## Supplemental Table S2 for "Major histocompatibility complex class I-restricted protection against murine cytomegalovirus requires missing-self recognition by the natural killer cell inhibitory Ly49 receptors"

A complete list of all guides with no or minimal off-target sites are indicated as identified by CCTop (see Methods). Specifically, the individual sgRNA sites only within the MCMV genome was interrogated for all CRISPR/Cas9 target sites. Any alternate cleavage sites are indicated. The two sgRNA guide sequences used in our experiments are identified as T303 (*m06* ) and T18561 (*m152* ) (see also **Table S3**) and is highlighted in yellow within this file.

<table>
<tr><td>PAM:</td><td>NGG</td><td></td><td></td><td></td><td></td><td></td><td></td><td></td><td></td><td></td><td></td></tr>
<tr><td>Target site length:</td><td>20</td><td></td><td></td><td></td><td></td><td></td><td></td><td></td><td></td><td></td><td></td></tr>
<tr><td>Core length:</td><td>12</td><td></td><td></td><td></td><td></td><td></td><td></td><td></td><td></td><td></td><td></td></tr>
<tr><td>Core MM:</td><td>2</td><td></td><td></td><td></td><td></td><td></td><td></td><td></td><td></td><td></td><td></td></tr>
<tr><td>Total MM:</td><td>4</td><td></td><td></td><td></td><td></td><td></td><td></td><td></td><td></td><td></td><td></td></tr>
<tr><td>T1</td><td>GCTTGATTCGTTCCCCCCCCCGG</td><td>1000</td><td></td><td></td><td></td><td></td><td></td><td></td><td></td><td></td><td></td></tr>
<tr><td>Oligo fwd</td><td>CACCGCTTGATTCGTTCCCCCCCC</td><td></td><td></td><td></td><td></td><td></td><td></td><td></td><td></td><td></td><td></td></tr>
<tr><td>Oligo rev</td><td>AAACGGGGGGGGAACGAATCAAGC</td><td></td><td></td><td></td><td></td><td></td><td></td><td></td><td></td><td></td><td></td></tr>
<tr><td>Chromosome</td><td>start</td><td>end</td><td>strand</td><td>MM</td><td>target_seq</td><td>PAM</td><td>alignment</td><td>distance</td><td>position</td><td>gene name</td><td>gene id</td></tr>
<tr><td>GU305914.1</td><td>20</td><td>42</td><td>+</td><td>0</td><td>GCTTGATTCGTTCCCCCCCC</td><td>CGG</td><td>||||||||[||||||||||||]PAM</td><td>NA</td><td>Intergenic</td><td>NA</td><td>NA</td></tr>
<tr><td>T2</td><td>CCCGGCCGTCTGAGTGCGCGCGG</td><td>1000</td><td></td><td></td><td></td><td></td><td></td><td></td><td></td><td></td><td></td></tr>
<tr><td>Oligo fwd</td><td>CACCCCCGGCCGTCTGAGTGCGCG</td><td></td><td></td><td></td><td></td><td></td><td></td><td></td><td></td><td></td><td></td></tr>
<tr><td>Oligo rev</td><td>AAACCGCGCACTCAGACGGCCGGG</td><td></td><td></td><td></td><td></td><td></td><td></td><td></td><td></td><td></td><td></td></tr>
<tr><td>Chromosome</td><td>start</td><td>end</td><td>strand</td><td>MM</td><td>target_seq</td><td>PAM</td><td>alignment</td><td>distance</td><td>position</td><td>gene name</td><td>gene id</td></tr>
<tr><td>GU305914.1</td><td>230353</td><td>230375</td><td>+</td><td>0</td><td>CCCGGCCGTCTGAGTGCGCG</td><td>CGG</td><td>||||||||[||||||||||||]PAM</td><td>NA</td><td>Intergenic</td><td>NA</td><td>NA</td></tr>
<tr><td>GU305914.1</td><td>38</td><td>60</td><td>+</td><td>0</td><td>CCCGGCCGTCTGAGTGCGCG</td><td>CGG</td><td>||||||||[||||||||||||]PAM</td><td>NA</td><td>Intergenic</td><td>NA</td><td>NA</td></tr>
<tr><td>GU305914.1</td><td>149055</td><td>149077</td><td>+</td><td>4</td><td>ACCGGCAGTCCGAGTCCGCG</td><td>CGG</td><td>-|||||-|[||-||||-||||]PAM</td><td>NA</td><td>Intergenic</td><td>NA</td><td>NA</td></tr>
<tr><td>T3</td><td>CCGGCCGTCTGAGTGCGCGCGGG</td><td>1000</td><td></td><td></td><td></td><td></td><td></td><td></td><td></td><td></td><td></td></tr>
<tr><td>Oligo fwd</td><td>CACCCCGGCCGTCTGAGTGCGCGC</td><td></td><td></td><td></td><td></td><td></td><td></td><td></td><td></td><td></td><td></td></tr>
<tr><td>Oligo rev</td><td>AAACGCGCGCACTCAGACGGCCGG</td><td></td><td></td><td></td><td></td><td></td><td></td><td></td><td></td><td></td><td></td></tr>
<tr><td>Chromosome</td><td>start</td><td>end</td><td>strand</td><td>MM</td><td>target_seq</td><td>PAM</td><td>alignment</td><td>distance</td><td>position</td><td>gene name</td><td>gene id</td></tr>
<tr><td>GU305914.1</td><td>230354</td><td>230376</td><td>+</td><td>0</td><td>CCGGCCGTCTGAGTGCGCGC</td><td>GGG</td><td>||||||||[||||||||||||]PAM</td><td>NA</td><td>Intergenic</td><td>NA</td><td>NA</td></tr>
<tr><td>GU305914.1</td><td>39</td><td>61</td><td>+</td><td>0</td><td>CCGGCCGTCTGAGTGCGCGC</td><td>GGG</td><td>||||||||[||||||||||||]PAM</td><td>NA</td><td>Intergenic</td><td>NA</td><td>NA</td></tr>
<tr><td>T4</td><td>TCTGAGTGCGCGCGGGCCCCCGG</td><td>1000</td><td></td><td></td><td></td><td></td><td></td><td></td><td></td><td></td><td></td></tr>
<tr><td>Oligo fwd</td><td>CACCTCTGAGTGCGCGCGGGCCCC</td><td></td><td></td><td></td><td></td><td></td><td></td><td></td><td></td><td></td><td></td></tr>
<tr><td>Oligo rev</td><td>AAACGGGGCCCGCGCGCACTCAGA</td><td></td><td></td><td></td><td></td><td></td><td></td><td></td><td></td><td></td><td></td></tr>
<tr><td>Chromosome</td><td>start</td><td>end</td><td>strand</td><td>MM</td><td>target_seq</td><td>PAM</td><td>alignment</td><td>distance</td><td>position</td><td>gene name</td><td>gene id</td></tr>
<tr><td>GU305914.1</td><td>46</td><td>68</td><td>+</td><td>0</td><td>TCTGAGTGCGCGCGGGCCCC</td><td>CGG</td><td>||||||||[||||||||||||]PAM</td><td>NA</td><td>Intergenic</td><td>NA</td><td>NA</td></tr>
<tr><td>T5</td><td>CTGAGTGCGCGCGGGCCCCCGGG</td><td>1000</td><td></td><td></td><td></td><td></td><td></td><td></td><td></td><td></td><td></td></tr>
<tr><td>Oligo fwd</td><td>CACCCTGAGTGCGCGCGGGCCCCC</td><td></td><td></td><td></td><td></td><td></td><td></td><td></td><td></td><td></td><td></td></tr>
<tr><td>Oligo rev</td><td>AAACGGGGGCCCGCGCGCACTCAG</td><td></td><td></td><td></td><td></td><td></td><td></td><td></td><td></td><td></td><td></td></tr>
<tr><td>Chromosome</td><td>start</td><td>end</td><td>strand</td><td>MM</td><td>target_seq</td><td>PAM</td><td>alignment</td><td>distance</td><td>position</td><td>gene name</td><td>gene id</td></tr>
<tr><td>GU305914.1</td><td>47</td><td>69</td><td>+</td><td>0</td><td>CTGAGTGCGCGCGGGCCCCC</td><td>GGG</td><td>||||||||[||||||||||||]PAM</td><td>NA</td><td>Intergenic</td><td>NA</td><td>NA</td></tr>
<tr><td>T6</td><td>TGAGTGCGCGCGGGCCCCCGGGG</td><td>1000</td><td></td><td></td><td></td><td></td><td></td><td></td><td></td><td></td><td></td></tr>
<tr><td>Oligo fwd</td><td>CACCTGAGTGCGCGCGGGCCCCCG</td><td></td><td></td><td></td><td></td><td></td><td></td><td></td><td></td><td></td><td></td></tr>
<tr><td>Oligo rev</td><td>AAACCGGGGGCCCGCGCGCACTCA</td><td></td><td></td><td></td><td></td><td></td><td></td><td></td><td></td><td></td><td></td></tr>
<tr><td>Chromosome</td><td>start</td><td>end</td><td>strand</td><td>MM</td><td>target_seq</td><td>PAM</td><td>alignment</td><td>distance</td><td>position</td><td>gene name</td><td>gene id</td></tr>
<tr><td>GU305914.1</td><td>48</td><td>70</td><td>+</td><td>0</td><td>TGAGTGCGCGCGGGCCCCCG</td><td>GGG</td><td>||||||||[||||||||||||]PAM</td><td>NA</td><td>Intergenic</td><td>NA</td><td>NA</td></tr>
<tr><td>T7</td><td>GAGTGCGCGCGGGCCCCCGGGGG</td><td>1000</td><td></td><td></td><td></td><td></td><td></td><td></td><td></td><td></td><td></td></tr>
<tr><td>Oligo fwd</td><td>CACCGAGTGCGCGCGGGCCCCCGG</td><td></td><td></td><td></td><td></td><td></td><td></td><td></td><td></td><td></td><td></td></tr>
<tr><td>Oligo rev</td><td>AAACCCGGGGGCCCGCGCGCACTC</td><td></td><td></td><td></td><td></td><td></td><td></td><td></td><td></td><td></td><td></td></tr>
<tr><td>Chromosome</td><td>start</td><td>end</td><td>strand</td><td>MM</td><td>target_seq</td><td>PAM</td><td>alignment</td><td>distance</td><td>position</td><td>gene name</td><td>gene id</td></tr>
<tr><td>GU305914.1</td><td>49</td><td>71</td><td>+</td><td>0</td><td>GAGTGCGCGCGGGCCCCCGG</td><td>GGG</td><td>||||||||[||||||||||||]PAM</td><td>NA</td><td>Intergenic</td><td>NA</td><td>NA</td></tr>
<tr><td>T8</td><td>AGTGCGCGCGGGCCCCCGGGGGG</td><td>1000</td><td></td><td></td><td></td><td></td><td></td><td></td><td></td><td></td><td></td></tr>
<tr><td>Oligo fwd</td><td>CACCAGTGCGCGCGGGCCCCCGGG</td><td></td><td></td><td></td><td></td><td></td><td></td><td></td><td></td><td></td><td></td></tr>
<tr><td>Oligo rev</td><td>AAACCCCGGGGGCCCGCGCGCACT</td><td></td><td></td><td></td><td></td><td></td><td></td><td></td><td></td><td></td><td></td></tr>
<tr><td>Chromosome</td><td>start</td><td>end</td><td>strand</td><td>MM</td><td>target_seq</td><td>PAM</td><td>alignment</td><td>distance</td><td>position</td><td>gene name</td><td>gene id</td></tr>
<tr><td>GU305914.1</td><td>50</td><td>72</td><td>+</td><td>0</td><td>AGTGCGCGCGGGCCCCCGGG</td><td>GGG</td><td>||||||||[||||||||||||]PAM</td><td>NA</td><td>Intergenic</td><td>NA</td><td>NA</td></tr>
<tr><td>T9</td><td>GTGCGCGCGGGCCCCCGGGGGGG</td><td>1000</td><td></td><td></td><td></td><td></td><td></td><td></td><td></td><td></td><td></td></tr>
<tr><td>Oligo fwd</td><td>CACCGTGCGCGCGGGCCCCCGGGG</td><td></td><td></td><td></td><td></td><td></td><td></td><td></td><td></td><td></td><td></td></tr>
<tr><td>Oligo rev</td><td>AAACCCCCGGGGGCCCGCGCGCAC</td><td></td><td></td><td></td><td></td><td></td><td></td><td></td><td></td><td></td><td></td></tr>
<tr><td>Chromosome</td><td>start</td><td>end</td><td>strand</td><td>MM</td><td>target_seq</td><td>PAM</td><td>alignment</td><td>distance</td><td>position</td><td>gene name</td><td>gene id</td></tr>
<tr><td>GU305914.1</td><td>51</td><td>73</td><td>+</td><td>0</td><td>GTGCGCGCGGGCCCCCGGGG</td><td>GGG</td><td>||||||||[||||||||||||]PAM</td><td>NA</td><td>Intergenic</td><td>NA</td><td>NA</td></tr>
<tr><td>T10</td><td>CCCCCGGGGGGGTATTTTGATGG</td><td>1000</td><td></td><td></td><td></td><td></td><td></td><td></td><td></td><td></td><td></td></tr>
<tr><td>Oligo fwd</td><td>CACCCCCCCGGGGGGGTATTTTGA</td><td></td><td></td><td></td><td></td><td></td><td></td><td></td><td></td><td></td><td></td></tr>
<tr><td>Oligo rev</td><td>AAACTCAAAATACCCCCCCGGGGG</td><td></td><td></td><td></td><td></td><td></td><td></td><td></td><td></td><td></td><td></td></tr>
<tr><td>Chromosome</td><td>start</td><td>end</td><td>strand</td><td>MM</td><td>target_seq</td><td>PAM</td><td>alignment</td><td>distance</td><td>position</td><td>gene name</td><td>gene id</td></tr>
<tr><td>GU305914.1</td><td>62</td><td>84</td><td>+</td><td>0</td><td>CCCCCGGGGGGGTATTTTGA</td><td>TGG</td><td>||||||||[||||||||||||]PAM</td><td>NA</td><td>Intergenic</td><td>NA</td><td>NA</td></tr>
<tr><td>T11</td><td>CCCCGGGGGGGTATTTTGATGGG</td><td>1000</td><td></td><td></td><td></td><td></td><td></td><td></td><td></td><td></td><td></td></tr>
<tr><td>Oligo fwd</td><td>CACCCCCCGGGGGGGTATTTTGAT</td><td></td><td></td><td></td><td></td><td></td><td></td><td></td><td></td><td></td><td></td></tr>
<tr><td>Oligo rev</td><td>AAACATCAAAATACCCCCCCGGGG</td><td></td><td></td><td></td><td></td><td></td><td></td><td></td><td></td><td></td><td></td></tr>
<tr><td>Chromosome</td><td>start</td><td>end</td><td>strand</td><td>MM</td><td>target_seq</td><td>PAM</td><td>alignment</td><td>distance</td><td>position</td><td>gene name</td><td>gene id</td></tr>
<tr><td>GU305914.1</td><td>63</td><td>85</td><td>+</td><td>0</td><td>CCCCGGGGGGGTATTTTGAT</td><td>GGG</td><td>||||||||[||||||||||||]PAM</td><td>NA</td><td>Intergenic</td><td>NA</td><td>NA</td></tr>
<tr><td>T12</td><td>CCCGGGGGGGTATTTTGATGGGG</td><td>1000</td><td></td><td></td><td></td><td></td><td></td><td></td><td></td><td></td><td></td></tr>
<tr><td>Oligo fwd</td><td>CACCCCCGGGGGGGTATTTTGATG</td><td></td><td></td><td></td><td></td><td></td><td></td><td></td><td></td><td></td><td></td></tr>
<tr><td>Oligo rev</td><td>AAACCATCAAAATACCCCCCCGGG</td><td></td><td></td><td></td><td></td><td></td><td></td><td></td><td></td><td></td><td></td></tr>
<tr><td>Chromosome</td><td>start</td><td>end</td><td>strand</td><td>MM</td><td>target_seq</td><td>PAM</td><td>alignment</td><td>distance</td><td>position</td><td>gene name</td><td>gene id</td></tr>
<tr><td>GU305914.1</td><td>64</td><td>86</td><td>+</td><td>0</td><td>CCCGGGGGGGTATTTTGATG</td><td>GGG</td><td>||||||||[||||||||||||]PAM</td><td>NA</td><td>Intergenic</td><td>NA</td><td>NA</td></tr>
<tr><td>T13</td><td>CCGGGGGGGTATTTTGATGGGGG</td><td>1000</td><td></td><td></td><td></td><td></td><td></td><td></td><td></td><td></td><td></td></tr>
<tr><td>Oligo fwd</td><td>CACCCCGGGGGGGTATTTTGATGG</td><td></td><td></td><td></td><td></td><td></td><td></td><td></td><td></td><td></td><td></td></tr>
<tr><td>Oligo rev</td><td>AAACCCATCAAAATACCCCCCCGG</td><td></td><td></td><td></td><td></td><td></td><td></td><td></td><td></td><td></td><td></td></tr>
</table>

<table>
<tr><td>Chromosome</td><td>start</td><td>end</td><td>strand</td><td>MM</td><td>target_seq</td><td>PAM</td><td>alignment</td><td>distance</td><td>position</td><td>gene name</td><td>gene id</td></tr>
<tr><td>GU305914.1</td><td>65</td><td>87</td><td>+</td><td>0</td><td>CCGGGGGGGTATTTTGATGG</td><td>GGG</td><td>||||||||[||||||||||||]PAM</td><td>NA</td><td>Intergenic</td><td>NA</td><td>NA</td></tr>
<tr><td></td><td></td><td></td><td></td><td></td><td></td><td></td><td></td><td></td><td></td><td></td><td></td></tr>
<tr><td>T14</td><td>CGGGGGGGTATTTTGATGGGGGG</td><td>1000</td><td></td><td></td><td></td><td></td><td></td><td></td><td></td><td></td><td></td></tr>
<tr><td>Oligo fwd</td><td>CACCCGGGGGGGTATTTTGATGGG</td><td></td><td></td><td></td><td></td><td></td><td></td><td></td><td></td><td></td><td></td></tr>
<tr><td>Oligo rev</td><td>AAACCCCATCAAAATACCCCCCCG</td><td></td><td></td><td></td><td></td><td></td><td></td><td></td><td></td><td></td><td></td></tr>
<tr><td>Chromosome</td><td>start</td><td>end</td><td>strand</td><td>MM</td><td>target_seq</td><td>PAM</td><td>alignment</td><td>distance</td><td>position</td><td>gene name</td><td>gene id</td></tr>
<tr><td>GU305914.1</td><td>66</td><td>88</td><td>+</td><td>0</td><td>CGGGGGGGTATTTTGATGGG</td><td>GGG</td><td>||||||||[||||||||||||]PAM</td><td>NA</td><td>Intergenic</td><td>NA</td><td>NA</td></tr>
<tr><td></td><td></td><td></td><td></td><td></td><td></td><td></td><td></td><td></td><td></td><td></td><td></td></tr>
<tr><td>T15</td><td>GGGGGGGTATTTTGATGGGGGGG</td><td>1000</td><td></td><td></td><td></td><td></td><td></td><td></td><td></td><td></td><td></td></tr>
<tr><td>Oligo fwd</td><td>CACCGGGGGGGTATTTTGATGGGG</td><td></td><td></td><td></td><td></td><td></td><td></td><td></td><td></td><td></td><td></td></tr>
<tr><td>Oligo rev</td><td>AAACCCCCATCAAAATACCCCCCC</td><td></td><td></td><td></td><td></td><td></td><td></td><td></td><td></td><td></td><td></td></tr>
<tr><td>Chromosome</td><td>start</td><td>end</td><td>strand</td><td>MM</td><td>target_seq</td><td>PAM</td><td>alignment</td><td>distance</td><td>position</td><td>gene name</td><td>gene id</td></tr>
<tr><td>GU305914.1</td><td>67</td><td>89</td><td>+</td><td>0</td><td>GGGGGGGTATTTTGATGGGG</td><td>GGG</td><td>||||||||[||||||||||||]PAM</td><td>NA</td><td>Intergenic</td><td>NA</td><td>NA</td></tr>
<tr><td></td><td></td><td></td><td></td><td></td><td></td><td></td><td></td><td></td><td></td><td></td><td></td></tr>
<tr><td>T16</td><td>GCTATATTTTTGTCCTCTAGAGG</td><td>1000</td><td></td><td></td><td></td><td></td><td></td><td></td><td></td><td></td><td></td></tr>
<tr><td>Oligo fwd</td><td>CACCGCTATATTTTTGTCCTCTAG</td><td></td><td></td><td></td><td></td><td></td><td></td><td></td><td></td><td></td><td></td></tr>
<tr><td>Oligo rev</td><td>AAACCTAGAGGACAAAAATATAGC</td><td></td><td></td><td></td><td></td><td></td><td></td><td></td><td></td><td></td><td></td></tr>
<tr><td>Chromosome</td><td>start</td><td>end</td><td>strand</td><td>MM</td><td>target_seq</td><td>PAM</td><td>alignment</td><td>distance</td><td>position</td><td>gene name</td><td>gene id</td></tr>
<tr><td>GU305914.1</td><td>89</td><td>111</td><td>+</td><td>0</td><td>GCTATATTTTTGTCCTCTAG</td><td>AGG</td><td>||||||||[||||||||||||]PAM</td><td>NA</td><td>Intergenic</td><td>NA</td><td>NA</td></tr>
<tr><td></td><td></td><td></td><td></td><td></td><td></td><td></td><td></td><td></td><td></td><td></td><td></td></tr>
<tr><td>T17</td><td>CTATATTTTTGTCCTCTAGAGGG</td><td>1000</td><td></td><td></td><td></td><td></td><td></td><td></td><td></td><td></td><td></td></tr>
<tr><td>Oligo fwd</td><td>CACCCTATATTTTTGTCCTCTAGA</td><td></td><td></td><td></td><td></td><td></td><td></td><td></td><td></td><td></td><td></td></tr>
<tr><td>Oligo rev</td><td>AAACTCTAGAGGACAAAAATATAG</td><td></td><td></td><td></td><td></td><td></td><td></td><td></td><td></td><td></td><td></td></tr>
<tr><td>Chromosome</td><td>start</td><td>end</td><td>strand</td><td>MM</td><td>target_seq</td><td>PAM</td><td>alignment</td><td>distance</td><td>position</td><td>gene name</td><td>gene id</td></tr>
<tr><td>GU305914.1</td><td>90</td><td>112</td><td>+</td><td>0</td><td>CTATATTTTTGTCCTCTAGA</td><td>GGG</td><td>||||||||[||||||||||||]PAM</td><td>NA</td><td>Intergenic</td><td>NA</td><td>NA</td></tr>
<tr><td></td><td></td><td></td><td></td><td></td><td></td><td></td><td></td><td></td><td></td><td></td><td></td></tr>
<tr><td>T18</td><td>AGAGGGCGCTGTTGCATCGATGG</td><td>1000</td><td></td><td></td><td></td><td></td><td></td><td></td><td></td><td></td><td></td></tr>
<tr><td>Oligo fwd</td><td>CACCAGAGGGCGCTGTTGCATCGA</td><td></td><td></td><td></td><td></td><td></td><td></td><td></td><td></td><td></td><td></td></tr>
<tr><td>Oligo rev</td><td>AAACTCGATGCAACAGCGCCCTCT</td><td></td><td></td><td></td><td></td><td></td><td></td><td></td><td></td><td></td><td></td></tr>
<tr><td>Chromosome</td><td>start</td><td>end</td><td>strand</td><td>MM</td><td>target_seq</td><td>PAM</td><td>alignment</td><td>distance</td><td>position</td><td>gene name</td><td>gene id</td></tr>
<tr><td>GU305914.1</td><td>107</td><td>129</td><td>+</td><td>0</td><td>AGAGGGCGCTGTTGCATCGA</td><td>TGG</td><td>||||||||[||||||||||||]PAM</td><td>NA</td><td>Intergenic</td><td>NA</td><td>NA</td></tr>
<tr><td></td><td></td><td></td><td></td><td></td><td></td><td></td><td></td><td></td><td></td><td></td><td></td></tr>
<tr><td>T19</td><td>CTGTTGCATCGATGGTCGTGAGG</td><td>1000</td><td></td><td></td><td></td><td></td><td></td><td></td><td></td><td></td><td></td></tr>
<tr><td>Oligo fwd</td><td>CACCCTGTTGCATCGATGGTCGTG</td><td></td><td></td><td></td><td></td><td></td><td></td><td></td><td></td><td></td><td></td></tr>
<tr><td>Oligo rev</td><td>AAACCACGACCATCGATGCAACAG</td><td></td><td></td><td></td><td></td><td></td><td></td><td></td><td></td><td></td><td></td></tr>
<tr><td>Chromosome</td><td>start</td><td>end</td><td>strand</td><td>MM</td><td>target_seq</td><td>PAM</td><td>alignment</td><td>distance</td><td>position</td><td>gene name</td><td>gene id</td></tr>
<tr><td>GU305914.1</td><td>115</td><td>137</td><td>+</td><td>0</td><td>CTGTTGCATCGATGGTCGTG</td><td>AGG</td><td>||||||||[||||||||||||]PAM</td><td>NA</td><td>Intergenic</td><td>NA</td><td>NA</td></tr>
<tr><td>GU305914.1</td><td>205998</td><td>206020</td><td>-</td><td>4</td><td>ATGTTGAATCGAAGGTCGGG</td><td>TGG</td><td>-|||||-|[||||-|||||-|]PAM</td><td>NA</td><td>Intergenic</td><td>NA</td><td>NA</td></tr>
<tr><td></td><td></td><td></td><td></td><td></td><td></td><td></td><td></td><td></td><td></td><td></td><td></td></tr>
<tr><td>T20</td><td>TGTTGCATCGATGGTCGTGAGGG</td><td>1000</td><td></td><td></td><td></td><td></td><td></td><td></td><td></td><td></td><td></td></tr>
<tr><td>Oligo fwd</td><td>CACCTGTTGCATCGATGGTCGTGA</td><td></td><td></td><td></td><td></td><td></td><td></td><td></td><td></td><td></td><td></td></tr>
<tr><td>Oligo rev</td><td>AAACTCACGACCATCGATGCAACA</td><td></td><td></td><td></td><td></td><td></td><td></td><td></td><td></td><td></td><td></td></tr>
<tr><td>Chromosome</td><td>start</td><td>end</td><td>strand</td><td>MM</td><td>target_seq</td><td>PAM</td><td>alignment</td><td>distance</td><td>position</td><td>gene name</td><td>gene id</td></tr>
<tr><td>GU305914.1</td><td>116</td><td>138</td><td>+</td><td>0</td><td>TGTTGCATCGATGGTCGTGA</td><td>GGG</td><td>||||||||[||||||||||||]PAM</td><td>NA</td><td>Intergenic</td><td>NA</td><td>NA</td></tr>
<tr><td></td><td></td><td></td><td></td><td></td><td></td><td></td><td></td><td></td><td></td><td></td><td></td></tr>
<tr><td>T21</td><td>GTGCTCCTGAGAGTGCGATGAGG</td><td>1000</td><td></td><td></td><td></td><td></td><td></td><td></td><td></td><td></td><td></td></tr>
<tr><td>Oligo fwd</td><td>CACCGTGCTCCTGAGAGTGCGATG</td><td></td><td></td><td></td><td></td><td></td><td></td><td></td><td></td><td></td><td></td></tr>
<tr><td>Oligo rev</td><td>AAACCATCGCACTCTCAGGAGCAC</td><td></td><td></td><td></td><td></td><td></td><td></td><td></td><td></td><td></td><td></td></tr>
<tr><td>Chromosome</td><td>start</td><td>end</td><td>strand</td><td>MM</td><td>target_seq</td><td>PAM</td><td>alignment</td><td>distance</td><td>position</td><td>gene name</td><td>gene id</td></tr>
<tr><td>GU305914.1</td><td>138</td><td>160</td><td>+</td><td>0</td><td>GTGCTCCTGAGAGTGCGATG</td><td>AGG</td><td>||||||||[||||||||||||]PAM</td><td>NA</td><td>Intergenic</td><td>NA</td><td>NA</td></tr>
<tr><td></td><td></td><td></td><td></td><td></td><td></td><td></td><td></td><td></td><td></td><td></td><td></td></tr>
<tr><td>T22</td><td>CTCCTGAGAGTGCGATGAGGTGG</td><td>1000</td><td></td><td></td><td></td><td></td><td></td><td></td><td></td><td></td><td></td></tr>
<tr><td>Oligo fwd</td><td>CACCCTCCTGAGAGTGCGATGAGG</td><td></td><td></td><td></td><td></td><td></td><td></td><td></td><td></td><td></td><td></td></tr>
<tr><td>Oligo rev</td><td>AAACCCTCATCGCACTCTCAGGAG</td><td></td><td></td><td></td><td></td><td></td><td></td><td></td><td></td><td></td><td></td></tr>
<tr><td>Chromosome</td><td>start</td><td>end</td><td>strand</td><td>MM</td><td>target_seq</td><td>PAM</td><td>alignment</td><td>distance</td><td>position</td><td>gene name</td><td>gene id</td></tr>
<tr><td>GU305914.1</td><td>141</td><td>163</td><td>+</td><td>0</td><td>CTCCTGAGAGTGCGATGAGG</td><td>TGG</td><td>||||||||[||||||||||||]PAM</td><td>NA</td><td>Intergenic</td><td>NA</td><td>NA</td></tr>
<tr><td></td><td></td><td></td><td></td><td></td><td></td><td></td><td></td><td></td><td></td><td></td><td></td></tr>
<tr><td>T23</td><td>TGGCTGCGCGCACGTCACCGAGG</td><td>1000</td><td></td><td></td><td></td><td></td><td></td><td></td><td></td><td></td><td></td></tr>
<tr><td>Oligo fwd</td><td>CACCTGGCTGCGCGCACGTCACCG</td><td></td><td></td><td></td><td></td><td></td><td></td><td></td><td></td><td></td><td></td></tr>
<tr><td>Oligo rev</td><td>AAACCGGTGACGTGCGCGCAGCCA</td><td></td><td></td><td></td><td></td><td></td><td></td><td></td><td></td><td></td><td></td></tr>
<tr><td>Chromosome</td><td>start</td><td>end</td><td>strand</td><td>MM</td><td>target_seq</td><td>PAM</td><td>alignment</td><td>distance</td><td>position</td><td>gene name</td><td>gene id</td></tr>
<tr><td>GU305914.1</td><td>161</td><td>183</td><td>+</td><td>0</td><td>TGGCTGCGCGCACGTCACCG</td><td>AGG</td><td>||||||||[||||||||||||]PAM</td><td>NA</td><td>Intergenic</td><td>NA</td><td>NA</td></tr>
<tr><td></td><td></td><td></td><td></td><td></td><td></td><td></td><td></td><td></td><td></td><td></td><td></td></tr>
<tr><td>T24</td><td>GGCTGCGCGCACGTCACCGAGGG</td><td>1000</td><td></td><td></td><td></td><td></td><td></td><td></td><td></td><td></td><td></td></tr>
<tr><td>Oligo fwd</td><td>CACCGGCTGCGCGCACGTCACCGA</td><td></td><td></td><td></td><td></td><td></td><td></td><td></td><td></td><td></td><td></td></tr>
<tr><td>Oligo rev</td><td>AAACTCGGTGACGTGCGCGCAGCC</td><td></td><td></td><td></td><td></td><td></td><td></td><td></td><td></td><td></td><td></td></tr>
<tr><td>Chromosome</td><td>start</td><td>end</td><td>strand</td><td>MM</td><td>target_seq</td><td>PAM</td><td>alignment</td><td>distance</td><td>position</td><td>gene name</td><td>gene id</td></tr>
<tr><td>GU305914.1</td><td>162</td><td>184</td><td>+</td><td>0</td><td>GGCTGCGCGCACGTCACCGA</td><td>GGG</td><td>||||||||[||||||||||||]PAM</td><td>NA</td><td>Intergenic</td><td>NA</td><td>NA</td></tr>
<tr><td></td><td></td><td></td><td></td><td></td><td></td><td></td><td></td><td></td><td></td><td></td><td></td></tr>
<tr><td>T25</td><td>CGCGCACGTCACCGAGGGCGTGG</td><td>1000</td><td></td><td></td><td></td><td></td><td></td><td></td><td></td><td></td><td></td></tr>
<tr><td>Oligo fwd</td><td>CACCCGCGCACGTCACCGAGGGCG</td><td></td><td></td><td></td><td></td><td></td><td></td><td></td><td></td><td></td><td></td></tr>
<tr><td>Oligo rev</td><td>AAACCGCCCTCGGTGACGTGCGCG</td><td></td><td></td><td></td><td></td><td></td><td></td><td></td><td></td><td></td><td></td></tr>
<tr><td>Chromosome</td><td>start</td><td>end</td><td>strand</td><td>MM</td><td>target_seq</td><td>PAM</td><td>alignment</td><td>distance</td><td>position</td><td>gene name</td><td>gene id</td></tr>
<tr><td>GU305914.1</td><td>167</td><td>189</td><td>+</td><td>0</td><td>CGCGCACGTCACCGAGGGCG</td><td>TGG</td><td>||||||||[||||||||||||]PAM</td><td>NA</td><td>Intergenic</td><td>NA</td><td>NA</td></tr>
<tr><td></td><td></td><td></td><td></td><td></td><td></td><td></td><td></td><td></td><td></td><td></td><td></td></tr>
<tr><td>T26</td><td>TCCGAGAAGTTCCCCTTTTCTGG</td><td>1000</td><td></td><td></td><td></td><td></td><td></td><td></td><td></td><td></td><td></td></tr>
<tr><td>Oligo fwd</td><td>CACCTCCGAGAAGTTCCCCTTTTC</td><td></td><td></td><td></td><td></td><td></td><td></td><td></td><td></td><td></td><td></td></tr>
<tr><td>Oligo rev</td><td>AAACGAAAAGGGGAACTTCTCGGA</td><td></td><td></td><td></td><td></td><td></td><td></td><td></td><td></td><td></td><td></td></tr>
<tr><td>Chromosome</td><td>start</td><td>end</td><td>strand</td><td>MM</td><td>target_seq</td><td>PAM</td><td>alignment</td><td>distance</td><td>position</td><td>gene name</td><td>gene id</td></tr>
<tr><td>GU305914.1</td><td>223</td><td>245</td><td>+</td><td>0</td><td>TCCGAGAAGTTCCCCTTTTC</td><td>TGG</td><td>||||||||[||||||||||||]PAM</td><td>NA</td><td>Intergenic</td><td>NA</td><td>NA</td></tr>
<tr><td></td><td></td><td></td><td></td><td></td><td></td><td></td><td></td><td></td><td></td><td></td><td></td></tr>
<tr><td>T27</td><td>CCGAGAAGTTCCCCTTTTCTGGG</td><td>1000</td><td></td><td></td><td></td><td></td><td></td><td></td><td></td><td></td><td></td></tr>
<tr><td>Oligo fwd</td><td>CACCCCGAGAAGTTCCCCTTTTCT</td><td></td><td></td><td></td><td></td><td></td><td></td><td></td><td></td><td></td><td></td></tr>
<tr><td>Oligo rev</td><td>AAACAGAAAAGGGGAACTTCTCGG</td><td></td><td></td><td></td><td></td><td></td><td></td><td></td><td></td><td></td><td></td></tr>
<tr><td>Chromosome</td><td>start</td><td>end</td><td>strand</td><td>MM</td><td>target_seq</td><td>PAM</td><td>alignment</td><td>distance</td><td>position</td><td>gene name</td><td>gene id</td></tr>
<tr><td>GU305914.1</td><td>224</td><td>246</td><td>+</td><td>0</td><td>CCGAGAAGTTCCCCTTTTCT</td><td>GGG</td><td>||||||||[||||||||||||]PAM</td><td>NA</td><td>Intergenic</td><td>NA</td><td>NA</td></tr>
<tr><td></td><td></td><td></td><td></td><td></td><td></td><td></td><td></td><td></td><td></td><td></td><td></td></tr>
<tr><td>T28</td><td>CCCCTTTTCTGGGAAGATTCTGG</td><td>1000</td><td></td><td></td><td></td><td></td><td></td><td></td><td></td><td></td><td></td></tr>
</table>

<table>
<tr><td>Oligo fwd</td><td>CACCCCCCTTTTCTGGGAAGATTC</td><td></td><td></td><td></td><td></td><td></td><td></td><td></td><td></td><td></td><td></td></tr>
<tr><td>Oligo rev</td><td>AAACGAATCTTCCCAGAAAAGGGG</td><td></td><td></td><td></td><td></td><td></td><td></td><td></td><td></td><td></td><td></td></tr>
<tr><td>Chromosome</td><td>start</td><td>end</td><td>strand</td><td>MM</td><td>target_seq</td><td>PAM</td><td>alignment</td><td>distance</td><td>position</td><td>gene name</td><td>gene id</td></tr>
<tr><td>GU305914.1</td><td>234</td><td>256</td><td>+</td><td>0</td><td>CCCCTTTTCTGGGAAGATTC</td><td>TGG</td><td>||||||||[||||||||||||]PAM</td><td>NA</td><td>Intergenic</td><td>NA</td><td>NA</td></tr>
<tr><td>T29</td><td>CTTTTCTGGGAAGATTCTGGCGG</td><td>1000</td><td></td><td></td><td></td><td></td><td></td><td></td><td></td><td></td><td></td></tr>
<tr><td>Oligo fwd</td><td>CACCCTTTTCTGGGAAGATTCTGG</td><td></td><td></td><td></td><td></td><td></td><td></td><td></td><td></td><td></td><td></td></tr>
<tr><td>Oligo rev</td><td>AAACCCAGAATCTTCCCAGAAAAG</td><td></td><td></td><td></td><td></td><td></td><td></td><td></td><td></td><td></td><td></td></tr>
<tr><td>Chromosome</td><td>start</td><td>end</td><td>strand</td><td>MM</td><td>target_seq</td><td>PAM</td><td>alignment</td><td>distance</td><td>position</td><td>gene name</td><td>gene id</td></tr>
<tr><td>GU305914.1</td><td>237</td><td>259</td><td>+</td><td>0</td><td>CTTTTCTGGGAAGATTCTGG</td><td>CGG</td><td>||||||||[||||||||||||]PAM</td><td>NA</td><td>Intergenic</td><td>NA</td><td>NA</td></tr>
<tr><td>T30</td><td>TTTTCTGGGAAGATTCTGGCGGG</td><td>1000</td><td></td><td></td><td></td><td></td><td></td><td></td><td></td><td></td><td></td></tr>
<tr><td>Oligo fwd</td><td>CACCTTTTCTGGGAAGATTCTGGC</td><td></td><td></td><td></td><td></td><td></td><td></td><td></td><td></td><td></td><td></td></tr>
<tr><td>Oligo rev</td><td>AAACGCCAGAATCTTCCCAGAAAA</td><td></td><td></td><td></td><td></td><td></td><td></td><td></td><td></td><td></td><td></td></tr>
<tr><td>Chromosome</td><td>start</td><td>end</td><td>strand</td><td>MM</td><td>target_seq</td><td>PAM</td><td>alignment</td><td>distance</td><td>position</td><td>gene name</td><td>gene id</td></tr>
<tr><td>GU305914.1</td><td>238</td><td>260</td><td>+</td><td>0</td><td>TTTTCTGGGAAGATTCTGGC</td><td>GGG</td><td>||||||||[||||||||||||]PAM</td><td>NA</td><td>Intergenic</td><td>NA</td><td>NA</td></tr>
<tr><td>T31</td><td>TGGCGGGAACTTCCCCTTTTGG</td><td>1000</td><td></td><td></td><td></td><td></td><td></td><td></td><td></td><td></td><td></td></tr>
<tr><td>Oligo fwd</td><td>CACCTGGCGGGAACTTCCCCTTTT</td><td></td><td></td><td></td><td></td><td></td><td></td><td></td><td></td><td></td><td></td></tr>
<tr><td>Oligo rev</td><td>AAACAAAAGGGGAAGTTCCCGCCA</td><td></td><td></td><td></td><td></td><td></td><td></td><td></td><td></td><td></td><td></td></tr>
<tr><td>Chromosome</td><td>start</td><td>end</td><td>strand</td><td>MM</td><td>target_seq</td><td>PAM</td><td>alignment</td><td>distance</td><td>position</td><td>gene name</td><td>gene id</td></tr>
<tr><td>GU305914.1</td><td>254</td><td>276</td><td>+</td><td>0</td><td>TGGCGGGAACTTCCCCTTTT</td><td>TGG</td><td>||||||||[||||||||||||]PAM</td><td>NA</td><td>Intergenic</td><td>NA</td><td>NA</td></tr>
<tr><td>T32</td><td>TGGAGCGCGTTCTGCTAGTTAGG</td><td>1000</td><td></td><td></td><td></td><td></td><td></td><td></td><td></td><td></td><td></td></tr>
<tr><td>Oligo fwd</td><td>CACCTGGAGCGCGTTCTGCTAGTT</td><td></td><td></td><td></td><td></td><td></td><td></td><td></td><td></td><td></td><td></td></tr>
<tr><td>Oligo rev</td><td>AAACAACTAGCAGAACGCGCTCCA</td><td></td><td></td><td></td><td></td><td></td><td></td><td></td><td></td><td></td><td></td></tr>
<tr><td>Chromosome</td><td>start</td><td>end</td><td>strand</td><td>MM</td><td>target_seq</td><td>PAM</td><td>alignment</td><td>distance</td><td>position</td><td>gene name</td><td>gene id</td></tr>
<tr><td>GU305914.1</td><td>274</td><td>296</td><td>+</td><td>0</td><td>TGGAGCGCGTTCTGCTAGTT</td><td>AGG</td><td>||||||||[||||||||||||]PAM</td><td>NA</td><td>Intergenic</td><td>NA</td><td>NA</td></tr>
<tr><td>T33</td><td>TTTCGTTTTCCCGTAAGCGCCGG</td><td>1000</td><td></td><td></td><td></td><td></td><td></td><td></td><td></td><td></td><td></td></tr>
<tr><td>Oligo fwd</td><td>CACCTTTCGTTTTCCCGTAAGCGC</td><td></td><td></td><td></td><td></td><td></td><td></td><td></td><td></td><td></td><td></td></tr>
<tr><td>Oligo rev</td><td>AAACGCGCTTACGGGAAAACGAAA</td><td></td><td></td><td></td><td></td><td></td><td></td><td></td><td></td><td></td><td></td></tr>
<tr><td>Chromosome</td><td>start</td><td>end</td><td>strand</td><td>MM</td><td>target_seq</td><td>PAM</td><td>alignment</td><td>distance</td><td>position</td><td>gene name</td><td>gene id</td></tr>
<tr><td>GU305914.1</td><td>297</td><td>319</td><td>+</td><td>0</td><td>TTTCGTTTTCCCGTAAGCGC</td><td>CGG</td><td>||||||||[||||||||||||]PAM</td><td>NA</td><td>Intergenic</td><td>NA</td><td>NA</td></tr>
<tr><td>T34</td><td>TTCGTTTTCCCGTAAGCGCCGGG</td><td>1000</td><td></td><td></td><td></td><td></td><td></td><td></td><td></td><td></td><td></td></tr>
<tr><td>Oligo fwd</td><td>CACCTTCGTTTTCCCGTAAGCGCC</td><td></td><td></td><td></td><td></td><td></td><td></td><td></td><td></td><td></td><td></td></tr>
<tr><td>Oligo rev</td><td>AAACGGCGCTTACGGGAAAACGAA</td><td></td><td></td><td></td><td></td><td></td><td></td><td></td><td></td><td></td><td></td></tr>
<tr><td>Chromosome</td><td>start</td><td>end</td><td>strand</td><td>MM</td><td>target_seq</td><td>PAM</td><td>alignment</td><td>distance</td><td>position</td><td>gene name</td><td>gene id</td></tr>
<tr><td>GU305914.1</td><td>298</td><td>320</td><td>+</td><td>0</td><td>TTCGTTTTCCCGTAAGCGCC</td><td>GGG</td><td>||||||||[||||||||||||]PAM</td><td>NA</td><td>Intergenic</td><td>NA</td><td>NA</td></tr>
<tr><td>T35</td><td>ACGCAGTTTTTGTGTCGTTTCGG</td><td>1000</td><td></td><td></td><td></td><td></td><td></td><td></td><td></td><td></td><td></td></tr>
<tr><td>Oligo fwd</td><td>CACCACGCAGTTTTTGTGTCGTTT</td><td></td><td></td><td></td><td></td><td></td><td></td><td></td><td></td><td></td><td></td></tr>
<tr><td>Oligo rev</td><td>AAACAAACGACACAAAAACTGCGT</td><td></td><td></td><td></td><td></td><td></td><td></td><td></td><td></td><td></td><td></td></tr>
<tr><td>Chromosome</td><td>start</td><td>end</td><td>strand</td><td>MM</td><td>target_seq</td><td>PAM</td><td>alignment</td><td>distance</td><td>position</td><td>gene name</td><td>gene id</td></tr>
<tr><td>GU305914.1</td><td>347</td><td>369</td><td>+</td><td>0</td><td>ACGCAGTTTTTGTGTCGTTT</td><td>CGG</td><td>||||||||[||||||||||||]PAM</td><td>NA</td><td>Intergenic</td><td>NA</td><td>NA</td></tr>
<tr><td>T36</td><td>CGCAGTTTTTGTGTCGTTTCGGG</td><td>1000</td><td></td><td></td><td></td><td></td><td></td><td></td><td></td><td></td><td></td></tr>
<tr><td>Oligo fwd</td><td>CACCCGCAGTTTTTGTGTCGTTTC</td><td></td><td></td><td></td><td></td><td></td><td></td><td></td><td></td><td></td><td></td></tr>
<tr><td>Oligo rev</td><td>AAACGAAACGACACAAAAACTGCG</td><td></td><td></td><td></td><td></td><td></td><td></td><td></td><td></td><td></td><td></td></tr>
<tr><td>Chromosome</td><td>start</td><td>end</td><td>strand</td><td>MM</td><td>target_seq</td><td>PAM</td><td>alignment</td><td>distance</td><td>position</td><td>gene name</td><td>gene id</td></tr>
<tr><td>GU305914.1</td><td>348</td><td>370</td><td>+</td><td>0</td><td>CGCAGTTTTTGTGTCGTTTC</td><td>GGG</td><td>||||||||[||||||||||||]PAM</td><td>NA</td><td>Intergenic</td><td>NA</td><td>NA</td></tr>
<tr><td>T37</td><td>GTTTTTGCGCTAGAACGCATAGG</td><td>1000</td><td></td><td></td><td></td><td></td><td></td><td></td><td></td><td></td><td></td></tr>
<tr><td>Oligo fwd</td><td>CACCGTTTTTGCGCTAGAACGCAT</td><td></td><td></td><td></td><td></td><td></td><td></td><td></td><td></td><td></td><td></td></tr>
<tr><td>Oligo rev</td><td>AAACATGCGTTCTAGCGCAAAAAC</td><td></td><td></td><td></td><td></td><td></td><td></td><td></td><td></td><td></td><td></td></tr>
<tr><td>Chromosome</td><td>start</td><td>end</td><td>strand</td><td>MM</td><td>target_seq</td><td>PAM</td><td>alignment</td><td>distance</td><td>position</td><td>gene name</td><td>gene id</td></tr>
<tr><td>GU305914.1</td><td>406</td><td>428</td><td>+</td><td>0</td><td>GTTTTTGCGCTAGAACGCAT</td><td>AGG</td><td>||||||||[||||||||||||]PAM</td><td>NA</td><td>Intergenic</td><td>NA</td><td>NA</td></tr>
<tr><td>T38</td><td>TAGGTTCTAATGCTACCACGCGG</td><td>1000</td><td></td><td></td><td></td><td></td><td></td><td></td><td></td><td></td><td></td></tr>
<tr><td>Oligo fwd</td><td>CACCTAGGTTCTAATGCTACCACG</td><td></td><td></td><td></td><td></td><td></td><td></td><td></td><td></td><td></td><td></td></tr>
<tr><td>Oligo rev</td><td>AAACCGTGGTAGCATTAGAACCTA</td><td></td><td></td><td></td><td></td><td></td><td></td><td></td><td></td><td></td><td></td></tr>
<tr><td>Chromosome</td><td>start</td><td>end</td><td>strand</td><td>MM</td><td>target_seq</td><td>PAM</td><td>alignment</td><td>distance</td><td>position</td><td>gene name</td><td>gene id</td></tr>
<tr><td>GU305914.1</td><td>425</td><td>447</td><td>+</td><td>0</td><td>TAGGTTCTAATGCTACCACG</td><td>CGG</td><td>||||||||[||||||||||||]PAM</td><td>NA</td><td>Intergenic</td><td>NA</td><td>NA</td></tr>
<tr><td>T39</td><td>TCACGCGCACGTGTTAGCATAGG</td><td>1000</td><td></td><td></td><td></td><td></td><td></td><td></td><td></td><td></td><td></td></tr>
<tr><td>Oligo fwd</td><td>CACCTCACGCGCACGTGTTAGCAT</td><td></td><td></td><td></td><td></td><td></td><td></td><td></td><td></td><td></td><td></td></tr>
<tr><td>Oligo rev</td><td>AAACATGCTAACACGTGCGCGTGA</td><td></td><td></td><td></td><td></td><td></td><td></td><td></td><td></td><td></td><td></td></tr>
<tr><td>Chromosome</td><td>start</td><td>end</td><td>strand</td><td>MM</td><td>target_seq</td><td>PAM</td><td>alignment</td><td>distance</td><td>position</td><td>gene name</td><td>gene id</td></tr>
<tr><td>GU305914.1</td><td>466</td><td>488</td><td>+</td><td>0</td><td>TCACGCGCACGTGTTAGCAT</td><td>AGG</td><td>||||||||[||||||||||||]PAM</td><td>NA</td><td>Intergenic</td><td>NA</td><td>NA</td></tr>
<tr><td>T40</td><td>CGCTCGCCTGAGTCAGCCTCCGG</td><td>1000</td><td></td><td></td><td></td><td></td><td></td><td></td><td></td><td></td><td></td></tr>
<tr><td>Oligo fwd</td><td>CACCCGCTCGCCTGAGTCAGCCTC</td><td></td><td></td><td></td><td></td><td></td><td></td><td></td><td></td><td></td><td></td></tr>
<tr><td>Oligo rev</td><td>AAACGAGGCTGACTCAGGCGAGCG</td><td></td><td></td><td></td><td></td><td></td><td></td><td></td><td></td><td></td><td></td></tr>
<tr><td>Chromosome</td><td>start</td><td>end</td><td>strand</td><td>MM</td><td>target_seq</td><td>PAM</td><td>alignment</td><td>distance</td><td>position</td><td>gene name</td><td>gene id</td></tr>
<tr><td>GU305914.1</td><td>501</td><td>523</td><td>+</td><td>0</td><td>CGCTCGCCTGAGTCAGCCTC</td><td>CGG</td><td>||||||||[||||||||||||]PAM</td><td>NA</td><td>Intergenic</td><td>NA</td><td>NA</td></tr>
<tr><td>T41</td><td>GCTCGCCTGAGTCAGCCTCCGGG</td><td>1000</td><td></td><td></td><td></td><td></td><td></td><td></td><td></td><td></td><td></td></tr>
<tr><td>Oligo fwd</td><td>CACCGCTCGCCTGAGTCAGCCTCC</td><td></td><td></td><td></td><td></td><td></td><td></td><td></td><td></td><td></td><td></td></tr>
<tr><td>Oligo rev</td><td>AAACGGAGGCTGACTCAGGCGAGC</td><td></td><td></td><td></td><td></td><td></td><td></td><td></td><td></td><td></td><td></td></tr>
<tr><td>Chromosome</td><td>start</td><td>end</td><td>strand</td><td>MM</td><td>target_seq</td><td>PAM</td><td>alignment</td><td>distance</td><td>position</td><td>gene name</td><td>gene id</td></tr>
<tr><td>GU305914.1</td><td>502</td><td>524</td><td>+</td><td>0</td><td>GCTCGCCTGAGTCAGCCTCC</td><td>GGG</td><td>||||||||[||||||||||||]PAM</td><td>NA</td><td>Intergenic</td><td>NA</td><td>NA</td></tr>
<tr><td>T42</td><td>CGGGCCGCGCGCCGCGTCCGCGG</td><td>1000</td><td></td><td></td><td></td><td></td><td></td><td></td><td></td><td></td><td></td></tr>
<tr><td>Oligo fwd</td><td>CACCCGGGCCGCGCGCCGCGTCCG</td><td></td><td></td><td></td><td></td><td></td><td></td><td></td><td></td><td></td><td></td></tr>
<tr><td>Oligo rev</td><td>AAACCGGACGCGGCGCGCGGCCCG</td><td></td><td></td><td></td><td></td><td></td><td></td><td></td><td></td><td></td><td></td></tr>
<tr><td>Chromosome</td><td>start</td><td>end</td><td>strand</td><td>MM</td><td>target_seq</td><td>PAM</td><td>alignment</td><td>distance</td><td>position</td><td>gene name</td><td>gene id</td></tr>
<tr><td>GU305914.1</td><td>521</td><td>543</td><td>+</td><td>0</td><td>CGGGCCGCGCGCCGCGTCCG</td><td>CGG</td><td>||||||||[||||||||||||]PAM</td><td>NA</td><td>Intergenic</td><td>NA</td><td>NA</td></tr>
<tr><td>GU305914.1</td><td>123893</td><td>123915</td><td>-</td><td>3</td><td>CGGGCAGCGCGCCACGTGCG</td><td>AGG</td><td>|||||-||[|||||-|||-||]PAM</td><td>NA</td><td>Intergenic</td><td>NA</td><td>NA</td></tr>
</table>

| | | | | | | | | | | | |
|---|---|---|---|---|---|---|---|---|---|---|---|
| T43 | GGGCCGCGCGCCGCGTCCGCGGG | 1000 | | | | | | | | | |
| Oligo fwd | CACCGGGCCGCGCGCCGCGTCCGC | | | | | | | | | | |
| Oligo rev | AAACGCGGACGCGGCGCGCGGCCC | | | | | | | | | | |
| Chromosome | start | end | strand | MM | target_seq | PAM | alignment | distance | position | gene name | gene id |
| GU305914.1 | 522 | 544 | + | 0 | GGGCCGCGCGCCGCGTCCGC | GGG | \|\|\|\|\|\|\|\|[\|\|\|\|\|\|\|\|\|\|\|\|]PAM | NA | Intergenic | NA | NA |
| T44 | CGCGCGCCGCGTCCGCGGGAAGG | 1000 | | | | | | | | | |
| Oligo fwd | CACCCGCGCGCCGCGTCCGCGGGA | | | | | | | | | | |
| Oligo rev | AAACTCCCGCGGACGCGGCGCGCG | | | | | | | | | | |
| Chromosome | start | end | strand | MM | target_seq | PAM | alignment | distance | position | gene name | gene id |
| GU305914.1 | 526 | 548 | + | 0 | CGCGCGCCGCGTCCGCGGGA | AGG | \|\|\|\|\|\|\|\|[\|\|\|\|\|\|\|\|\|\|\|\|]PAM | NA | Intergenic | NA | NA |
| T45 | GCGCCGCGTCCGCGGGAAGGCGG | 1000 | | | | | | | | | |
| Oligo fwd | CACCGCGCCGCGTCCGCGGGAAGG | | | | | | | | | | |
| Oligo rev | AAACCCTTCCCGCGGACGCGGCGC | | | | | | | | | | |
| Chromosome | start | end | strand | MM | target_seq | PAM | alignment | distance | position | gene name | gene id |
| GU305914.1 | 529 | 551 | + | 0 | GCGCCGCGTCCGCGGGAAGG | CGG | \|\|\|\|\|\|\|\|[\|\|\|\|\|\|\|\|\|\|\|\|]PAM | NA | Intergenic | NA | NA |
| T46 | CGCCGCGTCCGCGGGAAGGCGGG | 1000 | | | | | | | | | |
| Oligo fwd | CACCCGCCGCGTCCGCGGGAAGGC | | | | | | | | | | |
| Oligo rev | AAACGCCTTCCCGCGGACGCGGCG | | | | | | | | | | |
| Chromosome | start | end | strand | MM | target_seq | PAM | alignment | distance | position | gene name | gene id |
| GU305914.1 | 530 | 552 | + | 0 | CGCCGCGTCCGCGGGAAGGC | GGG | \|\|\|\|\|\|\|\|[\|\|\|\|\|\|\|\|\|\|\|\|]PAM | NA | Intergenic | NA | NA |
| T47 | GCCGCGTCCGCGGGAAGGCGGGG | 1000 | | | | | | | | | |
| Oligo fwd | CACCGCCGCGTCCGCGGGAAGGCG | | | | | | | | | | |
| Oligo rev | AAACCGCCTTCCCGCGGACGCGGC | | | | | | | | | | |
| Chromosome | start | end | strand | MM | target_seq | PAM | alignment | distance | position | gene name | gene id |
| GU305914.1 | 531 | 553 | + | 0 | GCCGCGTCCGCGGGAAGGCG | GGG | \|\|\|\|\|\|\|\|[\|\|\|\|\|\|\|\|\|\|\|\|]PAM | NA | Intergenic | NA | NA |
| T48 | GTCCGCGGGAAGGCGGGGTGAGG | 1000 | | | | | | | | | |
| Oligo fwd | CACCGTCCGCGGGAAGGCGGGGTG | | | | | | | | | | |
| Oligo rev | AAACCACCCCGCCTTCCCGCGGAC | | | | | | | | | | |
| Chromosome | start | end | strand | MM | target_seq | PAM | alignment | distance | position | gene name | gene id |
| GU305914.1 | 536 | 558 | + | 0 | GTCCGCGGGAAGGCGGGGTG | AGG | \|\|\|\|\|\|\|\|[\|\|\|\|\|\|\|\|\|\|\|\|]PAM | NA | Intergenic | NA | NA |
| T49 | TGTGAAGCTTCCGCGACGAGCGG | 1000 | | | | | | | | | |
| Oligo fwd | CACCTGTGAAGCTTCCGCGACGAG | | | | | | | | | | |
| Oligo rev | AAACCTCGTCGCGGAAGCTTCACA | | | | | | | | | | |
| Chromosome | start | end | strand | MM | target_seq | PAM | alignment | distance | position | gene name | gene id |
| GU305914.1 | 572 | 594 | + | 0 | TGTGAAGCTTCCGCGACGAG | CGG | \|\|\|\|\|\|\|\|[\|\|\|\|\|\|\|\|\|\|\|\|]PAM | NA | Intergenic | NA | NA |
| T50 | CTTCCGCGACGAGCGGCGCCCGG | 1000 | | | | | | | | | |
| Oligo fwd | CACCCTTCCGCGACGAGCGGCGCC | | | | | | | | | | |
| Oligo rev | AAACGGCGCCGCTCGTCGCGGAAG | | | | | | | | | | |
| Chromosome | start | end | strand | MM | target_seq | PAM | alignment | distance | position | gene name | gene id |
| GU305914.1 | 579 | 601 | + | 0 | CTTCCGCGACGAGCGGCGCC | CGG | \|\|\|\|\|\|\|\|[\|\|\|\|\|\|\|\|\|\|\|\|]PAM | NA | Intergenic | NA | NA |
| T51 | GCGACGAGCGGCGCCCGGCCCGG | 1000 | | | | | | | | | |
| Oligo fwd | CACCGCGACGAGCGGCGCCCGGCC | | | | | | | | | | |
| Oligo rev | AAACGGCCGGGCGCCGCTCGTCGC | | | | | | | | | | |
| Chromosome | start | end | strand | MM | target_seq | PAM | alignment | distance | position | gene name | gene id |
| GU305914.1 | 584 | 606 | + | 0 | GCGACGAGCGGCGCCCGGCC | CGG | \|\|\|\|\|\|\|\|[\|\|\|\|\|\|\|\|\|\|\|\|]PAM | NA | Intergenic | NA | NA |
| T52 | GAGCGCCGCCGTATGTTTCGAGG | 1000 | | | | | | | | | |
| Oligo fwd | CACCGAGCGCCGCCGTATGTTTCG | | | | | | | | | | |
| Oligo rev | AAACCGAAACATACGGCGGCGCTC | | | | | | | | | | |
| Chromosome | start | end | strand | MM | target_seq | PAM | alignment | distance | position | gene name | gene id |
| GU305914.1 | 631 | 653 | + | 0 | GAGCGCCGCCGTATGTTTCG | AGG | \|\|\|\|\|\|\|\|[\|\|\|\|\|\|\|\|\|\|\|\|]PAM | NA | Intergenic | NA | NA |
| T53 | ACAGTCGCTTCACCGCTCTTCGG | 1000 | | | | | | | | | |
| Oligo fwd | CACCACAGTCGCTTCACCGCTCTT | | | | | | | | | | |
| Oligo rev | AAACAAGAGCGGTGAAGCGACTGT | | | | | | | | | | |
| Chromosome | start | end | strand | MM | target_seq | PAM | alignment | distance | position | gene name | gene id |
| GU305914.1 | 656 | 678 | + | 0 | ACAGTCGCTTCACCGCTCTT | CGG | \|\|\|\|\|\|\|\|[\|\|\|\|\|\|\|\|\|\|\|\|]PAM | NA | Intergenic | NA | NA |
| GU305914.1 | 603 | 625 | - | 4 | ACGCTCGCTTCACGGCTCTC | CGG | \|\|--\|\|\|\|[\|\|\|\|\|-\|\|\|\|\|-]PAM | NA | Intergenic | NA | NA |
| T54 | CTTCACCGCTCTTCGGACCCCGG | 1000 | | | | | | | | | |
| Oligo fwd | CACCCTTCACCGCTCTTCGGACCC | | | | | | | | | | |
| Oligo rev | AAACGGGTCCGAAGAGCGGTGAAG | | | | | | | | | | |
| Chromosome | start | end | strand | MM | target_seq | PAM | alignment | distance | position | gene name | gene id |
| GU305914.1 | 663 | 685 | + | 0 | CTTCACCGCTCTTCGGACCC | CGG | \|\|\|\|\|\|\|\|[\|\|\|\|\|\|\|\|\|\|\|\|]PAM | NA | Intergenic | NA | NA |
| T55 | TTCACCGCTCTTCGGACCCCGGG | 1000 | | | | | | | | | |
| Oligo fwd | CACCTTCACCGCTCTTCGGACCCC | | | | | | | | | | |
| Oligo rev | AAACGGGGTCCGAAGAGCGGTGAA | | | | | | | | | | |
| Chromosome | start | end | strand | MM | target_seq | PAM | alignment | distance | position | gene name | gene id |
| GU305914.1 | 664 | 686 | + | 0 | TTCACCGCTCTTCGGACCCC | GGG | \|\|\|\|\|\|\|\|[\|\|\|\|\|\|\|\|\|\|\|\|]PAM | NA | Intergenic | NA | NA |
| T56 | TCACCGCTCTTCGGACCCCGGGG | 1000 | | | | | | | | | |
| Oligo fwd | CACCTCACCGCTCTTCGGACCCCG | | | | | | | | | | |
| Oligo rev | AAACCGGGGTCCGAAGAGCGGTGA | | | | | | | | | | |
| Chromosome | start | end | strand | MM | target_seq | PAM | alignment | distance | position | gene name | gene id |
| GU305914.1 | 665 | 687 | + | 0 | TCACCGCTCTTCGGACCCCG | GGG | \|\|\|\|\|\|\|\|[\|\|\|\|\|\|\|\|\|\|\|\|]PAM | NA | Intergenic | NA | NA |
| T57 | CCGCTCTTCGGACCCCGGGGTGG | 1000 | | | | | | | | | |
| Oligo fwd | CACCCCGCTCTTCGGACCCCGGGG | | | | | | | | | | |
| Oligo rev | AAACCCCCGGGGTCCGAAGAGCGG | | | | | | | | | | |

<table>
<tr><td>Chromosome</td><td>start</td><td>end</td><td>strand</td><td>MM</td><td>target_seq</td><td>PAM</td><td>alignment</td><td>distance</td><td>position</td><td>gene name</td><td>gene id</td></tr>
<tr><td>GU305914.1</td><td>668</td><td>690</td><td>+</td><td>0</td><td>CCGCTCTTCGGACCCCGGGG</td><td>TGG</td><td>||||||||[||||||||||||]PAM</td><td>NA</td><td>Intergenic</td><td>NA</td><td>NA</td></tr>
<tr><td></td><td></td><td></td><td></td><td></td><td></td><td></td><td></td><td></td><td></td><td></td><td></td></tr>
<tr><td>T58</td><td>CGCTCTTCGGACCCCGGGGTGGG</td><td>1000</td><td></td><td></td><td></td><td></td><td></td><td></td><td></td><td></td><td></td></tr>
<tr><td>Oligo fwd</td><td>CACCCGCTCTTCGGACCCCGGGGT</td><td></td><td></td><td></td><td></td><td></td><td></td><td></td><td></td><td></td><td></td></tr>
<tr><td>Oligo rev</td><td>AAACACCCCGGGGTCCGAAGAGCG</td><td></td><td></td><td></td><td></td><td></td><td></td><td></td><td></td><td></td><td></td></tr>
<tr><td>Chromosome</td><td>start</td><td>end</td><td>strand</td><td>MM</td><td>target_seq</td><td>PAM</td><td>alignment</td><td>distance</td><td>position</td><td>gene name</td><td>gene id</td></tr>
<tr><td>GU305914.1</td><td>669</td><td>691</td><td>+</td><td>0</td><td>CGCTCTTCGGACCCCGGGGT</td><td>GGG</td><td>||||||||[||||||||||||]PAM</td><td>NA</td><td>Intergenic</td><td>NA</td><td>NA</td></tr>
<tr><td></td><td></td><td></td><td></td><td></td><td></td><td></td><td></td><td></td><td></td><td></td><td></td></tr>
<tr><td>T59</td><td>GCTCTTCGGACCCCGGGGTGGGG</td><td>1000</td><td></td><td></td><td></td><td></td><td></td><td></td><td></td><td></td><td></td></tr>
<tr><td>Oligo fwd</td><td>CACCGCTCTTCGGACCCCGGGGTG</td><td></td><td></td><td></td><td></td><td></td><td></td><td></td><td></td><td></td><td></td></tr>
<tr><td>Oligo rev</td><td>AAACCACCCCGGGGTCCGAAGAGC</td><td></td><td></td><td></td><td></td><td></td><td></td><td></td><td></td><td></td><td></td></tr>
<tr><td>Chromosome</td><td>start</td><td>end</td><td>strand</td><td>MM</td><td>target_seq</td><td>PAM</td><td>alignment</td><td>distance</td><td>position</td><td>gene name</td><td>gene id</td></tr>
<tr><td>GU305914.1</td><td>670</td><td>692</td><td>+</td><td>0</td><td>GCTCTTCGGACCCCGGGGTG</td><td>GGG</td><td>||||||||[||||||||||||]PAM</td><td>NA</td><td>Intergenic</td><td>NA</td><td>NA</td></tr>
<tr><td></td><td></td><td></td><td></td><td></td><td></td><td></td><td></td><td></td><td></td><td></td><td></td></tr>
<tr><td>T60</td><td>CTCTTCGGACCCCGGGGTGGGGG</td><td>1000</td><td></td><td></td><td></td><td></td><td></td><td></td><td></td><td></td><td></td></tr>
<tr><td>Oligo fwd</td><td>CACCCTCTTCGGACCCCGGGGTGG</td><td></td><td></td><td></td><td></td><td></td><td></td><td></td><td></td><td></td><td></td></tr>
<tr><td>Oligo rev</td><td>AAACCCACCCCGGGGTCCGAAGAG</td><td></td><td></td><td></td><td></td><td></td><td></td><td></td><td></td><td></td><td></td></tr>
<tr><td>Chromosome</td><td>start</td><td>end</td><td>strand</td><td>MM</td><td>target_seq</td><td>PAM</td><td>alignment</td><td>distance</td><td>position</td><td>gene name</td><td>gene id</td></tr>
<tr><td>GU305914.1</td><td>671</td><td>693</td><td>+</td><td>0</td><td>CTCTTCGGACCCCGGGGTGG</td><td>GGG</td><td>||||||||[||||||||||||]PAM</td><td>NA</td><td>Intergenic</td><td>NA</td><td>NA</td></tr>
<tr><td></td><td></td><td></td><td></td><td></td><td></td><td></td><td></td><td></td><td></td><td></td><td></td></tr>
<tr><td>T61</td><td>TCTTCGGACCCCGGGGTGGGGGG</td><td>1000</td><td></td><td></td><td></td><td></td><td></td><td></td><td></td><td></td><td></td></tr>
<tr><td>Oligo fwd</td><td>CACCTCTTCGGACCCCGGGGTGGG</td><td></td><td></td><td></td><td></td><td></td><td></td><td></td><td></td><td></td><td></td></tr>
<tr><td>Oligo rev</td><td>AAACCCCACCCCGGGGTCCGAAGA</td><td></td><td></td><td></td><td></td><td></td><td></td><td></td><td></td><td></td><td></td></tr>
<tr><td>Chromosome</td><td>start</td><td>end</td><td>strand</td><td>MM</td><td>target_seq</td><td>PAM</td><td>alignment</td><td>distance</td><td>position</td><td>gene name</td><td>gene id</td></tr>
<tr><td>GU305914.1</td><td>672</td><td>694</td><td>+</td><td>0</td><td>TCTTCGGACCCCGGGGTGGG</td><td>GGG</td><td>||||||||[||||||||||||]PAM</td><td>NA</td><td>Intergenic</td><td>NA</td><td>NA</td></tr>
<tr><td></td><td></td><td></td><td></td><td></td><td></td><td></td><td></td><td></td><td></td><td></td><td></td></tr>
<tr><td>T62</td><td>CTTCGGACCCCGGGGTGGGGGGG</td><td>1000</td><td></td><td></td><td></td><td></td><td></td><td></td><td></td><td></td><td></td></tr>
<tr><td>Oligo fwd</td><td>CACCCTTCGGACCCCGGGGTGGGG</td><td></td><td></td><td></td><td></td><td></td><td></td><td></td><td></td><td></td><td></td></tr>
<tr><td>Oligo rev</td><td>AAACCCCCACCCCGGGGTCCGAAG</td><td></td><td></td><td></td><td></td><td></td><td></td><td></td><td></td><td></td><td></td></tr>
<tr><td>Chromosome</td><td>start</td><td>end</td><td>strand</td><td>MM</td><td>target_seq</td><td>PAM</td><td>alignment</td><td>distance</td><td>position</td><td>gene name</td><td>gene id</td></tr>
<tr><td>GU305914.1</td><td>673</td><td>695</td><td>+</td><td>0</td><td>CTTCGGACCCCGGGGTGGGG</td><td>GGG</td><td>||||||||[||||||||||||]PAM</td><td>NA</td><td>Intergenic</td><td>NA</td><td>NA</td></tr>
<tr><td></td><td></td><td></td><td></td><td></td><td></td><td></td><td></td><td></td><td></td><td></td><td></td></tr>
<tr><td>T63</td><td>CGGACCCCGGGGTGGGGGGGCGG</td><td>1000</td><td></td><td></td><td></td><td></td><td></td><td></td><td></td><td></td><td></td></tr>
<tr><td>Oligo fwd</td><td>CACCCGGACCCCGGGGTGGGGGGG</td><td></td><td></td><td></td><td></td><td></td><td></td><td></td><td></td><td></td><td></td></tr>
<tr><td>Oligo rev</td><td>AAACCCCCCCCACCCCGGGGTCCG</td><td></td><td></td><td></td><td></td><td></td><td></td><td></td><td></td><td></td><td></td></tr>
<tr><td>Chromosome</td><td>start</td><td>end</td><td>strand</td><td>MM</td><td>target_seq</td><td>PAM</td><td>alignment</td><td>distance</td><td>position</td><td>gene name</td><td>gene id</td></tr>
<tr><td>GU305914.1</td><td>676</td><td>698</td><td>+</td><td>0</td><td>CGGACCCCGGGGTGGGGGGG</td><td>CGG</td><td>||||||||[||||||||||||]PAM</td><td>NA</td><td>Intergenic</td><td>NA</td><td>NA</td></tr>
<tr><td></td><td></td><td></td><td></td><td></td><td></td><td></td><td></td><td></td><td></td><td></td><td></td></tr>
<tr><td>T64</td><td>GGACCCCGGGGTGGGGGGGCGGG</td><td>1000</td><td></td><td></td><td></td><td></td><td></td><td></td><td></td><td></td><td></td></tr>
<tr><td>Oligo fwd</td><td>CACCGGACCCCGGGGTGGGGGGGC</td><td></td><td></td><td></td><td></td><td></td><td></td><td></td><td></td><td></td><td></td></tr>
<tr><td>Oligo rev</td><td>AAACGCCCCCCCACCCCGGGGTCC</td><td></td><td></td><td></td><td></td><td></td><td></td><td></td><td></td><td></td><td></td></tr>
<tr><td>Chromosome</td><td>start</td><td>end</td><td>strand</td><td>MM</td><td>target_seq</td><td>PAM</td><td>alignment</td><td>distance</td><td>position</td><td>gene name</td><td>gene id</td></tr>
<tr><td>GU305914.1</td><td>677</td><td>699</td><td>+</td><td>0</td><td>GGACCCCGGGGTGGGGGGGC</td><td>GGG</td><td>||||||||[||||||||||||]PAM</td><td>NA</td><td>Intergenic</td><td>NA</td><td>NA</td></tr>
<tr><td></td><td></td><td></td><td></td><td></td><td></td><td></td><td></td><td></td><td></td><td></td><td></td></tr>
<tr><td>T65</td><td>GGGGGGGCGGGTATAATACCCGG</td><td>1000</td><td></td><td></td><td></td><td></td><td></td><td></td><td></td><td></td><td></td></tr>
<tr><td>Oligo fwd</td><td>CACCGGGGGGGCGGGTATAATACC</td><td></td><td></td><td></td><td></td><td></td><td></td><td></td><td></td><td></td><td></td></tr>
<tr><td>Oligo rev</td><td>AAACGGTATTATACCCGCCCCCCC</td><td></td><td></td><td></td><td></td><td></td><td></td><td></td><td></td><td></td><td></td></tr>
<tr><td>Chromosome</td><td>start</td><td>end</td><td>strand</td><td>MM</td><td>target_seq</td><td>PAM</td><td>alignment</td><td>distance</td><td>position</td><td>gene name</td><td>gene id</td></tr>
<tr><td>GU305914.1</td><td>689</td><td>711</td><td>+</td><td>0</td><td>GGGGGGGCGGGTATAATACC</td><td>CGG</td><td>||||||||[||||||||||||]PAM</td><td>NA</td><td>Intergenic</td><td>NA</td><td>NA</td></tr>
<tr><td></td><td></td><td></td><td></td><td></td><td></td><td></td><td></td><td></td><td></td><td></td><td></td></tr>
<tr><td>T66</td><td>GGGGGGCGGGTATAATACCCGGG</td><td>1000</td><td></td><td></td><td></td><td></td><td></td><td></td><td></td><td></td><td></td></tr>
<tr><td>Oligo fwd</td><td>CACCGGGGGGCGGGTATAATACCC</td><td></td><td></td><td></td><td></td><td></td><td></td><td></td><td></td><td></td><td></td></tr>
<tr><td>Oligo rev</td><td>AAACGGGTATTATACCCGCCCCCC</td><td></td><td></td><td></td><td></td><td></td><td></td><td></td><td></td><td></td><td></td></tr>
<tr><td>Chromosome</td><td>start</td><td>end</td><td>strand</td><td>MM</td><td>target_seq</td><td>PAM</td><td>alignment</td><td>distance</td><td>position</td><td>gene name</td><td>gene id</td></tr>
<tr><td>GU305914.1</td><td>690</td><td>712</td><td>+</td><td>0</td><td>GGGGGGCGGGTATAATACCC</td><td>GGG</td><td>||||||||[||||||||||||]PAM</td><td>NA</td><td>Intergenic</td><td>NA</td><td>NA</td></tr>
<tr><td></td><td></td><td></td><td></td><td></td><td></td><td></td><td></td><td></td><td></td><td></td><td></td></tr>
<tr><td>T67</td><td>GGGGGCGGGTATAATACCCGGGG</td><td>1000</td><td></td><td></td><td></td><td></td><td></td><td></td><td></td><td></td><td></td></tr>
<tr><td>Oligo fwd</td><td>CACCGGGGGCGGGTATAATACCCG</td><td></td><td></td><td></td><td></td><td></td><td></td><td></td><td></td><td></td><td></td></tr>
<tr><td>Oligo rev</td><td>AAACCGGGTATTATACCCGCCCCC</td><td></td><td></td><td></td><td></td><td></td><td></td><td></td><td></td><td></td><td></td></tr>
<tr><td>Chromosome</td><td>start</td><td>end</td><td>strand</td><td>MM</td><td>target_seq</td><td>PAM</td><td>alignment</td><td>distance</td><td>position</td><td>gene name</td><td>gene id</td></tr>
<tr><td>GU305914.1</td><td>691</td><td>713</td><td>+</td><td>0</td><td>GGGGGCGGGTATAATACCCG</td><td>GGG</td><td>||||||||[||||||||||||]PAM</td><td>NA</td><td>Intergenic</td><td>NA</td><td>NA</td></tr>
<tr><td></td><td></td><td></td><td></td><td></td><td></td><td></td><td></td><td></td><td></td><td></td><td></td></tr>
<tr><td>T68</td><td>GTATAATACCCGGGGAGTATTGG</td><td>1000</td><td></td><td></td><td></td><td></td><td></td><td></td><td></td><td></td><td></td></tr>
<tr><td>Oligo fwd</td><td>CACCGTATAATACCCGGGGAGTAT</td><td></td><td></td><td></td><td></td><td></td><td></td><td></td><td></td><td></td><td></td></tr>
<tr><td>Oligo rev</td><td>AAACATACTCCCCGGGTATTATAC</td><td></td><td></td><td></td><td></td><td></td><td></td><td></td><td></td><td></td><td></td></tr>
<tr><td>Chromosome</td><td>start</td><td>end</td><td>strand</td><td>MM</td><td>target_seq</td><td>PAM</td><td>alignment</td><td>distance</td><td>position</td><td>gene name</td><td>gene id</td></tr>
<tr><td>GU305914.1</td><td>699</td><td>721</td><td>+</td><td>0</td><td>GTATAATACCCGGGGAGTAT</td><td>TGG</td><td>||||||||[||||||||||||]PAM</td><td>NA</td><td>Intergenic</td><td>NA</td><td>NA</td></tr>
<tr><td></td><td></td><td></td><td></td><td></td><td></td><td></td><td></td><td></td><td></td><td></td><td></td></tr>
<tr><td>T69</td><td>TAATACCCGGGGAGTATTGGCGG</td><td>1000</td><td></td><td></td><td></td><td></td><td></td><td></td><td></td><td></td><td></td></tr>
<tr><td>Oligo fwd</td><td>CACCTAATACCCGGGGAGTATTGG</td><td></td><td></td><td></td><td></td><td></td><td></td><td></td><td></td><td></td><td></td></tr>
<tr><td>Oligo rev</td><td>AAACCCAATACTCCCCGGGTATTA</td><td></td><td></td><td></td><td></td><td></td><td></td><td></td><td></td><td></td><td></td></tr>
<tr><td>Chromosome</td><td>start</td><td>end</td><td>strand</td><td>MM</td><td>target_seq</td><td>PAM</td><td>alignment</td><td>distance</td><td>position</td><td>gene name</td><td>gene id</td></tr>
<tr><td>GU305914.1</td><td>702</td><td>724</td><td>+</td><td>0</td><td>TAATACCCGGGGAGTATTGG</td><td>CGG</td><td>||||||||[||||||||||||]PAM</td><td>NA</td><td>Intergenic</td><td>NA</td><td>NA</td></tr>
<tr><td></td><td></td><td></td><td></td><td></td><td></td><td></td><td></td><td></td><td></td><td></td><td></td></tr>
<tr><td>T70</td><td>AATACCCGGGGAGTATTGGCGGG</td><td>1000</td><td></td><td></td><td></td><td></td><td></td><td></td><td></td><td></td><td></td></tr>
<tr><td>Oligo fwd</td><td>CACCAATACCCGGGGAGTATTGGC</td><td></td><td></td><td></td><td></td><td></td><td></td><td></td><td></td><td></td><td></td></tr>
<tr><td>Oligo rev</td><td>AAACGCCAATACTCCCCGGGTATT</td><td></td><td></td><td></td><td></td><td></td><td></td><td></td><td></td><td></td><td></td></tr>
<tr><td>Chromosome</td><td>start</td><td>end</td><td>strand</td><td>MM</td><td>target_seq</td><td>PAM</td><td>alignment</td><td>distance</td><td>position</td><td>gene name</td><td>gene id</td></tr>
<tr><td>GU305914.1</td><td>703</td><td>725</td><td>+</td><td>0</td><td>AATACCCGGGGAGTATTGGC</td><td>GGG</td><td>||||||||[||||||||||||]PAM</td><td>NA</td><td>Intergenic</td><td>NA</td><td>NA</td></tr>
<tr><td></td><td></td><td></td><td></td><td></td><td></td><td></td><td></td><td></td><td></td><td></td><td></td></tr>
<tr><td>T71</td><td>ATACCCGGGGAGTATTGGCGGGG</td><td>1000</td><td></td><td></td><td></td><td></td><td></td><td></td><td></td><td></td><td></td></tr>
<tr><td>Oligo fwd</td><td>CACCATACCCGGGGAGTATTGGCG</td><td></td><td></td><td></td><td></td><td></td><td></td><td></td><td></td><td></td><td></td></tr>
<tr><td>Oligo rev</td><td>AAACCGCCAATACTCCCCGGGTAT</td><td></td><td></td><td></td><td></td><td></td><td></td><td></td><td></td><td></td><td></td></tr>
<tr><td>Chromosome</td><td>start</td><td>end</td><td>strand</td><td>MM</td><td>target_seq</td><td>PAM</td><td>alignment</td><td>distance</td><td>position</td><td>gene name</td><td>gene id</td></tr>
<tr><td>GU305914.1</td><td>704</td><td>726</td><td>+</td><td>0</td><td>ATACCCGGGGAGTATTGGCG</td><td>GGG</td><td>||||||||[||||||||||||]PAM</td><td>NA</td><td>Intergenic</td><td>NA</td><td>NA</td></tr>
<tr><td></td><td></td><td></td><td></td><td></td><td></td><td></td><td></td><td></td><td></td><td></td><td></td></tr>
<tr><td>T72</td><td>GGCGGGGCACCCCCCCCCGCGG</td><td>1000</td><td></td><td></td><td></td><td></td><td></td><td></td><td></td><td></td><td></td></tr>
<tr><td>Oligo fwd</td><td>CACCGGCGGGGCACCCCCCCCCG</td><td></td><td></td><td></td><td></td><td></td><td></td><td></td><td></td><td></td><td></td></tr>
</table>

<table>
<tr><td>Oligo rev</td><td>AAACCGGGGGGGGGGTGCCCCGCC</td><td></td><td></td><td></td><td></td><td></td><td></td><td></td><td></td><td></td><td></td></tr>
<tr><td>Chromosome</td><td>start</td><td>end</td><td>strand</td><td>MM</td><td>target_seq</td><td>PAM</td><td>alignment</td><td>distance</td><td>position</td><td>gene name</td><td>gene id</td></tr>
<tr><td>GU305914.1</td><td>720</td><td>742</td><td>+</td><td>0</td><td>GGCGGGGCACCCCCCCCCCG</td><td>CGG</td><td>||||||||[||||||||||||]PAM</td><td>NA</td><td>Intergenic</td><td>NA</td><td>NA</td></tr>
<tr><td>GU305914.1</td><td>68479</td><td>68501</td><td>+</td><td>4</td><td>GGCGCAGCACCCCATCCCCG</td><td>GGG</td><td>||||--||[|||||--|||||]PAM</td><td>NA</td><td>Intergenic</td><td>NA</td><td>NA</td></tr>
<tr><td></td><td></td><td></td><td></td><td></td><td></td><td></td><td></td><td></td><td></td><td></td><td></td></tr>
<tr><td>T73</td><td>GCGGGGCACCCCCCCCCCGCGGG</td><td>1000</td><td></td><td></td><td></td><td></td><td></td><td></td><td></td><td></td><td></td></tr>
<tr><td>Oligo fwd</td><td>CACCGCGGGGCACCCCCCCCCCGC</td><td></td><td></td><td></td><td></td><td></td><td></td><td></td><td></td><td></td><td></td></tr>
<tr><td>Oligo rev</td><td>AAACGCGGGGGGGGGGTGCCCCGC</td><td></td><td></td><td></td><td></td><td></td><td></td><td></td><td></td><td></td><td></td></tr>
<tr><td>Chromosome</td><td>start</td><td>end</td><td>strand</td><td>MM</td><td>target_seq</td><td>PAM</td><td>alignment</td><td>distance</td><td>position</td><td>gene name</td><td>gene id</td></tr>
<tr><td>GU305914.1</td><td>721</td><td>743</td><td>+</td><td>0</td><td>GCGGGGCACCCCCCCCCCGC</td><td>GGG</td><td>||||||||[||||||||||||]PAM</td><td>NA</td><td>Intergenic</td><td>NA</td><td>NA</td></tr>
<tr><td></td><td></td><td></td><td></td><td></td><td></td><td></td><td></td><td></td><td></td><td></td><td></td></tr>
<tr><td>T74</td><td>CGGGGCACCCCCCCCCCGCGGGG</td><td>1000</td><td></td><td></td><td></td><td></td><td></td><td></td><td></td><td></td><td></td></tr>
<tr><td>Oligo fwd</td><td>CACCCGGGGCACCCCCCCCCCGCG</td><td></td><td></td><td></td><td></td><td></td><td></td><td></td><td></td><td></td><td></td></tr>
<tr><td>Oligo rev</td><td>AAACCGCGGGGGGGGGGTGCCCCG</td><td></td><td></td><td></td><td></td><td></td><td></td><td></td><td></td><td></td><td></td></tr>
<tr><td>Chromosome</td><td>start</td><td>end</td><td>strand</td><td>MM</td><td>target_seq</td><td>PAM</td><td>alignment</td><td>distance</td><td>position</td><td>gene name</td><td>gene id</td></tr>
<tr><td>GU305914.1</td><td>722</td><td>744</td><td>+</td><td>0</td><td>CGGGGCACCCCCCCCCCGCG</td><td>GGG</td><td>||||||||[||||||||||||]PAM</td><td>NA</td><td>Intergenic</td><td>NA</td><td>NA</td></tr>
<tr><td></td><td></td><td></td><td></td><td></td><td></td><td></td><td></td><td></td><td></td><td></td><td></td></tr>
<tr><td>T75</td><td>ACCCCCCCCCCGCGGGGTTTTGG</td><td>1000</td><td></td><td></td><td></td><td></td><td></td><td></td><td></td><td></td><td></td></tr>
<tr><td>Oligo fwd</td><td>CACCACCCCCCCCCCGCGGGGTTT</td><td></td><td></td><td></td><td></td><td></td><td></td><td></td><td></td><td></td><td></td></tr>
<tr><td>Oligo rev</td><td>AAACAAACCCCGCGGGGGGGGGGT</td><td></td><td></td><td></td><td></td><td></td><td></td><td></td><td></td><td></td><td></td></tr>
<tr><td>Chromosome</td><td>start</td><td>end</td><td>strand</td><td>MM</td><td>target_seq</td><td>PAM</td><td>alignment</td><td>distance</td><td>position</td><td>gene name</td><td>gene id</td></tr>
<tr><td>GU305914.1</td><td>728</td><td>750</td><td>+</td><td>0</td><td>ACCCCCCCCCCGCGGGGTTT</td><td>TGG</td><td>||||||||[||||||||||||]PAM</td><td>NA</td><td>Intergenic</td><td>NA</td><td>NA</td></tr>
<tr><td></td><td></td><td></td><td></td><td></td><td></td><td></td><td></td><td></td><td></td><td></td><td></td></tr>
<tr><td>T76</td><td>CCCCCCCCCCGCGGGGTTTTGGG</td><td>1000</td><td></td><td></td><td></td><td></td><td></td><td></td><td></td><td></td><td></td></tr>
<tr><td>Oligo fwd</td><td>CACCCCCCCCCCCCGCGGGGTTTT</td><td></td><td></td><td></td><td></td><td></td><td></td><td></td><td></td><td></td><td></td></tr>
<tr><td>Oligo rev</td><td>AAACAAAACCCCGCGGGGGGGGGG</td><td></td><td></td><td></td><td></td><td></td><td></td><td></td><td></td><td></td><td></td></tr>
<tr><td>Chromosome</td><td>start</td><td>end</td><td>strand</td><td>MM</td><td>target_seq</td><td>PAM</td><td>alignment</td><td>distance</td><td>position</td><td>gene name</td><td>gene id</td></tr>
<tr><td>GU305914.1</td><td>729</td><td>751</td><td>+</td><td>0</td><td>CCCCCCCCCCGCGGGGTTTT</td><td>GGG</td><td>||||||||[||||||||||||]PAM</td><td>NA</td><td>Intergenic</td><td>NA</td><td>NA</td></tr>
<tr><td></td><td></td><td></td><td></td><td></td><td></td><td></td><td></td><td></td><td></td><td></td><td></td></tr>
<tr><td>T77</td><td>CCCCCCGCGGGGTTTTGGGTCGG</td><td>1000</td><td></td><td></td><td></td><td></td><td></td><td></td><td></td><td></td><td></td></tr>
<tr><td>Oligo fwd</td><td>CACCCCCCCCGCGGGGTTTTGGGT</td><td></td><td></td><td></td><td></td><td></td><td></td><td></td><td></td><td></td><td></td></tr>
<tr><td>Oligo rev</td><td>AAACACCCAAAACCCCGCGGGGGG</td><td></td><td></td><td></td><td></td><td></td><td></td><td></td><td></td><td></td><td></td></tr>
<tr><td>Chromosome</td><td>start</td><td>end</td><td>strand</td><td>MM</td><td>target_seq</td><td>PAM</td><td>alignment</td><td>distance</td><td>position</td><td>gene name</td><td>gene id</td></tr>
<tr><td>GU305914.1</td><td>733</td><td>755</td><td>+</td><td>0</td><td>CCCCCCGCGGGGTTTTGGGT</td><td>CGG</td><td>||||||||[||||||||||||]PAM</td><td>NA</td><td>Intergenic</td><td>NA</td><td>NA</td></tr>
<tr><td></td><td></td><td></td><td></td><td></td><td></td><td></td><td></td><td></td><td></td><td></td><td></td></tr>
<tr><td>T78</td><td>CCCCCGCGGGGTTTTGGGTCGGG</td><td>1000</td><td></td><td></td><td></td><td></td><td></td><td></td><td></td><td></td><td></td></tr>
<tr><td>Oligo fwd</td><td>CACCCCCCCGCGGGGTTTTGGGTC</td><td></td><td></td><td></td><td></td><td></td><td></td><td></td><td></td><td></td><td></td></tr>
<tr><td>Oligo rev</td><td>AAACGACCCAAAACCCCGCGGGGG</td><td></td><td></td><td></td><td></td><td></td><td></td><td></td><td></td><td></td><td></td></tr>
<tr><td>Chromosome</td><td>start</td><td>end</td><td>strand</td><td>MM</td><td>target_seq</td><td>PAM</td><td>alignment</td><td>distance</td><td>position</td><td>gene name</td><td>gene id</td></tr>
<tr><td>GU305914.1</td><td>734</td><td>756</td><td>+</td><td>0</td><td>CCCCCGCGGGGTTTTGGGTC</td><td>GGG</td><td>||||||||[||||||||||||]PAM</td><td>NA</td><td>Intergenic</td><td>NA</td><td>NA</td></tr>
<tr><td></td><td></td><td></td><td></td><td></td><td></td><td></td><td></td><td></td><td></td><td></td><td></td></tr>
<tr><td>T79</td><td>CCCCGCGGGGTTTTGGGTCGGGG</td><td>1000</td><td></td><td></td><td></td><td></td><td></td><td></td><td></td><td></td><td></td></tr>
<tr><td>Oligo fwd</td><td>CACCCCCCGCGGGGTTTTGGGTCG</td><td></td><td></td><td></td><td></td><td></td><td></td><td></td><td></td><td></td><td></td></tr>
<tr><td>Oligo rev</td><td>AAACCGACCCAAAACCCCGCGGGG</td><td></td><td></td><td></td><td></td><td></td><td></td><td></td><td></td><td></td><td></td></tr>
<tr><td>Chromosome</td><td>start</td><td>end</td><td>strand</td><td>MM</td><td>target_seq</td><td>PAM</td><td>alignment</td><td>distance</td><td>position</td><td>gene name</td><td>gene id</td></tr>
<tr><td>GU305914.1</td><td>735</td><td>757</td><td>+</td><td>0</td><td>CCCCGCGGGGTTTTGGGTCG</td><td>GGG</td><td>||||||||[||||||||||||]PAM</td><td>NA</td><td>Intergenic</td><td>NA</td><td>NA</td></tr>
<tr><td></td><td></td><td></td><td></td><td></td><td></td><td></td><td></td><td></td><td></td><td></td><td></td></tr>
<tr><td>T80</td><td>CCCGCGGGGTTTTGGGTCGGGGG</td><td>1000</td><td></td><td></td><td></td><td></td><td></td><td></td><td></td><td></td><td></td></tr>
<tr><td>Oligo fwd</td><td>CACCCCCGCGGGGTTTTGGGTCGG</td><td></td><td></td><td></td><td></td><td></td><td></td><td></td><td></td><td></td><td></td></tr>
<tr><td>Oligo rev</td><td>AAACCCGACCCAAAACCCCGCGGG</td><td></td><td></td><td></td><td></td><td></td><td></td><td></td><td></td><td></td><td></td></tr>
<tr><td>Chromosome</td><td>start</td><td>end</td><td>strand</td><td>MM</td><td>target_seq</td><td>PAM</td><td>alignment</td><td>distance</td><td>position</td><td>gene name</td><td>gene id</td></tr>
<tr><td>GU305914.1</td><td>736</td><td>758</td><td>+</td><td>0</td><td>CCCGCGGGGTTTTGGGTCGG</td><td>GGG</td><td>||||||||[||||||||||||]PAM</td><td>NA</td><td>Intergenic</td><td>NA</td><td>NA</td></tr>
<tr><td></td><td></td><td></td><td></td><td></td><td></td><td></td><td></td><td></td><td></td><td></td><td></td></tr>
<tr><td>T81</td><td>GAATGTCGAGCGTCTCTCAGCGG</td><td>1000</td><td></td><td></td><td></td><td></td><td></td><td></td><td></td><td></td><td></td></tr>
<tr><td>Oligo fwd</td><td>CACCGAATGTCGAGCGTCTCTCAG</td><td></td><td></td><td></td><td></td><td></td><td></td><td></td><td></td><td></td><td></td></tr>
<tr><td>Oligo rev</td><td>AAACCTGAGAGACGCTCGACATTC</td><td></td><td></td><td></td><td></td><td></td><td></td><td></td><td></td><td></td><td></td></tr>
<tr><td>Chromosome</td><td>start</td><td>end</td><td>strand</td><td>MM</td><td>target_seq</td><td>PAM</td><td>alignment</td><td>distance</td><td>position</td><td>gene name</td><td>gene id</td></tr>
<tr><td>GU305914.1</td><td>763</td><td>785</td><td>+</td><td>0</td><td>GAATGTCGAGCGTCTCTCAG</td><td>CGG</td><td>||||||||[||||||||||||]PAM</td><td>NA</td><td>Intergenic</td><td>NA</td><td>NA</td></tr>
<tr><td></td><td></td><td></td><td></td><td></td><td></td><td></td><td></td><td></td><td></td><td></td><td></td></tr>
<tr><td>T82</td><td>GCGTCTCTCAGCGGTGAGTCTGG</td><td>1000</td><td></td><td></td><td></td><td></td><td></td><td></td><td></td><td></td><td></td></tr>
<tr><td>Oligo fwd</td><td>CACCGCGTCTCTCAGCGGTGAGTC</td><td></td><td></td><td></td><td></td><td></td><td></td><td></td><td></td><td></td><td></td></tr>
<tr><td>Oligo rev</td><td>AAACGACTCACCGCTGAGAGACGC</td><td></td><td></td><td></td><td></td><td></td><td></td><td></td><td></td><td></td><td></td></tr>
<tr><td>Chromosome</td><td>start</td><td>end</td><td>strand</td><td>MM</td><td>target_seq</td><td>PAM</td><td>alignment</td><td>distance</td><td>position</td><td>gene name</td><td>gene id</td></tr>
<tr><td>GU305914.1</td><td>772</td><td>794</td><td>+</td><td>0</td><td>GCGTCTCTCAGCGGTGAGTC</td><td>TGG</td><td>||||||||[||||||||||||]PAM</td><td>NA</td><td>Intergenic</td><td>NA</td><td>NA</td></tr>
<tr><td></td><td></td><td></td><td></td><td></td><td></td><td></td><td></td><td></td><td></td><td></td><td></td></tr>
<tr><td>T83</td><td>CGTCTCTCAGCGGTGAGTCTGGG</td><td>1000</td><td></td><td></td><td></td><td></td><td></td><td></td><td></td><td></td><td></td></tr>
<tr><td>Oligo fwd</td><td>CACCCGTCTCTCAGCGGTGAGTCT</td><td></td><td></td><td></td><td></td><td></td><td></td><td></td><td></td><td></td><td></td></tr>
<tr><td>Oligo rev</td><td>AAACAGACTCACCGCTGAGAGACG</td><td></td><td></td><td></td><td></td><td></td><td></td><td></td><td></td><td></td><td></td></tr>
<tr><td>Chromosome</td><td>start</td><td>end</td><td>strand</td><td>MM</td><td>target_seq</td><td>PAM</td><td>alignment</td><td>distance</td><td>position</td><td>gene name</td><td>gene id</td></tr>
<tr><td>GU305914.1</td><td>773</td><td>795</td><td>+</td><td>0</td><td>CGTCTCTCAGCGGTGAGTCT</td><td>GGG</td><td>||||||||[||||||||||||]PAM</td><td>NA</td><td>Intergenic</td><td>NA</td><td>NA</td></tr>
<tr><td></td><td></td><td></td><td></td><td></td><td></td><td></td><td></td><td></td><td></td><td></td><td></td></tr>
<tr><td>T84</td><td>AGTCTGGGTAGCGCAGTCGAAGG</td><td>1000</td><td></td><td></td><td></td><td></td><td></td><td></td><td></td><td></td><td></td></tr>
<tr><td>Oligo fwd</td><td>CACCAGTCTGGGTAGCGCAGTCGA</td><td></td><td></td><td></td><td></td><td></td><td></td><td></td><td></td><td></td><td></td></tr>
<tr><td>Oligo rev</td><td>AAACTCGACTGCGCTACCCAGACT</td><td></td><td></td><td></td><td></td><td></td><td></td><td></td><td></td><td></td><td></td></tr>
<tr><td>Chromosome</td><td>start</td><td>end</td><td>strand</td><td>MM</td><td>target_seq</td><td>PAM</td><td>alignment</td><td>distance</td><td>position</td><td>gene name</td><td>gene id</td></tr>
<tr><td>GU305914.1</td><td>788</td><td>810</td><td>+</td><td>0</td><td>AGTCTGGGTAGCGCAGTCGA</td><td>AGG</td><td>||||||||[||||||||||||]PAM</td><td>NA</td><td>Intergenic</td><td>NA</td><td>NA</td></tr>
<tr><td></td><td></td><td></td><td></td><td></td><td></td><td></td><td></td><td></td><td></td><td></td><td></td></tr>
<tr><td>T85</td><td>TCGAAGGAAACCGTGTCGAACGG</td><td>1000</td><td></td><td></td><td></td><td></td><td></td><td></td><td></td><td></td><td></td></tr>
<tr><td>Oligo fwd</td><td>CACCTCGAAGGAAACCGTGTCGAA</td><td></td><td></td><td></td><td></td><td></td><td></td><td></td><td></td><td></td><td></td></tr>
<tr><td>Oligo rev</td><td>AAACTTCGACACGGTTTCCTTCGA</td><td></td><td></td><td></td><td></td><td></td><td></td><td></td><td></td><td></td><td></td></tr>
<tr><td>Chromosome</td><td>start</td><td>end</td><td>strand</td><td>MM</td><td>target_seq</td><td>PAM</td><td>alignment</td><td>distance</td><td>position</td><td>gene name</td><td>gene id</td></tr>
<tr><td>GU305914.1</td><td>804</td><td>826</td><td>+</td><td>0</td><td>TCGAAGGAAACCGTGTCGAA</td><td>CGG</td><td>||||||||[||||||||||||]PAM</td><td>NA</td><td>Intergenic</td><td>NA</td><td>NA</td></tr>
<tr><td></td><td></td><td></td><td></td><td></td><td></td><td></td><td></td><td></td><td></td><td></td><td></td></tr>
<tr><td>T86</td><td>CGTGTGAGCTATCCTCCCCGAGG</td><td>1000</td><td></td><td></td><td></td><td></td><td></td><td></td><td></td><td></td><td></td></tr>
<tr><td>Oligo fwd</td><td>CACCCGTGTGAGCTATCCTCCCCG</td><td></td><td></td><td></td><td></td><td></td><td></td><td></td><td></td><td></td><td></td></tr>
<tr><td>Oligo rev</td><td>AAACCGGGGAGGATAGCTCACACG</td><td></td><td></td><td></td><td></td><td></td><td></td><td></td><td></td><td></td><td></td></tr>
<tr><td>Chromosome</td><td>start</td><td>end</td><td>strand</td><td>MM</td><td>target_seq</td><td>PAM</td><td>alignment</td><td>distance</td><td>position</td><td>gene name</td><td>gene id</td></tr>
<tr><td>GU305914.1</td><td>900</td><td>922</td><td>+</td><td>0</td><td>CGTGTGAGCTATCCTCCCCG</td><td>AGG</td><td>||||||||[||||||||||||]PAM</td><td>NA</td><td>Intergenic</td><td>NA</td><td>NA</td></tr>
</table>

<table>
<tr><td>T87</td><td>CCCGAGGATCCCACCCAAGTAGG</td><td></td><td>1000</td><td></td><td></td><td></td><td></td><td></td><td></td><td></td><td></td><td></td></tr>
<tr><td>Oligo fwd</td><td>CACCCCCGAGGATCCCACCCAAGT</td><td></td><td></td><td></td><td></td><td></td><td></td><td></td><td></td><td></td><td></td><td></td></tr>
<tr><td>Oligo rev</td><td>AAACACTTGGGTGGGATCCTCGGG</td><td></td><td></td><td></td><td></td><td></td><td></td><td></td><td></td><td></td><td></td><td></td></tr>
<tr><td>Chromosome</td><td>start</td><td></td><td>end</td><td>strand</td><td>MM</td><td>target_seq</td><td>PAM</td><td>alignment</td><td>distance</td><td>position</td><td>gene name</td><td>gene id</td></tr>
<tr><td>GU305914.1</td><td></td><td>916</td><td>938</td><td>+</td><td>0</td><td>CCCGAGGATCCCACCCAAGT</td><td>AGG</td><td>||||||||[||||||||||||]PAM</td><td>NA</td><td>Intergenic</td><td>NA</td><td>NA</td></tr>
<tr><td>T88</td><td>CGTAATTCCCTCCCTCCGTCCGG</td><td></td><td>1000</td><td></td><td></td><td></td><td></td><td></td><td></td><td></td><td></td><td></td></tr>
<tr><td>Oligo fwd</td><td>CACCCGTAATTCCCTCCCTCCGTC</td><td></td><td></td><td></td><td></td><td></td><td></td><td></td><td></td><td></td><td></td><td></td></tr>
<tr><td>Oligo rev</td><td>AAACGACGGAGGGAGGGAATTACG</td><td></td><td></td><td></td><td></td><td></td><td></td><td></td><td></td><td></td><td></td><td></td></tr>
<tr><td>Chromosome</td><td>start</td><td></td><td>end</td><td>strand</td><td>MM</td><td>target_seq</td><td>PAM</td><td>alignment</td><td>distance</td><td>position</td><td>gene name</td><td>gene id</td></tr>
<tr><td>GU305914.1</td><td></td><td>950</td><td>972</td><td>+</td><td>0</td><td>CGTAATTCCCTCCCTCCGTC</td><td>CGG</td><td>||||||||[||||||||||||]PAM</td><td>NA</td><td>Intergenic</td><td>NA</td><td>NA</td></tr>
<tr><td>T89</td><td>TGATTCCCCCTAGCTGACAGAGG</td><td></td><td>1000</td><td></td><td></td><td></td><td></td><td></td><td></td><td></td><td></td><td></td></tr>
<tr><td>Oligo fwd</td><td>CACCTGATTCCCCCTAGCTGACAG</td><td></td><td></td><td></td><td></td><td></td><td></td><td></td><td></td><td></td><td></td><td></td></tr>
<tr><td>Oligo rev</td><td>AAACCTGTCAGCTAGGGGGAATCA</td><td></td><td></td><td></td><td></td><td></td><td></td><td></td><td></td><td></td><td></td><td></td></tr>
<tr><td>Chromosome</td><td>start</td><td></td><td>end</td><td>strand</td><td>MM</td><td>target_seq</td><td>PAM</td><td>alignment</td><td>distance</td><td>position</td><td>gene name</td><td>gene id</td></tr>
<tr><td>GU305914.1</td><td></td><td>988</td><td>1010</td><td>+</td><td>0</td><td>TGATTCCCCCTAGCTGACAG</td><td>AGG</td><td>||||||||[||||||||||||]PAM</td><td>NA</td><td>Intergenic</td><td>NA</td><td>NA</td></tr>
<tr><td>T90</td><td>AGAGGTGATAAATTACACCACGG</td><td></td><td>1000</td><td></td><td></td><td></td><td></td><td></td><td></td><td></td><td></td><td></td></tr>
<tr><td>Oligo fwd</td><td>CACCAGAGGTGATAAATTACACCA</td><td></td><td></td><td></td><td></td><td></td><td></td><td></td><td></td><td></td><td></td><td></td></tr>
<tr><td>Oligo rev</td><td>AAACTGGTGTAATTTATCACCTCT</td><td></td><td></td><td></td><td></td><td></td><td></td><td></td><td></td><td></td><td></td><td></td></tr>
<tr><td>Chromosome</td><td>start</td><td></td><td>end</td><td>strand</td><td>MM</td><td>target_seq</td><td>PAM</td><td>alignment</td><td>distance</td><td>position</td><td>gene name</td><td>gene id</td></tr>
<tr><td>GU305914.1</td><td></td><td>1006</td><td>1028</td><td>+</td><td>0</td><td>AGAGGTGATAAATTACACCA</td><td>CGG</td><td>||||||||[||||||||||||]PAM</td><td>NA</td><td>Intergenic</td><td>NA</td><td>NA</td></tr>
<tr><td>T91</td><td>TAAATTACACCACGGTGCCATGG</td><td></td><td>1000</td><td></td><td></td><td></td><td></td><td></td><td></td><td></td><td></td><td></td></tr>
<tr><td>Oligo fwd</td><td>CACCTAAATTACACCACGGTGCCA</td><td></td><td></td><td></td><td></td><td></td><td></td><td></td><td></td><td></td><td></td><td></td></tr>
<tr><td>Oligo rev</td><td>AAACTGGCACCGTGGTGTAATTTA</td><td></td><td></td><td></td><td></td><td></td><td></td><td></td><td></td><td></td><td></td><td></td></tr>
<tr><td>Chromosome</td><td>start</td><td></td><td>end</td><td>strand</td><td>MM</td><td>target_seq</td><td>PAM</td><td>alignment</td><td>distance</td><td>position</td><td>gene name</td><td>gene id</td></tr>
<tr><td>GU305914.1</td><td></td><td>1014</td><td>1036</td><td>+</td><td>0</td><td>TAAATTACACCACGGTGCCA</td><td>TGG</td><td>||||||||[||||||||||||]PAM</td><td>NA</td><td>Intergenic</td><td>NA</td><td>NA</td></tr>
<tr><td>T92</td><td>GGTGCCATGGCAGCCGCTGTCGG</td><td></td><td>1000</td><td></td><td></td><td></td><td></td><td></td><td></td><td></td><td></td><td></td></tr>
<tr><td>Oligo fwd</td><td>CACCGGTGCCATGGCAGCCGCTGT</td><td></td><td></td><td></td><td></td><td></td><td></td><td></td><td></td><td></td><td></td><td></td></tr>
<tr><td>Oligo rev</td><td>AAACACAGCGGCTGCCATGGCACC</td><td></td><td></td><td></td><td></td><td></td><td></td><td></td><td></td><td></td><td></td><td></td></tr>
<tr><td>Chromosome</td><td>start</td><td></td><td>end</td><td>strand</td><td>MM</td><td>target_seq</td><td>PAM</td><td>alignment</td><td>distance</td><td>position</td><td>gene name</td><td>gene id</td></tr>
<tr><td>GU305914.1</td><td></td><td>1027</td><td>1049</td><td>+</td><td>0</td><td>GGTGCCATGGCAGCCGCTGT</td><td>CGG</td><td>||||||||[||||||||||||]PAM</td><td>NA</td><td>Intergenic</td><td>NA</td><td>NA</td></tr>
<tr><td>T93</td><td>GTGCCATGGCAGCCGCTGTCGGG</td><td></td><td>1000</td><td></td><td></td><td></td><td></td><td></td><td></td><td></td><td></td><td></td></tr>
<tr><td>Oligo fwd</td><td>CACCGTGCCATGGCAGCCGCTGTC</td><td></td><td></td><td></td><td></td><td></td><td></td><td></td><td></td><td></td><td></td><td></td></tr>
<tr><td>Oligo rev</td><td>AAACGACAGCGGCTGCCATGGCAC</td><td></td><td></td><td></td><td></td><td></td><td></td><td></td><td></td><td></td><td></td><td></td></tr>
<tr><td>Chromosome</td><td>start</td><td></td><td>end</td><td>strand</td><td>MM</td><td>target_seq</td><td>PAM</td><td>alignment</td><td>distance</td><td>position</td><td>gene name</td><td>gene id</td></tr>
<tr><td>GU305914.1</td><td></td><td>1028</td><td>1050</td><td>+</td><td>0</td><td>GTGCCATGGCAGCCGCTGTC</td><td>GGG</td><td>||||||||[||||||||||||]PAM</td><td>NA</td><td>Intergenic</td><td>NA</td><td>NA</td></tr>
<tr><td>T94</td><td>GGGAGACGAGCGTGCCCCGCAGG</td><td></td><td>1000</td><td></td><td></td><td></td><td></td><td></td><td></td><td></td><td></td><td></td></tr>
<tr><td>Oligo fwd</td><td>CACCGGGAGACGAGCGTGCCCCGC</td><td></td><td></td><td></td><td></td><td></td><td></td><td></td><td></td><td></td><td></td><td></td></tr>
<tr><td>Oligo rev</td><td>AAACGCGGGGCACGCTCGTCTCCC</td><td></td><td></td><td></td><td></td><td></td><td></td><td></td><td></td><td></td><td></td><td></td></tr>
<tr><td>Chromosome</td><td>start</td><td></td><td>end</td><td>strand</td><td>MM</td><td>target_seq</td><td>PAM</td><td>alignment</td><td>distance</td><td>position</td><td>gene name</td><td>gene id</td></tr>
<tr><td>GU305914.1</td><td></td><td>1048</td><td>1070</td><td>+</td><td>0</td><td>GGGAGACGAGCGTGCCCCGC</td><td>AGG</td><td>||||||||[||||||||||||]PAM</td><td>NA</td><td>Intergenic</td><td>NA</td><td>NA</td></tr>
<tr><td>T95</td><td>GACGAGCGTGCCCCGCAGGCAGG</td><td></td><td>1000</td><td></td><td></td><td></td><td></td><td></td><td></td><td></td><td></td><td></td></tr>
<tr><td>Oligo fwd</td><td>CACCGACGAGCGTGCCCCGCAGGC</td><td></td><td></td><td></td><td></td><td></td><td></td><td></td><td></td><td></td><td></td><td></td></tr>
<tr><td>Oligo rev</td><td>AAACGCCTGCGGGGCACGCTCGTC</td><td></td><td></td><td></td><td></td><td></td><td></td><td></td><td></td><td></td><td></td><td></td></tr>
<tr><td>Chromosome</td><td>start</td><td></td><td>end</td><td>strand</td><td>MM</td><td>target_seq</td><td>PAM</td><td>alignment</td><td>distance</td><td>position</td><td>gene name</td><td>gene id</td></tr>
<tr><td>GU305914.1</td><td></td><td>1052</td><td>1074</td><td>+</td><td>0</td><td>GACGAGCGTGCCCCGCAGGC</td><td>AGG</td><td>||||||||[||||||||||||]PAM</td><td>NA</td><td>Intergenic</td><td>NA</td><td>NA</td></tr>
<tr><td>T96</td><td>GGCGCAGCGACGCAGTCATAAGG</td><td></td><td>1000</td><td></td><td></td><td></td><td></td><td></td><td></td><td></td><td></td><td></td></tr>
<tr><td>Oligo fwd</td><td>CACCGGCGCAGCGACGCAGTCATA</td><td></td><td></td><td></td><td></td><td></td><td></td><td></td><td></td><td></td><td></td><td></td></tr>
<tr><td>Oligo rev</td><td>AAACTATGACTGCGTCGCTGCGCC</td><td></td><td></td><td></td><td></td><td></td><td></td><td></td><td></td><td></td><td></td><td></td></tr>
<tr><td>Chromosome</td><td>start</td><td></td><td>end</td><td>strand</td><td>MM</td><td>target_seq</td><td>PAM</td><td>alignment</td><td>distance</td><td>position</td><td>gene name</td><td>gene id</td></tr>
<tr><td>GU305914.1</td><td></td><td>1073</td><td>1095</td><td>+</td><td>0</td><td>GGCGCAGCGACGCAGTCATA</td><td>AGG</td><td>||||||||[||||||||||||]PAM</td><td>NA</td><td>Intergenic</td><td>NA</td><td>NA</td></tr>
<tr><td>T97</td><td>GCGCAGCGACGCAGTCATAAGGG</td><td></td><td>1000</td><td></td><td></td><td></td><td></td><td></td><td></td><td></td><td></td><td></td></tr>
<tr><td>Oligo fwd</td><td>CACCGCGCAGCGACGCAGTCATAA</td><td></td><td></td><td></td><td></td><td></td><td></td><td></td><td></td><td></td><td></td><td></td></tr>
<tr><td>Oligo rev</td><td>AAACTTATGACTGCGTCGCTGCGC</td><td></td><td></td><td></td><td></td><td></td><td></td><td></td><td></td><td></td><td></td><td></td></tr>
<tr><td>Chromosome</td><td>start</td><td></td><td>end</td><td>strand</td><td>MM</td><td>target_seq</td><td>PAM</td><td>alignment</td><td>distance</td><td>position</td><td>gene name</td><td>gene id</td></tr>
<tr><td>GU305914.1</td><td></td><td>1074</td><td>1096</td><td>+</td><td>0</td><td>GCGCAGCGACGCAGTCATAA</td><td>GGG</td><td>||||||||[||||||||||||]PAM</td><td>NA</td><td>Intergenic</td><td>NA</td><td>NA</td></tr>
<tr><td>T98</td><td>GATCTGAGCGTCTCGAGATGAGG</td><td></td><td>1000</td><td></td><td></td><td></td><td></td><td></td><td></td><td></td><td></td><td></td></tr>
<tr><td>Oligo fwd</td><td>CACCGATCTGAGCGTCTCGAGATG</td><td></td><td></td><td></td><td></td><td></td><td></td><td></td><td></td><td></td><td></td><td></td></tr>
<tr><td>Oligo rev</td><td>AAACCATCTCGAGACGCTCAGATC</td><td></td><td></td><td></td><td></td><td></td><td></td><td></td><td></td><td></td><td></td><td></td></tr>
<tr><td>Chromosome</td><td>start</td><td></td><td>end</td><td>strand</td><td>MM</td><td>target_seq</td><td>PAM</td><td>alignment</td><td>distance</td><td>position</td><td>gene name</td><td>gene id</td></tr>
<tr><td>GU305914.1</td><td></td><td>1103</td><td>1125</td><td>+</td><td>0</td><td>GATCTGAGCGTCTCGAGATG</td><td>AGG</td><td>||||||||[||||||||||||]PAM</td><td>NA</td><td>Intergenic</td><td>NA</td><td>NA</td></tr>
<tr><td>T99</td><td>AGCGTCTCGAGATGAGGCACAGG</td><td></td><td>1000</td><td></td><td></td><td></td><td></td><td></td><td></td><td></td><td></td><td></td></tr>
<tr><td>Oligo fwd</td><td>CACCAGCGTCTCGAGATGAGGCAC</td><td></td><td></td><td></td><td></td><td></td><td></td><td></td><td></td><td></td><td></td><td></td></tr>
<tr><td>Oligo rev</td><td>AAACGTGCCTCATCTCGAGACGCT</td><td></td><td></td><td></td><td></td><td></td><td></td><td></td><td></td><td></td><td></td><td></td></tr>
<tr><td>Chromosome</td><td>start</td><td></td><td>end</td><td>strand</td><td>MM</td><td>target_seq</td><td>PAM</td><td>alignment</td><td>distance</td><td>position</td><td>gene name</td><td>gene id</td></tr>
<tr><td>GU305914.1</td><td></td><td>1109</td><td>1131</td><td>+</td><td>0</td><td>AGCGTCTCGAGATGAGGCAC</td><td>AGG</td><td>||||||||[||||||||||||]PAM</td><td>NA</td><td>Intergenic</td><td>NA</td><td>NA</td></tr>
<tr><td>T100</td><td>AGATGAGGCACAGGTCGAGTTGG</td><td></td><td>1000</td><td></td><td></td><td></td><td></td><td></td><td></td><td></td><td></td><td></td></tr>
<tr><td>Oligo fwd</td><td>CACCAGATGAGGCACAGGTCGAGT</td><td></td><td></td><td></td><td></td><td></td><td></td><td></td><td></td><td></td><td></td><td></td></tr>
<tr><td>Oligo rev</td><td>AAACACTCGACCTGTGCCTCATCT</td><td></td><td></td><td></td><td></td><td></td><td></td><td></td><td></td><td></td><td></td><td></td></tr>
<tr><td>Chromosome</td><td>start</td><td></td><td>end</td><td>strand</td><td>MM</td><td>target_seq</td><td>PAM</td><td>alignment</td><td>distance</td><td>position</td><td>gene name</td><td>gene id</td></tr>
<tr><td>GU305914.1</td><td></td><td>1118</td><td>1140</td><td>+</td><td>0</td><td>AGATGAGGCACAGGTCGAGT</td><td>TGG</td><td>||||||||[||||||||||||]PAM</td><td>NA</td><td>Intergenic</td><td>NA</td><td>NA</td></tr>
<tr><td>T101</td><td>TGGTACGCGTCTCGTCCCCGTGG</td><td></td><td>1000</td><td></td><td></td><td></td><td></td><td></td><td></td><td></td><td></td><td></td></tr>
<tr><td>Oligo fwd</td><td>CACCTGGTACGCGTCTCGTCCCCG</td><td></td><td></td><td></td><td></td><td></td><td></td><td></td><td></td><td></td><td></td><td></td></tr>
<tr><td>Oligo rev</td><td>AAACCGGGGACGAGACGCGTACCA</td><td></td><td></td><td></td><td></td><td></td><td></td><td></td><td></td><td></td><td></td><td></td></tr>
<tr><td>Chromosome</td><td>start</td><td></td><td>end</td><td>strand</td><td>MM</td><td>target_seq</td><td>PAM</td><td>alignment</td><td>distance</td><td>position</td><td>gene name</td><td>gene id</td></tr>
<tr><td>GU305914.1</td><td></td><td>1138</td><td>1160</td><td>+</td><td>0</td><td>TGGTACGCGTCTCGTCCCCG</td><td>TGG</td><td>||||||||[||||||||||||]PAM</td><td>NA</td><td>Intergenic</td><td>NA</td><td>NA</td></tr>
</table>

<table>
<tr><td>T102</td><td>CTCGTCCCCGTGGTGTCTGTCGG</td><td>1000</td><td></td><td></td><td></td><td></td><td></td><td></td><td></td><td></td><td></td></tr>
<tr><td>Oligo fwd</td><td>CACCCTCGTCCCCGTGGTGTCTGT</td><td></td><td></td><td></td><td></td><td></td><td></td><td></td><td></td><td></td><td></td></tr>
<tr><td>Oligo rev</td><td>AAACACAGACACCACGGGGACGAG</td><td></td><td></td><td></td><td></td><td></td><td></td><td></td><td></td><td></td><td></td></tr>
<tr><td>Chromosome</td><td>start</td><td>end</td><td>strand</td><td>MM</td><td>target_seq</td><td>PAM</td><td>alignment</td><td>distance</td><td>position</td><td>gene name</td><td>gene id</td></tr>
<tr><td>GU305914.1</td><td>1148</td><td>1170</td><td>+</td><td>0</td><td>CTCGTCCCCGTGGTGTCTGT</td><td>CGG</td><td>||||||||[||||||||||||]PAM</td><td>NA</td><td>Intergenic</td><td>NA</td><td>NA</td></tr>
<tr><td>GU305914.1</td><td>92323</td><td>92345</td><td>-</td><td>4</td><td>CTCGTCTGCGTAGTGTGTGT</td><td>CGG</td><td>||||||--[|||-||||-|||]PAM</td><td>NA</td><td>Intergenic</td><td>NA</td><td>NA</td></tr>
</table>

<table>
<tr><td>T103</td><td>GTTGTTGCTCGTGTGCGCCGCGG</td><td>1000</td><td></td><td></td><td></td><td></td><td></td><td></td><td></td><td></td><td></td></tr>
<tr><td>Oligo fwd</td><td>CACCGTTGTTGCTCGTGTGCGCCG</td><td></td><td></td><td></td><td></td><td></td><td></td><td></td><td></td><td></td><td></td></tr>
<tr><td>Oligo rev</td><td>AAACCGGCGCACACGAGCAACAAC</td><td></td><td></td><td></td><td></td><td></td><td></td><td></td><td></td><td></td><td></td></tr>
<tr><td>Chromosome</td><td>start</td><td>end</td><td>strand</td><td>MM</td><td>target_seq</td><td>PAM</td><td>alignment</td><td>distance</td><td>position</td><td>gene name</td><td>gene id</td></tr>
<tr><td>GU305914.1</td><td>1179</td><td>1201</td><td>+</td><td>0</td><td>GTTGTTGCTCGTGTGCGCCG</td><td>CGG</td><td>||||||||[||||||||||||]PAM</td><td>NA</td><td>Intergenic</td><td>NA</td><td>NA</td></tr>
<tr><td>GU305914.1</td><td>199339</td><td>199361</td><td>-</td><td>4</td><td>GTTGGTGTGTGTGTGCGCCG</td><td>GGG</td><td>||||-||-[--||||||||||]PAM</td><td>NA</td><td>Intergenic</td><td>NA</td><td>NA</td></tr>
</table>

<table>
<tr><td>T104</td><td>GTTGCTCGTGTGCGCCGCGGTGG</td><td>1000</td><td></td><td></td><td></td><td></td><td></td><td></td><td></td><td></td><td></td></tr>
<tr><td>Oligo fwd</td><td>CACCGTTGCTCGTGTGCGCCGCGG</td><td></td><td></td><td></td><td></td><td></td><td></td><td></td><td></td><td></td><td></td></tr>
<tr><td>Oligo rev</td><td>AAACCCGCGGCGCACACGAGCAAC</td><td></td><td></td><td></td><td></td><td></td><td></td><td></td><td></td><td></td><td></td></tr>
<tr><td>Chromosome</td><td>start</td><td>end</td><td>strand</td><td>MM</td><td>target_seq</td><td>PAM</td><td>alignment</td><td>distance</td><td>position</td><td>gene name</td><td>gene id</td></tr>
<tr><td>GU305914.1</td><td>1182</td><td>1204</td><td>+</td><td>0</td><td>GTTGCTCGTGTGCGCCGCGG</td><td>TGG</td><td>||||||||[||||||||||||]PAM</td><td>NA</td><td>Intergenic</td><td>NA</td><td>NA</td></tr>
</table>

<table>
<tr><td>T105</td><td>GGTGGTCCCCATTGTGTCGTCGG</td><td>1000</td><td></td><td></td><td></td><td></td><td></td><td></td><td></td><td></td><td></td></tr>
<tr><td>Oligo fwd</td><td>CACCGGTGGTCCCCATTGTGTCGT</td><td></td><td></td><td></td><td></td><td></td><td></td><td></td><td></td><td></td><td></td></tr>
<tr><td>Oligo rev</td><td>AAACACGACACAATGGGGACCACC</td><td></td><td></td><td></td><td></td><td></td><td></td><td></td><td></td><td></td><td></td></tr>
<tr><td>Chromosome</td><td>start</td><td>end</td><td>strand</td><td>MM</td><td>target_seq</td><td>PAM</td><td>alignment</td><td>distance</td><td>position</td><td>gene name</td><td>gene id</td></tr>
<tr><td>GU305914.1</td><td>1200</td><td>1222</td><td>+</td><td>0</td><td>GGTGGTCCCCATTGTGTCGT</td><td>CGG</td><td>||||||||[||||||||||||]PAM</td><td>NA</td><td>Intergenic</td><td>NA</td><td>NA</td></tr>
</table>

<table>
<tr><td>T106</td><td>TGTGTCGTCGGCGCCGCTGCCGG</td><td>1000</td><td></td><td></td><td></td><td></td><td></td><td></td><td></td><td></td><td></td></tr>
<tr><td>Oligo fwd</td><td>CACCTGTGTCGTCGGCGCCGCTGC</td><td></td><td></td><td></td><td></td><td></td><td></td><td></td><td></td><td></td><td></td></tr>
<tr><td>Oligo rev</td><td>AAACGCAGCGGCGCCGACGACACA</td><td></td><td></td><td></td><td></td><td></td><td></td><td></td><td></td><td></td><td></td></tr>
<tr><td>Chromosome</td><td>start</td><td>end</td><td>strand</td><td>MM</td><td>target_seq</td><td>PAM</td><td>alignment</td><td>distance</td><td>position</td><td>gene name</td><td>gene id</td></tr>
<tr><td>GU305914.1</td><td>1212</td><td>1234</td><td>+</td><td>0</td><td>TGTGTCGTCGGCGCCGCTGC</td><td>CGG</td><td>||||||||[||||||||||||]PAM</td><td>NA</td><td>Intergenic</td><td>NA</td><td>NA</td></tr>
<tr><td>GU305914.1</td><td>27311</td><td>27333</td><td>-</td><td>4</td><td>CGCGTCGTCGGCGGCGCCGC</td><td>CGG</td><td>-|-|||||[|||||-|||-||]PAM</td><td>NA</td><td>Intergenic</td><td>NA</td><td>NA</td></tr>
</table>

<table>
<tr><td>T107</td><td>GTGTCGTCGGCGCCGCTGCCGGG</td><td>1000</td><td></td><td></td><td></td><td></td><td></td><td></td><td></td><td></td><td></td></tr>
<tr><td>Oligo fwd</td><td>CACCGTGTCGTCGGCGCCGCTGCC</td><td></td><td></td><td></td><td></td><td></td><td></td><td></td><td></td><td></td><td></td></tr>
<tr><td>Oligo rev</td><td>AAACGGCAGCGGCGCCGACGACAC</td><td></td><td></td><td></td><td></td><td></td><td></td><td></td><td></td><td></td><td></td></tr>
<tr><td>Chromosome</td><td>start</td><td>end</td><td>strand</td><td>MM</td><td>target_seq</td><td>PAM</td><td>alignment</td><td>distance</td><td>position</td><td>gene name</td><td>gene id</td></tr>
<tr><td>GU305914.1</td><td>1213</td><td>1235</td><td>+</td><td>0</td><td>GTGTCGTCGGCGCCGCTGCC</td><td>GGG</td><td>||||||||[||||||||||||]PAM</td><td>NA</td><td>Intergenic</td><td>NA</td><td>NA</td></tr>
</table>

<table>
<tr><td>T108</td><td>TGTCGTCGGCGCCGCTGCCGGGG</td><td>1000</td><td></td><td></td><td></td><td></td><td></td><td></td><td></td><td></td><td></td></tr>
<tr><td>Oligo fwd</td><td>CACCTGTCGTCGGCGCCGCTGCCG</td><td></td><td></td><td></td><td></td><td></td><td></td><td></td><td></td><td></td><td></td></tr>
<tr><td>Oligo rev</td><td>AAACCGGCAGCGGCGCCGACGACA</td><td></td><td></td><td></td><td></td><td></td><td></td><td></td><td></td><td></td><td></td></tr>
<tr><td>Chromosome</td><td>start</td><td>end</td><td>strand</td><td>MM</td><td>target_seq</td><td>PAM</td><td>alignment</td><td>distance</td><td>position</td><td>gene name</td><td>gene id</td></tr>
<tr><td>GU305914.1</td><td>1214</td><td>1236</td><td>+</td><td>0</td><td>TGTCGTCGGCGCCGCTGCCG</td><td>GGG</td><td>||||||||[||||||||||||]PAM</td><td>NA</td><td>Intergenic</td><td>NA</td><td>NA</td></tr>
<tr><td>GU305914.1</td><td>153169</td><td>153191</td><td>+</td><td>3</td><td>TGTCGGCGGCGCTGCCGCCG</td><td>CGG</td><td>|||||-||[||||-||-||||]PAM</td><td>NA</td><td>Intergenic</td><td>NA</td><td>NA</td></tr>
</table>

<table>
<tr><td>T109</td><td>CTGCTACGCGAAGAACGCGACGG</td><td>1000</td><td></td><td></td><td></td><td></td><td></td><td></td><td></td><td></td><td></td></tr>
<tr><td>Oligo fwd</td><td>CACCCTGCTACGCGAAGAACGCGA</td><td></td><td></td><td></td><td></td><td></td><td></td><td></td><td></td><td></td><td></td></tr>
<tr><td>Oligo rev</td><td>AAACTCGCGTTCTTCGCGTAGCAG</td><td></td><td></td><td></td><td></td><td></td><td></td><td></td><td></td><td></td><td></td></tr>
<tr><td>Chromosome</td><td>start</td><td>end</td><td>strand</td><td>MM</td><td>target_seq</td><td>PAM</td><td>alignment</td><td>distance</td><td>position</td><td>gene name</td><td>gene id</td></tr>
<tr><td>GU305914.1</td><td>1242</td><td>1264</td><td>+</td><td>0</td><td>CTGCTACGCGAAGAACGCGA</td><td>CGG</td><td>||||||||[||||||||||||]PAM</td><td>NA</td><td>Intergenic</td><td>NA</td><td>NA</td></tr>
</table>

<table>
<tr><td>T110</td><td>CGAAGAACGCGACGGAGCGCAGG</td><td>1000</td><td></td><td></td><td></td><td></td><td></td><td></td><td></td><td></td><td></td></tr>
<tr><td>Oligo fwd</td><td>CACCCGAAGAACGCGACGGAGCGC</td><td></td><td></td><td></td><td></td><td></td><td></td><td></td><td></td><td></td><td></td></tr>
<tr><td>Oligo rev</td><td>AAACGCGCTCCGTCGCGTTCTTCG</td><td></td><td></td><td></td><td></td><td></td><td></td><td></td><td></td><td></td><td></td></tr>
<tr><td>Chromosome</td><td>start</td><td>end</td><td>strand</td><td>MM</td><td>target_seq</td><td>PAM</td><td>alignment</td><td>distance</td><td>position</td><td>gene name</td><td>gene id</td></tr>
<tr><td>GU305914.1</td><td>1250</td><td>1272</td><td>+</td><td>0</td><td>CGAAGAACGCGACGGAGCGC</td><td>AGG</td><td>||||||||[||||||||||||]PAM</td><td>NA</td><td>Intergenic</td><td>NA</td><td>NA</td></tr>
</table>

<table>
<tr><td>T111</td><td>CTAACATCGATCGCGATTACTGG</td><td>1000</td><td></td><td></td><td></td><td></td><td></td><td></td><td></td><td></td><td></td></tr>
<tr><td>Oligo fwd</td><td>CACCCTAACATCGATCGCGATTAC</td><td></td><td></td><td></td><td></td><td></td><td></td><td></td><td></td><td></td><td></td></tr>
<tr><td>Oligo rev</td><td>AAACGTAATCGCGATCGATGTTAG</td><td></td><td></td><td></td><td></td><td></td><td></td><td></td><td></td><td></td><td></td></tr>
<tr><td>Chromosome</td><td>start</td><td>end</td><td>strand</td><td>MM</td><td>target_seq</td><td>PAM</td><td>alignment</td><td>distance</td><td>position</td><td>gene name</td><td>gene id</td></tr>
<tr><td>GU305914.1</td><td>1277</td><td>1299</td><td>+</td><td>0</td><td>CTAACATCGATCGCGATTAC</td><td>TGG</td><td>||||||||[||||||||||||]PAM</td><td>NA</td><td>Intergenic</td><td>NA</td><td>NA</td></tr>
</table>

<table>
<tr><td>T112</td><td>ACATCGATCGCGATTACTGGAGG</td><td>1000</td><td></td><td></td><td></td><td></td><td></td><td></td><td></td><td></td><td></td></tr>
<tr><td>Oligo fwd</td><td>CACCACATCGATCGCGATTACTGG</td><td></td><td></td><td></td><td></td><td></td><td></td><td></td><td></td><td></td><td></td></tr>
<tr><td>Oligo rev</td><td>AAACCCAGTAATCGCGATCGATGT</td><td></td><td></td><td></td><td></td><td></td><td></td><td></td><td></td><td></td><td></td></tr>
<tr><td>Chromosome</td><td>start</td><td>end</td><td>strand</td><td>MM</td><td>target_seq</td><td>PAM</td><td>alignment</td><td>distance</td><td>position</td><td>gene name</td><td>gene id</td></tr>
<tr><td>GU305914.1</td><td>1280</td><td>1302</td><td>+</td><td>0</td><td>ACATCGATCGCGATTACTGG</td><td>AGG</td><td>||||||||[||||||||||||]PAM</td><td>NA</td><td>Intergenic</td><td>NA</td><td>NA</td></tr>
</table>

<table>
<tr><td>T113</td><td>CGATCGCGATTACTGGAGGCCGG</td><td>1000</td><td></td><td></td><td></td><td></td><td></td><td></td><td></td><td></td><td></td></tr>
<tr><td>Oligo fwd</td><td>CACCCGATCGCGATTACTGGAGGC</td><td></td><td></td><td></td><td></td><td></td><td></td><td></td><td></td><td></td><td></td></tr>
<tr><td>Oligo rev</td><td>AAACGCCTCCAGTAATCGCGATCG</td><td></td><td></td><td></td><td></td><td></td><td></td><td></td><td></td><td></td><td></td></tr>
<tr><td>Chromosome</td><td>start</td><td>end</td><td>strand</td><td>MM</td><td>target_seq</td><td>PAM</td><td>alignment</td><td>distance</td><td>position</td><td>gene name</td><td>gene id</td></tr>
<tr><td>GU305914.1</td><td>1284</td><td>1306</td><td>+</td><td>0</td><td>CGATCGCGATTACTGGAGGC</td><td>CGG</td><td>||||||||[||||||||||||]PAM</td><td>NA</td><td>Intergenic</td><td>NA</td><td>NA</td></tr>
</table>

<table>
<tr><td>T114</td><td>GATCGCGATTACTGGAGGCCGGG</td><td>1000</td><td></td><td></td><td></td><td></td><td></td><td></td><td></td><td></td><td></td></tr>
<tr><td>Oligo fwd</td><td>CACCGATCGCGATTACTGGAGGCC</td><td></td><td></td><td></td><td></td><td></td><td></td><td></td><td></td><td></td><td></td></tr>
<tr><td>Oligo rev</td><td>AAACGGCCTCCAGTAATCGCGATC</td><td></td><td></td><td></td><td></td><td></td><td></td><td></td><td></td><td></td><td></td></tr>
<tr><td>Chromosome</td><td>start</td><td>end</td><td>strand</td><td>MM</td><td>target_seq</td><td>PAM</td><td>alignment</td><td>distance</td><td>position</td><td>gene name</td><td>gene id</td></tr>
<tr><td>GU305914.1</td><td>1285</td><td>1307</td><td>+</td><td>0</td><td>GATCGCGATTACTGGAGGCC</td><td>GGG</td><td>||||||||[||||||||||||]PAM</td><td>NA</td><td>Intergenic</td><td>NA</td><td>NA</td></tr>
</table>

<table>
<tr><td>T115</td><td>TTACTGGAGGCCGGGCGTGTCGG</td><td>1000</td><td></td><td></td><td></td><td></td><td></td><td></td><td></td><td></td><td></td></tr>
<tr><td>Oligo fwd</td><td>CACCTTACTGGAGGCCGGGCGTGT</td><td></td><td></td><td></td><td></td><td></td><td></td><td></td><td></td><td></td><td></td></tr>
<tr><td>Oligo rev</td><td>AAACACACGCCCGGCCTCCAGTAA</td><td></td><td></td><td></td><td></td><td></td><td></td><td></td><td></td><td></td><td></td></tr>
<tr><td>Chromosome</td><td>start</td><td>end</td><td>strand</td><td>MM</td><td>target_seq</td><td>PAM</td><td>alignment</td><td>distance</td><td>position</td><td>gene name</td><td>gene id</td></tr>
<tr><td>GU305914.1</td><td>1293</td><td>1315</td><td>+</td><td>0</td><td>TTACTGGAGGCCGGGCGTGT</td><td>CGG</td><td>||||||||[||||||||||||]PAM</td><td>NA</td><td>Intergenic</td><td>NA</td><td>NA</td></tr>
</table>

<table>
<tr><td>T116</td><td>AGGCCGGGCGTGTCGGTGTTCGG</td><td>1000</td><td></td><td></td><td></td><td></td><td></td><td></td><td></td><td></td><td></td></tr>
<tr><td>Oligo fwd</td><td>CACCAGGCCGGGCGTGTCGGTGTT</td><td></td><td></td><td></td><td></td><td></td><td></td><td></td><td></td><td></td><td></td></tr>
<tr><td>Oligo rev</td><td>AAACAACACCGACACGCCCGGCCT</td><td></td><td></td><td></td><td></td><td></td><td></td><td></td><td></td><td></td><td></td></tr>
<tr><td>Chromosome</td><td>start</td><td>end</td><td>strand</td><td>MM</td><td>target_seq</td><td>PAM</td><td>alignment</td><td>distance</td><td>position</td><td>gene name</td><td>gene id</td></tr>
<tr><td>GU305914.1</td><td>1300</td><td>1322</td><td>+</td><td>0</td><td>AGGCCGGGCGTGTCGGTGTT</td><td>CGG</td><td>||||||||[||||||||||||]PAM</td><td>NA</td><td>Intergenic</td><td>NA</td><td>NA</td></tr>
<tr><td>T117</td><td>TTCGGTTGTCGCATGCCCAAAGG</td><td>1000</td><td></td><td></td><td></td><td></td><td></td><td></td><td></td><td></td><td></td></tr>
<tr><td>Oligo fwd</td><td>CACCTTCGGTTGTCGCATGCCCAA</td><td></td><td></td><td></td><td></td><td></td><td></td><td></td><td></td><td></td><td></td></tr>
<tr><td>Oligo rev</td><td>AAACTTGGGCATGCGACAACCGAA</td><td></td><td></td><td></td><td></td><td></td><td></td><td></td><td></td><td></td><td></td></tr>
<tr><td>Chromosome</td><td>start</td><td>end</td><td>strand</td><td>MM</td><td>target_seq</td><td>PAM</td><td>alignment</td><td>distance</td><td>position</td><td>gene name</td><td>gene id</td></tr>
<tr><td>GU305914.1</td><td>1318</td><td>1340</td><td>+</td><td>0</td><td>TTCGGTTGTCGCATGCCCAA</td><td>AGG</td><td>||||||||[||||||||||||]PAM</td><td>NA</td><td>Intergenic</td><td>NA</td><td>NA</td></tr>
<tr><td>T118</td><td>GCCCAAAGGAGTGTGTATCGAGG</td><td>1000</td><td></td><td></td><td></td><td></td><td></td><td></td><td></td><td></td><td></td></tr>
<tr><td>Oligo fwd</td><td>CACCGCCCAAAGGAGTGTGTATCG</td><td></td><td></td><td></td><td></td><td></td><td></td><td></td><td></td><td></td><td></td></tr>
<tr><td>Oligo rev</td><td>AAACCGATACACACTCCTTTGGGC</td><td></td><td></td><td></td><td></td><td></td><td></td><td></td><td></td><td></td><td></td></tr>
<tr><td>Chromosome</td><td>start</td><td>end</td><td>strand</td><td>MM</td><td>target_seq</td><td>PAM</td><td>alignment</td><td>distance</td><td>position</td><td>gene name</td><td>gene id</td></tr>
<tr><td>GU305914.1</td><td>1332</td><td>1354</td><td>+</td><td>0</td><td>GCCCAAAGGAGTGTGTATCG</td><td>AGG</td><td>||||||||[||||||||||||]PAM</td><td>NA</td><td>Intergenic</td><td>NA</td><td>NA</td></tr>
<tr><td>T119</td><td>CCCAAAGGAGTGTGTATCGAGGG</td><td>1000</td><td></td><td></td><td></td><td></td><td></td><td></td><td></td><td></td><td></td></tr>
<tr><td>Oligo fwd</td><td>CACCCCCAAAGGAGTGTGTATCGA</td><td></td><td></td><td></td><td></td><td></td><td></td><td></td><td></td><td></td><td></td></tr>
<tr><td>Oligo rev</td><td>AAACTCGATACACACTCCTTTGGG</td><td></td><td></td><td></td><td></td><td></td><td></td><td></td><td></td><td></td><td></td></tr>
<tr><td>Chromosome</td><td>start</td><td>end</td><td>strand</td><td>MM</td><td>target_seq</td><td>PAM</td><td>alignment</td><td>distance</td><td>position</td><td>gene name</td><td>gene id</td></tr>
<tr><td>GU305914.1</td><td>1333</td><td>1355</td><td>+</td><td>0</td><td>CCCAAAGGAGTGTGTATCGA</td><td>GGG</td><td>||||||||[||||||||||||]PAM</td><td>NA</td><td>Intergenic</td><td>NA</td><td>NA</td></tr>
<tr><td>T120</td><td>GAGTGTGTATCGAGGGCGAATGG</td><td>1000</td><td></td><td></td><td></td><td></td><td></td><td></td><td></td><td></td><td></td></tr>
<tr><td>Oligo fwd</td><td>CACCGAGTGTGTATCGAGGGCGAA</td><td></td><td></td><td></td><td></td><td></td><td></td><td></td><td></td><td></td><td></td></tr>
<tr><td>Oligo rev</td><td>AAACTTCGCCCTCGATACACACTC</td><td></td><td></td><td></td><td></td><td></td><td></td><td></td><td></td><td></td><td></td></tr>
<tr><td>Chromosome</td><td>start</td><td>end</td><td>strand</td><td>MM</td><td>target_seq</td><td>PAM</td><td>alignment</td><td>distance</td><td>position</td><td>gene name</td><td>gene id</td></tr>
<tr><td>GU305914.1</td><td>1340</td><td>1362</td><td>+</td><td>0</td><td>GAGTGTGTATCGAGGGCGAA</td><td>TGG</td><td>||||||||[||||||||||||]PAM</td><td>NA</td><td>Intergenic</td><td>NA</td><td>NA</td></tr>
<tr><td>T121</td><td>AGGGCGAATGGACGATCGAGTGG</td><td>1000</td><td></td><td></td><td></td><td></td><td></td><td></td><td></td><td></td><td></td></tr>
<tr><td>Oligo fwd</td><td>CACCAGGGCGAATGGACGATCGAG</td><td></td><td></td><td></td><td></td><td></td><td></td><td></td><td></td><td></td><td></td></tr>
<tr><td>Oligo rev</td><td>AAACCTCGATCGTCCATTCGCCCT</td><td></td><td></td><td></td><td></td><td></td><td></td><td></td><td></td><td></td><td></td></tr>
<tr><td>Chromosome</td><td>start</td><td>end</td><td>strand</td><td>MM</td><td>target_seq</td><td>PAM</td><td>alignment</td><td>distance</td><td>position</td><td>gene name</td><td>gene id</td></tr>
<tr><td>GU305914.1</td><td>1352</td><td>1374</td><td>+</td><td>0</td><td>AGGGCGAATGGACGATCGAG</td><td>TGG</td><td>||||||||[||||||||||||]PAM</td><td>NA</td><td>Intergenic</td><td>NA</td><td>NA</td></tr>
<tr><td>T122</td><td>GTGGTACATCCCCAGCCTACAGG</td><td>1000</td><td></td><td></td><td></td><td></td><td></td><td></td><td></td><td></td><td></td></tr>
<tr><td>Oligo fwd</td><td>CACCGTGGTACATCCCCAGCCTAC</td><td></td><td></td><td></td><td></td><td></td><td></td><td></td><td></td><td></td><td></td></tr>
<tr><td>Oligo rev</td><td>AAACGTAGGCTGGGGATGTACCAC</td><td></td><td></td><td></td><td></td><td></td><td></td><td></td><td></td><td></td><td></td></tr>
<tr><td>Chromosome</td><td>start</td><td>end</td><td>strand</td><td>MM</td><td>target_seq</td><td>PAM</td><td>alignment</td><td>distance</td><td>position</td><td>gene name</td><td>gene id</td></tr>
<tr><td>GU305914.1</td><td>1371</td><td>1393</td><td>+</td><td>0</td><td>GTGGTACATCCCCAGCCTAC</td><td>AGG</td><td>||||||||[||||||||||||]PAM</td><td>NA</td><td>Intergenic</td><td>NA</td><td>NA</td></tr>
<tr><td>T123</td><td>ACAGGCGTCCGTGATCAACCAGG</td><td>1000</td><td></td><td></td><td></td><td></td><td></td><td></td><td></td><td></td><td></td></tr>
<tr><td>Oligo fwd</td><td>CACCACAGGCGTCCGTGATCAACC</td><td></td><td></td><td></td><td></td><td></td><td></td><td></td><td></td><td></td><td></td></tr>
<tr><td>Oligo rev</td><td>AAACGGTTGATCACGGACGCCTGT</td><td></td><td></td><td></td><td></td><td></td><td></td><td></td><td></td><td></td><td></td></tr>
<tr><td>Chromosome</td><td>start</td><td>end</td><td>strand</td><td>MM</td><td>target_seq</td><td>PAM</td><td>alignment</td><td>distance</td><td>position</td><td>gene name</td><td>gene id</td></tr>
<tr><td>GU305914.1</td><td>1389</td><td>1411</td><td>+</td><td>0</td><td>ACAGGCGTCCGTGATCAACC</td><td>AGG</td><td>||||||||[||||||||||||]PAM</td><td>NA</td><td>Intergenic</td><td>NA</td><td>NA</td></tr>
<tr><td>T124</td><td>TCTTCTTTAAGTCACAGACGTGG</td><td>1000</td><td></td><td></td><td></td><td></td><td></td><td></td><td></td><td></td><td></td></tr>
<tr><td>Oligo fwd</td><td>CACCTCTTCTTTAAGTCACAGACG</td><td></td><td></td><td></td><td></td><td></td><td></td><td></td><td></td><td></td><td></td></tr>
<tr><td>Oligo rev</td><td>AAACCGTCTGTGACTTAAAGAAGA</td><td></td><td></td><td></td><td></td><td></td><td></td><td></td><td></td><td></td><td></td></tr>
<tr><td>Chromosome</td><td>start</td><td>end</td><td>strand</td><td>MM</td><td>target_seq</td><td>PAM</td><td>alignment</td><td>distance</td><td>position</td><td>gene name</td><td>gene id</td></tr>
<tr><td>GU305914.1</td><td>1412</td><td>1434</td><td>+</td><td>0</td><td>TCTTCTTTAAGTCACAGACG</td><td>TGG</td><td>||||||||[||||||||||||]PAM</td><td>NA</td><td>Intergenic</td><td>NA</td><td>NA</td></tr>
<tr><td>T125</td><td>CTTTAAGTCACAGACGTGGCTGG</td><td>1000</td><td></td><td></td><td></td><td></td><td></td><td></td><td></td><td></td><td></td></tr>
<tr><td>Oligo fwd</td><td>CACCCTTTAAGTCACAGACGTGGC</td><td></td><td></td><td></td><td></td><td></td><td></td><td></td><td></td><td></td><td></td></tr>
<tr><td>Oligo rev</td><td>AAACGCCACGTCTGTGACTTAAAG</td><td></td><td></td><td></td><td></td><td></td><td></td><td></td><td></td><td></td><td></td></tr>
<tr><td>Chromosome</td><td>start</td><td>end</td><td>strand</td><td>MM</td><td>target_seq</td><td>PAM</td><td>alignment</td><td>distance</td><td>position</td><td>gene name</td><td>gene id</td></tr>
<tr><td>GU305914.1</td><td>1416</td><td>1438</td><td>+</td><td>0</td><td>CTTTAAGTCACAGACGTGGC</td><td>TGG</td><td>||||||||[||||||||||||]PAM</td><td>NA</td><td>Intergenic</td><td>NA</td><td>NA</td></tr>
<tr><td>T126</td><td>TTTAAGTCACAGACGTGGCTGGG</td><td>1000</td><td></td><td></td><td></td><td></td><td></td><td></td><td></td><td></td><td></td></tr>
<tr><td>Oligo fwd</td><td>CACCTTTAAGTCACAGACGTGGCT</td><td></td><td></td><td></td><td></td><td></td><td></td><td></td><td></td><td></td><td></td></tr>
<tr><td>Oligo rev</td><td>AAACAGCCACGTCTGTGACTTAAA</td><td></td><td></td><td></td><td></td><td></td><td></td><td></td><td></td><td></td><td></td></tr>
<tr><td>Chromosome</td><td>start</td><td>end</td><td>strand</td><td>MM</td><td>target_seq</td><td>PAM</td><td>alignment</td><td>distance</td><td>position</td><td>gene name</td><td>gene id</td></tr>
<tr><td>GU305914.1</td><td>1417</td><td>1439</td><td>+</td><td>0</td><td>TTTAAGTCACAGACGTGGCT</td><td>GGG</td><td>||||||||[||||||||||||]PAM</td><td>NA</td><td>Intergenic</td><td>NA</td><td>NA</td></tr>
<tr><td>T127</td><td>ATCATACCGTCGTACGAGCGTGG</td><td>1000</td><td></td><td></td><td></td><td></td><td></td><td></td><td></td><td></td><td></td></tr>
<tr><td>Oligo fwd</td><td>CACCATCATACCGTCGTACGAGCG</td><td></td><td></td><td></td><td></td><td></td><td></td><td></td><td></td><td></td><td></td></tr>
<tr><td>Oligo rev</td><td>AAACCGCTCGTACGACGGTATGAT</td><td></td><td></td><td></td><td></td><td></td><td></td><td></td><td></td><td></td><td></td></tr>
<tr><td>Chromosome</td><td>start</td><td>end</td><td>strand</td><td>MM</td><td>target_seq</td><td>PAM</td><td>alignment</td><td>distance</td><td>position</td><td>gene name</td><td>gene id</td></tr>
<tr><td>GU305914.1</td><td>1456</td><td>1478</td><td>+</td><td>0</td><td>ATCATACCGTCGTACGAGCG</td><td>TGG</td><td>||||||||[||||||||||||]PAM</td><td>NA</td><td>Intergenic</td><td>NA</td><td>NA</td></tr>
<tr><td>T128</td><td>ACCGTCGTACGAGCGTGGTAAGG</td><td>1000</td><td></td><td></td><td></td><td></td><td></td><td></td><td></td><td></td><td></td></tr>
<tr><td>Oligo fwd</td><td>CACCACCGTCGTACGAGCGTGGTA</td><td></td><td></td><td></td><td></td><td></td><td></td><td></td><td></td><td></td><td></td></tr>
<tr><td>Oligo rev</td><td>AAACTACCACGCTCGTACGACGGT</td><td></td><td></td><td></td><td></td><td></td><td></td><td></td><td></td><td></td><td></td></tr>
<tr><td>Chromosome</td><td>start</td><td>end</td><td>strand</td><td>MM</td><td>target_seq</td><td>PAM</td><td>alignment</td><td>distance</td><td>position</td><td>gene name</td><td>gene id</td></tr>
<tr><td>GU305914.1</td><td>1461</td><td>1483</td><td>+</td><td>0</td><td>ACCGTCGTACGAGCGTGGTA</td><td>AGG</td><td>||||||||[||||||||||||]PAM</td><td>NA</td><td>Intergenic</td><td>NA</td><td>NA</td></tr>
<tr><td>T129</td><td>GTGGTAAGGAAGTGACCTGTAGG</td><td>1000</td><td></td><td></td><td></td><td></td><td></td><td></td><td></td><td></td><td></td></tr>
<tr><td>Oligo fwd</td><td>CACCGTGGTAAGGAAGTGACCTGT</td><td></td><td></td><td></td><td></td><td></td><td></td><td></td><td></td><td></td><td></td></tr>
<tr><td>Oligo rev</td><td>AAACACAGGTCACTTCCTTACCAC</td><td></td><td></td><td></td><td></td><td></td><td></td><td></td><td></td><td></td><td></td></tr>
<tr><td>Chromosome</td><td>start</td><td>end</td><td>strand</td><td>MM</td><td>target_seq</td><td>PAM</td><td>alignment</td><td>distance</td><td>position</td><td>gene name</td><td>gene id</td></tr>
<tr><td>GU305914.1</td><td>1475</td><td>1497</td><td>+</td><td>0</td><td>GTGGTAAGGAAGTGACCTGT</td><td>AGG</td><td>||||||||[||||||||||||]PAM</td><td>NA</td><td>Intergenic</td><td>NA</td><td>NA</td></tr>
<tr><td>T130</td><td>AAGGAAGTGACCTGTAGGCAAGG</td><td>1000</td><td></td><td></td><td></td><td></td><td></td><td></td><td></td><td></td><td></td></tr>
<tr><td>Oligo fwd</td><td>CACCAAGGAAGTGACCTGTAGGCA</td><td></td><td></td><td></td><td></td><td></td><td></td><td></td><td></td><td></td><td></td></tr>
<tr><td>Oligo rev</td><td>AAACTGCCTACAGGTCACTTCCTT</td><td></td><td></td><td></td><td></td><td></td><td></td><td></td><td></td><td></td><td></td></tr>
<tr><td>Chromosome</td><td>start</td><td>end</td><td>strand</td><td>MM</td><td>target_seq</td><td>PAM</td><td>alignment</td><td>distance</td><td>position</td><td>gene name</td><td>gene id</td></tr>
<tr><td>GU305914.1</td><td>1480</td><td>1502</td><td>+</td><td>0</td><td>AAGGAAGTGACCTGTAGGCA</td><td>AGG</td><td>||||||||[||||||||||||]PAM</td><td>NA</td><td>Intergenic</td><td>NA</td><td>NA</td></tr>
</table>

<table>
<tr><td>T131</td><td>TTCTGCGTCGACAGAGCAGAAGG</td><td>1000</td><td></td><td></td><td></td><td></td><td></td><td></td><td></td><td></td><td></td></tr>
<tr><td>Oligo fwd</td><td>CACCTTCTGCGTCGACAGAGCAGA</td><td></td><td></td><td></td><td></td><td></td><td></td><td></td><td></td><td></td><td></td></tr>
<tr><td>Oligo rev</td><td>AAACTCTGCTCTGTCGACGCAGAA</td><td></td><td></td><td></td><td></td><td></td><td></td><td></td><td></td><td></td><td></td></tr>
<tr><td>Chromosome</td><td>start</td><td>end</td><td>strand</td><td>MM</td><td>target_seq</td><td>PAM</td><td>alignment</td><td>distance</td><td>position</td><td>gene name</td><td>gene id</td></tr>
<tr><td>GU305914.1</td><td>1504</td><td>1526</td><td>+</td><td>0</td><td>TTCTGCGTCGACAGAGCAGA</td><td>AGG</td><td>||||||||[||||||||||||]PAM</td><td>NA</td><td>Intergenic</td><td>NA</td><td>NA</td></tr>
<tr><td>T132</td><td>TCATCACCGACAACAACACACGG</td><td>1000</td><td></td><td></td><td></td><td></td><td></td><td></td><td></td><td></td><td></td></tr>
<tr><td>Oligo fwd</td><td>CACCTCATCACCGACAACAACACA</td><td></td><td></td><td></td><td></td><td></td><td></td><td></td><td></td><td></td><td></td></tr>
<tr><td>Oligo rev</td><td>AAACTGTGTTGTTGTCGGTGATGA</td><td></td><td></td><td></td><td></td><td></td><td></td><td></td><td></td><td></td><td></td></tr>
<tr><td>Chromosome</td><td>start</td><td>end</td><td>strand</td><td>MM</td><td>target_seq</td><td>PAM</td><td>alignment</td><td>distance</td><td>position</td><td>gene name</td><td>gene id</td></tr>
<tr><td>GU305914.1</td><td>1535</td><td>1557</td><td>+</td><td>0</td><td>TCATCACCGACAACAACACA</td><td>CGG</td><td>||||||||[||||||||||||]PAM</td><td>NA</td><td>Intergenic</td><td>NA</td><td>NA</td></tr>
<tr><td>T133</td><td>ACAACACACGGAAAGAAGAGTGG</td><td>1000</td><td></td><td></td><td></td><td></td><td></td><td></td><td></td><td></td><td></td></tr>
<tr><td>Oligo fwd</td><td>CACCACAACACACGGAAAGAAGAG</td><td></td><td></td><td></td><td></td><td></td><td></td><td></td><td></td><td></td><td></td></tr>
<tr><td>Oligo rev</td><td>AAACCTCTTCTTTCCGTGTGTTGT</td><td></td><td></td><td></td><td></td><td></td><td></td><td></td><td></td><td></td><td></td></tr>
<tr><td>Chromosome</td><td>start</td><td>end</td><td>strand</td><td>MM</td><td>target_seq</td><td>PAM</td><td>alignment</td><td>distance</td><td>position</td><td>gene name</td><td>gene id</td></tr>
<tr><td>GU305914.1</td><td>1547</td><td>1569</td><td>+</td><td>0</td><td>ACAACACACGGAAAGAAGAG</td><td>TGG</td><td>||||||||[||||||||||||]PAM</td><td>NA</td><td>Intergenic</td><td>NA</td><td>NA</td></tr>
<tr><td>T134</td><td>CAACACACGGAAAGAAGAGTGGG</td><td>1000</td><td></td><td></td><td></td><td></td><td></td><td></td><td></td><td></td><td></td></tr>
<tr><td>Oligo fwd</td><td>CACCCAACACACGGAAAGAAGAGT</td><td></td><td></td><td></td><td></td><td></td><td></td><td></td><td></td><td></td><td></td></tr>
<tr><td>Oligo rev</td><td>AAACACTCTTCTTTCCGTGTGTTG</td><td></td><td></td><td></td><td></td><td></td><td></td><td></td><td></td><td></td><td></td></tr>
<tr><td>Chromosome</td><td>start</td><td>end</td><td>strand</td><td>MM</td><td>target_seq</td><td>PAM</td><td>alignment</td><td>distance</td><td>position</td><td>gene name</td><td>gene id</td></tr>
<tr><td>GU305914.1</td><td>1548</td><td>1570</td><td>+</td><td>0</td><td>CAACACACGGAAAGAAGAGT</td><td>GGG</td><td>||||||||[||||||||||||]PAM</td><td>NA</td><td>Intergenic</td><td>NA</td><td>NA</td></tr>
<tr><td>T135</td><td>ACGCAAGCCTACACGCGACGTGG</td><td>1000</td><td></td><td></td><td></td><td></td><td></td><td></td><td></td><td></td><td></td></tr>
<tr><td>Oligo fwd</td><td>CACCACGCAAGCCTACACGCGACG</td><td></td><td></td><td></td><td></td><td></td><td></td><td></td><td></td><td></td><td></td></tr>
<tr><td>Oligo rev</td><td>AAACCGTCGCGTGTAGGCTTGCGT</td><td></td><td></td><td></td><td></td><td></td><td></td><td></td><td></td><td></td><td></td></tr>
<tr><td>Chromosome</td><td>start</td><td>end</td><td>strand</td><td>MM</td><td>target_seq</td><td>PAM</td><td>alignment</td><td>distance</td><td>position</td><td>gene name</td><td>gene id</td></tr>
<tr><td>GU305914.1</td><td>1572</td><td>1594</td><td>+</td><td>0</td><td>ACGCAAGCCTACACGCGACG</td><td>TGG</td><td>||||||||[||||||||||||]PAM</td><td>NA</td><td>Intergenic</td><td>NA</td><td>NA</td></tr>
<tr><td>T136</td><td>CGACGTGGTGTGTAAGCTGACGG</td><td>1000</td><td></td><td></td><td></td><td></td><td></td><td></td><td></td><td></td><td></td></tr>
<tr><td>Oligo fwd</td><td>CACCCGACGTGGTGTGTAAGCTGA</td><td></td><td></td><td></td><td></td><td></td><td></td><td></td><td></td><td></td><td></td></tr>
<tr><td>Oligo rev</td><td>AAACTCAGCTTACACACCACGTCG</td><td></td><td></td><td></td><td></td><td></td><td></td><td></td><td></td><td></td><td></td></tr>
<tr><td>Chromosome</td><td>start</td><td>end</td><td>strand</td><td>MM</td><td>target_seq</td><td>PAM</td><td>alignment</td><td>distance</td><td>position</td><td>gene name</td><td>gene id</td></tr>
<tr><td>GU305914.1</td><td>1587</td><td>1609</td><td>+</td><td>0</td><td>CGACGTGGTGTGTAAGCTGA</td><td>CGG</td><td>||||||||[||||||||||||]PAM</td><td>NA</td><td>Intergenic</td><td>NA</td><td>NA</td></tr>
<tr><td>T137</td><td>GTATTCTCGCACAACATACGAGG</td><td>1000</td><td></td><td></td><td></td><td></td><td></td><td></td><td></td><td></td><td></td></tr>
<tr><td>Oligo fwd</td><td>CACCGTATTCTCGCACAACATACG</td><td></td><td></td><td></td><td></td><td></td><td></td><td></td><td></td><td></td><td></td></tr>
<tr><td>Oligo rev</td><td>AAACCGTATGTTGTGCGAGAATAC</td><td></td><td></td><td></td><td></td><td></td><td></td><td></td><td></td><td></td><td></td></tr>
<tr><td>Chromosome</td><td>start</td><td>end</td><td>strand</td><td>MM</td><td>target_seq</td><td>PAM</td><td>alignment</td><td>distance</td><td>position</td><td>gene name</td><td>gene id</td></tr>
<tr><td>GU305914.1</td><td>1638</td><td>1660</td><td>+</td><td>0</td><td>GTATTCTCGCACAACATACG</td><td>AGG</td><td>||||||||[||||||||||||]PAM</td><td>NA</td><td>Intergenic</td><td>NA</td><td>NA</td></tr>
<tr><td>T138</td><td>ACAACATACGAGGAATGTAACGG</td><td>1000</td><td></td><td></td><td></td><td></td><td></td><td></td><td></td><td></td><td></td></tr>
<tr><td>Oligo fwd</td><td>CACCACAACATACGAGGAATGTAA</td><td></td><td></td><td></td><td></td><td></td><td></td><td></td><td></td><td></td><td></td></tr>
<tr><td>Oligo rev</td><td>AAACTTACATTCCTCGTATGTTGT</td><td></td><td></td><td></td><td></td><td></td><td></td><td></td><td></td><td></td><td></td></tr>
<tr><td>Chromosome</td><td>start</td><td>end</td><td>strand</td><td>MM</td><td>target_seq</td><td>PAM</td><td>alignment</td><td>distance</td><td>position</td><td>gene name</td><td>gene id</td></tr>
<tr><td>GU305914.1</td><td>1648</td><td>1670</td><td>+</td><td>0</td><td>ACAACATACGAGGAATGTAA</td><td>CGG</td><td>||||||||[||||||||||||]PAM</td><td>NA</td><td>Intergenic</td><td>NA</td><td>NA</td></tr>
<tr><td>T139</td><td>CAACATACGAGGAATGTAACGGG</td><td>1000</td><td></td><td></td><td></td><td></td><td></td><td></td><td></td><td></td><td></td></tr>
<tr><td>Oligo fwd</td><td>CACCCAACATACGAGGAATGTAAC</td><td></td><td></td><td></td><td></td><td></td><td></td><td></td><td></td><td></td><td></td></tr>
<tr><td>Oligo rev</td><td>AAACGTTACATTCCTCGTATGTTG</td><td></td><td></td><td></td><td></td><td></td><td></td><td></td><td></td><td></td><td></td></tr>
<tr><td>Chromosome</td><td>start</td><td>end</td><td>strand</td><td>MM</td><td>target_seq</td><td>PAM</td><td>alignment</td><td>distance</td><td>position</td><td>gene name</td><td>gene id</td></tr>
<tr><td>GU305914.1</td><td>1649</td><td>1671</td><td>+</td><td>0</td><td>CAACATACGAGGAATGTAAC</td><td>GGG</td><td>||||||||[||||||||||||]PAM</td><td>NA</td><td>Intergenic</td><td>NA</td><td>NA</td></tr>
<tr><td>T140</td><td>CGAGGAATGTAACGGGACTCTGG</td><td>1000</td><td></td><td></td><td></td><td></td><td></td><td></td><td></td><td></td><td></td></tr>
<tr><td>Oligo fwd</td><td>CACCCGAGGAATGTAACGGGACTC</td><td></td><td></td><td></td><td></td><td></td><td></td><td></td><td></td><td></td><td></td></tr>
<tr><td>Oligo rev</td><td>AAACGAGTCCCGTTACATTCCTCG</td><td></td><td></td><td></td><td></td><td></td><td></td><td></td><td></td><td></td><td></td></tr>
<tr><td>Chromosome</td><td>start</td><td>end</td><td>strand</td><td>MM</td><td>target_seq</td><td>PAM</td><td>alignment</td><td>distance</td><td>position</td><td>gene name</td><td>gene id</td></tr>
<tr><td>GU305914.1</td><td>1656</td><td>1678</td><td>+</td><td>0</td><td>CGAGGAATGTAACGGGACTC</td><td>TGG</td><td>||||||||[||||||||||||]PAM</td><td>NA</td><td>Intergenic</td><td>NA</td><td>NA</td></tr>
<tr><td>T141</td><td>GGAATGTAACGGGACTCTGGAGG</td><td>1000</td><td></td><td></td><td></td><td></td><td></td><td></td><td></td><td></td><td></td></tr>
<tr><td>Oligo fwd</td><td>CACCGGAATGTAACGGGACTCTGG</td><td></td><td></td><td></td><td></td><td></td><td></td><td></td><td></td><td></td><td></td></tr>
<tr><td>Oligo rev</td><td>AAACCCAGAGTCCCGTTACATTCC</td><td></td><td></td><td></td><td></td><td></td><td></td><td></td><td></td><td></td><td></td></tr>
<tr><td>Chromosome</td><td>start</td><td>end</td><td>strand</td><td>MM</td><td>target_seq</td><td>PAM</td><td>alignment</td><td>distance</td><td>position</td><td>gene name</td><td>gene id</td></tr>
<tr><td>GU305914.1</td><td>1659</td><td>1681</td><td>+</td><td>0</td><td>GGAATGTAACGGGACTCTGG</td><td>AGG</td><td>||||||||[||||||||||||]PAM</td><td>NA</td><td>Intergenic</td><td>NA</td><td>NA</td></tr>
<tr><td>T142</td><td>GGAGGATTATCTAAGTTTACCGG</td><td>1000</td><td></td><td></td><td></td><td></td><td></td><td></td><td></td><td></td><td></td></tr>
<tr><td>Oligo fwd</td><td>CACCGGAGGATTATCTAAGTTTAC</td><td></td><td></td><td></td><td></td><td></td><td></td><td></td><td></td><td></td><td></td></tr>
<tr><td>Oligo rev</td><td>AAACGTAAACTTAGATAATCCTCC</td><td></td><td></td><td></td><td></td><td></td><td></td><td></td><td></td><td></td><td></td></tr>
<tr><td>Chromosome</td><td>start</td><td>end</td><td>strand</td><td>MM</td><td>target_seq</td><td>PAM</td><td>alignment</td><td>distance</td><td>position</td><td>gene name</td><td>gene id</td></tr>
<tr><td>GU305914.1</td><td>1677</td><td>1699</td><td>+</td><td>0</td><td>GGAGGATTATCTAAGTTTAC</td><td>CGG</td><td>||||||||[||||||||||||]PAM</td><td>NA</td><td>Intergenic</td><td>NA</td><td>NA</td></tr>
<tr><td>T143</td><td>GAACATATACGACATCTCGAAGG</td><td>1000</td><td></td><td></td><td></td><td></td><td></td><td></td><td></td><td></td><td></td></tr>
<tr><td>Oligo fwd</td><td>CACCGAACATATACGACATCTCGA</td><td></td><td></td><td></td><td></td><td></td><td></td><td></td><td></td><td></td><td></td></tr>
<tr><td>Oligo rev</td><td>AAACTCGAGATGTCGTATATGTTC</td><td></td><td></td><td></td><td></td><td></td><td></td><td></td><td></td><td></td><td></td></tr>
<tr><td>Chromosome</td><td>start</td><td>end</td><td>strand</td><td>MM</td><td>target_seq</td><td>PAM</td><td>alignment</td><td>distance</td><td>position</td><td>gene name</td><td>gene id</td></tr>
<tr><td>GU305914.1</td><td>1707</td><td>1729</td><td>+</td><td>0</td><td>GAACATATACGACATCTCGA</td><td>AGG</td><td>||||||||[||||||||||||]PAM</td><td>NA</td><td>Intergenic</td><td>NA</td><td>NA</td></tr>
<tr><td>T144</td><td>GCCGTCCCACCCAAAGTACCTGG</td><td>1000</td><td></td><td></td><td></td><td></td><td></td><td></td><td></td><td></td><td></td></tr>
<tr><td>Oligo fwd</td><td>CACCGCCGTCCCACCCAAAGTACC</td><td></td><td></td><td></td><td></td><td></td><td></td><td></td><td></td><td></td><td></td></tr>
<tr><td>Oligo rev</td><td>AAACGGTACTTTGGGTGGGACGGC</td><td></td><td></td><td></td><td></td><td></td><td></td><td></td><td></td><td></td><td></td></tr>
<tr><td>Chromosome</td><td>start</td><td>end</td><td>strand</td><td>MM</td><td>target_seq</td><td>PAM</td><td>alignment</td><td>distance</td><td>position</td><td>gene name</td><td>gene id</td></tr>
<tr><td>GU305914.1</td><td>1762</td><td>1784</td><td>+</td><td>0</td><td>GCCGTCCCACCCAAAGTACC</td><td>TGG</td><td>||||||||[||||||||||||]PAM</td><td>NA</td><td>Intergenic</td><td>NA</td><td>NA</td></tr>
<tr><td>T145</td><td>CCCACCCAAAGTACCTGGAAAGG</td><td>1000</td><td></td><td></td><td></td><td></td><td></td><td></td><td></td><td></td><td></td></tr>
<tr><td>Oligo fwd</td><td>CACCCCCACCCAAAGTACCTGGAA</td><td></td><td></td><td></td><td></td><td></td><td></td><td></td><td></td><td></td><td></td></tr>
<tr><td>Oligo rev</td><td>AAACTTCCAGGTACTTTGGGTGGG</td><td></td><td></td><td></td><td></td><td></td><td></td><td></td><td></td><td></td><td></td></tr>
<tr><td>Chromosome</td><td>start</td><td>end</td><td>strand</td><td>MM</td><td>target_seq</td><td>PAM</td><td>alignment</td><td>distance</td><td>position</td><td>gene name</td><td>gene id</td></tr>
</table>

GU305914.1 1767 1789 + 0 CCCACCCAAAGTACCTGGAA AGG ||||||||[||||||||||||]PAM NA Intergenic NA NA

T146 CAAAGTACCTGGAAAGGCCGAGG 1000
Oligo fwd CACCCAAAGTACCTGGAAAGGCCG
Oligo rev AAACCGGCCTTTCCAGGTACTTTG
Chromosome start end strand MM target_seq PAM alignment distance position gene name gene id
GU305914.1 1773 1795 + 0 CAAAGTACCTGGAAAGGCCG AGG ||||||||[||||||||||||]PAM NA Intergenic NA NA

T147 CCGCCAGACGAGTCGTGTATCGG 1000
Oligo fwd CACCCCGCCAGACGAGTCGTGTAT
Oligo rev AAACATACACGACTCGTCTGGCGG
Chromosome start end strand MM target_seq PAM alignment distance position gene name gene id
GU305914.1 1801 1823 + 0 CCGCCAGACGAGTCGTGTAT CGG ||||||||[||||||||||||]PAM NA Intergenic NA NA

T148 TGTATCGGATGCGACAATCCCGG 1000
Oligo fwd CACCTGTATCGGATGCGACAATCC
Oligo rev AAACGGATTGTCGCATCCGATACA
Chromosome start end strand MM target_seq PAM alignment distance position gene name gene id
GU305914.1 1816 1838 + 0 TGTATCGGATGCGACAATCC CGG ||||||||[||||||||||||]PAM NA Intergenic NA NA

T149 CGACAATCCCGGTCTGAACGCGG 1000
Oligo fwd CACCCGACAATCCCGGTCTGAACG
Oligo rev AAACCGTTCAGACCGGGATTGTCG
Chromosome start end strand MM target_seq PAM alignment distance position gene name gene id
GU305914.1 1827 1849 + 0 CGACAATCCCGGTCTGAACG CGG ||||||||[||||||||||||]PAM NA Intergenic NA NA

T150 CAATCCCGGTCTGAACGCGGCGG 1000
Oligo fwd CACCCAATCCCGGTCTGAACGCGG
Oligo rev AAACCCGCGTTCAGACCGGGATTG
Chromosome start end strand MM target_seq PAM alignment distance position gene name gene id
GU305914.1 1830 1852 + 0 CAATCCCGGTCTGAACGCGG CGG ||||||||[||||||||||||]PAM NA Intergenic NA NA

T151 TCTGAACGCGGCGGCGATCGCGG 1000
Oligo fwd CACCTCTGAACGCGGCGGCGATCG
Oligo rev AAACCGATCGCCGCCGCGTTCAGA
Chromosome start end strand MM target_seq PAM alignment distance position gene name gene id
GU305914.1 1839 1861 + 0 TCTGAACGCGGCGGCGATCG CGG ||||||||[||||||||||||]PAM NA Intergenic NA NA

T152 GGCGGCGATCGCGGTTCCCGTGG 1000
Oligo fwd CACCGGCGGCGATCGCGGTTCCCG
Oligo rev AAACCGGGAACCGCGATCGCCGCC
Chromosome start end strand MM target_seq PAM alignment distance position gene name gene id
GU305914.1 1848 1870 + 0 GGCGGCGATCGCGGTTCCCG TGG ||||||||[||||||||||||]PAM NA Intergenic NA NA

T153 GATCGCGGTTCCCGTGGTGACGG 1000
Oligo fwd CACCGATCGCGGTTCCCGTGGTGA
Oligo rev AAACTCACCACGGGAACCGCGATC
Chromosome start end strand MM target_seq PAM alignment distance position gene name gene id
GU305914.1 1854 1876 + 0 GATCGCGGTTCCCGTGGTGA CGG ||||||||[||||||||||||]PAM NA Intergenic NA NA

T154 CGTGGTGACGGTGATCGTGTTGG 1000
Oligo fwd CACCCGTGGTGACGGTGATCGTGT
Oligo rev AAACACACGATCACCGTCACCACG
Chromosome start end strand MM target_seq PAM alignment distance position gene name gene id
GU305914.1 1866 1888 + 0 CGTGGTGACGGTGATCGTGT TGG ||||||||[||||||||||||]PAM NA Intergenic NA NA

T155 ACGGTGATCGTGTTGGTGTCCGG 1000
Oligo fwd CACCACGGTGATCGTGTTGGTGTC
Oligo rev AAACGACACCAACACGATCACCGT
Chromosome start end strand MM target_seq PAM alignment distance position gene name gene id
GU305914.1 1873 1895 + 0 ACGGTGATCGTGTTGGTGTC CGG ||||||||[||||||||||||]PAM NA Intergenic NA NA

T156 ATCGTGTTGGTGTCCGGAATTGG 1000
Oligo fwd CACCATCGTGTTGGTGTCCGGAAT
Oligo rev AAACATTCCGGACACCAACACGAT
Chromosome start end strand MM target_seq PAM alignment distance position gene name gene id
GU305914.1 1879 1901 + 0 ATCGTGTTGGTGTCCGGAAT TGG ||||||||[||||||||||||]PAM NA Intergenic NA NA

T157 TCGTGTTGGTGTCCGGAATTGGG 1000
Oligo fwd CACCTCGTGTTGGTGTCCGGAATT
Oligo rev AAACAATTCCGGACACCAACACGA
Chromosome start end strand MM target_seq PAM alignment distance position gene name gene id
GU305914.1 1880 1902 + 0 TCGTGTTGGTGTCCGGAATT GGG ||||||||[||||||||||||]PAM NA Intergenic NA NA

T158 TGGGTATCTGTGCCGTTCCACGG 1000
Oligo fwd CACCTGGGTATCTGTGCCGTTCCA
Oligo rev AAACTGGAACGGCACAGATACCCA
Chromosome start end strand MM target_seq PAM alignment distance position gene name gene id
GU305914.1 1899 1921 + 0 TGGGTATCTGTGCCGTTCCA CGG ||||||||[||||||||||||]PAM NA Intergenic NA NA

T159 TGTGCCGTTCCACGGAGTCTCGG 1000
Oligo fwd CACCTGTGCCGTTCCACGGAGTCT
Oligo rev AAACAGACTCCGTGGAACGGCACA
Chromosome start end strand MM target_seq PAM alignment distance position gene name gene id
GU305914.1 1907 1929 + 0 TGTGCCGTTCCACGGAGTCT CGG ||||||||[||||||||||||]PAM NA Intergenic NA NA

T160 GTTCCACGGAGTCTCGGCAGCGG 1000
Oligo fwd CACCGTTCCACGGAGTCTCGGCAG
Oligo rev AAACCTGCCGAGACTCCGTGGAAC

<table>
<tr><td>Chromosome</td><td>start</td><td>end</td><td>strand</td><td>MM</td><td>target_seq</td><td>PAM</td><td>alignment</td><td>distance</td><td>position</td><td>gene name</td><td>gene id</td></tr>
<tr><td>GU305914.1</td><td>1913</td><td>1935</td><td>+</td><td>0</td><td>GTTCCACGGAGTCTCGGCAG</td><td>CGG</td><td>||||||||[||||||||||||]PAM</td><td>NA</td><td>Intergenic</td><td>NA</td><td>NA</td></tr>
<tr><td></td><td></td><td></td><td></td><td></td><td></td><td></td><td></td><td></td><td></td><td></td><td></td></tr>
<tr><td>T161</td><td>GGAGTCTCGGCAGCGGACGTTGG</td><td>1000</td><td></td><td></td><td></td><td></td><td></td><td></td><td></td><td></td><td></td></tr>
<tr><td>Oligo fwd</td><td>CACCGGAGTCTCGGCAGCGGACGT</td><td></td><td></td><td></td><td></td><td></td><td></td><td></td><td></td><td></td><td></td></tr>
<tr><td>Oligo rev</td><td>AAACACGTCCGCTGCCGAGACTCC</td><td></td><td></td><td></td><td></td><td></td><td></td><td></td><td></td><td></td><td></td></tr>
<tr><td>Chromosome</td><td>start</td><td>end</td><td>strand</td><td>MM</td><td>target_seq</td><td>PAM</td><td>alignment</td><td>distance</td><td>position</td><td>gene name</td><td>gene id</td></tr>
<tr><td>GU305914.1</td><td>1920</td><td>1942</td><td>+</td><td>0</td><td>GGAGTCTCGGCAGCGGACGT</td><td>TGG</td><td>||||||||[||||||||||||]PAM</td><td>NA</td><td>Intergenic</td><td>NA</td><td>NA</td></tr>
<tr><td></td><td></td><td></td><td></td><td></td><td></td><td></td><td></td><td></td><td></td><td></td><td></td></tr>
<tr><td>T162</td><td>TGGAGTTGTATCGAGATCTCTGG</td><td>1000</td><td></td><td></td><td></td><td></td><td></td><td></td><td></td><td></td><td></td></tr>
<tr><td>Oligo fwd</td><td>CACCTGGAGTTGTATCGAGATCTC</td><td></td><td></td><td></td><td></td><td></td><td></td><td></td><td></td><td></td><td></td></tr>
<tr><td>Oligo rev</td><td>AAACGAGATCTCGATACAACTCCA</td><td></td><td></td><td></td><td></td><td></td><td></td><td></td><td></td><td></td><td></td></tr>
<tr><td>Chromosome</td><td>start</td><td>end</td><td>strand</td><td>MM</td><td>target_seq</td><td>PAM</td><td>alignment</td><td>distance</td><td>position</td><td>gene name</td><td>gene id</td></tr>
<tr><td>GU305914.1</td><td>1940</td><td>1962</td><td>+</td><td>0</td><td>TGGAGTTGTATCGAGATCTC</td><td>TGG</td><td>||||||||[||||||||||||]PAM</td><td>NA</td><td>Intergenic</td><td>NA</td><td>NA</td></tr>
<tr><td></td><td></td><td></td><td></td><td></td><td></td><td></td><td></td><td></td><td></td><td></td><td></td></tr>
<tr><td>T163</td><td>GAGATCTCTGGTCCAGTCTGCGG</td><td>1000</td><td></td><td></td><td></td><td></td><td></td><td></td><td></td><td></td><td></td></tr>
<tr><td>Oligo fwd</td><td>CACCGAGATCTCTGGTCCAGTCTG</td><td></td><td></td><td></td><td></td><td></td><td></td><td></td><td></td><td></td><td></td></tr>
<tr><td>Oligo rev</td><td>AAACCAGACTGGACCAGAGATCTC</td><td></td><td></td><td></td><td></td><td></td><td></td><td></td><td></td><td></td><td></td></tr>
<tr><td>Chromosome</td><td>start</td><td>end</td><td>strand</td><td>MM</td><td>target_seq</td><td>PAM</td><td>alignment</td><td>distance</td><td>position</td><td>gene name</td><td>gene id</td></tr>
<tr><td>GU305914.1</td><td>1952</td><td>1974</td><td>+</td><td>0</td><td>GAGATCTCTGGTCCAGTCTG</td><td>CGG</td><td>||||||||[||||||||||||]PAM</td><td>NA</td><td>Intergenic</td><td>NA</td><td>NA</td></tr>
<tr><td></td><td></td><td></td><td></td><td></td><td></td><td></td><td></td><td></td><td></td><td></td><td></td></tr>
<tr><td>T164</td><td>CTGCGGCGTCGTCTCCACCGCGG</td><td>1000</td><td></td><td></td><td></td><td></td><td></td><td></td><td></td><td></td><td></td></tr>
<tr><td>Oligo fwd</td><td>CACCCTGCGGCGTCGTCTCCACCG</td><td></td><td></td><td></td><td></td><td></td><td></td><td></td><td></td><td></td><td></td></tr>
<tr><td>Oligo rev</td><td>AAACCGGTGGAGACGACGCCGCAG</td><td></td><td></td><td></td><td></td><td></td><td></td><td></td><td></td><td></td><td></td></tr>
<tr><td>Chromosome</td><td>start</td><td>end</td><td>strand</td><td>MM</td><td>target_seq</td><td>PAM</td><td>alignment</td><td>distance</td><td>position</td><td>gene name</td><td>gene id</td></tr>
<tr><td>GU305914.1</td><td>1969</td><td>1991</td><td>+</td><td>0</td><td>CTGCGGCGTCGTCTCCACCG</td><td>CGG</td><td>||||||||[||||||||||||]PAM</td><td>NA</td><td>Intergenic</td><td>NA</td><td>NA</td></tr>
<tr><td></td><td></td><td></td><td></td><td></td><td></td><td></td><td></td><td></td><td></td><td></td><td></td></tr>
<tr><td>T165</td><td>CGCGGAGACTACGCTCGAGACGG</td><td>1000</td><td></td><td></td><td></td><td></td><td></td><td></td><td></td><td></td><td></td></tr>
<tr><td>Oligo fwd</td><td>CACCCGCGGAGACTACGCTCGAGA</td><td></td><td></td><td></td><td></td><td></td><td></td><td></td><td></td><td></td><td></td></tr>
<tr><td>Oligo rev</td><td>AAACTCTCGAGCGTAGTCTCCGCG</td><td></td><td></td><td></td><td></td><td></td><td></td><td></td><td></td><td></td><td></td></tr>
<tr><td>Chromosome</td><td>start</td><td>end</td><td>strand</td><td>MM</td><td>target_seq</td><td>PAM</td><td>alignment</td><td>distance</td><td>position</td><td>gene name</td><td>gene id</td></tr>
<tr><td>GU305914.1</td><td>1987</td><td>2009</td><td>+</td><td>0</td><td>CGCGGAGACTACGCTCGAGA</td><td>CGG</td><td>||||||||[||||||||||||]PAM</td><td>NA</td><td>Intergenic</td><td>NA</td><td>NA</td></tr>
<tr><td></td><td></td><td></td><td></td><td></td><td></td><td></td><td></td><td></td><td></td><td></td><td></td></tr>
<tr><td>T166</td><td>ACTACGCTCGAGACGGTTGAAGG</td><td>1000</td><td></td><td></td><td></td><td></td><td></td><td></td><td></td><td></td><td></td></tr>
<tr><td>Oligo fwd</td><td>CACCACTACGCTCGAGACGGTTGA</td><td></td><td></td><td></td><td></td><td></td><td></td><td></td><td></td><td></td><td></td></tr>
<tr><td>Oligo rev</td><td>AAACTCAACCGTCTCGAGCGTAGT</td><td></td><td></td><td></td><td></td><td></td><td></td><td></td><td></td><td></td><td></td></tr>
<tr><td>Chromosome</td><td>start</td><td>end</td><td>strand</td><td>MM</td><td>target_seq</td><td>PAM</td><td>alignment</td><td>distance</td><td>position</td><td>gene name</td><td>gene id</td></tr>
<tr><td>GU305914.1</td><td>1994</td><td>2016</td><td>+</td><td>0</td><td>ACTACGCTCGAGACGGTTGA</td><td>AGG</td><td>||||||||[||||||||||||]PAM</td><td>NA</td><td>Intergenic</td><td>NA</td><td>NA</td></tr>
<tr><td></td><td></td><td></td><td></td><td></td><td></td><td></td><td></td><td></td><td></td><td></td><td></td></tr>
<tr><td>T167</td><td>CTACGCTCGAGACGGTTGAAGGG</td><td>1000</td><td></td><td></td><td></td><td></td><td></td><td></td><td></td><td></td><td></td></tr>
<tr><td>Oligo fwd</td><td>CACCCTACGCTCGAGACGGTTGAA</td><td></td><td></td><td></td><td></td><td></td><td></td><td></td><td></td><td></td><td></td></tr>
<tr><td>Oligo rev</td><td>AAACTTCAACCGTCTCGAGCGTAG</td><td></td><td></td><td></td><td></td><td></td><td></td><td></td><td></td><td></td><td></td></tr>
<tr><td>Chromosome</td><td>start</td><td>end</td><td>strand</td><td>MM</td><td>target_seq</td><td>PAM</td><td>alignment</td><td>distance</td><td>position</td><td>gene name</td><td>gene id</td></tr>
<tr><td>GU305914.1</td><td>1995</td><td>2017</td><td>+</td><td>0</td><td>CTACGCTCGAGACGGTTGAA</td><td>GGG</td><td>||||||||[||||||||||||]PAM</td><td>NA</td><td>Intergenic</td><td>NA</td><td>NA</td></tr>
<tr><td></td><td></td><td></td><td></td><td></td><td></td><td></td><td></td><td></td><td></td><td></td><td></td></tr>
<tr><td>T168</td><td>CGCTCGAGACGGTTGAAGGGTGG</td><td>1000</td><td></td><td></td><td></td><td></td><td></td><td></td><td></td><td></td><td></td></tr>
<tr><td>Oligo fwd</td><td>CACCCGCTCGAGACGGTTGAAGGG</td><td></td><td></td><td></td><td></td><td></td><td></td><td></td><td></td><td></td><td></td></tr>
<tr><td>Oligo rev</td><td>AAACCCCTTCAACCGTCTCGAGCG</td><td></td><td></td><td></td><td></td><td></td><td></td><td></td><td></td><td></td><td></td></tr>
<tr><td>Chromosome</td><td>start</td><td>end</td><td>strand</td><td>MM</td><td>target_seq</td><td>PAM</td><td>alignment</td><td>distance</td><td>position</td><td>gene name</td><td>gene id</td></tr>
<tr><td>GU305914.1</td><td>1998</td><td>2020</td><td>+</td><td>0</td><td>CGCTCGAGACGGTTGAAGGG</td><td>TGG</td><td>||||||||[||||||||||||]PAM</td><td>NA</td><td>Intergenic</td><td>NA</td><td>NA</td></tr>
<tr><td></td><td></td><td></td><td></td><td></td><td></td><td></td><td></td><td></td><td></td><td></td><td></td></tr>
<tr><td>T169</td><td>GACGTTATTCTCCTCTTGCGAGG</td><td>1000</td><td></td><td></td><td></td><td></td><td></td><td></td><td></td><td></td><td></td></tr>
<tr><td>Oligo fwd</td><td>CACCGACGTTATTCTCCTCTTGCG</td><td></td><td></td><td></td><td></td><td></td><td></td><td></td><td></td><td></td><td></td></tr>
<tr><td>Oligo rev</td><td>AAACCGCAAGAGGAGAATAACGTC</td><td></td><td></td><td></td><td></td><td></td><td></td><td></td><td></td><td></td><td></td></tr>
<tr><td>Chromosome</td><td>start</td><td>end</td><td>strand</td><td>MM</td><td>target_seq</td><td>PAM</td><td>alignment</td><td>distance</td><td>position</td><td>gene name</td><td>gene id</td></tr>
<tr><td>GU305914.1</td><td>2087</td><td>2109</td><td>+</td><td>0</td><td>GACGTTATTCTCCTCTTGCG</td><td>AGG</td><td>||||||||[||||||||||||]PAM</td><td>NA</td><td>Intergenic</td><td>NA</td><td>NA</td></tr>
<tr><td></td><td></td><td></td><td></td><td></td><td></td><td></td><td></td><td></td><td></td><td></td><td></td></tr>
<tr><td>T170</td><td>CTGCGCGCGTATGCCGAGCGTGG</td><td>1000</td><td></td><td></td><td></td><td></td><td></td><td></td><td></td><td></td><td></td></tr>
<tr><td>Oligo fwd</td><td>CACCCTGCGCGCGTATGCCGAGCG</td><td></td><td></td><td></td><td></td><td></td><td></td><td></td><td></td><td></td><td></td></tr>
<tr><td>Oligo rev</td><td>AAACCGCTCGGCATACGCGCGCAG</td><td></td><td></td><td></td><td></td><td></td><td></td><td></td><td></td><td></td><td></td></tr>
<tr><td>Chromosome</td><td>start</td><td>end</td><td>strand</td><td>MM</td><td>target_seq</td><td>PAM</td><td>alignment</td><td>distance</td><td>position</td><td>gene name</td><td>gene id</td></tr>
<tr><td>GU305914.1</td><td>2132</td><td>2154</td><td>+</td><td>0</td><td>CTGCGCGCGTATGCCGAGCG</td><td>TGG</td><td>||||||||[||||||||||||]PAM</td><td>NA</td><td>Intergenic</td><td>NA</td><td>NA</td></tr>
<tr><td></td><td></td><td></td><td></td><td></td><td></td><td></td><td></td><td></td><td></td><td></td><td></td></tr>
<tr><td>T171</td><td>TGCGCGCGTATGCCGAGCGTGGG</td><td>1000</td><td></td><td></td><td></td><td></td><td></td><td></td><td></td><td></td><td></td></tr>
<tr><td>Oligo fwd</td><td>CACCTGCGCGCGTATGCCGAGCGT</td><td></td><td></td><td></td><td></td><td></td><td></td><td></td><td></td><td></td><td></td></tr>
<tr><td>Oligo rev</td><td>AAACACGCTCGGCATACGCGCGCA</td><td></td><td></td><td></td><td></td><td></td><td></td><td></td><td></td><td></td><td></td></tr>
<tr><td>Chromosome</td><td>start</td><td>end</td><td>strand</td><td>MM</td><td>target_seq</td><td>PAM</td><td>alignment</td><td>distance</td><td>position</td><td>gene name</td><td>gene id</td></tr>
<tr><td>GU305914.1</td><td>2133</td><td>2155</td><td>+</td><td>0</td><td>TGCGCGCGTATGCCGAGCGT</td><td>GGG</td><td>||||||||[||||||||||||]PAM</td><td>NA</td><td>Intergenic</td><td>NA</td><td>NA</td></tr>
<tr><td></td><td></td><td></td><td></td><td></td><td></td><td></td><td></td><td></td><td></td><td></td><td></td></tr>
<tr><td>T172</td><td>CGAGCGTGGGAAGATGATGTCGG</td><td>1000</td><td></td><td></td><td></td><td></td><td></td><td></td><td></td><td></td><td></td></tr>
<tr><td>Oligo fwd</td><td>CACCCGAGCGTGGGAAGATGATGT</td><td></td><td></td><td></td><td></td><td></td><td></td><td></td><td></td><td></td><td></td></tr>
<tr><td>Oligo rev</td><td>AAACACATCATCTTCCCACGCTCG</td><td></td><td></td><td></td><td></td><td></td><td></td><td></td><td></td><td></td><td></td></tr>
<tr><td>Chromosome</td><td>start</td><td>end</td><td>strand</td><td>MM</td><td>target_seq</td><td>PAM</td><td>alignment</td><td>distance</td><td>position</td><td>gene name</td><td>gene id</td></tr>
<tr><td>GU305914.1</td><td>2146</td><td>2168</td><td>+</td><td>0</td><td>CGAGCGTGGGAAGATGATGT</td><td>CGG</td><td>||||||||[||||||||||||]PAM</td><td>NA</td><td>Intergenic</td><td>NA</td><td>NA</td></tr>
<tr><td></td><td></td><td></td><td></td><td></td><td></td><td></td><td></td><td></td><td></td><td></td><td></td></tr>
<tr><td>T173</td><td>AAGATGATGTCGGCCTTTATAGG</td><td>1000</td><td></td><td></td><td></td><td></td><td></td><td></td><td></td><td></td><td></td></tr>
<tr><td>Oligo fwd</td><td>CACCAAGATGATGTCGGCCTTTAT</td><td></td><td></td><td></td><td></td><td></td><td></td><td></td><td></td><td></td><td></td></tr>
<tr><td>Oligo rev</td><td>AAACATAAAGGCCGACATCATCTT</td><td></td><td></td><td></td><td></td><td></td><td></td><td></td><td></td><td></td><td></td></tr>
<tr><td>Chromosome</td><td>start</td><td>end</td><td>strand</td><td>MM</td><td>target_seq</td><td>PAM</td><td>alignment</td><td>distance</td><td>position</td><td>gene name</td><td>gene id</td></tr>
<tr><td>GU305914.1</td><td>2156</td><td>2178</td><td>+</td><td>0</td><td>AAGATGATGTCGGCCTTTAT</td><td>AGG</td><td>||||||||[||||||||||||]PAM</td><td>NA</td><td>Intergenic</td><td>NA</td><td>NA</td></tr>
<tr><td></td><td></td><td></td><td></td><td></td><td></td><td></td><td></td><td></td><td></td><td></td><td></td></tr>
<tr><td>T174</td><td>AGATGATGTCGGCCTTTATAGGG</td><td>1000</td><td></td><td></td><td></td><td></td><td></td><td></td><td></td><td></td><td></td></tr>
<tr><td>Oligo fwd</td><td>CACCAGATGATGTCGGCCTTTATA</td><td></td><td></td><td></td><td></td><td></td><td></td><td></td><td></td><td></td><td></td></tr>
<tr><td>Oligo rev</td><td>AAACTATAAAGGCCGACATCATCT</td><td></td><td></td><td></td><td></td><td></td><td></td><td></td><td></td><td></td><td></td></tr>
<tr><td>Chromosome</td><td>start</td><td>end</td><td>strand</td><td>MM</td><td>target_seq</td><td>PAM</td><td>alignment</td><td>distance</td><td>position</td><td>gene name</td><td>gene id</td></tr>
<tr><td>GU305914.1</td><td>2157</td><td>2179</td><td>+</td><td>0</td><td>AGATGATGTCGGCCTTTATA</td><td>GGG</td><td>||||||||[||||||||||||]PAM</td><td>NA</td><td>Intergenic</td><td>NA</td><td>NA</td></tr>
<tr><td></td><td></td><td></td><td></td><td></td><td></td><td></td><td></td><td></td><td></td><td></td><td></td></tr>
<tr><td>T175</td><td>CTACTCATTCACTCATCGCACGG</td><td>1000</td><td></td><td></td><td></td><td></td><td></td><td></td><td></td><td></td><td></td></tr>
<tr><td>Oligo fwd</td><td>CACCCTACTCATTCACTCATCGCA</td><td></td><td></td><td></td><td></td><td></td><td></td><td></td><td></td><td></td><td></td></tr>
</table>

<table>
<tr><td>Oligo rev</td><td>AAACTGCGATGAGTGAATGAGTAG</td><td></td><td></td><td></td><td></td><td></td><td></td><td></td><td></td><td></td><td></td></tr>
<tr><td>Chromosome</td><td>start</td><td>end</td><td>strand</td><td>MM</td><td>target_seq</td><td>PAM</td><td>alignment</td><td>distance</td><td>position</td><td>gene name</td><td>gene id</td></tr>
<tr><td>GU305914.1</td><td>2200</td><td>2222</td><td>+</td><td>0</td><td>CTACTCATTCACTCATCGCA</td><td>CGG</td><td>||||||||[||||||||||||]PAM</td><td>NA</td><td>Intergenic</td><td>NA</td><td>NA</td></tr>
<tr><td></td><td></td><td></td><td></td><td></td><td></td><td></td><td></td><td></td><td></td><td></td><td></td></tr>
<tr><td>T176</td><td>ATCATCAGCTCTCAGTGCTCCGG</td><td>1000</td><td></td><td></td><td></td><td></td><td></td><td></td><td></td><td></td><td></td></tr>
<tr><td>Oligo fwd</td><td>CACCATCATCAGCTCTCAGTGCTC</td><td></td><td></td><td></td><td></td><td></td><td></td><td></td><td></td><td></td><td></td></tr>
<tr><td>Oligo rev</td><td>AAACGAGCACTGAGAGCTGATGAT</td><td></td><td></td><td></td><td></td><td></td><td></td><td></td><td></td><td></td><td></td></tr>
<tr><td>Chromosome</td><td>start</td><td>end</td><td>strand</td><td>MM</td><td>target_seq</td><td>PAM</td><td>alignment</td><td>distance</td><td>position</td><td>gene name</td><td>gene id</td></tr>
<tr><td>GU305914.1</td><td>2232</td><td>2254</td><td>+</td><td>0</td><td>ATCATCAGCTCTCAGTGCTC</td><td>CGG</td><td>||||||||[||||||||||||]PAM</td><td>NA</td><td>Intergenic</td><td>NA</td><td>NA</td></tr>
<tr><td></td><td></td><td></td><td></td><td></td><td></td><td></td><td></td><td></td><td></td><td></td><td></td></tr>
<tr><td>T177</td><td>TAGAGAAGAAACATGACGATCGG</td><td>1000</td><td></td><td></td><td></td><td></td><td></td><td></td><td></td><td></td><td></td></tr>
<tr><td>Oligo fwd</td><td>CACCTAGAGAAGAAACATGACGAT</td><td></td><td></td><td></td><td></td><td></td><td></td><td></td><td></td><td></td><td></td></tr>
<tr><td>Oligo rev</td><td>AAACATCGTCATGTTTCTTCTCTA</td><td></td><td></td><td></td><td></td><td></td><td></td><td></td><td></td><td></td><td></td></tr>
<tr><td>Chromosome</td><td>start</td><td>end</td><td>strand</td><td>MM</td><td>target_seq</td><td>PAM</td><td>alignment</td><td>distance</td><td>position</td><td>gene name</td><td>gene id</td></tr>
<tr><td>GU305914.1</td><td>2258</td><td>2280</td><td>+</td><td>0</td><td>TAGAGAAGAAACATGACGAT</td><td>CGG</td><td>||||||||[||||||||||||]PAM</td><td>NA</td><td>Intergenic</td><td>NA</td><td>NA</td></tr>
<tr><td></td><td></td><td></td><td></td><td></td><td></td><td></td><td></td><td></td><td></td><td></td><td></td></tr>
<tr><td>T178</td><td>ATGTACGTCTCCGTTTGACACGG</td><td>1000</td><td></td><td></td><td></td><td></td><td></td><td></td><td></td><td></td><td></td></tr>
<tr><td>Oligo fwd</td><td>CACCATGTACGTCTCCGTTTGACA</td><td></td><td></td><td></td><td></td><td></td><td></td><td></td><td></td><td></td><td></td></tr>
<tr><td>Oligo rev</td><td>AAACTGTCAAACGGAGACGTACAT</td><td></td><td></td><td></td><td></td><td></td><td></td><td></td><td></td><td></td><td></td></tr>
<tr><td>Chromosome</td><td>start</td><td>end</td><td>strand</td><td>MM</td><td>target_seq</td><td>PAM</td><td>alignment</td><td>distance</td><td>position</td><td>gene name</td><td>gene id</td></tr>
<tr><td>GU305914.1</td><td>2304</td><td>2326</td><td>+</td><td>0</td><td>ATGTACGTCTCCGTTTGACA</td><td>CGG</td><td>||||||||[||||||||||||]PAM</td><td>NA</td><td>Intergenic</td><td>NA</td><td>NA</td></tr>
<tr><td></td><td></td><td></td><td></td><td></td><td></td><td></td><td></td><td></td><td></td><td></td><td></td></tr>
<tr><td>T179</td><td>GCTCAAGAATGTCCCAATTACGG</td><td>1000</td><td></td><td></td><td></td><td></td><td></td><td></td><td></td><td></td><td></td></tr>
<tr><td>Oligo fwd</td><td>CACCGCTCAAGAATGTCCCAATTA</td><td></td><td></td><td></td><td></td><td></td><td></td><td></td><td></td><td></td><td></td></tr>
<tr><td>Oligo rev</td><td>AAACTAATTGGGACATTCTTGAGC</td><td></td><td></td><td></td><td></td><td></td><td></td><td></td><td></td><td></td><td></td></tr>
<tr><td>Chromosome</td><td>start</td><td>end</td><td>strand</td><td>MM</td><td>target_seq</td><td>PAM</td><td>alignment</td><td>distance</td><td>position</td><td>gene name</td><td>gene id</td></tr>
<tr><td>GU305914.1</td><td>2366</td><td>2388</td><td>+</td><td>0</td><td>GCTCAAGAATGTCCCAATTA</td><td>CGG</td><td>||||||||[||||||||||||]PAM</td><td>NA</td><td>Intergenic</td><td>NA</td><td>NA</td></tr>
<tr><td></td><td></td><td></td><td></td><td></td><td></td><td></td><td></td><td></td><td></td><td></td><td></td></tr>
<tr><td>T180</td><td>CAAGAATGTCCCAATTACGGTGG</td><td>1000</td><td></td><td></td><td></td><td></td><td></td><td></td><td></td><td></td><td></td></tr>
<tr><td>Oligo fwd</td><td>CACCCAAGAATGTCCCAATTACGG</td><td></td><td></td><td></td><td></td><td></td><td></td><td></td><td></td><td></td><td></td></tr>
<tr><td>Oligo rev</td><td>AAACCCGTAATTGGGACATTCTTG</td><td></td><td></td><td></td><td></td><td></td><td></td><td></td><td></td><td></td><td></td></tr>
<tr><td>Chromosome</td><td>start</td><td>end</td><td>strand</td><td>MM</td><td>target_seq</td><td>PAM</td><td>alignment</td><td>distance</td><td>position</td><td>gene name</td><td>gene id</td></tr>
<tr><td>GU305914.1</td><td>2369</td><td>2391</td><td>+</td><td>0</td><td>CAAGAATGTCCCAATTACGG</td><td>TGG</td><td>||||||||[||||||||||||]PAM</td><td>NA</td><td>Intergenic</td><td>NA</td><td>NA</td></tr>
<tr><td></td><td></td><td></td><td></td><td></td><td></td><td></td><td></td><td></td><td></td><td></td><td></td></tr>
<tr><td>T181</td><td>AAAAGACAGATGACAGCGTTTGG</td><td>1000</td><td></td><td></td><td></td><td></td><td></td><td></td><td></td><td></td><td></td></tr>
<tr><td>Oligo fwd</td><td>CACCAAAAGACAGATGACAGCGTT</td><td></td><td></td><td></td><td></td><td></td><td></td><td></td><td></td><td></td><td></td></tr>
<tr><td>Oligo rev</td><td>AAACAACGCTGTCATCTGTCTTTT</td><td></td><td></td><td></td><td></td><td></td><td></td><td></td><td></td><td></td><td></td></tr>
<tr><td>Chromosome</td><td>start</td><td>end</td><td>strand</td><td>MM</td><td>target_seq</td><td>PAM</td><td>alignment</td><td>distance</td><td>position</td><td>gene name</td><td>gene id</td></tr>
<tr><td>GU305914.1</td><td>2394</td><td>2416</td><td>+</td><td>0</td><td>AAAAGACAGATGACAGCGTT</td><td>TGG</td><td>||||||||[||||||||||||]PAM</td><td>NA</td><td>Intergenic</td><td>NA</td><td>NA</td></tr>
<tr><td></td><td></td><td></td><td></td><td></td><td></td><td></td><td></td><td></td><td></td><td></td><td></td></tr>
<tr><td>T182</td><td>TGGAAGTACCGTAAAGACCTCGG</td><td>1000</td><td></td><td></td><td></td><td></td><td></td><td></td><td></td><td></td><td></td></tr>
<tr><td>Oligo fwd</td><td>CACCTGGAAGTACCGTAAAGACCT</td><td></td><td></td><td></td><td></td><td></td><td></td><td></td><td></td><td></td><td></td></tr>
<tr><td>Oligo rev</td><td>AAACAGGTCTTTACGGTACTTCCA</td><td></td><td></td><td></td><td></td><td></td><td></td><td></td><td></td><td></td><td></td></tr>
<tr><td>Chromosome</td><td>start</td><td>end</td><td>strand</td><td>MM</td><td>target_seq</td><td>PAM</td><td>alignment</td><td>distance</td><td>position</td><td>gene name</td><td>gene id</td></tr>
<tr><td>GU305914.1</td><td>2414</td><td>2436</td><td>+</td><td>0</td><td>TGGAAGTACCGTAAAGACCT</td><td>CGG</td><td>||||||||[||||||||||||]PAM</td><td>NA</td><td>Intergenic</td><td>NA</td><td>NA</td></tr>
<tr><td></td><td></td><td></td><td></td><td></td><td></td><td></td><td></td><td></td><td></td><td></td><td></td></tr>
<tr><td>T183</td><td>CATCACCTTGTACGAAAATAAGG</td><td>1000</td><td></td><td></td><td></td><td></td><td></td><td></td><td></td><td></td><td></td></tr>
<tr><td>Oligo fwd</td><td>CACCCATCACCTTGTACGAAAATA</td><td></td><td></td><td></td><td></td><td></td><td></td><td></td><td></td><td></td><td></td></tr>
<tr><td>Oligo rev</td><td>AAACTATTTTCGTACAAGGTGATG</td><td></td><td></td><td></td><td></td><td></td><td></td><td></td><td></td><td></td><td></td></tr>
<tr><td>Chromosome</td><td>start</td><td>end</td><td>strand</td><td>MM</td><td>target_seq</td><td>PAM</td><td>alignment</td><td>distance</td><td>position</td><td>gene name</td><td>gene id</td></tr>
<tr><td>GU305914.1</td><td>2461</td><td>2483</td><td>+</td><td>0</td><td>CATCACCTTGTACGAAAATA</td><td>AGG</td><td>||||||||[||||||||||||]PAM</td><td>NA</td><td>Intergenic</td><td>NA</td><td>NA</td></tr>
<tr><td></td><td></td><td></td><td></td><td></td><td></td><td></td><td></td><td></td><td></td><td></td><td></td></tr>
<tr><td>T184</td><td>ATCACCTTGTACGAAAATAAGGG</td><td>1000</td><td></td><td></td><td></td><td></td><td></td><td></td><td></td><td></td><td></td></tr>
<tr><td>Oligo fwd</td><td>CACCATCACCTTGTACGAAAATAA</td><td></td><td></td><td></td><td></td><td></td><td></td><td></td><td></td><td></td><td></td></tr>
<tr><td>Oligo rev</td><td>AAACTTATTTTCGTACAAGGTGAT</td><td></td><td></td><td></td><td></td><td></td><td></td><td></td><td></td><td></td><td></td></tr>
<tr><td>Chromosome</td><td>start</td><td>end</td><td>strand</td><td>MM</td><td>target_seq</td><td>PAM</td><td>alignment</td><td>distance</td><td>position</td><td>gene name</td><td>gene id</td></tr>
<tr><td>GU305914.1</td><td>2462</td><td>2484</td><td>+</td><td>0</td><td>ATCACCTTGTACGAAAATAA</td><td>GGG</td><td>||||||||[||||||||||||]PAM</td><td>NA</td><td>Intergenic</td><td>NA</td><td>NA</td></tr>
<tr><td></td><td></td><td></td><td></td><td></td><td></td><td></td><td></td><td></td><td></td><td></td><td></td></tr>
<tr><td>T185</td><td>CGAAAATAAGGGTATTAAGACGG</td><td>1000</td><td></td><td></td><td></td><td></td><td></td><td></td><td></td><td></td><td></td></tr>
<tr><td>Oligo fwd</td><td>CACCCGAAAATAAGGGTATTAAGA</td><td></td><td></td><td></td><td></td><td></td><td></td><td></td><td></td><td></td><td></td></tr>
<tr><td>Oligo rev</td><td>AAACTCTTAATACCCTTATTTTCG</td><td></td><td></td><td></td><td></td><td></td><td></td><td></td><td></td><td></td><td></td></tr>
<tr><td>Chromosome</td><td>start</td><td>end</td><td>strand</td><td>MM</td><td>target_seq</td><td>PAM</td><td>alignment</td><td>distance</td><td>position</td><td>gene name</td><td>gene id</td></tr>
<tr><td>GU305914.1</td><td>2473</td><td>2495</td><td>+</td><td>0</td><td>CGAAAATAAGGGTATTAAGA</td><td>CGG</td><td>||||||||[||||||||||||]PAM</td><td>NA</td><td>Intergenic</td><td>NA</td><td>NA</td></tr>
<tr><td></td><td></td><td></td><td></td><td></td><td></td><td></td><td></td><td></td><td></td><td></td><td></td></tr>
<tr><td>T186</td><td>ACGGCATATTTCATTCTTCCCGG</td><td>1000</td><td></td><td></td><td></td><td></td><td></td><td></td><td></td><td></td><td></td></tr>
<tr><td>Oligo fwd</td><td>CACCACGGCATATTTCATTCTTCC</td><td></td><td></td><td></td><td></td><td></td><td></td><td></td><td></td><td></td><td></td></tr>
<tr><td>Oligo rev</td><td>AAACGGAAGAATGAAATATGCCGT</td><td></td><td></td><td></td><td></td><td></td><td></td><td></td><td></td><td></td><td></td></tr>
<tr><td>Chromosome</td><td>start</td><td>end</td><td>strand</td><td>MM</td><td>target_seq</td><td>PAM</td><td>alignment</td><td>distance</td><td>position</td><td>gene name</td><td>gene id</td></tr>
<tr><td>GU305914.1</td><td>2492</td><td>2514</td><td>+</td><td>0</td><td>ACGGCATATTTCATTCTTCC</td><td>CGG</td><td>||||||||[||||||||||||]PAM</td><td>NA</td><td>Intergenic</td><td>NA</td><td>NA</td></tr>
<tr><td></td><td></td><td></td><td></td><td></td><td></td><td></td><td></td><td></td><td></td><td></td><td></td></tr>
<tr><td>T187</td><td>CAGTCTATGTCAACGCTTCATGG</td><td>1000</td><td></td><td></td><td></td><td></td><td></td><td></td><td></td><td></td><td></td></tr>
<tr><td>Oligo fwd</td><td>CACCCAGTCTATGTCAACGCTTCA</td><td></td><td></td><td></td><td></td><td></td><td></td><td></td><td></td><td></td><td></td></tr>
<tr><td>Oligo rev</td><td>AAACTGAAGCGTTGACATAGACTG</td><td></td><td></td><td></td><td></td><td></td><td></td><td></td><td></td><td></td><td></td></tr>
<tr><td>Chromosome</td><td>start</td><td>end</td><td>strand</td><td>MM</td><td>target_seq</td><td>PAM</td><td>alignment</td><td>distance</td><td>position</td><td>gene name</td><td>gene id</td></tr>
<tr><td>GU305914.1</td><td>2523</td><td>2545</td><td>+</td><td>0</td><td>CAGTCTATGTCAACGCTTCA</td><td>TGG</td><td>||||||||[||||||||||||]PAM</td><td>NA</td><td>Intergenic</td><td>NA</td><td>NA</td></tr>
<tr><td></td><td></td><td></td><td></td><td></td><td></td><td></td><td></td><td></td><td></td><td></td><td></td></tr>
<tr><td>T188</td><td>ACGCTTCATGGACCTTTGAATGG</td><td>1000</td><td></td><td></td><td></td><td></td><td></td><td></td><td></td><td></td><td></td></tr>
<tr><td>Oligo fwd</td><td>CACCACGCTTCATGGACCTTTGAA</td><td></td><td></td><td></td><td></td><td></td><td></td><td></td><td></td><td></td><td></td></tr>
<tr><td>Oligo rev</td><td>AAACTTCAAAGGTCCATGAAGCGT</td><td></td><td></td><td></td><td></td><td></td><td></td><td></td><td></td><td></td><td></td></tr>
<tr><td>Chromosome</td><td>start</td><td>end</td><td>strand</td><td>MM</td><td>target_seq</td><td>PAM</td><td>alignment</td><td>distance</td><td>position</td><td>gene name</td><td>gene id</td></tr>
<tr><td>GU305914.1</td><td>2535</td><td>2557</td><td>+</td><td>0</td><td>ACGCTTCATGGACCTTTGAA</td><td>TGG</td><td>||||||||[||||||||||||]PAM</td><td>NA</td><td>Intergenic</td><td>NA</td><td>NA</td></tr>
<tr><td>GU305914.1</td><td>3493</td><td>3515</td><td>+</td><td>3</td><td>ATGCTTCGTGGACCTTGGAA</td><td>TGG</td><td>|-|||||-[||||||||-|||]PAM</td><td>NA</td><td>Intergenic</td><td>NA</td><td>NA</td></tr>
<tr><td></td><td></td><td></td><td></td><td></td><td></td><td></td><td></td><td></td><td></td><td></td><td></td></tr>
<tr><td>T189</td><td>CATCAAACCGCTCTTCGCCTCGG</td><td>1000</td><td></td><td></td><td></td><td></td><td></td><td></td><td></td><td></td><td></td></tr>
<tr><td>Oligo fwd</td><td>CACCCATCAAACCGCTCTTCGCCT</td><td></td><td></td><td></td><td></td><td></td><td></td><td></td><td></td><td></td><td></td></tr>
<tr><td>Oligo rev</td><td>AAACAGGCGAAGAGCGGTTTGATG</td><td></td><td></td><td></td><td></td><td></td><td></td><td></td><td></td><td></td><td></td></tr>
<tr><td>Chromosome</td><td>start</td><td>end</td><td>strand</td><td>MM</td><td>target_seq</td><td>PAM</td><td>alignment</td><td>distance</td><td>position</td><td>gene name</td><td>gene id</td></tr>
<tr><td>GU305914.1</td><td>2560</td><td>2582</td><td>+</td><td>0</td><td>CATCAAACCGCTCTTCGCCT</td><td>CGG</td><td>||||||||[||||||||||||]PAM</td><td>NA</td><td>Intergenic</td><td>NA</td><td>NA</td></tr>
</table>

| | | | | | | | | | | | | |
|---|---|---|---|---|---|---|---|---|---|---|---|---|
| T190 | TTTTACCTCGACGACCACTTCGG | | 1000 | | | | | | | | | |
| Oligo fwd | CACCTTTTACCTCGACGACCACTT | | | | | | | | | | | |
| Oligo rev | AAACAAGTGGTCGTCGAGGTAAAA | | | | | | | | | | | |
| Chromosome | start | | end | strand | MM | target_seq | PAM | alignment | distance | position | gene name | gene id |
| GU305914.1 | | 2599 | 2621 | + | 0 | TTTTACCTCGACGACCACTT | CGG | \|\|\|\|\|\|\|\|[\|\|\|\|\|\|\|\|\|\|\|\|]PAM | NA | Intergenic | NA | NA |
| T191 | TTCTTACTATGATATGCATTCGG | | 1000 | | | | | | | | | |
| Oligo fwd | CACCTTCTTACTATGATATGCATT | | | | | | | | | | | |
| Oligo rev | AAACAATGCATATCATAGTAAGAA | | | | | | | | | | | |
| Chromosome | start | | end | strand | MM | target_seq | PAM | alignment | distance | position | gene name | gene id |
| GU305914.1 | | 2656 | 2678 | + | 0 | TTCTTACTATGATATGCATT | CGG | \|\|\|\|\|\|\|\|[\|\|\|\|\|\|\|\|\|\|\|\|]PAM | NA | Intergenic | NA | NA |
| T192 | TCTTACTATGATATGCATTCGGG | | 1000 | | | | | | | | | |
| Oligo fwd | CACCTCTTACTATGATATGCATTC | | | | | | | | | | | |
| Oligo rev | AAACGAATGCATATCATAGTAAGA | | | | | | | | | | | |
| Chromosome | start | | end | strand | MM | target_seq | PAM | alignment | distance | position | gene name | gene id |
| GU305914.1 | | 2657 | 2679 | + | 0 | TCTTACTATGATATGCATTC | GGG | \|\|\|\|\|\|\|\|[\|\|\|\|\|\|\|\|\|\|\|\|]PAM | NA | Intergenic | NA | NA |
| T193 | GGGTAGACTGAAGAAGACCGTGG | | 1000 | | | | | | | | | |
| Oligo fwd | CACCGGGTAGACTGAAGAAGACCG | | | | | | | | | | | |
| Oligo rev | AAACCGGTCTTCTTCAGTCTACCC | | | | | | | | | | | |
| Chromosome | start | | end | strand | MM | target_seq | PAM | alignment | distance | position | gene name | gene id |
| GU305914.1 | | 2677 | 2699 | + | 0 | GGGTAGACTGAAGAAGACCG | TGG | \|\|\|\|\|\|\|\|[\|\|\|\|\|\|\|\|\|\|\|\|]PAM | NA | Intergenic | NA | NA |
| T194 | GGTAGACTGAAGAAGACCGTGGG | | 1000 | | | | | | | | | |
| Oligo fwd | CACCGGTAGACTGAAGAAGACCGT | | | | | | | | | | | |
| Oligo rev | AAACACGGTCTTCTTCAGTCTACC | | | | | | | | | | | |
| Chromosome | start | | end | strand | MM | target_seq | PAM | alignment | distance | position | gene name | gene id |
| GU305914.1 | | 2678 | 2700 | + | 0 | GGTAGACTGAAGAAGACCGT | GGG | \|\|\|\|\|\|\|\|[\|\|\|\|\|\|\|\|\|\|\|\|]PAM | NA | Intergenic | NA | NA |
| T195 | AAGACCGTGGGATTAAAAGACGG | | 1000 | | | | | | | | | |
| Oligo fwd | CACCAAGACCGTGGGATTAAAAGA | | | | | | | | | | | |
| Oligo rev | AAACTCTTTTAATCCCACGGTCTT | | | | | | | | | | | |
| Chromosome | start | | end | strand | MM | target_seq | PAM | alignment | distance | position | gene name | gene id |
| GU305914.1 | | 2690 | 2712 | + | 0 | AAGACCGTGGGATTAAAAGA | CGG | \|\|\|\|\|\|\|\|[\|\|\|\|\|\|\|\|\|\|\|\|]PAM | NA | Intergenic | NA | NA |
| T196 | GGTTTCAAAGCCGATCATTCAGG | | 1000 | | | | | | | | | |
| Oligo fwd | CACCGGTTTCAAAGCCGATCATTC | | | | | | | | | | | |
| Oligo rev | AAACGAATGATCGGCTTTGAAACC | | | | | | | | | | | |
| Chromosome | start | | end | strand | MM | target_seq | PAM | alignment | distance | position | gene name | gene id |
| GU305914.1 | | 2711 | 2733 | + | 0 | GGTTTCAAAGCCGATCATTC | AGG | \|\|\|\|\|\|\|\|[\|\|\|\|\|\|\|\|\|\|\|\|]PAM | NA | Intergenic | NA | NA |
| T197 | CATCTTTCAGAACGCGAGCGTGG | | 1000 | | | | | | | | | |
| Oligo fwd | CACCCATCTTTCAGAACGCGAGCG | | | | | | | | | | | |
| Oligo rev | AAACCGCTCGCGTTCTGAAAGATG | | | | | | | | | | | |
| Chromosome | start | | end | strand | MM | target_seq | PAM | alignment | distance | position | gene name | gene id |
| GU305914.1 | | 2743 | 2765 | + | 0 | CATCTTTCAGAACGCGAGCG | TGG | \|\|\|\|\|\|\|\|[\|\|\|\|\|\|\|\|\|\|\|\|]PAM | NA | Intergenic | NA | NA |
| GU305914.1 | | 136753 | 136775 | + | 3 | CATCTACCAGAACGCGAGTG | TGG | \|\|\|\|\|--\|[\|\|\|\|\|\|\|\|\|\|-\|]PAM | NA | Intergenic | NA | NA |
| T198 | TTTCAGAACGCGAGCGTGGCAGG | | 1000 | | | | | | | | | |
| Oligo fwd | CACCTTTCAGAACGCGAGCGTGGC | | | | | | | | | | | |
| Oligo rev | AAACGCCACGCTCGCGTTCTGAAA | | | | | | | | | | | |
| Chromosome | start | | end | strand | MM | target_seq | PAM | alignment | distance | position | gene name | gene id |
| GU305914.1 | | 2747 | 2769 | + | 0 | TTTCAGAACGCGAGCGTGGC | AGG | \|\|\|\|\|\|\|\|[\|\|\|\|\|\|\|\|\|\|\|\|]PAM | NA | Intergenic | NA | NA |
| T199 | GAGCGTGGCAGGAAACATCACGG | | 1000 | | | | | | | | | |
| Oligo fwd | CACCGAGCGTGGCAGGAAACATCA | | | | | | | | | | | |
| Oligo rev | AAACTGATGTTTCCTGCCACGCTC | | | | | | | | | | | |
| Chromosome | start | | end | strand | MM | target_seq | PAM | alignment | distance | position | gene name | gene id |
| GU305914.1 | | 2758 | 2780 | + | 0 | GAGCGTGGCAGGAAACATCA | CGG | \|\|\|\|\|\|\|\|[\|\|\|\|\|\|\|\|\|\|\|\|]PAM | NA | Intergenic | NA | NA |
| T200 | ACGATACCATATGTGTACTAAGG | | 1000 | | | | | | | | | |
| Oligo fwd | CACCACGATACCATATGTGTACTA | | | | | | | | | | | |
| Oligo rev | AAACTAGTACACATATGGTATCGT | | | | | | | | | | | |
| Chromosome | start | | end | strand | MM | target_seq | PAM | alignment | distance | position | gene name | gene id |
| GU305914.1 | | 2784 | 2806 | + | 0 | ACGATACCATATGTGTACTA | AGG | \|\|\|\|\|\|\|\|[\|\|\|\|\|\|\|\|\|\|\|\|]PAM | NA | Intergenic | NA | NA |
| T201 | GTGTACTAAGGCTGTGTCTCTGG | | 1000 | | | | | | | | | |
| Oligo fwd | CACCGTGTACTAAGGCTGTGTCTC | | | | | | | | | | | |
| Oligo rev | AAACGAGACACAGCCTTAGTACAC | | | | | | | | | | | |
| Chromosome | start | | end | strand | MM | target_seq | PAM | alignment | distance | position | gene name | gene id |
| GU305914.1 | | 2796 | 2818 | + | 0 | GTGTACTAAGGCTGTGTCTC | TGG | \|\|\|\|\|\|\|\|[\|\|\|\|\|\|\|\|\|\|\|\|]PAM | NA | Intergenic | NA | NA |
| T202 | GCTGTGTCTCTGGACGACTAAGG | | 1000 | | | | | | | | | |
| Oligo fwd | CACCGCTGTGTCTCTGGACGACTA | | | | | | | | | | | |
| Oligo rev | AAACTAGTCGTCCAGAGACACAGC | | | | | | | | | | | |
| Chromosome | start | | end | strand | MM | target_seq | PAM | alignment | distance | position | gene name | gene id |
| GU305914.1 | | 2806 | 2828 | + | 0 | GCTGTGTCTCTGGACGACTA | AGG | \|\|\|\|\|\|\|\|[\|\|\|\|\|\|\|\|\|\|\|\|]PAM | NA | Intergenic | NA | NA |
| T203 | ACGATAAGCATAGATGATGCCGG | | 1000 | | | | | | | | | |
| Oligo fwd | CACCACGATAAGCATAGATGATGC | | | | | | | | | | | |
| Oligo rev | AAACGCATCATCTATGCTTATCGT | | | | | | | | | | | |
| Chromosome | start | | end | strand | MM | target_seq | PAM | alignment | distance | position | gene name | gene id |
| GU305914.1 | | 2834 | 2856 | + | 0 | ACGATAAGCATAGATGATGC | CGG | \|\|\|\|\|\|\|\|[\|\|\|\|\|\|\|\|\|\|\|\|]PAM | NA | Intergenic | NA | NA |
| T204 | CCAACATACGACGAGATTCGTGG | | 1000 | | | | | | | | | |
| Oligo fwd | CACCCCAACATACGACGAGATTCG | | | | | | | | | | | |
| Oligo rev | AAACCGAATCTCGTCGTATGTTGG | | | | | | | | | | | |
| Chromosome | start | | end | strand | MM | target_seq | PAM | alignment | distance | position | gene name | gene id |

<table>
<tr><td>GU305914.1</td><td></td><td>2891</td><td>2913</td><td>+</td><td>0</td><td>CCAACATACGACGAGATTCG</td><td>TGG</td><td>||||||||[||||||||||||]PAM</td><td>NA</td><td>Intergenic</td><td>NA</td><td>NA</td></tr>
<tr><td>T205</td><td>GTCCAGAGATCGAATGACGCTGG</td><td></td><td>1000</td><td></td><td></td><td></td><td></td><td></td><td></td><td></td><td></td><td></td></tr>
<tr><td>Oligo fwd</td><td>CACCGTCCAGAGATCGAATGACGC</td><td></td><td></td><td></td><td></td><td></td><td></td><td></td><td></td><td></td><td></td><td></td></tr>
<tr><td>Oligo rev</td><td>AAACGCGTCATTCGATCTCTGGAC</td><td></td><td></td><td></td><td></td><td></td><td></td><td></td><td></td><td></td><td></td><td></td></tr>
<tr><td>Chromosome</td><td>start</td><td></td><td>end</td><td>strand</td><td>MM</td><td>target_seq</td><td>PAM</td><td>alignment</td><td>distance</td><td>position</td><td>gene name</td><td>gene id</td></tr>
<tr><td>GU305914.1</td><td></td><td>2966</td><td>2988</td><td>+</td><td>0</td><td>GTCCAGAGATCGAATGACGC</td><td>TGG</td><td>||||||||[||||||||||||]PAM</td><td>NA</td><td>Intergenic</td><td>NA</td><td>NA</td></tr>
<tr><td>T206</td><td>GAGATCGAATGACGCTGGTCTGG</td><td></td><td>1000</td><td></td><td></td><td></td><td></td><td></td><td></td><td></td><td></td><td></td></tr>
<tr><td>Oligo fwd</td><td>CACCGAGATCGAATGACGCTGGTC</td><td></td><td></td><td></td><td></td><td></td><td></td><td></td><td></td><td></td><td></td><td></td></tr>
<tr><td>Oligo rev</td><td>AAACGACCAGCGTCATTCGATCTC</td><td></td><td></td><td></td><td></td><td></td><td></td><td></td><td></td><td></td><td></td><td></td></tr>
<tr><td>Chromosome</td><td>start</td><td></td><td>end</td><td>strand</td><td>MM</td><td>target_seq</td><td>PAM</td><td>alignment</td><td>distance</td><td>position</td><td>gene name</td><td>gene id</td></tr>
<tr><td>GU305914.1</td><td></td><td>2971</td><td>2993</td><td>+</td><td>0</td><td>GAGATCGAATGACGCTGGTC</td><td>TGG</td><td>||||||||[||||||||||||]PAM</td><td>NA</td><td>Intergenic</td><td>NA</td><td>NA</td></tr>
<tr><td>T207</td><td>GAATGACGCTGGTCTGGTCGCGG</td><td></td><td>1000</td><td></td><td></td><td></td><td></td><td></td><td></td><td></td><td></td><td></td></tr>
<tr><td>Oligo fwd</td><td>CACCGAATGACGCTGGTCTGGTCG</td><td></td><td></td><td></td><td></td><td></td><td></td><td></td><td></td><td></td><td></td><td></td></tr>
<tr><td>Oligo rev</td><td>AAACCGACCAGACCAGCGTCATTC</td><td></td><td></td><td></td><td></td><td></td><td></td><td></td><td></td><td></td><td></td><td></td></tr>
<tr><td>Chromosome</td><td>start</td><td></td><td>end</td><td>strand</td><td>MM</td><td>target_seq</td><td>PAM</td><td>alignment</td><td>distance</td><td>position</td><td>gene name</td><td>gene id</td></tr>
<tr><td>GU305914.1</td><td></td><td>2977</td><td>2999</td><td>+</td><td>0</td><td>GAATGACGCTGGTCTGGTCG</td><td>CGG</td><td>||||||||[||||||||||||]PAM</td><td>NA</td><td>Intergenic</td><td>NA</td><td>NA</td></tr>
<tr><td>T208</td><td>CTTCGTCGCCGCGCTCTTGTTGG</td><td></td><td>1000</td><td></td><td></td><td></td><td></td><td></td><td></td><td></td><td></td><td></td></tr>
<tr><td>Oligo fwd</td><td>CACCCTTCGTCGCCGCGCTCTTGT</td><td></td><td></td><td></td><td></td><td></td><td></td><td></td><td></td><td></td><td></td><td></td></tr>
<tr><td>Oligo rev</td><td>AAACACAAGAGCGCGGCGACGAAG</td><td></td><td></td><td></td><td></td><td></td><td></td><td></td><td></td><td></td><td></td><td></td></tr>
<tr><td>Chromosome</td><td>start</td><td></td><td>end</td><td>strand</td><td>MM</td><td>target_seq</td><td>PAM</td><td>alignment</td><td>distance</td><td>position</td><td>gene name</td><td>gene id</td></tr>
<tr><td>GU305914.1</td><td></td><td>3025</td><td>3047</td><td>+</td><td>0</td><td>CTTCGTCGCCGCGCTCTTGT</td><td>TGG</td><td>||||||||[||||||||||||]PAM</td><td>NA</td><td>Intergenic</td><td>NA</td><td>NA</td></tr>
<tr><td>T209</td><td>TTCGTCGCCGCGCTCTTGTTGGG</td><td></td><td>1000</td><td></td><td></td><td></td><td></td><td></td><td></td><td></td><td></td><td></td></tr>
<tr><td>Oligo fwd</td><td>CACCTTCGTCGCCGCGCTCTTGTT</td><td></td><td></td><td></td><td></td><td></td><td></td><td></td><td></td><td></td><td></td><td></td></tr>
<tr><td>Oligo rev</td><td>AAACAACAAGAGCGCGGCGACGAA</td><td></td><td></td><td></td><td></td><td></td><td></td><td></td><td></td><td></td><td></td><td></td></tr>
<tr><td>Chromosome</td><td>start</td><td></td><td>end</td><td>strand</td><td>MM</td><td>target_seq</td><td>PAM</td><td>alignment</td><td>distance</td><td>position</td><td>gene name</td><td>gene id</td></tr>
<tr><td>GU305914.1</td><td></td><td>3026</td><td>3048</td><td>+</td><td>0</td><td>TTCGTCGCCGCGCTCTTGTT</td><td>GGG</td><td>||||||||[||||||||||||]PAM</td><td>NA</td><td>Intergenic</td><td>NA</td><td>NA</td></tr>
<tr><td>T210</td><td>CATTCGTTATTCCAGAGTGACGG</td><td></td><td>1000</td><td></td><td></td><td></td><td></td><td></td><td></td><td></td><td></td><td></td></tr>
<tr><td>Oligo fwd</td><td>CACCCATTCGTTATTCCAGAGTGA</td><td></td><td></td><td></td><td></td><td></td><td></td><td></td><td></td><td></td><td></td><td></td></tr>
<tr><td>Oligo rev</td><td>AAACTCACTCTGGAATAACGAATG</td><td></td><td></td><td></td><td></td><td></td><td></td><td></td><td></td><td></td><td></td><td></td></tr>
<tr><td>Chromosome</td><td>start</td><td></td><td>end</td><td>strand</td><td>MM</td><td>target_seq</td><td>PAM</td><td>alignment</td><td>distance</td><td>position</td><td>gene name</td><td>gene id</td></tr>
<tr><td>GU305914.1</td><td></td><td>3234</td><td>3256</td><td>+</td><td>0</td><td>CATTCGTTATTCCAGAGTGA</td><td>CGG</td><td>||||||||[||||||||||||]PAM</td><td>NA</td><td>Intergenic</td><td>NA</td><td>NA</td></tr>
<tr><td>T211</td><td>GCAAGATGTCTCTCACACATCGG</td><td></td><td>1000</td><td></td><td></td><td></td><td></td><td></td><td></td><td></td><td></td><td></td></tr>
<tr><td>Oligo fwd</td><td>CACCGCAAGATGTCTCTCACACAT</td><td></td><td></td><td></td><td></td><td></td><td></td><td></td><td></td><td></td><td></td><td></td></tr>
<tr><td>Oligo rev</td><td>AAACATGTGTGAGAGACATCTTGC</td><td></td><td></td><td></td><td></td><td></td><td></td><td></td><td></td><td></td><td></td><td></td></tr>
<tr><td>Chromosome</td><td>start</td><td></td><td>end</td><td>strand</td><td>MM</td><td>target_seq</td><td>PAM</td><td>alignment</td><td>distance</td><td>position</td><td>gene name</td><td>gene id</td></tr>
<tr><td>GU305914.1</td><td></td><td>3262</td><td>3284</td><td>+</td><td>0</td><td>GCAAGATGTCTCTCACACAT</td><td>CGG</td><td>||||||||[||||||||||||]PAM</td><td>NA</td><td>Intergenic</td><td>NA</td><td>NA</td></tr>
<tr><td>T212</td><td>TCTCACACATCGGCCGTTGTTGG</td><td></td><td>1000</td><td></td><td></td><td></td><td></td><td></td><td></td><td></td><td></td><td></td></tr>
<tr><td>Oligo fwd</td><td>CACCTCTCACACATCGGCCGTTGT</td><td></td><td></td><td></td><td></td><td></td><td></td><td></td><td></td><td></td><td></td><td></td></tr>
<tr><td>Oligo rev</td><td>AAACACAACGGCCGATGTGTGAGA</td><td></td><td></td><td></td><td></td><td></td><td></td><td></td><td></td><td></td><td></td><td></td></tr>
<tr><td>Chromosome</td><td>start</td><td></td><td>end</td><td>strand</td><td>MM</td><td>target_seq</td><td>PAM</td><td>alignment</td><td>distance</td><td>position</td><td>gene name</td><td>gene id</td></tr>
<tr><td>GU305914.1</td><td></td><td>3272</td><td>3294</td><td>+</td><td>0</td><td>TCTCACACATCGGCCGTTGT</td><td>TGG</td><td>||||||||[||||||||||||]PAM</td><td>NA</td><td>Intergenic</td><td>NA</td><td>NA</td></tr>
<tr><td>T213</td><td>GTTGGTGATAGTCATTGTCTCGG</td><td></td><td>1000</td><td></td><td></td><td></td><td></td><td></td><td></td><td></td><td></td><td></td></tr>
<tr><td>Oligo fwd</td><td>CACCGTTGGTGATAGTCATTGTCT</td><td></td><td></td><td></td><td></td><td></td><td></td><td></td><td></td><td></td><td></td><td></td></tr>
<tr><td>Oligo rev</td><td>AAACAGACAATGACTATCACCAAC</td><td></td><td></td><td></td><td></td><td></td><td></td><td></td><td></td><td></td><td></td><td></td></tr>
<tr><td>Chromosome</td><td>start</td><td></td><td>end</td><td>strand</td><td>MM</td><td>target_seq</td><td>PAM</td><td>alignment</td><td>distance</td><td>position</td><td>gene name</td><td>gene id</td></tr>
<tr><td>GU305914.1</td><td></td><td>3290</td><td>3312</td><td>+</td><td>0</td><td>GTTGGTGATAGTCATTGTCT</td><td>CGG</td><td>||||||||[||||||||||||]PAM</td><td>NA</td><td>Intergenic</td><td>NA</td><td>NA</td></tr>
<tr><td>T214</td><td>TCGGTTGTCATCTCGTTCGTCGG</td><td></td><td>1000</td><td></td><td></td><td></td><td></td><td></td><td></td><td></td><td></td><td></td></tr>
<tr><td>Oligo fwd</td><td>CACCTCGGTTGTCATCTCGTTCGT</td><td></td><td></td><td></td><td></td><td></td><td></td><td></td><td></td><td></td><td></td><td></td></tr>
<tr><td>Oligo rev</td><td>AAACACGAACGAGATGACAACCGA</td><td></td><td></td><td></td><td></td><td></td><td></td><td></td><td></td><td></td><td></td><td></td></tr>
<tr><td>Chromosome</td><td>start</td><td></td><td>end</td><td>strand</td><td>MM</td><td>target_seq</td><td>PAM</td><td>alignment</td><td>distance</td><td>position</td><td>gene name</td><td>gene id</td></tr>
<tr><td>GU305914.1</td><td></td><td>3309</td><td>3331</td><td>+</td><td>0</td><td>TCGGTTGTCATCTCGTTCGT</td><td>CGG</td><td>||||||||[||||||||||||]PAM</td><td>NA</td><td>Intergenic</td><td>NA</td><td>NA</td></tr>
<tr><td>T215</td><td>CGGTTGTCATCTCGTTCGTCGGG</td><td></td><td>1000</td><td></td><td></td><td></td><td></td><td></td><td></td><td></td><td></td><td></td></tr>
<tr><td>Oligo fwd</td><td>CACCCGGTTGTCATCTCGTTCGTC</td><td></td><td></td><td></td><td></td><td></td><td></td><td></td><td></td><td></td><td></td><td></td></tr>
<tr><td>Oligo rev</td><td>AAACGACGAACGAGATGACAACCG</td><td></td><td></td><td></td><td></td><td></td><td></td><td></td><td></td><td></td><td></td><td></td></tr>
<tr><td>Chromosome</td><td>start</td><td></td><td>end</td><td>strand</td><td>MM</td><td>target_seq</td><td>PAM</td><td>alignment</td><td>distance</td><td>position</td><td>gene name</td><td>gene id</td></tr>
<tr><td>GU305914.1</td><td></td><td>3310</td><td>3332</td><td>+</td><td>0</td><td>CGGTTGTCATCTCGTTCGTC</td><td>GGG</td><td>||||||||[||||||||||||]PAM</td><td>NA</td><td>Intergenic</td><td>NA</td><td>NA</td></tr>
<tr><td>T216</td><td>CGATCAATGTAAAAAAGATGAGG</td><td></td><td>1000</td><td></td><td></td><td></td><td></td><td></td><td></td><td></td><td></td><td></td></tr>
<tr><td>Oligo fwd</td><td>CACCCGATCAATGTAAAAAAGATG</td><td></td><td></td><td></td><td></td><td></td><td></td><td></td><td></td><td></td><td></td><td></td></tr>
<tr><td>Oligo rev</td><td>AAACCATCTTTTTTACATTGATCG</td><td></td><td></td><td></td><td></td><td></td><td></td><td></td><td></td><td></td><td></td><td></td></tr>
<tr><td>Chromosome</td><td>start</td><td></td><td>end</td><td>strand</td><td>MM</td><td>target_seq</td><td>PAM</td><td>alignment</td><td>distance</td><td>position</td><td>gene name</td><td>gene id</td></tr>
<tr><td>GU305914.1</td><td></td><td>3344</td><td>3366</td><td>+</td><td>0</td><td>CGATCAATGTAAAAAAGATG</td><td>AGG</td><td>||||||||[||||||||||||]PAM</td><td>NA</td><td>Intergenic</td><td>NA</td><td>NA</td></tr>
<tr><td>T217</td><td>ATGCAGTATCGACATCCTCTCGG</td><td></td><td>1000</td><td></td><td></td><td></td><td></td><td></td><td></td><td></td><td></td><td></td></tr>
<tr><td>Oligo fwd</td><td>CACCATGCAGTATCGACATCCTCT</td><td></td><td></td><td></td><td></td><td></td><td></td><td></td><td></td><td></td><td></td><td></td></tr>
<tr><td>Oligo rev</td><td>AAACAGAGGATGTCGATACTGCAT</td><td></td><td></td><td></td><td></td><td></td><td></td><td></td><td></td><td></td><td></td><td></td></tr>
<tr><td>Chromosome</td><td>start</td><td></td><td>end</td><td>strand</td><td>MM</td><td>target_seq</td><td>PAM</td><td>alignment</td><td>distance</td><td>position</td><td>gene name</td><td>gene id</td></tr>
<tr><td>GU305914.1</td><td></td><td>3384</td><td>3406</td><td>+</td><td>0</td><td>ATGCAGTATCGACATCCTCT</td><td>CGG</td><td>||||||||[||||||||||||]PAM</td><td>NA</td><td>Intergenic</td><td>NA</td><td>NA</td></tr>
<tr><td>T218</td><td>CCTCTCGGTTGCTACTTCAAAGG</td><td></td><td>1000</td><td></td><td></td><td></td><td></td><td></td><td></td><td></td><td></td><td></td></tr>
<tr><td>Oligo fwd</td><td>CACCCCTCTCGGTTGCTACTTCAA</td><td></td><td></td><td></td><td></td><td></td><td></td><td></td><td></td><td></td><td></td><td></td></tr>
<tr><td>Oligo rev</td><td>AAACTTGAAGTAGCAACCGAGAGG</td><td></td><td></td><td></td><td></td><td></td><td></td><td></td><td></td><td></td><td></td><td></td></tr>
<tr><td>Chromosome</td><td>start</td><td></td><td>end</td><td>strand</td><td>MM</td><td>target_seq</td><td>PAM</td><td>alignment</td><td>distance</td><td>position</td><td>gene name</td><td>gene id</td></tr>
<tr><td>GU305914.1</td><td></td><td>3399</td><td>3421</td><td>+</td><td>0</td><td>CCTCTCGGTTGCTACTTCAA</td><td>AGG</td><td>||||||||[||||||||||||]PAM</td><td>NA</td><td>Intergenic</td><td>NA</td><td>NA</td></tr>
<tr><td>T219</td><td>ACCAACCAAGATACCGTCAAAGG</td><td></td><td>1000</td><td></td><td></td><td></td><td></td><td></td><td></td><td></td><td></td><td></td></tr>
<tr><td>Oligo fwd</td><td>CACCACCAACCAAGATACCGTCAA</td><td></td><td></td><td></td><td></td><td></td><td></td><td></td><td></td><td></td><td></td><td></td></tr>
<tr><td>Oligo rev</td><td>AAACTTGACGGTATCTTGGTTGGT</td><td></td><td></td><td></td><td></td><td></td><td></td><td></td><td></td><td></td><td></td><td></td></tr>
</table>

<table>
<tr><td>Chromosome</td><td>start</td><td></td><td>end</td><td>strand</td><td>MM</td><td>target_seq</td><td>PAM</td><td>alignment</td><td>distance</td><td>position</td><td>gene name</td><td>gene id</td></tr>
<tr><td>GU305914.1</td><td></td><td>3428</td><td>3450</td><td>+</td><td>0</td><td>ACCAACCAAGATACCGTCAA</td><td>AGG</td><td>||||||||[||||||||||||]PAM</td><td>NA</td><td>Intergenic</td><td>NA</td><td>NA</td></tr>
<tr><td></td><td></td><td></td><td></td><td></td><td></td><td></td><td></td><td></td><td></td><td></td><td></td><td></td></tr>
<tr><td>T220</td><td>CCAACCAAGATACCGTCAAAGGG</td><td></td><td>1000</td><td></td><td></td><td></td><td></td><td></td><td></td><td></td><td></td><td></td></tr>
<tr><td>Oligo fwd</td><td>CACCCCAACCAAGATACCGTCAAA</td><td></td><td></td><td></td><td></td><td></td><td></td><td></td><td></td><td></td><td></td><td></td></tr>
<tr><td>Oligo rev</td><td>AAACTTTGACGGTATCTTGGTTGG</td><td></td><td></td><td></td><td></td><td></td><td></td><td></td><td></td><td></td><td></td><td></td></tr>
<tr><td>Chromosome</td><td>start</td><td></td><td>end</td><td>strand</td><td>MM</td><td>target_seq</td><td>PAM</td><td>alignment</td><td>distance</td><td>position</td><td>gene name</td><td>gene id</td></tr>
<tr><td>GU305914.1</td><td></td><td>3429</td><td>3451</td><td>+</td><td>0</td><td>CCAACCAAGATACCGTCAAA</td><td>GGG</td><td>||||||||[||||||||||||]PAM</td><td>NA</td><td>Intergenic</td><td>NA</td><td>NA</td></tr>
<tr><td></td><td></td><td></td><td></td><td></td><td></td><td></td><td></td><td></td><td></td><td></td><td></td><td></td></tr>
<tr><td>T221</td><td>ACGTAAAAGTGAATGCTTCGTGG</td><td></td><td>1000</td><td></td><td></td><td></td><td></td><td></td><td></td><td></td><td></td><td></td></tr>
<tr><td>Oligo fwd</td><td>CACCACGTAAAAGTGAATGCTTCG</td><td></td><td></td><td></td><td></td><td></td><td></td><td></td><td></td><td></td><td></td><td></td></tr>
<tr><td>Oligo rev</td><td>AAACCGAAGCATTCACTTTTACGT</td><td></td><td></td><td></td><td></td><td></td><td></td><td></td><td></td><td></td><td></td><td></td></tr>
<tr><td>Chromosome</td><td>start</td><td></td><td>end</td><td>strand</td><td>MM</td><td>target_seq</td><td>PAM</td><td>alignment</td><td>distance</td><td>position</td><td>gene name</td><td>gene id</td></tr>
<tr><td>GU305914.1</td><td></td><td>3481</td><td>3503</td><td>+</td><td>0</td><td>ACGTAAAAGTGAATGCTTCG</td><td>TGG</td><td>||||||||[||||||||||||]PAM</td><td>NA</td><td>Intergenic</td><td>NA</td><td>NA</td></tr>
<tr><td></td><td></td><td></td><td></td><td></td><td></td><td></td><td></td><td></td><td></td><td></td><td></td><td></td></tr>
<tr><td>T222</td><td>AGTGAATGCTTCGTGGACCTTGG</td><td></td><td>1000</td><td></td><td></td><td></td><td></td><td></td><td></td><td></td><td></td><td></td></tr>
<tr><td>Oligo fwd</td><td>CACCAGTGAATGCTTCGTGGACCT</td><td></td><td></td><td></td><td></td><td></td><td></td><td></td><td></td><td></td><td></td><td></td></tr>
<tr><td>Oligo rev</td><td>AAACAGGTCCACGAAGCATTCACT</td><td></td><td></td><td></td><td></td><td></td><td></td><td></td><td></td><td></td><td></td><td></td></tr>
<tr><td>Chromosome</td><td>start</td><td></td><td>end</td><td>strand</td><td>MM</td><td>target_seq</td><td>PAM</td><td>alignment</td><td>distance</td><td>position</td><td>gene name</td><td>gene id</td></tr>
<tr><td>GU305914.1</td><td></td><td>3488</td><td>3510</td><td>+</td><td>0</td><td>AGTGAATGCTTCGTGGACCT</td><td>TGG</td><td>||||||||[||||||||||||]PAM</td><td>NA</td><td>Intergenic</td><td>NA</td><td>NA</td></tr>
<tr><td></td><td></td><td></td><td></td><td></td><td></td><td></td><td></td><td></td><td></td><td></td><td></td><td></td></tr>
<tr><td>T223</td><td>ATGCTTCGTGGACCTTGGAATGG</td><td></td><td>1000</td><td></td><td></td><td></td><td></td><td></td><td></td><td></td><td></td><td></td></tr>
<tr><td>Oligo fwd</td><td>CACCATGCTTCGTGGACCTTGGAA</td><td></td><td></td><td></td><td></td><td></td><td></td><td></td><td></td><td></td><td></td><td></td></tr>
<tr><td>Oligo rev</td><td>AAACTTCCAAGGTCCACGAAGCAT</td><td></td><td></td><td></td><td></td><td></td><td></td><td></td><td></td><td></td><td></td><td></td></tr>
<tr><td>Chromosome</td><td>start</td><td></td><td>end</td><td>strand</td><td>MM</td><td>target_seq</td><td>PAM</td><td>alignment</td><td>distance</td><td>position</td><td>gene name</td><td>gene id</td></tr>
<tr><td>GU305914.1</td><td></td><td>3493</td><td>3515</td><td>+</td><td>0</td><td>ATGCTTCGTGGACCTTGGAA</td><td>TGG</td><td>||||||||[||||||||||||]PAM</td><td>NA</td><td>Intergenic</td><td>NA</td><td>NA</td></tr>
<tr><td>GU305914.1</td><td></td><td>2535</td><td>2557</td><td>+</td><td>3</td><td>ACGCTTCATGGACCTTTGAA</td><td>TGG</td><td>|-|||||-[||||||||-|||]PAM</td><td>NA</td><td>Intergenic</td><td>NA</td><td>NA</td></tr>
<tr><td></td><td></td><td></td><td></td><td></td><td></td><td></td><td></td><td></td><td></td><td></td><td></td><td></td></tr>
<tr><td>T224</td><td>TGCTTCGTGGACCTTGGAATGGG</td><td></td><td>1000</td><td></td><td></td><td></td><td></td><td></td><td></td><td></td><td></td><td></td></tr>
<tr><td>Oligo fwd</td><td>CACCTGCTTCGTGGACCTTGGAAT</td><td></td><td></td><td></td><td></td><td></td><td></td><td></td><td></td><td></td><td></td><td></td></tr>
<tr><td>Oligo rev</td><td>AAACATTCCAAGGTCCACGAAGCA</td><td></td><td></td><td></td><td></td><td></td><td></td><td></td><td></td><td></td><td></td><td></td></tr>
<tr><td>Chromosome</td><td>start</td><td></td><td>end</td><td>strand</td><td>MM</td><td>target_seq</td><td>PAM</td><td>alignment</td><td>distance</td><td>position</td><td>gene name</td><td>gene id</td></tr>
<tr><td>GU305914.1</td><td></td><td>3494</td><td>3516</td><td>+</td><td>0</td><td>TGCTTCGTGGACCTTGGAAT</td><td>GGG</td><td>||||||||[||||||||||||]PAM</td><td>NA</td><td>Intergenic</td><td>NA</td><td>NA</td></tr>
<tr><td></td><td></td><td></td><td></td><td></td><td></td><td></td><td></td><td></td><td></td><td></td><td></td><td></td></tr>
<tr><td>T225</td><td>CGTCGTCAAGCTCCAGAATTCGG</td><td></td><td>1000</td><td></td><td></td><td></td><td></td><td></td><td></td><td></td><td></td><td></td></tr>
<tr><td>Oligo fwd</td><td>CACCCGTCGTCAAGCTCCAGAATT</td><td></td><td></td><td></td><td></td><td></td><td></td><td></td><td></td><td></td><td></td><td></td></tr>
<tr><td>Oligo rev</td><td>AAACAATTCTGGAGCTTGACGACG</td><td></td><td></td><td></td><td></td><td></td><td></td><td></td><td></td><td></td><td></td><td></td></tr>
<tr><td>Chromosome</td><td>start</td><td></td><td>end</td><td>strand</td><td>MM</td><td>target_seq</td><td>PAM</td><td>alignment</td><td>distance</td><td>position</td><td>gene name</td><td>gene id</td></tr>
<tr><td>GU305914.1</td><td></td><td>3518</td><td>3540</td><td>+</td><td>0</td><td>CGTCGTCAAGCTCCAGAATT</td><td>CGG</td><td>||||||||[||||||||||||]PAM</td><td>NA</td><td>Intergenic</td><td>NA</td><td>NA</td></tr>
<tr><td></td><td></td><td></td><td></td><td></td><td></td><td></td><td></td><td></td><td></td><td></td><td></td><td></td></tr>
<tr><td>T226</td><td>TTGAATTTCACTCCCATACACGG</td><td></td><td>1000</td><td></td><td></td><td></td><td></td><td></td><td></td><td></td><td></td><td></td></tr>
<tr><td>Oligo fwd</td><td>CACCTTGAATTTCACTCCCATACA</td><td></td><td></td><td></td><td></td><td></td><td></td><td></td><td></td><td></td><td></td><td></td></tr>
<tr><td>Oligo rev</td><td>AAACTGTATGGGAGTGAAATTCAA</td><td></td><td></td><td></td><td></td><td></td><td></td><td></td><td></td><td></td><td></td><td></td></tr>
<tr><td>Chromosome</td><td>start</td><td></td><td>end</td><td>strand</td><td>MM</td><td>target_seq</td><td>PAM</td><td>alignment</td><td>distance</td><td>position</td><td>gene name</td><td>gene id</td></tr>
<tr><td>GU305914.1</td><td></td><td>3603</td><td>3625</td><td>+</td><td>0</td><td>TTGAATTTCACTCCCATACA</td><td>CGG</td><td>||||||||[||||||||||||]PAM</td><td>NA</td><td>Intergenic</td><td>NA</td><td>NA</td></tr>
<tr><td></td><td></td><td></td><td></td><td></td><td></td><td></td><td></td><td></td><td></td><td></td><td></td><td></td></tr>
<tr><td>T227</td><td>GAATAACCTGTTGACAGTCAAGG</td><td></td><td>1000</td><td></td><td></td><td></td><td></td><td></td><td></td><td></td><td></td><td></td></tr>
<tr><td>Oligo fwd</td><td>CACCGAATAACCTGTTGACAGTCA</td><td></td><td></td><td></td><td></td><td></td><td></td><td></td><td></td><td></td><td></td><td></td></tr>
<tr><td>Oligo rev</td><td>AAACTGACTGTCAACAGGTTATTC</td><td></td><td></td><td></td><td></td><td></td><td></td><td></td><td></td><td></td><td></td><td></td></tr>
<tr><td>Chromosome</td><td>start</td><td></td><td>end</td><td>strand</td><td>MM</td><td>target_seq</td><td>PAM</td><td>alignment</td><td>distance</td><td>position</td><td>gene name</td><td>gene id</td></tr>
<tr><td>GU305914.1</td><td></td><td>3635</td><td>3657</td><td>+</td><td>0</td><td>GAATAACCTGTTGACAGTCA</td><td>AGG</td><td>||||||||[||||||||||||]PAM</td><td>NA</td><td>Intergenic</td><td>NA</td><td>NA</td></tr>
<tr><td></td><td></td><td></td><td></td><td></td><td></td><td></td><td></td><td></td><td></td><td></td><td></td><td></td></tr>
<tr><td>T228</td><td>AACCTGTTGACAGTCAAGGACGG</td><td></td><td>1000</td><td></td><td></td><td></td><td></td><td></td><td></td><td></td><td></td><td></td></tr>
<tr><td>Oligo fwd</td><td>CACCAACCTGTTGACAGTCAAGGA</td><td></td><td></td><td></td><td></td><td></td><td></td><td></td><td></td><td></td><td></td><td></td></tr>
<tr><td>Oligo rev</td><td>AAACTCCTTGACTGTCAACAGGTT</td><td></td><td></td><td></td><td></td><td></td><td></td><td></td><td></td><td></td><td></td><td></td></tr>
<tr><td>Chromosome</td><td>start</td><td></td><td>end</td><td>strand</td><td>MM</td><td>target_seq</td><td>PAM</td><td>alignment</td><td>distance</td><td>position</td><td>gene name</td><td>gene id</td></tr>
<tr><td>GU305914.1</td><td></td><td>3639</td><td>3661</td><td>+</td><td>0</td><td>AACCTGTTGACAGTCAAGGA</td><td>CGG</td><td>||||||||[||||||||||||]PAM</td><td>NA</td><td>Intergenic</td><td>NA</td><td>NA</td></tr>
<tr><td></td><td></td><td></td><td></td><td></td><td></td><td></td><td></td><td></td><td></td><td></td><td></td><td></td></tr>
<tr><td>T229</td><td>GTCGATAACAGCACTGATAACGG</td><td></td><td>1000</td><td></td><td></td><td></td><td></td><td></td><td></td><td></td><td></td><td></td></tr>
<tr><td>Oligo fwd</td><td>CACCGTCGATAACAGCACTGATAA</td><td></td><td></td><td></td><td></td><td></td><td></td><td></td><td></td><td></td><td></td><td></td></tr>
<tr><td>Oligo rev</td><td>AAACTTATCAGTGCTGTTATCGAC</td><td></td><td></td><td></td><td></td><td></td><td></td><td></td><td></td><td></td><td></td><td></td></tr>
<tr><td>Chromosome</td><td>start</td><td></td><td>end</td><td>strand</td><td>MM</td><td>target_seq</td><td>PAM</td><td>alignment</td><td>distance</td><td>position</td><td>gene name</td><td>gene id</td></tr>
<tr><td>GU305914.1</td><td></td><td>3669</td><td>3691</td><td>+</td><td>0</td><td>GTCGATAACAGCACTGATAA</td><td>CGG</td><td>||||||||[||||||||||||]PAM</td><td>NA</td><td>Intergenic</td><td>NA</td><td>NA</td></tr>
<tr><td></td><td></td><td></td><td></td><td></td><td></td><td></td><td></td><td></td><td></td><td></td><td></td><td></td></tr>
<tr><td>T230</td><td>AACAGCACTGATAACGGCAACGG</td><td></td><td>1000</td><td></td><td></td><td></td><td></td><td></td><td></td><td></td><td></td><td></td></tr>
<tr><td>Oligo fwd</td><td>CACCAACAGCACTGATAACGGCAA</td><td></td><td></td><td></td><td></td><td></td><td></td><td></td><td></td><td></td><td></td><td></td></tr>
<tr><td>Oligo rev</td><td>AAACTTGCCGTTATCAGTGCTGTT</td><td></td><td></td><td></td><td></td><td></td><td></td><td></td><td></td><td></td><td></td><td></td></tr>
<tr><td>Chromosome</td><td>start</td><td></td><td>end</td><td>strand</td><td>MM</td><td>target_seq</td><td>PAM</td><td>alignment</td><td>distance</td><td>position</td><td>gene name</td><td>gene id</td></tr>
<tr><td>GU305914.1</td><td></td><td>3675</td><td>3697</td><td>+</td><td>0</td><td>AACAGCACTGATAACGGCAA</td><td>CGG</td><td>||||||||[||||||||||||]PAM</td><td>NA</td><td>Intergenic</td><td>NA</td><td>NA</td></tr>
<tr><td></td><td></td><td></td><td></td><td></td><td></td><td></td><td></td><td></td><td></td><td></td><td></td><td></td></tr>
<tr><td>T231</td><td>AGCACTGATAACGGCAACGGTGG</td><td></td><td>1000</td><td></td><td></td><td></td><td></td><td></td><td></td><td></td><td></td><td></td></tr>
<tr><td>Oligo fwd</td><td>CACCAGCACTGATAACGGCAACGG</td><td></td><td></td><td></td><td></td><td></td><td></td><td></td><td></td><td></td><td></td><td></td></tr>
<tr><td>Oligo rev</td><td>AAACCCGTTGCCGTTATCAGTGCT</td><td></td><td></td><td></td><td></td><td></td><td></td><td></td><td></td><td></td><td></td><td></td></tr>
<tr><td>Chromosome</td><td>start</td><td></td><td>end</td><td>strand</td><td>MM</td><td>target_seq</td><td>PAM</td><td>alignment</td><td>distance</td><td>position</td><td>gene name</td><td>gene id</td></tr>
<tr><td>GU305914.1</td><td></td><td>3678</td><td>3700</td><td>+</td><td>0</td><td>AGCACTGATAACGGCAACGG</td><td>TGG</td><td>||||||||[||||||||||||]PAM</td><td>NA</td><td>Intergenic</td><td>NA</td><td>NA</td></tr>
<tr><td></td><td></td><td></td><td></td><td></td><td></td><td></td><td></td><td></td><td></td><td></td><td></td><td></td></tr>
<tr><td>T232</td><td>GTCTATTCGAACGCCAGCACAGG</td><td></td><td>1000</td><td></td><td></td><td></td><td></td><td></td><td></td><td></td><td></td><td></td></tr>
<tr><td>Oligo fwd</td><td>CACCGTCTATTCGAACGCCAGCAC</td><td></td><td></td><td></td><td></td><td></td><td></td><td></td><td></td><td></td><td></td><td></td></tr>
<tr><td>Oligo rev</td><td>AAACGTGCTGGCGTTCGAATAGAC</td><td></td><td></td><td></td><td></td><td></td><td></td><td></td><td></td><td></td><td></td><td></td></tr>
<tr><td>Chromosome</td><td>start</td><td></td><td>end</td><td>strand</td><td>MM</td><td>target_seq</td><td>PAM</td><td>alignment</td><td>distance</td><td>position</td><td>gene name</td><td>gene id</td></tr>
<tr><td>GU305914.1</td><td></td><td>3711</td><td>3733</td><td>+</td><td>0</td><td>GTCTATTCGAACGCCAGCAC</td><td>AGG</td><td>||||||||[||||||||||||]PAM</td><td>NA</td><td>Intergenic</td><td>NA</td><td>NA</td></tr>
<tr><td></td><td></td><td></td><td></td><td></td><td></td><td></td><td></td><td></td><td></td><td></td><td></td><td></td></tr>
<tr><td>T233</td><td>ATAGTGTCAGATGTCGTCTGAGG</td><td></td><td>1000</td><td></td><td></td><td></td><td></td><td></td><td></td><td></td><td></td><td></td></tr>
<tr><td>Oligo fwd</td><td>CACCATAGTGTCAGATGTCGTCTG</td><td></td><td></td><td></td><td></td><td></td><td></td><td></td><td></td><td></td><td></td><td></td></tr>
<tr><td>Oligo rev</td><td>AAACCAGACGACATCTGACACTAT</td><td></td><td></td><td></td><td></td><td></td><td></td><td></td><td></td><td></td><td></td><td></td></tr>
<tr><td>Chromosome</td><td>start</td><td></td><td>end</td><td>strand</td><td>MM</td><td>target_seq</td><td>PAM</td><td>alignment</td><td>distance</td><td>position</td><td>gene name</td><td>gene id</td></tr>
<tr><td>GU305914.1</td><td></td><td>3742</td><td>3764</td><td>+</td><td>0</td><td>ATAGTGTCAGATGTCGTCTG</td><td>AGG</td><td>||||||||[||||||||||||]PAM</td><td>NA</td><td>Intergenic</td><td>NA</td><td>NA</td></tr>
<tr><td></td><td></td><td></td><td></td><td></td><td></td><td></td><td></td><td></td><td></td><td></td><td></td><td></td></tr>
<tr><td>T234</td><td>GTCGTCTGAGGATGTGTCCGTGG</td><td></td><td>1000</td><td></td><td></td><td></td><td></td><td></td><td></td><td></td><td></td><td></td></tr>
</table>

<table>
<tr><td>Oligo fwd</td><td>CACCGTCGTCTGAGGATGTGTCCG</td><td></td><td></td><td></td><td></td><td></td><td></td><td></td><td></td><td></td><td></td></tr>
<tr><td>Oligo rev</td><td>AAACCGGACACATCCTCAGACGAC</td><td></td><td></td><td></td><td></td><td></td><td></td><td></td><td></td><td></td><td></td></tr>
<tr><td>Chromosome</td><td>start</td><td>end</td><td>strand</td><td>MM</td><td>target_seq</td><td>PAM</td><td>alignment</td><td>distance</td><td>position</td><td>gene name</td><td>gene id</td></tr>
<tr><td>GU305914.1</td><td>3754</td><td>3776</td><td>+</td><td>0</td><td>GTCGTCTGAGGATGTGTCCG</td><td>TGG</td><td>||||||||[||||||||||||]PAM</td><td>NA</td><td>Intergenic</td><td>NA</td><td>NA</td></tr>
<tr><td>T235</td><td>TAACAGCAAAATGACAGCTCCGG</td><td>1000</td><td></td><td></td><td></td><td></td><td></td><td></td><td></td><td></td><td></td></tr>
<tr><td>Oligo fwd</td><td>CACCTAACAGCAAAATGACAGCTC</td><td></td><td></td><td></td><td></td><td></td><td></td><td></td><td></td><td></td><td></td></tr>
<tr><td>Oligo rev</td><td>AAACGAGCTGTCATTTTGCTGTTA</td><td></td><td></td><td></td><td></td><td></td><td></td><td></td><td></td><td></td><td></td></tr>
<tr><td>Chromosome</td><td>start</td><td>end</td><td>strand</td><td>MM</td><td>target_seq</td><td>PAM</td><td>alignment</td><td>distance</td><td>position</td><td>gene name</td><td>gene id</td></tr>
<tr><td>GU305914.1</td><td>3782</td><td>3804</td><td>+</td><td>0</td><td>TAACAGCAAAATGACAGCTC</td><td>CGG</td><td>||||||||[||||||||||||]PAM</td><td>NA</td><td>Intergenic</td><td>NA</td><td>NA</td></tr>
<tr><td>T236</td><td>CAAAATGACAGCTCCGGATGAGG</td><td>1000</td><td></td><td></td><td></td><td></td><td></td><td></td><td></td><td></td><td></td></tr>
<tr><td>Oligo fwd</td><td>CACCCAAAATGACAGCTCCGGATG</td><td></td><td></td><td></td><td></td><td></td><td></td><td></td><td></td><td></td><td></td></tr>
<tr><td>Oligo rev</td><td>AAACCATCCGGAGCTGTCATTTTG</td><td></td><td></td><td></td><td></td><td></td><td></td><td></td><td></td><td></td><td></td></tr>
<tr><td>Chromosome</td><td>start</td><td>end</td><td>strand</td><td>MM</td><td>target_seq</td><td>PAM</td><td>alignment</td><td>distance</td><td>position</td><td>gene name</td><td>gene id</td></tr>
<tr><td>GU305914.1</td><td>3788</td><td>3810</td><td>+</td><td>0</td><td>CAAAATGACAGCTCCGGATG</td><td>AGG</td><td>||||||||[||||||||||||]PAM</td><td>NA</td><td>Intergenic</td><td>NA</td><td>NA</td></tr>
<tr><td>T237</td><td>CTCCGGATGAGGAGATGCTGCGG</td><td>1000</td><td></td><td></td><td></td><td></td><td></td><td></td><td></td><td></td><td></td></tr>
<tr><td>Oligo fwd</td><td>CACCCTCCGGATGAGGAGATGCTG</td><td></td><td></td><td></td><td></td><td></td><td></td><td></td><td></td><td></td><td></td></tr>
<tr><td>Oligo rev</td><td>AAACCAGCATCTCCTCATCCGGAG</td><td></td><td></td><td></td><td></td><td></td><td></td><td></td><td></td><td></td><td></td></tr>
<tr><td>Chromosome</td><td>start</td><td>end</td><td>strand</td><td>MM</td><td>target_seq</td><td>PAM</td><td>alignment</td><td>distance</td><td>position</td><td>gene name</td><td>gene id</td></tr>
<tr><td>GU305914.1</td><td>3799</td><td>3821</td><td>+</td><td>0</td><td>CTCCGGATGAGGAGATGCTG</td><td>CGG</td><td>||||||||[||||||||||||]PAM</td><td>NA</td><td>Intergenic</td><td>NA</td><td>NA</td></tr>
<tr><td>T238</td><td>GATGCTGCGGAAGATGAGCGAGG</td><td>1000</td><td></td><td></td><td></td><td></td><td></td><td></td><td></td><td></td><td></td></tr>
<tr><td>Oligo fwd</td><td>CACCGATGCTGCGGAAGATGAGCG</td><td></td><td></td><td></td><td></td><td></td><td></td><td></td><td></td><td></td><td></td></tr>
<tr><td>Oligo rev</td><td>AAACCGCTCATCTTCCGCAGCATC</td><td></td><td></td><td></td><td></td><td></td><td></td><td></td><td></td><td></td><td></td></tr>
<tr><td>Chromosome</td><td>start</td><td>end</td><td>strand</td><td>MM</td><td>target_seq</td><td>PAM</td><td>alignment</td><td>distance</td><td>position</td><td>gene name</td><td>gene id</td></tr>
<tr><td>GU305914.1</td><td>3812</td><td>3834</td><td>+</td><td>0</td><td>GATGCTGCGGAAGATGAGCG</td><td>AGG</td><td>||||||||[||||||||||||]PAM</td><td>NA</td><td>Intergenic</td><td>NA</td><td>NA</td></tr>
<tr><td>GU305914.1</td><td>147164</td><td>147186</td><td>-</td><td>4</td><td>CACGCGGCGGAAGATGCGCG</td><td>GGG</td><td>-|-||-||[||||||||-|||]PAM</td><td>NA</td><td>Intergenic</td><td>NA</td><td>NA</td></tr>
<tr><td>T239</td><td>GTACTCAACCTCCCAAACTACGG</td><td>1000</td><td></td><td></td><td></td><td></td><td></td><td></td><td></td><td></td><td></td></tr>
<tr><td>Oligo fwd</td><td>CACCGTACTCAACCTCCCAAACTA</td><td></td><td></td><td></td><td></td><td></td><td></td><td></td><td></td><td></td><td></td></tr>
<tr><td>Oligo rev</td><td>AAACTAGTTTGGGAGGTTGAGTAC</td><td></td><td></td><td></td><td></td><td></td><td></td><td></td><td></td><td></td><td></td></tr>
<tr><td>Chromosome</td><td>start</td><td>end</td><td>strand</td><td>MM</td><td>target_seq</td><td>PAM</td><td>alignment</td><td>distance</td><td>position</td><td>gene name</td><td>gene id</td></tr>
<tr><td>GU305914.1</td><td>3834</td><td>3856</td><td>+</td><td>0</td><td>GTACTCAACCTCCCAAACTA</td><td>CGG</td><td>||||||||[||||||||||||]PAM</td><td>NA</td><td>Intergenic</td><td>NA</td><td>NA</td></tr>
<tr><td>T240</td><td>CAGATCTGACCCCTCCAAGACGG</td><td>1000</td><td></td><td></td><td></td><td></td><td></td><td></td><td></td><td></td><td></td></tr>
<tr><td>Oligo fwd</td><td>CACCCAGATCTGACCCCTCCAAGA</td><td></td><td></td><td></td><td></td><td></td><td></td><td></td><td></td><td></td><td></td></tr>
<tr><td>Oligo rev</td><td>AAACTCTTGGAGGGGTCAGATCTG</td><td></td><td></td><td></td><td></td><td></td><td></td><td></td><td></td><td></td><td></td></tr>
<tr><td>Chromosome</td><td>start</td><td>end</td><td>strand</td><td>MM</td><td>target_seq</td><td>PAM</td><td>alignment</td><td>distance</td><td>position</td><td>gene name</td><td>gene id</td></tr>
<tr><td>GU305914.1</td><td>3862</td><td>3884</td><td>+</td><td>0</td><td>CAGATCTGACCCCTCCAAGA</td><td>CGG</td><td>||||||||[||||||||||||]PAM</td><td>NA</td><td>Intergenic</td><td>NA</td><td>NA</td></tr>
<tr><td>T241</td><td>AGATCTGACCCCTCCAAGACGGG</td><td>1000</td><td></td><td></td><td></td><td></td><td></td><td></td><td></td><td></td><td></td></tr>
<tr><td>Oligo fwd</td><td>CACCAGATCTGACCCCTCCAAGAC</td><td></td><td></td><td></td><td></td><td></td><td></td><td></td><td></td><td></td><td></td></tr>
<tr><td>Oligo rev</td><td>AAACGTCTTGGAGGGGTCAGATCT</td><td></td><td></td><td></td><td></td><td></td><td></td><td></td><td></td><td></td><td></td></tr>
<tr><td>Chromosome</td><td>start</td><td>end</td><td>strand</td><td>MM</td><td>target_seq</td><td>PAM</td><td>alignment</td><td>distance</td><td>position</td><td>gene name</td><td>gene id</td></tr>
<tr><td>GU305914.1</td><td>3863</td><td>3885</td><td>+</td><td>0</td><td>AGATCTGACCCCTCCAAGAC</td><td>GGG</td><td>||||||||[||||||||||||]PAM</td><td>NA</td><td>Intergenic</td><td>NA</td><td>NA</td></tr>
<tr><td>T242</td><td>ATAGTGACTCTTCTTTTTGTCGG</td><td>1000</td><td></td><td></td><td></td><td></td><td></td><td></td><td></td><td></td><td></td></tr>
<tr><td>Oligo fwd</td><td>CACCATAGTGACTCTTCTTTTTGT</td><td></td><td></td><td></td><td></td><td></td><td></td><td></td><td></td><td></td><td></td></tr>
<tr><td>Oligo rev</td><td>AAACACAAAAAGAAGAGTCACTAT</td><td></td><td></td><td></td><td></td><td></td><td></td><td></td><td></td><td></td><td></td></tr>
<tr><td>Chromosome</td><td>start</td><td>end</td><td>strand</td><td>MM</td><td>target_seq</td><td>PAM</td><td>alignment</td><td>distance</td><td>position</td><td>gene name</td><td>gene id</td></tr>
<tr><td>GU305914.1</td><td>3942</td><td>3964</td><td>+</td><td>0</td><td>ATAGTGACTCTTCTTTTTGT</td><td>CGG</td><td>||||||||[||||||||||||]PAM</td><td>NA</td><td>Intergenic</td><td>NA</td><td>NA</td></tr>
<tr><td>T243</td><td>CTCGTACTGATTTACTTGTACGG</td><td>1000</td><td></td><td></td><td></td><td></td><td></td><td></td><td></td><td></td><td></td></tr>
<tr><td>Oligo fwd</td><td>CACCCTCGTACTGATTTACTTGTA</td><td></td><td></td><td></td><td></td><td></td><td></td><td></td><td></td><td></td><td></td></tr>
<tr><td>Oligo rev</td><td>AAACTACAAGTAAATCAGTACGAG</td><td></td><td></td><td></td><td></td><td></td><td></td><td></td><td></td><td></td><td></td></tr>
<tr><td>Chromosome</td><td>start</td><td>end</td><td>strand</td><td>MM</td><td>target_seq</td><td>PAM</td><td>alignment</td><td>distance</td><td>position</td><td>gene name</td><td>gene id</td></tr>
<tr><td>GU305914.1</td><td>3972</td><td>3994</td><td>+</td><td>0</td><td>CTCGTACTGATTTACTTGTA</td><td>CGG</td><td>||||||||[||||||||||||]PAM</td><td>NA</td><td>Intergenic</td><td>NA</td><td>NA</td></tr>
<tr><td>T244</td><td>CCCTTGAGTAAACCATTTTCCGG</td><td>1000</td><td></td><td></td><td></td><td></td><td></td><td></td><td></td><td></td><td></td></tr>
<tr><td>Oligo fwd</td><td>CACCCCCTTGAGTAAACCATTTTC</td><td></td><td></td><td></td><td></td><td></td><td></td><td></td><td></td><td></td><td></td></tr>
<tr><td>Oligo rev</td><td>AAACGAAAATGGTTTACTCAAGGG</td><td></td><td></td><td></td><td></td><td></td><td></td><td></td><td></td><td></td><td></td></tr>
<tr><td>Chromosome</td><td>start</td><td>end</td><td>strand</td><td>MM</td><td>target_seq</td><td>PAM</td><td>alignment</td><td>distance</td><td>position</td><td>gene name</td><td>gene id</td></tr>
<tr><td>GU305914.1</td><td>4082</td><td>4104</td><td>+</td><td>0</td><td>CCCTTGAGTAAACCATTTTC</td><td>CGG</td><td>||||||||[||||||||||||]PAM</td><td>NA</td><td>Intergenic</td><td>NA</td><td>NA</td></tr>
<tr><td>T245</td><td>AACGTAGATATCCTCATACATGG</td><td>1000</td><td></td><td></td><td></td><td></td><td></td><td></td><td></td><td></td><td></td></tr>
<tr><td>Oligo fwd</td><td>CACCAACGTAGATATCCTCATACA</td><td></td><td></td><td></td><td></td><td></td><td></td><td></td><td></td><td></td><td></td></tr>
<tr><td>Oligo rev</td><td>AAACTGTATGAGGATATCTACGTT</td><td></td><td></td><td></td><td></td><td></td><td></td><td></td><td></td><td></td><td></td></tr>
<tr><td>Chromosome</td><td>start</td><td>end</td><td>strand</td><td>MM</td><td>target_seq</td><td>PAM</td><td>alignment</td><td>distance</td><td>position</td><td>gene name</td><td>gene id</td></tr>
<tr><td>GU305914.1</td><td>4133</td><td>4155</td><td>+</td><td>0</td><td>AACGTAGATATCCTCATACA</td><td>TGG</td><td>||||||||[||||||||||||]PAM</td><td>NA</td><td>Intergenic</td><td>NA</td><td>NA</td></tr>
<tr><td>T246</td><td>CTCATACATGGTCTAGAATCAGG</td><td>1000</td><td></td><td></td><td></td><td></td><td></td><td></td><td></td><td></td><td></td></tr>
<tr><td>Oligo fwd</td><td>CACCCTCATACATGGTCTAGAATC</td><td></td><td></td><td></td><td></td><td></td><td></td><td></td><td></td><td></td><td></td></tr>
<tr><td>Oligo rev</td><td>AAACGATTCTAGACCATGTATGAG</td><td></td><td></td><td></td><td></td><td></td><td></td><td></td><td></td><td></td><td></td></tr>
<tr><td>Chromosome</td><td>start</td><td>end</td><td>strand</td><td>MM</td><td>target_seq</td><td>PAM</td><td>alignment</td><td>distance</td><td>position</td><td>gene name</td><td>gene id</td></tr>
<tr><td>GU305914.1</td><td>4145</td><td>4167</td><td>+</td><td>0</td><td>CTCATACATGGTCTAGAATC</td><td>AGG</td><td>||||||||[||||||||||||]PAM</td><td>NA</td><td>Intergenic</td><td>NA</td><td>NA</td></tr>
<tr><td>T247</td><td>ATCAGGAGTTATCAGTCATGTGG</td><td>1000</td><td></td><td></td><td></td><td></td><td></td><td></td><td></td><td></td><td></td></tr>
<tr><td>Oligo fwd</td><td>CACCATCAGGAGTTATCAGTCATG</td><td></td><td></td><td></td><td></td><td></td><td></td><td></td><td></td><td></td><td></td></tr>
<tr><td>Oligo rev</td><td>AAACCATGACTGATAACTCCTGAT</td><td></td><td></td><td></td><td></td><td></td><td></td><td></td><td></td><td></td><td></td></tr>
<tr><td>Chromosome</td><td>start</td><td>end</td><td>strand</td><td>MM</td><td>target_seq</td><td>PAM</td><td>alignment</td><td>distance</td><td>position</td><td>gene name</td><td>gene id</td></tr>
<tr><td>GU305914.1</td><td>4162</td><td>4184</td><td>+</td><td>0</td><td>ATCAGGAGTTATCAGTCATG</td><td>TGG</td><td>||||||||[||||||||||||]PAM</td><td>NA</td><td>Intergenic</td><td>NA</td><td>NA</td></tr>
<tr><td>T248</td><td>TGTGTCCATGTACCGTCATGTGG</td><td>1000</td><td></td><td></td><td></td><td></td><td></td><td></td><td></td><td></td><td></td></tr>
<tr><td>Oligo fwd</td><td>CACCTGTGTCCATGTACCGTCATG</td><td></td><td></td><td></td><td></td><td></td><td></td><td></td><td></td><td></td><td></td></tr>
<tr><td>Oligo rev</td><td>AAACCATGACGGTACATGGACACA</td><td></td><td></td><td></td><td></td><td></td><td></td><td></td><td></td><td></td><td></td></tr>
<tr><td>Chromosome</td><td>start</td><td>end</td><td>strand</td><td>MM</td><td>target_seq</td><td>PAM</td><td>alignment</td><td>distance</td><td>position</td><td>gene name</td><td>gene id</td></tr>
<tr><td>GU305914.1</td><td>4192</td><td>4214</td><td>+</td><td>0</td><td>TGTGTCCATGTACCGTCATG</td><td>TGG</td><td>||||||||[||||||||||||]PAM</td><td>NA</td><td>Intergenic</td><td>NA</td><td>NA</td></tr>
</table>

T249 ATGTACCGTCATGTGGCGCTCGG 1000
Oligo fwd CACCATGTACCGTCATGTGGCGCT
Oligo rev AAACAGCGCCACATGACGGTACAT
Chromosome start end strand MM target_seq PAM alignment distance position gene name gene id
GU305914.1 4199 4221 + 0 ATGTACCGTCATGTGGCGCT CGG ||||||||[||||||||||||]PAM NA Intergenic NA NA

T250 TGTTTTTACTGTTCTCATCGTGG 1000
Oligo fwd CACCTGTTTTTACTGTTCTCATCG
Oligo rev AAACCGATGAGAACAGTAAAAACA
Chromosome start end strand MM target_seq PAM alignment distance position gene name gene id
GU305914.1 4228 4250 + 0 TGTTTTTACTGTTCTCATCG TGG ||||||||[||||||||||||]PAM NA Intergenic NA NA

T251 TCATCGTGGCTAACGACGTCCGG 1000
Oligo fwd CACCTCATCGTGGCTAACGACGTC
Oligo rev AAACGACGTCGTTAGCCACGATGA
Chromosome start end strand MM target_seq PAM alignment distance position gene name gene id
GU305914.1 4242 4264 + 0 TCATCGTGGCTAACGACGTC CGG ||||||||[||||||||||||]PAM NA Intergenic NA NA

T252 CCACCACAATCACCCAAGTCCGG 1000
Oligo fwd CACCCCACCACAATCACCCAAGTC
Oligo rev AAACGACTTGGGTGATTGTGGTGG
Chromosome start end strand MM target_seq PAM alignment distance position gene name gene id
GU305914.1 4290 4312 + 0 CCACCACAATCACCCAAGTC CGG ||||||||[||||||||||||]PAM NA Intergenic NA NA

T253 CCAAGTCCGGAATAACAACATGG 1000
Oligo fwd CACCCCAAGTCCGGAATAACAACA
Oligo rev AAACTGTTGTTATTCCGGACTTGG
Chromosome start end strand MM target_seq PAM alignment distance position gene name gene id
GU305914.1 4303 4325 + 0 CCAAGTCCGGAATAACAACA TGG ||||||||[||||||||||||]PAM NA Intergenic NA NA

T254 TCCGGAATAACAACATGGATAGG 1000
Oligo fwd CACCTCCGGAATAACAACATGGAT
Oligo rev AAACATCCATGTTGTTATTCCGGA
Chromosome start end strand MM target_seq PAM alignment distance position gene name gene id
GU305914.1 4308 4330 + 0 TCCGGAATAACAACATGGAT AGG ||||||||[||||||||||||]PAM NA Intergenic NA NA

T255 GGAATAACAACATGGATAGGAGG 1000
Oligo fwd CACCGGAATAACAACATGGATAGG
Oligo rev AAACCCTATCCATGTTGTTATTCC
Chromosome start end strand MM target_seq PAM alignment distance position gene name gene id
GU305914.1 4311 4333 + 0 GGAATAACAACATGGATAGG AGG ||||||||[||||||||||||]PAM NA Intergenic NA NA

T256 TAGGAGGATCATCCAGAACACGG 1000
Oligo fwd CACCTAGGAGGATCATCCAGAACA
Oligo rev AAACTGTTCTGGATGATCCTCCTA
Chromosome start end strand MM target_seq PAM alignment distance position gene name gene id
GU305914.1 4327 4349 + 0 TAGGAGGATCATCCAGAACA CGG ||||||||[||||||||||||]PAM NA Intergenic NA NA

T257 AAATATAAAACCATAGCTTTAGG 1000
Oligo fwd CACCAAATATAAAACCATAGCTTT
Oligo rev AAACAAAGCTATGGTTTTATATTT
Chromosome start end strand MM target_seq PAM alignment distance position gene name gene id
GU305914.1 4371 4393 + 0 AAATATAAAACCATAGCTTT AGG ||||||||[||||||||||||]PAM NA Intergenic NA NA

T258 TATAAAACCATAGCTTTAGGTGG 1000
Oligo fwd CACCTATAAAACCATAGCTTTAGG
Oligo rev AAACCCTAAAGCTATGGTTTTATA
Chromosome start end strand MM target_seq PAM alignment distance position gene name gene id
GU305914.1 4374 4396 + 0 TATAAAACCATAGCTTTAGG TGG ||||||||[||||||||||||]PAM NA Intergenic NA NA

T259 GTGGCGACAAACTTTCACCCTGG 1000
Oligo fwd CACCGTGGCGACAAACTTTCACCC
Oligo rev AAACGGGTGAAAGTTTGTCGCCAC
Chromosome start end strand MM target_seq PAM alignment distance position gene name gene id
GU305914.1 4393 4415 + 0 GTGGCGACAAACTTTCACCC TGG ||||||||[||||||||||||]PAM NA Intergenic NA NA

T260 AAGCAAGTGCTTCAACTCAAAGG 1000
Oligo fwd CACCAAGCAAGTGCTTCAACTCAA
Oligo rev AAACTTGAGTTGAAGCACTTGCTT
Chromosome start end strand MM target_seq PAM alignment distance position gene name gene id
GU305914.1 4424 4446 + 0 AAGCAAGTGCTTCAACTCAA AGG ||||||||[||||||||||||]PAM NA Intergenic NA NA

T261 TCAACTCAAAGGACGCCACGCGG 1000
Oligo fwd CACCTCAACTCAAAGGACGCCACG
Oligo rev AAACCGTGGCGTCCTTTGAGTTGA
Chromosome start end strand MM target_seq PAM alignment distance position gene name gene id
GU305914.1 4435 4457 + 0 TCAACTCAAAGGACGCCACG CGG ||||||||[||||||||||||]PAM NA Intergenic NA NA

T262 AAGTCTGTAGATTCAAAGACTGG 1000
Oligo fwd CACCAAGTCTGTAGATTCAAAGAC
Oligo rev AAACGTCTTTGAATCTACAGACTT
Chromosome start end strand MM target_seq PAM alignment distance position gene name gene id
GU305914.1 4467 4489 + 0 AAGTCTGTAGATTCAAAGAC TGG ||||||||[||||||||||||]PAM NA Intergenic NA NA

T263 GACGTTCACGTGCATGCCTACGG 1000
Oligo fwd CACCGACGTTCACGTGCATGCCTA
Oligo rev AAACTAGGCATGCACGTGAACGTC
Chromosome start end strand MM target_seq PAM alignment distance position gene name gene id

<table>
<tr><td>GU305914.1</td><td></td><td>4496</td><td>4518</td><td>+</td><td>0</td><td>GACGTTCACGTGCATGCCTA</td><td>CGG</td><td>||||||||[||||||||||||]PAM</td><td>NA</td><td>Intergenic</td><td>NA</td><td>NA</td></tr>
<tr><td>T264</td><td>ACGTCTACGCAAACGCTTCATGG</td><td></td><td>1000</td><td></td><td></td><td></td><td></td><td></td><td></td><td></td><td></td><td></td></tr>
<tr><td>Oligo fwd</td><td>CACCACGTCTACGCAAACGCTTCA</td><td></td><td></td><td></td><td></td><td></td><td></td><td></td><td></td><td></td><td></td><td></td></tr>
<tr><td>Oligo rev</td><td>AAACTGAAGCGTTTGCGTAGACGT</td><td></td><td></td><td></td><td></td><td></td><td></td><td></td><td></td><td></td><td></td><td></td></tr>
<tr><td>Chromosome</td><td>start</td><td></td><td>end</td><td>strand</td><td>MM</td><td>target_seq</td><td>PAM</td><td>alignment</td><td>distance</td><td>position</td><td>gene name</td><td>gene id</td></tr>
<tr><td>GU305914.1</td><td></td><td>4522</td><td>4544</td><td>+</td><td>0</td><td>ACGTCTACGCAAACGCTTCA</td><td>TGG</td><td>||||||||[||||||||||||]PAM</td><td>NA</td><td>Intergenic</td><td>NA</td><td>NA</td></tr>
<tr><td>T265</td><td>ACGCTTCATGGCGCATAGAATGG</td><td></td><td>1000</td><td></td><td></td><td></td><td></td><td></td><td></td><td></td><td></td><td></td></tr>
<tr><td>Oligo fwd</td><td>CACCACGCTTCATGGCGCATAGAA</td><td></td><td></td><td></td><td></td><td></td><td></td><td></td><td></td><td></td><td></td><td></td></tr>
<tr><td>Oligo rev</td><td>AAACTTCTATGCGCCATGAAGCGT</td><td></td><td></td><td></td><td></td><td></td><td></td><td></td><td></td><td></td><td></td><td></td></tr>
<tr><td>Chromosome</td><td>start</td><td></td><td>end</td><td>strand</td><td>MM</td><td>target_seq</td><td>PAM</td><td>alignment</td><td>distance</td><td>position</td><td>gene name</td><td>gene id</td></tr>
<tr><td>GU305914.1</td><td></td><td>4534</td><td>4556</td><td>+</td><td>0</td><td>ACGCTTCATGGCGCATAGAA</td><td>TGG</td><td>||||||||[||||||||||||]PAM</td><td>NA</td><td>Intergenic</td><td>NA</td><td>NA</td></tr>
<tr><td>T266</td><td>CGCTTCATGGCGCATAGAATGGG</td><td></td><td>1000</td><td></td><td></td><td></td><td></td><td></td><td></td><td></td><td></td><td></td></tr>
<tr><td>Oligo fwd</td><td>CACCCGCTTCATGGCGCATAGAAT</td><td></td><td></td><td></td><td></td><td></td><td></td><td></td><td></td><td></td><td></td><td></td></tr>
<tr><td>Oligo rev</td><td>AAACATTCTATGCGCCATGAAGCG</td><td></td><td></td><td></td><td></td><td></td><td></td><td></td><td></td><td></td><td></td><td></td></tr>
<tr><td>Chromosome</td><td>start</td><td></td><td>end</td><td>strand</td><td>MM</td><td>target_seq</td><td>PAM</td><td>alignment</td><td>distance</td><td>position</td><td>gene name</td><td>gene id</td></tr>
<tr><td>GU305914.1</td><td></td><td>4535</td><td>4557</td><td>+</td><td>0</td><td>CGCTTCATGGCGCATAGAAT</td><td>GGG</td><td>||||||||[||||||||||||]PAM</td><td>NA</td><td>Intergenic</td><td>NA</td><td>NA</td></tr>
<tr><td>T267</td><td>ATGGCGCATAGAATGGGTCCTGG</td><td></td><td>1000</td><td></td><td></td><td></td><td></td><td></td><td></td><td></td><td></td><td></td></tr>
<tr><td>Oligo fwd</td><td>CACCATGGCGCATAGAATGGGTCC</td><td></td><td></td><td></td><td></td><td></td><td></td><td></td><td></td><td></td><td></td><td></td></tr>
<tr><td>Oligo rev</td><td>AAACGGACCCATTCTATGCGCCAT</td><td></td><td></td><td></td><td></td><td></td><td></td><td></td><td></td><td></td><td></td><td></td></tr>
<tr><td>Chromosome</td><td>start</td><td></td><td>end</td><td>strand</td><td>MM</td><td>target_seq</td><td>PAM</td><td>alignment</td><td>distance</td><td>position</td><td>gene name</td><td>gene id</td></tr>
<tr><td>GU305914.1</td><td></td><td>4541</td><td>4563</td><td>+</td><td>0</td><td>ATGGCGCATAGAATGGGTCC</td><td>TGG</td><td>||||||||[||||||||||||]PAM</td><td>NA</td><td>Intergenic</td><td>NA</td><td>NA</td></tr>
<tr><td>T268</td><td>TGGCGCATAGAATGGGTCCTGGG</td><td></td><td>1000</td><td></td><td></td><td></td><td></td><td></td><td></td><td></td><td></td><td></td></tr>
<tr><td>Oligo fwd</td><td>CACCTGGCGCATAGAATGGGTCCT</td><td></td><td></td><td></td><td></td><td></td><td></td><td></td><td></td><td></td><td></td><td></td></tr>
<tr><td>Oligo rev</td><td>AAACAGGACCCATTCTATGCGCCA</td><td></td><td></td><td></td><td></td><td></td><td></td><td></td><td></td><td></td><td></td><td></td></tr>
<tr><td>Chromosome</td><td>start</td><td></td><td>end</td><td>strand</td><td>MM</td><td>target_seq</td><td>PAM</td><td>alignment</td><td>distance</td><td>position</td><td>gene name</td><td>gene id</td></tr>
<tr><td>GU305914.1</td><td></td><td>4542</td><td>4564</td><td>+</td><td>0</td><td>TGGCGCATAGAATGGGTCCT</td><td>GGG</td><td>||||||||[||||||||||||]PAM</td><td>NA</td><td>Intergenic</td><td>NA</td><td>NA</td></tr>
<tr><td>T269</td><td>ATAGAATGGGTCCTGGGAAATGG</td><td></td><td>1000</td><td></td><td></td><td></td><td></td><td></td><td></td><td></td><td></td><td></td></tr>
<tr><td>Oligo fwd</td><td>CACCATAGAATGGGTCCTGGGAAA</td><td></td><td></td><td></td><td></td><td></td><td></td><td></td><td></td><td></td><td></td><td></td></tr>
<tr><td>Oligo rev</td><td>AAACTTTCCCAGGACCCATTCTAT</td><td></td><td></td><td></td><td></td><td></td><td></td><td></td><td></td><td></td><td></td><td></td></tr>
<tr><td>Chromosome</td><td>start</td><td></td><td>end</td><td>strand</td><td>MM</td><td>target_seq</td><td>PAM</td><td>alignment</td><td>distance</td><td>position</td><td>gene name</td><td>gene id</td></tr>
<tr><td>GU305914.1</td><td></td><td>4548</td><td>4570</td><td>+</td><td>0</td><td>ATAGAATGGGTCCTGGGAAA</td><td>TGG</td><td>||||||||[||||||||||||]PAM</td><td>NA</td><td>Intergenic</td><td>NA</td><td>NA</td></tr>
<tr><td>T270</td><td>TAGAATGGGTCCTGGGAAATGGG</td><td></td><td>1000</td><td></td><td></td><td></td><td></td><td></td><td></td><td></td><td></td><td></td></tr>
<tr><td>Oligo fwd</td><td>CACCTAGAATGGGTCCTGGGAAAT</td><td></td><td></td><td></td><td></td><td></td><td></td><td></td><td></td><td></td><td></td><td></td></tr>
<tr><td>Oligo rev</td><td>AAACATTTCCCAGGACCCATTCTA</td><td></td><td></td><td></td><td></td><td></td><td></td><td></td><td></td><td></td><td></td><td></td></tr>
<tr><td>Chromosome</td><td>start</td><td></td><td>end</td><td>strand</td><td>MM</td><td>target_seq</td><td>PAM</td><td>alignment</td><td>distance</td><td>position</td><td>gene name</td><td>gene id</td></tr>
<tr><td>GU305914.1</td><td></td><td>4549</td><td>4571</td><td>+</td><td>0</td><td>TAGAATGGGTCCTGGGAAAT</td><td>GGG</td><td>||||||||[||||||||||||]PAM</td><td>NA</td><td>Intergenic</td><td>NA</td><td>NA</td></tr>
<tr><td>T271</td><td>TCCTGGGAAATGGGACAACTTGG</td><td></td><td>1000</td><td></td><td></td><td></td><td></td><td></td><td></td><td></td><td></td><td></td></tr>
<tr><td>Oligo fwd</td><td>CACCTCCTGGGAAATGGGACAACT</td><td></td><td></td><td></td><td></td><td></td><td></td><td></td><td></td><td></td><td></td><td></td></tr>
<tr><td>Oligo rev</td><td>AAACAGTTGTCCCATTTCCCAGGA</td><td></td><td></td><td></td><td></td><td></td><td></td><td></td><td></td><td></td><td></td><td></td></tr>
<tr><td>Chromosome</td><td>start</td><td></td><td>end</td><td>strand</td><td>MM</td><td>target_seq</td><td>PAM</td><td>alignment</td><td>distance</td><td>position</td><td>gene name</td><td>gene id</td></tr>
<tr><td>GU305914.1</td><td></td><td>4558</td><td>4580</td><td>+</td><td>0</td><td>TCCTGGGAAATGGGACAACT</td><td>TGG</td><td>||||||||[||||||||||||]PAM</td><td>NA</td><td>Intergenic</td><td>NA</td><td>NA</td></tr>
<tr><td>T272</td><td>TGGGACAACTTGGTTAACACCGG</td><td></td><td>1000</td><td></td><td></td><td></td><td></td><td></td><td></td><td></td><td></td><td></td></tr>
<tr><td>Oligo fwd</td><td>CACCTGGGACAACTTGGTTAACAC</td><td></td><td></td><td></td><td></td><td></td><td></td><td></td><td></td><td></td><td></td><td></td></tr>
<tr><td>Oligo rev</td><td>AAACGTGTTAACCAAGTTGTCCCA</td><td></td><td></td><td></td><td></td><td></td><td></td><td></td><td></td><td></td><td></td><td></td></tr>
<tr><td>Chromosome</td><td>start</td><td></td><td>end</td><td>strand</td><td>MM</td><td>target_seq</td><td>PAM</td><td>alignment</td><td>distance</td><td>position</td><td>gene name</td><td>gene id</td></tr>
<tr><td>GU305914.1</td><td></td><td>4568</td><td>4590</td><td>+</td><td>0</td><td>TGGGACAACTTGGTTAACAC</td><td>CGG</td><td>||||||||[||||||||||||]PAM</td><td>NA</td><td>Intergenic</td><td>NA</td><td>NA</td></tr>
<tr><td>T273</td><td>CGTTCGCCACCAACGTTCAACGG</td><td></td><td>1000</td><td></td><td></td><td></td><td></td><td></td><td></td><td></td><td></td><td></td></tr>
<tr><td>Oligo fwd</td><td>CACCCGTTCGCCACCAACGTTCAA</td><td></td><td></td><td></td><td></td><td></td><td></td><td></td><td></td><td></td><td></td><td></td></tr>
<tr><td>Oligo rev</td><td>AAACTTGAACGTTGGTGGCGAACG</td><td></td><td></td><td></td><td></td><td></td><td></td><td></td><td></td><td></td><td></td><td></td></tr>
<tr><td>Chromosome</td><td>start</td><td></td><td>end</td><td>strand</td><td>MM</td><td>target_seq</td><td>PAM</td><td>alignment</td><td>distance</td><td>position</td><td>gene name</td><td>gene id</td></tr>
<tr><td>GU305914.1</td><td></td><td>4614</td><td>4636</td><td>+</td><td>0</td><td>CGTTCGCCACCAACGTTCAA</td><td>CGG</td><td>||||||||[||||||||||||]PAM</td><td>NA</td><td>Intergenic</td><td>NA</td><td>NA</td></tr>
<tr><td>T274</td><td>CGTCGCTTACTCGACTAGATTGG</td><td></td><td>1000</td><td></td><td></td><td></td><td></td><td></td><td></td><td></td><td></td><td></td></tr>
<tr><td>Oligo fwd</td><td>CACCCGTCGCTTACTCGACTAGAT</td><td></td><td></td><td></td><td></td><td></td><td></td><td></td><td></td><td></td><td></td><td></td></tr>
<tr><td>Oligo rev</td><td>AAACATCTAGTCGAGTAAGCGACG</td><td></td><td></td><td></td><td></td><td></td><td></td><td></td><td></td><td></td><td></td><td></td></tr>
<tr><td>Chromosome</td><td>start</td><td></td><td>end</td><td>strand</td><td>MM</td><td>target_seq</td><td>PAM</td><td>alignment</td><td>distance</td><td>position</td><td>gene name</td><td>gene id</td></tr>
<tr><td>GU305914.1</td><td></td><td>4646</td><td>4668</td><td>+</td><td>0</td><td>CGTCGCTTACTCGACTAGAT</td><td>TGG</td><td>||||||||[||||||||||||]PAM</td><td>NA</td><td>Intergenic</td><td>NA</td><td>NA</td></tr>
<tr><td>T275</td><td>AATCCGAACCCAAATAAAACAGG</td><td></td><td>1000</td><td></td><td></td><td></td><td></td><td></td><td></td><td></td><td></td><td></td></tr>
<tr><td>Oligo fwd</td><td>CACCAATCCGAACCCAAATAAAAC</td><td></td><td></td><td></td><td></td><td></td><td></td><td></td><td></td><td></td><td></td><td></td></tr>
<tr><td>Oligo rev</td><td>AAACGTTTTATTTGGGTTCGGATT</td><td></td><td></td><td></td><td></td><td></td><td></td><td></td><td></td><td></td><td></td><td></td></tr>
<tr><td>Chromosome</td><td>start</td><td></td><td>end</td><td>strand</td><td>MM</td><td>target_seq</td><td>PAM</td><td>alignment</td><td>distance</td><td>position</td><td>gene name</td><td>gene id</td></tr>
<tr><td>GU305914.1</td><td></td><td>4689</td><td>4711</td><td>+</td><td>0</td><td>AATCCGAACCCAAATAAAAC</td><td>AGG</td><td>||||||||[||||||||||||]PAM</td><td>NA</td><td>Intergenic</td><td>NA</td><td>NA</td></tr>
<tr><td>T276</td><td>GGATTGCGCGTCGATAAGTCCGG</td><td></td><td>1000</td><td></td><td></td><td></td><td></td><td></td><td></td><td></td><td></td><td></td></tr>
<tr><td>Oligo fwd</td><td>CACCGGATTGCGCGTCGATAAGTC</td><td></td><td></td><td></td><td></td><td></td><td></td><td></td><td></td><td></td><td></td><td></td></tr>
<tr><td>Oligo rev</td><td>AAACGACTTATCGACGCGCAATCC</td><td></td><td></td><td></td><td></td><td></td><td></td><td></td><td></td><td></td><td></td><td></td></tr>
<tr><td>Chromosome</td><td>start</td><td></td><td>end</td><td>strand</td><td>MM</td><td>target_seq</td><td>PAM</td><td>alignment</td><td>distance</td><td>position</td><td>gene name</td><td>gene id</td></tr>
<tr><td>GU305914.1</td><td></td><td>4710</td><td>4732</td><td>+</td><td>0</td><td>GGATTGCGCGTCGATAAGTC</td><td>CGG</td><td>||||||||[||||||||||||]PAM</td><td>NA</td><td>Intergenic</td><td>NA</td><td>NA</td></tr>
<tr><td>T277</td><td>CGTACGAGCCGATCTCGTGAAGG</td><td></td><td>1000</td><td></td><td></td><td></td><td></td><td></td><td></td><td></td><td></td><td></td></tr>
<tr><td>Oligo fwd</td><td>CACCCGTACGAGCCGATCTCGTGA</td><td></td><td></td><td></td><td></td><td></td><td></td><td></td><td></td><td></td><td></td><td></td></tr>
<tr><td>Oligo rev</td><td>AAACTCACGAGATCGGCTCGTACG</td><td></td><td></td><td></td><td></td><td></td><td></td><td></td><td></td><td></td><td></td><td></td></tr>
<tr><td>Chromosome</td><td>start</td><td></td><td>end</td><td>strand</td><td>MM</td><td>target_seq</td><td>PAM</td><td>alignment</td><td>distance</td><td>position</td><td>gene name</td><td>gene id</td></tr>
<tr><td>GU305914.1</td><td></td><td>4751</td><td>4773</td><td>+</td><td>0</td><td>CGTACGAGCCGATCTCGTGA</td><td>AGG</td><td>||||||||[||||||||||||]PAM</td><td>NA</td><td>Intergenic</td><td>NA</td><td>NA</td></tr>
<tr><td>T278</td><td>GTACATTGTCAGTATGCTTGTGG</td><td></td><td>1000</td><td></td><td></td><td></td><td></td><td></td><td></td><td></td><td></td><td></td></tr>
<tr><td>Oligo fwd</td><td>CACCGTACATTGTCAGTATGCTTG</td><td></td><td></td><td></td><td></td><td></td><td></td><td></td><td></td><td></td><td></td><td></td></tr>
<tr><td>Oligo rev</td><td>AAACCAAGCATACTGACAATGTAC</td><td></td><td></td><td></td><td></td><td></td><td></td><td></td><td></td><td></td><td></td><td></td></tr>
</table>

Chromosome start end strand MM target_seq PAM alignment distance position gene name gene id
GU305914.1 4792 4814 + 0 GTACATTGTCAGTATGCTTG TGG ||||||||[||||||||||||]PAM NA Intergenic NA NA

T279 CAGTATGCTTGTGGACAGACAGG 1000
Oligo fwd CACCCAGTATGCTTGTGGACAGAC
Oligo rev AAACGTCTGTCCACAAGCATACTG
Chromosome start end strand MM target_seq PAM alignment distance position gene name gene id
GU305914.1 4801 4823 + 0 CAGTATGCTTGTGGACAGAC AGG ||||||||[||||||||||||]PAM NA Intergenic NA NA

T280 TCTGTATATAACACCTCCGATGG 1000
Oligo fwd CACCTCTGTATATAACACCTCCGA
Oligo rev AAACTCGGAGGTGTTATATACAGA
Chromosome start end strand MM target_seq PAM alignment distance position gene name gene id
GU305914.1 4827 4849 + 0 TCTGTATATAACACCTCCGA TGG ||||||||[||||||||||||]PAM NA Intergenic NA NA

T281 CTGTATATAACACCTCCGATGGG 1000
Oligo fwd CACCCTGTATATAACACCTCCGAT
Oligo rev AAACATCGGAGGTGTTATATACAG
Chromosome start end strand MM target_seq PAM alignment distance position gene name gene id
GU305914.1 4828 4850 + 0 CTGTATATAACACCTCCGAT GGG ||||||||[||||||||||||]PAM NA Intergenic NA NA

T282 GATGACTTCGTCTATCCACTCGG 1000
Oligo fwd CACCGATGACTTCGTCTATCCACT
Oligo rev AAACAGTGGATAGACGAAGTCATC
Chromosome start end strand MM target_seq PAM alignment distance position gene name gene id
GU305914.1 4917 4939 + 0 GATGACTTCGTCTATCCACT CGG ||||||||[||||||||||||]PAM NA Intergenic NA NA

T283 ATGACTTCGTCTATCCACTCGGG 1000
Oligo fwd CACCATGACTTCGTCTATCCACTC
Oligo rev AAACGAGTGGATAGACGAAGTCAT
Chromosome start end strand MM target_seq PAM alignment distance position gene name gene id
GU305914.1 4918 4940 + 0 ATGACTTCGTCTATCCACTC GGG ||||||||[||||||||||||]PAM NA Intergenic NA NA

T284 CTATCCACTCGGGTATCCAATGG 1000
Oligo fwd CACCCTATCCACTCGGGTATCCAA
Oligo rev AAACTTGGATACCCGAGTGGATAG
Chromosome start end strand MM target_seq PAM alignment distance position gene name gene id
GU305914.1 4928 4950 + 0 CTATCCACTCGGGTATCCAA TGG ||||||||[||||||||||||]PAM NA Intergenic NA NA

T285 CAAAGAAGAGCGCGTATAGTTGG 1000
Oligo fwd CACCCAAAGAAGAGCGCGTATAGT
Oligo rev AAACACTATACGCGCTCTTCTTTG
Chromosome start end strand MM target_seq PAM alignment distance position gene name gene id
GU305914.1 4957 4979 + 0 CAAAGAAGAGCGCGTATAGT TGG ||||||||[||||||||||||]PAM NA Intergenic NA NA

T286 ATGTCGATGAGTGTATTTTTTGG 1000
Oligo fwd CACCATGTCGATGAGTGTATTTTT
Oligo rev AAACAAAAATACACTCATCGACAT
Chromosome start end strand MM target_seq PAM alignment distance position gene name gene id
GU305914.1 5004 5026 + 0 ATGTCGATGAGTGTATTTTT TGG ||||||||[||||||||||||]PAM NA Intergenic NA NA

T287 GTATATATCGTGCTAATCATCGG 1000
Oligo fwd CACCGTATATATCGTGCTAATCAT
Oligo rev AAACATGATTAGCACGATATATAC
Chromosome start end strand MM target_seq PAM alignment distance position gene name gene id
GU305914.1 5037 5059 + 0 GTATATATCGTGCTAATCAT CGG ||||||||[||||||||||||]PAM NA Intergenic NA NA

T288 GACGTTGCTGTCGAATACCGAGG 1000
Oligo fwd CACCGACGTTGCTGTCGAATACCG
Oligo rev AAACCGGTATTCGACAGCAACGTC
Chromosome start end strand MM target_seq PAM alignment distance position gene name gene id
GU305914.1 5115 5137 + 0 GACGTTGCTGTCGAATACCG AGG ||||||||[||||||||||||]PAM NA Intergenic NA NA

T289 ACGTTGCTGTCGAATACCGAGGG 1000
Oligo fwd CACCACGTTGCTGTCGAATACCGA
Oligo rev AAACTCGGTATTCGACAGCAACGT
Chromosome start end strand MM target_seq PAM alignment distance position gene name gene id
GU305914.1 5116 5138 + 0 ACGTTGCTGTCGAATACCGA GGG ||||||||[||||||||||||]PAM NA Intergenic NA NA

T290 TGTAGTTGTTTCTTCCGATCTGG 1000
Oligo fwd CACCTGTAGTTGTTTCTTCCGATC
Oligo rev AAACGATCGGAAGAAACAACTACA
Chromosome start end strand MM target_seq PAM alignment distance position gene name gene id
GU305914.1 5156 5178 + 0 TGTAGTTGTTTCTTCCGATC TGG ||||||||[||||||||||||]PAM NA Intergenic NA NA

T291 CATCGAGATCTTAGACTAAGCGG 1000
Oligo fwd CACCCATCGAGATCTTAGACTAAG
Oligo rev AAACCTTAGTCTAAGATCTCGATG
Chromosome start end strand MM target_seq PAM alignment distance position gene name gene id
GU305914.1 5183 5205 + 0 CATCGAGATCTTAGACTAAG CGG ||||||||[||||||||||||]PAM NA Intergenic NA NA

T292 ACTAAGCGGCTCATAAATCGCGG 1000
Oligo fwd CACCACTAAGCGGCTCATAAATCG
Oligo rev AAACCGATTTATGAGCCGCTTAGT
Chromosome start end strand MM target_seq PAM alignment distance position gene name gene id
GU305914.1 5197 5219 + 0 ACTAAGCGGCTCATAAATCG CGG ||||||||[||||||||||||]PAM NA Intergenic NA NA

T293 GGCTCATAAATCGCGGCGACCGG 1000
Oligo fwd CACCGGCTCATAAATCGCGGCGAC

<table>
<tr><td>Oligo rev</td><td>AAACGTCGCCGCGATTTATGAGCC</td><td></td><td></td><td></td><td></td><td></td><td></td><td></td><td></td><td></td><td></td><td></td></tr>
<tr><td>Chromosome</td><td>start</td><td></td><td>end</td><td>strand</td><td>MM</td><td>target_seq</td><td>PAM</td><td>alignment</td><td>distance</td><td>position</td><td>gene name</td><td>gene id</td></tr>
<tr><td>GU305914.1</td><td></td><td>5204</td><td>5226</td><td>+</td><td>0</td><td>GGCTCATAAATCGCGGCGAC</td><td>CGG</td><td>||||||||[||||||||||||]PAM</td><td>NA</td><td>Intergenic</td><td>NA</td><td>NA</td></tr>
<tr><td>T294</td><td>ATCACGCAAACATATATAAATGG</td><td></td><td>1000</td><td></td><td></td><td></td><td></td><td></td><td></td><td></td><td></td><td></td></tr>
<tr><td>Oligo fwd</td><td>CACCATCACGCAAACATATATAAA</td><td></td><td></td><td></td><td></td><td></td><td></td><td></td><td></td><td></td><td></td><td></td></tr>
<tr><td>Oligo rev</td><td>AAACTTTATATATGTTTGCGTGAT</td><td></td><td></td><td></td><td></td><td></td><td></td><td></td><td></td><td></td><td></td><td></td></tr>
<tr><td>Chromosome</td><td>start</td><td></td><td>end</td><td>strand</td><td>MM</td><td>target_seq</td><td>PAM</td><td>alignment</td><td>distance</td><td>position</td><td>gene name</td><td>gene id</td></tr>
<tr><td>GU305914.1</td><td></td><td>5252</td><td>5274</td><td>+</td><td>0</td><td>ATCACGCAAACATATATAAA</td><td>TGG</td><td>||||||||[||||||||||||]PAM</td><td>NA</td><td>Intergenic</td><td>NA</td><td>NA</td></tr>
<tr><td>T295</td><td>GAAGCCAATCTATGCCCAGTTGG</td><td></td><td>1000</td><td></td><td></td><td></td><td></td><td></td><td></td><td></td><td></td><td></td></tr>
<tr><td>Oligo fwd</td><td>CACCGAAGCCAATCTATGCCCAGT</td><td></td><td></td><td></td><td></td><td></td><td></td><td></td><td></td><td></td><td></td><td></td></tr>
<tr><td>Oligo rev</td><td>AAACACTGGGCATAGATTGGCTTC</td><td></td><td></td><td></td><td></td><td></td><td></td><td></td><td></td><td></td><td></td><td></td></tr>
<tr><td>Chromosome</td><td>start</td><td></td><td>end</td><td>strand</td><td>MM</td><td>target_seq</td><td>PAM</td><td>alignment</td><td>distance</td><td>position</td><td>gene name</td><td>gene id</td></tr>
<tr><td>GU305914.1</td><td></td><td>5280</td><td>5302</td><td>+</td><td>0</td><td>GAAGCCAATCTATGCCCAGT</td><td>TGG</td><td>||||||||[||||||||||||]PAM</td><td>NA</td><td>Intergenic</td><td>NA</td><td>NA</td></tr>
<tr><td>T296</td><td>TTGGAGCGATACGTTGACAATGG</td><td></td><td>1000</td><td></td><td></td><td></td><td></td><td></td><td></td><td></td><td></td><td></td></tr>
<tr><td>Oligo fwd</td><td>CACCTTGGAGCGATACGTTGACAA</td><td></td><td></td><td></td><td></td><td></td><td></td><td></td><td></td><td></td><td></td><td></td></tr>
<tr><td>Oligo rev</td><td>AAACTTGTCAACGTATCGCTCCAA</td><td></td><td></td><td></td><td></td><td></td><td></td><td></td><td></td><td></td><td></td><td></td></tr>
<tr><td>Chromosome</td><td>start</td><td></td><td>end</td><td>strand</td><td>MM</td><td>target_seq</td><td>PAM</td><td>alignment</td><td>distance</td><td>position</td><td>gene name</td><td>gene id</td></tr>
<tr><td>GU305914.1</td><td></td><td>5299</td><td>5321</td><td>+</td><td>0</td><td>TTGGAGCGATACGTTGACAA</td><td>TGG</td><td>||||||||[||||||||||||]PAM</td><td>NA</td><td>Intergenic</td><td>NA</td><td>NA</td></tr>
<tr><td>T297</td><td>ACAATGGACACAACTGCGAGAGG</td><td></td><td>1000</td><td></td><td></td><td></td><td></td><td></td><td></td><td></td><td></td><td></td></tr>
<tr><td>Oligo fwd</td><td>CACCACAATGGACACAACTGCGAG</td><td></td><td></td><td></td><td></td><td></td><td></td><td></td><td></td><td></td><td></td><td></td></tr>
<tr><td>Oligo rev</td><td>AAACCTCGCAGTTGTGTCCATTGT</td><td></td><td></td><td></td><td></td><td></td><td></td><td></td><td></td><td></td><td></td><td></td></tr>
<tr><td>Chromosome</td><td>start</td><td></td><td>end</td><td>strand</td><td>MM</td><td>target_seq</td><td>PAM</td><td>alignment</td><td>distance</td><td>position</td><td>gene name</td><td>gene id</td></tr>
<tr><td>GU305914.1</td><td></td><td>5315</td><td>5337</td><td>+</td><td>0</td><td>ACAATGGACACAACTGCGAG</td><td>AGG</td><td>||||||||[||||||||||||]PAM</td><td>NA</td><td>Intergenic</td><td>NA</td><td>NA</td></tr>
<tr><td>T298</td><td>CAACTGCGAGAGGATCTACCCGG</td><td></td><td>1000</td><td></td><td></td><td></td><td></td><td></td><td></td><td></td><td></td><td></td></tr>
<tr><td>Oligo fwd</td><td>CACCCAACTGCGAGAGGATCTACC</td><td></td><td></td><td></td><td></td><td></td><td></td><td></td><td></td><td></td><td></td><td></td></tr>
<tr><td>Oligo rev</td><td>AAACGGTAGATCCTCTCGCAGTTG</td><td></td><td></td><td></td><td></td><td></td><td></td><td></td><td></td><td></td><td></td><td></td></tr>
<tr><td>Chromosome</td><td>start</td><td></td><td>end</td><td>strand</td><td>MM</td><td>target_seq</td><td>PAM</td><td>alignment</td><td>distance</td><td>position</td><td>gene name</td><td>gene id</td></tr>
<tr><td>GU305914.1</td><td></td><td>5325</td><td>5347</td><td>+</td><td>0</td><td>CAACTGCGAGAGGATCTACC</td><td>CGG</td><td>||||||||[||||||||||||]PAM</td><td>NA</td><td>Intergenic</td><td>NA</td><td>NA</td></tr>
<tr><td>T299</td><td>AGGATCTACCCGGTCGCGTCCGG</td><td></td><td>1000</td><td></td><td></td><td></td><td></td><td></td><td></td><td></td><td></td><td></td></tr>
<tr><td>Oligo fwd</td><td>CACCAGGATCTACCCGGTCGCGTC</td><td></td><td></td><td></td><td></td><td></td><td></td><td></td><td></td><td></td><td></td><td></td></tr>
<tr><td>Oligo rev</td><td>AAACGACGCGACCGGGTAGATCCT</td><td></td><td></td><td></td><td></td><td></td><td></td><td></td><td></td><td></td><td></td><td></td></tr>
<tr><td>Chromosome</td><td>start</td><td></td><td>end</td><td>strand</td><td>MM</td><td>target_seq</td><td>PAM</td><td>alignment</td><td>distance</td><td>position</td><td>gene name</td><td>gene id</td></tr>
<tr><td>GU305914.1</td><td></td><td>5335</td><td>5357</td><td>+</td><td>0</td><td>AGGATCTACCCGGTCGCGTC</td><td>CGG</td><td>||||||||[||||||||||||]PAM</td><td>NA</td><td>Intergenic</td><td>NA</td><td>NA</td></tr>
<tr><td>T300</td><td>GTCGCGTCCGGTACAGCTCTTGG</td><td></td><td>1000</td><td></td><td></td><td></td><td></td><td></td><td></td><td></td><td></td><td></td></tr>
<tr><td>Oligo fwd</td><td>CACCGTCGCGTCCGGTACAGCTCT</td><td></td><td></td><td></td><td></td><td></td><td></td><td></td><td></td><td></td><td></td><td></td></tr>
<tr><td>Oligo rev</td><td>AAACAGAGCTGTACCGGACGCGAC</td><td></td><td></td><td></td><td></td><td></td><td></td><td></td><td></td><td></td><td></td><td></td></tr>
<tr><td>Chromosome</td><td>start</td><td></td><td>end</td><td>strand</td><td>MM</td><td>target_seq</td><td>PAM</td><td>alignment</td><td>distance</td><td>position</td><td>gene name</td><td>gene id</td></tr>
<tr><td>GU305914.1</td><td></td><td>5347</td><td>5369</td><td>+</td><td>0</td><td>GTCGCGTCCGGTACAGCTCT</td><td>TGG</td><td>||||||||[||||||||||||]PAM</td><td>NA</td><td>Intergenic</td><td>NA</td><td>NA</td></tr>
<tr><td>T301</td><td>AGCCTCGATGACTTTCCAGATGG</td><td></td><td>1000</td><td></td><td></td><td></td><td></td><td></td><td></td><td></td><td></td><td></td></tr>
<tr><td>Oligo fwd</td><td>CACCAGCCTCGATGACTTTCCAGA</td><td></td><td></td><td></td><td></td><td></td><td></td><td></td><td></td><td></td><td></td><td></td></tr>
<tr><td>Oligo rev</td><td>AAACTCTGGAAAGTCATCGAGGCT</td><td></td><td></td><td></td><td></td><td></td><td></td><td></td><td></td><td></td><td></td><td></td></tr>
<tr><td>Chromosome</td><td>start</td><td></td><td>end</td><td>strand</td><td>MM</td><td>target_seq</td><td>PAM</td><td>alignment</td><td>distance</td><td>position</td><td>gene name</td><td>gene id</td></tr>
<tr><td>GU305914.1</td><td></td><td>5386</td><td>5408</td><td>+</td><td>0</td><td>AGCCTCGATGACTTTCCAGA</td><td>TGG</td><td>||||||||[||||||||||||]PAM</td><td>NA</td><td>Intergenic</td><td>NA</td><td>NA</td></tr>
<tr><td>T302</td><td>GCCTCGATGACTTTCCAGATGGG</td><td></td><td>1000</td><td></td><td></td><td></td><td></td><td></td><td></td><td></td><td></td><td></td></tr>
<tr><td>Oligo fwd</td><td>CACCGCCTCGATGACTTTCCAGAT</td><td></td><td></td><td></td><td></td><td></td><td></td><td></td><td></td><td></td><td></td><td></td></tr>
<tr><td>Oligo rev</td><td>AAACATCTGGAAAGTCATCGAGGC</td><td></td><td></td><td></td><td></td><td></td><td></td><td></td><td></td><td></td><td></td><td></td></tr>
<tr><td>Chromosome</td><td>start</td><td></td><td>end</td><td>strand</td><td>MM</td><td>target_seq</td><td>PAM</td><td>alignment</td><td>distance</td><td>position</td><td>gene name</td><td>gene id</td></tr>
<tr><td>GU305914.1</td><td></td><td>5387</td><td>5409</td><td>+</td><td>0</td><td>GCCTCGATGACTTTCCAGAT</td><td>GGG</td><td>||||||||[||||||||||||]PAM</td><td>NA</td><td>Intergenic</td><td>NA</td><td>NA</td></tr>
<tr><td>T303</td><td>AGAGTCTTACGTTAAGACAGAGG</td><td></td><td>1000</td><td></td><td></td><td></td><td></td><td></td><td></td><td></td><td></td><td></td></tr>
<tr><td>Oligo fwd</td><td>CACCAGAGTCTTACGTTAAGACAG</td><td></td><td></td><td></td><td></td><td></td><td></td><td></td><td></td><td></td><td></td><td></td></tr>
<tr><td>Oligo rev</td><td>AAACCTGTCTTAACGTAAGACTCT</td><td></td><td></td><td></td><td></td><td></td><td></td><td></td><td></td><td></td><td></td><td></td></tr>
<tr><td>Chromosome</td><td>start</td><td></td><td>end</td><td>strand</td><td>MM</td><td>target_seq</td><td>PAM</td><td>alignment</td><td>distance</td><td>position</td><td>gene name</td><td>gene id</td></tr>
<tr><td>GU305914.1</td><td></td><td>5515</td><td>5537</td><td>+</td><td>0</td><td>AGAGTCTTACGTTAAGACAG</td><td>AGG</td><td>||||||||[||||||||||||]PAM</td><td>NA</td><td>Intergenic</td><td>NA</td><td>NA</td></tr>
<tr><td>T304</td><td>TAAGACAGAGGAACCTATCGTGG</td><td></td><td>1000</td><td></td><td></td><td></td><td></td><td></td><td></td><td></td><td></td><td></td></tr>
<tr><td>Oligo fwd</td><td>CACCTAAGACAGAGGAACCTATCG</td><td></td><td></td><td></td><td></td><td></td><td></td><td></td><td></td><td></td><td></td><td></td></tr>
<tr><td>Oligo rev</td><td>AAACCGATAGGTTCCTCTGTCTTA</td><td></td><td></td><td></td><td></td><td></td><td></td><td></td><td></td><td></td><td></td><td></td></tr>
<tr><td>Chromosome</td><td>start</td><td></td><td>end</td><td>strand</td><td>MM</td><td>target_seq</td><td>PAM</td><td>alignment</td><td>distance</td><td>position</td><td>gene name</td><td>gene id</td></tr>
<tr><td>GU305914.1</td><td></td><td>5527</td><td>5549</td><td>+</td><td>0</td><td>TAAGACAGAGGAACCTATCG</td><td>TGG</td><td>||||||||[||||||||||||]PAM</td><td>NA</td><td>Intergenic</td><td>NA</td><td>NA</td></tr>
<tr><td>T305</td><td>AAGACAGAGGAACCTATCGTGGG</td><td></td><td>1000</td><td></td><td></td><td></td><td></td><td></td><td></td><td></td><td></td><td></td></tr>
<tr><td>Oligo fwd</td><td>CACCAAGACAGAGGAACCTATCGT</td><td></td><td></td><td></td><td></td><td></td><td></td><td></td><td></td><td></td><td></td><td></td></tr>
<tr><td>Oligo rev</td><td>AAACACGATAGGTTCCTCTGTCTT</td><td></td><td></td><td></td><td></td><td></td><td></td><td></td><td></td><td></td><td></td><td></td></tr>
<tr><td>Chromosome</td><td>start</td><td></td><td>end</td><td>strand</td><td>MM</td><td>target_seq</td><td>PAM</td><td>alignment</td><td>distance</td><td>position</td><td>gene name</td><td>gene id</td></tr>
<tr><td>GU305914.1</td><td></td><td>5528</td><td>5550</td><td>+</td><td>0</td><td>AAGACAGAGGAACCTATCGT</td><td>GGG</td><td>||||||||[||||||||||||]PAM</td><td>NA</td><td>Intergenic</td><td>NA</td><td>NA</td></tr>
<tr><td>T306</td><td>AGACAGAGGAACCTATCGTGGGG</td><td></td><td>1000</td><td></td><td></td><td></td><td></td><td></td><td></td><td></td><td></td><td></td></tr>
<tr><td>Oligo fwd</td><td>CACCAGACAGAGGAACCTATCGTG</td><td></td><td></td><td></td><td></td><td></td><td></td><td></td><td></td><td></td><td></td><td></td></tr>
<tr><td>Oligo rev</td><td>AAACCACGATAGGTTCCTCTGTCT</td><td></td><td></td><td></td><td></td><td></td><td></td><td></td><td></td><td></td><td></td><td></td></tr>
<tr><td>Chromosome</td><td>start</td><td></td><td>end</td><td>strand</td><td>MM</td><td>target_seq</td><td>PAM</td><td>alignment</td><td>distance</td><td>position</td><td>gene name</td><td>gene id</td></tr>
<tr><td>GU305914.1</td><td></td><td>5529</td><td>5551</td><td>+</td><td>0</td><td>AGACAGAGGAACCTATCGTG</td><td>GGG</td><td>||||||||[||||||||||||]PAM</td><td>NA</td><td>Intergenic</td><td>NA</td><td>NA</td></tr>
<tr><td>T307</td><td>CACATTCAAATGTTTCAAGAAGG</td><td></td><td>1000</td><td></td><td></td><td></td><td></td><td></td><td></td><td></td><td></td><td></td></tr>
<tr><td>Oligo fwd</td><td>CACCCACATTCAAATGTTTCAAGA</td><td></td><td></td><td></td><td></td><td></td><td></td><td></td><td></td><td></td><td></td><td></td></tr>
<tr><td>Oligo rev</td><td>AAACTCTTGAAACATTTGAATGTG</td><td></td><td></td><td></td><td></td><td></td><td></td><td></td><td></td><td></td><td></td><td></td></tr>
<tr><td>Chromosome</td><td>start</td><td></td><td>end</td><td>strand</td><td>MM</td><td>target_seq</td><td>PAM</td><td>alignment</td><td>distance</td><td>position</td><td>gene name</td><td>gene id</td></tr>
<tr><td>GU305914.1</td><td></td><td>5611</td><td>5633</td><td>+</td><td>0</td><td>CACATTCAAATGTTTCAAGA</td><td>AGG</td><td>||||||||[||||||||||||]PAM</td><td>NA</td><td>Intergenic</td><td>NA</td><td>NA</td></tr>
<tr><td>T308</td><td>TTCAAATGTTTCAAGAAGGACGG</td><td></td><td>1000</td><td></td><td></td><td></td><td></td><td></td><td></td><td></td><td></td><td></td></tr>
</table>

<table>
<tr><td>Oligo fwd</td><td>CACCTTCAAATGTTTCAAGAAGGA</td><td></td><td></td><td></td><td></td><td></td><td></td><td></td><td></td><td></td><td></td></tr>
<tr><td>Oligo rev</td><td>AAACTCCTTCTTGAAACATTTGAA</td><td></td><td></td><td></td><td></td><td></td><td></td><td></td><td></td><td></td><td></td></tr>
<tr><td>Chromosome</td><td>start</td><td>end</td><td>strand</td><td>MM</td><td>target_seq</td><td>PAM</td><td>alignment</td><td>distance</td><td>position</td><td>gene name</td><td>gene id</td></tr>
<tr><td>GU305914.1</td><td>5615</td><td>5637</td><td>+</td><td>0</td><td>TTCAAATGTTTCAAGAAGGA</td><td>CGG</td><td>||||||||[||||||||||||]PAM</td><td>NA</td><td>Intergenic</td><td>NA</td><td>NA</td></tr>
<tr><td></td><td></td><td></td><td></td><td></td><td></td><td></td><td></td><td></td><td></td><td></td><td></td></tr>
<tr><td>T309</td><td>TCACTCGATATGCAGACGACCGG</td><td>1000</td><td></td><td></td><td></td><td></td><td></td><td></td><td></td><td></td><td></td></tr>
<tr><td>Oligo fwd</td><td>CACCTCACTCGATATGCAGACGAC</td><td></td><td></td><td></td><td></td><td></td><td></td><td></td><td></td><td></td><td></td></tr>
<tr><td>Oligo rev</td><td>AAACGTCGTCTGCATATCGAGTGA</td><td></td><td></td><td></td><td></td><td></td><td></td><td></td><td></td><td></td><td></td></tr>
<tr><td>Chromosome</td><td>start</td><td>end</td><td>strand</td><td>MM</td><td>target_seq</td><td>PAM</td><td>alignment</td><td>distance</td><td>position</td><td>gene name</td><td>gene id</td></tr>
<tr><td>GU305914.1</td><td>5648</td><td>5670</td><td>+</td><td>0</td><td>TCACTCGATATGCAGACGAC</td><td>CGG</td><td>||||||||[||||||||||||]PAM</td><td>NA</td><td>Intergenic</td><td>NA</td><td>NA</td></tr>
<tr><td></td><td></td><td></td><td></td><td></td><td></td><td></td><td></td><td></td><td></td><td></td><td></td></tr>
<tr><td>T310</td><td>ACGTTTACGTGAATGCCACGTGG</td><td>1000</td><td></td><td></td><td></td><td></td><td></td><td></td><td></td><td></td><td></td></tr>
<tr><td>Oligo fwd</td><td>CACCACGTTTACGTGAATGCCACG</td><td></td><td></td><td></td><td></td><td></td><td></td><td></td><td></td><td></td><td></td></tr>
<tr><td>Oligo rev</td><td>AAACCGTGGCATTCACGTAAACGT</td><td></td><td></td><td></td><td></td><td></td><td></td><td></td><td></td><td></td><td></td></tr>
<tr><td>Chromosome</td><td>start</td><td>end</td><td>strand</td><td>MM</td><td>target_seq</td><td>PAM</td><td>alignment</td><td>distance</td><td>position</td><td>gene name</td><td>gene id</td></tr>
<tr><td>GU305914.1</td><td>5703</td><td>5725</td><td>+</td><td>0</td><td>ACGTTTACGTGAATGCCACG</td><td>TGG</td><td>||||||||[||||||||||||]PAM</td><td>NA</td><td>Intergenic</td><td>NA</td><td>NA</td></tr>
<tr><td></td><td></td><td></td><td></td><td></td><td></td><td></td><td></td><td></td><td></td><td></td><td></td></tr>
<tr><td>T311</td><td>ATGCCACGTGGTACGTACACTGG</td><td>1000</td><td></td><td></td><td></td><td></td><td></td><td></td><td></td><td></td><td></td></tr>
<tr><td>Oligo fwd</td><td>CACCATGCCACGTGGTACGTACAC</td><td></td><td></td><td></td><td></td><td></td><td></td><td></td><td></td><td></td><td></td></tr>
<tr><td>Oligo rev</td><td>AAACGTGTACGTACCACGTGGCAT</td><td></td><td></td><td></td><td></td><td></td><td></td><td></td><td></td><td></td><td></td></tr>
<tr><td>Chromosome</td><td>start</td><td>end</td><td>strand</td><td>MM</td><td>target_seq</td><td>PAM</td><td>alignment</td><td>distance</td><td>position</td><td>gene name</td><td>gene id</td></tr>
<tr><td>GU305914.1</td><td>5715</td><td>5737</td><td>+</td><td>0</td><td>ATGCCACGTGGTACGTACAC</td><td>TGG</td><td>||||||||[||||||||||||]PAM</td><td>NA</td><td>Intergenic</td><td>NA</td><td>NA</td></tr>
<tr><td></td><td></td><td></td><td></td><td></td><td></td><td></td><td></td><td></td><td></td><td></td><td></td></tr>
<tr><td>T312</td><td>TGCCACGTGGTACGTACACTGGG</td><td>1000</td><td></td><td></td><td></td><td></td><td></td><td></td><td></td><td></td><td></td></tr>
<tr><td>Oligo fwd</td><td>CACCTGCCACGTGGTACGTACACT</td><td></td><td></td><td></td><td></td><td></td><td></td><td></td><td></td><td></td><td></td></tr>
<tr><td>Oligo rev</td><td>AAACAGTGTACGTACCACGTGGCA</td><td></td><td></td><td></td><td></td><td></td><td></td><td></td><td></td><td></td><td></td></tr>
<tr><td>Chromosome</td><td>start</td><td>end</td><td>strand</td><td>MM</td><td>target_seq</td><td>PAM</td><td>alignment</td><td>distance</td><td>position</td><td>gene name</td><td>gene id</td></tr>
<tr><td>GU305914.1</td><td>5716</td><td>5738</td><td>+</td><td>0</td><td>TGCCACGTGGTACGTACACT</td><td>GGG</td><td>||||||||[||||||||||||]PAM</td><td>NA</td><td>Intergenic</td><td>NA</td><td>NA</td></tr>
<tr><td></td><td></td><td></td><td></td><td></td><td></td><td></td><td></td><td></td><td></td><td></td><td></td></tr>
<tr><td>T313</td><td>TGGTACGTACACTGGGTCGTCGG</td><td>1000</td><td></td><td></td><td></td><td></td><td></td><td></td><td></td><td></td><td></td></tr>
<tr><td>Oligo fwd</td><td>CACCTGGTACGTACACTGGGTCGT</td><td></td><td></td><td></td><td></td><td></td><td></td><td></td><td></td><td></td><td></td></tr>
<tr><td>Oligo rev</td><td>AAACACGACCCAGTGTACGTACCA</td><td></td><td></td><td></td><td></td><td></td><td></td><td></td><td></td><td></td><td></td></tr>
<tr><td>Chromosome</td><td>start</td><td>end</td><td>strand</td><td>MM</td><td>target_seq</td><td>PAM</td><td>alignment</td><td>distance</td><td>position</td><td>gene name</td><td>gene id</td></tr>
<tr><td>GU305914.1</td><td>5723</td><td>5745</td><td>+</td><td>0</td><td>TGGTACGTACACTGGGTCGT</td><td>CGG</td><td>||||||||[||||||||||||]PAM</td><td>NA</td><td>Intergenic</td><td>NA</td><td>NA</td></tr>
<tr><td></td><td></td><td></td><td></td><td></td><td></td><td></td><td></td><td></td><td></td><td></td><td></td></tr>
<tr><td>T314</td><td>GGTACGTACACTGGGTCGTCGGG</td><td>1000</td><td></td><td></td><td></td><td></td><td></td><td></td><td></td><td></td><td></td></tr>
<tr><td>Oligo fwd</td><td>CACCGGTACGTACACTGGGTCGTC</td><td></td><td></td><td></td><td></td><td></td><td></td><td></td><td></td><td></td><td></td></tr>
<tr><td>Oligo rev</td><td>AAACGACGACCCAGTGTACGTACC</td><td></td><td></td><td></td><td></td><td></td><td></td><td></td><td></td><td></td><td></td></tr>
<tr><td>Chromosome</td><td>start</td><td>end</td><td>strand</td><td>MM</td><td>target_seq</td><td>PAM</td><td>alignment</td><td>distance</td><td>position</td><td>gene name</td><td>gene id</td></tr>
<tr><td>GU305914.1</td><td>5724</td><td>5746</td><td>+</td><td>0</td><td>GGTACGTACACTGGGTCGTC</td><td>GGG</td><td>||||||||[||||||||||||]PAM</td><td>NA</td><td>Intergenic</td><td>NA</td><td>NA</td></tr>
<tr><td></td><td></td><td></td><td></td><td></td><td></td><td></td><td></td><td></td><td></td><td></td><td></td></tr>
<tr><td>T315</td><td>CGTCGGGAAGATCGCAGCCTCGG</td><td>1000</td><td></td><td></td><td></td><td></td><td></td><td></td><td></td><td></td><td></td></tr>
<tr><td>Oligo fwd</td><td>CACCCGTCGGGAAGATCGCAGCCT</td><td></td><td></td><td></td><td></td><td></td><td></td><td></td><td></td><td></td><td></td></tr>
<tr><td>Oligo rev</td><td>AAACAGGCTGCGATCTTCCCGACG</td><td></td><td></td><td></td><td></td><td></td><td></td><td></td><td></td><td></td><td></td></tr>
<tr><td>Chromosome</td><td>start</td><td>end</td><td>strand</td><td>MM</td><td>target_seq</td><td>PAM</td><td>alignment</td><td>distance</td><td>position</td><td>gene name</td><td>gene id</td></tr>
<tr><td>GU305914.1</td><td>5740</td><td>5762</td><td>+</td><td>0</td><td>CGTCGGGAAGATCGCAGCCT</td><td>CGG</td><td>||||||||[||||||||||||]PAM</td><td>NA</td><td>Intergenic</td><td>NA</td><td>NA</td></tr>
<tr><td></td><td></td><td></td><td></td><td></td><td></td><td></td><td></td><td></td><td></td><td></td><td></td></tr>
<tr><td>T316</td><td>AACGAGCTCACCGCCTACGTTGG</td><td>1000</td><td></td><td></td><td></td><td></td><td></td><td></td><td></td><td></td><td></td></tr>
<tr><td>Oligo fwd</td><td>CACCAACGAGCTCACCGCCTACGT</td><td></td><td></td><td></td><td></td><td></td><td></td><td></td><td></td><td></td><td></td></tr>
<tr><td>Oligo rev</td><td>AAACACGTAGGCGGTGAGCTCGTT</td><td></td><td></td><td></td><td></td><td></td><td></td><td></td><td></td><td></td><td></td></tr>
<tr><td>Chromosome</td><td>start</td><td>end</td><td>strand</td><td>MM</td><td>target_seq</td><td>PAM</td><td>alignment</td><td>distance</td><td>position</td><td>gene name</td><td>gene id</td></tr>
<tr><td>GU305914.1</td><td>5791</td><td>5813</td><td>+</td><td>0</td><td>AACGAGCTCACCGCCTACGT</td><td>TGG</td><td>||||||||[||||||||||||]PAM</td><td>NA</td><td>Intergenic</td><td>NA</td><td>NA</td></tr>
<tr><td></td><td></td><td></td><td></td><td></td><td></td><td></td><td></td><td></td><td></td><td></td><td></td></tr>
<tr><td>T317</td><td>AGCTCACCGCCTACGTTGGATGG</td><td>1000</td><td></td><td></td><td></td><td></td><td></td><td></td><td></td><td></td><td></td></tr>
<tr><td>Oligo fwd</td><td>CACCAGCTCACCGCCTACGTTGGA</td><td></td><td></td><td></td><td></td><td></td><td></td><td></td><td></td><td></td><td></td></tr>
<tr><td>Oligo rev</td><td>AAACTCCAACGTAGGCGGTGAGCT</td><td></td><td></td><td></td><td></td><td></td><td></td><td></td><td></td><td></td><td></td></tr>
<tr><td>Chromosome</td><td>start</td><td>end</td><td>strand</td><td>MM</td><td>target_seq</td><td>PAM</td><td>alignment</td><td>distance</td><td>position</td><td>gene name</td><td>gene id</td></tr>
<tr><td>GU305914.1</td><td>5795</td><td>5817</td><td>+</td><td>0</td><td>AGCTCACCGCCTACGTTGGA</td><td>TGG</td><td>||||||||[||||||||||||]PAM</td><td>NA</td><td>Intergenic</td><td>NA</td><td>NA</td></tr>
<tr><td></td><td></td><td></td><td></td><td></td><td></td><td></td><td></td><td></td><td></td><td></td><td></td></tr>
<tr><td>T318</td><td>ACATGCACCCTTTTTATCGACGG</td><td>1000</td><td></td><td></td><td></td><td></td><td></td><td></td><td></td><td></td><td></td></tr>
<tr><td>Oligo fwd</td><td>CACCACATGCACCCTTTTTATCGA</td><td></td><td></td><td></td><td></td><td></td><td></td><td></td><td></td><td></td><td></td></tr>
<tr><td>Oligo rev</td><td>AAACTCGATAAAAAGGGTGCATGT</td><td></td><td></td><td></td><td></td><td></td><td></td><td></td><td></td><td></td><td></td></tr>
<tr><td>Chromosome</td><td>start</td><td>end</td><td>strand</td><td>MM</td><td>target_seq</td><td>PAM</td><td>alignment</td><td>distance</td><td>position</td><td>gene name</td><td>gene id</td></tr>
<tr><td>GU305914.1</td><td>5820</td><td>5842</td><td>+</td><td>0</td><td>ACATGCACCCTTTTTATCGA</td><td>CGG</td><td>||||||||[||||||||||||]PAM</td><td>NA</td><td>Intergenic</td><td>NA</td><td>NA</td></tr>
<tr><td></td><td></td><td></td><td></td><td></td><td></td><td></td><td></td><td></td><td></td><td></td><td></td></tr>
<tr><td>T319</td><td>AAAATCGTAACCGCCGCGAACGG</td><td>1000</td><td></td><td></td><td></td><td></td><td></td><td></td><td></td><td></td><td></td></tr>
<tr><td>Oligo fwd</td><td>CACCAAAATCGTAACCGCCGCGAA</td><td></td><td></td><td></td><td></td><td></td><td></td><td></td><td></td><td></td><td></td></tr>
<tr><td>Oligo rev</td><td>AAACTTCGCGGCGGTTACGATTTT</td><td></td><td></td><td></td><td></td><td></td><td></td><td></td><td></td><td></td><td></td></tr>
<tr><td>Chromosome</td><td>start</td><td>end</td><td>strand</td><td>MM</td><td>target_seq</td><td>PAM</td><td>alignment</td><td>distance</td><td>position</td><td>gene name</td><td>gene id</td></tr>
<tr><td>GU305914.1</td><td>5843</td><td>5865</td><td>+</td><td>0</td><td>AAAATCGTAACCGCCGCGAA</td><td>CGG</td><td>||||||||[||||||||||||]PAM</td><td>NA</td><td>Intergenic</td><td>NA</td><td>NA</td></tr>
<tr><td></td><td></td><td></td><td></td><td></td><td></td><td></td><td></td><td></td><td></td><td></td><td></td></tr>
<tr><td>T320</td><td>TTCAAAGTCGACGAGAAGACTGG</td><td>1000</td><td></td><td></td><td></td><td></td><td></td><td></td><td></td><td></td><td></td></tr>
<tr><td>Oligo fwd</td><td>CACCTTCAAAGTCGACGAGAAGAC</td><td></td><td></td><td></td><td></td><td></td><td></td><td></td><td></td><td></td><td></td></tr>
<tr><td>Oligo rev</td><td>AAACGTCTTCTCGTCGACTTTGAA</td><td></td><td></td><td></td><td></td><td></td><td></td><td></td><td></td><td></td><td></td></tr>
<tr><td>Chromosome</td><td>start</td><td>end</td><td>strand</td><td>MM</td><td>target_seq</td><td>PAM</td><td>alignment</td><td>distance</td><td>position</td><td>gene name</td><td>gene id</td></tr>
<tr><td>GU305914.1</td><td>5867</td><td>5889</td><td>+</td><td>0</td><td>TTCAAAGTCGACGAGAAGAC</td><td>TGG</td><td>||||||||[||||||||||||]PAM</td><td>NA</td><td>Intergenic</td><td>NA</td><td>NA</td></tr>
<tr><td></td><td></td><td></td><td></td><td></td><td></td><td></td><td></td><td></td><td></td><td></td><td></td></tr>
<tr><td>T321</td><td>CGAGCCTCGCAGACAGCGTCCGG</td><td>1000</td><td></td><td></td><td></td><td></td><td></td><td></td><td></td><td></td><td></td></tr>
<tr><td>Oligo fwd</td><td>CACCCGAGCCTCGCAGACAGCGTC</td><td></td><td></td><td></td><td></td><td></td><td></td><td></td><td></td><td></td><td></td></tr>
<tr><td>Oligo rev</td><td>AAACGACGCTGTCTGCGAGGCTCG</td><td></td><td></td><td></td><td></td><td></td><td></td><td></td><td></td><td></td><td></td></tr>
<tr><td>Chromosome</td><td>start</td><td>end</td><td>strand</td><td>MM</td><td>target_seq</td><td>PAM</td><td>alignment</td><td>distance</td><td>position</td><td>gene name</td><td>gene id</td></tr>
<tr><td>GU305914.1</td><td>5916</td><td>5938</td><td>+</td><td>0</td><td>CGAGCCTCGCAGACAGCGTC</td><td>CGG</td><td>||||||||[||||||||||||]PAM</td><td>NA</td><td>Intergenic</td><td>NA</td><td>NA</td></tr>
<tr><td>GU305914.1</td><td>138101</td><td>138123</td><td>+</td><td>4</td><td>CGATCGCCGCCGACAGCGTC</td><td>GGG</td><td>|||-|--|[||-|||||||||]PAM</td><td>NA</td><td>Intergenic</td><td>NA</td><td>NA</td></tr>
<tr><td></td><td></td><td></td><td></td><td></td><td></td><td></td><td></td><td></td><td></td><td></td><td></td></tr>
<tr><td>T322</td><td>GTCGCCTGATAGTGTGTCTATGG</td><td>1000</td><td></td><td></td><td></td><td></td><td></td><td></td><td></td><td></td><td></td></tr>
<tr><td>Oligo fwd</td><td>CACCGTCGCCTGATAGTGTGTCTA</td><td></td><td></td><td></td><td></td><td></td><td></td><td></td><td></td><td></td><td></td></tr>
<tr><td>Oligo rev</td><td>AAACTAGACACACTATCAGGCGAC</td><td></td><td></td><td></td><td></td><td></td><td></td><td></td><td></td><td></td><td></td></tr>
<tr><td>Chromosome</td><td>start</td><td>end</td><td>strand</td><td>MM</td><td>target_seq</td><td>PAM</td><td>alignment</td><td>distance</td><td>position</td><td>gene name</td><td>gene id</td></tr>
<tr><td>GU305914.1</td><td>5940</td><td>5962</td><td>+</td><td>0</td><td>GTCGCCTGATAGTGTGTCTA</td><td>TGG</td><td>||||||||[||||||||||||]PAM</td><td>NA</td><td>Intergenic</td><td>NA</td><td>NA</td></tr>
</table>

T323 TGACAGCATCAGCGACCTGCCGG 1000
Oligo fwd CACCTGACAGCATCAGCGACCTGC
Oligo rev AAACGCAGGTCGCTGATGCTGTCA
Chromosome start end strand MM target_seq PAM alignment distance position gene name gene id
GU305914.1 5971 5993 + 0 TGACAGCATCAGCGACCTGC CGG ||||||||[||||||||||||]PAM NA Intergenic NA NA

T324 CCGCAGATGAAGAACATGAGCGG 1000
Oligo fwd CACCCCGCAGATGAAGAACATGAG
Oligo rev AAACCTCATGTTCTTCATCTGCGG
Chromosome start end strand MM target_seq PAM alignment distance position gene name gene id
GU305914.1 6002 6024 + 0 CCGCAGATGAAGAACATGAG CGG ||||||||[||||||||||||]PAM NA Intergenic NA NA

T325 ATCAAACTCCCAGATTACAGCGG 1000
Oligo fwd CACCATCAAACTCCCAGATTACAG
Oligo rev AAACCTGTAATCTGGGAGTTTGAT
Chromosome start end strand MM target_seq PAM alignment distance position gene name gene id
GU305914.1 6029 6051 + 0 ATCAAACTCCCAGATTACAG CGG ||||||||[||||||||||||]PAM NA Intergenic NA NA

T326 ACTCCCAGATTACAGCGGTCCGG 1000
Oligo fwd CACCACTCCCAGATTACAGCGGTC
Oligo rev AAACGACCGCTGTAATCTGGGAGT
Chromosome start end strand MM target_seq PAM alignment distance position gene name gene id
GU305914.1 6034 6056 + 0 ACTCCCAGATTACAGCGGTC CGG ||||||||[||||||||||||]PAM NA Intergenic NA NA

T327 CCGGACACGTTGCTCACAGACGG 1000
Oligo fwd CACCCCGGACACGTTGCTCACAGA
Oligo rev AAACTCTGTGAGCAACGTGTCCGG
Chromosome start end strand MM target_seq PAM alignment distance position gene name gene id
GU305914.1 6053 6075 + 0 CCGGACACGTTGCTCACAGA CGG ||||||||[||||||||||||]PAM NA Intergenic NA NA

T328 TACCATTCGACTACGCCGCGTGG 1000
Oligo fwd CACCTACCATTCGACTACGCCGCG
Oligo rev AAACCGCGGCGTAGTCGAATGGTA
Chromosome start end strand MM target_seq PAM alignment distance position gene name gene id
GU305914.1 6078 6100 + 0 TACCATTCGACTACGCCGCG TGG ||||||||[||||||||||||]PAM NA Intergenic NA NA

T329 CCGCGTGGCGTCAGAGAATGCGG 1000
Oligo fwd CACCCCGCGTGGCGTCAGAGAATG
Oligo rev AAACCATTCTCTGACGCCACGCGG
Chromosome start end strand MM target_seq PAM alignment distance position gene name gene id
GU305914.1 6093 6115 + 0 CCGCGTGGCGTCAGAGAATG CGG ||||||||[||||||||||||]PAM NA Intergenic NA NA

T330 GTGGCGTCAGAGAATGCGGACGG 1000
Oligo fwd CACCGTGGCGTCAGAGAATGCGGA
Oligo rev AAACTCCGCATTCTCTGACGCCAC
Chromosome start end strand MM target_seq PAM alignment distance position gene name gene id
GU305914.1 6097 6119 + 0 GTGGCGTCAGAGAATGCGGA CGG ||||||||[||||||||||||]PAM NA Intergenic NA NA

T331 TCAGAGAATGCGGACGGAGATGG 1000
Oligo fwd CACCTCAGAGAATGCGGACGGAGA
Oligo rev AAACTCTCCGTCCGCATTCTCTGA
Chromosome start end strand MM target_seq PAM alignment distance position gene name gene id
GU305914.1 6103 6125 + 0 TCAGAGAATGCGGACGGAGA TGG ||||||||[||||||||||||]PAM NA Intergenic NA NA

T332 GAGAATGCGGACGGAGATGGAGG 1000
Oligo fwd CACCGAGAATGCGGACGGAGATGG
Oligo rev AAACCCATCTCCGTCCGCATTCTC
Chromosome start end strand MM target_seq PAM alignment distance position gene name gene id
GU305914.1 6106 6128 + 0 GAGAATGCGGACGGAGATGG AGG ||||||||[||||||||||||]PAM NA Intergenic NA NA

T333 CCGTGATCGCCTCGCTACTCTGG 1000
Oligo fwd CACCCCGTGATCGCCTCGCTACTC
Oligo rev AAACGAGTAGCGAGGCGATCACGG
Chromosome start end strand MM target_seq PAM alignment distance position gene name gene id
GU305914.1 6168 6190 + 0 CCGTGATCGCCTCGCTACTC TGG ||||||||[||||||||||||]PAM NA Intergenic NA NA

T334 TCGCTACTCTGGCTCGCTGTCGG 1000
Oligo fwd CACCTCGCTACTCTGGCTCGCTGT
Oligo rev AAACACAGCGAGCCAGAGTAGCGA
Chromosome start end strand MM target_seq PAM alignment distance position gene name gene id
GU305914.1 6179 6201 + 0 TCGCTACTCTGGCTCGCTGT CGG ||||||||[||||||||||||]PAM NA Intergenic NA NA

T335 GCTATGTTGTTCTACACCTACGG 1000
Oligo fwd CACCGCTATGTTGTTCTACACCTA
Oligo rev AAACTAGGTGTAGAACAACATAGC
Chromosome start end strand MM target_seq PAM alignment distance position gene name gene id
GU305914.1 6203 6225 + 0 GCTATGTTGTTCTACACCTA CGG ||||||||[||||||||||||]PAM NA Intergenic NA NA

T336 GCTAGACTGCTCAAGCGATACGG 1000
Oligo fwd CACCGCTAGACTGCTCAAGCGATA
Oligo rev AAACTATCGCTTGAGCAGTCTAGC
Chromosome start end strand MM target_seq PAM alignment distance position gene name gene id
GU305914.1 6239 6261 + 0 GCTAGACTGCTCAAGCGATA CGG ||||||||[||||||||||||]PAM NA Intergenic NA NA

T337 GCGATACGGCAAGCAGCTCGCGG 1000
Oligo fwd CACCGCGATACGGCAAGCAGCTCG
Oligo rev AAACCGAGCTGCTTGCCGTATCGC
Chromosome start end strand MM target_seq PAM alignment distance position gene name gene id

| | | | | | | | | | | | | |
|---|---|---|---|---|---|---|---|---|---|---|---|---|
| GU305914.1 | | 6253 | 6275 | + | 0 | GCGATACGGCAAGCAGCTCG | CGG | \|\|\|\|\|\|\|\|[\|\|\|\|\|\|\|\|\|\|\|\|]PAM | NA | Intergenic | NA | NA |
| T338 | CGCGGCCGTCCGCATCCCCGCGG | | 1000 | | | | | | | | | |
| Oligo fwd | CACCCGCGGCCGTCCGCATCCCCG | | | | | | | | | | | |
| Oligo rev | AAACCGGGGATGCGGACGGCCGCG | | | | | | | | | | | |
| Chromosome | start | | end | strand | MM | target_seq | PAM | alignment | distance | position | gene name | gene id |
| GU305914.1 | | 6271 | 6293 | + | 0 | CGCGGCCGTCCGCATCCCCG | CGG | \|\|\|\|\|\|\|\|[\|\|\|\|\|\|\|\|\|\|\|\|]PAM | NA | Intergenic | NA | NA |
| T339 | GCCGTCCGCATCCCCGCGGACGG | | 1000 | | | | | | | | | |
| Oligo fwd | CACCGCCGTCCGCATCCCCGCGGA | | | | | | | | | | | |
| Oligo rev | AAACTCCGCGGGGATGCGGACGGC | | | | | | | | | | | |
| Chromosome | start | | end | strand | MM | target_seq | PAM | alignment | distance | position | gene name | gene id |
| GU305914.1 | | 6275 | 6297 | + | 0 | GCCGTCCGCATCCCCGCGGA | CGG | \|\|\|\|\|\|\|\|[\|\|\|\|\|\|\|\|\|\|\|\|]PAM | NA | Intergenic | NA | NA |
| T340 | CCTTGCTTACCAAATAAAAATGG | | 1000 | | | | | | | | | |
| Oligo fwd | CACCCCTTGCTTACCAAATAAAAA | | | | | | | | | | | |
| Oligo rev | AAACTTTTTATTTGGTAAGCAAGG | | | | | | | | | | | |
| Chromosome | start | | end | strand | MM | target_seq | PAM | alignment | distance | position | gene name | gene id |
| GU305914.1 | | 6321 | 6343 | + | 0 | CCTTGCTTACCAAATAAAAA | TGG | \|\|\|\|\|\|\|\|[\|\|\|\|\|\|\|\|\|\|\|\|]PAM | NA | Intergenic | NA | NA |
| T341 | GTTTTAGTACCTACACCAAACGG | | 1000 | | | | | | | | | |
| Oligo fwd | CACCGTTTTAGTACCTACACCAAA | | | | | | | | | | | |
| Oligo rev | AAACTTTGGTGTAGGTACTAAAAC | | | | | | | | | | | |
| Chromosome | start | | end | strand | MM | target_seq | PAM | alignment | distance | position | gene name | gene id |
| GU305914.1 | | 6349 | 6371 | + | 0 | GTTTTAGTACCTACACCAAA | CGG | \|\|\|\|\|\|\|\|[\|\|\|\|\|\|\|\|\|\|\|\|]PAM | NA | Intergenic | NA | NA |
| T342 | TGTTCTGTCTGCGTCGTCCGTGG | | 1000 | | | | | | | | | |
| Oligo fwd | CACCTGTTCTGTCTGCGTCGTCCG | | | | | | | | | | | |
| Oligo rev | AAACCGGACGACGCAGACAGAACA | | | | | | | | | | | |
| Chromosome | start | | end | strand | MM | target_seq | PAM | alignment | distance | position | gene name | gene id |
| GU305914.1 | | 6383 | 6405 | + | 0 | TGTTCTGTCTGCGTCGTCCG | TGG | \|\|\|\|\|\|\|\|[\|\|\|\|\|\|\|\|\|\|\|\|]PAM | NA | Intergenic | NA | NA |
| T343 | ACACTCTTGTCTGAGTCTTTTGG | | 1000 | | | | | | | | | |
| Oligo fwd | CACCACACTCTTGTCTGAGTCTTT | | | | | | | | | | | |
| Oligo rev | AAACAAAGACTCAGACAAGAGTGT | | | | | | | | | | | |
| Chromosome | start | | end | strand | MM | target_seq | PAM | alignment | distance | position | gene name | gene id |
| GU305914.1 | | 6414 | 6436 | + | 0 | ACACTCTTGTCTGAGTCTTT | TGG | \|\|\|\|\|\|\|\|[\|\|\|\|\|\|\|\|\|\|\|\|]PAM | NA | Intergenic | NA | NA |
| T344 | TCTGAGTCTTTTGGCGTCGTCGG | | 1000 | | | | | | | | | |
| Oligo fwd | CACCTCTGAGTCTTTTGGCGTCGT | | | | | | | | | | | |
| Oligo rev | AAACACGACGCCAAAAGACTCAGA | | | | | | | | | | | |
| Chromosome | start | | end | strand | MM | target_seq | PAM | alignment | distance | position | gene name | gene id |
| GU305914.1 | | 6423 | 6445 | + | 0 | TCTGAGTCTTTTGGCGTCGT | CGG | \|\|\|\|\|\|\|\|[\|\|\|\|\|\|\|\|\|\|\|\|]PAM | NA | Intergenic | NA | NA |
| T345 | CTGAGTCTTTTGGCGTCGTCGGG | | 1000 | | | | | | | | | |
| Oligo fwd | CACCCTGAGTCTTTTGGCGTCGTC | | | | | | | | | | | |
| Oligo rev | AAACGACGACGCCAAAAGACTCAG | | | | | | | | | | | |
| Chromosome | start | | end | strand | MM | target_seq | PAM | alignment | distance | position | gene name | gene id |
| GU305914.1 | | 6424 | 6446 | + | 0 | CTGAGTCTTTTGGCGTCGTC | GGG | \|\|\|\|\|\|\|\|[\|\|\|\|\|\|\|\|\|\|\|\|]PAM | NA | Intergenic | NA | NA |
| T346 | GTCTATCGAGTGAGAGAATGAGG | | 1000 | | | | | | | | | |
| Oligo fwd | CACCGTCTATCGAGTGAGAGAATG | | | | | | | | | | | |
| Oligo rev | AAACCATTCTCTCACTCGATAGAC | | | | | | | | | | | |
| Chromosome | start | | end | strand | MM | target_seq | PAM | alignment | distance | position | gene name | gene id |
| GU305914.1 | | 6446 | 6468 | + | 0 | GTCTATCGAGTGAGAGAATG | AGG | \|\|\|\|\|\|\|\|[\|\|\|\|\|\|\|\|\|\|\|\|]PAM | NA | Intergenic | NA | NA |
| T347 | TCTATCGAGTGAGAGAATGAGGG | | 1000 | | | | | | | | | |
| Oligo fwd | CACCTCTATCGAGTGAGAGAATGA | | | | | | | | | | | |
| Oligo rev | AAACTCATTCTCTCACTCGATAGA | | | | | | | | | | | |
| Chromosome | start | | end | strand | MM | target_seq | PAM | alignment | distance | position | gene name | gene id |
| GU305914.1 | | 6447 | 6469 | + | 0 | TCTATCGAGTGAGAGAATGA | GGG | \|\|\|\|\|\|\|\|[\|\|\|\|\|\|\|\|\|\|\|\|]PAM | NA | Intergenic | NA | NA |
| T348 | CTATCGAGTGAGAGAATGAGGGG | | 1000 | | | | | | | | | |
| Oligo fwd | CACCCTATCGAGTGAGAGAATGAG | | | | | | | | | | | |
| Oligo rev | AAACCTCATTCTCTCACTCGATAG | | | | | | | | | | | |
| Chromosome | start | | end | strand | MM | target_seq | PAM | alignment | distance | position | gene name | gene id |
| GU305914.1 | | 6448 | 6470 | + | 0 | CTATCGAGTGAGAGAATGAG | GGG | \|\|\|\|\|\|\|\|[\|\|\|\|\|\|\|\|\|\|\|\|]PAM | NA | Intergenic | NA | NA |
| T349 | TGAGAGAATGAGGGGATATACGG | | 1000 | | | | | | | | | |
| Oligo fwd | CACCTGAGAGAATGAGGGGATATA | | | | | | | | | | | |
| Oligo rev | AAACTATATCCCCTCATTCTCTCA | | | | | | | | | | | |
| Chromosome | start | | end | strand | MM | target_seq | PAM | alignment | distance | position | gene name | gene id |
| GU305914.1 | | 6456 | 6478 | + | 0 | TGAGAGAATGAGGGGATATA | CGG | \|\|\|\|\|\|\|\|[\|\|\|\|\|\|\|\|\|\|\|\|]PAM | NA | Intergenic | NA | NA |
| T350 | GAGAGAATGAGGGGATATACGGG | | 1000 | | | | | | | | | |
| Oligo fwd | CACCGAGAGAATGAGGGGATATAC | | | | | | | | | | | |
| Oligo rev | AAACGTATATCCCCTCATTCTCTC | | | | | | | | | | | |
| Chromosome | start | | end | strand | MM | target_seq | PAM | alignment | distance | position | gene name | gene id |
| GU305914.1 | | 6457 | 6479 | + | 0 | GAGAGAATGAGGGGATATAC | GGG | \|\|\|\|\|\|\|\|[\|\|\|\|\|\|\|\|\|\|\|\|]PAM | NA | Intergenic | NA | NA |
| T351 | GGGATATACGGGCGATACGATGG | | 1000 | | | | | | | | | |
| Oligo fwd | CACCGGGATATACGGGCGATACGA | | | | | | | | | | | |
| Oligo rev | AAACTCGTATCGCCCGTATATCCC | | | | | | | | | | | |
| Chromosome | start | | end | strand | MM | target_seq | PAM | alignment | distance | position | gene name | gene id |
| GU305914.1 | | 6468 | 6490 | + | 0 | GGGATATACGGGCGATACGA | TGG | \|\|\|\|\|\|\|\|[\|\|\|\|\|\|\|\|\|\|\|\|]PAM | NA | Intergenic | NA | NA |
| T352 | GGGCGATACGATGGTGTTAATGG | | 1000 | | | | | | | | | |
| Oligo fwd | CACCGGGCGATACGATGGTGTTAA | | | | | | | | | | | |
| Oligo rev | AAACTTAACACCATCGTATCGCCC | | | | | | | | | | | |

<table>
<tr><td>Chromosome</td><td>start</td><td></td><td>end</td><td>strand</td><td>MM</td><td>target_seq</td><td>PAM</td><td>alignment</td><td>distance</td><td>position</td><td>gene name</td><td>gene id</td></tr>
<tr><td>GU305914.1</td><td></td><td>6477</td><td>6499</td><td>+</td><td>0</td><td>GGGCGATACGATGGTGTTAA</td><td>TGG</td><td>||||||||[||||||||||||]PAM</td><td>NA</td><td>Intergenic</td><td>NA</td><td>NA</td></tr>
<tr><td></td><td></td><td></td><td></td><td></td><td></td><td></td><td></td><td></td><td></td><td></td><td></td><td></td></tr>
<tr><td>T353</td><td>TGTTAATGGTGTTCTGCGTATGG</td><td></td><td>1000</td><td></td><td></td><td></td><td></td><td></td><td></td><td></td><td></td><td></td></tr>
<tr><td>Oligo fwd</td><td>CACCTGTTAATGGTGTTCTGCGTA</td><td></td><td></td><td></td><td></td><td></td><td></td><td></td><td></td><td></td><td></td><td></td></tr>
<tr><td>Oligo rev</td><td>AAACTACGCAGAACACCATTAACA</td><td></td><td></td><td></td><td></td><td></td><td></td><td></td><td></td><td></td><td></td><td></td></tr>
<tr><td>Chromosome</td><td>start</td><td></td><td>end</td><td>strand</td><td>MM</td><td>target_seq</td><td>PAM</td><td>alignment</td><td>distance</td><td>position</td><td>gene name</td><td>gene id</td></tr>
<tr><td>GU305914.1</td><td></td><td>6491</td><td>6513</td><td>+</td><td>0</td><td>TGTTAATGGTGTTCTGCGTA</td><td>TGG</td><td>||||||||[||||||||||||]PAM</td><td>NA</td><td>Intergenic</td><td>NA</td><td>NA</td></tr>
<tr><td></td><td></td><td></td><td></td><td></td><td></td><td></td><td></td><td></td><td></td><td></td><td></td><td></td></tr>
<tr><td>T354</td><td>ATGGCTCGCGTACGTGTTCTCGG</td><td></td><td>1000</td><td></td><td></td><td></td><td></td><td></td><td></td><td></td><td></td><td></td></tr>
<tr><td>Oligo fwd</td><td>CACCATGGCTCGCGTACGTGTTCT</td><td></td><td></td><td></td><td></td><td></td><td></td><td></td><td></td><td></td><td></td><td></td></tr>
<tr><td>Oligo rev</td><td>AAACAGAACACGTACGCGAGCCAT</td><td></td><td></td><td></td><td></td><td></td><td></td><td></td><td></td><td></td><td></td><td></td></tr>
<tr><td>Chromosome</td><td>start</td><td></td><td>end</td><td>strand</td><td>MM</td><td>target_seq</td><td>PAM</td><td>alignment</td><td>distance</td><td>position</td><td>gene name</td><td>gene id</td></tr>
<tr><td>GU305914.1</td><td></td><td>6510</td><td>6532</td><td>+</td><td>0</td><td>ATGGCTCGCGTACGTGTTCT</td><td>CGG</td><td>||||||||[||||||||||||]PAM</td><td>NA</td><td>Intergenic</td><td>NA</td><td>NA</td></tr>
<tr><td></td><td></td><td></td><td></td><td></td><td></td><td></td><td></td><td></td><td></td><td></td><td></td><td></td></tr>
<tr><td>T355</td><td>GCTCGCGTACGTGTTCTCGGTGG</td><td></td><td>1000</td><td></td><td></td><td></td><td></td><td></td><td></td><td></td><td></td><td></td></tr>
<tr><td>Oligo fwd</td><td>CACCGCTCGCGTACGTGTTCTCGG</td><td></td><td></td><td></td><td></td><td></td><td></td><td></td><td></td><td></td><td></td><td></td></tr>
<tr><td>Oligo rev</td><td>AAACCCGAGAACACGTACGCGAGC</td><td></td><td></td><td></td><td></td><td></td><td></td><td></td><td></td><td></td><td></td><td></td></tr>
<tr><td>Chromosome</td><td>start</td><td></td><td>end</td><td>strand</td><td>MM</td><td>target_seq</td><td>PAM</td><td>alignment</td><td>distance</td><td>position</td><td>gene name</td><td>gene id</td></tr>
<tr><td>GU305914.1</td><td></td><td>6513</td><td>6535</td><td>+</td><td>0</td><td>GCTCGCGTACGTGTTCTCGG</td><td>TGG</td><td>||||||||[||||||||||||]PAM</td><td>NA</td><td>Intergenic</td><td>NA</td><td>NA</td></tr>
<tr><td></td><td></td><td></td><td></td><td></td><td></td><td></td><td></td><td></td><td></td><td></td><td></td><td></td></tr>
<tr><td>T356</td><td>CGTGTTCTCGGTGGAATCTAAGG</td><td></td><td>1000</td><td></td><td></td><td></td><td></td><td></td><td></td><td></td><td></td><td></td></tr>
<tr><td>Oligo fwd</td><td>CACCCGTGTTCTCGGTGGAATCTA</td><td></td><td></td><td></td><td></td><td></td><td></td><td></td><td></td><td></td><td></td><td></td></tr>
<tr><td>Oligo rev</td><td>AAACTAGATTCCACCGAGAACACG</td><td></td><td></td><td></td><td></td><td></td><td></td><td></td><td></td><td></td><td></td><td></td></tr>
<tr><td>Chromosome</td><td>start</td><td></td><td>end</td><td>strand</td><td>MM</td><td>target_seq</td><td>PAM</td><td>alignment</td><td>distance</td><td>position</td><td>gene name</td><td>gene id</td></tr>
<tr><td>GU305914.1</td><td></td><td>6522</td><td>6544</td><td>+</td><td>0</td><td>CGTGTTCTCGGTGGAATCTA</td><td>AGG</td><td>||||||||[||||||||||||]PAM</td><td>NA</td><td>Intergenic</td><td>NA</td><td>NA</td></tr>
<tr><td></td><td></td><td></td><td></td><td></td><td></td><td></td><td></td><td></td><td></td><td></td><td></td><td></td></tr>
<tr><td>T357</td><td>TCGGTGGAATCTAAGGCTAACGG</td><td></td><td>1000</td><td></td><td></td><td></td><td></td><td></td><td></td><td></td><td></td><td></td></tr>
<tr><td>Oligo fwd</td><td>CACCTCGGTGGAATCTAAGGCTAA</td><td></td><td></td><td></td><td></td><td></td><td></td><td></td><td></td><td></td><td></td><td></td></tr>
<tr><td>Oligo rev</td><td>AAACTTAGCCTTAGATTCCACCGA</td><td></td><td></td><td></td><td></td><td></td><td></td><td></td><td></td><td></td><td></td><td></td></tr>
<tr><td>Chromosome</td><td>start</td><td></td><td>end</td><td>strand</td><td>MM</td><td>target_seq</td><td>PAM</td><td>alignment</td><td>distance</td><td>position</td><td>gene name</td><td>gene id</td></tr>
<tr><td>GU305914.1</td><td></td><td>6529</td><td>6551</td><td>+</td><td>0</td><td>TCGGTGGAATCTAAGGCTAA</td><td>CGG</td><td>||||||||[||||||||||||]PAM</td><td>NA</td><td>Intergenic</td><td>NA</td><td>NA</td></tr>
<tr><td></td><td></td><td></td><td></td><td></td><td></td><td></td><td></td><td></td><td></td><td></td><td></td><td></td></tr>
<tr><td>T358</td><td>AACTACAGTAACTTAACACAAGG</td><td></td><td>1000</td><td></td><td></td><td></td><td></td><td></td><td></td><td></td><td></td><td></td></tr>
<tr><td>Oligo fwd</td><td>CACCAACTACAGTAACTTAACACA</td><td></td><td></td><td></td><td></td><td></td><td></td><td></td><td></td><td></td><td></td><td></td></tr>
<tr><td>Oligo rev</td><td>AAACTGTGTTAAGTTACTGTAGTT</td><td></td><td></td><td></td><td></td><td></td><td></td><td></td><td></td><td></td><td></td><td></td></tr>
<tr><td>Chromosome</td><td>start</td><td></td><td>end</td><td>strand</td><td>MM</td><td>target_seq</td><td>PAM</td><td>alignment</td><td>distance</td><td>position</td><td>gene name</td><td>gene id</td></tr>
<tr><td>GU305914.1</td><td></td><td>6574</td><td>6596</td><td>+</td><td>0</td><td>AACTACAGTAACTTAACACA</td><td>AGG</td><td>||||||||[||||||||||||]PAM</td><td>NA</td><td>Intergenic</td><td>NA</td><td>NA</td></tr>
<tr><td></td><td></td><td></td><td></td><td></td><td></td><td></td><td></td><td></td><td></td><td></td><td></td><td></td></tr>
<tr><td>T359</td><td>AGGTCAATGTGTGACAAGAAAGG</td><td></td><td>1000</td><td></td><td></td><td></td><td></td><td></td><td></td><td></td><td></td><td></td></tr>
<tr><td>Oligo fwd</td><td>CACCAGGTCAATGTGTGACAAGAA</td><td></td><td></td><td></td><td></td><td></td><td></td><td></td><td></td><td></td><td></td><td></td></tr>
<tr><td>Oligo rev</td><td>AAACTTCTTGTCACACATTGACCT</td><td></td><td></td><td></td><td></td><td></td><td></td><td></td><td></td><td></td><td></td><td></td></tr>
<tr><td>Chromosome</td><td>start</td><td></td><td>end</td><td>strand</td><td>MM</td><td>target_seq</td><td>PAM</td><td>alignment</td><td>distance</td><td>position</td><td>gene name</td><td>gene id</td></tr>
<tr><td>GU305914.1</td><td></td><td>6594</td><td>6616</td><td>+</td><td>0</td><td>AGGTCAATGTGTGACAAGAA</td><td>AGG</td><td>||||||||[||||||||||||]PAM</td><td>NA</td><td>Intergenic</td><td>NA</td><td>NA</td></tr>
<tr><td></td><td></td><td></td><td></td><td></td><td></td><td></td><td></td><td></td><td></td><td></td><td></td><td></td></tr>
<tr><td>T360</td><td>GGTCAATGTGTGACAAGAAAGGG</td><td></td><td>1000</td><td></td><td></td><td></td><td></td><td></td><td></td><td></td><td></td><td></td></tr>
<tr><td>Oligo fwd</td><td>CACCGGTCAATGTGTGACAAGAAA</td><td></td><td></td><td></td><td></td><td></td><td></td><td></td><td></td><td></td><td></td><td></td></tr>
<tr><td>Oligo rev</td><td>AAACTTTCTTGTCACACATTGACC</td><td></td><td></td><td></td><td></td><td></td><td></td><td></td><td></td><td></td><td></td><td></td></tr>
<tr><td>Chromosome</td><td>start</td><td></td><td>end</td><td>strand</td><td>MM</td><td>target_seq</td><td>PAM</td><td>alignment</td><td>distance</td><td>position</td><td>gene name</td><td>gene id</td></tr>
<tr><td>GU305914.1</td><td></td><td>6595</td><td>6617</td><td>+</td><td>0</td><td>GGTCAATGTGTGACAAGAAA</td><td>GGG</td><td>||||||||[||||||||||||]PAM</td><td>NA</td><td>Intergenic</td><td>NA</td><td>NA</td></tr>
<tr><td></td><td></td><td></td><td></td><td></td><td></td><td></td><td></td><td></td><td></td><td></td><td></td><td></td></tr>
<tr><td>T361</td><td>AAAGGGTGTCAAGAACGATACGG</td><td></td><td>1000</td><td></td><td></td><td></td><td></td><td></td><td></td><td></td><td></td><td></td></tr>
<tr><td>Oligo fwd</td><td>CACCAAAGGGTGTCAAGAACGATA</td><td></td><td></td><td></td><td></td><td></td><td></td><td></td><td></td><td></td><td></td><td></td></tr>
<tr><td>Oligo rev</td><td>AAACTATCGTTCTTGACACCCTTT</td><td></td><td></td><td></td><td></td><td></td><td></td><td></td><td></td><td></td><td></td><td></td></tr>
<tr><td>Chromosome</td><td>start</td><td></td><td>end</td><td>strand</td><td>MM</td><td>target_seq</td><td>PAM</td><td>alignment</td><td>distance</td><td>position</td><td>gene name</td><td>gene id</td></tr>
<tr><td>GU305914.1</td><td></td><td>6612</td><td>6634</td><td>+</td><td>0</td><td>AAAGGGTGTCAAGAACGATA</td><td>CGG</td><td>||||||||[||||||||||||]PAM</td><td>NA</td><td>Intergenic</td><td>NA</td><td>NA</td></tr>
<tr><td></td><td></td><td></td><td></td><td></td><td></td><td></td><td></td><td></td><td></td><td></td><td></td><td></td></tr>
<tr><td>T362</td><td>GTCAAGAACGATACGGAAGAAGG</td><td></td><td>1000</td><td></td><td></td><td></td><td></td><td></td><td></td><td></td><td></td><td></td></tr>
<tr><td>Oligo fwd</td><td>CACCGTCAAGAACGATACGGAAGA</td><td></td><td></td><td></td><td></td><td></td><td></td><td></td><td></td><td></td><td></td><td></td></tr>
<tr><td>Oligo rev</td><td>AAACTCTTCCGTATCGTTCTTGAC</td><td></td><td></td><td></td><td></td><td></td><td></td><td></td><td></td><td></td><td></td><td></td></tr>
<tr><td>Chromosome</td><td>start</td><td></td><td>end</td><td>strand</td><td>MM</td><td>target_seq</td><td>PAM</td><td>alignment</td><td>distance</td><td>position</td><td>gene name</td><td>gene id</td></tr>
<tr><td>GU305914.1</td><td></td><td>6619</td><td>6641</td><td>+</td><td>0</td><td>GTCAAGAACGATACGGAAGA</td><td>AGG</td><td>||||||||[||||||||||||]PAM</td><td>NA</td><td>Intergenic</td><td>NA</td><td>NA</td></tr>
<tr><td></td><td></td><td></td><td></td><td></td><td></td><td></td><td></td><td></td><td></td><td></td><td></td><td></td></tr>
<tr><td>T363</td><td>AGGATCGTACAGTTGTCTAACGG</td><td></td><td>1000</td><td></td><td></td><td></td><td></td><td></td><td></td><td></td><td></td><td></td></tr>
<tr><td>Oligo fwd</td><td>CACCAGGATCGTACAGTTGTCTAA</td><td></td><td></td><td></td><td></td><td></td><td></td><td></td><td></td><td></td><td></td><td></td></tr>
<tr><td>Oligo rev</td><td>AAACTTAGACAACTGTACGATCCT</td><td></td><td></td><td></td><td></td><td></td><td></td><td></td><td></td><td></td><td></td><td></td></tr>
<tr><td>Chromosome</td><td>start</td><td></td><td>end</td><td>strand</td><td>MM</td><td>target_seq</td><td>PAM</td><td>alignment</td><td>distance</td><td>position</td><td>gene name</td><td>gene id</td></tr>
<tr><td>GU305914.1</td><td></td><td>6639</td><td>6661</td><td>+</td><td>0</td><td>AGGATCGTACAGTTGTCTAA</td><td>CGG</td><td>||||||||[||||||||||||]PAM</td><td>NA</td><td>Intergenic</td><td>NA</td><td>NA</td></tr>
<tr><td></td><td></td><td></td><td></td><td></td><td></td><td></td><td></td><td></td><td></td><td></td><td></td><td></td></tr>
<tr><td>T364</td><td>CCATCTGTATTCCCGCATCGTGG</td><td></td><td>1000</td><td></td><td></td><td></td><td></td><td></td><td></td><td></td><td></td><td></td></tr>
<tr><td>Oligo fwd</td><td>CACCCCATCTGTATTCCCGCATCG</td><td></td><td></td><td></td><td></td><td></td><td></td><td></td><td></td><td></td><td></td><td></td></tr>
<tr><td>Oligo rev</td><td>AAACCGATGCGGGAATACAGATGG</td><td></td><td></td><td></td><td></td><td></td><td></td><td></td><td></td><td></td><td></td><td></td></tr>
<tr><td>Chromosome</td><td>start</td><td></td><td>end</td><td>strand</td><td>MM</td><td>target_seq</td><td>PAM</td><td>alignment</td><td>distance</td><td>position</td><td>gene name</td><td>gene id</td></tr>
<tr><td>GU305914.1</td><td></td><td>6665</td><td>6687</td><td>+</td><td>0</td><td>CCATCTGTATTCCCGCATCG</td><td>TGG</td><td>||||||||[||||||||||||]PAM</td><td>NA</td><td>Intergenic</td><td>NA</td><td>NA</td></tr>
<tr><td>GU305914.1</td><td></td><td>7658</td><td>7680</td><td>+</td><td>4</td><td>ATATCTGCATTCCGGCATCG</td><td>TGG</td><td>--|||||-[|||||-||||||]PAM</td><td>NA</td><td>Intergenic</td><td>NA</td><td>NA</td></tr>
<tr><td></td><td></td><td></td><td></td><td></td><td></td><td></td><td></td><td></td><td></td><td></td><td></td><td></td></tr>
<tr><td>T365</td><td>CATCTGTATTCCCGCATCGTGGG</td><td></td><td>1000</td><td></td><td></td><td></td><td></td><td></td><td></td><td></td><td></td><td></td></tr>
<tr><td>Oligo fwd</td><td>CACCCATCTGTATTCCCGCATCGT</td><td></td><td></td><td></td><td></td><td></td><td></td><td></td><td></td><td></td><td></td><td></td></tr>
<tr><td>Oligo rev</td><td>AAACACGATGCGGGAATACAGATG</td><td></td><td></td><td></td><td></td><td></td><td></td><td></td><td></td><td></td><td></td><td></td></tr>
<tr><td>Chromosome</td><td>start</td><td></td><td>end</td><td>strand</td><td>MM</td><td>target_seq</td><td>PAM</td><td>alignment</td><td>distance</td><td>position</td><td>gene name</td><td>gene id</td></tr>
<tr><td>GU305914.1</td><td></td><td>6666</td><td>6688</td><td>+</td><td>0</td><td>CATCTGTATTCCCGCATCGT</td><td>GGG</td><td>||||||||[||||||||||||]PAM</td><td>NA</td><td>Intergenic</td><td>NA</td><td>NA</td></tr>
<tr><td></td><td></td><td></td><td></td><td></td><td></td><td></td><td></td><td></td><td></td><td></td><td></td><td></td></tr>
<tr><td>T366</td><td>TGTATTCCCGCATCGTGGGAAGG</td><td></td><td>1000</td><td></td><td></td><td></td><td></td><td></td><td></td><td></td><td></td><td></td></tr>
<tr><td>Oligo fwd</td><td>CACCTGTATTCCCGCATCGTGGGA</td><td></td><td></td><td></td><td></td><td></td><td></td><td></td><td></td><td></td><td></td><td></td></tr>
<tr><td>Oligo rev</td><td>AAACTCCCACGATGCGGGAATACA</td><td></td><td></td><td></td><td></td><td></td><td></td><td></td><td></td><td></td><td></td><td></td></tr>
<tr><td>Chromosome</td><td>start</td><td></td><td>end</td><td>strand</td><td>MM</td><td>target_seq</td><td>PAM</td><td>alignment</td><td>distance</td><td>position</td><td>gene name</td><td>gene id</td></tr>
<tr><td>GU305914.1</td><td></td><td>6670</td><td>6692</td><td>+</td><td>0</td><td>TGTATTCCCGCATCGTGGGA</td><td>AGG</td><td>||||||||[||||||||||||]PAM</td><td>NA</td><td>Intergenic</td><td>NA</td><td>NA</td></tr>
<tr><td></td><td></td><td></td><td></td><td></td><td></td><td></td><td></td><td></td><td></td><td></td><td></td><td></td></tr>
<tr><td>T367</td><td>CCGCATCGTGGGAAGGTACGTGG</td><td></td><td>1000</td><td></td><td></td><td></td><td></td><td></td><td></td><td></td><td></td><td></td></tr>
</table>

| | | | | | | | | | | | | |
|---|---|---|---|---|---|---|---|---|---|---|---|---|
| Oligo fwd | CACCCCGCATCGTGGGAAGGTACG | | | | | | | | | | | |
| Oligo rev | AAACCGTACCTTCCCACGATGCGG | | | | | | | | | | | |
| Chromosome | start | | end | strand | MM | target_seq | PAM | alignment | distance | position | gene name | gene id |
| GU305914.1 | | 6677 | 6699 | + | 0 | CCGCATCGTGGGAAGGTACG | TGG | \|\|\|\|\|\|\|\|[\|\|\|\|\|\|\|\|\|\|\|\|]PAM | NA | Intergenic | NA | NA |
| T368 | ATCGTGGGAAGGTACGTGGCTGG | | 1000 | | | | | | | | | |
| Oligo fwd | CACCATCGTGGGAAGGTACGTGGC | | | | | | | | | | | |
| Oligo rev | AAACGCCACGTACCTTCCCACGAT | | | | | | | | | | | |
| Chromosome | start | | end | strand | MM | target_seq | PAM | alignment | distance | position | gene name | gene id |
| GU305914.1 | | 6681 | 6703 | + | 0 | ATCGTGGGAAGGTACGTGGC | TGG | \|\|\|\|\|\|\|\|[\|\|\|\|\|\|\|\|\|\|\|\|]PAM | NA | Intergenic | NA | NA |
| T369 | GTGGGAAGGTACGTGGCTGGTGG | | 1000 | | | | | | | | | |
| Oligo fwd | CACCGTGGGAAGGTACGTGGCTGG | | | | | | | | | | | |
| Oligo rev | AAACCCAGCCACGTACCTTCCCAC | | | | | | | | | | | |
| Chromosome | start | | end | strand | MM | target_seq | PAM | alignment | distance | position | gene name | gene id |
| GU305914.1 | | 6684 | 6706 | + | 0 | GTGGGAAGGTACGTGGCTGG | TGG | \|\|\|\|\|\|\|\|[\|\|\|\|\|\|\|\|\|\|\|\|]PAM | NA | Intergenic | NA | NA |
| T370 | GAAGGTACGTGGCTGGTGGACGG | | 1000 | | | | | | | | | |
| Oligo fwd | CACCGAAGGTACGTGGCTGGTGGA | | | | | | | | | | | |
| Oligo rev | AAACTCCACCAGCCACGTACCTTC | | | | | | | | | | | |
| Chromosome | start | | end | strand | MM | target_seq | PAM | alignment | distance | position | gene name | gene id |
| GU305914.1 | | 6688 | 6710 | + | 0 | GAAGGTACGTGGCTGGTGGA | CGG | \|\|\|\|\|\|\|\|[\|\|\|\|\|\|\|\|\|\|\|\|]PAM | NA | Intergenic | NA | NA |
| T371 | AAGGTACGTGGCTGGTGGACGGG | | 1000 | | | | | | | | | |
| Oligo fwd | CACCAAGGTACGTGGCTGGTGGAC | | | | | | | | | | | |
| Oligo rev | AAACGTCCACCAGCCACGTACCTT | | | | | | | | | | | |
| Chromosome | start | | end | strand | MM | target_seq | PAM | alignment | distance | position | gene name | gene id |
| GU305914.1 | | 6689 | 6711 | + | 0 | AAGGTACGTGGCTGGTGGAC | GGG | \|\|\|\|\|\|\|\|[\|\|\|\|\|\|\|\|\|\|\|\|]PAM | NA | Intergenic | NA | NA |
| T372 | GTACGTGGCTGGTGGACGGGTGG | | 1000 | | | | | | | | | |
| Oligo fwd | CACCGTACGTGGCTGGTGGACGGG | | | | | | | | | | | |
| Oligo rev | AAACCCCGTCCACCAGCCACGTAC | | | | | | | | | | | |
| Chromosome | start | | end | strand | MM | target_seq | PAM | alignment | distance | position | gene name | gene id |
| GU305914.1 | | 6692 | 6714 | + | 0 | GTACGTGGCTGGTGGACGGG | TGG | \|\|\|\|\|\|\|\|[\|\|\|\|\|\|\|\|\|\|\|\|]PAM | NA | Intergenic | NA | NA |
| T373 | GACGGGTGGCCGTTACCGTTCGG | | 1000 | | | | | | | | | |
| Oligo fwd | CACCGACGGGTGGCCGTTACCGTT | | | | | | | | | | | |
| Oligo rev | AAACAACGGTAACGGCCACCCGTC | | | | | | | | | | | |
| Chromosome | start | | end | strand | MM | target_seq | PAM | alignment | distance | position | gene name | gene id |
| GU305914.1 | | 6706 | 6728 | + | 0 | GACGGGTGGCCGTTACCGTT | CGG | \|\|\|\|\|\|\|\|[\|\|\|\|\|\|\|\|\|\|\|\|]PAM | NA | Intergenic | NA | NA |
| T374 | TATACACTAGATCCCACGTCAGG | | 1000 | | | | | | | | | |
| Oligo fwd | CACCTATACACTAGATCCCACGTC | | | | | | | | | | | |
| Oligo rev | AAACGACGTGGGATCTAGTGTATA | | | | | | | | | | | |
| Chromosome | start | | end | strand | MM | target_seq | PAM | alignment | distance | position | gene name | gene id |
| GU305914.1 | | 6829 | 6851 | + | 0 | TATACACTAGATCCCACGTC | AGG | \|\|\|\|\|\|\|\|[\|\|\|\|\|\|\|\|\|\|\|\|]PAM | NA | Intergenic | NA | NA |
| T375 | ATACACTAGATCCCACGTCAGGG | | 1000 | | | | | | | | | |
| Oligo fwd | CACCATACACTAGATCCCACGTCA | | | | | | | | | | | |
| Oligo rev | AAACTGACGTGGGATCTAGTGTAT | | | | | | | | | | | |
| Chromosome | start | | end | strand | MM | target_seq | PAM | alignment | distance | position | gene name | gene id |
| GU305914.1 | | 6830 | 6852 | + | 0 | ATACACTAGATCCCACGTCA | GGG | \|\|\|\|\|\|\|\|[\|\|\|\|\|\|\|\|\|\|\|\|]PAM | NA | Intergenic | NA | NA |
| T376 | ACGATTTGTCGACAGACCGACGG | | 1000 | | | | | | | | | |
| Oligo fwd | CACCACGATTTGTCGACAGACCGA | | | | | | | | | | | |
| Oligo rev | AAACTCGGTCTGTCGACAAATCGT | | | | | | | | | | | |
| Chromosome | start | | end | strand | MM | target_seq | PAM | alignment | distance | position | gene name | gene id |
| GU305914.1 | | 6907 | 6929 | + | 0 | ACGATTTGTCGACAGACCGA | CGG | \|\|\|\|\|\|\|\|[\|\|\|\|\|\|\|\|\|\|\|\|]PAM | NA | Intergenic | NA | NA |
| T377 | TGTCGACAGACCGACGGCTACGG | | 1000 | | | | | | | | | |
| Oligo fwd | CACCTGTCGACAGACCGACGGCTA | | | | | | | | | | | |
| Oligo rev | AAACTAGCCGTCGGTCTGTCGACA | | | | | | | | | | | |
| Chromosome | start | | end | strand | MM | target_seq | PAM | alignment | distance | position | gene name | gene id |
| GU305914.1 | | 6913 | 6935 | + | 0 | TGTCGACAGACCGACGGCTA | CGG | \|\|\|\|\|\|\|\|[\|\|\|\|\|\|\|\|\|\|\|\|]PAM | NA | Intergenic | NA | NA |
| T378 | GACCTAAACCTCCACCAAGACGG | | 1000 | | | | | | | | | |
| Oligo fwd | CACCGACCTAAACCTCCACCAAGA | | | | | | | | | | | |
| Oligo rev | AAACTCTTGGTGGAGGTTTAGGTC | | | | | | | | | | | |
| Chromosome | start | | end | strand | MM | target_seq | PAM | alignment | distance | position | gene name | gene id |
| GU305914.1 | | 6935 | 6957 | + | 0 | GACCTAAACCTCCACCAAGA | CGG | \|\|\|\|\|\|\|\|[\|\|\|\|\|\|\|\|\|\|\|\|]PAM | NA | Intergenic | NA | NA |
| T379 | GCCCAAGAAACGCACGACGACGG | | 1000 | | | | | | | | | |
| Oligo fwd | CACCGCCCAAGAAACGCACGACGA | | | | | | | | | | | |
| Oligo rev | AAACTCGTCGTGCGTTTCTTGGGC | | | | | | | | | | | |
| Chromosome | start | | end | strand | MM | target_seq | PAM | alignment | distance | position | gene name | gene id |
| GU305914.1 | | 6957 | 6979 | + | 0 | GCCCAAGAAACGCACGACGA | CGG | \|\|\|\|\|\|\|\|[\|\|\|\|\|\|\|\|\|\|\|\|]PAM | NA | Intergenic | NA | NA |
| T380 | GAAAACTCATCACCACACGCCGG | | 1000 | | | | | | | | | |
| Oligo fwd | CACCGAAAACTCATCACCACACGC | | | | | | | | | | | |
| Oligo rev | AAACGCGTGTGGTGATGAGTTTTC | | | | | | | | | | | |
| Chromosome | start | | end | strand | MM | target_seq | PAM | alignment | distance | position | gene name | gene id |
| GU305914.1 | | 7037 | 7059 | + | 0 | GAAAACTCATCACCACACGC | CGG | \|\|\|\|\|\|\|\|[\|\|\|\|\|\|\|\|\|\|\|\|]PAM | NA | Intergenic | NA | NA |
| T381 | AAAACTCATCACCACACGCCGGG | | 1000 | | | | | | | | | |
| Oligo fwd | CACCAAAACTCATCACCACACGCC | | | | | | | | | | | |
| Oligo rev | AAACGGCGTGTGGTGATGAGTTTT | | | | | | | | | | | |
| Chromosome | start | | end | strand | MM | target_seq | PAM | alignment | distance | position | gene name | gene id |
| GU305914.1 | | 7038 | 7060 | + | 0 | AAAACTCATCACCACACGCC | GGG | \|\|\|\|\|\|\|\|[\|\|\|\|\|\|\|\|\|\|\|\|]PAM | NA | Intergenic | NA | NA |

<table>
<tr><td>T382</td><td>AACACCGACGACGAGACGACCGG</td><td></td><td>1000</td><td></td><td></td><td></td><td></td><td></td><td></td><td></td><td></td><td></td></tr>
<tr><td>Oligo fwd</td><td>CACCAACACCGACGACGAGACGAC</td><td></td><td></td><td></td><td></td><td></td><td></td><td></td><td></td><td></td><td></td><td></td></tr>
<tr><td>Oligo rev</td><td>AAACGTCGTCTCGTCGTCGGTGTT</td><td></td><td></td><td></td><td></td><td></td><td></td><td></td><td></td><td></td><td></td><td></td></tr>
<tr><td>Chromosome</td><td>start</td><td></td><td>end</td><td>strand</td><td>MM</td><td>target_seq</td><td>PAM</td><td>alignment</td><td>distance</td><td>position</td><td>gene name</td><td>gene id</td></tr>
<tr><td>GU305914.1</td><td></td><td>7135</td><td>7157</td><td>+</td><td>0</td><td>AACACCGACGACGAGACGAC</td><td>CGG</td><td>||||||||[||||||||||||]PAM</td><td>NA</td><td>Intergenic</td><td>NA</td><td>NA</td></tr>
<tr><td>T383</td><td>ACACCGACGACGAGACGACCGGG</td><td></td><td>1000</td><td></td><td></td><td></td><td></td><td></td><td></td><td></td><td></td><td></td></tr>
<tr><td>Oligo fwd</td><td>CACCACACCGACGACGAGACGACC</td><td></td><td></td><td></td><td></td><td></td><td></td><td></td><td></td><td></td><td></td><td></td></tr>
<tr><td>Oligo rev</td><td>AAACGGTCGTCTCGTCGTCGGTGT</td><td></td><td></td><td></td><td></td><td></td><td></td><td></td><td></td><td></td><td></td><td></td></tr>
<tr><td>Chromosome</td><td>start</td><td></td><td>end</td><td>strand</td><td>MM</td><td>target_seq</td><td>PAM</td><td>alignment</td><td>distance</td><td>position</td><td>gene name</td><td>gene id</td></tr>
<tr><td>GU305914.1</td><td></td><td>7136</td><td>7158</td><td>+</td><td>0</td><td>ACACCGACGACGAGACGACC</td><td>GGG</td><td>||||||||[||||||||||||]PAM</td><td>NA</td><td>Intergenic</td><td>NA</td><td>NA</td></tr>
<tr><td>T384</td><td>CGACGAGACGACCGGGTCTATGG</td><td></td><td>1000</td><td></td><td></td><td></td><td></td><td></td><td></td><td></td><td></td><td></td></tr>
<tr><td>Oligo fwd</td><td>CACCCGACGAGACGACCGGGTCTA</td><td></td><td></td><td></td><td></td><td></td><td></td><td></td><td></td><td></td><td></td><td></td></tr>
<tr><td>Oligo rev</td><td>AAACTAGACCCGGTCGTCTCGTCG</td><td></td><td></td><td></td><td></td><td></td><td></td><td></td><td></td><td></td><td></td><td></td></tr>
<tr><td>Chromosome</td><td>start</td><td></td><td>end</td><td>strand</td><td>MM</td><td>target_seq</td><td>PAM</td><td>alignment</td><td>distance</td><td>position</td><td>gene name</td><td>gene id</td></tr>
<tr><td>GU305914.1</td><td></td><td>7143</td><td>7165</td><td>+</td><td>0</td><td>CGACGAGACGACCGGGTCTA</td><td>TGG</td><td>||||||||[||||||||||||]PAM</td><td>NA</td><td>Intergenic</td><td>NA</td><td>NA</td></tr>
<tr><td>T385</td><td>AGACGACCGGGTCTATGGAACGG</td><td></td><td>1000</td><td></td><td></td><td></td><td></td><td></td><td></td><td></td><td></td><td></td></tr>
<tr><td>Oligo fwd</td><td>CACCAGACGACCGGGTCTATGGAA</td><td></td><td></td><td></td><td></td><td></td><td></td><td></td><td></td><td></td><td></td><td></td></tr>
<tr><td>Oligo rev</td><td>AAACTTCCATAGACCCGGTCGTCT</td><td></td><td></td><td></td><td></td><td></td><td></td><td></td><td></td><td></td><td></td><td></td></tr>
<tr><td>Chromosome</td><td>start</td><td></td><td>end</td><td>strand</td><td>MM</td><td>target_seq</td><td>PAM</td><td>alignment</td><td>distance</td><td>position</td><td>gene name</td><td>gene id</td></tr>
<tr><td>GU305914.1</td><td></td><td>7148</td><td>7170</td><td>+</td><td>0</td><td>AGACGACCGGGTCTATGGAA</td><td>CGG</td><td>||||||||[||||||||||||]PAM</td><td>NA</td><td>Intergenic</td><td>NA</td><td>NA</td></tr>
<tr><td>T386</td><td>ATGCTCGTCTGCTTCTCGTTCGG</td><td></td><td>1000</td><td></td><td></td><td></td><td></td><td></td><td></td><td></td><td></td><td></td></tr>
<tr><td>Oligo fwd</td><td>CACCATGCTCGTCTGCTTCTCGTT</td><td></td><td></td><td></td><td></td><td></td><td></td><td></td><td></td><td></td><td></td><td></td></tr>
<tr><td>Oligo rev</td><td>AAACAACGAGAAGCAGACGAGCAT</td><td></td><td></td><td></td><td></td><td></td><td></td><td></td><td></td><td></td><td></td><td></td></tr>
<tr><td>Chromosome</td><td>start</td><td></td><td>end</td><td>strand</td><td>MM</td><td>target_seq</td><td>PAM</td><td>alignment</td><td>distance</td><td>position</td><td>gene name</td><td>gene id</td></tr>
<tr><td>GU305914.1</td><td></td><td>7201</td><td>7223</td><td>+</td><td>0</td><td>ATGCTCGTCTGCTTCTCGTT</td><td>CGG</td><td>||||||||[||||||||||||]PAM</td><td>NA</td><td>Intergenic</td><td>NA</td><td>NA</td></tr>
<tr><td>T387</td><td>GGTGCGTGCGTGCTGCTAGTAGG</td><td></td><td>1000</td><td></td><td></td><td></td><td></td><td></td><td></td><td></td><td></td><td></td></tr>
<tr><td>Oligo fwd</td><td>CACCGGTGCGTGCGTGCTGCTAGT</td><td></td><td></td><td></td><td></td><td></td><td></td><td></td><td></td><td></td><td></td><td></td></tr>
<tr><td>Oligo rev</td><td>AAACACTAGCAGCACGCACGCACC</td><td></td><td></td><td></td><td></td><td></td><td></td><td></td><td></td><td></td><td></td><td></td></tr>
<tr><td>Chromosome</td><td>start</td><td></td><td>end</td><td>strand</td><td>MM</td><td>target_seq</td><td>PAM</td><td>alignment</td><td>distance</td><td>position</td><td>gene name</td><td>gene id</td></tr>
<tr><td>GU305914.1</td><td></td><td>7222</td><td>7244</td><td>+</td><td>0</td><td>GGTGCGTGCGTGCTGCTAGT</td><td>AGG</td><td>||||||||[||||||||||||]PAM</td><td>NA</td><td>Intergenic</td><td>NA</td><td>NA</td></tr>
<tr><td>T388</td><td>GTGCGTGCTGCTAGTAGGCTTGG</td><td></td><td>1000</td><td></td><td></td><td></td><td></td><td></td><td></td><td></td><td></td><td></td></tr>
<tr><td>Oligo fwd</td><td>CACCGTGCGTGCTGCTAGTAGGCT</td><td></td><td></td><td></td><td></td><td></td><td></td><td></td><td></td><td></td><td></td><td></td></tr>
<tr><td>Oligo rev</td><td>AAACAGCCTACTAGCAGCACGCAC</td><td></td><td></td><td></td><td></td><td></td><td></td><td></td><td></td><td></td><td></td><td></td></tr>
<tr><td>Chromosome</td><td>start</td><td></td><td>end</td><td>strand</td><td>MM</td><td>target_seq</td><td>PAM</td><td>alignment</td><td>distance</td><td>position</td><td>gene name</td><td>gene id</td></tr>
<tr><td>GU305914.1</td><td></td><td>7227</td><td>7249</td><td>+</td><td>0</td><td>GTGCGTGCTGCTAGTAGGCT</td><td>TGG</td><td>||||||||[||||||||||||]PAM</td><td>NA</td><td>Intergenic</td><td>NA</td><td>NA</td></tr>
<tr><td>T389</td><td>GTAGGCTTGGCGTATTATTACGG</td><td></td><td>1000</td><td></td><td></td><td></td><td></td><td></td><td></td><td></td><td></td><td></td></tr>
<tr><td>Oligo fwd</td><td>CACCGTAGGCTTGGCGTATTATTA</td><td></td><td></td><td></td><td></td><td></td><td></td><td></td><td></td><td></td><td></td><td></td></tr>
<tr><td>Oligo rev</td><td>AAACTAATAATACGCCAAGCCTAC</td><td></td><td></td><td></td><td></td><td></td><td></td><td></td><td></td><td></td><td></td><td></td></tr>
<tr><td>Chromosome</td><td>start</td><td></td><td>end</td><td>strand</td><td>MM</td><td>target_seq</td><td>PAM</td><td>alignment</td><td>distance</td><td>position</td><td>gene name</td><td>gene id</td></tr>
<tr><td>GU305914.1</td><td></td><td>7240</td><td>7262</td><td>+</td><td>0</td><td>GTAGGCTTGGCGTATTATTA</td><td>CGG</td><td>||||||||[||||||||||||]PAM</td><td>NA</td><td>Intergenic</td><td>NA</td><td>NA</td></tr>
<tr><td>T390</td><td>GACACGCAGTCACACGCTCCCGG</td><td></td><td>1000</td><td></td><td></td><td></td><td></td><td></td><td></td><td></td><td></td><td></td></tr>
<tr><td>Oligo fwd</td><td>CACCGACACGCAGTCACACGCTCC</td><td></td><td></td><td></td><td></td><td></td><td></td><td></td><td></td><td></td><td></td><td></td></tr>
<tr><td>Oligo rev</td><td>AAACGGAGCGTGTGACTGCGTGTC</td><td></td><td></td><td></td><td></td><td></td><td></td><td></td><td></td><td></td><td></td><td></td></tr>
<tr><td>Chromosome</td><td>start</td><td></td><td>end</td><td>strand</td><td>MM</td><td>target_seq</td><td>PAM</td><td>alignment</td><td>distance</td><td>position</td><td>gene name</td><td>gene id</td></tr>
<tr><td>GU305914.1</td><td></td><td>7273</td><td>7295</td><td>+</td><td>0</td><td>GACACGCAGTCACACGCTCC</td><td>CGG</td><td>||||||||[||||||||||||]PAM</td><td>NA</td><td>Intergenic</td><td>NA</td><td>NA</td></tr>
<tr><td>T391</td><td>CAGTCACACGCTCCCGGTCACGG</td><td></td><td>1000</td><td></td><td></td><td></td><td></td><td></td><td></td><td></td><td></td><td></td></tr>
<tr><td>Oligo fwd</td><td>CACCCAGTCACACGCTCCCGGTCA</td><td></td><td></td><td></td><td></td><td></td><td></td><td></td><td></td><td></td><td></td><td></td></tr>
<tr><td>Oligo rev</td><td>AAACTGACCGGGAGCGTGTGACTG</td><td></td><td></td><td></td><td></td><td></td><td></td><td></td><td></td><td></td><td></td><td></td></tr>
<tr><td>Chromosome</td><td>start</td><td></td><td>end</td><td>strand</td><td>MM</td><td>target_seq</td><td>PAM</td><td>alignment</td><td>distance</td><td>position</td><td>gene name</td><td>gene id</td></tr>
<tr><td>GU305914.1</td><td></td><td>7279</td><td>7301</td><td>+</td><td>0</td><td>CAGTCACACGCTCCCGGTCA</td><td>CGG</td><td>||||||||[||||||||||||]PAM</td><td>NA</td><td>Intergenic</td><td>NA</td><td>NA</td></tr>
<tr><td>T392</td><td>CACGGATCCGAACCACGTTCCGG</td><td></td><td>1000</td><td></td><td></td><td></td><td></td><td></td><td></td><td></td><td></td><td></td></tr>
<tr><td>Oligo fwd</td><td>CACCCACGGATCCGAACCACGTTC</td><td></td><td></td><td></td><td></td><td></td><td></td><td></td><td></td><td></td><td></td><td></td></tr>
<tr><td>Oligo rev</td><td>AAACGAACGTGGTTCGGATCCGTG</td><td></td><td></td><td></td><td></td><td></td><td></td><td></td><td></td><td></td><td></td><td></td></tr>
<tr><td>Chromosome</td><td>start</td><td></td><td>end</td><td>strand</td><td>MM</td><td>target_seq</td><td>PAM</td><td>alignment</td><td>distance</td><td>position</td><td>gene name</td><td>gene id</td></tr>
<tr><td>GU305914.1</td><td></td><td>7297</td><td>7319</td><td>+</td><td>0</td><td>CACGGATCCGAACCACGTTC</td><td>CGG</td><td>||||||||[||||||||||||]PAM</td><td>NA</td><td>Intergenic</td><td>NA</td><td>NA</td></tr>
<tr><td>T393</td><td>ATCTGAACCTCTACGTCCGCCGG</td><td></td><td>1000</td><td></td><td></td><td></td><td></td><td></td><td></td><td></td><td></td><td></td></tr>
<tr><td>Oligo fwd</td><td>CACCATCTGAACCTCTACGTCCGC</td><td></td><td></td><td></td><td></td><td></td><td></td><td></td><td></td><td></td><td></td><td></td></tr>
<tr><td>Oligo rev</td><td>AAACGCGGACGTAGAGGTTCAGAT</td><td></td><td></td><td></td><td></td><td></td><td></td><td></td><td></td><td></td><td></td><td></td></tr>
<tr><td>Chromosome</td><td>start</td><td></td><td>end</td><td>strand</td><td>MM</td><td>target_seq</td><td>PAM</td><td>alignment</td><td>distance</td><td>position</td><td>gene name</td><td>gene id</td></tr>
<tr><td>GU305914.1</td><td></td><td>7323</td><td>7345</td><td>+</td><td>0</td><td>ATCTGAACCTCTACGTCCGC</td><td>CGG</td><td>||||||||[||||||||||||]PAM</td><td>NA</td><td>Intergenic</td><td>NA</td><td>NA</td></tr>
<tr><td>T394</td><td>CCTGTCACCGCCCACACCTGCGG</td><td></td><td>1000</td><td></td><td></td><td></td><td></td><td></td><td></td><td></td><td></td><td></td></tr>
<tr><td>Oligo fwd</td><td>CACCCCTGTCACCGCCCACACCTG</td><td></td><td></td><td></td><td></td><td></td><td></td><td></td><td></td><td></td><td></td><td></td></tr>
<tr><td>Oligo rev</td><td>AAACCAGGTGTGGGCGGTGACAGG</td><td></td><td></td><td></td><td></td><td></td><td></td><td></td><td></td><td></td><td></td><td></td></tr>
<tr><td>Chromosome</td><td>start</td><td></td><td>end</td><td>strand</td><td>MM</td><td>target_seq</td><td>PAM</td><td>alignment</td><td>distance</td><td>position</td><td>gene name</td><td>gene id</td></tr>
<tr><td>GU305914.1</td><td></td><td>7374</td><td>7396</td><td>+</td><td>0</td><td>CCTGTCACCGCCCACACCTG</td><td>CGG</td><td>||||||||[||||||||||||]PAM</td><td>NA</td><td>Intergenic</td><td>NA</td><td>NA</td></tr>
<tr><td>T395</td><td>CACCTGCGGTGAGTATGTAAAGG</td><td></td><td>1000</td><td></td><td></td><td></td><td></td><td></td><td></td><td></td><td></td><td></td></tr>
<tr><td>Oligo fwd</td><td>CACCCACCTGCGGTGAGTATGTAA</td><td></td><td></td><td></td><td></td><td></td><td></td><td></td><td></td><td></td><td></td><td></td></tr>
<tr><td>Oligo rev</td><td>AAACTTACATACTCACCGCAGGTG</td><td></td><td></td><td></td><td></td><td></td><td></td><td></td><td></td><td></td><td></td><td></td></tr>
<tr><td>Chromosome</td><td>start</td><td></td><td>end</td><td>strand</td><td>MM</td><td>target_seq</td><td>PAM</td><td>alignment</td><td>distance</td><td>position</td><td>gene name</td><td>gene id</td></tr>
<tr><td>GU305914.1</td><td></td><td>7388</td><td>7410</td><td>+</td><td>0</td><td>CACCTGCGGTGAGTATGTAA</td><td>AGG</td><td>||||||||[||||||||||||]PAM</td><td>NA</td><td>Intergenic</td><td>NA</td><td>NA</td></tr>
<tr><td>T396</td><td>ACCTGCGGTGAGTATGTAAAGGG</td><td></td><td>1000</td><td></td><td></td><td></td><td></td><td></td><td></td><td></td><td></td><td></td></tr>
<tr><td>Oligo fwd</td><td>CACCACCTGCGGTGAGTATGTAAA</td><td></td><td></td><td></td><td></td><td></td><td></td><td></td><td></td><td></td><td></td><td></td></tr>
<tr><td>Oligo rev</td><td>AAACTTTACATACTCACCGCAGGT</td><td></td><td></td><td></td><td></td><td></td><td></td><td></td><td></td><td></td><td></td><td></td></tr>
<tr><td>Chromosome</td><td>start</td><td></td><td>end</td><td>strand</td><td>MM</td><td>target_seq</td><td>PAM</td><td>alignment</td><td>distance</td><td>position</td><td>gene name</td><td>gene id</td></tr>
<tr><td>GU305914.1</td><td></td><td>7389</td><td>7411</td><td>+</td><td>0</td><td>ACCTGCGGTGAGTATGTAAA</td><td>GGG</td><td>||||||||[||||||||||||]PAM</td><td>NA</td><td>Intergenic</td><td>NA</td><td>NA</td></tr>
</table>

<table>
<tr><td>T397</td><td>TGTAAAGGGCCGACCGCCGCAGG</td><td>1000</td><td></td><td></td><td></td><td></td><td></td><td></td><td></td><td></td><td></td></tr>
<tr><td>Oligo fwd</td><td>CACCTGTAAAGGGCCGACCGCCGC</td><td></td><td></td><td></td><td></td><td></td><td></td><td></td><td></td><td></td><td></td></tr>
<tr><td>Oligo rev</td><td>AAACGCGGCGGTCGGCCCTTTACA</td><td></td><td></td><td></td><td></td><td></td><td></td><td></td><td></td><td></td><td></td></tr>
<tr><td>Chromosome</td><td>start</td><td>end</td><td>strand</td><td>MM</td><td>target_seq</td><td>PAM</td><td>alignment</td><td>distance</td><td>position</td><td>gene name</td><td>gene id</td></tr>
<tr><td>GU305914.1</td><td>7403</td><td>7425</td><td>+</td><td>0</td><td>TGTAAAGGGCCGACCGCCGC</td><td>AGG</td><td>||||||||[||||||||||||]PAM</td><td>NA</td><td>Intergenic</td><td>NA</td><td>NA</td></tr>
</table>

<table>
<tr><td>T398</td><td>GCCGACCGCCGCAGGTGCGACGG</td><td>1000</td><td></td><td></td><td></td><td></td><td></td><td></td><td></td><td></td><td></td></tr>
<tr><td>Oligo fwd</td><td>CACCGCCGACCGCCGCAGGTGCGA</td><td></td><td></td><td></td><td></td><td></td><td></td><td></td><td></td><td></td><td></td></tr>
<tr><td>Oligo rev</td><td>AAACTCGCACCTGCGGCGGTCGGC</td><td></td><td></td><td></td><td></td><td></td><td></td><td></td><td></td><td></td><td></td></tr>
<tr><td>Chromosome</td><td>start</td><td>end</td><td>strand</td><td>MM</td><td>target_seq</td><td>PAM</td><td>alignment</td><td>distance</td><td>position</td><td>gene name</td><td>gene id</td></tr>
<tr><td>GU305914.1</td><td>7411</td><td>7433</td><td>+</td><td>0</td><td>GCCGACCGCCGCAGGTGCGA</td><td>CGG</td><td>||||||||[||||||||||||]PAM</td><td>NA</td><td>Intergenic</td><td>NA</td><td>NA</td></tr>
</table>

<table>
<tr><td>T399</td><td>CGGTACGTATCAGATACACCTGG</td><td>1000</td><td></td><td></td><td></td><td></td><td></td><td></td><td></td><td></td><td></td></tr>
<tr><td>Oligo fwd</td><td>CACCCGGTACGTATCAGATACACC</td><td></td><td></td><td></td><td></td><td></td><td></td><td></td><td></td><td></td><td></td></tr>
<tr><td>Oligo rev</td><td>AAACGGTGTATCTGATACGTACCG</td><td></td><td></td><td></td><td></td><td></td><td></td><td></td><td></td><td></td><td></td></tr>
<tr><td>Chromosome</td><td>start</td><td>end</td><td>strand</td><td>MM</td><td>target_seq</td><td>PAM</td><td>alignment</td><td>distance</td><td>position</td><td>gene name</td><td>gene id</td></tr>
<tr><td>GU305914.1</td><td>7431</td><td>7453</td><td>+</td><td>0</td><td>CGGTACGTATCAGATACACC</td><td>TGG</td><td>||||||||[||||||||||||]PAM</td><td>NA</td><td>Intergenic</td><td>NA</td><td>NA</td></tr>
</table>

<table>
<tr><td>T400</td><td>AGTCATCGTAACAGATTCCGTGG</td><td>1000</td><td></td><td></td><td></td><td></td><td></td><td></td><td></td><td></td><td></td></tr>
<tr><td>Oligo fwd</td><td>CACCAGTCATCGTAACAGATTCCG</td><td></td><td></td><td></td><td></td><td></td><td></td><td></td><td></td><td></td><td></td></tr>
<tr><td>Oligo rev</td><td>AAACCGGAATCTGTTACGATGACT</td><td></td><td></td><td></td><td></td><td></td><td></td><td></td><td></td><td></td><td></td></tr>
<tr><td>Chromosome</td><td>start</td><td>end</td><td>strand</td><td>MM</td><td>target_seq</td><td>PAM</td><td>alignment</td><td>distance</td><td>position</td><td>gene name</td><td>gene id</td></tr>
<tr><td>GU305914.1</td><td>7500</td><td>7522</td><td>+</td><td>0</td><td>AGTCATCGTAACAGATTCCG</td><td>TGG</td><td>||||||||[||||||||||||]PAM</td><td>NA</td><td>Intergenic</td><td>NA</td><td>NA</td></tr>
</table>

<table>
<tr><td>T401</td><td>GCACATTCCAATCTACCACAAGG</td><td>1000</td><td></td><td></td><td></td><td></td><td></td><td></td><td></td><td></td><td></td></tr>
<tr><td>Oligo fwd</td><td>CACCGCACATTCCAATCTACCACA</td><td></td><td></td><td></td><td></td><td></td><td></td><td></td><td></td><td></td><td></td></tr>
<tr><td>Oligo rev</td><td>AAACTGTGGTAGATTGGAATGTGC</td><td></td><td></td><td></td><td></td><td></td><td></td><td></td><td></td><td></td><td></td></tr>
<tr><td>Chromosome</td><td>start</td><td>end</td><td>strand</td><td>MM</td><td>target_seq</td><td>PAM</td><td>alignment</td><td>distance</td><td>position</td><td>gene name</td><td>gene id</td></tr>
<tr><td>GU305914.1</td><td>7549</td><td>7571</td><td>+</td><td>0</td><td>GCACATTCCAATCTACCACA</td><td>AGG</td><td>||||||||[||||||||||||]PAM</td><td>NA</td><td>Intergenic</td><td>NA</td><td>NA</td></tr>
</table>

<table>
<tr><td>T402</td><td>GTACGTAACGACAGTAACCTAGG</td><td>1000</td><td></td><td></td><td></td><td></td><td></td><td></td><td></td><td></td><td></td></tr>
<tr><td>Oligo fwd</td><td>CACCGTACGTAACGACAGTAACCT</td><td></td><td></td><td></td><td></td><td></td><td></td><td></td><td></td><td></td><td></td></tr>
<tr><td>Oligo rev</td><td>AAACAGGTTACTGTCGTTACGTAC</td><td></td><td></td><td></td><td></td><td></td><td></td><td></td><td></td><td></td><td></td></tr>
<tr><td>Chromosome</td><td>start</td><td>end</td><td>strand</td><td>MM</td><td>target_seq</td><td>PAM</td><td>alignment</td><td>distance</td><td>position</td><td>gene name</td><td>gene id</td></tr>
<tr><td>GU305914.1</td><td>7600</td><td>7622</td><td>+</td><td>0</td><td>GTACGTAACGACAGTAACCT</td><td>AGG</td><td>||||||||[||||||||||||]PAM</td><td>NA</td><td>Intergenic</td><td>NA</td><td>NA</td></tr>
</table>

<table>
<tr><td>T403</td><td>TAACGACAGTAACCTAGGTGTGG</td><td>1000</td><td></td><td></td><td></td><td></td><td></td><td></td><td></td><td></td><td></td></tr>
<tr><td>Oligo fwd</td><td>CACCTAACGACAGTAACCTAGGTG</td><td></td><td></td><td></td><td></td><td></td><td></td><td></td><td></td><td></td><td></td></tr>
<tr><td>Oligo rev</td><td>AAACCACCTAGGTTACTGTCGTTA</td><td></td><td></td><td></td><td></td><td></td><td></td><td></td><td></td><td></td><td></td></tr>
<tr><td>Chromosome</td><td>start</td><td>end</td><td>strand</td><td>MM</td><td>target_seq</td><td>PAM</td><td>alignment</td><td>distance</td><td>position</td><td>gene name</td><td>gene id</td></tr>
<tr><td>GU305914.1</td><td>7605</td><td>7627</td><td>+</td><td>0</td><td>TAACGACAGTAACCTAGGTG</td><td>TGG</td><td>||||||||[||||||||||||]PAM</td><td>NA</td><td>Intergenic</td><td>NA</td><td>NA</td></tr>
</table>

<table>
<tr><td>T404</td><td>GACAGTAACCTAGGTGTGGACGG</td><td>1000</td><td></td><td></td><td></td><td></td><td></td><td></td><td></td><td></td><td></td></tr>
<tr><td>Oligo fwd</td><td>CACCGACAGTAACCTAGGTGTGGA</td><td></td><td></td><td></td><td></td><td></td><td></td><td></td><td></td><td></td><td></td></tr>
<tr><td>Oligo rev</td><td>AAACTCCACACCTAGGTTACTGTC</td><td></td><td></td><td></td><td></td><td></td><td></td><td></td><td></td><td></td><td></td></tr>
<tr><td>Chromosome</td><td>start</td><td>end</td><td>strand</td><td>MM</td><td>target_seq</td><td>PAM</td><td>alignment</td><td>distance</td><td>position</td><td>gene name</td><td>gene id</td></tr>
<tr><td>GU305914.1</td><td>7609</td><td>7631</td><td>+</td><td>0</td><td>GACAGTAACCTAGGTGTGGA</td><td>CGG</td><td>||||||||[||||||||||||]PAM</td><td>NA</td><td>Intergenic</td><td>NA</td><td>NA</td></tr>
</table>

<table>
<tr><td>T405</td><td>GTAACCTAGGTGTGGACGGTTGG</td><td>1000</td><td></td><td></td><td></td><td></td><td></td><td></td><td></td><td></td><td></td></tr>
<tr><td>Oligo fwd</td><td>CACCGTAACCTAGGTGTGGACGGT</td><td></td><td></td><td></td><td></td><td></td><td></td><td></td><td></td><td></td><td></td></tr>
<tr><td>Oligo rev</td><td>AAACACCGTCCACACCTAGGTTAC</td><td></td><td></td><td></td><td></td><td></td><td></td><td></td><td></td><td></td><td></td></tr>
<tr><td>Chromosome</td><td>start</td><td>end</td><td>strand</td><td>MM</td><td>target_seq</td><td>PAM</td><td>alignment</td><td>distance</td><td>position</td><td>gene name</td><td>gene id</td></tr>
<tr><td>GU305914.1</td><td>7613</td><td>7635</td><td>+</td><td>0</td><td>GTAACCTAGGTGTGGACGGT</td><td>TGG</td><td>||||||||[||||||||||||]PAM</td><td>NA</td><td>Intergenic</td><td>NA</td><td>NA</td></tr>
</table>

<table>
<tr><td>T406</td><td>TAACCTAGGTGTGGACGGTTGGG</td><td>1000</td><td></td><td></td><td></td><td></td><td></td><td></td><td></td><td></td><td></td></tr>
<tr><td>Oligo fwd</td><td>CACCTAACCTAGGTGTGGACGGTT</td><td></td><td></td><td></td><td></td><td></td><td></td><td></td><td></td><td></td><td></td></tr>
<tr><td>Oligo rev</td><td>AAACAACCGTCCACACCTAGGTTA</td><td></td><td></td><td></td><td></td><td></td><td></td><td></td><td></td><td></td><td></td></tr>
<tr><td>Chromosome</td><td>start</td><td>end</td><td>strand</td><td>MM</td><td>target_seq</td><td>PAM</td><td>alignment</td><td>distance</td><td>position</td><td>gene name</td><td>gene id</td></tr>
<tr><td>GU305914.1</td><td>7614</td><td>7636</td><td>+</td><td>0</td><td>TAACCTAGGTGTGGACGGTT</td><td>GGG</td><td>||||||||[||||||||||||]PAM</td><td>NA</td><td>Intergenic</td><td>NA</td><td>NA</td></tr>
</table>

<table>
<tr><td>T407</td><td>AACCTAGGTGTGGACGGTTGGGG</td><td>1000</td><td></td><td></td><td></td><td></td><td></td><td></td><td></td><td></td><td></td></tr>
<tr><td>Oligo fwd</td><td>CACCAACCTAGGTGTGGACGGTTG</td><td></td><td></td><td></td><td></td><td></td><td></td><td></td><td></td><td></td><td></td></tr>
<tr><td>Oligo rev</td><td>AAACCAACCGTCCACACCTAGGTT</td><td></td><td></td><td></td><td></td><td></td><td></td><td></td><td></td><td></td><td></td></tr>
<tr><td>Chromosome</td><td>start</td><td>end</td><td>strand</td><td>MM</td><td>target_seq</td><td>PAM</td><td>alignment</td><td>distance</td><td>position</td><td>gene name</td><td>gene id</td></tr>
<tr><td>GU305914.1</td><td>7615</td><td>7637</td><td>+</td><td>0</td><td>AACCTAGGTGTGGACGGTTG</td><td>GGG</td><td>||||||||[||||||||||||]PAM</td><td>NA</td><td>Intergenic</td><td>NA</td><td>NA</td></tr>
</table>

<table>
<tr><td>T408</td><td>TGGGGATACGAATGTCCACTCGG</td><td>1000</td><td></td><td></td><td></td><td></td><td></td><td></td><td></td><td></td><td></td></tr>
<tr><td>Oligo fwd</td><td>CACCTGGGGATACGAATGTCCACT</td><td></td><td></td><td></td><td></td><td></td><td></td><td></td><td></td><td></td><td></td></tr>
<tr><td>Oligo rev</td><td>AAACAGTGGACATTCGTATCCCCA</td><td></td><td></td><td></td><td></td><td></td><td></td><td></td><td></td><td></td><td></td></tr>
<tr><td>Chromosome</td><td>start</td><td>end</td><td>strand</td><td>MM</td><td>target_seq</td><td>PAM</td><td>alignment</td><td>distance</td><td>position</td><td>gene name</td><td>gene id</td></tr>
<tr><td>GU305914.1</td><td>7633</td><td>7655</td><td>+</td><td>0</td><td>TGGGGATACGAATGTCCACT</td><td>CGG</td><td>||||||||[||||||||||||]PAM</td><td>NA</td><td>Intergenic</td><td>NA</td><td>NA</td></tr>
</table>

<table>
<tr><td>T409</td><td>ACTCGGTGATATCTGCATTCCGG</td><td>1000</td><td></td><td></td><td></td><td></td><td></td><td></td><td></td><td></td><td></td></tr>
<tr><td>Oligo fwd</td><td>CACCACTCGGTGATATCTGCATTC</td><td></td><td></td><td></td><td></td><td></td><td></td><td></td><td></td><td></td><td></td></tr>
<tr><td>Oligo rev</td><td>AAACGAATGCAGATATCACCGAGT</td><td></td><td></td><td></td><td></td><td></td><td></td><td></td><td></td><td></td><td></td></tr>
<tr><td>Chromosome</td><td>start</td><td>end</td><td>strand</td><td>MM</td><td>target_seq</td><td>PAM</td><td>alignment</td><td>distance</td><td>position</td><td>gene name</td><td>gene id</td></tr>
<tr><td>GU305914.1</td><td>7650</td><td>7672</td><td>+</td><td>0</td><td>ACTCGGTGATATCTGCATTC</td><td>CGG</td><td>||||||||[||||||||||||]PAM</td><td>NA</td><td>Intergenic</td><td>NA</td><td>NA</td></tr>
</table>

<table>
<tr><td>T410</td><td>ATATCTGCATTCCGGCATCGTGG</td><td>1000</td><td></td><td></td><td></td><td></td><td></td><td></td><td></td><td></td><td></td></tr>
<tr><td>Oligo fwd</td><td>CACCATATCTGCATTCCGGCATCG</td><td></td><td></td><td></td><td></td><td></td><td></td><td></td><td></td><td></td><td></td></tr>
<tr><td>Oligo rev</td><td>AAACCGATGCCGGAATGCAGATAT</td><td></td><td></td><td></td><td></td><td></td><td></td><td></td><td></td><td></td><td></td></tr>
<tr><td>Chromosome</td><td>start</td><td>end</td><td>strand</td><td>MM</td><td>target_seq</td><td>PAM</td><td>alignment</td><td>distance</td><td>position</td><td>gene name</td><td>gene id</td></tr>
<tr><td>GU305914.1</td><td>7658</td><td>7680</td><td>+</td><td>0</td><td>ATATCTGCATTCCGGCATCG</td><td>TGG</td><td>||||||||[||||||||||||]PAM</td><td>NA</td><td>Intergenic</td><td>NA</td><td>NA</td></tr>
<tr><td>GU305914.1</td><td>6665</td><td>6687</td><td>+</td><td>4</td><td>CCATCTGTATTCCCGCATCG</td><td>TGG</td><td>--|||||-[|||||-||||||]PAM</td><td>NA</td><td>Intergenic</td><td>NA</td><td>NA</td></tr>
</table>

<table>
<tr><td>T411</td><td>CTGCATTCCGGCATCGTGGCAGG</td><td>1000</td></tr>
<tr><td>Oligo fwd</td><td>CACCCTGCATTCCGGCATCGTGGC</td><td></td></tr>
<tr><td>Oligo rev</td><td>AAACGCCACGATGCCGGAATGCAG</td><td></td></tr>
</table>

<table>
<tr><td>Chromosome</td><td>start</td><td>end</td><td>strand</td><td>MM</td><td>target_seq</td><td>PAM</td><td>alignment</td><td>distance</td><td>position</td><td>gene name</td><td>gene id</td></tr>
<tr><td>GU305914.1</td><td>7662</td><td>7684</td><td>+</td><td>0</td><td>CTGCATTCCGGCATCGTGGC</td><td>AGG</td><td>||||||||[||||||||||||]PAM</td><td>NA</td><td>Intergenic</td><td>NA</td><td>NA</td></tr>
<tr><td>T412</td><td>CGGCATCGTGGCAGGTTAAATGG</td><td>1000</td><td></td><td></td><td></td><td></td><td></td><td></td><td></td><td></td><td></td></tr>
<tr><td>Oligo fwd</td><td>CACCCGGCATCGTGGCAGGTTAAA</td><td></td><td></td><td></td><td></td><td></td><td></td><td></td><td></td><td></td><td></td></tr>
<tr><td>Oligo rev</td><td>AAACTTTAACCTGCCACGATGCCG</td><td></td><td></td><td></td><td></td><td></td><td></td><td></td><td></td><td></td><td></td></tr>
<tr><td>Chromosome</td><td>start</td><td>end</td><td>strand</td><td>MM</td><td>target_seq</td><td>PAM</td><td>alignment</td><td>distance</td><td>position</td><td>gene name</td><td>gene id</td></tr>
<tr><td>GU305914.1</td><td>7670</td><td>7692</td><td>+</td><td>0</td><td>CGGCATCGTGGCAGGTTAAA</td><td>TGG</td><td>||||||||[||||||||||||]PAM</td><td>NA</td><td>Intergenic</td><td>NA</td><td>NA</td></tr>
<tr><td>T413</td><td>TGGCAGGTTAAATGGTATATCGG</td><td>1000</td><td></td><td></td><td></td><td></td><td></td><td></td><td></td><td></td><td></td></tr>
<tr><td>Oligo fwd</td><td>CACCTGGCAGGTTAAATGGTATAT</td><td></td><td></td><td></td><td></td><td></td><td></td><td></td><td></td><td></td><td></td></tr>
<tr><td>Oligo rev</td><td>AAACATATACCATTTAACCTGCCA</td><td></td><td></td><td></td><td></td><td></td><td></td><td></td><td></td><td></td><td></td></tr>
<tr><td>Chromosome</td><td>start</td><td>end</td><td>strand</td><td>MM</td><td>target_seq</td><td>PAM</td><td>alignment</td><td>distance</td><td>position</td><td>gene name</td><td>gene id</td></tr>
<tr><td>GU305914.1</td><td>7678</td><td>7700</td><td>+</td><td>0</td><td>TGGCAGGTTAAATGGTATAT</td><td>CGG</td><td>||||||||[||||||||||||]PAM</td><td>NA</td><td>Intergenic</td><td>NA</td><td>NA</td></tr>
<tr><td>T414</td><td>GGCAGGTTAAATGGTATATCGGG</td><td>1000</td><td></td><td></td><td></td><td></td><td></td><td></td><td></td><td></td><td></td></tr>
<tr><td>Oligo fwd</td><td>CACCGGCAGGTTAAATGGTATATC</td><td></td><td></td><td></td><td></td><td></td><td></td><td></td><td></td><td></td><td></td></tr>
<tr><td>Oligo rev</td><td>AAACGATATACCATTTAACCTGCC</td><td></td><td></td><td></td><td></td><td></td><td></td><td></td><td></td><td></td><td></td></tr>
<tr><td>Chromosome</td><td>start</td><td>end</td><td>strand</td><td>MM</td><td>target_seq</td><td>PAM</td><td>alignment</td><td>distance</td><td>position</td><td>gene name</td><td>gene id</td></tr>
<tr><td>GU305914.1</td><td>7679</td><td>7701</td><td>+</td><td>0</td><td>GGCAGGTTAAATGGTATATC</td><td>GGG</td><td>||||||||[||||||||||||]PAM</td><td>NA</td><td>Intergenic</td><td>NA</td><td>NA</td></tr>
<tr><td>T415</td><td>GTATATCGGGCTTTACCAAATGG</td><td>1000</td><td></td><td></td><td></td><td></td><td></td><td></td><td></td><td></td><td></td></tr>
<tr><td>Oligo fwd</td><td>CACCGTATATCGGGCTTTACCAAA</td><td></td><td></td><td></td><td></td><td></td><td></td><td></td><td></td><td></td><td></td></tr>
<tr><td>Oligo rev</td><td>AAACTTTGGTAAAGCCCGATATAC</td><td></td><td></td><td></td><td></td><td></td><td></td><td></td><td></td><td></td><td></td></tr>
<tr><td>Chromosome</td><td>start</td><td>end</td><td>strand</td><td>MM</td><td>target_seq</td><td>PAM</td><td>alignment</td><td>distance</td><td>position</td><td>gene name</td><td>gene id</td></tr>
<tr><td>GU305914.1</td><td>7692</td><td>7714</td><td>+</td><td>0</td><td>GTATATCGGGCTTTACCAAA</td><td>TGG</td><td>||||||||[||||||||||||]PAM</td><td>NA</td><td>Intergenic</td><td>NA</td><td>NA</td></tr>
<tr><td>T416</td><td>CGGGCTTTACCAAATGGATACGG</td><td>1000</td><td></td><td></td><td></td><td></td><td></td><td></td><td></td><td></td><td></td></tr>
<tr><td>Oligo fwd</td><td>CACCCGGGCTTTACCAAATGGATA</td><td></td><td></td><td></td><td></td><td></td><td></td><td></td><td></td><td></td><td></td></tr>
<tr><td>Oligo rev</td><td>AAACTATCCATTTGGTAAAGCCCG</td><td></td><td></td><td></td><td></td><td></td><td></td><td></td><td></td><td></td><td></td></tr>
<tr><td>Chromosome</td><td>start</td><td>end</td><td>strand</td><td>MM</td><td>target_seq</td><td>PAM</td><td>alignment</td><td>distance</td><td>position</td><td>gene name</td><td>gene id</td></tr>
<tr><td>GU305914.1</td><td>7698</td><td>7720</td><td>+</td><td>0</td><td>CGGGCTTTACCAAATGGATA</td><td>CGG</td><td>||||||||[||||||||||||]PAM</td><td>NA</td><td>Intergenic</td><td>NA</td><td>NA</td></tr>
<tr><td>T417</td><td>GGGCTTTACCAAATGGATACGGG</td><td>1000</td><td></td><td></td><td></td><td></td><td></td><td></td><td></td><td></td><td></td></tr>
<tr><td>Oligo fwd</td><td>CACCGGGCTTTACCAAATGGATAC</td><td></td><td></td><td></td><td></td><td></td><td></td><td></td><td></td><td></td><td></td></tr>
<tr><td>Oligo rev</td><td>AAACGTATCCATTTGGTAAAGCCC</td><td></td><td></td><td></td><td></td><td></td><td></td><td></td><td></td><td></td><td></td></tr>
<tr><td>Chromosome</td><td>start</td><td>end</td><td>strand</td><td>MM</td><td>target_seq</td><td>PAM</td><td>alignment</td><td>distance</td><td>position</td><td>gene name</td><td>gene id</td></tr>
<tr><td>GU305914.1</td><td>7699</td><td>7721</td><td>+</td><td>0</td><td>GGGCTTTACCAAATGGATAC</td><td>GGG</td><td>||||||||[||||||||||||]PAM</td><td>NA</td><td>Intergenic</td><td>NA</td><td>NA</td></tr>
<tr><td>T418</td><td>CTTTACCAAATGGATACGGGCGG</td><td>1000</td><td></td><td></td><td></td><td></td><td></td><td></td><td></td><td></td><td></td></tr>
<tr><td>Oligo fwd</td><td>CACCCTTTACCAAATGGATACGGG</td><td></td><td></td><td></td><td></td><td></td><td></td><td></td><td></td><td></td><td></td></tr>
<tr><td>Oligo rev</td><td>AAACCCCGTATCCATTTGGTAAAG</td><td></td><td></td><td></td><td></td><td></td><td></td><td></td><td></td><td></td><td></td></tr>
<tr><td>Chromosome</td><td>start</td><td>end</td><td>strand</td><td>MM</td><td>target_seq</td><td>PAM</td><td>alignment</td><td>distance</td><td>position</td><td>gene name</td><td>gene id</td></tr>
<tr><td>GU305914.1</td><td>7702</td><td>7724</td><td>+</td><td>0</td><td>CTTTACCAAATGGATACGGG</td><td>CGG</td><td>||||||||[||||||||||||]PAM</td><td>NA</td><td>Intergenic</td><td>NA</td><td>NA</td></tr>
<tr><td>T419</td><td>ACAAAACCACCTACGTTTCAAGG</td><td>1000</td><td></td><td></td><td></td><td></td><td></td><td></td><td></td><td></td><td></td></tr>
<tr><td>Oligo fwd</td><td>CACCACAAAACCACCTACGTTTCA</td><td></td><td></td><td></td><td></td><td></td><td></td><td></td><td></td><td></td><td></td></tr>
<tr><td>Oligo rev</td><td>AAACTGAAACGTAGGTGGTTTTGT</td><td></td><td></td><td></td><td></td><td></td><td></td><td></td><td></td><td></td><td></td></tr>
<tr><td>Chromosome</td><td>start</td><td>end</td><td>strand</td><td>MM</td><td>target_seq</td><td>PAM</td><td>alignment</td><td>distance</td><td>position</td><td>gene name</td><td>gene id</td></tr>
<tr><td>GU305914.1</td><td>7744</td><td>7766</td><td>+</td><td>0</td><td>ACAAAACCACCTACGTTTCA</td><td>AGG</td><td>||||||||[||||||||||||]PAM</td><td>NA</td><td>Intergenic</td><td>NA</td><td>NA</td></tr>
<tr><td>T420</td><td>TGAAAGATAAAAACAACGCACGG</td><td>1000</td><td></td><td></td><td></td><td></td><td></td><td></td><td></td><td></td><td></td></tr>
<tr><td>Oligo fwd</td><td>CACCTGAAAGATAAAAACAACGCA</td><td></td><td></td><td></td><td></td><td></td><td></td><td></td><td></td><td></td><td></td></tr>
<tr><td>Oligo rev</td><td>AAACTGCGTTGTTTTTATCTTTCA</td><td></td><td></td><td></td><td></td><td></td><td></td><td></td><td></td><td></td><td></td></tr>
<tr><td>Chromosome</td><td>start</td><td>end</td><td>strand</td><td>MM</td><td>target_seq</td><td>PAM</td><td>alignment</td><td>distance</td><td>position</td><td>gene name</td><td>gene id</td></tr>
<tr><td>GU305914.1</td><td>7781</td><td>7803</td><td>+</td><td>0</td><td>TGAAAGATAAAAACAACGCA</td><td>CGG</td><td>||||||||[||||||||||||]PAM</td><td>NA</td><td>Intergenic</td><td>NA</td><td>NA</td></tr>
<tr><td>T421</td><td>AAAAACAACGCACGGCCAGATGG</td><td>1000</td><td></td><td></td><td></td><td></td><td></td><td></td><td></td><td></td><td></td></tr>
<tr><td>Oligo fwd</td><td>CACCAAAAACAACGCACGGCCAGA</td><td></td><td></td><td></td><td></td><td></td><td></td><td></td><td></td><td></td><td></td></tr>
<tr><td>Oligo rev</td><td>AAACTCTGGCCGTGCGTTGTTTTT</td><td></td><td></td><td></td><td></td><td></td><td></td><td></td><td></td><td></td><td></td></tr>
<tr><td>Chromosome</td><td>start</td><td>end</td><td>strand</td><td>MM</td><td>target_seq</td><td>PAM</td><td>alignment</td><td>distance</td><td>position</td><td>gene name</td><td>gene id</td></tr>
<tr><td>GU305914.1</td><td>7789</td><td>7811</td><td>+</td><td>0</td><td>AAAAACAACGCACGGCCAGA</td><td>TGG</td><td>||||||||[||||||||||||]PAM</td><td>NA</td><td>Intergenic</td><td>NA</td><td>NA</td></tr>
<tr><td>T422</td><td>AAAACAACGCACGGCCAGATGGG</td><td>1000</td><td></td><td></td><td></td><td></td><td></td><td></td><td></td><td></td><td></td></tr>
<tr><td>Oligo fwd</td><td>CACCAAAACAACGCACGGCCAGAT</td><td></td><td></td><td></td><td></td><td></td><td></td><td></td><td></td><td></td><td></td></tr>
<tr><td>Oligo rev</td><td>AAACATCTGGCCGTGCGTTGTTTT</td><td></td><td></td><td></td><td></td><td></td><td></td><td></td><td></td><td></td><td></td></tr>
<tr><td>Chromosome</td><td>start</td><td>end</td><td>strand</td><td>MM</td><td>target_seq</td><td>PAM</td><td>alignment</td><td>distance</td><td>position</td><td>gene name</td><td>gene id</td></tr>
<tr><td>GU305914.1</td><td>7790</td><td>7812</td><td>+</td><td>0</td><td>AAAACAACGCACGGCCAGAT</td><td>GGG</td><td>||||||||[||||||||||||]PAM</td><td>NA</td><td>Intergenic</td><td>NA</td><td>NA</td></tr>
<tr><td>T423</td><td>TTCGATATAAAACACGAAGACGG</td><td>1000</td><td></td><td></td><td></td><td></td><td></td><td></td><td></td><td></td><td></td></tr>
<tr><td>Oligo fwd</td><td>CACCTTCGATATAAAACACGAAGA</td><td></td><td></td><td></td><td></td><td></td><td></td><td></td><td></td><td></td><td></td></tr>
<tr><td>Oligo rev</td><td>AAACTCTTCGTGTTTTATATCGAA</td><td></td><td></td><td></td><td></td><td></td><td></td><td></td><td></td><td></td><td></td></tr>
<tr><td>Chromosome</td><td>start</td><td>end</td><td>strand</td><td>MM</td><td>target_seq</td><td>PAM</td><td>alignment</td><td>distance</td><td>position</td><td>gene name</td><td>gene id</td></tr>
<tr><td>GU305914.1</td><td>7813</td><td>7835</td><td>+</td><td>0</td><td>TTCGATATAAAACACGAAGA</td><td>CGG</td><td>||||||||[||||||||||||]PAM</td><td>NA</td><td>Intergenic</td><td>NA</td><td>NA</td></tr>
<tr><td>T424</td><td>CGGTCGTCTCATCGTTACTGTGG</td><td>1000</td><td></td><td></td><td></td><td></td><td></td><td></td><td></td><td></td><td></td></tr>
<tr><td>Oligo fwd</td><td>CACCCGGTCGTCTCATCGTTACTG</td><td></td><td></td><td></td><td></td><td></td><td></td><td></td><td></td><td></td><td></td></tr>
<tr><td>Oligo rev</td><td>AAACCAGTAACGATGAGACGACCG</td><td></td><td></td><td></td><td></td><td></td><td></td><td></td><td></td><td></td><td></td></tr>
<tr><td>Chromosome</td><td>start</td><td>end</td><td>strand</td><td>MM</td><td>target_seq</td><td>PAM</td><td>alignment</td><td>distance</td><td>position</td><td>gene name</td><td>gene id</td></tr>
<tr><td>GU305914.1</td><td>7833</td><td>7855</td><td>+</td><td>0</td><td>CGGTCGTCTCATCGTTACTG</td><td>TGG</td><td>||||||||[||||||||||||]PAM</td><td>NA</td><td>Intergenic</td><td>NA</td><td>NA</td></tr>
<tr><td>T425</td><td>CTCATCGTTACTGTGGATGCAGG</td><td>1000</td><td></td><td></td><td></td><td></td><td></td><td></td><td></td><td></td><td></td></tr>
<tr><td>Oligo fwd</td><td>CACCCTCATCGTTACTGTGGATGC</td><td></td><td></td><td></td><td></td><td></td><td></td><td></td><td></td><td></td><td></td></tr>
<tr><td>Oligo rev</td><td>AAACGCATCCACAGTAACGATGAG</td><td></td><td></td><td></td><td></td><td></td><td></td><td></td><td></td><td></td><td></td></tr>
<tr><td>Chromosome</td><td>start</td><td>end</td><td>strand</td><td>MM</td><td>target_seq</td><td>PAM</td><td>alignment</td><td>distance</td><td>position</td><td>gene name</td><td>gene id</td></tr>
<tr><td>GU305914.1</td><td>7840</td><td>7862</td><td>+</td><td>0</td><td>CTCATCGTTACTGTGGATGC</td><td>AGG</td><td>||||||||[||||||||||||]PAM</td><td>NA</td><td>Intergenic</td><td>NA</td><td>NA</td></tr>
<tr><td>T426</td><td>TCGTTACTGTGGATGCAGGTCGG</td><td>1000</td><td></td><td></td><td></td><td></td><td></td><td></td><td></td><td></td><td></td></tr>
<tr><td>Oligo fwd</td><td>CACCTCGTTACTGTGGATGCAGGT</td><td></td><td></td><td></td><td></td><td></td><td></td><td></td><td></td><td></td><td></td></tr>
</table>

<table>
<tr><td>Oligo rev</td><td>AAACACCTGCATCCACAGTAACGA</td><td></td><td></td><td></td><td></td><td></td><td></td><td></td><td></td><td></td><td></td><td></td></tr>
<tr><td>Chromosome</td><td>start</td><td></td><td>end</td><td>strand</td><td>MM</td><td>target_seq</td><td>PAM</td><td>alignment</td><td>distance</td><td>position</td><td>gene name</td><td>gene id</td></tr>
<tr><td>GU305914.1</td><td></td><td>7844</td><td>7866</td><td>+</td><td>0</td><td>TCGTTACTGTGGATGCAGGT</td><td>CGG</td><td>||||||||[||||||||||||]PAM</td><td>NA</td><td>Intergenic</td><td>NA</td><td>NA</td></tr>
<tr><td>T427</td><td>TGCAGGTCGGCCATTCACTGTGG</td><td></td><td>1000</td><td></td><td></td><td></td><td></td><td></td><td></td><td></td><td></td><td></td></tr>
<tr><td>Oligo fwd</td><td>CACCTGCAGGTCGGCCATTCACTG</td><td></td><td></td><td></td><td></td><td></td><td></td><td></td><td></td><td></td><td></td><td></td></tr>
<tr><td>Oligo rev</td><td>AAACCAGTGAATGGCCGACCTGCA</td><td></td><td></td><td></td><td></td><td></td><td></td><td></td><td></td><td></td><td></td><td></td></tr>
<tr><td>Chromosome</td><td>start</td><td></td><td>end</td><td>strand</td><td>MM</td><td>target_seq</td><td>PAM</td><td>alignment</td><td>distance</td><td>position</td><td>gene name</td><td>gene id</td></tr>
<tr><td>GU305914.1</td><td></td><td>7857</td><td>7879</td><td>+</td><td>0</td><td>TGCAGGTCGGCCATTCACTG</td><td>TGG</td><td>||||||||[||||||||||||]PAM</td><td>NA</td><td>Intergenic</td><td>NA</td><td>NA</td></tr>
<tr><td>T428</td><td>GCAGGTCGGCCATTCACTGTGGG</td><td></td><td>1000</td><td></td><td></td><td></td><td></td><td></td><td></td><td></td><td></td><td></td></tr>
<tr><td>Oligo fwd</td><td>CACCGCAGGTCGGCCATTCACTGT</td><td></td><td></td><td></td><td></td><td></td><td></td><td></td><td></td><td></td><td></td><td></td></tr>
<tr><td>Oligo rev</td><td>AAACACAGTGAATGGCCGACCTGC</td><td></td><td></td><td></td><td></td><td></td><td></td><td></td><td></td><td></td><td></td><td></td></tr>
<tr><td>Chromosome</td><td>start</td><td></td><td>end</td><td>strand</td><td>MM</td><td>target_seq</td><td>PAM</td><td>alignment</td><td>distance</td><td>position</td><td>gene name</td><td>gene id</td></tr>
<tr><td>GU305914.1</td><td></td><td>7858</td><td>7880</td><td>+</td><td>0</td><td>GCAGGTCGGCCATTCACTGT</td><td>GGG</td><td>||||||||[||||||||||||]PAM</td><td>NA</td><td>Intergenic</td><td>NA</td><td>NA</td></tr>
<tr><td>T429</td><td>GCCATTCACTGTGGGCTGTAAGG</td><td></td><td>1000</td><td></td><td></td><td></td><td></td><td></td><td></td><td></td><td></td><td></td></tr>
<tr><td>Oligo fwd</td><td>CACCGCCATTCACTGTGGGCTGTA</td><td></td><td></td><td></td><td></td><td></td><td></td><td></td><td></td><td></td><td></td><td></td></tr>
<tr><td>Oligo rev</td><td>AAACTACAGCCCACAGTGAATGGC</td><td></td><td></td><td></td><td></td><td></td><td></td><td></td><td></td><td></td><td></td><td></td></tr>
<tr><td>Chromosome</td><td>start</td><td></td><td>end</td><td>strand</td><td>MM</td><td>target_seq</td><td>PAM</td><td>alignment</td><td>distance</td><td>position</td><td>gene name</td><td>gene id</td></tr>
<tr><td>GU305914.1</td><td></td><td>7866</td><td>7888</td><td>+</td><td>0</td><td>GCCATTCACTGTGGGCTGTA</td><td>AGG</td><td>||||||||[||||||||||||]PAM</td><td>NA</td><td>Intergenic</td><td>NA</td><td>NA</td></tr>
<tr><td>T430</td><td>TGTAAGAGAGAACCCGATTATGG</td><td></td><td>1000</td><td></td><td></td><td></td><td></td><td></td><td></td><td></td><td></td><td></td></tr>
<tr><td>Oligo fwd</td><td>CACCTGTAAGAGAGAACCCGATTA</td><td></td><td></td><td></td><td></td><td></td><td></td><td></td><td></td><td></td><td></td><td></td></tr>
<tr><td>Oligo rev</td><td>AAACTAATCGGGTTCTCTCTTACA</td><td></td><td></td><td></td><td></td><td></td><td></td><td></td><td></td><td></td><td></td><td></td></tr>
<tr><td>Chromosome</td><td>start</td><td></td><td>end</td><td>strand</td><td>MM</td><td>target_seq</td><td>PAM</td><td>alignment</td><td>distance</td><td>position</td><td>gene name</td><td>gene id</td></tr>
<tr><td>GU305914.1</td><td></td><td>7897</td><td>7919</td><td>+</td><td>0</td><td>TGTAAGAGAGAACCCGATTA</td><td>TGG</td><td>||||||||[||||||||||||]PAM</td><td>NA</td><td>Intergenic</td><td>NA</td><td>NA</td></tr>
<tr><td>T431</td><td>CGATTATGGTCCAGCGACTGAGG</td><td></td><td>1000</td><td></td><td></td><td></td><td></td><td></td><td></td><td></td><td></td><td></td></tr>
<tr><td>Oligo fwd</td><td>CACCCGATTATGGTCCAGCGACTG</td><td></td><td></td><td></td><td></td><td></td><td></td><td></td><td></td><td></td><td></td><td></td></tr>
<tr><td>Oligo rev</td><td>AAACCAGTCGCTGGACCATAATCG</td><td></td><td></td><td></td><td></td><td></td><td></td><td></td><td></td><td></td><td></td><td></td></tr>
<tr><td>Chromosome</td><td>start</td><td></td><td>end</td><td>strand</td><td>MM</td><td>target_seq</td><td>PAM</td><td>alignment</td><td>distance</td><td>position</td><td>gene name</td><td>gene id</td></tr>
<tr><td>GU305914.1</td><td></td><td>7911</td><td>7933</td><td>+</td><td>0</td><td>CGATTATGGTCCAGCGACTG</td><td>AGG</td><td>||||||||[||||||||||||]PAM</td><td>NA</td><td>Intergenic</td><td>NA</td><td>NA</td></tr>
<tr><td>T432</td><td>TGCCGATTCCCACCACACAACGG</td><td></td><td>1000</td><td></td><td></td><td></td><td></td><td></td><td></td><td></td><td></td><td></td></tr>
<tr><td>Oligo fwd</td><td>CACCTGCCGATTCCCACCACACAA</td><td></td><td></td><td></td><td></td><td></td><td></td><td></td><td></td><td></td><td></td><td></td></tr>
<tr><td>Oligo rev</td><td>AAACTTGTGTGGTGGGAATCGGCA</td><td></td><td></td><td></td><td></td><td></td><td></td><td></td><td></td><td></td><td></td><td></td></tr>
<tr><td>Chromosome</td><td>start</td><td></td><td>end</td><td>strand</td><td>MM</td><td>target_seq</td><td>PAM</td><td>alignment</td><td>distance</td><td>position</td><td>gene name</td><td>gene id</td></tr>
<tr><td>GU305914.1</td><td></td><td>7970</td><td>7992</td><td>+</td><td>0</td><td>TGCCGATTCCCACCACACAA</td><td>CGG</td><td>||||||||[||||||||||||]PAM</td><td>NA</td><td>Intergenic</td><td>NA</td><td>NA</td></tr>
<tr><td>T433</td><td>TACTACCAGACCCACGACCACGG</td><td></td><td>1000</td><td></td><td></td><td></td><td></td><td></td><td></td><td></td><td></td><td></td></tr>
<tr><td>Oligo fwd</td><td>CACCTACTACCAGACCCACGACCA</td><td></td><td></td><td></td><td></td><td></td><td></td><td></td><td></td><td></td><td></td><td></td></tr>
<tr><td>Oligo rev</td><td>AAACTGGTCGTGGGTCTGGTAGTA</td><td></td><td></td><td></td><td></td><td></td><td></td><td></td><td></td><td></td><td></td><td></td></tr>
<tr><td>Chromosome</td><td>start</td><td></td><td>end</td><td>strand</td><td>MM</td><td>target_seq</td><td>PAM</td><td>alignment</td><td>distance</td><td>position</td><td>gene name</td><td>gene id</td></tr>
<tr><td>GU305914.1</td><td></td><td>8031</td><td>8053</td><td>+</td><td>0</td><td>TACTACCAGACCCACGACCA</td><td>CGG</td><td>||||||||[||||||||||||]PAM</td><td>NA</td><td>Intergenic</td><td>NA</td><td>NA</td></tr>
<tr><td>GU305914.1</td><td></td><td>8091</td><td>8113</td><td>+</td><td>1</td><td>TACTACCAGACCAACGACCA</td><td>CGG</td><td>||||||||[||||-|||||||]PAM</td><td>NA</td><td>Intergenic</td><td>NA</td><td>NA</td></tr>
<tr><td>GU305914.1</td><td></td><td>8061</td><td>8083</td><td>+</td><td>2</td><td>GACTACCAGACCAACGACCA</td><td>CGG</td><td>-|||||||[||||-|||||||]PAM</td><td>NA</td><td>Intergenic</td><td>NA</td><td>NA</td></tr>
<tr><td>T434</td><td>GACTACCAGACCAACGACCACGG</td><td></td><td>1000</td><td></td><td></td><td></td><td></td><td></td><td></td><td></td><td></td><td></td></tr>
<tr><td>Oligo fwd</td><td>CACCGACTACCAGACCAACGACCA</td><td></td><td></td><td></td><td></td><td></td><td></td><td></td><td></td><td></td><td></td><td></td></tr>
<tr><td>Oligo rev</td><td>AAACTGGTCGTTGGTCTGGTAGTC</td><td></td><td></td><td></td><td></td><td></td><td></td><td></td><td></td><td></td><td></td><td></td></tr>
<tr><td>Chromosome</td><td>start</td><td></td><td>end</td><td>strand</td><td>MM</td><td>target_seq</td><td>PAM</td><td>alignment</td><td>distance</td><td>position</td><td>gene name</td><td>gene id</td></tr>
<tr><td>GU305914.1</td><td></td><td>8061</td><td>8083</td><td>+</td><td>0</td><td>GACTACCAGACCAACGACCA</td><td>CGG</td><td>||||||||[||||||||||||]PAM</td><td>NA</td><td>Intergenic</td><td>NA</td><td>NA</td></tr>
<tr><td>GU305914.1</td><td></td><td>8091</td><td>8113</td><td>+</td><td>1</td><td>TACTACCAGACCAACGACCA</td><td>CGG</td><td>-|||||||[||||||||||||]PAM</td><td>NA</td><td>Intergenic</td><td>NA</td><td>NA</td></tr>
<tr><td>GU305914.1</td><td></td><td>8031</td><td>8053</td><td>+</td><td>2</td><td>TACTACCAGACCCACGACCA</td><td>CGG</td><td>-|||||||[||||-|||||||]PAM</td><td>NA</td><td>Intergenic</td><td>NA</td><td>NA</td></tr>
<tr><td>T435</td><td>TACTACCAGACCAACGACCACGG</td><td></td><td>1000</td><td></td><td></td><td></td><td></td><td></td><td></td><td></td><td></td><td></td></tr>
<tr><td>Oligo fwd</td><td>CACCTACTACCAGACCAACGACCA</td><td></td><td></td><td></td><td></td><td></td><td></td><td></td><td></td><td></td><td></td><td></td></tr>
<tr><td>Oligo rev</td><td>AAACTGGTCGTTGGTCTGGTAGTA</td><td></td><td></td><td></td><td></td><td></td><td></td><td></td><td></td><td></td><td></td><td></td></tr>
<tr><td>Chromosome</td><td>start</td><td></td><td>end</td><td>strand</td><td>MM</td><td>target_seq</td><td>PAM</td><td>alignment</td><td>distance</td><td>position</td><td>gene name</td><td>gene id</td></tr>
<tr><td>GU305914.1</td><td></td><td>8091</td><td>8113</td><td>+</td><td>0</td><td>TACTACCAGACCAACGACCA</td><td>CGG</td><td>||||||||[||||||||||||]PAM</td><td>NA</td><td>Intergenic</td><td>NA</td><td>NA</td></tr>
<tr><td>GU305914.1</td><td></td><td>8061</td><td>8083</td><td>+</td><td>1</td><td>GACTACCAGACCAACGACCA</td><td>CGG</td><td>-|||||||[||||||||||||]PAM</td><td>NA</td><td>Intergenic</td><td>NA</td><td>NA</td></tr>
<tr><td>GU305914.1</td><td></td><td>8031</td><td>8053</td><td>+</td><td>1</td><td>TACTACCAGACCCACGACCA</td><td>CGG</td><td>||||||||[||||-|||||||]PAM</td><td>NA</td><td>Intergenic</td><td>NA</td><td>NA</td></tr>
<tr><td>T436</td><td>TACCCAGAAACCAGTCCCACCGG</td><td></td><td>1000</td><td></td><td></td><td></td><td></td><td></td><td></td><td></td><td></td><td></td></tr>
<tr><td>Oligo fwd</td><td>CACCTACCCAGAAACCAGTCCCAC</td><td></td><td></td><td></td><td></td><td></td><td></td><td></td><td></td><td></td><td></td><td></td></tr>
<tr><td>Oligo rev</td><td>AAACGTGGGACTGGTTTCTGGGTA</td><td></td><td></td><td></td><td></td><td></td><td></td><td></td><td></td><td></td><td></td><td></td></tr>
<tr><td>Chromosome</td><td>start</td><td></td><td>end</td><td>strand</td><td>MM</td><td>target_seq</td><td>PAM</td><td>alignment</td><td>distance</td><td>position</td><td>gene name</td><td>gene id</td></tr>
<tr><td>GU305914.1</td><td></td><td>8145</td><td>8167</td><td>+</td><td>0</td><td>TACCCAGAAACCAGTCCCAC</td><td>CGG</td><td>||||||||[||||||||||||]PAM</td><td>NA</td><td>Intergenic</td><td>NA</td><td>NA</td></tr>
<tr><td>T437</td><td>AAAACATACCTCGATCACCCAGG</td><td></td><td>1000</td><td></td><td></td><td></td><td></td><td></td><td></td><td></td><td></td><td></td></tr>
<tr><td>Oligo fwd</td><td>CACCAAAACATACCTCGATCACCC</td><td></td><td></td><td></td><td></td><td></td><td></td><td></td><td></td><td></td><td></td><td></td></tr>
<tr><td>Oligo rev</td><td>AAACGGGTGATCGAGGTATGTTTT</td><td></td><td></td><td></td><td></td><td></td><td></td><td></td><td></td><td></td><td></td><td></td></tr>
<tr><td>Chromosome</td><td>start</td><td></td><td>end</td><td>strand</td><td>MM</td><td>target_seq</td><td>PAM</td><td>alignment</td><td>distance</td><td>position</td><td>gene name</td><td>gene id</td></tr>
<tr><td>GU305914.1</td><td></td><td>8190</td><td>8212</td><td>+</td><td>0</td><td>AAAACATACCTCGATCACCC</td><td>AGG</td><td>||||||||[||||||||||||]PAM</td><td>NA</td><td>Intergenic</td><td>NA</td><td>NA</td></tr>
<tr><td>T438</td><td>AGCCCCACAACTCTTTTTCCCGG</td><td></td><td>1000</td><td></td><td></td><td></td><td></td><td></td><td></td><td></td><td></td><td></td></tr>
<tr><td>Oligo fwd</td><td>CACCAGCCCCACAACTCTTTTTCC</td><td></td><td></td><td></td><td></td><td></td><td></td><td></td><td></td><td></td><td></td><td></td></tr>
<tr><td>Oligo rev</td><td>AAACGGAAAAAGAGTTGTGGGGCT</td><td></td><td></td><td></td><td></td><td></td><td></td><td></td><td></td><td></td><td></td><td></td></tr>
<tr><td>Chromosome</td><td>start</td><td></td><td>end</td><td>strand</td><td>MM</td><td>target_seq</td><td>PAM</td><td>alignment</td><td>distance</td><td>position</td><td>gene name</td><td>gene id</td></tr>
<tr><td>GU305914.1</td><td></td><td>8236</td><td>8258</td><td>+</td><td>0</td><td>AGCCCCACAACTCTTTTTCC</td><td>CGG</td><td>||||||||[||||||||||||]PAM</td><td>NA</td><td>Intergenic</td><td>NA</td><td>NA</td></tr>
<tr><td>T439</td><td>AGAGATGCAGACTTCGCATACGG</td><td></td><td>1000</td><td></td><td></td><td></td><td></td><td></td><td></td><td></td><td></td><td></td></tr>
<tr><td>Oligo fwd</td><td>CACCAGAGATGCAGACTTCGCATA</td><td></td><td></td><td></td><td></td><td></td><td></td><td></td><td></td><td></td><td></td><td></td></tr>
<tr><td>Oligo rev</td><td>AAACTATGCGAAGTCTGCATCTCT</td><td></td><td></td><td></td><td></td><td></td><td></td><td></td><td></td><td></td><td></td><td></td></tr>
<tr><td>Chromosome</td><td>start</td><td></td><td>end</td><td>strand</td><td>MM</td><td>target_seq</td><td>PAM</td><td>alignment</td><td>distance</td><td>position</td><td>gene name</td><td>gene id</td></tr>
<tr><td>GU305914.1</td><td></td><td>8295</td><td>8317</td><td>+</td><td>0</td><td>AGAGATGCAGACTTCGCATA</td><td>CGG</td><td>||||||||[||||||||||||]PAM</td><td>NA</td><td>Intergenic</td><td>NA</td><td>NA</td></tr>
<tr><td>T440</td><td>TACGGTTACGATCACCGTGTCGG</td><td></td><td>1000</td><td></td><td></td><td></td><td></td><td></td><td></td><td></td><td></td><td></td></tr>
</table>

<table>
<tr><td>Oligo fwd</td><td>CACCTACGGTTACGATCACCGTGT</td><td></td><td></td><td></td><td></td><td></td><td></td><td></td><td></td><td></td><td></td><td></td></tr>
<tr><td>Oligo rev</td><td>AAACACACGGTGATCGTAACCGTA</td><td></td><td></td><td></td><td></td><td></td><td></td><td></td><td></td><td></td><td></td><td></td></tr>
<tr><td>Chromosome</td><td>start</td><td></td><td>end</td><td>strand</td><td>MM</td><td>target_seq</td><td>PAM</td><td>alignment</td><td>distance</td><td>position</td><td>gene name</td><td>gene id</td></tr>
<tr><td>GU305914.1</td><td></td><td>8313</td><td>8335</td><td>+</td><td>0</td><td>TACGGTTACGATCACCGTGT</td><td>CGG</td><td>||||||||[||||||||||||]PAM</td><td>NA</td><td>Intergenic</td><td>NA</td><td>NA</td></tr>
<tr><td></td><td></td><td></td><td></td><td></td><td></td><td></td><td></td><td></td><td></td><td></td><td></td><td></td></tr>
<tr><td>T441</td><td>TACGATCACCGTGTCGGTAGCGG</td><td></td><td>1000</td><td></td><td></td><td></td><td></td><td></td><td></td><td></td><td></td><td></td></tr>
<tr><td>Oligo fwd</td><td>CACCTACGATCACCGTGTCGGTAG</td><td></td><td></td><td></td><td></td><td></td><td></td><td></td><td></td><td></td><td></td><td></td></tr>
<tr><td>Oligo rev</td><td>AAACCTACCGACACGGTGATCGTA</td><td></td><td></td><td></td><td></td><td></td><td></td><td></td><td></td><td></td><td></td><td></td></tr>
<tr><td>Chromosome</td><td>start</td><td></td><td>end</td><td>strand</td><td>MM</td><td>target_seq</td><td>PAM</td><td>alignment</td><td>distance</td><td>position</td><td>gene name</td><td>gene id</td></tr>
<tr><td>GU305914.1</td><td></td><td>8319</td><td>8341</td><td>+</td><td>0</td><td>TACGATCACCGTGTCGGTAG</td><td>CGG</td><td>||||||||[||||||||||||]PAM</td><td>NA</td><td>Intergenic</td><td>NA</td><td>NA</td></tr>
<tr><td>GU305914.1</td><td></td><td>9312</td><td>9334</td><td>+</td><td>4</td><td>CGCGATCGCCGTGTCGCTAG</td><td>TGG</td><td>--|||||-[||||||||-|||]PAM</td><td>NA</td><td>Intergenic</td><td>NA</td><td>NA</td></tr>
<tr><td></td><td></td><td></td><td></td><td></td><td></td><td></td><td></td><td></td><td></td><td></td><td></td><td></td></tr>
<tr><td>T442</td><td>TTAGCTGTGTTGTATCACTACGG</td><td></td><td>1000</td><td></td><td></td><td></td><td></td><td></td><td></td><td></td><td></td><td></td></tr>
<tr><td>Oligo fwd</td><td>CACCTTAGCTGTGTTGTATCACTA</td><td></td><td></td><td></td><td></td><td></td><td></td><td></td><td></td><td></td><td></td><td></td></tr>
<tr><td>Oligo rev</td><td>AAACTAGTGATACAACACAGCTAA</td><td></td><td></td><td></td><td></td><td></td><td></td><td></td><td></td><td></td><td></td><td></td></tr>
<tr><td>Chromosome</td><td>start</td><td></td><td>end</td><td>strand</td><td>MM</td><td>target_seq</td><td>PAM</td><td>alignment</td><td>distance</td><td>position</td><td>gene name</td><td>gene id</td></tr>
<tr><td>GU305914.1</td><td></td><td>8371</td><td>8393</td><td>+</td><td>0</td><td>TTAGCTGTGTTGTATCACTA</td><td>CGG</td><td>||||||||[||||||||||||]PAM</td><td>NA</td><td>Intergenic</td><td>NA</td><td>NA</td></tr>
<tr><td></td><td></td><td></td><td></td><td></td><td></td><td></td><td></td><td></td><td></td><td></td><td></td><td></td></tr>
<tr><td>T443</td><td>GTTGTATCACTACGGCTTTCTGG</td><td></td><td>1000</td><td></td><td></td><td></td><td></td><td></td><td></td><td></td><td></td><td></td></tr>
<tr><td>Oligo fwd</td><td>CACCGTTGTATCACTACGGCTTTC</td><td></td><td></td><td></td><td></td><td></td><td></td><td></td><td></td><td></td><td></td><td></td></tr>
<tr><td>Oligo rev</td><td>AAACGAAAGCCGTAGTGATACAAC</td><td></td><td></td><td></td><td></td><td></td><td></td><td></td><td></td><td></td><td></td><td></td></tr>
<tr><td>Chromosome</td><td>start</td><td></td><td>end</td><td>strand</td><td>MM</td><td>target_seq</td><td>PAM</td><td>alignment</td><td>distance</td><td>position</td><td>gene name</td><td>gene id</td></tr>
<tr><td>GU305914.1</td><td></td><td>8379</td><td>8401</td><td>+</td><td>0</td><td>GTTGTATCACTACGGCTTTC</td><td>TGG</td><td>||||||||[||||||||||||]PAM</td><td>NA</td><td>Intergenic</td><td>NA</td><td>NA</td></tr>
<tr><td>GU305914.1</td><td></td><td>9369</td><td>9391</td><td>+</td><td>2</td><td>ATTGTATCACTACGGCTTTT</td><td>TGG</td><td>-|||||||[|||||||||||-]PAM</td><td>NA</td><td>Intergenic</td><td>NA</td><td>NA</td></tr>
<tr><td></td><td></td><td></td><td></td><td></td><td></td><td></td><td></td><td></td><td></td><td></td><td></td><td></td></tr>
<tr><td>T444</td><td>TTGTATCACTACGGCTTTCTGGG</td><td></td><td>1000</td><td></td><td></td><td></td><td></td><td></td><td></td><td></td><td></td><td></td></tr>
<tr><td>Oligo fwd</td><td>CACCTTGTATCACTACGGCTTTCT</td><td></td><td></td><td></td><td></td><td></td><td></td><td></td><td></td><td></td><td></td><td></td></tr>
<tr><td>Oligo rev</td><td>AAACAGAAAGCCGTAGTGATACAA</td><td></td><td></td><td></td><td></td><td></td><td></td><td></td><td></td><td></td><td></td><td></td></tr>
<tr><td>Chromosome</td><td>start</td><td></td><td>end</td><td>strand</td><td>MM</td><td>target_seq</td><td>PAM</td><td>alignment</td><td>distance</td><td>position</td><td>gene name</td><td>gene id</td></tr>
<tr><td>GU305914.1</td><td></td><td>8380</td><td>8402</td><td>+</td><td>0</td><td>TTGTATCACTACGGCTTTCT</td><td>GGG</td><td>||||||||[||||||||||||]PAM</td><td>NA</td><td>Intergenic</td><td>NA</td><td>NA</td></tr>
<tr><td>GU305914.1</td><td></td><td>9370</td><td>9392</td><td>+</td><td>1</td><td>TTGTATCACTACGGCTTTTT</td><td>GGG</td><td>||||||||[||||||||||-|]PAM</td><td>NA</td><td>Intergenic</td><td>NA</td><td>NA</td></tr>
<tr><td></td><td></td><td></td><td></td><td></td><td></td><td></td><td></td><td></td><td></td><td></td><td></td><td></td></tr>
<tr><td>T445</td><td>TGTATCACTACGGCTTTCTGGGG</td><td></td><td>1000</td><td></td><td></td><td></td><td></td><td></td><td></td><td></td><td></td><td></td></tr>
<tr><td>Oligo fwd</td><td>CACCTGTATCACTACGGCTTTCTG</td><td></td><td></td><td></td><td></td><td></td><td></td><td></td><td></td><td></td><td></td><td></td></tr>
<tr><td>Oligo rev</td><td>AAACCAGAAAGCCGTAGTGATACA</td><td></td><td></td><td></td><td></td><td></td><td></td><td></td><td></td><td></td><td></td><td></td></tr>
<tr><td>Chromosome</td><td>start</td><td></td><td>end</td><td>strand</td><td>MM</td><td>target_seq</td><td>PAM</td><td>alignment</td><td>distance</td><td>position</td><td>gene name</td><td>gene id</td></tr>
<tr><td>GU305914.1</td><td></td><td>8381</td><td>8403</td><td>+</td><td>0</td><td>TGTATCACTACGGCTTTCTG</td><td>GGG</td><td>||||||||[||||||||||||]PAM</td><td>NA</td><td>Intergenic</td><td>NA</td><td>NA</td></tr>
<tr><td>GU305914.1</td><td></td><td>9371</td><td>9393</td><td>+</td><td>1</td><td>TGTATCACTACGGCTTTTTG</td><td>GGG</td><td>||||||||[|||||||||-||]PAM</td><td>NA</td><td>Intergenic</td><td>NA</td><td>NA</td></tr>
<tr><td></td><td></td><td></td><td></td><td></td><td></td><td></td><td></td><td></td><td></td><td></td><td></td><td></td></tr>
<tr><td>T446</td><td>CTACGGCTTTCTGGGGAGATCGG</td><td></td><td>1000</td><td></td><td></td><td></td><td></td><td></td><td></td><td></td><td></td><td></td></tr>
<tr><td>Oligo fwd</td><td>CACCCTACGGCTTTCTGGGGAGAT</td><td></td><td></td><td></td><td></td><td></td><td></td><td></td><td></td><td></td><td></td><td></td></tr>
<tr><td>Oligo rev</td><td>AAACATCTCCCCAGAAAGCCGTAG</td><td></td><td></td><td></td><td></td><td></td><td></td><td></td><td></td><td></td><td></td><td></td></tr>
<tr><td>Chromosome</td><td>start</td><td></td><td>end</td><td>strand</td><td>MM</td><td>target_seq</td><td>PAM</td><td>alignment</td><td>distance</td><td>position</td><td>gene name</td><td>gene id</td></tr>
<tr><td>GU305914.1</td><td></td><td>8388</td><td>8410</td><td>+</td><td>0</td><td>CTACGGCTTTCTGGGGAGAT</td><td>CGG</td><td>||||||||[||||||||||||]PAM</td><td>NA</td><td>Intergenic</td><td>NA</td><td>NA</td></tr>
<tr><td>GU305914.1</td><td></td><td>9378</td><td>9400</td><td>+</td><td>2</td><td>CTACGGCTTTTTGGGGAAAT</td><td>GGG</td><td>||||||||[||-||||||-||]PAM</td><td>NA</td><td>Intergenic</td><td>NA</td><td>NA</td></tr>
<tr><td></td><td></td><td></td><td></td><td></td><td></td><td></td><td></td><td></td><td></td><td></td><td></td><td></td></tr>
<tr><td>T447</td><td>TTAATCTGCTTACGTACACCTGG</td><td></td><td>1000</td><td></td><td></td><td></td><td></td><td></td><td></td><td></td><td></td><td></td></tr>
<tr><td>Oligo fwd</td><td>CACCTTAATCTGCTTACGTACACC</td><td></td><td></td><td></td><td></td><td></td><td></td><td></td><td></td><td></td><td></td><td></td></tr>
<tr><td>Oligo rev</td><td>AAACGGTGTACGTAAGCAGATTAA</td><td></td><td></td><td></td><td></td><td></td><td></td><td></td><td></td><td></td><td></td><td></td></tr>
<tr><td>Chromosome</td><td>start</td><td></td><td>end</td><td>strand</td><td>MM</td><td>target_seq</td><td>PAM</td><td>alignment</td><td>distance</td><td>position</td><td>gene name</td><td>gene id</td></tr>
<tr><td>GU305914.1</td><td></td><td>8526</td><td>8548</td><td>+</td><td>0</td><td>TTAATCTGCTTACGTACACC</td><td>TGG</td><td>||||||||[||||||||||||]PAM</td><td>NA</td><td>Intergenic</td><td>NA</td><td>NA</td></tr>
<tr><td></td><td></td><td></td><td></td><td></td><td></td><td></td><td></td><td></td><td></td><td></td><td></td><td></td></tr>
<tr><td>T448</td><td>TGTCCTGAGTATGTAAATACCGG</td><td></td><td>1000</td><td></td><td></td><td></td><td></td><td></td><td></td><td></td><td></td><td></td></tr>
<tr><td>Oligo fwd</td><td>CACCTGTCCTGAGTATGTAAATAC</td><td></td><td></td><td></td><td></td><td></td><td></td><td></td><td></td><td></td><td></td><td></td></tr>
<tr><td>Oligo rev</td><td>AAACGTATTTACATACTCAGGACA</td><td></td><td></td><td></td><td></td><td></td><td></td><td></td><td></td><td></td><td></td><td></td></tr>
<tr><td>Chromosome</td><td>start</td><td></td><td>end</td><td>strand</td><td>MM</td><td>target_seq</td><td>PAM</td><td>alignment</td><td>distance</td><td>position</td><td>gene name</td><td>gene id</td></tr>
<tr><td>GU305914.1</td><td></td><td>8562</td><td>8584</td><td>+</td><td>0</td><td>TGTCCTGAGTATGTAAATAC</td><td>CGG</td><td>||||||||[||||||||||||]PAM</td><td>NA</td><td>Intergenic</td><td>NA</td><td>NA</td></tr>
<tr><td></td><td></td><td></td><td></td><td></td><td></td><td></td><td></td><td></td><td></td><td></td><td></td><td></td></tr>
<tr><td>T449</td><td>TACCGGCGCACCTGTTGCGCCGG</td><td></td><td>1000</td><td></td><td></td><td></td><td></td><td></td><td></td><td></td><td></td><td></td></tr>
<tr><td>Oligo fwd</td><td>CACCTACCGGCGCACCTGTTGCGC</td><td></td><td></td><td></td><td></td><td></td><td></td><td></td><td></td><td></td><td></td><td></td></tr>
<tr><td>Oligo rev</td><td>AAACGCGCAACAGGTGCGCCGGTA</td><td></td><td></td><td></td><td></td><td></td><td></td><td></td><td></td><td></td><td></td><td></td></tr>
<tr><td>Chromosome</td><td>start</td><td></td><td>end</td><td>strand</td><td>MM</td><td>target_seq</td><td>PAM</td><td>alignment</td><td>distance</td><td>position</td><td>gene name</td><td>gene id</td></tr>
<tr><td>GU305914.1</td><td></td><td>8579</td><td>8601</td><td>+</td><td>0</td><td>TACCGGCGCACCTGTTGCGC</td><td>CGG</td><td>||||||||[||||||||||||]PAM</td><td>NA</td><td>Intergenic</td><td>NA</td><td>NA</td></tr>
<tr><td></td><td></td><td></td><td></td><td></td><td></td><td></td><td></td><td></td><td></td><td></td><td></td><td></td></tr>
<tr><td>T450</td><td>GAAATCGCTGATCACTTCTGCGG</td><td></td><td>1000</td><td></td><td></td><td></td><td></td><td></td><td></td><td></td><td></td><td></td></tr>
<tr><td>Oligo fwd</td><td>CACCGAAATCGCTGATCACTTCTG</td><td></td><td></td><td></td><td></td><td></td><td></td><td></td><td></td><td></td><td></td><td></td></tr>
<tr><td>Oligo rev</td><td>AAACCAGAAGTGATCAGCGATTTC</td><td></td><td></td><td></td><td></td><td></td><td></td><td></td><td></td><td></td><td></td><td></td></tr>
<tr><td>Chromosome</td><td>start</td><td></td><td>end</td><td>strand</td><td>MM</td><td>target_seq</td><td>PAM</td><td>alignment</td><td>distance</td><td>position</td><td>gene name</td><td>gene id</td></tr>
<tr><td>GU305914.1</td><td></td><td>8694</td><td>8716</td><td>+</td><td>0</td><td>GAAATCGCTGATCACTTCTG</td><td>CGG</td><td>||||||||[||||||||||||]PAM</td><td>NA</td><td>Intergenic</td><td>NA</td><td>NA</td></tr>
<tr><td></td><td></td><td></td><td></td><td></td><td></td><td></td><td></td><td></td><td></td><td></td><td></td><td></td></tr>
<tr><td>T451</td><td>CGGCGACGTGCAATTCAGTTCGG</td><td></td><td>1000</td><td></td><td></td><td></td><td></td><td></td><td></td><td></td><td></td><td></td></tr>
<tr><td>Oligo fwd</td><td>CACCCGGCGACGTGCAATTCAGTT</td><td></td><td></td><td></td><td></td><td></td><td></td><td></td><td></td><td></td><td></td><td></td></tr>
<tr><td>Oligo rev</td><td>AAACAACTGAATTGCACGTCGCCG</td><td></td><td></td><td></td><td></td><td></td><td></td><td></td><td></td><td></td><td></td><td></td></tr>
<tr><td>Chromosome</td><td>start</td><td></td><td>end</td><td>strand</td><td>MM</td><td>target_seq</td><td>PAM</td><td>alignment</td><td>distance</td><td>position</td><td>gene name</td><td>gene id</td></tr>
<tr><td>GU305914.1</td><td></td><td>8714</td><td>8736</td><td>+</td><td>0</td><td>CGGCGACGTGCAATTCAGTT</td><td>CGG</td><td>||||||||[||||||||||||]PAM</td><td>NA</td><td>Intergenic</td><td>NA</td><td>NA</td></tr>
<tr><td></td><td></td><td></td><td></td><td></td><td></td><td></td><td></td><td></td><td></td><td></td><td></td><td></td></tr>
<tr><td>T452</td><td>GGCGACGTGCAATTCAGTTCGGG</td><td></td><td>1000</td><td></td><td></td><td></td><td></td><td></td><td></td><td></td><td></td><td></td></tr>
<tr><td>Oligo fwd</td><td>CACCGGCGACGTGCAATTCAGTTC</td><td></td><td></td><td></td><td></td><td></td><td></td><td></td><td></td><td></td><td></td><td></td></tr>
<tr><td>Oligo rev</td><td>AAACGAACTGAATTGCACGTCGCC</td><td></td><td></td><td></td><td></td><td></td><td></td><td></td><td></td><td></td><td></td><td></td></tr>
<tr><td>Chromosome</td><td>start</td><td></td><td>end</td><td>strand</td><td>MM</td><td>target_seq</td><td>PAM</td><td>alignment</td><td>distance</td><td>position</td><td>gene name</td><td>gene id</td></tr>
<tr><td>GU305914.1</td><td></td><td>8715</td><td>8737</td><td>+</td><td>0</td><td>GGCGACGTGCAATTCAGTTC</td><td>GGG</td><td>||||||||[||||||||||||]PAM</td><td>NA</td><td>Intergenic</td><td>NA</td><td>NA</td></tr>
<tr><td></td><td></td><td></td><td></td><td></td><td></td><td></td><td></td><td></td><td></td><td></td><td></td><td></td></tr>
<tr><td>T453</td><td>GTGCAATTCAGTTCGGGATCTGG</td><td></td><td>1000</td><td></td><td></td><td></td><td></td><td></td><td></td><td></td><td></td><td></td></tr>
<tr><td>Oligo fwd</td><td>CACCGTGCAATTCAGTTCGGGATC</td><td></td><td></td><td></td><td></td><td></td><td></td><td></td><td></td><td></td><td></td><td></td></tr>
<tr><td>Oligo rev</td><td>AAACGATCCCGAACTGAATTGCAC</td><td></td><td></td><td></td><td></td><td></td><td></td><td></td><td></td><td></td><td></td><td></td></tr>
<tr><td>Chromosome</td><td>start</td><td></td><td>end</td><td>strand</td><td>MM</td><td>target_seq</td><td>PAM</td><td>alignment</td><td>distance</td><td>position</td><td>gene name</td><td>gene id</td></tr>
<tr><td>GU305914.1</td><td></td><td>8721</td><td>8743</td><td>+</td><td>0</td><td>GTGCAATTCAGTTCGGGATC</td><td>TGG</td><td>||||||||[||||||||||||]PAM</td><td>NA</td><td>Intergenic</td><td>NA</td><td>NA</td></tr>
<tr><td></td><td></td><td></td><td></td><td></td><td></td><td></td><td></td><td></td><td></td><td></td><td></td><td></td></tr>
<tr><td>T454</td><td>GTTCGGGATCTGGCTTTTTCTGG</td><td></td><td>1000</td><td></td><td></td><td></td><td></td><td></td><td></td><td></td><td></td><td></td></tr>
</table>

<table>
<tr><td>Oligo fwd</td><td>CACCGTTCGGGATCTGGCTTTTTC</td><td></td><td></td><td></td><td></td><td></td><td></td><td></td><td></td><td></td><td></td><td></td></tr>
<tr><td>Oligo rev</td><td>AAACGAAAAAGCCAGATCCCGAAC</td><td></td><td></td><td></td><td></td><td></td><td></td><td></td><td></td><td></td><td></td><td></td></tr>
<tr><td>Chromosome</td><td>start</td><td></td><td>end</td><td>strand</td><td>MM</td><td>target_seq</td><td>PAM</td><td>alignment</td><td>distance</td><td>position</td><td>gene name</td><td>gene id</td></tr>
<tr><td>GU305914.1</td><td></td><td>8731</td><td>8753</td><td>+</td><td>0</td><td>GTTCGGGATCTGGCTTTTTC</td><td>TGG</td><td>||||||||[||||||||||||]PAM</td><td>NA</td><td>Intergenic</td><td>NA</td><td>NA</td></tr>
<tr><td></td><td></td><td></td><td></td><td></td><td></td><td></td><td></td><td></td><td></td><td></td><td></td><td></td></tr>
<tr><td>T455</td><td>TTTTCTGGCACCGTCACACCAGG</td><td></td><td>1000</td><td></td><td></td><td></td><td></td><td></td><td></td><td></td><td></td><td></td></tr>
<tr><td>Oligo fwd</td><td>CACCTTTTCTGGCACCGTCACACC</td><td></td><td></td><td></td><td></td><td></td><td></td><td></td><td></td><td></td><td></td><td></td></tr>
<tr><td>Oligo rev</td><td>AAACGGTGTGACGGTGCCAGAAAA</td><td></td><td></td><td></td><td></td><td></td><td></td><td></td><td></td><td></td><td></td><td></td></tr>
<tr><td>Chromosome</td><td>start</td><td></td><td>end</td><td>strand</td><td>MM</td><td>target_seq</td><td>PAM</td><td>alignment</td><td>distance</td><td>position</td><td>gene name</td><td>gene id</td></tr>
<tr><td>GU305914.1</td><td></td><td>8746</td><td>8768</td><td>+</td><td>0</td><td>TTTTCTGGCACCGTCACACC</td><td>AGG</td><td>||||||||[||||||||||||]PAM</td><td>NA</td><td>Intergenic</td><td>NA</td><td>NA</td></tr>
<tr><td></td><td></td><td></td><td></td><td></td><td></td><td></td><td></td><td></td><td></td><td></td><td></td><td></td></tr>
<tr><td>T456</td><td>TTTCTGGCACCGTCACACCAGGG</td><td></td><td>1000</td><td></td><td></td><td></td><td></td><td></td><td></td><td></td><td></td><td></td></tr>
<tr><td>Oligo fwd</td><td>CACCTTTCTGGCACCGTCACACCA</td><td></td><td></td><td></td><td></td><td></td><td></td><td></td><td></td><td></td><td></td><td></td></tr>
<tr><td>Oligo rev</td><td>AAACTGGTGTGACGGTGCCAGAAA</td><td></td><td></td><td></td><td></td><td></td><td></td><td></td><td></td><td></td><td></td><td></td></tr>
<tr><td>Chromosome</td><td>start</td><td></td><td>end</td><td>strand</td><td>MM</td><td>target_seq</td><td>PAM</td><td>alignment</td><td>distance</td><td>position</td><td>gene name</td><td>gene id</td></tr>
<tr><td>GU305914.1</td><td></td><td>8747</td><td>8769</td><td>+</td><td>0</td><td>TTTCTGGCACCGTCACACCA</td><td>GGG</td><td>||||||||[||||||||||||]PAM</td><td>NA</td><td>Intergenic</td><td>NA</td><td>NA</td></tr>
<tr><td></td><td></td><td></td><td></td><td></td><td></td><td></td><td></td><td></td><td></td><td></td><td></td><td></td></tr>
<tr><td>T457</td><td>GTCACTCTCAAAGATATGTATGG</td><td></td><td>1000</td><td></td><td></td><td></td><td></td><td></td><td></td><td></td><td></td><td></td></tr>
<tr><td>Oligo fwd</td><td>CACCGTCACTCTCAAAGATATGTA</td><td></td><td></td><td></td><td></td><td></td><td></td><td></td><td></td><td></td><td></td><td></td></tr>
<tr><td>Oligo rev</td><td>AAACTACATATCTTTGAGAGTGAC</td><td></td><td></td><td></td><td></td><td></td><td></td><td></td><td></td><td></td><td></td><td></td></tr>
<tr><td>Chromosome</td><td>start</td><td></td><td>end</td><td>strand</td><td>MM</td><td>target_seq</td><td>PAM</td><td>alignment</td><td>distance</td><td>position</td><td>gene name</td><td>gene id</td></tr>
<tr><td>GU305914.1</td><td></td><td>8788</td><td>8810</td><td>+</td><td>0</td><td>GTCACTCTCAAAGATATGTA</td><td>TGG</td><td>||||||||[||||||||||||]PAM</td><td>NA</td><td>Intergenic</td><td>NA</td><td>NA</td></tr>
<tr><td></td><td></td><td></td><td></td><td></td><td></td><td></td><td></td><td></td><td></td><td></td><td></td><td></td></tr>
<tr><td>T458</td><td>CTCAAAGATATGTATGGATTTGG</td><td></td><td>1000</td><td></td><td></td><td></td><td></td><td></td><td></td><td></td><td></td><td></td></tr>
<tr><td>Oligo fwd</td><td>CACCCTCAAAGATATGTATGGATT</td><td></td><td></td><td></td><td></td><td></td><td></td><td></td><td></td><td></td><td></td><td></td></tr>
<tr><td>Oligo rev</td><td>AAACAATCCATACATATCTTTGAG</td><td></td><td></td><td></td><td></td><td></td><td></td><td></td><td></td><td></td><td></td><td></td></tr>
<tr><td>Chromosome</td><td>start</td><td></td><td>end</td><td>strand</td><td>MM</td><td>target_seq</td><td>PAM</td><td>alignment</td><td>distance</td><td>position</td><td>gene name</td><td>gene id</td></tr>
<tr><td>GU305914.1</td><td></td><td>8794</td><td>8816</td><td>+</td><td>0</td><td>CTCAAAGATATGTATGGATT</td><td>TGG</td><td>||||||||[||||||||||||]PAM</td><td>NA</td><td>Intergenic</td><td>NA</td><td>NA</td></tr>
<tr><td></td><td></td><td></td><td></td><td></td><td></td><td></td><td></td><td></td><td></td><td></td><td></td><td></td></tr>
<tr><td>T459</td><td>TCAAAGATATGTATGGATTTGGG</td><td></td><td>1000</td><td></td><td></td><td></td><td></td><td></td><td></td><td></td><td></td><td></td></tr>
<tr><td>Oligo fwd</td><td>CACCTCAAAGATATGTATGGATTT</td><td></td><td></td><td></td><td></td><td></td><td></td><td></td><td></td><td></td><td></td><td></td></tr>
<tr><td>Oligo rev</td><td>AAACAAATCCATACATATCTTTGA</td><td></td><td></td><td></td><td></td><td></td><td></td><td></td><td></td><td></td><td></td><td></td></tr>
<tr><td>Chromosome</td><td>start</td><td></td><td>end</td><td>strand</td><td>MM</td><td>target_seq</td><td>PAM</td><td>alignment</td><td>distance</td><td>position</td><td>gene name</td><td>gene id</td></tr>
<tr><td>GU305914.1</td><td></td><td>8795</td><td>8817</td><td>+</td><td>0</td><td>TCAAAGATATGTATGGATTT</td><td>GGG</td><td>||||||||[||||||||||||]PAM</td><td>NA</td><td>Intergenic</td><td>NA</td><td>NA</td></tr>
<tr><td></td><td></td><td></td><td></td><td></td><td></td><td></td><td></td><td></td><td></td><td></td><td></td><td></td></tr>
<tr><td>T460</td><td>GATTTGGGACTAGTTCCGTTTGG</td><td></td><td>1000</td><td></td><td></td><td></td><td></td><td></td><td></td><td></td><td></td><td></td></tr>
<tr><td>Oligo fwd</td><td>CACCGATTTGGGACTAGTTCCGTT</td><td></td><td></td><td></td><td></td><td></td><td></td><td></td><td></td><td></td><td></td><td></td></tr>
<tr><td>Oligo rev</td><td>AAACAACGGAACTAGTCCCAAATC</td><td></td><td></td><td></td><td></td><td></td><td></td><td></td><td></td><td></td><td></td><td></td></tr>
<tr><td>Chromosome</td><td>start</td><td></td><td>end</td><td>strand</td><td>MM</td><td>target_seq</td><td>PAM</td><td>alignment</td><td>distance</td><td>position</td><td>gene name</td><td>gene id</td></tr>
<tr><td>GU305914.1</td><td></td><td>8810</td><td>8832</td><td>+</td><td>0</td><td>GATTTGGGACTAGTTCCGTT</td><td>TGG</td><td>||||||||[||||||||||||]PAM</td><td>NA</td><td>Intergenic</td><td>NA</td><td>NA</td></tr>
<tr><td></td><td></td><td></td><td></td><td></td><td></td><td></td><td></td><td></td><td></td><td></td><td></td><td></td></tr>
<tr><td>T461</td><td>ATTTGGGACTAGTTCCGTTTGGG</td><td></td><td>1000</td><td></td><td></td><td></td><td></td><td></td><td></td><td></td><td></td><td></td></tr>
<tr><td>Oligo fwd</td><td>CACCATTTGGGACTAGTTCCGTTT</td><td></td><td></td><td></td><td></td><td></td><td></td><td></td><td></td><td></td><td></td><td></td></tr>
<tr><td>Oligo rev</td><td>AAACAAACGGAACTAGTCCCAAAT</td><td></td><td></td><td></td><td></td><td></td><td></td><td></td><td></td><td></td><td></td><td></td></tr>
<tr><td>Chromosome</td><td>start</td><td></td><td>end</td><td>strand</td><td>MM</td><td>target_seq</td><td>PAM</td><td>alignment</td><td>distance</td><td>position</td><td>gene name</td><td>gene id</td></tr>
<tr><td>GU305914.1</td><td></td><td>8811</td><td>8833</td><td>+</td><td>0</td><td>ATTTGGGACTAGTTCCGTTT</td><td>GGG</td><td>||||||||[||||||||||||]PAM</td><td>NA</td><td>Intergenic</td><td>NA</td><td>NA</td></tr>
<tr><td></td><td></td><td></td><td></td><td></td><td></td><td></td><td></td><td></td><td></td><td></td><td></td><td></td></tr>
<tr><td>T462</td><td>CATTTTGTATTAATGCAACATGG</td><td></td><td>1000</td><td></td><td></td><td></td><td></td><td></td><td></td><td></td><td></td><td></td></tr>
<tr><td>Oligo fwd</td><td>CACCCATTTTGTATTAATGCAACA</td><td></td><td></td><td></td><td></td><td></td><td></td><td></td><td></td><td></td><td></td><td></td></tr>
<tr><td>Oligo rev</td><td>AAACTGTTGCATTAATACAAAATG</td><td></td><td></td><td></td><td></td><td></td><td></td><td></td><td></td><td></td><td></td><td></td></tr>
<tr><td>Chromosome</td><td>start</td><td></td><td>end</td><td>strand</td><td>MM</td><td>target_seq</td><td>PAM</td><td>alignment</td><td>distance</td><td>position</td><td>gene name</td><td>gene id</td></tr>
<tr><td>GU305914.1</td><td></td><td>8855</td><td>8877</td><td>+</td><td>0</td><td>CATTTTGTATTAATGCAACA</td><td>TGG</td><td>||||||||[||||||||||||]PAM</td><td>NA</td><td>Intergenic</td><td>NA</td><td>NA</td></tr>
<tr><td></td><td></td><td></td><td></td><td></td><td></td><td></td><td></td><td></td><td></td><td></td><td></td><td></td></tr>
<tr><td>T463</td><td>TTTGTATTAATGCAACATGGAGG</td><td></td><td>1000</td><td></td><td></td><td></td><td></td><td></td><td></td><td></td><td></td><td></td></tr>
<tr><td>Oligo fwd</td><td>CACCTTTGTATTAATGCAACATGG</td><td></td><td></td><td></td><td></td><td></td><td></td><td></td><td></td><td></td><td></td><td></td></tr>
<tr><td>Oligo rev</td><td>AAACCCATGTTGCATTAATACAAA</td><td></td><td></td><td></td><td></td><td></td><td></td><td></td><td></td><td></td><td></td><td></td></tr>
<tr><td>Chromosome</td><td>start</td><td></td><td>end</td><td>strand</td><td>MM</td><td>target_seq</td><td>PAM</td><td>alignment</td><td>distance</td><td>position</td><td>gene name</td><td>gene id</td></tr>
<tr><td>GU305914.1</td><td></td><td>8858</td><td>8880</td><td>+</td><td>0</td><td>TTTGTATTAATGCAACATGG</td><td>AGG</td><td>||||||||[||||||||||||]PAM</td><td>NA</td><td>Intergenic</td><td>NA</td><td>NA</td></tr>
<tr><td></td><td></td><td></td><td></td><td></td><td></td><td></td><td></td><td></td><td></td><td></td><td></td><td></td></tr>
<tr><td>T464</td><td>ATGCAACATGGAGGTTAGAATGG</td><td></td><td>1000</td><td></td><td></td><td></td><td></td><td></td><td></td><td></td><td></td><td></td></tr>
<tr><td>Oligo fwd</td><td>CACCATGCAACATGGAGGTTAGAA</td><td></td><td></td><td></td><td></td><td></td><td></td><td></td><td></td><td></td><td></td><td></td></tr>
<tr><td>Oligo rev</td><td>AAACTTCTAACCTCCATGTTGCAT</td><td></td><td></td><td></td><td></td><td></td><td></td><td></td><td></td><td></td><td></td><td></td></tr>
<tr><td>Chromosome</td><td>start</td><td></td><td>end</td><td>strand</td><td>MM</td><td>target_seq</td><td>PAM</td><td>alignment</td><td>distance</td><td>position</td><td>gene name</td><td>gene id</td></tr>
<tr><td>GU305914.1</td><td></td><td>8867</td><td>8889</td><td>+</td><td>0</td><td>ATGCAACATGGAGGTTAGAA</td><td>TGG</td><td>||||||||[||||||||||||]PAM</td><td>NA</td><td>Intergenic</td><td>NA</td><td>NA</td></tr>
<tr><td></td><td></td><td></td><td></td><td></td><td></td><td></td><td></td><td></td><td></td><td></td><td></td><td></td></tr>
<tr><td>T465</td><td>GACTACCAGACCACCTCCGACGG</td><td></td><td>1000</td><td></td><td></td><td></td><td></td><td></td><td></td><td></td><td></td><td></td></tr>
<tr><td>Oligo fwd</td><td>CACCGACTACCAGACCACCTCCGA</td><td></td><td></td><td></td><td></td><td></td><td></td><td></td><td></td><td></td><td></td><td></td></tr>
<tr><td>Oligo rev</td><td>AAACTCGGAGGTGGTCTGGTAGTC</td><td></td><td></td><td></td><td></td><td></td><td></td><td></td><td></td><td></td><td></td><td></td></tr>
<tr><td>Chromosome</td><td>start</td><td></td><td>end</td><td>strand</td><td>MM</td><td>target_seq</td><td>PAM</td><td>alignment</td><td>distance</td><td>position</td><td>gene name</td><td>gene id</td></tr>
<tr><td>GU305914.1</td><td></td><td>8982</td><td>9004</td><td>+</td><td>0</td><td>GACTACCAGACCACCTCCGA</td><td>CGG</td><td>||||||||[||||||||||||]PAM</td><td>NA</td><td>Intergenic</td><td>NA</td><td>NA</td></tr>
<tr><td>GU305914.1</td><td></td><td>9971</td><td>9993</td><td>+</td><td>4</td><td>GACGACGAAACCACCTCCAA</td><td>AGG</td><td>|||-||-|[-|||||||||-|]PAM</td><td>NA</td><td>Intergenic</td><td>NA</td><td>NA</td></tr>
<tr><td></td><td></td><td></td><td></td><td></td><td></td><td></td><td></td><td></td><td></td><td></td><td></td><td></td></tr>
<tr><td>T466</td><td>ACTACCAGACCACCTCCGACGGG</td><td></td><td>1000</td><td></td><td></td><td></td><td></td><td></td><td></td><td></td><td></td><td></td></tr>
<tr><td>Oligo fwd</td><td>CACCACTACCAGACCACCTCCGAC</td><td></td><td></td><td></td><td></td><td></td><td></td><td></td><td></td><td></td><td></td><td></td></tr>
<tr><td>Oligo rev</td><td>AAACGTCGGAGGTGGTCTGGTAGT</td><td></td><td></td><td></td><td></td><td></td><td></td><td></td><td></td><td></td><td></td><td></td></tr>
<tr><td>Chromosome</td><td>start</td><td></td><td>end</td><td>strand</td><td>MM</td><td>target_seq</td><td>PAM</td><td>alignment</td><td>distance</td><td>position</td><td>gene name</td><td>gene id</td></tr>
<tr><td>GU305914.1</td><td></td><td>8983</td><td>9005</td><td>+</td><td>0</td><td>ACTACCAGACCACCTCCGAC</td><td>GGG</td><td>||||||||[||||||||||||]PAM</td><td>NA</td><td>Intergenic</td><td>NA</td><td>NA</td></tr>
<tr><td></td><td></td><td></td><td></td><td></td><td></td><td></td><td></td><td></td><td></td><td></td><td></td><td></td></tr>
<tr><td>T467</td><td>ACGGGTTTCGAATTAAATTCCGG</td><td></td><td>1000</td><td></td><td></td><td></td><td></td><td></td><td></td><td></td><td></td><td></td></tr>
<tr><td>Oligo fwd</td><td>CACCACGGGTTTCGAATTAAATTC</td><td></td><td></td><td></td><td></td><td></td><td></td><td></td><td></td><td></td><td></td><td></td></tr>
<tr><td>Oligo rev</td><td>AAACGAATTTAATTCGAAACCCGT</td><td></td><td></td><td></td><td></td><td></td><td></td><td></td><td></td><td></td><td></td><td></td></tr>
<tr><td>Chromosome</td><td>start</td><td></td><td>end</td><td>strand</td><td>MM</td><td>target_seq</td><td>PAM</td><td>alignment</td><td>distance</td><td>position</td><td>gene name</td><td>gene id</td></tr>
<tr><td>GU305914.1</td><td></td><td>9001</td><td>9023</td><td>+</td><td>0</td><td>ACGGGTTTCGAATTAAATTC</td><td>CGG</td><td>||||||||[||||||||||||]PAM</td><td>NA</td><td>Intergenic</td><td>NA</td><td>NA</td></tr>
<tr><td></td><td></td><td></td><td></td><td></td><td></td><td></td><td></td><td></td><td></td><td></td><td></td><td></td></tr>
<tr><td>T468</td><td>TTCGAATTAAATTCCGGTAACGG</td><td></td><td>1000</td><td></td><td></td><td></td><td></td><td></td><td></td><td></td><td></td><td></td></tr>
<tr><td>Oligo fwd</td><td>CACCTTCGAATTAAATTCCGGTAA</td><td></td><td></td><td></td><td></td><td></td><td></td><td></td><td></td><td></td><td></td><td></td></tr>
<tr><td>Oligo rev</td><td>AAACTTACCGGAATTTAATTCGAA</td><td></td><td></td><td></td><td></td><td></td><td></td><td></td><td></td><td></td><td></td><td></td></tr>
<tr><td>Chromosome</td><td>start</td><td></td><td>end</td><td>strand</td><td>MM</td><td>target_seq</td><td>PAM</td><td>alignment</td><td>distance</td><td>position</td><td>gene name</td><td>gene id</td></tr>
<tr><td>GU305914.1</td><td></td><td>9007</td><td>9029</td><td>+</td><td>0</td><td>TTCGAATTAAATTCCGGTAA</td><td>CGG</td><td>||||||||[||||||||||||]PAM</td><td>NA</td><td>Intergenic</td><td>NA</td><td>NA</td></tr>
</table>

T469 CTCTACATCTATCCTAGCGCCGG 1000
Oligo fwd CACCCTCTACATCTATCCTAGCGC
Oligo rev AAACGCGCTAGGATAGATGTAGAG
Chromosome start end strand MM target_seq PAM alignment distance position gene name gene id
GU305914.1 9034 9056 + 0 CTCTACATCTATCCTAGCGC CGG ||||||||[||||||||||||]PAM NA Intergenic NA NA

T470 CAGTTTCGATCTATCATGTACGG 1000
Oligo fwd CACCCAGTTTCGATCTATCATGTA
Oligo rev AAACTACATGATAGATCGAAACTG
Chromosome start end strand MM target_seq PAM alignment distance position gene name gene id
GU305914.1 9060 9082 + 0 CAGTTTCGATCTATCATGTA CGG ||||||||[||||||||||||]PAM NA Intergenic NA NA

T471 AGTTTCGATCTATCATGTACGGG 1000
Oligo fwd CACCAGTTTCGATCTATCATGTAC
Oligo rev AAACGTACATGATAGATCGAAACT
Chromosome start end strand MM target_seq PAM alignment distance position gene name gene id
GU305914.1 9061 9083 + 0 AGTTTCGATCTATCATGTAC GGG ||||||||[||||||||||||]PAM NA Intergenic NA NA

T472 GTTTCGATCTATCATGTACGGGG 1000
Oligo fwd CACCGTTTCGATCTATCATGTACG
Oligo rev AAACCGTACATGATAGATCGAAAC
Chromosome start end strand MM target_seq PAM alignment distance position gene name gene id
GU305914.1 9062 9084 + 0 GTTTCGATCTATCATGTACG GGG ||||||||[||||||||||||]PAM NA Intergenic NA NA

T473 CGCGATCGAGCACCGAGACACGG 1000
Oligo fwd CACCCGCGATCGAGCACCGAGACA
Oligo rev AAACTGTCTCGGTGCTCGATCGCG
Chromosome start end strand MM target_seq PAM alignment distance position gene name gene id
GU305914.1 9137 9159 + 0 CGCGATCGAGCACCGAGACA CGG ||||||||[||||||||||||]PAM NA Intergenic NA NA

T474 GAGACACGGACCACACGAAGCGG 1000
Oligo fwd CACCGAGACACGGACCACACGAAG
Oligo rev AAACCTTCGTGTGGTCCGTGTCTC
Chromosome start end strand MM target_seq PAM alignment distance position gene name gene id
GU305914.1 9151 9173 + 0 GAGACACGGACCACACGAAG CGG ||||||||[||||||||||||]PAM NA Intergenic NA NA

T475 ATCCCAGTCCGTCACCCCGCAGG 1000
Oligo fwd CACCATCCCAGTCCGTCACCCCGC
Oligo rev AAACGCGGGGTGACGGACTGGGAT
Chromosome start end strand MM target_seq PAM alignment distance position gene name gene id
GU305914.1 9210 9232 + 0 ATCCCAGTCCGTCACCCCGC AGG ||||||||[||||||||||||]PAM NA Intergenic NA NA
GU305914.1 10184 10206 + 3 ATCCCAGCCCGTCACCCGAC AGG |||||||-[|||||||||--|]PAM NA Intergenic NA NA

T476 CGCGATCGCCGTGTCGCTAGTGG 1000
Oligo fwd CACCCGCGATCGCCGTGTCGCTAG
Oligo rev AAACCTAGCGACACGGCGATCGCG
Chromosome start end strand MM target_seq PAM alignment distance position gene name gene id
GU305914.1 9312 9334 + 0 CGCGATCGCCGTGTCGCTAG TGG ||||||||[||||||||||||]PAM NA Intergenic NA NA
GU305914.1 8319 8341 + 4 TACGATCACCGTGTCGGTAG CGG --|||||-[||||||||-|||]PAM NA Intergenic NA NA

T477 AGTAGTTACCGTGATAGTGCTGG 1000
Oligo fwd CACCAGTAGTTACCGTGATAGTGC
Oligo rev AAACGCACTATCACGGTAACTACT
Chromosome start end strand MM target_seq PAM alignment distance position gene name gene id
GU305914.1 9342 9364 + 0 AGTAGTTACCGTGATAGTGC TGG ||||||||[||||||||||||]PAM NA Intergenic NA NA

T478 GTAGTTACCGTGATAGTGCTGGG 1000
Oligo fwd CACCGTAGTTACCGTGATAGTGCT
Oligo rev AAACAGCACTATCACGGTAACTAC
Chromosome start end strand MM target_seq PAM alignment distance position gene name gene id
GU305914.1 9343 9365 + 0 GTAGTTACCGTGATAGTGCT GGG ||||||||[||||||||||||]PAM NA Intergenic NA NA

T479 CTGGGTTTATTGTATCACTACGG 1000
Oligo fwd CACCCTGGGTTTATTGTATCACTA
Oligo rev AAACTAGTGATACAATAAACCCAG
Chromosome start end strand MM target_seq PAM alignment distance position gene name gene id
GU305914.1 9361 9383 + 0 CTGGGTTTATTGTATCACTA CGG ||||||||[||||||||||||]PAM NA Intergenic NA NA
GU305914.1 10347 10369 + 1 CTGGGTTTCTTGTATCACTA CGG ||||||||[-|||||||||||]PAM NA Intergenic NA NA

T480 ATTGTATCACTACGGCTTTTTGG 1000
Oligo fwd CACCATTGTATCACTACGGCTTTT
Oligo rev AAACAAAAGCCGTAGTGATACAAT
Chromosome start end strand MM target_seq PAM alignment distance position gene name gene id
GU305914.1 9369 9391 + 0 ATTGTATCACTACGGCTTTT TGG ||||||||[||||||||||||]PAM NA Intergenic NA NA
GU305914.1 8379 8401 + 2 GTTGTATCACTACGGCTTTC TGG -|||||||[|||||||||||-]PAM NA Intergenic NA NA

T481 TTGTATCACTACGGCTTTTTGGG 1000
Oligo fwd CACCTTGTATCACTACGGCTTTTT
Oligo rev AAACAAAAAGCCGTAGTGATACAA
Chromosome start end strand MM target_seq PAM alignment distance position gene name gene id
GU305914.1 9370 9392 + 0 TTGTATCACTACGGCTTTTT GGG ||||||||[||||||||||||]PAM NA Intergenic NA NA
GU305914.1 8380 8402 + 1 TTGTATCACTACGGCTTTCT GGG ||||||||[||||||||||-|]PAM NA Intergenic NA NA

T482 TGTATCACTACGGCTTTTTGGGG 1000
Oligo fwd CACCTGTATCACTACGGCTTTTTG
Oligo rev AAACCAAAAAGCCGTAGTGATACA
Chromosome start end strand MM target_seq PAM alignment distance position gene name gene id
GU305914.1 9371 9393 + 0 TGTATCACTACGGCTTTTTG GGG ||||||||[||||||||||||]PAM NA Intergenic NA NA

| | | | | | | | | | | | |
|---|---|---|---|---|---|---|---|---|---|---|---|
| GU305914.1 | 8381 | 8403 | + | 1 | TGTATCACTACGGCTTTCTG | GGG | \|\|\|\|\|\|\|\|[\|\|\|\|\|\|\|\|\|-\|\|]PAM | NA | Intergenic | NA | NA |
| T483 | ACTACGGCTTTTTGGGGAAATGG | 1000 | | | | | | | | | |
| Oligo fwd | CACCACTACGGCTTTTTGGGGAAA | | | | | | | | | | |
| Oligo rev | AAACTTTCCCCAAAAAGCCGTAGT | | | | | | | | | | |
| Chromosome | start | end | strand | MM | target_seq | PAM | alignment | distance | position | gene name | gene id |
| GU305914.1 | 9377 | 9399 | + | 0 | ACTACGGCTTTTTGGGGAAA | TGG | \|\|\|\|\|\|\|\|[\|\|\|\|\|\|\|\|\|\|\|\|]PAM | NA | Intergenic | NA | NA |
| T484 | CTACGGCTTTTTGGGGAAATGGG | 1000 | | | | | | | | | |
| Oligo fwd | CACCCTACGGCTTTTTGGGGAAAT | | | | | | | | | | |
| Oligo rev | AAACATTTCCCCAAAAAGCCGTAG | | | | | | | | | | |
| Chromosome | start | end | strand | MM | target_seq | PAM | alignment | distance | position | gene name | gene id |
| GU305914.1 | 9378 | 9400 | + | 0 | CTACGGCTTTTTGGGGAAAT | GGG | \|\|\|\|\|\|\|\|[\|\|\|\|\|\|\|\|\|\|\|\|]PAM | NA | Intergenic | NA | NA |
| GU305914.1 | 8388 | 8410 | + | 2 | CTACGGCTTTCTGGGGAGAT | CGG | \|\|\|\|\|\|\|\|[\|\|-\|\|\|\|\|\|-\|\|]PAM | NA | Intergenic | NA | NA |
| T485 | CCGTTTCCGCGTCCGTTTCATGG | 1000 | | | | | | | | | |
| Oligo fwd | CACCCCGTTTCCGCGTCCGTTTCA | | | | | | | | | | |
| Oligo rev | AAACTGAAACGGACGCGGAAACGG | | | | | | | | | | |
| Chromosome | start | end | strand | MM | target_seq | PAM | alignment | distance | position | gene name | gene id |
| GU305914.1 | 9423 | 9445 | + | 0 | CCGTTTCCGCGTCCGTTTCA | TGG | \|\|\|\|\|\|\|\|[\|\|\|\|\|\|\|\|\|\|\|\|]PAM | NA | Intergenic | NA | NA |
| T486 | CGTTTCCGCGTCCGTTTCATGGG | 1000 | | | | | | | | | |
| Oligo fwd | CACCCGTTTCCGCGTCCGTTTCAT | | | | | | | | | | |
| Oligo rev | AAACATGAAACGGACGCGGAAACG | | | | | | | | | | |
| Chromosome | start | end | strand | MM | target_seq | PAM | alignment | distance | position | gene name | gene id |
| GU305914.1 | 9424 | 9446 | + | 0 | CGTTTCCGCGTCCGTTTCAT | GGG | \|\|\|\|\|\|\|\|[\|\|\|\|\|\|\|\|\|\|\|\|]PAM | NA | Intergenic | NA | NA |
| T487 | AACCGAGCCTCTCCGTTCAACGG | 1000 | | | | | | | | | |
| Oligo fwd | CACCAACCGAGCCTCTCCGTTCAA | | | | | | | | | | |
| Oligo rev | AAACTTGAACGGAGAGGCTCGGTT | | | | | | | | | | |
| Chromosome | start | end | strand | MM | target_seq | PAM | alignment | distance | position | gene name | gene id |
| GU305914.1 | 9474 | 9496 | + | 0 | AACCGAGCCTCTCCGTTCAA | CGG | \|\|\|\|\|\|\|\|[\|\|\|\|\|\|\|\|\|\|\|\|]PAM | NA | Intergenic | NA | NA |
| T488 | CCACATTTTCTACGTACACCTGG | 1000 | | | | | | | | | |
| Oligo fwd | CACCCCACATTTTCTACGTACACC | | | | | | | | | | |
| Oligo rev | AAACGGTGTACGTAGAAAATGTGG | | | | | | | | | | |
| Chromosome | start | end | strand | MM | target_seq | PAM | alignment | distance | position | gene name | gene id |
| GU305914.1 | 9519 | 9541 | + | 0 | CCACATTTTCTACGTACACC | TGG | \|\|\|\|\|\|\|\|[\|\|\|\|\|\|\|\|\|\|\|\|]PAM | NA | Intergenic | NA | NA |
| T489 | CGCGACGTATTCACTAACACCGG | 1000 | | | | | | | | | |
| Oligo fwd | CACCCGCGACGTATTCACTAACAC | | | | | | | | | | |
| Oligo rev | AAACGTGTTAGTGAATACGTCGCG | | | | | | | | | | |
| Chromosome | start | end | strand | MM | target_seq | PAM | alignment | distance | position | gene name | gene id |
| GU305914.1 | 9591 | 9613 | + | 0 | CGCGACGTATTCACTAACAC | CGG | \|\|\|\|\|\|\|\|[\|\|\|\|\|\|\|\|\|\|\|\|]PAM | NA | Intergenic | NA | NA |
| T490 | GCGACGTATTCACTAACACCGGG | 1000 | | | | | | | | | |
| Oligo fwd | CACCGCGACGTATTCACTAACACC | | | | | | | | | | |
| Oligo rev | AAACGGTGTTAGTGAATACGTCGC | | | | | | | | | | |
| Chromosome | start | end | strand | MM | target_seq | PAM | alignment | distance | position | gene name | gene id |
| GU305914.1 | 9592 | 9614 | + | 0 | GCGACGTATTCACTAACACC | GGG | \|\|\|\|\|\|\|\|[\|\|\|\|\|\|\|\|\|\|\|\|]PAM | NA | Intergenic | NA | NA |
| T491 | TCGTCTGTTGCAACTCTCTCTGG | 1000 | | | | | | | | | |
| Oligo fwd | CACCTCGTCTGTTGCAACTCTCTC | | | | | | | | | | |
| Oligo rev | AAACGAGAGAGTTGCAACAGACGA | | | | | | | | | | |
| Chromosome | start | end | strand | MM | target_seq | PAM | alignment | distance | position | gene name | gene id |
| GU305914.1 | 9641 | 9663 | + | 0 | TCGTCTGTTGCAACTCTCTC | TGG | \|\|\|\|\|\|\|\|[\|\|\|\|\|\|\|\|\|\|\|\|]PAM | NA | Intergenic | NA | NA |
| T492 | CGCGTCGCGAAAATGTGATCCGG | 1000 | | | | | | | | | |
| Oligo fwd | CACCCGCGTCGCGAAAATGTGATC | | | | | | | | | | |
| Oligo rev | AAACGATCACATTTTCGCGACGCG | | | | | | | | | | |
| Chromosome | start | end | strand | MM | target_seq | PAM | alignment | distance | position | gene name | gene id |
| GU305914.1 | 9698 | 9720 | + | 0 | CGCGTCGCGAAAATGTGATC | CGG | \|\|\|\|\|\|\|\|[\|\|\|\|\|\|\|\|\|\|\|\|]PAM | NA | Intergenic | NA | NA |
| T493 | TCTCGAAATCTTACTCCTCCAGG | 1000 | | | | | | | | | |
| Oligo fwd | CACCTCTCGAAATCTTACTCCTCC | | | | | | | | | | |
| Oligo rev | AAACGGAGGAGTAAGATTTCGAGA | | | | | | | | | | |
| Chromosome | start | end | strand | MM | target_seq | PAM | alignment | distance | position | gene name | gene id |
| GU305914.1 | 9724 | 9746 | + | 0 | TCTCGAAATCTTACTCCTCC | AGG | \|\|\|\|\|\|\|\|[\|\|\|\|\|\|\|\|\|\|\|\|]PAM | NA | Intergenic | NA | NA |
| T494 | TACTCCTCCAGGACACCGCCAGG | 1000 | | | | | | | | | |
| Oligo fwd | CACCTACTCCTCCAGGACACCGCC | | | | | | | | | | |
| Oligo rev | AAACGGCGGTGTCCTGGAGGAGTA | | | | | | | | | | |
| Chromosome | start | end | strand | MM | target_seq | PAM | alignment | distance | position | gene name | gene id |
| GU305914.1 | 9735 | 9757 | + | 0 | TACTCCTCCAGGACACCGCC | AGG | \|\|\|\|\|\|\|\|[\|\|\|\|\|\|\|\|\|\|\|\|]PAM | NA | Intergenic | NA | NA |
| T495 | ACACCGCCAGGTACCTGTACAGG | 1000 | | | | | | | | | |
| Oligo fwd | CACCACACCGCCAGGTACCTGTAC | | | | | | | | | | |
| Oligo rev | AAACGTACAGGTACCTGGCGGTGT | | | | | | | | | | |
| Chromosome | start | end | strand | MM | target_seq | PAM | alignment | distance | position | gene name | gene id |
| GU305914.1 | 9747 | 9769 | + | 0 | ACACCGCCAGGTACCTGTAC | AGG | \|\|\|\|\|\|\|\|[\|\|\|\|\|\|\|\|\|\|\|\|]PAM | NA | Intergenic | NA | NA |
| T496 | AAAGTCACAGTACTCAAACTTGG | 1000 | | | | | | | | | |
| Oligo fwd | CACCAAAGTCACAGTACTCAAACT | | | | | | | | | | |
| Oligo rev | AAACAGTTTGAGTACTGTGACTTT | | | | | | | | | | |
| Chromosome | start | end | strand | MM | target_seq | PAM | alignment | distance | position | gene name | gene id |
| GU305914.1 | 9799 | 9821 | + | 0 | AAAGTCACAGTACTCAAACT | TGG | \|\|\|\|\|\|\|\|[\|\|\|\|\|\|\|\|\|\|\|\|]PAM | NA | Intergenic | NA | NA |
| T497 | ATACGAATGTCAGTTGCCTGAGG | 1000 | | | | | | | | | |
| Oligo fwd | CACCATACGAATGTCAGTTGCCTG | | | | | | | | | | |

| | | | | | | | | | | | | |
|---|---|---|---|---|---|---|---|---|---|---|---|---|
| Oligo rev | AAACCAGGCAACTGACATTCGTAT | | | | | | | | | | | |
| Chromosome | start | | end | strand | MM | target_seq | PAM | alignment | distance | position | gene name | gene id |
| GU305914.1 | | 9824 | 9846 | + | 0 | ATACGAATGTCAGTTGCCTG | AGG | \|\|\|\|\|\|\|\|[\|\|\|\|\|\|\|\|\|\|\|\|]PAM | NA | Intergenic | NA | NA |
| T498 | AGGCCTGTATTGAAGCAACATGG | | 1000 | | | | | | | | | |
| Oligo fwd | CACCAGGCCTGTATTGAAGCAACA | | | | | | | | | | | |
| Oligo rev | AAACTGTTGCTTCAATACAGGCCT | | | | | | | | | | | |
| Chromosome | start | | end | strand | MM | target_seq | PAM | alignment | distance | position | gene name | gene id |
| GU305914.1 | | 9844 | 9866 | + | 0 | AGGCCTGTATTGAAGCAACA | TGG | \|\|\|\|\|\|\|\|[\|\|\|\|\|\|\|\|\|\|\|\|]PAM | NA | Intergenic | NA | NA |
| T499 | CTGTATTGAAGCAACATGGCAGG | | 1000 | | | | | | | | | |
| Oligo fwd | CACCCTGTATTGAAGCAACATGGC | | | | | | | | | | | |
| Oligo rev | AAACGCCATGTTGCTTCAATACAG | | | | | | | | | | | |
| Chromosome | start | | end | strand | MM | target_seq | PAM | alignment | distance | position | gene name | gene id |
| GU305914.1 | | 9848 | 9870 | + | 0 | CTGTATTGAAGCAACATGGC | AGG | \|\|\|\|\|\|\|\|[\|\|\|\|\|\|\|\|\|\|\|\|]PAM | NA | Intergenic | NA | NA |
| T500 | AAGCAACATGGCAGGTAAAATGG | | 1000 | | | | | | | | | |
| Oligo fwd | CACCAAGCAACATGGCAGGTAAAA | | | | | | | | | | | |
| Oligo rev | AAACTTTTACCTGCCATGTTGCTT | | | | | | | | | | | |
| Chromosome | start | | end | strand | MM | target_seq | PAM | alignment | distance | position | gene name | gene id |
| GU305914.1 | | 9856 | 9878 | + | 0 | AAGCAACATGGCAGGTAAAA | TGG | \|\|\|\|\|\|\|\|[\|\|\|\|\|\|\|\|\|\|\|\|]PAM | NA | Intergenic | NA | NA |
| T501 | AGGTAAAATGGACCGTTAATCGG | | 1000 | | | | | | | | | |
| Oligo fwd | CACCAGGTAAAATGGACCGTTAAT | | | | | | | | | | | |
| Oligo rev | AAACATTAACGGTCCATTTTACCT | | | | | | | | | | | |
| Chromosome | start | | end | strand | MM | target_seq | PAM | alignment | distance | position | gene name | gene id |
| GU305914.1 | | 9868 | 9890 | + | 0 | AGGTAAAATGGACCGTTAAT | CGG | \|\|\|\|\|\|\|\|[\|\|\|\|\|\|\|\|\|\|\|\|]PAM | NA | Intergenic | NA | NA |
| T502 | GACGACGAAACCACCTCCAAAGG | | 1000 | | | | | | | | | |
| Oligo fwd | CACCGACGACGAAACCACCTCCAA | | | | | | | | | | | |
| Oligo rev | AAACTTGGAGGTGGTTTCGTCGTC | | | | | | | | | | | |
| Chromosome | start | | end | strand | MM | target_seq | PAM | alignment | distance | position | gene name | gene id |
| GU305914.1 | | 9971 | 9993 | + | 0 | GACGACGAAACCACCTCCAA | AGG | \|\|\|\|\|\|\|\|[\|\|\|\|\|\|\|\|\|\|\|\|]PAM | NA | Intergenic | NA | NA |
| GU305914.1 | | 8982 | 9004 | + | 4 | GACTACCAGACCACCTCCGA | CGG | \|\|\|-\|\|-\|[-\|\|\|\|\|\|\|\|\|-\|]PAM | NA | Intergenic | NA | NA |
| T503 | ACGACGAAACCACCTCCAAAGGG | | 1000 | | | | | | | | | |
| Oligo fwd | CACCACGACGAAACCACCTCCAAA | | | | | | | | | | | |
| Oligo rev | AAACTTTGGAGGTGGTTTCGTCGT | | | | | | | | | | | |
| Chromosome | start | | end | strand | MM | target_seq | PAM | alignment | distance | position | gene name | gene id |
| GU305914.1 | | 9972 | 9994 | + | 0 | ACGACGAAACCACCTCCAAA | GGG | \|\|\|\|\|\|\|\|[\|\|\|\|\|\|\|\|\|\|\|\|]PAM | NA | Intergenic | NA | NA |
| T504 | CGACGAAACCACCTCCAAAGGGG | | 1000 | | | | | | | | | |
| Oligo fwd | CACCCGACGAAACCACCTCCAAAG | | | | | | | | | | | |
| Oligo rev | AAACCTTTGGAGGTGGTTTCGTCG | | | | | | | | | | | |
| Chromosome | start | | end | strand | MM | target_seq | PAM | alignment | distance | position | gene name | gene id |
| GU305914.1 | | 9973 | 9995 | + | 0 | CGACGAAACCACCTCCAAAG | GGG | \|\|\|\|\|\|\|\|[\|\|\|\|\|\|\|\|\|\|\|\|]PAM | NA | Intergenic | NA | NA |
| T505 | ACCACCTCCAAAGGGGTTCACGG | | 1000 | | | | | | | | | |
| Oligo fwd | CACCACCACCTCCAAAGGGGTTCA | | | | | | | | | | | |
| Oligo rev | AAACTGAACCCCTTTGGAGGTGGT | | | | | | | | | | | |
| Chromosome | start | | end | strand | MM | target_seq | PAM | alignment | distance | position | gene name | gene id |
| GU305914.1 | | 9980 | 10002 | + | 0 | ACCACCTCCAAAGGGGTTCA | CGG | \|\|\|\|\|\|\|\|[\|\|\|\|\|\|\|\|\|\|\|\|]PAM | NA | Intergenic | NA | NA |
| T506 | TTCACGGTGAATCCGCATGACGG | | 1000 | | | | | | | | | |
| Oligo fwd | CACCTTCACGGTGAATCCGCATGA | | | | | | | | | | | |
| Oligo rev | AAACTCATGCGGATTCACCGTGAA | | | | | | | | | | | |
| Chromosome | start | | end | strand | MM | target_seq | PAM | alignment | distance | position | gene name | gene id |
| GU305914.1 | | 9996 | 10018 | + | 0 | TTCACGGTGAATCCGCATGA | CGG | \|\|\|\|\|\|\|\|[\|\|\|\|\|\|\|\|\|\|\|\|]PAM | NA | Intergenic | NA | NA |
| T507 | GTATCTCGTTTCGTCCCCAGAGG | | 1000 | | | | | | | | | |
| Oligo fwd | CACCGTATCTCGTTTCGTCCCCAG | | | | | | | | | | | |
| Oligo rev | AAACCTGGGGACGAAACGAGATAC | | | | | | | | | | | |
| Chromosome | start | | end | strand | MM | target_seq | PAM | alignment | distance | position | gene name | gene id |
| GU305914.1 | | 10025 | 10047 | + | 0 | GTATCTCGTTTCGTCCCCAG | AGG | \|\|\|\|\|\|\|\|[\|\|\|\|\|\|\|\|\|\|\|\|]PAM | NA | Intergenic | NA | NA |
| T508 | TTCTGTACATCTACAGTGTGAGG | | 1000 | | | | | | | | | |
| Oligo fwd | CACCTTCTGTACATCTACAGTGTG | | | | | | | | | | | |
| Oligo rev | AAACCACACTGTAGATGTACAGAA | | | | | | | | | | | |
| Chromosome | start | | end | strand | MM | target_seq | PAM | alignment | distance | position | gene name | gene id |
| GU305914.1 | | 10055 | 10077 | + | 0 | TTCTGTACATCTACAGTGTG | AGG | \|\|\|\|\|\|\|\|[\|\|\|\|\|\|\|\|\|\|\|\|]PAM | NA | Intergenic | NA | NA |
| T509 | TCTGTACATCTACAGTGTGAGGG | | 1000 | | | | | | | | | |
| Oligo fwd | CACCTCTGTACATCTACAGTGTGA | | | | | | | | | | | |
| Oligo rev | AAACTCACACTGTAGATGTACAGA | | | | | | | | | | | |
| Chromosome | start | | end | strand | MM | target_seq | PAM | alignment | distance | position | gene name | gene id |
| GU305914.1 | | 10056 | 10078 | + | 0 | TCTGTACATCTACAGTGTGA | GGG | \|\|\|\|\|\|\|\|[\|\|\|\|\|\|\|\|\|\|\|\|]PAM | NA | Intergenic | NA | NA |
| T510 | CTGTACATCTACAGTGTGAGGGG | | 1000 | | | | | | | | | |
| Oligo fwd | CACCCTGTACATCTACAGTGTGAG | | | | | | | | | | | |
| Oligo rev | AAACCTCACACTGTAGATGTACAG | | | | | | | | | | | |
| Chromosome | start | | end | strand | MM | target_seq | PAM | alignment | distance | position | gene name | gene id |
| GU305914.1 | | 10057 | 10079 | + | 0 | CTGTACATCTACAGTGTGAG | GGG | \|\|\|\|\|\|\|\|[\|\|\|\|\|\|\|\|\|\|\|\|]PAM | NA | Intergenic | NA | NA |
| T511 | ATCCCAGCCCGTCACCCGACAGG | | 1000 | | | | | | | | | |
| Oligo fwd | CACCATCCCAGCCCGTCACCCGAC | | | | | | | | | | | |
| Oligo rev | AAACGTCGGGTGACGGGCTGGGAT | | | | | | | | | | | |
| Chromosome | start | | end | strand | MM | target_seq | PAM | alignment | distance | position | gene name | gene id |
| GU305914.1 | | 10184 | 10206 | + | 0 | ATCCCAGCCCGTCACCCGAC | AGG | \|\|\|\|\|\|\|\|[\|\|\|\|\|\|\|\|\|\|\|\|]PAM | NA | Intergenic | NA | NA |
| GU305914.1 | | 9210 | 9232 | + | 3 | ATCCCAGTCCGTCACCCCGC | AGG | \|\|\|\|\|\|\|-[\|\|\|\|\|\|\|\|\|--\|]PAM | NA | Intergenic | NA | NA |

<table>
<tr><td>T512</td><td>AATGCAGACCTCGAATACGACGG</td><td>1000</td><td></td><td></td><td></td><td></td><td></td><td></td><td></td><td></td><td></td></tr>
<tr><td>Oligo fwd</td><td>CACCAATGCAGACCTCGAATACGA</td><td></td><td></td><td></td><td></td><td></td><td></td><td></td><td></td><td></td><td></td></tr>
<tr><td>Oligo rev</td><td>AAACTCGTATTCGAGGTCTGCATT</td><td></td><td></td><td></td><td></td><td></td><td></td><td></td><td></td><td></td><td></td></tr>
<tr><td>Chromosome</td><td>start</td><td>end</td><td>strand</td><td>MM</td><td>target_seq</td><td>PAM</td><td>alignment</td><td>distance</td><td>position</td><td>gene name</td><td>gene id</td></tr>
<tr><td>GU305914.1</td><td>10274</td><td>10296</td><td>+</td><td>0</td><td>AATGCAGACCTCGAATACGA</td><td>CGG</td><td>||||||||[||||||||||||]PAM</td><td>NA</td><td>Intergenic</td><td>NA</td><td>NA</td></tr>
<tr><td></td><td></td><td></td><td></td><td></td><td></td><td></td><td></td><td></td><td></td><td></td><td></td></tr>
<tr><td>T513</td><td>CGCGATCGTCGTGCCGTTAATGG</td><td>1000</td><td></td><td></td><td></td><td></td><td></td><td></td><td></td><td></td><td></td></tr>
<tr><td>Oligo fwd</td><td>CACCCGCGATCGTCGTGCCGTTAA</td><td></td><td></td><td></td><td></td><td></td><td></td><td></td><td></td><td></td><td></td></tr>
<tr><td>Oligo rev</td><td>AAACTTAACGGCACGACGATCGCG</td><td></td><td></td><td></td><td></td><td></td><td></td><td></td><td></td><td></td><td></td></tr>
<tr><td>Chromosome</td><td>start</td><td>end</td><td>strand</td><td>MM</td><td>target_seq</td><td>PAM</td><td>alignment</td><td>distance</td><td>position</td><td>gene name</td><td>gene id</td></tr>
<tr><td>GU305914.1</td><td>10298</td><td>10320</td><td>+</td><td>0</td><td>CGCGATCGTCGTGCCGTTAA</td><td>TGG</td><td>||||||||[||||||||||||]PAM</td><td>NA</td><td>Intergenic</td><td>NA</td><td>NA</td></tr>
<tr><td></td><td></td><td></td><td></td><td></td><td></td><td></td><td></td><td></td><td></td><td></td><td></td></tr>
<tr><td>T514</td><td>AGTAGCGACTGTGATCGTACTGG</td><td>1000</td><td></td><td></td><td></td><td></td><td></td><td></td><td></td><td></td><td></td></tr>
<tr><td>Oligo fwd</td><td>CACCAGTAGCGACTGTGATCGTAC</td><td></td><td></td><td></td><td></td><td></td><td></td><td></td><td></td><td></td><td></td></tr>
<tr><td>Oligo rev</td><td>AAACGTACGATCACAGTCGCTACT</td><td></td><td></td><td></td><td></td><td></td><td></td><td></td><td></td><td></td><td></td></tr>
<tr><td>Chromosome</td><td>start</td><td>end</td><td>strand</td><td>MM</td><td>target_seq</td><td>PAM</td><td>alignment</td><td>distance</td><td>position</td><td>gene name</td><td>gene id</td></tr>
<tr><td>GU305914.1</td><td>10328</td><td>10350</td><td>+</td><td>0</td><td>AGTAGCGACTGTGATCGTAC</td><td>TGG</td><td>||||||||[||||||||||||]PAM</td><td>NA</td><td>Intergenic</td><td>NA</td><td>NA</td></tr>
<tr><td></td><td></td><td></td><td></td><td></td><td></td><td></td><td></td><td></td><td></td><td></td><td></td></tr>
<tr><td>T515</td><td>GTAGCGACTGTGATCGTACTGGG</td><td>1000</td><td></td><td></td><td></td><td></td><td></td><td></td><td></td><td></td><td></td></tr>
<tr><td>Oligo fwd</td><td>CACCGTAGCGACTGTGATCGTACT</td><td></td><td></td><td></td><td></td><td></td><td></td><td></td><td></td><td></td><td></td></tr>
<tr><td>Oligo rev</td><td>AAACAGTACGATCACAGTCGCTAC</td><td></td><td></td><td></td><td></td><td></td><td></td><td></td><td></td><td></td><td></td></tr>
<tr><td>Chromosome</td><td>start</td><td>end</td><td>strand</td><td>MM</td><td>target_seq</td><td>PAM</td><td>alignment</td><td>distance</td><td>position</td><td>gene name</td><td>gene id</td></tr>
<tr><td>GU305914.1</td><td>10329</td><td>10351</td><td>+</td><td>0</td><td>GTAGCGACTGTGATCGTACT</td><td>GGG</td><td>||||||||[||||||||||||]PAM</td><td>NA</td><td>Intergenic</td><td>NA</td><td>NA</td></tr>
<tr><td></td><td></td><td></td><td></td><td></td><td></td><td></td><td></td><td></td><td></td><td></td><td></td></tr>
<tr><td>T516</td><td>CTGGGTTTCTTGTATCACTACGG</td><td>1000</td><td></td><td></td><td></td><td></td><td></td><td></td><td></td><td></td><td></td></tr>
<tr><td>Oligo fwd</td><td>CACCCTGGGTTTCTTGTATCACTA</td><td></td><td></td><td></td><td></td><td></td><td></td><td></td><td></td><td></td><td></td></tr>
<tr><td>Oligo rev</td><td>AAACTAGTGATACAAGAAACCCAG</td><td></td><td></td><td></td><td></td><td></td><td></td><td></td><td></td><td></td><td></td></tr>
<tr><td>Chromosome</td><td>start</td><td>end</td><td>strand</td><td>MM</td><td>target_seq</td><td>PAM</td><td>alignment</td><td>distance</td><td>position</td><td>gene name</td><td>gene id</td></tr>
<tr><td>GU305914.1</td><td>10347</td><td>10369</td><td>+</td><td>0</td><td>CTGGGTTTCTTGTATCACTA</td><td>CGG</td><td>||||||||[||||||||||||]PAM</td><td>NA</td><td>Intergenic</td><td>NA</td><td>NA</td></tr>
<tr><td>GU305914.1</td><td>9361</td><td>9383</td><td>+</td><td>1</td><td>CTGGGTTTATTGTATCACTA</td><td>CGG</td><td>||||||||[-|||||||||||]PAM</td><td>NA</td><td>Intergenic</td><td>NA</td><td>NA</td></tr>
<tr><td></td><td></td><td></td><td></td><td></td><td></td><td></td><td></td><td></td><td></td><td></td><td></td></tr>
<tr><td>T517</td><td>ACTACGGTTTCCTGAGAAAATGG</td><td>1000</td><td></td><td></td><td></td><td></td><td></td><td></td><td></td><td></td><td></td></tr>
<tr><td>Oligo fwd</td><td>CACCACTACGGTTTCCTGAGAAAA</td><td></td><td></td><td></td><td></td><td></td><td></td><td></td><td></td><td></td><td></td></tr>
<tr><td>Oligo rev</td><td>AAACTTTTCTCAGGAAACCGTAGT</td><td></td><td></td><td></td><td></td><td></td><td></td><td></td><td></td><td></td><td></td></tr>
<tr><td>Chromosome</td><td>start</td><td>end</td><td>strand</td><td>MM</td><td>target_seq</td><td>PAM</td><td>alignment</td><td>distance</td><td>position</td><td>gene name</td><td>gene id</td></tr>
<tr><td>GU305914.1</td><td>10363</td><td>10385</td><td>+</td><td>0</td><td>ACTACGGTTTCCTGAGAAAA</td><td>TGG</td><td>||||||||[||||||||||||]PAM</td><td>NA</td><td>Intergenic</td><td>NA</td><td>NA</td></tr>
<tr><td></td><td></td><td></td><td></td><td></td><td></td><td></td><td></td><td></td><td></td><td></td><td></td></tr>
<tr><td>T518</td><td>CTACGGTTTCCTGAGAAAATGGG</td><td>1000</td><td></td><td></td><td></td><td></td><td></td><td></td><td></td><td></td><td></td></tr>
<tr><td>Oligo fwd</td><td>CACCCTACGGTTTCCTGAGAAAAT</td><td></td><td></td><td></td><td></td><td></td><td></td><td></td><td></td><td></td><td></td></tr>
<tr><td>Oligo rev</td><td>AAACATTTTCTCAGGAAACCGTAG</td><td></td><td></td><td></td><td></td><td></td><td></td><td></td><td></td><td></td><td></td></tr>
<tr><td>Chromosome</td><td>start</td><td>end</td><td>strand</td><td>MM</td><td>target_seq</td><td>PAM</td><td>alignment</td><td>distance</td><td>position</td><td>gene name</td><td>gene id</td></tr>
<tr><td>GU305914.1</td><td>10364</td><td>10386</td><td>+</td><td>0</td><td>CTACGGTTTCCTGAGAAAAT</td><td>GGG</td><td>||||||||[||||||||||||]PAM</td><td>NA</td><td>Intergenic</td><td>NA</td><td>NA</td></tr>
<tr><td></td><td></td><td></td><td></td><td></td><td></td><td></td><td></td><td></td><td></td><td></td><td></td></tr>
<tr><td>T519</td><td>TCGTACCTCGCGCGTGTGCGCGG</td><td>1000</td><td></td><td></td><td></td><td></td><td></td><td></td><td></td><td></td><td></td></tr>
<tr><td>Oligo fwd</td><td>CACCTCGTACCTCGCGCGTGTGCG</td><td></td><td></td><td></td><td></td><td></td><td></td><td></td><td></td><td></td><td></td></tr>
<tr><td>Oligo rev</td><td>AAACCGCACACGCGCGAGGTACGA</td><td></td><td></td><td></td><td></td><td></td><td></td><td></td><td></td><td></td><td></td></tr>
<tr><td>Chromosome</td><td>start</td><td>end</td><td>strand</td><td>MM</td><td>target_seq</td><td>PAM</td><td>alignment</td><td>distance</td><td>position</td><td>gene name</td><td>gene id</td></tr>
<tr><td>GU305914.1</td><td>10398</td><td>10420</td><td>+</td><td>0</td><td>TCGTACCTCGCGCGTGTGCG</td><td>CGG</td><td>||||||||[||||||||||||]PAM</td><td>NA</td><td>Intergenic</td><td>NA</td><td>NA</td></tr>
<tr><td></td><td></td><td></td><td></td><td></td><td></td><td></td><td></td><td></td><td></td><td></td><td></td></tr>
<tr><td>T520</td><td>CAGTGAATAAAACGCTGTTTCGG</td><td>1000</td><td></td><td></td><td></td><td></td><td></td><td></td><td></td><td></td><td></td></tr>
<tr><td>Oligo fwd</td><td>CACCCAGTGAATAAAACGCTGTTT</td><td></td><td></td><td></td><td></td><td></td><td></td><td></td><td></td><td></td><td></td></tr>
<tr><td>Oligo rev</td><td>AAACAAACAGCGTTTTATTCACTG</td><td></td><td></td><td></td><td></td><td></td><td></td><td></td><td></td><td></td><td></td></tr>
<tr><td>Chromosome</td><td>start</td><td>end</td><td>strand</td><td>MM</td><td>target_seq</td><td>PAM</td><td>alignment</td><td>distance</td><td>position</td><td>gene name</td><td>gene id</td></tr>
<tr><td>GU305914.1</td><td>10510</td><td>10532</td><td>+</td><td>0</td><td>CAGTGAATAAAACGCTGTTT</td><td>CGG</td><td>||||||||[||||||||||||]PAM</td><td>NA</td><td>Intergenic</td><td>NA</td><td>NA</td></tr>
<tr><td></td><td></td><td></td><td></td><td></td><td></td><td></td><td></td><td></td><td></td><td></td><td></td></tr>
<tr><td>T521</td><td>TTTTCCGCCTCCGTTTACCTAGG</td><td>1000</td><td></td><td></td><td></td><td></td><td></td><td></td><td></td><td></td><td></td></tr>
<tr><td>Oligo fwd</td><td>CACCTTTTCCGCCTCCGTTTACCT</td><td></td><td></td><td></td><td></td><td></td><td></td><td></td><td></td><td></td><td></td></tr>
<tr><td>Oligo rev</td><td>AAACAGGTAAACGGAGGCGGAAAA</td><td></td><td></td><td></td><td></td><td></td><td></td><td></td><td></td><td></td><td></td></tr>
<tr><td>Chromosome</td><td>start</td><td>end</td><td>strand</td><td>MM</td><td>target_seq</td><td>PAM</td><td>alignment</td><td>distance</td><td>position</td><td>gene name</td><td>gene id</td></tr>
<tr><td>GU305914.1</td><td>10563</td><td>10585</td><td>+</td><td>0</td><td>TTTTCCGCCTCCGTTTACCT</td><td>AGG</td><td>||||||||[||||||||||||]PAM</td><td>NA</td><td>Intergenic</td><td>NA</td><td>NA</td></tr>
<tr><td></td><td></td><td></td><td></td><td></td><td></td><td></td><td></td><td></td><td></td><td></td><td></td></tr>
<tr><td>T522</td><td>GTAGATCCCGCCCACGTACCTGG</td><td>1000</td><td></td><td></td><td></td><td></td><td></td><td></td><td></td><td></td><td></td></tr>
<tr><td>Oligo fwd</td><td>CACCGTAGATCCCGCCCACGTACC</td><td></td><td></td><td></td><td></td><td></td><td></td><td></td><td></td><td></td><td></td></tr>
<tr><td>Oligo rev</td><td>AAACGGTACGTGGGCGGGATCTAC</td><td></td><td></td><td></td><td></td><td></td><td></td><td></td><td></td><td></td><td></td></tr>
<tr><td>Chromosome</td><td>start</td><td>end</td><td>strand</td><td>MM</td><td>target_seq</td><td>PAM</td><td>alignment</td><td>distance</td><td>position</td><td>gene name</td><td>gene id</td></tr>
<tr><td>GU305914.1</td><td>10611</td><td>10633</td><td>+</td><td>0</td><td>GTAGATCCCGCCCACGTACC</td><td>TGG</td><td>||||||||[||||||||||||]PAM</td><td>NA</td><td>Intergenic</td><td>NA</td><td>NA</td></tr>
<tr><td></td><td></td><td></td><td></td><td></td><td></td><td></td><td></td><td></td><td></td><td></td><td></td></tr>
<tr><td>T523</td><td>ACCTGGCGTATATGTAGCTGCGG</td><td>1000</td><td></td><td></td><td></td><td></td><td></td><td></td><td></td><td></td><td></td></tr>
<tr><td>Oligo fwd</td><td>CACCACCTGGCGTATATGTAGCTG</td><td></td><td></td><td></td><td></td><td></td><td></td><td></td><td></td><td></td><td></td></tr>
<tr><td>Oligo rev</td><td>AAACCAGCTACATATACGCCAGGT</td><td></td><td></td><td></td><td></td><td></td><td></td><td></td><td></td><td></td><td></td></tr>
<tr><td>Chromosome</td><td>start</td><td>end</td><td>strand</td><td>MM</td><td>target_seq</td><td>PAM</td><td>alignment</td><td>distance</td><td>position</td><td>gene name</td><td>gene id</td></tr>
<tr><td>GU305914.1</td><td>10628</td><td>10650</td><td>+</td><td>0</td><td>ACCTGGCGTATATGTAGCTG</td><td>CGG</td><td>||||||||[||||||||||||]PAM</td><td>NA</td><td>Intergenic</td><td>NA</td><td>NA</td></tr>
<tr><td></td><td></td><td></td><td></td><td></td><td></td><td></td><td></td><td></td><td></td><td></td><td></td></tr>
<tr><td>T524</td><td>CTGCGGAAAACACACCTGTACGG</td><td>1000</td><td></td><td></td><td></td><td></td><td></td><td></td><td></td><td></td><td></td></tr>
<tr><td>Oligo fwd</td><td>CACCCTGCGGAAAACACACCTGTA</td><td></td><td></td><td></td><td></td><td></td><td></td><td></td><td></td><td></td><td></td></tr>
<tr><td>Oligo rev</td><td>AAACTACAGGTGTGTTTTCCGCAG</td><td></td><td></td><td></td><td></td><td></td><td></td><td></td><td></td><td></td><td></td></tr>
<tr><td>Chromosome</td><td>start</td><td>end</td><td>strand</td><td>MM</td><td>target_seq</td><td>PAM</td><td>alignment</td><td>distance</td><td>position</td><td>gene name</td><td>gene id</td></tr>
<tr><td>GU305914.1</td><td>10645</td><td>10667</td><td>+</td><td>0</td><td>CTGCGGAAAACACACCTGTA</td><td>CGG</td><td>||||||||[||||||||||||]PAM</td><td>NA</td><td>Intergenic</td><td>NA</td><td>NA</td></tr>
<tr><td></td><td></td><td></td><td></td><td></td><td></td><td></td><td></td><td></td><td></td><td></td><td></td></tr>
<tr><td>T525</td><td>GGTAGATCACACTCTGCTCAAGG</td><td>1000</td><td></td><td></td><td></td><td></td><td></td><td></td><td></td><td></td><td></td></tr>
<tr><td>Oligo fwd</td><td>CACCGGTAGATCACACTCTGCTCA</td><td></td><td></td><td></td><td></td><td></td><td></td><td></td><td></td><td></td><td></td></tr>
<tr><td>Oligo rev</td><td>AAACTGAGCAGAGTGTGATCTACC</td><td></td><td></td><td></td><td></td><td></td><td></td><td></td><td></td><td></td><td></td></tr>
<tr><td>Chromosome</td><td>start</td><td>end</td><td>strand</td><td>MM</td><td>target_seq</td><td>PAM</td><td>alignment</td><td>distance</td><td>position</td><td>gene name</td><td>gene id</td></tr>
<tr><td>GU305914.1</td><td>10666</td><td>10688</td><td>+</td><td>0</td><td>GGTAGATCACACTCTGCTCA</td><td>AGG</td><td>||||||||[||||||||||||]PAM</td><td>NA</td><td>Intergenic</td><td>NA</td><td>NA</td></tr>
<tr><td>GU305914.1</td><td>74399</td><td>74421</td><td>+</td><td>4</td><td>TGTAGATGACACTCTTCTCC</td><td>TGG</td><td>-||||||-[|||||||-|||-]PAM</td><td>NA</td><td>Intergenic</td><td>NA</td><td>NA</td></tr>
<tr><td></td><td></td><td></td><td></td><td></td><td></td><td></td><td></td><td></td><td></td><td></td><td></td></tr>
<tr><td>T526</td><td>GATCACACTCTGCTCAAGGCAGG</td><td>1000</td><td></td><td></td><td></td><td></td><td></td><td></td><td></td><td></td><td></td></tr>
<tr><td>Oligo fwd</td><td>CACCGATCACACTCTGCTCAAGGC</td><td></td><td></td><td></td><td></td><td></td><td></td><td></td><td></td><td></td><td></td></tr>
</table>

| | | | | | | | | | | | |
|---|---|---|---|---|---|---|---|---|---|---|---|
| Oligo rev | AAACGCCTTGAGCAGAGTGTGATC | | | | | | | | | | |
| Chromosome | start | end | strand | MM | target_seq | PAM | alignment | distance | position | gene name | gene id |
| GU305914.1 | 10670 | 10692 | + | 0 | GATCACACTCTGCTCAAGGC | AGG | \|\|\|\|\|\|\|\|[\|\|\|\|\|\|\|\|\|\|\|\|]PAM | NA | Intergenic | NA | NA |
| T527 | GATCCCGTACAGCACCGAGATGG | 1000 | | | | | | | | | |
| Oligo fwd | CACCGATCCCGTACAGCACCGAGA | | | | | | | | | | |
| Oligo rev | AAACTCTCGGTGCTGTACGGGATC | | | | | | | | | | |
| Chromosome | start | end | strand | MM | target_seq | PAM | alignment | distance | position | gene name | gene id |
| GU305914.1 | 10696 | 10718 | + | 0 | GATCCCGTACAGCACCGAGA | TGG | \|\|\|\|\|\|\|\|[\|\|\|\|\|\|\|\|\|\|\|\|]PAM | NA | Intergenic | NA | NA |
| GU305914.1 | 33729 | 33751 | + | 4 | GTAGCCGTACAGCAGCGAGA | TGG | \|---\|\|\|\|[\|\|\|\|\|\|-\|\|\|\|\|]PAM | NA | Intergenic | NA | NA |
| T528 | CGAGATGGCTCACCGCTACCTGG | 1000 | | | | | | | | | |
| Oligo fwd | CACCCGAGATGGCTCACCGCTACC | | | | | | | | | | |
| Oligo rev | AAACGGTAGCGGTGAGCCATCTCG | | | | | | | | | | |
| Chromosome | start | end | strand | MM | target_seq | PAM | alignment | distance | position | gene name | gene id |
| GU305914.1 | 10711 | 10733 | + | 0 | CGAGATGGCTCACCGCTACC | TGG | \|\|\|\|\|\|\|\|[\|\|\|\|\|\|\|\|\|\|\|\|]PAM | NA | Intergenic | NA | NA |
| T529 | GAGATGGCTCACCGCTACCTGGG | 1000 | | | | | | | | | |
| Oligo fwd | CACCGAGATGGCTCACCGCTACCT | | | | | | | | | | |
| Oligo rev | AAACAGGTAGCGGTGAGCCATCTC | | | | | | | | | | |
| Chromosome | start | end | strand | MM | target_seq | PAM | alignment | distance | position | gene name | gene id |
| GU305914.1 | 10712 | 10734 | + | 0 | GAGATGGCTCACCGCTACCT | GGG | \|\|\|\|\|\|\|\|[\|\|\|\|\|\|\|\|\|\|\|\|]PAM | NA | Intergenic | NA | NA |
| T530 | AGATGGCTCACCGCTACCTGGGG | 1000 | | | | | | | | | |
| Oligo fwd | CACCAGATGGCTCACCGCTACCTG | | | | | | | | | | |
| Oligo rev | AAACCAGGTAGCGGTGAGCCATCT | | | | | | | | | | |
| Chromosome | start | end | strand | MM | target_seq | PAM | alignment | distance | position | gene name | gene id |
| GU305914.1 | 10713 | 10735 | + | 0 | AGATGGCTCACCGCTACCTG | GGG | \|\|\|\|\|\|\|\|[\|\|\|\|\|\|\|\|\|\|\|\|]PAM | NA | Intergenic | NA | NA |
| T531 | GTCGATATCGCGAGTTACACCGG | 1000 | | | | | | | | | |
| Oligo fwd | CACCGTCGATATCGCGAGTTACAC | | | | | | | | | | |
| Oligo rev | AAACGTGTAACTCGCGATATCGAC | | | | | | | | | | |
| Chromosome | start | end | strand | MM | target_seq | PAM | alignment | distance | position | gene name | gene id |
| GU305914.1 | 10774 | 10796 | + | 0 | GTCGATATCGCGAGTTACAC | CGG | \|\|\|\|\|\|\|\|[\|\|\|\|\|\|\|\|\|\|\|\|]PAM | NA | Intergenic | NA | NA |
| T532 | GATATCGCGAGTTACACCGGCGG | 1000 | | | | | | | | | |
| Oligo fwd | CACCGATATCGCGAGTTACACCGG | | | | | | | | | | |
| Oligo rev | AAACCCGGTGTAACTCGCGATATC | | | | | | | | | | |
| Chromosome | start | end | strand | MM | target_seq | PAM | alignment | distance | position | gene name | gene id |
| GU305914.1 | 10777 | 10799 | + | 0 | GATATCGCGAGTTACACCGG | CGG | \|\|\|\|\|\|\|\|[\|\|\|\|\|\|\|\|\|\|\|\|]PAM | NA | Intergenic | NA | NA |
| T533 | CTGTACGAAGTTTTCATATCCGG | 1000 | | | | | | | | | |
| Oligo fwd | CACCCTGTACGAAGTTTTCATATC | | | | | | | | | | |
| Oligo rev | AAACGATATGAAAACTTCGTACAG | | | | | | | | | | |
| Chromosome | start | end | strand | MM | target_seq | PAM | alignment | distance | position | gene name | gene id |
| GU305914.1 | 10828 | 10850 | + | 0 | CTGTACGAAGTTTTCATATC | CGG | \|\|\|\|\|\|\|\|[\|\|\|\|\|\|\|\|\|\|\|\|]PAM | NA | Intergenic | NA | NA |
| T534 | TGTACGAAGTTTTCATATCCGGG | 1000 | | | | | | | | | |
| Oligo fwd | CACCTGTACGAAGTTTTCATATCC | | | | | | | | | | |
| Oligo rev | AAACGGATATGAAAACTTCGTACA | | | | | | | | | | |
| Chromosome | start | end | strand | MM | target_seq | PAM | alignment | distance | position | gene name | gene id |
| GU305914.1 | 10829 | 10851 | + | 0 | TGTACGAAGTTTTCATATCC | GGG | \|\|\|\|\|\|\|\|[\|\|\|\|\|\|\|\|\|\|\|\|]PAM | NA | Intergenic | NA | NA |
| T535 | TTCATATCCGGGCGTCAGTCCGG | 1000 | | | | | | | | | |
| Oligo fwd | CACCTTCATATCCGGGCGTCAGTC | | | | | | | | | | |
| Oligo rev | AAACGACTGACGCCCGGATATGAA | | | | | | | | | | |
| Chromosome | start | end | strand | MM | target_seq | PAM | alignment | distance | position | gene name | gene id |
| GU305914.1 | 10840 | 10862 | + | 0 | TTCATATCCGGGCGTCAGTC | CGG | \|\|\|\|\|\|\|\|[\|\|\|\|\|\|\|\|\|\|\|\|]PAM | NA | Intergenic | NA | NA |
| T536 | TCATATCCGGGCGTCAGTCCGGG | 1000 | | | | | | | | | |
| Oligo fwd | CACCTCATATCCGGGCGTCAGTCC | | | | | | | | | | |
| Oligo rev | AAACGGACTGACGCCCGGATATGA | | | | | | | | | | |
| Chromosome | start | end | strand | MM | target_seq | PAM | alignment | distance | position | gene name | gene id |
| GU305914.1 | 10841 | 10863 | + | 0 | TCATATCCGGGCGTCAGTCC | GGG | \|\|\|\|\|\|\|\|[\|\|\|\|\|\|\|\|\|\|\|\|]PAM | NA | Intergenic | NA | NA |
| T537 | CGGGATACTGTACAGAAAGACGG | 1000 | | | | | | | | | |
| Oligo fwd | CACCCGGGATACTGTACAGAAAGA | | | | | | | | | | |
| Oligo rev | AAACTCTTTCTGTACAGTATCCCG | | | | | | | | | | |
| Chromosome | start | end | strand | MM | target_seq | PAM | alignment | distance | position | gene name | gene id |
| GU305914.1 | 10860 | 10882 | + | 0 | CGGGATACTGTACAGAAAGA | CGG | \|\|\|\|\|\|\|\|[\|\|\|\|\|\|\|\|\|\|\|\|]PAM | NA | Intergenic | NA | NA |
| T538 | GGGATACTGTACAGAAAGACGGG | 1000 | | | | | | | | | |
| Oligo fwd | CACCGGGATACTGTACAGAAAGAC | | | | | | | | | | |
| Oligo rev | AAACGTCTTTCTGTACAGTATCCC | | | | | | | | | | |
| Chromosome | start | end | strand | MM | target_seq | PAM | alignment | distance | position | gene name | gene id |
| GU305914.1 | 10861 | 10883 | + | 0 | GGGATACTGTACAGAAAGAC | GGG | \|\|\|\|\|\|\|\|[\|\|\|\|\|\|\|\|\|\|\|\|]PAM | NA | Intergenic | NA | NA |
| T539 | TTGATCACTAAGTTTAAAAACGG | 1000 | | | | | | | | | |
| Oligo fwd | CACCTTGATCACTAAGTTTAAAAA | | | | | | | | | | |
| Oligo rev | AAACTTTTTAAACTTAGTGATCAA | | | | | | | | | | |
| Chromosome | start | end | strand | MM | target_seq | PAM | alignment | distance | position | gene name | gene id |
| GU305914.1 | 10892 | 10914 | + | 0 | TTGATCACTAAGTTTAAAAA | CGG | \|\|\|\|\|\|\|\|[\|\|\|\|\|\|\|\|\|\|\|\|]PAM | NA | Intergenic | NA | NA |
| T540 | TAAGTTTAAAAACGGCACGAAGG | 1000 | | | | | | | | | |
| Oligo fwd | CACCTAAGTTTAAAAACGGCACGA | | | | | | | | | | |
| Oligo rev | AAACTCGTGCCGTTTTTAAACTTA | | | | | | | | | | |
| Chromosome | start | end | strand | MM | target_seq | PAM | alignment | distance | position | gene name | gene id |
| GU305914.1 | 10900 | 10922 | + | 0 | TAAGTTTAAAAACGGCACGA | AGG | \|\|\|\|\|\|\|\|[\|\|\|\|\|\|\|\|\|\|\|\|]PAM | NA | Intergenic | NA | NA |

<table>
<tr><td>T541</td><td>TGTGTCAACGCGCGAATGTCCGG</td><td></td><td>1000</td></tr>
<tr><td>Oligo fwd</td><td>CACCTGTGTCAACGCGCGAATGTC</td></tr>
<tr><td>Oligo rev</td><td>AAACGACATTCGCGCGTTGACACA</td></tr>
<tr><td>Chromosome</td><td>start</td><td></td><td>end</td><td>strand</td><td>MM</td><td>target_seq</td><td>PAM</td><td>alignment</td><td>distance</td><td>position</td><td>gene name</td><td>gene id</td></tr>
<tr><td>GU305914.1</td><td></td><td>10952</td><td>10974</td><td>+</td><td>0</td><td>TGTGTCAACGCGCGAATGTC</td><td>CGG</td><td>||||||||[||||||||||||]PAM</td><td>NA</td><td>Intergenic</td><td>NA</td><td>NA</td></tr>
<tr><td>T542</td><td>ACGCGCGAATGTCCGGTGTCTGG</td><td></td><td>1000</td></tr>
<tr><td>Oligo fwd</td><td>CACCACGCGCGAATGTCCGGTGTC</td></tr>
<tr><td>Oligo rev</td><td>AAACGACACCGGACATTCGCGCGT</td></tr>
<tr><td>Chromosome</td><td>start</td><td></td><td>end</td><td>strand</td><td>MM</td><td>target_seq</td><td>PAM</td><td>alignment</td><td>distance</td><td>position</td><td>gene name</td><td>gene id</td></tr>
<tr><td>GU305914.1</td><td></td><td>10959</td><td>10981</td><td>+</td><td>0</td><td>ACGCGCGAATGTCCGGTGTC</td><td>TGG</td><td>||||||||[||||||||||||]PAM</td><td>NA</td><td>Intergenic</td><td>NA</td><td>NA</td></tr>
<tr><td>T543</td><td>TGCGTAAATAATAGCGCCATCGG</td><td></td><td>1000</td></tr>
<tr><td>Oligo fwd</td><td>CACCTGCGTAAATAATAGCGCCAT</td></tr>
<tr><td>Oligo rev</td><td>AAACATGGCGCTATTATTTACGCA</td></tr>
<tr><td>Chromosome</td><td>start</td><td></td><td>end</td><td>strand</td><td>MM</td><td>target_seq</td><td>PAM</td><td>alignment</td><td>distance</td><td>position</td><td>gene name</td><td>gene id</td></tr>
<tr><td>GU305914.1</td><td></td><td>10982</td><td>11004</td><td>+</td><td>0</td><td>TGCGTAAATAATAGCGCCAT</td><td>CGG</td><td>||||||||[||||||||||||]PAM</td><td>NA</td><td>Intergenic</td><td>NA</td><td>NA</td></tr>
<tr><td>T544</td><td>CATACTCCTCCTATGTTTCATGG</td><td></td><td>1000</td></tr>
<tr><td>Oligo fwd</td><td>CACCCATACTCCTCCTATGTTTCA</td></tr>
<tr><td>Oligo rev</td><td>AAACTGAAACATAGGAGGAGTATG</td></tr>
<tr><td>Chromosome</td><td>start</td><td></td><td>end</td><td>strand</td><td>MM</td><td>target_seq</td><td>PAM</td><td>alignment</td><td>distance</td><td>position</td><td>gene name</td><td>gene id</td></tr>
<tr><td>GU305914.1</td><td></td><td>11030</td><td>11052</td><td>+</td><td>0</td><td>CATACTCCTCCTATGTTTCA</td><td>TGG</td><td>||||||||[||||||||||||]PAM</td><td>NA</td><td>Intergenic</td><td>NA</td><td>NA</td></tr>
<tr><td>T545</td><td>ACACATCACCGCCGATTATCTGG</td><td></td><td>1000</td></tr>
<tr><td>Oligo fwd</td><td>CACCACACATCACCGCCGATTATC</td></tr>
<tr><td>Oligo rev</td><td>AAACGATAATCGGCGGTGATGTGT</td></tr>
<tr><td>Chromosome</td><td>start</td><td></td><td>end</td><td>strand</td><td>MM</td><td>target_seq</td><td>PAM</td><td>alignment</td><td>distance</td><td>position</td><td>gene name</td><td>gene id</td></tr>
<tr><td>GU305914.1</td><td></td><td>11063</td><td>11085</td><td>+</td><td>0</td><td>ACACATCACCGCCGATTATC</td><td>TGG</td><td>||||||||[||||||||||||]PAM</td><td>NA</td><td>Intergenic</td><td>NA</td><td>NA</td></tr>
<tr><td>T546</td><td>ACCGCCGATTATCTGGTTTATGG</td><td></td><td>1000</td></tr>
<tr><td>Oligo fwd</td><td>CACCACCGCCGATTATCTGGTTTA</td></tr>
<tr><td>Oligo rev</td><td>AAACTAAACCAGATAATCGGCGGT</td></tr>
<tr><td>Chromosome</td><td>start</td><td></td><td>end</td><td>strand</td><td>MM</td><td>target_seq</td><td>PAM</td><td>alignment</td><td>distance</td><td>position</td><td>gene name</td><td>gene id</td></tr>
<tr><td>GU305914.1</td><td></td><td>11070</td><td>11092</td><td>+</td><td>0</td><td>ACCGCCGATTATCTGGTTTA</td><td>TGG</td><td>||||||||[||||||||||||]PAM</td><td>NA</td><td>Intergenic</td><td>NA</td><td>NA</td></tr>
<tr><td>T547</td><td>CCGCCGATTATCTGGTTTATGGG</td><td></td><td>1000</td></tr>
<tr><td>Oligo fwd</td><td>CACCCCGCCGATTATCTGGTTTAT</td></tr>
<tr><td>Oligo rev</td><td>AAACATAAACCAGATAATCGGCGG</td></tr>
<tr><td>Chromosome</td><td>start</td><td></td><td>end</td><td>strand</td><td>MM</td><td>target_seq</td><td>PAM</td><td>alignment</td><td>distance</td><td>position</td><td>gene name</td><td>gene id</td></tr>
<tr><td>GU305914.1</td><td></td><td>11071</td><td>11093</td><td>+</td><td>0</td><td>CCGCCGATTATCTGGTTTAT</td><td>GGG</td><td>||||||||[||||||||||||]PAM</td><td>NA</td><td>Intergenic</td><td>NA</td><td>NA</td></tr>
<tr><td>T548</td><td>TTATGGGTCGATTATCAAACCGG</td><td></td><td>1000</td></tr>
<tr><td>Oligo fwd</td><td>CACCTTATGGGTCGATTATCAAAC</td></tr>
<tr><td>Oligo rev</td><td>AAACGTTTGATAATCGACCCATAA</td></tr>
<tr><td>Chromosome</td><td>start</td><td></td><td>end</td><td>strand</td><td>MM</td><td>target_seq</td><td>PAM</td><td>alignment</td><td>distance</td><td>position</td><td>gene name</td><td>gene id</td></tr>
<tr><td>GU305914.1</td><td></td><td>11087</td><td>11109</td><td>+</td><td>0</td><td>TTATGGGTCGATTATCAAAC</td><td>CGG</td><td>||||||||[||||||||||||]PAM</td><td>NA</td><td>Intergenic</td><td>NA</td><td>NA</td></tr>
<tr><td>T549</td><td>CCCTAACGCCACGAAGAAACCGG</td><td></td><td>1000</td></tr>
<tr><td>Oligo fwd</td><td>CACCCCCTAACGCCACGAAGAAAC</td></tr>
<tr><td>Oligo rev</td><td>AAACGTTTCTTCGTGGCGTTAGGG</td></tr>
<tr><td>Chromosome</td><td>start</td><td></td><td>end</td><td>strand</td><td>MM</td><td>target_seq</td><td>PAM</td><td>alignment</td><td>distance</td><td>position</td><td>gene name</td><td>gene id</td></tr>
<tr><td>GU305914.1</td><td></td><td>11125</td><td>11147</td><td>+</td><td>0</td><td>CCCTAACGCCACGAAGAAAC</td><td>CGG</td><td>||||||||[||||||||||||]PAM</td><td>NA</td><td>Intergenic</td><td>NA</td><td>NA</td></tr>
<tr><td>T550</td><td>ACTACTACACGATCGTCGAGAGG</td><td></td><td>1000</td></tr>
<tr><td>Oligo fwd</td><td>CACCACTACTACACGATCGTCGAG</td></tr>
<tr><td>Oligo rev</td><td>AAACCTCGACGATCGTGTAGTAGT</td></tr>
<tr><td>Chromosome</td><td>start</td><td></td><td>end</td><td>strand</td><td>MM</td><td>target_seq</td><td>PAM</td><td>alignment</td><td>distance</td><td>position</td><td>gene name</td><td>gene id</td></tr>
<tr><td>GU305914.1</td><td></td><td>11207</td><td>11229</td><td>+</td><td>0</td><td>ACTACTACACGATCGTCGAG</td><td>AGG</td><td>||||||||[||||||||||||]PAM</td><td>NA</td><td>Intergenic</td><td>NA</td><td>NA</td></tr>
<tr><td>T551</td><td>ATCGTCGAGAGGAGTATCGACGG</td><td></td><td>1000</td></tr>
<tr><td>Oligo fwd</td><td>CACCATCGTCGAGAGGAGTATCGA</td></tr>
<tr><td>Oligo rev</td><td>AAACTCGATACTCCTCTCGACGAT</td></tr>
<tr><td>Chromosome</td><td>start</td><td></td><td>end</td><td>strand</td><td>MM</td><td>target_seq</td><td>PAM</td><td>alignment</td><td>distance</td><td>position</td><td>gene name</td><td>gene id</td></tr>
<tr><td>GU305914.1</td><td></td><td>11218</td><td>11240</td><td>+</td><td>0</td><td>ATCGTCGAGAGGAGTATCGA</td><td>CGG</td><td>||||||||[||||||||||||]PAM</td><td>NA</td><td>Intergenic</td><td>NA</td><td>NA</td></tr>
<tr><td>T552</td><td>TCGTCGAGAGGAGTATCGACGGG</td><td></td><td>1000</td></tr>
<tr><td>Oligo fwd</td><td>CACCTCGTCGAGAGGAGTATCGAC</td></tr>
<tr><td>Oligo rev</td><td>AAACGTCGATACTCCTCTCGACGA</td></tr>
<tr><td>Chromosome</td><td>start</td><td></td><td>end</td><td>strand</td><td>MM</td><td>target_seq</td><td>PAM</td><td>alignment</td><td>distance</td><td>position</td><td>gene name</td><td>gene id</td></tr>
<tr><td>GU305914.1</td><td></td><td>11219</td><td>11241</td><td>+</td><td>0</td><td>TCGTCGAGAGGAGTATCGAC</td><td>GGG</td><td>||||||||[||||||||||||]PAM</td><td>NA</td><td>Intergenic</td><td>NA</td><td>NA</td></tr>
<tr><td>T553</td><td>CGTCGAGAGGAGTATCGACGGGG</td><td></td><td>1000</td></tr>
<tr><td>Oligo fwd</td><td>CACCCGTCGAGAGGAGTATCGACG</td></tr>
<tr><td>Oligo rev</td><td>AAACCGTCGATACTCCTCTCGACG</td></tr>
<tr><td>Chromosome</td><td>start</td><td></td><td>end</td><td>strand</td><td>MM</td><td>target_seq</td><td>PAM</td><td>alignment</td><td>distance</td><td>position</td><td>gene name</td><td>gene id</td></tr>
<tr><td>GU305914.1</td><td></td><td>11220</td><td>11242</td><td>+</td><td>0</td><td>CGTCGAGAGGAGTATCGACG</td><td>GGG</td><td>||||||||[||||||||||||]PAM</td><td>NA</td><td>Intergenic</td><td>NA</td><td>NA</td></tr>
<tr><td>T554</td><td>TCGACGGGGATTTCGACGACAGG</td><td></td><td>1000</td></tr>
<tr><td>Oligo fwd</td><td>CACCTCGACGGGGATTTCGACGAC</td></tr>
<tr><td>Oligo rev</td><td>AAACGTCGTCGAAATCCCCGTCGA</td></tr>
<tr><td>Chromosome</td><td>start</td><td></td><td>end</td><td>strand</td><td>MM</td><td>target_seq</td><td>PAM</td><td>alignment</td><td>distance</td><td>position</td><td>gene name</td><td>gene id</td></tr>
<tr><td>GU305914.1</td><td></td><td>11234</td><td>11256</td><td>+</td><td>0</td><td>TCGACGGGGATTTCGACGAC</td><td>AGG</td><td>||||||||[||||||||||||]PAM</td><td>NA</td><td>Intergenic</td><td>NA</td><td>NA</td></tr>
<tr><td>T555</td><td>GGATTTCGACGACAGGACTATGG</td><td></td><td>1000</td></tr>
<tr><td>Oligo fwd</td><td>CACCGGATTTCGACGACAGGACTA</td></tr>
<tr><td>Oligo rev</td><td>AAACTAGTCCTGTCGTCGAAATCC</td></tr>
<tr><td>Chromosome</td><td>start</td><td></td><td>end</td><td>strand</td><td>MM</td><td>target_seq</td><td>PAM</td><td>alignment</td><td>distance</td><td>position</td><td>gene name</td><td>gene id</td></tr>
<tr><td>GU305914.1</td><td></td><td>11241</td><td>11263</td><td>+</td><td>0</td><td>GGATTTCGACGACAGGACTA</td><td>TGG</td><td>||||||||[||||||||||||]PAM</td><td>NA</td><td>Intergenic</td><td>NA</td><td>NA</td></tr>
</table>

<table>
<tr><td>T556</td><td>CGTCTTTCCTTTTGTATCCGCGG</td><td>1000</td><td></td><td></td><td></td><td></td><td></td><td></td><td></td><td></td><td></td></tr>
<tr><td>Oligo fwd</td><td>CACCCGTCTTTCCTTTTGTATCCG</td><td></td><td></td><td></td><td></td><td></td><td></td><td></td><td></td><td></td><td></td></tr>
<tr><td>Oligo rev</td><td>AAACCGGATACAAAAGGAAAGACG</td><td></td><td></td><td></td><td></td><td></td><td></td><td></td><td></td><td></td><td></td></tr>
<tr><td>Chromosome</td><td>start</td><td>end</td><td>strand</td><td>MM</td><td>target_seq</td><td>PAM</td><td>alignment</td><td>distance</td><td>position</td><td>gene name</td><td>gene id</td></tr>
<tr><td>GU305914.1</td><td>11281</td><td>11303</td><td>+</td><td>0</td><td>CGTCTTTCCTTTTGTATCCG</td><td>CGG</td><td>||||||||[||||||||||||]PAM</td><td>NA</td><td>Intergenic</td><td>NA</td><td>NA</td></tr>
<tr><td>T557</td><td>GTCTTTCCTTTTGTATCCGCGGG</td><td>1000</td><td></td><td></td><td></td><td></td><td></td><td></td><td></td><td></td><td></td></tr>
<tr><td>Oligo fwd</td><td>CACCGTCTTTCCTTTTGTATCCGC</td><td></td><td></td><td></td><td></td><td></td><td></td><td></td><td></td><td></td><td></td></tr>
<tr><td>Oligo rev</td><td>AAACGCGGATACAAAAGGAAAGAC</td><td></td><td></td><td></td><td></td><td></td><td></td><td></td><td></td><td></td><td></td></tr>
<tr><td>Chromosome</td><td>start</td><td>end</td><td>strand</td><td>MM</td><td>target_seq</td><td>PAM</td><td>alignment</td><td>distance</td><td>position</td><td>gene name</td><td>gene id</td></tr>
<tr><td>GU305914.1</td><td>11282</td><td>11304</td><td>+</td><td>0</td><td>GTCTTTCCTTTTGTATCCGC</td><td>GGG</td><td>||||||||[||||||||||||]PAM</td><td>NA</td><td>Intergenic</td><td>NA</td><td>NA</td></tr>
<tr><td>T558</td><td>GTATCCGCGGGTGAAAACTTCGG</td><td>1000</td><td></td><td></td><td></td><td></td><td></td><td></td><td></td><td></td><td></td></tr>
<tr><td>Oligo fwd</td><td>CACCGTATCCGCGGGTGAAAACTT</td><td></td><td></td><td></td><td></td><td></td><td></td><td></td><td></td><td></td><td></td></tr>
<tr><td>Oligo rev</td><td>AAACAAGTTTTCACCCGCGGATAC</td><td></td><td></td><td></td><td></td><td></td><td></td><td></td><td></td><td></td><td></td></tr>
<tr><td>Chromosome</td><td>start</td><td>end</td><td>strand</td><td>MM</td><td>target_seq</td><td>PAM</td><td>alignment</td><td>distance</td><td>position</td><td>gene name</td><td>gene id</td></tr>
<tr><td>GU305914.1</td><td>11294</td><td>11316</td><td>+</td><td>0</td><td>GTATCCGCGGGTGAAAACTT</td><td>CGG</td><td>||||||||[||||||||||||]PAM</td><td>NA</td><td>Intergenic</td><td>NA</td><td>NA</td></tr>
<tr><td>T559</td><td>TATCCGCGGGTGAAAACTTCGGG</td><td>1000</td><td></td><td></td><td></td><td></td><td></td><td></td><td></td><td></td><td></td></tr>
<tr><td>Oligo fwd</td><td>CACCTATCCGCGGGTGAAAACTTC</td><td></td><td></td><td></td><td></td><td></td><td></td><td></td><td></td><td></td><td></td></tr>
<tr><td>Oligo rev</td><td>AAACGAAGTTTTCACCCGCGGATA</td><td></td><td></td><td></td><td></td><td></td><td></td><td></td><td></td><td></td><td></td></tr>
<tr><td>Chromosome</td><td>start</td><td>end</td><td>strand</td><td>MM</td><td>target_seq</td><td>PAM</td><td>alignment</td><td>distance</td><td>position</td><td>gene name</td><td>gene id</td></tr>
<tr><td>GU305914.1</td><td>11295</td><td>11317</td><td>+</td><td>0</td><td>TATCCGCGGGTGAAAACTTC</td><td>GGG</td><td>||||||||[||||||||||||]PAM</td><td>NA</td><td>Intergenic</td><td>NA</td><td>NA</td></tr>
<tr><td>T560</td><td>ATCCGCGGGTGAAAACTTCGGGG</td><td>1000</td><td></td><td></td><td></td><td></td><td></td><td></td><td></td><td></td><td></td></tr>
<tr><td>Oligo fwd</td><td>CACCATCCGCGGGTGAAAACTTCG</td><td></td><td></td><td></td><td></td><td></td><td></td><td></td><td></td><td></td><td></td></tr>
<tr><td>Oligo rev</td><td>AAACCGAAGTTTTCACCCGCGGAT</td><td></td><td></td><td></td><td></td><td></td><td></td><td></td><td></td><td></td><td></td></tr>
<tr><td>Chromosome</td><td>start</td><td>end</td><td>strand</td><td>MM</td><td>target_seq</td><td>PAM</td><td>alignment</td><td>distance</td><td>position</td><td>gene name</td><td>gene id</td></tr>
<tr><td>GU305914.1</td><td>11296</td><td>11318</td><td>+</td><td>0</td><td>ATCCGCGGGTGAAAACTTCG</td><td>GGG</td><td>||||||||[||||||||||||]PAM</td><td>NA</td><td>Intergenic</td><td>NA</td><td>NA</td></tr>
<tr><td>T561</td><td>TCGCCACAGACAGCAAGTAGCGG</td><td>1000</td><td></td><td></td><td></td><td></td><td></td><td></td><td></td><td></td><td></td></tr>
<tr><td>Oligo fwd</td><td>CACCTCGCCACAGACAGCAAGTAG</td><td></td><td></td><td></td><td></td><td></td><td></td><td></td><td></td><td></td><td></td></tr>
<tr><td>Oligo rev</td><td>AAACCTACTTGCTGTCTGTGGCGA</td><td></td><td></td><td></td><td></td><td></td><td></td><td></td><td></td><td></td><td></td></tr>
<tr><td>Chromosome</td><td>start</td><td>end</td><td>strand</td><td>MM</td><td>target_seq</td><td>PAM</td><td>alignment</td><td>distance</td><td>position</td><td>gene name</td><td>gene id</td></tr>
<tr><td>GU305914.1</td><td>11327</td><td>11349</td><td>+</td><td>0</td><td>TCGCCACAGACAGCAAGTAG</td><td>CGG</td><td>||||||||[||||||||||||]PAM</td><td>NA</td><td>Intergenic</td><td>NA</td><td>NA</td></tr>
<tr><td>T562</td><td>ATCGTAATAGTCATCTGCGTCGG</td><td>1000</td><td></td><td></td><td></td><td></td><td></td><td></td><td></td><td></td><td></td></tr>
<tr><td>Oligo fwd</td><td>CACCATCGTAATAGTCATCTGCGT</td><td></td><td></td><td></td><td></td><td></td><td></td><td></td><td></td><td></td><td></td></tr>
<tr><td>Oligo rev</td><td>AAACACGCAGATGACTATTACGAT</td><td></td><td></td><td></td><td></td><td></td><td></td><td></td><td></td><td></td><td></td></tr>
<tr><td>Chromosome</td><td>start</td><td>end</td><td>strand</td><td>MM</td><td>target_seq</td><td>PAM</td><td>alignment</td><td>distance</td><td>position</td><td>gene name</td><td>gene id</td></tr>
<tr><td>GU305914.1</td><td>11369</td><td>11391</td><td>+</td><td>0</td><td>ATCGTAATAGTCATCTGCGT</td><td>CGG</td><td>||||||||[||||||||||||]PAM</td><td>NA</td><td>Intergenic</td><td>NA</td><td>NA</td></tr>
<tr><td>T563</td><td>CTGCGTCGGATTAGTCGTTCTGG</td><td>1000</td><td></td><td></td><td></td><td></td><td></td><td></td><td></td><td></td><td></td></tr>
<tr><td>Oligo fwd</td><td>CACCCTGCGTCGGATTAGTCGTTC</td><td></td><td></td><td></td><td></td><td></td><td></td><td></td><td></td><td></td><td></td></tr>
<tr><td>Oligo rev</td><td>AAACGAACGACTAATCCGACGCAG</td><td></td><td></td><td></td><td></td><td></td><td></td><td></td><td></td><td></td><td></td></tr>
<tr><td>Chromosome</td><td>start</td><td>end</td><td>strand</td><td>MM</td><td>target_seq</td><td>PAM</td><td>alignment</td><td>distance</td><td>position</td><td>gene name</td><td>gene id</td></tr>
<tr><td>GU305914.1</td><td>11383</td><td>11405</td><td>+</td><td>0</td><td>CTGCGTCGGATTAGTCGTTC</td><td>TGG</td><td>||||||||[||||||||||||]PAM</td><td>NA</td><td>Intergenic</td><td>NA</td><td>NA</td></tr>
<tr><td>T564</td><td>CGTCGGATTAGTCGTTCTGGCGG</td><td>1000</td><td></td><td></td><td></td><td></td><td></td><td></td><td></td><td></td><td></td></tr>
<tr><td>Oligo fwd</td><td>CACCCGTCGGATTAGTCGTTCTGG</td><td></td><td></td><td></td><td></td><td></td><td></td><td></td><td></td><td></td><td></td></tr>
<tr><td>Oligo rev</td><td>AAACCCAGAACGACTAATCCGACG</td><td></td><td></td><td></td><td></td><td></td><td></td><td></td><td></td><td></td><td></td></tr>
<tr><td>Chromosome</td><td>start</td><td>end</td><td>strand</td><td>MM</td><td>target_seq</td><td>PAM</td><td>alignment</td><td>distance</td><td>position</td><td>gene name</td><td>gene id</td></tr>
<tr><td>GU305914.1</td><td>11386</td><td>11408</td><td>+</td><td>0</td><td>CGTCGGATTAGTCGTTCTGG</td><td>CGG</td><td>||||||||[||||||||||||]PAM</td><td>NA</td><td>Intergenic</td><td>NA</td><td>NA</td></tr>
<tr><td>T565</td><td>CTGGCGGTGTTGTATTATCGCGG</td><td>1000</td><td></td><td></td><td></td><td></td><td></td><td></td><td></td><td></td><td></td></tr>
<tr><td>Oligo fwd</td><td>CACCCTGGCGGTGTTGTATTATCG</td><td></td><td></td><td></td><td></td><td></td><td></td><td></td><td></td><td></td><td></td></tr>
<tr><td>Oligo rev</td><td>AAACCGATAATACAACACCGCCAG</td><td></td><td></td><td></td><td></td><td></td><td></td><td></td><td></td><td></td><td></td></tr>
<tr><td>Chromosome</td><td>start</td><td>end</td><td>strand</td><td>MM</td><td>target_seq</td><td>PAM</td><td>alignment</td><td>distance</td><td>position</td><td>gene name</td><td>gene id</td></tr>
<tr><td>GU305914.1</td><td>11402</td><td>11424</td><td>+</td><td>0</td><td>CTGGCGGTGTTGTATTATCG</td><td>CGG</td><td>||||||||[||||||||||||]PAM</td><td>NA</td><td>Intergenic</td><td>NA</td><td>NA</td></tr>
<tr><td>T566</td><td>TTGTATTATCGCGGTTACATCGG</td><td>1000</td><td></td><td></td><td></td><td></td><td></td><td></td><td></td><td></td><td></td></tr>
<tr><td>Oligo fwd</td><td>CACCTTGTATTATCGCGGTTACAT</td><td></td><td></td><td></td><td></td><td></td><td></td><td></td><td></td><td></td><td></td></tr>
<tr><td>Oligo rev</td><td>AAACATGTAACCGCGATAATACAA</td><td></td><td></td><td></td><td></td><td></td><td></td><td></td><td></td><td></td><td></td></tr>
<tr><td>Chromosome</td><td>start</td><td>end</td><td>strand</td><td>MM</td><td>target_seq</td><td>PAM</td><td>alignment</td><td>distance</td><td>position</td><td>gene name</td><td>gene id</td></tr>
<tr><td>GU305914.1</td><td>11411</td><td>11433</td><td>+</td><td>0</td><td>TTGTATTATCGCGGTTACAT</td><td>CGG</td><td>||||||||[||||||||||||]PAM</td><td>NA</td><td>Intergenic</td><td>NA</td><td>NA</td></tr>
<tr><td>T567</td><td>ATAAGCAGTATTTGTATCCCCGG</td><td>1000</td><td></td><td></td><td></td><td></td><td></td><td></td><td></td><td></td><td></td></tr>
<tr><td>Oligo fwd</td><td>CACCATAAGCAGTATTTGTATCCC</td><td></td><td></td><td></td><td></td><td></td><td></td><td></td><td></td><td></td><td></td></tr>
<tr><td>Oligo rev</td><td>AAACGGGATACAAATACTGCTTAT</td><td></td><td></td><td></td><td></td><td></td><td></td><td></td><td></td><td></td><td></td></tr>
<tr><td>Chromosome</td><td>start</td><td>end</td><td>strand</td><td>MM</td><td>target_seq</td><td>PAM</td><td>alignment</td><td>distance</td><td>position</td><td>gene name</td><td>gene id</td></tr>
<tr><td>GU305914.1</td><td>11611</td><td>11633</td><td>+</td><td>0</td><td>ATAAGCAGTATTTGTATCCC</td><td>CGG</td><td>||||||||[||||||||||||]PAM</td><td>NA</td><td>Intergenic</td><td>NA</td><td>NA</td></tr>
<tr><td>T568</td><td>TGTTCATGTCCACTTTTAGATGG</td><td>1000</td><td></td><td></td><td></td><td></td><td></td><td></td><td></td><td></td><td></td></tr>
<tr><td>Oligo fwd</td><td>CACCTGTTCATGTCCACTTTTAGA</td><td></td><td></td><td></td><td></td><td></td><td></td><td></td><td></td><td></td><td></td></tr>
<tr><td>Oligo rev</td><td>AAACTCTAAAAGTGGACATGAACA</td><td></td><td></td><td></td><td></td><td></td><td></td><td></td><td></td><td></td><td></td></tr>
<tr><td>Chromosome</td><td>start</td><td>end</td><td>strand</td><td>MM</td><td>target_seq</td><td>PAM</td><td>alignment</td><td>distance</td><td>position</td><td>gene name</td><td>gene id</td></tr>
<tr><td>GU305914.1</td><td>11681</td><td>11703</td><td>+</td><td>0</td><td>TGTTCATGTCCACTTTTAGA</td><td>TGG</td><td>||||||||[||||||||||||]PAM</td><td>NA</td><td>Intergenic</td><td>NA</td><td>NA</td></tr>
<tr><td>T569</td><td>TCCTCACATTACACTCAACTCGG</td><td>1000</td><td></td><td></td><td></td><td></td><td></td><td></td><td></td><td></td><td></td></tr>
<tr><td>Oligo fwd</td><td>CACCTCCTCACATTACACTCAACT</td><td></td><td></td><td></td><td></td><td></td><td></td><td></td><td></td><td></td><td></td></tr>
<tr><td>Oligo rev</td><td>AAACAGTTGAGTGTAATGTGAGGA</td><td></td><td></td><td></td><td></td><td></td><td></td><td></td><td></td><td></td><td></td></tr>
<tr><td>Chromosome</td><td>start</td><td>end</td><td>strand</td><td>MM</td><td>target_seq</td><td>PAM</td><td>alignment</td><td>distance</td><td>position</td><td>gene name</td><td>gene id</td></tr>
<tr><td>GU305914.1</td><td>11735</td><td>11757</td><td>+</td><td>0</td><td>TCCTCACATTACACTCAACT</td><td>CGG</td><td>||||||||[||||||||||||]PAM</td><td>NA</td><td>Intergenic</td><td>NA</td><td>NA</td></tr>
<tr><td>T570</td><td>GATAACGTTACCGCTAGCACAGG</td><td>1000</td><td></td><td></td><td></td><td></td><td></td><td></td><td></td><td></td><td></td></tr>
<tr><td>Oligo fwd</td><td>CACCGATAACGTTACCGCTAGCAC</td><td></td><td></td><td></td><td></td><td></td><td></td><td></td><td></td><td></td><td></td></tr>
<tr><td>Oligo rev</td><td>AAACGTGCTAGCGGTAACGTTATC</td><td></td><td></td><td></td><td></td><td></td><td></td><td></td><td></td><td></td><td></td></tr>
<tr><td>Chromosome</td><td>start</td><td>end</td><td>strand</td><td>MM</td><td>target_seq</td><td>PAM</td><td>alignment</td><td>distance</td><td>position</td><td>gene name</td><td>gene id</td></tr>
</table>

| | | | | | | | | | | | | |
|---|---|---|---|---|---|---|---|---|---|---|---|---|
| GU305914.1 | | 11841 | 11863 | + | 0 | GATAACGTTACCGCTAGCAC | AGG | \|\|\|\|\|\|\|\|[\|\|\|\|\|\|\|\|\|\|\|\|]PAM | NA | Intergenic | NA | NA |
| T571 | AGACGTCCTCCTCCCGAACCAGG | | 1000 | | | | | | | | | |
| Oligo fwd | CACCAGACGTCCTCCTCCCGAACC | | | | | | | | | | | |
| Oligo rev | AAACGGTTCGGGAGGAGGACGTCT | | | | | | | | | | | |
| Chromosome | start | | end | strand | MM | target_seq | PAM | alignment | distance | position | gene name | gene id |
| GU305914.1 | | 11926 | 11948 | + | 0 | AGACGTCCTCCTCCCGAACC | AGG | \|\|\|\|\|\|\|\|[\|\|\|\|\|\|\|\|\|\|\|\|]PAM | NA | Intergenic | NA | NA |
| T572 | GGAAAATGCATGTCGCTGTTAGG | | 1000 | | | | | | | | | |
| Oligo fwd | CACCGGAAAATGCATGTCGCTGTT | | | | | | | | | | | |
| Oligo rev | AAACAACAGCGACATGCATTTTCC | | | | | | | | | | | |
| Chromosome | start | | end | strand | MM | target_seq | PAM | alignment | distance | position | gene name | gene id |
| GU305914.1 | | 11947 | 11969 | + | 0 | GGAAAATGCATGTCGCTGTT | AGG | \|\|\|\|\|\|\|\|[\|\|\|\|\|\|\|\|\|\|\|\|]PAM | NA | Intergenic | NA | NA |
| T573 | ACAAGCCGCGAATAAACAAGTGG | | 1000 | | | | | | | | | |
| Oligo fwd | CACCACAAGCCGCGAATAAACAAG | | | | | | | | | | | |
| Oligo rev | AAACCTTGTTTATTCGCGGCTTGT | | | | | | | | | | | |
| Chromosome | start | | end | strand | MM | target_seq | PAM | alignment | distance | position | gene name | gene id |
| GU305914.1 | | 11988 | 12010 | + | 0 | ACAAGCCGCGAATAAACAAG | TGG | \|\|\|\|\|\|\|\|[\|\|\|\|\|\|\|\|\|\|\|\|]PAM | NA | Intergenic | NA | NA |
| T574 | ACCTTTGCATGAACGCAACTTGG | | 1000 | | | | | | | | | |
| Oligo fwd | CACCACCTTTGCATGAACGCAACT | | | | | | | | | | | |
| Oligo rev | AAACAGTTGCGTTCATGCAAAGGT | | | | | | | | | | | |
| Chromosome | start | | end | strand | MM | target_seq | PAM | alignment | distance | position | gene name | gene id |
| GU305914.1 | | 12059 | 12081 | + | 0 | ACCTTTGCATGAACGCAACT | TGG | \|\|\|\|\|\|\|\|[\|\|\|\|\|\|\|\|\|\|\|\|]PAM | NA | Intergenic | NA | NA |
| T575 | TGCATGAACGCAACTTGGTCCGG | | 1000 | | | | | | | | | |
| Oligo fwd | CACCTGCATGAACGCAACTTGGTC | | | | | | | | | | | |
| Oligo rev | AAACGACCAAGTTGCGTTCATGCA | | | | | | | | | | | |
| Chromosome | start | | end | strand | MM | target_seq | PAM | alignment | distance | position | gene name | gene id |
| GU305914.1 | | 12064 | 12086 | + | 0 | TGCATGAACGCAACTTGGTC | CGG | \|\|\|\|\|\|\|\|[\|\|\|\|\|\|\|\|\|\|\|\|]PAM | NA | Intergenic | NA | NA |
| T576 | TCGATTGTTTGTACGCACACAGG | | 1000 | | | | | | | | | |
| Oligo fwd | CACCTCGATTGTTTGTACGCACAC | | | | | | | | | | | |
| Oligo rev | AAACGTGTGCGTACAAACAATCGA | | | | | | | | | | | |
| Chromosome | start | | end | strand | MM | target_seq | PAM | alignment | distance | position | gene name | gene id |
| GU305914.1 | | 12087 | 12109 | + | 0 | TCGATTGTTTGTACGCACAC | AGG | \|\|\|\|\|\|\|\|[\|\|\|\|\|\|\|\|\|\|\|\|]PAM | NA | Intergenic | NA | NA |
| T577 | GACAACAGAGAAATAATATTCGG | | 1000 | | | | | | | | | |
| Oligo fwd | CACCGACAACAGAGAAATAATATT | | | | | | | | | | | |
| Oligo rev | AAACAATATTATTTCTCTGTTGTC | | | | | | | | | | | |
| Chromosome | start | | end | strand | MM | target_seq | PAM | alignment | distance | position | gene name | gene id |
| GU305914.1 | | 12109 | 12131 | + | 0 | GACAACAGAGAAATAATATT | CGG | \|\|\|\|\|\|\|\|[\|\|\|\|\|\|\|\|\|\|\|\|]PAM | NA | Intergenic | NA | NA |
| T578 | TCTTCTCCTACCACGTTAACCGG | | 1000 | | | | | | | | | |
| Oligo fwd | CACCTCTTCTCCTACCACGTTAAC | | | | | | | | | | | |
| Oligo rev | AAACGTTAACGTGGTAGGAGAAGA | | | | | | | | | | | |
| Chromosome | start | | end | strand | MM | target_seq | PAM | alignment | distance | position | gene name | gene id |
| GU305914.1 | | 12157 | 12179 | + | 0 | TCTTCTCCTACCACGTTAAC | CGG | \|\|\|\|\|\|\|\|[\|\|\|\|\|\|\|\|\|\|\|\|]PAM | NA | Intergenic | NA | NA |
| T579 | GAATATCTACCTTAAAACCATGG | | 1000 | | | | | | | | | |
| Oligo fwd | CACCGAATATCTACCTTAAAACCA | | | | | | | | | | | |
| Oligo rev | AAACTGGTTTTAAGGTAGATATTC | | | | | | | | | | | |
| Chromosome | start | | end | strand | MM | target_seq | PAM | alignment | distance | position | gene name | gene id |
| GU305914.1 | | 12179 | 12201 | + | 0 | GAATATCTACCTTAAAACCA | TGG | \|\|\|\|\|\|\|\|[\|\|\|\|\|\|\|\|\|\|\|\|]PAM | NA | Intergenic | NA | NA |
| T580 | GGCCTTTATCATCAGACTTGTGG | | 1000 | | | | | | | | | |
| Oligo fwd | CACCGGCCTTTATCATCAGACTTG | | | | | | | | | | | |
| Oligo rev | AAACCAAGTCTGATGATAAAGGCC | | | | | | | | | | | |
| Chromosome | start | | end | strand | MM | target_seq | PAM | alignment | distance | position | gene name | gene id |
| GU305914.1 | | 12200 | 12222 | + | 0 | GGCCTTTATCATCAGACTTG | TGG | \|\|\|\|\|\|\|\|[\|\|\|\|\|\|\|\|\|\|\|\|]PAM | NA | Intergenic | NA | NA |
| T581 | TCAGACTTGTGGATCCATTCAGG | | 1000 | | | | | | | | | |
| Oligo fwd | CACCTCAGACTTGTGGATCCATTC | | | | | | | | | | | |
| Oligo rev | AAACGAATGGATCCACAAGTCTGA | | | | | | | | | | | |
| Chromosome | start | | end | strand | MM | target_seq | PAM | alignment | distance | position | gene name | gene id |
| GU305914.1 | | 12211 | 12233 | + | 0 | TCAGACTTGTGGATCCATTC | AGG | \|\|\|\|\|\|\|\|[\|\|\|\|\|\|\|\|\|\|\|\|]PAM | NA | Intergenic | NA | NA |
| T582 | TTGTGGATCCATTCAGGTACTGG | | 1000 | | | | | | | | | |
| Oligo fwd | CACCTTGTGGATCCATTCAGGTAC | | | | | | | | | | | |
| Oligo rev | AAACGTACCTGAATGGATCCACAA | | | | | | | | | | | |
| Chromosome | start | | end | strand | MM | target_seq | PAM | alignment | distance | position | gene name | gene id |
| GU305914.1 | | 12217 | 12239 | + | 0 | TTGTGGATCCATTCAGGTAC | TGG | \|\|\|\|\|\|\|\|[\|\|\|\|\|\|\|\|\|\|\|\|]PAM | NA | Intergenic | NA | NA |
| T583 | CAGAAAAAATGCGCATATCCCGG | | 1000 | | | | | | | | | |
| Oligo fwd | CACCCAGAAAAAATGCGCATATCC | | | | | | | | | | | |
| Oligo rev | AAACGGATATGCGCATTTTTTCTG | | | | | | | | | | | |
| Chromosome | start | | end | strand | MM | target_seq | PAM | alignment | distance | position | gene name | gene id |
| GU305914.1 | | 12306 | 12328 | + | 0 | CAGAAAAAATGCGCATATCC | CGG | \|\|\|\|\|\|\|\|[\|\|\|\|\|\|\|\|\|\|\|\|]PAM | NA | Intergenic | NA | NA |
| T584 | CAACGTGCTAACTTTGCTTTCGG | | 1000 | | | | | | | | | |
| Oligo fwd | CACCCAACGTGCTAACTTTGCTTT | | | | | | | | | | | |
| Oligo rev | AAACAAAGCAAAGTTAGCACGTTG | | | | | | | | | | | |
| Chromosome | start | | end | strand | MM | target_seq | PAM | alignment | distance | position | gene name | gene id |
| GU305914.1 | | 12331 | 12353 | + | 0 | CAACGTGCTAACTTTGCTTT | CGG | \|\|\|\|\|\|\|\|[\|\|\|\|\|\|\|\|\|\|\|\|]PAM | NA | Intergenic | NA | NA |
| T585 | TAACTTTGCTTTCGGAGTCACGG | | 1000 | | | | | | | | | |
| Oligo fwd | CACCTAACTTTGCTTTCGGAGTCA | | | | | | | | | | | |
| Oligo rev | AAACTGACTCCGAAAGCAAAGTTA | | | | | | | | | | | |

Chromosome start end strand MM target_seq PAM alignment distance position gene name gene id
GU305914.1 12339 12361 + 0 TAACTTTGCTTTCGGAGTCA CGG ||||||||[||||||||||||]PAM NA Intergenic NA NA

T586 TCGACACGAACAGTAGATAACGG 1000
Oligo fwd CACCTCGACACGAACAGTAGATAA
Oligo rev AAACTTATCTACTGTTCGTGTCGA
Chromosome start end strand MM target_seq PAM alignment distance position gene name gene id
GU305914.1 12385 12407 + 0 TCGACACGAACAGTAGATAA CGG ||||||||[||||||||||||]PAM NA Intergenic NA NA

T587 CTCGTATCTGCTATCGTCGTAGG 1000
Oligo fwd CACCCTCGTATCTGCTATCGTCGT
Oligo rev AAACACGACGATAGCAGATACGAG
Chromosome start end strand MM target_seq PAM alignment distance position gene name gene id
GU305914.1 12427 12449 + 0 CTCGTATCTGCTATCGTCGT AGG ||||||||[||||||||||||]PAM NA Intergenic NA NA

T588 GCAGCGTTGTATCGCCGCAGAGG 1000
Oligo fwd CACCGCAGCGTTGTATCGCCGCAG
Oligo rev AAACCTGCGGCGATACAACGCTGC
Chromosome start end strand MM target_seq PAM alignment distance position gene name gene id
GU305914.1 12463 12485 + 0 GCAGCGTTGTATCGCCGCAG AGG ||||||||[||||||||||||]PAM NA Intergenic NA NA

T589 TTTCGACCGCTTCAAATGATTGG 1000
Oligo fwd CACCTTTCGACCGCTTCAAATGAT
Oligo rev AAACATCATTTGAAGCGGTCGAAA
Chromosome start end strand MM target_seq PAM alignment distance position gene name gene id
GU305914.1 12486 12508 + 0 TTTCGACCGCTTCAAATGAT TGG ||||||||[||||||||||||]PAM NA Intergenic NA NA

T590 TTCGACCGCTTCAAATGATTGGG 1000
Oligo fwd CACCTTCGACCGCTTCAAATGATT
Oligo rev AAACAATCATTTGAAGCGGTCGAA
Chromosome start end strand MM target_seq PAM alignment distance position gene name gene id
GU305914.1 12487 12509 + 0 TTCGACCGCTTCAAATGATT GGG ||||||||[||||||||||||]PAM NA Intergenic NA NA

T591 CATTCAGCTACATACAGACAAGG 1000
Oligo fwd CACCCATTCAGCTACATACAGACA
Oligo rev AAACTGTCTGTATGTAGCTGAATG
Chromosome start end strand MM target_seq PAM alignment distance position gene name gene id
GU305914.1 12573 12595 + 0 CATTCAGCTACATACAGACA AGG ||||||||[||||||||||||]PAM NA Intergenic NA NA

T592 ATTCAGCTACATACAGACAAGGG 1000
Oligo fwd CACCATTCAGCTACATACAGACAA
Oligo rev AAACTTGTCTGTATGTAGCTGAAT
Chromosome start end strand MM target_seq PAM alignment distance position gene name gene id
GU305914.1 12574 12596 + 0 ATTCAGCTACATACAGACAA GGG ||||||||[||||||||||||]PAM NA Intergenic NA NA

T593 ACATACAGACAAGGGACATGCGG 1000
Oligo fwd CACCACATACAGACAAGGGACATG
Oligo rev AAACCATGTCCCTTGTCTGTATGT
Chromosome start end strand MM target_seq PAM alignment distance position gene name gene id
GU305914.1 12582 12604 + 0 ACATACAGACAAGGGACATG CGG ||||||||[||||||||||||]PAM NA Intergenic NA NA

T594 GATGTCTTTGTCGAAGTCAACGG 1000
Oligo fwd CACCGATGTCTTTGTCGAAGTCAA
Oligo rev AAACTTGACTTCGACAAAGACATC
Chromosome start end strand MM target_seq PAM alignment distance position gene name gene id
GU305914.1 12607 12629 + 0 GATGTCTTTGTCGAAGTCAA CGG ||||||||[||||||||||||]PAM NA Intergenic NA NA

T595 CAGAGCGCGTCGAGACGTCGCGG 1000
Oligo fwd CACCCAGAGCGCGTCGAGACGTCG
Oligo rev AAACCGACGTCTCGACGCGCTCTG
Chromosome start end strand MM target_seq PAM alignment distance position gene name gene id
GU305914.1 12632 12654 + 0 CAGAGCGCGTCGAGACGTCG CGG ||||||||[||||||||||||]PAM NA Intergenic NA NA

T596 CGAGACGTCGCGGTTCGACACGG 1000
Oligo fwd CACCCGAGACGTCGCGGTTCGACA
Oligo rev AAACTGTCGAACCGCGACGTCTCG
Chromosome start end strand MM target_seq PAM alignment distance position gene name gene id
GU305914.1 12642 12664 + 0 CGAGACGTCGCGGTTCGACA CGG ||||||||[||||||||||||]PAM NA Intergenic NA NA

T597 ACACGGCGAACCGCACCAGCCGG 1000
Oligo fwd CACCACACGGCGAACCGCACCAGC
Oligo rev AAACGCTGGTGCGGTTCGCCGTGT
Chromosome start end strand MM target_seq PAM alignment distance position gene name gene id
GU305914.1 12659 12681 + 0 ACACGGCGAACCGCACCAGC CGG ||||||||[||||||||||||]PAM NA Intergenic NA NA

T598 TCTGCATCGACGACGAACGTCGG 1000
Oligo fwd CACCTCTGCATCGACGACGAACGT
Oligo rev AAACACGTTCGTCGTCGATGCAGA
Chromosome start end strand MM target_seq PAM alignment distance position gene name gene id
GU305914.1 12701 12723 + 0 TCTGCATCGACGACGAACGT CGG ||||||||[||||||||||||]PAM NA Intergenic NA NA

T599 ACGAACGTCGGCAAGAGATGCGG 1000
Oligo fwd CACCACGAACGTCGGCAAGAGATG
Oligo rev AAACCATCTCTTGCCGACGTTCGT
Chromosome start end strand MM target_seq PAM alignment distance position gene name gene id
GU305914.1 12713 12735 + 0 ACGAACGTCGGCAAGAGATG CGG ||||||||[||||||||||||]PAM NA Intergenic NA NA

T600 GCAAGAGATGCGGCGATAAACGG 1000
Oligo fwd CACCGCAAGAGATGCGGCGATAAA

Oligo rev AAACTTTATCGCCGCATCTCTTGC
Chromosome start end strand MM target_seq PAM alignment distance position gene name gene id
GU305914.1 12723 12745 + 0 GCAAGAGATGCGGCGATAAA CGG ||||||||[||||||||||||]PAM NA Intergenic NA NA

T601 TAAACGGCCACGTAAGAGCACGG 1000
Oligo fwd CACCTAAACGGCCACGTAAGAGCA
Oligo rev AAACTGCTCTTACGTGGCCGTTTA
Chromosome start end strand MM target_seq PAM alignment distance position gene name gene id
GU305914.1 12739 12761 + 0 TAAACGGCCACGTAAGAGCA CGG ||||||||[||||||||||||]PAM NA Intergenic NA NA

T602 CCAACGACCATTTATTTGAATGG 1000
Oligo fwd CACCCCAACGACCATTTATTTGAA
Oligo rev AAACTTCAAATAAATGGTCGTTGG
Chromosome start end strand MM target_seq PAM alignment distance position gene name gene id
GU305914.1 12800 12822 + 0 CCAACGACCATTTATTTGAA TGG ||||||||[||||||||||||]PAM NA Intergenic NA NA

T603 ACCATTTATTTGAATGGTGACGG 1000
Oligo fwd CACCACCATTTATTTGAATGGTGA
Oligo rev AAACTCACCATTCAAATAAATGGT
Chromosome start end strand MM target_seq PAM alignment distance position gene name gene id
GU305914.1 12806 12828 + 0 ACCATTTATTTGAATGGTGA CGG ||||||||[||||||||||||]PAM NA Intergenic NA NA

T604 AGCACACACGAGAACGAGCGCGG 1000
Oligo fwd CACCAGCACACACGAGAACGAGCG
Oligo rev AAACCGCTCGTTCTCGTGTGTGCT
Chromosome start end strand MM target_seq PAM alignment distance position gene name gene id
GU305914.1 12880 12902 + 0 AGCACACACGAGAACGAGCG CGG ||||||||[||||||||||||]PAM NA Intergenic NA NA
GU305914.1 12416 12438 - 4 AGCAGATACGAGCACGATCG CGG ||||-|-|[||||-||||-||]PAM NA Intergenic NA NA

T605 TCTATCATCGTCCTGACGATAGG 1000
Oligo fwd CACCTCTATCATCGTCCTGACGAT
Oligo rev AAACATCGTCAGGACGATGATAGA
Chromosome start end strand MM target_seq PAM alignment distance position gene name gene id
GU305914.1 12911 12933 + 0 TCTATCATCGTCCTGACGAT AGG ||||||||[||||||||||||]PAM NA Intergenic NA NA

T606 CTATCATCGTCCTGACGATAGGG 1000
Oligo fwd CACCCTATCATCGTCCTGACGATA
Oligo rev AAACTATCGTCAGGACGATGATAG
Chromosome start end strand MM target_seq PAM alignment distance position gene name gene id
GU305914.1 12912 12934 + 0 CTATCATCGTCCTGACGATA GGG ||||||||[||||||||||||]PAM NA Intergenic NA NA

T607 CCTGACGATAGGGCTCCTGCTGG 1000
Oligo fwd CACCCCTGACGATAGGGCTCCTGC
Oligo rev AAACGCAGGAGCCCTATCGTCAGG
Chromosome start end strand MM target_seq PAM alignment distance position gene name gene id
GU305914.1 12922 12944 + 0 CCTGACGATAGGGCTCCTGC TGG ||||||||[||||||||||||]PAM NA Intergenic NA NA

T608 CTGACGATAGGGCTCCTGCTGGG 1000
Oligo fwd CACCCTGACGATAGGGCTCCTGCT
Oligo rev AAACAGCAGGAGCCCTATCGTCAG
Chromosome start end strand MM target_seq PAM alignment distance position gene name gene id
GU305914.1 12923 12945 + 0 CTGACGATAGGGCTCCTGCT GGG ||||||||[||||||||||||]PAM NA Intergenic NA NA

T609 GCCGCGACGATCTATTCTTTCGG 1000
Oligo fwd CACCGCCGCGACGATCTATTCTTT
Oligo rev AAACAAAGAATAGATCGTCGCGGC
Chromosome start end strand MM target_seq PAM alignment distance position gene name gene id
GU305914.1 12956 12978 + 0 GCCGCGACGATCTATTCTTT CGG ||||||||[||||||||||||]PAM NA Intergenic NA NA

T610 TTATCAATTATACCAGAGTTTGG 1000
Oligo fwd CACCTTATCAATTATACCAGAGTT
Oligo rev AAACAACTCTGGTATAATTGATAA
Chromosome start end strand MM target_seq PAM alignment distance position gene name gene id
GU305914.1 13045 13067 + 0 TTATCAATTATACCAGAGTT TGG ||||||||[||||||||||||]PAM NA Intergenic NA NA

T611 TTAATCTGAGATGCGTCGTCTGG 1000
Oligo fwd CACCTTAATCTGAGATGCGTCGTC
Oligo rev AAACGACGACGCATCTCAGATTAA
Chromosome start end strand MM target_seq PAM alignment distance position gene name gene id
GU305914.1 13075 13097 + 0 TTAATCTGAGATGCGTCGTC TGG ||||||||[||||||||||||]PAM NA Intergenic NA NA

T612 TAATCTGAGATGCGTCGTCTGGG 1000
Oligo fwd CACCTAATCTGAGATGCGTCGTCT
Oligo rev AAACAGACGACGCATCTCAGATTA
Chromosome start end strand MM target_seq PAM alignment distance position gene name gene id
GU305914.1 13076 13098 + 0 TAATCTGAGATGCGTCGTCT GGG ||||||||[||||||||||||]PAM NA Intergenic NA NA

T613 TTTTTTCGATAATCTTTACGAGG 1000
Oligo fwd CACCTTTTTTCGATAATCTTTACG
Oligo rev AAACCGTAAAGATTATCGAAAAAA
Chromosome start end strand MM target_seq PAM alignment distance position gene name gene id
GU305914.1 13195 13217 + 0 TTTTTTCGATAATCTTTACG AGG ||||||||[||||||||||||]PAM NA Intergenic NA NA

T614 TCGACGAGAAAGATACCAACAGG 1000
Oligo fwd CACCTCGACGAGAAAGATACCAAC
Oligo rev AAACGTTGGTATCTTTCTCGTCGA
Chromosome start end strand MM target_seq PAM alignment distance position gene name gene id
GU305914.1 13230 13252 + 0 TCGACGAGAAAGATACCAAC AGG ||||||||[||||||||||||]PAM NA Intergenic NA NA

<table>
<tr><td>T615</td><td>CGACGAGAAAGATACCAACAGGG</td><td>1000</td><td></td><td></td><td></td><td></td><td></td><td></td><td></td><td></td><td></td></tr>
<tr><td>Oligo fwd</td><td>CACCCGACGAGAAAGATACCAACA</td><td></td><td></td><td></td><td></td><td></td><td></td><td></td><td></td><td></td><td></td></tr>
<tr><td>Oligo rev</td><td>AAACTGTTGGTATCTTTCTCGTCG</td><td></td><td></td><td></td><td></td><td></td><td></td><td></td><td></td><td></td><td></td></tr>
<tr><td>Chromosome</td><td>start</td><td>end</td><td>strand</td><td>MM</td><td>target_seq</td><td>PAM</td><td>alignment</td><td>distance</td><td>position</td><td>gene name</td><td>gene id</td></tr>
<tr><td>GU305914.1</td><td>13231</td><td>13253</td><td>+</td><td>0</td><td>CGACGAGAAAGATACCAACA</td><td>GGG</td><td>||||||||[||||||||||||]PAM</td><td>NA</td><td>Intergenic</td><td>NA</td><td>NA</td></tr>
<tr><td>T616</td><td>ACTACTCATGTCAATTCAAATGG</td><td>1000</td><td></td><td></td><td></td><td></td><td></td><td></td><td></td><td></td><td></td></tr>
<tr><td>Oligo fwd</td><td>CACCACTACTCATGTCAATTCAAA</td><td></td><td></td><td></td><td></td><td></td><td></td><td></td><td></td><td></td><td></td></tr>
<tr><td>Oligo rev</td><td>AAACTTTGAATTGACATGAGTAGT</td><td></td><td></td><td></td><td></td><td></td><td></td><td></td><td></td><td></td><td></td></tr>
<tr><td>Chromosome</td><td>start</td><td>end</td><td>strand</td><td>MM</td><td>target_seq</td><td>PAM</td><td>alignment</td><td>distance</td><td>position</td><td>gene name</td><td>gene id</td></tr>
<tr><td>GU305914.1</td><td>13257</td><td>13279</td><td>+</td><td>0</td><td>ACTACTCATGTCAATTCAAA</td><td>TGG</td><td>||||||||[||||||||||||]PAM</td><td>NA</td><td>Intergenic</td><td>NA</td><td>NA</td></tr>
<tr><td>T617</td><td>CAAATGGAACATCTGTACCCAGG</td><td>1000</td><td></td><td></td><td></td><td></td><td></td><td></td><td></td><td></td><td></td></tr>
<tr><td>Oligo fwd</td><td>CACCCAAATGGAACATCTGTACCC</td><td></td><td></td><td></td><td></td><td></td><td></td><td></td><td></td><td></td><td></td></tr>
<tr><td>Oligo rev</td><td>AAACGGGTACAGATGTTCCATTTG</td><td></td><td></td><td></td><td></td><td></td><td></td><td></td><td></td><td></td><td></td></tr>
<tr><td>Chromosome</td><td>start</td><td>end</td><td>strand</td><td>MM</td><td>target_seq</td><td>PAM</td><td>alignment</td><td>distance</td><td>position</td><td>gene name</td><td>gene id</td></tr>
<tr><td>GU305914.1</td><td>13273</td><td>13295</td><td>+</td><td>0</td><td>CAAATGGAACATCTGTACCC</td><td>AGG</td><td>||||||||[||||||||||||]PAM</td><td>NA</td><td>Intergenic</td><td>NA</td><td>NA</td></tr>
<tr><td>T618</td><td>ACATCTGTACCCAGGCAACGTGG</td><td>1000</td><td></td><td></td><td></td><td></td><td></td><td></td><td></td><td></td><td></td></tr>
<tr><td>Oligo fwd</td><td>CACCACATCTGTACCCAGGCAACG</td><td></td><td></td><td></td><td></td><td></td><td></td><td></td><td></td><td></td><td></td></tr>
<tr><td>Oligo rev</td><td>AAACCGTTGCCTGGGTACAGATGT</td><td></td><td></td><td></td><td></td><td></td><td></td><td></td><td></td><td></td><td></td></tr>
<tr><td>Chromosome</td><td>start</td><td>end</td><td>strand</td><td>MM</td><td>target_seq</td><td>PAM</td><td>alignment</td><td>distance</td><td>position</td><td>gene name</td><td>gene id</td></tr>
<tr><td>GU305914.1</td><td>13281</td><td>13303</td><td>+</td><td>0</td><td>ACATCTGTACCCAGGCAACG</td><td>TGG</td><td>||||||||[||||||||||||]PAM</td><td>NA</td><td>Intergenic</td><td>NA</td><td>NA</td></tr>
<tr><td>T619</td><td>TGTACCCAGGCAACGTGGTACGG</td><td>1000</td><td></td><td></td><td></td><td></td><td></td><td></td><td></td><td></td><td></td></tr>
<tr><td>Oligo fwd</td><td>CACCTGTACCCAGGCAACGTGGTA</td><td></td><td></td><td></td><td></td><td></td><td></td><td></td><td></td><td></td><td></td></tr>
<tr><td>Oligo rev</td><td>AAACTACCACGTTGCCTGGGTACA</td><td></td><td></td><td></td><td></td><td></td><td></td><td></td><td></td><td></td><td></td></tr>
<tr><td>Chromosome</td><td>start</td><td>end</td><td>strand</td><td>MM</td><td>target_seq</td><td>PAM</td><td>alignment</td><td>distance</td><td>position</td><td>gene name</td><td>gene id</td></tr>
<tr><td>GU305914.1</td><td>13286</td><td>13308</td><td>+</td><td>0</td><td>TGTACCCAGGCAACGTGGTA</td><td>CGG</td><td>||||||||[||||||||||||]PAM</td><td>NA</td><td>Intergenic</td><td>NA</td><td>NA</td></tr>
<tr><td>T620</td><td>GTACCCAGGCAACGTGGTACGGG</td><td>1000</td><td></td><td></td><td></td><td></td><td></td><td></td><td></td><td></td><td></td></tr>
<tr><td>Oligo fwd</td><td>CACCGTACCCAGGCAACGTGGTAC</td><td></td><td></td><td></td><td></td><td></td><td></td><td></td><td></td><td></td><td></td></tr>
<tr><td>Oligo rev</td><td>AAACGTACCACGTTGCCTGGGTAC</td><td></td><td></td><td></td><td></td><td></td><td></td><td></td><td></td><td></td><td></td></tr>
<tr><td>Chromosome</td><td>start</td><td>end</td><td>strand</td><td>MM</td><td>target_seq</td><td>PAM</td><td>alignment</td><td>distance</td><td>position</td><td>gene name</td><td>gene id</td></tr>
<tr><td>GU305914.1</td><td>13287</td><td>13309</td><td>+</td><td>0</td><td>GTACCCAGGCAACGTGGTAC</td><td>GGG</td><td>||||||||[||||||||||||]PAM</td><td>NA</td><td>Intergenic</td><td>NA</td><td>NA</td></tr>
<tr><td>T621</td><td>AGGCAACGTGGTACGGGACGTGG</td><td>1000</td><td></td><td></td><td></td><td></td><td></td><td></td><td></td><td></td><td></td></tr>
<tr><td>Oligo fwd</td><td>CACCAGGCAACGTGGTACGGGACG</td><td></td><td></td><td></td><td></td><td></td><td></td><td></td><td></td><td></td><td></td></tr>
<tr><td>Oligo rev</td><td>AAACCGTCCCGTACCACGTTGCCT</td><td></td><td></td><td></td><td></td><td></td><td></td><td></td><td></td><td></td><td></td></tr>
<tr><td>Chromosome</td><td>start</td><td>end</td><td>strand</td><td>MM</td><td>target_seq</td><td>PAM</td><td>alignment</td><td>distance</td><td>position</td><td>gene name</td><td>gene id</td></tr>
<tr><td>GU305914.1</td><td>13293</td><td>13315</td><td>+</td><td>0</td><td>AGGCAACGTGGTACGGGACG</td><td>TGG</td><td>||||||||[||||||||||||]PAM</td><td>NA</td><td>Intergenic</td><td>NA</td><td>NA</td></tr>
<tr><td>T622</td><td>GATACGAATGTCATTCGATGAGG</td><td>1000</td><td></td><td></td><td></td><td></td><td></td><td></td><td></td><td></td><td></td></tr>
<tr><td>Oligo fwd</td><td>CACCGATACGAATGTCATTCGATG</td><td></td><td></td><td></td><td></td><td></td><td></td><td></td><td></td><td></td><td></td></tr>
<tr><td>Oligo rev</td><td>AAACCATCGAATGACATTCGTATC</td><td></td><td></td><td></td><td></td><td></td><td></td><td></td><td></td><td></td><td></td></tr>
<tr><td>Chromosome</td><td>start</td><td>end</td><td>strand</td><td>MM</td><td>target_seq</td><td>PAM</td><td>alignment</td><td>distance</td><td>position</td><td>gene name</td><td>gene id</td></tr>
<tr><td>GU305914.1</td><td>13315</td><td>13337</td><td>+</td><td>0</td><td>GATACGAATGTCATTCGATG</td><td>AGG</td><td>||||||||[||||||||||||]PAM</td><td>NA</td><td>Intergenic</td><td>NA</td><td>NA</td></tr>
<tr><td>T623</td><td>ACATCTCCACCAACATTAGTCGG</td><td>1000</td><td></td><td></td><td></td><td></td><td></td><td></td><td></td><td></td><td></td></tr>
<tr><td>Oligo fwd</td><td>CACCACATCTCCACCAACATTAGT</td><td></td><td></td><td></td><td></td><td></td><td></td><td></td><td></td><td></td><td></td></tr>
<tr><td>Oligo rev</td><td>AAACACTAATGTTGGTGGAGATGT</td><td></td><td></td><td></td><td></td><td></td><td></td><td></td><td></td><td></td><td></td></tr>
<tr><td>Chromosome</td><td>start</td><td>end</td><td>strand</td><td>MM</td><td>target_seq</td><td>PAM</td><td>alignment</td><td>distance</td><td>position</td><td>gene name</td><td>gene id</td></tr>
<tr><td>GU305914.1</td><td>13391</td><td>13413</td><td>+</td><td>0</td><td>ACATCTCCACCAACATTAGT</td><td>CGG</td><td>||||||||[||||||||||||]PAM</td><td>NA</td><td>Intergenic</td><td>NA</td><td>NA</td></tr>
<tr><td>T624</td><td>CACCAACATTAGTCGGCTTCTGG</td><td>1000</td><td></td><td></td><td></td><td></td><td></td><td></td><td></td><td></td><td></td></tr>
<tr><td>Oligo fwd</td><td>CACCCACCAACATTAGTCGGCTTC</td><td></td><td></td><td></td><td></td><td></td><td></td><td></td><td></td><td></td><td></td></tr>
<tr><td>Oligo rev</td><td>AAACGAAGCCGACTAATGTTGGTG</td><td></td><td></td><td></td><td></td><td></td><td></td><td></td><td></td><td></td><td></td></tr>
<tr><td>Chromosome</td><td>start</td><td>end</td><td>strand</td><td>MM</td><td>target_seq</td><td>PAM</td><td>alignment</td><td>distance</td><td>position</td><td>gene name</td><td>gene id</td></tr>
<tr><td>GU305914.1</td><td>13398</td><td>13420</td><td>+</td><td>0</td><td>CACCAACATTAGTCGGCTTC</td><td>TGG</td><td>||||||||[||||||||||||]PAM</td><td>NA</td><td>Intergenic</td><td>NA</td><td>NA</td></tr>
<tr><td>T625</td><td>CCTTTCACATAACAGTTCTGCGG</td><td>1000</td><td></td><td></td><td></td><td></td><td></td><td></td><td></td><td></td><td></td></tr>
<tr><td>Oligo fwd</td><td>CACCCCTTTCACATAACAGTTCTG</td><td></td><td></td><td></td><td></td><td></td><td></td><td></td><td></td><td></td><td></td></tr>
<tr><td>Oligo rev</td><td>AAACCAGAACTGTTATGTGAAAGG</td><td></td><td></td><td></td><td></td><td></td><td></td><td></td><td></td><td></td><td></td></tr>
<tr><td>Chromosome</td><td>start</td><td>end</td><td>strand</td><td>MM</td><td>target_seq</td><td>PAM</td><td>alignment</td><td>distance</td><td>position</td><td>gene name</td><td>gene id</td></tr>
<tr><td>GU305914.1</td><td>13450</td><td>13472</td><td>+</td><td>0</td><td>CCTTTCACATAACAGTTCTG</td><td>CGG</td><td>||||||||[||||||||||||]PAM</td><td>NA</td><td>Intergenic</td><td>NA</td><td>NA</td></tr>
<tr><td>T626</td><td>ATAACAGTTCTGCGGAGATATGG</td><td>1000</td><td></td><td></td><td></td><td></td><td></td><td></td><td></td><td></td><td></td></tr>
<tr><td>Oligo fwd</td><td>CACCATAACAGTTCTGCGGAGATA</td><td></td><td></td><td></td><td></td><td></td><td></td><td></td><td></td><td></td><td></td></tr>
<tr><td>Oligo rev</td><td>AAACTATCTCCGCAGAACTGTTAT</td><td></td><td></td><td></td><td></td><td></td><td></td><td></td><td></td><td></td><td></td></tr>
<tr><td>Chromosome</td><td>start</td><td>end</td><td>strand</td><td>MM</td><td>target_seq</td><td>PAM</td><td>alignment</td><td>distance</td><td>position</td><td>gene name</td><td>gene id</td></tr>
<tr><td>GU305914.1</td><td>13458</td><td>13480</td><td>+</td><td>0</td><td>ATAACAGTTCTGCGGAGATA</td><td>TGG</td><td>||||||||[||||||||||||]PAM</td><td>NA</td><td>Intergenic</td><td>NA</td><td>NA</td></tr>
<tr><td>T627</td><td>ACAAGTATCATCAAACCAATTGG</td><td>1000</td><td></td><td></td><td></td><td></td><td></td><td></td><td></td><td></td><td></td></tr>
<tr><td>Oligo fwd</td><td>CACCACAAGTATCATCAAACCAAT</td><td></td><td></td><td></td><td></td><td></td><td></td><td></td><td></td><td></td><td></td></tr>
<tr><td>Oligo rev</td><td>AAACATTGGTTTGATGATACTTGT</td><td></td><td></td><td></td><td></td><td></td><td></td><td></td><td></td><td></td><td></td></tr>
<tr><td>Chromosome</td><td>start</td><td>end</td><td>strand</td><td>MM</td><td>target_seq</td><td>PAM</td><td>alignment</td><td>distance</td><td>position</td><td>gene name</td><td>gene id</td></tr>
<tr><td>GU305914.1</td><td>13516</td><td>13538</td><td>+</td><td>0</td><td>ACAAGTATCATCAAACCAAT</td><td>TGG</td><td>||||||||[||||||||||||]PAM</td><td>NA</td><td>Intergenic</td><td>NA</td><td>NA</td></tr>
<tr><td>T628</td><td>CAAGTATCATCAAACCAATTGGG</td><td>1000</td><td></td><td></td><td></td><td></td><td></td><td></td><td></td><td></td><td></td></tr>
<tr><td>Oligo fwd</td><td>CACCCAAGTATCATCAAACCAATT</td><td></td><td></td><td></td><td></td><td></td><td></td><td></td><td></td><td></td><td></td></tr>
<tr><td>Oligo rev</td><td>AAACAATTGGTTTGATGATACTTG</td><td></td><td></td><td></td><td></td><td></td><td></td><td></td><td></td><td></td><td></td></tr>
<tr><td>Chromosome</td><td>start</td><td>end</td><td>strand</td><td>MM</td><td>target_seq</td><td>PAM</td><td>alignment</td><td>distance</td><td>position</td><td>gene name</td><td>gene id</td></tr>
<tr><td>GU305914.1</td><td>13517</td><td>13539</td><td>+</td><td>0</td><td>CAAGTATCATCAAACCAATT</td><td>GGG</td><td>||||||||[||||||||||||]PAM</td><td>NA</td><td>Intergenic</td><td>NA</td><td>NA</td></tr>
<tr><td>T629</td><td>GGACTAATCTGTCAACTGACCGG</td><td>1000</td><td></td><td></td><td></td><td></td><td></td><td></td><td></td><td></td><td></td></tr>
<tr><td>Oligo fwd</td><td>CACCGGACTAATCTGTCAACTGAC</td><td></td><td></td><td></td><td></td><td></td><td></td><td></td><td></td><td></td><td></td></tr>
<tr><td>Oligo rev</td><td>AAACGTCAGTTGACAGATTAGTCC</td><td></td><td></td><td></td><td></td><td></td><td></td><td></td><td></td><td></td><td></td></tr>
<tr><td>Chromosome</td><td>start</td><td>end</td><td>strand</td><td>MM</td><td>target_seq</td><td>PAM</td><td>alignment</td><td>distance</td><td>position</td><td>gene name</td><td>gene id</td></tr>
<tr><td>GU305914.1</td><td>13538</td><td>13560</td><td>+</td><td>0</td><td>GGACTAATCTGTCAACTGAC</td><td>CGG</td><td>||||||||[||||||||||||]PAM</td><td>NA</td><td>Intergenic</td><td>NA</td><td>NA</td></tr>
</table>

T630 CGGCTGTCTAAACTCGAAGATGG 1000
Oligo fwd CACCCGGCTGTCTAAACTCGAAGA
Oligo rev AAACTCTTCGAGTTTAGACAGCCG
Chromosome start end strand MM target_seq PAM alignment distance position gene name gene id
GU305914.1 13558 13580 + 0 CGGCTGTCTAAACTCGAAGA TGG ||||||||[||||||||||||]PAM NA Intergenic NA NA

T631 CGAGAAAATCGAGACCCAGACGG 1000
Oligo fwd CACCCGAGAAAATCGAGACCCAGA
Oligo rev AAACTCTGGGTCTCGATTTTCTCG
Chromosome start end strand MM target_seq PAM alignment distance position gene name gene id
GU305914.1 13606 13628 + 0 CGAGAAAATCGAGACCCAGA CGG ||||||||[||||||||||||]PAM NA Intergenic NA NA

T632 ACCCAGACGGCAATTGCGATCGG 1000
Oligo fwd CACCACCCAGACGGCAATTGCGAT
Oligo rev AAACATCGCAATTGCCGTCTGGGT
Chromosome start end strand MM target_seq PAM alignment distance position gene name gene id
GU305914.1 13619 13641 + 0 ACCCAGACGGCAATTGCGAT CGG ||||||||[||||||||||||]PAM NA Intergenic NA NA

T633 AATTGCGATCGGTAAACCCGCGG 1000
Oligo fwd CACCAATTGCGATCGGTAAACCCG
Oligo rev AAACCGGGTTTACCGATCGCAATT
Chromosome start end strand MM target_seq PAM alignment distance position gene name gene id
GU305914.1 13630 13652 + 0 AATTGCGATCGGTAAACCCG CGG ||||||||[||||||||||||]PAM NA Intergenic NA NA

T634 CGCGGATAAACAAAGTGAGTTGG 1000
Oligo fwd CACCCGCGGATAAACAAAGTGAGT
Oligo rev AAACACTCACTTTGTTTATCCGCG
Chromosome start end strand MM target_seq PAM alignment distance position gene name gene id
GU305914.1 13648 13670 + 0 CGCGGATAAACAAAGTGAGT TGG ||||||||[||||||||||||]PAM NA Intergenic NA NA

T635 ATGATCAGACCTACAACAAGAGG 1000
Oligo fwd CACCATGATCAGACCTACAACAAG
Oligo rev AAACCTTGTTGTAGGTCTGATCAT
Chromosome start end strand MM target_seq PAM alignment distance position gene name gene id
GU305914.1 13745 13767 + 0 ATGATCAGACCTACAACAAG AGG ||||||||[||||||||||||]PAM NA Intergenic NA NA

T636 ACCTACAACAAGAGGAACATCGG 1000
Oligo fwd CACCACCTACAACAAGAGGAACAT
Oligo rev AAACATGTTCCTCTTGTTGTAGGT
Chromosome start end strand MM target_seq PAM alignment distance position gene name gene id
GU305914.1 13753 13775 + 0 ACCTACAACAAGAGGAACAT CGG ||||||||[||||||||||||]PAM NA Intergenic NA NA

T637 CAAGAGGAACATCGGTTTCGAGG 1000
Oligo fwd CACCCAAGAGGAACATCGGTTTCG
Oligo rev AAACCGAAACCGATGTTCCTCTTG
Chromosome start end strand MM target_seq PAM alignment distance position gene name gene id
GU305914.1 13761 13783 + 0 CAAGAGGAACATCGGTTTCG AGG ||||||||[||||||||||||]PAM NA Intergenic NA NA

T638 GCCCGAGATCACCGTGTCGTCGG 1000
Oligo fwd CACCGCCCGAGATCACCGTGTCGT
Oligo rev AAACACGACACGGTGATCTCGGGC
Chromosome start end strand MM target_seq PAM alignment distance position gene name gene id
GU305914.1 13786 13808 + 0 GCCCGAGATCACCGTGTCGT CGG ||||||||[||||||||||||]PAM NA Intergenic NA NA

T639 CGCCGCGCCTTTACGTCACGCGG 1000
Oligo fwd CACCCGCCGCGCCTTTACGTCACG
Oligo rev AAACCGTGACGTAAAGGCGCGGCG
Chromosome start end strand MM target_seq PAM alignment distance position gene name gene id
GU305914.1 13828 13850 + 0 CGCCGCGCCTTTACGTCACG CGG ||||||||[||||||||||||]PAM NA Intergenic NA NA

T640 CGCGGCACAAGCGTACGTAACGG 1000
Oligo fwd CACCCGCGGCACAAGCGTACGTAA
Oligo rev AAACTTACGTACGCTTGTGCCGCG
Chromosome start end strand MM target_seq PAM alignment distance position gene name gene id
GU305914.1 13846 13868 + 0 CGCGGCACAAGCGTACGTAA CGG ||||||||[||||||||||||]PAM NA Intergenic NA NA

T641 GCGTACGTAACGGTCGTCTTAGG 1000
Oligo fwd CACCGCGTACGTAACGGTCGTCTT
Oligo rev AAACAAGACGACCGTTACGTACGC
Chromosome start end strand MM target_seq PAM alignment distance position gene name gene id
GU305914.1 13856 13878 + 0 GCGTACGTAACGGTCGTCTT AGG ||||||||[||||||||||||]PAM NA Intergenic NA NA

T642 CGTACGTAACGGTCGTCTTAGGG 1000
Oligo fwd CACCCGTACGTAACGGTCGTCTTA
Oligo rev AAACTAAGACGACCGTTACGTACG
Chromosome start end strand MM target_seq PAM alignment distance position gene name gene id
GU305914.1 13857 13879 + 0 CGTACGTAACGGTCGTCTTA GGG ||||||||[||||||||||||]PAM NA Intergenic NA NA

T643 GTACGTAACGGTCGTCTTAGGGG 1000
Oligo fwd CACCGTACGTAACGGTCGTCTTAG
Oligo rev AAACCTAAGACGACCGTTACGTAC
Chromosome start end strand MM target_seq PAM alignment distance position gene name gene id
GU305914.1 13858 13880 + 0 GTACGTAACGGTCGTCTTAG GGG ||||||||[||||||||||||]PAM NA Intergenic NA NA

T644 GGTCGTCTTAGGGGTAATCGCGG 1000
Oligo fwd CACCGGTCGTCTTAGGGGTAATCG
Oligo rev AAACCGATTACCCCTAAGACGACC
Chromosome start end strand MM target_seq PAM alignment distance position gene name gene id

GU305914.1 13867 13889 + 0 GGTCGTCTTAGGGGTAATCG CGG ||||||||[||||||||||||]PAM NA Intergenic NA NA

T645 GTCGTCTTAGGGGTAATCGCGGG 1000
Oligo fwd CACCGTCGTCTTAGGGGTAATCGC
Oligo rev AAACGCGATTACCCCTAAGACGAC
Chromosome start end strand MM target_seq PAM alignment distance position gene name gene id
GU305914.1 13868 13890 + 0 GTCGTCTTAGGGGTAATCGC GGG ||||||||[||||||||||||]PAM NA Intergenic NA NA

T646 TAATCGCGGGTCTCTTCTTGTGG 1000
Oligo fwd CACCTAATCGCGGGTCTCTTCTTG
Oligo rev AAACCAAGAAGAGACCCGCGATTA
Chromosome start end strand MM target_seq PAM alignment distance position gene name gene id
GU305914.1 13881 13903 + 0 TAATCGCGGGTCTCTTCTTG TGG ||||||||[||||||||||||]PAM NA Intergenic NA NA

T647 TGTGATCGCCGATCCCCGTCAGG 1000
Oligo fwd CACCTGTGATCGCCGATCCCCGTC
Oligo rev AAACGACGGGGATCGGCGATCACA
Chromosome start end strand MM target_seq PAM alignment distance position gene name gene id
GU305914.1 13924 13946 + 0 TGTGATCGCCGATCCCCGTC AGG ||||||||[||||||||||||]PAM NA Intergenic NA NA

T648 AGGATCAGCGCACGATCGACCGG 1000
Oligo fwd CACCAGGATCAGCGCACGATCGAC
Oligo rev AAACGTCGATCGTGCGCTGATCCT
Chromosome start end strand MM target_seq PAM alignment distance position gene name gene id
GU305914.1 13944 13966 + 0 AGGATCAGCGCACGATCGAC CGG ||||||||[||||||||||||]PAM NA Intergenic NA NA

T649 TCAGCATTCGTCATAAGCTGTGG 1000
Oligo fwd CACCTCAGCATTCGTCATAAGCTG
Oligo rev AAACCAGCTTATGACGAATGCTGA
Chromosome start end strand MM target_seq PAM alignment distance position gene name gene id
GU305914.1 13975 13997 + 0 TCAGCATTCGTCATAAGCTG TGG ||||||||[||||||||||||]PAM NA Intergenic NA NA

T650 AAGAAGATGTACCGTCACGTCGG 1000
Oligo fwd CACCAAGAAGATGTACCGTCACGT
Oligo rev AAACACGTGACGGTACATCTTCTT
Chromosome start end strand MM target_seq PAM alignment distance position gene name gene id
GU305914.1 14079 14101 + 0 AAGAAGATGTACCGTCACGT CGG ||||||||[||||||||||||]PAM NA Intergenic NA NA

T651 AACTGCCAAAGCCCGTCTTCCGG 1000
Oligo fwd CACCAACTGCCAAAGCCCGTCTTC
Oligo rev AAACGAAGACGGGCTTTGGCAGTT
Chromosome start end strand MM target_seq PAM alignment distance position gene name gene id
GU305914.1 14157 14179 + 0 AACTGCCAAAGCCCGTCTTC CGG ||||||||[||||||||||||]PAM NA Intergenic NA NA

T652 CCAAAGCCCGTCTTCCGGTATGG 1000
Oligo fwd CACCCCAAAGCCCGTCTTCCGGTA
Oligo rev AAACTACCGGAAGACGGGCTTTGG
Chromosome start end strand MM target_seq PAM alignment distance position gene name gene id
GU305914.1 14162 14184 + 0 CCAAAGCCCGTCTTCCGGTA TGG ||||||||[||||||||||||]PAM NA Intergenic NA NA

T653 TCCGGTATGGATTTTACTCCAGG 1000
Oligo fwd CACCTCCGGTATGGATTTTACTCC
Oligo rev AAACGGAGTAAAATCCATACCGGA
Chromosome start end strand MM target_seq PAM alignment distance position gene name gene id
GU305914.1 14175 14197 + 0 TCCGGTATGGATTTTACTCC AGG ||||||||[||||||||||||]PAM NA Intergenic NA NA

T654 TTACTCCAGGTATGTGTGTACGG 1000
Oligo fwd CACCTTACTCCAGGTATGTGTGTA
Oligo rev AAACTACACACATACCTGGAGTAA
Chromosome start end strand MM target_seq PAM alignment distance position gene name gene id
GU305914.1 14188 14210 + 0 TTACTCCAGGTATGTGTGTA CGG ||||||||[||||||||||||]PAM NA Intergenic NA NA

T655 TACTCCAGGTATGTGTGTACGGG 1000
Oligo fwd CACCTACTCCAGGTATGTGTGTAC
Oligo rev AAACGTACACACATACCTGGAGTA
Chromosome start end strand MM target_seq PAM alignment distance position gene name gene id
GU305914.1 14189 14211 + 0 TACTCCAGGTATGTGTGTAC GGG ||||||||[||||||||||||]PAM NA Intergenic NA NA

T656 GTTGCAAGCGCATCACGCAACGG 1000
Oligo fwd CACCGTTGCAAGCGCATCACGCAA
Oligo rev AAACTTGCGTGATGCGCTTGCAAC
Chromosome start end strand MM target_seq PAM alignment distance position gene name gene id
GU305914.1 14226 14248 + 0 GTTGCAAGCGCATCACGCAA CGG ||||||||[||||||||||||]PAM NA Intergenic NA NA

T657 CTGTATGTATTCCTGCCATATGG 1000
Oligo fwd CACCCTGTATGTATTCCTGCCATA
Oligo rev AAACTATGGCAGGAATACATACAG
Chromosome start end strand MM target_seq PAM alignment distance position gene name gene id
GU305914.1 14278 14300 + 0 CTGTATGTATTCCTGCCATA TGG ||||||||[||||||||||||]PAM NA Intergenic NA NA

T658 CTGCCATATGGTACGCAACATGG 1000
Oligo fwd CACCCTGCCATATGGTACGCAACA
Oligo rev AAACTGTTGCGTACCATATGGCAG
Chromosome start end strand MM target_seq PAM alignment distance position gene name gene id
GU305914.1 14290 14312 + 0 CTGCCATATGGTACGCAACA TGG ||||||||[||||||||||||]PAM NA Intergenic NA NA

T659 AAGAACTTCTACATCTGTTCCGG 1000
Oligo fwd CACCAAGAACTTCTACATCTGTTC
Oligo rev AAACGAACAGATGTAGAAGTTCTT

<table>
<tr><td>Chromosome</td><td>start</td><td></td><td>end</td><td>strand</td><td>MM</td><td>target_seq</td><td>PAM</td><td>alignment</td><td>distance</td><td>position</td><td>gene name</td><td>gene id</td></tr>
<tr><td>GU305914.1</td><td></td><td>14318</td><td>14340</td><td>+</td><td>0</td><td>AAGAACTTCTACATCTGTTC</td><td>CGG</td><td>||||||||[||||||||||||]PAM</td><td>NA</td><td>Intergenic</td><td>NA</td><td>NA</td></tr>
<tr><td></td><td></td><td></td><td></td><td></td><td></td><td></td><td></td><td></td><td></td><td></td><td></td><td></td></tr>
<tr><td>T660</td><td>CATCACTTCGATTTCTGATACGG</td><td></td><td>1000</td><td></td><td></td><td></td><td></td><td></td><td></td><td></td><td></td><td></td></tr>
<tr><td>Oligo fwd</td><td>CACCCATCACTTCGATTTCTGATA</td><td></td><td></td><td></td><td></td><td></td><td></td><td></td><td></td><td></td><td></td><td></td></tr>
<tr><td>Oligo rev</td><td>AAACTATCAGAAATCGAAGTGATG</td><td></td><td></td><td></td><td></td><td></td><td></td><td></td><td></td><td></td><td></td><td></td></tr>
<tr><td>Chromosome</td><td>start</td><td></td><td>end</td><td>strand</td><td>MM</td><td>target_seq</td><td>PAM</td><td>alignment</td><td>distance</td><td>position</td><td>gene name</td><td>gene id</td></tr>
<tr><td>GU305914.1</td><td></td><td>14342</td><td>14364</td><td>+</td><td>0</td><td>CATCACTTCGATTTCTGATA</td><td>CGG</td><td>||||||||[||||||||||||]PAM</td><td>NA</td><td>Intergenic</td><td>NA</td><td>NA</td></tr>
<tr><td></td><td></td><td></td><td></td><td></td><td></td><td></td><td></td><td></td><td></td><td></td><td></td><td></td></tr>
<tr><td>T661</td><td>CACTTCGATTTCTGATACGGAGG</td><td></td><td>1000</td><td></td><td></td><td></td><td></td><td></td><td></td><td></td><td></td><td></td></tr>
<tr><td>Oligo fwd</td><td>CACCCACTTCGATTTCTGATACGG</td><td></td><td></td><td></td><td></td><td></td><td></td><td></td><td></td><td></td><td></td><td></td></tr>
<tr><td>Oligo rev</td><td>AAACCCGTATCAGAAATCGAAGTG</td><td></td><td></td><td></td><td></td><td></td><td></td><td></td><td></td><td></td><td></td><td></td></tr>
<tr><td>Chromosome</td><td>start</td><td></td><td>end</td><td>strand</td><td>MM</td><td>target_seq</td><td>PAM</td><td>alignment</td><td>distance</td><td>position</td><td>gene name</td><td>gene id</td></tr>
<tr><td>GU305914.1</td><td></td><td>14345</td><td>14367</td><td>+</td><td>0</td><td>CACTTCGATTTCTGATACGG</td><td>AGG</td><td>||||||||[||||||||||||]PAM</td><td>NA</td><td>Intergenic</td><td>NA</td><td>NA</td></tr>
<tr><td></td><td></td><td></td><td></td><td></td><td></td><td></td><td></td><td></td><td></td><td></td><td></td><td></td></tr>
<tr><td>T662</td><td>TGATACGGAGGATGAGTATTCGG</td><td></td><td>1000</td><td></td><td></td><td></td><td></td><td></td><td></td><td></td><td></td><td></td></tr>
<tr><td>Oligo fwd</td><td>CACCTGATACGGAGGATGAGTATT</td><td></td><td></td><td></td><td></td><td></td><td></td><td></td><td></td><td></td><td></td><td></td></tr>
<tr><td>Oligo rev</td><td>AAACAATACTCATCCTCCGTATCA</td><td></td><td></td><td></td><td></td><td></td><td></td><td></td><td></td><td></td><td></td><td></td></tr>
<tr><td>Chromosome</td><td>start</td><td></td><td>end</td><td>strand</td><td>MM</td><td>target_seq</td><td>PAM</td><td>alignment</td><td>distance</td><td>position</td><td>gene name</td><td>gene id</td></tr>
<tr><td>GU305914.1</td><td></td><td>14357</td><td>14379</td><td>+</td><td>0</td><td>TGATACGGAGGATGAGTATT</td><td>CGG</td><td>||||||||[||||||||||||]PAM</td><td>NA</td><td>Intergenic</td><td>NA</td><td>NA</td></tr>
<tr><td></td><td></td><td></td><td></td><td></td><td></td><td></td><td></td><td></td><td></td><td></td><td></td><td></td></tr>
<tr><td>T663</td><td>GATACGGAGGATGAGTATTCGGG</td><td></td><td>1000</td><td></td><td></td><td></td><td></td><td></td><td></td><td></td><td></td><td></td></tr>
<tr><td>Oligo fwd</td><td>CACCGATACGGAGGATGAGTATTC</td><td></td><td></td><td></td><td></td><td></td><td></td><td></td><td></td><td></td><td></td><td></td></tr>
<tr><td>Oligo rev</td><td>AAACGAATACTCATCCTCCGTATC</td><td></td><td></td><td></td><td></td><td></td><td></td><td></td><td></td><td></td><td></td><td></td></tr>
<tr><td>Chromosome</td><td>start</td><td></td><td>end</td><td>strand</td><td>MM</td><td>target_seq</td><td>PAM</td><td>alignment</td><td>distance</td><td>position</td><td>gene name</td><td>gene id</td></tr>
<tr><td>GU305914.1</td><td></td><td>14358</td><td>14380</td><td>+</td><td>0</td><td>GATACGGAGGATGAGTATTC</td><td>GGG</td><td>||||||||[||||||||||||]PAM</td><td>NA</td><td>Intergenic</td><td>NA</td><td>NA</td></tr>
<tr><td></td><td></td><td></td><td></td><td></td><td></td><td></td><td></td><td></td><td></td><td></td><td></td><td></td></tr>
<tr><td>T664</td><td>GGAGGATGAGTATTCGGGATCGG</td><td></td><td>1000</td><td></td><td></td><td></td><td></td><td></td><td></td><td></td><td></td><td></td></tr>
<tr><td>Oligo fwd</td><td>CACCGGAGGATGAGTATTCGGGAT</td><td></td><td></td><td></td><td></td><td></td><td></td><td></td><td></td><td></td><td></td><td></td></tr>
<tr><td>Oligo rev</td><td>AAACATCCCGAATACTCATCCTCC</td><td></td><td></td><td></td><td></td><td></td><td></td><td></td><td></td><td></td><td></td><td></td></tr>
<tr><td>Chromosome</td><td>start</td><td></td><td>end</td><td>strand</td><td>MM</td><td>target_seq</td><td>PAM</td><td>alignment</td><td>distance</td><td>position</td><td>gene name</td><td>gene id</td></tr>
<tr><td>GU305914.1</td><td></td><td>14363</td><td>14385</td><td>+</td><td>0</td><td>GGAGGATGAGTATTCGGGAT</td><td>CGG</td><td>||||||||[||||||||||||]PAM</td><td>NA</td><td>Intergenic</td><td>NA</td><td>NA</td></tr>
<tr><td></td><td></td><td></td><td></td><td></td><td></td><td></td><td></td><td></td><td></td><td></td><td></td><td></td></tr>
<tr><td>T665</td><td>GAGGATGAGTATTCGGGATCGGG</td><td></td><td>1000</td><td></td><td></td><td></td><td></td><td></td><td></td><td></td><td></td><td></td></tr>
<tr><td>Oligo fwd</td><td>CACCGAGGATGAGTATTCGGGATC</td><td></td><td></td><td></td><td></td><td></td><td></td><td></td><td></td><td></td><td></td><td></td></tr>
<tr><td>Oligo rev</td><td>AAACGATCCCGAATACTCATCCTC</td><td></td><td></td><td></td><td></td><td></td><td></td><td></td><td></td><td></td><td></td><td></td></tr>
<tr><td>Chromosome</td><td>start</td><td></td><td>end</td><td>strand</td><td>MM</td><td>target_seq</td><td>PAM</td><td>alignment</td><td>distance</td><td>position</td><td>gene name</td><td>gene id</td></tr>
<tr><td>GU305914.1</td><td></td><td>14364</td><td>14386</td><td>+</td><td>0</td><td>GAGGATGAGTATTCGGGATC</td><td>GGG</td><td>||||||||[||||||||||||]PAM</td><td>NA</td><td>Intergenic</td><td>NA</td><td>NA</td></tr>
<tr><td></td><td></td><td></td><td></td><td></td><td></td><td></td><td></td><td></td><td></td><td></td><td></td><td></td></tr>
<tr><td>T666</td><td>CGGGCGCACCACCAAAACATCGG</td><td></td><td>1000</td><td></td><td></td><td></td><td></td><td></td><td></td><td></td><td></td><td></td></tr>
<tr><td>Oligo fwd</td><td>CACCCGGGCGCACCACCAAAACAT</td><td></td><td></td><td></td><td></td><td></td><td></td><td></td><td></td><td></td><td></td><td></td></tr>
<tr><td>Oligo rev</td><td>AAACATGTTTTGGTGGTGCGCCCG</td><td></td><td></td><td></td><td></td><td></td><td></td><td></td><td></td><td></td><td></td><td></td></tr>
<tr><td>Chromosome</td><td>start</td><td></td><td>end</td><td>strand</td><td>MM</td><td>target_seq</td><td>PAM</td><td>alignment</td><td>distance</td><td>position</td><td>gene name</td><td>gene id</td></tr>
<tr><td>GU305914.1</td><td></td><td>14383</td><td>14405</td><td>+</td><td>0</td><td>CGGGCGCACCACCAAAACAT</td><td>CGG</td><td>||||||||[||||||||||||]PAM</td><td>NA</td><td>Intergenic</td><td>NA</td><td>NA</td></tr>
<tr><td></td><td></td><td></td><td></td><td></td><td></td><td></td><td></td><td></td><td></td><td></td><td></td><td></td></tr>
<tr><td>T667</td><td>CATCGGCAACCAAGATCGTATGG</td><td></td><td>1000</td><td></td><td></td><td></td><td></td><td></td><td></td><td></td><td></td><td></td></tr>
<tr><td>Oligo fwd</td><td>CACCCATCGGCAACCAAGATCGTA</td><td></td><td></td><td></td><td></td><td></td><td></td><td></td><td></td><td></td><td></td><td></td></tr>
<tr><td>Oligo rev</td><td>AAACTACGATCTTGGTTGCCGATG</td><td></td><td></td><td></td><td></td><td></td><td></td><td></td><td></td><td></td><td></td><td></td></tr>
<tr><td>Chromosome</td><td>start</td><td></td><td>end</td><td>strand</td><td>MM</td><td>target_seq</td><td>PAM</td><td>alignment</td><td>distance</td><td>position</td><td>gene name</td><td>gene id</td></tr>
<tr><td>GU305914.1</td><td></td><td>14400</td><td>14422</td><td>+</td><td>0</td><td>CATCGGCAACCAAGATCGTA</td><td>TGG</td><td>||||||||[||||||||||||]PAM</td><td>NA</td><td>Intergenic</td><td>NA</td><td>NA</td></tr>
<tr><td></td><td></td><td></td><td></td><td></td><td></td><td></td><td></td><td></td><td></td><td></td><td></td><td></td></tr>
<tr><td>T668</td><td>AAGATCGTATGGCGCATATTTGG</td><td></td><td>1000</td><td></td><td></td><td></td><td></td><td></td><td></td><td></td><td></td><td></td></tr>
<tr><td>Oligo fwd</td><td>CACCAAGATCGTATGGCGCATATT</td><td></td><td></td><td></td><td></td><td></td><td></td><td></td><td></td><td></td><td></td><td></td></tr>
<tr><td>Oligo rev</td><td>AAACAATATGCGCCATACGATCTT</td><td></td><td></td><td></td><td></td><td></td><td></td><td></td><td></td><td></td><td></td><td></td></tr>
<tr><td>Chromosome</td><td>start</td><td></td><td>end</td><td>strand</td><td>MM</td><td>target_seq</td><td>PAM</td><td>alignment</td><td>distance</td><td>position</td><td>gene name</td><td>gene id</td></tr>
<tr><td>GU305914.1</td><td></td><td>14411</td><td>14433</td><td>+</td><td>0</td><td>AAGATCGTATGGCGCATATT</td><td>TGG</td><td>||||||||[||||||||||||]PAM</td><td>NA</td><td>Intergenic</td><td>NA</td><td>NA</td></tr>
<tr><td></td><td></td><td></td><td></td><td></td><td></td><td></td><td></td><td></td><td></td><td></td><td></td><td></td></tr>
<tr><td>T669</td><td>ACCTCAATACCGTATTACACTGG</td><td></td><td>1000</td><td></td><td></td><td></td><td></td><td></td><td></td><td></td><td></td><td></td></tr>
<tr><td>Oligo fwd</td><td>CACCACCTCAATACCGTATTACAC</td><td></td><td></td><td></td><td></td><td></td><td></td><td></td><td></td><td></td><td></td><td></td></tr>
<tr><td>Oligo rev</td><td>AAACGTGTAATACGGTATTGAGGT</td><td></td><td></td><td></td><td></td><td></td><td></td><td></td><td></td><td></td><td></td><td></td></tr>
<tr><td>Chromosome</td><td>start</td><td></td><td>end</td><td>strand</td><td>MM</td><td>target_seq</td><td>PAM</td><td>alignment</td><td>distance</td><td>position</td><td>gene name</td><td>gene id</td></tr>
<tr><td>GU305914.1</td><td></td><td>14484</td><td>14506</td><td>+</td><td>0</td><td>ACCTCAATACCGTATTACAC</td><td>TGG</td><td>||||||||[||||||||||||]PAM</td><td>NA</td><td>Intergenic</td><td>NA</td><td>NA</td></tr>
<tr><td></td><td></td><td></td><td></td><td></td><td></td><td></td><td></td><td></td><td></td><td></td><td></td><td></td></tr>
<tr><td>T670</td><td>TACCGTATTACACTGGCACTTGG</td><td></td><td>1000</td><td></td><td></td><td></td><td></td><td></td><td></td><td></td><td></td><td></td></tr>
<tr><td>Oligo fwd</td><td>CACCTACCGTATTACACTGGCACT</td><td></td><td></td><td></td><td></td><td></td><td></td><td></td><td></td><td></td><td></td><td></td></tr>
<tr><td>Oligo rev</td><td>AAACAGTGCCAGTGTAATACGGTA</td><td></td><td></td><td></td><td></td><td></td><td></td><td></td><td></td><td></td><td></td><td></td></tr>
<tr><td>Chromosome</td><td>start</td><td></td><td>end</td><td>strand</td><td>MM</td><td>target_seq</td><td>PAM</td><td>alignment</td><td>distance</td><td>position</td><td>gene name</td><td>gene id</td></tr>
<tr><td>GU305914.1</td><td></td><td>14491</td><td>14513</td><td>+</td><td>0</td><td>TACCGTATTACACTGGCACT</td><td>TGG</td><td>||||||||[||||||||||||]PAM</td><td>NA</td><td>Intergenic</td><td>NA</td><td>NA</td></tr>
<tr><td></td><td></td><td></td><td></td><td></td><td></td><td></td><td></td><td></td><td></td><td></td><td></td><td></td></tr>
<tr><td>T671</td><td>TGGCACTTGGAATAACGATTCGG</td><td></td><td>1000</td><td></td><td></td><td></td><td></td><td></td><td></td><td></td><td></td><td></td></tr>
<tr><td>Oligo fwd</td><td>CACCTGGCACTTGGAATAACGATT</td><td></td><td></td><td></td><td></td><td></td><td></td><td></td><td></td><td></td><td></td><td></td></tr>
<tr><td>Oligo rev</td><td>AAACAATCGTTATTCCAAGTGCCA</td><td></td><td></td><td></td><td></td><td></td><td></td><td></td><td></td><td></td><td></td><td></td></tr>
<tr><td>Chromosome</td><td>start</td><td></td><td>end</td><td>strand</td><td>MM</td><td>target_seq</td><td>PAM</td><td>alignment</td><td>distance</td><td>position</td><td>gene name</td><td>gene id</td></tr>
<tr><td>GU305914.1</td><td></td><td>14504</td><td>14526</td><td>+</td><td>0</td><td>TGGCACTTGGAATAACGATT</td><td>CGG</td><td>||||||||[||||||||||||]PAM</td><td>NA</td><td>Intergenic</td><td>NA</td><td>NA</td></tr>
<tr><td></td><td></td><td></td><td></td><td></td><td></td><td></td><td></td><td></td><td></td><td></td><td></td><td></td></tr>
<tr><td>T672</td><td>CATGTGAATCAGAGCCTCCTAGG</td><td></td><td>1000</td><td></td><td></td><td></td><td></td><td></td><td></td><td></td><td></td><td></td></tr>
<tr><td>Oligo fwd</td><td>CACCCATGTGAATCAGAGCCTCCT</td><td></td><td></td><td></td><td></td><td></td><td></td><td></td><td></td><td></td><td></td><td></td></tr>
<tr><td>Oligo rev</td><td>AAACAGGAGGCTCTGATTCACATG</td><td></td><td></td><td></td><td></td><td></td><td></td><td></td><td></td><td></td><td></td><td></td></tr>
<tr><td>Chromosome</td><td>start</td><td></td><td>end</td><td>strand</td><td>MM</td><td>target_seq</td><td>PAM</td><td>alignment</td><td>distance</td><td>position</td><td>gene name</td><td>gene id</td></tr>
<tr><td>GU305914.1</td><td></td><td>14538</td><td>14560</td><td>+</td><td>0</td><td>CATGTGAATCAGAGCCTCCT</td><td>AGG</td><td>||||||||[||||||||||||]PAM</td><td>NA</td><td>Intergenic</td><td>NA</td><td>NA</td></tr>
<tr><td></td><td></td><td></td><td></td><td></td><td></td><td></td><td></td><td></td><td></td><td></td><td></td><td></td></tr>
<tr><td>T673</td><td>ATGTGAATCAGAGCCTCCTAGGG</td><td></td><td>1000</td><td></td><td></td><td></td><td></td><td></td><td></td><td></td><td></td><td></td></tr>
<tr><td>Oligo fwd</td><td>CACCATGTGAATCAGAGCCTCCTA</td><td></td><td></td><td></td><td></td><td></td><td></td><td></td><td></td><td></td><td></td><td></td></tr>
<tr><td>Oligo rev</td><td>AAACTAGGAGGCTCTGATTCACAT</td><td></td><td></td><td></td><td></td><td></td><td></td><td></td><td></td><td></td><td></td><td></td></tr>
<tr><td>Chromosome</td><td>start</td><td></td><td>end</td><td>strand</td><td>MM</td><td>target_seq</td><td>PAM</td><td>alignment</td><td>distance</td><td>position</td><td>gene name</td><td>gene id</td></tr>
<tr><td>GU305914.1</td><td></td><td>14539</td><td>14561</td><td>+</td><td>0</td><td>ATGTGAATCAGAGCCTCCTA</td><td>GGG</td><td>||||||||[||||||||||||]PAM</td><td>NA</td><td>Intergenic</td><td>NA</td><td>NA</td></tr>
<tr><td></td><td></td><td></td><td></td><td></td><td></td><td></td><td></td><td></td><td></td><td></td><td></td><td></td></tr>
<tr><td>T674</td><td>ATACACTTCAATCAACAAAATGG</td><td></td><td>1000</td><td></td><td></td><td></td><td></td><td></td><td></td><td></td><td></td><td></td></tr>
<tr><td>Oligo fwd</td><td>CACCATACACTTCAATCAACAAAA</td><td></td><td></td><td></td><td></td><td></td><td></td><td></td><td></td><td></td><td></td><td></td></tr>
</table>

<table>
<tr><td>Oligo rev</td><td>AAACTTTTGTTGATTGAAGTGTAT</td><td></td><td></td><td></td><td></td><td></td><td></td><td></td><td></td><td></td><td></td><td></td></tr>
<tr><td>Chromosome</td><td>start</td><td></td><td>end</td><td>strand</td><td>MM</td><td>target_seq</td><td>PAM</td><td>alignment</td><td>distance</td><td>position</td><td>gene name</td><td>gene id</td></tr>
<tr><td>GU305914.1</td><td></td><td>14562</td><td>14584</td><td>+</td><td>0</td><td>ATACACTTCAATCAACAAAA</td><td>TGG</td><td>||||||||[||||||||||||]PAM</td><td>NA</td><td>Intergenic</td><td>NA</td><td>NA</td></tr>
<tr><td>T675</td><td>ATATCATAATTTTGAATGTGAGG</td><td></td><td>1000</td><td></td><td></td><td></td><td></td><td></td><td></td><td></td><td></td><td></td></tr>
<tr><td>Oligo fwd</td><td>CACCATATCATAATTTTGAATGTG</td><td></td><td></td><td></td><td></td><td></td><td></td><td></td><td></td><td></td><td></td><td></td></tr>
<tr><td>Oligo rev</td><td>AAACCACATTCAAAATTATGATAT</td><td></td><td></td><td></td><td></td><td></td><td></td><td></td><td></td><td></td><td></td><td></td></tr>
<tr><td>Chromosome</td><td>start</td><td></td><td>end</td><td>strand</td><td>MM</td><td>target_seq</td><td>PAM</td><td>alignment</td><td>distance</td><td>position</td><td>gene name</td><td>gene id</td></tr>
<tr><td>GU305914.1</td><td></td><td>14618</td><td>14640</td><td>+</td><td>0</td><td>ATATCATAATTTTGAATGTG</td><td>AGG</td><td>||||||||[||||||||||||]PAM</td><td>NA</td><td>Intergenic</td><td>NA</td><td>NA</td></tr>
<tr><td>T676</td><td>CCGTCGACTCCACCACCCAACGG</td><td></td><td>1000</td><td></td><td></td><td></td><td></td><td></td><td></td><td></td><td></td><td></td></tr>
<tr><td>Oligo fwd</td><td>CACCCCGTCGACTCCACCACCCAA</td><td></td><td></td><td></td><td></td><td></td><td></td><td></td><td></td><td></td><td></td><td></td></tr>
<tr><td>Oligo rev</td><td>AAACTTGGGTGGTGGAGTCGACGG</td><td></td><td></td><td></td><td></td><td></td><td></td><td></td><td></td><td></td><td></td><td></td></tr>
<tr><td>Chromosome</td><td>start</td><td></td><td>end</td><td>strand</td><td>MM</td><td>target_seq</td><td>PAM</td><td>alignment</td><td>distance</td><td>position</td><td>gene name</td><td>gene id</td></tr>
<tr><td>GU305914.1</td><td></td><td>14691</td><td>14713</td><td>+</td><td>0</td><td>CCGTCGACTCCACCACCCAA</td><td>CGG</td><td>||||||||[||||||||||||]PAM</td><td>NA</td><td>Intergenic</td><td>NA</td><td>NA</td></tr>
<tr><td>T677</td><td>AGAATCGCAAACGACATCAAAGG</td><td></td><td>1000</td><td></td><td></td><td></td><td></td><td></td><td></td><td></td><td></td><td></td></tr>
<tr><td>Oligo fwd</td><td>CACCAGAATCGCAAACGACATCAA</td><td></td><td></td><td></td><td></td><td></td><td></td><td></td><td></td><td></td><td></td><td></td></tr>
<tr><td>Oligo rev</td><td>AAACTTGATGTCGTTTGCGATTCT</td><td></td><td></td><td></td><td></td><td></td><td></td><td></td><td></td><td></td><td></td><td></td></tr>
<tr><td>Chromosome</td><td>start</td><td></td><td>end</td><td>strand</td><td>MM</td><td>target_seq</td><td>PAM</td><td>alignment</td><td>distance</td><td>position</td><td>gene name</td><td>gene id</td></tr>
<tr><td>GU305914.1</td><td></td><td>14762</td><td>14784</td><td>+</td><td>0</td><td>AGAATCGCAAACGACATCAA</td><td>AGG</td><td>||||||||[||||||||||||]PAM</td><td>NA</td><td>Intergenic</td><td>NA</td><td>NA</td></tr>
<tr><td>T678</td><td>CAAGATATATTCGAATACAACGG</td><td></td><td>1000</td><td></td><td></td><td></td><td></td><td></td><td></td><td></td><td></td><td></td></tr>
<tr><td>Oligo fwd</td><td>CACCCAAGATATATTCGAATACAA</td><td></td><td></td><td></td><td></td><td></td><td></td><td></td><td></td><td></td><td></td><td></td></tr>
<tr><td>Oligo rev</td><td>AAACTTGTATTCGAATATATCTTG</td><td></td><td></td><td></td><td></td><td></td><td></td><td></td><td></td><td></td><td></td><td></td></tr>
<tr><td>Chromosome</td><td>start</td><td></td><td>end</td><td>strand</td><td>MM</td><td>target_seq</td><td>PAM</td><td>alignment</td><td>distance</td><td>position</td><td>gene name</td><td>gene id</td></tr>
<tr><td>GU305914.1</td><td></td><td>14813</td><td>14835</td><td>+</td><td>0</td><td>CAAGATATATTCGAATACAA</td><td>CGG</td><td>||||||||[||||||||||||]PAM</td><td>NA</td><td>Intergenic</td><td>NA</td><td>NA</td></tr>
<tr><td>T679</td><td>GAATTCGTTTTTTTACATTCTGG</td><td></td><td>1000</td><td></td><td></td><td></td><td></td><td></td><td></td><td></td><td></td><td></td></tr>
<tr><td>Oligo fwd</td><td>CACCGAATTCGTTTTTTTACATTC</td><td></td><td></td><td></td><td></td><td></td><td></td><td></td><td></td><td></td><td></td><td></td></tr>
<tr><td>Oligo rev</td><td>AAACGAATGTAAAAAAACGAATTC</td><td></td><td></td><td></td><td></td><td></td><td></td><td></td><td></td><td></td><td></td><td></td></tr>
<tr><td>Chromosome</td><td>start</td><td></td><td>end</td><td>strand</td><td>MM</td><td>target_seq</td><td>PAM</td><td>alignment</td><td>distance</td><td>position</td><td>gene name</td><td>gene id</td></tr>
<tr><td>GU305914.1</td><td></td><td>14964</td><td>14986</td><td>+</td><td>0</td><td>GAATTCGTTTTTTTACATTC</td><td>TGG</td><td>||||||||[||||||||||||]PAM</td><td>NA</td><td>Intergenic</td><td>NA</td><td>NA</td></tr>
<tr><td>T680</td><td>TCGTCGCAATGAGCCTCCTATGG</td><td></td><td>1000</td><td></td><td></td><td></td><td></td><td></td><td></td><td></td><td></td><td></td></tr>
<tr><td>Oligo fwd</td><td>CACCTCGTCGCAATGAGCCTCCTA</td><td></td><td></td><td></td><td></td><td></td><td></td><td></td><td></td><td></td><td></td><td></td></tr>
<tr><td>Oligo rev</td><td>AAACTAGGAGGCTCATTGCGACGA</td><td></td><td></td><td></td><td></td><td></td><td></td><td></td><td></td><td></td><td></td><td></td></tr>
<tr><td>Chromosome</td><td>start</td><td></td><td>end</td><td>strand</td><td>MM</td><td>target_seq</td><td>PAM</td><td>alignment</td><td>distance</td><td>position</td><td>gene name</td><td>gene id</td></tr>
<tr><td>GU305914.1</td><td></td><td>15025</td><td>15047</td><td>+</td><td>0</td><td>TCGTCGCAATGAGCCTCCTA</td><td>TGG</td><td>||||||||[||||||||||||]PAM</td><td>NA</td><td>Intergenic</td><td>NA</td><td>NA</td></tr>
<tr><td>T681</td><td>TTTACAGAGATTTTTATGTACGG</td><td></td><td>1000</td><td></td><td></td><td></td><td></td><td></td><td></td><td></td><td></td><td></td></tr>
<tr><td>Oligo fwd</td><td>CACCTTTACAGAGATTTTTATGTA</td><td></td><td></td><td></td><td></td><td></td><td></td><td></td><td></td><td></td><td></td><td></td></tr>
<tr><td>Oligo rev</td><td>AAACTACATAAAAATCTCTGTAAA</td><td></td><td></td><td></td><td></td><td></td><td></td><td></td><td></td><td></td><td></td><td></td></tr>
<tr><td>Chromosome</td><td>start</td><td></td><td>end</td><td>strand</td><td>MM</td><td>target_seq</td><td>PAM</td><td>alignment</td><td>distance</td><td>position</td><td>gene name</td><td>gene id</td></tr>
<tr><td>GU305914.1</td><td></td><td>15127</td><td>15149</td><td>+</td><td>0</td><td>TTTACAGAGATTTTTATGTA</td><td>CGG</td><td>||||||||[||||||||||||]PAM</td><td>NA</td><td>Intergenic</td><td>NA</td><td>NA</td></tr>
<tr><td>T682</td><td>ATACGACCTTGATATCGCACTGG</td><td></td><td>1000</td><td></td><td></td><td></td><td></td><td></td><td></td><td></td><td></td><td></td></tr>
<tr><td>Oligo fwd</td><td>CACCATACGACCTTGATATCGCAC</td><td></td><td></td><td></td><td></td><td></td><td></td><td></td><td></td><td></td><td></td><td></td></tr>
<tr><td>Oligo rev</td><td>AAACGTGCGATATCAAGGTCGTAT</td><td></td><td></td><td></td><td></td><td></td><td></td><td></td><td></td><td></td><td></td><td></td></tr>
<tr><td>Chromosome</td><td>start</td><td></td><td>end</td><td>strand</td><td>MM</td><td>target_seq</td><td>PAM</td><td>alignment</td><td>distance</td><td>position</td><td>gene name</td><td>gene id</td></tr>
<tr><td>GU305914.1</td><td></td><td>15166</td><td>15188</td><td>+</td><td>0</td><td>ATACGACCTTGATATCGCAC</td><td>TGG</td><td>||||||||[||||||||||||]PAM</td><td>NA</td><td>Intergenic</td><td>NA</td><td>NA</td></tr>
<tr><td>T683</td><td>ACTGGACGCTTGTAAAGAACAGG</td><td></td><td>1000</td><td></td><td></td><td></td><td></td><td></td><td></td><td></td><td></td><td></td></tr>
<tr><td>Oligo fwd</td><td>CACCACTGGACGCTTGTAAAGAAC</td><td></td><td></td><td></td><td></td><td></td><td></td><td></td><td></td><td></td><td></td><td></td></tr>
<tr><td>Oligo rev</td><td>AAACGTTCTTTACAAGCGTCCAGT</td><td></td><td></td><td></td><td></td><td></td><td></td><td></td><td></td><td></td><td></td><td></td></tr>
<tr><td>Chromosome</td><td>start</td><td></td><td>end</td><td>strand</td><td>MM</td><td>target_seq</td><td>PAM</td><td>alignment</td><td>distance</td><td>position</td><td>gene name</td><td>gene id</td></tr>
<tr><td>GU305914.1</td><td></td><td>15184</td><td>15206</td><td>+</td><td>0</td><td>ACTGGACGCTTGTAAAGAAC</td><td>AGG</td><td>||||||||[||||||||||||]PAM</td><td>NA</td><td>Intergenic</td><td>NA</td><td>NA</td></tr>
<tr><td>T684</td><td>GACGCTTGTAAAGAACAGGATGG</td><td></td><td>1000</td><td></td><td></td><td></td><td></td><td></td><td></td><td></td><td></td><td></td></tr>
<tr><td>Oligo fwd</td><td>CACCGACGCTTGTAAAGAACAGGA</td><td></td><td></td><td></td><td></td><td></td><td></td><td></td><td></td><td></td><td></td><td></td></tr>
<tr><td>Oligo rev</td><td>AAACTCCTGTTCTTTACAAGCGTC</td><td></td><td></td><td></td><td></td><td></td><td></td><td></td><td></td><td></td><td></td><td></td></tr>
<tr><td>Chromosome</td><td>start</td><td></td><td>end</td><td>strand</td><td>MM</td><td>target_seq</td><td>PAM</td><td>alignment</td><td>distance</td><td>position</td><td>gene name</td><td>gene id</td></tr>
<tr><td>GU305914.1</td><td></td><td>15188</td><td>15210</td><td>+</td><td>0</td><td>GACGCTTGTAAAGAACAGGA</td><td>TGG</td><td>||||||||[||||||||||||]PAM</td><td>NA</td><td>Intergenic</td><td>NA</td><td>NA</td></tr>
<tr><td>T685</td><td>GCTTGTAAAGAACAGGATGGAGG</td><td></td><td>1000</td><td></td><td></td><td></td><td></td><td></td><td></td><td></td><td></td><td></td></tr>
<tr><td>Oligo fwd</td><td>CACCGCTTGTAAAGAACAGGATGG</td><td></td><td></td><td></td><td></td><td></td><td></td><td></td><td></td><td></td><td></td><td></td></tr>
<tr><td>Oligo rev</td><td>AAACCCATCCTGTTCTTTACAAGC</td><td></td><td></td><td></td><td></td><td></td><td></td><td></td><td></td><td></td><td></td><td></td></tr>
<tr><td>Chromosome</td><td>start</td><td></td><td>end</td><td>strand</td><td>MM</td><td>target_seq</td><td>PAM</td><td>alignment</td><td>distance</td><td>position</td><td>gene name</td><td>gene id</td></tr>
<tr><td>GU305914.1</td><td></td><td>15191</td><td>15213</td><td>+</td><td>0</td><td>GCTTGTAAAGAACAGGATGG</td><td>AGG</td><td>||||||||[||||||||||||]PAM</td><td>NA</td><td>Intergenic</td><td>NA</td><td>NA</td></tr>
<tr><td>T686</td><td>CTGTTTGCGCATCTGCATCGTGG</td><td></td><td>1000</td><td></td><td></td><td></td><td></td><td></td><td></td><td></td><td></td><td></td></tr>
<tr><td>Oligo fwd</td><td>CACCCTGTTTGCGCATCTGCATCG</td><td></td><td></td><td></td><td></td><td></td><td></td><td></td><td></td><td></td><td></td><td></td></tr>
<tr><td>Oligo rev</td><td>AAACCGATGCAGATGCGCAAACAG</td><td></td><td></td><td></td><td></td><td></td><td></td><td></td><td></td><td></td><td></td><td></td></tr>
<tr><td>Chromosome</td><td>start</td><td></td><td>end</td><td>strand</td><td>MM</td><td>target_seq</td><td>PAM</td><td>alignment</td><td>distance</td><td>position</td><td>gene name</td><td>gene id</td></tr>
<tr><td>GU305914.1</td><td></td><td>15270</td><td>15292</td><td>+</td><td>0</td><td>CTGTTTGCGCATCTGCATCG</td><td>TGG</td><td>||||||||[||||||||||||]PAM</td><td>NA</td><td>Intergenic</td><td>NA</td><td>NA</td></tr>
<tr><td>T687</td><td>CTGCATCGTGGTACGCAACGTGG</td><td></td><td>1000</td><td></td><td></td><td></td><td></td><td></td><td></td><td></td><td></td><td></td></tr>
<tr><td>Oligo fwd</td><td>CACCCTGCATCGTGGTACGCAACG</td><td></td><td></td><td></td><td></td><td></td><td></td><td></td><td></td><td></td><td></td><td></td></tr>
<tr><td>Oligo rev</td><td>AAACCGTTGCGTACCACGATGCAG</td><td></td><td></td><td></td><td></td><td></td><td></td><td></td><td></td><td></td><td></td><td></td></tr>
<tr><td>Chromosome</td><td>start</td><td></td><td>end</td><td>strand</td><td>MM</td><td>target_seq</td><td>PAM</td><td>alignment</td><td>distance</td><td>position</td><td>gene name</td><td>gene id</td></tr>
<tr><td>GU305914.1</td><td></td><td>15282</td><td>15304</td><td>+</td><td>0</td><td>CTGCATCGTGGTACGCAACG</td><td>TGG</td><td>||||||||[||||||||||||]PAM</td><td>NA</td><td>Intergenic</td><td>NA</td><td>NA</td></tr>
<tr><td>T688</td><td>TGCATCGTGGTACGCAACGTGGG</td><td></td><td>1000</td><td></td><td></td><td></td><td></td><td></td><td></td><td></td><td></td><td></td></tr>
<tr><td>Oligo fwd</td><td>CACCTGCATCGTGGTACGCAACGT</td><td></td><td></td><td></td><td></td><td></td><td></td><td></td><td></td><td></td><td></td><td></td></tr>
<tr><td>Oligo rev</td><td>AAACACGTTGCGTACCACGATGCA</td><td></td><td></td><td></td><td></td><td></td><td></td><td></td><td></td><td></td><td></td><td></td></tr>
<tr><td>Chromosome</td><td>start</td><td></td><td>end</td><td>strand</td><td>MM</td><td>target_seq</td><td>PAM</td><td>alignment</td><td>distance</td><td>position</td><td>gene name</td><td>gene id</td></tr>
<tr><td>GU305914.1</td><td></td><td>15283</td><td>15305</td><td>+</td><td>0</td><td>TGCATCGTGGTACGCAACGT</td><td>GGG</td><td>||||||||[||||||||||||]PAM</td><td>NA</td><td>Intergenic</td><td>NA</td><td>NA</td></tr>
<tr><td>T689</td><td>ACGTGGGTTGCATCTTCATCTGG</td><td></td><td>1000</td><td></td><td></td><td></td><td></td><td></td><td></td><td></td><td></td><td></td></tr>
</table>

<table>
<tr><td>Oligo fwd</td><td>CACCACGTGGGTTGCATCTTCATC</td><td></td><td></td><td></td><td></td><td></td><td></td><td></td><td></td><td></td><td></td></tr>
<tr><td>Oligo rev</td><td>AAACGATGAAGATGCAACCCACGT</td><td></td><td></td><td></td><td></td><td></td><td></td><td></td><td></td><td></td><td></td></tr>
<tr><td>Chromosome</td><td>start</td><td>end</td><td>strand</td><td>MM</td><td>target_seq</td><td>PAM</td><td>alignment</td><td>distance</td><td>position</td><td>gene name</td><td>gene id</td></tr>
<tr><td>GU305914.1</td><td>15299</td><td>15321</td><td>+</td><td>0</td><td>ACGTGGGTTGCATCTTCATC</td><td>TGG</td><td>||||||||[||||||||||||]PAM</td><td>NA</td><td>Intergenic</td><td>NA</td><td>NA</td></tr>
<tr><td>T690</td><td>TCTGGCACACAATACGTTATCGG</td><td>1000</td><td></td><td></td><td></td><td></td><td></td><td></td><td></td><td></td><td></td></tr>
<tr><td>Oligo fwd</td><td>CACCTCTGGCACACAATACGTTAT</td><td></td><td></td><td></td><td></td><td></td><td></td><td></td><td></td><td></td><td></td></tr>
<tr><td>Oligo rev</td><td>AAACATAACGTATTGTGTGCCAGA</td><td></td><td></td><td></td><td></td><td></td><td></td><td></td><td></td><td></td><td></td></tr>
<tr><td>Chromosome</td><td>start</td><td>end</td><td>strand</td><td>MM</td><td>target_seq</td><td>PAM</td><td>alignment</td><td>distance</td><td>position</td><td>gene name</td><td>gene id</td></tr>
<tr><td>GU305914.1</td><td>15317</td><td>15339</td><td>+</td><td>0</td><td>TCTGGCACACAATACGTTAT</td><td>CGG</td><td>||||||||[||||||||||||]PAM</td><td>NA</td><td>Intergenic</td><td>NA</td><td>NA</td></tr>
<tr><td>T691</td><td>TATACTTCTTCGTCAACATCCGG</td><td>1000</td><td></td><td></td><td></td><td></td><td></td><td></td><td></td><td></td><td></td></tr>
<tr><td>Oligo fwd</td><td>CACCTATACTTCTTCGTCAACATC</td><td></td><td></td><td></td><td></td><td></td><td></td><td></td><td></td><td></td><td></td></tr>
<tr><td>Oligo rev</td><td>AAACGATGTTGACGAAGAAGTATA</td><td></td><td></td><td></td><td></td><td></td><td></td><td></td><td></td><td></td><td></td></tr>
<tr><td>Chromosome</td><td>start</td><td>end</td><td>strand</td><td>MM</td><td>target_seq</td><td>PAM</td><td>alignment</td><td>distance</td><td>position</td><td>gene name</td><td>gene id</td></tr>
<tr><td>GU305914.1</td><td>15350</td><td>15372</td><td>+</td><td>0</td><td>TATACTTCTTCGTCAACATC</td><td>CGG</td><td>||||||||[||||||||||||]PAM</td><td>NA</td><td>Intergenic</td><td>NA</td><td>NA</td></tr>
<tr><td>T692</td><td>GCGCGACATTCACCGACCTGTGG</td><td>1000</td><td></td><td></td><td></td><td></td><td></td><td></td><td></td><td></td><td></td></tr>
<tr><td>Oligo fwd</td><td>CACCGCGCGACATTCACCGACCTG</td><td></td><td></td><td></td><td></td><td></td><td></td><td></td><td></td><td></td><td></td></tr>
<tr><td>Oligo rev</td><td>AAACCAGGTCGGTGAATGTCGCGC</td><td></td><td></td><td></td><td></td><td></td><td></td><td></td><td></td><td></td><td></td></tr>
<tr><td>Chromosome</td><td>start</td><td>end</td><td>strand</td><td>MM</td><td>target_seq</td><td>PAM</td><td>alignment</td><td>distance</td><td>position</td><td>gene name</td><td>gene id</td></tr>
<tr><td>GU305914.1</td><td>15372</td><td>15394</td><td>+</td><td>0</td><td>GCGCGACATTCACCGACCTG</td><td>TGG</td><td>||||||||[||||||||||||]PAM</td><td>NA</td><td>Intergenic</td><td>NA</td><td>NA</td></tr>
<tr><td>T693</td><td>CGACCTGTGGACAAATTCTTCGG</td><td>1000</td><td></td><td></td><td></td><td></td><td></td><td></td><td></td><td></td><td></td></tr>
<tr><td>Oligo fwd</td><td>CACCCGACCTGTGGACAAATTCTT</td><td></td><td></td><td></td><td></td><td></td><td></td><td></td><td></td><td></td><td></td></tr>
<tr><td>Oligo rev</td><td>AAACAAGAATTTGTCCACAGGTCG</td><td></td><td></td><td></td><td></td><td></td><td></td><td></td><td></td><td></td><td></td></tr>
<tr><td>Chromosome</td><td>start</td><td>end</td><td>strand</td><td>MM</td><td>target_seq</td><td>PAM</td><td>alignment</td><td>distance</td><td>position</td><td>gene name</td><td>gene id</td></tr>
<tr><td>GU305914.1</td><td>15385</td><td>15407</td><td>+</td><td>0</td><td>CGACCTGTGGACAAATTCTT</td><td>CGG</td><td>||||||||[||||||||||||]PAM</td><td>NA</td><td>Intergenic</td><td>NA</td><td>NA</td></tr>
<tr><td>T694</td><td>CTCAAACTCGACAACAAAACCGG</td><td>1000</td><td></td><td></td><td></td><td></td><td></td><td></td><td></td><td></td><td></td></tr>
<tr><td>Oligo fwd</td><td>CACCCTCAAACTCGACAACAAAAC</td><td></td><td></td><td></td><td></td><td></td><td></td><td></td><td></td><td></td><td></td></tr>
<tr><td>Oligo rev</td><td>AAACGTTTTGTTGTCGAGTTTGAG</td><td></td><td></td><td></td><td></td><td></td><td></td><td></td><td></td><td></td><td></td></tr>
<tr><td>Chromosome</td><td>start</td><td>end</td><td>strand</td><td>MM</td><td>target_seq</td><td>PAM</td><td>alignment</td><td>distance</td><td>position</td><td>gene name</td><td>gene id</td></tr>
<tr><td>GU305914.1</td><td>15437</td><td>15459</td><td>+</td><td>0</td><td>CTCAAACTCGACAACAAAAC</td><td>CGG</td><td>||||||||[||||||||||||]PAM</td><td>NA</td><td>Intergenic</td><td>NA</td><td>NA</td></tr>
<tr><td>T695</td><td>TCTCCTTCATGTTCGAGCAGTGG</td><td>1000</td><td></td><td></td><td></td><td></td><td></td><td></td><td></td><td></td><td></td></tr>
<tr><td>Oligo fwd</td><td>CACCTCTCCTTCATGTTCGAGCAG</td><td></td><td></td><td></td><td></td><td></td><td></td><td></td><td></td><td></td><td></td></tr>
<tr><td>Oligo rev</td><td>AAACCTGCTCGAACATGAAGGAGA</td><td></td><td></td><td></td><td></td><td></td><td></td><td></td><td></td><td></td><td></td></tr>
<tr><td>Chromosome</td><td>start</td><td>end</td><td>strand</td><td>MM</td><td>target_seq</td><td>PAM</td><td>alignment</td><td>distance</td><td>position</td><td>gene name</td><td>gene id</td></tr>
<tr><td>GU305914.1</td><td>15473</td><td>15495</td><td>+</td><td>0</td><td>TCTCCTTCATGTTCGAGCAG</td><td>TGG</td><td>||||||||[||||||||||||]PAM</td><td>NA</td><td>Intergenic</td><td>NA</td><td>NA</td></tr>
<tr><td>T696</td><td>TATTGTACGCCATCGAAATAAGG</td><td>1000</td><td></td><td></td><td></td><td></td><td></td><td></td><td></td><td></td><td></td></tr>
<tr><td>Oligo fwd</td><td>CACCTATTGTACGCCATCGAAATA</td><td></td><td></td><td></td><td></td><td></td><td></td><td></td><td></td><td></td><td></td></tr>
<tr><td>Oligo rev</td><td>AAACTATTTCGATGGCGTACAATA</td><td></td><td></td><td></td><td></td><td></td><td></td><td></td><td></td><td></td><td></td></tr>
<tr><td>Chromosome</td><td>start</td><td>end</td><td>strand</td><td>MM</td><td>target_seq</td><td>PAM</td><td>alignment</td><td>distance</td><td>position</td><td>gene name</td><td>gene id</td></tr>
<tr><td>GU305914.1</td><td>15721</td><td>15743</td><td>+</td><td>0</td><td>TATTGTACGCCATCGAAATA</td><td>AGG</td><td>||||||||[||||||||||||]PAM</td><td>NA</td><td>Intergenic</td><td>NA</td><td>NA</td></tr>
<tr><td>T697</td><td>ATTGTACGCCATCGAAATAAGGG</td><td>1000</td><td></td><td></td><td></td><td></td><td></td><td></td><td></td><td></td><td></td></tr>
<tr><td>Oligo fwd</td><td>CACCATTGTACGCCATCGAAATAA</td><td></td><td></td><td></td><td></td><td></td><td></td><td></td><td></td><td></td><td></td></tr>
<tr><td>Oligo rev</td><td>AAACTTATTTCGATGGCGTACAAT</td><td></td><td></td><td></td><td></td><td></td><td></td><td></td><td></td><td></td><td></td></tr>
<tr><td>Chromosome</td><td>start</td><td>end</td><td>strand</td><td>MM</td><td>target_seq</td><td>PAM</td><td>alignment</td><td>distance</td><td>position</td><td>gene name</td><td>gene id</td></tr>
<tr><td>GU305914.1</td><td>15722</td><td>15744</td><td>+</td><td>0</td><td>ATTGTACGCCATCGAAATAA</td><td>GGG</td><td>||||||||[||||||||||||]PAM</td><td>NA</td><td>Intergenic</td><td>NA</td><td>NA</td></tr>
<tr><td>T698</td><td>TTGTACGCCATCGAAATAAGGGG</td><td>1000</td><td></td><td></td><td></td><td></td><td></td><td></td><td></td><td></td><td></td></tr>
<tr><td>Oligo fwd</td><td>CACCTTGTACGCCATCGAAATAAG</td><td></td><td></td><td></td><td></td><td></td><td></td><td></td><td></td><td></td><td></td></tr>
<tr><td>Oligo rev</td><td>AAACCTTATTTCGATGGCGTACAA</td><td></td><td></td><td></td><td></td><td></td><td></td><td></td><td></td><td></td><td></td></tr>
<tr><td>Chromosome</td><td>start</td><td>end</td><td>strand</td><td>MM</td><td>target_seq</td><td>PAM</td><td>alignment</td><td>distance</td><td>position</td><td>gene name</td><td>gene id</td></tr>
<tr><td>GU305914.1</td><td>15723</td><td>15745</td><td>+</td><td>0</td><td>TTGTACGCCATCGAAATAAG</td><td>GGG</td><td>||||||||[||||||||||||]PAM</td><td>NA</td><td>Intergenic</td><td>NA</td><td>NA</td></tr>
<tr><td>T699</td><td>AAATAAGGGGAGCTCAGACTTGG</td><td>1000</td><td></td><td></td><td></td><td></td><td></td><td></td><td></td><td></td><td></td></tr>
<tr><td>Oligo fwd</td><td>CACCAAATAAGGGGAGCTCAGACT</td><td></td><td></td><td></td><td></td><td></td><td></td><td></td><td></td><td></td><td></td></tr>
<tr><td>Oligo rev</td><td>AAACAGTCTGAGCTCCCCTTATTT</td><td></td><td></td><td></td><td></td><td></td><td></td><td></td><td></td><td></td><td></td></tr>
<tr><td>Chromosome</td><td>start</td><td>end</td><td>strand</td><td>MM</td><td>target_seq</td><td>PAM</td><td>alignment</td><td>distance</td><td>position</td><td>gene name</td><td>gene id</td></tr>
<tr><td>GU305914.1</td><td>15736</td><td>15758</td><td>+</td><td>0</td><td>AAATAAGGGGAGCTCAGACT</td><td>TGG</td><td>||||||||[||||||||||||]PAM</td><td>NA</td><td>Intergenic</td><td>NA</td><td>NA</td></tr>
<tr><td>T700</td><td>AATAAGGGGAGCTCAGACTTGGG</td><td>1000</td><td></td><td></td><td></td><td></td><td></td><td></td><td></td><td></td><td></td></tr>
<tr><td>Oligo fwd</td><td>CACCAATAAGGGGAGCTCAGACTT</td><td></td><td></td><td></td><td></td><td></td><td></td><td></td><td></td><td></td><td></td></tr>
<tr><td>Oligo rev</td><td>AAACAAGTCTGAGCTCCCCTTATT</td><td></td><td></td><td></td><td></td><td></td><td></td><td></td><td></td><td></td><td></td></tr>
<tr><td>Chromosome</td><td>start</td><td>end</td><td>strand</td><td>MM</td><td>target_seq</td><td>PAM</td><td>alignment</td><td>distance</td><td>position</td><td>gene name</td><td>gene id</td></tr>
<tr><td>GU305914.1</td><td>15737</td><td>15759</td><td>+</td><td>0</td><td>AATAAGGGGAGCTCAGACTT</td><td>GGG</td><td>||||||||[||||||||||||]PAM</td><td>NA</td><td>Intergenic</td><td>NA</td><td>NA</td></tr>
<tr><td>T701</td><td>ATAAGGGGAGCTCAGACTTGGGG</td><td>1000</td><td></td><td></td><td></td><td></td><td></td><td></td><td></td><td></td><td></td></tr>
<tr><td>Oligo fwd</td><td>CACCATAAGGGGAGCTCAGACTTG</td><td></td><td></td><td></td><td></td><td></td><td></td><td></td><td></td><td></td><td></td></tr>
<tr><td>Oligo rev</td><td>AAACCAAGTCTGAGCTCCCCTTAT</td><td></td><td></td><td></td><td></td><td></td><td></td><td></td><td></td><td></td><td></td></tr>
<tr><td>Chromosome</td><td>start</td><td>end</td><td>strand</td><td>MM</td><td>target_seq</td><td>PAM</td><td>alignment</td><td>distance</td><td>position</td><td>gene name</td><td>gene id</td></tr>
<tr><td>GU305914.1</td><td>15738</td><td>15760</td><td>+</td><td>0</td><td>ATAAGGGGAGCTCAGACTTG</td><td>GGG</td><td>||||||||[||||||||||||]PAM</td><td>NA</td><td>Intergenic</td><td>NA</td><td>NA</td></tr>
<tr><td>T702</td><td>CTTGGGGCAGACGCTGAAATCGG</td><td>1000</td><td></td><td></td><td></td><td></td><td></td><td></td><td></td><td></td><td></td></tr>
<tr><td>Oligo fwd</td><td>CACCCTTGGGGCAGACGCTGAAAT</td><td></td><td></td><td></td><td></td><td></td><td></td><td></td><td></td><td></td><td></td></tr>
<tr><td>Oligo rev</td><td>AAACATTTCAGCGTCTGCCCCAAG</td><td></td><td></td><td></td><td></td><td></td><td></td><td></td><td></td><td></td><td></td></tr>
<tr><td>Chromosome</td><td>start</td><td>end</td><td>strand</td><td>MM</td><td>target_seq</td><td>PAM</td><td>alignment</td><td>distance</td><td>position</td><td>gene name</td><td>gene id</td></tr>
<tr><td>GU305914.1</td><td>15754</td><td>15776</td><td>+</td><td>0</td><td>CTTGGGGCAGACGCTGAAAT</td><td>CGG</td><td>||||||||[||||||||||||]PAM</td><td>NA</td><td>Intergenic</td><td>NA</td><td>NA</td></tr>
<tr><td>T703</td><td>GGGGCAGACGCTGAAATCGGTGG</td><td>1000</td><td></td><td></td><td></td><td></td><td></td><td></td><td></td><td></td><td></td></tr>
<tr><td>Oligo fwd</td><td>CACCGGGGCAGACGCTGAAATCGG</td><td></td><td></td><td></td><td></td><td></td><td></td><td></td><td></td><td></td><td></td></tr>
<tr><td>Oligo rev</td><td>AAACCCGATTTCAGCGTCTGCCCC</td><td></td><td></td><td></td><td></td><td></td><td></td><td></td><td></td><td></td><td></td></tr>
<tr><td>Chromosome</td><td>start</td><td>end</td><td>strand</td><td>MM</td><td>target_seq</td><td>PAM</td><td>alignment</td><td>distance</td><td>position</td><td>gene name</td><td>gene id</td></tr>
<tr><td>GU305914.1</td><td>15757</td><td>15779</td><td>+</td><td>0</td><td>GGGGCAGACGCTGAAATCGG</td><td>TGG</td><td>||||||||[||||||||||||]PAM</td><td>NA</td><td>Intergenic</td><td>NA</td><td>NA</td></tr>
</table>

<table>
<tr><td>T704</td><td>CTATTCCTCTGTAACGTGAGCGG</td><td></td><td>1000</td><td></td><td></td><td></td><td></td><td></td><td></td><td></td><td></td><td></td></tr>
<tr><td>Oligo fwd</td><td>CACCCTATTCCTCTGTAACGTGAG</td><td></td><td></td><td></td><td></td><td></td><td></td><td></td><td></td><td></td><td></td><td></td></tr>
<tr><td>Oligo rev</td><td>AAACCTCACGTTACAGAGGAATAG</td><td></td><td></td><td></td><td></td><td></td><td></td><td></td><td></td><td></td><td></td><td></td></tr>
<tr><td>Chromosome</td><td>start</td><td></td><td>end</td><td>strand</td><td>MM</td><td>target_seq</td><td>PAM</td><td>alignment</td><td>distance</td><td>position</td><td>gene name</td><td>gene id</td></tr>
<tr><td>GU305914.1</td><td></td><td>15790</td><td>15812</td><td>+</td><td>0</td><td>CTATTCCTCTGTAACGTGAG</td><td>CGG</td><td>||||||||[||||||||||||]PAM</td><td>NA</td><td>Intergenic</td><td>NA</td><td>NA</td></tr>
<tr><td>T705</td><td>TATTCCTCTGTAACGTGAGCGGG</td><td></td><td>1000</td><td></td><td></td><td></td><td></td><td></td><td></td><td></td><td></td><td></td></tr>
<tr><td>Oligo fwd</td><td>CACCTATTCCTCTGTAACGTGAGC</td><td></td><td></td><td></td><td></td><td></td><td></td><td></td><td></td><td></td><td></td><td></td></tr>
<tr><td>Oligo rev</td><td>AAACGCTCACGTTACAGAGGAATA</td><td></td><td></td><td></td><td></td><td></td><td></td><td></td><td></td><td></td><td></td><td></td></tr>
<tr><td>Chromosome</td><td>start</td><td></td><td>end</td><td>strand</td><td>MM</td><td>target_seq</td><td>PAM</td><td>alignment</td><td>distance</td><td>position</td><td>gene name</td><td>gene id</td></tr>
<tr><td>GU305914.1</td><td></td><td>15791</td><td>15813</td><td>+</td><td>0</td><td>TATTCCTCTGTAACGTGAGC</td><td>GGG</td><td>||||||||[||||||||||||]PAM</td><td>NA</td><td>Intergenic</td><td>NA</td><td>NA</td></tr>
<tr><td>T706</td><td>TGTTGCCTATGCATAAGTTCAGG</td><td></td><td>1000</td><td></td><td></td><td></td><td></td><td></td><td></td><td></td><td></td><td></td></tr>
<tr><td>Oligo fwd</td><td>CACCTGTTGCCTATGCATAAGTTC</td><td></td><td></td><td></td><td></td><td></td><td></td><td></td><td></td><td></td><td></td><td></td></tr>
<tr><td>Oligo rev</td><td>AAACGAACTTATGCATAGGCAACA</td><td></td><td></td><td></td><td></td><td></td><td></td><td></td><td></td><td></td><td></td><td></td></tr>
<tr><td>Chromosome</td><td>start</td><td></td><td>end</td><td>strand</td><td>MM</td><td>target_seq</td><td>PAM</td><td>alignment</td><td>distance</td><td>position</td><td>gene name</td><td>gene id</td></tr>
<tr><td>GU305914.1</td><td></td><td>15859</td><td>15881</td><td>+</td><td>0</td><td>TGTTGCCTATGCATAAGTTC</td><td>AGG</td><td>||||||||[||||||||||||]PAM</td><td>NA</td><td>Intergenic</td><td>NA</td><td>NA</td></tr>
<tr><td>T707</td><td>GATCAAACAGATAGCAAAAATGG</td><td></td><td>1000</td><td></td><td></td><td></td><td></td><td></td><td></td><td></td><td></td><td></td></tr>
<tr><td>Oligo fwd</td><td>CACCGATCAAACAGATAGCAAAAA</td><td></td><td></td><td></td><td></td><td></td><td></td><td></td><td></td><td></td><td></td><td></td></tr>
<tr><td>Oligo rev</td><td>AAACTTTTTGCTATCTGTTTGATC</td><td></td><td></td><td></td><td></td><td></td><td></td><td></td><td></td><td></td><td></td><td></td></tr>
<tr><td>Chromosome</td><td>start</td><td></td><td>end</td><td>strand</td><td>MM</td><td>target_seq</td><td>PAM</td><td>alignment</td><td>distance</td><td>position</td><td>gene name</td><td>gene id</td></tr>
<tr><td>GU305914.1</td><td></td><td>15881</td><td>15903</td><td>+</td><td>0</td><td>GATCAAACAGATAGCAAAAA</td><td>TGG</td><td>||||||||[||||||||||||]PAM</td><td>NA</td><td>Intergenic</td><td>NA</td><td>NA</td></tr>
<tr><td>T708</td><td>ATCAAACAGATAGCAAAAATGGG</td><td></td><td>1000</td><td></td><td></td><td></td><td></td><td></td><td></td><td></td><td></td><td></td></tr>
<tr><td>Oligo fwd</td><td>CACCATCAAACAGATAGCAAAAAT</td><td></td><td></td><td></td><td></td><td></td><td></td><td></td><td></td><td></td><td></td><td></td></tr>
<tr><td>Oligo rev</td><td>AAACATTTTTGCTATCTGTTTGAT</td><td></td><td></td><td></td><td></td><td></td><td></td><td></td><td></td><td></td><td></td><td></td></tr>
<tr><td>Chromosome</td><td>start</td><td></td><td>end</td><td>strand</td><td>MM</td><td>target_seq</td><td>PAM</td><td>alignment</td><td>distance</td><td>position</td><td>gene name</td><td>gene id</td></tr>
<tr><td>GU305914.1</td><td></td><td>15882</td><td>15904</td><td>+</td><td>0</td><td>ATCAAACAGATAGCAAAAAT</td><td>GGG</td><td>||||||||[||||||||||||]PAM</td><td>NA</td><td>Intergenic</td><td>NA</td><td>NA</td></tr>
<tr><td>T709</td><td>TGGGTAAGAATATCAGCGCGAGG</td><td></td><td>1000</td><td></td><td></td><td></td><td></td><td></td><td></td><td></td><td></td><td></td></tr>
<tr><td>Oligo fwd</td><td>CACCTGGGTAAGAATATCAGCGCG</td><td></td><td></td><td></td><td></td><td></td><td></td><td></td><td></td><td></td><td></td><td></td></tr>
<tr><td>Oligo rev</td><td>AAACCGCGCTGATATTCTTACCCA</td><td></td><td></td><td></td><td></td><td></td><td></td><td></td><td></td><td></td><td></td><td></td></tr>
<tr><td>Chromosome</td><td>start</td><td></td><td>end</td><td>strand</td><td>MM</td><td>target_seq</td><td>PAM</td><td>alignment</td><td>distance</td><td>position</td><td>gene name</td><td>gene id</td></tr>
<tr><td>GU305914.1</td><td></td><td>15901</td><td>15923</td><td>+</td><td>0</td><td>TGGGTAAGAATATCAGCGCG</td><td>AGG</td><td>||||||||[||||||||||||]PAM</td><td>NA</td><td>Intergenic</td><td>NA</td><td>NA</td></tr>
<tr><td>T710</td><td>CGAGGTTTAGAATTTGTCCGAGG</td><td></td><td>1000</td><td></td><td></td><td></td><td></td><td></td><td></td><td></td><td></td><td></td></tr>
<tr><td>Oligo fwd</td><td>CACCCGAGGTTTAGAATTTGTCCG</td><td></td><td></td><td></td><td></td><td></td><td></td><td></td><td></td><td></td><td></td><td></td></tr>
<tr><td>Oligo rev</td><td>AAACCGGACAAATTCTAAACCTCG</td><td></td><td></td><td></td><td></td><td></td><td></td><td></td><td></td><td></td><td></td><td></td></tr>
<tr><td>Chromosome</td><td>start</td><td></td><td>end</td><td>strand</td><td>MM</td><td>target_seq</td><td>PAM</td><td>alignment</td><td>distance</td><td>position</td><td>gene name</td><td>gene id</td></tr>
<tr><td>GU305914.1</td><td></td><td>15919</td><td>15941</td><td>+</td><td>0</td><td>CGAGGTTTAGAATTTGTCCG</td><td>AGG</td><td>||||||||[||||||||||||]PAM</td><td>NA</td><td>Intergenic</td><td>NA</td><td>NA</td></tr>
<tr><td>T711</td><td>ACTTCGTGTCGAAGAACCTTCGG</td><td></td><td>1000</td><td></td><td></td><td></td><td></td><td></td><td></td><td></td><td></td><td></td></tr>
<tr><td>Oligo fwd</td><td>CACCACTTCGTGTCGAAGAACCTT</td><td></td><td></td><td></td><td></td><td></td><td></td><td></td><td></td><td></td><td></td><td></td></tr>
<tr><td>Oligo rev</td><td>AAACAAGGTTCTTCGACACGAAGT</td><td></td><td></td><td></td><td></td><td></td><td></td><td></td><td></td><td></td><td></td><td></td></tr>
<tr><td>Chromosome</td><td>start</td><td></td><td>end</td><td>strand</td><td>MM</td><td>target_seq</td><td>PAM</td><td>alignment</td><td>distance</td><td>position</td><td>gene name</td><td>gene id</td></tr>
<tr><td>GU305914.1</td><td></td><td>15982</td><td>16004</td><td>+</td><td>0</td><td>ACTTCGTGTCGAAGAACCTT</td><td>CGG</td><td>||||||||[||||||||||||]PAM</td><td>NA</td><td>Intergenic</td><td>NA</td><td>NA</td></tr>
<tr><td>T712</td><td>TGAACTCGCAGCCACGCACATGG</td><td></td><td>1000</td><td></td><td></td><td></td><td></td><td></td><td></td><td></td><td></td><td></td></tr>
<tr><td>Oligo fwd</td><td>CACCTGAACTCGCAGCCACGCACA</td><td></td><td></td><td></td><td></td><td></td><td></td><td></td><td></td><td></td><td></td><td></td></tr>
<tr><td>Oligo rev</td><td>AAACTGTGCGTGGCTGCGAGTTCA</td><td></td><td></td><td></td><td></td><td></td><td></td><td></td><td></td><td></td><td></td><td></td></tr>
<tr><td>Chromosome</td><td>start</td><td></td><td>end</td><td>strand</td><td>MM</td><td>target_seq</td><td>PAM</td><td>alignment</td><td>distance</td><td>position</td><td>gene name</td><td>gene id</td></tr>
<tr><td>GU305914.1</td><td></td><td>16024</td><td>16046</td><td>+</td><td>0</td><td>TGAACTCGCAGCCACGCACA</td><td>TGG</td><td>||||||||[||||||||||||]PAM</td><td>NA</td><td>Intergenic</td><td>NA</td><td>NA</td></tr>
<tr><td>T713</td><td>CTCGCAGCCACGCACATGGCCGG</td><td></td><td>1000</td><td></td><td></td><td></td><td></td><td></td><td></td><td></td><td></td><td></td></tr>
<tr><td>Oligo fwd</td><td>CACCCTCGCAGCCACGCACATGGC</td><td></td><td></td><td></td><td></td><td></td><td></td><td></td><td></td><td></td><td></td><td></td></tr>
<tr><td>Oligo rev</td><td>AAACGCCATGTGCGTGGCTGCGAG</td><td></td><td></td><td></td><td></td><td></td><td></td><td></td><td></td><td></td><td></td><td></td></tr>
<tr><td>Chromosome</td><td>start</td><td></td><td>end</td><td>strand</td><td>MM</td><td>target_seq</td><td>PAM</td><td>alignment</td><td>distance</td><td>position</td><td>gene name</td><td>gene id</td></tr>
<tr><td>GU305914.1</td><td></td><td>16028</td><td>16050</td><td>+</td><td>0</td><td>CTCGCAGCCACGCACATGGC</td><td>CGG</td><td>||||||||[||||||||||||]PAM</td><td>NA</td><td>Intergenic</td><td>NA</td><td>NA</td></tr>
<tr><td>T714</td><td>TCGCAGCCACGCACATGGCCGGG</td><td></td><td>1000</td><td></td><td></td><td></td><td></td><td></td><td></td><td></td><td></td><td></td></tr>
<tr><td>Oligo fwd</td><td>CACCTCGCAGCCACGCACATGGCC</td><td></td><td></td><td></td><td></td><td></td><td></td><td></td><td></td><td></td><td></td><td></td></tr>
<tr><td>Oligo rev</td><td>AAACGGCCATGTGCGTGGCTGCGA</td><td></td><td></td><td></td><td></td><td></td><td></td><td></td><td></td><td></td><td></td><td></td></tr>
<tr><td>Chromosome</td><td>start</td><td></td><td>end</td><td>strand</td><td>MM</td><td>target_seq</td><td>PAM</td><td>alignment</td><td>distance</td><td>position</td><td>gene name</td><td>gene id</td></tr>
<tr><td>GU305914.1</td><td></td><td>16029</td><td>16051</td><td>+</td><td>0</td><td>TCGCAGCCACGCACATGGCC</td><td>GGG</td><td>||||||||[||||||||||||]PAM</td><td>NA</td><td>Intergenic</td><td>NA</td><td>NA</td></tr>
<tr><td>T715</td><td>CCACGCACATGGCCGGGTGTTGG</td><td></td><td>1000</td><td></td><td></td><td></td><td></td><td></td><td></td><td></td><td></td><td></td></tr>
<tr><td>Oligo fwd</td><td>CACCCCACGCACATGGCCGGGTGT</td><td></td><td></td><td></td><td></td><td></td><td></td><td></td><td></td><td></td><td></td><td></td></tr>
<tr><td>Oligo rev</td><td>AAACACACCCGGCCATGTGCGTGG</td><td></td><td></td><td></td><td></td><td></td><td></td><td></td><td></td><td></td><td></td><td></td></tr>
<tr><td>Chromosome</td><td>start</td><td></td><td>end</td><td>strand</td><td>MM</td><td>target_seq</td><td>PAM</td><td>alignment</td><td>distance</td><td>position</td><td>gene name</td><td>gene id</td></tr>
<tr><td>GU305914.1</td><td></td><td>16035</td><td>16057</td><td>+</td><td>0</td><td>CCACGCACATGGCCGGGTGT</td><td>TGG</td><td>||||||||[||||||||||||]PAM</td><td>NA</td><td>Intergenic</td><td>NA</td><td>NA</td></tr>
<tr><td>T716</td><td>TTGGCAGTTCATAGCTCCAGCGG</td><td></td><td>1000</td><td></td><td></td><td></td><td></td><td></td><td></td><td></td><td></td><td></td></tr>
<tr><td>Oligo fwd</td><td>CACCTTGGCAGTTCATAGCTCCAG</td><td></td><td></td><td></td><td></td><td></td><td></td><td></td><td></td><td></td><td></td><td></td></tr>
<tr><td>Oligo rev</td><td>AAACCTGGAGCTATGAACTGCCAA</td><td></td><td></td><td></td><td></td><td></td><td></td><td></td><td></td><td></td><td></td><td></td></tr>
<tr><td>Chromosome</td><td>start</td><td></td><td>end</td><td>strand</td><td>MM</td><td>target_seq</td><td>PAM</td><td>alignment</td><td>distance</td><td>position</td><td>gene name</td><td>gene id</td></tr>
<tr><td>GU305914.1</td><td></td><td>16054</td><td>16076</td><td>+</td><td>0</td><td>TTGGCAGTTCATAGCTCCAG</td><td>CGG</td><td>||||||||[||||||||||||]PAM</td><td>NA</td><td>Intergenic</td><td>NA</td><td>NA</td></tr>
<tr><td>T717</td><td>CATAGCTCCAGCGGCCGAACAGG</td><td></td><td>1000</td><td></td><td></td><td></td><td></td><td></td><td></td><td></td><td></td><td></td></tr>
<tr><td>Oligo fwd</td><td>CACCCATAGCTCCAGCGGCCGAAC</td><td></td><td></td><td></td><td></td><td></td><td></td><td></td><td></td><td></td><td></td><td></td></tr>
<tr><td>Oligo rev</td><td>AAACGTTCGGCCGCTGGAGCTATG</td><td></td><td></td><td></td><td></td><td></td><td></td><td></td><td></td><td></td><td></td><td></td></tr>
<tr><td>Chromosome</td><td>start</td><td></td><td>end</td><td>strand</td><td>MM</td><td>target_seq</td><td>PAM</td><td>alignment</td><td>distance</td><td>position</td><td>gene name</td><td>gene id</td></tr>
<tr><td>GU305914.1</td><td></td><td>16063</td><td>16085</td><td>+</td><td>0</td><td>CATAGCTCCAGCGGCCGAAC</td><td>AGG</td><td>||||||||[||||||||||||]PAM</td><td>NA</td><td>Intergenic</td><td>NA</td><td>NA</td></tr>
<tr><td>T718</td><td>ATAGCTCCAGCGGCCGAACAGGG</td><td></td><td>1000</td><td></td><td></td><td></td><td></td><td></td><td></td><td></td><td></td><td></td></tr>
<tr><td>Oligo fwd</td><td>CACCATAGCTCCAGCGGCCGAACA</td><td></td><td></td><td></td><td></td><td></td><td></td><td></td><td></td><td></td><td></td><td></td></tr>
<tr><td>Oligo rev</td><td>AAACTGTTCGGCCGCTGGAGCTAT</td><td></td><td></td><td></td><td></td><td></td><td></td><td></td><td></td><td></td><td></td><td></td></tr>
<tr><td>Chromosome</td><td>start</td><td></td><td>end</td><td>strand</td><td>MM</td><td>target_seq</td><td>PAM</td><td>alignment</td><td>distance</td><td>position</td><td>gene name</td><td>gene id</td></tr>
<tr><td>GU305914.1</td><td></td><td>16064</td><td>16086</td><td>+</td><td>0</td><td>ATAGCTCCAGCGGCCGAACA</td><td>GGG</td><td>||||||||[||||||||||||]PAM</td><td>NA</td><td>Intergenic</td><td>NA</td><td>NA</td></tr>
</table>

| | | | | | | | | | | | |
|---|---|---|---|---|---|---|---|---|---|---|---|
| T719 | GAACAGGGTCTGAAAATCGCAGG | 1000 | | | | | | | | | |
| Oligo fwd | CACCGAACAGGGTCTGAAAATCGC | | | | | | | | | | |
| Oligo rev | AAACGCGATTTTCAGACCCTGTTC | | | | | | | | | | |
| Chromosome | start | end | strand | MM | target_seq | PAM | alignment | distance | position | gene name | gene id |
| GU305914.1 | 16079 | 16101 | + | 0 | GAACAGGGTCTGAAAATCGC | AGG | ||||||||[||||||||||||]PAM | NA | Intergenic | NA | NA |
| T720 | GCAGGTCGAGTTCGTTTCGTTGG | 1000 | | | | | | | | | |
| Oligo fwd | CACCGCAGGTCGAGTTCGTTTCGT | | | | | | | | | | |
| Oligo rev | AAACACGAAACGAACTCGACCTGC | | | | | | | | | | |
| Chromosome | start | end | strand | MM | target_seq | PAM | alignment | distance | position | gene name | gene id |
| GU305914.1 | 16097 | 16119 | + | 0 | GCAGGTCGAGTTCGTTTCGT | TGG | ||||||||[||||||||||||]PAM | NA | Intergenic | NA | NA |
| T721 | GTTGACCGTGCTCGACACGCCGG | 1000 | | | | | | | | | |
| Oligo fwd | CACCGTTGACCGTGCTCGACACGC | | | | | | | | | | |
| Oligo rev | AAACGCGTGTCGAGCACGGTCAAC | | | | | | | | | | |
| Chromosome | start | end | strand | MM | target_seq | PAM | alignment | distance | position | gene name | gene id |
| GU305914.1 | 16130 | 16152 | + | 0 | GTTGACCGTGCTCGACACGC | CGG | ||||||||[||||||||||||]PAM | NA | Intergenic | NA | NA |
| T722 | GACCGTGCTCGACACGCCGGTGG | 1000 | | | | | | | | | |
| Oligo fwd | CACCGACCGTGCTCGACACGCCGG | | | | | | | | | | |
| Oligo rev | AAACCCGGCGTGTCGAGCACGGTC | | | | | | | | | | |
| Chromosome | start | end | strand | MM | target_seq | PAM | alignment | distance | position | gene name | gene id |
| GU305914.1 | 16133 | 16155 | + | 0 | GACCGTGCTCGACACGCCGG | TGG | ||||||||[||||||||||||]PAM | NA | Intergenic | NA | NA |
| T723 | GCCGGTGGTGTAGTTGTATCTGG | 1000 | | | | | | | | | |
| Oligo fwd | CACCGCCGGTGGTGTAGTTGTATC | | | | | | | | | | |
| Oligo rev | AAACGATACAACTACACCACCGGC | | | | | | | | | | |
| Chromosome | start | end | strand | MM | target_seq | PAM | alignment | distance | position | gene name | gene id |
| GU305914.1 | 16148 | 16170 | + | 0 | GCCGGTGGTGTAGTTGTATC | TGG | ||||||||[||||||||||||]PAM | NA | Intergenic | NA | NA |
| T724 | TATCTGGTATACATTCCGTAAGG | 1000 | | | | | | | | | |
| Oligo fwd | CACCTATCTGGTATACATTCCGTA | | | | | | | | | | |
| Oligo rev | AAACTACGGAATGTATACCAGATA | | | | | | | | | | |
| Chromosome | start | end | strand | MM | target_seq | PAM | alignment | distance | position | gene name | gene id |
| GU305914.1 | 16164 | 16186 | + | 0 | TATCTGGTATACATTCCGTA | AGG | ||||||||[||||||||||||]PAM | NA | Intergenic | NA | NA |
| T725 | ATCTGGTATACATTCCGTAAGGG | 1000 | | | | | | | | | |
| Oligo fwd | CACCATCTGGTATACATTCCGTAA | | | | | | | | | | |
| Oligo rev | AAACTTACGGAATGTATACCAGAT | | | | | | | | | | |
| Chromosome | start | end | strand | MM | target_seq | PAM | alignment | distance | position | gene name | gene id |
| GU305914.1 | 16165 | 16187 | + | 0 | ATCTGGTATACATTCCGTAA | GGG | ||||||||[||||||||||||]PAM | NA | Intergenic | NA | NA |
| T726 | TCTGGTATACATTCCGTAAGGGG | 1000 | | | | | | | | | |
| Oligo fwd | CACCTCTGGTATACATTCCGTAAG | | | | | | | | | | |
| Oligo rev | AAACCTTACGGAATGTATACCAGA | | | | | | | | | | |
| Chromosome | start | end | strand | MM | target_seq | PAM | alignment | distance | position | gene name | gene id |
| GU305914.1 | 16166 | 16188 | + | 0 | TCTGGTATACATTCCGTAAG | GGG | ||||||||[||||||||||||]PAM | NA | Intergenic | NA | NA |
| T727 | CATACCCGCCATGAATCTAGCGG | 1000 | | | | | | | | | |
| Oligo fwd | CACCCATACCCGCCATGAATCTAG | | | | | | | | | | |
| Oligo rev | AAACCTAGATTCATGGCGGGTATG | | | | | | | | | | |
| Chromosome | start | end | strand | MM | target_seq | PAM | alignment | distance | position | gene name | gene id |
| GU305914.1 | 16193 | 16215 | + | 0 | CATACCCGCCATGAATCTAG | CGG | ||||||||[||||||||||||]PAM | NA | Intergenic | NA | NA |
| T728 | ATACCCGCCATGAATCTAGCGGG | 1000 | | | | | | | | | |
| Oligo fwd | CACCATACCCGCCATGAATCTAGC | | | | | | | | | | |
| Oligo rev | AAACGCTAGATTCATGGCGGGTAT | | | | | | | | | | |
| Chromosome | start | end | strand | MM | target_seq | PAM | alignment | distance | position | gene name | gene id |
| GU305914.1 | 16194 | 16216 | + | 0 | ATACCCGCCATGAATCTAGC | GGG | ||||||||[||||||||||||]PAM | NA | Intergenic | NA | NA |
| T729 | GCCATGAATCTAGCGGGTGATGG | 1000 | | | | | | | | | |
| Oligo fwd | CACCGCCATGAATCTAGCGGGTGA | | | | | | | | | | |
| Oligo rev | AAACTCACCCGCTAGATTCATGGC | | | | | | | | | | |
| Chromosome | start | end | strand | MM | target_seq | PAM | alignment | distance | position | gene name | gene id |
| GU305914.1 | 16200 | 16222 | + | 0 | GCCATGAATCTAGCGGGTGA | TGG | ||||||||[||||||||||||]PAM | NA | Intergenic | NA | NA |
| T730 | TGACTCAATCCGACATCTGACGG | 1000 | | | | | | | | | |
| Oligo fwd | CACCTGACTCAATCCGACATCTGA | | | | | | | | | | |
| Oligo rev | AAACTCAGATGTCGGATTGAGTCA | | | | | | | | | | |
| Chromosome | start | end | strand | MM | target_seq | PAM | alignment | distance | position | gene name | gene id |
| GU305914.1 | 16223 | 16245 | + | 0 | TGACTCAATCCGACATCTGA | CGG | ||||||||[||||||||||||]PAM | NA | Intergenic | NA | NA |
| T731 | GTTAGCATATCTCCGCATCGAGG | 1000 | | | | | | | | | |
| Oligo fwd | CACCGTTAGCATATCTCCGCATCG | | | | | | | | | | |
| Oligo rev | AAACCGATGCGGAGATATGCTAAC | | | | | | | | | | |
| Chromosome | start | end | strand | MM | target_seq | PAM | alignment | distance | position | gene name | gene id |
| GU305914.1 | 16271 | 16293 | + | 0 | GTTAGCATATCTCCGCATCG | AGG | ||||||||[||||||||||||]PAM | NA | Intergenic | NA | NA |
| T732 | CACAGACCTCCTTCCACCGTTGG | 1000 | | | | | | | | | |
| Oligo fwd | CACCCACAGACCTCCTTCCACCGT | | | | | | | | | | |
| Oligo rev | AAACACGGTGGAAGGAGGTCTGTG | | | | | | | | | | |
| Chromosome | start | end | strand | MM | target_seq | PAM | alignment | distance | position | gene name | gene id |
| GU305914.1 | 16309 | 16331 | + | 0 | CACAGACCTCCTTCCACCGT | TGG | ||||||||[||||||||||||]PAM | NA | Intergenic | NA | NA |
| T733 | CCTTCCACCGTTGGTATATGTGG | 1000 | | | | | | | | | |
| Oligo fwd | CACCCCTTCCACCGTTGGTATATG | | | | | | | | | | |
| Oligo rev | AAACCATATACCAACGGTGGAAGG | | | | | | | | | | |
| Chromosome | start | end | strand | MM | target_seq | PAM | alignment | distance | position | gene name | gene id |

GU305914.1 16318 16340 + 0 CCTTCCACCGTTGGTATATG TGG ||||||||[||||||||||||]PAM NA Intergenic NA NA

T734 GATACTGAACGCCTACCCCACGG 1000
Oligo fwd CACCGATACTGAACGCCTACCCCA
Oligo rev AAACTGGGGTAGGCGTTCAGTATC
Chromosome start end strand MM target_seq PAM alignment distance position gene name gene id
GU305914.1 16357 16379 + 0 GATACTGAACGCCTACCCCA CGG ||||||||[||||||||||||]PAM NA Intergenic NA NA

T735 ATACTGAACGCCTACCCCACGGG 1000
Oligo fwd CACCATACTGAACGCCTACCCCAC
Oligo rev AAACGTGGGGTAGGCGTTCAGTAT
Chromosome start end strand MM target_seq PAM alignment distance position gene name gene id
GU305914.1 16358 16380 + 0 ATACTGAACGCCTACCCCAC GGG ||||||||[||||||||||||]PAM NA Intergenic NA NA

T736 AACGCCTACCCCACGGGCCACGG 1000
Oligo fwd CACCAACGCCTACCCCACGGGCCA
Oligo rev AAACTGGCCCGTGGGGTAGGCGTT
Chromosome start end strand MM target_seq PAM alignment distance position gene name gene id
GU305914.1 16364 16386 + 0 AACGCCTACCCCACGGGCCA CGG ||||||||[||||||||||||]PAM NA Intergenic NA NA

T737 AGTTTTCCAACGTCTCCCACCGG 1000
Oligo fwd CACCAGTTTTCCAACGTCTCCCAC
Oligo rev AAACGTGGGAGACGTTGGAAAACT
Chromosome start end strand MM target_seq PAM alignment distance position gene name gene id
GU305914.1 16395 16417 + 0 AGTTTTCCAACGTCTCCCAC CGG ||||||||[||||||||||||]PAM NA Intergenic NA NA

T738 GTTTTCCAACGTCTCCCACCGGG 1000
Oligo fwd CACCGTTTTCCAACGTCTCCCACC
Oligo rev AAACGGTGGGAGACGTTGGAAAAC
Chromosome start end strand MM target_seq PAM alignment distance position gene name gene id
GU305914.1 16396 16418 + 0 GTTTTCCAACGTCTCCCACC GGG ||||||||[||||||||||||]PAM NA Intergenic NA NA

T739 CGTCTCCCACCGGGTACGTATGG 1000
Oligo fwd CACCCGTCTCCCACCGGGTACGTA
Oligo rev AAACTACGTACCCGGTGGGAGACG
Chromosome start end strand MM target_seq PAM alignment distance position gene name gene id
GU305914.1 16405 16427 + 0 CGTCTCCCACCGGGTACGTA TGG ||||||||[||||||||||||]PAM NA Intergenic NA NA

T740 ACGTATGGTTTCTGTAATTACGG 1000
Oligo fwd CACCACGTATGGTTTCTGTAATTA
Oligo rev AAACTAATTACAGAAACCATACGT
Chromosome start end strand MM target_seq PAM alignment distance position gene name gene id
GU305914.1 16420 16442 + 0 ACGTATGGTTTCTGTAATTA CGG ||||||||[||||||||||||]PAM NA Intergenic NA NA

T741 CACGACATCTGACCGTCACGTGG 1000
Oligo fwd CACCCACGACATCTGACCGTCACG
Oligo rev AAACCGTGACGGTCAGATGTCGTG
Chromosome start end strand MM target_seq PAM alignment distance position gene name gene id
GU305914.1 16483 16505 + 0 CACGACATCTGACCGTCACG TGG ||||||||[||||||||||||]PAM NA Intergenic NA NA

T742 TGACCGTCACGTGGCTCACGCGG 1000
Oligo fwd CACCTGACCGTCACGTGGCTCACG
Oligo rev AAACCGTGAGCCACGTGACGGTCA
Chromosome start end strand MM target_seq PAM alignment distance position gene name gene id
GU305914.1 16492 16514 + 0 TGACCGTCACGTGGCTCACG CGG ||||||||[||||||||||||]PAM NA Intergenic NA NA

T743 CCGATGTGAGTAAAATCTTGCGG 1000
Oligo fwd CACCCCGATGTGAGTAAAATCTTG
Oligo rev AAACCAAGATTTTACTCACATCGG
Chromosome start end strand MM target_seq PAM alignment distance position gene name gene id
GU305914.1 16530 16552 + 0 CCGATGTGAGTAAAATCTTG CGG ||||||||[||||||||||||]PAM NA Intergenic NA NA

T744 ACTTCTTTACATGTTCCGTCTGG 1000
Oligo fwd CACCACTTCTTTACATGTTCCGTC
Oligo rev AAACGACGGAACATGTAAAGAAGT
Chromosome start end strand MM target_seq PAM alignment distance position gene name gene id
GU305914.1 16591 16613 + 0 ACTTCTTTACATGTTCCGTC TGG ||||||||[||||||||||||]PAM NA Intergenic NA NA

T745 CTTCTTTACATGTTCCGTCTGGG 1000
Oligo fwd CACCCTTCTTTACATGTTCCGTCT
Oligo rev AAACAGACGGAACATGTAAAGAAG
Chromosome start end strand MM target_seq PAM alignment distance position gene name gene id
GU305914.1 16592 16614 + 0 CTTCTTTACATGTTCCGTCT GGG ||||||||[||||||||||||]PAM NA Intergenic NA NA

T746 CCATCTAACGTCGTTGTTGATGG 1000
Oligo fwd CACCCCATCTAACGTCGTTGTTGA
Oligo rev AAACTCAACAACGACGTTAGATGG
Chromosome start end strand MM target_seq PAM alignment distance position gene name gene id
GU305914.1 16658 16680 + 0 CCATCTAACGTCGTTGTTGA TGG ||||||||[||||||||||||]PAM NA Intergenic NA NA

T747 TCGTTGTTGATGGTACCGTTCGG 1000
Oligo fwd CACCTCGTTGTTGATGGTACCGTT
Oligo rev AAACAACGGTACCATCAACAACGA
Chromosome start end strand MM target_seq PAM alignment distance position gene name gene id
GU305914.1 16668 16690 + 0 TCGTTGTTGATGGTACCGTT CGG ||||||||[||||||||||||]PAM NA Intergenic NA NA

T748 CGGCAAGACTTCGAACAGAGCGG 1000
Oligo fwd CACCCGGCAAGACTTCGAACAGAG
Oligo rev AAACCTCTGTTCGAAGTCTTGCCG

<table>
<tr><td>Chromosome</td><td>start</td><td></td><td>end</td><td>strand</td><td>MM</td><td>target_seq</td><td>PAM</td><td>alignment</td><td>distance</td><td>position</td><td>gene name</td><td>gene id</td></tr>
<tr><td>GU305914.1</td><td></td><td>16688</td><td>16710</td><td>+</td><td>0</td><td>CGGCAAGACTTCGAACAGAG</td><td>CGG</td><td>||||||||[||||||||||||]PAM</td><td>NA</td><td>Intergenic</td><td>NA</td><td>NA</td></tr>
<tr><td>T749</td><td>CAAGACTTCGAACAGAGCGGCGG</td><td></td><td>1000</td><td></td><td></td><td></td><td></td><td></td><td></td><td></td><td></td><td></td></tr>
<tr><td>Oligo fwd</td><td>CACCCAAGACTTCGAACAGAGCGG</td><td></td><td></td><td></td><td></td><td></td><td></td><td></td><td></td><td></td><td></td><td></td></tr>
<tr><td>Oligo rev</td><td>AAACCCGCTCTGTTCGAAGTCTTG</td><td></td><td></td><td></td><td></td><td></td><td></td><td></td><td></td><td></td><td></td><td></td></tr>
<tr><td>Chromosome</td><td>start</td><td></td><td>end</td><td>strand</td><td>MM</td><td>target_seq</td><td>PAM</td><td>alignment</td><td>distance</td><td>position</td><td>gene name</td><td>gene id</td></tr>
<tr><td>GU305914.1</td><td></td><td>16691</td><td>16713</td><td>+</td><td>0</td><td>CAAGACTTCGAACAGAGCGG</td><td>CGG</td><td>||||||||[||||||||||||]PAM</td><td>NA</td><td>Intergenic</td><td>NA</td><td>NA</td></tr>
<tr><td>T750</td><td>GACTTCGAACAGAGCGGCGGTGG</td><td></td><td>1000</td><td></td><td></td><td></td><td></td><td></td><td></td><td></td><td></td><td></td></tr>
<tr><td>Oligo fwd</td><td>CACCGACTTCGAACAGAGCGGCGG</td><td></td><td></td><td></td><td></td><td></td><td></td><td></td><td></td><td></td><td></td><td></td></tr>
<tr><td>Oligo rev</td><td>AAACCCGCCGCTCTGTTCGAAGTC</td><td></td><td></td><td></td><td></td><td></td><td></td><td></td><td></td><td></td><td></td><td></td></tr>
<tr><td>Chromosome</td><td>start</td><td></td><td>end</td><td>strand</td><td>MM</td><td>target_seq</td><td>PAM</td><td>alignment</td><td>distance</td><td>position</td><td>gene name</td><td>gene id</td></tr>
<tr><td>GU305914.1</td><td></td><td>16694</td><td>16716</td><td>+</td><td>0</td><td>GACTTCGAACAGAGCGGCGG</td><td>TGG</td><td>||||||||[||||||||||||]PAM</td><td>NA</td><td>Intergenic</td><td>NA</td><td>NA</td></tr>
<tr><td>T751</td><td>GCTCGTGTTGAGAACCATGCCGG</td><td></td><td>1000</td><td></td><td></td><td></td><td></td><td></td><td></td><td></td><td></td><td></td></tr>
<tr><td>Oligo fwd</td><td>CACCGCTCGTGTTGAGAACCATGC</td><td></td><td></td><td></td><td></td><td></td><td></td><td></td><td></td><td></td><td></td><td></td></tr>
<tr><td>Oligo rev</td><td>AAACGCATGGTTCTCAACACGAGC</td><td></td><td></td><td></td><td></td><td></td><td></td><td></td><td></td><td></td><td></td><td></td></tr>
<tr><td>Chromosome</td><td>start</td><td></td><td>end</td><td>strand</td><td>MM</td><td>target_seq</td><td>PAM</td><td>alignment</td><td>distance</td><td>position</td><td>gene name</td><td>gene id</td></tr>
<tr><td>GU305914.1</td><td></td><td>16745</td><td>16767</td><td>+</td><td>0</td><td>GCTCGTGTTGAGAACCATGC</td><td>CGG</td><td>||||||||[||||||||||||]PAM</td><td>NA</td><td>Intergenic</td><td>NA</td><td>NA</td></tr>
<tr><td>T752</td><td>TCCGCACTCGCTCCGCACCGTGG</td><td></td><td>1000</td><td></td><td></td><td></td><td></td><td></td><td></td><td></td><td></td><td></td></tr>
<tr><td>Oligo fwd</td><td>CACCTCCGCACTCGCTCCGCACCG</td><td></td><td></td><td></td><td></td><td></td><td></td><td></td><td></td><td></td><td></td><td></td></tr>
<tr><td>Oligo rev</td><td>AAACCGGTGCGGAGCGAGTGCGGA</td><td></td><td></td><td></td><td></td><td></td><td></td><td></td><td></td><td></td><td></td><td></td></tr>
<tr><td>Chromosome</td><td>start</td><td></td><td>end</td><td>strand</td><td>MM</td><td>target_seq</td><td>PAM</td><td>alignment</td><td>distance</td><td>position</td><td>gene name</td><td>gene id</td></tr>
<tr><td>GU305914.1</td><td></td><td>16793</td><td>16815</td><td>+</td><td>0</td><td>TCCGCACTCGCTCCGCACCG</td><td>TGG</td><td>||||||||[||||||||||||]PAM</td><td>NA</td><td>Intergenic</td><td>NA</td><td>NA</td></tr>
<tr><td>T753</td><td>GCACTCGCTCCGCACCGTGGCGG</td><td></td><td>1000</td><td></td><td></td><td></td><td></td><td></td><td></td><td></td><td></td><td></td></tr>
<tr><td>Oligo fwd</td><td>CACCGCACTCGCTCCGCACCGTGG</td><td></td><td></td><td></td><td></td><td></td><td></td><td></td><td></td><td></td><td></td><td></td></tr>
<tr><td>Oligo rev</td><td>AAACCCACGGTGCGGAGCGAGTGC</td><td></td><td></td><td></td><td></td><td></td><td></td><td></td><td></td><td></td><td></td><td></td></tr>
<tr><td>Chromosome</td><td>start</td><td></td><td>end</td><td>strand</td><td>MM</td><td>target_seq</td><td>PAM</td><td>alignment</td><td>distance</td><td>position</td><td>gene name</td><td>gene id</td></tr>
<tr><td>GU305914.1</td><td></td><td>16796</td><td>16818</td><td>+</td><td>0</td><td>GCACTCGCTCCGCACCGTGG</td><td>CGG</td><td>||||||||[||||||||||||]PAM</td><td>NA</td><td>Intergenic</td><td>NA</td><td>NA</td></tr>
<tr><td>T754</td><td>TTCATCAGCTAACAGATCATCGG</td><td></td><td>1000</td><td></td><td></td><td></td><td></td><td></td><td></td><td></td><td></td><td></td></tr>
<tr><td>Oligo fwd</td><td>CACCTTCATCAGCTAACAGATCAT</td><td></td><td></td><td></td><td></td><td></td><td></td><td></td><td></td><td></td><td></td><td></td></tr>
<tr><td>Oligo rev</td><td>AAACATGATCTGTTAGCTGATGAA</td><td></td><td></td><td></td><td></td><td></td><td></td><td></td><td></td><td></td><td></td><td></td></tr>
<tr><td>Chromosome</td><td>start</td><td></td><td>end</td><td>strand</td><td>MM</td><td>target_seq</td><td>PAM</td><td>alignment</td><td>distance</td><td>position</td><td>gene name</td><td>gene id</td></tr>
<tr><td>GU305914.1</td><td></td><td>16826</td><td>16848</td><td>+</td><td>0</td><td>TTCATCAGCTAACAGATCAT</td><td>CGG</td><td>||||||||[||||||||||||]PAM</td><td>NA</td><td>Intergenic</td><td>NA</td><td>NA</td></tr>
<tr><td>T755</td><td>TCATCAGCTAACAGATCATCGGG</td><td></td><td>1000</td><td></td><td></td><td></td><td></td><td></td><td></td><td></td><td></td><td></td></tr>
<tr><td>Oligo fwd</td><td>CACCTCATCAGCTAACAGATCATC</td><td></td><td></td><td></td><td></td><td></td><td></td><td></td><td></td><td></td><td></td><td></td></tr>
<tr><td>Oligo rev</td><td>AAACGATGATCTGTTAGCTGATGA</td><td></td><td></td><td></td><td></td><td></td><td></td><td></td><td></td><td></td><td></td><td></td></tr>
<tr><td>Chromosome</td><td>start</td><td></td><td>end</td><td>strand</td><td>MM</td><td>target_seq</td><td>PAM</td><td>alignment</td><td>distance</td><td>position</td><td>gene name</td><td>gene id</td></tr>
<tr><td>GU305914.1</td><td></td><td>16827</td><td>16849</td><td>+</td><td>0</td><td>TCATCAGCTAACAGATCATC</td><td>GGG</td><td>||||||||[||||||||||||]PAM</td><td>NA</td><td>Intergenic</td><td>NA</td><td>NA</td></tr>
<tr><td>T756</td><td>GCCGCAGTTCTGCGCACGCACGG</td><td></td><td>1000</td><td></td><td></td><td></td><td></td><td></td><td></td><td></td><td></td><td></td></tr>
<tr><td>Oligo fwd</td><td>CACCGCCGCAGTTCTGCGCACGCA</td><td></td><td></td><td></td><td></td><td></td><td></td><td></td><td></td><td></td><td></td><td></td></tr>
<tr><td>Oligo rev</td><td>AAACTGCGTGCGCAGAACTGCGGC</td><td></td><td></td><td></td><td></td><td></td><td></td><td></td><td></td><td></td><td></td><td></td></tr>
<tr><td>Chromosome</td><td>start</td><td></td><td>end</td><td>strand</td><td>MM</td><td>target_seq</td><td>PAM</td><td>alignment</td><td>distance</td><td>position</td><td>gene name</td><td>gene id</td></tr>
<tr><td>GU305914.1</td><td></td><td>16862</td><td>16884</td><td>+</td><td>0</td><td>GCCGCAGTTCTGCGCACGCA</td><td>CGG</td><td>||||||||[||||||||||||]PAM</td><td>NA</td><td>Intergenic</td><td>NA</td><td>NA</td></tr>
<tr><td>T757</td><td>AGTTCTGCGCACGCACGGCGAGG</td><td></td><td>1000</td><td></td><td></td><td></td><td></td><td></td><td></td><td></td><td></td><td></td></tr>
<tr><td>Oligo fwd</td><td>CACCAGTTCTGCGCACGCACGGCG</td><td></td><td></td><td></td><td></td><td></td><td></td><td></td><td></td><td></td><td></td><td></td></tr>
<tr><td>Oligo rev</td><td>AAACCGCCGTGCGTGCGCAGAACT</td><td></td><td></td><td></td><td></td><td></td><td></td><td></td><td></td><td></td><td></td><td></td></tr>
<tr><td>Chromosome</td><td>start</td><td></td><td>end</td><td>strand</td><td>MM</td><td>target_seq</td><td>PAM</td><td>alignment</td><td>distance</td><td>position</td><td>gene name</td><td>gene id</td></tr>
<tr><td>GU305914.1</td><td></td><td>16867</td><td>16889</td><td>+</td><td>0</td><td>AGTTCTGCGCACGCACGGCG</td><td>AGG</td><td>||||||||[||||||||||||]PAM</td><td>NA</td><td>Intergenic</td><td>NA</td><td>NA</td></tr>
<tr><td>T758</td><td>GTTCTGCGCACGCACGGCGAGGG</td><td></td><td>1000</td><td></td><td></td><td></td><td></td><td></td><td></td><td></td><td></td><td></td></tr>
<tr><td>Oligo fwd</td><td>CACCGTTCTGCGCACGCACGGCGA</td><td></td><td></td><td></td><td></td><td></td><td></td><td></td><td></td><td></td><td></td><td></td></tr>
<tr><td>Oligo rev</td><td>AAACTCGCCGTGCGTGCGCAGAAC</td><td></td><td></td><td></td><td></td><td></td><td></td><td></td><td></td><td></td><td></td><td></td></tr>
<tr><td>Chromosome</td><td>start</td><td></td><td>end</td><td>strand</td><td>MM</td><td>target_seq</td><td>PAM</td><td>alignment</td><td>distance</td><td>position</td><td>gene name</td><td>gene id</td></tr>
<tr><td>GU305914.1</td><td></td><td>16868</td><td>16890</td><td>+</td><td>0</td><td>GTTCTGCGCACGCACGGCGA</td><td>GGG</td><td>||||||||[||||||||||||]PAM</td><td>NA</td><td>Intergenic</td><td>NA</td><td>NA</td></tr>
<tr><td>T759</td><td>CCTCCCGTCCCCATTCTCGAAGG</td><td></td><td>1000</td><td></td><td></td><td></td><td></td><td></td><td></td><td></td><td></td><td></td></tr>
<tr><td>Oligo fwd</td><td>CACCCCTCCCGTCCCCATTCTCGA</td><td></td><td></td><td></td><td></td><td></td><td></td><td></td><td></td><td></td><td></td><td></td></tr>
<tr><td>Oligo rev</td><td>AAACTCGAGAATGGGGACGGGAGG</td><td></td><td></td><td></td><td></td><td></td><td></td><td></td><td></td><td></td><td></td><td></td></tr>
<tr><td>Chromosome</td><td>start</td><td></td><td>end</td><td>strand</td><td>MM</td><td>target_seq</td><td>PAM</td><td>alignment</td><td>distance</td><td>position</td><td>gene name</td><td>gene id</td></tr>
<tr><td>GU305914.1</td><td></td><td>16938</td><td>16960</td><td>+</td><td>0</td><td>CCTCCCGTCCCCATTCTCGA</td><td>AGG</td><td>||||||||[||||||||||||]PAM</td><td>NA</td><td>Intergenic</td><td>NA</td><td>NA</td></tr>
<tr><td>T760</td><td>CCCGTCCCCATTCTCGAAGGCGG</td><td></td><td>1000</td><td></td><td></td><td></td><td></td><td></td><td></td><td></td><td></td><td></td></tr>
<tr><td>Oligo fwd</td><td>CACCCCCGTCCCCATTCTCGAAGG</td><td></td><td></td><td></td><td></td><td></td><td></td><td></td><td></td><td></td><td></td><td></td></tr>
<tr><td>Oligo rev</td><td>AAACCCTTCGAGAATGGGGACGGG</td><td></td><td></td><td></td><td></td><td></td><td></td><td></td><td></td><td></td><td></td><td></td></tr>
<tr><td>Chromosome</td><td>start</td><td></td><td>end</td><td>strand</td><td>MM</td><td>target_seq</td><td>PAM</td><td>alignment</td><td>distance</td><td>position</td><td>gene name</td><td>gene id</td></tr>
<tr><td>GU305914.1</td><td></td><td>16941</td><td>16963</td><td>+</td><td>0</td><td>CCCGTCCCCATTCTCGAAGG</td><td>CGG</td><td>||||||||[||||||||||||]PAM</td><td>NA</td><td>Intergenic</td><td>NA</td><td>NA</td></tr>
<tr><td>T761</td><td>TCGAAGGCGGACGTTCTGATCGG</td><td></td><td>1000</td><td></td><td></td><td></td><td></td><td></td><td></td><td></td><td></td><td></td></tr>
<tr><td>Oligo fwd</td><td>CACCTCGAAGGCGGACGTTCTGAT</td><td></td><td></td><td></td><td></td><td></td><td></td><td></td><td></td><td></td><td></td><td></td></tr>
<tr><td>Oligo rev</td><td>AAACATCAGAACGTCCGCCTTCGA</td><td></td><td></td><td></td><td></td><td></td><td></td><td></td><td></td><td></td><td></td><td></td></tr>
<tr><td>Chromosome</td><td>start</td><td></td><td>end</td><td>strand</td><td>MM</td><td>target_seq</td><td>PAM</td><td>alignment</td><td>distance</td><td>position</td><td>gene name</td><td>gene id</td></tr>
<tr><td>GU305914.1</td><td></td><td>16954</td><td>16976</td><td>+</td><td>0</td><td>TCGAAGGCGGACGTTCTGAT</td><td>CGG</td><td>||||||||[||||||||||||]PAM</td><td>NA</td><td>Intergenic</td><td>NA</td><td>NA</td></tr>
<tr><td>T762</td><td>CGATTTCAAAGAAAGCAGACAGG</td><td></td><td>1000</td><td></td><td></td><td></td><td></td><td></td><td></td><td></td><td></td><td></td></tr>
<tr><td>Oligo fwd</td><td>CACCCGATTTCAAAGAAAGCAGAC</td><td></td><td></td><td></td><td></td><td></td><td></td><td></td><td></td><td></td><td></td><td></td></tr>
<tr><td>Oligo rev</td><td>AAACGTCTGCTTTCTTTGAAATCG</td><td></td><td></td><td></td><td></td><td></td><td></td><td></td><td></td><td></td><td></td><td></td></tr>
<tr><td>Chromosome</td><td>start</td><td></td><td>end</td><td>strand</td><td>MM</td><td>target_seq</td><td>PAM</td><td>alignment</td><td>distance</td><td>position</td><td>gene name</td><td>gene id</td></tr>
<tr><td>GU305914.1</td><td></td><td>16982</td><td>17004</td><td>+</td><td>0</td><td>CGATTTCAAAGAAAGCAGAC</td><td>AGG</td><td>||||||||[||||||||||||]PAM</td><td>NA</td><td>Intergenic</td><td>NA</td><td>NA</td></tr>
<tr><td>T763</td><td>TTGAATACGATTGTTTTTATTGG</td><td></td><td>1000</td><td></td><td></td><td></td><td></td><td></td><td></td><td></td><td></td><td></td></tr>
<tr><td>Oligo fwd</td><td>CACCTTGAATACGATTGTTTTTAT</td><td></td><td></td><td></td><td></td><td></td><td></td><td></td><td></td><td></td><td></td><td></td></tr>
</table>

Oligo rev AAACATAAAAACAATCGTATTCAA
Chromosome start end strand MM target_seq PAM alignment distance position gene name gene id
GU305914.1 17021 17043 + 0 TTGAATACGATTGTTTTTAT TGG ||||||||[||||||||||||]PAM NA Intergenic NA NA

T764 CCTCAATCATCCCACCAGAGAGG 1000
Oligo fwd CACCCCTCAATCATCCCACCAGAG
Oligo rev AAACCTCTGGTGGGATGATTGAGG
Chromosome start end strand MM target_seq PAM alignment distance position gene name gene id
GU305914.1 17069 17091 + 0 CCTCAATCATCCCACCAGAG AGG ||||||||[||||||||||||]PAM NA Intergenic NA NA

T765 GCTAACGACCTCGACGACCCTGG 1000
Oligo fwd CACCGCTAACGACCTCGACGACCC
Oligo rev AAACGGGTCGTCGAGGTCGTTAGC
Chromosome start end strand MM target_seq PAM alignment distance position gene name gene id
GU305914.1 17143 17165 + 0 GCTAACGACCTCGACGACCC TGG ||||||||[||||||||||||]PAM NA Intergenic NA NA

T766 GCAACTCCCGTCATGCCATCAGG 1000
Oligo fwd CACCGCAACTCCCGTCATGCCATC
Oligo rev AAACGATGGCATGACGGGAGTTGC
Chromosome start end strand MM target_seq PAM alignment distance position gene name gene id
GU305914.1 17165 17187 + 0 GCAACTCCCGTCATGCCATC AGG ||||||||[||||||||||||]PAM NA Intergenic NA NA

T767 CGTCATGCCATCAGGTACGACGG 1000
Oligo fwd CACCCGTCATGCCATCAGGTACGA
Oligo rev AAACTCGTACCTGATGGCATGACG
Chromosome start end strand MM target_seq PAM alignment distance position gene name gene id
GU305914.1 17173 17195 + 0 CGTCATGCCATCAGGTACGA CGG ||||||||[||||||||||||]PAM NA Intergenic NA NA

T768 GGTACGACGGCATCCCCCAAAGG 1000
Oligo fwd CACCGGTACGACGGCATCCCCCAA
Oligo rev AAACTTGGGGGATGCCGTCGTACC
Chromosome start end strand MM target_seq PAM alignment distance position gene name gene id
GU305914.1 17186 17208 + 0 GGTACGACGGCATCCCCCAA AGG ||||||||[||||||||||||]PAM NA Intergenic NA NA

T769 AAAGGATCGTCCTCGTCCCCAGG 1000
Oligo fwd CACCAAAGGATCGTCCTCGTCCCC
Oligo rev AAACGGGGACGAGGACGATCCTTT
Chromosome start end strand MM target_seq PAM alignment distance position gene name gene id
GU305914.1 17204 17226 + 0 AAAGGATCGTCCTCGTCCCC AGG ||||||||[||||||||||||]PAM NA Intergenic NA NA

T770 GCAGCGCCAAAGCGTCCGAAAGG 1000
Oligo fwd CACCGCAGCGCCAAAGCGTCCGAA
Oligo rev AAACTTCGGACGCTTTGGCGCTGC
Chromosome start end strand MM target_seq PAM alignment distance position gene name gene id
GU305914.1 17235 17257 + 0 GCAGCGCCAAAGCGTCCGAA AGG ||||||||[||||||||||||]PAM NA Intergenic NA NA

T771 CAGCGCCAAAGCGTCCGAAAGGG 1000
Oligo fwd CACCCAGCGCCAAAGCGTCCGAAA
Oligo rev AAACTTTCGGACGCTTTGGCGCTG
Chromosome start end strand MM target_seq PAM alignment distance position gene name gene id
GU305914.1 17236 17258 + 0 CAGCGCCAAAGCGTCCGAAA GGG ||||||||[||||||||||||]PAM NA Intergenic NA NA

T772 GACCTCCTCTTCTTCACCGCCGG 1000
Oligo fwd CACCGACCTCCTCTTCTTCACCGC
Oligo rev AAACGCGGTGAAGAAGAGGAGGTC
Chromosome start end strand MM target_seq PAM alignment distance position gene name gene id
GU305914.1 17281 17303 + 0 GACCTCCTCTTCTTCACCGC CGG ||||||||[||||||||||||]PAM NA Intergenic NA NA

T773 CTCCTCTTCTTCACCGCCGGAGG 1000
Oligo fwd CACCCTCCTCTTCTTCACCGCCGG
Oligo rev AAACCCGGCGGTGAAGAAGAGGAG
Chromosome start end strand MM target_seq PAM alignment distance position gene name gene id
GU305914.1 17284 17306 + 0 CTCCTCTTCTTCACCGCCGG AGG ||||||||[||||||||||||]PAM NA Intergenic NA NA

T774 CTCTTCTTCACCGCCGGAGGTGG 1000
Oligo fwd CACCCTCTTCTTCACCGCCGGAGG
Oligo rev AAACCCTCCGGCGGTGAAGAAGAG
Chromosome start end strand MM target_seq PAM alignment distance position gene name gene id
GU305914.1 17287 17309 + 0 CTCTTCTTCACCGCCGGAGG TGG ||||||||[||||||||||||]PAM NA Intergenic NA NA

T775 TTCACCGCCGGAGGTGGTGTCGG 1000
Oligo fwd CACCTTCACCGCCGGAGGTGGTGT
Oligo rev AAACACACCACCTCCGGCGGTGAA
Chromosome start end strand MM target_seq PAM alignment distance position gene name gene id
GU305914.1 17293 17315 + 0 TTCACCGCCGGAGGTGGTGT CGG ||||||||[||||||||||||]PAM NA Intergenic NA NA

T776 TCACCGCCGGAGGTGGTGTCGGG 1000
Oligo fwd CACCTCACCGCCGGAGGTGGTGTC
Oligo rev AAACGACACCACCTCCGGCGGTGA
Chromosome start end strand MM target_seq PAM alignment distance position gene name gene id
GU305914.1 17294 17316 + 0 TCACCGCCGGAGGTGGTGTC GGG ||||||||[||||||||||||]PAM NA Intergenic NA NA

T777 GCCGGAGGTGGTGTCGGGTGCGG 1000
Oligo fwd CACCGCCGGAGGTGGTGTCGGGTG
Oligo rev AAACCACCCGACACCACCTCCGGC
Chromosome start end strand MM target_seq PAM alignment distance position gene name gene id
GU305914.1 17299 17321 + 0 GCCGGAGGTGGTGTCGGGTG CGG ||||||||[||||||||||||]PAM NA Intergenic NA NA

T778 GGAGGTGGTGTCGGGTGCGGCGG 1000

<table>
<tr><td>Oligo fwd</td><td>CACCGGAGGTGGTGTCGGGTGCGG</td><td></td><td></td><td></td><td></td><td></td><td></td><td></td><td></td><td></td><td></td><td></td></tr>
<tr><td>Oligo rev</td><td>AAACCCGCACCCGACACCACCTCC</td><td></td><td></td><td></td><td></td><td></td><td></td><td></td><td></td><td></td><td></td><td></td></tr>
<tr><td>Chromosome</td><td>start</td><td></td><td>end</td><td>strand</td><td>MM</td><td>target_seq</td><td>PAM</td><td>alignment</td><td>distance</td><td>position</td><td>gene name</td><td>gene id</td></tr>
<tr><td>GU305914.1</td><td></td><td>17302</td><td>17324</td><td>+</td><td>0</td><td>GGAGGTGGTGTCGGGTGCGG</td><td>CGG</td><td>||||||||[||||||||||||]PAM</td><td>NA</td><td>Intergenic</td><td>NA</td><td>NA</td></tr>
<tr><td></td><td></td><td></td><td></td><td></td><td></td><td></td><td></td><td></td><td></td><td></td><td></td><td></td></tr>
<tr><td>T779</td><td>GTCGGGTGCGGCGGAGACAACGG</td><td></td><td>1000</td><td></td><td></td><td></td><td></td><td></td><td></td><td></td><td></td><td></td></tr>
<tr><td>Oligo fwd</td><td>CACCGTCGGGTGCGGCGGAGACAA</td><td></td><td></td><td></td><td></td><td></td><td></td><td></td><td></td><td></td><td></td><td></td></tr>
<tr><td>Oligo rev</td><td>AAACTTGTCTCCGCCGCACCCGAC</td><td></td><td></td><td></td><td></td><td></td><td></td><td></td><td></td><td></td><td></td><td></td></tr>
<tr><td>Chromosome</td><td>start</td><td></td><td>end</td><td>strand</td><td>MM</td><td>target_seq</td><td>PAM</td><td>alignment</td><td>distance</td><td>position</td><td>gene name</td><td>gene id</td></tr>
<tr><td>GU305914.1</td><td></td><td>17311</td><td>17333</td><td>+</td><td>0</td><td>GTCGGGTGCGGCGGAGACAA</td><td>CGG</td><td>||||||||[||||||||||||]PAM</td><td>NA</td><td>Intergenic</td><td>NA</td><td>NA</td></tr>
<tr><td></td><td></td><td></td><td></td><td></td><td></td><td></td><td></td><td></td><td></td><td></td><td></td><td></td></tr>
<tr><td>T780</td><td>TGCGGCGGAGACAACGGCCTCGG</td><td></td><td>1000</td><td></td><td></td><td></td><td></td><td></td><td></td><td></td><td></td><td></td></tr>
<tr><td>Oligo fwd</td><td>CACCTGCGGCGGAGACAACGGCCT</td><td></td><td></td><td></td><td></td><td></td><td></td><td></td><td></td><td></td><td></td><td></td></tr>
<tr><td>Oligo rev</td><td>AAACAGGCCGTTGTCTCCGCCGCA</td><td></td><td></td><td></td><td></td><td></td><td></td><td></td><td></td><td></td><td></td><td></td></tr>
<tr><td>Chromosome</td><td>start</td><td></td><td>end</td><td>strand</td><td>MM</td><td>target_seq</td><td>PAM</td><td>alignment</td><td>distance</td><td>position</td><td>gene name</td><td>gene id</td></tr>
<tr><td>GU305914.1</td><td></td><td>17317</td><td>17339</td><td>+</td><td>0</td><td>TGCGGCGGAGACAACGGCCT</td><td>CGG</td><td>||||||||[||||||||||||]PAM</td><td>NA</td><td>Intergenic</td><td>NA</td><td>NA</td></tr>
<tr><td></td><td></td><td></td><td></td><td></td><td></td><td></td><td></td><td></td><td></td><td></td><td></td><td></td></tr>
<tr><td>T781</td><td>CTCGGAGCCGCCTTCGAGTCCGG</td><td></td><td>1000</td><td></td><td></td><td></td><td></td><td></td><td></td><td></td><td></td><td></td></tr>
<tr><td>Oligo fwd</td><td>CACCCTCGGAGCCGCCTTCGAGTC</td><td></td><td></td><td></td><td></td><td></td><td></td><td></td><td></td><td></td><td></td><td></td></tr>
<tr><td>Oligo rev</td><td>AAACGACTCGAAGGCGGCTCCGAG</td><td></td><td></td><td></td><td></td><td></td><td></td><td></td><td></td><td></td><td></td><td></td></tr>
<tr><td>Chromosome</td><td>start</td><td></td><td>end</td><td>strand</td><td>MM</td><td>target_seq</td><td>PAM</td><td>alignment</td><td>distance</td><td>position</td><td>gene name</td><td>gene id</td></tr>
<tr><td>GU305914.1</td><td></td><td>17335</td><td>17357</td><td>+</td><td>0</td><td>CTCGGAGCCGCCTTCGAGTC</td><td>CGG</td><td>||||||||[||||||||||||]PAM</td><td>NA</td><td>Intergenic</td><td>NA</td><td>NA</td></tr>
<tr><td></td><td></td><td></td><td></td><td></td><td></td><td></td><td></td><td></td><td></td><td></td><td></td><td></td></tr>
<tr><td>T782</td><td>GCCTTCGAGTCCGGTGTCCTCGG</td><td></td><td>1000</td><td></td><td></td><td></td><td></td><td></td><td></td><td></td><td></td><td></td></tr>
<tr><td>Oligo fwd</td><td>CACCGCCTTCGAGTCCGGTGTCCT</td><td></td><td></td><td></td><td></td><td></td><td></td><td></td><td></td><td></td><td></td><td></td></tr>
<tr><td>Oligo rev</td><td>AAACAGGACACCGGACTCGAAGGC</td><td></td><td></td><td></td><td></td><td></td><td></td><td></td><td></td><td></td><td></td><td></td></tr>
<tr><td>Chromosome</td><td>start</td><td></td><td>end</td><td>strand</td><td>MM</td><td>target_seq</td><td>PAM</td><td>alignment</td><td>distance</td><td>position</td><td>gene name</td><td>gene id</td></tr>
<tr><td>GU305914.1</td><td></td><td>17344</td><td>17366</td><td>+</td><td>0</td><td>GCCTTCGAGTCCGGTGTCCT</td><td>CGG</td><td>||||||||[||||||||||||]PAM</td><td>NA</td><td>Intergenic</td><td>NA</td><td>NA</td></tr>
<tr><td></td><td></td><td></td><td></td><td></td><td></td><td></td><td></td><td></td><td></td><td></td><td></td><td></td></tr>
<tr><td>T783</td><td>ATGCCGCGAGAGACTGTTGTTGG</td><td></td><td>1000</td><td></td><td></td><td></td><td></td><td></td><td></td><td></td><td></td><td></td></tr>
<tr><td>Oligo fwd</td><td>CACCATGCCGCGAGAGACTGTTGT</td><td></td><td></td><td></td><td></td><td></td><td></td><td></td><td></td><td></td><td></td><td></td></tr>
<tr><td>Oligo rev</td><td>AAACACAACAGTCTCTCGCGGCAT</td><td></td><td></td><td></td><td></td><td></td><td></td><td></td><td></td><td></td><td></td><td></td></tr>
<tr><td>Chromosome</td><td>start</td><td></td><td>end</td><td>strand</td><td>MM</td><td>target_seq</td><td>PAM</td><td>alignment</td><td>distance</td><td>position</td><td>gene name</td><td>gene id</td></tr>
<tr><td>GU305914.1</td><td></td><td>17371</td><td>17393</td><td>+</td><td>0</td><td>ATGCCGCGAGAGACTGTTGT</td><td>TGG</td><td>||||||||[||||||||||||]PAM</td><td>NA</td><td>Intergenic</td><td>NA</td><td>NA</td></tr>
<tr><td></td><td></td><td></td><td></td><td></td><td></td><td></td><td></td><td></td><td></td><td></td><td></td><td></td></tr>
<tr><td>T784</td><td>TGCCGCGAGAGACTGTTGTTGGG</td><td></td><td>1000</td><td></td><td></td><td></td><td></td><td></td><td></td><td></td><td></td><td></td></tr>
<tr><td>Oligo fwd</td><td>CACCTGCCGCGAGAGACTGTTGTT</td><td></td><td></td><td></td><td></td><td></td><td></td><td></td><td></td><td></td><td></td><td></td></tr>
<tr><td>Oligo rev</td><td>AAACAACAACAGTCTCTCGCGGCA</td><td></td><td></td><td></td><td></td><td></td><td></td><td></td><td></td><td></td><td></td><td></td></tr>
<tr><td>Chromosome</td><td>start</td><td></td><td>end</td><td>strand</td><td>MM</td><td>target_seq</td><td>PAM</td><td>alignment</td><td>distance</td><td>position</td><td>gene name</td><td>gene id</td></tr>
<tr><td>GU305914.1</td><td></td><td>17372</td><td>17394</td><td>+</td><td>0</td><td>TGCCGCGAGAGACTGTTGTT</td><td>GGG</td><td>||||||||[||||||||||||]PAM</td><td>NA</td><td>Intergenic</td><td>NA</td><td>NA</td></tr>
<tr><td></td><td></td><td></td><td></td><td></td><td></td><td></td><td></td><td></td><td></td><td></td><td></td><td></td></tr>
<tr><td>T785</td><td>GCCGCGAGAGACTGTTGTTGGGG</td><td></td><td>1000</td><td></td><td></td><td></td><td></td><td></td><td></td><td></td><td></td><td></td></tr>
<tr><td>Oligo fwd</td><td>CACCGCCGCGAGAGACTGTTGTTG</td><td></td><td></td><td></td><td></td><td></td><td></td><td></td><td></td><td></td><td></td><td></td></tr>
<tr><td>Oligo rev</td><td>AAACCAACAACAGTCTCTCGCGGC</td><td></td><td></td><td></td><td></td><td></td><td></td><td></td><td></td><td></td><td></td><td></td></tr>
<tr><td>Chromosome</td><td>start</td><td></td><td>end</td><td>strand</td><td>MM</td><td>target_seq</td><td>PAM</td><td>alignment</td><td>distance</td><td>position</td><td>gene name</td><td>gene id</td></tr>
<tr><td>GU305914.1</td><td></td><td>17373</td><td>17395</td><td>+</td><td>0</td><td>GCCGCGAGAGACTGTTGTTG</td><td>GGG</td><td>||||||||[||||||||||||]PAM</td><td>NA</td><td>Intergenic</td><td>NA</td><td>NA</td></tr>
<tr><td></td><td></td><td></td><td></td><td></td><td></td><td></td><td></td><td></td><td></td><td></td><td></td><td></td></tr>
<tr><td>T786</td><td>GGGTTGACGAGCTGTTGCTGTGG</td><td></td><td>1000</td><td></td><td></td><td></td><td></td><td></td><td></td><td></td><td></td><td></td></tr>
<tr><td>Oligo fwd</td><td>CACCGGGTTGACGAGCTGTTGCTG</td><td></td><td></td><td></td><td></td><td></td><td></td><td></td><td></td><td></td><td></td><td></td></tr>
<tr><td>Oligo rev</td><td>AAACCAGCAACAGCTCGTCAACCC</td><td></td><td></td><td></td><td></td><td></td><td></td><td></td><td></td><td></td><td></td><td></td></tr>
<tr><td>Chromosome</td><td>start</td><td></td><td>end</td><td>strand</td><td>MM</td><td>target_seq</td><td>PAM</td><td>alignment</td><td>distance</td><td>position</td><td>gene name</td><td>gene id</td></tr>
<tr><td>GU305914.1</td><td></td><td>17393</td><td>17415</td><td>+</td><td>0</td><td>GGGTTGACGAGCTGTTGCTG</td><td>TGG</td><td>||||||||[||||||||||||]PAM</td><td>NA</td><td>Intergenic</td><td>NA</td><td>NA</td></tr>
<tr><td></td><td></td><td></td><td></td><td></td><td></td><td></td><td></td><td></td><td></td><td></td><td></td><td></td></tr>
<tr><td>T787</td><td>GAGCTGTTGCTGTGGTGTCGAGG</td><td></td><td>1000</td><td></td><td></td><td></td><td></td><td></td><td></td><td></td><td></td><td></td></tr>
<tr><td>Oligo fwd</td><td>CACCGAGCTGTTGCTGTGGTGTCG</td><td></td><td></td><td></td><td></td><td></td><td></td><td></td><td></td><td></td><td></td><td></td></tr>
<tr><td>Oligo rev</td><td>AAACCGACACCACAGCAACAGCTC</td><td></td><td></td><td></td><td></td><td></td><td></td><td></td><td></td><td></td><td></td><td></td></tr>
<tr><td>Chromosome</td><td>start</td><td></td><td>end</td><td>strand</td><td>MM</td><td>target_seq</td><td>PAM</td><td>alignment</td><td>distance</td><td>position</td><td>gene name</td><td>gene id</td></tr>
<tr><td>GU305914.1</td><td></td><td>17401</td><td>17423</td><td>+</td><td>0</td><td>GAGCTGTTGCTGTGGTGTCG</td><td>AGG</td><td>||||||||[||||||||||||]PAM</td><td>NA</td><td>Intergenic</td><td>NA</td><td>NA</td></tr>
<tr><td></td><td></td><td></td><td></td><td></td><td></td><td></td><td></td><td></td><td></td><td></td><td></td><td></td></tr>
<tr><td>T788</td><td>AGCTGTTGCTGTGGTGTCGAGGG</td><td></td><td>1000</td><td></td><td></td><td></td><td></td><td></td><td></td><td></td><td></td><td></td></tr>
<tr><td>Oligo fwd</td><td>CACCAGCTGTTGCTGTGGTGTCGA</td><td></td><td></td><td></td><td></td><td></td><td></td><td></td><td></td><td></td><td></td><td></td></tr>
<tr><td>Oligo rev</td><td>AAACTCGACACCACAGCAACAGCT</td><td></td><td></td><td></td><td></td><td></td><td></td><td></td><td></td><td></td><td></td><td></td></tr>
<tr><td>Chromosome</td><td>start</td><td></td><td>end</td><td>strand</td><td>MM</td><td>target_seq</td><td>PAM</td><td>alignment</td><td>distance</td><td>position</td><td>gene name</td><td>gene id</td></tr>
<tr><td>GU305914.1</td><td></td><td>17402</td><td>17424</td><td>+</td><td>0</td><td>AGCTGTTGCTGTGGTGTCGA</td><td>GGG</td><td>||||||||[||||||||||||]PAM</td><td>NA</td><td>Intergenic</td><td>NA</td><td>NA</td></tr>
<tr><td></td><td></td><td></td><td></td><td></td><td></td><td></td><td></td><td></td><td></td><td></td><td></td><td></td></tr>
<tr><td>T789</td><td>GTTGCTGTGGTGTCGAGGGTCGG</td><td></td><td>1000</td><td></td><td></td><td></td><td></td><td></td><td></td><td></td><td></td><td></td></tr>
<tr><td>Oligo fwd</td><td>CACCGTTGCTGTGGTGTCGAGGGT</td><td></td><td></td><td></td><td></td><td></td><td></td><td></td><td></td><td></td><td></td><td></td></tr>
<tr><td>Oligo rev</td><td>AAACACCCTCGACACCACAGCAAC</td><td></td><td></td><td></td><td></td><td></td><td></td><td></td><td></td><td></td><td></td><td></td></tr>
<tr><td>Chromosome</td><td>start</td><td></td><td>end</td><td>strand</td><td>MM</td><td>target_seq</td><td>PAM</td><td>alignment</td><td>distance</td><td>position</td><td>gene name</td><td>gene id</td></tr>
<tr><td>GU305914.1</td><td></td><td>17406</td><td>17428</td><td>+</td><td>0</td><td>GTTGCTGTGGTGTCGAGGGT</td><td>CGG</td><td>||||||||[||||||||||||]PAM</td><td>NA</td><td>Intergenic</td><td>NA</td><td>NA</td></tr>
<tr><td></td><td></td><td></td><td></td><td></td><td></td><td></td><td></td><td></td><td></td><td></td><td></td><td></td></tr>
<tr><td>T790</td><td>TGTGGTGTCGAGGGTCGGAGAGG</td><td></td><td>1000</td><td></td><td></td><td></td><td></td><td></td><td></td><td></td><td></td><td></td></tr>
<tr><td>Oligo fwd</td><td>CACCTGTGGTGTCGAGGGTCGGAG</td><td></td><td></td><td></td><td></td><td></td><td></td><td></td><td></td><td></td><td></td><td></td></tr>
<tr><td>Oligo rev</td><td>AAACCTCCGACCCTCGACACCACA</td><td></td><td></td><td></td><td></td><td></td><td></td><td></td><td></td><td></td><td></td><td></td></tr>
<tr><td>Chromosome</td><td>start</td><td></td><td>end</td><td>strand</td><td>MM</td><td>target_seq</td><td>PAM</td><td>alignment</td><td>distance</td><td>position</td><td>gene name</td><td>gene id</td></tr>
<tr><td>GU305914.1</td><td></td><td>17411</td><td>17433</td><td>+</td><td>0</td><td>TGTGGTGTCGAGGGTCGGAG</td><td>AGG</td><td>||||||||[||||||||||||]PAM</td><td>NA</td><td>Intergenic</td><td>NA</td><td>NA</td></tr>
<tr><td></td><td></td><td></td><td></td><td></td><td></td><td></td><td></td><td></td><td></td><td></td><td></td><td></td></tr>
<tr><td>T791</td><td>GTCGAGGGTCGGAGAGGCGAAGG</td><td></td><td>1000</td><td></td><td></td><td></td><td></td><td></td><td></td><td></td><td></td><td></td></tr>
<tr><td>Oligo fwd</td><td>CACCGTCGAGGGTCGGAGAGGCGA</td><td></td><td></td><td></td><td></td><td></td><td></td><td></td><td></td><td></td><td></td><td></td></tr>
<tr><td>Oligo rev</td><td>AAACTCGCCTCTCCGACCCTCGAC</td><td></td><td></td><td></td><td></td><td></td><td></td><td></td><td></td><td></td><td></td><td></td></tr>
<tr><td>Chromosome</td><td>start</td><td></td><td>end</td><td>strand</td><td>MM</td><td>target_seq</td><td>PAM</td><td>alignment</td><td>distance</td><td>position</td><td>gene name</td><td>gene id</td></tr>
<tr><td>GU305914.1</td><td></td><td>17417</td><td>17439</td><td>+</td><td>0</td><td>GTCGAGGGTCGGAGAGGCGA</td><td>AGG</td><td>||||||||[||||||||||||]PAM</td><td>NA</td><td>Intergenic</td><td>NA</td><td>NA</td></tr>
<tr><td></td><td></td><td></td><td></td><td></td><td></td><td></td><td></td><td></td><td></td><td></td><td></td><td></td></tr>
<tr><td>T792</td><td>GAGGGTCGGAGAGGCGAAGGAGG</td><td></td><td>1000</td><td></td><td></td><td></td><td></td><td></td><td></td><td></td><td></td><td></td></tr>
<tr><td>Oligo fwd</td><td>CACCGAGGGTCGGAGAGGCGAAGG</td><td></td><td></td><td></td><td></td><td></td><td></td><td></td><td></td><td></td><td></td><td></td></tr>
<tr><td>Oligo rev</td><td>AAACCCTTCGCCTCTCCGACCCTC</td><td></td><td></td><td></td><td></td><td></td><td></td><td></td><td></td><td></td><td></td><td></td></tr>
<tr><td>Chromosome</td><td>start</td><td></td><td>end</td><td>strand</td><td>MM</td><td>target_seq</td><td>PAM</td><td>alignment</td><td>distance</td><td>position</td><td>gene name</td><td>gene id</td></tr>
<tr><td>GU305914.1</td><td></td><td>17420</td><td>17442</td><td>+</td><td>0</td><td>GAGGGTCGGAGAGGCGAAGG</td><td>AGG</td><td>||||||||[||||||||||||]PAM</td><td>NA</td><td>Intergenic</td><td>NA</td><td>NA</td></tr>
</table>

<table>
<tr><td>T793</td><td>AGAGGCGAAGGAGGCAAGAGCGG</td><td>1000</td><td></td><td></td><td></td><td></td><td></td><td></td><td></td><td></td><td></td></tr>
<tr><td>Oligo fwd</td><td>CACCAGAGGCGAAGGAGGCAAGAG</td><td></td><td></td><td></td><td></td><td></td><td></td><td></td><td></td><td></td><td></td></tr>
<tr><td>Oligo rev</td><td>AAACCTCTTGCCTCCTTCGCCTCT</td><td></td><td></td><td></td><td></td><td></td><td></td><td></td><td></td><td></td><td></td></tr>
<tr><td>Chromosome</td><td>start</td><td>end</td><td>strand</td><td>MM</td><td>target_seq</td><td>PAM</td><td>alignment</td><td>distance</td><td>position</td><td>gene name</td><td>gene id</td></tr>
<tr><td>GU305914.1</td><td>17429</td><td>17451</td><td>+</td><td>0</td><td>AGAGGCGAAGGAGGCAAGAG</td><td>CGG</td><td>||||||||[||||||||||||]PAM</td><td>NA</td><td>Intergenic</td><td>NA</td><td>NA</td></tr>
<tr><td>T794</td><td>AGGAGGCAAGAGCGGTCGTTTGG</td><td>1000</td><td></td><td></td><td></td><td></td><td></td><td></td><td></td><td></td><td></td></tr>
<tr><td>Oligo fwd</td><td>CACCAGGAGGCAAGAGCGGTCGTT</td><td></td><td></td><td></td><td></td><td></td><td></td><td></td><td></td><td></td><td></td></tr>
<tr><td>Oligo rev</td><td>AAACAACGACCGCTCTTGCCTCCT</td><td></td><td></td><td></td><td></td><td></td><td></td><td></td><td></td><td></td><td></td></tr>
<tr><td>Chromosome</td><td>start</td><td>end</td><td>strand</td><td>MM</td><td>target_seq</td><td>PAM</td><td>alignment</td><td>distance</td><td>position</td><td>gene name</td><td>gene id</td></tr>
<tr><td>GU305914.1</td><td>17437</td><td>17459</td><td>+</td><td>0</td><td>AGGAGGCAAGAGCGGTCGTT</td><td>TGG</td><td>||||||||[||||||||||||]PAM</td><td>NA</td><td>Intergenic</td><td>NA</td><td>NA</td></tr>
<tr><td>T795</td><td>AGGCAAGAGCGGTCGTTTGGCGG</td><td>1000</td><td></td><td></td><td></td><td></td><td></td><td></td><td></td><td></td><td></td></tr>
<tr><td>Oligo fwd</td><td>CACCAGGCAAGAGCGGTCGTTTGG</td><td></td><td></td><td></td><td></td><td></td><td></td><td></td><td></td><td></td><td></td></tr>
<tr><td>Oligo rev</td><td>AAACCCAAACGACCGCTCTTGCCT</td><td></td><td></td><td></td><td></td><td></td><td></td><td></td><td></td><td></td><td></td></tr>
<tr><td>Chromosome</td><td>start</td><td>end</td><td>strand</td><td>MM</td><td>target_seq</td><td>PAM</td><td>alignment</td><td>distance</td><td>position</td><td>gene name</td><td>gene id</td></tr>
<tr><td>GU305914.1</td><td>17440</td><td>17462</td><td>+</td><td>0</td><td>AGGCAAGAGCGGTCGTTTGG</td><td>CGG</td><td>||||||||[||||||||||||]PAM</td><td>NA</td><td>Intergenic</td><td>NA</td><td>NA</td></tr>
<tr><td>T796</td><td>CGGTCGTTTGGCGGCCTCTCTGG</td><td>1000</td><td></td><td></td><td></td><td></td><td></td><td></td><td></td><td></td><td></td></tr>
<tr><td>Oligo fwd</td><td>CACCCGGTCGTTTGGCGGCCTCTC</td><td></td><td></td><td></td><td></td><td></td><td></td><td></td><td></td><td></td><td></td></tr>
<tr><td>Oligo rev</td><td>AAACGAGAGGCCGCCAAACGACCG</td><td></td><td></td><td></td><td></td><td></td><td></td><td></td><td></td><td></td><td></td></tr>
<tr><td>Chromosome</td><td>start</td><td>end</td><td>strand</td><td>MM</td><td>target_seq</td><td>PAM</td><td>alignment</td><td>distance</td><td>position</td><td>gene name</td><td>gene id</td></tr>
<tr><td>GU305914.1</td><td>17449</td><td>17471</td><td>+</td><td>0</td><td>CGGTCGTTTGGCGGCCTCTC</td><td>TGG</td><td>||||||||[||||||||||||]PAM</td><td>NA</td><td>Intergenic</td><td>NA</td><td>NA</td></tr>
<tr><td>T797</td><td>CAAGTTTCTCTGCACAAAACCGG</td><td>1000</td><td></td><td></td><td></td><td></td><td></td><td></td><td></td><td></td><td></td></tr>
<tr><td>Oligo fwd</td><td>CACCCAAGTTTCTCTGCACAAAAC</td><td></td><td></td><td></td><td></td><td></td><td></td><td></td><td></td><td></td><td></td></tr>
<tr><td>Oligo rev</td><td>AAACGTTTTGTGCAGAGAAACTTG</td><td></td><td></td><td></td><td></td><td></td><td></td><td></td><td></td><td></td><td></td></tr>
<tr><td>Chromosome</td><td>start</td><td>end</td><td>strand</td><td>MM</td><td>target_seq</td><td>PAM</td><td>alignment</td><td>distance</td><td>position</td><td>gene name</td><td>gene id</td></tr>
<tr><td>GU305914.1</td><td>17494</td><td>17516</td><td>+</td><td>0</td><td>CAAGTTTCTCTGCACAAAAC</td><td>CGG</td><td>||||||||[||||||||||||]PAM</td><td>NA</td><td>Intergenic</td><td>NA</td><td>NA</td></tr>
<tr><td>T798</td><td>AAGTTTCTCTGCACAAAACCGGG</td><td>1000</td><td></td><td></td><td></td><td></td><td></td><td></td><td></td><td></td><td></td></tr>
<tr><td>Oligo fwd</td><td>CACCAAGTTTCTCTGCACAAAACC</td><td></td><td></td><td></td><td></td><td></td><td></td><td></td><td></td><td></td><td></td></tr>
<tr><td>Oligo rev</td><td>AAACGGTTTTGTGCAGAGAAACTT</td><td></td><td></td><td></td><td></td><td></td><td></td><td></td><td></td><td></td><td></td></tr>
<tr><td>Chromosome</td><td>start</td><td>end</td><td>strand</td><td>MM</td><td>target_seq</td><td>PAM</td><td>alignment</td><td>distance</td><td>position</td><td>gene name</td><td>gene id</td></tr>
<tr><td>GU305914.1</td><td>17495</td><td>17517</td><td>+</td><td>0</td><td>AAGTTTCTCTGCACAAAACC</td><td>GGG</td><td>||||||||[||||||||||||]PAM</td><td>NA</td><td>Intergenic</td><td>NA</td><td>NA</td></tr>
<tr><td>T799</td><td>ACCGGGTCCGCGAGTATCTTTGG</td><td>1000</td><td></td><td></td><td></td><td></td><td></td><td></td><td></td><td></td><td></td></tr>
<tr><td>Oligo fwd</td><td>CACCACCGGGTCCGCGAGTATCTT</td><td></td><td></td><td></td><td></td><td></td><td></td><td></td><td></td><td></td><td></td></tr>
<tr><td>Oligo rev</td><td>AAACAAGATACTCGCGGACCCGGT</td><td></td><td></td><td></td><td></td><td></td><td></td><td></td><td></td><td></td><td></td></tr>
<tr><td>Chromosome</td><td>start</td><td>end</td><td>strand</td><td>MM</td><td>target_seq</td><td>PAM</td><td>alignment</td><td>distance</td><td>position</td><td>gene name</td><td>gene id</td></tr>
<tr><td>GU305914.1</td><td>17512</td><td>17534</td><td>+</td><td>0</td><td>ACCGGGTCCGCGAGTATCTT</td><td>TGG</td><td>||||||||[||||||||||||]PAM</td><td>NA</td><td>Intergenic</td><td>NA</td><td>NA</td></tr>
<tr><td>T800</td><td>CCGGGTCCGCGAGTATCTTTGGG</td><td>1000</td><td></td><td></td><td></td><td></td><td></td><td></td><td></td><td></td><td></td></tr>
<tr><td>Oligo fwd</td><td>CACCCCGGGTCCGCGAGTATCTTT</td><td></td><td></td><td></td><td></td><td></td><td></td><td></td><td></td><td></td><td></td></tr>
<tr><td>Oligo rev</td><td>AAACAAAGATACTCGCGGACCCGG</td><td></td><td></td><td></td><td></td><td></td><td></td><td></td><td></td><td></td><td></td></tr>
<tr><td>Chromosome</td><td>start</td><td>end</td><td>strand</td><td>MM</td><td>target_seq</td><td>PAM</td><td>alignment</td><td>distance</td><td>position</td><td>gene name</td><td>gene id</td></tr>
<tr><td>GU305914.1</td><td>17513</td><td>17535</td><td>+</td><td>0</td><td>CCGGGTCCGCGAGTATCTTT</td><td>GGG</td><td>||||||||[||||||||||||]PAM</td><td>NA</td><td>Intergenic</td><td>NA</td><td>NA</td></tr>
<tr><td>T801</td><td>AGTATCTTTGGGTTCGTCTTCGG</td><td>1000</td><td></td><td></td><td></td><td></td><td></td><td></td><td></td><td></td><td></td></tr>
<tr><td>Oligo fwd</td><td>CACCAGTATCTTTGGGTTCGTCTT</td><td></td><td></td><td></td><td></td><td></td><td></td><td></td><td></td><td></td><td></td></tr>
<tr><td>Oligo rev</td><td>AAACAAGACGAACCCAAAGATACT</td><td></td><td></td><td></td><td></td><td></td><td></td><td></td><td></td><td></td><td></td></tr>
<tr><td>Chromosome</td><td>start</td><td>end</td><td>strand</td><td>MM</td><td>target_seq</td><td>PAM</td><td>alignment</td><td>distance</td><td>position</td><td>gene name</td><td>gene id</td></tr>
<tr><td>GU305914.1</td><td>17524</td><td>17546</td><td>+</td><td>0</td><td>AGTATCTTTGGGTTCGTCTT</td><td>CGG</td><td>||||||||[||||||||||||]PAM</td><td>NA</td><td>Intergenic</td><td>NA</td><td>NA</td></tr>
<tr><td>T802</td><td>GTCTTCGGCCGTCATCGCAGCGG</td><td>1000</td><td></td><td></td><td></td><td></td><td></td><td></td><td></td><td></td><td></td></tr>
<tr><td>Oligo fwd</td><td>CACCGTCTTCGGCCGTCATCGCAG</td><td></td><td></td><td></td><td></td><td></td><td></td><td></td><td></td><td></td><td></td></tr>
<tr><td>Oligo rev</td><td>AAACCTGCGATGACGGCCGAAGAC</td><td></td><td></td><td></td><td></td><td></td><td></td><td></td><td></td><td></td><td></td></tr>
<tr><td>Chromosome</td><td>start</td><td>end</td><td>strand</td><td>MM</td><td>target_seq</td><td>PAM</td><td>alignment</td><td>distance</td><td>position</td><td>gene name</td><td>gene id</td></tr>
<tr><td>GU305914.1</td><td>17539</td><td>17561</td><td>+</td><td>0</td><td>GTCTTCGGCCGTCATCGCAG</td><td>CGG</td><td>||||||||[||||||||||||]PAM</td><td>NA</td><td>Intergenic</td><td>NA</td><td>NA</td></tr>
<tr><td>T803</td><td>TCTTCGGCCGTCATCGCAGCGGG</td><td>1000</td><td></td><td></td><td></td><td></td><td></td><td></td><td></td><td></td><td></td></tr>
<tr><td>Oligo fwd</td><td>CACCTCTTCGGCCGTCATCGCAGC</td><td></td><td></td><td></td><td></td><td></td><td></td><td></td><td></td><td></td><td></td></tr>
<tr><td>Oligo rev</td><td>AAACGCTGCGATGACGGCCGAAGA</td><td></td><td></td><td></td><td></td><td></td><td></td><td></td><td></td><td></td><td></td></tr>
<tr><td>Chromosome</td><td>start</td><td>end</td><td>strand</td><td>MM</td><td>target_seq</td><td>PAM</td><td>alignment</td><td>distance</td><td>position</td><td>gene name</td><td>gene id</td></tr>
<tr><td>GU305914.1</td><td>17540</td><td>17562</td><td>+</td><td>0</td><td>TCTTCGGCCGTCATCGCAGC</td><td>GGG</td><td>||||||||[||||||||||||]PAM</td><td>NA</td><td>Intergenic</td><td>NA</td><td>NA</td></tr>
<tr><td>T804</td><td>CGTCATCGCAGCGGGCGATTTGG</td><td>1000</td><td></td><td></td><td></td><td></td><td></td><td></td><td></td><td></td><td></td></tr>
<tr><td>Oligo fwd</td><td>CACCCGTCATCGCAGCGGGCGATT</td><td></td><td></td><td></td><td></td><td></td><td></td><td></td><td></td><td></td><td></td></tr>
<tr><td>Oligo rev</td><td>AAACAATCGCCCGCTGCGATGACG</td><td></td><td></td><td></td><td></td><td></td><td></td><td></td><td></td><td></td><td></td></tr>
<tr><td>Chromosome</td><td>start</td><td>end</td><td>strand</td><td>MM</td><td>target_seq</td><td>PAM</td><td>alignment</td><td>distance</td><td>position</td><td>gene name</td><td>gene id</td></tr>
<tr><td>GU305914.1</td><td>17548</td><td>17570</td><td>+</td><td>0</td><td>CGTCATCGCAGCGGGCGATT</td><td>TGG</td><td>||||||||[||||||||||||]PAM</td><td>NA</td><td>Intergenic</td><td>NA</td><td>NA</td></tr>
<tr><td>T805</td><td>GCGCGCTCGTTCCCCTTGCTCGG</td><td>1000</td><td></td><td></td><td></td><td></td><td></td><td></td><td></td><td></td><td></td></tr>
<tr><td>Oligo fwd</td><td>CACCGCGCGCTCGTTCCCCTTGCT</td><td></td><td></td><td></td><td></td><td></td><td></td><td></td><td></td><td></td><td></td></tr>
<tr><td>Oligo rev</td><td>AAACAGCAAGGGGAACGAGCGCGC</td><td></td><td></td><td></td><td></td><td></td><td></td><td></td><td></td><td></td><td></td></tr>
<tr><td>Chromosome</td><td>start</td><td>end</td><td>strand</td><td>MM</td><td>target_seq</td><td>PAM</td><td>alignment</td><td>distance</td><td>position</td><td>gene name</td><td>gene id</td></tr>
<tr><td>GU305914.1</td><td>17570</td><td>17592</td><td>+</td><td>0</td><td>GCGCGCTCGTTCCCCTTGCT</td><td>CGG</td><td>||||||||[||||||||||||]PAM</td><td>NA</td><td>Intergenic</td><td>NA</td><td>NA</td></tr>
<tr><td>T806</td><td>CGCGCTCGTTCCCCTTGCTCGGG</td><td>1000</td><td></td><td></td><td></td><td></td><td></td><td></td><td></td><td></td><td></td></tr>
<tr><td>Oligo fwd</td><td>CACCCGCGCTCGTTCCCCTTGCTC</td><td></td><td></td><td></td><td></td><td></td><td></td><td></td><td></td><td></td><td></td></tr>
<tr><td>Oligo rev</td><td>AAACGAGCAAGGGGAACGAGCGCG</td><td></td><td></td><td></td><td></td><td></td><td></td><td></td><td></td><td></td><td></td></tr>
<tr><td>Chromosome</td><td>start</td><td>end</td><td>strand</td><td>MM</td><td>target_seq</td><td>PAM</td><td>alignment</td><td>distance</td><td>position</td><td>gene name</td><td>gene id</td></tr>
<tr><td>GU305914.1</td><td>17571</td><td>17593</td><td>+</td><td>0</td><td>CGCGCTCGTTCCCCTTGCTC</td><td>GGG</td><td>||||||||[||||||||||||]PAM</td><td>NA</td><td>Intergenic</td><td>NA</td><td>NA</td></tr>
<tr><td>T807</td><td>CCTCATCGACCGATATTTTGTGG</td><td>1000</td><td></td><td></td><td></td><td></td><td></td><td></td><td></td><td></td><td></td></tr>
<tr><td>Oligo fwd</td><td>CACCCCTCATCGACCGATATTTTG</td><td></td><td></td><td></td><td></td><td></td><td></td><td></td><td></td><td></td><td></td></tr>
<tr><td>Oligo rev</td><td>AAACCAAAATATCGGTCGATGAGG</td><td></td><td></td><td></td><td></td><td></td><td></td><td></td><td></td><td></td><td></td></tr>
<tr><td>Chromosome</td><td>start</td><td>end</td><td>strand</td><td>MM</td><td>target_seq</td><td>PAM</td><td>alignment</td><td>distance</td><td>position</td><td>gene name</td><td>gene id</td></tr>
<tr><td>GU305914.1</td><td>17622</td><td>17644</td><td>+</td><td>0</td><td>CCTCATCGACCGATATTTTG</td><td>TGG</td><td>||||||||[||||||||||||]PAM</td><td>NA</td><td>Intergenic</td><td>NA</td><td>NA</td></tr>
</table>

| | | | | | | | | | | | |
|---|---|---|---|---|---|---|---|---|---|---|---|
| T808 | CGACCGATATTTTGTGGTCGCGG | 1000 | | | | | | | | | |
| Oligo fwd | CACCCGACCGATATTTTGTGGTCG | | | | | | | | | | |
| Oligo rev | AAACCGACCACAAAATATCGGTCG | | | | | | | | | | |
| Chromosome | start | end | strand | MM | target_seq | PAM | alignment | distance | position | gene name | gene id |
| GU305914.1 | 17628 | 17650 | + | 0 | CGACCGATATTTTGTGGTCG | CGG | \|\|\|\|\|\|\|\|[\|\|\|\|\|\|\|\|\|\|\|\|]PAM | NA | Intergenic | NA | NA |

| | | | | | | | | | | | |
|---|---|---|---|---|---|---|---|---|---|---|---|
| T809 | TGGTCGCGGCGCGTCTTGCGAGG | 1000 | | | | | | | | | |
| Oligo fwd | CACCTGGTCGCGGCGCGTCTTGCG | | | | | | | | | | |
| Oligo rev | AAACCGCAAGACGCGCCGCGACCA | | | | | | | | | | |
| Chromosome | start | end | strand | MM | target_seq | PAM | alignment | distance | position | gene name | gene id |
| GU305914.1 | 17642 | 17664 | + | 0 | TGGTCGCGGCGCGTCTTGCG | AGG | \|\|\|\|\|\|\|\|[\|\|\|\|\|\|\|\|\|\|\|\|]PAM | NA | Intergenic | NA | NA |

| | | | | | | | | | | | |
|---|---|---|---|---|---|---|---|---|---|---|---|
| T810 | GCGCGTCTTGCGAGGAAGATCGG | 1000 | | | | | | | | | |
| Oligo fwd | CACCGCGCGTCTTGCGAGGAAGAT | | | | | | | | | | |
| Oligo rev | AAACATCTTCCTCGCAAGACGCGC | | | | | | | | | | |
| Chromosome | start | end | strand | MM | target_seq | PAM | alignment | distance | position | gene name | gene id |
| GU305914.1 | 17650 | 17672 | + | 0 | GCGCGTCTTGCGAGGAAGAT | CGG | \|\|\|\|\|\|\|\|[\|\|\|\|\|\|\|\|\|\|\|\|]PAM | NA | Intergenic | NA | NA |

| | | | | | | | | | | | |
|---|---|---|---|---|---|---|---|---|---|---|---|
| T811 | CTTGCGAGGAAGATCGGCACCGG | 1000 | | | | | | | | | |
| Oligo fwd | CACCCTTGCGAGGAAGATCGGCAC | | | | | | | | | | |
| Oligo rev | AAACGTGCCGATCTTCCTCGCAAG | | | | | | | | | | |
| Chromosome | start | end | strand | MM | target_seq | PAM | alignment | distance | position | gene name | gene id |
| GU305914.1 | 17656 | 17678 | + | 0 | CTTGCGAGGAAGATCGGCAC | CGG | \|\|\|\|\|\|\|\|[\|\|\|\|\|\|\|\|\|\|\|\|]PAM | NA | Intergenic | NA | NA |

| | | | | | | | | | | | |
|---|---|---|---|---|---|---|---|---|---|---|---|
| T812 | AGGAAGATCGGCACCGGAGTCGG | 1000 | | | | | | | | | |
| Oligo fwd | CACCAGGAAGATCGGCACCGGAGT | | | | | | | | | | |
| Oligo rev | AAACACTCCGGTGCCGATCTTCCT | | | | | | | | | | |
| Chromosome | start | end | strand | MM | target_seq | PAM | alignment | distance | position | gene name | gene id |
| GU305914.1 | 17662 | 17684 | + | 0 | AGGAAGATCGGCACCGGAGT | CGG | \|\|\|\|\|\|\|\|[\|\|\|\|\|\|\|\|\|\|\|\|]PAM | NA | Intergenic | NA | NA |

| | | | | | | | | | | | |
|---|---|---|---|---|---|---|---|---|---|---|---|
| T813 | CGGAGTCGGATGACGACCGACGG | 1000 | | | | | | | | | |
| Oligo fwd | CACCCGGAGTCGGATGACGACCGA | | | | | | | | | | |
| Oligo rev | AAACTCGGTCGTCATCCGACTCCG | | | | | | | | | | |
| Chromosome | start | end | strand | MM | target_seq | PAM | alignment | distance | position | gene name | gene id |
| GU305914.1 | 17676 | 17698 | + | 0 | CGGAGTCGGATGACGACCGA | CGG | \|\|\|\|\|\|\|\|[\|\|\|\|\|\|\|\|\|\|\|\|]PAM | NA | Intergenic | NA | NA |

| | | | | | | | | | | | |
|---|---|---|---|---|---|---|---|---|---|---|---|
| T814 | CGTCCGTCGCGCCATCGCCGTGG | 1000 | | | | | | | | | |
| Oligo fwd | CACCCGTCCGTCGCGCCATCGCCG | | | | | | | | | | |
| Oligo rev | AAACCGGCGATGGCGCGACGGACG | | | | | | | | | | |
| Chromosome | start | end | strand | MM | target_seq | PAM | alignment | distance | position | gene name | gene id |
| GU305914.1 | 17700 | 17722 | + | 0 | CGTCCGTCGCGCCATCGCCG | TGG | \|\|\|\|\|\|\|\|[\|\|\|\|\|\|\|\|\|\|\|\|]PAM | NA | Intergenic | NA | NA |

| | | | | | | | | | | | |
|---|---|---|---|---|---|---|---|---|---|---|---|
| T815 | TCGCTTTGTTGTTGCGCCTCAGG | 1000 | | | | | | | | | |
| Oligo fwd | CACCTCGCTTTGTTGTTGCGCCTC | | | | | | | | | | |
| Oligo rev | AAACGAGGCGCAACAACAAAGCGA | | | | | | | | | | |
| Chromosome | start | end | strand | MM | target_seq | PAM | alignment | distance | position | gene name | gene id |
| GU305914.1 | 17732 | 17754 | + | 0 | TCGCTTTGTTGTTGCGCCTC | AGG | \|\|\|\|\|\|\|\|[\|\|\|\|\|\|\|\|\|\|\|\|]PAM | NA | Intergenic | NA | NA |

| | | | | | | | | | | | |
|---|---|---|---|---|---|---|---|---|---|---|---|
| T816 | TTGTTGCGCCTCAGGCGATCTGG | 1000 | | | | | | | | | |
| Oligo fwd | CACCTTGTTGCGCCTCAGGCGATC | | | | | | | | | | |
| Oligo rev | AAACGATCGCCTGAGGCGCAACAA | | | | | | | | | | |
| Chromosome | start | end | strand | MM | target_seq | PAM | alignment | distance | position | gene name | gene id |
| GU305914.1 | 17740 | 17762 | + | 0 | TTGTTGCGCCTCAGGCGATC | TGG | \|\|\|\|\|\|\|\|[\|\|\|\|\|\|\|\|\|\|\|\|]PAM | NA | Intergenic | NA | NA |

| | | | | | | | | | | | |
|---|---|---|---|---|---|---|---|---|---|---|---|
| T817 | TGTTGCGCCTCAGGCGATCTGGG | 1000 | | | | | | | | | |
| Oligo fwd | CACCTGTTGCGCCTCAGGCGATCT | | | | | | | | | | |
| Oligo rev | AAACAGATCGCCTGAGGCGCAACA | | | | | | | | | | |
| Chromosome | start | end | strand | MM | target_seq | PAM | alignment | distance | position | gene name | gene id |
| GU305914.1 | 17741 | 17763 | + | 0 | TGTTGCGCCTCAGGCGATCT | GGG | \|\|\|\|\|\|\|\|[\|\|\|\|\|\|\|\|\|\|\|\|]PAM | NA | Intergenic | NA | NA |

| | | | | | | | | | | | |
|---|---|---|---|---|---|---|---|---|---|---|---|
| T818 | TGCGCCTCAGGCGATCTGGGAGG | 1000 | | | | | | | | | |
| Oligo fwd | CACCTGCGCCTCAGGCGATCTGGG | | | | | | | | | | |
| Oligo rev | AAACCCCAGATCGCCTGAGGCGCA | | | | | | | | | | |
| Chromosome | start | end | strand | MM | target_seq | PAM | alignment | distance | position | gene name | gene id |
| GU305914.1 | 17744 | 17766 | + | 0 | TGCGCCTCAGGCGATCTGGG | AGG | \|\|\|\|\|\|\|\|[\|\|\|\|\|\|\|\|\|\|\|\|]PAM | NA | Intergenic | NA | NA |

| | | | | | | | | | | | |
|---|---|---|---|---|---|---|---|---|---|---|---|
| T819 | CTGGGAGGCTTTTTCTTCCTCGG | 1000 | | | | | | | | | |
| Oligo fwd | CACCCTGGGAGGCTTTTTCTTCCT | | | | | | | | | | |
| Oligo rev | AAACAGGAAGAAAAAGCCTCCCAG | | | | | | | | | | |
| Chromosome | start | end | strand | MM | target_seq | PAM | alignment | distance | position | gene name | gene id |
| GU305914.1 | 17759 | 17781 | + | 0 | CTGGGAGGCTTTTTCTTCCT | CGG | \|\|\|\|\|\|\|\|[\|\|\|\|\|\|\|\|\|\|\|\|]PAM | NA | Intergenic | NA | NA |

| | | | | | | | | | | | |
|---|---|---|---|---|---|---|---|---|---|---|---|
| T820 | GGAGGCTTTTTCTTCCTCGGAGG | 1000 | | | | | | | | | |
| Oligo fwd | CACCGGAGGCTTTTTCTTCCTCGG | | | | | | | | | | |
| Oligo rev | AAACCCGAGGAAGAAAAAGCCTCC | | | | | | | | | | |
| Chromosome | start | end | strand | MM | target_seq | PAM | alignment | distance | position | gene name | gene id |
| GU305914.1 | 17762 | 17784 | + | 0 | GGAGGCTTTTTCTTCCTCGG | AGG | \|\|\|\|\|\|\|\|[\|\|\|\|\|\|\|\|\|\|\|\|]PAM | NA | Intergenic | NA | NA |

| | | | | | | | | | | | |
|---|---|---|---|---|---|---|---|---|---|---|---|
| T821 | TTCCTCGGAGGAGTGTGCAGCGG | 1000 | | | | | | | | | |
| Oligo fwd | CACCTTCCTCGGAGGAGTGTGCAG | | | | | | | | | | |
| Oligo rev | AAACCTGCACACTCCTCCGAGGAA | | | | | | | | | | |
| Chromosome | start | end | strand | MM | target_seq | PAM | alignment | distance | position | gene name | gene id |
| GU305914.1 | 17774 | 17796 | + | 0 | TTCCTCGGAGGAGTGTGCAG | CGG | \|\|\|\|\|\|\|\|[\|\|\|\|\|\|\|\|\|\|\|\|]PAM | NA | Intergenic | NA | NA |

| | | | | | | | | | | | |
|---|---|---|---|---|---|---|---|---|---|---|---|
| T822 | GTGTGCAGCGGCTTAGACAGCGG | 1000 | | | | | | | | | |
| Oligo fwd | CACCGTGTGCAGCGGCTTAGACAG | | | | | | | | | | |
| Oligo rev | AAACCTGTCTAAGCCGCTGCACAC | | | | | | | | | | |
| Chromosome | start | end | strand | MM | target_seq | PAM | alignment | distance | position | gene name | gene id |

GU305914.1 17786 17808 + 0 GTGTGCAGCGGCTTAGACAG CGG ||||||||[||||||||||||]PAM NA Intergenic NA NA

T823 TGTGCAGCGGCTTAGACAGCGGG 1000
Oligo fwd CACCTGTGCAGCGGCTTAGACAGC
Oligo rev AAACGCTGTCTAAGCCGCTGCACA
Chromosome start end strand MM target_seq PAM alignment distance position gene name gene id
GU305914.1 17787 17809 + 0 TGTGCAGCGGCTTAGACAGC GGG ||||||||[||||||||||||]PAM NA Intergenic NA NA

T824 GCAGCGGCTTAGACAGCGGGTGG 1000
Oligo fwd CACCGCAGCGGCTTAGACAGCGGG
Oligo rev AAACCCCGCTGTCTAAGCCGCTGC
Chromosome start end strand MM target_seq PAM alignment distance position gene name gene id
GU305914.1 17790 17812 + 0 GCAGCGGCTTAGACAGCGGG TGG ||||||||[||||||||||||]PAM NA Intergenic NA NA

T825 CAGCGGGTGGCCGTCGCGTTTGG 1000
Oligo fwd CACCCAGCGGGTGGCCGTCGCGTT
Oligo rev AAACAACGCGACGGCCACCCGCTG
Chromosome start end strand MM target_seq PAM alignment distance position gene name gene id
GU305914.1 17803 17825 + 0 CAGCGGGTGGCCGTCGCGTT TGG ||||||||[||||||||||||]PAM NA Intergenic NA NA

T826 GTGATCACCCCGTTTTCCACAGG 1000
Oligo fwd CACCGTGATCACCCCGTTTTCCAC
Oligo rev AAACGTGGAAAACGGGGTGATCAC
Chromosome start end strand MM target_seq PAM alignment distance position gene name gene id
GU305914.1 17837 17859 + 0 GTGATCACCCCGTTTTCCAC AGG ||||||||[||||||||||||]PAM NA Intergenic NA NA

T827 CACCCCGTTTTCCACAGGTCTGG 1000
Oligo fwd CACCCACCCCGTTTTCCACAGGTC
Oligo rev AAACGACCTGTGGAAAACGGGGTG
Chromosome start end strand MM target_seq PAM alignment distance position gene name gene id
GU305914.1 17842 17864 + 0 CACCCCGTTTTCCACAGGTC TGG ||||||||[||||||||||||]PAM NA Intergenic NA NA

T828 ACCCCGTTTTCCACAGGTCTGGG 1000
Oligo fwd CACCACCCCGTTTTCCACAGGTCT
Oligo rev AAACAGACCTGTGGAAAACGGGGT
Chromosome start end strand MM target_seq PAM alignment distance position gene name gene id
GU305914.1 17843 17865 + 0 ACCCCGTTTTCCACAGGTCT GGG ||||||||[||||||||||||]PAM NA Intergenic NA NA

T829 AGTCGAGCAGCCGAGCTTTCAGG 1000
Oligo fwd CACCAGTCGAGCAGCCGAGCTTTC
Oligo rev AAACGAAAGCTCGGCTGCTCGACT
Chromosome start end strand MM target_seq PAM alignment distance position gene name gene id
GU305914.1 17876 17898 + 0 AGTCGAGCAGCCGAGCTTTC AGG ||||||||[||||||||||||]PAM NA Intergenic NA NA

T830 CGAGCTTTCAGGAAAAGTAGCGG 1000
Oligo fwd CACCCGAGCTTTCAGGAAAAGTAG
Oligo rev AAACCTACTTTTCCTGAAAGCTCG
Chromosome start end strand MM target_seq PAM alignment distance position gene name gene id
GU305914.1 17887 17909 + 0 CGAGCTTTCAGGAAAAGTAG CGG ||||||||[||||||||||||]PAM NA Intergenic NA NA

T831 GCTTTCAGGAAAAGTAGCGGCGG 1000
Oligo fwd CACCGCTTTCAGGAAAAGTAGCGG
Oligo rev AAACCCGCTACTTTTCCTGAAAGC
Chromosome start end strand MM target_seq PAM alignment distance position gene name gene id
GU305914.1 17890 17912 + 0 GCTTTCAGGAAAAGTAGCGG CGG ||||||||[||||||||||||]PAM NA Intergenic NA NA

T832 TCAGGAAAAGTAGCGGCGGACGG 1000
Oligo fwd CACCTCAGGAAAAGTAGCGGCGGA
Oligo rev AAACTCCGCCGCTACTTTTCCTGA
Chromosome start end strand MM target_seq PAM alignment distance position gene name gene id
GU305914.1 17894 17916 + 0 TCAGGAAAAGTAGCGGCGGA CGG ||||||||[||||||||||||]PAM NA Intergenic NA NA

T833 GTCGAGAGACTGTCGCCTCCTGG 1000
Oligo fwd CACCGTCGAGAGACTGTCGCCTCC
Oligo rev AAACGGAGGCGACAGTCTCTCGAC
Chromosome start end strand MM target_seq PAM alignment distance position gene name gene id
GU305914.1 17923 17945 + 0 GTCGAGAGACTGTCGCCTCC TGG ||||||||[||||||||||||]PAM NA Intergenic NA NA

T834 GACTGTCGCCTCCTGGATATCGG 1000
Oligo fwd CACCGACTGTCGCCTCCTGGATAT
Oligo rev AAACATATCCAGGAGGCGACAGTC
Chromosome start end strand MM target_seq PAM alignment distance position gene name gene id
GU305914.1 17930 17952 + 0 GACTGTCGCCTCCTGGATAT CGG ||||||||[||||||||||||]PAM NA Intergenic NA NA

T835 ACTGTCGCCTCCTGGATATCGGG 1000
Oligo fwd CACCACTGTCGCCTCCTGGATATC
Oligo rev AAACGATATCCAGGAGGCGACAGT
Chromosome start end strand MM target_seq PAM alignment distance position gene name gene id
GU305914.1 17931 17953 + 0 ACTGTCGCCTCCTGGATATC GGG ||||||||[||||||||||||]PAM NA Intergenic NA NA

T836 CTGTCGCCTCCTGGATATCGGGG 1000
Oligo fwd CACCCTGTCGCCTCCTGGATATCG
Oligo rev AAACCGATATCCAGGAGGCGACAG
Chromosome start end strand MM target_seq PAM alignment distance position gene name gene id
GU305914.1 17932 17954 + 0 CTGTCGCCTCCTGGATATCG GGG ||||||||[||||||||||||]PAM NA Intergenic NA NA

T837 CGCCTCCTGGATATCGGGGCAGG 1000
Oligo fwd CACCCGCCTCCTGGATATCGGGGC
Oligo rev AAACGCCCCGATATCCAGGAGGCG

<table>
<tr><td>Chromosome</td><td>start</td><td>end</td><td>strand</td><td>MM</td><td>target_seq</td><td>PAM</td><td>alignment</td><td>distance</td><td>position</td><td>gene name</td><td>gene id</td></tr>
<tr><td>GU305914.1</td><td>17936</td><td>17958</td><td>+</td><td>0</td><td>CGCCTCCTGGATATCGGGGC</td><td>AGG</td><td>||||||||[||||||||||||]PAM</td><td>NA</td><td>Intergenic</td><td>NA</td><td>NA</td></tr>
<tr><td></td><td></td><td></td><td></td><td></td><td></td><td></td><td></td><td></td><td></td><td></td><td></td></tr>
<tr><td>T838</td><td>GGATATCGGGGCAGGTACGATGG</td><td>1000</td><td></td><td></td><td></td><td></td><td></td><td></td><td></td><td></td><td></td></tr>
<tr><td>Oligo fwd</td><td>CACCGGATATCGGGGCAGGTACGA</td><td></td><td></td><td></td><td></td><td></td><td></td><td></td><td></td><td></td><td></td></tr>
<tr><td>Oligo rev</td><td>AAACTCGTACCTGCCCCGATATCC</td><td></td><td></td><td></td><td></td><td></td><td></td><td></td><td></td><td></td><td></td></tr>
<tr><td>Chromosome</td><td>start</td><td>end</td><td>strand</td><td>MM</td><td>target_seq</td><td>PAM</td><td>alignment</td><td>distance</td><td>position</td><td>gene name</td><td>gene id</td></tr>
<tr><td>GU305914.1</td><td>17944</td><td>17966</td><td>+</td><td>0</td><td>GGATATCGGGGCAGGTACGA</td><td>TGG</td><td>||||||||[||||||||||||]PAM</td><td>NA</td><td>Intergenic</td><td>NA</td><td>NA</td></tr>
<tr><td></td><td></td><td></td><td></td><td></td><td></td><td></td><td></td><td></td><td></td><td></td><td></td></tr>
<tr><td>T839</td><td>GATATCGGGGCAGGTACGATGGG</td><td>1000</td><td></td><td></td><td></td><td></td><td></td><td></td><td></td><td></td><td></td></tr>
<tr><td>Oligo fwd</td><td>CACCGATATCGGGGCAGGTACGAT</td><td></td><td></td><td></td><td></td><td></td><td></td><td></td><td></td><td></td><td></td></tr>
<tr><td>Oligo rev</td><td>AAACATCGTACCTGCCCCGATATC</td><td></td><td></td><td></td><td></td><td></td><td></td><td></td><td></td><td></td><td></td></tr>
<tr><td>Chromosome</td><td>start</td><td>end</td><td>strand</td><td>MM</td><td>target_seq</td><td>PAM</td><td>alignment</td><td>distance</td><td>position</td><td>gene name</td><td>gene id</td></tr>
<tr><td>GU305914.1</td><td>17945</td><td>17967</td><td>+</td><td>0</td><td>GATATCGGGGCAGGTACGAT</td><td>GGG</td><td>||||||||[||||||||||||]PAM</td><td>NA</td><td>Intergenic</td><td>NA</td><td>NA</td></tr>
<tr><td></td><td></td><td></td><td></td><td></td><td></td><td></td><td></td><td></td><td></td><td></td><td></td></tr>
<tr><td>T840</td><td>ATATCGGGGCAGGTACGATGGGG</td><td>1000</td><td></td><td></td><td></td><td></td><td></td><td></td><td></td><td></td><td></td></tr>
<tr><td>Oligo fwd</td><td>CACCATATCGGGGCAGGTACGATG</td><td></td><td></td><td></td><td></td><td></td><td></td><td></td><td></td><td></td><td></td></tr>
<tr><td>Oligo rev</td><td>AAACCATCGTACCTGCCCCGATAT</td><td></td><td></td><td></td><td></td><td></td><td></td><td></td><td></td><td></td><td></td></tr>
<tr><td>Chromosome</td><td>start</td><td>end</td><td>strand</td><td>MM</td><td>target_seq</td><td>PAM</td><td>alignment</td><td>distance</td><td>position</td><td>gene name</td><td>gene id</td></tr>
<tr><td>GU305914.1</td><td>17946</td><td>17968</td><td>+</td><td>0</td><td>ATATCGGGGCAGGTACGATG</td><td>GGG</td><td>||||||||[||||||||||||]PAM</td><td>NA</td><td>Intergenic</td><td>NA</td><td>NA</td></tr>
<tr><td></td><td></td><td></td><td></td><td></td><td></td><td></td><td></td><td></td><td></td><td></td><td></td></tr>
<tr><td>T841</td><td>GGCAGGTACGATGGGGTCGCAGG</td><td>1000</td><td></td><td></td><td></td><td></td><td></td><td></td><td></td><td></td><td></td></tr>
<tr><td>Oligo fwd</td><td>CACCGGCAGGTACGATGGGGTCGC</td><td></td><td></td><td></td><td></td><td></td><td></td><td></td><td></td><td></td><td></td></tr>
<tr><td>Oligo rev</td><td>AAACGCGACCCCATCGTACCTGCC</td><td></td><td></td><td></td><td></td><td></td><td></td><td></td><td></td><td></td><td></td></tr>
<tr><td>Chromosome</td><td>start</td><td>end</td><td>strand</td><td>MM</td><td>target_seq</td><td>PAM</td><td>alignment</td><td>distance</td><td>position</td><td>gene name</td><td>gene id</td></tr>
<tr><td>GU305914.1</td><td>17953</td><td>17975</td><td>+</td><td>0</td><td>GGCAGGTACGATGGGGTCGC</td><td>AGG</td><td>||||||||[||||||||||||]PAM</td><td>NA</td><td>Intergenic</td><td>NA</td><td>NA</td></tr>
<tr><td></td><td></td><td></td><td></td><td></td><td></td><td></td><td></td><td></td><td></td><td></td><td></td></tr>
<tr><td>T842</td><td>GGGGTCGCAGGTCGCCGTGACGG</td><td>1000</td><td></td><td></td><td></td><td></td><td></td><td></td><td></td><td></td><td></td></tr>
<tr><td>Oligo fwd</td><td>CACCGGGGTCGCAGGTCGCCGTGA</td><td></td><td></td><td></td><td></td><td></td><td></td><td></td><td></td><td></td><td></td></tr>
<tr><td>Oligo rev</td><td>AAACTCACGGCGACCTGCGACCCC</td><td></td><td></td><td></td><td></td><td></td><td></td><td></td><td></td><td></td><td></td></tr>
<tr><td>Chromosome</td><td>start</td><td>end</td><td>strand</td><td>MM</td><td>target_seq</td><td>PAM</td><td>alignment</td><td>distance</td><td>position</td><td>gene name</td><td>gene id</td></tr>
<tr><td>GU305914.1</td><td>17965</td><td>17987</td><td>+</td><td>0</td><td>GGGGTCGCAGGTCGCCGTGA</td><td>CGG</td><td>||||||||[||||||||||||]PAM</td><td>NA</td><td>Intergenic</td><td>NA</td><td>NA</td></tr>
<tr><td></td><td></td><td></td><td></td><td></td><td></td><td></td><td></td><td></td><td></td><td></td><td></td></tr>
<tr><td>T843</td><td>CCGTCTGCCGCGCGACCTCTTGG</td><td>1000</td><td></td><td></td><td></td><td></td><td></td><td></td><td></td><td></td><td></td></tr>
<tr><td>Oligo fwd</td><td>CACCCCGTCTGCCGCGCGACCTCT</td><td></td><td></td><td></td><td></td><td></td><td></td><td></td><td></td><td></td><td></td></tr>
<tr><td>Oligo rev</td><td>AAACAGAGGTCGCGCGGCAGACGG</td><td></td><td></td><td></td><td></td><td></td><td></td><td></td><td></td><td></td><td></td></tr>
<tr><td>Chromosome</td><td>start</td><td>end</td><td>strand</td><td>MM</td><td>target_seq</td><td>PAM</td><td>alignment</td><td>distance</td><td>position</td><td>gene name</td><td>gene id</td></tr>
<tr><td>GU305914.1</td><td>17992</td><td>18014</td><td>+</td><td>0</td><td>CCGTCTGCCGCGCGACCTCT</td><td>TGG</td><td>||||||||[||||||||||||]PAM</td><td>NA</td><td>Intergenic</td><td>NA</td><td>NA</td></tr>
<tr><td></td><td></td><td></td><td></td><td></td><td></td><td></td><td></td><td></td><td></td><td></td><td></td></tr>
<tr><td>T844</td><td>CGTCTGCCGCGCGACCTCTTGGG</td><td>1000</td><td></td><td></td><td></td><td></td><td></td><td></td><td></td><td></td><td></td></tr>
<tr><td>Oligo fwd</td><td>CACCCGTCTGCCGCGCGACCTCTT</td><td></td><td></td><td></td><td></td><td></td><td></td><td></td><td></td><td></td><td></td></tr>
<tr><td>Oligo rev</td><td>AAACAAGAGGTCGCGCGGCAGACG</td><td></td><td></td><td></td><td></td><td></td><td></td><td></td><td></td><td></td><td></td></tr>
<tr><td>Chromosome</td><td>start</td><td>end</td><td>strand</td><td>MM</td><td>target_seq</td><td>PAM</td><td>alignment</td><td>distance</td><td>position</td><td>gene name</td><td>gene id</td></tr>
<tr><td>GU305914.1</td><td>17993</td><td>18015</td><td>+</td><td>0</td><td>CGTCTGCCGCGCGACCTCTT</td><td>GGG</td><td>||||||||[||||||||||||]PAM</td><td>NA</td><td>Intergenic</td><td>NA</td><td>NA</td></tr>
<tr><td></td><td></td><td></td><td></td><td></td><td></td><td></td><td></td><td></td><td></td><td></td><td></td></tr>
<tr><td>T845</td><td>CCGCGCGACCTCTTGGGTGTCGG</td><td>1000</td><td></td><td></td><td></td><td></td><td></td><td></td><td></td><td></td><td></td></tr>
<tr><td>Oligo fwd</td><td>CACCCCGCGCGACCTCTTGGGTGT</td><td></td><td></td><td></td><td></td><td></td><td></td><td></td><td></td><td></td><td></td></tr>
<tr><td>Oligo rev</td><td>AAACACACCCAAGAGGTCGCGCGG</td><td></td><td></td><td></td><td></td><td></td><td></td><td></td><td></td><td></td><td></td></tr>
<tr><td>Chromosome</td><td>start</td><td>end</td><td>strand</td><td>MM</td><td>target_seq</td><td>PAM</td><td>alignment</td><td>distance</td><td>position</td><td>gene name</td><td>gene id</td></tr>
<tr><td>GU305914.1</td><td>17999</td><td>18021</td><td>+</td><td>0</td><td>CCGCGCGACCTCTTGGGTGT</td><td>CGG</td><td>||||||||[||||||||||||]PAM</td><td>NA</td><td>Intergenic</td><td>NA</td><td>NA</td></tr>
<tr><td></td><td></td><td></td><td></td><td></td><td></td><td></td><td></td><td></td><td></td><td></td><td></td></tr>
<tr><td>T846</td><td>CGCGCGACCTCTTGGGTGTCGGG</td><td>1000</td><td></td><td></td><td></td><td></td><td></td><td></td><td></td><td></td><td></td></tr>
<tr><td>Oligo fwd</td><td>CACCCGCGCGACCTCTTGGGTGTC</td><td></td><td></td><td></td><td></td><td></td><td></td><td></td><td></td><td></td><td></td></tr>
<tr><td>Oligo rev</td><td>AAACGACACCCAAGAGGTCGCGCG</td><td></td><td></td><td></td><td></td><td></td><td></td><td></td><td></td><td></td><td></td></tr>
<tr><td>Chromosome</td><td>start</td><td>end</td><td>strand</td><td>MM</td><td>target_seq</td><td>PAM</td><td>alignment</td><td>distance</td><td>position</td><td>gene name</td><td>gene id</td></tr>
<tr><td>GU305914.1</td><td>18000</td><td>18022</td><td>+</td><td>0</td><td>CGCGCGACCTCTTGGGTGTC</td><td>GGG</td><td>||||||||[||||||||||||]PAM</td><td>NA</td><td>Intergenic</td><td>NA</td><td>NA</td></tr>
<tr><td>GU305914.1</td><td>202678</td><td>202700</td><td>-</td><td>4</td><td>TGCGCGATGTCTTGGGTGAC</td><td>CGG</td><td>-||||||-[-|||||||||-|]PAM</td><td>NA</td><td>Intergenic</td><td>NA</td><td>NA</td></tr>
<tr><td></td><td></td><td></td><td></td><td></td><td></td><td></td><td></td><td></td><td></td><td></td><td></td></tr>
<tr><td>T847</td><td>GCGCGACCTCTTGGGTGTCGGGG</td><td>1000</td><td></td><td></td><td></td><td></td><td></td><td></td><td></td><td></td><td></td></tr>
<tr><td>Oligo fwd</td><td>CACCGCGCGACCTCTTGGGTGTCG</td><td></td><td></td><td></td><td></td><td></td><td></td><td></td><td></td><td></td><td></td></tr>
<tr><td>Oligo rev</td><td>AAACCGACACCCAAGAGGTCGCGC</td><td></td><td></td><td></td><td></td><td></td><td></td><td></td><td></td><td></td><td></td></tr>
<tr><td>Chromosome</td><td>start</td><td>end</td><td>strand</td><td>MM</td><td>target_seq</td><td>PAM</td><td>alignment</td><td>distance</td><td>position</td><td>gene name</td><td>gene id</td></tr>
<tr><td>GU305914.1</td><td>18001</td><td>18023</td><td>+</td><td>0</td><td>GCGCGACCTCTTGGGTGTCG</td><td>GGG</td><td>||||||||[||||||||||||]PAM</td><td>NA</td><td>Intergenic</td><td>NA</td><td>NA</td></tr>
<tr><td></td><td></td><td></td><td></td><td></td><td></td><td></td><td></td><td></td><td></td><td></td><td></td></tr>
<tr><td>T848</td><td>CCTCTTGGGTGTCGGGGAGCTGG</td><td>1000</td><td></td><td></td><td></td><td></td><td></td><td></td><td></td><td></td><td></td></tr>
<tr><td>Oligo fwd</td><td>CACCCCTCTTGGGTGTCGGGGAGC</td><td></td><td></td><td></td><td></td><td></td><td></td><td></td><td></td><td></td><td></td></tr>
<tr><td>Oligo rev</td><td>AAACGCTCCCCGACACCCAAGAGG</td><td></td><td></td><td></td><td></td><td></td><td></td><td></td><td></td><td></td><td></td></tr>
<tr><td>Chromosome</td><td>start</td><td>end</td><td>strand</td><td>MM</td><td>target_seq</td><td>PAM</td><td>alignment</td><td>distance</td><td>position</td><td>gene name</td><td>gene id</td></tr>
<tr><td>GU305914.1</td><td>18007</td><td>18029</td><td>+</td><td>0</td><td>CCTCTTGGGTGTCGGGGAGC</td><td>TGG</td><td>||||||||[||||||||||||]PAM</td><td>NA</td><td>Intergenic</td><td>NA</td><td>NA</td></tr>
<tr><td></td><td></td><td></td><td></td><td></td><td></td><td></td><td></td><td></td><td></td><td></td><td></td></tr>
<tr><td>T849</td><td>CTTGGGTGTCGGGGAGCTGGAGG</td><td>1000</td><td></td><td></td><td></td><td></td><td></td><td></td><td></td><td></td><td></td></tr>
<tr><td>Oligo fwd</td><td>CACCCTTGGGTGTCGGGGAGCTGG</td><td></td><td></td><td></td><td></td><td></td><td></td><td></td><td></td><td></td><td></td></tr>
<tr><td>Oligo rev</td><td>AAACCCAGCTCCCCGACACCCAAG</td><td></td><td></td><td></td><td></td><td></td><td></td><td></td><td></td><td></td><td></td></tr>
<tr><td>Chromosome</td><td>start</td><td>end</td><td>strand</td><td>MM</td><td>target_seq</td><td>PAM</td><td>alignment</td><td>distance</td><td>position</td><td>gene name</td><td>gene id</td></tr>
<tr><td>GU305914.1</td><td>18010</td><td>18032</td><td>+</td><td>0</td><td>CTTGGGTGTCGGGGAGCTGG</td><td>AGG</td><td>||||||||[||||||||||||]PAM</td><td>NA</td><td>Intergenic</td><td>NA</td><td>NA</td></tr>
<tr><td></td><td></td><td></td><td></td><td></td><td></td><td></td><td></td><td></td><td></td><td></td><td></td></tr>
<tr><td>T850</td><td>TTGGGTGTCGGGGAGCTGGAGGG</td><td>1000</td><td></td><td></td><td></td><td></td><td></td><td></td><td></td><td></td><td></td></tr>
<tr><td>Oligo fwd</td><td>CACCTTGGGTGTCGGGGAGCTGGA</td><td></td><td></td><td></td><td></td><td></td><td></td><td></td><td></td><td></td><td></td></tr>
<tr><td>Oligo rev</td><td>AAACTCCAGCTCCCCGACACCCAA</td><td></td><td></td><td></td><td></td><td></td><td></td><td></td><td></td><td></td><td></td></tr>
<tr><td>Chromosome</td><td>start</td><td>end</td><td>strand</td><td>MM</td><td>target_seq</td><td>PAM</td><td>alignment</td><td>distance</td><td>position</td><td>gene name</td><td>gene id</td></tr>
<tr><td>GU305914.1</td><td>18011</td><td>18033</td><td>+</td><td>0</td><td>TTGGGTGTCGGGGAGCTGGA</td><td>GGG</td><td>||||||||[||||||||||||]PAM</td><td>NA</td><td>Intergenic</td><td>NA</td><td>NA</td></tr>
<tr><td></td><td></td><td></td><td></td><td></td><td></td><td></td><td></td><td></td><td></td><td></td><td></td></tr>
<tr><td>T851</td><td>GGAGGGTGTCGTAGTCGTAGAGG</td><td>1000</td><td></td><td></td><td></td><td></td><td></td><td></td><td></td><td></td><td></td></tr>
<tr><td>Oligo fwd</td><td>CACCGGAGGGTGTCGTAGTCGTAG</td><td></td><td></td><td></td><td></td><td></td><td></td><td></td><td></td><td></td><td></td></tr>
<tr><td>Oligo rev</td><td>AAACCTACGACTACGACACCCTCC</td><td></td><td></td><td></td><td></td><td></td><td></td><td></td><td></td><td></td><td></td></tr>
<tr><td>Chromosome</td><td>start</td><td>end</td><td>strand</td><td>MM</td><td>target_seq</td><td>PAM</td><td>alignment</td><td>distance</td><td>position</td><td>gene name</td><td>gene id</td></tr>
<tr><td>GU305914.1</td><td>18028</td><td>18050</td><td>+</td><td>0</td><td>GGAGGGTGTCGTAGTCGTAG</td><td>AGG</td><td>||||||||[||||||||||||]PAM</td><td>NA</td><td>Intergenic</td><td>NA</td><td>NA</td></tr>
<tr><td></td><td></td><td></td><td></td><td></td><td></td><td></td><td></td><td></td><td></td><td></td><td></td></tr>
<tr><td>T852</td><td>GTGTCGTAGTCGTAGAGGTCTGG</td><td>1000</td><td></td><td></td><td></td><td></td><td></td><td></td><td></td><td></td><td></td></tr>
</table>

<table>
<tr><td>Oligo fwd</td><td>CACCGTGTCGTAGTCGTAGAGGTC</td><td></td><td></td><td></td><td></td><td></td><td></td><td></td><td></td><td></td><td></td><td></td></tr>
<tr><td>Oligo rev</td><td>AAACGACCTCTACGACTACGACAC</td><td></td><td></td><td></td><td></td><td></td><td></td><td></td><td></td><td></td><td></td><td></td></tr>
<tr><td>Chromosome</td><td>start</td><td></td><td>end</td><td>strand</td><td>MM</td><td>target_seq</td><td>PAM</td><td>alignment</td><td>distance</td><td>position</td><td>gene name</td><td>gene id</td></tr>
<tr><td>GU305914.1</td><td></td><td>18033</td><td>18055</td><td>+</td><td>0</td><td>GTGTCGTAGTCGTAGAGGTC</td><td>TGG</td><td>||||||||[||||||||||||]PAM</td><td>NA</td><td>Intergenic</td><td>NA</td><td>NA</td></tr>
<tr><td>GU305914.1</td><td></td><td>60152</td><td>60174</td><td>+</td><td>4</td><td>TCGTCGTCGTCGAAGAGGTC</td><td>GGG</td><td>--|||||-[||||-|||||||]PAM</td><td>NA</td><td>Intergenic</td><td>NA</td><td>NA</td></tr>
<tr><td></td><td></td><td></td><td></td><td></td><td></td><td></td><td></td><td></td><td></td><td></td><td></td><td></td></tr>
<tr><td>T853</td><td>GTCGTAGAGGTCTGGTTCGTCGG</td><td></td><td>1000</td><td></td><td></td><td></td><td></td><td></td><td></td><td></td><td></td><td></td></tr>
<tr><td>Oligo fwd</td><td>CACCGTCGTAGAGGTCTGGTTCGT</td><td></td><td></td><td></td><td></td><td></td><td></td><td></td><td></td><td></td><td></td><td></td></tr>
<tr><td>Oligo rev</td><td>AAACACGAACCAGACCTCTACGAC</td><td></td><td></td><td></td><td></td><td></td><td></td><td></td><td></td><td></td><td></td><td></td></tr>
<tr><td>Chromosome</td><td>start</td><td></td><td>end</td><td>strand</td><td>MM</td><td>target_seq</td><td>PAM</td><td>alignment</td><td>distance</td><td>position</td><td>gene name</td><td>gene id</td></tr>
<tr><td>GU305914.1</td><td></td><td>18041</td><td>18063</td><td>+</td><td>0</td><td>GTCGTAGAGGTCTGGTTCGT</td><td>CGG</td><td>||||||||[||||||||||||]PAM</td><td>NA</td><td>Intergenic</td><td>NA</td><td>NA</td></tr>
<tr><td></td><td></td><td></td><td></td><td></td><td></td><td></td><td></td><td></td><td></td><td></td><td></td><td></td></tr>
<tr><td>T854</td><td>TCGTAGAGGTCTGGTTCGTCGGG</td><td></td><td>1000</td><td></td><td></td><td></td><td></td><td></td><td></td><td></td><td></td><td></td></tr>
<tr><td>Oligo fwd</td><td>CACCTCGTAGAGGTCTGGTTCGTC</td><td></td><td></td><td></td><td></td><td></td><td></td><td></td><td></td><td></td><td></td><td></td></tr>
<tr><td>Oligo rev</td><td>AAACGACGAACCAGACCTCTACGA</td><td></td><td></td><td></td><td></td><td></td><td></td><td></td><td></td><td></td><td></td><td></td></tr>
<tr><td>Chromosome</td><td>start</td><td></td><td>end</td><td>strand</td><td>MM</td><td>target_seq</td><td>PAM</td><td>alignment</td><td>distance</td><td>position</td><td>gene name</td><td>gene id</td></tr>
<tr><td>GU305914.1</td><td></td><td>18042</td><td>18064</td><td>+</td><td>0</td><td>TCGTAGAGGTCTGGTTCGTC</td><td>GGG</td><td>||||||||[||||||||||||]PAM</td><td>NA</td><td>Intergenic</td><td>NA</td><td>NA</td></tr>
<tr><td></td><td></td><td></td><td></td><td></td><td></td><td></td><td></td><td></td><td></td><td></td><td></td><td></td></tr>
<tr><td>T855</td><td>CTGGTTCGTCGGGCTGCGCGAGG</td><td></td><td>1000</td><td></td><td></td><td></td><td></td><td></td><td></td><td></td><td></td><td></td></tr>
<tr><td>Oligo fwd</td><td>CACCCTGGTTCGTCGGGCTGCGCG</td><td></td><td></td><td></td><td></td><td></td><td></td><td></td><td></td><td></td><td></td><td></td></tr>
<tr><td>Oligo rev</td><td>AAACCGCGCAGCCCGACGAACCAG</td><td></td><td></td><td></td><td></td><td></td><td></td><td></td><td></td><td></td><td></td><td></td></tr>
<tr><td>Chromosome</td><td>start</td><td></td><td>end</td><td>strand</td><td>MM</td><td>target_seq</td><td>PAM</td><td>alignment</td><td>distance</td><td>position</td><td>gene name</td><td>gene id</td></tr>
<tr><td>GU305914.1</td><td></td><td>18052</td><td>18074</td><td>+</td><td>0</td><td>CTGGTTCGTCGGGCTGCGCG</td><td>AGG</td><td>||||||||[||||||||||||]PAM</td><td>NA</td><td>Intergenic</td><td>NA</td><td>NA</td></tr>
<tr><td></td><td></td><td></td><td></td><td></td><td></td><td></td><td></td><td></td><td></td><td></td><td></td><td></td></tr>
<tr><td>T856</td><td>GTCGAGTCTGATCTCGACGATGG</td><td></td><td>1000</td><td></td><td></td><td></td><td></td><td></td><td></td><td></td><td></td><td></td></tr>
<tr><td>Oligo fwd</td><td>CACCGTCGAGTCTGATCTCGACGA</td><td></td><td></td><td></td><td></td><td></td><td></td><td></td><td></td><td></td><td></td><td></td></tr>
<tr><td>Oligo rev</td><td>AAACTCGTCGAGATCAGACTCGAC</td><td></td><td></td><td></td><td></td><td></td><td></td><td></td><td></td><td></td><td></td><td></td></tr>
<tr><td>Chromosome</td><td>start</td><td></td><td>end</td><td>strand</td><td>MM</td><td>target_seq</td><td>PAM</td><td>alignment</td><td>distance</td><td>position</td><td>gene name</td><td>gene id</td></tr>
<tr><td>GU305914.1</td><td></td><td>18074</td><td>18096</td><td>+</td><td>0</td><td>GTCGAGTCTGATCTCGACGA</td><td>TGG</td><td>||||||||[||||||||||||]PAM</td><td>NA</td><td>Intergenic</td><td>NA</td><td>NA</td></tr>
<tr><td></td><td></td><td></td><td></td><td></td><td></td><td></td><td></td><td></td><td></td><td></td><td></td><td></td></tr>
<tr><td>T857</td><td>GAGTCTGATCTCGACGATGGCGG</td><td></td><td>1000</td><td></td><td></td><td></td><td></td><td></td><td></td><td></td><td></td><td></td></tr>
<tr><td>Oligo fwd</td><td>CACCGAGTCTGATCTCGACGATGG</td><td></td><td></td><td></td><td></td><td></td><td></td><td></td><td></td><td></td><td></td><td></td></tr>
<tr><td>Oligo rev</td><td>AAACCCATCGTCGAGATCAGACTC</td><td></td><td></td><td></td><td></td><td></td><td></td><td></td><td></td><td></td><td></td><td></td></tr>
<tr><td>Chromosome</td><td>start</td><td></td><td>end</td><td>strand</td><td>MM</td><td>target_seq</td><td>PAM</td><td>alignment</td><td>distance</td><td>position</td><td>gene name</td><td>gene id</td></tr>
<tr><td>GU305914.1</td><td></td><td>18077</td><td>18099</td><td>+</td><td>0</td><td>GAGTCTGATCTCGACGATGG</td><td>CGG</td><td>||||||||[||||||||||||]PAM</td><td>NA</td><td>Intergenic</td><td>NA</td><td>NA</td></tr>
<tr><td></td><td></td><td></td><td></td><td></td><td></td><td></td><td></td><td></td><td></td><td></td><td></td><td></td></tr>
<tr><td>T858</td><td>CAGCGACCGCCGCAGCCTCGTGG</td><td></td><td>1000</td><td></td><td></td><td></td><td></td><td></td><td></td><td></td><td></td><td></td></tr>
<tr><td>Oligo fwd</td><td>CACCCAGCGACCGCCGCAGCCTCG</td><td></td><td></td><td></td><td></td><td></td><td></td><td></td><td></td><td></td><td></td><td></td></tr>
<tr><td>Oligo rev</td><td>AAACCGAGGCTGCGGCGGTCGCTG</td><td></td><td></td><td></td><td></td><td></td><td></td><td></td><td></td><td></td><td></td><td></td></tr>
<tr><td>Chromosome</td><td>start</td><td></td><td>end</td><td>strand</td><td>MM</td><td>target_seq</td><td>PAM</td><td>alignment</td><td>distance</td><td>position</td><td>gene name</td><td>gene id</td></tr>
<tr><td>GU305914.1</td><td></td><td>18100</td><td>18122</td><td>+</td><td>0</td><td>CAGCGACCGCCGCAGCCTCG</td><td>TGG</td><td>||||||||[||||||||||||]PAM</td><td>NA</td><td>Intergenic</td><td>NA</td><td>NA</td></tr>
<tr><td></td><td></td><td></td><td></td><td></td><td></td><td></td><td></td><td></td><td></td><td></td><td></td><td></td></tr>
<tr><td>T859</td><td>GACCGCCGCAGCCTCGTGGCCGG</td><td></td><td>1000</td><td></td><td></td><td></td><td></td><td></td><td></td><td></td><td></td><td></td></tr>
<tr><td>Oligo fwd</td><td>CACCGACCGCCGCAGCCTCGTGGC</td><td></td><td></td><td></td><td></td><td></td><td></td><td></td><td></td><td></td><td></td><td></td></tr>
<tr><td>Oligo rev</td><td>AAACGCCACGAGGCTGCGGCGGTC</td><td></td><td></td><td></td><td></td><td></td><td></td><td></td><td></td><td></td><td></td><td></td></tr>
<tr><td>Chromosome</td><td>start</td><td></td><td>end</td><td>strand</td><td>MM</td><td>target_seq</td><td>PAM</td><td>alignment</td><td>distance</td><td>position</td><td>gene name</td><td>gene id</td></tr>
<tr><td>GU305914.1</td><td></td><td>18104</td><td>18126</td><td>+</td><td>0</td><td>GACCGCCGCAGCCTCGTGGC</td><td>CGG</td><td>||||||||[||||||||||||]PAM</td><td>NA</td><td>Intergenic</td><td>NA</td><td>NA</td></tr>
<tr><td></td><td></td><td></td><td></td><td></td><td></td><td></td><td></td><td></td><td></td><td></td><td></td><td></td></tr>
<tr><td>T860</td><td>CGTGGCCGGCATCTCTGTCGCGG</td><td></td><td>1000</td><td></td><td></td><td></td><td></td><td></td><td></td><td></td><td></td><td></td></tr>
<tr><td>Oligo fwd</td><td>CACCCGTGGCCGGCATCTCTGTCG</td><td></td><td></td><td></td><td></td><td></td><td></td><td></td><td></td><td></td><td></td><td></td></tr>
<tr><td>Oligo rev</td><td>AAACCGACAGAGATGCCGGCCACG</td><td></td><td></td><td></td><td></td><td></td><td></td><td></td><td></td><td></td><td></td><td></td></tr>
<tr><td>Chromosome</td><td>start</td><td></td><td>end</td><td>strand</td><td>MM</td><td>target_seq</td><td>PAM</td><td>alignment</td><td>distance</td><td>position</td><td>gene name</td><td>gene id</td></tr>
<tr><td>GU305914.1</td><td></td><td>18118</td><td>18140</td><td>+</td><td>0</td><td>CGTGGCCGGCATCTCTGTCG</td><td>CGG</td><td>||||||||[||||||||||||]PAM</td><td>NA</td><td>Intergenic</td><td>NA</td><td>NA</td></tr>
<tr><td></td><td></td><td></td><td></td><td></td><td></td><td></td><td></td><td></td><td></td><td></td><td></td><td></td></tr>
<tr><td>T861</td><td>CTCTGTCGCGGAAGAACCTCCGG</td><td></td><td>1000</td><td></td><td></td><td></td><td></td><td></td><td></td><td></td><td></td><td></td></tr>
<tr><td>Oligo fwd</td><td>CACCCTCTGTCGCGGAAGAACCTC</td><td></td><td></td><td></td><td></td><td></td><td></td><td></td><td></td><td></td><td></td><td></td></tr>
<tr><td>Oligo rev</td><td>AAACGAGGTTCTTCCGCGACAGAG</td><td></td><td></td><td></td><td></td><td></td><td></td><td></td><td></td><td></td><td></td><td></td></tr>
<tr><td>Chromosome</td><td>start</td><td></td><td>end</td><td>strand</td><td>MM</td><td>target_seq</td><td>PAM</td><td>alignment</td><td>distance</td><td>position</td><td>gene name</td><td>gene id</td></tr>
<tr><td>GU305914.1</td><td></td><td>18130</td><td>18152</td><td>+</td><td>0</td><td>CTCTGTCGCGGAAGAACCTC</td><td>CGG</td><td>||||||||[||||||||||||]PAM</td><td>NA</td><td>Intergenic</td><td>NA</td><td>NA</td></tr>
<tr><td></td><td></td><td></td><td></td><td></td><td></td><td></td><td></td><td></td><td></td><td></td><td></td><td></td></tr>
<tr><td>T862</td><td>TCTGTCGCGGAAGAACCTCCGGG</td><td></td><td>1000</td><td></td><td></td><td></td><td></td><td></td><td></td><td></td><td></td><td></td></tr>
<tr><td>Oligo fwd</td><td>CACCTCTGTCGCGGAAGAACCTCC</td><td></td><td></td><td></td><td></td><td></td><td></td><td></td><td></td><td></td><td></td><td></td></tr>
<tr><td>Oligo rev</td><td>AAACGGAGGTTCTTCCGCGACAGA</td><td></td><td></td><td></td><td></td><td></td><td></td><td></td><td></td><td></td><td></td><td></td></tr>
<tr><td>Chromosome</td><td>start</td><td></td><td>end</td><td>strand</td><td>MM</td><td>target_seq</td><td>PAM</td><td>alignment</td><td>distance</td><td>position</td><td>gene name</td><td>gene id</td></tr>
<tr><td>GU305914.1</td><td></td><td>18131</td><td>18153</td><td>+</td><td>0</td><td>TCTGTCGCGGAAGAACCTCC</td><td>GGG</td><td>||||||||[||||||||||||]PAM</td><td>NA</td><td>Intergenic</td><td>NA</td><td>NA</td></tr>
<tr><td></td><td></td><td></td><td></td><td></td><td></td><td></td><td></td><td></td><td></td><td></td><td></td><td></td></tr>
<tr><td>T863</td><td>AAGAACCTCCGGGCCCGCCGCGG</td><td></td><td>1000</td><td></td><td></td><td></td><td></td><td></td><td></td><td></td><td></td><td></td></tr>
<tr><td>Oligo fwd</td><td>CACCAAGAACCTCCGGGCCCGCCG</td><td></td><td></td><td></td><td></td><td></td><td></td><td></td><td></td><td></td><td></td><td></td></tr>
<tr><td>Oligo rev</td><td>AAACCGGCGGGCCCGGAGGTTCTT</td><td></td><td></td><td></td><td></td><td></td><td></td><td></td><td></td><td></td><td></td><td></td></tr>
<tr><td>Chromosome</td><td>start</td><td></td><td>end</td><td>strand</td><td>MM</td><td>target_seq</td><td>PAM</td><td>alignment</td><td>distance</td><td>position</td><td>gene name</td><td>gene id</td></tr>
<tr><td>GU305914.1</td><td></td><td>18141</td><td>18163</td><td>+</td><td>0</td><td>AAGAACCTCCGGGCCCGCCG</td><td>CGG</td><td>||||||||[||||||||||||]PAM</td><td>NA</td><td>Intergenic</td><td>NA</td><td>NA</td></tr>
<tr><td></td><td></td><td></td><td></td><td></td><td></td><td></td><td></td><td></td><td></td><td></td><td></td><td></td></tr>
<tr><td>T864</td><td>AGAACCTCCGGGCCCGCCGCGGG</td><td></td><td>1000</td><td></td><td></td><td></td><td></td><td></td><td></td><td></td><td></td><td></td></tr>
<tr><td>Oligo fwd</td><td>CACCAGAACCTCCGGGCCCGCCGC</td><td></td><td></td><td></td><td></td><td></td><td></td><td></td><td></td><td></td><td></td><td></td></tr>
<tr><td>Oligo rev</td><td>AAACGCGGCGGGCCCGGAGGTTCT</td><td></td><td></td><td></td><td></td><td></td><td></td><td></td><td></td><td></td><td></td><td></td></tr>
<tr><td>Chromosome</td><td>start</td><td></td><td>end</td><td>strand</td><td>MM</td><td>target_seq</td><td>PAM</td><td>alignment</td><td>distance</td><td>position</td><td>gene name</td><td>gene id</td></tr>
<tr><td>GU305914.1</td><td></td><td>18142</td><td>18164</td><td>+</td><td>0</td><td>AGAACCTCCGGGCCCGCCGC</td><td>GGG</td><td>||||||||[||||||||||||]PAM</td><td>NA</td><td>Intergenic</td><td>NA</td><td>NA</td></tr>
<tr><td></td><td></td><td></td><td></td><td></td><td></td><td></td><td></td><td></td><td></td><td></td><td></td><td></td></tr>
<tr><td>T865</td><td>CCTCCGGGCCCGCCGCGGGACGG</td><td></td><td>1000</td><td></td><td></td><td></td><td></td><td></td><td></td><td></td><td></td><td></td></tr>
<tr><td>Oligo fwd</td><td>CACCCCTCCGGGCCCGCCGCGGGA</td><td></td><td></td><td></td><td></td><td></td><td></td><td></td><td></td><td></td><td></td><td></td></tr>
<tr><td>Oligo rev</td><td>AAACTCCCGCGGCGGGCCCGGAGG</td><td></td><td></td><td></td><td></td><td></td><td></td><td></td><td></td><td></td><td></td><td></td></tr>
<tr><td>Chromosome</td><td>start</td><td></td><td>end</td><td>strand</td><td>MM</td><td>target_seq</td><td>PAM</td><td>alignment</td><td>distance</td><td>position</td><td>gene name</td><td>gene id</td></tr>
<tr><td>GU305914.1</td><td></td><td>18146</td><td>18168</td><td>+</td><td>0</td><td>CCTCCGGGCCCGCCGCGGGA</td><td>CGG</td><td>||||||||[||||||||||||]PAM</td><td>NA</td><td>Intergenic</td><td>NA</td><td>NA</td></tr>
<tr><td></td><td></td><td></td><td></td><td></td><td></td><td></td><td></td><td></td><td></td><td></td><td></td><td></td></tr>
<tr><td>T866</td><td>CTCCGGGCCCGCCGCGGGACGGG</td><td></td><td>1000</td><td></td><td></td><td></td><td></td><td></td><td></td><td></td><td></td><td></td></tr>
<tr><td>Oligo fwd</td><td>CACCCTCCGGGCCCGCCGCGGGAC</td><td></td><td></td><td></td><td></td><td></td><td></td><td></td><td></td><td></td><td></td><td></td></tr>
<tr><td>Oligo rev</td><td>AAACGTCCCGCGGCGGGCCCGGAG</td><td></td><td></td><td></td><td></td><td></td><td></td><td></td><td></td><td></td><td></td><td></td></tr>
<tr><td>Chromosome</td><td>start</td><td></td><td>end</td><td>strand</td><td>MM</td><td>target_seq</td><td>PAM</td><td>alignment</td><td>distance</td><td>position</td><td>gene name</td><td>gene id</td></tr>
<tr><td>GU305914.1</td><td></td><td>18147</td><td>18169</td><td>+</td><td>0</td><td>CTCCGGGCCCGCCGCGGGAC</td><td>GGG</td><td>||||||||[||||||||||||]PAM</td><td>NA</td><td>Intergenic</td><td>NA</td><td>NA</td></tr>
</table>

T867 CCCGCCGCGGGACGGGATGTTGG 1000
Oligo fwd CACCCCCGCCGCGGGACGGGATGT
Oligo rev AAACACATCCCGTCCCGCGGCGGG
Chromosome start end strand MM target_seq PAM alignment distance position gene name gene id
GU305914.1 18154 18176 + 0 CCCGCCGCGGGACGGGATGT TGG ||||||||[||||||||||||]PAM NA Intergenic NA NA

T868 GCCGCGGGACGGGATGTTGGCGG 1000
Oligo fwd CACCGCCGCGGGACGGGATGTTGG
Oligo rev AAACCCAACATCCCGTCCCGCGGC
Chromosome start end strand MM target_seq PAM alignment distance position gene name gene id
GU305914.1 18157 18179 + 0 GCCGCGGGACGGGATGTTGG CGG ||||||||[||||||||||||]PAM NA Intergenic NA NA

T869 GATGTTGGCGGTATCGTCCGCGG 1000
Oligo fwd CACCGATGTTGGCGGTATCGTCCG
Oligo rev AAACCGGACGATACCGCCAACATC
Chromosome start end strand MM target_seq PAM alignment distance position gene name gene id
GU305914.1 18169 18191 + 0 GATGTTGGCGGTATCGTCCG CGG ||||||||[||||||||||||]PAM NA Intergenic NA NA

T870 ATGTTGGCGGTATCGTCCGCGGG 1000
Oligo fwd CACCATGTTGGCGGTATCGTCCGC
Oligo rev AAACGCGGACGATACCGCCAACAT
Chromosome start end strand MM target_seq PAM alignment distance position gene name gene id
GU305914.1 18170 18192 + 0 ATGTTGGCGGTATCGTCCGC GGG ||||||||[||||||||||||]PAM NA Intergenic NA NA

T871 TGTTGGCGGTATCGTCCGCGGGG 1000
Oligo fwd CACCTGTTGGCGGTATCGTCCGCG
Oligo rev AAACCGCGGACGATACCGCCAACA
Chromosome start end strand MM target_seq PAM alignment distance position gene name gene id
GU305914.1 18171 18193 + 0 TGTTGGCGGTATCGTCCGCG GGG ||||||||[||||||||||||]PAM NA Intergenic NA NA

T872 TATCGTCCGCGGGGCGTTCTCGG 1000
Oligo fwd CACCTATCGTCCGCGGGGCGTTCT
Oligo rev AAACAGAACGCCCCGCGGACGATA
Chromosome start end strand MM target_seq PAM alignment distance position gene name gene id
GU305914.1 18180 18202 + 0 TATCGTCCGCGGGGCGTTCT CGG ||||||||[||||||||||||]PAM NA Intergenic NA NA

T873 ATCGTCCGCGGGGCGTTCTCGGG 1000
Oligo fwd CACCATCGTCCGCGGGGCGTTCTC
Oligo rev AAACGAGAACGCCCCGCGGACGAT
Chromosome start end strand MM target_seq PAM alignment distance position gene name gene id
GU305914.1 18181 18203 + 0 ATCGTCCGCGGGGCGTTCTC GGG ||||||||[||||||||||||]PAM NA Intergenic NA NA

T874 TTCTCGGGACGAATCTGAGCTGG 1000
Oligo fwd CACCTTCTCGGGACGAATCTGAGC
Oligo rev AAACGCTCAGATTCGTCCCGAGAA
Chromosome start end strand MM target_seq PAM alignment distance position gene name gene id
GU305914.1 18196 18218 + 0 TTCTCGGGACGAATCTGAGC TGG ||||||||[||||||||||||]PAM NA Intergenic NA NA

T875 CGGGACGAATCTGAGCTGGACGG 1000
Oligo fwd CACCCGGGACGAATCTGAGCTGGA
Oligo rev AAACTCCAGCTCAGATTCGTCCCG
Chromosome start end strand MM target_seq PAM alignment distance position gene name gene id
GU305914.1 18200 18222 + 0 CGGGACGAATCTGAGCTGGA CGG ||||||||[||||||||||||]PAM NA Intergenic NA NA

T876 GCTCGCGACTGTCGACAAGTCGG 1000
Oligo fwd CACCGCTCGCGACTGTCGACAAGT
Oligo rev AAACACTTGTCGACAGTCGCGAGC
Chromosome start end strand MM target_seq PAM alignment distance position gene name gene id
GU305914.1 18222 18244 + 0 GCTCGCGACTGTCGACAAGT CGG ||||||||[||||||||||||]PAM NA Intergenic NA NA

T877 CTCTTCCTCCCGCGCCTCTCCGG 1000
Oligo fwd CACCCTCTTCCTCCCGCGCCTCTC
Oligo rev AAACGAGAGGCGCGGGAGGAAGAG
Chromosome start end strand MM target_seq PAM alignment distance position gene name gene id
GU305914.1 18245 18267 + 0 CTCTTCCTCCCGCGCCTCTC CGG ||||||||[||||||||||||]PAM NA Intergenic NA NA

T878 CTCCGGAGTCGTCTCCCGACTGG 1000
Oligo fwd CACCCTCCGGAGTCGTCTCCCGAC
Oligo rev AAACGTCGGGAGACGACTCCGGAG
Chromosome start end strand MM target_seq PAM alignment distance position gene name gene id
GU305914.1 18262 18284 + 0 CTCCGGAGTCGTCTCCCGAC TGG ||||||||[||||||||||||]PAM NA Intergenic NA NA
GU305914.1 77064 77086 - 4 CTCCAGCATCGTCTCCCGCC CGG ||||-|--[||||||||||-|]PAM NA Intergenic NA NA

T879 CGGAGTCGTCTCCCGACTGGAGG 1000
Oligo fwd CACCCGGAGTCGTCTCCCGACTGG
Oligo rev AAACCCAGTCGGGAGACGACTCCG
Chromosome start end strand MM target_seq PAM alignment distance position gene name gene id
GU305914.1 18265 18287 + 0 CGGAGTCGTCTCCCGACTGG AGG ||||||||[||||||||||||]PAM NA Intergenic NA NA

T880 TCCCGACTGGAGGCGTCGCTCGG 1000
Oligo fwd CACCTCCCGACTGGAGGCGTCGCT
Oligo rev AAACAGCGACGCCTCCAGTCGGGA
Chromosome start end strand MM target_seq PAM alignment distance position gene name gene id
GU305914.1 18275 18297 + 0 TCCCGACTGGAGGCGTCGCT CGG ||||||||[||||||||||||]PAM NA Intergenic NA NA

T881 TCGCTCGGAGTAGCGCCGATAGG 1000
Oligo fwd CACCTCGCTCGGAGTAGCGCCGAT
Oligo rev AAACATCGGCGCTACTCCGAGCGA

<table>
<tr><td>Chromosome</td><td>start</td><td></td><td>end</td><td>strand</td><td>MM</td><td>target_seq</td><td>PAM</td><td>alignment</td><td>distance</td><td>position</td><td>gene name</td><td>gene id</td></tr>
<tr><td>GU305914.1</td><td></td><td>18290</td><td>18312</td><td>+</td><td>0</td><td>TCGCTCGGAGTAGCGCCGAT</td><td>AGG</td><td>||||||||[||||||||||||]PAM</td><td>NA</td><td>Intergenic</td><td>NA</td><td>NA</td></tr>
<tr><td>T882</td><td>GGACGAAAGCCGACACGAGCCGG</td><td></td><td>1000</td><td></td><td></td><td></td><td></td><td></td><td></td><td></td><td></td><td></td></tr>
<tr><td>Oligo fwd</td><td>CACCGGACGAAAGCCGACACGAGC</td><td></td><td></td><td></td><td></td><td></td><td></td><td></td><td></td><td></td><td></td><td></td></tr>
<tr><td>Oligo rev</td><td>AAACGCTCGTGTCGGCTTTCGTCC</td><td></td><td></td><td></td><td></td><td></td><td></td><td></td><td></td><td></td><td></td><td></td></tr>
<tr><td>Chromosome</td><td>start</td><td></td><td>end</td><td>strand</td><td>MM</td><td>target_seq</td><td>PAM</td><td>alignment</td><td>distance</td><td>position</td><td>gene name</td><td>gene id</td></tr>
<tr><td>GU305914.1</td><td></td><td>18311</td><td>18333</td><td>+</td><td>0</td><td>GGACGAAAGCCGACACGAGC</td><td>CGG</td><td>||||||||[||||||||||||]PAM</td><td>NA</td><td>Intergenic</td><td>NA</td><td>NA</td></tr>
<tr><td>T883</td><td>GATCTCTTCTGATCGTTCGACGG</td><td></td><td>1000</td><td></td><td></td><td></td><td></td><td></td><td></td><td></td><td></td><td></td></tr>
<tr><td>Oligo fwd</td><td>CACCGATCTCTTCTGATCGTTCGA</td><td></td><td></td><td></td><td></td><td></td><td></td><td></td><td></td><td></td><td></td><td></td></tr>
<tr><td>Oligo rev</td><td>AAACTCGAACGATCAGAAGAGATC</td><td></td><td></td><td></td><td></td><td></td><td></td><td></td><td></td><td></td><td></td><td></td></tr>
<tr><td>Chromosome</td><td>start</td><td></td><td>end</td><td>strand</td><td>MM</td><td>target_seq</td><td>PAM</td><td>alignment</td><td>distance</td><td>position</td><td>gene name</td><td>gene id</td></tr>
<tr><td>GU305914.1</td><td></td><td>18353</td><td>18375</td><td>+</td><td>0</td><td>GATCTCTTCTGATCGTTCGA</td><td>CGG</td><td>||||||||[||||||||||||]PAM</td><td>NA</td><td>Intergenic</td><td>NA</td><td>NA</td></tr>
<tr><td>T884</td><td>CCATTTCGACTCCGAGTCCGTGG</td><td></td><td>1000</td><td></td><td></td><td></td><td></td><td></td><td></td><td></td><td></td><td></td></tr>
<tr><td>Oligo fwd</td><td>CACCCCATTTCGACTCCGAGTCCG</td><td></td><td></td><td></td><td></td><td></td><td></td><td></td><td></td><td></td><td></td><td></td></tr>
<tr><td>Oligo rev</td><td>AAACCGGACTCGGAGTCGAAATGG</td><td></td><td></td><td></td><td></td><td></td><td></td><td></td><td></td><td></td><td></td><td></td></tr>
<tr><td>Chromosome</td><td>start</td><td></td><td>end</td><td>strand</td><td>MM</td><td>target_seq</td><td>PAM</td><td>alignment</td><td>distance</td><td>position</td><td>gene name</td><td>gene id</td></tr>
<tr><td>GU305914.1</td><td></td><td>18409</td><td>18431</td><td>+</td><td>0</td><td>CCATTTCGACTCCGAGTCCG</td><td>TGG</td><td>||||||||[||||||||||||]PAM</td><td>NA</td><td>Intergenic</td><td>NA</td><td>NA</td></tr>
<tr><td>T885</td><td>TTTCGACTCCGAGTCCGTGGTGG</td><td></td><td>1000</td><td></td><td></td><td></td><td></td><td></td><td></td><td></td><td></td><td></td></tr>
<tr><td>Oligo fwd</td><td>CACCTTTCGACTCCGAGTCCGTGG</td><td></td><td></td><td></td><td></td><td></td><td></td><td></td><td></td><td></td><td></td><td></td></tr>
<tr><td>Oligo rev</td><td>AAACCCACGGACTCGGAGTCGAAA</td><td></td><td></td><td></td><td></td><td></td><td></td><td></td><td></td><td></td><td></td><td></td></tr>
<tr><td>Chromosome</td><td>start</td><td></td><td>end</td><td>strand</td><td>MM</td><td>target_seq</td><td>PAM</td><td>alignment</td><td>distance</td><td>position</td><td>gene name</td><td>gene id</td></tr>
<tr><td>GU305914.1</td><td></td><td>18412</td><td>18434</td><td>+</td><td>0</td><td>TTTCGACTCCGAGTCCGTGG</td><td>TGG</td><td>||||||||[||||||||||||]PAM</td><td>NA</td><td>Intergenic</td><td>NA</td><td>NA</td></tr>
<tr><td>T886</td><td>CGACTCCGAGTCCGTGGTGGTGG</td><td></td><td>1000</td><td></td><td></td><td></td><td></td><td></td><td></td><td></td><td></td><td></td></tr>
<tr><td>Oligo fwd</td><td>CACCCGACTCCGAGTCCGTGGTGG</td><td></td><td></td><td></td><td></td><td></td><td></td><td></td><td></td><td></td><td></td><td></td></tr>
<tr><td>Oligo rev</td><td>AAACCCACCACGGACTCGGAGTCG</td><td></td><td></td><td></td><td></td><td></td><td></td><td></td><td></td><td></td><td></td><td></td></tr>
<tr><td>Chromosome</td><td>start</td><td></td><td>end</td><td>strand</td><td>MM</td><td>target_seq</td><td>PAM</td><td>alignment</td><td>distance</td><td>position</td><td>gene name</td><td>gene id</td></tr>
<tr><td>GU305914.1</td><td></td><td>18415</td><td>18437</td><td>+</td><td>0</td><td>CGACTCCGAGTCCGTGGTGG</td><td>TGG</td><td>||||||||[||||||||||||]PAM</td><td>NA</td><td>Intergenic</td><td>NA</td><td>NA</td></tr>
<tr><td>T887</td><td>GGTGGTGGAATCTGAAGCGATGG</td><td></td><td>1000</td><td></td><td></td><td></td><td></td><td></td><td></td><td></td><td></td><td></td></tr>
<tr><td>Oligo fwd</td><td>CACCGGTGGTGGAATCTGAAGCGA</td><td></td><td></td><td></td><td></td><td></td><td></td><td></td><td></td><td></td><td></td><td></td></tr>
<tr><td>Oligo rev</td><td>AAACTCGCTTCAGATTCCACCACC</td><td></td><td></td><td></td><td></td><td></td><td></td><td></td><td></td><td></td><td></td><td></td></tr>
<tr><td>Chromosome</td><td>start</td><td></td><td>end</td><td>strand</td><td>MM</td><td>target_seq</td><td>PAM</td><td>alignment</td><td>distance</td><td>position</td><td>gene name</td><td>gene id</td></tr>
<tr><td>GU305914.1</td><td></td><td>18430</td><td>18452</td><td>+</td><td>0</td><td>GGTGGTGGAATCTGAAGCGA</td><td>TGG</td><td>||||||||[||||||||||||]PAM</td><td>NA</td><td>Intergenic</td><td>NA</td><td>NA</td></tr>
<tr><td>T888</td><td>ATGGTAGACGAGCGAGACCTCGG</td><td></td><td>1000</td><td></td><td></td><td></td><td></td><td></td><td></td><td></td><td></td><td></td></tr>
<tr><td>Oligo fwd</td><td>CACCATGGTAGACGAGCGAGACCT</td><td></td><td></td><td></td><td></td><td></td><td></td><td></td><td></td><td></td><td></td><td></td></tr>
<tr><td>Oligo rev</td><td>AAACAGGTCTCGCTCGTCTACCAT</td><td></td><td></td><td></td><td></td><td></td><td></td><td></td><td></td><td></td><td></td><td></td></tr>
<tr><td>Chromosome</td><td>start</td><td></td><td>end</td><td>strand</td><td>MM</td><td>target_seq</td><td>PAM</td><td>alignment</td><td>distance</td><td>position</td><td>gene name</td><td>gene id</td></tr>
<tr><td>GU305914.1</td><td></td><td>18449</td><td>18471</td><td>+</td><td>0</td><td>ATGGTAGACGAGCGAGACCT</td><td>CGG</td><td>||||||||[||||||||||||]PAM</td><td>NA</td><td>Intergenic</td><td>NA</td><td>NA</td></tr>
<tr><td>T889</td><td>GCGAGACCTCGGACAGCCGCCGG</td><td></td><td>1000</td><td></td><td></td><td></td><td></td><td></td><td></td><td></td><td></td><td></td></tr>
<tr><td>Oligo fwd</td><td>CACCGCGAGACCTCGGACAGCCGC</td><td></td><td></td><td></td><td></td><td></td><td></td><td></td><td></td><td></td><td></td><td></td></tr>
<tr><td>Oligo rev</td><td>AAACGCGGCTGTCCGAGGTCTCGC</td><td></td><td></td><td></td><td></td><td></td><td></td><td></td><td></td><td></td><td></td><td></td></tr>
<tr><td>Chromosome</td><td>start</td><td></td><td>end</td><td>strand</td><td>MM</td><td>target_seq</td><td>PAM</td><td>alignment</td><td>distance</td><td>position</td><td>gene name</td><td>gene id</td></tr>
<tr><td>GU305914.1</td><td></td><td>18460</td><td>18482</td><td>+</td><td>0</td><td>GCGAGACCTCGGACAGCCGC</td><td>CGG</td><td>||||||||[||||||||||||]PAM</td><td>NA</td><td>Intergenic</td><td>NA</td><td>NA</td></tr>
<tr><td>T890</td><td>CAGCCGCCGGATTCCTCGAACGG</td><td></td><td>1000</td><td></td><td></td><td></td><td></td><td></td><td></td><td></td><td></td><td></td></tr>
<tr><td>Oligo fwd</td><td>CACCCAGCCGCCGGATTCCTCGAA</td><td></td><td></td><td></td><td></td><td></td><td></td><td></td><td></td><td></td><td></td><td></td></tr>
<tr><td>Oligo rev</td><td>AAACTTCGAGGAATCCGGCGGCTG</td><td></td><td></td><td></td><td></td><td></td><td></td><td></td><td></td><td></td><td></td><td></td></tr>
<tr><td>Chromosome</td><td>start</td><td></td><td>end</td><td>strand</td><td>MM</td><td>target_seq</td><td>PAM</td><td>alignment</td><td>distance</td><td>position</td><td>gene name</td><td>gene id</td></tr>
<tr><td>GU305914.1</td><td></td><td>18473</td><td>18495</td><td>+</td><td>0</td><td>CAGCCGCCGGATTCCTCGAA</td><td>CGG</td><td>||||||||[||||||||||||]PAM</td><td>NA</td><td>Intergenic</td><td>NA</td><td>NA</td></tr>
<tr><td>T891</td><td>AGCCGCCGGATTCCTCGAACGGG</td><td></td><td>1000</td><td></td><td></td><td></td><td></td><td></td><td></td><td></td><td></td><td></td></tr>
<tr><td>Oligo fwd</td><td>CACCAGCCGCCGGATTCCTCGAAC</td><td></td><td></td><td></td><td></td><td></td><td></td><td></td><td></td><td></td><td></td><td></td></tr>
<tr><td>Oligo rev</td><td>AAACGTTCGAGGAATCCGGCGGCT</td><td></td><td></td><td></td><td></td><td></td><td></td><td></td><td></td><td></td><td></td><td></td></tr>
<tr><td>Chromosome</td><td>start</td><td></td><td>end</td><td>strand</td><td>MM</td><td>target_seq</td><td>PAM</td><td>alignment</td><td>distance</td><td>position</td><td>gene name</td><td>gene id</td></tr>
<tr><td>GU305914.1</td><td></td><td>18474</td><td>18496</td><td>+</td><td>0</td><td>AGCCGCCGGATTCCTCGAAC</td><td>GGG</td><td>||||||||[||||||||||||]PAM</td><td>NA</td><td>Intergenic</td><td>NA</td><td>NA</td></tr>
<tr><td>T892</td><td>TCCTCGAACGGGCCAGAAGAAGG</td><td></td><td>1000</td><td></td><td></td><td></td><td></td><td></td><td></td><td></td><td></td><td></td></tr>
<tr><td>Oligo fwd</td><td>CACCTCCTCGAACGGGCCAGAAGA</td><td></td><td></td><td></td><td></td><td></td><td></td><td></td><td></td><td></td><td></td><td></td></tr>
<tr><td>Oligo rev</td><td>AAACTCTTCTGGCCCGTTCGAGGA</td><td></td><td></td><td></td><td></td><td></td><td></td><td></td><td></td><td></td><td></td><td></td></tr>
<tr><td>Chromosome</td><td>start</td><td></td><td>end</td><td>strand</td><td>MM</td><td>target_seq</td><td>PAM</td><td>alignment</td><td>distance</td><td>position</td><td>gene name</td><td>gene id</td></tr>
<tr><td>GU305914.1</td><td></td><td>18485</td><td>18507</td><td>+</td><td>0</td><td>TCCTCGAACGGGCCAGAAGA</td><td>AGG</td><td>||||||||[||||||||||||]PAM</td><td>NA</td><td>Intergenic</td><td>NA</td><td>NA</td></tr>
<tr><td>T893</td><td>CCAGAAGAAGGTACCAACGACGG</td><td></td><td>1000</td><td></td><td></td><td></td><td></td><td></td><td></td><td></td><td></td><td></td></tr>
<tr><td>Oligo fwd</td><td>CACCCCAGAAGAAGGTACCAACGA</td><td></td><td></td><td></td><td></td><td></td><td></td><td></td><td></td><td></td><td></td><td></td></tr>
<tr><td>Oligo rev</td><td>AAACTCGTTGGTACCTTCTTCTGG</td><td></td><td></td><td></td><td></td><td></td><td></td><td></td><td></td><td></td><td></td><td></td></tr>
<tr><td>Chromosome</td><td>start</td><td></td><td>end</td><td>strand</td><td>MM</td><td>target_seq</td><td>PAM</td><td>alignment</td><td>distance</td><td>position</td><td>gene name</td><td>gene id</td></tr>
<tr><td>GU305914.1</td><td></td><td>18497</td><td>18519</td><td>+</td><td>0</td><td>CCAGAAGAAGGTACCAACGA</td><td>CGG</td><td>||||||||[||||||||||||]PAM</td><td>NA</td><td>Intergenic</td><td>NA</td><td>NA</td></tr>
<tr><td>T894</td><td>CAGAAGAAGGTACCAACGACGGG</td><td></td><td>1000</td><td></td><td></td><td></td><td></td><td></td><td></td><td></td><td></td><td></td></tr>
<tr><td>Oligo fwd</td><td>CACCCAGAAGAAGGTACCAACGAC</td><td></td><td></td><td></td><td></td><td></td><td></td><td></td><td></td><td></td><td></td><td></td></tr>
<tr><td>Oligo rev</td><td>AAACGTCGTTGGTACCTTCTTCTG</td><td></td><td></td><td></td><td></td><td></td><td></td><td></td><td></td><td></td><td></td><td></td></tr>
<tr><td>Chromosome</td><td>start</td><td></td><td>end</td><td>strand</td><td>MM</td><td>target_seq</td><td>PAM</td><td>alignment</td><td>distance</td><td>position</td><td>gene name</td><td>gene id</td></tr>
<tr><td>GU305914.1</td><td></td><td>18498</td><td>18520</td><td>+</td><td>0</td><td>CAGAAGAAGGTACCAACGAC</td><td>GGG</td><td>||||||||[||||||||||||]PAM</td><td>NA</td><td>Intergenic</td><td>NA</td><td>NA</td></tr>
<tr><td>T895</td><td>AGAAGAAGGTACCAACGACGGGG</td><td></td><td>1000</td><td></td><td></td><td></td><td></td><td></td><td></td><td></td><td></td><td></td></tr>
<tr><td>Oligo fwd</td><td>CACCAGAAGAAGGTACCAACGACG</td><td></td><td></td><td></td><td></td><td></td><td></td><td></td><td></td><td></td><td></td><td></td></tr>
<tr><td>Oligo rev</td><td>AAACCGTCGTTGGTACCTTCTTCT</td><td></td><td></td><td></td><td></td><td></td><td></td><td></td><td></td><td></td><td></td><td></td></tr>
<tr><td>Chromosome</td><td>start</td><td></td><td>end</td><td>strand</td><td>MM</td><td>target_seq</td><td>PAM</td><td>alignment</td><td>distance</td><td>position</td><td>gene name</td><td>gene id</td></tr>
<tr><td>GU305914.1</td><td></td><td>18499</td><td>18521</td><td>+</td><td>0</td><td>AGAAGAAGGTACCAACGACG</td><td>GGG</td><td>||||||||[||||||||||||]PAM</td><td>NA</td><td>Intergenic</td><td>NA</td><td>NA</td></tr>
<tr><td>T896</td><td>ACCCTCAGAGCCGAACCCGACGG</td><td></td><td>1000</td><td></td><td></td><td></td><td></td><td></td><td></td><td></td><td></td><td></td></tr>
<tr><td>Oligo fwd</td><td>CACCACCCTCAGAGCCGAACCCGA</td><td></td><td></td><td></td><td></td><td></td><td></td><td></td><td></td><td></td><td></td><td></td></tr>
</table>

<table>
<tr><td>Oligo rev</td><td>AAACTCGGGTTCGGCTCTGAGGGT</td><td></td><td></td><td></td><td></td><td></td><td></td><td></td><td></td><td></td><td></td><td></td></tr>
<tr><td>Chromosome</td><td>start</td><td></td><td>end</td><td>strand</td><td>MM</td><td>target_seq</td><td>PAM</td><td>alignment</td><td>distance</td><td>position</td><td>gene name</td><td>gene id</td></tr>
<tr><td>GU305914.1</td><td></td><td>18539</td><td>18561</td><td>+</td><td>0</td><td>ACCCTCAGAGCCGAACCCGA</td><td>CGG</td><td>||||||||[||||||||||||]PAM</td><td>NA</td><td>Intergenic</td><td>NA</td><td>NA</td></tr>
<tr><td></td><td></td><td></td><td></td><td></td><td></td><td></td><td></td><td></td><td></td><td></td><td></td><td></td></tr>
<tr><td>T897</td><td>CAGAGCCGAACCCGACGGTTCGG</td><td></td><td>1000</td><td></td><td></td><td></td><td></td><td></td><td></td><td></td><td></td><td></td></tr>
<tr><td>Oligo fwd</td><td>CACCCAGAGCCGAACCCGACGGTT</td><td></td><td></td><td></td><td></td><td></td><td></td><td></td><td></td><td></td><td></td><td></td></tr>
<tr><td>Oligo rev</td><td>AAACAACCGTCGGGTTCGGCTCTG</td><td></td><td></td><td></td><td></td><td></td><td></td><td></td><td></td><td></td><td></td><td></td></tr>
<tr><td>Chromosome</td><td>start</td><td></td><td>end</td><td>strand</td><td>MM</td><td>target_seq</td><td>PAM</td><td>alignment</td><td>distance</td><td>position</td><td>gene name</td><td>gene id</td></tr>
<tr><td>GU305914.1</td><td></td><td>18544</td><td>18566</td><td>+</td><td>0</td><td>CAGAGCCGAACCCGACGGTT</td><td>CGG</td><td>||||||||[||||||||||||]PAM</td><td>NA</td><td>Intergenic</td><td>NA</td><td>NA</td></tr>
<tr><td></td><td></td><td></td><td></td><td></td><td></td><td></td><td></td><td></td><td></td><td></td><td></td><td></td></tr>
<tr><td>T898</td><td>CTCGACGCCGACGACCTCTCCGG</td><td></td><td>1000</td><td></td><td></td><td></td><td></td><td></td><td></td><td></td><td></td><td></td></tr>
<tr><td>Oligo fwd</td><td>CACCCTCGACGCCGACGACCTCTC</td><td></td><td></td><td></td><td></td><td></td><td></td><td></td><td></td><td></td><td></td><td></td></tr>
<tr><td>Oligo rev</td><td>AAACGAGAGGTCGTCGGCGTCGAG</td><td></td><td></td><td></td><td></td><td></td><td></td><td></td><td></td><td></td><td></td><td></td></tr>
<tr><td>Chromosome</td><td>start</td><td></td><td>end</td><td>strand</td><td>MM</td><td>target_seq</td><td>PAM</td><td>alignment</td><td>distance</td><td>position</td><td>gene name</td><td>gene id</td></tr>
<tr><td>GU305914.1</td><td></td><td>18568</td><td>18590</td><td>+</td><td>0</td><td>CTCGACGCCGACGACCTCTC</td><td>CGG</td><td>||||||||[||||||||||||]PAM</td><td>NA</td><td>Intergenic</td><td>NA</td><td>NA</td></tr>
<tr><td></td><td></td><td></td><td></td><td></td><td></td><td></td><td></td><td></td><td></td><td></td><td></td><td></td></tr>
<tr><td>T899</td><td>CCTCTCCGGCGACTCCTCCGAGG</td><td></td><td>1000</td><td></td><td></td><td></td><td></td><td></td><td></td><td></td><td></td><td></td></tr>
<tr><td>Oligo fwd</td><td>CACCCCTCTCCGGCGACTCCTCCG</td><td></td><td></td><td></td><td></td><td></td><td></td><td></td><td></td><td></td><td></td><td></td></tr>
<tr><td>Oligo rev</td><td>AAACCGGAGGAGTCGCCGGAGAGG</td><td></td><td></td><td></td><td></td><td></td><td></td><td></td><td></td><td></td><td></td><td></td></tr>
<tr><td>Chromosome</td><td>start</td><td></td><td>end</td><td>strand</td><td>MM</td><td>target_seq</td><td>PAM</td><td>alignment</td><td>distance</td><td>position</td><td>gene name</td><td>gene id</td></tr>
<tr><td>GU305914.1</td><td></td><td>18582</td><td>18604</td><td>+</td><td>0</td><td>CCTCTCCGGCGACTCCTCCG</td><td>AGG</td><td>||||||||[||||||||||||]PAM</td><td>NA</td><td>Intergenic</td><td>NA</td><td>NA</td></tr>
<tr><td></td><td></td><td></td><td></td><td></td><td></td><td></td><td></td><td></td><td></td><td></td><td></td><td></td></tr>
<tr><td>T900</td><td>CCGGCGACTCCTCCGAGGAGCGG</td><td></td><td>1000</td><td></td><td></td><td></td><td></td><td></td><td></td><td></td><td></td><td></td></tr>
<tr><td>Oligo fwd</td><td>CACCCCGGCGACTCCTCCGAGGAG</td><td></td><td></td><td></td><td></td><td></td><td></td><td></td><td></td><td></td><td></td><td></td></tr>
<tr><td>Oligo rev</td><td>AAACCTCCTCGGAGGAGTCGCCGG</td><td></td><td></td><td></td><td></td><td></td><td></td><td></td><td></td><td></td><td></td><td></td></tr>
<tr><td>Chromosome</td><td>start</td><td></td><td>end</td><td>strand</td><td>MM</td><td>target_seq</td><td>PAM</td><td>alignment</td><td>distance</td><td>position</td><td>gene name</td><td>gene id</td></tr>
<tr><td>GU305914.1</td><td></td><td>18587</td><td>18609</td><td>+</td><td>0</td><td>CCGGCGACTCCTCCGAGGAG</td><td>CGG</td><td>||||||||[||||||||||||]PAM</td><td>NA</td><td>Intergenic</td><td>NA</td><td>NA</td></tr>
<tr><td></td><td></td><td></td><td></td><td></td><td></td><td></td><td></td><td></td><td></td><td></td><td></td><td></td></tr>
<tr><td>T901</td><td>GCGACTCCTCCGAGGAGCGGAGG</td><td></td><td>1000</td><td></td><td></td><td></td><td></td><td></td><td></td><td></td><td></td><td></td></tr>
<tr><td>Oligo fwd</td><td>CACCGCGACTCCTCCGAGGAGCGG</td><td></td><td></td><td></td><td></td><td></td><td></td><td></td><td></td><td></td><td></td><td></td></tr>
<tr><td>Oligo rev</td><td>AAACCCGCTCCTCGGAGGAGTCGC</td><td></td><td></td><td></td><td></td><td></td><td></td><td></td><td></td><td></td><td></td><td></td></tr>
<tr><td>Chromosome</td><td>start</td><td></td><td>end</td><td>strand</td><td>MM</td><td>target_seq</td><td>PAM</td><td>alignment</td><td>distance</td><td>position</td><td>gene name</td><td>gene id</td></tr>
<tr><td>GU305914.1</td><td></td><td>18590</td><td>18612</td><td>+</td><td>0</td><td>GCGACTCCTCCGAGGAGCGG</td><td>AGG</td><td>||||||||[||||||||||||]PAM</td><td>NA</td><td>Intergenic</td><td>NA</td><td>NA</td></tr>
<tr><td></td><td></td><td></td><td></td><td></td><td></td><td></td><td></td><td></td><td></td><td></td><td></td><td></td></tr>
<tr><td>T902</td><td>CGACTCCTCCGAGGAGCGGAGGG</td><td></td><td>1000</td><td></td><td></td><td></td><td></td><td></td><td></td><td></td><td></td><td></td></tr>
<tr><td>Oligo fwd</td><td>CACCCGACTCCTCCGAGGAGCGGA</td><td></td><td></td><td></td><td></td><td></td><td></td><td></td><td></td><td></td><td></td><td></td></tr>
<tr><td>Oligo rev</td><td>AAACTCCGCTCCTCGGAGGAGTCG</td><td></td><td></td><td></td><td></td><td></td><td></td><td></td><td></td><td></td><td></td><td></td></tr>
<tr><td>Chromosome</td><td>start</td><td></td><td>end</td><td>strand</td><td>MM</td><td>target_seq</td><td>PAM</td><td>alignment</td><td>distance</td><td>position</td><td>gene name</td><td>gene id</td></tr>
<tr><td>GU305914.1</td><td></td><td>18591</td><td>18613</td><td>+</td><td>0</td><td>CGACTCCTCCGAGGAGCGGA</td><td>GGG</td><td>||||||||[||||||||||||]PAM</td><td>NA</td><td>Intergenic</td><td>NA</td><td>NA</td></tr>
<tr><td></td><td></td><td></td><td></td><td></td><td></td><td></td><td></td><td></td><td></td><td></td><td></td><td></td></tr>
<tr><td>T903</td><td>GACTCCTCCGAGGAGCGGAGGGG</td><td></td><td>1000</td><td></td><td></td><td></td><td></td><td></td><td></td><td></td><td></td><td></td></tr>
<tr><td>Oligo fwd</td><td>CACCGACTCCTCCGAGGAGCGGAG</td><td></td><td></td><td></td><td></td><td></td><td></td><td></td><td></td><td></td><td></td><td></td></tr>
<tr><td>Oligo rev</td><td>AAACCTCCGCTCCTCGGAGGAGTC</td><td></td><td></td><td></td><td></td><td></td><td></td><td></td><td></td><td></td><td></td><td></td></tr>
<tr><td>Chromosome</td><td>start</td><td></td><td>end</td><td>strand</td><td>MM</td><td>target_seq</td><td>PAM</td><td>alignment</td><td>distance</td><td>position</td><td>gene name</td><td>gene id</td></tr>
<tr><td>GU305914.1</td><td></td><td>18592</td><td>18614</td><td>+</td><td>0</td><td>GACTCCTCCGAGGAGCGGAG</td><td>GGG</td><td>||||||||[||||||||||||]PAM</td><td>NA</td><td>Intergenic</td><td>NA</td><td>NA</td></tr>
<tr><td></td><td></td><td></td><td></td><td></td><td></td><td></td><td></td><td></td><td></td><td></td><td></td><td></td></tr>
<tr><td>T904</td><td>GAGGAGCGGAGGGGCCTCGTTGG</td><td></td><td>1000</td><td></td><td></td><td></td><td></td><td></td><td></td><td></td><td></td><td></td></tr>
<tr><td>Oligo fwd</td><td>CACCGAGGAGCGGAGGGGCCTCGT</td><td></td><td></td><td></td><td></td><td></td><td></td><td></td><td></td><td></td><td></td><td></td></tr>
<tr><td>Oligo rev</td><td>AAACACGAGGCCCCTCCGCTCCTC</td><td></td><td></td><td></td><td></td><td></td><td></td><td></td><td></td><td></td><td></td><td></td></tr>
<tr><td>Chromosome</td><td>start</td><td></td><td>end</td><td>strand</td><td>MM</td><td>target_seq</td><td>PAM</td><td>alignment</td><td>distance</td><td>position</td><td>gene name</td><td>gene id</td></tr>
<tr><td>GU305914.1</td><td></td><td>18601</td><td>18623</td><td>+</td><td>0</td><td>GAGGAGCGGAGGGGCCTCGT</td><td>TGG</td><td>||||||||[||||||||||||]PAM</td><td>NA</td><td>Intergenic</td><td>NA</td><td>NA</td></tr>
<tr><td></td><td></td><td></td><td></td><td></td><td></td><td></td><td></td><td></td><td></td><td></td><td></td><td></td></tr>
<tr><td>T905</td><td>TGGTGTTGTCGTCACCGCCCAGG</td><td></td><td>1000</td><td></td><td></td><td></td><td></td><td></td><td></td><td></td><td></td><td></td></tr>
<tr><td>Oligo fwd</td><td>CACCTGGTGTTGTCGTCACCGCCC</td><td></td><td></td><td></td><td></td><td></td><td></td><td></td><td></td><td></td><td></td><td></td></tr>
<tr><td>Oligo rev</td><td>AAACGGGCGGTGACGACAACACCA</td><td></td><td></td><td></td><td></td><td></td><td></td><td></td><td></td><td></td><td></td><td></td></tr>
<tr><td>Chromosome</td><td>start</td><td></td><td>end</td><td>strand</td><td>MM</td><td>target_seq</td><td>PAM</td><td>alignment</td><td>distance</td><td>position</td><td>gene name</td><td>gene id</td></tr>
<tr><td>GU305914.1</td><td></td><td>18621</td><td>18643</td><td>+</td><td>0</td><td>TGGTGTTGTCGTCACCGCCC</td><td>AGG</td><td>||||||||[||||||||||||]PAM</td><td>NA</td><td>Intergenic</td><td>NA</td><td>NA</td></tr>
<tr><td></td><td></td><td></td><td></td><td></td><td></td><td></td><td></td><td></td><td></td><td></td><td></td><td></td></tr>
<tr><td>T906</td><td>TCTTGTAGCATCTTCGCCAGCGG</td><td></td><td>1000</td><td></td><td></td><td></td><td></td><td></td><td></td><td></td><td></td><td></td></tr>
<tr><td>Oligo fwd</td><td>CACCTCTTGTAGCATCTTCGCCAG</td><td></td><td></td><td></td><td></td><td></td><td></td><td></td><td></td><td></td><td></td><td></td></tr>
<tr><td>Oligo rev</td><td>AAACCTGGCGAAGATGCTACAAGA</td><td></td><td></td><td></td><td></td><td></td><td></td><td></td><td></td><td></td><td></td><td></td></tr>
<tr><td>Chromosome</td><td>start</td><td></td><td>end</td><td>strand</td><td>MM</td><td>target_seq</td><td>PAM</td><td>alignment</td><td>distance</td><td>position</td><td>gene name</td><td>gene id</td></tr>
<tr><td>GU305914.1</td><td></td><td>18679</td><td>18701</td><td>+</td><td>0</td><td>TCTTGTAGCATCTTCGCCAG</td><td>CGG</td><td>||||||||[||||||||||||]PAM</td><td>NA</td><td>Intergenic</td><td>NA</td><td>NA</td></tr>
<tr><td></td><td></td><td></td><td></td><td></td><td></td><td></td><td></td><td></td><td></td><td></td><td></td><td></td></tr>
<tr><td>T907</td><td>AGCATCTTCGCCAGCGGTCCCGG</td><td></td><td>1000</td><td></td><td></td><td></td><td></td><td></td><td></td><td></td><td></td><td></td></tr>
<tr><td>Oligo fwd</td><td>CACCAGCATCTTCGCCAGCGGTCC</td><td></td><td></td><td></td><td></td><td></td><td></td><td></td><td></td><td></td><td></td><td></td></tr>
<tr><td>Oligo rev</td><td>AAACGGACCGCTGGCGAAGATGCT</td><td></td><td></td><td></td><td></td><td></td><td></td><td></td><td></td><td></td><td></td><td></td></tr>
<tr><td>Chromosome</td><td>start</td><td></td><td>end</td><td>strand</td><td>MM</td><td>target_seq</td><td>PAM</td><td>alignment</td><td>distance</td><td>position</td><td>gene name</td><td>gene id</td></tr>
<tr><td>GU305914.1</td><td></td><td>18685</td><td>18707</td><td>+</td><td>0</td><td>AGCATCTTCGCCAGCGGTCC</td><td>CGG</td><td>||||||||[||||||||||||]PAM</td><td>NA</td><td>Intergenic</td><td>NA</td><td>NA</td></tr>
<tr><td></td><td></td><td></td><td></td><td></td><td></td><td></td><td></td><td></td><td></td><td></td><td></td><td></td></tr>
<tr><td>T908</td><td>GCCCACCACCGAGTCGTCGAAGG</td><td></td><td>1000</td><td></td><td></td><td></td><td></td><td></td><td></td><td></td><td></td><td></td></tr>
<tr><td>Oligo fwd</td><td>CACCGCCCACCACCGAGTCGTCGA</td><td></td><td></td><td></td><td></td><td></td><td></td><td></td><td></td><td></td><td></td><td></td></tr>
<tr><td>Oligo rev</td><td>AAACTCGACGACTCGGTGGTGGGC</td><td></td><td></td><td></td><td></td><td></td><td></td><td></td><td></td><td></td><td></td><td></td></tr>
<tr><td>Chromosome</td><td>start</td><td></td><td>end</td><td>strand</td><td>MM</td><td>target_seq</td><td>PAM</td><td>alignment</td><td>distance</td><td>position</td><td>gene name</td><td>gene id</td></tr>
<tr><td>GU305914.1</td><td></td><td>18712</td><td>18734</td><td>+</td><td>0</td><td>GCCCACCACCGAGTCGTCGA</td><td>AGG</td><td>||||||||[||||||||||||]PAM</td><td>NA</td><td>Intergenic</td><td>NA</td><td>NA</td></tr>
<tr><td></td><td></td><td></td><td></td><td></td><td></td><td></td><td></td><td></td><td></td><td></td><td></td><td></td></tr>
<tr><td>T909</td><td>TCCAGCAGCAGTGACAGTAGCGG</td><td></td><td>1000</td><td></td><td></td><td></td><td></td><td></td><td></td><td></td><td></td><td></td></tr>
<tr><td>Oligo fwd</td><td>CACCTCCAGCAGCAGTGACAGTAG</td><td></td><td></td><td></td><td></td><td></td><td></td><td></td><td></td><td></td><td></td><td></td></tr>
<tr><td>Oligo rev</td><td>AAACCTACTGTCACTGCTGCTGGA</td><td></td><td></td><td></td><td></td><td></td><td></td><td></td><td></td><td></td><td></td><td></td></tr>
<tr><td>Chromosome</td><td>start</td><td></td><td>end</td><td>strand</td><td>MM</td><td>target_seq</td><td>PAM</td><td>alignment</td><td>distance</td><td>position</td><td>gene name</td><td>gene id</td></tr>
<tr><td>GU305914.1</td><td></td><td>18755</td><td>18777</td><td>+</td><td>0</td><td>TCCAGCAGCAGTGACAGTAG</td><td>CGG</td><td>||||||||[||||||||||||]PAM</td><td>NA</td><td>Intergenic</td><td>NA</td><td>NA</td></tr>
<tr><td></td><td></td><td></td><td></td><td></td><td></td><td></td><td></td><td></td><td></td><td></td><td></td><td></td></tr>
<tr><td>T910</td><td>CCAGCAGCAGTGACAGTAGCGGG</td><td></td><td>1000</td><td></td><td></td><td></td><td></td><td></td><td></td><td></td><td></td><td></td></tr>
<tr><td>Oligo fwd</td><td>CACCCCAGCAGCAGTGACAGTAGC</td><td></td><td></td><td></td><td></td><td></td><td></td><td></td><td></td><td></td><td></td><td></td></tr>
<tr><td>Oligo rev</td><td>AAACGCTACTGTCACTGCTGCTGG</td><td></td><td></td><td></td><td></td><td></td><td></td><td></td><td></td><td></td><td></td><td></td></tr>
<tr><td>Chromosome</td><td>start</td><td></td><td>end</td><td>strand</td><td>MM</td><td>target_seq</td><td>PAM</td><td>alignment</td><td>distance</td><td>position</td><td>gene name</td><td>gene id</td></tr>
<tr><td>GU305914.1</td><td></td><td>18756</td><td>18778</td><td>+</td><td>0</td><td>CCAGCAGCAGTGACAGTAGC</td><td>GGG</td><td>||||||||[||||||||||||]PAM</td><td>NA</td><td>Intergenic</td><td>NA</td><td>NA</td></tr>
<tr><td></td><td></td><td></td><td></td><td></td><td></td><td></td><td></td><td></td><td></td><td></td><td></td><td></td></tr>
<tr><td>T911</td><td>CAGCAGCAGTGACAGTAGCGGGG</td><td></td><td>1000</td><td></td><td></td><td></td><td></td><td></td><td></td><td></td><td></td><td></td></tr>
</table>

<table>
<tr><td>Oligo fwd</td><td>CACCCAGCAGCAGTGACAGTAGCG</td><td></td><td></td><td></td><td></td><td></td><td></td><td></td><td></td><td></td><td></td><td></td></tr>
<tr><td>Oligo rev</td><td>AAACCGCTACTGTCACTGCTGCTG</td><td></td><td></td><td></td><td></td><td></td><td></td><td></td><td></td><td></td><td></td><td></td></tr>
<tr><td>Chromosome</td><td>start</td><td></td><td>end</td><td>strand</td><td>MM</td><td>target_seq</td><td>PAM</td><td>alignment</td><td>distance</td><td>position</td><td>gene name</td><td>gene id</td></tr>
<tr><td>GU305914.1</td><td></td><td>18757</td><td>18779</td><td>+</td><td>0</td><td>CAGCAGCAGTGACAGTAGCG</td><td>GGG</td><td>||||||||[||||||||||||]PAM</td><td>NA</td><td>Intergenic</td><td>NA</td><td>NA</td></tr>
<tr><td>T912</td><td>CGAAAACGAGACGTTCGTCTTGG</td><td></td><td>1000</td><td></td><td></td><td></td><td></td><td></td><td></td><td></td><td></td><td></td></tr>
<tr><td>Oligo fwd</td><td>CACCCGAAAACGAGACGTTCGTCT</td><td></td><td></td><td></td><td></td><td></td><td></td><td></td><td></td><td></td><td></td><td></td></tr>
<tr><td>Oligo rev</td><td>AAACAGACGAACGTCTCGTTTTCG</td><td></td><td></td><td></td><td></td><td></td><td></td><td></td><td></td><td></td><td></td><td></td></tr>
<tr><td>Chromosome</td><td>start</td><td></td><td>end</td><td>strand</td><td>MM</td><td>target_seq</td><td>PAM</td><td>alignment</td><td>distance</td><td>position</td><td>gene name</td><td>gene id</td></tr>
<tr><td>GU305914.1</td><td></td><td>18784</td><td>18806</td><td>+</td><td>0</td><td>CGAAAACGAGACGTTCGTCT</td><td>TGG</td><td>||||||||[||||||||||||]PAM</td><td>NA</td><td>Intergenic</td><td>NA</td><td>NA</td></tr>
<tr><td>T913</td><td>GACGTTCGTCTTGGCGTCCGAGG</td><td></td><td>1000</td><td></td><td></td><td></td><td></td><td></td><td></td><td></td><td></td><td></td></tr>
<tr><td>Oligo fwd</td><td>CACCGACGTTCGTCTTGGCGTCCG</td><td></td><td></td><td></td><td></td><td></td><td></td><td></td><td></td><td></td><td></td><td></td></tr>
<tr><td>Oligo rev</td><td>AAACCGGACGCCAAGACGAACGTC</td><td></td><td></td><td></td><td></td><td></td><td></td><td></td><td></td><td></td><td></td><td></td></tr>
<tr><td>Chromosome</td><td>start</td><td></td><td>end</td><td>strand</td><td>MM</td><td>target_seq</td><td>PAM</td><td>alignment</td><td>distance</td><td>position</td><td>gene name</td><td>gene id</td></tr>
<tr><td>GU305914.1</td><td></td><td>18793</td><td>18815</td><td>+</td><td>0</td><td>GACGTTCGTCTTGGCGTCCG</td><td>AGG</td><td>||||||||[||||||||||||]PAM</td><td>NA</td><td>Intergenic</td><td>NA</td><td>NA</td></tr>
<tr><td>T914</td><td>GTCTTGGCGTCCGAGGAGATTGG</td><td></td><td>1000</td><td></td><td></td><td></td><td></td><td></td><td></td><td></td><td></td><td></td></tr>
<tr><td>Oligo fwd</td><td>CACCGTCTTGGCGTCCGAGGAGAT</td><td></td><td></td><td></td><td></td><td></td><td></td><td></td><td></td><td></td><td></td><td></td></tr>
<tr><td>Oligo rev</td><td>AAACATCTCCTCGGACGCCAAGAC</td><td></td><td></td><td></td><td></td><td></td><td></td><td></td><td></td><td></td><td></td><td></td></tr>
<tr><td>Chromosome</td><td>start</td><td></td><td>end</td><td>strand</td><td>MM</td><td>target_seq</td><td>PAM</td><td>alignment</td><td>distance</td><td>position</td><td>gene name</td><td>gene id</td></tr>
<tr><td>GU305914.1</td><td></td><td>18800</td><td>18822</td><td>+</td><td>0</td><td>GTCTTGGCGTCCGAGGAGAT</td><td>TGG</td><td>||||||||[||||||||||||]PAM</td><td>NA</td><td>Intergenic</td><td>NA</td><td>NA</td></tr>
<tr><td>T915</td><td>ACCTAAGAAAAGATTATCGCAGG</td><td></td><td>1000</td><td></td><td></td><td></td><td></td><td></td><td></td><td></td><td></td><td></td></tr>
<tr><td>Oligo fwd</td><td>CACCACCTAAGAAAAGATTATCGC</td><td></td><td></td><td></td><td></td><td></td><td></td><td></td><td></td><td></td><td></td><td></td></tr>
<tr><td>Oligo rev</td><td>AAACGCGATAATCTTTTCTTAGGT</td><td></td><td></td><td></td><td></td><td></td><td></td><td></td><td></td><td></td><td></td><td></td></tr>
<tr><td>Chromosome</td><td>start</td><td></td><td>end</td><td>strand</td><td>MM</td><td>target_seq</td><td>PAM</td><td>alignment</td><td>distance</td><td>position</td><td>gene name</td><td>gene id</td></tr>
<tr><td>GU305914.1</td><td></td><td>18826</td><td>18848</td><td>+</td><td>0</td><td>ACCTAAGAAAAGATTATCGC</td><td>AGG</td><td>||||||||[||||||||||||]PAM</td><td>NA</td><td>Intergenic</td><td>NA</td><td>NA</td></tr>
<tr><td>T916</td><td>GACTGACGAGTGCGACATCCCGG</td><td></td><td>1000</td><td></td><td></td><td></td><td></td><td></td><td></td><td></td><td></td><td></td></tr>
<tr><td>Oligo fwd</td><td>CACCGACTGACGAGTGCGACATCC</td><td></td><td></td><td></td><td></td><td></td><td></td><td></td><td></td><td></td><td></td><td></td></tr>
<tr><td>Oligo rev</td><td>AAACGGATGTCGCACTCGTCAGTC</td><td></td><td></td><td></td><td></td><td></td><td></td><td></td><td></td><td></td><td></td><td></td></tr>
<tr><td>Chromosome</td><td>start</td><td></td><td>end</td><td>strand</td><td>MM</td><td>target_seq</td><td>PAM</td><td>alignment</td><td>distance</td><td>position</td><td>gene name</td><td>gene id</td></tr>
<tr><td>GU305914.1</td><td></td><td>18874</td><td>18896</td><td>+</td><td>0</td><td>GACTGACGAGTGCGACATCC</td><td>CGG</td><td>||||||||[||||||||||||]PAM</td><td>NA</td><td>Intergenic</td><td>NA</td><td>NA</td></tr>
<tr><td>T917</td><td>CACGCCCGCTCCCCCCCGAACGG</td><td></td><td>1000</td><td></td><td></td><td></td><td></td><td></td><td></td><td></td><td></td><td></td></tr>
<tr><td>Oligo fwd</td><td>CACCCACGCCCGCTCCCCCCCGAA</td><td></td><td></td><td></td><td></td><td></td><td></td><td></td><td></td><td></td><td></td><td></td></tr>
<tr><td>Oligo rev</td><td>AAACTTCGGGGGGGAGCGGGCGTG</td><td></td><td></td><td></td><td></td><td></td><td></td><td></td><td></td><td></td><td></td><td></td></tr>
<tr><td>Chromosome</td><td>start</td><td></td><td>end</td><td>strand</td><td>MM</td><td>target_seq</td><td>PAM</td><td>alignment</td><td>distance</td><td>position</td><td>gene name</td><td>gene id</td></tr>
<tr><td>GU305914.1</td><td></td><td>18897</td><td>18919</td><td>+</td><td>0</td><td>CACGCCCGCTCCCCCCCGAA</td><td>CGG</td><td>||||||||[||||||||||||]PAM</td><td>NA</td><td>Intergenic</td><td>NA</td><td>NA</td></tr>
<tr><td>T918</td><td>CTCGATGTCGCGTGTGCTGCTGG</td><td></td><td>1000</td><td></td><td></td><td></td><td></td><td></td><td></td><td></td><td></td><td></td></tr>
<tr><td>Oligo fwd</td><td>CACCCTCGATGTCGCGTGTGCTGC</td><td></td><td></td><td></td><td></td><td></td><td></td><td></td><td></td><td></td><td></td><td></td></tr>
<tr><td>Oligo rev</td><td>AAACGCAGCACACGCGACATCGAG</td><td></td><td></td><td></td><td></td><td></td><td></td><td></td><td></td><td></td><td></td><td></td></tr>
<tr><td>Chromosome</td><td>start</td><td></td><td>end</td><td>strand</td><td>MM</td><td>target_seq</td><td>PAM</td><td>alignment</td><td>distance</td><td>position</td><td>gene name</td><td>gene id</td></tr>
<tr><td>GU305914.1</td><td></td><td>18933</td><td>18955</td><td>+</td><td>0</td><td>CTCGATGTCGCGTGTGCTGC</td><td>TGG</td><td>||||||||[||||||||||||]PAM</td><td>NA</td><td>Intergenic</td><td>NA</td><td>NA</td></tr>
<tr><td>T919</td><td>GCGTGTGCTGCTGGTACTGTTGG</td><td></td><td>1000</td><td></td><td></td><td></td><td></td><td></td><td></td><td></td><td></td><td></td></tr>
<tr><td>Oligo fwd</td><td>CACCGCGTGTGCTGCTGGTACTGT</td><td></td><td></td><td></td><td></td><td></td><td></td><td></td><td></td><td></td><td></td><td></td></tr>
<tr><td>Oligo rev</td><td>AAACACAGTACCAGCAGCACACGC</td><td></td><td></td><td></td><td></td><td></td><td></td><td></td><td></td><td></td><td></td><td></td></tr>
<tr><td>Chromosome</td><td>start</td><td></td><td>end</td><td>strand</td><td>MM</td><td>target_seq</td><td>PAM</td><td>alignment</td><td>distance</td><td>position</td><td>gene name</td><td>gene id</td></tr>
<tr><td>GU305914.1</td><td></td><td>18942</td><td>18964</td><td>+</td><td>0</td><td>GCGTGTGCTGCTGGTACTGT</td><td>TGG</td><td>||||||||[||||||||||||]PAM</td><td>NA</td><td>Intergenic</td><td>NA</td><td>NA</td></tr>
<tr><td>T920</td><td>GCATCTGTTGCTGCTGCACCTGG</td><td></td><td>1000</td><td></td><td></td><td></td><td></td><td></td><td></td><td></td><td></td><td></td></tr>
<tr><td>Oligo fwd</td><td>CACCGCATCTGTTGCTGCTGCACC</td><td></td><td></td><td></td><td></td><td></td><td></td><td></td><td></td><td></td><td></td><td></td></tr>
<tr><td>Oligo rev</td><td>AAACGGTGCAGCAGCAACAGATGC</td><td></td><td></td><td></td><td></td><td></td><td></td><td></td><td></td><td></td><td></td><td></td></tr>
<tr><td>Chromosome</td><td>start</td><td></td><td>end</td><td>strand</td><td>MM</td><td>target_seq</td><td>PAM</td><td>alignment</td><td>distance</td><td>position</td><td>gene name</td><td>gene id</td></tr>
<tr><td>GU305914.1</td><td></td><td>18969</td><td>18991</td><td>+</td><td>0</td><td>GCATCTGTTGCTGCTGCACC</td><td>TGG</td><td>||||||||[||||||||||||]PAM</td><td>NA</td><td>Intergenic</td><td>NA</td><td>NA</td></tr>
<tr><td>T921</td><td>CATCTGTTGCTGCTGCACCTGGG</td><td></td><td>1000</td><td></td><td></td><td></td><td></td><td></td><td></td><td></td><td></td><td></td></tr>
<tr><td>Oligo fwd</td><td>CACCCATCTGTTGCTGCTGCACCT</td><td></td><td></td><td></td><td></td><td></td><td></td><td></td><td></td><td></td><td></td><td></td></tr>
<tr><td>Oligo rev</td><td>AAACAGGTGCAGCAGCAACAGATG</td><td></td><td></td><td></td><td></td><td></td><td></td><td></td><td></td><td></td><td></td><td></td></tr>
<tr><td>Chromosome</td><td>start</td><td></td><td>end</td><td>strand</td><td>MM</td><td>target_seq</td><td>PAM</td><td>alignment</td><td>distance</td><td>position</td><td>gene name</td><td>gene id</td></tr>
<tr><td>GU305914.1</td><td></td><td>18970</td><td>18992</td><td>+</td><td>0</td><td>CATCTGTTGCTGCTGCACCT</td><td>GGG</td><td>||||||||[||||||||||||]PAM</td><td>NA</td><td>Intergenic</td><td>NA</td><td>NA</td></tr>
<tr><td>T922</td><td>GTTGCTGCTGCACCTGGGTGTGG</td><td></td><td>1000</td><td></td><td></td><td></td><td></td><td></td><td></td><td></td><td></td><td></td></tr>
<tr><td>Oligo fwd</td><td>CACCGTTGCTGCTGCACCTGGGTG</td><td></td><td></td><td></td><td></td><td></td><td></td><td></td><td></td><td></td><td></td><td></td></tr>
<tr><td>Oligo rev</td><td>AAACCACCCAGGTGCAGCAGCAAC</td><td></td><td></td><td></td><td></td><td></td><td></td><td></td><td></td><td></td><td></td><td></td></tr>
<tr><td>Chromosome</td><td>start</td><td></td><td>end</td><td>strand</td><td>MM</td><td>target_seq</td><td>PAM</td><td>alignment</td><td>distance</td><td>position</td><td>gene name</td><td>gene id</td></tr>
<tr><td>GU305914.1</td><td></td><td>18975</td><td>18997</td><td>+</td><td>0</td><td>GTTGCTGCTGCACCTGGGTG</td><td>TGG</td><td>||||||||[||||||||||||]PAM</td><td>NA</td><td>Intergenic</td><td>NA</td><td>NA</td></tr>
<tr><td>GU305914.1</td><td></td><td>168320</td><td>168342</td><td>+</td><td>4</td><td>GCTGGTGCTGCTCCTGGTTG</td><td>CGG</td><td>|-||-|||[|||-|||||-||]PAM</td><td>NA</td><td>Intergenic</td><td>NA</td><td>NA</td></tr>
<tr><td>T923</td><td>GCTGCTGCACCTGGGTGTGGTGG</td><td></td><td>1000</td><td></td><td></td><td></td><td></td><td></td><td></td><td></td><td></td><td></td></tr>
<tr><td>Oligo fwd</td><td>CACCGCTGCTGCACCTGGGTGTGG</td><td></td><td></td><td></td><td></td><td></td><td></td><td></td><td></td><td></td><td></td><td></td></tr>
<tr><td>Oligo rev</td><td>AAACCCACACCCAGGTGCAGCAGC</td><td></td><td></td><td></td><td></td><td></td><td></td><td></td><td></td><td></td><td></td><td></td></tr>
<tr><td>Chromosome</td><td>start</td><td></td><td>end</td><td>strand</td><td>MM</td><td>target_seq</td><td>PAM</td><td>alignment</td><td>distance</td><td>position</td><td>gene name</td><td>gene id</td></tr>
<tr><td>GU305914.1</td><td></td><td>18978</td><td>19000</td><td>+</td><td>0</td><td>GCTGCTGCACCTGGGTGTGG</td><td>TGG</td><td>||||||||[||||||||||||]PAM</td><td>NA</td><td>Intergenic</td><td>NA</td><td>NA</td></tr>
<tr><td>T924</td><td>TGCTGCTGTCCCTGCTGTCCCGG</td><td></td><td>1000</td><td></td><td></td><td></td><td></td><td></td><td></td><td></td><td></td><td></td></tr>
<tr><td>Oligo fwd</td><td>CACCTGCTGCTGTCCCTGCTGTCC</td><td></td><td></td><td></td><td></td><td></td><td></td><td></td><td></td><td></td><td></td><td></td></tr>
<tr><td>Oligo rev</td><td>AAACGGACAGCAGGGACAGCAGCA</td><td></td><td></td><td></td><td></td><td></td><td></td><td></td><td></td><td></td><td></td><td></td></tr>
<tr><td>Chromosome</td><td>start</td><td></td><td>end</td><td>strand</td><td>MM</td><td>target_seq</td><td>PAM</td><td>alignment</td><td>distance</td><td>position</td><td>gene name</td><td>gene id</td></tr>
<tr><td>GU305914.1</td><td></td><td>19007</td><td>19029</td><td>+</td><td>0</td><td>TGCTGCTGTCCCTGCTGTCC</td><td>CGG</td><td>||||||||[||||||||||||]PAM</td><td>NA</td><td>Intergenic</td><td>NA</td><td>NA</td></tr>
<tr><td>T925</td><td>AAACTGATCGACGACTCCGAAGG</td><td></td><td>1000</td><td></td><td></td><td></td><td></td><td></td><td></td><td></td><td></td><td></td></tr>
<tr><td>Oligo fwd</td><td>CACCAAACTGATCGACGACTCCGA</td><td></td><td></td><td></td><td></td><td></td><td></td><td></td><td></td><td></td><td></td><td></td></tr>
<tr><td>Oligo rev</td><td>AAACTCGGAGTCGTCGATCAGTTT</td><td></td><td></td><td></td><td></td><td></td><td></td><td></td><td></td><td></td><td></td><td></td></tr>
<tr><td>Chromosome</td><td>start</td><td></td><td>end</td><td>strand</td><td>MM</td><td>target_seq</td><td>PAM</td><td>alignment</td><td>distance</td><td>position</td><td>gene name</td><td>gene id</td></tr>
<tr><td>GU305914.1</td><td></td><td>19033</td><td>19055</td><td>+</td><td>0</td><td>AAACTGATCGACGACTCCGA</td><td>AGG</td><td>||||||||[||||||||||||]PAM</td><td>NA</td><td>Intergenic</td><td>NA</td><td>NA</td></tr>
</table>

| | | | | | | | | | | | |
|---|---|---|---|---|---|---|---|---|---|---|---|
| T926 | AACTGATCGACGACTCCGAAGGG | 1000 | | | | | | | | | |
| Oligo fwd | CACCAACTGATCGACGACTCCGAA | | | | | | | | | | |
| Oligo rev | AAACTTCGGAGTCGTCGATCAGTT | | | | | | | | | | |
| Chromosome | start | end | strand | MM | target_seq | PAM | alignment | distance | position | gene name | gene id |
| GU305914.1 | 19034 | 19056 | + | 0 | AACTGATCGACGACTCCGAA | GGG | \|\|\|\|\|\|\|\|[\|\|\|\|\|\|\|\|\|\|\|\|]PAM | NA | Intergenic | NA | NA |
| T927 | GATCGACGACTCCGAAGGGTTGG | 1000 | | | | | | | | | |
| Oligo fwd | CACCGATCGACGACTCCGAAGGGT | | | | | | | | | | |
| Oligo rev | AAACACCCTTCGGAGTCGTCGATC | | | | | | | | | | |
| Chromosome | start | end | strand | MM | target_seq | PAM | alignment | distance | position | gene name | gene id |
| GU305914.1 | 19038 | 19060 | + | 0 | GATCGACGACTCCGAAGGGT | TGG | \|\|\|\|\|\|\|\|[\|\|\|\|\|\|\|\|\|\|\|\|]PAM | NA | Intergenic | NA | NA |
| T928 | GACTCCGAAGGGTTGGACGACGG | 1000 | | | | | | | | | |
| Oligo fwd | CACCGACTCCGAAGGGTTGGACGA | | | | | | | | | | |
| Oligo rev | AAACTCGTCCAACCCTTCGGAGTC | | | | | | | | | | |
| Chromosome | start | end | strand | MM | target_seq | PAM | alignment | distance | position | gene name | gene id |
| GU305914.1 | 19045 | 19067 | + | 0 | GACTCCGAAGGGTTGGACGA | CGG | \|\|\|\|\|\|\|\|[\|\|\|\|\|\|\|\|\|\|\|\|]PAM | NA | Intergenic | NA | NA |
| T929 | TCCGAAGGGTTGGACGACGGCGG | 1000 | | | | | | | | | |
| Oligo fwd | CACCTCCGAAGGGTTGGACGACGG | | | | | | | | | | |
| Oligo rev | AAACCCGTCGTCCAACCCTTCGGA | | | | | | | | | | |
| Chromosome | start | end | strand | MM | target_seq | PAM | alignment | distance | position | gene name | gene id |
| GU305914.1 | 19048 | 19070 | + | 0 | TCCGAAGGGTTGGACGACGG | CGG | \|\|\|\|\|\|\|\|[\|\|\|\|\|\|\|\|\|\|\|\|]PAM | NA | Intergenic | NA | NA |
| T930 | GAAGGGTTGGACGACGGCGGCGG | 1000 | | | | | | | | | |
| Oligo fwd | CACCGAAGGGTTGGACGACGGCGG | | | | | | | | | | |
| Oligo rev | AAACCCGCCGTCGTCCAACCCTTC | | | | | | | | | | |
| Chromosome | start | end | strand | MM | target_seq | PAM | alignment | distance | position | gene name | gene id |
| GU305914.1 | 19051 | 19073 | + | 0 | GAAGGGTTGGACGACGGCGG | CGG | \|\|\|\|\|\|\|\|[\|\|\|\|\|\|\|\|\|\|\|\|]PAM | NA | Intergenic | NA | NA |
| T931 | GGGTTGGACGACGGCGGCGGTGG | 1000 | | | | | | | | | |
| Oligo fwd | CACCGGGTTGGACGACGGCGGCGG | | | | | | | | | | |
| Oligo rev | AAACCCGCCGCCGTCGTCCAACCC | | | | | | | | | | |
| Chromosome | start | end | strand | MM | target_seq | PAM | alignment | distance | position | gene name | gene id |
| GU305914.1 | 19054 | 19076 | + | 0 | GGGTTGGACGACGGCGGCGG | TGG | \|\|\|\|\|\|\|\|[\|\|\|\|\|\|\|\|\|\|\|\|]PAM | NA | Intergenic | NA | NA |
| GU305914.1 | 110629 | 110651 | - | 4 | GTGCAGGACGCCGGCGGCGG | AGG | \|-\|--\|\|\|[\|\|-\|\|\|\|\|\|\|\|\|]PAM | NA | Intergenic | NA | NA |
| GU305914.1 | 35650 | 35672 | + | 4 | GGGATCCACGGCGGCGGCGG | CGG | \|\|\|-\|--\|[\|\|-\|\|\|\|\|\|\|\|\|]PAM | NA | Intergenic | NA | NA |
| T932 | GGCCCACATCTGTGACGACGAGG | 1000 | | | | | | | | | |
| Oligo fwd | CACCGGCCCACATCTGTGACGACG | | | | | | | | | | |
| Oligo rev | AAACCGTCGTCACAGATGTGGGCC | | | | | | | | | | |
| Chromosome | start | end | strand | MM | target_seq | PAM | alignment | distance | position | gene name | gene id |
| GU305914.1 | 19075 | 19097 | + | 0 | GGCCCACATCTGTGACGACG | AGG | \|\|\|\|\|\|\|\|[\|\|\|\|\|\|\|\|\|\|\|\|]PAM | NA | Intergenic | NA | NA |
| GU305914.1 | 169375 | 169397 | - | 4 | GGCCCGCGTCTGTCAGGACG | TGG | \|\|\|\|\|-\|-[\|\|\|\|\|-\|-\|\|\|\|]PAM | NA | Intergenic | NA | NA |
| T933 | CCACATCTGTGACGACGAGGAGG | 1000 | | | | | | | | | |
| Oligo fwd | CACCCCACATCTGTGACGACGAGG | | | | | | | | | | |
| Oligo rev | AAACCCTCGTCGTCACAGATGTGG | | | | | | | | | | |
| Chromosome | start | end | strand | MM | target_seq | PAM | alignment | distance | position | gene name | gene id |
| GU305914.1 | 19078 | 19100 | + | 0 | CCACATCTGTGACGACGAGG | AGG | \|\|\|\|\|\|\|\|[\|\|\|\|\|\|\|\|\|\|\|\|]PAM | NA | Intergenic | NA | NA |
| T934 | ACGAGGAGGAGACCGCGCCCAGG | 1000 | | | | | | | | | |
| Oligo fwd | CACCACGAGGAGGAGACCGCGCCC | | | | | | | | | | |
| Oligo rev | AAACGGGCGCGGTCTCCTCCTCGT | | | | | | | | | | |
| Chromosome | start | end | strand | MM | target_seq | PAM | alignment | distance | position | gene name | gene id |
| GU305914.1 | 19092 | 19114 | + | 0 | ACGAGGAGGAGACCGCGCCC | AGG | \|\|\|\|\|\|\|\|[\|\|\|\|\|\|\|\|\|\|\|\|]PAM | NA | Intergenic | NA | NA |
| T935 | GAGGAGACCGCGCCCAGGTTCGG | 1000 | | | | | | | | | |
| Oligo fwd | CACCGAGGAGACCGCGCCCAGGTT | | | | | | | | | | |
| Oligo rev | AAACAACCTGGGCGCGGTCTCCTC | | | | | | | | | | |
| Chromosome | start | end | strand | MM | target_seq | PAM | alignment | distance | position | gene name | gene id |
| GU305914.1 | 19097 | 19119 | + | 0 | GAGGAGACCGCGCCCAGGTT | CGG | \|\|\|\|\|\|\|\|[\|\|\|\|\|\|\|\|\|\|\|\|]PAM | NA | Intergenic | NA | NA |
| T936 | AGGAGACCGCGCCCAGGTTCGGG | 1000 | | | | | | | | | |
| Oligo fwd | CACCAGGAGACCGCGCCCAGGTTC | | | | | | | | | | |
| Oligo rev | AAACGAACCTGGGCGCGGTCTCCT | | | | | | | | | | |
| Chromosome | start | end | strand | MM | target_seq | PAM | alignment | distance | position | gene name | gene id |
| GU305914.1 | 19098 | 19120 | + | 0 | AGGAGACCGCGCCCAGGTTC | GGG | \|\|\|\|\|\|\|\|[\|\|\|\|\|\|\|\|\|\|\|\|]PAM | NA | Intergenic | NA | NA |
| T937 | GGTCACTGACCGAGATGATCCGG | 1000 | | | | | | | | | |
| Oligo fwd | CACCGGTCACTGACCGAGATGATC | | | | | | | | | | |
| Oligo rev | AAACGATCATCTCGGTCAGTGACC | | | | | | | | | | |
| Chromosome | start | end | strand | MM | target_seq | PAM | alignment | distance | position | gene name | gene id |
| GU305914.1 | 19119 | 19141 | + | 0 | GGTCACTGACCGAGATGATC | CGG | \|\|\|\|\|\|\|\|[\|\|\|\|\|\|\|\|\|\|\|\|]PAM | NA | Intergenic | NA | NA |
| T938 | GAGATGATCCGGCCACGAGACGG | 1000 | | | | | | | | | |
| Oligo fwd | CACCGAGATGATCCGGCCACGAGA | | | | | | | | | | |
| Oligo rev | AAACTCTCGTGGCCGGATCATCTC | | | | | | | | | | |
| Chromosome | start | end | strand | MM | target_seq | PAM | alignment | distance | position | gene name | gene id |
| GU305914.1 | 19130 | 19152 | + | 0 | GAGATGATCCGGCCACGAGA | CGG | \|\|\|\|\|\|\|\|[\|\|\|\|\|\|\|\|\|\|\|\|]PAM | NA | Intergenic | NA | NA |
| T939 | GATCCGGCCACGAGACGGTCCGG | 1000 | | | | | | | | | |
| Oligo fwd | CACCGATCCGGCCACGAGACGGTC | | | | | | | | | | |
| Oligo rev | AAACGACCGTCTCGTGGCCGGATC | | | | | | | | | | |
| Chromosome | start | end | strand | MM | target_seq | PAM | alignment | distance | position | gene name | gene id |
| GU305914.1 | 19135 | 19157 | + | 0 | GATCCGGCCACGAGACGGTC | CGG | \|\|\|\|\|\|\|\|[\|\|\|\|\|\|\|\|\|\|\|\|]PAM | NA | Intergenic | NA | NA |
| T940 | CGGTCCGGAAGAAGATGCCGAGG | 1000 | | | | | | | | | |

<table>
<tr><td>Oligo fwd</td><td>CACCCGGTCCGGAAGAAGATGCCG</td><td></td><td></td><td></td><td></td><td></td><td></td><td></td><td></td><td></td><td></td></tr>
<tr><td>Oligo rev</td><td>AAACCGGCATCTTCTTCCGGACCG</td><td></td><td></td><td></td><td></td><td></td><td></td><td></td><td></td><td></td><td></td></tr>
<tr><td>Chromosome</td><td>start</td><td>end</td><td>strand</td><td>MM</td><td>target_seq</td><td>PAM</td><td>alignment</td><td>distance</td><td>position</td><td>gene name</td><td>gene id</td></tr>
<tr><td>GU305914.1</td><td>19150</td><td>19172</td><td>+</td><td>0</td><td>CGGTCCGGAAGAAGATGCCG</td><td>AGG</td><td>||||||||[||||||||||||]PAM</td><td>NA</td><td>Intergenic</td><td>NA</td><td>NA</td></tr>
<tr><td></td><td></td><td></td><td></td><td></td><td></td><td></td><td></td><td></td><td></td><td></td><td></td></tr>
<tr><td>T941</td><td>GCCGATGCCGCTTTCTGCCGAGG</td><td>1000</td><td></td><td></td><td></td><td></td><td></td><td></td><td></td><td></td><td></td></tr>
<tr><td>Oligo fwd</td><td>CACCGCCGATGCCGCTTTCTGCCG</td><td></td><td></td><td></td><td></td><td></td><td></td><td></td><td></td><td></td><td></td></tr>
<tr><td>Oligo rev</td><td>AAACCGGCAGAAAGCGGCATCGGC</td><td></td><td></td><td></td><td></td><td></td><td></td><td></td><td></td><td></td><td></td></tr>
<tr><td>Chromosome</td><td>start</td><td>end</td><td>strand</td><td>MM</td><td>target_seq</td><td>PAM</td><td>alignment</td><td>distance</td><td>position</td><td>gene name</td><td>gene id</td></tr>
<tr><td>GU305914.1</td><td>19187</td><td>19209</td><td>+</td><td>0</td><td>GCCGATGCCGCTTTCTGCCG</td><td>AGG</td><td>||||||||[||||||||||||]PAM</td><td>NA</td><td>Intergenic</td><td>NA</td><td>NA</td></tr>
<tr><td></td><td></td><td></td><td></td><td></td><td></td><td></td><td></td><td></td><td></td><td></td><td></td></tr>
<tr><td>T942</td><td>GATGCCGCTTTCTGCCGAGGCGG</td><td>1000</td><td></td><td></td><td></td><td></td><td></td><td></td><td></td><td></td><td></td></tr>
<tr><td>Oligo fwd</td><td>CACCGATGCCGCTTTCTGCCGAGG</td><td></td><td></td><td></td><td></td><td></td><td></td><td></td><td></td><td></td><td></td></tr>
<tr><td>Oligo rev</td><td>AAACCCTCGGCAGAAAGCGGCATC</td><td></td><td></td><td></td><td></td><td></td><td></td><td></td><td></td><td></td><td></td></tr>
<tr><td>Chromosome</td><td>start</td><td>end</td><td>strand</td><td>MM</td><td>target_seq</td><td>PAM</td><td>alignment</td><td>distance</td><td>position</td><td>gene name</td><td>gene id</td></tr>
<tr><td>GU305914.1</td><td>19190</td><td>19212</td><td>+</td><td>0</td><td>GATGCCGCTTTCTGCCGAGG</td><td>CGG</td><td>||||||||[||||||||||||]PAM</td><td>NA</td><td>Intergenic</td><td>NA</td><td>NA</td></tr>
<tr><td></td><td></td><td></td><td></td><td></td><td></td><td></td><td></td><td></td><td></td><td></td><td></td></tr>
<tr><td>T943</td><td>ATGCCGCTTTCTGCCGAGGCGGG</td><td>1000</td><td></td><td></td><td></td><td></td><td></td><td></td><td></td><td></td><td></td></tr>
<tr><td>Oligo fwd</td><td>CACCATGCCGCTTTCTGCCGAGGC</td><td></td><td></td><td></td><td></td><td></td><td></td><td></td><td></td><td></td><td></td></tr>
<tr><td>Oligo rev</td><td>AAACGCCTCGGCAGAAAGCGGCAT</td><td></td><td></td><td></td><td></td><td></td><td></td><td></td><td></td><td></td><td></td></tr>
<tr><td>Chromosome</td><td>start</td><td>end</td><td>strand</td><td>MM</td><td>target_seq</td><td>PAM</td><td>alignment</td><td>distance</td><td>position</td><td>gene name</td><td>gene id</td></tr>
<tr><td>GU305914.1</td><td>19191</td><td>19213</td><td>+</td><td>0</td><td>ATGCCGCTTTCTGCCGAGGC</td><td>GGG</td><td>||||||||[||||||||||||]PAM</td><td>NA</td><td>Intergenic</td><td>NA</td><td>NA</td></tr>
<tr><td></td><td></td><td></td><td></td><td></td><td></td><td></td><td></td><td></td><td></td><td></td><td></td></tr>
<tr><td>T944</td><td>CTGCCGAGGCGGGACACCTCCGG</td><td>1000</td><td></td><td></td><td></td><td></td><td></td><td></td><td></td><td></td><td></td></tr>
<tr><td>Oligo fwd</td><td>CACCCTGCCGAGGCGGGACACCTC</td><td></td><td></td><td></td><td></td><td></td><td></td><td></td><td></td><td></td><td></td></tr>
<tr><td>Oligo rev</td><td>AAACGAGGTGTCCCGCCTCGGCAG</td><td></td><td></td><td></td><td></td><td></td><td></td><td></td><td></td><td></td><td></td></tr>
<tr><td>Chromosome</td><td>start</td><td>end</td><td>strand</td><td>MM</td><td>target_seq</td><td>PAM</td><td>alignment</td><td>distance</td><td>position</td><td>gene name</td><td>gene id</td></tr>
<tr><td>GU305914.1</td><td>19201</td><td>19223</td><td>+</td><td>0</td><td>CTGCCGAGGCGGGACACCTC</td><td>CGG</td><td>||||||||[||||||||||||]PAM</td><td>NA</td><td>Intergenic</td><td>NA</td><td>NA</td></tr>
<tr><td></td><td></td><td></td><td></td><td></td><td></td><td></td><td></td><td></td><td></td><td></td><td></td></tr>
<tr><td>T945</td><td>TCCGGCCGCTCTGCGCGAGCCGG</td><td>1000</td><td></td><td></td><td></td><td></td><td></td><td></td><td></td><td></td><td></td></tr>
<tr><td>Oligo fwd</td><td>CACCTCCGGCCGCTCTGCGCGAGC</td><td></td><td></td><td></td><td></td><td></td><td></td><td></td><td></td><td></td><td></td></tr>
<tr><td>Oligo rev</td><td>AAACGCTCGCGCAGAGCGGCCGGA</td><td></td><td></td><td></td><td></td><td></td><td></td><td></td><td></td><td></td><td></td></tr>
<tr><td>Chromosome</td><td>start</td><td>end</td><td>strand</td><td>MM</td><td>target_seq</td><td>PAM</td><td>alignment</td><td>distance</td><td>position</td><td>gene name</td><td>gene id</td></tr>
<tr><td>GU305914.1</td><td>19219</td><td>19241</td><td>+</td><td>0</td><td>TCCGGCCGCTCTGCGCGAGC</td><td>CGG</td><td>||||||||[||||||||||||]PAM</td><td>NA</td><td>Intergenic</td><td>NA</td><td>NA</td></tr>
<tr><td></td><td></td><td></td><td></td><td></td><td></td><td></td><td></td><td></td><td></td><td></td><td></td></tr>
<tr><td>T946</td><td>GGCCGCTCTGCGCGAGCCGGTGG</td><td>1000</td><td></td><td></td><td></td><td></td><td></td><td></td><td></td><td></td><td></td></tr>
<tr><td>Oligo fwd</td><td>CACCGGCCGCTCTGCGCGAGCCGG</td><td></td><td></td><td></td><td></td><td></td><td></td><td></td><td></td><td></td><td></td></tr>
<tr><td>Oligo rev</td><td>AAACCCGGCTCGCGCAGAGCGGCC</td><td></td><td></td><td></td><td></td><td></td><td></td><td></td><td></td><td></td><td></td></tr>
<tr><td>Chromosome</td><td>start</td><td>end</td><td>strand</td><td>MM</td><td>target_seq</td><td>PAM</td><td>alignment</td><td>distance</td><td>position</td><td>gene name</td><td>gene id</td></tr>
<tr><td>GU305914.1</td><td>19222</td><td>19244</td><td>+</td><td>0</td><td>GGCCGCTCTGCGCGAGCCGG</td><td>TGG</td><td>||||||||[||||||||||||]PAM</td><td>NA</td><td>Intergenic</td><td>NA</td><td>NA</td></tr>
<tr><td></td><td></td><td></td><td></td><td></td><td></td><td></td><td></td><td></td><td></td><td></td><td></td></tr>
<tr><td>T947</td><td>GAGCCGGTGGTCGTCTCCACAGG</td><td>1000</td><td></td><td></td><td></td><td></td><td></td><td></td><td></td><td></td><td></td></tr>
<tr><td>Oligo fwd</td><td>CACCGAGCCGGTGGTCGTCTCCAC</td><td></td><td></td><td></td><td></td><td></td><td></td><td></td><td></td><td></td><td></td></tr>
<tr><td>Oligo rev</td><td>AAACGTGGAGACGACCACCGGCTC</td><td></td><td></td><td></td><td></td><td></td><td></td><td></td><td></td><td></td><td></td></tr>
<tr><td>Chromosome</td><td>start</td><td>end</td><td>strand</td><td>MM</td><td>target_seq</td><td>PAM</td><td>alignment</td><td>distance</td><td>position</td><td>gene name</td><td>gene id</td></tr>
<tr><td>GU305914.1</td><td>19235</td><td>19257</td><td>+</td><td>0</td><td>GAGCCGGTGGTCGTCTCCAC</td><td>AGG</td><td>||||||||[||||||||||||]PAM</td><td>NA</td><td>Intergenic</td><td>NA</td><td>NA</td></tr>
<tr><td></td><td></td><td></td><td></td><td></td><td></td><td></td><td></td><td></td><td></td><td></td><td></td></tr>
<tr><td>T948</td><td>CTTTTTCTTTCCTCGCGCGACGG</td><td>1000</td><td></td><td></td><td></td><td></td><td></td><td></td><td></td><td></td><td></td></tr>
<tr><td>Oligo fwd</td><td>CACCCTTTTTCTTTCCTCGCGCGA</td><td></td><td></td><td></td><td></td><td></td><td></td><td></td><td></td><td></td><td></td></tr>
<tr><td>Oligo rev</td><td>AAACTCGCGCGAGGAAAGAAAAAG</td><td></td><td></td><td></td><td></td><td></td><td></td><td></td><td></td><td></td><td></td></tr>
<tr><td>Chromosome</td><td>start</td><td>end</td><td>strand</td><td>MM</td><td>target_seq</td><td>PAM</td><td>alignment</td><td>distance</td><td>position</td><td>gene name</td><td>gene id</td></tr>
<tr><td>GU305914.1</td><td>19291</td><td>19313</td><td>+</td><td>0</td><td>CTTTTTCTTTCCTCGCGCGA</td><td>CGG</td><td>||||||||[||||||||||||]PAM</td><td>NA</td><td>Intergenic</td><td>NA</td><td>NA</td></tr>
<tr><td></td><td></td><td></td><td></td><td></td><td></td><td></td><td></td><td></td><td></td><td></td><td></td></tr>
<tr><td>T949</td><td>TTTTTCTTTCCTCGCGCGACGGG</td><td>1000</td><td></td><td></td><td></td><td></td><td></td><td></td><td></td><td></td><td></td></tr>
<tr><td>Oligo fwd</td><td>CACCTTTTTCTTTCCTCGCGCGAC</td><td></td><td></td><td></td><td></td><td></td><td></td><td></td><td></td><td></td><td></td></tr>
<tr><td>Oligo rev</td><td>AAACGTCGCGCGAGGAAAGAAAAA</td><td></td><td></td><td></td><td></td><td></td><td></td><td></td><td></td><td></td><td></td></tr>
<tr><td>Chromosome</td><td>start</td><td>end</td><td>strand</td><td>MM</td><td>target_seq</td><td>PAM</td><td>alignment</td><td>distance</td><td>position</td><td>gene name</td><td>gene id</td></tr>
<tr><td>GU305914.1</td><td>19292</td><td>19314</td><td>+</td><td>0</td><td>TTTTTCTTTCCTCGCGCGAC</td><td>GGG</td><td>||||||||[||||||||||||]PAM</td><td>NA</td><td>Intergenic</td><td>NA</td><td>NA</td></tr>
<tr><td></td><td></td><td></td><td></td><td></td><td></td><td></td><td></td><td></td><td></td><td></td><td></td></tr>
<tr><td>T950</td><td>TTTTCTTTCCTCGCGCGACGGGG</td><td>1000</td><td></td><td></td><td></td><td></td><td></td><td></td><td></td><td></td><td></td></tr>
<tr><td>Oligo fwd</td><td>CACCTTTTCTTTCCTCGCGCGACG</td><td></td><td></td><td></td><td></td><td></td><td></td><td></td><td></td><td></td><td></td></tr>
<tr><td>Oligo rev</td><td>AAACCGTCGCGCGAGGAAAGAAAA</td><td></td><td></td><td></td><td></td><td></td><td></td><td></td><td></td><td></td><td></td></tr>
<tr><td>Chromosome</td><td>start</td><td>end</td><td>strand</td><td>MM</td><td>target_seq</td><td>PAM</td><td>alignment</td><td>distance</td><td>position</td><td>gene name</td><td>gene id</td></tr>
<tr><td>GU305914.1</td><td>19293</td><td>19315</td><td>+</td><td>0</td><td>TTTTCTTTCCTCGCGCGACG</td><td>GGG</td><td>||||||||[||||||||||||]PAM</td><td>NA</td><td>Intergenic</td><td>NA</td><td>NA</td></tr>
<tr><td></td><td></td><td></td><td></td><td></td><td></td><td></td><td></td><td></td><td></td><td></td><td></td></tr>
<tr><td>T951</td><td>TTTCTTTCCTCGCGCGACGGGGG</td><td>1000</td><td></td><td></td><td></td><td></td><td></td><td></td><td></td><td></td><td></td></tr>
<tr><td>Oligo fwd</td><td>CACCTTTCTTTCCTCGCGCGACGG</td><td></td><td></td><td></td><td></td><td></td><td></td><td></td><td></td><td></td><td></td></tr>
<tr><td>Oligo rev</td><td>AAACCCGTCGCGCGAGGAAAGAAA</td><td></td><td></td><td></td><td></td><td></td><td></td><td></td><td></td><td></td><td></td></tr>
<tr><td>Chromosome</td><td>start</td><td>end</td><td>strand</td><td>MM</td><td>target_seq</td><td>PAM</td><td>alignment</td><td>distance</td><td>position</td><td>gene name</td><td>gene id</td></tr>
<tr><td>GU305914.1</td><td>19294</td><td>19316</td><td>+</td><td>0</td><td>TTTCTTTCCTCGCGCGACGG</td><td>GGG</td><td>||||||||[||||||||||||]PAM</td><td>NA</td><td>Intergenic</td><td>NA</td><td>NA</td></tr>
<tr><td></td><td></td><td></td><td></td><td></td><td></td><td></td><td></td><td></td><td></td><td></td><td></td></tr>
<tr><td>T952</td><td>TCCTCGCGCGACGGGGGATCTGG</td><td>1000</td><td></td><td></td><td></td><td></td><td></td><td></td><td></td><td></td><td></td></tr>
<tr><td>Oligo fwd</td><td>CACCTCCTCGCGCGACGGGGGATC</td><td></td><td></td><td></td><td></td><td></td><td></td><td></td><td></td><td></td><td></td></tr>
<tr><td>Oligo rev</td><td>AAACGATCCCCCGTCGCGCGAGGA</td><td></td><td></td><td></td><td></td><td></td><td></td><td></td><td></td><td></td><td></td></tr>
<tr><td>Chromosome</td><td>start</td><td>end</td><td>strand</td><td>MM</td><td>target_seq</td><td>PAM</td><td>alignment</td><td>distance</td><td>position</td><td>gene name</td><td>gene id</td></tr>
<tr><td>GU305914.1</td><td>19300</td><td>19322</td><td>+</td><td>0</td><td>TCCTCGCGCGACGGGGGATC</td><td>TGG</td><td>||||||||[||||||||||||]PAM</td><td>NA</td><td>Intergenic</td><td>NA</td><td>NA</td></tr>
<tr><td></td><td></td><td></td><td></td><td></td><td></td><td></td><td></td><td></td><td></td><td></td><td></td></tr>
<tr><td>T953</td><td>GACGGGGGATCTGGACCGTCCGG</td><td>1000</td><td></td><td></td><td></td><td></td><td></td><td></td><td></td><td></td><td></td></tr>
<tr><td>Oligo fwd</td><td>CACCGACGGGGGATCTGGACCGTC</td><td></td><td></td><td></td><td></td><td></td><td></td><td></td><td></td><td></td><td></td></tr>
<tr><td>Oligo rev</td><td>AAACGACGGTCCAGATCCCCCGTC</td><td></td><td></td><td></td><td></td><td></td><td></td><td></td><td></td><td></td><td></td></tr>
<tr><td>Chromosome</td><td>start</td><td>end</td><td>strand</td><td>MM</td><td>target_seq</td><td>PAM</td><td>alignment</td><td>distance</td><td>position</td><td>gene name</td><td>gene id</td></tr>
<tr><td>GU305914.1</td><td>19309</td><td>19331</td><td>+</td><td>0</td><td>GACGGGGGATCTGGACCGTC</td><td>CGG</td><td>||||||||[||||||||||||]PAM</td><td>NA</td><td>Intergenic</td><td>NA</td><td>NA</td></tr>
<tr><td></td><td></td><td></td><td></td><td></td><td></td><td></td><td></td><td></td><td></td><td></td><td></td></tr>
<tr><td>T954</td><td>GGATCTGGACCGTCCGGCGACGG</td><td>1000</td><td></td><td></td><td></td><td></td><td></td><td></td><td></td><td></td><td></td></tr>
<tr><td>Oligo fwd</td><td>CACCGGATCTGGACCGTCCGGCGA</td><td></td><td></td><td></td><td></td><td></td><td></td><td></td><td></td><td></td><td></td></tr>
<tr><td>Oligo rev</td><td>AAACTCGCCGGACGGTCCAGATCC</td><td></td><td></td><td></td><td></td><td></td><td></td><td></td><td></td><td></td><td></td></tr>
<tr><td>Chromosome</td><td>start</td><td>end</td><td>strand</td><td>MM</td><td>target_seq</td><td>PAM</td><td>alignment</td><td>distance</td><td>position</td><td>gene name</td><td>gene id</td></tr>
<tr><td>GU305914.1</td><td>19315</td><td>19337</td><td>+</td><td>0</td><td>GGATCTGGACCGTCCGGCGA</td><td>CGG</td><td>||||||||[||||||||||||]PAM</td><td>NA</td><td>Intergenic</td><td>NA</td><td>NA</td></tr>
<tr><td>GU305914.1</td><td>26912</td><td>26934</td><td>-</td><td>4</td><td>GCCTCTGGACCGGCCGGCGG</td><td>TGG</td><td>|--|||||[||||-||||||-]PAM</td><td>NA</td><td>Intergenic</td><td>NA</td><td>NA</td></tr>
</table>

<table>
<tr><td>T955</td><td>GATCTGGACCGTCCGGCGACGGG</td><td>1000</td><td></td><td></td><td></td><td></td><td></td><td></td><td></td><td></td><td></td></tr>
<tr><td>Oligo fwd</td><td>CACCGATCTGGACCGTCCGGCGAC</td><td></td><td></td><td></td><td></td><td></td><td></td><td></td><td></td><td></td><td></td></tr>
<tr><td>Oligo rev</td><td>AAACGTCGCCGGACGGTCCAGATC</td><td></td><td></td><td></td><td></td><td></td><td></td><td></td><td></td><td></td><td></td></tr>
<tr><td>Chromosome</td><td>start</td><td>end</td><td>strand</td><td>MM</td><td>target_seq</td><td>PAM</td><td>alignment</td><td>distance</td><td>position</td><td>gene name</td><td>gene id</td></tr>
<tr><td>GU305914.1</td><td>19316</td><td>19338</td><td>+</td><td>0</td><td>GATCTGGACCGTCCGGCGAC</td><td>GGG</td><td>||||||||[||||||||||||]PAM</td><td>NA</td><td>Intergenic</td><td>NA</td><td>NA</td></tr>
<tr><td>T956</td><td>GCGACGGGCGTCATCCTACCCGG</td><td>1000</td><td></td><td></td><td></td><td></td><td></td><td></td><td></td><td></td><td></td></tr>
<tr><td>Oligo fwd</td><td>CACCGCGACGGGCGTCATCCTACC</td><td></td><td></td><td></td><td></td><td></td><td></td><td></td><td></td><td></td><td></td></tr>
<tr><td>Oligo rev</td><td>AAACGGTAGGATGACGCCCGTCGC</td><td></td><td></td><td></td><td></td><td></td><td></td><td></td><td></td><td></td><td></td></tr>
<tr><td>Chromosome</td><td>start</td><td>end</td><td>strand</td><td>MM</td><td>target_seq</td><td>PAM</td><td>alignment</td><td>distance</td><td>position</td><td>gene name</td><td>gene id</td></tr>
<tr><td>GU305914.1</td><td>19331</td><td>19353</td><td>+</td><td>0</td><td>GCGACGGGCGTCATCCTACC</td><td>CGG</td><td>||||||||[||||||||||||]PAM</td><td>NA</td><td>Intergenic</td><td>NA</td><td>NA</td></tr>
<tr><td>T957</td><td>ACGGGCGTCATCCTACCCGGAGG</td><td>1000</td><td></td><td></td><td></td><td></td><td></td><td></td><td></td><td></td><td></td></tr>
<tr><td>Oligo fwd</td><td>CACCACGGGCGTCATCCTACCCGG</td><td></td><td></td><td></td><td></td><td></td><td></td><td></td><td></td><td></td><td></td></tr>
<tr><td>Oligo rev</td><td>AAACCCGGGTAGGATGACGCCCGT</td><td></td><td></td><td></td><td></td><td></td><td></td><td></td><td></td><td></td><td></td></tr>
<tr><td>Chromosome</td><td>start</td><td>end</td><td>strand</td><td>MM</td><td>target_seq</td><td>PAM</td><td>alignment</td><td>distance</td><td>position</td><td>gene name</td><td>gene id</td></tr>
<tr><td>GU305914.1</td><td>19334</td><td>19356</td><td>+</td><td>0</td><td>ACGGGCGTCATCCTACCCGG</td><td>AGG</td><td>||||||||[||||||||||||]PAM</td><td>NA</td><td>Intergenic</td><td>NA</td><td>NA</td></tr>
<tr><td>T958</td><td>GGAGGACGCACGCTACATCGCGG</td><td>1000</td><td></td><td></td><td></td><td></td><td></td><td></td><td></td><td></td><td></td></tr>
<tr><td>Oligo fwd</td><td>CACCGGAGGACGCACGCTACATCG</td><td></td><td></td><td></td><td></td><td></td><td></td><td></td><td></td><td></td><td></td></tr>
<tr><td>Oligo rev</td><td>AAACCGATGTAGCGTGCGTCCTCC</td><td></td><td></td><td></td><td></td><td></td><td></td><td></td><td></td><td></td><td></td></tr>
<tr><td>Chromosome</td><td>start</td><td>end</td><td>strand</td><td>MM</td><td>target_seq</td><td>PAM</td><td>alignment</td><td>distance</td><td>position</td><td>gene name</td><td>gene id</td></tr>
<tr><td>GU305914.1</td><td>19352</td><td>19374</td><td>+</td><td>0</td><td>GGAGGACGCACGCTACATCG</td><td>CGG</td><td>||||||||[||||||||||||]PAM</td><td>NA</td><td>Intergenic</td><td>NA</td><td>NA</td></tr>
<tr><td>T959</td><td>GGACGCACGCTACATCGCGGAGG</td><td>1000</td><td></td><td></td><td></td><td></td><td></td><td></td><td></td><td></td><td></td></tr>
<tr><td>Oligo fwd</td><td>CACCGGACGCACGCTACATCGCGG</td><td></td><td></td><td></td><td></td><td></td><td></td><td></td><td></td><td></td><td></td></tr>
<tr><td>Oligo rev</td><td>AAACCCGCGATGTAGCGTGCGTCC</td><td></td><td></td><td></td><td></td><td></td><td></td><td></td><td></td><td></td><td></td></tr>
<tr><td>Chromosome</td><td>start</td><td>end</td><td>strand</td><td>MM</td><td>target_seq</td><td>PAM</td><td>alignment</td><td>distance</td><td>position</td><td>gene name</td><td>gene id</td></tr>
<tr><td>GU305914.1</td><td>19355</td><td>19377</td><td>+</td><td>0</td><td>GGACGCACGCTACATCGCGG</td><td>AGG</td><td>||||||||[||||||||||||]PAM</td><td>NA</td><td>Intergenic</td><td>NA</td><td>NA</td></tr>
<tr><td>T960</td><td>GCTACATCGCGGAGGTCTCTCGG</td><td>1000</td><td></td><td></td><td></td><td></td><td></td><td></td><td></td><td></td><td></td></tr>
<tr><td>Oligo fwd</td><td>CACCGCTACATCGCGGAGGTCTCT</td><td></td><td></td><td></td><td></td><td></td><td></td><td></td><td></td><td></td><td></td></tr>
<tr><td>Oligo rev</td><td>AAACAGAGACCTCCGCGATGTAGC</td><td></td><td></td><td></td><td></td><td></td><td></td><td></td><td></td><td></td><td></td></tr>
<tr><td>Chromosome</td><td>start</td><td>end</td><td>strand</td><td>MM</td><td>target_seq</td><td>PAM</td><td>alignment</td><td>distance</td><td>position</td><td>gene name</td><td>gene id</td></tr>
<tr><td>GU305914.1</td><td>19363</td><td>19385</td><td>+</td><td>0</td><td>GCTACATCGCGGAGGTCTCT</td><td>CGG</td><td>||||||||[||||||||||||]PAM</td><td>NA</td><td>Intergenic</td><td>NA</td><td>NA</td></tr>
<tr><td>T961</td><td>CTACATCGCGGAGGTCTCTCGGG</td><td>1000</td><td></td><td></td><td></td><td></td><td></td><td></td><td></td><td></td><td></td></tr>
<tr><td>Oligo fwd</td><td>CACCCTACATCGCGGAGGTCTCTC</td><td></td><td></td><td></td><td></td><td></td><td></td><td></td><td></td><td></td><td></td></tr>
<tr><td>Oligo rev</td><td>AAACGAGAGACCTCCGCGATGTAG</td><td></td><td></td><td></td><td></td><td></td><td></td><td></td><td></td><td></td><td></td></tr>
<tr><td>Chromosome</td><td>start</td><td>end</td><td>strand</td><td>MM</td><td>target_seq</td><td>PAM</td><td>alignment</td><td>distance</td><td>position</td><td>gene name</td><td>gene id</td></tr>
<tr><td>GU305914.1</td><td>19364</td><td>19386</td><td>+</td><td>0</td><td>CTACATCGCGGAGGTCTCTC</td><td>GGG</td><td>||||||||[||||||||||||]PAM</td><td>NA</td><td>Intergenic</td><td>NA</td><td>NA</td></tr>
<tr><td>T962</td><td>CATCGCGGAGGTCTCTCGGGAGG</td><td>1000</td><td></td><td></td><td></td><td></td><td></td><td></td><td></td><td></td><td></td></tr>
<tr><td>Oligo fwd</td><td>CACCCATCGCGGAGGTCTCTCGGG</td><td></td><td></td><td></td><td></td><td></td><td></td><td></td><td></td><td></td><td></td></tr>
<tr><td>Oligo rev</td><td>AAACCCCGAGAGACCTCCGCGATG</td><td></td><td></td><td></td><td></td><td></td><td></td><td></td><td></td><td></td><td></td></tr>
<tr><td>Chromosome</td><td>start</td><td>end</td><td>strand</td><td>MM</td><td>target_seq</td><td>PAM</td><td>alignment</td><td>distance</td><td>position</td><td>gene name</td><td>gene id</td></tr>
<tr><td>GU305914.1</td><td>19367</td><td>19389</td><td>+</td><td>0</td><td>CATCGCGGAGGTCTCTCGGG</td><td>AGG</td><td>||||||||[||||||||||||]PAM</td><td>NA</td><td>Intergenic</td><td>NA</td><td>NA</td></tr>
<tr><td>T963</td><td>AGGTCTCTCGGGAGGCGTCTCGG</td><td>1000</td><td></td><td></td><td></td><td></td><td></td><td></td><td></td><td></td><td></td></tr>
<tr><td>Oligo fwd</td><td>CACCAGGTCTCTCGGGAGGCGTCT</td><td></td><td></td><td></td><td></td><td></td><td></td><td></td><td></td><td></td><td></td></tr>
<tr><td>Oligo rev</td><td>AAACAGACGCCTCCCGAGAGACCT</td><td></td><td></td><td></td><td></td><td></td><td></td><td></td><td></td><td></td><td></td></tr>
<tr><td>Chromosome</td><td>start</td><td>end</td><td>strand</td><td>MM</td><td>target_seq</td><td>PAM</td><td>alignment</td><td>distance</td><td>position</td><td>gene name</td><td>gene id</td></tr>
<tr><td>GU305914.1</td><td>19375</td><td>19397</td><td>+</td><td>0</td><td>AGGTCTCTCGGGAGGCGTCT</td><td>CGG</td><td>||||||||[||||||||||||]PAM</td><td>NA</td><td>Intergenic</td><td>NA</td><td>NA</td></tr>
<tr><td>T964</td><td>AGGCGTCTCGGTCTCGCGCTCGG</td><td>1000</td><td></td><td></td><td></td><td></td><td></td><td></td><td></td><td></td><td></td></tr>
<tr><td>Oligo fwd</td><td>CACCAGGCGTCTCGGTCTCGCGCT</td><td></td><td></td><td></td><td></td><td></td><td></td><td></td><td></td><td></td><td></td></tr>
<tr><td>Oligo rev</td><td>AAACAGCGCGAGACCGAGACGCCT</td><td></td><td></td><td></td><td></td><td></td><td></td><td></td><td></td><td></td><td></td></tr>
<tr><td>Chromosome</td><td>start</td><td>end</td><td>strand</td><td>MM</td><td>target_seq</td><td>PAM</td><td>alignment</td><td>distance</td><td>position</td><td>gene name</td><td>gene id</td></tr>
<tr><td>GU305914.1</td><td>19387</td><td>19409</td><td>+</td><td>0</td><td>AGGCGTCTCGGTCTCGCGCT</td><td>CGG</td><td>||||||||[||||||||||||]PAM</td><td>NA</td><td>Intergenic</td><td>NA</td><td>NA</td></tr>
<tr><td>T965</td><td>GGCGTCTCGGTCTCGCGCTCGGG</td><td>1000</td><td></td><td></td><td></td><td></td><td></td><td></td><td></td><td></td><td></td></tr>
<tr><td>Oligo fwd</td><td>CACCGGCGTCTCGGTCTCGCGCTC</td><td></td><td></td><td></td><td></td><td></td><td></td><td></td><td></td><td></td><td></td></tr>
<tr><td>Oligo rev</td><td>AAACGAGCGCGAGACCGAGACGCC</td><td></td><td></td><td></td><td></td><td></td><td></td><td></td><td></td><td></td><td></td></tr>
<tr><td>Chromosome</td><td>start</td><td>end</td><td>strand</td><td>MM</td><td>target_seq</td><td>PAM</td><td>alignment</td><td>distance</td><td>position</td><td>gene name</td><td>gene id</td></tr>
<tr><td>GU305914.1</td><td>19388</td><td>19410</td><td>+</td><td>0</td><td>GGCGTCTCGGTCTCGCGCTC</td><td>GGG</td><td>||||||||[||||||||||||]PAM</td><td>NA</td><td>Intergenic</td><td>NA</td><td>NA</td></tr>
<tr><td>T966</td><td>CTCGGTCTCGCGCTCGGGCCTGG</td><td>1000</td><td></td><td></td><td></td><td></td><td></td><td></td><td></td><td></td><td></td></tr>
<tr><td>Oligo fwd</td><td>CACCCTCGGTCTCGCGCTCGGGCC</td><td></td><td></td><td></td><td></td><td></td><td></td><td></td><td></td><td></td><td></td></tr>
<tr><td>Oligo rev</td><td>AAACGGCCCGAGCGCGAGACCGAG</td><td></td><td></td><td></td><td></td><td></td><td></td><td></td><td></td><td></td><td></td></tr>
<tr><td>Chromosome</td><td>start</td><td>end</td><td>strand</td><td>MM</td><td>target_seq</td><td>PAM</td><td>alignment</td><td>distance</td><td>position</td><td>gene name</td><td>gene id</td></tr>
<tr><td>GU305914.1</td><td>19393</td><td>19415</td><td>+</td><td>0</td><td>CTCGGTCTCGCGCTCGGGCC</td><td>TGG</td><td>||||||||[||||||||||||]PAM</td><td>NA</td><td>Intergenic</td><td>NA</td><td>NA</td></tr>
<tr><td>T967</td><td>TCGGTCTCGCGCTCGGGCCTGGG</td><td>1000</td><td></td><td></td><td></td><td></td><td></td><td></td><td></td><td></td><td></td></tr>
<tr><td>Oligo fwd</td><td>CACCTCGGTCTCGCGCTCGGGCCT</td><td></td><td></td><td></td><td></td><td></td><td></td><td></td><td></td><td></td><td></td></tr>
<tr><td>Oligo rev</td><td>AAACAGGCCCGAGCGCGAGACCGA</td><td></td><td></td><td></td><td></td><td></td><td></td><td></td><td></td><td></td><td></td></tr>
<tr><td>Chromosome</td><td>start</td><td>end</td><td>strand</td><td>MM</td><td>target_seq</td><td>PAM</td><td>alignment</td><td>distance</td><td>position</td><td>gene name</td><td>gene id</td></tr>
<tr><td>GU305914.1</td><td>19394</td><td>19416</td><td>+</td><td>0</td><td>TCGGTCTCGCGCTCGGGCCT</td><td>GGG</td><td>||||||||[||||||||||||]PAM</td><td>NA</td><td>Intergenic</td><td>NA</td><td>NA</td></tr>
<tr><td>T968</td><td>GCGCTCGGGCCTGGGTAGTATGG</td><td>1000</td><td></td><td></td><td></td><td></td><td></td><td></td><td></td><td></td><td></td></tr>
<tr><td>Oligo fwd</td><td>CACCGCGCTCGGGCCTGGGTAGTA</td><td></td><td></td><td></td><td></td><td></td><td></td><td></td><td></td><td></td><td></td></tr>
<tr><td>Oligo rev</td><td>AAACTACTACCCAGGCCCGAGCGC</td><td></td><td></td><td></td><td></td><td></td><td></td><td></td><td></td><td></td><td></td></tr>
<tr><td>Chromosome</td><td>start</td><td>end</td><td>strand</td><td>MM</td><td>target_seq</td><td>PAM</td><td>alignment</td><td>distance</td><td>position</td><td>gene name</td><td>gene id</td></tr>
<tr><td>GU305914.1</td><td>19402</td><td>19424</td><td>+</td><td>0</td><td>GCGCTCGGGCCTGGGTAGTA</td><td>TGG</td><td>||||||||[||||||||||||]PAM</td><td>NA</td><td>Intergenic</td><td>NA</td><td>NA</td></tr>
<tr><td>T969</td><td>CGCTCGGGCCTGGGTAGTATGGG</td><td>1000</td><td></td><td></td><td></td><td></td><td></td><td></td><td></td><td></td><td></td></tr>
<tr><td>Oligo fwd</td><td>CACCCGCTCGGGCCTGGGTAGTAT</td><td></td><td></td><td></td><td></td><td></td><td></td><td></td><td></td><td></td><td></td></tr>
<tr><td>Oligo rev</td><td>AAACATACTACCCAGGCCCGAGCG</td><td></td><td></td><td></td><td></td><td></td><td></td><td></td><td></td><td></td><td></td></tr>
<tr><td>Chromosome</td><td>start</td><td>end</td><td>strand</td><td>MM</td><td>target_seq</td><td>PAM</td><td>alignment</td><td>distance</td><td>position</td><td>gene name</td><td>gene id</td></tr>
</table>

<table>
<tr><td>GU305914.1</td><td>19403</td><td>19425</td><td>+</td><td>0</td><td>CGCTCGGGCCTGGGTAGTAT</td><td>GGG</td><td>||||||||[||||||||||||]PAM</td><td>NA</td><td>Intergenic</td><td>NA</td><td>NA</td></tr>
</table>

<table>
<tr><td>T970</td><td>GCTCGGGCCTGGGTAGTATGGGG</td><td>1000</td></tr>
<tr><td>Oligo fwd</td><td>CACCGCTCGGGCCTGGGTAGTATG</td><td></td></tr>
<tr><td>Oligo rev</td><td>AAACCATACTACCCAGGCCCGAGC</td><td></td></tr>
</table>

<table>
<tr><td>Chromosome</td><td>start</td><td>end</td><td>strand</td><td>MM</td><td>target_seq</td><td>PAM</td><td>alignment</td><td>distance</td><td>position</td><td>gene name</td><td>gene id</td></tr>
<tr><td>GU305914.1</td><td>19404</td><td>19426</td><td>+</td><td>0</td><td>GCTCGGGCCTGGGTAGTATG</td><td>GGG</td><td>||||||||[||||||||||||]PAM</td><td>NA</td><td>Intergenic</td><td>NA</td><td>NA</td></tr>
</table>

<table>
<tr><td>T971</td><td>AGTATGGGGACGTTACCCGCCGG</td><td>1000</td></tr>
<tr><td>Oligo fwd</td><td>CACCAGTATGGGGACGTTACCCGC</td><td></td></tr>
<tr><td>Oligo rev</td><td>AAACGCGGGTAACGTCCCCATACT</td><td></td></tr>
</table>

<table>
<tr><td>Chromosome</td><td>start</td><td>end</td><td>strand</td><td>MM</td><td>target_seq</td><td>PAM</td><td>alignment</td><td>distance</td><td>position</td><td>gene name</td><td>gene id</td></tr>
<tr><td>GU305914.1</td><td>19418</td><td>19440</td><td>+</td><td>0</td><td>AGTATGGGGACGTTACCCGC</td><td>CGG</td><td>||||||||[||||||||||||]PAM</td><td>NA</td><td>Intergenic</td><td>NA</td><td>NA</td></tr>
</table>

<table>
<tr><td>T972</td><td>GGGACGTTACCCGCCGGCGTCGG</td><td>1000</td></tr>
<tr><td>Oligo fwd</td><td>CACCGGGACGTTACCCGCCGGCGT</td><td></td></tr>
<tr><td>Oligo rev</td><td>AAACACGCCGGCGGGTAACGTCCC</td><td></td></tr>
</table>

<table>
<tr><td>Chromosome</td><td>start</td><td>end</td><td>strand</td><td>MM</td><td>target_seq</td><td>PAM</td><td>alignment</td><td>distance</td><td>position</td><td>gene name</td><td>gene id</td></tr>
<tr><td>GU305914.1</td><td>19424</td><td>19446</td><td>+</td><td>0</td><td>GGGACGTTACCCGCCGGCGT</td><td>CGG</td><td>||||||||[||||||||||||]PAM</td><td>NA</td><td>Intergenic</td><td>NA</td><td>NA</td></tr>
</table>

<table>
<tr><td>T973</td><td>ACGTTACCCGCCGGCGTCGGCGG</td><td>1000</td></tr>
<tr><td>Oligo fwd</td><td>CACCACGTTACCCGCCGGCGTCGG</td><td></td></tr>
<tr><td>Oligo rev</td><td>AAACCCGACGCCGGCGGGTAACGT</td><td></td></tr>
</table>

<table>
<tr><td>Chromosome</td><td>start</td><td>end</td><td>strand</td><td>MM</td><td>target_seq</td><td>PAM</td><td>alignment</td><td>distance</td><td>position</td><td>gene name</td><td>gene id</td></tr>
<tr><td>GU305914.1</td><td>19427</td><td>19449</td><td>+</td><td>0</td><td>ACGTTACCCGCCGGCGTCGG</td><td>CGG</td><td>||||||||[||||||||||||]PAM</td><td>NA</td><td>Intergenic</td><td>NA</td><td>NA</td></tr>
</table>

<table>
<tr><td>T974</td><td>TTACCCGCCGGCGTCGGCGGAGG</td><td>1000</td></tr>
<tr><td>Oligo fwd</td><td>CACCTTACCCGCCGGCGTCGGCGG</td><td></td></tr>
<tr><td>Oligo rev</td><td>AAACCCGCCGACGCCGGCGGGTAA</td><td></td></tr>
</table>

<table>
<tr><td>Chromosome</td><td>start</td><td>end</td><td>strand</td><td>MM</td><td>target_seq</td><td>PAM</td><td>alignment</td><td>distance</td><td>position</td><td>gene name</td><td>gene id</td></tr>
<tr><td>GU305914.1</td><td>19430</td><td>19452</td><td>+</td><td>0</td><td>TTACCCGCCGGCGTCGGCGG</td><td>AGG</td><td>||||||||[||||||||||||]PAM</td><td>NA</td><td>Intergenic</td><td>NA</td><td>NA</td></tr>
</table>

<table>
<tr><td>T975</td><td>GCCGGCGTCGGCGGAGGTGCCGG</td><td>1000</td></tr>
<tr><td>Oligo fwd</td><td>CACCGCCGGCGTCGGCGGAGGTGC</td><td></td></tr>
<tr><td>Oligo rev</td><td>AAACGCACCTCCGCCGACGCCGGC</td><td></td></tr>
</table>

<table>
<tr><td>Chromosome</td><td>start</td><td>end</td><td>strand</td><td>MM</td><td>target_seq</td><td>PAM</td><td>alignment</td><td>distance</td><td>position</td><td>gene name</td><td>gene id</td></tr>
<tr><td>GU305914.1</td><td>19436</td><td>19458</td><td>+</td><td>0</td><td>GCCGGCGTCGGCGGAGGTGC</td><td>CGG</td><td>||||||||[||||||||||||]PAM</td><td>NA</td><td>Intergenic</td><td>NA</td><td>NA</td></tr>
<tr><td>GU305914.1</td><td>152550</td><td>152572</td><td>+</td><td>4</td><td>TCCTGCGGCGGCGGCGGTGC</td><td>CGG</td><td>-||-|||-[||||||-|||||]PAM</td><td>NA</td><td>Intergenic</td><td>NA</td><td>NA</td></tr>
<tr><td>GU305914.1</td><td>161748</td><td>161770</td><td>-</td><td>4</td><td>GGAGGCGGCGGCGGAGGTGA</td><td>CGG</td><td>|--||||-[|||||||||||-]PAM</td><td>NA</td><td>Intergenic</td><td>NA</td><td>NA</td></tr>
<tr><td>GU305914.1</td><td>111175</td><td>111197</td><td>-</td><td>4</td><td>GCCGACGACGGCGAAGGTGG</td><td>AGG</td><td>||||-||-[|||||-|||||-]PAM</td><td>NA</td><td>Intergenic</td><td>NA</td><td>NA</td></tr>
<tr><td>GU305914.1</td><td>90259</td><td>90281</td><td>-</td><td>4</td><td>GCGGGCGACGGCGGAGGCGG</td><td>CGG</td><td>||-||||-[|||||||||-|-]PAM</td><td>NA</td><td>Intergenic</td><td>NA</td><td>NA</td></tr>
</table>

<table>
<tr><td>T976</td><td>GGCGTCGGCGGAGGTGCCGGAGG</td><td>1000</td></tr>
<tr><td>Oligo fwd</td><td>CACCGGCGTCGGCGGAGGTGCCGG</td><td></td></tr>
<tr><td>Oligo rev</td><td>AAACCCGGCACCTCCGCCGACGCC</td><td></td></tr>
</table>

<table>
<tr><td>Chromosome</td><td>start</td><td>end</td><td>strand</td><td>MM</td><td>target_seq</td><td>PAM</td><td>alignment</td><td>distance</td><td>position</td><td>gene name</td><td>gene id</td></tr>
<tr><td>GU305914.1</td><td>19439</td><td>19461</td><td>+</td><td>0</td><td>GGCGTCGGCGGAGGTGCCGG</td><td>AGG</td><td>||||||||[||||||||||||]PAM</td><td>NA</td><td>Intergenic</td><td>NA</td><td>NA</td></tr>
<tr><td>GU305914.1</td><td>152553</td><td>152575</td><td>+</td><td>3</td><td>TGCGGCGGCGGCGGTGCCGG</td><td>CGG</td><td>-|||-|||[|||-||||||||]PAM</td><td>NA</td><td>Intergenic</td><td>NA</td><td>NA</td></tr>
<tr><td>GU305914.1</td><td>90250</td><td>90272</td><td>-</td><td>4</td><td>GGCGGAGGCGGCGGAGCCGG</td><td>CGG</td><td>||||--||[|||-||-|||||]PAM</td><td>NA</td><td>Intergenic</td><td>NA</td><td>NA</td></tr>
</table>

<table>
<tr><td>T977</td><td>TGCCGGAGGCACCGCGCTGCTGG</td><td>1000</td></tr>
<tr><td>Oligo fwd</td><td>CACCTGCCGGAGGCACCGCGCTGC</td><td></td></tr>
<tr><td>Oligo rev</td><td>AAACGCAGCGCGGTGCCTCCGGCA</td><td></td></tr>
</table>

<table>
<tr><td>Chromosome</td><td>start</td><td>end</td><td>strand</td><td>MM</td><td>target_seq</td><td>PAM</td><td>alignment</td><td>distance</td><td>position</td><td>gene name</td><td>gene id</td></tr>
<tr><td>GU305914.1</td><td>19453</td><td>19475</td><td>+</td><td>0</td><td>TGCCGGAGGCACCGCGCTGC</td><td>TGG</td><td>||||||||[||||||||||||]PAM</td><td>NA</td><td>Intergenic</td><td>NA</td><td>NA</td></tr>
</table>

<table>
<tr><td>T978</td><td>CGCGCTGCTGGCGATGAGCGTGG</td><td>1000</td></tr>
<tr><td>Oligo fwd</td><td>CACCCGCGCTGCTGGCGATGAGCG</td><td></td></tr>
<tr><td>Oligo rev</td><td>AAACCGCTCATCGCCAGCAGCGCG</td><td></td></tr>
</table>

<table>
<tr><td>Chromosome</td><td>start</td><td>end</td><td>strand</td><td>MM</td><td>target_seq</td><td>PAM</td><td>alignment</td><td>distance</td><td>position</td><td>gene name</td><td>gene id</td></tr>
<tr><td>GU305914.1</td><td>19465</td><td>19487</td><td>+</td><td>0</td><td>CGCGCTGCTGGCGATGAGCG</td><td>TGG</td><td>||||||||[||||||||||||]PAM</td><td>NA</td><td>Intergenic</td><td>NA</td><td>NA</td></tr>
</table>

<table>
<tr><td>T979</td><td>GCTGCTGGCGATGAGCGTGGTGG</td><td>1000</td></tr>
<tr><td>Oligo fwd</td><td>CACCGCTGCTGGCGATGAGCGTGG</td><td></td></tr>
<tr><td>Oligo rev</td><td>AAACCCACGCTCATCGCCAGCAGC</td><td></td></tr>
</table>

<table>
<tr><td>Chromosome</td><td>start</td><td>end</td><td>strand</td><td>MM</td><td>target_seq</td><td>PAM</td><td>alignment</td><td>distance</td><td>position</td><td>gene name</td><td>gene id</td></tr>
<tr><td>GU305914.1</td><td>19468</td><td>19490</td><td>+</td><td>0</td><td>GCTGCTGGCGATGAGCGTGG</td><td>TGG</td><td>||||||||[||||||||||||]PAM</td><td>NA</td><td>Intergenic</td><td>NA</td><td>NA</td></tr>
</table>

<table>
<tr><td>T980</td><td>CTGCTGGCGATGAGCGTGGTGGG</td><td>1000</td></tr>
<tr><td>Oligo fwd</td><td>CACCCTGCTGGCGATGAGCGTGGT</td><td></td></tr>
<tr><td>Oligo rev</td><td>AAACACCACGCTCATCGCCAGCAG</td><td></td></tr>
</table>

<table>
<tr><td>Chromosome</td><td>start</td><td>end</td><td>strand</td><td>MM</td><td>target_seq</td><td>PAM</td><td>alignment</td><td>distance</td><td>position</td><td>gene name</td><td>gene id</td></tr>
<tr><td>GU305914.1</td><td>19469</td><td>19491</td><td>+</td><td>0</td><td>CTGCTGGCGATGAGCGTGGT</td><td>GGG</td><td>||||||||[||||||||||||]PAM</td><td>NA</td><td>Intergenic</td><td>NA</td><td>NA</td></tr>
</table>

<table>
<tr><td>T981</td><td>TGCTGGCGATGAGCGTGGTGGGG</td><td>1000</td></tr>
<tr><td>Oligo fwd</td><td>CACCTGCTGGCGATGAGCGTGGTG</td><td></td></tr>
<tr><td>Oligo rev</td><td>AAACCACCACGCTCATCGCCAGCA</td><td></td></tr>
</table>

<table>
<tr><td>Chromosome</td><td>start</td><td>end</td><td>strand</td><td>MM</td><td>target_seq</td><td>PAM</td><td>alignment</td><td>distance</td><td>position</td><td>gene name</td><td>gene id</td></tr>
<tr><td>GU305914.1</td><td>19470</td><td>19492</td><td>+</td><td>0</td><td>TGCTGGCGATGAGCGTGGTG</td><td>GGG</td><td>||||||||[||||||||||||]PAM</td><td>NA</td><td>Intergenic</td><td>NA</td><td>NA</td></tr>
</table>

<table>
<tr><td>T982</td><td>GGCGATGAGCGTGGTGGGGACGG</td><td>1000</td></tr>
<tr><td>Oligo fwd</td><td>CACCGGCGATGAGCGTGGTGGGGA</td><td></td></tr>
<tr><td>Oligo rev</td><td>AAACTCCCCACCACGCTCATCGCC</td><td></td></tr>
</table>

<table>
<tr><td>Chromosome</td><td>start</td><td>end</td><td>strand</td><td>MM</td><td>target_seq</td><td>PAM</td><td>alignment</td><td>distance</td><td>position</td><td>gene name</td><td>gene id</td></tr>
<tr><td>GU305914.1</td><td>19474</td><td>19496</td><td>+</td><td>0</td><td>GGCGATGAGCGTGGTGGGGA</td><td>CGG</td><td>||||||||[||||||||||||]PAM</td><td>NA</td><td>Intergenic</td><td>NA</td><td>NA</td></tr>
<tr><td>GU305914.1</td><td>124295</td><td>124317</td><td>-</td><td>4</td><td>CGCGCTGACCGTGGTGCGGA</td><td>CGG</td><td>-|||-|||[-|||||||-|||]PAM</td><td>NA</td><td>Intergenic</td><td>NA</td><td>NA</td></tr>
<tr><td>GU305914.1</td><td>79906</td><td>79928</td><td>+</td><td>4</td><td>GGGCATGAGCGTGTTGGTGA</td><td>CGG</td><td>||--||||[|||||-|||-||]PAM</td><td>NA</td><td>Intergenic</td><td>NA</td><td>NA</td></tr>
</table>

<table>
<tr><td>T983</td><td>GCGATGAGCGTGGTGGGGACGGG</td><td>1000</td></tr>
</table>

| | | | | | | | | | | | |
|---|---|---|---|---|---|---|---|---|---|---|---|
| Oligo fwd | CACCGCGATGAGCGTGGTGGGGAC | | | | | | | | | | |
| Oligo rev | AAACGTCCCCACCACGCTCATCGC | | | | | | | | | | |
| Chromosome | start | end | strand | MM | target_seq | PAM | alignment | distance | position | gene name | gene id |
| GU305914.1 | 19475 | 19497 | + | 0 | GCGATGAGCGTGGTGGGGAC | GGG | \|\|\|\|\|\|\|\|[\|\|\|\|\|\|\|\|\|\|\|\|]PAM | NA | Intergenic | NA | NA |
| T984 | AGCGTGGTGGGGACGGGAGTCGG | 1000 | | | | | | | | | |
| Oligo fwd | CACCAGCGTGGTGGGGACGGGAGT | | | | | | | | | | |
| Oligo rev | AAACACTCCCGTCCCCACCACGCT | | | | | | | | | | |
| Chromosome | start | end | strand | MM | target_seq | PAM | alignment | distance | position | gene name | gene id |
| GU305914.1 | 19481 | 19503 | + | 0 | AGCGTGGTGGGGACGGGAGT | CGG | \|\|\|\|\|\|\|\|[\|\|\|\|\|\|\|\|\|\|\|\|]PAM | NA | Intergenic | NA | NA |
| T985 | GTGGTGGGGACGGGAGTCGGTGG | 1000 | | | | | | | | | |
| Oligo fwd | CACCGTGGTGGGGACGGGAGTCGG | | | | | | | | | | |
| Oligo rev | AAACCCGACTCCCGTCCCCACCAC | | | | | | | | | | |
| Chromosome | start | end | strand | MM | target_seq | PAM | alignment | distance | position | gene name | gene id |
| GU305914.1 | 19484 | 19506 | + | 0 | GTGGTGGGGACGGGAGTCGG | TGG | \|\|\|\|\|\|\|\|[\|\|\|\|\|\|\|\|\|\|\|\|]PAM | NA | Intergenic | NA | NA |
| T986 | TGGAGCGCCCATCGTCGCGATGG | 1000 | | | | | | | | | |
| Oligo fwd | CACCTGGAGCGCCCATCGTCGCGA | | | | | | | | | | |
| Oligo rev | AAACTCGCGACGATGGGCGCTCCA | | | | | | | | | | |
| Chromosome | start | end | strand | MM | target_seq | PAM | alignment | distance | position | gene name | gene id |
| GU305914.1 | 19504 | 19526 | + | 0 | TGGAGCGCCCATCGTCGCGA | TGG | \|\|\|\|\|\|\|\|[\|\|\|\|\|\|\|\|\|\|\|\|]PAM | NA | Intergenic | NA | NA |
| T987 | GGAGCGCCCATCGTCGCGATGGG | 1000 | | | | | | | | | |
| Oligo fwd | CACCGGAGCGCCCATCGTCGCGAT | | | | | | | | | | |
| Oligo rev | AAACATCGCGACGATGGGCGCTCC | | | | | | | | | | |
| Chromosome | start | end | strand | MM | target_seq | PAM | alignment | distance | position | gene name | gene id |
| GU305914.1 | 19505 | 19527 | + | 0 | GGAGCGCCCATCGTCGCGAT | GGG | \|\|\|\|\|\|\|\|[\|\|\|\|\|\|\|\|\|\|\|\|]PAM | NA | Intergenic | NA | NA |
| T988 | GAGCGCCCATCGTCGCGATGGGG | 1000 | | | | | | | | | |
| Oligo fwd | CACCGAGCGCCCATCGTCGCGATG | | | | | | | | | | |
| Oligo rev | AAACCATCGCGACGATGGGCGCTC | | | | | | | | | | |
| Chromosome | start | end | strand | MM | target_seq | PAM | alignment | distance | position | gene name | gene id |
| GU305914.1 | 19506 | 19528 | + | 0 | GAGCGCCCATCGTCGCGATG | GGG | \|\|\|\|\|\|\|\|[\|\|\|\|\|\|\|\|\|\|\|\|]PAM | NA | Intergenic | NA | NA |
| T989 | GCCCATCGTCGCGATGGGGACGG | 1000 | | | | | | | | | |
| Oligo fwd | CACCGCCCATCGTCGCGATGGGGA | | | | | | | | | | |
| Oligo rev | AAACTCCCCATCGCGACGATGGGC | | | | | | | | | | |
| Chromosome | start | end | strand | MM | target_seq | PAM | alignment | distance | position | gene name | gene id |
| GU305914.1 | 19510 | 19532 | + | 0 | GCCCATCGTCGCGATGGGGA | CGG | \|\|\|\|\|\|\|\|[\|\|\|\|\|\|\|\|\|\|\|\|]PAM | NA | Intergenic | NA | NA |
| T990 | CATCGTCGCGATGGGGACGGCGG | 1000 | | | | | | | | | |
| Oligo fwd | CACCCATCGTCGCGATGGGGACGG | | | | | | | | | | |
| Oligo rev | AAACCCGTCCCCATCGCGACGATG | | | | | | | | | | |
| Chromosome | start | end | strand | MM | target_seq | PAM | alignment | distance | position | gene name | gene id |
| GU305914.1 | 19513 | 19535 | + | 0 | CATCGTCGCGATGGGGACGG | CGG | \|\|\|\|\|\|\|\|[\|\|\|\|\|\|\|\|\|\|\|\|]PAM | NA | Intergenic | NA | NA |
| GU305914.1 | 76594 | 76616 | + | 4 | CATCTTCTCGTTGAGGACGG | CGG | \|\|\|\|-\|\|-[\|\|-\|\|-\|\|\|\|\|\|]PAM | NA | Intergenic | NA | NA |
| GU305914.1 | 91746 | 91768 | + | 4 | CAGCGGCGCGATGACGACGG | GGG | \|\|-\|\|-\|\|[\|\|\|\|\|--\|\|\|\|\|]PAM | NA | Intergenic | NA | NA |
| T991 | ACGGCGGAACCGCTGCCGAACGG | 1000 | | | | | | | | | |
| Oligo fwd | CACCACGGCGGAACCGCTGCCGAA | | | | | | | | | | |
| Oligo rev | AAACTTCGGCAGCGGTTCCGCCGT | | | | | | | | | | |
| Chromosome | start | end | strand | MM | target_seq | PAM | alignment | distance | position | gene name | gene id |
| GU305914.1 | 19529 | 19551 | + | 0 | ACGGCGGAACCGCTGCCGAA | CGG | \|\|\|\|\|\|\|\|[\|\|\|\|\|\|\|\|\|\|\|\|]PAM | NA | Intergenic | NA | NA |
| GU305914.1 | 69470 | 69492 | + | 4 | CTGGCGGAACAGCTGCCGAC | GGG | --\|\|\|\|\|\|[\|\|-\|\|\|\|\|\|\|\|-]PAM | NA | Intergenic | NA | NA |
| T992 | AACGGCGATACGATTCCGAACGG | 1000 | | | | | | | | | |
| Oligo fwd | CACCAACGGCGATACGATTCCGAA | | | | | | | | | | |
| Oligo rev | AAACTTCGGAATCGTATCGCCGTT | | | | | | | | | | |
| Chromosome | start | end | strand | MM | target_seq | PAM | alignment | distance | position | gene name | gene id |
| GU305914.1 | 19547 | 19569 | + | 0 | AACGGCGATACGATTCCGAA | CGG | \|\|\|\|\|\|\|\|[\|\|\|\|\|\|\|\|\|\|\|\|]PAM | NA | Intergenic | NA | NA |
| T993 | GCGATACGATTCCGAACGGCAGG | 1000 | | | | | | | | | |
| Oligo fwd | CACCGCGATACGATTCCGAACGGC | | | | | | | | | | |
| Oligo rev | AAACGCCGTTCGGAATCGTATCGC | | | | | | | | | | |
| Chromosome | start | end | strand | MM | target_seq | PAM | alignment | distance | position | gene name | gene id |
| GU305914.1 | 19551 | 19573 | + | 0 | GCGATACGATTCCGAACGGC | AGG | \|\|\|\|\|\|\|\|[\|\|\|\|\|\|\|\|\|\|\|\|]PAM | NA | Intergenic | NA | NA |
| T994 | GAACGGCAGGATGAACTGCGTGG | 1000 | | | | | | | | | |
| Oligo fwd | CACCGAACGGCAGGATGAACTGCG | | | | | | | | | | |
| Oligo rev | AAACCGCAGTTCATCCTGCCGTTC | | | | | | | | | | |
| Chromosome | start | end | strand | MM | target_seq | PAM | alignment | distance | position | gene name | gene id |
| GU305914.1 | 19564 | 19586 | + | 0 | GAACGGCAGGATGAACTGCG | TGG | \|\|\|\|\|\|\|\|[\|\|\|\|\|\|\|\|\|\|\|\|]PAM | NA | Intergenic | NA | NA |
| T995 | AACGGCAGGATGAACTGCGTGGG | 1000 | | | | | | | | | |
| Oligo fwd | CACCAACGGCAGGATGAACTGCGT | | | | | | | | | | |
| Oligo rev | AAACACGCAGTTCATCCTGCCGTT | | | | | | | | | | |
| Chromosome | start | end | strand | MM | target_seq | PAM | alignment | distance | position | gene name | gene id |
| GU305914.1 | 19565 | 19587 | + | 0 | AACGGCAGGATGAACTGCGT | GGG | \|\|\|\|\|\|\|\|[\|\|\|\|\|\|\|\|\|\|\|\|]PAM | NA | Intergenic | NA | NA |
| T996 | AGGATGAACTGCGTGGGTGTCGG | 1000 | | | | | | | | | |
| Oligo fwd | CACCAGGATGAACTGCGTGGGTGT | | | | | | | | | | |
| Oligo rev | AAACACACCCACGCAGTTCATCCT | | | | | | | | | | |
| Chromosome | start | end | strand | MM | target_seq | PAM | alignment | distance | position | gene name | gene id |
| GU305914.1 | 19571 | 19593 | + | 0 | AGGATGAACTGCGTGGGTGT | CGG | \|\|\|\|\|\|\|\|[\|\|\|\|\|\|\|\|\|\|\|\|]PAM | NA | Intergenic | NA | NA |
| T997 | ATGAACTGCGTGGGTGTCGGTGG | 1000 | | | | | | | | | |
| Oligo fwd | CACCATGAACTGCGTGGGTGTCGG | | | | | | | | | | |
| Oligo rev | AAACCCGACACCCACGCAGTTCAT | | | | | | | | | | |

| | | | | | | | | | | | |
|---|---|---|---|---|---|---|---|---|---|---|---|
| Chromosome | start | end | strand | MM | target_seq | PAM | alignment | distance | position | gene name | gene id |
| GU305914.1 | 19574 | 19596 | + | 0 | ATGAACTGCGTGGGTGTCGG | TGG | \|\|\|\|\|\|\|\|[\|\|\|\|\|\|\|\|\|\|\|\|]PAM | NA | Intergenic | NA | NA |
| T998 | AACTGCGTGGGTGTCGGTGGCGG | 1000 | | | | | | | | | |
| Oligo fwd | CACCAACTGCGTGGGTGTCGGTGG | | | | | | | | | | |
| Oligo rev | AAACCCACCGACACCCACGCAGTT | | | | | | | | | | |
| Chromosome | start | end | strand | MM | target_seq | PAM | alignment | distance | position | gene name | gene id |
| GU305914.1 | 19577 | 19599 | + | 0 | AACTGCGTGGGTGTCGGTGG | CGG | \|\|\|\|\|\|\|\|[\|\|\|\|\|\|\|\|\|\|\|\|]PAM | NA | Intergenic | NA | NA |
| T999 | TGCGTGGGTGTCGGTGGCGGCGG | 1000 | | | | | | | | | |
| Oligo fwd | CACCTGCGTGGGTGTCGGTGGCGG | | | | | | | | | | |
| Oligo rev | AAACCCGCCACCGACACCCACGCA | | | | | | | | | | |
| Chromosome | start | end | strand | MM | target_seq | PAM | alignment | distance | position | gene name | gene id |
| GU305914.1 | 19580 | 19602 | + | 0 | TGCGTGGGTGTCGGTGGCGG | CGG | \|\|\|\|\|\|\|\|[\|\|\|\|\|\|\|\|\|\|\|\|]PAM | NA | Intergenic | NA | NA |
| T1000 | GTCGGTGGCGGCGGTAGCGTCGG | 1000 | | | | | | | | | |
| Oligo fwd | CACCGTCGGTGGCGGCGGTAGCGT | | | | | | | | | | |
| Oligo rev | AAACACGCTACCGCCGCCACCGAC | | | | | | | | | | |
| Chromosome | start | end | strand | MM | target_seq | PAM | alignment | distance | position | gene name | gene id |
| GU305914.1 | 19589 | 19611 | + | 0 | GTCGGTGGCGGCGGTAGCGT | CGG | \|\|\|\|\|\|\|\|[\|\|\|\|\|\|\|\|\|\|\|\|]PAM | NA | Intergenic | NA | NA |
| GU305914.1 | 62551 | 62573 | + | 4 | AGCGGCCGCGGCGGTAGCGT | TGG | --\|\|\|--\|[\|\|\|\|\|\|\|\|\|\|\|\|]PAM | NA | Intergenic | NA | NA |
| T1001 | CGGCTGCGAAAAATCGTAGACGG | 1000 | | | | | | | | | |
| Oligo fwd | CACCCGGCTGCGAAAAATCGTAGA | | | | | | | | | | |
| Oligo rev | AAACTCTACGATTTTTCGCAGCCG | | | | | | | | | | |
| Chromosome | start | end | strand | MM | target_seq | PAM | alignment | distance | position | gene name | gene id |
| GU305914.1 | 19609 | 19631 | + | 0 | CGGCTGCGAAAAATCGTAGA | CGG | \|\|\|\|\|\|\|\|[\|\|\|\|\|\|\|\|\|\|\|\|]PAM | NA | Intergenic | NA | NA |
| T1002 | AATCGTAGACGGACGTATCGAGG | 1000 | | | | | | | | | |
| Oligo fwd | CACCAATCGTAGACGGACGTATCG | | | | | | | | | | |
| Oligo rev | AAACCGATACGTCCGTCTACGATT | | | | | | | | | | |
| Chromosome | start | end | strand | MM | target_seq | PAM | alignment | distance | position | gene name | gene id |
| GU305914.1 | 19620 | 19642 | + | 0 | AATCGTAGACGGACGTATCG | AGG | \|\|\|\|\|\|\|\|[\|\|\|\|\|\|\|\|\|\|\|\|]PAM | NA | Intergenic | NA | NA |
| T1003 | ATCGTAGACGGACGTATCGAGGG | 1000 | | | | | | | | | |
| Oligo fwd | CACCATCGTAGACGGACGTATCGA | | | | | | | | | | |
| Oligo rev | AAACTCGATACGTCCGTCTACGAT | | | | | | | | | | |
| Chromosome | start | end | strand | MM | target_seq | PAM | alignment | distance | position | gene name | gene id |
| GU305914.1 | 19621 | 19643 | + | 0 | ATCGTAGACGGACGTATCGA | GGG | \|\|\|\|\|\|\|\|[\|\|\|\|\|\|\|\|\|\|\|\|]PAM | NA | Intergenic | NA | NA |
| T1004 | TCGTAGACGGACGTATCGAGGGG | 1000 | | | | | | | | | |
| Oligo fwd | CACCTCGTAGACGGACGTATCGAG | | | | | | | | | | |
| Oligo rev | AAACCTCGATACGTCCGTCTACGA | | | | | | | | | | |
| Chromosome | start | end | strand | MM | target_seq | PAM | alignment | distance | position | gene name | gene id |
| GU305914.1 | 19622 | 19644 | + | 0 | TCGTAGACGGACGTATCGAG | GGG | \|\|\|\|\|\|\|\|[\|\|\|\|\|\|\|\|\|\|\|\|]PAM | NA | Intergenic | NA | NA |
| T1005 | ACGGACGTATCGAGGGGTAAAGG | 1000 | | | | | | | | | |
| Oligo fwd | CACCACGGACGTATCGAGGGGTAA | | | | | | | | | | |
| Oligo rev | AAACTTACCCCTCGATACGTCCGT | | | | | | | | | | |
| Chromosome | start | end | strand | MM | target_seq | PAM | alignment | distance | position | gene name | gene id |
| GU305914.1 | 19628 | 19650 | + | 0 | ACGGACGTATCGAGGGGTAA | AGG | \|\|\|\|\|\|\|\|[\|\|\|\|\|\|\|\|\|\|\|\|]PAM | NA | Intergenic | NA | NA |
| T1006 | GAGGGGTAAAGGATTGATCGTGG | 1000 | | | | | | | | | |
| Oligo fwd | CACCGAGGGGTAAAGGATTGATCG | | | | | | | | | | |
| Oligo rev | AAACCGATCAATCCTTTACCCCTC | | | | | | | | | | |
| Chromosome | start | end | strand | MM | target_seq | PAM | alignment | distance | position | gene name | gene id |
| GU305914.1 | 19639 | 19661 | + | 0 | GAGGGGTAAAGGATTGATCG | TGG | \|\|\|\|\|\|\|\|[\|\|\|\|\|\|\|\|\|\|\|\|]PAM | NA | Intergenic | NA | NA |
| T1007 | AGGGGTAAAGGATTGATCGTGGG | 1000 | | | | | | | | | |
| Oligo fwd | CACCAGGGGTAAAGGATTGATCGT | | | | | | | | | | |
| Oligo rev | AAACACGATCAATCCTTTACCCCT | | | | | | | | | | |
| Chromosome | start | end | strand | MM | target_seq | PAM | alignment | distance | position | gene name | gene id |
| GU305914.1 | 19640 | 19662 | + | 0 | AGGGGTAAAGGATTGATCGT | GGG | \|\|\|\|\|\|\|\|[\|\|\|\|\|\|\|\|\|\|\|\|]PAM | NA | Intergenic | NA | NA |
| T1008 | TAAAGGATTGATCGTGGGATAGG | 1000 | | | | | | | | | |
| Oligo fwd | CACCTAAAGGATTGATCGTGGGAT | | | | | | | | | | |
| Oligo rev | AAACATCCCACGATCAATCCTTTA | | | | | | | | | | |
| Chromosome | start | end | strand | MM | target_seq | PAM | alignment | distance | position | gene name | gene id |
| GU305914.1 | 19645 | 19667 | + | 0 | TAAAGGATTGATCGTGGGAT | AGG | \|\|\|\|\|\|\|\|[\|\|\|\|\|\|\|\|\|\|\|\|]PAM | NA | Intergenic | NA | NA |
| T1009 | ATTGATCGTGGGATAGGCTGCGG | 1000 | | | | | | | | | |
| Oligo fwd | CACCATTGATCGTGGGATAGGCTG | | | | | | | | | | |
| Oligo rev | AAACCAGCCTATCCCACGATCAAT | | | | | | | | | | |
| Chromosome | start | end | strand | MM | target_seq | PAM | alignment | distance | position | gene name | gene id |
| GU305914.1 | 19651 | 19673 | + | 0 | ATTGATCGTGGGATAGGCTG | CGG | \|\|\|\|\|\|\|\|[\|\|\|\|\|\|\|\|\|\|\|\|]PAM | NA | Intergenic | NA | NA |
| T1010 | TTGATCGTGGGATAGGCTGCGGG | 1000 | | | | | | | | | |
| Oligo fwd | CACCTTGATCGTGGGATAGGCTGC | | | | | | | | | | |
| Oligo rev | AAACGCAGCCTATCCCACGATCAA | | | | | | | | | | |
| Chromosome | start | end | strand | MM | target_seq | PAM | alignment | distance | position | gene name | gene id |
| GU305914.1 | 19652 | 19674 | + | 0 | TTGATCGTGGGATAGGCTGC | GGG | \|\|\|\|\|\|\|\|[\|\|\|\|\|\|\|\|\|\|\|\|]PAM | NA | Intergenic | NA | NA |
| T1011 | GGATAGGCTGCGGGCAGCCAAGG | 1000 | | | | | | | | | |
| Oligo fwd | CACCGGATAGGCTGCGGGCAGCCA | | | | | | | | | | |
| Oligo rev | AAACTGGCTGCCCGCAGCCTATCC | | | | | | | | | | |
| Chromosome | start | end | strand | MM | target_seq | PAM | alignment | distance | position | gene name | gene id |
| GU305914.1 | 19661 | 19683 | + | 0 | GGATAGGCTGCGGGCAGCCA | AGG | \|\|\|\|\|\|\|\|[\|\|\|\|\|\|\|\|\|\|\|\|]PAM | NA | Intergenic | NA | NA |
| T1012 | GATAGGCTGCGGGCAGCCAAGGG | 1000 | | | | | | | | | |

| | | | | | | | | | | | |
|---|---|---|---|---|---|---|---|---|---|---|---|
| Oligo fwd | CACCGATAGGCTGCGGGCAGCCAA | | | | | | | | | | |
| Oligo rev | AAACTTGGCTGCCCGCAGCCTATC | | | | | | | | | | |
| Chromosome | start | end | strand | MM | target_seq | PAM | alignment | distance | position | gene name | gene id |
| GU305914.1 | 19662 | 19684 | + | 0 | GATAGGCTGCGGGCAGCCAA | GGG | \|\|\|\|\|\|\|\|[\|\|\|\|\|\|\|\|\|\|\|\|]PAM | NA | Intergenic | NA | NA |
| T1013 | GGCTGCGGGCAGCCAAGGGAAGG | 1000 | | | | | | | | | |
| Oligo fwd | CACCGGCTGCGGGCAGCCAAGGGA | | | | | | | | | | |
| Oligo rev | AAACTCCCTTGGCTGCCCGCAGCC | | | | | | | | | | |
| Chromosome | start | end | strand | MM | target_seq | PAM | alignment | distance | position | gene name | gene id |
| GU305914.1 | 19666 | 19688 | + | 0 | GGCTGCGGGCAGCCAAGGGA | AGG | \|\|\|\|\|\|\|\|[\|\|\|\|\|\|\|\|\|\|\|\|]PAM | NA | Intergenic | NA | NA |
| T1014 | GCTGCGGGCAGCCAAGGGAAGGG | 1000 | | | | | | | | | |
| Oligo fwd | CACCGCTGCGGGCAGCCAAGGGAA | | | | | | | | | | |
| Oligo rev | AAACTTCCCTTGGCTGCCCGCAGC | | | | | | | | | | |
| Chromosome | start | end | strand | MM | target_seq | PAM | alignment | distance | position | gene name | gene id |
| GU305914.1 | 19667 | 19689 | + | 0 | GCTGCGGGCAGCCAAGGGAA | GGG | \|\|\|\|\|\|\|\|[\|\|\|\|\|\|\|\|\|\|\|\|]PAM | NA | Intergenic | NA | NA |
| T1015 | CGGGCAGCCAAGGGAAGGGTTGG | 1000 | | | | | | | | | |
| Oligo fwd | CACCCGGGCAGCCAAGGGAAGGGT | | | | | | | | | | |
| Oligo rev | AAACACCCTTCCCTTGGCTGCCCG | | | | | | | | | | |
| Chromosome | start | end | strand | MM | target_seq | PAM | alignment | distance | position | gene name | gene id |
| GU305914.1 | 19671 | 19693 | + | 0 | CGGGCAGCCAAGGGAAGGGT | TGG | \|\|\|\|\|\|\|\|[\|\|\|\|\|\|\|\|\|\|\|\|]PAM | NA | Intergenic | NA | NA |
| T1016 | CAGCCAAGGGAAGGGTTGGAAGG | 1000 | | | | | | | | | |
| Oligo fwd | CACCCAGCCAAGGGAAGGGTTGGA | | | | | | | | | | |
| Oligo rev | AAACTCCAACCCTTCCCTTGGCTG | | | | | | | | | | |
| Chromosome | start | end | strand | MM | target_seq | PAM | alignment | distance | position | gene name | gene id |
| GU305914.1 | 19675 | 19697 | + | 0 | CAGCCAAGGGAAGGGTTGGA | AGG | \|\|\|\|\|\|\|\|[\|\|\|\|\|\|\|\|\|\|\|\|]PAM | NA | Intergenic | NA | NA |
| T1017 | CCAAGGGAAGGGTTGGAAGGTGG | 1000 | | | | | | | | | |
| Oligo fwd | CACCCCAAGGGAAGGGTTGGAAGG | | | | | | | | | | |
| Oligo rev | AAACCCTTCCAACCCTTCCCTTGG | | | | | | | | | | |
| Chromosome | start | end | strand | MM | target_seq | PAM | alignment | distance | position | gene name | gene id |
| GU305914.1 | 19678 | 19700 | + | 0 | CCAAGGGAAGGGTTGGAAGG | TGG | \|\|\|\|\|\|\|\|[\|\|\|\|\|\|\|\|\|\|\|\|]PAM | NA | Intergenic | NA | NA |
| T1018 | GGTTGGAAGGTGGCGTATGCCGG | 1000 | | | | | | | | | |
| Oligo fwd | CACCGGTTGGAAGGTGGCGTATGC | | | | | | | | | | |
| Oligo rev | AAACGCATACGCCACCTTCCAACC | | | | | | | | | | |
| Chromosome | start | end | strand | MM | target_seq | PAM | alignment | distance | position | gene name | gene id |
| GU305914.1 | 19688 | 19710 | + | 0 | GGTTGGAAGGTGGCGTATGC | CGG | \|\|\|\|\|\|\|\|[\|\|\|\|\|\|\|\|\|\|\|\|]PAM | NA | Intergenic | NA | NA |
| T1019 | GTTGGAAGGTGGCGTATGCCGGG | 1000 | | | | | | | | | |
| Oligo fwd | CACCGTTGGAAGGTGGCGTATGCC | | | | | | | | | | |
| Oligo rev | AAACGGCATACGCCACCTTCCAAC | | | | | | | | | | |
| Chromosome | start | end | strand | MM | target_seq | PAM | alignment | distance | position | gene name | gene id |
| GU305914.1 | 19689 | 19711 | + | 0 | GTTGGAAGGTGGCGTATGCC | GGG | \|\|\|\|\|\|\|\|[\|\|\|\|\|\|\|\|\|\|\|\|]PAM | NA | Intergenic | NA | NA |
| T1020 | GAAGGTGGCGTATGCCGGGCCGG | 1000 | | | | | | | | | |
| Oligo fwd | CACCGAAGGTGGCGTATGCCGGGC | | | | | | | | | | |
| Oligo rev | AAACGCCCGGCATACGCCACCTTC | | | | | | | | | | |
| Chromosome | start | end | strand | MM | target_seq | PAM | alignment | distance | position | gene name | gene id |
| GU305914.1 | 19693 | 19715 | + | 0 | GAAGGTGGCGTATGCCGGGC | CGG | \|\|\|\|\|\|\|\|[\|\|\|\|\|\|\|\|\|\|\|\|]PAM | NA | Intergenic | NA | NA |
| T1021 | AAGGTGGCGTATGCCGGGCCGGG | 1000 | | | | | | | | | |
| Oligo fwd | CACCAAGGTGGCGTATGCCGGGCC | | | | | | | | | | |
| Oligo rev | AAACGGCCCGGCATACGCCACCTT | | | | | | | | | | |
| Chromosome | start | end | strand | MM | target_seq | PAM | alignment | distance | position | gene name | gene id |
| GU305914.1 | 19694 | 19716 | + | 0 | AAGGTGGCGTATGCCGGGCC | GGG | \|\|\|\|\|\|\|\|[\|\|\|\|\|\|\|\|\|\|\|\|]PAM | NA | Intergenic | NA | NA |
| T1022 | GCGTATGCCGGGCCGGGTAACGG | 1000 | | | | | | | | | |
| Oligo fwd | CACCGCGTATGCCGGGCCGGGTAA | | | | | | | | | | |
| Oligo rev | AAACTTACCCGGCCCGGCATACGC | | | | | | | | | | |
| Chromosome | start | end | strand | MM | target_seq | PAM | alignment | distance | position | gene name | gene id |
| GU305914.1 | 19700 | 19722 | + | 0 | GCGTATGCCGGGCCGGGTAA | CGG | \|\|\|\|\|\|\|\|[\|\|\|\|\|\|\|\|\|\|\|\|]PAM | NA | Intergenic | NA | NA |
| T1023 | GCCGGGTAACGGCTGAAACGCGG | 1000 | | | | | | | | | |
| Oligo fwd | CACCGCCGGGTAACGGCTGAAACG | | | | | | | | | | |
| Oligo rev | AAACCGTTTCAGCCGTTACCCGGC | | | | | | | | | | |
| Chromosome | start | end | strand | MM | target_seq | PAM | alignment | distance | position | gene name | gene id |
| GU305914.1 | 19711 | 19733 | + | 0 | GCCGGGTAACGGCTGAAACG | CGG | \|\|\|\|\|\|\|\|[\|\|\|\|\|\|\|\|\|\|\|\|]PAM | NA | Intergenic | NA | NA |
| T1024 | GGGTAACGGCTGAAACGCGGAGG | 1000 | | | | | | | | | |
| Oligo fwd | CACCGGGTAACGGCTGAAACGCGG | | | | | | | | | | |
| Oligo rev | AAACCCGCGTTTCAGCCGTTACCC | | | | | | | | | | |
| Chromosome | start | end | strand | MM | target_seq | PAM | alignment | distance | position | gene name | gene id |
| GU305914.1 | 19714 | 19736 | + | 0 | GGGTAACGGCTGAAACGCGG | AGG | \|\|\|\|\|\|\|\|[\|\|\|\|\|\|\|\|\|\|\|\|]PAM | NA | Intergenic | NA | NA |
| T1025 | GGCTGAAACGCGGAGGCGTGCGG | 1000 | | | | | | | | | |
| Oligo fwd | CACCGGCTGAAACGCGGAGGCGTG | | | | | | | | | | |
| Oligo rev | AAACCACGCCTCCGCGTTTCAGCC | | | | | | | | | | |
| Chromosome | start | end | strand | MM | target_seq | PAM | alignment | distance | position | gene name | gene id |
| GU305914.1 | 19721 | 19743 | + | 0 | GGCTGAAACGCGGAGGCGTG | CGG | \|\|\|\|\|\|\|\|[\|\|\|\|\|\|\|\|\|\|\|\|]PAM | NA | Intergenic | NA | NA |
| T1026 | AGGCGTGCGGCCATCCGAGATGG | 1000 | | | | | | | | | |
| Oligo fwd | CACCAGGCGTGCGGCCATCCGAGA | | | | | | | | | | |
| Oligo rev | AAACTCTCGGATGGCCGCACGCCT | | | | | | | | | | |
| Chromosome | start | end | strand | MM | target_seq | PAM | alignment | distance | position | gene name | gene id |
| GU305914.1 | 19734 | 19756 | + | 0 | AGGCGTGCGGCCATCCGAGA | TGG | \|\|\|\|\|\|\|\|[\|\|\|\|\|\|\|\|\|\|\|\|]PAM | NA | Intergenic | NA | NA |

<table>
<tr><td>T1027</td><td>GGCGTGCGGCCATCCGAGATGGG</td><td>1000</td></tr>
<tr><td>Oligo fwd</td><td>CACCGGCGTGCGGCCATCCGAGAT</td></tr>
<tr><td>Oligo rev</td><td>AAACATCTCGGATGGCCGCACGCC</td></tr>
<tr><td>Chromosome</td><td>start</td><td>end</td><td>strand</td><td>MM</td><td>target_seq</td><td>PAM</td><td>alignment</td><td>distance</td><td>position</td><td>gene name</td><td>gene id</td></tr>
<tr><td>GU305914.1</td><td>19735</td><td>19757</td><td>+</td><td>0</td><td>GGCGTGCGGCCATCCGAGAT</td><td>GGG</td><td>||||||||[||||||||||||]PAM</td><td>NA</td><td>Intergenic</td><td>NA</td><td>NA</td></tr>
<tr><td>T1028</td><td>GCGTGCGGCCATCCGAGATGGGG</td><td>1000</td></tr>
<tr><td>Oligo fwd</td><td>CACCGCGTGCGGCCATCCGAGATG</td></tr>
<tr><td>Oligo rev</td><td>AAACCATCTCGGATGGCCGCACGC</td></tr>
<tr><td>Chromosome</td><td>start</td><td>end</td><td>strand</td><td>MM</td><td>target_seq</td><td>PAM</td><td>alignment</td><td>distance</td><td>position</td><td>gene name</td><td>gene id</td></tr>
<tr><td>GU305914.1</td><td>19736</td><td>19758</td><td>+</td><td>0</td><td>GCGTGCGGCCATCCGAGATG</td><td>GGG</td><td>||||||||[||||||||||||]PAM</td><td>NA</td><td>Intergenic</td><td>NA</td><td>NA</td></tr>
<tr><td>T1029</td><td>CGAGATGGGGCGCGTATCCCAGG</td><td>1000</td></tr>
<tr><td>Oligo fwd</td><td>CACCCGAGATGGGGCGCGTATCCC</td></tr>
<tr><td>Oligo rev</td><td>AAACGGGATACGCGCCCCATCTCG</td></tr>
<tr><td>Chromosome</td><td>start</td><td>end</td><td>strand</td><td>MM</td><td>target_seq</td><td>PAM</td><td>alignment</td><td>distance</td><td>position</td><td>gene name</td><td>gene id</td></tr>
<tr><td>GU305914.1</td><td>19749</td><td>19771</td><td>+</td><td>0</td><td>CGAGATGGGGCGCGTATCCC</td><td>AGG</td><td>||||||||[||||||||||||]PAM</td><td>NA</td><td>Intergenic</td><td>NA</td><td>NA</td></tr>
<tr><td>T1030</td><td>GGGCGCGTATCCCAGGAAGACGG</td><td>1000</td></tr>
<tr><td>Oligo fwd</td><td>CACCGGGCGCGTATCCCAGGAAGA</td></tr>
<tr><td>Oligo rev</td><td>AAACTCTTCCTGGGATACGCGCCC</td></tr>
<tr><td>Chromosome</td><td>start</td><td>end</td><td>strand</td><td>MM</td><td>target_seq</td><td>PAM</td><td>alignment</td><td>distance</td><td>position</td><td>gene name</td><td>gene id</td></tr>
<tr><td>GU305914.1</td><td>19756</td><td>19778</td><td>+</td><td>0</td><td>GGGCGCGTATCCCAGGAAGA</td><td>CGG</td><td>||||||||[||||||||||||]PAM</td><td>NA</td><td>Intergenic</td><td>NA</td><td>NA</td></tr>
<tr><td>T1031</td><td>GGCGCGTATCCCAGGAAGACGGG</td><td>1000</td></tr>
<tr><td>Oligo fwd</td><td>CACCGGCGCGTATCCCAGGAAGAC</td></tr>
<tr><td>Oligo rev</td><td>AAACGTCTTCCTGGGATACGCGCC</td></tr>
<tr><td>Chromosome</td><td>start</td><td>end</td><td>strand</td><td>MM</td><td>target_seq</td><td>PAM</td><td>alignment</td><td>distance</td><td>position</td><td>gene name</td><td>gene id</td></tr>
<tr><td>GU305914.1</td><td>19757</td><td>19779</td><td>+</td><td>0</td><td>GGCGCGTATCCCAGGAAGAC</td><td>GGG</td><td>||||||||[||||||||||||]PAM</td><td>NA</td><td>Intergenic</td><td>NA</td><td>NA</td></tr>
<tr><td>T1032</td><td>GCGCGTATCCCAGGAAGACGGGG</td><td>1000</td></tr>
<tr><td>Oligo fwd</td><td>CACCGCGCGTATCCCAGGAAGACG</td></tr>
<tr><td>Oligo rev</td><td>AAACCGTCTTCCTGGGATACGCGC</td></tr>
<tr><td>Chromosome</td><td>start</td><td>end</td><td>strand</td><td>MM</td><td>target_seq</td><td>PAM</td><td>alignment</td><td>distance</td><td>position</td><td>gene name</td><td>gene id</td></tr>
<tr><td>GU305914.1</td><td>19758</td><td>19780</td><td>+</td><td>0</td><td>GCGCGTATCCCAGGAAGACG</td><td>GGG</td><td>||||||||[||||||||||||]PAM</td><td>NA</td><td>Intergenic</td><td>NA</td><td>NA</td></tr>
<tr><td>T1033</td><td>CGCGTATCCCAGGAAGACGGGGG</td><td>1000</td></tr>
<tr><td>Oligo fwd</td><td>CACCCGCGTATCCCAGGAAGACGG</td></tr>
<tr><td>Oligo rev</td><td>AAACCCGTCTTCCTGGGATACGCG</td></tr>
<tr><td>Chromosome</td><td>start</td><td>end</td><td>strand</td><td>MM</td><td>target_seq</td><td>PAM</td><td>alignment</td><td>distance</td><td>position</td><td>gene name</td><td>gene id</td></tr>
<tr><td>GU305914.1</td><td>19759</td><td>19781</td><td>+</td><td>0</td><td>CGCGTATCCCAGGAAGACGG</td><td>GGG</td><td>||||||||[||||||||||||]PAM</td><td>NA</td><td>Intergenic</td><td>NA</td><td>NA</td></tr>
<tr><td>T1034</td><td>CCCAGGAAGACGGGGGCGTGCGG</td><td>1000</td></tr>
<tr><td>Oligo fwd</td><td>CACCCCCAGGAAGACGGGGGCGTG</td></tr>
<tr><td>Oligo rev</td><td>AAACCACGCCCCCGTCTTCCTGGG</td></tr>
<tr><td>Chromosome</td><td>start</td><td>end</td><td>strand</td><td>MM</td><td>target_seq</td><td>PAM</td><td>alignment</td><td>distance</td><td>position</td><td>gene name</td><td>gene id</td></tr>
<tr><td>GU305914.1</td><td>19766</td><td>19788</td><td>+</td><td>0</td><td>CCCAGGAAGACGGGGGCGTG</td><td>CGG</td><td>||||||||[||||||||||||]PAM</td><td>NA</td><td>Intergenic</td><td>NA</td><td>NA</td></tr>
<tr><td>T1035</td><td>CCAGGAAGACGGGGGCGTGCGGG</td><td>1000</td></tr>
<tr><td>Oligo fwd</td><td>CACCCCAGGAAGACGGGGGCGTGC</td></tr>
<tr><td>Oligo rev</td><td>AAACGCACGCCCCCGTCTTCCTGG</td></tr>
<tr><td>Chromosome</td><td>start</td><td>end</td><td>strand</td><td>MM</td><td>target_seq</td><td>PAM</td><td>alignment</td><td>distance</td><td>position</td><td>gene name</td><td>gene id</td></tr>
<tr><td>GU305914.1</td><td>19767</td><td>19789</td><td>+</td><td>0</td><td>CCAGGAAGACGGGGGCGTGC</td><td>GGG</td><td>||||||||[||||||||||||]PAM</td><td>NA</td><td>Intergenic</td><td>NA</td><td>NA</td></tr>
<tr><td>T1036</td><td>CAGGAAGACGGGGGCGTGCGGGG</td><td>1000</td></tr>
<tr><td>Oligo fwd</td><td>CACCCAGGAAGACGGGGGCGTGCG</td></tr>
<tr><td>Oligo rev</td><td>AAACCGCACGCCCCCGTCTTCCTG</td></tr>
<tr><td>Chromosome</td><td>start</td><td>end</td><td>strand</td><td>MM</td><td>target_seq</td><td>PAM</td><td>alignment</td><td>distance</td><td>position</td><td>gene name</td><td>gene id</td></tr>
<tr><td>GU305914.1</td><td>19768</td><td>19790</td><td>+</td><td>0</td><td>CAGGAAGACGGGGGCGTGCG</td><td>GGG</td><td>||||||||[||||||||||||]PAM</td><td>NA</td><td>Intergenic</td><td>NA</td><td>NA</td></tr>
<tr><td>T1037</td><td>AGGAAGACGGGGGCGTGCGGGGG</td><td>1000</td></tr>
<tr><td>Oligo fwd</td><td>CACCAGGAAGACGGGGGCGTGCGG</td></tr>
<tr><td>Oligo rev</td><td>AAACCCGCACGCCCCCGTCTTCCT</td></tr>
<tr><td>Chromosome</td><td>start</td><td>end</td><td>strand</td><td>MM</td><td>target_seq</td><td>PAM</td><td>alignment</td><td>distance</td><td>position</td><td>gene name</td><td>gene id</td></tr>
<tr><td>GU305914.1</td><td>19769</td><td>19791</td><td>+</td><td>0</td><td>AGGAAGACGGGGGCGTGCGG</td><td>GGG</td><td>||||||||[||||||||||||]PAM</td><td>NA</td><td>Intergenic</td><td>NA</td><td>NA</td></tr>
<tr><td>T1038</td><td>GACGGGGGCGTGCGGGGGTGTGG</td><td>1000</td></tr>
<tr><td>Oligo fwd</td><td>CACCGACGGGGGCGTGCGGGGGTG</td></tr>
<tr><td>Oligo rev</td><td>AAACCACCCCCGCACGCCCCCGTC</td></tr>
<tr><td>Chromosome</td><td>start</td><td>end</td><td>strand</td><td>MM</td><td>target_seq</td><td>PAM</td><td>alignment</td><td>distance</td><td>position</td><td>gene name</td><td>gene id</td></tr>
<tr><td>GU305914.1</td><td>19774</td><td>19796</td><td>+</td><td>0</td><td>GACGGGGGCGTGCGGGGGTG</td><td>TGG</td><td>||||||||[||||||||||||]PAM</td><td>NA</td><td>Intergenic</td><td>NA</td><td>NA</td></tr>
<tr><td>T1039</td><td>ACGGGGGCGTGCGGGGGTGTGGG</td><td>1000</td></tr>
<tr><td>Oligo fwd</td><td>CACCACGGGGGCGTGCGGGGGTGT</td></tr>
<tr><td>Oligo rev</td><td>AAACACACCCCCGCACGCCCCCGT</td></tr>
<tr><td>Chromosome</td><td>start</td><td>end</td><td>strand</td><td>MM</td><td>target_seq</td><td>PAM</td><td>alignment</td><td>distance</td><td>position</td><td>gene name</td><td>gene id</td></tr>
<tr><td>GU305914.1</td><td>19775</td><td>19797</td><td>+</td><td>0</td><td>ACGGGGGCGTGCGGGGGTGT</td><td>GGG</td><td>||||||||[||||||||||||]PAM</td><td>NA</td><td>Intergenic</td><td>NA</td><td>NA</td></tr>
<tr><td>T1040</td><td>GCGTGCGGGGGTGTGGGCGACGG</td><td>1000</td></tr>
<tr><td>Oligo fwd</td><td>CACCGCGTGCGGGGGTGTGGGCGA</td></tr>
<tr><td>Oligo rev</td><td>AAACTCGCCCACACCCCCGCACGC</td></tr>
<tr><td>Chromosome</td><td>start</td><td>end</td><td>strand</td><td>MM</td><td>target_seq</td><td>PAM</td><td>alignment</td><td>distance</td><td>position</td><td>gene name</td><td>gene id</td></tr>
<tr><td>GU305914.1</td><td>19781</td><td>19803</td><td>+</td><td>0</td><td>GCGTGCGGGGGTGTGGGCGA</td><td>CGG</td><td>||||||||[||||||||||||]PAM</td><td>NA</td><td>Intergenic</td><td>NA</td><td>NA</td></tr>
<tr><td>T1041</td><td>GGGGGTGTGGGCGACGGAGCCGG</td><td>1000</td></tr>
<tr><td>Oligo fwd</td><td>CACCGGGGGTGTGGGCGACGGAGC</td></tr>
<tr><td>Oligo rev</td><td>AAACGCTCCGTCGCCCACACCCCC</td></tr>
<tr><td>Chromosome</td><td>start</td><td>end</td><td>strand</td><td>MM</td><td>target_seq</td><td>PAM</td><td>alignment</td><td>distance</td><td>position</td><td>gene name</td><td>gene id</td></tr>
<tr><td>GU305914.1</td><td>19787</td><td>19809</td><td>+</td><td>0</td><td>GGGGGTGTGGGCGACGGAGC</td><td>CGG</td><td>||||||||[||||||||||||]PAM</td><td>NA</td><td>Intergenic</td><td>NA</td><td>NA</td></tr>
</table>

T1042 AGTCAAAAGAGTCGTCCACGAGG 1000
Oligo fwd CACCAGTCAAAAGAGTCGTCCACG
Oligo rev AAACCGTGGACGACTCTTTTGACT
Chromosome start end strand MM target_seq PAM alignment distance position gene name gene id
GU305914.1 19810 19832 + 0 AGTCAAAAGAGTCGTCCACG AGG ||||||||[||||||||||||]PAM NA Intergenic NA NA

T1043 AAAAGAGTCGTCCACGAGGTAGG 1000
Oligo fwd CACCAAAAGAGTCGTCCACGAGGT
Oligo rev AAACACCTCGTGGACGACTCTTTT
Chromosome start end strand MM target_seq PAM alignment distance position gene name gene id
GU305914.1 19814 19836 + 0 AAAAGAGTCGTCCACGAGGT AGG ||||||||[||||||||||||]PAM NA Intergenic NA NA

T1044 GTCGTCCACGAGGTAGGAGCAGG 1000
Oligo fwd CACCGTCGTCCACGAGGTAGGAGC
Oligo rev AAACGCTCCTACCTCGTGGACGAC
Chromosome start end strand MM target_seq PAM alignment distance position gene name gene id
GU305914.1 19820 19842 + 0 GTCGTCCACGAGGTAGGAGC AGG ||||||||[||||||||||||]PAM NA Intergenic NA NA

T1045 GGAGCAGGTGATTGTATCTGTGG 1000
Oligo fwd CACCGGAGCAGGTGATTGTATCTG
Oligo rev AAACCAGATACAATCACCTGCTCC
Chromosome start end strand MM target_seq PAM alignment distance position gene name gene id
GU305914.1 19835 19857 + 0 GGAGCAGGTGATTGTATCTG TGG ||||||||[||||||||||||]PAM NA Intergenic NA NA

T1046 GAGCAGGTGATTGTATCTGTGGG 1000
Oligo fwd CACCGAGCAGGTGATTGTATCTGT
Oligo rev AAACACAGATACAATCACCTGCTC
Chromosome start end strand MM target_seq PAM alignment distance position gene name gene id
GU305914.1 19836 19858 + 0 GAGCAGGTGATTGTATCTGT GGG ||||||||[||||||||||||]PAM NA Intergenic NA NA

T1047 GGTGATTGTATCTGTGGGAGCGG 1000
Oligo fwd CACCGGTGATTGTATCTGTGGGAG
Oligo rev AAACCTCCCACAGATACAATCACC
Chromosome start end strand MM target_seq PAM alignment distance position gene name gene id
GU305914.1 19841 19863 + 0 GGTGATTGTATCTGTGGGAG CGG ||||||||[||||||||||||]PAM NA Intergenic NA NA

T1048 GATTGTATCTGTGGGAGCGGCGG 1000
Oligo fwd CACCGATTGTATCTGTGGGAGCGG
Oligo rev AAACCCGCTCCCACAGATACAATC
Chromosome start end strand MM target_seq PAM alignment distance position gene name gene id
GU305914.1 19844 19866 + 0 GATTGTATCTGTGGGAGCGG CGG ||||||||[||||||||||||]PAM NA Intergenic NA NA

T1049 ATCTGTGGGAGCGGCGGTGTCGG 1000
Oligo fwd CACCATCTGTGGGAGCGGCGGTGT
Oligo rev AAACACACCGCCGCTCCCACAGAT
Chromosome start end strand MM target_seq PAM alignment distance position gene name gene id
GU305914.1 19850 19872 + 0 ATCTGTGGGAGCGGCGGTGT CGG ||||||||[||||||||||||]PAM NA Intergenic NA NA

T1050 TCTGTGGGAGCGGCGGTGTCGGG 1000
Oligo fwd CACCTCTGTGGGAGCGGCGGTGTC
Oligo rev AAACGACACCGCCGCTCCCACAGA
Chromosome start end strand MM target_seq PAM alignment distance position gene name gene id
GU305914.1 19851 19873 + 0 TCTGTGGGAGCGGCGGTGTC GGG ||||||||[||||||||||||]PAM NA Intergenic NA NA

T1051 GTCGGGACGAAGCTGCCGAATGG 1000
Oligo fwd CACCGTCGGGACGAAGCTGCCGAA
Oligo rev AAACTTCGGCAGCTTCGTCCCGAC
Chromosome start end strand MM target_seq PAM alignment distance position gene name gene id
GU305914.1 19868 19890 + 0 GTCGGGACGAAGCTGCCGAA TGG ||||||||[||||||||||||]PAM NA Intergenic NA NA

T1052 GATCCACCCCGTCGCCGCCGAGG 1000
Oligo fwd CACCGATCCACCCCGTCGCCGCCG
Oligo rev AAACCGGCGGCGACGGGGTGGATC
Chromosome start end strand MM target_seq PAM alignment distance position gene name gene id
GU305914.1 19906 19928 + 0 GATCCACCCCGTCGCCGCCG AGG ||||||||[||||||||||||]PAM NA Intergenic NA NA

T1053 CGCCGCCGAGGAATCGAAGACGG 1000
Oligo fwd CACCCGCCGCCGAGGAATCGAAGA
Oligo rev AAACTCTTCGATTCCTCGGCGGCG
Chromosome start end strand MM target_seq PAM alignment distance position gene name gene id
GU305914.1 19918 19940 + 0 CGCCGCCGAGGAATCGAAGA CGG ||||||||[||||||||||||]PAM NA Intergenic NA NA

T1054 CGAGGAATCGAAGACGGTAGCGG 1000
Oligo fwd CACCCGAGGAATCGAAGACGGTAG
Oligo rev AAACCTACCGTCTTCGATTCCTCG
Chromosome start end strand MM target_seq PAM alignment distance position gene name gene id
GU305914.1 19924 19946 + 0 CGAGGAATCGAAGACGGTAG CGG ||||||||[||||||||||||]PAM NA Intergenic NA NA

T1055 GGAATCGAAGACGGTAGCGGTGG 1000
Oligo fwd CACCGGAATCGAAGACGGTAGCGG
Oligo rev AAACCCGCTACCGTCTTCGATTCC
Chromosome start end strand MM target_seq PAM alignment distance position gene name gene id
GU305914.1 19927 19949 + 0 GGAATCGAAGACGGTAGCGG TGG ||||||||[||||||||||||]PAM NA Intergenic NA NA

T1056 ATCGAAGACGGTAGCGGTGGTGG 1000
Oligo fwd CACCATCGAAGACGGTAGCGGTGG
Oligo rev AAACCCACCGCTACCGTCTTCGAT
Chromosome start end strand MM target_seq PAM alignment distance position gene name gene id

<table>
<tr><td>GU305914.1</td><td></td><td>19930</td><td>19952</td><td>+</td><td>0</td><td>ATCGAAGACGGTAGCGGTGG</td><td>TGG</td><td>||||||||[||||||||||||]PAM</td><td>NA</td><td>Intergenic</td><td>NA</td><td>NA</td></tr>
<tr><td colspan="13"></td></tr>
<tr><td>T1057</td><td>GAAGACGGTAGCGGTGGTGGCGG</td><td></td><td>1000</td><td></td><td></td><td></td><td></td><td></td><td></td><td></td><td></td><td></td></tr>
<tr><td>Oligo fwd</td><td>CACCGAAGACGGTAGCGGTGGTGG</td><td colspan="11"></td></tr>
<tr><td>Oligo rev</td><td>AAACCCACCACCGCTACCGTCTTC</td><td colspan="11"></td></tr>
<tr><td>Chromosome</td><td>start</td><td></td><td>end</td><td>strand</td><td>MM</td><td>target_seq</td><td>PAM</td><td>alignment</td><td>distance</td><td>position</td><td>gene name</td><td>gene id</td></tr>
<tr><td>GU305914.1</td><td></td><td>19933</td><td>19955</td><td>+</td><td>0</td><td>GAAGACGGTAGCGGTGGTGG</td><td>CGG</td><td>||||||||[||||||||||||]PAM</td><td>NA</td><td>Intergenic</td><td>NA</td><td>NA</td></tr>
<tr><td colspan="13"></td></tr>
<tr><td>T1058</td><td>GACGGTAGCGGTGGTGGCGGTGG</td><td></td><td>1000</td><td colspan="9"></td></tr>
<tr><td>Oligo fwd</td><td>CACCGACGGTAGCGGTGGTGGCGG</td><td colspan="11"></td></tr>
<tr><td>Oligo rev</td><td>AAACCCGCCACCACCGCTACCGTC</td><td colspan="11"></td></tr>
<tr><td>Chromosome</td><td>start</td><td></td><td>end</td><td>strand</td><td>MM</td><td>target_seq</td><td>PAM</td><td>alignment</td><td>distance</td><td>position</td><td>gene name</td><td>gene id</td></tr>
<tr><td>GU305914.1</td><td></td><td>19936</td><td>19958</td><td>+</td><td>0</td><td>GACGGTAGCGGTGGTGGCGG</td><td>TGG</td><td>||||||||[||||||||||||]PAM</td><td>NA</td><td>Intergenic</td><td>NA</td><td>NA</td></tr>
<tr><td>GU305914.1</td><td></td><td>114661</td><td>114683</td><td>-</td><td>3</td><td>GACGGCGGTGGTGGTGGCGG</td><td>CGG</td><td>|||||--|[-|||||||||||]PAM</td><td>NA</td><td>Intergenic</td><td>NA</td><td>NA</td></tr>
<tr><td>GU305914.1</td><td></td><td>62443</td><td>62465</td><td>+</td><td>4</td><td>GGCGGCGGGGGTGGTGGCGG</td><td>CGG</td><td>|-|||--|[-|||||||||||]PAM</td><td>NA</td><td>Intergenic</td><td>NA</td><td>NA</td></tr>
<tr><td>GU305914.1</td><td></td><td>48156</td><td>48178</td><td>+</td><td>4</td><td>GGCGGCGGCGGTGGCGGCGG</td><td>CGG</td><td>|-|||--|[||||||-|||||]PAM</td><td>NA</td><td>Intergenic</td><td>NA</td><td>NA</td></tr>
<tr><td>GU305914.1</td><td></td><td>114658</td><td>114680</td><td>-</td><td>4</td><td>GGCGGTGGTGGTGGCGGCGG</td><td>CGG</td><td>|-||||-|[-|||||-|||||]PAM</td><td>NA</td><td>Intergenic</td><td>NA</td><td>NA</td></tr>
<tr><td>GU305914.1</td><td></td><td>154404</td><td>154426</td><td>+</td><td>4</td><td>GGCGGTGGCGGCGGCGGCGG</td><td>CGG</td><td>|-||||-|[|||-||-|||||]PAM</td><td>NA</td><td>Intergenic</td><td>NA</td><td>NA</td></tr>
<tr><td>GU305914.1</td><td></td><td>58126</td><td>58148</td><td>-</td><td>4</td><td>GGCGGTGGCGGCGGCGGCGG</td><td>CGG</td><td>|-||||-|[|||-||-|||||]PAM</td><td>NA</td><td>Intergenic</td><td>NA</td><td>NA</td></tr>
<tr><td>GU305914.1</td><td></td><td>45258</td><td>45280</td><td>-</td><td>4</td><td>AACGGTGGCGGTGGCGGCAG</td><td>CGG</td><td>-|||||-|[||||||-|||-|]PAM</td><td>NA</td><td>Intergenic</td><td>NA</td><td>NA</td></tr>
<tr><td colspan="13"></td></tr>
<tr><td>T1059</td><td>GGTGGTGGCGGTGGTTACGACGG</td><td></td><td>1000</td><td colspan="9"></td></tr>
<tr><td>Oligo fwd</td><td>CACCGGTGGTGGCGGTGGTTACGA</td><td colspan="11"></td></tr>
<tr><td>Oligo rev</td><td>AAACTCGTAACCACCGCCACCACC</td><td colspan="11"></td></tr>
<tr><td>Chromosome</td><td>start</td><td></td><td>end</td><td>strand</td><td>MM</td><td>target_seq</td><td>PAM</td><td>alignment</td><td>distance</td><td>position</td><td>gene name</td><td>gene id</td></tr>
<tr><td>GU305914.1</td><td></td><td>19945</td><td>19967</td><td>+</td><td>0</td><td>GGTGGTGGCGGTGGTTACGA</td><td>CGG</td><td>||||||||[||||||||||||]PAM</td><td>NA</td><td>Intergenic</td><td>NA</td><td>NA</td></tr>
<tr><td colspan="13"></td></tr>
<tr><td>T1060</td><td>GTGGTTACGACGGACACCGTCGG</td><td></td><td>1000</td><td colspan="9"></td></tr>
<tr><td>Oligo fwd</td><td>CACCGTGGTTACGACGGACACCGT</td><td colspan="11"></td></tr>
<tr><td>Oligo rev</td><td>AAACACGGTGTCCGTCGTAACCAC</td><td colspan="11"></td></tr>
<tr><td>Chromosome</td><td>start</td><td></td><td>end</td><td>strand</td><td>MM</td><td>target_seq</td><td>PAM</td><td>alignment</td><td>distance</td><td>position</td><td>gene name</td><td>gene id</td></tr>
<tr><td>GU305914.1</td><td></td><td>19955</td><td>19977</td><td>+</td><td>0</td><td>GTGGTTACGACGGACACCGT</td><td>CGG</td><td>||||||||[||||||||||||]PAM</td><td>NA</td><td>Intergenic</td><td>NA</td><td>NA</td></tr>
<tr><td colspan="13"></td></tr>
<tr><td>T1061</td><td>GACGCTGTTGCCTTTGAGATCGG</td><td></td><td>1000</td><td colspan="9"></td></tr>
<tr><td>Oligo fwd</td><td>CACCGACGCTGTTGCCTTTGAGAT</td><td colspan="11"></td></tr>
<tr><td>Oligo rev</td><td>AAACATCTCAAAGGCAACAGCGTC</td><td colspan="11"></td></tr>
<tr><td>Chromosome</td><td>start</td><td></td><td>end</td><td>strand</td><td>MM</td><td>target_seq</td><td>PAM</td><td>alignment</td><td>distance</td><td>position</td><td>gene name</td><td>gene id</td></tr>
<tr><td>GU305914.1</td><td></td><td>20043</td><td>20065</td><td>+</td><td>0</td><td>GACGCTGTTGCCTTTGAGAT</td><td>CGG</td><td>||||||||[||||||||||||]PAM</td><td>NA</td><td>Intergenic</td><td>NA</td><td>NA</td></tr>
<tr><td colspan="13"></td></tr>
<tr><td>T1062</td><td>ACGCTGTTGCCTTTGAGATCGGG</td><td></td><td>1000</td><td colspan="9"></td></tr>
<tr><td>Oligo fwd</td><td>CACCACGCTGTTGCCTTTGAGATC</td><td colspan="11"></td></tr>
<tr><td>Oligo rev</td><td>AAACGATCTCAAAGGCAACAGCGT</td><td colspan="11"></td></tr>
<tr><td>Chromosome</td><td>start</td><td></td><td>end</td><td>strand</td><td>MM</td><td>target_seq</td><td>PAM</td><td>alignment</td><td>distance</td><td>position</td><td>gene name</td><td>gene id</td></tr>
<tr><td>GU305914.1</td><td></td><td>20044</td><td>20066</td><td>+</td><td>0</td><td>ACGCTGTTGCCTTTGAGATC</td><td>GGG</td><td>||||||||[||||||||||||]PAM</td><td>NA</td><td>Intergenic</td><td>NA</td><td>NA</td></tr>
<tr><td colspan="13"></td></tr>
<tr><td>T1063</td><td>CTGTTGCCTTTGAGATCGGGCGG</td><td></td><td>1000</td><td colspan="9"></td></tr>
<tr><td>Oligo fwd</td><td>CACCCTGTTGCCTTTGAGATCGGG</td><td colspan="11"></td></tr>
<tr><td>Oligo rev</td><td>AAACCCCGATCTCAAAGGCAACAG</td><td colspan="11"></td></tr>
<tr><td>Chromosome</td><td>start</td><td></td><td>end</td><td>strand</td><td>MM</td><td>target_seq</td><td>PAM</td><td>alignment</td><td>distance</td><td>position</td><td>gene name</td><td>gene id</td></tr>
<tr><td>GU305914.1</td><td></td><td>20047</td><td>20069</td><td>+</td><td>0</td><td>CTGTTGCCTTTGAGATCGGG</td><td>CGG</td><td>||||||||[||||||||||||]PAM</td><td>NA</td><td>Intergenic</td><td>NA</td><td>NA</td></tr>
<tr><td colspan="13"></td></tr>
<tr><td>T1064</td><td>GTTGAATTATTCCCACCTACCGG</td><td></td><td>1000</td><td colspan="9"></td></tr>
<tr><td>Oligo fwd</td><td>CACCGTTGAATTATTCCCACCTAC</td><td colspan="11"></td></tr>
<tr><td>Oligo rev</td><td>AAACGTAGGTGGGAATAATTCAAC</td><td colspan="11"></td></tr>
<tr><td>Chromosome</td><td>start</td><td></td><td>end</td><td>strand</td><td>MM</td><td>target_seq</td><td>PAM</td><td>alignment</td><td>distance</td><td>position</td><td>gene name</td><td>gene id</td></tr>
<tr><td>GU305914.1</td><td></td><td>20069</td><td>20091</td><td>+</td><td>0</td><td>GTTGAATTATTCCCACCTAC</td><td>CGG</td><td>||||||||[||||||||||||]PAM</td><td>NA</td><td>Intergenic</td><td>NA</td><td>NA</td></tr>
<tr><td colspan="13"></td></tr>
<tr><td>T1065</td><td>CCCACCTACCGGCTCTGCCGCGG</td><td></td><td>1000</td><td colspan="9"></td></tr>
<tr><td>Oligo fwd</td><td>CACCCCCACCTACCGGCTCTGCCG</td><td colspan="11"></td></tr>
<tr><td>Oligo rev</td><td>AAACCGGCAGAGCCGGTAGGTGGG</td><td colspan="11"></td></tr>
<tr><td>Chromosome</td><td>start</td><td></td><td>end</td><td>strand</td><td>MM</td><td>target_seq</td><td>PAM</td><td>alignment</td><td>distance</td><td>position</td><td>gene name</td><td>gene id</td></tr>
<tr><td>GU305914.1</td><td></td><td>20080</td><td>20102</td><td>+</td><td>0</td><td>CCCACCTACCGGCTCTGCCG</td><td>CGG</td><td>||||||||[||||||||||||]PAM</td><td>NA</td><td>Intergenic</td><td>NA</td><td>NA</td></tr>
<tr><td colspan="13"></td></tr>
<tr><td>T1066</td><td>GCCGCGGCACTGCCGTCTGACGG</td><td></td><td>1000</td><td colspan="9"></td></tr>
<tr><td>Oligo fwd</td><td>CACCGCCGCGGCACTGCCGTCTGA</td><td colspan="11"></td></tr>
<tr><td>Oligo rev</td><td>AAACTCAGACGGCAGTGCCGCGGC</td><td colspan="11"></td></tr>
<tr><td>Chromosome</td><td>start</td><td></td><td>end</td><td>strand</td><td>MM</td><td>target_seq</td><td>PAM</td><td>alignment</td><td>distance</td><td>position</td><td>gene name</td><td>gene id</td></tr>
<tr><td>GU305914.1</td><td></td><td>20096</td><td>20118</td><td>+</td><td>0</td><td>GCCGCGGCACTGCCGTCTGA</td><td>CGG</td><td>||||||||[||||||||||||]PAM</td><td>NA</td><td>Intergenic</td><td>NA</td><td>NA</td></tr>
<tr><td colspan="13"></td></tr>
<tr><td>T1067</td><td>GCCGTCTGACGGCATATCGACGG</td><td></td><td>1000</td><td colspan="9"></td></tr>
<tr><td>Oligo fwd</td><td>CACCGCCGTCTGACGGCATATCGA</td><td colspan="11"></td></tr>
<tr><td>Oligo rev</td><td>AAACTCGATATGCCGTCAGACGGC</td><td colspan="11"></td></tr>
<tr><td>Chromosome</td><td>start</td><td></td><td>end</td><td>strand</td><td>MM</td><td>target_seq</td><td>PAM</td><td>alignment</td><td>distance</td><td>position</td><td>gene name</td><td>gene id</td></tr>
<tr><td>GU305914.1</td><td></td><td>20107</td><td>20129</td><td>+</td><td>0</td><td>GCCGTCTGACGGCATATCGA</td><td>CGG</td><td>||||||||[||||||||||||]PAM</td><td>NA</td><td>Intergenic</td><td>NA</td><td>NA</td></tr>
<tr><td colspan="13"></td></tr>
<tr><td>T1068</td><td>CCGTCTGACGGCATATCGACGGG</td><td></td><td>1000</td><td colspan="9"></td></tr>
<tr><td>Oligo fwd</td><td>CACCCCGTCTGACGGCATATCGAC</td><td colspan="11"></td></tr>
<tr><td>Oligo rev</td><td>AAACGTCGATATGCCGTCAGACGG</td><td colspan="11"></td></tr>
<tr><td>Chromosome</td><td>start</td><td></td><td>end</td><td>strand</td><td>MM</td><td>target_seq</td><td>PAM</td><td>alignment</td><td>distance</td><td>position</td><td>gene name</td><td>gene id</td></tr>
<tr><td>GU305914.1</td><td></td><td>20108</td><td>20130</td><td>+</td><td>0</td><td>CCGTCTGACGGCATATCGAC</td><td>GGG</td><td>||||||||[||||||||||||]PAM</td><td>NA</td><td>Intergenic</td><td>NA</td><td>NA</td></tr>
<tr><td colspan="13"></td></tr>
<tr><td>T1069</td><td>CGGCATATCGACGGGAACGACGG</td><td></td><td>1000</td><td colspan="9"></td></tr>
<tr><td>Oligo fwd</td><td>CACCCGGCATATCGACGGGAACGA</td><td colspan="11"></td></tr>
<tr><td>Oligo rev</td><td>AAACTCGTTCCCGTCGATATGCCG</td><td colspan="11"></td></tr>
<tr><td>Chromosome</td><td>start</td><td></td><td>end</td><td>strand</td><td>MM</td><td>target_seq</td><td>PAM</td><td>alignment</td><td>distance</td><td>position</td><td>gene name</td><td>gene id</td></tr>
<tr><td>GU305914.1</td><td></td><td>20116</td><td>20138</td><td>+</td><td>0</td><td>CGGCATATCGACGGGAACGA</td><td>CGG</td><td>||||||||[||||||||||||]PAM</td><td>NA</td><td>Intergenic</td><td>NA</td><td>NA</td></tr>
<tr><td colspan="13"></td></tr>
<tr><td>T1070</td><td>GGCATATCGACGGGAACGACGGG</td><td></td><td>1000</td><td colspan="9"></td></tr>
<tr><td>Oligo fwd</td><td>CACCGGCATATCGACGGGAACGAC</td><td colspan="11"></td></tr>
</table>

<table>
<tr><td>Oligo rev</td><td>AAACGTCGTTCCCGTCGATATGCC</td></tr>
<tr><td>Chromosome</td><td>start</td><td></td><td>end</td><td>strand</td><td>MM</td><td>target_seq</td><td>PAM</td><td>alignment</td><td>distance</td><td>position</td><td>gene name</td><td>gene id</td></tr>
<tr><td>GU305914.1</td><td></td><td>20117</td><td>20139</td><td>+</td><td>0</td><td>GGCATATCGACGGGAACGAC</td><td>GGG</td><td>||||||||[||||||||||||]PAM</td><td>NA</td><td>Intergenic</td><td>NA</td><td>NA</td></tr>
<tr><td>T1071</td><td>CGACCCTCGAACCTCTCGTTCGG</td><td></td><td>1000</td></tr>
<tr><td>Oligo fwd</td><td>CACCCGACCCTCGAACCTCTCGTT</td></tr>
<tr><td>Oligo rev</td><td>AAACAACGAGAGGTTCGAGGGTCG</td></tr>
<tr><td>Chromosome</td><td>start</td><td></td><td>end</td><td>strand</td><td>MM</td><td>target_seq</td><td>PAM</td><td>alignment</td><td>distance</td><td>position</td><td>gene name</td><td>gene id</td></tr>
<tr><td>GU305914.1</td><td></td><td>20148</td><td>20170</td><td>+</td><td>0</td><td>CGACCCTCGAACCTCTCGTT</td><td>CGG</td><td>||||||||[||||||||||||]PAM</td><td>NA</td><td>Intergenic</td><td>NA</td><td>NA</td></tr>
<tr><td>T1072</td><td>GACCCTCGAACCTCTCGTTCGGG</td><td></td><td>1000</td></tr>
<tr><td>Oligo fwd</td><td>CACCGACCCTCGAACCTCTCGTTC</td></tr>
<tr><td>Oligo rev</td><td>AAACGAACGAGAGGTTCGAGGGTC</td></tr>
<tr><td>Chromosome</td><td>start</td><td></td><td>end</td><td>strand</td><td>MM</td><td>target_seq</td><td>PAM</td><td>alignment</td><td>distance</td><td>position</td><td>gene name</td><td>gene id</td></tr>
<tr><td>GU305914.1</td><td></td><td>20149</td><td>20171</td><td>+</td><td>0</td><td>GACCCTCGAACCTCTCGTTC</td><td>GGG</td><td>||||||||[||||||||||||]PAM</td><td>NA</td><td>Intergenic</td><td>NA</td><td>NA</td></tr>
<tr><td>T1073</td><td>CGTGCCCAGTGTCAGCCATGCGG</td><td></td><td>1000</td></tr>
<tr><td>Oligo fwd</td><td>CACCCGTGCCCAGTGTCAGCCATG</td></tr>
<tr><td>Oligo rev</td><td>AAACCATGGCTGACACTGGGCACG</td></tr>
<tr><td>Chromosome</td><td>start</td><td></td><td>end</td><td>strand</td><td>MM</td><td>target_seq</td><td>PAM</td><td>alignment</td><td>distance</td><td>position</td><td>gene name</td><td>gene id</td></tr>
<tr><td>GU305914.1</td><td></td><td>20175</td><td>20197</td><td>+</td><td>0</td><td>CGTGCCCAGTGTCAGCCATG</td><td>CGG</td><td>||||||||[||||||||||||]PAM</td><td>NA</td><td>Intergenic</td><td>NA</td><td>NA</td></tr>
<tr><td>T1074</td><td>GCCCAGTGTCAGCCATGCGGCGG</td><td></td><td>1000</td></tr>
<tr><td>Oligo fwd</td><td>CACCGCCCAGTGTCAGCCATGCGG</td></tr>
<tr><td>Oligo rev</td><td>AAACCCGCATGGCTGACACTGGGC</td></tr>
<tr><td>Chromosome</td><td>start</td><td></td><td>end</td><td>strand</td><td>MM</td><td>target_seq</td><td>PAM</td><td>alignment</td><td>distance</td><td>position</td><td>gene name</td><td>gene id</td></tr>
<tr><td>GU305914.1</td><td></td><td>20178</td><td>20200</td><td>+</td><td>0</td><td>GCCCAGTGTCAGCCATGCGG</td><td>CGG</td><td>||||||||[||||||||||||]PAM</td><td>NA</td><td>Intergenic</td><td>NA</td><td>NA</td></tr>
<tr><td>T1075</td><td>CAGCCATGCGGCGGACGTCTCGG</td><td></td><td>1000</td></tr>
<tr><td>Oligo fwd</td><td>CACCCAGCCATGCGGCGGACGTCT</td></tr>
<tr><td>Oligo rev</td><td>AAACAGACGTCCGCCGCATGGCTG</td></tr>
<tr><td>Chromosome</td><td>start</td><td></td><td>end</td><td>strand</td><td>MM</td><td>target_seq</td><td>PAM</td><td>alignment</td><td>distance</td><td>position</td><td>gene name</td><td>gene id</td></tr>
<tr><td>GU305914.1</td><td></td><td>20187</td><td>20209</td><td>+</td><td>0</td><td>CAGCCATGCGGCGGACGTCT</td><td>CGG</td><td>||||||||[||||||||||||]PAM</td><td>NA</td><td>Intergenic</td><td>NA</td><td>NA</td></tr>
<tr><td>T1076</td><td>TGTATGTCGCTCTGAAACTCAGG</td><td></td><td>1000</td></tr>
<tr><td>Oligo fwd</td><td>CACCTGTATGTCGCTCTGAAACTC</td></tr>
<tr><td>Oligo rev</td><td>AAACGAGTTTCAGAGCGACATACA</td></tr>
<tr><td>Chromosome</td><td>start</td><td></td><td>end</td><td>strand</td><td>MM</td><td>target_seq</td><td>PAM</td><td>alignment</td><td>distance</td><td>position</td><td>gene name</td><td>gene id</td></tr>
<tr><td>GU305914.1</td><td></td><td>20245</td><td>20267</td><td>+</td><td>0</td><td>TGTATGTCGCTCTGAAACTC</td><td>AGG</td><td>||||||||[||||||||||||]PAM</td><td>NA</td><td>Intergenic</td><td>NA</td><td>NA</td></tr>
<tr><td>T1077</td><td>AAACTCAGGACCGACAGCAGTGG</td><td></td><td>1000</td></tr>
<tr><td>Oligo fwd</td><td>CACCAAACTCAGGACCGACAGCAG</td></tr>
<tr><td>Oligo rev</td><td>AAACCTGCTGTCGGTCCTGAGTTT</td></tr>
<tr><td>Chromosome</td><td>start</td><td></td><td>end</td><td>strand</td><td>MM</td><td>target_seq</td><td>PAM</td><td>alignment</td><td>distance</td><td>position</td><td>gene name</td><td>gene id</td></tr>
<tr><td>GU305914.1</td><td></td><td>20259</td><td>20281</td><td>+</td><td>0</td><td>AAACTCAGGACCGACAGCAG</td><td>TGG</td><td>||||||||[||||||||||||]PAM</td><td>NA</td><td>Intergenic</td><td>NA</td><td>NA</td></tr>
<tr><td>T1078</td><td>ACTGACGTTTACTCCGACTCAGG</td><td></td><td>1000</td></tr>
<tr><td>Oligo fwd</td><td>CACCACTGACGTTTACTCCGACTC</td></tr>
<tr><td>Oligo rev</td><td>AAACGAGTCGGAGTAAACGTCAGT</td></tr>
<tr><td>Chromosome</td><td>start</td><td></td><td>end</td><td>strand</td><td>MM</td><td>target_seq</td><td>PAM</td><td>alignment</td><td>distance</td><td>position</td><td>gene name</td><td>gene id</td></tr>
<tr><td>GU305914.1</td><td></td><td>20313</td><td>20335</td><td>+</td><td>0</td><td>ACTGACGTTTACTCCGACTC</td><td>AGG</td><td>||||||||[||||||||||||]PAM</td><td>NA</td><td>Intergenic</td><td>NA</td><td>NA</td></tr>
<tr><td>T1079</td><td>CCCCTACCATTATGTTTCCCCGG</td><td></td><td>1000</td></tr>
<tr><td>Oligo fwd</td><td>CACCCCCCTACCATTATGTTTCCC</td></tr>
<tr><td>Oligo rev</td><td>AAACGGGAAACATAATGGTAGGGG</td></tr>
<tr><td>Chromosome</td><td>start</td><td></td><td>end</td><td>strand</td><td>MM</td><td>target_seq</td><td>PAM</td><td>alignment</td><td>distance</td><td>position</td><td>gene name</td><td>gene id</td></tr>
<tr><td>GU305914.1</td><td></td><td>20361</td><td>20383</td><td>+</td><td>0</td><td>CCCCTACCATTATGTTTCCC</td><td>CGG</td><td>||||||||[||||||||||||]PAM</td><td>NA</td><td>Intergenic</td><td>NA</td><td>NA</td></tr>
<tr><td>T1080</td><td>AAGAAACCAGAAGTGAGACACGG</td><td></td><td>1000</td></tr>
<tr><td>Oligo fwd</td><td>CACCAAGAAACCAGAAGTGAGACA</td></tr>
<tr><td>Oligo rev</td><td>AAACTGTCTCACTTCTGGTTTCTT</td></tr>
<tr><td>Chromosome</td><td>start</td><td></td><td>end</td><td>strand</td><td>MM</td><td>target_seq</td><td>PAM</td><td>alignment</td><td>distance</td><td>position</td><td>gene name</td><td>gene id</td></tr>
<tr><td>GU305914.1</td><td></td><td>20402</td><td>20424</td><td>+</td><td>0</td><td>AAGAAACCAGAAGTGAGACA</td><td>CGG</td><td>||||||||[||||||||||||]PAM</td><td>NA</td><td>Intergenic</td><td>NA</td><td>NA</td></tr>
<tr><td>T1081</td><td>CAGAAGTGAGACACGGACATCGG</td><td></td><td>1000</td></tr>
<tr><td>Oligo fwd</td><td>CACCCAGAAGTGAGACACGGACAT</td></tr>
<tr><td>Oligo rev</td><td>AAACATGTCCGTGTCTCACTTCTG</td></tr>
<tr><td>Chromosome</td><td>start</td><td></td><td>end</td><td>strand</td><td>MM</td><td>target_seq</td><td>PAM</td><td>alignment</td><td>distance</td><td>position</td><td>gene name</td><td>gene id</td></tr>
<tr><td>GU305914.1</td><td></td><td>20409</td><td>20431</td><td>+</td><td>0</td><td>CAGAAGTGAGACACGGACAT</td><td>CGG</td><td>||||||||[||||||||||||]PAM</td><td>NA</td><td>Intergenic</td><td>NA</td><td>NA</td></tr>
<tr><td>T1082</td><td>ACACGGACATCGGACTCGAATGG</td><td></td><td>1000</td></tr>
<tr><td>Oligo fwd</td><td>CACCACACGGACATCGGACTCGAA</td></tr>
<tr><td>Oligo rev</td><td>AAACTTCGAGTCCGATGTCCGTGT</td></tr>
<tr><td>Chromosome</td><td>start</td><td></td><td>end</td><td>strand</td><td>MM</td><td>target_seq</td><td>PAM</td><td>alignment</td><td>distance</td><td>position</td><td>gene name</td><td>gene id</td></tr>
<tr><td>GU305914.1</td><td></td><td>20419</td><td>20441</td><td>+</td><td>0</td><td>ACACGGACATCGGACTCGAA</td><td>TGG</td><td>||||||||[||||||||||||]PAM</td><td>NA</td><td>Intergenic</td><td>NA</td><td>NA</td></tr>
<tr><td>T1083</td><td>GTATGATTTTTTATTACAAAAGG</td><td></td><td>1000</td></tr>
<tr><td>Oligo fwd</td><td>CACCGTATGATTTTTTATTACAAA</td></tr>
<tr><td>Oligo rev</td><td>AAACTTTGTAATAAAAAATCATAC</td></tr>
<tr><td>Chromosome</td><td>start</td><td></td><td>end</td><td>strand</td><td>MM</td><td>target_seq</td><td>PAM</td><td>alignment</td><td>distance</td><td>position</td><td>gene name</td><td>gene id</td></tr>
<tr><td>GU305914.1</td><td></td><td>20441</td><td>20463</td><td>+</td><td>0</td><td>GTATGATTTTTTATTACAAA</td><td>AGG</td><td>||||||||[||||||||||||]PAM</td><td>NA</td><td>Intergenic</td><td>NA</td><td>NA</td></tr>
<tr><td>T1084</td><td>CCGCACGCCCGCCAGCCAGCCGG</td><td></td><td>1000</td></tr>
<tr><td>Oligo fwd</td><td>CACCCCGCACGCCCGCCAGCCAGC</td></tr>
<tr><td>Oligo rev</td><td>AAACGCTGGCTGGCGGGCGTGCGG</td></tr>
<tr><td>Chromosome</td><td>start</td><td></td><td>end</td><td>strand</td><td>MM</td><td>target_seq</td><td>PAM</td><td>alignment</td><td>distance</td><td>position</td><td>gene name</td><td>gene id</td></tr>
<tr><td>GU305914.1</td><td></td><td>20474</td><td>20496</td><td>+</td><td>0</td><td>CCGCACGCCCGCCAGCCAGC</td><td>CGG</td><td>||||||||[||||||||||||]PAM</td><td>NA</td><td>Intergenic</td><td>NA</td><td>NA</td></tr>
<tr><td>GU305914.1</td><td></td><td>20478</td><td>20500</td><td>+</td><td>4</td><td>ACGCCCGCCAGCCAGCCGGC</td><td>CGG</td><td>-|||-|||[|-|||||||-||]PAM</td><td>NA</td><td>Intergenic</td><td>NA</td><td>NA</td></tr>
</table>

<table>
<tr><td>T1085</td><td>ACGCCCGCCAGCCAGCCGGCCGG</td><td>1000</td><td></td><td></td><td></td><td></td><td></td><td></td><td></td><td></td><td></td></tr>
<tr><td>Oligo fwd</td><td>CACCACGCCCGCCAGCCAGCCGGC</td><td></td><td></td><td></td><td></td><td></td><td></td><td></td><td></td><td></td><td></td></tr>
<tr><td>Oligo rev</td><td>AAACGCCGGCTGGCTGGCGGGCGT</td><td></td><td></td><td></td><td></td><td></td><td></td><td></td><td></td><td></td><td></td></tr>
<tr><td>Chromosome</td><td>start</td><td>end</td><td>strand</td><td>MM</td><td>target_seq</td><td>PAM</td><td>alignment</td><td>distance</td><td>position</td><td>gene name</td><td>gene id</td></tr>
<tr><td>GU305914.1</td><td>20478</td><td>20500</td><td>+</td><td>0</td><td>ACGCCCGCCAGCCAGCCGGC</td><td>CGG</td><td>||||||||[||||||||||||]PAM</td><td>NA</td><td>Intergenic</td><td>NA</td><td>NA</td></tr>
<tr><td>GU305914.1</td><td>20474</td><td>20496</td><td>+</td><td>4</td><td>CCGCACGCCCGCCAGCCAGC</td><td>CGG</td><td>-|||-|||[|-|||||||-||]PAM</td><td>NA</td><td>Intergenic</td><td>NA</td><td>NA</td></tr>
<tr><td></td><td></td><td></td><td></td><td></td><td></td><td></td><td></td><td></td><td></td><td></td><td></td></tr>
<tr><td>T1086</td><td>CGCCCGCCAGCCAGCCGGCCGGG</td><td>1000</td><td></td><td></td><td></td><td></td><td></td><td></td><td></td><td></td><td></td></tr>
<tr><td>Oligo fwd</td><td>CACCCGCCCGCCAGCCAGCCGGCC</td><td></td><td></td><td></td><td></td><td></td><td></td><td></td><td></td><td></td><td></td></tr>
<tr><td>Oligo rev</td><td>AAACGGCCGGCTGGCTGGCGGGCG</td><td></td><td></td><td></td><td></td><td></td><td></td><td></td><td></td><td></td><td></td></tr>
<tr><td>Chromosome</td><td>start</td><td>end</td><td>strand</td><td>MM</td><td>target_seq</td><td>PAM</td><td>alignment</td><td>distance</td><td>position</td><td>gene name</td><td>gene id</td></tr>
<tr><td>GU305914.1</td><td>20479</td><td>20501</td><td>+</td><td>0</td><td>CGCCCGCCAGCCAGCCGGCC</td><td>GGG</td><td>||||||||[||||||||||||]PAM</td><td>NA</td><td>Intergenic</td><td>NA</td><td>NA</td></tr>
<tr><td></td><td></td><td></td><td></td><td></td><td></td><td></td><td></td><td></td><td></td><td></td><td></td></tr>
<tr><td>T1087</td><td>CCGGGCCTCCTTCAGTCCCGCGG</td><td>1000</td><td></td><td></td><td></td><td></td><td></td><td></td><td></td><td></td><td></td></tr>
<tr><td>Oligo fwd</td><td>CACCCCGGGCCTCCTTCAGTCCCG</td><td></td><td></td><td></td><td></td><td></td><td></td><td></td><td></td><td></td><td></td></tr>
<tr><td>Oligo rev</td><td>AAACCGGGACTGAAGGAGGCCCGG</td><td></td><td></td><td></td><td></td><td></td><td></td><td></td><td></td><td></td><td></td></tr>
<tr><td>Chromosome</td><td>start</td><td>end</td><td>strand</td><td>MM</td><td>target_seq</td><td>PAM</td><td>alignment</td><td>distance</td><td>position</td><td>gene name</td><td>gene id</td></tr>
<tr><td>GU305914.1</td><td>20497</td><td>20519</td><td>+</td><td>0</td><td>CCGGGCCTCCTTCAGTCCCG</td><td>CGG</td><td>||||||||[||||||||||||]PAM</td><td>NA</td><td>Intergenic</td><td>NA</td><td>NA</td></tr>
<tr><td></td><td></td><td></td><td></td><td></td><td></td><td></td><td></td><td></td><td></td><td></td><td></td></tr>
<tr><td>T1088</td><td>GCCTCCTTCAGTCCCGCGGTCGG</td><td>1000</td><td></td><td></td><td></td><td></td><td></td><td></td><td></td><td></td><td></td></tr>
<tr><td>Oligo fwd</td><td>CACCGCCTCCTTCAGTCCCGCGGT</td><td></td><td></td><td></td><td></td><td></td><td></td><td></td><td></td><td></td><td></td></tr>
<tr><td>Oligo rev</td><td>AAACACCGCGGGACTGAAGGAGGC</td><td></td><td></td><td></td><td></td><td></td><td></td><td></td><td></td><td></td><td></td></tr>
<tr><td>Chromosome</td><td>start</td><td>end</td><td>strand</td><td>MM</td><td>target_seq</td><td>PAM</td><td>alignment</td><td>distance</td><td>position</td><td>gene name</td><td>gene id</td></tr>
<tr><td>GU305914.1</td><td>20501</td><td>20523</td><td>+</td><td>0</td><td>GCCTCCTTCAGTCCCGCGGT</td><td>CGG</td><td>||||||||[||||||||||||]PAM</td><td>NA</td><td>Intergenic</td><td>NA</td><td>NA</td></tr>
<tr><td></td><td></td><td></td><td></td><td></td><td></td><td></td><td></td><td></td><td></td><td></td><td></td></tr>
<tr><td>T1089</td><td>CCTCCTTCAGTCCCGCGGTCGGG</td><td>1000</td><td></td><td></td><td></td><td></td><td></td><td></td><td></td><td></td><td></td></tr>
<tr><td>Oligo fwd</td><td>CACCCCTCCTTCAGTCCCGCGGTC</td><td></td><td></td><td></td><td></td><td></td><td></td><td></td><td></td><td></td><td></td></tr>
<tr><td>Oligo rev</td><td>AAACGACCGCGGGACTGAAGGAGG</td><td></td><td></td><td></td><td></td><td></td><td></td><td></td><td></td><td></td><td></td></tr>
<tr><td>Chromosome</td><td>start</td><td>end</td><td>strand</td><td>MM</td><td>target_seq</td><td>PAM</td><td>alignment</td><td>distance</td><td>position</td><td>gene name</td><td>gene id</td></tr>
<tr><td>GU305914.1</td><td>20502</td><td>20524</td><td>+</td><td>0</td><td>CCTCCTTCAGTCCCGCGGTC</td><td>GGG</td><td>||||||||[||||||||||||]PAM</td><td>NA</td><td>Intergenic</td><td>NA</td><td>NA</td></tr>
<tr><td></td><td></td><td></td><td></td><td></td><td></td><td></td><td></td><td></td><td></td><td></td><td></td></tr>
<tr><td>T1090</td><td>GGTCGGGATGTTCTAGTTAGAGG</td><td>1000</td><td></td><td></td><td></td><td></td><td></td><td></td><td></td><td></td><td></td></tr>
<tr><td>Oligo fwd</td><td>CACCGGTCGGGATGTTCTAGTTAG</td><td></td><td></td><td></td><td></td><td></td><td></td><td></td><td></td><td></td><td></td></tr>
<tr><td>Oligo rev</td><td>AAACCTAACTAGAACATCCCGACC</td><td></td><td></td><td></td><td></td><td></td><td></td><td></td><td></td><td></td><td></td></tr>
<tr><td>Chromosome</td><td>start</td><td>end</td><td>strand</td><td>MM</td><td>target_seq</td><td>PAM</td><td>alignment</td><td>distance</td><td>position</td><td>gene name</td><td>gene id</td></tr>
<tr><td>GU305914.1</td><td>20518</td><td>20540</td><td>+</td><td>0</td><td>GGTCGGGATGTTCTAGTTAG</td><td>AGG</td><td>||||||||[||||||||||||]PAM</td><td>NA</td><td>Intergenic</td><td>NA</td><td>NA</td></tr>
<tr><td></td><td></td><td></td><td></td><td></td><td></td><td></td><td></td><td></td><td></td><td></td><td></td></tr>
<tr><td>T1091</td><td>ATGTTCTAGTTAGAGGAGAAAGG</td><td>1000</td><td></td><td></td><td></td><td></td><td></td><td></td><td></td><td></td><td></td></tr>
<tr><td>Oligo fwd</td><td>CACCATGTTCTAGTTAGAGGAGAA</td><td></td><td></td><td></td><td></td><td></td><td></td><td></td><td></td><td></td><td></td></tr>
<tr><td>Oligo rev</td><td>AAACTTCTCCTCTAACTAGAACAT</td><td></td><td></td><td></td><td></td><td></td><td></td><td></td><td></td><td></td><td></td></tr>
<tr><td>Chromosome</td><td>start</td><td>end</td><td>strand</td><td>MM</td><td>target_seq</td><td>PAM</td><td>alignment</td><td>distance</td><td>position</td><td>gene name</td><td>gene id</td></tr>
<tr><td>GU305914.1</td><td>20525</td><td>20547</td><td>+</td><td>0</td><td>ATGTTCTAGTTAGAGGAGAA</td><td>AGG</td><td>||||||||[||||||||||||]PAM</td><td>NA</td><td>Intergenic</td><td>NA</td><td>NA</td></tr>
<tr><td></td><td></td><td></td><td></td><td></td><td></td><td></td><td></td><td></td><td></td><td></td><td></td></tr>
<tr><td>T1092</td><td>TGTTCTAGTTAGAGGAGAAAGGG</td><td>1000</td><td></td><td></td><td></td><td></td><td></td><td></td><td></td><td></td><td></td></tr>
<tr><td>Oligo fwd</td><td>CACCTGTTCTAGTTAGAGGAGAAA</td><td></td><td></td><td></td><td></td><td></td><td></td><td></td><td></td><td></td><td></td></tr>
<tr><td>Oligo rev</td><td>AAACTTTCTCCTCTAACTAGAACA</td><td></td><td></td><td></td><td></td><td></td><td></td><td></td><td></td><td></td><td></td></tr>
<tr><td>Chromosome</td><td>start</td><td>end</td><td>strand</td><td>MM</td><td>target_seq</td><td>PAM</td><td>alignment</td><td>distance</td><td>position</td><td>gene name</td><td>gene id</td></tr>
<tr><td>GU305914.1</td><td>20526</td><td>20548</td><td>+</td><td>0</td><td>TGTTCTAGTTAGAGGAGAAA</td><td>GGG</td><td>||||||||[||||||||||||]PAM</td><td>NA</td><td>Intergenic</td><td>NA</td><td>NA</td></tr>
<tr><td></td><td></td><td></td><td></td><td></td><td></td><td></td><td></td><td></td><td></td><td></td><td></td></tr>
<tr><td>T1093</td><td>GGAAGAGAAAGCAAGCAGAAAGG</td><td>1000</td><td></td><td></td><td></td><td></td><td></td><td></td><td></td><td></td><td></td></tr>
<tr><td>Oligo fwd</td><td>CACCGGAAGAGAAAGCAAGCAGAA</td><td></td><td></td><td></td><td></td><td></td><td></td><td></td><td></td><td></td><td></td></tr>
<tr><td>Oligo rev</td><td>AAACTTCTGCTTGCTTTCTCTTCC</td><td></td><td></td><td></td><td></td><td></td><td></td><td></td><td></td><td></td><td></td></tr>
<tr><td>Chromosome</td><td>start</td><td>end</td><td>strand</td><td>MM</td><td>target_seq</td><td>PAM</td><td>alignment</td><td>distance</td><td>position</td><td>gene name</td><td>gene id</td></tr>
<tr><td>GU305914.1</td><td>20547</td><td>20569</td><td>+</td><td>0</td><td>GGAAGAGAAAGCAAGCAGAA</td><td>AGG</td><td>||||||||[||||||||||||]PAM</td><td>NA</td><td>Intergenic</td><td>NA</td><td>NA</td></tr>
<tr><td></td><td></td><td></td><td></td><td></td><td></td><td></td><td></td><td></td><td></td><td></td><td></td></tr>
<tr><td>T1094</td><td>GAAGAGAAAGCAAGCAGAAAGGG</td><td>1000</td><td></td><td></td><td></td><td></td><td></td><td></td><td></td><td></td><td></td></tr>
<tr><td>Oligo fwd</td><td>CACCGAAGAGAAAGCAAGCAGAAA</td><td></td><td></td><td></td><td></td><td></td><td></td><td></td><td></td><td></td><td></td></tr>
<tr><td>Oligo rev</td><td>AAACTTTCTGCTTGCTTTCTCTTC</td><td></td><td></td><td></td><td></td><td></td><td></td><td></td><td></td><td></td><td></td></tr>
<tr><td>Chromosome</td><td>start</td><td>end</td><td>strand</td><td>MM</td><td>target_seq</td><td>PAM</td><td>alignment</td><td>distance</td><td>position</td><td>gene name</td><td>gene id</td></tr>
<tr><td>GU305914.1</td><td>20548</td><td>20570</td><td>+</td><td>0</td><td>GAAGAGAAAGCAAGCAGAAA</td><td>GGG</td><td>||||||||[||||||||||||]PAM</td><td>NA</td><td>Intergenic</td><td>NA</td><td>NA</td></tr>
<tr><td></td><td></td><td></td><td></td><td></td><td></td><td></td><td></td><td></td><td></td><td></td><td></td></tr>
<tr><td>T1095</td><td>GAGAAAGCAAGCAGAAAGGGAGG</td><td>1000</td><td></td><td></td><td></td><td></td><td></td><td></td><td></td><td></td><td></td></tr>
<tr><td>Oligo fwd</td><td>CACCGAGAAAGCAAGCAGAAAGGG</td><td></td><td></td><td></td><td></td><td></td><td></td><td></td><td></td><td></td><td></td></tr>
<tr><td>Oligo rev</td><td>AAACCCCTTTCTGCTTGCTTTCTC</td><td></td><td></td><td></td><td></td><td></td><td></td><td></td><td></td><td></td><td></td></tr>
<tr><td>Chromosome</td><td>start</td><td>end</td><td>strand</td><td>MM</td><td>target_seq</td><td>PAM</td><td>alignment</td><td>distance</td><td>position</td><td>gene name</td><td>gene id</td></tr>
<tr><td>GU305914.1</td><td>20551</td><td>20573</td><td>+</td><td>0</td><td>GAGAAAGCAAGCAGAAAGGG</td><td>AGG</td><td>||||||||[||||||||||||]PAM</td><td>NA</td><td>Intergenic</td><td>NA</td><td>NA</td></tr>
<tr><td></td><td></td><td></td><td></td><td></td><td></td><td></td><td></td><td></td><td></td><td></td><td></td></tr>
<tr><td>T1096</td><td>AAGCAAGCAGAAAGGGAGGAAGG</td><td>1000</td><td></td><td></td><td></td><td></td><td></td><td></td><td></td><td></td><td></td></tr>
<tr><td>Oligo fwd</td><td>CACCAAGCAAGCAGAAAGGGAGGA</td><td></td><td></td><td></td><td></td><td></td><td></td><td></td><td></td><td></td><td></td></tr>
<tr><td>Oligo rev</td><td>AAACTCCTCCCTTTCTGCTTGCTT</td><td></td><td></td><td></td><td></td><td></td><td></td><td></td><td></td><td></td><td></td></tr>
<tr><td>Chromosome</td><td>start</td><td>end</td><td>strand</td><td>MM</td><td>target_seq</td><td>PAM</td><td>alignment</td><td>distance</td><td>position</td><td>gene name</td><td>gene id</td></tr>
<tr><td>GU305914.1</td><td>20555</td><td>20577</td><td>+</td><td>0</td><td>AAGCAAGCAGAAAGGGAGGA</td><td>AGG</td><td>||||||||[||||||||||||]PAM</td><td>NA</td><td>Intergenic</td><td>NA</td><td>NA</td></tr>
<tr><td></td><td></td><td></td><td></td><td></td><td></td><td></td><td></td><td></td><td></td><td></td><td></td></tr>
<tr><td>T1097</td><td>AGCAAGCAGAAAGGGAGGAAGGG</td><td>1000</td><td></td><td></td><td></td><td></td><td></td><td></td><td></td><td></td><td></td></tr>
<tr><td>Oligo fwd</td><td>CACCAGCAAGCAGAAAGGGAGGAA</td><td></td><td></td><td></td><td></td><td></td><td></td><td></td><td></td><td></td><td></td></tr>
<tr><td>Oligo rev</td><td>AAACTTCCTCCCTTTCTGCTTGCT</td><td></td><td></td><td></td><td></td><td></td><td></td><td></td><td></td><td></td><td></td></tr>
<tr><td>Chromosome</td><td>start</td><td>end</td><td>strand</td><td>MM</td><td>target_seq</td><td>PAM</td><td>alignment</td><td>distance</td><td>position</td><td>gene name</td><td>gene id</td></tr>
<tr><td>GU305914.1</td><td>20556</td><td>20578</td><td>+</td><td>0</td><td>AGCAAGCAGAAAGGGAGGAA</td><td>GGG</td><td>||||||||[||||||||||||]PAM</td><td>NA</td><td>Intergenic</td><td>NA</td><td>NA</td></tr>
<tr><td></td><td></td><td></td><td></td><td></td><td></td><td></td><td></td><td></td><td></td><td></td><td></td></tr>
<tr><td>T1098</td><td>GTCAGTCGCGTAACTCTTGTTGG</td><td>1000</td><td></td><td></td><td></td><td></td><td></td><td></td><td></td><td></td><td></td></tr>
<tr><td>Oligo fwd</td><td>CACCGTCAGTCGCGTAACTCTTGT</td><td></td><td></td><td></td><td></td><td></td><td></td><td></td><td></td><td></td><td></td></tr>
<tr><td>Oligo rev</td><td>AAACACAAGAGTTACGCGACTGAC</td><td></td><td></td><td></td><td></td><td></td><td></td><td></td><td></td><td></td><td></td></tr>
<tr><td>Chromosome</td><td>start</td><td>end</td><td>strand</td><td>MM</td><td>target_seq</td><td>PAM</td><td>alignment</td><td>distance</td><td>position</td><td>gene name</td><td>gene id</td></tr>
<tr><td>GU305914.1</td><td>20578</td><td>20600</td><td>+</td><td>0</td><td>GTCAGTCGCGTAACTCTTGT</td><td>TGG</td><td>||||||||[||||||||||||]PAM</td><td>NA</td><td>Intergenic</td><td>NA</td><td>NA</td></tr>
<tr><td></td><td></td><td></td><td></td><td></td><td></td><td></td><td></td><td></td><td></td><td></td><td></td></tr>
<tr><td>T1099</td><td>GCGTAACTCTTGTTGGTCCTTGG</td><td>1000</td><td></td><td></td><td></td><td></td><td></td><td></td><td></td><td></td><td></td></tr>
<tr><td>Oligo fwd</td><td>CACCGCGTAACTCTTGTTGGTCCT</td><td></td><td></td><td></td><td></td><td></td><td></td><td></td><td></td><td></td><td></td></tr>
<tr><td>Oligo rev</td><td>AAACAGGACCAACAAGAGTTACGC</td><td></td><td></td><td></td><td></td><td></td><td></td><td></td><td></td><td></td><td></td></tr>
<tr><td>Chromosome</td><td>start</td><td>end</td><td>strand</td><td>MM</td><td>target_seq</td><td>PAM</td><td>alignment</td><td>distance</td><td>position</td><td>gene name</td><td>gene id</td></tr>
</table>

| | | | | | | | | | | | |
|---|---|---|---|---|---|---|---|---|---|---|---|
| GU305914.1 | 20585 | 20607 | + | 0 | GCGTAACTCTTGTTGGTCCT | TGG | \|\|\|\|\|\|\|\|[\|\|\|\|\|\|\|\|\|\|\|\|]PAM | NA | Intergenic | NA | NA |
| T1100 | CGTAACTCTTGTTGGTCCTTGGG | 1000 | | | | | | | | | |
| Oligo fwd | CACCCGTAACTCTTGTTGGTCCTT | | | | | | | | | | |
| Oligo rev | AAACAAGGACCAACAAGAGTTACG | | | | | | | | | | |
| Chromosome | start | end | strand | MM | target_seq | PAM | alignment | distance | position | gene name | gene id |
| GU305914.1 | 20586 | 20608 | + | 0 | CGTAACTCTTGTTGGTCCTT | GGG | \|\|\|\|\|\|\|\|[\|\|\|\|\|\|\|\|\|\|\|\|]PAM | NA | Intergenic | NA | NA |
| T1101 | CTTGGGCGACTGCTCCTCGACGG | 1000 | | | | | | | | | |
| Oligo fwd | CACCCTTGGGCGACTGCTCCTCGA | | | | | | | | | | |
| Oligo rev | AAACTCGAGGAGCAGTCGCCCAAG | | | | | | | | | | |
| Chromosome | start | end | strand | MM | target_seq | PAM | alignment | distance | position | gene name | gene id |
| GU305914.1 | 20603 | 20625 | + | 0 | CTTGGGCGACTGCTCCTCGA | CGG | \|\|\|\|\|\|\|\|[\|\|\|\|\|\|\|\|\|\|\|\|]PAM | NA | Intergenic | NA | NA |
| T1102 | CGGCCATCTGTTCCTCTTCCAGG | 1000 | | | | | | | | | |
| Oligo fwd | CACCCGGCCATCTGTTCCTCTTCC | | | | | | | | | | |
| Oligo rev | AAACGGAAGAGGAACAGATGGCCG | | | | | | | | | | |
| Chromosome | start | end | strand | MM | target_seq | PAM | alignment | distance | position | gene name | gene id |
| GU305914.1 | 20623 | 20645 | + | 0 | CGGCCATCTGTTCCTCTTCC | AGG | \|\|\|\|\|\|\|\|[\|\|\|\|\|\|\|\|\|\|\|\|]PAM | NA | Intergenic | NA | NA |
| T1103 | CGTATCGCCGTCTCCGTCGCAGG | 1000 | | | | | | | | | |
| Oligo fwd | CACCCGTATCGCCGTCTCCGTCGC | | | | | | | | | | |
| Oligo rev | AAACGCGACGGAGACGGCGATACG | | | | | | | | | | |
| Chromosome | start | end | strand | MM | target_seq | PAM | alignment | distance | position | gene name | gene id |
| GU305914.1 | 20660 | 20682 | + | 0 | CGTATCGCCGTCTCCGTCGC | AGG | \|\|\|\|\|\|\|\|[\|\|\|\|\|\|\|\|\|\|\|\|]PAM | NA | Intergenic | NA | NA |
| T1104 | GCCGTCTCCGTCGCAGGCCTCGG | 1000 | | | | | | | | | |
| Oligo fwd | CACCGCCGTCTCCGTCGCAGGCCT | | | | | | | | | | |
| Oligo rev | AAACAGGCCTGCGACGGAGACGGC | | | | | | | | | | |
| Chromosome | start | end | strand | MM | target_seq | PAM | alignment | distance | position | gene name | gene id |
| GU305914.1 | 20666 | 20688 | + | 0 | GCCGTCTCCGTCGCAGGCCT | CGG | \|\|\|\|\|\|\|\|[\|\|\|\|\|\|\|\|\|\|\|\|]PAM | NA | Intergenic | NA | NA |
| T1105 | CGCGTCTTCCACCACGCCGCTGG | 1000 | | | | | | | | | |
| Oligo fwd | CACCCGCGTCTTCCACCACGCCGC | | | | | | | | | | |
| Oligo rev | AAACGCGGCGTGGTGGAAGACGCG | | | | | | | | | | |
| Chromosome | start | end | strand | MM | target_seq | PAM | alignment | distance | position | gene name | gene id |
| GU305914.1 | 20693 | 20715 | + | 0 | CGCGTCTTCCACCACGCCGC | TGG | \|\|\|\|\|\|\|\|[\|\|\|\|\|\|\|\|\|\|\|\|]PAM | NA | Intergenic | NA | NA |
| T1106 | TTCCACCACGCCGCTGGCACCGG | 1000 | | | | | | | | | |
| Oligo fwd | CACCTTCCACCACGCCGCTGGCAC | | | | | | | | | | |
| Oligo rev | AAACGTGCCAGCGGCGTGGTGGAA | | | | | | | | | | |
| Chromosome | start | end | strand | MM | target_seq | PAM | alignment | distance | position | gene name | gene id |
| GU305914.1 | 20699 | 20721 | + | 0 | TTCCACCACGCCGCTGGCAC | CGG | \|\|\|\|\|\|\|\|[\|\|\|\|\|\|\|\|\|\|\|\|]PAM | NA | Intergenic | NA | NA |
| T1107 | TCCACCACGCCGCTGGCACCGGG | 1000 | | | | | | | | | |
| Oligo fwd | CACCTCCACCACGCCGCTGGCACC | | | | | | | | | | |
| Oligo rev | AAACGGTGCCAGCGGCGTGGTGGA | | | | | | | | | | |
| Chromosome | start | end | strand | MM | target_seq | PAM | alignment | distance | position | gene name | gene id |
| GU305914.1 | 20700 | 20722 | + | 0 | TCCACCACGCCGCTGGCACC | GGG | \|\|\|\|\|\|\|\|[\|\|\|\|\|\|\|\|\|\|\|\|]PAM | NA | Intergenic | NA | NA |
| T1108 | CAGTCCCCAGTAGCACACGACGG | 1000 | | | | | | | | | |
| Oligo fwd | CACCCAGTCCCCAGTAGCACACGA | | | | | | | | | | |
| Oligo rev | AAACTCGTGTGCTACTGGGGACTG | | | | | | | | | | |
| Chromosome | start | end | strand | MM | target_seq | PAM | alignment | distance | position | gene name | gene id |
| GU305914.1 | 20723 | 20745 | + | 0 | CAGTCCCCAGTAGCACACGA | CGG | \|\|\|\|\|\|\|\|[\|\|\|\|\|\|\|\|\|\|\|\|]PAM | NA | Intergenic | NA | NA |
| T1109 | GTAGCACACGACGGCACCGACGG | 1000 | | | | | | | | | |
| Oligo fwd | CACCGTAGCACACGACGGCACCGA | | | | | | | | | | |
| Oligo rev | AAACTCGGTGCCGTCGTGTGCTAC | | | | | | | | | | |
| Chromosome | start | end | strand | MM | target_seq | PAM | alignment | distance | position | gene name | gene id |
| GU305914.1 | 20732 | 20754 | + | 0 | GTAGCACACGACGGCACCGA | CGG | \|\|\|\|\|\|\|\|[\|\|\|\|\|\|\|\|\|\|\|\|]PAM | NA | Intergenic | NA | NA |
| T1110 | GACGGCACCGACGGCGACGACGG | 1000 | | | | | | | | | |
| Oligo fwd | CACCGACGGCACCGACGGCGACGA | | | | | | | | | | |
| Oligo rev | AAACTCGTCGCCGTCGGTGCCGTC | | | | | | | | | | |
| Chromosome | start | end | strand | MM | target_seq | PAM | alignment | distance | position | gene name | gene id |
| GU305914.1 | 20741 | 20763 | + | 0 | GACGGCACCGACGGCGACGA | CGG | \|\|\|\|\|\|\|\|[\|\|\|\|\|\|\|\|\|\|\|\|]PAM | NA | Intergenic | NA | NA |
| GU305914.1 | 117379 | 117401 | + | 3 | AACGGCACCGACAGCGCCGA | CGG | -\|\|\|\|\|\|\|[\|\|\|\|-\|\|\|-\|\|\|]PAM | NA | Intergenic | NA | NA |
| GU305914.1 | 20750 | 20772 | + | 4 | GACGGCGACGACGGCGATCA | CGG | \|\|\|\|\|\|--[\|\|\|\|\|\|\|\|\|--\|]PAM | NA | Intergenic | NA | NA |
| T1111 | GACGGCGACGACGGCGATCACGG | 1000 | | | | | | | | | |
| Oligo fwd | CACCGACGGCGACGACGGCGATCA | | | | | | | | | | |
| Oligo rev | AAACTGATCGCCGTCGTCGCCGTC | | | | | | | | | | |
| Chromosome | start | end | strand | MM | target_seq | PAM | alignment | distance | position | gene name | gene id |
| GU305914.1 | 20750 | 20772 | + | 0 | GACGGCGACGACGGCGATCA | CGG | \|\|\|\|\|\|\|\|[\|\|\|\|\|\|\|\|\|\|\|\|]PAM | NA | Intergenic | NA | NA |
| GU305914.1 | 20741 | 20763 | + | 4 | GACGGCACCGACGGCGACGA | CGG | \|\|\|\|\|\|--[\|\|\|\|\|\|\|\|\|--\|]PAM | NA | Intergenic | NA | NA |
| T1112 | CGACGACGGCGATCACGGCGAGG | 1000 | | | | | | | | | |
| Oligo fwd | CACCCGACGACGGCGATCACGGCG | | | | | | | | | | |
| Oligo rev | AAACCGCCGTGATCGCCGTCGTCG | | | | | | | | | | |
| Chromosome | start | end | strand | MM | target_seq | PAM | alignment | distance | position | gene name | gene id |
| GU305914.1 | 20755 | 20777 | + | 0 | CGACGACGGCGATCACGGCG | AGG | \|\|\|\|\|\|\|\|[\|\|\|\|\|\|\|\|\|\|\|\|]PAM | NA | Intergenic | NA | NA |
| T1113 | GACGACGGCGATCACGGCGAGGG | 1000 | | | | | | | | | |
| Oligo fwd | CACCGACGACGGCGATCACGGCGA | | | | | | | | | | |
| Oligo rev | AAACTCGCCGTGATCGCCGTCGTC | | | | | | | | | | |
| Chromosome | start | end | strand | MM | target_seq | PAM | alignment | distance | position | gene name | gene id |
| GU305914.1 | 20756 | 20778 | + | 0 | GACGACGGCGATCACGGCGA | GGG | \|\|\|\|\|\|\|\|[\|\|\|\|\|\|\|\|\|\|\|\|]PAM | NA | Intergenic | NA | NA |
| GU305914.1 | 17711 | 17733 | - | 3 | GACGACGACGGCCACGGCGA | TGG | \|\|\|\|\|\|\|-[\|\|--\|\|\|\|\|\|\|\|]PAM | NA | Intergenic | NA | NA |

| | | | | | | | | | | | |
|---|---|---|---|---|---|---|---|---|---|---|---|
| GU305914.1 | | 96936 | 96958 | + | 4 | GGCGAGGGCGAGCAGGGCGA | CGG | \|-\|\|\|-\|\|[\|\|\|-\|\|-\|\|\|\|\|]PAM | NA | Intergenic | NA | NA |
| GU305914.1 | | 215568 | 215590 | - | 4 | GACAACGACGATTACTGCGA | TGG | \|\|\|-\|\|\|-[\|\|\|\|-\|\|-\|\|\|\|]PAM | NA | Intergenic | NA | NA |
| GU305914.1 | | 40442 | 40464 | - | 4 | GAAGACGACGATGACGGCGG | AGG | \|\|-\|\|\|\|-[\|\|\|\|-\|\|\|\|\|\|-]PAM | NA | Intergenic | NA | NA |
| T1114 | CGATCACGGCGAGGGTGAAGAGG | | 1000 | | | | | | | | | |
| Oligo fwd | CACCCGATCACGGCGAGGGTGAAG | | | | | | | | | | | |
| Oligo rev | AAACCTTCACCCTCGCCGTGATCG | | | | | | | | | | | |
| Chromosome | start | | end | strand | MM | target_seq | PAM | alignment | distance | position | gene name | gene id |
| GU305914.1 | | 20764 | 20786 | + | 0 | CGATCACGGCGAGGGTGAAG | AGG | \|\|\|\|\|\|\|\|[\|\|\|\|\|\|\|\|\|\|\|\|]PAM | NA | Intergenic | NA | NA |
| T1115 | GATCACGGCGAGGGTGAAGAGGG | | 1000 | | | | | | | | | |
| Oligo fwd | CACCGATCACGGCGAGGGTGAAGA | | | | | | | | | | | |
| Oligo rev | AAACTCTTCACCCTCGCCGTGATC | | | | | | | | | | | |
| Chromosome | start | | end | strand | MM | target_seq | PAM | alignment | distance | position | gene name | gene id |
| GU305914.1 | | 20765 | 20787 | + | 0 | GATCACGGCGAGGGTGAAGA | GGG | \|\|\|\|\|\|\|\|[\|\|\|\|\|\|\|\|\|\|\|\|]PAM | NA | Intergenic | NA | NA |
| T1116 | GGCGAGGGTGAAGAGGGTGATGG | | 1000 | | | | | | | | | |
| Oligo fwd | CACCGGCGAGGGTGAAGAGGGTGA | | | | | | | | | | | |
| Oligo rev | AAACTCACCCTCTTCACCCTCGCC | | | | | | | | | | | |
| Chromosome | start | | end | strand | MM | target_seq | PAM | alignment | distance | position | gene name | gene id |
| GU305914.1 | | 20771 | 20793 | + | 0 | GGCGAGGGTGAAGAGGGTGA | TGG | \|\|\|\|\|\|\|\|[\|\|\|\|\|\|\|\|\|\|\|\|]PAM | NA | Intergenic | NA | NA |
| T1117 | TTCCAACATCGTCCTTCTCGAGG | | 1000 | | | | | | | | | |
| Oligo fwd | CACCTTCCAACATCGTCCTTCTCG | | | | | | | | | | | |
| Oligo rev | AAACCGAGAAGGACGATGTTGGAA | | | | | | | | | | | |
| Chromosome | start | | end | strand | MM | target_seq | PAM | alignment | distance | position | gene name | gene id |
| GU305914.1 | | 20833 | 20855 | + | 0 | TTCCAACATCGTCCTTCTCG | AGG | \|\|\|\|\|\|\|\|[\|\|\|\|\|\|\|\|\|\|\|\|]PAM | NA | Intergenic | NA | NA |
| T1118 | TCCAACATCGTCCTTCTCGAGGG | | 1000 | | | | | | | | | |
| Oligo fwd | CACCTCCAACATCGTCCTTCTCGA | | | | | | | | | | | |
| Oligo rev | AAACTCGAGAAGGACGATGTTGGA | | | | | | | | | | | |
| Chromosome | start | | end | strand | MM | target_seq | PAM | alignment | distance | position | gene name | gene id |
| GU305914.1 | | 20834 | 20856 | + | 0 | TCCAACATCGTCCTTCTCGA | GGG | \|\|\|\|\|\|\|\|[\|\|\|\|\|\|\|\|\|\|\|\|]PAM | NA | Intergenic | NA | NA |
| T1119 | GTCCTTCTCGAGGGTCGTCGAGG | | 1000 | | | | | | | | | |
| Oligo fwd | CACCGTCCTTCTCGAGGGTCGTCG | | | | | | | | | | | |
| Oligo rev | AAACCGACGACCCTCGAGAAGGAC | | | | | | | | | | | |
| Chromosome | start | | end | strand | MM | target_seq | PAM | alignment | distance | position | gene name | gene id |
| GU305914.1 | | 20843 | 20865 | + | 0 | GTCCTTCTCGAGGGTCGTCG | AGG | \|\|\|\|\|\|\|\|[\|\|\|\|\|\|\|\|\|\|\|\|]PAM | NA | Intergenic | NA | NA |
| T1120 | AGGGTCGTCGAGGAAAGCGCCGG | | 1000 | | | | | | | | | |
| Oligo fwd | CACCAGGGTCGTCGAGGAAAGCGC | | | | | | | | | | | |
| Oligo rev | AAACGCGCTTTCCTCGACGACCCT | | | | | | | | | | | |
| Chromosome | start | | end | strand | MM | target_seq | PAM | alignment | distance | position | gene name | gene id |
| GU305914.1 | | 20853 | 20875 | + | 0 | AGGGTCGTCGAGGAAAGCGC | CGG | \|\|\|\|\|\|\|\|[\|\|\|\|\|\|\|\|\|\|\|\|]PAM | NA | Intergenic | NA | NA |
| T1121 | GGAAAGCGCCGGTCCGCGAGCGG | | 1000 | | | | | | | | | |
| Oligo fwd | CACCGGAAAGCGCCGGTCCGCGAG | | | | | | | | | | | |
| Oligo rev | AAACCTCGCGGACCGGCGCTTTCC | | | | | | | | | | | |
| Chromosome | start | | end | strand | MM | target_seq | PAM | alignment | distance | position | gene name | gene id |
| GU305914.1 | | 20864 | 20886 | + | 0 | GGAAAGCGCCGGTCCGCGAG | CGG | \|\|\|\|\|\|\|\|[\|\|\|\|\|\|\|\|\|\|\|\|]PAM | NA | Intergenic | NA | NA |
| T1122 | AGCGCCGGTCCGCGAGCGGTAGG | | 1000 | | | | | | | | | |
| Oligo fwd | CACCAGCGCCGGTCCGCGAGCGGT | | | | | | | | | | | |
| Oligo rev | AAACACCGCTCGCGGACCGGCGCT | | | | | | | | | | | |
| Chromosome | start | | end | strand | MM | target_seq | PAM | alignment | distance | position | gene name | gene id |
| GU305914.1 | | 20868 | 20890 | + | 0 | AGCGCCGGTCCGCGAGCGGT | AGG | \|\|\|\|\|\|\|\|[\|\|\|\|\|\|\|\|\|\|\|\|]PAM | NA | Intergenic | NA | NA |
| T1123 | TCCGCGAGCGGTAGGTAGAGAGG | | 1000 | | | | | | | | | |
| Oligo fwd | CACCTCCGCGAGCGGTAGGTAGAG | | | | | | | | | | | |
| Oligo rev | AAACCTCTACCTACCGCTCGCGGA | | | | | | | | | | | |
| Chromosome | start | | end | strand | MM | target_seq | PAM | alignment | distance | position | gene name | gene id |
| GU305914.1 | | 20876 | 20898 | + | 0 | TCCGCGAGCGGTAGGTAGAG | AGG | \|\|\|\|\|\|\|\|[\|\|\|\|\|\|\|\|\|\|\|\|]PAM | NA | Intergenic | NA | NA |
| T1124 | AGGTAGAGAGGCGAGCGACACGG | | 1000 | | | | | | | | | |
| Oligo fwd | CACCAGGTAGAGAGGCGAGCGACA | | | | | | | | | | | |
| Oligo rev | AAACTGTCGCTCGCCTCTCTACCT | | | | | | | | | | | |
| Chromosome | start | | end | strand | MM | target_seq | PAM | alignment | distance | position | gene name | gene id |
| GU305914.1 | | 20888 | 20910 | + | 0 | AGGTAGAGAGGCGAGCGACA | CGG | \|\|\|\|\|\|\|\|[\|\|\|\|\|\|\|\|\|\|\|\|]PAM | NA | Intergenic | NA | NA |
| T1125 | AGAGGCGAGCGACACGGCCGCGG | | 1000 | | | | | | | | | |
| Oligo fwd | CACCAGAGGCGAGCGACACGGCCG | | | | | | | | | | | |
| Oligo rev | AAACCGGCCGTGTCGCTCGCCTCT | | | | | | | | | | | |
| Chromosome | start | | end | strand | MM | target_seq | PAM | alignment | distance | position | gene name | gene id |
| GU305914.1 | | 20894 | 20916 | + | 0 | AGAGGCGAGCGACACGGCCG | CGG | \|\|\|\|\|\|\|\|[\|\|\|\|\|\|\|\|\|\|\|\|]PAM | NA | Intergenic | NA | NA |
| T1126 | GAGCGACACGGCCGCGGCGCTGG | | 1000 | | | | | | | | | |
| Oligo fwd | CACCGAGCGACACGGCCGCGGCGC | | | | | | | | | | | |
| Oligo rev | AAACGCGCCGCGGCCGTGTCGCTC | | | | | | | | | | | |
| Chromosome | start | | end | strand | MM | target_seq | PAM | alignment | distance | position | gene name | gene id |
| GU305914.1 | | 20900 | 20922 | + | 0 | GAGCGACACGGCCGCGGCGC | TGG | \|\|\|\|\|\|\|\|[\|\|\|\|\|\|\|\|\|\|\|\|]PAM | NA | Intergenic | NA | NA |
| T1127 | AGCGACACGGCCGCGGCGCTGGG | | 1000 | | | | | | | | | |
| Oligo fwd | CACCAGCGACACGGCCGCGGCGCT | | | | | | | | | | | |
| Oligo rev | AAACAGCGCCGCGGCCGTGTCGCT | | | | | | | | | | | |
| Chromosome | start | | end | strand | MM | target_seq | PAM | alignment | distance | position | gene name | gene id |
| GU305914.1 | | 20901 | 20923 | + | 0 | AGCGACACGGCCGCGGCGCT | GGG | \|\|\|\|\|\|\|\|[\|\|\|\|\|\|\|\|\|\|\|\|]PAM | NA | Intergenic | NA | NA |
| T1128 | GCGACACGGCCGCGGCGCTGGGG | | 1000 | | | | | | | | | |

Oligo fwd CACCGCGACACGGCCGCGGCGCTG
Oligo rev AAACCAGCGCCGCGGCCGTGTCGC
Chromosome start end strand MM target_seq PAM alignment distance position gene name gene id
GU305914.1 20902 20924 + 0 GCGACACGGCCGCGGCGCTG GGG ||||||||[||||||||||||]PAM NA Intergenic NA NA

T1129 TCGCCTCGTTGTCCGCTTTGAGG 1000
Oligo fwd CACCTCGCCTCGTTGTCCGCTTTG
Oligo rev AAACCAAAGCGGACAACGAGGCGA
Chromosome start end strand MM target_seq PAM alignment distance position gene name gene id
GU305914.1 20938 20960 + 0 TCGCCTCGTTGTCCGCTTTG AGG ||||||||[||||||||||||]PAM NA Intergenic NA NA

T1130 CCTCGTTGTCCGCTTTGAGGCGG 1000
Oligo fwd CACCCCTCGTTGTCCGCTTTGAGG
Oligo rev AAACCCTCAAAGCGGACAACGAGG
Chromosome start end strand MM target_seq PAM alignment distance position gene name gene id
GU305914.1 20941 20963 + 0 CCTCGTTGTCCGCTTTGAGG CGG ||||||||[||||||||||||]PAM NA Intergenic NA NA

T1131 CTCGTTGTCCGCTTTGAGGCGGG 1000
Oligo fwd CACCCTCGTTGTCCGCTTTGAGGC
Oligo rev AAACGCCTCAAAGCGGACAACGAG
Chromosome start end strand MM target_seq PAM alignment distance position gene name gene id
GU305914.1 20942 20964 + 0 CTCGTTGTCCGCTTTGAGGC GGG ||||||||[||||||||||||]PAM NA Intergenic NA NA

T1132 CCGCTTTGAGGCGGGCGCACTGG 1000
Oligo fwd CACCCCGCTTTGAGGCGGGCGCAC
Oligo rev AAACGTGCGCCCGCCTCAAAGCGG
Chromosome start end strand MM target_seq PAM alignment distance position gene name gene id
GU305914.1 20950 20972 + 0 CCGCTTTGAGGCGGGCGCAC TGG ||||||||[||||||||||||]PAM NA Intergenic NA NA

T1133 CTCGACCACTGTGAAGACGCAGG 1000
Oligo fwd CACCCTCGACCACTGTGAAGACGC
Oligo rev AAACGCGTCTTCACAGTGGTCGAG
Chromosome start end strand MM target_seq PAM alignment distance position gene name gene id
GU305914.1 20975 20997 + 0 CTCGACCACTGTGAAGACGC AGG ||||||||[||||||||||||]PAM NA Intergenic NA NA

T1134 CTGTGAAGACGCAGGAGTAGCGG 1000
Oligo fwd CACCCTGTGAAGACGCAGGAGTAG
Oligo rev AAACCTACTCCTGCGTCTTCACAG
Chromosome start end strand MM target_seq PAM alignment distance position gene name gene id
GU305914.1 20983 21005 + 0 CTGTGAAGACGCAGGAGTAG CGG ||||||||[||||||||||||]PAM NA Intergenic NA NA

T1135 GTAGCGGCGAACCCCGTCGATGG 1000
Oligo fwd CACCGTAGCGGCGAACCCCGTCGA
Oligo rev AAACTCGACGGGGTTCGCCGCTAC
Chromosome start end strand MM target_seq PAM alignment distance position gene name gene id
GU305914.1 20999 21021 + 0 GTAGCGGCGAACCCCGTCGA TGG ||||||||[||||||||||||]PAM NA Intergenic NA NA

T1136 GGCGAACCCCGTCGATGGTGAGG 1000
Oligo fwd CACCGGCGAACCCCGTCGATGGTG
Oligo rev AAACCACCATCGACGGGGTTCGCC
Chromosome start end strand MM target_seq PAM alignment distance position gene name gene id
GU305914.1 21004 21026 + 0 GGCGAACCCCGTCGATGGTG AGG ||||||||[||||||||||||]PAM NA Intergenic NA NA

T1137 CGTCGATGGTGAGGAGACCTCGG 1000
Oligo fwd CACCCGTCGATGGTGAGGAGACCT
Oligo rev AAACAGGTCTCCTCACCATCGACG
Chromosome start end strand MM target_seq PAM alignment distance position gene name gene id
GU305914.1 21013 21035 + 0 CGTCGATGGTGAGGAGACCT CGG ||||||||[||||||||||||]PAM NA Intergenic NA NA

T1138 GAGGAGACCTCGGTATACGCCGG 1000
Oligo fwd CACCGAGGAGACCTCGGTATACGC
Oligo rev AAACGCGTATACCGAGGTCTCCTC
Chromosome start end strand MM target_seq PAM alignment distance position gene name gene id
GU305914.1 21023 21045 + 0 GAGGAGACCTCGGTATACGC CGG ||||||||[||||||||||||]PAM NA Intergenic NA NA

T1139 CTTTCTAACCACGAATCTGAAGG 1000
Oligo fwd CACCCTTTCTAACCACGAATCTGA
Oligo rev AAACTCAGATTCGTGGTTAGAAAG
Chromosome start end strand MM target_seq PAM alignment distance position gene name gene id
GU305914.1 21047 21069 + 0 CTTTCTAACCACGAATCTGA AGG ||||||||[||||||||||||]PAM NA Intergenic NA NA

T1140 TAACCACGAATCTGAAGGTGAGG 1000
Oligo fwd CACCTAACCACGAATCTGAAGGTG
Oligo rev AAACCACCTTCAGATTCGTGGTTA
Chromosome start end strand MM target_seq PAM alignment distance position gene name gene id
GU305914.1 21052 21074 + 0 TAACCACGAATCTGAAGGTG AGG ||||||||[||||||||||||]PAM NA Intergenic NA NA

T1141 TCTCGCCACACCGTGTCCGTAGG 1000
Oligo fwd CACCTCTCGCCACACCGTGTCCGT
Oligo rev AAACACGGACACGGTGTGGCGAGA
Chromosome start end strand MM target_seq PAM alignment distance position gene name gene id
GU305914.1 21087 21109 + 0 TCTCGCCACACCGTGTCCGT AGG ||||||||[||||||||||||]PAM NA Intergenic NA NA

T1142 CTCGCCACACCGTGTCCGTAGGG 1000
Oligo fwd CACCCTCGCCACACCGTGTCCGTA
Oligo rev AAACTACGGACACGGTGTGGCGAG
Chromosome start end strand MM target_seq PAM alignment distance position gene name gene id
GU305914.1 21088 21110 + 0 CTCGCCACACCGTGTCCGTA GGG ||||||||[||||||||||||]PAM NA Intergenic NA NA

<table>
<tr><td>T1143</td><td>TCGCCACACCGTGTCCGTAGGGG</td><td>1000</td></tr>
<tr><td>Oligo fwd</td><td>CACCTCGCCACACCGTGTCCGTAG</td></tr>
<tr><td>Oligo rev</td><td>AAACCTACGGACACGGTGTGGCGA</td></tr>
<tr><td>Chromosome</td><td>start</td><td>end</td><td>strand</td><td>MM</td><td>target_seq</td><td>PAM</td><td>alignment</td><td>distance</td><td>position</td><td>gene name</td><td>gene id</td></tr>
<tr><td>GU305914.1</td><td>21089</td><td>21111</td><td>+</td><td>0</td><td>TCGCCACACCGTGTCCGTAG</td><td>GGG</td><td>||||||||[||||||||||||]PAM</td><td>NA</td><td>Intergenic</td><td>NA</td><td>NA</td></tr>
<tr><td>T1144</td><td>GGACATGACGTACGCTCGATCGG</td><td>1000</td></tr>
<tr><td>Oligo fwd</td><td>CACCGGACATGACGTACGCTCGAT</td></tr>
<tr><td>Oligo rev</td><td>AAACATCGAGCGTACGTCATGTCC</td></tr>
<tr><td>Chromosome</td><td>start</td><td>end</td><td>strand</td><td>MM</td><td>target_seq</td><td>PAM</td><td>alignment</td><td>distance</td><td>position</td><td>gene name</td><td>gene id</td></tr>
<tr><td>GU305914.1</td><td>21110</td><td>21132</td><td>+</td><td>0</td><td>GGACATGACGTACGCTCGAT</td><td>CGG</td><td>||||||||[||||||||||||]PAM</td><td>NA</td><td>Intergenic</td><td>NA</td><td>NA</td></tr>
<tr><td>T1145</td><td>CACCCTCCGTTCCGTGTATGAGG</td><td>1000</td></tr>
<tr><td>Oligo fwd</td><td>CACCCACCCTCCGTTCCGTGTATG</td></tr>
<tr><td>Oligo rev</td><td>AAACCATACACGGAACGGAGGGTG</td></tr>
<tr><td>Chromosome</td><td>start</td><td>end</td><td>strand</td><td>MM</td><td>target_seq</td><td>PAM</td><td>alignment</td><td>distance</td><td>position</td><td>gene name</td><td>gene id</td></tr>
<tr><td>GU305914.1</td><td>21133</td><td>21155</td><td>+</td><td>0</td><td>CACCCTCCGTTCCGTGTATG</td><td>AGG</td><td>||||||||[||||||||||||]PAM</td><td>NA</td><td>Intergenic</td><td>NA</td><td>NA</td></tr>
<tr><td>T1146</td><td>CCTGTTAAACAACACAGCCCCGG</td><td>1000</td></tr>
<tr><td>Oligo fwd</td><td>CACCCCTGTTAAACAACACAGCCC</td></tr>
<tr><td>Oligo rev</td><td>AAACGGGCTGTGTTGTTTAACAGG</td></tr>
<tr><td>Chromosome</td><td>start</td><td>end</td><td>strand</td><td>MM</td><td>target_seq</td><td>PAM</td><td>alignment</td><td>distance</td><td>position</td><td>gene name</td><td>gene id</td></tr>
<tr><td>GU305914.1</td><td>21167</td><td>21189</td><td>+</td><td>0</td><td>CCTGTTAAACAACACAGCCC</td><td>CGG</td><td>||||||||[||||||||||||]PAM</td><td>NA</td><td>Intergenic</td><td>NA</td><td>NA</td></tr>
<tr><td>T1147</td><td>TAAACAACACAGCCCCGGCGAGG</td><td>1000</td></tr>
<tr><td>Oligo fwd</td><td>CACCTAAACAACACAGCCCCGGCG</td></tr>
<tr><td>Oligo rev</td><td>AAACCGCCGGGGCTGTGTTGTTTA</td></tr>
<tr><td>Chromosome</td><td>start</td><td>end</td><td>strand</td><td>MM</td><td>target_seq</td><td>PAM</td><td>alignment</td><td>distance</td><td>position</td><td>gene name</td><td>gene id</td></tr>
<tr><td>GU305914.1</td><td>21172</td><td>21194</td><td>+</td><td>0</td><td>TAAACAACACAGCCCCGGCG</td><td>AGG</td><td>||||||||[||||||||||||]PAM</td><td>NA</td><td>Intergenic</td><td>NA</td><td>NA</td></tr>
<tr><td>T1148</td><td>GCCCCGGCGAGGATCCGCAACGG</td><td>1000</td></tr>
<tr><td>Oligo fwd</td><td>CACCGCCCCGGCGAGGATCCGCAA</td></tr>
<tr><td>Oligo rev</td><td>AAACTTGCGGATCCTCGCCGGGGC</td></tr>
<tr><td>Chromosome</td><td>start</td><td>end</td><td>strand</td><td>MM</td><td>target_seq</td><td>PAM</td><td>alignment</td><td>distance</td><td>position</td><td>gene name</td><td>gene id</td></tr>
<tr><td>GU305914.1</td><td>21183</td><td>21205</td><td>+</td><td>0</td><td>GCCCCGGCGAGGATCCGCAA</td><td>CGG</td><td>||||||||[||||||||||||]PAM</td><td>NA</td><td>Intergenic</td><td>NA</td><td>NA</td></tr>
<tr><td>T1149</td><td>TCCATTATCTCCCGTGCGATCGG</td><td>1000</td></tr>
<tr><td>Oligo fwd</td><td>CACCTCCATTATCTCCCGTGCGAT</td></tr>
<tr><td>Oligo rev</td><td>AAACATCGCACGGGAGATAATGGA</td></tr>
<tr><td>Chromosome</td><td>start</td><td>end</td><td>strand</td><td>MM</td><td>target_seq</td><td>PAM</td><td>alignment</td><td>distance</td><td>position</td><td>gene name</td><td>gene id</td></tr>
<tr><td>GU305914.1</td><td>21209</td><td>21231</td><td>+</td><td>0</td><td>TCCATTATCTCCCGTGCGAT</td><td>CGG</td><td>||||||||[||||||||||||]PAM</td><td>NA</td><td>Intergenic</td><td>NA</td><td>NA</td></tr>
<tr><td>T1150</td><td>CCATTATCTCCCGTGCGATCGGG</td><td>1000</td></tr>
<tr><td>Oligo fwd</td><td>CACCCCATTATCTCCCGTGCGATC</td></tr>
<tr><td>Oligo rev</td><td>AAACGATCGCACGGGAGATAATGG</td></tr>
<tr><td>Chromosome</td><td>start</td><td>end</td><td>strand</td><td>MM</td><td>target_seq</td><td>PAM</td><td>alignment</td><td>distance</td><td>position</td><td>gene name</td><td>gene id</td></tr>
<tr><td>GU305914.1</td><td>21210</td><td>21232</td><td>+</td><td>0</td><td>CCATTATCTCCCGTGCGATC</td><td>GGG</td><td>||||||||[||||||||||||]PAM</td><td>NA</td><td>Intergenic</td><td>NA</td><td>NA</td></tr>
<tr><td>T1151</td><td>CATTATCTCCCGTGCGATCGGGG</td><td>1000</td></tr>
<tr><td>Oligo fwd</td><td>CACCCATTATCTCCCGTGCGATCG</td></tr>
<tr><td>Oligo rev</td><td>AAACCGATCGCACGGGAGATAATG</td></tr>
<tr><td>Chromosome</td><td>start</td><td>end</td><td>strand</td><td>MM</td><td>target_seq</td><td>PAM</td><td>alignment</td><td>distance</td><td>position</td><td>gene name</td><td>gene id</td></tr>
<tr><td>GU305914.1</td><td>21211</td><td>21233</td><td>+</td><td>0</td><td>CATTATCTCCCGTGCGATCG</td><td>GGG</td><td>||||||||[||||||||||||]PAM</td><td>NA</td><td>Intergenic</td><td>NA</td><td>NA</td></tr>
<tr><td>T1152</td><td>ATTATCTCCCGTGCGATCGGGGG</td><td>1000</td></tr>
<tr><td>Oligo fwd</td><td>CACCATTATCTCCCGTGCGATCGG</td></tr>
<tr><td>Oligo rev</td><td>AAACCCGATCGCACGGGAGATAAT</td></tr>
<tr><td>Chromosome</td><td>start</td><td>end</td><td>strand</td><td>MM</td><td>target_seq</td><td>PAM</td><td>alignment</td><td>distance</td><td>position</td><td>gene name</td><td>gene id</td></tr>
<tr><td>GU305914.1</td><td>21212</td><td>21234</td><td>+</td><td>0</td><td>ATTATCTCCCGTGCGATCGG</td><td>GGG</td><td>||||||||[||||||||||||]PAM</td><td>NA</td><td>Intergenic</td><td>NA</td><td>NA</td></tr>
<tr><td>T1153</td><td>CGTGCGATCGGGGGAATCGAAGG</td><td>1000</td></tr>
<tr><td>Oligo fwd</td><td>CACCCGTGCGATCGGGGGAATCGA</td></tr>
<tr><td>Oligo rev</td><td>AAACTCGATTCCCCCGATCGCACG</td></tr>
<tr><td>Chromosome</td><td>start</td><td>end</td><td>strand</td><td>MM</td><td>target_seq</td><td>PAM</td><td>alignment</td><td>distance</td><td>position</td><td>gene name</td><td>gene id</td></tr>
<tr><td>GU305914.1</td><td>21221</td><td>21243</td><td>+</td><td>0</td><td>CGTGCGATCGGGGGAATCGA</td><td>AGG</td><td>||||||||[||||||||||||]PAM</td><td>NA</td><td>Intergenic</td><td>NA</td><td>NA</td></tr>
<tr><td>T1154</td><td>TCCAGTTGAAAATCCCGATGCGG</td><td>1000</td></tr>
<tr><td>Oligo fwd</td><td>CACCTCCAGTTGAAAATCCCGATG</td></tr>
<tr><td>Oligo rev</td><td>AAACCATCGGGATTTTCAACTGGA</td></tr>
<tr><td>Chromosome</td><td>start</td><td>end</td><td>strand</td><td>MM</td><td>target_seq</td><td>PAM</td><td>alignment</td><td>distance</td><td>position</td><td>gene name</td><td>gene id</td></tr>
<tr><td>GU305914.1</td><td>21244</td><td>21266</td><td>+</td><td>0</td><td>TCCAGTTGAAAATCCCGATG</td><td>CGG</td><td>||||||||[||||||||||||]PAM</td><td>NA</td><td>Intergenic</td><td>NA</td><td>NA</td></tr>
<tr><td>T1155</td><td>CCGATGCGGTAATAGACCAGAGG</td><td>1000</td></tr>
<tr><td>Oligo fwd</td><td>CACCCCGATGCGGTAATAGACCAG</td></tr>
<tr><td>Oligo rev</td><td>AAACCTGGTCTATTACCGCATCGG</td></tr>
<tr><td>Chromosome</td><td>start</td><td>end</td><td>strand</td><td>MM</td><td>target_seq</td><td>PAM</td><td>alignment</td><td>distance</td><td>position</td><td>gene name</td><td>gene id</td></tr>
<tr><td>GU305914.1</td><td>21258</td><td>21280</td><td>+</td><td>0</td><td>CCGATGCGGTAATAGACCAG</td><td>AGG</td><td>||||||||[||||||||||||]PAM</td><td>NA</td><td>Intergenic</td><td>NA</td><td>NA</td></tr>
<tr><td>T1156</td><td>GACCAGAGGAATGTCGATGATGG</td><td>1000</td></tr>
<tr><td>Oligo fwd</td><td>CACCGACCAGAGGAATGTCGATGA</td></tr>
<tr><td>Oligo rev</td><td>AAACTCATCGACATTCCTCTGGTC</td></tr>
<tr><td>Chromosome</td><td>start</td><td>end</td><td>strand</td><td>MM</td><td>target_seq</td><td>PAM</td><td>alignment</td><td>distance</td><td>position</td><td>gene name</td><td>gene id</td></tr>
<tr><td>GU305914.1</td><td>21272</td><td>21294</td><td>+</td><td>0</td><td>GACCAGAGGAATGTCGATGA</td><td>TGG</td><td>||||||||[||||||||||||]PAM</td><td>NA</td><td>Intergenic</td><td>NA</td><td>NA</td></tr>
<tr><td>T1157</td><td>AGGAATGTCGATGATGGACTCGG</td><td>1000</td></tr>
<tr><td>Oligo fwd</td><td>CACCAGGAATGTCGATGATGGACT</td></tr>
<tr><td>Oligo rev</td><td>AAACAGTCCATCATCGACATTCCT</td></tr>
<tr><td>Chromosome</td><td>start</td><td>end</td><td>strand</td><td>MM</td><td>target_seq</td><td>PAM</td><td>alignment</td><td>distance</td><td>position</td><td>gene name</td><td>gene id</td></tr>
<tr><td>GU305914.1</td><td>21278</td><td>21300</td><td>+</td><td>0</td><td>AGGAATGTCGATGATGGACT</td><td>CGG</td><td>||||||||[||||||||||||]PAM</td><td>NA</td><td>Intergenic</td><td>NA</td><td>NA</td></tr>
</table>

T1158 TCGATGATGGACTCGGTGTACGG 1000
Oligo fwd CACCTCGATGATGGACTCGGTGTA
Oligo rev AAACTACACCGAGTCCATCATCGA
Chromosome start end strand MM target_seq PAM alignment distance position gene name gene id
GU305914.1 21285 21307 + 0 TCGATGATGGACTCGGTGTA CGG ||||||||[||||||||||||]PAM NA Intergenic NA NA

T1159 GGACTCGGTGTACGGCGTCATGG 1000
Oligo fwd CACCGGACTCGGTGTACGGCGTCA
Oligo rev AAACTGACGCCGTACACCGAGTCC
Chromosome start end strand MM target_seq PAM alignment distance position gene name gene id
GU305914.1 21293 21315 + 0 GGACTCGGTGTACGGCGTCA TGG ||||||||[||||||||||||]PAM NA Intergenic NA NA
GU305914.1 48079 48101 - 4 GGACCCGCAGTACGTCGTCA GGG ||||-||-[-|||||-|||||]PAM NA Intergenic NA NA

T1160 ACATCCGTGCACGACCGCCTCGG 1000
Oligo fwd CACCACATCCGTGCACGACCGCCT
Oligo rev AAACAGGCGGTCGTGCACGGATGT
Chromosome start end strand MM target_seq PAM alignment distance position gene name gene id
GU305914.1 21326 21348 + 0 ACATCCGTGCACGACCGCCT CGG ||||||||[||||||||||||]PAM NA Intergenic NA NA

T1161 CACGACCGCCTCGGCGTGCGCGG 1000
Oligo fwd CACCCACGACCGCCTCGGCGTGCG
Oligo rev AAACCGCACGCCGAGGCGGTCGTG
Chromosome start end strand MM target_seq PAM alignment distance position gene name gene id
GU305914.1 21335 21357 + 0 CACGACCGCCTCGGCGTGCG CGG ||||||||[||||||||||||]PAM NA Intergenic NA NA
GU305914.1 134237 134259 - 3 CACTACCGCCCCCGCGTGCG TGG |||-||||[||-|-|||||||]PAM NA Intergenic NA NA

T1162 GACCGCCTCGGCGTGCGCGGCGG 1000
Oligo fwd CACCGACCGCCTCGGCGTGCGCGG
Oligo rev AAACCCGCGCACGCCGAGGCGGTC
Chromosome start end strand MM target_seq PAM alignment distance position gene name gene id
GU305914.1 21338 21360 + 0 GACCGCCTCGGCGTGCGCGG CGG ||||||||[||||||||||||]PAM NA Intergenic NA NA
GU305914.1 26973 26995 + 4 GACCGCTCCGGCGGCCGCGG CGG ||||||--[|||||--|||||]PAM NA Intergenic NA NA

T1163 GGCGGTCAACATCTTTTCTTTGG 1000
Oligo fwd CACCGGCGGTCAACATCTTTTCTT
Oligo rev AAACAAGAAAAGATGTTGACCGCC
Chromosome start end strand MM target_seq PAM alignment distance position gene name gene id
GU305914.1 21356 21378 + 0 GGCGGTCAACATCTTTTCTT TGG ||||||||[||||||||||||]PAM NA Intergenic NA NA

T1164 TGCTGTAGTCGACCAACACCTGG 1000
Oligo fwd CACCTGCTGTAGTCGACCAACACC
Oligo rev AAACGGTGTTGGTCGACTACAGCA
Chromosome start end strand MM target_seq PAM alignment distance position gene name gene id
GU305914.1 21382 21404 + 0 TGCTGTAGTCGACCAACACC TGG ||||||||[||||||||||||]PAM NA Intergenic NA NA

T1165 TACTCACGTTGCCGTTGTGCAGG 1000
Oligo fwd CACCTACTCACGTTGCCGTTGTGC
Oligo rev AAACGCACAACGGCAACGTGAGTA
Chromosome start end strand MM target_seq PAM alignment distance position gene name gene id
GU305914.1 21409 21431 + 0 TACTCACGTTGCCGTTGTGC AGG ||||||||[||||||||||||]PAM NA Intergenic NA NA

T1166 GTTGTGCAGGTCCGAGAACATGG 1000
Oligo fwd CACCGTTGTGCAGGTCCGAGAACA
Oligo rev AAACTGTTCTCGGACCTGCACAAC
Chromosome start end strand MM target_seq PAM alignment distance position gene name gene id
GU305914.1 21422 21444 + 0 GTTGTGCAGGTCCGAGAACA TGG ||||||||[||||||||||||]PAM NA Intergenic NA NA

T1167 GTGCAGGTCCGAGAACATGGCGG 1000
Oligo fwd CACCGTGCAGGTCCGAGAACATGG
Oligo rev AAACCCATGTTCTCGGACCTGCAC
Chromosome start end strand MM target_seq PAM alignment distance position gene name gene id
GU305914.1 21425 21447 + 0 GTGCAGGTCCGAGAACATGG CGG ||||||||[||||||||||||]PAM NA Intergenic NA NA

T1168 GAACATGGCGGTGAAGTAGACGG 1000
Oligo fwd CACCGAACATGGCGGTGAAGTAGA
Oligo rev AAACTCTACTTCACCGCCATGTTC
Chromosome start end strand MM target_seq PAM alignment distance position gene name gene id
GU305914.1 21437 21459 + 0 GAACATGGCGGTGAAGTAGA CGG ||||||||[||||||||||||]PAM NA Intergenic NA NA

T1169 GACGGCGCAGCGATACATTCCGG 1000
Oligo fwd CACCGACGGCGCAGCGATACATTC
Oligo rev AAACGAATGTATCGCTGCGCCGTC
Chromosome start end strand MM target_seq PAM alignment distance position gene name gene id
GU305914.1 21455 21477 + 0 GACGGCGCAGCGATACATTC CGG ||||||||[||||||||||||]PAM NA Intergenic NA NA

T1170 ATACATTCCGGCCAGATCGTTGG 1000
Oligo fwd CACCATACATTCCGGCCAGATCGT
Oligo rev AAACACGATCTGGCCGGAATGTAT
Chromosome start end strand MM target_seq PAM alignment distance position gene name gene id
GU305914.1 21467 21489 + 0 ATACATTCCGGCCAGATCGT TGG ||||||||[||||||||||||]PAM NA Intergenic NA NA

T1171 TTTTCCGAAGTTCCACACGCAGG 1000
Oligo fwd CACCTTTTCCGAAGTTCCACACGC
Oligo rev AAACGCGTGTGGAACTTCGGAAAA
Chromosome start end strand MM target_seq PAM alignment distance position gene name gene id
GU305914.1 21494 21516 + 0 TTTTCCGAAGTTCCACACGC AGG ||||||||[||||||||||||]PAM NA Intergenic NA NA

T1172 CCTCTTGTCGCGCTGTCCGTAGG 1000

| | | | | | | | | | | | | |
|---|---|---|---|---|---|---|---|---|---|---|---|---|
| Oligo fwd | CACCCCTCTTGTCGCGCTGTCCGT | | | | | | | | | | | |
| Oligo rev | AAACACGGACAGCGCGACAAGAGG | | | | | | | | | | | |
| Chromosome | start | | end | strand | MM | target_seq | PAM | alignment | distance | position | gene name | gene id |
| GU305914.1 | | 21539 | 21561 | + | 0 | CCTCTTGTCGCGCTGTCCGT | AGG | \|\|\|\|\|\|\|\|[\|\|\|\|\|\|\|\|\|\|\|\|]PAM | NA | Intergenic | NA | NA |
| T1173 | CTCTTGTCGCGCTGTCCGTAGGG | | 1000 | | | | | | | | | |
| Oligo fwd | CACCCTCTTGTCGCGCTGTCCGTA | | | | | | | | | | | |
| Oligo rev | AAACTACGGACAGCGCGACAAGAG | | | | | | | | | | | |
| Chromosome | start | | end | strand | MM | target_seq | PAM | alignment | distance | position | gene name | gene id |
| GU305914.1 | | 21540 | 21562 | + | 0 | CTCTTGTCGCGCTGTCCGTA | GGG | \|\|\|\|\|\|\|\|[\|\|\|\|\|\|\|\|\|\|\|\|]PAM | NA | Intergenic | NA | NA |
| T1174 | TCTTGTCGCGCTGTCCGTAGGGG | | 1000 | | | | | | | | | |
| Oligo fwd | CACCTCTTGTCGCGCTGTCCGTAG | | | | | | | | | | | |
| Oligo rev | AAACCTACGGACAGCGCGACAAGA | | | | | | | | | | | |
| Chromosome | start | | end | strand | MM | target_seq | PAM | alignment | distance | position | gene name | gene id |
| GU305914.1 | | 21541 | 21563 | + | 0 | TCTTGTCGCGCTGTCCGTAG | GGG | \|\|\|\|\|\|\|\|[\|\|\|\|\|\|\|\|\|\|\|\|]PAM | NA | Intergenic | NA | NA |
| T1175 | CCCTGCGTATGTTGCAACGCTGG | | 1000 | | | | | | | | | |
| Oligo fwd | CACCCCCTGCGTATGTTGCAACGC | | | | | | | | | | | |
| Oligo rev | AAACGCGTTGCAACATACGCAGGG | | | | | | | | | | | |
| Chromosome | start | | end | strand | MM | target_seq | PAM | alignment | distance | position | gene name | gene id |
| GU305914.1 | | 21589 | 21611 | + | 0 | CCCTGCGTATGTTGCAACGC | TGG | \|\|\|\|\|\|\|\|[\|\|\|\|\|\|\|\|\|\|\|\|]PAM | NA | Intergenic | NA | NA |
| T1176 | GGTCGTTCCATTCCCTGTGCAGG | | 1000 | | | | | | | | | |
| Oligo fwd | CACCGGTCGTTCCATTCCCTGTGC | | | | | | | | | | | |
| Oligo rev | AAACGCACAGGGAATGGAACGACC | | | | | | | | | | | |
| Chromosome | start | | end | strand | MM | target_seq | PAM | alignment | distance | position | gene name | gene id |
| GU305914.1 | | 21610 | 21632 | + | 0 | GGTCGTTCCATTCCCTGTGC | AGG | \|\|\|\|\|\|\|\|[\|\|\|\|\|\|\|\|\|\|\|\|]PAM | NA | Intergenic | NA | NA |
| T1177 | CCTGCAACTCGTAGCCGCTCTGG | | 1000 | | | | | | | | | |
| Oligo fwd | CACCCCTGCAACTCGTAGCCGCTC | | | | | | | | | | | |
| Oligo rev | AAACGAGCGGCTACGAGTTGCAGG | | | | | | | | | | | |
| Chromosome | start | | end | strand | MM | target_seq | PAM | alignment | distance | position | gene name | gene id |
| GU305914.1 | | 21634 | 21656 | + | 0 | CCTGCAACTCGTAGCCGCTC | TGG | \|\|\|\|\|\|\|\|[\|\|\|\|\|\|\|\|\|\|\|\|]PAM | NA | Intergenic | NA | NA |
| T1178 | CTGCAACTCGTAGCCGCTCTGGG | | 1000 | | | | | | | | | |
| Oligo fwd | CACCCTGCAACTCGTAGCCGCTCT | | | | | | | | | | | |
| Oligo rev | AAACAGAGCGGCTACGAGTTGCAG | | | | | | | | | | | |
| Chromosome | start | | end | strand | MM | target_seq | PAM | alignment | distance | position | gene name | gene id |
| GU305914.1 | | 21635 | 21657 | + | 0 | CTGCAACTCGTAGCCGCTCT | GGG | \|\|\|\|\|\|\|\|[\|\|\|\|\|\|\|\|\|\|\|\|]PAM | NA | Intergenic | NA | NA |
| T1179 | CAACTCGTAGCCGCTCTGGGAGG | | 1000 | | | | | | | | | |
| Oligo fwd | CACCCAACTCGTAGCCGCTCTGGG | | | | | | | | | | | |
| Oligo rev | AAACCCCAGAGCGGCTACGAGTTG | | | | | | | | | | | |
| Chromosome | start | | end | strand | MM | target_seq | PAM | alignment | distance | position | gene name | gene id |
| GU305914.1 | | 21638 | 21660 | + | 0 | CAACTCGTAGCCGCTCTGGG | AGG | \|\|\|\|\|\|\|\|[\|\|\|\|\|\|\|\|\|\|\|\|]PAM | NA | Intergenic | NA | NA |
| T1180 | TCACGTCCAAGACCTGTCCGCGG | | 1000 | | | | | | | | | |
| Oligo fwd | CACCTCACGTCCAAGACCTGTCCG | | | | | | | | | | | |
| Oligo rev | AAACCGGACAGGTCTTGGACGTGA | | | | | | | | | | | |
| Chromosome | start | | end | strand | MM | target_seq | PAM | alignment | distance | position | gene name | gene id |
| GU305914.1 | | 21673 | 21695 | + | 0 | TCACGTCCAAGACCTGTCCG | CGG | \|\|\|\|\|\|\|\|[\|\|\|\|\|\|\|\|\|\|\|\|]PAM | NA | Intergenic | NA | NA |
| T1181 | CACGTCCAAGACCTGTCCGCGGG | | 1000 | | | | | | | | | |
| Oligo fwd | CACCCACGTCCAAGACCTGTCCGC | | | | | | | | | | | |
| Oligo rev | AAACGCGGACAGGTCTTGGACGTG | | | | | | | | | | | |
| Chromosome | start | | end | strand | MM | target_seq | PAM | alignment | distance | position | gene name | gene id |
| GU305914.1 | | 21674 | 21696 | + | 0 | CACGTCCAAGACCTGTCCGC | GGG | \|\|\|\|\|\|\|\|[\|\|\|\|\|\|\|\|\|\|\|\|]PAM | NA | Intergenic | NA | NA |
| T1182 | CCTGTCCGCGGGTCTTGAAACGG | | 1000 | | | | | | | | | |
| Oligo fwd | CACCCCTGTCCGCGGGTCTTGAAA | | | | | | | | | | | |
| Oligo rev | AAACTTTCAAGACCCGCGGACAGG | | | | | | | | | | | |
| Chromosome | start | | end | strand | MM | target_seq | PAM | alignment | distance | position | gene name | gene id |
| GU305914.1 | | 21685 | 21707 | + | 0 | CCTGTCCGCGGGTCTTGAAA | CGG | \|\|\|\|\|\|\|\|[\|\|\|\|\|\|\|\|\|\|\|\|]PAM | NA | Intergenic | NA | NA |
| T1183 | TCCGCGGGTCTTGAAACGGCAGG | | 1000 | | | | | | | | | |
| Oligo fwd | CACCTCCGCGGGTCTTGAAACGGC | | | | | | | | | | | |
| Oligo rev | AAACGCCGTTTCAAGACCCGCGGA | | | | | | | | | | | |
| Chromosome | start | | end | strand | MM | target_seq | PAM | alignment | distance | position | gene name | gene id |
| GU305914.1 | | 21689 | 21711 | + | 0 | TCCGCGGGTCTTGAAACGGC | AGG | \|\|\|\|\|\|\|\|[\|\|\|\|\|\|\|\|\|\|\|\|]PAM | NA | Intergenic | NA | NA |
| T1184 | GTAGCGACGCAGTCGCCGTCCGG | | 1000 | | | | | | | | | |
| Oligo fwd | CACCGTAGCGACGCAGTCGCCGTC | | | | | | | | | | | |
| Oligo rev | AAACGACGGCGACTGCGTCGCTAC | | | | | | | | | | | |
| Chromosome | start | | end | strand | MM | target_seq | PAM | alignment | distance | position | gene name | gene id |
| GU305914.1 | | 21711 | 21733 | + | 0 | GTAGCGACGCAGTCGCCGTC | CGG | \|\|\|\|\|\|\|\|[\|\|\|\|\|\|\|\|\|\|\|\|]PAM | NA | Intergenic | NA | NA |
| T1185 | GTCGCCGTCCGGACAATCGAAGG | | 1000 | | | | | | | | | |
| Oligo fwd | CACCGTCGCCGTCCGGACAATCGA | | | | | | | | | | | |
| Oligo rev | AAACTCGATTGTCCGGACGGCGAC | | | | | | | | | | | |
| Chromosome | start | | end | strand | MM | target_seq | PAM | alignment | distance | position | gene name | gene id |
| GU305914.1 | | 21722 | 21744 | + | 0 | GTCGCCGTCCGGACAATCGA | AGG | \|\|\|\|\|\|\|\|[\|\|\|\|\|\|\|\|\|\|\|\|]PAM | NA | Intergenic | NA | NA |
| T1186 | CGTCCGGACAATCGAAGGTCAGG | | 1000 | | | | | | | | | |
| Oligo fwd | CACCCGTCCGGACAATCGAAGGTC | | | | | | | | | | | |
| Oligo rev | AAACGACCTTCGATTGTCCGGACG | | | | | | | | | | | |
| Chromosome | start | | end | strand | MM | target_seq | PAM | alignment | distance | position | gene name | gene id |
| GU305914.1 | | 21727 | 21749 | + | 0 | CGTCCGGACAATCGAAGGTC | AGG | \|\|\|\|\|\|\|\|[\|\|\|\|\|\|\|\|\|\|\|\|]PAM | NA | Intergenic | NA | NA |

<table>
<tr><td>T1187</td><td>CGGACAATCGAAGGTCAGGAAGG</td><td></td><td>1000</td><td></td><td></td><td></td><td></td><td></td><td></td><td></td><td></td><td></td></tr>
<tr><td>Oligo fwd</td><td>CACCCGGACAATCGAAGGTCAGGA</td><td></td><td></td><td></td><td></td><td></td><td></td><td></td><td></td><td></td><td></td><td></td></tr>
<tr><td>Oligo rev</td><td>AAACTCCTGACCTTCGATTGTCCG</td><td></td><td></td><td></td><td></td><td></td><td></td><td></td><td></td><td></td><td></td><td></td></tr>
<tr><td>Chromosome</td><td>start</td><td></td><td>end</td><td>strand</td><td>MM</td><td>target_seq</td><td>PAM</td><td>alignment</td><td>distance</td><td>position</td><td>gene name</td><td>gene id</td></tr>
<tr><td>GU305914.1</td><td></td><td>21731</td><td>21753</td><td>+</td><td>0</td><td>CGGACAATCGAAGGTCAGGA</td><td>AGG</td><td>||||||||[||||||||||||]PAM</td><td>NA</td><td>Intergenic</td><td>NA</td><td>NA</td></tr>
<tr><td>T1188</td><td>AGGTTTCGCTGCCGCTATCTTGG</td><td></td><td>1000</td><td></td><td></td><td></td><td></td><td></td><td></td><td></td><td></td><td></td></tr>
<tr><td>Oligo fwd</td><td>CACCAGGTTTCGCTGCCGCTATCT</td><td></td><td></td><td></td><td></td><td></td><td></td><td></td><td></td><td></td><td></td><td></td></tr>
<tr><td>Oligo rev</td><td>AAACAGATAGCGGCAGCGAAACCT</td><td></td><td></td><td></td><td></td><td></td><td></td><td></td><td></td><td></td><td></td><td></td></tr>
<tr><td>Chromosome</td><td>start</td><td></td><td>end</td><td>strand</td><td>MM</td><td>target_seq</td><td>PAM</td><td>alignment</td><td>distance</td><td>position</td><td>gene name</td><td>gene id</td></tr>
<tr><td>GU305914.1</td><td></td><td>21751</td><td>21773</td><td>+</td><td>0</td><td>AGGTTTCGCTGCCGCTATCT</td><td>TGG</td><td>||||||||[||||||||||||]PAM</td><td>NA</td><td>Intergenic</td><td>NA</td><td>NA</td></tr>
<tr><td>T1189</td><td>GTACGTCGTCAGCGCCTCCTCGG</td><td></td><td>1000</td><td></td><td></td><td></td><td></td><td></td><td></td><td></td><td></td><td></td></tr>
<tr><td>Oligo fwd</td><td>CACCGTACGTCGTCAGCGCCTCCT</td><td></td><td></td><td></td><td></td><td></td><td></td><td></td><td></td><td></td><td></td><td></td></tr>
<tr><td>Oligo rev</td><td>AAACAGGAGGCGCTGACGACGTAC</td><td></td><td></td><td></td><td></td><td></td><td></td><td></td><td></td><td></td><td></td><td></td></tr>
<tr><td>Chromosome</td><td>start</td><td></td><td>end</td><td>strand</td><td>MM</td><td>target_seq</td><td>PAM</td><td>alignment</td><td>distance</td><td>position</td><td>gene name</td><td>gene id</td></tr>
<tr><td>GU305914.1</td><td></td><td>21773</td><td>21795</td><td>+</td><td>0</td><td>GTACGTCGTCAGCGCCTCCT</td><td>CGG</td><td>||||||||[||||||||||||]PAM</td><td>NA</td><td>Intergenic</td><td>NA</td><td>NA</td></tr>
<tr><td>T1190</td><td>GGCGTCGTAGAAAGAAGCGTTGG</td><td></td><td>1000</td><td></td><td></td><td></td><td></td><td></td><td></td><td></td><td></td><td></td></tr>
<tr><td>Oligo fwd</td><td>CACCGGCGTCGTAGAAAGAAGCGT</td><td></td><td></td><td></td><td></td><td></td><td></td><td></td><td></td><td></td><td></td><td></td></tr>
<tr><td>Oligo rev</td><td>AAACACGCTTCTTTCTACGACGCC</td><td></td><td></td><td></td><td></td><td></td><td></td><td></td><td></td><td></td><td></td><td></td></tr>
<tr><td>Chromosome</td><td>start</td><td></td><td>end</td><td>strand</td><td>MM</td><td>target_seq</td><td>PAM</td><td>alignment</td><td>distance</td><td>position</td><td>gene name</td><td>gene id</td></tr>
<tr><td>GU305914.1</td><td></td><td>21794</td><td>21816</td><td>+</td><td>0</td><td>GGCGTCGTAGAAAGAAGCGT</td><td>TGG</td><td>||||||||[||||||||||||]PAM</td><td>NA</td><td>Intergenic</td><td>NA</td><td>NA</td></tr>
<tr><td>T1191</td><td>GCGTCGTAGAAAGAAGCGTTGGG</td><td></td><td>1000</td><td></td><td></td><td></td><td></td><td></td><td></td><td></td><td></td><td></td></tr>
<tr><td>Oligo fwd</td><td>CACCGCGTCGTAGAAAGAAGCGTT</td><td></td><td></td><td></td><td></td><td></td><td></td><td></td><td></td><td></td><td></td><td></td></tr>
<tr><td>Oligo rev</td><td>AAACAACGCTTCTTTCTACGACGC</td><td></td><td></td><td></td><td></td><td></td><td></td><td></td><td></td><td></td><td></td><td></td></tr>
<tr><td>Chromosome</td><td>start</td><td></td><td>end</td><td>strand</td><td>MM</td><td>target_seq</td><td>PAM</td><td>alignment</td><td>distance</td><td>position</td><td>gene name</td><td>gene id</td></tr>
<tr><td>GU305914.1</td><td></td><td>21795</td><td>21817</td><td>+</td><td>0</td><td>GCGTCGTAGAAAGAAGCGTT</td><td>GGG</td><td>||||||||[||||||||||||]PAM</td><td>NA</td><td>Intergenic</td><td>NA</td><td>NA</td></tr>
<tr><td>T1192</td><td>AAAGAAGCGTTGGGTTCATCCGG</td><td></td><td>1000</td><td></td><td></td><td></td><td></td><td></td><td></td><td></td><td></td><td></td></tr>
<tr><td>Oligo fwd</td><td>CACCAAAGAAGCGTTGGGTTCATC</td><td></td><td></td><td></td><td></td><td></td><td></td><td></td><td></td><td></td><td></td><td></td></tr>
<tr><td>Oligo rev</td><td>AAACGATGAACCCAACGCTTCTTT</td><td></td><td></td><td></td><td></td><td></td><td></td><td></td><td></td><td></td><td></td><td></td></tr>
<tr><td>Chromosome</td><td>start</td><td></td><td>end</td><td>strand</td><td>MM</td><td>target_seq</td><td>PAM</td><td>alignment</td><td>distance</td><td>position</td><td>gene name</td><td>gene id</td></tr>
<tr><td>GU305914.1</td><td></td><td>21804</td><td>21826</td><td>+</td><td>0</td><td>AAAGAAGCGTTGGGTTCATC</td><td>CGG</td><td>||||||||[||||||||||||]PAM</td><td>NA</td><td>Intergenic</td><td>NA</td><td>NA</td></tr>
<tr><td>T1193</td><td>AGCGTTGGGTTCATCCGGCGTGG</td><td></td><td>1000</td><td></td><td></td><td></td><td></td><td></td><td></td><td></td><td></td><td></td></tr>
<tr><td>Oligo fwd</td><td>CACCAGCGTTGGGTTCATCCGGCG</td><td></td><td></td><td></td><td></td><td></td><td></td><td></td><td></td><td></td><td></td><td></td></tr>
<tr><td>Oligo rev</td><td>AAACCGCCGGATGAACCCAACGCT</td><td></td><td></td><td></td><td></td><td></td><td></td><td></td><td></td><td></td><td></td><td></td></tr>
<tr><td>Chromosome</td><td>start</td><td></td><td>end</td><td>strand</td><td>MM</td><td>target_seq</td><td>PAM</td><td>alignment</td><td>distance</td><td>position</td><td>gene name</td><td>gene id</td></tr>
<tr><td>GU305914.1</td><td></td><td>21809</td><td>21831</td><td>+</td><td>0</td><td>AGCGTTGGGTTCATCCGGCG</td><td>TGG</td><td>||||||||[||||||||||||]PAM</td><td>NA</td><td>Intergenic</td><td>NA</td><td>NA</td></tr>
<tr><td>T1194</td><td>CTCTTGAATCCTCGCCCGCCCGG</td><td></td><td>1000</td><td></td><td></td><td></td><td></td><td></td><td></td><td></td><td></td><td></td></tr>
<tr><td>Oligo fwd</td><td>CACCCTCTTGAATCCTCGCCCGCC</td><td></td><td></td><td></td><td></td><td></td><td></td><td></td><td></td><td></td><td></td><td></td></tr>
<tr><td>Oligo rev</td><td>AAACGGCGGGCGAGGATTCAAGAG</td><td></td><td></td><td></td><td></td><td></td><td></td><td></td><td></td><td></td><td></td><td></td></tr>
<tr><td>Chromosome</td><td>start</td><td></td><td>end</td><td>strand</td><td>MM</td><td>target_seq</td><td>PAM</td><td>alignment</td><td>distance</td><td>position</td><td>gene name</td><td>gene id</td></tr>
<tr><td>GU305914.1</td><td></td><td>21843</td><td>21865</td><td>+</td><td>0</td><td>CTCTTGAATCCTCGCCCGCC</td><td>CGG</td><td>||||||||[||||||||||||]PAM</td><td>NA</td><td>Intergenic</td><td>NA</td><td>NA</td></tr>
<tr><td>T1195</td><td>TCTTGAATCCTCGCCCGCCCGGG</td><td></td><td>1000</td><td></td><td></td><td></td><td></td><td></td><td></td><td></td><td></td><td></td></tr>
<tr><td>Oligo fwd</td><td>CACCTCTTGAATCCTCGCCCGCCC</td><td></td><td></td><td></td><td></td><td></td><td></td><td></td><td></td><td></td><td></td><td></td></tr>
<tr><td>Oligo rev</td><td>AAACGGGCGGGCGAGGATTCAAGA</td><td></td><td></td><td></td><td></td><td></td><td></td><td></td><td></td><td></td><td></td><td></td></tr>
<tr><td>Chromosome</td><td>start</td><td></td><td>end</td><td>strand</td><td>MM</td><td>target_seq</td><td>PAM</td><td>alignment</td><td>distance</td><td>position</td><td>gene name</td><td>gene id</td></tr>
<tr><td>GU305914.1</td><td></td><td>21844</td><td>21866</td><td>+</td><td>0</td><td>TCTTGAATCCTCGCCCGCCC</td><td>GGG</td><td>||||||||[||||||||||||]PAM</td><td>NA</td><td>Intergenic</td><td>NA</td><td>NA</td></tr>
<tr><td>T1196</td><td>CTTGAATCCTCGCCCGCCCGGGG</td><td></td><td>1000</td><td></td><td></td><td></td><td></td><td></td><td></td><td></td><td></td><td></td></tr>
<tr><td>Oligo fwd</td><td>CACCCTTGAATCCTCGCCCGCCCG</td><td></td><td></td><td></td><td></td><td></td><td></td><td></td><td></td><td></td><td></td><td></td></tr>
<tr><td>Oligo rev</td><td>AAACCGGGCGGGCGAGGATTCAAG</td><td></td><td></td><td></td><td></td><td></td><td></td><td></td><td></td><td></td><td></td><td></td></tr>
<tr><td>Chromosome</td><td>start</td><td></td><td>end</td><td>strand</td><td>MM</td><td>target_seq</td><td>PAM</td><td>alignment</td><td>distance</td><td>position</td><td>gene name</td><td>gene id</td></tr>
<tr><td>GU305914.1</td><td></td><td>21845</td><td>21867</td><td>+</td><td>0</td><td>CTTGAATCCTCGCCCGCCCG</td><td>GGG</td><td>||||||||[||||||||||||]PAM</td><td>NA</td><td>Intergenic</td><td>NA</td><td>NA</td></tr>
<tr><td>T1197</td><td>TTGAATCCTCGCCCGCCCGGGGG</td><td></td><td>1000</td><td></td><td></td><td></td><td></td><td></td><td></td><td></td><td></td><td></td></tr>
<tr><td>Oligo fwd</td><td>CACCTTGAATCCTCGCCCGCCCGG</td><td></td><td></td><td></td><td></td><td></td><td></td><td></td><td></td><td></td><td></td><td></td></tr>
<tr><td>Oligo rev</td><td>AAACCCGGGCGGGCGAGGATTCAA</td><td></td><td></td><td></td><td></td><td></td><td></td><td></td><td></td><td></td><td></td><td></td></tr>
<tr><td>Chromosome</td><td>start</td><td></td><td>end</td><td>strand</td><td>MM</td><td>target_seq</td><td>PAM</td><td>alignment</td><td>distance</td><td>position</td><td>gene name</td><td>gene id</td></tr>
<tr><td>GU305914.1</td><td></td><td>21846</td><td>21868</td><td>+</td><td>0</td><td>TTGAATCCTCGCCCGCCCGG</td><td>GGG</td><td>||||||||[||||||||||||]PAM</td><td>NA</td><td>Intergenic</td><td>NA</td><td>NA</td></tr>
<tr><td>T1198</td><td>CGCCCGCCCGGGGGATACCGCGG</td><td></td><td>1000</td><td></td><td></td><td></td><td></td><td></td><td></td><td></td><td></td><td></td></tr>
<tr><td>Oligo fwd</td><td>CACCCGCCCGCCCGGGGGATACCG</td><td></td><td></td><td></td><td></td><td></td><td></td><td></td><td></td><td></td><td></td><td></td></tr>
<tr><td>Oligo rev</td><td>AAACCGGTATCCCCCGGGCGGGCG</td><td></td><td></td><td></td><td></td><td></td><td></td><td></td><td></td><td></td><td></td><td></td></tr>
<tr><td>Chromosome</td><td>start</td><td></td><td>end</td><td>strand</td><td>MM</td><td>target_seq</td><td>PAM</td><td>alignment</td><td>distance</td><td>position</td><td>gene name</td><td>gene id</td></tr>
<tr><td>GU305914.1</td><td></td><td>21855</td><td>21877</td><td>+</td><td>0</td><td>CGCCCGCCCGGGGGATACCG</td><td>CGG</td><td>||||||||[||||||||||||]PAM</td><td>NA</td><td>Intergenic</td><td>NA</td><td>NA</td></tr>
<tr><td>T1199</td><td>ATACCGCGGCTTATCGAACTCGG</td><td></td><td>1000</td><td></td><td></td><td></td><td></td><td></td><td></td><td></td><td></td><td></td></tr>
<tr><td>Oligo fwd</td><td>CACCATACCGCGGCTTATCGAACT</td><td></td><td></td><td></td><td></td><td></td><td></td><td></td><td></td><td></td><td></td><td></td></tr>
<tr><td>Oligo rev</td><td>AAACAGTTCGATAAGCCGCGGTAT</td><td></td><td></td><td></td><td></td><td></td><td></td><td></td><td></td><td></td><td></td><td></td></tr>
<tr><td>Chromosome</td><td>start</td><td></td><td>end</td><td>strand</td><td>MM</td><td>target_seq</td><td>PAM</td><td>alignment</td><td>distance</td><td>position</td><td>gene name</td><td>gene id</td></tr>
<tr><td>GU305914.1</td><td></td><td>21869</td><td>21891</td><td>+</td><td>0</td><td>ATACCGCGGCTTATCGAACT</td><td>CGG</td><td>||||||||[||||||||||||]PAM</td><td>NA</td><td>Intergenic</td><td>NA</td><td>NA</td></tr>
<tr><td>T1200</td><td>CGCGGCTTATCGAACTCGGAAGG</td><td></td><td>1000</td><td></td><td></td><td></td><td></td><td></td><td></td><td></td><td></td><td></td></tr>
<tr><td>Oligo fwd</td><td>CACCCGCGGCTTATCGAACTCGGA</td><td></td><td></td><td></td><td></td><td></td><td></td><td></td><td></td><td></td><td></td><td></td></tr>
<tr><td>Oligo rev</td><td>AAACTCCGAGTTCGATAAGCCGCG</td><td></td><td></td><td></td><td></td><td></td><td></td><td></td><td></td><td></td><td></td><td></td></tr>
<tr><td>Chromosome</td><td>start</td><td></td><td>end</td><td>strand</td><td>MM</td><td>target_seq</td><td>PAM</td><td>alignment</td><td>distance</td><td>position</td><td>gene name</td><td>gene id</td></tr>
<tr><td>GU305914.1</td><td></td><td>21873</td><td>21895</td><td>+</td><td>0</td><td>CGCGGCTTATCGAACTCGGA</td><td>AGG</td><td>||||||||[||||||||||||]PAM</td><td>NA</td><td>Intergenic</td><td>NA</td><td>NA</td></tr>
<tr><td>T1201</td><td>GCGGCTTATCGAACTCGGAAGGG</td><td></td><td>1000</td><td></td><td></td><td></td><td></td><td></td><td></td><td></td><td></td><td></td></tr>
<tr><td>Oligo fwd</td><td>CACCGCGGCTTATCGAACTCGGAA</td><td></td><td></td><td></td><td></td><td></td><td></td><td></td><td></td><td></td><td></td><td></td></tr>
<tr><td>Oligo rev</td><td>AAACTTCCGAGTTCGATAAGCCGC</td><td></td><td></td><td></td><td></td><td></td><td></td><td></td><td></td><td></td><td></td><td></td></tr>
<tr><td>Chromosome</td><td>start</td><td></td><td>end</td><td>strand</td><td>MM</td><td>target_seq</td><td>PAM</td><td>alignment</td><td>distance</td><td>position</td><td>gene name</td><td>gene id</td></tr>
<tr><td>GU305914.1</td><td></td><td>21874</td><td>21896</td><td>+</td><td>0</td><td>GCGGCTTATCGAACTCGGAA</td><td>GGG</td><td>||||||||[||||||||||||]PAM</td><td>NA</td><td>Intergenic</td><td>NA</td><td>NA</td></tr>
</table>

| | | | | | | | | | | | |
|---|---|---|---|---|---|---|---|---|---|---|---|
| T1202 | CATTCCCCCACTCCACCACTTGG | 1000 | | | | | | | | | |
| Oligo fwd | CACCCATTCCCCCACTCCACCACT | | | | | | | | | | |
| Oligo rev | AAACAGTGGTGGAGTGGGGGAATG | | | | | | | | | | |
| Chromosome | start | end | strand | MM | target_seq | PAM | alignment | distance | position | gene name | gene id |
| GU305914.1 | 21905 | 21927 | + | 0 | CATTCCCCCACTCCACCACT | TGG | \|\|\|\|\|\|\|\|[\|\|\|\|\|\|\|\|\|\|\|\|]PAM | NA | Intergenic | NA | NA |
| T1203 | TCCCCCACTCCACCACTTGGAGG | 1000 | | | | | | | | | |
| Oligo fwd | CACCTCCCCCACTCCACCACTTGG | | | | | | | | | | |
| Oligo rev | AAACCCAAGTGGTGGAGTGGGGGA | | | | | | | | | | |
| Chromosome | start | end | strand | MM | target_seq | PAM | alignment | distance | position | gene name | gene id |
| GU305914.1 | 21908 | 21930 | + | 0 | TCCCCCACTCCACCACTTGG | AGG | \|\|\|\|\|\|\|\|[\|\|\|\|\|\|\|\|\|\|\|\|]PAM | NA | Intergenic | NA | NA |
| T1204 | CTGATGCTGATGCTGATGCGCGG | 1000 | | | | | | | | | |
| Oligo fwd | CACCCTGATGCTGATGCTGATGCG | | | | | | | | | | |
| Oligo rev | AAACCGCATCAGCATCAGCATCAG | | | | | | | | | | |
| Chromosome | start | end | strand | MM | target_seq | PAM | alignment | distance | position | gene name | gene id |
| GU305914.1 | 21959 | 21981 | + | 0 | CTGATGCTGATGCTGATGCG | CGG | \|\|\|\|\|\|\|\|[\|\|\|\|\|\|\|\|\|\|\|\|]PAM | NA | Intergenic | NA | NA |
| T1205 | GCTGATGCTGATGCGCGGCCAGG | 1000 | | | | | | | | | |
| Oligo fwd | CACCGCTGATGCTGATGCGCGGCC | | | | | | | | | | |
| Oligo rev | AAACGGCCGCGCATCAGCATCAGC | | | | | | | | | | |
| Chromosome | start | end | strand | MM | target_seq | PAM | alignment | distance | position | gene name | gene id |
| GU305914.1 | 21964 | 21986 | + | 0 | GCTGATGCTGATGCGCGGCC | AGG | \|\|\|\|\|\|\|\|[\|\|\|\|\|\|\|\|\|\|\|\|]PAM | NA | Intergenic | NA | NA |
| T1206 | CTGATGCTGATGCGCGGCCAGGG | 1000 | | | | | | | | | |
| Oligo fwd | CACCCTGATGCTGATGCGCGGCCA | | | | | | | | | | |
| Oligo rev | AAACTGGCCGCGCATCAGCATCAG | | | | | | | | | | |
| Chromosome | start | end | strand | MM | target_seq | PAM | alignment | distance | position | gene name | gene id |
| GU305914.1 | 21965 | 21987 | + | 0 | CTGATGCTGATGCGCGGCCA | GGG | \|\|\|\|\|\|\|\|[\|\|\|\|\|\|\|\|\|\|\|\|]PAM | NA | Intergenic | NA | NA |
| T1207 | TGCGCGGCCAGGGCCGTCGTCGG | 1000 | | | | | | | | | |
| Oligo fwd | CACCTGCGCGGCCAGGGCCGTCGT | | | | | | | | | | |
| Oligo rev | AAACACGACGGCCCTGGCCGCGCA | | | | | | | | | | |
| Chromosome | start | end | strand | MM | target_seq | PAM | alignment | distance | position | gene name | gene id |
| GU305914.1 | 21975 | 21997 | + | 0 | TGCGCGGCCAGGGCCGTCGT | CGG | \|\|\|\|\|\|\|\|[\|\|\|\|\|\|\|\|\|\|\|\|]PAM | NA | Intergenic | NA | NA |
| T1208 | CGGCCAGGGCCGTCGTCGGCAGG | 1000 | | | | | | | | | |
| Oligo fwd | CACCCGGCCAGGGCCGTCGTCGGC | | | | | | | | | | |
| Oligo rev | AAACGCCGACGACGGCCCTGGCCG | | | | | | | | | | |
| Chromosome | start | end | strand | MM | target_seq | PAM | alignment | distance | position | gene name | gene id |
| GU305914.1 | 21979 | 22001 | + | 0 | CGGCCAGGGCCGTCGTCGGC | AGG | \|\|\|\|\|\|\|\|[\|\|\|\|\|\|\|\|\|\|\|\|]PAM | NA | Intergenic | NA | NA |
| T1209 | CAGGGCCGTCGTCGGCAGGACGG | 1000 | | | | | | | | | |
| Oligo fwd | CACCCAGGGCCGTCGTCGGCAGGA | | | | | | | | | | |
| Oligo rev | AAACTCCTGCCGACGACGGCCCTG | | | | | | | | | | |
| Chromosome | start | end | strand | MM | target_seq | PAM | alignment | distance | position | gene name | gene id |
| GU305914.1 | 21983 | 22005 | + | 0 | CAGGGCCGTCGTCGGCAGGA | CGG | \|\|\|\|\|\|\|\|[\|\|\|\|\|\|\|\|\|\|\|\|]PAM | NA | Intergenic | NA | NA |
| GU305914.1 | 115383 | 115405 | + | 4 | CGGCACCGTCGTCGCCAGGA | TGG | \|-\|--\|\|\|[\|\|\|\|\|\|-\|\|\|\|\|]PAM | NA | Intergenic | NA | NA |
| T1210 | CCGTCGTCGGCAGGACGGTGAGG | 1000 | | | | | | | | | |
| Oligo fwd | CACCCCGTCGTCGGCAGGACGGTG | | | | | | | | | | |
| Oligo rev | AAACCACCGTCCTGCCGACGACGG | | | | | | | | | | |
| Chromosome | start | end | strand | MM | target_seq | PAM | alignment | distance | position | gene name | gene id |
| GU305914.1 | 21988 | 22010 | + | 0 | CCGTCGTCGGCAGGACGGTG | AGG | \|\|\|\|\|\|\|\|[\|\|\|\|\|\|\|\|\|\|\|\|]PAM | NA | Intergenic | NA | NA |
| T1211 | CGTCGTCGGCAGGACGGTGAGGG | 1000 | | | | | | | | | |
| Oligo fwd | CACCCGTCGTCGGCAGGACGGTGA | | | | | | | | | | |
| Oligo rev | AAACTCACCGTCCTGCCGACGACG | | | | | | | | | | |
| Chromosome | start | end | strand | MM | target_seq | PAM | alignment | distance | position | gene name | gene id |
| GU305914.1 | 21989 | 22011 | + | 0 | CGTCGTCGGCAGGACGGTGA | GGG | \|\|\|\|\|\|\|\|[\|\|\|\|\|\|\|\|\|\|\|\|]PAM | NA | Intergenic | NA | NA |
| T1212 | CAGGACGGTGAGGGCCGCCACGG | 1000 | | | | | | | | | |
| Oligo fwd | CACCCAGGACGGTGAGGGCCGCCA | | | | | | | | | | |
| Oligo rev | AAACTGGCGGCCCTCACCGTCCTG | | | | | | | | | | |
| Chromosome | start | end | strand | MM | target_seq | PAM | alignment | distance | position | gene name | gene id |
| GU305914.1 | 21998 | 22020 | + | 0 | CAGGACGGTGAGGGCCGCCA | CGG | \|\|\|\|\|\|\|\|[\|\|\|\|\|\|\|\|\|\|\|\|]PAM | NA | Intergenic | NA | NA |
| T1213 | GGCCGCCACGGCCACGCCGACGG | 1000 | | | | | | | | | |
| Oligo fwd | CACCGGCCGCCACGGCCACGCCGA | | | | | | | | | | |
| Oligo rev | AAACTCGGCGTGGCCGTGGCGGCC | | | | | | | | | | |
| Chromosome | start | end | strand | MM | target_seq | PAM | alignment | distance | position | gene name | gene id |
| GU305914.1 | 22010 | 22032 | + | 0 | GGCCGCCACGGCCACGCCGA | CGG | \|\|\|\|\|\|\|\|[\|\|\|\|\|\|\|\|\|\|\|\|]PAM | NA | Intergenic | NA | NA |
| T1214 | ACGCCGACGGTCATCGCCGCCGG | 1000 | | | | | | | | | |
| Oligo fwd | CACCACGCCGACGGTCATCGCCGC | | | | | | | | | | |
| Oligo rev | AAACGCGGCGATGACCGTCGGCGT | | | | | | | | | | |
| Chromosome | start | end | strand | MM | target_seq | PAM | alignment | distance | position | gene name | gene id |
| GU305914.1 | 22023 | 22045 | + | 0 | ACGCCGACGGTCATCGCCGC | CGG | \|\|\|\|\|\|\|\|[\|\|\|\|\|\|\|\|\|\|\|\|]PAM | NA | Intergenic | NA | NA |
| T1215 | CATCGCCGCCGGCACTAACCCGG | 1000 | | | | | | | | | |
| Oligo fwd | CACCCATCGCCGCCGGCACTAACC | | | | | | | | | | |
| Oligo rev | AAACGGTTAGTGCCGGCGGCGATG | | | | | | | | | | |
| Chromosome | start | end | strand | MM | target_seq | PAM | alignment | distance | position | gene name | gene id |
| GU305914.1 | 22034 | 22056 | + | 0 | CATCGCCGCCGGCACTAACC | CGG | \|\|\|\|\|\|\|\|[\|\|\|\|\|\|\|\|\|\|\|\|]PAM | NA | Intergenic | NA | NA |
| T1216 | CGCCGGCACTAACCCGGCCACGG | 1000 | | | | | | | | | |
| Oligo fwd | CACCCGCCGGCACTAACCCGGCCA | | | | | | | | | | |
| Oligo rev | AAACTGGCCGGGTTAGTGCCGGCG | | | | | | | | | | |

<table>
<tr><td>Chromosome</td><td>start</td><td>end</td><td>strand</td><td>MM</td><td>target_seq</td><td>PAM</td><td>alignment</td><td>distance</td><td>position</td><td>gene name</td><td>gene id</td></tr>
<tr><td>GU305914.1</td><td>22040</td><td>22062</td><td>+</td><td>0</td><td>CGCCGGCACTAACCCGGCCA</td><td>CGG</td><td>||||||||[||||||||||||]PAM</td><td>NA</td><td>Intergenic</td><td>NA</td><td>NA</td></tr>
<tr><td>T1217</td><td>CACTAACCCGGCCACGGCCCCGG</td><td>1000</td><td></td><td></td><td></td><td></td><td></td><td></td><td></td><td></td><td></td></tr>
<tr><td>Oligo fwd</td><td>CACCCACTAACCCGGCCACGGCCC</td><td></td><td></td><td></td><td></td><td></td><td></td><td></td><td></td><td></td><td></td></tr>
<tr><td>Oligo rev</td><td>AAACGGGCCGTGGCCGGGTTAGTG</td><td></td><td></td><td></td><td></td><td></td><td></td><td></td><td></td><td></td><td></td></tr>
<tr><td>Chromosome</td><td>start</td><td>end</td><td>strand</td><td>MM</td><td>target_seq</td><td>PAM</td><td>alignment</td><td>distance</td><td>position</td><td>gene name</td><td>gene id</td></tr>
<tr><td>GU305914.1</td><td>22046</td><td>22068</td><td>+</td><td>0</td><td>CACTAACCCGGCCACGGCCC</td><td>CGG</td><td>||||||||[||||||||||||]PAM</td><td>NA</td><td>Intergenic</td><td>NA</td><td>NA</td></tr>
<tr><td>T1218</td><td>ACTAACCCGGCCACGGCCCCGGG</td><td>1000</td><td></td><td></td><td></td><td></td><td></td><td></td><td></td><td></td><td></td></tr>
<tr><td>Oligo fwd</td><td>CACCACTAACCCGGCCACGGCCCC</td><td></td><td></td><td></td><td></td><td></td><td></td><td></td><td></td><td></td><td></td></tr>
<tr><td>Oligo rev</td><td>AAACGGGGCCGTGGCCGGGTTAGT</td><td></td><td></td><td></td><td></td><td></td><td></td><td></td><td></td><td></td><td></td></tr>
<tr><td>Chromosome</td><td>start</td><td>end</td><td>strand</td><td>MM</td><td>target_seq</td><td>PAM</td><td>alignment</td><td>distance</td><td>position</td><td>gene name</td><td>gene id</td></tr>
<tr><td>GU305914.1</td><td>22047</td><td>22069</td><td>+</td><td>0</td><td>ACTAACCCGGCCACGGCCCC</td><td>GGG</td><td>||||||||[||||||||||||]PAM</td><td>NA</td><td>Intergenic</td><td>NA</td><td>NA</td></tr>
<tr><td>T1219</td><td>ACCCGGCCACGGCCCCGGGCAGG</td><td>1000</td><td></td><td></td><td></td><td></td><td></td><td></td><td></td><td></td><td></td></tr>
<tr><td>Oligo fwd</td><td>CACCACCCGGCCACGGCCCCGGGC</td><td></td><td></td><td></td><td></td><td></td><td></td><td></td><td></td><td></td><td></td></tr>
<tr><td>Oligo rev</td><td>AAACGCCCGGGGCCGTGGCCGGGT</td><td></td><td></td><td></td><td></td><td></td><td></td><td></td><td></td><td></td><td></td></tr>
<tr><td>Chromosome</td><td>start</td><td>end</td><td>strand</td><td>MM</td><td>target_seq</td><td>PAM</td><td>alignment</td><td>distance</td><td>position</td><td>gene name</td><td>gene id</td></tr>
<tr><td>GU305914.1</td><td>22051</td><td>22073</td><td>+</td><td>0</td><td>ACCCGGCCACGGCCCCGGGC</td><td>AGG</td><td>||||||||[||||||||||||]PAM</td><td>NA</td><td>Intergenic</td><td>NA</td><td>NA</td></tr>
<tr><td>T1220</td><td>CCACGGCCCCGGGCAGGCCGCGG</td><td>1000</td><td></td><td></td><td></td><td></td><td></td><td></td><td></td><td></td><td></td></tr>
<tr><td>Oligo fwd</td><td>CACCCCACGGCCCCGGGCAGGCCG</td><td></td><td></td><td></td><td></td><td></td><td></td><td></td><td></td><td></td><td></td></tr>
<tr><td>Oligo rev</td><td>AAACCGGCCTGCCCGGGGCCGTGG</td><td></td><td></td><td></td><td></td><td></td><td></td><td></td><td></td><td></td><td></td></tr>
<tr><td>Chromosome</td><td>start</td><td>end</td><td>strand</td><td>MM</td><td>target_seq</td><td>PAM</td><td>alignment</td><td>distance</td><td>position</td><td>gene name</td><td>gene id</td></tr>
<tr><td>GU305914.1</td><td>22057</td><td>22079</td><td>+</td><td>0</td><td>CCACGGCCCCGGGCAGGCCG</td><td>CGG</td><td>||||||||[||||||||||||]PAM</td><td>NA</td><td>Intergenic</td><td>NA</td><td>NA</td></tr>
<tr><td>T1221</td><td>GGCAGGCCGCGGTATCCCTCCGG</td><td>1000</td><td></td><td></td><td></td><td></td><td></td><td></td><td></td><td></td><td></td></tr>
<tr><td>Oligo fwd</td><td>CACCGGCAGGCCGCGGTATCCCTC</td><td></td><td></td><td></td><td></td><td></td><td></td><td></td><td></td><td></td><td></td></tr>
<tr><td>Oligo rev</td><td>AAACGAGGGATACCGCGGCCTGCC</td><td></td><td></td><td></td><td></td><td></td><td></td><td></td><td></td><td></td><td></td></tr>
<tr><td>Chromosome</td><td>start</td><td>end</td><td>strand</td><td>MM</td><td>target_seq</td><td>PAM</td><td>alignment</td><td>distance</td><td>position</td><td>gene name</td><td>gene id</td></tr>
<tr><td>GU305914.1</td><td>22068</td><td>22090</td><td>+</td><td>0</td><td>GGCAGGCCGCGGTATCCCTC</td><td>CGG</td><td>||||||||[||||||||||||]PAM</td><td>NA</td><td>Intergenic</td><td>NA</td><td>NA</td></tr>
<tr><td>GU305914.1</td><td>21861</td><td>21883</td><td>-</td><td>4</td><td>GATAAGCCGCGGTATCCCCC</td><td>GGG</td><td>|--|-|||[||||||||||-|]PAM</td><td>NA</td><td>Intergenic</td><td>NA</td><td>NA</td></tr>
<tr><td>T1222</td><td>GGCCGCGGTATCCCTCCGGTCGG</td><td>1000</td><td></td><td></td><td></td><td></td><td></td><td></td><td></td><td></td><td></td></tr>
<tr><td>Oligo fwd</td><td>CACCGGCCGCGGTATCCCTCCGGT</td><td></td><td></td><td></td><td></td><td></td><td></td><td></td><td></td><td></td><td></td></tr>
<tr><td>Oligo rev</td><td>AAACACCGGAGGGATACCGCGGCC</td><td></td><td></td><td></td><td></td><td></td><td></td><td></td><td></td><td></td><td></td></tr>
<tr><td>Chromosome</td><td>start</td><td>end</td><td>strand</td><td>MM</td><td>target_seq</td><td>PAM</td><td>alignment</td><td>distance</td><td>position</td><td>gene name</td><td>gene id</td></tr>
<tr><td>GU305914.1</td><td>22072</td><td>22094</td><td>+</td><td>0</td><td>GGCCGCGGTATCCCTCCGGT</td><td>CGG</td><td>||||||||[||||||||||||]PAM</td><td>NA</td><td>Intergenic</td><td>NA</td><td>NA</td></tr>
<tr><td>T1223</td><td>GGTATCCCTCCGGTCGGTCGCGG</td><td>1000</td><td></td><td></td><td></td><td></td><td></td><td></td><td></td><td></td><td></td></tr>
<tr><td>Oligo fwd</td><td>CACCGGTATCCCTCCGGTCGGTCG</td><td></td><td></td><td></td><td></td><td></td><td></td><td></td><td></td><td></td><td></td></tr>
<tr><td>Oligo rev</td><td>AAACCGACCGACCGGAGGGATACC</td><td></td><td></td><td></td><td></td><td></td><td></td><td></td><td></td><td></td><td></td></tr>
<tr><td>Chromosome</td><td>start</td><td>end</td><td>strand</td><td>MM</td><td>target_seq</td><td>PAM</td><td>alignment</td><td>distance</td><td>position</td><td>gene name</td><td>gene id</td></tr>
<tr><td>GU305914.1</td><td>22078</td><td>22100</td><td>+</td><td>0</td><td>GGTATCCCTCCGGTCGGTCG</td><td>CGG</td><td>||||||||[||||||||||||]PAM</td><td>NA</td><td>Intergenic</td><td>NA</td><td>NA</td></tr>
<tr><td>T1224</td><td>ATCCCTCCGGTCGGTCGCGGCGG</td><td>1000</td><td></td><td></td><td></td><td></td><td></td><td></td><td></td><td></td><td></td></tr>
<tr><td>Oligo fwd</td><td>CACCATCCCTCCGGTCGGTCGCGG</td><td></td><td></td><td></td><td></td><td></td><td></td><td></td><td></td><td></td><td></td></tr>
<tr><td>Oligo rev</td><td>AAACCCGCGACCGACCGGAGGGAT</td><td></td><td></td><td></td><td></td><td></td><td></td><td></td><td></td><td></td><td></td></tr>
<tr><td>Chromosome</td><td>start</td><td>end</td><td>strand</td><td>MM</td><td>target_seq</td><td>PAM</td><td>alignment</td><td>distance</td><td>position</td><td>gene name</td><td>gene id</td></tr>
<tr><td>GU305914.1</td><td>22081</td><td>22103</td><td>+</td><td>0</td><td>ATCCCTCCGGTCGGTCGCGG</td><td>CGG</td><td>||||||||[||||||||||||]PAM</td><td>NA</td><td>Intergenic</td><td>NA</td><td>NA</td></tr>
<tr><td>GU305914.1</td><td>154395</td><td>154417</td><td>+</td><td>4</td><td>ATCGCCCCGGGCGGTGGCGG</td><td>CGG</td><td>|||-|-||[||-||||-||||]PAM</td><td>NA</td><td>Intergenic</td><td>NA</td><td>NA</td></tr>
<tr><td>T1225</td><td>GAAGATCATATGCGCGTCGACGG</td><td>1000</td><td></td><td></td><td></td><td></td><td></td><td></td><td></td><td></td><td></td></tr>
<tr><td>Oligo fwd</td><td>CACCGAAGATCATATGCGCGTCGA</td><td></td><td></td><td></td><td></td><td></td><td></td><td></td><td></td><td></td><td></td></tr>
<tr><td>Oligo rev</td><td>AAACTCGACGCGCATATGATCTTC</td><td></td><td></td><td></td><td></td><td></td><td></td><td></td><td></td><td></td><td></td></tr>
<tr><td>Chromosome</td><td>start</td><td>end</td><td>strand</td><td>MM</td><td>target_seq</td><td>PAM</td><td>alignment</td><td>distance</td><td>position</td><td>gene name</td><td>gene id</td></tr>
<tr><td>GU305914.1</td><td>22121</td><td>22143</td><td>+</td><td>0</td><td>GAAGATCATATGCGCGTCGA</td><td>CGG</td><td>||||||||[||||||||||||]PAM</td><td>NA</td><td>Intergenic</td><td>NA</td><td>NA</td></tr>
<tr><td>T1226</td><td>AAGATCATATGCGCGTCGACGGG</td><td>1000</td><td></td><td></td><td></td><td></td><td></td><td></td><td></td><td></td><td></td></tr>
<tr><td>Oligo fwd</td><td>CACCAAGATCATATGCGCGTCGAC</td><td></td><td></td><td></td><td></td><td></td><td></td><td></td><td></td><td></td><td></td></tr>
<tr><td>Oligo rev</td><td>AAACGTCGACGCGCATATGATCTT</td><td></td><td></td><td></td><td></td><td></td><td></td><td></td><td></td><td></td><td></td></tr>
<tr><td>Chromosome</td><td>start</td><td>end</td><td>strand</td><td>MM</td><td>target_seq</td><td>PAM</td><td>alignment</td><td>distance</td><td>position</td><td>gene name</td><td>gene id</td></tr>
<tr><td>GU305914.1</td><td>22122</td><td>22144</td><td>+</td><td>0</td><td>AAGATCATATGCGCGTCGAC</td><td>GGG</td><td>||||||||[||||||||||||]PAM</td><td>NA</td><td>Intergenic</td><td>NA</td><td>NA</td></tr>
<tr><td>T1227</td><td>AGATCATATGCGCGTCGACGGGG</td><td>1000</td><td></td><td></td><td></td><td></td><td></td><td></td><td></td><td></td><td></td></tr>
<tr><td>Oligo fwd</td><td>CACCAGATCATATGCGCGTCGACG</td><td></td><td></td><td></td><td></td><td></td><td></td><td></td><td></td><td></td><td></td></tr>
<tr><td>Oligo rev</td><td>AAACCGTCGACGCGCATATGATCT</td><td></td><td></td><td></td><td></td><td></td><td></td><td></td><td></td><td></td><td></td></tr>
<tr><td>Chromosome</td><td>start</td><td>end</td><td>strand</td><td>MM</td><td>target_seq</td><td>PAM</td><td>alignment</td><td>distance</td><td>position</td><td>gene name</td><td>gene id</td></tr>
<tr><td>GU305914.1</td><td>22123</td><td>22145</td><td>+</td><td>0</td><td>AGATCATATGCGCGTCGACG</td><td>GGG</td><td>||||||||[||||||||||||]PAM</td><td>NA</td><td>Intergenic</td><td>NA</td><td>NA</td></tr>
<tr><td>T1228</td><td>TTTCGAACCGTTGAAAATGCCGG</td><td>1000</td><td></td><td></td><td></td><td></td><td></td><td></td><td></td><td></td><td></td></tr>
<tr><td>Oligo fwd</td><td>CACCTTTCGAACCGTTGAAAATGC</td><td></td><td></td><td></td><td></td><td></td><td></td><td></td><td></td><td></td><td></td></tr>
<tr><td>Oligo rev</td><td>AAACGCATTTTCAACGGTTCGAAA</td><td></td><td></td><td></td><td></td><td></td><td></td><td></td><td></td><td></td><td></td></tr>
<tr><td>Chromosome</td><td>start</td><td>end</td><td>strand</td><td>MM</td><td>target_seq</td><td>PAM</td><td>alignment</td><td>distance</td><td>position</td><td>gene name</td><td>gene id</td></tr>
<tr><td>GU305914.1</td><td>22147</td><td>22169</td><td>+</td><td>0</td><td>TTTCGAACCGTTGAAAATGC</td><td>CGG</td><td>||||||||[||||||||||||]PAM</td><td>NA</td><td>Intergenic</td><td>NA</td><td>NA</td></tr>
<tr><td>T1229</td><td>AACCGTACCGCGATTCCCACAGG</td><td>1000</td><td></td><td></td><td></td><td></td><td></td><td></td><td></td><td></td><td></td></tr>
<tr><td>Oligo fwd</td><td>CACCAACCGTACCGCGATTCCCAC</td><td></td><td></td><td></td><td></td><td></td><td></td><td></td><td></td><td></td><td></td></tr>
<tr><td>Oligo rev</td><td>AAACGTGGGAATCGCGGTACGGTT</td><td></td><td></td><td></td><td></td><td></td><td></td><td></td><td></td><td></td><td></td></tr>
<tr><td>Chromosome</td><td>start</td><td>end</td><td>strand</td><td>MM</td><td>target_seq</td><td>PAM</td><td>alignment</td><td>distance</td><td>position</td><td>gene name</td><td>gene id</td></tr>
<tr><td>GU305914.1</td><td>22180</td><td>22202</td><td>+</td><td>0</td><td>AACCGTACCGCGATTCCCAC</td><td>AGG</td><td>||||||||[||||||||||||]PAM</td><td>NA</td><td>Intergenic</td><td>NA</td><td>NA</td></tr>
<tr><td>T1230</td><td>ACCGTACCGCGATTCCCACAGGG</td><td>1000</td><td></td><td></td><td></td><td></td><td></td><td></td><td></td><td></td><td></td></tr>
<tr><td>Oligo fwd</td><td>CACCACCGTACCGCGATTCCCACA</td><td></td><td></td><td></td><td></td><td></td><td></td><td></td><td></td><td></td><td></td></tr>
<tr><td>Oligo rev</td><td>AAACTGTGGGAATCGCGGTACGGT</td><td></td><td></td><td></td><td></td><td></td><td></td><td></td><td></td><td></td><td></td></tr>
<tr><td>Chromosome</td><td>start</td><td>end</td><td>strand</td><td>MM</td><td>target_seq</td><td>PAM</td><td>alignment</td><td>distance</td><td>position</td><td>gene name</td><td>gene id</td></tr>
<tr><td>GU305914.1</td><td>22181</td><td>22203</td><td>+</td><td>0</td><td>ACCGTACCGCGATTCCCACA</td><td>GGG</td><td>||||||||[||||||||||||]PAM</td><td>NA</td><td>Intergenic</td><td>NA</td><td>NA</td></tr>
</table>

<table>
<tr><td>T1231</td><td>CGTCACGATCTTGAAAATGACGG</td><td></td><td>1000</td><td></td><td></td><td></td><td></td><td></td><td></td><td></td><td></td><td></td></tr>
<tr><td>Oligo fwd</td><td>CACCCGTCACGATCTTGAAAATGA</td><td></td><td></td><td></td><td></td><td></td><td></td><td></td><td></td><td></td><td></td><td></td></tr>
<tr><td>Oligo rev</td><td>AAACTCATTTTCAAGATCGTGACG</td><td></td><td></td><td></td><td></td><td></td><td></td><td></td><td></td><td></td><td></td><td></td></tr>
<tr><td>Chromosome</td><td>start</td><td></td><td>end</td><td>strand</td><td>MM</td><td>target_seq</td><td>PAM</td><td>alignment</td><td>distance</td><td>position</td><td>gene name</td><td>gene id</td></tr>
<tr><td>GU305914.1</td><td></td><td>22205</td><td>22227</td><td>+</td><td>0</td><td>CGTCACGATCTTGAAAATGA</td><td>CGG</td><td>||||||||[||||||||||||]PAM</td><td>NA</td><td>Intergenic</td><td>NA</td><td>NA</td></tr>
<tr><td>T1232</td><td>GACGGCGAACAGAATTCCGACGG</td><td></td><td>1000</td><td></td><td></td><td></td><td></td><td></td><td></td><td></td><td></td><td></td></tr>
<tr><td>Oligo fwd</td><td>CACCGACGGCGAACAGAATTCCGA</td><td></td><td></td><td></td><td></td><td></td><td></td><td></td><td></td><td></td><td></td><td></td></tr>
<tr><td>Oligo rev</td><td>AAACTCGGAATTCTGTTCGCCGTC</td><td></td><td></td><td></td><td></td><td></td><td></td><td></td><td></td><td></td><td></td><td></td></tr>
<tr><td>Chromosome</td><td>start</td><td></td><td>end</td><td>strand</td><td>MM</td><td>target_seq</td><td>PAM</td><td>alignment</td><td>distance</td><td>position</td><td>gene name</td><td>gene id</td></tr>
<tr><td>GU305914.1</td><td></td><td>22223</td><td>22245</td><td>+</td><td>0</td><td>GACGGCGAACAGAATTCCGA</td><td>CGG</td><td>||||||||[||||||||||||]PAM</td><td>NA</td><td>Intergenic</td><td>NA</td><td>NA</td></tr>
<tr><td>T1233</td><td>CGCGATGACCGTCATCGTGCCGG</td><td></td><td>1000</td><td></td><td></td><td></td><td></td><td></td><td></td><td></td><td></td><td></td></tr>
<tr><td>Oligo fwd</td><td>CACCCGCGATGACCGTCATCGTGC</td><td></td><td></td><td></td><td></td><td></td><td></td><td></td><td></td><td></td><td></td><td></td></tr>
<tr><td>Oligo rev</td><td>AAACGCACGATGACGGTCATCGCG</td><td></td><td></td><td></td><td></td><td></td><td></td><td></td><td></td><td></td><td></td><td></td></tr>
<tr><td>Chromosome</td><td>start</td><td></td><td>end</td><td>strand</td><td>MM</td><td>target_seq</td><td>PAM</td><td>alignment</td><td>distance</td><td>position</td><td>gene name</td><td>gene id</td></tr>
<tr><td>GU305914.1</td><td></td><td>22247</td><td>22269</td><td>+</td><td>0</td><td>CGCGATGACCGTCATCGTGC</td><td>CGG</td><td>||||||||[||||||||||||]PAM</td><td>NA</td><td>Intergenic</td><td>NA</td><td>NA</td></tr>
<tr><td>T1234</td><td>CATCGTGCCGGCGCCCAACATGG</td><td></td><td>1000</td><td></td><td></td><td></td><td></td><td></td><td></td><td></td><td></td><td></td></tr>
<tr><td>Oligo fwd</td><td>CACCCATCGTGCCGGCGCCCAACA</td><td></td><td></td><td></td><td></td><td></td><td></td><td></td><td></td><td></td><td></td><td></td></tr>
<tr><td>Oligo rev</td><td>AAACTGTTGGGCGCCGGCACGATG</td><td></td><td></td><td></td><td></td><td></td><td></td><td></td><td></td><td></td><td></td><td></td></tr>
<tr><td>Chromosome</td><td>start</td><td></td><td>end</td><td>strand</td><td>MM</td><td>target_seq</td><td>PAM</td><td>alignment</td><td>distance</td><td>position</td><td>gene name</td><td>gene id</td></tr>
<tr><td>GU305914.1</td><td></td><td>22259</td><td>22281</td><td>+</td><td>0</td><td>CATCGTGCCGGCGCCCAACA</td><td>TGG</td><td>||||||||[||||||||||||]PAM</td><td>NA</td><td>Intergenic</td><td>NA</td><td>NA</td></tr>
<tr><td>T1235</td><td>ACATGGTCCCGACGAGAAACCGG</td><td></td><td>1000</td><td></td><td></td><td></td><td></td><td></td><td></td><td></td><td></td><td></td></tr>
<tr><td>Oligo fwd</td><td>CACCACATGGTCCCGACGAGAAAC</td><td></td><td></td><td></td><td></td><td></td><td></td><td></td><td></td><td></td><td></td><td></td></tr>
<tr><td>Oligo rev</td><td>AAACGTTTCTCGTCGGGACCATGT</td><td></td><td></td><td></td><td></td><td></td><td></td><td></td><td></td><td></td><td></td><td></td></tr>
<tr><td>Chromosome</td><td>start</td><td></td><td>end</td><td>strand</td><td>MM</td><td>target_seq</td><td>PAM</td><td>alignment</td><td>distance</td><td>position</td><td>gene name</td><td>gene id</td></tr>
<tr><td>GU305914.1</td><td></td><td>22276</td><td>22298</td><td>+</td><td>0</td><td>ACATGGTCCCGACGAGAAAC</td><td>CGG</td><td>||||||||[||||||||||||]PAM</td><td>NA</td><td>Intergenic</td><td>NA</td><td>NA</td></tr>
<tr><td>T1236</td><td>CCCGACGAGAAACCGGAACCCGG</td><td></td><td>1000</td><td></td><td></td><td></td><td></td><td></td><td></td><td></td><td></td><td></td></tr>
<tr><td>Oligo fwd</td><td>CACCCCCGACGAGAAACCGGAACC</td><td></td><td></td><td></td><td></td><td></td><td></td><td></td><td></td><td></td><td></td><td></td></tr>
<tr><td>Oligo rev</td><td>AAACGGTTCCGGTTTCTCGTCGGG</td><td></td><td></td><td></td><td></td><td></td><td></td><td></td><td></td><td></td><td></td><td></td></tr>
<tr><td>Chromosome</td><td>start</td><td></td><td>end</td><td>strand</td><td>MM</td><td>target_seq</td><td>PAM</td><td>alignment</td><td>distance</td><td>position</td><td>gene name</td><td>gene id</td></tr>
<tr><td>GU305914.1</td><td></td><td>22283</td><td>22305</td><td>+</td><td>0</td><td>CCCGACGAGAAACCGGAACC</td><td>CGG</td><td>||||||||[||||||||||||]PAM</td><td>NA</td><td>Intergenic</td><td>NA</td><td>NA</td></tr>
<tr><td>T1237</td><td>CCGACGAGAAACCGGAACCCGGG</td><td></td><td>1000</td><td></td><td></td><td></td><td></td><td></td><td></td><td></td><td></td><td></td></tr>
<tr><td>Oligo fwd</td><td>CACCCCGACGAGAAACCGGAACCC</td><td></td><td></td><td></td><td></td><td></td><td></td><td></td><td></td><td></td><td></td><td></td></tr>
<tr><td>Oligo rev</td><td>AAACGGGTTCCGGTTTCTCGTCGG</td><td></td><td></td><td></td><td></td><td></td><td></td><td></td><td></td><td></td><td></td><td></td></tr>
<tr><td>Chromosome</td><td>start</td><td></td><td>end</td><td>strand</td><td>MM</td><td>target_seq</td><td>PAM</td><td>alignment</td><td>distance</td><td>position</td><td>gene name</td><td>gene id</td></tr>
<tr><td>GU305914.1</td><td></td><td>22284</td><td>22306</td><td>+</td><td>0</td><td>CCGACGAGAAACCGGAACCC</td><td>GGG</td><td>||||||||[||||||||||||]PAM</td><td>NA</td><td>Intergenic</td><td>NA</td><td>NA</td></tr>
<tr><td>T1238</td><td>CCGGGAGAGCTCTCGAATCTCGG</td><td></td><td>1000</td><td></td><td></td><td></td><td></td><td></td><td></td><td></td><td></td><td></td></tr>
<tr><td>Oligo fwd</td><td>CACCCCGGGAGAGCTCTCGAATCT</td><td></td><td></td><td></td><td></td><td></td><td></td><td></td><td></td><td></td><td></td><td></td></tr>
<tr><td>Oligo rev</td><td>AAACAGATTCGAGAGCTCTCCCGG</td><td></td><td></td><td></td><td></td><td></td><td></td><td></td><td></td><td></td><td></td><td></td></tr>
<tr><td>Chromosome</td><td>start</td><td></td><td>end</td><td>strand</td><td>MM</td><td>target_seq</td><td>PAM</td><td>alignment</td><td>distance</td><td>position</td><td>gene name</td><td>gene id</td></tr>
<tr><td>GU305914.1</td><td></td><td>22302</td><td>22324</td><td>+</td><td>0</td><td>CCGGGAGAGCTCTCGAATCT</td><td>CGG</td><td>||||||||[||||||||||||]PAM</td><td>NA</td><td>Intergenic</td><td>NA</td><td>NA</td></tr>
<tr><td>T1239</td><td>GTTTTTCCCAGCGACATCAGCGG</td><td></td><td>1000</td><td></td><td></td><td></td><td></td><td></td><td></td><td></td><td></td><td></td></tr>
<tr><td>Oligo fwd</td><td>CACCGTTTTTCCCAGCGACATCAG</td><td></td><td></td><td></td><td></td><td></td><td></td><td></td><td></td><td></td><td></td><td></td></tr>
<tr><td>Oligo rev</td><td>AAACCTGATGTCGCTGGGAAAAAC</td><td></td><td></td><td></td><td></td><td></td><td></td><td></td><td></td><td></td><td></td><td></td></tr>
<tr><td>Chromosome</td><td>start</td><td></td><td>end</td><td>strand</td><td>MM</td><td>target_seq</td><td>PAM</td><td>alignment</td><td>distance</td><td>position</td><td>gene name</td><td>gene id</td></tr>
<tr><td>GU305914.1</td><td></td><td>22334</td><td>22356</td><td>+</td><td>0</td><td>GTTTTTCCCAGCGACATCAG</td><td>CGG</td><td>||||||||[||||||||||||]PAM</td><td>NA</td><td>Intergenic</td><td>NA</td><td>NA</td></tr>
<tr><td>T1240</td><td>TTTTTCCCAGCGACATCAGCGGG</td><td></td><td>1000</td><td></td><td></td><td></td><td></td><td></td><td></td><td></td><td></td><td></td></tr>
<tr><td>Oligo fwd</td><td>CACCTTTTTCCCAGCGACATCAGC</td><td></td><td></td><td></td><td></td><td></td><td></td><td></td><td></td><td></td><td></td><td></td></tr>
<tr><td>Oligo rev</td><td>AAACGCTGATGTCGCTGGGAAAAA</td><td></td><td></td><td></td><td></td><td></td><td></td><td></td><td></td><td></td><td></td><td></td></tr>
<tr><td>Chromosome</td><td>start</td><td></td><td>end</td><td>strand</td><td>MM</td><td>target_seq</td><td>PAM</td><td>alignment</td><td>distance</td><td>position</td><td>gene name</td><td>gene id</td></tr>
<tr><td>GU305914.1</td><td></td><td>22335</td><td>22357</td><td>+</td><td>0</td><td>TTTTTCCCAGCGACATCAGC</td><td>GGG</td><td>||||||||[||||||||||||]PAM</td><td>NA</td><td>Intergenic</td><td>NA</td><td>NA</td></tr>
<tr><td>T1241</td><td>TTTTCCCAGCGACATCAGCGGGG</td><td></td><td>1000</td><td></td><td></td><td></td><td></td><td></td><td></td><td></td><td></td><td></td></tr>
<tr><td>Oligo fwd</td><td>CACCTTTTCCCAGCGACATCAGCG</td><td></td><td></td><td></td><td></td><td></td><td></td><td></td><td></td><td></td><td></td><td></td></tr>
<tr><td>Oligo rev</td><td>AAACCGCTGATGTCGCTGGGAAAA</td><td></td><td></td><td></td><td></td><td></td><td></td><td></td><td></td><td></td><td></td><td></td></tr>
<tr><td>Chromosome</td><td>start</td><td></td><td>end</td><td>strand</td><td>MM</td><td>target_seq</td><td>PAM</td><td>alignment</td><td>distance</td><td>position</td><td>gene name</td><td>gene id</td></tr>
<tr><td>GU305914.1</td><td></td><td>22336</td><td>22358</td><td>+</td><td>0</td><td>TTTTCCCAGCGACATCAGCG</td><td>GGG</td><td>||||||||[||||||||||||]PAM</td><td>NA</td><td>Intergenic</td><td>NA</td><td>NA</td></tr>
<tr><td>T1242</td><td>GTAGTCCCACCCGAGAGATGCGG</td><td></td><td>1000</td><td></td><td></td><td></td><td></td><td></td><td></td><td></td><td></td><td></td></tr>
<tr><td>Oligo fwd</td><td>CACCGTAGTCCCACCCGAGAGATG</td><td></td><td></td><td></td><td></td><td></td><td></td><td></td><td></td><td></td><td></td><td></td></tr>
<tr><td>Oligo rev</td><td>AAACCATCTCTCGGGTGGGACTAC</td><td></td><td></td><td></td><td></td><td></td><td></td><td></td><td></td><td></td><td></td><td></td></tr>
<tr><td>Chromosome</td><td>start</td><td></td><td>end</td><td>strand</td><td>MM</td><td>target_seq</td><td>PAM</td><td>alignment</td><td>distance</td><td>position</td><td>gene name</td><td>gene id</td></tr>
<tr><td>GU305914.1</td><td></td><td>22381</td><td>22403</td><td>+</td><td>0</td><td>GTAGTCCCACCCGAGAGATG</td><td>CGG</td><td>||||||||[||||||||||||]PAM</td><td>NA</td><td>Intergenic</td><td>NA</td><td>NA</td></tr>
<tr><td>T1243</td><td>GCGGTTCGACAAGATGTCCATGG</td><td></td><td>1000</td><td></td><td></td><td></td><td></td><td></td><td></td><td></td><td></td><td></td></tr>
<tr><td>Oligo fwd</td><td>CACCGCGGTTCGACAAGATGTCCA</td><td></td><td></td><td></td><td></td><td></td><td></td><td></td><td></td><td></td><td></td><td></td></tr>
<tr><td>Oligo rev</td><td>AAACTGGACATCTTGTCGAACCGC</td><td></td><td></td><td></td><td></td><td></td><td></td><td></td><td></td><td></td><td></td><td></td></tr>
<tr><td>Chromosome</td><td>start</td><td></td><td>end</td><td>strand</td><td>MM</td><td>target_seq</td><td>PAM</td><td>alignment</td><td>distance</td><td>position</td><td>gene name</td><td>gene id</td></tr>
<tr><td>GU305914.1</td><td></td><td>22400</td><td>22422</td><td>+</td><td>0</td><td>GCGGTTCGACAAGATGTCCA</td><td>TGG</td><td>||||||||[||||||||||||]PAM</td><td>NA</td><td>Intergenic</td><td>NA</td><td>NA</td></tr>
<tr><td>T1244</td><td>CGGTTCGACAAGATGTCCATGGG</td><td></td><td>1000</td><td></td><td></td><td></td><td></td><td></td><td></td><td></td><td></td><td></td></tr>
<tr><td>Oligo fwd</td><td>CACCCGGTTCGACAAGATGTCCAT</td><td></td><td></td><td></td><td></td><td></td><td></td><td></td><td></td><td></td><td></td><td></td></tr>
<tr><td>Oligo rev</td><td>AAACATGGACATCTTGTCGAACCG</td><td></td><td></td><td></td><td></td><td></td><td></td><td></td><td></td><td></td><td></td><td></td></tr>
<tr><td>Chromosome</td><td>start</td><td></td><td>end</td><td>strand</td><td>MM</td><td>target_seq</td><td>PAM</td><td>alignment</td><td>distance</td><td>position</td><td>gene name</td><td>gene id</td></tr>
<tr><td>GU305914.1</td><td></td><td>22401</td><td>22423</td><td>+</td><td>0</td><td>CGGTTCGACAAGATGTCCAT</td><td>GGG</td><td>||||||||[||||||||||||]PAM</td><td>NA</td><td>Intergenic</td><td>NA</td><td>NA</td></tr>
<tr><td>T1245</td><td>GGGAAAGATGCCGAGCGCCACGG</td><td></td><td>1000</td><td></td><td></td><td></td><td></td><td></td><td></td><td></td><td></td><td></td></tr>
<tr><td>Oligo fwd</td><td>CACCGGGAAAGATGCCGAGCGCCA</td><td></td><td></td><td></td><td></td><td></td><td></td><td></td><td></td><td></td><td></td><td></td></tr>
<tr><td>Oligo rev</td><td>AAACTGGCGCTCGGCATCTTTCCC</td><td></td><td></td><td></td><td></td><td></td><td></td><td></td><td></td><td></td><td></td><td></td></tr>
<tr><td>Chromosome</td><td>start</td><td></td><td>end</td><td>strand</td><td>MM</td><td>target_seq</td><td>PAM</td><td>alignment</td><td>distance</td><td>position</td><td>gene name</td><td>gene id</td></tr>
<tr><td>GU305914.1</td><td></td><td>22421</td><td>22443</td><td>+</td><td>0</td><td>GGGAAAGATGCCGAGCGCCA</td><td>CGG</td><td>||||||||[||||||||||||]PAM</td><td>NA</td><td>Intergenic</td><td>NA</td><td>NA</td></tr>
</table>

<table>
<tr><td>T1246</td><td>CGTGCTCTCTCACTGTCAAACGG</td><td></td><td>1000</td><td></td><td></td><td></td><td></td><td></td><td></td><td></td><td></td><td></td></tr>
<tr><td>Oligo fwd</td><td>CACCCGTGCTCTCTCACTGTCAAA</td><td></td><td></td><td></td><td></td><td></td><td></td><td></td><td></td><td></td><td></td><td></td></tr>
<tr><td>Oligo rev</td><td>AAACTTTGACAGTGAGAGAGCACG</td><td></td><td></td><td></td><td></td><td></td><td></td><td></td><td></td><td></td><td></td><td></td></tr>
<tr><td>Chromosome</td><td>start</td><td></td><td>end</td><td>strand</td><td>MM</td><td>target_seq</td><td>PAM</td><td>alignment</td><td>distance</td><td>position</td><td>gene name</td><td>gene id</td></tr>
<tr><td>GU305914.1</td><td></td><td>22459</td><td>22481</td><td>+</td><td>0</td><td>CGTGCTCTCTCACTGTCAAA</td><td>CGG</td><td>||||||||[||||||||||||]PAM</td><td>NA</td><td>Intergenic</td><td>NA</td><td>NA</td></tr>
<tr><td>T1247</td><td>TCACTGTCAAACGGCAAGACAGG</td><td></td><td>1000</td><td></td><td></td><td></td><td></td><td></td><td></td><td></td><td></td><td></td></tr>
<tr><td>Oligo fwd</td><td>CACCTCACTGTCAAACGGCAAGAC</td><td></td><td></td><td></td><td></td><td></td><td></td><td></td><td></td><td></td><td></td><td></td></tr>
<tr><td>Oligo rev</td><td>AAACGTCTTGCCGTTTGACAGTGA</td><td></td><td></td><td></td><td></td><td></td><td></td><td></td><td></td><td></td><td></td><td></td></tr>
<tr><td>Chromosome</td><td>start</td><td></td><td>end</td><td>strand</td><td>MM</td><td>target_seq</td><td>PAM</td><td>alignment</td><td>distance</td><td>position</td><td>gene name</td><td>gene id</td></tr>
<tr><td>GU305914.1</td><td></td><td>22468</td><td>22490</td><td>+</td><td>0</td><td>TCACTGTCAAACGGCAAGAC</td><td>AGG</td><td>||||||||[||||||||||||]PAM</td><td>NA</td><td>Intergenic</td><td>NA</td><td>NA</td></tr>
<tr><td>T1248</td><td>TCAAACGGCAAGACAGGTCCTGG</td><td></td><td>1000</td><td></td><td></td><td></td><td></td><td></td><td></td><td></td><td></td><td></td></tr>
<tr><td>Oligo fwd</td><td>CACCTCAAACGGCAAGACAGGTCC</td><td></td><td></td><td></td><td></td><td></td><td></td><td></td><td></td><td></td><td></td><td></td></tr>
<tr><td>Oligo rev</td><td>AAACGGACCTGTCTTGCCGTTTGA</td><td></td><td></td><td></td><td></td><td></td><td></td><td></td><td></td><td></td><td></td><td></td></tr>
<tr><td>Chromosome</td><td>start</td><td></td><td>end</td><td>strand</td><td>MM</td><td>target_seq</td><td>PAM</td><td>alignment</td><td>distance</td><td>position</td><td>gene name</td><td>gene id</td></tr>
<tr><td>GU305914.1</td><td></td><td>22474</td><td>22496</td><td>+</td><td>0</td><td>TCAAACGGCAAGACAGGTCC</td><td>TGG</td><td>||||||||[||||||||||||]PAM</td><td>NA</td><td>Intergenic</td><td>NA</td><td>NA</td></tr>
<tr><td>T1249</td><td>TTCTTGTAGTCGACGTCCTTTGG</td><td></td><td>1000</td><td></td><td></td><td></td><td></td><td></td><td></td><td></td><td></td><td></td></tr>
<tr><td>Oligo fwd</td><td>CACCTTCTTGTAGTCGACGTCCTT</td><td></td><td></td><td></td><td></td><td></td><td></td><td></td><td></td><td></td><td></td><td></td></tr>
<tr><td>Oligo rev</td><td>AAACAAGGACGTCGACTACAAGAA</td><td></td><td></td><td></td><td></td><td></td><td></td><td></td><td></td><td></td><td></td><td></td></tr>
<tr><td>Chromosome</td><td>start</td><td></td><td>end</td><td>strand</td><td>MM</td><td>target_seq</td><td>PAM</td><td>alignment</td><td>distance</td><td>position</td><td>gene name</td><td>gene id</td></tr>
<tr><td>GU305914.1</td><td></td><td>22500</td><td>22522</td><td>+</td><td>0</td><td>TTCTTGTAGTCGACGTCCTT</td><td>TGG</td><td>||||||||[||||||||||||]PAM</td><td>NA</td><td>Intergenic</td><td>NA</td><td>NA</td></tr>
<tr><td>T1250</td><td>TTGTAGTCGACGTCCTTTGGAGG</td><td></td><td>1000</td><td></td><td></td><td></td><td></td><td></td><td></td><td></td><td></td><td></td></tr>
<tr><td>Oligo fwd</td><td>CACCTTGTAGTCGACGTCCTTTGG</td><td></td><td></td><td></td><td></td><td></td><td></td><td></td><td></td><td></td><td></td><td></td></tr>
<tr><td>Oligo rev</td><td>AAACCCAAAGGACGTCGACTACAA</td><td></td><td></td><td></td><td></td><td></td><td></td><td></td><td></td><td></td><td></td><td></td></tr>
<tr><td>Chromosome</td><td>start</td><td></td><td>end</td><td>strand</td><td>MM</td><td>target_seq</td><td>PAM</td><td>alignment</td><td>distance</td><td>position</td><td>gene name</td><td>gene id</td></tr>
<tr><td>GU305914.1</td><td></td><td>22503</td><td>22525</td><td>+</td><td>0</td><td>TTGTAGTCGACGTCCTTTGG</td><td>AGG</td><td>||||||||[||||||||||||]PAM</td><td>NA</td><td>Intergenic</td><td>NA</td><td>NA</td></tr>
<tr><td>T1251</td><td>TGTAGTCGACGTCCTTTGGAGGG</td><td></td><td>1000</td><td></td><td></td><td></td><td></td><td></td><td></td><td></td><td></td><td></td></tr>
<tr><td>Oligo fwd</td><td>CACCTGTAGTCGACGTCCTTTGGA</td><td></td><td></td><td></td><td></td><td></td><td></td><td></td><td></td><td></td><td></td><td></td></tr>
<tr><td>Oligo rev</td><td>AAACTCCAAAGGACGTCGACTACA</td><td></td><td></td><td></td><td></td><td></td><td></td><td></td><td></td><td></td><td></td><td></td></tr>
<tr><td>Chromosome</td><td>start</td><td></td><td>end</td><td>strand</td><td>MM</td><td>target_seq</td><td>PAM</td><td>alignment</td><td>distance</td><td>position</td><td>gene name</td><td>gene id</td></tr>
<tr><td>GU305914.1</td><td></td><td>22504</td><td>22526</td><td>+</td><td>0</td><td>TGTAGTCGACGTCCTTTGGA</td><td>GGG</td><td>||||||||[||||||||||||]PAM</td><td>NA</td><td>Intergenic</td><td>NA</td><td>NA</td></tr>
<tr><td>T1252</td><td>GTGCGACTTTAAGCCCCACCCGG</td><td></td><td>1000</td><td></td><td></td><td></td><td></td><td></td><td></td><td></td><td></td><td></td></tr>
<tr><td>Oligo fwd</td><td>CACCGTGCGACTTTAAGCCCCACC</td><td></td><td></td><td></td><td></td><td></td><td></td><td></td><td></td><td></td><td></td><td></td></tr>
<tr><td>Oligo rev</td><td>AAACGGTGGGGCTTAAAGTCGCAC</td><td></td><td></td><td></td><td></td><td></td><td></td><td></td><td></td><td></td><td></td><td></td></tr>
<tr><td>Chromosome</td><td>start</td><td></td><td>end</td><td>strand</td><td>MM</td><td>target_seq</td><td>PAM</td><td>alignment</td><td>distance</td><td>position</td><td>gene name</td><td>gene id</td></tr>
<tr><td>GU305914.1</td><td></td><td>22528</td><td>22550</td><td>+</td><td>0</td><td>GTGCGACTTTAAGCCCCACC</td><td>CGG</td><td>||||||||[||||||||||||]PAM</td><td>NA</td><td>Intergenic</td><td>NA</td><td>NA</td></tr>
<tr><td>T1253</td><td>TTTAAGCCCCACCCGGACAACGG</td><td></td><td>1000</td><td></td><td></td><td></td><td></td><td></td><td></td><td></td><td></td><td></td></tr>
<tr><td>Oligo fwd</td><td>CACCTTTAAGCCCCACCCGGACAA</td><td></td><td></td><td></td><td></td><td></td><td></td><td></td><td></td><td></td><td></td><td></td></tr>
<tr><td>Oligo rev</td><td>AAACTTGTCCGGGTGGGGCTTAAA</td><td></td><td></td><td></td><td></td><td></td><td></td><td></td><td></td><td></td><td></td><td></td></tr>
<tr><td>Chromosome</td><td>start</td><td></td><td>end</td><td>strand</td><td>MM</td><td>target_seq</td><td>PAM</td><td>alignment</td><td>distance</td><td>position</td><td>gene name</td><td>gene id</td></tr>
<tr><td>GU305914.1</td><td></td><td>22535</td><td>22557</td><td>+</td><td>0</td><td>TTTAAGCCCCACCCGGACAA</td><td>CGG</td><td>||||||||[||||||||||||]PAM</td><td>NA</td><td>Intergenic</td><td>NA</td><td>NA</td></tr>
<tr><td>T1254</td><td>GACAACGGTATTCCCCGTTCCGG</td><td></td><td>1000</td><td></td><td></td><td></td><td></td><td></td><td></td><td></td><td></td><td></td></tr>
<tr><td>Oligo fwd</td><td>CACCGACAACGGTATTCCCCGTTC</td><td></td><td></td><td></td><td></td><td></td><td></td><td></td><td></td><td></td><td></td><td></td></tr>
<tr><td>Oligo rev</td><td>AAACGAACGGGGAATACCGTTGTC</td><td></td><td></td><td></td><td></td><td></td><td></td><td></td><td></td><td></td><td></td><td></td></tr>
<tr><td>Chromosome</td><td>start</td><td></td><td>end</td><td>strand</td><td>MM</td><td>target_seq</td><td>PAM</td><td>alignment</td><td>distance</td><td>position</td><td>gene name</td><td>gene id</td></tr>
<tr><td>GU305914.1</td><td></td><td>22550</td><td>22572</td><td>+</td><td>0</td><td>GACAACGGTATTCCCCGTTC</td><td>CGG</td><td>||||||||[||||||||||||]PAM</td><td>NA</td><td>Intergenic</td><td>NA</td><td>NA</td></tr>
<tr><td>T1255</td><td>CCCACCTCAACCCCAGCGCACGG</td><td></td><td>1000</td><td></td><td></td><td></td><td></td><td></td><td></td><td></td><td></td><td></td></tr>
<tr><td>Oligo fwd</td><td>CACCCCCACCTCAACCCCAGCGCA</td><td></td><td></td><td></td><td></td><td></td><td></td><td></td><td></td><td></td><td></td><td></td></tr>
<tr><td>Oligo rev</td><td>AAACTGCGCTGGGGTTGAGGTGGG</td><td></td><td></td><td></td><td></td><td></td><td></td><td></td><td></td><td></td><td></td><td></td></tr>
<tr><td>Chromosome</td><td>start</td><td></td><td>end</td><td>strand</td><td>MM</td><td>target_seq</td><td>PAM</td><td>alignment</td><td>distance</td><td>position</td><td>gene name</td><td>gene id</td></tr>
<tr><td>GU305914.1</td><td></td><td>22575</td><td>22597</td><td>+</td><td>0</td><td>CCCACCTCAACCCCAGCGCA</td><td>CGG</td><td>||||||||[||||||||||||]PAM</td><td>NA</td><td>Intergenic</td><td>NA</td><td>NA</td></tr>
<tr><td>T1256</td><td>GCACGGCTCACCCCAACTCGTGG</td><td></td><td>1000</td><td></td><td></td><td></td><td></td><td></td><td></td><td></td><td></td><td></td></tr>
<tr><td>Oligo fwd</td><td>CACCGCACGGCTCACCCCAACTCG</td><td></td><td></td><td></td><td></td><td></td><td></td><td></td><td></td><td></td><td></td><td></td></tr>
<tr><td>Oligo rev</td><td>AAACCGAGTTGGGGTGAGCCGTGC</td><td></td><td></td><td></td><td></td><td></td><td></td><td></td><td></td><td></td><td></td><td></td></tr>
<tr><td>Chromosome</td><td>start</td><td></td><td>end</td><td>strand</td><td>MM</td><td>target_seq</td><td>PAM</td><td>alignment</td><td>distance</td><td>position</td><td>gene name</td><td>gene id</td></tr>
<tr><td>GU305914.1</td><td></td><td>22592</td><td>22614</td><td>+</td><td>0</td><td>GCACGGCTCACCCCAACTCG</td><td>TGG</td><td>||||||||[||||||||||||]PAM</td><td>NA</td><td>Intergenic</td><td>NA</td><td>NA</td></tr>
<tr><td>T1257</td><td>CTCGTGGTGTAATTCGTGCCCGG</td><td></td><td>1000</td><td></td><td></td><td></td><td></td><td></td><td></td><td></td><td></td><td></td></tr>
<tr><td>Oligo fwd</td><td>CACCCTCGTGGTGTAATTCGTGCC</td><td></td><td></td><td></td><td></td><td></td><td></td><td></td><td></td><td></td><td></td><td></td></tr>
<tr><td>Oligo rev</td><td>AAACGGCACGAATTACACCACGAG</td><td></td><td></td><td></td><td></td><td></td><td></td><td></td><td></td><td></td><td></td><td></td></tr>
<tr><td>Chromosome</td><td>start</td><td></td><td>end</td><td>strand</td><td>MM</td><td>target_seq</td><td>PAM</td><td>alignment</td><td>distance</td><td>position</td><td>gene name</td><td>gene id</td></tr>
<tr><td>GU305914.1</td><td></td><td>22608</td><td>22630</td><td>+</td><td>0</td><td>CTCGTGGTGTAATTCGTGCC</td><td>CGG</td><td>||||||||[||||||||||||]PAM</td><td>NA</td><td>Intergenic</td><td>NA</td><td>NA</td></tr>
<tr><td>T1258</td><td>GTGTAATTCGTGCCCGGTAAAGG</td><td></td><td>1000</td><td></td><td></td><td></td><td></td><td></td><td></td><td></td><td></td><td></td></tr>
<tr><td>Oligo fwd</td><td>CACCGTGTAATTCGTGCCCGGTAA</td><td></td><td></td><td></td><td></td><td></td><td></td><td></td><td></td><td></td><td></td><td></td></tr>
<tr><td>Oligo rev</td><td>AAACTTACCGGGCACGAATTACAC</td><td></td><td></td><td></td><td></td><td></td><td></td><td></td><td></td><td></td><td></td><td></td></tr>
<tr><td>Chromosome</td><td>start</td><td></td><td>end</td><td>strand</td><td>MM</td><td>target_seq</td><td>PAM</td><td>alignment</td><td>distance</td><td>position</td><td>gene name</td><td>gene id</td></tr>
<tr><td>GU305914.1</td><td></td><td>22614</td><td>22636</td><td>+</td><td>0</td><td>GTGTAATTCGTGCCCGGTAA</td><td>AGG</td><td>||||||||[||||||||||||]PAM</td><td>NA</td><td>Intergenic</td><td>NA</td><td>NA</td></tr>
<tr><td>T1259</td><td>TGTAATTCGTGCCCGGTAAAGGG</td><td></td><td>1000</td><td></td><td></td><td></td><td></td><td></td><td></td><td></td><td></td><td></td></tr>
<tr><td>Oligo fwd</td><td>CACCTGTAATTCGTGCCCGGTAAA</td><td></td><td></td><td></td><td></td><td></td><td></td><td></td><td></td><td></td><td></td><td></td></tr>
<tr><td>Oligo rev</td><td>AAACTTTACCGGGCACGAATTACA</td><td></td><td></td><td></td><td></td><td></td><td></td><td></td><td></td><td></td><td></td><td></td></tr>
<tr><td>Chromosome</td><td>start</td><td></td><td>end</td><td>strand</td><td>MM</td><td>target_seq</td><td>PAM</td><td>alignment</td><td>distance</td><td>position</td><td>gene name</td><td>gene id</td></tr>
<tr><td>GU305914.1</td><td></td><td>22615</td><td>22637</td><td>+</td><td>0</td><td>TGTAATTCGTGCCCGGTAAA</td><td>GGG</td><td>||||||||[||||||||||||]PAM</td><td>NA</td><td>Intergenic</td><td>NA</td><td>NA</td></tr>
<tr><td>T1260</td><td>GTAATTCGTGCCCGGTAAAGGGG</td><td></td><td>1000</td><td></td><td></td><td></td><td></td><td></td><td></td><td></td><td></td><td></td></tr>
<tr><td>Oligo fwd</td><td>CACCGTAATTCGTGCCCGGTAAAG</td><td></td><td></td><td></td><td></td><td></td><td></td><td></td><td></td><td></td><td></td><td></td></tr>
<tr><td>Oligo rev</td><td>AAACCTTTACCGGGCACGAATTAC</td><td></td><td></td><td></td><td></td><td></td><td></td><td></td><td></td><td></td><td></td><td></td></tr>
<tr><td>Chromosome</td><td>start</td><td></td><td>end</td><td>strand</td><td>MM</td><td>target_seq</td><td>PAM</td><td>alignment</td><td>distance</td><td>position</td><td>gene name</td><td>gene id</td></tr>
</table>

GU305914.1 22616 22638 + 0 GTAATTCGTGCCCGGTAAAG GGG ||||||||[||||||||||||]PAM NA Intergenic NA NA

T1261 GGTAAAGGGGTTTTCATGACAGG 1000
Oligo fwd CACCGGTAAAGGGGTTTTCATGAC
Oligo rev AAACGTCATGAAAACCCCTTTACC
Chromosome start end strand MM target_seq PAM alignment distance position gene name gene id
GU305914.1 22629 22651 + 0 GGTAAAGGGGTTTTCATGAC AGG ||||||||[||||||||||||]PAM NA Intergenic NA NA

T1262 TTCATGACAGGTATCCGATACGG 1000
Oligo fwd CACCTTCATGACAGGTATCCGATA
Oligo rev AAACTATCGGATACCTGTCATGAA
Chromosome start end strand MM target_seq PAM alignment distance position gene name gene id
GU305914.1 22641 22663 + 0 TTCATGACAGGTATCCGATA CGG ||||||||[||||||||||||]PAM NA Intergenic NA NA

T1263 CAGGTATCCGATACGGACCCAGG 1000
Oligo fwd CACCCAGGTATCCGATACGGACCC
Oligo rev AAACGGGTCCGTATCGGATACCTG
Chromosome start end strand MM target_seq PAM alignment distance position gene name gene id
GU305914.1 22648 22670 + 0 CAGGTATCCGATACGGACCC AGG ||||||||[||||||||||||]PAM NA Intergenic NA NA

T1264 CATTCTGCTGTACGACCTGCCGG 1000
Oligo fwd CACCCATTCTGCTGTACGACCTGC
Oligo rev AAACGCAGGTCGTACAGCAGAATG
Chromosome start end strand MM target_seq PAM alignment distance position gene name gene id
GU305914.1 22673 22695 + 0 CATTCTGCTGTACGACCTGC CGG ||||||||[||||||||||||]PAM NA Intergenic NA NA
GU305914.1 90872 90894 - 4 CGATCTGCTGGACAACCTGC GGG |--|||||[||-||-||||||]PAM NA Intergenic NA NA

T1265 GCTGTACGACCTGCCGGTCGCGG 1000
Oligo fwd CACCGCTGTACGACCTGCCGGTCG
Oligo rev AAACCGACCGGCAGGTCGTACAGC
Chromosome start end strand MM target_seq PAM alignment distance position gene name gene id
GU305914.1 22679 22701 + 0 GCTGTACGACCTGCCGGTCG CGG ||||||||[||||||||||||]PAM NA Intergenic NA NA

T1266 CCACCTCCCAGATCAAATCCAGG 1000
Oligo fwd CACCCCACCTCCCAGATCAAATCC
Oligo rev AAACGGATTTGATCTGGGAGGTGG
Chromosome start end strand MM target_seq PAM alignment distance position gene name gene id
GU305914.1 22714 22736 + 0 CCACCTCCCAGATCAAATCC AGG ||||||||[||||||||||||]PAM NA Intergenic NA NA

T1267 CACCTCCCAGATCAAATCCAGGG 1000
Oligo fwd CACCCACCTCCCAGATCAAATCCA
Oligo rev AAACTGGATTTGATCTGGGAGGTG
Chromosome start end strand MM target_seq PAM alignment distance position gene name gene id
GU305914.1 22715 22737 + 0 CACCTCCCAGATCAAATCCA GGG ||||||||[||||||||||||]PAM NA Intergenic NA NA

T1268 TCCACCGTCCCGAAACCGTCTGG 1000
Oligo fwd CACCTCCACCGTCCCGAAACCGTC
Oligo rev AAACGACGGTTTCGGGACGGTGGA
Chromosome start end strand MM target_seq PAM alignment distance position gene name gene id
GU305914.1 22738 22760 + 0 TCCACCGTCCCGAAACCGTC TGG ||||||||[||||||||||||]PAM NA Intergenic NA NA

T1269 ACCATCGTCGTCCTCACGACCGG 1000
Oligo fwd CACCACCATCGTCGTCCTCACGAC
Oligo rev AAACGTCGTGAGGACGACGATGGT
Chromosome start end strand MM target_seq PAM alignment distance position gene name gene id
GU305914.1 22805 22827 + 0 ACCATCGTCGTCCTCACGAC CGG ||||||||[||||||||||||]PAM NA Intergenic NA NA

T1270 CCACGCTCAAAACGTGCGCGAGG 1000
Oligo fwd CACCCCACGCTCAAAACGTGCGCG
Oligo rev AAACCGCGCACGTTTTGAGCGTGG
Chromosome start end strand MM target_seq PAM alignment distance position gene name gene id
GU305914.1 22861 22883 + 0 CCACGCTCAAAACGTGCGCG AGG ||||||||[||||||||||||]PAM NA Intergenic NA NA

T1271 CAAAACGTGCGCGAGGTTCACGG 1000
Oligo fwd CACCCAAAACGTGCGCGAGGTTCA
Oligo rev AAACTGAACCTCGCGCACGTTTTG
Chromosome start end strand MM target_seq PAM alignment distance position gene name gene id
GU305914.1 22868 22890 + 0 CAAAACGTGCGCGAGGTTCA CGG ||||||||[||||||||||||]PAM NA Intergenic NA NA

T1272 GCTGTCGCAGCCTCCCAACGCGG 1000
Oligo fwd CACCGCTGTCGCAGCCTCCCAACG
Oligo rev AAACCGTTGGGAGGCTGCGACAGC
Chromosome start end strand MM target_seq PAM alignment distance position gene name gene id
GU305914.1 22913 22935 + 0 GCTGTCGCAGCCTCCCAACG CGG ||||||||[||||||||||||]PAM NA Intergenic NA NA

T1273 CAGCCTCCCAACGCGGTCAGTGG 1000
Oligo fwd CACCCAGCCTCCCAACGCGGTCAG
Oligo rev AAACCTGACCGCGTTGGGAGGCTG
Chromosome start end strand MM target_seq PAM alignment distance position gene name gene id
GU305914.1 22920 22942 + 0 CAGCCTCCCAACGCGGTCAG TGG ||||||||[||||||||||||]PAM NA Intergenic NA NA

T1274 CCTCCCAACGCGGTCAGTGGCGG 1000
Oligo fwd CACCCCTCCCAACGCGGTCAGTGG
Oligo rev AAACCCACTGACCGCGTTGGGAGG
Chromosome start end strand MM target_seq PAM alignment distance position gene name gene id
GU305914.1 22923 22945 + 0 CCTCCCAACGCGGTCAGTGG CGG ||||||||[||||||||||||]PAM NA Intergenic NA NA

T1275 CCCAACGCGGTCAGTGGCGGCGG 1000
Oligo fwd CACCCCCAACGCGGTCAGTGGCGG

| | | | | | | | | | | | |
|---|---|---|---|---|---|---|---|---|---|---|---|
| Oligo rev | AAACCCGCCACTGACCGCGTTGGG | | | | | | | | | | |
| Chromosome | start | end | strand | MM | target_seq | PAM | alignment | distance | position | gene name | gene id |
| GU305914.1 | 22926 | 22948 | + | 0 | CCCAACGCGGTCAGTGGCGG | CGG | ||||||||[||||||||||||]PAM | NA | Intergenic | NA | NA |
| GU305914.1 | 58135 | 58157 | - | 4 | CGCAAGGCGGGCGGTGGCGG | CGG | |-|||-||[||-|-|||||||]PAM | NA | Intergenic | NA | NA |
| T1276 | ACGCGGTCAGTGGCGGCGGTCGG | 1000 | | | | | | | | | |
| Oligo fwd | CACCACGCGGTCAGTGGCGGCGGT | | | | | | | | | | |
| Oligo rev | AAACACCGCCGCCACTGACCGCGT | | | | | | | | | | |
| Chromosome | start | end | strand | MM | target_seq | PAM | alignment | distance | position | gene name | gene id |
| GU305914.1 | 22930 | 22952 | + | 0 | ACGCGGTCAGTGGCGGCGGT | CGG | ||||||||[||||||||||||]PAM | NA | Intergenic | NA | NA |
| T1277 | CGGTCAGTGGCGGCGGTCGGCGG | 1000 | | | | | | | | | |
| Oligo fwd | CACCCGGTCAGTGGCGGCGGTCGG | | | | | | | | | | |
| Oligo rev | AAACCCGACCGCCGCCACTGACCG | | | | | | | | | | |
| Chromosome | start | end | strand | MM | target_seq | PAM | alignment | distance | position | gene name | gene id |
| GU305914.1 | 22933 | 22955 | + | 0 | CGGTCAGTGGCGGCGGTCGG | CGG | ||||||||[||||||||||||]PAM | NA | Intergenic | NA | NA |
| GU305914.1 | 128859 | 128881 | + | 4 | CGGTCGTTGGCGGAGGTCAG | GGG | |||||--|[|||||-||||-|]PAM | NA | Intergenic | NA | NA |
| T1278 | CTCGTCGTCAGATTTGTCGCCGG | 1000 | | | | | | | | | |
| Oligo fwd | CACCCTCGTCGTCAGATTTGTCGC | | | | | | | | | | |
| Oligo rev | AAACGCGACAAATCTGACGACGAG | | | | | | | | | | |
| Chromosome | start | end | strand | MM | target_seq | PAM | alignment | distance | position | gene name | gene id |
| GU305914.1 | 22962 | 22984 | + | 0 | CTCGTCGTCAGATTTGTCGC | CGG | ||||||||[||||||||||||]PAM | NA | Intergenic | NA | NA |
| T1279 | CAGATTTGTCGCCGGCATCGAGG | 1000 | | | | | | | | | |
| Oligo fwd | CACCCAGATTTGTCGCCGGCATCG | | | | | | | | | | |
| Oligo rev | AAACCGATGCCGGCGACAAATCTG | | | | | | | | | | |
| Chromosome | start | end | strand | MM | target_seq | PAM | alignment | distance | position | gene name | gene id |
| GU305914.1 | 22970 | 22992 | + | 0 | CAGATTTGTCGCCGGCATCG | AGG | ||||||||[||||||||||||]PAM | NA | Intergenic | NA | NA |
| T1280 | ATTTGTCGCCGGCATCGAGGCGG | 1000 | | | | | | | | | |
| Oligo fwd | CACCATTTGTCGCCGGCATCGAGG | | | | | | | | | | |
| Oligo rev | AAACCCTCGATGCCGGCGACAAAT | | | | | | | | | | |
| Chromosome | start | end | strand | MM | target_seq | PAM | alignment | distance | position | gene name | gene id |
| GU305914.1 | 22973 | 22995 | + | 0 | ATTTGTCGCCGGCATCGAGG | CGG | ||||||||[||||||||||||]PAM | NA | Intergenic | NA | NA |
| T1281 | CACACTCGAGCCATAGCGAACGG | 1000 | | | | | | | | | |
| Oligo fwd | CACCCACACTCGAGCCATAGCGAA | | | | | | | | | | |
| Oligo rev | AAACTTCGCTATGGCTCGAGTGTG | | | | | | | | | | |
| Chromosome | start | end | strand | MM | target_seq | PAM | alignment | distance | position | gene name | gene id |
| GU305914.1 | 23031 | 23053 | + | 0 | CACACTCGAGCCATAGCGAA | CGG | ||||||||[||||||||||||]PAM | NA | Intergenic | NA | NA |
| T1282 | GAACGGTTCGAGACGAGATACGG | 1000 | | | | | | | | | |
| Oligo fwd | CACCGAACGGTTCGAGACGAGATA | | | | | | | | | | |
| Oligo rev | AAACTATCTCGTCTCGAACCGTTC | | | | | | | | | | |
| Chromosome | start | end | strand | MM | target_seq | PAM | alignment | distance | position | gene name | gene id |
| GU305914.1 | 23048 | 23070 | + | 0 | GAACGGTTCGAGACGAGATA | CGG | ||||||||[||||||||||||]PAM | NA | Intergenic | NA | NA |
| T1283 | CGGTTCGAGACGAGATACGGAGG | 1000 | | | | | | | | | |
| Oligo fwd | CACCCGGTTCGAGACGAGATACGG | | | | | | | | | | |
| Oligo rev | AAACCCGTATCTCGTCTCGAACCG | | | | | | | | | | |
| Chromosome | start | end | strand | MM | target_seq | PAM | alignment | distance | position | gene name | gene id |
| GU305914.1 | 23051 | 23073 | + | 0 | CGGTTCGAGACGAGATACGG | AGG | ||||||||[||||||||||||]PAM | NA | Intergenic | NA | NA |
| T1284 | GGTTCGAGACGAGATACGGAGGG | 1000 | | | | | | | | | |
| Oligo fwd | CACCGGTTCGAGACGAGATACGGA | | | | | | | | | | |
| Oligo rev | AAACTCCGTATCTCGTCTCGAACC | | | | | | | | | | |
| Chromosome | start | end | strand | MM | target_seq | PAM | alignment | distance | position | gene name | gene id |
| GU305914.1 | 23052 | 23074 | + | 0 | GGTTCGAGACGAGATACGGA | GGG | ||||||||[||||||||||||]PAM | NA | Intergenic | NA | NA |
| T1285 | GTTCGAGACGAGATACGGAGGGG | 1000 | | | | | | | | | |
| Oligo fwd | CACCGTTCGAGACGAGATACGGAG | | | | | | | | | | |
| Oligo rev | AAACCTCCGTATCTCGTCTCGAAC | | | | | | | | | | |
| Chromosome | start | end | strand | MM | target_seq | PAM | alignment | distance | position | gene name | gene id |
| GU305914.1 | 23053 | 23075 | + | 0 | GTTCGAGACGAGATACGGAG | GGG | ||||||||[||||||||||||]PAM | NA | Intergenic | NA | NA |
| T1286 | AGACGAGATACGGAGGGGTCCGG | 1000 | | | | | | | | | |
| Oligo fwd | CACCAGACGAGATACGGAGGGGTC | | | | | | | | | | |
| Oligo rev | AAACGACCCCTCCGTATCTCGTCT | | | | | | | | | | |
| Chromosome | start | end | strand | MM | target_seq | PAM | alignment | distance | position | gene name | gene id |
| GU305914.1 | 23058 | 23080 | + | 0 | AGACGAGATACGGAGGGGTC | CGG | ||||||||[||||||||||||]PAM | NA | Intergenic | NA | NA |
| T1287 | GTCCGGAGACCTCAAAATTCTGG | 1000 | | | | | | | | | |
| Oligo fwd | CACCGTCCGGAGACCTCAAAATTC | | | | | | | | | | |
| Oligo rev | AAACGAATTTTGAGGTCTCCGGAC | | | | | | | | | | |
| Chromosome | start | end | strand | MM | target_seq | PAM | alignment | distance | position | gene name | gene id |
| GU305914.1 | 23075 | 23097 | + | 0 | GTCCGGAGACCTCAAAATTC | TGG | ||||||||[||||||||||||]PAM | NA | Intergenic | NA | NA |
| T1288 | TTCTGGTGACATGTTTGTCTTGG | 1000 | | | | | | | | | |
| Oligo fwd | CACCTTCTGGTGACATGTTTGTCT | | | | | | | | | | |
| Oligo rev | AAACAGACAAACATGTCACCAGAA | | | | | | | | | | |
| Chromosome | start | end | strand | MM | target_seq | PAM | alignment | distance | position | gene name | gene id |
| GU305914.1 | 23092 | 23114 | + | 0 | TTCTGGTGACATGTTTGTCT | TGG | ||||||||[||||||||||||]PAM | NA | Intergenic | NA | NA |
| T1289 | TCTGGTGACATGTTTGTCTTGGG | 1000 | | | | | | | | | |
| Oligo fwd | CACCTCTGGTGACATGTTTGTCTT | | | | | | | | | | |
| Oligo rev | AAACAAGACAAACATGTCACCAGA | | | | | | | | | | |
| Chromosome | start | end | strand | MM | target_seq | PAM | alignment | distance | position | gene name | gene id |
| GU305914.1 | 23093 | 23115 | + | 0 | TCTGGTGACATGTTTGTCTT | GGG | ||||||||[||||||||||||]PAM | NA | Intergenic | NA | NA |

<table>
<tr><td>T1290</td><td>TTGTCAGTCATCGATCGTCACGG</td><td>1000</td></tr>
<tr><td>Oligo fwd</td><td>CACCTTGTCAGTCATCGATCGTCA</td></tr>
<tr><td>Oligo rev</td><td>AAACTGACGATCGATGACTGACAA</td></tr>
<tr><td>Chromosome</td><td>start</td><td>end</td><td>strand</td><td>MM</td><td>target_seq</td><td>PAM</td><td>alignment</td><td>distance</td><td>position</td><td>gene name</td><td>gene id</td></tr>
<tr><td>GU305914.1</td><td>23140</td><td>23162</td><td>+</td><td>0</td><td>TTGTCAGTCATCGATCGTCA</td><td>CGG</td><td>||||||||[||||||||||||]PAM</td><td>NA</td><td>Intergenic</td><td>NA</td><td>NA</td></tr>
<tr><td>T1291</td><td>GTCATCGATCGTCACGGTCCCGG</td><td>1000</td></tr>
<tr><td>Oligo fwd</td><td>CACCGTCATCGATCGTCACGGTCC</td></tr>
<tr><td>Oligo rev</td><td>AAACGGACCGTGACGATCGATGAC</td></tr>
<tr><td>Chromosome</td><td>start</td><td>end</td><td>strand</td><td>MM</td><td>target_seq</td><td>PAM</td><td>alignment</td><td>distance</td><td>position</td><td>gene name</td><td>gene id</td></tr>
<tr><td>GU305914.1</td><td>23146</td><td>23168</td><td>+</td><td>0</td><td>GTCATCGATCGTCACGGTCC</td><td>CGG</td><td>||||||||[||||||||||||]PAM</td><td>NA</td><td>Intergenic</td><td>NA</td><td>NA</td></tr>
<tr><td>T1292</td><td>TCATCGATCGTCACGGTCCCGGG</td><td>1000</td></tr>
<tr><td>Oligo fwd</td><td>CACCTCATCGATCGTCACGGTCCC</td></tr>
<tr><td>Oligo rev</td><td>AAACGGGACCGTGACGATCGATGA</td></tr>
<tr><td>Chromosome</td><td>start</td><td>end</td><td>strand</td><td>MM</td><td>target_seq</td><td>PAM</td><td>alignment</td><td>distance</td><td>position</td><td>gene name</td><td>gene id</td></tr>
<tr><td>GU305914.1</td><td>23147</td><td>23169</td><td>+</td><td>0</td><td>TCATCGATCGTCACGGTCCC</td><td>GGG</td><td>||||||||[||||||||||||]PAM</td><td>NA</td><td>Intergenic</td><td>NA</td><td>NA</td></tr>
<tr><td>T1293</td><td>CCCGGGAGAATCCCACTCACCGG</td><td>1000</td></tr>
<tr><td>Oligo fwd</td><td>CACCCCCGGGAGAATCCCACTCAC</td></tr>
<tr><td>Oligo rev</td><td>AAACGTGAGTGGGATTCTCCCGGG</td></tr>
<tr><td>Chromosome</td><td>start</td><td>end</td><td>strand</td><td>MM</td><td>target_seq</td><td>PAM</td><td>alignment</td><td>distance</td><td>position</td><td>gene name</td><td>gene id</td></tr>
<tr><td>GU305914.1</td><td>23164</td><td>23186</td><td>+</td><td>0</td><td>CCCGGGAGAATCCCACTCAC</td><td>CGG</td><td>||||||||[||||||||||||]PAM</td><td>NA</td><td>Intergenic</td><td>NA</td><td>NA</td></tr>
<tr><td>T1294</td><td>GTAACCAGCGATTCTCGAGCAGG</td><td>1000</td></tr>
<tr><td>Oligo fwd</td><td>CACCGTAACCAGCGATTCTCGAGC</td></tr>
<tr><td>Oligo rev</td><td>AAACGCTCGAGAATCGCTGGTTAC</td></tr>
<tr><td>Chromosome</td><td>start</td><td>end</td><td>strand</td><td>MM</td><td>target_seq</td><td>PAM</td><td>alignment</td><td>distance</td><td>position</td><td>gene name</td><td>gene id</td></tr>
<tr><td>GU305914.1</td><td>23193</td><td>23215</td><td>+</td><td>0</td><td>GTAACCAGCGATTCTCGAGC</td><td>AGG</td><td>||||||||[||||||||||||]PAM</td><td>NA</td><td>Intergenic</td><td>NA</td><td>NA</td></tr>
<tr><td>T1295</td><td>CCAGCGATTCTCGAGCAGGTCGG</td><td>1000</td></tr>
<tr><td>Oligo fwd</td><td>CACCCCAGCGATTCTCGAGCAGGT</td></tr>
<tr><td>Oligo rev</td><td>AAACACCTGCTCGAGAATCGCTGG</td></tr>
<tr><td>Chromosome</td><td>start</td><td>end</td><td>strand</td><td>MM</td><td>target_seq</td><td>PAM</td><td>alignment</td><td>distance</td><td>position</td><td>gene name</td><td>gene id</td></tr>
<tr><td>GU305914.1</td><td>23197</td><td>23219</td><td>+</td><td>0</td><td>CCAGCGATTCTCGAGCAGGT</td><td>CGG</td><td>||||||||[||||||||||||]PAM</td><td>NA</td><td>Intergenic</td><td>NA</td><td>NA</td></tr>
<tr><td>T1296</td><td>ATCCCGCCCGAATTCGCAGCAGG</td><td>1000</td></tr>
<tr><td>Oligo fwd</td><td>CACCATCCCGCCCGAATTCGCAGC</td></tr>
<tr><td>Oligo rev</td><td>AAACGCTGCGAATTCGGGCGGGAT</td></tr>
<tr><td>Chromosome</td><td>start</td><td>end</td><td>strand</td><td>MM</td><td>target_seq</td><td>PAM</td><td>alignment</td><td>distance</td><td>position</td><td>gene name</td><td>gene id</td></tr>
<tr><td>GU305914.1</td><td>23228</td><td>23250</td><td>+</td><td>0</td><td>ATCCCGCCCGAATTCGCAGC</td><td>AGG</td><td>||||||||[||||||||||||]PAM</td><td>NA</td><td>Intergenic</td><td>NA</td><td>NA</td></tr>
<tr><td>T1297</td><td>CGACACCCTCCCTCCGCCAACGG</td><td>1000</td></tr>
<tr><td>Oligo fwd</td><td>CACCCGACACCCTCCCTCCGCCAA</td></tr>
<tr><td>Oligo rev</td><td>AAACTTGGCGGAGGGAGGGTGTCG</td></tr>
<tr><td>Chromosome</td><td>start</td><td>end</td><td>strand</td><td>MM</td><td>target_seq</td><td>PAM</td><td>alignment</td><td>distance</td><td>position</td><td>gene name</td><td>gene id</td></tr>
<tr><td>GU305914.1</td><td>23267</td><td>23289</td><td>+</td><td>0</td><td>CGACACCCTCCCTCCGCCAA</td><td>CGG</td><td>||||||||[||||||||||||]PAM</td><td>NA</td><td>Intergenic</td><td>NA</td><td>NA</td></tr>
<tr><td>T1298</td><td>CACCCTCCCTCCGCCAACGGCGG</td><td>1000</td></tr>
<tr><td>Oligo fwd</td><td>CACCCACCCTCCCTCCGCCAACGG</td></tr>
<tr><td>Oligo rev</td><td>AAACCCGTTGGCGGAGGGAGGGTG</td></tr>
<tr><td>Chromosome</td><td>start</td><td>end</td><td>strand</td><td>MM</td><td>target_seq</td><td>PAM</td><td>alignment</td><td>distance</td><td>position</td><td>gene name</td><td>gene id</td></tr>
<tr><td>GU305914.1</td><td>23270</td><td>23292</td><td>+</td><td>0</td><td>CACCCTCCCTCCGCCAACGG</td><td>CGG</td><td>||||||||[||||||||||||]PAM</td><td>NA</td><td>Intergenic</td><td>NA</td><td>NA</td></tr>
<tr><td>T1299</td><td>TTGACAAACGAGCGAAACTCAGG</td><td>1000</td></tr>
<tr><td>Oligo fwd</td><td>CACCTTGACAAACGAGCGAAACTC</td></tr>
<tr><td>Oligo rev</td><td>AAACGAGTTTCGCTCGTTTGTCAA</td></tr>
<tr><td>Chromosome</td><td>start</td><td>end</td><td>strand</td><td>MM</td><td>target_seq</td><td>PAM</td><td>alignment</td><td>distance</td><td>position</td><td>gene name</td><td>gene id</td></tr>
<tr><td>GU305914.1</td><td>23293</td><td>23315</td><td>+</td><td>0</td><td>TTGACAAACGAGCGAAACTC</td><td>AGG</td><td>||||||||[||||||||||||]PAM</td><td>NA</td><td>Intergenic</td><td>NA</td><td>NA</td></tr>
<tr><td>T1300</td><td>ACGAGCGAAACTCAGGCAGATGG</td><td>1000</td></tr>
<tr><td>Oligo fwd</td><td>CACCACGAGCGAAACTCAGGCAGA</td></tr>
<tr><td>Oligo rev</td><td>AAACTCTGCCTGAGTTTCGCTCGT</td></tr>
<tr><td>Chromosome</td><td>start</td><td>end</td><td>strand</td><td>MM</td><td>target_seq</td><td>PAM</td><td>alignment</td><td>distance</td><td>position</td><td>gene name</td><td>gene id</td></tr>
<tr><td>GU305914.1</td><td>23300</td><td>23322</td><td>+</td><td>0</td><td>ACGAGCGAAACTCAGGCAGA</td><td>TGG</td><td>||||||||[||||||||||||]PAM</td><td>NA</td><td>Intergenic</td><td>NA</td><td>NA</td></tr>
<tr><td>T1301</td><td>AGATGGTATCGATTTAACAGAGG</td><td>1000</td></tr>
<tr><td>Oligo fwd</td><td>CACCAGATGGTATCGATTTAACAG</td></tr>
<tr><td>Oligo rev</td><td>AAACCTGTTAAATCGATACCATCT</td></tr>
<tr><td>Chromosome</td><td>start</td><td>end</td><td>strand</td><td>MM</td><td>target_seq</td><td>PAM</td><td>alignment</td><td>distance</td><td>position</td><td>gene name</td><td>gene id</td></tr>
<tr><td>GU305914.1</td><td>23317</td><td>23339</td><td>+</td><td>0</td><td>AGATGGTATCGATTTAACAG</td><td>AGG</td><td>||||||||[||||||||||||]PAM</td><td>NA</td><td>Intergenic</td><td>NA</td><td>NA</td></tr>
<tr><td>T1302</td><td>ACCGTCCATTTACATGACAGCGG</td><td>1000</td></tr>
<tr><td>Oligo fwd</td><td>CACCACCGTCCATTTACATGACAG</td></tr>
<tr><td>Oligo rev</td><td>AAACCTGTCATGTAAATGGACGGT</td></tr>
<tr><td>Chromosome</td><td>start</td><td>end</td><td>strand</td><td>MM</td><td>target_seq</td><td>PAM</td><td>alignment</td><td>distance</td><td>position</td><td>gene name</td><td>gene id</td></tr>
<tr><td>GU305914.1</td><td>23363</td><td>23385</td><td>+</td><td>0</td><td>ACCGTCCATTTACATGACAG</td><td>CGG</td><td>||||||||[||||||||||||]PAM</td><td>NA</td><td>Intergenic</td><td>NA</td><td>NA</td></tr>
<tr><td>T1303</td><td>CCGTCCATTTACATGACAGCGGG</td><td>1000</td></tr>
<tr><td>Oligo fwd</td><td>CACCCCGTCCATTTACATGACAGC</td></tr>
<tr><td>Oligo rev</td><td>AAACGCTGTCATGTAAATGGACGG</td></tr>
<tr><td>Chromosome</td><td>start</td><td>end</td><td>strand</td><td>MM</td><td>target_seq</td><td>PAM</td><td>alignment</td><td>distance</td><td>position</td><td>gene name</td><td>gene id</td></tr>
<tr><td>GU305914.1</td><td>23364</td><td>23386</td><td>+</td><td>0</td><td>CCGTCCATTTACATGACAGC</td><td>GGG</td><td>||||||||[||||||||||||]PAM</td><td>NA</td><td>Intergenic</td><td>NA</td><td>NA</td></tr>
<tr><td>T1304</td><td>CAAGCGACACCACCCCTCTCCGG</td><td>1000</td></tr>
<tr><td>Oligo fwd</td><td>CACCCAAGCGACACCACCCCTCTC</td></tr>
<tr><td>Oligo rev</td><td>AAACGAGAGGGGTGGTGTCGCTTG</td></tr>
<tr><td>Chromosome</td><td>start</td><td>end</td><td>strand</td><td>MM</td><td>target_seq</td><td>PAM</td><td>alignment</td><td>distance</td><td>position</td><td>gene name</td><td>gene id</td></tr>
</table>

GU305914.1 23388 23410 + 0 CAAGCGACACCACCCCTCTC CGG ||||||||[||||||||||||]PAM NA Intergenic NA NA

T1305 AAACAGACAGATAGACAGACAGG 1000
Oligo fwd CACCAAACAGACAGATAGACAGAC
Oligo rev AAACGTCTGTCTATCTGTCTGTTT
Chromosome start end strand MM target_seq PAM alignment distance position gene name gene id
GU305914.1 23417 23439 + 0 AAACAGACAGATAGACAGAC AGG ||||||||[||||||||||||]PAM NA Intergenic NA NA

T1306 CAGGCTCAGTCTCATACCGTCGG 1000
Oligo fwd CACCCAGGCTCAGTCTCATACCGT
Oligo rev AAACACGGTATGAGACTGAGCCTG
Chromosome start end strand MM target_seq PAM alignment distance position gene name gene id
GU305914.1 23436 23458 + 0 CAGGCTCAGTCTCATACCGT CGG ||||||||[||||||||||||]PAM NA Intergenic NA NA

T1307 GCCATCCCCCCGCTTGACCGAGG 1000
Oligo fwd CACCGCCATCCCCCCGCTTGACCG
Oligo rev AAACCGGTCAAGCGGGGGGATGGC
Chromosome start end strand MM target_seq PAM alignment distance position gene name gene id
GU305914.1 23458 23480 + 0 GCCATCCCCCCGCTTGACCG AGG ||||||||[||||||||||||]PAM NA Intergenic NA NA

T1308 CGCAGTAAACTACCACCGATAGG 1000
Oligo fwd CACCCGCAGTAAACTACCACCGAT
Oligo rev AAACATCGGTGGTAGTTTACTGCG
Chromosome start end strand MM target_seq PAM alignment distance position gene name gene id
GU305914.1 23496 23518 + 0 CGCAGTAAACTACCACCGAT AGG ||||||||[||||||||||||]PAM NA Intergenic NA NA

T1309 GCAGTAAACTACCACCGATAGGG 1000
Oligo fwd CACCGCAGTAAACTACCACCGATA
Oligo rev AAACTATCGGTGGTAGTTTACTGC
Chromosome start end strand MM target_seq PAM alignment distance position gene name gene id
GU305914.1 23497 23519 + 0 GCAGTAAACTACCACCGATA GGG ||||||||[||||||||||||]PAM NA Intergenic NA NA

T1310 CAGTAAACTACCACCGATAGGGG 1000
Oligo fwd CACCCAGTAAACTACCACCGATAG
Oligo rev AAACCTATCGGTGGTAGTTTACTG
Chromosome start end strand MM target_seq PAM alignment distance position gene name gene id
GU305914.1 23498 23520 + 0 CAGTAAACTACCACCGATAG GGG ||||||||[||||||||||||]PAM NA Intergenic NA NA

T1311 GATAGGGGACACGTTCAAGCCGG 1000
Oligo fwd CACCGATAGGGGACACGTTCAAGC
Oligo rev AAACGCTTGAACGTGTCCCCTATC
Chromosome start end strand MM target_seq PAM alignment distance position gene name gene id
GU305914.1 23513 23535 + 0 GATAGGGGACACGTTCAAGC CGG ||||||||[||||||||||||]PAM NA Intergenic NA NA

T1312 ATAGGGGACACGTTCAAGCCGGG 1000
Oligo fwd CACCATAGGGGACACGTTCAAGCC
Oligo rev AAACGGCTTGAACGTGTCCCCTAT
Chromosome start end strand MM target_seq PAM alignment distance position gene name gene id
GU305914.1 23514 23536 + 0 ATAGGGGACACGTTCAAGCC GGG ||||||||[||||||||||||]PAM NA Intergenic NA NA

T1313 TAGGGGACACGTTCAAGCCGGGG 1000
Oligo fwd CACCTAGGGGACACGTTCAAGCCG
Oligo rev AAACCGGCTTGAACGTGTCCCCTA
Chromosome start end strand MM target_seq PAM alignment distance position gene name gene id
GU305914.1 23515 23537 + 0 TAGGGGACACGTTCAAGCCG GGG ||||||||[||||||||||||]PAM NA Intergenic NA NA

T1314 GCGCCTCGATCGACGAGCGTCGG 1000
Oligo fwd CACCGCGCCTCGATCGACGAGCGT
Oligo rev AAACACGCTCGTCGATCGAGGCGC
Chromosome start end strand MM target_seq PAM alignment distance position gene name gene id
GU305914.1 23540 23562 + 0 GCGCCTCGATCGACGAGCGT CGG ||||||||[||||||||||||]PAM NA Intergenic NA NA
GU305914.1 99540 99562 + 4 CCTCCTCGCCCGACGAGCGT TGG -|-|||||[--||||||||||]PAM NA Intergenic NA NA

T1315 CGAGCGTCGGACAAAGAAACCGG 1000
Oligo fwd CACCCGAGCGTCGGACAAAGAAAC
Oligo rev AAACGTTTCTTTGTCCGACGCTCG
Chromosome start end strand MM target_seq PAM alignment distance position gene name gene id
GU305914.1 23553 23575 + 0 CGAGCGTCGGACAAAGAAAC CGG ||||||||[||||||||||||]PAM NA Intergenic NA NA

T1316 GAGCGTCGGACAAAGAAACCGGG 1000
Oligo fwd CACCGAGCGTCGGACAAAGAAACC
Oligo rev AAACGGTTTCTTTGTCCGACGCTC
Chromosome start end strand MM target_seq PAM alignment distance position gene name gene id
GU305914.1 23554 23576 + 0 GAGCGTCGGACAAAGAAACC GGG ||||||||[||||||||||||]PAM NA Intergenic NA NA

T1317 GAAACCGGGAGAAGAAGATGCGG 1000
Oligo fwd CACCGAAACCGGGAGAAGAAGATG
Oligo rev AAACCATCTTCTTCTCCCGGTTTC
Chromosome start end strand MM target_seq PAM alignment distance position gene name gene id
GU305914.1 23568 23590 + 0 GAAACCGGGAGAAGAAGATG CGG ||||||||[||||||||||||]PAM NA Intergenic NA NA

T1318 AAACCGGGAGAAGAAGATGCGGG 1000
Oligo fwd CACCAAACCGGGAGAAGAAGATGC
Oligo rev AAACGCATCTTCTTCTCCCGGTTT
Chromosome start end strand MM target_seq PAM alignment distance position gene name gene id
GU305914.1 23569 23591 + 0 AAACCGGGAGAAGAAGATGC GGG ||||||||[||||||||||||]PAM NA Intergenic NA NA

T1319 CCGGGAGAAGAAGATGCGGGCGG 1000
Oligo fwd CACCCCGGGAGAAGAAGATGCGGG

| | | | | | | | | | | | |
|---|---|---|---|---|---|---|---|---|---|---|---|
| Oligo rev | AAACCCCGCATCTTCTTCTCCCGG | | | | | | | | | | |
| Chromosome | start | end | strand | MM | target_seq | PAM | alignment | distance | position | gene name | gene id |
| GU305914.1 | 23572 | 23594 | + | 0 | CCGGGAGAAGAAGATGCGGG | CGG | ||||||||[||||||||||||]PAM | NA | Intergenic | NA | NA |
| T1320 | CGGGAGAAGAAGATGCGGGCGGG | 1000 | | | | | | | | | |
| Oligo fwd | CACCCGGGAGAAGAAGATGCGGGC | | | | | | | | | | |
| Oligo rev | AAACGCCCGCATCTTCTTCTCCCG | | | | | | | | | | |
| Chromosome | start | end | strand | MM | target_seq | PAM | alignment | distance | position | gene name | gene id |
| GU305914.1 | 23573 | 23595 | + | 0 | CGGGAGAAGAAGATGCGGGC | GGG | ||||||||[||||||||||||]PAM | NA | Intergenic | NA | NA |
| T1321 | GAGAAGAAGATGCGGGCGGGCGG | 1000 | | | | | | | | | |
| Oligo fwd | CACCGAGAAGAAGATGCGGGCGGG | | | | | | | | | | |
| Oligo rev | AAACCCCGCCCGCATCTTCTTCTC | | | | | | | | | | |
| Chromosome | start | end | strand | MM | target_seq | PAM | alignment | distance | position | gene name | gene id |
| GU305914.1 | 23576 | 23598 | + | 0 | GAGAAGAAGATGCGGGCGGG | CGG | ||||||||[||||||||||||]PAM | NA | Intergenic | NA | NA |
| T1322 | AGAAGAAGATGCGGGCGGGCGGG | 1000 | | | | | | | | | |
| Oligo fwd | CACCAGAAGAAGATGCGGGCGGGC | | | | | | | | | | |
| Oligo rev | AAACGCCCGCCCGCATCTTCTTCT | | | | | | | | | | |
| Chromosome | start | end | strand | MM | target_seq | PAM | alignment | distance | position | gene name | gene id |
| GU305914.1 | 23577 | 23599 | + | 0 | AGAAGAAGATGCGGGCGGGC | GGG | ||||||||[||||||||||||]PAM | NA | Intergenic | NA | NA |
| T1323 | GAAGATGCGGGCGGGCGGGCAGG | 1000 | | | | | | | | | |
| Oligo fwd | CACCGAAGATGCGGGCGGGCGGGC | | | | | | | | | | |
| Oligo rev | AAACGCCCGCCCGCCCGCATCTTC | | | | | | | | | | |
| Chromosome | start | end | strand | MM | target_seq | PAM | alignment | distance | position | gene name | gene id |
| GU305914.1 | 23581 | 23603 | + | 0 | GAAGATGCGGGCGGGCGGGC | AGG | ||||||||[||||||||||||]PAM | NA | Intergenic | NA | NA |
| T1324 | ATGCGGGCGGGCGGGCAGGCAGG | 1000 | | | | | | | | | |
| Oligo fwd | CACCATGCGGGCGGGCGGGCAGGC | | | | | | | | | | |
| Oligo rev | AAACGCCTGCCCGCCCGCCCGCAT | | | | | | | | | | |
| Chromosome | start | end | strand | MM | target_seq | PAM | alignment | distance | position | gene name | gene id |
| GU305914.1 | 23585 | 23607 | + | 0 | ATGCGGGCGGGCGGGCAGGC | AGG | ||||||||[||||||||||||]PAM | NA | Intergenic | NA | NA |
| GU305914.1 | 23589 | 23611 | + | 3 | GGGCGGGCGGGCAGGCAGGC | GGG | --||||||[||||-|||||||]PAM | NA | Intergenic | NA | NA |
| GU305914.1 | 23629 | 23651 | + | 4 | AGGCGGTGAGGCGGGCAGGC | GGG | |-||||--[-|||||||||||]PAM | NA | Intergenic | NA | NA |
| T1325 | CGGGCGGGCGGGCAGGCAGGCGG | 1000 | | | | | | | | | |
| Oligo fwd | CACCCGGGCGGGCGGGCAGGCAGG | | | | | | | | | | |
| Oligo rev | AAACCCTGCCTGCCCGCCCGCCCG | | | | | | | | | | |
| Chromosome | start | end | strand | MM | target_seq | PAM | alignment | distance | position | gene name | gene id |
| GU305914.1 | 23588 | 23610 | + | 0 | CGGGCGGGCGGGCAGGCAGG | CGG | ||||||||[||||||||||||]PAM | NA | Intergenic | NA | NA |
| GU305914.1 | 23632 | 23654 | + | 4 | CGGTGAGGCGGGCAGGCGGG | TGG | |||---||[|||||||||-||]PAM | NA | Intergenic | NA | NA |
| T1326 | GGGCGGGCGGGCAGGCAGGCGGG | 1000 | | | | | | | | | |
| Oligo fwd | CACCGGGCGGGCGGGCAGGCAGGC | | | | | | | | | | |
| Oligo rev | AAACGCCTGCCTGCCCGCCCGCCC | | | | | | | | | | |
| Chromosome | start | end | strand | MM | target_seq | PAM | alignment | distance | position | gene name | gene id |
| GU305914.1 | 23589 | 23611 | + | 0 | GGGCGGGCGGGCAGGCAGGC | GGG | ||||||||[||||||||||||]PAM | NA | Intergenic | NA | NA |
| GU305914.1 | 23585 | 23607 | + | 3 | ATGCGGGCGGGCGGGCAGGC | AGG | --||||||[||||-|||||||]PAM | NA | Intergenic | NA | NA |
| GU305914.1 | 23593 | 23615 | + | 2 | GGGCGGGCAGGCAGGCGGGC | TGG | ||||||||[-|||||||-|||]PAM | NA | Intergenic | NA | NA |
| T1327 | GGGCGGGCAGGCAGGCGGGCTGG | 1000 | | | | | | | | | |
| Oligo fwd | CACCGGGCGGGCAGGCAGGCGGGC | | | | | | | | | | |
| Oligo rev | AAACGCCCGCCTGCCTGCCCGCCC | | | | | | | | | | |
| Chromosome | start | end | strand | MM | target_seq | PAM | alignment | distance | position | gene name | gene id |
| GU305914.1 | 23593 | 23615 | + | 0 | GGGCGGGCAGGCAGGCGGGC | TGG | ||||||||[||||||||||||]PAM | NA | Intergenic | NA | NA |
| GU305914.1 | 23589 | 23611 | + | 2 | GGGCGGGCGGGCAGGCAGGC | GGG | ||||||||[-|||||||-|||]PAM | NA | Intergenic | NA | NA |
| T1328 | AGGCAGGCGGGCTGGTGAACAGG | 1000 | | | | | | | | | |
| Oligo fwd | CACCAGGCAGGCGGGCTGGTGAAC | | | | | | | | | | |
| Oligo rev | AAACGTTCACCAGCCCGCCTGCCT | | | | | | | | | | |
| Chromosome | start | end | strand | MM | target_seq | PAM | alignment | distance | position | gene name | gene id |
| GU305914.1 | 23601 | 23623 | + | 0 | AGGCAGGCGGGCTGGTGAAC | AGG | ||||||||[||||||||||||]PAM | NA | Intergenic | NA | NA |
| T1329 | AGGCGGGCTGGTGAACAGGAAGG | 1000 | | | | | | | | | |
| Oligo fwd | CACCAGGCGGGCTGGTGAACAGGA | | | | | | | | | | |
| Oligo rev | AAACTCCTGTTCACCAGCCCGCCT | | | | | | | | | | |
| Chromosome | start | end | strand | MM | target_seq | PAM | alignment | distance | position | gene name | gene id |
| GU305914.1 | 23605 | 23627 | + | 0 | AGGCGGGCTGGTGAACAGGA | AGG | ||||||||[||||||||||||]PAM | NA | Intergenic | NA | NA |
| T1330 | GGGCTGGTGAACAGGAAGGCAGG | 1000 | | | | | | | | | |
| Oligo fwd | CACCGGGCTGGTGAACAGGAAGGC | | | | | | | | | | |
| Oligo rev | AAACGCCTTCCTGTTCACCAGCCC | | | | | | | | | | |
| Chromosome | start | end | strand | MM | target_seq | PAM | alignment | distance | position | gene name | gene id |
| GU305914.1 | 23609 | 23631 | + | 0 | GGGCTGGTGAACAGGAAGGC | AGG | ||||||||[||||||||||||]PAM | NA | Intergenic | NA | NA |
| T1331 | CTGGTGAACAGGAAGGCAGGCGG | 1000 | | | | | | | | | |
| Oligo fwd | CACCCTGGTGAACAGGAAGGCAGG | | | | | | | | | | |
| Oligo rev | AAACCCTGCCTTCCTGTTCACCAG | | | | | | | | | | |
| Chromosome | start | end | strand | MM | target_seq | PAM | alignment | distance | position | gene name | gene id |
| GU305914.1 | 23612 | 23634 | + | 0 | CTGGTGAACAGGAAGGCAGG | CGG | ||||||||[||||||||||||]PAM | NA | Intergenic | NA | NA |
| T1332 | GAACAGGAAGGCAGGCGGTGAGG | 1000 | | | | | | | | | |
| Oligo fwd | CACCGAACAGGAAGGCAGGCGGTG | | | | | | | | | | |
| Oligo rev | AAACCACCGCCTGCCTTCCTGTTC | | | | | | | | | | |
| Chromosome | start | end | strand | MM | target_seq | PAM | alignment | distance | position | gene name | gene id |
| GU305914.1 | 23617 | 23639 | + | 0 | GAACAGGAAGGCAGGCGGTG | AGG | ||||||||[||||||||||||]PAM | NA | Intergenic | NA | NA |
| T1333 | CAGGAAGGCAGGCGGTGAGGCGG | 1000 | | | | | | | | | |

| | | | | | | | | | | | |
|---|---|---|---|---|---|---|---|---|---|---|---|
| Oligo fwd | CACCCAGGAAGGCAGGCGGTGAGG | | | | | | | | | | |
| Oligo rev | AAACCCTCACCGCCTGCCTTCCTG | | | | | | | | | | |
| Chromosome | start | end | strand | MM | target_seq | PAM | alignment | distance | position | gene name | gene id |
| GU305914.1 | 23620 | 23642 | + | 0 | CAGGAAGGCAGGCGGTGAGG | CGG | \|\|\|\|\|\|\|\|[\|\|\|\|\|\|\|\|\|\|\|\|]PAM | NA | Intergenic | NA | NA |
| T1334 | AGGAAGGCAGGCGGTGAGGCGGG | 1000 | | | | | | | | | |
| Oligo fwd | CACCAGGAAGGCAGGCGGTGAGGC | | | | | | | | | | |
| Oligo rev | AAACGCCTCACCGCCTGCCTTCCT | | | | | | | | | | |
| Chromosome | start | end | strand | MM | target_seq | PAM | alignment | distance | position | gene name | gene id |
| GU305914.1 | 23621 | 23643 | + | 0 | AGGAAGGCAGGCGGTGAGGC | GGG | \|\|\|\|\|\|\|\|[\|\|\|\|\|\|\|\|\|\|\|\|]PAM | NA | Intergenic | NA | NA |
| T1335 | AGGCAGGCGGTGAGGCGGGCAGG | 1000 | | | | | | | | | |
| Oligo fwd | CACCAGGCAGGCGGTGAGGCGGGC | | | | | | | | | | |
| Oligo rev | AAACGCCCGCCTCACCGCCTGCCT | | | | | | | | | | |
| Chromosome | start | end | strand | MM | target_seq | PAM | alignment | distance | position | gene name | gene id |
| GU305914.1 | 23625 | 23647 | + | 0 | AGGCAGGCGGTGAGGCGGGC | AGG | \|\|\|\|\|\|\|\|[\|\|\|\|\|\|\|\|\|\|\|\|]PAM | NA | Intergenic | NA | NA |
| T1336 | CAGGCGGTGAGGCGGGCAGGCGG | 1000 | | | | | | | | | |
| Oligo fwd | CACCCAGGCGGTGAGGCGGGCAGG | | | | | | | | | | |
| Oligo rev | AAACCCTGCCCGCCTCACCGCCTG | | | | | | | | | | |
| Chromosome | start | end | strand | MM | target_seq | PAM | alignment | distance | position | gene name | gene id |
| GU305914.1 | 23628 | 23650 | + | 0 | CAGGCGGTGAGGCGGGCAGG | CGG | \|\|\|\|\|\|\|\|[\|\|\|\|\|\|\|\|\|\|\|\|]PAM | NA | Intergenic | NA | NA |
| T1337 | AGGCGGTGAGGCGGGCAGGCGGG | 1000 | | | | | | | | | |
| Oligo fwd | CACCAGGCGGTGAGGCGGGCAGGC | | | | | | | | | | |
| Oligo rev | AAACGCCTGCCCGCCTCACCGCCT | | | | | | | | | | |
| Chromosome | start | end | strand | MM | target_seq | PAM | alignment | distance | position | gene name | gene id |
| GU305914.1 | 23629 | 23651 | + | 0 | AGGCGGTGAGGCGGGCAGGC | GGG | \|\|\|\|\|\|\|\|[\|\|\|\|\|\|\|\|\|\|\|\|]PAM | NA | Intergenic | NA | NA |
| GU305914.1 | 23585 | 23607 | + | 4 | ATGCGGGCGGGCGGGCAGGC | AGG | \|-\|\|\|\|--[-\|\|\|\|\|\|\|\|\|\|\|]PAM | NA | Intergenic | NA | NA |
| T1338 | CGGTGAGGCGGGCAGGCGGGTGG | 1000 | | | | | | | | | |
| Oligo fwd | CACCCGGTGAGGCGGGCAGGCGGG | | | | | | | | | | |
| Oligo rev | AAACCCCGCCTGCCCGCCTCACCG | | | | | | | | | | |
| Chromosome | start | end | strand | MM | target_seq | PAM | alignment | distance | position | gene name | gene id |
| GU305914.1 | 23632 | 23654 | + | 0 | CGGTGAGGCGGGCAGGCGGG | TGG | \|\|\|\|\|\|\|\|[\|\|\|\|\|\|\|\|\|\|\|\|]PAM | NA | Intergenic | NA | NA |
| GU305914.1 | 23588 | 23610 | + | 4 | CGGGCGGGCGGGCAGGCAGG | CGG | \|\|\|---\|\|[\|\|\|\|\|\|\|\|\|-\|\|]PAM | NA | Intergenic | NA | NA |
| T1339 | GAGGCGGGCAGGCGGGTGGACGG | 1000 | | | | | | | | | |
| Oligo fwd | CACCGAGGCGGGCAGGCGGGTGGA | | | | | | | | | | |
| Oligo rev | AAACTCCACCCGCCTGCCCGCCTC | | | | | | | | | | |
| Chromosome | start | end | strand | MM | target_seq | PAM | alignment | distance | position | gene name | gene id |
| GU305914.1 | 23636 | 23658 | + | 0 | GAGGCGGGCAGGCGGGTGGA | CGG | \|\|\|\|\|\|\|\|[\|\|\|\|\|\|\|\|\|\|\|\|]PAM | NA | Intergenic | NA | NA |
| T1340 | AGGCGGGCAGGCGGGTGGACGGG | 1000 | | | | | | | | | |
| Oligo fwd | CACCAGGCGGGCAGGCGGGTGGAC | | | | | | | | | | |
| Oligo rev | AAACGTCCACCCGCCTGCCCGCCT | | | | | | | | | | |
| Chromosome | start | end | strand | MM | target_seq | PAM | alignment | distance | position | gene name | gene id |
| GU305914.1 | 23637 | 23659 | + | 0 | AGGCGGGCAGGCGGGTGGAC | GGG | \|\|\|\|\|\|\|\|[\|\|\|\|\|\|\|\|\|\|\|\|]PAM | NA | Intergenic | NA | NA |
| T1341 | CGGGCAGGCGGGTGGACGGGCGG | 1000 | | | | | | | | | |
| Oligo fwd | CACCCGGGCAGGCGGGTGGACGGG | | | | | | | | | | |
| Oligo rev | AAACCCCGTCCACCCGCCTGCCCG | | | | | | | | | | |
| Chromosome | start | end | strand | MM | target_seq | PAM | alignment | distance | position | gene name | gene id |
| GU305914.1 | 23640 | 23662 | + | 0 | CGGGCAGGCGGGTGGACGGG | CGG | \|\|\|\|\|\|\|\|[\|\|\|\|\|\|\|\|\|\|\|\|]PAM | NA | Intergenic | NA | NA |
| T1342 | CAGGCGGGTGGACGGGCGGCTGG | 1000 | | | | | | | | | |
| Oligo fwd | CACCCAGGCGGGTGGACGGGCGGC | | | | | | | | | | |
| Oligo rev | AAACGCCGCCCGTCCACCCGCCTG | | | | | | | | | | |
| Chromosome | start | end | strand | MM | target_seq | PAM | alignment | distance | position | gene name | gene id |
| GU305914.1 | 23644 | 23666 | + | 0 | CAGGCGGGTGGACGGGCGGC | TGG | \|\|\|\|\|\|\|\|[\|\|\|\|\|\|\|\|\|\|\|\|]PAM | NA | Intergenic | NA | NA |
| T1343 | GGTGGACGGGCGGCTGGCCTCGG | 1000 | | | | | | | | | |
| Oligo fwd | CACCGGTGGACGGGCGGCTGGCCT | | | | | | | | | | |
| Oligo rev | AAACAGGCCAGCCGCCCGTCCACC | | | | | | | | | | |
| Chromosome | start | end | strand | MM | target_seq | PAM | alignment | distance | position | gene name | gene id |
| GU305914.1 | 23650 | 23672 | + | 0 | GGTGGACGGGCGGCTGGCCT | CGG | \|\|\|\|\|\|\|\|[\|\|\|\|\|\|\|\|\|\|\|\|]PAM | NA | Intergenic | NA | NA |
| T1344 | GTGGACGGGCGGCTGGCCTCGGG | 1000 | | | | | | | | | |
| Oligo fwd | CACCGTGGACGGGCGGCTGGCCTC | | | | | | | | | | |
| Oligo rev | AAACGAGGCCAGCCGCCCGTCCAC | | | | | | | | | | |
| Chromosome | start | end | strand | MM | target_seq | PAM | alignment | distance | position | gene name | gene id |
| GU305914.1 | 23651 | 23673 | + | 0 | GTGGACGGGCGGCTGGCCTC | GGG | \|\|\|\|\|\|\|\|[\|\|\|\|\|\|\|\|\|\|\|\|]PAM | NA | Intergenic | NA | NA |
| T1345 | GGCTGGCCTCGGGCCGACGCAGG | 1000 | | | | | | | | | |
| Oligo fwd | CACCGGCTGGCCTCGGGCCGACGC | | | | | | | | | | |
| Oligo rev | AAACGCGTCGGCCCGAGGCCAGCC | | | | | | | | | | |
| Chromosome | start | end | strand | MM | target_seq | PAM | alignment | distance | position | gene name | gene id |
| GU305914.1 | 23661 | 23683 | + | 0 | GGCTGGCCTCGGGCCGACGC | AGG | \|\|\|\|\|\|\|\|[\|\|\|\|\|\|\|\|\|\|\|\|]PAM | NA | Intergenic | NA | NA |
| T1346 | GAGCCCGTTCACAATACACACGG | 1000 | | | | | | | | | |
| Oligo fwd | CACCGAGCCCGTTCACAATACACA | | | | | | | | | | |
| Oligo rev | AAACTGTGTATTGTGAACGGGCTC | | | | | | | | | | |
| Chromosome | start | end | strand | MM | target_seq | PAM | alignment | distance | position | gene name | gene id |
| GU305914.1 | 23683 | 23705 | + | 0 | GAGCCCGTTCACAATACACA | CGG | \|\|\|\|\|\|\|\|[\|\|\|\|\|\|\|\|\|\|\|\|]PAM | NA | Intergenic | NA | NA |
| T1347 | ACGGTTCCCCGTCCGTACCGAGG | 1000 | | | | | | | | | |
| Oligo fwd | CACCACGGTTCCCCGTCCGTACCG | | | | | | | | | | |
| Oligo rev | AAACCGGTACGGACGGGGAACCGT | | | | | | | | | | |
| Chromosome | start | end | strand | MM | target_seq | PAM | alignment | distance | position | gene name | gene id |

<table>
<tr><td>GU305914.1</td><td>23702</td><td>23724</td><td>+</td><td>0</td><td>ACGGTTCCCCGTCCGTACCG</td><td>AGG</td><td>||||||||[||||||||||||]PAM</td><td>NA</td><td>Intergenic</td><td>NA</td><td>NA</td></tr>
<tr><td>T1348</td><td>ACCCGCCCGACAAGCAGTCGCGG</td><td>1000</td><td></td><td></td><td></td><td></td><td></td><td></td><td></td><td></td><td></td></tr>
<tr><td>Oligo fwd</td><td>CACCACCCGCCCGACAAGCAGTCG</td><td></td><td></td><td></td><td></td><td></td><td></td><td></td><td></td><td></td><td></td></tr>
<tr><td>Oligo rev</td><td>AAACCGACTGCTTGTCGGGCGGGT</td><td></td><td></td><td></td><td></td><td></td><td></td><td></td><td></td><td></td><td></td></tr>
<tr><td>Chromosome</td><td>start</td><td>end</td><td>strand</td><td>MM</td><td>target_seq</td><td>PAM</td><td>alignment</td><td>distance</td><td>position</td><td>gene name</td><td>gene id</td></tr>
<tr><td>GU305914.1</td><td>23731</td><td>23753</td><td>+</td><td>0</td><td>ACCCGCCCGACAAGCAGTCG</td><td>CGG</td><td>||||||||[||||||||||||]PAM</td><td>NA</td><td>Intergenic</td><td>NA</td><td>NA</td></tr>
<tr><td>T1349</td><td>ACAAGCAGTCGCGGTCCGTCCGG</td><td>1000</td><td></td><td></td><td></td><td></td><td></td><td></td><td></td><td></td><td></td></tr>
<tr><td>Oligo fwd</td><td>CACCACAAGCAGTCGCGGTCCGTC</td><td></td><td></td><td></td><td></td><td></td><td></td><td></td><td></td><td></td><td></td></tr>
<tr><td>Oligo rev</td><td>AAACGACGGACCGCGACTGCTTGT</td><td></td><td></td><td></td><td></td><td></td><td></td><td></td><td></td><td></td><td></td></tr>
<tr><td>Chromosome</td><td>start</td><td>end</td><td>strand</td><td>MM</td><td>target_seq</td><td>PAM</td><td>alignment</td><td>distance</td><td>position</td><td>gene name</td><td>gene id</td></tr>
<tr><td>GU305914.1</td><td>23740</td><td>23762</td><td>+</td><td>0</td><td>ACAAGCAGTCGCGGTCCGTC</td><td>CGG</td><td>||||||||[||||||||||||]PAM</td><td>NA</td><td>Intergenic</td><td>NA</td><td>NA</td></tr>
<tr><td>T1350</td><td>ATCCGATCACCTCCTTTTGATGG</td><td>1000</td><td></td><td></td><td></td><td></td><td></td><td></td><td></td><td></td><td></td></tr>
<tr><td>Oligo fwd</td><td>CACCATCCGATCACCTCCTTTTGA</td><td></td><td></td><td></td><td></td><td></td><td></td><td></td><td></td><td></td><td></td></tr>
<tr><td>Oligo rev</td><td>AAACTCAAAAGGAGGTGATCGGAT</td><td></td><td></td><td></td><td></td><td></td><td></td><td></td><td></td><td></td><td></td></tr>
<tr><td>Chromosome</td><td>start</td><td>end</td><td>strand</td><td>MM</td><td>target_seq</td><td>PAM</td><td>alignment</td><td>distance</td><td>position</td><td>gene name</td><td>gene id</td></tr>
<tr><td>GU305914.1</td><td>23771</td><td>23793</td><td>+</td><td>0</td><td>ATCCGATCACCTCCTTTTGA</td><td>TGG</td><td>||||||||[||||||||||||]PAM</td><td>NA</td><td>Intergenic</td><td>NA</td><td>NA</td></tr>
<tr><td>T1351</td><td>CTTTCTCATGCTGTTCCTCCCGG</td><td>1000</td><td></td><td></td><td></td><td></td><td></td><td></td><td></td><td></td><td></td></tr>
<tr><td>Oligo fwd</td><td>CACCCTTTCTCATGCTGTTCCTCC</td><td></td><td></td><td></td><td></td><td></td><td></td><td></td><td></td><td></td><td></td></tr>
<tr><td>Oligo rev</td><td>AAACGGAGGAACAGCATGAGAAAG</td><td></td><td></td><td></td><td></td><td></td><td></td><td></td><td></td><td></td><td></td></tr>
<tr><td>Chromosome</td><td>start</td><td>end</td><td>strand</td><td>MM</td><td>target_seq</td><td>PAM</td><td>alignment</td><td>distance</td><td>position</td><td>gene name</td><td>gene id</td></tr>
<tr><td>GU305914.1</td><td>23816</td><td>23838</td><td>+</td><td>0</td><td>CTTTCTCATGCTGTTCCTCC</td><td>CGG</td><td>||||||||[||||||||||||]PAM</td><td>NA</td><td>Intergenic</td><td>NA</td><td>NA</td></tr>
<tr><td>T1352</td><td>CCTCCCGGTTTTCTTCTCCCAGG</td><td>1000</td><td></td><td></td><td></td><td></td><td></td><td></td><td></td><td></td><td></td></tr>
<tr><td>Oligo fwd</td><td>CACCCCTCCCGGTTTTCTTCTCCC</td><td></td><td></td><td></td><td></td><td></td><td></td><td></td><td></td><td></td><td></td></tr>
<tr><td>Oligo rev</td><td>AAACGGGAGAAGAAAACCGGGAGG</td><td></td><td></td><td></td><td></td><td></td><td></td><td></td><td></td><td></td><td></td></tr>
<tr><td>Chromosome</td><td>start</td><td>end</td><td>strand</td><td>MM</td><td>target_seq</td><td>PAM</td><td>alignment</td><td>distance</td><td>position</td><td>gene name</td><td>gene id</td></tr>
<tr><td>GU305914.1</td><td>23831</td><td>23853</td><td>+</td><td>0</td><td>CCTCCCGGTTTTCTTCTCCC</td><td>AGG</td><td>||||||||[||||||||||||]PAM</td><td>NA</td><td>Intergenic</td><td>NA</td><td>NA</td></tr>
<tr><td>T1353</td><td>CCGGTTTTCTTCTCCCAGGTTGG</td><td>1000</td><td></td><td></td><td></td><td></td><td></td><td></td><td></td><td></td><td></td></tr>
<tr><td>Oligo fwd</td><td>CACCCCGGTTTTCTTCTCCCAGGT</td><td></td><td></td><td></td><td></td><td></td><td></td><td></td><td></td><td></td><td></td></tr>
<tr><td>Oligo rev</td><td>AAACACCTGGGAGAAGAAAACCGG</td><td></td><td></td><td></td><td></td><td></td><td></td><td></td><td></td><td></td><td></td></tr>
<tr><td>Chromosome</td><td>start</td><td>end</td><td>strand</td><td>MM</td><td>target_seq</td><td>PAM</td><td>alignment</td><td>distance</td><td>position</td><td>gene name</td><td>gene id</td></tr>
<tr><td>GU305914.1</td><td>23835</td><td>23857</td><td>+</td><td>0</td><td>CCGGTTTTCTTCTCCCAGGT</td><td>TGG</td><td>||||||||[||||||||||||]PAM</td><td>NA</td><td>Intergenic</td><td>NA</td><td>NA</td></tr>
<tr><td>T1354</td><td>CAGGTTGGTTCGCTACCATGCGG</td><td>1000</td><td></td><td></td><td></td><td></td><td></td><td></td><td></td><td></td><td></td></tr>
<tr><td>Oligo fwd</td><td>CACCCAGGTTGGTTCGCTACCATG</td><td></td><td></td><td></td><td></td><td></td><td></td><td></td><td></td><td></td><td></td></tr>
<tr><td>Oligo rev</td><td>AAACCATGGTAGCGAACCAACCTG</td><td></td><td></td><td></td><td></td><td></td><td></td><td></td><td></td><td></td><td></td></tr>
<tr><td>Chromosome</td><td>start</td><td>end</td><td>strand</td><td>MM</td><td>target_seq</td><td>PAM</td><td>alignment</td><td>distance</td><td>position</td><td>gene name</td><td>gene id</td></tr>
<tr><td>GU305914.1</td><td>23850</td><td>23872</td><td>+</td><td>0</td><td>CAGGTTGGTTCGCTACCATG</td><td>CGG</td><td>||||||||[||||||||||||]PAM</td><td>NA</td><td>Intergenic</td><td>NA</td><td>NA</td></tr>
<tr><td>T1355</td><td>GCTACCATGCGGCACCCCATTGG</td><td>1000</td><td></td><td></td><td></td><td></td><td></td><td></td><td></td><td></td><td></td></tr>
<tr><td>Oligo fwd</td><td>CACCGCTACCATGCGGCACCCCAT</td><td></td><td></td><td></td><td></td><td></td><td></td><td></td><td></td><td></td><td></td></tr>
<tr><td>Oligo rev</td><td>AAACATGGGGTGCCGCATGGTAGC</td><td></td><td></td><td></td><td></td><td></td><td></td><td></td><td></td><td></td><td></td></tr>
<tr><td>Chromosome</td><td>start</td><td>end</td><td>strand</td><td>MM</td><td>target_seq</td><td>PAM</td><td>alignment</td><td>distance</td><td>position</td><td>gene name</td><td>gene id</td></tr>
<tr><td>GU305914.1</td><td>23861</td><td>23883</td><td>+</td><td>0</td><td>GCTACCATGCGGCACCCCAT</td><td>TGG</td><td>||||||||[||||||||||||]PAM</td><td>NA</td><td>Intergenic</td><td>NA</td><td>NA</td></tr>
<tr><td>T1356</td><td>CGGCACCCCATTGGTCGCCCTGG</td><td>1000</td><td></td><td></td><td></td><td></td><td></td><td></td><td></td><td></td><td></td></tr>
<tr><td>Oligo fwd</td><td>CACCCGGCACCCCATTGGTCGCCC</td><td></td><td></td><td></td><td></td><td></td><td></td><td></td><td></td><td></td><td></td></tr>
<tr><td>Oligo rev</td><td>AAACGGGCGACCAATGGGGTGCCG</td><td></td><td></td><td></td><td></td><td></td><td></td><td></td><td></td><td></td><td></td></tr>
<tr><td>Chromosome</td><td>start</td><td>end</td><td>strand</td><td>MM</td><td>target_seq</td><td>PAM</td><td>alignment</td><td>distance</td><td>position</td><td>gene name</td><td>gene id</td></tr>
<tr><td>GU305914.1</td><td>23870</td><td>23892</td><td>+</td><td>0</td><td>CGGCACCCCATTGGTCGCCC</td><td>TGG</td><td>||||||||[||||||||||||]PAM</td><td>NA</td><td>Intergenic</td><td>NA</td><td>NA</td></tr>
<tr><td>T1357</td><td>CCTGGAGTGTAAGAAGTTGTTGG</td><td>1000</td><td></td><td></td><td></td><td></td><td></td><td></td><td></td><td></td><td></td></tr>
<tr><td>Oligo fwd</td><td>CACCCCTGGAGTGTAAGAAGTTGT</td><td></td><td></td><td></td><td></td><td></td><td></td><td></td><td></td><td></td><td></td></tr>
<tr><td>Oligo rev</td><td>AAACACAACTTCTTACACTCCAGG</td><td></td><td></td><td></td><td></td><td></td><td></td><td></td><td></td><td></td><td></td></tr>
<tr><td>Chromosome</td><td>start</td><td>end</td><td>strand</td><td>MM</td><td>target_seq</td><td>PAM</td><td>alignment</td><td>distance</td><td>position</td><td>gene name</td><td>gene id</td></tr>
<tr><td>GU305914.1</td><td>23888</td><td>23910</td><td>+</td><td>0</td><td>CCTGGAGTGTAAGAAGTTGT</td><td>TGG</td><td>||||||||[||||||||||||]PAM</td><td>NA</td><td>Intergenic</td><td>NA</td><td>NA</td></tr>
<tr><td>T1358</td><td>CTGGAGTGTAAGAAGTTGTTGGG</td><td>1000</td><td></td><td></td><td></td><td></td><td></td><td></td><td></td><td></td><td></td></tr>
<tr><td>Oligo fwd</td><td>CACCCTGGAGTGTAAGAAGTTGTT</td><td></td><td></td><td></td><td></td><td></td><td></td><td></td><td></td><td></td><td></td></tr>
<tr><td>Oligo rev</td><td>AAACAACAACTTCTTACACTCCAG</td><td></td><td></td><td></td><td></td><td></td><td></td><td></td><td></td><td></td><td></td></tr>
<tr><td>Chromosome</td><td>start</td><td>end</td><td>strand</td><td>MM</td><td>target_seq</td><td>PAM</td><td>alignment</td><td>distance</td><td>position</td><td>gene name</td><td>gene id</td></tr>
<tr><td>GU305914.1</td><td>23889</td><td>23911</td><td>+</td><td>0</td><td>CTGGAGTGTAAGAAGTTGTT</td><td>GGG</td><td>||||||||[||||||||||||]PAM</td><td>NA</td><td>Intergenic</td><td>NA</td><td>NA</td></tr>
<tr><td>T1359</td><td>TAAGAAGTTGTTGGGACACTCGG</td><td>1000</td><td></td><td></td><td></td><td></td><td></td><td></td><td></td><td></td><td></td></tr>
<tr><td>Oligo fwd</td><td>CACCTAAGAAGTTGTTGGGACACT</td><td></td><td></td><td></td><td></td><td></td><td></td><td></td><td></td><td></td><td></td></tr>
<tr><td>Oligo rev</td><td>AAACAGTGTCCCAACAACTTCTTA</td><td></td><td></td><td></td><td></td><td></td><td></td><td></td><td></td><td></td><td></td></tr>
<tr><td>Chromosome</td><td>start</td><td>end</td><td>strand</td><td>MM</td><td>target_seq</td><td>PAM</td><td>alignment</td><td>distance</td><td>position</td><td>gene name</td><td>gene id</td></tr>
<tr><td>GU305914.1</td><td>23897</td><td>23919</td><td>+</td><td>0</td><td>TAAGAAGTTGTTGGGACACT</td><td>CGG</td><td>||||||||[||||||||||||]PAM</td><td>NA</td><td>Intergenic</td><td>NA</td><td>NA</td></tr>
<tr><td>T1360</td><td>GTTGGGACACTCGGTCTTCTTGG</td><td>1000</td><td></td><td></td><td></td><td></td><td></td><td></td><td></td><td></td><td></td></tr>
<tr><td>Oligo fwd</td><td>CACCGTTGGGACACTCGGTCTTCT</td><td></td><td></td><td></td><td></td><td></td><td></td><td></td><td></td><td></td><td></td></tr>
<tr><td>Oligo rev</td><td>AAACAGAAGACCGAGTGTCCCAAC</td><td></td><td></td><td></td><td></td><td></td><td></td><td></td><td></td><td></td><td></td></tr>
<tr><td>Chromosome</td><td>start</td><td>end</td><td>strand</td><td>MM</td><td>target_seq</td><td>PAM</td><td>alignment</td><td>distance</td><td>position</td><td>gene name</td><td>gene id</td></tr>
<tr><td>GU305914.1</td><td>23906</td><td>23928</td><td>+</td><td>0</td><td>GTTGGGACACTCGGTCTTCT</td><td>TGG</td><td>||||||||[||||||||||||]PAM</td><td>NA</td><td>Intergenic</td><td>NA</td><td>NA</td></tr>
<tr><td>T1361</td><td>CTCGGTCTTCTTGGTGAGCATGG</td><td>1000</td><td></td><td></td><td></td><td></td><td></td><td></td><td></td><td></td><td></td></tr>
<tr><td>Oligo fwd</td><td>CACCCTCGGTCTTCTTGGTGAGCA</td><td></td><td></td><td></td><td></td><td></td><td></td><td></td><td></td><td></td><td></td></tr>
<tr><td>Oligo rev</td><td>AAACTGCTCACCAAGAAGACCGAG</td><td></td><td></td><td></td><td></td><td></td><td></td><td></td><td></td><td></td><td></td></tr>
<tr><td>Chromosome</td><td>start</td><td>end</td><td>strand</td><td>MM</td><td>target_seq</td><td>PAM</td><td>alignment</td><td>distance</td><td>position</td><td>gene name</td><td>gene id</td></tr>
<tr><td>GU305914.1</td><td>23915</td><td>23937</td><td>+</td><td>0</td><td>CTCGGTCTTCTTGGTGAGCA</td><td>TGG</td><td>||||||||[||||||||||||]PAM</td><td>NA</td><td>Intergenic</td><td>NA</td><td>NA</td></tr>
<tr><td>T1362</td><td>TTGGTGAGCATGGCCTCGAGCGG</td><td>1000</td><td></td><td></td><td></td><td></td><td></td><td></td><td></td><td></td><td></td></tr>
<tr><td>Oligo fwd</td><td>CACCTTGGTGAGCATGGCCTCGAG</td><td></td><td></td><td></td><td></td><td></td><td></td><td></td><td></td><td></td><td></td></tr>
<tr><td>Oligo rev</td><td>AAACCTCGAGGCCATGCTCACCAA</td><td></td><td></td><td></td><td></td><td></td><td></td><td></td><td></td><td></td><td></td></tr>
</table>

| Chromosome | start | end | strand | MM | target_seq | PAM | alignment | distance | position | gene name | gene id |
|---|---|---|---|---|---|---|---|---|---|---|---|
| GU305914.1 | 23925 | 23947 | + | 0 | TTGGTGAGCATGGCCTCGAG | CGG | \|\|\|\|\|\|\|\|[\|\|\|\|\|\|\|\|\|\|\|\|]PAM | NA | Intergenic | NA | NA |
| T1363 | AGTACCTGTTCTTGTGATAGCGG | 1000 | | | | | | | | | |
| Oligo fwd | CACCAGTACCTGTTCTTGTGATAG | | | | | | | | | | |
| Oligo rev | AAACCTATCACAAGAACAGGTACT | | | | | | | | | | |
| Chromosome | start | end | strand | MM | target_seq | PAM | alignment | distance | position | gene name | gene id |
| GU305914.1 | 23959 | 23981 | + | 0 | AGTACCTGTTCTTGTGATAG | CGG | \|\|\|\|\|\|\|\|[\|\|\|\|\|\|\|\|\|\|\|\|]PAM | NA | Intergenic | NA | NA |
| T1364 | GACGAACTCAAAGTAGTCCTCGG | 1000 | | | | | | | | | |
| Oligo fwd | CACCGACGAACTCAAAGTAGTCCT | | | | | | | | | | |
| Oligo rev | AAACAGGACTACTTTGAGTTCGTC | | | | | | | | | | |
| Chromosome | start | end | strand | MM | target_seq | PAM | alignment | distance | position | gene name | gene id |
| GU305914.1 | 23996 | 24018 | + | 0 | GACGAACTCAAAGTAGTCCT | CGG | \|\|\|\|\|\|\|\|[\|\|\|\|\|\|\|\|\|\|\|\|]PAM | NA | Intergenic | NA | NA |
| T1365 | GAACCACGTAGACGCGACAGAGG | 1000 | | | | | | | | | |
| Oligo fwd | CACCGAACCACGTAGACGCGACAG | | | | | | | | | | |
| Oligo rev | AAACCTGTCGCGTCTACGTGGTTC | | | | | | | | | | |
| Chromosome | start | end | strand | MM | target_seq | PAM | alignment | distance | position | gene name | gene id |
| GU305914.1 | 24046 | 24068 | + | 0 | GAACCACGTAGACGCGACAG | AGG | \|\|\|\|\|\|\|\|[\|\|\|\|\|\|\|\|\|\|\|\|]PAM | NA | Intergenic | NA | NA |
| T1366 | GTAGACGCGACAGAGGTCGTCGG | 1000 | | | | | | | | | |
| Oligo fwd | CACCGTAGACGCGACAGAGGTCGT | | | | | | | | | | |
| Oligo rev | AAACACGACCTCTGTCGCGTCTAC | | | | | | | | | | |
| Chromosome | start | end | strand | MM | target_seq | PAM | alignment | distance | position | gene name | gene id |
| GU305914.1 | 24053 | 24075 | + | 0 | GTAGACGCGACAGAGGTCGT | CGG | \|\|\|\|\|\|\|\|[\|\|\|\|\|\|\|\|\|\|\|\|]PAM | NA | Intergenic | NA | NA |
| GU305914.1 | 151810 | 151832 | - | 4 | GTTGCTGCGGCAGAGGTCGT | CGG | \|\|-\|--\|\|[\|-\|\|\|\|\|\|\|\|\|\|]PAM | NA | Intergenic | NA | NA |
| T1367 | CGCGACAGAGGTCGTCGGTGAGG | 1000 | | | | | | | | | |
| Oligo fwd | CACCCGCGACAGAGGTCGTCGGTG | | | | | | | | | | |
| Oligo rev | AAACCACCGACGACCTCTGTCGCG | | | | | | | | | | |
| Chromosome | start | end | strand | MM | target_seq | PAM | alignment | distance | position | gene name | gene id |
| GU305914.1 | 24058 | 24080 | + | 0 | CGCGACAGAGGTCGTCGGTG | AGG | \|\|\|\|\|\|\|\|[\|\|\|\|\|\|\|\|\|\|\|\|]PAM | NA | Intergenic | NA | NA |
| T1368 | ACAGAGGTCGTCGGTGAGGATGG | 1000 | | | | | | | | | |
| Oligo fwd | CACCACAGAGGTCGTCGGTGAGGA | | | | | | | | | | |
| Oligo rev | AAACTCCTCACCGACGACCTCTGT | | | | | | | | | | |
| Chromosome | start | end | strand | MM | target_seq | PAM | alignment | distance | position | gene name | gene id |
| GU305914.1 | 24062 | 24084 | + | 0 | ACAGAGGTCGTCGGTGAGGA | TGG | \|\|\|\|\|\|\|\|[\|\|\|\|\|\|\|\|\|\|\|\|]PAM | NA | Intergenic | NA | NA |
| T1369 | TCGTCGGTGAGGATGGTGAGCGG | 1000 | | | | | | | | | |
| Oligo fwd | CACCTCGTCGGTGAGGATGGTGAG | | | | | | | | | | |
| Oligo rev | AAACCTCACCATCCTCACCGACGA | | | | | | | | | | |
| Chromosome | start | end | strand | MM | target_seq | PAM | alignment | distance | position | gene name | gene id |
| GU305914.1 | 24069 | 24091 | + | 0 | TCGTCGGTGAGGATGGTGAG | CGG | \|\|\|\|\|\|\|\|[\|\|\|\|\|\|\|\|\|\|\|\|]PAM | NA | Intergenic | NA | NA |
| T1370 | TCGGTGAGGATGGTGAGCGGCGG | 1000 | | | | | | | | | |
| Oligo fwd | CACCTCGGTGAGGATGGTGAGCGG | | | | | | | | | | |
| Oligo rev | AAACCCGCTCACCATCCTCACCGA | | | | | | | | | | |
| Chromosome | start | end | strand | MM | target_seq | PAM | alignment | distance | position | gene name | gene id |
| GU305914.1 | 24072 | 24094 | + | 0 | TCGGTGAGGATGGTGAGCGG | CGG | \|\|\|\|\|\|\|\|[\|\|\|\|\|\|\|\|\|\|\|\|]PAM | NA | Intergenic | NA | NA |
| T1371 | GTGAGCGGCGGCAGCATGAGCGG | 1000 | | | | | | | | | |
| Oligo fwd | CACCGTGAGCGGCGGCAGCATGAG | | | | | | | | | | |
| Oligo rev | AAACCTCATGCTGCCGCCGCTCAC | | | | | | | | | | |
| Chromosome | start | end | strand | MM | target_seq | PAM | alignment | distance | position | gene name | gene id |
| GU305914.1 | 24084 | 24106 | + | 0 | GTGAGCGGCGGCAGCATGAG | CGG | \|\|\|\|\|\|\|\|[\|\|\|\|\|\|\|\|\|\|\|\|]PAM | NA | Intergenic | NA | NA |
| T1372 | GCGGCTCGATGATGATGCCGAGG | 1000 | | | | | | | | | |
| Oligo fwd | CACCGCGGCTCGATGATGATGCCG | | | | | | | | | | |
| Oligo rev | AAACCGGCATCATCATCGAGCCGC | | | | | | | | | | |
| Chromosome | start | end | strand | MM | target_seq | PAM | alignment | distance | position | gene name | gene id |
| GU305914.1 | 24103 | 24125 | + | 0 | GCGGCTCGATGATGATGCCG | AGG | \|\|\|\|\|\|\|\|[\|\|\|\|\|\|\|\|\|\|\|\|]PAM | NA | Intergenic | NA | NA |
| T1373 | GATGATGCCGAGGACGTCCATGG | 1000 | | | | | | | | | |
| Oligo fwd | CACCGATGATGCCGAGGACGTCCA | | | | | | | | | | |
| Oligo rev | AAACTGGACGTCCTCGGCATCATC | | | | | | | | | | |
| Chromosome | start | end | strand | MM | target_seq | PAM | alignment | distance | position | gene name | gene id |
| GU305914.1 | 24113 | 24135 | + | 0 | GATGATGCCGAGGACGTCCA | TGG | \|\|\|\|\|\|\|\|[\|\|\|\|\|\|\|\|\|\|\|\|]PAM | NA | Intergenic | NA | NA |
| T1374 | GACGTCCATGGCGCCGACGTCGG | 1000 | | | | | | | | | |
| Oligo fwd | CACCGACGTCCATGGCGCCGACGT | | | | | | | | | | |
| Oligo rev | AAACACGTCGGCGCCATGGACGTC | | | | | | | | | | |
| Chromosome | start | end | strand | MM | target_seq | PAM | alignment | distance | position | gene name | gene id |
| GU305914.1 | 24125 | 24147 | + | 0 | GACGTCCATGGCGCCGACGT | CGG | \|\|\|\|\|\|\|\|[\|\|\|\|\|\|\|\|\|\|\|\|]PAM | NA | Intergenic | NA | NA |
| GU305914.1 | 118564 | 118586 | + | 2 | GACGTCCATGGCGTCGGCGT | CGG | \|\|\|\|\|\|\|\|[\|\|\|\|\|-\|\|-\|\|\|]PAM | NA | Intergenic | NA | NA |
| T1375 | CATGGCGCCGACGTCGGCCTCGG | 1000 | | | | | | | | | |
| Oligo fwd | CACCCATGGCGCCGACGTCGGCCT | | | | | | | | | | |
| Oligo rev | AAACAGGCCGACGTCGGCGCCATG | | | | | | | | | | |
| Chromosome | start | end | strand | MM | target_seq | PAM | alignment | distance | position | gene name | gene id |
| GU305914.1 | 24131 | 24153 | + | 0 | CATGGCGCCGACGTCGGCCT | CGG | \|\|\|\|\|\|\|\|[\|\|\|\|\|\|\|\|\|\|\|\|]PAM | NA | Intergenic | NA | NA |
| T1376 | GGCGCCGACGTCGGCCTCGGCGG | 1000 | | | | | | | | | |
| Oligo fwd | CACCGGCGCCGACGTCGGCCTCGG | | | | | | | | | | |
| Oligo rev | AAACCCGAGGCCGACGTCGGCGCC | | | | | | | | | | |
| Chromosome | start | end | strand | MM | target_seq | PAM | alignment | distance | position | gene name | gene id |
| GU305914.1 | 24134 | 24156 | + | 0 | GGCGCCGACGTCGGCCTCGG | CGG | \|\|\|\|\|\|\|\|[\|\|\|\|\|\|\|\|\|\|\|\|]PAM | NA | Intergenic | NA | NA |
| GU305914.1 | 128973 | 128995 | - | 4 | GCTGCCGACGGCTGCCTCGG | CGG | \|--\|\|\|\|\|[\|\|-\|-\|\|\|\|\|\|\|]PAM | NA | Intergenic | NA | NA |

| | | | | | | | | | | | |
|---|---|---|---|---|---|---|---|---|---|---|---|
| GU305914.1 | 57831 | 57853 | + | 4 | GCCGCCGCCGTCGCTCTCGG | TGG | \|-\|\|\|\|\|-[\|\|\|\|\|--\|\|\|\|\|]PAM | NA | Intergenic | NA | NA |
| GU305914.1 | 24884 | 24906 | + | 3 | GGCGGCGACGGCGGCCTCGA | CGG | \|\|\|\|-\|\|\|[\|\|-\|\|\|\|\|\|\|\|-]PAM | NA | Intergenic | NA | NA |
| T1377 | CCTCGGCGGCCCATCCGTCAAGG | 1000 | | | | | | | | | |
| Oligo fwd | CACCCCTCGGCGGCCCATCCGTCA | | | | | | | | | | |
| Oligo rev | AAACTGACGGATGGGCCGCCGAGG | | | | | | | | | | |
| Chromosome | start | end | strand | MM | target_seq | PAM | alignment | distance | position | gene name | gene id |
| GU305914.1 | 24148 | 24170 | + | 0 | CCTCGGCGGCCCATCCGTCA | AGG | \|\|\|\|\|\|\|\|[\|\|\|\|\|\|\|\|\|\|\|\|]PAM | NA | Intergenic | NA | NA |
| T1378 | CTCGGCGGCCCATCCGTCAAGGG | 1000 | | | | | | | | | |
| Oligo fwd | CACCCTCGGCGGCCCATCCGTCAA | | | | | | | | | | |
| Oligo rev | AAACTTGACGGATGGGCCGCCGAG | | | | | | | | | | |
| Chromosome | start | end | strand | MM | target_seq | PAM | alignment | distance | position | gene name | gene id |
| GU305914.1 | 24149 | 24171 | + | 0 | CTCGGCGGCCCATCCGTCAA | GGG | \|\|\|\|\|\|\|\|[\|\|\|\|\|\|\|\|\|\|\|\|]PAM | NA | Intergenic | NA | NA |
| T1379 | TCCCACGAAGAACTCGTCACAGG | 1000 | | | | | | | | | |
| Oligo fwd | CACCTCCCACGAAGAACTCGTCAC | | | | | | | | | | |
| Oligo rev | AAACGTGACGAGTTCTTCGTGGGA | | | | | | | | | | |
| Chromosome | start | end | strand | MM | target_seq | PAM | alignment | distance | position | gene name | gene id |
| GU305914.1 | 24185 | 24207 | + | 0 | TCCCACGAAGAACTCGTCAC | AGG | \|\|\|\|\|\|\|\|[\|\|\|\|\|\|\|\|\|\|\|\|]PAM | NA | Intergenic | NA | NA |
| T1380 | CTCGTCACAGGCGCTGAGTTCGG | 1000 | | | | | | | | | |
| Oligo fwd | CACCCTCGTCACAGGCGCTGAGTT | | | | | | | | | | |
| Oligo rev | AAACAACTCAGCGCCTGTGACGAG | | | | | | | | | | |
| Chromosome | start | end | strand | MM | target_seq | PAM | alignment | distance | position | gene name | gene id |
| GU305914.1 | 24197 | 24219 | + | 0 | CTCGTCACAGGCGCTGAGTT | CGG | \|\|\|\|\|\|\|\|[\|\|\|\|\|\|\|\|\|\|\|\|]PAM | NA | Intergenic | NA | NA |
| T1381 | ATTTATCCCCCAAGAAGACGCGG | 1000 | | | | | | | | | |
| Oligo fwd | CACCATTTATCCCCCAAGAAGACG | | | | | | | | | | |
| Oligo rev | AAACCGTCTTCTTGGGGGATAAAT | | | | | | | | | | |
| Chromosome | start | end | strand | MM | target_seq | PAM | alignment | distance | position | gene name | gene id |
| GU305914.1 | 24220 | 24242 | + | 0 | ATTTATCCCCCAAGAAGACG | CGG | \|\|\|\|\|\|\|\|[\|\|\|\|\|\|\|\|\|\|\|\|]PAM | NA | Intergenic | NA | NA |
| T1382 | ATCCCCCAAGAAGACGCGGTTGG | 1000 | | | | | | | | | |
| Oligo fwd | CACCATCCCCCAAGAAGACGCGGT | | | | | | | | | | |
| Oligo rev | AAACACCGCGTCTTCTTGGGGGAT | | | | | | | | | | |
| Chromosome | start | end | strand | MM | target_seq | PAM | alignment | distance | position | gene name | gene id |
| GU305914.1 | 24224 | 24246 | + | 0 | ATCCCCCAAGAAGACGCGGT | TGG | \|\|\|\|\|\|\|\|[\|\|\|\|\|\|\|\|\|\|\|\|]PAM | NA | Intergenic | NA | NA |
| T1383 | CCAAGAAGACGCGGTTGGCGAGG | 1000 | | | | | | | | | |
| Oligo fwd | CACCCCAAGAAGACGCGGTTGGCG | | | | | | | | | | |
| Oligo rev | AAACCGCCAACCGCGTCTTCTTGG | | | | | | | | | | |
| Chromosome | start | end | strand | MM | target_seq | PAM | alignment | distance | position | gene name | gene id |
| GU305914.1 | 24229 | 24251 | + | 0 | CCAAGAAGACGCGGTTGGCG | AGG | \|\|\|\|\|\|\|\|[\|\|\|\|\|\|\|\|\|\|\|\|]PAM | NA | Intergenic | NA | NA |
| T1384 | GGCGAGGTTTCGCTCGCGCCAGG | 1000 | | | | | | | | | |
| Oligo fwd | CACCGGCGAGGTTTCGCTCGCGCC | | | | | | | | | | |
| Oligo rev | AAACGGCGCGAGCGAAACCTCGCC | | | | | | | | | | |
| Chromosome | start | end | strand | MM | target_seq | PAM | alignment | distance | position | gene name | gene id |
| GU305914.1 | 24245 | 24267 | + | 0 | GGCGAGGTTTCGCTCGCGCC | AGG | \|\|\|\|\|\|\|\|[\|\|\|\|\|\|\|\|\|\|\|\|]PAM | NA | Intergenic | NA | NA |
| T1385 | GGTTTCGCTCGCGCCAGGACAGG | 1000 | | | | | | | | | |
| Oligo fwd | CACCGGTTTCGCTCGCGCCAGGAC | | | | | | | | | | |
| Oligo rev | AAACGTCCTGGCGCGAGCGAAACC | | | | | | | | | | |
| Chromosome | start | end | strand | MM | target_seq | PAM | alignment | distance | position | gene name | gene id |
| GU305914.1 | 24250 | 24272 | + | 0 | GGTTTCGCTCGCGCCAGGAC | AGG | \|\|\|\|\|\|\|\|[\|\|\|\|\|\|\|\|\|\|\|\|]PAM | NA | Intergenic | NA | NA |
| T1386 | CCAGGACAGGATGTCGTTGTTGG | 1000 | | | | | | | | | |
| Oligo fwd | CACCCCAGGACAGGATGTCGTTGT | | | | | | | | | | |
| Oligo rev | AAACACAACGACATCCTGTCCTGG | | | | | | | | | | |
| Chromosome | start | end | strand | MM | target_seq | PAM | alignment | distance | position | gene name | gene id |
| GU305914.1 | 24263 | 24285 | + | 0 | CCAGGACAGGATGTCGTTGT | TGG | \|\|\|\|\|\|\|\|[\|\|\|\|\|\|\|\|\|\|\|\|]PAM | NA | Intergenic | NA | NA |
| T1387 | ACAGGATGTCGTTGTTGGTGCGG | 1000 | | | | | | | | | |
| Oligo fwd | CACCACAGGATGTCGTTGTTGGTG | | | | | | | | | | |
| Oligo rev | AAACCACCAACAACGACATCCTGT | | | | | | | | | | |
| Chromosome | start | end | strand | MM | target_seq | PAM | alignment | distance | position | gene name | gene id |
| GU305914.1 | 24268 | 24290 | + | 0 | ACAGGATGTCGTTGTTGGTG | CGG | \|\|\|\|\|\|\|\|[\|\|\|\|\|\|\|\|\|\|\|\|]PAM | NA | Intergenic | NA | NA |
| T1388 | GTCGTTGTTGGTGCGGCACTCGG | 1000 | | | | | | | | | |
| Oligo fwd | CACCGTCGTTGTTGGTGCGGCACT | | | | | | | | | | |
| Oligo rev | AAACAGTGCCGCACCAACAACGAC | | | | | | | | | | |
| Chromosome | start | end | strand | MM | target_seq | PAM | alignment | distance | position | gene name | gene id |
| GU305914.1 | 24275 | 24297 | + | 0 | GTCGTTGTTGGTGCGGCACT | CGG | \|\|\|\|\|\|\|\|[\|\|\|\|\|\|\|\|\|\|\|\|]PAM | NA | Intergenic | NA | NA |
| T1389 | TGCGGCACTCGGTGAGCTCACGG | 1000 | | | | | | | | | |
| Oligo fwd | CACCTGCGGCACTCGGTGAGCTCA | | | | | | | | | | |
| Oligo rev | AAACTGAGCTCACCGAGTGCCGCA | | | | | | | | | | |
| Chromosome | start | end | strand | MM | target_seq | PAM | alignment | distance | position | gene name | gene id |
| GU305914.1 | 24286 | 24308 | + | 0 | TGCGGCACTCGGTGAGCTCA | CGG | \|\|\|\|\|\|\|\|[\|\|\|\|\|\|\|\|\|\|\|\|]PAM | NA | Intergenic | NA | NA |
| T1390 | CTCGGTGAGCTCACGGCTCGCGG | 1000 | | | | | | | | | |
| Oligo fwd | CACCCTCGGTGAGCTCACGGCTCG | | | | | | | | | | |
| Oligo rev | AAACCGAGCCGTGAGCTCACCGAG | | | | | | | | | | |
| Chromosome | start | end | strand | MM | target_seq | PAM | alignment | distance | position | gene name | gene id |
| GU305914.1 | 24293 | 24315 | + | 0 | CTCGGTGAGCTCACGGCTCG | CGG | \|\|\|\|\|\|\|\|[\|\|\|\|\|\|\|\|\|\|\|\|]PAM | NA | Intergenic | NA | NA |
| T1391 | CGCGGCCCCGCGACAGATGATGG | 1000 | | | | | | | | | |
| Oligo fwd | CACCCGCGGCCCCGCGACAGATGA | | | | | | | | | | |

Oligo rev AAACTCATCTGTCGCGGGGCCGCG
Chromosome start end strand MM target_seq PAM alignment distance position gene name gene id
GU305914.1 24311 24333 + 0 CGCGGCCCCGCGACAGATGA TGG ||||||||[||||||||||||]PAM NA Intergenic NA NA
GU305914.1 75898 75920 + 4 CCTGGCACGGCGACAGATGA GGG |--|||-|[-|||||||||||]PAM NA Intergenic NA NA

T1392 GCGGCCCCGCGACAGATGATGGG 1000
Oligo fwd CACCGCGGCCCCGCGACAGATGAT
Oligo rev AAACATCATCTGTCGCGGGGCCGC
Chromosome start end strand MM target_seq PAM alignment distance position gene name gene id
GU305914.1 24312 24334 + 0 GCGGCCCCGCGACAGATGAT GGG ||||||||[||||||||||||]PAM NA Intergenic NA NA

T1393 CGGCCCCGCGACAGATGATGGGG 1000
Oligo fwd CACCCGGCCCCGCGACAGATGATG
Oligo rev AAACCATCATCTGTCGCGGGGCCG
Chromosome start end strand MM target_seq PAM alignment distance position gene name gene id
GU305914.1 24313 24335 + 0 CGGCCCCGCGACAGATGATG GGG ||||||||[||||||||||||]PAM NA Intergenic NA NA

T1394 CAGATGATGGGGCGACGCACCGG 1000
Oligo fwd CACCCAGATGATGGGGCGACGCAC
Oligo rev AAACGTGCGTCGCCCCATCATCTG
Chromosome start end strand MM target_seq PAM alignment distance position gene name gene id
GU305914.1 24324 24346 + 0 CAGATGATGGGGCGACGCAC CGG ||||||||[||||||||||||]PAM NA Intergenic NA NA

T1395 TGATGGGGCGACGCACCGGCAGG 1000
Oligo fwd CACCTGATGGGGCGACGCACCGGC
Oligo rev AAACGCCGGTGCGTCGCCCCATCA
Chromosome start end strand MM target_seq PAM alignment distance position gene name gene id
GU305914.1 24328 24350 + 0 TGATGGGGCGACGCACCGGC AGG ||||||||[||||||||||||]PAM NA Intergenic NA NA

T1396 GACGATCTCGCGCATGCTGTCGG 1000
Oligo fwd CACCGACGATCTCGCGCATGCTGT
Oligo rev AAACACAGCATGCGCGAGATCGTC
Chromosome start end strand MM target_seq PAM alignment distance position gene name gene id
GU305914.1 24374 24396 + 0 GACGATCTCGCGCATGCTGT CGG ||||||||[||||||||||||]PAM NA Intergenic NA NA

T1397 CGCGCATGCTGTCGGCGACGTGG 1000
Oligo fwd CACCCGCGCATGCTGTCGGCGACG
Oligo rev AAACCGTCGCCGACAGCATGCGCG
Chromosome start end strand MM target_seq PAM alignment distance position gene name gene id
GU305914.1 24382 24404 + 0 CGCGCATGCTGTCGGCGACG TGG ||||||||[||||||||||||]PAM NA Intergenic NA NA

T1398 GCGCATGCTGTCGGCGACGTGGG 1000
Oligo fwd CACCGCGCATGCTGTCGGCGACGT
Oligo rev AAACACGTCGCCGACAGCATGCGC
Chromosome start end strand MM target_seq PAM alignment distance position gene name gene id
GU305914.1 24383 24405 + 0 GCGCATGCTGTCGGCGACGT GGG ||||||||[||||||||||||]PAM NA Intergenic NA NA

T1399 GCTGTCGGCGACGTGGGCAAAGG 1000
Oligo fwd CACCGCTGTCGGCGACGTGGGCAA
Oligo rev AAACTTGCCCACGTCGCCGACAGC
Chromosome start end strand MM target_seq PAM alignment distance position gene name gene id
GU305914.1 24389 24411 + 0 GCTGTCGGCGACGTGGGCAA AGG ||||||||[||||||||||||]PAM NA Intergenic NA NA

T1400 CCTCGCTGCGCGTGAGAAGCAGG 1000
Oligo fwd CACCCCTCGCTGCGCGTGAGAAGC
Oligo rev AAACGCTTCTCACGCGCAGCGAGG
Chromosome start end strand MM target_seq PAM alignment distance position gene name gene id
GU305914.1 24412 24434 + 0 CCTCGCTGCGCGTGAGAAGC AGG ||||||||[||||||||||||]PAM NA Intergenic NA NA

T1401 GCTGCTGCCGACGTAGACCTCGG 1000
Oligo fwd CACCGCTGCTGCCGACGTAGACCT
Oligo rev AAACAGGTCTACGTCGGCAGCAGC
Chromosome start end strand MM target_seq PAM alignment distance position gene name gene id
GU305914.1 24452 24474 + 0 GCTGCTGCCGACGTAGACCT CGG ||||||||[||||||||||||]PAM NA Intergenic NA NA

T1402 CTGCTGCCGACGTAGACCTCGGG 1000
Oligo fwd CACCCTGCTGCCGACGTAGACCTC
Oligo rev AAACGAGGTCTACGTCGGCAGCAG
Chromosome start end strand MM target_seq PAM alignment distance position gene name gene id
GU305914.1 24453 24475 + 0 CTGCTGCCGACGTAGACCTC GGG ||||||||[||||||||||||]PAM NA Intergenic NA NA

T1403 CTGCCGACGTAGACCTCGGGAGG 1000
Oligo fwd CACCCTGCCGACGTAGACCTCGGG
Oligo rev AAACCCCGAGGTCTACGTCGGCAG
Chromosome start end strand MM target_seq PAM alignment distance position gene name gene id
GU305914.1 24456 24478 + 0 CTGCCGACGTAGACCTCGGG AGG ||||||||[||||||||||||]PAM NA Intergenic NA NA

T1404 CGACGTAGACCTCGGGAGGCTGG 1000
Oligo fwd CACCCGACGTAGACCTCGGGAGGC
Oligo rev AAACGCCTCCCGAGGTCTACGTCG
Chromosome start end strand MM target_seq PAM alignment distance position gene name gene id
GU305914.1 24460 24482 + 0 CGACGTAGACCTCGGGAGGC TGG ||||||||[||||||||||||]PAM NA Intergenic NA NA

T1405 CGTAGACCTCGGGAGGCTGGCGG 1000
Oligo fwd CACCCGTAGACCTCGGGAGGCTGG
Oligo rev AAACCCAGCCTCCCGAGGTCTACG
Chromosome start end strand MM target_seq PAM alignment distance position gene name gene id
GU305914.1 24463 24485 + 0 CGTAGACCTCGGGAGGCTGG CGG ||||||||[||||||||||||]PAM NA Intergenic NA NA

<table>
<tr><td>T1406</td><td>CCTCGGGAGGCTGGCGGTAGTGG</td><td>1000</td><td></td><td></td><td></td><td></td><td></td><td></td><td></td><td></td><td></td></tr>
<tr><td>Oligo fwd</td><td>CACCCCTCGGGAGGCTGGCGGTAG</td><td></td><td></td><td></td><td></td><td></td><td></td><td></td><td></td><td></td><td></td></tr>
<tr><td>Oligo rev</td><td>AAACCTACCGCCAGCCTCCCGAGG</td><td></td><td></td><td></td><td></td><td></td><td></td><td></td><td></td><td></td><td></td></tr>
<tr><td>Chromosome</td><td>start</td><td>end</td><td>strand</td><td>MM</td><td>target_seq</td><td>PAM</td><td>alignment</td><td>distance</td><td>position</td><td>gene name</td><td>gene id</td></tr>
<tr><td>GU305914.1</td><td>24469</td><td>24491</td><td>+</td><td>0</td><td>CCTCGGGAGGCTGGCGGTAG</td><td>TGG</td><td>||||||||[||||||||||||]PAM</td><td>NA</td><td>Intergenic</td><td>NA</td><td>NA</td></tr>
<tr><td></td><td></td><td></td><td></td><td></td><td></td><td></td><td></td><td></td><td></td><td></td><td></td></tr>
<tr><td>T1407</td><td>CTCGGGAGGCTGGCGGTAGTGGG</td><td>1000</td><td></td><td></td><td></td><td></td><td></td><td></td><td></td><td></td><td></td></tr>
<tr><td>Oligo fwd</td><td>CACCCTCGGGAGGCTGGCGGTAGT</td><td></td><td></td><td></td><td></td><td></td><td></td><td></td><td></td><td></td><td></td></tr>
<tr><td>Oligo rev</td><td>AAACACTACCGCCAGCCTCCCGAG</td><td></td><td></td><td></td><td></td><td></td><td></td><td></td><td></td><td></td><td></td></tr>
<tr><td>Chromosome</td><td>start</td><td>end</td><td>strand</td><td>MM</td><td>target_seq</td><td>PAM</td><td>alignment</td><td>distance</td><td>position</td><td>gene name</td><td>gene id</td></tr>
<tr><td>GU305914.1</td><td>24470</td><td>24492</td><td>+</td><td>0</td><td>CTCGGGAGGCTGGCGGTAGT</td><td>GGG</td><td>||||||||[||||||||||||]PAM</td><td>NA</td><td>Intergenic</td><td>NA</td><td>NA</td></tr>
<tr><td></td><td></td><td></td><td></td><td></td><td></td><td></td><td></td><td></td><td></td><td></td><td></td></tr>
<tr><td>T1408</td><td>TCGGGAGGCTGGCGGTAGTGGGG</td><td>1000</td><td></td><td></td><td></td><td></td><td></td><td></td><td></td><td></td><td></td></tr>
<tr><td>Oligo fwd</td><td>CACCTCGGGAGGCTGGCGGTAGTG</td><td></td><td></td><td></td><td></td><td></td><td></td><td></td><td></td><td></td><td></td></tr>
<tr><td>Oligo rev</td><td>AAACCACTACCGCCAGCCTCCCGA</td><td></td><td></td><td></td><td></td><td></td><td></td><td></td><td></td><td></td><td></td></tr>
<tr><td>Chromosome</td><td>start</td><td>end</td><td>strand</td><td>MM</td><td>target_seq</td><td>PAM</td><td>alignment</td><td>distance</td><td>position</td><td>gene name</td><td>gene id</td></tr>
<tr><td>GU305914.1</td><td>24471</td><td>24493</td><td>+</td><td>0</td><td>TCGGGAGGCTGGCGGTAGTG</td><td>GGG</td><td>||||||||[||||||||||||]PAM</td><td>NA</td><td>Intergenic</td><td>NA</td><td>NA</td></tr>
<tr><td></td><td></td><td></td><td></td><td></td><td></td><td></td><td></td><td></td><td></td><td></td><td></td></tr>
<tr><td>T1409</td><td>CTGGCGGTAGTGGGGCAGCTCGG</td><td>1000</td><td></td><td></td><td></td><td></td><td></td><td></td><td></td><td></td><td></td></tr>
<tr><td>Oligo fwd</td><td>CACCCTGGCGGTAGTGGGGCAGCT</td><td></td><td></td><td></td><td></td><td></td><td></td><td></td><td></td><td></td><td></td></tr>
<tr><td>Oligo rev</td><td>AAACAGCTGCCCCACTACCGCCAG</td><td></td><td></td><td></td><td></td><td></td><td></td><td></td><td></td><td></td><td></td></tr>
<tr><td>Chromosome</td><td>start</td><td>end</td><td>strand</td><td>MM</td><td>target_seq</td><td>PAM</td><td>alignment</td><td>distance</td><td>position</td><td>gene name</td><td>gene id</td></tr>
<tr><td>GU305914.1</td><td>24479</td><td>24501</td><td>+</td><td>0</td><td>CTGGCGGTAGTGGGGCAGCT</td><td>CGG</td><td>||||||||[||||||||||||]PAM</td><td>NA</td><td>Intergenic</td><td>NA</td><td>NA</td></tr>
<tr><td></td><td></td><td></td><td></td><td></td><td></td><td></td><td></td><td></td><td></td><td></td><td></td></tr>
<tr><td>T1410</td><td>GTAGTGGGGCAGCTCGGACATGG</td><td>1000</td><td></td><td></td><td></td><td></td><td></td><td></td><td></td><td></td><td></td></tr>
<tr><td>Oligo fwd</td><td>CACCGTAGTGGGGCAGCTCGGACA</td><td></td><td></td><td></td><td></td><td></td><td></td><td></td><td></td><td></td><td></td></tr>
<tr><td>Oligo rev</td><td>AAACTGTCCGAGCTGCCCCACTAC</td><td></td><td></td><td></td><td></td><td></td><td></td><td></td><td></td><td></td><td></td></tr>
<tr><td>Chromosome</td><td>start</td><td>end</td><td>strand</td><td>MM</td><td>target_seq</td><td>PAM</td><td>alignment</td><td>distance</td><td>position</td><td>gene name</td><td>gene id</td></tr>
<tr><td>GU305914.1</td><td>24485</td><td>24507</td><td>+</td><td>0</td><td>GTAGTGGGGCAGCTCGGACA</td><td>TGG</td><td>||||||||[||||||||||||]PAM</td><td>NA</td><td>Intergenic</td><td>NA</td><td>NA</td></tr>
<tr><td></td><td></td><td></td><td></td><td></td><td></td><td></td><td></td><td></td><td></td><td></td><td></td></tr>
<tr><td>T1411</td><td>TAGTGGGGCAGCTCGGACATGGG</td><td>1000</td><td></td><td></td><td></td><td></td><td></td><td></td><td></td><td></td><td></td></tr>
<tr><td>Oligo fwd</td><td>CACCTAGTGGGGCAGCTCGGACAT</td><td></td><td></td><td></td><td></td><td></td><td></td><td></td><td></td><td></td><td></td></tr>
<tr><td>Oligo rev</td><td>AAACATGTCCGAGCTGCCCCACTA</td><td></td><td></td><td></td><td></td><td></td><td></td><td></td><td></td><td></td><td></td></tr>
<tr><td>Chromosome</td><td>start</td><td>end</td><td>strand</td><td>MM</td><td>target_seq</td><td>PAM</td><td>alignment</td><td>distance</td><td>position</td><td>gene name</td><td>gene id</td></tr>
<tr><td>GU305914.1</td><td>24486</td><td>24508</td><td>+</td><td>0</td><td>TAGTGGGGCAGCTCGGACAT</td><td>GGG</td><td>||||||||[||||||||||||]PAM</td><td>NA</td><td>Intergenic</td><td>NA</td><td>NA</td></tr>
<tr><td></td><td></td><td></td><td></td><td></td><td></td><td></td><td></td><td></td><td></td><td></td><td></td></tr>
<tr><td>T1412</td><td>CTCGGACATGGGCGCGTAGATGG</td><td>1000</td><td></td><td></td><td></td><td></td><td></td><td></td><td></td><td></td><td></td></tr>
<tr><td>Oligo fwd</td><td>CACCCTCGGACATGGGCGCGTAGA</td><td></td><td></td><td></td><td></td><td></td><td></td><td></td><td></td><td></td><td></td></tr>
<tr><td>Oligo rev</td><td>AAACTCTACGCGCCCATGTCCGAG</td><td></td><td></td><td></td><td></td><td></td><td></td><td></td><td></td><td></td><td></td></tr>
<tr><td>Chromosome</td><td>start</td><td>end</td><td>strand</td><td>MM</td><td>target_seq</td><td>PAM</td><td>alignment</td><td>distance</td><td>position</td><td>gene name</td><td>gene id</td></tr>
<tr><td>GU305914.1</td><td>24497</td><td>24519</td><td>+</td><td>0</td><td>CTCGGACATGGGCGCGTAGA</td><td>TGG</td><td>||||||||[||||||||||||]PAM</td><td>NA</td><td>Intergenic</td><td>NA</td><td>NA</td></tr>
<tr><td></td><td></td><td></td><td></td><td></td><td></td><td></td><td></td><td></td><td></td><td></td><td></td></tr>
<tr><td>T1413</td><td>CGCGTAGATGGCACCGCCGCCGG</td><td>1000</td><td></td><td></td><td></td><td></td><td></td><td></td><td></td><td></td><td></td></tr>
<tr><td>Oligo fwd</td><td>CACCCGCGTAGATGGCACCGCCGC</td><td></td><td></td><td></td><td></td><td></td><td></td><td></td><td></td><td></td><td></td></tr>
<tr><td>Oligo rev</td><td>AAACGCGGCGGTGCCATCTACGCG</td><td></td><td></td><td></td><td></td><td></td><td></td><td></td><td></td><td></td><td></td></tr>
<tr><td>Chromosome</td><td>start</td><td>end</td><td>strand</td><td>MM</td><td>target_seq</td><td>PAM</td><td>alignment</td><td>distance</td><td>position</td><td>gene name</td><td>gene id</td></tr>
<tr><td>GU305914.1</td><td>24509</td><td>24531</td><td>+</td><td>0</td><td>CGCGTAGATGGCACCGCCGC</td><td>CGG</td><td>||||||||[||||||||||||]PAM</td><td>NA</td><td>Intergenic</td><td>NA</td><td>NA</td></tr>
<tr><td></td><td></td><td></td><td></td><td></td><td></td><td></td><td></td><td></td><td></td><td></td><td></td></tr>
<tr><td>T1414</td><td>CCGCCGCCGGCCCTGAGCGCCGG</td><td>1000</td><td></td><td></td><td></td><td></td><td></td><td></td><td></td><td></td><td></td></tr>
<tr><td>Oligo fwd</td><td>CACCCCGCCGCCGGCCCTGAGCGC</td><td></td><td></td><td></td><td></td><td></td><td></td><td></td><td></td><td></td><td></td></tr>
<tr><td>Oligo rev</td><td>AAACGCGCTCAGGGCCGGCGGCGG</td><td></td><td></td><td></td><td></td><td></td><td></td><td></td><td></td><td></td><td></td></tr>
<tr><td>Chromosome</td><td>start</td><td>end</td><td>strand</td><td>MM</td><td>target_seq</td><td>PAM</td><td>alignment</td><td>distance</td><td>position</td><td>gene name</td><td>gene id</td></tr>
<tr><td>GU305914.1</td><td>24522</td><td>24544</td><td>+</td><td>0</td><td>CCGCCGCCGGCCCTGAGCGC</td><td>CGG</td><td>||||||||[||||||||||||]PAM</td><td>NA</td><td>Intergenic</td><td>NA</td><td>NA</td></tr>
<tr><td></td><td></td><td></td><td></td><td></td><td></td><td></td><td></td><td></td><td></td><td></td><td></td></tr>
<tr><td>T1415</td><td>CCTGAGCGCCGGCCCCAAGATGG</td><td>1000</td><td></td><td></td><td></td><td></td><td></td><td></td><td></td><td></td><td></td></tr>
<tr><td>Oligo fwd</td><td>CACCCCTGAGCGCCGGCCCCAAGA</td><td></td><td></td><td></td><td></td><td></td><td></td><td></td><td></td><td></td><td></td></tr>
<tr><td>Oligo rev</td><td>AAACTCTTGGGGCCGGCGCTCAGG</td><td></td><td></td><td></td><td></td><td></td><td></td><td></td><td></td><td></td><td></td></tr>
<tr><td>Chromosome</td><td>start</td><td>end</td><td>strand</td><td>MM</td><td>target_seq</td><td>PAM</td><td>alignment</td><td>distance</td><td>position</td><td>gene name</td><td>gene id</td></tr>
<tr><td>GU305914.1</td><td>24533</td><td>24555</td><td>+</td><td>0</td><td>CCTGAGCGCCGGCCCCAAGA</td><td>TGG</td><td>||||||||[||||||||||||]PAM</td><td>NA</td><td>Intergenic</td><td>NA</td><td>NA</td></tr>
<tr><td></td><td></td><td></td><td></td><td></td><td></td><td></td><td></td><td></td><td></td><td></td><td></td></tr>
<tr><td>T1416</td><td>GAGCGCCGGCCCCAAGATGGCGG</td><td>1000</td><td></td><td></td><td></td><td></td><td></td><td></td><td></td><td></td><td></td></tr>
<tr><td>Oligo fwd</td><td>CACCGAGCGCCGGCCCCAAGATGG</td><td></td><td></td><td></td><td></td><td></td><td></td><td></td><td></td><td></td><td></td></tr>
<tr><td>Oligo rev</td><td>AAACCCATCTTGGGGCCGGCGCTC</td><td></td><td></td><td></td><td></td><td></td><td></td><td></td><td></td><td></td><td></td></tr>
<tr><td>Chromosome</td><td>start</td><td>end</td><td>strand</td><td>MM</td><td>target_seq</td><td>PAM</td><td>alignment</td><td>distance</td><td>position</td><td>gene name</td><td>gene id</td></tr>
<tr><td>GU305914.1</td><td>24536</td><td>24558</td><td>+</td><td>0</td><td>GAGCGCCGGCCCCAAGATGG</td><td>CGG</td><td>||||||||[||||||||||||]PAM</td><td>NA</td><td>Intergenic</td><td>NA</td><td>NA</td></tr>
<tr><td></td><td></td><td></td><td></td><td></td><td></td><td></td><td></td><td></td><td></td><td></td><td></td></tr>
<tr><td>T1417</td><td>CGCCGGCCCCAAGATGGCGGCGG</td><td>1000</td><td></td><td></td><td></td><td></td><td></td><td></td><td></td><td></td><td></td></tr>
<tr><td>Oligo fwd</td><td>CACCCGCCGGCCCCAAGATGGCGG</td><td></td><td></td><td></td><td></td><td></td><td></td><td></td><td></td><td></td><td></td></tr>
<tr><td>Oligo rev</td><td>AAACCCGCCATCTTGGGGCCGGCG</td><td></td><td></td><td></td><td></td><td></td><td></td><td></td><td></td><td></td><td></td></tr>
<tr><td>Chromosome</td><td>start</td><td>end</td><td>strand</td><td>MM</td><td>target_seq</td><td>PAM</td><td>alignment</td><td>distance</td><td>position</td><td>gene name</td><td>gene id</td></tr>
<tr><td>GU305914.1</td><td>24539</td><td>24561</td><td>+</td><td>0</td><td>CGCCGGCCCCAAGATGGCGG</td><td>CGG</td><td>||||||||[||||||||||||]PAM</td><td>NA</td><td>Intergenic</td><td>NA</td><td>NA</td></tr>
<tr><td></td><td></td><td></td><td></td><td></td><td></td><td></td><td></td><td></td><td></td><td></td><td></td></tr>
<tr><td>T1418</td><td>GATGGCGGCGGCAGCGACGACGG</td><td>1000</td><td></td><td></td><td></td><td></td><td></td><td></td><td></td><td></td><td></td></tr>
<tr><td>Oligo fwd</td><td>CACCGATGGCGGCGGCAGCGACGA</td><td></td><td></td><td></td><td></td><td></td><td></td><td></td><td></td><td></td><td></td></tr>
<tr><td>Oligo rev</td><td>AAACTCGTCGCTGCCGCCGCCATC</td><td></td><td></td><td></td><td></td><td></td><td></td><td></td><td></td><td></td><td></td></tr>
<tr><td>Chromosome</td><td>start</td><td>end</td><td>strand</td><td>MM</td><td>target_seq</td><td>PAM</td><td>alignment</td><td>distance</td><td>position</td><td>gene name</td><td>gene id</td></tr>
<tr><td>GU305914.1</td><td>24551</td><td>24573</td><td>+</td><td>0</td><td>GATGGCGGCGGCAGCGACGA</td><td>CGG</td><td>||||||||[||||||||||||]PAM</td><td>NA</td><td>Intergenic</td><td>NA</td><td>NA</td></tr>
<tr><td>GU305914.1</td><td>113918</td><td>113940</td><td>-</td><td>3</td><td>GTCGGCGGCGGCAGCGGCGA</td><td>CGG</td><td>|--|||||[||||||||-|||]PAM</td><td>NA</td><td>Intergenic</td><td>NA</td><td>NA</td></tr>
<tr><td></td><td></td><td></td><td></td><td></td><td></td><td></td><td></td><td></td><td></td><td></td><td></td></tr>
<tr><td>T1419</td><td>GGCGGCAGCGACGACGGCAGCGG</td><td>1000</td><td></td><td></td><td></td><td></td><td></td><td></td><td></td><td></td><td></td></tr>
<tr><td>Oligo fwd</td><td>CACCGGCGGCAGCGACGACGGCAG</td><td></td><td></td><td></td><td></td><td></td><td></td><td></td><td></td><td></td><td></td></tr>
<tr><td>Oligo rev</td><td>AAACCTGCCGTCGTCGCTGCCGCC</td><td></td><td></td><td></td><td></td><td></td><td></td><td></td><td></td><td></td><td></td></tr>
<tr><td>Chromosome</td><td>start</td><td>end</td><td>strand</td><td>MM</td><td>target_seq</td><td>PAM</td><td>alignment</td><td>distance</td><td>position</td><td>gene name</td><td>gene id</td></tr>
<tr><td>GU305914.1</td><td>24557</td><td>24579</td><td>+</td><td>0</td><td>GGCGGCAGCGACGACGGCAG</td><td>CGG</td><td>||||||||[||||||||||||]PAM</td><td>NA</td><td>Intergenic</td><td>NA</td><td>NA</td></tr>
<tr><td>GU305914.1</td><td>30006</td><td>30028</td><td>+</td><td>3</td><td>GGCGGCGGCGGCGGCGGCAG</td><td>TGG</td><td>||||||-|[||-||-||||||]PAM</td><td>NA</td><td>Intergenic</td><td>NA</td><td>NA</td></tr>
<tr><td>GU305914.1</td><td>149476</td><td>149498</td><td>-</td><td>3</td><td>GGCGGCGGCGGCGGCGGCAG</td><td>CGG</td><td>||||||-|[||-||-||||||]PAM</td><td>NA</td><td>Intergenic</td><td>NA</td><td>NA</td></tr>
<tr><td>GU305914.1</td><td>163902</td><td>163924</td><td>+</td><td>4</td><td>AGCGGCGGCGGCGGCGGCAG</td><td>CGG</td><td>-|||||-|[||-||-||||||]PAM</td><td>NA</td><td>Intergenic</td><td>NA</td><td>NA</td></tr>
<tr><td></td><td></td><td></td><td></td><td></td><td></td><td></td><td></td><td></td><td></td><td></td><td></td></tr>
<tr><td>T1420</td><td>AGCGGAGACGAGCCCCTCGTCGG</td><td>1000</td><td></td><td></td><td></td><td></td><td></td><td></td><td></td><td></td><td></td></tr>
</table>

| | | | | | | | | | | | |
|---|---|---|---|---|---|---|---|---|---|---|---|
| Oligo fwd | CACCAGCGGAGACGAGCCCCTCGT | | | | | | | | | | |
| Oligo rev | AAACACGAGGGGCTCGTCTCCGCT | | | | | | | | | | |
| Chromosome | start | end | strand | MM | target_seq | PAM | alignment | distance | position | gene name | gene id |
| GU305914.1 | 24575 | 24597 | + | 0 | AGCGGAGACGAGCCCCTCGT | CGG | \|\|\|\|\|\|\|\|[\|\|\|\|\|\|\|\|\|\|\|\|]PAM | NA | Intergenic | NA | NA |
| T1421 | GCGGAGACGAGCCCCTCGTCGGG | 1000 | | | | | | | | | |
| Oligo fwd | CACCGCGGAGACGAGCCCCTCGTC | | | | | | | | | | |
| Oligo rev | AAACGACGAGGGGCTCGTCTCCGC | | | | | | | | | | |
| Chromosome | start | end | strand | MM | target_seq | PAM | alignment | distance | position | gene name | gene id |
| GU305914.1 | 24576 | 24598 | + | 0 | GCGGAGACGAGCCCCTCGTC | GGG | \|\|\|\|\|\|\|\|[\|\|\|\|\|\|\|\|\|\|\|\|]PAM | NA | Intergenic | NA | NA |
| T1422 | AGCCCCTCGTCGGGATCGCGCGG | 1000 | | | | | | | | | |
| Oligo fwd | CACCAGCCCCTCGTCGGGATCGCG | | | | | | | | | | |
| Oligo rev | AAACCGCGATCCCGACGAGGGGCT | | | | | | | | | | |
| Chromosome | start | end | strand | MM | target_seq | PAM | alignment | distance | position | gene name | gene id |
| GU305914.1 | 24585 | 24607 | + | 0 | AGCCCCTCGTCGGGATCGCG | CGG | \|\|\|\|\|\|\|\|[\|\|\|\|\|\|\|\|\|\|\|\|]PAM | NA | Intergenic | NA | NA |
| T1423 | GTCGCCGCCACCGCCGCCGCCGG | 1000 | | | | | | | | | |
| Oligo fwd | CACCGTCGCCGCCACCGCCGCCGC | | | | | | | | | | |
| Oligo rev | AAACGCGGCGGCGGTGGCGGCGAC | | | | | | | | | | |
| Chromosome | start | end | strand | MM | target_seq | PAM | alignment | distance | position | gene name | gene id |
| GU305914.1 | 24620 | 24642 | + | 0 | GTCGCCGCCACCGCCGCCGC | CGG | \|\|\|\|\|\|\|\|[\|\|\|\|\|\|\|\|\|\|\|\|]PAM | NA | Intergenic | NA | NA |
| GU305914.1 | 164964 | 164986 | + | 4 | GCTGCTGCCGCCGCCGCCGC | CGG | \|--\|\|-\|\|[\|-\|\|\|\|\|\|\|\|\|\|]PAM | NA | Intergenic | NA | NA |
| GU305914.1 | 152551 | 152573 | - | 4 | GCCGGCACCGCCGCCGCCGC | AGG | \|-\|\|-\|-\|[\|-\|\|\|\|\|\|\|\|\|\|]PAM | NA | Intergenic | NA | NA |
| GU305914.1 | 48151 | 48173 | - | 4 | GCCGCCACCGCCGCCGCCAC | AGG | \|-\|\|\|\|-\|[\|-\|\|\|\|\|\|\|\|-\|]PAM | NA | Intergenic | NA | NA |
| T1424 | TGTTGCGAAAGCGCCAGTTGAGG | 1000 | | | | | | | | | |
| Oligo fwd | CACCTGTTGCGAAAGCGCCAGTTG | | | | | | | | | | |
| Oligo rev | AAACCAACTGGCGCTTTCGCAACA | | | | | | | | | | |
| Chromosome | start | end | strand | MM | target_seq | PAM | alignment | distance | position | gene name | gene id |
| GU305914.1 | 24643 | 24665 | + | 0 | TGTTGCGAAAGCGCCAGTTG | AGG | \|\|\|\|\|\|\|\|[\|\|\|\|\|\|\|\|\|\|\|\|]PAM | NA | Intergenic | NA | NA |
| T1425 | TGCGAAAGCGCCAGTTGAGGAGG | 1000 | | | | | | | | | |
| Oligo fwd | CACCTGCGAAAGCGCCAGTTGAGG | | | | | | | | | | |
| Oligo rev | AAACCCTCAACTGGCGCTTTCGCA | | | | | | | | | | |
| Chromosome | start | end | strand | MM | target_seq | PAM | alignment | distance | position | gene name | gene id |
| GU305914.1 | 24646 | 24668 | + | 0 | TGCGAAAGCGCCAGTTGAGG | AGG | \|\|\|\|\|\|\|\|[\|\|\|\|\|\|\|\|\|\|\|\|]PAM | NA | Intergenic | NA | NA |
| T1426 | CCAGTTGAGGAGGCGCGATCTGG | 1000 | | | | | | | | | |
| Oligo fwd | CACCCCAGTTGAGGAGGCGCGATC | | | | | | | | | | |
| Oligo rev | AAACGATCGCGCCTCCTCAACTGG | | | | | | | | | | |
| Chromosome | start | end | strand | MM | target_seq | PAM | alignment | distance | position | gene name | gene id |
| GU305914.1 | 24656 | 24678 | + | 0 | CCAGTTGAGGAGGCGCGATC | TGG | \|\|\|\|\|\|\|\|[\|\|\|\|\|\|\|\|\|\|\|\|]PAM | NA | Intergenic | NA | NA |
| T1427 | GGAGGCGCGATCTGGTGTTCTGG | 1000 | | | | | | | | | |
| Oligo fwd | CACCGGAGGCGCGATCTGGTGTTC | | | | | | | | | | |
| Oligo rev | AAACGAACACCAGATCGCGCCTCC | | | | | | | | | | |
| Chromosome | start | end | strand | MM | target_seq | PAM | alignment | distance | position | gene name | gene id |
| GU305914.1 | 24664 | 24686 | + | 0 | GGAGGCGCGATCTGGTGTTC | TGG | \|\|\|\|\|\|\|\|[\|\|\|\|\|\|\|\|\|\|\|\|]PAM | NA | Intergenic | NA | NA |
| T1428 | TGGTGTTCTGGCGACCCTCGAGG | 1000 | | | | | | | | | |
| Oligo fwd | CACCTGGTGTTCTGGCGACCCTCG | | | | | | | | | | |
| Oligo rev | AAACCGAGGGTCGCCAGAACACCA | | | | | | | | | | |
| Chromosome | start | end | strand | MM | target_seq | PAM | alignment | distance | position | gene name | gene id |
| GU305914.1 | 24676 | 24698 | + | 0 | TGGTGTTCTGGCGACCCTCG | AGG | \|\|\|\|\|\|\|\|[\|\|\|\|\|\|\|\|\|\|\|\|]PAM | NA | Intergenic | NA | NA |
| T1429 | GCGACCCTCGAGGTTAGAATCGG | 1000 | | | | | | | | | |
| Oligo fwd | CACCGCGACCCTCGAGGTTAGAAT | | | | | | | | | | |
| Oligo rev | AAACATTCTAACCTCGAGGGTCGC | | | | | | | | | | |
| Chromosome | start | end | strand | MM | target_seq | PAM | alignment | distance | position | gene name | gene id |
| GU305914.1 | 24686 | 24708 | + | 0 | GCGACCCTCGAGGTTAGAAT | CGG | \|\|\|\|\|\|\|\|[\|\|\|\|\|\|\|\|\|\|\|\|]PAM | NA | Intergenic | NA | NA |
| T1430 | ACCCTCGAGGTTAGAATCGGAGG | 1000 | | | | | | | | | |
| Oligo fwd | CACCACCCTCGAGGTTAGAATCGG | | | | | | | | | | |
| Oligo rev | AAACCCGATTCTAACCTCGAGGGT | | | | | | | | | | |
| Chromosome | start | end | strand | MM | target_seq | PAM | alignment | distance | position | gene name | gene id |
| GU305914.1 | 24689 | 24711 | + | 0 | ACCCTCGAGGTTAGAATCGG | AGG | \|\|\|\|\|\|\|\|[\|\|\|\|\|\|\|\|\|\|\|\|]PAM | NA | Intergenic | NA | NA |
| T1431 | CTCGTGACAGCAGAGAAGAGCGG | 1000 | | | | | | | | | |
| Oligo fwd | CACCCTCGTGACAGCAGAGAAGAG | | | | | | | | | | |
| Oligo rev | AAACCTCTTCTCTGCTGTCACGAG | | | | | | | | | | |
| Chromosome | start | end | strand | MM | target_seq | PAM | alignment | distance | position | gene name | gene id |
| GU305914.1 | 24740 | 24762 | + | 0 | CTCGTGACAGCAGAGAAGAG | CGG | \|\|\|\|\|\|\|\|[\|\|\|\|\|\|\|\|\|\|\|\|]PAM | NA | Intergenic | NA | NA |
| T1432 | GCAGAGAAGAGCGGCGAGTCTGG | 1000 | | | | | | | | | |
| Oligo fwd | CACCGCAGAGAAGAGCGGCGAGTC | | | | | | | | | | |
| Oligo rev | AAACGACTCGCCGCTCTTCTCTGC | | | | | | | | | | |
| Chromosome | start | end | strand | MM | target_seq | PAM | alignment | distance | position | gene name | gene id |
| GU305914.1 | 24749 | 24771 | + | 0 | GCAGAGAAGAGCGGCGAGTC | TGG | \|\|\|\|\|\|\|\|[\|\|\|\|\|\|\|\|\|\|\|\|]PAM | NA | Intergenic | NA | NA |
| T1433 | GGCGAGTCTGGCGCACTCGTTGG | 1000 | | | | | | | | | |
| Oligo fwd | CACCGGCGAGTCTGGCGCACTCGT | | | | | | | | | | |
| Oligo rev | AAACACGAGTGCGCCAGACTCGCC | | | | | | | | | | |
| Chromosome | start | end | strand | MM | target_seq | PAM | alignment | distance | position | gene name | gene id |
| GU305914.1 | 24761 | 24783 | + | 0 | GGCGAGTCTGGCGCACTCGT | TGG | \|\|\|\|\|\|\|\|[\|\|\|\|\|\|\|\|\|\|\|\|]PAM | NA | Intergenic | NA | NA |
| T1434 | GCGAGTCTGGCGCACTCGTTGGG | 1000 | | | | | | | | | |
| Oligo fwd | CACCGCGAGTCTGGCGCACTCGTT | | | | | | | | | | |
| Oligo rev | AAACAACGAGTGCGCCAGACTCGC | | | | | | | | | | |

Chromosome start end strand MM target_seq PAM alignment distance position gene name gene id
GU305914.1 24762 24784 + 0 GCGAGTCTGGCGCACTCGTT GGG ||||||||[||||||||||||]PAM NA Intergenic NA NA

T1435 AGTCTGGCGCACTCGTTGGGCGG 1000
Oligo fwd CACCAGTCTGGCGCACTCGTTGGG
Oligo rev AAACCCCAACGAGTGCGCCAGACT
Chromosome start end strand MM target_seq PAM alignment distance position gene name gene id
GU305914.1 24765 24787 + 0 AGTCTGGCGCACTCGTTGGG CGG ||||||||[||||||||||||]PAM NA Intergenic NA NA

T1436 CTCGTTGGGCGGCCCCCCGTTGG 1000
Oligo fwd CACCCTCGTTGGGCGGCCCCCCGT
Oligo rev AAACACGGGGGGCCGCCCAACGAG
Chromosome start end strand MM target_seq PAM alignment distance position gene name gene id
GU305914.1 24776 24798 + 0 CTCGTTGGGCGGCCCCCCGT TGG ||||||||[||||||||||||]PAM NA Intergenic NA NA

T1437 TTGGGCGGCCCCCCGTTGGCCGG 1000
Oligo fwd CACCTTGGGCGGCCCCCCGTTGGC
Oligo rev AAACGCCAACGGGGGGCCGCCCAA
Chromosome start end strand MM target_seq PAM alignment distance position gene name gene id
GU305914.1 24780 24802 + 0 TTGGGCGGCCCCCCGTTGGC CGG ||||||||[||||||||||||]PAM NA Intergenic NA NA

T1438 CGGCCCCCCGTTGGCCGGCGCGG 1000
Oligo fwd CACCCGGCCCCCCGTTGGCCGGCG
Oligo rev AAACCGCCGGCCAACGGGGGGCCG
Chromosome start end strand MM target_seq PAM alignment distance position gene name gene id
GU305914.1 24785 24807 + 0 CGGCCCCCCGTTGGCCGGCG CGG ||||||||[||||||||||||]PAM NA Intergenic NA NA

T1439 GGCCCCCCGTTGGCCGGCGCGGG 1000
Oligo fwd CACCGGCCCCCCGTTGGCCGGCGC
Oligo rev AAACGCGCCGGCCAACGGGGGGCC
Chromosome start end strand MM target_seq PAM alignment distance position gene name gene id
GU305914.1 24786 24808 + 0 GGCCCCCCGTTGGCCGGCGC GGG ||||||||[||||||||||||]PAM NA Intergenic NA NA

T1440 ATGCAGAGCGAGAGCCCTTTCGG 1000
Oligo fwd CACCATGCAGAGCGAGAGCCCTTT
Oligo rev AAACAAAGGGCTCTCGCTCTGCAT
Chromosome start end strand MM target_seq PAM alignment distance position gene name gene id
GU305914.1 24831 24853 + 0 ATGCAGAGCGAGAGCCCTTT CGG ||||||||[||||||||||||]PAM NA Intergenic NA NA

T1441 TGCAGAGCGAGAGCCCTTTCGGG 1000
Oligo fwd CACCTGCAGAGCGAGAGCCCTTTC
Oligo rev AAACGAAAGGGCTCTCGCTCTGCA
Chromosome start end strand MM target_seq PAM alignment distance position gene name gene id
GU305914.1 24832 24854 + 0 TGCAGAGCGAGAGCCCTTTC GGG ||||||||[||||||||||||]PAM NA Intergenic NA NA

T1442 AGAGCGAGAGCCCTTTCGGGTGG 1000
Oligo fwd CACCAGAGCGAGAGCCCTTTCGGG
Oligo rev AAACCCCGAAAGGGCTCTCGCTCT
Chromosome start end strand MM target_seq PAM alignment distance position gene name gene id
GU305914.1 24835 24857 + 0 AGAGCGAGAGCCCTTTCGGG TGG ||||||||[||||||||||||]PAM NA Intergenic NA NA

T1443 GCGAGAGCCCTTTCGGGTGGTGG 1000
Oligo fwd CACCGCGAGAGCCCTTTCGGGTGG
Oligo rev AAACCCACCCGAAAGGGCTCTCGC
Chromosome start end strand MM target_seq PAM alignment distance position gene name gene id
GU305914.1 24838 24860 + 0 GCGAGAGCCCTTTCGGGTGG TGG ||||||||[||||||||||||]PAM NA Intergenic NA NA

T1444 TCGGGTGGTGGAGACGCACGCGG 1000
Oligo fwd CACCTCGGGTGGTGGAGACGCACG
Oligo rev AAACCGTGCGTCTCCACCACCCGA
Chromosome start end strand MM target_seq PAM alignment distance position gene name gene id
GU305914.1 24850 24872 + 0 TCGGGTGGTGGAGACGCACG CGG ||||||||[||||||||||||]PAM NA Intergenic NA NA

T1445 ACGCACGCGGCGCCCGATGTTGG 1000
Oligo fwd CACCACGCACGCGGCGCCCGATGT
Oligo rev AAACACATCGGGCGCCGCGTGCGT
Chromosome start end strand MM target_seq PAM alignment distance position gene name gene id
GU305914.1 24863 24885 + 0 ACGCACGCGGCGCCCGATGT TGG ||||||||[||||||||||||]PAM NA Intergenic NA NA

T1446 CACGCGGCGCCCGATGTTGGCGG 1000
Oligo fwd CACCCACGCGGCGCCCGATGTTGG
Oligo rev AAACCCAACATCGGGCGCCGCGTG
Chromosome start end strand MM target_seq PAM alignment distance position gene name gene id
GU305914.1 24866 24888 + 0 CACGCGGCGCCCGATGTTGG CGG ||||||||[||||||||||||]PAM NA Intergenic NA NA

T1447 GCGCCCGATGTTGGCGGCGACGG 1000
Oligo fwd CACCGCGCCCGATGTTGGCGGCGA
Oligo rev AAACTCGCCGCCAACATCGGGCGC
Chromosome start end strand MM target_seq PAM alignment distance position gene name gene id
GU305914.1 24872 24894 + 0 GCGCCCGATGTTGGCGGCGA CGG ||||||||[||||||||||||]PAM NA Intergenic NA NA
GU305914.1 170536 170558 - 4 GTGGCCGATGGTGGCGGCGG CGG |-|-||||[||-||||||||-]PAM NA Intergenic NA NA

T1448 CCCGATGTTGGCGGCGACGGCGG 1000
Oligo fwd CACCCCCGATGTTGGCGGCGACGG
Oligo rev AAACCCGTCGCCGCCAACATCGGG
Chromosome start end strand MM target_seq PAM alignment distance position gene name gene id
GU305914.1 24875 24897 + 0 CCCGATGTTGGCGGCGACGG CGG ||||||||[||||||||||||]PAM NA Intergenic NA NA
GU305914.1 29997 30019 + 4 ACCGGTGCTGGCGGCGGCGG CGG -|||-||-[||||||||-|||]PAM NA Intergenic NA NA

<table>
<tr><td>T1449</td><td>GGCGGCGACGGCGGCCTCGACGG</td><td>1000</td><td></td><td></td><td></td><td></td><td></td><td></td><td></td><td></td><td></td></tr>
<tr><td>Oligo fwd</td><td>CACCGGCGGCGACGGCGGCCTCGA</td><td></td><td></td><td></td><td></td><td></td><td></td><td></td><td></td><td></td><td></td></tr>
<tr><td>Oligo rev</td><td>AAACTCGAGGCCGCCGTCGCCGCC</td><td></td><td></td><td></td><td></td><td></td><td></td><td></td><td></td><td></td><td></td></tr>
<tr><td>Chromosome</td><td>start</td><td>end</td><td>strand</td><td>MM</td><td>target_seq</td><td>PAM</td><td>alignment</td><td>distance</td><td>position</td><td>gene name</td><td>gene id</td></tr>
<tr><td>GU305914.1</td><td>24884</td><td>24906</td><td>+</td><td>0</td><td>GGCGGCGACGGCGGCCTCGA</td><td>CGG</td><td>||||||||[||||||||||||]PAM</td><td>NA</td><td>Intergenic</td><td>NA</td><td>NA</td></tr>
<tr><td>GU305914.1</td><td>24134</td><td>24156</td><td>+</td><td>3</td><td>GGCGCCGACGTCGGCCTCGG</td><td>CGG</td><td>||||-|||[||-||||||||-]PAM</td><td>NA</td><td>Intergenic</td><td>NA</td><td>NA</td></tr>
<tr><td>GU305914.1</td><td>110125</td><td>110147</td><td>-</td><td>4</td><td>GGACGCGACGGCGGCCGCGG</td><td>CGG</td><td>||--||||[||||||||-||-]PAM</td><td>NA</td><td>Intergenic</td><td>NA</td><td>NA</td></tr>
<tr><td></td><td></td><td></td><td></td><td></td><td></td><td></td><td></td><td></td><td></td><td></td><td></td></tr>
<tr><td>T1450</td><td>GGCGACGGCGGCCTCGACGGTGG</td><td>1000</td><td></td><td></td><td></td><td></td><td></td><td></td><td></td><td></td><td></td></tr>
<tr><td>Oligo fwd</td><td>CACCGGCGACGGCGGCCTCGACGG</td><td></td><td></td><td></td><td></td><td></td><td></td><td></td><td></td><td></td><td></td></tr>
<tr><td>Oligo rev</td><td>AAACCCGTCGAGGCCGCCGTCGCC</td><td></td><td></td><td></td><td></td><td></td><td></td><td></td><td></td><td></td><td></td></tr>
<tr><td>Chromosome</td><td>start</td><td>end</td><td>strand</td><td>MM</td><td>target_seq</td><td>PAM</td><td>alignment</td><td>distance</td><td>position</td><td>gene name</td><td>gene id</td></tr>
<tr><td>GU305914.1</td><td>24887</td><td>24909</td><td>+</td><td>0</td><td>GGCGACGGCGGCCTCGACGG</td><td>TGG</td><td>||||||||[||||||||||||]PAM</td><td>NA</td><td>Intergenic</td><td>NA</td><td>NA</td></tr>
<tr><td></td><td></td><td></td><td></td><td></td><td></td><td></td><td></td><td></td><td></td><td></td><td></td></tr>
<tr><td>T1451</td><td>AAGATGCTCTCGAGAGCTTCCGG</td><td>1000</td><td></td><td></td><td></td><td></td><td></td><td></td><td></td><td></td><td></td></tr>
<tr><td>Oligo fwd</td><td>CACCAAGATGCTCTCGAGAGCTTC</td><td></td><td></td><td></td><td></td><td></td><td></td><td></td><td></td><td></td><td></td></tr>
<tr><td>Oligo rev</td><td>AAACGAAGCTCTCGAGAGCATCTT</td><td></td><td></td><td></td><td></td><td></td><td></td><td></td><td></td><td></td><td></td></tr>
<tr><td>Chromosome</td><td>start</td><td>end</td><td>strand</td><td>MM</td><td>target_seq</td><td>PAM</td><td>alignment</td><td>distance</td><td>position</td><td>gene name</td><td>gene id</td></tr>
<tr><td>GU305914.1</td><td>24921</td><td>24943</td><td>+</td><td>0</td><td>AAGATGCTCTCGAGAGCTTC</td><td>CGG</td><td>||||||||[||||||||||||]PAM</td><td>NA</td><td>Intergenic</td><td>NA</td><td>NA</td></tr>
<tr><td></td><td></td><td></td><td></td><td></td><td></td><td></td><td></td><td></td><td></td><td></td><td></td></tr>
<tr><td>T1452</td><td>ATGCTCTCGAGAGCTTCCGGAGG</td><td>1000</td><td></td><td></td><td></td><td></td><td></td><td></td><td></td><td></td><td></td></tr>
<tr><td>Oligo fwd</td><td>CACCATGCTCTCGAGAGCTTCCGG</td><td></td><td></td><td></td><td></td><td></td><td></td><td></td><td></td><td></td><td></td></tr>
<tr><td>Oligo rev</td><td>AAACCCGGAAGCTCTCGAGAGCAT</td><td></td><td></td><td></td><td></td><td></td><td></td><td></td><td></td><td></td><td></td></tr>
<tr><td>Chromosome</td><td>start</td><td>end</td><td>strand</td><td>MM</td><td>target_seq</td><td>PAM</td><td>alignment</td><td>distance</td><td>position</td><td>gene name</td><td>gene id</td></tr>
<tr><td>GU305914.1</td><td>24924</td><td>24946</td><td>+</td><td>0</td><td>ATGCTCTCGAGAGCTTCCGG</td><td>AGG</td><td>||||||||[||||||||||||]PAM</td><td>NA</td><td>Intergenic</td><td>NA</td><td>NA</td></tr>
<tr><td></td><td></td><td></td><td></td><td></td><td></td><td></td><td></td><td></td><td></td><td></td><td></td></tr>
<tr><td>T1453</td><td>GTTTTTCGAGCTCGAGACGATGG</td><td>1000</td><td></td><td></td><td></td><td></td><td></td><td></td><td></td><td></td><td></td></tr>
<tr><td>Oligo fwd</td><td>CACCGTTTTTCGAGCTCGAGACGA</td><td></td><td></td><td></td><td></td><td></td><td></td><td></td><td></td><td></td><td></td></tr>
<tr><td>Oligo rev</td><td>AAACTCGTCTCGAGCTCGAAAAAC</td><td></td><td></td><td></td><td></td><td></td><td></td><td></td><td></td><td></td><td></td></tr>
<tr><td>Chromosome</td><td>start</td><td>end</td><td>strand</td><td>MM</td><td>target_seq</td><td>PAM</td><td>alignment</td><td>distance</td><td>position</td><td>gene name</td><td>gene id</td></tr>
<tr><td>GU305914.1</td><td>24970</td><td>24992</td><td>+</td><td>0</td><td>GTTTTTCGAGCTCGAGACGA</td><td>TGG</td><td>||||||||[||||||||||||]PAM</td><td>NA</td><td>Intergenic</td><td>NA</td><td>NA</td></tr>
<tr><td></td><td></td><td></td><td></td><td></td><td></td><td></td><td></td><td></td><td></td><td></td><td></td></tr>
<tr><td>T1454</td><td>TTTTTCGAGCTCGAGACGATGGG</td><td>1000</td><td></td><td></td><td></td><td></td><td></td><td></td><td></td><td></td><td></td></tr>
<tr><td>Oligo fwd</td><td>CACCTTTTTCGAGCTCGAGACGAT</td><td></td><td></td><td></td><td></td><td></td><td></td><td></td><td></td><td></td><td></td></tr>
<tr><td>Oligo rev</td><td>AAACATCGTCTCGAGCTCGAAAAA</td><td></td><td></td><td></td><td></td><td></td><td></td><td></td><td></td><td></td><td></td></tr>
<tr><td>Chromosome</td><td>start</td><td>end</td><td>strand</td><td>MM</td><td>target_seq</td><td>PAM</td><td>alignment</td><td>distance</td><td>position</td><td>gene name</td><td>gene id</td></tr>
<tr><td>GU305914.1</td><td>24971</td><td>24993</td><td>+</td><td>0</td><td>TTTTTCGAGCTCGAGACGAT</td><td>GGG</td><td>||||||||[||||||||||||]PAM</td><td>NA</td><td>Intergenic</td><td>NA</td><td>NA</td></tr>
<tr><td></td><td></td><td></td><td></td><td></td><td></td><td></td><td></td><td></td><td></td><td></td><td></td></tr>
<tr><td>T1455</td><td>TCGAGCTCGAGACGATGGGTCGG</td><td>1000</td><td></td><td></td><td></td><td></td><td></td><td></td><td></td><td></td><td></td></tr>
<tr><td>Oligo fwd</td><td>CACCTCGAGCTCGAGACGATGGGT</td><td></td><td></td><td></td><td></td><td></td><td></td><td></td><td></td><td></td><td></td></tr>
<tr><td>Oligo rev</td><td>AAACACCCATCGTCTCGAGCTCGA</td><td></td><td></td><td></td><td></td><td></td><td></td><td></td><td></td><td></td><td></td></tr>
<tr><td>Chromosome</td><td>start</td><td>end</td><td>strand</td><td>MM</td><td>target_seq</td><td>PAM</td><td>alignment</td><td>distance</td><td>position</td><td>gene name</td><td>gene id</td></tr>
<tr><td>GU305914.1</td><td>24975</td><td>24997</td><td>+</td><td>0</td><td>TCGAGCTCGAGACGATGGGT</td><td>CGG</td><td>||||||||[||||||||||||]PAM</td><td>NA</td><td>Intergenic</td><td>NA</td><td>NA</td></tr>
<tr><td></td><td></td><td></td><td></td><td></td><td></td><td></td><td></td><td></td><td></td><td></td><td></td></tr>
<tr><td>T1456</td><td>TCGAGACGATGGGTCGGTCGTGG</td><td>1000</td><td></td><td></td><td></td><td></td><td></td><td></td><td></td><td></td><td></td></tr>
<tr><td>Oligo fwd</td><td>CACCTCGAGACGATGGGTCGGTCG</td><td></td><td></td><td></td><td></td><td></td><td></td><td></td><td></td><td></td><td></td></tr>
<tr><td>Oligo rev</td><td>AAACCGACCGACCCATCGTCTCGA</td><td></td><td></td><td></td><td></td><td></td><td></td><td></td><td></td><td></td><td></td></tr>
<tr><td>Chromosome</td><td>start</td><td>end</td><td>strand</td><td>MM</td><td>target_seq</td><td>PAM</td><td>alignment</td><td>distance</td><td>position</td><td>gene name</td><td>gene id</td></tr>
<tr><td>GU305914.1</td><td>24981</td><td>25003</td><td>+</td><td>0</td><td>TCGAGACGATGGGTCGGTCG</td><td>TGG</td><td>||||||||[||||||||||||]PAM</td><td>NA</td><td>Intergenic</td><td>NA</td><td>NA</td></tr>
<tr><td></td><td></td><td></td><td></td><td></td><td></td><td></td><td></td><td></td><td></td><td></td><td></td></tr>
<tr><td>T1457</td><td>CGAGACGATGGGTCGGTCGTGGG</td><td>1000</td><td></td><td></td><td></td><td></td><td></td><td></td><td></td><td></td><td></td></tr>
<tr><td>Oligo fwd</td><td>CACCCGAGACGATGGGTCGGTCGT</td><td></td><td></td><td></td><td></td><td></td><td></td><td></td><td></td><td></td><td></td></tr>
<tr><td>Oligo rev</td><td>AAACACGACCGACCCATCGTCTCG</td><td></td><td></td><td></td><td></td><td></td><td></td><td></td><td></td><td></td><td></td></tr>
<tr><td>Chromosome</td><td>start</td><td>end</td><td>strand</td><td>MM</td><td>target_seq</td><td>PAM</td><td>alignment</td><td>distance</td><td>position</td><td>gene name</td><td>gene id</td></tr>
<tr><td>GU305914.1</td><td>24982</td><td>25004</td><td>+</td><td>0</td><td>CGAGACGATGGGTCGGTCGT</td><td>GGG</td><td>||||||||[||||||||||||]PAM</td><td>NA</td><td>Intergenic</td><td>NA</td><td>NA</td></tr>
<tr><td>GU305914.1</td><td>213545</td><td>213567</td><td>+</td><td>4</td><td>GTGGACGATGGTTCGGTCGT</td><td>CGG</td><td>---|||||[|||-||||||||]PAM</td><td>NA</td><td>Intergenic</td><td>NA</td><td>NA</td></tr>
<tr><td></td><td></td><td></td><td></td><td></td><td></td><td></td><td></td><td></td><td></td><td></td><td></td></tr>
<tr><td>T1458</td><td>GACGATGGGTCGGTCGTGGGTGG</td><td>1000</td><td></td><td></td><td></td><td></td><td></td><td></td><td></td><td></td><td></td></tr>
<tr><td>Oligo fwd</td><td>CACCGACGATGGGTCGGTCGTGGG</td><td></td><td></td><td></td><td></td><td></td><td></td><td></td><td></td><td></td><td></td></tr>
<tr><td>Oligo rev</td><td>AAACCCCACGACCGACCCATCGTC</td><td></td><td></td><td></td><td></td><td></td><td></td><td></td><td></td><td></td><td></td></tr>
<tr><td>Chromosome</td><td>start</td><td>end</td><td>strand</td><td>MM</td><td>target_seq</td><td>PAM</td><td>alignment</td><td>distance</td><td>position</td><td>gene name</td><td>gene id</td></tr>
<tr><td>GU305914.1</td><td>24985</td><td>25007</td><td>+</td><td>0</td><td>GACGATGGGTCGGTCGTGGG</td><td>TGG</td><td>||||||||[||||||||||||]PAM</td><td>NA</td><td>Intergenic</td><td>NA</td><td>NA</td></tr>
<tr><td></td><td></td><td></td><td></td><td></td><td></td><td></td><td></td><td></td><td></td><td></td><td></td></tr>
<tr><td>T1459</td><td>ACGATGGGTCGGTCGTGGGTGGG</td><td>1000</td><td></td><td></td><td></td><td></td><td></td><td></td><td></td><td></td><td></td></tr>
<tr><td>Oligo fwd</td><td>CACCACGATGGGTCGGTCGTGGGT</td><td></td><td></td><td></td><td></td><td></td><td></td><td></td><td></td><td></td><td></td></tr>
<tr><td>Oligo rev</td><td>AAACACCCACGACCGACCCATCGT</td><td></td><td></td><td></td><td></td><td></td><td></td><td></td><td></td><td></td><td></td></tr>
<tr><td>Chromosome</td><td>start</td><td>end</td><td>strand</td><td>MM</td><td>target_seq</td><td>PAM</td><td>alignment</td><td>distance</td><td>position</td><td>gene name</td><td>gene id</td></tr>
<tr><td>GU305914.1</td><td>24986</td><td>25008</td><td>+</td><td>0</td><td>ACGATGGGTCGGTCGTGGGT</td><td>GGG</td><td>||||||||[||||||||||||]PAM</td><td>NA</td><td>Intergenic</td><td>NA</td><td>NA</td></tr>
<tr><td></td><td></td><td></td><td></td><td></td><td></td><td></td><td></td><td></td><td></td><td></td><td></td></tr>
<tr><td>T1460</td><td>GGTCGTGGGTGGGCGTCCCTCGG</td><td>1000</td><td></td><td></td><td></td><td></td><td></td><td></td><td></td><td></td><td></td></tr>
<tr><td>Oligo fwd</td><td>CACCGGTCGTGGGTGGGCGTCCCT</td><td></td><td></td><td></td><td></td><td></td><td></td><td></td><td></td><td></td><td></td></tr>
<tr><td>Oligo rev</td><td>AAACAGGGACGCCCACCCACGACC</td><td></td><td></td><td></td><td></td><td></td><td></td><td></td><td></td><td></td><td></td></tr>
<tr><td>Chromosome</td><td>start</td><td>end</td><td>strand</td><td>MM</td><td>target_seq</td><td>PAM</td><td>alignment</td><td>distance</td><td>position</td><td>gene name</td><td>gene id</td></tr>
<tr><td>GU305914.1</td><td>24996</td><td>25018</td><td>+</td><td>0</td><td>GGTCGTGGGTGGGCGTCCCT</td><td>CGG</td><td>||||||||[||||||||||||]PAM</td><td>NA</td><td>Intergenic</td><td>NA</td><td>NA</td></tr>
<tr><td></td><td></td><td></td><td></td><td></td><td></td><td></td><td></td><td></td><td></td><td></td><td></td></tr>
<tr><td>T1461</td><td>GTCCCTCGGCCTTTTGTGTGTGG</td><td>1000</td><td></td><td></td><td></td><td></td><td></td><td></td><td></td><td></td><td></td></tr>
<tr><td>Oligo fwd</td><td>CACCGTCCCTCGGCCTTTTGTGTG</td><td></td><td></td><td></td><td></td><td></td><td></td><td></td><td></td><td></td><td></td></tr>
<tr><td>Oligo rev</td><td>AAACCACACAAAAGGCCGAGGGAC</td><td></td><td></td><td></td><td></td><td></td><td></td><td></td><td></td><td></td><td></td></tr>
<tr><td>Chromosome</td><td>start</td><td>end</td><td>strand</td><td>MM</td><td>target_seq</td><td>PAM</td><td>alignment</td><td>distance</td><td>position</td><td>gene name</td><td>gene id</td></tr>
<tr><td>GU305914.1</td><td>25010</td><td>25032</td><td>+</td><td>0</td><td>GTCCCTCGGCCTTTTGTGTG</td><td>TGG</td><td>||||||||[||||||||||||]PAM</td><td>NA</td><td>Intergenic</td><td>NA</td><td>NA</td></tr>
<tr><td></td><td></td><td></td><td></td><td></td><td></td><td></td><td></td><td></td><td></td><td></td><td></td></tr>
<tr><td>T1462</td><td>TCCCTCGGCCTTTTGTGTGTGGG</td><td>1000</td><td></td><td></td><td></td><td></td><td></td><td></td><td></td><td></td><td></td></tr>
<tr><td>Oligo fwd</td><td>CACCTCCCTCGGCCTTTTGTGTGT</td><td></td><td></td><td></td><td></td><td></td><td></td><td></td><td></td><td></td><td></td></tr>
<tr><td>Oligo rev</td><td>AAACACACACAAAAGGCCGAGGGA</td><td></td><td></td><td></td><td></td><td></td><td></td><td></td><td></td><td></td><td></td></tr>
<tr><td>Chromosome</td><td>start</td><td>end</td><td>strand</td><td>MM</td><td>target_seq</td><td>PAM</td><td>alignment</td><td>distance</td><td>position</td><td>gene name</td><td>gene id</td></tr>
<tr><td>GU305914.1</td><td>25011</td><td>25033</td><td>+</td><td>0</td><td>TCCCTCGGCCTTTTGTGTGT</td><td>GGG</td><td>||||||||[||||||||||||]PAM</td><td>NA</td><td>Intergenic</td><td>NA</td><td>NA</td></tr>
<tr><td></td><td></td><td></td><td></td><td></td><td></td><td></td><td></td><td></td><td></td><td></td><td></td></tr>
<tr><td>T1463</td><td>CTTTTGTGTGTGGGCGTCCGTGG</td><td>1000</td><td></td><td></td><td></td><td></td><td></td><td></td><td></td><td></td><td></td></tr>
<tr><td>Oligo fwd</td><td>CACCCTTTTGTGTGTGGGCGTCCG</td><td></td><td></td><td></td><td></td><td></td><td></td><td></td><td></td><td></td><td></td></tr>
</table>

Oligo rev AAACCGGACGCCCACACACAAAAG
Chromosome start end strand MM target_seq PAM alignment distance position gene name gene id
GU305914.1 25020 25042 + 0 CTTTTGTGTGTGGGCGTCCG TGG ||||||||[||||||||||||]PAM NA Intergenic NA NA

T1464 TTTTGTGTGTGGGCGTCCGTGGG 1000
Oligo fwd CACCTTTTGTGTGTGGGCGTCCGT
Oligo rev AAACACGGACGCCCACACACAAAA
Chromosome start end strand MM target_seq PAM alignment distance position gene name gene id
GU305914.1 25021 25043 + 0 TTTTGTGTGTGGGCGTCCGT GGG ||||||||[||||||||||||]PAM NA Intergenic NA NA

T1465 TGTGTGGGCGTCCGTGGGCGCGG 1000
Oligo fwd CACCTGTGTGGGCGTCCGTGGGCG
Oligo rev AAACCGCCCACGGACGCCCACACA
Chromosome start end strand MM target_seq PAM alignment distance position gene name gene id
GU305914.1 25026 25048 + 0 TGTGTGGGCGTCCGTGGGCG CGG ||||||||[||||||||||||]PAM NA Intergenic NA NA

T1466 GGGCGTCCGTGGGCGCGGCAAGG 1000
Oligo fwd CACCGGGCGTCCGTGGGCGCGGCA
Oligo rev AAACTGCCGCGCCCACGGACGCCC
Chromosome start end strand MM target_seq PAM alignment distance position gene name gene id
GU305914.1 25031 25053 + 0 GGGCGTCCGTGGGCGCGGCA AGG ||||||||[||||||||||||]PAM NA Intergenic NA NA

T1467 GGCGTCCGTGGGCGCGGCAAGGG 1000
Oligo fwd CACCGGCGTCCGTGGGCGCGGCAA
Oligo rev AAACTTGCCGCGCCCACGGACGCC
Chromosome start end strand MM target_seq PAM alignment distance position gene name gene id
GU305914.1 25032 25054 + 0 GGCGTCCGTGGGCGCGGCAA GGG ||||||||[||||||||||||]PAM NA Intergenic NA NA

T1468 GCGTCCGTGGGCGCGGCAAGGGG 1000
Oligo fwd CACCGCGTCCGTGGGCGCGGCAAG
Oligo rev AAACCTTGCCGCGCCCACGGACGC
Chromosome start end strand MM target_seq PAM alignment distance position gene name gene id
GU305914.1 25033 25055 + 0 GCGTCCGTGGGCGCGGCAAG GGG ||||||||[||||||||||||]PAM NA Intergenic NA NA

T1469 ATCGACCGATCGACAGAGAAAGG 1000
Oligo fwd CACCATCGACCGATCGACAGAGAA
Oligo rev AAACTTCTCTGTCGATCGGTCGAT
Chromosome start end strand MM target_seq PAM alignment distance position gene name gene id
GU305914.1 25101 25123 + 0 ATCGACCGATCGACAGAGAA AGG ||||||||[||||||||||||]PAM NA Intergenic NA NA

T1470 GACCGATCGACAGAGAAAGGAGG 1000
Oligo fwd CACCGACCGATCGACAGAGAAAGG
Oligo rev AAACCCTTTCTCTGTCGATCGGTC
Chromosome start end strand MM target_seq PAM alignment distance position gene name gene id
GU305914.1 25104 25126 + 0 GACCGATCGACAGAGAAAGG AGG ||||||||[||||||||||||]PAM NA Intergenic NA NA

T1471 GACAGAGAAAGGAGGAAAAGTGG 1000
Oligo fwd CACCGACAGAGAAAGGAGGAAAAG
Oligo rev AAACCTTTTCCTCCTTTCTCTGTC
Chromosome start end strand MM target_seq PAM alignment distance position gene name gene id
GU305914.1 25112 25134 + 0 GACAGAGAAAGGAGGAAAAG TGG ||||||||[||||||||||||]PAM NA Intergenic NA NA

T1472 ACAGAGAAAGGAGGAAAAGTGGG 1000
Oligo fwd CACCACAGAGAAAGGAGGAAAAGT
Oligo rev AAACACTTTTCCTCCTTTCTCTGT
Chromosome start end strand MM target_seq PAM alignment distance position gene name gene id
GU305914.1 25113 25135 + 0 ACAGAGAAAGGAGGAAAAGT GGG ||||||||[||||||||||||]PAM NA Intergenic NA NA

T1473 AAAGGAGGAAAAGTGGGTCACGG 1000
Oligo fwd CACCAAAGGAGGAAAAGTGGGTCA
Oligo rev AAACTGACCCACTTTTCCTCCTTT
Chromosome start end strand MM target_seq PAM alignment distance position gene name gene id
GU305914.1 25119 25141 + 0 AAAGGAGGAAAAGTGGGTCA CGG ||||||||[||||||||||||]PAM NA Intergenic NA NA

T1474 GGAAAGACTAGAACTCGCTGAGG 1000
Oligo fwd CACCGGAAAGACTAGAACTCGCTG
Oligo rev AAACCAGCGAGTTCTAGTCTTTCC
Chromosome start end strand MM target_seq PAM alignment distance position gene name gene id
GU305914.1 25140 25162 + 0 GGAAAGACTAGAACTCGCTG AGG ||||||||[||||||||||||]PAM NA Intergenic NA NA

T1475 GACTAGAACTCGCTGAGGCGCGG 1000
Oligo fwd CACCGACTAGAACTCGCTGAGGCG
Oligo rev AAACCGCCTCAGCGAGTTCTAGTC
Chromosome start end strand MM target_seq PAM alignment distance position gene name gene id
GU305914.1 25145 25167 + 0 GACTAGAACTCGCTGAGGCG CGG ||||||||[||||||||||||]PAM NA Intergenic NA NA

T1476 ACTAGAACTCGCTGAGGCGCGGG 1000
Oligo fwd CACCACTAGAACTCGCTGAGGCGC
Oligo rev AAACGCGCCTCAGCGAGTTCTAGT
Chromosome start end strand MM target_seq PAM alignment distance position gene name gene id
GU305914.1 25146 25168 + 0 ACTAGAACTCGCTGAGGCGC GGG ||||||||[||||||||||||]PAM NA Intergenic NA NA

T1477 CTAGAACTCGCTGAGGCGCGGGG 1000
Oligo fwd CACCCTAGAACTCGCTGAGGCGCG
Oligo rev AAACCGCGCCTCAGCGAGTTCTAG
Chromosome start end strand MM target_seq PAM alignment distance position gene name gene id
GU305914.1 25147 25169 + 0 CTAGAACTCGCTGAGGCGCG GGG ||||||||[||||||||||||]PAM NA Intergenic NA NA

T1478 TAGAACTCGCTGAGGCGCGGGGG 1000

Oligo fwd CACCTAGAACTCGCTGAGGCGCGG
Oligo rev AAACCCGCGCCTCAGCGAGTTCTA
Chromosome start end strand MM target_seq PAM alignment distance position gene name gene id
GU305914.1 25148 25170 + 0 TAGAACTCGCTGAGGCGCGG GGG ||||||||[||||||||||||]PAM NA Intergenic NA NA

T1479 AACTCGCTGAGGCGCGGGGGAGG 1000
Oligo fwd CACCAACTCGCTGAGGCGCGGGGG
Oligo rev AAACCCCCCGCGCCTCAGCGAGTT
Chromosome start end strand MM target_seq PAM alignment distance position gene name gene id
GU305914.1 25151 25173 + 0 AACTCGCTGAGGCGCGGGGG AGG ||||||||[||||||||||||]PAM NA Intergenic NA NA

T1480 AGGCCTCTCACAGTTCCTCCCGG 1000
Oligo fwd CACCAGGCCTCTCACAGTTCCTCC
Oligo rev AAACGGAGGAACTGTGAGAGGCCT
Chromosome start end strand MM target_seq PAM alignment distance position gene name gene id
GU305914.1 25171 25193 + 0 AGGCCTCTCACAGTTCCTCC CGG ||||||||[||||||||||||]PAM NA Intergenic NA NA

T1481 CGGAGTCCCGCTCGCTGATGAGG 1000
Oligo fwd CACCCGGAGTCCCGCTCGCTGATG
Oligo rev AAACCATCAGCGAGCGGGACTCCG
Chromosome start end strand MM target_seq PAM alignment distance position gene name gene id
GU305914.1 25191 25213 + 0 CGGAGTCCCGCTCGCTGATG AGG ||||||||[||||||||||||]PAM NA Intergenic NA NA

T1482 ACGTCGATTCTCAAAGTACCTGG 1000
Oligo fwd CACCACGTCGATTCTCAAAGTACC
Oligo rev AAACGGTACTTTGAGAATCGACGT
Chromosome start end strand MM target_seq PAM alignment distance position gene name gene id
GU305914.1 25219 25241 + 0 ACGTCGATTCTCAAAGTACC TGG ||||||||[||||||||||||]PAM NA Intergenic NA NA

T1483 TCGATTCTCAAAGTACCTGGTGG 1000
Oligo fwd CACCTCGATTCTCAAAGTACCTGG
Oligo rev AAACCCAGGTACTTTGAGAATCGA
Chromosome start end strand MM target_seq PAM alignment distance position gene name gene id
GU305914.1 25222 25244 + 0 TCGATTCTCAAAGTACCTGG TGG ||||||||[||||||||||||]PAM NA Intergenic NA NA

T1484 GCGCCGATGCCGACGAAGCCCGG 1000
Oligo fwd CACCGCGCCGATGCCGACGAAGCC
Oligo rev AAACGGCTTCGTCGGCATCGGCGC
Chromosome start end strand MM target_seq PAM alignment distance position gene name gene id
GU305914.1 25244 25266 + 0 GCGCCGATGCCGACGAAGCC CGG ||||||||[||||||||||||]PAM NA Intergenic NA NA

T1485 CGCCGATGCCGACGAAGCCCGGG 1000
Oligo fwd CACCCGCCGATGCCGACGAAGCCC
Oligo rev AAACGGGCTTCGTCGGCATCGGCG
Chromosome start end strand MM target_seq PAM alignment distance position gene name gene id
GU305914.1 25245 25267 + 0 CGCCGATGCCGACGAAGCCC GGG ||||||||[||||||||||||]PAM NA Intergenic NA NA

T1486 GACGAAGCCCGGGATCGAGTCGG 1000
Oligo fwd CACCGACGAAGCCCGGGATCGAGT
Oligo rev AAACACTCGATCCCGGGCTTCGTC
Chromosome start end strand MM target_seq PAM alignment distance position gene name gene id
GU305914.1 25255 25277 + 0 GACGAAGCCCGGGATCGAGT CGG ||||||||[||||||||||||]PAM NA Intergenic NA NA

T1487 GAGATAGTAAGAGCTGTCCCCGG 1000
Oligo fwd CACCGAGATAGTAAGAGCTGTCCC
Oligo rev AAACGGGACAGCTCTTACTATCTC
Chromosome start end strand MM target_seq PAM alignment distance position gene name gene id
GU305914.1 25282 25304 + 0 GAGATAGTAAGAGCTGTCCC CGG ||||||||[||||||||||||]PAM NA Intergenic NA NA

T1488 AAGAGCTGTCCCCGGCCTGTCGG 1000
Oligo fwd CACCAAGAGCTGTCCCCGGCCTGT
Oligo rev AAACACAGGCCGGGGACAGCTCTT
Chromosome start end strand MM target_seq PAM alignment distance position gene name gene id
GU305914.1 25290 25312 + 0 AAGAGCTGTCCCCGGCCTGT CGG ||||||||[||||||||||||]PAM NA Intergenic NA NA

T1489 AGAGCTGTCCCCGGCCTGTCGGG 1000
Oligo fwd CACCAGAGCTGTCCCCGGCCTGTC
Oligo rev AAACGACAGGCCGGGGACAGCTCT
Chromosome start end strand MM target_seq PAM alignment distance position gene name gene id
GU305914.1 25291 25313 + 0 AGAGCTGTCCCCGGCCTGTC GGG ||||||||[||||||||||||]PAM NA Intergenic NA NA

T1490 CTGTCGGGCGTACACCCTACCGG 1000
Oligo fwd CACCCTGTCGGGCGTACACCCTAC
Oligo rev AAACGTAGGGTGTACGCCCGACAG
Chromosome start end strand MM target_seq PAM alignment distance position gene name gene id
GU305914.1 25306 25328 + 0 CTGTCGGGCGTACACCCTAC CGG ||||||||[||||||||||||]PAM NA Intergenic NA NA

T1491 TCGGGCGTACACCCTACCGGAGG 1000
Oligo fwd CACCTCGGGCGTACACCCTACCGG
Oligo rev AAACCCGGTAGGGTGTACGCCCGA
Chromosome start end strand MM target_seq PAM alignment distance position gene name gene id
GU305914.1 25309 25331 + 0 TCGGGCGTACACCCTACCGG AGG ||||||||[||||||||||||]PAM NA Intergenic NA NA

T1492 CGGGCGTACACCCTACCGGAGGG 1000
Oligo fwd CACCCGGGCGTACACCCTACCGGA
Oligo rev AAACTCCGGTAGGGTGTACGCCCG
Chromosome start end strand MM target_seq PAM alignment distance position gene name gene id
GU305914.1 25310 25332 + 0 CGGGCGTACACCCTACCGGA GGG ||||||||[||||||||||||]PAM NA Intergenic NA NA

| | | | | | | | | | | | | |
|---|---|---|---|---|---|---|---|---|---|---|---|---|
| T1493 | CCCACGAACACAGCGATCCCAGG | | 1000 | | | | | | | | | |
| Oligo fwd | CACCCCCACGAACACAGCGATCCC | | | | | | | | | | | |
| Oligo rev | AAACGGGATCGCTGTGTTCGTGGG | | | | | | | | | | | |
| Chromosome | start | | end | strand | MM | target_seq | PAM | alignment | distance | position | gene name | gene id |
| GU305914.1 | | 25337 | 25359 | + | 0 | CCCACGAACACAGCGATCCC | AGG | \|\|\|\|\|\|\|\|[\|\|\|\|\|\|\|\|\|\|\|\|]PAM | NA | Intergenic | NA | NA |
| T1494 | CCACGAACACAGCGATCCCAGGG | | 1000 | | | | | | | | | |
| Oligo fwd | CACCCCACGAACACAGCGATCCCA | | | | | | | | | | | |
| Oligo rev | AAACTGGGATCGCTGTGTTCGTGG | | | | | | | | | | | |
| Chromosome | start | | end | strand | MM | target_seq | PAM | alignment | distance | position | gene name | gene id |
| GU305914.1 | | 25338 | 25360 | + | 0 | CCACGAACACAGCGATCCCA | GGG | \|\|\|\|\|\|\|\|[\|\|\|\|\|\|\|\|\|\|\|\|]PAM | NA | Intergenic | NA | NA |
| T1495 | ACAGCGATCCCAGGGTCCTGCGG | | 1000 | | | | | | | | | |
| Oligo fwd | CACCACAGCGATCCCAGGGTCCTG | | | | | | | | | | | |
| Oligo rev | AAACCAGGACCCTGGGATCGCTGT | | | | | | | | | | | |
| Chromosome | start | | end | strand | MM | target_seq | PAM | alignment | distance | position | gene name | gene id |
| GU305914.1 | | 25346 | 25368 | + | 0 | ACAGCGATCCCAGGGTCCTG | CGG | \|\|\|\|\|\|\|\|[\|\|\|\|\|\|\|\|\|\|\|\|]PAM | NA | Intergenic | NA | NA |
| T1496 | CCCTTTTTGACTCGACCAAGCGG | | 1000 | | | | | | | | | |
| Oligo fwd | CACCCCCTTTTTGACTCGACCAAG | | | | | | | | | | | |
| Oligo rev | AAACCTTGGTCGAGTCAAAAAGGG | | | | | | | | | | | |
| Chromosome | start | | end | strand | MM | target_seq | PAM | alignment | distance | position | gene name | gene id |
| GU305914.1 | | 25373 | 25395 | + | 0 | CCCTTTTTGACTCGACCAAG | CGG | \|\|\|\|\|\|\|\|[\|\|\|\|\|\|\|\|\|\|\|\|]PAM | NA | Intergenic | NA | NA |
| T1497 | TTTGACTCGACCAAGCGGAACGG | | 1000 | | | | | | | | | |
| Oligo fwd | CACCTTTGACTCGACCAAGCGGAA | | | | | | | | | | | |
| Oligo rev | AAACTTCCGCTTGGTCGAGTCAAA | | | | | | | | | | | |
| Chromosome | start | | end | strand | MM | target_seq | PAM | alignment | distance | position | gene name | gene id |
| GU305914.1 | | 25378 | 25400 | + | 0 | TTTGACTCGACCAAGCGGAA | CGG | \|\|\|\|\|\|\|\|[\|\|\|\|\|\|\|\|\|\|\|\|]PAM | NA | Intergenic | NA | NA |
| T1498 | ACGGTGCTATATTCTCCCAGTGG | | 1000 | | | | | | | | | |
| Oligo fwd | CACCACGGTGCTATATTCTCCCAG | | | | | | | | | | | |
| Oligo rev | AAACCTGGGAGAATATAGCACCGT | | | | | | | | | | | |
| Chromosome | start | | end | strand | MM | target_seq | PAM | alignment | distance | position | gene name | gene id |
| GU305914.1 | | 25397 | 25419 | + | 0 | ACGGTGCTATATTCTCCCAG | TGG | \|\|\|\|\|\|\|\|[\|\|\|\|\|\|\|\|\|\|\|\|]PAM | NA | Intergenic | NA | NA |
| T1499 | CGGTGCTATATTCTCCCAGTGGG | | 1000 | | | | | | | | | |
| Oligo fwd | CACCCGGTGCTATATTCTCCCAGT | | | | | | | | | | | |
| Oligo rev | AAACACTGGGAGAATATAGCACCG | | | | | | | | | | | |
| Chromosome | start | | end | strand | MM | target_seq | PAM | alignment | distance | position | gene name | gene id |
| GU305914.1 | | 25398 | 25420 | + | 0 | CGGTGCTATATTCTCCCAGT | GGG | \|\|\|\|\|\|\|\|[\|\|\|\|\|\|\|\|\|\|\|\|]PAM | NA | Intergenic | NA | NA |
| T1500 | GGTGCTATATTCTCCCAGTGGGG | | 1000 | | | | | | | | | |
| Oligo fwd | CACCGGTGCTATATTCTCCCAGTG | | | | | | | | | | | |
| Oligo rev | AAACCACTGGGAGAATATAGCACC | | | | | | | | | | | |
| Chromosome | start | | end | strand | MM | target_seq | PAM | alignment | distance | position | gene name | gene id |
| GU305914.1 | | 25399 | 25421 | + | 0 | GGTGCTATATTCTCCCAGTG | GGG | \|\|\|\|\|\|\|\|[\|\|\|\|\|\|\|\|\|\|\|\|]PAM | NA | Intergenic | NA | NA |
| T1501 | GTGCTATATTCTCCCAGTGGGGG | | 1000 | | | | | | | | | |
| Oligo fwd | CACCGTGCTATATTCTCCCAGTGG | | | | | | | | | | | |
| Oligo rev | AAACCCACTGGGAGAATATAGCAC | | | | | | | | | | | |
| Chromosome | start | | end | strand | MM | target_seq | PAM | alignment | distance | position | gene name | gene id |
| GU305914.1 | | 25400 | 25422 | + | 0 | GTGCTATATTCTCCCAGTGG | GGG | \|\|\|\|\|\|\|\|[\|\|\|\|\|\|\|\|\|\|\|\|]PAM | NA | Intergenic | NA | NA |
| T1502 | CTATATTCTCCCAGTGGGGGCGG | | 1000 | | | | | | | | | |
| Oligo fwd | CACCCTATATTCTCCCAGTGGGGG | | | | | | | | | | | |
| Oligo rev | AAACCCCCCACTGGGAGAATATAG | | | | | | | | | | | |
| Chromosome | start | | end | strand | MM | target_seq | PAM | alignment | distance | position | gene name | gene id |
| GU305914.1 | | 25403 | 25425 | + | 0 | CTATATTCTCCCAGTGGGGG | CGG | \|\|\|\|\|\|\|\|[\|\|\|\|\|\|\|\|\|\|\|\|]PAM | NA | Intergenic | NA | NA |
| T1503 | TTCTCCCAGTGGGGGCGGACAGG | | 1000 | | | | | | | | | |
| Oligo fwd | CACCTTCTCCCAGTGGGGGCGGAC | | | | | | | | | | | |
| Oligo rev | AAACGTCCGCCCCCACTGGGAGAA | | | | | | | | | | | |
| Chromosome | start | | end | strand | MM | target_seq | PAM | alignment | distance | position | gene name | gene id |
| GU305914.1 | | 25408 | 25430 | + | 0 | TTCTCCCAGTGGGGGCGGAC | AGG | \|\|\|\|\|\|\|\|[\|\|\|\|\|\|\|\|\|\|\|\|]PAM | NA | Intergenic | NA | NA |
| T1504 | TCTCCCAGTGGGGGCGGACAGGG | | 1000 | | | | | | | | | |
| Oligo fwd | CACCTCTCCCAGTGGGGGCGGACA | | | | | | | | | | | |
| Oligo rev | AAACTGTCCGCCCCCACTGGGAGA | | | | | | | | | | | |
| Chromosome | start | | end | strand | MM | target_seq | PAM | alignment | distance | position | gene name | gene id |
| GU305914.1 | | 25409 | 25431 | + | 0 | TCTCCCAGTGGGGGCGGACA | GGG | \|\|\|\|\|\|\|\|[\|\|\|\|\|\|\|\|\|\|\|\|]PAM | NA | Intergenic | NA | NA |
| T1505 | GCACCTCCCGCCGCCCGACGAGG | | 1000 | | | | | | | | | |
| Oligo fwd | CACCGCACCTCCCGCCGCCCGACG | | | | | | | | | | | |
| Oligo rev | AAACCGTCGGGCGGCGGGAGGTGC | | | | | | | | | | | |
| Chromosome | start | | end | strand | MM | target_seq | PAM | alignment | distance | position | gene name | gene id |
| GU305914.1 | | 25503 | 25525 | + | 0 | GCACCTCCCGCCGCCCGACG | AGG | \|\|\|\|\|\|\|\|[\|\|\|\|\|\|\|\|\|\|\|\|]PAM | NA | Intergenic | NA | NA |
| T1506 | CCGCCGCCCGACGAGGCGTCTGG | | 1000 | | | | | | | | | |
| Oligo fwd | CACCCCGCCGCCCGACGAGGCGTC | | | | | | | | | | | |
| Oligo rev | AAACGACGCCTCGTCGGGCGGCGG | | | | | | | | | | | |
| Chromosome | start | | end | strand | MM | target_seq | PAM | alignment | distance | position | gene name | gene id |
| GU305914.1 | | 25510 | 25532 | + | 0 | CCGCCGCCCGACGAGGCGTC | TGG | \|\|\|\|\|\|\|\|[\|\|\|\|\|\|\|\|\|\|\|\|]PAM | NA | Intergenic | NA | NA |
| T1507 | GAGGCGTCTGGCGACCACGACGG | | 1000 | | | | | | | | | |
| Oligo fwd | CACCGAGGCGTCTGGCGACCACGA | | | | | | | | | | | |
| Oligo rev | AAACTCGTGGTCGCCAGACGCCTC | | | | | | | | | | | |
| Chromosome | start | | end | strand | MM | target_seq | PAM | alignment | distance | position | gene name | gene id |
| GU305914.1 | | 25522 | 25544 | + | 0 | GAGGCGTCTGGCGACCACGA | CGG | \|\|\|\|\|\|\|\|[\|\|\|\|\|\|\|\|\|\|\|\|]PAM | NA | Intergenic | NA | NA |

<table>
<tr><td>T1508</td><td>GCGCACGACAAAGCGACAGACGG</td><td>1000</td><td></td><td></td><td></td><td></td><td></td><td></td><td></td><td></td><td></td></tr>
<tr><td>Oligo fwd</td><td>CACCGCGCACGACAAAGCGACAGA</td><td></td><td></td><td></td><td></td><td></td><td></td><td></td><td></td><td></td><td></td></tr>
<tr><td>Oligo rev</td><td>AAACTCTGTCGCTTTGTCGTGCGC</td><td></td><td></td><td></td><td></td><td></td><td></td><td></td><td></td><td></td><td></td></tr>
<tr><td>Chromosome</td><td>start</td><td>end</td><td>strand</td><td>MM</td><td>target_seq</td><td>PAM</td><td>alignment</td><td>distance</td><td>position</td><td>gene name</td><td>gene id</td></tr>
<tr><td>GU305914.1</td><td>25558</td><td>25580</td><td>+</td><td>0</td><td>GCGCACGACAAAGCGACAGA</td><td>CGG</td><td>||||||||[||||||||||||]PAM</td><td>NA</td><td>Intergenic</td><td>NA</td><td>NA</td></tr>
<tr><td>T1509</td><td>GACAAAGCGACAGACGGCGTCGG</td><td>1000</td><td></td><td></td><td></td><td></td><td></td><td></td><td></td><td></td><td></td></tr>
<tr><td>Oligo fwd</td><td>CACCGACAAAGCGACAGACGGCGT</td><td></td><td></td><td></td><td></td><td></td><td></td><td></td><td></td><td></td><td></td></tr>
<tr><td>Oligo rev</td><td>AAACACGCCGTCTGTCGCTTTGTC</td><td></td><td></td><td></td><td></td><td></td><td></td><td></td><td></td><td></td><td></td></tr>
<tr><td>Chromosome</td><td>start</td><td>end</td><td>strand</td><td>MM</td><td>target_seq</td><td>PAM</td><td>alignment</td><td>distance</td><td>position</td><td>gene name</td><td>gene id</td></tr>
<tr><td>GU305914.1</td><td>25564</td><td>25586</td><td>+</td><td>0</td><td>GACAAAGCGACAGACGGCGT</td><td>CGG</td><td>||||||||[||||||||||||]PAM</td><td>NA</td><td>Intergenic</td><td>NA</td><td>NA</td></tr>
<tr><td>T1510</td><td>GCGACAGACGGCGTCGGCGTCGG</td><td>1000</td><td></td><td></td><td></td><td></td><td></td><td></td><td></td><td></td><td></td></tr>
<tr><td>Oligo fwd</td><td>CACCGCGACAGACGGCGTCGGCGT</td><td></td><td></td><td></td><td></td><td></td><td></td><td></td><td></td><td></td><td></td></tr>
<tr><td>Oligo rev</td><td>AAACACGCCGACGCCGTCTGTCGC</td><td></td><td></td><td></td><td></td><td></td><td></td><td></td><td></td><td></td><td></td></tr>
<tr><td>Chromosome</td><td>start</td><td>end</td><td>strand</td><td>MM</td><td>target_seq</td><td>PAM</td><td>alignment</td><td>distance</td><td>position</td><td>gene name</td><td>gene id</td></tr>
<tr><td>GU305914.1</td><td>25570</td><td>25592</td><td>+</td><td>0</td><td>GCGACAGACGGCGTCGGCGT</td><td>CGG</td><td>||||||||[||||||||||||]PAM</td><td>NA</td><td>Intergenic</td><td>NA</td><td>NA</td></tr>
<tr><td>T1511</td><td>CGCGCCTGATCATCATCCAGAGG</td><td>1000</td><td></td><td></td><td></td><td></td><td></td><td></td><td></td><td></td><td></td></tr>
<tr><td>Oligo fwd</td><td>CACCCGCGCCTGATCATCATCCAG</td><td></td><td></td><td></td><td></td><td></td><td></td><td></td><td></td><td></td><td></td></tr>
<tr><td>Oligo rev</td><td>AAACCTGGATGATGATCAGGCGCG</td><td></td><td></td><td></td><td></td><td></td><td></td><td></td><td></td><td></td><td></td></tr>
<tr><td>Chromosome</td><td>start</td><td>end</td><td>strand</td><td>MM</td><td>target_seq</td><td>PAM</td><td>alignment</td><td>distance</td><td>position</td><td>gene name</td><td>gene id</td></tr>
<tr><td>GU305914.1</td><td>25596</td><td>25618</td><td>+</td><td>0</td><td>CGCGCCTGATCATCATCCAG</td><td>AGG</td><td>||||||||[||||||||||||]PAM</td><td>NA</td><td>Intergenic</td><td>NA</td><td>NA</td></tr>
<tr><td>T1512</td><td>GCCTGATCATCATCCAGAGGCGG</td><td>1000</td><td></td><td></td><td></td><td></td><td></td><td></td><td></td><td></td><td></td></tr>
<tr><td>Oligo fwd</td><td>CACCGCCTGATCATCATCCAGAGG</td><td></td><td></td><td></td><td></td><td></td><td></td><td></td><td></td><td></td><td></td></tr>
<tr><td>Oligo rev</td><td>AAACCCTCTGGATGATGATCAGGC</td><td></td><td></td><td></td><td></td><td></td><td></td><td></td><td></td><td></td><td></td></tr>
<tr><td>Chromosome</td><td>start</td><td>end</td><td>strand</td><td>MM</td><td>target_seq</td><td>PAM</td><td>alignment</td><td>distance</td><td>position</td><td>gene name</td><td>gene id</td></tr>
<tr><td>GU305914.1</td><td>25599</td><td>25621</td><td>+</td><td>0</td><td>GCCTGATCATCATCCAGAGG</td><td>CGG</td><td>||||||||[||||||||||||]PAM</td><td>NA</td><td>Intergenic</td><td>NA</td><td>NA</td></tr>
<tr><td>T1513</td><td>TCCAGAGGCGGCTGAGCAGCAGG</td><td>1000</td><td></td><td></td><td></td><td></td><td></td><td></td><td></td><td></td><td></td></tr>
<tr><td>Oligo fwd</td><td>CACCTCCAGAGGCGGCTGAGCAGC</td><td></td><td></td><td></td><td></td><td></td><td></td><td></td><td></td><td></td><td></td></tr>
<tr><td>Oligo rev</td><td>AAACGCTGCTCAGCCGCCTCTGGA</td><td></td><td></td><td></td><td></td><td></td><td></td><td></td><td></td><td></td><td></td></tr>
<tr><td>Chromosome</td><td>start</td><td>end</td><td>strand</td><td>MM</td><td>target_seq</td><td>PAM</td><td>alignment</td><td>distance</td><td>position</td><td>gene name</td><td>gene id</td></tr>
<tr><td>GU305914.1</td><td>25611</td><td>25633</td><td>+</td><td>0</td><td>TCCAGAGGCGGCTGAGCAGC</td><td>AGG</td><td>||||||||[||||||||||||]PAM</td><td>NA</td><td>Intergenic</td><td>NA</td><td>NA</td></tr>
<tr><td>T1514</td><td>GAGCAGCAGGCCGATCTCGATGG</td><td>1000</td><td></td><td></td><td></td><td></td><td></td><td></td><td></td><td></td><td></td></tr>
<tr><td>Oligo fwd</td><td>CACCGAGCAGCAGGCCGATCTCGA</td><td></td><td></td><td></td><td></td><td></td><td></td><td></td><td></td><td></td><td></td></tr>
<tr><td>Oligo rev</td><td>AAACTCGAGATCGGCCTGCTGCTC</td><td></td><td></td><td></td><td></td><td></td><td></td><td></td><td></td><td></td><td></td></tr>
<tr><td>Chromosome</td><td>start</td><td>end</td><td>strand</td><td>MM</td><td>target_seq</td><td>PAM</td><td>alignment</td><td>distance</td><td>position</td><td>gene name</td><td>gene id</td></tr>
<tr><td>GU305914.1</td><td>25624</td><td>25646</td><td>+</td><td>0</td><td>GAGCAGCAGGCCGATCTCGA</td><td>TGG</td><td>||||||||[||||||||||||]PAM</td><td>NA</td><td>Intergenic</td><td>NA</td><td>NA</td></tr>
<tr><td>T1515</td><td>AGCAGCAGGCCGATCTCGATGGG</td><td>1000</td><td></td><td></td><td></td><td></td><td></td><td></td><td></td><td></td><td></td></tr>
<tr><td>Oligo fwd</td><td>CACCAGCAGCAGGCCGATCTCGAT</td><td></td><td></td><td></td><td></td><td></td><td></td><td></td><td></td><td></td><td></td></tr>
<tr><td>Oligo rev</td><td>AAACATCGAGATCGGCCTGCTGCT</td><td></td><td></td><td></td><td></td><td></td><td></td><td></td><td></td><td></td><td></td></tr>
<tr><td>Chromosome</td><td>start</td><td>end</td><td>strand</td><td>MM</td><td>target_seq</td><td>PAM</td><td>alignment</td><td>distance</td><td>position</td><td>gene name</td><td>gene id</td></tr>
<tr><td>GU305914.1</td><td>25625</td><td>25647</td><td>+</td><td>0</td><td>AGCAGCAGGCCGATCTCGAT</td><td>GGG</td><td>||||||||[||||||||||||]PAM</td><td>NA</td><td>Intergenic</td><td>NA</td><td>NA</td></tr>
<tr><td>T1516</td><td>GCAGCAGGCCGATCTCGATGGGG</td><td>1000</td><td></td><td></td><td></td><td></td><td></td><td></td><td></td><td></td><td></td></tr>
<tr><td>Oligo fwd</td><td>CACCGCAGCAGGCCGATCTCGATG</td><td></td><td></td><td></td><td></td><td></td><td></td><td></td><td></td><td></td><td></td></tr>
<tr><td>Oligo rev</td><td>AAACCATCGAGATCGGCCTGCTGC</td><td></td><td></td><td></td><td></td><td></td><td></td><td></td><td></td><td></td><td></td></tr>
<tr><td>Chromosome</td><td>start</td><td>end</td><td>strand</td><td>MM</td><td>target_seq</td><td>PAM</td><td>alignment</td><td>distance</td><td>position</td><td>gene name</td><td>gene id</td></tr>
<tr><td>GU305914.1</td><td>25626</td><td>25648</td><td>+</td><td>0</td><td>GCAGCAGGCCGATCTCGATG</td><td>GGG</td><td>||||||||[||||||||||||]PAM</td><td>NA</td><td>Intergenic</td><td>NA</td><td>NA</td></tr>
<tr><td>GU305914.1</td><td>91127</td><td>91149</td><td>+</td><td>4</td><td>GCTGCAGACCGGCCTCGATG</td><td>AGG</td><td>||-||||-[|||--|||||||]PAM</td><td>NA</td><td>Intergenic</td><td>NA</td><td>NA</td></tr>
<tr><td>T1517</td><td>GGGCCTGCCGTGCAGTGAATCGG</td><td>1000</td><td></td><td></td><td></td><td></td><td></td><td></td><td></td><td></td><td></td></tr>
<tr><td>Oligo fwd</td><td>CACCGGGCCTGCCGTGCAGTGAAT</td><td></td><td></td><td></td><td></td><td></td><td></td><td></td><td></td><td></td><td></td></tr>
<tr><td>Oligo rev</td><td>AAACATTCACTGCACGGCAGGCCC</td><td></td><td></td><td></td><td></td><td></td><td></td><td></td><td></td><td></td><td></td></tr>
<tr><td>Chromosome</td><td>start</td><td>end</td><td>strand</td><td>MM</td><td>target_seq</td><td>PAM</td><td>alignment</td><td>distance</td><td>position</td><td>gene name</td><td>gene id</td></tr>
<tr><td>GU305914.1</td><td>25646</td><td>25668</td><td>+</td><td>0</td><td>GGGCCTGCCGTGCAGTGAAT</td><td>CGG</td><td>||||||||[||||||||||||]PAM</td><td>NA</td><td>Intergenic</td><td>NA</td><td>NA</td></tr>
<tr><td>T1518</td><td>AAAATATCTGAACCCGATGACGG</td><td>1000</td><td></td><td></td><td></td><td></td><td></td><td></td><td></td><td></td><td></td></tr>
<tr><td>Oligo fwd</td><td>CACCAAAATATCTGAACCCGATGA</td><td></td><td></td><td></td><td></td><td></td><td></td><td></td><td></td><td></td><td></td></tr>
<tr><td>Oligo rev</td><td>AAACTCATCGGGTTCAGATATTTT</td><td></td><td></td><td></td><td></td><td></td><td></td><td></td><td></td><td></td><td></td></tr>
<tr><td>Chromosome</td><td>start</td><td>end</td><td>strand</td><td>MM</td><td>target_seq</td><td>PAM</td><td>alignment</td><td>distance</td><td>position</td><td>gene name</td><td>gene id</td></tr>
<tr><td>GU305914.1</td><td>25672</td><td>25694</td><td>+</td><td>0</td><td>AAAATATCTGAACCCGATGA</td><td>CGG</td><td>||||||||[||||||||||||]PAM</td><td>NA</td><td>Intergenic</td><td>NA</td><td>NA</td></tr>
<tr><td>T1519</td><td>CCGATGACGGAGAACTGCTCCGG</td><td>1000</td><td></td><td></td><td></td><td></td><td></td><td></td><td></td><td></td><td></td></tr>
<tr><td>Oligo fwd</td><td>CACCCCGATGACGGAGAACTGCTC</td><td></td><td></td><td></td><td></td><td></td><td></td><td></td><td></td><td></td><td></td></tr>
<tr><td>Oligo rev</td><td>AAACGAGCAGTTCTCCGTCATCGG</td><td></td><td></td><td></td><td></td><td></td><td></td><td></td><td></td><td></td><td></td></tr>
<tr><td>Chromosome</td><td>start</td><td>end</td><td>strand</td><td>MM</td><td>target_seq</td><td>PAM</td><td>alignment</td><td>distance</td><td>position</td><td>gene name</td><td>gene id</td></tr>
<tr><td>GU305914.1</td><td>25685</td><td>25707</td><td>+</td><td>0</td><td>CCGATGACGGAGAACTGCTC</td><td>CGG</td><td>||||||||[||||||||||||]PAM</td><td>NA</td><td>Intergenic</td><td>NA</td><td>NA</td></tr>
<tr><td>T1520</td><td>GACGGAGAACTGCTCCGGCGAGG</td><td>1000</td><td></td><td></td><td></td><td></td><td></td><td></td><td></td><td></td><td></td></tr>
<tr><td>Oligo fwd</td><td>CACCGACGGAGAACTGCTCCGGCG</td><td></td><td></td><td></td><td></td><td></td><td></td><td></td><td></td><td></td><td></td></tr>
<tr><td>Oligo rev</td><td>AAACCGCCGGAGCAGTTCTCCGTC</td><td></td><td></td><td></td><td></td><td></td><td></td><td></td><td></td><td></td><td></td></tr>
<tr><td>Chromosome</td><td>start</td><td>end</td><td>strand</td><td>MM</td><td>target_seq</td><td>PAM</td><td>alignment</td><td>distance</td><td>position</td><td>gene name</td><td>gene id</td></tr>
<tr><td>GU305914.1</td><td>25690</td><td>25712</td><td>+</td><td>0</td><td>GACGGAGAACTGCTCCGGCG</td><td>AGG</td><td>||||||||[||||||||||||]PAM</td><td>NA</td><td>Intergenic</td><td>NA</td><td>NA</td></tr>
<tr><td>T1521</td><td>GGAGAACTGCTCCGGCGAGGCGG</td><td>1000</td><td></td><td></td><td></td><td></td><td></td><td></td><td></td><td></td><td></td></tr>
<tr><td>Oligo fwd</td><td>CACCGGAGAACTGCTCCGGCGAGG</td><td></td><td></td><td></td><td></td><td></td><td></td><td></td><td></td><td></td><td></td></tr>
<tr><td>Oligo rev</td><td>AAACCCTCGCCGGAGCAGTTCTCC</td><td></td><td></td><td></td><td></td><td></td><td></td><td></td><td></td><td></td><td></td></tr>
<tr><td>Chromosome</td><td>start</td><td>end</td><td>strand</td><td>MM</td><td>target_seq</td><td>PAM</td><td>alignment</td><td>distance</td><td>position</td><td>gene name</td><td>gene id</td></tr>
<tr><td>GU305914.1</td><td>25693</td><td>25715</td><td>+</td><td>0</td><td>GGAGAACTGCTCCGGCGAGG</td><td>CGG</td><td>||||||||[||||||||||||]PAM</td><td>NA</td><td>Intergenic</td><td>NA</td><td>NA</td></tr>
<tr><td>T1522</td><td>ACTGCTCCGGCGAGGCGGCGAGG</td><td>1000</td><td></td><td></td><td></td><td></td><td></td><td></td><td></td><td></td><td></td></tr>
<tr><td>Oligo fwd</td><td>CACCACTGCTCCGGCGAGGCGGCG</td><td></td><td></td><td></td><td></td><td></td><td></td><td></td><td></td><td></td><td></td></tr>
<tr><td>Oligo rev</td><td>AAACCGCCGCCTCGCCGGAGCAGT</td><td></td><td></td><td></td><td></td><td></td><td></td><td></td><td></td><td></td><td></td></tr>
</table>

| | | | | | | | | | | | |
|---|---|---|---|---|---|---|---|---|---|---|---|
| Chromosome | start | end | strand | MM | target_seq | PAM | alignment | distance | position | gene name | gene id |
| GU305914.1 | 25698 | 25720 | + | 0 | ACTGCTCCGGCGAGGCGGCG | AGG | \|\|\|\|\|\|\|\|[\|\|\|\|\|\|\|\|\|\|\|\|]PAM | NA | Intergenic | NA | NA |
| T1523 | CGAGGCGGCGAGGTAGTAGATGG | 1000 | | | | | | | | | |
| Oligo fwd | CACCCGAGGCGGCGAGGTAGTAGA | | | | | | | | | | |
| Oligo rev | AAACTCTACTACCTCGCCGCCTCG | | | | | | | | | | |
| Chromosome | start | end | strand | MM | target_seq | PAM | alignment | distance | position | gene name | gene id |
| GU305914.1 | 25708 | 25730 | + | 0 | CGAGGCGGCGAGGTAGTAGA | TGG | \|\|\|\|\|\|\|\|[\|\|\|\|\|\|\|\|\|\|\|\|]PAM | NA | Intergenic | NA | NA |
| T1524 | AGGTAGTAGATGGCGTCGTCCGG | 1000 | | | | | | | | | |
| Oligo fwd | CACCAGGTAGTAGATGGCGTCGTC | | | | | | | | | | |
| Oligo rev | AAACGACGACGCCATCTACTACCT | | | | | | | | | | |
| Chromosome | start | end | strand | MM | target_seq | PAM | alignment | distance | position | gene name | gene id |
| GU305914.1 | 25718 | 25740 | + | 0 | AGGTAGTAGATGGCGTCGTC | CGG | \|\|\|\|\|\|\|\|[\|\|\|\|\|\|\|\|\|\|\|\|]PAM | NA | Intergenic | NA | NA |
| T1525 | GGTAGTAGATGGCGTCGTCCGGG | 1000 | | | | | | | | | |
| Oligo fwd | CACCGGTAGTAGATGGCGTCGTCC | | | | | | | | | | |
| Oligo rev | AAACGGACGACGCCATCTACTACC | | | | | | | | | | |
| Chromosome | start | end | strand | MM | target_seq | PAM | alignment | distance | position | gene name | gene id |
| GU305914.1 | 25719 | 25741 | + | 0 | GGTAGTAGATGGCGTCGTCC | GGG | \|\|\|\|\|\|\|\|[\|\|\|\|\|\|\|\|\|\|\|\|]PAM | NA | Intergenic | NA | NA |
| T1526 | GTAGTAGATGGCGTCGTCCGGGG | 1000 | | | | | | | | | |
| Oligo fwd | CACCGTAGTAGATGGCGTCGTCCG | | | | | | | | | | |
| Oligo rev | AAACCGGACGACGCCATCTACTAC | | | | | | | | | | |
| Chromosome | start | end | strand | MM | target_seq | PAM | alignment | distance | position | gene name | gene id |
| GU305914.1 | 25720 | 25742 | + | 0 | GTAGTAGATGGCGTCGTCCG | GGG | \|\|\|\|\|\|\|\|[\|\|\|\|\|\|\|\|\|\|\|\|]PAM | NA | Intergenic | NA | NA |
| T1527 | TAGTAGATGGCGTCGTCCGGGGG | 1000 | | | | | | | | | |
| Oligo fwd | CACCTAGTAGATGGCGTCGTCCGG | | | | | | | | | | |
| Oligo rev | AAACCCGGACGACGCCATCTACTA | | | | | | | | | | |
| Chromosome | start | end | strand | MM | target_seq | PAM | alignment | distance | position | gene name | gene id |
| GU305914.1 | 25721 | 25743 | + | 0 | TAGTAGATGGCGTCGTCCGG | GGG | \|\|\|\|\|\|\|\|[\|\|\|\|\|\|\|\|\|\|\|\|]PAM | NA | Intergenic | NA | NA |
| T1528 | GCCCGCGATAACAGTAGAAACGG | 1000 | | | | | | | | | |
| Oligo fwd | CACCGCCCGCGATAACAGTAGAAA | | | | | | | | | | |
| Oligo rev | AAACTTTCTACTGTTATCGCGGGC | | | | | | | | | | |
| Chromosome | start | end | strand | MM | target_seq | PAM | alignment | distance | position | gene name | gene id |
| GU305914.1 | 25743 | 25765 | + | 0 | GCCCGCGATAACAGTAGAAA | CGG | \|\|\|\|\|\|\|\|[\|\|\|\|\|\|\|\|\|\|\|\|]PAM | NA | Intergenic | NA | NA |
| T1529 | AACGGCCGAAGACGTCGACGAGG | 1000 | | | | | | | | | |
| Oligo fwd | CACCAACGGCCGAAGACGTCGACG | | | | | | | | | | |
| Oligo rev | AAACCGTCGACGTCTTCGGCCGTT | | | | | | | | | | |
| Chromosome | start | end | strand | MM | target_seq | PAM | alignment | distance | position | gene name | gene id |
| GU305914.1 | 25761 | 25783 | + | 0 | AACGGCCGAAGACGTCGACG | AGG | \|\|\|\|\|\|\|\|[\|\|\|\|\|\|\|\|\|\|\|\|]PAM | NA | Intergenic | NA | NA |
| T1530 | CGAAGACGTCGACGAGGACCAGG | 1000 | | | | | | | | | |
| Oligo fwd | CACCCGAAGACGTCGACGAGGACC | | | | | | | | | | |
| Oligo rev | AAACGGTCCTCGTCGACGTCTTCG | | | | | | | | | | |
| Chromosome | start | end | strand | MM | target_seq | PAM | alignment | distance | position | gene name | gene id |
| GU305914.1 | 25767 | 25789 | + | 0 | CGAAGACGTCGACGAGGACC | AGG | \|\|\|\|\|\|\|\|[\|\|\|\|\|\|\|\|\|\|\|\|]PAM | NA | Intergenic | NA | NA |
| T1531 | GAAGACGTCGACGAGGACCAGGG | 1000 | | | | | | | | | |
| Oligo fwd | CACCGAAGACGTCGACGAGGACCA | | | | | | | | | | |
| Oligo rev | AAACTGGTCCTCGTCGACGTCTTC | | | | | | | | | | |
| Chromosome | start | end | strand | MM | target_seq | PAM | alignment | distance | position | gene name | gene id |
| GU305914.1 | 25768 | 25790 | + | 0 | GAAGACGTCGACGAGGACCA | GGG | \|\|\|\|\|\|\|\|[\|\|\|\|\|\|\|\|\|\|\|\|]PAM | NA | Intergenic | NA | NA |
| GU305914.1 | 65115 | 65137 | + | 3 | GAAGACGACGACGAGGAGGA | GGG | \|\|\|\|\|\|\|-[\|\|\|\|\|\|\|\|\|--\|]PAM | NA | Intergenic | NA | NA |
| T1532 | GACGAGGACCAGGGTGTCGTCGG | 1000 | | | | | | | | | |
| Oligo fwd | CACCGACGAGGACCAGGGTGTCGT | | | | | | | | | | |
| Oligo rev | AAACACGACACCCTGGTCCTCGTC | | | | | | | | | | |
| Chromosome | start | end | strand | MM | target_seq | PAM | alignment | distance | position | gene name | gene id |
| GU305914.1 | 25777 | 25799 | + | 0 | GACGAGGACCAGGGTGTCGT | CGG | \|\|\|\|\|\|\|\|[\|\|\|\|\|\|\|\|\|\|\|\|]PAM | NA | Intergenic | NA | NA |
| GU305914.1 | 146852 | 146874 | - | 4 | GACGGCGTCCAGGGTGTCGA | CGG | \|\|\|\|--\|-[\|\|\|\|\|\|\|\|\|\|\|-]PAM | NA | Intergenic | NA | NA |
| T1533 | GTCGGCGACCTTGAGTCCGCTGG | 1000 | | | | | | | | | |
| Oligo fwd | CACCGTCGGCGACCTTGAGTCCGC | | | | | | | | | | |
| Oligo rev | AAACGCGGACTCAAGGTCGCCGAC | | | | | | | | | | |
| Chromosome | start | end | strand | MM | target_seq | PAM | alignment | distance | position | gene name | gene id |
| GU305914.1 | 25795 | 25817 | + | 0 | GTCGGCGACCTTGAGTCCGC | TGG | \|\|\|\|\|\|\|\|[\|\|\|\|\|\|\|\|\|\|\|\|]PAM | NA | Intergenic | NA | NA |
| T1534 | GAGTCCGCTGGCATAGCCGATGG | 1000 | | | | | | | | | |
| Oligo fwd | CACCGAGTCCGCTGGCATAGCCGA | | | | | | | | | | |
| Oligo rev | AAACTCGGCTATGCCAGCGGACTC | | | | | | | | | | |
| Chromosome | start | end | strand | MM | target_seq | PAM | alignment | distance | position | gene name | gene id |
| GU305914.1 | 25807 | 25829 | + | 0 | GAGTCCGCTGGCATAGCCGA | TGG | \|\|\|\|\|\|\|\|[\|\|\|\|\|\|\|\|\|\|\|\|]PAM | NA | Intergenic | NA | NA |
| T1535 | AGTCCGCTGGCATAGCCGATGGG | 1000 | | | | | | | | | |
| Oligo fwd | CACCAGTCCGCTGGCATAGCCGAT | | | | | | | | | | |
| Oligo rev | AAACATCGGCTATGCCAGCGGACT | | | | | | | | | | |
| Chromosome | start | end | strand | MM | target_seq | PAM | alignment | distance | position | gene name | gene id |
| GU305914.1 | 25808 | 25830 | + | 0 | AGTCCGCTGGCATAGCCGAT | GGG | \|\|\|\|\|\|\|\|[\|\|\|\|\|\|\|\|\|\|\|\|]PAM | NA | Intergenic | NA | NA |
| T1536 | AGCAAAAGTATCTGAGTACGCGG | 1000 | | | | | | | | | |
| Oligo fwd | CACCAGCAAAAGTATCTGAGTACG | | | | | | | | | | |
| Oligo rev | AAACCGTACTCAGATACTTTTGCT | | | | | | | | | | |
| Chromosome | start | end | strand | MM | target_seq | PAM | alignment | distance | position | gene name | gene id |
| GU305914.1 | 25848 | 25870 | + | 0 | AGCAAAAGTATCTGAGTACG | CGG | \|\|\|\|\|\|\|\|[\|\|\|\|\|\|\|\|\|\|\|\|]PAM | NA | Intergenic | NA | NA |

<table>
<tr><td>T1537</td><td>CGGTTGCGACGTCCAGACTGCGG</td><td>1000</td></tr>
<tr><td>Oligo fwd</td><td>CACCCGGTTGCGACGTCCAGACTG</td></tr>
<tr><td>Oligo rev</td><td>AAACCAGTCTGGACGTCGCAACCG</td></tr>
<tr><td>Chromosome</td><td>start</td><td>end</td><td>strand</td><td>MM</td><td>target_seq</td><td>PAM</td><td>alignment</td><td>distance</td><td>position</td><td>gene name</td><td>gene id</td></tr>
<tr><td>GU305914.1</td><td>25868</td><td>25890</td><td>+</td><td>0</td><td>CGGTTGCGACGTCCAGACTG</td><td>CGG</td><td>||||||||[||||||||||||]PAM</td><td>NA</td><td>Intergenic</td><td>NA</td><td>NA</td></tr>
<tr><td>T1538</td><td>CAGACTGCGGCTCCCAACCCTGG</td><td>1000</td></tr>
<tr><td>Oligo fwd</td><td>CACCCAGACTGCGGCTCCCAACCC</td></tr>
<tr><td>Oligo rev</td><td>AAACGGGTTGGGAGCCGCAGTCTG</td></tr>
<tr><td>Chromosome</td><td>start</td><td>end</td><td>strand</td><td>MM</td><td>target_seq</td><td>PAM</td><td>alignment</td><td>distance</td><td>position</td><td>gene name</td><td>gene id</td></tr>
<tr><td>GU305914.1</td><td>25881</td><td>25903</td><td>+</td><td>0</td><td>CAGACTGCGGCTCCCAACCC</td><td>TGG</td><td>||||||||[||||||||||||]PAM</td><td>NA</td><td>Intergenic</td><td>NA</td><td>NA</td></tr>
<tr><td>T1539</td><td>CGGCTCCCAACCCTGGATATCGG</td><td>1000</td></tr>
<tr><td>Oligo fwd</td><td>CACCCGGCTCCCAACCCTGGATAT</td></tr>
<tr><td>Oligo rev</td><td>AAACATATCCAGGGTTGGGAGCCG</td></tr>
<tr><td>Chromosome</td><td>start</td><td>end</td><td>strand</td><td>MM</td><td>target_seq</td><td>PAM</td><td>alignment</td><td>distance</td><td>position</td><td>gene name</td><td>gene id</td></tr>
<tr><td>GU305914.1</td><td>25888</td><td>25910</td><td>+</td><td>0</td><td>CGGCTCCCAACCCTGGATAT</td><td>CGG</td><td>||||||||[||||||||||||]PAM</td><td>NA</td><td>Intergenic</td><td>NA</td><td>NA</td></tr>
<tr><td>T1540</td><td>GGCTCCCAACCCTGGATATCGGG</td><td>1000</td></tr>
<tr><td>Oligo fwd</td><td>CACCGGCTCCCAACCCTGGATATC</td></tr>
<tr><td>Oligo rev</td><td>AAACGATATCCAGGGTTGGGAGCC</td></tr>
<tr><td>Chromosome</td><td>start</td><td>end</td><td>strand</td><td>MM</td><td>target_seq</td><td>PAM</td><td>alignment</td><td>distance</td><td>position</td><td>gene name</td><td>gene id</td></tr>
<tr><td>GU305914.1</td><td>25889</td><td>25911</td><td>+</td><td>0</td><td>GGCTCCCAACCCTGGATATC</td><td>GGG</td><td>||||||||[||||||||||||]PAM</td><td>NA</td><td>Intergenic</td><td>NA</td><td>NA</td></tr>
<tr><td>T1541</td><td>GCTCCCAACCCTGGATATCGGGG</td><td>1000</td></tr>
<tr><td>Oligo fwd</td><td>CACCGCTCCCAACCCTGGATATCG</td></tr>
<tr><td>Oligo rev</td><td>AAACCGATATCCAGGGTTGGGAGC</td></tr>
<tr><td>Chromosome</td><td>start</td><td>end</td><td>strand</td><td>MM</td><td>target_seq</td><td>PAM</td><td>alignment</td><td>distance</td><td>position</td><td>gene name</td><td>gene id</td></tr>
<tr><td>GU305914.1</td><td>25890</td><td>25912</td><td>+</td><td>0</td><td>GCTCCCAACCCTGGATATCG</td><td>GGG</td><td>||||||||[||||||||||||]PAM</td><td>NA</td><td>Intergenic</td><td>NA</td><td>NA</td></tr>
<tr><td>T1542</td><td>ACCCTGGATATCGGGGACGAAGG</td><td>1000</td></tr>
<tr><td>Oligo fwd</td><td>CACCACCCTGGATATCGGGGACGA</td></tr>
<tr><td>Oligo rev</td><td>AAACTCGTCCCCGATATCCAGGGT</td></tr>
<tr><td>Chromosome</td><td>start</td><td>end</td><td>strand</td><td>MM</td><td>target_seq</td><td>PAM</td><td>alignment</td><td>distance</td><td>position</td><td>gene name</td><td>gene id</td></tr>
<tr><td>GU305914.1</td><td>25897</td><td>25919</td><td>+</td><td>0</td><td>ACCCTGGATATCGGGGACGA</td><td>AGG</td><td>||||||||[||||||||||||]PAM</td><td>NA</td><td>Intergenic</td><td>NA</td><td>NA</td></tr>
<tr><td>T1543</td><td>ATCGGGGACGAAGGTTATCACGG</td><td>1000</td></tr>
<tr><td>Oligo fwd</td><td>CACCATCGGGGACGAAGGTTATCA</td></tr>
<tr><td>Oligo rev</td><td>AAACTGATAACCTTCGTCCCCGAT</td></tr>
<tr><td>Chromosome</td><td>start</td><td>end</td><td>strand</td><td>MM</td><td>target_seq</td><td>PAM</td><td>alignment</td><td>distance</td><td>position</td><td>gene name</td><td>gene id</td></tr>
<tr><td>GU305914.1</td><td>25906</td><td>25928</td><td>+</td><td>0</td><td>ATCGGGGACGAAGGTTATCA</td><td>CGG</td><td>||||||||[||||||||||||]PAM</td><td>NA</td><td>Intergenic</td><td>NA</td><td>NA</td></tr>
<tr><td>T1544</td><td>GACGAAGGTTATCACGGCCCCGG</td><td>1000</td></tr>
<tr><td>Oligo fwd</td><td>CACCGACGAAGGTTATCACGGCCC</td></tr>
<tr><td>Oligo rev</td><td>AAACGGGCCGTGATAACCTTCGTC</td></tr>
<tr><td>Chromosome</td><td>start</td><td>end</td><td>strand</td><td>MM</td><td>target_seq</td><td>PAM</td><td>alignment</td><td>distance</td><td>position</td><td>gene name</td><td>gene id</td></tr>
<tr><td>GU305914.1</td><td>25912</td><td>25934</td><td>+</td><td>0</td><td>GACGAAGGTTATCACGGCCC</td><td>CGG</td><td>||||||||[||||||||||||]PAM</td><td>NA</td><td>Intergenic</td><td>NA</td><td>NA</td></tr>
<tr><td>T1545</td><td>GAAGGTTATCACGGCCCCGGCGG</td><td>1000</td></tr>
<tr><td>Oligo fwd</td><td>CACCGAAGGTTATCACGGCCCCGG</td></tr>
<tr><td>Oligo rev</td><td>AAACCCGGGGCCGTGATAACCTTC</td></tr>
<tr><td>Chromosome</td><td>start</td><td>end</td><td>strand</td><td>MM</td><td>target_seq</td><td>PAM</td><td>alignment</td><td>distance</td><td>position</td><td>gene name</td><td>gene id</td></tr>
<tr><td>GU305914.1</td><td>25915</td><td>25937</td><td>+</td><td>0</td><td>GAAGGTTATCACGGCCCCGG</td><td>CGG</td><td>||||||||[||||||||||||]PAM</td><td>NA</td><td>Intergenic</td><td>NA</td><td>NA</td></tr>
<tr><td>T1546</td><td>AAGGTTATCACGGCCCCGGCGGG</td><td>1000</td></tr>
<tr><td>Oligo fwd</td><td>CACCAAGGTTATCACGGCCCCGGC</td></tr>
<tr><td>Oligo rev</td><td>AAACGCCGGGGCCGTGATAACCTT</td></tr>
<tr><td>Chromosome</td><td>start</td><td>end</td><td>strand</td><td>MM</td><td>target_seq</td><td>PAM</td><td>alignment</td><td>distance</td><td>position</td><td>gene name</td><td>gene id</td></tr>
<tr><td>GU305914.1</td><td>25916</td><td>25938</td><td>+</td><td>0</td><td>AAGGTTATCACGGCCCCGGC</td><td>GGG</td><td>||||||||[||||||||||||]PAM</td><td>NA</td><td>Intergenic</td><td>NA</td><td>NA</td></tr>
<tr><td>T1547</td><td>TATCACGGCCCCGGCGGGCCAGG</td><td>1000</td></tr>
<tr><td>Oligo fwd</td><td>CACCTATCACGGCCCCGGCGGGCC</td></tr>
<tr><td>Oligo rev</td><td>AAACGGCCCGCCGGGGCCGTGATA</td></tr>
<tr><td>Chromosome</td><td>start</td><td>end</td><td>strand</td><td>MM</td><td>target_seq</td><td>PAM</td><td>alignment</td><td>distance</td><td>position</td><td>gene name</td><td>gene id</td></tr>
<tr><td>GU305914.1</td><td>25921</td><td>25943</td><td>+</td><td>0</td><td>TATCACGGCCCCGGCGGGCC</td><td>AGG</td><td>||||||||[||||||||||||]PAM</td><td>NA</td><td>Intergenic</td><td>NA</td><td>NA</td></tr>
<tr><td>T1548</td><td>CGGCGGGCCAGGCCATGCGCAGG</td><td>1000</td></tr>
<tr><td>Oligo fwd</td><td>CACCCGGCGGGCCAGGCCATGCGC</td></tr>
<tr><td>Oligo rev</td><td>AAACGCGCATGGCCTGGCCCGCCG</td></tr>
<tr><td>Chromosome</td><td>start</td><td>end</td><td>strand</td><td>MM</td><td>target_seq</td><td>PAM</td><td>alignment</td><td>distance</td><td>position</td><td>gene name</td><td>gene id</td></tr>
<tr><td>GU305914.1</td><td>25932</td><td>25954</td><td>+</td><td>0</td><td>CGGCGGGCCAGGCCATGCGC</td><td>AGG</td><td>||||||||[||||||||||||]PAM</td><td>NA</td><td>Intergenic</td><td>NA</td><td>NA</td></tr>
<tr><td>T1549</td><td>GGCCATGCGCAGGTTCCCGCCGG</td><td>1000</td></tr>
<tr><td>Oligo fwd</td><td>CACCGGCCATGCGCAGGTTCCCGC</td></tr>
<tr><td>Oligo rev</td><td>AAACGCGGGAACCTGCGCATGGCC</td></tr>
<tr><td>Chromosome</td><td>start</td><td>end</td><td>strand</td><td>MM</td><td>target_seq</td><td>PAM</td><td>alignment</td><td>distance</td><td>position</td><td>gene name</td><td>gene id</td></tr>
<tr><td>GU305914.1</td><td>25942</td><td>25964</td><td>+</td><td>0</td><td>GGCCATGCGCAGGTTCCCGC</td><td>CGG</td><td>||||||||[||||||||||||]PAM</td><td>NA</td><td>Intergenic</td><td>NA</td><td>NA</td></tr>
<tr><td>T1550</td><td>GCAGGTTCCCGCCGGCGAAGAGG</td><td>1000</td></tr>
<tr><td>Oligo fwd</td><td>CACCGCAGGTTCCCGCCGGCGAAG</td></tr>
<tr><td>Oligo rev</td><td>AAACCTTCGCCGGCGGGAACCTGC</td></tr>
<tr><td>Chromosome</td><td>start</td><td>end</td><td>strand</td><td>MM</td><td>target_seq</td><td>PAM</td><td>alignment</td><td>distance</td><td>position</td><td>gene name</td><td>gene id</td></tr>
<tr><td>GU305914.1</td><td>25950</td><td>25972</td><td>+</td><td>0</td><td>GCAGGTTCCCGCCGGCGAAG</td><td>AGG</td><td>||||||||[||||||||||||]PAM</td><td>NA</td><td>Intergenic</td><td>NA</td><td>NA</td></tr>
<tr><td>T1551</td><td>AGAGTTTCTCTCTGAGCTCTAGG</td><td>1000</td></tr>
<tr><td>Oligo fwd</td><td>CACCAGAGTTTCTCTCTGAGCTCT</td></tr>
<tr><td>Oligo rev</td><td>AAACAGAGCTCAGAGAGAAACTCT</td></tr>
<tr><td>Chromosome</td><td>start</td><td>end</td><td>strand</td><td>MM</td><td>target_seq</td><td>PAM</td><td>alignment</td><td>distance</td><td>position</td><td>gene name</td><td>gene id</td></tr>
<tr><td>GU305914.1</td><td>25974</td><td>25996</td><td>+</td><td>0</td><td>AGAGTTTCTCTCTGAGCTCT</td><td>AGG</td><td>||||||||[||||||||||||]PAM</td><td>NA</td><td>Intergenic</td><td>NA</td><td>NA</td></tr>
</table>

T1552 GAGTTTCTCTCTGAGCTCTAGGG 1000
Oligo fwd CACCGAGTTTCTCTCTGAGCTCTA
Oligo rev AAACTAGAGCTCAGAGAGAAACTC
Chromosome start end strand MM target_seq PAM alignment distance position gene name gene id
GU305914.1 25975 25997 + 0 GAGTTTCTCTCTGAGCTCTA GGG ||||||||[||||||||||||]PAM NA Intergenic NA NA

T1553 CTCTGAGCTCTAGGGATCCCCGG 1000
Oligo fwd CACCCTCTGAGCTCTAGGGATCCC
Oligo rev AAACGGGATCCCTAGAGCTCAGAG
Chromosome start end strand MM target_seq PAM alignment distance position gene name gene id
GU305914.1 25983 26005 + 0 CTCTGAGCTCTAGGGATCCC CGG ||||||||[||||||||||||]PAM NA Intergenic NA NA

T1554 GAGCTCTAGGGATCCCCGGCCGG 1000
Oligo fwd CACCGAGCTCTAGGGATCCCCGGC
Oligo rev AAACGCCGGGGATCCCTAGAGCTC
Chromosome start end strand MM target_seq PAM alignment distance position gene name gene id
GU305914.1 25987 26009 + 0 GAGCTCTAGGGATCCCCGGC CGG ||||||||[||||||||||||]PAM NA Intergenic NA NA

T1555 CTCTAGGGATCCCCGGCCGGCGG 1000
Oligo fwd CACCCTCTAGGGATCCCCGGCCGG
Oligo rev AAACCCGGCCGGGGATCCCTAGAG
Chromosome start end strand MM target_seq PAM alignment distance position gene name gene id
GU305914.1 25990 26012 + 0 CTCTAGGGATCCCCGGCCGG CGG ||||||||[||||||||||||]PAM NA Intergenic NA NA

T1556 GTACAGCACGTCGCTGACCGTGG 1000
Oligo fwd CACCGTACAGCACGTCGCTGACCG
Oligo rev AAACCGGTCAGCGACGTGCTGTAC
Chromosome start end strand MM target_seq PAM alignment distance position gene name gene id
GU305914.1 26026 26048 + 0 GTACAGCACGTCGCTGACCG TGG ||||||||[||||||||||||]PAM NA Intergenic NA NA

T1557 CACGTCGCTGACCGTGGCCTCGG 1000
Oligo fwd CACCCACGTCGCTGACCGTGGCCT
Oligo rev AAACAGGCCACGGTCAGCGACGTG
Chromosome start end strand MM target_seq PAM alignment distance position gene name gene id
GU305914.1 26032 26054 + 0 CACGTCGCTGACCGTGGCCT CGG ||||||||[||||||||||||]PAM NA Intergenic NA NA

T1558 GCCTCGGCGTCGTCCACCAGCGG 1000
Oligo fwd CACCGCCTCGGCGTCGTCCACCAG
Oligo rev AAACCTGGTGGACGACGCCGAGGC
Chromosome start end strand MM target_seq PAM alignment distance position gene name gene id
GU305914.1 26048 26070 + 0 GCCTCGGCGTCGTCCACCAG CGG ||||||||[||||||||||||]PAM NA Intergenic NA NA

T1559 GTCCACCAGCGGCTCGCACGCGG 1000
Oligo fwd CACCGTCCACCAGCGGCTCGCACG
Oligo rev AAACCGTGCGAGCCGCTGGTGGAC
Chromosome start end strand MM target_seq PAM alignment distance position gene name gene id
GU305914.1 26059 26081 + 0 GTCCACCAGCGGCTCGCACG CGG ||||||||[||||||||||||]PAM NA Intergenic NA NA

T1560 ATCCGAACCCTGCCCAGAGATGG 1000
Oligo fwd CACCATCCGAACCCTGCCCAGAGA
Oligo rev AAACTCTCTGGGCAGGGTTCGGAT
Chromosome start end strand MM target_seq PAM alignment distance position gene name gene id
GU305914.1 26125 26147 + 0 ATCCGAACCCTGCCCAGAGA TGG ||||||||[||||||||||||]PAM NA Intergenic NA NA

T1561 CTATATAAAATCACGAGAGCCGG 1000
Oligo fwd CACCCTATATAAAATCACGAGAGC
Oligo rev AAACGCTCTCGTGATTTTATATAG
Chromosome start end strand MM target_seq PAM alignment distance position gene name gene id
GU305914.1 26158 26180 + 0 CTATATAAAATCACGAGAGC CGG ||||||||[||||||||||||]PAM NA Intergenic NA NA

T1562 TATATAAAATCACGAGAGCCGGG 1000
Oligo fwd CACCTATATAAAATCACGAGAGCC
Oligo rev AAACGGCTCTCGTGATTTTATATA
Chromosome start end strand MM target_seq PAM alignment distance position gene name gene id
GU305914.1 26159 26181 + 0 TATATAAAATCACGAGAGCC GGG ||||||||[||||||||||||]PAM NA Intergenic NA NA

T1563 ATATAAAATCACGAGAGCCGGGG 1000
Oligo fwd CACCATATAAAATCACGAGAGCCG
Oligo rev AAACCGGCTCTCGTGATTTTATAT
Chromosome start end strand MM target_seq PAM alignment distance position gene name gene id
GU305914.1 26160 26182 + 0 ATATAAAATCACGAGAGCCG GGG ||||||||[||||||||||||]PAM NA Intergenic NA NA

T1564 AGCCGTCCTATTCCCCGTGCCGG 1000
Oligo fwd CACCAGCCGTCCTATTCCCCGTGC
Oligo rev AAACGCACGGGGAATAGGACGGCT
Chromosome start end strand MM target_seq PAM alignment distance position gene name gene id
GU305914.1 26193 26215 + 0 AGCCGTCCTATTCCCCGTGC CGG ||||||||[||||||||||||]PAM NA Intergenic NA NA

T1565 GATGAACCGTCGATCCTCCAAGG 1000
Oligo fwd CACCGATGAACCGTCGATCCTCCA
Oligo rev AAACTGGAGGATCGACGGTTCATC
Chromosome start end strand MM target_seq PAM alignment distance position gene name gene id
GU305914.1 26268 26290 + 0 GATGAACCGTCGATCCTCCA AGG ||||||||[||||||||||||]PAM NA Intergenic NA NA

T1566 GTTCGTCACCGACGACTCGTCGG 1000
Oligo fwd CACCGTTCGTCACCGACGACTCGT
Oligo rev AAACACGAGTCGTCGGTGACGAAC
Chromosome start end strand MM target_seq PAM alignment distance position gene name gene id

<table>
<tr><td>GU305914.1</td><td>26301</td><td>26323</td><td>+</td><td>0</td><td>GTTCGTCACCGACGACTCGT</td><td>CGG</td><td>||||||||[||||||||||||]PAM</td><td>NA</td><td>Intergenic</td><td>NA</td><td>NA</td></tr>
<tr><td></td><td></td><td></td><td></td><td></td><td></td><td></td><td></td><td></td><td></td><td></td><td></td></tr>
<tr><td>T1567</td><td>CGACGACGATGTGATGATCATGG</td><td>1000</td><td></td><td></td><td></td><td></td><td></td><td></td><td></td><td></td><td></td></tr>
<tr><td>Oligo fwd</td><td>CACCCGACGACGATGTGATGATCA</td><td></td><td></td><td></td><td></td><td></td><td></td><td></td><td></td><td></td><td></td></tr>
<tr><td>Oligo rev</td><td>AAACTGATCATCACATCGTCGTCG</td><td></td><td></td><td></td><td></td><td></td><td></td><td></td><td></td><td></td><td></td></tr>
<tr><td>Chromosome</td><td>start</td><td>end</td><td>strand</td><td>MM</td><td>target_seq</td><td>PAM</td><td>alignment</td><td>distance</td><td>position</td><td>gene name</td><td>gene id</td></tr>
<tr><td>GU305914.1</td><td>26331</td><td>26353</td><td>+</td><td>0</td><td>CGACGACGATGTGATGATCA</td><td>TGG</td><td>||||||||[||||||||||||]PAM</td><td>NA</td><td>Intergenic</td><td>NA</td><td>NA</td></tr>
<tr><td></td><td></td><td></td><td></td><td></td><td></td><td></td><td></td><td></td><td></td><td></td><td></td></tr>
<tr><td>T1568</td><td>TGTGATGATCATGGACCCCCCGG</td><td>1000</td><td></td><td></td><td></td><td></td><td></td><td></td><td></td><td></td><td></td></tr>
<tr><td>Oligo fwd</td><td>CACCTGTGATGATCATGGACCCCC</td><td></td><td></td><td></td><td></td><td></td><td></td><td></td><td></td><td></td><td></td></tr>
<tr><td>Oligo rev</td><td>AAACGGGGGTCCATGATCATCACA</td><td></td><td></td><td></td><td></td><td></td><td></td><td></td><td></td><td></td><td></td></tr>
<tr><td>Chromosome</td><td>start</td><td>end</td><td>strand</td><td>MM</td><td>target_seq</td><td>PAM</td><td>alignment</td><td>distance</td><td>position</td><td>gene name</td><td>gene id</td></tr>
<tr><td>GU305914.1</td><td>26340</td><td>26362</td><td>+</td><td>0</td><td>TGTGATGATCATGGACCCCC</td><td>CGG</td><td>||||||||[||||||||||||]PAM</td><td>NA</td><td>Intergenic</td><td>NA</td><td>NA</td></tr>
<tr><td></td><td></td><td></td><td></td><td></td><td></td><td></td><td></td><td></td><td></td><td></td><td></td></tr>
<tr><td>T1569</td><td>GACGACGTCGTCGTCGTCATCGG</td><td>1000</td><td></td><td></td><td></td><td></td><td></td><td></td><td></td><td></td><td></td></tr>
<tr><td>Oligo fwd</td><td>CACCGACGACGTCGTCGTCGTCAT</td><td></td><td></td><td></td><td></td><td></td><td></td><td></td><td></td><td></td><td></td></tr>
<tr><td>Oligo rev</td><td>AAACATGACGACGACGACGTCGTC</td><td></td><td></td><td></td><td></td><td></td><td></td><td></td><td></td><td></td><td></td></tr>
<tr><td>Chromosome</td><td>start</td><td>end</td><td>strand</td><td>MM</td><td>target_seq</td><td>PAM</td><td>alignment</td><td>distance</td><td>position</td><td>gene name</td><td>gene id</td></tr>
<tr><td>GU305914.1</td><td>26364</td><td>26386</td><td>+</td><td>0</td><td>GACGACGTCGTCGTCGTCAT</td><td>CGG</td><td>||||||||[||||||||||||]PAM</td><td>NA</td><td>Intergenic</td><td>NA</td><td>NA</td></tr>
<tr><td></td><td></td><td></td><td></td><td></td><td></td><td></td><td></td><td></td><td></td><td></td><td></td></tr>
<tr><td>T1570</td><td>GTCGTCGTCGTCATCGGCGCTGG</td><td>1000</td><td></td><td></td><td></td><td></td><td></td><td></td><td></td><td></td><td></td></tr>
<tr><td>Oligo fwd</td><td>CACCGTCGTCGTCGTCATCGGCGC</td><td></td><td></td><td></td><td></td><td></td><td></td><td></td><td></td><td></td><td></td></tr>
<tr><td>Oligo rev</td><td>AAACGCGCCGATGACGACGACGAC</td><td></td><td></td><td></td><td></td><td></td><td></td><td></td><td></td><td></td><td></td></tr>
<tr><td>Chromosome</td><td>start</td><td>end</td><td>strand</td><td>MM</td><td>target_seq</td><td>PAM</td><td>alignment</td><td>distance</td><td>position</td><td>gene name</td><td>gene id</td></tr>
<tr><td>GU305914.1</td><td>26370</td><td>26392</td><td>+</td><td>0</td><td>GTCGTCGTCGTCATCGGCGC</td><td>TGG</td><td>||||||||[||||||||||||]PAM</td><td>NA</td><td>Intergenic</td><td>NA</td><td>NA</td></tr>
<tr><td></td><td></td><td></td><td></td><td></td><td></td><td></td><td></td><td></td><td></td><td></td><td></td></tr>
<tr><td>T1571</td><td>GTCGTCATCGGCGCTGGCGACGG</td><td>1000</td><td></td><td></td><td></td><td></td><td></td><td></td><td></td><td></td><td></td></tr>
<tr><td>Oligo fwd</td><td>CACCGTCGTCATCGGCGCTGGCGA</td><td></td><td></td><td></td><td></td><td></td><td></td><td></td><td></td><td></td><td></td></tr>
<tr><td>Oligo rev</td><td>AAACTCGCCAGCGCCGATGACGAC</td><td></td><td></td><td></td><td></td><td></td><td></td><td></td><td></td><td></td><td></td></tr>
<tr><td>Chromosome</td><td>start</td><td>end</td><td>strand</td><td>MM</td><td>target_seq</td><td>PAM</td><td>alignment</td><td>distance</td><td>position</td><td>gene name</td><td>gene id</td></tr>
<tr><td>GU305914.1</td><td>26376</td><td>26398</td><td>+</td><td>0</td><td>GTCGTCATCGGCGCTGGCGA</td><td>CGG</td><td>||||||||[||||||||||||]PAM</td><td>NA</td><td>Intergenic</td><td>NA</td><td>NA</td></tr>
<tr><td></td><td></td><td></td><td></td><td></td><td></td><td></td><td></td><td></td><td></td><td></td><td></td></tr>
<tr><td>T1572</td><td>GTCATCGGCGCTGGCGACGGCGG</td><td>1000</td><td></td><td></td><td></td><td></td><td></td><td></td><td></td><td></td><td></td></tr>
<tr><td>Oligo fwd</td><td>CACCGTCATCGGCGCTGGCGACGG</td><td></td><td></td><td></td><td></td><td></td><td></td><td></td><td></td><td></td><td></td></tr>
<tr><td>Oligo rev</td><td>AAACCCGTCGCCAGCGCCGATGAC</td><td></td><td></td><td></td><td></td><td></td><td></td><td></td><td></td><td></td><td></td></tr>
<tr><td>Chromosome</td><td>start</td><td>end</td><td>strand</td><td>MM</td><td>target_seq</td><td>PAM</td><td>alignment</td><td>distance</td><td>position</td><td>gene name</td><td>gene id</td></tr>
<tr><td>GU305914.1</td><td>26379</td><td>26401</td><td>+</td><td>0</td><td>GTCATCGGCGCTGGCGACGG</td><td>CGG</td><td>||||||||[||||||||||||]PAM</td><td>NA</td><td>Intergenic</td><td>NA</td><td>NA</td></tr>
<tr><td>GU305914.1</td><td>127139</td><td>127161</td><td>-</td><td>4</td><td>GGGAACGGCGTTGGCGACGG</td><td>CGG</td><td>|--|-|||[||-|||||||||]PAM</td><td>NA</td><td>Intergenic</td><td>NA</td><td>NA</td></tr>
<tr><td></td><td></td><td></td><td></td><td></td><td></td><td></td><td></td><td></td><td></td><td></td><td></td></tr>
<tr><td>T1573</td><td>ATCGGCGCTGGCGACGGCGGCGG</td><td>1000</td><td></td><td></td><td></td><td></td><td></td><td></td><td></td><td></td><td></td></tr>
<tr><td>Oligo fwd</td><td>CACCATCGGCGCTGGCGACGGCGG</td><td></td><td></td><td></td><td></td><td></td><td></td><td></td><td></td><td></td><td></td></tr>
<tr><td>Oligo rev</td><td>AAACCCGCCGTCGCCAGCGCCGAT</td><td></td><td></td><td></td><td></td><td></td><td></td><td></td><td></td><td></td><td></td></tr>
<tr><td>Chromosome</td><td>start</td><td>end</td><td>strand</td><td>MM</td><td>target_seq</td><td>PAM</td><td>alignment</td><td>distance</td><td>position</td><td>gene name</td><td>gene id</td></tr>
<tr><td>GU305914.1</td><td>26382</td><td>26404</td><td>+</td><td>0</td><td>ATCGGCGCTGGCGACGGCGG</td><td>CGG</td><td>||||||||[||||||||||||]PAM</td><td>NA</td><td>Intergenic</td><td>NA</td><td>NA</td></tr>
<tr><td>GU305914.1</td><td>29997</td><td>30019</td><td>+</td><td>3</td><td>ACCGGTGCTGGCGGCGGCGG</td><td>CGG</td><td>|-|||-||[|||||-||||||]PAM</td><td>NA</td><td>Intergenic</td><td>NA</td><td>NA</td></tr>
<tr><td>GU305914.1</td><td>48159</td><td>48181</td><td>+</td><td>4</td><td>GGCGGCGGTGGCGGCGGCGG</td><td>TGG</td><td>--|||||-[|||||-||||||]PAM</td><td>NA</td><td>Intergenic</td><td>NA</td><td>NA</td></tr>
<tr><td>GU305914.1</td><td>91257</td><td>91279</td><td>+</td><td>3</td><td>ATCGTCGCAGGCGACGGCCG</td><td>CGG</td><td>||||-|||[-|||||||||-|]PAM</td><td>NA</td><td>Intergenic</td><td>NA</td><td>NA</td></tr>
<tr><td></td><td></td><td></td><td></td><td></td><td></td><td></td><td></td><td></td><td></td><td></td><td></td></tr>
<tr><td>T1574</td><td>TCGGCGCTGGCGACGGCGGCGGG</td><td>1000</td><td></td><td></td><td></td><td></td><td></td><td></td><td></td><td></td><td></td></tr>
<tr><td>Oligo fwd</td><td>CACCTCGGCGCTGGCGACGGCGGC</td><td></td><td></td><td></td><td></td><td></td><td></td><td></td><td></td><td></td><td></td></tr>
<tr><td>Oligo rev</td><td>AAACGCCGCCGTCGCCAGCGCCGA</td><td></td><td></td><td></td><td></td><td></td><td></td><td></td><td></td><td></td><td></td></tr>
<tr><td>Chromosome</td><td>start</td><td>end</td><td>strand</td><td>MM</td><td>target_seq</td><td>PAM</td><td>alignment</td><td>distance</td><td>position</td><td>gene name</td><td>gene id</td></tr>
<tr><td>GU305914.1</td><td>26383</td><td>26405</td><td>+</td><td>0</td><td>TCGGCGCTGGCGACGGCGGC</td><td>GGG</td><td>||||||||[||||||||||||]PAM</td><td>NA</td><td>Intergenic</td><td>NA</td><td>NA</td></tr>
<tr><td>GU305914.1</td><td>48160</td><td>48182</td><td>+</td><td>4</td><td>GCGGCGGTGGCGGCGGCGGT</td><td>GGG</td><td>-|||||-|[||||-||||||-]PAM</td><td>NA</td><td>Intergenic</td><td>NA</td><td>NA</td></tr>
<tr><td></td><td></td><td></td><td></td><td></td><td></td><td></td><td></td><td></td><td></td><td></td><td></td></tr>
<tr><td>T1575</td><td>CGGCGCTGGCGACGGCGGCGGGG</td><td>1000</td><td></td><td></td><td></td><td></td><td></td><td></td><td></td><td></td><td></td></tr>
<tr><td>Oligo fwd</td><td>CACCCGGCGCTGGCGACGGCGGCG</td><td></td><td></td><td></td><td></td><td></td><td></td><td></td><td></td><td></td><td></td></tr>
<tr><td>Oligo rev</td><td>AAACCGCCGCCGTCGCCAGCGCCG</td><td></td><td></td><td></td><td></td><td></td><td></td><td></td><td></td><td></td><td></td></tr>
<tr><td>Chromosome</td><td>start</td><td>end</td><td>strand</td><td>MM</td><td>target_seq</td><td>PAM</td><td>alignment</td><td>distance</td><td>position</td><td>gene name</td><td>gene id</td></tr>
<tr><td>GU305914.1</td><td>26384</td><td>26406</td><td>+</td><td>0</td><td>CGGCGCTGGCGACGGCGGCG</td><td>GGG</td><td>||||||||[||||||||||||]PAM</td><td>NA</td><td>Intergenic</td><td>NA</td><td>NA</td></tr>
<tr><td>GU305914.1</td><td>91844</td><td>91866</td><td>+</td><td>4</td><td>CGGCGGGAGCGACGGGGGCG</td><td>AGG</td><td>|||||---[|||||||-||||]PAM</td><td>NA</td><td>Intergenic</td><td>NA</td><td>NA</td></tr>
<tr><td></td><td></td><td></td><td></td><td></td><td></td><td></td><td></td><td></td><td></td><td></td><td></td></tr>
<tr><td>T1576</td><td>TGGCGACGGCGGCGGGGATACGG</td><td>1000</td><td></td><td></td><td></td><td></td><td></td><td></td><td></td><td></td><td></td></tr>
<tr><td>Oligo fwd</td><td>CACCTGGCGACGGCGGCGGGGATA</td><td></td><td></td><td></td><td></td><td></td><td></td><td></td><td></td><td></td><td></td></tr>
<tr><td>Oligo rev</td><td>AAACTATCCCCGCCGCCGTCGCCA</td><td></td><td></td><td></td><td></td><td></td><td></td><td></td><td></td><td></td><td></td></tr>
<tr><td>Chromosome</td><td>start</td><td>end</td><td>strand</td><td>MM</td><td>target_seq</td><td>PAM</td><td>alignment</td><td>distance</td><td>position</td><td>gene name</td><td>gene id</td></tr>
<tr><td>GU305914.1</td><td>26390</td><td>26412</td><td>+</td><td>0</td><td>TGGCGACGGCGGCGGGGATA</td><td>CGG</td><td>||||||||[||||||||||||]PAM</td><td>NA</td><td>Intergenic</td><td>NA</td><td>NA</td></tr>
<tr><td></td><td></td><td></td><td></td><td></td><td></td><td></td><td></td><td></td><td></td><td></td><td></td></tr>
<tr><td>T1577</td><td>GGCGACGGCGGCGGGGATACGGG</td><td>1000</td><td></td><td></td><td></td><td></td><td></td><td></td><td></td><td></td><td></td></tr>
<tr><td>Oligo fwd</td><td>CACCGGCGACGGCGGCGGGGATAC</td><td></td><td></td><td></td><td></td><td></td><td></td><td></td><td></td><td></td><td></td></tr>
<tr><td>Oligo rev</td><td>AAACGTATCCCCGCCGCCGTCGCC</td><td></td><td></td><td></td><td></td><td></td><td></td><td></td><td></td><td></td><td></td></tr>
<tr><td>Chromosome</td><td>start</td><td>end</td><td>strand</td><td>MM</td><td>target_seq</td><td>PAM</td><td>alignment</td><td>distance</td><td>position</td><td>gene name</td><td>gene id</td></tr>
<tr><td>GU305914.1</td><td>26391</td><td>26413</td><td>+</td><td>0</td><td>GGCGACGGCGGCGGGGATAC</td><td>GGG</td><td>||||||||[||||||||||||]PAM</td><td>NA</td><td>Intergenic</td><td>NA</td><td>NA</td></tr>
<tr><td></td><td></td><td></td><td></td><td></td><td></td><td></td><td></td><td></td><td></td><td></td><td></td></tr>
<tr><td>T1578</td><td>GCGACGGCGGCGGGGATACGGGG</td><td>1000</td><td></td><td></td><td></td><td></td><td></td><td></td><td></td><td></td><td></td></tr>
<tr><td>Oligo fwd</td><td>CACCGCGACGGCGGCGGGGATACG</td><td></td><td></td><td></td><td></td><td></td><td></td><td></td><td></td><td></td><td></td></tr>
<tr><td>Oligo rev</td><td>AAACCGTATCCCCGCCGCCGTCGC</td><td></td><td></td><td></td><td></td><td></td><td></td><td></td><td></td><td></td><td></td></tr>
<tr><td>Chromosome</td><td>start</td><td>end</td><td>strand</td><td>MM</td><td>target_seq</td><td>PAM</td><td>alignment</td><td>distance</td><td>position</td><td>gene name</td><td>gene id</td></tr>
<tr><td>GU305914.1</td><td>26392</td><td>26414</td><td>+</td><td>0</td><td>GCGACGGCGGCGGGGATACG</td><td>GGG</td><td>||||||||[||||||||||||]PAM</td><td>NA</td><td>Intergenic</td><td>NA</td><td>NA</td></tr>
<tr><td>GU305914.1</td><td>199022</td><td>199044</td><td>-</td><td>4</td><td>GCGAGGGTTGCGAGGATACG</td><td>GGG</td><td>||||-||-[-|||-|||||||]PAM</td><td>NA</td><td>Intergenic</td><td>NA</td><td>NA</td></tr>
<tr><td></td><td></td><td></td><td></td><td></td><td></td><td></td><td></td><td></td><td></td><td></td><td></td></tr>
<tr><td>T1579</td><td>CCCCGCCTCCAACTCCGCCACGG</td><td>1000</td><td></td><td></td><td></td><td></td><td></td><td></td><td></td><td></td><td></td></tr>
<tr><td>Oligo fwd</td><td>CACCCCCCGCCTCCAACTCCGCCA</td><td></td><td></td><td></td><td></td><td></td><td></td><td></td><td></td><td></td><td></td></tr>
<tr><td>Oligo rev</td><td>AAACTGGCGGAGTTGGAGGCGGGG</td><td></td><td></td><td></td><td></td><td></td><td></td><td></td><td></td><td></td><td></td></tr>
<tr><td>Chromosome</td><td>start</td><td>end</td><td>strand</td><td>MM</td><td>target_seq</td><td>PAM</td><td>alignment</td><td>distance</td><td>position</td><td>gene name</td><td>gene id</td></tr>
<tr><td>GU305914.1</td><td>26427</td><td>26449</td><td>+</td><td>0</td><td>CCCCGCCTCCAACTCCGCCA</td><td>CGG</td><td>||||||||[||||||||||||]PAM</td><td>NA</td><td>Intergenic</td><td>NA</td><td>NA</td></tr>
<tr><td></td><td></td><td></td><td></td><td></td><td></td><td></td><td></td><td></td><td></td><td></td><td></td></tr>
<tr><td>T1580</td><td>CGCCTCCAACTCCGCCACGGCGG</td><td>1000</td><td></td><td></td><td></td><td></td><td></td><td></td><td></td><td></td><td></td></tr>
<tr><td>Oligo fwd</td><td>CACCCGCCTCCAACTCCGCCACGG</td><td></td><td></td><td></td><td></td><td></td><td></td><td></td><td></td><td></td><td></td></tr>
</table>

<table>
<tr><td>Oligo rev</td><td>AAACCCGTGGCGGAGTTGGAGGCG</td><td></td><td></td><td></td><td></td><td></td><td></td><td></td><td></td><td></td><td></td></tr>
<tr><td>Chromosome</td><td>start</td><td>end</td><td>strand</td><td>MM</td><td>target_seq</td><td>PAM</td><td>alignment</td><td>distance</td><td>position</td><td>gene name</td><td>gene id</td></tr>
<tr><td>GU305914.1</td><td>26430</td><td>26452</td><td>+</td><td>0</td><td>CGCCTCCAACTCCGCCACGG</td><td>CGG</td><td>||||||||[||||||||||||]PAM</td><td>NA</td><td>Intergenic</td><td>NA</td><td>NA</td></tr>
<tr><td></td><td></td><td></td><td></td><td></td><td></td><td></td><td></td><td></td><td></td><td></td><td></td></tr>
<tr><td>T1581</td><td>CTCCAACTCCGCCACGGCGGCGG</td><td>1000</td><td></td><td></td><td></td><td></td><td></td><td></td><td></td><td></td><td></td></tr>
<tr><td>Oligo fwd</td><td>CACCCTCCAACTCCGCCACGGCGG</td><td></td><td></td><td></td><td></td><td></td><td></td><td></td><td></td><td></td><td></td></tr>
<tr><td>Oligo rev</td><td>AAACCCGCCGTGGCGGAGTTGGAG</td><td></td><td></td><td></td><td></td><td></td><td></td><td></td><td></td><td></td><td></td></tr>
<tr><td>Chromosome</td><td>start</td><td>end</td><td>strand</td><td>MM</td><td>target_seq</td><td>PAM</td><td>alignment</td><td>distance</td><td>position</td><td>gene name</td><td>gene id</td></tr>
<tr><td>GU305914.1</td><td>26433</td><td>26455</td><td>+</td><td>0</td><td>CTCCAACTCCGCCACGGCGG</td><td>CGG</td><td>||||||||[||||||||||||]PAM</td><td>NA</td><td>Intergenic</td><td>NA</td><td>NA</td></tr>
<tr><td></td><td></td><td></td><td></td><td></td><td></td><td></td><td></td><td></td><td></td><td></td><td></td></tr>
<tr><td>T1582</td><td>CAACTCCGCCACGGCGGCGGCGG</td><td>1000</td><td></td><td></td><td></td><td></td><td></td><td></td><td></td><td></td><td></td></tr>
<tr><td>Oligo fwd</td><td>CACCCAACTCCGCCACGGCGGCGG</td><td></td><td></td><td></td><td></td><td></td><td></td><td></td><td></td><td></td><td></td></tr>
<tr><td>Oligo rev</td><td>AAACCCGCCGCCGTGGCGGAGTTG</td><td></td><td></td><td></td><td></td><td></td><td></td><td></td><td></td><td></td><td></td></tr>
<tr><td>Chromosome</td><td>start</td><td>end</td><td>strand</td><td>MM</td><td>target_seq</td><td>PAM</td><td>alignment</td><td>distance</td><td>position</td><td>gene name</td><td>gene id</td></tr>
<tr><td>GU305914.1</td><td>26436</td><td>26458</td><td>+</td><td>0</td><td>CAACTCCGCCACGGCGGCGG</td><td>CGG</td><td>||||||||[||||||||||||]PAM</td><td>NA</td><td>Intergenic</td><td>NA</td><td>NA</td></tr>
<tr><td>GU305914.1</td><td>56767</td><td>56789</td><td>+</td><td>4</td><td>CTTCTCCGCCGCAGCGGCGG</td><td>CGG</td><td>|--|||||[||-|-|||||||]PAM</td><td>NA</td><td>Intergenic</td><td>NA</td><td>NA</td></tr>
<tr><td></td><td></td><td></td><td></td><td></td><td></td><td></td><td></td><td></td><td></td><td></td><td></td></tr>
<tr><td>T1583</td><td>GCCATTGAGCATCCCGTCTGAGG</td><td>1000</td><td></td><td></td><td></td><td></td><td></td><td></td><td></td><td></td><td></td></tr>
<tr><td>Oligo fwd</td><td>CACCGCCATTGAGCATCCCGTCTG</td><td></td><td></td><td></td><td></td><td></td><td></td><td></td><td></td><td></td><td></td></tr>
<tr><td>Oligo rev</td><td>AAACCAGACGGGATGCTCAATGGC</td><td></td><td></td><td></td><td></td><td></td><td></td><td></td><td></td><td></td><td></td></tr>
<tr><td>Chromosome</td><td>start</td><td>end</td><td>strand</td><td>MM</td><td>target_seq</td><td>PAM</td><td>alignment</td><td>distance</td><td>position</td><td>gene name</td><td>gene id</td></tr>
<tr><td>GU305914.1</td><td>26478</td><td>26500</td><td>+</td><td>0</td><td>GCCATTGAGCATCCCGTCTG</td><td>AGG</td><td>||||||||[||||||||||||]PAM</td><td>NA</td><td>Intergenic</td><td>NA</td><td>NA</td></tr>
<tr><td></td><td></td><td></td><td></td><td></td><td></td><td></td><td></td><td></td><td></td><td></td><td></td></tr>
<tr><td>T1584</td><td>ATCCCGTCTGAGGAACTAAACGG</td><td>1000</td><td></td><td></td><td></td><td></td><td></td><td></td><td></td><td></td><td></td></tr>
<tr><td>Oligo fwd</td><td>CACCATCCCGTCTGAGGAACTAAA</td><td></td><td></td><td></td><td></td><td></td><td></td><td></td><td></td><td></td><td></td></tr>
<tr><td>Oligo rev</td><td>AAACTTTAGTTCCTCAGACGGGAT</td><td></td><td></td><td></td><td></td><td></td><td></td><td></td><td></td><td></td><td></td></tr>
<tr><td>Chromosome</td><td>start</td><td>end</td><td>strand</td><td>MM</td><td>target_seq</td><td>PAM</td><td>alignment</td><td>distance</td><td>position</td><td>gene name</td><td>gene id</td></tr>
<tr><td>GU305914.1</td><td>26488</td><td>26510</td><td>+</td><td>0</td><td>ATCCCGTCTGAGGAACTAAA</td><td>CGG</td><td>||||||||[||||||||||||]PAM</td><td>NA</td><td>Intergenic</td><td>NA</td><td>NA</td></tr>
<tr><td></td><td></td><td></td><td></td><td></td><td></td><td></td><td></td><td></td><td></td><td></td><td></td></tr>
<tr><td>T1585</td><td>TGAGGAACTAAACGGCGAAGAGG</td><td>1000</td><td></td><td></td><td></td><td></td><td></td><td></td><td></td><td></td><td></td></tr>
<tr><td>Oligo fwd</td><td>CACCTGAGGAACTAAACGGCGAAG</td><td></td><td></td><td></td><td></td><td></td><td></td><td></td><td></td><td></td><td></td></tr>
<tr><td>Oligo rev</td><td>AAACCTTCGCCGTTTAGTTCCTCA</td><td></td><td></td><td></td><td></td><td></td><td></td><td></td><td></td><td></td><td></td></tr>
<tr><td>Chromosome</td><td>start</td><td>end</td><td>strand</td><td>MM</td><td>target_seq</td><td>PAM</td><td>alignment</td><td>distance</td><td>position</td><td>gene name</td><td>gene id</td></tr>
<tr><td>GU305914.1</td><td>26496</td><td>26518</td><td>+</td><td>0</td><td>TGAGGAACTAAACGGCGAAG</td><td>AGG</td><td>||||||||[||||||||||||]PAM</td><td>NA</td><td>Intergenic</td><td>NA</td><td>NA</td></tr>
<tr><td></td><td></td><td></td><td></td><td></td><td></td><td></td><td></td><td></td><td></td><td></td><td></td></tr>
<tr><td>T1586</td><td>GGAACTAAACGGCGAAGAGGAGG</td><td>1000</td><td></td><td></td><td></td><td></td><td></td><td></td><td></td><td></td><td></td></tr>
<tr><td>Oligo fwd</td><td>CACCGGAACTAAACGGCGAAGAGG</td><td></td><td></td><td></td><td></td><td></td><td></td><td></td><td></td><td></td><td></td></tr>
<tr><td>Oligo rev</td><td>AAACCCTCTTCGCCGTTTAGTTCC</td><td></td><td></td><td></td><td></td><td></td><td></td><td></td><td></td><td></td><td></td></tr>
<tr><td>Chromosome</td><td>start</td><td>end</td><td>strand</td><td>MM</td><td>target_seq</td><td>PAM</td><td>alignment</td><td>distance</td><td>position</td><td>gene name</td><td>gene id</td></tr>
<tr><td>GU305914.1</td><td>26499</td><td>26521</td><td>+</td><td>0</td><td>GGAACTAAACGGCGAAGAGG</td><td>AGG</td><td>||||||||[||||||||||||]PAM</td><td>NA</td><td>Intergenic</td><td>NA</td><td>NA</td></tr>
<tr><td></td><td></td><td></td><td></td><td></td><td></td><td></td><td></td><td></td><td></td><td></td><td></td></tr>
<tr><td>T1587</td><td>GAAGAAGACATGTCACGCGACGG</td><td>1000</td><td></td><td></td><td></td><td></td><td></td><td></td><td></td><td></td><td></td></tr>
<tr><td>Oligo fwd</td><td>CACCGAAGAAGACATGTCACGCGA</td><td></td><td></td><td></td><td></td><td></td><td></td><td></td><td></td><td></td><td></td></tr>
<tr><td>Oligo rev</td><td>AAACTCGCGTGACATGTCTTCTTC</td><td></td><td></td><td></td><td></td><td></td><td></td><td></td><td></td><td></td><td></td></tr>
<tr><td>Chromosome</td><td>start</td><td>end</td><td>strand</td><td>MM</td><td>target_seq</td><td>PAM</td><td>alignment</td><td>distance</td><td>position</td><td>gene name</td><td>gene id</td></tr>
<tr><td>GU305914.1</td><td>26530</td><td>26552</td><td>+</td><td>0</td><td>GAAGAAGACATGTCACGCGA</td><td>CGG</td><td>||||||||[||||||||||||]PAM</td><td>NA</td><td>Intergenic</td><td>NA</td><td>NA</td></tr>
<tr><td></td><td></td><td></td><td></td><td></td><td></td><td></td><td></td><td></td><td></td><td></td><td></td></tr>
<tr><td>T1588</td><td>TGTCACGCGACGGACCCCGCCGG</td><td>1000</td><td></td><td></td><td></td><td></td><td></td><td></td><td></td><td></td><td></td></tr>
<tr><td>Oligo fwd</td><td>CACCTGTCACGCGACGGACCCCGC</td><td></td><td></td><td></td><td></td><td></td><td></td><td></td><td></td><td></td><td></td></tr>
<tr><td>Oligo rev</td><td>AAACGCGGGGTCCGTCGCGTGACA</td><td></td><td></td><td></td><td></td><td></td><td></td><td></td><td></td><td></td><td></td></tr>
<tr><td>Chromosome</td><td>start</td><td>end</td><td>strand</td><td>MM</td><td>target_seq</td><td>PAM</td><td>alignment</td><td>distance</td><td>position</td><td>gene name</td><td>gene id</td></tr>
<tr><td>GU305914.1</td><td>26540</td><td>26562</td><td>+</td><td>0</td><td>TGTCACGCGACGGACCCCGC</td><td>CGG</td><td>||||||||[||||||||||||]PAM</td><td>NA</td><td>Intergenic</td><td>NA</td><td>NA</td></tr>
<tr><td></td><td></td><td></td><td></td><td></td><td></td><td></td><td></td><td></td><td></td><td></td><td></td></tr>
<tr><td>T1589</td><td>CGGACCCCGCCGGCACAGCCAGG</td><td>1000</td><td></td><td></td><td></td><td></td><td></td><td></td><td></td><td></td><td></td></tr>
<tr><td>Oligo fwd</td><td>CACCCGGACCCCGCCGGCACAGCC</td><td></td><td></td><td></td><td></td><td></td><td></td><td></td><td></td><td></td><td></td></tr>
<tr><td>Oligo rev</td><td>AAACGGCTGTGCCGGCGGGGTCCG</td><td></td><td></td><td></td><td></td><td></td><td></td><td></td><td></td><td></td><td></td></tr>
<tr><td>Chromosome</td><td>start</td><td>end</td><td>strand</td><td>MM</td><td>target_seq</td><td>PAM</td><td>alignment</td><td>distance</td><td>position</td><td>gene name</td><td>gene id</td></tr>
<tr><td>GU305914.1</td><td>26550</td><td>26572</td><td>+</td><td>0</td><td>CGGACCCCGCCGGCACAGCC</td><td>AGG</td><td>||||||||[||||||||||||]PAM</td><td>NA</td><td>Intergenic</td><td>NA</td><td>NA</td></tr>
<tr><td></td><td></td><td></td><td></td><td></td><td></td><td></td><td></td><td></td><td></td><td></td><td></td></tr>
<tr><td>T1590</td><td>GGACGACGACTTTACTTATGCGG</td><td>1000</td><td></td><td></td><td></td><td></td><td></td><td></td><td></td><td></td><td></td></tr>
<tr><td>Oligo fwd</td><td>CACCGGACGACGACTTTACTTATG</td><td></td><td></td><td></td><td></td><td></td><td></td><td></td><td></td><td></td><td></td></tr>
<tr><td>Oligo rev</td><td>AAACCATAAGTAAAGTCGTCGTCC</td><td></td><td></td><td></td><td></td><td></td><td></td><td></td><td></td><td></td><td></td></tr>
<tr><td>Chromosome</td><td>start</td><td>end</td><td>strand</td><td>MM</td><td>target_seq</td><td>PAM</td><td>alignment</td><td>distance</td><td>position</td><td>gene name</td><td>gene id</td></tr>
<tr><td>GU305914.1</td><td>26571</td><td>26593</td><td>+</td><td>0</td><td>GGACGACGACTTTACTTATG</td><td>CGG</td><td>||||||||[||||||||||||]PAM</td><td>NA</td><td>Intergenic</td><td>NA</td><td>NA</td></tr>
<tr><td></td><td></td><td></td><td></td><td></td><td></td><td></td><td></td><td></td><td></td><td></td><td></td></tr>
<tr><td>T1591</td><td>CGACTTTACTTATGCGGACCCGG</td><td>1000</td><td></td><td></td><td></td><td></td><td></td><td></td><td></td><td></td><td></td></tr>
<tr><td>Oligo fwd</td><td>CACCCGACTTTACTTATGCGGACC</td><td></td><td></td><td></td><td></td><td></td><td></td><td></td><td></td><td></td><td></td></tr>
<tr><td>Oligo rev</td><td>AAACGGTCCGCATAAGTAAAGTCG</td><td></td><td></td><td></td><td></td><td></td><td></td><td></td><td></td><td></td><td></td></tr>
<tr><td>Chromosome</td><td>start</td><td>end</td><td>strand</td><td>MM</td><td>target_seq</td><td>PAM</td><td>alignment</td><td>distance</td><td>position</td><td>gene name</td><td>gene id</td></tr>
<tr><td>GU305914.1</td><td>26577</td><td>26599</td><td>+</td><td>0</td><td>CGACTTTACTTATGCGGACC</td><td>CGG</td><td>||||||||[||||||||||||]PAM</td><td>NA</td><td>Intergenic</td><td>NA</td><td>NA</td></tr>
<tr><td></td><td></td><td></td><td></td><td></td><td></td><td></td><td></td><td></td><td></td><td></td><td></td></tr>
<tr><td>T1592</td><td>ACCCGGCCGATGTGAGACTGCGG</td><td>1000</td><td></td><td></td><td></td><td></td><td></td><td></td><td></td><td></td><td></td></tr>
<tr><td>Oligo fwd</td><td>CACCACCCGGCCGATGTGAGACTG</td><td></td><td></td><td></td><td></td><td></td><td></td><td></td><td></td><td></td><td></td></tr>
<tr><td>Oligo rev</td><td>AAACCAGTCTCACATCGGCCGGGT</td><td></td><td></td><td></td><td></td><td></td><td></td><td></td><td></td><td></td><td></td></tr>
<tr><td>Chromosome</td><td>start</td><td>end</td><td>strand</td><td>MM</td><td>target_seq</td><td>PAM</td><td>alignment</td><td>distance</td><td>position</td><td>gene name</td><td>gene id</td></tr>
<tr><td>GU305914.1</td><td>26594</td><td>26616</td><td>+</td><td>0</td><td>ACCCGGCCGATGTGAGACTG</td><td>CGG</td><td>||||||||[||||||||||||]PAM</td><td>NA</td><td>Intergenic</td><td>NA</td><td>NA</td></tr>
<tr><td></td><td></td><td></td><td></td><td></td><td></td><td></td><td></td><td></td><td></td><td></td><td></td></tr>
<tr><td>T1593</td><td>CCCGGCCGATGTGAGACTGCGGG</td><td>1000</td><td></td><td></td><td></td><td></td><td></td><td></td><td></td><td></td><td></td></tr>
<tr><td>Oligo fwd</td><td>CACCCCCGGCCGATGTGAGACTGC</td><td></td><td></td><td></td><td></td><td></td><td></td><td></td><td></td><td></td><td></td></tr>
<tr><td>Oligo rev</td><td>AAACGCAGTCTCACATCGGCCGGG</td><td></td><td></td><td></td><td></td><td></td><td></td><td></td><td></td><td></td><td></td></tr>
<tr><td>Chromosome</td><td>start</td><td>end</td><td>strand</td><td>MM</td><td>target_seq</td><td>PAM</td><td>alignment</td><td>distance</td><td>position</td><td>gene name</td><td>gene id</td></tr>
<tr><td>GU305914.1</td><td>26595</td><td>26617</td><td>+</td><td>0</td><td>CCCGGCCGATGTGAGACTGC</td><td>GGG</td><td>||||||||[||||||||||||]PAM</td><td>NA</td><td>Intergenic</td><td>NA</td><td>NA</td></tr>
<tr><td></td><td></td><td></td><td></td><td></td><td></td><td></td><td></td><td></td><td></td><td></td><td></td></tr>
<tr><td>T1594</td><td>TGTGAGACTGCGGGCGATGATGG</td><td>1000</td><td></td><td></td><td></td><td></td><td></td><td></td><td></td><td></td><td></td></tr>
<tr><td>Oligo fwd</td><td>CACCTGTGAGACTGCGGGCGATGA</td><td></td><td></td><td></td><td></td><td></td><td></td><td></td><td></td><td></td><td></td></tr>
<tr><td>Oligo rev</td><td>AAACTCATCGCCCGCAGTCTCACA</td><td></td><td></td><td></td><td></td><td></td><td></td><td></td><td></td><td></td><td></td></tr>
<tr><td>Chromosome</td><td>start</td><td>end</td><td>strand</td><td>MM</td><td>target_seq</td><td>PAM</td><td>alignment</td><td>distance</td><td>position</td><td>gene name</td><td>gene id</td></tr>
<tr><td>GU305914.1</td><td>26604</td><td>26626</td><td>+</td><td>0</td><td>TGTGAGACTGCGGGCGATGA</td><td>TGG</td><td>||||||||[||||||||||||]PAM</td><td>NA</td><td>Intergenic</td><td>NA</td><td>NA</td></tr>
</table>

<table>
<tr><td>T1595</td><td>GTGAGACTGCGGGCGATGATGGG</td><td>1000</td></tr>
<tr><td>Oligo fwd</td><td>CACCGTGAGACTGCGGGCGATGAT</td></tr>
<tr><td>Oligo rev</td><td>AAACATCATCGCCCGCAGTCTCAC</td></tr>
<tr><td>Chromosome</td><td>start</td><td>end</td><td>strand</td><td>MM</td><td>target_seq</td><td>PAM</td><td>alignment</td><td>distance</td><td>position</td><td>gene name</td><td>gene id</td></tr>
<tr><td>GU305914.1</td><td>26605</td><td>26627</td><td>+</td><td>0</td><td>GTGAGACTGCGGGCGATGAT</td><td>GGG</td><td>||||||||[||||||||||||]PAM</td><td>NA</td><td>Intergenic</td><td>NA</td><td>NA</td></tr>
<tr><td></td></tr>
<tr><td>T1596</td><td>TGCGGGCGATGATGGGCAATAGG</td><td>1000</td></tr>
<tr><td>Oligo fwd</td><td>CACCTGCGGGCGATGATGGGCAAT</td></tr>
<tr><td>Oligo rev</td><td>AAACATTGCCCATCATCGCCCGCA</td></tr>
<tr><td>Chromosome</td><td>start</td><td>end</td><td>strand</td><td>MM</td><td>target_seq</td><td>PAM</td><td>alignment</td><td>distance</td><td>position</td><td>gene name</td><td>gene id</td></tr>
<tr><td>GU305914.1</td><td>26612</td><td>26634</td><td>+</td><td>0</td><td>TGCGGGCGATGATGGGCAAT</td><td>AGG</td><td>||||||||[||||||||||||]PAM</td><td>NA</td><td>Intergenic</td><td>NA</td><td>NA</td></tr>
<tr><td></td></tr>
<tr><td>T1597</td><td>GCGATGATGGGCAATAGGTACGG</td><td>1000</td></tr>
<tr><td>Oligo fwd</td><td>CACCGCGATGATGGGCAATAGGTA</td></tr>
<tr><td>Oligo rev</td><td>AAACTACCTATTGCCCATCATCGC</td></tr>
<tr><td>Chromosome</td><td>start</td><td>end</td><td>strand</td><td>MM</td><td>target_seq</td><td>PAM</td><td>alignment</td><td>distance</td><td>position</td><td>gene name</td><td>gene id</td></tr>
<tr><td>GU305914.1</td><td>26617</td><td>26639</td><td>+</td><td>0</td><td>GCGATGATGGGCAATAGGTA</td><td>CGG</td><td>||||||||[||||||||||||]PAM</td><td>NA</td><td>Intergenic</td><td>NA</td><td>NA</td></tr>
<tr><td></td></tr>
<tr><td>T1598</td><td>ATGATGGGCAATAGGTACGGCGG</td><td>1000</td></tr>
<tr><td>Oligo fwd</td><td>CACCATGATGGGCAATAGGTACGG</td></tr>
<tr><td>Oligo rev</td><td>AAACCCGTACCTATTGCCCATCAT</td></tr>
<tr><td>Chromosome</td><td>start</td><td>end</td><td>strand</td><td>MM</td><td>target_seq</td><td>PAM</td><td>alignment</td><td>distance</td><td>position</td><td>gene name</td><td>gene id</td></tr>
<tr><td>GU305914.1</td><td>26620</td><td>26642</td><td>+</td><td>0</td><td>ATGATGGGCAATAGGTACGG</td><td>CGG</td><td>||||||||[||||||||||||]PAM</td><td>NA</td><td>Intergenic</td><td>NA</td><td>NA</td></tr>
<tr><td></td></tr>
<tr><td>T1599</td><td>ATAGGTACGGCGGACAGAGTCGG</td><td>1000</td></tr>
<tr><td>Oligo fwd</td><td>CACCATAGGTACGGCGGACAGAGT</td></tr>
<tr><td>Oligo rev</td><td>AAACACTCTGTCCGCCGTACCTAT</td></tr>
<tr><td>Chromosome</td><td>start</td><td>end</td><td>strand</td><td>MM</td><td>target_seq</td><td>PAM</td><td>alignment</td><td>distance</td><td>position</td><td>gene name</td><td>gene id</td></tr>
<tr><td>GU305914.1</td><td>26630</td><td>26652</td><td>+</td><td>0</td><td>ATAGGTACGGCGGACAGAGT</td><td>CGG</td><td>||||||||[||||||||||||]PAM</td><td>NA</td><td>Intergenic</td><td>NA</td><td>NA</td></tr>
<tr><td></td></tr>
<tr><td>T1600</td><td>CGGCGGACAGAGTCGGAGTGCGG</td><td>1000</td></tr>
<tr><td>Oligo fwd</td><td>CACCCGGCGGACAGAGTCGGAGTG</td></tr>
<tr><td>Oligo rev</td><td>AAACCACTCCGACTCTGTCCGCCG</td></tr>
<tr><td>Chromosome</td><td>start</td><td>end</td><td>strand</td><td>MM</td><td>target_seq</td><td>PAM</td><td>alignment</td><td>distance</td><td>position</td><td>gene name</td><td>gene id</td></tr>
<tr><td>GU305914.1</td><td>26637</td><td>26659</td><td>+</td><td>0</td><td>CGGCGGACAGAGTCGGAGTG</td><td>CGG</td><td>||||||||[||||||||||||]PAM</td><td>NA</td><td>Intergenic</td><td>NA</td><td>NA</td></tr>
<tr><td></td></tr>
<tr><td>T1601</td><td>GAGTCGGAGTGCGGCGACAGCGG</td><td>1000</td></tr>
<tr><td>Oligo fwd</td><td>CACCGAGTCGGAGTGCGGCGACAG</td></tr>
<tr><td>Oligo rev</td><td>AAACCTGTCGCCGCACTCCGACTC</td></tr>
<tr><td>Chromosome</td><td>start</td><td>end</td><td>strand</td><td>MM</td><td>target_seq</td><td>PAM</td><td>alignment</td><td>distance</td><td>position</td><td>gene name</td><td>gene id</td></tr>
<tr><td>GU305914.1</td><td>26646</td><td>26668</td><td>+</td><td>0</td><td>GAGTCGGAGTGCGGCGACAG</td><td>CGG</td><td>||||||||[||||||||||||]PAM</td><td>NA</td><td>Intergenic</td><td>NA</td><td>NA</td></tr>
<tr><td></td></tr>
<tr><td>T1602</td><td>AGTCGGAGTGCGGCGACAGCGGG</td><td>1000</td></tr>
<tr><td>Oligo fwd</td><td>CACCAGTCGGAGTGCGGCGACAGC</td></tr>
<tr><td>Oligo rev</td><td>AAACGCTGTCGCCGCACTCCGACT</td></tr>
<tr><td>Chromosome</td><td>start</td><td>end</td><td>strand</td><td>MM</td><td>target_seq</td><td>PAM</td><td>alignment</td><td>distance</td><td>position</td><td>gene name</td><td>gene id</td></tr>
<tr><td>GU305914.1</td><td>26647</td><td>26669</td><td>+</td><td>0</td><td>AGTCGGAGTGCGGCGACAGC</td><td>GGG</td><td>||||||||[||||||||||||]PAM</td><td>NA</td><td>Intergenic</td><td>NA</td><td>NA</td></tr>
<tr><td>GU305914.1</td><td>132081</td><td>132103</td><td>+</td><td>4</td><td>TCTCGGCGTGCGCCGACAGC</td><td>AGG</td><td>--||||-|[||||-|||||||]PAM</td><td>NA</td><td>Intergenic</td><td>NA</td><td>NA</td></tr>
<tr><td></td></tr>
<tr><td>T1603</td><td>CGACAGCGGGAGCAGCGTCCCGG</td><td>1000</td></tr>
<tr><td>Oligo fwd</td><td>CACCCGACAGCGGGAGCAGCGTCC</td></tr>
<tr><td>Oligo rev</td><td>AAACGGACGCTGCTCCCGCTGTCG</td></tr>
<tr><td>Chromosome</td><td>start</td><td>end</td><td>strand</td><td>MM</td><td>target_seq</td><td>PAM</td><td>alignment</td><td>distance</td><td>position</td><td>gene name</td><td>gene id</td></tr>
<tr><td>GU305914.1</td><td>26660</td><td>26682</td><td>+</td><td>0</td><td>CGACAGCGGGAGCAGCGTCC</td><td>CGG</td><td>||||||||[||||||||||||]PAM</td><td>NA</td><td>Intergenic</td><td>NA</td><td>NA</td></tr>
<tr><td></td></tr>
<tr><td>T1604</td><td>GCAGCGTCCCGGAATGATAGTGG</td><td>1000</td></tr>
<tr><td>Oligo fwd</td><td>CACCGCAGCGTCCCGGAATGATAG</td></tr>
<tr><td>Oligo rev</td><td>AAACCTATCATTCCGGGACGCTGC</td></tr>
<tr><td>Chromosome</td><td>start</td><td>end</td><td>strand</td><td>MM</td><td>target_seq</td><td>PAM</td><td>alignment</td><td>distance</td><td>position</td><td>gene name</td><td>gene id</td></tr>
<tr><td>GU305914.1</td><td>26671</td><td>26693</td><td>+</td><td>0</td><td>GCAGCGTCCCGGAATGATAG</td><td>TGG</td><td>||||||||[||||||||||||]PAM</td><td>NA</td><td>Intergenic</td><td>NA</td><td>NA</td></tr>
<tr><td></td></tr>
<tr><td>T1605</td><td>GCCCGTGACCCTTTTTGATGAGG</td><td>1000</td></tr>
<tr><td>Oligo fwd</td><td>CACCGCCCGTGACCCTTTTTGATG</td></tr>
<tr><td>Oligo rev</td><td>AAACCATCAAAAAGGGTCACGGGC</td></tr>
<tr><td>Chromosome</td><td>start</td><td>end</td><td>strand</td><td>MM</td><td>target_seq</td><td>PAM</td><td>alignment</td><td>distance</td><td>position</td><td>gene name</td><td>gene id</td></tr>
<tr><td>GU305914.1</td><td>26703</td><td>26725</td><td>+</td><td>0</td><td>GCCCGTGACCCTTTTTGATG</td><td>AGG</td><td>||||||||[||||||||||||]PAM</td><td>NA</td><td>Intergenic</td><td>NA</td><td>NA</td></tr>
<tr><td></td></tr>
<tr><td>T1606</td><td>GTGACCCTTTTTGATGAGGACGG</td><td>1000</td></tr>
<tr><td>Oligo fwd</td><td>CACCGTGACCCTTTTTGATGAGGA</td></tr>
<tr><td>Oligo rev</td><td>AAACTCCTCATCAAAAAGGGTCAC</td></tr>
<tr><td>Chromosome</td><td>start</td><td>end</td><td>strand</td><td>MM</td><td>target_seq</td><td>PAM</td><td>alignment</td><td>distance</td><td>position</td><td>gene name</td><td>gene id</td></tr>
<tr><td>GU305914.1</td><td>26707</td><td>26729</td><td>+</td><td>0</td><td>GTGACCCTTTTTGATGAGGA</td><td>CGG</td><td>||||||||[||||||||||||]PAM</td><td>NA</td><td>Intergenic</td><td>NA</td><td>NA</td></tr>
<tr><td></td></tr>
<tr><td>T1607</td><td>CGATTCGCGACACGTCGTCGTGG</td><td>1000</td></tr>
<tr><td>Oligo fwd</td><td>CACCCGATTCGCGACACGTCGTCG</td></tr>
<tr><td>Oligo rev</td><td>AAACCGACGACGTGTCGCGAATCG</td></tr>
<tr><td>Chromosome</td><td>start</td><td>end</td><td>strand</td><td>MM</td><td>target_seq</td><td>PAM</td><td>alignment</td><td>distance</td><td>position</td><td>gene name</td><td>gene id</td></tr>
<tr><td>GU305914.1</td><td>26766</td><td>26788</td><td>+</td><td>0</td><td>CGATTCGCGACACGTCGTCG</td><td>TGG</td><td>||||||||[||||||||||||]PAM</td><td>NA</td><td>Intergenic</td><td>NA</td><td>NA</td></tr>
<tr><td>GU305914.1</td><td>79977</td><td>79999</td><td>-</td><td>2</td><td>GGATCCGCGACACGTCGTCG</td><td>AGG</td><td>-|||-|||[||||||||||||]PAM</td><td>NA</td><td>Intergenic</td><td>NA</td><td>NA</td></tr>
<tr><td></td></tr>
<tr><td>T1608</td><td>GTCGTGGACGACGACGACGACGG</td><td>1000</td></tr>
<tr><td>Oligo fwd</td><td>CACCGTCGTGGACGACGACGACGA</td></tr>
<tr><td>Oligo rev</td><td>AAACTCGTCGTCGTCGTCCACGAC</td></tr>
<tr><td>Chromosome</td><td>start</td><td>end</td><td>strand</td><td>MM</td><td>target_seq</td><td>PAM</td><td>alignment</td><td>distance</td><td>position</td><td>gene name</td><td>gene id</td></tr>
<tr><td>GU305914.1</td><td>26782</td><td>26804</td><td>+</td><td>0</td><td>GTCGTGGACGACGACGACGA</td><td>CGG</td><td>||||||||[||||||||||||]PAM</td><td>NA</td><td>Intergenic</td><td>NA</td><td>NA</td></tr>
<tr><td></td></tr>
<tr><td>T1609</td><td>GGAGATTATGATTCACACTACGG</td><td>1000</td></tr>
<tr><td>Oligo fwd</td><td>CACCGGAGATTATGATTCACACTA</td></tr>
<tr><td>Oligo rev</td><td>AAACTAGTGTGAATCATAATCTCC</td></tr>
</table>

<table>
<tr><td>Chromosome</td><td>start</td><td>end</td><td>strand</td><td>MM</td><td>target_seq</td><td>PAM</td><td>alignment</td><td>distance</td><td>position</td><td>gene name</td><td>gene id</td></tr>
<tr><td>GU305914.1</td><td>26803</td><td>26825</td><td>+</td><td>0</td><td>GGAGATTATGATTCACACTA</td><td>CGG</td><td>||||||||[||||||||||||]PAM</td><td>NA</td><td>Intergenic</td><td>NA</td><td>NA</td></tr>
<tr><td>T1610</td><td>TCGACCTTCGACCAAAAAATCGG</td><td>1000</td><td></td><td></td><td></td><td></td><td></td><td></td><td></td><td></td><td></td></tr>
<tr><td>Oligo fwd</td><td>CACCTCGACCTTCGACCAAAAAAT</td><td></td><td></td><td></td><td></td><td></td><td></td><td></td><td></td><td></td><td></td></tr>
<tr><td>Oligo rev</td><td>AAACATTTTTTGGTCGAAGGTCGA</td><td></td><td></td><td></td><td></td><td></td><td></td><td></td><td></td><td></td><td></td></tr>
<tr><td>Chromosome</td><td>start</td><td>end</td><td>strand</td><td>MM</td><td>target_seq</td><td>PAM</td><td>alignment</td><td>distance</td><td>position</td><td>gene name</td><td>gene id</td></tr>
<tr><td>GU305914.1</td><td>26871</td><td>26893</td><td>+</td><td>0</td><td>TCGACCTTCGACCAAAAAAT</td><td>CGG</td><td>||||||||[||||||||||||]PAM</td><td>NA</td><td>Intergenic</td><td>NA</td><td>NA</td></tr>
<tr><td>T1611</td><td>ACCTTCGACCAAAAAATCGGCGG</td><td>1000</td><td></td><td></td><td></td><td></td><td></td><td></td><td></td><td></td><td></td></tr>
<tr><td>Oligo fwd</td><td>CACCACCTTCGACCAAAAAATCGG</td><td></td><td></td><td></td><td></td><td></td><td></td><td></td><td></td><td></td><td></td></tr>
<tr><td>Oligo rev</td><td>AAACCCGATTTTTTGGTCGAAGGT</td><td></td><td></td><td></td><td></td><td></td><td></td><td></td><td></td><td></td><td></td></tr>
<tr><td>Chromosome</td><td>start</td><td>end</td><td>strand</td><td>MM</td><td>target_seq</td><td>PAM</td><td>alignment</td><td>distance</td><td>position</td><td>gene name</td><td>gene id</td></tr>
<tr><td>GU305914.1</td><td>26874</td><td>26896</td><td>+</td><td>0</td><td>ACCTTCGACCAAAAAATCGG</td><td>CGG</td><td>||||||||[||||||||||||]PAM</td><td>NA</td><td>Intergenic</td><td>NA</td><td>NA</td></tr>
<tr><td>T1612</td><td>TTCGACCAAAAAATCGGCGGAGG</td><td>1000</td><td></td><td></td><td></td><td></td><td></td><td></td><td></td><td></td><td></td></tr>
<tr><td>Oligo fwd</td><td>CACCTTCGACCAAAAAATCGGCGG</td><td></td><td></td><td></td><td></td><td></td><td></td><td></td><td></td><td></td><td></td></tr>
<tr><td>Oligo rev</td><td>AAACCCGCCGATTTTTTGGTCGAA</td><td></td><td></td><td></td><td></td><td></td><td></td><td></td><td></td><td></td><td></td></tr>
<tr><td>Chromosome</td><td>start</td><td>end</td><td>strand</td><td>MM</td><td>target_seq</td><td>PAM</td><td>alignment</td><td>distance</td><td>position</td><td>gene name</td><td>gene id</td></tr>
<tr><td>GU305914.1</td><td>26877</td><td>26899</td><td>+</td><td>0</td><td>TTCGACCAAAAAATCGGCGG</td><td>AGG</td><td>||||||||[||||||||||||]PAM</td><td>NA</td><td>Intergenic</td><td>NA</td><td>NA</td></tr>
<tr><td>T1613</td><td>GAAAAGCAATCTTCCACCGCCGG</td><td>1000</td><td></td><td></td><td></td><td></td><td></td><td></td><td></td><td></td><td></td></tr>
<tr><td>Oligo fwd</td><td>CACCGAAAAGCAATCTTCCACCGC</td><td></td><td></td><td></td><td></td><td></td><td></td><td></td><td></td><td></td><td></td></tr>
<tr><td>Oligo rev</td><td>AAACGCGGTGGAAGATTGCTTTTC</td><td></td><td></td><td></td><td></td><td></td><td></td><td></td><td></td><td></td><td></td></tr>
<tr><td>Chromosome</td><td>start</td><td>end</td><td>strand</td><td>MM</td><td>target_seq</td><td>PAM</td><td>alignment</td><td>distance</td><td>position</td><td>gene name</td><td>gene id</td></tr>
<tr><td>GU305914.1</td><td>26899</td><td>26921</td><td>+</td><td>0</td><td>GAAAAGCAATCTTCCACCGC</td><td>CGG</td><td>||||||||[||||||||||||]PAM</td><td>NA</td><td>Intergenic</td><td>NA</td><td>NA</td></tr>
<tr><td>T1614</td><td>AGCAATCTTCCACCGCCGGCCGG</td><td>1000</td><td></td><td></td><td></td><td></td><td></td><td></td><td></td><td></td><td></td></tr>
<tr><td>Oligo fwd</td><td>CACCAGCAATCTTCCACCGCCGGC</td><td></td><td></td><td></td><td></td><td></td><td></td><td></td><td></td><td></td><td></td></tr>
<tr><td>Oligo rev</td><td>AAACGCCGGCGGTGGAAGATTGCT</td><td></td><td></td><td></td><td></td><td></td><td></td><td></td><td></td><td></td><td></td></tr>
<tr><td>Chromosome</td><td>start</td><td>end</td><td>strand</td><td>MM</td><td>target_seq</td><td>PAM</td><td>alignment</td><td>distance</td><td>position</td><td>gene name</td><td>gene id</td></tr>
<tr><td>GU305914.1</td><td>26903</td><td>26925</td><td>+</td><td>0</td><td>AGCAATCTTCCACCGCCGGC</td><td>CGG</td><td>||||||||[||||||||||||]PAM</td><td>NA</td><td>Intergenic</td><td>NA</td><td>NA</td></tr>
<tr><td>T1615</td><td>TCCACCGCCGGCCGGTCCAGAGG</td><td>1000</td><td></td><td></td><td></td><td></td><td></td><td></td><td></td><td></td><td></td></tr>
<tr><td>Oligo fwd</td><td>CACCTCCACCGCCGGCCGGTCCAG</td><td></td><td></td><td></td><td></td><td></td><td></td><td></td><td></td><td></td><td></td></tr>
<tr><td>Oligo rev</td><td>AAACCTGGACCGGCCGGCGGTGGA</td><td></td><td></td><td></td><td></td><td></td><td></td><td></td><td></td><td></td><td></td></tr>
<tr><td>Chromosome</td><td>start</td><td>end</td><td>strand</td><td>MM</td><td>target_seq</td><td>PAM</td><td>alignment</td><td>distance</td><td>position</td><td>gene name</td><td>gene id</td></tr>
<tr><td>GU305914.1</td><td>26911</td><td>26933</td><td>+</td><td>0</td><td>TCCACCGCCGGCCGGTCCAG</td><td>AGG</td><td>||||||||[||||||||||||]PAM</td><td>NA</td><td>Intergenic</td><td>NA</td><td>NA</td></tr>
<tr><td>T1616</td><td>GAGGCCGCTCGACCGCACGCCGG</td><td>1000</td><td></td><td></td><td></td><td></td><td></td><td></td><td></td><td></td><td></td></tr>
<tr><td>Oligo fwd</td><td>CACCGAGGCCGCTCGACCGCACGC</td><td></td><td></td><td></td><td></td><td></td><td></td><td></td><td></td><td></td><td></td></tr>
<tr><td>Oligo rev</td><td>AAACGCGTGCGGTCGAGCGGCCTC</td><td></td><td></td><td></td><td></td><td></td><td></td><td></td><td></td><td></td><td></td></tr>
<tr><td>Chromosome</td><td>start</td><td>end</td><td>strand</td><td>MM</td><td>target_seq</td><td>PAM</td><td>alignment</td><td>distance</td><td>position</td><td>gene name</td><td>gene id</td></tr>
<tr><td>GU305914.1</td><td>26930</td><td>26952</td><td>+</td><td>0</td><td>GAGGCCGCTCGACCGCACGC</td><td>CGG</td><td>||||||||[||||||||||||]PAM</td><td>NA</td><td>Intergenic</td><td>NA</td><td>NA</td></tr>
<tr><td>T1617</td><td>CGCACGCCGGACCCCGAAGAAGG</td><td>1000</td><td></td><td></td><td></td><td></td><td></td><td></td><td></td><td></td><td></td></tr>
<tr><td>Oligo fwd</td><td>CACCCGCACGCCGGACCCCGAAGA</td><td></td><td></td><td></td><td></td><td></td><td></td><td></td><td></td><td></td><td></td></tr>
<tr><td>Oligo rev</td><td>AAACTCTTCGGGGTCCGGCGTGCG</td><td></td><td></td><td></td><td></td><td></td><td></td><td></td><td></td><td></td><td></td></tr>
<tr><td>Chromosome</td><td>start</td><td>end</td><td>strand</td><td>MM</td><td>target_seq</td><td>PAM</td><td>alignment</td><td>distance</td><td>position</td><td>gene name</td><td>gene id</td></tr>
<tr><td>GU305914.1</td><td>26943</td><td>26965</td><td>+</td><td>0</td><td>CGCACGCCGGACCCCGAAGA</td><td>AGG</td><td>||||||||[||||||||||||]PAM</td><td>NA</td><td>Intergenic</td><td>NA</td><td>NA</td></tr>
<tr><td>T1618</td><td>GAAGGCTCAAGAGACCGCTCCGG</td><td>1000</td><td></td><td></td><td></td><td></td><td></td><td></td><td></td><td></td><td></td></tr>
<tr><td>Oligo fwd</td><td>CACCGAAGGCTCAAGAGACCGCTC</td><td></td><td></td><td></td><td></td><td></td><td></td><td></td><td></td><td></td><td></td></tr>
<tr><td>Oligo rev</td><td>AAACGAGCGGTCTCTTGAGCCTTC</td><td></td><td></td><td></td><td></td><td></td><td></td><td></td><td></td><td></td><td></td></tr>
<tr><td>Chromosome</td><td>start</td><td>end</td><td>strand</td><td>MM</td><td>target_seq</td><td>PAM</td><td>alignment</td><td>distance</td><td>position</td><td>gene name</td><td>gene id</td></tr>
<tr><td>GU305914.1</td><td>26961</td><td>26983</td><td>+</td><td>0</td><td>GAAGGCTCAAGAGACCGCTC</td><td>CGG</td><td>||||||||[||||||||||||]PAM</td><td>NA</td><td>Intergenic</td><td>NA</td><td>NA</td></tr>
<tr><td>T1619</td><td>GGCTCAAGAGACCGCTCCGGCGG</td><td>1000</td><td></td><td></td><td></td><td></td><td></td><td></td><td></td><td></td><td></td></tr>
<tr><td>Oligo fwd</td><td>CACCGGCTCAAGAGACCGCTCCGG</td><td></td><td></td><td></td><td></td><td></td><td></td><td></td><td></td><td></td><td></td></tr>
<tr><td>Oligo rev</td><td>AAACCCGGAGCGGTCTCTTGAGCC</td><td></td><td></td><td></td><td></td><td></td><td></td><td></td><td></td><td></td><td></td></tr>
<tr><td>Chromosome</td><td>start</td><td>end</td><td>strand</td><td>MM</td><td>target_seq</td><td>PAM</td><td>alignment</td><td>distance</td><td>position</td><td>gene name</td><td>gene id</td></tr>
<tr><td>GU305914.1</td><td>26964</td><td>26986</td><td>+</td><td>0</td><td>GGCTCAAGAGACCGCTCCGG</td><td>CGG</td><td>||||||||[||||||||||||]PAM</td><td>NA</td><td>Intergenic</td><td>NA</td><td>NA</td></tr>
<tr><td>T1620</td><td>AGAGACCGCTCCGGCGGCCGCGG</td><td>1000</td><td></td><td></td><td></td><td></td><td></td><td></td><td></td><td></td><td></td></tr>
<tr><td>Oligo fwd</td><td>CACCAGAGACCGCTCCGGCGGCCG</td><td></td><td></td><td></td><td></td><td></td><td></td><td></td><td></td><td></td><td></td></tr>
<tr><td>Oligo rev</td><td>AAACCGGCCGCCGGAGCGGTCTCT</td><td></td><td></td><td></td><td></td><td></td><td></td><td></td><td></td><td></td><td></td></tr>
<tr><td>Chromosome</td><td>start</td><td>end</td><td>strand</td><td>MM</td><td>target_seq</td><td>PAM</td><td>alignment</td><td>distance</td><td>position</td><td>gene name</td><td>gene id</td></tr>
<tr><td>GU305914.1</td><td>26970</td><td>26992</td><td>+</td><td>0</td><td>AGAGACCGCTCCGGCGGCCG</td><td>CGG</td><td>||||||||[||||||||||||]PAM</td><td>NA</td><td>Intergenic</td><td>NA</td><td>NA</td></tr>
<tr><td>T1621</td><td>GACCGCTCCGGCGGCCGCGGCGG</td><td>1000</td><td></td><td></td><td></td><td></td><td></td><td></td><td></td><td></td><td></td></tr>
<tr><td>Oligo fwd</td><td>CACCGACCGCTCCGGCGGCCGCGG</td><td></td><td></td><td></td><td></td><td></td><td></td><td></td><td></td><td></td><td></td></tr>
<tr><td>Oligo rev</td><td>AAACCCGCGGCCGCCGGAGCGGTC</td><td></td><td></td><td></td><td></td><td></td><td></td><td></td><td></td><td></td><td></td></tr>
<tr><td>Chromosome</td><td>start</td><td>end</td><td>strand</td><td>MM</td><td>target_seq</td><td>PAM</td><td>alignment</td><td>distance</td><td>position</td><td>gene name</td><td>gene id</td></tr>
<tr><td>GU305914.1</td><td>26973</td><td>26995</td><td>+</td><td>0</td><td>GACCGCTCCGGCGGCCGCGG</td><td>CGG</td><td>||||||||[||||||||||||]PAM</td><td>NA</td><td>Intergenic</td><td>NA</td><td>NA</td></tr>
<tr><td>GU305914.1</td><td>110125</td><td>110147</td><td>-</td><td>4</td><td>GGACGCGACGGCGGCCGCGG</td><td>CGG</td><td>|--|||--[||||||||||||]PAM</td><td>NA</td><td>Intergenic</td><td>NA</td><td>NA</td></tr>
<tr><td>GU305914.1</td><td>21338</td><td>21360</td><td>+</td><td>4</td><td>GACCGCCTCGGCGTGCGCGG</td><td>CGG</td><td>||||||--[|||||--|||||]PAM</td><td>NA</td><td>Intergenic</td><td>NA</td><td>NA</td></tr>
<tr><td>T1622</td><td>ACCGCTCCGGCGGCCGCGGCGGG</td><td>1000</td><td></td><td></td><td></td><td></td><td></td><td></td><td></td><td></td><td></td></tr>
<tr><td>Oligo fwd</td><td>CACCACCGCTCCGGCGGCCGCGGC</td><td></td><td></td><td></td><td></td><td></td><td></td><td></td><td></td><td></td><td></td></tr>
<tr><td>Oligo rev</td><td>AAACGCCGCGGCCGCCGGAGCGGT</td><td></td><td></td><td></td><td></td><td></td><td></td><td></td><td></td><td></td><td></td></tr>
<tr><td>Chromosome</td><td>start</td><td>end</td><td>strand</td><td>MM</td><td>target_seq</td><td>PAM</td><td>alignment</td><td>distance</td><td>position</td><td>gene name</td><td>gene id</td></tr>
<tr><td>GU305914.1</td><td>26974</td><td>26996</td><td>+</td><td>0</td><td>ACCGCTCCGGCGGCCGCGGC</td><td>GGG</td><td>||||||||[||||||||||||]PAM</td><td>NA</td><td>Intergenic</td><td>NA</td><td>NA</td></tr>
<tr><td>T1623</td><td>CCGGCGGCCGCGGCGGGCTCCGG</td><td>1000</td><td></td><td></td><td></td><td></td><td></td><td></td><td></td><td></td><td></td></tr>
<tr><td>Oligo fwd</td><td>CACCCCGGCGGCCGCGGCGGGCTC</td><td></td><td></td><td></td><td></td><td></td><td></td><td></td><td></td><td></td><td></td></tr>
<tr><td>Oligo rev</td><td>AAACGAGCCCGCCGCGGCCGCCGG</td><td></td><td></td><td></td><td></td><td></td><td></td><td></td><td></td><td></td><td></td></tr>
<tr><td>Chromosome</td><td>start</td><td>end</td><td>strand</td><td>MM</td><td>target_seq</td><td>PAM</td><td>alignment</td><td>distance</td><td>position</td><td>gene name</td><td>gene id</td></tr>
<tr><td>GU305914.1</td><td>26980</td><td>27002</td><td>+</td><td>0</td><td>CCGGCGGCCGCGGCGGGCTC</td><td>CGG</td><td>||||||||[||||||||||||]PAM</td><td>NA</td><td>Intergenic</td><td>NA</td><td>NA</td></tr>
</table>

<table>
<tr><td>T1624</td><td>TCCCCCACCTCCGCCCGTACAGG</td><td>1000</td><td></td><td></td><td></td><td></td><td></td><td></td><td></td><td></td><td></td></tr>
<tr><td>Oligo fwd</td><td>CACCTCCCCCACCTCCGCCCGTAC</td><td></td><td></td><td></td><td></td><td></td><td></td><td></td><td></td><td></td><td></td></tr>
<tr><td>Oligo rev</td><td>AAACGTACGGGCGGAGGTGGGGGA</td><td></td><td></td><td></td><td></td><td></td><td></td><td></td><td></td><td></td><td></td></tr>
<tr><td>Chromosome</td><td>start</td><td>end</td><td>strand</td><td>MM</td><td>target_seq</td><td>PAM</td><td>alignment</td><td>distance</td><td>position</td><td>gene name</td><td>gene id</td></tr>
<tr><td>GU305914.1</td><td>27081</td><td>27103</td><td>+</td><td>0</td><td>TCCCCCACCTCCGCCCGTAC</td><td>AGG</td><td>||||||||[||||||||||||]PAM</td><td>NA</td><td>Intergenic</td><td>NA</td><td>NA</td></tr>
</table>

<table>
<tr><td>T1625</td><td>CTGTGTCGCGTCCTCTCCCCAGG</td><td>1000</td><td></td><td></td><td></td><td></td><td></td><td></td><td></td><td></td><td></td></tr>
<tr><td>Oligo fwd</td><td>CACCCTGTGTCGCGTCCTCTCCCC</td><td></td><td></td><td></td><td></td><td></td><td></td><td></td><td></td><td></td><td></td></tr>
<tr><td>Oligo rev</td><td>AAACGGGGAGAGGACGCGACACAG</td><td></td><td></td><td></td><td></td><td></td><td></td><td></td><td></td><td></td><td></td></tr>
<tr><td>Chromosome</td><td>start</td><td>end</td><td>strand</td><td>MM</td><td>target_seq</td><td>PAM</td><td>alignment</td><td>distance</td><td>position</td><td>gene name</td><td>gene id</td></tr>
<tr><td>GU305914.1</td><td>27107</td><td>27129</td><td>+</td><td>0</td><td>CTGTGTCGCGTCCTCTCCCC</td><td>AGG</td><td>||||||||[||||||||||||]PAM</td><td>NA</td><td>Intergenic</td><td>NA</td><td>NA</td></tr>
</table>

<table>
<tr><td>T1626</td><td>CGACGATGACAACGACGAGCCGG</td><td>1000</td><td></td><td></td><td></td><td></td><td></td><td></td><td></td><td></td><td></td></tr>
<tr><td>Oligo fwd</td><td>CACCCGACGATGACAACGACGAGC</td><td></td><td></td><td></td><td></td><td></td><td></td><td></td><td></td><td></td><td></td></tr>
<tr><td>Oligo rev</td><td>AAACGCTCGTCGTTGTCATCGTCG</td><td></td><td></td><td></td><td></td><td></td><td></td><td></td><td></td><td></td><td></td></tr>
<tr><td>Chromosome</td><td>start</td><td>end</td><td>strand</td><td>MM</td><td>target_seq</td><td>PAM</td><td>alignment</td><td>distance</td><td>position</td><td>gene name</td><td>gene id</td></tr>
<tr><td>GU305914.1</td><td>27156</td><td>27178</td><td>+</td><td>0</td><td>CGACGATGACAACGACGAGC</td><td>CGG</td><td>||||||||[||||||||||||]PAM</td><td>NA</td><td>Intergenic</td><td>NA</td><td>NA</td></tr>
</table>

<table>
<tr><td>T1627</td><td>GACGATGACAACGACGAGCCGGG</td><td>1000</td><td></td><td></td><td></td><td></td><td></td><td></td><td></td><td></td><td></td></tr>
<tr><td>Oligo fwd</td><td>CACCGACGATGACAACGACGAGCC</td><td></td><td></td><td></td><td></td><td></td><td></td><td></td><td></td><td></td><td></td></tr>
<tr><td>Oligo rev</td><td>AAACGGCTCGTCGTTGTCATCGTC</td><td></td><td></td><td></td><td></td><td></td><td></td><td></td><td></td><td></td><td></td></tr>
<tr><td>Chromosome</td><td>start</td><td>end</td><td>strand</td><td>MM</td><td>target_seq</td><td>PAM</td><td>alignment</td><td>distance</td><td>position</td><td>gene name</td><td>gene id</td></tr>
<tr><td>GU305914.1</td><td>27157</td><td>27179</td><td>+</td><td>0</td><td>GACGATGACAACGACGAGCC</td><td>GGG</td><td>||||||||[||||||||||||]PAM</td><td>NA</td><td>Intergenic</td><td>NA</td><td>NA</td></tr>
</table>

<table>
<tr><td>T1628</td><td>GGTCCGAGCAACACACGCCGCGG</td><td>1000</td><td></td><td></td><td></td><td></td><td></td><td></td><td></td><td></td><td></td></tr>
<tr><td>Oligo fwd</td><td>CACCGGTCCGAGCAACACACGCCG</td><td></td><td></td><td></td><td></td><td></td><td></td><td></td><td></td><td></td><td></td></tr>
<tr><td>Oligo rev</td><td>AAACCGGCGTGTGTTGCTCGGACC</td><td></td><td></td><td></td><td></td><td></td><td></td><td></td><td></td><td></td><td></td></tr>
<tr><td>Chromosome</td><td>start</td><td>end</td><td>strand</td><td>MM</td><td>target_seq</td><td>PAM</td><td>alignment</td><td>distance</td><td>position</td><td>gene name</td><td>gene id</td></tr>
<tr><td>GU305914.1</td><td>27178</td><td>27200</td><td>+</td><td>0</td><td>GGTCCGAGCAACACACGCCG</td><td>CGG</td><td>||||||||[||||||||||||]PAM</td><td>NA</td><td>Intergenic</td><td>NA</td><td>NA</td></tr>
<tr><td>GU305914.1</td><td>226273</td><td>226295</td><td>-</td><td>3</td><td>CGTACGAGCAACACCCGCCG</td><td>CGG</td><td>-||-||||[||||||-|||||]PAM</td><td>NA</td><td>Intergenic</td><td>NA</td><td>NA</td></tr>
</table>

<table>
<tr><td>T1629</td><td>CTGCCGTCGTGTCGATCACACGG</td><td>1000</td><td></td><td></td><td></td><td></td><td></td><td></td><td></td><td></td><td></td></tr>
<tr><td>Oligo fwd</td><td>CACCCTGCCGTCGTGTCGATCACA</td><td></td><td></td><td></td><td></td><td></td><td></td><td></td><td></td><td></td><td></td></tr>
<tr><td>Oligo rev</td><td>AAACTGTGATCGACACGACGGCAG</td><td></td><td></td><td></td><td></td><td></td><td></td><td></td><td></td><td></td><td></td></tr>
<tr><td>Chromosome</td><td>start</td><td>end</td><td>strand</td><td>MM</td><td>target_seq</td><td>PAM</td><td>alignment</td><td>distance</td><td>position</td><td>gene name</td><td>gene id</td></tr>
<tr><td>GU305914.1</td><td>27210</td><td>27232</td><td>+</td><td>0</td><td>CTGCCGTCGTGTCGATCACA</td><td>CGG</td><td>||||||||[||||||||||||]PAM</td><td>NA</td><td>Intergenic</td><td>NA</td><td>NA</td></tr>
</table>

<table>
<tr><td>T1630</td><td>CCCGATATCCGCCACCGCCATGG</td><td>1000</td><td></td><td></td><td></td><td></td><td></td><td></td><td></td><td></td><td></td></tr>
<tr><td>Oligo fwd</td><td>CACCCCCGATATCCGCCACCGCCA</td><td></td><td></td><td></td><td></td><td></td><td></td><td></td><td></td><td></td><td></td></tr>
<tr><td>Oligo rev</td><td>AAACTGGCGGTGGCGGATATCGGG</td><td></td><td></td><td></td><td></td><td></td><td></td><td></td><td></td><td></td><td></td></tr>
<tr><td>Chromosome</td><td>start</td><td>end</td><td>strand</td><td>MM</td><td>target_seq</td><td>PAM</td><td>alignment</td><td>distance</td><td>position</td><td>gene name</td><td>gene id</td></tr>
<tr><td>GU305914.1</td><td>27255</td><td>27277</td><td>+</td><td>0</td><td>CCCGATATCCGCCACCGCCA</td><td>TGG</td><td>||||||||[||||||||||||]PAM</td><td>NA</td><td>Intergenic</td><td>NA</td><td>NA</td></tr>
</table>

<table>
<tr><td>T1631</td><td>GATCGATATCGAAGATGACGAGG</td><td>1000</td><td></td><td></td><td></td><td></td><td></td><td></td><td></td><td></td><td></td></tr>
<tr><td>Oligo fwd</td><td>CACCGATCGATATCGAAGATGACG</td><td></td><td></td><td></td><td></td><td></td><td></td><td></td><td></td><td></td><td></td></tr>
<tr><td>Oligo rev</td><td>AAACCGTCATCTTCGATATCGATC</td><td></td><td></td><td></td><td></td><td></td><td></td><td></td><td></td><td></td><td></td></tr>
<tr><td>Chromosome</td><td>start</td><td>end</td><td>strand</td><td>MM</td><td>target_seq</td><td>PAM</td><td>alignment</td><td>distance</td><td>position</td><td>gene name</td><td>gene id</td></tr>
<tr><td>GU305914.1</td><td>27279</td><td>27301</td><td>+</td><td>0</td><td>GATCGATATCGAAGATGACG</td><td>AGG</td><td>||||||||[||||||||||||]PAM</td><td>NA</td><td>Intergenic</td><td>NA</td><td>NA</td></tr>
</table>

<table>
<tr><td>T1632</td><td>GATGACGAGGACGAAGAGACCGG</td><td>1000</td><td></td><td></td><td></td><td></td><td></td><td></td><td></td><td></td><td></td></tr>
<tr><td>Oligo fwd</td><td>CACCGATGACGAGGACGAAGAGAC</td><td></td><td></td><td></td><td></td><td></td><td></td><td></td><td></td><td></td><td></td></tr>
<tr><td>Oligo rev</td><td>AAACGTCTCTTCGTCCTCGTCATC</td><td></td><td></td><td></td><td></td><td></td><td></td><td></td><td></td><td></td><td></td></tr>
<tr><td>Chromosome</td><td>start</td><td>end</td><td>strand</td><td>MM</td><td>target_seq</td><td>PAM</td><td>alignment</td><td>distance</td><td>position</td><td>gene name</td><td>gene id</td></tr>
<tr><td>GU305914.1</td><td>27292</td><td>27314</td><td>+</td><td>0</td><td>GATGACGAGGACGAAGAGAC</td><td>CGG</td><td>||||||||[||||||||||||]PAM</td><td>NA</td><td>Intergenic</td><td>NA</td><td>NA</td></tr>
</table>

<table>
<tr><td>T1633</td><td>GACGAGGACGAAGAGACCGGCGG</td><td>1000</td><td></td><td></td><td></td><td></td><td></td><td></td><td></td><td></td><td></td></tr>
<tr><td>Oligo fwd</td><td>CACCGACGAGGACGAAGAGACCGG</td><td></td><td></td><td></td><td></td><td></td><td></td><td></td><td></td><td></td><td></td></tr>
<tr><td>Oligo rev</td><td>AAACCCGGTCTCTTCGTCCTCGTC</td><td></td><td></td><td></td><td></td><td></td><td></td><td></td><td></td><td></td><td></td></tr>
<tr><td>Chromosome</td><td>start</td><td>end</td><td>strand</td><td>MM</td><td>target_seq</td><td>PAM</td><td>alignment</td><td>distance</td><td>position</td><td>gene name</td><td>gene id</td></tr>
<tr><td>GU305914.1</td><td>27295</td><td>27317</td><td>+</td><td>0</td><td>GACGAGGACGAAGAGACCGG</td><td>CGG</td><td>||||||||[||||||||||||]PAM</td><td>NA</td><td>Intergenic</td><td>NA</td><td>NA</td></tr>
</table>

<table>
<tr><td>T1634</td><td>CGAAGATGATGATGAAGAAGAGG</td><td>1000</td><td></td><td></td><td></td><td></td><td></td><td></td><td></td><td></td><td></td></tr>
<tr><td>Oligo fwd</td><td>CACCCGAAGATGATGATGAAGAAG</td><td></td><td></td><td></td><td></td><td></td><td></td><td></td><td></td><td></td><td></td></tr>
<tr><td>Oligo rev</td><td>AAACCTTCTTCATCATCATCTTCG</td><td></td><td></td><td></td><td></td><td></td><td></td><td></td><td></td><td></td><td></td></tr>
<tr><td>Chromosome</td><td>start</td><td>end</td><td>strand</td><td>MM</td><td>target_seq</td><td>PAM</td><td>alignment</td><td>distance</td><td>position</td><td>gene name</td><td>gene id</td></tr>
<tr><td>GU305914.1</td><td>27345</td><td>27367</td><td>+</td><td>0</td><td>CGAAGATGATGATGAAGAAG</td><td>AGG</td><td>||||||||[||||||||||||]PAM</td><td>NA</td><td>Intergenic</td><td>NA</td><td>NA</td></tr>
<tr><td>GU305914.1</td><td>168589</td><td>168611</td><td>+</td><td>4</td><td>TGAATATGAGGATGATGAAG</td><td>AGG</td><td>-|||-|||[|-|||||-||||]PAM</td><td>NA</td><td>Intergenic</td><td>NA</td><td>NA</td></tr>
<tr><td>GU305914.1</td><td>118515</td><td>118537</td><td>-</td><td>4</td><td>CGATGAGGACGATGATGAAG</td><td>AGG</td><td>|||-||-|[|-|||||-||||]PAM</td><td>NA</td><td>Intergenic</td><td>NA</td><td>NA</td></tr>
<tr><td>GU305914.1</td><td>118512</td><td>118534</td><td>-</td><td>4</td><td>TGAGGACGATGATGAAGAGG</td><td>AGG</td><td>-||-||-|[||||||||||-|]PAM</td><td>NA</td><td>Intergenic</td><td>NA</td><td>NA</td></tr>
<tr><td>GU305914.1</td><td>65090</td><td>65112</td><td>+</td><td>4</td><td>CGAAAAGGAGGATGAAGAGG</td><td>AGG</td><td>||||-|-|[|-||||||||-|]PAM</td><td>NA</td><td>Intergenic</td><td>NA</td><td>NA</td></tr>
</table>

<table>
<tr><td>T1635</td><td>AGAAGAGGAGAACGATTGCGAGG</td><td>1000</td><td></td><td></td><td></td><td></td><td></td><td></td><td></td><td></td><td></td></tr>
<tr><td>Oligo fwd</td><td>CACCAGAAGAGGAGAACGATTGCG</td><td></td><td></td><td></td><td></td><td></td><td></td><td></td><td></td><td></td><td></td></tr>
<tr><td>Oligo rev</td><td>AAACCGCAATCGTTCTCCTCTTCT</td><td></td><td></td><td></td><td></td><td></td><td></td><td></td><td></td><td></td><td></td></tr>
<tr><td>Chromosome</td><td>start</td><td>end</td><td>strand</td><td>MM</td><td>target_seq</td><td>PAM</td><td>alignment</td><td>distance</td><td>position</td><td>gene name</td><td>gene id</td></tr>
<tr><td>GU305914.1</td><td>27360</td><td>27382</td><td>+</td><td>0</td><td>AGAAGAGGAGAACGATTGCG</td><td>AGG</td><td>||||||||[||||||||||||]PAM</td><td>NA</td><td>Intergenic</td><td>NA</td><td>NA</td></tr>
</table>

<table>
<tr><td>T1636</td><td>GATTGCGAGGAGATCTGCGACGG</td><td>1000</td><td></td><td></td><td></td><td></td><td></td><td></td><td></td><td></td><td></td></tr>
<tr><td>Oligo fwd</td><td>CACCGATTGCGAGGAGATCTGCGA</td><td></td><td></td><td></td><td></td><td></td><td></td><td></td><td></td><td></td><td></td></tr>
<tr><td>Oligo rev</td><td>AAACTCGCAGATCTCCTCGCAATC</td><td></td><td></td><td></td><td></td><td></td><td></td><td></td><td></td><td></td><td></td></tr>
<tr><td>Chromosome</td><td>start</td><td>end</td><td>strand</td><td>MM</td><td>target_seq</td><td>PAM</td><td>alignment</td><td>distance</td><td>position</td><td>gene name</td><td>gene id</td></tr>
<tr><td>GU305914.1</td><td>27373</td><td>27395</td><td>+</td><td>0</td><td>GATTGCGAGGAGATCTGCGA</td><td>CGG</td><td>||||||||[||||||||||||]PAM</td><td>NA</td><td>Intergenic</td><td>NA</td><td>NA</td></tr>
<tr><td>GU305914.1</td><td>86021</td><td>86043</td><td>-</td><td>4</td><td>GACGGCGATGAGATCGGCGA</td><td>CGG</td><td>||--||||[-||||||-||||]PAM</td><td>NA</td><td>Intergenic</td><td>NA</td><td>NA</td></tr>
</table>

<table>
<tr><td>T1637</td><td>ATTGCGAGGAGATCTGCGACGGG</td><td>1000</td><td></td><td></td><td></td><td></td><td></td><td></td><td></td><td></td><td></td></tr>
<tr><td>Oligo fwd</td><td>CACCATTGCGAGGAGATCTGCGAC</td><td></td><td></td><td></td><td></td><td></td><td></td><td></td><td></td><td></td><td></td></tr>
<tr><td>Oligo rev</td><td>AAACGTCGCAGATCTCCTCGCAAT</td><td></td><td></td><td></td><td></td><td></td><td></td><td></td><td></td><td></td><td></td></tr>
<tr><td>Chromosome</td><td>start</td><td>end</td><td>strand</td><td>MM</td><td>target_seq</td><td>PAM</td><td>alignment</td><td>distance</td><td>position</td><td>gene name</td><td>gene id</td></tr>
<tr><td>GU305914.1</td><td>27374</td><td>27396</td><td>+</td><td>0</td><td>ATTGCGAGGAGATCTGCGAC</td><td>GGG</td><td>||||||||[||||||||||||]PAM</td><td>NA</td><td>Intergenic</td><td>NA</td><td>NA</td></tr>
</table>

| | | | | | | | | | | | |
|---|---|---|---|---|---|---|---|---|---|---|---|
| GU305914.1 | 86020 | 86042 | - | 4 | ACGGCGATGAGATCGGCGAC | GGG | \|--\|\|\|\|-[\|\|\|\|\|\|-\|\|\|\|\|]PAM | NA | Intergenic | NA | NA |
| T1638 | TTGCGAGGAGATCTGCGACGGGG | 1000 | | | | | | | | | |
| Oligo fwd | CACCTTGCGAGGAGATCTGCGACG | | | | | | | | | | |
| Oligo rev | AAACCGTCGCAGATCTCCTCGCAA | | | | | | | | | | |
| Chromosome | start | end | strand | MM | target_seq | PAM | alignment | distance | position | gene name | gene id |
| GU305914.1 | 27375 | 27397 | + | 0 | TTGCGAGGAGATCTGCGACG | GGG | \|\|\|\|\|\|\|\|[\|\|\|\|\|\|\|\|\|\|\|\|]PAM | NA | Intergenic | NA | NA |
| GU305914.1 | 153197 | 153219 | + | 4 | CGACGAGGAGATCTCCGACG | TGG | ---\|\|\|\|\|[\|\|\|\|\|\|-\|\|\|\|\|]PAM | NA | Intergenic | NA | NA |
| T1639 | CTGCGACGGGGAAGAAGAGCCGG | 1000 | | | | | | | | | |
| Oligo fwd | CACCCTGCGACGGGGAAGAAGAGC | | | | | | | | | | |
| Oligo rev | AAACGCTCTTCTTCCCCGTCGCAG | | | | | | | | | | |
| Chromosome | start | end | strand | MM | target_seq | PAM | alignment | distance | position | gene name | gene id |
| GU305914.1 | 27387 | 27409 | + | 0 | CTGCGACGGGGAAGAAGAGC | CGG | \|\|\|\|\|\|\|\|[\|\|\|\|\|\|\|\|\|\|\|\|]PAM | NA | Intergenic | NA | NA |
| GU305914.1 | 17283 | 17305 | - | 4 | CTCCGGCGGTGAAGAAGAGG | AGG | \|\|-\|\|-\|\|[\|-\|\|\|\|\|\|\|\|\|-]PAM | NA | Intergenic | NA | NA |
| T1640 | GGCAGCAGCAGCAGCAGCAGCGG | 1000 | | | | | | | | | |
| Oligo fwd | CACCGGCAGCAGCAGCAGCAGCAG | | | | | | | | | | |
| Oligo rev | AAACCTGCTGCTGCTGCTGCTGCC | | | | | | | | | | |
| Chromosome | start | end | strand | MM | target_seq | PAM | alignment | distance | position | gene name | gene id |
| GU305914.1 | 27408 | 27430 | + | 0 | GGCAGCAGCAGCAGCAGCAG | CGG | \|\|\|\|\|\|\|\|[\|\|\|\|\|\|\|\|\|\|\|\|]PAM | NA | Intergenic | NA | NA |
| GU305914.1 | 71741 | 71763 | + | 4 | GGCCTCCTCAGCAGCAGCAG | CGG | \|\|\|--\|--[\|\|\|\|\|\|\|\|\|\|\|\|]PAM | NA | Intergenic | NA | NA |
| GU305914.1 | 73813 | 73835 | - | 4 | GGGAGTAGCGGCGGCAGCAG | TGG | \|\|-\|\|-\|\|[\|-\|\|-\|\|\|\|\|\|\|]PAM | NA | Intergenic | NA | NA |
| GU305914.1 | 163908 | 163930 | + | 4 | GGCGGCGGCGGCAGCGGCAG | CGG | \|\|\|-\|\|-\|[\|-\|\|\|\|\|-\|\|\|\|]PAM | NA | Intergenic | NA | NA |
| GU305914.1 | 27411 | 27433 | + | 2 | AGCAGCAGCAGCAGCAGCGG | CGG | -\|\|\|\|\|\|\|[\|\|\|\|\|\|\|\|\|\|-\|]PAM | NA | Intergenic | NA | NA |
| GU305914.1 | 164959 | 164981 | - | 4 | GGCGGCGGCGGCAGCAGCTG | CGG | \|\|\|-\|\|-\|[\|-\|\|\|\|\|\|\|\|-\|]PAM | NA | Intergenic | NA | NA |
| GU305914.1 | 226682 | 226704 | - | 4 | GTCAGCAACAGCAGAAGAAG | TGG | \|-\|\|\|\|\|-[\|\|\|\|\|\|-\|\|-\|\|]PAM | NA | Intergenic | NA | NA |
| GU305914.1 | 163911 | 163933 | + | 4 | GGCGGCGGCAGCGGCAGCGG | AGG | \|\|\|-\|\|-\|[\|\|\|\|-\|\|\|\|\|-\|]PAM | NA | Intergenic | NA | NA |
| GU305914.1 | 71747 | 71769 | + | 4 | CTCAGCAGCAGCAGCGGCGG | CGG | --\|\|\|\|\|\|[\|\|\|\|\|\|\|-\|\|-\|]PAM | NA | Intergenic | NA | NA |
| T1641 | AGCAGCAGCAGCAGCAGCGGCGG | 1000 | | | | | | | | | |
| Oligo fwd | CACCAGCAGCAGCAGCAGCAGCGG | | | | | | | | | | |
| Oligo rev | AAACCCGCTGCTGCTGCTGCTGCT | | | | | | | | | | |
| Chromosome | start | end | strand | MM | target_seq | PAM | alignment | distance | position | gene name | gene id |
| GU305914.1 | 27411 | 27433 | + | 0 | AGCAGCAGCAGCAGCAGCGG | CGG | \|\|\|\|\|\|\|\|[\|\|\|\|\|\|\|\|\|\|\|\|]PAM | NA | Intergenic | NA | NA |
| GU305914.1 | 71744 | 71766 | + | 4 | CTCCTCAGCAGCAGCAGCGG | CGG | --\|--\|\|\|[\|\|\|\|\|\|\|\|\|\|\|\|]PAM | NA | Intergenic | NA | NA |
| GU305914.1 | 163911 | 163933 | + | 4 | GGCGGCGGCAGCGGCAGCGG | AGG | -\|\|-\|\|-\|[\|\|\|\|-\|\|\|\|\|\|\|]PAM | NA | Intergenic | NA | NA |
| GU305914.1 | 73798 | 73820 | - | 4 | AGCAGTGGCGCCAGCAGCGG | CGG | \|\|\|\|\|--\|[\|--\|\|\|\|\|\|\|\|\|]PAM | NA | Intergenic | NA | NA |
| GU305914.1 | 71747 | 71769 | + | 3 | CTCAGCAGCAGCAGCGGCGG | CGG | --\|\|\|\|\|\|[\|\|\|\|\|\|\|-\|\|\|\|]PAM | NA | Intergenic | NA | NA |
| GU305914.1 | 27408 | 27430 | + | 2 | GGCAGCAGCAGCAGCAGCAG | CGG | -\|\|\|\|\|\|\|[\|\|\|\|\|\|\|\|\|\|-\|]PAM | NA | Intergenic | NA | NA |
| GU305914.1 | 62100 | 62122 | - | 3 | ACCACCAGCAGCAGCAGCGA | TGG | \|-\|\|-\|\|\|[\|\|\|\|\|\|\|\|\|\|\|-]PAM | NA | Intergenic | NA | NA |
| GU305914.1 | 128721 | 128743 | - | 4 | GGCGGCAGCAGCAGCGGCGC | TGG | -\|\|-\|\|\|\|[\|\|\|\|\|\|\|-\|\|\|-]PAM | NA | Intergenic | NA | NA |
| T1642 | CCGCGCATCACGCGCGAGCTCGG | 1000 | | | | | | | | | |
| Oligo fwd | CACCCCGCGCATCACGCGCGAGCT | | | | | | | | | | |
| Oligo rev | AAACAGCTCGCGCGTGATGCGCGG | | | | | | | | | | |
| Chromosome | start | end | strand | MM | target_seq | PAM | alignment | distance | position | gene name | gene id |
| GU305914.1 | 27487 | 27509 | + | 0 | CCGCGCATCACGCGCGAGCT | CGG | \|\|\|\|\|\|\|\|[\|\|\|\|\|\|\|\|\|\|\|\|]PAM | NA | Intergenic | NA | NA |
| T1643 | CGCGCATCACGCGCGAGCTCGGG | 1000 | | | | | | | | | |
| Oligo fwd | CACCCGCGCATCACGCGCGAGCTC | | | | | | | | | | |
| Oligo rev | AAACGAGCTCGCGCGTGATGCGCG | | | | | | | | | | |
| Chromosome | start | end | strand | MM | target_seq | PAM | alignment | distance | position | gene name | gene id |
| GU305914.1 | 27488 | 27510 | + | 0 | CGCGCATCACGCGCGAGCTC | GGG | \|\|\|\|\|\|\|\|[\|\|\|\|\|\|\|\|\|\|\|\|]PAM | NA | Intergenic | NA | NA |
| T1644 | CGCGAGCTCGGGTTCCTGCCCGG | 1000 | | | | | | | | | |
| Oligo fwd | CACCCGCGAGCTCGGGTTCCTGCC | | | | | | | | | | |
| Oligo rev | AAACGGCAGGAACCCGAGCTCGCG | | | | | | | | | | |
| Chromosome | start | end | strand | MM | target_seq | PAM | alignment | distance | position | gene name | gene id |
| GU305914.1 | 27499 | 27521 | + | 0 | CGCGAGCTCGGGTTCCTGCC | CGG | \|\|\|\|\|\|\|\|[\|\|\|\|\|\|\|\|\|\|\|\|]PAM | NA | Intergenic | NA | NA |
| T1645 | TTCCTGCCCGGTGTAGTGAGCGG | 1000 | | | | | | | | | |
| Oligo fwd | CACCTTCCTGCCCGGTGTAGTGAG | | | | | | | | | | |
| Oligo rev | AAACCTCACTACACCGGGCAGGAA | | | | | | | | | | |
| Chromosome | start | end | strand | MM | target_seq | PAM | alignment | distance | position | gene name | gene id |
| GU305914.1 | 27511 | 27533 | + | 0 | TTCCTGCCCGGTGTAGTGAG | CGG | \|\|\|\|\|\|\|\|[\|\|\|\|\|\|\|\|\|\|\|\|]PAM | NA | Intergenic | NA | NA |
| T1646 | GCCCGGTGTAGTGAGCGGTCAGG | 1000 | | | | | | | | | |
| Oligo fwd | CACCGCCCGGTGTAGTGAGCGGTC | | | | | | | | | | |
| Oligo rev | AAACGACCGCTCACTACACCGGGC | | | | | | | | | | |
| Chromosome | start | end | strand | MM | target_seq | PAM | alignment | distance | position | gene name | gene id |
| GU305914.1 | 27516 | 27538 | + | 0 | GCCCGGTGTAGTGAGCGGTC | AGG | \|\|\|\|\|\|\|\|[\|\|\|\|\|\|\|\|\|\|\|\|]PAM | NA | Intergenic | NA | NA |
| T1647 | TCAGGACGCCAGATTCATCGCGG | 1000 | | | | | | | | | |
| Oligo fwd | CACCTCAGGACGCCAGATTCATCG | | | | | | | | | | |
| Oligo rev | AAACCGATGAATCTGGCGTCCTGA | | | | | | | | | | |
| Chromosome | start | end | strand | MM | target_seq | PAM | alignment | distance | position | gene name | gene id |
| GU305914.1 | 27534 | 27556 | + | 0 | TCAGGACGCCAGATTCATCG | CGG | \|\|\|\|\|\|\|\|[\|\|\|\|\|\|\|\|\|\|\|\|]PAM | NA | Intergenic | NA | NA |
| T1648 | CCACCACTCGCACGCGCCGCAGG | 1000 | | | | | | | | | |
| Oligo fwd | CACCCCACCACTCGCACGCGCCGC | | | | | | | | | | |
| Oligo rev | AAACGCGGCGCGTGCGAGTGGTGG | | | | | | | | | | |
| Chromosome | start | end | strand | MM | target_seq | PAM | alignment | distance | position | gene name | gene id |
| GU305914.1 | 27564 | 27586 | + | 0 | CCACCACTCGCACGCGCCGC | AGG | \|\|\|\|\|\|\|\|[\|\|\|\|\|\|\|\|\|\|\|\|]PAM | NA | Intergenic | NA | NA |
| T1649 | CCTAGACGCGCTATCTGAGCCGG | 1000 | | | | | | | | | |
| Oligo fwd | CACCCCTAGACGCGCTATCTGAGC | | | | | | | | | | |
| Oligo rev | AAACGCTCAGATAGCGCGTCTAGG | | | | | | | | | | |
| Chromosome | start | end | strand | MM | target_seq | PAM | alignment | distance | position | gene name | gene id |

GU305914.1 27627 27649 + 0 CCTAGACGCGCTATCTGAGC CGG ||||||||[||||||||||||]PAM NA Intergenic NA NA

T1650 TGAGCCGGTCCTGACCAAGAAGG 1000
Oligo fwd CACCTGAGCCGGTCCTGACCAAGA
Oligo rev AAACTCTTGGTCAGGACCGGCTCA
Chromosome start end strand MM target_seq PAM alignment distance position gene name gene id
GU305914.1 27642 27664 + 0 TGAGCCGGTCCTGACCAAGA AGG ||||||||[||||||||||||]PAM NA Intergenic NA NA

T1651 CAAGAAGGCGCTGCGCTGCGCGG 1000
Oligo fwd CACCCAAGAAGGCGCTGCGCTGCG
Oligo rev AAACCGCAGCGCAGCGCCTTCTTG
Chromosome start end strand MM target_seq PAM alignment distance position gene name gene id
GU305914.1 27657 27679 + 0 CAAGAAGGCGCTGCGCTGCG CGG ||||||||[||||||||||||]PAM NA Intergenic NA NA

T1652 AAGAAGGCGCTGCGCTGCGCGGG 1000
Oligo fwd CACCAAGAAGGCGCTGCGCTGCGC
Oligo rev AAACGCGCAGCGCAGCGCCTTCTT
Chromosome start end strand MM target_seq PAM alignment distance position gene name gene id
GU305914.1 27658 27680 + 0 AAGAAGGCGCTGCGCTGCGC GGG ||||||||[||||||||||||]PAM NA Intergenic NA NA

T1653 AGAAGGCGCTGCGCTGCGCGGGG 1000
Oligo fwd CACCAGAAGGCGCTGCGCTGCGCG
Oligo rev AAACCGCGCAGCGCAGCGCCTTCT
Chromosome start end strand MM target_seq PAM alignment distance position gene name gene id
GU305914.1 27659 27681 + 0 AGAAGGCGCTGCGCTGCGCG GGG ||||||||[||||||||||||]PAM NA Intergenic NA NA

T1654 GAAGGCGCTGCGCTGCGCGGGGG 1000
Oligo fwd CACCGAAGGCGCTGCGCTGCGCGG
Oligo rev AAACCCGCGCAGCGCAGCGCCTTC
Chromosome start end strand MM target_seq PAM alignment distance position gene name gene id
GU305914.1 27660 27682 + 0 GAAGGCGCTGCGCTGCGCGG GGG ||||||||[||||||||||||]PAM NA Intergenic NA NA

T1655 TGCGCTGCGCGGGGGTGCTGCGG 1000
Oligo fwd CACCTGCGCTGCGCGGGGGTGCTG
Oligo rev AAACCAGCACCCCCGCGCAGCGCA
Chromosome start end strand MM target_seq PAM alignment distance position gene name gene id
GU305914.1 27668 27690 + 0 TGCGCTGCGCGGGGGTGCTG CGG ||||||||[||||||||||||]PAM NA Intergenic NA NA

T1656 CGTCATCAAGCTAGCCATCCTGG 1000
Oligo fwd CACCCGTCATCAAGCTAGCCATCC
Oligo rev AAACGGATGGCTAGCTTGATGACG
Chromosome start end strand MM target_seq PAM alignment distance position gene name gene id
GU305914.1 27693 27715 + 0 CGTCATCAAGCTAGCCATCC TGG ||||||||[||||||||||||]PAM NA Intergenic NA NA

T1657 CTGGTGAATTACTACTGCGTAGG 1000
Oligo fwd CACCCTGGTGAATTACTACTGCGT
Oligo rev AAACACGCAGTAGTAATTCACCAG
Chromosome start end strand MM target_seq PAM alignment distance position gene name gene id
GU305914.1 27712 27734 + 0 CTGGTGAATTACTACTGCGT AGG ||||||||[||||||||||||]PAM NA Intergenic NA NA

T1658 TGGTGAATTACTACTGCGTAGGG 1000
Oligo fwd CACCTGGTGAATTACTACTGCGTA
Oligo rev AAACTACGCAGTAGTAATTCACCA
Chromosome start end strand MM target_seq PAM alignment distance position gene name gene id
GU305914.1 27713 27735 + 0 TGGTGAATTACTACTGCGTA GGG ||||||||[||||||||||||]PAM NA Intergenic NA NA

T1659 AATTACTACTGCGTAGGGATCGG 1000
Oligo fwd CACCAATTACTACTGCGTAGGGAT
Oligo rev AAACATCCCTACGCAGTAGTAATT
Chromosome start end strand MM target_seq PAM alignment distance position gene name gene id
GU305914.1 27718 27740 + 0 AATTACTACTGCGTAGGGAT CGG ||||||||[||||||||||||]PAM NA Intergenic NA NA

T1660 ATTACTACTGCGTAGGGATCGGG 1000
Oligo fwd CACCATTACTACTGCGTAGGGATC
Oligo rev AAACGATCCCTACGCAGTAGTAAT
Chromosome start end strand MM target_seq PAM alignment distance position gene name gene id
GU305914.1 27719 27741 + 0 ATTACTACTGCGTAGGGATC GGG ||||||||[||||||||||||]PAM NA Intergenic NA NA

T1661 CTGCGTAGGGATCGGGCGTCTGG 1000
Oligo fwd CACCCTGCGTAGGGATCGGGCGTC
Oligo rev AAACGACGCCCGATCCCTACGCAG
Chromosome start end strand MM target_seq PAM alignment distance position gene name gene id
GU305914.1 27726 27748 + 0 CTGCGTAGGGATCGGGCGTC TGG ||||||||[||||||||||||]PAM NA Intergenic NA NA

T1662 CGAGACGCTGCGCCGCCGTCTGG 1000
Oligo fwd CACCCGAGACGCTGCGCCGCCGTC
Oligo rev AAACGACGGCGGCGCAGCGTCTCG
Chromosome start end strand MM target_seq PAM alignment distance position gene name gene id
GU305914.1 27795 27817 + 0 CGAGACGCTGCGCCGCCGTC TGG ||||||||[||||||||||||]PAM NA Intergenic NA NA

T1663 GACGCTGCGCCGCCGTCTGGAGG 1000
Oligo fwd CACCGACGCTGCGCCGCCGTCTGG
Oligo rev AAACCCAGACGGCGGCGCAGCGTC
Chromosome start end strand MM target_seq PAM alignment distance position gene name gene id
GU305914.1 27798 27820 + 0 GACGCTGCGCCGCCGTCTGG AGG ||||||||[||||||||||||]PAM NA Intergenic NA NA

T1664 ACGCTGCGCCGCCGTCTGGAGGG 1000
Oligo fwd CACCACGCTGCGCCGCCGTCTGGA
Oligo rev AAACTCCAGACGGCGGCGCAGCGT

<table>
<tr><td>Chromosome</td><td>start</td><td>end</td><td>strand</td><td>MM</td><td>target_seq</td><td>PAM</td><td>alignment</td><td>distance</td><td>position</td><td>gene name</td><td>gene id</td></tr>
<tr><td>GU305914.1</td><td>27799</td><td>27821</td><td>+</td><td>0</td><td>ACGCTGCGCCGCCGTCTGGA</td><td>GGG</td><td>||||||||[||||||||||||]PAM</td><td>NA</td><td>Intergenic</td><td>NA</td><td>NA</td></tr>
<tr><td></td><td></td><td></td><td></td><td></td><td></td><td></td><td></td><td></td><td></td><td></td><td></td></tr>
<tr><td>T1665</td><td>CCAGCCCTCCGATGTGCCTGCGG</td><td>1000</td><td></td><td></td><td></td><td></td><td></td><td></td><td></td><td></td><td></td></tr>
<tr><td>Oligo fwd</td><td>CACCCCAGCCCTCCGATGTGCCTG</td><td></td><td></td><td></td><td></td><td></td><td></td><td></td><td></td><td></td><td></td></tr>
<tr><td>Oligo rev</td><td>AAACCAGGCACATCGGAGGGCTGG</td><td></td><td></td><td></td><td></td><td></td><td></td><td></td><td></td><td></td><td></td></tr>
<tr><td>Chromosome</td><td>start</td><td>end</td><td>strand</td><td>MM</td><td>target_seq</td><td>PAM</td><td>alignment</td><td>distance</td><td>position</td><td>gene name</td><td>gene id</td></tr>
<tr><td>GU305914.1</td><td>27845</td><td>27867</td><td>+</td><td>0</td><td>CCAGCCCTCCGATGTGCCTG</td><td>CGG</td><td>||||||||[||||||||||||]PAM</td><td>NA</td><td>Intergenic</td><td>NA</td><td>NA</td></tr>
<tr><td></td><td></td><td></td><td></td><td></td><td></td><td></td><td></td><td></td><td></td><td></td><td></td></tr>
<tr><td>T1666</td><td>CAGCCCTCCGATGTGCCTGCGGG</td><td>1000</td><td></td><td></td><td></td><td></td><td></td><td></td><td></td><td></td><td></td></tr>
<tr><td>Oligo fwd</td><td>CACCCAGCCCTCCGATGTGCCTGC</td><td></td><td></td><td></td><td></td><td></td><td></td><td></td><td></td><td></td><td></td></tr>
<tr><td>Oligo rev</td><td>AAACGCAGGCACATCGGAGGGCTG</td><td></td><td></td><td></td><td></td><td></td><td></td><td></td><td></td><td></td><td></td></tr>
<tr><td>Chromosome</td><td>start</td><td>end</td><td>strand</td><td>MM</td><td>target_seq</td><td>PAM</td><td>alignment</td><td>distance</td><td>position</td><td>gene name</td><td>gene id</td></tr>
<tr><td>GU305914.1</td><td>27846</td><td>27868</td><td>+</td><td>0</td><td>CAGCCCTCCGATGTGCCTGC</td><td>GGG</td><td>||||||||[||||||||||||]PAM</td><td>NA</td><td>Intergenic</td><td>NA</td><td>NA</td></tr>
<tr><td></td><td></td><td></td><td></td><td></td><td></td><td></td><td></td><td></td><td></td><td></td><td></td></tr>
<tr><td>T1667</td><td>CCGATGTGCCTGCGGGTGCTCGG</td><td>1000</td><td></td><td></td><td></td><td></td><td></td><td></td><td></td><td></td><td></td></tr>
<tr><td>Oligo fwd</td><td>CACCCCGATGTGCCTGCGGGTGCT</td><td></td><td></td><td></td><td></td><td></td><td></td><td></td><td></td><td></td><td></td></tr>
<tr><td>Oligo rev</td><td>AAACAGCACCCGCAGGCACATCGG</td><td></td><td></td><td></td><td></td><td></td><td></td><td></td><td></td><td></td><td></td></tr>
<tr><td>Chromosome</td><td>start</td><td>end</td><td>strand</td><td>MM</td><td>target_seq</td><td>PAM</td><td>alignment</td><td>distance</td><td>position</td><td>gene name</td><td>gene id</td></tr>
<tr><td>GU305914.1</td><td>27853</td><td>27875</td><td>+</td><td>0</td><td>CCGATGTGCCTGCGGGTGCT</td><td>CGG</td><td>||||||||[||||||||||||]PAM</td><td>NA</td><td>Intergenic</td><td>NA</td><td>NA</td></tr>
<tr><td></td><td></td><td></td><td></td><td></td><td></td><td></td><td></td><td></td><td></td><td></td><td></td></tr>
<tr><td>T1668</td><td>CGGCCGCCTGAACATCACCGCGG</td><td>1000</td><td></td><td></td><td></td><td></td><td></td><td></td><td></td><td></td><td></td></tr>
<tr><td>Oligo fwd</td><td>CACCCGGCCGCCTGAACATCACCG</td><td></td><td></td><td></td><td></td><td></td><td></td><td></td><td></td><td></td><td></td></tr>
<tr><td>Oligo rev</td><td>AAACCGGTGATGTTCAGGCGGCCG</td><td></td><td></td><td></td><td></td><td></td><td></td><td></td><td></td><td></td><td></td></tr>
<tr><td>Chromosome</td><td>start</td><td>end</td><td>strand</td><td>MM</td><td>target_seq</td><td>PAM</td><td>alignment</td><td>distance</td><td>position</td><td>gene name</td><td>gene id</td></tr>
<tr><td>GU305914.1</td><td>27873</td><td>27895</td><td>+</td><td>0</td><td>CGGCCGCCTGAACATCACCG</td><td>CGG</td><td>||||||||[||||||||||||]PAM</td><td>NA</td><td>Intergenic</td><td>NA</td><td>NA</td></tr>
<tr><td></td><td></td><td></td><td></td><td></td><td></td><td></td><td></td><td></td><td></td><td></td><td></td></tr>
<tr><td>T1669</td><td>CATCACCGCGGCGCAGCACAAGG</td><td>1000</td><td></td><td></td><td></td><td></td><td></td><td></td><td></td><td></td><td></td></tr>
<tr><td>Oligo fwd</td><td>CACCCATCACCGCGGCGCAGCACA</td><td></td><td></td><td></td><td></td><td></td><td></td><td></td><td></td><td></td><td></td></tr>
<tr><td>Oligo rev</td><td>AAACTGTGCTGCGCCGCGGTGATG</td><td></td><td></td><td></td><td></td><td></td><td></td><td></td><td></td><td></td><td></td></tr>
<tr><td>Chromosome</td><td>start</td><td>end</td><td>strand</td><td>MM</td><td>target_seq</td><td>PAM</td><td>alignment</td><td>distance</td><td>position</td><td>gene name</td><td>gene id</td></tr>
<tr><td>GU305914.1</td><td>27885</td><td>27907</td><td>+</td><td>0</td><td>CATCACCGCGGCGCAGCACA</td><td>AGG</td><td>||||||||[||||||||||||]PAM</td><td>NA</td><td>Intergenic</td><td>NA</td><td>NA</td></tr>
<tr><td></td><td></td><td></td><td></td><td></td><td></td><td></td><td></td><td></td><td></td><td></td><td></td></tr>
<tr><td>T1670</td><td>CCAACTGATGAAACCGATGCAGG</td><td>1000</td><td></td><td></td><td></td><td></td><td></td><td></td><td></td><td></td><td></td></tr>
<tr><td>Oligo fwd</td><td>CACCCCAACTGATGAAACCGATGC</td><td></td><td></td><td></td><td></td><td></td><td></td><td></td><td></td><td></td><td></td></tr>
<tr><td>Oligo rev</td><td>AAACGCATCGGTTTCATCAGTTGG</td><td></td><td></td><td></td><td></td><td></td><td></td><td></td><td></td><td></td><td></td></tr>
<tr><td>Chromosome</td><td>start</td><td>end</td><td>strand</td><td>MM</td><td>target_seq</td><td>PAM</td><td>alignment</td><td>distance</td><td>position</td><td>gene name</td><td>gene id</td></tr>
<tr><td>GU305914.1</td><td>27927</td><td>27949</td><td>+</td><td>0</td><td>CCAACTGATGAAACCGATGC</td><td>AGG</td><td>||||||||[||||||||||||]PAM</td><td>NA</td><td>Intergenic</td><td>NA</td><td>NA</td></tr>
<tr><td></td><td></td><td></td><td></td><td></td><td></td><td></td><td></td><td></td><td></td><td></td><td></td></tr>
<tr><td>T1671</td><td>TCCCCCGCGTTTCACCCGCGAGG</td><td>1000</td><td></td><td></td><td></td><td></td><td></td><td></td><td></td><td></td><td></td></tr>
<tr><td>Oligo fwd</td><td>CACCTCCCCCGCGTTTCACCCGCG</td><td></td><td></td><td></td><td></td><td></td><td></td><td></td><td></td><td></td><td></td></tr>
<tr><td>Oligo rev</td><td>AAACCGCGGGTGAAACGCGGGGGA</td><td></td><td></td><td></td><td></td><td></td><td></td><td></td><td></td><td></td><td></td></tr>
<tr><td>Chromosome</td><td>start</td><td>end</td><td>strand</td><td>MM</td><td>target_seq</td><td>PAM</td><td>alignment</td><td>distance</td><td>position</td><td>gene name</td><td>gene id</td></tr>
<tr><td>GU305914.1</td><td>28017</td><td>28039</td><td>+</td><td>0</td><td>TCCCCCGCGTTTCACCCGCG</td><td>AGG</td><td>||||||||[||||||||||||]PAM</td><td>NA</td><td>Intergenic</td><td>NA</td><td>NA</td></tr>
<tr><td></td><td></td><td></td><td></td><td></td><td></td><td></td><td></td><td></td><td></td><td></td><td></td></tr>
<tr><td>T1672</td><td>CCCCCGCGTTTCACCCGCGAGGG</td><td>1000</td><td></td><td></td><td></td><td></td><td></td><td></td><td></td><td></td><td></td></tr>
<tr><td>Oligo fwd</td><td>CACCCCCCCGCGTTTCACCCGCGA</td><td></td><td></td><td></td><td></td><td></td><td></td><td></td><td></td><td></td><td></td></tr>
<tr><td>Oligo rev</td><td>AAACTCGCGGGTGAAACGCGGGGG</td><td></td><td></td><td></td><td></td><td></td><td></td><td></td><td></td><td></td><td></td></tr>
<tr><td>Chromosome</td><td>start</td><td>end</td><td>strand</td><td>MM</td><td>target_seq</td><td>PAM</td><td>alignment</td><td>distance</td><td>position</td><td>gene name</td><td>gene id</td></tr>
<tr><td>GU305914.1</td><td>28018</td><td>28040</td><td>+</td><td>0</td><td>CCCCCGCGTTTCACCCGCGA</td><td>GGG</td><td>||||||||[||||||||||||]PAM</td><td>NA</td><td>Intergenic</td><td>NA</td><td>NA</td></tr>
<tr><td></td><td></td><td></td><td></td><td></td><td></td><td></td><td></td><td></td><td></td><td></td><td></td></tr>
<tr><td>T1673</td><td>CCCCGCGTTTCACCCGCGAGGGG</td><td>1000</td><td></td><td></td><td></td><td></td><td></td><td></td><td></td><td></td><td></td></tr>
<tr><td>Oligo fwd</td><td>CACCCCCCGCGTTTCACCCGCGAG</td><td></td><td></td><td></td><td></td><td></td><td></td><td></td><td></td><td></td><td></td></tr>
<tr><td>Oligo rev</td><td>AAACCTCGCGGGTGAAACGCGGGG</td><td></td><td></td><td></td><td></td><td></td><td></td><td></td><td></td><td></td><td></td></tr>
<tr><td>Chromosome</td><td>start</td><td>end</td><td>strand</td><td>MM</td><td>target_seq</td><td>PAM</td><td>alignment</td><td>distance</td><td>position</td><td>gene name</td><td>gene id</td></tr>
<tr><td>GU305914.1</td><td>28019</td><td>28041</td><td>+</td><td>0</td><td>CCCCGCGTTTCACCCGCGAG</td><td>GGG</td><td>||||||||[||||||||||||]PAM</td><td>NA</td><td>Intergenic</td><td>NA</td><td>NA</td></tr>
<tr><td></td><td></td><td></td><td></td><td></td><td></td><td></td><td></td><td></td><td></td><td></td><td></td></tr>
<tr><td>T1674</td><td>CCCGCGTTTCACCCGCGAGGGGG</td><td>1000</td><td></td><td></td><td></td><td></td><td></td><td></td><td></td><td></td><td></td></tr>
<tr><td>Oligo fwd</td><td>CACCCCCGCGTTTCACCCGCGAGG</td><td></td><td></td><td></td><td></td><td></td><td></td><td></td><td></td><td></td><td></td></tr>
<tr><td>Oligo rev</td><td>AAACCCTCGCGGGTGAAACGCGGG</td><td></td><td></td><td></td><td></td><td></td><td></td><td></td><td></td><td></td><td></td></tr>
<tr><td>Chromosome</td><td>start</td><td>end</td><td>strand</td><td>MM</td><td>target_seq</td><td>PAM</td><td>alignment</td><td>distance</td><td>position</td><td>gene name</td><td>gene id</td></tr>
<tr><td>GU305914.1</td><td>28020</td><td>28042</td><td>+</td><td>0</td><td>CCCGCGTTTCACCCGCGAGG</td><td>GGG</td><td>||||||||[||||||||||||]PAM</td><td>NA</td><td>Intergenic</td><td>NA</td><td>NA</td></tr>
<tr><td></td><td></td><td></td><td></td><td></td><td></td><td></td><td></td><td></td><td></td><td></td><td></td></tr>
<tr><td>T1675</td><td>GGGCGAAGACCCTGTACATGCGG</td><td>1000</td><td></td><td></td><td></td><td></td><td></td><td></td><td></td><td></td><td></td></tr>
<tr><td>Oligo fwd</td><td>CACCGGGCGAAGACCCTGTACATG</td><td></td><td></td><td></td><td></td><td></td><td></td><td></td><td></td><td></td><td></td></tr>
<tr><td>Oligo rev</td><td>AAACCATGTACAGGGTCTTCGCCC</td><td></td><td></td><td></td><td></td><td></td><td></td><td></td><td></td><td></td><td></td></tr>
<tr><td>Chromosome</td><td>start</td><td>end</td><td>strand</td><td>MM</td><td>target_seq</td><td>PAM</td><td>alignment</td><td>distance</td><td>position</td><td>gene name</td><td>gene id</td></tr>
<tr><td>GU305914.1</td><td>28040</td><td>28062</td><td>+</td><td>0</td><td>GGGCGAAGACCCTGTACATG</td><td>CGG</td><td>||||||||[||||||||||||]PAM</td><td>NA</td><td>Intergenic</td><td>NA</td><td>NA</td></tr>
<tr><td></td><td></td><td></td><td></td><td></td><td></td><td></td><td></td><td></td><td></td><td></td><td></td></tr>
<tr><td>T1676</td><td>CAAGATCCTGAACAGCGACGAGG</td><td>1000</td><td></td><td></td><td></td><td></td><td></td><td></td><td></td><td></td><td></td></tr>
<tr><td>Oligo fwd</td><td>CACCCAAGATCCTGAACAGCGACG</td><td></td><td></td><td></td><td></td><td></td><td></td><td></td><td></td><td></td><td></td></tr>
<tr><td>Oligo rev</td><td>AAACCGTCGCTGTTCAGGATCTTG</td><td></td><td></td><td></td><td></td><td></td><td></td><td></td><td></td><td></td><td></td></tr>
<tr><td>Chromosome</td><td>start</td><td>end</td><td>strand</td><td>MM</td><td>target_seq</td><td>PAM</td><td>alignment</td><td>distance</td><td>position</td><td>gene name</td><td>gene id</td></tr>
<tr><td>GU305914.1</td><td>28068</td><td>28090</td><td>+</td><td>0</td><td>CAAGATCCTGAACAGCGACG</td><td>AGG</td><td>||||||||[||||||||||||]PAM</td><td>NA</td><td>Intergenic</td><td>NA</td><td>NA</td></tr>
<tr><td></td><td></td><td></td><td></td><td></td><td></td><td></td><td></td><td></td><td></td><td></td><td></td></tr>
<tr><td>T1677</td><td>GATCCTGAACAGCGACGAGGAGG</td><td>1000</td><td></td><td></td><td></td><td></td><td></td><td></td><td></td><td></td><td></td></tr>
<tr><td>Oligo fwd</td><td>CACCGATCCTGAACAGCGACGAGG</td><td></td><td></td><td></td><td></td><td></td><td></td><td></td><td></td><td></td><td></td></tr>
<tr><td>Oligo rev</td><td>AAACCCTCGTCGCTGTTCAGGATC</td><td></td><td></td><td></td><td></td><td></td><td></td><td></td><td></td><td></td><td></td></tr>
<tr><td>Chromosome</td><td>start</td><td>end</td><td>strand</td><td>MM</td><td>target_seq</td><td>PAM</td><td>alignment</td><td>distance</td><td>position</td><td>gene name</td><td>gene id</td></tr>
<tr><td>GU305914.1</td><td>28071</td><td>28093</td><td>+</td><td>0</td><td>GATCCTGAACAGCGACGAGG</td><td>AGG</td><td>||||||||[||||||||||||]PAM</td><td>NA</td><td>Intergenic</td><td>NA</td><td>NA</td></tr>
<tr><td></td><td></td><td></td><td></td><td></td><td></td><td></td><td></td><td></td><td></td><td></td><td></td></tr>
<tr><td>T1678</td><td>CCCTGAATCCGCATCCCACACGG</td><td>1000</td><td></td><td></td><td></td><td></td><td></td><td></td><td></td><td></td><td></td></tr>
<tr><td>Oligo fwd</td><td>CACCCCCTGAATCCGCATCCCACA</td><td></td><td></td><td></td><td></td><td></td><td></td><td></td><td></td><td></td><td></td></tr>
<tr><td>Oligo rev</td><td>AAACTGTGGGATGCGGATTCAGGG</td><td></td><td></td><td></td><td></td><td></td><td></td><td></td><td></td><td></td><td></td></tr>
<tr><td>Chromosome</td><td>start</td><td>end</td><td>strand</td><td>MM</td><td>target_seq</td><td>PAM</td><td>alignment</td><td>distance</td><td>position</td><td>gene name</td><td>gene id</td></tr>
<tr><td>GU305914.1</td><td>28118</td><td>28140</td><td>+</td><td>0</td><td>CCCTGAATCCGCATCCCACA</td><td>CGG</td><td>||||||||[||||||||||||]PAM</td><td>NA</td><td>Intergenic</td><td>NA</td><td>NA</td></tr>
<tr><td></td><td></td><td></td><td></td><td></td><td></td><td></td><td></td><td></td><td></td><td></td><td></td></tr>
<tr><td>T1679</td><td>CCTGAATCCGCATCCCACACGGG</td><td>1000</td><td></td><td></td><td></td><td></td><td></td><td></td><td></td><td></td><td></td></tr>
<tr><td>Oligo fwd</td><td>CACCCCTGAATCCGCATCCCACAC</td><td></td><td></td><td></td><td></td><td></td><td></td><td></td><td></td><td></td><td></td></tr>
</table>

Oligo rev AAACGTGTGGGATGCGGATTCAGG
Chromosome start end strand MM target_seq PAM alignment distance position gene name gene id
GU305914.1 28119 28141 + 0 CCTGAATCCGCATCCCACAC GGG ||||||||[||||||||||||]PAM NA Intergenic NA NA

T1680 CGCGATCTTTTGCCTGTCGCTGG 1000
Oligo fwd CACCCGCGATCTTTTGCCTGTCGC
Oligo rev AAACGCGACAGGCAAAAGATCGCG
Chromosome start end strand MM target_seq PAM alignment distance position gene name gene id
GU305914.1 28158 28180 + 0 CGCGATCTTTTGCCTGTCGC TGG ||||||||[||||||||||||]PAM NA Intergenic NA NA

T1681 GCGATCTTTTGCCTGTCGCTGGG 1000
Oligo fwd CACCGCGATCTTTTGCCTGTCGCT
Oligo rev AAACAGCGACAGGCAAAAGATCGC
Chromosome start end strand MM target_seq PAM alignment distance position gene name gene id
GU305914.1 28159 28181 + 0 GCGATCTTTTGCCTGTCGCT GGG ||||||||[||||||||||||]PAM NA Intergenic NA NA

T1682 CGCGCGCTCGAAGAACTGCGCGG 1000
Oligo fwd CACCCGCGCGCTCGAAGAACTGCG
Oligo rev AAACCGCAGTTCTTCGAGCGCGCG
Chromosome start end strand MM target_seq PAM alignment distance position gene name gene id
GU305914.1 28204 28226 + 0 CGCGCGCTCGAAGAACTGCG CGG ||||||||[||||||||||||]PAM NA Intergenic NA NA

T1683 AATGATCAGATGCCAGTTCGAGG 1000
Oligo fwd CACCAATGATCAGATGCCAGTTCG
Oligo rev AAACCGAACTGGCATCTGATCATT
Chromosome start end strand MM target_seq PAM alignment distance position gene name gene id
GU305914.1 28227 28249 + 0 AATGATCAGATGCCAGTTCG AGG ||||||||[||||||||||||]PAM NA Intergenic NA NA

T1684 CCTGACCGAGACCCTGTACGCGG 1000
Oligo fwd CACCCCTGACCGAGACCCTGTACG
Oligo rev AAACCGTACAGGGTCTCGGTCAGG
Chromosome start end strand MM target_seq PAM alignment distance position gene name gene id
GU305914.1 28251 28273 + 0 CCTGACCGAGACCCTGTACG CGG ||||||||[||||||||||||]PAM NA Intergenic NA NA

T1685 CTACCGCGTGCTGTGCTCCGAGG 1000
Oligo fwd CACCCTACCGCGTGCTGTGCTCCG
Oligo rev AAACCGGAGCACAGCACGCGGTAG
Chromosome start end strand MM target_seq PAM alignment distance position gene name gene id
GU305914.1 28305 28327 + 0 CTACCGCGTGCTGTGCTCCG AGG ||||||||[||||||||||||]PAM NA Intergenic NA NA

T1686 CCGCGTGCTGTGCTCCGAGGTGG 1000
Oligo fwd CACCCCGCGTGCTGTGCTCCGAGG
Oligo rev AAACCCTCGGAGCACAGCACGCGG
Chromosome start end strand MM target_seq PAM alignment distance position gene name gene id
GU305914.1 28308 28330 + 0 CCGCGTGCTGTGCTCCGAGG TGG ||||||||[||||||||||||]PAM NA Intergenic NA NA

T1687 ATCACGTCGCCCCGTGAAGACGG 1000
Oligo fwd CACCATCACGTCGCCCCGTGAAGA
Oligo rev AAACTCTTCACGGGGCGACGTGAT
Chromosome start end strand MM target_seq PAM alignment distance position gene name gene id
GU305914.1 28339 28361 + 0 ATCACGTCGCCCCGTGAAGA CGG ||||||||[||||||||||||]PAM NA Intergenic NA NA

T1688 GTCGCCCCGTGAAGACGGACAGG 1000
Oligo fwd CACCGTCGCCCCGTGAAGACGGAC
Oligo rev AAACGTCCGTCTTCACGGGGCGAC
Chromosome start end strand MM target_seq PAM alignment distance position gene name gene id
GU305914.1 28344 28366 + 0 GTCGCCCCGTGAAGACGGAC AGG ||||||||[||||||||||||]PAM NA Intergenic NA NA

T1689 TCGCCCCGTGAAGACGGACAGGG 1000
Oligo fwd CACCTCGCCCCGTGAAGACGGACA
Oligo rev AAACTGTCCGTCTTCACGGGGCGA
Chromosome start end strand MM target_seq PAM alignment distance position gene name gene id
GU305914.1 28345 28367 + 0 TCGCCCCGTGAAGACGGACA GGG ||||||||[||||||||||||]PAM NA Intergenic NA NA

T1690 GCGCGCCGCTGCTACAACGAAGG 1000
Oligo fwd CACCGCGCGCCGCTGCTACAACGA
Oligo rev AAACTCGTTGTAGCAGCGGCGCGC
Chromosome start end strand MM target_seq PAM alignment distance position gene name gene id
GU305914.1 28402 28424 + 0 GCGCGCCGCTGCTACAACGA AGG ||||||||[||||||||||||]PAM NA Intergenic NA NA

T1691 CAAGTATCTGATCAAGTGCGCGG 1000
Oligo fwd CACCCAAGTATCTGATCAAGTGCG
Oligo rev AAACCGCACTTGATCAGATACTTG
Chromosome start end strand MM target_seq PAM alignment distance position gene name gene id
GU305914.1 28449 28471 + 0 CAAGTATCTGATCAAGTGCG CGG ||||||||[||||||||||||]PAM NA Intergenic NA NA

T1692 TCTGATCAAGTGCGCGGCCATGG 1000
Oligo fwd CACCTCTGATCAAGTGCGCGGCCA
Oligo rev AAACTGGCCGCGCACTTGATCAGA
Chromosome start end strand MM target_seq PAM alignment distance position gene name gene id
GU305914.1 28455 28477 + 0 TCTGATCAAGTGCGCGGCCA TGG ||||||||[||||||||||||]PAM NA Intergenic NA NA

T1693 CAAGTGCGCGGCCATGGAAGAGG 1000
Oligo fwd CACCCAAGTGCGCGGCCATGGAAG
Oligo rev AAACCTTCCATGGCCGCGCACTTG
Chromosome start end strand MM target_seq PAM alignment distance position gene name gene id
GU305914.1 28461 28483 + 0 CAAGTGCGCGGCCATGGAAG AGG ||||||||[||||||||||||]PAM NA Intergenic NA NA

T1694 TGCGCGGCCATGGAAGAGGCAGG 1000

<table>
<tr><td>Oligo fwd</td><td>CACCTGCGCGGCCATGGAAGAGGC</td></tr>
<tr><td>Oligo rev</td><td>AAACGCCTCTTCCATGGCCGCGCA</td></tr>
<tr><td>Chromosome</td><td>start</td><td></td><td>end</td><td>strand</td><td>MM</td><td>target_seq</td><td>PAM</td><td>alignment</td><td>distance</td><td>position</td><td>gene name</td><td>gene id</td></tr>
<tr><td>GU305914.1</td><td></td><td>28465</td><td>28487</td><td>+</td><td>0</td><td>TGCGCGGCCATGGAAGAGGC</td><td>AGG</td><td>||||||||[||||||||||||]PAM</td><td>NA</td><td>Intergenic</td><td>NA</td><td>NA</td></tr>
<tr><td></td></tr>
<tr><td>T1695</td><td>ATGGAAGAGGCAGGCTTCGAAGG</td><td></td><td>1000</td></tr>
<tr><td>Oligo fwd</td><td>CACCATGGAAGAGGCAGGCTTCGA</td></tr>
<tr><td>Oligo rev</td><td>AAACTCGAAGCCTGCCTCTTCCAT</td></tr>
<tr><td>Chromosome</td><td>start</td><td></td><td>end</td><td>strand</td><td>MM</td><td>target_seq</td><td>PAM</td><td>alignment</td><td>distance</td><td>position</td><td>gene name</td><td>gene id</td></tr>
<tr><td>GU305914.1</td><td></td><td>28474</td><td>28496</td><td>+</td><td>0</td><td>ATGGAAGAGGCAGGCTTCGA</td><td>AGG</td><td>||||||||[||||||||||||]PAM</td><td>NA</td><td>Intergenic</td><td>NA</td><td>NA</td></tr>
<tr><td></td></tr>
<tr><td>T1696</td><td>CGAAGGCTACTCGCTCGAGTCGG</td><td></td><td>1000</td></tr>
<tr><td>Oligo fwd</td><td>CACCCGAAGGCTACTCGCTCGAGT</td></tr>
<tr><td>Oligo rev</td><td>AAACACTCGAGCGAGTAGCCTTCG</td></tr>
<tr><td>Chromosome</td><td>start</td><td></td><td>end</td><td>strand</td><td>MM</td><td>target_seq</td><td>PAM</td><td>alignment</td><td>distance</td><td>position</td><td>gene name</td><td>gene id</td></tr>
<tr><td>GU305914.1</td><td></td><td>28491</td><td>28513</td><td>+</td><td>0</td><td>CGAAGGCTACTCGCTCGAGT</td><td>CGG</td><td>||||||||[||||||||||||]PAM</td><td>NA</td><td>Intergenic</td><td>NA</td><td>NA</td></tr>
<tr><td></td></tr>
<tr><td>T1697</td><td>CGAGTCGGCCGCGAGATCTCTGG</td><td></td><td>1000</td></tr>
<tr><td>Oligo fwd</td><td>CACCCGAGTCGGCCGCGAGATCTC</td></tr>
<tr><td>Oligo rev</td><td>AAACGAGATCTCGCGGCCGACTCG</td></tr>
<tr><td>Chromosome</td><td>start</td><td></td><td>end</td><td>strand</td><td>MM</td><td>target_seq</td><td>PAM</td><td>alignment</td><td>distance</td><td>position</td><td>gene name</td><td>gene id</td></tr>
<tr><td>GU305914.1</td><td></td><td>28506</td><td>28528</td><td>+</td><td>0</td><td>CGAGTCGGCCGCGAGATCTC</td><td>TGG</td><td>||||||||[||||||||||||]PAM</td><td>NA</td><td>Intergenic</td><td>NA</td><td>NA</td></tr>
<tr><td></td></tr>
<tr><td>T1698</td><td>ATACACTTCGTGCGCGTCGACGG</td><td></td><td>1000</td></tr>
<tr><td>Oligo fwd</td><td>CACCATACACTTCGTGCGCGTCGA</td></tr>
<tr><td>Oligo rev</td><td>AAACTCGACGCGCACGAAGTGTAT</td></tr>
<tr><td>Chromosome</td><td>start</td><td></td><td>end</td><td>strand</td><td>MM</td><td>target_seq</td><td>PAM</td><td>alignment</td><td>distance</td><td>position</td><td>gene name</td><td>gene id</td></tr>
<tr><td>GU305914.1</td><td></td><td>28594</td><td>28616</td><td>+</td><td>0</td><td>ATACACTTCGTGCGCGTCGA</td><td>CGG</td><td>||||||||[||||||||||||]PAM</td><td>NA</td><td>Intergenic</td><td>NA</td><td>NA</td></tr>
<tr><td></td></tr>
<tr><td>T1699</td><td>TACACTTCGTGCGCGTCGACGGG</td><td></td><td>1000</td></tr>
<tr><td>Oligo fwd</td><td>CACCTACACTTCGTGCGCGTCGAC</td></tr>
<tr><td>Oligo rev</td><td>AAACGTCGACGCGCACGAAGTGTA</td></tr>
<tr><td>Chromosome</td><td>start</td><td></td><td>end</td><td>strand</td><td>MM</td><td>target_seq</td><td>PAM</td><td>alignment</td><td>distance</td><td>position</td><td>gene name</td><td>gene id</td></tr>
<tr><td>GU305914.1</td><td></td><td>28595</td><td>28617</td><td>+</td><td>0</td><td>TACACTTCGTGCGCGTCGAC</td><td>GGG</td><td>||||||||[||||||||||||]PAM</td><td>NA</td><td>Intergenic</td><td>NA</td><td>NA</td></tr>
<tr><td></td></tr>
<tr><td>T1700</td><td>ATCATCCTCTCCTCCTCCCCCGG</td><td></td><td>1000</td></tr>
<tr><td>Oligo fwd</td><td>CACCATCATCCTCTCCTCCTCCCC</td></tr>
<tr><td>Oligo rev</td><td>AAACGGGGAGGAGGAGAGGATGAT</td></tr>
<tr><td>Chromosome</td><td>start</td><td></td><td>end</td><td>strand</td><td>MM</td><td>target_seq</td><td>PAM</td><td>alignment</td><td>distance</td><td>position</td><td>gene name</td><td>gene id</td></tr>
<tr><td>GU305914.1</td><td></td><td>28749</td><td>28771</td><td>+</td><td>0</td><td>ATCATCCTCTCCTCCTCCCC</td><td>CGG</td><td>||||||||[||||||||||||]PAM</td><td>NA</td><td>Intergenic</td><td>NA</td><td>NA</td></tr>
<tr><td></td></tr>
<tr><td>T1701</td><td>TCTCCTCCTCCCCCGGCCGCCGG</td><td></td><td>1000</td></tr>
<tr><td>Oligo fwd</td><td>CACCTCTCCTCCTCCCCCGGCCGC</td></tr>
<tr><td>Oligo rev</td><td>AAACGCGGCCGGGGGAGGAGGAGA</td></tr>
<tr><td>Chromosome</td><td>start</td><td></td><td>end</td><td>strand</td><td>MM</td><td>target_seq</td><td>PAM</td><td>alignment</td><td>distance</td><td>position</td><td>gene name</td><td>gene id</td></tr>
<tr><td>GU305914.1</td><td></td><td>28756</td><td>28778</td><td>+</td><td>0</td><td>TCTCCTCCTCCCCCGGCCGC</td><td>CGG</td><td>||||||||[||||||||||||]PAM</td><td>NA</td><td>Intergenic</td><td>NA</td><td>NA</td></tr>
<tr><td></td></tr>
<tr><td>T1702</td><td>CCTCCTCCCCCGGCCGCCGGAGG</td><td></td><td>1000</td></tr>
<tr><td>Oligo fwd</td><td>CACCCCTCCTCCCCCGGCCGCCGG</td></tr>
<tr><td>Oligo rev</td><td>AAACCCGGCGGCCGGGGGAGGAGG</td></tr>
<tr><td>Chromosome</td><td>start</td><td></td><td>end</td><td>strand</td><td>MM</td><td>target_seq</td><td>PAM</td><td>alignment</td><td>distance</td><td>position</td><td>gene name</td><td>gene id</td></tr>
<tr><td>GU305914.1</td><td></td><td>28759</td><td>28781</td><td>+</td><td>0</td><td>CCTCCTCCCCCGGCCGCCGG</td><td>AGG</td><td>||||||||[||||||||||||]PAM</td><td>NA</td><td>Intergenic</td><td>NA</td><td>NA</td></tr>
<tr><td></td></tr>
<tr><td>T1703</td><td>CTCCTCCCCCGGCCGCCGGAGGG</td><td></td><td>1000</td></tr>
<tr><td>Oligo fwd</td><td>CACCCTCCTCCCCCGGCCGCCGGA</td></tr>
<tr><td>Oligo rev</td><td>AAACTCCGGCGGCCGGGGGAGGAG</td></tr>
<tr><td>Chromosome</td><td>start</td><td></td><td>end</td><td>strand</td><td>MM</td><td>target_seq</td><td>PAM</td><td>alignment</td><td>distance</td><td>position</td><td>gene name</td><td>gene id</td></tr>
<tr><td>GU305914.1</td><td></td><td>28760</td><td>28782</td><td>+</td><td>0</td><td>CTCCTCCCCCGGCCGCCGGA</td><td>GGG</td><td>||||||||[||||||||||||]PAM</td><td>NA</td><td>Intergenic</td><td>NA</td><td>NA</td></tr>
<tr><td></td></tr>
<tr><td>T1704</td><td>GTGTCTGACTTTATTGATATCGG</td><td></td><td>1000</td></tr>
<tr><td>Oligo fwd</td><td>CACCGTGTCTGACTTTATTGATAT</td></tr>
<tr><td>Oligo rev</td><td>AAACATATCAATAAAGTCAGACAC</td></tr>
<tr><td>Chromosome</td><td>start</td><td></td><td>end</td><td>strand</td><td>MM</td><td>target_seq</td><td>PAM</td><td>alignment</td><td>distance</td><td>position</td><td>gene name</td><td>gene id</td></tr>
<tr><td>GU305914.1</td><td></td><td>29000</td><td>29022</td><td>+</td><td>0</td><td>GTGTCTGACTTTATTGATAT</td><td>CGG</td><td>||||||||[||||||||||||]PAM</td><td>NA</td><td>Intergenic</td><td>NA</td><td>NA</td></tr>
<tr><td></td></tr>
<tr><td>T1705</td><td>GGCATTGTTGTCGTCGTTTTCGG</td><td></td><td>1000</td></tr>
<tr><td>Oligo fwd</td><td>CACCGGCATTGTTGTCGTCGTTTT</td></tr>
<tr><td>Oligo rev</td><td>AAACAAAACGACGACAACAATGCC</td></tr>
<tr><td>Chromosome</td><td>start</td><td></td><td>end</td><td>strand</td><td>MM</td><td>target_seq</td><td>PAM</td><td>alignment</td><td>distance</td><td>position</td><td>gene name</td><td>gene id</td></tr>
<tr><td>GU305914.1</td><td></td><td>29021</td><td>29043</td><td>+</td><td>0</td><td>GGCATTGTTGTCGTCGTTTT</td><td>CGG</td><td>||||||||[||||||||||||]PAM</td><td>NA</td><td>Intergenic</td><td>NA</td><td>NA</td></tr>
<tr><td></td></tr>
<tr><td>T1706</td><td>TCCCTTAGCCAGCAGATTGAAGG</td><td></td><td>1000</td></tr>
<tr><td>Oligo fwd</td><td>CACCTCCCTTAGCCAGCAGATTGA</td></tr>
<tr><td>Oligo rev</td><td>AAACTCAATCTGCTGGCTAAGGGA</td></tr>
<tr><td>Chromosome</td><td>start</td><td></td><td>end</td><td>strand</td><td>MM</td><td>target_seq</td><td>PAM</td><td>alignment</td><td>distance</td><td>position</td><td>gene name</td><td>gene id</td></tr>
<tr><td>GU305914.1</td><td></td><td>29048</td><td>29070</td><td>+</td><td>0</td><td>TCCCTTAGCCAGCAGATTGA</td><td>AGG</td><td>||||||||[||||||||||||]PAM</td><td>NA</td><td>Intergenic</td><td>NA</td><td>NA</td></tr>
<tr><td></td></tr>
<tr><td>T1707</td><td>GCAGATTGAAGGACCGACCGAGG</td><td></td><td>1000</td></tr>
<tr><td>Oligo fwd</td><td>CACCGCAGATTGAAGGACCGACCG</td></tr>
<tr><td>Oligo rev</td><td>AAACCGGTCGGTCCTTCAATCTGC</td></tr>
<tr><td>Chromosome</td><td>start</td><td></td><td>end</td><td>strand</td><td>MM</td><td>target_seq</td><td>PAM</td><td>alignment</td><td>distance</td><td>position</td><td>gene name</td><td>gene id</td></tr>
<tr><td>GU305914.1</td><td></td><td>29059</td><td>29081</td><td>+</td><td>0</td><td>GCAGATTGAAGGACCGACCG</td><td>AGG</td><td>||||||||[||||||||||||]PAM</td><td>NA</td><td>Intergenic</td><td>NA</td><td>NA</td></tr>
<tr><td></td></tr>
<tr><td>T1708</td><td>GGACCGACCGAGGCAACGTAAGG</td><td></td><td>1000</td></tr>
<tr><td>Oligo fwd</td><td>CACCGGACCGACCGAGGCAACGTA</td></tr>
<tr><td>Oligo rev</td><td>AAACTACGTTGCCTCGGTCGGTCC</td></tr>
<tr><td>Chromosome</td><td>start</td><td></td><td>end</td><td>strand</td><td>MM</td><td>target_seq</td><td>PAM</td><td>alignment</td><td>distance</td><td>position</td><td>gene name</td><td>gene id</td></tr>
<tr><td>GU305914.1</td><td></td><td>29069</td><td>29091</td><td>+</td><td>0</td><td>GGACCGACCGAGGCAACGTA</td><td>AGG</td><td>||||||||[||||||||||||]PAM</td><td>NA</td><td>Intergenic</td><td>NA</td><td>NA</td></tr>
</table>

T1709 TCTTGTCCTGTGAAGAACACCGG 1000
Oligo fwd CACCTCTTGTCCTGTGAAGAACAC
Oligo rev AAACGTGTTCTTCACAGGACAAGA
Chromosome start end strand MM target_seq PAM alignment distance position gene name gene id
GU305914.1 29112 29134 + 0 TCTTGTCCTGTGAAGAACAC CGG ||||||||[||||||||||||]PAM NA Intergenic NA NA

T1710 TGAAGAACACCGGACTGACGTGG 1000
Oligo fwd CACCTGAAGAACACCGGACTGACG
Oligo rev AAACCGTCAGTCCGGTGTTCTTCA
Chromosome start end strand MM target_seq PAM alignment distance position gene name gene id
GU305914.1 29122 29144 + 0 TGAAGAACACCGGACTGACG TGG ||||||||[||||||||||||]PAM NA Intergenic NA NA

T1711 GAAGAACACCGGACTGACGTGGG 1000
Oligo fwd CACCGAAGAACACCGGACTGACGT
Oligo rev AAACACGTCAGTCCGGTGTTCTTC
Chromosome start end strand MM target_seq PAM alignment distance position gene name gene id
GU305914.1 29123 29145 + 0 GAAGAACACCGGACTGACGT GGG ||||||||[||||||||||||]PAM NA Intergenic NA NA

T1712 AAGAACACCGGACTGACGTGGGG 1000
Oligo fwd CACCAAGAACACCGGACTGACGTG
Oligo rev AAACCACGTCAGTCCGGTGTTCTT
Chromosome start end strand MM target_seq PAM alignment distance position gene name gene id
GU305914.1 29124 29146 + 0 AAGAACACCGGACTGACGTG GGG ||||||||[||||||||||||]PAM NA Intergenic NA NA

T1713 CGGACTGACGTGGGGACACTTGG 1000
Oligo fwd CACCCGGACTGACGTGGGGACACT
Oligo rev AAACAGTGTCCCCACGTCAGTCCG
Chromosome start end strand MM target_seq PAM alignment distance position gene name gene id
GU305914.1 29132 29154 + 0 CGGACTGACGTGGGGACACT TGG ||||||||[||||||||||||]PAM NA Intergenic NA NA

T1714 GGACTGACGTGGGGACACTTGGG 1000
Oligo fwd CACCGGACTGACGTGGGGACACTT
Oligo rev AAACAAGTGTCCCCACGTCAGTCC
Chromosome start end strand MM target_seq PAM alignment distance position gene name gene id
GU305914.1 29133 29155 + 0 GGACTGACGTGGGGACACTT GGG ||||||||[||||||||||||]PAM NA Intergenic NA NA

T1715 CTGACGTGGGGACACTTGGGAGG 1000
Oligo fwd CACCCTGACGTGGGGACACTTGGG
Oligo rev AAACCCCAAGTGTCCCCACGTCAG
Chromosome start end strand MM target_seq PAM alignment distance position gene name gene id
GU305914.1 29136 29158 + 0 CTGACGTGGGGACACTTGGG AGG ||||||||[||||||||||||]PAM NA Intergenic NA NA

T1716 CGTGGGGACACTTGGGAGGCCGG 1000
Oligo fwd CACCCGTGGGGACACTTGGGAGGC
Oligo rev AAACGCCTCCCAAGTGTCCCCACG
Chromosome start end strand MM target_seq PAM alignment distance position gene name gene id
GU305914.1 29140 29162 + 0 CGTGGGGACACTTGGGAGGC CGG ||||||||[||||||||||||]PAM NA Intergenic NA NA

T1717 AGGCCGGTCGCCCAGCGCGACGG 1000
Oligo fwd CACCAGGCCGGTCGCCCAGCGCGA
Oligo rev AAACTCGCGCTGGGCGACCGGCCT
Chromosome start end strand MM target_seq PAM alignment distance position gene name gene id
GU305914.1 29156 29178 + 0 AGGCCGGTCGCCCAGCGCGA CGG ||||||||[||||||||||||]PAM NA Intergenic NA NA

T1718 GGCCGGTCGCCCAGCGCGACGGG 1000
Oligo fwd CACCGGCCGGTCGCCCAGCGCGAC
Oligo rev AAACGTCGCGCTGGGCGACCGGCC
Chromosome start end strand MM target_seq PAM alignment distance position gene name gene id
GU305914.1 29157 29179 + 0 GGCCGGTCGCCCAGCGCGAC GGG ||||||||[||||||||||||]PAM NA Intergenic NA NA

T1719 GATAGTTCTTTTTGATTCCCCGG 1000
Oligo fwd CACCGATAGTTCTTTTTGATTCCC
Oligo rev AAACGGGAATCAAAAAGAACTATC
Chromosome start end strand MM target_seq PAM alignment distance position gene name gene id
GU305914.1 29197 29219 + 0 GATAGTTCTTTTTGATTCCC CGG ||||||||[||||||||||||]PAM NA Intergenic NA NA

T1720 CATGACGCCCTCGCAGATCTTGG 1000
Oligo fwd CACCCATGACGCCCTCGCAGATCT
Oligo rev AAACAGATCTGCGAGGGCGTCATG
Chromosome start end strand MM target_seq PAM alignment distance position gene name gene id
GU305914.1 29240 29262 + 0 CATGACGCCCTCGCAGATCT TGG ||||||||[||||||||||||]PAM NA Intergenic NA NA

T1721 CTTGGTGATCTCGTCGTCCCAGG 1000
Oligo fwd CACCCTTGGTGATCTCGTCGTCCC
Oligo rev AAACGGGACGACGAGATCACCAAG
Chromosome start end strand MM target_seq PAM alignment distance position gene name gene id
GU305914.1 29258 29280 + 0 CTTGGTGATCTCGTCGTCCC AGG ||||||||[||||||||||||]PAM NA Intergenic NA NA

T1722 TTGGTGATCTCGTCGTCCCAGGG 1000
Oligo fwd CACCTTGGTGATCTCGTCGTCCCA
Oligo rev AAACTGGGACGACGAGATCACCAA
Chromosome start end strand MM target_seq PAM alignment distance position gene name gene id
GU305914.1 29259 29281 + 0 TTGGTGATCTCGTCGTCCCA GGG ||||||||[||||||||||||]PAM NA Intergenic NA NA

T1723 TGGTGATCTCGTCGTCCCAGGGG 1000
Oligo fwd CACCTGGTGATCTCGTCGTCCCAG
Oligo rev AAACCTGGGACGACGAGATCACCA
Chromosome start end strand MM target_seq PAM alignment distance position gene name gene id
GU305914.1 29260 29282 + 0 TGGTGATCTCGTCGTCCCAG GGG ||||||||[||||||||||||]PAM NA Intergenic NA NA

| | | | | | | | | | | | |
|---|---|---|---|---|---|---|---|---|---|---|---|
| T1724 | CTCGTCGTCCCAGGGGTTGTAGG | 1000 | | | | | | | | | |
| Oligo fwd | CACCCTCGTCGTCCCAGGGGTTGT | | | | | | | | | | |
| Oligo rev | AAACACAACCCCTGGGACGACGAG | | | | | | | | | | |
| Chromosome | start | end | strand | MM | target_seq | PAM | alignment | distance | position | gene name | gene id |
| GU305914.1 | 29267 | 29289 | + | 0 | CTCGTCGTCCCAGGGGTTGT | AGG | \|\|\|\|\|\|\|\|[\|\|\|\|\|\|\|\|\|\|\|\|]PAM | NA | Intergenic | NA | NA |
| T1725 | CCAGGGGTTGTAGGAAAAGATGG | 1000 | | | | | | | | | |
| Oligo fwd | CACCCCAGGGGTTGTAGGAAAAGA | | | | | | | | | | |
| Oligo rev | AAACTCTTTTCCTACAACCCCTGG | | | | | | | | | | |
| Chromosome | start | end | strand | MM | target_seq | PAM | alignment | distance | position | gene name | gene id |
| GU305914.1 | 29276 | 29298 | + | 0 | CCAGGGGTTGTAGGAAAAGA | TGG | \|\|\|\|\|\|\|\|[\|\|\|\|\|\|\|\|\|\|\|\|]PAM | NA | Intergenic | NA | NA |
| T1726 | GATGGCACCTTTCTCGTTGACGG | 1000 | | | | | | | | | |
| Oligo fwd | CACCGATGGCACCTTTCTCGTTGA | | | | | | | | | | |
| Oligo rev | AAACTCAACGAGAAAGGTGCCATC | | | | | | | | | | |
| Chromosome | start | end | strand | MM | target_seq | PAM | alignment | distance | position | gene name | gene id |
| GU305914.1 | 29294 | 29316 | + | 0 | GATGGCACCTTTCTCGTTGA | CGG | \|\|\|\|\|\|\|\|[\|\|\|\|\|\|\|\|\|\|\|\|]PAM | NA | Intergenic | NA | NA |
| T1727 | CTTTCTCGTTGACGGCCAAGAGG | 1000 | | | | | | | | | |
| Oligo fwd | CACCCTTTCTCGTTGACGGCCAAG | | | | | | | | | | |
| Oligo rev | AAACCTTGGCCGTCAACGAGAAAG | | | | | | | | | | |
| Chromosome | start | end | strand | MM | target_seq | PAM | alignment | distance | position | gene name | gene id |
| GU305914.1 | 29302 | 29324 | + | 0 | CTTTCTCGTTGACGGCCAAG | AGG | \|\|\|\|\|\|\|\|[\|\|\|\|\|\|\|\|\|\|\|\|]PAM | NA | Intergenic | NA | NA |
| T1728 | TGACGGCCAAGAGGACCTCGCGG | 1000 | | | | | | | | | |
| Oligo fwd | CACCTGACGGCCAAGAGGACCTCG | | | | | | | | | | |
| Oligo rev | AAACCGAGGTCCTCTTGGCCGTCA | | | | | | | | | | |
| Chromosome | start | end | strand | MM | target_seq | PAM | alignment | distance | position | gene name | gene id |
| GU305914.1 | 29311 | 29333 | + | 0 | TGACGGCCAAGAGGACCTCG | CGG | \|\|\|\|\|\|\|\|[\|\|\|\|\|\|\|\|\|\|\|\|]PAM | NA | Intergenic | NA | NA |
| T1729 | CAAGAGGACCTCGCGGAAGACGG | 1000 | | | | | | | | | |
| Oligo fwd | CACCCAAGAGGACCTCGCGGAAGA | | | | | | | | | | |
| Oligo rev | AAACTCTTCCGCGAGGTCCTCTTG | | | | | | | | | | |
| Chromosome | start | end | strand | MM | target_seq | PAM | alignment | distance | position | gene name | gene id |
| GU305914.1 | 29318 | 29340 | + | 0 | CAAGAGGACCTCGCGGAAGA | CGG | \|\|\|\|\|\|\|\|[\|\|\|\|\|\|\|\|\|\|\|\|]PAM | NA | Intergenic | NA | NA |
| T1730 | AAGAGGACCTCGCGGAAGACGGG | 1000 | | | | | | | | | |
| Oligo fwd | CACCAAGAGGACCTCGCGGAAGAC | | | | | | | | | | |
| Oligo rev | AAACGTCTTCCGCGAGGTCCTCTT | | | | | | | | | | |
| Chromosome | start | end | strand | MM | target_seq | PAM | alignment | distance | position | gene name | gene id |
| GU305914.1 | 29319 | 29341 | + | 0 | AAGAGGACCTCGCGGAAGAC | GGG | \|\|\|\|\|\|\|\|[\|\|\|\|\|\|\|\|\|\|\|\|]PAM | NA | Intergenic | NA | NA |
| T1731 | AGAGGACCTCGCGGAAGACGGGG | 1000 | | | | | | | | | |
| Oligo fwd | CACCAGAGGACCTCGCGGAAGACG | | | | | | | | | | |
| Oligo rev | AAACCGTCTTCCGCGAGGTCCTCT | | | | | | | | | | |
| Chromosome | start | end | strand | MM | target_seq | PAM | alignment | distance | position | gene name | gene id |
| GU305914.1 | 29320 | 29342 | + | 0 | AGAGGACCTCGCGGAAGACG | GGG | \|\|\|\|\|\|\|\|[\|\|\|\|\|\|\|\|\|\|\|\|]PAM | NA | Intergenic | NA | NA |
| T1732 | GTCCAAGTAACGTCCGATGACGG | 1000 | | | | | | | | | |
| Oligo fwd | CACCGTCCAAGTAACGTCCGATGA | | | | | | | | | | |
| Oligo rev | AAACTCATCGGACGTTACTTGGAC | | | | | | | | | | |
| Chromosome | start | end | strand | MM | target_seq | PAM | alignment | distance | position | gene name | gene id |
| GU305914.1 | 29342 | 29364 | + | 0 | GTCCAAGTAACGTCCGATGA | CGG | \|\|\|\|\|\|\|\|[\|\|\|\|\|\|\|\|\|\|\|\|]PAM | NA | Intergenic | NA | NA |
| T1733 | GACGGCCGCGAAGTAGACCATGG | 1000 | | | | | | | | | |
| Oligo fwd | CACCGACGGCCGCGAAGTAGACCA | | | | | | | | | | |
| Oligo rev | AAACTGGTCTACTTCGCGGCCGTC | | | | | | | | | | |
| Chromosome | start | end | strand | MM | target_seq | PAM | alignment | distance | position | gene name | gene id |
| GU305914.1 | 29360 | 29382 | + | 0 | GACGGCCGCGAAGTAGACCA | TGG | \|\|\|\|\|\|\|\|[\|\|\|\|\|\|\|\|\|\|\|\|]PAM | NA | Intergenic | NA | NA |
| T1734 | CGTGACGAACTTGCGTCGTCTGG | 1000 | | | | | | | | | |
| Oligo fwd | CACCCGTGACGAACTTGCGTCGTC | | | | | | | | | | |
| Oligo rev | AAACGACGACGCAAGTTCGTCACG | | | | | | | | | | |
| Chromosome | start | end | strand | MM | target_seq | PAM | alignment | distance | position | gene name | gene id |
| GU305914.1 | 29399 | 29421 | + | 0 | CGTGACGAACTTGCGTCGTC | TGG | \|\|\|\|\|\|\|\|[\|\|\|\|\|\|\|\|\|\|\|\|]PAM | NA | Intergenic | NA | NA |
| T1735 | GTGACGAACTTGCGTCGTCTGGG | 1000 | | | | | | | | | |
| Oligo fwd | CACCGTGACGAACTTGCGTCGTCT | | | | | | | | | | |
| Oligo rev | AAACAGACGACGCAAGTTCGTCAC | | | | | | | | | | |
| Chromosome | start | end | strand | MM | target_seq | PAM | alignment | distance | position | gene name | gene id |
| GU305914.1 | 29400 | 29422 | + | 0 | GTGACGAACTTGCGTCGTCT | GGG | \|\|\|\|\|\|\|\|[\|\|\|\|\|\|\|\|\|\|\|\|]PAM | NA | Intergenic | NA | NA |
| T1736 | ACGAACTTGCGTCGTCTGGGTGG | 1000 | | | | | | | | | |
| Oligo fwd | CACCACGAACTTGCGTCGTCTGGG | | | | | | | | | | |
| Oligo rev | AAACCCCAGACGACGCAAGTTCGT | | | | | | | | | | |
| Chromosome | start | end | strand | MM | target_seq | PAM | alignment | distance | position | gene name | gene id |
| GU305914.1 | 29403 | 29425 | + | 0 | ACGAACTTGCGTCGTCTGGG | TGG | \|\|\|\|\|\|\|\|[\|\|\|\|\|\|\|\|\|\|\|\|]PAM | NA | Intergenic | NA | NA |
| T1737 | CGAACTTGCGTCGTCTGGGTGGG | 1000 | | | | | | | | | |
| Oligo fwd | CACCCGAACTTGCGTCGTCTGGGT | | | | | | | | | | |
| Oligo rev | AAACACCCAGACGACGCAAGTTCG | | | | | | | | | | |
| Chromosome | start | end | strand | MM | target_seq | PAM | alignment | distance | position | gene name | gene id |
| GU305914.1 | 29404 | 29426 | + | 0 | CGAACTTGCGTCGTCTGGGT | GGG | \|\|\|\|\|\|\|\|[\|\|\|\|\|\|\|\|\|\|\|\|]PAM | NA | Intergenic | NA | NA |
| T1738 | CTTGCGTCGTCTGGGTGGGATGG | 1000 | | | | | | | | | |
| Oligo fwd | CACCCTTGCGTCGTCTGGGTGGGA | | | | | | | | | | |
| Oligo rev | AAACTCCCACCCAGACGACGCAAG | | | | | | | | | | |
| Chromosome | start | end | strand | MM | target_seq | PAM | alignment | distance | position | gene name | gene id |

<table>
<tr><td>GU305914.1</td><td>29408</td><td>29430</td><td>+</td><td>0</td><td>CTTGCGTCGTCTGGGTGGGA</td><td>TGG</td><td>||||||||[||||||||||||]PAM</td><td>NA</td><td>Intergenic</td><td>NA</td><td>NA</td></tr>
<tr><td></td><td></td><td></td><td></td><td></td><td></td><td></td><td></td><td></td><td></td><td></td><td></td></tr>
<tr><td>T1739</td><td>GGGTGGGATGGTTTTGATGTAGG</td><td>1000</td><td></td><td></td><td></td><td></td><td></td><td></td><td></td><td></td><td></td></tr>
<tr><td>Oligo fwd</td><td>CACCGGGTGGGATGGTTTTGATGT</td><td></td><td></td><td></td><td></td><td></td><td></td><td></td><td></td><td></td><td></td></tr>
<tr><td>Oligo rev</td><td>AAACACATCAAAACCATCCCACCC</td><td></td><td></td><td></td><td></td><td></td><td></td><td></td><td></td><td></td><td></td></tr>
<tr><td>Chromosome</td><td>start</td><td>end</td><td>strand</td><td>MM</td><td>target_seq</td><td>PAM</td><td>alignment</td><td>distance</td><td>position</td><td>gene name</td><td>gene id</td></tr>
<tr><td>GU305914.1</td><td>29420</td><td>29442</td><td>+</td><td>0</td><td>GGGTGGGATGGTTTTGATGT</td><td>AGG</td><td>||||||||[||||||||||||]PAM</td><td>NA</td><td>Intergenic</td><td>NA</td><td>NA</td></tr>
<tr><td></td><td></td><td></td><td></td><td></td><td></td><td></td><td></td><td></td><td></td><td></td><td></td></tr>
<tr><td>T1740</td><td>GATGGTTTTGATGTAGGCGCCGG</td><td>1000</td><td></td><td></td><td></td><td></td><td></td><td></td><td></td><td></td><td></td></tr>
<tr><td>Oligo fwd</td><td>CACCGATGGTTTTGATGTAGGCGC</td><td></td><td></td><td></td><td></td><td></td><td></td><td></td><td></td><td></td><td></td></tr>
<tr><td>Oligo rev</td><td>AAACGCGCCTACATCAAAACCATC</td><td></td><td></td><td></td><td></td><td></td><td></td><td></td><td></td><td></td><td></td></tr>
<tr><td>Chromosome</td><td>start</td><td>end</td><td>strand</td><td>MM</td><td>target_seq</td><td>PAM</td><td>alignment</td><td>distance</td><td>position</td><td>gene name</td><td>gene id</td></tr>
<tr><td>GU305914.1</td><td>29426</td><td>29448</td><td>+</td><td>0</td><td>GATGGTTTTGATGTAGGCGC</td><td>CGG</td><td>||||||||[||||||||||||]PAM</td><td>NA</td><td>Intergenic</td><td>NA</td><td>NA</td></tr>
<tr><td></td><td></td><td></td><td></td><td></td><td></td><td></td><td></td><td></td><td></td><td></td><td></td></tr>
<tr><td>T1741</td><td>GGTTTTGATGTAGGCGCCGGTGG</td><td>1000</td><td></td><td></td><td></td><td></td><td></td><td></td><td></td><td></td><td></td></tr>
<tr><td>Oligo fwd</td><td>CACCGGTTTTGATGTAGGCGCCGG</td><td></td><td></td><td></td><td></td><td></td><td></td><td></td><td></td><td></td><td></td></tr>
<tr><td>Oligo rev</td><td>AAACCCGGCGCCTACATCAAAACC</td><td></td><td></td><td></td><td></td><td></td><td></td><td></td><td></td><td></td><td></td></tr>
<tr><td>Chromosome</td><td>start</td><td>end</td><td>strand</td><td>MM</td><td>target_seq</td><td>PAM</td><td>alignment</td><td>distance</td><td>position</td><td>gene name</td><td>gene id</td></tr>
<tr><td>GU305914.1</td><td>29429</td><td>29451</td><td>+</td><td>0</td><td>GGTTTTGATGTAGGCGCCGG</td><td>TGG</td><td>||||||||[||||||||||||]PAM</td><td>NA</td><td>Intergenic</td><td>NA</td><td>NA</td></tr>
<tr><td></td><td></td><td></td><td></td><td></td><td></td><td></td><td></td><td></td><td></td><td></td><td></td></tr>
<tr><td>T1742</td><td>GTTTTGATGTAGGCGCCGGTGGG</td><td>1000</td><td></td><td></td><td></td><td></td><td></td><td></td><td></td><td></td><td></td></tr>
<tr><td>Oligo fwd</td><td>CACCGTTTTGATGTAGGCGCCGGT</td><td></td><td></td><td></td><td></td><td></td><td></td><td></td><td></td><td></td><td></td></tr>
<tr><td>Oligo rev</td><td>AAACACCGGCGCCTACATCAAAAC</td><td></td><td></td><td></td><td></td><td></td><td></td><td></td><td></td><td></td><td></td></tr>
<tr><td>Chromosome</td><td>start</td><td>end</td><td>strand</td><td>MM</td><td>target_seq</td><td>PAM</td><td>alignment</td><td>distance</td><td>position</td><td>gene name</td><td>gene id</td></tr>
<tr><td>GU305914.1</td><td>29430</td><td>29452</td><td>+</td><td>0</td><td>GTTTTGATGTAGGCGCCGGT</td><td>GGG</td><td>||||||||[||||||||||||]PAM</td><td>NA</td><td>Intergenic</td><td>NA</td><td>NA</td></tr>
<tr><td></td><td></td><td></td><td></td><td></td><td></td><td></td><td></td><td></td><td></td><td></td><td></td></tr>
<tr><td>T1743</td><td>GATGTAGGCGCCGGTGGGCCAGG</td><td>1000</td><td></td><td></td><td></td><td></td><td></td><td></td><td></td><td></td><td></td></tr>
<tr><td>Oligo fwd</td><td>CACCGATGTAGGCGCCGGTGGGCC</td><td></td><td></td><td></td><td></td><td></td><td></td><td></td><td></td><td></td><td></td></tr>
<tr><td>Oligo rev</td><td>AAACGGCCCACCGGCGCCTACATC</td><td></td><td></td><td></td><td></td><td></td><td></td><td></td><td></td><td></td><td></td></tr>
<tr><td>Chromosome</td><td>start</td><td>end</td><td>strand</td><td>MM</td><td>target_seq</td><td>PAM</td><td>alignment</td><td>distance</td><td>position</td><td>gene name</td><td>gene id</td></tr>
<tr><td>GU305914.1</td><td>29435</td><td>29457</td><td>+</td><td>0</td><td>GATGTAGGCGCCGGTGGGCC</td><td>AGG</td><td>||||||||[||||||||||||]PAM</td><td>NA</td><td>Intergenic</td><td>NA</td><td>NA</td></tr>
<tr><td></td><td></td><td></td><td></td><td></td><td></td><td></td><td></td><td></td><td></td><td></td><td></td></tr>
<tr><td>T1744</td><td>ATGTAGGCGCCGGTGGGCCAGGG</td><td>1000</td><td></td><td></td><td></td><td></td><td></td><td></td><td></td><td></td><td></td></tr>
<tr><td>Oligo fwd</td><td>CACCATGTAGGCGCCGGTGGGCCA</td><td></td><td></td><td></td><td></td><td></td><td></td><td></td><td></td><td></td><td></td></tr>
<tr><td>Oligo rev</td><td>AAACTGGCCCACCGGCGCCTACAT</td><td></td><td></td><td></td><td></td><td></td><td></td><td></td><td></td><td></td><td></td></tr>
<tr><td>Chromosome</td><td>start</td><td>end</td><td>strand</td><td>MM</td><td>target_seq</td><td>PAM</td><td>alignment</td><td>distance</td><td>position</td><td>gene name</td><td>gene id</td></tr>
<tr><td>GU305914.1</td><td>29436</td><td>29458</td><td>+</td><td>0</td><td>ATGTAGGCGCCGGTGGGCCA</td><td>GGG</td><td>||||||||[||||||||||||]PAM</td><td>NA</td><td>Intergenic</td><td>NA</td><td>NA</td></tr>
<tr><td></td><td></td><td></td><td></td><td></td><td></td><td></td><td></td><td></td><td></td><td></td><td></td></tr>
<tr><td>T1745</td><td>AGGCGCCGGTGGGCCAGGGCAGG</td><td>1000</td><td></td><td></td><td></td><td></td><td></td><td></td><td></td><td></td><td></td></tr>
<tr><td>Oligo fwd</td><td>CACCAGGCGCCGGTGGGCCAGGGC</td><td></td><td></td><td></td><td></td><td></td><td></td><td></td><td></td><td></td><td></td></tr>
<tr><td>Oligo rev</td><td>AAACGCCCTGGCCCACCGGCGCCT</td><td></td><td></td><td></td><td></td><td></td><td></td><td></td><td></td><td></td><td></td></tr>
<tr><td>Chromosome</td><td>start</td><td>end</td><td>strand</td><td>MM</td><td>target_seq</td><td>PAM</td><td>alignment</td><td>distance</td><td>position</td><td>gene name</td><td>gene id</td></tr>
<tr><td>GU305914.1</td><td>29440</td><td>29462</td><td>+</td><td>0</td><td>AGGCGCCGGTGGGCCAGGGC</td><td>AGG</td><td>||||||||[||||||||||||]PAM</td><td>NA</td><td>Intergenic</td><td>NA</td><td>NA</td></tr>
<tr><td></td><td></td><td></td><td></td><td></td><td></td><td></td><td></td><td></td><td></td><td></td><td></td></tr>
<tr><td>T1746</td><td>GGCGCCGGTGGGCCAGGGCAGGG</td><td>1000</td><td></td><td></td><td></td><td></td><td></td><td></td><td></td><td></td><td></td></tr>
<tr><td>Oligo fwd</td><td>CACCGGCGCCGGTGGGCCAGGGCA</td><td></td><td></td><td></td><td></td><td></td><td></td><td></td><td></td><td></td><td></td></tr>
<tr><td>Oligo rev</td><td>AAACTGCCCTGGCCCACCGGCGCC</td><td></td><td></td><td></td><td></td><td></td><td></td><td></td><td></td><td></td><td></td></tr>
<tr><td>Chromosome</td><td>start</td><td>end</td><td>strand</td><td>MM</td><td>target_seq</td><td>PAM</td><td>alignment</td><td>distance</td><td>position</td><td>gene name</td><td>gene id</td></tr>
<tr><td>GU305914.1</td><td>29441</td><td>29463</td><td>+</td><td>0</td><td>GGCGCCGGTGGGCCAGGGCA</td><td>GGG</td><td>||||||||[||||||||||||]PAM</td><td>NA</td><td>Intergenic</td><td>NA</td><td>NA</td></tr>
<tr><td></td><td></td><td></td><td></td><td></td><td></td><td></td><td></td><td></td><td></td><td></td><td></td></tr>
<tr><td>T1747</td><td>GGGCCAGGGCAGGGAGAAGACGG</td><td>1000</td><td></td><td></td><td></td><td></td><td></td><td></td><td></td><td></td><td></td></tr>
<tr><td>Oligo fwd</td><td>CACCGGGCCAGGGCAGGGAGAAGA</td><td></td><td></td><td></td><td></td><td></td><td></td><td></td><td></td><td></td><td></td></tr>
<tr><td>Oligo rev</td><td>AAACTCTTCTCCCTGCCCTGGCCC</td><td></td><td></td><td></td><td></td><td></td><td></td><td></td><td></td><td></td><td></td></tr>
<tr><td>Chromosome</td><td>start</td><td>end</td><td>strand</td><td>MM</td><td>target_seq</td><td>PAM</td><td>alignment</td><td>distance</td><td>position</td><td>gene name</td><td>gene id</td></tr>
<tr><td>GU305914.1</td><td>29450</td><td>29472</td><td>+</td><td>0</td><td>GGGCCAGGGCAGGGAGAAGA</td><td>CGG</td><td>||||||||[||||||||||||]PAM</td><td>NA</td><td>Intergenic</td><td>NA</td><td>NA</td></tr>
<tr><td>GU305914.1</td><td>146711</td><td>146733</td><td>-</td><td>3</td><td>GCGCCAGGGCACGGAGTAGA</td><td>CGG</td><td>|-||||||[|||-||||-|||]PAM</td><td>NA</td><td>Intergenic</td><td>NA</td><td>NA</td></tr>
<tr><td></td><td></td><td></td><td></td><td></td><td></td><td></td><td></td><td></td><td></td><td></td><td></td></tr>
<tr><td>T1748</td><td>GGGCAGGGAGAAGACGGTGCCGG</td><td>1000</td><td></td><td></td><td></td><td></td><td></td><td></td><td></td><td></td><td></td></tr>
<tr><td>Oligo fwd</td><td>CACCGGGCAGGGAGAAGACGGTGC</td><td></td><td></td><td></td><td></td><td></td><td></td><td></td><td></td><td></td><td></td></tr>
<tr><td>Oligo rev</td><td>AAACGCACCGTCTTCTCCCTGCCC</td><td></td><td></td><td></td><td></td><td></td><td></td><td></td><td></td><td></td><td></td></tr>
<tr><td>Chromosome</td><td>start</td><td>end</td><td>strand</td><td>MM</td><td>target_seq</td><td>PAM</td><td>alignment</td><td>distance</td><td>position</td><td>gene name</td><td>gene id</td></tr>
<tr><td>GU305914.1</td><td>29456</td><td>29478</td><td>+</td><td>0</td><td>GGGCAGGGAGAAGACGGTGC</td><td>CGG</td><td>||||||||[||||||||||||]PAM</td><td>NA</td><td>Intergenic</td><td>NA</td><td>NA</td></tr>
<tr><td></td><td></td><td></td><td></td><td></td><td></td><td></td><td></td><td></td><td></td><td></td><td></td></tr>
<tr><td>T1749</td><td>GGCAGGGAGAAGACGGTGCCGGG</td><td>1000</td><td></td><td></td><td></td><td></td><td></td><td></td><td></td><td></td><td></td></tr>
<tr><td>Oligo fwd</td><td>CACCGGCAGGGAGAAGACGGTGCC</td><td></td><td></td><td></td><td></td><td></td><td></td><td></td><td></td><td></td><td></td></tr>
<tr><td>Oligo rev</td><td>AAACGGCACCGTCTTCTCCCTGCC</td><td></td><td></td><td></td><td></td><td></td><td></td><td></td><td></td><td></td><td></td></tr>
<tr><td>Chromosome</td><td>start</td><td>end</td><td>strand</td><td>MM</td><td>target_seq</td><td>PAM</td><td>alignment</td><td>distance</td><td>position</td><td>gene name</td><td>gene id</td></tr>
<tr><td>GU305914.1</td><td>29457</td><td>29479</td><td>+</td><td>0</td><td>GGCAGGGAGAAGACGGTGCC</td><td>GGG</td><td>||||||||[||||||||||||]PAM</td><td>NA</td><td>Intergenic</td><td>NA</td><td>NA</td></tr>
<tr><td></td><td></td><td></td><td></td><td></td><td></td><td></td><td></td><td></td><td></td><td></td><td></td></tr>
<tr><td>T1750</td><td>GCAGGGAGAAGACGGTGCCGGGG</td><td>1000</td><td></td><td></td><td></td><td></td><td></td><td></td><td></td><td></td><td></td></tr>
<tr><td>Oligo fwd</td><td>CACCGCAGGGAGAAGACGGTGCCG</td><td></td><td></td><td></td><td></td><td></td><td></td><td></td><td></td><td></td><td></td></tr>
<tr><td>Oligo rev</td><td>AAACCGGCACCGTCTTCTCCCTGC</td><td></td><td></td><td></td><td></td><td></td><td></td><td></td><td></td><td></td><td></td></tr>
<tr><td>Chromosome</td><td>start</td><td>end</td><td>strand</td><td>MM</td><td>target_seq</td><td>PAM</td><td>alignment</td><td>distance</td><td>position</td><td>gene name</td><td>gene id</td></tr>
<tr><td>GU305914.1</td><td>29458</td><td>29480</td><td>+</td><td>0</td><td>GCAGGGAGAAGACGGTGCCG</td><td>GGG</td><td>||||||||[||||||||||||]PAM</td><td>NA</td><td>Intergenic</td><td>NA</td><td>NA</td></tr>
<tr><td></td><td></td><td></td><td></td><td></td><td></td><td></td><td></td><td></td><td></td><td></td><td></td></tr>
<tr><td>T1751</td><td>GGAGAAGACGGTGCCGGGGTAGG</td><td>1000</td><td></td><td></td><td></td><td></td><td></td><td></td><td></td><td></td><td></td></tr>
<tr><td>Oligo fwd</td><td>CACCGGAGAAGACGGTGCCGGGGT</td><td></td><td></td><td></td><td></td><td></td><td></td><td></td><td></td><td></td><td></td></tr>
<tr><td>Oligo rev</td><td>AAACACCCCGGCACCGTCTTCTCC</td><td></td><td></td><td></td><td></td><td></td><td></td><td></td><td></td><td></td><td></td></tr>
<tr><td>Chromosome</td><td>start</td><td>end</td><td>strand</td><td>MM</td><td>target_seq</td><td>PAM</td><td>alignment</td><td>distance</td><td>position</td><td>gene name</td><td>gene id</td></tr>
<tr><td>GU305914.1</td><td>29462</td><td>29484</td><td>+</td><td>0</td><td>GGAGAAGACGGTGCCGGGGT</td><td>AGG</td><td>||||||||[||||||||||||]PAM</td><td>NA</td><td>Intergenic</td><td>NA</td><td>NA</td></tr>
<tr><td>GU305914.1</td><td>108777</td><td>108799</td><td>-</td><td>4</td><td>GGAGCGGATGGTGCCGGTGT</td><td>CGG</td><td>||||--||[-||||||||-||]PAM</td><td>NA</td><td>Intergenic</td><td>NA</td><td>NA</td></tr>
<tr><td></td><td></td><td></td><td></td><td></td><td></td><td></td><td></td><td></td><td></td><td></td><td></td></tr>
<tr><td>T1752</td><td>GAAGACGGTGCCGGGGTAGGTGG</td><td>1000</td><td></td><td></td><td></td><td></td><td></td><td></td><td></td><td></td><td></td></tr>
<tr><td>Oligo fwd</td><td>CACCGAAGACGGTGCCGGGGTAGG</td><td></td><td></td><td></td><td></td><td></td><td></td><td></td><td></td><td></td><td></td></tr>
<tr><td>Oligo rev</td><td>AAACCCTACCCCGGCACCGTCTTC</td><td></td><td></td><td></td><td></td><td></td><td></td><td></td><td></td><td></td><td></td></tr>
<tr><td>Chromosome</td><td>start</td><td>end</td><td>strand</td><td>MM</td><td>target_seq</td><td>PAM</td><td>alignment</td><td>distance</td><td>position</td><td>gene name</td><td>gene id</td></tr>
<tr><td>GU305914.1</td><td>29465</td><td>29487</td><td>+</td><td>0</td><td>GAAGACGGTGCCGGGGTAGG</td><td>TGG</td><td>||||||||[||||||||||||]PAM</td><td>NA</td><td>Intergenic</td><td>NA</td><td>NA</td></tr>
<tr><td></td><td></td><td></td><td></td><td></td><td></td><td></td><td></td><td></td><td></td><td></td><td></td></tr>
<tr><td>T1753</td><td>GACGGTGCCGGGGTAGGTGGCGG</td><td>1000</td><td></td><td></td><td></td><td></td><td></td><td></td><td></td><td></td><td></td></tr>
</table>

<table>
<tr><td>Oligo fwd</td><td>CACCGACGGTGCCGGGGTAGGTGG</td><td></td><td></td><td></td><td></td><td></td><td></td><td></td><td></td><td></td><td></td></tr>
<tr><td>Oligo rev</td><td>AAACCCACCTACCCCGGCACCGTC</td><td></td><td></td><td></td><td></td><td></td><td></td><td></td><td></td><td></td><td></td></tr>
<tr><td>Chromosome</td><td>start</td><td>end</td><td>strand</td><td>MM</td><td>target_seq</td><td>PAM</td><td>alignment</td><td>distance</td><td>position</td><td>gene name</td><td>gene id</td></tr>
<tr><td>GU305914.1</td><td>29468</td><td>29490</td><td>+</td><td>0</td><td>GACGGTGCCGGGGTAGGTGG</td><td>CGG</td><td>||||||||[||||||||||||]PAM</td><td>NA</td><td>Intergenic</td><td>NA</td><td>NA</td></tr>
<tr><td>T1754</td><td>GGGGTAGGTGGCGGAGAACTTGG</td><td>1000</td><td></td><td></td><td></td><td></td><td></td><td></td><td></td><td></td><td></td></tr>
<tr><td>Oligo fwd</td><td>CACCGGGGTAGGTGGCGGAGAACT</td><td></td><td></td><td></td><td></td><td></td><td></td><td></td><td></td><td></td><td></td></tr>
<tr><td>Oligo rev</td><td>AAACAGTTCTCCGCCACCTACCCC</td><td></td><td></td><td></td><td></td><td></td><td></td><td></td><td></td><td></td><td></td></tr>
<tr><td>Chromosome</td><td>start</td><td>end</td><td>strand</td><td>MM</td><td>target_seq</td><td>PAM</td><td>alignment</td><td>distance</td><td>position</td><td>gene name</td><td>gene id</td></tr>
<tr><td>GU305914.1</td><td>29477</td><td>29499</td><td>+</td><td>0</td><td>GGGGTAGGTGGCGGAGAACT</td><td>TGG</td><td>||||||||[||||||||||||]PAM</td><td>NA</td><td>Intergenic</td><td>NA</td><td>NA</td></tr>
<tr><td>T1755</td><td>GAACTTGGCGATCGAGAGCGCGG</td><td>1000</td><td></td><td></td><td></td><td></td><td></td><td></td><td></td><td></td><td></td></tr>
<tr><td>Oligo fwd</td><td>CACCGAACTTGGCGATCGAGAGCG</td><td></td><td></td><td></td><td></td><td></td><td></td><td></td><td></td><td></td><td></td></tr>
<tr><td>Oligo rev</td><td>AAACCGCTCTCGATCGCCAAGTTC</td><td></td><td></td><td></td><td></td><td></td><td></td><td></td><td></td><td></td><td></td></tr>
<tr><td>Chromosome</td><td>start</td><td>end</td><td>strand</td><td>MM</td><td>target_seq</td><td>PAM</td><td>alignment</td><td>distance</td><td>position</td><td>gene name</td><td>gene id</td></tr>
<tr><td>GU305914.1</td><td>29492</td><td>29514</td><td>+</td><td>0</td><td>GAACTTGGCGATCGAGAGCG</td><td>CGG</td><td>||||||||[||||||||||||]PAM</td><td>NA</td><td>Intergenic</td><td>NA</td><td>NA</td></tr>
<tr><td>T1756</td><td>GATCGAGAGCGCGGTGCTCTTGG</td><td>1000</td><td></td><td></td><td></td><td></td><td></td><td></td><td></td><td></td><td></td></tr>
<tr><td>Oligo fwd</td><td>CACCGATCGAGAGCGCGGTGCTCT</td><td></td><td></td><td></td><td></td><td></td><td></td><td></td><td></td><td></td><td></td></tr>
<tr><td>Oligo rev</td><td>AAACAGAGCACCGCGCTCTCGATC</td><td></td><td></td><td></td><td></td><td></td><td></td><td></td><td></td><td></td><td></td></tr>
<tr><td>Chromosome</td><td>start</td><td>end</td><td>strand</td><td>MM</td><td>target_seq</td><td>PAM</td><td>alignment</td><td>distance</td><td>position</td><td>gene name</td><td>gene id</td></tr>
<tr><td>GU305914.1</td><td>29501</td><td>29523</td><td>+</td><td>0</td><td>GATCGAGAGCGCGGTGCTCT</td><td>TGG</td><td>||||||||[||||||||||||]PAM</td><td>NA</td><td>Intergenic</td><td>NA</td><td>NA</td></tr>
<tr><td>T1757</td><td>AGAGCGCGGTGCTCTTGGTCAGG</td><td>1000</td><td></td><td></td><td></td><td></td><td></td><td></td><td></td><td></td><td></td></tr>
<tr><td>Oligo fwd</td><td>CACCAGAGCGCGGTGCTCTTGGTC</td><td></td><td></td><td></td><td></td><td></td><td></td><td></td><td></td><td></td><td></td></tr>
<tr><td>Oligo rev</td><td>AAACGACCAAGAGCACCGCGCTCT</td><td></td><td></td><td></td><td></td><td></td><td></td><td></td><td></td><td></td><td></td></tr>
<tr><td>Chromosome</td><td>start</td><td>end</td><td>strand</td><td>MM</td><td>target_seq</td><td>PAM</td><td>alignment</td><td>distance</td><td>position</td><td>gene name</td><td>gene id</td></tr>
<tr><td>GU305914.1</td><td>29506</td><td>29528</td><td>+</td><td>0</td><td>AGAGCGCGGTGCTCTTGGTC</td><td>AGG</td><td>||||||||[||||||||||||]PAM</td><td>NA</td><td>Intergenic</td><td>NA</td><td>NA</td></tr>
<tr><td>T1758</td><td>TGCTCTTGGTCAGGCGTTCCAGG</td><td>1000</td><td></td><td></td><td></td><td></td><td></td><td></td><td></td><td></td><td></td></tr>
<tr><td>Oligo fwd</td><td>CACCTGCTCTTGGTCAGGCGTTCC</td><td></td><td></td><td></td><td></td><td></td><td></td><td></td><td></td><td></td><td></td></tr>
<tr><td>Oligo rev</td><td>AAACGGAACGCCTGACCAAGAGCA</td><td></td><td></td><td></td><td></td><td></td><td></td><td></td><td></td><td></td><td></td></tr>
<tr><td>Chromosome</td><td>start</td><td>end</td><td>strand</td><td>MM</td><td>target_seq</td><td>PAM</td><td>alignment</td><td>distance</td><td>position</td><td>gene name</td><td>gene id</td></tr>
<tr><td>GU305914.1</td><td>29515</td><td>29537</td><td>+</td><td>0</td><td>TGCTCTTGGTCAGGCGTTCC</td><td>AGG</td><td>||||||||[||||||||||||]PAM</td><td>NA</td><td>Intergenic</td><td>NA</td><td>NA</td></tr>
<tr><td>T1759</td><td>GGTCAGGCGTTCCAGGTACTCGG</td><td>1000</td><td></td><td></td><td></td><td></td><td></td><td></td><td></td><td></td><td></td></tr>
<tr><td>Oligo fwd</td><td>CACCGGTCAGGCGTTCCAGGTACT</td><td></td><td></td><td></td><td></td><td></td><td></td><td></td><td></td><td></td><td></td></tr>
<tr><td>Oligo rev</td><td>AAACAGTACCTGGAACGCCTGACC</td><td></td><td></td><td></td><td></td><td></td><td></td><td></td><td></td><td></td><td></td></tr>
<tr><td>Chromosome</td><td>start</td><td>end</td><td>strand</td><td>MM</td><td>target_seq</td><td>PAM</td><td>alignment</td><td>distance</td><td>position</td><td>gene name</td><td>gene id</td></tr>
<tr><td>GU305914.1</td><td>29522</td><td>29544</td><td>+</td><td>0</td><td>GGTCAGGCGTTCCAGGTACT</td><td>CGG</td><td>||||||||[||||||||||||]PAM</td><td>NA</td><td>Intergenic</td><td>NA</td><td>NA</td></tr>
<tr><td>T1760</td><td>TTCCAGGTACTCGGTGACGCCGG</td><td>1000</td><td></td><td></td><td></td><td></td><td></td><td></td><td></td><td></td><td></td></tr>
<tr><td>Oligo fwd</td><td>CACCTTCCAGGTACTCGGTGACGC</td><td></td><td></td><td></td><td></td><td></td><td></td><td></td><td></td><td></td><td></td></tr>
<tr><td>Oligo rev</td><td>AAACGCGTCACCGAGTACCTGGAA</td><td></td><td></td><td></td><td></td><td></td><td></td><td></td><td></td><td></td><td></td></tr>
<tr><td>Chromosome</td><td>start</td><td>end</td><td>strand</td><td>MM</td><td>target_seq</td><td>PAM</td><td>alignment</td><td>distance</td><td>position</td><td>gene name</td><td>gene id</td></tr>
<tr><td>GU305914.1</td><td>29531</td><td>29553</td><td>+</td><td>0</td><td>TTCCAGGTACTCGGTGACGC</td><td>CGG</td><td>||||||||[||||||||||||]PAM</td><td>NA</td><td>Intergenic</td><td>NA</td><td>NA</td></tr>
<tr><td>T1761</td><td>CGTCTCTATCTCCCGATAGATGG</td><td>1000</td><td></td><td></td><td></td><td></td><td></td><td></td><td></td><td></td><td></td></tr>
<tr><td>Oligo fwd</td><td>CACCCGTCTCTATCTCCCGATAGA</td><td></td><td></td><td></td><td></td><td></td><td></td><td></td><td></td><td></td><td></td></tr>
<tr><td>Oligo rev</td><td>AAACTCTATCGGGAGATAGAGACG</td><td></td><td></td><td></td><td></td><td></td><td></td><td></td><td></td><td></td><td></td></tr>
<tr><td>Chromosome</td><td>start</td><td>end</td><td>strand</td><td>MM</td><td>target_seq</td><td>PAM</td><td>alignment</td><td>distance</td><td>position</td><td>gene name</td><td>gene id</td></tr>
<tr><td>GU305914.1</td><td>29567</td><td>29589</td><td>+</td><td>0</td><td>CGTCTCTATCTCCCGATAGA</td><td>TGG</td><td>||||||||[||||||||||||]PAM</td><td>NA</td><td>Intergenic</td><td>NA</td><td>NA</td></tr>
<tr><td>T1762</td><td>GTCTCTATCTCCCGATAGATGGG</td><td>1000</td><td></td><td></td><td></td><td></td><td></td><td></td><td></td><td></td><td></td></tr>
<tr><td>Oligo fwd</td><td>CACCGTCTCTATCTCCCGATAGAT</td><td></td><td></td><td></td><td></td><td></td><td></td><td></td><td></td><td></td><td></td></tr>
<tr><td>Oligo rev</td><td>AAACATCTATCGGGAGATAGAGAC</td><td></td><td></td><td></td><td></td><td></td><td></td><td></td><td></td><td></td><td></td></tr>
<tr><td>Chromosome</td><td>start</td><td>end</td><td>strand</td><td>MM</td><td>target_seq</td><td>PAM</td><td>alignment</td><td>distance</td><td>position</td><td>gene name</td><td>gene id</td></tr>
<tr><td>GU305914.1</td><td>29568</td><td>29590</td><td>+</td><td>0</td><td>GTCTCTATCTCCCGATAGAT</td><td>GGG</td><td>||||||||[||||||||||||]PAM</td><td>NA</td><td>Intergenic</td><td>NA</td><td>NA</td></tr>
<tr><td>T1763</td><td>TCTCTATCTCCCGATAGATGGGG</td><td>1000</td><td></td><td></td><td></td><td></td><td></td><td></td><td></td><td></td><td></td></tr>
<tr><td>Oligo fwd</td><td>CACCTCTCTATCTCCCGATAGATG</td><td></td><td></td><td></td><td></td><td></td><td></td><td></td><td></td><td></td><td></td></tr>
<tr><td>Oligo rev</td><td>AAACCATCTATCGGGAGATAGAGA</td><td></td><td></td><td></td><td></td><td></td><td></td><td></td><td></td><td></td><td></td></tr>
<tr><td>Chromosome</td><td>start</td><td>end</td><td>strand</td><td>MM</td><td>target_seq</td><td>PAM</td><td>alignment</td><td>distance</td><td>position</td><td>gene name</td><td>gene id</td></tr>
<tr><td>GU305914.1</td><td>29569</td><td>29591</td><td>+</td><td>0</td><td>TCTCTATCTCCCGATAGATG</td><td>GGG</td><td>||||||||[||||||||||||]PAM</td><td>NA</td><td>Intergenic</td><td>NA</td><td>NA</td></tr>
<tr><td>T1764</td><td>CCCCTCTTGATGAAGCCCTCCGG</td><td>1000</td><td></td><td></td><td></td><td></td><td></td><td></td><td></td><td></td><td></td></tr>
<tr><td>Oligo fwd</td><td>CACCCCCCTCTTGATGAAGCCCTC</td><td></td><td></td><td></td><td></td><td></td><td></td><td></td><td></td><td></td><td></td></tr>
<tr><td>Oligo rev</td><td>AAACGAGGGCTTCATCAAGAGGGG</td><td></td><td></td><td></td><td></td><td></td><td></td><td></td><td></td><td></td><td></td></tr>
<tr><td>Chromosome</td><td>start</td><td>end</td><td>strand</td><td>MM</td><td>target_seq</td><td>PAM</td><td>alignment</td><td>distance</td><td>position</td><td>gene name</td><td>gene id</td></tr>
<tr><td>GU305914.1</td><td>29607</td><td>29629</td><td>+</td><td>0</td><td>CCCCTCTTGATGAAGCCCTC</td><td>CGG</td><td>||||||||[||||||||||||]PAM</td><td>NA</td><td>Intergenic</td><td>NA</td><td>NA</td></tr>
<tr><td>T1765</td><td>CCCTCTTGATGAAGCCCTCCGGG</td><td>1000</td><td></td><td></td><td></td><td></td><td></td><td></td><td></td><td></td><td></td></tr>
<tr><td>Oligo fwd</td><td>CACCCCCTCTTGATGAAGCCCTCC</td><td></td><td></td><td></td><td></td><td></td><td></td><td></td><td></td><td></td><td></td></tr>
<tr><td>Oligo rev</td><td>AAACGGAGGGCTTCATCAAGAGGG</td><td></td><td></td><td></td><td></td><td></td><td></td><td></td><td></td><td></td><td></td></tr>
<tr><td>Chromosome</td><td>start</td><td>end</td><td>strand</td><td>MM</td><td>target_seq</td><td>PAM</td><td>alignment</td><td>distance</td><td>position</td><td>gene name</td><td>gene id</td></tr>
<tr><td>GU305914.1</td><td>29608</td><td>29630</td><td>+</td><td>0</td><td>CCCTCTTGATGAAGCCCTCC</td><td>GGG</td><td>||||||||[||||||||||||]PAM</td><td>NA</td><td>Intergenic</td><td>NA</td><td>NA</td></tr>
<tr><td>T1766</td><td>CCTCTTGATGAAGCCCTCCGGGG</td><td>1000</td><td></td><td></td><td></td><td></td><td></td><td></td><td></td><td></td><td></td></tr>
<tr><td>Oligo fwd</td><td>CACCCCTCTTGATGAAGCCCTCCG</td><td></td><td></td><td></td><td></td><td></td><td></td><td></td><td></td><td></td><td></td></tr>
<tr><td>Oligo rev</td><td>AAACCGGAGGGCTTCATCAAGAGG</td><td></td><td></td><td></td><td></td><td></td><td></td><td></td><td></td><td></td><td></td></tr>
<tr><td>Chromosome</td><td>start</td><td>end</td><td>strand</td><td>MM</td><td>target_seq</td><td>PAM</td><td>alignment</td><td>distance</td><td>position</td><td>gene name</td><td>gene id</td></tr>
<tr><td>GU305914.1</td><td>29609</td><td>29631</td><td>+</td><td>0</td><td>CCTCTTGATGAAGCCCTCCG</td><td>GGG</td><td>||||||||[||||||||||||]PAM</td><td>NA</td><td>Intergenic</td><td>NA</td><td>NA</td></tr>
<tr><td>T1767</td><td>GGAGTCCCCGACCAGATAGATGG</td><td>1000</td><td></td><td></td><td></td><td></td><td></td><td></td><td></td><td></td><td></td></tr>
<tr><td>Oligo fwd</td><td>CACCGGAGTCCCCGACCAGATAGA</td><td></td><td></td><td></td><td></td><td></td><td></td><td></td><td></td><td></td><td></td></tr>
<tr><td>Oligo rev</td><td>AAACTCTATCTGGTCGGGGACTCC</td><td></td><td></td><td></td><td></td><td></td><td></td><td></td><td></td><td></td><td></td></tr>
<tr><td>Chromosome</td><td>start</td><td>end</td><td>strand</td><td>MM</td><td>target_seq</td><td>PAM</td><td>alignment</td><td>distance</td><td>position</td><td>gene name</td><td>gene id</td></tr>
<tr><td>GU305914.1</td><td>29630</td><td>29652</td><td>+</td><td>0</td><td>GGAGTCCCCGACCAGATAGA</td><td>TGG</td><td>||||||||[||||||||||||]PAM</td><td>NA</td><td>Intergenic</td><td>NA</td><td>NA</td></tr>
</table>

<table>
<tr><td>T1768</td><td>ATAGATGGCGTCGTCGACGTCGG</td><td>1000</td><td></td><td></td><td></td><td></td><td></td><td></td><td></td><td></td><td></td></tr>
<tr><td>Oligo fwd</td><td>CACCATAGATGGCGTCGTCGACGT</td><td></td><td></td><td></td><td></td><td></td><td></td><td></td><td></td><td></td><td></td></tr>
<tr><td>Oligo rev</td><td>AAACACGTCGACGACGCCATCTAT</td><td></td><td></td><td></td><td></td><td></td><td></td><td></td><td></td><td></td><td></td></tr>
<tr><td>Chromosome</td><td>start</td><td>end</td><td>strand</td><td>MM</td><td>target_seq</td><td>PAM</td><td>alignment</td><td>distance</td><td>position</td><td>gene name</td><td>gene id</td></tr>
<tr><td>GU305914.1</td><td>29645</td><td>29667</td><td>+</td><td>0</td><td>ATAGATGGCGTCGTCGACGT</td><td>CGG</td><td>||||||||[||||||||||||]PAM</td><td>NA</td><td>Intergenic</td><td>NA</td><td>NA</td></tr>
<tr><td>T1769</td><td>TAGATGGCGTCGTCGACGTCGGG</td><td>1000</td><td></td><td></td><td></td><td></td><td></td><td></td><td></td><td></td><td></td></tr>
<tr><td>Oligo fwd</td><td>CACCTAGATGGCGTCGTCGACGTC</td><td></td><td></td><td></td><td></td><td></td><td></td><td></td><td></td><td></td><td></td></tr>
<tr><td>Oligo rev</td><td>AAACGACGTCGACGACGCCATCTA</td><td></td><td></td><td></td><td></td><td></td><td></td><td></td><td></td><td></td><td></td></tr>
<tr><td>Chromosome</td><td>start</td><td>end</td><td>strand</td><td>MM</td><td>target_seq</td><td>PAM</td><td>alignment</td><td>distance</td><td>position</td><td>gene name</td><td>gene id</td></tr>
<tr><td>GU305914.1</td><td>29646</td><td>29668</td><td>+</td><td>0</td><td>TAGATGGCGTCGTCGACGTC</td><td>GGG</td><td>||||||||[||||||||||||]PAM</td><td>NA</td><td>Intergenic</td><td>NA</td><td>NA</td></tr>
<tr><td>T1770</td><td>GTAGACGCGTCCCTCCTCGCCGG</td><td>1000</td><td></td><td></td><td></td><td></td><td></td><td></td><td></td><td></td><td></td></tr>
<tr><td>Oligo fwd</td><td>CACCGTAGACGCGTCCCTCCTCGC</td><td></td><td></td><td></td><td></td><td></td><td></td><td></td><td></td><td></td><td></td></tr>
<tr><td>Oligo rev</td><td>AAACGCGAGGAGGGACGCGTCTAC</td><td></td><td></td><td></td><td></td><td></td><td></td><td></td><td></td><td></td><td></td></tr>
<tr><td>Chromosome</td><td>start</td><td>end</td><td>strand</td><td>MM</td><td>target_seq</td><td>PAM</td><td>alignment</td><td>distance</td><td>position</td><td>gene name</td><td>gene id</td></tr>
<tr><td>GU305914.1</td><td>29678</td><td>29700</td><td>+</td><td>0</td><td>GTAGACGCGTCCCTCCTCGC</td><td>CGG</td><td>||||||||[||||||||||||]PAM</td><td>NA</td><td>Intergenic</td><td>NA</td><td>NA</td></tr>
<tr><td>T1771</td><td>GTCCCTCCTCGCCGGCGAAGAGG</td><td>1000</td><td></td><td></td><td></td><td></td><td></td><td></td><td></td><td></td><td></td></tr>
<tr><td>Oligo fwd</td><td>CACCGTCCCTCCTCGCCGGCGAAG</td><td></td><td></td><td></td><td></td><td></td><td></td><td></td><td></td><td></td><td></td></tr>
<tr><td>Oligo rev</td><td>AAACCTTCGCCGGCGAGGAGGGAC</td><td></td><td></td><td></td><td></td><td></td><td></td><td></td><td></td><td></td><td></td></tr>
<tr><td>Chromosome</td><td>start</td><td>end</td><td>strand</td><td>MM</td><td>target_seq</td><td>PAM</td><td>alignment</td><td>distance</td><td>position</td><td>gene name</td><td>gene id</td></tr>
<tr><td>GU305914.1</td><td>29686</td><td>29708</td><td>+</td><td>0</td><td>GTCCCTCCTCGCCGGCGAAG</td><td>AGG</td><td>||||||||[||||||||||||]PAM</td><td>NA</td><td>Intergenic</td><td>NA</td><td>NA</td></tr>
<tr><td>T1772</td><td>CGCCGGCGAAGAGGACGCCCAGG</td><td>1000</td><td></td><td></td><td></td><td></td><td></td><td></td><td></td><td></td><td></td></tr>
<tr><td>Oligo fwd</td><td>CACCCGCCGGCGAAGAGGACGCCC</td><td></td><td></td><td></td><td></td><td></td><td></td><td></td><td></td><td></td><td></td></tr>
<tr><td>Oligo rev</td><td>AAACGGGCGTCCTCTTCGCCGGCG</td><td></td><td></td><td></td><td></td><td></td><td></td><td></td><td></td><td></td><td></td></tr>
<tr><td>Chromosome</td><td>start</td><td>end</td><td>strand</td><td>MM</td><td>target_seq</td><td>PAM</td><td>alignment</td><td>distance</td><td>position</td><td>gene name</td><td>gene id</td></tr>
<tr><td>GU305914.1</td><td>29695</td><td>29717</td><td>+</td><td>0</td><td>CGCCGGCGAAGAGGACGCCC</td><td>AGG</td><td>||||||||[||||||||||||]PAM</td><td>NA</td><td>Intergenic</td><td>NA</td><td>NA</td></tr>
<tr><td>T1773</td><td>GTCCTCGTTCTTGTCATCGCCGG</td><td>1000</td><td></td><td></td><td></td><td></td><td></td><td></td><td></td><td></td><td></td></tr>
<tr><td>Oligo fwd</td><td>CACCGTCCTCGTTCTTGTCATCGC</td><td></td><td></td><td></td><td></td><td></td><td></td><td></td><td></td><td></td><td></td></tr>
<tr><td>Oligo rev</td><td>AAACGCGATGACAAGAACGAGGAC</td><td></td><td></td><td></td><td></td><td></td><td></td><td></td><td></td><td></td><td></td></tr>
<tr><td>Chromosome</td><td>start</td><td>end</td><td>strand</td><td>MM</td><td>target_seq</td><td>PAM</td><td>alignment</td><td>distance</td><td>position</td><td>gene name</td><td>gene id</td></tr>
<tr><td>GU305914.1</td><td>29717</td><td>29739</td><td>+</td><td>0</td><td>GTCCTCGTTCTTGTCATCGC</td><td>CGG</td><td>||||||||[||||||||||||]PAM</td><td>NA</td><td>Intergenic</td><td>NA</td><td>NA</td></tr>
<tr><td>T1774</td><td>TCTTGTCATCGCCGGTCGAGCGG</td><td>1000</td><td></td><td></td><td></td><td></td><td></td><td></td><td></td><td></td><td></td></tr>
<tr><td>Oligo fwd</td><td>CACCTCTTGTCATCGCCGGTCGAG</td><td></td><td></td><td></td><td></td><td></td><td></td><td></td><td></td><td></td><td></td></tr>
<tr><td>Oligo rev</td><td>AAACCTCGACCGGCGATGACAAGA</td><td></td><td></td><td></td><td></td><td></td><td></td><td></td><td></td><td></td><td></td></tr>
<tr><td>Chromosome</td><td>start</td><td>end</td><td>strand</td><td>MM</td><td>target_seq</td><td>PAM</td><td>alignment</td><td>distance</td><td>position</td><td>gene name</td><td>gene id</td></tr>
<tr><td>GU305914.1</td><td>29725</td><td>29747</td><td>+</td><td>0</td><td>TCTTGTCATCGCCGGTCGAG</td><td>CGG</td><td>||||||||[||||||||||||]PAM</td><td>NA</td><td>Intergenic</td><td>NA</td><td>NA</td></tr>
<tr><td>T1775</td><td>TGTCATCGCCGGTCGAGCGGCGG</td><td>1000</td><td></td><td></td><td></td><td></td><td></td><td></td><td></td><td></td><td></td></tr>
<tr><td>Oligo fwd</td><td>CACCTGTCATCGCCGGTCGAGCGG</td><td></td><td></td><td></td><td></td><td></td><td></td><td></td><td></td><td></td><td></td></tr>
<tr><td>Oligo rev</td><td>AAACCCGCTCGACCGGCGATGACA</td><td></td><td></td><td></td><td></td><td></td><td></td><td></td><td></td><td></td><td></td></tr>
<tr><td>Chromosome</td><td>start</td><td>end</td><td>strand</td><td>MM</td><td>target_seq</td><td>PAM</td><td>alignment</td><td>distance</td><td>position</td><td>gene name</td><td>gene id</td></tr>
<tr><td>GU305914.1</td><td>29728</td><td>29750</td><td>+</td><td>0</td><td>TGTCATCGCCGGTCGAGCGG</td><td>CGG</td><td>||||||||[||||||||||||]PAM</td><td>NA</td><td>Intergenic</td><td>NA</td><td>NA</td></tr>
<tr><td>T1776</td><td>TCGAGCGGCGGACGCCGCCCAGG</td><td>1000</td><td></td><td></td><td></td><td></td><td></td><td></td><td></td><td></td><td></td></tr>
<tr><td>Oligo fwd</td><td>CACCTCGAGCGGCGGACGCCGCCC</td><td></td><td></td><td></td><td></td><td></td><td></td><td></td><td></td><td></td><td></td></tr>
<tr><td>Oligo rev</td><td>AAACGGGCGGCGTCCGCCGCTCGA</td><td></td><td></td><td></td><td></td><td></td><td></td><td></td><td></td><td></td><td></td></tr>
<tr><td>Chromosome</td><td>start</td><td>end</td><td>strand</td><td>MM</td><td>target_seq</td><td>PAM</td><td>alignment</td><td>distance</td><td>position</td><td>gene name</td><td>gene id</td></tr>
<tr><td>GU305914.1</td><td>29740</td><td>29762</td><td>+</td><td>0</td><td>TCGAGCGGCGGACGCCGCCC</td><td>AGG</td><td>||||||||[||||||||||||]PAM</td><td>NA</td><td>Intergenic</td><td>NA</td><td>NA</td></tr>
<tr><td>GU305914.1</td><td>34722</td><td>34744</td><td>+</td><td>4</td><td>TCAAGACGCGGGCGCCGCCC</td><td>AGG</td><td>||-||--|[|||-||||||||]PAM</td><td>NA</td><td>Intergenic</td><td>NA</td><td>NA</td></tr>
<tr><td>T1777</td><td>CCCAGGACGATGAGATCCTCCGG</td><td>1000</td><td></td><td></td><td></td><td></td><td></td><td></td><td></td><td></td><td></td></tr>
<tr><td>Oligo fwd</td><td>CACCCCCAGGACGATGAGATCCTC</td><td></td><td></td><td></td><td></td><td></td><td></td><td></td><td></td><td></td><td></td></tr>
<tr><td>Oligo rev</td><td>AAACGAGGATCTCATCGTCCTGGG</td><td></td><td></td><td></td><td></td><td></td><td></td><td></td><td></td><td></td><td></td></tr>
<tr><td>Chromosome</td><td>start</td><td>end</td><td>strand</td><td>MM</td><td>target_seq</td><td>PAM</td><td>alignment</td><td>distance</td><td>position</td><td>gene name</td><td>gene id</td></tr>
<tr><td>GU305914.1</td><td>29757</td><td>29779</td><td>+</td><td>0</td><td>CCCAGGACGATGAGATCCTC</td><td>CGG</td><td>||||||||[||||||||||||]PAM</td><td>NA</td><td>Intergenic</td><td>NA</td><td>NA</td></tr>
<tr><td>T1778</td><td>TGAGATCCTCCGGACAGCAGAGG</td><td>1000</td><td></td><td></td><td></td><td></td><td></td><td></td><td></td><td></td><td></td></tr>
<tr><td>Oligo fwd</td><td>CACCTGAGATCCTCCGGACAGCAG</td><td></td><td></td><td></td><td></td><td></td><td></td><td></td><td></td><td></td><td></td></tr>
<tr><td>Oligo rev</td><td>AAACCTGCTGTCCGGAGGATCTCA</td><td></td><td></td><td></td><td></td><td></td><td></td><td></td><td></td><td></td><td></td></tr>
<tr><td>Chromosome</td><td>start</td><td>end</td><td>strand</td><td>MM</td><td>target_seq</td><td>PAM</td><td>alignment</td><td>distance</td><td>position</td><td>gene name</td><td>gene id</td></tr>
<tr><td>GU305914.1</td><td>29767</td><td>29789</td><td>+</td><td>0</td><td>TGAGATCCTCCGGACAGCAG</td><td>AGG</td><td>||||||||[||||||||||||]PAM</td><td>NA</td><td>Intergenic</td><td>NA</td><td>NA</td></tr>
<tr><td>T1779</td><td>GAGCCGACTGATCTCCCCGACGG</td><td>1000</td><td></td><td></td><td></td><td></td><td></td><td></td><td></td><td></td><td></td></tr>
<tr><td>Oligo fwd</td><td>CACCGAGCCGACTGATCTCCCCGA</td><td></td><td></td><td></td><td></td><td></td><td></td><td></td><td></td><td></td><td></td></tr>
<tr><td>Oligo rev</td><td>AAACTCGGGGAGATCAGTCGGCTC</td><td></td><td></td><td></td><td></td><td></td><td></td><td></td><td></td><td></td><td></td></tr>
<tr><td>Chromosome</td><td>start</td><td>end</td><td>strand</td><td>MM</td><td>target_seq</td><td>PAM</td><td>alignment</td><td>distance</td><td>position</td><td>gene name</td><td>gene id</td></tr>
<tr><td>GU305914.1</td><td>29801</td><td>29823</td><td>+</td><td>0</td><td>GAGCCGACTGATCTCCCCGA</td><td>CGG</td><td>||||||||[||||||||||||]PAM</td><td>NA</td><td>Intergenic</td><td>NA</td><td>NA</td></tr>
<tr><td>T1780</td><td>AGCCGACTGATCTCCCCGACGGG</td><td>1000</td><td></td><td></td><td></td><td></td><td></td><td></td><td></td><td></td><td></td></tr>
<tr><td>Oligo fwd</td><td>CACCAGCCGACTGATCTCCCCGAC</td><td></td><td></td><td></td><td></td><td></td><td></td><td></td><td></td><td></td><td></td></tr>
<tr><td>Oligo rev</td><td>AAACGTCGGGGAGATCAGTCGGCT</td><td></td><td></td><td></td><td></td><td></td><td></td><td></td><td></td><td></td><td></td></tr>
<tr><td>Chromosome</td><td>start</td><td>end</td><td>strand</td><td>MM</td><td>target_seq</td><td>PAM</td><td>alignment</td><td>distance</td><td>position</td><td>gene name</td><td>gene id</td></tr>
<tr><td>GU305914.1</td><td>29802</td><td>29824</td><td>+</td><td>0</td><td>AGCCGACTGATCTCCCCGAC</td><td>GGG</td><td>||||||||[||||||||||||]PAM</td><td>NA</td><td>Intergenic</td><td>NA</td><td>NA</td></tr>
<tr><td>T1781</td><td>TGATCTCCCCGACGGGCATGCGG</td><td>1000</td><td></td><td></td><td></td><td></td><td></td><td></td><td></td><td></td><td></td></tr>
<tr><td>Oligo fwd</td><td>CACCTGATCTCCCCGACGGGCATG</td><td></td><td></td><td></td><td></td><td></td><td></td><td></td><td></td><td></td><td></td></tr>
<tr><td>Oligo rev</td><td>AAACCATGCCCGTCGGGGAGATCA</td><td></td><td></td><td></td><td></td><td></td><td></td><td></td><td></td><td></td><td></td></tr>
<tr><td>Chromosome</td><td>start</td><td>end</td><td>strand</td><td>MM</td><td>target_seq</td><td>PAM</td><td>alignment</td><td>distance</td><td>position</td><td>gene name</td><td>gene id</td></tr>
<tr><td>GU305914.1</td><td>29809</td><td>29831</td><td>+</td><td>0</td><td>TGATCTCCCCGACGGGCATG</td><td>CGG</td><td>||||||||[||||||||||||]PAM</td><td>NA</td><td>Intergenic</td><td>NA</td><td>NA</td></tr>
<tr><td>T1782</td><td>CCCGACGGGCATGCGGAGTTCGG</td><td>1000</td><td></td><td></td><td></td><td></td><td></td><td></td><td></td><td></td><td></td></tr>
<tr><td>Oligo fwd</td><td>CACCCCCGACGGGCATGCGGAGTT</td><td></td><td></td><td></td><td></td><td></td><td></td><td></td><td></td><td></td><td></td></tr>
<tr><td>Oligo rev</td><td>AAACAACTCCGCATGCCCGTCGGG</td><td></td><td></td><td></td><td></td><td></td><td></td><td></td><td></td><td></td><td></td></tr>
<tr><td>Chromosome</td><td>start</td><td>end</td><td>strand</td><td>MM</td><td>target_seq</td><td>PAM</td><td>alignment</td><td>distance</td><td>position</td><td>gene name</td><td>gene id</td></tr>
</table>

<table>
<tr><td>GU305914.1</td><td>29816</td><td>29838</td><td>+</td><td>0</td><td>CCCGACGGGCATGCGGAGTT</td><td>CGG</td><td>||||||||[||||||||||||]PAM</td><td>NA</td><td>Intergenic</td><td>NA</td><td>NA</td></tr>
<tr><td></td><td></td><td></td><td></td><td></td><td></td><td></td><td></td><td></td><td></td><td></td><td></td></tr>
<tr><td>T1783</td><td>GGGCATGCGGAGTTCGGCCTCGG</td><td>1000</td><td></td><td></td><td></td><td></td><td></td><td></td><td></td><td></td><td></td></tr>
<tr><td>Oligo fwd</td><td>CACCGGGCATGCGGAGTTCGGCCT</td><td></td><td></td><td></td><td></td><td></td><td></td><td></td><td></td><td></td><td></td></tr>
<tr><td>Oligo rev</td><td>AAACAGGCCGAACTCCGCATGCCC</td><td></td><td></td><td></td><td></td><td></td><td></td><td></td><td></td><td></td><td></td></tr>
<tr><td>Chromosome</td><td>start</td><td>end</td><td>strand</td><td>MM</td><td>target_seq</td><td>PAM</td><td>alignment</td><td>distance</td><td>position</td><td>gene name</td><td>gene id</td></tr>
<tr><td>GU305914.1</td><td>29822</td><td>29844</td><td>+</td><td>0</td><td>GGGCATGCGGAGTTCGGCCT</td><td>CGG</td><td>||||||||[||||||||||||]PAM</td><td>NA</td><td>Intergenic</td><td>NA</td><td>NA</td></tr>
<tr><td></td><td></td><td></td><td></td><td></td><td></td><td></td><td></td><td></td><td></td><td></td><td></td></tr>
<tr><td>T1784</td><td>CAACCAGAGTGTGTAATTCTTGG</td><td>1000</td><td></td><td></td><td></td><td></td><td></td><td></td><td></td><td></td><td></td></tr>
<tr><td>Oligo fwd</td><td>CACCCAACCAGAGTGTGTAATTCT</td><td></td><td></td><td></td><td></td><td></td><td></td><td></td><td></td><td></td><td></td></tr>
<tr><td>Oligo rev</td><td>AAACAGAATTACACACTCTGGTTG</td><td></td><td></td><td></td><td></td><td></td><td></td><td></td><td></td><td></td><td></td></tr>
<tr><td>Chromosome</td><td>start</td><td>end</td><td>strand</td><td>MM</td><td>target_seq</td><td>PAM</td><td>alignment</td><td>distance</td><td>position</td><td>gene name</td><td>gene id</td></tr>
<tr><td>GU305914.1</td><td>29849</td><td>29871</td><td>+</td><td>0</td><td>CAACCAGAGTGTGTAATTCT</td><td>TGG</td><td>||||||||[||||||||||||]PAM</td><td>NA</td><td>Intergenic</td><td>NA</td><td>NA</td></tr>
<tr><td></td><td></td><td></td><td></td><td></td><td></td><td></td><td></td><td></td><td></td><td></td><td></td></tr>
<tr><td>T1785</td><td>AACCAGAGTGTGTAATTCTTGGG</td><td>1000</td><td></td><td></td><td></td><td></td><td></td><td></td><td></td><td></td><td></td></tr>
<tr><td>Oligo fwd</td><td>CACCAACCAGAGTGTGTAATTCTT</td><td></td><td></td><td></td><td></td><td></td><td></td><td></td><td></td><td></td><td></td></tr>
<tr><td>Oligo rev</td><td>AAACAAGAATTACACACTCTGGTT</td><td></td><td></td><td></td><td></td><td></td><td></td><td></td><td></td><td></td><td></td></tr>
<tr><td>Chromosome</td><td>start</td><td>end</td><td>strand</td><td>MM</td><td>target_seq</td><td>PAM</td><td>alignment</td><td>distance</td><td>position</td><td>gene name</td><td>gene id</td></tr>
<tr><td>GU305914.1</td><td>29850</td><td>29872</td><td>+</td><td>0</td><td>AACCAGAGTGTGTAATTCTT</td><td>GGG</td><td>||||||||[||||||||||||]PAM</td><td>NA</td><td>Intergenic</td><td>NA</td><td>NA</td></tr>
<tr><td></td><td></td><td></td><td></td><td></td><td></td><td></td><td></td><td></td><td></td><td></td><td></td></tr>
<tr><td>T1786</td><td>GAGTGTGTAATTCTTGGGCCAGG</td><td>1000</td><td></td><td></td><td></td><td></td><td></td><td></td><td></td><td></td><td></td></tr>
<tr><td>Oligo fwd</td><td>CACCGAGTGTGTAATTCTTGGGCC</td><td></td><td></td><td></td><td></td><td></td><td></td><td></td><td></td><td></td><td></td></tr>
<tr><td>Oligo rev</td><td>AAACGGCCCAAGAATTACACACTC</td><td></td><td></td><td></td><td></td><td></td><td></td><td></td><td></td><td></td><td></td></tr>
<tr><td>Chromosome</td><td>start</td><td>end</td><td>strand</td><td>MM</td><td>target_seq</td><td>PAM</td><td>alignment</td><td>distance</td><td>position</td><td>gene name</td><td>gene id</td></tr>
<tr><td>GU305914.1</td><td>29855</td><td>29877</td><td>+</td><td>0</td><td>GAGTGTGTAATTCTTGGGCC</td><td>AGG</td><td>||||||||[||||||||||||]PAM</td><td>NA</td><td>Intergenic</td><td>NA</td><td>NA</td></tr>
<tr><td></td><td></td><td></td><td></td><td></td><td></td><td></td><td></td><td></td><td></td><td></td><td></td></tr>
<tr><td>T1787</td><td>TAATTCTTGGGCCAGGTCAGCGG</td><td>1000</td><td></td><td></td><td></td><td></td><td></td><td></td><td></td><td></td><td></td></tr>
<tr><td>Oligo fwd</td><td>CACCTAATTCTTGGGCCAGGTCAG</td><td></td><td></td><td></td><td></td><td></td><td></td><td></td><td></td><td></td><td></td></tr>
<tr><td>Oligo rev</td><td>AAACCTGACCTGGCCCAAGAATTA</td><td></td><td></td><td></td><td></td><td></td><td></td><td></td><td></td><td></td><td></td></tr>
<tr><td>Chromosome</td><td>start</td><td>end</td><td>strand</td><td>MM</td><td>target_seq</td><td>PAM</td><td>alignment</td><td>distance</td><td>position</td><td>gene name</td><td>gene id</td></tr>
<tr><td>GU305914.1</td><td>29862</td><td>29884</td><td>+</td><td>0</td><td>TAATTCTTGGGCCAGGTCAG</td><td>CGG</td><td>||||||||[||||||||||||]PAM</td><td>NA</td><td>Intergenic</td><td>NA</td><td>NA</td></tr>
<tr><td></td><td></td><td></td><td></td><td></td><td></td><td></td><td></td><td></td><td></td><td></td><td></td></tr>
<tr><td>T1788</td><td>GCGGCATCCGCCATCCCTTGCGG</td><td>1000</td><td></td><td></td><td></td><td></td><td></td><td></td><td></td><td></td><td></td></tr>
<tr><td>Oligo fwd</td><td>CACCGCGGCATCCGCCATCCCTTG</td><td></td><td></td><td></td><td></td><td></td><td></td><td></td><td></td><td></td><td></td></tr>
<tr><td>Oligo rev</td><td>AAACCAAGGGATGGCGGATGCCGC</td><td></td><td></td><td></td><td></td><td></td><td></td><td></td><td></td><td></td><td></td></tr>
<tr><td>Chromosome</td><td>start</td><td>end</td><td>strand</td><td>MM</td><td>target_seq</td><td>PAM</td><td>alignment</td><td>distance</td><td>position</td><td>gene name</td><td>gene id</td></tr>
<tr><td>GU305914.1</td><td>29881</td><td>29903</td><td>+</td><td>0</td><td>GCGGCATCCGCCATCCCTTG</td><td>CGG</td><td>||||||||[||||||||||||]PAM</td><td>NA</td><td>Intergenic</td><td>NA</td><td>NA</td></tr>
<tr><td></td><td></td><td></td><td></td><td></td><td></td><td></td><td></td><td></td><td></td><td></td><td></td></tr>
<tr><td>T1789</td><td>TGCGGATCTTGACGAACTCCCGG</td><td>1000</td><td></td><td></td><td></td><td></td><td></td><td></td><td></td><td></td><td></td></tr>
<tr><td>Oligo fwd</td><td>CACCTGCGGATCTTGACGAACTCC</td><td></td><td></td><td></td><td></td><td></td><td></td><td></td><td></td><td></td><td></td></tr>
<tr><td>Oligo rev</td><td>AAACGGAGTTCGTCAAGATCCGCA</td><td></td><td></td><td></td><td></td><td></td><td></td><td></td><td></td><td></td><td></td></tr>
<tr><td>Chromosome</td><td>start</td><td>end</td><td>strand</td><td>MM</td><td>target_seq</td><td>PAM</td><td>alignment</td><td>distance</td><td>position</td><td>gene name</td><td>gene id</td></tr>
<tr><td>GU305914.1</td><td>29899</td><td>29921</td><td>+</td><td>0</td><td>TGCGGATCTTGACGAACTCC</td><td>CGG</td><td>||||||||[||||||||||||]PAM</td><td>NA</td><td>Intergenic</td><td>NA</td><td>NA</td></tr>
<tr><td></td><td></td><td></td><td></td><td></td><td></td><td></td><td></td><td></td><td></td><td></td><td></td></tr>
<tr><td>T1790</td><td>GACGAACTCCCGGCAGAGATCGG</td><td>1000</td><td></td><td></td><td></td><td></td><td></td><td></td><td></td><td></td><td></td></tr>
<tr><td>Oligo fwd</td><td>CACCGACGAACTCCCGGCAGAGAT</td><td></td><td></td><td></td><td></td><td></td><td></td><td></td><td></td><td></td><td></td></tr>
<tr><td>Oligo rev</td><td>AAACATCTCTGCCGGGAGTTCGTC</td><td></td><td></td><td></td><td></td><td></td><td></td><td></td><td></td><td></td><td></td></tr>
<tr><td>Chromosome</td><td>start</td><td>end</td><td>strand</td><td>MM</td><td>target_seq</td><td>PAM</td><td>alignment</td><td>distance</td><td>position</td><td>gene name</td><td>gene id</td></tr>
<tr><td>GU305914.1</td><td>29909</td><td>29931</td><td>+</td><td>0</td><td>GACGAACTCCCGGCAGAGAT</td><td>CGG</td><td>||||||||[||||||||||||]PAM</td><td>NA</td><td>Intergenic</td><td>NA</td><td>NA</td></tr>
<tr><td></td><td></td><td></td><td></td><td></td><td></td><td></td><td></td><td></td><td></td><td></td><td></td></tr>
<tr><td>T1791</td><td>ACGAACTCCCGGCAGAGATCGGG</td><td>1000</td><td></td><td></td><td></td><td></td><td></td><td></td><td></td><td></td><td></td></tr>
<tr><td>Oligo fwd</td><td>CACCACGAACTCCCGGCAGAGATC</td><td></td><td></td><td></td><td></td><td></td><td></td><td></td><td></td><td></td><td></td></tr>
<tr><td>Oligo rev</td><td>AAACGATCTCTGCCGGGAGTTCGT</td><td></td><td></td><td></td><td></td><td></td><td></td><td></td><td></td><td></td><td></td></tr>
<tr><td>Chromosome</td><td>start</td><td>end</td><td>strand</td><td>MM</td><td>target_seq</td><td>PAM</td><td>alignment</td><td>distance</td><td>position</td><td>gene name</td><td>gene id</td></tr>
<tr><td>GU305914.1</td><td>29910</td><td>29932</td><td>+</td><td>0</td><td>ACGAACTCCCGGCAGAGATC</td><td>GGG</td><td>||||||||[||||||||||||]PAM</td><td>NA</td><td>Intergenic</td><td>NA</td><td>NA</td></tr>
<tr><td></td><td></td><td></td><td></td><td></td><td></td><td></td><td></td><td></td><td></td><td></td><td></td></tr>
<tr><td>T1792</td><td>GCAGAGATCGGGATTGTTGACGG</td><td>1000</td><td></td><td></td><td></td><td></td><td></td><td></td><td></td><td></td><td></td></tr>
<tr><td>Oligo fwd</td><td>CACCGCAGAGATCGGGATTGTTGA</td><td></td><td></td><td></td><td></td><td></td><td></td><td></td><td></td><td></td><td></td></tr>
<tr><td>Oligo rev</td><td>AAACTCAACAATCCCGATCTCTGC</td><td></td><td></td><td></td><td></td><td></td><td></td><td></td><td></td><td></td><td></td></tr>
<tr><td>Chromosome</td><td>start</td><td>end</td><td>strand</td><td>MM</td><td>target_seq</td><td>PAM</td><td>alignment</td><td>distance</td><td>position</td><td>gene name</td><td>gene id</td></tr>
<tr><td>GU305914.1</td><td>29921</td><td>29943</td><td>+</td><td>0</td><td>GCAGAGATCGGGATTGTTGA</td><td>CGG</td><td>||||||||[||||||||||||]PAM</td><td>NA</td><td>Intergenic</td><td>NA</td><td>NA</td></tr>
<tr><td></td><td></td><td></td><td></td><td></td><td></td><td></td><td></td><td></td><td></td><td></td><td></td></tr>
<tr><td>T1793</td><td>GACGGCCCGCGCGAAGTCCTCGG</td><td>1000</td><td></td><td></td><td></td><td></td><td></td><td></td><td></td><td></td><td></td></tr>
<tr><td>Oligo fwd</td><td>CACCGACGGCCCGCGCGAAGTCCT</td><td></td><td></td><td></td><td></td><td></td><td></td><td></td><td></td><td></td><td></td></tr>
<tr><td>Oligo rev</td><td>AAACAGGACTTCGCGCGGGCCGTC</td><td></td><td></td><td></td><td></td><td></td><td></td><td></td><td></td><td></td><td></td></tr>
<tr><td>Chromosome</td><td>start</td><td>end</td><td>strand</td><td>MM</td><td>target_seq</td><td>PAM</td><td>alignment</td><td>distance</td><td>position</td><td>gene name</td><td>gene id</td></tr>
<tr><td>GU305914.1</td><td>29939</td><td>29961</td><td>+</td><td>0</td><td>GACGGCCCGCGCGAAGTCCT</td><td>CGG</td><td>||||||||[||||||||||||]PAM</td><td>NA</td><td>Intergenic</td><td>NA</td><td>NA</td></tr>
<tr><td>GU305914.1</td><td>65755</td><td>65777</td><td>-</td><td>4</td><td>GCACGCCCGCGCGATGTCCT</td><td>CGG</td><td>|---||||[||||||-|||||]PAM</td><td>NA</td><td>Intergenic</td><td>NA</td><td>NA</td></tr>
<tr><td></td><td></td><td></td><td></td><td></td><td></td><td></td><td></td><td></td><td></td><td></td><td></td></tr>
<tr><td>T1794</td><td>ACGGCCCGCGCGAAGTCCTCGGG</td><td>1000</td><td></td><td></td><td></td><td></td><td></td><td></td><td></td><td></td><td></td></tr>
<tr><td>Oligo fwd</td><td>CACCACGGCCCGCGCGAAGTCCTC</td><td></td><td></td><td></td><td></td><td></td><td></td><td></td><td></td><td></td><td></td></tr>
<tr><td>Oligo rev</td><td>AAACGAGGACTTCGCGCGGGCCGT</td><td></td><td></td><td></td><td></td><td></td><td></td><td></td><td></td><td></td><td></td></tr>
<tr><td>Chromosome</td><td>start</td><td>end</td><td>strand</td><td>MM</td><td>target_seq</td><td>PAM</td><td>alignment</td><td>distance</td><td>position</td><td>gene name</td><td>gene id</td></tr>
<tr><td>GU305914.1</td><td>29940</td><td>29962</td><td>+</td><td>0</td><td>ACGGCCCGCGCGAAGTCCTC</td><td>GGG</td><td>||||||||[||||||||||||]PAM</td><td>NA</td><td>Intergenic</td><td>NA</td><td>NA</td></tr>
<tr><td>GU305914.1</td><td>65754</td><td>65776</td><td>-</td><td>4</td><td>CACGCCCGCGCGATGTCCTC</td><td>GGG</td><td>---|||||[|||||-||||||]PAM</td><td>NA</td><td>Intergenic</td><td>NA</td><td>NA</td></tr>
<tr><td></td><td></td><td></td><td></td><td></td><td></td><td></td><td></td><td></td><td></td><td></td><td></td></tr>
<tr><td>T1795</td><td>TCGGGCTCGATGATCCGCTGAGG</td><td>1000</td><td></td><td></td><td></td><td></td><td></td><td></td><td></td><td></td><td></td></tr>
<tr><td>Oligo fwd</td><td>CACCTCGGGCTCGATGATCCGCTG</td><td></td><td></td><td></td><td></td><td></td><td></td><td></td><td></td><td></td><td></td></tr>
<tr><td>Oligo rev</td><td>AAACCAGCGGATCATCGAGCCCGA</td><td></td><td></td><td></td><td></td><td></td><td></td><td></td><td></td><td></td><td></td></tr>
<tr><td>Chromosome</td><td>start</td><td>end</td><td>strand</td><td>MM</td><td>target_seq</td><td>PAM</td><td>alignment</td><td>distance</td><td>position</td><td>gene name</td><td>gene id</td></tr>
<tr><td>GU305914.1</td><td>29958</td><td>29980</td><td>+</td><td>0</td><td>TCGGGCTCGATGATCCGCTG</td><td>AGG</td><td>||||||||[||||||||||||]PAM</td><td>NA</td><td>Intergenic</td><td>NA</td><td>NA</td></tr>
<tr><td></td><td></td><td></td><td></td><td></td><td></td><td></td><td></td><td></td><td></td><td></td><td></td></tr>
<tr><td>T1796</td><td>GGCTCGATGATCCGCTGAGGAGG</td><td>1000</td><td></td><td></td><td></td><td></td><td></td><td></td><td></td><td></td><td></td></tr>
<tr><td>Oligo fwd</td><td>CACCGGCTCGATGATCCGCTGAGG</td><td></td><td></td><td></td><td></td><td></td><td></td><td></td><td></td><td></td><td></td></tr>
<tr><td>Oligo rev</td><td>AAACCCTCAGCGGATCATCGAGCC</td><td></td><td></td><td></td><td></td><td></td><td></td><td></td><td></td><td></td><td></td></tr>
<tr><td>Chromosome</td><td>start</td><td>end</td><td>strand</td><td>MM</td><td>target_seq</td><td>PAM</td><td>alignment</td><td>distance</td><td>position</td><td>gene name</td><td>gene id</td></tr>
<tr><td>GU305914.1</td><td>29961</td><td>29983</td><td>+</td><td>0</td><td>GGCTCGATGATCCGCTGAGG</td><td>AGG</td><td>||||||||[||||||||||||]PAM</td><td>NA</td><td>Intergenic</td><td>NA</td><td>NA</td></tr>
<tr><td></td><td></td><td></td><td></td><td></td><td></td><td></td><td></td><td></td><td></td><td></td><td></td></tr>
<tr><td>T1797</td><td>CCGCTGAGGAGGAGACTTTTCGG</td><td>1000</td><td></td><td></td><td></td><td></td><td></td><td></td><td></td><td></td><td></td></tr>
</table>

<table>
<tr><td>Oligo fwd</td><td>CACCCCGCTGAGGAGGAGACTTTT</td><td></td><td></td><td></td><td></td><td></td><td></td><td></td><td></td><td></td><td></td></tr>
<tr><td>Oligo rev</td><td>AAACAAAAGTCTCCTCCTCAGCGG</td><td></td><td></td><td></td><td></td><td></td><td></td><td></td><td></td><td></td><td></td></tr>
<tr><td>Chromosome</td><td>start</td><td>end</td><td>strand</td><td>MM</td><td>target_seq</td><td>PAM</td><td>alignment</td><td>distance</td><td>position</td><td>gene name</td><td>gene id</td></tr>
<tr><td>GU305914.1</td><td>29972</td><td>29994</td><td>+</td><td>0</td><td>CCGCTGAGGAGGAGACTTTT</td><td>CGG</td><td>||||||||[||||||||||||]PAM</td><td>NA</td><td>Intergenic</td><td>NA</td><td>NA</td></tr>
<tr><td></td><td></td><td></td><td></td><td></td><td></td><td></td><td></td><td></td><td></td><td></td><td></td></tr>
<tr><td>T1798</td><td>CGCTGAGGAGGAGACTTTTCGGG</td><td>1000</td><td></td><td></td><td></td><td></td><td></td><td></td><td></td><td></td><td></td></tr>
<tr><td>Oligo fwd</td><td>CACCCGCTGAGGAGGAGACTTTTC</td><td></td><td></td><td></td><td></td><td></td><td></td><td></td><td></td><td></td><td></td></tr>
<tr><td>Oligo rev</td><td>AAACGAAAAGTCTCCTCCTCAGCG</td><td></td><td></td><td></td><td></td><td></td><td></td><td></td><td></td><td></td><td></td></tr>
<tr><td>Chromosome</td><td>start</td><td>end</td><td>strand</td><td>MM</td><td>target_seq</td><td>PAM</td><td>alignment</td><td>distance</td><td>position</td><td>gene name</td><td>gene id</td></tr>
<tr><td>GU305914.1</td><td>29973</td><td>29995</td><td>+</td><td>0</td><td>CGCTGAGGAGGAGACTTTTC</td><td>GGG</td><td>||||||||[||||||||||||]PAM</td><td>NA</td><td>Intergenic</td><td>NA</td><td>NA</td></tr>
<tr><td></td><td></td><td></td><td></td><td></td><td></td><td></td><td></td><td></td><td></td><td></td><td></td></tr>
<tr><td>T1799</td><td>GCTGAGGAGGAGACTTTTCGGGG</td><td>1000</td><td></td><td></td><td></td><td></td><td></td><td></td><td></td><td></td><td></td></tr>
<tr><td>Oligo fwd</td><td>CACCGCTGAGGAGGAGACTTTTCG</td><td></td><td></td><td></td><td></td><td></td><td></td><td></td><td></td><td></td><td></td></tr>
<tr><td>Oligo rev</td><td>AAACCGAAAAGTCTCCTCCTCAGC</td><td></td><td></td><td></td><td></td><td></td><td></td><td></td><td></td><td></td><td></td></tr>
<tr><td>Chromosome</td><td>start</td><td>end</td><td>strand</td><td>MM</td><td>target_seq</td><td>PAM</td><td>alignment</td><td>distance</td><td>position</td><td>gene name</td><td>gene id</td></tr>
<tr><td>GU305914.1</td><td>29974</td><td>29996</td><td>+</td><td>0</td><td>GCTGAGGAGGAGACTTTTCG</td><td>GGG</td><td>||||||||[||||||||||||]PAM</td><td>NA</td><td>Intergenic</td><td>NA</td><td>NA</td></tr>
<tr><td></td><td></td><td></td><td></td><td></td><td></td><td></td><td></td><td></td><td></td><td></td><td></td></tr>
<tr><td>T1800</td><td>GGAGGAGACTTTTCGGGGACCGG</td><td>1000</td><td></td><td></td><td></td><td></td><td></td><td></td><td></td><td></td><td></td></tr>
<tr><td>Oligo fwd</td><td>CACCGGAGGAGACTTTTCGGGGAC</td><td></td><td></td><td></td><td></td><td></td><td></td><td></td><td></td><td></td><td></td></tr>
<tr><td>Oligo rev</td><td>AAACGTCCCCGAAAAGTCTCCTCC</td><td></td><td></td><td></td><td></td><td></td><td></td><td></td><td></td><td></td><td></td></tr>
<tr><td>Chromosome</td><td>start</td><td>end</td><td>strand</td><td>MM</td><td>target_seq</td><td>PAM</td><td>alignment</td><td>distance</td><td>position</td><td>gene name</td><td>gene id</td></tr>
<tr><td>GU305914.1</td><td>29979</td><td>30001</td><td>+</td><td>0</td><td>GGAGGAGACTTTTCGGGGAC</td><td>CGG</td><td>||||||||[||||||||||||]PAM</td><td>NA</td><td>Intergenic</td><td>NA</td><td>NA</td></tr>
<tr><td></td><td></td><td></td><td></td><td></td><td></td><td></td><td></td><td></td><td></td><td></td><td></td></tr>
<tr><td>T1801</td><td>GACTTTTCGGGGACCGGTGCTGG</td><td>1000</td><td></td><td></td><td></td><td></td><td></td><td></td><td></td><td></td><td></td></tr>
<tr><td>Oligo fwd</td><td>CACCGACTTTTCGGGGACCGGTGC</td><td></td><td></td><td></td><td></td><td></td><td></td><td></td><td></td><td></td><td></td></tr>
<tr><td>Oligo rev</td><td>AAACGCACCGGTCCCCGAAAAGTC</td><td></td><td></td><td></td><td></td><td></td><td></td><td></td><td></td><td></td><td></td></tr>
<tr><td>Chromosome</td><td>start</td><td>end</td><td>strand</td><td>MM</td><td>target_seq</td><td>PAM</td><td>alignment</td><td>distance</td><td>position</td><td>gene name</td><td>gene id</td></tr>
<tr><td>GU305914.1</td><td>29985</td><td>30007</td><td>+</td><td>0</td><td>GACTTTTCGGGGACCGGTGC</td><td>TGG</td><td>||||||||[||||||||||||]PAM</td><td>NA</td><td>Intergenic</td><td>NA</td><td>NA</td></tr>
<tr><td></td><td></td><td></td><td></td><td></td><td></td><td></td><td></td><td></td><td></td><td></td><td></td></tr>
<tr><td>T1802</td><td>TTTTCGGGGACCGGTGCTGGCGG</td><td>1000</td><td></td><td></td><td></td><td></td><td></td><td></td><td></td><td></td><td></td></tr>
<tr><td>Oligo fwd</td><td>CACCTTTTCGGGGACCGGTGCTGG</td><td></td><td></td><td></td><td></td><td></td><td></td><td></td><td></td><td></td><td></td></tr>
<tr><td>Oligo rev</td><td>AAACCCAGCACCGGTCCCCGAAAA</td><td></td><td></td><td></td><td></td><td></td><td></td><td></td><td></td><td></td><td></td></tr>
<tr><td>Chromosome</td><td>start</td><td>end</td><td>strand</td><td>MM</td><td>target_seq</td><td>PAM</td><td>alignment</td><td>distance</td><td>position</td><td>gene name</td><td>gene id</td></tr>
<tr><td>GU305914.1</td><td>29988</td><td>30010</td><td>+</td><td>0</td><td>TTTTCGGGGACCGGTGCTGG</td><td>CGG</td><td>||||||||[||||||||||||]PAM</td><td>NA</td><td>Intergenic</td><td>NA</td><td>NA</td></tr>
<tr><td></td><td></td><td></td><td></td><td></td><td></td><td></td><td></td><td></td><td></td><td></td><td></td></tr>
<tr><td>T1803</td><td>TCGGGGACCGGTGCTGGCGGCGG</td><td>1000</td><td></td><td></td><td></td><td></td><td></td><td></td><td></td><td></td><td></td></tr>
<tr><td>Oligo fwd</td><td>CACCTCGGGGACCGGTGCTGGCGG</td><td></td><td></td><td></td><td></td><td></td><td></td><td></td><td></td><td></td><td></td></tr>
<tr><td>Oligo rev</td><td>AAACCCGCCAGCACCGGTCCCCGA</td><td></td><td></td><td></td><td></td><td></td><td></td><td></td><td></td><td></td><td></td></tr>
<tr><td>Chromosome</td><td>start</td><td>end</td><td>strand</td><td>MM</td><td>target_seq</td><td>PAM</td><td>alignment</td><td>distance</td><td>position</td><td>gene name</td><td>gene id</td></tr>
<tr><td>GU305914.1</td><td>29991</td><td>30013</td><td>+</td><td>0</td><td>TCGGGGACCGGTGCTGGCGG</td><td>CGG</td><td>||||||||[||||||||||||]PAM</td><td>NA</td><td>Intergenic</td><td>NA</td><td>NA</td></tr>
<tr><td>GU305914.1</td><td>88498</td><td>88520</td><td>-</td><td>4</td><td>GCGGGCTCCGCTGCTGGCGG</td><td>TGG</td><td>-||||--|[||-|||||||||]PAM</td><td>NA</td><td>Intergenic</td><td>NA</td><td>NA</td></tr>
<tr><td></td><td></td><td></td><td></td><td></td><td></td><td></td><td></td><td></td><td></td><td></td><td></td></tr>
<tr><td>T1804</td><td>GGGACCGGTGCTGGCGGCGGCGG</td><td>1000</td><td></td><td></td><td></td><td></td><td></td><td></td><td></td><td></td><td></td></tr>
<tr><td>Oligo fwd</td><td>CACCGGGACCGGTGCTGGCGGCGG</td><td></td><td></td><td></td><td></td><td></td><td></td><td></td><td></td><td></td><td></td></tr>
<tr><td>Oligo rev</td><td>AAACCCGCCGCCAGCACCGGTCCC</td><td></td><td></td><td></td><td></td><td></td><td></td><td></td><td></td><td></td><td></td></tr>
<tr><td>Chromosome</td><td>start</td><td>end</td><td>strand</td><td>MM</td><td>target_seq</td><td>PAM</td><td>alignment</td><td>distance</td><td>position</td><td>gene name</td><td>gene id</td></tr>
<tr><td>GU305914.1</td><td>29994</td><td>30016</td><td>+</td><td>0</td><td>GGGACCGGTGCTGGCGGCGG</td><td>CGG</td><td>||||||||[||||||||||||]PAM</td><td>NA</td><td>Intergenic</td><td>NA</td><td>NA</td></tr>
<tr><td>GU305914.1</td><td>170536</td><td>170558</td><td>-</td><td>4</td><td>GTGGCCGATGGTGGCGGCGG</td><td>CGG</td><td>|-|-|||-[||-|||||||||]PAM</td><td>NA</td><td>Intergenic</td><td>NA</td><td>NA</td></tr>
<tr><td>GU305914.1</td><td>48147</td><td>48169</td><td>+</td><td>4</td><td>CGGACCTGTGGCGGCGGCGG</td><td>TGG</td><td>-|||||-|[||--||||||||]PAM</td><td>NA</td><td>Intergenic</td><td>NA</td><td>NA</td></tr>
<tr><td>GU305914.1</td><td>88495</td><td>88517</td><td>-</td><td>4</td><td>GGCTCCGCTGCTGGCGGTGG</td><td>CGG</td><td>||--|||-[|||||||||-||]PAM</td><td>NA</td><td>Intergenic</td><td>NA</td><td>NA</td></tr>
<tr><td></td><td></td><td></td><td></td><td></td><td></td><td></td><td></td><td></td><td></td><td></td><td></td></tr>
<tr><td>T1805</td><td>ACCGGTGCTGGCGGCGGCGGCGG</td><td>1000</td><td></td><td></td><td></td><td></td><td></td><td></td><td></td><td></td><td></td></tr>
<tr><td>Oligo fwd</td><td>CACCACCGGTGCTGGCGGCGGCGG</td><td></td><td></td><td></td><td></td><td></td><td></td><td></td><td></td><td></td><td></td></tr>
<tr><td>Oligo rev</td><td>AAACCCGCCGCCGCCAGCACCGGT</td><td></td><td></td><td></td><td></td><td></td><td></td><td></td><td></td><td></td><td></td></tr>
<tr><td>Chromosome</td><td>start</td><td>end</td><td>strand</td><td>MM</td><td>target_seq</td><td>PAM</td><td>alignment</td><td>distance</td><td>position</td><td>gene name</td><td>gene id</td></tr>
<tr><td>GU305914.1</td><td>29997</td><td>30019</td><td>+</td><td>0</td><td>ACCGGTGCTGGCGGCGGCGG</td><td>CGG</td><td>||||||||[||||||||||||]PAM</td><td>NA</td><td>Intergenic</td><td>NA</td><td>NA</td></tr>
<tr><td>GU305914.1</td><td>48159</td><td>48181</td><td>+</td><td>4</td><td>GGCGGCGGTGGCGGCGGCGG</td><td>TGG</td><td>--|||-|-[||||||||||||]PAM</td><td>NA</td><td>Intergenic</td><td>NA</td><td>NA</td></tr>
<tr><td>GU305914.1</td><td>154401</td><td>154423</td><td>+</td><td>4</td><td>CCGGGCGGTGGCGGCGGCGG</td><td>CGG</td><td>-|-||-|-[||||||||||||]PAM</td><td>NA</td><td>Intergenic</td><td>NA</td><td>NA</td></tr>
<tr><td>GU305914.1</td><td>58129</td><td>58151</td><td>-</td><td>4</td><td>GCGGGCGGTGGCGGCGGCGG</td><td>CGG</td><td>-|-||-|-[||||||||||||]PAM</td><td>NA</td><td>Intergenic</td><td>NA</td><td>NA</td></tr>
<tr><td>GU305914.1</td><td>154404</td><td>154426</td><td>+</td><td>4</td><td>GGCGGTGGCGGCGGCGGCGG</td><td>CGG</td><td>--|||||-[-|||||||||||]PAM</td><td>NA</td><td>Intergenic</td><td>NA</td><td>NA</td></tr>
<tr><td>GU305914.1</td><td>58126</td><td>58148</td><td>-</td><td>4</td><td>GGCGGTGGCGGCGGCGGCGG</td><td>CGG</td><td>--|||||-[-|||||||||||]PAM</td><td>NA</td><td>Intergenic</td><td>NA</td><td>NA</td></tr>
<tr><td>GU305914.1</td><td>114658</td><td>114680</td><td>-</td><td>4</td><td>GGCGGTGGTGGTGGCGGCGG</td><td>CGG</td><td>--|||||-[|||-||||||||]PAM</td><td>NA</td><td>Intergenic</td><td>NA</td><td>NA</td></tr>
<tr><td>GU305914.1</td><td>26382</td><td>26404</td><td>+</td><td>3</td><td>ATCGGCGCTGGCGACGGCGG</td><td>CGG</td><td>|-|||-||[|||||-||||||]PAM</td><td>NA</td><td>Intergenic</td><td>NA</td><td>NA</td></tr>
<tr><td>GU305914.1</td><td>168817</td><td>168839</td><td>+</td><td>4</td><td>GGCGGTGGTGGCAGCGGCGG</td><td>CGG</td><td>--|||||-[||||-|||||||]PAM</td><td>NA</td><td>Intergenic</td><td>NA</td><td>NA</td></tr>
<tr><td>GU305914.1</td><td>52716</td><td>52738</td><td>+</td><td>4</td><td>GCCGCCGCTGGCGGTGGCGG</td><td>TGG</td><td>-|||--||[||||||-|||||]PAM</td><td>NA</td><td>Intergenic</td><td>NA</td><td>NA</td></tr>
<tr><td>GU305914.1</td><td>24875</td><td>24897</td><td>+</td><td>4</td><td>CCCGATGTTGGCGGCGACGG</td><td>CGG</td><td>-|||-||-[||||||||-|||]PAM</td><td>NA</td><td>Intergenic</td><td>NA</td><td>NA</td></tr>
<tr><td>GU305914.1</td><td>48150</td><td>48172</td><td>+</td><td>4</td><td>ACCTGTGGCGGCGGCGGTGG</td><td>CGG</td><td>|||-|||-[-||||||||-||]PAM</td><td>NA</td><td>Intergenic</td><td>NA</td><td>NA</td></tr>
<tr><td></td><td></td><td></td><td></td><td></td><td></td><td></td><td></td><td></td><td></td><td></td><td></td></tr>
<tr><td>T1806</td><td>GGTGCTGGCGGCGGCGGCGGCGG</td><td>1000</td><td></td><td></td><td></td><td></td><td></td><td></td><td></td><td></td><td></td></tr>
<tr><td>Oligo fwd</td><td>CACCGGTGCTGGCGGCGGCGGCGG</td><td></td><td></td><td></td><td></td><td></td><td></td><td></td><td></td><td></td><td></td></tr>
<tr><td>Oligo rev</td><td>AAACCCGCCGCCGCCGCCAGCACC</td><td></td><td></td><td></td><td></td><td></td><td></td><td></td><td></td><td></td><td></td></tr>
<tr><td>Chromosome</td><td>start</td><td>end</td><td>strand</td><td>MM</td><td>target_seq</td><td>PAM</td><td>alignment</td><td>distance</td><td>position</td><td>gene name</td><td>gene id</td></tr>
<tr><td>GU305914.1</td><td>30000</td><td>30022</td><td>+</td><td>0</td><td>GGTGCTGGCGGCGGCGGCGG</td><td>CGG</td><td>||||||||[||||||||||||]PAM</td><td>NA</td><td>Intergenic</td><td>NA</td><td>NA</td></tr>
<tr><td>GU305914.1</td><td>154404</td><td>154426</td><td>+</td><td>2</td><td>GGCGGTGGCGGCGGCGGCGG</td><td>CGG</td><td>||-|-|||[||||||||||||]PAM</td><td>NA</td><td>Intergenic</td><td>NA</td><td>NA</td></tr>
<tr><td>GU305914.1</td><td>58126</td><td>58148</td><td>-</td><td>2</td><td>GGCGGTGGCGGCGGCGGCGG</td><td>CGG</td><td>||-|-|||[||||||||||||]PAM</td><td>NA</td><td>Intergenic</td><td>NA</td><td>NA</td></tr>
<tr><td>GU305914.1</td><td>154407</td><td>154429</td><td>+</td><td>2</td><td>GGTGGCGGCGGCGGCGGCGG</td><td>CGG</td><td>||||--||[||||||||||||]PAM</td><td>NA</td><td>Intergenic</td><td>NA</td><td>NA</td></tr>
<tr><td>GU305914.1</td><td>58123</td><td>58145</td><td>-</td><td>2</td><td>GGTGGCGGCGGCGGCGGCGG</td><td>AGG</td><td>||||--||[||||||||||||]PAM</td><td>NA</td><td>Intergenic</td><td>NA</td><td>NA</td></tr>
<tr><td>GU305914.1</td><td>149482</td><td>149504</td><td>-</td><td>3</td><td>GGCGACGGCGGCGGCGGCGG</td><td>CGG</td><td>||-|--||[||||||||||||]PAM</td><td>NA</td><td>Intergenic</td><td>NA</td><td>NA</td></tr>
<tr><td>GU305914.1</td><td>168820</td><td>168842</td><td>+</td><td>2</td><td>GGTGGTGGCAGCGGCGGCGG</td><td>CGG</td><td>||||-|||[|-||||||||||]PAM</td><td>NA</td><td>Intergenic</td><td>NA</td><td>NA</td></tr>
<tr><td>GU305914.1</td><td>67742</td><td>67764</td><td>+</td><td>4</td><td>GGGAGCGGCGGCGGCGGCGG</td><td>CGG</td><td>||----||[||||||||||||]PAM</td><td>NA</td><td>Intergenic</td><td>NA</td><td>NA</td></tr>
<tr><td>GU305914.1</td><td>163896</td><td>163918</td><td>+</td><td>4</td><td>GGATCGAGCGGCGGCGGCGG</td><td>CGG</td><td>||--|--|[||||||||||||]PAM</td><td>NA</td><td>Intergenic</td><td>NA</td><td>NA</td></tr>
<tr><td>GU305914.1</td><td>149488</td><td>149510</td><td>-</td><td>3</td><td>GGAGCAGGCGACGGCGGCGG</td><td>CGG</td><td>||-||-||[||-|||||||||]PAM</td><td>NA</td><td>Intergenic</td><td>NA</td><td>NA</td></tr>
<tr><td>GU305914.1</td><td>48159</td><td>48181</td><td>+</td><td>4</td><td>GGCGGCGGTGGCGGCGGCGG</td><td>TGG</td><td>||-|--||[-|||||||||||]PAM</td><td>NA</td><td>Intergenic</td><td>NA</td><td>NA</td></tr>
<tr><td>GU305914.1</td><td>176454</td><td>176476</td><td>+</td><td>4</td><td>GACGATGGCGACGGCGGCGG</td><td>AGG</td><td>|--|-|||[||-|||||||||]PAM</td><td>NA</td><td>Intergenic</td><td>NA</td><td>NA</td></tr>
<tr><td>GU305914.1</td><td>164971</td><td>164993</td><td>-</td><td>4</td><td>GGTGGGACCGGCGGCGGCGG</td><td>CGG</td><td>||||----[||||||||||||]PAM</td><td>NA</td><td>Intergenic</td><td>NA</td><td>NA</td></tr>
<tr><td>GU305914.1</td><td>73792</td><td>73814</td><td>-</td><td>4</td><td>GGCGCCAGCAGCGGCGGCGG</td><td>AGG</td><td>||-||--|[|-||||||||||]PAM</td><td>NA</td><td>Intergenic</td><td>NA</td><td>NA</td></tr>
<tr><td>GU305914.1</td><td>48156</td><td>48178</td><td>+</td><td>4</td><td>GGCGGCGGCGGTGGCGGCGG</td><td>CGG</td><td>||-|--||[|||-||||||||]PAM</td><td>NA</td><td>Intergenic</td><td>NA</td><td>NA</td></tr>
<tr><td>GU305914.1</td><td>114658</td><td>114680</td><td>-</td><td>4</td><td>GGCGGTGGTGGTGGCGGCGG</td><td>CGG</td><td>||-|-|||[-||-||||||||]PAM</td><td>NA</td><td>Intergenic</td><td>NA</td><td>NA</td></tr>
<tr><td>GU305914.1</td><td>168817</td><td>168839</td><td>+</td><td>4</td><td>GGCGGTGGTGGCAGCGGCGG</td><td>CGG</td><td>||-|-|||[-|||-|||||||]PAM</td><td>NA</td><td>Intergenic</td><td>NA</td><td>NA</td></tr>
</table>

| | | | | | | | | | | | |
|---|---|---|---|---|---|---|---|---|---|---|---|
| GU305914.1 | 48153 | 48175 | + | 4 | TGTGGCGGCGGCGGTGGCGG | CGG | -|||--||[||||||-|||||]PAM | NA | Intergenic | NA | NA |
| GU305914.1 | 62672 | 62694 | - | 4 | GCTGCCGCCGGCGGGGGCGG | CGG | |-|||-|-[||||||-|||||]PAM | NA | Intergenic | NA | NA |
| GU305914.1 | 149292 | 149314 | + | 4 | GGAGCGGGCGACGACGGCGG | CGG | ||-||-||[||-||-||||||]PAM | NA | Intergenic | NA | NA |
| GU305914.1 | 45255 | 45277 | - | 4 | GGTGGCGGTGGCGGCAGCGG | CGG | ||||--||[-||||||-||||]PAM | NA | Intergenic | NA | NA |
| GU305914.1 | 62449 | 62471 | + | 4 | GGGGGTGGTGGCGGCGGGGG | TGG | ||-|-|||[-||||||||-||]PAM | NA | Intergenic | NA | NA |
| GU305914.1 | 52719 | 52741 | + | 4 | GCCGCTGGCGGTGGCGGTGG | CGG | |--|||||[|||-|||||-||]PAM | NA | Intergenic | NA | NA |
| GU305914.1 | 30006 | 30028 | + | 4 | GGCGGCGGCGGCGGCGGCAG | TGG | ||-|--||[||||||||||-|]PAM | NA | Intergenic | NA | NA |
| GU305914.1 | 149476 | 149498 | - | 4 | GGCGGCGGCGGCGGCGGCAG | CGG | ||-|--||[||||||||||-|]PAM | NA | Intergenic | NA | NA |
| GU305914.1 | 62675 | 62697 | - | 4 | GCTGCTGCCGCCGGCGGGGG | CGG | |-|||||-[||-||||||-||]PAM | NA | Intergenic | NA | NA |
| GU305914.1 | 74673 | 74695 | + | 4 | GGTGCCGTCGGTGGCGGTGG | CGG | |||||-|-[|||-|||||-||]PAM | NA | Intergenic | NA | NA |
| GU305914.1 | 62452 | 62474 | + | 3 | GGTGGTGGCGGCGGGGGTGG | CGG | ||||-|||[||||||-||-||]PAM | NA | Intergenic | NA | NA |
| GU305914.1 | 62437 | 62459 | + | 4 | GGCGGTGGCGGCGGGGGTGG | TGG | ||-|-|||[||||||-||-||]PAM | NA | Intergenic | NA | NA |
| GU305914.1 | 170530 | 170552 | - | 4 | GATGGTGGCGGCGGCGGCCA | CGG | |-||-|||[||||||||||--]PAM | NA | Intergenic | NA | NA |

| T1807 | GGCGGCGGCGGCGGCGGCAGTGG | 1000 | | | | | | | | | |
|---|---|---|---|---|---|---|---|---|---|---|---|
| Oligo fwd | CACCGGCGGCGGCGGCGGCGGCAG | | | | | | | | | | |
| Oligo rev | AAACCTGCCGCCGCCGCCGCCGCC | | | | | | | | | | |
| Chromosome | start | end | strand | MM | target_seq | PAM | alignment | distance | position | gene name | gene id |
| GU305914.1 | 30006 | 30028 | + | 0 | GGCGGCGGCGGCGGCGGCAG | TGG | ||||||||[||||||||||||]PAM | NA | Intergenic | NA | NA |
| GU305914.1 | 149476 | 149498 | - | 0 | GGCGGCGGCGGCGGCGGCAG | CGG | ||||||||[||||||||||||]PAM | NA | Intergenic | NA | NA |
| GU305914.1 | 163902 | 163924 | + | 1 | AGCGGCGGCGGCGGCGGCAG | CGG | -|||||||[||||||||||||]PAM | NA | Intergenic | NA | NA |
| GU305914.1 | 113924 | 113946 | - | 2 | GTCGGCGTCGGCGGCGGCAG | CGG | |-|||||-[||||||||||||]PAM | NA | Intergenic | NA | NA |
| GU305914.1 | 163908 | 163930 | + | 1 | GGCGGCGGCGGCAGCGGCAG | CGG | ||||||||[||||-|||||||]PAM | NA | Intergenic | NA | NA |
| GU305914.1 | 166410 | 166432 | + | 3 | GACGTCGGCGCCGGCGGCAG | CGG | |-||-|||[||-|||||||||]PAM | NA | Intergenic | NA | NA |
| GU305914.1 | 45258 | 45280 | - | 4 | AACGGTGGCGGTGGCGGCAG | CGG | --|||-||[|||-||||||||]PAM | NA | Intergenic | NA | NA |
| GU305914.1 | 65721 | 65743 | - | 4 | TCCTGCGGCGGCAGCGGCAG | CGG | --|-||||[||||-|||||||]PAM | NA | Intergenic | NA | NA |
| GU305914.1 | 62152 | 62174 | + | 4 | AGGGGCGACGGCAGCGGCAG | CGG | -|-||||-[||||-|||||||]PAM | NA | Intergenic | NA | NA |
| GU305914.1 | 168829 | 168851 | + | 2 | AGCGGCGGCGGCGGCGACAG | CGG | -|||||||[||||||||-|||]PAM | NA | Intergenic | NA | NA |
| GU305914.1 | 24557 | 24579 | + | 3 | GGCGGCAGCGACGACGGCAG | CGG | ||||||-|[||-||-||||||]PAM | NA | Intergenic | NA | NA |
| GU305914.1 | 163920 | 163942 | + | 4 | AGCGGCAGCGGAGGCAGCAG | CGG | -|||||-|[|||-|||-||||]PAM | NA | Intergenic | NA | NA |
| GU305914.1 | 154407 | 154429 | + | 2 | GGTGGCGGCGGCGGCGGCGG | CGG | ||-|||||[||||||||||-|]PAM | NA | Intergenic | NA | NA |
| GU305914.1 | 58123 | 58145 | - | 2 | GGTGGCGGCGGCGGCGGCGG | AGG | ||-|||||[||||||||||-|]PAM | NA | Intergenic | NA | NA |
| GU305914.1 | 128727 | 128749 | - | 4 | GGTGGAGGCGGCAGCAGCAG | CGG | ||-||-||[||||-||-||||]PAM | NA | Intergenic | NA | NA |
| GU305914.1 | 149482 | 149504 | - | 2 | GGCGACGGCGGCGGCGGCGG | CGG | ||||-|||[||||||||||-|]PAM | NA | Intergenic | NA | NA |
| GU305914.1 | 154404 | 154426 | + | 2 | GGCGGTGGCGGCGGCGGCGG | CGG | |||||-||[||||||||||-|]PAM | NA | Intergenic | NA | NA |
| GU305914.1 | 58126 | 58148 | - | 2 | GGCGGTGGCGGCGGCGGCGG | CGG | |||||-||[||||||||||-|]PAM | NA | Intergenic | NA | NA |
| GU305914.1 | 67742 | 67764 | + | 3 | GGGAGCGGCGGCGGCGGCGG | CGG | ||--||||[||||||||||-|]PAM | NA | Intergenic | NA | NA |
| GU305914.1 | 48159 | 48181 | + | 2 | GGCGGCGGTGGCGGCGGCGG | TGG | ||||||||[-|||||||||-|]PAM | NA | Intergenic | NA | NA |
| GU305914.1 | 56773 | 56795 | + | 4 | CGCCGCAGCGGCGGCGGCGG | CGG | -||-||-|[||||||||||-|]PAM | NA | Intergenic | NA | NA |
| GU305914.1 | 30000 | 30022 | + | 4 | GGTGCTGGCGGCGGCGGCGG | CGG | ||-|--||[||||||||||-|]PAM | NA | Intergenic | NA | NA |
| GU305914.1 | 149485 | 149507 | - | 4 | GCAGGCGACGGCGGCGGCGG | CGG | |--||||-[||||||||||-|]PAM | NA | Intergenic | NA | NA |
| GU305914.1 | 58129 | 58151 | - | 4 | GCGGGCGGTGGCGGCGGCGG | CGG | |--|||||[-|||||||||-|]PAM | NA | Intergenic | NA | NA |
| GU305914.1 | 48156 | 48178 | + | 2 | GGCGGCGGCGGTGGCGGCGG | CGG | ||||||||[|||-||||||-|]PAM | NA | Intergenic | NA | NA |
| GU305914.1 | 161754 | 161776 | - | 4 | GGCCGAGGAGGCGGCGGCGG | AGG | |||-|-||[-|||||||||-|]PAM | NA | Intergenic | NA | NA |
| GU305914.1 | 168820 | 168842 | + | 4 | GGTGGTGGCAGCGGCGGCGG | CGG | ||-||-||[|-||||||||-|]PAM | NA | Intergenic | NA | NA |
| GU305914.1 | 73792 | 73814 | - | 4 | GGCGCCAGCAGCGGCGGCGG | AGG | ||||-|-|[|-||||||||-|]PAM | NA | Intergenic | NA | NA |
| GU305914.1 | 71753 | 71775 | + | 4 | AGCAGCAGCGGCGGCGGCAA | CGG | -||-||-|[|||||||||||-]PAM | NA | Intergenic | NA | NA |
| GU305914.1 | 48153 | 48175 | + | 4 | TGTGGCGGCGGCGGTGGCGG | CGG | -|-|||||[||||||-|||-|]PAM | NA | Intergenic | NA | NA |

| T1808 | GGCGGCAGTGGTATCGCCAGAGG | 1000 | | | | | | | | | |
|---|---|---|---|---|---|---|---|---|---|---|---|
| Oligo fwd | CACCGGCGGCAGTGGTATCGCCAG | | | | | | | | | | |
| Oligo rev | AAACCTGGCGATACCACTGCCGCC | | | | | | | | | | |
| Chromosome | start | end | strand | MM | target_seq | PAM | alignment | distance | position | gene name | gene id |
| GU305914.1 | 30018 | 30040 | + | 0 | GGCGGCAGTGGTATCGCCAG | AGG | ||||||||[||||||||||||]PAM | NA | Intergenic | NA | NA |

| T1809 | AGAGGCTGCGCGACCGTCACCGG | 1000 | | | | | | | | | |
|---|---|---|---|---|---|---|---|---|---|---|---|
| Oligo fwd | CACCAGAGGCTGCGCGACCGTCAC | | | | | | | | | | |
| Oligo rev | AAACGTGACGGTCGCGCAGCCTCT | | | | | | | | | | |
| Chromosome | start | end | strand | MM | target_seq | PAM | alignment | distance | position | gene name | gene id |
| GU305914.1 | 30036 | 30058 | + | 0 | AGAGGCTGCGCGACCGTCAC | CGG | ||||||||[||||||||||||]PAM | NA | Intergenic | NA | NA |

| T1810 | CTGCGCGACCGTCACCGGCGCGG | 1000 | | | | | | | | | |
|---|---|---|---|---|---|---|---|---|---|---|---|
| Oligo fwd | CACCCTGCGCGACCGTCACCGGCG | | | | | | | | | | |
| Oligo rev | AAACCGCCGGTGACGGTCGCGCAG | | | | | | | | | | |
| Chromosome | start | end | strand | MM | target_seq | PAM | alignment | distance | position | gene name | gene id |
| GU305914.1 | 30041 | 30063 | + | 0 | CTGCGCGACCGTCACCGGCG | CGG | ||||||||[||||||||||||]PAM | NA | Intergenic | NA | NA |
| GU305914.1 | 132327 | 132349 | - | 3 | CCGCGCGTCCGGCACCGGCG | TGG | |-|||||-[|||-||||||||]PAM | NA | Intergenic | NA | NA |

| T1811 | CGCGACCGTCACCGGCGCGGTGG | 1000 | | | | | | | | | |
|---|---|---|---|---|---|---|---|---|---|---|---|
| Oligo fwd | CACCCGCGACCGTCACCGGCGCGG | | | | | | | | | | |
| Oligo rev | AAACCCGCGCCGGTGACGGTCGCG | | | | | | | | | | |
| Chromosome | start | end | strand | MM | target_seq | PAM | alignment | distance | position | gene name | gene id |
| GU305914.1 | 30044 | 30066 | + | 0 | CGCGACCGTCACCGGCGCGG | TGG | ||||||||[||||||||||||]PAM | NA | Intergenic | NA | NA |

| T1812 | CGTCACCGGCGCGGTGGTGACGG | 1000 | | | | | | | | | |
|---|---|---|---|---|---|---|---|---|---|---|---|
| Oligo fwd | CACCCGTCACCGGCGCGGTGGTGA | | | | | | | | | | |
| Oligo rev | AAACTCACCACCGCGCCGGTGACG | | | | | | | | | | |
| Chromosome | start | end | strand | MM | target_seq | PAM | alignment | distance | position | gene name | gene id |
| GU305914.1 | 30050 | 30072 | + | 0 | CGTCACCGGCGCGGTGGTGA | CGG | ||||||||[||||||||||||]PAM | NA | Intergenic | NA | NA |

| T1813 | GTCACCGGCGCGGTGGTGACGGG | 1000 | | | | | | | | | |
|---|---|---|---|---|---|---|---|---|---|---|---|
| Oligo fwd | CACCGTCACCGGCGCGGTGGTGAC | | | | | | | | | | |
| Oligo rev | AAACGTCACCACCGCGCCGGTGAC | | | | | | | | | | |
| Chromosome | start | end | strand | MM | target_seq | PAM | alignment | distance | position | gene name | gene id |
| GU305914.1 | 30051 | 30073 | + | 0 | GTCACCGGCGCGGTGGTGAC | GGG | ||||||||[||||||||||||]PAM | NA | Intergenic | NA | NA |

| T1814 | GGTGGTGACGGGCACGAACGAGG | 1000 |
|---|---|---|
| Oligo fwd | CACCGGTGGTGACGGGCACGAACG | |
| Oligo rev | AAACCGTTCGTGCCCGTCACCACC | |

| | | | | | | | | | | | |
|---|---|---|---|---|---|---|---|---|---|---|---|
| Chromosome | start | end | strand | MM | target_seq | PAM | alignment | distance | position | gene name | gene id |
| GU305914.1 | 30062 | 30084 | + | 0 | GGTGGTGACGGGCACGAACG | AGG | \|\|\|\|\|\|\|\|[\|\|\|\|\|\|\|\|\|\|\|\|]PAM | NA | Intergenic | NA | NA |
| T1815 | ACGGGCACGAACGAGGACAGCGG | 1000 | | | | | | | | | |
| Oligo fwd | CACCACGGGCACGAACGAGGACAG | | | | | | | | | | |
| Oligo rev | AAACCTGTCCTCGTTCGTGCCCGT | | | | | | | | | | |
| Chromosome | start | end | strand | MM | target_seq | PAM | alignment | distance | position | gene name | gene id |
| GU305914.1 | 30069 | 30091 | + | 0 | ACGGGCACGAACGAGGACAG | CGG | \|\|\|\|\|\|\|\|[\|\|\|\|\|\|\|\|\|\|\|\|]PAM | NA | Intergenic | NA | NA |
| T1816 | AGCGGCAGCGTTATCGTTGATGG | 1000 | | | | | | | | | |
| Oligo fwd | CACCAGCGGCAGCGTTATCGTTGA | | | | | | | | | | |
| Oligo rev | AAACTCAACGATAACGCTGCCGCT | | | | | | | | | | |
| Chromosome | start | end | strand | MM | target_seq | PAM | alignment | distance | position | gene name | gene id |
| GU305914.1 | 30087 | 30109 | + | 0 | AGCGGCAGCGTTATCGTTGA | TGG | \|\|\|\|\|\|\|\|[\|\|\|\|\|\|\|\|\|\|\|\|]PAM | NA | Intergenic | NA | NA |
| T1817 | GGCAGCGTTATCGTTGATGGCGG | 1000 | | | | | | | | | |
| Oligo fwd | CACCGGCAGCGTTATCGTTGATGG | | | | | | | | | | |
| Oligo rev | AAACCCATCAACGATAACGCTGCC | | | | | | | | | | |
| Chromosome | start | end | strand | MM | target_seq | PAM | alignment | distance | position | gene name | gene id |
| GU305914.1 | 30090 | 30112 | + | 0 | GGCAGCGTTATCGTTGATGG | CGG | \|\|\|\|\|\|\|\|[\|\|\|\|\|\|\|\|\|\|\|\|]PAM | NA | Intergenic | NA | NA |
| T1818 | AGCGTTATCGTTGATGGCGGCGG | 1000 | | | | | | | | | |
| Oligo fwd | CACCAGCGTTATCGTTGATGGCGG | | | | | | | | | | |
| Oligo rev | AAACCCGCCATCAACGATAACGCT | | | | | | | | | | |
| Chromosome | start | end | strand | MM | target_seq | PAM | alignment | distance | position | gene name | gene id |
| GU305914.1 | 30093 | 30115 | + | 0 | AGCGTTATCGTTGATGGCGG | CGG | \|\|\|\|\|\|\|\|[\|\|\|\|\|\|\|\|\|\|\|\|]PAM | NA | Intergenic | NA | NA |
| T1819 | GTTATCGTTGATGGCGGCGGTGG | 1000 | | | | | | | | | |
| Oligo fwd | CACCGTTATCGTTGATGGCGGCGG | | | | | | | | | | |
| Oligo rev | AAACCCGCCGCCATCAACGATAAC | | | | | | | | | | |
| Chromosome | start | end | strand | MM | target_seq | PAM | alignment | distance | position | gene name | gene id |
| GU305914.1 | 30096 | 30118 | + | 0 | GTTATCGTTGATGGCGGCGG | TGG | \|\|\|\|\|\|\|\|[\|\|\|\|\|\|\|\|\|\|\|\|]PAM | NA | Intergenic | NA | NA |
| T1820 | TGATGGCGGCGGTGGAAGCGAGG | 1000 | | | | | | | | | |
| Oligo fwd | CACCTGATGGCGGCGGTGGAAGCG | | | | | | | | | | |
| Oligo rev | AAACCGCTTCCACCGCCGCCATCA | | | | | | | | | | |
| Chromosome | start | end | strand | MM | target_seq | PAM | alignment | distance | position | gene name | gene id |
| GU305914.1 | 30104 | 30126 | + | 0 | TGATGGCGGCGGTGGAAGCG | AGG | \|\|\|\|\|\|\|\|[\|\|\|\|\|\|\|\|\|\|\|\|]PAM | NA | Intergenic | NA | NA |
| T1821 | GGCGGTGGAAGCGAGGCCAGCGG | 1000 | | | | | | | | | |
| Oligo fwd | CACCGGCGGTGGAAGCGAGGCCAG | | | | | | | | | | |
| Oligo rev | AAACCTGGCCTCGCTTCCACCGCC | | | | | | | | | | |
| Chromosome | start | end | strand | MM | target_seq | PAM | alignment | distance | position | gene name | gene id |
| GU305914.1 | 30111 | 30133 | + | 0 | GGCGGTGGAAGCGAGGCCAG | CGG | \|\|\|\|\|\|\|\|[\|\|\|\|\|\|\|\|\|\|\|\|]PAM | NA | Intergenic | NA | NA |
| GU305914.1 | 92348 | 92370 | + | 4 | TGCGGTCGAAGCGTGGTCAG | CGG | -\|\|\|\|\|-\|[\|\|\|\|\|-\|\|-\|\|\|]PAM | NA | Intergenic | NA | NA |
| T1822 | GGTAGCGAGAGCTGCAGCCCCGG | 1000 | | | | | | | | | |
| Oligo fwd | CACCGGTAGCGAGAGCTGCAGCCC | | | | | | | | | | |
| Oligo rev | AAACGGGCTGCAGCTCTCGCTACC | | | | | | | | | | |
| Chromosome | start | end | strand | MM | target_seq | PAM | alignment | distance | position | gene name | gene id |
| GU305914.1 | 30132 | 30154 | + | 0 | GGTAGCGAGAGCTGCAGCCC | CGG | \|\|\|\|\|\|\|\|[\|\|\|\|\|\|\|\|\|\|\|\|]PAM | NA | Intergenic | NA | NA |
| T1823 | AGCGAGAGCTGCAGCCCCGGCGG | 1000 | | | | | | | | | |
| Oligo fwd | CACCAGCGAGAGCTGCAGCCCCGG | | | | | | | | | | |
| Oligo rev | AAACCCGGGGCTGCAGCTCTCGCT | | | | | | | | | | |
| Chromosome | start | end | strand | MM | target_seq | PAM | alignment | distance | position | gene name | gene id |
| GU305914.1 | 30135 | 30157 | + | 0 | AGCGAGAGCTGCAGCCCCGG | CGG | \|\|\|\|\|\|\|\|[\|\|\|\|\|\|\|\|\|\|\|\|]PAM | NA | Intergenic | NA | NA |
| T1824 | GAGCTGCAGCCCCGGCGGCCTGG | 1000 | | | | | | | | | |
| Oligo fwd | CACCGAGCTGCAGCCCCGGCGGCC | | | | | | | | | | |
| Oligo rev | AAACGGCCGCCGGGGCTGCAGCTC | | | | | | | | | | |
| Chromosome | start | end | strand | MM | target_seq | PAM | alignment | distance | position | gene name | gene id |
| GU305914.1 | 30140 | 30162 | + | 0 | GAGCTGCAGCCCCGGCGGCC | TGG | \|\|\|\|\|\|\|\|[\|\|\|\|\|\|\|\|\|\|\|\|]PAM | NA | Intergenic | NA | NA |
| T1825 | AGCTGCAGCCCCGGCGGCCTGGG | 1000 | | | | | | | | | |
| Oligo fwd | CACCAGCTGCAGCCCCGGCGGCCT | | | | | | | | | | |
| Oligo rev | AAACAGGCCGCCGGGGCTGCAGCT | | | | | | | | | | |
| Chromosome | start | end | strand | MM | target_seq | PAM | alignment | distance | position | gene name | gene id |
| GU305914.1 | 30141 | 30163 | + | 0 | AGCTGCAGCCCCGGCGGCCT | GGG | \|\|\|\|\|\|\|\|[\|\|\|\|\|\|\|\|\|\|\|\|]PAM | NA | Intergenic | NA | NA |
| T1826 | GCTGCAGCCCCGGCGGCCTGGGG | 1000 | | | | | | | | | |
| Oligo fwd | CACCGCTGCAGCCCCGGCGGCCTG | | | | | | | | | | |
| Oligo rev | AAACCAGGCCGCCGGGGCTGCAGC | | | | | | | | | | |
| Chromosome | start | end | strand | MM | target_seq | PAM | alignment | distance | position | gene name | gene id |
| GU305914.1 | 30142 | 30164 | + | 0 | GCTGCAGCCCCGGCGGCCTG | GGG | \|\|\|\|\|\|\|\|[\|\|\|\|\|\|\|\|\|\|\|\|]PAM | NA | Intergenic | NA | NA |
| T1827 | GCAGCCCCGGCGGCCTGGGGAGG | 1000 | | | | | | | | | |
| Oligo fwd | CACCGCAGCCCCGGCGGCCTGGGG | | | | | | | | | | |
| Oligo rev | AAACCCCCAGGCCGCCGGGGCTGC | | | | | | | | | | |
| Chromosome | start | end | strand | MM | target_seq | PAM | alignment | distance | position | gene name | gene id |
| GU305914.1 | 30145 | 30167 | + | 0 | GCAGCCCCGGCGGCCTGGGG | AGG | \|\|\|\|\|\|\|\|[\|\|\|\|\|\|\|\|\|\|\|\|]PAM | NA | Intergenic | NA | NA |
| T1828 | CAGCCCCGGCGGCCTGGGGAGGG | 1000 | | | | | | | | | |
| Oligo fwd | CACCCAGCCCCGGCGGCCTGGGGA | | | | | | | | | | |
| Oligo rev | AAACTCCCCAGGCCGCCGGGGCTG | | | | | | | | | | |
| Chromosome | start | end | strand | MM | target_seq | PAM | alignment | distance | position | gene name | gene id |
| GU305914.1 | 30146 | 30168 | + | 0 | CAGCCCCGGCGGCCTGGGGA | GGG | \|\|\|\|\|\|\|\|[\|\|\|\|\|\|\|\|\|\|\|\|]PAM | NA | Intergenic | NA | NA |
| T1829 | CCCCGGCGGCCTGGGGAGGGCGG | 1000 | | | | | | | | | |

<table>
<tr><td>Oligo fwd</td><td>CACCCCCCGGCGGCCTGGGGAGGG</td><td></td><td></td><td></td><td></td><td></td><td></td><td></td><td></td><td></td><td></td></tr>
<tr><td>Oligo rev</td><td>AAACCCCTCCCCAGGCCGCCGGGG</td><td></td><td></td><td></td><td></td><td></td><td></td><td></td><td></td><td></td><td></td></tr>
<tr><td>Chromosome</td><td>start</td><td>end</td><td>strand</td><td>MM</td><td>target_seq</td><td>PAM</td><td>alignment</td><td>distance</td><td>position</td><td>gene name</td><td>gene id</td></tr>
<tr><td>GU305914.1</td><td>30149</td><td>30171</td><td>+</td><td>0</td><td>CCCCGGCGGCCTGGGGAGGG</td><td>CGG</td><td>||||||||[||||||||||||]PAM</td><td>NA</td><td>Intergenic</td><td>NA</td><td>NA</td></tr>
<tr><td>T1830</td><td>CCCGGCGGCCTGGGGAGGGCGGG</td><td>1000</td><td></td><td></td><td></td><td></td><td></td><td></td><td></td><td></td><td></td></tr>
<tr><td>Oligo fwd</td><td>CACCCCCGGCGGCCTGGGGAGGGC</td><td></td><td></td><td></td><td></td><td></td><td></td><td></td><td></td><td></td><td></td></tr>
<tr><td>Oligo rev</td><td>AAACGCCCTCCCCAGGCCGCCGGG</td><td></td><td></td><td></td><td></td><td></td><td></td><td></td><td></td><td></td><td></td></tr>
<tr><td>Chromosome</td><td>start</td><td>end</td><td>strand</td><td>MM</td><td>target_seq</td><td>PAM</td><td>alignment</td><td>distance</td><td>position</td><td>gene name</td><td>gene id</td></tr>
<tr><td>GU305914.1</td><td>30150</td><td>30172</td><td>+</td><td>0</td><td>CCCGGCGGCCTGGGGAGGGC</td><td>GGG</td><td>||||||||[||||||||||||]PAM</td><td>NA</td><td>Intergenic</td><td>NA</td><td>NA</td></tr>
<tr><td>T1831</td><td>GGCGGCCTGGGGAGGGCGGGTGG</td><td>1000</td><td></td><td></td><td></td><td></td><td></td><td></td><td></td><td></td><td></td></tr>
<tr><td>Oligo fwd</td><td>CACCGGCGGCCTGGGGAGGGCGGG</td><td></td><td></td><td></td><td></td><td></td><td></td><td></td><td></td><td></td><td></td></tr>
<tr><td>Oligo rev</td><td>AAACCCCGCCCTCCCCAGGCCGCC</td><td></td><td></td><td></td><td></td><td></td><td></td><td></td><td></td><td></td><td></td></tr>
<tr><td>Chromosome</td><td>start</td><td>end</td><td>strand</td><td>MM</td><td>target_seq</td><td>PAM</td><td>alignment</td><td>distance</td><td>position</td><td>gene name</td><td>gene id</td></tr>
<tr><td>GU305914.1</td><td>30153</td><td>30175</td><td>+</td><td>0</td><td>GGCGGCCTGGGGAGGGCGGG</td><td>TGG</td><td>||||||||[||||||||||||]PAM</td><td>NA</td><td>Intergenic</td><td>NA</td><td>NA</td></tr>
<tr><td>T1832</td><td>GGCCTGGGGAGGGCGGGTGGCGG</td><td>1000</td><td></td><td></td><td></td><td></td><td></td><td></td><td></td><td></td><td></td></tr>
<tr><td>Oligo fwd</td><td>CACCGGCCTGGGGAGGGCGGGTGG</td><td></td><td></td><td></td><td></td><td></td><td></td><td></td><td></td><td></td><td></td></tr>
<tr><td>Oligo rev</td><td>AAACCCACCCGCCCTCCCCAGGCC</td><td></td><td></td><td></td><td></td><td></td><td></td><td></td><td></td><td></td><td></td></tr>
<tr><td>Chromosome</td><td>start</td><td>end</td><td>strand</td><td>MM</td><td>target_seq</td><td>PAM</td><td>alignment</td><td>distance</td><td>position</td><td>gene name</td><td>gene id</td></tr>
<tr><td>GU305914.1</td><td>30156</td><td>30178</td><td>+</td><td>0</td><td>GGCCTGGGGAGGGCGGGTGG</td><td>CGG</td><td>||||||||[||||||||||||]PAM</td><td>NA</td><td>Intergenic</td><td>NA</td><td>NA</td></tr>
<tr><td>T1833</td><td>AGGGCGGGTGGCGGAGCGACCGG</td><td>1000</td><td></td><td></td><td></td><td></td><td></td><td></td><td></td><td></td><td></td></tr>
<tr><td>Oligo fwd</td><td>CACCAGGGCGGGTGGCGGAGCGAC</td><td></td><td></td><td></td><td></td><td></td><td></td><td></td><td></td><td></td><td></td></tr>
<tr><td>Oligo rev</td><td>AAACGTCGCTCCGCCACCCGCCCT</td><td></td><td></td><td></td><td></td><td></td><td></td><td></td><td></td><td></td><td></td></tr>
<tr><td>Chromosome</td><td>start</td><td>end</td><td>strand</td><td>MM</td><td>target_seq</td><td>PAM</td><td>alignment</td><td>distance</td><td>position</td><td>gene name</td><td>gene id</td></tr>
<tr><td>GU305914.1</td><td>30165</td><td>30187</td><td>+</td><td>0</td><td>AGGGCGGGTGGCGGAGCGAC</td><td>CGG</td><td>||||||||[||||||||||||]PAM</td><td>NA</td><td>Intergenic</td><td>NA</td><td>NA</td></tr>
<tr><td>T1834</td><td>GGAGCGACCGGACCCAGCCGCGG</td><td>1000</td><td></td><td></td><td></td><td></td><td></td><td></td><td></td><td></td><td></td></tr>
<tr><td>Oligo fwd</td><td>CACCGGAGCGACCGGACCCAGCCG</td><td></td><td></td><td></td><td></td><td></td><td></td><td></td><td></td><td></td><td></td></tr>
<tr><td>Oligo rev</td><td>AAACCGGCTGGGTCCGGTCGCTCC</td><td></td><td></td><td></td><td></td><td></td><td></td><td></td><td></td><td></td><td></td></tr>
<tr><td>Chromosome</td><td>start</td><td>end</td><td>strand</td><td>MM</td><td>target_seq</td><td>PAM</td><td>alignment</td><td>distance</td><td>position</td><td>gene name</td><td>gene id</td></tr>
<tr><td>GU305914.1</td><td>30177</td><td>30199</td><td>+</td><td>0</td><td>GGAGCGACCGGACCCAGCCG</td><td>CGG</td><td>||||||||[||||||||||||]PAM</td><td>NA</td><td>Intergenic</td><td>NA</td><td>NA</td></tr>
<tr><td>T1835</td><td>GAGCGACCGGACCCAGCCGCGGG</td><td>1000</td><td></td><td></td><td></td><td></td><td></td><td></td><td></td><td></td><td></td></tr>
<tr><td>Oligo fwd</td><td>CACCGAGCGACCGGACCCAGCCGC</td><td></td><td></td><td></td><td></td><td></td><td></td><td></td><td></td><td></td><td></td></tr>
<tr><td>Oligo rev</td><td>AAACGCGGCTGGGTCCGGTCGCTC</td><td></td><td></td><td></td><td></td><td></td><td></td><td></td><td></td><td></td><td></td></tr>
<tr><td>Chromosome</td><td>start</td><td>end</td><td>strand</td><td>MM</td><td>target_seq</td><td>PAM</td><td>alignment</td><td>distance</td><td>position</td><td>gene name</td><td>gene id</td></tr>
<tr><td>GU305914.1</td><td>30178</td><td>30200</td><td>+</td><td>0</td><td>GAGCGACCGGACCCAGCCGC</td><td>GGG</td><td>||||||||[||||||||||||]PAM</td><td>NA</td><td>Intergenic</td><td>NA</td><td>NA</td></tr>
<tr><td>T1836</td><td>GACCCAGCCGCGGGATGTAGCGG</td><td>1000</td><td></td><td></td><td></td><td></td><td></td><td></td><td></td><td></td><td></td></tr>
<tr><td>Oligo fwd</td><td>CACCGACCCAGCCGCGGGATGTAG</td><td></td><td></td><td></td><td></td><td></td><td></td><td></td><td></td><td></td><td></td></tr>
<tr><td>Oligo rev</td><td>AAACCTACATCCCGCGGCTGGGTC</td><td></td><td></td><td></td><td></td><td></td><td></td><td></td><td></td><td></td><td></td></tr>
<tr><td>Chromosome</td><td>start</td><td>end</td><td>strand</td><td>MM</td><td>target_seq</td><td>PAM</td><td>alignment</td><td>distance</td><td>position</td><td>gene name</td><td>gene id</td></tr>
<tr><td>GU305914.1</td><td>30187</td><td>30209</td><td>+</td><td>0</td><td>GACCCAGCCGCGGGATGTAG</td><td>CGG</td><td>||||||||[||||||||||||]PAM</td><td>NA</td><td>Intergenic</td><td>NA</td><td>NA</td></tr>
<tr><td>T1837</td><td>CTGTCGTCCCCACATCTGCTCGG</td><td>1000</td><td></td><td></td><td></td><td></td><td></td><td></td><td></td><td></td><td></td></tr>
<tr><td>Oligo fwd</td><td>CACCCTGTCGTCCCCACATCTGCT</td><td></td><td></td><td></td><td></td><td></td><td></td><td></td><td></td><td></td><td></td></tr>
<tr><td>Oligo rev</td><td>AAACAGCAGATGTGGGGACGACAG</td><td></td><td></td><td></td><td></td><td></td><td></td><td></td><td></td><td></td><td></td></tr>
<tr><td>Chromosome</td><td>start</td><td>end</td><td>strand</td><td>MM</td><td>target_seq</td><td>PAM</td><td>alignment</td><td>distance</td><td>position</td><td>gene name</td><td>gene id</td></tr>
<tr><td>GU305914.1</td><td>30266</td><td>30288</td><td>+</td><td>0</td><td>CTGTCGTCCCCACATCTGCT</td><td>CGG</td><td>||||||||[||||||||||||]PAM</td><td>NA</td><td>Intergenic</td><td>NA</td><td>NA</td></tr>
<tr><td>T1838</td><td>GAGTCTGAGCAGCTTTGACACGG</td><td>1000</td><td></td><td></td><td></td><td></td><td></td><td></td><td></td><td></td><td></td></tr>
<tr><td>Oligo fwd</td><td>CACCGAGTCTGAGCAGCTTTGACA</td><td></td><td></td><td></td><td></td><td></td><td></td><td></td><td></td><td></td><td></td></tr>
<tr><td>Oligo rev</td><td>AAACTGTCAAAGCTGCTCAGACTC</td><td></td><td></td><td></td><td></td><td></td><td></td><td></td><td></td><td></td><td></td></tr>
<tr><td>Chromosome</td><td>start</td><td>end</td><td>strand</td><td>MM</td><td>target_seq</td><td>PAM</td><td>alignment</td><td>distance</td><td>position</td><td>gene name</td><td>gene id</td></tr>
<tr><td>GU305914.1</td><td>30355</td><td>30377</td><td>+</td><td>0</td><td>GAGTCTGAGCAGCTTTGACA</td><td>CGG</td><td>||||||||[||||||||||||]PAM</td><td>NA</td><td>Intergenic</td><td>NA</td><td>NA</td></tr>
<tr><td>T1839</td><td>CTTTGACACGGCGTGATGAATGG</td><td>1000</td><td></td><td></td><td></td><td></td><td></td><td></td><td></td><td></td><td></td></tr>
<tr><td>Oligo fwd</td><td>CACCCTTTGACACGGCGTGATGAA</td><td></td><td></td><td></td><td></td><td></td><td></td><td></td><td></td><td></td><td></td></tr>
<tr><td>Oligo rev</td><td>AAACTTCATCACGCCGTGTCAAAG</td><td></td><td></td><td></td><td></td><td></td><td></td><td></td><td></td><td></td><td></td></tr>
<tr><td>Chromosome</td><td>start</td><td>end</td><td>strand</td><td>MM</td><td>target_seq</td><td>PAM</td><td>alignment</td><td>distance</td><td>position</td><td>gene name</td><td>gene id</td></tr>
<tr><td>GU305914.1</td><td>30367</td><td>30389</td><td>+</td><td>0</td><td>CTTTGACACGGCGTGATGAA</td><td>TGG</td><td>||||||||[||||||||||||]PAM</td><td>NA</td><td>Intergenic</td><td>NA</td><td>NA</td></tr>
<tr><td>T1840</td><td>AATGGCATAGACCGCGTGATCGG</td><td>1000</td><td></td><td></td><td></td><td></td><td></td><td></td><td></td><td></td><td></td></tr>
<tr><td>Oligo fwd</td><td>CACCAATGGCATAGACCGCGTGAT</td><td></td><td></td><td></td><td></td><td></td><td></td><td></td><td></td><td></td><td></td></tr>
<tr><td>Oligo rev</td><td>AAACATCACGCGGTCTATGCCATT</td><td></td><td></td><td></td><td></td><td></td><td></td><td></td><td></td><td></td><td></td></tr>
<tr><td>Chromosome</td><td>start</td><td>end</td><td>strand</td><td>MM</td><td>target_seq</td><td>PAM</td><td>alignment</td><td>distance</td><td>position</td><td>gene name</td><td>gene id</td></tr>
<tr><td>GU305914.1</td><td>30385</td><td>30407</td><td>+</td><td>0</td><td>AATGGCATAGACCGCGTGAT</td><td>CGG</td><td>||||||||[||||||||||||]PAM</td><td>NA</td><td>Intergenic</td><td>NA</td><td>NA</td></tr>
<tr><td>T1841</td><td>GTGATCGGCCCCAATGAAAACGG</td><td>1000</td><td></td><td></td><td></td><td></td><td></td><td></td><td></td><td></td><td></td></tr>
<tr><td>Oligo fwd</td><td>CACCGTGATCGGCCCCAATGAAAA</td><td></td><td></td><td></td><td></td><td></td><td></td><td></td><td></td><td></td><td></td></tr>
<tr><td>Oligo rev</td><td>AAACTTTTCATTGGGGCCGATCAC</td><td></td><td></td><td></td><td></td><td></td><td></td><td></td><td></td><td></td><td></td></tr>
<tr><td>Chromosome</td><td>start</td><td>end</td><td>strand</td><td>MM</td><td>target_seq</td><td>PAM</td><td>alignment</td><td>distance</td><td>position</td><td>gene name</td><td>gene id</td></tr>
<tr><td>GU305914.1</td><td>30400</td><td>30422</td><td>+</td><td>0</td><td>GTGATCGGCCCCAATGAAAA</td><td>CGG</td><td>||||||||[||||||||||||]PAM</td><td>NA</td><td>Intergenic</td><td>NA</td><td>NA</td></tr>
<tr><td>T1842</td><td>CCAATGAAAACGGCCGTACGCGG</td><td>1000</td><td></td><td></td><td></td><td></td><td></td><td></td><td></td><td></td><td></td></tr>
<tr><td>Oligo fwd</td><td>CACCCCAATGAAAACGGCCGTACG</td><td></td><td></td><td></td><td></td><td></td><td></td><td></td><td></td><td></td><td></td></tr>
<tr><td>Oligo rev</td><td>AAACCGTACGGCCGTTTTCATTGG</td><td></td><td></td><td></td><td></td><td></td><td></td><td></td><td></td><td></td><td></td></tr>
<tr><td>Chromosome</td><td>start</td><td>end</td><td>strand</td><td>MM</td><td>target_seq</td><td>PAM</td><td>alignment</td><td>distance</td><td>position</td><td>gene name</td><td>gene id</td></tr>
<tr><td>GU305914.1</td><td>30410</td><td>30432</td><td>+</td><td>0</td><td>CCAATGAAAACGGCCGTACG</td><td>CGG</td><td>||||||||[||||||||||||]PAM</td><td>NA</td><td>Intergenic</td><td>NA</td><td>NA</td></tr>
<tr><td>T1843</td><td>GAAAACGGCCGTACGCGGTCTGG</td><td>1000</td><td></td><td></td><td></td><td></td><td></td><td></td><td></td><td></td><td></td></tr>
<tr><td>Oligo fwd</td><td>CACCGAAAACGGCCGTACGCGGTC</td><td></td><td></td><td></td><td></td><td></td><td></td><td></td><td></td><td></td><td></td></tr>
<tr><td>Oligo rev</td><td>AAACGACCGCGTACGGCCGTTTTC</td><td></td><td></td><td></td><td></td><td></td><td></td><td></td><td></td><td></td><td></td></tr>
<tr><td>Chromosome</td><td>start</td><td>end</td><td>strand</td><td>MM</td><td>target_seq</td><td>PAM</td><td>alignment</td><td>distance</td><td>position</td><td>gene name</td><td>gene id</td></tr>
<tr><td>GU305914.1</td><td>30415</td><td>30437</td><td>+</td><td>0</td><td>GAAAACGGCCGTACGCGGTC</td><td>TGG</td><td>||||||||[||||||||||||]PAM</td><td>NA</td><td>Intergenic</td><td>NA</td><td>NA</td></tr>
</table>

<table>
<tr><td>T1844</td><td>CAGTCGTAAAGCGTCGATGCAGG</td><td>1000</td><td></td><td></td><td></td><td></td><td></td><td></td><td></td><td></td><td></td></tr>
<tr><td>Oligo fwd</td><td>CACCCAGTCGTAAAGCGTCGATGC</td><td></td><td></td><td></td><td></td><td></td><td></td><td></td><td></td><td></td><td></td></tr>
<tr><td>Oligo rev</td><td>AAACGCATCGACGCTTTACGACTG</td><td></td><td></td><td></td><td></td><td></td><td></td><td></td><td></td><td></td><td></td></tr>
<tr><td>Chromosome</td><td>start</td><td>end</td><td>strand</td><td>MM</td><td>target_seq</td><td>PAM</td><td>alignment</td><td>distance</td><td>position</td><td>gene name</td><td>gene id</td></tr>
<tr><td>GU305914.1</td><td>30481</td><td>30503</td><td>+</td><td>0</td><td>CAGTCGTAAAGCGTCGATGC</td><td>AGG</td><td>||||||||[||||||||||||]PAM</td><td>NA</td><td>Intergenic</td><td>NA</td><td>NA</td></tr>
<tr><td>T1845</td><td>AGTCGTAAAGCGTCGATGCAGGG</td><td>1000</td><td></td><td></td><td></td><td></td><td></td><td></td><td></td><td></td><td></td></tr>
<tr><td>Oligo fwd</td><td>CACCAGTCGTAAAGCGTCGATGCA</td><td></td><td></td><td></td><td></td><td></td><td></td><td></td><td></td><td></td><td></td></tr>
<tr><td>Oligo rev</td><td>AAACTGCATCGACGCTTTACGACT</td><td></td><td></td><td></td><td></td><td></td><td></td><td></td><td></td><td></td><td></td></tr>
<tr><td>Chromosome</td><td>start</td><td>end</td><td>strand</td><td>MM</td><td>target_seq</td><td>PAM</td><td>alignment</td><td>distance</td><td>position</td><td>gene name</td><td>gene id</td></tr>
<tr><td>GU305914.1</td><td>30482</td><td>30504</td><td>+</td><td>0</td><td>AGTCGTAAAGCGTCGATGCA</td><td>GGG</td><td>||||||||[||||||||||||]PAM</td><td>NA</td><td>Intergenic</td><td>NA</td><td>NA</td></tr>
<tr><td>T1846</td><td>GTCGTAAAGCGTCGATGCAGGGG</td><td>1000</td><td></td><td></td><td></td><td></td><td></td><td></td><td></td><td></td><td></td></tr>
<tr><td>Oligo fwd</td><td>CACCGTCGTAAAGCGTCGATGCAG</td><td></td><td></td><td></td><td></td><td></td><td></td><td></td><td></td><td></td><td></td></tr>
<tr><td>Oligo rev</td><td>AAACCTGCATCGACGCTTTACGAC</td><td></td><td></td><td></td><td></td><td></td><td></td><td></td><td></td><td></td><td></td></tr>
<tr><td>Chromosome</td><td>start</td><td>end</td><td>strand</td><td>MM</td><td>target_seq</td><td>PAM</td><td>alignment</td><td>distance</td><td>position</td><td>gene name</td><td>gene id</td></tr>
<tr><td>GU305914.1</td><td>30483</td><td>30505</td><td>+</td><td>0</td><td>GTCGTAAAGCGTCGATGCAG</td><td>GGG</td><td>||||||||[||||||||||||]PAM</td><td>NA</td><td>Intergenic</td><td>NA</td><td>NA</td></tr>
<tr><td>T1847</td><td>GGGGCTTTGCTCTTTGTTTAAGG</td><td>1000</td><td></td><td></td><td></td><td></td><td></td><td></td><td></td><td></td><td></td></tr>
<tr><td>Oligo fwd</td><td>CACCGGGGCTTTGCTCTTTGTTTA</td><td></td><td></td><td></td><td></td><td></td><td></td><td></td><td></td><td></td><td></td></tr>
<tr><td>Oligo rev</td><td>AAACTAAACAAAGAGCAAAGCCCC</td><td></td><td></td><td></td><td></td><td></td><td></td><td></td><td></td><td></td><td></td></tr>
<tr><td>Chromosome</td><td>start</td><td>end</td><td>strand</td><td>MM</td><td>target_seq</td><td>PAM</td><td>alignment</td><td>distance</td><td>position</td><td>gene name</td><td>gene id</td></tr>
<tr><td>GU305914.1</td><td>30502</td><td>30524</td><td>+</td><td>0</td><td>GGGGCTTTGCTCTTTGTTTA</td><td>AGG</td><td>||||||||[||||||||||||]PAM</td><td>NA</td><td>Intergenic</td><td>NA</td><td>NA</td></tr>
<tr><td>T1848</td><td>TTGCTCTTTGTTTAAGGATGAGG</td><td>1000</td><td></td><td></td><td></td><td></td><td></td><td></td><td></td><td></td><td></td></tr>
<tr><td>Oligo fwd</td><td>CACCTTGCTCTTTGTTTAAGGATG</td><td></td><td></td><td></td><td></td><td></td><td></td><td></td><td></td><td></td><td></td></tr>
<tr><td>Oligo rev</td><td>AAACCATCCTTAAACAAAGAGCAA</td><td></td><td></td><td></td><td></td><td></td><td></td><td></td><td></td><td></td><td></td></tr>
<tr><td>Chromosome</td><td>start</td><td>end</td><td>strand</td><td>MM</td><td>target_seq</td><td>PAM</td><td>alignment</td><td>distance</td><td>position</td><td>gene name</td><td>gene id</td></tr>
<tr><td>GU305914.1</td><td>30508</td><td>30530</td><td>+</td><td>0</td><td>TTGCTCTTTGTTTAAGGATG</td><td>AGG</td><td>||||||||[||||||||||||]PAM</td><td>NA</td><td>Intergenic</td><td>NA</td><td>NA</td></tr>
<tr><td>T1849</td><td>TTTGTTTAAGGATGAGGCGACGG</td><td>1000</td><td></td><td></td><td></td><td></td><td></td><td></td><td></td><td></td><td></td></tr>
<tr><td>Oligo fwd</td><td>CACCTTTGTTTAAGGATGAGGCGA</td><td></td><td></td><td></td><td></td><td></td><td></td><td></td><td></td><td></td><td></td></tr>
<tr><td>Oligo rev</td><td>AAACTCGCCTCATCCTTAAACAAA</td><td></td><td></td><td></td><td></td><td></td><td></td><td></td><td></td><td></td><td></td></tr>
<tr><td>Chromosome</td><td>start</td><td>end</td><td>strand</td><td>MM</td><td>target_seq</td><td>PAM</td><td>alignment</td><td>distance</td><td>position</td><td>gene name</td><td>gene id</td></tr>
<tr><td>GU305914.1</td><td>30514</td><td>30536</td><td>+</td><td>0</td><td>TTTGTTTAAGGATGAGGCGA</td><td>CGG</td><td>||||||||[||||||||||||]PAM</td><td>NA</td><td>Intergenic</td><td>NA</td><td>NA</td></tr>
<tr><td>T1850</td><td>TAAGGATGAGGCGACGGCGTTGG</td><td>1000</td><td></td><td></td><td></td><td></td><td></td><td></td><td></td><td></td><td></td></tr>
<tr><td>Oligo fwd</td><td>CACCTAAGGATGAGGCGACGGCGT</td><td></td><td></td><td></td><td></td><td></td><td></td><td></td><td></td><td></td><td></td></tr>
<tr><td>Oligo rev</td><td>AAACACGCCGTCGCCTCATCCTTA</td><td></td><td></td><td></td><td></td><td></td><td></td><td></td><td></td><td></td><td></td></tr>
<tr><td>Chromosome</td><td>start</td><td>end</td><td>strand</td><td>MM</td><td>target_seq</td><td>PAM</td><td>alignment</td><td>distance</td><td>position</td><td>gene name</td><td>gene id</td></tr>
<tr><td>GU305914.1</td><td>30520</td><td>30542</td><td>+</td><td>0</td><td>TAAGGATGAGGCGACGGCGT</td><td>TGG</td><td>||||||||[||||||||||||]PAM</td><td>NA</td><td>Intergenic</td><td>NA</td><td>NA</td></tr>
<tr><td>T1851</td><td>GCGACGGCGTTGGATGATTCCGG</td><td>1000</td><td></td><td></td><td></td><td></td><td></td><td></td><td></td><td></td><td></td></tr>
<tr><td>Oligo fwd</td><td>CACCGCGACGGCGTTGGATGATTC</td><td></td><td></td><td></td><td></td><td></td><td></td><td></td><td></td><td></td><td></td></tr>
<tr><td>Oligo rev</td><td>AAACGAATCATCCAACGCCGTCGC</td><td></td><td></td><td></td><td></td><td></td><td></td><td></td><td></td><td></td><td></td></tr>
<tr><td>Chromosome</td><td>start</td><td>end</td><td>strand</td><td>MM</td><td>target_seq</td><td>PAM</td><td>alignment</td><td>distance</td><td>position</td><td>gene name</td><td>gene id</td></tr>
<tr><td>GU305914.1</td><td>30530</td><td>30552</td><td>+</td><td>0</td><td>GCGACGGCGTTGGATGATTC</td><td>CGG</td><td>||||||||[||||||||||||]PAM</td><td>NA</td><td>Intergenic</td><td>NA</td><td>NA</td></tr>
<tr><td>T1852</td><td>CGACGGCGTTGGATGATTCCGGG</td><td>1000</td><td></td><td></td><td></td><td></td><td></td><td></td><td></td><td></td><td></td></tr>
<tr><td>Oligo fwd</td><td>CACCCGACGGCGTTGGATGATTCC</td><td></td><td></td><td></td><td></td><td></td><td></td><td></td><td></td><td></td><td></td></tr>
<tr><td>Oligo rev</td><td>AAACGGAATCATCCAACGCCGTCG</td><td></td><td></td><td></td><td></td><td></td><td></td><td></td><td></td><td></td><td></td></tr>
<tr><td>Chromosome</td><td>start</td><td>end</td><td>strand</td><td>MM</td><td>target_seq</td><td>PAM</td><td>alignment</td><td>distance</td><td>position</td><td>gene name</td><td>gene id</td></tr>
<tr><td>GU305914.1</td><td>30531</td><td>30553</td><td>+</td><td>0</td><td>CGACGGCGTTGGATGATTCC</td><td>GGG</td><td>||||||||[||||||||||||]PAM</td><td>NA</td><td>Intergenic</td><td>NA</td><td>NA</td></tr>
<tr><td>T1853</td><td>GACGGCGTTGGATGATTCCGGGG</td><td>1000</td><td></td><td></td><td></td><td></td><td></td><td></td><td></td><td></td><td></td></tr>
<tr><td>Oligo fwd</td><td>CACCGACGGCGTTGGATGATTCCG</td><td></td><td></td><td></td><td></td><td></td><td></td><td></td><td></td><td></td><td></td></tr>
<tr><td>Oligo rev</td><td>AAACCGGAATCATCCAACGCCGTC</td><td></td><td></td><td></td><td></td><td></td><td></td><td></td><td></td><td></td><td></td></tr>
<tr><td>Chromosome</td><td>start</td><td>end</td><td>strand</td><td>MM</td><td>target_seq</td><td>PAM</td><td>alignment</td><td>distance</td><td>position</td><td>gene name</td><td>gene id</td></tr>
<tr><td>GU305914.1</td><td>30532</td><td>30554</td><td>+</td><td>0</td><td>GACGGCGTTGGATGATTCCG</td><td>GGG</td><td>||||||||[||||||||||||]PAM</td><td>NA</td><td>Intergenic</td><td>NA</td><td>NA</td></tr>
<tr><td>T1854</td><td>ACGGCGTTGGATGATTCCGGGGG</td><td>1000</td><td></td><td></td><td></td><td></td><td></td><td></td><td></td><td></td><td></td></tr>
<tr><td>Oligo fwd</td><td>CACCACGGCGTTGGATGATTCCGG</td><td></td><td></td><td></td><td></td><td></td><td></td><td></td><td></td><td></td><td></td></tr>
<tr><td>Oligo rev</td><td>AAACCCGGAATCATCCAACGCCGT</td><td></td><td></td><td></td><td></td><td></td><td></td><td></td><td></td><td></td><td></td></tr>
<tr><td>Chromosome</td><td>start</td><td>end</td><td>strand</td><td>MM</td><td>target_seq</td><td>PAM</td><td>alignment</td><td>distance</td><td>position</td><td>gene name</td><td>gene id</td></tr>
<tr><td>GU305914.1</td><td>30533</td><td>30555</td><td>+</td><td>0</td><td>ACGGCGTTGGATGATTCCGG</td><td>GGG</td><td>||||||||[||||||||||||]PAM</td><td>NA</td><td>Intergenic</td><td>NA</td><td>NA</td></tr>
<tr><td>T1855</td><td>AAAACGAAGTCGTAATCTTCTGG</td><td>1000</td><td></td><td></td><td></td><td></td><td></td><td></td><td></td><td></td><td></td></tr>
<tr><td>Oligo fwd</td><td>CACCAAAACGAAGTCGTAATCTTC</td><td></td><td></td><td></td><td></td><td></td><td></td><td></td><td></td><td></td><td></td></tr>
<tr><td>Oligo rev</td><td>AAACGAAGATTACGACTTCGTTTT</td><td></td><td></td><td></td><td></td><td></td><td></td><td></td><td></td><td></td><td></td></tr>
<tr><td>Chromosome</td><td>start</td><td>end</td><td>strand</td><td>MM</td><td>target_seq</td><td>PAM</td><td>alignment</td><td>distance</td><td>position</td><td>gene name</td><td>gene id</td></tr>
<tr><td>GU305914.1</td><td>30563</td><td>30585</td><td>+</td><td>0</td><td>AAAACGAAGTCGTAATCTTC</td><td>TGG</td><td>||||||||[||||||||||||]PAM</td><td>NA</td><td>Intergenic</td><td>NA</td><td>NA</td></tr>
<tr><td>T1856</td><td>CGTAATCTTCTGGCCACTGTAGG</td><td>1000</td><td></td><td></td><td></td><td></td><td></td><td></td><td></td><td></td><td></td></tr>
<tr><td>Oligo fwd</td><td>CACCCGTAATCTTCTGGCCACTGT</td><td></td><td></td><td></td><td></td><td></td><td></td><td></td><td></td><td></td><td></td></tr>
<tr><td>Oligo rev</td><td>AAACACAGTGGCCAGAAGATTACG</td><td></td><td></td><td></td><td></td><td></td><td></td><td></td><td></td><td></td><td></td></tr>
<tr><td>Chromosome</td><td>start</td><td>end</td><td>strand</td><td>MM</td><td>target_seq</td><td>PAM</td><td>alignment</td><td>distance</td><td>position</td><td>gene name</td><td>gene id</td></tr>
<tr><td>GU305914.1</td><td>30573</td><td>30595</td><td>+</td><td>0</td><td>CGTAATCTTCTGGCCACTGT</td><td>AGG</td><td>||||||||[||||||||||||]PAM</td><td>NA</td><td>Intergenic</td><td>NA</td><td>NA</td></tr>
<tr><td>T1857</td><td>GTAATCTTCTGGCCACTGTAGGG</td><td>1000</td><td></td><td></td><td></td><td></td><td></td><td></td><td></td><td></td><td></td></tr>
<tr><td>Oligo fwd</td><td>CACCGTAATCTTCTGGCCACTGTA</td><td></td><td></td><td></td><td></td><td></td><td></td><td></td><td></td><td></td><td></td></tr>
<tr><td>Oligo rev</td><td>AAACTACAGTGGCCAGAAGATTAC</td><td></td><td></td><td></td><td></td><td></td><td></td><td></td><td></td><td></td><td></td></tr>
<tr><td>Chromosome</td><td>start</td><td>end</td><td>strand</td><td>MM</td><td>target_seq</td><td>PAM</td><td>alignment</td><td>distance</td><td>position</td><td>gene name</td><td>gene id</td></tr>
<tr><td>GU305914.1</td><td>30574</td><td>30596</td><td>+</td><td>0</td><td>GTAATCTTCTGGCCACTGTA</td><td>GGG</td><td>||||||||[||||||||||||]PAM</td><td>NA</td><td>Intergenic</td><td>NA</td><td>NA</td></tr>
<tr><td>T1858</td><td>TACCAGCCCCCGTTCCGCCGCGG</td><td>1000</td><td></td><td></td><td></td><td></td><td></td><td></td><td></td><td></td><td></td></tr>
<tr><td>Oligo fwd</td><td>CACCTACCAGCCCCCGTTCCGCCG</td><td></td><td></td><td></td><td></td><td></td><td></td><td></td><td></td><td></td><td></td></tr>
<tr><td>Oligo rev</td><td>AAACCGGCGGAACGGGGGCTGGTA</td><td></td><td></td><td></td><td></td><td></td><td></td><td></td><td></td><td></td><td></td></tr>
<tr><td>Chromosome</td><td>start</td><td>end</td><td>strand</td><td>MM</td><td>target_seq</td><td>PAM</td><td>alignment</td><td>distance</td><td>position</td><td>gene name</td><td>gene id</td></tr>
<tr><td>GU305914.1</td><td>30601</td><td>30623</td><td>+</td><td>0</td><td>TACCAGCCCCCGTTCCGCCG</td><td>CGG</td><td>||||||||[||||||||||||]PAM</td><td>NA</td><td>Intergenic</td><td>NA</td><td>NA</td></tr>
</table>

| | | | | | | | | | | | |
|---|---|---|---|---|---|---|---|---|---|---|---|
| T1859 | AGCGCCTCGTCCGCGTTCCGAGG | 1000 | | | | | | | | | |
| Oligo fwd | CACCAGCGCCTCGTCCGCGTTCCG | | | | | | | | | | |
| Oligo rev | AAACCGGAACGCGGACGAGGCGCT | | | | | | | | | | |
| Chromosome | start | end | strand | MM | target_seq | PAM | alignment | distance | position | gene name | gene id |
| GU305914.1 | 30626 | 30648 | + | 0 | AGCGCCTCGTCCGCGTTCCG | AGG | ||||||||[||||||||||||]PAM | NA | Intergenic | NA | NA |

| | | | | | | | | | | | |
|---|---|---|---|---|---|---|---|---|---|---|---|
| T1860 | ACGCGTGTCCGTCCTTCACCCGG | 1000 | | | | | | | | | |
| Oligo fwd | CACCACGCGTGTCCGTCCTTCACC | | | | | | | | | | |
| Oligo rev | AAACGGTGAAGGACGGACACGCGT | | | | | | | | | | |
| Chromosome | start | end | strand | MM | target_seq | PAM | alignment | distance | position | gene name | gene id |
| GU305914.1 | 30654 | 30676 | + | 0 | ACGCGTGTCCGTCCTTCACC | CGG | ||||||||[||||||||||||]PAM | NA | Intergenic | NA | NA |

| | | | | | | | | | | | |
|---|---|---|---|---|---|---|---|---|---|---|---|
| T1861 | GATTGCATCCTCGACACAGCAGG | 1000 | | | | | | | | | |
| Oligo fwd | CACCGATTGCATCCTCGACACAGC | | | | | | | | | | |
| Oligo rev | AAACGCTGTGTCGAGGATGCAATC | | | | | | | | | | |
| Chromosome | start | end | strand | MM | target_seq | PAM | alignment | distance | position | gene name | gene id |
| GU305914.1 | 30676 | 30698 | + | 0 | GATTGCATCCTCGACACAGC | AGG | ||||||||[||||||||||||]PAM | NA | Intergenic | NA | NA |

| | | | | | | | | | | | |
|---|---|---|---|---|---|---|---|---|---|---|---|
| T1862 | ATTGCATCCTCGACACAGCAGGG | 1000 | | | | | | | | | |
| Oligo fwd | CACCATTGCATCCTCGACACAGCA | | | | | | | | | | |
| Oligo rev | AAACTGCTGTGTCGAGGATGCAAT | | | | | | | | | | |
| Chromosome | start | end | strand | MM | target_seq | PAM | alignment | distance | position | gene name | gene id |
| GU305914.1 | 30677 | 30699 | + | 0 | ATTGCATCCTCGACACAGCA | GGG | ||||||||[||||||||||||]PAM | NA | Intergenic | NA | NA |

| | | | | | | | | | | | |
|---|---|---|---|---|---|---|---|---|---|---|---|
| T1863 | CACGAAGCCGATCATAGTGTTGG | 1000 | | | | | | | | | |
| Oligo fwd | CACCCACGAAGCCGATCATAGTGT | | | | | | | | | | |
| Oligo rev | AAACACACTATGATCGGCTTCGTG | | | | | | | | | | |
| Chromosome | start | end | strand | MM | target_seq | PAM | alignment | distance | position | gene name | gene id |
| GU305914.1 | 30739 | 30761 | + | 0 | CACGAAGCCGATCATAGTGT | TGG | ||||||||[||||||||||||]PAM | NA | Intergenic | NA | NA |

| | | | | | | | | | | | |
|---|---|---|---|---|---|---|---|---|---|---|---|
| T1864 | AAGCCTATATAGTCCCCCGTAGG | 1000 | | | | | | | | | |
| Oligo fwd | CACCAAGCCTATATAGTCCCCCGT | | | | | | | | | | |
| Oligo rev | AAACACGGGGGACTATATAGGCTT | | | | | | | | | | |
| Chromosome | start | end | strand | MM | target_seq | PAM | alignment | distance | position | gene name | gene id |
| GU305914.1 | 30763 | 30785 | + | 0 | AAGCCTATATAGTCCCCCGT | AGG | ||||||||[||||||||||||]PAM | NA | Intergenic | NA | NA |

| | | | | | | | | | | | |
|---|---|---|---|---|---|---|---|---|---|---|---|
| T1865 | AGCCTATATAGTCCCCCGTAGGG | 1000 | | | | | | | | | |
| Oligo fwd | CACCAGCCTATATAGTCCCCCGTA | | | | | | | | | | |
| Oligo rev | AAACTACGGGGGACTATATAGGCT | | | | | | | | | | |
| Chromosome | start | end | strand | MM | target_seq | PAM | alignment | distance | position | gene name | gene id |
| GU305914.1 | 30764 | 30786 | + | 0 | AGCCTATATAGTCCCCCGTA | GGG | ||||||||[||||||||||||]PAM | NA | Intergenic | NA | NA |

| | | | | | | | | | | | |
|---|---|---|---|---|---|---|---|---|---|---|---|
| T1866 | TATAGTCCCCCGTAGGGCGCCGG | 1000 | | | | | | | | | |
| Oligo fwd | CACCTATAGTCCCCCGTAGGGCGC | | | | | | | | | | |
| Oligo rev | AAACGCGCCCTACGGGGGACTATA | | | | | | | | | | |
| Chromosome | start | end | strand | MM | target_seq | PAM | alignment | distance | position | gene name | gene id |
| GU305914.1 | 30770 | 30792 | + | 0 | TATAGTCCCCCGTAGGGCGC | CGG | ||||||||[||||||||||||]PAM | NA | Intergenic | NA | NA |

| | | | | | | | | | | | |
|---|---|---|---|---|---|---|---|---|---|---|---|
| T1867 | ATAGTCCCCCGTAGGGCGCCGGG | 1000 | | | | | | | | | |
| Oligo fwd | CACCATAGTCCCCCGTAGGGCGCC | | | | | | | | | | |
| Oligo rev | AAACGGCGCCCTACGGGGGACTAT | | | | | | | | | | |
| Chromosome | start | end | strand | MM | target_seq | PAM | alignment | distance | position | gene name | gene id |
| GU305914.1 | 30771 | 30793 | + | 0 | ATAGTCCCCCGTAGGGCGCC | GGG | ||||||||[||||||||||||]PAM | NA | Intergenic | NA | NA |

| | | | | | | | | | | | |
|---|---|---|---|---|---|---|---|---|---|---|---|
| T1868 | TAGATGAAGACCGCGCTCGACGG | 1000 | | | | | | | | | |
| Oligo fwd | CACCTAGATGAAGACCGCGCTCGA | | | | | | | | | | |
| Oligo rev | AAACTCGAGCGCGGTCTTCATCTA | | | | | | | | | | |
| Chromosome | start | end | strand | MM | target_seq | PAM | alignment | distance | position | gene name | gene id |
| GU305914.1 | 30797 | 30819 | + | 0 | TAGATGAAGACCGCGCTCGA | CGG | ||||||||[||||||||||||]PAM | NA | Intergenic | NA | NA |

| | | | | | | | | | | | |
|---|---|---|---|---|---|---|---|---|---|---|---|
| T1869 | TGCACCTGTCGTCACCGTCGCGG | 1000 | | | | | | | | | |
| Oligo fwd | CACCTGCACCTGTCGTCACCGTCG | | | | | | | | | | |
| Oligo rev | AAACCGACGGTGACGACAGGTGCA | | | | | | | | | | |
| Chromosome | start | end | strand | MM | target_seq | PAM | alignment | distance | position | gene name | gene id |
| GU305914.1 | 30834 | 30856 | + | 0 | TGCACCTGTCGTCACCGTCG | CGG | ||||||||[||||||||||||]PAM | NA | Intergenic | NA | NA |

| | | | | | | | | | | | |
|---|---|---|---|---|---|---|---|---|---|---|---|
| T1870 | GACGACGACCAGCCTGTCGACGG | 1000 | | | | | | | | | |
| Oligo fwd | CACCGACGACGACCAGCCTGTCGA | | | | | | | | | | |
| Oligo rev | AAACTCGACAGGCTGGTCGTCGTC | | | | | | | | | | |
| Chromosome | start | end | strand | MM | target_seq | PAM | alignment | distance | position | gene name | gene id |
| GU305914.1 | 30886 | 30908 | + | 0 | GACGACGACCAGCCTGTCGA | CGG | ||||||||[||||||||||||]PAM | NA | Intergenic | NA | NA |
| GU305914.1 | 146852 | 146874 | - | 4 | GACGGCGTCCAGGGTGTCGA | CGG | ||||-||-[||||--||||||]PAM | NA | Intergenic | NA | NA |

| | | | | | | | | | | | |
|---|---|---|---|---|---|---|---|---|---|---|---|
| T1871 | GACGACCAGCCTGTCGACGGCGG | 1000 | | | | | | | | | |
| Oligo fwd | CACCGACGACCAGCCTGTCGACGG | | | | | | | | | | |
| Oligo rev | AAACCCGTCGACAGGCTGGTCGTC | | | | | | | | | | |
| Chromosome | start | end | strand | MM | target_seq | PAM | alignment | distance | position | gene name | gene id |
| GU305914.1 | 30889 | 30911 | + | 0 | GACGACCAGCCTGTCGACGG | CGG | ||||||||[||||||||||||]PAM | NA | Intergenic | NA | NA |

| | | | | | | | | | | | |
|---|---|---|---|---|---|---|---|---|---|---|---|
| T1872 | ACGACCAGCCTGTCGACGGCGGG | 1000 | | | | | | | | | |
| Oligo fwd | CACCACGACCAGCCTGTCGACGGC | | | | | | | | | | |
| Oligo rev | AAACGCCGTCGACAGGCTGGTCGT | | | | | | | | | | |
| Chromosome | start | end | strand | MM | target_seq | PAM | alignment | distance | position | gene name | gene id |
| GU305914.1 | 30890 | 30912 | + | 0 | ACGACCAGCCTGTCGACGGC | GGG | ||||||||[||||||||||||]PAM | NA | Intergenic | NA | NA |

| | | |
|---|---|---|
| T1873 | CGGCGGGTTCGAGCGCCCGCAGG | 1000 |
| Oligo fwd | CACCCGGCGGGTTCGAGCGCCCGC | |
| Oligo rev | AAACGCGGGCGCTCGAACCCGCCG | |

<table>
<tr><td>Chromosome</td><td>start</td><td>end</td><td>strand</td><td>MM</td><td>target_seq</td><td>PAM</td><td>alignment</td><td>distance</td><td>position</td><td>gene name</td><td>gene id</td></tr>
<tr><td>GU305914.1</td><td>30906</td><td>30928</td><td>+</td><td>0</td><td>CGGCGGGTTCGAGCGCCCGC</td><td>AGG</td><td>||||||||[||||||||||||]PAM</td><td>NA</td><td>Intergenic</td><td>NA</td><td>NA</td></tr>
<tr><td>T1874</td><td>GGCGGGTTCGAGCGCCCGCAGGG</td><td>1000</td><td></td><td></td><td></td><td></td><td></td><td></td><td></td><td></td><td></td></tr>
<tr><td>Oligo fwd</td><td>CACCGGCGGGTTCGAGCGCCCGCA</td><td></td><td></td><td></td><td></td><td></td><td></td><td></td><td></td><td></td><td></td></tr>
<tr><td>Oligo rev</td><td>AAACTGCGGGCGCTCGAACCCGCC</td><td></td><td></td><td></td><td></td><td></td><td></td><td></td><td></td><td></td><td></td></tr>
<tr><td>Chromosome</td><td>start</td><td>end</td><td>strand</td><td>MM</td><td>target_seq</td><td>PAM</td><td>alignment</td><td>distance</td><td>position</td><td>gene name</td><td>gene id</td></tr>
<tr><td>GU305914.1</td><td>30907</td><td>30929</td><td>+</td><td>0</td><td>GGCGGGTTCGAGCGCCCGCA</td><td>GGG</td><td>||||||||[||||||||||||]PAM</td><td>NA</td><td>Intergenic</td><td>NA</td><td>NA</td></tr>
<tr><td>T1875</td><td>TTCGAGCGCCCGCAGGGCGTCGG</td><td>1000</td><td></td><td></td><td></td><td></td><td></td><td></td><td></td><td></td><td></td></tr>
<tr><td>Oligo fwd</td><td>CACCTTCGAGCGCCCGCAGGGCGT</td><td></td><td></td><td></td><td></td><td></td><td></td><td></td><td></td><td></td><td></td></tr>
<tr><td>Oligo rev</td><td>AAACACGCCCTGCGGGCGCTCGAA</td><td></td><td></td><td></td><td></td><td></td><td></td><td></td><td></td><td></td><td></td></tr>
<tr><td>Chromosome</td><td>start</td><td>end</td><td>strand</td><td>MM</td><td>target_seq</td><td>PAM</td><td>alignment</td><td>distance</td><td>position</td><td>gene name</td><td>gene id</td></tr>
<tr><td>GU305914.1</td><td>30913</td><td>30935</td><td>+</td><td>0</td><td>TTCGAGCGCCCGCAGGGCGT</td><td>CGG</td><td>||||||||[||||||||||||]PAM</td><td>NA</td><td>Intergenic</td><td>NA</td><td>NA</td></tr>
<tr><td>T1876</td><td>TCGAGCGCCCGCAGGGCGTCGGG</td><td>1000</td><td></td><td></td><td></td><td></td><td></td><td></td><td></td><td></td><td></td></tr>
<tr><td>Oligo fwd</td><td>CACCTCGAGCGCCCGCAGGGCGTC</td><td></td><td></td><td></td><td></td><td></td><td></td><td></td><td></td><td></td><td></td></tr>
<tr><td>Oligo rev</td><td>AAACGACGCCCTGCGGGCGCTCGA</td><td></td><td></td><td></td><td></td><td></td><td></td><td></td><td></td><td></td><td></td></tr>
<tr><td>Chromosome</td><td>start</td><td>end</td><td>strand</td><td>MM</td><td>target_seq</td><td>PAM</td><td>alignment</td><td>distance</td><td>position</td><td>gene name</td><td>gene id</td></tr>
<tr><td>GU305914.1</td><td>30914</td><td>30936</td><td>+</td><td>0</td><td>TCGAGCGCCCGCAGGGCGTC</td><td>GGG</td><td>||||||||[||||||||||||]PAM</td><td>NA</td><td>Intergenic</td><td>NA</td><td>NA</td></tr>
<tr><td>T1877</td><td>CGAGCGCCCGCAGGGCGTCGGGG</td><td>1000</td><td></td><td></td><td></td><td></td><td></td><td></td><td></td><td></td><td></td></tr>
<tr><td>Oligo fwd</td><td>CACCCGAGCGCCCGCAGGGCGTCG</td><td></td><td></td><td></td><td></td><td></td><td></td><td></td><td></td><td></td><td></td></tr>
<tr><td>Oligo rev</td><td>AAACCGACGCCCTGCGGGCGCTCG</td><td></td><td></td><td></td><td></td><td></td><td></td><td></td><td></td><td></td><td></td></tr>
<tr><td>Chromosome</td><td>start</td><td>end</td><td>strand</td><td>MM</td><td>target_seq</td><td>PAM</td><td>alignment</td><td>distance</td><td>position</td><td>gene name</td><td>gene id</td></tr>
<tr><td>GU305914.1</td><td>30915</td><td>30937</td><td>+</td><td>0</td><td>CGAGCGCCCGCAGGGCGTCG</td><td>GGG</td><td>||||||||[||||||||||||]PAM</td><td>NA</td><td>Intergenic</td><td>NA</td><td>NA</td></tr>
<tr><td>T1878</td><td>CTCGTGTCCGTCGACATCGCCGG</td><td>1000</td><td></td><td></td><td></td><td></td><td></td><td></td><td></td><td></td><td></td></tr>
<tr><td>Oligo fwd</td><td>CACCCTCGTGTCCGTCGACATCGC</td><td></td><td></td><td></td><td></td><td></td><td></td><td></td><td></td><td></td><td></td></tr>
<tr><td>Oligo rev</td><td>AAACGCGATGTCGACGGACACGAG</td><td></td><td></td><td></td><td></td><td></td><td></td><td></td><td></td><td></td><td></td></tr>
<tr><td>Chromosome</td><td>start</td><td>end</td><td>strand</td><td>MM</td><td>target_seq</td><td>PAM</td><td>alignment</td><td>distance</td><td>position</td><td>gene name</td><td>gene id</td></tr>
<tr><td>GU305914.1</td><td>30943</td><td>30965</td><td>+</td><td>0</td><td>CTCGTGTCCGTCGACATCGC</td><td>CGG</td><td>||||||||[||||||||||||]PAM</td><td>NA</td><td>Intergenic</td><td>NA</td><td>NA</td></tr>
<tr><td>T1879</td><td>TCGTGTCCGTCGACATCGCCGGG</td><td>1000</td><td></td><td></td><td></td><td></td><td></td><td></td><td></td><td></td><td></td></tr>
<tr><td>Oligo fwd</td><td>CACCTCGTGTCCGTCGACATCGCC</td><td></td><td></td><td></td><td></td><td></td><td></td><td></td><td></td><td></td><td></td></tr>
<tr><td>Oligo rev</td><td>AAACGGCGATGTCGACGGACACGA</td><td></td><td></td><td></td><td></td><td></td><td></td><td></td><td></td><td></td><td></td></tr>
<tr><td>Chromosome</td><td>start</td><td>end</td><td>strand</td><td>MM</td><td>target_seq</td><td>PAM</td><td>alignment</td><td>distance</td><td>position</td><td>gene name</td><td>gene id</td></tr>
<tr><td>GU305914.1</td><td>30944</td><td>30966</td><td>+</td><td>0</td><td>TCGTGTCCGTCGACATCGCC</td><td>GGG</td><td>||||||||[||||||||||||]PAM</td><td>NA</td><td>Intergenic</td><td>NA</td><td>NA</td></tr>
<tr><td>T1880</td><td>CGACATCGCCGGGAGCGCAGAGG</td><td>1000</td><td></td><td></td><td></td><td></td><td></td><td></td><td></td><td></td><td></td></tr>
<tr><td>Oligo fwd</td><td>CACCCGACATCGCCGGGAGCGCAG</td><td></td><td></td><td></td><td></td><td></td><td></td><td></td><td></td><td></td><td></td></tr>
<tr><td>Oligo rev</td><td>AAACCTGCGCTCCCGGCGATGTCG</td><td></td><td></td><td></td><td></td><td></td><td></td><td></td><td></td><td></td><td></td></tr>
<tr><td>Chromosome</td><td>start</td><td>end</td><td>strand</td><td>MM</td><td>target_seq</td><td>PAM</td><td>alignment</td><td>distance</td><td>position</td><td>gene name</td><td>gene id</td></tr>
<tr><td>GU305914.1</td><td>30954</td><td>30976</td><td>+</td><td>0</td><td>CGACATCGCCGGGAGCGCAG</td><td>AGG</td><td>||||||||[||||||||||||]PAM</td><td>NA</td><td>Intergenic</td><td>NA</td><td>NA</td></tr>
<tr><td>T1881</td><td>AGCGCAGAGGATGAGATACTTGG</td><td>1000</td><td></td><td></td><td></td><td></td><td></td><td></td><td></td><td></td><td></td></tr>
<tr><td>Oligo fwd</td><td>CACCAGCGCAGAGGATGAGATACT</td><td></td><td></td><td></td><td></td><td></td><td></td><td></td><td></td><td></td><td></td></tr>
<tr><td>Oligo rev</td><td>AAACAGTATCTCATCCTCTGCGCT</td><td></td><td></td><td></td><td></td><td></td><td></td><td></td><td></td><td></td><td></td></tr>
<tr><td>Chromosome</td><td>start</td><td>end</td><td>strand</td><td>MM</td><td>target_seq</td><td>PAM</td><td>alignment</td><td>distance</td><td>position</td><td>gene name</td><td>gene id</td></tr>
<tr><td>GU305914.1</td><td>30967</td><td>30989</td><td>+</td><td>0</td><td>AGCGCAGAGGATGAGATACT</td><td>TGG</td><td>||||||||[||||||||||||]PAM</td><td>NA</td><td>Intergenic</td><td>NA</td><td>NA</td></tr>
<tr><td>T1882</td><td>GAGATACTTGGCGCACTTTTCGG</td><td>1000</td><td></td><td></td><td></td><td></td><td></td><td></td><td></td><td></td><td></td></tr>
<tr><td>Oligo fwd</td><td>CACCGAGATACTTGGCGCACTTTT</td><td></td><td></td><td></td><td></td><td></td><td></td><td></td><td></td><td></td><td></td></tr>
<tr><td>Oligo rev</td><td>AAACAAAAGTGCGCCAAGTATCTC</td><td></td><td></td><td></td><td></td><td></td><td></td><td></td><td></td><td></td><td></td></tr>
<tr><td>Chromosome</td><td>start</td><td>end</td><td>strand</td><td>MM</td><td>target_seq</td><td>PAM</td><td>alignment</td><td>distance</td><td>position</td><td>gene name</td><td>gene id</td></tr>
<tr><td>GU305914.1</td><td>30979</td><td>31001</td><td>+</td><td>0</td><td>GAGATACTTGGCGCACTTTT</td><td>CGG</td><td>||||||||[||||||||||||]PAM</td><td>NA</td><td>Intergenic</td><td>NA</td><td>NA</td></tr>
<tr><td>T1883</td><td>AGATACTTGGCGCACTTTTCGGG</td><td>1000</td><td></td><td></td><td></td><td></td><td></td><td></td><td></td><td></td><td></td></tr>
<tr><td>Oligo fwd</td><td>CACCAGATACTTGGCGCACTTTTC</td><td></td><td></td><td></td><td></td><td></td><td></td><td></td><td></td><td></td><td></td></tr>
<tr><td>Oligo rev</td><td>AAACGAAAAGTGCGCCAAGTATCT</td><td></td><td></td><td></td><td></td><td></td><td></td><td></td><td></td><td></td><td></td></tr>
<tr><td>Chromosome</td><td>start</td><td>end</td><td>strand</td><td>MM</td><td>target_seq</td><td>PAM</td><td>alignment</td><td>distance</td><td>position</td><td>gene name</td><td>gene id</td></tr>
<tr><td>GU305914.1</td><td>30980</td><td>31002</td><td>+</td><td>0</td><td>AGATACTTGGCGCACTTTTC</td><td>GGG</td><td>||||||||[||||||||||||]PAM</td><td>NA</td><td>Intergenic</td><td>NA</td><td>NA</td></tr>
<tr><td>T1884</td><td>GATACTTGGCGCACTTTTCGGGG</td><td>1000</td><td></td><td></td><td></td><td></td><td></td><td></td><td></td><td></td><td></td></tr>
<tr><td>Oligo fwd</td><td>CACCGATACTTGGCGCACTTTTCG</td><td></td><td></td><td></td><td></td><td></td><td></td><td></td><td></td><td></td><td></td></tr>
<tr><td>Oligo rev</td><td>AAACCGAAAAGTGCGCCAAGTATC</td><td></td><td></td><td></td><td></td><td></td><td></td><td></td><td></td><td></td><td></td></tr>
<tr><td>Chromosome</td><td>start</td><td>end</td><td>strand</td><td>MM</td><td>target_seq</td><td>PAM</td><td>alignment</td><td>distance</td><td>position</td><td>gene name</td><td>gene id</td></tr>
<tr><td>GU305914.1</td><td>30981</td><td>31003</td><td>+</td><td>0</td><td>GATACTTGGCGCACTTTTCG</td><td>GGG</td><td>||||||||[||||||||||||]PAM</td><td>NA</td><td>Intergenic</td><td>NA</td><td>NA</td></tr>
<tr><td>T1885</td><td>CACTTTTCGGGGAAGACGAGTGG</td><td>1000</td><td></td><td></td><td></td><td></td><td></td><td></td><td></td><td></td><td></td></tr>
<tr><td>Oligo fwd</td><td>CACCCACTTTTCGGGGAAGACGAG</td><td></td><td></td><td></td><td></td><td></td><td></td><td></td><td></td><td></td><td></td></tr>
<tr><td>Oligo rev</td><td>AAACCTCGTCTTCCCCGAAAAGTG</td><td></td><td></td><td></td><td></td><td></td><td></td><td></td><td></td><td></td><td></td></tr>
<tr><td>Chromosome</td><td>start</td><td>end</td><td>strand</td><td>MM</td><td>target_seq</td><td>PAM</td><td>alignment</td><td>distance</td><td>position</td><td>gene name</td><td>gene id</td></tr>
<tr><td>GU305914.1</td><td>30992</td><td>31014</td><td>+</td><td>0</td><td>CACTTTTCGGGGAAGACGAG</td><td>TGG</td><td>||||||||[||||||||||||]PAM</td><td>NA</td><td>Intergenic</td><td>NA</td><td>NA</td></tr>
<tr><td>T1886</td><td>ACTTTTCGGGGAAGACGAGTGGG</td><td>1000</td><td></td><td></td><td></td><td></td><td></td><td></td><td></td><td></td><td></td></tr>
<tr><td>Oligo fwd</td><td>CACCACTTTTCGGGGAAGACGAGT</td><td></td><td></td><td></td><td></td><td></td><td></td><td></td><td></td><td></td><td></td></tr>
<tr><td>Oligo rev</td><td>AAACACTCGTCTTCCCCGAAAAGT</td><td></td><td></td><td></td><td></td><td></td><td></td><td></td><td></td><td></td><td></td></tr>
<tr><td>Chromosome</td><td>start</td><td>end</td><td>strand</td><td>MM</td><td>target_seq</td><td>PAM</td><td>alignment</td><td>distance</td><td>position</td><td>gene name</td><td>gene id</td></tr>
<tr><td>GU305914.1</td><td>30993</td><td>31015</td><td>+</td><td>0</td><td>ACTTTTCGGGGAAGACGAGT</td><td>GGG</td><td>||||||||[||||||||||||]PAM</td><td>NA</td><td>Intergenic</td><td>NA</td><td>NA</td></tr>
<tr><td>T1887</td><td>GTGGGATGCGCTCGCCGCGTCGG</td><td>1000</td><td></td><td></td><td></td><td></td><td></td><td></td><td></td><td></td><td></td></tr>
<tr><td>Oligo fwd</td><td>CACCGTGGGATGCGCTCGCCGCGT</td><td></td><td></td><td></td><td></td><td></td><td></td><td></td><td></td><td></td><td></td></tr>
<tr><td>Oligo rev</td><td>AAACACGCGGCGAGCGCATCCCAC</td><td></td><td></td><td></td><td></td><td></td><td></td><td></td><td></td><td></td><td></td></tr>
<tr><td>Chromosome</td><td>start</td><td>end</td><td>strand</td><td>MM</td><td>target_seq</td><td>PAM</td><td>alignment</td><td>distance</td><td>position</td><td>gene name</td><td>gene id</td></tr>
<tr><td>GU305914.1</td><td>31011</td><td>31033</td><td>+</td><td>0</td><td>GTGGGATGCGCTCGCCGCGT</td><td>CGG</td><td>||||||||[||||||||||||]PAM</td><td>NA</td><td>Intergenic</td><td>NA</td><td>NA</td></tr>
<tr><td>T1888</td><td>TGGGATGCGCTCGCCGCGTCGGG</td><td>1000</td><td></td><td></td><td></td><td></td><td></td><td></td><td></td><td></td><td></td></tr>
<tr><td>Oligo fwd</td><td>CACCTGGGATGCGCTCGCCGCGTC</td><td></td><td></td><td></td><td></td><td></td><td></td><td></td><td></td><td></td><td></td></tr>
</table>

Oligo rev AAACGACGCGGCGAGCGCATCCCA
Chromosome start end strand MM target_seq PAM alignment distance position gene name gene id
GU305914.1 31012 31034 + 0 TGGGATGCGCTCGCCGCGTC GGG ||||||||[||||||||||||]PAM NA Intergenic NA NA

T1889 CCACTGTTCGAGCTTGCCCTCGG 1000
Oligo fwd CACCCCACTGTTCGAGCTTGCCCT
Oligo rev AAACAGGGCAAGCTCGAACAGTGG
Chromosome start end strand MM target_seq PAM alignment distance position gene name gene id
GU305914.1 31042 31064 + 0 CCACTGTTCGAGCTTGCCCT CGG ||||||||[||||||||||||]PAM NA Intergenic NA NA

T1890 TTCGAGCTTGCCCTCGGCCGCGG 1000
Oligo fwd CACCTTCGAGCTTGCCCTCGGCCG
Oligo rev AAACCGGCCGAGGGCAAGCTCGAA
Chromosome start end strand MM target_seq PAM alignment distance position gene name gene id
GU305914.1 31048 31070 + 0 TTCGAGCTTGCCCTCGGCCG CGG ||||||||[||||||||||||]PAM NA Intergenic NA NA

T1891 CCGCGGCCCTGCCGAGCGCCAGG 1000
Oligo fwd CACCCCGCGGCCCTGCCGAGCGCC
Oligo rev AAACGGCGCTCGGCAGGGCCGCGG
Chromosome start end strand MM target_seq PAM alignment distance position gene name gene id
GU305914.1 31065 31087 + 0 CCGCGGCCCTGCCGAGCGCC AGG ||||||||[||||||||||||]PAM NA Intergenic NA NA

T1892 GAGCGCCAGGAGTCTGTTGACGG 1000
Oligo fwd CACCGAGCGCCAGGAGTCTGTTGA
Oligo rev AAACTCAACAGACTCCTGGCGCTC
Chromosome start end strand MM target_seq PAM alignment distance position gene name gene id
GU305914.1 31078 31100 + 0 GAGCGCCAGGAGTCTGTTGA CGG ||||||||[||||||||||||]PAM NA Intergenic NA NA

T1893 GACGGCGTACGCCTTCCTCATGG 1000
Oligo fwd CACCGACGGCGTACGCCTTCCTCA
Oligo rev AAACTGAGGAAGGCGTACGCCGTC
Chromosome start end strand MM target_seq PAM alignment distance position gene name gene id
GU305914.1 31096 31118 + 0 GACGGCGTACGCCTTCCTCA TGG ||||||||[||||||||||||]PAM NA Intergenic NA NA

T1894 GGCGTACGCCTTCCTCATGGCGG 1000
Oligo fwd CACCGGCGTACGCCTTCCTCATGG
Oligo rev AAACCCATGAGGAAGGCGTACGCC
Chromosome start end strand MM target_seq PAM alignment distance position gene name gene id
GU305914.1 31099 31121 + 0 GGCGTACGCCTTCCTCATGG CGG ||||||||[||||||||||||]PAM NA Intergenic NA NA

T1895 GTACGCCTTCCTCATGGCGGCGG 1000
Oligo fwd CACCGTACGCCTTCCTCATGGCGG
Oligo rev AAACCCGCCATGAGGAAGGCGTAC
Chromosome start end strand MM target_seq PAM alignment distance position gene name gene id
GU305914.1 31102 31124 + 0 GTACGCCTTCCTCATGGCGG CGG ||||||||[||||||||||||]PAM NA Intergenic NA NA

T1896 CTTCCTCATGGCGGCGGCCTTGG 1000
Oligo fwd CACCCTTCCTCATGGCGGCGGCCT
Oligo rev AAACAGGCCGCCGCCATGAGGAAG
Chromosome start end strand MM target_seq PAM alignment distance position gene name gene id
GU305914.1 31108 31130 + 0 CTTCCTCATGGCGGCGGCCT TGG ||||||||[||||||||||||]PAM NA Intergenic NA NA

T1897 TGGCGGCGGCCTTGGCCGCCAGG 1000
Oligo fwd CACCTGGCGGCGGCCTTGGCCGCC
Oligo rev AAACGGCGGCCAAGGCCGCCGCCA
Chromosome start end strand MM target_seq PAM alignment distance position gene name gene id
GU305914.1 31116 31138 + 0 TGGCGGCGGCCTTGGCCGCC AGG ||||||||[||||||||||||]PAM NA Intergenic NA NA

T1898 GGCGGCGGCCTTGGCCGCCAGGG 1000
Oligo fwd CACCGGCGGCGGCCTTGGCCGCCA
Oligo rev AAACTGGCGGCCAAGGCCGCCGCC
Chromosome start end strand MM target_seq PAM alignment distance position gene name gene id
GU305914.1 31117 31139 + 0 GGCGGCGGCCTTGGCCGCCA GGG ||||||||[||||||||||||]PAM NA Intergenic NA NA

T1899 GGCGGCCTTGGCCGCCAGGGTGG 1000
Oligo fwd CACCGGCGGCCTTGGCCGCCAGGG
Oligo rev AAACCCCTGGCGGCCAAGGCCGCC
Chromosome start end strand MM target_seq PAM alignment distance position gene name gene id
GU305914.1 31120 31142 + 0 GGCGGCCTTGGCCGCCAGGG TGG ||||||||[||||||||||||]PAM NA Intergenic NA NA

T1900 CCTTGGCCGCCAGGGTGGAGCGG 1000
Oligo fwd CACCCCTTGGCCGCCAGGGTGGAG
Oligo rev AAACCTCCACCCTGGCGGCCAAGG
Chromosome start end strand MM target_seq PAM alignment distance position gene name gene id
GU305914.1 31125 31147 + 0 CCTTGGCCGCCAGGGTGGAG CGG ||||||||[||||||||||||]PAM NA Intergenic NA NA

T1901 CTTGGCCGCCAGGGTGGAGCGGG 1000
Oligo fwd CACCCTTGGCCGCCAGGGTGGAGC
Oligo rev AAACGCTCCACCCTGGCGGCCAAG
Chromosome start end strand MM target_seq PAM alignment distance position gene name gene id
GU305914.1 31126 31148 + 0 CTTGGCCGCCAGGGTGGAGC GGG ||||||||[||||||||||||]PAM NA Intergenic NA NA

T1902 GCCGCCAGGGTGGAGCGGGCAGG 1000
Oligo fwd CACCGCCGCCAGGGTGGAGCGGGC
Oligo rev AAACGCCCGCTCCACCCTGGCGGC
Chromosome start end strand MM target_seq PAM alignment distance position gene name gene id
GU305914.1 31130 31152 + 0 GCCGCCAGGGTGGAGCGGGC AGG ||||||||[||||||||||||]PAM NA Intergenic NA NA

T1903 CACCGTCTCCGTCTCCGACTCGG 1000

<table>
<tr><td>Oligo fwd</td><td>CACCCACCGTCTCCGTCTCCGACT</td><td></td><td></td><td></td><td></td><td></td><td></td><td></td><td></td><td></td><td></td></tr>
<tr><td>Oligo rev</td><td>AAACAGTCGGAGACGGAGACGGTG</td><td></td><td></td><td></td><td></td><td></td><td></td><td></td><td></td><td></td><td></td></tr>
<tr><td>Chromosome</td><td>start</td><td>end</td><td>strand</td><td>MM</td><td>target_seq</td><td>PAM</td><td>alignment</td><td>distance</td><td>position</td><td>gene name</td><td>gene id</td></tr>
<tr><td>GU305914.1</td><td>31162</td><td>31184</td><td>+</td><td>0</td><td>CACCGTCTCCGTCTCCGACT</td><td>CGG</td><td>||||||||[||||||||||||]PAM</td><td>NA</td><td>Intergenic</td><td>NA</td><td>NA</td></tr>
<tr><td>GU305914.1</td><td>122850</td><td>122872</td><td>-</td><td>3</td><td>CCTCGTCTCCGTCTCGGACT</td><td>GGG</td><td>|--|||||[|||||||-||||]PAM</td><td>NA</td><td>Intergenic</td><td>NA</td><td>NA</td></tr>
<tr><td></td><td></td><td></td><td></td><td></td><td></td><td></td><td></td><td></td><td></td><td></td><td></td></tr>
<tr><td>T1904</td><td>ACCGTCTCCGTCTCCGACTCGGG</td><td>1000</td><td></td><td></td><td></td><td></td><td></td><td></td><td></td><td></td><td></td></tr>
<tr><td>Oligo fwd</td><td>CACCACCGTCTCCGTCTCCGACTC</td><td></td><td></td><td></td><td></td><td></td><td></td><td></td><td></td><td></td><td></td></tr>
<tr><td>Oligo rev</td><td>AAACGAGTCGGAGACGGAGACGGT</td><td></td><td></td><td></td><td></td><td></td><td></td><td></td><td></td><td></td><td></td></tr>
<tr><td>Chromosome</td><td>start</td><td>end</td><td>strand</td><td>MM</td><td>target_seq</td><td>PAM</td><td>alignment</td><td>distance</td><td>position</td><td>gene name</td><td>gene id</td></tr>
<tr><td>GU305914.1</td><td>31163</td><td>31185</td><td>+</td><td>0</td><td>ACCGTCTCCGTCTCCGACTC</td><td>GGG</td><td>||||||||[||||||||||||]PAM</td><td>NA</td><td>Intergenic</td><td>NA</td><td>NA</td></tr>
<tr><td></td><td></td><td></td><td></td><td></td><td></td><td></td><td></td><td></td><td></td><td></td><td></td></tr>
<tr><td>T1905</td><td>CTCGGGCATCGACACCGCCGCGG</td><td>1000</td><td></td><td></td><td></td><td></td><td></td><td></td><td></td><td></td><td></td></tr>
<tr><td>Oligo fwd</td><td>CACCCTCGGGCATCGACACCGCCG</td><td></td><td></td><td></td><td></td><td></td><td></td><td></td><td></td><td></td><td></td></tr>
<tr><td>Oligo rev</td><td>AAACCGGCGGTGTCGATGCCCGAG</td><td></td><td></td><td></td><td></td><td></td><td></td><td></td><td></td><td></td><td></td></tr>
<tr><td>Chromosome</td><td>start</td><td>end</td><td>strand</td><td>MM</td><td>target_seq</td><td>PAM</td><td>alignment</td><td>distance</td><td>position</td><td>gene name</td><td>gene id</td></tr>
<tr><td>GU305914.1</td><td>31180</td><td>31202</td><td>+</td><td>0</td><td>CTCGGGCATCGACACCGCCG</td><td>CGG</td><td>||||||||[||||||||||||]PAM</td><td>NA</td><td>Intergenic</td><td>NA</td><td>NA</td></tr>
<tr><td></td><td></td><td></td><td></td><td></td><td></td><td></td><td></td><td></td><td></td><td></td><td></td></tr>
<tr><td>T1906</td><td>GGCATCGACACCGCCGCGGCCGG</td><td>1000</td><td></td><td></td><td></td><td></td><td></td><td></td><td></td><td></td><td></td></tr>
<tr><td>Oligo fwd</td><td>CACCGGCATCGACACCGCCGCGGC</td><td></td><td></td><td></td><td></td><td></td><td></td><td></td><td></td><td></td><td></td></tr>
<tr><td>Oligo rev</td><td>AAACGCCGCGGCGGTGTCGATGCC</td><td></td><td></td><td></td><td></td><td></td><td></td><td></td><td></td><td></td><td></td></tr>
<tr><td>Chromosome</td><td>start</td><td>end</td><td>strand</td><td>MM</td><td>target_seq</td><td>PAM</td><td>alignment</td><td>distance</td><td>position</td><td>gene name</td><td>gene id</td></tr>
<tr><td>GU305914.1</td><td>31184</td><td>31206</td><td>+</td><td>0</td><td>GGCATCGACACCGCCGCGGC</td><td>CGG</td><td>||||||||[||||||||||||]PAM</td><td>NA</td><td>Intergenic</td><td>NA</td><td>NA</td></tr>
<tr><td></td><td></td><td></td><td></td><td></td><td></td><td></td><td></td><td></td><td></td><td></td><td></td></tr>
<tr><td>T1907</td><td>CGCCGCGGCCGGAGCTAGCATGG</td><td>1000</td><td></td><td></td><td></td><td></td><td></td><td></td><td></td><td></td><td></td></tr>
<tr><td>Oligo fwd</td><td>CACCCGCCGCGGCCGGAGCTAGCA</td><td></td><td></td><td></td><td></td><td></td><td></td><td></td><td></td><td></td><td></td></tr>
<tr><td>Oligo rev</td><td>AAACTGCTAGCTCCGGCCGCGGCG</td><td></td><td></td><td></td><td></td><td></td><td></td><td></td><td></td><td></td><td></td></tr>
<tr><td>Chromosome</td><td>start</td><td>end</td><td>strand</td><td>MM</td><td>target_seq</td><td>PAM</td><td>alignment</td><td>distance</td><td>position</td><td>gene name</td><td>gene id</td></tr>
<tr><td>GU305914.1</td><td>31195</td><td>31217</td><td>+</td><td>0</td><td>CGCCGCGGCCGGAGCTAGCA</td><td>TGG</td><td>||||||||[||||||||||||]PAM</td><td>NA</td><td>Intergenic</td><td>NA</td><td>NA</td></tr>
<tr><td></td><td></td><td></td><td></td><td></td><td></td><td></td><td></td><td></td><td></td><td></td><td></td></tr>
<tr><td>T1908</td><td>CGGCCGGAGCTAGCATGGTCTGG</td><td>1000</td><td></td><td></td><td></td><td></td><td></td><td></td><td></td><td></td><td></td></tr>
<tr><td>Oligo fwd</td><td>CACCCGGCCGGAGCTAGCATGGTC</td><td></td><td></td><td></td><td></td><td></td><td></td><td></td><td></td><td></td><td></td></tr>
<tr><td>Oligo rev</td><td>AAACGACCATGCTAGCTCCGGCCG</td><td></td><td></td><td></td><td></td><td></td><td></td><td></td><td></td><td></td><td></td></tr>
<tr><td>Chromosome</td><td>start</td><td>end</td><td>strand</td><td>MM</td><td>target_seq</td><td>PAM</td><td>alignment</td><td>distance</td><td>position</td><td>gene name</td><td>gene id</td></tr>
<tr><td>GU305914.1</td><td>31200</td><td>31222</td><td>+</td><td>0</td><td>CGGCCGGAGCTAGCATGGTC</td><td>TGG</td><td>||||||||[||||||||||||]PAM</td><td>NA</td><td>Intergenic</td><td>NA</td><td>NA</td></tr>
<tr><td></td><td></td><td></td><td></td><td></td><td></td><td></td><td></td><td></td><td></td><td></td><td></td></tr>
<tr><td>T1909</td><td>ACCGTGTCCGATCGTCTCTGCGG</td><td>1000</td><td></td><td></td><td></td><td></td><td></td><td></td><td></td><td></td><td></td></tr>
<tr><td>Oligo fwd</td><td>CACCACCGTGTCCGATCGTCTCTG</td><td></td><td></td><td></td><td></td><td></td><td></td><td></td><td></td><td></td><td></td></tr>
<tr><td>Oligo rev</td><td>AAACCAGAGACGATCGGACACGGT</td><td></td><td></td><td></td><td></td><td></td><td></td><td></td><td></td><td></td><td></td></tr>
<tr><td>Chromosome</td><td>start</td><td>end</td><td>strand</td><td>MM</td><td>target_seq</td><td>PAM</td><td>alignment</td><td>distance</td><td>position</td><td>gene name</td><td>gene id</td></tr>
<tr><td>GU305914.1</td><td>31283</td><td>31305</td><td>+</td><td>0</td><td>ACCGTGTCCGATCGTCTCTG</td><td>CGG</td><td>||||||||[||||||||||||]PAM</td><td>NA</td><td>Intergenic</td><td>NA</td><td>NA</td></tr>
<tr><td></td><td></td><td></td><td></td><td></td><td></td><td></td><td></td><td></td><td></td><td></td><td></td></tr>
<tr><td>T1910</td><td>GATCGTCTCTGCGGTCGCCACGG</td><td>1000</td><td></td><td></td><td></td><td></td><td></td><td></td><td></td><td></td><td></td></tr>
<tr><td>Oligo fwd</td><td>CACCGATCGTCTCTGCGGTCGCCA</td><td></td><td></td><td></td><td></td><td></td><td></td><td></td><td></td><td></td><td></td></tr>
<tr><td>Oligo rev</td><td>AAACTGGCGACCGCAGAGACGATC</td><td></td><td></td><td></td><td></td><td></td><td></td><td></td><td></td><td></td><td></td></tr>
<tr><td>Chromosome</td><td>start</td><td>end</td><td>strand</td><td>MM</td><td>target_seq</td><td>PAM</td><td>alignment</td><td>distance</td><td>position</td><td>gene name</td><td>gene id</td></tr>
<tr><td>GU305914.1</td><td>31292</td><td>31314</td><td>+</td><td>0</td><td>GATCGTCTCTGCGGTCGCCA</td><td>CGG</td><td>||||||||[||||||||||||]PAM</td><td>NA</td><td>Intergenic</td><td>NA</td><td>NA</td></tr>
<tr><td></td><td></td><td></td><td></td><td></td><td></td><td></td><td></td><td></td><td></td><td></td><td></td></tr>
<tr><td>T1911</td><td>GTCTCTGCGGTCGCCACGGACGG</td><td>1000</td><td></td><td></td><td></td><td></td><td></td><td></td><td></td><td></td><td></td></tr>
<tr><td>Oligo fwd</td><td>CACCGTCTCTGCGGTCGCCACGGA</td><td></td><td></td><td></td><td></td><td></td><td></td><td></td><td></td><td></td><td></td></tr>
<tr><td>Oligo rev</td><td>AAACTCCGTGGCGACCGCAGAGAC</td><td></td><td></td><td></td><td></td><td></td><td></td><td></td><td></td><td></td><td></td></tr>
<tr><td>Chromosome</td><td>start</td><td>end</td><td>strand</td><td>MM</td><td>target_seq</td><td>PAM</td><td>alignment</td><td>distance</td><td>position</td><td>gene name</td><td>gene id</td></tr>
<tr><td>GU305914.1</td><td>31296</td><td>31318</td><td>+</td><td>0</td><td>GTCTCTGCGGTCGCCACGGA</td><td>CGG</td><td>||||||||[||||||||||||]PAM</td><td>NA</td><td>Intergenic</td><td>NA</td><td>NA</td></tr>
<tr><td></td><td></td><td></td><td></td><td></td><td></td><td></td><td></td><td></td><td></td><td></td><td></td></tr>
<tr><td>T1912</td><td>TGCGGTCGCCACGGACGGAGAGG</td><td>1000</td><td></td><td></td><td></td><td></td><td></td><td></td><td></td><td></td><td></td></tr>
<tr><td>Oligo fwd</td><td>CACCTGCGGTCGCCACGGACGGAG</td><td></td><td></td><td></td><td></td><td></td><td></td><td></td><td></td><td></td><td></td></tr>
<tr><td>Oligo rev</td><td>AAACCTCCGTCCGTGGCGACCGCA</td><td></td><td></td><td></td><td></td><td></td><td></td><td></td><td></td><td></td><td></td></tr>
<tr><td>Chromosome</td><td>start</td><td>end</td><td>strand</td><td>MM</td><td>target_seq</td><td>PAM</td><td>alignment</td><td>distance</td><td>position</td><td>gene name</td><td>gene id</td></tr>
<tr><td>GU305914.1</td><td>31301</td><td>31323</td><td>+</td><td>0</td><td>TGCGGTCGCCACGGACGGAG</td><td>AGG</td><td>||||||||[||||||||||||]PAM</td><td>NA</td><td>Intergenic</td><td>NA</td><td>NA</td></tr>
<tr><td></td><td></td><td></td><td></td><td></td><td></td><td></td><td></td><td></td><td></td><td></td><td></td></tr>
<tr><td>T1913</td><td>GGTCGCCACGGACGGAGAGGAGG</td><td>1000</td><td></td><td></td><td></td><td></td><td></td><td></td><td></td><td></td><td></td></tr>
<tr><td>Oligo fwd</td><td>CACCGGTCGCCACGGACGGAGAGG</td><td></td><td></td><td></td><td></td><td></td><td></td><td></td><td></td><td></td><td></td></tr>
<tr><td>Oligo rev</td><td>AAACCCTCTCCGTCCGTGGCGACC</td><td></td><td></td><td></td><td></td><td></td><td></td><td></td><td></td><td></td><td></td></tr>
<tr><td>Chromosome</td><td>start</td><td>end</td><td>strand</td><td>MM</td><td>target_seq</td><td>PAM</td><td>alignment</td><td>distance</td><td>position</td><td>gene name</td><td>gene id</td></tr>
<tr><td>GU305914.1</td><td>31304</td><td>31326</td><td>+</td><td>0</td><td>GGTCGCCACGGACGGAGAGG</td><td>AGG</td><td>||||||||[||||||||||||]PAM</td><td>NA</td><td>Intergenic</td><td>NA</td><td>NA</td></tr>
<tr><td></td><td></td><td></td><td></td><td></td><td></td><td></td><td></td><td></td><td></td><td></td><td></td></tr>
<tr><td>T1914</td><td>ACGGACGGAGAGGAGGACGTCGG</td><td>1000</td><td></td><td></td><td></td><td></td><td></td><td></td><td></td><td></td><td></td></tr>
<tr><td>Oligo fwd</td><td>CACCACGGACGGAGAGGAGGACGT</td><td></td><td></td><td></td><td></td><td></td><td></td><td></td><td></td><td></td><td></td></tr>
<tr><td>Oligo rev</td><td>AAACACGTCCTCCTCTCCGTCCGT</td><td></td><td></td><td></td><td></td><td></td><td></td><td></td><td></td><td></td><td></td></tr>
<tr><td>Chromosome</td><td>start</td><td>end</td><td>strand</td><td>MM</td><td>target_seq</td><td>PAM</td><td>alignment</td><td>distance</td><td>position</td><td>gene name</td><td>gene id</td></tr>
<tr><td>GU305914.1</td><td>31311</td><td>31333</td><td>+</td><td>0</td><td>ACGGACGGAGAGGAGGACGT</td><td>CGG</td><td>||||||||[||||||||||||]PAM</td><td>NA</td><td>Intergenic</td><td>NA</td><td>NA</td></tr>
<tr><td></td><td></td><td></td><td></td><td></td><td></td><td></td><td></td><td></td><td></td><td></td><td></td></tr>
<tr><td>T1915</td><td>GACGGAGAGGAGGACGTCGGCGG</td><td>1000</td><td></td><td></td><td></td><td></td><td></td><td></td><td></td><td></td><td></td></tr>
<tr><td>Oligo fwd</td><td>CACCGACGGAGAGGAGGACGTCGG</td><td></td><td></td><td></td><td></td><td></td><td></td><td></td><td></td><td></td><td></td></tr>
<tr><td>Oligo rev</td><td>AAACCCGACGTCCTCCTCTCCGTC</td><td></td><td></td><td></td><td></td><td></td><td></td><td></td><td></td><td></td><td></td></tr>
<tr><td>Chromosome</td><td>start</td><td>end</td><td>strand</td><td>MM</td><td>target_seq</td><td>PAM</td><td>alignment</td><td>distance</td><td>position</td><td>gene name</td><td>gene id</td></tr>
<tr><td>GU305914.1</td><td>31314</td><td>31336</td><td>+</td><td>0</td><td>GACGGAGAGGAGGACGTCGG</td><td>CGG</td><td>||||||||[||||||||||||]PAM</td><td>NA</td><td>Intergenic</td><td>NA</td><td>NA</td></tr>
<tr><td></td><td></td><td></td><td></td><td></td><td></td><td></td><td></td><td></td><td></td><td></td><td></td></tr>
<tr><td>T1916</td><td>GCGGCGTCGTCGTTCACATGTGG</td><td>1000</td><td></td><td></td><td></td><td></td><td></td><td></td><td></td><td></td><td></td></tr>
<tr><td>Oligo fwd</td><td>CACCGCGGCGTCGTCGTTCACATG</td><td></td><td></td><td></td><td></td><td></td><td></td><td></td><td></td><td></td><td></td></tr>
<tr><td>Oligo rev</td><td>AAACCATGTGAACGACGACGCCGC</td><td></td><td></td><td></td><td></td><td></td><td></td><td></td><td></td><td></td><td></td></tr>
<tr><td>Chromosome</td><td>start</td><td>end</td><td>strand</td><td>MM</td><td>target_seq</td><td>PAM</td><td>alignment</td><td>distance</td><td>position</td><td>gene name</td><td>gene id</td></tr>
<tr><td>GU305914.1</td><td>31333</td><td>31355</td><td>+</td><td>0</td><td>GCGGCGTCGTCGTTCACATG</td><td>TGG</td><td>||||||||[||||||||||||]PAM</td><td>NA</td><td>Intergenic</td><td>NA</td><td>NA</td></tr>
<tr><td></td><td></td><td></td><td></td><td></td><td></td><td></td><td></td><td></td><td></td><td></td><td></td></tr>
<tr><td>T1917</td><td>CGGCGTCGTCGTTCACATGTGGG</td><td>1000</td><td></td><td></td><td></td><td></td><td></td><td></td><td></td><td></td><td></td></tr>
<tr><td>Oligo fwd</td><td>CACCCGGCGTCGTCGTTCACATGT</td><td></td><td></td><td></td><td></td><td></td><td></td><td></td><td></td><td></td><td></td></tr>
<tr><td>Oligo rev</td><td>AAACACATGTGAACGACGACGCCG</td><td></td><td></td><td></td><td></td><td></td><td></td><td></td><td></td><td></td><td></td></tr>
<tr><td>Chromosome</td><td>start</td><td>end</td><td>strand</td><td>MM</td><td>target_seq</td><td>PAM</td><td>alignment</td><td>distance</td><td>position</td><td>gene name</td><td>gene id</td></tr>
<tr><td>GU305914.1</td><td>31334</td><td>31356</td><td>+</td><td>0</td><td>CGGCGTCGTCGTTCACATGT</td><td>GGG</td><td>||||||||[||||||||||||]PAM</td><td>NA</td><td>Intergenic</td><td>NA</td><td>NA</td></tr>
</table>

<table>
<tr><td>T1918</td><td>TTCACATGTGGGACAGAGCCAGG</td><td>1000</td><td></td><td></td><td></td><td></td><td></td><td></td><td></td><td></td><td></td></tr>
<tr><td>Oligo fwd</td><td>CACCTTCACATGTGGGACAGAGCC</td><td></td><td></td><td></td><td></td><td></td><td></td><td></td><td></td><td></td><td></td></tr>
<tr><td>Oligo rev</td><td>AAACGGCTCTGTCCCACATGTGAA</td><td></td><td></td><td></td><td></td><td></td><td></td><td></td><td></td><td></td><td></td></tr>
<tr><td>Chromosome</td><td>start</td><td>end</td><td>strand</td><td>MM</td><td>target_seq</td><td>PAM</td><td>alignment</td><td>distance</td><td>position</td><td>gene name</td><td>gene id</td></tr>
<tr><td>GU305914.1</td><td>31345</td><td>31367</td><td>+</td><td>0</td><td>TTCACATGTGGGACAGAGCC</td><td>AGG</td><td>||||||||[||||||||||||]PAM</td><td>NA</td><td>Intergenic</td><td>NA</td><td>NA</td></tr>
<tr><td>T1919</td><td>TCACATGTGGGACAGAGCCAGGG</td><td>1000</td><td></td><td></td><td></td><td></td><td></td><td></td><td></td><td></td><td></td></tr>
<tr><td>Oligo fwd</td><td>CACCTCACATGTGGGACAGAGCCA</td><td></td><td></td><td></td><td></td><td></td><td></td><td></td><td></td><td></td><td></td></tr>
<tr><td>Oligo rev</td><td>AAACTGGCTCTGTCCCACATGTGA</td><td></td><td></td><td></td><td></td><td></td><td></td><td></td><td></td><td></td><td></td></tr>
<tr><td>Chromosome</td><td>start</td><td>end</td><td>strand</td><td>MM</td><td>target_seq</td><td>PAM</td><td>alignment</td><td>distance</td><td>position</td><td>gene name</td><td>gene id</td></tr>
<tr><td>GU305914.1</td><td>31346</td><td>31368</td><td>+</td><td>0</td><td>TCACATGTGGGACAGAGCCA</td><td>GGG</td><td>||||||||[||||||||||||]PAM</td><td>NA</td><td>Intergenic</td><td>NA</td><td>NA</td></tr>
<tr><td>T1920</td><td>GCCGCCTTAAACACCCCGTCCGG</td><td>1000</td><td></td><td></td><td></td><td></td><td></td><td></td><td></td><td></td><td></td></tr>
<tr><td>Oligo fwd</td><td>CACCGCCGCCTTAAACACCCCGTC</td><td></td><td></td><td></td><td></td><td></td><td></td><td></td><td></td><td></td><td></td></tr>
<tr><td>Oligo rev</td><td>AAACGACGGGGTGTTTAAGGCGGC</td><td></td><td></td><td></td><td></td><td></td><td></td><td></td><td></td><td></td><td></td></tr>
<tr><td>Chromosome</td><td>start</td><td>end</td><td>strand</td><td>MM</td><td>target_seq</td><td>PAM</td><td>alignment</td><td>distance</td><td>position</td><td>gene name</td><td>gene id</td></tr>
<tr><td>GU305914.1</td><td>31368</td><td>31390</td><td>+</td><td>0</td><td>GCCGCCTTAAACACCCCGTC</td><td>CGG</td><td>||||||||[||||||||||||]PAM</td><td>NA</td><td>Intergenic</td><td>NA</td><td>NA</td></tr>
<tr><td>T1921</td><td>CACCCCGTCCGGAAAATCCGCGG</td><td>1000</td><td></td><td></td><td></td><td></td><td></td><td></td><td></td><td></td><td></td></tr>
<tr><td>Oligo fwd</td><td>CACCCACCCCGTCCGGAAAATCCG</td><td></td><td></td><td></td><td></td><td></td><td></td><td></td><td></td><td></td><td></td></tr>
<tr><td>Oligo rev</td><td>AAACCGGATTTTCCGGACGGGGTG</td><td></td><td></td><td></td><td></td><td></td><td></td><td></td><td></td><td></td><td></td></tr>
<tr><td>Chromosome</td><td>start</td><td>end</td><td>strand</td><td>MM</td><td>target_seq</td><td>PAM</td><td>alignment</td><td>distance</td><td>position</td><td>gene name</td><td>gene id</td></tr>
<tr><td>GU305914.1</td><td>31379</td><td>31401</td><td>+</td><td>0</td><td>CACCCCGTCCGGAAAATCCG</td><td>CGG</td><td>||||||||[||||||||||||]PAM</td><td>NA</td><td>Intergenic</td><td>NA</td><td>NA</td></tr>
<tr><td>T1922</td><td>ACCCCGTCCGGAAAATCCGCGGG</td><td>1000</td><td></td><td></td><td></td><td></td><td></td><td></td><td></td><td></td><td></td></tr>
<tr><td>Oligo fwd</td><td>CACCACCCCGTCCGGAAAATCCGC</td><td></td><td></td><td></td><td></td><td></td><td></td><td></td><td></td><td></td><td></td></tr>
<tr><td>Oligo rev</td><td>AAACGCGGATTTTCCGGACGGGGT</td><td></td><td></td><td></td><td></td><td></td><td></td><td></td><td></td><td></td><td></td></tr>
<tr><td>Chromosome</td><td>start</td><td>end</td><td>strand</td><td>MM</td><td>target_seq</td><td>PAM</td><td>alignment</td><td>distance</td><td>position</td><td>gene name</td><td>gene id</td></tr>
<tr><td>GU305914.1</td><td>31380</td><td>31402</td><td>+</td><td>0</td><td>ACCCCGTCCGGAAAATCCGC</td><td>GGG</td><td>||||||||[||||||||||||]PAM</td><td>NA</td><td>Intergenic</td><td>NA</td><td>NA</td></tr>
<tr><td>T1923</td><td>CGTCCGGAAAATCCGCGGGACGG</td><td>1000</td><td></td><td></td><td></td><td></td><td></td><td></td><td></td><td></td><td></td></tr>
<tr><td>Oligo fwd</td><td>CACCCGTCCGGAAAATCCGCGGGA</td><td></td><td></td><td></td><td></td><td></td><td></td><td></td><td></td><td></td><td></td></tr>
<tr><td>Oligo rev</td><td>AAACTCCCGCGGATTTTCCGGACG</td><td></td><td></td><td></td><td></td><td></td><td></td><td></td><td></td><td></td><td></td></tr>
<tr><td>Chromosome</td><td>start</td><td>end</td><td>strand</td><td>MM</td><td>target_seq</td><td>PAM</td><td>alignment</td><td>distance</td><td>position</td><td>gene name</td><td>gene id</td></tr>
<tr><td>GU305914.1</td><td>31384</td><td>31406</td><td>+</td><td>0</td><td>CGTCCGGAAAATCCGCGGGA</td><td>CGG</td><td>||||||||[||||||||||||]PAM</td><td>NA</td><td>Intergenic</td><td>NA</td><td>NA</td></tr>
<tr><td>T1924</td><td>TCCGCGGGACGGAGTCTGATCGG</td><td>1000</td><td></td><td></td><td></td><td></td><td></td><td></td><td></td><td></td><td></td></tr>
<tr><td>Oligo fwd</td><td>CACCTCCGCGGGACGGAGTCTGAT</td><td></td><td></td><td></td><td></td><td></td><td></td><td></td><td></td><td></td><td></td></tr>
<tr><td>Oligo rev</td><td>AAACATCAGACTCCGTCCCGCGGA</td><td></td><td></td><td></td><td></td><td></td><td></td><td></td><td></td><td></td><td></td></tr>
<tr><td>Chromosome</td><td>start</td><td>end</td><td>strand</td><td>MM</td><td>target_seq</td><td>PAM</td><td>alignment</td><td>distance</td><td>position</td><td>gene name</td><td>gene id</td></tr>
<tr><td>GU305914.1</td><td>31395</td><td>31417</td><td>+</td><td>0</td><td>TCCGCGGGACGGAGTCTGAT</td><td>CGG</td><td>||||||||[||||||||||||]PAM</td><td>NA</td><td>Intergenic</td><td>NA</td><td>NA</td></tr>
<tr><td>T1925</td><td>CGGGACGGAGTCTGATCGGCCGG</td><td>1000</td><td></td><td></td><td></td><td></td><td></td><td></td><td></td><td></td><td></td></tr>
<tr><td>Oligo fwd</td><td>CACCCGGGACGGAGTCTGATCGGC</td><td></td><td></td><td></td><td></td><td></td><td></td><td></td><td></td><td></td><td></td></tr>
<tr><td>Oligo rev</td><td>AAACGCCGATCAGACTCCGTCCCG</td><td></td><td></td><td></td><td></td><td></td><td></td><td></td><td></td><td></td><td></td></tr>
<tr><td>Chromosome</td><td>start</td><td>end</td><td>strand</td><td>MM</td><td>target_seq</td><td>PAM</td><td>alignment</td><td>distance</td><td>position</td><td>gene name</td><td>gene id</td></tr>
<tr><td>GU305914.1</td><td>31399</td><td>31421</td><td>+</td><td>0</td><td>CGGGACGGAGTCTGATCGGC</td><td>CGG</td><td>||||||||[||||||||||||]PAM</td><td>NA</td><td>Intergenic</td><td>NA</td><td>NA</td></tr>
<tr><td>T1926</td><td>GACGGAGTCTGATCGGCCGGCGG</td><td>1000</td><td></td><td></td><td></td><td></td><td></td><td></td><td></td><td></td><td></td></tr>
<tr><td>Oligo fwd</td><td>CACCGACGGAGTCTGATCGGCCGG</td><td></td><td></td><td></td><td></td><td></td><td></td><td></td><td></td><td></td><td></td></tr>
<tr><td>Oligo rev</td><td>AAACCCGGCCGATCAGACTCCGTC</td><td></td><td></td><td></td><td></td><td></td><td></td><td></td><td></td><td></td><td></td></tr>
<tr><td>Chromosome</td><td>start</td><td>end</td><td>strand</td><td>MM</td><td>target_seq</td><td>PAM</td><td>alignment</td><td>distance</td><td>position</td><td>gene name</td><td>gene id</td></tr>
<tr><td>GU305914.1</td><td>31402</td><td>31424</td><td>+</td><td>0</td><td>GACGGAGTCTGATCGGCCGG</td><td>CGG</td><td>||||||||[||||||||||||]PAM</td><td>NA</td><td>Intergenic</td><td>NA</td><td>NA</td></tr>
<tr><td>T1927</td><td>GATCGGCCGGCGGATCCTCTCGG</td><td>1000</td><td></td><td></td><td></td><td></td><td></td><td></td><td></td><td></td><td></td></tr>
<tr><td>Oligo fwd</td><td>CACCGATCGGCCGGCGGATCCTCT</td><td></td><td></td><td></td><td></td><td></td><td></td><td></td><td></td><td></td><td></td></tr>
<tr><td>Oligo rev</td><td>AAACAGAGGATCCGCCGGCCGATC</td><td></td><td></td><td></td><td></td><td></td><td></td><td></td><td></td><td></td><td></td></tr>
<tr><td>Chromosome</td><td>start</td><td>end</td><td>strand</td><td>MM</td><td>target_seq</td><td>PAM</td><td>alignment</td><td>distance</td><td>position</td><td>gene name</td><td>gene id</td></tr>
<tr><td>GU305914.1</td><td>31412</td><td>31434</td><td>+</td><td>0</td><td>GATCGGCCGGCGGATCCTCT</td><td>CGG</td><td>||||||||[||||||||||||]PAM</td><td>NA</td><td>Intergenic</td><td>NA</td><td>NA</td></tr>
<tr><td>T1928</td><td>GGCCGGCGGATCCTCTCGGCCGG</td><td>1000</td><td></td><td></td><td></td><td></td><td></td><td></td><td></td><td></td><td></td></tr>
<tr><td>Oligo fwd</td><td>CACCGGCCGGCGGATCCTCTCGGC</td><td></td><td></td><td></td><td></td><td></td><td></td><td></td><td></td><td></td><td></td></tr>
<tr><td>Oligo rev</td><td>AAACGCCGAGAGGATCCGCCGGCC</td><td></td><td></td><td></td><td></td><td></td><td></td><td></td><td></td><td></td><td></td></tr>
<tr><td>Chromosome</td><td>start</td><td>end</td><td>strand</td><td>MM</td><td>target_seq</td><td>PAM</td><td>alignment</td><td>distance</td><td>position</td><td>gene name</td><td>gene id</td></tr>
<tr><td>GU305914.1</td><td>31416</td><td>31438</td><td>+</td><td>0</td><td>GGCCGGCGGATCCTCTCGGC</td><td>CGG</td><td>||||||||[||||||||||||]PAM</td><td>NA</td><td>Intergenic</td><td>NA</td><td>NA</td></tr>
<tr><td>T1929</td><td>GCCGGCGGATCCTCTCGGCCGGG</td><td>1000</td><td></td><td></td><td></td><td></td><td></td><td></td><td></td><td></td><td></td></tr>
<tr><td>Oligo fwd</td><td>CACCGCCGGCGGATCCTCTCGGCC</td><td></td><td></td><td></td><td></td><td></td><td></td><td></td><td></td><td></td><td></td></tr>
<tr><td>Oligo rev</td><td>AAACGGCCGAGAGGATCCGCCGGC</td><td></td><td></td><td></td><td></td><td></td><td></td><td></td><td></td><td></td><td></td></tr>
<tr><td>Chromosome</td><td>start</td><td>end</td><td>strand</td><td>MM</td><td>target_seq</td><td>PAM</td><td>alignment</td><td>distance</td><td>position</td><td>gene name</td><td>gene id</td></tr>
<tr><td>GU305914.1</td><td>31417</td><td>31439</td><td>+</td><td>0</td><td>GCCGGCGGATCCTCTCGGCC</td><td>GGG</td><td>||||||||[||||||||||||]PAM</td><td>NA</td><td>Intergenic</td><td>NA</td><td>NA</td></tr>
<tr><td>T1930</td><td>GATCCTCTCGGCCGGGACCCTGG</td><td>1000</td><td></td><td></td><td></td><td></td><td></td><td></td><td></td><td></td><td></td></tr>
<tr><td>Oligo fwd</td><td>CACCGATCCTCTCGGCCGGGACCC</td><td></td><td></td><td></td><td></td><td></td><td></td><td></td><td></td><td></td><td></td></tr>
<tr><td>Oligo rev</td><td>AAACGGGTCCCGGCCGAGAGGATC</td><td></td><td></td><td></td><td></td><td></td><td></td><td></td><td></td><td></td><td></td></tr>
<tr><td>Chromosome</td><td>start</td><td>end</td><td>strand</td><td>MM</td><td>target_seq</td><td>PAM</td><td>alignment</td><td>distance</td><td>position</td><td>gene name</td><td>gene id</td></tr>
<tr><td>GU305914.1</td><td>31424</td><td>31446</td><td>+</td><td>0</td><td>GATCCTCTCGGCCGGGACCC</td><td>TGG</td><td>||||||||[||||||||||||]PAM</td><td>NA</td><td>Intergenic</td><td>NA</td><td>NA</td></tr>
<tr><td>T1931</td><td>GAGTACGCCTCCGAGCTCGACGG</td><td>1000</td><td></td><td></td><td></td><td></td><td></td><td></td><td></td><td></td><td></td></tr>
<tr><td>Oligo fwd</td><td>CACCGAGTACGCCTCCGAGCTCGA</td><td></td><td></td><td></td><td></td><td></td><td></td><td></td><td></td><td></td><td></td></tr>
<tr><td>Oligo rev</td><td>AAACTCGAGCTCGGAGGCGTACTC</td><td></td><td></td><td></td><td></td><td></td><td></td><td></td><td></td><td></td><td></td></tr>
<tr><td>Chromosome</td><td>start</td><td>end</td><td>strand</td><td>MM</td><td>target_seq</td><td>PAM</td><td>alignment</td><td>distance</td><td>position</td><td>gene name</td><td>gene id</td></tr>
<tr><td>GU305914.1</td><td>31451</td><td>31473</td><td>+</td><td>0</td><td>GAGTACGCCTCCGAGCTCGA</td><td>CGG</td><td>||||||||[||||||||||||]PAM</td><td>NA</td><td>Intergenic</td><td>NA</td><td>NA</td></tr>
<tr><td>T1932</td><td>AGTACGCCTCCGAGCTCGACGGG</td><td>1000</td><td></td><td></td><td></td><td></td><td></td><td></td><td></td><td></td><td></td></tr>
<tr><td>Oligo fwd</td><td>CACCAGTACGCCTCCGAGCTCGAC</td><td></td><td></td><td></td><td></td><td></td><td></td><td></td><td></td><td></td><td></td></tr>
<tr><td>Oligo rev</td><td>AAACGTCGAGCTCGGAGGCGTACT</td><td></td><td></td><td></td><td></td><td></td><td></td><td></td><td></td><td></td><td></td></tr>
<tr><td>Chromosome</td><td>start</td><td>end</td><td>strand</td><td>MM</td><td>target_seq</td><td>PAM</td><td>alignment</td><td>distance</td><td>position</td><td>gene name</td><td>gene id</td></tr>
</table>

GU305914.1 31452 31474 + 0 AGTACGCCTCCGAGCTCGAC GGG ||||||||[||||||||||||]PAM NA Intergenic NA NA

T1933 CGAGCTCGACGGGCAGCCACTGG 1000
Oligo fwd CACCCGAGCTCGACGGGCAGCCAC
Oligo rev AAACGTGGCTGCCCGTCGAGCTCG
Chromosome start end strand MM target_seq PAM alignment distance position gene name gene id
GU305914.1 31462 31484 + 0 CGAGCTCGACGGGCAGCCAC TGG ||||||||[||||||||||||]PAM NA Intergenic NA NA

T1934 GACGGGCAGCCACTGGCTGAAGG 1000
Oligo fwd CACCGACGGGCAGCCACTGGCTGA
Oligo rev AAACTCAGCCAGTGGCTGCCCGTC
Chromosome start end strand MM target_seq PAM alignment distance position gene name gene id
GU305914.1 31469 31491 + 0 GACGGGCAGCCACTGGCTGA AGG ||||||||[||||||||||||]PAM NA Intergenic NA NA

T1935 GGCAGCCACTGGCTGAAGGACGG 1000
Oligo fwd CACCGGCAGCCACTGGCTGAAGGA
Oligo rev AAACTCCTTCAGCCAGTGGCTGCC
Chromosome start end strand MM target_seq PAM alignment distance position gene name gene id
GU305914.1 31473 31495 + 0 GGCAGCCACTGGCTGAAGGA CGG ||||||||[||||||||||||]PAM NA Intergenic NA NA

T1936 AGCCACTGGCTGAAGGACGGCGG 1000
Oligo fwd CACCAGCCACTGGCTGAAGGACGG
Oligo rev AAACCCGTCCTTCAGCCAGTGGCT
Chromosome start end strand MM target_seq PAM alignment distance position gene name gene id
GU305914.1 31476 31498 + 0 AGCCACTGGCTGAAGGACGG CGG ||||||||[||||||||||||]PAM NA Intergenic NA NA

T1937 GCTGAAGGACGGCGGTAGCCAGG 1000
Oligo fwd CACCGCTGAAGGACGGCGGTAGCC
Oligo rev AAACGGCTACCGCCGTCCTTCAGC
Chromosome start end strand MM target_seq PAM alignment distance position gene name gene id
GU305914.1 31484 31506 + 0 GCTGAAGGACGGCGGTAGCC AGG ||||||||[||||||||||||]PAM NA Intergenic NA NA

T1938 CTGAAGGACGGCGGTAGCCAGGG 1000
Oligo fwd CACCCTGAAGGACGGCGGTAGCCA
Oligo rev AAACTGGCTACCGCCGTCCTTCAG
Chromosome start end strand MM target_seq PAM alignment distance position gene name gene id
GU305914.1 31485 31507 + 0 CTGAAGGACGGCGGTAGCCA GGG ||||||||[||||||||||||]PAM NA Intergenic NA NA

T1939 GGCGGTAGCCAGGGAGACCGCGG 1000
Oligo fwd CACCGGCGGTAGCCAGGGAGACCG
Oligo rev AAACCGGTCTCCCTGGCTACCGCC
Chromosome start end strand MM target_seq PAM alignment distance position gene name gene id
GU305914.1 31494 31516 + 0 GGCGGTAGCCAGGGAGACCG CGG ||||||||[||||||||||||]PAM NA Intergenic NA NA
GU305914.1 70978 71000 + 4 GGCGATCGCCAGGAAGATCG AGG ||||-|-|[|||||-|||-||]PAM NA Intergenic NA NA

T1940 GAGACCGCGGCTCGTTACACTGG 1000
Oligo fwd CACCGAGACCGCGGCTCGTTACAC
Oligo rev AAACGTGTAACGAGCCGCGGTCTC
Chromosome start end strand MM target_seq PAM alignment distance position gene name gene id
GU305914.1 31507 31529 + 0 GAGACCGCGGCTCGTTACAC TGG ||||||||[||||||||||||]PAM NA Intergenic NA NA

T1941 TGGCAGATCTCGACGATCTGCGG 1000
Oligo fwd CACCTGGCAGATCTCGACGATCTG
Oligo rev AAACCAGATCGTCGAGATCTGCCA
Chromosome start end strand MM target_seq PAM alignment distance position gene name gene id
GU305914.1 31527 31549 + 0 TGGCAGATCTCGACGATCTG CGG ||||||||[||||||||||||]PAM NA Intergenic NA NA

T1942 AGATCTCGACGATCTGCGGTAGG 1000
Oligo fwd CACCAGATCTCGACGATCTGCGGT
Oligo rev AAACACCGCAGATCGTCGAGATCT
Chromosome start end strand MM target_seq PAM alignment distance position gene name gene id
GU305914.1 31531 31553 + 0 AGATCTCGACGATCTGCGGT AGG ||||||||[||||||||||||]PAM NA Intergenic NA NA

T1943 GCGGTAGGACGCCCTCACGCAGG 1000
Oligo fwd CACCGCGGTAGGACGCCCTCACGC
Oligo rev AAACGCGTGAGGGCGTCCTACCGC
Chromosome start end strand MM target_seq PAM alignment distance position gene name gene id
GU305914.1 31546 31568 + 0 GCGGTAGGACGCCCTCACGC AGG ||||||||[||||||||||||]PAM NA Intergenic NA NA

T1944 CACGCAGGTCGCGCTGTACCTGG 1000
Oligo fwd CACCCACGCAGGTCGCGCTGTACC
Oligo rev AAACGGTACAGCGCGACCTGCGTG
Chromosome start end strand MM target_seq PAM alignment distance position gene name gene id
GU305914.1 31561 31583 + 0 CACGCAGGTCGCGCTGTACC TGG ||||||||[||||||||||||]PAM NA Intergenic NA NA

T1945 CGTCCGAGATGAGCTGCGTGAGG 1000
Oligo fwd CACCCGTCCGAGATGAGCTGCGTG
Oligo rev AAACCACGCAGCTCATCTCGGACG
Chromosome start end strand MM target_seq PAM alignment distance position gene name gene id
GU305914.1 31603 31625 + 0 CGTCCGAGATGAGCTGCGTG AGG ||||||||[||||||||||||]PAM NA Intergenic NA NA

T1946 CGAGACGTACATGATGAAGCAGG 1000
Oligo fwd CACCCGAGACGTACATGATGAAGC
Oligo rev AAACGCTTCATCATGTACGTCTCG
Chromosome start end strand MM target_seq PAM alignment distance position gene name gene id
GU305914.1 31631 31653 + 0 CGAGACGTACATGATGAAGC AGG ||||||||[||||||||||||]PAM NA Intergenic NA NA

T1947 GCAGGCATCGTTGATCTCGTAGG 1000
Oligo fwd CACCGCAGGCATCGTTGATCTCGT

<table>
<tr><td>Oligo rev</td><td>AAACACGAGATCAACGATGCCTGC</td><td></td><td></td><td></td><td></td><td></td><td></td><td></td><td></td><td></td><td></td></tr>
<tr><td>Chromosome</td><td>start</td><td>end</td><td>strand</td><td>MM</td><td>target_seq</td><td>PAM</td><td>alignment</td><td>distance</td><td>position</td><td>gene name</td><td>gene id</td></tr>
<tr><td>GU305914.1</td><td>31649</td><td>31671</td><td>+</td><td>0</td><td>GCAGGCATCGTTGATCTCGT</td><td>AGG</td><td>||||||||[||||||||||||]PAM</td><td>NA</td><td>Intergenic</td><td>NA</td><td>NA</td></tr>
<tr><td></td><td></td><td></td><td></td><td></td><td></td><td></td><td></td><td></td><td></td><td></td><td></td></tr>
<tr><td>T1948</td><td>TGATCTCGTAGGTGAAGACCTGG</td><td>1000</td><td></td><td></td><td></td><td></td><td></td><td></td><td></td><td></td><td></td></tr>
<tr><td>Oligo fwd</td><td>CACCTGATCTCGTAGGTGAAGACC</td><td></td><td></td><td></td><td></td><td></td><td></td><td></td><td></td><td></td><td></td></tr>
<tr><td>Oligo rev</td><td>AAACGGTCTTCACCTACGAGATCA</td><td></td><td></td><td></td><td></td><td></td><td></td><td></td><td></td><td></td><td></td></tr>
<tr><td>Chromosome</td><td>start</td><td>end</td><td>strand</td><td>MM</td><td>target_seq</td><td>PAM</td><td>alignment</td><td>distance</td><td>position</td><td>gene name</td><td>gene id</td></tr>
<tr><td>GU305914.1</td><td>31660</td><td>31682</td><td>+</td><td>0</td><td>TGATCTCGTAGGTGAAGACC</td><td>TGG</td><td>||||||||[||||||||||||]PAM</td><td>NA</td><td>Intergenic</td><td>NA</td><td>NA</td></tr>
<tr><td>GU305914.1</td><td>197232</td><td>197254</td><td>+</td><td>4</td><td>GAATCTCGTAGACGAAGACC</td><td>CGG</td><td>--||||||[|||--|||||||]PAM</td><td>NA</td><td>Intergenic</td><td>NA</td><td>NA</td></tr>
<tr><td></td><td></td><td></td><td></td><td></td><td></td><td></td><td></td><td></td><td></td><td></td><td></td></tr>
<tr><td>T1949</td><td>GTAGGTGAAGACCTGGCCCACGG</td><td>1000</td><td></td><td></td><td></td><td></td><td></td><td></td><td></td><td></td><td></td></tr>
<tr><td>Oligo fwd</td><td>CACCGTAGGTGAAGACCTGGCCCA</td><td></td><td></td><td></td><td></td><td></td><td></td><td></td><td></td><td></td><td></td></tr>
<tr><td>Oligo rev</td><td>AAACTGGGCCAGGTCTTCACCTAC</td><td></td><td></td><td></td><td></td><td></td><td></td><td></td><td></td><td></td><td></td></tr>
<tr><td>Chromosome</td><td>start</td><td>end</td><td>strand</td><td>MM</td><td>target_seq</td><td>PAM</td><td>alignment</td><td>distance</td><td>position</td><td>gene name</td><td>gene id</td></tr>
<tr><td>GU305914.1</td><td>31667</td><td>31689</td><td>+</td><td>0</td><td>GTAGGTGAAGACCTGGCCCA</td><td>CGG</td><td>||||||||[||||||||||||]PAM</td><td>NA</td><td>Intergenic</td><td>NA</td><td>NA</td></tr>
<tr><td></td><td></td><td></td><td></td><td></td><td></td><td></td><td></td><td></td><td></td><td></td><td></td></tr>
<tr><td>T1950</td><td>CGGCGTTGACGCCGACCACCCGG</td><td>1000</td><td></td><td></td><td></td><td></td><td></td><td></td><td></td><td></td><td></td></tr>
<tr><td>Oligo fwd</td><td>CACCCGGCGTTGACGCCGACCACC</td><td></td><td></td><td></td><td></td><td></td><td></td><td></td><td></td><td></td><td></td></tr>
<tr><td>Oligo rev</td><td>AAACGGTGGTCGGCGTCAACGCCG</td><td></td><td></td><td></td><td></td><td></td><td></td><td></td><td></td><td></td><td></td></tr>
<tr><td>Chromosome</td><td>start</td><td>end</td><td>strand</td><td>MM</td><td>target_seq</td><td>PAM</td><td>alignment</td><td>distance</td><td>position</td><td>gene name</td><td>gene id</td></tr>
<tr><td>GU305914.1</td><td>31687</td><td>31709</td><td>+</td><td>0</td><td>CGGCGTTGACGCCGACCACC</td><td>CGG</td><td>||||||||[||||||||||||]PAM</td><td>NA</td><td>Intergenic</td><td>NA</td><td>NA</td></tr>
<tr><td></td><td></td><td></td><td></td><td></td><td></td><td></td><td></td><td></td><td></td><td></td><td></td></tr>
<tr><td>T1951</td><td>CGTTGACGCCGACCACCCGGCGG</td><td>1000</td><td></td><td></td><td></td><td></td><td></td><td></td><td></td><td></td><td></td></tr>
<tr><td>Oligo fwd</td><td>CACCCGTTGACGCCGACCACCCGG</td><td></td><td></td><td></td><td></td><td></td><td></td><td></td><td></td><td></td><td></td></tr>
<tr><td>Oligo rev</td><td>AAACCCGGGTGGTCGGCGTCAACG</td><td></td><td></td><td></td><td></td><td></td><td></td><td></td><td></td><td></td><td></td></tr>
<tr><td>Chromosome</td><td>start</td><td>end</td><td>strand</td><td>MM</td><td>target_seq</td><td>PAM</td><td>alignment</td><td>distance</td><td>position</td><td>gene name</td><td>gene id</td></tr>
<tr><td>GU305914.1</td><td>31690</td><td>31712</td><td>+</td><td>0</td><td>CGTTGACGCCGACCACCCGG</td><td>CGG</td><td>||||||||[||||||||||||]PAM</td><td>NA</td><td>Intergenic</td><td>NA</td><td>NA</td></tr>
<tr><td></td><td></td><td></td><td></td><td></td><td></td><td></td><td></td><td></td><td></td><td></td><td></td></tr>
<tr><td>T1952</td><td>CACCCGGCGGTCCCAAGCGAAGG</td><td>1000</td><td></td><td></td><td></td><td></td><td></td><td></td><td></td><td></td><td></td></tr>
<tr><td>Oligo fwd</td><td>CACCCACCCGGCGGTCCCAAGCGA</td><td></td><td></td><td></td><td></td><td></td><td></td><td></td><td></td><td></td><td></td></tr>
<tr><td>Oligo rev</td><td>AAACTCGCTTGGGACCGCCGGGTG</td><td></td><td></td><td></td><td></td><td></td><td></td><td></td><td></td><td></td><td></td></tr>
<tr><td>Chromosome</td><td>start</td><td>end</td><td>strand</td><td>MM</td><td>target_seq</td><td>PAM</td><td>alignment</td><td>distance</td><td>position</td><td>gene name</td><td>gene id</td></tr>
<tr><td>GU305914.1</td><td>31703</td><td>31725</td><td>+</td><td>0</td><td>CACCCGGCGGTCCCAAGCGA</td><td>AGG</td><td>||||||||[||||||||||||]PAM</td><td>NA</td><td>Intergenic</td><td>NA</td><td>NA</td></tr>
<tr><td></td><td></td><td></td><td></td><td></td><td></td><td></td><td></td><td></td><td></td><td></td><td></td></tr>
<tr><td>T1953</td><td>GGTCCCAAGCGAAGGTGCGTCGG</td><td>1000</td><td></td><td></td><td></td><td></td><td></td><td></td><td></td><td></td><td></td></tr>
<tr><td>Oligo fwd</td><td>CACCGGTCCCAAGCGAAGGTGCGT</td><td></td><td></td><td></td><td></td><td></td><td></td><td></td><td></td><td></td><td></td></tr>
<tr><td>Oligo rev</td><td>AAACACGCACCTTCGCTTGGGACC</td><td></td><td></td><td></td><td></td><td></td><td></td><td></td><td></td><td></td><td></td></tr>
<tr><td>Chromosome</td><td>start</td><td>end</td><td>strand</td><td>MM</td><td>target_seq</td><td>PAM</td><td>alignment</td><td>distance</td><td>position</td><td>gene name</td><td>gene id</td></tr>
<tr><td>GU305914.1</td><td>31711</td><td>31733</td><td>+</td><td>0</td><td>GGTCCCAAGCGAAGGTGCGT</td><td>CGG</td><td>||||||||[||||||||||||]PAM</td><td>NA</td><td>Intergenic</td><td>NA</td><td>NA</td></tr>
<tr><td></td><td></td><td></td><td></td><td></td><td></td><td></td><td></td><td></td><td></td><td></td><td></td></tr>
<tr><td>T1954</td><td>GTGCCCGAGCGTGCCGATGACGG</td><td>1000</td><td></td><td></td><td></td><td></td><td></td><td></td><td></td><td></td><td></td></tr>
<tr><td>Oligo fwd</td><td>CACCGTGCCCGAGCGTGCCGATGA</td><td></td><td></td><td></td><td></td><td></td><td></td><td></td><td></td><td></td><td></td></tr>
<tr><td>Oligo rev</td><td>AAACTCATCGGCACGCTCGGGCAC</td><td></td><td></td><td></td><td></td><td></td><td></td><td></td><td></td><td></td><td></td></tr>
<tr><td>Chromosome</td><td>start</td><td>end</td><td>strand</td><td>MM</td><td>target_seq</td><td>PAM</td><td>alignment</td><td>distance</td><td>position</td><td>gene name</td><td>gene id</td></tr>
<tr><td>GU305914.1</td><td>31733</td><td>31755</td><td>+</td><td>0</td><td>GTGCCCGAGCGTGCCGATGA</td><td>CGG</td><td>||||||||[||||||||||||]PAM</td><td>NA</td><td>Intergenic</td><td>NA</td><td>NA</td></tr>
<tr><td></td><td></td><td></td><td></td><td></td><td></td><td></td><td></td><td></td><td></td><td></td><td></td></tr>
<tr><td>T1955</td><td>GTGAGCTCCCCGCACATGAGCGG</td><td>1000</td><td></td><td></td><td></td><td></td><td></td><td></td><td></td><td></td><td></td></tr>
<tr><td>Oligo fwd</td><td>CACCGTGAGCTCCCCGCACATGAG</td><td></td><td></td><td></td><td></td><td></td><td></td><td></td><td></td><td></td><td></td></tr>
<tr><td>Oligo rev</td><td>AAACCTCATGTGCGGGGAGCTCAC</td><td></td><td></td><td></td><td></td><td></td><td></td><td></td><td></td><td></td><td></td></tr>
<tr><td>Chromosome</td><td>start</td><td>end</td><td>strand</td><td>MM</td><td>target_seq</td><td>PAM</td><td>alignment</td><td>distance</td><td>position</td><td>gene name</td><td>gene id</td></tr>
<tr><td>GU305914.1</td><td>31755</td><td>31777</td><td>+</td><td>0</td><td>GTGAGCTCCCCGCACATGAG</td><td>CGG</td><td>||||||||[||||||||||||]PAM</td><td>NA</td><td>Intergenic</td><td>NA</td><td>NA</td></tr>
<tr><td></td><td></td><td></td><td></td><td></td><td></td><td></td><td></td><td></td><td></td><td></td><td></td></tr>
<tr><td>T1956</td><td>TGAGCTCCCCGCACATGAGCGGG</td><td>1000</td><td></td><td></td><td></td><td></td><td></td><td></td><td></td><td></td><td></td></tr>
<tr><td>Oligo fwd</td><td>CACCTGAGCTCCCCGCACATGAGC</td><td></td><td></td><td></td><td></td><td></td><td></td><td></td><td></td><td></td><td></td></tr>
<tr><td>Oligo rev</td><td>AAACGCTCATGTGCGGGGAGCTCA</td><td></td><td></td><td></td><td></td><td></td><td></td><td></td><td></td><td></td><td></td></tr>
<tr><td>Chromosome</td><td>start</td><td>end</td><td>strand</td><td>MM</td><td>target_seq</td><td>PAM</td><td>alignment</td><td>distance</td><td>position</td><td>gene name</td><td>gene id</td></tr>
<tr><td>GU305914.1</td><td>31756</td><td>31778</td><td>+</td><td>0</td><td>TGAGCTCCCCGCACATGAGC</td><td>GGG</td><td>||||||||[||||||||||||]PAM</td><td>NA</td><td>Intergenic</td><td>NA</td><td>NA</td></tr>
<tr><td></td><td></td><td></td><td></td><td></td><td></td><td></td><td></td><td></td><td></td><td></td><td></td></tr>
<tr><td>T1957</td><td>GCGGGTGCATCCGCCAGAAATGG</td><td>1000</td><td></td><td></td><td></td><td></td><td></td><td></td><td></td><td></td><td></td></tr>
<tr><td>Oligo fwd</td><td>CACCGCGGGTGCATCCGCCAGAAA</td><td></td><td></td><td></td><td></td><td></td><td></td><td></td><td></td><td></td><td></td></tr>
<tr><td>Oligo rev</td><td>AAACTTTCTGGCGGATGCACCCGC</td><td></td><td></td><td></td><td></td><td></td><td></td><td></td><td></td><td></td><td></td></tr>
<tr><td>Chromosome</td><td>start</td><td>end</td><td>strand</td><td>MM</td><td>target_seq</td><td>PAM</td><td>alignment</td><td>distance</td><td>position</td><td>gene name</td><td>gene id</td></tr>
<tr><td>GU305914.1</td><td>31774</td><td>31796</td><td>+</td><td>0</td><td>GCGGGTGCATCCGCCAGAAA</td><td>TGG</td><td>||||||||[||||||||||||]PAM</td><td>NA</td><td>Intergenic</td><td>NA</td><td>NA</td></tr>
<tr><td></td><td></td><td></td><td></td><td></td><td></td><td></td><td></td><td></td><td></td><td></td><td></td></tr>
<tr><td>T1958</td><td>CCAGAAATGGAGATCGCACGAGG</td><td>1000</td><td></td><td></td><td></td><td></td><td></td><td></td><td></td><td></td><td></td></tr>
<tr><td>Oligo fwd</td><td>CACCCCAGAAATGGAGATCGCACG</td><td></td><td></td><td></td><td></td><td></td><td></td><td></td><td></td><td></td><td></td></tr>
<tr><td>Oligo rev</td><td>AAACCGTGCGATCTCCATTTCTGG</td><td></td><td></td><td></td><td></td><td></td><td></td><td></td><td></td><td></td><td></td></tr>
<tr><td>Chromosome</td><td>start</td><td>end</td><td>strand</td><td>MM</td><td>target_seq</td><td>PAM</td><td>alignment</td><td>distance</td><td>position</td><td>gene name</td><td>gene id</td></tr>
<tr><td>GU305914.1</td><td>31787</td><td>31809</td><td>+</td><td>0</td><td>CCAGAAATGGAGATCGCACG</td><td>AGG</td><td>||||||||[||||||||||||]PAM</td><td>NA</td><td>Intergenic</td><td>NA</td><td>NA</td></tr>
<tr><td></td><td></td><td></td><td></td><td></td><td></td><td></td><td></td><td></td><td></td><td></td><td></td></tr>
<tr><td>T1959</td><td>GAGATCGCACGAGGTGAGTATGG</td><td>1000</td><td></td><td></td><td></td><td></td><td></td><td></td><td></td><td></td><td></td></tr>
<tr><td>Oligo fwd</td><td>CACCGAGATCGCACGAGGTGAGTA</td><td></td><td></td><td></td><td></td><td></td><td></td><td></td><td></td><td></td><td></td></tr>
<tr><td>Oligo rev</td><td>AAACTACTCACCTCGTGCGATCTC</td><td></td><td></td><td></td><td></td><td></td><td></td><td></td><td></td><td></td><td></td></tr>
<tr><td>Chromosome</td><td>start</td><td>end</td><td>strand</td><td>MM</td><td>target_seq</td><td>PAM</td><td>alignment</td><td>distance</td><td>position</td><td>gene name</td><td>gene id</td></tr>
<tr><td>GU305914.1</td><td>31796</td><td>31818</td><td>+</td><td>0</td><td>GAGATCGCACGAGGTGAGTA</td><td>TGG</td><td>||||||||[||||||||||||]PAM</td><td>NA</td><td>Intergenic</td><td>NA</td><td>NA</td></tr>
<tr><td></td><td></td><td></td><td></td><td></td><td></td><td></td><td></td><td></td><td></td><td></td><td></td></tr>
<tr><td>T1960</td><td>GATGTAAAAGATGTACCCCGTGG</td><td>1000</td><td></td><td></td><td></td><td></td><td></td><td></td><td></td><td></td><td></td></tr>
<tr><td>Oligo fwd</td><td>CACCGATGTAAAAGATGTACCCCG</td><td></td><td></td><td></td><td></td><td></td><td></td><td></td><td></td><td></td><td></td></tr>
<tr><td>Oligo rev</td><td>AAACCGGGGTACATCTTTTACATC</td><td></td><td></td><td></td><td></td><td></td><td></td><td></td><td></td><td></td><td></td></tr>
<tr><td>Chromosome</td><td>start</td><td>end</td><td>strand</td><td>MM</td><td>target_seq</td><td>PAM</td><td>alignment</td><td>distance</td><td>position</td><td>gene name</td><td>gene id</td></tr>
<tr><td>GU305914.1</td><td>31835</td><td>31857</td><td>+</td><td>0</td><td>GATGTAAAAGATGTACCCCG</td><td>TGG</td><td>||||||||[||||||||||||]PAM</td><td>NA</td><td>Intergenic</td><td>NA</td><td>NA</td></tr>
<tr><td></td><td></td><td></td><td></td><td></td><td></td><td></td><td></td><td></td><td></td><td></td><td></td></tr>
<tr><td>T1961</td><td>ATGTAAAAGATGTACCCCGTGGG</td><td>1000</td><td></td><td></td><td></td><td></td><td></td><td></td><td></td><td></td><td></td></tr>
<tr><td>Oligo fwd</td><td>CACCATGTAAAAGATGTACCCCGT</td><td></td><td></td><td></td><td></td><td></td><td></td><td></td><td></td><td></td><td></td></tr>
<tr><td>Oligo rev</td><td>AAACACGGGGTACATCTTTTACAT</td><td></td><td></td><td></td><td></td><td></td><td></td><td></td><td></td><td></td><td></td></tr>
<tr><td>Chromosome</td><td>start</td><td>end</td><td>strand</td><td>MM</td><td>target_seq</td><td>PAM</td><td>alignment</td><td>distance</td><td>position</td><td>gene name</td><td>gene id</td></tr>
<tr><td>GU305914.1</td><td>31836</td><td>31858</td><td>+</td><td>0</td><td>ATGTAAAAGATGTACCCCGT</td><td>GGG</td><td>||||||||[||||||||||||]PAM</td><td>NA</td><td>Intergenic</td><td>NA</td><td>NA</td></tr>
</table>

<table>
<tr><td>T1962</td><td>TGTAAAAGATGTACCCCGTGGGG</td><td>1000</td><td></td><td></td><td></td><td></td><td></td><td></td><td></td><td></td><td></td></tr>
<tr><td>Oligo fwd</td><td>CACCTGTAAAAGATGTACCCCGTG</td><td></td><td></td><td></td><td></td><td></td><td></td><td></td><td></td><td></td><td></td></tr>
<tr><td>Oligo rev</td><td>AAACCACGGGGTACATCTTTTACA</td><td></td><td></td><td></td><td></td><td></td><td></td><td></td><td></td><td></td><td></td></tr>
<tr><td>Chromosome</td><td>start</td><td>end</td><td>strand</td><td>MM</td><td>target_seq</td><td>PAM</td><td>alignment</td><td>distance</td><td>position</td><td>gene name</td><td>gene id</td></tr>
<tr><td>GU305914.1</td><td>31837</td><td>31859</td><td>+</td><td>0</td><td>TGTAAAAGATGTACCCCGTG</td><td>GGG</td><td>||||||||[||||||||||||]PAM</td><td>NA</td><td>Intergenic</td><td>NA</td><td>NA</td></tr>
<tr><td>T1963</td><td>AAGATGTACCCCGTGGGGAAAGG</td><td>1000</td><td></td><td></td><td></td><td></td><td></td><td></td><td></td><td></td><td></td></tr>
<tr><td>Oligo fwd</td><td>CACCAAGATGTACCCCGTGGGGAA</td><td></td><td></td><td></td><td></td><td></td><td></td><td></td><td></td><td></td><td></td></tr>
<tr><td>Oligo rev</td><td>AAACTTCCCCACGGGGTACATCTT</td><td></td><td></td><td></td><td></td><td></td><td></td><td></td><td></td><td></td><td></td></tr>
<tr><td>Chromosome</td><td>start</td><td>end</td><td>strand</td><td>MM</td><td>target_seq</td><td>PAM</td><td>alignment</td><td>distance</td><td>position</td><td>gene name</td><td>gene id</td></tr>
<tr><td>GU305914.1</td><td>31842</td><td>31864</td><td>+</td><td>0</td><td>AAGATGTACCCCGTGGGGAA</td><td>AGG</td><td>||||||||[||||||||||||]PAM</td><td>NA</td><td>Intergenic</td><td>NA</td><td>NA</td></tr>
<tr><td>T1964</td><td>AGATGTACCCCGTGGGGAAAGGG</td><td>1000</td><td></td><td></td><td></td><td></td><td></td><td></td><td></td><td></td><td></td></tr>
<tr><td>Oligo fwd</td><td>CACCAGATGTACCCCGTGGGGAAA</td><td></td><td></td><td></td><td></td><td></td><td></td><td></td><td></td><td></td><td></td></tr>
<tr><td>Oligo rev</td><td>AAACTTTCCCCACGGGGTACATCT</td><td></td><td></td><td></td><td></td><td></td><td></td><td></td><td></td><td></td><td></td></tr>
<tr><td>Chromosome</td><td>start</td><td>end</td><td>strand</td><td>MM</td><td>target_seq</td><td>PAM</td><td>alignment</td><td>distance</td><td>position</td><td>gene name</td><td>gene id</td></tr>
<tr><td>GU305914.1</td><td>31843</td><td>31865</td><td>+</td><td>0</td><td>AGATGTACCCCGTGGGGAAA</td><td>GGG</td><td>||||||||[||||||||||||]PAM</td><td>NA</td><td>Intergenic</td><td>NA</td><td>NA</td></tr>
<tr><td>T1965</td><td>GTCTCTGTACACATCCCCTCAGG</td><td>1000</td><td></td><td></td><td></td><td></td><td></td><td></td><td></td><td></td><td></td></tr>
<tr><td>Oligo fwd</td><td>CACCGTCTCTGTACACATCCCCTC</td><td></td><td></td><td></td><td></td><td></td><td></td><td></td><td></td><td></td><td></td></tr>
<tr><td>Oligo rev</td><td>AAACGAGGGGATGTGTACAGAGAC</td><td></td><td></td><td></td><td></td><td></td><td></td><td></td><td></td><td></td><td></td></tr>
<tr><td>Chromosome</td><td>start</td><td>end</td><td>strand</td><td>MM</td><td>target_seq</td><td>PAM</td><td>alignment</td><td>distance</td><td>position</td><td>gene name</td><td>gene id</td></tr>
<tr><td>GU305914.1</td><td>31888</td><td>31910</td><td>+</td><td>0</td><td>GTCTCTGTACACATCCCCTC</td><td>AGG</td><td>||||||||[||||||||||||]PAM</td><td>NA</td><td>Intergenic</td><td>NA</td><td>NA</td></tr>
<tr><td>T1966</td><td>CCTCAGGCATTCGCCACCCATGG</td><td>1000</td><td></td><td></td><td></td><td></td><td></td><td></td><td></td><td></td><td></td></tr>
<tr><td>Oligo fwd</td><td>CACCCCTCAGGCATTCGCCACCCA</td><td></td><td></td><td></td><td></td><td></td><td></td><td></td><td></td><td></td><td></td></tr>
<tr><td>Oligo rev</td><td>AAACTGGGTGGCGAATGCCTGAGG</td><td></td><td></td><td></td><td></td><td></td><td></td><td></td><td></td><td></td><td></td></tr>
<tr><td>Chromosome</td><td>start</td><td>end</td><td>strand</td><td>MM</td><td>target_seq</td><td>PAM</td><td>alignment</td><td>distance</td><td>position</td><td>gene name</td><td>gene id</td></tr>
<tr><td>GU305914.1</td><td>31904</td><td>31926</td><td>+</td><td>0</td><td>CCTCAGGCATTCGCCACCCA</td><td>TGG</td><td>||||||||[||||||||||||]PAM</td><td>NA</td><td>Intergenic</td><td>NA</td><td>NA</td></tr>
<tr><td>T1967</td><td>CAGGCATTCGCCACCCATGGCGG</td><td>1000</td><td></td><td></td><td></td><td></td><td></td><td></td><td></td><td></td><td></td></tr>
<tr><td>Oligo fwd</td><td>CACCCAGGCATTCGCCACCCATGG</td><td></td><td></td><td></td><td></td><td></td><td></td><td></td><td></td><td></td><td></td></tr>
<tr><td>Oligo rev</td><td>AAACCCATGGGTGGCGAATGCCTG</td><td></td><td></td><td></td><td></td><td></td><td></td><td></td><td></td><td></td><td></td></tr>
<tr><td>Chromosome</td><td>start</td><td>end</td><td>strand</td><td>MM</td><td>target_seq</td><td>PAM</td><td>alignment</td><td>distance</td><td>position</td><td>gene name</td><td>gene id</td></tr>
<tr><td>GU305914.1</td><td>31907</td><td>31929</td><td>+</td><td>0</td><td>CAGGCATTCGCCACCCATGG</td><td>CGG</td><td>||||||||[||||||||||||]PAM</td><td>NA</td><td>Intergenic</td><td>NA</td><td>NA</td></tr>
<tr><td>T1968</td><td>GCATTCGCCACCCATGGCGGCGG</td><td>1000</td><td></td><td></td><td></td><td></td><td></td><td></td><td></td><td></td><td></td></tr>
<tr><td>Oligo fwd</td><td>CACCGCATTCGCCACCCATGGCGG</td><td></td><td></td><td></td><td></td><td></td><td></td><td></td><td></td><td></td><td></td></tr>
<tr><td>Oligo rev</td><td>AAACCCGCCATGGGTGGCGAATGC</td><td></td><td></td><td></td><td></td><td></td><td></td><td></td><td></td><td></td><td></td></tr>
<tr><td>Chromosome</td><td>start</td><td>end</td><td>strand</td><td>MM</td><td>target_seq</td><td>PAM</td><td>alignment</td><td>distance</td><td>position</td><td>gene name</td><td>gene id</td></tr>
<tr><td>GU305914.1</td><td>31910</td><td>31932</td><td>+</td><td>0</td><td>GCATTCGCCACCCATGGCGG</td><td>CGG</td><td>||||||||[||||||||||||]PAM</td><td>NA</td><td>Intergenic</td><td>NA</td><td>NA</td></tr>
<tr><td>T1969</td><td>CCAGCTCGCAGACCTCGTTCTGG</td><td>1000</td><td></td><td></td><td></td><td></td><td></td><td></td><td></td><td></td><td></td></tr>
<tr><td>Oligo fwd</td><td>CACCCCAGCTCGCAGACCTCGTTC</td><td></td><td></td><td></td><td></td><td></td><td></td><td></td><td></td><td></td><td></td></tr>
<tr><td>Oligo rev</td><td>AAACGAACGAGGTCTGCGAGCTGG</td><td></td><td></td><td></td><td></td><td></td><td></td><td></td><td></td><td></td><td></td></tr>
<tr><td>Chromosome</td><td>start</td><td>end</td><td>strand</td><td>MM</td><td>target_seq</td><td>PAM</td><td>alignment</td><td>distance</td><td>position</td><td>gene name</td><td>gene id</td></tr>
<tr><td>GU305914.1</td><td>31939</td><td>31961</td><td>+</td><td>0</td><td>CCAGCTCGCAGACCTCGTTC</td><td>TGG</td><td>||||||||[||||||||||||]PAM</td><td>NA</td><td>Intergenic</td><td>NA</td><td>NA</td></tr>
<tr><td>T1970</td><td>CAGCTCGCAGACCTCGTTCTGGG</td><td>1000</td><td></td><td></td><td></td><td></td><td></td><td></td><td></td><td></td><td></td></tr>
<tr><td>Oligo fwd</td><td>CACCCAGCTCGCAGACCTCGTTCT</td><td></td><td></td><td></td><td></td><td></td><td></td><td></td><td></td><td></td><td></td></tr>
<tr><td>Oligo rev</td><td>AAACAGAACGAGGTCTGCGAGCTG</td><td></td><td></td><td></td><td></td><td></td><td></td><td></td><td></td><td></td><td></td></tr>
<tr><td>Chromosome</td><td>start</td><td>end</td><td>strand</td><td>MM</td><td>target_seq</td><td>PAM</td><td>alignment</td><td>distance</td><td>position</td><td>gene name</td><td>gene id</td></tr>
<tr><td>GU305914.1</td><td>31940</td><td>31962</td><td>+</td><td>0</td><td>CAGCTCGCAGACCTCGTTCT</td><td>GGG</td><td>||||||||[||||||||||||]PAM</td><td>NA</td><td>Intergenic</td><td>NA</td><td>NA</td></tr>
<tr><td>T1971</td><td>CTCGTTCTGGGACGCGCCGTAGG</td><td>1000</td><td></td><td></td><td></td><td></td><td></td><td></td><td></td><td></td><td></td></tr>
<tr><td>Oligo fwd</td><td>CACCCTCGTTCTGGGACGCGCCGT</td><td></td><td></td><td></td><td></td><td></td><td></td><td></td><td></td><td></td><td></td></tr>
<tr><td>Oligo rev</td><td>AAACACGGCGCGTCCCAGAACGAG</td><td></td><td></td><td></td><td></td><td></td><td></td><td></td><td></td><td></td><td></td></tr>
<tr><td>Chromosome</td><td>start</td><td>end</td><td>strand</td><td>MM</td><td>target_seq</td><td>PAM</td><td>alignment</td><td>distance</td><td>position</td><td>gene name</td><td>gene id</td></tr>
<tr><td>GU305914.1</td><td>31952</td><td>31974</td><td>+</td><td>0</td><td>CTCGTTCTGGGACGCGCCGT</td><td>AGG</td><td>||||||||[||||||||||||]PAM</td><td>NA</td><td>Intergenic</td><td>NA</td><td>NA</td></tr>
<tr><td>T1972</td><td>TCGTTCTGGGACGCGCCGTAGGG</td><td>1000</td><td></td><td></td><td></td><td></td><td></td><td></td><td></td><td></td><td></td></tr>
<tr><td>Oligo fwd</td><td>CACCTCGTTCTGGGACGCGCCGTA</td><td></td><td></td><td></td><td></td><td></td><td></td><td></td><td></td><td></td><td></td></tr>
<tr><td>Oligo rev</td><td>AAACTACGGCGCGTCCCAGAACGA</td><td></td><td></td><td></td><td></td><td></td><td></td><td></td><td></td><td></td><td></td></tr>
<tr><td>Chromosome</td><td>start</td><td>end</td><td>strand</td><td>MM</td><td>target_seq</td><td>PAM</td><td>alignment</td><td>distance</td><td>position</td><td>gene name</td><td>gene id</td></tr>
<tr><td>GU305914.1</td><td>31953</td><td>31975</td><td>+</td><td>0</td><td>TCGTTCTGGGACGCGCCGTA</td><td>GGG</td><td>||||||||[||||||||||||]PAM</td><td>NA</td><td>Intergenic</td><td>NA</td><td>NA</td></tr>
<tr><td>T1973</td><td>CGTCCGCAACTCCATCCCCTTGG</td><td>1000</td><td></td><td></td><td></td><td></td><td></td><td></td><td></td><td></td><td></td></tr>
<tr><td>Oligo fwd</td><td>CACCCGTCCGCAACTCCATCCCCT</td><td></td><td></td><td></td><td></td><td></td><td></td><td></td><td></td><td></td><td></td></tr>
<tr><td>Oligo rev</td><td>AAACAGGGGATGGAGTTGCGGACG</td><td></td><td></td><td></td><td></td><td></td><td></td><td></td><td></td><td></td><td></td></tr>
<tr><td>Chromosome</td><td>start</td><td>end</td><td>strand</td><td>MM</td><td>target_seq</td><td>PAM</td><td>alignment</td><td>distance</td><td>position</td><td>gene name</td><td>gene id</td></tr>
<tr><td>GU305914.1</td><td>32009</td><td>32031</td><td>+</td><td>0</td><td>CGTCCGCAACTCCATCCCCT</td><td>TGG</td><td>||||||||[||||||||||||]PAM</td><td>NA</td><td>Intergenic</td><td>NA</td><td>NA</td></tr>
<tr><td>T1974</td><td>CTCCATCCCCTTGGAACCGCCGG</td><td>1000</td><td></td><td></td><td></td><td></td><td></td><td></td><td></td><td></td><td></td></tr>
<tr><td>Oligo fwd</td><td>CACCCTCCATCCCCTTGGAACCGC</td><td></td><td></td><td></td><td></td><td></td><td></td><td></td><td></td><td></td><td></td></tr>
<tr><td>Oligo rev</td><td>AAACGCGGTTCCAAGGGGATGGAG</td><td></td><td></td><td></td><td></td><td></td><td></td><td></td><td></td><td></td><td></td></tr>
<tr><td>Chromosome</td><td>start</td><td>end</td><td>strand</td><td>MM</td><td>target_seq</td><td>PAM</td><td>alignment</td><td>distance</td><td>position</td><td>gene name</td><td>gene id</td></tr>
<tr><td>GU305914.1</td><td>32018</td><td>32040</td><td>+</td><td>0</td><td>CTCCATCCCCTTGGAACCGC</td><td>CGG</td><td>||||||||[||||||||||||]PAM</td><td>NA</td><td>Intergenic</td><td>NA</td><td>NA</td></tr>
<tr><td>T1975</td><td>GGCGACCGCGACCGCCTCCGAGG</td><td>1000</td><td></td><td></td><td></td><td></td><td></td><td></td><td></td><td></td><td></td></tr>
<tr><td>Oligo fwd</td><td>CACCGGCGACCGCGACCGCCTCCG</td><td></td><td></td><td></td><td></td><td></td><td></td><td></td><td></td><td></td><td></td></tr>
<tr><td>Oligo rev</td><td>AAACCGGAGGCGGTCGCGGTCGCC</td><td></td><td></td><td></td><td></td><td></td><td></td><td></td><td></td><td></td><td></td></tr>
<tr><td>Chromosome</td><td>start</td><td>end</td><td>strand</td><td>MM</td><td>target_seq</td><td>PAM</td><td>alignment</td><td>distance</td><td>position</td><td>gene name</td><td>gene id</td></tr>
<tr><td>GU305914.1</td><td>32039</td><td>32061</td><td>+</td><td>0</td><td>GGCGACCGCGACCGCCTCCG</td><td>AGG</td><td>||||||||[||||||||||||]PAM</td><td>NA</td><td>Intergenic</td><td>NA</td><td>NA</td></tr>
<tr><td>T1976</td><td>GTGTCCCGTCCTGCCCACTCTGG</td><td>1000</td><td></td><td></td><td></td><td></td><td></td><td></td><td></td><td></td><td></td></tr>
<tr><td>Oligo fwd</td><td>CACCGTGTCCCGTCCTGCCCACTC</td><td></td><td></td><td></td><td></td><td></td><td></td><td></td><td></td><td></td><td></td></tr>
<tr><td>Oligo rev</td><td>AAACGAGTGGGCAGGACGGGACAC</td><td></td><td></td><td></td><td></td><td></td><td></td><td></td><td></td><td></td><td></td></tr>
<tr><td>Chromosome</td><td>start</td><td>end</td><td>strand</td><td>MM</td><td>target_seq</td><td>PAM</td><td>alignment</td><td>distance</td><td>position</td><td>gene name</td><td>gene id</td></tr>
<tr><td>GU305914.1</td><td>32072</td><td>32094</td><td>+</td><td>0</td><td>GTGTCCCGTCCTGCCCACTC</td><td>TGG</td><td>||||||||[||||||||||||]PAM</td><td>NA</td><td>Intergenic</td><td>NA</td><td>NA</td></tr>
</table>

<table>
<tr><td>T1977</td><td>CCGACACAGACAGACCCTCACGG</td><td>1000</td><td></td><td></td><td></td><td></td><td></td><td></td><td></td><td></td><td></td></tr>
<tr><td>Oligo fwd</td><td>CACCCCGACACAGACAGACCCTCA</td><td></td><td></td><td></td><td></td><td></td><td></td><td></td><td></td><td></td><td></td></tr>
<tr><td>Oligo rev</td><td>AAACTGAGGGTCTGTCTGTGTCGG</td><td></td><td></td><td></td><td></td><td></td><td></td><td></td><td></td><td></td><td></td></tr>
<tr><td>Chromosome</td><td>start</td><td>end</td><td>strand</td><td>MM</td><td>target_seq</td><td>PAM</td><td>alignment</td><td>distance</td><td>position</td><td>gene name</td><td>gene id</td></tr>
<tr><td>GU305914.1</td><td>32103</td><td>32125</td><td>+</td><td>0</td><td>CCGACACAGACAGACCCTCA</td><td>CGG</td><td>||||||||[||||||||||||]PAM</td><td>NA</td><td>Intergenic</td><td>NA</td><td>NA</td></tr>
<tr><td>T1978</td><td>CGAAACTTTATAATAATAACCGG</td><td>1000</td><td></td><td></td><td></td><td></td><td></td><td></td><td></td><td></td><td></td></tr>
<tr><td>Oligo fwd</td><td>CACCCGAAACTTTATAATAATAAC</td><td></td><td></td><td></td><td></td><td></td><td></td><td></td><td></td><td></td><td></td></tr>
<tr><td>Oligo rev</td><td>AAACGTTATTATTATAAAGTTTCG</td><td></td><td></td><td></td><td></td><td></td><td></td><td></td><td></td><td></td><td></td></tr>
<tr><td>Chromosome</td><td>start</td><td>end</td><td>strand</td><td>MM</td><td>target_seq</td><td>PAM</td><td>alignment</td><td>distance</td><td>position</td><td>gene name</td><td>gene id</td></tr>
<tr><td>GU305914.1</td><td>32142</td><td>32164</td><td>+</td><td>0</td><td>CGAAACTTTATAATAATAAC</td><td>CGG</td><td>||||||||[||||||||||||]PAM</td><td>NA</td><td>Intergenic</td><td>NA</td><td>NA</td></tr>
<tr><td>T1979</td><td>GTGACTTAACACCGCCCACTCGG</td><td>1000</td><td></td><td></td><td></td><td></td><td></td><td></td><td></td><td></td><td></td></tr>
<tr><td>Oligo fwd</td><td>CACCGTGACTTAACACCGCCCACT</td><td></td><td></td><td></td><td></td><td></td><td></td><td></td><td></td><td></td><td></td></tr>
<tr><td>Oligo rev</td><td>AAACAGTGGGCGGTGTTAAGTCAC</td><td></td><td></td><td></td><td></td><td></td><td></td><td></td><td></td><td></td><td></td></tr>
<tr><td>Chromosome</td><td>start</td><td>end</td><td>strand</td><td>MM</td><td>target_seq</td><td>PAM</td><td>alignment</td><td>distance</td><td>position</td><td>gene name</td><td>gene id</td></tr>
<tr><td>GU305914.1</td><td>32170</td><td>32192</td><td>+</td><td>0</td><td>GTGACTTAACACCGCCCACT</td><td>CGG</td><td>||||||||[||||||||||||]PAM</td><td>NA</td><td>Intergenic</td><td>NA</td><td>NA</td></tr>
<tr><td>T1980</td><td>CACCCGCTCCACCACAAACTCGG</td><td>1000</td><td></td><td></td><td></td><td></td><td></td><td></td><td></td><td></td><td></td></tr>
<tr><td>Oligo fwd</td><td>CACCCACCCGCTCCACCACAAACT</td><td></td><td></td><td></td><td></td><td></td><td></td><td></td><td></td><td></td><td></td></tr>
<tr><td>Oligo rev</td><td>AAACAGTTTGTGGTGGAGCGGGTG</td><td></td><td></td><td></td><td></td><td></td><td></td><td></td><td></td><td></td><td></td></tr>
<tr><td>Chromosome</td><td>start</td><td>end</td><td>strand</td><td>MM</td><td>target_seq</td><td>PAM</td><td>alignment</td><td>distance</td><td>position</td><td>gene name</td><td>gene id</td></tr>
<tr><td>GU305914.1</td><td>32250</td><td>32272</td><td>+</td><td>0</td><td>CACCCGCTCCACCACAAACT</td><td>CGG</td><td>||||||||[||||||||||||]PAM</td><td>NA</td><td>Intergenic</td><td>NA</td><td>NA</td></tr>
<tr><td>T1981</td><td>CACCACAAACTCGGAGCCGTCGG</td><td>1000</td><td></td><td></td><td></td><td></td><td></td><td></td><td></td><td></td><td></td></tr>
<tr><td>Oligo fwd</td><td>CACCCACCACAAACTCGGAGCCGT</td><td></td><td></td><td></td><td></td><td></td><td></td><td></td><td></td><td></td><td></td></tr>
<tr><td>Oligo rev</td><td>AAACACGGCTCCGAGTTTGTGGTG</td><td></td><td></td><td></td><td></td><td></td><td></td><td></td><td></td><td></td><td></td></tr>
<tr><td>Chromosome</td><td>start</td><td>end</td><td>strand</td><td>MM</td><td>target_seq</td><td>PAM</td><td>alignment</td><td>distance</td><td>position</td><td>gene name</td><td>gene id</td></tr>
<tr><td>GU305914.1</td><td>32259</td><td>32281</td><td>+</td><td>0</td><td>CACCACAAACTCGGAGCCGT</td><td>CGG</td><td>||||||||[||||||||||||]PAM</td><td>NA</td><td>Intergenic</td><td>NA</td><td>NA</td></tr>
<tr><td>T1982</td><td>GTCGGACTCGCCGCCATCATCGG</td><td>1000</td><td></td><td></td><td></td><td></td><td></td><td></td><td></td><td></td><td></td></tr>
<tr><td>Oligo fwd</td><td>CACCGTCGGACTCGCCGCCATCAT</td><td></td><td></td><td></td><td></td><td></td><td></td><td></td><td></td><td></td><td></td></tr>
<tr><td>Oligo rev</td><td>AAACATGATGGCGGCGAGTCCGAC</td><td></td><td></td><td></td><td></td><td></td><td></td><td></td><td></td><td></td><td></td></tr>
<tr><td>Chromosome</td><td>start</td><td>end</td><td>strand</td><td>MM</td><td>target_seq</td><td>PAM</td><td>alignment</td><td>distance</td><td>position</td><td>gene name</td><td>gene id</td></tr>
<tr><td>GU305914.1</td><td>32277</td><td>32299</td><td>+</td><td>0</td><td>GTCGGACTCGCCGCCATCAT</td><td>CGG</td><td>||||||||[||||||||||||]PAM</td><td>NA</td><td>Intergenic</td><td>NA</td><td>NA</td></tr>
<tr><td>T1983</td><td>GACTCGCCGCCATCATCGGACGG</td><td>1000</td><td></td><td></td><td></td><td></td><td></td><td></td><td></td><td></td><td></td></tr>
<tr><td>Oligo fwd</td><td>CACCGACTCGCCGCCATCATCGGA</td><td></td><td></td><td></td><td></td><td></td><td></td><td></td><td></td><td></td><td></td></tr>
<tr><td>Oligo rev</td><td>AAACTCCGATGATGGCGGCGAGTC</td><td></td><td></td><td></td><td></td><td></td><td></td><td></td><td></td><td></td><td></td></tr>
<tr><td>Chromosome</td><td>start</td><td>end</td><td>strand</td><td>MM</td><td>target_seq</td><td>PAM</td><td>alignment</td><td>distance</td><td>position</td><td>gene name</td><td>gene id</td></tr>
<tr><td>GU305914.1</td><td>32281</td><td>32303</td><td>+</td><td>0</td><td>GACTCGCCGCCATCATCGGA</td><td>CGG</td><td>||||||||[||||||||||||]PAM</td><td>NA</td><td>Intergenic</td><td>NA</td><td>NA</td></tr>
<tr><td>T1984</td><td>GTGCCCGCTGTCCGCGTCCTCGG</td><td>1000</td><td></td><td></td><td></td><td></td><td></td><td></td><td></td><td></td><td></td></tr>
<tr><td>Oligo fwd</td><td>CACCGTGCCCGCTGTCCGCGTCCT</td><td></td><td></td><td></td><td></td><td></td><td></td><td></td><td></td><td></td><td></td></tr>
<tr><td>Oligo rev</td><td>AAACAGGACGCGGACAGCGGGCAC</td><td></td><td></td><td></td><td></td><td></td><td></td><td></td><td></td><td></td><td></td></tr>
<tr><td>Chromosome</td><td>start</td><td>end</td><td>strand</td><td>MM</td><td>target_seq</td><td>PAM</td><td>alignment</td><td>distance</td><td>position</td><td>gene name</td><td>gene id</td></tr>
<tr><td>GU305914.1</td><td>32310</td><td>32332</td><td>+</td><td>0</td><td>GTGCCCGCTGTCCGCGTCCT</td><td>CGG</td><td>||||||||[||||||||||||]PAM</td><td>NA</td><td>Intergenic</td><td>NA</td><td>NA</td></tr>
<tr><td>T1985</td><td>CGCGTCCTCGGTCTCCTCGTCGG</td><td>1000</td><td></td><td></td><td></td><td></td><td></td><td></td><td></td><td></td><td></td></tr>
<tr><td>Oligo fwd</td><td>CACCCGCGTCCTCGGTCTCCTCGT</td><td></td><td></td><td></td><td></td><td></td><td></td><td></td><td></td><td></td><td></td></tr>
<tr><td>Oligo rev</td><td>AAACACGAGGAGACCGAGGACGCG</td><td></td><td></td><td></td><td></td><td></td><td></td><td></td><td></td><td></td><td></td></tr>
<tr><td>Chromosome</td><td>start</td><td>end</td><td>strand</td><td>MM</td><td>target_seq</td><td>PAM</td><td>alignment</td><td>distance</td><td>position</td><td>gene name</td><td>gene id</td></tr>
<tr><td>GU305914.1</td><td>32322</td><td>32344</td><td>+</td><td>0</td><td>CGCGTCCTCGGTCTCCTCGT</td><td>CGG</td><td>||||||||[||||||||||||]PAM</td><td>NA</td><td>Intergenic</td><td>NA</td><td>NA</td></tr>
<tr><td>GU305914.1</td><td>201600</td><td>201622</td><td>+</td><td>4</td><td>CGCCACCGCGGTCTCCTCCT</td><td>CGG</td><td>|||--||-[||||||||||-|]PAM</td><td>NA</td><td>Intergenic</td><td>NA</td><td>NA</td></tr>
<tr><td>T1986</td><td>GAGCTCGACGAGCTCCCGCTTGG</td><td>1000</td><td></td><td></td><td></td><td></td><td></td><td></td><td></td><td></td><td></td></tr>
<tr><td>Oligo fwd</td><td>CACCGAGCTCGACGAGCTCCCGCT</td><td></td><td></td><td></td><td></td><td></td><td></td><td></td><td></td><td></td><td></td></tr>
<tr><td>Oligo rev</td><td>AAACAGCGGGAGCTCGTCGAGCTC</td><td></td><td></td><td></td><td></td><td></td><td></td><td></td><td></td><td></td><td></td></tr>
<tr><td>Chromosome</td><td>start</td><td>end</td><td>strand</td><td>MM</td><td>target_seq</td><td>PAM</td><td>alignment</td><td>distance</td><td>position</td><td>gene name</td><td>gene id</td></tr>
<tr><td>GU305914.1</td><td>32373</td><td>32395</td><td>+</td><td>0</td><td>GAGCTCGACGAGCTCCCGCT</td><td>TGG</td><td>||||||||[||||||||||||]PAM</td><td>NA</td><td>Intergenic</td><td>NA</td><td>NA</td></tr>
<tr><td>T1987</td><td>AGCTCGACGAGCTCCCGCTTGGG</td><td>1000</td><td></td><td></td><td></td><td></td><td></td><td></td><td></td><td></td><td></td></tr>
<tr><td>Oligo fwd</td><td>CACCAGCTCGACGAGCTCCCGCTT</td><td></td><td></td><td></td><td></td><td></td><td></td><td></td><td></td><td></td><td></td></tr>
<tr><td>Oligo rev</td><td>AAACAAGCGGGAGCTCGTCGAGCT</td><td></td><td></td><td></td><td></td><td></td><td></td><td></td><td></td><td></td><td></td></tr>
<tr><td>Chromosome</td><td>start</td><td>end</td><td>strand</td><td>MM</td><td>target_seq</td><td>PAM</td><td>alignment</td><td>distance</td><td>position</td><td>gene name</td><td>gene id</td></tr>
<tr><td>GU305914.1</td><td>32374</td><td>32396</td><td>+</td><td>0</td><td>AGCTCGACGAGCTCCCGCTT</td><td>GGG</td><td>||||||||[||||||||||||]PAM</td><td>NA</td><td>Intergenic</td><td>NA</td><td>NA</td></tr>
<tr><td>T1988</td><td>GCTCGACGAGCTCCCGCTTGGGG</td><td>1000</td><td></td><td></td><td></td><td></td><td></td><td></td><td></td><td></td><td></td></tr>
<tr><td>Oligo fwd</td><td>CACCGCTCGACGAGCTCCCGCTTG</td><td></td><td></td><td></td><td></td><td></td><td></td><td></td><td></td><td></td><td></td></tr>
<tr><td>Oligo rev</td><td>AAACCAAGCGGGAGCTCGTCGAGC</td><td></td><td></td><td></td><td></td><td></td><td></td><td></td><td></td><td></td><td></td></tr>
<tr><td>Chromosome</td><td>start</td><td>end</td><td>strand</td><td>MM</td><td>target_seq</td><td>PAM</td><td>alignment</td><td>distance</td><td>position</td><td>gene name</td><td>gene id</td></tr>
<tr><td>GU305914.1</td><td>32375</td><td>32397</td><td>+</td><td>0</td><td>GCTCGACGAGCTCCCGCTTG</td><td>GGG</td><td>||||||||[||||||||||||]PAM</td><td>NA</td><td>Intergenic</td><td>NA</td><td>NA</td></tr>
<tr><td>T1989</td><td>CAGCATGTCGACGTACTCGAAGG</td><td>1000</td><td></td><td></td><td></td><td></td><td></td><td></td><td></td><td></td><td></td></tr>
<tr><td>Oligo fwd</td><td>CACCCAGCATGTCGACGTACTCGA</td><td></td><td></td><td></td><td></td><td></td><td></td><td></td><td></td><td></td><td></td></tr>
<tr><td>Oligo rev</td><td>AAACTCGAGTACGTCGACATGCTG</td><td></td><td></td><td></td><td></td><td></td><td></td><td></td><td></td><td></td><td></td></tr>
<tr><td>Chromosome</td><td>start</td><td>end</td><td>strand</td><td>MM</td><td>target_seq</td><td>PAM</td><td>alignment</td><td>distance</td><td>position</td><td>gene name</td><td>gene id</td></tr>
<tr><td>GU305914.1</td><td>32454</td><td>32476</td><td>+</td><td>0</td><td>CAGCATGTCGACGTACTCGA</td><td>AGG</td><td>||||||||[||||||||||||]PAM</td><td>NA</td><td>Intergenic</td><td>NA</td><td>NA</td></tr>
<tr><td>T1990</td><td>AGCGCAGCCGCATGCCCACCAGG</td><td>1000</td><td></td><td></td><td></td><td></td><td></td><td></td><td></td><td></td><td></td></tr>
<tr><td>Oligo fwd</td><td>CACCAGCGCAGCCGCATGCCCACC</td><td></td><td></td><td></td><td></td><td></td><td></td><td></td><td></td><td></td><td></td></tr>
<tr><td>Oligo rev</td><td>AAACGGTGGGCATGCGGCTGCGCT</td><td></td><td></td><td></td><td></td><td></td><td></td><td></td><td></td><td></td><td></td></tr>
<tr><td>Chromosome</td><td>start</td><td>end</td><td>strand</td><td>MM</td><td>target_seq</td><td>PAM</td><td>alignment</td><td>distance</td><td>position</td><td>gene name</td><td>gene id</td></tr>
<tr><td>GU305914.1</td><td>32486</td><td>32508</td><td>+</td><td>0</td><td>AGCGCAGCCGCATGCCCACC</td><td>AGG</td><td>||||||||[||||||||||||]PAM</td><td>NA</td><td>Intergenic</td><td>NA</td><td>NA</td></tr>
<tr><td>T1991</td><td>CCGCATGCCCACCAGGTCGTCGG</td><td>1000</td><td></td><td></td><td></td><td></td><td></td><td></td><td></td><td></td><td></td></tr>
<tr><td>Oligo fwd</td><td>CACCCCGCATGCCCACCAGGTCGT</td><td></td><td></td><td></td><td></td><td></td><td></td><td></td><td></td><td></td><td></td></tr>
<tr><td>Oligo rev</td><td>AAACACGACCTGGTGGGCATGCGG</td><td></td><td></td><td></td><td></td><td></td><td></td><td></td><td></td><td></td><td></td></tr>
</table>

| | | | | | | | | | | | | |
|---|---|---|---|---|---|---|---|---|---|---|---|---|
| Chromosome | start | | end | strand | MM | target_seq | PAM | alignment | distance | position | gene name | gene id |
| GU305914.1 | | 32493 | 32515 | + | 0 | CCGCATGCCCACCAGGTCGT | CGG | \|\|\|\|\|\|\|\|[\|\|\|\|\|\|\|\|\|\|\|\|]PAM | NA | Intergenic | NA | NA |
| T1992 | CCAGTGTCCCTTGACGATCTCGG | | 1000 | | | | | | | | | |
| Oligo fwd | CACCCCAGTGTCCCTTGACGATCT | | | | | | | | | | | |
| Oligo rev | AAACAGATCGTCAAGGGACACTGG | | | | | | | | | | | |
| Chromosome | start | | end | strand | MM | target_seq | PAM | alignment | distance | position | gene name | gene id |
| GU305914.1 | | 32520 | 32542 | + | 0 | CCAGTGTCCCTTGACGATCT | CGG | \|\|\|\|\|\|\|\|[\|\|\|\|\|\|\|\|\|\|\|\|]PAM | NA | Intergenic | NA | NA |
| T1993 | TGACGATCTCGGTGTCACTGAGG | | 1000 | | | | | | | | | |
| Oligo fwd | CACCTGACGATCTCGGTGTCACTG | | | | | | | | | | | |
| Oligo rev | AAACCAGTGACACCGAGATCGTCA | | | | | | | | | | | |
| Chromosome | start | | end | strand | MM | target_seq | PAM | alignment | distance | position | gene name | gene id |
| GU305914.1 | | 32531 | 32553 | + | 0 | TGACGATCTCGGTGTCACTG | AGG | \|\|\|\|\|\|\|\|[\|\|\|\|\|\|\|\|\|\|\|\|]PAM | NA | Intergenic | NA | NA |
| T1994 | GACGATCTCGGTGTCACTGAGGG | | 1000 | | | | | | | | | |
| Oligo fwd | CACCGACGATCTCGGTGTCACTGA | | | | | | | | | | | |
| Oligo rev | AAACTCAGTGACACCGAGATCGTC | | | | | | | | | | | |
| Chromosome | start | | end | strand | MM | target_seq | PAM | alignment | distance | position | gene name | gene id |
| GU305914.1 | | 32532 | 32554 | + | 0 | GACGATCTCGGTGTCACTGA | GGG | \|\|\|\|\|\|\|\|[\|\|\|\|\|\|\|\|\|\|\|\|]PAM | NA | Intergenic | NA | NA |
| T1995 | CTCGGTGTCACTGAGGGTGACGG | | 1000 | | | | | | | | | |
| Oligo fwd | CACCCTCGGTGTCACTGAGGGTGA | | | | | | | | | | | |
| Oligo rev | AAACTCACCCTCAGTGACACCGAG | | | | | | | | | | | |
| Chromosome | start | | end | strand | MM | target_seq | PAM | alignment | distance | position | gene name | gene id |
| GU305914.1 | | 32538 | 32560 | + | 0 | CTCGGTGTCACTGAGGGTGA | CGG | \|\|\|\|\|\|\|\|[\|\|\|\|\|\|\|\|\|\|\|\|]PAM | NA | Intergenic | NA | NA |
| T1996 | GACGGCGCCCATCCTGATGACGG | | 1000 | | | | | | | | | |
| Oligo fwd | CACCGACGGCGCCCATCCTGATGA | | | | | | | | | | | |
| Oligo rev | AAACTCATCAGGATGGGCGCCGTC | | | | | | | | | | | |
| Chromosome | start | | end | strand | MM | target_seq | PAM | alignment | distance | position | gene name | gene id |
| GU305914.1 | | 32556 | 32578 | + | 0 | GACGGCGCCCATCCTGATGA | CGG | \|\|\|\|\|\|\|\|[\|\|\|\|\|\|\|\|\|\|\|\|]PAM | NA | Intergenic | NA | NA |
| GU305914.1 | | 49741 | 49763 | + | 4 | GCAGGCGTCCCTCCTGATGA | CGG | \|--\|\|\|\|-[\|\|-\|\|\|\|\|\|\|\|\|]PAM | NA | Intergenic | NA | NA |
| T1997 | TGATGACGGCCCCGTCGCGCCGG | | 1000 | | | | | | | | | |
| Oligo fwd | CACCTGATGACGGCCCCGTCGCGC | | | | | | | | | | | |
| Oligo rev | AAACGCGCGACGGGGCCGTCATCA | | | | | | | | | | | |
| Chromosome | start | | end | strand | MM | target_seq | PAM | alignment | distance | position | gene name | gene id |
| GU305914.1 | | 32570 | 32592 | + | 0 | TGATGACGGCCCCGTCGCGC | CGG | \|\|\|\|\|\|\|\|[\|\|\|\|\|\|\|\|\|\|\|\|]PAM | NA | Intergenic | NA | NA |
| T1998 | TGACGGCCCCGTCGCGCCGGCGG | | 1000 | | | | | | | | | |
| Oligo fwd | CACCTGACGGCCCCGTCGCGCCGG | | | | | | | | | | | |
| Oligo rev | AAACCCGGCGCGACGGGGCCGTCA | | | | | | | | | | | |
| Chromosome | start | | end | strand | MM | target_seq | PAM | alignment | distance | position | gene name | gene id |
| GU305914.1 | | 32573 | 32595 | + | 0 | TGACGGCCCCGTCGCGCCGG | CGG | \|\|\|\|\|\|\|\|[\|\|\|\|\|\|\|\|\|\|\|\|]PAM | NA | Intergenic | NA | NA |
| T1999 | TCCGCCGACAAACATGTAGTAGG | | 1000 | | | | | | | | | |
| Oligo fwd | CACCTCCGCCGACAAACATGTAGT | | | | | | | | | | | |
| Oligo rev | AAACACTACATGTTTGTCGGCGGA | | | | | | | | | | | |
| Chromosome | start | | end | strand | MM | target_seq | PAM | alignment | distance | position | gene name | gene id |
| GU305914.1 | | 32598 | 32620 | + | 0 | TCCGCCGACAAACATGTAGT | AGG | \|\|\|\|\|\|\|\|[\|\|\|\|\|\|\|\|\|\|\|\|]PAM | NA | Intergenic | NA | NA |
| T2000 | CCGCCGACAAACATGTAGTAGGG | | 1000 | | | | | | | | | |
| Oligo fwd | CACCCCGCCGACAAACATGTAGTA | | | | | | | | | | | |
| Oligo rev | AAACTACTACATGTTTGTCGGCGG | | | | | | | | | | | |
| Chromosome | start | | end | strand | MM | target_seq | PAM | alignment | distance | position | gene name | gene id |
| GU305914.1 | | 32599 | 32621 | + | 0 | CCGCCGACAAACATGTAGTA | GGG | \|\|\|\|\|\|\|\|[\|\|\|\|\|\|\|\|\|\|\|\|]PAM | NA | Intergenic | NA | NA |
| T2001 | CGCCGACAAACATGTAGTAGGGG | | 1000 | | | | | | | | | |
| Oligo fwd | CACCCGCCGACAAACATGTAGTAG | | | | | | | | | | | |
| Oligo rev | AAACCTACTACATGTTTGTCGGCG | | | | | | | | | | | |
| Chromosome | start | | end | strand | MM | target_seq | PAM | alignment | distance | position | gene name | gene id |
| GU305914.1 | | 32600 | 32622 | + | 0 | CGCCGACAAACATGTAGTAG | GGG | \|\|\|\|\|\|\|\|[\|\|\|\|\|\|\|\|\|\|\|\|]PAM | NA | Intergenic | NA | NA |
| T2002 | GAGCCTGTCGCCGTAGAAGCCGG | | 1000 | | | | | | | | | |
| Oligo fwd | CACCGAGCCTGTCGCCGTAGAAGC | | | | | | | | | | | |
| Oligo rev | AAACGCTTCTACGGCGACAGGCTC | | | | | | | | | | | |
| Chromosome | start | | end | strand | MM | target_seq | PAM | alignment | distance | position | gene name | gene id |
| GU305914.1 | | 32625 | 32647 | + | 0 | GAGCCTGTCGCCGTAGAAGC | CGG | \|\|\|\|\|\|\|\|[\|\|\|\|\|\|\|\|\|\|\|\|]PAM | NA | Intergenic | NA | NA |
| GU305914.1 | | 82417 | 82439 | + | 4 | GAGGCCGTAGCCGTAGAAGG | AGG | \|\|\|-\|-\|\|[-\|\|\|\|\|\|\|\|\|\|-]PAM | NA | Intergenic | NA | NA |
| T2003 | TCGCCGTAGAAGCCGGCCAGCGG | | 1000 | | | | | | | | | |
| Oligo fwd | CACCTCGCCGTAGAAGCCGGCCAG | | | | | | | | | | | |
| Oligo rev | AAACCTGGCCGGCTTCTACGGCGA | | | | | | | | | | | |
| Chromosome | start | | end | strand | MM | target_seq | PAM | alignment | distance | position | gene name | gene id |
| GU305914.1 | | 32632 | 32654 | + | 0 | TCGCCGTAGAAGCCGGCCAG | CGG | \|\|\|\|\|\|\|\|[\|\|\|\|\|\|\|\|\|\|\|\|]PAM | NA | Intergenic | NA | NA |
| T2004 | CGCCGTAGAAGCCGGCCAGCGGG | | 1000 | | | | | | | | | |
| Oligo fwd | CACCCGCCGTAGAAGCCGGCCAGC | | | | | | | | | | | |
| Oligo rev | AAACGCTGGCCGGCTTCTACGGCG | | | | | | | | | | | |
| Chromosome | start | | end | strand | MM | target_seq | PAM | alignment | distance | position | gene name | gene id |
| GU305914.1 | | 32633 | 32655 | + | 0 | CGCCGTAGAAGCCGGCCAGC | GGG | \|\|\|\|\|\|\|\|[\|\|\|\|\|\|\|\|\|\|\|\|]PAM | NA | Intergenic | NA | NA |
| T2005 | GTACATCCTCTTGATGAAGAAGG | | 1000 | | | | | | | | | |
| Oligo fwd | CACCGTACATCCTCTTGATGAAGA | | | | | | | | | | | |
| Oligo rev | AAACTCTTCATCAAGAGGATGTAC | | | | | | | | | | | |
| Chromosome | start | | end | strand | MM | target_seq | PAM | alignment | distance | position | gene name | gene id |
| GU305914.1 | | 32664 | 32686 | + | 0 | GTACATCCTCTTGATGAAGA | AGG | \|\|\|\|\|\|\|\|[\|\|\|\|\|\|\|\|\|\|\|\|]PAM | NA | Intergenic | NA | NA |

<table>
<tr><td>T2006</td><td>TCTTGATGAAGAAGGTGTGCAGG</td><td>1000</td></tr>
<tr><td>Oligo fwd</td><td>CACCTCTTGATGAAGAAGGTGTGC</td></tr>
<tr><td>Oligo rev</td><td>AAACGCACACCTTCTTCATCAAGA</td></tr>
<tr><td>Chromosome</td><td>start</td><td>end</td><td>strand</td><td>MM</td><td>target_seq</td><td>PAM</td><td>alignment</td><td>distance</td><td>position</td><td>gene name</td><td>gene id</td></tr>
<tr><td>GU305914.1</td><td>32672</td><td>32694</td><td>+</td><td>0</td><td>TCTTGATGAAGAAGGTGTGC</td><td>AGG</td><td>||||||||[||||||||||||]PAM</td><td>NA</td><td>Intergenic</td><td>NA</td><td>NA</td></tr>
<tr><td>T2007</td><td>TGATGAAGAAGGTGTGCAGGAGG</td><td>1000</td></tr>
<tr><td>Oligo fwd</td><td>CACCTGATGAAGAAGGTGTGCAGG</td></tr>
<tr><td>Oligo rev</td><td>AAACCCTGCACACCTTCTTCATCA</td></tr>
<tr><td>Chromosome</td><td>start</td><td>end</td><td>strand</td><td>MM</td><td>target_seq</td><td>PAM</td><td>alignment</td><td>distance</td><td>position</td><td>gene name</td><td>gene id</td></tr>
<tr><td>GU305914.1</td><td>32675</td><td>32697</td><td>+</td><td>0</td><td>TGATGAAGAAGGTGTGCAGG</td><td>AGG</td><td>||||||||[||||||||||||]PAM</td><td>NA</td><td>Intergenic</td><td>NA</td><td>NA</td></tr>
<tr><td>T2008</td><td>TGTGCAGGAGGAGTTCCTCGCGG</td><td>1000</td></tr>
<tr><td>Oligo fwd</td><td>CACCTGTGCAGGAGGAGTTCCTCG</td></tr>
<tr><td>Oligo rev</td><td>AAACCGAGGAACTCCTCCTGCACA</td></tr>
<tr><td>Chromosome</td><td>start</td><td>end</td><td>strand</td><td>MM</td><td>target_seq</td><td>PAM</td><td>alignment</td><td>distance</td><td>position</td><td>gene name</td><td>gene id</td></tr>
<tr><td>GU305914.1</td><td>32687</td><td>32709</td><td>+</td><td>0</td><td>TGTGCAGGAGGAGTTCCTCG</td><td>CGG</td><td>||||||||[||||||||||||]PAM</td><td>NA</td><td>Intergenic</td><td>NA</td><td>NA</td></tr>
<tr><td>T2009</td><td>GTGCAGGAGGAGTTCCTCGCGGG</td><td>1000</td></tr>
<tr><td>Oligo fwd</td><td>CACCGTGCAGGAGGAGTTCCTCGC</td></tr>
<tr><td>Oligo rev</td><td>AAACGCGAGGAACTCCTCCTGCAC</td></tr>
<tr><td>Chromosome</td><td>start</td><td>end</td><td>strand</td><td>MM</td><td>target_seq</td><td>PAM</td><td>alignment</td><td>distance</td><td>position</td><td>gene name</td><td>gene id</td></tr>
<tr><td>GU305914.1</td><td>32688</td><td>32710</td><td>+</td><td>0</td><td>GTGCAGGAGGAGTTCCTCGC</td><td>GGG</td><td>||||||||[||||||||||||]PAM</td><td>NA</td><td>Intergenic</td><td>NA</td><td>NA</td></tr>
<tr><td>T2010</td><td>GGAGGAGTTCCTCGCGGGCGCGG</td><td>1000</td></tr>
<tr><td>Oligo fwd</td><td>CACCGGAGGAGTTCCTCGCGGGCG</td></tr>
<tr><td>Oligo rev</td><td>AAACCGCCCGCGAGGAACTCCTCC</td></tr>
<tr><td>Chromosome</td><td>start</td><td>end</td><td>strand</td><td>MM</td><td>target_seq</td><td>PAM</td><td>alignment</td><td>distance</td><td>position</td><td>gene name</td><td>gene id</td></tr>
<tr><td>GU305914.1</td><td>32693</td><td>32715</td><td>+</td><td>0</td><td>GGAGGAGTTCCTCGCGGGCG</td><td>CGG</td><td>||||||||[||||||||||||]PAM</td><td>NA</td><td>Intergenic</td><td>NA</td><td>NA</td></tr>
<tr><td>T2011</td><td>CGGTTCTCGCGCAGCGCCTTCGG</td><td>1000</td></tr>
<tr><td>Oligo fwd</td><td>CACCCGGTTCTCGCGCAGCGCCTT</td></tr>
<tr><td>Oligo rev</td><td>AAACAAGGCGCTGCGCGAGAACCG</td></tr>
<tr><td>Chromosome</td><td>start</td><td>end</td><td>strand</td><td>MM</td><td>target_seq</td><td>PAM</td><td>alignment</td><td>distance</td><td>position</td><td>gene name</td><td>gene id</td></tr>
<tr><td>GU305914.1</td><td>32713</td><td>32735</td><td>+</td><td>0</td><td>CGGTTCTCGCGCAGCGCCTT</td><td>CGG</td><td>||||||||[||||||||||||]PAM</td><td>NA</td><td>Intergenic</td><td>NA</td><td>NA</td></tr>
<tr><td>T2012</td><td>GGTTCTCGCGCAGCGCCTTCGGG</td><td>1000</td></tr>
<tr><td>Oligo fwd</td><td>CACCGGTTCTCGCGCAGCGCCTTC</td></tr>
<tr><td>Oligo rev</td><td>AAACGAAGGCGCTGCGCGAGAACC</td></tr>
<tr><td>Chromosome</td><td>start</td><td>end</td><td>strand</td><td>MM</td><td>target_seq</td><td>PAM</td><td>alignment</td><td>distance</td><td>position</td><td>gene name</td><td>gene id</td></tr>
<tr><td>GU305914.1</td><td>32714</td><td>32736</td><td>+</td><td>0</td><td>GGTTCTCGCGCAGCGCCTTC</td><td>GGG</td><td>||||||||[||||||||||||]PAM</td><td>NA</td><td>Intergenic</td><td>NA</td><td>NA</td></tr>
<tr><td>T2013</td><td>CGCGCAGCGCCTTCGGGACGCGG</td><td>1000</td></tr>
<tr><td>Oligo fwd</td><td>CACCCGCGCAGCGCCTTCGGGACG</td></tr>
<tr><td>Oligo rev</td><td>AAACCGTCCCGAAGGCGCTGCGCG</td></tr>
<tr><td>Chromosome</td><td>start</td><td>end</td><td>strand</td><td>MM</td><td>target_seq</td><td>PAM</td><td>alignment</td><td>distance</td><td>position</td><td>gene name</td><td>gene id</td></tr>
<tr><td>GU305914.1</td><td>32720</td><td>32742</td><td>+</td><td>0</td><td>CGCGCAGCGCCTTCGGGACG</td><td>CGG</td><td>||||||||[||||||||||||]PAM</td><td>NA</td><td>Intergenic</td><td>NA</td><td>NA</td></tr>
<tr><td>T2014</td><td>GGACGCGGAACGCATCCAAGAGG</td><td>1000</td></tr>
<tr><td>Oligo fwd</td><td>CACCGGACGCGGAACGCATCCAAG</td></tr>
<tr><td>Oligo rev</td><td>AAACCTTGGATGCGTTCCGCGTCC</td></tr>
<tr><td>Chromosome</td><td>start</td><td>end</td><td>strand</td><td>MM</td><td>target_seq</td><td>PAM</td><td>alignment</td><td>distance</td><td>position</td><td>gene name</td><td>gene id</td></tr>
<tr><td>GU305914.1</td><td>32735</td><td>32757</td><td>+</td><td>0</td><td>GGACGCGGAACGCATCCAAG</td><td>AGG</td><td>||||||||[||||||||||||]PAM</td><td>NA</td><td>Intergenic</td><td>NA</td><td>NA</td></tr>
<tr><td>T2015</td><td>ATCCAAGAGGTCCATGAACTTGG</td><td>1000</td></tr>
<tr><td>Oligo fwd</td><td>CACCATCCAAGAGGTCCATGAACT</td></tr>
<tr><td>Oligo rev</td><td>AAACAGTTCATGGACCTCTTGGAT</td></tr>
<tr><td>Chromosome</td><td>start</td><td>end</td><td>strand</td><td>MM</td><td>target_seq</td><td>PAM</td><td>alignment</td><td>distance</td><td>position</td><td>gene name</td><td>gene id</td></tr>
<tr><td>GU305914.1</td><td>32748</td><td>32770</td><td>+</td><td>0</td><td>ATCCAAGAGGTCCATGAACT</td><td>TGG</td><td>||||||||[||||||||||||]PAM</td><td>NA</td><td>Intergenic</td><td>NA</td><td>NA</td></tr>
<tr><td>T2016</td><td>CAAGAGGTCCATGAACTTGGCGG</td><td>1000</td></tr>
<tr><td>Oligo fwd</td><td>CACCCAAGAGGTCCATGAACTTGG</td></tr>
<tr><td>Oligo rev</td><td>AAACCCAAGTTCATGGACCTCTTG</td></tr>
<tr><td>Chromosome</td><td>start</td><td>end</td><td>strand</td><td>MM</td><td>target_seq</td><td>PAM</td><td>alignment</td><td>distance</td><td>position</td><td>gene name</td><td>gene id</td></tr>
<tr><td>GU305914.1</td><td>32751</td><td>32773</td><td>+</td><td>0</td><td>CAAGAGGTCCATGAACTTGG</td><td>CGG</td><td>||||||||[||||||||||||]PAM</td><td>NA</td><td>Intergenic</td><td>NA</td><td>NA</td></tr>
<tr><td>T2017</td><td>TGCGCTCGTAGCGCTGCTCGAGG</td><td>1000</td></tr>
<tr><td>Oligo fwd</td><td>CACCTGCGCTCGTAGCGCTGCTCG</td></tr>
<tr><td>Oligo rev</td><td>AAACCGAGCAGCGCTACGAGCGCA</td></tr>
<tr><td>Chromosome</td><td>start</td><td>end</td><td>strand</td><td>MM</td><td>target_seq</td><td>PAM</td><td>alignment</td><td>distance</td><td>position</td><td>gene name</td><td>gene id</td></tr>
<tr><td>GU305914.1</td><td>32795</td><td>32817</td><td>+</td><td>0</td><td>TGCGCTCGTAGCGCTGCTCG</td><td>AGG</td><td>||||||||[||||||||||||]PAM</td><td>NA</td><td>Intergenic</td><td>NA</td><td>NA</td></tr>
<tr><td>T2018</td><td>GCTCGAGGAACATGCGCTCGAGG</td><td>1000</td></tr>
<tr><td>Oligo fwd</td><td>CACCGCTCGAGGAACATGCGCTCG</td></tr>
<tr><td>Oligo rev</td><td>AAACCGAGCGCATGTTCCTCGAGC</td></tr>
<tr><td>Chromosome</td><td>start</td><td>end</td><td>strand</td><td>MM</td><td>target_seq</td><td>PAM</td><td>alignment</td><td>distance</td><td>position</td><td>gene name</td><td>gene id</td></tr>
<tr><td>GU305914.1</td><td>32810</td><td>32832</td><td>+</td><td>0</td><td>GCTCGAGGAACATGCGCTCG</td><td>AGG</td><td>||||||||[||||||||||||]PAM</td><td>NA</td><td>Intergenic</td><td>NA</td><td>NA</td></tr>
<tr><td>T2019</td><td>CTCGAGGAACATGCGCTCGAGGG</td><td>1000</td></tr>
<tr><td>Oligo fwd</td><td>CACCCTCGAGGAACATGCGCTCGA</td></tr>
<tr><td>Oligo rev</td><td>AAACTCGAGCGCATGTTCCTCGAG</td></tr>
<tr><td>Chromosome</td><td>start</td><td>end</td><td>strand</td><td>MM</td><td>target_seq</td><td>PAM</td><td>alignment</td><td>distance</td><td>position</td><td>gene name</td><td>gene id</td></tr>
<tr><td>GU305914.1</td><td>32811</td><td>32833</td><td>+</td><td>0</td><td>CTCGAGGAACATGCGCTCGA</td><td>GGG</td><td>||||||||[||||||||||||]PAM</td><td>NA</td><td>Intergenic</td><td>NA</td><td>NA</td></tr>
<tr><td>T2020</td><td>GTCGTCGTCCTCGTAGCCGTCGG</td><td>1000</td></tr>
<tr><td>Oligo fwd</td><td>CACCGTCGTCGTCCTCGTAGCCGT</td></tr>
<tr><td>Oligo rev</td><td>AAACACGGCTACGAGGACGACGAC</td></tr>
<tr><td>Chromosome</td><td>start</td><td>end</td><td>strand</td><td>MM</td><td>target_seq</td><td>PAM</td><td>alignment</td><td>distance</td><td>position</td><td>gene name</td><td>gene id</td></tr>
<tr><td>GU305914.1</td><td>32841</td><td>32863</td><td>+</td><td>0</td><td>GTCGTCGTCCTCGTAGCCGT</td><td>CGG</td><td>||||||||[||||||||||||]PAM</td><td>NA</td><td>Intergenic</td><td>NA</td><td>NA</td></tr>
</table>

| | | | | | | | | | | | |
|---|---|---|---|---|---|---|---|---|---|---|---|
| T2021 | TCGTCGTCCTCGTAGCCGTCGGG | 1000 | | | | | | | | | |
| Oligo fwd | CACCTCGTCGTCCTCGTAGCCGTC | | | | | | | | | | |
| Oligo rev | AAACGACGGCTACGAGGACGACGA | | | | | | | | | | |
| Chromosome | start | end | strand | MM | target_seq | PAM | alignment | distance | position | gene name | gene id |
| GU305914.1 | 32842 | 32864 | + | 0 | TCGTCGTCCTCGTAGCCGTC | GGG | \|\|\|\|\|\|\|\|[\|\|\|\|\|\|\|\|\|\|\|\|]PAM | NA | Intergenic | NA | NA |
| T2022 | GTCGGGCCGCTCGTCCGAGAAGG | 1000 | | | | | | | | | |
| Oligo fwd | CACCGTCGGGCCGCTCGTCCGAGA | | | | | | | | | | |
| Oligo rev | AAACTCTCGGACGAGCGGCCCGAC | | | | | | | | | | |
| Chromosome | start | end | strand | MM | target_seq | PAM | alignment | distance | position | gene name | gene id |
| GU305914.1 | 32859 | 32881 | + | 0 | GTCGGGCCGCTCGTCCGAGA | AGG | \|\|\|\|\|\|\|\|[\|\|\|\|\|\|\|\|\|\|\|\|]PAM | NA | Intergenic | NA | NA |
| T2023 | GCTCGTCCGAGAAGGTGAGCCGG | 1000 | | | | | | | | | |
| Oligo fwd | CACCGCTCGTCCGAGAAGGTGAGC | | | | | | | | | | |
| Oligo rev | AAACGCTCACCTTCTCGGACGAGC | | | | | | | | | | |
| Chromosome | start | end | strand | MM | target_seq | PAM | alignment | distance | position | gene name | gene id |
| GU305914.1 | 32867 | 32889 | + | 0 | GCTCGTCCGAGAAGGTGAGC | CGG | \|\|\|\|\|\|\|\|[\|\|\|\|\|\|\|\|\|\|\|\|]PAM | NA | Intergenic | NA | NA |
| T2024 | GGTGAGCCGGTCCAGCGCGTTGG | 1000 | | | | | | | | | |
| Oligo fwd | CACCGGTGAGCCGGTCCAGCGCGT | | | | | | | | | | |
| Oligo rev | AAACACGCGCTGGACCGGCTCACC | | | | | | | | | | |
| Chromosome | start | end | strand | MM | target_seq | PAM | alignment | distance | position | gene name | gene id |
| GU305914.1 | 32880 | 32902 | + | 0 | GGTGAGCCGGTCCAGCGCGT | TGG | \|\|\|\|\|\|\|\|[\|\|\|\|\|\|\|\|\|\|\|\|]PAM | NA | Intergenic | NA | NA |
| T2025 | CCGGTCCAGCGCGTTGGCGATGG | 1000 | | | | | | | | | |
| Oligo fwd | CACCCCGGTCCAGCGCGTTGGCGA | | | | | | | | | | |
| Oligo rev | AAACTCGCCAACGCGCTGGACCGG | | | | | | | | | | |
| Chromosome | start | end | strand | MM | target_seq | PAM | alignment | distance | position | gene name | gene id |
| GU305914.1 | 32886 | 32908 | + | 0 | CCGGTCCAGCGCGTTGGCGA | TGG | \|\|\|\|\|\|\|\|[\|\|\|\|\|\|\|\|\|\|\|\|]PAM | NA | Intergenic | NA | NA |
| T2026 | CCAGCGCGTTGGCGATGGACAGG | 1000 | | | | | | | | | |
| Oligo fwd | CACCCCAGCGCGTTGGCGATGGAC | | | | | | | | | | |
| Oligo rev | AAACGTCCATCGCCAACGCGCTGG | | | | | | | | | | |
| Chromosome | start | end | strand | MM | target_seq | PAM | alignment | distance | position | gene name | gene id |
| GU305914.1 | 32891 | 32913 | + | 0 | CCAGCGCGTTGGCGATGGAC | AGG | \|\|\|\|\|\|\|\|[\|\|\|\|\|\|\|\|\|\|\|\|]PAM | NA | Intergenic | NA | NA |
| T2027 | CGTCCGAGATGCCGTCCTGTCGG | 1000 | | | | | | | | | |
| Oligo fwd | CACCCGTCCGAGATGCCGTCCTGT | | | | | | | | | | |
| Oligo rev | AAACACAGGACGGCATCTCGGACG | | | | | | | | | | |
| Chromosome | start | end | strand | MM | target_seq | PAM | alignment | distance | position | gene name | gene id |
| GU305914.1 | 32924 | 32946 | + | 0 | CGTCCGAGATGCCGTCCTGT | CGG | \|\|\|\|\|\|\|\|[\|\|\|\|\|\|\|\|\|\|\|\|]PAM | NA | Intergenic | NA | NA |
| T2028 | GTCCGAGATGCCGTCCTGTCGGG | 1000 | | | | | | | | | |
| Oligo fwd | CACCGTCCGAGATGCCGTCCTGTC | | | | | | | | | | |
| Oligo rev | AAACGACAGGACGGCATCTCGGAC | | | | | | | | | | |
| Chromosome | start | end | strand | MM | target_seq | PAM | alignment | distance | position | gene name | gene id |
| GU305914.1 | 32925 | 32947 | + | 0 | GTCCGAGATGCCGTCCTGTC | GGG | \|\|\|\|\|\|\|\|[\|\|\|\|\|\|\|\|\|\|\|\|]PAM | NA | Intergenic | NA | NA |
| T2029 | TCCGAGATGCCGTCCTGTCGGGG | 1000 | | | | | | | | | |
| Oligo fwd | CACCTCCGAGATGCCGTCCTGTCG | | | | | | | | | | |
| Oligo rev | AAACCGACAGGACGGCATCTCGGA | | | | | | | | | | |
| Chromosome | start | end | strand | MM | target_seq | PAM | alignment | distance | position | gene name | gene id |
| GU305914.1 | 32926 | 32948 | + | 0 | TCCGAGATGCCGTCCTGTCG | GGG | \|\|\|\|\|\|\|\|[\|\|\|\|\|\|\|\|\|\|\|\|]PAM | NA | Intergenic | NA | NA |
| T2030 | CCGAGATGCCGTCCTGTCGGGGG | 1000 | | | | | | | | | |
| Oligo fwd | CACCCCGAGATGCCGTCCTGTCGG | | | | | | | | | | |
| Oligo rev | AAACCCGACAGGACGGCATCTCGG | | | | | | | | | | |
| Chromosome | start | end | strand | MM | target_seq | PAM | alignment | distance | position | gene name | gene id |
| GU305914.1 | 32927 | 32949 | + | 0 | CCGAGATGCCGTCCTGTCGG | GGG | \|\|\|\|\|\|\|\|[\|\|\|\|\|\|\|\|\|\|\|\|]PAM | NA | Intergenic | NA | NA |
| T2031 | GATGCCGTCCTGTCGGGGGTCGG | 1000 | | | | | | | | | |
| Oligo fwd | CACCGATGCCGTCCTGTCGGGGGT | | | | | | | | | | |
| Oligo rev | AAACACCCCCGACAGGACGGCATC | | | | | | | | | | |
| Chromosome | start | end | strand | MM | target_seq | PAM | alignment | distance | position | gene name | gene id |
| GU305914.1 | 32931 | 32953 | + | 0 | GATGCCGTCCTGTCGGGGGT | CGG | \|\|\|\|\|\|\|\|[\|\|\|\|\|\|\|\|\|\|\|\|]PAM | NA | Intergenic | NA | NA |
| T2032 | GCCGTCCTGTCGGGGGTCGGAGG | 1000 | | | | | | | | | |
| Oligo fwd | CACCGCCGTCCTGTCGGGGGTCGG | | | | | | | | | | |
| Oligo rev | AAACCCGACCCCCGACAGGACGGC | | | | | | | | | | |
| Chromosome | start | end | strand | MM | target_seq | PAM | alignment | distance | position | gene name | gene id |
| GU305914.1 | 32934 | 32956 | + | 0 | GCCGTCCTGTCGGGGGTCGG | AGG | \|\|\|\|\|\|\|\|[\|\|\|\|\|\|\|\|\|\|\|\|]PAM | NA | Intergenic | NA | NA |
| T2033 | GGGGTCGGAGGCGACGAAGACGG | 1000 | | | | | | | | | |
| Oligo fwd | CACCGGGGTCGGAGGCGACGAAGA | | | | | | | | | | |
| Oligo rev | AAACTCTTCGTCGCCTCCGACCCC | | | | | | | | | | |
| Chromosome | start | end | strand | MM | target_seq | PAM | alignment | distance | position | gene name | gene id |
| GU305914.1 | 32946 | 32968 | + | 0 | GGGGTCGGAGGCGACGAAGA | CGG | \|\|\|\|\|\|\|\|[\|\|\|\|\|\|\|\|\|\|\|\|]PAM | NA | Intergenic | NA | NA |
| T2034 | GGGTCGGAGGCGACGAAGACGGG | 1000 | | | | | | | | | |
| Oligo fwd | CACCGGGTCGGAGGCGACGAAGAC | | | | | | | | | | |
| Oligo rev | AAACGTCTTCGTCGCCTCCGACCC | | | | | | | | | | |
| Chromosome | start | end | strand | MM | target_seq | PAM | alignment | distance | position | gene name | gene id |
| GU305914.1 | 32947 | 32969 | + | 0 | GGGTCGGAGGCGACGAAGAC | GGG | \|\|\|\|\|\|\|\|[\|\|\|\|\|\|\|\|\|\|\|\|]PAM | NA | Intergenic | NA | NA |
| T2035 | CGGAGGCGACGAAGACGGGCCGG | 1000 | | | | | | | | | |
| Oligo fwd | CACCCGGAGGCGACGAAGACGGGC | | | | | | | | | | |
| Oligo rev | AAACGCCCGTCTTCGTCGCCTCCG | | | | | | | | | | |
| Chromosome | start | end | strand | MM | target_seq | PAM | alignment | distance | position | gene name | gene id |

<table>
<tr><td>GU305914.1</td><td></td><td>32951</td><td>32973</td><td>+</td><td>0</td><td>CGGAGGCGACGAAGACGGGC</td><td>CGG</td><td>||||||||[||||||||||||]PAM</td><td>NA</td><td>Intergenic</td><td>NA</td><td>NA</td></tr>
<tr><td>GU305914.1</td><td></td><td>68313</td><td>68335</td><td>+</td><td>4</td><td>CGGCGGTGACGAGGACGGAC</td><td>CGG</td><td>|||-||-|[||||-|||||-|]PAM</td><td>NA</td><td>Intergenic</td><td>NA</td><td>NA</td></tr>
<tr><td>T2036</td><td>GGCGACGAAGACGGGCCGGTCGG</td><td></td><td>1000</td><td></td><td></td><td></td><td></td><td></td><td></td><td></td><td></td><td></td></tr>
<tr><td>Oligo fwd</td><td>CACCGGCGACGAAGACGGGCCGGT</td><td></td><td></td><td></td><td></td><td></td><td></td><td></td><td></td><td></td><td></td><td></td></tr>
<tr><td>Oligo rev</td><td>AAACACCGGCCCGTCTTCGTCGCC</td><td></td><td></td><td></td><td></td><td></td><td></td><td></td><td></td><td></td><td></td><td></td></tr>
<tr><td>Chromosome</td><td>start</td><td></td><td>end</td><td>strand</td><td>MM</td><td>target_seq</td><td>PAM</td><td>alignment</td><td>distance</td><td>position</td><td>gene name</td><td>gene id</td></tr>
<tr><td>GU305914.1</td><td></td><td>32955</td><td>32977</td><td>+</td><td>0</td><td>GGCGACGAAGACGGGCCGGT</td><td>CGG</td><td>||||||||[||||||||||||]PAM</td><td>NA</td><td>Intergenic</td><td>NA</td><td>NA</td></tr>
<tr><td>T2037</td><td>CGGGCCGGTCGGCTCCCGCGCGG</td><td></td><td>1000</td><td></td><td></td><td></td><td></td><td></td><td></td><td></td><td></td><td></td></tr>
<tr><td>Oligo fwd</td><td>CACCCGGGCCGGTCGGCTCCCGCG</td><td></td><td></td><td></td><td></td><td></td><td></td><td></td><td></td><td></td><td></td><td></td></tr>
<tr><td>Oligo rev</td><td>AAACCGCGGGAGCCGACCGGCCCG</td><td></td><td></td><td></td><td></td><td></td><td></td><td></td><td></td><td></td><td></td><td></td></tr>
<tr><td>Chromosome</td><td>start</td><td></td><td>end</td><td>strand</td><td>MM</td><td>target_seq</td><td>PAM</td><td>alignment</td><td>distance</td><td>position</td><td>gene name</td><td>gene id</td></tr>
<tr><td>GU305914.1</td><td></td><td>32966</td><td>32988</td><td>+</td><td>0</td><td>CGGGCCGGTCGGCTCCCGCG</td><td>CGG</td><td>||||||||[||||||||||||]PAM</td><td>NA</td><td>Intergenic</td><td>NA</td><td>NA</td></tr>
<tr><td>T2038</td><td>GGTCGGCTCCCGCGCGGATGCGG</td><td></td><td>1000</td><td></td><td></td><td></td><td></td><td></td><td></td><td></td><td></td><td></td></tr>
<tr><td>Oligo fwd</td><td>CACCGGTCGGCTCCCGCGCGGATG</td><td></td><td></td><td></td><td></td><td></td><td></td><td></td><td></td><td></td><td></td><td></td></tr>
<tr><td>Oligo rev</td><td>AAACCATCCGCGCGGGAGCCGACC</td><td></td><td></td><td></td><td></td><td></td><td></td><td></td><td></td><td></td><td></td><td></td></tr>
<tr><td>Chromosome</td><td>start</td><td></td><td>end</td><td>strand</td><td>MM</td><td>target_seq</td><td>PAM</td><td>alignment</td><td>distance</td><td>position</td><td>gene name</td><td>gene id</td></tr>
<tr><td>GU305914.1</td><td></td><td>32972</td><td>32994</td><td>+</td><td>0</td><td>GGTCGGCTCCCGCGCGGATG</td><td>CGG</td><td>||||||||[||||||||||||]PAM</td><td>NA</td><td>Intergenic</td><td>NA</td><td>NA</td></tr>
<tr><td>T2039</td><td>GTCGGCTCCCGCGCGGATGCGGG</td><td></td><td>1000</td><td></td><td></td><td></td><td></td><td></td><td></td><td></td><td></td><td></td></tr>
<tr><td>Oligo fwd</td><td>CACCGTCGGCTCCCGCGCGGATGC</td><td></td><td></td><td></td><td></td><td></td><td></td><td></td><td></td><td></td><td></td><td></td></tr>
<tr><td>Oligo rev</td><td>AAACGCATCCGCGCGGGAGCCGAC</td><td></td><td></td><td></td><td></td><td></td><td></td><td></td><td></td><td></td><td></td><td></td></tr>
<tr><td>Chromosome</td><td>start</td><td></td><td>end</td><td>strand</td><td>MM</td><td>target_seq</td><td>PAM</td><td>alignment</td><td>distance</td><td>position</td><td>gene name</td><td>gene id</td></tr>
<tr><td>GU305914.1</td><td></td><td>32973</td><td>32995</td><td>+</td><td>0</td><td>GTCGGCTCCCGCGCGGATGC</td><td>GGG</td><td>||||||||[||||||||||||]PAM</td><td>NA</td><td>Intergenic</td><td>NA</td><td>NA</td></tr>
<tr><td>T2040</td><td>GGATGCGGGCGTCGACGTCGCGG</td><td></td><td>1000</td><td></td><td></td><td></td><td></td><td></td><td></td><td></td><td></td><td></td></tr>
<tr><td>Oligo fwd</td><td>CACCGGATGCGGGCGTCGACGTCG</td><td></td><td></td><td></td><td></td><td></td><td></td><td></td><td></td><td></td><td></td><td></td></tr>
<tr><td>Oligo rev</td><td>AAACCGACGTCGACGCCCGCATCC</td><td></td><td></td><td></td><td></td><td></td><td></td><td></td><td></td><td></td><td></td><td></td></tr>
<tr><td>Chromosome</td><td>start</td><td></td><td>end</td><td>strand</td><td>MM</td><td>target_seq</td><td>PAM</td><td>alignment</td><td>distance</td><td>position</td><td>gene name</td><td>gene id</td></tr>
<tr><td>GU305914.1</td><td></td><td>32987</td><td>33009</td><td>+</td><td>0</td><td>GGATGCGGGCGTCGACGTCG</td><td>CGG</td><td>||||||||[||||||||||||]PAM</td><td>NA</td><td>Intergenic</td><td>NA</td><td>NA</td></tr>
<tr><td>T2041</td><td>TGCGGGCGTCGACGTCGCGGCGG</td><td></td><td>1000</td><td></td><td></td><td></td><td></td><td></td><td></td><td></td><td></td><td></td></tr>
<tr><td>Oligo fwd</td><td>CACCTGCGGGCGTCGACGTCGCGG</td><td></td><td></td><td></td><td></td><td></td><td></td><td></td><td></td><td></td><td></td><td></td></tr>
<tr><td>Oligo rev</td><td>AAACCCGCGACGTCGACGCCCGCA</td><td></td><td></td><td></td><td></td><td></td><td></td><td></td><td></td><td></td><td></td><td></td></tr>
<tr><td>Chromosome</td><td>start</td><td></td><td>end</td><td>strand</td><td>MM</td><td>target_seq</td><td>PAM</td><td>alignment</td><td>distance</td><td>position</td><td>gene name</td><td>gene id</td></tr>
<tr><td>GU305914.1</td><td></td><td>32990</td><td>33012</td><td>+</td><td>0</td><td>TGCGGGCGTCGACGTCGCGG</td><td>CGG</td><td>||||||||[||||||||||||]PAM</td><td>NA</td><td>Intergenic</td><td>NA</td><td>NA</td></tr>
<tr><td>T2042</td><td>GACGTCGCGGCGGAGCATGACGG</td><td></td><td>1000</td><td></td><td></td><td></td><td></td><td></td><td></td><td></td><td></td><td></td></tr>
<tr><td>Oligo fwd</td><td>CACCGACGTCGCGGCGGAGCATGA</td><td></td><td></td><td></td><td></td><td></td><td></td><td></td><td></td><td></td><td></td><td></td></tr>
<tr><td>Oligo rev</td><td>AAACTCATGCTCCGCCGCGACGTC</td><td></td><td></td><td></td><td></td><td></td><td></td><td></td><td></td><td></td><td></td><td></td></tr>
<tr><td>Chromosome</td><td>start</td><td></td><td>end</td><td>strand</td><td>MM</td><td>target_seq</td><td>PAM</td><td>alignment</td><td>distance</td><td>position</td><td>gene name</td><td>gene id</td></tr>
<tr><td>GU305914.1</td><td></td><td>33000</td><td>33022</td><td>+</td><td>0</td><td>GACGTCGCGGCGGAGCATGA</td><td>CGG</td><td>||||||||[||||||||||||]PAM</td><td>NA</td><td>Intergenic</td><td>NA</td><td>NA</td></tr>
<tr><td>T2043</td><td>GTCGCGGCGGAGCATGACGGAGG</td><td></td><td>1000</td><td></td><td></td><td></td><td></td><td></td><td></td><td></td><td></td><td></td></tr>
<tr><td>Oligo fwd</td><td>CACCGTCGCGGCGGAGCATGACGG</td><td></td><td></td><td></td><td></td><td></td><td></td><td></td><td></td><td></td><td></td><td></td></tr>
<tr><td>Oligo rev</td><td>AAACCCGTCATGCTCCGCCGCGAC</td><td></td><td></td><td></td><td></td><td></td><td></td><td></td><td></td><td></td><td></td><td></td></tr>
<tr><td>Chromosome</td><td>start</td><td></td><td>end</td><td>strand</td><td>MM</td><td>target_seq</td><td>PAM</td><td>alignment</td><td>distance</td><td>position</td><td>gene name</td><td>gene id</td></tr>
<tr><td>GU305914.1</td><td></td><td>33003</td><td>33025</td><td>+</td><td>0</td><td>GTCGCGGCGGAGCATGACGG</td><td>AGG</td><td>||||||||[||||||||||||]PAM</td><td>NA</td><td>Intergenic</td><td>NA</td><td>NA</td></tr>
<tr><td>T2044</td><td>GGAGCATGACGGAGGCCTCGCGG</td><td></td><td>1000</td><td></td><td></td><td></td><td></td><td></td><td></td><td></td><td></td><td></td></tr>
<tr><td>Oligo fwd</td><td>CACCGGAGCATGACGGAGGCCTCG</td><td></td><td></td><td></td><td></td><td></td><td></td><td></td><td></td><td></td><td></td><td></td></tr>
<tr><td>Oligo rev</td><td>AAACCGAGGCCTCCGTCATGCTCC</td><td></td><td></td><td></td><td></td><td></td><td></td><td></td><td></td><td></td><td></td><td></td></tr>
<tr><td>Chromosome</td><td>start</td><td></td><td>end</td><td>strand</td><td>MM</td><td>target_seq</td><td>PAM</td><td>alignment</td><td>distance</td><td>position</td><td>gene name</td><td>gene id</td></tr>
<tr><td>GU305914.1</td><td></td><td>33011</td><td>33033</td><td>+</td><td>0</td><td>GGAGCATGACGGAGGCCTCG</td><td>CGG</td><td>||||||||[||||||||||||]PAM</td><td>NA</td><td>Intergenic</td><td>NA</td><td>NA</td></tr>
<tr><td>T2045</td><td>TGACGGAGGCCTCGCGGACCAGG</td><td></td><td>1000</td><td></td><td></td><td></td><td></td><td></td><td></td><td></td><td></td><td></td></tr>
<tr><td>Oligo fwd</td><td>CACCTGACGGAGGCCTCGCGGACC</td><td></td><td></td><td></td><td></td><td></td><td></td><td></td><td></td><td></td><td></td><td></td></tr>
<tr><td>Oligo rev</td><td>AAACGGTCCGCGAGGCCTCCGTCA</td><td></td><td></td><td></td><td></td><td></td><td></td><td></td><td></td><td></td><td></td><td></td></tr>
<tr><td>Chromosome</td><td>start</td><td></td><td>end</td><td>strand</td><td>MM</td><td>target_seq</td><td>PAM</td><td>alignment</td><td>distance</td><td>position</td><td>gene name</td><td>gene id</td></tr>
<tr><td>GU305914.1</td><td></td><td>33017</td><td>33039</td><td>+</td><td>0</td><td>TGACGGAGGCCTCGCGGACC</td><td>AGG</td><td>||||||||[||||||||||||]PAM</td><td>NA</td><td>Intergenic</td><td>NA</td><td>NA</td></tr>
<tr><td>T2046</td><td>GGAGGCCTCGCGGACCAGGCCGG</td><td></td><td>1000</td><td></td><td></td><td></td><td></td><td></td><td></td><td></td><td></td><td></td></tr>
<tr><td>Oligo fwd</td><td>CACCGGAGGCCTCGCGGACCAGGC</td><td></td><td></td><td></td><td></td><td></td><td></td><td></td><td></td><td></td><td></td><td></td></tr>
<tr><td>Oligo rev</td><td>AAACGCCTGGTCCGCGAGGCCTCC</td><td></td><td></td><td></td><td></td><td></td><td></td><td></td><td></td><td></td><td></td><td></td></tr>
<tr><td>Chromosome</td><td>start</td><td></td><td>end</td><td>strand</td><td>MM</td><td>target_seq</td><td>PAM</td><td>alignment</td><td>distance</td><td>position</td><td>gene name</td><td>gene id</td></tr>
<tr><td>GU305914.1</td><td></td><td>33021</td><td>33043</td><td>+</td><td>0</td><td>GGAGGCCTCGCGGACCAGGC</td><td>CGG</td><td>||||||||[||||||||||||]PAM</td><td>NA</td><td>Intergenic</td><td>NA</td><td>NA</td></tr>
<tr><td>T2047</td><td>CGCGGACCAGGCCGGTGACGAGG</td><td></td><td>1000</td><td></td><td></td><td></td><td></td><td></td><td></td><td></td><td></td><td></td></tr>
<tr><td>Oligo fwd</td><td>CACCCGCGGACCAGGCCGGTGACG</td><td></td><td></td><td></td><td></td><td></td><td></td><td></td><td></td><td></td><td></td><td></td></tr>
<tr><td>Oligo rev</td><td>AAACCGTCACCGGCCTGGTCCGCG</td><td></td><td></td><td></td><td></td><td></td><td></td><td></td><td></td><td></td><td></td><td></td></tr>
<tr><td>Chromosome</td><td>start</td><td></td><td>end</td><td>strand</td><td>MM</td><td>target_seq</td><td>PAM</td><td>alignment</td><td>distance</td><td>position</td><td>gene name</td><td>gene id</td></tr>
<tr><td>GU305914.1</td><td></td><td>33029</td><td>33051</td><td>+</td><td>0</td><td>CGCGGACCAGGCCGGTGACG</td><td>AGG</td><td>||||||||[||||||||||||]PAM</td><td>NA</td><td>Intergenic</td><td>NA</td><td>NA</td></tr>
<tr><td>T2048</td><td>GCTCGCTACGCACTACGTCGCGG</td><td></td><td>1000</td><td></td><td></td><td></td><td></td><td></td><td></td><td></td><td></td><td></td></tr>
<tr><td>Oligo fwd</td><td>CACCGCTCGCTACGCACTACGTCG</td><td></td><td></td><td></td><td></td><td></td><td></td><td></td><td></td><td></td><td></td><td></td></tr>
<tr><td>Oligo rev</td><td>AAACCGACGTAGTGCGTAGCGAGC</td><td></td><td></td><td></td><td></td><td></td><td></td><td></td><td></td><td></td><td></td><td></td></tr>
<tr><td>Chromosome</td><td>start</td><td></td><td>end</td><td>strand</td><td>MM</td><td>target_seq</td><td>PAM</td><td>alignment</td><td>distance</td><td>position</td><td>gene name</td><td>gene id</td></tr>
<tr><td>GU305914.1</td><td></td><td>33068</td><td>33090</td><td>+</td><td>0</td><td>GCTCGCTACGCACTACGTCG</td><td>CGG</td><td>||||||||[||||||||||||]PAM</td><td>NA</td><td>Intergenic</td><td>NA</td><td>NA</td></tr>
<tr><td>T2049</td><td>GCACTACGTCGCGGACGCAGCGG</td><td></td><td>1000</td><td></td><td></td><td></td><td></td><td></td><td></td><td></td><td></td><td></td></tr>
<tr><td>Oligo fwd</td><td>CACCGCACTACGTCGCGGACGCAG</td><td></td><td></td><td></td><td></td><td></td><td></td><td></td><td></td><td></td><td></td><td></td></tr>
<tr><td>Oligo rev</td><td>AAACCTGCGTCCGCGACGTAGTGC</td><td></td><td></td><td></td><td></td><td></td><td></td><td></td><td></td><td></td><td></td><td></td></tr>
<tr><td>Chromosome</td><td>start</td><td></td><td>end</td><td>strand</td><td>MM</td><td>target_seq</td><td>PAM</td><td>alignment</td><td>distance</td><td>position</td><td>gene name</td><td>gene id</td></tr>
<tr><td>GU305914.1</td><td></td><td>33077</td><td>33099</td><td>+</td><td>0</td><td>GCACTACGTCGCGGACGCAG</td><td>CGG</td><td>||||||||[||||||||||||]PAM</td><td>NA</td><td>Intergenic</td><td>NA</td><td>NA</td></tr>
<tr><td>T2050</td><td>GAGCATGAAGTAGACGTTGTTGG</td><td></td><td>1000</td><td></td><td></td><td></td><td></td><td></td><td></td><td></td><td></td><td></td></tr>
<tr><td>Oligo fwd</td><td>CACCGAGCATGAAGTAGACGTTGT</td><td></td><td></td><td></td><td></td><td></td><td></td><td></td><td></td><td></td><td></td><td></td></tr>
</table>

<table>
<tr><td>Oligo rev</td><td>AAACACAACGTCTACTTCATGCTC</td><td></td><td></td><td></td><td></td><td></td><td></td><td></td><td></td><td></td><td></td></tr>
<tr><td>Chromosome</td><td>start</td><td>end</td><td>strand</td><td>MM</td><td>target_seq</td><td>PAM</td><td>alignment</td><td>distance</td><td>position</td><td>gene name</td><td>gene id</td></tr>
<tr><td>GU305914.1</td><td>33108</td><td>33130</td><td>+</td><td>0</td><td>GAGCATGAAGTAGACGTTGT</td><td>TGG</td><td>||||||||[||||||||||||]PAM</td><td>NA</td><td>Intergenic</td><td>NA</td><td>NA</td></tr>
<tr><td></td><td></td><td></td><td></td><td></td><td></td><td></td><td></td><td></td><td></td><td></td><td></td></tr>
<tr><td>T2051</td><td>ATCGATGTCCATGCAGTCGACGG</td><td>1000</td><td></td><td></td><td></td><td></td><td></td><td></td><td></td><td></td><td></td></tr>
<tr><td>Oligo fwd</td><td>CACCATCGATGTCCATGCAGTCGA</td><td></td><td></td><td></td><td></td><td></td><td></td><td></td><td></td><td></td><td></td></tr>
<tr><td>Oligo rev</td><td>AAACTCGACTGCATGGACATCGAT</td><td></td><td></td><td></td><td></td><td></td><td></td><td></td><td></td><td></td><td></td></tr>
<tr><td>Chromosome</td><td>start</td><td>end</td><td>strand</td><td>MM</td><td>target_seq</td><td>PAM</td><td>alignment</td><td>distance</td><td>position</td><td>gene name</td><td>gene id</td></tr>
<tr><td>GU305914.1</td><td>33153</td><td>33175</td><td>+</td><td>0</td><td>ATCGATGTCCATGCAGTCGA</td><td>CGG</td><td>||||||||[||||||||||||]PAM</td><td>NA</td><td>Intergenic</td><td>NA</td><td>NA</td></tr>
<tr><td></td><td></td><td></td><td></td><td></td><td></td><td></td><td></td><td></td><td></td><td></td><td></td></tr>
<tr><td>T2052</td><td>CATGCAGTCGACGGCCGTGAAGG</td><td>1000</td><td></td><td></td><td></td><td></td><td></td><td></td><td></td><td></td><td></td></tr>
<tr><td>Oligo fwd</td><td>CACCCATGCAGTCGACGGCCGTGA</td><td></td><td></td><td></td><td></td><td></td><td></td><td></td><td></td><td></td><td></td></tr>
<tr><td>Oligo rev</td><td>AAACTCACGGCCGTCGACTGCATG</td><td></td><td></td><td></td><td></td><td></td><td></td><td></td><td></td><td></td><td></td></tr>
<tr><td>Chromosome</td><td>start</td><td>end</td><td>strand</td><td>MM</td><td>target_seq</td><td>PAM</td><td>alignment</td><td>distance</td><td>position</td><td>gene name</td><td>gene id</td></tr>
<tr><td>GU305914.1</td><td>33162</td><td>33184</td><td>+</td><td>0</td><td>CATGCAGTCGACGGCCGTGA</td><td>AGG</td><td>||||||||[||||||||||||]PAM</td><td>NA</td><td>Intergenic</td><td>NA</td><td>NA</td></tr>
<tr><td></td><td></td><td></td><td></td><td></td><td></td><td></td><td></td><td></td><td></td><td></td><td></td></tr>
<tr><td>T2053</td><td>ATGCAGTCGACGGCCGTGAAGGG</td><td>1000</td><td></td><td></td><td></td><td></td><td></td><td></td><td></td><td></td><td></td></tr>
<tr><td>Oligo fwd</td><td>CACCATGCAGTCGACGGCCGTGAA</td><td></td><td></td><td></td><td></td><td></td><td></td><td></td><td></td><td></td><td></td></tr>
<tr><td>Oligo rev</td><td>AAACTTCACGGCCGTCGACTGCAT</td><td></td><td></td><td></td><td></td><td></td><td></td><td></td><td></td><td></td><td></td></tr>
<tr><td>Chromosome</td><td>start</td><td>end</td><td>strand</td><td>MM</td><td>target_seq</td><td>PAM</td><td>alignment</td><td>distance</td><td>position</td><td>gene name</td><td>gene id</td></tr>
<tr><td>GU305914.1</td><td>33163</td><td>33185</td><td>+</td><td>0</td><td>ATGCAGTCGACGGCCGTGAA</td><td>GGG</td><td>||||||||[||||||||||||]PAM</td><td>NA</td><td>Intergenic</td><td>NA</td><td>NA</td></tr>
<tr><td></td><td></td><td></td><td></td><td></td><td></td><td></td><td></td><td></td><td></td><td></td><td></td></tr>
<tr><td>T2054</td><td>AGTCGACGGCCGTGAAGGGCAGG</td><td>1000</td><td></td><td></td><td></td><td></td><td></td><td></td><td></td><td></td><td></td></tr>
<tr><td>Oligo fwd</td><td>CACCAGTCGACGGCCGTGAAGGGC</td><td></td><td></td><td></td><td></td><td></td><td></td><td></td><td></td><td></td><td></td></tr>
<tr><td>Oligo rev</td><td>AAACGCCCTTCACGGCCGTCGACT</td><td></td><td></td><td></td><td></td><td></td><td></td><td></td><td></td><td></td><td></td></tr>
<tr><td>Chromosome</td><td>start</td><td>end</td><td>strand</td><td>MM</td><td>target_seq</td><td>PAM</td><td>alignment</td><td>distance</td><td>position</td><td>gene name</td><td>gene id</td></tr>
<tr><td>GU305914.1</td><td>33167</td><td>33189</td><td>+</td><td>0</td><td>AGTCGACGGCCGTGAAGGGC</td><td>AGG</td><td>||||||||[||||||||||||]PAM</td><td>NA</td><td>Intergenic</td><td>NA</td><td>NA</td></tr>
<tr><td></td><td></td><td></td><td></td><td></td><td></td><td></td><td></td><td></td><td></td><td></td><td></td></tr>
<tr><td>T2055</td><td>GTCGACGGCCGTGAAGGGCAGGG</td><td>1000</td><td></td><td></td><td></td><td></td><td></td><td></td><td></td><td></td><td></td></tr>
<tr><td>Oligo fwd</td><td>CACCGTCGACGGCCGTGAAGGGCA</td><td></td><td></td><td></td><td></td><td></td><td></td><td></td><td></td><td></td><td></td></tr>
<tr><td>Oligo rev</td><td>AAACTGCCCTTCACGGCCGTCGAC</td><td></td><td></td><td></td><td></td><td></td><td></td><td></td><td></td><td></td><td></td></tr>
<tr><td>Chromosome</td><td>start</td><td>end</td><td>strand</td><td>MM</td><td>target_seq</td><td>PAM</td><td>alignment</td><td>distance</td><td>position</td><td>gene name</td><td>gene id</td></tr>
<tr><td>GU305914.1</td><td>33168</td><td>33190</td><td>+</td><td>0</td><td>GTCGACGGCCGTGAAGGGCA</td><td>GGG</td><td>||||||||[||||||||||||]PAM</td><td>NA</td><td>Intergenic</td><td>NA</td><td>NA</td></tr>
<tr><td></td><td></td><td></td><td></td><td></td><td></td><td></td><td></td><td></td><td></td><td></td><td></td></tr>
<tr><td>T2056</td><td>TCGACGGCCGTGAAGGGCAGGGG</td><td>1000</td><td></td><td></td><td></td><td></td><td></td><td></td><td></td><td></td><td></td></tr>
<tr><td>Oligo fwd</td><td>CACCTCGACGGCCGTGAAGGGCAG</td><td></td><td></td><td></td><td></td><td></td><td></td><td></td><td></td><td></td><td></td></tr>
<tr><td>Oligo rev</td><td>AAACCTGCCCTTCACGGCCGTCGA</td><td></td><td></td><td></td><td></td><td></td><td></td><td></td><td></td><td></td><td></td></tr>
<tr><td>Chromosome</td><td>start</td><td>end</td><td>strand</td><td>MM</td><td>target_seq</td><td>PAM</td><td>alignment</td><td>distance</td><td>position</td><td>gene name</td><td>gene id</td></tr>
<tr><td>GU305914.1</td><td>33169</td><td>33191</td><td>+</td><td>0</td><td>TCGACGGCCGTGAAGGGCAG</td><td>GGG</td><td>||||||||[||||||||||||]PAM</td><td>NA</td><td>Intergenic</td><td>NA</td><td>NA</td></tr>
<tr><td></td><td></td><td></td><td></td><td></td><td></td><td></td><td></td><td></td><td></td><td></td><td></td></tr>
<tr><td>T2057</td><td>CGACGGCCGTGAAGGGCAGGGGG</td><td>1000</td><td></td><td></td><td></td><td></td><td></td><td></td><td></td><td></td><td></td></tr>
<tr><td>Oligo fwd</td><td>CACCCGACGGCCGTGAAGGGCAGG</td><td></td><td></td><td></td><td></td><td></td><td></td><td></td><td></td><td></td><td></td></tr>
<tr><td>Oligo rev</td><td>AAACCCTGCCCTTCACGGCCGTCG</td><td></td><td></td><td></td><td></td><td></td><td></td><td></td><td></td><td></td><td></td></tr>
<tr><td>Chromosome</td><td>start</td><td>end</td><td>strand</td><td>MM</td><td>target_seq</td><td>PAM</td><td>alignment</td><td>distance</td><td>position</td><td>gene name</td><td>gene id</td></tr>
<tr><td>GU305914.1</td><td>33170</td><td>33192</td><td>+</td><td>0</td><td>CGACGGCCGTGAAGGGCAGG</td><td>GGG</td><td>||||||||[||||||||||||]PAM</td><td>NA</td><td>Intergenic</td><td>NA</td><td>NA</td></tr>
<tr><td></td><td></td><td></td><td></td><td></td><td></td><td></td><td></td><td></td><td></td><td></td><td></td></tr>
<tr><td>T2058</td><td>ATGCCGTTGACGCGCGCGAGCGG</td><td>1000</td><td></td><td></td><td></td><td></td><td></td><td></td><td></td><td></td><td></td></tr>
<tr><td>Oligo fwd</td><td>CACCATGCCGTTGACGCGCGCGAG</td><td></td><td></td><td></td><td></td><td></td><td></td><td></td><td></td><td></td><td></td></tr>
<tr><td>Oligo rev</td><td>AAACCTCGCGCGCGTCAACGGCAT</td><td></td><td></td><td></td><td></td><td></td><td></td><td></td><td></td><td></td><td></td></tr>
<tr><td>Chromosome</td><td>start</td><td>end</td><td>strand</td><td>MM</td><td>target_seq</td><td>PAM</td><td>alignment</td><td>distance</td><td>position</td><td>gene name</td><td>gene id</td></tr>
<tr><td>GU305914.1</td><td>33193</td><td>33215</td><td>+</td><td>0</td><td>ATGCCGTTGACGCGCGCGAG</td><td>CGG</td><td>||||||||[||||||||||||]PAM</td><td>NA</td><td>Intergenic</td><td>NA</td><td>NA</td></tr>
<tr><td></td><td></td><td></td><td></td><td></td><td></td><td></td><td></td><td></td><td></td><td></td><td></td></tr>
<tr><td>T2059</td><td>GTTGACGCGCGCGAGCGGACTGG</td><td>1000</td><td></td><td></td><td></td><td></td><td></td><td></td><td></td><td></td><td></td></tr>
<tr><td>Oligo fwd</td><td>CACCGTTGACGCGCGCGAGCGGAC</td><td></td><td></td><td></td><td></td><td></td><td></td><td></td><td></td><td></td><td></td></tr>
<tr><td>Oligo rev</td><td>AAACGTCCGCTCGCGCGCGTCAAC</td><td></td><td></td><td></td><td></td><td></td><td></td><td></td><td></td><td></td><td></td></tr>
<tr><td>Chromosome</td><td>start</td><td>end</td><td>strand</td><td>MM</td><td>target_seq</td><td>PAM</td><td>alignment</td><td>distance</td><td>position</td><td>gene name</td><td>gene id</td></tr>
<tr><td>GU305914.1</td><td>33198</td><td>33220</td><td>+</td><td>0</td><td>GTTGACGCGCGCGAGCGGAC</td><td>TGG</td><td>||||||||[||||||||||||]PAM</td><td>NA</td><td>Intergenic</td><td>NA</td><td>NA</td></tr>
<tr><td></td><td></td><td></td><td></td><td></td><td></td><td></td><td></td><td></td><td></td><td></td><td></td></tr>
<tr><td>T2060</td><td>GACGCGCGCGAGCGGACTGGAGG</td><td>1000</td><td></td><td></td><td></td><td></td><td></td><td></td><td></td><td></td><td></td></tr>
<tr><td>Oligo fwd</td><td>CACCGACGCGCGCGAGCGGACTGG</td><td></td><td></td><td></td><td></td><td></td><td></td><td></td><td></td><td></td><td></td></tr>
<tr><td>Oligo rev</td><td>AAACCCAGTCCGCTCGCGCGCGTC</td><td></td><td></td><td></td><td></td><td></td><td></td><td></td><td></td><td></td><td></td></tr>
<tr><td>Chromosome</td><td>start</td><td>end</td><td>strand</td><td>MM</td><td>target_seq</td><td>PAM</td><td>alignment</td><td>distance</td><td>position</td><td>gene name</td><td>gene id</td></tr>
<tr><td>GU305914.1</td><td>33201</td><td>33223</td><td>+</td><td>0</td><td>GACGCGCGCGAGCGGACTGG</td><td>AGG</td><td>||||||||[||||||||||||]PAM</td><td>NA</td><td>Intergenic</td><td>NA</td><td>NA</td></tr>
<tr><td></td><td></td><td></td><td></td><td></td><td></td><td></td><td></td><td></td><td></td><td></td><td></td></tr>
<tr><td>T2061</td><td>CGAAGTGCACGTCGTACATGAGG</td><td>1000</td><td></td><td></td><td></td><td></td><td></td><td></td><td></td><td></td><td></td></tr>
<tr><td>Oligo fwd</td><td>CACCCGAAGTGCACGTCGTACATG</td><td></td><td></td><td></td><td></td><td></td><td></td><td></td><td></td><td></td><td></td></tr>
<tr><td>Oligo rev</td><td>AAACCATGTACGACGTGCACTTCG</td><td></td><td></td><td></td><td></td><td></td><td></td><td></td><td></td><td></td><td></td></tr>
<tr><td>Chromosome</td><td>start</td><td>end</td><td>strand</td><td>MM</td><td>target_seq</td><td>PAM</td><td>alignment</td><td>distance</td><td>position</td><td>gene name</td><td>gene id</td></tr>
<tr><td>GU305914.1</td><td>33227</td><td>33249</td><td>+</td><td>0</td><td>CGAAGTGCACGTCGTACATG</td><td>AGG</td><td>||||||||[||||||||||||]PAM</td><td>NA</td><td>Intergenic</td><td>NA</td><td>NA</td></tr>
<tr><td></td><td></td><td></td><td></td><td></td><td></td><td></td><td></td><td></td><td></td><td></td><td></td></tr>
<tr><td>T2062</td><td>GCACGTCGTACATGAGGTCGCGG</td><td>1000</td><td></td><td></td><td></td><td></td><td></td><td></td><td></td><td></td><td></td></tr>
<tr><td>Oligo fwd</td><td>CACCGCACGTCGTACATGAGGTCG</td><td></td><td></td><td></td><td></td><td></td><td></td><td></td><td></td><td></td><td></td></tr>
<tr><td>Oligo rev</td><td>AAACCGACCTCATGTACGACGTGC</td><td></td><td></td><td></td><td></td><td></td><td></td><td></td><td></td><td></td><td></td></tr>
<tr><td>Chromosome</td><td>start</td><td>end</td><td>strand</td><td>MM</td><td>target_seq</td><td>PAM</td><td>alignment</td><td>distance</td><td>position</td><td>gene name</td><td>gene id</td></tr>
<tr><td>GU305914.1</td><td>33233</td><td>33255</td><td>+</td><td>0</td><td>GCACGTCGTACATGAGGTCG</td><td>CGG</td><td>||||||||[||||||||||||]PAM</td><td>NA</td><td>Intergenic</td><td>NA</td><td>NA</td></tr>
<tr><td></td><td></td><td></td><td></td><td></td><td></td><td></td><td></td><td></td><td></td><td></td><td></td></tr>
<tr><td>T2063</td><td>CACGTCGTACATGAGGTCGCGGG</td><td>1000</td><td></td><td></td><td></td><td></td><td></td><td></td><td></td><td></td><td></td></tr>
<tr><td>Oligo fwd</td><td>CACCCACGTCGTACATGAGGTCGC</td><td></td><td></td><td></td><td></td><td></td><td></td><td></td><td></td><td></td><td></td></tr>
<tr><td>Oligo rev</td><td>AAACGCGACCTCATGTACGACGTG</td><td></td><td></td><td></td><td></td><td></td><td></td><td></td><td></td><td></td><td></td></tr>
<tr><td>Chromosome</td><td>start</td><td>end</td><td>strand</td><td>MM</td><td>target_seq</td><td>PAM</td><td>alignment</td><td>distance</td><td>position</td><td>gene name</td><td>gene id</td></tr>
<tr><td>GU305914.1</td><td>33234</td><td>33256</td><td>+</td><td>0</td><td>CACGTCGTACATGAGGTCGC</td><td>GGG</td><td>||||||||[||||||||||||]PAM</td><td>NA</td><td>Intergenic</td><td>NA</td><td>NA</td></tr>
<tr><td></td><td></td><td></td><td></td><td></td><td></td><td></td><td></td><td></td><td></td><td></td><td></td></tr>
<tr><td>T2064</td><td>CGTACATGAGGTCGCGGGCGAGG</td><td>1000</td><td></td><td></td><td></td><td></td><td></td><td></td><td></td><td></td><td></td></tr>
<tr><td>Oligo fwd</td><td>CACCCGTACATGAGGTCGCGGGCG</td><td></td><td></td><td></td><td></td><td></td><td></td><td></td><td></td><td></td><td></td></tr>
<tr><td>Oligo rev</td><td>AAACCGCCCGCGACCTCATGTACG</td><td></td><td></td><td></td><td></td><td></td><td></td><td></td><td></td><td></td><td></td></tr>
<tr><td>Chromosome</td><td>start</td><td>end</td><td>strand</td><td>MM</td><td>target_seq</td><td>PAM</td><td>alignment</td><td>distance</td><td>position</td><td>gene name</td><td>gene id</td></tr>
<tr><td>GU305914.1</td><td>33239</td><td>33261</td><td>+</td><td>0</td><td>CGTACATGAGGTCGCGGGCG</td><td>AGG</td><td>||||||||[||||||||||||]PAM</td><td>NA</td><td>Intergenic</td><td>NA</td><td>NA</td></tr>
<tr><td></td><td></td><td></td><td></td><td></td><td></td><td></td><td></td><td></td><td></td><td></td><td></td></tr>
<tr><td>T2065</td><td>GGGCGAGGTGTCCGCAGACGAGG</td><td>1000</td><td></td><td></td><td></td><td></td><td></td><td></td><td></td><td></td><td></td></tr>
</table>

Oligo fwd CACCGGGCGAGGTGTCCGCAGACG
Oligo rev AAACCGTCTGCGGACACCTCGCCC
Chromosome start end strand MM target_seq PAM alignment distance position gene name gene id
GU305914.1 33254 33276 + 0 GGGCGAGGTGTCCGCAGACG AGG ||||||||[||||||||||||]PAM NA Intergenic NA NA

T2066 GGCGAGGTGTCCGCAGACGAGGG 1000
Oligo fwd CACCGGCGAGGTGTCCGCAGACGA
Oligo rev AAACTCGTCTGCGGACACCTCGCC
Chromosome start end strand MM target_seq PAM alignment distance position gene name gene id
GU305914.1 33255 33277 + 0 GGCGAGGTGTCCGCAGACGA GGG ||||||||[||||||||||||]PAM NA Intergenic NA NA

T2067 GGTGTCCGCAGACGAGGGTCTGG 1000
Oligo fwd CACCGGTGTCCGCAGACGAGGGTC
Oligo rev AAACGACCCTCGTCTGCGGACACC
Chromosome start end strand MM target_seq PAM alignment distance position gene name gene id
GU305914.1 33260 33282 + 0 GGTGTCCGCAGACGAGGGTC TGG ||||||||[||||||||||||]PAM NA Intergenic NA NA

T2068 GGGTCTGGTGTTCGTCGCTGAGG 1000
Oligo fwd CACCGGGTCTGGTGTTCGTCGCTG
Oligo rev AAACCAGCGACGAACACCAGACCC
Chromosome start end strand MM target_seq PAM alignment distance position gene name gene id
GU305914.1 33275 33297 + 0 GGGTCTGGTGTTCGTCGCTG AGG ||||||||[||||||||||||]PAM NA Intergenic NA NA

T2069 TCTGGTGTTCGTCGCTGAGGTGG 1000
Oligo fwd CACCTCTGGTGTTCGTCGCTGAGG
Oligo rev AAACCCTCAGCGACGAACACCAGA
Chromosome start end strand MM target_seq PAM alignment distance position gene name gene id
GU305914.1 33278 33300 + 0 TCTGGTGTTCGTCGCTGAGG TGG ||||||||[||||||||||||]PAM NA Intergenic NA NA

T2070 GGTGTTCGTCGCTGAGGTGGCGG 1000
Oligo fwd CACCGGTGTTCGTCGCTGAGGTGG
Oligo rev AAACCCACCTCAGCGACGAACACC
Chromosome start end strand MM target_seq PAM alignment distance position gene name gene id
GU305914.1 33281 33303 + 0 GGTGTTCGTCGCTGAGGTGG CGG ||||||||[||||||||||||]PAM NA Intergenic NA NA

T2071 GTCGCTGAGGTGGCGGATCTTGG 1000
Oligo fwd CACCGTCGCTGAGGTGGCGGATCT
Oligo rev AAACAGATCCGCCACCTCAGCGAC
Chromosome start end strand MM target_seq PAM alignment distance position gene name gene id
GU305914.1 33288 33310 + 0 GTCGCTGAGGTGGCGGATCT TGG ||||||||[||||||||||||]PAM NA Intergenic NA NA

T2072 TCGCTGAGGTGGCGGATCTTGGG 1000
Oligo fwd CACCTCGCTGAGGTGGCGGATCTT
Oligo rev AAACAAGATCCGCCACCTCAGCGA
Chromosome start end strand MM target_seq PAM alignment distance position gene name gene id
GU305914.1 33289 33311 + 0 TCGCTGAGGTGGCGGATCTT GGG ||||||||[||||||||||||]PAM NA Intergenic NA NA

T2073 TGAGGTGGCGGATCTTGGGCAGG 1000
Oligo fwd CACCTGAGGTGGCGGATCTTGGGC
Oligo rev AAACGCCCAAGATCCGCCACCTCA
Chromosome start end strand MM target_seq PAM alignment distance position gene name gene id
GU305914.1 33293 33315 + 0 TGAGGTGGCGGATCTTGGGC AGG ||||||||[||||||||||||]PAM NA Intergenic NA NA

T2074 CAGGCGCAGCACGCCGATGTTGG 1000
Oligo fwd CACCCAGGCGCAGCACGCCGATGT
Oligo rev AAACACATCGGCGTGCTGCGCCTG
Chromosome start end strand MM target_seq PAM alignment distance position gene name gene id
GU305914.1 33312 33334 + 0 CAGGCGCAGCACGCCGATGT TGG ||||||||[||||||||||||]PAM NA Intergenic NA NA

T2075 AGGCGCAGCACGCCGATGTTGGG 1000
Oligo fwd CACCAGGCGCAGCACGCCGATGTT
Oligo rev AAACAACATCGGCGTGCTGCGCCT
Chromosome start end strand MM target_seq PAM alignment distance position gene name gene id
GU305914.1 33313 33335 + 0 AGGCGCAGCACGCCGATGTT GGG ||||||||[||||||||||||]PAM NA Intergenic NA NA

T2076 GGCGCAGCACGCCGATGTTGGGG 1000
Oligo fwd CACCGGCGCAGCACGCCGATGTTG
Oligo rev AAACCAACATCGGCGTGCTGCGCC
Chromosome start end strand MM target_seq PAM alignment distance position gene name gene id
GU305914.1 33314 33336 + 0 GGCGCAGCACGCCGATGTTG GGG ||||||||[||||||||||||]PAM NA Intergenic NA NA

T2077 CAGCACGCCGATGTTGGGGTAGG 1000
Oligo fwd CACCCAGCACGCCGATGTTGGGGT
Oligo rev AAACACCCCAACATCGGCGTGCTG
Chromosome start end strand MM target_seq PAM alignment distance position gene name gene id
GU305914.1 33318 33340 + 0 CAGCACGCCGATGTTGGGGT AGG ||||||||[||||||||||||]PAM NA Intergenic NA NA

T2078 CACGCCGATGTTGGGGTAGGCGG 1000
Oligo fwd CACCCACGCCGATGTTGGGGTAGG
Oligo rev AAACCCTACCCCAACATCGGCGTG
Chromosome start end strand MM target_seq PAM alignment distance position gene name gene id
GU305914.1 33321 33343 + 0 CACGCCGATGTTGGGGTAGG CGG ||||||||[||||||||||||]PAM NA Intergenic NA NA

T2079 GAACGTAATTCTCCTCGAAGAGG 1000
Oligo fwd CACCGAACGTAATTCTCCTCGAAG
Oligo rev AAACCTTCGAGGAGAATTACGTTC
Chromosome start end strand MM target_seq PAM alignment distance position gene name gene id
GU305914.1 33350 33372 + 0 GAACGTAATTCTCCTCGAAG AGG ||||||||[||||||||||||]PAM NA Intergenic NA NA

<table>
<tr><td>T2080</td><td>CGTAATTCTCCTCGAAGAGGCGG</td><td></td><td>1000</td><td></td><td></td><td></td><td></td><td></td><td></td><td></td><td></td><td></td></tr>
<tr><td>Oligo fwd</td><td>CACCCGTAATTCTCCTCGAAGAGG</td><td></td><td></td><td></td><td></td><td></td><td></td><td></td><td></td><td></td><td></td><td></td></tr>
<tr><td>Oligo rev</td><td>AAACCCTCTTCGAGGAGAATTACG</td><td></td><td></td><td></td><td></td><td></td><td></td><td></td><td></td><td></td><td></td><td></td></tr>
<tr><td>Chromosome</td><td>start</td><td></td><td>end</td><td>strand</td><td>MM</td><td>target_seq</td><td>PAM</td><td>alignment</td><td>distance</td><td>position</td><td>gene name</td><td>gene id</td></tr>
<tr><td>GU305914.1</td><td></td><td>33353</td><td>33375</td><td>+</td><td>0</td><td>CGTAATTCTCCTCGAAGAGG</td><td>CGG</td><td>||||||||[||||||||||||]PAM</td><td>NA</td><td>Intergenic</td><td>NA</td><td>NA</td></tr>
<tr><td>T2081</td><td>CTCGAAGAGGCGGATCCCGTCGG</td><td></td><td>1000</td><td></td><td></td><td></td><td></td><td></td><td></td><td></td><td></td><td></td></tr>
<tr><td>Oligo fwd</td><td>CACCCTCGAAGAGGCGGATCCCGT</td><td></td><td></td><td></td><td></td><td></td><td></td><td></td><td></td><td></td><td></td><td></td></tr>
<tr><td>Oligo rev</td><td>AAACACGGGATCCGCCTCTTCGAG</td><td></td><td></td><td></td><td></td><td></td><td></td><td></td><td></td><td></td><td></td><td></td></tr>
<tr><td>Chromosome</td><td>start</td><td></td><td>end</td><td>strand</td><td>MM</td><td>target_seq</td><td>PAM</td><td>alignment</td><td>distance</td><td>position</td><td>gene name</td><td>gene id</td></tr>
<tr><td>GU305914.1</td><td></td><td>33363</td><td>33385</td><td>+</td><td>0</td><td>CTCGAAGAGGCGGATCCCGT</td><td>CGG</td><td>||||||||[||||||||||||]PAM</td><td>NA</td><td>Intergenic</td><td>NA</td><td>NA</td></tr>
<tr><td>T2082</td><td>TCGAAGAGGCGGATCCCGTCGGG</td><td></td><td>1000</td><td></td><td></td><td></td><td></td><td></td><td></td><td></td><td></td><td></td></tr>
<tr><td>Oligo fwd</td><td>CACCTCGAAGAGGCGGATCCCGTC</td><td></td><td></td><td></td><td></td><td></td><td></td><td></td><td></td><td></td><td></td><td></td></tr>
<tr><td>Oligo rev</td><td>AAACGACGGGATCCGCCTCTTCGA</td><td></td><td></td><td></td><td></td><td></td><td></td><td></td><td></td><td></td><td></td><td></td></tr>
<tr><td>Chromosome</td><td>start</td><td></td><td>end</td><td>strand</td><td>MM</td><td>target_seq</td><td>PAM</td><td>alignment</td><td>distance</td><td>position</td><td>gene name</td><td>gene id</td></tr>
<tr><td>GU305914.1</td><td></td><td>33364</td><td>33386</td><td>+</td><td>0</td><td>TCGAAGAGGCGGATCCCGTC</td><td>GGG</td><td>||||||||[||||||||||||]PAM</td><td>NA</td><td>Intergenic</td><td>NA</td><td>NA</td></tr>
<tr><td>T2083</td><td>CGAAGAGGCGGATCCCGTCGGGG</td><td></td><td>1000</td><td></td><td></td><td></td><td></td><td></td><td></td><td></td><td></td><td></td></tr>
<tr><td>Oligo fwd</td><td>CACCCGAAGAGGCGGATCCCGTCG</td><td></td><td></td><td></td><td></td><td></td><td></td><td></td><td></td><td></td><td></td><td></td></tr>
<tr><td>Oligo rev</td><td>AAACCGACGGGATCCGCCTCTTCG</td><td></td><td></td><td></td><td></td><td></td><td></td><td></td><td></td><td></td><td></td><td></td></tr>
<tr><td>Chromosome</td><td>start</td><td></td><td>end</td><td>strand</td><td>MM</td><td>target_seq</td><td>PAM</td><td>alignment</td><td>distance</td><td>position</td><td>gene name</td><td>gene id</td></tr>
<tr><td>GU305914.1</td><td></td><td>33365</td><td>33387</td><td>+</td><td>0</td><td>CGAAGAGGCGGATCCCGTCG</td><td>GGG</td><td>||||||||[||||||||||||]PAM</td><td>NA</td><td>Intergenic</td><td>NA</td><td>NA</td></tr>
<tr><td>T2084</td><td>TCCCGTCGGGGTTCCGACGCCGG</td><td></td><td>1000</td><td></td><td></td><td></td><td></td><td></td><td></td><td></td><td></td><td></td></tr>
<tr><td>Oligo fwd</td><td>CACCTCCCGTCGGGGTTCCGACGC</td><td></td><td></td><td></td><td></td><td></td><td></td><td></td><td></td><td></td><td></td><td></td></tr>
<tr><td>Oligo rev</td><td>AAACGCGTCGGAACCCCGACGGGA</td><td></td><td></td><td></td><td></td><td></td><td></td><td></td><td></td><td></td><td></td><td></td></tr>
<tr><td>Chromosome</td><td>start</td><td></td><td>end</td><td>strand</td><td>MM</td><td>target_seq</td><td>PAM</td><td>alignment</td><td>distance</td><td>position</td><td>gene name</td><td>gene id</td></tr>
<tr><td>GU305914.1</td><td></td><td>33377</td><td>33399</td><td>+</td><td>0</td><td>TCCCGTCGGGGTTCCGACGC</td><td>CGG</td><td>||||||||[||||||||||||]PAM</td><td>NA</td><td>Intergenic</td><td>NA</td><td>NA</td></tr>
<tr><td>T2085</td><td>TCGTGCCGATCTTCTGCGCGCGG</td><td></td><td>1000</td><td></td><td></td><td></td><td></td><td></td><td></td><td></td><td></td><td></td></tr>
<tr><td>Oligo fwd</td><td>CACCTCGTGCCGATCTTCTGCGCG</td><td></td><td></td><td></td><td></td><td></td><td></td><td></td><td></td><td></td><td></td><td></td></tr>
<tr><td>Oligo rev</td><td>AAACCGCGCAGAAGATCGGCACGA</td><td></td><td></td><td></td><td></td><td></td><td></td><td></td><td></td><td></td><td></td><td></td></tr>
<tr><td>Chromosome</td><td>start</td><td></td><td>end</td><td>strand</td><td>MM</td><td>target_seq</td><td>PAM</td><td>alignment</td><td>distance</td><td>position</td><td>gene name</td><td>gene id</td></tr>
<tr><td>GU305914.1</td><td></td><td>33404</td><td>33426</td><td>+</td><td>0</td><td>TCGTGCCGATCTTCTGCGCG</td><td>CGG</td><td>||||||||[||||||||||||]PAM</td><td>NA</td><td>Intergenic</td><td>NA</td><td>NA</td></tr>
<tr><td>T2086</td><td>TCTGCGCGCGGAGCCTCTCGAGG</td><td></td><td>1000</td><td></td><td></td><td></td><td></td><td></td><td></td><td></td><td></td><td></td></tr>
<tr><td>Oligo fwd</td><td>CACCTCTGCGCGCGGAGCCTCTCG</td><td></td><td></td><td></td><td></td><td></td><td></td><td></td><td></td><td></td><td></td><td></td></tr>
<tr><td>Oligo rev</td><td>AAACCGAGAGGCTCCGCGCGCAGA</td><td></td><td></td><td></td><td></td><td></td><td></td><td></td><td></td><td></td><td></td><td></td></tr>
<tr><td>Chromosome</td><td>start</td><td></td><td>end</td><td>strand</td><td>MM</td><td>target_seq</td><td>PAM</td><td>alignment</td><td>distance</td><td>position</td><td>gene name</td><td>gene id</td></tr>
<tr><td>GU305914.1</td><td></td><td>33416</td><td>33438</td><td>+</td><td>0</td><td>TCTGCGCGCGGAGCCTCTCG</td><td>AGG</td><td>||||||||[||||||||||||]PAM</td><td>NA</td><td>Intergenic</td><td>NA</td><td>NA</td></tr>
<tr><td>T2087</td><td>GCCTCTCGAGGCGCTCGCCCCGG</td><td></td><td>1000</td><td></td><td></td><td></td><td></td><td></td><td></td><td></td><td></td><td></td></tr>
<tr><td>Oligo fwd</td><td>CACCGCCTCTCGAGGCGCTCGCCC</td><td></td><td></td><td></td><td></td><td></td><td></td><td></td><td></td><td></td><td></td><td></td></tr>
<tr><td>Oligo rev</td><td>AAACGGGCGAGCGCCTCGAGAGGC</td><td></td><td></td><td></td><td></td><td></td><td></td><td></td><td></td><td></td><td></td><td></td></tr>
<tr><td>Chromosome</td><td>start</td><td></td><td>end</td><td>strand</td><td>MM</td><td>target_seq</td><td>PAM</td><td>alignment</td><td>distance</td><td>position</td><td>gene name</td><td>gene id</td></tr>
<tr><td>GU305914.1</td><td></td><td>33428</td><td>33450</td><td>+</td><td>0</td><td>GCCTCTCGAGGCGCTCGCCC</td><td>CGG</td><td>||||||||[||||||||||||]PAM</td><td>NA</td><td>Intergenic</td><td>NA</td><td>NA</td></tr>
<tr><td>T2088</td><td>TTGATCGTAGATGACCTCCATGG</td><td></td><td>1000</td><td></td><td></td><td></td><td></td><td></td><td></td><td></td><td></td><td></td></tr>
<tr><td>Oligo fwd</td><td>CACCTTGATCGTAGATGACCTCCA</td><td></td><td></td><td></td><td></td><td></td><td></td><td></td><td></td><td></td><td></td><td></td></tr>
<tr><td>Oligo rev</td><td>AAACTGGAGGTCATCTACGATCAA</td><td></td><td></td><td></td><td></td><td></td><td></td><td></td><td></td><td></td><td></td><td></td></tr>
<tr><td>Chromosome</td><td>start</td><td></td><td>end</td><td>strand</td><td>MM</td><td>target_seq</td><td>PAM</td><td>alignment</td><td>distance</td><td>position</td><td>gene name</td><td>gene id</td></tr>
<tr><td>GU305914.1</td><td></td><td>33480</td><td>33502</td><td>+</td><td>0</td><td>TTGATCGTAGATGACCTCCA</td><td>TGG</td><td>||||||||[||||||||||||]PAM</td><td>NA</td><td>Intergenic</td><td>NA</td><td>NA</td></tr>
<tr><td>T2089</td><td>GTCGAGCAGACAGCTCTCGAAGG</td><td></td><td>1000</td><td></td><td></td><td></td><td></td><td></td><td></td><td></td><td></td><td></td></tr>
<tr><td>Oligo fwd</td><td>CACCGTCGAGCAGACAGCTCTCGA</td><td></td><td></td><td></td><td></td><td></td><td></td><td></td><td></td><td></td><td></td><td></td></tr>
<tr><td>Oligo rev</td><td>AAACTCGAGAGCTGTCTGCTCGAC</td><td></td><td></td><td></td><td></td><td></td><td></td><td></td><td></td><td></td><td></td><td></td></tr>
<tr><td>Chromosome</td><td>start</td><td></td><td>end</td><td>strand</td><td>MM</td><td>target_seq</td><td>PAM</td><td>alignment</td><td>distance</td><td>position</td><td>gene name</td><td>gene id</td></tr>
<tr><td>GU305914.1</td><td></td><td>33528</td><td>33550</td><td>+</td><td>0</td><td>GTCGAGCAGACAGCTCTCGA</td><td>AGG</td><td>||||||||[||||||||||||]PAM</td><td>NA</td><td>Intergenic</td><td>NA</td><td>NA</td></tr>
<tr><td>T2090</td><td>CAGACAGCTCTCGAAGGAGAAGG</td><td></td><td>1000</td><td></td><td></td><td></td><td></td><td></td><td></td><td></td><td></td><td></td></tr>
<tr><td>Oligo fwd</td><td>CACCCAGACAGCTCTCGAAGGAGA</td><td></td><td></td><td></td><td></td><td></td><td></td><td></td><td></td><td></td><td></td><td></td></tr>
<tr><td>Oligo rev</td><td>AAACTCTCCTTCGAGAGCTGTCTG</td><td></td><td></td><td></td><td></td><td></td><td></td><td></td><td></td><td></td><td></td><td></td></tr>
<tr><td>Chromosome</td><td>start</td><td></td><td>end</td><td>strand</td><td>MM</td><td>target_seq</td><td>PAM</td><td>alignment</td><td>distance</td><td>position</td><td>gene name</td><td>gene id</td></tr>
<tr><td>GU305914.1</td><td></td><td>33534</td><td>33556</td><td>+</td><td>0</td><td>CAGACAGCTCTCGAAGGAGA</td><td>AGG</td><td>||||||||[||||||||||||]PAM</td><td>NA</td><td>Intergenic</td><td>NA</td><td>NA</td></tr>
<tr><td>T2091</td><td>AGACAGCTCTCGAAGGAGAAGGG</td><td></td><td>1000</td><td></td><td></td><td></td><td></td><td></td><td></td><td></td><td></td><td></td></tr>
<tr><td>Oligo fwd</td><td>CACCAGACAGCTCTCGAAGGAGAA</td><td></td><td></td><td></td><td></td><td></td><td></td><td></td><td></td><td></td><td></td><td></td></tr>
<tr><td>Oligo rev</td><td>AAACTTCTCCTTCGAGAGCTGTCT</td><td></td><td></td><td></td><td></td><td></td><td></td><td></td><td></td><td></td><td></td><td></td></tr>
<tr><td>Chromosome</td><td>start</td><td></td><td>end</td><td>strand</td><td>MM</td><td>target_seq</td><td>PAM</td><td>alignment</td><td>distance</td><td>position</td><td>gene name</td><td>gene id</td></tr>
<tr><td>GU305914.1</td><td></td><td>33535</td><td>33557</td><td>+</td><td>0</td><td>AGACAGCTCTCGAAGGAGAA</td><td>GGG</td><td>||||||||[||||||||||||]PAM</td><td>NA</td><td>Intergenic</td><td>NA</td><td>NA</td></tr>
<tr><td>T2092</td><td>GACAGCTCTCGAAGGAGAAGGGG</td><td></td><td>1000</td><td></td><td></td><td></td><td></td><td></td><td></td><td></td><td></td><td></td></tr>
<tr><td>Oligo fwd</td><td>CACCGACAGCTCTCGAAGGAGAAG</td><td></td><td></td><td></td><td></td><td></td><td></td><td></td><td></td><td></td><td></td><td></td></tr>
<tr><td>Oligo rev</td><td>AAACCTTCTCCTTCGAGAGCTGTC</td><td></td><td></td><td></td><td></td><td></td><td></td><td></td><td></td><td></td><td></td><td></td></tr>
<tr><td>Chromosome</td><td>start</td><td></td><td>end</td><td>strand</td><td>MM</td><td>target_seq</td><td>PAM</td><td>alignment</td><td>distance</td><td>position</td><td>gene name</td><td>gene id</td></tr>
<tr><td>GU305914.1</td><td></td><td>33536</td><td>33558</td><td>+</td><td>0</td><td>GACAGCTCTCGAAGGAGAAG</td><td>GGG</td><td>||||||||[||||||||||||]PAM</td><td>NA</td><td>Intergenic</td><td>NA</td><td>NA</td></tr>
<tr><td>T2093</td><td>AGAAGGGGATACCGAGTTGCTGG</td><td></td><td>1000</td><td></td><td></td><td></td><td></td><td></td><td></td><td></td><td></td><td></td></tr>
<tr><td>Oligo fwd</td><td>CACCAGAAGGGGATACCGAGTTGC</td><td></td><td></td><td></td><td></td><td></td><td></td><td></td><td></td><td></td><td></td><td></td></tr>
<tr><td>Oligo rev</td><td>AAACGCAACTCGGTATCCCCTTCT</td><td></td><td></td><td></td><td></td><td></td><td></td><td></td><td></td><td></td><td></td><td></td></tr>
<tr><td>Chromosome</td><td>start</td><td></td><td>end</td><td>strand</td><td>MM</td><td>target_seq</td><td>PAM</td><td>alignment</td><td>distance</td><td>position</td><td>gene name</td><td>gene id</td></tr>
<tr><td>GU305914.1</td><td></td><td>33551</td><td>33573</td><td>+</td><td>0</td><td>AGAAGGGGATACCGAGTTGC</td><td>TGG</td><td>||||||||[||||||||||||]PAM</td><td>NA</td><td>Intergenic</td><td>NA</td><td>NA</td></tr>
<tr><td>T2094</td><td>GAGTTGCTGGCAGATGCGCTCGG</td><td></td><td>1000</td><td></td><td></td><td></td><td></td><td></td><td></td><td></td><td></td><td></td></tr>
<tr><td>Oligo fwd</td><td>CACCGAGTTGCTGGCAGATGCGCT</td><td></td><td></td><td></td><td></td><td></td><td></td><td></td><td></td><td></td><td></td><td></td></tr>
<tr><td>Oligo rev</td><td>AAACAGCGCATCTGCCAGCAACTC</td><td></td><td></td><td></td><td></td><td></td><td></td><td></td><td></td><td></td><td></td><td></td></tr>
<tr><td>Chromosome</td><td>start</td><td></td><td>end</td><td>strand</td><td>MM</td><td>target_seq</td><td>PAM</td><td>alignment</td><td>distance</td><td>position</td><td>gene name</td><td>gene id</td></tr>
<tr><td>GU305914.1</td><td></td><td>33564</td><td>33586</td><td>+</td><td>0</td><td>GAGTTGCTGGCAGATGCGCT</td><td>CGG</td><td>||||||||[||||||||||||]PAM</td><td>NA</td><td>Intergenic</td><td>NA</td><td>NA</td></tr>
</table>

<table>
<tr><td>T2095</td><td>AGTTGCTGGCAGATGCGCTCGGG</td><td>1000</td><td></td><td></td><td></td><td></td><td></td><td></td><td></td><td></td><td></td></tr>
<tr><td>Oligo fwd</td><td>CACCAGTTGCTGGCAGATGCGCTC</td><td></td><td></td><td></td><td></td><td></td><td></td><td></td><td></td><td></td><td></td></tr>
<tr><td>Oligo rev</td><td>AAACGAGCGCATCTGCCAGCAACT</td><td></td><td></td><td></td><td></td><td></td><td></td><td></td><td></td><td></td><td></td></tr>
<tr><td>Chromosome</td><td>start</td><td>end</td><td>strand</td><td>MM</td><td>target_seq</td><td>PAM</td><td>alignment</td><td>distance</td><td>position</td><td>gene name</td><td>gene id</td></tr>
<tr><td>GU305914.1</td><td>33565</td><td>33587</td><td>+</td><td>0</td><td>AGTTGCTGGCAGATGCGCTC</td><td>GGG</td><td>||||||||[||||||||||||]PAM</td><td>NA</td><td>Intergenic</td><td>NA</td><td>NA</td></tr>
<tr><td>T2096</td><td>GCTGGCAGATGCGCTCGGGCAGG</td><td>1000</td><td></td><td></td><td></td><td></td><td></td><td></td><td></td><td></td><td></td></tr>
<tr><td>Oligo fwd</td><td>CACCGCTGGCAGATGCGCTCGGGC</td><td></td><td></td><td></td><td></td><td></td><td></td><td></td><td></td><td></td><td></td></tr>
<tr><td>Oligo rev</td><td>AAACGCCCGAGCGCATCTGCCAGC</td><td></td><td></td><td></td><td></td><td></td><td></td><td></td><td></td><td></td><td></td></tr>
<tr><td>Chromosome</td><td>start</td><td>end</td><td>strand</td><td>MM</td><td>target_seq</td><td>PAM</td><td>alignment</td><td>distance</td><td>position</td><td>gene name</td><td>gene id</td></tr>
<tr><td>GU305914.1</td><td>33569</td><td>33591</td><td>+</td><td>0</td><td>GCTGGCAGATGCGCTCGGGC</td><td>AGG</td><td>||||||||[||||||||||||]PAM</td><td>NA</td><td>Intergenic</td><td>NA</td><td>NA</td></tr>
<tr><td>T2097</td><td>GGAGACGCTGCACGATCTCGAGG</td><td>1000</td><td></td><td></td><td></td><td></td><td></td><td></td><td></td><td></td><td></td></tr>
<tr><td>Oligo fwd</td><td>CACCGGAGACGCTGCACGATCTCG</td><td></td><td></td><td></td><td></td><td></td><td></td><td></td><td></td><td></td><td></td></tr>
<tr><td>Oligo rev</td><td>AAACCGAGATCGTGCAGCGTCTCC</td><td></td><td></td><td></td><td></td><td></td><td></td><td></td><td></td><td></td><td></td></tr>
<tr><td>Chromosome</td><td>start</td><td>end</td><td>strand</td><td>MM</td><td>target_seq</td><td>PAM</td><td>alignment</td><td>distance</td><td>position</td><td>gene name</td><td>gene id</td></tr>
<tr><td>GU305914.1</td><td>33590</td><td>33612</td><td>+</td><td>0</td><td>GGAGACGCTGCACGATCTCG</td><td>AGG</td><td>||||||||[||||||||||||]PAM</td><td>NA</td><td>Intergenic</td><td>NA</td><td>NA</td></tr>
<tr><td>T2098</td><td>ACGATCTCGAGGTCGTGTCCCGG</td><td>1000</td><td></td><td></td><td></td><td></td><td></td><td></td><td></td><td></td><td></td></tr>
<tr><td>Oligo fwd</td><td>CACCACGATCTCGAGGTCGTGTCC</td><td></td><td></td><td></td><td></td><td></td><td></td><td></td><td></td><td></td><td></td></tr>
<tr><td>Oligo rev</td><td>AAACGGACACGACCTCGAGATCGT</td><td></td><td></td><td></td><td></td><td></td><td></td><td></td><td></td><td></td><td></td></tr>
<tr><td>Chromosome</td><td>start</td><td>end</td><td>strand</td><td>MM</td><td>target_seq</td><td>PAM</td><td>alignment</td><td>distance</td><td>position</td><td>gene name</td><td>gene id</td></tr>
<tr><td>GU305914.1</td><td>33601</td><td>33623</td><td>+</td><td>0</td><td>ACGATCTCGAGGTCGTGTCC</td><td>CGG</td><td>||||||||[||||||||||||]PAM</td><td>NA</td><td>Intergenic</td><td>NA</td><td>NA</td></tr>
<tr><td>T2099</td><td>CGATCTCGAGGTCGTGTCCCGGG</td><td>1000</td><td></td><td></td><td></td><td></td><td></td><td></td><td></td><td></td><td></td></tr>
<tr><td>Oligo fwd</td><td>CACCCGATCTCGAGGTCGTGTCCC</td><td></td><td></td><td></td><td></td><td></td><td></td><td></td><td></td><td></td><td></td></tr>
<tr><td>Oligo rev</td><td>AAACGGGACACGACCTCGAGATCG</td><td></td><td></td><td></td><td></td><td></td><td></td><td></td><td></td><td></td><td></td></tr>
<tr><td>Chromosome</td><td>start</td><td>end</td><td>strand</td><td>MM</td><td>target_seq</td><td>PAM</td><td>alignment</td><td>distance</td><td>position</td><td>gene name</td><td>gene id</td></tr>
<tr><td>GU305914.1</td><td>33602</td><td>33624</td><td>+</td><td>0</td><td>CGATCTCGAGGTCGTGTCCC</td><td>GGG</td><td>||||||||[||||||||||||]PAM</td><td>NA</td><td>Intergenic</td><td>NA</td><td>NA</td></tr>
<tr><td>T2100</td><td>CGAGGTCGTGTCCCGGGATGCGG</td><td>1000</td><td></td><td></td><td></td><td></td><td></td><td></td><td></td><td></td><td></td></tr>
<tr><td>Oligo fwd</td><td>CACCCGAGGTCGTGTCCCGGGATG</td><td></td><td></td><td></td><td></td><td></td><td></td><td></td><td></td><td></td><td></td></tr>
<tr><td>Oligo rev</td><td>AAACCATCCCGGGACACGACCTCG</td><td></td><td></td><td></td><td></td><td></td><td></td><td></td><td></td><td></td><td></td></tr>
<tr><td>Chromosome</td><td>start</td><td>end</td><td>strand</td><td>MM</td><td>target_seq</td><td>PAM</td><td>alignment</td><td>distance</td><td>position</td><td>gene name</td><td>gene id</td></tr>
<tr><td>GU305914.1</td><td>33608</td><td>33630</td><td>+</td><td>0</td><td>CGAGGTCGTGTCCCGGGATG</td><td>CGG</td><td>||||||||[||||||||||||]PAM</td><td>NA</td><td>Intergenic</td><td>NA</td><td>NA</td></tr>
<tr><td>T2101</td><td>GGTCGTGTCCCGGGATGCGGAGG</td><td>1000</td><td></td><td></td><td></td><td></td><td></td><td></td><td></td><td></td><td></td></tr>
<tr><td>Oligo fwd</td><td>CACCGGTCGTGTCCCGGGATGCGG</td><td></td><td></td><td></td><td></td><td></td><td></td><td></td><td></td><td></td><td></td></tr>
<tr><td>Oligo rev</td><td>AAACCCGCATCCCGGGACACGACC</td><td></td><td></td><td></td><td></td><td></td><td></td><td></td><td></td><td></td><td></td></tr>
<tr><td>Chromosome</td><td>start</td><td>end</td><td>strand</td><td>MM</td><td>target_seq</td><td>PAM</td><td>alignment</td><td>distance</td><td>position</td><td>gene name</td><td>gene id</td></tr>
<tr><td>GU305914.1</td><td>33611</td><td>33633</td><td>+</td><td>0</td><td>GGTCGTGTCCCGGGATGCGG</td><td>AGG</td><td>||||||||[||||||||||||]PAM</td><td>NA</td><td>Intergenic</td><td>NA</td><td>NA</td></tr>
<tr><td>T2102</td><td>GTGTCCCGGGATGCGGAGGCTGG</td><td>1000</td><td></td><td></td><td></td><td></td><td></td><td></td><td></td><td></td><td></td></tr>
<tr><td>Oligo fwd</td><td>CACCGTGTCCCGGGATGCGGAGGC</td><td></td><td></td><td></td><td></td><td></td><td></td><td></td><td></td><td></td><td></td></tr>
<tr><td>Oligo rev</td><td>AAACGCCTCCGCATCCCGGGACAC</td><td></td><td></td><td></td><td></td><td></td><td></td><td></td><td></td><td></td><td></td></tr>
<tr><td>Chromosome</td><td>start</td><td>end</td><td>strand</td><td>MM</td><td>target_seq</td><td>PAM</td><td>alignment</td><td>distance</td><td>position</td><td>gene name</td><td>gene id</td></tr>
<tr><td>GU305914.1</td><td>33615</td><td>33637</td><td>+</td><td>0</td><td>GTGTCCCGGGATGCGGAGGC</td><td>TGG</td><td>||||||||[||||||||||||]PAM</td><td>NA</td><td>Intergenic</td><td>NA</td><td>NA</td></tr>
<tr><td>T2103</td><td>GGATGCGGAGGCTGGACATGAGG</td><td>1000</td><td></td><td></td><td></td><td></td><td></td><td></td><td></td><td></td><td></td></tr>
<tr><td>Oligo fwd</td><td>CACCGGATGCGGAGGCTGGACATG</td><td></td><td></td><td></td><td></td><td></td><td></td><td></td><td></td><td></td><td></td></tr>
<tr><td>Oligo rev</td><td>AAACCATGTCCAGCCTCCGCATCC</td><td></td><td></td><td></td><td></td><td></td><td></td><td></td><td></td><td></td><td></td></tr>
<tr><td>Chromosome</td><td>start</td><td>end</td><td>strand</td><td>MM</td><td>target_seq</td><td>PAM</td><td>alignment</td><td>distance</td><td>position</td><td>gene name</td><td>gene id</td></tr>
<tr><td>GU305914.1</td><td>33623</td><td>33645</td><td>+</td><td>0</td><td>GGATGCGGAGGCTGGACATG</td><td>AGG</td><td>||||||||[||||||||||||]PAM</td><td>NA</td><td>Intergenic</td><td>NA</td><td>NA</td></tr>
<tr><td>T2104</td><td>TGTTGCGCATCCTGTCGTGCAGG</td><td>1000</td><td></td><td></td><td></td><td></td><td></td><td></td><td></td><td></td><td></td></tr>
<tr><td>Oligo fwd</td><td>CACCTGTTGCGCATCCTGTCGTGC</td><td></td><td></td><td></td><td></td><td></td><td></td><td></td><td></td><td></td><td></td></tr>
<tr><td>Oligo rev</td><td>AAACGCACGACAGGATGCGCAACA</td><td></td><td></td><td></td><td></td><td></td><td></td><td></td><td></td><td></td><td></td></tr>
<tr><td>Chromosome</td><td>start</td><td>end</td><td>strand</td><td>MM</td><td>target_seq</td><td>PAM</td><td>alignment</td><td>distance</td><td>position</td><td>gene name</td><td>gene id</td></tr>
<tr><td>GU305914.1</td><td>33650</td><td>33672</td><td>+</td><td>0</td><td>TGTTGCGCATCCTGTCGTGC</td><td>AGG</td><td>||||||||[||||||||||||]PAM</td><td>NA</td><td>Intergenic</td><td>NA</td><td>NA</td></tr>
<tr><td>T2105</td><td>CGATGAGCTCGTTGACGAGCCGG</td><td>1000</td><td></td><td></td><td></td><td></td><td></td><td></td><td></td><td></td><td></td></tr>
<tr><td>Oligo fwd</td><td>CACCCGATGAGCTCGTTGACGAGC</td><td></td><td></td><td></td><td></td><td></td><td></td><td></td><td></td><td></td><td></td></tr>
<tr><td>Oligo rev</td><td>AAACGCTCGTCAACGAGCTCATCG</td><td></td><td></td><td></td><td></td><td></td><td></td><td></td><td></td><td></td><td></td></tr>
<tr><td>Chromosome</td><td>start</td><td>end</td><td>strand</td><td>MM</td><td>target_seq</td><td>PAM</td><td>alignment</td><td>distance</td><td>position</td><td>gene name</td><td>gene id</td></tr>
<tr><td>GU305914.1</td><td>33674</td><td>33696</td><td>+</td><td>0</td><td>CGATGAGCTCGTTGACGAGC</td><td>CGG</td><td>||||||||[||||||||||||]PAM</td><td>NA</td><td>Intergenic</td><td>NA</td><td>NA</td></tr>
<tr><td>T2106</td><td>TGAGCTCGTTGACGAGCCGGCGG</td><td>1000</td><td></td><td></td><td></td><td></td><td></td><td></td><td></td><td></td><td></td></tr>
<tr><td>Oligo fwd</td><td>CACCTGAGCTCGTTGACGAGCCGG</td><td></td><td></td><td></td><td></td><td></td><td></td><td></td><td></td><td></td><td></td></tr>
<tr><td>Oligo rev</td><td>AAACCCGGCTCGTCAACGAGCTCA</td><td></td><td></td><td></td><td></td><td></td><td></td><td></td><td></td><td></td><td></td></tr>
<tr><td>Chromosome</td><td>start</td><td>end</td><td>strand</td><td>MM</td><td>target_seq</td><td>PAM</td><td>alignment</td><td>distance</td><td>position</td><td>gene name</td><td>gene id</td></tr>
<tr><td>GU305914.1</td><td>33677</td><td>33699</td><td>+</td><td>0</td><td>TGAGCTCGTTGACGAGCCGG</td><td>CGG</td><td>||||||||[||||||||||||]PAM</td><td>NA</td><td>Intergenic</td><td>NA</td><td>NA</td></tr>
<tr><td>T2107</td><td>TGACGAGCCGGCGGAGATAGCGG</td><td>1000</td><td></td><td></td><td></td><td></td><td></td><td></td><td></td><td></td><td></td></tr>
<tr><td>Oligo fwd</td><td>CACCTGACGAGCCGGCGGAGATAG</td><td></td><td></td><td></td><td></td><td></td><td></td><td></td><td></td><td></td><td></td></tr>
<tr><td>Oligo rev</td><td>AAACCTATCTCCGCCGGCTCGTCA</td><td></td><td></td><td></td><td></td><td></td><td></td><td></td><td></td><td></td><td></td></tr>
<tr><td>Chromosome</td><td>start</td><td>end</td><td>strand</td><td>MM</td><td>target_seq</td><td>PAM</td><td>alignment</td><td>distance</td><td>position</td><td>gene name</td><td>gene id</td></tr>
<tr><td>GU305914.1</td><td>33686</td><td>33708</td><td>+</td><td>0</td><td>TGACGAGCCGGCGGAGATAG</td><td>CGG</td><td>||||||||[||||||||||||]PAM</td><td>NA</td><td>Intergenic</td><td>NA</td><td>NA</td></tr>
<tr><td>T2108</td><td>GACGAGCCGGCGGAGATAGCGGG</td><td>1000</td><td></td><td></td><td></td><td></td><td></td><td></td><td></td><td></td><td></td></tr>
<tr><td>Oligo fwd</td><td>CACCGACGAGCCGGCGGAGATAGC</td><td></td><td></td><td></td><td></td><td></td><td></td><td></td><td></td><td></td><td></td></tr>
<tr><td>Oligo rev</td><td>AAACGCTATCTCCGCCGGCTCGTC</td><td></td><td></td><td></td><td></td><td></td><td></td><td></td><td></td><td></td><td></td></tr>
<tr><td>Chromosome</td><td>start</td><td>end</td><td>strand</td><td>MM</td><td>target_seq</td><td>PAM</td><td>alignment</td><td>distance</td><td>position</td><td>gene name</td><td>gene id</td></tr>
<tr><td>GU305914.1</td><td>33687</td><td>33709</td><td>+</td><td>0</td><td>GACGAGCCGGCGGAGATAGC</td><td>GGG</td><td>||||||||[||||||||||||]PAM</td><td>NA</td><td>Intergenic</td><td>NA</td><td>NA</td></tr>
<tr><td>T2109</td><td>CCGGCGGAGATAGCGGGACATGG</td><td>1000</td><td></td><td></td><td></td><td></td><td></td><td></td><td></td><td></td><td></td></tr>
<tr><td>Oligo fwd</td><td>CACCCCGGCGGAGATAGCGGGACA</td><td></td><td></td><td></td><td></td><td></td><td></td><td></td><td></td><td></td><td></td></tr>
<tr><td>Oligo rev</td><td>AAACTGTCCCGCTATCTCCGCCGG</td><td></td><td></td><td></td><td></td><td></td><td></td><td></td><td></td><td></td><td></td></tr>
<tr><td>Chromosome</td><td>start</td><td>end</td><td>strand</td><td>MM</td><td>target_seq</td><td>PAM</td><td>alignment</td><td>distance</td><td>position</td><td>gene name</td><td>gene id</td></tr>
</table>

<table>
<tr><td>GU305914.1</td><td></td><td>33693</td><td>33715</td><td>+</td><td>0</td><td>CCGGCGGAGATAGCGGGACA</td><td>TGG</td><td>||||||||[||||||||||||]PAM</td><td>NA</td><td>Intergenic</td><td>NA</td><td>NA</td></tr>
<tr><td>T2110</td><td>GCGGAGATAGCGGGACATGGAGG</td><td></td><td>1000</td><td></td><td></td><td></td><td></td><td></td><td></td><td></td><td></td><td></td></tr>
<tr><td>Oligo fwd</td><td>CACCGCGGAGATAGCGGGACATGG</td><td></td><td></td><td></td><td></td><td></td><td></td><td></td><td></td><td></td><td></td><td></td></tr>
<tr><td>Oligo rev</td><td>AAACCCATGTCCCGCTATCTCCGC</td><td></td><td></td><td></td><td></td><td></td><td></td><td></td><td></td><td></td><td></td><td></td></tr>
<tr><td>Chromosome</td><td>start</td><td></td><td>end</td><td>strand</td><td>MM</td><td>target_seq</td><td>PAM</td><td>alignment</td><td>distance</td><td>position</td><td>gene name</td><td>gene id</td></tr>
<tr><td>GU305914.1</td><td></td><td>33696</td><td>33718</td><td>+</td><td>0</td><td>GCGGAGATAGCGGGACATGG</td><td>AGG</td><td>||||||||[||||||||||||]PAM</td><td>NA</td><td>Intergenic</td><td>NA</td><td>NA</td></tr>
<tr><td>T2111</td><td>GTAGCCGTACAGCAGCGAGATGG</td><td></td><td>1000</td><td></td><td></td><td></td><td></td><td></td><td></td><td></td><td></td><td></td></tr>
<tr><td>Oligo fwd</td><td>CACCGTAGCCGTACAGCAGCGAGA</td><td></td><td></td><td></td><td></td><td></td><td></td><td></td><td></td><td></td><td></td><td></td></tr>
<tr><td>Oligo rev</td><td>AAACTCTCGCTGCTGTACGGCTAC</td><td></td><td></td><td></td><td></td><td></td><td></td><td></td><td></td><td></td><td></td><td></td></tr>
<tr><td>Chromosome</td><td>start</td><td></td><td>end</td><td>strand</td><td>MM</td><td>target_seq</td><td>PAM</td><td>alignment</td><td>distance</td><td>position</td><td>gene name</td><td>gene id</td></tr>
<tr><td>GU305914.1</td><td></td><td>33729</td><td>33751</td><td>+</td><td>0</td><td>GTAGCCGTACAGCAGCGAGA</td><td>TGG</td><td>||||||||[||||||||||||]PAM</td><td>NA</td><td>Intergenic</td><td>NA</td><td>NA</td></tr>
<tr><td>GU305914.1</td><td></td><td>10696</td><td>10718</td><td>+</td><td>4</td><td>GATCCCGTACAGCACCGAGA</td><td>TGG</td><td>|---||||[||||||-|||||]PAM</td><td>NA</td><td>Intergenic</td><td>NA</td><td>NA</td></tr>
<tr><td>T2112</td><td>CAGCGAGATGGCGTACTGTATGG</td><td></td><td>1000</td><td></td><td></td><td></td><td></td><td></td><td></td><td></td><td></td><td></td></tr>
<tr><td>Oligo fwd</td><td>CACCCAGCGAGATGGCGTACTGTA</td><td></td><td></td><td></td><td></td><td></td><td></td><td></td><td></td><td></td><td></td><td></td></tr>
<tr><td>Oligo rev</td><td>AAACTACAGTACGCCATCTCGCTG</td><td></td><td></td><td></td><td></td><td></td><td></td><td></td><td></td><td></td><td></td><td></td></tr>
<tr><td>Chromosome</td><td>start</td><td></td><td>end</td><td>strand</td><td>MM</td><td>target_seq</td><td>PAM</td><td>alignment</td><td>distance</td><td>position</td><td>gene name</td><td>gene id</td></tr>
<tr><td>GU305914.1</td><td></td><td>33741</td><td>33763</td><td>+</td><td>0</td><td>CAGCGAGATGGCGTACTGTA</td><td>TGG</td><td>||||||||[||||||||||||]PAM</td><td>NA</td><td>Intergenic</td><td>NA</td><td>NA</td></tr>
<tr><td>T2113</td><td>GATGGCGTACTGTATGGCGACGG</td><td></td><td>1000</td><td></td><td></td><td></td><td></td><td></td><td></td><td></td><td></td><td></td></tr>
<tr><td>Oligo fwd</td><td>CACCGATGGCGTACTGTATGGCGA</td><td></td><td></td><td></td><td></td><td></td><td></td><td></td><td></td><td></td><td></td><td></td></tr>
<tr><td>Oligo rev</td><td>AAACTCGCCATACAGTACGCCATC</td><td></td><td></td><td></td><td></td><td></td><td></td><td></td><td></td><td></td><td></td><td></td></tr>
<tr><td>Chromosome</td><td>start</td><td></td><td>end</td><td>strand</td><td>MM</td><td>target_seq</td><td>PAM</td><td>alignment</td><td>distance</td><td>position</td><td>gene name</td><td>gene id</td></tr>
<tr><td>GU305914.1</td><td></td><td>33747</td><td>33769</td><td>+</td><td>0</td><td>GATGGCGTACTGTATGGCGA</td><td>CGG</td><td>||||||||[||||||||||||]PAM</td><td>NA</td><td>Intergenic</td><td>NA</td><td>NA</td></tr>
<tr><td>T2114</td><td>GGCGTACTGTATGGCGACGGTGG</td><td></td><td>1000</td><td></td><td></td><td></td><td></td><td></td><td></td><td></td><td></td><td></td></tr>
<tr><td>Oligo fwd</td><td>CACCGGCGTACTGTATGGCGACGG</td><td></td><td></td><td></td><td></td><td></td><td></td><td></td><td></td><td></td><td></td><td></td></tr>
<tr><td>Oligo rev</td><td>AAACCCGTCGCCATACAGTACGCC</td><td></td><td></td><td></td><td></td><td></td><td></td><td></td><td></td><td></td><td></td><td></td></tr>
<tr><td>Chromosome</td><td>start</td><td></td><td>end</td><td>strand</td><td>MM</td><td>target_seq</td><td>PAM</td><td>alignment</td><td>distance</td><td>position</td><td>gene name</td><td>gene id</td></tr>
<tr><td>GU305914.1</td><td></td><td>33750</td><td>33772</td><td>+</td><td>0</td><td>GGCGTACTGTATGGCGACGG</td><td>TGG</td><td>||||||||[||||||||||||]PAM</td><td>NA</td><td>Intergenic</td><td>NA</td><td>NA</td></tr>
<tr><td>T2115</td><td>ACTGTATGGCGACGGTGGCTCGG</td><td></td><td>1000</td><td></td><td></td><td></td><td></td><td></td><td></td><td></td><td></td><td></td></tr>
<tr><td>Oligo fwd</td><td>CACCACTGTATGGCGACGGTGGCT</td><td></td><td></td><td></td><td></td><td></td><td></td><td></td><td></td><td></td><td></td><td></td></tr>
<tr><td>Oligo rev</td><td>AAACAGCCACCGTCGCCATACAGT</td><td></td><td></td><td></td><td></td><td></td><td></td><td></td><td></td><td></td><td></td><td></td></tr>
<tr><td>Chromosome</td><td>start</td><td></td><td>end</td><td>strand</td><td>MM</td><td>target_seq</td><td>PAM</td><td>alignment</td><td>distance</td><td>position</td><td>gene name</td><td>gene id</td></tr>
<tr><td>GU305914.1</td><td></td><td>33755</td><td>33777</td><td>+</td><td>0</td><td>ACTGTATGGCGACGGTGGCT</td><td>CGG</td><td>||||||||[||||||||||||]PAM</td><td>NA</td><td>Intergenic</td><td>NA</td><td>NA</td></tr>
<tr><td>T2116</td><td>GGCTCGGCCCAGCGCCTTGCAGG</td><td></td><td>1000</td><td></td><td></td><td></td><td></td><td></td><td></td><td></td><td></td><td></td></tr>
<tr><td>Oligo fwd</td><td>CACCGGCTCGGCCCAGCGCCTTGC</td><td></td><td></td><td></td><td></td><td></td><td></td><td></td><td></td><td></td><td></td><td></td></tr>
<tr><td>Oligo rev</td><td>AAACGCAAGGCGCTGGGCCGAGCC</td><td></td><td></td><td></td><td></td><td></td><td></td><td></td><td></td><td></td><td></td><td></td></tr>
<tr><td>Chromosome</td><td>start</td><td></td><td>end</td><td>strand</td><td>MM</td><td>target_seq</td><td>PAM</td><td>alignment</td><td>distance</td><td>position</td><td>gene name</td><td>gene id</td></tr>
<tr><td>GU305914.1</td><td></td><td>33771</td><td>33793</td><td>+</td><td>0</td><td>GGCTCGGCCCAGCGCCTTGC</td><td>AGG</td><td>||||||||[||||||||||||]PAM</td><td>NA</td><td>Intergenic</td><td>NA</td><td>NA</td></tr>
<tr><td>T2117</td><td>GCCCAGCGCCTTGCAGGAGTCGG</td><td></td><td>1000</td><td></td><td></td><td></td><td></td><td></td><td></td><td></td><td></td><td></td></tr>
<tr><td>Oligo fwd</td><td>CACCGCCCAGCGCCTTGCAGGAGT</td><td></td><td></td><td></td><td></td><td></td><td></td><td></td><td></td><td></td><td></td><td></td></tr>
<tr><td>Oligo rev</td><td>AAACACTCCTGCAAGGCGCTGGGC</td><td></td><td></td><td></td><td></td><td></td><td></td><td></td><td></td><td></td><td></td><td></td></tr>
<tr><td>Chromosome</td><td>start</td><td></td><td>end</td><td>strand</td><td>MM</td><td>target_seq</td><td>PAM</td><td>alignment</td><td>distance</td><td>position</td><td>gene name</td><td>gene id</td></tr>
<tr><td>GU305914.1</td><td></td><td>33777</td><td>33799</td><td>+</td><td>0</td><td>GCCCAGCGCCTTGCAGGAGT</td><td>CGG</td><td>||||||||[||||||||||||]PAM</td><td>NA</td><td>Intergenic</td><td>NA</td><td>NA</td></tr>
<tr><td>T2118</td><td>CAGCGCCTTGCAGGAGTCGGAGG</td><td></td><td>1000</td><td></td><td></td><td></td><td></td><td></td><td></td><td></td><td></td><td></td></tr>
<tr><td>Oligo fwd</td><td>CACCCAGCGCCTTGCAGGAGTCGG</td><td></td><td></td><td></td><td></td><td></td><td></td><td></td><td></td><td></td><td></td><td></td></tr>
<tr><td>Oligo rev</td><td>AAACCCGACTCCTGCAAGGCGCTG</td><td></td><td></td><td></td><td></td><td></td><td></td><td></td><td></td><td></td><td></td><td></td></tr>
<tr><td>Chromosome</td><td>start</td><td></td><td>end</td><td>strand</td><td>MM</td><td>target_seq</td><td>PAM</td><td>alignment</td><td>distance</td><td>position</td><td>gene name</td><td>gene id</td></tr>
<tr><td>GU305914.1</td><td></td><td>33780</td><td>33802</td><td>+</td><td>0</td><td>CAGCGCCTTGCAGGAGTCGG</td><td>AGG</td><td>||||||||[||||||||||||]PAM</td><td>NA</td><td>Intergenic</td><td>NA</td><td>NA</td></tr>
<tr><td>T2119</td><td>CTTGCAGGAGTCGGAGGCCTCGG</td><td></td><td>1000</td><td></td><td></td><td></td><td></td><td></td><td></td><td></td><td></td><td></td></tr>
<tr><td>Oligo fwd</td><td>CACCCTTGCAGGAGTCGGAGGCCT</td><td></td><td></td><td></td><td></td><td></td><td></td><td></td><td></td><td></td><td></td><td></td></tr>
<tr><td>Oligo rev</td><td>AAACAGGCCTCCGACTCCTGCAAG</td><td></td><td></td><td></td><td></td><td></td><td></td><td></td><td></td><td></td><td></td><td></td></tr>
<tr><td>Chromosome</td><td>start</td><td></td><td>end</td><td>strand</td><td>MM</td><td>target_seq</td><td>PAM</td><td>alignment</td><td>distance</td><td>position</td><td>gene name</td><td>gene id</td></tr>
<tr><td>GU305914.1</td><td></td><td>33786</td><td>33808</td><td>+</td><td>0</td><td>CTTGCAGGAGTCGGAGGCCT</td><td>CGG</td><td>||||||||[||||||||||||]PAM</td><td>NA</td><td>Intergenic</td><td>NA</td><td>NA</td></tr>
<tr><td>T2120</td><td>TTGCAGGAGTCGGAGGCCTCGGG</td><td></td><td>1000</td><td></td><td></td><td></td><td></td><td></td><td></td><td></td><td></td><td></td></tr>
<tr><td>Oligo fwd</td><td>CACCTTGCAGGAGTCGGAGGCCTC</td><td></td><td></td><td></td><td></td><td></td><td></td><td></td><td></td><td></td><td></td><td></td></tr>
<tr><td>Oligo rev</td><td>AAACGAGGCCTCCGACTCCTGCAA</td><td></td><td></td><td></td><td></td><td></td><td></td><td></td><td></td><td></td><td></td><td></td></tr>
<tr><td>Chromosome</td><td>start</td><td></td><td>end</td><td>strand</td><td>MM</td><td>target_seq</td><td>PAM</td><td>alignment</td><td>distance</td><td>position</td><td>gene name</td><td>gene id</td></tr>
<tr><td>GU305914.1</td><td></td><td>33787</td><td>33809</td><td>+</td><td>0</td><td>TTGCAGGAGTCGGAGGCCTC</td><td>GGG</td><td>||||||||[||||||||||||]PAM</td><td>NA</td><td>Intergenic</td><td>NA</td><td>NA</td></tr>
<tr><td>T2121</td><td>GCCTCCATTCGAGATCGAACAGG</td><td></td><td>1000</td><td></td><td></td><td></td><td></td><td></td><td></td><td></td><td></td><td></td></tr>
<tr><td>Oligo fwd</td><td>CACCGCCTCCATTCGAGATCGAAC</td><td></td><td></td><td></td><td></td><td></td><td></td><td></td><td></td><td></td><td></td><td></td></tr>
<tr><td>Oligo rev</td><td>AAACGTTCGATCTCGAATGGAGGC</td><td></td><td></td><td></td><td></td><td></td><td></td><td></td><td></td><td></td><td></td><td></td></tr>
<tr><td>Chromosome</td><td>start</td><td></td><td>end</td><td>strand</td><td>MM</td><td>target_seq</td><td>PAM</td><td>alignment</td><td>distance</td><td>position</td><td>gene name</td><td>gene id</td></tr>
<tr><td>GU305914.1</td><td></td><td>33809</td><td>33831</td><td>+</td><td>0</td><td>GCCTCCATTCGAGATCGAAC</td><td>AGG</td><td>||||||||[||||||||||||]PAM</td><td>NA</td><td>Intergenic</td><td>NA</td><td>NA</td></tr>
<tr><td>T2122</td><td>TTCGAGATCGAACAGGATCTTGG</td><td></td><td>1000</td><td></td><td></td><td></td><td></td><td></td><td></td><td></td><td></td><td></td></tr>
<tr><td>Oligo fwd</td><td>CACCTTCGAGATCGAACAGGATCT</td><td></td><td></td><td></td><td></td><td></td><td></td><td></td><td></td><td></td><td></td><td></td></tr>
<tr><td>Oligo rev</td><td>AAACAGATCCTGTTCGATCTCGAA</td><td></td><td></td><td></td><td></td><td></td><td></td><td></td><td></td><td></td><td></td><td></td></tr>
<tr><td>Chromosome</td><td>start</td><td></td><td>end</td><td>strand</td><td>MM</td><td>target_seq</td><td>PAM</td><td>alignment</td><td>distance</td><td>position</td><td>gene name</td><td>gene id</td></tr>
<tr><td>GU305914.1</td><td></td><td>33816</td><td>33838</td><td>+</td><td>0</td><td>TTCGAGATCGAACAGGATCT</td><td>TGG</td><td>||||||||[||||||||||||]PAM</td><td>NA</td><td>Intergenic</td><td>NA</td><td>NA</td></tr>
<tr><td>T2123</td><td>CTTGGCGATCCCGTCGCACGAGG</td><td></td><td>1000</td><td></td><td></td><td></td><td></td><td></td><td></td><td></td><td></td><td></td></tr>
<tr><td>Oligo fwd</td><td>CACCCTTGGCGATCCCGTCGCACG</td><td></td><td></td><td></td><td></td><td></td><td></td><td></td><td></td><td></td><td></td><td></td></tr>
<tr><td>Oligo rev</td><td>AAACCGTGCGACGGGATCGCCAAG</td><td></td><td></td><td></td><td></td><td></td><td></td><td></td><td></td><td></td><td></td><td></td></tr>
<tr><td>Chromosome</td><td>start</td><td></td><td>end</td><td>strand</td><td>MM</td><td>target_seq</td><td>PAM</td><td>alignment</td><td>distance</td><td>position</td><td>gene name</td><td>gene id</td></tr>
<tr><td>GU305914.1</td><td></td><td>33834</td><td>33856</td><td>+</td><td>0</td><td>CTTGGCGATCCCGTCGCACG</td><td>AGG</td><td>||||||||[||||||||||||]PAM</td><td>NA</td><td>Intergenic</td><td>NA</td><td>NA</td></tr>
<tr><td>T2124</td><td>TTGGCGATCCCGTCGCACGAGGG</td><td></td><td>1000</td><td></td><td></td><td></td><td></td><td></td><td></td><td></td><td></td><td></td></tr>
<tr><td>Oligo fwd</td><td>CACCTTGGCGATCCCGTCGCACGA</td><td></td><td></td><td></td><td></td><td></td><td></td><td></td><td></td><td></td><td></td><td></td></tr>
</table>

<table>
<tr><td>Oligo rev</td><td>AAACTCGTGCGACGGGATCGCCAA</td><td></td><td></td><td></td><td></td><td></td><td></td><td></td><td></td><td></td><td></td><td></td></tr>
<tr><td>Chromosome</td><td>start</td><td></td><td>end</td><td>strand</td><td>MM</td><td>target_seq</td><td>PAM</td><td>alignment</td><td>distance</td><td>position</td><td>gene name</td><td>gene id</td></tr>
<tr><td>GU305914.1</td><td></td><td>33835</td><td>33857</td><td>+</td><td>0</td><td>TTGGCGATCCCGTCGCACGA</td><td>GGG</td><td>||||||||[||||||||||||]PAM</td><td>NA</td><td>Intergenic</td><td>NA</td><td>NA</td></tr>
<tr><td>T2125</td><td>TGGCGATCCCGTCGCACGAGGGG</td><td></td><td>1000</td><td></td><td></td><td></td><td></td><td></td><td></td><td></td><td></td><td></td></tr>
<tr><td>Oligo fwd</td><td>CACCTGGCGATCCCGTCGCACGAG</td><td></td><td></td><td></td><td></td><td></td><td></td><td></td><td></td><td></td><td></td><td></td></tr>
<tr><td>Oligo rev</td><td>AAACCTCGTGCGACGGGATCGCCA</td><td></td><td></td><td></td><td></td><td></td><td></td><td></td><td></td><td></td><td></td><td></td></tr>
<tr><td>Chromosome</td><td>start</td><td></td><td>end</td><td>strand</td><td>MM</td><td>target_seq</td><td>PAM</td><td>alignment</td><td>distance</td><td>position</td><td>gene name</td><td>gene id</td></tr>
<tr><td>GU305914.1</td><td></td><td>33836</td><td>33858</td><td>+</td><td>0</td><td>TGGCGATCCCGTCGCACGAG</td><td>GGG</td><td>||||||||[||||||||||||]PAM</td><td>NA</td><td>Intergenic</td><td>NA</td><td>NA</td></tr>
<tr><td>T2126</td><td>GATCCCGTCGCACGAGGGGAAGG</td><td></td><td>1000</td><td></td><td></td><td></td><td></td><td></td><td></td><td></td><td></td><td></td></tr>
<tr><td>Oligo fwd</td><td>CACCGATCCCGTCGCACGAGGGGA</td><td></td><td></td><td></td><td></td><td></td><td></td><td></td><td></td><td></td><td></td><td></td></tr>
<tr><td>Oligo rev</td><td>AAACTCCCCTCGTGCGACGGGATC</td><td></td><td></td><td></td><td></td><td></td><td></td><td></td><td></td><td></td><td></td><td></td></tr>
<tr><td>Chromosome</td><td>start</td><td></td><td>end</td><td>strand</td><td>MM</td><td>target_seq</td><td>PAM</td><td>alignment</td><td>distance</td><td>position</td><td>gene name</td><td>gene id</td></tr>
<tr><td>GU305914.1</td><td></td><td>33840</td><td>33862</td><td>+</td><td>0</td><td>GATCCCGTCGCACGAGGGGA</td><td>AGG</td><td>||||||||[||||||||||||]PAM</td><td>NA</td><td>Intergenic</td><td>NA</td><td>NA</td></tr>
<tr><td>T2127</td><td>GTCGCACGAGGGGAAGGAGAAGG</td><td></td><td>1000</td><td></td><td></td><td></td><td></td><td></td><td></td><td></td><td></td><td></td></tr>
<tr><td>Oligo fwd</td><td>CACCGTCGCACGAGGGGAAGGAGA</td><td></td><td></td><td></td><td></td><td></td><td></td><td></td><td></td><td></td><td></td><td></td></tr>
<tr><td>Oligo rev</td><td>AAACTCTCCTTCCCCTCGTGCGAC</td><td></td><td></td><td></td><td></td><td></td><td></td><td></td><td></td><td></td><td></td><td></td></tr>
<tr><td>Chromosome</td><td>start</td><td></td><td>end</td><td>strand</td><td>MM</td><td>target_seq</td><td>PAM</td><td>alignment</td><td>distance</td><td>position</td><td>gene name</td><td>gene id</td></tr>
<tr><td>GU305914.1</td><td></td><td>33846</td><td>33868</td><td>+</td><td>0</td><td>GTCGCACGAGGGGAAGGAGA</td><td>AGG</td><td>||||||||[||||||||||||]PAM</td><td>NA</td><td>Intergenic</td><td>NA</td><td>NA</td></tr>
<tr><td>T2128</td><td>TCGCACGAGGGGAAGGAGAAGGG</td><td></td><td>1000</td><td></td><td></td><td></td><td></td><td></td><td></td><td></td><td></td><td></td></tr>
<tr><td>Oligo fwd</td><td>CACCTCGCACGAGGGGAAGGAGAA</td><td></td><td></td><td></td><td></td><td></td><td></td><td></td><td></td><td></td><td></td><td></td></tr>
<tr><td>Oligo rev</td><td>AAACTTCTCCTTCCCCTCGTGCGA</td><td></td><td></td><td></td><td></td><td></td><td></td><td></td><td></td><td></td><td></td><td></td></tr>
<tr><td>Chromosome</td><td>start</td><td></td><td>end</td><td>strand</td><td>MM</td><td>target_seq</td><td>PAM</td><td>alignment</td><td>distance</td><td>position</td><td>gene name</td><td>gene id</td></tr>
<tr><td>GU305914.1</td><td></td><td>33847</td><td>33869</td><td>+</td><td>0</td><td>TCGCACGAGGGGAAGGAGAA</td><td>GGG</td><td>||||||||[||||||||||||]PAM</td><td>NA</td><td>Intergenic</td><td>NA</td><td>NA</td></tr>
<tr><td>T2129</td><td>CGCACGAGGGGAAGGAGAAGGGG</td><td></td><td>1000</td><td></td><td></td><td></td><td></td><td></td><td></td><td></td><td></td><td></td></tr>
<tr><td>Oligo fwd</td><td>CACCCGCACGAGGGGAAGGAGAAG</td><td></td><td></td><td></td><td></td><td></td><td></td><td></td><td></td><td></td><td></td><td></td></tr>
<tr><td>Oligo rev</td><td>AAACCTTCTCCTTCCCCTCGTGCG</td><td></td><td></td><td></td><td></td><td></td><td></td><td></td><td></td><td></td><td></td><td></td></tr>
<tr><td>Chromosome</td><td>start</td><td></td><td>end</td><td>strand</td><td>MM</td><td>target_seq</td><td>PAM</td><td>alignment</td><td>distance</td><td>position</td><td>gene name</td><td>gene id</td></tr>
<tr><td>GU305914.1</td><td></td><td>33848</td><td>33870</td><td>+</td><td>0</td><td>CGCACGAGGGGAAGGAGAAG</td><td>GGG</td><td>||||||||[||||||||||||]PAM</td><td>NA</td><td>Intergenic</td><td>NA</td><td>NA</td></tr>
<tr><td>T2130</td><td>CGAGGGGAAGGAGAAGGGGTTGG</td><td></td><td>1000</td><td></td><td></td><td></td><td></td><td></td><td></td><td></td><td></td><td></td></tr>
<tr><td>Oligo fwd</td><td>CACCCGAGGGGAAGGAGAAGGGGT</td><td></td><td></td><td></td><td></td><td></td><td></td><td></td><td></td><td></td><td></td><td></td></tr>
<tr><td>Oligo rev</td><td>AAACACCCCTTCTCCTTCCCCTCG</td><td></td><td></td><td></td><td></td><td></td><td></td><td></td><td></td><td></td><td></td><td></td></tr>
<tr><td>Chromosome</td><td>start</td><td></td><td>end</td><td>strand</td><td>MM</td><td>target_seq</td><td>PAM</td><td>alignment</td><td>distance</td><td>position</td><td>gene name</td><td>gene id</td></tr>
<tr><td>GU305914.1</td><td></td><td>33852</td><td>33874</td><td>+</td><td>0</td><td>CGAGGGGAAGGAGAAGGGGT</td><td>TGG</td><td>||||||||[||||||||||||]PAM</td><td>NA</td><td>Intergenic</td><td>NA</td><td>NA</td></tr>
<tr><td>GU305914.1</td><td></td><td>164995</td><td>165017</td><td>+</td><td>3</td><td>CGCGGGGAAGGAGCAGGAGT</td><td>AGG</td><td>||-|||||[|||||-|||-||]PAM</td><td>NA</td><td>Intergenic</td><td>NA</td><td>NA</td></tr>
<tr><td>T2131</td><td>GAAGGAGAAGGGGTTGGCGAAGG</td><td></td><td>1000</td><td></td><td></td><td></td><td></td><td></td><td></td><td></td><td></td><td></td></tr>
<tr><td>Oligo fwd</td><td>CACCGAAGGAGAAGGGGTTGGCGA</td><td></td><td></td><td></td><td></td><td></td><td></td><td></td><td></td><td></td><td></td><td></td></tr>
<tr><td>Oligo rev</td><td>AAACTCGCCAACCCCTTCTCCTTC</td><td></td><td></td><td></td><td></td><td></td><td></td><td></td><td></td><td></td><td></td><td></td></tr>
<tr><td>Chromosome</td><td>start</td><td></td><td>end</td><td>strand</td><td>MM</td><td>target_seq</td><td>PAM</td><td>alignment</td><td>distance</td><td>position</td><td>gene name</td><td>gene id</td></tr>
<tr><td>GU305914.1</td><td></td><td>33858</td><td>33880</td><td>+</td><td>0</td><td>GAAGGAGAAGGGGTTGGCGA</td><td>AGG</td><td>||||||||[||||||||||||]PAM</td><td>NA</td><td>Intergenic</td><td>NA</td><td>NA</td></tr>
<tr><td>T2132</td><td>AAGGAGAAGGGGTTGGCGAAGGG</td><td></td><td>1000</td><td></td><td></td><td></td><td></td><td></td><td></td><td></td><td></td><td></td></tr>
<tr><td>Oligo fwd</td><td>CACCAAGGAGAAGGGGTTGGCGAA</td><td></td><td></td><td></td><td></td><td></td><td></td><td></td><td></td><td></td><td></td><td></td></tr>
<tr><td>Oligo rev</td><td>AAACTTCGCCAACCCCTTCTCCTT</td><td></td><td></td><td></td><td></td><td></td><td></td><td></td><td></td><td></td><td></td><td></td></tr>
<tr><td>Chromosome</td><td>start</td><td></td><td>end</td><td>strand</td><td>MM</td><td>target_seq</td><td>PAM</td><td>alignment</td><td>distance</td><td>position</td><td>gene name</td><td>gene id</td></tr>
<tr><td>GU305914.1</td><td></td><td>33859</td><td>33881</td><td>+</td><td>0</td><td>AAGGAGAAGGGGTTGGCGAA</td><td>GGG</td><td>||||||||[||||||||||||]PAM</td><td>NA</td><td>Intergenic</td><td>NA</td><td>NA</td></tr>
<tr><td>T2133</td><td>AGGAGAAGGGGTTGGCGAAGGGG</td><td></td><td>1000</td><td></td><td></td><td></td><td></td><td></td><td></td><td></td><td></td><td></td></tr>
<tr><td>Oligo fwd</td><td>CACCAGGAGAAGGGGTTGGCGAAG</td><td></td><td></td><td></td><td></td><td></td><td></td><td></td><td></td><td></td><td></td><td></td></tr>
<tr><td>Oligo rev</td><td>AAACCTTCGCCAACCCCTTCTCCT</td><td></td><td></td><td></td><td></td><td></td><td></td><td></td><td></td><td></td><td></td><td></td></tr>
<tr><td>Chromosome</td><td>start</td><td></td><td>end</td><td>strand</td><td>MM</td><td>target_seq</td><td>PAM</td><td>alignment</td><td>distance</td><td>position</td><td>gene name</td><td>gene id</td></tr>
<tr><td>GU305914.1</td><td></td><td>33860</td><td>33882</td><td>+</td><td>0</td><td>AGGAGAAGGGGTTGGCGAAG</td><td>GGG</td><td>||||||||[||||||||||||]PAM</td><td>NA</td><td>Intergenic</td><td>NA</td><td>NA</td></tr>
<tr><td>T2134</td><td>GGACGAACTCGATCTCCCTGAGG</td><td></td><td>1000</td><td></td><td></td><td></td><td></td><td></td><td></td><td></td><td></td><td></td></tr>
<tr><td>Oligo fwd</td><td>CACCGGACGAACTCGATCTCCCTG</td><td></td><td></td><td></td><td></td><td></td><td></td><td></td><td></td><td></td><td></td><td></td></tr>
<tr><td>Oligo rev</td><td>AAACCAGGGAGATCGAGTTCGTCC</td><td></td><td></td><td></td><td></td><td></td><td></td><td></td><td></td><td></td><td></td><td></td></tr>
<tr><td>Chromosome</td><td>start</td><td></td><td>end</td><td>strand</td><td>MM</td><td>target_seq</td><td>PAM</td><td>alignment</td><td>distance</td><td>position</td><td>gene name</td><td>gene id</td></tr>
<tr><td>GU305914.1</td><td></td><td>33881</td><td>33903</td><td>+</td><td>0</td><td>GGACGAACTCGATCTCCCTG</td><td>AGG</td><td>||||||||[||||||||||||]PAM</td><td>NA</td><td>Intergenic</td><td>NA</td><td>NA</td></tr>
<tr><td>T2135</td><td>CTCCCTGAGGAGCCCCCAGATGG</td><td></td><td>1000</td><td></td><td></td><td></td><td></td><td></td><td></td><td></td><td></td><td></td></tr>
<tr><td>Oligo fwd</td><td>CACCCTCCCTGAGGAGCCCCCAGA</td><td></td><td></td><td></td><td></td><td></td><td></td><td></td><td></td><td></td><td></td><td></td></tr>
<tr><td>Oligo rev</td><td>AAACTCTGGGGGCTCCTCAGGGAG</td><td></td><td></td><td></td><td></td><td></td><td></td><td></td><td></td><td></td><td></td><td></td></tr>
<tr><td>Chromosome</td><td>start</td><td></td><td>end</td><td>strand</td><td>MM</td><td>target_seq</td><td>PAM</td><td>alignment</td><td>distance</td><td>position</td><td>gene name</td><td>gene id</td></tr>
<tr><td>GU305914.1</td><td></td><td>33894</td><td>33916</td><td>+</td><td>0</td><td>CTCCCTGAGGAGCCCCCAGA</td><td>TGG</td><td>||||||||[||||||||||||]PAM</td><td>NA</td><td>Intergenic</td><td>NA</td><td>NA</td></tr>
<tr><td>T2136</td><td>CCTGAGGAGCCCCCAGATGGCGG</td><td></td><td>1000</td><td></td><td></td><td></td><td></td><td></td><td></td><td></td><td></td><td></td></tr>
<tr><td>Oligo fwd</td><td>CACCCCTGAGGAGCCCCCAGATGG</td><td></td><td></td><td></td><td></td><td></td><td></td><td></td><td></td><td></td><td></td><td></td></tr>
<tr><td>Oligo rev</td><td>AAACCCATCTGGGGGCTCCTCAGG</td><td></td><td></td><td></td><td></td><td></td><td></td><td></td><td></td><td></td><td></td><td></td></tr>
<tr><td>Chromosome</td><td>start</td><td></td><td>end</td><td>strand</td><td>MM</td><td>target_seq</td><td>PAM</td><td>alignment</td><td>distance</td><td>position</td><td>gene name</td><td>gene id</td></tr>
<tr><td>GU305914.1</td><td></td><td>33897</td><td>33919</td><td>+</td><td>0</td><td>CCTGAGGAGCCCCCAGATGG</td><td>CGG</td><td>||||||||[||||||||||||]PAM</td><td>NA</td><td>Intergenic</td><td>NA</td><td>NA</td></tr>
<tr><td>T2137</td><td>CCCCCAGATGGCGGAGTCCCAGG</td><td></td><td>1000</td><td></td><td></td><td></td><td></td><td></td><td></td><td></td><td></td><td></td></tr>
<tr><td>Oligo fwd</td><td>CACCCCCCCAGATGGCGGAGTCCC</td><td></td><td></td><td></td><td></td><td></td><td></td><td></td><td></td><td></td><td></td><td></td></tr>
<tr><td>Oligo rev</td><td>AAACGGGACTCCGCCATCTGGGGG</td><td></td><td></td><td></td><td></td><td></td><td></td><td></td><td></td><td></td><td></td><td></td></tr>
<tr><td>Chromosome</td><td>start</td><td></td><td>end</td><td>strand</td><td>MM</td><td>target_seq</td><td>PAM</td><td>alignment</td><td>distance</td><td>position</td><td>gene name</td><td>gene id</td></tr>
<tr><td>GU305914.1</td><td></td><td>33906</td><td>33928</td><td>+</td><td>0</td><td>CCCCCAGATGGCGGAGTCCC</td><td>AGG</td><td>||||||||[||||||||||||]PAM</td><td>NA</td><td>Intergenic</td><td>NA</td><td>NA</td></tr>
<tr><td>T2138</td><td>AGATGGCGGAGTCCCAGGCGAGG</td><td></td><td>1000</td><td></td><td></td><td></td><td></td><td></td><td></td><td></td><td></td><td></td></tr>
<tr><td>Oligo fwd</td><td>CACCAGATGGCGGAGTCCCAGGCG</td><td></td><td></td><td></td><td></td><td></td><td></td><td></td><td></td><td></td><td></td><td></td></tr>
<tr><td>Oligo rev</td><td>AAACCGCCTGGGACTCCGCCATCT</td><td></td><td></td><td></td><td></td><td></td><td></td><td></td><td></td><td></td><td></td><td></td></tr>
<tr><td>Chromosome</td><td>start</td><td></td><td>end</td><td>strand</td><td>MM</td><td>target_seq</td><td>PAM</td><td>alignment</td><td>distance</td><td>position</td><td>gene name</td><td>gene id</td></tr>
<tr><td>GU305914.1</td><td></td><td>33911</td><td>33933</td><td>+</td><td>0</td><td>AGATGGCGGAGTCCCAGGCG</td><td>AGG</td><td>||||||||[||||||||||||]PAM</td><td>NA</td><td>Intergenic</td><td>NA</td><td>NA</td></tr>
</table>

<table>
<tr><td>T2139</td><td>GATGGCGGAGTCCCAGGCGAGGG</td><td></td><td>1000</td><td></td><td></td><td></td><td></td><td></td><td></td><td></td><td></td><td></td></tr>
<tr><td>Oligo fwd</td><td>CACCGATGGCGGAGTCCCAGGCGA</td><td></td><td></td><td></td><td></td><td></td><td></td><td></td><td></td><td></td><td></td><td></td></tr>
<tr><td>Oligo rev</td><td>AAACTCGCCTGGGACTCCGCCATC</td><td></td><td></td><td></td><td></td><td></td><td></td><td></td><td></td><td></td><td></td><td></td></tr>
<tr><td>Chromosome</td><td>start</td><td></td><td>end</td><td>strand</td><td>MM</td><td>target_seq</td><td>PAM</td><td>alignment</td><td>distance</td><td>position</td><td>gene name</td><td>gene id</td></tr>
<tr><td>GU305914.1</td><td></td><td>33912</td><td>33934</td><td>+</td><td>0</td><td>GATGGCGGAGTCCCAGGCGA</td><td>GGG</td><td>||||||||[||||||||||||]PAM</td><td>NA</td><td>Intergenic</td><td>NA</td><td>NA</td></tr>
<tr><td>T2140</td><td>CCCAGGCGAGGGTGAGACCGCGG</td><td></td><td>1000</td><td></td><td></td><td></td><td></td><td></td><td></td><td></td><td></td><td></td></tr>
<tr><td>Oligo fwd</td><td>CACCCCCAGGCGAGGGTGAGACCG</td><td></td><td></td><td></td><td></td><td></td><td></td><td></td><td></td><td></td><td></td><td></td></tr>
<tr><td>Oligo rev</td><td>AAACCGGTCTCACCCTCGCCTGGG</td><td></td><td></td><td></td><td></td><td></td><td></td><td></td><td></td><td></td><td></td><td></td></tr>
<tr><td>Chromosome</td><td>start</td><td></td><td>end</td><td>strand</td><td>MM</td><td>target_seq</td><td>PAM</td><td>alignment</td><td>distance</td><td>position</td><td>gene name</td><td>gene id</td></tr>
<tr><td>GU305914.1</td><td></td><td>33923</td><td>33945</td><td>+</td><td>0</td><td>CCCAGGCGAGGGTGAGACCG</td><td>CGG</td><td>||||||||[||||||||||||]PAM</td><td>NA</td><td>Intergenic</td><td>NA</td><td>NA</td></tr>
<tr><td>T2141</td><td>GCGGATGCCGTACCCGCCGTTGG</td><td></td><td>1000</td><td></td><td></td><td></td><td></td><td></td><td></td><td></td><td></td><td></td></tr>
<tr><td>Oligo fwd</td><td>CACCGCGGATGCCGTACCCGCCGT</td><td></td><td></td><td></td><td></td><td></td><td></td><td></td><td></td><td></td><td></td><td></td></tr>
<tr><td>Oligo rev</td><td>AAACACGGCGGGTACGGCATCCGC</td><td></td><td></td><td></td><td></td><td></td><td></td><td></td><td></td><td></td><td></td><td></td></tr>
<tr><td>Chromosome</td><td>start</td><td></td><td>end</td><td>strand</td><td>MM</td><td>target_seq</td><td>PAM</td><td>alignment</td><td>distance</td><td>position</td><td>gene name</td><td>gene id</td></tr>
<tr><td>GU305914.1</td><td></td><td>33942</td><td>33964</td><td>+</td><td>0</td><td>GCGGATGCCGTACCCGCCGT</td><td>TGG</td><td>||||||||[||||||||||||]PAM</td><td>NA</td><td>Intergenic</td><td>NA</td><td>NA</td></tr>
<tr><td>T2142</td><td>GATGCCGTACCCGCCGTTGGCGG</td><td></td><td>1000</td><td></td><td></td><td></td><td></td><td></td><td></td><td></td><td></td><td></td></tr>
<tr><td>Oligo fwd</td><td>CACCGATGCCGTACCCGCCGTTGG</td><td></td><td></td><td></td><td></td><td></td><td></td><td></td><td></td><td></td><td></td><td></td></tr>
<tr><td>Oligo rev</td><td>AAACCCAACGGCGGGTACGGCATC</td><td></td><td></td><td></td><td></td><td></td><td></td><td></td><td></td><td></td><td></td><td></td></tr>
<tr><td>Chromosome</td><td>start</td><td></td><td>end</td><td>strand</td><td>MM</td><td>target_seq</td><td>PAM</td><td>alignment</td><td>distance</td><td>position</td><td>gene name</td><td>gene id</td></tr>
<tr><td>GU305914.1</td><td></td><td>33945</td><td>33967</td><td>+</td><td>0</td><td>GATGCCGTACCCGCCGTTGG</td><td>CGG</td><td>||||||||[||||||||||||]PAM</td><td>NA</td><td>Intergenic</td><td>NA</td><td>NA</td></tr>
<tr><td>T2143</td><td>GCCGTACCCGCCGTTGGCGGTGG</td><td></td><td>1000</td><td></td><td></td><td></td><td></td><td></td><td></td><td></td><td></td><td></td></tr>
<tr><td>Oligo fwd</td><td>CACCGCCGTACCCGCCGTTGGCGG</td><td></td><td></td><td></td><td></td><td></td><td></td><td></td><td></td><td></td><td></td><td></td></tr>
<tr><td>Oligo rev</td><td>AAACCCGCCAACGGCGGGTACGGC</td><td></td><td></td><td></td><td></td><td></td><td></td><td></td><td></td><td></td><td></td><td></td></tr>
<tr><td>Chromosome</td><td>start</td><td></td><td>end</td><td>strand</td><td>MM</td><td>target_seq</td><td>PAM</td><td>alignment</td><td>distance</td><td>position</td><td>gene name</td><td>gene id</td></tr>
<tr><td>GU305914.1</td><td></td><td>33948</td><td>33970</td><td>+</td><td>0</td><td>GCCGTACCCGCCGTTGGCGG</td><td>TGG</td><td>||||||||[||||||||||||]PAM</td><td>NA</td><td>Intergenic</td><td>NA</td><td>NA</td></tr>
<tr><td>T2144</td><td>TGGCGAGCCGCGCGTTCTTCAGG</td><td></td><td>1000</td><td></td><td></td><td></td><td></td><td></td><td></td><td></td><td></td><td></td></tr>
<tr><td>Oligo fwd</td><td>CACCTGGCGAGCCGCGCGTTCTTC</td><td></td><td></td><td></td><td></td><td></td><td></td><td></td><td></td><td></td><td></td><td></td></tr>
<tr><td>Oligo rev</td><td>AAACGAAGAACGCGCGGCTCGCCA</td><td></td><td></td><td></td><td></td><td></td><td></td><td></td><td></td><td></td><td></td><td></td></tr>
<tr><td>Chromosome</td><td>start</td><td></td><td>end</td><td>strand</td><td>MM</td><td>target_seq</td><td>PAM</td><td>alignment</td><td>distance</td><td>position</td><td>gene name</td><td>gene id</td></tr>
<tr><td>GU305914.1</td><td></td><td>33968</td><td>33990</td><td>+</td><td>0</td><td>TGGCGAGCCGCGCGTTCTTC</td><td>AGG</td><td>||||||||[||||||||||||]PAM</td><td>NA</td><td>Intergenic</td><td>NA</td><td>NA</td></tr>
<tr><td>T2145</td><td>CTTGAGTTCGATGTGATTAAAGG</td><td></td><td>1000</td><td></td><td></td><td></td><td></td><td></td><td></td><td></td><td></td><td></td></tr>
<tr><td>Oligo fwd</td><td>CACCCTTGAGTTCGATGTGATTAA</td><td></td><td></td><td></td><td></td><td></td><td></td><td></td><td></td><td></td><td></td><td></td></tr>
<tr><td>Oligo rev</td><td>AAACTTAATCACATCGAACTCAAG</td><td></td><td></td><td></td><td></td><td></td><td></td><td></td><td></td><td></td><td></td><td></td></tr>
<tr><td>Chromosome</td><td>start</td><td></td><td>end</td><td>strand</td><td>MM</td><td>target_seq</td><td>PAM</td><td>alignment</td><td>distance</td><td>position</td><td>gene name</td><td>gene id</td></tr>
<tr><td>GU305914.1</td><td></td><td>33999</td><td>34021</td><td>+</td><td>0</td><td>CTTGAGTTCGATGTGATTAA</td><td>AGG</td><td>||||||||[||||||||||||]PAM</td><td>NA</td><td>Intergenic</td><td>NA</td><td>NA</td></tr>
<tr><td>T2146</td><td>TTGAGTTCGATGTGATTAAAGGG</td><td></td><td>1000</td><td></td><td></td><td></td><td></td><td></td><td></td><td></td><td></td><td></td></tr>
<tr><td>Oligo fwd</td><td>CACCTTGAGTTCGATGTGATTAAA</td><td></td><td></td><td></td><td></td><td></td><td></td><td></td><td></td><td></td><td></td><td></td></tr>
<tr><td>Oligo rev</td><td>AAACTTTAATCACATCGAACTCAA</td><td></td><td></td><td></td><td></td><td></td><td></td><td></td><td></td><td></td><td></td><td></td></tr>
<tr><td>Chromosome</td><td>start</td><td></td><td>end</td><td>strand</td><td>MM</td><td>target_seq</td><td>PAM</td><td>alignment</td><td>distance</td><td>position</td><td>gene name</td><td>gene id</td></tr>
<tr><td>GU305914.1</td><td></td><td>34000</td><td>34022</td><td>+</td><td>0</td><td>TTGAGTTCGATGTGATTAAA</td><td>GGG</td><td>||||||||[||||||||||||]PAM</td><td>NA</td><td>Intergenic</td><td>NA</td><td>NA</td></tr>
<tr><td>T2147</td><td>TGTGATTAAAGGGCTTATTGAGG</td><td></td><td>1000</td><td></td><td></td><td></td><td></td><td></td><td></td><td></td><td></td><td></td></tr>
<tr><td>Oligo fwd</td><td>CACCTGTGATTAAAGGGCTTATTG</td><td></td><td></td><td></td><td></td><td></td><td></td><td></td><td></td><td></td><td></td><td></td></tr>
<tr><td>Oligo rev</td><td>AAACCAATAAGCCCTTTAATCACA</td><td></td><td></td><td></td><td></td><td></td><td></td><td></td><td></td><td></td><td></td><td></td></tr>
<tr><td>Chromosome</td><td>start</td><td></td><td>end</td><td>strand</td><td>MM</td><td>target_seq</td><td>PAM</td><td>alignment</td><td>distance</td><td>position</td><td>gene name</td><td>gene id</td></tr>
<tr><td>GU305914.1</td><td></td><td>34010</td><td>34032</td><td>+</td><td>0</td><td>TGTGATTAAAGGGCTTATTG</td><td>AGG</td><td>||||||||[||||||||||||]PAM</td><td>NA</td><td>Intergenic</td><td>NA</td><td>NA</td></tr>
<tr><td>T2148</td><td>GATTAAAGGGCTTATTGAGGCGG</td><td></td><td>1000</td><td></td><td></td><td></td><td></td><td></td><td></td><td></td><td></td><td></td></tr>
<tr><td>Oligo fwd</td><td>CACCGATTAAAGGGCTTATTGAGG</td><td></td><td></td><td></td><td></td><td></td><td></td><td></td><td></td><td></td><td></td><td></td></tr>
<tr><td>Oligo rev</td><td>AAACCCTCAATAAGCCCTTTAATC</td><td></td><td></td><td></td><td></td><td></td><td></td><td></td><td></td><td></td><td></td><td></td></tr>
<tr><td>Chromosome</td><td>start</td><td></td><td>end</td><td>strand</td><td>MM</td><td>target_seq</td><td>PAM</td><td>alignment</td><td>distance</td><td>position</td><td>gene name</td><td>gene id</td></tr>
<tr><td>GU305914.1</td><td></td><td>34013</td><td>34035</td><td>+</td><td>0</td><td>GATTAAAGGGCTTATTGAGG</td><td>CGG</td><td>||||||||[||||||||||||]PAM</td><td>NA</td><td>Intergenic</td><td>NA</td><td>NA</td></tr>
<tr><td>T2149</td><td>TATTGAGGCGGATGAAACTGCGG</td><td></td><td>1000</td><td></td><td></td><td></td><td></td><td></td><td></td><td></td><td></td><td></td></tr>
<tr><td>Oligo fwd</td><td>CACCTATTGAGGCGGATGAAACTG</td><td></td><td></td><td></td><td></td><td></td><td></td><td></td><td></td><td></td><td></td><td></td></tr>
<tr><td>Oligo rev</td><td>AAACCAGTTTCATCCGCCTCAATA</td><td></td><td></td><td></td><td></td><td></td><td></td><td></td><td></td><td></td><td></td><td></td></tr>
<tr><td>Chromosome</td><td>start</td><td></td><td>end</td><td>strand</td><td>MM</td><td>target_seq</td><td>PAM</td><td>alignment</td><td>distance</td><td>position</td><td>gene name</td><td>gene id</td></tr>
<tr><td>GU305914.1</td><td></td><td>34025</td><td>34047</td><td>+</td><td>0</td><td>TATTGAGGCGGATGAAACTG</td><td>CGG</td><td>||||||||[||||||||||||]PAM</td><td>NA</td><td>Intergenic</td><td>NA</td><td>NA</td></tr>
<tr><td>T2150</td><td>GCGGATGAAACTGCGGATCTCGG</td><td></td><td>1000</td><td></td><td></td><td></td><td></td><td></td><td></td><td></td><td></td><td></td></tr>
<tr><td>Oligo fwd</td><td>CACCGCGGATGAAACTGCGGATCT</td><td></td><td></td><td></td><td></td><td></td><td></td><td></td><td></td><td></td><td></td><td></td></tr>
<tr><td>Oligo rev</td><td>AAACAGATCCGCAGTTTCATCCGC</td><td></td><td></td><td></td><td></td><td></td><td></td><td></td><td></td><td></td><td></td><td></td></tr>
<tr><td>Chromosome</td><td>start</td><td></td><td>end</td><td>strand</td><td>MM</td><td>target_seq</td><td>PAM</td><td>alignment</td><td>distance</td><td>position</td><td>gene name</td><td>gene id</td></tr>
<tr><td>GU305914.1</td><td></td><td>34032</td><td>34054</td><td>+</td><td>0</td><td>GCGGATGAAACTGCGGATCT</td><td>CGG</td><td>||||||||[||||||||||||]PAM</td><td>NA</td><td>Intergenic</td><td>NA</td><td>NA</td></tr>
<tr><td>T2151</td><td>GATGAAACTGCGGATCTCGGTGG</td><td></td><td>1000</td><td></td><td></td><td></td><td></td><td></td><td></td><td></td><td></td><td></td></tr>
<tr><td>Oligo fwd</td><td>CACCGATGAAACTGCGGATCTCGG</td><td></td><td></td><td></td><td></td><td></td><td></td><td></td><td></td><td></td><td></td><td></td></tr>
<tr><td>Oligo rev</td><td>AAACCCGAGATCCGCAGTTTCATC</td><td></td><td></td><td></td><td></td><td></td><td></td><td></td><td></td><td></td><td></td><td></td></tr>
<tr><td>Chromosome</td><td>start</td><td></td><td>end</td><td>strand</td><td>MM</td><td>target_seq</td><td>PAM</td><td>alignment</td><td>distance</td><td>position</td><td>gene name</td><td>gene id</td></tr>
<tr><td>GU305914.1</td><td></td><td>34035</td><td>34057</td><td>+</td><td>0</td><td>GATGAAACTGCGGATCTCGG</td><td>TGG</td><td>||||||||[||||||||||||]PAM</td><td>NA</td><td>Intergenic</td><td>NA</td><td>NA</td></tr>
<tr><td>T2152</td><td>GGTGGCCTTGCCCGACTCCCTGG</td><td></td><td>1000</td><td></td><td></td><td></td><td></td><td></td><td></td><td></td><td></td><td></td></tr>
<tr><td>Oligo fwd</td><td>CACCGGTGGCCTTGCCCGACTCCC</td><td></td><td></td><td></td><td></td><td></td><td></td><td></td><td></td><td></td><td></td><td></td></tr>
<tr><td>Oligo rev</td><td>AAACGGGAGTCGGGCAAGGCCACC</td><td></td><td></td><td></td><td></td><td></td><td></td><td></td><td></td><td></td><td></td><td></td></tr>
<tr><td>Chromosome</td><td>start</td><td></td><td>end</td><td>strand</td><td>MM</td><td>target_seq</td><td>PAM</td><td>alignment</td><td>distance</td><td>position</td><td>gene name</td><td>gene id</td></tr>
<tr><td>GU305914.1</td><td></td><td>34053</td><td>34075</td><td>+</td><td>0</td><td>GGTGGCCTTGCCCGACTCCC</td><td>TGG</td><td>||||||||[||||||||||||]PAM</td><td>NA</td><td>Intergenic</td><td>NA</td><td>NA</td></tr>
<tr><td>T2153</td><td>CTTGCCCGACTCCCTGGACCAGG</td><td></td><td>1000</td><td></td><td></td><td></td><td></td><td></td><td></td><td></td><td></td><td></td></tr>
<tr><td>Oligo fwd</td><td>CACCCTTGCCCGACTCCCTGGACC</td><td></td><td></td><td></td><td></td><td></td><td></td><td></td><td></td><td></td><td></td><td></td></tr>
<tr><td>Oligo rev</td><td>AAACGGTCCAGGGAGTCGGGCAAG</td><td></td><td></td><td></td><td></td><td></td><td></td><td></td><td></td><td></td><td></td><td></td></tr>
<tr><td>Chromosome</td><td>start</td><td></td><td>end</td><td>strand</td><td>MM</td><td>target_seq</td><td>PAM</td><td>alignment</td><td>distance</td><td>position</td><td>gene name</td><td>gene id</td></tr>
<tr><td>GU305914.1</td><td></td><td>34059</td><td>34081</td><td>+</td><td>0</td><td>CTTGCCCGACTCCCTGGACC</td><td>AGG</td><td>||||||||[||||||||||||]PAM</td><td>NA</td><td>Intergenic</td><td>NA</td><td>NA</td></tr>
</table>

| | | | | | | | | | | | |
|---|---|---|---|---|---|---|---|---|---|---|---|
| T2154 | CATGACGAAGCCGAAGCGCGAGG | 1000 | | | | | | | | | |
| Oligo fwd | CACCCATGACGAAGCCGAAGCGCG | | | | | | | | | | |
| Oligo rev | AAACCGCGCTTCGGCTTCGTCATG | | | | | | | | | | |
| Chromosome | start | end | strand | MM | target_seq | PAM | alignment | distance | position | gene name | gene id |
| GU305914.1 | 34086 | 34108 | + | 0 | CATGACGAAGCCGAAGCGCG | AGG | \|\|\|\|\|\|\|\|[\|\|\|\|\|\|\|\|\|\|\|\|]PAM | NA | Intergenic | NA | NA |
| T2155 | CCTGCTCGTAGTTCCTGAGCTGG | 1000 | | | | | | | | | |
| Oligo fwd | CACCCCTGCTCGTAGTTCCTGAGC | | | | | | | | | | |
| Oligo rev | AAACGCTCAGGAACTACGAGCAGG | | | | | | | | | | |
| Chromosome | start | end | strand | MM | target_seq | PAM | alignment | distance | position | gene name | gene id |
| GU305914.1 | 34109 | 34131 | + | 0 | CCTGCTCGTAGTTCCTGAGC | TGG | \|\|\|\|\|\|\|\|[\|\|\|\|\|\|\|\|\|\|\|\|]PAM | NA | Intergenic | NA | NA |
| T2156 | CGTAGTTCCTGAGCTGGACCAGG | 1000 | | | | | | | | | |
| Oligo fwd | CACCCGTAGTTCCTGAGCTGGACC | | | | | | | | | | |
| Oligo rev | AAACGGTCCAGCTCAGGAACTACG | | | | | | | | | | |
| Chromosome | start | end | strand | MM | target_seq | PAM | alignment | distance | position | gene name | gene id |
| GU305914.1 | 34115 | 34137 | + | 0 | CGTAGTTCCTGAGCTGGACC | AGG | \|\|\|\|\|\|\|\|[\|\|\|\|\|\|\|\|\|\|\|\|]PAM | NA | Intergenic | NA | NA |
| T2157 | CCTGAGCTGGACCAGGTGATCGG | 1000 | | | | | | | | | |
| Oligo fwd | CACCCCTGAGCTGGACCAGGTGAT | | | | | | | | | | |
| Oligo rev | AAACATCACCTGGTCCAGCTCAGG | | | | | | | | | | |
| Chromosome | start | end | strand | MM | target_seq | PAM | alignment | distance | position | gene name | gene id |
| GU305914.1 | 34122 | 34144 | + | 0 | CCTGAGCTGGACCAGGTGAT | CGG | \|\|\|\|\|\|\|\|[\|\|\|\|\|\|\|\|\|\|\|\|]PAM | NA | Intergenic | NA | NA |
| T2158 | GGTGATCGGCGATAAAGCGCCGG | 1000 | | | | | | | | | |
| Oligo fwd | CACCGGTGATCGGCGATAAAGCGC | | | | | | | | | | |
| Oligo rev | AAACGCGCTTTATCGCCGATCACC | | | | | | | | | | |
| Chromosome | start | end | strand | MM | target_seq | PAM | alignment | distance | position | gene name | gene id |
| GU305914.1 | 34136 | 34158 | + | 0 | GGTGATCGGCGATAAAGCGC | CGG | \|\|\|\|\|\|\|\|[\|\|\|\|\|\|\|\|\|\|\|\|]PAM | NA | Intergenic | NA | NA |
| T2159 | GAACTGCGTGTCGTCGACCTCGG | 1000 | | | | | | | | | |
| Oligo fwd | CACCGAACTGCGTGTCGTCGACCT | | | | | | | | | | |
| Oligo rev | AAACAGGTCGACGACACGCAGTTC | | | | | | | | | | |
| Chromosome | start | end | strand | MM | target_seq | PAM | alignment | distance | position | gene name | gene id |
| GU305914.1 | 34200 | 34222 | + | 0 | GAACTGCGTGTCGTCGACCT | CGG | \|\|\|\|\|\|\|\|[\|\|\|\|\|\|\|\|\|\|\|\|]PAM | NA | Intergenic | NA | NA |
| T2160 | GTCGTCGACCTCGGAAGACTCGG | 1000 | | | | | | | | | |
| Oligo fwd | CACCGTCGTCGACCTCGGAAGACT | | | | | | | | | | |
| Oligo rev | AAACAGTCTTCCGAGGTCGACGAC | | | | | | | | | | |
| Chromosome | start | end | strand | MM | target_seq | PAM | alignment | distance | position | gene name | gene id |
| GU305914.1 | 34209 | 34231 | + | 0 | GTCGTCGACCTCGGAAGACT | CGG | \|\|\|\|\|\|\|\|[\|\|\|\|\|\|\|\|\|\|\|\|]PAM | NA | Intergenic | NA | NA |
| T2161 | TCGACCTCGGAAGACTCGGACGG | 1000 | | | | | | | | | |
| Oligo fwd | CACCTCGACCTCGGAAGACTCGGA | | | | | | | | | | |
| Oligo rev | AAACTCCGAGTCTTCCGAGGTCGA | | | | | | | | | | |
| Chromosome | start | end | strand | MM | target_seq | PAM | alignment | distance | position | gene name | gene id |
| GU305914.1 | 34213 | 34235 | + | 0 | TCGACCTCGGAAGACTCGGA | CGG | \|\|\|\|\|\|\|\|[\|\|\|\|\|\|\|\|\|\|\|\|]PAM | NA | Intergenic | NA | NA |
| T2162 | CGACCTCGGAAGACTCGGACGGG | 1000 | | | | | | | | | |
| Oligo fwd | CACCCGACCTCGGAAGACTCGGAC | | | | | | | | | | |
| Oligo rev | AAACGTCCGAGTCTTCCGAGGTCG | | | | | | | | | | |
| Chromosome | start | end | strand | MM | target_seq | PAM | alignment | distance | position | gene name | gene id |
| GU305914.1 | 34214 | 34236 | + | 0 | CGACCTCGGAAGACTCGGAC | GGG | \|\|\|\|\|\|\|\|[\|\|\|\|\|\|\|\|\|\|\|\|]PAM | NA | Intergenic | NA | NA |
| T2163 | CCCACCCGACCCTTCAGCGCAGG | 1000 | | | | | | | | | |
| Oligo fwd | CACCCCCACCCGACCCTTCAGCGC | | | | | | | | | | |
| Oligo rev | AAACGCGCTGAAGGGTCGGGTGGG | | | | | | | | | | |
| Chromosome | start | end | strand | MM | target_seq | PAM | alignment | distance | position | gene name | gene id |
| GU305914.1 | 34257 | 34279 | + | 0 | CCCACCCGACCCTTCAGCGC | AGG | \|\|\|\|\|\|\|\|[\|\|\|\|\|\|\|\|\|\|\|\|]PAM | NA | Intergenic | NA | NA |
| T2164 | CCACCCGACCCTTCAGCGCAGGG | 1000 | | | | | | | | | |
| Oligo fwd | CACCCCACCCGACCCTTCAGCGCA | | | | | | | | | | |
| Oligo rev | AAACTGCGCTGAAGGGTCGGGTGG | | | | | | | | | | |
| Chromosome | start | end | strand | MM | target_seq | PAM | alignment | distance | position | gene name | gene id |
| GU305914.1 | 34258 | 34280 | + | 0 | CCACCCGACCCTTCAGCGCA | GGG | \|\|\|\|\|\|\|\|[\|\|\|\|\|\|\|\|\|\|\|\|]PAM | NA | Intergenic | NA | NA |
| T2165 | ACCCTTCAGCGCAGGGCCCGCGG | 1000 | | | | | | | | | |
| Oligo fwd | CACCACCCTTCAGCGCAGGGCCCG | | | | | | | | | | |
| Oligo rev | AAACCGGGCCCTGCGCTGAAGGGT | | | | | | | | | | |
| Chromosome | start | end | strand | MM | target_seq | PAM | alignment | distance | position | gene name | gene id |
| GU305914.1 | 34265 | 34287 | + | 0 | ACCCTTCAGCGCAGGGCCCG | CGG | \|\|\|\|\|\|\|\|[\|\|\|\|\|\|\|\|\|\|\|\|]PAM | NA | Intergenic | NA | NA |
| T2166 | GCAGGGCCCGCGGTCCGCCATGG | 1000 | | | | | | | | | |
| Oligo fwd | CACCGCAGGGCCCGCGGTCCGCCA | | | | | | | | | | |
| Oligo rev | AAACTGGCGGACCGCGGGCCCTGC | | | | | | | | | | |
| Chromosome | start | end | strand | MM | target_seq | PAM | alignment | distance | position | gene name | gene id |
| GU305914.1 | 34275 | 34297 | + | 0 | GCAGGGCCCGCGGTCCGCCA | TGG | \|\|\|\|\|\|\|\|[\|\|\|\|\|\|\|\|\|\|\|\|]PAM | NA | Intergenic | NA | NA |
| T2167 | CCGCCATGGTCAGCGCGACACGG | 1000 | | | | | | | | | |
| Oligo fwd | CACCCCGCCATGGTCAGCGCGACA | | | | | | | | | | |
| Oligo rev | AAACTGTCGCGCTGACCATGGCGG | | | | | | | | | | |
| Chromosome | start | end | strand | MM | target_seq | PAM | alignment | distance | position | gene name | gene id |
| GU305914.1 | 34289 | 34311 | + | 0 | CCGCCATGGTCAGCGCGACA | CGG | \|\|\|\|\|\|\|\|[\|\|\|\|\|\|\|\|\|\|\|\|]PAM | NA | Intergenic | NA | NA |
| T2168 | GCACGCCACCCTCCGCCGCTCGG | 1000 | | | | | | | | | |
| Oligo fwd | CACCGCACGCCACCCTCCGCCGCT | | | | | | | | | | |
| Oligo rev | AAACAGCGGCGGAGGGTGGCGTGC | | | | | | | | | | |
| Chromosome | start | end | strand | MM | target_seq | PAM | alignment | distance | position | gene name | gene id |

<table>
<tr><td>GU305914.1</td><td>34395</td><td>34417</td><td>+</td><td>0</td><td>GCACGCCACCCTCCGCCGCT</td><td>CGG</td><td>||||||||[||||||||||||]PAM</td><td>NA</td><td>Intergenic</td><td>NA</td><td>NA</td></tr>
<tr><td></td><td></td><td></td><td></td><td></td><td></td><td></td><td></td><td></td><td></td><td></td><td></td></tr>
<tr><td>T2169</td><td>ACCCTCCGCCGCTCGGCTGCCGG</td><td>1000</td><td></td><td></td><td></td><td></td><td></td><td></td><td></td><td></td><td></td></tr>
<tr><td>Oligo fwd</td><td>CACCACCCTCCGCCGCTCGGCTGC</td><td></td><td></td><td></td><td></td><td></td><td></td><td></td><td></td><td></td><td></td></tr>
<tr><td>Oligo rev</td><td>AAACGCAGCCGAGCGGCGGAGGGT</td><td></td><td></td><td></td><td></td><td></td><td></td><td></td><td></td><td></td><td></td></tr>
<tr><td>Chromosome</td><td>start</td><td>end</td><td>strand</td><td>MM</td><td>target_seq</td><td>PAM</td><td>alignment</td><td>distance</td><td>position</td><td>gene name</td><td>gene id</td></tr>
<tr><td>GU305914.1</td><td>34402</td><td>34424</td><td>+</td><td>0</td><td>ACCCTCCGCCGCTCGGCTGC</td><td>CGG</td><td>||||||||[||||||||||||]PAM</td><td>NA</td><td>Intergenic</td><td>NA</td><td>NA</td></tr>
<tr><td></td><td></td><td></td><td></td><td></td><td></td><td></td><td></td><td></td><td></td><td></td><td></td></tr>
<tr><td>T2170</td><td>CCCTCCGCCGCTCGGCTGCCGGG</td><td>1000</td><td></td><td></td><td></td><td></td><td></td><td></td><td></td><td></td><td></td></tr>
<tr><td>Oligo fwd</td><td>CACCCCCTCCGCCGCTCGGCTGCC</td><td></td><td></td><td></td><td></td><td></td><td></td><td></td><td></td><td></td><td></td></tr>
<tr><td>Oligo rev</td><td>AAACGGCAGCCGAGCGGCGGAGGG</td><td></td><td></td><td></td><td></td><td></td><td></td><td></td><td></td><td></td><td></td></tr>
<tr><td>Chromosome</td><td>start</td><td>end</td><td>strand</td><td>MM</td><td>target_seq</td><td>PAM</td><td>alignment</td><td>distance</td><td>position</td><td>gene name</td><td>gene id</td></tr>
<tr><td>GU305914.1</td><td>34403</td><td>34425</td><td>+</td><td>0</td><td>CCCTCCGCCGCTCGGCTGCC</td><td>GGG</td><td>||||||||[||||||||||||]PAM</td><td>NA</td><td>Intergenic</td><td>NA</td><td>NA</td></tr>
<tr><td></td><td></td><td></td><td></td><td></td><td></td><td></td><td></td><td></td><td></td><td></td><td></td></tr>
<tr><td>T2171</td><td>TCCGCCGCTCGGCTGCCGGGAGG</td><td>1000</td><td></td><td></td><td></td><td></td><td></td><td></td><td></td><td></td><td></td></tr>
<tr><td>Oligo fwd</td><td>CACCTCCGCCGCTCGGCTGCCGGG</td><td></td><td></td><td></td><td></td><td></td><td></td><td></td><td></td><td></td><td></td></tr>
<tr><td>Oligo rev</td><td>AAACCCCGGCAGCCGAGCGGCGGA</td><td></td><td></td><td></td><td></td><td></td><td></td><td></td><td></td><td></td><td></td></tr>
<tr><td>Chromosome</td><td>start</td><td>end</td><td>strand</td><td>MM</td><td>target_seq</td><td>PAM</td><td>alignment</td><td>distance</td><td>position</td><td>gene name</td><td>gene id</td></tr>
<tr><td>GU305914.1</td><td>34406</td><td>34428</td><td>+</td><td>0</td><td>TCCGCCGCTCGGCTGCCGGG</td><td>AGG</td><td>||||||||[||||||||||||]PAM</td><td>NA</td><td>Intergenic</td><td>NA</td><td>NA</td></tr>
<tr><td></td><td></td><td></td><td></td><td></td><td></td><td></td><td></td><td></td><td></td><td></td><td></td></tr>
<tr><td>T2172</td><td>CGCTCGGCTGCCGGGAGGCGAGG</td><td>1000</td><td></td><td></td><td></td><td></td><td></td><td></td><td></td><td></td><td></td></tr>
<tr><td>Oligo fwd</td><td>CACCCGCTCGGCTGCCGGGAGGCG</td><td></td><td></td><td></td><td></td><td></td><td></td><td></td><td></td><td></td><td></td></tr>
<tr><td>Oligo rev</td><td>AAACCGCCTCCCGGCAGCCGAGCG</td><td></td><td></td><td></td><td></td><td></td><td></td><td></td><td></td><td></td><td></td></tr>
<tr><td>Chromosome</td><td>start</td><td>end</td><td>strand</td><td>MM</td><td>target_seq</td><td>PAM</td><td>alignment</td><td>distance</td><td>position</td><td>gene name</td><td>gene id</td></tr>
<tr><td>GU305914.1</td><td>34411</td><td>34433</td><td>+</td><td>0</td><td>CGCTCGGCTGCCGGGAGGCG</td><td>AGG</td><td>||||||||[||||||||||||]PAM</td><td>NA</td><td>Intergenic</td><td>NA</td><td>NA</td></tr>
<tr><td></td><td></td><td></td><td></td><td></td><td></td><td></td><td></td><td></td><td></td><td></td><td></td></tr>
<tr><td>T2173</td><td>GCCGGGAGGCGAGGCGAAACTGG</td><td>1000</td><td></td><td></td><td></td><td></td><td></td><td></td><td></td><td></td><td></td></tr>
<tr><td>Oligo fwd</td><td>CACCGCCGGGAGGCGAGGCGAAAC</td><td></td><td></td><td></td><td></td><td></td><td></td><td></td><td></td><td></td><td></td></tr>
<tr><td>Oligo rev</td><td>AAACGTTTCGCCTCGCCTCCCGGC</td><td></td><td></td><td></td><td></td><td></td><td></td><td></td><td></td><td></td><td></td></tr>
<tr><td>Chromosome</td><td>start</td><td>end</td><td>strand</td><td>MM</td><td>target_seq</td><td>PAM</td><td>alignment</td><td>distance</td><td>position</td><td>gene name</td><td>gene id</td></tr>
<tr><td>GU305914.1</td><td>34420</td><td>34442</td><td>+</td><td>0</td><td>GCCGGGAGGCGAGGCGAAAC</td><td>TGG</td><td>||||||||[||||||||||||]PAM</td><td>NA</td><td>Intergenic</td><td>NA</td><td>NA</td></tr>
<tr><td></td><td></td><td></td><td></td><td></td><td></td><td></td><td></td><td></td><td></td><td></td><td></td></tr>
<tr><td>T2174</td><td>GGGAGGCGAGGCGAAACTGGCGG</td><td>1000</td><td></td><td></td><td></td><td></td><td></td><td></td><td></td><td></td><td></td></tr>
<tr><td>Oligo fwd</td><td>CACCGGGAGGCGAGGCGAAACTGG</td><td></td><td></td><td></td><td></td><td></td><td></td><td></td><td></td><td></td><td></td></tr>
<tr><td>Oligo rev</td><td>AAACCCAGTTTCGCCTCGCCTCCC</td><td></td><td></td><td></td><td></td><td></td><td></td><td></td><td></td><td></td><td></td></tr>
<tr><td>Chromosome</td><td>start</td><td>end</td><td>strand</td><td>MM</td><td>target_seq</td><td>PAM</td><td>alignment</td><td>distance</td><td>position</td><td>gene name</td><td>gene id</td></tr>
<tr><td>GU305914.1</td><td>34423</td><td>34445</td><td>+</td><td>0</td><td>GGGAGGCGAGGCGAAACTGG</td><td>CGG</td><td>||||||||[||||||||||||]PAM</td><td>NA</td><td>Intergenic</td><td>NA</td><td>NA</td></tr>
<tr><td></td><td></td><td></td><td></td><td></td><td></td><td></td><td></td><td></td><td></td><td></td><td></td></tr>
<tr><td>T2175</td><td>GGAGGCGAGGCGAAACTGGCGGG</td><td>1000</td><td></td><td></td><td></td><td></td><td></td><td></td><td></td><td></td><td></td></tr>
<tr><td>Oligo fwd</td><td>CACCGGAGGCGAGGCGAAACTGGC</td><td></td><td></td><td></td><td></td><td></td><td></td><td></td><td></td><td></td><td></td></tr>
<tr><td>Oligo rev</td><td>AAACGCCAGTTTCGCCTCGCCTCC</td><td></td><td></td><td></td><td></td><td></td><td></td><td></td><td></td><td></td><td></td></tr>
<tr><td>Chromosome</td><td>start</td><td>end</td><td>strand</td><td>MM</td><td>target_seq</td><td>PAM</td><td>alignment</td><td>distance</td><td>position</td><td>gene name</td><td>gene id</td></tr>
<tr><td>GU305914.1</td><td>34424</td><td>34446</td><td>+</td><td>0</td><td>GGAGGCGAGGCGAAACTGGC</td><td>GGG</td><td>||||||||[||||||||||||]PAM</td><td>NA</td><td>Intergenic</td><td>NA</td><td>NA</td></tr>
<tr><td></td><td></td><td></td><td></td><td></td><td></td><td></td><td></td><td></td><td></td><td></td><td></td></tr>
<tr><td>T2176</td><td>TGGCGGGATAACTGCAAGAGAGG</td><td>1000</td><td></td><td></td><td></td><td></td><td></td><td></td><td></td><td></td><td></td></tr>
<tr><td>Oligo fwd</td><td>CACCTGGCGGGATAACTGCAAGAG</td><td></td><td></td><td></td><td></td><td></td><td></td><td></td><td></td><td></td><td></td></tr>
<tr><td>Oligo rev</td><td>AAACCTCTTGCAGTTATCCCGCCA</td><td></td><td></td><td></td><td></td><td></td><td></td><td></td><td></td><td></td><td></td></tr>
<tr><td>Chromosome</td><td>start</td><td>end</td><td>strand</td><td>MM</td><td>target_seq</td><td>PAM</td><td>alignment</td><td>distance</td><td>position</td><td>gene name</td><td>gene id</td></tr>
<tr><td>GU305914.1</td><td>34440</td><td>34462</td><td>+</td><td>0</td><td>TGGCGGGATAACTGCAAGAG</td><td>AGG</td><td>||||||||[||||||||||||]PAM</td><td>NA</td><td>Intergenic</td><td>NA</td><td>NA</td></tr>
<tr><td></td><td></td><td></td><td></td><td></td><td></td><td></td><td></td><td></td><td></td><td></td><td></td></tr>
<tr><td>T2177</td><td>GGCGGGATAACTGCAAGAGAGGG</td><td>1000</td><td></td><td></td><td></td><td></td><td></td><td></td><td></td><td></td><td></td></tr>
<tr><td>Oligo fwd</td><td>CACCGGCGGGATAACTGCAAGAGA</td><td></td><td></td><td></td><td></td><td></td><td></td><td></td><td></td><td></td><td></td></tr>
<tr><td>Oligo rev</td><td>AAACTCTCTTGCAGTTATCCCGCC</td><td></td><td></td><td></td><td></td><td></td><td></td><td></td><td></td><td></td><td></td></tr>
<tr><td>Chromosome</td><td>start</td><td>end</td><td>strand</td><td>MM</td><td>target_seq</td><td>PAM</td><td>alignment</td><td>distance</td><td>position</td><td>gene name</td><td>gene id</td></tr>
<tr><td>GU305914.1</td><td>34441</td><td>34463</td><td>+</td><td>0</td><td>GGCGGGATAACTGCAAGAGA</td><td>GGG</td><td>||||||||[||||||||||||]PAM</td><td>NA</td><td>Intergenic</td><td>NA</td><td>NA</td></tr>
<tr><td></td><td></td><td></td><td></td><td></td><td></td><td></td><td></td><td></td><td></td><td></td><td></td></tr>
<tr><td>T2178</td><td>GCGGGATAACTGCAAGAGAGGGG</td><td>1000</td><td></td><td></td><td></td><td></td><td></td><td></td><td></td><td></td><td></td></tr>
<tr><td>Oligo fwd</td><td>CACCGCGGGATAACTGCAAGAGAG</td><td></td><td></td><td></td><td></td><td></td><td></td><td></td><td></td><td></td><td></td></tr>
<tr><td>Oligo rev</td><td>AAACCTCTCTTGCAGTTATCCCGC</td><td></td><td></td><td></td><td></td><td></td><td></td><td></td><td></td><td></td><td></td></tr>
<tr><td>Chromosome</td><td>start</td><td>end</td><td>strand</td><td>MM</td><td>target_seq</td><td>PAM</td><td>alignment</td><td>distance</td><td>position</td><td>gene name</td><td>gene id</td></tr>
<tr><td>GU305914.1</td><td>34442</td><td>34464</td><td>+</td><td>0</td><td>GCGGGATAACTGCAAGAGAG</td><td>GGG</td><td>||||||||[||||||||||||]PAM</td><td>NA</td><td>Intergenic</td><td>NA</td><td>NA</td></tr>
<tr><td></td><td></td><td></td><td></td><td></td><td></td><td></td><td></td><td></td><td></td><td></td><td></td></tr>
<tr><td>T2179</td><td>ACTGCAAGAGAGGGGAAAAGCGG</td><td>1000</td><td></td><td></td><td></td><td></td><td></td><td></td><td></td><td></td><td></td></tr>
<tr><td>Oligo fwd</td><td>CACCACTGCAAGAGAGGGGAAAAG</td><td></td><td></td><td></td><td></td><td></td><td></td><td></td><td></td><td></td><td></td></tr>
<tr><td>Oligo rev</td><td>AAACCTTTTCCCCTCTCTTGCAGT</td><td></td><td></td><td></td><td></td><td></td><td></td><td></td><td></td><td></td><td></td></tr>
<tr><td>Chromosome</td><td>start</td><td>end</td><td>strand</td><td>MM</td><td>target_seq</td><td>PAM</td><td>alignment</td><td>distance</td><td>position</td><td>gene name</td><td>gene id</td></tr>
<tr><td>GU305914.1</td><td>34450</td><td>34472</td><td>+</td><td>0</td><td>ACTGCAAGAGAGGGGAAAAG</td><td>CGG</td><td>||||||||[||||||||||||]PAM</td><td>NA</td><td>Intergenic</td><td>NA</td><td>NA</td></tr>
<tr><td></td><td></td><td></td><td></td><td></td><td></td><td></td><td></td><td></td><td></td><td></td><td></td></tr>
<tr><td>T2180</td><td>TCCCAGCCGTCACCCGAGCGTGG</td><td>1000</td><td></td><td></td><td></td><td></td><td></td><td></td><td></td><td></td><td></td></tr>
<tr><td>Oligo fwd</td><td>CACCTCCCAGCCGTCACCCGAGCG</td><td></td><td></td><td></td><td></td><td></td><td></td><td></td><td></td><td></td><td></td></tr>
<tr><td>Oligo rev</td><td>AAACCGCTCGGGTGACGGCTGGGA</td><td></td><td></td><td></td><td></td><td></td><td></td><td></td><td></td><td></td><td></td></tr>
<tr><td>Chromosome</td><td>start</td><td>end</td><td>strand</td><td>MM</td><td>target_seq</td><td>PAM</td><td>alignment</td><td>distance</td><td>position</td><td>gene name</td><td>gene id</td></tr>
<tr><td>GU305914.1</td><td>34477</td><td>34499</td><td>+</td><td>0</td><td>TCCCAGCCGTCACCCGAGCG</td><td>TGG</td><td>||||||||[||||||||||||]PAM</td><td>NA</td><td>Intergenic</td><td>NA</td><td>NA</td></tr>
<tr><td></td><td></td><td></td><td></td><td></td><td></td><td></td><td></td><td></td><td></td><td></td><td></td></tr>
<tr><td>T2181</td><td>CCCAGCCGTCACCCGAGCGTGGG</td><td>1000</td><td></td><td></td><td></td><td></td><td></td><td></td><td></td><td></td><td></td></tr>
<tr><td>Oligo fwd</td><td>CACCCCCAGCCGTCACCCGAGCGT</td><td></td><td></td><td></td><td></td><td></td><td></td><td></td><td></td><td></td><td></td></tr>
<tr><td>Oligo rev</td><td>AAACACGCTCGGGTGACGGCTGGG</td><td></td><td></td><td></td><td></td><td></td><td></td><td></td><td></td><td></td><td></td></tr>
<tr><td>Chromosome</td><td>start</td><td>end</td><td>strand</td><td>MM</td><td>target_seq</td><td>PAM</td><td>alignment</td><td>distance</td><td>position</td><td>gene name</td><td>gene id</td></tr>
<tr><td>GU305914.1</td><td>34478</td><td>34500</td><td>+</td><td>0</td><td>CCCAGCCGTCACCCGAGCGT</td><td>GGG</td><td>||||||||[||||||||||||]PAM</td><td>NA</td><td>Intergenic</td><td>NA</td><td>NA</td></tr>
<tr><td></td><td></td><td></td><td></td><td></td><td></td><td></td><td></td><td></td><td></td><td></td><td></td></tr>
<tr><td>T2182</td><td>TCACCCGAGCGTGGGCACGCAGG</td><td>1000</td><td></td><td></td><td></td><td></td><td></td><td></td><td></td><td></td><td></td></tr>
<tr><td>Oligo fwd</td><td>CACCTCACCCGAGCGTGGGCACGC</td><td></td><td></td><td></td><td></td><td></td><td></td><td></td><td></td><td></td><td></td></tr>
<tr><td>Oligo rev</td><td>AAACGCGTGCCCACGCTCGGGTGA</td><td></td><td></td><td></td><td></td><td></td><td></td><td></td><td></td><td></td><td></td></tr>
<tr><td>Chromosome</td><td>start</td><td>end</td><td>strand</td><td>MM</td><td>target_seq</td><td>PAM</td><td>alignment</td><td>distance</td><td>position</td><td>gene name</td><td>gene id</td></tr>
<tr><td>GU305914.1</td><td>34486</td><td>34508</td><td>+</td><td>0</td><td>TCACCCGAGCGTGGGCACGC</td><td>AGG</td><td>||||||||[||||||||||||]PAM</td><td>NA</td><td>Intergenic</td><td>NA</td><td>NA</td></tr>
<tr><td></td><td></td><td></td><td></td><td></td><td></td><td></td><td></td><td></td><td></td><td></td><td></td></tr>
<tr><td>T2183</td><td>GAGCGTGGGCACGCAGGTCTCGG</td><td>1000</td><td></td><td></td><td></td><td></td><td></td><td></td><td></td><td></td><td></td></tr>
<tr><td>Oligo fwd</td><td>CACCGAGCGTGGGCACGCAGGTCT</td><td></td><td></td><td></td><td></td><td></td><td></td><td></td><td></td><td></td><td></td></tr>
<tr><td>Oligo rev</td><td>AAACAGACCTGCGTGCCCACGCTC</td><td></td><td></td><td></td><td></td><td></td><td></td><td></td><td></td><td></td><td></td></tr>
</table>

Chromosome start end strand MM target_seq PAM alignment distance position gene name gene id
GU305914.1 34492 34514 + 0 GAGCGTGGGCACGCAGGTCT CGG ||||||||[||||||||||||]PAM NA Intergenic NA NA

T2184 CACGCAGGTCTCGGCTATCCCGG 1000
Oligo fwd CACCCACGCAGGTCTCGGCTATCC
Oligo rev AAACGGATAGCCGAGACCTGCGTG
Chromosome start end strand MM target_seq PAM alignment distance position gene name gene id
GU305914.1 34501 34523 + 0 CACGCAGGTCTCGGCTATCC CGG ||||||||[||||||||||||]PAM NA Intergenic NA NA

T2185 CAGGTCTCGGCTATCCCGGACGG 1000
Oligo fwd CACCCAGGTCTCGGCTATCCCGGA
Oligo rev AAACTCCGGGATAGCCGAGACCTG
Chromosome start end strand MM target_seq PAM alignment distance position gene name gene id
GU305914.1 34505 34527 + 0 CAGGTCTCGGCTATCCCGGA CGG ||||||||[||||||||||||]PAM NA Intergenic NA NA

T2186 AGCCCGCAACGCCAACCACTTGG 1000
Oligo fwd CACCAGCCCGCAACGCCAACCACT
Oligo rev AAACAGTGGTTGGCGTTGCGGGCT
Chromosome start end strand MM target_seq PAM alignment distance position gene name gene id
GU305914.1 34528 34550 + 0 AGCCCGCAACGCCAACCACT TGG ||||||||[||||||||||||]PAM NA Intergenic NA NA

T2187 CGCCAACCACTTGGCGAAGTCGG 1000
Oligo fwd CACCCGCCAACCACTTGGCGAAGT
Oligo rev AAACACTTCGCCAAGTGGTTGGCG
Chromosome start end strand MM target_seq PAM alignment distance position gene name gene id
GU305914.1 34537 34559 + 0 CGCCAACCACTTGGCGAAGT CGG ||||||||[||||||||||||]PAM NA Intergenic NA NA

T2188 CTTGGCGAAGTCGGCACCGCCGG 1000
Oligo fwd CACCCTTGGCGAAGTCGGCACCGC
Oligo rev AAACGCGGTGCCGACTTCGCCAAG
Chromosome start end strand MM target_seq PAM alignment distance position gene name gene id
GU305914.1 34546 34568 + 0 CTTGGCGAAGTCGGCACCGC CGG ||||||||[||||||||||||]PAM NA Intergenic NA NA

T2189 TTGGCGAAGTCGGCACCGCCGGG 1000
Oligo fwd CACCTTGGCGAAGTCGGCACCGCC
Oligo rev AAACGGCGGTGCCGACTTCGCCAA
Chromosome start end strand MM target_seq PAM alignment distance position gene name gene id
GU305914.1 34547 34569 + 0 TTGGCGAAGTCGGCACCGCC GGG ||||||||[||||||||||||]PAM NA Intergenic NA NA

T2190 GCGCTCCGCGACGCCGTTCCTGG 1000
Oligo fwd CACCGCGCTCCGCGACGCCGTTCC
Oligo rev AAACGGAACGGCGTCGCGGAGCGC
Chromosome start end strand MM target_seq PAM alignment distance position gene name gene id
GU305914.1 34624 34646 + 0 GCGCTCCGCGACGCCGTTCC TGG ||||||||[||||||||||||]PAM NA Intergenic NA NA

T2191 CGTTCCTGGCGAACACCCTCCGG 1000
Oligo fwd CACCCGTTCCTGGCGAACACCCTC
Oligo rev AAACGAGGGTGTTCGCCAGGAACG
Chromosome start end strand MM target_seq PAM alignment distance position gene name gene id
GU305914.1 34638 34660 + 0 CGTTCCTGGCGAACACCCTC CGG ||||||||[||||||||||||]PAM NA Intergenic NA NA

T2192 CCTGGCGAACACCCTCCGGTAGG 1000
Oligo fwd CACCCCTGGCGAACACCCTCCGGT
Oligo rev AAACACCGGAGGGTGTTCGCCAGG
Chromosome start end strand MM target_seq PAM alignment distance position gene name gene id
GU305914.1 34642 34664 + 0 CCTGGCGAACACCCTCCGGT AGG ||||||||[||||||||||||]PAM NA Intergenic NA NA

T2193 CTGGCGAACACCCTCCGGTAGGG 1000
Oligo fwd CACCCTGGCGAACACCCTCCGGTA
Oligo rev AAACTACCGGAGGGTGTTCGCCAG
Chromosome start end strand MM target_seq PAM alignment distance position gene name gene id
GU305914.1 34643 34665 + 0 CTGGCGAACACCCTCCGGTA GGG ||||||||[||||||||||||]PAM NA Intergenic NA NA

T2194 ACACCCTCCGGTAGGGCTGCAGG 1000
Oligo fwd CACCACACCCTCCGGTAGGGCTGC
Oligo rev AAACGCAGCCCTACCGGAGGGTGT
Chromosome start end strand MM target_seq PAM alignment distance position gene name gene id
GU305914.1 34650 34672 + 0 ACACCCTCCGGTAGGGCTGC AGG ||||||||[||||||||||||]PAM NA Intergenic NA NA

T2195 GCTGCAGGTTGATCCTGAGCCGG 1000
Oligo fwd CACCGCTGCAGGTTGATCCTGAGC
Oligo rev AAACGCTCAGGATCAACCTGCAGC
Chromosome start end strand MM target_seq PAM alignment distance position gene name gene id
GU305914.1 34665 34687 + 0 GCTGCAGGTTGATCCTGAGC CGG ||||||||[||||||||||||]PAM NA Intergenic NA NA

T2196 GGTTGATCCTGAGCCGGTCCCGG 1000
Oligo fwd CACCGGTTGATCCTGAGCCGGTCC
Oligo rev AAACGGACCGGCTCAGGATCAACC
Chromosome start end strand MM target_seq PAM alignment distance position gene name gene id
GU305914.1 34671 34693 + 0 GGTTGATCCTGAGCCGGTCC CGG ||||||||[||||||||||||]PAM NA Intergenic NA NA

T2197 TGATCCTGAGCCGGTCCCGGAGG 1000
Oligo fwd CACCTGATCCTGAGCCGGTCCCGG
Oligo rev AAACCCGGGACCGGCTCAGGATCA
Chromosome start end strand MM target_seq PAM alignment distance position gene name gene id
GU305914.1 34674 34696 + 0 TGATCCTGAGCCGGTCCCGG AGG ||||||||[||||||||||||]PAM NA Intergenic NA NA

T2198 GAGGAATCCCGCTATCGTGTTGG 1000
Oligo fwd CACCGAGGAATCCCGCTATCGTGT

| | | | | | | | | | | | |
|---|---|---|---|---|---|---|---|---|---|---|---|
| Oligo rev | AAACACACGATAGCGGGATTCCTC | | | | | | | | | | |
| Chromosome | start | end | strand | MM | target_seq | PAM | alignment | distance | position | gene name | gene id |
| GU305914.1 | 34693 | 34715 | + | 0 | GAGGAATCCCGCTATCGTGT | TGG | \|\|\|\|\|\|\|\|[\|\|\|\|\|\|\|\|\|\|\|\|]PAM | NA | Intergenic | NA | NA |
| T2199 | TGTTGGCCACCTTCAAGACGCGG | 1000 | | | | | | | | | |
| Oligo fwd | CACCTGTTGGCCACCTTCAAGACG | | | | | | | | | | |
| Oligo rev | AAACCGTCTTGAAGGTGGCCAACA | | | | | | | | | | |
| Chromosome | start | end | strand | MM | target_seq | PAM | alignment | distance | position | gene name | gene id |
| GU305914.1 | 34710 | 34732 | + | 0 | TGTTGGCCACCTTCAAGACG | CGG | \|\|\|\|\|\|\|\|[\|\|\|\|\|\|\|\|\|\|\|\|]PAM | NA | Intergenic | NA | NA |
| T2200 | GTTGGCCACCTTCAAGACGCGGG | 1000 | | | | | | | | | |
| Oligo fwd | CACCGTTGGCCACCTTCAAGACGC | | | | | | | | | | |
| Oligo rev | AAACGCGTCTTGAAGGTGGCCAAC | | | | | | | | | | |
| Chromosome | start | end | strand | MM | target_seq | PAM | alignment | distance | position | gene name | gene id |
| GU305914.1 | 34711 | 34733 | + | 0 | GTTGGCCACCTTCAAGACGC | GGG | \|\|\|\|\|\|\|\|[\|\|\|\|\|\|\|\|\|\|\|\|]PAM | NA | Intergenic | NA | NA |
| T2201 | TCAAGACGCGGGCGCCGCCCAGG | 1000 | | | | | | | | | |
| Oligo fwd | CACCTCAAGACGCGGGCGCCGCCC | | | | | | | | | | |
| Oligo rev | AAACGGGCGGCGCCCGCGTCTTGA | | | | | | | | | | |
| Chromosome | start | end | strand | MM | target_seq | PAM | alignment | distance | position | gene name | gene id |
| GU305914.1 | 34722 | 34744 | + | 0 | TCAAGACGCGGGCGCCGCCC | AGG | \|\|\|\|\|\|\|\|[\|\|\|\|\|\|\|\|\|\|\|\|]PAM | NA | Intergenic | NA | NA |
| GU305914.1 | 29740 | 29762 | + | 4 | TCGAGCGGCGGACGCCGCCC | AGG | \|\|-\|\|--\|[\|\|\|-\|\|\|\|\|\|\|\|]PAM | NA | Intergenic | NA | NA |
| T2202 | CAGCACGATGAACTGCATCTCGG | 1000 | | | | | | | | | |
| Oligo fwd | CACCCAGCACGATGAACTGCATCT | | | | | | | | | | |
| Oligo rev | AAACAGATGCAGTTCATCGTGCTG | | | | | | | | | | |
| Chromosome | start | end | strand | MM | target_seq | PAM | alignment | distance | position | gene name | gene id |
| GU305914.1 | 34774 | 34796 | + | 0 | CAGCACGATGAACTGCATCT | CGG | \|\|\|\|\|\|\|\|[\|\|\|\|\|\|\|\|\|\|\|\|]PAM | NA | Intergenic | NA | NA |
| T2203 | CGATGACGCGATAGCCCGCCCGG | 1000 | | | | | | | | | |
| Oligo fwd | CACCCGATGACGCGATAGCCCGCC | | | | | | | | | | |
| Oligo rev | AAACGGCGGGCTATCGCGTCATCG | | | | | | | | | | |
| Chromosome | start | end | strand | MM | target_seq | PAM | alignment | distance | position | gene name | gene id |
| GU305914.1 | 34803 | 34825 | + | 0 | CGATGACGCGATAGCCCGCC | CGG | \|\|\|\|\|\|\|\|[\|\|\|\|\|\|\|\|\|\|\|\|]PAM | NA | Intergenic | NA | NA |
| T2204 | AAGATAGAAGAACACCCGTCCGG | 1000 | | | | | | | | | |
| Oligo fwd | CACCAAGATAGAAGAACACCCGTC | | | | | | | | | | |
| Oligo rev | AAACGACGGGTGTTCTTCTATCTT | | | | | | | | | | |
| Chromosome | start | end | strand | MM | target_seq | PAM | alignment | distance | position | gene name | gene id |
| GU305914.1 | 34828 | 34850 | + | 0 | AAGATAGAAGAACACCCGTC | CGG | \|\|\|\|\|\|\|\|[\|\|\|\|\|\|\|\|\|\|\|\|]PAM | NA | Intergenic | NA | NA |
| T2205 | AGATAGAAGAACACCCGTCCGGG | 1000 | | | | | | | | | |
| Oligo fwd | CACCAGATAGAAGAACACCCGTCC | | | | | | | | | | |
| Oligo rev | AAACGGACGGGTGTTCTTCTATCT | | | | | | | | | | |
| Chromosome | start | end | strand | MM | target_seq | PAM | alignment | distance | position | gene name | gene id |
| GU305914.1 | 34829 | 34851 | + | 0 | AGATAGAAGAACACCCGTCC | GGG | \|\|\|\|\|\|\|\|[\|\|\|\|\|\|\|\|\|\|\|\|]PAM | NA | Intergenic | NA | NA |
| T2206 | ACACCCGTCCGGGCACCACCTGG | 1000 | | | | | | | | | |
| Oligo fwd | CACCACACCCGTCCGGGCACCACC | | | | | | | | | | |
| Oligo rev | AAACGGTGGTGCCCGGACGGGTGT | | | | | | | | | | |
| Chromosome | start | end | strand | MM | target_seq | PAM | alignment | distance | position | gene name | gene id |
| GU305914.1 | 34839 | 34861 | + | 0 | ACACCCGTCCGGGCACCACC | TGG | \|\|\|\|\|\|\|\|[\|\|\|\|\|\|\|\|\|\|\|\|]PAM | NA | Intergenic | NA | NA |
| T2207 | AGAACTCGCCCATAAACCTGAGG | 1000 | | | | | | | | | |
| Oligo fwd | CACCAGAACTCGCCCATAAACCTG | | | | | | | | | | |
| Oligo rev | AAACCAGGTTTATGGGCGAGTTCT | | | | | | | | | | |
| Chromosome | start | end | strand | MM | target_seq | PAM | alignment | distance | position | gene name | gene id |
| GU305914.1 | 34905 | 34927 | + | 0 | AGAACTCGCCCATAAACCTG | AGG | \|\|\|\|\|\|\|\|[\|\|\|\|\|\|\|\|\|\|\|\|]PAM | NA | Intergenic | NA | NA |
| T2208 | GCCCATAAACCTGAGGTTACTGG | 1000 | | | | | | | | | |
| Oligo fwd | CACCGCCCATAAACCTGAGGTTAC | | | | | | | | | | |
| Oligo rev | AAACGTAACCTCAGGTTTATGGGC | | | | | | | | | | |
| Chromosome | start | end | strand | MM | target_seq | PAM | alignment | distance | position | gene name | gene id |
| GU305914.1 | 34912 | 34934 | + | 0 | GCCCATAAACCTGAGGTTAC | TGG | \|\|\|\|\|\|\|\|[\|\|\|\|\|\|\|\|\|\|\|\|]PAM | NA | Intergenic | NA | NA |
| T2209 | TAAACCTGAGGTTACTGGCCTGG | 1000 | | | | | | | | | |
| Oligo fwd | CACCTAAACCTGAGGTTACTGGCC | | | | | | | | | | |
| Oligo rev | AAACGGCCAGTAACCTCAGGTTTA | | | | | | | | | | |
| Chromosome | start | end | strand | MM | target_seq | PAM | alignment | distance | position | gene name | gene id |
| GU305914.1 | 34917 | 34939 | + | 0 | TAAACCTGAGGTTACTGGCC | TGG | \|\|\|\|\|\|\|\|[\|\|\|\|\|\|\|\|\|\|\|\|]PAM | NA | Intergenic | NA | NA |
| T2210 | ACCTGAGGTTACTGGCCTGGCGG | 1000 | | | | | | | | | |
| Oligo fwd | CACCACCTGAGGTTACTGGCCTGG | | | | | | | | | | |
| Oligo rev | AAACCCAGGCCAGTAACCTCAGGT | | | | | | | | | | |
| Chromosome | start | end | strand | MM | target_seq | PAM | alignment | distance | position | gene name | gene id |
| GU305914.1 | 34920 | 34942 | + | 0 | ACCTGAGGTTACTGGCCTGG | CGG | \|\|\|\|\|\|\|\|[\|\|\|\|\|\|\|\|\|\|\|\|]PAM | NA | Intergenic | NA | NA |
| T2211 | CTGGCGGAAAGTAGACAGCCCGG | 1000 | | | | | | | | | |
| Oligo fwd | CACCCTGGCGGAAAGTAGACAGCC | | | | | | | | | | |
| Oligo rev | AAACGGCTGTCTACTTTCCGCCAG | | | | | | | | | | |
| Chromosome | start | end | strand | MM | target_seq | PAM | alignment | distance | position | gene name | gene id |
| GU305914.1 | 34936 | 34958 | + | 0 | CTGGCGGAAAGTAGACAGCC | CGG | \|\|\|\|\|\|\|\|[\|\|\|\|\|\|\|\|\|\|\|\|]PAM | NA | Intergenic | NA | NA |
| T2212 | GGAAAGTAGACAGCCCGGTGCGG | 1000 | | | | | | | | | |
| Oligo fwd | CACCGGAAAGTAGACAGCCCGGTG | | | | | | | | | | |
| Oligo rev | AAACCACCGGGCTGTCTACTTTCC | | | | | | | | | | |
| Chromosome | start | end | strand | MM | target_seq | PAM | alignment | distance | position | gene name | gene id |
| GU305914.1 | 34941 | 34963 | + | 0 | GGAAAGTAGACAGCCCGGTG | CGG | \|\|\|\|\|\|\|\|[\|\|\|\|\|\|\|\|\|\|\|\|]PAM | NA | Intergenic | NA | NA |

<table>
<tr><td>T2213</td><td>TAGACAGCCCGGTGCGGTCGCGG</td><td></td><td>1000</td><td></td><td></td><td></td><td></td><td></td><td></td><td></td><td></td><td></td></tr>
<tr><td>Oligo fwd</td><td>CACCTAGACAGCCCGGTGCGGTCG</td><td></td><td></td><td></td><td></td><td></td><td></td><td></td><td></td><td></td><td></td><td></td></tr>
<tr><td>Oligo rev</td><td>AAACCGACCGCACCGGGCTGTCTA</td><td></td><td></td><td></td><td></td><td></td><td></td><td></td><td></td><td></td><td></td><td></td></tr>
<tr><td>Chromosome</td><td>start</td><td></td><td>end</td><td>strand</td><td>MM</td><td>target_seq</td><td>PAM</td><td>alignment</td><td>distance</td><td>position</td><td>gene name</td><td>gene id</td></tr>
<tr><td>GU305914.1</td><td></td><td>34947</td><td>34969</td><td>+</td><td>0</td><td>TAGACAGCCCGGTGCGGTCG</td><td>CGG</td><td>||||||||[||||||||||||]PAM</td><td>NA</td><td>Intergenic</td><td>NA</td><td>NA</td></tr>
<tr><td>T2214</td><td>ATTCTATCGATTCGAGATCCCGG</td><td></td><td>1000</td><td></td><td></td><td></td><td></td><td></td><td></td><td></td><td></td><td></td></tr>
<tr><td>Oligo fwd</td><td>CACCATTCTATCGATTCGAGATCC</td><td></td><td></td><td></td><td></td><td></td><td></td><td></td><td></td><td></td><td></td><td></td></tr>
<tr><td>Oligo rev</td><td>AAACGGATCTCGAATCGATAGAAT</td><td></td><td></td><td></td><td></td><td></td><td></td><td></td><td></td><td></td><td></td><td></td></tr>
<tr><td>Chromosome</td><td>start</td><td></td><td>end</td><td>strand</td><td>MM</td><td>target_seq</td><td>PAM</td><td>alignment</td><td>distance</td><td>position</td><td>gene name</td><td>gene id</td></tr>
<tr><td>GU305914.1</td><td></td><td>34977</td><td>34999</td><td>+</td><td>0</td><td>ATTCTATCGATTCGAGATCC</td><td>CGG</td><td>||||||||[||||||||||||]PAM</td><td>NA</td><td>Intergenic</td><td>NA</td><td>NA</td></tr>
<tr><td>T2215</td><td>CGATTCGAGATCCCGGAGCTCGG</td><td></td><td>1000</td><td></td><td></td><td></td><td></td><td></td><td></td><td></td><td></td><td></td></tr>
<tr><td>Oligo fwd</td><td>CACCCGATTCGAGATCCCGGAGCT</td><td></td><td></td><td></td><td></td><td></td><td></td><td></td><td></td><td></td><td></td><td></td></tr>
<tr><td>Oligo rev</td><td>AAACAGCTCCGGGATCTCGAATCG</td><td></td><td></td><td></td><td></td><td></td><td></td><td></td><td></td><td></td><td></td><td></td></tr>
<tr><td>Chromosome</td><td>start</td><td></td><td>end</td><td>strand</td><td>MM</td><td>target_seq</td><td>PAM</td><td>alignment</td><td>distance</td><td>position</td><td>gene name</td><td>gene id</td></tr>
<tr><td>GU305914.1</td><td></td><td>34984</td><td>35006</td><td>+</td><td>0</td><td>CGATTCGAGATCCCGGAGCT</td><td>CGG</td><td>||||||||[||||||||||||]PAM</td><td>NA</td><td>Intergenic</td><td>NA</td><td>NA</td></tr>
<tr><td>T2216</td><td>CGAGATCCCGGAGCTCGGTCAGG</td><td></td><td>1000</td><td></td><td></td><td></td><td></td><td></td><td></td><td></td><td></td><td></td></tr>
<tr><td>Oligo fwd</td><td>CACCCGAGATCCCGGAGCTCGGTC</td><td></td><td></td><td></td><td></td><td></td><td></td><td></td><td></td><td></td><td></td><td></td></tr>
<tr><td>Oligo rev</td><td>AAACGACCGAGCTCCGGGATCTCG</td><td></td><td></td><td></td><td></td><td></td><td></td><td></td><td></td><td></td><td></td><td></td></tr>
<tr><td>Chromosome</td><td>start</td><td></td><td>end</td><td>strand</td><td>MM</td><td>target_seq</td><td>PAM</td><td>alignment</td><td>distance</td><td>position</td><td>gene name</td><td>gene id</td></tr>
<tr><td>GU305914.1</td><td></td><td>34989</td><td>35011</td><td>+</td><td>0</td><td>CGAGATCCCGGAGCTCGGTC</td><td>AGG</td><td>||||||||[||||||||||||]PAM</td><td>NA</td><td>Intergenic</td><td>NA</td><td>NA</td></tr>
<tr><td>T2217</td><td>GAGATCCCGGAGCTCGGTCAGGG</td><td></td><td>1000</td><td></td><td></td><td></td><td></td><td></td><td></td><td></td><td></td><td></td></tr>
<tr><td>Oligo fwd</td><td>CACCGAGATCCCGGAGCTCGGTCA</td><td></td><td></td><td></td><td></td><td></td><td></td><td></td><td></td><td></td><td></td><td></td></tr>
<tr><td>Oligo rev</td><td>AAACTGACCGAGCTCCGGGATCTC</td><td></td><td></td><td></td><td></td><td></td><td></td><td></td><td></td><td></td><td></td><td></td></tr>
<tr><td>Chromosome</td><td>start</td><td></td><td>end</td><td>strand</td><td>MM</td><td>target_seq</td><td>PAM</td><td>alignment</td><td>distance</td><td>position</td><td>gene name</td><td>gene id</td></tr>
<tr><td>GU305914.1</td><td></td><td>34990</td><td>35012</td><td>+</td><td>0</td><td>GAGATCCCGGAGCTCGGTCA</td><td>GGG</td><td>||||||||[||||||||||||]PAM</td><td>NA</td><td>Intergenic</td><td>NA</td><td>NA</td></tr>
<tr><td>T2218</td><td>AGATCCCGGAGCTCGGTCAGGGG</td><td></td><td>1000</td><td></td><td></td><td></td><td></td><td></td><td></td><td></td><td></td><td></td></tr>
<tr><td>Oligo fwd</td><td>CACCAGATCCCGGAGCTCGGTCAG</td><td></td><td></td><td></td><td></td><td></td><td></td><td></td><td></td><td></td><td></td><td></td></tr>
<tr><td>Oligo rev</td><td>AAACCTGACCGAGCTCCGGGATCT</td><td></td><td></td><td></td><td></td><td></td><td></td><td></td><td></td><td></td><td></td><td></td></tr>
<tr><td>Chromosome</td><td>start</td><td></td><td>end</td><td>strand</td><td>MM</td><td>target_seq</td><td>PAM</td><td>alignment</td><td>distance</td><td>position</td><td>gene name</td><td>gene id</td></tr>
<tr><td>GU305914.1</td><td></td><td>34991</td><td>35013</td><td>+</td><td>0</td><td>AGATCCCGGAGCTCGGTCAG</td><td>GGG</td><td>||||||||[||||||||||||]PAM</td><td>NA</td><td>Intergenic</td><td>NA</td><td>NA</td></tr>
<tr><td>T2219</td><td>CTCGGTCAGGGGCAGTAAACAGG</td><td></td><td>1000</td><td></td><td></td><td></td><td></td><td></td><td></td><td></td><td></td><td></td></tr>
<tr><td>Oligo fwd</td><td>CACCCTCGGTCAGGGGCAGTAAAC</td><td></td><td></td><td></td><td></td><td></td><td></td><td></td><td></td><td></td><td></td><td></td></tr>
<tr><td>Oligo rev</td><td>AAACGTTTACTGCCCCTGACCGAG</td><td></td><td></td><td></td><td></td><td></td><td></td><td></td><td></td><td></td><td></td><td></td></tr>
<tr><td>Chromosome</td><td>start</td><td></td><td>end</td><td>strand</td><td>MM</td><td>target_seq</td><td>PAM</td><td>alignment</td><td>distance</td><td>position</td><td>gene name</td><td>gene id</td></tr>
<tr><td>GU305914.1</td><td></td><td>35002</td><td>35024</td><td>+</td><td>0</td><td>CTCGGTCAGGGGCAGTAAAC</td><td>AGG</td><td>||||||||[||||||||||||]PAM</td><td>NA</td><td>Intergenic</td><td>NA</td><td>NA</td></tr>
<tr><td>T2220</td><td>GTCAGGGGCAGTAAACAGGAAGG</td><td></td><td>1000</td><td></td><td></td><td></td><td></td><td></td><td></td><td></td><td></td><td></td></tr>
<tr><td>Oligo fwd</td><td>CACCGTCAGGGGCAGTAAACAGGA</td><td></td><td></td><td></td><td></td><td></td><td></td><td></td><td></td><td></td><td></td><td></td></tr>
<tr><td>Oligo rev</td><td>AAACTCCTGTTTACTGCCCCTGAC</td><td></td><td></td><td></td><td></td><td></td><td></td><td></td><td></td><td></td><td></td><td></td></tr>
<tr><td>Chromosome</td><td>start</td><td></td><td>end</td><td>strand</td><td>MM</td><td>target_seq</td><td>PAM</td><td>alignment</td><td>distance</td><td>position</td><td>gene name</td><td>gene id</td></tr>
<tr><td>GU305914.1</td><td></td><td>35006</td><td>35028</td><td>+</td><td>0</td><td>GTCAGGGGCAGTAAACAGGA</td><td>AGG</td><td>||||||||[||||||||||||]PAM</td><td>NA</td><td>Intergenic</td><td>NA</td><td>NA</td></tr>
<tr><td>T2221</td><td>TCGACGTCGTCGCCCGTCGTTGG</td><td></td><td>1000</td><td></td><td></td><td></td><td></td><td></td><td></td><td></td><td></td><td></td></tr>
<tr><td>Oligo fwd</td><td>CACCTCGACGTCGTCGCCCGTCGT</td><td></td><td></td><td></td><td></td><td></td><td></td><td></td><td></td><td></td><td></td><td></td></tr>
<tr><td>Oligo rev</td><td>AAACACGACGGGCGACGACGTCGA</td><td></td><td></td><td></td><td></td><td></td><td></td><td></td><td></td><td></td><td></td><td></td></tr>
<tr><td>Chromosome</td><td>start</td><td></td><td>end</td><td>strand</td><td>MM</td><td>target_seq</td><td>PAM</td><td>alignment</td><td>distance</td><td>position</td><td>gene name</td><td>gene id</td></tr>
<tr><td>GU305914.1</td><td></td><td>35033</td><td>35055</td><td>+</td><td>0</td><td>TCGACGTCGTCGCCCGTCGT</td><td>TGG</td><td>||||||||[||||||||||||]PAM</td><td>NA</td><td>Intergenic</td><td>NA</td><td>NA</td></tr>
<tr><td>T2222</td><td>ACGTCGTCGCCCGTCGTTGGCGG</td><td></td><td>1000</td><td></td><td></td><td></td><td></td><td></td><td></td><td></td><td></td><td></td></tr>
<tr><td>Oligo fwd</td><td>CACCACGTCGTCGCCCGTCGTTGG</td><td></td><td></td><td></td><td></td><td></td><td></td><td></td><td></td><td></td><td></td><td></td></tr>
<tr><td>Oligo rev</td><td>AAACCCAACGACGGGCGACGACGT</td><td></td><td></td><td></td><td></td><td></td><td></td><td></td><td></td><td></td><td></td><td></td></tr>
<tr><td>Chromosome</td><td>start</td><td></td><td>end</td><td>strand</td><td>MM</td><td>target_seq</td><td>PAM</td><td>alignment</td><td>distance</td><td>position</td><td>gene name</td><td>gene id</td></tr>
<tr><td>GU305914.1</td><td></td><td>35036</td><td>35058</td><td>+</td><td>0</td><td>ACGTCGTCGCCCGTCGTTGG</td><td>CGG</td><td>||||||||[||||||||||||]PAM</td><td>NA</td><td>Intergenic</td><td>NA</td><td>NA</td></tr>
<tr><td>T2223</td><td>CGAACATCTCATATATGTGCCGG</td><td></td><td>1000</td><td></td><td></td><td></td><td></td><td></td><td></td><td></td><td></td><td></td></tr>
<tr><td>Oligo fwd</td><td>CACCCGAACATCTCATATATGTGC</td><td></td><td></td><td></td><td></td><td></td><td></td><td></td><td></td><td></td><td></td><td></td></tr>
<tr><td>Oligo rev</td><td>AAACGCACATATATGAGATGTTCG</td><td></td><td></td><td></td><td></td><td></td><td></td><td></td><td></td><td></td><td></td><td></td></tr>
<tr><td>Chromosome</td><td>start</td><td></td><td>end</td><td>strand</td><td>MM</td><td>target_seq</td><td>PAM</td><td>alignment</td><td>distance</td><td>position</td><td>gene name</td><td>gene id</td></tr>
<tr><td>GU305914.1</td><td></td><td>35064</td><td>35086</td><td>+</td><td>0</td><td>CGAACATCTCATATATGTGC</td><td>CGG</td><td>||||||||[||||||||||||]PAM</td><td>NA</td><td>Intergenic</td><td>NA</td><td>NA</td></tr>
<tr><td>T2224</td><td>AAGATCCACCAGACCCCGTTCGG</td><td></td><td>1000</td><td></td><td></td><td></td><td></td><td></td><td></td><td></td><td></td><td></td></tr>
<tr><td>Oligo fwd</td><td>CACCAAGATCCACCAGACCCCGTT</td><td></td><td></td><td></td><td></td><td></td><td></td><td></td><td></td><td></td><td></td><td></td></tr>
<tr><td>Oligo rev</td><td>AAACAACGGGGTCTGGTGGATCTT</td><td></td><td></td><td></td><td></td><td></td><td></td><td></td><td></td><td></td><td></td><td></td></tr>
<tr><td>Chromosome</td><td>start</td><td></td><td>end</td><td>strand</td><td>MM</td><td>target_seq</td><td>PAM</td><td>alignment</td><td>distance</td><td>position</td><td>gene name</td><td>gene id</td></tr>
<tr><td>GU305914.1</td><td></td><td>35104</td><td>35126</td><td>+</td><td>0</td><td>AAGATCCACCAGACCCCGTT</td><td>CGG</td><td>||||||||[||||||||||||]PAM</td><td>NA</td><td>Intergenic</td><td>NA</td><td>NA</td></tr>
<tr><td>T2225</td><td>GGAGACGATATAGAGCAGATCGG</td><td></td><td>1000</td><td></td><td></td><td></td><td></td><td></td><td></td><td></td><td></td><td></td></tr>
<tr><td>Oligo fwd</td><td>CACCGGAGACGATATAGAGCAGAT</td><td></td><td></td><td></td><td></td><td></td><td></td><td></td><td></td><td></td><td></td><td></td></tr>
<tr><td>Oligo rev</td><td>AAACATCTGCTCTATATCGTCTCC</td><td></td><td></td><td></td><td></td><td></td><td></td><td></td><td></td><td></td><td></td><td></td></tr>
<tr><td>Chromosome</td><td>start</td><td></td><td>end</td><td>strand</td><td>MM</td><td>target_seq</td><td>PAM</td><td>alignment</td><td>distance</td><td>position</td><td>gene name</td><td>gene id</td></tr>
<tr><td>GU305914.1</td><td></td><td>35125</td><td>35147</td><td>+</td><td>0</td><td>GGAGACGATATAGAGCAGAT</td><td>CGG</td><td>||||||||[||||||||||||]PAM</td><td>NA</td><td>Intergenic</td><td>NA</td><td>NA</td></tr>
<tr><td>T2226</td><td>TATAGAGCAGATCGGTGTCCTGG</td><td></td><td>1000</td><td></td><td></td><td></td><td></td><td></td><td></td><td></td><td></td><td></td></tr>
<tr><td>Oligo fwd</td><td>CACCTATAGAGCAGATCGGTGTCC</td><td></td><td></td><td></td><td></td><td></td><td></td><td></td><td></td><td></td><td></td><td></td></tr>
<tr><td>Oligo rev</td><td>AAACGGACACCGATCTGCTCTATA</td><td></td><td></td><td></td><td></td><td></td><td></td><td></td><td></td><td></td><td></td><td></td></tr>
<tr><td>Chromosome</td><td>start</td><td></td><td>end</td><td>strand</td><td>MM</td><td>target_seq</td><td>PAM</td><td>alignment</td><td>distance</td><td>position</td><td>gene name</td><td>gene id</td></tr>
<tr><td>GU305914.1</td><td></td><td>35133</td><td>35155</td><td>+</td><td>0</td><td>TATAGAGCAGATCGGTGTCC</td><td>TGG</td><td>||||||||[||||||||||||]PAM</td><td>NA</td><td>Intergenic</td><td>NA</td><td>NA</td></tr>
<tr><td>T2227</td><td>CACGTAGCAACGATCTCCGCCGG</td><td></td><td>1000</td><td></td><td></td><td></td><td></td><td></td><td></td><td></td><td></td><td></td></tr>
<tr><td>Oligo fwd</td><td>CACCCACGTAGCAACGATCTCCGC</td><td></td><td></td><td></td><td></td><td></td><td></td><td></td><td></td><td></td><td></td><td></td></tr>
<tr><td>Oligo rev</td><td>AAACGCGGAGATCGTTGCTACGTG</td><td></td><td></td><td></td><td></td><td></td><td></td><td></td><td></td><td></td><td></td><td></td></tr>
<tr><td>Chromosome</td><td>start</td><td></td><td>end</td><td>strand</td><td>MM</td><td>target_seq</td><td>PAM</td><td>alignment</td><td>distance</td><td>position</td><td>gene name</td><td>gene id</td></tr>
<tr><td>GU305914.1</td><td></td><td>35161</td><td>35183</td><td>+</td><td>0</td><td>CACGTAGCAACGATCTCCGC</td><td>CGG</td><td>||||||||[||||||||||||]PAM</td><td>NA</td><td>Intergenic</td><td>NA</td><td>NA</td></tr>
</table>

T2228 GATCAGATCGCCGTGACTCCTGG 1000
Oligo fwd CACCGATCAGATCGCCGTGACTCC
Oligo rev AAACGGAGTCACGGCGATCTGATC
Chromosome start end strand MM target_seq PAM alignment distance position gene name gene id
GU305914.1 35194 35216 + 0 GATCAGATCGCCGTGACTCC TGG ||||||||[||||||||||||]PAM NA Intergenic NA NA

T2229 GAAAACGATCCGACCGACGACGG 1000
Oligo fwd CACCGAAAACGATCCGACCGACGA
Oligo rev AAACTCGTCGGTCGGATCGTTTTC
Chromosome start end strand MM target_seq PAM alignment distance position gene name gene id
GU305914.1 35221 35243 + 0 GAAAACGATCCGACCGACGA CGG ||||||||[||||||||||||]PAM NA Intergenic NA NA

T2230 CCGACCGACGACGGTCATCTCGG 1000
Oligo fwd CACCCCGACCGACGACGGTCATCT
Oligo rev AAACAGATGACCGTCGTCGGTCGG
Chromosome start end strand MM target_seq PAM alignment distance position gene name gene id
GU305914.1 35230 35252 + 0 CCGACCGACGACGGTCATCT CGG ||||||||[||||||||||||]PAM NA Intergenic NA NA

T2231 ACCGACGACGGTCATCTCGGAGG 1000
Oligo fwd CACCACCGACGACGGTCATCTCGG
Oligo rev AAACCCGAGATGACCGTCGTCGGT
Chromosome start end strand MM target_seq PAM alignment distance position gene name gene id
GU305914.1 35233 35255 + 0 ACCGACGACGGTCATCTCGG AGG ||||||||[||||||||||||]PAM NA Intergenic NA NA

T2232 CGGAGGACCGCTCGCTACAGCGG 1000
Oligo fwd CACCCGGAGGACCGCTCGCTACAG
Oligo rev AAACCTGTAGCGAGCGGTCCTCCG
Chromosome start end strand MM target_seq PAM alignment distance position gene name gene id
GU305914.1 35250 35272 + 0 CGGAGGACCGCTCGCTACAG CGG ||||||||[||||||||||||]PAM NA Intergenic NA NA

T2233 CAGCTTTTCTACGTCGAACGTGG 1000
Oligo fwd CACCCAGCTTTTCTACGTCGAACG
Oligo rev AAACCGTTCGACGTAGAAAAGCTG
Chromosome start end strand MM target_seq PAM alignment distance position gene name gene id
GU305914.1 35281 35303 + 0 CAGCTTTTCTACGTCGAACG TGG ||||||||[||||||||||||]PAM NA Intergenic NA NA

T2234 AGCTTTTCTACGTCGAACGTGGG 1000
Oligo fwd CACCAGCTTTTCTACGTCGAACGT
Oligo rev AAACACGTTCGACGTAGAAAAGCT
Chromosome start end strand MM target_seq PAM alignment distance position gene name gene id
GU305914.1 35282 35304 + 0 AGCTTTTCTACGTCGAACGT GGG ||||||||[||||||||||||]PAM NA Intergenic NA NA

T2235 GAACGTGGGACAGATGTTATCGG 1000
Oligo fwd CACCGAACGTGGGACAGATGTTAT
Oligo rev AAACATAACATCTGTCCCACGTTC
Chromosome start end strand MM target_seq PAM alignment distance position gene name gene id
GU305914.1 35296 35318 + 0 GAACGTGGGACAGATGTTAT CGG ||||||||[||||||||||||]PAM NA Intergenic NA NA

T2236 GCAACGTGATATCCGCCATGTGG 1000
Oligo fwd CACCGCAACGTGATATCCGCCATG
Oligo rev AAACCATGGCGGATATCACGTTGC
Chromosome start end strand MM target_seq PAM alignment distance position gene name gene id
GU305914.1 35331 35353 + 0 GCAACGTGATATCCGCCATG TGG ||||||||[||||||||||||]PAM NA Intergenic NA NA

T2237 GTCCAGCCCGAAACGCCGTCCGG 1000
Oligo fwd CACCGTCCAGCCCGAAACGCCGTC
Oligo rev AAACGACGGCGTTTCGGGCTGGAC
Chromosome start end strand MM target_seq PAM alignment distance position gene name gene id
GU305914.1 35353 35375 + 0 GTCCAGCCCGAAACGCCGTC CGG ||||||||[||||||||||||]PAM NA Intergenic NA NA

T2238 GAAAGCCTTCACCTCGTCGAAGG 1000
Oligo fwd CACCGAAAGCCTTCACCTCGTCGA
Oligo rev AAACTCGACGAGGTGAAGGCTTTC
Chromosome start end strand MM target_seq PAM alignment distance position gene name gene id
GU305914.1 35389 35411 + 0 GAAAGCCTTCACCTCGTCGA AGG ||||||||[||||||||||||]PAM NA Intergenic NA NA

T2239 GGAGCTGCCATTGCACGCCCTGG 1000
Oligo fwd CACCGGAGCTGCCATTGCACGCCC
Oligo rev AAACGGGCGTGCAATGGCAGCTCC
Chromosome start end strand MM target_seq PAM alignment distance position gene name gene id
GU305914.1 35410 35432 + 0 GGAGCTGCCATTGCACGCCC TGG ||||||||[||||||||||||]PAM NA Intergenic NA NA

T2240 GCAACAGCTCCCCCCTCGCGTGG 1000
Oligo fwd CACCGCAACAGCTCCCCCCTCGCG
Oligo rev AAACCGCGAGGGGGGAGCTGTTGC
Chromosome start end strand MM target_seq PAM alignment distance position gene name gene id
GU305914.1 35439 35461 + 0 GCAACAGCTCCCCCCTCGCG TGG ||||||||[||||||||||||]PAM NA Intergenic NA NA

T2241 CGCGTGGAAGTTGTCGACGCTGG 1000
Oligo fwd CACCCGCGTGGAAGTTGTCGACGC
Oligo rev AAACGCGTCGACAACTTCCACGCG
Chromosome start end strand MM target_seq PAM alignment distance position gene name gene id
GU305914.1 35455 35477 + 0 CGCGTGGAAGTTGTCGACGC TGG ||||||||[||||||||||||]PAM NA Intergenic NA NA

T2242 GACGCTGGTACCGCGACTGCCGG 1000
Oligo fwd CACCGACGCTGGTACCGCGACTGC
Oligo rev AAACGCAGTCGCGGTACCAGCGTC
Chromosome start end strand MM target_seq PAM alignment distance position gene name gene id

<table>
<tr><td>GU305914.1</td><td></td><td>35470</td><td>35492</td><td>+</td><td>0</td><td>GACGCTGGTACCGCGACTGC</td><td>CGG</td><td>||||||||[||||||||||||]PAM</td><td>NA</td><td>Intergenic</td><td>NA</td><td>NA</td></tr>
<tr><td>T2243</td><td>ATCCAAGTCGACGCGCTGCGCGG</td><td></td><td>1000</td><td></td><td></td><td></td><td></td><td></td><td></td><td></td><td></td><td></td></tr>
<tr><td>Oligo fwd</td><td>CACCATCCAAGTCGACGCGCTGCG</td><td></td><td></td><td></td><td></td><td></td><td></td><td></td><td></td><td></td><td></td><td></td></tr>
<tr><td>Oligo rev</td><td>AAACCGCAGCGCGTCGACTTGGAT</td><td></td><td></td><td></td><td></td><td></td><td></td><td></td><td></td><td></td><td></td><td></td></tr>
<tr><td>Chromosome</td><td>start</td><td></td><td>end</td><td>strand</td><td>MM</td><td>target_seq</td><td>PAM</td><td>alignment</td><td>distance</td><td>position</td><td>gene name</td><td>gene id</td></tr>
<tr><td>GU305914.1</td><td></td><td>35494</td><td>35516</td><td>+</td><td>0</td><td>ATCCAAGTCGACGCGCTGCG</td><td>CGG</td><td>||||||||[||||||||||||]PAM</td><td>NA</td><td>Intergenic</td><td>NA</td><td>NA</td></tr>
<tr><td>T2244</td><td>GGTGCCCGTGACGACGTTCACGG</td><td></td><td>1000</td><td></td><td></td><td></td><td></td><td></td><td></td><td></td><td></td><td></td></tr>
<tr><td>Oligo fwd</td><td>CACCGGTGCCCGTGACGACGTTCA</td><td></td><td></td><td></td><td></td><td></td><td></td><td></td><td></td><td></td><td></td><td></td></tr>
<tr><td>Oligo rev</td><td>AAACTGAACGTCGTCACGGGCACC</td><td></td><td></td><td></td><td></td><td></td><td></td><td></td><td></td><td></td><td></td><td></td></tr>
<tr><td>Chromosome</td><td>start</td><td></td><td>end</td><td>strand</td><td>MM</td><td>target_seq</td><td>PAM</td><td>alignment</td><td>distance</td><td>position</td><td>gene name</td><td>gene id</td></tr>
<tr><td>GU305914.1</td><td></td><td>35515</td><td>35537</td><td>+</td><td>0</td><td>GGTGCCCGTGACGACGTTCA</td><td>CGG</td><td>||||||||[||||||||||||]PAM</td><td>NA</td><td>Intergenic</td><td>NA</td><td>NA</td></tr>
<tr><td>T2245</td><td>GTGCCCGTGACGACGTTCACGGG</td><td></td><td>1000</td><td></td><td></td><td></td><td></td><td></td><td></td><td></td><td></td><td></td></tr>
<tr><td>Oligo fwd</td><td>CACCGTGCCCGTGACGACGTTCAC</td><td></td><td></td><td></td><td></td><td></td><td></td><td></td><td></td><td></td><td></td><td></td></tr>
<tr><td>Oligo rev</td><td>AAACGTGAACGTCGTCACGGGCAC</td><td></td><td></td><td></td><td></td><td></td><td></td><td></td><td></td><td></td><td></td><td></td></tr>
<tr><td>Chromosome</td><td>start</td><td></td><td>end</td><td>strand</td><td>MM</td><td>target_seq</td><td>PAM</td><td>alignment</td><td>distance</td><td>position</td><td>gene name</td><td>gene id</td></tr>
<tr><td>GU305914.1</td><td></td><td>35516</td><td>35538</td><td>+</td><td>0</td><td>GTGCCCGTGACGACGTTCAC</td><td>GGG</td><td>||||||||[||||||||||||]PAM</td><td>NA</td><td>Intergenic</td><td>NA</td><td>NA</td></tr>
<tr><td>T2246</td><td>ACGACGTTCACGGGCACGACCGG</td><td></td><td>1000</td><td></td><td></td><td></td><td></td><td></td><td></td><td></td><td></td><td></td></tr>
<tr><td>Oligo fwd</td><td>CACCACGACGTTCACGGGCACGAC</td><td></td><td></td><td></td><td></td><td></td><td></td><td></td><td></td><td></td><td></td><td></td></tr>
<tr><td>Oligo rev</td><td>AAACGTCGTGCCCGTGAACGTCGT</td><td></td><td></td><td></td><td></td><td></td><td></td><td></td><td></td><td></td><td></td><td></td></tr>
<tr><td>Chromosome</td><td>start</td><td></td><td>end</td><td>strand</td><td>MM</td><td>target_seq</td><td>PAM</td><td>alignment</td><td>distance</td><td>position</td><td>gene name</td><td>gene id</td></tr>
<tr><td>GU305914.1</td><td></td><td>35525</td><td>35547</td><td>+</td><td>0</td><td>ACGACGTTCACGGGCACGAC</td><td>CGG</td><td>||||||||[||||||||||||]PAM</td><td>NA</td><td>Intergenic</td><td>NA</td><td>NA</td></tr>
<tr><td>T2247</td><td>CACGGGCACGACCGGCGTCTTGG</td><td></td><td>1000</td><td></td><td></td><td></td><td></td><td></td><td></td><td></td><td></td><td></td></tr>
<tr><td>Oligo fwd</td><td>CACCCACGGGCACGACCGGCGTCT</td><td></td><td></td><td></td><td></td><td></td><td></td><td></td><td></td><td></td><td></td><td></td></tr>
<tr><td>Oligo rev</td><td>AAACAGACGCCGGTCGTGCCCGTG</td><td></td><td></td><td></td><td></td><td></td><td></td><td></td><td></td><td></td><td></td><td></td></tr>
<tr><td>Chromosome</td><td>start</td><td></td><td>end</td><td>strand</td><td>MM</td><td>target_seq</td><td>PAM</td><td>alignment</td><td>distance</td><td>position</td><td>gene name</td><td>gene id</td></tr>
<tr><td>GU305914.1</td><td></td><td>35533</td><td>35555</td><td>+</td><td>0</td><td>CACGGGCACGACCGGCGTCT</td><td>TGG</td><td>||||||||[||||||||||||]PAM</td><td>NA</td><td>Intergenic</td><td>NA</td><td>NA</td></tr>
<tr><td>T2248</td><td>GGGCACGACCGGCGTCTTGGAGG</td><td></td><td>1000</td><td></td><td></td><td></td><td></td><td></td><td></td><td></td><td></td><td></td></tr>
<tr><td>Oligo fwd</td><td>CACCGGGCACGACCGGCGTCTTGG</td><td></td><td></td><td></td><td></td><td></td><td></td><td></td><td></td><td></td><td></td><td></td></tr>
<tr><td>Oligo rev</td><td>AAACCCAAGACGCCGGTCGTGCCC</td><td></td><td></td><td></td><td></td><td></td><td></td><td></td><td></td><td></td><td></td><td></td></tr>
<tr><td>Chromosome</td><td>start</td><td></td><td>end</td><td>strand</td><td>MM</td><td>target_seq</td><td>PAM</td><td>alignment</td><td>distance</td><td>position</td><td>gene name</td><td>gene id</td></tr>
<tr><td>GU305914.1</td><td></td><td>35536</td><td>35558</td><td>+</td><td>0</td><td>GGGCACGACCGGCGTCTTGG</td><td>AGG</td><td>||||||||[||||||||||||]PAM</td><td>NA</td><td>Intergenic</td><td>NA</td><td>NA</td></tr>
<tr><td>T2249</td><td>GACCGGCGTCTTGGAGGCGACGG</td><td></td><td>1000</td><td></td><td></td><td></td><td></td><td></td><td></td><td></td><td></td><td></td></tr>
<tr><td>Oligo fwd</td><td>CACCGACCGGCGTCTTGGAGGCGA</td><td></td><td></td><td></td><td></td><td></td><td></td><td></td><td></td><td></td><td></td><td></td></tr>
<tr><td>Oligo rev</td><td>AAACTCGCCTCCAAGACGCCGGTC</td><td></td><td></td><td></td><td></td><td></td><td></td><td></td><td></td><td></td><td></td><td></td></tr>
<tr><td>Chromosome</td><td>start</td><td></td><td>end</td><td>strand</td><td>MM</td><td>target_seq</td><td>PAM</td><td>alignment</td><td>distance</td><td>position</td><td>gene name</td><td>gene id</td></tr>
<tr><td>GU305914.1</td><td></td><td>35542</td><td>35564</td><td>+</td><td>0</td><td>GACCGGCGTCTTGGAGGCGA</td><td>CGG</td><td>||||||||[||||||||||||]PAM</td><td>NA</td><td>Intergenic</td><td>NA</td><td>NA</td></tr>
<tr><td>T2250</td><td>CGGCGTCTTGGAGGCGACGGAGG</td><td></td><td>1000</td><td></td><td></td><td></td><td></td><td></td><td></td><td></td><td></td><td></td></tr>
<tr><td>Oligo fwd</td><td>CACCCGGCGTCTTGGAGGCGACGG</td><td></td><td></td><td></td><td></td><td></td><td></td><td></td><td></td><td></td><td></td><td></td></tr>
<tr><td>Oligo rev</td><td>AAACCCGTCGCCTCCAAGACGCCG</td><td></td><td></td><td></td><td></td><td></td><td></td><td></td><td></td><td></td><td></td><td></td></tr>
<tr><td>Chromosome</td><td>start</td><td></td><td>end</td><td>strand</td><td>MM</td><td>target_seq</td><td>PAM</td><td>alignment</td><td>distance</td><td>position</td><td>gene name</td><td>gene id</td></tr>
<tr><td>GU305914.1</td><td></td><td>35545</td><td>35567</td><td>+</td><td>0</td><td>CGGCGTCTTGGAGGCGACGG</td><td>AGG</td><td>||||||||[||||||||||||]PAM</td><td>NA</td><td>Intergenic</td><td>NA</td><td>NA</td></tr>
<tr><td>T2251</td><td>CGTCTTGGAGGCGACGGAGGTGG</td><td></td><td>1000</td><td></td><td></td><td></td><td></td><td></td><td></td><td></td><td></td><td></td></tr>
<tr><td>Oligo fwd</td><td>CACCCGTCTTGGAGGCGACGGAGG</td><td></td><td></td><td></td><td></td><td></td><td></td><td></td><td></td><td></td><td></td><td></td></tr>
<tr><td>Oligo rev</td><td>AAACCCTCCGTCGCCTCCAAGACG</td><td></td><td></td><td></td><td></td><td></td><td></td><td></td><td></td><td></td><td></td><td></td></tr>
<tr><td>Chromosome</td><td>start</td><td></td><td>end</td><td>strand</td><td>MM</td><td>target_seq</td><td>PAM</td><td>alignment</td><td>distance</td><td>position</td><td>gene name</td><td>gene id</td></tr>
<tr><td>GU305914.1</td><td></td><td>35548</td><td>35570</td><td>+</td><td>0</td><td>CGTCTTGGAGGCGACGGAGG</td><td>TGG</td><td>||||||||[||||||||||||]PAM</td><td>NA</td><td>Intergenic</td><td>NA</td><td>NA</td></tr>
<tr><td>T2252</td><td>GTCTTGGAGGCGACGGAGGTGGG</td><td></td><td>1000</td><td></td><td></td><td></td><td></td><td></td><td></td><td></td><td></td><td></td></tr>
<tr><td>Oligo fwd</td><td>CACCGTCTTGGAGGCGACGGAGGT</td><td></td><td></td><td></td><td></td><td></td><td></td><td></td><td></td><td></td><td></td><td></td></tr>
<tr><td>Oligo rev</td><td>AAACACCTCCGTCGCCTCCAAGAC</td><td></td><td></td><td></td><td></td><td></td><td></td><td></td><td></td><td></td><td></td><td></td></tr>
<tr><td>Chromosome</td><td>start</td><td></td><td>end</td><td>strand</td><td>MM</td><td>target_seq</td><td>PAM</td><td>alignment</td><td>distance</td><td>position</td><td>gene name</td><td>gene id</td></tr>
<tr><td>GU305914.1</td><td></td><td>35549</td><td>35571</td><td>+</td><td>0</td><td>GTCTTGGAGGCGACGGAGGT</td><td>GGG</td><td>||||||||[||||||||||||]PAM</td><td>NA</td><td>Intergenic</td><td>NA</td><td>NA</td></tr>
<tr><td>T2253</td><td>TCTTGGAGGCGACGGAGGTGGGG</td><td></td><td>1000</td><td></td><td></td><td></td><td></td><td></td><td></td><td></td><td></td><td></td></tr>
<tr><td>Oligo fwd</td><td>CACCTCTTGGAGGCGACGGAGGTG</td><td></td><td></td><td></td><td></td><td></td><td></td><td></td><td></td><td></td><td></td><td></td></tr>
<tr><td>Oligo rev</td><td>AAACCACCTCCGTCGCCTCCAAGA</td><td></td><td></td><td></td><td></td><td></td><td></td><td></td><td></td><td></td><td></td><td></td></tr>
<tr><td>Chromosome</td><td>start</td><td></td><td>end</td><td>strand</td><td>MM</td><td>target_seq</td><td>PAM</td><td>alignment</td><td>distance</td><td>position</td><td>gene name</td><td>gene id</td></tr>
<tr><td>GU305914.1</td><td></td><td>35550</td><td>35572</td><td>+</td><td>0</td><td>TCTTGGAGGCGACGGAGGTG</td><td>GGG</td><td>||||||||[||||||||||||]PAM</td><td>NA</td><td>Intergenic</td><td>NA</td><td>NA</td></tr>
<tr><td>T2254</td><td>GGAGGCGACGGAGGTGGGGACGG</td><td></td><td>1000</td><td></td><td></td><td></td><td></td><td></td><td></td><td></td><td></td><td></td></tr>
<tr><td>Oligo fwd</td><td>CACCGGAGGCGACGGAGGTGGGGA</td><td></td><td></td><td></td><td></td><td></td><td></td><td></td><td></td><td></td><td></td><td></td></tr>
<tr><td>Oligo rev</td><td>AAACTCCCCACCTCCGTCGCCTCC</td><td></td><td></td><td></td><td></td><td></td><td></td><td></td><td></td><td></td><td></td><td></td></tr>
<tr><td>Chromosome</td><td>start</td><td></td><td>end</td><td>strand</td><td>MM</td><td>target_seq</td><td>PAM</td><td>alignment</td><td>distance</td><td>position</td><td>gene name</td><td>gene id</td></tr>
<tr><td>GU305914.1</td><td></td><td>35554</td><td>35576</td><td>+</td><td>0</td><td>GGAGGCGACGGAGGTGGGGA</td><td>CGG</td><td>||||||||[||||||||||||]PAM</td><td>NA</td><td>Intergenic</td><td>NA</td><td>NA</td></tr>
<tr><td>T2255</td><td>GAGGCGACGGAGGTGGGGACGGG</td><td></td><td>1000</td><td></td><td></td><td></td><td></td><td></td><td></td><td></td><td></td><td></td></tr>
<tr><td>Oligo fwd</td><td>CACCGAGGCGACGGAGGTGGGGAC</td><td></td><td></td><td></td><td></td><td></td><td></td><td></td><td></td><td></td><td></td><td></td></tr>
<tr><td>Oligo rev</td><td>AAACGTCCCCACCTCCGTCGCCTC</td><td></td><td></td><td></td><td></td><td></td><td></td><td></td><td></td><td></td><td></td><td></td></tr>
<tr><td>Chromosome</td><td>start</td><td></td><td>end</td><td>strand</td><td>MM</td><td>target_seq</td><td>PAM</td><td>alignment</td><td>distance</td><td>position</td><td>gene name</td><td>gene id</td></tr>
<tr><td>GU305914.1</td><td></td><td>35555</td><td>35577</td><td>+</td><td>0</td><td>GAGGCGACGGAGGTGGGGAC</td><td>GGG</td><td>||||||||[||||||||||||]PAM</td><td>NA</td><td>Intergenic</td><td>NA</td><td>NA</td></tr>
<tr><td>T2256</td><td>GACGGAGGTGGGGACGGGCACGG</td><td></td><td>1000</td><td></td><td></td><td></td><td></td><td></td><td></td><td></td><td></td><td></td></tr>
<tr><td>Oligo fwd</td><td>CACCGACGGAGGTGGGGACGGGCA</td><td></td><td></td><td></td><td></td><td></td><td></td><td></td><td></td><td></td><td></td><td></td></tr>
<tr><td>Oligo rev</td><td>AAACTGCCCGTCCCCACCTCCGTC</td><td></td><td></td><td></td><td></td><td></td><td></td><td></td><td></td><td></td><td></td><td></td></tr>
<tr><td>Chromosome</td><td>start</td><td></td><td>end</td><td>strand</td><td>MM</td><td>target_seq</td><td>PAM</td><td>alignment</td><td>distance</td><td>position</td><td>gene name</td><td>gene id</td></tr>
<tr><td>GU305914.1</td><td></td><td>35560</td><td>35582</td><td>+</td><td>0</td><td>GACGGAGGTGGGGACGGGCA</td><td>CGG</td><td>||||||||[||||||||||||]PAM</td><td>NA</td><td>Intergenic</td><td>NA</td><td>NA</td></tr>
<tr><td>T2257</td><td>GAGGTGGGGACGGGCACGGTCGG</td><td></td><td>1000</td><td></td><td></td><td></td><td></td><td></td><td></td><td></td><td></td><td></td></tr>
<tr><td>Oligo fwd</td><td>CACCGAGGTGGGGACGGGCACGGT</td><td></td><td></td><td></td><td></td><td></td><td></td><td></td><td></td><td></td><td></td><td></td></tr>
<tr><td>Oligo rev</td><td>AAACACCGTGCCCGTCCCCACCTC</td><td></td><td></td><td></td><td></td><td></td><td></td><td></td><td></td><td></td><td></td><td></td></tr>
</table>

<table>
<tr><td>Chromosome</td><td>start</td><td>end</td><td>strand</td><td>MM</td><td>target_seq</td><td>PAM</td><td>alignment</td><td>distance</td><td>position</td><td>gene name</td><td>gene id</td></tr>
<tr><td>GU305914.1</td><td>35564</td><td>35586</td><td>+</td><td>0</td><td>GAGGTGGGGACGGGCACGGT</td><td>CGG</td><td>||||||||[||||||||||||]PAM</td><td>NA</td><td>Intergenic</td><td>NA</td><td>NA</td></tr>
<tr><td>T2258</td><td>TGGGGACGGGCACGGTCGGTTGG</td><td>1000</td><td></td><td></td><td></td><td></td><td></td><td></td><td></td><td></td><td></td></tr>
<tr><td>Oligo fwd</td><td>CACCTGGGGACGGGCACGGTCGGT</td><td></td><td></td><td></td><td></td><td></td><td></td><td></td><td></td><td></td><td></td></tr>
<tr><td>Oligo rev</td><td>AAACACCGACCGTGCCCGTCCCCA</td><td></td><td></td><td></td><td></td><td></td><td></td><td></td><td></td><td></td><td></td></tr>
<tr><td>Chromosome</td><td>start</td><td>end</td><td>strand</td><td>MM</td><td>target_seq</td><td>PAM</td><td>alignment</td><td>distance</td><td>position</td><td>gene name</td><td>gene id</td></tr>
<tr><td>GU305914.1</td><td>35568</td><td>35590</td><td>+</td><td>0</td><td>TGGGGACGGGCACGGTCGGT</td><td>TGG</td><td>||||||||[||||||||||||]PAM</td><td>NA</td><td>Intergenic</td><td>NA</td><td>NA</td></tr>
<tr><td>T2259</td><td>TTGGATAACCATCTCCGAGAAGG</td><td>1000</td><td></td><td></td><td></td><td></td><td></td><td></td><td></td><td></td><td></td></tr>
<tr><td>Oligo fwd</td><td>CACCTTGGATAACCATCTCCGAGA</td><td></td><td></td><td></td><td></td><td></td><td></td><td></td><td></td><td></td><td></td></tr>
<tr><td>Oligo rev</td><td>AAACTCTCGGAGATGGTTATCCAA</td><td></td><td></td><td></td><td></td><td></td><td></td><td></td><td></td><td></td><td></td></tr>
<tr><td>Chromosome</td><td>start</td><td>end</td><td>strand</td><td>MM</td><td>target_seq</td><td>PAM</td><td>alignment</td><td>distance</td><td>position</td><td>gene name</td><td>gene id</td></tr>
<tr><td>GU305914.1</td><td>35587</td><td>35609</td><td>+</td><td>0</td><td>TTGGATAACCATCTCCGAGA</td><td>AGG</td><td>||||||||[||||||||||||]PAM</td><td>NA</td><td>Intergenic</td><td>NA</td><td>NA</td></tr>
<tr><td>T2260</td><td>GATAACCATCTCCGAGAAGGCGG</td><td>1000</td><td></td><td></td><td></td><td></td><td></td><td></td><td></td><td></td><td></td></tr>
<tr><td>Oligo fwd</td><td>CACCGATAACCATCTCCGAGAAGG</td><td></td><td></td><td></td><td></td><td></td><td></td><td></td><td></td><td></td><td></td></tr>
<tr><td>Oligo rev</td><td>AAACCCTTCTCGGAGATGGTTATC</td><td></td><td></td><td></td><td></td><td></td><td></td><td></td><td></td><td></td><td></td></tr>
<tr><td>Chromosome</td><td>start</td><td>end</td><td>strand</td><td>MM</td><td>target_seq</td><td>PAM</td><td>alignment</td><td>distance</td><td>position</td><td>gene name</td><td>gene id</td></tr>
<tr><td>GU305914.1</td><td>35590</td><td>35612</td><td>+</td><td>0</td><td>GATAACCATCTCCGAGAAGG</td><td>CGG</td><td>||||||||[||||||||||||]PAM</td><td>NA</td><td>Intergenic</td><td>NA</td><td>NA</td></tr>
<tr><td>T2261</td><td>ATCTCCGAGAAGGCGGCCACAGG</td><td>1000</td><td></td><td></td><td></td><td></td><td></td><td></td><td></td><td></td><td></td></tr>
<tr><td>Oligo fwd</td><td>CACCATCTCCGAGAAGGCGGCCAC</td><td></td><td></td><td></td><td></td><td></td><td></td><td></td><td></td><td></td><td></td></tr>
<tr><td>Oligo rev</td><td>AAACGTGGCCGCCTTCTCGGAGAT</td><td></td><td></td><td></td><td></td><td></td><td></td><td></td><td></td><td></td><td></td></tr>
<tr><td>Chromosome</td><td>start</td><td>end</td><td>strand</td><td>MM</td><td>target_seq</td><td>PAM</td><td>alignment</td><td>distance</td><td>position</td><td>gene name</td><td>gene id</td></tr>
<tr><td>GU305914.1</td><td>35597</td><td>35619</td><td>+</td><td>0</td><td>ATCTCCGAGAAGGCGGCCAC</td><td>AGG</td><td>||||||||[||||||||||||]PAM</td><td>NA</td><td>Intergenic</td><td>NA</td><td>NA</td></tr>
<tr><td>T2262</td><td>TCCGAGAAGGCGGCCACAGGCGG</td><td>1000</td><td></td><td></td><td></td><td></td><td></td><td></td><td></td><td></td><td></td></tr>
<tr><td>Oligo fwd</td><td>CACCTCCGAGAAGGCGGCCACAGG</td><td></td><td></td><td></td><td></td><td></td><td></td><td></td><td></td><td></td><td></td></tr>
<tr><td>Oligo rev</td><td>AAACCCTGTGGCCGCCTTCTCGGA</td><td></td><td></td><td></td><td></td><td></td><td></td><td></td><td></td><td></td><td></td></tr>
<tr><td>Chromosome</td><td>start</td><td>end</td><td>strand</td><td>MM</td><td>target_seq</td><td>PAM</td><td>alignment</td><td>distance</td><td>position</td><td>gene name</td><td>gene id</td></tr>
<tr><td>GU305914.1</td><td>35600</td><td>35622</td><td>+</td><td>0</td><td>TCCGAGAAGGCGGCCACAGG</td><td>CGG</td><td>||||||||[||||||||||||]PAM</td><td>NA</td><td>Intergenic</td><td>NA</td><td>NA</td></tr>
<tr><td>T2263</td><td>AAGGCGGCCACAGGCGGAATCGG</td><td>1000</td><td></td><td></td><td></td><td></td><td></td><td></td><td></td><td></td><td></td></tr>
<tr><td>Oligo fwd</td><td>CACCAAGGCGGCCACAGGCGGAAT</td><td></td><td></td><td></td><td></td><td></td><td></td><td></td><td></td><td></td><td></td></tr>
<tr><td>Oligo rev</td><td>AAACATTCCGCCTGTGGCCGCCTT</td><td></td><td></td><td></td><td></td><td></td><td></td><td></td><td></td><td></td><td></td></tr>
<tr><td>Chromosome</td><td>start</td><td>end</td><td>strand</td><td>MM</td><td>target_seq</td><td>PAM</td><td>alignment</td><td>distance</td><td>position</td><td>gene name</td><td>gene id</td></tr>
<tr><td>GU305914.1</td><td>35606</td><td>35628</td><td>+</td><td>0</td><td>AAGGCGGCCACAGGCGGAAT</td><td>CGG</td><td>||||||||[||||||||||||]PAM</td><td>NA</td><td>Intergenic</td><td>NA</td><td>NA</td></tr>
<tr><td>T2264</td><td>GCGGCCACAGGCGGAATCGGTGG</td><td>1000</td><td></td><td></td><td></td><td></td><td></td><td></td><td></td><td></td><td></td></tr>
<tr><td>Oligo fwd</td><td>CACCGCGGCCACAGGCGGAATCGG</td><td></td><td></td><td></td><td></td><td></td><td></td><td></td><td></td><td></td><td></td></tr>
<tr><td>Oligo rev</td><td>AAACCCGATTCCGCCTGTGGCCGC</td><td></td><td></td><td></td><td></td><td></td><td></td><td></td><td></td><td></td><td></td></tr>
<tr><td>Chromosome</td><td>start</td><td>end</td><td>strand</td><td>MM</td><td>target_seq</td><td>PAM</td><td>alignment</td><td>distance</td><td>position</td><td>gene name</td><td>gene id</td></tr>
<tr><td>GU305914.1</td><td>35609</td><td>35631</td><td>+</td><td>0</td><td>GCGGCCACAGGCGGAATCGG</td><td>TGG</td><td>||||||||[||||||||||||]PAM</td><td>NA</td><td>Intergenic</td><td>NA</td><td>NA</td></tr>
<tr><td>T2265</td><td>ACAGGCGGAATCGGTGGCGTCGG</td><td>1000</td><td></td><td></td><td></td><td></td><td></td><td></td><td></td><td></td><td></td></tr>
<tr><td>Oligo fwd</td><td>CACCACAGGCGGAATCGGTGGCGT</td><td></td><td></td><td></td><td></td><td></td><td></td><td></td><td></td><td></td><td></td></tr>
<tr><td>Oligo rev</td><td>AAACACGCCACCGATTCCGCCTGT</td><td></td><td></td><td></td><td></td><td></td><td></td><td></td><td></td><td></td><td></td></tr>
<tr><td>Chromosome</td><td>start</td><td>end</td><td>strand</td><td>MM</td><td>target_seq</td><td>PAM</td><td>alignment</td><td>distance</td><td>position</td><td>gene name</td><td>gene id</td></tr>
<tr><td>GU305914.1</td><td>35615</td><td>35637</td><td>+</td><td>0</td><td>ACAGGCGGAATCGGTGGCGT</td><td>CGG</td><td>||||||||[||||||||||||]PAM</td><td>NA</td><td>Intergenic</td><td>NA</td><td>NA</td></tr>
<tr><td>T2266</td><td>GGCGGAATCGGTGGCGTCGGCGG</td><td>1000</td><td></td><td></td><td></td><td></td><td></td><td></td><td></td><td></td><td></td></tr>
<tr><td>Oligo fwd</td><td>CACCGGCGGAATCGGTGGCGTCGG</td><td></td><td></td><td></td><td></td><td></td><td></td><td></td><td></td><td></td><td></td></tr>
<tr><td>Oligo rev</td><td>AAACCCGACGCCACCGATTCCGCC</td><td></td><td></td><td></td><td></td><td></td><td></td><td></td><td></td><td></td><td></td></tr>
<tr><td>Chromosome</td><td>start</td><td>end</td><td>strand</td><td>MM</td><td>target_seq</td><td>PAM</td><td>alignment</td><td>distance</td><td>position</td><td>gene name</td><td>gene id</td></tr>
<tr><td>GU305914.1</td><td>35618</td><td>35640</td><td>+</td><td>0</td><td>GGCGGAATCGGTGGCGTCGG</td><td>CGG</td><td>||||||||[||||||||||||]PAM</td><td>NA</td><td>Intergenic</td><td>NA</td><td>NA</td></tr>
<tr><td>GU305914.1</td><td>48156</td><td>48178</td><td>+</td><td>4</td><td>GGCGGCGGCGGTGGCGGCGG</td><td>CGG</td><td>|||||---[||||||||-|||]PAM</td><td>NA</td><td>Intergenic</td><td>NA</td><td>NA</td></tr>
<tr><td>T2267</td><td>ATCGGTGGCGTCGGCGGTGTCGG</td><td>1000</td><td></td><td></td><td></td><td></td><td></td><td></td><td></td><td></td><td></td></tr>
<tr><td>Oligo fwd</td><td>CACCATCGGTGGCGTCGGCGGTGT</td><td></td><td></td><td></td><td></td><td></td><td></td><td></td><td></td><td></td><td></td></tr>
<tr><td>Oligo rev</td><td>AAACACACCGCCGACGCCACCGAT</td><td></td><td></td><td></td><td></td><td></td><td></td><td></td><td></td><td></td><td></td></tr>
<tr><td>Chromosome</td><td>start</td><td>end</td><td>strand</td><td>MM</td><td>target_seq</td><td>PAM</td><td>alignment</td><td>distance</td><td>position</td><td>gene name</td><td>gene id</td></tr>
<tr><td>GU305914.1</td><td>35624</td><td>35646</td><td>+</td><td>0</td><td>ATCGGTGGCGTCGGCGGTGT</td><td>CGG</td><td>||||||||[||||||||||||]PAM</td><td>NA</td><td>Intergenic</td><td>NA</td><td>NA</td></tr>
<tr><td>GU305914.1</td><td>48150</td><td>48172</td><td>+</td><td>4</td><td>ACCTGTGGCGGCGGCGGTGG</td><td>CGG</td><td>|-|-||||[||-||||||||-]PAM</td><td>NA</td><td>Intergenic</td><td>NA</td><td>NA</td></tr>
<tr><td>T2268</td><td>TGGCGTCGGCGGTGTCGGTGCGG</td><td>1000</td><td></td><td></td><td></td><td></td><td></td><td></td><td></td><td></td><td></td></tr>
<tr><td>Oligo fwd</td><td>CACCTGGCGTCGGCGGTGTCGGTG</td><td></td><td></td><td></td><td></td><td></td><td></td><td></td><td></td><td></td><td></td></tr>
<tr><td>Oligo rev</td><td>AAACCACCGACACCGCCGACGCCA</td><td></td><td></td><td></td><td></td><td></td><td></td><td></td><td></td><td></td><td></td></tr>
<tr><td>Chromosome</td><td>start</td><td>end</td><td>strand</td><td>MM</td><td>target_seq</td><td>PAM</td><td>alignment</td><td>distance</td><td>position</td><td>gene name</td><td>gene id</td></tr>
<tr><td>GU305914.1</td><td>35629</td><td>35651</td><td>+</td><td>0</td><td>TGGCGTCGGCGGTGTCGGTG</td><td>CGG</td><td>||||||||[||||||||||||]PAM</td><td>NA</td><td>Intergenic</td><td>NA</td><td>NA</td></tr>
<tr><td>GU305914.1</td><td>152555</td><td>152577</td><td>+</td><td>4</td><td>CGGCGGCGGCGGTGCCGGCG</td><td>GGG</td><td>-||||-||[||||||-|||-|]PAM</td><td>NA</td><td>Intergenic</td><td>NA</td><td>NA</td></tr>
<tr><td>T2269</td><td>GGCGTCGGCGGTGTCGGTGCGGG</td><td>1000</td><td></td><td></td><td></td><td></td><td></td><td></td><td></td><td></td><td></td></tr>
<tr><td>Oligo fwd</td><td>CACCGGCGTCGGCGGTGTCGGTGC</td><td></td><td></td><td></td><td></td><td></td><td></td><td></td><td></td><td></td><td></td></tr>
<tr><td>Oligo rev</td><td>AAACGCACCGACACCGCCGACGCC</td><td></td><td></td><td></td><td></td><td></td><td></td><td></td><td></td><td></td><td></td></tr>
<tr><td>Chromosome</td><td>start</td><td>end</td><td>strand</td><td>MM</td><td>target_seq</td><td>PAM</td><td>alignment</td><td>distance</td><td>position</td><td>gene name</td><td>gene id</td></tr>
<tr><td>GU305914.1</td><td>35630</td><td>35652</td><td>+</td><td>0</td><td>GGCGTCGGCGGTGTCGGTGC</td><td>GGG</td><td>||||||||[||||||||||||]PAM</td><td>NA</td><td>Intergenic</td><td>NA</td><td>NA</td></tr>
<tr><td>T2270</td><td>CGGTGTCGGTGCGGGATCCACGG</td><td>1000</td><td></td><td></td><td></td><td></td><td></td><td></td><td></td><td></td><td></td></tr>
<tr><td>Oligo fwd</td><td>CACCCGGTGTCGGTGCGGGATCCA</td><td></td><td></td><td></td><td></td><td></td><td></td><td></td><td></td><td></td><td></td></tr>
<tr><td>Oligo rev</td><td>AAACTGGATCCCGCACCGACACCG</td><td></td><td></td><td></td><td></td><td></td><td></td><td></td><td></td><td></td><td></td></tr>
<tr><td>Chromosome</td><td>start</td><td>end</td><td>strand</td><td>MM</td><td>target_seq</td><td>PAM</td><td>alignment</td><td>distance</td><td>position</td><td>gene name</td><td>gene id</td></tr>
<tr><td>GU305914.1</td><td>35638</td><td>35660</td><td>+</td><td>0</td><td>CGGTGTCGGTGCGGGATCCA</td><td>CGG</td><td>||||||||[||||||||||||]PAM</td><td>NA</td><td>Intergenic</td><td>NA</td><td>NA</td></tr>
<tr><td>T2271</td><td>TGTCGGTGCGGGATCCACGGCGG</td><td>1000</td><td></td><td></td><td></td><td></td><td></td><td></td><td></td><td></td><td></td></tr>
<tr><td>Oligo fwd</td><td>CACCTGTCGGTGCGGGATCCACGG</td><td></td><td></td><td></td><td></td><td></td><td></td><td></td><td></td><td></td><td></td></tr>
<tr><td>Oligo rev</td><td>AAACCCGTGGATCCCGCACCGACA</td><td></td><td></td><td></td><td></td><td></td><td></td><td></td><td></td><td></td><td></td></tr>
<tr><td>Chromosome</td><td>start</td><td>end</td><td>strand</td><td>MM</td><td>target_seq</td><td>PAM</td><td>alignment</td><td>distance</td><td>position</td><td>gene name</td><td>gene id</td></tr>
<tr><td>GU305914.1</td><td>35641</td><td>35663</td><td>+</td><td>0</td><td>TGTCGGTGCGGGATCCACGG</td><td>CGG</td><td>||||||||[||||||||||||]PAM</td><td>NA</td><td>Intergenic</td><td>NA</td><td>NA</td></tr>
</table>

| | | | | | | | | | | | |
|---|---|---|---|---|---|---|---|---|---|---|---|
| T2272 | CGGTGCGGGATCCACGGCGGCGG | 1000 | | | | | | | | | |
| Oligo fwd | CACCCGGTGCGGGATCCACGGCGG | | | | | | | | | | |
| Oligo rev | AAACCCGCCGTGGATCCCGCACCG | | | | | | | | | | |
| Chromosome | start | end | strand | MM | target_seq | PAM | alignment | distance | position | gene name | gene id |
| GU305914.1 | 35644 | 35666 | + | 0 | CGGTGCGGGATCCACGGCGG | CGG | \|\|\|\|\|\|\|\|[\|\|\|\|\|\|\|\|\|\|\|\|]PAM | NA | Intergenic | NA | NA |
| T2273 | TGCGGGATCCACGGCGGCGGCGG | 1000 | | | | | | | | | |
| Oligo fwd | CACCTGCGGGATCCACGGCGGCGG | | | | | | | | | | |
| Oligo rev | AAACCCGCCGCCGTGGATCCCGCA | | | | | | | | | | |
| Chromosome | start | end | strand | MM | target_seq | PAM | alignment | distance | position | gene name | gene id |
| GU305914.1 | 35647 | 35669 | + | 0 | TGCGGGATCCACGGCGGCGG | CGG | \|\|\|\|\|\|\|\|[\|\|\|\|\|\|\|\|\|\|\|\|]PAM | NA | Intergenic | NA | NA |
| T2274 | GGGATCCACGGCGGCGGCGGCGG | 1000 | | | | | | | | | |
| Oligo fwd | CACCGGGATCCACGGCGGCGGCGG | | | | | | | | | | |
| Oligo rev | AAACCCGCCGCCGCCGTGGATCCC | | | | | | | | | | |
| Chromosome | start | end | strand | MM | target_seq | PAM | alignment | distance | position | gene name | gene id |
| GU305914.1 | 35650 | 35672 | + | 0 | GGGATCCACGGCGGCGGCGG | CGG | \|\|\|\|\|\|\|\|[\|\|\|\|\|\|\|\|\|\|\|\|]PAM | NA | Intergenic | NA | NA |
| GU305914.1 | 67742 | 67764 | + | 3 | GGGAGCGGCGGCGGCGGCGG | CGG | \|\|\|\|-\|--[\|\|\|\|\|\|\|\|\|\|\|\|]PAM | NA | Intergenic | NA | NA |
| GU305914.1 | 19054 | 19076 | + | 4 | GGGTTGGACGACGGCGGCGG | TGG | \|\|\|-\|--\|[\|\|-\|\|\|\|\|\|\|\|\|]PAM | NA | Intergenic | NA | NA |
| T2275 | ATCCACGGCGGCGGCGGCGGCGG | 1000 | | | | | | | | | |
| Oligo fwd | CACCATCCACGGCGGCGGCGGCGG | | | | | | | | | | |
| Oligo rev | AAACCCGCCGCCGCCGCCGTGGAT | | | | | | | | | | |
| Chromosome | start | end | strand | MM | target_seq | PAM | alignment | distance | position | gene name | gene id |
| GU305914.1 | 35653 | 35675 | + | 0 | ATCCACGGCGGCGGCGGCGG | CGG | \|\|\|\|\|\|\|\|[\|\|\|\|\|\|\|\|\|\|\|\|]PAM | NA | Intergenic | NA | NA |
| GU305914.1 | 149482 | 149504 | - | 3 | GGCGACGGCGGCGGCGGCGG | CGG | --\|-\|\|\|\|[\|\|\|\|\|\|\|\|\|\|\|\|]PAM | NA | Intergenic | NA | NA |
| GU305914.1 | 56773 | 56795 | + | 4 | CGCCGCAGCGGCGGCGGCGG | CGG | --\|\|-\|-\|[\|\|\|\|\|\|\|\|\|\|\|\|]PAM | NA | Intergenic | NA | NA |
| GU305914.1 | 62431 | 62453 | + | 3 | TTCGACGGCGGTGGCGGCGG | GGG | -\|\|-\|\|\|\|[\|\|\|-\|\|\|\|\|\|\|\|]PAM | NA | Intergenic | NA | NA |
| GU305914.1 | 56770 | 56792 | + | 4 | CTCCGCCGCAGCGGCGGCGG | CGG | -\|\|\|-\|-\|[\|-\|\|\|\|\|\|\|\|\|\|]PAM | NA | Intergenic | NA | NA |
| GU305914.1 | 98352 | 98374 | - | 4 | AGCCTAGGCGGCGGTGGCGG | CGG | \|-\|\|--\|\|[\|\|\|\|\|\|-\|\|\|\|\|]PAM | NA | Intergenic | NA | NA |
| GU305914.1 | 24624 | 24646 | - | 4 | AACACCGGCGGCGGCGGTGG | CGG | \|-\|--\|\|\|[\|\|\|\|\|\|\|\|\|-\|\|]PAM | NA | Intergenic | NA | NA |
| GU305914.1 | 163902 | 163924 | + | 4 | AGCGGCGGCGGCGGCGGCAG | CGG | \|-\|--\|\|\|[\|\|\|\|\|\|\|\|\|\|-\|]PAM | NA | Intergenic | NA | NA |
| T2276 | TCCACGGCGGCGGCGGCGGCGGG | 1000 | | | | | | | | | |
| Oligo fwd | CACCTCCACGGCGGCGGCGGCGGC | | | | | | | | | | |
| Oligo rev | AAACGCCGCCGCCGCCGCCGTGGA | | | | | | | | | | |
| Chromosome | start | end | strand | MM | target_seq | PAM | alignment | distance | position | gene name | gene id |
| GU305914.1 | 35654 | 35676 | + | 0 | TCCACGGCGGCGGCGGCGGC | GGG | \|\|\|\|\|\|\|\|[\|\|\|\|\|\|\|\|\|\|\|\|]PAM | NA | Intergenic | NA | NA |
| GU305914.1 | 67743 | 67765 | + | 4 | GGAGCGGCGGCGGCGGCGGC | GGG | ----\|\|\|\|[\|\|\|\|\|\|\|\|\|\|\|\|]PAM | NA | Intergenic | NA | NA |
| GU305914.1 | 149475 | 149497 | - | 4 | GCGGCGGCGGCGGCGGCAGC | GGG | -\|--\|\|\|\|[\|\|\|\|\|\|\|\|\|-\|\|]PAM | NA | Intergenic | NA | NA |
| GU305914.1 | 225977 | 225999 | - | 4 | GCGACGGCGGCGGCGGGAGC | CGG | -\|-\|\|\|\|\|[\|\|\|\|\|\|\|\|--\|\|]PAM | NA | Intergenic | NA | NA |
| T2277 | CCACGGCGGCGGCGGCGGCGGGG | 1000 | | | | | | | | | |
| Oligo fwd | CACCCCACGGCGGCGGCGGCGGCG | | | | | | | | | | |
| Oligo rev | AAACCGCCGCCGCCGCCGCCGTGG | | | | | | | | | | |
| Chromosome | start | end | strand | MM | target_seq | PAM | alignment | distance | position | gene name | gene id |
| GU305914.1 | 35655 | 35677 | + | 0 | CCACGGCGGCGGCGGCGGCG | GGG | \|\|\|\|\|\|\|\|[\|\|\|\|\|\|\|\|\|\|\|\|]PAM | NA | Intergenic | NA | NA |
| GU305914.1 | 159954 | 159976 | - | 4 | TATCGGCGGCGACGGCGGCG | AGG | ---\|\|\|\|\|[\|\|\|-\|\|\|\|\|\|\|\|]PAM | NA | Intergenic | NA | NA |
| GU305914.1 | 75558 | 75580 | + | 4 | CGAGGGCGATGGCGGCGGCG | AGG | \|-\|-\|\|\|\|[--\|\|\|\|\|\|\|\|\|\|]PAM | NA | Intergenic | NA | NA |
| GU305914.1 | 152555 | 152577 | + | 4 | CGGCGGCGGCGGTGCCGGCG | GGG | \|--\|\|\|\|\|[\|\|\|\|-\|-\|\|\|\|\|]PAM | NA | Intergenic | NA | NA |
| GU305914.1 | 58121 | 58143 | - | 4 | TGGCGGCGGCGGCGGCGGAG | GGG | ---\|\|\|\|\|[\|\|\|\|\|\|\|\|\|\|-\|]PAM | NA | Intergenic | NA | NA |
| T2278 | GGCGGCGGCGGGGATGTCATCGG | 1000 | | | | | | | | | |
| Oligo fwd | CACCGGCGGCGGCGGGGATGTCAT | | | | | | | | | | |
| Oligo rev | AAACATGACATCCCCGCCGCCGCC | | | | | | | | | | |
| Chromosome | start | end | strand | MM | target_seq | PAM | alignment | distance | position | gene name | gene id |
| GU305914.1 | 35665 | 35687 | + | 0 | GGCGGCGGCGGGGATGTCAT | CGG | \|\|\|\|\|\|\|\|[\|\|\|\|\|\|\|\|\|\|\|\|]PAM | NA | Intergenic | NA | NA |
| GU305914.1 | 122058 | 122080 | + | 4 | GGAGGTGGCGGGGATGTCTG | GGG | \|\|-\|\|-\|\|[\|\|\|\|\|\|\|\|\|\|--]PAM | NA | Intergenic | NA | NA |
| T2279 | GCGGGGATGTCATCGGCGACAGG | 1000 | | | | | | | | | |
| Oligo fwd | CACCGCGGGGATGTCATCGGCGAC | | | | | | | | | | |
| Oligo rev | AAACGTCGCCGATGACATCCCCGC | | | | | | | | | | |
| Chromosome | start | end | strand | MM | target_seq | PAM | alignment | distance | position | gene name | gene id |
| GU305914.1 | 35672 | 35694 | + | 0 | GCGGGGATGTCATCGGCGAC | AGG | \|\|\|\|\|\|\|\|[\|\|\|\|\|\|\|\|\|\|\|\|]PAM | NA | Intergenic | NA | NA |
| GU305914.1 | 86020 | 86042 | - | 4 | ACGGCGATGAGATCGGCGAC | GGG | -\|\|\|-\|\|\|[\|--\|\|\|\|\|\|\|\|\|]PAM | NA | Intergenic | NA | NA |
| T2280 | CGGGGATGTCATCGGCGACAGGG | 1000 | | | | | | | | | |
| Oligo fwd | CACCCGGGGATGTCATCGGCGACA | | | | | | | | | | |
| Oligo rev | AAACTGTCGCCGATGACATCCCCG | | | | | | | | | | |
| Chromosome | start | end | strand | MM | target_seq | PAM | alignment | distance | position | gene name | gene id |
| GU305914.1 | 35673 | 35695 | + | 0 | CGGGGATGTCATCGGCGACA | GGG | \|\|\|\|\|\|\|\|[\|\|\|\|\|\|\|\|\|\|\|\|]PAM | NA | Intergenic | NA | NA |
| T2281 | CCCCCTGTTCATCGTCGCGAAGG | 1000 | | | | | | | | | |
| Oligo fwd | CACCCCCCCTGTTCATCGTCGCGA | | | | | | | | | | |
| Oligo rev | AAACTCGCGACGATGAACAGGGGG | | | | | | | | | | |
| Chromosome | start | end | strand | MM | target_seq | PAM | alignment | distance | position | gene name | gene id |
| GU305914.1 | 35697 | 35719 | + | 0 | CCCCCTGTTCATCGTCGCGA | AGG | \|\|\|\|\|\|\|\|[\|\|\|\|\|\|\|\|\|\|\|\|]PAM | NA | Intergenic | NA | NA |
| T2282 | CGCGAAGGAACACCTCATCGAGG | 1000 | | | | | | | | | |
| Oligo fwd | CACCCGCGAAGGAACACCTCATCG | | | | | | | | | | |
| Oligo rev | AAACCGATGAGGTGTTCCTTCGCG | | | | | | | | | | |
| Chromosome | start | end | strand | MM | target_seq | PAM | alignment | distance | position | gene name | gene id |
| GU305914.1 | 35712 | 35734 | + | 0 | CGCGAAGGAACACCTCATCG | AGG | \|\|\|\|\|\|\|\|[\|\|\|\|\|\|\|\|\|\|\|\|]PAM | NA | Intergenic | NA | NA |
| T2283 | TCCGCATACCGACAGCTTCCAGG | 1000 | | | | | | | | | |
| Oligo fwd | CACCTCCGCATACCGACAGCTTCC | | | | | | | | | | |
| Oligo rev | AAACGGAAGCTGTCGGTATGCGGA | | | | | | | | | | |
| Chromosome | start | end | strand | MM | target_seq | PAM | alignment | distance | position | gene name | gene id |

<table>
<tr><td>GU305914.1</td><td>35751</td><td>35773</td><td>+</td><td>0</td><td>TCCGCATACCGACAGCTTCC</td><td>AGG</td><td>||||||||[||||||||||||]PAM</td><td>NA</td><td>Intergenic</td><td>NA</td><td>NA</td></tr>
<tr><td>T2284</td><td>CATATCGACTGATTTTTATCTGG</td><td>1000</td><td></td><td></td><td></td><td></td><td></td><td></td><td></td><td></td><td></td></tr>
<tr><td>Oligo fwd</td><td>CACCCATATCGACTGATTTTTATC</td><td></td><td></td><td></td><td></td><td></td><td></td><td></td><td></td><td></td><td></td></tr>
<tr><td>Oligo rev</td><td>AAACGATAAAAATCAGTCGATATG</td><td></td><td></td><td></td><td></td><td></td><td></td><td></td><td></td><td></td><td></td></tr>
<tr><td>Chromosome</td><td>start</td><td>end</td><td>strand</td><td>MM</td><td>target_seq</td><td>PAM</td><td>alignment</td><td>distance</td><td>position</td><td>gene name</td><td>gene id</td></tr>
<tr><td>GU305914.1</td><td>35776</td><td>35798</td><td>+</td><td>0</td><td>CATATCGACTGATTTTTATC</td><td>TGG</td><td>||||||||[||||||||||||]PAM</td><td>NA</td><td>Intergenic</td><td>NA</td><td>NA</td></tr>
<tr><td>T2285</td><td>ATATCGACTGATTTTTATCTGGG</td><td>1000</td><td></td><td></td><td></td><td></td><td></td><td></td><td></td><td></td><td></td></tr>
<tr><td>Oligo fwd</td><td>CACCATATCGACTGATTTTTATCT</td><td></td><td></td><td></td><td></td><td></td><td></td><td></td><td></td><td></td><td></td></tr>
<tr><td>Oligo rev</td><td>AAACAGATAAAAATCAGTCGATAT</td><td></td><td></td><td></td><td></td><td></td><td></td><td></td><td></td><td></td><td></td></tr>
<tr><td>Chromosome</td><td>start</td><td>end</td><td>strand</td><td>MM</td><td>target_seq</td><td>PAM</td><td>alignment</td><td>distance</td><td>position</td><td>gene name</td><td>gene id</td></tr>
<tr><td>GU305914.1</td><td>35777</td><td>35799</td><td>+</td><td>0</td><td>ATATCGACTGATTTTTATCT</td><td>GGG</td><td>||||||||[||||||||||||]PAM</td><td>NA</td><td>Intergenic</td><td>NA</td><td>NA</td></tr>
<tr><td>T2286</td><td>GACTGATTTTTATCTGGGATCGG</td><td>1000</td><td></td><td></td><td></td><td></td><td></td><td></td><td></td><td></td><td></td></tr>
<tr><td>Oligo fwd</td><td>CACCGACTGATTTTTATCTGGGAT</td><td></td><td></td><td></td><td></td><td></td><td></td><td></td><td></td><td></td><td></td></tr>
<tr><td>Oligo rev</td><td>AAACATCCCAGATAAAAATCAGTC</td><td></td><td></td><td></td><td></td><td></td><td></td><td></td><td></td><td></td><td></td></tr>
<tr><td>Chromosome</td><td>start</td><td>end</td><td>strand</td><td>MM</td><td>target_seq</td><td>PAM</td><td>alignment</td><td>distance</td><td>position</td><td>gene name</td><td>gene id</td></tr>
<tr><td>GU305914.1</td><td>35782</td><td>35804</td><td>+</td><td>0</td><td>GACTGATTTTTATCTGGGAT</td><td>CGG</td><td>||||||||[||||||||||||]PAM</td><td>NA</td><td>Intergenic</td><td>NA</td><td>NA</td></tr>
<tr><td>T2287</td><td>AAAAGAGATAGAAATGCACACGG</td><td>1000</td><td></td><td></td><td></td><td></td><td></td><td></td><td></td><td></td><td></td></tr>
<tr><td>Oligo fwd</td><td>CACCAAAAGAGATAGAAATGCACA</td><td></td><td></td><td></td><td></td><td></td><td></td><td></td><td></td><td></td><td></td></tr>
<tr><td>Oligo rev</td><td>AAACTGTGCATTTCTATCTCTTTT</td><td></td><td></td><td></td><td></td><td></td><td></td><td></td><td></td><td></td><td></td></tr>
<tr><td>Chromosome</td><td>start</td><td>end</td><td>strand</td><td>MM</td><td>target_seq</td><td>PAM</td><td>alignment</td><td>distance</td><td>position</td><td>gene name</td><td>gene id</td></tr>
<tr><td>GU305914.1</td><td>35815</td><td>35837</td><td>+</td><td>0</td><td>AAAAGAGATAGAAATGCACA</td><td>CGG</td><td>||||||||[||||||||||||]PAM</td><td>NA</td><td>Intergenic</td><td>NA</td><td>NA</td></tr>
<tr><td>T2288</td><td>AAAAAAAGCTTTTACCCGAATGG</td><td>1000</td><td></td><td></td><td></td><td></td><td></td><td></td><td></td><td></td><td></td></tr>
<tr><td>Oligo fwd</td><td>CACCAAAAAAAGCTTTTACCCGAA</td><td></td><td></td><td></td><td></td><td></td><td></td><td></td><td></td><td></td><td></td></tr>
<tr><td>Oligo rev</td><td>AAACTTCGGGTAAAAGCTTTTTTT</td><td></td><td></td><td></td><td></td><td></td><td></td><td></td><td></td><td></td><td></td></tr>
<tr><td>Chromosome</td><td>start</td><td>end</td><td>strand</td><td>MM</td><td>target_seq</td><td>PAM</td><td>alignment</td><td>distance</td><td>position</td><td>gene name</td><td>gene id</td></tr>
<tr><td>GU305914.1</td><td>35838</td><td>35860</td><td>+</td><td>0</td><td>AAAAAAAGCTTTTACCCGAA</td><td>TGG</td><td>||||||||[||||||||||||]PAM</td><td>NA</td><td>Intergenic</td><td>NA</td><td>NA</td></tr>
<tr><td>T2289</td><td>ACCTCCTCTACGATCTGAGCTGG</td><td>1000</td><td></td><td></td><td></td><td></td><td></td><td></td><td></td><td></td><td></td></tr>
<tr><td>Oligo fwd</td><td>CACCACCTCCTCTACGATCTGAGC</td><td></td><td></td><td></td><td></td><td></td><td></td><td></td><td></td><td></td><td></td></tr>
<tr><td>Oligo rev</td><td>AAACGCTCAGATCGTAGAGGAGGT</td><td></td><td></td><td></td><td></td><td></td><td></td><td></td><td></td><td></td><td></td></tr>
<tr><td>Chromosome</td><td>start</td><td>end</td><td>strand</td><td>MM</td><td>target_seq</td><td>PAM</td><td>alignment</td><td>distance</td><td>position</td><td>gene name</td><td>gene id</td></tr>
<tr><td>GU305914.1</td><td>35916</td><td>35938</td><td>+</td><td>0</td><td>ACCTCCTCTACGATCTGAGC</td><td>TGG</td><td>||||||||[||||||||||||]PAM</td><td>NA</td><td>Intergenic</td><td>NA</td><td>NA</td></tr>
<tr><td>T2290</td><td>CTACGATCTGAGCTGGCAAATGG</td><td>1000</td><td></td><td></td><td></td><td></td><td></td><td></td><td></td><td></td><td></td></tr>
<tr><td>Oligo fwd</td><td>CACCCTACGATCTGAGCTGGCAAA</td><td></td><td></td><td></td><td></td><td></td><td></td><td></td><td></td><td></td><td></td></tr>
<tr><td>Oligo rev</td><td>AAACTTTGCCAGCTCAGATCGTAG</td><td></td><td></td><td></td><td></td><td></td><td></td><td></td><td></td><td></td><td></td></tr>
<tr><td>Chromosome</td><td>start</td><td>end</td><td>strand</td><td>MM</td><td>target_seq</td><td>PAM</td><td>alignment</td><td>distance</td><td>position</td><td>gene name</td><td>gene id</td></tr>
<tr><td>GU305914.1</td><td>35923</td><td>35945</td><td>+</td><td>0</td><td>CTACGATCTGAGCTGGCAAA</td><td>TGG</td><td>||||||||[||||||||||||]PAM</td><td>NA</td><td>Intergenic</td><td>NA</td><td>NA</td></tr>
<tr><td>T2291</td><td>AAATGGCGAAACCTCCCGCATGG</td><td>1000</td><td></td><td></td><td></td><td></td><td></td><td></td><td></td><td></td><td></td></tr>
<tr><td>Oligo fwd</td><td>CACCAAATGGCGAAACCTCCCGCA</td><td></td><td></td><td></td><td></td><td></td><td></td><td></td><td></td><td></td><td></td></tr>
<tr><td>Oligo rev</td><td>AAACTGCGGGAGGTTTCGCCATTT</td><td></td><td></td><td></td><td></td><td></td><td></td><td></td><td></td><td></td><td></td></tr>
<tr><td>Chromosome</td><td>start</td><td>end</td><td>strand</td><td>MM</td><td>target_seq</td><td>PAM</td><td>alignment</td><td>distance</td><td>position</td><td>gene name</td><td>gene id</td></tr>
<tr><td>GU305914.1</td><td>35940</td><td>35962</td><td>+</td><td>0</td><td>AAATGGCGAAACCTCCCGCA</td><td>TGG</td><td>||||||||[||||||||||||]PAM</td><td>NA</td><td>Intergenic</td><td>NA</td><td>NA</td></tr>
<tr><td>T2292</td><td>CCTCCCGCATGGCGACCGTTCGG</td><td>1000</td><td></td><td></td><td></td><td></td><td></td><td></td><td></td><td></td><td></td></tr>
<tr><td>Oligo fwd</td><td>CACCCCTCCCGCATGGCGACCGTT</td><td></td><td></td><td></td><td></td><td></td><td></td><td></td><td></td><td></td><td></td></tr>
<tr><td>Oligo rev</td><td>AAACAACGGTCGCCATGCGGGAGG</td><td></td><td></td><td></td><td></td><td></td><td></td><td></td><td></td><td></td><td></td></tr>
<tr><td>Chromosome</td><td>start</td><td>end</td><td>strand</td><td>MM</td><td>target_seq</td><td>PAM</td><td>alignment</td><td>distance</td><td>position</td><td>gene name</td><td>gene id</td></tr>
<tr><td>GU305914.1</td><td>35951</td><td>35973</td><td>+</td><td>0</td><td>CCTCCCGCATGGCGACCGTT</td><td>CGG</td><td>||||||||[||||||||||||]PAM</td><td>NA</td><td>Intergenic</td><td>NA</td><td>NA</td></tr>
<tr><td>T2293</td><td>GACCGTTCGGTGCTCTGTCGCGG</td><td>1000</td><td></td><td></td><td></td><td></td><td></td><td></td><td></td><td></td><td></td></tr>
<tr><td>Oligo fwd</td><td>CACCGACCGTTCGGTGCTCTGTCG</td><td></td><td></td><td></td><td></td><td></td><td></td><td></td><td></td><td></td><td></td></tr>
<tr><td>Oligo rev</td><td>AAACCGACAGAGCACCGAACGGTC</td><td></td><td></td><td></td><td></td><td></td><td></td><td></td><td></td><td></td><td></td></tr>
<tr><td>Chromosome</td><td>start</td><td>end</td><td>strand</td><td>MM</td><td>target_seq</td><td>PAM</td><td>alignment</td><td>distance</td><td>position</td><td>gene name</td><td>gene id</td></tr>
<tr><td>GU305914.1</td><td>35964</td><td>35986</td><td>+</td><td>0</td><td>GACCGTTCGGTGCTCTGTCG</td><td>CGG</td><td>||||||||[||||||||||||]PAM</td><td>NA</td><td>Intergenic</td><td>NA</td><td>NA</td></tr>
<tr><td>T2294</td><td>CGGTGCTCTGTCGCGGCCGCAGG</td><td>1000</td><td></td><td></td><td></td><td></td><td></td><td></td><td></td><td></td><td></td></tr>
<tr><td>Oligo fwd</td><td>CACCCGGTGCTCTGTCGCGGCCGC</td><td></td><td></td><td></td><td></td><td></td><td></td><td></td><td></td><td></td><td></td></tr>
<tr><td>Oligo rev</td><td>AAACGCGGCCGCGACAGAGCACCG</td><td></td><td></td><td></td><td></td><td></td><td></td><td></td><td></td><td></td><td></td></tr>
<tr><td>Chromosome</td><td>start</td><td>end</td><td>strand</td><td>MM</td><td>target_seq</td><td>PAM</td><td>alignment</td><td>distance</td><td>position</td><td>gene name</td><td>gene id</td></tr>
<tr><td>GU305914.1</td><td>35971</td><td>35993</td><td>+</td><td>0</td><td>CGGTGCTCTGTCGCGGCCGC</td><td>AGG</td><td>||||||||[||||||||||||]PAM</td><td>NA</td><td>Intergenic</td><td>NA</td><td>NA</td></tr>
<tr><td>T2295</td><td>CTGTCGCGGCCGCAGGTCGTCGG</td><td>1000</td><td></td><td></td><td></td><td></td><td></td><td></td><td></td><td></td><td></td></tr>
<tr><td>Oligo fwd</td><td>CACCCTGTCGCGGCCGCAGGTCGT</td><td></td><td></td><td></td><td></td><td></td><td></td><td></td><td></td><td></td><td></td></tr>
<tr><td>Oligo rev</td><td>AAACACGACCTGCGGCCGCGACAG</td><td></td><td></td><td></td><td></td><td></td><td></td><td></td><td></td><td></td><td></td></tr>
<tr><td>Chromosome</td><td>start</td><td>end</td><td>strand</td><td>MM</td><td>target_seq</td><td>PAM</td><td>alignment</td><td>distance</td><td>position</td><td>gene name</td><td>gene id</td></tr>
<tr><td>GU305914.1</td><td>35978</td><td>36000</td><td>+</td><td>0</td><td>CTGTCGCGGCCGCAGGTCGT</td><td>CGG</td><td>||||||||[||||||||||||]PAM</td><td>NA</td><td>Intergenic</td><td>NA</td><td>NA</td></tr>
<tr><td>T2296</td><td>CTCCTGCCTCACTTACTAGATGG</td><td>1000</td><td></td><td></td><td></td><td></td><td></td><td></td><td></td><td></td><td></td></tr>
<tr><td>Oligo fwd</td><td>CACCCTCCTGCCTCACTTACTAGA</td><td></td><td></td><td></td><td></td><td></td><td></td><td></td><td></td><td></td><td></td></tr>
<tr><td>Oligo rev</td><td>AAACTCTAGTAAGTGAGGCAGGAG</td><td></td><td></td><td></td><td></td><td></td><td></td><td></td><td></td><td></td><td></td></tr>
<tr><td>Chromosome</td><td>start</td><td>end</td><td>strand</td><td>MM</td><td>target_seq</td><td>PAM</td><td>alignment</td><td>distance</td><td>position</td><td>gene name</td><td>gene id</td></tr>
<tr><td>GU305914.1</td><td>36014</td><td>36036</td><td>+</td><td>0</td><td>CTCCTGCCTCACTTACTAGA</td><td>TGG</td><td>||||||||[||||||||||||]PAM</td><td>NA</td><td>Intergenic</td><td>NA</td><td>NA</td></tr>
<tr><td>T2297</td><td>CTGCCTCACTTACTAGATGGTGG</td><td>1000</td><td></td><td></td><td></td><td></td><td></td><td></td><td></td><td></td><td></td></tr>
<tr><td>Oligo fwd</td><td>CACCCTGCCTCACTTACTAGATGG</td><td></td><td></td><td></td><td></td><td></td><td></td><td></td><td></td><td></td><td></td></tr>
<tr><td>Oligo rev</td><td>AAACCCATCTAGTAAGTGAGGCAG</td><td></td><td></td><td></td><td></td><td></td><td></td><td></td><td></td><td></td><td></td></tr>
<tr><td>Chromosome</td><td>start</td><td>end</td><td>strand</td><td>MM</td><td>target_seq</td><td>PAM</td><td>alignment</td><td>distance</td><td>position</td><td>gene name</td><td>gene id</td></tr>
<tr><td>GU305914.1</td><td>36017</td><td>36039</td><td>+</td><td>0</td><td>CTGCCTCACTTACTAGATGG</td><td>TGG</td><td>||||||||[||||||||||||]PAM</td><td>NA</td><td>Intergenic</td><td>NA</td><td>NA</td></tr>
<tr><td>T2298</td><td>CTCCTGCTCGATTTTCTGATCGG</td><td>1000</td><td></td><td></td><td></td><td></td><td></td><td></td><td></td><td></td><td></td></tr>
<tr><td>Oligo fwd</td><td>CACCCTCCTGCTCGATTTTCTGAT</td><td></td><td></td><td></td><td></td><td></td><td></td><td></td><td></td><td></td><td></td></tr>
<tr><td>Oligo rev</td><td>AAACATCAGAAAATCGAGCAGGAG</td><td></td><td></td><td></td><td></td><td></td><td></td><td></td><td></td><td></td><td></td></tr>
</table>

<table>
<tr><td>Chromosome</td><td>start</td><td></td><td>end</td><td>strand</td><td>MM</td><td>target_seq</td><td>PAM</td><td>alignment</td><td>distance</td><td>position</td><td>gene name</td><td>gene id</td></tr>
<tr><td>GU305914.1</td><td></td><td>36045</td><td>36067</td><td>+</td><td>0</td><td>CTCCTGCTCGATTTTCTGAT</td><td>CGG</td><td>||||||||[||||||||||||]PAM</td><td>NA</td><td>Intergenic</td><td>NA</td><td>NA</td></tr>
<tr><td>T2299</td><td>TCCTGCTCGATTTTCTGATCGGG</td><td>1000</td><td colspan="10"></td></tr>
<tr><td>Oligo fwd</td><td>CACCTCCTGCTCGATTTTCTGATC</td><td colspan="11"></td></tr>
<tr><td>Oligo rev</td><td>AAACGATCAGAAAATCGAGCAGGA</td><td colspan="11"></td></tr>
<tr><td>Chromosome</td><td>start</td><td></td><td>end</td><td>strand</td><td>MM</td><td>target_seq</td><td>PAM</td><td>alignment</td><td>distance</td><td>position</td><td>gene name</td><td>gene id</td></tr>
<tr><td>GU305914.1</td><td></td><td>36046</td><td>36068</td><td>+</td><td>0</td><td>TCCTGCTCGATTTTCTGATC</td><td>GGG</td><td>||||||||[||||||||||||]PAM</td><td>NA</td><td>Intergenic</td><td>NA</td><td>NA</td></tr>
<tr><td>T2300</td><td>CTCGATTTTCTGATCGGGACAGG</td><td>1000</td><td colspan="10"></td></tr>
<tr><td>Oligo fwd</td><td>CACCCTCGATTTTCTGATCGGGAC</td><td colspan="11"></td></tr>
<tr><td>Oligo rev</td><td>AAACGTCCCGATCAGAAAATCGAG</td><td colspan="11"></td></tr>
<tr><td>Chromosome</td><td>start</td><td></td><td>end</td><td>strand</td><td>MM</td><td>target_seq</td><td>PAM</td><td>alignment</td><td>distance</td><td>position</td><td>gene name</td><td>gene id</td></tr>
<tr><td>GU305914.1</td><td></td><td>36051</td><td>36073</td><td>+</td><td>0</td><td>CTCGATTTTCTGATCGGGAC</td><td>AGG</td><td>||||||||[||||||||||||]PAM</td><td>NA</td><td>Intergenic</td><td>NA</td><td>NA</td></tr>
<tr><td>T2301</td><td>GATTTTCTGATCGGGACAGGAGG</td><td>1000</td><td colspan="10"></td></tr>
<tr><td>Oligo fwd</td><td>CACCGATTTTCTGATCGGGACAGG</td><td colspan="11"></td></tr>
<tr><td>Oligo rev</td><td>AAACCCTGTCCCGATCAGAAAATC</td><td colspan="11"></td></tr>
<tr><td>Chromosome</td><td>start</td><td></td><td>end</td><td>strand</td><td>MM</td><td>target_seq</td><td>PAM</td><td>alignment</td><td>distance</td><td>position</td><td>gene name</td><td>gene id</td></tr>
<tr><td>GU305914.1</td><td></td><td>36054</td><td>36076</td><td>+</td><td>0</td><td>GATTTTCTGATCGGGACAGG</td><td>AGG</td><td>||||||||[||||||||||||]PAM</td><td>NA</td><td>Intergenic</td><td>NA</td><td>NA</td></tr>
<tr><td>T2302</td><td>TTTCTGATCGGGACAGGAGGTGG</td><td>1000</td><td colspan="10"></td></tr>
<tr><td>Oligo fwd</td><td>CACCTTTCTGATCGGGACAGGAGG</td><td colspan="11"></td></tr>
<tr><td>Oligo rev</td><td>AAACCCTCCTGTCCCGATCAGAAA</td><td colspan="11"></td></tr>
<tr><td>Chromosome</td><td>start</td><td></td><td>end</td><td>strand</td><td>MM</td><td>target_seq</td><td>PAM</td><td>alignment</td><td>distance</td><td>position</td><td>gene name</td><td>gene id</td></tr>
<tr><td>GU305914.1</td><td></td><td>36057</td><td>36079</td><td>+</td><td>0</td><td>TTTCTGATCGGGACAGGAGG</td><td>TGG</td><td>||||||||[||||||||||||]PAM</td><td>NA</td><td>Intergenic</td><td>NA</td><td>NA</td></tr>
<tr><td>T2303</td><td>GGTTGTCTCTTCTCGCCGATAGG</td><td>1000</td><td colspan="10"></td></tr>
<tr><td>Oligo fwd</td><td>CACCGGTTGTCTCTTCTCGCCGAT</td><td colspan="11"></td></tr>
<tr><td>Oligo rev</td><td>AAACATCGGCGAGAAGAGACAACC</td><td colspan="11"></td></tr>
<tr><td>Chromosome</td><td>start</td><td></td><td>end</td><td>strand</td><td>MM</td><td>target_seq</td><td>PAM</td><td>alignment</td><td>distance</td><td>position</td><td>gene name</td><td>gene id</td></tr>
<tr><td>GU305914.1</td><td></td><td>36078</td><td>36100</td><td>+</td><td>0</td><td>GGTTGTCTCTTCTCGCCGAT</td><td>AGG</td><td>||||||||[||||||||||||]PAM</td><td>NA</td><td>Intergenic</td><td>NA</td><td>NA</td></tr>
<tr><td>T2304</td><td>GTTGTCTCTTCTCGCCGATAGGG</td><td>1000</td><td colspan="10"></td></tr>
<tr><td>Oligo fwd</td><td>CACCGTTGTCTCTTCTCGCCGATA</td><td colspan="11"></td></tr>
<tr><td>Oligo rev</td><td>AAACTATCGGCGAGAAGAGACAAC</td><td colspan="11"></td></tr>
<tr><td>Chromosome</td><td>start</td><td></td><td>end</td><td>strand</td><td>MM</td><td>target_seq</td><td>PAM</td><td>alignment</td><td>distance</td><td>position</td><td>gene name</td><td>gene id</td></tr>
<tr><td>GU305914.1</td><td></td><td>36079</td><td>36101</td><td>+</td><td>0</td><td>GTTGTCTCTTCTCGCCGATA</td><td>GGG</td><td>||||||||[||||||||||||]PAM</td><td>NA</td><td>Intergenic</td><td>NA</td><td>NA</td></tr>
<tr><td>T2305</td><td>GGGACACTCCTCACGAACAGAGG</td><td>1000</td><td colspan="10"></td></tr>
<tr><td>Oligo fwd</td><td>CACCGGGACACTCCTCACGAACAG</td><td colspan="11"></td></tr>
<tr><td>Oligo rev</td><td>AAACCTGTTCGTGAGGAGTGTCCC</td><td colspan="11"></td></tr>
<tr><td>Chromosome</td><td>start</td><td></td><td>end</td><td>strand</td><td>MM</td><td>target_seq</td><td>PAM</td><td>alignment</td><td>distance</td><td>position</td><td>gene name</td><td>gene id</td></tr>
<tr><td>GU305914.1</td><td></td><td>36099</td><td>36121</td><td>+</td><td>0</td><td>GGGACACTCCTCACGAACAG</td><td>AGG</td><td>||||||||[||||||||||||]PAM</td><td>NA</td><td>Intergenic</td><td>NA</td><td>NA</td></tr>
<tr><td>T2306</td><td>ACTCCTCACGAACAGAGGAGCGG</td><td>1000</td><td colspan="10"></td></tr>
<tr><td>Oligo fwd</td><td>CACCACTCCTCACGAACAGAGGAG</td><td colspan="11"></td></tr>
<tr><td>Oligo rev</td><td>AAACCTCCTCTGTTCGTGAGGAGT</td><td colspan="11"></td></tr>
<tr><td>Chromosome</td><td>start</td><td></td><td>end</td><td>strand</td><td>MM</td><td>target_seq</td><td>PAM</td><td>alignment</td><td>distance</td><td>position</td><td>gene name</td><td>gene id</td></tr>
<tr><td>GU305914.1</td><td></td><td>36104</td><td>36126</td><td>+</td><td>0</td><td>ACTCCTCACGAACAGAGGAG</td><td>CGG</td><td>||||||||[||||||||||||]PAM</td><td>NA</td><td>Intergenic</td><td>NA</td><td>NA</td></tr>
<tr><td>T2307</td><td>GAACAGAGGAGCGGCGTTAGTGG</td><td>1000</td><td colspan="10"></td></tr>
<tr><td>Oligo fwd</td><td>CACCGAACAGAGGAGCGGCGTTAG</td><td colspan="11"></td></tr>
<tr><td>Oligo rev</td><td>AAACCTAACGCCGCTCCTCTGTTC</td><td colspan="11"></td></tr>
<tr><td>Chromosome</td><td>start</td><td></td><td>end</td><td>strand</td><td>MM</td><td>target_seq</td><td>PAM</td><td>alignment</td><td>distance</td><td>position</td><td>gene name</td><td>gene id</td></tr>
<tr><td>GU305914.1</td><td></td><td>36113</td><td>36135</td><td>+</td><td>0</td><td>GAACAGAGGAGCGGCGTTAG</td><td>TGG</td><td>||||||||[||||||||||||]PAM</td><td>NA</td><td>Intergenic</td><td>NA</td><td>NA</td></tr>
<tr><td>T2308</td><td>GGCGTTAGTGGAAACAACCGTGG</td><td>1000</td><td colspan="10"></td></tr>
<tr><td>Oligo fwd</td><td>CACCGGCGTTAGTGGAAACAACCG</td><td colspan="11"></td></tr>
<tr><td>Oligo rev</td><td>AAACCGGTTGTTTCCACTAACGCC</td><td colspan="11"></td></tr>
<tr><td>Chromosome</td><td>start</td><td></td><td>end</td><td>strand</td><td>MM</td><td>target_seq</td><td>PAM</td><td>alignment</td><td>distance</td><td>position</td><td>gene name</td><td>gene id</td></tr>
<tr><td>GU305914.1</td><td></td><td>36125</td><td>36147</td><td>+</td><td>0</td><td>GGCGTTAGTGGAAACAACCG</td><td>TGG</td><td>||||||||[||||||||||||]PAM</td><td>NA</td><td>Intergenic</td><td>NA</td><td>NA</td></tr>
<tr><td>T2309</td><td>GTTAGTGGAAACAACCGTGGCGG</td><td>1000</td><td colspan="10"></td></tr>
<tr><td>Oligo fwd</td><td>CACCGTTAGTGGAAACAACCGTGG</td><td colspan="11"></td></tr>
<tr><td>Oligo rev</td><td>AAACCCACGGTTGTTTCCACTAAC</td><td colspan="11"></td></tr>
<tr><td>Chromosome</td><td>start</td><td></td><td>end</td><td>strand</td><td>MM</td><td>target_seq</td><td>PAM</td><td>alignment</td><td>distance</td><td>position</td><td>gene name</td><td>gene id</td></tr>
<tr><td>GU305914.1</td><td></td><td>36128</td><td>36150</td><td>+</td><td>0</td><td>GTTAGTGGAAACAACCGTGG</td><td>CGG</td><td>||||||||[||||||||||||]PAM</td><td>NA</td><td>Intergenic</td><td>NA</td><td>NA</td></tr>
<tr><td>T2310</td><td>GGAAACAACCGTGGCGGCAACGG</td><td>1000</td><td colspan="10"></td></tr>
<tr><td>Oligo fwd</td><td>CACCGGAAACAACCGTGGCGGCAA</td><td colspan="11"></td></tr>
<tr><td>Oligo rev</td><td>AAACTTGCCGCCACGGTTGTTTCC</td><td colspan="11"></td></tr>
<tr><td>Chromosome</td><td>start</td><td></td><td>end</td><td>strand</td><td>MM</td><td>target_seq</td><td>PAM</td><td>alignment</td><td>distance</td><td>position</td><td>gene name</td><td>gene id</td></tr>
<tr><td>GU305914.1</td><td></td><td>36134</td><td>36156</td><td>+</td><td>0</td><td>GGAAACAACCGTGGCGGCAA</td><td>CGG</td><td>||||||||[||||||||||||]PAM</td><td>NA</td><td>Intergenic</td><td>NA</td><td>NA</td></tr>
<tr><td>T2311</td><td>GAAACAACCGTGGCGGCAACGGG</td><td>1000</td><td colspan="10"></td></tr>
<tr><td>Oligo fwd</td><td>CACCGAAACAACCGTGGCGGCAAC</td><td colspan="11"></td></tr>
<tr><td>Oligo rev</td><td>AAACGTTGCCGCCACGGTTGTTTC</td><td colspan="11"></td></tr>
<tr><td>Chromosome</td><td>start</td><td></td><td>end</td><td>strand</td><td>MM</td><td>target_seq</td><td>PAM</td><td>alignment</td><td>distance</td><td>position</td><td>gene name</td><td>gene id</td></tr>
<tr><td>GU305914.1</td><td></td><td>36135</td><td>36157</td><td>+</td><td>0</td><td>GAAACAACCGTGGCGGCAAC</td><td>GGG</td><td>||||||||[||||||||||||]PAM</td><td>NA</td><td>Intergenic</td><td>NA</td><td>NA</td></tr>
<tr><td>T2312</td><td>AAACAACCGTGGCGGCAACGGGG</td><td>1000</td><td colspan="10"></td></tr>
<tr><td>Oligo fwd</td><td>CACCAAACAACCGTGGCGGCAACG</td><td colspan="11"></td></tr>
<tr><td>Oligo rev</td><td>AAACCGTTGCCGCCACGGTTGTTT</td><td colspan="11"></td></tr>
<tr><td>Chromosome</td><td>start</td><td></td><td>end</td><td>strand</td><td>MM</td><td>target_seq</td><td>PAM</td><td>alignment</td><td>distance</td><td>position</td><td>gene name</td><td>gene id</td></tr>
<tr><td>GU305914.1</td><td></td><td>36136</td><td>36158</td><td>+</td><td>0</td><td>AAACAACCGTGGCGGCAACG</td><td>GGG</td><td>||||||||[||||||||||||]PAM</td><td>NA</td><td>Intergenic</td><td>NA</td><td>NA</td></tr>
<tr><td>T2313</td><td>GGCGGCAACGGGGCCGCTGCCGG</td><td>1000</td><td colspan="10"></td></tr>
<tr><td>Oligo fwd</td><td>CACCGGCGGCAACGGGGCCGCTGC</td><td colspan="11"></td></tr>
</table>

<table>
<tr><td>Oligo rev</td><td>AAACGCAGCGGCCCCGTTGCCGCC</td><td></td><td></td><td></td><td></td><td></td><td></td><td></td><td></td><td></td><td></td></tr>
<tr><td>Chromosome</td><td>start</td><td>end</td><td>strand</td><td>MM</td><td>target_seq</td><td>PAM</td><td>alignment</td><td>distance</td><td>position</td><td>gene name</td><td>gene id</td></tr>
<tr><td>GU305914.1</td><td>36146</td><td>36168</td><td>+</td><td>0</td><td>GGCGGCAACGGGGCCGCTGC</td><td>CGG</td><td>||||||||[||||||||||||]PAM</td><td>NA</td><td>Intergenic</td><td>NA</td><td>NA</td></tr>
<tr><td>T2314</td><td>GGCAACGGGGCCGCTGCCGGCGG</td><td>1000</td><td></td><td></td><td></td><td></td><td></td><td></td><td></td><td></td><td></td></tr>
<tr><td>Oligo fwd</td><td>CACCGGCAACGGGGCCGCTGCCGG</td><td></td><td></td><td></td><td></td><td></td><td></td><td></td><td></td><td></td><td></td></tr>
<tr><td>Oligo rev</td><td>AAACCCGGCAGCGGCCCCGTTGCC</td><td></td><td></td><td></td><td></td><td></td><td></td><td></td><td></td><td></td><td></td></tr>
<tr><td>Chromosome</td><td>start</td><td>end</td><td>strand</td><td>MM</td><td>target_seq</td><td>PAM</td><td>alignment</td><td>distance</td><td>position</td><td>gene name</td><td>gene id</td></tr>
<tr><td>GU305914.1</td><td>36149</td><td>36171</td><td>+</td><td>0</td><td>GGCAACGGGGCCGCTGCCGG</td><td>CGG</td><td>||||||||[||||||||||||]PAM</td><td>NA</td><td>Intergenic</td><td>NA</td><td>NA</td></tr>
<tr><td>T2315</td><td>AACGGGGCCGCTGCCGGCGGCGG</td><td>1000</td><td></td><td></td><td></td><td></td><td></td><td></td><td></td><td></td><td></td></tr>
<tr><td>Oligo fwd</td><td>CACCAACGGGGCCGCTGCCGGCGG</td><td></td><td></td><td></td><td></td><td></td><td></td><td></td><td></td><td></td><td></td></tr>
<tr><td>Oligo rev</td><td>AAACCCGCCGGCAGCGGCCCCGTT</td><td></td><td></td><td></td><td></td><td></td><td></td><td></td><td></td><td></td><td></td></tr>
<tr><td>Chromosome</td><td>start</td><td>end</td><td>strand</td><td>MM</td><td>target_seq</td><td>PAM</td><td>alignment</td><td>distance</td><td>position</td><td>gene name</td><td>gene id</td></tr>
<tr><td>GU305914.1</td><td>36152</td><td>36174</td><td>+</td><td>0</td><td>AACGGGGCCGCTGCCGGCGG</td><td>CGG</td><td>||||||||[||||||||||||]PAM</td><td>NA</td><td>Intergenic</td><td>NA</td><td>NA</td></tr>
<tr><td>T2316</td><td>GCCGCTGCCGGCGGCGGTTGTGG</td><td>1000</td><td></td><td></td><td></td><td></td><td></td><td></td><td></td><td></td><td></td></tr>
<tr><td>Oligo fwd</td><td>CACCGCCGCTGCCGGCGGCGGTTG</td><td></td><td></td><td></td><td></td><td></td><td></td><td></td><td></td><td></td><td></td></tr>
<tr><td>Oligo rev</td><td>AAACCAACCGCCGCCGGCAGCGGC</td><td></td><td></td><td></td><td></td><td></td><td></td><td></td><td></td><td></td><td></td></tr>
<tr><td>Chromosome</td><td>start</td><td>end</td><td>strand</td><td>MM</td><td>target_seq</td><td>PAM</td><td>alignment</td><td>distance</td><td>position</td><td>gene name</td><td>gene id</td></tr>
<tr><td>GU305914.1</td><td>36158</td><td>36180</td><td>+</td><td>0</td><td>GCCGCTGCCGGCGGCGGTTG</td><td>TGG</td><td>||||||||[||||||||||||]PAM</td><td>NA</td><td>Intergenic</td><td>NA</td><td>NA</td></tr>
<tr><td>GU305914.1</td><td>52719</td><td>52741</td><td>+</td><td>3</td><td>GCCGCTGGCGGTGGCGGTGG</td><td>CGG</td><td>|||||||-[|||-||||||-|]PAM</td><td>NA</td><td>Intergenic</td><td>NA</td><td>NA</td></tr>
<tr><td>T2317</td><td>GCTGCCGGCGGCGGTTGTGGTGG</td><td>1000</td><td></td><td></td><td></td><td></td><td></td><td></td><td></td><td></td><td></td></tr>
<tr><td>Oligo fwd</td><td>CACCGCTGCCGGCGGCGGTTGTGG</td><td></td><td></td><td></td><td></td><td></td><td></td><td></td><td></td><td></td><td></td></tr>
<tr><td>Oligo rev</td><td>AAACCCACAACCGCCGCCGGCAGC</td><td></td><td></td><td></td><td></td><td></td><td></td><td></td><td></td><td></td><td></td></tr>
<tr><td>Chromosome</td><td>start</td><td>end</td><td>strand</td><td>MM</td><td>target_seq</td><td>PAM</td><td>alignment</td><td>distance</td><td>position</td><td>gene name</td><td>gene id</td></tr>
<tr><td>GU305914.1</td><td>36161</td><td>36183</td><td>+</td><td>0</td><td>GCTGCCGGCGGCGGTTGTGG</td><td>TGG</td><td>||||||||[||||||||||||]PAM</td><td>NA</td><td>Intergenic</td><td>NA</td><td>NA</td></tr>
<tr><td>GU305914.1</td><td>225498</td><td>225520</td><td>+</td><td>4</td><td>GCCGAGGGCGGCGGTGGTGG</td><td>TGG</td><td>||-|--||[|||||||-||||]PAM</td><td>NA</td><td>Intergenic</td><td>NA</td><td>NA</td></tr>
<tr><td>GU305914.1</td><td>62440</td><td>62462</td><td>+</td><td>4</td><td>GGTGGCGGCGGGGGTGGTGG</td><td>CGG</td><td>|-||-|||[|||-|||-||||]PAM</td><td>NA</td><td>Intergenic</td><td>NA</td><td>NA</td></tr>
<tr><td>T2318</td><td>GGCGGTTGTGGTGGTTGTGTTGG</td><td>1000</td><td></td><td></td><td></td><td></td><td></td><td></td><td></td><td></td><td></td></tr>
<tr><td>Oligo fwd</td><td>CACCGGCGGTTGTGGTGGTTGTGT</td><td></td><td></td><td></td><td></td><td></td><td></td><td></td><td></td><td></td><td></td></tr>
<tr><td>Oligo rev</td><td>AAACACACAACCACCACAACCGCC</td><td></td><td></td><td></td><td></td><td></td><td></td><td></td><td></td><td></td><td></td></tr>
<tr><td>Chromosome</td><td>start</td><td>end</td><td>strand</td><td>MM</td><td>target_seq</td><td>PAM</td><td>alignment</td><td>distance</td><td>position</td><td>gene name</td><td>gene id</td></tr>
<tr><td>GU305914.1</td><td>36170</td><td>36192</td><td>+</td><td>0</td><td>GGCGGTTGTGGTGGTTGTGT</td><td>TGG</td><td>||||||||[||||||||||||]PAM</td><td>NA</td><td>Intergenic</td><td>NA</td><td>NA</td></tr>
<tr><td>T2319</td><td>GGTTGTGGTGGTTGTGTTGGTGG</td><td>1000</td><td></td><td></td><td></td><td></td><td></td><td></td><td></td><td></td><td></td></tr>
<tr><td>Oligo fwd</td><td>CACCGGTTGTGGTGGTTGTGTTGG</td><td></td><td></td><td></td><td></td><td></td><td></td><td></td><td></td><td></td><td></td></tr>
<tr><td>Oligo rev</td><td>AAACCCAACACAACCACCACAACC</td><td></td><td></td><td></td><td></td><td></td><td></td><td></td><td></td><td></td><td></td></tr>
<tr><td>Chromosome</td><td>start</td><td>end</td><td>strand</td><td>MM</td><td>target_seq</td><td>PAM</td><td>alignment</td><td>distance</td><td>position</td><td>gene name</td><td>gene id</td></tr>
<tr><td>GU305914.1</td><td>36173</td><td>36195</td><td>+</td><td>0</td><td>GGTTGTGGTGGTTGTGTTGG</td><td>TGG</td><td>||||||||[||||||||||||]PAM</td><td>NA</td><td>Intergenic</td><td>NA</td><td>NA</td></tr>
<tr><td>T2320</td><td>GGTGGTTGTGTTGGTGGATGCGG</td><td>1000</td><td></td><td></td><td></td><td></td><td></td><td></td><td></td><td></td><td></td></tr>
<tr><td>Oligo fwd</td><td>CACCGGTGGTTGTGTTGGTGGATG</td><td></td><td></td><td></td><td></td><td></td><td></td><td></td><td></td><td></td><td></td></tr>
<tr><td>Oligo rev</td><td>AAACCATCCACCAACACAACCACC</td><td></td><td></td><td></td><td></td><td></td><td></td><td></td><td></td><td></td><td></td></tr>
<tr><td>Chromosome</td><td>start</td><td>end</td><td>strand</td><td>MM</td><td>target_seq</td><td>PAM</td><td>alignment</td><td>distance</td><td>position</td><td>gene name</td><td>gene id</td></tr>
<tr><td>GU305914.1</td><td>36179</td><td>36201</td><td>+</td><td>0</td><td>GGTGGTTGTGTTGGTGGATG</td><td>CGG</td><td>||||||||[||||||||||||]PAM</td><td>NA</td><td>Intergenic</td><td>NA</td><td>NA</td></tr>
<tr><td>T2321</td><td>GGTTGTGTTGGTGGATGCGGCGG</td><td>1000</td><td></td><td></td><td></td><td></td><td></td><td></td><td></td><td></td><td></td></tr>
<tr><td>Oligo fwd</td><td>CACCGGTTGTGTTGGTGGATGCGG</td><td></td><td></td><td></td><td></td><td></td><td></td><td></td><td></td><td></td><td></td></tr>
<tr><td>Oligo rev</td><td>AAACCCGCATCCACCAACACAACC</td><td></td><td></td><td></td><td></td><td></td><td></td><td></td><td></td><td></td><td></td></tr>
<tr><td>Chromosome</td><td>start</td><td>end</td><td>strand</td><td>MM</td><td>target_seq</td><td>PAM</td><td>alignment</td><td>distance</td><td>position</td><td>gene name</td><td>gene id</td></tr>
<tr><td>GU305914.1</td><td>36182</td><td>36204</td><td>+</td><td>0</td><td>GGTTGTGTTGGTGGATGCGG</td><td>CGG</td><td>||||||||[||||||||||||]PAM</td><td>NA</td><td>Intergenic</td><td>NA</td><td>NA</td></tr>
<tr><td>T2322</td><td>GTTGTGTTGGTGGATGCGGCGGG</td><td>1000</td><td></td><td></td><td></td><td></td><td></td><td></td><td></td><td></td><td></td></tr>
<tr><td>Oligo fwd</td><td>CACCGTTGTGTTGGTGGATGCGGC</td><td></td><td></td><td></td><td></td><td></td><td></td><td></td><td></td><td></td><td></td></tr>
<tr><td>Oligo rev</td><td>AAACGCCGCATCCACCAACACAAC</td><td></td><td></td><td></td><td></td><td></td><td></td><td></td><td></td><td></td><td></td></tr>
<tr><td>Chromosome</td><td>start</td><td>end</td><td>strand</td><td>MM</td><td>target_seq</td><td>PAM</td><td>alignment</td><td>distance</td><td>position</td><td>gene name</td><td>gene id</td></tr>
<tr><td>GU305914.1</td><td>36183</td><td>36205</td><td>+</td><td>0</td><td>GTTGTGTTGGTGGATGCGGC</td><td>GGG</td><td>||||||||[||||||||||||]PAM</td><td>NA</td><td>Intergenic</td><td>NA</td><td>NA</td></tr>
<tr><td>T2323</td><td>GGATGCGGCGGGAGTCGCCGCGG</td><td>1000</td><td></td><td></td><td></td><td></td><td></td><td></td><td></td><td></td><td></td></tr>
<tr><td>Oligo fwd</td><td>CACCGGATGCGGCGGGAGTCGCCG</td><td></td><td></td><td></td><td></td><td></td><td></td><td></td><td></td><td></td><td></td></tr>
<tr><td>Oligo rev</td><td>AAACCGGCGACTCCCGCCGCATCC</td><td></td><td></td><td></td><td></td><td></td><td></td><td></td><td></td><td></td><td></td></tr>
<tr><td>Chromosome</td><td>start</td><td>end</td><td>strand</td><td>MM</td><td>target_seq</td><td>PAM</td><td>alignment</td><td>distance</td><td>position</td><td>gene name</td><td>gene id</td></tr>
<tr><td>GU305914.1</td><td>36194</td><td>36216</td><td>+</td><td>0</td><td>GGATGCGGCGGGAGTCGCCG</td><td>CGG</td><td>||||||||[||||||||||||]PAM</td><td>NA</td><td>Intergenic</td><td>NA</td><td>NA</td></tr>
<tr><td>T2324</td><td>AGTCGCCGCGGACGAAGCGACGG</td><td>1000</td><td></td><td></td><td></td><td></td><td></td><td></td><td></td><td></td><td></td></tr>
<tr><td>Oligo fwd</td><td>CACCAGTCGCCGCGGACGAAGCGA</td><td></td><td></td><td></td><td></td><td></td><td></td><td></td><td></td><td></td><td></td></tr>
<tr><td>Oligo rev</td><td>AAACTCGCTTCGTCCGCGGCGACT</td><td></td><td></td><td></td><td></td><td></td><td></td><td></td><td></td><td></td><td></td></tr>
<tr><td>Chromosome</td><td>start</td><td>end</td><td>strand</td><td>MM</td><td>target_seq</td><td>PAM</td><td>alignment</td><td>distance</td><td>position</td><td>gene name</td><td>gene id</td></tr>
<tr><td>GU305914.1</td><td>36206</td><td>36228</td><td>+</td><td>0</td><td>AGTCGCCGCGGACGAAGCGA</td><td>CGG</td><td>||||||||[||||||||||||]PAM</td><td>NA</td><td>Intergenic</td><td>NA</td><td>NA</td></tr>
<tr><td>GU305914.1</td><td>44717</td><td>44739</td><td>+</td><td>4</td><td>CGTCGTCGGCGACGAAGCGA</td><td>TGG</td><td>-||||-||[--||||||||||]PAM</td><td>NA</td><td>Intergenic</td><td>NA</td><td>NA</td></tr>
<tr><td>T2325</td><td>GCGGACGAAGCGACGGCAGCAGG</td><td>1000</td><td></td><td></td><td></td><td></td><td></td><td></td><td></td><td></td><td></td></tr>
<tr><td>Oligo fwd</td><td>CACCGCGGACGAAGCGACGGCAGC</td><td></td><td></td><td></td><td></td><td></td><td></td><td></td><td></td><td></td><td></td></tr>
<tr><td>Oligo rev</td><td>AAACGCTGCCGTCGCTTCGTCCGC</td><td></td><td></td><td></td><td></td><td></td><td></td><td></td><td></td><td></td><td></td></tr>
<tr><td>Chromosome</td><td>start</td><td>end</td><td>strand</td><td>MM</td><td>target_seq</td><td>PAM</td><td>alignment</td><td>distance</td><td>position</td><td>gene name</td><td>gene id</td></tr>
<tr><td>GU305914.1</td><td>36213</td><td>36235</td><td>+</td><td>0</td><td>GCGGACGAAGCGACGGCAGC</td><td>AGG</td><td>||||||||[||||||||||||]PAM</td><td>NA</td><td>Intergenic</td><td>NA</td><td>NA</td></tr>
<tr><td>T2326</td><td>CGAAGCGACGGCAGCAGGAATGG</td><td>1000</td><td></td><td></td><td></td><td></td><td></td><td></td><td></td><td></td><td></td></tr>
<tr><td>Oligo fwd</td><td>CACCCGAAGCGACGGCAGCAGGAA</td><td></td><td></td><td></td><td></td><td></td><td></td><td></td><td></td><td></td><td></td></tr>
<tr><td>Oligo rev</td><td>AAACTTCCTGCTGCCGTCGCTTCG</td><td></td><td></td><td></td><td></td><td></td><td></td><td></td><td></td><td></td><td></td></tr>
<tr><td>Chromosome</td><td>start</td><td>end</td><td>strand</td><td>MM</td><td>target_seq</td><td>PAM</td><td>alignment</td><td>distance</td><td>position</td><td>gene name</td><td>gene id</td></tr>
<tr><td>GU305914.1</td><td>36218</td><td>36240</td><td>+</td><td>0</td><td>CGAAGCGACGGCAGCAGGAA</td><td>TGG</td><td>||||||||[||||||||||||]PAM</td><td>NA</td><td>Intergenic</td><td>NA</td><td>NA</td></tr>
<tr><td>T2327</td><td>AGCGACGGCAGCAGGAATGGTGG</td><td>1000</td><td></td><td></td><td></td><td></td><td></td><td></td><td></td><td></td><td></td></tr>
<tr><td>Oligo fwd</td><td>CACCAGCGACGGCAGCAGGAATGG</td><td></td><td></td><td></td><td></td><td></td><td></td><td></td><td></td><td></td><td></td></tr>
<tr><td>Oligo rev</td><td>AAACCCATTCCTGCTGCCGTCGCT</td><td></td><td></td><td></td><td></td><td></td><td></td><td></td><td></td><td></td><td></td></tr>
</table>

Chromosome start end strand MM target_seq PAM alignment distance position gene name gene id
GU305914.1 36221 36243 + 0 AGCGACGGCAGCAGGAATGG TGG ||||||||[||||||||||||]PAM NA Intergenic NA NA

T2328 GACGGCAGCAGGAATGGTGGCGG 1000
Oligo fwd CACCGACGGCAGCAGGAATGGTGG
Oligo rev AAACCCACCATTCCTGCTGCCGTC
Chromosome start end strand MM target_seq PAM alignment distance position gene name gene id
GU305914.1 36224 36246 + 0 GACGGCAGCAGGAATGGTGG CGG ||||||||[||||||||||||]PAM NA Intergenic NA NA

T2329 TGGCGGTTACAGCCGCACCGTGG 1000
Oligo fwd CACCTGGCGGTTACAGCCGCACCG
Oligo rev AAACCGGTGCGGCTGTAACCGCCA
Chromosome start end strand MM target_seq PAM alignment distance position gene name gene id
GU305914.1 36241 36263 + 0 TGGCGGTTACAGCCGCACCG TGG ||||||||[||||||||||||]PAM NA Intergenic NA NA

T2330 GTCGCCCTTTCTTTTGCGCTTGG 1000
Oligo fwd CACCGTCGCCCTTTCTTTTGCGCT
Oligo rev AAACAGCGCAAAAGAAAGGGCGAC
Chromosome start end strand MM target_seq PAM alignment distance position gene name gene id
GU305914.1 36263 36285 + 0 GTCGCCCTTTCTTTTGCGCT TGG ||||||||[||||||||||||]PAM NA Intergenic NA NA

T2331 TTTGCGCTTGGCCTTCCGCTCGG 1000
Oligo fwd CACCTTTGCGCTTGGCCTTCCGCT
Oligo rev AAACAGCGGAAGGCCAAGCGCAAA
Chromosome start end strand MM target_seq PAM alignment distance position gene name gene id
GU305914.1 36275 36297 + 0 TTTGCGCTTGGCCTTCCGCT CGG ||||||||[||||||||||||]PAM NA Intergenic NA NA

T2332 GCTTGGCCTTCCGCTCGGAGCGG 1000
Oligo fwd CACCGCTTGGCCTTCCGCTCGGAG
Oligo rev AAACCTCCGAGCGGAAGGCCAAGC
Chromosome start end strand MM target_seq PAM alignment distance position gene name gene id
GU305914.1 36280 36302 + 0 GCTTGGCCTTCCGCTCGGAG CGG ||||||||[||||||||||||]PAM NA Intergenic NA NA

T2333 CTCGGAGCGGCGCTTTTCGCTGG 1000
Oligo fwd CACCCTCGGAGCGGCGCTTTTCGC
Oligo rev AAACGCGAAAAGCGCCGCTCCGAG
Chromosome start end strand MM target_seq PAM alignment distance position gene name gene id
GU305914.1 36293 36315 + 0 CTCGGAGCGGCGCTTTTCGC TGG ||||||||[||||||||||||]PAM NA Intergenic NA NA

T2334 ACGCGAGCATTTTATTCATAAGG 1000
Oligo fwd CACCACGCGAGCATTTTATTCATA
Oligo rev AAACTATGAATAAAATGCTCGCGT
Chromosome start end strand MM target_seq PAM alignment distance position gene name gene id
GU305914.1 36319 36341 + 0 ACGCGAGCATTTTATTCATA AGG ||||||||[||||||||||||]PAM NA Intergenic NA NA

T2335 CGAGCATTTTATTCATAAGGAGG 1000
Oligo fwd CACCCGAGCATTTTATTCATAAGG
Oligo rev AAACCCTTATGAATAAAATGCTCG
Chromosome start end strand MM target_seq PAM alignment distance position gene name gene id
GU305914.1 36322 36344 + 0 CGAGCATTTTATTCATAAGG AGG ||||||||[||||||||||||]PAM NA Intergenic NA NA

T2336 CATCCTCTCGATATCTTCGAAGG 1000
Oligo fwd CACCCATCCTCTCGATATCTTCGA
Oligo rev AAACTCGAAGATATCGAGAGGATG
Chromosome start end strand MM target_seq PAM alignment distance position gene name gene id
GU305914.1 36347 36369 + 0 CATCCTCTCGATATCTTCGA AGG ||||||||[||||||||||||]PAM NA Intergenic NA NA

T2337 ATTCATGTCCGAGCCACGACAGG 1000
Oligo fwd CACCATTCATGTCCGAGCCACGAC
Oligo rev AAACGTCGTGGCTCGGACATGAAT
Chromosome start end strand MM target_seq PAM alignment distance position gene name gene id
GU305914.1 36383 36405 + 0 ATTCATGTCCGAGCCACGAC AGG ||||||||[||||||||||||]PAM NA Intergenic NA NA

T2338 CTCTTCACTAGACGCTAGTATGG 1000
Oligo fwd CACCCTCTTCACTAGACGCTAGTA
Oligo rev AAACTACTAGCGTCTAGTGAAGAG
Chromosome start end strand MM target_seq PAM alignment distance position gene name gene id
GU305914.1 36458 36480 + 0 CTCTTCACTAGACGCTAGTA TGG ||||||||[||||||||||||]PAM NA Intergenic NA NA

T2339 CACTAGACGCTAGTATGGAATGG 1000
Oligo fwd CACCCACTAGACGCTAGTATGGAA
Oligo rev AAACTTCCATACTAGCGTCTAGTG
Chromosome start end strand MM target_seq PAM alignment distance position gene name gene id
GU305914.1 36463 36485 + 0 CACTAGACGCTAGTATGGAA TGG ||||||||[||||||||||||]PAM NA Intergenic NA NA

T2340 CACGACCTTCATGTCGCTAACGG 1000
Oligo fwd CACCCACGACCTTCATGTCGCTAA
Oligo rev AAACTTAGCGACATGAAGGTCGTG
Chromosome start end strand MM target_seq PAM alignment distance position gene name gene id
GU305914.1 36566 36588 + 0 CACGACCTTCATGTCGCTAA CGG ||||||||[||||||||||||]PAM NA Intergenic NA NA

T2341 ACGACCTTCATGTCGCTAACGGG 1000
Oligo fwd CACCACGACCTTCATGTCGCTAAC
Oligo rev AAACGTTAGCGACATGAAGGTCGT
Chromosome start end strand MM target_seq PAM alignment distance position gene name gene id
GU305914.1 36567 36589 + 0 ACGACCTTCATGTCGCTAAC GGG ||||||||[||||||||||||]PAM NA Intergenic NA NA

T2342 GTCGCTAACGGGCATCCGTCCGG 1000
Oligo fwd CACCGTCGCTAACGGGCATCCGTC

<table>
<tr><td>Oligo rev</td><td>AAACGACGGATGCCCGTTAGCGAC</td><td></td><td></td><td></td><td></td><td></td><td></td><td></td><td></td><td></td><td></td></tr>
<tr><td>Chromosome</td><td>start</td><td>end</td><td>strand</td><td>MM</td><td>target_seq</td><td>PAM</td><td>alignment</td><td>distance</td><td>position</td><td>gene name</td><td>gene id</td></tr>
<tr><td>GU305914.1</td><td>36578</td><td>36600</td><td>+</td><td>0</td><td>GTCGCTAACGGGCATCCGTC</td><td>CGG</td><td>||||||||[||||||||||||]PAM</td><td>NA</td><td>Intergenic</td><td>NA</td><td>NA</td></tr>
<tr><td>T2343</td><td>GAAGTTGACCATGAGATCAAAGG</td><td>1000</td><td></td><td></td><td></td><td></td><td></td><td></td><td></td><td></td><td></td></tr>
<tr><td>Oligo fwd</td><td>CACCGAAGTTGACCATGAGATCAA</td><td></td><td></td><td></td><td></td><td></td><td></td><td></td><td></td><td></td><td></td></tr>
<tr><td>Oligo rev</td><td>AAACTTGATCTCATGGTCAACTTC</td><td></td><td></td><td></td><td></td><td></td><td></td><td></td><td></td><td></td><td></td></tr>
<tr><td>Chromosome</td><td>start</td><td>end</td><td>strand</td><td>MM</td><td>target_seq</td><td>PAM</td><td>alignment</td><td>distance</td><td>position</td><td>gene name</td><td>gene id</td></tr>
<tr><td>GU305914.1</td><td>36608</td><td>36630</td><td>+</td><td>0</td><td>GAAGTTGACCATGAGATCAA</td><td>AGG</td><td>||||||||[||||||||||||]PAM</td><td>NA</td><td>Intergenic</td><td>NA</td><td>NA</td></tr>
<tr><td>T2344</td><td>CACGCCCCCCAGATCAACGTCGG</td><td>1000</td><td></td><td></td><td></td><td></td><td></td><td></td><td></td><td></td><td></td></tr>
<tr><td>Oligo fwd</td><td>CACCCACGCCCCCCAGATCAACGT</td><td></td><td></td><td></td><td></td><td></td><td></td><td></td><td></td><td></td><td></td></tr>
<tr><td>Oligo rev</td><td>AAACACGTTGATCTGGGGGGCGTG</td><td></td><td></td><td></td><td></td><td></td><td></td><td></td><td></td><td></td><td></td></tr>
<tr><td>Chromosome</td><td>start</td><td>end</td><td>strand</td><td>MM</td><td>target_seq</td><td>PAM</td><td>alignment</td><td>distance</td><td>position</td><td>gene name</td><td>gene id</td></tr>
<tr><td>GU305914.1</td><td>36676</td><td>36698</td><td>+</td><td>0</td><td>CACGCCCCCCAGATCAACGT</td><td>CGG</td><td>||||||||[||||||||||||]PAM</td><td>NA</td><td>Intergenic</td><td>NA</td><td>NA</td></tr>
<tr><td>T2345</td><td>GCATATATAGACACAAGCCCCGG</td><td>1000</td><td></td><td></td><td></td><td></td><td></td><td></td><td></td><td></td><td></td></tr>
<tr><td>Oligo fwd</td><td>CACCGCATATATAGACACAAGCCC</td><td></td><td></td><td></td><td></td><td></td><td></td><td></td><td></td><td></td><td></td></tr>
<tr><td>Oligo rev</td><td>AAACGGGCTTGTGTCTATATATGC</td><td></td><td></td><td></td><td></td><td></td><td></td><td></td><td></td><td></td><td></td></tr>
<tr><td>Chromosome</td><td>start</td><td>end</td><td>strand</td><td>MM</td><td>target_seq</td><td>PAM</td><td>alignment</td><td>distance</td><td>position</td><td>gene name</td><td>gene id</td></tr>
<tr><td>GU305914.1</td><td>36706</td><td>36728</td><td>+</td><td>0</td><td>GCATATATAGACACAAGCCC</td><td>CGG</td><td>||||||||[||||||||||||]PAM</td><td>NA</td><td>Intergenic</td><td>NA</td><td>NA</td></tr>
<tr><td>T2346</td><td>ACCTATTTTATACGCGAGCGTGG</td><td>1000</td><td></td><td></td><td></td><td></td><td></td><td></td><td></td><td></td><td></td></tr>
<tr><td>Oligo fwd</td><td>CACCACCTATTTTATACGCGAGCG</td><td></td><td></td><td></td><td></td><td></td><td></td><td></td><td></td><td></td><td></td></tr>
<tr><td>Oligo rev</td><td>AAACCGCTCGCGTATAAAATAGGT</td><td></td><td></td><td></td><td></td><td></td><td></td><td></td><td></td><td></td><td></td></tr>
<tr><td>Chromosome</td><td>start</td><td>end</td><td>strand</td><td>MM</td><td>target_seq</td><td>PAM</td><td>alignment</td><td>distance</td><td>position</td><td>gene name</td><td>gene id</td></tr>
<tr><td>GU305914.1</td><td>36774</td><td>36796</td><td>+</td><td>0</td><td>ACCTATTTTATACGCGAGCG</td><td>TGG</td><td>||||||||[||||||||||||]PAM</td><td>NA</td><td>Intergenic</td><td>NA</td><td>NA</td></tr>
<tr><td>T2347</td><td>TATTTTATACGCGAGCGTGGAGG</td><td>1000</td><td></td><td></td><td></td><td></td><td></td><td></td><td></td><td></td><td></td></tr>
<tr><td>Oligo fwd</td><td>CACCTATTTTATACGCGAGCGTGG</td><td></td><td></td><td></td><td></td><td></td><td></td><td></td><td></td><td></td><td></td></tr>
<tr><td>Oligo rev</td><td>AAACCCACGCTCGCGTATAAAATA</td><td></td><td></td><td></td><td></td><td></td><td></td><td></td><td></td><td></td><td></td></tr>
<tr><td>Chromosome</td><td>start</td><td>end</td><td>strand</td><td>MM</td><td>target_seq</td><td>PAM</td><td>alignment</td><td>distance</td><td>position</td><td>gene name</td><td>gene id</td></tr>
<tr><td>GU305914.1</td><td>36777</td><td>36799</td><td>+</td><td>0</td><td>TATTTTATACGCGAGCGTGG</td><td>AGG</td><td>||||||||[||||||||||||]PAM</td><td>NA</td><td>Intergenic</td><td>NA</td><td>NA</td></tr>
<tr><td>T2348</td><td>TTATACGCGAGCGTGGAGGACGG</td><td>1000</td><td></td><td></td><td></td><td></td><td></td><td></td><td></td><td></td><td></td></tr>
<tr><td>Oligo fwd</td><td>CACCTTATACGCGAGCGTGGAGGA</td><td></td><td></td><td></td><td></td><td></td><td></td><td></td><td></td><td></td><td></td></tr>
<tr><td>Oligo rev</td><td>AAACTCCTCCACGCTCGCGTATAA</td><td></td><td></td><td></td><td></td><td></td><td></td><td></td><td></td><td></td><td></td></tr>
<tr><td>Chromosome</td><td>start</td><td>end</td><td>strand</td><td>MM</td><td>target_seq</td><td>PAM</td><td>alignment</td><td>distance</td><td>position</td><td>gene name</td><td>gene id</td></tr>
<tr><td>GU305914.1</td><td>36781</td><td>36803</td><td>+</td><td>0</td><td>TTATACGCGAGCGTGGAGGA</td><td>CGG</td><td>||||||||[||||||||||||]PAM</td><td>NA</td><td>Intergenic</td><td>NA</td><td>NA</td></tr>
<tr><td>T2349</td><td>GCGAGCGTGGAGGACGGCGCAGG</td><td>1000</td><td></td><td></td><td></td><td></td><td></td><td></td><td></td><td></td><td></td></tr>
<tr><td>Oligo fwd</td><td>CACCGCGAGCGTGGAGGACGGCGC</td><td></td><td></td><td></td><td></td><td></td><td></td><td></td><td></td><td></td><td></td></tr>
<tr><td>Oligo rev</td><td>AAACGCGCCGTCCTCCACGCTCGC</td><td></td><td></td><td></td><td></td><td></td><td></td><td></td><td></td><td></td><td></td></tr>
<tr><td>Chromosome</td><td>start</td><td>end</td><td>strand</td><td>MM</td><td>target_seq</td><td>PAM</td><td>alignment</td><td>distance</td><td>position</td><td>gene name</td><td>gene id</td></tr>
<tr><td>GU305914.1</td><td>36787</td><td>36809</td><td>+</td><td>0</td><td>GCGAGCGTGGAGGACGGCGC</td><td>AGG</td><td>||||||||[||||||||||||]PAM</td><td>NA</td><td>Intergenic</td><td>NA</td><td>NA</td></tr>
<tr><td>T2350</td><td>AATTCTCCTAACGAGAACAGAGG</td><td>1000</td><td></td><td></td><td></td><td></td><td></td><td></td><td></td><td></td><td></td></tr>
<tr><td>Oligo fwd</td><td>CACCAATTCTCCTAACGAGAACAG</td><td></td><td></td><td></td><td></td><td></td><td></td><td></td><td></td><td></td><td></td></tr>
<tr><td>Oligo rev</td><td>AAACCTGTTCTCGTTAGGAGAATT</td><td></td><td></td><td></td><td></td><td></td><td></td><td></td><td></td><td></td><td></td></tr>
<tr><td>Chromosome</td><td>start</td><td>end</td><td>strand</td><td>MM</td><td>target_seq</td><td>PAM</td><td>alignment</td><td>distance</td><td>position</td><td>gene name</td><td>gene id</td></tr>
<tr><td>GU305914.1</td><td>36861</td><td>36883</td><td>+</td><td>0</td><td>AATTCTCCTAACGAGAACAG</td><td>AGG</td><td>||||||||[||||||||||||]PAM</td><td>NA</td><td>Intergenic</td><td>NA</td><td>NA</td></tr>
<tr><td>T2351</td><td>ATTCTCCTAACGAGAACAGAGGG</td><td>1000</td><td></td><td></td><td></td><td></td><td></td><td></td><td></td><td></td><td></td></tr>
<tr><td>Oligo fwd</td><td>CACCATTCTCCTAACGAGAACAGA</td><td></td><td></td><td></td><td></td><td></td><td></td><td></td><td></td><td></td><td></td></tr>
<tr><td>Oligo rev</td><td>AAACTCTGTTCTCGTTAGGAGAAT</td><td></td><td></td><td></td><td></td><td></td><td></td><td></td><td></td><td></td><td></td></tr>
<tr><td>Chromosome</td><td>start</td><td>end</td><td>strand</td><td>MM</td><td>target_seq</td><td>PAM</td><td>alignment</td><td>distance</td><td>position</td><td>gene name</td><td>gene id</td></tr>
<tr><td>GU305914.1</td><td>36862</td><td>36884</td><td>+</td><td>0</td><td>ATTCTCCTAACGAGAACAGA</td><td>GGG</td><td>||||||||[||||||||||||]PAM</td><td>NA</td><td>Intergenic</td><td>NA</td><td>NA</td></tr>
<tr><td>T2352</td><td>TCCTAACGAGAACAGAGGGATGG</td><td>1000</td><td></td><td></td><td></td><td></td><td></td><td></td><td></td><td></td><td></td></tr>
<tr><td>Oligo fwd</td><td>CACCTCCTAACGAGAACAGAGGGA</td><td></td><td></td><td></td><td></td><td></td><td></td><td></td><td></td><td></td><td></td></tr>
<tr><td>Oligo rev</td><td>AAACTCCCTCTGTTCTCGTTAGGA</td><td></td><td></td><td></td><td></td><td></td><td></td><td></td><td></td><td></td><td></td></tr>
<tr><td>Chromosome</td><td>start</td><td>end</td><td>strand</td><td>MM</td><td>target_seq</td><td>PAM</td><td>alignment</td><td>distance</td><td>position</td><td>gene name</td><td>gene id</td></tr>
<tr><td>GU305914.1</td><td>36866</td><td>36888</td><td>+</td><td>0</td><td>TCCTAACGAGAACAGAGGGA</td><td>TGG</td><td>||||||||[||||||||||||]PAM</td><td>NA</td><td>Intergenic</td><td>NA</td><td>NA</td></tr>
<tr><td>T2353</td><td>GCGCATCGCCGCCCAGCAGCAGG</td><td>1000</td><td></td><td></td><td></td><td></td><td></td><td></td><td></td><td></td><td></td></tr>
<tr><td>Oligo fwd</td><td>CACCGCGCATCGCCGCCCAGCAGC</td><td></td><td></td><td></td><td></td><td></td><td></td><td></td><td></td><td></td><td></td></tr>
<tr><td>Oligo rev</td><td>AAACGCTGCTGGGCGGCGATGCGC</td><td></td><td></td><td></td><td></td><td></td><td></td><td></td><td></td><td></td><td></td></tr>
<tr><td>Chromosome</td><td>start</td><td>end</td><td>strand</td><td>MM</td><td>target_seq</td><td>PAM</td><td>alignment</td><td>distance</td><td>position</td><td>gene name</td><td>gene id</td></tr>
<tr><td>GU305914.1</td><td>36907</td><td>36929</td><td>+</td><td>0</td><td>GCGCATCGCCGCCCAGCAGC</td><td>AGG</td><td>||||||||[||||||||||||]PAM</td><td>NA</td><td>Intergenic</td><td>NA</td><td>NA</td></tr>
<tr><td>T2354</td><td>CAGCAGCAGGCGTCCTGACGCGG</td><td>1000</td><td></td><td></td><td></td><td></td><td></td><td></td><td></td><td></td><td></td></tr>
<tr><td>Oligo fwd</td><td>CACCCAGCAGCAGGCGTCCTGACG</td><td></td><td></td><td></td><td></td><td></td><td></td><td></td><td></td><td></td><td></td></tr>
<tr><td>Oligo rev</td><td>AAACCGTCAGGACGCCTGCTGCTG</td><td></td><td></td><td></td><td></td><td></td><td></td><td></td><td></td><td></td><td></td></tr>
<tr><td>Chromosome</td><td>start</td><td>end</td><td>strand</td><td>MM</td><td>target_seq</td><td>PAM</td><td>alignment</td><td>distance</td><td>position</td><td>gene name</td><td>gene id</td></tr>
<tr><td>GU305914.1</td><td>36920</td><td>36942</td><td>+</td><td>0</td><td>CAGCAGCAGGCGTCCTGACG</td><td>CGG</td><td>||||||||[||||||||||||]PAM</td><td>NA</td><td>Intergenic</td><td>NA</td><td>NA</td></tr>
<tr><td>T2355</td><td>GACTGAGCACGACATCCTCGAGG</td><td>1000</td><td></td><td></td><td></td><td></td><td></td><td></td><td></td><td></td><td></td></tr>
<tr><td>Oligo fwd</td><td>CACCGACTGAGCACGACATCCTCG</td><td></td><td></td><td></td><td></td><td></td><td></td><td></td><td></td><td></td><td></td></tr>
<tr><td>Oligo rev</td><td>AAACCGAGGATGTCGTGCTCAGTC</td><td></td><td></td><td></td><td></td><td></td><td></td><td></td><td></td><td></td><td></td></tr>
<tr><td>Chromosome</td><td>start</td><td>end</td><td>strand</td><td>MM</td><td>target_seq</td><td>PAM</td><td>alignment</td><td>distance</td><td>position</td><td>gene name</td><td>gene id</td></tr>
<tr><td>GU305914.1</td><td>36955</td><td>36977</td><td>+</td><td>0</td><td>GACTGAGCACGACATCCTCG</td><td>AGG</td><td>||||||||[||||||||||||]PAM</td><td>NA</td><td>Intergenic</td><td>NA</td><td>NA</td></tr>
<tr><td>T2356</td><td>ACTGAGCACGACATCCTCGAGGG</td><td>1000</td><td></td><td></td><td></td><td></td><td></td><td></td><td></td><td></td><td></td></tr>
<tr><td>Oligo fwd</td><td>CACCACTGAGCACGACATCCTCGA</td><td></td><td></td><td></td><td></td><td></td><td></td><td></td><td></td><td></td><td></td></tr>
<tr><td>Oligo rev</td><td>AAACTCGAGGATGTCGTGCTCAGT</td><td></td><td></td><td></td><td></td><td></td><td></td><td></td><td></td><td></td><td></td></tr>
<tr><td>Chromosome</td><td>start</td><td>end</td><td>strand</td><td>MM</td><td>target_seq</td><td>PAM</td><td>alignment</td><td>distance</td><td>position</td><td>gene name</td><td>gene id</td></tr>
<tr><td>GU305914.1</td><td>36956</td><td>36978</td><td>+</td><td>0</td><td>ACTGAGCACGACATCCTCGA</td><td>GGG</td><td>||||||||[||||||||||||]PAM</td><td>NA</td><td>Intergenic</td><td>NA</td><td>NA</td></tr>
<tr><td>T2357</td><td>CATCCTCGAGGGAGAGATCCAGG</td><td>1000</td><td></td><td></td><td></td><td></td><td></td><td></td><td></td><td></td><td></td></tr>
</table>

<table>
<tr><td>Oligo fwd</td><td>CACCCATCCTCGAGGGAGAGATCC</td><td></td><td></td><td></td><td></td><td></td><td></td><td></td><td></td><td></td><td></td></tr>
<tr><td>Oligo rev</td><td>AAACGGATCTCTCCCTCGAGGATG</td><td></td><td></td><td></td><td></td><td></td><td></td><td></td><td></td><td></td><td></td></tr>
<tr><td>Chromosome</td><td>start</td><td>end</td><td>strand</td><td>MM</td><td>target_seq</td><td>PAM</td><td>alignment</td><td>distance</td><td>position</td><td>gene name</td><td>gene id</td></tr>
<tr><td>GU305914.1</td><td>36967</td><td>36989</td><td>+</td><td>0</td><td>CATCCTCGAGGGAGAGATCC</td><td>AGG</td><td>||||||||[||||||||||||]PAM</td><td>NA</td><td>Intergenic</td><td>NA</td><td>NA</td></tr>
<tr><td></td><td></td><td></td><td></td><td></td><td></td><td></td><td></td><td></td><td></td><td></td><td></td></tr>
<tr><td>T2358</td><td>AGACAAATCGCGAAGCGTTTCGG</td><td>1000</td><td></td><td></td><td></td><td></td><td></td><td></td><td></td><td></td><td></td></tr>
<tr><td>Oligo fwd</td><td>CACCAGACAAATCGCGAAGCGTTT</td><td></td><td></td><td></td><td></td><td></td><td></td><td></td><td></td><td></td><td></td></tr>
<tr><td>Oligo rev</td><td>AAACAAACGCTTCGCGATTTGTCT</td><td></td><td></td><td></td><td></td><td></td><td></td><td></td><td></td><td></td><td></td></tr>
<tr><td>Chromosome</td><td>start</td><td>end</td><td>strand</td><td>MM</td><td>target_seq</td><td>PAM</td><td>alignment</td><td>distance</td><td>position</td><td>gene name</td><td>gene id</td></tr>
<tr><td>GU305914.1</td><td>37019</td><td>37041</td><td>+</td><td>0</td><td>AGACAAATCGCGAAGCGTTT</td><td>CGG</td><td>||||||||[||||||||||||]PAM</td><td>NA</td><td>Intergenic</td><td>NA</td><td>NA</td></tr>
<tr><td></td><td></td><td></td><td></td><td></td><td></td><td></td><td></td><td></td><td></td><td></td><td></td></tr>
<tr><td>T2359</td><td>AACCCCCCGAGAGCGCAGATCGG</td><td>1000</td><td></td><td></td><td></td><td></td><td></td><td></td><td></td><td></td><td></td></tr>
<tr><td>Oligo fwd</td><td>CACCAACCCCCCGAGAGCGCAGAT</td><td></td><td></td><td></td><td></td><td></td><td></td><td></td><td></td><td></td><td></td></tr>
<tr><td>Oligo rev</td><td>AAACATCTGCGCTCTCGGGGGGTT</td><td></td><td></td><td></td><td></td><td></td><td></td><td></td><td></td><td></td><td></td></tr>
<tr><td>Chromosome</td><td>start</td><td>end</td><td>strand</td><td>MM</td><td>target_seq</td><td>PAM</td><td>alignment</td><td>distance</td><td>position</td><td>gene name</td><td>gene id</td></tr>
<tr><td>GU305914.1</td><td>37118</td><td>37140</td><td>+</td><td>0</td><td>AACCCCCCGAGAGCGCAGAT</td><td>CGG</td><td>||||||||[||||||||||||]PAM</td><td>NA</td><td>Intergenic</td><td>NA</td><td>NA</td></tr>
<tr><td></td><td></td><td></td><td></td><td></td><td></td><td></td><td></td><td></td><td></td><td></td><td></td></tr>
<tr><td>T2360</td><td>GGTCAGCGCCATCAAAGAGCCGG</td><td>1000</td><td></td><td></td><td></td><td></td><td></td><td></td><td></td><td></td><td></td></tr>
<tr><td>Oligo fwd</td><td>CACCGGTCAGCGCCATCAAAGAGC</td><td></td><td></td><td></td><td></td><td></td><td></td><td></td><td></td><td></td><td></td></tr>
<tr><td>Oligo rev</td><td>AAACGCTCTTTGATGGCGCTGACC</td><td></td><td></td><td></td><td></td><td></td><td></td><td></td><td></td><td></td><td></td></tr>
<tr><td>Chromosome</td><td>start</td><td>end</td><td>strand</td><td>MM</td><td>target_seq</td><td>PAM</td><td>alignment</td><td>distance</td><td>position</td><td>gene name</td><td>gene id</td></tr>
<tr><td>GU305914.1</td><td>37139</td><td>37161</td><td>+</td><td>0</td><td>GGTCAGCGCCATCAAAGAGC</td><td>CGG</td><td>||||||||[||||||||||||]PAM</td><td>NA</td><td>Intergenic</td><td>NA</td><td>NA</td></tr>
<tr><td></td><td></td><td></td><td></td><td></td><td></td><td></td><td></td><td></td><td></td><td></td><td></td></tr>
<tr><td>T2361</td><td>CGCAGTTGCAGTGTCAACATCGG</td><td>1000</td><td></td><td></td><td></td><td></td><td></td><td></td><td></td><td></td><td></td></tr>
<tr><td>Oligo fwd</td><td>CACCCGCAGTTGCAGTGTCAACAT</td><td></td><td></td><td></td><td></td><td></td><td></td><td></td><td></td><td></td><td></td></tr>
<tr><td>Oligo rev</td><td>AAACATGTTGACACTGCAACTGCG</td><td></td><td></td><td></td><td></td><td></td><td></td><td></td><td></td><td></td><td></td></tr>
<tr><td>Chromosome</td><td>start</td><td>end</td><td>strand</td><td>MM</td><td>target_seq</td><td>PAM</td><td>alignment</td><td>distance</td><td>position</td><td>gene name</td><td>gene id</td></tr>
<tr><td>GU305914.1</td><td>37179</td><td>37201</td><td>+</td><td>0</td><td>CGCAGTTGCAGTGTCAACAT</td><td>CGG</td><td>||||||||[||||||||||||]PAM</td><td>NA</td><td>Intergenic</td><td>NA</td><td>NA</td></tr>
<tr><td></td><td></td><td></td><td></td><td></td><td></td><td></td><td></td><td></td><td></td><td></td><td></td></tr>
<tr><td>T2362</td><td>AGTTGCAGTGTCAACATCGGCGG</td><td>1000</td><td></td><td></td><td></td><td></td><td></td><td></td><td></td><td></td><td></td></tr>
<tr><td>Oligo fwd</td><td>CACCAGTTGCAGTGTCAACATCGG</td><td></td><td></td><td></td><td></td><td></td><td></td><td></td><td></td><td></td><td></td></tr>
<tr><td>Oligo rev</td><td>AAACCCGATGTTGACACTGCAACT</td><td></td><td></td><td></td><td></td><td></td><td></td><td></td><td></td><td></td><td></td></tr>
<tr><td>Chromosome</td><td>start</td><td>end</td><td>strand</td><td>MM</td><td>target_seq</td><td>PAM</td><td>alignment</td><td>distance</td><td>position</td><td>gene name</td><td>gene id</td></tr>
<tr><td>GU305914.1</td><td>37182</td><td>37204</td><td>+</td><td>0</td><td>AGTTGCAGTGTCAACATCGG</td><td>CGG</td><td>||||||||[||||||||||||]PAM</td><td>NA</td><td>Intergenic</td><td>NA</td><td>NA</td></tr>
<tr><td></td><td></td><td></td><td></td><td></td><td></td><td></td><td></td><td></td><td></td><td></td><td></td></tr>
<tr><td>T2363</td><td>TCCGCCGCCGTCACCGAGAGCGG</td><td>1000</td><td></td><td></td><td></td><td></td><td></td><td></td><td></td><td></td><td></td></tr>
<tr><td>Oligo fwd</td><td>CACCTCCGCCGCCGTCACCGAGAG</td><td></td><td></td><td></td><td></td><td></td><td></td><td></td><td></td><td></td><td></td></tr>
<tr><td>Oligo rev</td><td>AAACCTCTCGGTGACGGCGGCGGA</td><td></td><td></td><td></td><td></td><td></td><td></td><td></td><td></td><td></td><td></td></tr>
<tr><td>Chromosome</td><td>start</td><td>end</td><td>strand</td><td>MM</td><td>target_seq</td><td>PAM</td><td>alignment</td><td>distance</td><td>position</td><td>gene name</td><td>gene id</td></tr>
<tr><td>GU305914.1</td><td>37221</td><td>37243</td><td>+</td><td>0</td><td>TCCGCCGCCGTCACCGAGAG</td><td>CGG</td><td>||||||||[||||||||||||]PAM</td><td>NA</td><td>Intergenic</td><td>NA</td><td>NA</td></tr>
<tr><td></td><td></td><td></td><td></td><td></td><td></td><td></td><td></td><td></td><td></td><td></td><td></td></tr>
<tr><td>T2364</td><td>AGACGCCGCCGTCTCCATGCAGG</td><td>1000</td><td></td><td></td><td></td><td></td><td></td><td></td><td></td><td></td><td></td></tr>
<tr><td>Oligo fwd</td><td>CACCAGACGCCGCCGTCTCCATGC</td><td></td><td></td><td></td><td></td><td></td><td></td><td></td><td></td><td></td><td></td></tr>
<tr><td>Oligo rev</td><td>AAACGCATGGAGACGGCGGCGTCT</td><td></td><td></td><td></td><td></td><td></td><td></td><td></td><td></td><td></td><td></td></tr>
<tr><td>Chromosome</td><td>start</td><td>end</td><td>strand</td><td>MM</td><td>target_seq</td><td>PAM</td><td>alignment</td><td>distance</td><td>position</td><td>gene name</td><td>gene id</td></tr>
<tr><td>GU305914.1</td><td>37265</td><td>37287</td><td>+</td><td>0</td><td>AGACGCCGCCGTCTCCATGC</td><td>AGG</td><td>||||||||[||||||||||||]PAM</td><td>NA</td><td>Intergenic</td><td>NA</td><td>NA</td></tr>
<tr><td></td><td></td><td></td><td></td><td></td><td></td><td></td><td></td><td></td><td></td><td></td><td></td></tr>
<tr><td>T2365</td><td>GCAGGCGAAGCAGCGAAAACCGG</td><td>1000</td><td></td><td></td><td></td><td></td><td></td><td></td><td></td><td></td><td></td></tr>
<tr><td>Oligo fwd</td><td>CACCGCAGGCGAAGCAGCGAAAAC</td><td></td><td></td><td></td><td></td><td></td><td></td><td></td><td></td><td></td><td></td></tr>
<tr><td>Oligo rev</td><td>AAACGTTTTCGCTGCTTCGCCTGC</td><td></td><td></td><td></td><td></td><td></td><td></td><td></td><td></td><td></td><td></td></tr>
<tr><td>Chromosome</td><td>start</td><td>end</td><td>strand</td><td>MM</td><td>target_seq</td><td>PAM</td><td>alignment</td><td>distance</td><td>position</td><td>gene name</td><td>gene id</td></tr>
<tr><td>GU305914.1</td><td>37283</td><td>37305</td><td>+</td><td>0</td><td>GCAGGCGAAGCAGCGAAAAC</td><td>CGG</td><td>||||||||[||||||||||||]PAM</td><td>NA</td><td>Intergenic</td><td>NA</td><td>NA</td></tr>
<tr><td></td><td></td><td></td><td></td><td></td><td></td><td></td><td></td><td></td><td></td><td></td><td></td></tr>
<tr><td>T2366</td><td>CCAGAGAAGAGCGCCAGTTTCGG</td><td>1000</td><td></td><td></td><td></td><td></td><td></td><td></td><td></td><td></td><td></td></tr>
<tr><td>Oligo fwd</td><td>CACCCCAGAGAAGAGCGCCAGTTT</td><td></td><td></td><td></td><td></td><td></td><td></td><td></td><td></td><td></td><td></td></tr>
<tr><td>Oligo rev</td><td>AAACAAACTGGCGCTCTTCTCTGG</td><td></td><td></td><td></td><td></td><td></td><td></td><td></td><td></td><td></td><td></td></tr>
<tr><td>Chromosome</td><td>start</td><td>end</td><td>strand</td><td>MM</td><td>target_seq</td><td>PAM</td><td>alignment</td><td>distance</td><td>position</td><td>gene name</td><td>gene id</td></tr>
<tr><td>GU305914.1</td><td>37320</td><td>37342</td><td>+</td><td>0</td><td>CCAGAGAAGAGCGCCAGTTT</td><td>CGG</td><td>||||||||[||||||||||||]PAM</td><td>NA</td><td>Intergenic</td><td>NA</td><td>NA</td></tr>
<tr><td></td><td></td><td></td><td></td><td></td><td></td><td></td><td></td><td></td><td></td><td></td><td></td></tr>
<tr><td>T2367</td><td>GAGAAGAGCGCCAGTTTCGGCGG</td><td>1000</td><td></td><td></td><td></td><td></td><td></td><td></td><td></td><td></td><td></td></tr>
<tr><td>Oligo fwd</td><td>CACCGAGAAGAGCGCCAGTTTCGG</td><td></td><td></td><td></td><td></td><td></td><td></td><td></td><td></td><td></td><td></td></tr>
<tr><td>Oligo rev</td><td>AAACCCGAAACTGGCGCTCTTCTC</td><td></td><td></td><td></td><td></td><td></td><td></td><td></td><td></td><td></td><td></td></tr>
<tr><td>Chromosome</td><td>start</td><td>end</td><td>strand</td><td>MM</td><td>target_seq</td><td>PAM</td><td>alignment</td><td>distance</td><td>position</td><td>gene name</td><td>gene id</td></tr>
<tr><td>GU305914.1</td><td>37323</td><td>37345</td><td>+</td><td>0</td><td>GAGAAGAGCGCCAGTTTCGG</td><td>CGG</td><td>||||||||[||||||||||||]PAM</td><td>NA</td><td>Intergenic</td><td>NA</td><td>NA</td></tr>
<tr><td></td><td></td><td></td><td></td><td></td><td></td><td></td><td></td><td></td><td></td><td></td><td></td></tr>
<tr><td>T2368</td><td>TCGGCGGCCTCAAGAGCCCCCGG</td><td>1000</td><td></td><td></td><td></td><td></td><td></td><td></td><td></td><td></td><td></td></tr>
<tr><td>Oligo fwd</td><td>CACCTCGGCGGCCTCAAGAGCCCC</td><td></td><td></td><td></td><td></td><td></td><td></td><td></td><td></td><td></td><td></td></tr>
<tr><td>Oligo rev</td><td>AAACGGGGCTCTTGAGGCCGCCGA</td><td></td><td></td><td></td><td></td><td></td><td></td><td></td><td></td><td></td><td></td></tr>
<tr><td>Chromosome</td><td>start</td><td>end</td><td>strand</td><td>MM</td><td>target_seq</td><td>PAM</td><td>alignment</td><td>distance</td><td>position</td><td>gene name</td><td>gene id</td></tr>
<tr><td>GU305914.1</td><td>37339</td><td>37361</td><td>+</td><td>0</td><td>TCGGCGGCCTCAAGAGCCCC</td><td>CGG</td><td>||||||||[||||||||||||]PAM</td><td>NA</td><td>Intergenic</td><td>NA</td><td>NA</td></tr>
<tr><td></td><td></td><td></td><td></td><td></td><td></td><td></td><td></td><td></td><td></td><td></td><td></td></tr>
<tr><td>T2369</td><td>CGGCCTCAAGAGCCCCCGGTCGG</td><td>1000</td><td></td><td></td><td></td><td></td><td></td><td></td><td></td><td></td><td></td></tr>
<tr><td>Oligo fwd</td><td>CACCCGGCCTCAAGAGCCCCCGGT</td><td></td><td></td><td></td><td></td><td></td><td></td><td></td><td></td><td></td><td></td></tr>
<tr><td>Oligo rev</td><td>AAACACCGGGGGCTCTTGAGGCCG</td><td></td><td></td><td></td><td></td><td></td><td></td><td></td><td></td><td></td><td></td></tr>
<tr><td>Chromosome</td><td>start</td><td>end</td><td>strand</td><td>MM</td><td>target_seq</td><td>PAM</td><td>alignment</td><td>distance</td><td>position</td><td>gene name</td><td>gene id</td></tr>
<tr><td>GU305914.1</td><td>37343</td><td>37365</td><td>+</td><td>0</td><td>CGGCCTCAAGAGCCCCCGGT</td><td>CGG</td><td>||||||||[||||||||||||]PAM</td><td>NA</td><td>Intergenic</td><td>NA</td><td>NA</td></tr>
<tr><td></td><td></td><td></td><td></td><td></td><td></td><td></td><td></td><td></td><td></td><td></td><td></td></tr>
<tr><td>T2370</td><td>CGGCACCAGCTTCCGTCATCCGG</td><td>1000</td><td></td><td></td><td></td><td></td><td></td><td></td><td></td><td></td><td></td></tr>
<tr><td>Oligo fwd</td><td>CACCCGGCACCAGCTTCCGTCATC</td><td></td><td></td><td></td><td></td><td></td><td></td><td></td><td></td><td></td><td></td></tr>
<tr><td>Oligo rev</td><td>AAACGATGACGGAAGCTGGTGCCG</td><td></td><td></td><td></td><td></td><td></td><td></td><td></td><td></td><td></td><td></td></tr>
<tr><td>Chromosome</td><td>start</td><td>end</td><td>strand</td><td>MM</td><td>target_seq</td><td>PAM</td><td>alignment</td><td>distance</td><td>position</td><td>gene name</td><td>gene id</td></tr>
<tr><td>GU305914.1</td><td>37363</td><td>37385</td><td>+</td><td>0</td><td>CGGCACCAGCTTCCGTCATC</td><td>CGG</td><td>||||||||[||||||||||||]PAM</td><td>NA</td><td>Intergenic</td><td>NA</td><td>NA</td></tr>
<tr><td></td><td></td><td></td><td></td><td></td><td></td><td></td><td></td><td></td><td></td><td></td><td></td></tr>
<tr><td>T2371</td><td>TCCGGAGCAGCCCACCGCAGAGG</td><td>1000</td><td></td><td></td><td></td><td></td><td></td><td></td><td></td><td></td><td></td></tr>
<tr><td>Oligo fwd</td><td>CACCTCCGGAGCAGCCCACCGCAG</td><td></td><td></td><td></td><td></td><td></td><td></td><td></td><td></td><td></td><td></td></tr>
<tr><td>Oligo rev</td><td>AAACCTGCGGTGGGCTGCTCCGGA</td><td></td><td></td><td></td><td></td><td></td><td></td><td></td><td></td><td></td><td></td></tr>
<tr><td>Chromosome</td><td>start</td><td>end</td><td>strand</td><td>MM</td><td>target_seq</td><td>PAM</td><td>alignment</td><td>distance</td><td>position</td><td>gene name</td><td>gene id</td></tr>
<tr><td>GU305914.1</td><td>37381</td><td>37403</td><td>+</td><td>0</td><td>TCCGGAGCAGCCCACCGCAG</td><td>AGG</td><td>||||||||[||||||||||||]PAM</td><td>NA</td><td>Intergenic</td><td>NA</td><td>NA</td></tr>
</table>

<table>
<tr><td>T2372</td><td>CATCCTCGATCCGCATCATGCGG</td><td>1000</td></tr>
<tr><td>Oligo fwd</td><td>CACCCATCCTCGATCCGCATCATG</td></tr>
<tr><td>Oligo rev</td><td>AAACCATGATGCGGATCGAGGATG</td></tr>
<tr><td>Chromosome</td><td>start</td><td>end</td><td>strand</td><td>MM</td><td>target_seq</td><td>PAM</td><td>alignment</td><td>distance</td><td>position</td><td>gene name</td><td>gene id</td></tr>
<tr><td>GU305914.1</td><td>37463</td><td>37485</td><td>+</td><td>0</td><td>CATCCTCGATCCGCATCATG</td><td>CGG</td><td>||||||||[||||||||||||]PAM</td><td>NA</td><td>Intergenic</td><td>NA</td><td>NA</td></tr>
<tr><td></td></tr>
<tr><td>T2373</td><td>TTCTCCGAAGCTCATCCCGCAGG</td><td>1000</td></tr>
<tr><td>Oligo fwd</td><td>CACCTTCTCCGAAGCTCATCCCGC</td></tr>
<tr><td>Oligo rev</td><td>AAACGCGGGATGAGCTTCGGAGAA</td></tr>
<tr><td>Chromosome</td><td>start</td><td>end</td><td>strand</td><td>MM</td><td>target_seq</td><td>PAM</td><td>alignment</td><td>distance</td><td>position</td><td>gene name</td><td>gene id</td></tr>
<tr><td>GU305914.1</td><td>37509</td><td>37531</td><td>+</td><td>0</td><td>TTCTCCGAAGCTCATCCCGC</td><td>AGG</td><td>||||||||[||||||||||||]PAM</td><td>NA</td><td>Intergenic</td><td>NA</td><td>NA</td></tr>
<tr><td></td></tr>
<tr><td>T2374</td><td>TCTCCGAAGCTCATCCCGCAGGG</td><td>1000</td></tr>
<tr><td>Oligo fwd</td><td>CACCTCTCCGAAGCTCATCCCGCA</td></tr>
<tr><td>Oligo rev</td><td>AAACTGCGGGATGAGCTTCGGAGA</td></tr>
<tr><td>Chromosome</td><td>start</td><td>end</td><td>strand</td><td>MM</td><td>target_seq</td><td>PAM</td><td>alignment</td><td>distance</td><td>position</td><td>gene name</td><td>gene id</td></tr>
<tr><td>GU305914.1</td><td>37510</td><td>37532</td><td>+</td><td>0</td><td>TCTCCGAAGCTCATCCCGCA</td><td>GGG</td><td>||||||||[||||||||||||]PAM</td><td>NA</td><td>Intergenic</td><td>NA</td><td>NA</td></tr>
<tr><td></td></tr>
<tr><td>T2375</td><td>GCTCATCCCGCAGGGCTGTGCGG</td><td>1000</td></tr>
<tr><td>Oligo fwd</td><td>CACCGCTCATCCCGCAGGGCTGTG</td></tr>
<tr><td>Oligo rev</td><td>AAACCACAGCCCTGCGGGATGAGC</td></tr>
<tr><td>Chromosome</td><td>start</td><td>end</td><td>strand</td><td>MM</td><td>target_seq</td><td>PAM</td><td>alignment</td><td>distance</td><td>position</td><td>gene name</td><td>gene id</td></tr>
<tr><td>GU305914.1</td><td>37518</td><td>37540</td><td>+</td><td>0</td><td>GCTCATCCCGCAGGGCTGTG</td><td>CGG</td><td>||||||||[||||||||||||]PAM</td><td>NA</td><td>Intergenic</td><td>NA</td><td>NA</td></tr>
<tr><td></td></tr>
<tr><td>T2376</td><td>CGCAGGGCTGTGCGGACCCCTGG</td><td>1000</td></tr>
<tr><td>Oligo fwd</td><td>CACCCGCAGGGCTGTGCGGACCCC</td></tr>
<tr><td>Oligo rev</td><td>AAACGGGGTCCGCACAGCCCTGCG</td></tr>
<tr><td>Chromosome</td><td>start</td><td>end</td><td>strand</td><td>MM</td><td>target_seq</td><td>PAM</td><td>alignment</td><td>distance</td><td>position</td><td>gene name</td><td>gene id</td></tr>
<tr><td>GU305914.1</td><td>37526</td><td>37548</td><td>+</td><td>0</td><td>CGCAGGGCTGTGCGGACCCC</td><td>TGG</td><td>||||||||[||||||||||||]PAM</td><td>NA</td><td>Intergenic</td><td>NA</td><td>NA</td></tr>
<tr><td></td></tr>
<tr><td>T2377</td><td>AGGGCTGTGCGGACCCCTGGCGG</td><td>1000</td></tr>
<tr><td>Oligo fwd</td><td>CACCAGGGCTGTGCGGACCCCTGG</td></tr>
<tr><td>Oligo rev</td><td>AAACCCAGGGGTCCGCACAGCCCT</td></tr>
<tr><td>Chromosome</td><td>start</td><td>end</td><td>strand</td><td>MM</td><td>target_seq</td><td>PAM</td><td>alignment</td><td>distance</td><td>position</td><td>gene name</td><td>gene id</td></tr>
<tr><td>GU305914.1</td><td>37529</td><td>37551</td><td>+</td><td>0</td><td>AGGGCTGTGCGGACCCCTGG</td><td>CGG</td><td>||||||||[||||||||||||]PAM</td><td>NA</td><td>Intergenic</td><td>NA</td><td>NA</td></tr>
<tr><td></td></tr>
<tr><td>T2378</td><td>CCTGGCGGCTCTCCGCCAAGCGG</td><td>1000</td></tr>
<tr><td>Oligo fwd</td><td>CACCCCTGGCGGCTCTCCGCCAAG</td></tr>
<tr><td>Oligo rev</td><td>AAACCTTGGCGGAGAGCCGCCAGG</td></tr>
<tr><td>Chromosome</td><td>start</td><td>end</td><td>strand</td><td>MM</td><td>target_seq</td><td>PAM</td><td>alignment</td><td>distance</td><td>position</td><td>gene name</td><td>gene id</td></tr>
<tr><td>GU305914.1</td><td>37544</td><td>37566</td><td>+</td><td>0</td><td>CCTGGCGGCTCTCCGCCAAG</td><td>CGG</td><td>||||||||[||||||||||||]PAM</td><td>NA</td><td>Intergenic</td><td>NA</td><td>NA</td></tr>
<tr><td></td></tr>
<tr><td>T2379</td><td>GGCGGCTCTCCGCCAAGCGGTGG</td><td>1000</td></tr>
<tr><td>Oligo fwd</td><td>CACCGGCGGCTCTCCGCCAAGCGG</td></tr>
<tr><td>Oligo rev</td><td>AAACCCGCTTGGCGGAGAGCCGCC</td></tr>
<tr><td>Chromosome</td><td>start</td><td>end</td><td>strand</td><td>MM</td><td>target_seq</td><td>PAM</td><td>alignment</td><td>distance</td><td>position</td><td>gene name</td><td>gene id</td></tr>
<tr><td>GU305914.1</td><td>37547</td><td>37569</td><td>+</td><td>0</td><td>GGCGGCTCTCCGCCAAGCGG</td><td>TGG</td><td>||||||||[||||||||||||]PAM</td><td>NA</td><td>Intergenic</td><td>NA</td><td>NA</td></tr>
<tr><td></td></tr>
<tr><td>T2380</td><td>CAACAGCCCTCCATCATCGTCGG</td><td>1000</td></tr>
<tr><td>Oligo fwd</td><td>CACCCAACAGCCCTCCATCATCGT</td></tr>
<tr><td>Oligo rev</td><td>AAACACGATGATGGAGGGCTGTTG</td></tr>
<tr><td>Chromosome</td><td>start</td><td>end</td><td>strand</td><td>MM</td><td>target_seq</td><td>PAM</td><td>alignment</td><td>distance</td><td>position</td><td>gene name</td><td>gene id</td></tr>
<tr><td>GU305914.1</td><td>37581</td><td>37603</td><td>+</td><td>0</td><td>CAACAGCCCTCCATCATCGT</td><td>CGG</td><td>||||||||[||||||||||||]PAM</td><td>NA</td><td>Intergenic</td><td>NA</td><td>NA</td></tr>
<tr><td></td></tr>
<tr><td>T2381</td><td>AACAGCCCTCCATCATCGTCGGG</td><td>1000</td></tr>
<tr><td>Oligo fwd</td><td>CACCAACAGCCCTCCATCATCGTC</td></tr>
<tr><td>Oligo rev</td><td>AAACGACGATGATGGAGGGCTGTT</td></tr>
<tr><td>Chromosome</td><td>start</td><td>end</td><td>strand</td><td>MM</td><td>target_seq</td><td>PAM</td><td>alignment</td><td>distance</td><td>position</td><td>gene name</td><td>gene id</td></tr>
<tr><td>GU305914.1</td><td>37582</td><td>37604</td><td>+</td><td>0</td><td>AACAGCCCTCCATCATCGTC</td><td>GGG</td><td>||||||||[||||||||||||]PAM</td><td>NA</td><td>Intergenic</td><td>NA</td><td>NA</td></tr>
<tr><td></td></tr>
<tr><td>T2382</td><td>ACAGCCCTCCATCATCGTCGGGG</td><td>1000</td></tr>
<tr><td>Oligo fwd</td><td>CACCACAGCCCTCCATCATCGTCG</td></tr>
<tr><td>Oligo rev</td><td>AAACCGACGATGATGGAGGGCTGT</td></tr>
<tr><td>Chromosome</td><td>start</td><td>end</td><td>strand</td><td>MM</td><td>target_seq</td><td>PAM</td><td>alignment</td><td>distance</td><td>position</td><td>gene name</td><td>gene id</td></tr>
<tr><td>GU305914.1</td><td>37583</td><td>37605</td><td>+</td><td>0</td><td>ACAGCCCTCCATCATCGTCG</td><td>GGG</td><td>||||||||[||||||||||||]PAM</td><td>NA</td><td>Intergenic</td><td>NA</td><td>NA</td></tr>
<tr><td></td></tr>
<tr><td>T2383</td><td>CGTCGGGGCACCCAGTTCGTCGG</td><td>1000</td></tr>
<tr><td>Oligo fwd</td><td>CACCCGTCGGGGCACCCAGTTCGT</td></tr>
<tr><td>Oligo rev</td><td>AAACACGAACTGGGTGCCCCGACG</td></tr>
<tr><td>Chromosome</td><td>start</td><td>end</td><td>strand</td><td>MM</td><td>target_seq</td><td>PAM</td><td>alignment</td><td>distance</td><td>position</td><td>gene name</td><td>gene id</td></tr>
<tr><td>GU305914.1</td><td>37598</td><td>37620</td><td>+</td><td>0</td><td>CGTCGGGGCACCCAGTTCGT</td><td>CGG</td><td>||||||||[||||||||||||]PAM</td><td>NA</td><td>Intergenic</td><td>NA</td><td>NA</td></tr>
<tr><td></td></tr>
<tr><td>T2384</td><td>CGGGGCACCCAGTTCGTCGGTGG</td><td>1000</td></tr>
<tr><td>Oligo fwd</td><td>CACCCGGGGCACCCAGTTCGTCGG</td></tr>
<tr><td>Oligo rev</td><td>AAACCCGACGAACTGGGTGCCCCG</td></tr>
<tr><td>Chromosome</td><td>start</td><td>end</td><td>strand</td><td>MM</td><td>target_seq</td><td>PAM</td><td>alignment</td><td>distance</td><td>position</td><td>gene name</td><td>gene id</td></tr>
<tr><td>GU305914.1</td><td>37601</td><td>37623</td><td>+</td><td>0</td><td>CGGGGCACCCAGTTCGTCGG</td><td>TGG</td><td>||||||||[||||||||||||]PAM</td><td>NA</td><td>Intergenic</td><td>NA</td><td>NA</td></tr>
<tr><td></td></tr>
<tr><td>T2385</td><td>TCGTCGGTGGCGCTCAATGCCGG</td><td>1000</td></tr>
<tr><td>Oligo fwd</td><td>CACCTCGTCGGTGGCGCTCAATGC</td></tr>
<tr><td>Oligo rev</td><td>AAACGCATTGAGCGCCACCGACGA</td></tr>
<tr><td>Chromosome</td><td>start</td><td>end</td><td>strand</td><td>MM</td><td>target_seq</td><td>PAM</td><td>alignment</td><td>distance</td><td>position</td><td>gene name</td><td>gene id</td></tr>
<tr><td>GU305914.1</td><td>37614</td><td>37636</td><td>+</td><td>0</td><td>TCGTCGGTGGCGCTCAATGC</td><td>CGG</td><td>||||||||[||||||||||||]PAM</td><td>NA</td><td>Intergenic</td><td>NA</td><td>NA</td></tr>
<tr><td></td></tr>
<tr><td>T2386</td><td>TGCCGGAGATTTCCATGCTCAGG</td><td>1000</td></tr>
<tr><td>Oligo fwd</td><td>CACCTGCCGGAGATTTCCATGCTC</td></tr>
<tr><td>Oligo rev</td><td>AAACGAGCATGGAAATCTCCGGCA</td></tr>
<tr><td>Chromosome</td><td>start</td><td>end</td><td>strand</td><td>MM</td><td>target_seq</td><td>PAM</td><td>alignment</td><td>distance</td><td>position</td><td>gene name</td><td>gene id</td></tr>
<tr><td>GU305914.1</td><td>37631</td><td>37653</td><td>+</td><td>0</td><td>TGCCGGAGATTTCCATGCTC</td><td>AGG</td><td>||||||||[||||||||||||]PAM</td><td>NA</td><td>Intergenic</td><td>NA</td><td>NA</td></tr>
</table>

<table>
<tr><td>T2387</td><td>GCTCAGGTCGTCATGCTGTACGG</td><td>1000</td><td></td><td></td><td></td><td></td><td></td><td></td><td></td><td></td><td></td></tr>
<tr><td>Oligo fwd</td><td>CACCGCTCAGGTCGTCATGCTGTA</td><td></td><td></td><td></td><td></td><td></td><td></td><td></td><td></td><td></td><td></td></tr>
<tr><td>Oligo rev</td><td>AAACTACAGCATGACGACCTGAGC</td><td></td><td></td><td></td><td></td><td></td><td></td><td></td><td></td><td></td><td></td></tr>
<tr><td>Chromosome</td><td>start</td><td>end</td><td>strand</td><td>MM</td><td>target_seq</td><td>PAM</td><td>alignment</td><td>distance</td><td>position</td><td>gene name</td><td>gene id</td></tr>
<tr><td>GU305914.1</td><td>37647</td><td>37669</td><td>+</td><td>0</td><td>GCTCAGGTCGTCATGCTGTA</td><td>CGG</td><td>||||||||[||||||||||||]PAM</td><td>NA</td><td>Intergenic</td><td>NA</td><td>NA</td></tr>
<tr><td>T2388</td><td>CGTCATGCTGTACGGCTCACAGG</td><td>1000</td><td></td><td></td><td></td><td></td><td></td><td></td><td></td><td></td><td></td></tr>
<tr><td>Oligo fwd</td><td>CACCCGTCATGCTGTACGGCTCAC</td><td></td><td></td><td></td><td></td><td></td><td></td><td></td><td></td><td></td><td></td></tr>
<tr><td>Oligo rev</td><td>AAACGTGAGCCGTACAGCATGACG</td><td></td><td></td><td></td><td></td><td></td><td></td><td></td><td></td><td></td><td></td></tr>
<tr><td>Chromosome</td><td>start</td><td>end</td><td>strand</td><td>MM</td><td>target_seq</td><td>PAM</td><td>alignment</td><td>distance</td><td>position</td><td>gene name</td><td>gene id</td></tr>
<tr><td>GU305914.1</td><td>37655</td><td>37677</td><td>+</td><td>0</td><td>CGTCATGCTGTACGGCTCAC</td><td>AGG</td><td>||||||||[||||||||||||]PAM</td><td>NA</td><td>Intergenic</td><td>NA</td><td>NA</td></tr>
<tr><td>T2389</td><td>CGGCTCACAGGCCACCGTCCTGG</td><td>1000</td><td></td><td></td><td></td><td></td><td></td><td></td><td></td><td></td><td></td></tr>
<tr><td>Oligo fwd</td><td>CACCCGGCTCACAGGCCACCGTCC</td><td></td><td></td><td></td><td></td><td></td><td></td><td></td><td></td><td></td><td></td></tr>
<tr><td>Oligo rev</td><td>AAACGGACGGTGGCCTGTGAGCCG</td><td></td><td></td><td></td><td></td><td></td><td></td><td></td><td></td><td></td><td></td></tr>
<tr><td>Chromosome</td><td>start</td><td>end</td><td>strand</td><td>MM</td><td>target_seq</td><td>PAM</td><td>alignment</td><td>distance</td><td>position</td><td>gene name</td><td>gene id</td></tr>
<tr><td>GU305914.1</td><td>37667</td><td>37689</td><td>+</td><td>0</td><td>CGGCTCACAGGCCACCGTCC</td><td>TGG</td><td>||||||||[||||||||||||]PAM</td><td>NA</td><td>Intergenic</td><td>NA</td><td>NA</td></tr>
<tr><td>T2390</td><td>ACAGGCCACCGTCCTGGACACGG</td><td>1000</td><td></td><td></td><td></td><td></td><td></td><td></td><td></td><td></td><td></td></tr>
<tr><td>Oligo fwd</td><td>CACCACAGGCCACCGTCCTGGACA</td><td></td><td></td><td></td><td></td><td></td><td></td><td></td><td></td><td></td><td></td></tr>
<tr><td>Oligo rev</td><td>AAACTGTCCAGGACGGTGGCCTGT</td><td></td><td></td><td></td><td></td><td></td><td></td><td></td><td></td><td></td><td></td></tr>
<tr><td>Chromosome</td><td>start</td><td>end</td><td>strand</td><td>MM</td><td>target_seq</td><td>PAM</td><td>alignment</td><td>distance</td><td>position</td><td>gene name</td><td>gene id</td></tr>
<tr><td>GU305914.1</td><td>37673</td><td>37695</td><td>+</td><td>0</td><td>ACAGGCCACCGTCCTGGACA</td><td>CGG</td><td>||||||||[||||||||||||]PAM</td><td>NA</td><td>Intergenic</td><td>NA</td><td>NA</td></tr>
<tr><td>T2391</td><td>CTCGATCGACGCGTCGTGCCCGG</td><td>1000</td><td></td><td></td><td></td><td></td><td></td><td></td><td></td><td></td><td></td></tr>
<tr><td>Oligo fwd</td><td>CACCCTCGATCGACGCGTCGTGCC</td><td></td><td></td><td></td><td></td><td></td><td></td><td></td><td></td><td></td><td></td></tr>
<tr><td>Oligo rev</td><td>AAACGGCACGACGCGTCGATCGAG</td><td></td><td></td><td></td><td></td><td></td><td></td><td></td><td></td><td></td><td></td></tr>
<tr><td>Chromosome</td><td>start</td><td>end</td><td>strand</td><td>MM</td><td>target_seq</td><td>PAM</td><td>alignment</td><td>distance</td><td>position</td><td>gene name</td><td>gene id</td></tr>
<tr><td>GU305914.1</td><td>37745</td><td>37767</td><td>+</td><td>0</td><td>CTCGATCGACGCGTCGTGCC</td><td>CGG</td><td>||||||||[||||||||||||]PAM</td><td>NA</td><td>Intergenic</td><td>NA</td><td>NA</td></tr>
<tr><td>T2392</td><td>CCCGGACACGCGCGTGAAGCTGG</td><td>1000</td><td></td><td></td><td></td><td></td><td></td><td></td><td></td><td></td><td></td></tr>
<tr><td>Oligo fwd</td><td>CACCCCCGGACACGCGCGTGAAGC</td><td></td><td></td><td></td><td></td><td></td><td></td><td></td><td></td><td></td><td></td></tr>
<tr><td>Oligo rev</td><td>AAACGCTTCACGCGCGTGTCCGGG</td><td></td><td></td><td></td><td></td><td></td><td></td><td></td><td></td><td></td><td></td></tr>
<tr><td>Chromosome</td><td>start</td><td>end</td><td>strand</td><td>MM</td><td>target_seq</td><td>PAM</td><td>alignment</td><td>distance</td><td>position</td><td>gene name</td><td>gene id</td></tr>
<tr><td>GU305914.1</td><td>37763</td><td>37785</td><td>+</td><td>0</td><td>CCCGGACACGCGCGTGAAGC</td><td>TGG</td><td>||||||||[||||||||||||]PAM</td><td>NA</td><td>Intergenic</td><td>NA</td><td>NA</td></tr>
<tr><td>T2393</td><td>AGCTGGATCGCACCGAGAGACGG</td><td>1000</td><td></td><td></td><td></td><td></td><td></td><td></td><td></td><td></td><td></td></tr>
<tr><td>Oligo fwd</td><td>CACCAGCTGGATCGCACCGAGAGA</td><td></td><td></td><td></td><td></td><td></td><td></td><td></td><td></td><td></td><td></td></tr>
<tr><td>Oligo rev</td><td>AAACTCTCTCGGTGCGATCCAGCT</td><td></td><td></td><td></td><td></td><td></td><td></td><td></td><td></td><td></td><td></td></tr>
<tr><td>Chromosome</td><td>start</td><td>end</td><td>strand</td><td>MM</td><td>target_seq</td><td>PAM</td><td>alignment</td><td>distance</td><td>position</td><td>gene name</td><td>gene id</td></tr>
<tr><td>GU305914.1</td><td>37780</td><td>37802</td><td>+</td><td>0</td><td>AGCTGGATCGCACCGAGAGA</td><td>CGG</td><td>||||||||[||||||||||||]PAM</td><td>NA</td><td>Intergenic</td><td>NA</td><td>NA</td></tr>
<tr><td>T2394</td><td>TTCCAACTACCGCTACCGACTGG</td><td>1000</td><td></td><td></td><td></td><td></td><td></td><td></td><td></td><td></td><td></td></tr>
<tr><td>Oligo fwd</td><td>CACCTTCCAACTACCGCTACCGAC</td><td></td><td></td><td></td><td></td><td></td><td></td><td></td><td></td><td></td><td></td></tr>
<tr><td>Oligo rev</td><td>AAACGTCGGTAGCGGTAGTTGGAA</td><td></td><td></td><td></td><td></td><td></td><td></td><td></td><td></td><td></td><td></td></tr>
<tr><td>Chromosome</td><td>start</td><td>end</td><td>strand</td><td>MM</td><td>target_seq</td><td>PAM</td><td>alignment</td><td>distance</td><td>position</td><td>gene name</td><td>gene id</td></tr>
<tr><td>GU305914.1</td><td>37808</td><td>37830</td><td>+</td><td>0</td><td>TTCCAACTACCGCTACCGAC</td><td>TGG</td><td>||||||||[||||||||||||]PAM</td><td>NA</td><td>Intergenic</td><td>NA</td><td>NA</td></tr>
<tr><td>T2395</td><td>CAGCGCCAGCCAGTACAACGAGG</td><td>1000</td><td></td><td></td><td></td><td></td><td></td><td></td><td></td><td></td><td></td></tr>
<tr><td>Oligo fwd</td><td>CACCCAGCGCCAGCCAGTACAACG</td><td></td><td></td><td></td><td></td><td></td><td></td><td></td><td></td><td></td><td></td></tr>
<tr><td>Oligo rev</td><td>AAACCGTTGTACTGGCTGGCGCTG</td><td></td><td></td><td></td><td></td><td></td><td></td><td></td><td></td><td></td><td></td></tr>
<tr><td>Chromosome</td><td>start</td><td>end</td><td>strand</td><td>MM</td><td>target_seq</td><td>PAM</td><td>alignment</td><td>distance</td><td>position</td><td>gene name</td><td>gene id</td></tr>
<tr><td>GU305914.1</td><td>37835</td><td>37857</td><td>+</td><td>0</td><td>CAGCGCCAGCCAGTACAACG</td><td>AGG</td><td>||||||||[||||||||||||]PAM</td><td>NA</td><td>Intergenic</td><td>NA</td><td>NA</td></tr>
<tr><td>T2396</td><td>CGCCAGCCAGTACAACGAGGTGG</td><td>1000</td><td></td><td></td><td></td><td></td><td></td><td></td><td></td><td></td><td></td></tr>
<tr><td>Oligo fwd</td><td>CACCCGCCAGCCAGTACAACGAGG</td><td></td><td></td><td></td><td></td><td></td><td></td><td></td><td></td><td></td><td></td></tr>
<tr><td>Oligo rev</td><td>AAACCCTCGTTGTACTGGCTGGCG</td><td></td><td></td><td></td><td></td><td></td><td></td><td></td><td></td><td></td><td></td></tr>
<tr><td>Chromosome</td><td>start</td><td>end</td><td>strand</td><td>MM</td><td>target_seq</td><td>PAM</td><td>alignment</td><td>distance</td><td>position</td><td>gene name</td><td>gene id</td></tr>
<tr><td>GU305914.1</td><td>37838</td><td>37860</td><td>+</td><td>0</td><td>CGCCAGCCAGTACAACGAGG</td><td>TGG</td><td>||||||||[||||||||||||]PAM</td><td>NA</td><td>Intergenic</td><td>NA</td><td>NA</td></tr>
<tr><td>T2397</td><td>GTACAACGAGGTGGTGCCGTCGG</td><td>1000</td><td></td><td></td><td></td><td></td><td></td><td></td><td></td><td></td><td></td></tr>
<tr><td>Oligo fwd</td><td>CACCGTACAACGAGGTGGTGCCGT</td><td></td><td></td><td></td><td></td><td></td><td></td><td></td><td></td><td></td><td></td></tr>
<tr><td>Oligo rev</td><td>AAACACGGCACCACCTCGTTGTAC</td><td></td><td></td><td></td><td></td><td></td><td></td><td></td><td></td><td></td><td></td></tr>
<tr><td>Chromosome</td><td>start</td><td>end</td><td>strand</td><td>MM</td><td>target_seq</td><td>PAM</td><td>alignment</td><td>distance</td><td>position</td><td>gene name</td><td>gene id</td></tr>
<tr><td>GU305914.1</td><td>37847</td><td>37869</td><td>+</td><td>0</td><td>GTACAACGAGGTGGTGCCGT</td><td>CGG</td><td>||||||||[||||||||||||]PAM</td><td>NA</td><td>Intergenic</td><td>NA</td><td>NA</td></tr>
<tr><td>T2398</td><td>CTGCGCGCTCGCTACTACAAAGG</td><td>1000</td><td></td><td></td><td></td><td></td><td></td><td></td><td></td><td></td><td></td></tr>
<tr><td>Oligo fwd</td><td>CACCCTGCGCGCTCGCTACTACAA</td><td></td><td></td><td></td><td></td><td></td><td></td><td></td><td></td><td></td><td></td></tr>
<tr><td>Oligo rev</td><td>AAACTTGTAGTAGCGAGCGCGCAG</td><td></td><td></td><td></td><td></td><td></td><td></td><td></td><td></td><td></td><td></td></tr>
<tr><td>Chromosome</td><td>start</td><td>end</td><td>strand</td><td>MM</td><td>target_seq</td><td>PAM</td><td>alignment</td><td>distance</td><td>position</td><td>gene name</td><td>gene id</td></tr>
<tr><td>GU305914.1</td><td>37872</td><td>37894</td><td>+</td><td>0</td><td>CTGCGCGCTCGCTACTACAA</td><td>AGG</td><td>||||||||[||||||||||||]PAM</td><td>NA</td><td>Intergenic</td><td>NA</td><td>NA</td></tr>
<tr><td>T2399</td><td>AGGCGCCATTCACTTCGCGACGG</td><td>1000</td><td></td><td></td><td></td><td></td><td></td><td></td><td></td><td></td><td></td></tr>
<tr><td>Oligo fwd</td><td>CACCAGGCGCCATTCACTTCGCGA</td><td></td><td></td><td></td><td></td><td></td><td></td><td></td><td></td><td></td><td></td></tr>
<tr><td>Oligo rev</td><td>AAACTCGCGAAGTGAATGGCGCCT</td><td></td><td></td><td></td><td></td><td></td><td></td><td></td><td></td><td></td><td></td></tr>
<tr><td>Chromosome</td><td>start</td><td>end</td><td>strand</td><td>MM</td><td>target_seq</td><td>PAM</td><td>alignment</td><td>distance</td><td>position</td><td>gene name</td><td>gene id</td></tr>
<tr><td>GU305914.1</td><td>37892</td><td>37914</td><td>+</td><td>0</td><td>AGGCGCCATTCACTTCGCGA</td><td>CGG</td><td>||||||||[||||||||||||]PAM</td><td>NA</td><td>Intergenic</td><td>NA</td><td>NA</td></tr>
<tr><td>T2400</td><td>CGCCATTCACTTCGCGACGGCGG</td><td>1000</td><td></td><td></td><td></td><td></td><td></td><td></td><td></td><td></td><td></td></tr>
<tr><td>Oligo fwd</td><td>CACCCGCCATTCACTTCGCGACGG</td><td></td><td></td><td></td><td></td><td></td><td></td><td></td><td></td><td></td><td></td></tr>
<tr><td>Oligo rev</td><td>AAACCCGTCGCGAAGTGAATGGCG</td><td></td><td></td><td></td><td></td><td></td><td></td><td></td><td></td><td></td><td></td></tr>
<tr><td>Chromosome</td><td>start</td><td>end</td><td>strand</td><td>MM</td><td>target_seq</td><td>PAM</td><td>alignment</td><td>distance</td><td>position</td><td>gene name</td><td>gene id</td></tr>
<tr><td>GU305914.1</td><td>37895</td><td>37917</td><td>+</td><td>0</td><td>CGCCATTCACTTCGCGACGG</td><td>CGG</td><td>||||||||[||||||||||||]PAM</td><td>NA</td><td>Intergenic</td><td>NA</td><td>NA</td></tr>
<tr><td>T2401</td><td>CACTTCGCGACGGCGGTGATCGG</td><td>1000</td><td></td><td></td><td></td><td></td><td></td><td></td><td></td><td></td><td></td></tr>
<tr><td>Oligo fwd</td><td>CACCCACTTCGCGACGGCGGTGAT</td><td></td><td></td><td></td><td></td><td></td><td></td><td></td><td></td><td></td><td></td></tr>
<tr><td>Oligo rev</td><td>AAACATCACCGCCGTCGCGAAGTG</td><td></td><td></td><td></td><td></td><td></td><td></td><td></td><td></td><td></td><td></td></tr>
<tr><td>Chromosome</td><td>start</td><td>end</td><td>strand</td><td>MM</td><td>target_seq</td><td>PAM</td><td>alignment</td><td>distance</td><td>position</td><td>gene name</td><td>gene id</td></tr>
</table>

| | | | | | | | | | | | |
|---|---|---|---|---|---|---|---|---|---|---|---|
| GU305914.1 | 37902 | 37924 | + | 0 | CACTTCGCGACGGCGGTGAT | CGG | \|\|\|\|\|\|\|\|[\|\|\|\|\|\|\|\|\|\|\|\|]PAM | NA | Intergenic | NA | NA |
| T2402 | CGGCACGATGTTCCGCACCTCGG | 1000 | | | | | | | | | |
| Oligo fwd | CACCCGGCACGATGTTCCGCACCT | | | | | | | | | | |
| Oligo rev | AAACAGGTGCGGAACATCGTGCCG | | | | | | | | | | |
| Chromosome | start | end | strand | MM | target_seq | PAM | alignment | distance | position | gene name | gene id |
| GU305914.1 | 37922 | 37944 | + | 0 | CGGCACGATGTTCCGCACCT | CGG | \|\|\|\|\|\|\|\|[\|\|\|\|\|\|\|\|\|\|\|\|]PAM | NA | Intergenic | NA | NA |
| T2403 | CGGACGAGCTGACGTACACCTGG | 1000 | | | | | | | | | |
| Oligo fwd | CACCCGGACGAGCTGACGTACACC | | | | | | | | | | |
| Oligo rev | AAACGGTGTACGTCAGCTCGTCCG | | | | | | | | | | |
| Chromosome | start | end | strand | MM | target_seq | PAM | alignment | distance | position | gene name | gene id |
| GU305914.1 | 37942 | 37964 | + | 0 | CGGACGAGCTGACGTACACC | TGG | \|\|\|\|\|\|\|\|[\|\|\|\|\|\|\|\|\|\|\|\|]PAM | NA | Intergenic | NA | NA |
| T2404 | GTACACCTGGACCACGCTGCTGG | 1000 | | | | | | | | | |
| Oligo fwd | CACCGTACACCTGGACCACGCTGC | | | | | | | | | | |
| Oligo rev | AAACGCAGCGTGGTCCAGGTGTAC | | | | | | | | | | |
| Chromosome | start | end | strand | MM | target_seq | PAM | alignment | distance | position | gene name | gene id |
| GU305914.1 | 37955 | 37977 | + | 0 | GTACACCTGGACCACGCTGC | TGG | \|\|\|\|\|\|\|\|[\|\|\|\|\|\|\|\|\|\|\|\|]PAM | NA | Intergenic | NA | NA |
| T2405 | CCCTCGCGTCTCGAACATCCCGG | 1000 | | | | | | | | | |
| Oligo fwd | CACCCCCTCGCGTCTCGAACATCC | | | | | | | | | | |
| Oligo rev | AAACGGATGTTCGAGACGCGAGGG | | | | | | | | | | |
| Chromosome | start | end | strand | MM | target_seq | PAM | alignment | distance | position | gene name | gene id |
| GU305914.1 | 37979 | 38001 | + | 0 | CCCTCGCGTCTCGAACATCC | CGG | \|\|\|\|\|\|\|\|[\|\|\|\|\|\|\|\|\|\|\|\|]PAM | NA | Intergenic | NA | NA |
| T2406 | CCTCGCGTCTCGAACATCCCGGG | 1000 | | | | | | | | | |
| Oligo fwd | CACCCCTCGCGTCTCGAACATCCC | | | | | | | | | | |
| Oligo rev | AAACGGGATGTTCGAGACGCGAGG | | | | | | | | | | |
| Chromosome | start | end | strand | MM | target_seq | PAM | alignment | distance | position | gene name | gene id |
| GU305914.1 | 37980 | 38002 | + | 0 | CCTCGCGTCTCGAACATCCC | GGG | \|\|\|\|\|\|\|\|[\|\|\|\|\|\|\|\|\|\|\|\|]PAM | NA | Intergenic | NA | NA |
| T2407 | CGCGTCTCGAACATCCCGGGCGG | 1000 | | | | | | | | | |
| Oligo fwd | CACCCGCGTCTCGAACATCCCGGG | | | | | | | | | | |
| Oligo rev | AAACCCCGGGATGTTCGAGACGCG | | | | | | | | | | |
| Chromosome | start | end | strand | MM | target_seq | PAM | alignment | distance | position | gene name | gene id |
| GU305914.1 | 37983 | 38005 | + | 0 | CGCGTCTCGAACATCCCGGG | CGG | \|\|\|\|\|\|\|\|[\|\|\|\|\|\|\|\|\|\|\|\|]PAM | NA | Intergenic | NA | NA |
| T2408 | GTTCCTGCTGTGCTCCCTGAAGG | 1000 | | | | | | | | | |
| Oligo fwd | CACCGTTCCTGCTGTGCTCCCTGA | | | | | | | | | | |
| Oligo rev | AAACTCAGGGAGCACAGCAGGAAC | | | | | | | | | | |
| Chromosome | start | end | strand | MM | target_seq | PAM | alignment | distance | position | gene name | gene id |
| GU305914.1 | 38009 | 38031 | + | 0 | GTTCCTGCTGTGCTCCCTGA | AGG | \|\|\|\|\|\|\|\|[\|\|\|\|\|\|\|\|\|\|\|\|]PAM | NA | Intergenic | NA | NA |
| T2409 | TTCCTGCTGTGCTCCCTGAAGGG | 1000 | | | | | | | | | |
| Oligo fwd | CACCTTCCTGCTGTGCTCCCTGAA | | | | | | | | | | |
| Oligo rev | AAACTTCAGGGAGCACAGCAGGAA | | | | | | | | | | |
| Chromosome | start | end | strand | MM | target_seq | PAM | alignment | distance | position | gene name | gene id |
| GU305914.1 | 38010 | 38032 | + | 0 | TTCCTGCTGTGCTCCCTGAA | GGG | \|\|\|\|\|\|\|\|[\|\|\|\|\|\|\|\|\|\|\|\|]PAM | NA | Intergenic | NA | NA |
| T2410 | CTCCCTGAAGGGAAACGCAGCGG | 1000 | | | | | | | | | |
| Oligo fwd | CACCCTCCCTGAAGGGAAACGCAG | | | | | | | | | | |
| Oligo rev | AAACCTGCGTTTCCCTTCAGGGAG | | | | | | | | | | |
| Chromosome | start | end | strand | MM | target_seq | PAM | alignment | distance | position | gene name | gene id |
| GU305914.1 | 38021 | 38043 | + | 0 | CTCCCTGAAGGGAAACGCAG | CGG | \|\|\|\|\|\|\|\|[\|\|\|\|\|\|\|\|\|\|\|\|]PAM | NA | Intergenic | NA | NA |
| T2411 | GTGTCAACCGACCGTGACCACGG | 1000 | | | | | | | | | |
| Oligo fwd | CACCGTGTCAACCGACCGTGACCA | | | | | | | | | | |
| Oligo rev | AAACTGGTCACGGTCGGTTGACAC | | | | | | | | | | |
| Chromosome | start | end | strand | MM | target_seq | PAM | alignment | distance | position | gene name | gene id |
| GU305914.1 | 38045 | 38067 | + | 0 | GTGTCAACCGACCGTGACCA | CGG | \|\|\|\|\|\|\|\|[\|\|\|\|\|\|\|\|\|\|\|\|]PAM | NA | Intergenic | NA | NA |
| T2412 | TGTCAACCGACCGTGACCACGGG | 1000 | | | | | | | | | |
| Oligo fwd | CACCTGTCAACCGACCGTGACCAC | | | | | | | | | | |
| Oligo rev | AAACGTGGTCACGGTCGGTTGACA | | | | | | | | | | |
| Chromosome | start | end | strand | MM | target_seq | PAM | alignment | distance | position | gene name | gene id |
| GU305914.1 | 38046 | 38068 | + | 0 | TGTCAACCGACCGTGACCAC | GGG | \|\|\|\|\|\|\|\|[\|\|\|\|\|\|\|\|\|\|\|\|]PAM | NA | Intergenic | NA | NA |
| T2413 | CAACCGACCGTGACCACGGGTGG | 1000 | | | | | | | | | |
| Oligo fwd | CACCCAACCGACCGTGACCACGGG | | | | | | | | | | |
| Oligo rev | AAACCCCGTGGTCACGGTCGGTTG | | | | | | | | | | |
| Chromosome | start | end | strand | MM | target_seq | PAM | alignment | distance | position | gene name | gene id |
| GU305914.1 | 38049 | 38071 | + | 0 | CAACCGACCGTGACCACGGG | TGG | \|\|\|\|\|\|\|\|[\|\|\|\|\|\|\|\|\|\|\|\|]PAM | NA | Intergenic | NA | NA |
| T2414 | TGGCCCGCACACCATCCTGCTGG | 1000 | | | | | | | | | |
| Oligo fwd | CACCTGGCCCGCACACCATCCTGC | | | | | | | | | | |
| Oligo rev | AAACGCAGGATGGTGTGCGGGCCA | | | | | | | | | | |
| Chromosome | start | end | strand | MM | target_seq | PAM | alignment | distance | position | gene name | gene id |
| GU305914.1 | 38069 | 38091 | + | 0 | TGGCCCGCACACCATCCTGC | TGG | \|\|\|\|\|\|\|\|[\|\|\|\|\|\|\|\|\|\|\|\|]PAM | NA | Intergenic | NA | NA |
| T2415 | CGACCCGTCCAACGTGCCGCCGG | 1000 | | | | | | | | | |
| Oligo fwd | CACCCGACCCGTCCAACGTGCCGC | | | | | | | | | | |
| Oligo rev | AAACGCGGCACGTTGGACGGGTCG | | | | | | | | | | |
| Chromosome | start | end | strand | MM | target_seq | PAM | alignment | distance | position | gene name | gene id |
| GU305914.1 | 38096 | 38118 | + | 0 | CGACCCGTCCAACGTGCCGC | CGG | \|\|\|\|\|\|\|\|[\|\|\|\|\|\|\|\|\|\|\|\|]PAM | NA | Intergenic | NA | NA |
| GU305914.1 | 198812 | 198834 | - | 4 | CGCCTCGTGCGACGTGCCGC | GGG | \|\|-\|-\|\|\|[-\|-\|\|\|\|\|\|\|\|\|]PAM | NA | Intergenic | NA | NA |
| T2416 | CCAACGTGCCGCCGGACAACAGG | 1000 | | | | | | | | | |
| Oligo fwd | CACCCCAACGTGCCGCCGGACAAC | | | | | | | | | | |

<table>
<tr><td>Oligo rev</td><td>AAACGTTGTCCGGCGGCACGTTGG</td><td></td><td></td><td></td><td></td><td></td><td></td><td></td><td></td><td></td><td></td></tr>
<tr><td>Chromosome</td><td>start</td><td>end</td><td>strand</td><td>MM</td><td>target_seq</td><td>PAM</td><td>alignment</td><td>distance</td><td>position</td><td>gene name</td><td>gene id</td></tr>
<tr><td>GU305914.1</td><td>38104</td><td>38126</td><td>+</td><td>0</td><td>CCAACGTGCCGCCGGACAAC</td><td>AGG</td><td>||||||||[||||||||||||]PAM</td><td>NA</td><td>Intergenic</td><td>NA</td><td>NA</td></tr>
<tr><td>T2417</td><td>CGCCGGACAACAGGCCCGTGAGG</td><td>1000</td><td></td><td></td><td></td><td></td><td></td><td></td><td></td><td></td><td></td></tr>
<tr><td>Oligo fwd</td><td>CACCCGCCGGACAACAGGCCCGTG</td><td></td><td></td><td></td><td></td><td></td><td></td><td></td><td></td><td></td><td></td></tr>
<tr><td>Oligo rev</td><td>AAACCACGGGCCTGTTGTCCGGCG</td><td></td><td></td><td></td><td></td><td></td><td></td><td></td><td></td><td></td><td></td></tr>
<tr><td>Chromosome</td><td>start</td><td>end</td><td>strand</td><td>MM</td><td>target_seq</td><td>PAM</td><td>alignment</td><td>distance</td><td>position</td><td>gene name</td><td>gene id</td></tr>
<tr><td>GU305914.1</td><td>38113</td><td>38135</td><td>+</td><td>0</td><td>CGCCGGACAACAGGCCCGTG</td><td>AGG</td><td>||||||||[||||||||||||]PAM</td><td>NA</td><td>Intergenic</td><td>NA</td><td>NA</td></tr>
<tr><td>T2418</td><td>GGACAACAGGCCCGTGAGGCTGG</td><td>1000</td><td></td><td></td><td></td><td></td><td></td><td></td><td></td><td></td><td></td></tr>
<tr><td>Oligo fwd</td><td>CACCGGACAACAGGCCCGTGAGGC</td><td></td><td></td><td></td><td></td><td></td><td></td><td></td><td></td><td></td><td></td></tr>
<tr><td>Oligo rev</td><td>AAACGCCTCACGGGCCTGTTGTCC</td><td></td><td></td><td></td><td></td><td></td><td></td><td></td><td></td><td></td><td></td></tr>
<tr><td>Chromosome</td><td>start</td><td>end</td><td>strand</td><td>MM</td><td>target_seq</td><td>PAM</td><td>alignment</td><td>distance</td><td>position</td><td>gene name</td><td>gene id</td></tr>
<tr><td>GU305914.1</td><td>38117</td><td>38139</td><td>+</td><td>0</td><td>GGACAACAGGCCCGTGAGGC</td><td>TGG</td><td>||||||||[||||||||||||]PAM</td><td>NA</td><td>Intergenic</td><td>NA</td><td>NA</td></tr>
<tr><td>T2419</td><td>GCCCGTGAGGCTGGAGCTGTCGG</td><td>1000</td><td></td><td></td><td></td><td></td><td></td><td></td><td></td><td></td><td></td></tr>
<tr><td>Oligo fwd</td><td>CACCGCCCGTGAGGCTGGAGCTGT</td><td></td><td></td><td></td><td></td><td></td><td></td><td></td><td></td><td></td><td></td></tr>
<tr><td>Oligo rev</td><td>AAACACAGCTCCAGCCTCACGGGC</td><td></td><td></td><td></td><td></td><td></td><td></td><td></td><td></td><td></td><td></td></tr>
<tr><td>Chromosome</td><td>start</td><td>end</td><td>strand</td><td>MM</td><td>target_seq</td><td>PAM</td><td>alignment</td><td>distance</td><td>position</td><td>gene name</td><td>gene id</td></tr>
<tr><td>GU305914.1</td><td>38126</td><td>38148</td><td>+</td><td>0</td><td>GCCCGTGAGGCTGGAGCTGT</td><td>CGG</td><td>||||||||[||||||||||||]PAM</td><td>NA</td><td>Intergenic</td><td>NA</td><td>NA</td></tr>
<tr><td>T2420</td><td>CTTCGTGCCGTACGTCGCGCCGG</td><td>1000</td><td></td><td></td><td></td><td></td><td></td><td></td><td></td><td></td><td></td></tr>
<tr><td>Oligo fwd</td><td>CACCCTTCGTGCCGTACGTCGCGC</td><td></td><td></td><td></td><td></td><td></td><td></td><td></td><td></td><td></td><td></td></tr>
<tr><td>Oligo rev</td><td>AAACGCGCGACGTACGGCACGAAG</td><td></td><td></td><td></td><td></td><td></td><td></td><td></td><td></td><td></td><td></td></tr>
<tr><td>Chromosome</td><td>start</td><td>end</td><td>strand</td><td>MM</td><td>target_seq</td><td>PAM</td><td>alignment</td><td>distance</td><td>position</td><td>gene name</td><td>gene id</td></tr>
<tr><td>GU305914.1</td><td>38156</td><td>38178</td><td>+</td><td>0</td><td>CTTCGTGCCGTACGTCGCGC</td><td>CGG</td><td>||||||||[||||||||||||]PAM</td><td>NA</td><td>Intergenic</td><td>NA</td><td>NA</td></tr>
<tr><td>T2421</td><td>CCGTACGTCGCGCCGGACTTCGG</td><td>1000</td><td></td><td></td><td></td><td></td><td></td><td></td><td></td><td></td><td></td></tr>
<tr><td>Oligo fwd</td><td>CACCCCGTACGTCGCGCCGGACTT</td><td></td><td></td><td></td><td></td><td></td><td></td><td></td><td></td><td></td><td></td></tr>
<tr><td>Oligo rev</td><td>AAACAAGTCCGGCGCGACGTACGG</td><td></td><td></td><td></td><td></td><td></td><td></td><td></td><td></td><td></td><td></td></tr>
<tr><td>Chromosome</td><td>start</td><td>end</td><td>strand</td><td>MM</td><td>target_seq</td><td>PAM</td><td>alignment</td><td>distance</td><td>position</td><td>gene name</td><td>gene id</td></tr>
<tr><td>GU305914.1</td><td>38163</td><td>38185</td><td>+</td><td>0</td><td>CCGTACGTCGCGCCGGACTT</td><td>CGG</td><td>||||||||[||||||||||||]PAM</td><td>NA</td><td>Intergenic</td><td>NA</td><td>NA</td></tr>
<tr><td>T2422</td><td>CGGCGTGCGCATGCTGACGCTGG</td><td>1000</td><td></td><td></td><td></td><td></td><td></td><td></td><td></td><td></td><td></td></tr>
<tr><td>Oligo fwd</td><td>CACCCGGCGTGCGCATGCTGACGC</td><td></td><td></td><td></td><td></td><td></td><td></td><td></td><td></td><td></td><td></td></tr>
<tr><td>Oligo rev</td><td>AAACGCGTCAGCATGCGCACGCCG</td><td></td><td></td><td></td><td></td><td></td><td></td><td></td><td></td><td></td><td></td></tr>
<tr><td>Chromosome</td><td>start</td><td>end</td><td>strand</td><td>MM</td><td>target_seq</td><td>PAM</td><td>alignment</td><td>distance</td><td>position</td><td>gene name</td><td>gene id</td></tr>
<tr><td>GU305914.1</td><td>38183</td><td>38205</td><td>+</td><td>0</td><td>CGGCGTGCGCATGCTGACGC</td><td>TGG</td><td>||||||||[||||||||||||]PAM</td><td>NA</td><td>Intergenic</td><td>NA</td><td>NA</td></tr>
<tr><td>T2423</td><td>CATGCTGACGCTGGAAGACGCGG</td><td>1000</td><td></td><td></td><td></td><td></td><td></td><td></td><td></td><td></td><td></td></tr>
<tr><td>Oligo fwd</td><td>CACCCATGCTGACGCTGGAAGACG</td><td></td><td></td><td></td><td></td><td></td><td></td><td></td><td></td><td></td><td></td></tr>
<tr><td>Oligo rev</td><td>AAACCGTCTTCCAGCGTCAGCATG</td><td></td><td></td><td></td><td></td><td></td><td></td><td></td><td></td><td></td><td></td></tr>
<tr><td>Chromosome</td><td>start</td><td>end</td><td>strand</td><td>MM</td><td>target_seq</td><td>PAM</td><td>alignment</td><td>distance</td><td>position</td><td>gene name</td><td>gene id</td></tr>
<tr><td>GU305914.1</td><td>38192</td><td>38214</td><td>+</td><td>0</td><td>CATGCTGACGCTGGAAGACG</td><td>CGG</td><td>||||||||[||||||||||||]PAM</td><td>NA</td><td>Intergenic</td><td>NA</td><td>NA</td></tr>
<tr><td>T2424</td><td>GACGCGGCGATCCGCGCCAGCGG</td><td>1000</td><td></td><td></td><td></td><td></td><td></td><td></td><td></td><td></td><td></td></tr>
<tr><td>Oligo fwd</td><td>CACCGACGCGGCGATCCGCGCCAG</td><td></td><td></td><td></td><td></td><td></td><td></td><td></td><td></td><td></td><td></td></tr>
<tr><td>Oligo rev</td><td>AAACCTGGCGCGGATCGCCGCGTC</td><td></td><td></td><td></td><td></td><td></td><td></td><td></td><td></td><td></td><td></td></tr>
<tr><td>Chromosome</td><td>start</td><td>end</td><td>strand</td><td>MM</td><td>target_seq</td><td>PAM</td><td>alignment</td><td>distance</td><td>position</td><td>gene name</td><td>gene id</td></tr>
<tr><td>GU305914.1</td><td>38208</td><td>38230</td><td>+</td><td>0</td><td>GACGCGGCGATCCGCGCCAG</td><td>CGG</td><td>||||||||[||||||||||||]PAM</td><td>NA</td><td>Intergenic</td><td>NA</td><td>NA</td></tr>
<tr><td>T2425</td><td>CGCGCCAGCGGCATCAGCTTCGG</td><td>1000</td><td></td><td></td><td></td><td></td><td></td><td></td><td></td><td></td><td></td></tr>
<tr><td>Oligo fwd</td><td>CACCCGCGCCAGCGGCATCAGCTT</td><td></td><td></td><td></td><td></td><td></td><td></td><td></td><td></td><td></td><td></td></tr>
<tr><td>Oligo rev</td><td>AAACAAGCTGATGCCGCTGGCGCG</td><td></td><td></td><td></td><td></td><td></td><td></td><td></td><td></td><td></td><td></td></tr>
<tr><td>Chromosome</td><td>start</td><td>end</td><td>strand</td><td>MM</td><td>target_seq</td><td>PAM</td><td>alignment</td><td>distance</td><td>position</td><td>gene name</td><td>gene id</td></tr>
<tr><td>GU305914.1</td><td>38220</td><td>38242</td><td>+</td><td>0</td><td>CGCGCCAGCGGCATCAGCTT</td><td>CGG</td><td>||||||||[||||||||||||]PAM</td><td>NA</td><td>Intergenic</td><td>NA</td><td>NA</td></tr>
<tr><td>T2426</td><td>TCGGCATCATGACCCCCGTGCGG</td><td>1000</td><td></td><td></td><td></td><td></td><td></td><td></td><td></td><td></td><td></td></tr>
<tr><td>Oligo fwd</td><td>CACCTCGGCATCATGACCCCCGTG</td><td></td><td></td><td></td><td></td><td></td><td></td><td></td><td></td><td></td><td></td></tr>
<tr><td>Oligo rev</td><td>AAACCACGGGGGTCATGATGCCGA</td><td></td><td></td><td></td><td></td><td></td><td></td><td></td><td></td><td></td><td></td></tr>
<tr><td>Chromosome</td><td>start</td><td>end</td><td>strand</td><td>MM</td><td>target_seq</td><td>PAM</td><td>alignment</td><td>distance</td><td>position</td><td>gene name</td><td>gene id</td></tr>
<tr><td>GU305914.1</td><td>38239</td><td>38261</td><td>+</td><td>0</td><td>TCGGCATCATGACCCCCGTG</td><td>CGG</td><td>||||||||[||||||||||||]PAM</td><td>NA</td><td>Intergenic</td><td>NA</td><td>NA</td></tr>
<tr><td>T2427</td><td>GCATCATGACCCCCGTGCGGAGG</td><td>1000</td><td></td><td></td><td></td><td></td><td></td><td></td><td></td><td></td><td></td></tr>
<tr><td>Oligo fwd</td><td>CACCGCATCATGACCCCCGTGCGG</td><td></td><td></td><td></td><td></td><td></td><td></td><td></td><td></td><td></td><td></td></tr>
<tr><td>Oligo rev</td><td>AAACCCGCACGGGGGTCATGATGC</td><td></td><td></td><td></td><td></td><td></td><td></td><td></td><td></td><td></td><td></td></tr>
<tr><td>Chromosome</td><td>start</td><td>end</td><td>strand</td><td>MM</td><td>target_seq</td><td>PAM</td><td>alignment</td><td>distance</td><td>position</td><td>gene name</td><td>gene id</td></tr>
<tr><td>GU305914.1</td><td>38242</td><td>38264</td><td>+</td><td>0</td><td>GCATCATGACCCCCGTGCGG</td><td>AGG</td><td>||||||||[||||||||||||]PAM</td><td>NA</td><td>Intergenic</td><td>NA</td><td>NA</td></tr>
<tr><td>GU305914.1</td><td>126467</td><td>126489</td><td>+</td><td>4</td><td>GCGTCTTCACGCCCGTGCGG</td><td>CGG</td><td>||-||-|-[||-|||||||||]PAM</td><td>NA</td><td>Intergenic</td><td>NA</td><td>NA</td></tr>
<tr><td>T2428</td><td>CATGACCCCCGTGCGGAGGACGG</td><td>1000</td><td></td><td></td><td></td><td></td><td></td><td></td><td></td><td></td><td></td></tr>
<tr><td>Oligo fwd</td><td>CACCCATGACCCCCGTGCGGAGGA</td><td></td><td></td><td></td><td></td><td></td><td></td><td></td><td></td><td></td><td></td></tr>
<tr><td>Oligo rev</td><td>AAACTCCTCCGCACGGGGGTCATG</td><td></td><td></td><td></td><td></td><td></td><td></td><td></td><td></td><td></td><td></td></tr>
<tr><td>Chromosome</td><td>start</td><td>end</td><td>strand</td><td>MM</td><td>target_seq</td><td>PAM</td><td>alignment</td><td>distance</td><td>position</td><td>gene name</td><td>gene id</td></tr>
<tr><td>GU305914.1</td><td>38246</td><td>38268</td><td>+</td><td>0</td><td>CATGACCCCCGTGCGGAGGA</td><td>CGG</td><td>||||||||[||||||||||||]PAM</td><td>NA</td><td>Intergenic</td><td>NA</td><td>NA</td></tr>
<tr><td>T2429</td><td>ATGACCCCCGTGCGGAGGACGGG</td><td>1000</td><td></td><td></td><td></td><td></td><td></td><td></td><td></td><td></td><td></td></tr>
<tr><td>Oligo fwd</td><td>CACCATGACCCCCGTGCGGAGGAC</td><td></td><td></td><td></td><td></td><td></td><td></td><td></td><td></td><td></td><td></td></tr>
<tr><td>Oligo rev</td><td>AAACGTCCTCCGCACGGGGGTCAT</td><td></td><td></td><td></td><td></td><td></td><td></td><td></td><td></td><td></td><td></td></tr>
<tr><td>Chromosome</td><td>start</td><td>end</td><td>strand</td><td>MM</td><td>target_seq</td><td>PAM</td><td>alignment</td><td>distance</td><td>position</td><td>gene name</td><td>gene id</td></tr>
<tr><td>GU305914.1</td><td>38247</td><td>38269</td><td>+</td><td>0</td><td>ATGACCCCCGTGCGGAGGAC</td><td>GGG</td><td>||||||||[||||||||||||]PAM</td><td>NA</td><td>Intergenic</td><td>NA</td><td>NA</td></tr>
<tr><td>T2430</td><td>ACCCCCGTGCGGAGGACGGGCGG</td><td>1000</td><td></td><td></td><td></td><td></td><td></td><td></td><td></td><td></td><td></td></tr>
<tr><td>Oligo fwd</td><td>CACCACCCCCGTGCGGAGGACGGG</td><td></td><td></td><td></td><td></td><td></td><td></td><td></td><td></td><td></td><td></td></tr>
<tr><td>Oligo rev</td><td>AAACCCCGTCCTCCGCACGGGGGT</td><td></td><td></td><td></td><td></td><td></td><td></td><td></td><td></td><td></td><td></td></tr>
<tr><td>Chromosome</td><td>start</td><td>end</td><td>strand</td><td>MM</td><td>target_seq</td><td>PAM</td><td>alignment</td><td>distance</td><td>position</td><td>gene name</td><td>gene id</td></tr>
<tr><td>GU305914.1</td><td>38250</td><td>38272</td><td>+</td><td>0</td><td>ACCCCCGTGCGGAGGACGGG</td><td>CGG</td><td>||||||||[||||||||||||]PAM</td><td>NA</td><td>Intergenic</td><td>NA</td><td>NA</td></tr>
</table>

T2431 CCCCCGTGCGGAGGACGGGCGGG 1000
Oligo fwd CACCCCCCCGTGCGGAGGACGGGC
Oligo rev AAACGCCCGTCCTCCGCACGGGGG
Chromosome start end strand MM target_seq PAM alignment distance position gene name gene id
GU305914.1 38251 38273 + 0 CCCCCGTGCGGAGGACGGGC GGG ||||||||[||||||||||||]PAM NA Intergenic NA NA

T2432 CCCCGTGCGGAGGACGGGCGGGG 1000
Oligo fwd CACCCCCCGTGCGGAGGACGGGCG
Oligo rev AAACCGCCCGTCCTCCGCACGGGG
Chromosome start end strand MM target_seq PAM alignment distance position gene name gene id
GU305914.1 38252 38274 + 0 CCCCGTGCGGAGGACGGGCG GGG ||||||||[||||||||||||]PAM NA Intergenic NA NA

T2433 AGGACGGGCGGGGTGTGTATCGG 1000
Oligo fwd CACCAGGACGGGCGGGGTGTGTAT
Oligo rev AAACATACACACCCCGCCCGTCCT
Chromosome start end strand MM target_seq PAM alignment distance position gene name gene id
GU305914.1 38262 38284 + 0 AGGACGGGCGGGGTGTGTAT CGG ||||||||[||||||||||||]PAM NA Intergenic NA NA

T2434 GGTGTGTATCGGACTGTTCTCGG 1000
Oligo fwd CACCGGTGTGTATCGGACTGTTCT
Oligo rev AAACAGAACAGTCCGATACACACC
Chromosome start end strand MM target_seq PAM alignment distance position gene name gene id
GU305914.1 38273 38295 + 0 GGTGTGTATCGGACTGTTCT CGG ||||||||[||||||||||||]PAM NA Intergenic NA NA

T2435 GACTGTTCTCGGCCCTGAGCTGG 1000
Oligo fwd CACCGACTGTTCTCGGCCCTGAGC
Oligo rev AAACGCTCAGGGCCGAGAACAGTC
Chromosome start end strand MM target_seq PAM alignment distance position gene name gene id
GU305914.1 38284 38306 + 0 GACTGTTCTCGGCCCTGAGC TGG ||||||||[||||||||||||]PAM NA Intergenic NA NA

T2436 CGCACCATCATGAACGAAAACGG 1000
Oligo fwd CACCCGCACCATCATGAACGAAAA
Oligo rev AAACTTTTCGTTCATGATGGTGCG
Chromosome start end strand MM target_seq PAM alignment distance position gene name gene id
GU305914.1 38313 38335 + 0 CGCACCATCATGAACGAAAA CGG ||||||||[||||||||||||]PAM NA Intergenic NA NA

T2437 CGACCACGAGCTACGTAGCCTGG 1000
Oligo fwd CACCCGACCACGAGCTACGTAGCC
Oligo rev AAACGGCTACGTAGCTCGTGGTCG
Chromosome start end strand MM target_seq PAM alignment distance position gene name gene id
GU305914.1 38341 38363 + 0 CGACCACGAGCTACGTAGCC TGG ||||||||[||||||||||||]PAM NA Intergenic NA NA

T2438 AGCCTGGTTCGAGTGCACCCAGG 1000
Oligo fwd CACCAGCCTGGTTCGAGTGCACCC
Oligo rev AAACGGGTGCACTCGAACCAGGCT
Chromosome start end strand MM target_seq PAM alignment distance position gene name gene id
GU305914.1 38357 38379 + 0 AGCCTGGTTCGAGTGCACCC AGG ||||||||[||||||||||||]PAM NA Intergenic NA NA

T2439 GCTACTCCCTATGCAAGTACTGG 1000
Oligo fwd CACCGCTACTCCCTATGCAAGTAC
Oligo rev AAACGTACTTGCATAGGGAGTAGC
Chromosome start end strand MM target_seq PAM alignment distance position gene name gene id
GU305914.1 38386 38408 + 0 GCTACTCCCTATGCAAGTAC TGG ||||||||[||||||||||||]PAM NA Intergenic NA NA

T2440 GCAAGTACTGGCGACCGTACAGG 1000
Oligo fwd CACCGCAAGTACTGGCGACCGTAC
Oligo rev AAACGTACGGTCGCCAGTACTTGC
Chromosome start end strand MM target_seq PAM alignment distance position gene name gene id
GU305914.1 38398 38420 + 0 GCAAGTACTGGCGACCGTAC AGG ||||||||[||||||||||||]PAM NA Intergenic NA NA

T2441 GTACTGGCGACCGTACAGGAAGG 1000
Oligo fwd CACCGTACTGGCGACCGTACAGGA
Oligo rev AAACTCCTGTACGGTCGCCAGTAC
Chromosome start end strand MM target_seq PAM alignment distance position gene name gene id
GU305914.1 38402 38424 + 0 GTACTGGCGACCGTACAGGA AGG ||||||||[||||||||||||]PAM NA Intergenic NA NA

T2442 AGGAAGGTGACCTACGAGTGCGG 1000
Oligo fwd CACCAGGAAGGTGACCTACGAGTG
Oligo rev AAACCACTCGTAGGTCACCTTCCT
Chromosome start end strand MM target_seq PAM alignment distance position gene name gene id
GU305914.1 38418 38440 + 0 AGGAAGGTGACCTACGAGTG CGG ||||||||[||||||||||||]PAM NA Intergenic NA NA

T2443 CTACGAGTGCGGCGATTTCGAGG 1000
Oligo fwd CACCCTACGAGTGCGGCGATTTCG
Oligo rev AAACCGAAATCGCCGCACTCGTAG
Chromosome start end strand MM target_seq PAM alignment distance position gene name gene id
GU305914.1 38429 38451 + 0 CTACGAGTGCGGCGATTTCG AGG ||||||||[||||||||||||]PAM NA Intergenic NA NA

T2444 GAGTGCGGCGATTTCGAGGCCGG 1000
Oligo fwd CACCGAGTGCGGCGATTTCGAGGC
Oligo rev AAACGCCTCGAAATCGCCGCACTC
Chromosome start end strand MM target_seq PAM alignment distance position gene name gene id
GU305914.1 38433 38455 + 0 GAGTGCGGCGATTTCGAGGC CGG ||||||||[||||||||||||]PAM NA Intergenic NA NA

T2445 CGTCGAGAGCATGTACGTCGTGG 1000
Oligo fwd CACCCGTCGAGAGCATGTACGTCG
Oligo rev AAACCGACGTACATGCTCTCGACG
Chromosome start end strand MM target_seq PAM alignment distance position gene name gene id
GU305914.1 38462 38484 + 0 CGTCGAGAGCATGTACGTCG TGG ||||||||[||||||||||||]PAM NA Intergenic NA NA

<table>
<tr><td>T2446</td><td>GAGAGCATGTACGTCGTGGACGG</td><td></td><td>1000</td><td></td><td></td><td></td><td></td><td></td><td></td><td></td><td></td><td></td></tr>
<tr><td>Oligo fwd</td><td>CACCGAGAGCATGTACGTCGTGGA</td><td></td><td></td><td></td><td></td><td></td><td></td><td></td><td></td><td></td><td></td><td></td></tr>
<tr><td>Oligo rev</td><td>AAACTCCACGACGTACATGCTCTC</td><td></td><td></td><td></td><td></td><td></td><td></td><td></td><td></td><td></td><td></td><td></td></tr>
<tr><td>Chromosome</td><td>start</td><td></td><td>end</td><td>strand</td><td>MM</td><td>target_seq</td><td>PAM</td><td>alignment</td><td>distance</td><td>position</td><td>gene name</td><td>gene id</td></tr>
<tr><td>GU305914.1</td><td></td><td>38466</td><td>38488</td><td>+</td><td>0</td><td>GAGAGCATGTACGTCGTGGA</td><td>CGG</td><td>||||||||[||||||||||||]PAM</td><td>NA</td><td>Intergenic</td><td>NA</td><td>NA</td></tr>
<tr><td>T2447</td><td>AGCATGTACGTCGTGGACGGCGG</td><td></td><td>1000</td><td></td><td></td><td></td><td></td><td></td><td></td><td></td><td></td><td></td></tr>
<tr><td>Oligo fwd</td><td>CACCAGCATGTACGTCGTGGACGG</td><td></td><td></td><td></td><td></td><td></td><td></td><td></td><td></td><td></td><td></td><td></td></tr>
<tr><td>Oligo rev</td><td>AAACCCGTCCACGACGTACATGCT</td><td></td><td></td><td></td><td></td><td></td><td></td><td></td><td></td><td></td><td></td><td></td></tr>
<tr><td>Chromosome</td><td>start</td><td></td><td>end</td><td>strand</td><td>MM</td><td>target_seq</td><td>PAM</td><td>alignment</td><td>distance</td><td>position</td><td>gene name</td><td>gene id</td></tr>
<tr><td>GU305914.1</td><td></td><td>38469</td><td>38491</td><td>+</td><td>0</td><td>AGCATGTACGTCGTGGACGG</td><td>CGG</td><td>||||||||[||||||||||||]PAM</td><td>NA</td><td>Intergenic</td><td>NA</td><td>NA</td></tr>
<tr><td>T2448</td><td>TGTACGTCGTGGACGGCGGCAGG</td><td></td><td>1000</td><td></td><td></td><td></td><td></td><td></td><td></td><td></td><td></td><td></td></tr>
<tr><td>Oligo fwd</td><td>CACCTGTACGTCGTGGACGGCGGC</td><td></td><td></td><td></td><td></td><td></td><td></td><td></td><td></td><td></td><td></td><td></td></tr>
<tr><td>Oligo rev</td><td>AAACGCCGCCGTCCACGACGTACA</td><td></td><td></td><td></td><td></td><td></td><td></td><td></td><td></td><td></td><td></td><td></td></tr>
<tr><td>Chromosome</td><td>start</td><td></td><td>end</td><td>strand</td><td>MM</td><td>target_seq</td><td>PAM</td><td>alignment</td><td>distance</td><td>position</td><td>gene name</td><td>gene id</td></tr>
<tr><td>GU305914.1</td><td></td><td>38473</td><td>38495</td><td>+</td><td>0</td><td>TGTACGTCGTGGACGGCGGC</td><td>AGG</td><td>||||||||[||||||||||||]PAM</td><td>NA</td><td>Intergenic</td><td>NA</td><td>NA</td></tr>
<tr><td>T2449</td><td>ACGTCGTGGACGGCGGCAGGAGG</td><td></td><td>1000</td><td></td><td></td><td></td><td></td><td></td><td></td><td></td><td></td><td></td></tr>
<tr><td>Oligo fwd</td><td>CACCACGTCGTGGACGGCGGCAGG</td><td></td><td></td><td></td><td></td><td></td><td></td><td></td><td></td><td></td><td></td><td></td></tr>
<tr><td>Oligo rev</td><td>AAACCCTGCCGCCGTCCACGACGT</td><td></td><td></td><td></td><td></td><td></td><td></td><td></td><td></td><td></td><td></td><td></td></tr>
<tr><td>Chromosome</td><td>start</td><td></td><td>end</td><td>strand</td><td>MM</td><td>target_seq</td><td>PAM</td><td>alignment</td><td>distance</td><td>position</td><td>gene name</td><td>gene id</td></tr>
<tr><td>GU305914.1</td><td></td><td>38476</td><td>38498</td><td>+</td><td>0</td><td>ACGTCGTGGACGGCGGCAGG</td><td>AGG</td><td>||||||||[||||||||||||]PAM</td><td>NA</td><td>Intergenic</td><td>NA</td><td>NA</td></tr>
<tr><td>T2450</td><td>GACGGCGGCAGGAGGCTCATCGG</td><td></td><td>1000</td><td></td><td></td><td></td><td></td><td></td><td></td><td></td><td></td><td></td></tr>
<tr><td>Oligo fwd</td><td>CACCGACGGCGGCAGGAGGCTCAT</td><td></td><td></td><td></td><td></td><td></td><td></td><td></td><td></td><td></td><td></td><td></td></tr>
<tr><td>Oligo rev</td><td>AAACATGAGCCTCCTGCCGCCGTC</td><td></td><td></td><td></td><td></td><td></td><td></td><td></td><td></td><td></td><td></td><td></td></tr>
<tr><td>Chromosome</td><td>start</td><td></td><td>end</td><td>strand</td><td>MM</td><td>target_seq</td><td>PAM</td><td>alignment</td><td>distance</td><td>position</td><td>gene name</td><td>gene id</td></tr>
<tr><td>GU305914.1</td><td></td><td>38484</td><td>38506</td><td>+</td><td>0</td><td>GACGGCGGCAGGAGGCTCAT</td><td>CGG</td><td>||||||||[||||||||||||]PAM</td><td>NA</td><td>Intergenic</td><td>NA</td><td>NA</td></tr>
<tr><td>T2451</td><td>CAGGAGGCTCATCGGCACACTGG</td><td></td><td>1000</td><td></td><td></td><td></td><td></td><td></td><td></td><td></td><td></td><td></td></tr>
<tr><td>Oligo fwd</td><td>CACCCAGGAGGCTCATCGGCACAC</td><td></td><td></td><td></td><td></td><td></td><td></td><td></td><td></td><td></td><td></td><td></td></tr>
<tr><td>Oligo rev</td><td>AAACGTGTGCCGATGAGCCTCCTG</td><td></td><td></td><td></td><td></td><td></td><td></td><td></td><td></td><td></td><td></td><td></td></tr>
<tr><td>Chromosome</td><td>start</td><td></td><td>end</td><td>strand</td><td>MM</td><td>target_seq</td><td>PAM</td><td>alignment</td><td>distance</td><td>position</td><td>gene name</td><td>gene id</td></tr>
<tr><td>GU305914.1</td><td></td><td>38492</td><td>38514</td><td>+</td><td>0</td><td>CAGGAGGCTCATCGGCACAC</td><td>TGG</td><td>||||||||[||||||||||||]PAM</td><td>NA</td><td>Intergenic</td><td>NA</td><td>NA</td></tr>
<tr><td>T2452</td><td>GAGGCTCATCGGCACACTGGTGG</td><td></td><td>1000</td><td></td><td></td><td></td><td></td><td></td><td></td><td></td><td></td><td></td></tr>
<tr><td>Oligo fwd</td><td>CACCGAGGCTCATCGGCACACTGG</td><td></td><td></td><td></td><td></td><td></td><td></td><td></td><td></td><td></td><td></td><td></td></tr>
<tr><td>Oligo rev</td><td>AAACCCAGTGTGCCGATGAGCCTC</td><td></td><td></td><td></td><td></td><td></td><td></td><td></td><td></td><td></td><td></td><td></td></tr>
<tr><td>Chromosome</td><td>start</td><td></td><td>end</td><td>strand</td><td>MM</td><td>target_seq</td><td>PAM</td><td>alignment</td><td>distance</td><td>position</td><td>gene name</td><td>gene id</td></tr>
<tr><td>GU305914.1</td><td></td><td>38495</td><td>38517</td><td>+</td><td>0</td><td>GAGGCTCATCGGCACACTGG</td><td>TGG</td><td>||||||||[||||||||||||]PAM</td><td>NA</td><td>Intergenic</td><td>NA</td><td>NA</td></tr>
<tr><td>T2453</td><td>GCTCATCGGCACACTGGTGGTGG</td><td></td><td>1000</td><td></td><td></td><td></td><td></td><td></td><td></td><td></td><td></td><td></td></tr>
<tr><td>Oligo fwd</td><td>CACCGCTCATCGGCACACTGGTGG</td><td></td><td></td><td></td><td></td><td></td><td></td><td></td><td></td><td></td><td></td><td></td></tr>
<tr><td>Oligo rev</td><td>AAACCCACCAGTGTGCCGATGAGC</td><td></td><td></td><td></td><td></td><td></td><td></td><td></td><td></td><td></td><td></td><td></td></tr>
<tr><td>Chromosome</td><td>start</td><td></td><td>end</td><td>strand</td><td>MM</td><td>target_seq</td><td>PAM</td><td>alignment</td><td>distance</td><td>position</td><td>gene name</td><td>gene id</td></tr>
<tr><td>GU305914.1</td><td></td><td>38498</td><td>38520</td><td>+</td><td>0</td><td>GCTCATCGGCACACTGGTGG</td><td>TGG</td><td>||||||||[||||||||||||]PAM</td><td>NA</td><td>Intergenic</td><td>NA</td><td>NA</td></tr>
<tr><td>T2454</td><td>CTCATCGGCACACTGGTGGTGGG</td><td></td><td>1000</td><td></td><td></td><td></td><td></td><td></td><td></td><td></td><td></td><td></td></tr>
<tr><td>Oligo fwd</td><td>CACCCTCATCGGCACACTGGTGGT</td><td></td><td></td><td></td><td></td><td></td><td></td><td></td><td></td><td></td><td></td><td></td></tr>
<tr><td>Oligo rev</td><td>AAACACCACCAGTGTGCCGATGAG</td><td></td><td></td><td></td><td></td><td></td><td></td><td></td><td></td><td></td><td></td><td></td></tr>
<tr><td>Chromosome</td><td>start</td><td></td><td>end</td><td>strand</td><td>MM</td><td>target_seq</td><td>PAM</td><td>alignment</td><td>distance</td><td>position</td><td>gene name</td><td>gene id</td></tr>
<tr><td>GU305914.1</td><td></td><td>38499</td><td>38521</td><td>+</td><td>0</td><td>CTCATCGGCACACTGGTGGT</td><td>GGG</td><td>||||||||[||||||||||||]PAM</td><td>NA</td><td>Intergenic</td><td>NA</td><td>NA</td></tr>
<tr><td>T2455</td><td>GAACACGCCGTCGCTGTCACCGG</td><td></td><td>1000</td><td></td><td></td><td></td><td></td><td></td><td></td><td></td><td></td><td></td></tr>
<tr><td>Oligo fwd</td><td>CACCGAACACGCCGTCGCTGTCAC</td><td></td><td></td><td></td><td></td><td></td><td></td><td></td><td></td><td></td><td></td><td></td></tr>
<tr><td>Oligo rev</td><td>AAACGTGACAGCGACGGCGTGTTC</td><td></td><td></td><td></td><td></td><td></td><td></td><td></td><td></td><td></td><td></td><td></td></tr>
<tr><td>Chromosome</td><td>start</td><td></td><td>end</td><td>strand</td><td>MM</td><td>target_seq</td><td>PAM</td><td>alignment</td><td>distance</td><td>position</td><td>gene name</td><td>gene id</td></tr>
<tr><td>GU305914.1</td><td></td><td>38537</td><td>38559</td><td>+</td><td>0</td><td>GAACACGCCGTCGCTGTCAC</td><td>CGG</td><td>||||||||[||||||||||||]PAM</td><td>NA</td><td>Intergenic</td><td>NA</td><td>NA</td></tr>
<tr><td>T2456</td><td>ACCCCACCCTGCACCTCACCTGG</td><td></td><td>1000</td><td></td><td></td><td></td><td></td><td></td><td></td><td></td><td></td><td></td></tr>
<tr><td>Oligo fwd</td><td>CACCACCCCACCCTGCACCTCACC</td><td></td><td></td><td></td><td></td><td></td><td></td><td></td><td></td><td></td><td></td><td></td></tr>
<tr><td>Oligo rev</td><td>AAACGGTGAGGTGCAGGGTGGGGT</td><td></td><td></td><td></td><td></td><td></td><td></td><td></td><td></td><td></td><td></td><td></td></tr>
<tr><td>Chromosome</td><td>start</td><td></td><td>end</td><td>strand</td><td>MM</td><td>target_seq</td><td>PAM</td><td>alignment</td><td>distance</td><td>position</td><td>gene name</td><td>gene id</td></tr>
<tr><td>GU305914.1</td><td></td><td>38641</td><td>38663</td><td>+</td><td>0</td><td>ACCCCACCCTGCACCTCACC</td><td>TGG</td><td>||||||||[||||||||||||]PAM</td><td>NA</td><td>Intergenic</td><td>NA</td><td>NA</td></tr>
<tr><td>T2457</td><td>CTACGATCTCTGCTTCAGAGAGG</td><td></td><td>1000</td><td></td><td></td><td></td><td></td><td></td><td></td><td></td><td></td><td></td></tr>
<tr><td>Oligo fwd</td><td>CACCCTACGATCTCTGCTTCAGAG</td><td></td><td></td><td></td><td></td><td></td><td></td><td></td><td></td><td></td><td></td><td></td></tr>
<tr><td>Oligo rev</td><td>AAACCTCTGAAGCAGAGATCGTAG</td><td></td><td></td><td></td><td></td><td></td><td></td><td></td><td></td><td></td><td></td><td></td></tr>
<tr><td>Chromosome</td><td>start</td><td></td><td>end</td><td>strand</td><td>MM</td><td>target_seq</td><td>PAM</td><td>alignment</td><td>distance</td><td>position</td><td>gene name</td><td>gene id</td></tr>
<tr><td>GU305914.1</td><td></td><td>38705</td><td>38727</td><td>+</td><td>0</td><td>CTACGATCTCTGCTTCAGAG</td><td>AGG</td><td>||||||||[||||||||||||]PAM</td><td>NA</td><td>Intergenic</td><td>NA</td><td>NA</td></tr>
<tr><td>T2458</td><td>TACGATCTCTGCTTCAGAGAGGG</td><td></td><td>1000</td><td></td><td></td><td></td><td></td><td></td><td></td><td></td><td></td><td></td></tr>
<tr><td>Oligo fwd</td><td>CACCTACGATCTCTGCTTCAGAGA</td><td></td><td></td><td></td><td></td><td></td><td></td><td></td><td></td><td></td><td></td><td></td></tr>
<tr><td>Oligo rev</td><td>AAACTCTCTGAAGCAGAGATCGTA</td><td></td><td></td><td></td><td></td><td></td><td></td><td></td><td></td><td></td><td></td><td></td></tr>
<tr><td>Chromosome</td><td>start</td><td></td><td>end</td><td>strand</td><td>MM</td><td>target_seq</td><td>PAM</td><td>alignment</td><td>distance</td><td>position</td><td>gene name</td><td>gene id</td></tr>
<tr><td>GU305914.1</td><td></td><td>38706</td><td>38728</td><td>+</td><td>0</td><td>TACGATCTCTGCTTCAGAGA</td><td>GGG</td><td>||||||||[||||||||||||]PAM</td><td>NA</td><td>Intergenic</td><td>NA</td><td>NA</td></tr>
<tr><td>T2459</td><td>GGGTAAGTGCGCCGTCTTCATGG</td><td></td><td>1000</td><td></td><td></td><td></td><td></td><td></td><td></td><td></td><td></td><td></td></tr>
<tr><td>Oligo fwd</td><td>CACCGGGTAAGTGCGCCGTCTTCA</td><td></td><td></td><td></td><td></td><td></td><td></td><td></td><td></td><td></td><td></td><td></td></tr>
<tr><td>Oligo rev</td><td>AAACTGAAGACGGCGCACTTACCC</td><td></td><td></td><td></td><td></td><td></td><td></td><td></td><td></td><td></td><td></td><td></td></tr>
<tr><td>Chromosome</td><td>start</td><td></td><td>end</td><td>strand</td><td>MM</td><td>target_seq</td><td>PAM</td><td>alignment</td><td>distance</td><td>position</td><td>gene name</td><td>gene id</td></tr>
<tr><td>GU305914.1</td><td></td><td>38726</td><td>38748</td><td>+</td><td>0</td><td>GGGTAAGTGCGCCGTCTTCA</td><td>TGG</td><td>||||||||[||||||||||||]PAM</td><td>NA</td><td>Intergenic</td><td>NA</td><td>NA</td></tr>
<tr><td>T2460</td><td>CATGGACATCAGATACGCGAAGG</td><td></td><td>1000</td><td></td><td></td><td></td><td></td><td></td><td></td><td></td><td></td><td></td></tr>
<tr><td>Oligo fwd</td><td>CACCCATGGACATCAGATACGCGA</td><td></td><td></td><td></td><td></td><td></td><td></td><td></td><td></td><td></td><td></td><td></td></tr>
<tr><td>Oligo rev</td><td>AAACTCGCGTATCTGATGTCCATG</td><td></td><td></td><td></td><td></td><td></td><td></td><td></td><td></td><td></td><td></td><td></td></tr>
<tr><td>Chromosome</td><td>start</td><td></td><td>end</td><td>strand</td><td>MM</td><td>target_seq</td><td>PAM</td><td>alignment</td><td>distance</td><td>position</td><td>gene name</td><td>gene id</td></tr>
</table>

<table>
<tr><td>GU305914.1</td><td>38744</td><td>38766</td><td>+</td><td>0</td><td>CATGGACATCAGATACGCGA</td><td>AGG</td><td>||||||||[||||||||||||]PAM</td><td>NA</td><td>Intergenic</td><td>NA</td><td>NA</td></tr>
<tr><td>T2461</td><td>GGACATCAGATACGCGAAGGCGG</td><td>1000</td><td></td><td></td><td></td><td></td><td></td><td></td><td></td><td></td><td></td></tr>
<tr><td>Oligo fwd</td><td>CACCGGACATCAGATACGCGAAGG</td><td></td><td></td><td></td><td></td><td></td><td></td><td></td><td></td><td></td><td></td></tr>
<tr><td>Oligo rev</td><td>AAACCCTTCGCGTATCTGATGTCC</td><td></td><td></td><td></td><td></td><td></td><td></td><td></td><td></td><td></td><td></td></tr>
<tr><td>Chromosome</td><td>start</td><td>end</td><td>strand</td><td>MM</td><td>target_seq</td><td>PAM</td><td>alignment</td><td>distance</td><td>position</td><td>gene name</td><td>gene id</td></tr>
<tr><td>GU305914.1</td><td>38747</td><td>38769</td><td>+</td><td>0</td><td>GGACATCAGATACGCGAAGG</td><td>CGG</td><td>||||||||[||||||||||||]PAM</td><td>NA</td><td>Intergenic</td><td>NA</td><td>NA</td></tr>
<tr><td>T2462</td><td>AACCGCTGCTTTTTCATCTCCGG</td><td>1000</td><td></td><td></td><td></td><td></td><td></td><td></td><td></td><td></td><td></td></tr>
<tr><td>Oligo fwd</td><td>CACCAACCGCTGCTTTTTCATCTC</td><td></td><td></td><td></td><td></td><td></td><td></td><td></td><td></td><td></td><td></td></tr>
<tr><td>Oligo rev</td><td>AAACGAGATGAAAAAGCAGCGGTT</td><td></td><td></td><td></td><td></td><td></td><td></td><td></td><td></td><td></td><td></td></tr>
<tr><td>Chromosome</td><td>start</td><td>end</td><td>strand</td><td>MM</td><td>target_seq</td><td>PAM</td><td>alignment</td><td>distance</td><td>position</td><td>gene name</td><td>gene id</td></tr>
<tr><td>GU305914.1</td><td>38775</td><td>38797</td><td>+</td><td>0</td><td>AACCGCTGCTTTTTCATCTC</td><td>CGG</td><td>||||||||[||||||||||||]PAM</td><td>NA</td><td>Intergenic</td><td>NA</td><td>NA</td></tr>
<tr><td>T2463</td><td>TTTTTCATCTCCGGCGTGCCCGG</td><td>1000</td><td></td><td></td><td></td><td></td><td></td><td></td><td></td><td></td><td></td></tr>
<tr><td>Oligo fwd</td><td>CACCTTTTTCATCTCCGGCGTGCC</td><td></td><td></td><td></td><td></td><td></td><td></td><td></td><td></td><td></td><td></td></tr>
<tr><td>Oligo rev</td><td>AAACGGCACGCCGGAGATGAAAAA</td><td></td><td></td><td></td><td></td><td></td><td></td><td></td><td></td><td></td><td></td></tr>
<tr><td>Chromosome</td><td>start</td><td>end</td><td>strand</td><td>MM</td><td>target_seq</td><td>PAM</td><td>alignment</td><td>distance</td><td>position</td><td>gene name</td><td>gene id</td></tr>
<tr><td>GU305914.1</td><td>38784</td><td>38806</td><td>+</td><td>0</td><td>TTTTTCATCTCCGGCGTGCC</td><td>CGG</td><td>||||||||[||||||||||||]PAM</td><td>NA</td><td>Intergenic</td><td>NA</td><td>NA</td></tr>
<tr><td>T2464</td><td>GGCGAGTACCGTTTCCACACCGG</td><td>1000</td><td></td><td></td><td></td><td></td><td></td><td></td><td></td><td></td><td></td></tr>
<tr><td>Oligo fwd</td><td>CACCGGCGAGTACCGTTTCCACAC</td><td></td><td></td><td></td><td></td><td></td><td></td><td></td><td></td><td></td><td></td></tr>
<tr><td>Oligo rev</td><td>AAACGTGTGGAAACGGTACTCGCC</td><td></td><td></td><td></td><td></td><td></td><td></td><td></td><td></td><td></td><td></td></tr>
<tr><td>Chromosome</td><td>start</td><td>end</td><td>strand</td><td>MM</td><td>target_seq</td><td>PAM</td><td>alignment</td><td>distance</td><td>position</td><td>gene name</td><td>gene id</td></tr>
<tr><td>GU305914.1</td><td>38805</td><td>38827</td><td>+</td><td>0</td><td>GGCGAGTACCGTTTCCACAC</td><td>CGG</td><td>||||||||[||||||||||||]PAM</td><td>NA</td><td>Intergenic</td><td>NA</td><td>NA</td></tr>
<tr><td>T2465</td><td>GCGAGTACCGTTTCCACACCGGG</td><td>1000</td><td></td><td></td><td></td><td></td><td></td><td></td><td></td><td></td><td></td></tr>
<tr><td>Oligo fwd</td><td>CACCGCGAGTACCGTTTCCACACC</td><td></td><td></td><td></td><td></td><td></td><td></td><td></td><td></td><td></td><td></td></tr>
<tr><td>Oligo rev</td><td>AAACGGTGTGGAAACGGTACTCGC</td><td></td><td></td><td></td><td></td><td></td><td></td><td></td><td></td><td></td><td></td></tr>
<tr><td>Chromosome</td><td>start</td><td>end</td><td>strand</td><td>MM</td><td>target_seq</td><td>PAM</td><td>alignment</td><td>distance</td><td>position</td><td>gene name</td><td>gene id</td></tr>
<tr><td>GU305914.1</td><td>38806</td><td>38828</td><td>+</td><td>0</td><td>GCGAGTACCGTTTCCACACC</td><td>GGG</td><td>||||||||[||||||||||||]PAM</td><td>NA</td><td>Intergenic</td><td>NA</td><td>NA</td></tr>
<tr><td>T2466</td><td>CCGTTTCCACACCGGGCTGACGG</td><td>1000</td><td></td><td></td><td></td><td></td><td></td><td></td><td></td><td></td><td></td></tr>
<tr><td>Oligo fwd</td><td>CACCCCGTTTCCACACCGGGCTGA</td><td></td><td></td><td></td><td></td><td></td><td></td><td></td><td></td><td></td><td></td></tr>
<tr><td>Oligo rev</td><td>AAACTCAGCCCGGTGTGGAAACGG</td><td></td><td></td><td></td><td></td><td></td><td></td><td></td><td></td><td></td><td></td></tr>
<tr><td>Chromosome</td><td>start</td><td>end</td><td>strand</td><td>MM</td><td>target_seq</td><td>PAM</td><td>alignment</td><td>distance</td><td>position</td><td>gene name</td><td>gene id</td></tr>
<tr><td>GU305914.1</td><td>38813</td><td>38835</td><td>+</td><td>0</td><td>CCGTTTCCACACCGGGCTGA</td><td>CGG</td><td>||||||||[||||||||||||]PAM</td><td>NA</td><td>Intergenic</td><td>NA</td><td>NA</td></tr>
<tr><td>T2467</td><td>TCCACACCGGGCTGACGGTCTGG</td><td>1000</td><td></td><td></td><td></td><td></td><td></td><td></td><td></td><td></td><td></td></tr>
<tr><td>Oligo fwd</td><td>CACCTCCACACCGGGCTGACGGTC</td><td></td><td></td><td></td><td></td><td></td><td></td><td></td><td></td><td></td><td></td></tr>
<tr><td>Oligo rev</td><td>AAACGACCGTCAGCCCGGTGTGGA</td><td></td><td></td><td></td><td></td><td></td><td></td><td></td><td></td><td></td><td></td></tr>
<tr><td>Chromosome</td><td>start</td><td>end</td><td>strand</td><td>MM</td><td>target_seq</td><td>PAM</td><td>alignment</td><td>distance</td><td>position</td><td>gene name</td><td>gene id</td></tr>
<tr><td>GU305914.1</td><td>38818</td><td>38840</td><td>+</td><td>0</td><td>TCCACACCGGGCTGACGGTC</td><td>TGG</td><td>||||||||[||||||||||||]PAM</td><td>NA</td><td>Intergenic</td><td>NA</td><td>NA</td></tr>
<tr><td>T2468</td><td>CCCAGCCTCCACATCTGCCGCGG</td><td>1000</td><td></td><td></td><td></td><td></td><td></td><td></td><td></td><td></td><td></td></tr>
<tr><td>Oligo fwd</td><td>CACCCCCAGCCTCCACATCTGCCG</td><td></td><td></td><td></td><td></td><td></td><td></td><td></td><td></td><td></td><td></td></tr>
<tr><td>Oligo rev</td><td>AAACCGGCAGATGTGGAGGCTGGG</td><td></td><td></td><td></td><td></td><td></td><td></td><td></td><td></td><td></td><td></td></tr>
<tr><td>Chromosome</td><td>start</td><td>end</td><td>strand</td><td>MM</td><td>target_seq</td><td>PAM</td><td>alignment</td><td>distance</td><td>position</td><td>gene name</td><td>gene id</td></tr>
<tr><td>GU305914.1</td><td>38883</td><td>38905</td><td>+</td><td>0</td><td>CCCAGCCTCCACATCTGCCG</td><td>CGG</td><td>||||||||[||||||||||||]PAM</td><td>NA</td><td>Intergenic</td><td>NA</td><td>NA</td></tr>
<tr><td>T2469</td><td>CCAGCCTCCACATCTGCCGCGGG</td><td>1000</td><td></td><td></td><td></td><td></td><td></td><td></td><td></td><td></td><td></td></tr>
<tr><td>Oligo fwd</td><td>CACCCCAGCCTCCACATCTGCCGC</td><td></td><td></td><td></td><td></td><td></td><td></td><td></td><td></td><td></td><td></td></tr>
<tr><td>Oligo rev</td><td>AAACGCGGCAGATGTGGAGGCTGG</td><td></td><td></td><td></td><td></td><td></td><td></td><td></td><td></td><td></td><td></td></tr>
<tr><td>Chromosome</td><td>start</td><td>end</td><td>strand</td><td>MM</td><td>target_seq</td><td>PAM</td><td>alignment</td><td>distance</td><td>position</td><td>gene name</td><td>gene id</td></tr>
<tr><td>GU305914.1</td><td>38884</td><td>38906</td><td>+</td><td>0</td><td>CCAGCCTCCACATCTGCCGC</td><td>GGG</td><td>||||||||[||||||||||||]PAM</td><td>NA</td><td>Intergenic</td><td>NA</td><td>NA</td></tr>
<tr><td>T2470</td><td>CTTCTTCATGCACTCCACAGAGG</td><td>1000</td><td></td><td></td><td></td><td></td><td></td><td></td><td></td><td></td><td></td></tr>
<tr><td>Oligo fwd</td><td>CACCCTTCTTCATGCACTCCACAG</td><td></td><td></td><td></td><td></td><td></td><td></td><td></td><td></td><td></td><td></td></tr>
<tr><td>Oligo rev</td><td>AAACCTGTGGAGTGCATGAAGAAG</td><td></td><td></td><td></td><td></td><td></td><td></td><td></td><td></td><td></td><td></td></tr>
<tr><td>Chromosome</td><td>start</td><td>end</td><td>strand</td><td>MM</td><td>target_seq</td><td>PAM</td><td>alignment</td><td>distance</td><td>position</td><td>gene name</td><td>gene id</td></tr>
<tr><td>GU305914.1</td><td>38924</td><td>38946</td><td>+</td><td>0</td><td>CTTCTTCATGCACTCCACAG</td><td>AGG</td><td>||||||||[||||||||||||]PAM</td><td>NA</td><td>Intergenic</td><td>NA</td><td>NA</td></tr>
<tr><td>T2471</td><td>TTCTTCATGCACTCCACAGAGGG</td><td>1000</td><td></td><td></td><td></td><td></td><td></td><td></td><td></td><td></td><td></td></tr>
<tr><td>Oligo fwd</td><td>CACCTTCTTCATGCACTCCACAGA</td><td></td><td></td><td></td><td></td><td></td><td></td><td></td><td></td><td></td><td></td></tr>
<tr><td>Oligo rev</td><td>AAACTCTGTGGAGTGCATGAAGAA</td><td></td><td></td><td></td><td></td><td></td><td></td><td></td><td></td><td></td><td></td></tr>
<tr><td>Chromosome</td><td>start</td><td>end</td><td>strand</td><td>MM</td><td>target_seq</td><td>PAM</td><td>alignment</td><td>distance</td><td>position</td><td>gene name</td><td>gene id</td></tr>
<tr><td>GU305914.1</td><td>38925</td><td>38947</td><td>+</td><td>0</td><td>TTCTTCATGCACTCCACAGA</td><td>GGG</td><td>||||||||[||||||||||||]PAM</td><td>NA</td><td>Intergenic</td><td>NA</td><td>NA</td></tr>
<tr><td>T2472</td><td>CTACCAACACAACGAACGCACGG</td><td>1000</td><td></td><td></td><td></td><td></td><td></td><td></td><td></td><td></td><td></td></tr>
<tr><td>Oligo fwd</td><td>CACCCTACCAACACAACGAACGCA</td><td></td><td></td><td></td><td></td><td></td><td></td><td></td><td></td><td></td><td></td></tr>
<tr><td>Oligo rev</td><td>AAACTGCGTTCGTTGTGTTGGTAG</td><td></td><td></td><td></td><td></td><td></td><td></td><td></td><td></td><td></td><td></td></tr>
<tr><td>Chromosome</td><td>start</td><td>end</td><td>strand</td><td>MM</td><td>target_seq</td><td>PAM</td><td>alignment</td><td>distance</td><td>position</td><td>gene name</td><td>gene id</td></tr>
<tr><td>GU305914.1</td><td>38954</td><td>38976</td><td>+</td><td>0</td><td>CTACCAACACAACGAACGCA</td><td>CGG</td><td>||||||||[||||||||||||]PAM</td><td>NA</td><td>Intergenic</td><td>NA</td><td>NA</td></tr>
<tr><td>T2473</td><td>GGCGACCCGAGTGATCAAGACGG</td><td>1000</td><td></td><td></td><td></td><td></td><td></td><td></td><td></td><td></td><td></td></tr>
<tr><td>Oligo fwd</td><td>CACCGGCGACCCGAGTGATCAAGA</td><td></td><td></td><td></td><td></td><td></td><td></td><td></td><td></td><td></td><td></td></tr>
<tr><td>Oligo rev</td><td>AAACTCTTGATCACTCGGGTCGCC</td><td></td><td></td><td></td><td></td><td></td><td></td><td></td><td></td><td></td><td></td></tr>
<tr><td>Chromosome</td><td>start</td><td>end</td><td>strand</td><td>MM</td><td>target_seq</td><td>PAM</td><td>alignment</td><td>distance</td><td>position</td><td>gene name</td><td>gene id</td></tr>
<tr><td>GU305914.1</td><td>38975</td><td>38997</td><td>+</td><td>0</td><td>GGCGACCCGAGTGATCAAGA</td><td>CGG</td><td>||||||||[||||||||||||]PAM</td><td>NA</td><td>Intergenic</td><td>NA</td><td>NA</td></tr>
<tr><td>T2474</td><td>GCGACCCGAGTGATCAAGACGGG</td><td>1000</td><td></td><td></td><td></td><td></td><td></td><td></td><td></td><td></td><td></td></tr>
<tr><td>Oligo fwd</td><td>CACCGCGACCCGAGTGATCAAGAC</td><td></td><td></td><td></td><td></td><td></td><td></td><td></td><td></td><td></td><td></td></tr>
<tr><td>Oligo rev</td><td>AAACGTCTTGATCACTCGGGTCGC</td><td></td><td></td><td></td><td></td><td></td><td></td><td></td><td></td><td></td><td></td></tr>
<tr><td>Chromosome</td><td>start</td><td>end</td><td>strand</td><td>MM</td><td>target_seq</td><td>PAM</td><td>alignment</td><td>distance</td><td>position</td><td>gene name</td><td>gene id</td></tr>
<tr><td>GU305914.1</td><td>38976</td><td>38998</td><td>+</td><td>0</td><td>GCGACCCGAGTGATCAAGAC</td><td>GGG</td><td>||||||||[||||||||||||]PAM</td><td>NA</td><td>Intergenic</td><td>NA</td><td>NA</td></tr>
<tr><td>T2475</td><td>AGACCGAATCAACCTTCACTTGG</td><td>1000</td><td></td><td></td><td></td><td></td><td></td><td></td><td></td><td></td><td></td></tr>
<tr><td>Oligo fwd</td><td>CACCAGACCGAATCAACCTTCACT</td><td></td><td></td><td></td><td></td><td></td><td></td><td></td><td></td><td></td><td></td></tr>
<tr><td>Oligo rev</td><td>AAACAGTGAAGGTTGATTCGGTCT</td><td></td><td></td><td></td><td></td><td></td><td></td><td></td><td></td><td></td><td></td></tr>
</table>

<table>
<tr><td>Chromosome</td><td>start</td><td>end</td><td>strand</td><td>MM</td><td>target_seq</td><td>PAM</td><td>alignment</td><td>distance</td><td>position</td><td>gene name</td><td>gene id</td></tr>
<tr><td>GU305914.1</td><td>38999</td><td>39021</td><td>+</td><td>0</td><td>AGACCGAATCAACCTTCACT</td><td>TGG</td><td>||||||||[||||||||||||]PAM</td><td>NA</td><td>Intergenic</td><td>NA</td><td>NA</td></tr>
<tr><td>T2476</td><td>GACCGAATCAACCTTCACTTGGG</td><td>1000</td><td></td><td></td><td></td><td></td><td></td><td></td><td></td><td></td><td></td></tr>
<tr><td>Oligo fwd</td><td>CACCGACCGAATCAACCTTCACTT</td><td></td><td></td><td></td><td></td><td></td><td></td><td></td><td></td><td></td><td></td></tr>
<tr><td>Oligo rev</td><td>AAACAAGTGAAGGTTGATTCGGTC</td><td></td><td></td><td></td><td></td><td></td><td></td><td></td><td></td><td></td><td></td></tr>
<tr><td>Chromosome</td><td>start</td><td>end</td><td>strand</td><td>MM</td><td>target_seq</td><td>PAM</td><td>alignment</td><td>distance</td><td>position</td><td>gene name</td><td>gene id</td></tr>
<tr><td>GU305914.1</td><td>39000</td><td>39022</td><td>+</td><td>0</td><td>GACCGAATCAACCTTCACTT</td><td>GGG</td><td>||||||||[||||||||||||]PAM</td><td>NA</td><td>Intergenic</td><td>NA</td><td>NA</td></tr>
<tr><td>T2477</td><td>TTATCAGAACTTTACAACCTCGG</td><td>1000</td><td></td><td></td><td></td><td></td><td></td><td></td><td></td><td></td><td></td></tr>
<tr><td>Oligo fwd</td><td>CACCTTATCAGAACTTTACAACCT</td><td></td><td></td><td></td><td></td><td></td><td></td><td></td><td></td><td></td><td></td></tr>
<tr><td>Oligo rev</td><td>AAACAGGTTGTAAAGTTCTGATAA</td><td></td><td></td><td></td><td></td><td></td><td></td><td></td><td></td><td></td><td></td></tr>
<tr><td>Chromosome</td><td>start</td><td>end</td><td>strand</td><td>MM</td><td>target_seq</td><td>PAM</td><td>alignment</td><td>distance</td><td>position</td><td>gene name</td><td>gene id</td></tr>
<tr><td>GU305914.1</td><td>39038</td><td>39060</td><td>+</td><td>0</td><td>TTATCAGAACTTTACAACCT</td><td>CGG</td><td>||||||||[||||||||||||]PAM</td><td>NA</td><td>Intergenic</td><td>NA</td><td>NA</td></tr>
<tr><td>T2478</td><td>AAAAATAATAAAAACCAACCAGG</td><td>1000</td><td></td><td></td><td></td><td></td><td></td><td></td><td></td><td></td><td></td></tr>
<tr><td>Oligo fwd</td><td>CACCAAAAATAATAAAAACCAACC</td><td></td><td></td><td></td><td></td><td></td><td></td><td></td><td></td><td></td><td></td></tr>
<tr><td>Oligo rev</td><td>AAACGGTTGGTTTTTATTATTTTT</td><td></td><td></td><td></td><td></td><td></td><td></td><td></td><td></td><td></td><td></td></tr>
<tr><td>Chromosome</td><td>start</td><td>end</td><td>strand</td><td>MM</td><td>target_seq</td><td>PAM</td><td>alignment</td><td>distance</td><td>position</td><td>gene name</td><td>gene id</td></tr>
<tr><td>GU305914.1</td><td>39238</td><td>39260</td><td>+</td><td>0</td><td>AAAAATAATAAAAACCAACC</td><td>AGG</td><td>||||||||[||||||||||||]PAM</td><td>NA</td><td>Intergenic</td><td>NA</td><td>NA</td></tr>
<tr><td>T2479</td><td>ACGACGACGATTTTTTTACCAGG</td><td>1000</td><td></td><td></td><td></td><td></td><td></td><td></td><td></td><td></td><td></td></tr>
<tr><td>Oligo fwd</td><td>CACCACGACGACGATTTTTTTACC</td><td></td><td></td><td></td><td></td><td></td><td></td><td></td><td></td><td></td><td></td></tr>
<tr><td>Oligo rev</td><td>AAACGGTAAAAAAATCGTCGTCGT</td><td></td><td></td><td></td><td></td><td></td><td></td><td></td><td></td><td></td><td></td></tr>
<tr><td>Chromosome</td><td>start</td><td>end</td><td>strand</td><td>MM</td><td>target_seq</td><td>PAM</td><td>alignment</td><td>distance</td><td>position</td><td>gene name</td><td>gene id</td></tr>
<tr><td>GU305914.1</td><td>39291</td><td>39313</td><td>+</td><td>0</td><td>ACGACGACGATTTTTTTACC</td><td>AGG</td><td>||||||||[||||||||||||]PAM</td><td>NA</td><td>Intergenic</td><td>NA</td><td>NA</td></tr>
<tr><td>T2480</td><td>GCCACCGTCCCCATCAGAACCGG</td><td>1000</td><td></td><td></td><td></td><td></td><td></td><td></td><td></td><td></td><td></td></tr>
<tr><td>Oligo fwd</td><td>CACCGCCACCGTCCCCATCAGAAC</td><td></td><td></td><td></td><td></td><td></td><td></td><td></td><td></td><td></td><td></td></tr>
<tr><td>Oligo rev</td><td>AAACGTTCTGATGGGGACGGTGGC</td><td></td><td></td><td></td><td></td><td></td><td></td><td></td><td></td><td></td><td></td></tr>
<tr><td>Chromosome</td><td>start</td><td>end</td><td>strand</td><td>MM</td><td>target_seq</td><td>PAM</td><td>alignment</td><td>distance</td><td>position</td><td>gene name</td><td>gene id</td></tr>
<tr><td>GU305914.1</td><td>39316</td><td>39338</td><td>+</td><td>0</td><td>GCCACCGTCCCCATCAGAAC</td><td>CGG</td><td>||||||||[||||||||||||]PAM</td><td>NA</td><td>Intergenic</td><td>NA</td><td>NA</td></tr>
<tr><td>T2481</td><td>TCCCCATCAGAACCGGCGAATGG</td><td>1000</td><td></td><td></td><td></td><td></td><td></td><td></td><td></td><td></td><td></td></tr>
<tr><td>Oligo fwd</td><td>CACCTCCCCATCAGAACCGGCGAA</td><td></td><td></td><td></td><td></td><td></td><td></td><td></td><td></td><td></td><td></td></tr>
<tr><td>Oligo rev</td><td>AAACTTCGCCGGTTCTGATGGGGA</td><td></td><td></td><td></td><td></td><td></td><td></td><td></td><td></td><td></td><td></td></tr>
<tr><td>Chromosome</td><td>start</td><td>end</td><td>strand</td><td>MM</td><td>target_seq</td><td>PAM</td><td>alignment</td><td>distance</td><td>position</td><td>gene name</td><td>gene id</td></tr>
<tr><td>GU305914.1</td><td>39323</td><td>39345</td><td>+</td><td>0</td><td>TCCCCATCAGAACCGGCGAA</td><td>TGG</td><td>||||||||[||||||||||||]PAM</td><td>NA</td><td>Intergenic</td><td>NA</td><td>NA</td></tr>
<tr><td>T2482</td><td>CCCCATCAGAACCGGCGAATGGG</td><td>1000</td><td></td><td></td><td></td><td></td><td></td><td></td><td></td><td></td><td></td></tr>
<tr><td>Oligo fwd</td><td>CACCCCCCATCAGAACCGGCGAAT</td><td></td><td></td><td></td><td></td><td></td><td></td><td></td><td></td><td></td><td></td></tr>
<tr><td>Oligo rev</td><td>AAACATTCGCCGGTTCTGATGGGG</td><td></td><td></td><td></td><td></td><td></td><td></td><td></td><td></td><td></td><td></td></tr>
<tr><td>Chromosome</td><td>start</td><td>end</td><td>strand</td><td>MM</td><td>target_seq</td><td>PAM</td><td>alignment</td><td>distance</td><td>position</td><td>gene name</td><td>gene id</td></tr>
<tr><td>GU305914.1</td><td>39324</td><td>39346</td><td>+</td><td>0</td><td>CCCCATCAGAACCGGCGAAT</td><td>GGG</td><td>||||||||[||||||||||||]PAM</td><td>NA</td><td>Intergenic</td><td>NA</td><td>NA</td></tr>
<tr><td>T2483</td><td>CCCATCAGAACCGGCGAATGGGG</td><td>1000</td><td></td><td></td><td></td><td></td><td></td><td></td><td></td><td></td><td></td></tr>
<tr><td>Oligo fwd</td><td>CACCCCCATCAGAACCGGCGAATG</td><td></td><td></td><td></td><td></td><td></td><td></td><td></td><td></td><td></td><td></td></tr>
<tr><td>Oligo rev</td><td>AAACCATTCGCCGGTTCTGATGGG</td><td></td><td></td><td></td><td></td><td></td><td></td><td></td><td></td><td></td><td></td></tr>
<tr><td>Chromosome</td><td>start</td><td>end</td><td>strand</td><td>MM</td><td>target_seq</td><td>PAM</td><td>alignment</td><td>distance</td><td>position</td><td>gene name</td><td>gene id</td></tr>
<tr><td>GU305914.1</td><td>39325</td><td>39347</td><td>+</td><td>0</td><td>CCCATCAGAACCGGCGAATG</td><td>GGG</td><td>||||||||[||||||||||||]PAM</td><td>NA</td><td>Intergenic</td><td>NA</td><td>NA</td></tr>
<tr><td>T2484</td><td>AGAACCGGCGAATGGGGAGCCGG</td><td>1000</td><td></td><td></td><td></td><td></td><td></td><td></td><td></td><td></td><td></td></tr>
<tr><td>Oligo fwd</td><td>CACCAGAACCGGCGAATGGGGAGC</td><td></td><td></td><td></td><td></td><td></td><td></td><td></td><td></td><td></td><td></td></tr>
<tr><td>Oligo rev</td><td>AAACGCTCCCCATTCGCCGGTTCT</td><td></td><td></td><td></td><td></td><td></td><td></td><td></td><td></td><td></td><td></td></tr>
<tr><td>Chromosome</td><td>start</td><td>end</td><td>strand</td><td>MM</td><td>target_seq</td><td>PAM</td><td>alignment</td><td>distance</td><td>position</td><td>gene name</td><td>gene id</td></tr>
<tr><td>GU305914.1</td><td>39331</td><td>39353</td><td>+</td><td>0</td><td>AGAACCGGCGAATGGGGAGC</td><td>CGG</td><td>||||||||[||||||||||||]PAM</td><td>NA</td><td>Intergenic</td><td>NA</td><td>NA</td></tr>
<tr><td>T2485</td><td>ATGGGGAGCCGGCGAACTGTTGG</td><td>1000</td><td></td><td></td><td></td><td></td><td></td><td></td><td></td><td></td><td></td></tr>
<tr><td>Oligo fwd</td><td>CACCATGGGGAGCCGGCGAACTGT</td><td></td><td></td><td></td><td></td><td></td><td></td><td></td><td></td><td></td><td></td></tr>
<tr><td>Oligo rev</td><td>AAACACAGTTCGCCGGCTCCCCAT</td><td></td><td></td><td></td><td></td><td></td><td></td><td></td><td></td><td></td><td></td></tr>
<tr><td>Chromosome</td><td>start</td><td>end</td><td>strand</td><td>MM</td><td>target_seq</td><td>PAM</td><td>alignment</td><td>distance</td><td>position</td><td>gene name</td><td>gene id</td></tr>
<tr><td>GU305914.1</td><td>39342</td><td>39364</td><td>+</td><td>0</td><td>ATGGGGAGCCGGCGAACTGT</td><td>TGG</td><td>||||||||[||||||||||||]PAM</td><td>NA</td><td>Intergenic</td><td>NA</td><td>NA</td></tr>
<tr><td>T2486</td><td>TGGGGAGCCGGCGAACTGTTGGG</td><td>1000</td><td></td><td></td><td></td><td></td><td></td><td></td><td></td><td></td><td></td></tr>
<tr><td>Oligo fwd</td><td>CACCTGGGGAGCCGGCGAACTGTT</td><td></td><td></td><td></td><td></td><td></td><td></td><td></td><td></td><td></td><td></td></tr>
<tr><td>Oligo rev</td><td>AAACAACAGTTCGCCGGCTCCCCA</td><td></td><td></td><td></td><td></td><td></td><td></td><td></td><td></td><td></td><td></td></tr>
<tr><td>Chromosome</td><td>start</td><td>end</td><td>strand</td><td>MM</td><td>target_seq</td><td>PAM</td><td>alignment</td><td>distance</td><td>position</td><td>gene name</td><td>gene id</td></tr>
<tr><td>GU305914.1</td><td>39343</td><td>39365</td><td>+</td><td>0</td><td>TGGGGAGCCGGCGAACTGTT</td><td>GGG</td><td>||||||||[||||||||||||]PAM</td><td>NA</td><td>Intergenic</td><td>NA</td><td>NA</td></tr>
<tr><td>T2487</td><td>CCGGCGAACTGTTGGGAGAGTGG</td><td>1000</td><td></td><td></td><td></td><td></td><td></td><td></td><td></td><td></td><td></td></tr>
<tr><td>Oligo fwd</td><td>CACCCCGGCGAACTGTTGGGAGAG</td><td></td><td></td><td></td><td></td><td></td><td></td><td></td><td></td><td></td><td></td></tr>
<tr><td>Oligo rev</td><td>AAACCTCTCCCAACAGTTCGCCGG</td><td></td><td></td><td></td><td></td><td></td><td></td><td></td><td></td><td></td><td></td></tr>
<tr><td>Chromosome</td><td>start</td><td>end</td><td>strand</td><td>MM</td><td>target_seq</td><td>PAM</td><td>alignment</td><td>distance</td><td>position</td><td>gene name</td><td>gene id</td></tr>
<tr><td>GU305914.1</td><td>39350</td><td>39372</td><td>+</td><td>0</td><td>CCGGCGAACTGTTGGGAGAG</td><td>TGG</td><td>||||||||[||||||||||||]PAM</td><td>NA</td><td>Intergenic</td><td>NA</td><td>NA</td></tr>
<tr><td>T2488</td><td>GCGAACTGTTGGGAGAGTGGTGG</td><td>1000</td><td></td><td></td><td></td><td></td><td></td><td></td><td></td><td></td><td></td></tr>
<tr><td>Oligo fwd</td><td>CACCGCGAACTGTTGGGAGAGTGG</td><td></td><td></td><td></td><td></td><td></td><td></td><td></td><td></td><td></td><td></td></tr>
<tr><td>Oligo rev</td><td>AAACCCACTCTCCCAACAGTTCGC</td><td></td><td></td><td></td><td></td><td></td><td></td><td></td><td></td><td></td><td></td></tr>
<tr><td>Chromosome</td><td>start</td><td>end</td><td>strand</td><td>MM</td><td>target_seq</td><td>PAM</td><td>alignment</td><td>distance</td><td>position</td><td>gene name</td><td>gene id</td></tr>
<tr><td>GU305914.1</td><td>39353</td><td>39375</td><td>+</td><td>0</td><td>GCGAACTGTTGGGAGAGTGG</td><td>TGG</td><td>||||||||[||||||||||||]PAM</td><td>NA</td><td>Intergenic</td><td>NA</td><td>NA</td></tr>
<tr><td>T2489</td><td>GTTGGGAGAGTGGTGGCTGTAGG</td><td>1000</td><td></td><td></td><td></td><td></td><td></td><td></td><td></td><td></td><td></td></tr>
<tr><td>Oligo fwd</td><td>CACCGTTGGGAGAGTGGTGGCTGT</td><td></td><td></td><td></td><td></td><td></td><td></td><td></td><td></td><td></td><td></td></tr>
<tr><td>Oligo rev</td><td>AAACACAGCCACCACTCTCCCAAC</td><td></td><td></td><td></td><td></td><td></td><td></td><td></td><td></td><td></td><td></td></tr>
<tr><td>Chromosome</td><td>start</td><td>end</td><td>strand</td><td>MM</td><td>target_seq</td><td>PAM</td><td>alignment</td><td>distance</td><td>position</td><td>gene name</td><td>gene id</td></tr>
<tr><td>GU305914.1</td><td>39360</td><td>39382</td><td>+</td><td>0</td><td>GTTGGGAGAGTGGTGGCTGT</td><td>AGG</td><td>||||||||[||||||||||||]PAM</td><td>NA</td><td>Intergenic</td><td>NA</td><td>NA</td></tr>
<tr><td>T2490</td><td>CTGTAGGCTACCACCGCCACCGG</td><td>1000</td><td></td><td></td><td></td><td></td><td></td><td></td><td></td><td></td><td></td></tr>
<tr><td>Oligo fwd</td><td>CACCCTGTAGGCTACCACCGCCAC</td><td></td><td></td><td></td><td></td><td></td><td></td><td></td><td></td><td></td><td></td></tr>
</table>

<table>
<tr><td>Oligo rev</td><td>AAACGTGGCGGTGGTAGCCTACAG</td><td></td><td></td><td></td><td></td><td></td><td></td><td></td><td></td><td></td><td></td><td></td></tr>
<tr><td>Chromosome</td><td>start</td><td></td><td>end</td><td>strand</td><td>MM</td><td>target_seq</td><td>PAM</td><td>alignment</td><td>distance</td><td>position</td><td>gene name</td><td>gene id</td></tr>
<tr><td>GU305914.1</td><td></td><td>39376</td><td>39398</td><td>+</td><td>0</td><td>CTGTAGGCTACCACCGCCAC</td><td>CGG</td><td>||||||||[||||||||||||]PAM</td><td>NA</td><td>Intergenic</td><td>NA</td><td>NA</td></tr>
<tr><td>T2491</td><td>CTATACCCTCCAGACCTTTACGG</td><td></td><td>1000</td><td></td><td></td><td></td><td></td><td></td><td></td><td></td><td></td><td></td></tr>
<tr><td>Oligo fwd</td><td>CACCCTATACCCTCCAGACCTTTA</td><td></td><td></td><td></td><td></td><td></td><td></td><td></td><td></td><td></td><td></td><td></td></tr>
<tr><td>Oligo rev</td><td>AAACTAAAGGTCTGGAGGGTATAG</td><td></td><td></td><td></td><td></td><td></td><td></td><td></td><td></td><td></td><td></td><td></td></tr>
<tr><td>Chromosome</td><td>start</td><td></td><td>end</td><td>strand</td><td>MM</td><td>target_seq</td><td>PAM</td><td>alignment</td><td>distance</td><td>position</td><td>gene name</td><td>gene id</td></tr>
<tr><td>GU305914.1</td><td></td><td>39408</td><td>39430</td><td>+</td><td>0</td><td>CTATACCCTCCAGACCTTTA</td><td>CGG</td><td>||||||||[||||||||||||]PAM</td><td>NA</td><td>Intergenic</td><td>NA</td><td>NA</td></tr>
<tr><td>T2492</td><td>TACCCTCCAGACCTTTACGGAGG</td><td></td><td>1000</td><td></td><td></td><td></td><td></td><td></td><td></td><td></td><td></td><td></td></tr>
<tr><td>Oligo fwd</td><td>CACCTACCCTCCAGACCTTTACGG</td><td></td><td></td><td></td><td></td><td></td><td></td><td></td><td></td><td></td><td></td><td></td></tr>
<tr><td>Oligo rev</td><td>AAACCCGTAAAGGTCTGGAGGGTA</td><td></td><td></td><td></td><td></td><td></td><td></td><td></td><td></td><td></td><td></td><td></td></tr>
<tr><td>Chromosome</td><td>start</td><td></td><td>end</td><td>strand</td><td>MM</td><td>target_seq</td><td>PAM</td><td>alignment</td><td>distance</td><td>position</td><td>gene name</td><td>gene id</td></tr>
<tr><td>GU305914.1</td><td></td><td>39411</td><td>39433</td><td>+</td><td>0</td><td>TACCCTCCAGACCTTTACGG</td><td>AGG</td><td>||||||||[||||||||||||]PAM</td><td>NA</td><td>Intergenic</td><td>NA</td><td>NA</td></tr>
<tr><td>T2493</td><td>ACCACCTCCATCCTCCCCGACGG</td><td></td><td>1000</td><td></td><td></td><td></td><td></td><td></td><td></td><td></td><td></td><td></td></tr>
<tr><td>Oligo fwd</td><td>CACCACCACCTCCATCCTCCCCGA</td><td></td><td></td><td></td><td></td><td></td><td></td><td></td><td></td><td></td><td></td><td></td></tr>
<tr><td>Oligo rev</td><td>AAACTCGGGGAGGATGGAGGTGGT</td><td></td><td></td><td></td><td></td><td></td><td></td><td></td><td></td><td></td><td></td><td></td></tr>
<tr><td>Chromosome</td><td>start</td><td></td><td>end</td><td>strand</td><td>MM</td><td>target_seq</td><td>PAM</td><td>alignment</td><td>distance</td><td>position</td><td>gene name</td><td>gene id</td></tr>
<tr><td>GU305914.1</td><td></td><td>39445</td><td>39467</td><td>+</td><td>0</td><td>ACCACCTCCATCCTCCCCGA</td><td>CGG</td><td>||||||||[||||||||||||]PAM</td><td>NA</td><td>Intergenic</td><td>NA</td><td>NA</td></tr>
<tr><td>T2494</td><td>CCACCTCCATCCTCCCCGACGGG</td><td></td><td>1000</td><td></td><td></td><td></td><td></td><td></td><td></td><td></td><td></td><td></td></tr>
<tr><td>Oligo fwd</td><td>CACCCCACCTCCATCCTCCCCGAC</td><td></td><td></td><td></td><td></td><td></td><td></td><td></td><td></td><td></td><td></td><td></td></tr>
<tr><td>Oligo rev</td><td>AAACGTCGGGGAGGATGGAGGTGG</td><td></td><td></td><td></td><td></td><td></td><td></td><td></td><td></td><td></td><td></td><td></td></tr>
<tr><td>Chromosome</td><td>start</td><td></td><td>end</td><td>strand</td><td>MM</td><td>target_seq</td><td>PAM</td><td>alignment</td><td>distance</td><td>position</td><td>gene name</td><td>gene id</td></tr>
<tr><td>GU305914.1</td><td></td><td>39446</td><td>39468</td><td>+</td><td>0</td><td>CCACCTCCATCCTCCCCGAC</td><td>GGG</td><td>||||||||[||||||||||||]PAM</td><td>NA</td><td>Intergenic</td><td>NA</td><td>NA</td></tr>
<tr><td>GU305914.1</td><td></td><td>99827</td><td>99849</td><td>-</td><td>4</td><td>CCATCTTCATCGTCCCCGCC</td><td>GGG</td><td>|||-||-|[|||-||||||-|]PAM</td><td>NA</td><td>Intergenic</td><td>NA</td><td>NA</td></tr>
<tr><td>T2495</td><td>AATACTCGTCGTCTTTATTATGG</td><td></td><td>1000</td><td></td><td></td><td></td><td></td><td></td><td></td><td></td><td></td><td></td></tr>
<tr><td>Oligo fwd</td><td>CACCAATACTCGTCGTCTTTATTA</td><td></td><td></td><td></td><td></td><td></td><td></td><td></td><td></td><td></td><td></td><td></td></tr>
<tr><td>Oligo rev</td><td>AAACTAATAAAGACGACGAGTATT</td><td></td><td></td><td></td><td></td><td></td><td></td><td></td><td></td><td></td><td></td><td></td></tr>
<tr><td>Chromosome</td><td>start</td><td></td><td>end</td><td>strand</td><td>MM</td><td>target_seq</td><td>PAM</td><td>alignment</td><td>distance</td><td>position</td><td>gene name</td><td>gene id</td></tr>
<tr><td>GU305914.1</td><td></td><td>39495</td><td>39517</td><td>+</td><td>0</td><td>AATACTCGTCGTCTTTATTA</td><td>TGG</td><td>||||||||[||||||||||||]PAM</td><td>NA</td><td>Intergenic</td><td>NA</td><td>NA</td></tr>
<tr><td>T2496</td><td>ATACTCGTCGTCTTTATTATGGG</td><td></td><td>1000</td><td></td><td></td><td></td><td></td><td></td><td></td><td></td><td></td><td></td></tr>
<tr><td>Oligo fwd</td><td>CACCATACTCGTCGTCTTTATTAT</td><td></td><td></td><td></td><td></td><td></td><td></td><td></td><td></td><td></td><td></td><td></td></tr>
<tr><td>Oligo rev</td><td>AAACATAATAAAGACGACGAGTAT</td><td></td><td></td><td></td><td></td><td></td><td></td><td></td><td></td><td></td><td></td><td></td></tr>
<tr><td>Chromosome</td><td>start</td><td></td><td>end</td><td>strand</td><td>MM</td><td>target_seq</td><td>PAM</td><td>alignment</td><td>distance</td><td>position</td><td>gene name</td><td>gene id</td></tr>
<tr><td>GU305914.1</td><td></td><td>39496</td><td>39518</td><td>+</td><td>0</td><td>ATACTCGTCGTCTTTATTAT</td><td>GGG</td><td>||||||||[||||||||||||]PAM</td><td>NA</td><td>Intergenic</td><td>NA</td><td>NA</td></tr>
<tr><td>T2497</td><td>TACTCGTCGTCTTTATTATGGGG</td><td></td><td>1000</td><td></td><td></td><td></td><td></td><td></td><td></td><td></td><td></td><td></td></tr>
<tr><td>Oligo fwd</td><td>CACCTACTCGTCGTCTTTATTATG</td><td></td><td></td><td></td><td></td><td></td><td></td><td></td><td></td><td></td><td></td><td></td></tr>
<tr><td>Oligo rev</td><td>AAACCATAATAAAGACGACGAGTA</td><td></td><td></td><td></td><td></td><td></td><td></td><td></td><td></td><td></td><td></td><td></td></tr>
<tr><td>Chromosome</td><td>start</td><td></td><td>end</td><td>strand</td><td>MM</td><td>target_seq</td><td>PAM</td><td>alignment</td><td>distance</td><td>position</td><td>gene name</td><td>gene id</td></tr>
<tr><td>GU305914.1</td><td></td><td>39497</td><td>39519</td><td>+</td><td>0</td><td>TACTCGTCGTCTTTATTATG</td><td>GGG</td><td>||||||||[||||||||||||]PAM</td><td>NA</td><td>Intergenic</td><td>NA</td><td>NA</td></tr>
<tr><td>T2498</td><td>TCGTCGTCTTTATTATGGGGTGG</td><td></td><td>1000</td><td></td><td></td><td></td><td></td><td></td><td></td><td></td><td></td><td></td></tr>
<tr><td>Oligo fwd</td><td>CACCTCGTCGTCTTTATTATGGGG</td><td></td><td></td><td></td><td></td><td></td><td></td><td></td><td></td><td></td><td></td><td></td></tr>
<tr><td>Oligo rev</td><td>AAACCCCCATAATAAAGACGACGA</td><td></td><td></td><td></td><td></td><td></td><td></td><td></td><td></td><td></td><td></td><td></td></tr>
<tr><td>Chromosome</td><td>start</td><td></td><td>end</td><td>strand</td><td>MM</td><td>target_seq</td><td>PAM</td><td>alignment</td><td>distance</td><td>position</td><td>gene name</td><td>gene id</td></tr>
<tr><td>GU305914.1</td><td></td><td>39500</td><td>39522</td><td>+</td><td>0</td><td>TCGTCGTCTTTATTATGGGG</td><td>TGG</td><td>||||||||[||||||||||||]PAM</td><td>NA</td><td>Intergenic</td><td>NA</td><td>NA</td></tr>
<tr><td>T2499</td><td>TCGTCTTTATTATGGGGTGGTGG</td><td></td><td>1000</td><td></td><td></td><td></td><td></td><td></td><td></td><td></td><td></td><td></td></tr>
<tr><td>Oligo fwd</td><td>CACCTCGTCTTTATTATGGGGTGG</td><td></td><td></td><td></td><td></td><td></td><td></td><td></td><td></td><td></td><td></td><td></td></tr>
<tr><td>Oligo rev</td><td>AAACCCACCCCATAATAAAGACGA</td><td></td><td></td><td></td><td></td><td></td><td></td><td></td><td></td><td></td><td></td><td></td></tr>
<tr><td>Chromosome</td><td>start</td><td></td><td>end</td><td>strand</td><td>MM</td><td>target_seq</td><td>PAM</td><td>alignment</td><td>distance</td><td>position</td><td>gene name</td><td>gene id</td></tr>
<tr><td>GU305914.1</td><td></td><td>39503</td><td>39525</td><td>+</td><td>0</td><td>TCGTCTTTATTATGGGGTGG</td><td>TGG</td><td>||||||||[||||||||||||]PAM</td><td>NA</td><td>Intergenic</td><td>NA</td><td>NA</td></tr>
<tr><td>T2500</td><td>GGTCTCTGTTTATAATTCTCAGG</td><td></td><td>1000</td><td></td><td></td><td></td><td></td><td></td><td></td><td></td><td></td><td></td></tr>
<tr><td>Oligo fwd</td><td>CACCGGTCTCTGTTTATAATTCTC</td><td></td><td></td><td></td><td></td><td></td><td></td><td></td><td></td><td></td><td></td><td></td></tr>
<tr><td>Oligo rev</td><td>AAACGAGAATTATAAACAGAGACC</td><td></td><td></td><td></td><td></td><td></td><td></td><td></td><td></td><td></td><td></td><td></td></tr>
<tr><td>Chromosome</td><td>start</td><td></td><td>end</td><td>strand</td><td>MM</td><td>target_seq</td><td>PAM</td><td>alignment</td><td>distance</td><td>position</td><td>gene name</td><td>gene id</td></tr>
<tr><td>GU305914.1</td><td></td><td>39524</td><td>39546</td><td>+</td><td>0</td><td>GGTCTCTGTTTATAATTCTC</td><td>AGG</td><td>||||||||[||||||||||||]PAM</td><td>NA</td><td>Intergenic</td><td>NA</td><td>NA</td></tr>
<tr><td>T2501</td><td>TTTATAATTCTCAGGAAAAGTGG</td><td></td><td>1000</td><td></td><td></td><td></td><td></td><td></td><td></td><td></td><td></td><td></td></tr>
<tr><td>Oligo fwd</td><td>CACCTTTATAATTCTCAGGAAAAG</td><td></td><td></td><td></td><td></td><td></td><td></td><td></td><td></td><td></td><td></td><td></td></tr>
<tr><td>Oligo rev</td><td>AAACCTTTTCCTGAGAATTATAAA</td><td></td><td></td><td></td><td></td><td></td><td></td><td></td><td></td><td></td><td></td><td></td></tr>
<tr><td>Chromosome</td><td>start</td><td></td><td>end</td><td>strand</td><td>MM</td><td>target_seq</td><td>PAM</td><td>alignment</td><td>distance</td><td>position</td><td>gene name</td><td>gene id</td></tr>
<tr><td>GU305914.1</td><td></td><td>39532</td><td>39554</td><td>+</td><td>0</td><td>TTTATAATTCTCAGGAAAAG</td><td>TGG</td><td>||||||||[||||||||||||]PAM</td><td>NA</td><td>Intergenic</td><td>NA</td><td>NA</td></tr>
<tr><td>T2502</td><td>TTATAATTCTCAGGAAAAGTGGG</td><td></td><td>1000</td><td></td><td></td><td></td><td></td><td></td><td></td><td></td><td></td><td></td></tr>
<tr><td>Oligo fwd</td><td>CACCTTATAATTCTCAGGAAAAGT</td><td></td><td></td><td></td><td></td><td></td><td></td><td></td><td></td><td></td><td></td><td></td></tr>
<tr><td>Oligo rev</td><td>AAACACTTTTCCTGAGAATTATAA</td><td></td><td></td><td></td><td></td><td></td><td></td><td></td><td></td><td></td><td></td><td></td></tr>
<tr><td>Chromosome</td><td>start</td><td></td><td>end</td><td>strand</td><td>MM</td><td>target_seq</td><td>PAM</td><td>alignment</td><td>distance</td><td>position</td><td>gene name</td><td>gene id</td></tr>
<tr><td>GU305914.1</td><td></td><td>39533</td><td>39555</td><td>+</td><td>0</td><td>TTATAATTCTCAGGAAAAGT</td><td>GGG</td><td>||||||||[||||||||||||]PAM</td><td>NA</td><td>Intergenic</td><td>NA</td><td>NA</td></tr>
<tr><td>T2503</td><td>TATAATTCTCAGGAAAAGTGGGG</td><td></td><td>1000</td><td></td><td></td><td></td><td></td><td></td><td></td><td></td><td></td><td></td></tr>
<tr><td>Oligo fwd</td><td>CACCTATAATTCTCAGGAAAAGTG</td><td></td><td></td><td></td><td></td><td></td><td></td><td></td><td></td><td></td><td></td><td></td></tr>
<tr><td>Oligo rev</td><td>AAACCACTTTTCCTGAGAATTATA</td><td></td><td></td><td></td><td></td><td></td><td></td><td></td><td></td><td></td><td></td><td></td></tr>
<tr><td>Chromosome</td><td>start</td><td></td><td>end</td><td>strand</td><td>MM</td><td>target_seq</td><td>PAM</td><td>alignment</td><td>distance</td><td>position</td><td>gene name</td><td>gene id</td></tr>
<tr><td>GU305914.1</td><td></td><td>39534</td><td>39556</td><td>+</td><td>0</td><td>TATAATTCTCAGGAAAAGTG</td><td>GGG</td><td>||||||||[||||||||||||]PAM</td><td>NA</td><td>Intergenic</td><td>NA</td><td>NA</td></tr>
<tr><td>T2504</td><td>AGGAAAAGTGGGGTGCTTCTTGG</td><td></td><td>1000</td><td></td><td></td><td></td><td></td><td></td><td></td><td></td><td></td><td></td></tr>
<tr><td>Oligo fwd</td><td>CACCAGGAAAAGTGGGGTGCTTCT</td><td></td><td></td><td></td><td></td><td></td><td></td><td></td><td></td><td></td><td></td><td></td></tr>
<tr><td>Oligo rev</td><td>AAACAGAAGCACCCCACTTTTCCT</td><td></td><td></td><td></td><td></td><td></td><td></td><td></td><td></td><td></td><td></td><td></td></tr>
<tr><td>Chromosome</td><td>start</td><td></td><td>end</td><td>strand</td><td>MM</td><td>target_seq</td><td>PAM</td><td>alignment</td><td>distance</td><td>position</td><td>gene name</td><td>gene id</td></tr>
<tr><td>GU305914.1</td><td></td><td>39544</td><td>39566</td><td>+</td><td>0</td><td>AGGAAAAGTGGGGTGCTTCT</td><td>TGG</td><td>||||||||[||||||||||||]PAM</td><td>NA</td><td>Intergenic</td><td>NA</td><td>NA</td></tr>
</table>

<table>
<tr><td>T2505</td><td>TGCTTCTTGGTCCTGCTCTTTGG</td><td>1000</td><td></td><td></td><td></td><td></td><td></td><td></td><td></td><td></td><td></td></tr>
<tr><td>Oligo fwd</td><td>CACCTGCTTCTTGGTCCTGCTCTT</td><td></td><td></td><td></td><td></td><td></td><td></td><td></td><td></td><td></td><td></td></tr>
<tr><td>Oligo rev</td><td>AAACAAGAGCAGGACCAAGAAGCA</td><td></td><td></td><td></td><td></td><td></td><td></td><td></td><td></td><td></td><td></td></tr>
<tr><td>Chromosome</td><td>start</td><td>end</td><td>strand</td><td>MM</td><td>target_seq</td><td>PAM</td><td>alignment</td><td>distance</td><td>position</td><td>gene name</td><td>gene id</td></tr>
<tr><td>GU305914.1</td><td>39557</td><td>39579</td><td>+</td><td>0</td><td>TGCTTCTTGGTCCTGCTCTT</td><td>TGG</td><td>||||||||[||||||||||||]PAM</td><td>NA</td><td>Intergenic</td><td>NA</td><td>NA</td></tr>
<tr><td>T2506</td><td>TCTTGCCGTATCTGCTGTCCTGG</td><td>1000</td><td></td><td></td><td></td><td></td><td></td><td></td><td></td><td></td><td></td></tr>
<tr><td>Oligo fwd</td><td>CACCTCTTGCCGTATCTGCTGTCC</td><td></td><td></td><td></td><td></td><td></td><td></td><td></td><td></td><td></td><td></td></tr>
<tr><td>Oligo rev</td><td>AAACGGACAGCAGATACGGCAAGA</td><td></td><td></td><td></td><td></td><td></td><td></td><td></td><td></td><td></td><td></td></tr>
<tr><td>Chromosome</td><td>start</td><td>end</td><td>strand</td><td>MM</td><td>target_seq</td><td>PAM</td><td>alignment</td><td>distance</td><td>position</td><td>gene name</td><td>gene id</td></tr>
<tr><td>GU305914.1</td><td>39582</td><td>39604</td><td>+</td><td>0</td><td>TCTTGCCGTATCTGCTGTCC</td><td>TGG</td><td>||||||||[||||||||||||]PAM</td><td>NA</td><td>Intergenic</td><td>NA</td><td>NA</td></tr>
<tr><td>T2507</td><td>GTCTTCGTCCGCGAGAACCCCGG</td><td>1000</td><td></td><td></td><td></td><td></td><td></td><td></td><td></td><td></td><td></td></tr>
<tr><td>Oligo fwd</td><td>CACCGTCTTCGTCCGCGAGAACCC</td><td></td><td></td><td></td><td></td><td></td><td></td><td></td><td></td><td></td><td></td></tr>
<tr><td>Oligo rev</td><td>AAACGGGTTCTCGCGGACGAAGAC</td><td></td><td></td><td></td><td></td><td></td><td></td><td></td><td></td><td></td><td></td></tr>
<tr><td>Chromosome</td><td>start</td><td>end</td><td>strand</td><td>MM</td><td>target_seq</td><td>PAM</td><td>alignment</td><td>distance</td><td>position</td><td>gene name</td><td>gene id</td></tr>
<tr><td>GU305914.1</td><td>39607</td><td>39629</td><td>+</td><td>0</td><td>GTCTTCGTCCGCGAGAACCC</td><td>CGG</td><td>||||||||[||||||||||||]PAM</td><td>NA</td><td>Intergenic</td><td>NA</td><td>NA</td></tr>
<tr><td>T2508</td><td>GCGAGAACCCCGGAGAGATCCGG</td><td>1000</td><td></td><td></td><td></td><td></td><td></td><td></td><td></td><td></td><td></td></tr>
<tr><td>Oligo fwd</td><td>CACCGCGAGAACCCCGGAGAGATC</td><td></td><td></td><td></td><td></td><td></td><td></td><td></td><td></td><td></td><td></td></tr>
<tr><td>Oligo rev</td><td>AAACGATCTCTCCGGGGTTCTCGC</td><td></td><td></td><td></td><td></td><td></td><td></td><td></td><td></td><td></td><td></td></tr>
<tr><td>Chromosome</td><td>start</td><td>end</td><td>strand</td><td>MM</td><td>target_seq</td><td>PAM</td><td>alignment</td><td>distance</td><td>position</td><td>gene name</td><td>gene id</td></tr>
<tr><td>GU305914.1</td><td>39617</td><td>39639</td><td>+</td><td>0</td><td>GCGAGAACCCCGGAGAGATC</td><td>CGG</td><td>||||||||[||||||||||||]PAM</td><td>NA</td><td>Intergenic</td><td>NA</td><td>NA</td></tr>
<tr><td>T2509</td><td>CCCGGAGAGATCCGGCAGATCGG</td><td>1000</td><td></td><td></td><td></td><td></td><td></td><td></td><td></td><td></td><td></td></tr>
<tr><td>Oligo fwd</td><td>CACCCCCGGAGAGATCCGGCAGAT</td><td></td><td></td><td></td><td></td><td></td><td></td><td></td><td></td><td></td><td></td></tr>
<tr><td>Oligo rev</td><td>AAACATCTGCCGGATCTCTCCGGG</td><td></td><td></td><td></td><td></td><td></td><td></td><td></td><td></td><td></td><td></td></tr>
<tr><td>Chromosome</td><td>start</td><td>end</td><td>strand</td><td>MM</td><td>target_seq</td><td>PAM</td><td>alignment</td><td>distance</td><td>position</td><td>gene name</td><td>gene id</td></tr>
<tr><td>GU305914.1</td><td>39625</td><td>39647</td><td>+</td><td>0</td><td>CCCGGAGAGATCCGGCAGAT</td><td>CGG</td><td>||||||||[||||||||||||]PAM</td><td>NA</td><td>Intergenic</td><td>NA</td><td>NA</td></tr>
<tr><td>T2510</td><td>CCGGAGAGATCCGGCAGATCGGG</td><td>1000</td><td></td><td></td><td></td><td></td><td></td><td></td><td></td><td></td><td></td></tr>
<tr><td>Oligo fwd</td><td>CACCCCGGAGAGATCCGGCAGATC</td><td></td><td></td><td></td><td></td><td></td><td></td><td></td><td></td><td></td><td></td></tr>
<tr><td>Oligo rev</td><td>AAACGATCTGCCGGATCTCTCCGG</td><td></td><td></td><td></td><td></td><td></td><td></td><td></td><td></td><td></td><td></td></tr>
<tr><td>Chromosome</td><td>start</td><td>end</td><td>strand</td><td>MM</td><td>target_seq</td><td>PAM</td><td>alignment</td><td>distance</td><td>position</td><td>gene name</td><td>gene id</td></tr>
<tr><td>GU305914.1</td><td>39626</td><td>39648</td><td>+</td><td>0</td><td>CCGGAGAGATCCGGCAGATC</td><td>GGG</td><td>||||||||[||||||||||||]PAM</td><td>NA</td><td>Intergenic</td><td>NA</td><td>NA</td></tr>
<tr><td>T2511</td><td>CGGAGAGATCCGGCAGATCGGGG</td><td>1000</td><td></td><td></td><td></td><td></td><td></td><td></td><td></td><td></td><td></td></tr>
<tr><td>Oligo fwd</td><td>CACCCGGAGAGATCCGGCAGATCG</td><td></td><td></td><td></td><td></td><td></td><td></td><td></td><td></td><td></td><td></td></tr>
<tr><td>Oligo rev</td><td>AAACCGATCTGCCGGATCTCTCCG</td><td></td><td></td><td></td><td></td><td></td><td></td><td></td><td></td><td></td><td></td></tr>
<tr><td>Chromosome</td><td>start</td><td>end</td><td>strand</td><td>MM</td><td>target_seq</td><td>PAM</td><td>alignment</td><td>distance</td><td>position</td><td>gene name</td><td>gene id</td></tr>
<tr><td>GU305914.1</td><td>39627</td><td>39649</td><td>+</td><td>0</td><td>CGGAGAGATCCGGCAGATCG</td><td>GGG</td><td>||||||||[||||||||||||]PAM</td><td>NA</td><td>Intergenic</td><td>NA</td><td>NA</td></tr>
<tr><td>T2512</td><td>GGAGAGATCCGGCAGATCGGGGG</td><td>1000</td><td></td><td></td><td></td><td></td><td></td><td></td><td></td><td></td><td></td></tr>
<tr><td>Oligo fwd</td><td>CACCGGAGAGATCCGGCAGATCGG</td><td></td><td></td><td></td><td></td><td></td><td></td><td></td><td></td><td></td><td></td></tr>
<tr><td>Oligo rev</td><td>AAACCCGATCTGCCGGATCTCTCC</td><td></td><td></td><td></td><td></td><td></td><td></td><td></td><td></td><td></td><td></td></tr>
<tr><td>Chromosome</td><td>start</td><td>end</td><td>strand</td><td>MM</td><td>target_seq</td><td>PAM</td><td>alignment</td><td>distance</td><td>position</td><td>gene name</td><td>gene id</td></tr>
<tr><td>GU305914.1</td><td>39628</td><td>39650</td><td>+</td><td>0</td><td>GGAGAGATCCGGCAGATCGG</td><td>GGG</td><td>||||||||[||||||||||||]PAM</td><td>NA</td><td>Intergenic</td><td>NA</td><td>NA</td></tr>
<tr><td>T2513</td><td>GAGAGATCCGGCAGATCGGGGGG</td><td>1000</td><td></td><td></td><td></td><td></td><td></td><td></td><td></td><td></td><td></td></tr>
<tr><td>Oligo fwd</td><td>CACCGAGAGATCCGGCAGATCGGG</td><td></td><td></td><td></td><td></td><td></td><td></td><td></td><td></td><td></td><td></td></tr>
<tr><td>Oligo rev</td><td>AAACCCCGATCTGCCGGATCTCTC</td><td></td><td></td><td></td><td></td><td></td><td></td><td></td><td></td><td></td><td></td></tr>
<tr><td>Chromosome</td><td>start</td><td>end</td><td>strand</td><td>MM</td><td>target_seq</td><td>PAM</td><td>alignment</td><td>distance</td><td>position</td><td>gene name</td><td>gene id</td></tr>
<tr><td>GU305914.1</td><td>39629</td><td>39651</td><td>+</td><td>0</td><td>GAGAGATCCGGCAGATCGGG</td><td>GGG</td><td>||||||||[||||||||||||]PAM</td><td>NA</td><td>Intergenic</td><td>NA</td><td>NA</td></tr>
<tr><td>T2514</td><td>AGAGATCCGGCAGATCGGGGGGG</td><td>1000</td><td></td><td></td><td></td><td></td><td></td><td></td><td></td><td></td><td></td></tr>
<tr><td>Oligo fwd</td><td>CACCAGAGATCCGGCAGATCGGGG</td><td></td><td></td><td></td><td></td><td></td><td></td><td></td><td></td><td></td><td></td></tr>
<tr><td>Oligo rev</td><td>AAACCCCCGATCTGCCGGATCTCT</td><td></td><td></td><td></td><td></td><td></td><td></td><td></td><td></td><td></td><td></td></tr>
<tr><td>Chromosome</td><td>start</td><td>end</td><td>strand</td><td>MM</td><td>target_seq</td><td>PAM</td><td>alignment</td><td>distance</td><td>position</td><td>gene name</td><td>gene id</td></tr>
<tr><td>GU305914.1</td><td>39630</td><td>39652</td><td>+</td><td>0</td><td>AGAGATCCGGCAGATCGGGG</td><td>GGG</td><td>||||||||[||||||||||||]PAM</td><td>NA</td><td>Intergenic</td><td>NA</td><td>NA</td></tr>
<tr><td>T2515</td><td>GGGGAGAGTCAAAGTGCTACCGG</td><td>1000</td><td></td><td></td><td></td><td></td><td></td><td></td><td></td><td></td><td></td></tr>
<tr><td>Oligo fwd</td><td>CACCGGGGAGAGTCAAAGTGCTAC</td><td></td><td></td><td></td><td></td><td></td><td></td><td></td><td></td><td></td><td></td></tr>
<tr><td>Oligo rev</td><td>AAACGTAGCACTTTGACTCTCCCC</td><td></td><td></td><td></td><td></td><td></td><td></td><td></td><td></td><td></td><td></td></tr>
<tr><td>Chromosome</td><td>start</td><td>end</td><td>strand</td><td>MM</td><td>target_seq</td><td>PAM</td><td>alignment</td><td>distance</td><td>position</td><td>gene name</td><td>gene id</td></tr>
<tr><td>GU305914.1</td><td>39649</td><td>39671</td><td>+</td><td>0</td><td>GGGGAGAGTCAAAGTGCTAC</td><td>CGG</td><td>||||||||[||||||||||||]PAM</td><td>NA</td><td>Intergenic</td><td>NA</td><td>NA</td></tr>
<tr><td>T2516</td><td>GGGAGAGTCAAAGTGCTACCGGG</td><td>1000</td><td></td><td></td><td></td><td></td><td></td><td></td><td></td><td></td><td></td></tr>
<tr><td>Oligo fwd</td><td>CACCGGGAGAGTCAAAGTGCTACC</td><td></td><td></td><td></td><td></td><td></td><td></td><td></td><td></td><td></td><td></td></tr>
<tr><td>Oligo rev</td><td>AAACGGTAGCACTTTGACTCTCCC</td><td></td><td></td><td></td><td></td><td></td><td></td><td></td><td></td><td></td><td></td></tr>
<tr><td>Chromosome</td><td>start</td><td>end</td><td>strand</td><td>MM</td><td>target_seq</td><td>PAM</td><td>alignment</td><td>distance</td><td>position</td><td>gene name</td><td>gene id</td></tr>
<tr><td>GU305914.1</td><td>39650</td><td>39672</td><td>+</td><td>0</td><td>GGGAGAGTCAAAGTGCTACC</td><td>GGG</td><td>||||||||[||||||||||||]PAM</td><td>NA</td><td>Intergenic</td><td>NA</td><td>NA</td></tr>
<tr><td>T2517</td><td>AAGTGCTACCGGGCCCTCCGAGG</td><td>1000</td><td></td><td></td><td></td><td></td><td></td><td></td><td></td><td></td><td></td></tr>
<tr><td>Oligo fwd</td><td>CACCAAGTGCTACCGGGCCCTCCG</td><td></td><td></td><td></td><td></td><td></td><td></td><td></td><td></td><td></td><td></td></tr>
<tr><td>Oligo rev</td><td>AAACCGGAGGGCCCGGTAGCACTT</td><td></td><td></td><td></td><td></td><td></td><td></td><td></td><td></td><td></td><td></td></tr>
<tr><td>Chromosome</td><td>start</td><td>end</td><td>strand</td><td>MM</td><td>target_seq</td><td>PAM</td><td>alignment</td><td>distance</td><td>position</td><td>gene name</td><td>gene id</td></tr>
<tr><td>GU305914.1</td><td>39660</td><td>39682</td><td>+</td><td>0</td><td>AAGTGCTACCGGGCCCTCCG</td><td>AGG</td><td>||||||||[||||||||||||]PAM</td><td>NA</td><td>Intergenic</td><td>NA</td><td>NA</td></tr>
<tr><td>T2518</td><td>AGTGCTACCGGGCCCTCCGAGGG</td><td>1000</td><td></td><td></td><td></td><td></td><td></td><td></td><td></td><td></td><td></td></tr>
<tr><td>Oligo fwd</td><td>CACCAGTGCTACCGGGCCCTCCGA</td><td></td><td></td><td></td><td></td><td></td><td></td><td></td><td></td><td></td><td></td></tr>
<tr><td>Oligo rev</td><td>AAACTCGGAGGGCCCGGTAGCACT</td><td></td><td></td><td></td><td></td><td></td><td></td><td></td><td></td><td></td><td></td></tr>
<tr><td>Chromosome</td><td>start</td><td>end</td><td>strand</td><td>MM</td><td>target_seq</td><td>PAM</td><td>alignment</td><td>distance</td><td>position</td><td>gene name</td><td>gene id</td></tr>
<tr><td>GU305914.1</td><td>39661</td><td>39683</td><td>+</td><td>0</td><td>AGTGCTACCGGGCCCTCCGA</td><td>GGG</td><td>||||||||[||||||||||||]PAM</td><td>NA</td><td>Intergenic</td><td>NA</td><td>NA</td></tr>
<tr><td>T2519</td><td>GTGCTACCGGGCCCTCCGAGGGG</td><td>1000</td><td></td><td></td><td></td><td></td><td></td><td></td><td></td><td></td><td></td></tr>
<tr><td>Oligo fwd</td><td>CACCGTGCTACCGGGCCCTCCGAG</td><td></td><td></td><td></td><td></td><td></td><td></td><td></td><td></td><td></td><td></td></tr>
<tr><td>Oligo rev</td><td>AAACCTCGGAGGGCCCGGTAGCAC</td><td></td><td></td><td></td><td></td><td></td><td></td><td></td><td></td><td></td><td></td></tr>
<tr><td>Chromosome</td><td>start</td><td>end</td><td>strand</td><td>MM</td><td>target_seq</td><td>PAM</td><td>alignment</td><td>distance</td><td>position</td><td>gene name</td><td>gene id</td></tr>
<tr><td>GU305914.1</td><td>39662</td><td>39684</td><td>+</td><td>0</td><td>GTGCTACCGGGCCCTCCGAG</td><td>GGG</td><td>||||||||[||||||||||||]PAM</td><td>NA</td><td>Intergenic</td><td>NA</td><td>NA</td></tr>
</table>

<table>
<tr><td>T2520</td><td>CTACCGGGCCCTCCGAGGGGAGG</td><td>1000</td><td></td><td></td><td></td><td></td><td></td><td></td><td></td><td></td><td></td></tr>
<tr><td>Oligo fwd</td><td>CACCCTACCGGGCCCTCCGAGGGG</td><td></td><td></td><td></td><td></td><td></td><td></td><td></td><td></td><td></td><td></td></tr>
<tr><td>Oligo rev</td><td>AAACCCCCTCGGAGGGCCCGGTAG</td><td></td><td></td><td></td><td></td><td></td><td></td><td></td><td></td><td></td><td></td></tr>
<tr><td>Chromosome</td><td>start</td><td>end</td><td>strand</td><td>MM</td><td>target_seq</td><td>PAM</td><td>alignment</td><td>distance</td><td>position</td><td>gene name</td><td>gene id</td></tr>
<tr><td>GU305914.1</td><td>39665</td><td>39687</td><td>+</td><td>0</td><td>CTACCGGGCCCTCCGAGGGG</td><td>AGG</td><td>||||||||[||||||||||||]PAM</td><td>NA</td><td>Intergenic</td><td>NA</td><td>NA</td></tr>
<tr><td>T2521</td><td>CCGGGCCCTCCGAGGGGAGGAGG</td><td>1000</td><td></td><td></td><td></td><td></td><td></td><td></td><td></td><td></td><td></td></tr>
<tr><td>Oligo fwd</td><td>CACCCCGGGCCCTCCGAGGGGAGG</td><td></td><td></td><td></td><td></td><td></td><td></td><td></td><td></td><td></td><td></td></tr>
<tr><td>Oligo rev</td><td>AAACCCTCCCCTCGGAGGGCCCGG</td><td></td><td></td><td></td><td></td><td></td><td></td><td></td><td></td><td></td><td></td></tr>
<tr><td>Chromosome</td><td>start</td><td>end</td><td>strand</td><td>MM</td><td>target_seq</td><td>PAM</td><td>alignment</td><td>distance</td><td>position</td><td>gene name</td><td>gene id</td></tr>
<tr><td>GU305914.1</td><td>39668</td><td>39690</td><td>+</td><td>0</td><td>CCGGGCCCTCCGAGGGGAGG</td><td>AGG</td><td>||||||||[||||||||||||]PAM</td><td>NA</td><td>Intergenic</td><td>NA</td><td>NA</td></tr>
<tr><td>T2522</td><td>GCCCTCCGAGGGGAGGAGGAAGG</td><td>1000</td><td></td><td></td><td></td><td></td><td></td><td></td><td></td><td></td><td></td></tr>
<tr><td>Oligo fwd</td><td>CACCGCCCTCCGAGGGGAGGAGGA</td><td></td><td></td><td></td><td></td><td></td><td></td><td></td><td></td><td></td><td></td></tr>
<tr><td>Oligo rev</td><td>AAACTCCTCCTCCCCTCGGAGGGC</td><td></td><td></td><td></td><td></td><td></td><td></td><td></td><td></td><td></td><td></td></tr>
<tr><td>Chromosome</td><td>start</td><td>end</td><td>strand</td><td>MM</td><td>target_seq</td><td>PAM</td><td>alignment</td><td>distance</td><td>position</td><td>gene name</td><td>gene id</td></tr>
<tr><td>GU305914.1</td><td>39672</td><td>39694</td><td>+</td><td>0</td><td>GCCCTCCGAGGGGAGGAGGA</td><td>AGG</td><td>||||||||[||||||||||||]PAM</td><td>NA</td><td>Intergenic</td><td>NA</td><td>NA</td></tr>
<tr><td>GU305914.1</td><td>78501</td><td>78523</td><td>+</td><td>4</td><td>GCCGACCGAGAAGAGGAGGA</td><td>AGG</td><td>|||--|||[||--||||||||]PAM</td><td>NA</td><td>Intergenic</td><td>NA</td><td>NA</td></tr>
<tr><td>T2523</td><td>GAGGGGAGGAGGAAGGAGCACGG</td><td>1000</td><td></td><td></td><td></td><td></td><td></td><td></td><td></td><td></td><td></td></tr>
<tr><td>Oligo fwd</td><td>CACCGAGGGGAGGAGGAAGGAGCA</td><td></td><td></td><td></td><td></td><td></td><td></td><td></td><td></td><td></td><td></td></tr>
<tr><td>Oligo rev</td><td>AAACTGCTCCTTCCTCCTCCCCTC</td><td></td><td></td><td></td><td></td><td></td><td></td><td></td><td></td><td></td><td></td></tr>
<tr><td>Chromosome</td><td>start</td><td>end</td><td>strand</td><td>MM</td><td>target_seq</td><td>PAM</td><td>alignment</td><td>distance</td><td>position</td><td>gene name</td><td>gene id</td></tr>
<tr><td>GU305914.1</td><td>39679</td><td>39701</td><td>+</td><td>0</td><td>GAGGGGAGGAGGAAGGAGCA</td><td>CGG</td><td>||||||||[||||||||||||]PAM</td><td>NA</td><td>Intergenic</td><td>NA</td><td>NA</td></tr>
<tr><td>T2524</td><td>GAAGGAGCACGGACGCGCCTTGG</td><td>1000</td><td></td><td></td><td></td><td></td><td></td><td></td><td></td><td></td><td></td></tr>
<tr><td>Oligo fwd</td><td>CACCGAAGGAGCACGGACGCGCCT</td><td></td><td></td><td></td><td></td><td></td><td></td><td></td><td></td><td></td><td></td></tr>
<tr><td>Oligo rev</td><td>AAACAGGCGCGTCCGTGCTCCTTC</td><td></td><td></td><td></td><td></td><td></td><td></td><td></td><td></td><td></td><td></td></tr>
<tr><td>Chromosome</td><td>start</td><td>end</td><td>strand</td><td>MM</td><td>target_seq</td><td>PAM</td><td>alignment</td><td>distance</td><td>position</td><td>gene name</td><td>gene id</td></tr>
<tr><td>GU305914.1</td><td>39690</td><td>39712</td><td>+</td><td>0</td><td>GAAGGAGCACGGACGCGCCT</td><td>TGG</td><td>||||||||[||||||||||||]PAM</td><td>NA</td><td>Intergenic</td><td>NA</td><td>NA</td></tr>
<tr><td>T2525</td><td>AAGGAGCACGGACGCGCCTTGGG</td><td>1000</td><td></td><td></td><td></td><td></td><td></td><td></td><td></td><td></td><td></td></tr>
<tr><td>Oligo fwd</td><td>CACCAAGGAGCACGGACGCGCCTT</td><td></td><td></td><td></td><td></td><td></td><td></td><td></td><td></td><td></td><td></td></tr>
<tr><td>Oligo rev</td><td>AAACAAGGCGCGTCCGTGCTCCTT</td><td></td><td></td><td></td><td></td><td></td><td></td><td></td><td></td><td></td><td></td></tr>
<tr><td>Chromosome</td><td>start</td><td>end</td><td>strand</td><td>MM</td><td>target_seq</td><td>PAM</td><td>alignment</td><td>distance</td><td>position</td><td>gene name</td><td>gene id</td></tr>
<tr><td>GU305914.1</td><td>39691</td><td>39713</td><td>+</td><td>0</td><td>AAGGAGCACGGACGCGCCTT</td><td>GGG</td><td>||||||||[||||||||||||]PAM</td><td>NA</td><td>Intergenic</td><td>NA</td><td>NA</td></tr>
<tr><td>T2526</td><td>GACGCGCCTTGGGTCGTGAAAGG</td><td>1000</td><td></td><td></td><td></td><td></td><td></td><td></td><td></td><td></td><td></td></tr>
<tr><td>Oligo fwd</td><td>CACCGACGCGCCTTGGGTCGTGAA</td><td></td><td></td><td></td><td></td><td></td><td></td><td></td><td></td><td></td><td></td></tr>
<tr><td>Oligo rev</td><td>AAACTTCACGACCCAAGGCGCGTC</td><td></td><td></td><td></td><td></td><td></td><td></td><td></td><td></td><td></td><td></td></tr>
<tr><td>Chromosome</td><td>start</td><td>end</td><td>strand</td><td>MM</td><td>target_seq</td><td>PAM</td><td>alignment</td><td>distance</td><td>position</td><td>gene name</td><td>gene id</td></tr>
<tr><td>GU305914.1</td><td>39701</td><td>39723</td><td>+</td><td>0</td><td>GACGCGCCTTGGGTCGTGAA</td><td>AGG</td><td>||||||||[||||||||||||]PAM</td><td>NA</td><td>Intergenic</td><td>NA</td><td>NA</td></tr>
<tr><td>T2527</td><td>GTCGTCGAATCCCTTCGACGAGG</td><td>1000</td><td></td><td></td><td></td><td></td><td></td><td></td><td></td><td></td><td></td></tr>
<tr><td>Oligo fwd</td><td>CACCGTCGTCGAATCCCTTCGACG</td><td></td><td></td><td></td><td></td><td></td><td></td><td></td><td></td><td></td><td></td></tr>
<tr><td>Oligo rev</td><td>AAACCGTCGAAGGGATTCGACGAC</td><td></td><td></td><td></td><td></td><td></td><td></td><td></td><td></td><td></td><td></td></tr>
<tr><td>Chromosome</td><td>start</td><td>end</td><td>strand</td><td>MM</td><td>target_seq</td><td>PAM</td><td>alignment</td><td>distance</td><td>position</td><td>gene name</td><td>gene id</td></tr>
<tr><td>GU305914.1</td><td>39727</td><td>39749</td><td>+</td><td>0</td><td>GTCGTCGAATCCCTTCGACG</td><td>AGG</td><td>||||||||[||||||||||||]PAM</td><td>NA</td><td>Intergenic</td><td>NA</td><td>NA</td></tr>
<tr><td>T2528</td><td>GTCGAATCCCTTCGACGAGGAGG</td><td>1000</td><td></td><td></td><td></td><td></td><td></td><td></td><td></td><td></td><td></td></tr>
<tr><td>Oligo fwd</td><td>CACCGTCGAATCCCTTCGACGAGG</td><td></td><td></td><td></td><td></td><td></td><td></td><td></td><td></td><td></td><td></td></tr>
<tr><td>Oligo rev</td><td>AAACCCTCGTCGAAGGGATTCGAC</td><td></td><td></td><td></td><td></td><td></td><td></td><td></td><td></td><td></td><td></td></tr>
<tr><td>Chromosome</td><td>start</td><td>end</td><td>strand</td><td>MM</td><td>target_seq</td><td>PAM</td><td>alignment</td><td>distance</td><td>position</td><td>gene name</td><td>gene id</td></tr>
<tr><td>GU305914.1</td><td>39730</td><td>39752</td><td>+</td><td>0</td><td>GTCGAATCCCTTCGACGAGG</td><td>AGG</td><td>||||||||[||||||||||||]PAM</td><td>NA</td><td>Intergenic</td><td>NA</td><td>NA</td></tr>
<tr><td>T2529</td><td>GAATCCCTTCGACGAGGAGGTGG</td><td>1000</td><td></td><td></td><td></td><td></td><td></td><td></td><td></td><td></td><td></td></tr>
<tr><td>Oligo fwd</td><td>CACCGAATCCCTTCGACGAGGAGG</td><td></td><td></td><td></td><td></td><td></td><td></td><td></td><td></td><td></td><td></td></tr>
<tr><td>Oligo rev</td><td>AAACCCTCCTCGTCGAAGGGATTC</td><td></td><td></td><td></td><td></td><td></td><td></td><td></td><td></td><td></td><td></td></tr>
<tr><td>Chromosome</td><td>start</td><td>end</td><td>strand</td><td>MM</td><td>target_seq</td><td>PAM</td><td>alignment</td><td>distance</td><td>position</td><td>gene name</td><td>gene id</td></tr>
<tr><td>GU305914.1</td><td>39733</td><td>39755</td><td>+</td><td>0</td><td>GAATCCCTTCGACGAGGAGG</td><td>TGG</td><td>||||||||[||||||||||||]PAM</td><td>NA</td><td>Intergenic</td><td>NA</td><td>NA</td></tr>
<tr><td>T2530</td><td>GTATCTCCGAACGCCGCGACCGG</td><td>1000</td><td></td><td></td><td></td><td></td><td></td><td></td><td></td><td></td><td></td></tr>
<tr><td>Oligo fwd</td><td>CACCGTATCTCCGAACGCCGCGAC</td><td></td><td></td><td></td><td></td><td></td><td></td><td></td><td></td><td></td><td></td></tr>
<tr><td>Oligo rev</td><td>AAACGTCGCGGCGTTCGGAGATAC</td><td></td><td></td><td></td><td></td><td></td><td></td><td></td><td></td><td></td><td></td></tr>
<tr><td>Chromosome</td><td>start</td><td>end</td><td>strand</td><td>MM</td><td>target_seq</td><td>PAM</td><td>alignment</td><td>distance</td><td>position</td><td>gene name</td><td>gene id</td></tr>
<tr><td>GU305914.1</td><td>39764</td><td>39786</td><td>+</td><td>0</td><td>GTATCTCCGAACGCCGCGAC</td><td>CGG</td><td>||||||||[||||||||||||]PAM</td><td>NA</td><td>Intergenic</td><td>NA</td><td>NA</td></tr>
<tr><td>T2531</td><td>CGCGACCGGATCGCCGACGAAGG</td><td>1000</td><td></td><td></td><td></td><td></td><td></td><td></td><td></td><td></td><td></td></tr>
<tr><td>Oligo fwd</td><td>CACCCGCGACCGGATCGCCGACGA</td><td></td><td></td><td></td><td></td><td></td><td></td><td></td><td></td><td></td><td></td></tr>
<tr><td>Oligo rev</td><td>AAACTCGTCGGCGATCCGGTCGCG</td><td></td><td></td><td></td><td></td><td></td><td></td><td></td><td></td><td></td><td></td></tr>
<tr><td>Chromosome</td><td>start</td><td>end</td><td>strand</td><td>MM</td><td>target_seq</td><td>PAM</td><td>alignment</td><td>distance</td><td>position</td><td>gene name</td><td>gene id</td></tr>
<tr><td>GU305914.1</td><td>39778</td><td>39800</td><td>+</td><td>0</td><td>CGCGACCGGATCGCCGACGA</td><td>AGG</td><td>||||||||[||||||||||||]PAM</td><td>NA</td><td>Intergenic</td><td>NA</td><td>NA</td></tr>
<tr><td>T2532</td><td>ACGAAGGCGACTTTCCGCAGCGG</td><td>1000</td><td></td><td></td><td></td><td></td><td></td><td></td><td></td><td></td><td></td></tr>
<tr><td>Oligo fwd</td><td>CACCACGAAGGCGACTTTCCGCAG</td><td></td><td></td><td></td><td></td><td></td><td></td><td></td><td></td><td></td><td></td></tr>
<tr><td>Oligo rev</td><td>AAACCTGCGGAAAGTCGCCTTCGT</td><td></td><td></td><td></td><td></td><td></td><td></td><td></td><td></td><td></td><td></td></tr>
<tr><td>Chromosome</td><td>start</td><td>end</td><td>strand</td><td>MM</td><td>target_seq</td><td>PAM</td><td>alignment</td><td>distance</td><td>position</td><td>gene name</td><td>gene id</td></tr>
<tr><td>GU305914.1</td><td>39794</td><td>39816</td><td>+</td><td>0</td><td>ACGAAGGCGACTTTCCGCAG</td><td>CGG</td><td>||||||||[||||||||||||]PAM</td><td>NA</td><td>Intergenic</td><td>NA</td><td>NA</td></tr>
<tr><td>T2533</td><td>GGCGACTTTCCGCAGCGGCTCGG</td><td>1000</td><td></td><td></td><td></td><td></td><td></td><td></td><td></td><td></td><td></td></tr>
<tr><td>Oligo fwd</td><td>CACCGGCGACTTTCCGCAGCGGCT</td><td></td><td></td><td></td><td></td><td></td><td></td><td></td><td></td><td></td><td></td></tr>
<tr><td>Oligo rev</td><td>AAACAGCCGCTGCGGAAAGTCGCC</td><td></td><td></td><td></td><td></td><td></td><td></td><td></td><td></td><td></td><td></td></tr>
<tr><td>Chromosome</td><td>start</td><td>end</td><td>strand</td><td>MM</td><td>target_seq</td><td>PAM</td><td>alignment</td><td>distance</td><td>position</td><td>gene name</td><td>gene id</td></tr>
<tr><td>GU305914.1</td><td>39799</td><td>39821</td><td>+</td><td>0</td><td>GGCGACTTTCCGCAGCGGCT</td><td>CGG</td><td>||||||||[||||||||||||]PAM</td><td>NA</td><td>Intergenic</td><td>NA</td><td>NA</td></tr>
<tr><td>T2534</td><td>CTTTCCGCAGCGGCTCGGTCCGG</td><td>1000</td><td></td><td></td><td></td><td></td><td></td><td></td><td></td><td></td><td></td></tr>
<tr><td>Oligo fwd</td><td>CACCCTTTCCGCAGCGGCTCGGTC</td><td></td><td></td><td></td><td></td><td></td><td></td><td></td><td></td><td></td><td></td></tr>
<tr><td>Oligo rev</td><td>AAACGACCGAGCCGCTGCGGAAAG</td><td></td><td></td><td></td><td></td><td></td><td></td><td></td><td></td><td></td><td></td></tr>
</table>

<table>
<tr><td>Chromosome</td><td>start</td><td>end</td><td>strand</td><td>MM</td><td>target_seq</td><td>PAM</td><td>alignment</td><td>distance</td><td>position</td><td>gene name</td><td>gene id</td></tr>
<tr><td>GU305914.1</td><td>39804</td><td>39826</td><td>+</td><td>0</td><td>CTTTCCGCAGCGGCTCGGTC</td><td>CGG</td><td>||||||||[||||||||||||]PAM</td><td>NA</td><td>Intergenic</td><td>NA</td><td>NA</td></tr>
<tr><td>T2535</td><td>TTTCCGCAGCGGCTCGGTCCGGG</td><td>1000</td><td></td><td></td><td></td><td></td><td></td><td></td><td></td><td></td><td></td></tr>
<tr><td>Oligo fwd</td><td>CACCTTTCCGCAGCGGCTCGGTCC</td><td></td><td></td><td></td><td></td><td></td><td></td><td></td><td></td><td></td><td></td></tr>
<tr><td>Oligo rev</td><td>AAACGGACCGAGCCGCTGCGGAAA</td><td></td><td></td><td></td><td></td><td></td><td></td><td></td><td></td><td></td><td></td></tr>
<tr><td>Chromosome</td><td>start</td><td>end</td><td>strand</td><td>MM</td><td>target_seq</td><td>PAM</td><td>alignment</td><td>distance</td><td>position</td><td>gene name</td><td>gene id</td></tr>
<tr><td>GU305914.1</td><td>39805</td><td>39827</td><td>+</td><td>0</td><td>TTTCCGCAGCGGCTCGGTCC</td><td>GGG</td><td>||||||||[||||||||||||]PAM</td><td>NA</td><td>Intergenic</td><td>NA</td><td>NA</td></tr>
<tr><td>T2536</td><td>TTCCGCAGCGGCTCGGTCCGGGG</td><td>1000</td><td></td><td></td><td></td><td></td><td></td><td></td><td></td><td></td><td></td></tr>
<tr><td>Oligo fwd</td><td>CACCTTCCGCAGCGGCTCGGTCCG</td><td></td><td></td><td></td><td></td><td></td><td></td><td></td><td></td><td></td><td></td></tr>
<tr><td>Oligo rev</td><td>AAACCGGACCGAGCCGCTGCGGAA</td><td></td><td></td><td></td><td></td><td></td><td></td><td></td><td></td><td></td><td></td></tr>
<tr><td>Chromosome</td><td>start</td><td>end</td><td>strand</td><td>MM</td><td>target_seq</td><td>PAM</td><td>alignment</td><td>distance</td><td>position</td><td>gene name</td><td>gene id</td></tr>
<tr><td>GU305914.1</td><td>39806</td><td>39828</td><td>+</td><td>0</td><td>TTCCGCAGCGGCTCGGTCCG</td><td>GGG</td><td>||||||||[||||||||||||]PAM</td><td>NA</td><td>Intergenic</td><td>NA</td><td>NA</td></tr>
<tr><td>T2537</td><td>GCAGCGGCTCGGTCCGGGGAAGG</td><td>1000</td><td></td><td></td><td></td><td></td><td></td><td></td><td></td><td></td><td></td></tr>
<tr><td>Oligo fwd</td><td>CACCGCAGCGGCTCGGTCCGGGGA</td><td></td><td></td><td></td><td></td><td></td><td></td><td></td><td></td><td></td><td></td></tr>
<tr><td>Oligo rev</td><td>AAACTCCCCGGACCGAGCCGCTGC</td><td></td><td></td><td></td><td></td><td></td><td></td><td></td><td></td><td></td><td></td></tr>
<tr><td>Chromosome</td><td>start</td><td>end</td><td>strand</td><td>MM</td><td>target_seq</td><td>PAM</td><td>alignment</td><td>distance</td><td>position</td><td>gene name</td><td>gene id</td></tr>
<tr><td>GU305914.1</td><td>39810</td><td>39832</td><td>+</td><td>0</td><td>GCAGCGGCTCGGTCCGGGGA</td><td>AGG</td><td>||||||||[||||||||||||]PAM</td><td>NA</td><td>Intergenic</td><td>NA</td><td>NA</td></tr>
<tr><td>T2538</td><td>GATTGTGCTCAGATCCGAGTCGG</td><td>1000</td><td></td><td></td><td></td><td></td><td></td><td></td><td></td><td></td><td></td></tr>
<tr><td>Oligo fwd</td><td>CACCGATTGTGCTCAGATCCGAGT</td><td></td><td></td><td></td><td></td><td></td><td></td><td></td><td></td><td></td><td></td></tr>
<tr><td>Oligo rev</td><td>AAACACTCGGATCTGAGCACAATC</td><td></td><td></td><td></td><td></td><td></td><td></td><td></td><td></td><td></td><td></td></tr>
<tr><td>Chromosome</td><td>start</td><td>end</td><td>strand</td><td>MM</td><td>target_seq</td><td>PAM</td><td>alignment</td><td>distance</td><td>position</td><td>gene name</td><td>gene id</td></tr>
<tr><td>GU305914.1</td><td>39832</td><td>39854</td><td>+</td><td>0</td><td>GATTGTGCTCAGATCCGAGT</td><td>CGG</td><td>||||||||[||||||||||||]PAM</td><td>NA</td><td>Intergenic</td><td>NA</td><td>NA</td></tr>
<tr><td>T2539</td><td>CGTTTCCCCGACGCCGTTGACGG</td><td>1000</td><td></td><td></td><td></td><td></td><td></td><td></td><td></td><td></td><td></td></tr>
<tr><td>Oligo fwd</td><td>CACCCGTTTCCCCGACGCCGTTGA</td><td></td><td></td><td></td><td></td><td></td><td></td><td></td><td></td><td></td><td></td></tr>
<tr><td>Oligo rev</td><td>AAACTCAACGGCGTCGGGGAAACG</td><td></td><td></td><td></td><td></td><td></td><td></td><td></td><td></td><td></td><td></td></tr>
<tr><td>Chromosome</td><td>start</td><td>end</td><td>strand</td><td>MM</td><td>target_seq</td><td>PAM</td><td>alignment</td><td>distance</td><td>position</td><td>gene name</td><td>gene id</td></tr>
<tr><td>GU305914.1</td><td>40010</td><td>40032</td><td>+</td><td>0</td><td>CGTTTCCCCGACGCCGTTGA</td><td>CGG</td><td>||||||||[||||||||||||]PAM</td><td>NA</td><td>Intergenic</td><td>NA</td><td>NA</td></tr>
<tr><td>T2540</td><td>AACGCTATTACTTCCCGAGACGG</td><td>1000</td><td></td><td></td><td></td><td></td><td></td><td></td><td></td><td></td><td></td></tr>
<tr><td>Oligo fwd</td><td>CACCAACGCTATTACTTCCCGAGA</td><td></td><td></td><td></td><td></td><td></td><td></td><td></td><td></td><td></td><td></td></tr>
<tr><td>Oligo rev</td><td>AAACTCTCGGGAAGTAATAGCGTT</td><td></td><td></td><td></td><td></td><td></td><td></td><td></td><td></td><td></td><td></td></tr>
<tr><td>Chromosome</td><td>start</td><td>end</td><td>strand</td><td>MM</td><td>target_seq</td><td>PAM</td><td>alignment</td><td>distance</td><td>position</td><td>gene name</td><td>gene id</td></tr>
<tr><td>GU305914.1</td><td>40051</td><td>40073</td><td>+</td><td>0</td><td>AACGCTATTACTTCCCGAGA</td><td>CGG</td><td>||||||||[||||||||||||]PAM</td><td>NA</td><td>Intergenic</td><td>NA</td><td>NA</td></tr>
<tr><td>T2541</td><td>ACGCTATTACTTCCCGAGACGGG</td><td>1000</td><td></td><td></td><td></td><td></td><td></td><td></td><td></td><td></td><td></td></tr>
<tr><td>Oligo fwd</td><td>CACCACGCTATTACTTCCCGAGAC</td><td></td><td></td><td></td><td></td><td></td><td></td><td></td><td></td><td></td><td></td></tr>
<tr><td>Oligo rev</td><td>AAACGTCTCGGGAAGTAATAGCGT</td><td></td><td></td><td></td><td></td><td></td><td></td><td></td><td></td><td></td><td></td></tr>
<tr><td>Chromosome</td><td>start</td><td>end</td><td>strand</td><td>MM</td><td>target_seq</td><td>PAM</td><td>alignment</td><td>distance</td><td>position</td><td>gene name</td><td>gene id</td></tr>
<tr><td>GU305914.1</td><td>40052</td><td>40074</td><td>+</td><td>0</td><td>ACGCTATTACTTCCCGAGAC</td><td>GGG</td><td>||||||||[||||||||||||]PAM</td><td>NA</td><td>Intergenic</td><td>NA</td><td>NA</td></tr>
<tr><td>T2542</td><td>CTATTACTTCCCGAGACGGGTGG</td><td>1000</td><td></td><td></td><td></td><td></td><td></td><td></td><td></td><td></td><td></td></tr>
<tr><td>Oligo fwd</td><td>CACCCTATTACTTCCCGAGACGGG</td><td></td><td></td><td></td><td></td><td></td><td></td><td></td><td></td><td></td><td></td></tr>
<tr><td>Oligo rev</td><td>AAACCCCGTCTCGGGAAGTAATAG</td><td></td><td></td><td></td><td></td><td></td><td></td><td></td><td></td><td></td><td></td></tr>
<tr><td>Chromosome</td><td>start</td><td>end</td><td>strand</td><td>MM</td><td>target_seq</td><td>PAM</td><td>alignment</td><td>distance</td><td>position</td><td>gene name</td><td>gene id</td></tr>
<tr><td>GU305914.1</td><td>40055</td><td>40077</td><td>+</td><td>0</td><td>CTATTACTTCCCGAGACGGG</td><td>TGG</td><td>||||||||[||||||||||||]PAM</td><td>NA</td><td>Intergenic</td><td>NA</td><td>NA</td></tr>
<tr><td>T2543</td><td>GGTGGCGTGACATTCTTCTTAGG</td><td>1000</td><td></td><td></td><td></td><td></td><td></td><td></td><td></td><td></td><td></td></tr>
<tr><td>Oligo fwd</td><td>CACCGGTGGCGTGACATTCTTCTT</td><td></td><td></td><td></td><td></td><td></td><td></td><td></td><td></td><td></td><td></td></tr>
<tr><td>Oligo rev</td><td>AAACAAGAAGAATGTCACGCCACC</td><td></td><td></td><td></td><td></td><td></td><td></td><td></td><td></td><td></td><td></td></tr>
<tr><td>Chromosome</td><td>start</td><td>end</td><td>strand</td><td>MM</td><td>target_seq</td><td>PAM</td><td>alignment</td><td>distance</td><td>position</td><td>gene name</td><td>gene id</td></tr>
<tr><td>GU305914.1</td><td>40073</td><td>40095</td><td>+</td><td>0</td><td>GGTGGCGTGACATTCTTCTT</td><td>AGG</td><td>||||||||[||||||||||||]PAM</td><td>NA</td><td>Intergenic</td><td>NA</td><td>NA</td></tr>
<tr><td>T2544</td><td>GGCGTGACATTCTTCTTAGGCGG</td><td>1000</td><td></td><td></td><td></td><td></td><td></td><td></td><td></td><td></td><td></td></tr>
<tr><td>Oligo fwd</td><td>CACCGGCGTGACATTCTTCTTAGG</td><td></td><td></td><td></td><td></td><td></td><td></td><td></td><td></td><td></td><td></td></tr>
<tr><td>Oligo rev</td><td>AAACCCTAAGAAGAATGTCACGCC</td><td></td><td></td><td></td><td></td><td></td><td></td><td></td><td></td><td></td><td></td></tr>
<tr><td>Chromosome</td><td>start</td><td>end</td><td>strand</td><td>MM</td><td>target_seq</td><td>PAM</td><td>alignment</td><td>distance</td><td>position</td><td>gene name</td><td>gene id</td></tr>
<tr><td>GU305914.1</td><td>40076</td><td>40098</td><td>+</td><td>0</td><td>GGCGTGACATTCTTCTTAGG</td><td>CGG</td><td>||||||||[||||||||||||]PAM</td><td>NA</td><td>Intergenic</td><td>NA</td><td>NA</td></tr>
<tr><td>T2545</td><td>GTGACATTCTTCTTAGGCGGAGG</td><td>1000</td><td></td><td></td><td></td><td></td><td></td><td></td><td></td><td></td><td></td></tr>
<tr><td>Oligo fwd</td><td>CACCGTGACATTCTTCTTAGGCGG</td><td></td><td></td><td></td><td></td><td></td><td></td><td></td><td></td><td></td><td></td></tr>
<tr><td>Oligo rev</td><td>AAACCCGCCTAAGAAGAATGTCAC</td><td></td><td></td><td></td><td></td><td></td><td></td><td></td><td></td><td></td><td></td></tr>
<tr><td>Chromosome</td><td>start</td><td>end</td><td>strand</td><td>MM</td><td>target_seq</td><td>PAM</td><td>alignment</td><td>distance</td><td>position</td><td>gene name</td><td>gene id</td></tr>
<tr><td>GU305914.1</td><td>40079</td><td>40101</td><td>+</td><td>0</td><td>GTGACATTCTTCTTAGGCGG</td><td>AGG</td><td>||||||||[||||||||||||]PAM</td><td>NA</td><td>Intergenic</td><td>NA</td><td>NA</td></tr>
<tr><td>T2546</td><td>TGACATTCTTCTTAGGCGGAGGG</td><td>1000</td><td></td><td></td><td></td><td></td><td></td><td></td><td></td><td></td><td></td></tr>
<tr><td>Oligo fwd</td><td>CACCTGACATTCTTCTTAGGCGGA</td><td></td><td></td><td></td><td></td><td></td><td></td><td></td><td></td><td></td><td></td></tr>
<tr><td>Oligo rev</td><td>AAACTCCGCCTAAGAAGAATGTCA</td><td></td><td></td><td></td><td></td><td></td><td></td><td></td><td></td><td></td><td></td></tr>
<tr><td>Chromosome</td><td>start</td><td>end</td><td>strand</td><td>MM</td><td>target_seq</td><td>PAM</td><td>alignment</td><td>distance</td><td>position</td><td>gene name</td><td>gene id</td></tr>
<tr><td>GU305914.1</td><td>40080</td><td>40102</td><td>+</td><td>0</td><td>TGACATTCTTCTTAGGCGGA</td><td>GGG</td><td>||||||||[||||||||||||]PAM</td><td>NA</td><td>Intergenic</td><td>NA</td><td>NA</td></tr>
<tr><td>T2547</td><td>CTTCTTATTACCACCGTCGTCGG</td><td>1000</td><td></td><td></td><td></td><td></td><td></td><td></td><td></td><td></td><td></td></tr>
<tr><td>Oligo fwd</td><td>CACCCTTCTTATTACCACCGTCGT</td><td></td><td></td><td></td><td></td><td></td><td></td><td></td><td></td><td></td><td></td></tr>
<tr><td>Oligo rev</td><td>AAACACGACGGTGGTAATAAGAAG</td><td></td><td></td><td></td><td></td><td></td><td></td><td></td><td></td><td></td><td></td></tr>
<tr><td>Chromosome</td><td>start</td><td>end</td><td>strand</td><td>MM</td><td>target_seq</td><td>PAM</td><td>alignment</td><td>distance</td><td>position</td><td>gene name</td><td>gene id</td></tr>
<tr><td>GU305914.1</td><td>40105</td><td>40127</td><td>+</td><td>0</td><td>CTTCTTATTACCACCGTCGT</td><td>CGG</td><td>||||||||[||||||||||||]PAM</td><td>NA</td><td>Intergenic</td><td>NA</td><td>NA</td></tr>
<tr><td>T2548</td><td>CGTCGTCGGAGCTTGCCTTTCGG</td><td>1000</td><td></td><td></td><td></td><td></td><td></td><td></td><td></td><td></td><td></td></tr>
<tr><td>Oligo fwd</td><td>CACCCGTCGTCGGAGCTTGCCTTT</td><td></td><td></td><td></td><td></td><td></td><td></td><td></td><td></td><td></td><td></td></tr>
<tr><td>Oligo rev</td><td>AAACAAAGGCAAGCTCCGACGACG</td><td></td><td></td><td></td><td></td><td></td><td></td><td></td><td></td><td></td><td></td></tr>
<tr><td>Chromosome</td><td>start</td><td>end</td><td>strand</td><td>MM</td><td>target_seq</td><td>PAM</td><td>alignment</td><td>distance</td><td>position</td><td>gene name</td><td>gene id</td></tr>
<tr><td>GU305914.1</td><td>40119</td><td>40141</td><td>+</td><td>0</td><td>CGTCGTCGGAGCTTGCCTTT</td><td>CGG</td><td>||||||||[||||||||||||]PAM</td><td>NA</td><td>Intergenic</td><td>NA</td><td>NA</td></tr>
<tr><td>T2549</td><td>GCTTGCCTTTCGGAGTTCCTTGG</td><td>1000</td><td></td><td></td><td></td><td></td><td></td><td></td><td></td><td></td><td></td></tr>
<tr><td>Oligo fwd</td><td>CACCGCTTGCCTTTCGGAGTTCCT</td><td></td><td></td><td></td><td></td><td></td><td></td><td></td><td></td><td></td><td></td></tr>
</table>

<table>
<tr><td>Oligo rev</td><td>AAACAGGAACTCCGAAAGGCAAGC</td><td></td><td></td><td></td><td></td><td></td><td></td><td></td><td></td><td></td><td></td></tr>
<tr><td>Chromosome</td><td>start</td><td>end</td><td>strand</td><td>MM</td><td>target_seq</td><td>PAM</td><td>alignment</td><td>distance</td><td>position</td><td>gene name</td><td>gene id</td></tr>
<tr><td>GU305914.1</td><td>40129</td><td>40151</td><td>+</td><td>0</td><td>GCTTGCCTTTCGGAGTTCCT</td><td>TGG</td><td>||||||||[||||||||||||]PAM</td><td>NA</td><td>Intergenic</td><td>NA</td><td>NA</td></tr>
<tr><td>T2550</td><td>CTTGCCTTTCGGAGTTCCTTGGG</td><td>1000</td><td></td><td></td><td></td><td></td><td></td><td></td><td></td><td></td><td></td></tr>
<tr><td>Oligo fwd</td><td>CACCCTTGCCTTTCGGAGTTCCTT</td><td></td><td></td><td></td><td></td><td></td><td></td><td></td><td></td><td></td><td></td></tr>
<tr><td>Oligo rev</td><td>AAACAAGGAACTCCGAAAGGCAAG</td><td></td><td></td><td></td><td></td><td></td><td></td><td></td><td></td><td></td><td></td></tr>
<tr><td>Chromosome</td><td>start</td><td>end</td><td>strand</td><td>MM</td><td>target_seq</td><td>PAM</td><td>alignment</td><td>distance</td><td>position</td><td>gene name</td><td>gene id</td></tr>
<tr><td>GU305914.1</td><td>40130</td><td>40152</td><td>+</td><td>0</td><td>CTTGCCTTTCGGAGTTCCTT</td><td>GGG</td><td>||||||||[||||||||||||]PAM</td><td>NA</td><td>Intergenic</td><td>NA</td><td>NA</td></tr>
<tr><td>T2551</td><td>GCCTTTCGGAGTTCCTTGGGCGG</td><td>1000</td><td></td><td></td><td></td><td></td><td></td><td></td><td></td><td></td><td></td></tr>
<tr><td>Oligo fwd</td><td>CACCGCCTTTCGGAGTTCCTTGGG</td><td></td><td></td><td></td><td></td><td></td><td></td><td></td><td></td><td></td><td></td></tr>
<tr><td>Oligo rev</td><td>AAACCCCAAGGAACTCCGAAAGGC</td><td></td><td></td><td></td><td></td><td></td><td></td><td></td><td></td><td></td><td></td></tr>
<tr><td>Chromosome</td><td>start</td><td>end</td><td>strand</td><td>MM</td><td>target_seq</td><td>PAM</td><td>alignment</td><td>distance</td><td>position</td><td>gene name</td><td>gene id</td></tr>
<tr><td>GU305914.1</td><td>40133</td><td>40155</td><td>+</td><td>0</td><td>GCCTTTCGGAGTTCCTTGGG</td><td>CGG</td><td>||||||||[||||||||||||]PAM</td><td>NA</td><td>Intergenic</td><td>NA</td><td>NA</td></tr>
<tr><td>T2552</td><td>TTTCGGAGTTCCTTGGGCGGTGG</td><td>1000</td><td></td><td></td><td></td><td></td><td></td><td></td><td></td><td></td><td></td></tr>
<tr><td>Oligo fwd</td><td>CACCTTTCGGAGTTCCTTGGGCGG</td><td></td><td></td><td></td><td></td><td></td><td></td><td></td><td></td><td></td><td></td></tr>
<tr><td>Oligo rev</td><td>AAACCCGCCCAAGGAACTCCGAAA</td><td></td><td></td><td></td><td></td><td></td><td></td><td></td><td></td><td></td><td></td></tr>
<tr><td>Chromosome</td><td>start</td><td>end</td><td>strand</td><td>MM</td><td>target_seq</td><td>PAM</td><td>alignment</td><td>distance</td><td>position</td><td>gene name</td><td>gene id</td></tr>
<tr><td>GU305914.1</td><td>40136</td><td>40158</td><td>+</td><td>0</td><td>TTTCGGAGTTCCTTGGGCGG</td><td>TGG</td><td>||||||||[||||||||||||]PAM</td><td>NA</td><td>Intergenic</td><td>NA</td><td>NA</td></tr>
<tr><td>T2553</td><td>GCCGTGTTCGTCGATCCAGACGG</td><td>1000</td><td></td><td></td><td></td><td></td><td></td><td></td><td></td><td></td><td></td></tr>
<tr><td>Oligo fwd</td><td>CACCGCCGTGTTCGTCGATCCAGA</td><td></td><td></td><td></td><td></td><td></td><td></td><td></td><td></td><td></td><td></td></tr>
<tr><td>Oligo rev</td><td>AAACTCTGGATCGACGAACACGGC</td><td></td><td></td><td></td><td></td><td></td><td></td><td></td><td></td><td></td><td></td></tr>
<tr><td>Chromosome</td><td>start</td><td>end</td><td>strand</td><td>MM</td><td>target_seq</td><td>PAM</td><td>alignment</td><td>distance</td><td>position</td><td>gene name</td><td>gene id</td></tr>
<tr><td>GU305914.1</td><td>40174</td><td>40196</td><td>+</td><td>0</td><td>GCCGTGTTCGTCGATCCAGA</td><td>CGG</td><td>||||||||[||||||||||||]PAM</td><td>NA</td><td>Intergenic</td><td>NA</td><td>NA</td></tr>
<tr><td>T2554</td><td>GTGTTCGTCGATCCAGACGGCGG</td><td>1000</td><td></td><td></td><td></td><td></td><td></td><td></td><td></td><td></td><td></td></tr>
<tr><td>Oligo fwd</td><td>CACCGTGTTCGTCGATCCAGACGG</td><td></td><td></td><td></td><td></td><td></td><td></td><td></td><td></td><td></td><td></td></tr>
<tr><td>Oligo rev</td><td>AAACCCGTCTGGATCGACGAACAC</td><td></td><td></td><td></td><td></td><td></td><td></td><td></td><td></td><td></td><td></td></tr>
<tr><td>Chromosome</td><td>start</td><td>end</td><td>strand</td><td>MM</td><td>target_seq</td><td>PAM</td><td>alignment</td><td>distance</td><td>position</td><td>gene name</td><td>gene id</td></tr>
<tr><td>GU305914.1</td><td>40177</td><td>40199</td><td>+</td><td>0</td><td>GTGTTCGTCGATCCAGACGG</td><td>CGG</td><td>||||||||[||||||||||||]PAM</td><td>NA</td><td>Intergenic</td><td>NA</td><td>NA</td></tr>
<tr><td>T2555</td><td>CTCGTTGTCCACCATTATCCCGG</td><td>1000</td><td></td><td></td><td></td><td></td><td></td><td></td><td></td><td></td><td></td></tr>
<tr><td>Oligo fwd</td><td>CACCCTCGTTGTCCACCATTATCC</td><td></td><td></td><td></td><td></td><td></td><td></td><td></td><td></td><td></td><td></td></tr>
<tr><td>Oligo rev</td><td>AAACGGATAATGGTGGACAACGAG</td><td></td><td></td><td></td><td></td><td></td><td></td><td></td><td></td><td></td><td></td></tr>
<tr><td>Chromosome</td><td>start</td><td>end</td><td>strand</td><td>MM</td><td>target_seq</td><td>PAM</td><td>alignment</td><td>distance</td><td>position</td><td>gene name</td><td>gene id</td></tr>
<tr><td>GU305914.1</td><td>40201</td><td>40223</td><td>+</td><td>0</td><td>CTCGTTGTCCACCATTATCC</td><td>CGG</td><td>||||||||[||||||||||||]PAM</td><td>NA</td><td>Intergenic</td><td>NA</td><td>NA</td></tr>
<tr><td>T2556</td><td>TTGTCCACCATTATCCCGGTCGG</td><td>1000</td><td></td><td></td><td></td><td></td><td></td><td></td><td></td><td></td><td></td></tr>
<tr><td>Oligo fwd</td><td>CACCTTGTCCACCATTATCCCGGT</td><td></td><td></td><td></td><td></td><td></td><td></td><td></td><td></td><td></td><td></td></tr>
<tr><td>Oligo rev</td><td>AAACACCGGGATAATGGTGGACAA</td><td></td><td></td><td></td><td></td><td></td><td></td><td></td><td></td><td></td><td></td></tr>
<tr><td>Chromosome</td><td>start</td><td>end</td><td>strand</td><td>MM</td><td>target_seq</td><td>PAM</td><td>alignment</td><td>distance</td><td>position</td><td>gene name</td><td>gene id</td></tr>
<tr><td>GU305914.1</td><td>40205</td><td>40227</td><td>+</td><td>0</td><td>TTGTCCACCATTATCCCGGT</td><td>CGG</td><td>||||||||[||||||||||||]PAM</td><td>NA</td><td>Intergenic</td><td>NA</td><td>NA</td></tr>
<tr><td>T2557</td><td>CCCAAGACACCGCTCGCTATCGG</td><td>1000</td><td></td><td></td><td></td><td></td><td></td><td></td><td></td><td></td><td></td></tr>
<tr><td>Oligo fwd</td><td>CACCCCCAAGACACCGCTCGCTAT</td><td></td><td></td><td></td><td></td><td></td><td></td><td></td><td></td><td></td><td></td></tr>
<tr><td>Oligo rev</td><td>AAACATAGCGAGCGGTGTCTTGGG</td><td></td><td></td><td></td><td></td><td></td><td></td><td></td><td></td><td></td><td></td></tr>
<tr><td>Chromosome</td><td>start</td><td>end</td><td>strand</td><td>MM</td><td>target_seq</td><td>PAM</td><td>alignment</td><td>distance</td><td>position</td><td>gene name</td><td>gene id</td></tr>
<tr><td>GU305914.1</td><td>40267</td><td>40289</td><td>+</td><td>0</td><td>CCCAAGACACCGCTCGCTAT</td><td>CGG</td><td>||||||||[||||||||||||]PAM</td><td>NA</td><td>Intergenic</td><td>NA</td><td>NA</td></tr>
<tr><td>T2558</td><td>CCAAGACACCGCTCGCTATCGGG</td><td>1000</td><td></td><td></td><td></td><td></td><td></td><td></td><td></td><td></td><td></td></tr>
<tr><td>Oligo fwd</td><td>CACCCCAAGACACCGCTCGCTATC</td><td></td><td></td><td></td><td></td><td></td><td></td><td></td><td></td><td></td><td></td></tr>
<tr><td>Oligo rev</td><td>AAACGATAGCGAGCGGTGTCTTGG</td><td></td><td></td><td></td><td></td><td></td><td></td><td></td><td></td><td></td><td></td></tr>
<tr><td>Chromosome</td><td>start</td><td>end</td><td>strand</td><td>MM</td><td>target_seq</td><td>PAM</td><td>alignment</td><td>distance</td><td>position</td><td>gene name</td><td>gene id</td></tr>
<tr><td>GU305914.1</td><td>40268</td><td>40290</td><td>+</td><td>0</td><td>CCAAGACACCGCTCGCTATC</td><td>GGG</td><td>||||||||[||||||||||||]PAM</td><td>NA</td><td>Intergenic</td><td>NA</td><td>NA</td></tr>
<tr><td>T2559</td><td>GAGACACGTGTTTGCGATGCCGG</td><td>1000</td><td></td><td></td><td></td><td></td><td></td><td></td><td></td><td></td><td></td></tr>
<tr><td>Oligo fwd</td><td>CACCGAGACACGTGTTTGCGATGC</td><td></td><td></td><td></td><td></td><td></td><td></td><td></td><td></td><td></td><td></td></tr>
<tr><td>Oligo rev</td><td>AAACGCATCGCAAACACGTGTCTC</td><td></td><td></td><td></td><td></td><td></td><td></td><td></td><td></td><td></td><td></td></tr>
<tr><td>Chromosome</td><td>start</td><td>end</td><td>strand</td><td>MM</td><td>target_seq</td><td>PAM</td><td>alignment</td><td>distance</td><td>position</td><td>gene name</td><td>gene id</td></tr>
<tr><td>GU305914.1</td><td>40348</td><td>40370</td><td>+</td><td>0</td><td>GAGACACGTGTTTGCGATGC</td><td>CGG</td><td>||||||||[||||||||||||]PAM</td><td>NA</td><td>Intergenic</td><td>NA</td><td>NA</td></tr>
<tr><td>T2560</td><td>TTGCGATGCCGGTGTCGTCGTGG</td><td>1000</td><td></td><td></td><td></td><td></td><td></td><td></td><td></td><td></td><td></td></tr>
<tr><td>Oligo fwd</td><td>CACCTTGCGATGCCGGTGTCGTCG</td><td></td><td></td><td></td><td></td><td></td><td></td><td></td><td></td><td></td><td></td></tr>
<tr><td>Oligo rev</td><td>AAACCGACGACACCGGCATCGCAA</td><td></td><td></td><td></td><td></td><td></td><td></td><td></td><td></td><td></td><td></td></tr>
<tr><td>Chromosome</td><td>start</td><td>end</td><td>strand</td><td>MM</td><td>target_seq</td><td>PAM</td><td>alignment</td><td>distance</td><td>position</td><td>gene name</td><td>gene id</td></tr>
<tr><td>GU305914.1</td><td>40359</td><td>40381</td><td>+</td><td>0</td><td>TTGCGATGCCGGTGTCGTCG</td><td>TGG</td><td>||||||||[||||||||||||]PAM</td><td>NA</td><td>Intergenic</td><td>NA</td><td>NA</td></tr>
<tr><td>T2561</td><td>CGATGCCGGTGTCGTCGTGGAGG</td><td>1000</td><td></td><td></td><td></td><td></td><td></td><td></td><td></td><td></td><td></td></tr>
<tr><td>Oligo fwd</td><td>CACCCGATGCCGGTGTCGTCGTGG</td><td></td><td></td><td></td><td></td><td></td><td></td><td></td><td></td><td></td><td></td></tr>
<tr><td>Oligo rev</td><td>AAACCCACGACGACACCGGCATCG</td><td></td><td></td><td></td><td></td><td></td><td></td><td></td><td></td><td></td><td></td></tr>
<tr><td>Chromosome</td><td>start</td><td>end</td><td>strand</td><td>MM</td><td>target_seq</td><td>PAM</td><td>alignment</td><td>distance</td><td>position</td><td>gene name</td><td>gene id</td></tr>
<tr><td>GU305914.1</td><td>40362</td><td>40384</td><td>+</td><td>0</td><td>CGATGCCGGTGTCGTCGTGG</td><td>AGG</td><td>||||||||[||||||||||||]PAM</td><td>NA</td><td>Intergenic</td><td>NA</td><td>NA</td></tr>
<tr><td>T2562</td><td>GATGCCGGTGTCGTCGTGGAGGG</td><td>1000</td><td></td><td></td><td></td><td></td><td></td><td></td><td></td><td></td><td></td></tr>
<tr><td>Oligo fwd</td><td>CACCGATGCCGGTGTCGTCGTGGA</td><td></td><td></td><td></td><td></td><td></td><td></td><td></td><td></td><td></td><td></td></tr>
<tr><td>Oligo rev</td><td>AAACTCCACGACGACACCGGCATC</td><td></td><td></td><td></td><td></td><td></td><td></td><td></td><td></td><td></td><td></td></tr>
<tr><td>Chromosome</td><td>start</td><td>end</td><td>strand</td><td>MM</td><td>target_seq</td><td>PAM</td><td>alignment</td><td>distance</td><td>position</td><td>gene name</td><td>gene id</td></tr>
<tr><td>GU305914.1</td><td>40363</td><td>40385</td><td>+</td><td>0</td><td>GATGCCGGTGTCGTCGTGGA</td><td>GGG</td><td>||||||||[||||||||||||]PAM</td><td>NA</td><td>Intergenic</td><td>NA</td><td>NA</td></tr>
<tr><td>T2563</td><td>TCGTCGTGGAGGGACATCCCCGG</td><td>1000</td><td></td><td></td><td></td><td></td><td></td><td></td><td></td><td></td><td></td></tr>
<tr><td>Oligo fwd</td><td>CACCTCGTCGTGGAGGGACATCCC</td><td></td><td></td><td></td><td></td><td></td><td></td><td></td><td></td><td></td><td></td></tr>
<tr><td>Oligo rev</td><td>AAACGGGATGTCCCTCCACGACGA</td><td></td><td></td><td></td><td></td><td></td><td></td><td></td><td></td><td></td><td></td></tr>
<tr><td>Chromosome</td><td>start</td><td>end</td><td>strand</td><td>MM</td><td>target_seq</td><td>PAM</td><td>alignment</td><td>distance</td><td>position</td><td>gene name</td><td>gene id</td></tr>
<tr><td>GU305914.1</td><td>40373</td><td>40395</td><td>+</td><td>0</td><td>TCGTCGTGGAGGGACATCCC</td><td>CGG</td><td>||||||||[||||||||||||]PAM</td><td>NA</td><td>Intergenic</td><td>NA</td><td>NA</td></tr>
<tr><td>T2564</td><td>GTGGAGGGACATCCCCGGATCGG</td><td>1000</td><td></td><td></td><td></td><td></td><td></td><td></td><td></td><td></td><td></td></tr>
</table>

<table>
<tr><td>Oligo fwd</td><td>CACCGTGGAGGGACATCCCCGGAT</td><td></td><td></td><td></td><td></td><td></td><td></td><td></td><td></td><td></td><td></td><td></td></tr>
<tr><td>Oligo rev</td><td>AAACATCCGGGGATGTCCCTCCAC</td><td></td><td></td><td></td><td></td><td></td><td></td><td></td><td></td><td></td><td></td><td></td></tr>
<tr><td>Chromosome</td><td>start</td><td></td><td>end</td><td>strand</td><td>MM</td><td>target_seq</td><td>PAM</td><td>alignment</td><td>distance</td><td>position</td><td>gene name</td><td>gene id</td></tr>
<tr><td>GU305914.1</td><td></td><td>40378</td><td>40400</td><td>+</td><td>0</td><td>GTGGAGGGACATCCCCGGAT</td><td>CGG</td><td>||||||||[||||||||||||]PAM</td><td>NA</td><td>Intergenic</td><td>NA</td><td>NA</td></tr>
<tr><td>T2565</td><td>TGGAGGGACATCCCCGGATCGGG</td><td></td><td>1000</td><td></td><td></td><td></td><td></td><td></td><td></td><td></td><td></td><td></td></tr>
<tr><td>Oligo fwd</td><td>CACCTGGAGGGACATCCCCGGATC</td><td></td><td></td><td></td><td></td><td></td><td></td><td></td><td></td><td></td><td></td><td></td></tr>
<tr><td>Oligo rev</td><td>AAACGATCCGGGGATGTCCCTCCA</td><td></td><td></td><td></td><td></td><td></td><td></td><td></td><td></td><td></td><td></td><td></td></tr>
<tr><td>Chromosome</td><td>start</td><td></td><td>end</td><td>strand</td><td>MM</td><td>target_seq</td><td>PAM</td><td>alignment</td><td>distance</td><td>position</td><td>gene name</td><td>gene id</td></tr>
<tr><td>GU305914.1</td><td></td><td>40379</td><td>40401</td><td>+</td><td>0</td><td>TGGAGGGACATCCCCGGATC</td><td>GGG</td><td>||||||||[||||||||||||]PAM</td><td>NA</td><td>Intergenic</td><td>NA</td><td>NA</td></tr>
<tr><td>T2566</td><td>CGACTCGTCGTCATGTCCTCCGG</td><td></td><td>1000</td><td></td><td></td><td></td><td></td><td></td><td></td><td></td><td></td><td></td></tr>
<tr><td>Oligo fwd</td><td>CACCCGACTCGTCGTCATGTCCTC</td><td></td><td></td><td></td><td></td><td></td><td></td><td></td><td></td><td></td><td></td><td></td></tr>
<tr><td>Oligo rev</td><td>AAACGAGGACATGACGACGAGTCG</td><td></td><td></td><td></td><td></td><td></td><td></td><td></td><td></td><td></td><td></td><td></td></tr>
<tr><td>Chromosome</td><td>start</td><td></td><td>end</td><td>strand</td><td>MM</td><td>target_seq</td><td>PAM</td><td>alignment</td><td>distance</td><td>position</td><td>gene name</td><td>gene id</td></tr>
<tr><td>GU305914.1</td><td></td><td>40417</td><td>40439</td><td>+</td><td>0</td><td>CGACTCGTCGTCATGTCCTC</td><td>CGG</td><td>||||||||[||||||||||||]PAM</td><td>NA</td><td>Intergenic</td><td>NA</td><td>NA</td></tr>
<tr><td>T2567</td><td>GACTCGTCGTCATGTCCTCCGGG</td><td></td><td>1000</td><td></td><td></td><td></td><td></td><td></td><td></td><td></td><td></td><td></td></tr>
<tr><td>Oligo fwd</td><td>CACCGACTCGTCGTCATGTCCTCC</td><td></td><td></td><td></td><td></td><td></td><td></td><td></td><td></td><td></td><td></td><td></td></tr>
<tr><td>Oligo rev</td><td>AAACGGAGGACATGACGACGAGTC</td><td></td><td></td><td></td><td></td><td></td><td></td><td></td><td></td><td></td><td></td><td></td></tr>
<tr><td>Chromosome</td><td>start</td><td></td><td>end</td><td>strand</td><td>MM</td><td>target_seq</td><td>PAM</td><td>alignment</td><td>distance</td><td>position</td><td>gene name</td><td>gene id</td></tr>
<tr><td>GU305914.1</td><td></td><td>40418</td><td>40440</td><td>+</td><td>0</td><td>GACTCGTCGTCATGTCCTCC</td><td>GGG</td><td>||||||||[||||||||||||]PAM</td><td>NA</td><td>Intergenic</td><td>NA</td><td>NA</td></tr>
<tr><td>T2568</td><td>CGTCGTCTTCTTCATCTTCGAGG</td><td></td><td>1000</td><td></td><td></td><td></td><td></td><td></td><td></td><td></td><td></td><td></td></tr>
<tr><td>Oligo fwd</td><td>CACCCGTCGTCTTCTTCATCTTCG</td><td></td><td></td><td></td><td></td><td></td><td></td><td></td><td></td><td></td><td></td><td></td></tr>
<tr><td>Oligo rev</td><td>AAACCGAAGATGAAGAAGACGACG</td><td></td><td></td><td></td><td></td><td></td><td></td><td></td><td></td><td></td><td></td><td></td></tr>
<tr><td>Chromosome</td><td>start</td><td></td><td>end</td><td>strand</td><td>MM</td><td>target_seq</td><td>PAM</td><td>alignment</td><td>distance</td><td>position</td><td>gene name</td><td>gene id</td></tr>
<tr><td>GU305914.1</td><td></td><td>40455</td><td>40477</td><td>+</td><td>0</td><td>CGTCGTCTTCTTCATCTTCG</td><td>AGG</td><td>||||||||[||||||||||||]PAM</td><td>NA</td><td>Intergenic</td><td>NA</td><td>NA</td></tr>
<tr><td>T2569</td><td>CTTCTTCATCTTCGAGGTCCAGG</td><td></td><td>1000</td><td></td><td></td><td></td><td></td><td></td><td></td><td></td><td></td><td></td></tr>
<tr><td>Oligo fwd</td><td>CACCCTTCTTCATCTTCGAGGTCC</td><td></td><td></td><td></td><td></td><td></td><td></td><td></td><td></td><td></td><td></td><td></td></tr>
<tr><td>Oligo rev</td><td>AAACGGACCTCGAAGATGAAGAAG</td><td></td><td></td><td></td><td></td><td></td><td></td><td></td><td></td><td></td><td></td><td></td></tr>
<tr><td>Chromosome</td><td>start</td><td></td><td>end</td><td>strand</td><td>MM</td><td>target_seq</td><td>PAM</td><td>alignment</td><td>distance</td><td>position</td><td>gene name</td><td>gene id</td></tr>
<tr><td>GU305914.1</td><td></td><td>40461</td><td>40483</td><td>+</td><td>0</td><td>CTTCTTCATCTTCGAGGTCC</td><td>AGG</td><td>||||||||[||||||||||||]PAM</td><td>NA</td><td>Intergenic</td><td>NA</td><td>NA</td></tr>
<tr><td>GU305914.1</td><td></td><td>43816</td><td>43838</td><td>-</td><td>4</td><td>ATTCATCATCTTGTAGGTCC</td><td>CGG</td><td>-|||-|||[||||--||||||]PAM</td><td>NA</td><td>Intergenic</td><td>NA</td><td>NA</td></tr>
<tr><td>T2570</td><td>TTCGAGGTCCAGGTCCGAGCCGG</td><td></td><td>1000</td><td></td><td></td><td></td><td></td><td></td><td></td><td></td><td></td><td></td></tr>
<tr><td>Oligo fwd</td><td>CACCTTCGAGGTCCAGGTCCGAGC</td><td></td><td></td><td></td><td></td><td></td><td></td><td></td><td></td><td></td><td></td><td></td></tr>
<tr><td>Oligo rev</td><td>AAACGCTCGGACCTGGACCTCGAA</td><td></td><td></td><td></td><td></td><td></td><td></td><td></td><td></td><td></td><td></td><td></td></tr>
<tr><td>Chromosome</td><td>start</td><td></td><td>end</td><td>strand</td><td>MM</td><td>target_seq</td><td>PAM</td><td>alignment</td><td>distance</td><td>position</td><td>gene name</td><td>gene id</td></tr>
<tr><td>GU305914.1</td><td></td><td>40471</td><td>40493</td><td>+</td><td>0</td><td>TTCGAGGTCCAGGTCCGAGC</td><td>CGG</td><td>||||||||[||||||||||||]PAM</td><td>NA</td><td>Intergenic</td><td>NA</td><td>NA</td></tr>
<tr><td>T2571</td><td>ATCTCCCTCGCTCTCGAGCTCGG</td><td></td><td>1000</td><td></td><td></td><td></td><td></td><td></td><td></td><td></td><td></td><td></td></tr>
<tr><td>Oligo fwd</td><td>CACCATCTCCCTCGCTCTCGAGCT</td><td></td><td></td><td></td><td></td><td></td><td></td><td></td><td></td><td></td><td></td><td></td></tr>
<tr><td>Oligo rev</td><td>AAACAGCTCGAGAGCGAGGGAGAT</td><td></td><td></td><td></td><td></td><td></td><td></td><td></td><td></td><td></td><td></td><td></td></tr>
<tr><td>Chromosome</td><td>start</td><td></td><td>end</td><td>strand</td><td>MM</td><td>target_seq</td><td>PAM</td><td>alignment</td><td>distance</td><td>position</td><td>gene name</td><td>gene id</td></tr>
<tr><td>GU305914.1</td><td></td><td>40504</td><td>40526</td><td>+</td><td>0</td><td>ATCTCCCTCGCTCTCGAGCT</td><td>CGG</td><td>||||||||[||||||||||||]PAM</td><td>NA</td><td>Intergenic</td><td>NA</td><td>NA</td></tr>
<tr><td>T2572</td><td>CTCGGACTCTAACCCCCAGTCGG</td><td></td><td>1000</td><td></td><td></td><td></td><td></td><td></td><td></td><td></td><td></td><td></td></tr>
<tr><td>Oligo fwd</td><td>CACCCTCGGACTCTAACCCCCAGT</td><td></td><td></td><td></td><td></td><td></td><td></td><td></td><td></td><td></td><td></td><td></td></tr>
<tr><td>Oligo rev</td><td>AAACACTGGGGGTTAGAGTCCGAG</td><td></td><td></td><td></td><td></td><td></td><td></td><td></td><td></td><td></td><td></td><td></td></tr>
<tr><td>Chromosome</td><td>start</td><td></td><td>end</td><td>strand</td><td>MM</td><td>target_seq</td><td>PAM</td><td>alignment</td><td>distance</td><td>position</td><td>gene name</td><td>gene id</td></tr>
<tr><td>GU305914.1</td><td></td><td>40522</td><td>40544</td><td>+</td><td>0</td><td>CTCGGACTCTAACCCCCAGT</td><td>CGG</td><td>||||||||[||||||||||||]PAM</td><td>NA</td><td>Intergenic</td><td>NA</td><td>NA</td></tr>
<tr><td>T2573</td><td>TCGGACTCTAACCCCCAGTCGGG</td><td></td><td>1000</td><td></td><td></td><td></td><td></td><td></td><td></td><td></td><td></td><td></td></tr>
<tr><td>Oligo fwd</td><td>CACCTCGGACTCTAACCCCCAGTC</td><td></td><td></td><td></td><td></td><td></td><td></td><td></td><td></td><td></td><td></td><td></td></tr>
<tr><td>Oligo rev</td><td>AAACGACTGGGGGTTAGAGTCCGA</td><td></td><td></td><td></td><td></td><td></td><td></td><td></td><td></td><td></td><td></td><td></td></tr>
<tr><td>Chromosome</td><td>start</td><td></td><td>end</td><td>strand</td><td>MM</td><td>target_seq</td><td>PAM</td><td>alignment</td><td>distance</td><td>position</td><td>gene name</td><td>gene id</td></tr>
<tr><td>GU305914.1</td><td></td><td>40523</td><td>40545</td><td>+</td><td>0</td><td>TCGGACTCTAACCCCCAGTC</td><td>GGG</td><td>||||||||[||||||||||||]PAM</td><td>NA</td><td>Intergenic</td><td>NA</td><td>NA</td></tr>
<tr><td>T2574</td><td>ACCTAAGCGCGTTTCTGAAGAGG</td><td></td><td>1000</td><td></td><td></td><td></td><td></td><td></td><td></td><td></td><td></td><td></td></tr>
<tr><td>Oligo fwd</td><td>CACCACCTAAGCGCGTTTCTGAAG</td><td></td><td></td><td></td><td></td><td></td><td></td><td></td><td></td><td></td><td></td><td></td></tr>
<tr><td>Oligo rev</td><td>AAACCTTCAGAAACGCGCTTAGGT</td><td></td><td></td><td></td><td></td><td></td><td></td><td></td><td></td><td></td><td></td><td></td></tr>
<tr><td>Chromosome</td><td>start</td><td></td><td>end</td><td>strand</td><td>MM</td><td>target_seq</td><td>PAM</td><td>alignment</td><td>distance</td><td>position</td><td>gene name</td><td>gene id</td></tr>
<tr><td>GU305914.1</td><td></td><td>40554</td><td>40576</td><td>+</td><td>0</td><td>ACCTAAGCGCGTTTCTGAAG</td><td>AGG</td><td>||||||||[||||||||||||]PAM</td><td>NA</td><td>Intergenic</td><td>NA</td><td>NA</td></tr>
<tr><td>T2575</td><td>TAAGCGCGTTTCTGAAGAGGAGG</td><td></td><td>1000</td><td></td><td></td><td></td><td></td><td></td><td></td><td></td><td></td><td></td></tr>
<tr><td>Oligo fwd</td><td>CACCTAAGCGCGTTTCTGAAGAGG</td><td></td><td></td><td></td><td></td><td></td><td></td><td></td><td></td><td></td><td></td><td></td></tr>
<tr><td>Oligo rev</td><td>AAACCCTCTTCAGAAACGCGCTTA</td><td></td><td></td><td></td><td></td><td></td><td></td><td></td><td></td><td></td><td></td><td></td></tr>
<tr><td>Chromosome</td><td>start</td><td></td><td>end</td><td>strand</td><td>MM</td><td>target_seq</td><td>PAM</td><td>alignment</td><td>distance</td><td>position</td><td>gene name</td><td>gene id</td></tr>
<tr><td>GU305914.1</td><td></td><td>40557</td><td>40579</td><td>+</td><td>0</td><td>TAAGCGCGTTTCTGAAGAGG</td><td>AGG</td><td>||||||||[||||||||||||]PAM</td><td>NA</td><td>Intergenic</td><td>NA</td><td>NA</td></tr>
<tr><td>T2576</td><td>CCGCCGCAGTAGATTATACGCGG</td><td></td><td>1000</td><td></td><td></td><td></td><td></td><td></td><td></td><td></td><td></td><td></td></tr>
<tr><td>Oligo fwd</td><td>CACCCCGCCGCAGTAGATTATACG</td><td></td><td></td><td></td><td></td><td></td><td></td><td></td><td></td><td></td><td></td><td></td></tr>
<tr><td>Oligo rev</td><td>AAACCGTATAATCTACTGCGGCGG</td><td></td><td></td><td></td><td></td><td></td><td></td><td></td><td></td><td></td><td></td><td></td></tr>
<tr><td>Chromosome</td><td>start</td><td></td><td>end</td><td>strand</td><td>MM</td><td>target_seq</td><td>PAM</td><td>alignment</td><td>distance</td><td>position</td><td>gene name</td><td>gene id</td></tr>
<tr><td>GU305914.1</td><td></td><td>40594</td><td>40616</td><td>+</td><td>0</td><td>CCGCCGCAGTAGATTATACG</td><td>CGG</td><td>||||||||[||||||||||||]PAM</td><td>NA</td><td>Intergenic</td><td>NA</td><td>NA</td></tr>
<tr><td>T2577</td><td>ATACGCGGTCGTACCGTCGACGG</td><td></td><td>1000</td><td></td><td></td><td></td><td></td><td></td><td></td><td></td><td></td><td></td></tr>
<tr><td>Oligo fwd</td><td>CACCATACGCGGTCGTACCGTCGA</td><td></td><td></td><td></td><td></td><td></td><td></td><td></td><td></td><td></td><td></td><td></td></tr>
<tr><td>Oligo rev</td><td>AAACTCGACGGTACGACCGCGTAT</td><td></td><td></td><td></td><td></td><td></td><td></td><td></td><td></td><td></td><td></td><td></td></tr>
<tr><td>Chromosome</td><td>start</td><td></td><td>end</td><td>strand</td><td>MM</td><td>target_seq</td><td>PAM</td><td>alignment</td><td>distance</td><td>position</td><td>gene name</td><td>gene id</td></tr>
<tr><td>GU305914.1</td><td></td><td>40609</td><td>40631</td><td>+</td><td>0</td><td>ATACGCGGTCGTACCGTCGA</td><td>CGG</td><td>||||||||[||||||||||||]PAM</td><td>NA</td><td>Intergenic</td><td>NA</td><td>NA</td></tr>
<tr><td>T2578</td><td>TACGCGGTCGTACCGTCGACGGG</td><td></td><td>1000</td><td></td><td></td><td></td><td></td><td></td><td></td><td></td><td></td><td></td></tr>
<tr><td>Oligo fwd</td><td>CACCTACGCGGTCGTACCGTCGAC</td><td></td><td></td><td></td><td></td><td></td><td></td><td></td><td></td><td></td><td></td><td></td></tr>
<tr><td>Oligo rev</td><td>AAACGTCGACGGTACGACCGCGTA</td><td></td><td></td><td></td><td></td><td></td><td></td><td></td><td></td><td></td><td></td><td></td></tr>
<tr><td>Chromosome</td><td>start</td><td></td><td>end</td><td>strand</td><td>MM</td><td>target_seq</td><td>PAM</td><td>alignment</td><td>distance</td><td>position</td><td>gene name</td><td>gene id</td></tr>
<tr><td>GU305914.1</td><td></td><td>40610</td><td>40632</td><td>+</td><td>0</td><td>TACGCGGTCGTACCGTCGAC</td><td>GGG</td><td>||||||||[||||||||||||]PAM</td><td>NA</td><td>Intergenic</td><td>NA</td><td>NA</td></tr>
</table>

<table>
<tr><td>T2579</td><td>TCGAGCGTAGACTCGAGCGAAGG</td><td>1000</td><td></td><td></td><td></td><td></td><td></td><td></td><td></td><td></td><td></td></tr>
<tr><td>Oligo fwd</td><td>CACCTCGAGCGTAGACTCGAGCGA</td><td></td><td></td><td></td><td></td><td></td><td></td><td></td><td></td><td></td><td></td></tr>
<tr><td>Oligo rev</td><td>AAACTCGCTCGAGTCTACGCTCGA</td><td></td><td></td><td></td><td></td><td></td><td></td><td></td><td></td><td></td><td></td></tr>
<tr><td>Chromosome</td><td>start</td><td>end</td><td>strand</td><td>MM</td><td>target_seq</td><td>PAM</td><td>alignment</td><td>distance</td><td>position</td><td>gene name</td><td>gene id</td></tr>
<tr><td>GU305914.1</td><td>40648</td><td>40670</td><td>+</td><td>0</td><td>TCGAGCGTAGACTCGAGCGA</td><td>AGG</td><td>||||||||[||||||||||||]PAM</td><td>NA</td><td>Intergenic</td><td>NA</td><td>NA</td></tr>
<tr><td>T2580</td><td>GCGTAGACTCGAGCGAAGGCCGG</td><td>1000</td><td></td><td></td><td></td><td></td><td></td><td></td><td></td><td></td><td></td></tr>
<tr><td>Oligo fwd</td><td>CACCGCGTAGACTCGAGCGAAGGC</td><td></td><td></td><td></td><td></td><td></td><td></td><td></td><td></td><td></td><td></td></tr>
<tr><td>Oligo rev</td><td>AAACGCCTTCGCTCGAGTCTACGC</td><td></td><td></td><td></td><td></td><td></td><td></td><td></td><td></td><td></td><td></td></tr>
<tr><td>Chromosome</td><td>start</td><td>end</td><td>strand</td><td>MM</td><td>target_seq</td><td>PAM</td><td>alignment</td><td>distance</td><td>position</td><td>gene name</td><td>gene id</td></tr>
<tr><td>GU305914.1</td><td>40652</td><td>40674</td><td>+</td><td>0</td><td>GCGTAGACTCGAGCGAAGGC</td><td>CGG</td><td>||||||||[||||||||||||]PAM</td><td>NA</td><td>Intergenic</td><td>NA</td><td>NA</td></tr>
<tr><td>T2581</td><td>TGATGCGATCCAGACACATGTGG</td><td>1000</td><td></td><td></td><td></td><td></td><td></td><td></td><td></td><td></td><td></td></tr>
<tr><td>Oligo fwd</td><td>CACCTGATGCGATCCAGACACATG</td><td></td><td></td><td></td><td></td><td></td><td></td><td></td><td></td><td></td><td></td></tr>
<tr><td>Oligo rev</td><td>AAACCATGTGTCTGGATCGCATCA</td><td></td><td></td><td></td><td></td><td></td><td></td><td></td><td></td><td></td><td></td></tr>
<tr><td>Chromosome</td><td>start</td><td>end</td><td>strand</td><td>MM</td><td>target_seq</td><td>PAM</td><td>alignment</td><td>distance</td><td>position</td><td>gene name</td><td>gene id</td></tr>
<tr><td>GU305914.1</td><td>40716</td><td>40738</td><td>+</td><td>0</td><td>TGATGCGATCCAGACACATG</td><td>TGG</td><td>||||||||[||||||||||||]PAM</td><td>NA</td><td>Intergenic</td><td>NA</td><td>NA</td></tr>
<tr><td>T2582</td><td>GATGCGATCCAGACACATGTGGG</td><td>1000</td><td></td><td></td><td></td><td></td><td></td><td></td><td></td><td></td><td></td></tr>
<tr><td>Oligo fwd</td><td>CACCGATGCGATCCAGACACATGT</td><td></td><td></td><td></td><td></td><td></td><td></td><td></td><td></td><td></td><td></td></tr>
<tr><td>Oligo rev</td><td>AAACACATGTGTCTGGATCGCATC</td><td></td><td></td><td></td><td></td><td></td><td></td><td></td><td></td><td></td><td></td></tr>
<tr><td>Chromosome</td><td>start</td><td>end</td><td>strand</td><td>MM</td><td>target_seq</td><td>PAM</td><td>alignment</td><td>distance</td><td>position</td><td>gene name</td><td>gene id</td></tr>
<tr><td>GU305914.1</td><td>40717</td><td>40739</td><td>+</td><td>0</td><td>GATGCGATCCAGACACATGT</td><td>GGG</td><td>||||||||[||||||||||||]PAM</td><td>NA</td><td>Intergenic</td><td>NA</td><td>NA</td></tr>
<tr><td>T2583</td><td>CGTCTGCCAAGACTTACTGATGG</td><td>1000</td><td></td><td></td><td></td><td></td><td></td><td></td><td></td><td></td><td></td></tr>
<tr><td>Oligo fwd</td><td>CACCCGTCTGCCAAGACTTACTGA</td><td></td><td></td><td></td><td></td><td></td><td></td><td></td><td></td><td></td><td></td></tr>
<tr><td>Oligo rev</td><td>AAACTCAGTAAGTCTTGGCAGACG</td><td></td><td></td><td></td><td></td><td></td><td></td><td></td><td></td><td></td><td></td></tr>
<tr><td>Chromosome</td><td>start</td><td>end</td><td>strand</td><td>MM</td><td>target_seq</td><td>PAM</td><td>alignment</td><td>distance</td><td>position</td><td>gene name</td><td>gene id</td></tr>
<tr><td>GU305914.1</td><td>40744</td><td>40766</td><td>+</td><td>0</td><td>CGTCTGCCAAGACTTACTGA</td><td>TGG</td><td>||||||||[||||||||||||]PAM</td><td>NA</td><td>Intergenic</td><td>NA</td><td>NA</td></tr>
<tr><td>T2584</td><td>CCAAGACTTACTGATGGCGTAGG</td><td>1000</td><td></td><td></td><td></td><td></td><td></td><td></td><td></td><td></td><td></td></tr>
<tr><td>Oligo fwd</td><td>CACCCCAAGACTTACTGATGGCGT</td><td></td><td></td><td></td><td></td><td></td><td></td><td></td><td></td><td></td><td></td></tr>
<tr><td>Oligo rev</td><td>AAACACGCCATCAGTAAGTCTTGG</td><td></td><td></td><td></td><td></td><td></td><td></td><td></td><td></td><td></td><td></td></tr>
<tr><td>Chromosome</td><td>start</td><td>end</td><td>strand</td><td>MM</td><td>target_seq</td><td>PAM</td><td>alignment</td><td>distance</td><td>position</td><td>gene name</td><td>gene id</td></tr>
<tr><td>GU305914.1</td><td>40750</td><td>40772</td><td>+</td><td>0</td><td>CCAAGACTTACTGATGGCGT</td><td>AGG</td><td>||||||||[||||||||||||]PAM</td><td>NA</td><td>Intergenic</td><td>NA</td><td>NA</td></tr>
<tr><td>T2585</td><td>ACTTACTGATGGCGTAGGCGAGG</td><td>1000</td><td></td><td></td><td></td><td></td><td></td><td></td><td></td><td></td><td></td></tr>
<tr><td>Oligo fwd</td><td>CACCACTTACTGATGGCGTAGGCG</td><td></td><td></td><td></td><td></td><td></td><td></td><td></td><td></td><td></td><td></td></tr>
<tr><td>Oligo rev</td><td>AAACCGCCTACGCCATCAGTAAGT</td><td></td><td></td><td></td><td></td><td></td><td></td><td></td><td></td><td></td><td></td></tr>
<tr><td>Chromosome</td><td>start</td><td>end</td><td>strand</td><td>MM</td><td>target_seq</td><td>PAM</td><td>alignment</td><td>distance</td><td>position</td><td>gene name</td><td>gene id</td></tr>
<tr><td>GU305914.1</td><td>40755</td><td>40777</td><td>+</td><td>0</td><td>ACTTACTGATGGCGTAGGCG</td><td>AGG</td><td>||||||||[||||||||||||]PAM</td><td>NA</td><td>Intergenic</td><td>NA</td><td>NA</td></tr>
<tr><td>T2586</td><td>TACTGATGGCGTAGGCGAGGCGG</td><td>1000</td><td></td><td></td><td></td><td></td><td></td><td></td><td></td><td></td><td></td></tr>
<tr><td>Oligo fwd</td><td>CACCTACTGATGGCGTAGGCGAGG</td><td></td><td></td><td></td><td></td><td></td><td></td><td></td><td></td><td></td><td></td></tr>
<tr><td>Oligo rev</td><td>AAACCCTCGCCTACGCCATCAGTA</td><td></td><td></td><td></td><td></td><td></td><td></td><td></td><td></td><td></td><td></td></tr>
<tr><td>Chromosome</td><td>start</td><td>end</td><td>strand</td><td>MM</td><td>target_seq</td><td>PAM</td><td>alignment</td><td>distance</td><td>position</td><td>gene name</td><td>gene id</td></tr>
<tr><td>GU305914.1</td><td>40758</td><td>40780</td><td>+</td><td>0</td><td>TACTGATGGCGTAGGCGAGG</td><td>CGG</td><td>||||||||[||||||||||||]PAM</td><td>NA</td><td>Intergenic</td><td>NA</td><td>NA</td></tr>
<tr><td>T2587</td><td>TGGCGTAGGCGAGGCGGCAGAGG</td><td>1000</td><td></td><td></td><td></td><td></td><td></td><td></td><td></td><td></td><td></td></tr>
<tr><td>Oligo fwd</td><td>CACCTGGCGTAGGCGAGGCGGCAG</td><td></td><td></td><td></td><td></td><td></td><td></td><td></td><td></td><td></td><td></td></tr>
<tr><td>Oligo rev</td><td>AAACCTGCCGCCTCGCCTACGCCA</td><td></td><td></td><td></td><td></td><td></td><td></td><td></td><td></td><td></td><td></td></tr>
<tr><td>Chromosome</td><td>start</td><td>end</td><td>strand</td><td>MM</td><td>target_seq</td><td>PAM</td><td>alignment</td><td>distance</td><td>position</td><td>gene name</td><td>gene id</td></tr>
<tr><td>GU305914.1</td><td>40764</td><td>40786</td><td>+</td><td>0</td><td>TGGCGTAGGCGAGGCGGCAG</td><td>AGG</td><td>||||||||[||||||||||||]PAM</td><td>NA</td><td>Intergenic</td><td>NA</td><td>NA</td></tr>
<tr><td>T2588</td><td>AGACGAGACGATTGTGCACGAGG</td><td>1000</td><td></td><td></td><td></td><td></td><td></td><td></td><td></td><td></td><td></td></tr>
<tr><td>Oligo fwd</td><td>CACCAGACGAGACGATTGTGCACG</td><td></td><td></td><td></td><td></td><td></td><td></td><td></td><td></td><td></td><td></td></tr>
<tr><td>Oligo rev</td><td>AAACCGTGCACAATCGTCTCGTCT</td><td></td><td></td><td></td><td></td><td></td><td></td><td></td><td></td><td></td><td></td></tr>
<tr><td>Chromosome</td><td>start</td><td>end</td><td>strand</td><td>MM</td><td>target_seq</td><td>PAM</td><td>alignment</td><td>distance</td><td>position</td><td>gene name</td><td>gene id</td></tr>
<tr><td>GU305914.1</td><td>40788</td><td>40810</td><td>+</td><td>0</td><td>AGACGAGACGATTGTGCACG</td><td>AGG</td><td>||||||||[||||||||||||]PAM</td><td>NA</td><td>Intergenic</td><td>NA</td><td>NA</td></tr>
<tr><td>T2589</td><td>GAGACGATTGTGCACGAGGACGG</td><td>1000</td><td></td><td></td><td></td><td></td><td></td><td></td><td></td><td></td><td></td></tr>
<tr><td>Oligo fwd</td><td>CACCGAGACGATTGTGCACGAGGA</td><td></td><td></td><td></td><td></td><td></td><td></td><td></td><td></td><td></td><td></td></tr>
<tr><td>Oligo rev</td><td>AAACTCCTCGTGCACAATCGTCTC</td><td></td><td></td><td></td><td></td><td></td><td></td><td></td><td></td><td></td><td></td></tr>
<tr><td>Chromosome</td><td>start</td><td>end</td><td>strand</td><td>MM</td><td>target_seq</td><td>PAM</td><td>alignment</td><td>distance</td><td>position</td><td>gene name</td><td>gene id</td></tr>
<tr><td>GU305914.1</td><td>40792</td><td>40814</td><td>+</td><td>0</td><td>GAGACGATTGTGCACGAGGA</td><td>CGG</td><td>||||||||[||||||||||||]PAM</td><td>NA</td><td>Intergenic</td><td>NA</td><td>NA</td></tr>
<tr><td>T2590</td><td>CGATTGTGCACGAGGACGGACGG</td><td>1000</td><td></td><td></td><td></td><td></td><td></td><td></td><td></td><td></td><td></td></tr>
<tr><td>Oligo fwd</td><td>CACCCGATTGTGCACGAGGACGGA</td><td></td><td></td><td></td><td></td><td></td><td></td><td></td><td></td><td></td><td></td></tr>
<tr><td>Oligo rev</td><td>AAACTCCGTCCTCGTGCACAATCG</td><td></td><td></td><td></td><td></td><td></td><td></td><td></td><td></td><td></td><td></td></tr>
<tr><td>Chromosome</td><td>start</td><td>end</td><td>strand</td><td>MM</td><td>target_seq</td><td>PAM</td><td>alignment</td><td>distance</td><td>position</td><td>gene name</td><td>gene id</td></tr>
<tr><td>GU305914.1</td><td>40796</td><td>40818</td><td>+</td><td>0</td><td>CGATTGTGCACGAGGACGGA</td><td>CGG</td><td>||||||||[||||||||||||]PAM</td><td>NA</td><td>Intergenic</td><td>NA</td><td>NA</td></tr>
<tr><td>T2591</td><td>ACGGACGGATCTTCGTTGAGAGG</td><td>1000</td><td></td><td></td><td></td><td></td><td></td><td></td><td></td><td></td><td></td></tr>
<tr><td>Oligo fwd</td><td>CACCACGGACGGATCTTCGTTGAG</td><td></td><td></td><td></td><td></td><td></td><td></td><td></td><td></td><td></td><td></td></tr>
<tr><td>Oligo rev</td><td>AAACCTCAACGAAGATCCGTCCGT</td><td></td><td></td><td></td><td></td><td></td><td></td><td></td><td></td><td></td><td></td></tr>
<tr><td>Chromosome</td><td>start</td><td>end</td><td>strand</td><td>MM</td><td>target_seq</td><td>PAM</td><td>alignment</td><td>distance</td><td>position</td><td>gene name</td><td>gene id</td></tr>
<tr><td>GU305914.1</td><td>40811</td><td>40833</td><td>+</td><td>0</td><td>ACGGACGGATCTTCGTTGAG</td><td>AGG</td><td>||||||||[||||||||||||]PAM</td><td>NA</td><td>Intergenic</td><td>NA</td><td>NA</td></tr>
<tr><td>T2592</td><td>CGGACGGATCTTCGTTGAGAGGG</td><td>1000</td><td></td><td></td><td></td><td></td><td></td><td></td><td></td><td></td><td></td></tr>
<tr><td>Oligo fwd</td><td>CACCCGGACGGATCTTCGTTGAGA</td><td></td><td></td><td></td><td></td><td></td><td></td><td></td><td></td><td></td><td></td></tr>
<tr><td>Oligo rev</td><td>AAACTCTCAACGAAGATCCGTCCG</td><td></td><td></td><td></td><td></td><td></td><td></td><td></td><td></td><td></td><td></td></tr>
<tr><td>Chromosome</td><td>start</td><td>end</td><td>strand</td><td>MM</td><td>target_seq</td><td>PAM</td><td>alignment</td><td>distance</td><td>position</td><td>gene name</td><td>gene id</td></tr>
<tr><td>GU305914.1</td><td>40812</td><td>40834</td><td>+</td><td>0</td><td>CGGACGGATCTTCGTTGAGA</td><td>GGG</td><td>||||||||[||||||||||||]PAM</td><td>NA</td><td>Intergenic</td><td>NA</td><td>NA</td></tr>
<tr><td>T2593</td><td>CGTCCAAGACTTCGTAGACGAGG</td><td>1000</td><td></td><td></td><td></td><td></td><td></td><td></td><td></td><td></td><td></td></tr>
<tr><td>Oligo fwd</td><td>CACCCGTCCAAGACTTCGTAGACG</td><td></td><td></td><td></td><td></td><td></td><td></td><td></td><td></td><td></td><td></td></tr>
<tr><td>Oligo rev</td><td>AAACCGTCTACGAAGTCTTGGACG</td><td></td><td></td><td></td><td></td><td></td><td></td><td></td><td></td><td></td><td></td></tr>
<tr><td>Chromosome</td><td>start</td><td>end</td><td>strand</td><td>MM</td><td>target_seq</td><td>PAM</td><td>alignment</td><td>distance</td><td>position</td><td>gene name</td><td>gene id</td></tr>
</table>

| | | | | | | | | | | | |
|---|---|---|---|---|---|---|---|---|---|---|---|
| GU305914.1 | 40842 | 40864 | + | 0 | CGTCCAAGACTTCGTAGACG | AGG | \|\|\|\|\|\|\|\|[\|\|\|\|\|\|\|\|\|\|\|\|]PAM | NA | Intergenic | NA | NA |
| T2594 | TTCGTAGACGAGGACGCGTTCGG | 1000 | | | | | | | | | |
| Oligo fwd | CACCTTCGTAGACGAGGACGCGTT | | | | | | | | | | |
| Oligo rev | AAACAACGCGTCCTCGTCTACGAA | | | | | | | | | | |
| Chromosome | start | end | strand | MM | target_seq | PAM | alignment | distance | position | gene name | gene id |
| GU305914.1 | 40852 | 40874 | + | 0 | TTCGTAGACGAGGACGCGTT | CGG | \|\|\|\|\|\|\|\|[\|\|\|\|\|\|\|\|\|\|\|\|]PAM | NA | Intergenic | NA | NA |
| T2595 | CGTCCCTGACGTTGTCCATCAGG | 1000 | | | | | | | | | |
| Oligo fwd | CACCCGTCCCTGACGTTGTCCATC | | | | | | | | | | |
| Oligo rev | AAACGATGGACAACGTCAGGGACG | | | | | | | | | | |
| Chromosome | start | end | strand | MM | target_seq | PAM | alignment | distance | position | gene name | gene id |
| GU305914.1 | 40878 | 40900 | + | 0 | CGTCCCTGACGTTGTCCATC | AGG | \|\|\|\|\|\|\|\|[\|\|\|\|\|\|\|\|\|\|\|\|]PAM | NA | Intergenic | NA | NA |
| T2596 | CCCATGATCACCGCACGCGCCGG | 1000 | | | | | | | | | |
| Oligo fwd | CACCCCCATGATCACCGCACGCGC | | | | | | | | | | |
| Oligo rev | AAACGCGCGTGCGGTGATCATGGG | | | | | | | | | | |
| Chromosome | start | end | strand | MM | target_seq | PAM | alignment | distance | position | gene name | gene id |
| GU305914.1 | 40934 | 40956 | + | 0 | CCCATGATCACCGCACGCGC | CGG | \|\|\|\|\|\|\|\|[\|\|\|\|\|\|\|\|\|\|\|\|]PAM | NA | Intergenic | NA | NA |
| T2597 | ACCGCACGCGCCGGCTTGTCCGG | 1000 | | | | | | | | | |
| Oligo fwd | CACCACCGCACGCGCCGGCTTGTC | | | | | | | | | | |
| Oligo rev | AAACGACAAGCCGGCGCGTGCGGT | | | | | | | | | | |
| Chromosome | start | end | strand | MM | target_seq | PAM | alignment | distance | position | gene name | gene id |
| GU305914.1 | 40943 | 40965 | + | 0 | ACCGCACGCGCCGGCTTGTC | CGG | \|\|\|\|\|\|\|\|[\|\|\|\|\|\|\|\|\|\|\|\|]PAM | NA | Intergenic | NA | NA |
| T2598 | CCGCACGCGCCGGCTTGTCCGGG | 1000 | | | | | | | | | |
| Oligo fwd | CACCCCGCACGCGCCGGCTTGTCC | | | | | | | | | | |
| Oligo rev | AAACGGACAAGCCGGCGCGTGCGG | | | | | | | | | | |
| Chromosome | start | end | strand | MM | target_seq | PAM | alignment | distance | position | gene name | gene id |
| GU305914.1 | 40944 | 40966 | + | 0 | CCGCACGCGCCGGCTTGTCC | GGG | \|\|\|\|\|\|\|\|[\|\|\|\|\|\|\|\|\|\|\|\|]PAM | NA | Intergenic | NA | NA |
| T2599 | GGCCCTGAGCACAGAGACACCGG | 1000 | | | | | | | | | |
| Oligo fwd | CACCGGCCCTGAGCACAGAGACAC | | | | | | | | | | |
| Oligo rev | AAACGTGTCTCTGTGCTCAGGGCC | | | | | | | | | | |
| Chromosome | start | end | strand | MM | target_seq | PAM | alignment | distance | position | gene name | gene id |
| GU305914.1 | 40965 | 40987 | + | 0 | GGCCCTGAGCACAGAGACAC | CGG | \|\|\|\|\|\|\|\|[\|\|\|\|\|\|\|\|\|\|\|\|]PAM | NA | Intergenic | NA | NA |
| T2600 | GCCCTGAGCACAGAGACACCGGG | 1000 | | | | | | | | | |
| Oligo fwd | CACCGCCCTGAGCACAGAGACACC | | | | | | | | | | |
| Oligo rev | AAACGGTGTCTCTGTGCTCAGGGC | | | | | | | | | | |
| Chromosome | start | end | strand | MM | target_seq | PAM | alignment | distance | position | gene name | gene id |
| GU305914.1 | 40966 | 40988 | + | 0 | GCCCTGAGCACAGAGACACC | GGG | \|\|\|\|\|\|\|\|[\|\|\|\|\|\|\|\|\|\|\|\|]PAM | NA | Intergenic | NA | NA |
| T2601 | GAGCACAGAGACACCGGGCGAGG | 1000 | | | | | | | | | |
| Oligo fwd | CACCGAGCACAGAGACACCGGGCG | | | | | | | | | | |
| Oligo rev | AAACCGCCCGGTGTCTCTGTGCTC | | | | | | | | | | |
| Chromosome | start | end | strand | MM | target_seq | PAM | alignment | distance | position | gene name | gene id |
| GU305914.1 | 40971 | 40993 | + | 0 | GAGCACAGAGACACCGGGCG | AGG | \|\|\|\|\|\|\|\|[\|\|\|\|\|\|\|\|\|\|\|\|]PAM | NA | Intergenic | NA | NA |
| T2602 | CGAGGTCCTCGAGCCCCCGCAGG | 1000 | | | | | | | | | |
| Oligo fwd | CACCCGAGGTCCTCGAGCCCCCGC | | | | | | | | | | |
| Oligo rev | AAACGCGGGGGCTCGAGGACCTCG | | | | | | | | | | |
| Chromosome | start | end | strand | MM | target_seq | PAM | alignment | distance | position | gene name | gene id |
| GU305914.1 | 40989 | 41011 | + | 0 | CGAGGTCCTCGAGCCCCCGC | AGG | \|\|\|\|\|\|\|\|[\|\|\|\|\|\|\|\|\|\|\|\|]PAM | NA | Intergenic | NA | NA |
| T2603 | CGCAGGAAGACGTCCACGCGAGG | 1000 | | | | | | | | | |
| Oligo fwd | CACCCGCAGGAAGACGTCCACGCG | | | | | | | | | | |
| Oligo rev | AAACCGCGTGGACGTCTTCCTGCG | | | | | | | | | | |
| Chromosome | start | end | strand | MM | target_seq | PAM | alignment | distance | position | gene name | gene id |
| GU305914.1 | 41006 | 41028 | + | 0 | CGCAGGAAGACGTCCACGCG | AGG | \|\|\|\|\|\|\|\|[\|\|\|\|\|\|\|\|\|\|\|\|]PAM | NA | Intergenic | NA | NA |
| T2604 | CACATAATCGTCGCCGTAGACGG | 1000 | | | | | | | | | |
| Oligo fwd | CACCCACATAATCGTCGCCGTAGA | | | | | | | | | | |
| Oligo rev | AAACTCTACGGCGACGATTATGTG | | | | | | | | | | |
| Chromosome | start | end | strand | MM | target_seq | PAM | alignment | distance | position | gene name | gene id |
| GU305914.1 | 41029 | 41051 | + | 0 | CACATAATCGTCGCCGTAGA | CGG | \|\|\|\|\|\|\|\|[\|\|\|\|\|\|\|\|\|\|\|\|]PAM | NA | Intergenic | NA | NA |
| T2605 | TCCGTCGAAGCCGCAGAGCTCGG | 1000 | | | | | | | | | |
| Oligo fwd | CACCTCCGTCGAAGCCGCAGAGCT | | | | | | | | | | |
| Oligo rev | AAACAGCTCTGCGGCTTCGACGGA | | | | | | | | | | |
| Chromosome | start | end | strand | MM | target_seq | PAM | alignment | distance | position | gene name | gene id |
| GU305914.1 | 41080 | 41102 | + | 0 | TCCGTCGAAGCCGCAGAGCT | CGG | \|\|\|\|\|\|\|\|[\|\|\|\|\|\|\|\|\|\|\|\|]PAM | NA | Intergenic | NA | NA |
| T2606 | CCGTCGAAGCCGCAGAGCTCGGG | 1000 | | | | | | | | | |
| Oligo fwd | CACCCCGTCGAAGCCGCAGAGCTC | | | | | | | | | | |
| Oligo rev | AAACGAGCTCTGCGGCTTCGACGG | | | | | | | | | | |
| Chromosome | start | end | strand | MM | target_seq | PAM | alignment | distance | position | gene name | gene id |
| GU305914.1 | 41081 | 41103 | + | 0 | CCGTCGAAGCCGCAGAGCTC | GGG | \|\|\|\|\|\|\|\|[\|\|\|\|\|\|\|\|\|\|\|\|]PAM | NA | Intergenic | NA | NA |
| T2607 | GCCGCAGAGCTCGGGCCGCGCGG | 1000 | | | | | | | | | |
| Oligo fwd | CACCGCCGCAGAGCTCGGGCCGCG | | | | | | | | | | |
| Oligo rev | AAACCGCGGCCCGAGCTCTGCGGC | | | | | | | | | | |
| Chromosome | start | end | strand | MM | target_seq | PAM | alignment | distance | position | gene name | gene id |
| GU305914.1 | 41089 | 41111 | + | 0 | GCCGCAGAGCTCGGGCCGCG | CGG | \|\|\|\|\|\|\|\|[\|\|\|\|\|\|\|\|\|\|\|\|]PAM | NA | Intergenic | NA | NA |
| T2608 | AATCTTGTCTGATCCTAGTGAGG | 1000 | | | | | | | | | |
| Oligo fwd | CACCAATCTTGTCTGATCCTAGTG | | | | | | | | | | |
| Oligo rev | AAACCACTAGGATCAGACAAGATT | | | | | | | | | | |

<table>
<tr><td>Chromosome</td><td>start</td><td>end</td><td>strand</td><td>MM</td><td>target_seq</td><td>PAM</td><td>alignment</td><td>distance</td><td>position</td><td>gene name</td><td>gene id</td></tr>
<tr><td>GU305914.1</td><td>41136</td><td>41158</td><td>+</td><td>0</td><td>AATCTTGTCTGATCCTAGTG</td><td>AGG</td><td>||||||||[||||||||||||]PAM</td><td>NA</td><td>Intergenic</td><td>NA</td><td>NA</td></tr>
<tr><td></td><td></td><td></td><td></td><td></td><td></td><td></td><td></td><td></td><td></td><td></td><td></td></tr>
<tr><td>T2609</td><td>TGTCTGATCCTAGTGAGGAGCGG</td><td>1000</td><td></td><td></td><td></td><td></td><td></td><td></td><td></td><td></td><td></td></tr>
<tr><td>Oligo fwd</td><td>CACCTGTCTGATCCTAGTGAGGAG</td><td></td><td></td><td></td><td></td><td></td><td></td><td></td><td></td><td></td><td></td></tr>
<tr><td>Oligo rev</td><td>AAACCTCCTCACTAGGATCAGACA</td><td></td><td></td><td></td><td></td><td></td><td></td><td></td><td></td><td></td><td></td></tr>
<tr><td>Chromosome</td><td>start</td><td>end</td><td>strand</td><td>MM</td><td>target_seq</td><td>PAM</td><td>alignment</td><td>distance</td><td>position</td><td>gene name</td><td>gene id</td></tr>
<tr><td>GU305914.1</td><td>41141</td><td>41163</td><td>+</td><td>0</td><td>TGTCTGATCCTAGTGAGGAG</td><td>CGG</td><td>||||||||[||||||||||||]PAM</td><td>NA</td><td>Intergenic</td><td>NA</td><td>NA</td></tr>
<tr><td></td><td></td><td></td><td></td><td></td><td></td><td></td><td></td><td></td><td></td><td></td><td></td></tr>
<tr><td>T2610</td><td>TGATCCTAGTGAGGAGCGGCAGG</td><td>1000</td><td></td><td></td><td></td><td></td><td></td><td></td><td></td><td></td><td></td></tr>
<tr><td>Oligo fwd</td><td>CACCTGATCCTAGTGAGGAGCGGC</td><td></td><td></td><td></td><td></td><td></td><td></td><td></td><td></td><td></td><td></td></tr>
<tr><td>Oligo rev</td><td>AAACGCCGCTCCTCACTAGGATCA</td><td></td><td></td><td></td><td></td><td></td><td></td><td></td><td></td><td></td><td></td></tr>
<tr><td>Chromosome</td><td>start</td><td>end</td><td>strand</td><td>MM</td><td>target_seq</td><td>PAM</td><td>alignment</td><td>distance</td><td>position</td><td>gene name</td><td>gene id</td></tr>
<tr><td>GU305914.1</td><td>41145</td><td>41167</td><td>+</td><td>0</td><td>TGATCCTAGTGAGGAGCGGC</td><td>AGG</td><td>||||||||[||||||||||||]PAM</td><td>NA</td><td>Intergenic</td><td>NA</td><td>NA</td></tr>
<tr><td></td><td></td><td></td><td></td><td></td><td></td><td></td><td></td><td></td><td></td><td></td><td></td></tr>
<tr><td>T2611</td><td>AGTGAGGAGCGGCAGGTGTCCGG</td><td>1000</td><td></td><td></td><td></td><td></td><td></td><td></td><td></td><td></td><td></td></tr>
<tr><td>Oligo fwd</td><td>CACCAGTGAGGAGCGGCAGGTGTC</td><td></td><td></td><td></td><td></td><td></td><td></td><td></td><td></td><td></td><td></td></tr>
<tr><td>Oligo rev</td><td>AAACGACACCTGCCGCTCCTCACT</td><td></td><td></td><td></td><td></td><td></td><td></td><td></td><td></td><td></td><td></td></tr>
<tr><td>Chromosome</td><td>start</td><td>end</td><td>strand</td><td>MM</td><td>target_seq</td><td>PAM</td><td>alignment</td><td>distance</td><td>position</td><td>gene name</td><td>gene id</td></tr>
<tr><td>GU305914.1</td><td>41152</td><td>41174</td><td>+</td><td>0</td><td>AGTGAGGAGCGGCAGGTGTC</td><td>CGG</td><td>||||||||[||||||||||||]PAM</td><td>NA</td><td>Intergenic</td><td>NA</td><td>NA</td></tr>
<tr><td></td><td></td><td></td><td></td><td></td><td></td><td></td><td></td><td></td><td></td><td></td><td></td></tr>
<tr><td>T2612</td><td>GTGAGGAGCGGCAGGTGTCCGGG</td><td>1000</td><td></td><td></td><td></td><td></td><td></td><td></td><td></td><td></td><td></td></tr>
<tr><td>Oligo fwd</td><td>CACCGTGAGGAGCGGCAGGTGTCC</td><td></td><td></td><td></td><td></td><td></td><td></td><td></td><td></td><td></td><td></td></tr>
<tr><td>Oligo rev</td><td>AAACGGACACCTGCCGCTCCTCAC</td><td></td><td></td><td></td><td></td><td></td><td></td><td></td><td></td><td></td><td></td></tr>
<tr><td>Chromosome</td><td>start</td><td>end</td><td>strand</td><td>MM</td><td>target_seq</td><td>PAM</td><td>alignment</td><td>distance</td><td>position</td><td>gene name</td><td>gene id</td></tr>
<tr><td>GU305914.1</td><td>41153</td><td>41175</td><td>+</td><td>0</td><td>GTGAGGAGCGGCAGGTGTCC</td><td>GGG</td><td>||||||||[||||||||||||]PAM</td><td>NA</td><td>Intergenic</td><td>NA</td><td>NA</td></tr>
<tr><td></td><td></td><td></td><td></td><td></td><td></td><td></td><td></td><td></td><td></td><td></td><td></td></tr>
<tr><td>T2613</td><td>GCGGCAGGTGTCCGGGTCGTCGG</td><td>1000</td><td></td><td></td><td></td><td></td><td></td><td></td><td></td><td></td><td></td></tr>
<tr><td>Oligo fwd</td><td>CACCGCGGCAGGTGTCCGGGTCGT</td><td></td><td></td><td></td><td></td><td></td><td></td><td></td><td></td><td></td><td></td></tr>
<tr><td>Oligo rev</td><td>AAACACGACCCGGACACCTGCCGC</td><td></td><td></td><td></td><td></td><td></td><td></td><td></td><td></td><td></td><td></td></tr>
<tr><td>Chromosome</td><td>start</td><td>end</td><td>strand</td><td>MM</td><td>target_seq</td><td>PAM</td><td>alignment</td><td>distance</td><td>position</td><td>gene name</td><td>gene id</td></tr>
<tr><td>GU305914.1</td><td>41160</td><td>41182</td><td>+</td><td>0</td><td>GCGGCAGGTGTCCGGGTCGT</td><td>CGG</td><td>||||||||[||||||||||||]PAM</td><td>NA</td><td>Intergenic</td><td>NA</td><td>NA</td></tr>
<tr><td></td><td></td><td></td><td></td><td></td><td></td><td></td><td></td><td></td><td></td><td></td><td></td></tr>
<tr><td>T2614</td><td>CGGCAGGTGTCCGGGTCGTCGGG</td><td>1000</td><td></td><td></td><td></td><td></td><td></td><td></td><td></td><td></td><td></td></tr>
<tr><td>Oligo fwd</td><td>CACCCGGCAGGTGTCCGGGTCGTC</td><td></td><td></td><td></td><td></td><td></td><td></td><td></td><td></td><td></td><td></td></tr>
<tr><td>Oligo rev</td><td>AAACGACGACCCGGACACCTGCCG</td><td></td><td></td><td></td><td></td><td></td><td></td><td></td><td></td><td></td><td></td></tr>
<tr><td>Chromosome</td><td>start</td><td>end</td><td>strand</td><td>MM</td><td>target_seq</td><td>PAM</td><td>alignment</td><td>distance</td><td>position</td><td>gene name</td><td>gene id</td></tr>
<tr><td>GU305914.1</td><td>41161</td><td>41183</td><td>+</td><td>0</td><td>CGGCAGGTGTCCGGGTCGTC</td><td>GGG</td><td>||||||||[||||||||||||]PAM</td><td>NA</td><td>Intergenic</td><td>NA</td><td>NA</td></tr>
<tr><td></td><td></td><td></td><td></td><td></td><td></td><td></td><td></td><td></td><td></td><td></td><td></td></tr>
<tr><td>T2615</td><td>GTCGGGCCGTGAGCTTCTCGTGG</td><td>1000</td><td></td><td></td><td></td><td></td><td></td><td></td><td></td><td></td><td></td></tr>
<tr><td>Oligo fwd</td><td>CACCGTCGGGCCGTGAGCTTCTCG</td><td></td><td></td><td></td><td></td><td></td><td></td><td></td><td></td><td></td><td></td></tr>
<tr><td>Oligo rev</td><td>AAACCGAGAAGCTCACGGCCCGAC</td><td></td><td></td><td></td><td></td><td></td><td></td><td></td><td></td><td></td><td></td></tr>
<tr><td>Chromosome</td><td>start</td><td>end</td><td>strand</td><td>MM</td><td>target_seq</td><td>PAM</td><td>alignment</td><td>distance</td><td>position</td><td>gene name</td><td>gene id</td></tr>
<tr><td>GU305914.1</td><td>41178</td><td>41200</td><td>+</td><td>0</td><td>GTCGGGCCGTGAGCTTCTCG</td><td>TGG</td><td>||||||||[||||||||||||]PAM</td><td>NA</td><td>Intergenic</td><td>NA</td><td>NA</td></tr>
<tr><td></td><td></td><td></td><td></td><td></td><td></td><td></td><td></td><td></td><td></td><td></td><td></td></tr>
<tr><td>T2616</td><td>TCTCGTGGTACCTGAGCCAGAGG</td><td>1000</td><td></td><td></td><td></td><td></td><td></td><td></td><td></td><td></td><td></td></tr>
<tr><td>Oligo fwd</td><td>CACCTCTCGTGGTACCTGAGCCAG</td><td></td><td></td><td></td><td></td><td></td><td></td><td></td><td></td><td></td><td></td></tr>
<tr><td>Oligo rev</td><td>AAACCTGGCTCAGGTACCACGAGA</td><td></td><td></td><td></td><td></td><td></td><td></td><td></td><td></td><td></td><td></td></tr>
<tr><td>Chromosome</td><td>start</td><td>end</td><td>strand</td><td>MM</td><td>target_seq</td><td>PAM</td><td>alignment</td><td>distance</td><td>position</td><td>gene name</td><td>gene id</td></tr>
<tr><td>GU305914.1</td><td>41193</td><td>41215</td><td>+</td><td>0</td><td>TCTCGTGGTACCTGAGCCAG</td><td>AGG</td><td>||||||||[||||||||||||]PAM</td><td>NA</td><td>Intergenic</td><td>NA</td><td>NA</td></tr>
<tr><td></td><td></td><td></td><td></td><td></td><td></td><td></td><td></td><td></td><td></td><td></td><td></td></tr>
<tr><td>T2617</td><td>GTTGTAGAGCTGCGTCTCCCTGG</td><td>1000</td><td></td><td></td><td></td><td></td><td></td><td></td><td></td><td></td><td></td></tr>
<tr><td>Oligo fwd</td><td>CACCGTTGTAGAGCTGCGTCTCCC</td><td></td><td></td><td></td><td></td><td></td><td></td><td></td><td></td><td></td><td></td></tr>
<tr><td>Oligo rev</td><td>AAACGGGAGACGCAGCTCTACAAC</td><td></td><td></td><td></td><td></td><td></td><td></td><td></td><td></td><td></td><td></td></tr>
<tr><td>Chromosome</td><td>start</td><td>end</td><td>strand</td><td>MM</td><td>target_seq</td><td>PAM</td><td>alignment</td><td>distance</td><td>position</td><td>gene name</td><td>gene id</td></tr>
<tr><td>GU305914.1</td><td>41224</td><td>41246</td><td>+</td><td>0</td><td>GTTGTAGAGCTGCGTCTCCC</td><td>TGG</td><td>||||||||[||||||||||||]PAM</td><td>NA</td><td>Intergenic</td><td>NA</td><td>NA</td></tr>
<tr><td></td><td></td><td></td><td></td><td></td><td></td><td></td><td></td><td></td><td></td><td></td><td></td></tr>
<tr><td>T2618</td><td>AGAGCTGCGTCTCCCTGGAAAGG</td><td>1000</td><td></td><td></td><td></td><td></td><td></td><td></td><td></td><td></td><td></td></tr>
<tr><td>Oligo fwd</td><td>CACCAGAGCTGCGTCTCCCTGGAA</td><td></td><td></td><td></td><td></td><td></td><td></td><td></td><td></td><td></td><td></td></tr>
<tr><td>Oligo rev</td><td>AAACTTCCAGGGAGACGCAGCTCT</td><td></td><td></td><td></td><td></td><td></td><td></td><td></td><td></td><td></td><td></td></tr>
<tr><td>Chromosome</td><td>start</td><td>end</td><td>strand</td><td>MM</td><td>target_seq</td><td>PAM</td><td>alignment</td><td>distance</td><td>position</td><td>gene name</td><td>gene id</td></tr>
<tr><td>GU305914.1</td><td>41229</td><td>41251</td><td>+</td><td>0</td><td>AGAGCTGCGTCTCCCTGGAA</td><td>AGG</td><td>||||||||[||||||||||||]PAM</td><td>NA</td><td>Intergenic</td><td>NA</td><td>NA</td></tr>
<tr><td></td><td></td><td></td><td></td><td></td><td></td><td></td><td></td><td></td><td></td><td></td><td></td></tr>
<tr><td>T2619</td><td>CTGCGTCTCCCTGGAAAGGTCGG</td><td>1000</td><td></td><td></td><td></td><td></td><td></td><td></td><td></td><td></td><td></td></tr>
<tr><td>Oligo fwd</td><td>CACCCTGCGTCTCCCTGGAAAGGT</td><td></td><td></td><td></td><td></td><td></td><td></td><td></td><td></td><td></td><td></td></tr>
<tr><td>Oligo rev</td><td>AAACACCTTTCCAGGGAGACGCAG</td><td></td><td></td><td></td><td></td><td></td><td></td><td></td><td></td><td></td><td></td></tr>
<tr><td>Chromosome</td><td>start</td><td>end</td><td>strand</td><td>MM</td><td>target_seq</td><td>PAM</td><td>alignment</td><td>distance</td><td>position</td><td>gene name</td><td>gene id</td></tr>
<tr><td>GU305914.1</td><td>41233</td><td>41255</td><td>+</td><td>0</td><td>CTGCGTCTCCCTGGAAAGGT</td><td>CGG</td><td>||||||||[||||||||||||]PAM</td><td>NA</td><td>Intergenic</td><td>NA</td><td>NA</td></tr>
<tr><td></td><td></td><td></td><td></td><td></td><td></td><td></td><td></td><td></td><td></td><td></td><td></td></tr>
<tr><td>T2620</td><td>CGACATTCCGCAATCACACGTGG</td><td>1000</td><td></td><td></td><td></td><td></td><td></td><td></td><td></td><td></td><td></td></tr>
<tr><td>Oligo fwd</td><td>CACCCGACATTCCGCAATCACACG</td><td></td><td></td><td></td><td></td><td></td><td></td><td></td><td></td><td></td><td></td></tr>
<tr><td>Oligo rev</td><td>AAACCGTGTGATTGCGGAATGTCG</td><td></td><td></td><td></td><td></td><td></td><td></td><td></td><td></td><td></td><td></td></tr>
<tr><td>Chromosome</td><td>start</td><td>end</td><td>strand</td><td>MM</td><td>target_seq</td><td>PAM</td><td>alignment</td><td>distance</td><td>position</td><td>gene name</td><td>gene id</td></tr>
<tr><td>GU305914.1</td><td>41261</td><td>41283</td><td>+</td><td>0</td><td>CGACATTCCGCAATCACACG</td><td>TGG</td><td>||||||||[||||||||||||]PAM</td><td>NA</td><td>Intergenic</td><td>NA</td><td>NA</td></tr>
<tr><td></td><td></td><td></td><td></td><td></td><td></td><td></td><td></td><td></td><td></td><td></td><td></td></tr>
<tr><td>T2621</td><td>GACATTCCGCAATCACACGTGGG</td><td>1000</td><td></td><td></td><td></td><td></td><td></td><td></td><td></td><td></td><td></td></tr>
<tr><td>Oligo fwd</td><td>CACCGACATTCCGCAATCACACGT</td><td></td><td></td><td></td><td></td><td></td><td></td><td></td><td></td><td></td><td></td></tr>
<tr><td>Oligo rev</td><td>AAACACGTGTGATTGCGGAATGTC</td><td></td><td></td><td></td><td></td><td></td><td></td><td></td><td></td><td></td><td></td></tr>
<tr><td>Chromosome</td><td>start</td><td>end</td><td>strand</td><td>MM</td><td>target_seq</td><td>PAM</td><td>alignment</td><td>distance</td><td>position</td><td>gene name</td><td>gene id</td></tr>
<tr><td>GU305914.1</td><td>41262</td><td>41284</td><td>+</td><td>0</td><td>GACATTCCGCAATCACACGT</td><td>GGG</td><td>||||||||[||||||||||||]PAM</td><td>NA</td><td>Intergenic</td><td>NA</td><td>NA</td></tr>
<tr><td></td><td></td><td></td><td></td><td></td><td></td><td></td><td></td><td></td><td></td><td></td><td></td></tr>
<tr><td>T2622</td><td>ATTCCGCAATCACACGTGGGTGG</td><td>1000</td><td></td><td></td><td></td><td></td><td></td><td></td><td></td><td></td><td></td></tr>
<tr><td>Oligo fwd</td><td>CACCATTCCGCAATCACACGTGGG</td><td></td><td></td><td></td><td></td><td></td><td></td><td></td><td></td><td></td><td></td></tr>
<tr><td>Oligo rev</td><td>AAACCCCACGTGTGATTGCGGAAT</td><td></td><td></td><td></td><td></td><td></td><td></td><td></td><td></td><td></td><td></td></tr>
<tr><td>Chromosome</td><td>start</td><td>end</td><td>strand</td><td>MM</td><td>target_seq</td><td>PAM</td><td>alignment</td><td>distance</td><td>position</td><td>gene name</td><td>gene id</td></tr>
<tr><td>GU305914.1</td><td>41265</td><td>41287</td><td>+</td><td>0</td><td>ATTCCGCAATCACACGTGGG</td><td>TGG</td><td>||||||||[||||||||||||]PAM</td><td>NA</td><td>Intergenic</td><td>NA</td><td>NA</td></tr>
<tr><td></td><td></td><td></td><td></td><td></td><td></td><td></td><td></td><td></td><td></td><td></td><td></td></tr>
<tr><td>T2623</td><td>TGGGTGGCGACAGAGATTAATGG</td><td>1000</td><td></td><td></td><td></td><td></td><td></td><td></td><td></td><td></td><td></td></tr>
<tr><td>Oligo fwd</td><td>CACCTGGGTGGCGACAGAGATTAA</td><td></td><td></td><td></td><td></td><td></td><td></td><td></td><td></td><td></td><td></td></tr>
</table>

<table>
<tr><td>Oligo rev</td><td>AAACTTAATCTCTGTCGCCACCCA</td><td></td><td></td><td></td><td></td><td></td><td></td><td></td><td></td><td></td><td></td></tr>
<tr><td>Chromosome</td><td>start</td><td>end</td><td>strand</td><td>MM</td><td>target_seq</td><td>PAM</td><td>alignment</td><td>distance</td><td>position</td><td>gene name</td><td>gene id</td></tr>
<tr><td>GU305914.1</td><td>41281</td><td>41303</td><td>+</td><td>0</td><td>TGGGTGGCGACAGAGATTAA</td><td>TGG</td><td>||||||||[||||||||||||]PAM</td><td>NA</td><td>Intergenic</td><td>NA</td><td>NA</td></tr>
<tr><td></td><td></td><td></td><td></td><td></td><td></td><td></td><td></td><td></td><td></td><td></td><td></td></tr>
<tr><td>T2624</td><td>AAACTTCTTAACCTTTCCAACGG</td><td>1000</td><td></td><td></td><td></td><td></td><td></td><td></td><td></td><td></td><td></td></tr>
<tr><td>Oligo fwd</td><td>CACCAAACTTCTTAACCTTTCCAA</td><td></td><td></td><td></td><td></td><td></td><td></td><td></td><td></td><td></td><td></td></tr>
<tr><td>Oligo rev</td><td>AAACTTGGAAAGGTTAAGAAGTTT</td><td></td><td></td><td></td><td></td><td></td><td></td><td></td><td></td><td></td><td></td></tr>
<tr><td>Chromosome</td><td>start</td><td>end</td><td>strand</td><td>MM</td><td>target_seq</td><td>PAM</td><td>alignment</td><td>distance</td><td>position</td><td>gene name</td><td>gene id</td></tr>
<tr><td>GU305914.1</td><td>41329</td><td>41351</td><td>+</td><td>0</td><td>AAACTTCTTAACCTTTCCAA</td><td>CGG</td><td>||||||||[||||||||||||]PAM</td><td>NA</td><td>Intergenic</td><td>NA</td><td>NA</td></tr>
<tr><td></td><td></td><td></td><td></td><td></td><td></td><td></td><td></td><td></td><td></td><td></td><td></td></tr>
<tr><td>T2625</td><td>AACCTTTCCAACGGAGCTCTTGG</td><td>1000</td><td></td><td></td><td></td><td></td><td></td><td></td><td></td><td></td><td></td></tr>
<tr><td>Oligo fwd</td><td>CACCAACCTTTCCAACGGAGCTCT</td><td></td><td></td><td></td><td></td><td></td><td></td><td></td><td></td><td></td><td></td></tr>
<tr><td>Oligo rev</td><td>AAACAGAGCTCCGTTGGAAAGGTT</td><td></td><td></td><td></td><td></td><td></td><td></td><td></td><td></td><td></td><td></td></tr>
<tr><td>Chromosome</td><td>start</td><td>end</td><td>strand</td><td>MM</td><td>target_seq</td><td>PAM</td><td>alignment</td><td>distance</td><td>position</td><td>gene name</td><td>gene id</td></tr>
<tr><td>GU305914.1</td><td>41338</td><td>41360</td><td>+</td><td>0</td><td>AACCTTTCCAACGGAGCTCT</td><td>TGG</td><td>||||||||[||||||||||||]PAM</td><td>NA</td><td>Intergenic</td><td>NA</td><td>NA</td></tr>
<tr><td></td><td></td><td></td><td></td><td></td><td></td><td></td><td></td><td></td><td></td><td></td><td></td></tr>
<tr><td>T2626</td><td>TGGAGAGCCTGACGAACCCGAGG</td><td>1000</td><td></td><td></td><td></td><td></td><td></td><td></td><td></td><td></td><td></td></tr>
<tr><td>Oligo fwd</td><td>CACCTGGAGAGCCTGACGAACCCG</td><td></td><td></td><td></td><td></td><td></td><td></td><td></td><td></td><td></td><td></td></tr>
<tr><td>Oligo rev</td><td>AAACCGGGTTCGTCAGGCTCTCCA</td><td></td><td></td><td></td><td></td><td></td><td></td><td></td><td></td><td></td><td></td></tr>
<tr><td>Chromosome</td><td>start</td><td>end</td><td>strand</td><td>MM</td><td>target_seq</td><td>PAM</td><td>alignment</td><td>distance</td><td>position</td><td>gene name</td><td>gene id</td></tr>
<tr><td>GU305914.1</td><td>41358</td><td>41380</td><td>+</td><td>0</td><td>TGGAGAGCCTGACGAACCCG</td><td>AGG</td><td>||||||||[||||||||||||]PAM</td><td>NA</td><td>Intergenic</td><td>NA</td><td>NA</td></tr>
<tr><td></td><td></td><td></td><td></td><td></td><td></td><td></td><td></td><td></td><td></td><td></td><td></td></tr>
<tr><td>T2627</td><td>GGAGAGCCTGACGAACCCGAGGG</td><td>1000</td><td></td><td></td><td></td><td></td><td></td><td></td><td></td><td></td><td></td></tr>
<tr><td>Oligo fwd</td><td>CACCGGAGAGCCTGACGAACCCGA</td><td></td><td></td><td></td><td></td><td></td><td></td><td></td><td></td><td></td><td></td></tr>
<tr><td>Oligo rev</td><td>AAACTCGGGTTCGTCAGGCTCTCC</td><td></td><td></td><td></td><td></td><td></td><td></td><td></td><td></td><td></td><td></td></tr>
<tr><td>Chromosome</td><td>start</td><td>end</td><td>strand</td><td>MM</td><td>target_seq</td><td>PAM</td><td>alignment</td><td>distance</td><td>position</td><td>gene name</td><td>gene id</td></tr>
<tr><td>GU305914.1</td><td>41359</td><td>41381</td><td>+</td><td>0</td><td>GGAGAGCCTGACGAACCCGA</td><td>GGG</td><td>||||||||[||||||||||||]PAM</td><td>NA</td><td>Intergenic</td><td>NA</td><td>NA</td></tr>
<tr><td></td><td></td><td></td><td></td><td></td><td></td><td></td><td></td><td></td><td></td><td></td><td></td></tr>
<tr><td>T2628</td><td>GAGCCTGACGAACCCGAGGGTGG</td><td>1000</td><td></td><td></td><td></td><td></td><td></td><td></td><td></td><td></td><td></td></tr>
<tr><td>Oligo fwd</td><td>CACCGAGCCTGACGAACCCGAGGG</td><td></td><td></td><td></td><td></td><td></td><td></td><td></td><td></td><td></td><td></td></tr>
<tr><td>Oligo rev</td><td>AAACCCCTCGGGTTCGTCAGGCTC</td><td></td><td></td><td></td><td></td><td></td><td></td><td></td><td></td><td></td><td></td></tr>
<tr><td>Chromosome</td><td>start</td><td>end</td><td>strand</td><td>MM</td><td>target_seq</td><td>PAM</td><td>alignment</td><td>distance</td><td>position</td><td>gene name</td><td>gene id</td></tr>
<tr><td>GU305914.1</td><td>41362</td><td>41384</td><td>+</td><td>0</td><td>GAGCCTGACGAACCCGAGGG</td><td>TGG</td><td>||||||||[||||||||||||]PAM</td><td>NA</td><td>Intergenic</td><td>NA</td><td>NA</td></tr>
<tr><td></td><td></td><td></td><td></td><td></td><td></td><td></td><td></td><td></td><td></td><td></td><td></td></tr>
<tr><td>T2629</td><td>CCCGAGGGTGGCCATCAGTTCGG</td><td>1000</td><td></td><td></td><td></td><td></td><td></td><td></td><td></td><td></td><td></td></tr>
<tr><td>Oligo fwd</td><td>CACCCCCGAGGGTGGCCATCAGTT</td><td></td><td></td><td></td><td></td><td></td><td></td><td></td><td></td><td></td><td></td></tr>
<tr><td>Oligo rev</td><td>AAACAACTGATGGCCACCCTCGGG</td><td></td><td></td><td></td><td></td><td></td><td></td><td></td><td></td><td></td><td></td></tr>
<tr><td>Chromosome</td><td>start</td><td>end</td><td>strand</td><td>MM</td><td>target_seq</td><td>PAM</td><td>alignment</td><td>distance</td><td>position</td><td>gene name</td><td>gene id</td></tr>
<tr><td>GU305914.1</td><td>41374</td><td>41396</td><td>+</td><td>0</td><td>CCCGAGGGTGGCCATCAGTT</td><td>CGG</td><td>||||||||[||||||||||||]PAM</td><td>NA</td><td>Intergenic</td><td>NA</td><td>NA</td></tr>
<tr><td></td><td></td><td></td><td></td><td></td><td></td><td></td><td></td><td></td><td></td><td></td><td></td></tr>
<tr><td>T2630</td><td>TCCCGCCGCGCGCCCTCGAGCGG</td><td>1000</td><td></td><td></td><td></td><td></td><td></td><td></td><td></td><td></td><td></td></tr>
<tr><td>Oligo fwd</td><td>CACCTCCCGCCGCGCGCCCTCGAG</td><td></td><td></td><td></td><td></td><td></td><td></td><td></td><td></td><td></td><td></td></tr>
<tr><td>Oligo rev</td><td>AAACCTCGAGGGCGCGCGGCGGGA</td><td></td><td></td><td></td><td></td><td></td><td></td><td></td><td></td><td></td><td></td></tr>
<tr><td>Chromosome</td><td>start</td><td>end</td><td>strand</td><td>MM</td><td>target_seq</td><td>PAM</td><td>alignment</td><td>distance</td><td>position</td><td>gene name</td><td>gene id</td></tr>
<tr><td>GU305914.1</td><td>41413</td><td>41435</td><td>+</td><td>0</td><td>TCCCGCCGCGCGCCCTCGAG</td><td>CGG</td><td>||||||||[||||||||||||]PAM</td><td>NA</td><td>Intergenic</td><td>NA</td><td>NA</td></tr>
<tr><td>GU305914.1</td><td>138665</td><td>138687</td><td>+</td><td>4</td><td>AACCGCCGTGCGTCCTCGAG</td><td>GGG</td><td>--||||||[-|||-|||||||]PAM</td><td>NA</td><td>Intergenic</td><td>NA</td><td>NA</td></tr>
<tr><td></td><td></td><td></td><td></td><td></td><td></td><td></td><td></td><td></td><td></td><td></td><td></td></tr>
<tr><td>T2631</td><td>TCGTTAAATACAGCGACTCGCGG</td><td>1000</td><td></td><td></td><td></td><td></td><td></td><td></td><td></td><td></td><td></td></tr>
<tr><td>Oligo fwd</td><td>CACCTCGTTAAATACAGCGACTCG</td><td></td><td></td><td></td><td></td><td></td><td></td><td></td><td></td><td></td><td></td></tr>
<tr><td>Oligo rev</td><td>AAACCGAGTCGCTGTATTTAACGA</td><td></td><td></td><td></td><td></td><td></td><td></td><td></td><td></td><td></td><td></td></tr>
<tr><td>Chromosome</td><td>start</td><td>end</td><td>strand</td><td>MM</td><td>target_seq</td><td>PAM</td><td>alignment</td><td>distance</td><td>position</td><td>gene name</td><td>gene id</td></tr>
<tr><td>GU305914.1</td><td>41439</td><td>41461</td><td>+</td><td>0</td><td>TCGTTAAATACAGCGACTCG</td><td>CGG</td><td>||||||||[||||||||||||]PAM</td><td>NA</td><td>Intergenic</td><td>NA</td><td>NA</td></tr>
<tr><td></td><td></td><td></td><td></td><td></td><td></td><td></td><td></td><td></td><td></td><td></td><td></td></tr>
<tr><td>T2632</td><td>AGCGACTCGCGGAAGTTCTCCGG</td><td>1000</td><td></td><td></td><td></td><td></td><td></td><td></td><td></td><td></td><td></td></tr>
<tr><td>Oligo fwd</td><td>CACCAGCGACTCGCGGAAGTTCTC</td><td></td><td></td><td></td><td></td><td></td><td></td><td></td><td></td><td></td><td></td></tr>
<tr><td>Oligo rev</td><td>AAACGAGAACTTCCGCGAGTCGCT</td><td></td><td></td><td></td><td></td><td></td><td></td><td></td><td></td><td></td><td></td></tr>
<tr><td>Chromosome</td><td>start</td><td>end</td><td>strand</td><td>MM</td><td>target_seq</td><td>PAM</td><td>alignment</td><td>distance</td><td>position</td><td>gene name</td><td>gene id</td></tr>
<tr><td>GU305914.1</td><td>41450</td><td>41472</td><td>+</td><td>0</td><td>AGCGACTCGCGGAAGTTCTC</td><td>CGG</td><td>||||||||[||||||||||||]PAM</td><td>NA</td><td>Intergenic</td><td>NA</td><td>NA</td></tr>
<tr><td></td><td></td><td></td><td></td><td></td><td></td><td></td><td></td><td></td><td></td><td></td><td></td></tr>
<tr><td>T2633</td><td>GGAAGTTCTCCGGCGACGACCGG</td><td>1000</td><td></td><td></td><td></td><td></td><td></td><td></td><td></td><td></td><td></td></tr>
<tr><td>Oligo fwd</td><td>CACCGGAAGTTCTCCGGCGACGAC</td><td></td><td></td><td></td><td></td><td></td><td></td><td></td><td></td><td></td><td></td></tr>
<tr><td>Oligo rev</td><td>AAACGTCGTCGCCGGAGAACTTCC</td><td></td><td></td><td></td><td></td><td></td><td></td><td></td><td></td><td></td><td></td></tr>
<tr><td>Chromosome</td><td>start</td><td>end</td><td>strand</td><td>MM</td><td>target_seq</td><td>PAM</td><td>alignment</td><td>distance</td><td>position</td><td>gene name</td><td>gene id</td></tr>
<tr><td>GU305914.1</td><td>41460</td><td>41482</td><td>+</td><td>0</td><td>GGAAGTTCTCCGGCGACGAC</td><td>CGG</td><td>||||||||[||||||||||||]PAM</td><td>NA</td><td>Intergenic</td><td>NA</td><td>NA</td></tr>
<tr><td></td><td></td><td></td><td></td><td></td><td></td><td></td><td></td><td></td><td></td><td></td><td></td></tr>
<tr><td>T2634</td><td>CGGCGACGACCGGCTCGAGATGG</td><td>1000</td><td></td><td></td><td></td><td></td><td></td><td></td><td></td><td></td><td></td></tr>
<tr><td>Oligo fwd</td><td>CACCCGGCGACGACCGGCTCGAGA</td><td></td><td></td><td></td><td></td><td></td><td></td><td></td><td></td><td></td><td></td></tr>
<tr><td>Oligo rev</td><td>AAACTCTCGAGCCGGTCGTCGCCG</td><td></td><td></td><td></td><td></td><td></td><td></td><td></td><td></td><td></td><td></td></tr>
<tr><td>Chromosome</td><td>start</td><td>end</td><td>strand</td><td>MM</td><td>target_seq</td><td>PAM</td><td>alignment</td><td>distance</td><td>position</td><td>gene name</td><td>gene id</td></tr>
<tr><td>GU305914.1</td><td>41470</td><td>41492</td><td>+</td><td>0</td><td>CGGCGACGACCGGCTCGAGA</td><td>TGG</td><td>||||||||[||||||||||||]PAM</td><td>NA</td><td>Intergenic</td><td>NA</td><td>NA</td></tr>
<tr><td></td><td></td><td></td><td></td><td></td><td></td><td></td><td></td><td></td><td></td><td></td><td></td></tr>
<tr><td>T2635</td><td>GCTCGAGATGGACGTCCTTTTGG</td><td>1000</td><td></td><td></td><td></td><td></td><td></td><td></td><td></td><td></td><td></td></tr>
<tr><td>Oligo fwd</td><td>CACCGCTCGAGATGGACGTCCTTT</td><td></td><td></td><td></td><td></td><td></td><td></td><td></td><td></td><td></td><td></td></tr>
<tr><td>Oligo rev</td><td>AAACAAAGGACGTCCATCTCGAGC</td><td></td><td></td><td></td><td></td><td></td><td></td><td></td><td></td><td></td><td></td></tr>
<tr><td>Chromosome</td><td>start</td><td>end</td><td>strand</td><td>MM</td><td>target_seq</td><td>PAM</td><td>alignment</td><td>distance</td><td>position</td><td>gene name</td><td>gene id</td></tr>
<tr><td>GU305914.1</td><td>41482</td><td>41504</td><td>+</td><td>0</td><td>GCTCGAGATGGACGTCCTTT</td><td>TGG</td><td>||||||||[||||||||||||]PAM</td><td>NA</td><td>Intergenic</td><td>NA</td><td>NA</td></tr>
<tr><td></td><td></td><td></td><td></td><td></td><td></td><td></td><td></td><td></td><td></td><td></td><td></td></tr>
<tr><td>T2636</td><td>CTCGAGATGGACGTCCTTTTGGG</td><td>1000</td><td></td><td></td><td></td><td></td><td></td><td></td><td></td><td></td><td></td></tr>
<tr><td>Oligo fwd</td><td>CACCCTCGAGATGGACGTCCTTTT</td><td></td><td></td><td></td><td></td><td></td><td></td><td></td><td></td><td></td><td></td></tr>
<tr><td>Oligo rev</td><td>AAACAAAAGGACGTCCATCTCGAG</td><td></td><td></td><td></td><td></td><td></td><td></td><td></td><td></td><td></td><td></td></tr>
<tr><td>Chromosome</td><td>start</td><td>end</td><td>strand</td><td>MM</td><td>target_seq</td><td>PAM</td><td>alignment</td><td>distance</td><td>position</td><td>gene name</td><td>gene id</td></tr>
<tr><td>GU305914.1</td><td>41483</td><td>41505</td><td>+</td><td>0</td><td>CTCGAGATGGACGTCCTTTT</td><td>GGG</td><td>||||||||[||||||||||||]PAM</td><td>NA</td><td>Intergenic</td><td>NA</td><td>NA</td></tr>
<tr><td></td><td></td><td></td><td></td><td></td><td></td><td></td><td></td><td></td><td></td><td></td><td></td></tr>
<tr><td>T2637</td><td>AGATGGACGTCCTTTTGGGCCGG</td><td>1000</td><td></td><td></td><td></td><td></td><td></td><td></td><td></td><td></td><td></td></tr>
<tr><td>Oligo fwd</td><td>CACCAGATGGACGTCCTTTTGGGC</td><td></td><td></td><td></td><td></td><td></td><td></td><td></td><td></td><td></td><td></td></tr>
<tr><td>Oligo rev</td><td>AAACGCCCAAAAGGACGTCCATCT</td><td></td><td></td><td></td><td></td><td></td><td></td><td></td><td></td><td></td><td></td></tr>
<tr><td>Chromosome</td><td>start</td><td>end</td><td>strand</td><td>MM</td><td>target_seq</td><td>PAM</td><td>alignment</td><td>distance</td><td>position</td><td>gene name</td><td>gene id</td></tr>
<tr><td>GU305914.1</td><td>41487</td><td>41509</td><td>+</td><td>0</td><td>AGATGGACGTCCTTTTGGGC</td><td>CGG</td><td>||||||||[||||||||||||]PAM</td><td>NA</td><td>Intergenic</td><td>NA</td><td>NA</td></tr>
</table>

<table>
<tr><td>T2638</td><td>GATGGACGTCCTTTTGGGCCGGG</td><td></td><td>1000</td></tr>
<tr><td>Oligo fwd</td><td>CACCGATGGACGTCCTTTTGGGCC</td></tr>
<tr><td>Oligo rev</td><td>AAACGGCCCAAAAGGACGTCCATC</td></tr>
<tr><td>Chromosome</td><td>start</td><td></td><td>end</td><td>strand</td><td>MM</td><td>target_seq</td><td>PAM</td><td>alignment</td><td>distance</td><td>position</td><td>gene name</td><td>gene id</td></tr>
<tr><td>GU305914.1</td><td></td><td>41488</td><td>41510</td><td>+</td><td>0</td><td>GATGGACGTCCTTTTGGGCC</td><td>GGG</td><td>||||||||[||||||||||||]PAM</td><td>NA</td><td>Intergenic</td><td>NA</td><td>NA</td></tr>
<tr><td>T2639</td><td>TTTGGGCCGGGACGAGACTATGG</td><td></td><td>1000</td></tr>
<tr><td>Oligo fwd</td><td>CACCTTTGGGCCGGGACGAGACTA</td></tr>
<tr><td>Oligo rev</td><td>AAACTAGTCTCGTCCCGGCCCAAA</td></tr>
<tr><td>Chromosome</td><td>start</td><td></td><td>end</td><td>strand</td><td>MM</td><td>target_seq</td><td>PAM</td><td>alignment</td><td>distance</td><td>position</td><td>gene name</td><td>gene id</td></tr>
<tr><td>GU305914.1</td><td></td><td>41500</td><td>41522</td><td>+</td><td>0</td><td>TTTGGGCCGGGACGAGACTA</td><td>TGG</td><td>||||||||[||||||||||||]PAM</td><td>NA</td><td>Intergenic</td><td>NA</td><td>NA</td></tr>
<tr><td>T2640</td><td>CTATGGTAAGCAGCGTGCGCCGG</td><td></td><td>1000</td></tr>
<tr><td>Oligo fwd</td><td>CACCCTATGGTAAGCAGCGTGCGC</td></tr>
<tr><td>Oligo rev</td><td>AAACGCGCACGCTGCTTACCATAG</td></tr>
<tr><td>Chromosome</td><td>start</td><td></td><td>end</td><td>strand</td><td>MM</td><td>target_seq</td><td>PAM</td><td>alignment</td><td>distance</td><td>position</td><td>gene name</td><td>gene id</td></tr>
<tr><td>GU305914.1</td><td></td><td>41517</td><td>41539</td><td>+</td><td>0</td><td>CTATGGTAAGCAGCGTGCGC</td><td>CGG</td><td>||||||||[||||||||||||]PAM</td><td>NA</td><td>Intergenic</td><td>NA</td><td>NA</td></tr>
<tr><td>T2641</td><td>TATGGTAAGCAGCGTGCGCCGGG</td><td></td><td>1000</td></tr>
<tr><td>Oligo fwd</td><td>CACCTATGGTAAGCAGCGTGCGCC</td></tr>
<tr><td>Oligo rev</td><td>AAACGGCGCACGCTGCTTACCATA</td></tr>
<tr><td>Chromosome</td><td>start</td><td></td><td>end</td><td>strand</td><td>MM</td><td>target_seq</td><td>PAM</td><td>alignment</td><td>distance</td><td>position</td><td>gene name</td><td>gene id</td></tr>
<tr><td>GU305914.1</td><td></td><td>41518</td><td>41540</td><td>+</td><td>0</td><td>TATGGTAAGCAGCGTGCGCC</td><td>GGG</td><td>||||||||[||||||||||||]PAM</td><td>NA</td><td>Intergenic</td><td>NA</td><td>NA</td></tr>
<tr><td>T2642</td><td>ATGGTAAGCAGCGTGCGCCGGGG</td><td></td><td>1000</td></tr>
<tr><td>Oligo fwd</td><td>CACCATGGTAAGCAGCGTGCGCCG</td></tr>
<tr><td>Oligo rev</td><td>AAACCGGCGCACGCTGCTTACCAT</td></tr>
<tr><td>Chromosome</td><td>start</td><td></td><td>end</td><td>strand</td><td>MM</td><td>target_seq</td><td>PAM</td><td>alignment</td><td>distance</td><td>position</td><td>gene name</td><td>gene id</td></tr>
<tr><td>GU305914.1</td><td></td><td>41519</td><td>41541</td><td>+</td><td>0</td><td>ATGGTAAGCAGCGTGCGCCG</td><td>GGG</td><td>||||||||[||||||||||||]PAM</td><td>NA</td><td>Intergenic</td><td>NA</td><td>NA</td></tr>
<tr><td>T2643</td><td>AGCAGCGTGCGCCGGGGCGTCGG</td><td></td><td>1000</td></tr>
<tr><td>Oligo fwd</td><td>CACCAGCAGCGTGCGCCGGGGCGT</td></tr>
<tr><td>Oligo rev</td><td>AAACACGCCCCGGCGCACGCTGCT</td></tr>
<tr><td>Chromosome</td><td>start</td><td></td><td>end</td><td>strand</td><td>MM</td><td>target_seq</td><td>PAM</td><td>alignment</td><td>distance</td><td>position</td><td>gene name</td><td>gene id</td></tr>
<tr><td>GU305914.1</td><td></td><td>41525</td><td>41547</td><td>+</td><td>0</td><td>AGCAGCGTGCGCCGGGGCGT</td><td>CGG</td><td>||||||||[||||||||||||]PAM</td><td>NA</td><td>Intergenic</td><td>NA</td><td>NA</td></tr>
<tr><td>T2644</td><td>GCAGCGTGCGCCGGGGCGTCGGG</td><td></td><td>1000</td></tr>
<tr><td>Oligo fwd</td><td>CACCGCAGCGTGCGCCGGGGCGTC</td></tr>
<tr><td>Oligo rev</td><td>AAACGACGCCCCGGCGCACGCTGC</td></tr>
<tr><td>Chromosome</td><td>start</td><td></td><td>end</td><td>strand</td><td>MM</td><td>target_seq</td><td>PAM</td><td>alignment</td><td>distance</td><td>position</td><td>gene name</td><td>gene id</td></tr>
<tr><td>GU305914.1</td><td></td><td>41526</td><td>41548</td><td>+</td><td>0</td><td>GCAGCGTGCGCCGGGGCGTC</td><td>GGG</td><td>||||||||[||||||||||||]PAM</td><td>NA</td><td>Intergenic</td><td>NA</td><td>NA</td></tr>
<tr><td>T2645</td><td>CGTGCGCCGGGGCGTCGGGACGG</td><td></td><td>1000</td></tr>
<tr><td>Oligo fwd</td><td>CACCCGTGCGCCGGGGCGTCGGGA</td></tr>
<tr><td>Oligo rev</td><td>AAACTCCCGACGCCCCGGCGCACG</td></tr>
<tr><td>Chromosome</td><td>start</td><td></td><td>end</td><td>strand</td><td>MM</td><td>target_seq</td><td>PAM</td><td>alignment</td><td>distance</td><td>position</td><td>gene name</td><td>gene id</td></tr>
<tr><td>GU305914.1</td><td></td><td>41530</td><td>41552</td><td>+</td><td>0</td><td>CGTGCGCCGGGGCGTCGGGA</td><td>CGG</td><td>||||||||[||||||||||||]PAM</td><td>NA</td><td>Intergenic</td><td>NA</td><td>NA</td></tr>
<tr><td>T2646</td><td>GTGCGCCGGGGCGTCGGGACGGG</td><td></td><td>1000</td></tr>
<tr><td>Oligo fwd</td><td>CACCGTGCGCCGGGGCGTCGGGAC</td></tr>
<tr><td>Oligo rev</td><td>AAACGTCCCGACGCCCCGGCGCAC</td></tr>
<tr><td>Chromosome</td><td>start</td><td></td><td>end</td><td>strand</td><td>MM</td><td>target_seq</td><td>PAM</td><td>alignment</td><td>distance</td><td>position</td><td>gene name</td><td>gene id</td></tr>
<tr><td>GU305914.1</td><td></td><td>41531</td><td>41553</td><td>+</td><td>0</td><td>GTGCGCCGGGGCGTCGGGAC</td><td>GGG</td><td>||||||||[||||||||||||]PAM</td><td>NA</td><td>Intergenic</td><td>NA</td><td>NA</td></tr>
<tr><td>T2647</td><td>TGCGCCGGGGCGTCGGGACGGGG</td><td></td><td>1000</td></tr>
<tr><td>Oligo fwd</td><td>CACCTGCGCCGGGGCGTCGGGACG</td></tr>
<tr><td>Oligo rev</td><td>AAACCGTCCCGACGCCCCGGCGCA</td></tr>
<tr><td>Chromosome</td><td>start</td><td></td><td>end</td><td>strand</td><td>MM</td><td>target_seq</td><td>PAM</td><td>alignment</td><td>distance</td><td>position</td><td>gene name</td><td>gene id</td></tr>
<tr><td>GU305914.1</td><td></td><td>41532</td><td>41554</td><td>+</td><td>0</td><td>TGCGCCGGGGCGTCGGGACG</td><td>GGG</td><td>||||||||[||||||||||||]PAM</td><td>NA</td><td>Intergenic</td><td>NA</td><td>NA</td></tr>
<tr><td>T2648</td><td>GCGTCGGGACGGGGAGCGCGCGG</td><td></td><td>1000</td></tr>
<tr><td>Oligo fwd</td><td>CACCGCGTCGGGACGGGGAGCGCG</td></tr>
<tr><td>Oligo rev</td><td>AAACCGCGCTCCCCGTCCCGACGC</td></tr>
<tr><td>Chromosome</td><td>start</td><td></td><td>end</td><td>strand</td><td>MM</td><td>target_seq</td><td>PAM</td><td>alignment</td><td>distance</td><td>position</td><td>gene name</td><td>gene id</td></tr>
<tr><td>GU305914.1</td><td></td><td>41541</td><td>41563</td><td>+</td><td>0</td><td>GCGTCGGGACGGGGAGCGCG</td><td>CGG</td><td>||||||||[||||||||||||]PAM</td><td>NA</td><td>Intergenic</td><td>NA</td><td>NA</td></tr>
<tr><td>T2649</td><td>CGTCGGGACGGGGAGCGCGCGGG</td><td></td><td>1000</td></tr>
<tr><td>Oligo fwd</td><td>CACCCGTCGGGACGGGGAGCGCGC</td></tr>
<tr><td>Oligo rev</td><td>AAACGCGCGCTCCCCGTCCCGACG</td></tr>
<tr><td>Chromosome</td><td>start</td><td></td><td>end</td><td>strand</td><td>MM</td><td>target_seq</td><td>PAM</td><td>alignment</td><td>distance</td><td>position</td><td>gene name</td><td>gene id</td></tr>
<tr><td>GU305914.1</td><td></td><td>41542</td><td>41564</td><td>+</td><td>0</td><td>CGTCGGGACGGGGAGCGCGC</td><td>GGG</td><td>||||||||[||||||||||||]PAM</td><td>NA</td><td>Intergenic</td><td>NA</td><td>NA</td></tr>
<tr><td>T2650</td><td>CGGGACGGGGAGCGCGCGGGCGG</td><td></td><td>1000</td></tr>
<tr><td>Oligo fwd</td><td>CACCCGGGACGGGGAGCGCGCGGG</td></tr>
<tr><td>Oligo rev</td><td>AAACCCCGCGCGCTCCCCGTCCCG</td></tr>
<tr><td>Chromosome</td><td>start</td><td></td><td>end</td><td>strand</td><td>MM</td><td>target_seq</td><td>PAM</td><td>alignment</td><td>distance</td><td>position</td><td>gene name</td><td>gene id</td></tr>
<tr><td>GU305914.1</td><td></td><td>41545</td><td>41567</td><td>+</td><td>0</td><td>CGGGACGGGGAGCGCGCGGG</td><td>CGG</td><td>||||||||[||||||||||||]PAM</td><td>NA</td><td>Intergenic</td><td>NA</td><td>NA</td></tr>
<tr><td>GU305914.1</td><td></td><td>58965</td><td>58987</td><td>+</td><td>4</td><td>CTGGACGATGAGCGCGGGGG</td><td>TGG</td><td>|-|||||-[-|||||||-|||]PAM</td><td>NA</td><td>Intergenic</td><td>NA</td><td>NA</td></tr>
<tr><td>T2651</td><td>GGGCGGTCCCGCCAGAGTCGCGG</td><td></td><td>1000</td></tr>
<tr><td>Oligo fwd</td><td>CACCGGGCGGTCCCGCCAGAGTCG</td></tr>
<tr><td>Oligo rev</td><td>AAACCGACTCTGGCGGGACCGCCC</td></tr>
<tr><td>Chromosome</td><td>start</td><td></td><td>end</td><td>strand</td><td>MM</td><td>target_seq</td><td>PAM</td><td>alignment</td><td>distance</td><td>position</td><td>gene name</td><td>gene id</td></tr>
<tr><td>GU305914.1</td><td></td><td>41562</td><td>41584</td><td>+</td><td>0</td><td>GGGCGGTCCCGCCAGAGTCG</td><td>CGG</td><td>||||||||[||||||||||||]PAM</td><td>NA</td><td>Intergenic</td><td>NA</td><td>NA</td></tr>
<tr><td>T2652</td><td>CGGCCTATCTGTCAAGAGAAAGG</td><td></td><td>1000</td></tr>
<tr><td>Oligo fwd</td><td>CACCCGGCCTATCTGTCAAGAGAA</td></tr>
<tr><td>Oligo rev</td><td>AAACTTCTCTTGACAGATAGGCCG</td></tr>
<tr><td>Chromosome</td><td>start</td><td></td><td>end</td><td>strand</td><td>MM</td><td>target_seq</td><td>PAM</td><td>alignment</td><td>distance</td><td>position</td><td>gene name</td><td>gene id</td></tr>
</table>

GU305914.1 41582 41604 + 0 CGGCCTATCTGTCAAGAGAA AGG ||||||||[||||||||||||]PAM NA Intergenic NA NA

T2653 CTATCTGTCAAGAGAAAGGCAGG 1000
Oligo fwd CACCCTATCTGTCAAGAGAAAGGC
Oligo rev AAACGCCTTTCTCTTGACAGATAG
Chromosome start end strand MM target_seq PAM alignment distance position gene name gene id
GU305914.1 41586 41608 + 0 CTATCTGTCAAGAGAAAGGC AGG ||||||||[||||||||||||]PAM NA Intergenic NA NA

T2654 GATTGATATCCGTCTATCTTCGG 1000
Oligo fwd CACCGATTGATATCCGTCTATCTT
Oligo rev AAACAAGATAGACGGATATCAATC
Chromosome start end strand MM target_seq PAM alignment distance position gene name gene id
GU305914.1 41614 41636 + 0 GATTGATATCCGTCTATCTT CGG ||||||||[||||||||||||]PAM NA Intergenic NA NA

T2655 ATCCGTCTATCTTCGGTGTCCGG 1000
Oligo fwd CACCATCCGTCTATCTTCGGTGTC
Oligo rev AAACGACACCGAAGATAGACGGAT
Chromosome start end strand MM target_seq PAM alignment distance position gene name gene id
GU305914.1 41621 41643 + 0 ATCCGTCTATCTTCGGTGTC CGG ||||||||[||||||||||||]PAM NA Intergenic NA NA

T2656 TATCTCGCCCTCTCGCCCGCAGG 1000
Oligo fwd CACCTATCTCGCCCTCTCGCCCGC
Oligo rev AAACGCGGGCGAGAGGGCGAGATA
Chromosome start end strand MM target_seq PAM alignment distance position gene name gene id
GU305914.1 41661 41683 + 0 TATCTCGCCCTCTCGCCCGC AGG ||||||||[||||||||||||]PAM NA Intergenic NA NA

T2657 CAACACGTGCGCCCCCTCGCTGG 1000
Oligo fwd CACCCAACACGTGCGCCCCCTCGC
Oligo rev AAACGCGAGGGGGCGCACGTGTTG
Chromosome start end strand MM target_seq PAM alignment distance position gene name gene id
GU305914.1 41709 41731 + 0 CAACACGTGCGCCCCCTCGC TGG ||||||||[||||||||||||]PAM NA Intergenic NA NA

T2658 AACACGTGCGCCCCCTCGCTGGG 1000
Oligo fwd CACCAACACGTGCGCCCCCTCGCT
Oligo rev AAACAGCGAGGGGGCGCACGTGTT
Chromosome start end strand MM target_seq PAM alignment distance position gene name gene id
GU305914.1 41710 41732 + 0 AACACGTGCGCCCCCTCGCT GGG ||||||||[||||||||||||]PAM NA Intergenic NA NA

T2659 ACACGTGCGCCCCCTCGCTGGGG 1000
Oligo fwd CACCACACGTGCGCCCCCTCGCTG
Oligo rev AAACCAGCGAGGGGGCGCACGTGT
Chromosome start end strand MM target_seq PAM alignment distance position gene name gene id
GU305914.1 41711 41733 + 0 ACACGTGCGCCCCCTCGCTG GGG ||||||||[||||||||||||]PAM NA Intergenic NA NA

T2660 CGCTGGGGCTGAGCATCGCGAGG 1000
Oligo fwd CACCCGCTGGGGCTGAGCATCGCG
Oligo rev AAACCGCGATGCTCAGCCCCAGCG
Chromosome start end strand MM target_seq PAM alignment distance position gene name gene id
GU305914.1 41726 41748 + 0 CGCTGGGGCTGAGCATCGCG AGG ||||||||[||||||||||||]PAM NA Intergenic NA NA

T2661 GCTGGGGCTGAGCATCGCGAGGG 1000
Oligo fwd CACCGCTGGGGCTGAGCATCGCGA
Oligo rev AAACTCGCGATGCTCAGCCCCAGC
Chromosome start end strand MM target_seq PAM alignment distance position gene name gene id
GU305914.1 41727 41749 + 0 GCTGGGGCTGAGCATCGCGA GGG ||||||||[||||||||||||]PAM NA Intergenic NA NA

T2662 GCTGAGCATCGCGAGGGACGCGG 1000
Oligo fwd CACCGCTGAGCATCGCGAGGGACG
Oligo rev AAACCGTCCCTCGCGATGCTCAGC
Chromosome start end strand MM target_seq PAM alignment distance position gene name gene id
GU305914.1 41733 41755 + 0 GCTGAGCATCGCGAGGGACG CGG ||||||||[||||||||||||]PAM NA Intergenic NA NA

T2663 CATCGCGAGGGACGCGGAGACGG 1000
Oligo fwd CACCCATCGCGAGGGACGCGGAGA
Oligo rev AAACTCTCCGCGTCCCTCGCGATG
Chromosome start end strand MM target_seq PAM alignment distance position gene name gene id
GU305914.1 41739 41761 + 0 CATCGCGAGGGACGCGGAGA CGG ||||||||[||||||||||||]PAM NA Intergenic NA NA

T2664 CGCGGAGACGGCCATCAACACGG 1000
Oligo fwd CACCCGCGGAGACGGCCATCAACA
Oligo rev AAACTGTTGATGGCCGTCTCCGCG
Chromosome start end strand MM target_seq PAM alignment distance position gene name gene id
GU305914.1 41751 41773 + 0 CGCGGAGACGGCCATCAACA CGG ||||||||[||||||||||||]PAM NA Intergenic NA NA

T2665 AACACGGTGATCATCCTGATCGG 1000
Oligo fwd CACCAACACGGTGATCATCCTGAT
Oligo rev AAACATCAGGATGATCACCGTGTT
Chromosome start end strand MM target_seq PAM alignment distance position gene name gene id
GU305914.1 41767 41789 + 0 AACACGGTGATCATCCTGAT CGG ||||||||[||||||||||||]PAM NA Intergenic NA NA

T2666 ACGGTGATCATCCTGATCGGCGG 1000
Oligo fwd CACCACGGTGATCATCCTGATCGG
Oligo rev AAACCCGATCAGGATGATCACCGT
Chromosome start end strand MM target_seq PAM alignment distance position gene name gene id
GU305914.1 41770 41792 + 0 ACGGTGATCATCCTGATCGG CGG ||||||||[||||||||||||]PAM NA Intergenic NA NA

T2667 CGGTGATCATCCTGATCGGCGGG 1000
Oligo fwd CACCCGGTGATCATCCTGATCGGC
Oligo rev AAACGCCGATCAGGATGATCACCG

<table>
<tr><td>Chromosome</td><td>start</td><td></td><td>end</td><td>strand</td><td>MM</td><td>target_seq</td><td>PAM</td><td>alignment</td><td>distance</td><td>position</td><td>gene name</td><td>gene id</td></tr>
<tr><td>GU305914.1</td><td></td><td>41771</td><td>41793</td><td>+</td><td>0</td><td>CGGTGATCATCCTGATCGGC</td><td>GGG</td><td>||||||||[||||||||||||]PAM</td><td>NA</td><td>Intergenic</td><td>NA</td><td>NA</td></tr>
<tr><td>T2668</td><td>CGGCGGGCCGATGAACTTCGTGG</td><td></td><td>1000</td></tr>
<tr><td>Oligo fwd</td><td>CACCCGGCGGGCCGATGAACTTCG</td></tr>
<tr><td>Oligo rev</td><td>AAACCGAAGTTCATCGGCCCGCCG</td></tr>
<tr><td>Chromosome</td><td>start</td><td></td><td>end</td><td>strand</td><td>MM</td><td>target_seq</td><td>PAM</td><td>alignment</td><td>distance</td><td>position</td><td>gene name</td><td>gene id</td></tr>
<tr><td>GU305914.1</td><td></td><td>41787</td><td>41809</td><td>+</td><td>0</td><td>CGGCGGGCCGATGAACTTCG</td><td>TGG</td><td>||||||||[||||||||||||]PAM</td><td>NA</td><td>Intergenic</td><td>NA</td><td>NA</td></tr>
<tr><td>T2669</td><td>GAACCGCATCTATCGCAGCTCGG</td><td></td><td>1000</td></tr>
<tr><td>Oligo fwd</td><td>CACCGAACCGCATCTATCGCAGCT</td></tr>
<tr><td>Oligo rev</td><td>AAACAGCTGCGATAGATGCGGTTC</td></tr>
<tr><td>Chromosome</td><td>start</td><td></td><td>end</td><td>strand</td><td>MM</td><td>target_seq</td><td>PAM</td><td>alignment</td><td>distance</td><td>position</td><td>gene name</td><td>gene id</td></tr>
<tr><td>GU305914.1</td><td></td><td>41832</td><td>41854</td><td>+</td><td>0</td><td>GAACCGCATCTATCGCAGCT</td><td>CGG</td><td>||||||||[||||||||||||]PAM</td><td>NA</td><td>Intergenic</td><td>NA</td><td>NA</td></tr>
<tr><td>T2670</td><td>GTACTTTGCGAACCTGTTGACGG</td><td></td><td>1000</td></tr>
<tr><td>Oligo fwd</td><td>CACCGTACTTTGCGAACCTGTTGA</td></tr>
<tr><td>Oligo rev</td><td>AAACTCAACAGGTTCGCAAAGTAC</td></tr>
<tr><td>Chromosome</td><td>start</td><td></td><td>end</td><td>strand</td><td>MM</td><td>target_seq</td><td>PAM</td><td>alignment</td><td>distance</td><td>position</td><td>gene name</td><td>gene id</td></tr>
<tr><td>GU305914.1</td><td></td><td>41880</td><td>41902</td><td>+</td><td>0</td><td>GTACTTTGCGAACCTGTTGA</td><td>CGG</td><td>||||||||[||||||||||||]PAM</td><td>NA</td><td>Intergenic</td><td>NA</td><td>NA</td></tr>
<tr><td>T2671</td><td>CGTTCCTGATCCTGAGCAATCGG</td><td></td><td>1000</td></tr>
<tr><td>Oligo fwd</td><td>CACCCGTTCCTGATCCTGAGCAAT</td></tr>
<tr><td>Oligo rev</td><td>AAACATTGCTCAGGATCAGGAACG</td></tr>
<tr><td>Chromosome</td><td>start</td><td></td><td>end</td><td>strand</td><td>MM</td><td>target_seq</td><td>PAM</td><td>alignment</td><td>distance</td><td>position</td><td>gene name</td><td>gene id</td></tr>
<tr><td>GU305914.1</td><td></td><td>41915</td><td>41937</td><td>+</td><td>0</td><td>CGTTCCTGATCCTGAGCAAT</td><td>CGG</td><td>||||||||[||||||||||||]PAM</td><td>NA</td><td>Intergenic</td><td>NA</td><td>NA</td></tr>
<tr><td>T2672</td><td>GTTCCTGATCCTGAGCAATCGGG</td><td></td><td>1000</td></tr>
<tr><td>Oligo fwd</td><td>CACCGTTCCTGATCCTGAGCAATC</td></tr>
<tr><td>Oligo rev</td><td>AAACGATTGCTCAGGATCAGGAAC</td></tr>
<tr><td>Chromosome</td><td>start</td><td></td><td>end</td><td>strand</td><td>MM</td><td>target_seq</td><td>PAM</td><td>alignment</td><td>distance</td><td>position</td><td>gene name</td><td>gene id</td></tr>
<tr><td>GU305914.1</td><td></td><td>41916</td><td>41938</td><td>+</td><td>0</td><td>GTTCCTGATCCTGAGCAATC</td><td>GGG</td><td>||||||||[||||||||||||]PAM</td><td>NA</td><td>Intergenic</td><td>NA</td><td>NA</td></tr>
<tr><td>T2673</td><td>TTCCTGATCCTGAGCAATCGGGG</td><td></td><td>1000</td></tr>
<tr><td>Oligo fwd</td><td>CACCTTCCTGATCCTGAGCAATCG</td></tr>
<tr><td>Oligo rev</td><td>AAACCGATTGCTCAGGATCAGGAA</td></tr>
<tr><td>Chromosome</td><td>start</td><td></td><td>end</td><td>strand</td><td>MM</td><td>target_seq</td><td>PAM</td><td>alignment</td><td>distance</td><td>position</td><td>gene name</td><td>gene id</td></tr>
<tr><td>GU305914.1</td><td></td><td>41917</td><td>41939</td><td>+</td><td>0</td><td>TTCCTGATCCTGAGCAATCG</td><td>GGG</td><td>||||||||[||||||||||||]PAM</td><td>NA</td><td>Intergenic</td><td>NA</td><td>NA</td></tr>
<tr><td>T2674</td><td>GGGTCAGATCAGCAGCTCCCCGG</td><td></td><td>1000</td></tr>
<tr><td>Oligo fwd</td><td>CACCGGGTCAGATCAGCAGCTCCC</td></tr>
<tr><td>Oligo rev</td><td>AAACGGGAGCTGCTGATCTGACCC</td></tr>
<tr><td>Chromosome</td><td>start</td><td></td><td>end</td><td>strand</td><td>MM</td><td>target_seq</td><td>PAM</td><td>alignment</td><td>distance</td><td>position</td><td>gene name</td><td>gene id</td></tr>
<tr><td>GU305914.1</td><td></td><td>41937</td><td>41959</td><td>+</td><td>0</td><td>GGGTCAGATCAGCAGCTCCC</td><td>CGG</td><td>||||||||[||||||||||||]PAM</td><td>NA</td><td>Intergenic</td><td>NA</td><td>NA</td></tr>
<tr><td>T2675</td><td>TCAGATCAGCAGCTCCCCGGAGG</td><td></td><td>1000</td></tr>
<tr><td>Oligo fwd</td><td>CACCTCAGATCAGCAGCTCCCCGG</td></tr>
<tr><td>Oligo rev</td><td>AAACCCGGGGAGCTGCTGATCTGA</td></tr>
<tr><td>Chromosome</td><td>start</td><td></td><td>end</td><td>strand</td><td>MM</td><td>target_seq</td><td>PAM</td><td>alignment</td><td>distance</td><td>position</td><td>gene name</td><td>gene id</td></tr>
<tr><td>GU305914.1</td><td></td><td>41940</td><td>41962</td><td>+</td><td>0</td><td>TCAGATCAGCAGCTCCCCGG</td><td>AGG</td><td>||||||||[||||||||||||]PAM</td><td>NA</td><td>Intergenic</td><td>NA</td><td>NA</td></tr>
<tr><td>T2676</td><td>CAGATCAGCAGCTCCCCGGAGGG</td><td></td><td>1000</td></tr>
<tr><td>Oligo fwd</td><td>CACCCAGATCAGCAGCTCCCCGGA</td></tr>
<tr><td>Oligo rev</td><td>AAACTCCGGGGAGCTGCTGATCTG</td></tr>
<tr><td>Chromosome</td><td>start</td><td></td><td>end</td><td>strand</td><td>MM</td><td>target_seq</td><td>PAM</td><td>alignment</td><td>distance</td><td>position</td><td>gene name</td><td>gene id</td></tr>
<tr><td>GU305914.1</td><td></td><td>41941</td><td>41963</td><td>+</td><td>0</td><td>CAGATCAGCAGCTCCCCGGA</td><td>GGG</td><td>||||||||[||||||||||||]PAM</td><td>NA</td><td>Intergenic</td><td>NA</td><td>NA</td></tr>
<tr><td>T2677</td><td>AGATCAGCAGCTCCCCGGAGGGG</td><td></td><td>1000</td></tr>
<tr><td>Oligo fwd</td><td>CACCAGATCAGCAGCTCCCCGGAG</td></tr>
<tr><td>Oligo rev</td><td>AAACCTCCGGGGAGCTGCTGATCT</td></tr>
<tr><td>Chromosome</td><td>start</td><td></td><td>end</td><td>strand</td><td>MM</td><td>target_seq</td><td>PAM</td><td>alignment</td><td>distance</td><td>position</td><td>gene name</td><td>gene id</td></tr>
<tr><td>GU305914.1</td><td></td><td>41942</td><td>41964</td><td>+</td><td>0</td><td>AGATCAGCAGCTCCCCGGAG</td><td>GGG</td><td>||||||||[||||||||||||]PAM</td><td>NA</td><td>Intergenic</td><td>NA</td><td>NA</td></tr>
<tr><td>T2678</td><td>CTCCCCGGAGGGGTGCAAGCTGG</td><td></td><td>1000</td></tr>
<tr><td>Oligo fwd</td><td>CACCCTCCCCGGAGGGGTGCAAGC</td></tr>
<tr><td>Oligo rev</td><td>AAACGCTTGCACCCCTCCGGGGAG</td></tr>
<tr><td>Chromosome</td><td>start</td><td></td><td>end</td><td>strand</td><td>MM</td><td>target_seq</td><td>PAM</td><td>alignment</td><td>distance</td><td>position</td><td>gene name</td><td>gene id</td></tr>
<tr><td>GU305914.1</td><td></td><td>41952</td><td>41974</td><td>+</td><td>0</td><td>CTCCCCGGAGGGGTGCAAGC</td><td>TGG</td><td>||||||||[||||||||||||]PAM</td><td>NA</td><td>Intergenic</td><td>NA</td><td>NA</td></tr>
<tr><td>T2679</td><td>CCCGGAGGGGTGCAAGCTGGTGG</td><td></td><td>1000</td></tr>
<tr><td>Oligo fwd</td><td>CACCCCCGGAGGGGTGCAAGCTGG</td></tr>
<tr><td>Oligo rev</td><td>AAACCCAGCTTGCACCCCTCCGGG</td></tr>
<tr><td>Chromosome</td><td>start</td><td></td><td>end</td><td>strand</td><td>MM</td><td>target_seq</td><td>PAM</td><td>alignment</td><td>distance</td><td>position</td><td>gene name</td><td>gene id</td></tr>
<tr><td>GU305914.1</td><td></td><td>41955</td><td>41977</td><td>+</td><td>0</td><td>CCCGGAGGGGTGCAAGCTGG</td><td>TGG</td><td>||||||||[||||||||||||]PAM</td><td>NA</td><td>Intergenic</td><td>NA</td><td>NA</td></tr>
<tr><td>T2680</td><td>GGAGGGGTGCAAGCTGGTGGCGG</td><td></td><td>1000</td></tr>
<tr><td>Oligo fwd</td><td>CACCGGAGGGGTGCAAGCTGGTGG</td></tr>
<tr><td>Oligo rev</td><td>AAACCCACCAGCTTGCACCCCTCC</td></tr>
<tr><td>Chromosome</td><td>start</td><td></td><td>end</td><td>strand</td><td>MM</td><td>target_seq</td><td>PAM</td><td>alignment</td><td>distance</td><td>position</td><td>gene name</td><td>gene id</td></tr>
<tr><td>GU305914.1</td><td></td><td>41958</td><td>41980</td><td>+</td><td>0</td><td>GGAGGGGTGCAAGCTGGTGG</td><td>CGG</td><td>||||||||[||||||||||||]PAM</td><td>NA</td><td>Intergenic</td><td>NA</td><td>NA</td></tr>
<tr><td>T2681</td><td>GACCTACTACGCGTCGTGCACGG</td><td></td><td>1000</td></tr>
<tr><td>Oligo fwd</td><td>CACCGACCTACTACGCGTCGTGCA</td></tr>
<tr><td>Oligo rev</td><td>AAACTGCACGACGCGTAGTAGGTC</td></tr>
<tr><td>Chromosome</td><td>start</td><td></td><td>end</td><td>strand</td><td>MM</td><td>target_seq</td><td>PAM</td><td>alignment</td><td>distance</td><td>position</td><td>gene name</td><td>gene id</td></tr>
<tr><td>GU305914.1</td><td></td><td>41982</td><td>42004</td><td>+</td><td>0</td><td>GACCTACTACGCGTCGTGCA</td><td>CGG</td><td>||||||||[||||||||||||]PAM</td><td>NA</td><td>Intergenic</td><td>NA</td><td>NA</td></tr>
<tr><td>T2682</td><td>TACTACGCGTCGTGCACGGCCGG</td><td></td><td>1000</td></tr>
<tr><td>Oligo fwd</td><td>CACCTACTACGCGTCGTGCACGGC</td></tr>
</table>

<table>
<tr><td>Oligo rev</td><td>AAACGCCGTGCACGACGCGTAGTA</td><td></td><td></td><td></td><td></td><td></td><td></td><td></td><td></td><td></td><td></td><td></td></tr>
<tr><td>Chromosome</td><td>start</td><td></td><td>end</td><td>strand</td><td>MM</td><td>target_seq</td><td>PAM</td><td>alignment</td><td>distance</td><td>position</td><td>gene name</td><td>gene id</td></tr>
<tr><td>GU305914.1</td><td></td><td>41986</td><td>42008</td><td>+</td><td>0</td><td>TACTACGCGTCGTGCACGGC</td><td>CGG</td><td>||||||||[||||||||||||]PAM</td><td>NA</td><td>Intergenic</td><td>NA</td><td>NA</td></tr>
<tr><td>T2683</td><td>ACTACGCGTCGTGCACGGCCGGG</td><td></td><td>1000</td><td></td><td></td><td></td><td></td><td></td><td></td><td></td><td></td><td></td></tr>
<tr><td>Oligo fwd</td><td>CACCACTACGCGTCGTGCACGGCC</td><td></td><td></td><td></td><td></td><td></td><td></td><td></td><td></td><td></td><td></td><td></td></tr>
<tr><td>Oligo rev</td><td>AAACGGCCGTGCACGACGCGTAGT</td><td></td><td></td><td></td><td></td><td></td><td></td><td></td><td></td><td></td><td></td><td></td></tr>
<tr><td>Chromosome</td><td>start</td><td></td><td>end</td><td>strand</td><td>MM</td><td>target_seq</td><td>PAM</td><td>alignment</td><td>distance</td><td>position</td><td>gene name</td><td>gene id</td></tr>
<tr><td>GU305914.1</td><td></td><td>41987</td><td>42009</td><td>+</td><td>0</td><td>ACTACGCGTCGTGCACGGCC</td><td>GGG</td><td>||||||||[||||||||||||]PAM</td><td>NA</td><td>Intergenic</td><td>NA</td><td>NA</td></tr>
<tr><td>T2684</td><td>CACGGCCGGGTTCTCGACGCTGG</td><td></td><td>1000</td><td></td><td></td><td></td><td></td><td></td><td></td><td></td><td></td><td></td></tr>
<tr><td>Oligo fwd</td><td>CACCCACGGCCGGGTTCTCGACGC</td><td></td><td></td><td></td><td></td><td></td><td></td><td></td><td></td><td></td><td></td><td></td></tr>
<tr><td>Oligo rev</td><td>AAACGCGTCGAGAACCCGGCCGTG</td><td></td><td></td><td></td><td></td><td></td><td></td><td></td><td></td><td></td><td></td><td></td></tr>
<tr><td>Chromosome</td><td>start</td><td></td><td>end</td><td>strand</td><td>MM</td><td>target_seq</td><td>PAM</td><td>alignment</td><td>distance</td><td>position</td><td>gene name</td><td>gene id</td></tr>
<tr><td>GU305914.1</td><td></td><td>42000</td><td>42022</td><td>+</td><td>0</td><td>CACGGCCGGGTTCTCGACGC</td><td>TGG</td><td>||||||||[||||||||||||]PAM</td><td>NA</td><td>Intergenic</td><td>NA</td><td>NA</td></tr>
<tr><td>T2685</td><td>CTCGACGCTGGCGCTGATCTCGG</td><td></td><td>1000</td><td></td><td></td><td></td><td></td><td></td><td></td><td></td><td></td><td></td></tr>
<tr><td>Oligo fwd</td><td>CACCCTCGACGCTGGCGCTGATCT</td><td></td><td></td><td></td><td></td><td></td><td></td><td></td><td></td><td></td><td></td><td></td></tr>
<tr><td>Oligo rev</td><td>AAACAGATCAGCGCCAGCGTCGAG</td><td></td><td></td><td></td><td></td><td></td><td></td><td></td><td></td><td></td><td></td><td></td></tr>
<tr><td>Chromosome</td><td>start</td><td></td><td>end</td><td>strand</td><td>MM</td><td>target_seq</td><td>PAM</td><td>alignment</td><td>distance</td><td>position</td><td>gene name</td><td>gene id</td></tr>
<tr><td>GU305914.1</td><td></td><td>42012</td><td>42034</td><td>+</td><td>0</td><td>CTCGACGCTGGCGCTGATCT</td><td>CGG</td><td>||||||||[||||||||||||]PAM</td><td>NA</td><td>Intergenic</td><td>NA</td><td>NA</td></tr>
<tr><td>T2686</td><td>TGGCGCTGATCTCGGTGAACAGG</td><td></td><td>1000</td><td></td><td></td><td></td><td></td><td></td><td></td><td></td><td></td><td></td></tr>
<tr><td>Oligo fwd</td><td>CACCTGGCGCTGATCTCGGTGAAC</td><td></td><td></td><td></td><td></td><td></td><td></td><td></td><td></td><td></td><td></td><td></td></tr>
<tr><td>Oligo rev</td><td>AAACGTTCACCGAGATCAGCGCCA</td><td></td><td></td><td></td><td></td><td></td><td></td><td></td><td></td><td></td><td></td><td></td></tr>
<tr><td>Chromosome</td><td>start</td><td></td><td>end</td><td>strand</td><td>MM</td><td>target_seq</td><td>PAM</td><td>alignment</td><td>distance</td><td>position</td><td>gene name</td><td>gene id</td></tr>
<tr><td>GU305914.1</td><td></td><td>42020</td><td>42042</td><td>+</td><td>0</td><td>TGGCGCTGATCTCGGTGAAC</td><td>AGG</td><td>||||||||[||||||||||||]PAM</td><td>NA</td><td>Intergenic</td><td>NA</td><td>NA</td></tr>
<tr><td>T2687</td><td>CCAGAGCACGAACAAGAACGCGG</td><td></td><td>1000</td><td></td><td></td><td></td><td></td><td></td><td></td><td></td><td></td><td></td></tr>
<tr><td>Oligo fwd</td><td>CACCCCAGAGCACGAACAAGAACG</td><td></td><td></td><td></td><td></td><td></td><td></td><td></td><td></td><td></td><td></td><td></td></tr>
<tr><td>Oligo rev</td><td>AAACCGTTCTTGTTCGTGCTCTGG</td><td></td><td></td><td></td><td></td><td></td><td></td><td></td><td></td><td></td><td></td><td></td></tr>
<tr><td>Chromosome</td><td>start</td><td></td><td>end</td><td>strand</td><td>MM</td><td>target_seq</td><td>PAM</td><td>alignment</td><td>distance</td><td>position</td><td>gene name</td><td>gene id</td></tr>
<tr><td>GU305914.1</td><td></td><td>42057</td><td>42079</td><td>+</td><td>0</td><td>CCAGAGCACGAACAAGAACG</td><td>CGG</td><td>||||||||[||||||||||||]PAM</td><td>NA</td><td>Intergenic</td><td>NA</td><td>NA</td></tr>
<tr><td>T2688</td><td>GAGCACGAACAAGAACGCGGCGG</td><td></td><td>1000</td><td></td><td></td><td></td><td></td><td></td><td></td><td></td><td></td><td></td></tr>
<tr><td>Oligo fwd</td><td>CACCGAGCACGAACAAGAACGCGG</td><td></td><td></td><td></td><td></td><td></td><td></td><td></td><td></td><td></td><td></td><td></td></tr>
<tr><td>Oligo rev</td><td>AAACCCGCGTTCTTGTTCGTGCTC</td><td></td><td></td><td></td><td></td><td></td><td></td><td></td><td></td><td></td><td></td><td></td></tr>
<tr><td>Chromosome</td><td>start</td><td></td><td>end</td><td>strand</td><td>MM</td><td>target_seq</td><td>PAM</td><td>alignment</td><td>distance</td><td>position</td><td>gene name</td><td>gene id</td></tr>
<tr><td>GU305914.1</td><td></td><td>42060</td><td>42082</td><td>+</td><td>0</td><td>GAGCACGAACAAGAACGCGG</td><td>CGG</td><td>||||||||[||||||||||||]PAM</td><td>NA</td><td>Intergenic</td><td>NA</td><td>NA</td></tr>
<tr><td>T2689</td><td>AGCACGAACAAGAACGCGGCGGG</td><td></td><td>1000</td><td></td><td></td><td></td><td></td><td></td><td></td><td></td><td></td><td></td></tr>
<tr><td>Oligo fwd</td><td>CACCAGCACGAACAAGAACGCGGC</td><td></td><td></td><td></td><td></td><td></td><td></td><td></td><td></td><td></td><td></td><td></td></tr>
<tr><td>Oligo rev</td><td>AAACGCCGCGTTCTTGTTCGTGCT</td><td></td><td></td><td></td><td></td><td></td><td></td><td></td><td></td><td></td><td></td><td></td></tr>
<tr><td>Chromosome</td><td>start</td><td></td><td>end</td><td>strand</td><td>MM</td><td>target_seq</td><td>PAM</td><td>alignment</td><td>distance</td><td>position</td><td>gene name</td><td>gene id</td></tr>
<tr><td>GU305914.1</td><td></td><td>42061</td><td>42083</td><td>+</td><td>0</td><td>AGCACGAACAAGAACGCGGC</td><td>GGG</td><td>||||||||[||||||||||||]PAM</td><td>NA</td><td>Intergenic</td><td>NA</td><td>NA</td></tr>
<tr><td>T2690</td><td>GCACGAACAAGAACGCGGCGGGG</td><td></td><td>1000</td><td></td><td></td><td></td><td></td><td></td><td></td><td></td><td></td><td></td></tr>
<tr><td>Oligo fwd</td><td>CACCGCACGAACAAGAACGCGGCG</td><td></td><td></td><td></td><td></td><td></td><td></td><td></td><td></td><td></td><td></td><td></td></tr>
<tr><td>Oligo rev</td><td>AAACCGCCGCGTTCTTGTTCGTGC</td><td></td><td></td><td></td><td></td><td></td><td></td><td></td><td></td><td></td><td></td><td></td></tr>
<tr><td>Chromosome</td><td>start</td><td></td><td>end</td><td>strand</td><td>MM</td><td>target_seq</td><td>PAM</td><td>alignment</td><td>distance</td><td>position</td><td>gene name</td><td>gene id</td></tr>
<tr><td>GU305914.1</td><td></td><td>42062</td><td>42084</td><td>+</td><td>0</td><td>GCACGAACAAGAACGCGGCG</td><td>GGG</td><td>||||||||[||||||||||||]PAM</td><td>NA</td><td>Intergenic</td><td>NA</td><td>NA</td></tr>
<tr><td>T2691</td><td>GGGTCGAAGAAGAAGACTTACGG</td><td></td><td>1000</td><td></td><td></td><td></td><td></td><td></td><td></td><td></td><td></td><td></td></tr>
<tr><td>Oligo fwd</td><td>CACCGGGTCGAAGAAGAAGACTTA</td><td></td><td></td><td></td><td></td><td></td><td></td><td></td><td></td><td></td><td></td><td></td></tr>
<tr><td>Oligo rev</td><td>AAACTAAGTCTTCTTCTTCGACCC</td><td></td><td></td><td></td><td></td><td></td><td></td><td></td><td></td><td></td><td></td><td></td></tr>
<tr><td>Chromosome</td><td>start</td><td></td><td>end</td><td>strand</td><td>MM</td><td>target_seq</td><td>PAM</td><td>alignment</td><td>distance</td><td>position</td><td>gene name</td><td>gene id</td></tr>
<tr><td>GU305914.1</td><td></td><td>42082</td><td>42104</td><td>+</td><td>0</td><td>GGGTCGAAGAAGAAGACTTA</td><td>CGG</td><td>||||||||[||||||||||||]PAM</td><td>NA</td><td>Intergenic</td><td>NA</td><td>NA</td></tr>
<tr><td>T2692</td><td>GAAGAAGACTTACGGCGTGCTGG</td><td></td><td>1000</td><td></td><td></td><td></td><td></td><td></td><td></td><td></td><td></td><td></td></tr>
<tr><td>Oligo fwd</td><td>CACCGAAGAAGACTTACGGCGTGC</td><td></td><td></td><td></td><td></td><td></td><td></td><td></td><td></td><td></td><td></td><td></td></tr>
<tr><td>Oligo rev</td><td>AAACGCACGCCGTAAGTCTTCTTC</td><td></td><td></td><td></td><td></td><td></td><td></td><td></td><td></td><td></td><td></td><td></td></tr>
<tr><td>Chromosome</td><td>start</td><td></td><td>end</td><td>strand</td><td>MM</td><td>target_seq</td><td>PAM</td><td>alignment</td><td>distance</td><td>position</td><td>gene name</td><td>gene id</td></tr>
<tr><td>GU305914.1</td><td></td><td>42090</td><td>42112</td><td>+</td><td>0</td><td>GAAGAAGACTTACGGCGTGC</td><td>TGG</td><td>||||||||[||||||||||||]PAM</td><td>NA</td><td>Intergenic</td><td>NA</td><td>NA</td></tr>
<tr><td>T2693</td><td>GACTTACGGCGTGCTGGCGCTGG</td><td></td><td>1000</td><td></td><td></td><td></td><td></td><td></td><td></td><td></td><td></td><td></td></tr>
<tr><td>Oligo fwd</td><td>CACCGACTTACGGCGTGCTGGCGC</td><td></td><td></td><td></td><td></td><td></td><td></td><td></td><td></td><td></td><td></td><td></td></tr>
<tr><td>Oligo rev</td><td>AAACGCGCCAGCACGCCGTAAGTC</td><td></td><td></td><td></td><td></td><td></td><td></td><td></td><td></td><td></td><td></td><td></td></tr>
<tr><td>Chromosome</td><td>start</td><td></td><td>end</td><td>strand</td><td>MM</td><td>target_seq</td><td>PAM</td><td>alignment</td><td>distance</td><td>position</td><td>gene name</td><td>gene id</td></tr>
<tr><td>GU305914.1</td><td></td><td>42096</td><td>42118</td><td>+</td><td>0</td><td>GACTTACGGCGTGCTGGCGC</td><td>TGG</td><td>||||||||[||||||||||||]PAM</td><td>NA</td><td>Intergenic</td><td>NA</td><td>NA</td></tr>
<tr><td>T2694</td><td>ACGGCGTGCTGGCGCTGGTGTGG</td><td></td><td>1000</td><td></td><td></td><td></td><td></td><td></td><td></td><td></td><td></td><td></td></tr>
<tr><td>Oligo fwd</td><td>CACCACGGCGTGCTGGCGCTGGTG</td><td></td><td></td><td></td><td></td><td></td><td></td><td></td><td></td><td></td><td></td><td></td></tr>
<tr><td>Oligo rev</td><td>AAACCACCAGCGCCAGCACGCCGT</td><td></td><td></td><td></td><td></td><td></td><td></td><td></td><td></td><td></td><td></td><td></td></tr>
<tr><td>Chromosome</td><td>start</td><td></td><td>end</td><td>strand</td><td>MM</td><td>target_seq</td><td>PAM</td><td>alignment</td><td>distance</td><td>position</td><td>gene name</td><td>gene id</td></tr>
<tr><td>GU305914.1</td><td></td><td>42101</td><td>42123</td><td>+</td><td>0</td><td>ACGGCGTGCTGGCGCTGGTG</td><td>TGG</td><td>||||||||[||||||||||||]PAM</td><td>NA</td><td>Intergenic</td><td>NA</td><td>NA</td></tr>
<tr><td>T2695</td><td>CTCGCTGATGTGCGCGTCGCCGG</td><td></td><td>1000</td><td></td><td></td><td></td><td></td><td></td><td></td><td></td><td></td><td></td></tr>
<tr><td>Oligo fwd</td><td>CACCCTCGCTGATGTGCGCGTCGC</td><td></td><td></td><td></td><td></td><td></td><td></td><td></td><td></td><td></td><td></td><td></td></tr>
<tr><td>Oligo rev</td><td>AAACGCGACGCGCACATCAGCGAG</td><td></td><td></td><td></td><td></td><td></td><td></td><td></td><td></td><td></td><td></td><td></td></tr>
<tr><td>Chromosome</td><td>start</td><td></td><td>end</td><td>strand</td><td>MM</td><td>target_seq</td><td>PAM</td><td>alignment</td><td>distance</td><td>position</td><td>gene name</td><td>gene id</td></tr>
<tr><td>GU305914.1</td><td></td><td>42129</td><td>42151</td><td>+</td><td>0</td><td>CTCGCTGATGTGCGCGTCGC</td><td>CGG</td><td>||||||||[||||||||||||]PAM</td><td>NA</td><td>Intergenic</td><td>NA</td><td>NA</td></tr>
<tr><td>T2696</td><td>GCCGACGTACGTGACCGTTTTGG</td><td></td><td>1000</td><td></td><td></td><td></td><td></td><td></td><td></td><td></td><td></td><td></td></tr>
<tr><td>Oligo fwd</td><td>CACCGCCGACGTACGTGACCGTTT</td><td></td><td></td><td></td><td></td><td></td><td></td><td></td><td></td><td></td><td></td><td></td></tr>
<tr><td>Oligo rev</td><td>AAACAAACGGTCACGTACGTCGGC</td><td></td><td></td><td></td><td></td><td></td><td></td><td></td><td></td><td></td><td></td><td></td></tr>
<tr><td>Chromosome</td><td>start</td><td></td><td>end</td><td>strand</td><td>MM</td><td>target_seq</td><td>PAM</td><td>alignment</td><td>distance</td><td>position</td><td>gene name</td><td>gene id</td></tr>
<tr><td>GU305914.1</td><td></td><td>42153</td><td>42175</td><td>+</td><td>0</td><td>GCCGACGTACGTGACCGTTT</td><td>TGG</td><td>||||||||[||||||||||||]PAM</td><td>NA</td><td>Intergenic</td><td>NA</td><td>NA</td></tr>
<tr><td>T2697</td><td>GTGACCGTTTTGGCGCACGACGG</td><td></td><td>1000</td><td></td><td></td><td></td><td></td><td></td><td></td><td></td><td></td><td></td></tr>
</table>

Oligo fwd CACCGTGACCGTTTTGGCGCACGA
Oligo rev AAACTCGTGCGCCAAAACGGTCAC
Chromosome start end strand MM target_seq PAM alignment distance position gene name gene id
GU305914.1 42163 42185 + 0 GTGACCGTTTTGGCGCACGA CGG ||||||||[||||||||||||]PAM NA Intergenic NA NA

T2698 TGACCGTTTTGGCGCACGACGGG 1000
Oligo fwd CACCTGACCGTTTTGGCGCACGAC
Oligo rev AAACGTCGTGCGCCAAAACGGTCA
Chromosome start end strand MM target_seq PAM alignment distance position gene name gene id
GU305914.1 42164 42186 + 0 TGACCGTTTTGGCGCACGAC GGG ||||||||[||||||||||||]PAM NA Intergenic NA NA

T2699 GACCGTTTTGGCGCACGACGGGG 1000
Oligo fwd CACCGACCGTTTTGGCGCACGACG
Oligo rev AAACCGTCGTGCGCCAAAACGGTC
Chromosome start end strand MM target_seq PAM alignment distance position gene name gene id
GU305914.1 42165 42187 + 0 GACCGTTTTGGCGCACGACG GGG ||||||||[||||||||||||]PAM NA Intergenic NA NA

T2700 GGCGCACGACGGGGACACGCCGG 1000
Oligo fwd CACCGGCGCACGACGGGGACACGC
Oligo rev AAACGCGTGTCCCCGTCGTGCGCC
Chromosome start end strand MM target_seq PAM alignment distance position gene name gene id
GU305914.1 42174 42196 + 0 GGCGCACGACGGGGACACGC CGG ||||||||[||||||||||||]PAM NA Intergenic NA NA

T2701 CATCTTCTTCAACTACGATCAGG 1000
Oligo fwd CACCCATCTTCTTCAACTACGATC
Oligo rev AAACGATCGTAGTTGAAGAAGATG
Chromosome start end strand MM target_seq PAM alignment distance position gene name gene id
GU305914.1 42219 42241 + 0 CATCTTCTTCAACTACGATC AGG ||||||||[||||||||||||]PAM NA Intergenic NA NA

T2702 CGATCAGGTCAAAACCGTGCTGG 1000
Oligo fwd CACCCGATCAGGTCAAAACCGTGC
Oligo rev AAACGCACGGTTTTGACCTGATCG
Chromosome start end strand MM target_seq PAM alignment distance position gene name gene id
GU305914.1 42234 42256 + 0 CGATCAGGTCAAAACCGTGC TGG ||||||||[||||||||||||]PAM NA Intergenic NA NA

T2703 AGATCCTCATTTGCATAGTATGG 1000
Oligo fwd CACCAGATCCTCATTTGCATAGTA
Oligo rev AAACTACTATGCAAATGAGGATCT
Chromosome start end strand MM target_seq PAM alignment distance position gene name gene id
GU305914.1 42266 42288 + 0 AGATCCTCATTTGCATAGTA TGG ||||||||[||||||||||||]PAM NA Intergenic NA NA

T2704 GATCCTCATTTGCATAGTATGGG 1000
Oligo fwd CACCGATCCTCATTTGCATAGTAT
Oligo rev AAACATACTATGCAAATGAGGATC
Chromosome start end strand MM target_seq PAM alignment distance position gene name gene id
GU305914.1 42267 42289 + 0 GATCCTCATTTGCATAGTAT GGG ||||||||[||||||||||||]PAM NA Intergenic NA NA

T2705 ATCCTCATTTGCATAGTATGGGG 1000
Oligo fwd CACCATCCTCATTTGCATAGTATG
Oligo rev AAACCATACTATGCAAATGAGGAT
Chromosome start end strand MM target_seq PAM alignment distance position gene name gene id
GU305914.1 42268 42290 + 0 ATCCTCATTTGCATAGTATG GGG ||||||||[||||||||||||]PAM NA Intergenic NA NA

T2706 AGTATGGGGCGTCATGCCAGTGG 1000
Oligo fwd CACCAGTATGGGGCGTCATGCCAG
Oligo rev AAACCTGGCATGACGCCCCATACT
Chromosome start end strand MM target_seq PAM alignment distance position gene name gene id
GU305914.1 42282 42304 + 0 AGTATGGGGCGTCATGCCAG TGG ||||||||[||||||||||||]PAM NA Intergenic NA NA

T2707 TGCCAGTGGTGATGATGACATGG 1000
Oligo fwd CACCTGCCAGTGGTGATGATGACA
Oligo rev AAACTGTCATCATCACCACTGGCA
Chromosome start end strand MM target_seq PAM alignment distance position gene name gene id
GU305914.1 42296 42318 + 0 TGCCAGTGGTGATGATGACA TGG ||||||||[||||||||||||]PAM NA Intergenic NA NA

T2708 TGAAACTGACCTCGTACCGCAGG 1000
Oligo fwd CACCTGAAACTGACCTCGTACCGC
Oligo rev AAACGCGGTACGAGGTCAGTTTCA
Chromosome start end strand MM target_seq PAM alignment distance position gene name gene id
GU305914.1 42344 42366 + 0 TGAAACTGACCTCGTACCGC AGG ||||||||[||||||||||||]PAM NA Intergenic NA NA

T2709 CCGCAGGCGCAGCCACACGCTGG 1000
Oligo fwd CACCCCGCAGGCGCAGCCACACGC
Oligo rev AAACGCGTGTGGCTGCGCCTGCGG
Chromosome start end strand MM target_seq PAM alignment distance position gene name gene id
GU305914.1 42360 42382 + 0 CCGCAGGCGCAGCCACACGC TGG ||||||||[||||||||||||]PAM NA Intergenic NA NA

T2710 AGTGCTGCAGACCCCGTTCGTGG 1000
Oligo fwd CACCAGTGCTGCAGACCCCGTTCG
Oligo rev AAACCGAACGGGGTCTGCAGCACT
Chromosome start end strand MM target_seq PAM alignment distance position gene name gene id
GU305914.1 42414 42436 + 0 AGTGCTGCAGACCCCGTTCG TGG ||||||||[||||||||||||]PAM NA Intergenic NA NA

T2711 GTGCTGCAGACCCCGTTCGTGGG 1000
Oligo fwd CACCGTGCTGCAGACCCCGTTCGT
Oligo rev AAACACGAACGGGGTCTGCAGCAC
Chromosome start end strand MM target_seq PAM alignment distance position gene name gene id
GU305914.1 42415 42437 + 0 GTGCTGCAGACCCCGTTCGT GGG ||||||||[||||||||||||]PAM NA Intergenic NA NA

<table>
<tr><td>T2712</td><td>ATAGTTACGCCGTGATCGAGTGG</td><td></td><td>1000</td><td></td><td></td><td></td><td></td><td></td><td></td><td></td><td></td><td></td></tr>
<tr><td>Oligo fwd</td><td>CACCATAGTTACGCCGTGATCGAG</td><td></td><td></td><td></td><td></td><td></td><td></td><td></td><td></td><td></td><td></td><td></td></tr>
<tr><td>Oligo rev</td><td>AAACCTCGATCACGGCGTAACTAT</td><td></td><td></td><td></td><td></td><td></td><td></td><td></td><td></td><td></td><td></td><td></td></tr>
<tr><td>Chromosome</td><td>start</td><td></td><td>end</td><td>strand</td><td>MM</td><td>target_seq</td><td>PAM</td><td>alignment</td><td>distance</td><td>position</td><td>gene name</td><td>gene id</td></tr>
<tr><td>GU305914.1</td><td></td><td>42452</td><td>42474</td><td>+</td><td>0</td><td>ATAGTTACGCCGTGATCGAG</td><td>TGG</td><td>||||||||[||||||||||||]PAM</td><td>NA</td><td>Intergenic</td><td>NA</td><td>NA</td></tr>
<tr><td>T2713</td><td>TAGTTACGCCGTGATCGAGTGGG</td><td></td><td>1000</td><td></td><td></td><td></td><td></td><td></td><td></td><td></td><td></td><td></td></tr>
<tr><td>Oligo fwd</td><td>CACCTAGTTACGCCGTGATCGAGT</td><td></td><td></td><td></td><td></td><td></td><td></td><td></td><td></td><td></td><td></td><td></td></tr>
<tr><td>Oligo rev</td><td>AAACACTCGATCACGGCGTAACTA</td><td></td><td></td><td></td><td></td><td></td><td></td><td></td><td></td><td></td><td></td><td></td></tr>
<tr><td>Chromosome</td><td>start</td><td></td><td>end</td><td>strand</td><td>MM</td><td>target_seq</td><td>PAM</td><td>alignment</td><td>distance</td><td>position</td><td>gene name</td><td>gene id</td></tr>
<tr><td>GU305914.1</td><td></td><td>42453</td><td>42475</td><td>+</td><td>0</td><td>TAGTTACGCCGTGATCGAGT</td><td>GGG</td><td>||||||||[||||||||||||]PAM</td><td>NA</td><td>Intergenic</td><td>NA</td><td>NA</td></tr>
<tr><td>T2714</td><td>GTCGATCAACAGCCGAGACGCGG</td><td></td><td>1000</td><td></td><td></td><td></td><td></td><td></td><td></td><td></td><td></td><td></td></tr>
<tr><td>Oligo fwd</td><td>CACCGTCGATCAACAGCCGAGACG</td><td></td><td></td><td></td><td></td><td></td><td></td><td></td><td></td><td></td><td></td><td></td></tr>
<tr><td>Oligo rev</td><td>AAACCGTCTCGGCTGTTGATCGAC</td><td></td><td></td><td></td><td></td><td></td><td></td><td></td><td></td><td></td><td></td><td></td></tr>
<tr><td>Chromosome</td><td>start</td><td></td><td>end</td><td>strand</td><td>MM</td><td>target_seq</td><td>PAM</td><td>alignment</td><td>distance</td><td>position</td><td>gene name</td><td>gene id</td></tr>
<tr><td>GU305914.1</td><td></td><td>42489</td><td>42511</td><td>+</td><td>0</td><td>GTCGATCAACAGCCGAGACG</td><td>CGG</td><td>||||||||[||||||||||||]PAM</td><td>NA</td><td>Intergenic</td><td>NA</td><td>NA</td></tr>
<tr><td>T2715</td><td>CCGAGACGCGGTCGCCATGCTGG</td><td></td><td>1000</td><td></td><td></td><td></td><td></td><td></td><td></td><td></td><td></td><td></td></tr>
<tr><td>Oligo fwd</td><td>CACCCCGAGACGCGGTCGCCATGC</td><td></td><td></td><td></td><td></td><td></td><td></td><td></td><td></td><td></td><td></td><td></td></tr>
<tr><td>Oligo rev</td><td>AAACGCATGGCGACCGCGTCTCGG</td><td></td><td></td><td></td><td></td><td></td><td></td><td></td><td></td><td></td><td></td><td></td></tr>
<tr><td>Chromosome</td><td>start</td><td></td><td>end</td><td>strand</td><td>MM</td><td>target_seq</td><td>PAM</td><td>alignment</td><td>distance</td><td>position</td><td>gene name</td><td>gene id</td></tr>
<tr><td>GU305914.1</td><td></td><td>42501</td><td>42523</td><td>+</td><td>0</td><td>CCGAGACGCGGTCGCCATGC</td><td>TGG</td><td>||||||||[||||||||||||]PAM</td><td>NA</td><td>Intergenic</td><td>NA</td><td>NA</td></tr>
<tr><td>T2716</td><td>GGTCGCCATGCTGGCGCGAGTGG</td><td></td><td>1000</td><td></td><td></td><td></td><td></td><td></td><td></td><td></td><td></td><td></td></tr>
<tr><td>Oligo fwd</td><td>CACCGGTCGCCATGCTGGCGCGAG</td><td></td><td></td><td></td><td></td><td></td><td></td><td></td><td></td><td></td><td></td><td></td></tr>
<tr><td>Oligo rev</td><td>AAACCTCGCGCCAGCATGGCGACC</td><td></td><td></td><td></td><td></td><td></td><td></td><td></td><td></td><td></td><td></td><td></td></tr>
<tr><td>Chromosome</td><td>start</td><td></td><td>end</td><td>strand</td><td>MM</td><td>target_seq</td><td>PAM</td><td>alignment</td><td>distance</td><td>position</td><td>gene name</td><td>gene id</td></tr>
<tr><td>GU305914.1</td><td></td><td>42510</td><td>42532</td><td>+</td><td>0</td><td>GGTCGCCATGCTGGCGCGAG</td><td>TGG</td><td>||||||||[||||||||||||]PAM</td><td>NA</td><td>Intergenic</td><td>NA</td><td>NA</td></tr>
<tr><td>T2717</td><td>CCCGTGCTGTACGCCTTCCTCGG</td><td></td><td>1000</td><td></td><td></td><td></td><td></td><td></td><td></td><td></td><td></td><td></td></tr>
<tr><td>Oligo fwd</td><td>CACCCCCGTGCTGTACGCCTTCCT</td><td></td><td></td><td></td><td></td><td></td><td></td><td></td><td></td><td></td><td></td><td></td></tr>
<tr><td>Oligo rev</td><td>AAACAGGAAGGCGTACAGCACGGG</td><td></td><td></td><td></td><td></td><td></td><td></td><td></td><td></td><td></td><td></td><td></td></tr>
<tr><td>Chromosome</td><td>start</td><td></td><td>end</td><td>strand</td><td>MM</td><td>target_seq</td><td>PAM</td><td>alignment</td><td>distance</td><td>position</td><td>gene name</td><td>gene id</td></tr>
<tr><td>GU305914.1</td><td></td><td>42559</td><td>42581</td><td>+</td><td>0</td><td>CCCGTGCTGTACGCCTTCCT</td><td>CGG</td><td>||||||||[||||||||||||]PAM</td><td>NA</td><td>Intergenic</td><td>NA</td><td>NA</td></tr>
<tr><td>T2718</td><td>CCGTGCTGTACGCCTTCCTCGGG</td><td></td><td>1000</td><td></td><td></td><td></td><td></td><td></td><td></td><td></td><td></td><td></td></tr>
<tr><td>Oligo fwd</td><td>CACCCCGTGCTGTACGCCTTCCTC</td><td></td><td></td><td></td><td></td><td></td><td></td><td></td><td></td><td></td><td></td><td></td></tr>
<tr><td>Oligo rev</td><td>AAACGAGGAAGGCGTACAGCACGG</td><td></td><td></td><td></td><td></td><td></td><td></td><td></td><td></td><td></td><td></td><td></td></tr>
<tr><td>Chromosome</td><td>start</td><td></td><td>end</td><td>strand</td><td>MM</td><td>target_seq</td><td>PAM</td><td>alignment</td><td>distance</td><td>position</td><td>gene name</td><td>gene id</td></tr>
<tr><td>GU305914.1</td><td></td><td>42560</td><td>42582</td><td>+</td><td>0</td><td>CCGTGCTGTACGCCTTCCTC</td><td>GGG</td><td>||||||||[||||||||||||]PAM</td><td>NA</td><td>Intergenic</td><td>NA</td><td>NA</td></tr>
<tr><td>T2719</td><td>TGCTGTACGCCTTCCTCGGGCGG</td><td></td><td>1000</td><td></td><td></td><td></td><td></td><td></td><td></td><td></td><td></td><td></td></tr>
<tr><td>Oligo fwd</td><td>CACCTGCTGTACGCCTTCCTCGGG</td><td></td><td></td><td></td><td></td><td></td><td></td><td></td><td></td><td></td><td></td><td></td></tr>
<tr><td>Oligo rev</td><td>AAACCCCGAGGAAGGCGTACAGCA</td><td></td><td></td><td></td><td></td><td></td><td></td><td></td><td></td><td></td><td></td><td></td></tr>
<tr><td>Chromosome</td><td>start</td><td></td><td>end</td><td>strand</td><td>MM</td><td>target_seq</td><td>PAM</td><td>alignment</td><td>distance</td><td>position</td><td>gene name</td><td>gene id</td></tr>
<tr><td>GU305914.1</td><td></td><td>42563</td><td>42585</td><td>+</td><td>0</td><td>TGCTGTACGCCTTCCTCGGG</td><td>CGG</td><td>||||||||[||||||||||||]PAM</td><td>NA</td><td>Intergenic</td><td>NA</td><td>NA</td></tr>
<tr><td>T2720</td><td>GCTGTACGCCTTCCTCGGGCGGG</td><td></td><td>1000</td><td></td><td></td><td></td><td></td><td></td><td></td><td></td><td></td><td></td></tr>
<tr><td>Oligo fwd</td><td>CACCGCTGTACGCCTTCCTCGGGC</td><td></td><td></td><td></td><td></td><td></td><td></td><td></td><td></td><td></td><td></td><td></td></tr>
<tr><td>Oligo rev</td><td>AAACGCCCGAGGAAGGCGTACAGC</td><td></td><td></td><td></td><td></td><td></td><td></td><td></td><td></td><td></td><td></td><td></td></tr>
<tr><td>Chromosome</td><td>start</td><td></td><td>end</td><td>strand</td><td>MM</td><td>target_seq</td><td>PAM</td><td>alignment</td><td>distance</td><td>position</td><td>gene name</td><td>gene id</td></tr>
<tr><td>GU305914.1</td><td></td><td>42564</td><td>42586</td><td>+</td><td>0</td><td>GCTGTACGCCTTCCTCGGGC</td><td>GGG</td><td>||||||||[||||||||||||]PAM</td><td>NA</td><td>Intergenic</td><td>NA</td><td>NA</td></tr>
<tr><td>T2721</td><td>CGCTTCATGCAATGCATAACTGG</td><td></td><td>1000</td><td></td><td></td><td></td><td></td><td></td><td></td><td></td><td></td><td></td></tr>
<tr><td>Oligo fwd</td><td>CACCCGCTTCATGCAATGCATAAC</td><td></td><td></td><td></td><td></td><td></td><td></td><td></td><td></td><td></td><td></td><td></td></tr>
<tr><td>Oligo rev</td><td>AAACGTTATGCATTGCATGAAGCG</td><td></td><td></td><td></td><td></td><td></td><td></td><td></td><td></td><td></td><td></td><td></td></tr>
<tr><td>Chromosome</td><td>start</td><td></td><td>end</td><td>strand</td><td>MM</td><td>target_seq</td><td>PAM</td><td>alignment</td><td>distance</td><td>position</td><td>gene name</td><td>gene id</td></tr>
<tr><td>GU305914.1</td><td></td><td>42598</td><td>42620</td><td>+</td><td>0</td><td>CGCTTCATGCAATGCATAAC</td><td>TGG</td><td>||||||||[||||||||||||]PAM</td><td>NA</td><td>Intergenic</td><td>NA</td><td>NA</td></tr>
<tr><td>T2722</td><td>TAACTGGCAAGCTCTTCAGTCGG</td><td></td><td>1000</td><td></td><td></td><td></td><td></td><td></td><td></td><td></td><td></td><td></td></tr>
<tr><td>Oligo fwd</td><td>CACCTAACTGGCAAGCTCTTCAGT</td><td></td><td></td><td></td><td></td><td></td><td></td><td></td><td></td><td></td><td></td><td></td></tr>
<tr><td>Oligo rev</td><td>AAACACTGAAGAGCTTGCCAGTTA</td><td></td><td></td><td></td><td></td><td></td><td></td><td></td><td></td><td></td><td></td><td></td></tr>
<tr><td>Chromosome</td><td>start</td><td></td><td>end</td><td>strand</td><td>MM</td><td>target_seq</td><td>PAM</td><td>alignment</td><td>distance</td><td>position</td><td>gene name</td><td>gene id</td></tr>
<tr><td>GU305914.1</td><td></td><td>42614</td><td>42636</td><td>+</td><td>0</td><td>TAACTGGCAAGCTCTTCAGT</td><td>CGG</td><td>||||||||[||||||||||||]PAM</td><td>NA</td><td>Intergenic</td><td>NA</td><td>NA</td></tr>
<tr><td>T2723</td><td>CAGTCGGCGCCGCATGCTTCAGG</td><td></td><td>1000</td><td></td><td></td><td></td><td></td><td></td><td></td><td></td><td></td><td></td></tr>
<tr><td>Oligo fwd</td><td>CACCCAGTCGGCGCCGCATGCTTC</td><td></td><td></td><td></td><td></td><td></td><td></td><td></td><td></td><td></td><td></td><td></td></tr>
<tr><td>Oligo rev</td><td>AAACGAAGCATGCGGCGCCGACTG</td><td></td><td></td><td></td><td></td><td></td><td></td><td></td><td></td><td></td><td></td><td></td></tr>
<tr><td>Chromosome</td><td>start</td><td></td><td>end</td><td>strand</td><td>MM</td><td>target_seq</td><td>PAM</td><td>alignment</td><td>distance</td><td>position</td><td>gene name</td><td>gene id</td></tr>
<tr><td>GU305914.1</td><td></td><td>42630</td><td>42652</td><td>+</td><td>0</td><td>CAGTCGGCGCCGCATGCTTC</td><td>AGG</td><td>||||||||[||||||||||||]PAM</td><td>NA</td><td>Intergenic</td><td>NA</td><td>NA</td></tr>
<tr><td>T2724</td><td>CGCATGCTTCAGGAGCGCGCTGG</td><td></td><td>1000</td><td></td><td></td><td></td><td></td><td></td><td></td><td></td><td></td><td></td></tr>
<tr><td>Oligo fwd</td><td>CACCCGCATGCTTCAGGAGCGCGC</td><td></td><td></td><td></td><td></td><td></td><td></td><td></td><td></td><td></td><td></td><td></td></tr>
<tr><td>Oligo rev</td><td>AAACGCGCGCTCCTGAAGCATGCG</td><td></td><td></td><td></td><td></td><td></td><td></td><td></td><td></td><td></td><td></td><td></td></tr>
<tr><td>Chromosome</td><td>start</td><td></td><td>end</td><td>strand</td><td>MM</td><td>target_seq</td><td>PAM</td><td>alignment</td><td>distance</td><td>position</td><td>gene name</td><td>gene id</td></tr>
<tr><td>GU305914.1</td><td></td><td>42640</td><td>42662</td><td>+</td><td>0</td><td>CGCATGCTTCAGGAGCGCGC</td><td>TGG</td><td>||||||||[||||||||||||]PAM</td><td>NA</td><td>Intergenic</td><td>NA</td><td>NA</td></tr>
<tr><td>T2725</td><td>GCATGCTTCAGGAGCGCGCTGGG</td><td></td><td>1000</td><td></td><td></td><td></td><td></td><td></td><td></td><td></td><td></td><td></td></tr>
<tr><td>Oligo fwd</td><td>CACCGCATGCTTCAGGAGCGCGCT</td><td></td><td></td><td></td><td></td><td></td><td></td><td></td><td></td><td></td><td></td><td></td></tr>
<tr><td>Oligo rev</td><td>AAACAGCGCGCTCCTGAAGCATGC</td><td></td><td></td><td></td><td></td><td></td><td></td><td></td><td></td><td></td><td></td><td></td></tr>
<tr><td>Chromosome</td><td>start</td><td></td><td>end</td><td>strand</td><td>MM</td><td>target_seq</td><td>PAM</td><td>alignment</td><td>distance</td><td>position</td><td>gene name</td><td>gene id</td></tr>
<tr><td>GU305914.1</td><td></td><td>42641</td><td>42663</td><td>+</td><td>0</td><td>GCATGCTTCAGGAGCGCGCT</td><td>GGG</td><td>||||||||[||||||||||||]PAM</td><td>NA</td><td>Intergenic</td><td>NA</td><td>NA</td></tr>
<tr><td>T2726</td><td>CATGCTTCAGGAGCGCGCTGGGG</td><td></td><td>1000</td><td></td><td></td><td></td><td></td><td></td><td></td><td></td><td></td><td></td></tr>
<tr><td>Oligo fwd</td><td>CACCCATGCTTCAGGAGCGCGCTG</td><td></td><td></td><td></td><td></td><td></td><td></td><td></td><td></td><td></td><td></td><td></td></tr>
<tr><td>Oligo rev</td><td>AAACCAGCGCGCTCCTGAAGCATG</td><td></td><td></td><td></td><td></td><td></td><td></td><td></td><td></td><td></td><td></td><td></td></tr>
<tr><td>Chromosome</td><td>start</td><td></td><td>end</td><td>strand</td><td>MM</td><td>target_seq</td><td>PAM</td><td>alignment</td><td>distance</td><td>position</td><td>gene name</td><td>gene id</td></tr>
<tr><td>GU305914.1</td><td></td><td>42642</td><td>42664</td><td>+</td><td>0</td><td>CATGCTTCAGGAGCGCGCTG</td><td>GGG</td><td>||||||||[||||||||||||]PAM</td><td>NA</td><td>Intergenic</td><td>NA</td><td>NA</td></tr>
</table>

<table>
<tr><td>T2727</td><td>ACACCGCGCCGCGCGCCAGCTGG</td><td>1000</td><td></td><td></td><td></td><td></td><td></td><td></td><td></td><td></td><td></td></tr>
<tr><td>Oligo fwd</td><td>CACCACACCGCGCCGCGCGCCAGC</td><td></td><td></td><td></td><td></td><td></td><td></td><td></td><td></td><td></td><td></td></tr>
<tr><td>Oligo rev</td><td>AAACGCTGGCGCGCGGCGCGGTGT</td><td></td><td></td><td></td><td></td><td></td><td></td><td></td><td></td><td></td><td></td></tr>
<tr><td>Chromosome</td><td>start</td><td>end</td><td>strand</td><td>MM</td><td>target_seq</td><td>PAM</td><td>alignment</td><td>distance</td><td>position</td><td>gene name</td><td>gene id</td></tr>
<tr><td>GU305914.1</td><td>42684</td><td>42706</td><td>+</td><td>0</td><td>ACACCGCGCCGCGCGCCAGC</td><td>TGG</td><td>||||||||[||||||||||||]PAM</td><td>NA</td><td>Intergenic</td><td>NA</td><td>NA</td></tr>
<tr><td>T2728</td><td>GCGCGCCAGCTGGCAAAGATCGG</td><td>1000</td><td></td><td></td><td></td><td></td><td></td><td></td><td></td><td></td><td></td></tr>
<tr><td>Oligo fwd</td><td>CACCGCGCGCCAGCTGGCAAAGAT</td><td></td><td></td><td></td><td></td><td></td><td></td><td></td><td></td><td></td><td></td></tr>
<tr><td>Oligo rev</td><td>AAACATCTTTGCCAGCTGGCGCGC</td><td></td><td></td><td></td><td></td><td></td><td></td><td></td><td></td><td></td><td></td></tr>
<tr><td>Chromosome</td><td>start</td><td>end</td><td>strand</td><td>MM</td><td>target_seq</td><td>PAM</td><td>alignment</td><td>distance</td><td>position</td><td>gene name</td><td>gene id</td></tr>
<tr><td>GU305914.1</td><td>42694</td><td>42716</td><td>+</td><td>0</td><td>GCGCGCCAGCTGGCAAAGAT</td><td>CGG</td><td>||||||||[||||||||||||]PAM</td><td>NA</td><td>Intergenic</td><td>NA</td><td>NA</td></tr>
<tr><td>T2729</td><td>CGCGCCAGCTGGCAAAGATCGGG</td><td>1000</td><td></td><td></td><td></td><td></td><td></td><td></td><td></td><td></td><td></td></tr>
<tr><td>Oligo fwd</td><td>CACCCGCGCCAGCTGGCAAAGATC</td><td></td><td></td><td></td><td></td><td></td><td></td><td></td><td></td><td></td><td></td></tr>
<tr><td>Oligo rev</td><td>AAACGATCTTTGCCAGCTGGCGCG</td><td></td><td></td><td></td><td></td><td></td><td></td><td></td><td></td><td></td><td></td></tr>
<tr><td>Chromosome</td><td>start</td><td>end</td><td>strand</td><td>MM</td><td>target_seq</td><td>PAM</td><td>alignment</td><td>distance</td><td>position</td><td>gene name</td><td>gene id</td></tr>
<tr><td>GU305914.1</td><td>42695</td><td>42717</td><td>+</td><td>0</td><td>CGCGCCAGCTGGCAAAGATC</td><td>GGG</td><td>||||||||[||||||||||||]PAM</td><td>NA</td><td>Intergenic</td><td>NA</td><td>NA</td></tr>
<tr><td>T2730</td><td>GACGCGAAGCTGCTCGCGTTCGG</td><td>1000</td><td></td><td></td><td></td><td></td><td></td><td></td><td></td><td></td><td></td></tr>
<tr><td>Oligo fwd</td><td>CACCGACGCGAAGCTGCTCGCGTT</td><td></td><td></td><td></td><td></td><td></td><td></td><td></td><td></td><td></td><td></td></tr>
<tr><td>Oligo rev</td><td>AAACAACGCGAGCAGCTTCGCGTC</td><td></td><td></td><td></td><td></td><td></td><td></td><td></td><td></td><td></td><td></td></tr>
<tr><td>Chromosome</td><td>start</td><td>end</td><td>strand</td><td>MM</td><td>target_seq</td><td>PAM</td><td>alignment</td><td>distance</td><td>position</td><td>gene name</td><td>gene id</td></tr>
<tr><td>GU305914.1</td><td>42723</td><td>42745</td><td>+</td><td>0</td><td>GACGCGAAGCTGCTCGCGTT</td><td>CGG</td><td>||||||||[||||||||||||]PAM</td><td>NA</td><td>Intergenic</td><td>NA</td><td>NA</td></tr>
<tr><td>T2731</td><td>GCTCGCGTTCGGAGCTGCAGAGG</td><td>1000</td><td></td><td></td><td></td><td></td><td></td><td></td><td></td><td></td><td></td></tr>
<tr><td>Oligo fwd</td><td>CACCGCTCGCGTTCGGAGCTGCAG</td><td></td><td></td><td></td><td></td><td></td><td></td><td></td><td></td><td></td><td></td></tr>
<tr><td>Oligo rev</td><td>AAACCTGCAGCTCCGAACGCGAGC</td><td></td><td></td><td></td><td></td><td></td><td></td><td></td><td></td><td></td><td></td></tr>
<tr><td>Chromosome</td><td>start</td><td>end</td><td>strand</td><td>MM</td><td>target_seq</td><td>PAM</td><td>alignment</td><td>distance</td><td>position</td><td>gene name</td><td>gene id</td></tr>
<tr><td>GU305914.1</td><td>42734</td><td>42756</td><td>+</td><td>0</td><td>GCTCGCGTTCGGAGCTGCAG</td><td>AGG</td><td>||||||||[||||||||||||]PAM</td><td>NA</td><td>Intergenic</td><td>NA</td><td>NA</td></tr>
<tr><td>T2732</td><td>GCGTTCGGAGCTGCAGAGGTCGG</td><td>1000</td><td></td><td></td><td></td><td></td><td></td><td></td><td></td><td></td><td></td></tr>
<tr><td>Oligo fwd</td><td>CACCGCGTTCGGAGCTGCAGAGGT</td><td></td><td></td><td></td><td></td><td></td><td></td><td></td><td></td><td></td><td></td></tr>
<tr><td>Oligo rev</td><td>AAACACCTCTGCAGCTCCGAACGC</td><td></td><td></td><td></td><td></td><td></td><td></td><td></td><td></td><td></td><td></td></tr>
<tr><td>Chromosome</td><td>start</td><td>end</td><td>strand</td><td>MM</td><td>target_seq</td><td>PAM</td><td>alignment</td><td>distance</td><td>position</td><td>gene name</td><td>gene id</td></tr>
<tr><td>GU305914.1</td><td>42738</td><td>42760</td><td>+</td><td>0</td><td>GCGTTCGGAGCTGCAGAGGT</td><td>CGG</td><td>||||||||[||||||||||||]PAM</td><td>NA</td><td>Intergenic</td><td>NA</td><td>NA</td></tr>
<tr><td>T2733</td><td>CCGCCCCAGTGATCGCAGCGCGG</td><td>1000</td><td></td><td></td><td></td><td></td><td></td><td></td><td></td><td></td><td></td></tr>
<tr><td>Oligo fwd</td><td>CACCCCGCCCCAGTGATCGCAGCG</td><td></td><td></td><td></td><td></td><td></td><td></td><td></td><td></td><td></td><td></td></tr>
<tr><td>Oligo rev</td><td>AAACCGCTGCGATCACTGGGGCGG</td><td></td><td></td><td></td><td></td><td></td><td></td><td></td><td></td><td></td><td></td></tr>
<tr><td>Chromosome</td><td>start</td><td>end</td><td>strand</td><td>MM</td><td>target_seq</td><td>PAM</td><td>alignment</td><td>distance</td><td>position</td><td>gene name</td><td>gene id</td></tr>
<tr><td>GU305914.1</td><td>42772</td><td>42794</td><td>+</td><td>0</td><td>CCGCCCCAGTGATCGCAGCG</td><td>CGG</td><td>||||||||[||||||||||||]PAM</td><td>NA</td><td>Intergenic</td><td>NA</td><td>NA</td></tr>
<tr><td>T2734</td><td>CCCCAGTGATCGCAGCGCGGCGG</td><td>1000</td><td></td><td></td><td></td><td></td><td></td><td></td><td></td><td></td><td></td></tr>
<tr><td>Oligo fwd</td><td>CACCCCCCAGTGATCGCAGCGCGG</td><td></td><td></td><td></td><td></td><td></td><td></td><td></td><td></td><td></td><td></td></tr>
<tr><td>Oligo rev</td><td>AAACCCGCGCTGCGATCACTGGGG</td><td></td><td></td><td></td><td></td><td></td><td></td><td></td><td></td><td></td><td></td></tr>
<tr><td>Chromosome</td><td>start</td><td>end</td><td>strand</td><td>MM</td><td>target_seq</td><td>PAM</td><td>alignment</td><td>distance</td><td>position</td><td>gene name</td><td>gene id</td></tr>
<tr><td>GU305914.1</td><td>42775</td><td>42797</td><td>+</td><td>0</td><td>CCCCAGTGATCGCAGCGCGG</td><td>CGG</td><td>||||||||[||||||||||||]PAM</td><td>NA</td><td>Intergenic</td><td>NA</td><td>NA</td></tr>
<tr><td>T2735</td><td>CTTCCTCATCCTCCCCCGTGCGG</td><td>1000</td><td></td><td></td><td></td><td></td><td></td><td></td><td></td><td></td><td></td></tr>
<tr><td>Oligo fwd</td><td>CACCCTTCCTCATCCTCCCCCGTG</td><td></td><td></td><td></td><td></td><td></td><td></td><td></td><td></td><td></td><td></td></tr>
<tr><td>Oligo rev</td><td>AAACCACGGGGGAGGATGAGGAAG</td><td></td><td></td><td></td><td></td><td></td><td></td><td></td><td></td><td></td><td></td></tr>
<tr><td>Chromosome</td><td>start</td><td>end</td><td>strand</td><td>MM</td><td>target_seq</td><td>PAM</td><td>alignment</td><td>distance</td><td>position</td><td>gene name</td><td>gene id</td></tr>
<tr><td>GU305914.1</td><td>42864</td><td>42886</td><td>+</td><td>0</td><td>CTTCCTCATCCTCCCCCGTG</td><td>CGG</td><td>||||||||[||||||||||||]PAM</td><td>NA</td><td>Intergenic</td><td>NA</td><td>NA</td></tr>
<tr><td>T2736</td><td>CCTCATCCTCCCCCGTGCGGAGG</td><td>1000</td><td></td><td></td><td></td><td></td><td></td><td></td><td></td><td></td><td></td></tr>
<tr><td>Oligo fwd</td><td>CACCCCTCATCCTCCCCCGTGCGG</td><td></td><td></td><td></td><td></td><td></td><td></td><td></td><td></td><td></td><td></td></tr>
<tr><td>Oligo rev</td><td>AAACCCGCACGGGGGAGGATGAGG</td><td></td><td></td><td></td><td></td><td></td><td></td><td></td><td></td><td></td><td></td></tr>
<tr><td>Chromosome</td><td>start</td><td>end</td><td>strand</td><td>MM</td><td>target_seq</td><td>PAM</td><td>alignment</td><td>distance</td><td>position</td><td>gene name</td><td>gene id</td></tr>
<tr><td>GU305914.1</td><td>42867</td><td>42889</td><td>+</td><td>0</td><td>CCTCATCCTCCCCCGTGCGG</td><td>AGG</td><td>||||||||[||||||||||||]PAM</td><td>NA</td><td>Intergenic</td><td>NA</td><td>NA</td></tr>
<tr><td>T2737</td><td>CTCATCCTCCCCCGTGCGGAGGG</td><td>1000</td><td></td><td></td><td></td><td></td><td></td><td></td><td></td><td></td><td></td></tr>
<tr><td>Oligo fwd</td><td>CACCCTCATCCTCCCCCGTGCGGA</td><td></td><td></td><td></td><td></td><td></td><td></td><td></td><td></td><td></td><td></td></tr>
<tr><td>Oligo rev</td><td>AAACTCCGCACGGGGGAGGATGAG</td><td></td><td></td><td></td><td></td><td></td><td></td><td></td><td></td><td></td><td></td></tr>
<tr><td>Chromosome</td><td>start</td><td>end</td><td>strand</td><td>MM</td><td>target_seq</td><td>PAM</td><td>alignment</td><td>distance</td><td>position</td><td>gene name</td><td>gene id</td></tr>
<tr><td>GU305914.1</td><td>42868</td><td>42890</td><td>+</td><td>0</td><td>CTCATCCTCCCCCGTGCGGA</td><td>GGG</td><td>||||||||[||||||||||||]PAM</td><td>NA</td><td>Intergenic</td><td>NA</td><td>NA</td></tr>
<tr><td>T2738</td><td>TCCTCCCCCGTGCGGAGGGTCGG</td><td>1000</td><td></td><td></td><td></td><td></td><td></td><td></td><td></td><td></td><td></td></tr>
<tr><td>Oligo fwd</td><td>CACCTCCTCCCCCGTGCGGAGGGT</td><td></td><td></td><td></td><td></td><td></td><td></td><td></td><td></td><td></td><td></td></tr>
<tr><td>Oligo rev</td><td>AAACACCCTCCGCACGGGGGAGGA</td><td></td><td></td><td></td><td></td><td></td><td></td><td></td><td></td><td></td><td></td></tr>
<tr><td>Chromosome</td><td>start</td><td>end</td><td>strand</td><td>MM</td><td>target_seq</td><td>PAM</td><td>alignment</td><td>distance</td><td>position</td><td>gene name</td><td>gene id</td></tr>
<tr><td>GU305914.1</td><td>42872</td><td>42894</td><td>+</td><td>0</td><td>TCCTCCCCCGTGCGGAGGGT</td><td>CGG</td><td>||||||||[||||||||||||]PAM</td><td>NA</td><td>Intergenic</td><td>NA</td><td>NA</td></tr>
<tr><td>T2739</td><td>GAGGGTCGGTCGAGACCCCTCGG</td><td>1000</td><td></td><td></td><td></td><td></td><td></td><td></td><td></td><td></td><td></td></tr>
<tr><td>Oligo fwd</td><td>CACCGAGGGTCGGTCGAGACCCCT</td><td></td><td></td><td></td><td></td><td></td><td></td><td></td><td></td><td></td><td></td></tr>
<tr><td>Oligo rev</td><td>AAACAGGGGTCTCGACCGACCCTC</td><td></td><td></td><td></td><td></td><td></td><td></td><td></td><td></td><td></td><td></td></tr>
<tr><td>Chromosome</td><td>start</td><td>end</td><td>strand</td><td>MM</td><td>target_seq</td><td>PAM</td><td>alignment</td><td>distance</td><td>position</td><td>gene name</td><td>gene id</td></tr>
<tr><td>GU305914.1</td><td>42886</td><td>42908</td><td>+</td><td>0</td><td>GAGGGTCGGTCGAGACCCCT</td><td>CGG</td><td>||||||||[||||||||||||]PAM</td><td>NA</td><td>Intergenic</td><td>NA</td><td>NA</td></tr>
<tr><td>T2740</td><td>GAGACCCCTCGGAGAGCGCGAGG</td><td>1000</td><td></td><td></td><td></td><td></td><td></td><td></td><td></td><td></td><td></td></tr>
<tr><td>Oligo fwd</td><td>CACCGAGACCCCTCGGAGAGCGCG</td><td></td><td></td><td></td><td></td><td></td><td></td><td></td><td></td><td></td><td></td></tr>
<tr><td>Oligo rev</td><td>AAACCGCGCTCTCCGAGGGGTCTC</td><td></td><td></td><td></td><td></td><td></td><td></td><td></td><td></td><td></td><td></td></tr>
<tr><td>Chromosome</td><td>start</td><td>end</td><td>strand</td><td>MM</td><td>target_seq</td><td>PAM</td><td>alignment</td><td>distance</td><td>position</td><td>gene name</td><td>gene id</td></tr>
<tr><td>GU305914.1</td><td>42897</td><td>42919</td><td>+</td><td>0</td><td>GAGACCCCTCGGAGAGCGCG</td><td>AGG</td><td>||||||||[||||||||||||]PAM</td><td>NA</td><td>Intergenic</td><td>NA</td><td>NA</td></tr>
<tr><td>T2741</td><td>AGACCCCTCGGAGAGCGCGAGGG</td><td>1000</td><td></td><td></td><td></td><td></td><td></td><td></td><td></td><td></td><td></td></tr>
<tr><td>Oligo fwd</td><td>CACCAGACCCCTCGGAGAGCGCGA</td><td></td><td></td><td></td><td></td><td></td><td></td><td></td><td></td><td></td><td></td></tr>
<tr><td>Oligo rev</td><td>AAACTCGCGCTCTCCGAGGGGTCT</td><td></td><td></td><td></td><td></td><td></td><td></td><td></td><td></td><td></td><td></td></tr>
<tr><td>Chromosome</td><td>start</td><td>end</td><td>strand</td><td>MM</td><td>target_seq</td><td>PAM</td><td>alignment</td><td>distance</td><td>position</td><td>gene name</td><td>gene id</td></tr>
</table>

<table>
<tr><td>GU305914.1</td><td></td><td>42898</td><td>42920</td><td>+</td><td>0</td><td>AGACCCCTCGGAGAGCGCGA</td><td>GGG</td><td>||||||||[||||||||||||]PAM</td><td>NA</td><td>Intergenic</td><td>NA</td><td>NA</td></tr>
<tr><td>T2742</td><td>TCTTTCGCAGAATACCGTGTAGG</td><td></td><td>1000</td><td></td><td></td><td></td><td></td><td></td><td></td><td></td><td></td><td></td></tr>
<tr><td>Oligo fwd</td><td>CACCTCTTTCGCAGAATACCGTGT</td><td></td><td></td><td></td><td></td><td></td><td></td><td></td><td></td><td></td><td></td><td></td></tr>
<tr><td>Oligo rev</td><td>AAACACACGGTATTCTGCGAAAGA</td><td></td><td></td><td></td><td></td><td></td><td></td><td></td><td></td><td></td><td></td><td></td></tr>
<tr><td>Chromosome</td><td>start</td><td></td><td>end</td><td>strand</td><td>MM</td><td>target_seq</td><td>PAM</td><td>alignment</td><td>distance</td><td>position</td><td>gene name</td><td>gene id</td></tr>
<tr><td>GU305914.1</td><td></td><td>42981</td><td>43003</td><td>+</td><td>0</td><td>TCTTTCGCAGAATACCGTGT</td><td>AGG</td><td>||||||||[||||||||||||]PAM</td><td>NA</td><td>Intergenic</td><td>NA</td><td>NA</td></tr>
<tr><td>T2743</td><td>CGCAGAATACCGTGTAGGAGTGG</td><td></td><td>1000</td><td></td><td></td><td></td><td></td><td></td><td></td><td></td><td></td><td></td></tr>
<tr><td>Oligo fwd</td><td>CACCCGCAGAATACCGTGTAGGAG</td><td></td><td></td><td></td><td></td><td></td><td></td><td></td><td></td><td></td><td></td><td></td></tr>
<tr><td>Oligo rev</td><td>AAACCTCCTACACGGTATTCTGCG</td><td></td><td></td><td></td><td></td><td></td><td></td><td></td><td></td><td></td><td></td><td></td></tr>
<tr><td>Chromosome</td><td>start</td><td></td><td>end</td><td>strand</td><td>MM</td><td>target_seq</td><td>PAM</td><td>alignment</td><td>distance</td><td>position</td><td>gene name</td><td>gene id</td></tr>
<tr><td>GU305914.1</td><td></td><td>42986</td><td>43008</td><td>+</td><td>0</td><td>CGCAGAATACCGTGTAGGAG</td><td>TGG</td><td>||||||||[||||||||||||]PAM</td><td>NA</td><td>Intergenic</td><td>NA</td><td>NA</td></tr>
<tr><td>T2744</td><td>AGAATACCGTGTAGGAGTGGCGG</td><td></td><td>1000</td><td></td><td></td><td></td><td></td><td></td><td></td><td></td><td></td><td></td></tr>
<tr><td>Oligo fwd</td><td>CACCAGAATACCGTGTAGGAGTGG</td><td></td><td></td><td></td><td></td><td></td><td></td><td></td><td></td><td></td><td></td><td></td></tr>
<tr><td>Oligo rev</td><td>AAACCCACTCCTACACGGTATTCT</td><td></td><td></td><td></td><td></td><td></td><td></td><td></td><td></td><td></td><td></td><td></td></tr>
<tr><td>Chromosome</td><td>start</td><td></td><td>end</td><td>strand</td><td>MM</td><td>target_seq</td><td>PAM</td><td>alignment</td><td>distance</td><td>position</td><td>gene name</td><td>gene id</td></tr>
<tr><td>GU305914.1</td><td></td><td>42989</td><td>43011</td><td>+</td><td>0</td><td>AGAATACCGTGTAGGAGTGG</td><td>CGG</td><td>||||||||[||||||||||||]PAM</td><td>NA</td><td>Intergenic</td><td>NA</td><td>NA</td></tr>
<tr><td>T2745</td><td>ACCGTGTAGGAGTGGCGGCCCGG</td><td></td><td>1000</td><td></td><td></td><td></td><td></td><td></td><td></td><td></td><td></td><td></td></tr>
<tr><td>Oligo fwd</td><td>CACCACCGTGTAGGAGTGGCGGCC</td><td></td><td></td><td></td><td></td><td></td><td></td><td></td><td></td><td></td><td></td><td></td></tr>
<tr><td>Oligo rev</td><td>AAACGGCCGCCACTCCTACACGGT</td><td></td><td></td><td></td><td></td><td></td><td></td><td></td><td></td><td></td><td></td><td></td></tr>
<tr><td>Chromosome</td><td>start</td><td></td><td>end</td><td>strand</td><td>MM</td><td>target_seq</td><td>PAM</td><td>alignment</td><td>distance</td><td>position</td><td>gene name</td><td>gene id</td></tr>
<tr><td>GU305914.1</td><td></td><td>42994</td><td>43016</td><td>+</td><td>0</td><td>ACCGTGTAGGAGTGGCGGCC</td><td>CGG</td><td>||||||||[||||||||||||]PAM</td><td>NA</td><td>Intergenic</td><td>NA</td><td>NA</td></tr>
<tr><td>T2746</td><td>CGGCCCGGCTAAAAGCGCAGCGG</td><td></td><td>1000</td><td></td><td></td><td></td><td></td><td></td><td></td><td></td><td></td><td></td></tr>
<tr><td>Oligo fwd</td><td>CACCCGGCCCGGCTAAAAGCGCAG</td><td></td><td></td><td></td><td></td><td></td><td></td><td></td><td></td><td></td><td></td><td></td></tr>
<tr><td>Oligo rev</td><td>AAACCTGCGCTTTTAGCCGGGCCG</td><td></td><td></td><td></td><td></td><td></td><td></td><td></td><td></td><td></td><td></td><td></td></tr>
<tr><td>Chromosome</td><td>start</td><td></td><td>end</td><td>strand</td><td>MM</td><td>target_seq</td><td>PAM</td><td>alignment</td><td>distance</td><td>position</td><td>gene name</td><td>gene id</td></tr>
<tr><td>GU305914.1</td><td></td><td>43009</td><td>43031</td><td>+</td><td>0</td><td>CGGCCCGGCTAAAAGCGCAG</td><td>CGG</td><td>||||||||[||||||||||||]PAM</td><td>NA</td><td>Intergenic</td><td>NA</td><td>NA</td></tr>
<tr><td>T2747</td><td>CTAACAGTTTGCTCGCTCGATGG</td><td></td><td>1000</td><td></td><td></td><td></td><td></td><td></td><td></td><td></td><td></td><td></td></tr>
<tr><td>Oligo fwd</td><td>CACCCTAACAGTTTGCTCGCTCGA</td><td></td><td></td><td></td><td></td><td></td><td></td><td></td><td></td><td></td><td></td><td></td></tr>
<tr><td>Oligo rev</td><td>AAACTCGAGCGAGCAAACTGTTAG</td><td></td><td></td><td></td><td></td><td></td><td></td><td></td><td></td><td></td><td></td><td></td></tr>
<tr><td>Chromosome</td><td>start</td><td></td><td>end</td><td>strand</td><td>MM</td><td>target_seq</td><td>PAM</td><td>alignment</td><td>distance</td><td>position</td><td>gene name</td><td>gene id</td></tr>
<tr><td>GU305914.1</td><td></td><td>43067</td><td>43089</td><td>+</td><td>0</td><td>CTAACAGTTTGCTCGCTCGA</td><td>TGG</td><td>||||||||[||||||||||||]PAM</td><td>NA</td><td>Intergenic</td><td>NA</td><td>NA</td></tr>
<tr><td>T2748</td><td>TCGCTCGATGGAAACCGCGTCGG</td><td></td><td>1000</td><td></td><td></td><td></td><td></td><td></td><td></td><td></td><td></td><td></td></tr>
<tr><td>Oligo fwd</td><td>CACCTCGCTCGATGGAAACCGCGT</td><td></td><td></td><td></td><td></td><td></td><td></td><td></td><td></td><td></td><td></td><td></td></tr>
<tr><td>Oligo rev</td><td>AAACACGCGGTTTCCATCGAGCGA</td><td></td><td></td><td></td><td></td><td></td><td></td><td></td><td></td><td></td><td></td><td></td></tr>
<tr><td>Chromosome</td><td>start</td><td></td><td>end</td><td>strand</td><td>MM</td><td>target_seq</td><td>PAM</td><td>alignment</td><td>distance</td><td>position</td><td>gene name</td><td>gene id</td></tr>
<tr><td>GU305914.1</td><td></td><td>43079</td><td>43101</td><td>+</td><td>0</td><td>TCGCTCGATGGAAACCGCGT</td><td>CGG</td><td>||||||||[||||||||||||]PAM</td><td>NA</td><td>Intergenic</td><td>NA</td><td>NA</td></tr>
<tr><td>T2749</td><td>GAAACCGCGTCGGCTTCTCGAGG</td><td></td><td>1000</td><td></td><td></td><td></td><td></td><td></td><td></td><td></td><td></td><td></td></tr>
<tr><td>Oligo fwd</td><td>CACCGAAACCGCGTCGGCTTCTCG</td><td></td><td></td><td></td><td></td><td></td><td></td><td></td><td></td><td></td><td></td><td></td></tr>
<tr><td>Oligo rev</td><td>AAACCGAGAAGCCGACGCGGTTTC</td><td></td><td></td><td></td><td></td><td></td><td></td><td></td><td></td><td></td><td></td><td></td></tr>
<tr><td>Chromosome</td><td>start</td><td></td><td>end</td><td>strand</td><td>MM</td><td>target_seq</td><td>PAM</td><td>alignment</td><td>distance</td><td>position</td><td>gene name</td><td>gene id</td></tr>
<tr><td>GU305914.1</td><td></td><td>43089</td><td>43111</td><td>+</td><td>0</td><td>GAAACCGCGTCGGCTTCTCG</td><td>AGG</td><td>||||||||[||||||||||||]PAM</td><td>NA</td><td>Intergenic</td><td>NA</td><td>NA</td></tr>
<tr><td>T2750</td><td>GCGTCGGCTTCTCGAGGCGTCGG</td><td></td><td>1000</td><td></td><td></td><td></td><td></td><td></td><td></td><td></td><td></td><td></td></tr>
<tr><td>Oligo fwd</td><td>CACCGCGTCGGCTTCTCGAGGCGT</td><td></td><td></td><td></td><td></td><td></td><td></td><td></td><td></td><td></td><td></td><td></td></tr>
<tr><td>Oligo rev</td><td>AAACACGCCTCGAGAAGCCGACGC</td><td></td><td></td><td></td><td></td><td></td><td></td><td></td><td></td><td></td><td></td><td></td></tr>
<tr><td>Chromosome</td><td>start</td><td></td><td>end</td><td>strand</td><td>MM</td><td>target_seq</td><td>PAM</td><td>alignment</td><td>distance</td><td>position</td><td>gene name</td><td>gene id</td></tr>
<tr><td>GU305914.1</td><td></td><td>43095</td><td>43117</td><td>+</td><td>0</td><td>GCGTCGGCTTCTCGAGGCGT</td><td>CGG</td><td>||||||||[||||||||||||]PAM</td><td>NA</td><td>Intergenic</td><td>NA</td><td>NA</td></tr>
<tr><td>T2751</td><td>GGCGCGACAGACCGCACGTTCGG</td><td></td><td>1000</td><td></td><td></td><td></td><td></td><td></td><td></td><td></td><td></td><td></td></tr>
<tr><td>Oligo fwd</td><td>CACCGGCGCGACAGACCGCACGTT</td><td></td><td></td><td></td><td></td><td></td><td></td><td></td><td></td><td></td><td></td><td></td></tr>
<tr><td>Oligo rev</td><td>AAACAACGTGCGGTCTGTCGCGCC</td><td></td><td></td><td></td><td></td><td></td><td></td><td></td><td></td><td></td><td></td><td></td></tr>
<tr><td>Chromosome</td><td>start</td><td></td><td>end</td><td>strand</td><td>MM</td><td>target_seq</td><td>PAM</td><td>alignment</td><td>distance</td><td>position</td><td>gene name</td><td>gene id</td></tr>
<tr><td>GU305914.1</td><td></td><td>43116</td><td>43138</td><td>+</td><td>0</td><td>GGCGCGACAGACCGCACGTT</td><td>CGG</td><td>||||||||[||||||||||||]PAM</td><td>NA</td><td>Intergenic</td><td>NA</td><td>NA</td></tr>
<tr><td>T2752</td><td>CGACAGACCGCACGTTCGGACGG</td><td></td><td>1000</td><td></td><td></td><td></td><td></td><td></td><td></td><td></td><td></td><td></td></tr>
<tr><td>Oligo fwd</td><td>CACCCGACAGACCGCACGTTCGGA</td><td></td><td></td><td></td><td></td><td></td><td></td><td></td><td></td><td></td><td></td><td></td></tr>
<tr><td>Oligo rev</td><td>AAACTCCGAACGTGCGGTCTGTCG</td><td></td><td></td><td></td><td></td><td></td><td></td><td></td><td></td><td></td><td></td><td></td></tr>
<tr><td>Chromosome</td><td>start</td><td></td><td>end</td><td>strand</td><td>MM</td><td>target_seq</td><td>PAM</td><td>alignment</td><td>distance</td><td>position</td><td>gene name</td><td>gene id</td></tr>
<tr><td>GU305914.1</td><td></td><td>43120</td><td>43142</td><td>+</td><td>0</td><td>CGACAGACCGCACGTTCGGA</td><td>CGG</td><td>||||||||[||||||||||||]PAM</td><td>NA</td><td>Intergenic</td><td>NA</td><td>NA</td></tr>
<tr><td>T2753</td><td>GACCGCACGTTCGGACGGTCCGG</td><td></td><td>1000</td><td></td><td></td><td></td><td></td><td></td><td></td><td></td><td></td><td></td></tr>
<tr><td>Oligo fwd</td><td>CACCGACCGCACGTTCGGACGGTC</td><td></td><td></td><td></td><td></td><td></td><td></td><td></td><td></td><td></td><td></td><td></td></tr>
<tr><td>Oligo rev</td><td>AAACGACCGTCCGAACGTGCGGTC</td><td></td><td></td><td></td><td></td><td></td><td></td><td></td><td></td><td></td><td></td><td></td></tr>
<tr><td>Chromosome</td><td>start</td><td></td><td>end</td><td>strand</td><td>MM</td><td>target_seq</td><td>PAM</td><td>alignment</td><td>distance</td><td>position</td><td>gene name</td><td>gene id</td></tr>
<tr><td>GU305914.1</td><td></td><td>43125</td><td>43147</td><td>+</td><td>0</td><td>GACCGCACGTTCGGACGGTC</td><td>CGG</td><td>||||||||[||||||||||||]PAM</td><td>NA</td><td>Intergenic</td><td>NA</td><td>NA</td></tr>
<tr><td>T2754</td><td>ACCGCACGTTCGGACGGTCCGGG</td><td></td><td>1000</td><td></td><td></td><td></td><td></td><td></td><td></td><td></td><td></td><td></td></tr>
<tr><td>Oligo fwd</td><td>CACCACCGCACGTTCGGACGGTCC</td><td></td><td></td><td></td><td></td><td></td><td></td><td></td><td></td><td></td><td></td><td></td></tr>
<tr><td>Oligo rev</td><td>AAACGGACCGTCCGAACGTGCGGT</td><td></td><td></td><td></td><td></td><td></td><td></td><td></td><td></td><td></td><td></td><td></td></tr>
<tr><td>Chromosome</td><td>start</td><td></td><td>end</td><td>strand</td><td>MM</td><td>target_seq</td><td>PAM</td><td>alignment</td><td>distance</td><td>position</td><td>gene name</td><td>gene id</td></tr>
<tr><td>GU305914.1</td><td></td><td>43126</td><td>43148</td><td>+</td><td>0</td><td>ACCGCACGTTCGGACGGTCC</td><td>GGG</td><td>||||||||[||||||||||||]PAM</td><td>NA</td><td>Intergenic</td><td>NA</td><td>NA</td></tr>
<tr><td>T2755</td><td>CGGTCCGGGCATCCGCGACCCGG</td><td></td><td>1000</td><td></td><td></td><td></td><td></td><td></td><td></td><td></td><td></td><td></td></tr>
<tr><td>Oligo fwd</td><td>CACCCGGTCCGGGCATCCGCGACC</td><td></td><td></td><td></td><td></td><td></td><td></td><td></td><td></td><td></td><td></td><td></td></tr>
<tr><td>Oligo rev</td><td>AAACGGTCGCGGATGCCCGGACCG</td><td></td><td></td><td></td><td></td><td></td><td></td><td></td><td></td><td></td><td></td><td></td></tr>
<tr><td>Chromosome</td><td>start</td><td></td><td>end</td><td>strand</td><td>MM</td><td>target_seq</td><td>PAM</td><td>alignment</td><td>distance</td><td>position</td><td>gene name</td><td>gene id</td></tr>
<tr><td>GU305914.1</td><td></td><td>43140</td><td>43162</td><td>+</td><td>0</td><td>CGGTCCGGGCATCCGCGACC</td><td>CGG</td><td>||||||||[||||||||||||]PAM</td><td>NA</td><td>Intergenic</td><td>NA</td><td>NA</td></tr>
<tr><td>T2756</td><td>TCCGCGCTCGCGAAGACCGCCGG</td><td></td><td>1000</td><td></td><td></td><td></td><td></td><td></td><td></td><td></td><td></td><td></td></tr>
<tr><td>Oligo fwd</td><td>CACCTCCGCGCTCGCGAAGACCGC</td><td></td><td></td><td></td><td></td><td></td><td></td><td></td><td></td><td></td><td></td><td></td></tr>
<tr><td>Oligo rev</td><td>AAACGCGGTCTTCGCGAGCGCGGA</td><td></td><td></td><td></td><td></td><td></td><td></td><td></td><td></td><td></td><td></td><td></td></tr>
</table>

<table>
<tr><td>Chromosome</td><td>start</td><td>end</td><td>strand</td><td>MM</td><td>target_seq</td><td>PAM</td><td>alignment</td><td>distance</td><td>position</td><td>gene name</td><td>gene id</td></tr>
<tr><td>GU305914.1</td><td>43169</td><td>43191</td><td>+</td><td>0</td><td>TCCGCGCTCGCGAAGACCGC</td><td>CGG</td><td>||||||||[||||||||||||]PAM</td><td>NA</td><td>Intergenic</td><td>NA</td><td>NA</td></tr>
<tr><td></td><td></td><td></td><td></td><td></td><td></td><td></td><td></td><td></td><td></td><td></td><td></td></tr>
<tr><td>T2757</td><td>CTCGCGAAGACCGCCGGCCGCGG</td><td>1000</td><td></td><td></td><td></td><td></td><td></td><td></td><td></td><td></td><td></td></tr>
<tr><td>Oligo fwd</td><td>CACCCTCGCGAAGACCGCCGGCCG</td><td></td><td></td><td></td><td></td><td></td><td></td><td></td><td></td><td></td><td></td></tr>
<tr><td>Oligo rev</td><td>AAACCGGCCGGCGGTCTTCGCGAG</td><td></td><td></td><td></td><td></td><td></td><td></td><td></td><td></td><td></td><td></td></tr>
<tr><td>Chromosome</td><td>start</td><td>end</td><td>strand</td><td>MM</td><td>target_seq</td><td>PAM</td><td>alignment</td><td>distance</td><td>position</td><td>gene name</td><td>gene id</td></tr>
<tr><td>GU305914.1</td><td>43175</td><td>43197</td><td>+</td><td>0</td><td>CTCGCGAAGACCGCCGGCCG</td><td>CGG</td><td>||||||||[||||||||||||]PAM</td><td>NA</td><td>Intergenic</td><td>NA</td><td>NA</td></tr>
<tr><td></td><td></td><td></td><td></td><td></td><td></td><td></td><td></td><td></td><td></td><td></td><td></td></tr>
<tr><td>T2758</td><td>GGCCGCGGAAGACACGCGCACGG</td><td>1000</td><td></td><td></td><td></td><td></td><td></td><td></td><td></td><td></td><td></td></tr>
<tr><td>Oligo fwd</td><td>CACCGGCCGCGGAAGACACGCGCA</td><td></td><td></td><td></td><td></td><td></td><td></td><td></td><td></td><td></td><td></td></tr>
<tr><td>Oligo rev</td><td>AAACTGCGCGTGTCTTCCGCGGCC</td><td></td><td></td><td></td><td></td><td></td><td></td><td></td><td></td><td></td><td></td></tr>
<tr><td>Chromosome</td><td>start</td><td>end</td><td>strand</td><td>MM</td><td>target_seq</td><td>PAM</td><td>alignment</td><td>distance</td><td>position</td><td>gene name</td><td>gene id</td></tr>
<tr><td>GU305914.1</td><td>43190</td><td>43212</td><td>+</td><td>0</td><td>GGCCGCGGAAGACACGCGCA</td><td>CGG</td><td>||||||||[||||||||||||]PAM</td><td>NA</td><td>Intergenic</td><td>NA</td><td>NA</td></tr>
<tr><td></td><td></td><td></td><td></td><td></td><td></td><td></td><td></td><td></td><td></td><td></td><td></td></tr>
<tr><td>T2759</td><td>GCGGAAGACACGCGCACGGACGG</td><td>1000</td><td></td><td></td><td></td><td></td><td></td><td></td><td></td><td></td><td></td></tr>
<tr><td>Oligo fwd</td><td>CACCGCGGAAGACACGCGCACGGA</td><td></td><td></td><td></td><td></td><td></td><td></td><td></td><td></td><td></td><td></td></tr>
<tr><td>Oligo rev</td><td>AAACTCCGTGCGCGTGTCTTCCGC</td><td></td><td></td><td></td><td></td><td></td><td></td><td></td><td></td><td></td><td></td></tr>
<tr><td>Chromosome</td><td>start</td><td>end</td><td>strand</td><td>MM</td><td>target_seq</td><td>PAM</td><td>alignment</td><td>distance</td><td>position</td><td>gene name</td><td>gene id</td></tr>
<tr><td>GU305914.1</td><td>43194</td><td>43216</td><td>+</td><td>0</td><td>GCGGAAGACACGCGCACGGA</td><td>CGG</td><td>||||||||[||||||||||||]PAM</td><td>NA</td><td>Intergenic</td><td>NA</td><td>NA</td></tr>
<tr><td></td><td></td><td></td><td></td><td></td><td></td><td></td><td></td><td></td><td></td><td></td><td></td></tr>
<tr><td>T2760</td><td>GAAGACACGCGCACGGACGGCGG</td><td>1000</td><td></td><td></td><td></td><td></td><td></td><td></td><td></td><td></td><td></td></tr>
<tr><td>Oligo fwd</td><td>CACCGAAGACACGCGCACGGACGG</td><td></td><td></td><td></td><td></td><td></td><td></td><td></td><td></td><td></td><td></td></tr>
<tr><td>Oligo rev</td><td>AAACCCGTCCGTGCGCGTGTCTTC</td><td></td><td></td><td></td><td></td><td></td><td></td><td></td><td></td><td></td><td></td></tr>
<tr><td>Chromosome</td><td>start</td><td>end</td><td>strand</td><td>MM</td><td>target_seq</td><td>PAM</td><td>alignment</td><td>distance</td><td>position</td><td>gene name</td><td>gene id</td></tr>
<tr><td>GU305914.1</td><td>43197</td><td>43219</td><td>+</td><td>0</td><td>GAAGACACGCGCACGGACGG</td><td>CGG</td><td>||||||||[||||||||||||]PAM</td><td>NA</td><td>Intergenic</td><td>NA</td><td>NA</td></tr>
<tr><td></td><td></td><td></td><td></td><td></td><td></td><td></td><td></td><td></td><td></td><td></td><td></td></tr>
<tr><td>T2761</td><td>AAGACACGCGCACGGACGGCGGG</td><td>1000</td><td></td><td></td><td></td><td></td><td></td><td></td><td></td><td></td><td></td></tr>
<tr><td>Oligo fwd</td><td>CACCAAGACACGCGCACGGACGGC</td><td></td><td></td><td></td><td></td><td></td><td></td><td></td><td></td><td></td><td></td></tr>
<tr><td>Oligo rev</td><td>AAACGCCGTCCGTGCGCGTGTCTT</td><td></td><td></td><td></td><td></td><td></td><td></td><td></td><td></td><td></td><td></td></tr>
<tr><td>Chromosome</td><td>start</td><td>end</td><td>strand</td><td>MM</td><td>target_seq</td><td>PAM</td><td>alignment</td><td>distance</td><td>position</td><td>gene name</td><td>gene id</td></tr>
<tr><td>GU305914.1</td><td>43198</td><td>43220</td><td>+</td><td>0</td><td>AAGACACGCGCACGGACGGC</td><td>GGG</td><td>||||||||[||||||||||||]PAM</td><td>NA</td><td>Intergenic</td><td>NA</td><td>NA</td></tr>
<tr><td></td><td></td><td></td><td></td><td></td><td></td><td></td><td></td><td></td><td></td><td></td><td></td></tr>
<tr><td>T2762</td><td>ACGCGCACGGACGGCGGGAACGG</td><td>1000</td><td></td><td></td><td></td><td></td><td></td><td></td><td></td><td></td><td></td></tr>
<tr><td>Oligo fwd</td><td>CACCACGCGCACGGACGGCGGGAA</td><td></td><td></td><td></td><td></td><td></td><td></td><td></td><td></td><td></td><td></td></tr>
<tr><td>Oligo rev</td><td>AAACTTCCCGCCGTCCGTGCGCGT</td><td></td><td></td><td></td><td></td><td></td><td></td><td></td><td></td><td></td><td></td></tr>
<tr><td>Chromosome</td><td>start</td><td>end</td><td>strand</td><td>MM</td><td>target_seq</td><td>PAM</td><td>alignment</td><td>distance</td><td>position</td><td>gene name</td><td>gene id</td></tr>
<tr><td>GU305914.1</td><td>43203</td><td>43225</td><td>+</td><td>0</td><td>ACGCGCACGGACGGCGGGAA</td><td>CGG</td><td>||||||||[||||||||||||]PAM</td><td>NA</td><td>Intergenic</td><td>NA</td><td>NA</td></tr>
<tr><td></td><td></td><td></td><td></td><td></td><td></td><td></td><td></td><td></td><td></td><td></td><td></td></tr>
<tr><td>T2763</td><td>ACCGACACAGACGACGCAAGAGG</td><td>1000</td><td></td><td></td><td></td><td></td><td></td><td></td><td></td><td></td><td></td></tr>
<tr><td>Oligo fwd</td><td>CACCACCGACACAGACGACGCAAG</td><td></td><td></td><td></td><td></td><td></td><td></td><td></td><td></td><td></td><td></td></tr>
<tr><td>Oligo rev</td><td>AAACCTTGCGTCGTCTGTGTCGGT</td><td></td><td></td><td></td><td></td><td></td><td></td><td></td><td></td><td></td><td></td></tr>
<tr><td>Chromosome</td><td>start</td><td>end</td><td>strand</td><td>MM</td><td>target_seq</td><td>PAM</td><td>alignment</td><td>distance</td><td>position</td><td>gene name</td><td>gene id</td></tr>
<tr><td>GU305914.1</td><td>43231</td><td>43253</td><td>+</td><td>0</td><td>ACCGACACAGACGACGCAAG</td><td>AGG</td><td>||||||||[||||||||||||]PAM</td><td>NA</td><td>Intergenic</td><td>NA</td><td>NA</td></tr>
<tr><td></td><td></td><td></td><td></td><td></td><td></td><td></td><td></td><td></td><td></td><td></td><td></td></tr>
<tr><td>T2764</td><td>TTCTGACAACAATAACAACGAGG</td><td>1000</td><td></td><td></td><td></td><td></td><td></td><td></td><td></td><td></td><td></td></tr>
<tr><td>Oligo fwd</td><td>CACCTTCTGACAACAATAACAACG</td><td></td><td></td><td></td><td></td><td></td><td></td><td></td><td></td><td></td><td></td></tr>
<tr><td>Oligo rev</td><td>AAACCGTTGTTATTGTTGTCAGAA</td><td></td><td></td><td></td><td></td><td></td><td></td><td></td><td></td><td></td><td></td></tr>
<tr><td>Chromosome</td><td>start</td><td>end</td><td>strand</td><td>MM</td><td>target_seq</td><td>PAM</td><td>alignment</td><td>distance</td><td>position</td><td>gene name</td><td>gene id</td></tr>
<tr><td>GU305914.1</td><td>43259</td><td>43281</td><td>+</td><td>0</td><td>TTCTGACAACAATAACAACG</td><td>AGG</td><td>||||||||[||||||||||||]PAM</td><td>NA</td><td>Intergenic</td><td>NA</td><td>NA</td></tr>
<tr><td></td><td></td><td></td><td></td><td></td><td></td><td></td><td></td><td></td><td></td><td></td><td></td></tr>
<tr><td>T2765</td><td>TCTGACAACAATAACAACGAGGG</td><td>1000</td><td></td><td></td><td></td><td></td><td></td><td></td><td></td><td></td><td></td></tr>
<tr><td>Oligo fwd</td><td>CACCTCTGACAACAATAACAACGA</td><td></td><td></td><td></td><td></td><td></td><td></td><td></td><td></td><td></td><td></td></tr>
<tr><td>Oligo rev</td><td>AAACTCGTTGTTATTGTTGTCAGA</td><td></td><td></td><td></td><td></td><td></td><td></td><td></td><td></td><td></td><td></td></tr>
<tr><td>Chromosome</td><td>start</td><td>end</td><td>strand</td><td>MM</td><td>target_seq</td><td>PAM</td><td>alignment</td><td>distance</td><td>position</td><td>gene name</td><td>gene id</td></tr>
<tr><td>GU305914.1</td><td>43260</td><td>43282</td><td>+</td><td>0</td><td>TCTGACAACAATAACAACGA</td><td>GGG</td><td>||||||||[||||||||||||]PAM</td><td>NA</td><td>Intergenic</td><td>NA</td><td>NA</td></tr>
<tr><td></td><td></td><td></td><td></td><td></td><td></td><td></td><td></td><td></td><td></td><td></td><td></td></tr>
<tr><td>T2766</td><td>CTGACAACAATAACAACGAGGGG</td><td>1000</td><td></td><td></td><td></td><td></td><td></td><td></td><td></td><td></td><td></td></tr>
<tr><td>Oligo fwd</td><td>CACCCTGACAACAATAACAACGAG</td><td></td><td></td><td></td><td></td><td></td><td></td><td></td><td></td><td></td><td></td></tr>
<tr><td>Oligo rev</td><td>AAACCTCGTTGTTATTGTTGTCAG</td><td></td><td></td><td></td><td></td><td></td><td></td><td></td><td></td><td></td><td></td></tr>
<tr><td>Chromosome</td><td>start</td><td>end</td><td>strand</td><td>MM</td><td>target_seq</td><td>PAM</td><td>alignment</td><td>distance</td><td>position</td><td>gene name</td><td>gene id</td></tr>
<tr><td>GU305914.1</td><td>43261</td><td>43283</td><td>+</td><td>0</td><td>CTGACAACAATAACAACGAG</td><td>GGG</td><td>||||||||[||||||||||||]PAM</td><td>NA</td><td>Intergenic</td><td>NA</td><td>NA</td></tr>
<tr><td></td><td></td><td></td><td></td><td></td><td></td><td></td><td></td><td></td><td></td><td></td><td></td></tr>
<tr><td>T2767</td><td>TGACAACAATAACAACGAGGGGG</td><td>1000</td><td></td><td></td><td></td><td></td><td></td><td></td><td></td><td></td><td></td></tr>
<tr><td>Oligo fwd</td><td>CACCTGACAACAATAACAACGAGG</td><td></td><td></td><td></td><td></td><td></td><td></td><td></td><td></td><td></td><td></td></tr>
<tr><td>Oligo rev</td><td>AAACCCTCGTTGTTATTGTTGTCA</td><td></td><td></td><td></td><td></td><td></td><td></td><td></td><td></td><td></td><td></td></tr>
<tr><td>Chromosome</td><td>start</td><td>end</td><td>strand</td><td>MM</td><td>target_seq</td><td>PAM</td><td>alignment</td><td>distance</td><td>position</td><td>gene name</td><td>gene id</td></tr>
<tr><td>GU305914.1</td><td>43262</td><td>43284</td><td>+</td><td>0</td><td>TGACAACAATAACAACGAGG</td><td>GGG</td><td>||||||||[||||||||||||]PAM</td><td>NA</td><td>Intergenic</td><td>NA</td><td>NA</td></tr>
<tr><td></td><td></td><td></td><td></td><td></td><td></td><td></td><td></td><td></td><td></td><td></td><td></td></tr>
<tr><td>T2768</td><td>ACAATAACAACGAGGGGGTCCGG</td><td>1000</td><td></td><td></td><td></td><td></td><td></td><td></td><td></td><td></td><td></td></tr>
<tr><td>Oligo fwd</td><td>CACCACAATAACAACGAGGGGGTC</td><td></td><td></td><td></td><td></td><td></td><td></td><td></td><td></td><td></td><td></td></tr>
<tr><td>Oligo rev</td><td>AAACGACCCCCTCGTTGTTATTGT</td><td></td><td></td><td></td><td></td><td></td><td></td><td></td><td></td><td></td><td></td></tr>
<tr><td>Chromosome</td><td>start</td><td>end</td><td>strand</td><td>MM</td><td>target_seq</td><td>PAM</td><td>alignment</td><td>distance</td><td>position</td><td>gene name</td><td>gene id</td></tr>
<tr><td>GU305914.1</td><td>43267</td><td>43289</td><td>+</td><td>0</td><td>ACAATAACAACGAGGGGGTC</td><td>CGG</td><td>||||||||[||||||||||||]PAM</td><td>NA</td><td>Intergenic</td><td>NA</td><td>NA</td></tr>
<tr><td></td><td></td><td></td><td></td><td></td><td></td><td></td><td></td><td></td><td></td><td></td><td></td></tr>
<tr><td>T2769</td><td>CAATAACAACGAGGGGGTCCGGG</td><td>1000</td><td></td><td></td><td></td><td></td><td></td><td></td><td></td><td></td><td></td></tr>
<tr><td>Oligo fwd</td><td>CACCCAATAACAACGAGGGGGTCC</td><td></td><td></td><td></td><td></td><td></td><td></td><td></td><td></td><td></td><td></td></tr>
<tr><td>Oligo rev</td><td>AAACGGACCCCCTCGTTGTTATTG</td><td></td><td></td><td></td><td></td><td></td><td></td><td></td><td></td><td></td><td></td></tr>
<tr><td>Chromosome</td><td>start</td><td>end</td><td>strand</td><td>MM</td><td>target_seq</td><td>PAM</td><td>alignment</td><td>distance</td><td>position</td><td>gene name</td><td>gene id</td></tr>
<tr><td>GU305914.1</td><td>43268</td><td>43290</td><td>+</td><td>0</td><td>CAATAACAACGAGGGGGTCC</td><td>GGG</td><td>||||||||[||||||||||||]PAM</td><td>NA</td><td>Intergenic</td><td>NA</td><td>NA</td></tr>
<tr><td></td><td></td><td></td><td></td><td></td><td></td><td></td><td></td><td></td><td></td><td></td><td></td></tr>
<tr><td>T2770</td><td>AATAACAACGAGGGGGTCCGGGG</td><td>1000</td><td></td><td></td><td></td><td></td><td></td><td></td><td></td><td></td><td></td></tr>
<tr><td>Oligo fwd</td><td>CACCAATAACAACGAGGGGGTCCG</td><td></td><td></td><td></td><td></td><td></td><td></td><td></td><td></td><td></td><td></td></tr>
<tr><td>Oligo rev</td><td>AAACCGGACCCCCTCGTTGTTATT</td><td></td><td></td><td></td><td></td><td></td><td></td><td></td><td></td><td></td><td></td></tr>
<tr><td>Chromosome</td><td>start</td><td>end</td><td>strand</td><td>MM</td><td>target_seq</td><td>PAM</td><td>alignment</td><td>distance</td><td>position</td><td>gene name</td><td>gene id</td></tr>
<tr><td>GU305914.1</td><td>43269</td><td>43291</td><td>+</td><td>0</td><td>AATAACAACGAGGGGGTCCG</td><td>GGG</td><td>||||||||[||||||||||||]PAM</td><td>NA</td><td>Intergenic</td><td>NA</td><td>NA</td></tr>
<tr><td></td><td></td><td></td><td></td><td></td><td></td><td></td><td></td><td></td><td></td><td></td><td></td></tr>
<tr><td>T2771</td><td>ACAAAAACGACGTCGTCATCTGG</td><td>1000</td><td></td><td></td><td></td><td></td><td></td><td></td><td></td><td></td><td></td></tr>
<tr><td>Oligo fwd</td><td>CACCACAAAAACGACGTCGTCATC</td><td></td><td></td><td></td><td></td><td></td><td></td><td></td><td></td><td></td><td></td></tr>
</table>

<table>
<tr><td>Oligo rev</td><td>AAACGATGACGACGTCGTTTTTGT</td><td></td><td></td><td></td><td></td><td></td><td></td><td></td><td></td><td></td><td></td></tr>
<tr><td>Chromosome</td><td>start</td><td>end</td><td>strand</td><td>MM</td><td>target_seq</td><td>PAM</td><td>alignment</td><td>distance</td><td>position</td><td>gene name</td><td>gene id</td></tr>
<tr><td>GU305914.1</td><td>43356</td><td>43378</td><td>+</td><td>0</td><td>ACAAAAACGACGTCGTCATC</td><td>TGG</td><td>||||||||[||||||||||||]PAM</td><td>NA</td><td>Intergenic</td><td>NA</td><td>NA</td></tr>
<tr><td>T2772</td><td>AACGACGTCGTCATCTGGCCCGG</td><td>1000</td><td></td><td></td><td></td><td></td><td></td><td></td><td></td><td></td><td></td></tr>
<tr><td>Oligo fwd</td><td>CACCAACGACGTCGTCATCTGGCC</td><td></td><td></td><td></td><td></td><td></td><td></td><td></td><td></td><td></td><td></td></tr>
<tr><td>Oligo rev</td><td>AAACGGCCAGATGACGACGTCGTT</td><td></td><td></td><td></td><td></td><td></td><td></td><td></td><td></td><td></td><td></td></tr>
<tr><td>Chromosome</td><td>start</td><td>end</td><td>strand</td><td>MM</td><td>target_seq</td><td>PAM</td><td>alignment</td><td>distance</td><td>position</td><td>gene name</td><td>gene id</td></tr>
<tr><td>GU305914.1</td><td>43361</td><td>43383</td><td>+</td><td>0</td><td>AACGACGTCGTCATCTGGCC</td><td>CGG</td><td>||||||||[||||||||||||]PAM</td><td>NA</td><td>Intergenic</td><td>NA</td><td>NA</td></tr>
<tr><td>T2773</td><td>GACGTCGTCATCTGGCCCGGCGG</td><td>1000</td><td></td><td></td><td></td><td></td><td></td><td></td><td></td><td></td><td></td></tr>
<tr><td>Oligo fwd</td><td>CACCGACGTCGTCATCTGGCCCGG</td><td></td><td></td><td></td><td></td><td></td><td></td><td></td><td></td><td></td><td></td></tr>
<tr><td>Oligo rev</td><td>AAACCCGGGCCAGATGACGACGTC</td><td></td><td></td><td></td><td></td><td></td><td></td><td></td><td></td><td></td><td></td></tr>
<tr><td>Chromosome</td><td>start</td><td>end</td><td>strand</td><td>MM</td><td>target_seq</td><td>PAM</td><td>alignment</td><td>distance</td><td>position</td><td>gene name</td><td>gene id</td></tr>
<tr><td>GU305914.1</td><td>43364</td><td>43386</td><td>+</td><td>0</td><td>GACGTCGTCATCTGGCCCGG</td><td>CGG</td><td>||||||||[||||||||||||]PAM</td><td>NA</td><td>Intergenic</td><td>NA</td><td>NA</td></tr>
<tr><td>T2774</td><td>ACGTCGTCATCTGGCCCGGCGGG</td><td>1000</td><td></td><td></td><td></td><td></td><td></td><td></td><td></td><td></td><td></td></tr>
<tr><td>Oligo fwd</td><td>CACCACGTCGTCATCTGGCCCGGC</td><td></td><td></td><td></td><td></td><td></td><td></td><td></td><td></td><td></td><td></td></tr>
<tr><td>Oligo rev</td><td>AAACGCCGGGCCAGATGACGACGT</td><td></td><td></td><td></td><td></td><td></td><td></td><td></td><td></td><td></td><td></td></tr>
<tr><td>Chromosome</td><td>start</td><td>end</td><td>strand</td><td>MM</td><td>target_seq</td><td>PAM</td><td>alignment</td><td>distance</td><td>position</td><td>gene name</td><td>gene id</td></tr>
<tr><td>GU305914.1</td><td>43365</td><td>43387</td><td>+</td><td>0</td><td>ACGTCGTCATCTGGCCCGGC</td><td>GGG</td><td>||||||||[||||||||||||]PAM</td><td>NA</td><td>Intergenic</td><td>NA</td><td>NA</td></tr>
<tr><td>T2775</td><td>CGTCGTCATCTGGCCCGGCGGGG</td><td>1000</td><td></td><td></td><td></td><td></td><td></td><td></td><td></td><td></td><td></td></tr>
<tr><td>Oligo fwd</td><td>CACCCGTCGTCATCTGGCCCGGCG</td><td></td><td></td><td></td><td></td><td></td><td></td><td></td><td></td><td></td><td></td></tr>
<tr><td>Oligo rev</td><td>AAACCGCCGGGCCAGATGACGACG</td><td></td><td></td><td></td><td></td><td></td><td></td><td></td><td></td><td></td><td></td></tr>
<tr><td>Chromosome</td><td>start</td><td>end</td><td>strand</td><td>MM</td><td>target_seq</td><td>PAM</td><td>alignment</td><td>distance</td><td>position</td><td>gene name</td><td>gene id</td></tr>
<tr><td>GU305914.1</td><td>43366</td><td>43388</td><td>+</td><td>0</td><td>CGTCGTCATCTGGCCCGGCG</td><td>GGG</td><td>||||||||[||||||||||||]PAM</td><td>NA</td><td>Intergenic</td><td>NA</td><td>NA</td></tr>
<tr><td>T2776</td><td>CGTCCGAGTCGACGACGTATCGG</td><td>1000</td><td></td><td></td><td></td><td></td><td></td><td></td><td></td><td></td><td></td></tr>
<tr><td>Oligo fwd</td><td>CACCCGTCCGAGTCGACGACGTAT</td><td></td><td></td><td></td><td></td><td></td><td></td><td></td><td></td><td></td><td></td></tr>
<tr><td>Oligo rev</td><td>AAACATACGTCGTCGACTCGGACG</td><td></td><td></td><td></td><td></td><td></td><td></td><td></td><td></td><td></td><td></td></tr>
<tr><td>Chromosome</td><td>start</td><td>end</td><td>strand</td><td>MM</td><td>target_seq</td><td>PAM</td><td>alignment</td><td>distance</td><td>position</td><td>gene name</td><td>gene id</td></tr>
<tr><td>GU305914.1</td><td>43417</td><td>43439</td><td>+</td><td>0</td><td>CGTCCGAGTCGACGACGTAT</td><td>CGG</td><td>||||||||[||||||||||||]PAM</td><td>NA</td><td>Intergenic</td><td>NA</td><td>NA</td></tr>
<tr><td>T2777</td><td>CCGAGTCGACGACGTATCGGCGG</td><td>1000</td><td></td><td></td><td></td><td></td><td></td><td></td><td></td><td></td><td></td></tr>
<tr><td>Oligo fwd</td><td>CACCCCGAGTCGACGACGTATCGG</td><td></td><td></td><td></td><td></td><td></td><td></td><td></td><td></td><td></td><td></td></tr>
<tr><td>Oligo rev</td><td>AAACCCGATACGTCGTCGACTCGG</td><td></td><td></td><td></td><td></td><td></td><td></td><td></td><td></td><td></td><td></td></tr>
<tr><td>Chromosome</td><td>start</td><td>end</td><td>strand</td><td>MM</td><td>target_seq</td><td>PAM</td><td>alignment</td><td>distance</td><td>position</td><td>gene name</td><td>gene id</td></tr>
<tr><td>GU305914.1</td><td>43420</td><td>43442</td><td>+</td><td>0</td><td>CCGAGTCGACGACGTATCGG</td><td>CGG</td><td>||||||||[||||||||||||]PAM</td><td>NA</td><td>Intergenic</td><td>NA</td><td>NA</td></tr>
<tr><td>T2778</td><td>TCCGCCGCAGCACGTGTCCTCGG</td><td>1000</td><td></td><td></td><td></td><td></td><td></td><td></td><td></td><td></td><td></td></tr>
<tr><td>Oligo fwd</td><td>CACCTCCGCCGCAGCACGTGTCCT</td><td></td><td></td><td></td><td></td><td></td><td></td><td></td><td></td><td></td><td></td></tr>
<tr><td>Oligo rev</td><td>AAACAGGACACGTGCTGCGGCGGA</td><td></td><td></td><td></td><td></td><td></td><td></td><td></td><td></td><td></td><td></td></tr>
<tr><td>Chromosome</td><td>start</td><td>end</td><td>strand</td><td>MM</td><td>target_seq</td><td>PAM</td><td>alignment</td><td>distance</td><td>position</td><td>gene name</td><td>gene id</td></tr>
<tr><td>GU305914.1</td><td>43517</td><td>43539</td><td>+</td><td>0</td><td>TCCGCCGCAGCACGTGTCCT</td><td>CGG</td><td>||||||||[||||||||||||]PAM</td><td>NA</td><td>Intergenic</td><td>NA</td><td>NA</td></tr>
<tr><td>T2779</td><td>GGCGCCGCCGATACATCACTTGG</td><td>1000</td><td></td><td></td><td></td><td></td><td></td><td></td><td></td><td></td><td></td></tr>
<tr><td>Oligo fwd</td><td>CACCGGCGCCGCCGATACATCACT</td><td></td><td></td><td></td><td></td><td></td><td></td><td></td><td></td><td></td><td></td></tr>
<tr><td>Oligo rev</td><td>AAACAGTGATGTATCGGCGGCGCC</td><td></td><td></td><td></td><td></td><td></td><td></td><td></td><td></td><td></td><td></td></tr>
<tr><td>Chromosome</td><td>start</td><td>end</td><td>strand</td><td>MM</td><td>target_seq</td><td>PAM</td><td>alignment</td><td>distance</td><td>position</td><td>gene name</td><td>gene id</td></tr>
<tr><td>GU305914.1</td><td>43538</td><td>43560</td><td>+</td><td>0</td><td>GGCGCCGCCGATACATCACT</td><td>TGG</td><td>||||||||[||||||||||||]PAM</td><td>NA</td><td>Intergenic</td><td>NA</td><td>NA</td></tr>
<tr><td>T2780</td><td>CGATACATCACTTGGAGAGACGG</td><td>1000</td><td></td><td></td><td></td><td></td><td></td><td></td><td></td><td></td><td></td></tr>
<tr><td>Oligo fwd</td><td>CACCCGATACATCACTTGGAGAGA</td><td></td><td></td><td></td><td></td><td></td><td></td><td></td><td></td><td></td><td></td></tr>
<tr><td>Oligo rev</td><td>AAACTCTCTCCAAGTGATGTATCG</td><td></td><td></td><td></td><td></td><td></td><td></td><td></td><td></td><td></td><td></td></tr>
<tr><td>Chromosome</td><td>start</td><td>end</td><td>strand</td><td>MM</td><td>target_seq</td><td>PAM</td><td>alignment</td><td>distance</td><td>position</td><td>gene name</td><td>gene id</td></tr>
<tr><td>GU305914.1</td><td>43546</td><td>43568</td><td>+</td><td>0</td><td>CGATACATCACTTGGAGAGA</td><td>CGG</td><td>||||||||[||||||||||||]PAM</td><td>NA</td><td>Intergenic</td><td>NA</td><td>NA</td></tr>
<tr><td>T2781</td><td>GGAGAGACGGAGATCCGACCCGG</td><td>1000</td><td></td><td></td><td></td><td></td><td></td><td></td><td></td><td></td><td></td></tr>
<tr><td>Oligo fwd</td><td>CACCGGAGAGACGGAGATCCGACC</td><td></td><td></td><td></td><td></td><td></td><td></td><td></td><td></td><td></td><td></td></tr>
<tr><td>Oligo rev</td><td>AAACGGTCGGATCTCCGTCTCTCC</td><td></td><td></td><td></td><td></td><td></td><td></td><td></td><td></td><td></td><td></td></tr>
<tr><td>Chromosome</td><td>start</td><td>end</td><td>strand</td><td>MM</td><td>target_seq</td><td>PAM</td><td>alignment</td><td>distance</td><td>position</td><td>gene name</td><td>gene id</td></tr>
<tr><td>GU305914.1</td><td>43559</td><td>43581</td><td>+</td><td>0</td><td>GGAGAGACGGAGATCCGACC</td><td>CGG</td><td>||||||||[||||||||||||]PAM</td><td>NA</td><td>Intergenic</td><td>NA</td><td>NA</td></tr>
<tr><td>T2782</td><td>GCAGCACCAGCAACAACAACAGG</td><td>1000</td><td></td><td></td><td></td><td></td><td></td><td></td><td></td><td></td><td></td></tr>
<tr><td>Oligo fwd</td><td>CACCGCAGCACCAGCAACAACAAC</td><td></td><td></td><td></td><td></td><td></td><td></td><td></td><td></td><td></td><td></td></tr>
<tr><td>Oligo rev</td><td>AAACGTTGTTGTTGCTGGTGCTGC</td><td></td><td></td><td></td><td></td><td></td><td></td><td></td><td></td><td></td><td></td></tr>
<tr><td>Chromosome</td><td>start</td><td>end</td><td>strand</td><td>MM</td><td>target_seq</td><td>PAM</td><td>alignment</td><td>distance</td><td>position</td><td>gene name</td><td>gene id</td></tr>
<tr><td>GU305914.1</td><td>43616</td><td>43638</td><td>+</td><td>0</td><td>GCAGCACCAGCAACAACAAC</td><td>AGG</td><td>||||||||[||||||||||||]PAM</td><td>NA</td><td>Intergenic</td><td>NA</td><td>NA</td></tr>
<tr><td>T2783</td><td>CAGCACCAGCAACAACAACAGGG</td><td>1000</td><td></td><td></td><td></td><td></td><td></td><td></td><td></td><td></td><td></td></tr>
<tr><td>Oligo fwd</td><td>CACCCAGCACCAGCAACAACAACA</td><td></td><td></td><td></td><td></td><td></td><td></td><td></td><td></td><td></td><td></td></tr>
<tr><td>Oligo rev</td><td>AAACTGTTGTTGTTGCTGGTGCTG</td><td></td><td></td><td></td><td></td><td></td><td></td><td></td><td></td><td></td><td></td></tr>
<tr><td>Chromosome</td><td>start</td><td>end</td><td>strand</td><td>MM</td><td>target_seq</td><td>PAM</td><td>alignment</td><td>distance</td><td>position</td><td>gene name</td><td>gene id</td></tr>
<tr><td>GU305914.1</td><td>43617</td><td>43639</td><td>+</td><td>0</td><td>CAGCACCAGCAACAACAACA</td><td>GGG</td><td>||||||||[||||||||||||]PAM</td><td>NA</td><td>Intergenic</td><td>NA</td><td>NA</td></tr>
<tr><td>T2784</td><td>CACCAGCAACAACAACAGGGCGG</td><td>1000</td><td></td><td></td><td></td><td></td><td></td><td></td><td></td><td></td><td></td></tr>
<tr><td>Oligo fwd</td><td>CACCCACCAGCAACAACAACAGGG</td><td></td><td></td><td></td><td></td><td></td><td></td><td></td><td></td><td></td><td></td></tr>
<tr><td>Oligo rev</td><td>AAACCCCTGTTGTTGTTGCTGGTG</td><td></td><td></td><td></td><td></td><td></td><td></td><td></td><td></td><td></td><td></td></tr>
<tr><td>Chromosome</td><td>start</td><td>end</td><td>strand</td><td>MM</td><td>target_seq</td><td>PAM</td><td>alignment</td><td>distance</td><td>position</td><td>gene name</td><td>gene id</td></tr>
<tr><td>GU305914.1</td><td>43620</td><td>43642</td><td>+</td><td>0</td><td>CACCAGCAACAACAACAGGG</td><td>CGG</td><td>||||||||[||||||||||||]PAM</td><td>NA</td><td>Intergenic</td><td>NA</td><td>NA</td></tr>
<tr><td>T2785</td><td>CAGCAACAACAACAGGGCGGCGG</td><td>1000</td><td></td><td></td><td></td><td></td><td></td><td></td><td></td><td></td><td></td></tr>
<tr><td>Oligo fwd</td><td>CACCCAGCAACAACAACAGGGCGG</td><td></td><td></td><td></td><td></td><td></td><td></td><td></td><td></td><td></td><td></td></tr>
<tr><td>Oligo rev</td><td>AAACCCGCCCTGTTGTTGTTGCTG</td><td></td><td></td><td></td><td></td><td></td><td></td><td></td><td></td><td></td><td></td></tr>
<tr><td>Chromosome</td><td>start</td><td>end</td><td>strand</td><td>MM</td><td>target_seq</td><td>PAM</td><td>alignment</td><td>distance</td><td>position</td><td>gene name</td><td>gene id</td></tr>
<tr><td>GU305914.1</td><td>43623</td><td>43645</td><td>+</td><td>0</td><td>CAGCAACAACAACAGGGCGG</td><td>CGG</td><td>||||||||[||||||||||||]PAM</td><td>NA</td><td>Intergenic</td><td>NA</td><td>NA</td></tr>
<tr><td>T2786</td><td>CAACAACAACAGGGCGGCGGCGG</td><td>1000</td><td></td><td></td><td></td><td></td><td></td><td></td><td></td><td></td><td></td></tr>
</table>

<table>
<tr><td>Oligo fwd</td><td>CACCCAACAACAACAGGGCGGCGG</td><td></td><td></td><td></td><td></td><td></td><td></td><td></td><td></td><td></td><td></td><td></td></tr>
<tr><td>Oligo rev</td><td>AAACCCGCCGCCCTGTTGTTGTTG</td><td></td><td></td><td></td><td></td><td></td><td></td><td></td><td></td><td></td><td></td><td></td></tr>
<tr><td>Chromosome</td><td>start</td><td></td><td>end</td><td>strand</td><td>MM</td><td>target_seq</td><td>PAM</td><td>alignment</td><td>distance</td><td>position</td><td>gene name</td><td>gene id</td></tr>
<tr><td>GU305914.1</td><td>43626</td><td></td><td>43648</td><td>+</td><td>0</td><td>CAACAACAACAGGGCGGCGG</td><td>CGG</td><td>||||||||[||||||||||||]PAM</td><td>NA</td><td>Intergenic</td><td>NA</td><td>NA</td></tr>
<tr><td>T2787</td><td>CAACAGGGCGGCGGCGGCAACGG</td><td>1000</td><td></td><td></td><td></td><td></td><td></td><td></td><td></td><td></td><td></td><td></td></tr>
<tr><td>Oligo fwd</td><td>CACCCAACAGGGCGGCGGCGGCAA</td><td></td><td></td><td></td><td></td><td></td><td></td><td></td><td></td><td></td><td></td><td></td></tr>
<tr><td>Oligo rev</td><td>AAACTTGCCGCCGCCGCCCTGTTG</td><td></td><td></td><td></td><td></td><td></td><td></td><td></td><td></td><td></td><td></td><td></td></tr>
<tr><td>Chromosome</td><td>start</td><td></td><td>end</td><td>strand</td><td>MM</td><td>target_seq</td><td>PAM</td><td>alignment</td><td>distance</td><td>position</td><td>gene name</td><td>gene id</td></tr>
<tr><td>GU305914.1</td><td>43632</td><td></td><td>43654</td><td>+</td><td>0</td><td>CAACAGGGCGGCGGCGGCAA</td><td>CGG</td><td>||||||||[||||||||||||]PAM</td><td>NA</td><td>Intergenic</td><td>NA</td><td>NA</td></tr>
<tr><td>GU305914.1</td><td>90321</td><td></td><td>90343</td><td>+</td><td>4</td><td>CAAGAGGTCGGCGTCGGCGA</td><td>AGG</td><td>|||-|||-[|||||-||||-|]PAM</td><td>NA</td><td>Intergenic</td><td>NA</td><td>NA</td></tr>
<tr><td>T2788</td><td>AGGGCGGCGGCGGCAACGGTAGG</td><td>1000</td><td></td><td></td><td></td><td></td><td></td><td></td><td></td><td></td><td></td><td></td></tr>
<tr><td>Oligo fwd</td><td>CACCAGGGCGGCGGCGGCAACGGT</td><td></td><td></td><td></td><td></td><td></td><td></td><td></td><td></td><td></td><td></td><td></td></tr>
<tr><td>Oligo rev</td><td>AAACACCGTTGCCGCCGCCGCCCT</td><td></td><td></td><td></td><td></td><td></td><td></td><td></td><td></td><td></td><td></td><td></td></tr>
<tr><td>Chromosome</td><td>start</td><td></td><td>end</td><td>strand</td><td>MM</td><td>target_seq</td><td>PAM</td><td>alignment</td><td>distance</td><td>position</td><td>gene name</td><td>gene id</td></tr>
<tr><td>GU305914.1</td><td>43636</td><td></td><td>43658</td><td>+</td><td>0</td><td>AGGGCGGCGGCGGCAACGGT</td><td>AGG</td><td>||||||||[||||||||||||]PAM</td><td>NA</td><td>Intergenic</td><td>NA</td><td>NA</td></tr>
<tr><td>T2789</td><td>GGCAACGGTAGGAAAGAGAGCGG</td><td>1000</td><td></td><td></td><td></td><td></td><td></td><td></td><td></td><td></td><td></td><td></td></tr>
<tr><td>Oligo fwd</td><td>CACCGGCAACGGTAGGAAAGAGAG</td><td></td><td></td><td></td><td></td><td></td><td></td><td></td><td></td><td></td><td></td><td></td></tr>
<tr><td>Oligo rev</td><td>AAACCTCTCTTTCCTACCGTTGCC</td><td></td><td></td><td></td><td></td><td></td><td></td><td></td><td></td><td></td><td></td><td></td></tr>
<tr><td>Chromosome</td><td>start</td><td></td><td>end</td><td>strand</td><td>MM</td><td>target_seq</td><td>PAM</td><td>alignment</td><td>distance</td><td>position</td><td>gene name</td><td>gene id</td></tr>
<tr><td>GU305914.1</td><td>43647</td><td></td><td>43669</td><td>+</td><td>0</td><td>GGCAACGGTAGGAAAGAGAG</td><td>CGG</td><td>||||||||[||||||||||||]PAM</td><td>NA</td><td>Intergenic</td><td>NA</td><td>NA</td></tr>
<tr><td>T2790</td><td>AACGGTAGGAAAGAGAGCGGCGG</td><td>1000</td><td></td><td></td><td></td><td></td><td></td><td></td><td></td><td></td><td></td><td></td></tr>
<tr><td>Oligo fwd</td><td>CACCAACGGTAGGAAAGAGAGCGG</td><td></td><td></td><td></td><td></td><td></td><td></td><td></td><td></td><td></td><td></td><td></td></tr>
<tr><td>Oligo rev</td><td>AAACCCGCTCTCTTTCCTACCGTT</td><td></td><td></td><td></td><td></td><td></td><td></td><td></td><td></td><td></td><td></td><td></td></tr>
<tr><td>Chromosome</td><td>start</td><td></td><td>end</td><td>strand</td><td>MM</td><td>target_seq</td><td>PAM</td><td>alignment</td><td>distance</td><td>position</td><td>gene name</td><td>gene id</td></tr>
<tr><td>GU305914.1</td><td>43650</td><td></td><td>43672</td><td>+</td><td>0</td><td>AACGGTAGGAAAGAGAGCGG</td><td>CGG</td><td>||||||||[||||||||||||]PAM</td><td>NA</td><td>Intergenic</td><td>NA</td><td>NA</td></tr>
<tr><td>T2791</td><td>GGTAGGAAAGAGAGCGGCGGCGG</td><td>1000</td><td></td><td></td><td></td><td></td><td></td><td></td><td></td><td></td><td></td><td></td></tr>
<tr><td>Oligo fwd</td><td>CACCGGTAGGAAAGAGAGCGGCGG</td><td></td><td></td><td></td><td></td><td></td><td></td><td></td><td></td><td></td><td></td><td></td></tr>
<tr><td>Oligo rev</td><td>AAACCCGCCGCTCTCTTTCCTACC</td><td></td><td></td><td></td><td></td><td></td><td></td><td></td><td></td><td></td><td></td><td></td></tr>
<tr><td>Chromosome</td><td>start</td><td></td><td>end</td><td>strand</td><td>MM</td><td>target_seq</td><td>PAM</td><td>alignment</td><td>distance</td><td>position</td><td>gene name</td><td>gene id</td></tr>
<tr><td>GU305914.1</td><td>43653</td><td></td><td>43675</td><td>+</td><td>0</td><td>GGTAGGAAAGAGAGCGGCGG</td><td>CGG</td><td>||||||||[||||||||||||]PAM</td><td>NA</td><td>Intergenic</td><td>NA</td><td>NA</td></tr>
<tr><td>T2792</td><td>CCGCGCCGTCGAAGATCGTGAGG</td><td>1000</td><td></td><td></td><td></td><td></td><td></td><td></td><td></td><td></td><td></td><td></td></tr>
<tr><td>Oligo fwd</td><td>CACCCCGCGCCGTCGAAGATCGTG</td><td></td><td></td><td></td><td></td><td></td><td></td><td></td><td></td><td></td><td></td><td></td></tr>
<tr><td>Oligo rev</td><td>AAACCACGATCTTCGACGGCGCGG</td><td></td><td></td><td></td><td></td><td></td><td></td><td></td><td></td><td></td><td></td><td></td></tr>
<tr><td>Chromosome</td><td>start</td><td></td><td>end</td><td>strand</td><td>MM</td><td>target_seq</td><td>PAM</td><td>alignment</td><td>distance</td><td>position</td><td>gene name</td><td>gene id</td></tr>
<tr><td>GU305914.1</td><td>43741</td><td></td><td>43763</td><td>+</td><td>0</td><td>CCGCGCCGTCGAAGATCGTG</td><td>AGG</td><td>||||||||[||||||||||||]PAM</td><td>NA</td><td>Intergenic</td><td>NA</td><td>NA</td></tr>
<tr><td>T2793</td><td>GAAGATCGTGAGGTACGAAGCGG</td><td>1000</td><td></td><td></td><td></td><td></td><td></td><td></td><td></td><td></td><td></td><td></td></tr>
<tr><td>Oligo fwd</td><td>CACCGAAGATCGTGAGGTACGAAG</td><td></td><td></td><td></td><td></td><td></td><td></td><td></td><td></td><td></td><td></td><td></td></tr>
<tr><td>Oligo rev</td><td>AAACCTTCGTACCTCACGATCTTC</td><td></td><td></td><td></td><td></td><td></td><td></td><td></td><td></td><td></td><td></td><td></td></tr>
<tr><td>Chromosome</td><td>start</td><td></td><td>end</td><td>strand</td><td>MM</td><td>target_seq</td><td>PAM</td><td>alignment</td><td>distance</td><td>position</td><td>gene name</td><td>gene id</td></tr>
<tr><td>GU305914.1</td><td>43751</td><td></td><td>43773</td><td>+</td><td>0</td><td>GAAGATCGTGAGGTACGAAG</td><td>CGG</td><td>||||||||[||||||||||||]PAM</td><td>NA</td><td>Intergenic</td><td>NA</td><td>NA</td></tr>
<tr><td>T2794</td><td>TGAGGTACGAAGCGGCGCACAGG</td><td>1000</td><td></td><td></td><td></td><td></td><td></td><td></td><td></td><td></td><td></td><td></td></tr>
<tr><td>Oligo fwd</td><td>CACCTGAGGTACGAAGCGGCGCAC</td><td></td><td></td><td></td><td></td><td></td><td></td><td></td><td></td><td></td><td></td><td></td></tr>
<tr><td>Oligo rev</td><td>AAACGTGCGCCGCTTCGTACCTCA</td><td></td><td></td><td></td><td></td><td></td><td></td><td></td><td></td><td></td><td></td><td></td></tr>
<tr><td>Chromosome</td><td>start</td><td></td><td>end</td><td>strand</td><td>MM</td><td>target_seq</td><td>PAM</td><td>alignment</td><td>distance</td><td>position</td><td>gene name</td><td>gene id</td></tr>
<tr><td>GU305914.1</td><td>43759</td><td></td><td>43781</td><td>+</td><td>0</td><td>TGAGGTACGAAGCGGCGCAC</td><td>AGG</td><td>||||||||[||||||||||||]PAM</td><td>NA</td><td>Intergenic</td><td>NA</td><td>NA</td></tr>
<tr><td>T2795</td><td>AGCGGCGCACAGGTACACGATGG</td><td>1000</td><td></td><td></td><td></td><td></td><td></td><td></td><td></td><td></td><td></td><td></td></tr>
<tr><td>Oligo fwd</td><td>CACCAGCGGCGCACAGGTACACGA</td><td></td><td></td><td></td><td></td><td></td><td></td><td></td><td></td><td></td><td></td><td></td></tr>
<tr><td>Oligo rev</td><td>AAACTCGTGTACCTGTGCGCCGCT</td><td></td><td></td><td></td><td></td><td></td><td></td><td></td><td></td><td></td><td></td><td></td></tr>
<tr><td>Chromosome</td><td>start</td><td></td><td>end</td><td>strand</td><td>MM</td><td>target_seq</td><td>PAM</td><td>alignment</td><td>distance</td><td>position</td><td>gene name</td><td>gene id</td></tr>
<tr><td>GU305914.1</td><td>43769</td><td></td><td>43791</td><td>+</td><td>0</td><td>AGCGGCGCACAGGTACACGA</td><td>TGG</td><td>||||||||[||||||||||||]PAM</td><td>NA</td><td>Intergenic</td><td>NA</td><td>NA</td></tr>
<tr><td>T2796</td><td>GCACAGGTACACGATGGCCGAGG</td><td>1000</td><td></td><td></td><td></td><td></td><td></td><td></td><td></td><td></td><td></td><td></td></tr>
<tr><td>Oligo fwd</td><td>CACCGCACAGGTACACGATGGCCG</td><td></td><td></td><td></td><td></td><td></td><td></td><td></td><td></td><td></td><td></td><td></td></tr>
<tr><td>Oligo rev</td><td>AAACCGGCCATCGTGTACCTGTGC</td><td></td><td></td><td></td><td></td><td></td><td></td><td></td><td></td><td></td><td></td><td></td></tr>
<tr><td>Chromosome</td><td>start</td><td></td><td>end</td><td>strand</td><td>MM</td><td>target_seq</td><td>PAM</td><td>alignment</td><td>distance</td><td>position</td><td>gene name</td><td>gene id</td></tr>
<tr><td>GU305914.1</td><td>43775</td><td></td><td>43797</td><td>+</td><td>0</td><td>GCACAGGTACACGATGGCCG</td><td>AGG</td><td>||||||||[||||||||||||]PAM</td><td>NA</td><td>Intergenic</td><td>NA</td><td>NA</td></tr>
<tr><td>T2797</td><td>CACAGGTACACGATGGCCGAGGG</td><td>1000</td><td></td><td></td><td></td><td></td><td></td><td></td><td></td><td></td><td></td><td></td></tr>
<tr><td>Oligo fwd</td><td>CACCCACAGGTACACGATGGCCGA</td><td></td><td></td><td></td><td></td><td></td><td></td><td></td><td></td><td></td><td></td><td></td></tr>
<tr><td>Oligo rev</td><td>AAACTCGGCCATCGTGTACCTGTG</td><td></td><td></td><td></td><td></td><td></td><td></td><td></td><td></td><td></td><td></td><td></td></tr>
<tr><td>Chromosome</td><td>start</td><td></td><td>end</td><td>strand</td><td>MM</td><td>target_seq</td><td>PAM</td><td>alignment</td><td>distance</td><td>position</td><td>gene name</td><td>gene id</td></tr>
<tr><td>GU305914.1</td><td>43776</td><td></td><td>43798</td><td>+</td><td>0</td><td>CACAGGTACACGATGGCCGA</td><td>GGG</td><td>||||||||[||||||||||||]PAM</td><td>NA</td><td>Intergenic</td><td>NA</td><td>NA</td></tr>
<tr><td>T2798</td><td>GGTTTCATCCGCTCCGTGTCCGG</td><td>1000</td><td></td><td></td><td></td><td></td><td></td><td></td><td></td><td></td><td></td><td></td></tr>
<tr><td>Oligo fwd</td><td>CACCGGTTTCATCCGCTCCGTGTC</td><td></td><td></td><td></td><td></td><td></td><td></td><td></td><td></td><td></td><td></td><td></td></tr>
<tr><td>Oligo rev</td><td>AAACGACACGGAGCGGATGAAACC</td><td></td><td></td><td></td><td></td><td></td><td></td><td></td><td></td><td></td><td></td><td></td></tr>
<tr><td>Chromosome</td><td>start</td><td></td><td>end</td><td>strand</td><td>MM</td><td>target_seq</td><td>PAM</td><td>alignment</td><td>distance</td><td>position</td><td>gene name</td><td>gene id</td></tr>
<tr><td>GU305914.1</td><td>43797</td><td></td><td>43819</td><td>+</td><td>0</td><td>GGTTTCATCCGCTCCGTGTC</td><td>CGG</td><td>||||||||[||||||||||||]PAM</td><td>NA</td><td>Intergenic</td><td>NA</td><td>NA</td></tr>
<tr><td>T2799</td><td>GTTTCATCCGCTCCGTGTCCGGG</td><td>1000</td><td></td><td></td><td></td><td></td><td></td><td></td><td></td><td></td><td></td><td></td></tr>
<tr><td>Oligo fwd</td><td>CACCGTTTCATCCGCTCCGTGTCC</td><td></td><td></td><td></td><td></td><td></td><td></td><td></td><td></td><td></td><td></td><td></td></tr>
<tr><td>Oligo rev</td><td>AAACGGACACGGAGCGGATGAAAC</td><td></td><td></td><td></td><td></td><td></td><td></td><td></td><td></td><td></td><td></td><td></td></tr>
<tr><td>Chromosome</td><td>start</td><td></td><td>end</td><td>strand</td><td>MM</td><td>target_seq</td><td>PAM</td><td>alignment</td><td>distance</td><td>position</td><td>gene name</td><td>gene id</td></tr>
<tr><td>GU305914.1</td><td>43798</td><td></td><td>43820</td><td>+</td><td>0</td><td>GTTTCATCCGCTCCGTGTCC</td><td>GGG</td><td>||||||||[||||||||||||]PAM</td><td>NA</td><td>Intergenic</td><td>NA</td><td>NA</td></tr>
<tr><td>T2800</td><td>CTACAAGATGATGAATCGAGCGG</td><td>1000</td><td></td><td></td><td></td><td></td><td></td><td></td><td></td><td></td><td></td><td></td></tr>
<tr><td>Oligo fwd</td><td>CACCCTACAAGATGATGAATCGAG</td><td></td><td></td><td></td><td></td><td></td><td></td><td></td><td></td><td></td><td></td><td></td></tr>
<tr><td>Oligo rev</td><td>AAACCTCGATTCATCATCTTGTAG</td><td></td><td></td><td></td><td></td><td></td><td></td><td></td><td></td><td></td><td></td><td></td></tr>
<tr><td>Chromosome</td><td>start</td><td></td><td>end</td><td>strand</td><td>MM</td><td>target_seq</td><td>PAM</td><td>alignment</td><td>distance</td><td>position</td><td>gene name</td><td>gene id</td></tr>
<tr><td>GU305914.1</td><td>43823</td><td></td><td>43845</td><td>+</td><td>0</td><td>CTACAAGATGATGAATCGAG</td><td>CGG</td><td>||||||||[||||||||||||]PAM</td><td>NA</td><td>Intergenic</td><td>NA</td><td>NA</td></tr>
</table>

<table>
<tr><td>T2801</td><td>CAAGATGATGAATCGAGCGGAGG</td><td>1000</td><td></td><td></td><td></td><td></td><td></td><td></td><td></td><td></td><td></td></tr>
<tr><td>Oligo fwd</td><td>CACCCAAGATGATGAATCGAGCGG</td><td></td><td></td><td></td><td></td><td></td><td></td><td></td><td></td><td></td><td></td></tr>
<tr><td>Oligo rev</td><td>AAACCCGCTCGATTCATCATCTTG</td><td></td><td></td><td></td><td></td><td></td><td></td><td></td><td></td><td></td><td></td></tr>
<tr><td>Chromosome</td><td>start</td><td>end</td><td>strand</td><td>MM</td><td>target_seq</td><td>PAM</td><td>alignment</td><td>distance</td><td>position</td><td>gene name</td><td>gene id</td></tr>
<tr><td>GU305914.1</td><td>43826</td><td>43848</td><td>+</td><td>0</td><td>CAAGATGATGAATCGAGCGG</td><td>AGG</td><td>||||||||[||||||||||||]PAM</td><td>NA</td><td>Intergenic</td><td>NA</td><td>NA</td></tr>
<tr><td>T2802</td><td>ATCGAGCGGAGGATCGCAAGAGG</td><td>1000</td><td></td><td></td><td></td><td></td><td></td><td></td><td></td><td></td><td></td></tr>
<tr><td>Oligo fwd</td><td>CACCATCGAGCGGAGGATCGCAAG</td><td></td><td></td><td></td><td></td><td></td><td></td><td></td><td></td><td></td><td></td></tr>
<tr><td>Oligo rev</td><td>AAACCTTGCGATCCTCCGCTCGAT</td><td></td><td></td><td></td><td></td><td></td><td></td><td></td><td></td><td></td><td></td></tr>
<tr><td>Chromosome</td><td>start</td><td>end</td><td>strand</td><td>MM</td><td>target_seq</td><td>PAM</td><td>alignment</td><td>distance</td><td>position</td><td>gene name</td><td>gene id</td></tr>
<tr><td>GU305914.1</td><td>43837</td><td>43859</td><td>+</td><td>0</td><td>ATCGAGCGGAGGATCGCAAG</td><td>AGG</td><td>||||||||[||||||||||||]PAM</td><td>NA</td><td>Intergenic</td><td>NA</td><td>NA</td></tr>
<tr><td>T2803</td><td>GTTCATCTTCTGCACGTCGTCGG</td><td>1000</td><td></td><td></td><td></td><td></td><td></td><td></td><td></td><td></td><td></td></tr>
<tr><td>Oligo fwd</td><td>CACCGTTCATCTTCTGCACGTCGT</td><td></td><td></td><td></td><td></td><td></td><td></td><td></td><td></td><td></td><td></td></tr>
<tr><td>Oligo rev</td><td>AAACACGACGTGCAGAAGATGAAC</td><td></td><td></td><td></td><td></td><td></td><td></td><td></td><td></td><td></td><td></td></tr>
<tr><td>Chromosome</td><td>start</td><td>end</td><td>strand</td><td>MM</td><td>target_seq</td><td>PAM</td><td>alignment</td><td>distance</td><td>position</td><td>gene name</td><td>gene id</td></tr>
<tr><td>GU305914.1</td><td>43913</td><td>43935</td><td>+</td><td>0</td><td>GTTCATCTTCTGCACGTCGT</td><td>CGG</td><td>||||||||[||||||||||||]PAM</td><td>NA</td><td>Intergenic</td><td>NA</td><td>NA</td></tr>
<tr><td>T2804</td><td>ATCTTCTGCACGTCGTCGGACGG</td><td>1000</td><td></td><td></td><td></td><td></td><td></td><td></td><td></td><td></td><td></td></tr>
<tr><td>Oligo fwd</td><td>CACCATCTTCTGCACGTCGTCGGA</td><td></td><td></td><td></td><td></td><td></td><td></td><td></td><td></td><td></td><td></td></tr>
<tr><td>Oligo rev</td><td>AAACTCCGACGACGTGCAGAAGAT</td><td></td><td></td><td></td><td></td><td></td><td></td><td></td><td></td><td></td><td></td></tr>
<tr><td>Chromosome</td><td>start</td><td>end</td><td>strand</td><td>MM</td><td>target_seq</td><td>PAM</td><td>alignment</td><td>distance</td><td>position</td><td>gene name</td><td>gene id</td></tr>
<tr><td>GU305914.1</td><td>43917</td><td>43939</td><td>+</td><td>0</td><td>ATCTTCTGCACGTCGTCGGA</td><td>CGG</td><td>||||||||[||||||||||||]PAM</td><td>NA</td><td>Intergenic</td><td>NA</td><td>NA</td></tr>
<tr><td>T2805</td><td>GTCGTCGGACGGAGACATCCTGG</td><td>1000</td><td></td><td></td><td></td><td></td><td></td><td></td><td></td><td></td><td></td></tr>
<tr><td>Oligo fwd</td><td>CACCGTCGTCGGACGGAGACATCC</td><td></td><td></td><td></td><td></td><td></td><td></td><td></td><td></td><td></td><td></td></tr>
<tr><td>Oligo rev</td><td>AAACGGATGTCTCCGTCCGACGAC</td><td></td><td></td><td></td><td></td><td></td><td></td><td></td><td></td><td></td><td></td></tr>
<tr><td>Chromosome</td><td>start</td><td>end</td><td>strand</td><td>MM</td><td>target_seq</td><td>PAM</td><td>alignment</td><td>distance</td><td>position</td><td>gene name</td><td>gene id</td></tr>
<tr><td>GU305914.1</td><td>43928</td><td>43950</td><td>+</td><td>0</td><td>GTCGTCGGACGGAGACATCC</td><td>TGG</td><td>||||||||[||||||||||||]PAM</td><td>NA</td><td>Intergenic</td><td>NA</td><td>NA</td></tr>
<tr><td>T2806</td><td>GGACGGAGACATCCTGGAGTCGG</td><td>1000</td><td></td><td></td><td></td><td></td><td></td><td></td><td></td><td></td><td></td></tr>
<tr><td>Oligo fwd</td><td>CACCGGACGGAGACATCCTGGAGT</td><td></td><td></td><td></td><td></td><td></td><td></td><td></td><td></td><td></td><td></td></tr>
<tr><td>Oligo rev</td><td>AAACACTCCAGGATGTCTCCGTCC</td><td></td><td></td><td></td><td></td><td></td><td></td><td></td><td></td><td></td><td></td></tr>
<tr><td>Chromosome</td><td>start</td><td>end</td><td>strand</td><td>MM</td><td>target_seq</td><td>PAM</td><td>alignment</td><td>distance</td><td>position</td><td>gene name</td><td>gene id</td></tr>
<tr><td>GU305914.1</td><td>43934</td><td>43956</td><td>+</td><td>0</td><td>GGACGGAGACATCCTGGAGT</td><td>CGG</td><td>||||||||[||||||||||||]PAM</td><td>NA</td><td>Intergenic</td><td>NA</td><td>NA</td></tr>
<tr><td>T2807</td><td>CCGCCACCAGAAACACATCACGG</td><td>1000</td><td></td><td></td><td></td><td></td><td></td><td></td><td></td><td></td><td></td></tr>
<tr><td>Oligo fwd</td><td>CACCCCGCCACCAGAAACACATCA</td><td></td><td></td><td></td><td></td><td></td><td></td><td></td><td></td><td></td><td></td></tr>
<tr><td>Oligo rev</td><td>AAACTGATGTGTTTCTGGTGGCGG</td><td></td><td></td><td></td><td></td><td></td><td></td><td></td><td></td><td></td><td></td></tr>
<tr><td>Chromosome</td><td>start</td><td>end</td><td>strand</td><td>MM</td><td>target_seq</td><td>PAM</td><td>alignment</td><td>distance</td><td>position</td><td>gene name</td><td>gene id</td></tr>
<tr><td>GU305914.1</td><td>43973</td><td>43995</td><td>+</td><td>0</td><td>CCGCCACCAGAAACACATCA</td><td>CGG</td><td>||||||||[||||||||||||]PAM</td><td>NA</td><td>Intergenic</td><td>NA</td><td>NA</td></tr>
<tr><td>T2808</td><td>CATCACGGATTTCAGCAAGCTGG</td><td>1000</td><td></td><td></td><td></td><td></td><td></td><td></td><td></td><td></td><td></td></tr>
<tr><td>Oligo fwd</td><td>CACCCATCACGGATTTCAGCAAGC</td><td></td><td></td><td></td><td></td><td></td><td></td><td></td><td></td><td></td><td></td></tr>
<tr><td>Oligo rev</td><td>AAACGCTTGCTGAAATCCGTGATG</td><td></td><td></td><td></td><td></td><td></td><td></td><td></td><td></td><td></td><td></td></tr>
<tr><td>Chromosome</td><td>start</td><td>end</td><td>strand</td><td>MM</td><td>target_seq</td><td>PAM</td><td>alignment</td><td>distance</td><td>position</td><td>gene name</td><td>gene id</td></tr>
<tr><td>GU305914.1</td><td>43988</td><td>44010</td><td>+</td><td>0</td><td>CATCACGGATTTCAGCAAGC</td><td>TGG</td><td>||||||||[||||||||||||]PAM</td><td>NA</td><td>Intergenic</td><td>NA</td><td>NA</td></tr>
<tr><td>T2809</td><td>AAGCTGGACCGCGCGATCCACGG</td><td>1000</td><td></td><td></td><td></td><td></td><td></td><td></td><td></td><td></td><td></td></tr>
<tr><td>Oligo fwd</td><td>CACCAAGCTGGACCGCGCGATCCA</td><td></td><td></td><td></td><td></td><td></td><td></td><td></td><td></td><td></td><td></td></tr>
<tr><td>Oligo rev</td><td>AAACTGGATCGCGCGGTCCAGCTT</td><td></td><td></td><td></td><td></td><td></td><td></td><td></td><td></td><td></td><td></td></tr>
<tr><td>Chromosome</td><td>start</td><td>end</td><td>strand</td><td>MM</td><td>target_seq</td><td>PAM</td><td>alignment</td><td>distance</td><td>position</td><td>gene name</td><td>gene id</td></tr>
<tr><td>GU305914.1</td><td>44004</td><td>44026</td><td>+</td><td>0</td><td>AAGCTGGACCGCGCGATCCA</td><td>CGG</td><td>||||||||[||||||||||||]PAM</td><td>NA</td><td>Intergenic</td><td>NA</td><td>NA</td></tr>
<tr><td>T2810</td><td>AGCTGGACCGCGCGATCCACGGG</td><td>1000</td><td></td><td></td><td></td><td></td><td></td><td></td><td></td><td></td><td></td></tr>
<tr><td>Oligo fwd</td><td>CACCAGCTGGACCGCGCGATCCAC</td><td></td><td></td><td></td><td></td><td></td><td></td><td></td><td></td><td></td><td></td></tr>
<tr><td>Oligo rev</td><td>AAACGTGGATCGCGCGGTCCAGCT</td><td></td><td></td><td></td><td></td><td></td><td></td><td></td><td></td><td></td><td></td></tr>
<tr><td>Chromosome</td><td>start</td><td>end</td><td>strand</td><td>MM</td><td>target_seq</td><td>PAM</td><td>alignment</td><td>distance</td><td>position</td><td>gene name</td><td>gene id</td></tr>
<tr><td>GU305914.1</td><td>44005</td><td>44027</td><td>+</td><td>0</td><td>AGCTGGACCGCGCGATCCAC</td><td>GGG</td><td>||||||||[||||||||||||]PAM</td><td>NA</td><td>Intergenic</td><td>NA</td><td>NA</td></tr>
<tr><td>T2811</td><td>GCTGGACCGCGCGATCCACGGGG</td><td>1000</td><td></td><td></td><td></td><td></td><td></td><td></td><td></td><td></td><td></td></tr>
<tr><td>Oligo fwd</td><td>CACCGCTGGACCGCGCGATCCACG</td><td></td><td></td><td></td><td></td><td></td><td></td><td></td><td></td><td></td><td></td></tr>
<tr><td>Oligo rev</td><td>AAACCGTGGATCGCGCGGTCCAGC</td><td></td><td></td><td></td><td></td><td></td><td></td><td></td><td></td><td></td><td></td></tr>
<tr><td>Chromosome</td><td>start</td><td>end</td><td>strand</td><td>MM</td><td>target_seq</td><td>PAM</td><td>alignment</td><td>distance</td><td>position</td><td>gene name</td><td>gene id</td></tr>
<tr><td>GU305914.1</td><td>44006</td><td>44028</td><td>+</td><td>0</td><td>GCTGGACCGCGCGATCCACG</td><td>GGG</td><td>||||||||[||||||||||||]PAM</td><td>NA</td><td>Intergenic</td><td>NA</td><td>NA</td></tr>
<tr><td>T2812</td><td>CGATCCACGGGGTGAGCTGCAGG</td><td>1000</td><td></td><td></td><td></td><td></td><td></td><td></td><td></td><td></td><td></td></tr>
<tr><td>Oligo fwd</td><td>CACCCGATCCACGGGGTGAGCTGC</td><td></td><td></td><td></td><td></td><td></td><td></td><td></td><td></td><td></td><td></td></tr>
<tr><td>Oligo rev</td><td>AAACGCAGCTCACCCCGTGGATCG</td><td></td><td></td><td></td><td></td><td></td><td></td><td></td><td></td><td></td><td></td></tr>
<tr><td>Chromosome</td><td>start</td><td>end</td><td>strand</td><td>MM</td><td>target_seq</td><td>PAM</td><td>alignment</td><td>distance</td><td>position</td><td>gene name</td><td>gene id</td></tr>
<tr><td>GU305914.1</td><td>44017</td><td>44039</td><td>+</td><td>0</td><td>CGATCCACGGGGTGAGCTGC</td><td>AGG</td><td>||||||||[||||||||||||]PAM</td><td>NA</td><td>Intergenic</td><td>NA</td><td>NA</td></tr>
<tr><td>T2813</td><td>CCCGAACCTGAGCGAACAGCAGG</td><td>1000</td><td></td><td></td><td></td><td></td><td></td><td></td><td></td><td></td><td></td></tr>
<tr><td>Oligo fwd</td><td>CACCCCCGAACCTGAGCGAACAGC</td><td></td><td></td><td></td><td></td><td></td><td></td><td></td><td></td><td></td><td></td></tr>
<tr><td>Oligo rev</td><td>AAACGCTGTTCGCTCAGGTTCGGG</td><td></td><td></td><td></td><td></td><td></td><td></td><td></td><td></td><td></td><td></td></tr>
<tr><td>Chromosome</td><td>start</td><td>end</td><td>strand</td><td>MM</td><td>target_seq</td><td>PAM</td><td>alignment</td><td>distance</td><td>position</td><td>gene name</td><td>gene id</td></tr>
<tr><td>GU305914.1</td><td>44060</td><td>44082</td><td>+</td><td>0</td><td>CCCGAACCTGAGCGAACAGC</td><td>AGG</td><td>||||||||[||||||||||||]PAM</td><td>NA</td><td>Intergenic</td><td>NA</td><td>NA</td></tr>
<tr><td>T2814</td><td>GAACCTGAGCGAACAGCAGGTGG</td><td>1000</td><td></td><td></td><td></td><td></td><td></td><td></td><td></td><td></td><td></td></tr>
<tr><td>Oligo fwd</td><td>CACCGAACCTGAGCGAACAGCAGG</td><td></td><td></td><td></td><td></td><td></td><td></td><td></td><td></td><td></td><td></td></tr>
<tr><td>Oligo rev</td><td>AAACCCTGCTGTTCGCTCAGGTTC</td><td></td><td></td><td></td><td></td><td></td><td></td><td></td><td></td><td></td><td></td></tr>
<tr><td>Chromosome</td><td>start</td><td>end</td><td>strand</td><td>MM</td><td>target_seq</td><td>PAM</td><td>alignment</td><td>distance</td><td>position</td><td>gene name</td><td>gene id</td></tr>
<tr><td>GU305914.1</td><td>44063</td><td>44085</td><td>+</td><td>0</td><td>GAACCTGAGCGAACAGCAGG</td><td>TGG</td><td>||||||||[||||||||||||]PAM</td><td>NA</td><td>Intergenic</td><td>NA</td><td>NA</td></tr>
<tr><td>T2815</td><td>CTTCAACCGCATCACGTACGTGG</td><td>1000</td><td></td><td></td><td></td><td></td><td></td><td></td><td></td><td></td><td></td></tr>
<tr><td>Oligo fwd</td><td>CACCCTTCAACCGCATCACGTACG</td><td></td><td></td><td></td><td></td><td></td><td></td><td></td><td></td><td></td><td></td></tr>
<tr><td>Oligo rev</td><td>AAACCGTACGTGATGCGGTTGAAG</td><td></td><td></td><td></td><td></td><td></td><td></td><td></td><td></td><td></td><td></td></tr>
<tr><td>Chromosome</td><td>start</td><td>end</td><td>strand</td><td>MM</td><td>target_seq</td><td>PAM</td><td>alignment</td><td>distance</td><td>position</td><td>gene name</td><td>gene id</td></tr>
</table>

GU305914.1 44120 44142 + 0 CTTCAACCGCATCACGTACG TGG ||||||||[||||||||||||]PAM NA Intergenic NA NA

T2816 CGTCAAACATTACTGCGACAAGG 1000
Oligo fwd CACCGTCAAACATTACTGCGACA
Oligo rev AAACTGTCGCAGTAATGTTTGACG
Chromosome start end strand MM target_seq PAM alignment distance position gene name gene id
GU305914.1 44147 44169 + 0 CGTCAAACATTACTGCGACA AGG ||||||||[||||||||||||]PAM NA Intergenic NA NA

T2817 ATTACTGCGACAAGGATTCCAGG 1000
Oligo fwd CACCATTACTGCGACAAGGATTCC
Oligo rev AAACGGAATCCTTGTCGCAGTAAT
Chromosome start end strand MM target_seq PAM alignment distance position gene name gene id
GU305914.1 44155 44177 + 0 ATTACTGCGACAAGGATTCC AGG ||||||||[||||||||||||]PAM NA Intergenic NA NA

T2818 TCCGCGACCAGCTCACCAAACGG 1000
Oligo fwd CACCTCCGCGACCAGCTCACCAAA
Oligo rev AAACTTTGGTGAGCTGGTCGCGGA
Chromosome start end strand MM target_seq PAM alignment distance position gene name gene id
GU305914.1 44191 44213 + 0 TCCGCGACCAGCTCACCAAA CGG ||||||||[||||||||||||]PAM NA Intergenic NA NA

T2819 CTCACCAAACGGTGCAGCGAAGG 1000
Oligo fwd CACCCTCACCAAACGGTGCAGCGA
Oligo rev AAACTCGCTGCACCGTTTGGTGAG
Chromosome start end strand MM target_seq PAM alignment distance position gene name gene id
GU305914.1 44202 44224 + 0 CTCACCAAACGGTGCAGCGA AGG ||||||||[||||||||||||]PAM NA Intergenic NA NA

T2820 GAAGGCTCGCGCCTCAACCTCGG 1000
Oligo fwd CACCGAAGGCTCGCGCCTCAACCT
Oligo rev AAACAGGTTGAGGCGCGAGCCTTC
Chromosome start end strand MM target_seq PAM alignment distance position gene name gene id
GU305914.1 44220 44242 + 0 GAAGGCTCGCGCCTCAACCT CGG ||||||||[||||||||||||]PAM NA Intergenic NA NA

T2821 AAGGCTCGCGCCTCAACCTCGGG 1000
Oligo fwd CACCAAGGCTCGCGCCTCAACCTC
Oligo rev AAACGAGGTTGAGGCGCGAGCCTT
Chromosome start end strand MM target_seq PAM alignment distance position gene name gene id
GU305914.1 44221 44243 + 0 AAGGCTCGCGCCTCAACCTC GGG ||||||||[||||||||||||]PAM NA Intergenic NA NA

T2822 GATACGTCGTTTCATCAGCTCGG 1000
Oligo fwd CACCGATACGTCGTTTCATCAGCT
Oligo rev AAACAGCTGATGAAACGACGTATC
Chromosome start end strand MM target_seq PAM alignment distance position gene name gene id
GU305914.1 44243 44265 + 0 GATACGTCGTTTCATCAGCT CGG ||||||||[||||||||||||]PAM NA Intergenic NA NA

T2823 TTTCATCAGCTCGGTGAACGTGG 1000
Oligo fwd CACCTTTCATCAGCTCGGTGAACG
Oligo rev AAACCGTTCACCGAGCTGATGAAA
Chromosome start end strand MM target_seq PAM alignment distance position gene name gene id
GU305914.1 44252 44274 + 0 TTTCATCAGCTCGGTGAACG TGG ||||||||[||||||||||||]PAM NA Intergenic NA NA

T2824 CCGAGACCTCTGCATGATCCTGG 1000
Oligo fwd CACCCCGAGACCTCTGCATGATCC
Oligo rev AAACGGATCATGCAGAGGTCTCGG
Chromosome start end strand MM target_seq PAM alignment distance position gene name gene id
GU305914.1 44282 44304 + 0 CCGAGACCTCTGCATGATCC TGG ||||||||[||||||||||||]PAM NA Intergenic NA NA

T2825 AGACCTCTGCATGATCCTGGTGG 1000
Oligo fwd CACCAGACCTCTGCATGATCCTGG
Oligo rev AAACCCAGGATCATGCAGAGGTCT
Chromosome start end strand MM target_seq PAM alignment distance position gene name gene id
GU305914.1 44285 44307 + 0 AGACCTCTGCATGATCCTGG TGG ||||||||[||||||||||||]PAM NA Intergenic NA NA

T2826 GACCTCTGCATGATCCTGGTGGG 1000
Oligo fwd CACCGACCTCTGCATGATCCTGGT
Oligo rev AAACACCAGGATCATGCAGAGGTC
Chromosome start end strand MM target_seq PAM alignment distance position gene name gene id
GU305914.1 44286 44308 + 0 GACCTCTGCATGATCCTGGT GGG ||||||||[||||||||||||]PAM NA Intergenic NA NA

T2827 ACCTCTGCATGATCCTGGTGGGG 1000
Oligo fwd CACCACCTCTGCATGATCCTGGTG
Oligo rev AAACCACCAGGATCATGCAGAGGT
Chromosome start end strand MM target_seq PAM alignment distance position gene name gene id
GU305914.1 44287 44309 + 0 ACCTCTGCATGATCCTGGTG GGG ||||||||[||||||||||||]PAM NA Intergenic NA NA

T2828 TGTGCCAGACGCCCCACATGTGG 1000
Oligo fwd CACCTGTGCCAGACGCCCCACATG
Oligo rev AAACCATGTGGGGCGTCTGGCACA
Chromosome start end strand MM target_seq PAM alignment distance position gene name gene id
GU305914.1 44314 44336 + 0 TGTGCCAGACGCCCCACATG TGG ||||||||[||||||||||||]PAM NA Intergenic NA NA

T2829 GTGCCAGACGCCCCACATGTGGG 1000
Oligo fwd CACCGTGCCAGACGCCCCACATGT
Oligo rev AAACACATGTGGGGCGTCTGGCAC
Chromosome start end strand MM target_seq PAM alignment distance position gene name gene id
GU305914.1 44315 44337 + 0 GTGCCAGACGCCCCACATGT GGG ||||||||[||||||||||||]PAM NA Intergenic NA NA

T2830 AGACGCCCCACATGTGGGCCCGG 1000
Oligo fwd CACCAGACGCCCCACATGTGGGCC
Oligo rev AAACGGCCCACATGTGGGGCGTCT

<table>
<tr><td>Chromosome</td><td>start</td><td>end</td><td>strand</td><td>MM</td><td>target_seq</td><td>PAM</td><td>alignment</td><td>distance</td><td>position</td><td>gene name</td><td>gene id</td></tr>
<tr><td>GU305914.1</td><td>44320</td><td>44342</td><td>+</td><td>0</td><td>AGACGCCCCACATGTGGGCC</td><td>CGG</td><td>||||||||[||||||||||||]PAM</td><td>NA</td><td>Intergenic</td><td>NA</td><td>NA</td></tr>
<tr><td></td><td></td><td></td><td></td><td></td><td></td><td></td><td></td><td></td><td></td><td></td><td></td></tr>
<tr><td>T2831</td><td>CCGACGAGAAGATCGACCTCAGG</td><td>1000</td><td></td><td></td><td></td><td></td><td></td><td></td><td></td><td></td><td></td></tr>
<tr><td>Oligo fwd</td><td>CACCCCGACGAGAAGATCGACCTC</td><td></td><td></td><td></td><td></td><td></td><td></td><td></td><td></td><td></td><td></td></tr>
<tr><td>Oligo rev</td><td>AAACGAGGTCGATCTTCTCGTCGG</td><td></td><td></td><td></td><td></td><td></td><td></td><td></td><td></td><td></td><td></td></tr>
<tr><td>Chromosome</td><td>start</td><td>end</td><td>strand</td><td>MM</td><td>target_seq</td><td>PAM</td><td>alignment</td><td>distance</td><td>position</td><td>gene name</td><td>gene id</td></tr>
<tr><td>GU305914.1</td><td>44404</td><td>44426</td><td>+</td><td>0</td><td>CCGACGAGAAGATCGACCTC</td><td>AGG</td><td>||||||||[||||||||||||]PAM</td><td>NA</td><td>Intergenic</td><td>NA</td><td>NA</td></tr>
<tr><td></td><td></td><td></td><td></td><td></td><td></td><td></td><td></td><td></td><td></td><td></td><td></td></tr>
<tr><td>T2832</td><td>CGACGAGAAGATCGACCTCAGGG</td><td>1000</td><td></td><td></td><td></td><td></td><td></td><td></td><td></td><td></td><td></td></tr>
<tr><td>Oligo fwd</td><td>CACCCGACGAGAAGATCGACCTCA</td><td></td><td></td><td></td><td></td><td></td><td></td><td></td><td></td><td></td><td></td></tr>
<tr><td>Oligo rev</td><td>AAACTGAGGTCGATCTTCTCGTCG</td><td></td><td></td><td></td><td></td><td></td><td></td><td></td><td></td><td></td><td></td></tr>
<tr><td>Chromosome</td><td>start</td><td>end</td><td>strand</td><td>MM</td><td>target_seq</td><td>PAM</td><td>alignment</td><td>distance</td><td>position</td><td>gene name</td><td>gene id</td></tr>
<tr><td>GU305914.1</td><td>44405</td><td>44427</td><td>+</td><td>0</td><td>CGACGAGAAGATCGACCTCA</td><td>GGG</td><td>||||||||[||||||||||||]PAM</td><td>NA</td><td>Intergenic</td><td>NA</td><td>NA</td></tr>
<tr><td></td><td></td><td></td><td></td><td></td><td></td><td></td><td></td><td></td><td></td><td></td><td></td></tr>
<tr><td>T2833</td><td>CGAGCTGCAGTACCACTCGACGG</td><td>1000</td><td></td><td></td><td></td><td></td><td></td><td></td><td></td><td></td><td></td></tr>
<tr><td>Oligo fwd</td><td>CACCCGAGCTGCAGTACCACTCGA</td><td></td><td></td><td></td><td></td><td></td><td></td><td></td><td></td><td></td><td></td></tr>
<tr><td>Oligo rev</td><td>AAACTCGAGTGGTACTGCAGCTCG</td><td></td><td></td><td></td><td></td><td></td><td></td><td></td><td></td><td></td><td></td></tr>
<tr><td>Chromosome</td><td>start</td><td>end</td><td>strand</td><td>MM</td><td>target_seq</td><td>PAM</td><td>alignment</td><td>distance</td><td>position</td><td>gene name</td><td>gene id</td></tr>
<tr><td>GU305914.1</td><td>44435</td><td>44457</td><td>+</td><td>0</td><td>CGAGCTGCAGTACCACTCGA</td><td>CGG</td><td>||||||||[||||||||||||]PAM</td><td>NA</td><td>Intergenic</td><td>NA</td><td>NA</td></tr>
<tr><td>GU305914.1</td><td>88974</td><td>88996</td><td>+</td><td>4</td><td>GGAGCTGGAGTACTTCTCGA</td><td>CGG</td><td>-||||||-[|||||--|||||]PAM</td><td>NA</td><td>Intergenic</td><td>NA</td><td>NA</td></tr>
<tr><td></td><td></td><td></td><td></td><td></td><td></td><td></td><td></td><td></td><td></td><td></td><td></td></tr>
<tr><td>T2834</td><td>GAGCTGCAGTACCACTCGACGGG</td><td>1000</td><td></td><td></td><td></td><td></td><td></td><td></td><td></td><td></td><td></td></tr>
<tr><td>Oligo fwd</td><td>CACCGAGCTGCAGTACCACTCGAC</td><td></td><td></td><td></td><td></td><td></td><td></td><td></td><td></td><td></td><td></td></tr>
<tr><td>Oligo rev</td><td>AAACGTCGAGTGGTACTGCAGCTC</td><td></td><td></td><td></td><td></td><td></td><td></td><td></td><td></td><td></td><td></td></tr>
<tr><td>Chromosome</td><td>start</td><td>end</td><td>strand</td><td>MM</td><td>target_seq</td><td>PAM</td><td>alignment</td><td>distance</td><td>position</td><td>gene name</td><td>gene id</td></tr>
<tr><td>GU305914.1</td><td>44436</td><td>44458</td><td>+</td><td>0</td><td>GAGCTGCAGTACCACTCGAC</td><td>GGG</td><td>||||||||[||||||||||||]PAM</td><td>NA</td><td>Intergenic</td><td>NA</td><td>NA</td></tr>
<tr><td></td><td></td><td></td><td></td><td></td><td></td><td></td><td></td><td></td><td></td><td></td><td></td></tr>
<tr><td>T2835</td><td>CGCGTTCGTCTTGAACGACGCGG</td><td>1000</td><td></td><td></td><td></td><td></td><td></td><td></td><td></td><td></td><td></td></tr>
<tr><td>Oligo fwd</td><td>CACCCGCGTTCGTCTTGAACGACG</td><td></td><td></td><td></td><td></td><td></td><td></td><td></td><td></td><td></td><td></td></tr>
<tr><td>Oligo rev</td><td>AAACCGTCGTTCAAGACGAACGCG</td><td></td><td></td><td></td><td></td><td></td><td></td><td></td><td></td><td></td><td></td></tr>
<tr><td>Chromosome</td><td>start</td><td>end</td><td>strand</td><td>MM</td><td>target_seq</td><td>PAM</td><td>alignment</td><td>distance</td><td>position</td><td>gene name</td><td>gene id</td></tr>
<tr><td>GU305914.1</td><td>44498</td><td>44520</td><td>+</td><td>0</td><td>CGCGTTCGTCTTGAACGACG</td><td>CGG</td><td>||||||||[||||||||||||]PAM</td><td>NA</td><td>Intergenic</td><td>NA</td><td>NA</td></tr>
<tr><td>GU305914.1</td><td>129385</td><td>129407</td><td>+</td><td>4</td><td>CACATTCGTCTCTAACGACG</td><td>CGG</td><td>|-|-||||[|||--|||||||]PAM</td><td>NA</td><td>Intergenic</td><td>NA</td><td>NA</td></tr>
<tr><td></td><td></td><td></td><td></td><td></td><td></td><td></td><td></td><td></td><td></td><td></td><td></td></tr>
<tr><td>T2836</td><td>GTTCGTCTTGAACGACGCGGTGG</td><td>1000</td><td></td><td></td><td></td><td></td><td></td><td></td><td></td><td></td><td></td></tr>
<tr><td>Oligo fwd</td><td>CACCGTTCGTCTTGAACGACGCGG</td><td></td><td></td><td></td><td></td><td></td><td></td><td></td><td></td><td></td><td></td></tr>
<tr><td>Oligo rev</td><td>AAACCCGCGTCGTTCAAGACGAAC</td><td></td><td></td><td></td><td></td><td></td><td></td><td></td><td></td><td></td><td></td></tr>
<tr><td>Chromosome</td><td>start</td><td>end</td><td>strand</td><td>MM</td><td>target_seq</td><td>PAM</td><td>alignment</td><td>distance</td><td>position</td><td>gene name</td><td>gene id</td></tr>
<tr><td>GU305914.1</td><td>44501</td><td>44523</td><td>+</td><td>0</td><td>GTTCGTCTTGAACGACGCGG</td><td>TGG</td><td>||||||||[||||||||||||]PAM</td><td>NA</td><td>Intergenic</td><td>NA</td><td>NA</td></tr>
<tr><td></td><td></td><td></td><td></td><td></td><td></td><td></td><td></td><td></td><td></td><td></td><td></td></tr>
<tr><td>T2837</td><td>TTCGTCTTGAACGACGCGGTGGG</td><td>1000</td><td></td><td></td><td></td><td></td><td></td><td></td><td></td><td></td><td></td></tr>
<tr><td>Oligo fwd</td><td>CACCTTCGTCTTGAACGACGCGGT</td><td></td><td></td><td></td><td></td><td></td><td></td><td></td><td></td><td></td><td></td></tr>
<tr><td>Oligo rev</td><td>AAACACCGCGTCGTTCAAGACGAA</td><td></td><td></td><td></td><td></td><td></td><td></td><td></td><td></td><td></td><td></td></tr>
<tr><td>Chromosome</td><td>start</td><td>end</td><td>strand</td><td>MM</td><td>target_seq</td><td>PAM</td><td>alignment</td><td>distance</td><td>position</td><td>gene name</td><td>gene id</td></tr>
<tr><td>GU305914.1</td><td>44502</td><td>44524</td><td>+</td><td>0</td><td>TTCGTCTTGAACGACGCGGT</td><td>GGG</td><td>||||||||[||||||||||||]PAM</td><td>NA</td><td>Intergenic</td><td>NA</td><td>NA</td></tr>
<tr><td></td><td></td><td></td><td></td><td></td><td></td><td></td><td></td><td></td><td></td><td></td><td></td></tr>
<tr><td>T2838</td><td>GAACGACGCGGTGGGCACAGTGG</td><td>1000</td><td></td><td></td><td></td><td></td><td></td><td></td><td></td><td></td><td></td></tr>
<tr><td>Oligo fwd</td><td>CACCGAACGACGCGGTGGGCACAG</td><td></td><td></td><td></td><td></td><td></td><td></td><td></td><td></td><td></td><td></td></tr>
<tr><td>Oligo rev</td><td>AAACCTGTGCCCACCGCGTCGTTC</td><td></td><td></td><td></td><td></td><td></td><td></td><td></td><td></td><td></td><td></td></tr>
<tr><td>Chromosome</td><td>start</td><td>end</td><td>strand</td><td>MM</td><td>target_seq</td><td>PAM</td><td>alignment</td><td>distance</td><td>position</td><td>gene name</td><td>gene id</td></tr>
<tr><td>GU305914.1</td><td>44510</td><td>44532</td><td>+</td><td>0</td><td>GAACGACGCGGTGGGCACAG</td><td>TGG</td><td>||||||||[||||||||||||]PAM</td><td>NA</td><td>Intergenic</td><td>NA</td><td>NA</td></tr>
<tr><td></td><td></td><td></td><td></td><td></td><td></td><td></td><td></td><td></td><td></td><td></td><td></td></tr>
<tr><td>T2839</td><td>CAACGTCAATCGTCTTCGTCCGG</td><td>1000</td><td></td><td></td><td></td><td></td><td></td><td></td><td></td><td></td><td></td></tr>
<tr><td>Oligo fwd</td><td>CACCCAACGTCAATCGTCTTCGTC</td><td></td><td></td><td></td><td></td><td></td><td></td><td></td><td></td><td></td><td></td></tr>
<tr><td>Oligo rev</td><td>AAACGACGAAGACGATTGACGTTG</td><td></td><td></td><td></td><td></td><td></td><td></td><td></td><td></td><td></td><td></td></tr>
<tr><td>Chromosome</td><td>start</td><td>end</td><td>strand</td><td>MM</td><td>target_seq</td><td>PAM</td><td>alignment</td><td>distance</td><td>position</td><td>gene name</td><td>gene id</td></tr>
<tr><td>GU305914.1</td><td>44547</td><td>44569</td><td>+</td><td>0</td><td>CAACGTCAATCGTCTTCGTC</td><td>CGG</td><td>||||||||[||||||||||||]PAM</td><td>NA</td><td>Intergenic</td><td>NA</td><td>NA</td></tr>
<tr><td></td><td></td><td></td><td></td><td></td><td></td><td></td><td></td><td></td><td></td><td></td><td></td></tr>
<tr><td>T2840</td><td>CGTCAATCGTCTTCGTCCGGCGG</td><td>1000</td><td></td><td></td><td></td><td></td><td></td><td></td><td></td><td></td><td></td></tr>
<tr><td>Oligo fwd</td><td>CACCCGTCAATCGTCTTCGTCCGG</td><td></td><td></td><td></td><td></td><td></td><td></td><td></td><td></td><td></td><td></td></tr>
<tr><td>Oligo rev</td><td>AAACCCGGACGAAGACGATTGACG</td><td></td><td></td><td></td><td></td><td></td><td></td><td></td><td></td><td></td><td></td></tr>
<tr><td>Chromosome</td><td>start</td><td>end</td><td>strand</td><td>MM</td><td>target_seq</td><td>PAM</td><td>alignment</td><td>distance</td><td>position</td><td>gene name</td><td>gene id</td></tr>
<tr><td>GU305914.1</td><td>44550</td><td>44572</td><td>+</td><td>0</td><td>CGTCAATCGTCTTCGTCCGG</td><td>CGG</td><td>||||||||[||||||||||||]PAM</td><td>NA</td><td>Intergenic</td><td>NA</td><td>NA</td></tr>
<tr><td></td><td></td><td></td><td></td><td></td><td></td><td></td><td></td><td></td><td></td><td></td><td></td></tr>
<tr><td>T2841</td><td>GTCCGGCGGCGCGATGTTACCGG</td><td>1000</td><td></td><td></td><td></td><td></td><td></td><td></td><td></td><td></td><td></td></tr>
<tr><td>Oligo fwd</td><td>CACCGTCCGGCGGCGCGATGTTAC</td><td></td><td></td><td></td><td></td><td></td><td></td><td></td><td></td><td></td><td></td></tr>
<tr><td>Oligo rev</td><td>AAACGTAACATCGCGCCGCCGGAC</td><td></td><td></td><td></td><td></td><td></td><td></td><td></td><td></td><td></td><td></td></tr>
<tr><td>Chromosome</td><td>start</td><td>end</td><td>strand</td><td>MM</td><td>target_seq</td><td>PAM</td><td>alignment</td><td>distance</td><td>position</td><td>gene name</td><td>gene id</td></tr>
<tr><td>GU305914.1</td><td>44564</td><td>44586</td><td>+</td><td>0</td><td>GTCCGGCGGCGCGATGTTAC</td><td>CGG</td><td>||||||||[||||||||||||]PAM</td><td>NA</td><td>Intergenic</td><td>NA</td><td>NA</td></tr>
<tr><td></td><td></td><td></td><td></td><td></td><td></td><td></td><td></td><td></td><td></td><td></td><td></td></tr>
<tr><td>T2842</td><td>TCCGGCGGCGCGATGTTACCGGG</td><td>1000</td><td></td><td></td><td></td><td></td><td></td><td></td><td></td><td></td><td></td></tr>
<tr><td>Oligo fwd</td><td>CACCTCCGGCGGCGCGATGTTACC</td><td></td><td></td><td></td><td></td><td></td><td></td><td></td><td></td><td></td><td></td></tr>
<tr><td>Oligo rev</td><td>AAACGGTAACATCGCGCCGCCGGA</td><td></td><td></td><td></td><td></td><td></td><td></td><td></td><td></td><td></td><td></td></tr>
<tr><td>Chromosome</td><td>start</td><td>end</td><td>strand</td><td>MM</td><td>target_seq</td><td>PAM</td><td>alignment</td><td>distance</td><td>position</td><td>gene name</td><td>gene id</td></tr>
<tr><td>GU305914.1</td><td>44565</td><td>44587</td><td>+</td><td>0</td><td>TCCGGCGGCGCGATGTTACC</td><td>GGG</td><td>||||||||[||||||||||||]PAM</td><td>NA</td><td>Intergenic</td><td>NA</td><td>NA</td></tr>
<tr><td></td><td></td><td></td><td></td><td></td><td></td><td></td><td></td><td></td><td></td><td></td><td></td></tr>
<tr><td>T2843</td><td>CCGGCGGCGCGATGTTACCGGGG</td><td>1000</td><td></td><td></td><td></td><td></td><td></td><td></td><td></td><td></td><td></td></tr>
<tr><td>Oligo fwd</td><td>CACCCCGGCGGCGCGATGTTACCG</td><td></td><td></td><td></td><td></td><td></td><td></td><td></td><td></td><td></td><td></td></tr>
<tr><td>Oligo rev</td><td>AAACCGGTAACATCGCGCCGCCGG</td><td></td><td></td><td></td><td></td><td></td><td></td><td></td><td></td><td></td><td></td></tr>
<tr><td>Chromosome</td><td>start</td><td>end</td><td>strand</td><td>MM</td><td>target_seq</td><td>PAM</td><td>alignment</td><td>distance</td><td>position</td><td>gene name</td><td>gene id</td></tr>
<tr><td>GU305914.1</td><td>44566</td><td>44588</td><td>+</td><td>0</td><td>CCGGCGGCGCGATGTTACCG</td><td>GGG</td><td>||||||||[||||||||||||]PAM</td><td>NA</td><td>Intergenic</td><td>NA</td><td>NA</td></tr>
<tr><td></td><td></td><td></td><td></td><td></td><td></td><td></td><td></td><td></td><td></td><td></td><td></td></tr>
<tr><td>T2844</td><td>CGGCGCGATGTTACCGGGGACGG</td><td>1000</td><td></td><td></td><td></td><td></td><td></td><td></td><td></td><td></td><td></td></tr>
<tr><td>Oligo fwd</td><td>CACCCGGCGCGATGTTACCGGGGA</td><td></td><td></td><td></td><td></td><td></td><td></td><td></td><td></td><td></td><td></td></tr>
<tr><td>Oligo rev</td><td>AAACTCCCCGGTAACATCGCGCCG</td><td></td><td></td><td></td><td></td><td></td><td></td><td></td><td></td><td></td><td></td></tr>
<tr><td>Chromosome</td><td>start</td><td>end</td><td>strand</td><td>MM</td><td>target_seq</td><td>PAM</td><td>alignment</td><td>distance</td><td>position</td><td>gene name</td><td>gene id</td></tr>
<tr><td>GU305914.1</td><td>44570</td><td>44592</td><td>+</td><td>0</td><td>CGGCGCGATGTTACCGGGGA</td><td>CGG</td><td>||||||||[||||||||||||]PAM</td><td>NA</td><td>Intergenic</td><td>NA</td><td>NA</td></tr>
</table>

<table>
<tr><td>T2845</td><td>CGCGATGTTACCGGGGACGGTGG</td><td>1000</td><td></td><td></td><td></td><td></td><td></td><td></td><td></td><td></td><td></td></tr>
<tr><td>Oligo fwd</td><td>CACCCGCGATGTTACCGGGGACGG</td><td></td><td></td><td></td><td></td><td></td><td></td><td></td><td></td><td></td><td></td></tr>
<tr><td>Oligo rev</td><td>AAACCCGTCCCCGGTAACATCGCG</td><td></td><td></td><td></td><td></td><td></td><td></td><td></td><td></td><td></td><td></td></tr>
<tr><td>Chromosome</td><td>start</td><td>end</td><td>strand</td><td>MM</td><td>target_seq</td><td>PAM</td><td>alignment</td><td>distance</td><td>position</td><td>gene name</td><td>gene id</td></tr>
<tr><td>GU305914.1</td><td>44573</td><td>44595</td><td>+</td><td>0</td><td>CGCGATGTTACCGGGGACGG</td><td>TGG</td><td>||||||||[||||||||||||]PAM</td><td>NA</td><td>Intergenic</td><td>NA</td><td>NA</td></tr>
<tr><td>T2846</td><td>GATGTTACCGGGGACGGTGGCGG</td><td>1000</td><td></td><td></td><td></td><td></td><td></td><td></td><td></td><td></td><td></td></tr>
<tr><td>Oligo fwd</td><td>CACCGATGTTACCGGGGACGGTGG</td><td></td><td></td><td></td><td></td><td></td><td></td><td></td><td></td><td></td><td></td></tr>
<tr><td>Oligo rev</td><td>AAACCCACCGTCCCCGGTAACATC</td><td></td><td></td><td></td><td></td><td></td><td></td><td></td><td></td><td></td><td></td></tr>
<tr><td>Chromosome</td><td>start</td><td>end</td><td>strand</td><td>MM</td><td>target_seq</td><td>PAM</td><td>alignment</td><td>distance</td><td>position</td><td>gene name</td><td>gene id</td></tr>
<tr><td>GU305914.1</td><td>44576</td><td>44598</td><td>+</td><td>0</td><td>GATGTTACCGGGGACGGTGG</td><td>CGG</td><td>||||||||[||||||||||||]PAM</td><td>NA</td><td>Intergenic</td><td>NA</td><td>NA</td></tr>
<tr><td>T2847</td><td>CCTCATCGCTTCGAAGACGACGG</td><td>1000</td><td></td><td></td><td></td><td></td><td></td><td></td><td></td><td></td><td></td></tr>
<tr><td>Oligo fwd</td><td>CACCCCTCATCGCTTCGAAGACGA</td><td></td><td></td><td></td><td></td><td></td><td></td><td></td><td></td><td></td><td></td></tr>
<tr><td>Oligo rev</td><td>AAACTCGTCTTCGAAGCGATGAGG</td><td></td><td></td><td></td><td></td><td></td><td></td><td></td><td></td><td></td><td></td></tr>
<tr><td>Chromosome</td><td>start</td><td>end</td><td>strand</td><td>MM</td><td>target_seq</td><td>PAM</td><td>alignment</td><td>distance</td><td>position</td><td>gene name</td><td>gene id</td></tr>
<tr><td>GU305914.1</td><td>44694</td><td>44716</td><td>+</td><td>0</td><td>CCTCATCGCTTCGAAGACGA</td><td>CGG</td><td>||||||||[||||||||||||]PAM</td><td>NA</td><td>Intergenic</td><td>NA</td><td>NA</td></tr>
<tr><td>T2848</td><td>TTCGAAGACGACGGCGTCGTCGG</td><td>1000</td><td></td><td></td><td></td><td></td><td></td><td></td><td></td><td></td><td></td></tr>
<tr><td>Oligo fwd</td><td>CACCTTCGAAGACGACGGCGTCGT</td><td></td><td></td><td></td><td></td><td></td><td></td><td></td><td></td><td></td><td></td></tr>
<tr><td>Oligo rev</td><td>AAACACGACGCCGTCGTCTTCGAA</td><td></td><td></td><td></td><td></td><td></td><td></td><td></td><td></td><td></td><td></td></tr>
<tr><td>Chromosome</td><td>start</td><td>end</td><td>strand</td><td>MM</td><td>target_seq</td><td>PAM</td><td>alignment</td><td>distance</td><td>position</td><td>gene name</td><td>gene id</td></tr>
<tr><td>GU305914.1</td><td>44703</td><td>44725</td><td>+</td><td>0</td><td>TTCGAAGACGACGGCGTCGT</td><td>CGG</td><td>||||||||[||||||||||||]PAM</td><td>NA</td><td>Intergenic</td><td>NA</td><td>NA</td></tr>
<tr><td>T2849</td><td>CGTCGTCGGCGACGAAGCGATGG</td><td>1000</td><td></td><td></td><td></td><td></td><td></td><td></td><td></td><td></td><td></td></tr>
<tr><td>Oligo fwd</td><td>CACCCGTCGTCGGCGACGAAGCGA</td><td></td><td></td><td></td><td></td><td></td><td></td><td></td><td></td><td></td><td></td></tr>
<tr><td>Oligo rev</td><td>AAACTCGCTTCGTCGCCGACGACG</td><td></td><td></td><td></td><td></td><td></td><td></td><td></td><td></td><td></td><td></td></tr>
<tr><td>Chromosome</td><td>start</td><td>end</td><td>strand</td><td>MM</td><td>target_seq</td><td>PAM</td><td>alignment</td><td>distance</td><td>position</td><td>gene name</td><td>gene id</td></tr>
<tr><td>GU305914.1</td><td>44717</td><td>44739</td><td>+</td><td>0</td><td>CGTCGTCGGCGACGAAGCGA</td><td>TGG</td><td>||||||||[||||||||||||]PAM</td><td>NA</td><td>Intergenic</td><td>NA</td><td>NA</td></tr>
<tr><td>GU305914.1</td><td>36206</td><td>36228</td><td>+</td><td>4</td><td>AGTCGCCGCGGACGAAGCGA</td><td>CGG</td><td>-||||-||[--||||||||||]PAM</td><td>NA</td><td>Intergenic</td><td>NA</td><td>NA</td></tr>
<tr><td>T2850</td><td>GCGATGGTCGTCGACCAGAGCGG</td><td>1000</td><td></td><td></td><td></td><td></td><td></td><td></td><td></td><td></td><td></td></tr>
<tr><td>Oligo fwd</td><td>CACCGCGATGGTCGTCGACCAGAG</td><td></td><td></td><td></td><td></td><td></td><td></td><td></td><td></td><td></td><td></td></tr>
<tr><td>Oligo rev</td><td>AAACCTCTGGTCGACGACCATCGC</td><td></td><td></td><td></td><td></td><td></td><td></td><td></td><td></td><td></td><td></td></tr>
<tr><td>Chromosome</td><td>start</td><td>end</td><td>strand</td><td>MM</td><td>target_seq</td><td>PAM</td><td>alignment</td><td>distance</td><td>position</td><td>gene name</td><td>gene id</td></tr>
<tr><td>GU305914.1</td><td>44733</td><td>44755</td><td>+</td><td>0</td><td>GCGATGGTCGTCGACCAGAG</td><td>CGG</td><td>||||||||[||||||||||||]PAM</td><td>NA</td><td>Intergenic</td><td>NA</td><td>NA</td></tr>
<tr><td>T2851</td><td>GTACATCTTCTACCCGCGCCCGG</td><td>1000</td><td></td><td></td><td></td><td></td><td></td><td></td><td></td><td></td><td></td></tr>
<tr><td>Oligo fwd</td><td>CACCGTACATCTTCTACCCGCGCC</td><td></td><td></td><td></td><td></td><td></td><td></td><td></td><td></td><td></td><td></td></tr>
<tr><td>Oligo rev</td><td>AAACGGCGCGGGTAGAAGATGTAC</td><td></td><td></td><td></td><td></td><td></td><td></td><td></td><td></td><td></td><td></td></tr>
<tr><td>Chromosome</td><td>start</td><td>end</td><td>strand</td><td>MM</td><td>target_seq</td><td>PAM</td><td>alignment</td><td>distance</td><td>position</td><td>gene name</td><td>gene id</td></tr>
<tr><td>GU305914.1</td><td>44813</td><td>44835</td><td>+</td><td>0</td><td>GTACATCTTCTACCCGCGCC</td><td>CGG</td><td>||||||||[||||||||||||]PAM</td><td>NA</td><td>Intergenic</td><td>NA</td><td>NA</td></tr>
<tr><td>T2852</td><td>TACATCTTCTACCCGCGCCCGGG</td><td>1000</td><td></td><td></td><td></td><td></td><td></td><td></td><td></td><td></td><td></td></tr>
<tr><td>Oligo fwd</td><td>CACCTACATCTTCTACCCGCGCCC</td><td></td><td></td><td></td><td></td><td></td><td></td><td></td><td></td><td></td><td></td></tr>
<tr><td>Oligo rev</td><td>AAACGGGCGCGGGTAGAAGATGTA</td><td></td><td></td><td></td><td></td><td></td><td></td><td></td><td></td><td></td><td></td></tr>
<tr><td>Chromosome</td><td>start</td><td>end</td><td>strand</td><td>MM</td><td>target_seq</td><td>PAM</td><td>alignment</td><td>distance</td><td>position</td><td>gene name</td><td>gene id</td></tr>
<tr><td>GU305914.1</td><td>44814</td><td>44836</td><td>+</td><td>0</td><td>TACATCTTCTACCCGCGCCC</td><td>GGG</td><td>||||||||[||||||||||||]PAM</td><td>NA</td><td>Intergenic</td><td>NA</td><td>NA</td></tr>
<tr><td>T2853</td><td>GTCCGACGTGTCCTCCGATACGG</td><td>1000</td><td></td><td></td><td></td><td></td><td></td><td></td><td></td><td></td><td></td></tr>
<tr><td>Oligo fwd</td><td>CACCGTCCGACGTGTCCTCCGATA</td><td></td><td></td><td></td><td></td><td></td><td></td><td></td><td></td><td></td><td></td></tr>
<tr><td>Oligo rev</td><td>AAACTATCGGAGGACACGTCGGAC</td><td></td><td></td><td></td><td></td><td></td><td></td><td></td><td></td><td></td><td></td></tr>
<tr><td>Chromosome</td><td>start</td><td>end</td><td>strand</td><td>MM</td><td>target_seq</td><td>PAM</td><td>alignment</td><td>distance</td><td>position</td><td>gene name</td><td>gene id</td></tr>
<tr><td>GU305914.1</td><td>44843</td><td>44865</td><td>+</td><td>0</td><td>GTCCGACGTGTCCTCCGATA</td><td>CGG</td><td>||||||||[||||||||||||]PAM</td><td>NA</td><td>Intergenic</td><td>NA</td><td>NA</td></tr>
<tr><td>T2854</td><td>TCCGCGCCACCACCGCGACACGG</td><td>1000</td><td></td><td></td><td></td><td></td><td></td><td></td><td></td><td></td><td></td></tr>
<tr><td>Oligo fwd</td><td>CACCTCCGCGCCACCACCGCGACA</td><td></td><td></td><td></td><td></td><td></td><td></td><td></td><td></td><td></td><td></td></tr>
<tr><td>Oligo rev</td><td>AAACTGTCGCGGTGGTGGCGCGGA</td><td></td><td></td><td></td><td></td><td></td><td></td><td></td><td></td><td></td><td></td></tr>
<tr><td>Chromosome</td><td>start</td><td>end</td><td>strand</td><td>MM</td><td>target_seq</td><td>PAM</td><td>alignment</td><td>distance</td><td>position</td><td>gene name</td><td>gene id</td></tr>
<tr><td>GU305914.1</td><td>44870</td><td>44892</td><td>+</td><td>0</td><td>TCCGCGCCACCACCGCGACA</td><td>CGG</td><td>||||||||[||||||||||||]PAM</td><td>NA</td><td>Intergenic</td><td>NA</td><td>NA</td></tr>
<tr><td>T2855</td><td>CCGCGACACGGACAGCGACACGG</td><td>1000</td><td></td><td></td><td></td><td></td><td></td><td></td><td></td><td></td><td></td></tr>
<tr><td>Oligo fwd</td><td>CACCCCGCGACACGGACAGCGACA</td><td></td><td></td><td></td><td></td><td></td><td></td><td></td><td></td><td></td><td></td></tr>
<tr><td>Oligo rev</td><td>AAACTGTCGCTGTCCGTGTCGCGG</td><td></td><td></td><td></td><td></td><td></td><td></td><td></td><td></td><td></td><td></td></tr>
<tr><td>Chromosome</td><td>start</td><td>end</td><td>strand</td><td>MM</td><td>target_seq</td><td>PAM</td><td>alignment</td><td>distance</td><td>position</td><td>gene name</td><td>gene id</td></tr>
<tr><td>GU305914.1</td><td>44882</td><td>44904</td><td>+</td><td>0</td><td>CCGCGACACGGACAGCGACA</td><td>CGG</td><td>||||||||[||||||||||||]PAM</td><td>NA</td><td>Intergenic</td><td>NA</td><td>NA</td></tr>
<tr><td>T2856</td><td>CGCGACACGGACAGCGACACGGG</td><td>1000</td><td></td><td></td><td></td><td></td><td></td><td></td><td></td><td></td><td></td></tr>
<tr><td>Oligo fwd</td><td>CACCCGCGACACGGACAGCGACAC</td><td></td><td></td><td></td><td></td><td></td><td></td><td></td><td></td><td></td><td></td></tr>
<tr><td>Oligo rev</td><td>AAACGTGTCGCTGTCCGTGTCGCG</td><td></td><td></td><td></td><td></td><td></td><td></td><td></td><td></td><td></td><td></td></tr>
<tr><td>Chromosome</td><td>start</td><td>end</td><td>strand</td><td>MM</td><td>target_seq</td><td>PAM</td><td>alignment</td><td>distance</td><td>position</td><td>gene name</td><td>gene id</td></tr>
<tr><td>GU305914.1</td><td>44883</td><td>44905</td><td>+</td><td>0</td><td>CGCGACACGGACAGCGACAC</td><td>GGG</td><td>||||||||[||||||||||||]PAM</td><td>NA</td><td>Intergenic</td><td>NA</td><td>NA</td></tr>
<tr><td>T2857</td><td>ACACGGACAGCGACACGGGCCGG</td><td>1000</td><td></td><td></td><td></td><td></td><td></td><td></td><td></td><td></td><td></td></tr>
<tr><td>Oligo fwd</td><td>CACCACACGGACAGCGACACGGGC</td><td></td><td></td><td></td><td></td><td></td><td></td><td></td><td></td><td></td><td></td></tr>
<tr><td>Oligo rev</td><td>AAACGCCCGTGTCGCTGTCCGTGT</td><td></td><td></td><td></td><td></td><td></td><td></td><td></td><td></td><td></td><td></td></tr>
<tr><td>Chromosome</td><td>start</td><td>end</td><td>strand</td><td>MM</td><td>target_seq</td><td>PAM</td><td>alignment</td><td>distance</td><td>position</td><td>gene name</td><td>gene id</td></tr>
<tr><td>GU305914.1</td><td>44887</td><td>44909</td><td>+</td><td>0</td><td>ACACGGACAGCGACACGGGC</td><td>CGG</td><td>||||||||[||||||||||||]PAM</td><td>NA</td><td>Intergenic</td><td>NA</td><td>NA</td></tr>
<tr><td>T2858</td><td>GCGTCGCCTCTCGCTCGAGCAGG</td><td>1000</td><td></td><td></td><td></td><td></td><td></td><td></td><td></td><td></td><td></td></tr>
<tr><td>Oligo fwd</td><td>CACCGCGTCGCCTCTCGCTCGAGC</td><td></td><td></td><td></td><td></td><td></td><td></td><td></td><td></td><td></td><td></td></tr>
<tr><td>Oligo rev</td><td>AAACGCTCGAGCGAGAGGCGACGC</td><td></td><td></td><td></td><td></td><td></td><td></td><td></td><td></td><td></td><td></td></tr>
<tr><td>Chromosome</td><td>start</td><td>end</td><td>strand</td><td>MM</td><td>target_seq</td><td>PAM</td><td>alignment</td><td>distance</td><td>position</td><td>gene name</td><td>gene id</td></tr>
<tr><td>GU305914.1</td><td>44914</td><td>44936</td><td>+</td><td>0</td><td>GCGTCGCCTCTCGCTCGAGC</td><td>AGG</td><td>||||||||[||||||||||||]PAM</td><td>NA</td><td>Intergenic</td><td>NA</td><td>NA</td></tr>
<tr><td>T2859</td><td>GAGCAGGACGCCTCCGTTCGCGG</td><td>1000</td><td></td><td></td><td></td><td></td><td></td><td></td><td></td><td></td><td></td></tr>
<tr><td>Oligo fwd</td><td>CACCGAGCAGGACGCCTCCGTTCG</td><td></td><td></td><td></td><td></td><td></td><td></td><td></td><td></td><td></td><td></td></tr>
<tr><td>Oligo rev</td><td>AAACCGAACGGAGGCGTCCTGCTC</td><td></td><td></td><td></td><td></td><td></td><td></td><td></td><td></td><td></td><td></td></tr>
<tr><td>Chromosome</td><td>start</td><td>end</td><td>strand</td><td>MM</td><td>target_seq</td><td>PAM</td><td>alignment</td><td>distance</td><td>position</td><td>gene name</td><td>gene id</td></tr>
</table>

<table>
<tr><td>GU305914.1</td><td>44930</td><td>44952</td><td>+</td><td>0</td><td>GAGCAGGACGCCTCCGTTCG</td><td>CGG</td><td>||||||||[||||||||||||]PAM</td><td>NA</td><td>Intergenic</td><td>NA</td><td>NA</td></tr>
<tr><td></td><td></td><td></td><td></td><td></td><td></td><td></td><td></td><td></td><td></td><td></td><td></td></tr>
<tr><td>T2860</td><td>CAGGACGCCTCCGTTCGCGGAGG</td><td>1000</td><td></td><td></td><td></td><td></td><td></td><td></td><td></td><td></td><td></td></tr>
<tr><td>Oligo fwd</td><td>CACCCAGGACGCCTCCGTTCGCGG</td><td></td><td></td><td></td><td></td><td></td><td></td><td></td><td></td><td></td><td></td></tr>
<tr><td>Oligo rev</td><td>AAACCCGCGAACGGAGGCGTCCTG</td><td></td><td></td><td></td><td></td><td></td><td></td><td></td><td></td><td></td><td></td></tr>
<tr><td>Chromosome</td><td>start</td><td>end</td><td>strand</td><td>MM</td><td>target_seq</td><td>PAM</td><td>alignment</td><td>distance</td><td>position</td><td>gene name</td><td>gene id</td></tr>
<tr><td>GU305914.1</td><td>44933</td><td>44955</td><td>+</td><td>0</td><td>CAGGACGCCTCCGTTCGCGG</td><td>AGG</td><td>||||||||[||||||||||||]PAM</td><td>NA</td><td>Intergenic</td><td>NA</td><td>NA</td></tr>
<tr><td></td><td></td><td></td><td></td><td></td><td></td><td></td><td></td><td></td><td></td><td></td><td></td></tr>
<tr><td>T2861</td><td>GGAGGCAGCAGAAGAGACCTCGG</td><td>1000</td><td></td><td></td><td></td><td></td><td></td><td></td><td></td><td></td><td></td></tr>
<tr><td>Oligo fwd</td><td>CACCGGAGGCAGCAGAAGAGACCT</td><td></td><td></td><td></td><td></td><td></td><td></td><td></td><td></td><td></td><td></td></tr>
<tr><td>Oligo rev</td><td>AAACAGGTCTCTTCTGCTGCCTCC</td><td></td><td></td><td></td><td></td><td></td><td></td><td></td><td></td><td></td><td></td></tr>
<tr><td>Chromosome</td><td>start</td><td>end</td><td>strand</td><td>MM</td><td>target_seq</td><td>PAM</td><td>alignment</td><td>distance</td><td>position</td><td>gene name</td><td>gene id</td></tr>
<tr><td>GU305914.1</td><td>44951</td><td>44973</td><td>+</td><td>0</td><td>GGAGGCAGCAGAAGAGACCT</td><td>CGG</td><td>||||||||[||||||||||||]PAM</td><td>NA</td><td>Intergenic</td><td>NA</td><td>NA</td></tr>
<tr><td></td><td></td><td></td><td></td><td></td><td></td><td></td><td></td><td></td><td></td><td></td><td></td></tr>
<tr><td>T2862</td><td>AGAAGAGACCTCGGATCTAGAGG</td><td>1000</td><td></td><td></td><td></td><td></td><td></td><td></td><td></td><td></td><td></td></tr>
<tr><td>Oligo fwd</td><td>CACCAGAAGAGACCTCGGATCTAG</td><td></td><td></td><td></td><td></td><td></td><td></td><td></td><td></td><td></td><td></td></tr>
<tr><td>Oligo rev</td><td>AAACCTAGATCCGAGGTCTCTTCT</td><td></td><td></td><td></td><td></td><td></td><td></td><td></td><td></td><td></td><td></td></tr>
<tr><td>Chromosome</td><td>start</td><td>end</td><td>strand</td><td>MM</td><td>target_seq</td><td>PAM</td><td>alignment</td><td>distance</td><td>position</td><td>gene name</td><td>gene id</td></tr>
<tr><td>GU305914.1</td><td>44960</td><td>44982</td><td>+</td><td>0</td><td>AGAAGAGACCTCGGATCTAG</td><td>AGG</td><td>||||||||[||||||||||||]PAM</td><td>NA</td><td>Intergenic</td><td>NA</td><td>NA</td></tr>
<tr><td></td><td></td><td></td><td></td><td></td><td></td><td></td><td></td><td></td><td></td><td></td><td></td></tr>
<tr><td>T2863</td><td>AGAGACCTCGGATCTAGAGGAGG</td><td>1000</td><td></td><td></td><td></td><td></td><td></td><td></td><td></td><td></td><td></td></tr>
<tr><td>Oligo fwd</td><td>CACCAGAGACCTCGGATCTAGAGG</td><td></td><td></td><td></td><td></td><td></td><td></td><td></td><td></td><td></td><td></td></tr>
<tr><td>Oligo rev</td><td>AAACCCTCTAGATCCGAGGTCTCT</td><td></td><td></td><td></td><td></td><td></td><td></td><td></td><td></td><td></td><td></td></tr>
<tr><td>Chromosome</td><td>start</td><td>end</td><td>strand</td><td>MM</td><td>target_seq</td><td>PAM</td><td>alignment</td><td>distance</td><td>position</td><td>gene name</td><td>gene id</td></tr>
<tr><td>GU305914.1</td><td>44963</td><td>44985</td><td>+</td><td>0</td><td>AGAGACCTCGGATCTAGAGG</td><td>AGG</td><td>||||||||[||||||||||||]PAM</td><td>NA</td><td>Intergenic</td><td>NA</td><td>NA</td></tr>
<tr><td></td><td></td><td></td><td></td><td></td><td></td><td></td><td></td><td></td><td></td><td></td><td></td></tr>
<tr><td>T2864</td><td>TCGGATCTAGAGGAGGAAGACGG</td><td>1000</td><td></td><td></td><td></td><td></td><td></td><td></td><td></td><td></td><td></td></tr>
<tr><td>Oligo fwd</td><td>CACCTCGGATCTAGAGGAGGAAGA</td><td></td><td></td><td></td><td></td><td></td><td></td><td></td><td></td><td></td><td></td></tr>
<tr><td>Oligo rev</td><td>AAACTCTTCCTCCTCTAGATCCGA</td><td></td><td></td><td></td><td></td><td></td><td></td><td></td><td></td><td></td><td></td></tr>
<tr><td>Chromosome</td><td>start</td><td>end</td><td>strand</td><td>MM</td><td>target_seq</td><td>PAM</td><td>alignment</td><td>distance</td><td>position</td><td>gene name</td><td>gene id</td></tr>
<tr><td>GU305914.1</td><td>44970</td><td>44992</td><td>+</td><td>0</td><td>TCGGATCTAGAGGAGGAAGA</td><td>CGG</td><td>||||||||[||||||||||||]PAM</td><td>NA</td><td>Intergenic</td><td>NA</td><td>NA</td></tr>
<tr><td></td><td></td><td></td><td></td><td></td><td></td><td></td><td></td><td></td><td></td><td></td><td></td></tr>
<tr><td>T2865</td><td>CGGATCTAGAGGAGGAAGACGGG</td><td>1000</td><td></td><td></td><td></td><td></td><td></td><td></td><td></td><td></td><td></td></tr>
<tr><td>Oligo fwd</td><td>CACCCGGATCTAGAGGAGGAAGAC</td><td></td><td></td><td></td><td></td><td></td><td></td><td></td><td></td><td></td><td></td></tr>
<tr><td>Oligo rev</td><td>AAACGTCTTCCTCCTCTAGATCCG</td><td></td><td></td><td></td><td></td><td></td><td></td><td></td><td></td><td></td><td></td></tr>
<tr><td>Chromosome</td><td>start</td><td>end</td><td>strand</td><td>MM</td><td>target_seq</td><td>PAM</td><td>alignment</td><td>distance</td><td>position</td><td>gene name</td><td>gene id</td></tr>
<tr><td>GU305914.1</td><td>44971</td><td>44993</td><td>+</td><td>0</td><td>CGGATCTAGAGGAGGAAGAC</td><td>GGG</td><td>||||||||[||||||||||||]PAM</td><td>NA</td><td>Intergenic</td><td>NA</td><td>NA</td></tr>
<tr><td></td><td></td><td></td><td></td><td></td><td></td><td></td><td></td><td></td><td></td><td></td><td></td></tr>
<tr><td>T2866</td><td>GGATCTAGAGGAGGAAGACGGGG</td><td>1000</td><td></td><td></td><td></td><td></td><td></td><td></td><td></td><td></td><td></td></tr>
<tr><td>Oligo fwd</td><td>CACCGGATCTAGAGGAGGAAGACG</td><td></td><td></td><td></td><td></td><td></td><td></td><td></td><td></td><td></td><td></td></tr>
<tr><td>Oligo rev</td><td>AAACCGTCTTCCTCCTCTAGATCC</td><td></td><td></td><td></td><td></td><td></td><td></td><td></td><td></td><td></td><td></td></tr>
<tr><td>Chromosome</td><td>start</td><td>end</td><td>strand</td><td>MM</td><td>target_seq</td><td>PAM</td><td>alignment</td><td>distance</td><td>position</td><td>gene name</td><td>gene id</td></tr>
<tr><td>GU305914.1</td><td>44972</td><td>44994</td><td>+</td><td>0</td><td>GGATCTAGAGGAGGAAGACG</td><td>GGG</td><td>||||||||[||||||||||||]PAM</td><td>NA</td><td>Intergenic</td><td>NA</td><td>NA</td></tr>
<tr><td></td><td></td><td></td><td></td><td></td><td></td><td></td><td></td><td></td><td></td><td></td><td></td></tr>
<tr><td>T2867</td><td>GGAGGAAGACGGGGAAGAGACGG</td><td>1000</td><td></td><td></td><td></td><td></td><td></td><td></td><td></td><td></td><td></td></tr>
<tr><td>Oligo fwd</td><td>CACCGGAGGAAGACGGGGAAGAGA</td><td></td><td></td><td></td><td></td><td></td><td></td><td></td><td></td><td></td><td></td></tr>
<tr><td>Oligo rev</td><td>AAACTCTCTTCCCCGTCTTCCTCC</td><td></td><td></td><td></td><td></td><td></td><td></td><td></td><td></td><td></td><td></td></tr>
<tr><td>Chromosome</td><td>start</td><td>end</td><td>strand</td><td>MM</td><td>target_seq</td><td>PAM</td><td>alignment</td><td>distance</td><td>position</td><td>gene name</td><td>gene id</td></tr>
<tr><td>GU305914.1</td><td>44981</td><td>45003</td><td>+</td><td>0</td><td>GGAGGAAGACGGGGAAGAGA</td><td>CGG</td><td>||||||||[||||||||||||]PAM</td><td>NA</td><td>Intergenic</td><td>NA</td><td>NA</td></tr>
<tr><td>GU305914.1</td><td>42858</td><td>42880</td><td>-</td><td>4</td><td>GGGGGAGGATGAGGAAGAGA</td><td>AGG</td><td>||-|||-|[|-|-||||||||]PAM</td><td>NA</td><td>Intergenic</td><td>NA</td><td>NA</td></tr>
<tr><td></td><td></td><td></td><td></td><td></td><td></td><td></td><td></td><td></td><td></td><td></td><td></td></tr>
<tr><td>T2868</td><td>GACAGCGACCCGTTCTCCCCCGG</td><td>1000</td><td></td><td></td><td></td><td></td><td></td><td></td><td></td><td></td><td></td></tr>
<tr><td>Oligo fwd</td><td>CACCGACAGCGACCCGTTCTCCCC</td><td></td><td></td><td></td><td></td><td></td><td></td><td></td><td></td><td></td><td></td></tr>
<tr><td>Oligo rev</td><td>AAACGGGGAGAACGGGTCGCTGTC</td><td></td><td></td><td></td><td></td><td></td><td></td><td></td><td></td><td></td><td></td></tr>
<tr><td>Chromosome</td><td>start</td><td>end</td><td>strand</td><td>MM</td><td>target_seq</td><td>PAM</td><td>alignment</td><td>distance</td><td>position</td><td>gene name</td><td>gene id</td></tr>
<tr><td>GU305914.1</td><td>45024</td><td>45046</td><td>+</td><td>0</td><td>GACAGCGACCCGTTCTCCCC</td><td>CGG</td><td>||||||||[||||||||||||]PAM</td><td>NA</td><td>Intergenic</td><td>NA</td><td>NA</td></tr>
<tr><td></td><td></td><td></td><td></td><td></td><td></td><td></td><td></td><td></td><td></td><td></td><td></td></tr>
<tr><td>T2869</td><td>GACCCGTTCTCCCCCGGCCGAGG</td><td>1000</td><td></td><td></td><td></td><td></td><td></td><td></td><td></td><td></td><td></td></tr>
<tr><td>Oligo fwd</td><td>CACCGACCCGTTCTCCCCCGGCCG</td><td></td><td></td><td></td><td></td><td></td><td></td><td></td><td></td><td></td><td></td></tr>
<tr><td>Oligo rev</td><td>AAACCGGCCGGGGGAGAACGGGTC</td><td></td><td></td><td></td><td></td><td></td><td></td><td></td><td></td><td></td><td></td></tr>
<tr><td>Chromosome</td><td>start</td><td>end</td><td>strand</td><td>MM</td><td>target_seq</td><td>PAM</td><td>alignment</td><td>distance</td><td>position</td><td>gene name</td><td>gene id</td></tr>
<tr><td>GU305914.1</td><td>45030</td><td>45052</td><td>+</td><td>0</td><td>GACCCGTTCTCCCCCGGCCG</td><td>AGG</td><td>||||||||[||||||||||||]PAM</td><td>NA</td><td>Intergenic</td><td>NA</td><td>NA</td></tr>
<tr><td></td><td></td><td></td><td></td><td></td><td></td><td></td><td></td><td></td><td></td><td></td><td></td></tr>
<tr><td>T2870</td><td>CCGTTCTCCCCCGGCCGAGGAGG</td><td>1000</td><td></td><td></td><td></td><td></td><td></td><td></td><td></td><td></td><td></td></tr>
<tr><td>Oligo fwd</td><td>CACCCCGTTCTCCCCCGGCCGAGG</td><td></td><td></td><td></td><td></td><td></td><td></td><td></td><td></td><td></td><td></td></tr>
<tr><td>Oligo rev</td><td>AAACCCTCGGCCGGGGGAGAACGG</td><td></td><td></td><td></td><td></td><td></td><td></td><td></td><td></td><td></td><td></td></tr>
<tr><td>Chromosome</td><td>start</td><td>end</td><td>strand</td><td>MM</td><td>target_seq</td><td>PAM</td><td>alignment</td><td>distance</td><td>position</td><td>gene name</td><td>gene id</td></tr>
<tr><td>GU305914.1</td><td>45033</td><td>45055</td><td>+</td><td>0</td><td>CCGTTCTCCCCCGGCCGAGG</td><td>AGG</td><td>||||||||[||||||||||||]PAM</td><td>NA</td><td>Intergenic</td><td>NA</td><td>NA</td></tr>
<tr><td></td><td></td><td></td><td></td><td></td><td></td><td></td><td></td><td></td><td></td><td></td><td></td></tr>
<tr><td>T2871</td><td>CCCGGCCGAGGAGGAGTAGTAGG</td><td>1000</td><td></td><td></td><td></td><td></td><td></td><td></td><td></td><td></td><td></td></tr>
<tr><td>Oligo fwd</td><td>CACCCCCGGCCGAGGAGGAGTAGT</td><td></td><td></td><td></td><td></td><td></td><td></td><td></td><td></td><td></td><td></td></tr>
<tr><td>Oligo rev</td><td>AAACACTACTCCTCCTCGGCCGGG</td><td></td><td></td><td></td><td></td><td></td><td></td><td></td><td></td><td></td><td></td></tr>
<tr><td>Chromosome</td><td>start</td><td>end</td><td>strand</td><td>MM</td><td>target_seq</td><td>PAM</td><td>alignment</td><td>distance</td><td>position</td><td>gene name</td><td>gene id</td></tr>
<tr><td>GU305914.1</td><td>45042</td><td>45064</td><td>+</td><td>0</td><td>CCCGGCCGAGGAGGAGTAGT</td><td>AGG</td><td>||||||||[||||||||||||]PAM</td><td>NA</td><td>Intergenic</td><td>NA</td><td>NA</td></tr>
<tr><td></td><td></td><td></td><td></td><td></td><td></td><td></td><td></td><td></td><td></td><td></td><td></td></tr>
<tr><td>T2872</td><td>GGCCGAGGAGGAGTAGTAGGAGG</td><td>1000</td><td></td><td></td><td></td><td></td><td></td><td></td><td></td><td></td><td></td></tr>
<tr><td>Oligo fwd</td><td>CACCGGCCGAGGAGGAGTAGTAGG</td><td></td><td></td><td></td><td></td><td></td><td></td><td></td><td></td><td></td><td></td></tr>
<tr><td>Oligo rev</td><td>AAACCCTACTACTCCTCCTCGGCC</td><td></td><td></td><td></td><td></td><td></td><td></td><td></td><td></td><td></td><td></td></tr>
<tr><td>Chromosome</td><td>start</td><td>end</td><td>strand</td><td>MM</td><td>target_seq</td><td>PAM</td><td>alignment</td><td>distance</td><td>position</td><td>gene name</td><td>gene id</td></tr>
<tr><td>GU305914.1</td><td>45045</td><td>45067</td><td>+</td><td>0</td><td>GGCCGAGGAGGAGTAGTAGG</td><td>AGG</td><td>||||||||[||||||||||||]PAM</td><td>NA</td><td>Intergenic</td><td>NA</td><td>NA</td></tr>
<tr><td></td><td></td><td></td><td></td><td></td><td></td><td></td><td></td><td></td><td></td><td></td><td></td></tr>
<tr><td>T2873</td><td>AGGAGTAGTAGGAGGAGAAGAGG</td><td>1000</td><td></td><td></td><td></td><td></td><td></td><td></td><td></td><td></td><td></td></tr>
<tr><td>Oligo fwd</td><td>CACCAGGAGTAGTAGGAGGAGAAG</td><td></td><td></td><td></td><td></td><td></td><td></td><td></td><td></td><td></td><td></td></tr>
<tr><td>Oligo rev</td><td>AAACCTTCTCCTCCTACTACTCCT</td><td></td><td></td><td></td><td></td><td></td><td></td><td></td><td></td><td></td><td></td></tr>
<tr><td>Chromosome</td><td>start</td><td>end</td><td>strand</td><td>MM</td><td>target_seq</td><td>PAM</td><td>alignment</td><td>distance</td><td>position</td><td>gene name</td><td>gene id</td></tr>
<tr><td>GU305914.1</td><td>45053</td><td>45075</td><td>+</td><td>0</td><td>AGGAGTAGTAGGAGGAGAAG</td><td>AGG</td><td>||||||||[||||||||||||]PAM</td><td>NA</td><td>Intergenic</td><td>NA</td><td>NA</td></tr>
<tr><td></td><td></td><td></td><td></td><td></td><td></td><td></td><td></td><td></td><td></td><td></td><td></td></tr>
<tr><td>T2874</td><td>AGTAGTAGGAGGAGAAGAGGAGG</td><td>1000</td><td></td><td></td><td></td><td></td><td></td><td></td><td></td><td></td><td></td></tr>
<tr><td>Oligo fwd</td><td>CACCAGTAGTAGGAGGAGAAGAGG</td><td></td><td></td><td></td><td></td><td></td><td></td><td></td><td></td><td></td><td></td></tr>
</table>

| | | | | | | | | | | | |
|---|---|---|---|---|---|---|---|---|---|---|---|
| Oligo rev | AAACCCTCTTCTCCTCCTACTACT | | | | | | | | | | |
| Chromosome | start | end | strand | MM | target_seq | PAM | alignment | distance | position | gene name | gene id |
| GU305914.1 | 45056 | 45078 | + | 0 | AGTAGTAGGAGGAGAAGAGG | AGG | \|\|\|\|\|\|\|\|[\|\|\|\|\|\|\|\|\|\|\|\|]PAM | NA | Intergenic | NA | NA |
| GU305914.1 | 45059 | 45081 | + | 3 | AGTAGGAGGAGAAGAGGAGG | AGG | \|\|\|\|\|-\|\|[\|\|\|-\|\|\|-\|\|\|\|]PAM | NA | Intergenic | NA | NA |
| GU305914.1 | 165009 | 165031 | + | 3 | AGGAGTAGGAGGGGAAGACG | AGG | \|\|-\|\|\|\|\|[\|\|\|\|-\|\|\|\|\|-\|]PAM | NA | Intergenic | NA | NA |
| T2875 | AGTAGGAGGAGAAGAGGAGGAGG | 1000 | | | | | | | | | |
| Oligo fwd | CACCAGTAGGAGGAGAAGAGGAGG | | | | | | | | | | |
| Oligo rev | AAACCCTCCTCTTCTCCTCCTACT | | | | | | | | | | |
| Chromosome | start | end | strand | MM | target_seq | PAM | alignment | distance | position | gene name | gene id |
| GU305914.1 | 45059 | 45081 | + | 0 | AGTAGGAGGAGAAGAGGAGG | AGG | \|\|\|\|\|\|\|\|[\|\|\|\|\|\|\|\|\|\|\|\|]PAM | NA | Intergenic | NA | NA |
| GU305914.1 | 45056 | 45078 | + | 3 | AGTAGTAGGAGGAGAAGAGG | AGG | \|\|\|\|\|-\|\|[\|\|\|-\|\|\|-\|\|\|\|]PAM | NA | Intergenic | NA | NA |
| GU305914.1 | 86793 | 86815 | - | 4 | GGCAGGAGGGGAAGAGGGGG | CGG | -\|-\|\|\|\|\|[\|-\|\|\|\|\|\|\|-\|\|]PAM | NA | Intergenic | NA | NA |
| T2876 | AGGAGAAGAGGAGGAGGAAGAGG | 1000 | | | | | | | | | |
| Oligo fwd | CACCAGGAGAAGAGGAGGAGGAAG | | | | | | | | | | |
| Oligo rev | AAACCTTCCTCCTCCTCTTCTCCT | | | | | | | | | | |
| Chromosome | start | end | strand | MM | target_seq | PAM | alignment | distance | position | gene name | gene id |
| GU305914.1 | 45065 | 45087 | + | 0 | AGGAGAAGAGGAGGAGGAAG | AGG | \|\|\|\|\|\|\|\|[\|\|\|\|\|\|\|\|\|\|\|\|]PAM | NA | Intergenic | NA | NA |
| GU305914.1 | 220274 | 220296 | - | 4 | AGAGGAAGATGAAGAGGAAG | AGG | \|\|--\|\|\|\|[\|-\|\|-\|\|\|\|\|\|\|]PAM | NA | Intergenic | NA | NA |
| GU305914.1 | 96415 | 96437 | - | 4 | GGGAGACGCGGAGGAAGAAG | AGG | -\|\|\|\|\|-\|[-\|\|\|\|\|\|-\|\|\|\|]PAM | NA | Intergenic | NA | NA |
| T2877 | AAGAGGAAGAGAACAACAAACGG | 1000 | | | | | | | | | |
| Oligo fwd | CACCAAGAGGAAGAGAACAACAAA | | | | | | | | | | |
| Oligo rev | AAACTTTGTTGTTCTCTTCCTCTT | | | | | | | | | | |
| Chromosome | start | end | strand | MM | target_seq | PAM | alignment | distance | position | gene name | gene id |
| GU305914.1 | 45082 | 45104 | + | 0 | AAGAGGAAGAGAACAACAAA | CGG | \|\|\|\|\|\|\|\|[\|\|\|\|\|\|\|\|\|\|\|\|]PAM | NA | Intergenic | NA | NA |
| T2878 | AGAGGAAGAGAACAACAAACGGG | 1000 | | | | | | | | | |
| Oligo fwd | CACCAGAGGAAGAGAACAACAAAC | | | | | | | | | | |
| Oligo rev | AAACGTTTGTTGTTCTCTTCCTCT | | | | | | | | | | |
| Chromosome | start | end | strand | MM | target_seq | PAM | alignment | distance | position | gene name | gene id |
| GU305914.1 | 45083 | 45105 | + | 0 | AGAGGAAGAGAACAACAAAC | GGG | \|\|\|\|\|\|\|\|[\|\|\|\|\|\|\|\|\|\|\|\|]PAM | NA | Intergenic | NA | NA |
| T2879 | TCGTCGAAATCGAGTTCGCGTGG | 1000 | | | | | | | | | |
| Oligo fwd | CACCTCGTCGAAATCGAGTTCGCG | | | | | | | | | | |
| Oligo rev | AAACCGCGAACTCGATTTCGACGA | | | | | | | | | | |
| Chromosome | start | end | strand | MM | target_seq | PAM | alignment | distance | position | gene name | gene id |
| GU305914.1 | 45111 | 45133 | + | 0 | TCGTCGAAATCGAGTTCGCG | TGG | \|\|\|\|\|\|\|\|[\|\|\|\|\|\|\|\|\|\|\|\|]PAM | NA | Intergenic | NA | NA |
| T2880 | TCGAAATCGAGTTCGCGTGGAGG | 1000 | | | | | | | | | |
| Oligo fwd | CACCTCGAAATCGAGTTCGCGTGG | | | | | | | | | | |
| Oligo rev | AAACCCACGCGAACTCGATTTCGA | | | | | | | | | | |
| Chromosome | start | end | strand | MM | target_seq | PAM | alignment | distance | position | gene name | gene id |
| GU305914.1 | 45114 | 45136 | + | 0 | TCGAAATCGAGTTCGCGTGG | AGG | \|\|\|\|\|\|\|\|[\|\|\|\|\|\|\|\|\|\|\|\|]PAM | NA | Intergenic | NA | NA |
| T2881 | AGGTAGTTCGACGTGCAGCTCGG | 1000 | | | | | | | | | |
| Oligo fwd | CACCAGGTAGTTCGACGTGCAGCT | | | | | | | | | | |
| Oligo rev | AAACAGCTGCACGTCGAACTACCT | | | | | | | | | | |
| Chromosome | start | end | strand | MM | target_seq | PAM | alignment | distance | position | gene name | gene id |
| GU305914.1 | 45134 | 45156 | + | 0 | AGGTAGTTCGACGTGCAGCT | CGG | \|\|\|\|\|\|\|\|[\|\|\|\|\|\|\|\|\|\|\|\|]PAM | NA | Intergenic | NA | NA |
| T2882 | GGTAGTTCGACGTGCAGCTCGGG | 1000 | | | | | | | | | |
| Oligo fwd | CACCGGTAGTTCGACGTGCAGCTC | | | | | | | | | | |
| Oligo rev | AAACGAGCTGCACGTCGAACTACC | | | | | | | | | | |
| Chromosome | start | end | strand | MM | target_seq | PAM | alignment | distance | position | gene name | gene id |
| GU305914.1 | 45135 | 45157 | + | 0 | GGTAGTTCGACGTGCAGCTC | GGG | \|\|\|\|\|\|\|\|[\|\|\|\|\|\|\|\|\|\|\|\|]PAM | NA | Intergenic | NA | NA |
| T2883 | GTAGTTCGACGTGCAGCTCGGGG | 1000 | | | | | | | | | |
| Oligo fwd | CACCGTAGTTCGACGTGCAGCTCG | | | | | | | | | | |
| Oligo rev | AAACCGAGCTGCACGTCGAACTAC | | | | | | | | | | |
| Chromosome | start | end | strand | MM | target_seq | PAM | alignment | distance | position | gene name | gene id |
| GU305914.1 | 45136 | 45158 | + | 0 | GTAGTTCGACGTGCAGCTCG | GGG | \|\|\|\|\|\|\|\|[\|\|\|\|\|\|\|\|\|\|\|\|]PAM | NA | Intergenic | NA | NA |
| T2884 | TGCAGCTCGGGGTCCTCCAGCGG | 1000 | | | | | | | | | |
| Oligo fwd | CACCTGCAGCTCGGGGTCCTCCAG | | | | | | | | | | |
| Oligo rev | AAACCTGGAGGACCCCGAGCTGCA | | | | | | | | | | |
| Chromosome | start | end | strand | MM | target_seq | PAM | alignment | distance | position | gene name | gene id |
| GU305914.1 | 45147 | 45169 | + | 0 | TGCAGCTCGGGGTCCTCCAG | CGG | \|\|\|\|\|\|\|\|[\|\|\|\|\|\|\|\|\|\|\|\|]PAM | NA | Intergenic | NA | NA |
| T2885 | AGCTCGGGGTCCTCCAGCGGCGG | 1000 | | | | | | | | | |
| Oligo fwd | CACCAGCTCGGGGTCCTCCAGCGG | | | | | | | | | | |
| Oligo rev | AAACCCGCTGGAGGACCCCGAGCT | | | | | | | | | | |
| Chromosome | start | end | strand | MM | target_seq | PAM | alignment | distance | position | gene name | gene id |
| GU305914.1 | 45150 | 45172 | + | 0 | AGCTCGGGGTCCTCCAGCGG | CGG | \|\|\|\|\|\|\|\|[\|\|\|\|\|\|\|\|\|\|\|\|]PAM | NA | Intergenic | NA | NA |
| T2886 | GGGTCCTCCAGCGGCGGCGAAGG | 1000 | | | | | | | | | |
| Oligo fwd | CACCGGGTCCTCCAGCGGCGGCGA | | | | | | | | | | |
| Oligo rev | AAACTCGCCGCCGCTGGAGGACCC | | | | | | | | | | |
| Chromosome | start | end | strand | MM | target_seq | PAM | alignment | distance | position | gene name | gene id |
| GU305914.1 | 45156 | 45178 | + | 0 | GGGTCCTCCAGCGGCGGCGA | AGG | \|\|\|\|\|\|\|\|[\|\|\|\|\|\|\|\|\|\|\|\|]PAM | NA | Intergenic | NA | NA |
| T2887 | CGGCGGCGAAGGCGACTCGTCGG | 1000 | | | | | | | | | |
| Oligo fwd | CACCCGGCGGCGAAGGCGACTCGT | | | | | | | | | | |
| Oligo rev | AAACACGAGTCGCCTTCGCCGCCG | | | | | | | | | | |
| Chromosome | start | end | strand | MM | target_seq | PAM | alignment | distance | position | gene name | gene id |
| GU305914.1 | 45167 | 45189 | + | 0 | CGGCGGCGAAGGCGACTCGT | CGG | \|\|\|\|\|\|\|\|[\|\|\|\|\|\|\|\|\|\|\|\|]PAM | NA | Intergenic | NA | NA |
| T2888 | CGACTCGTCGGACTCGTCCACGG | 1000 | | | | | | | | | |

<table>
<tr><td>Oligo fwd</td><td>CACCCGACTCGTCGGACTCGTCCA</td><td></td><td></td><td></td><td></td><td></td><td></td><td></td><td></td><td></td><td></td></tr>
<tr><td>Oligo rev</td><td>AAACTGGACGAGTCCGACGAGTCG</td><td></td><td></td><td></td><td></td><td></td><td></td><td></td><td></td><td></td><td></td></tr>
<tr><td>Chromosome</td><td>start</td><td>end</td><td>strand</td><td>MM</td><td>target_seq</td><td>PAM</td><td>alignment</td><td>distance</td><td>position</td><td>gene name</td><td>gene id</td></tr>
<tr><td>GU305914.1</td><td>45179</td><td>45201</td><td>+</td><td>0</td><td>CGACTCGTCGGACTCGTCCA</td><td>CGG</td><td>||||||||[||||||||||||]PAM</td><td>NA</td><td>Intergenic</td><td>NA</td><td>NA</td></tr>
<tr><td>GU305914.1</td><td>106460</td><td>106482</td><td>+</td><td>4</td><td>CACCTCGTCGTACTCGACCA</td><td>GGG</td><td>|--|||||[||-|||||-|||]PAM</td><td>NA</td><td>Intergenic</td><td>NA</td><td>NA</td></tr>
<tr><td></td><td></td><td></td><td></td><td></td><td></td><td></td><td></td><td></td><td></td><td></td><td></td></tr>
<tr><td>T2889</td><td>GACTCGTCGGACTCGTCCACGGG</td><td>1000</td><td></td><td></td><td></td><td></td><td></td><td></td><td></td><td></td><td></td></tr>
<tr><td>Oligo fwd</td><td>CACCGACTCGTCGGACTCGTCCAC</td><td></td><td></td><td></td><td></td><td></td><td></td><td></td><td></td><td></td><td></td></tr>
<tr><td>Oligo rev</td><td>AAACGTGGACGAGTCCGACGAGTC</td><td></td><td></td><td></td><td></td><td></td><td></td><td></td><td></td><td></td><td></td></tr>
<tr><td>Chromosome</td><td>start</td><td>end</td><td>strand</td><td>MM</td><td>target_seq</td><td>PAM</td><td>alignment</td><td>distance</td><td>position</td><td>gene name</td><td>gene id</td></tr>
<tr><td>GU305914.1</td><td>45180</td><td>45202</td><td>+</td><td>0</td><td>GACTCGTCGGACTCGTCCAC</td><td>GGG</td><td>||||||||[||||||||||||]PAM</td><td>NA</td><td>Intergenic</td><td>NA</td><td>NA</td></tr>
<tr><td></td><td></td><td></td><td></td><td></td><td></td><td></td><td></td><td></td><td></td><td></td><td></td></tr>
<tr><td>T2890</td><td>TCGTCGGACTCGTCCACGGGTGG</td><td>1000</td><td></td><td></td><td></td><td></td><td></td><td></td><td></td><td></td><td></td></tr>
<tr><td>Oligo fwd</td><td>CACCTCGTCGGACTCGTCCACGGG</td><td></td><td></td><td></td><td></td><td></td><td></td><td></td><td></td><td></td><td></td></tr>
<tr><td>Oligo rev</td><td>AAACCCCGTGGACGAGTCCGACGA</td><td></td><td></td><td></td><td></td><td></td><td></td><td></td><td></td><td></td><td></td></tr>
<tr><td>Chromosome</td><td>start</td><td>end</td><td>strand</td><td>MM</td><td>target_seq</td><td>PAM</td><td>alignment</td><td>distance</td><td>position</td><td>gene name</td><td>gene id</td></tr>
<tr><td>GU305914.1</td><td>45183</td><td>45205</td><td>+</td><td>0</td><td>TCGTCGGACTCGTCCACGGG</td><td>TGG</td><td>||||||||[||||||||||||]PAM</td><td>NA</td><td>Intergenic</td><td>NA</td><td>NA</td></tr>
<tr><td>GU305914.1</td><td>73070</td><td>73092</td><td>-</td><td>4</td><td>TCGTCGCCCTCGTCCTCGTG</td><td>GGG</td><td>||||||--[|||||||-||-|]PAM</td><td>NA</td><td>Intergenic</td><td>NA</td><td>NA</td></tr>
<tr><td></td><td></td><td></td><td></td><td></td><td></td><td></td><td></td><td></td><td></td><td></td><td></td></tr>
<tr><td>T2891</td><td>ACTCGTCCACGGGTGGTTCTCGG</td><td>1000</td><td></td><td></td><td></td><td></td><td></td><td></td><td></td><td></td><td></td></tr>
<tr><td>Oligo fwd</td><td>CACCACTCGTCCACGGGTGGTTCT</td><td></td><td></td><td></td><td></td><td></td><td></td><td></td><td></td><td></td><td></td></tr>
<tr><td>Oligo rev</td><td>AAACAGAACCACCCGTGGACGAGT</td><td></td><td></td><td></td><td></td><td></td><td></td><td></td><td></td><td></td><td></td></tr>
<tr><td>Chromosome</td><td>start</td><td>end</td><td>strand</td><td>MM</td><td>target_seq</td><td>PAM</td><td>alignment</td><td>distance</td><td>position</td><td>gene name</td><td>gene id</td></tr>
<tr><td>GU305914.1</td><td>45190</td><td>45212</td><td>+</td><td>0</td><td>ACTCGTCCACGGGTGGTTCT</td><td>CGG</td><td>||||||||[||||||||||||]PAM</td><td>NA</td><td>Intergenic</td><td>NA</td><td>NA</td></tr>
<tr><td></td><td></td><td></td><td></td><td></td><td></td><td></td><td></td><td></td><td></td><td></td><td></td></tr>
<tr><td>T2892</td><td>GGTGGTTCTCGGTCCCACACCGG</td><td>1000</td><td></td><td></td><td></td><td></td><td></td><td></td><td></td><td></td><td></td></tr>
<tr><td>Oligo fwd</td><td>CACCGGTGGTTCTCGGTCCCACAC</td><td></td><td></td><td></td><td></td><td></td><td></td><td></td><td></td><td></td><td></td></tr>
<tr><td>Oligo rev</td><td>AAACGTGTGGGACCGAGAACCACC</td><td></td><td></td><td></td><td></td><td></td><td></td><td></td><td></td><td></td><td></td></tr>
<tr><td>Chromosome</td><td>start</td><td>end</td><td>strand</td><td>MM</td><td>target_seq</td><td>PAM</td><td>alignment</td><td>distance</td><td>position</td><td>gene name</td><td>gene id</td></tr>
<tr><td>GU305914.1</td><td>45201</td><td>45223</td><td>+</td><td>0</td><td>GGTGGTTCTCGGTCCCACAC</td><td>CGG</td><td>||||||||[||||||||||||]PAM</td><td>NA</td><td>Intergenic</td><td>NA</td><td>NA</td></tr>
<tr><td></td><td></td><td></td><td></td><td></td><td></td><td></td><td></td><td></td><td></td><td></td><td></td></tr>
<tr><td>T2893</td><td>GTTCACCGACGTTATCCGACAGG</td><td>1000</td><td></td><td></td><td></td><td></td><td></td><td></td><td></td><td></td><td></td></tr>
<tr><td>Oligo fwd</td><td>CACCGTTCACCGACGTTATCCGAC</td><td></td><td></td><td></td><td></td><td></td><td></td><td></td><td></td><td></td><td></td></tr>
<tr><td>Oligo rev</td><td>AAACGTCGGATAACGTCGGTGAAC</td><td></td><td></td><td></td><td></td><td></td><td></td><td></td><td></td><td></td><td></td></tr>
<tr><td>Chromosome</td><td>start</td><td>end</td><td>strand</td><td>MM</td><td>target_seq</td><td>PAM</td><td>alignment</td><td>distance</td><td>position</td><td>gene name</td><td>gene id</td></tr>
<tr><td>GU305914.1</td><td>45223</td><td>45245</td><td>+</td><td>0</td><td>GTTCACCGACGTTATCCGAC</td><td>AGG</td><td>||||||||[||||||||||||]PAM</td><td>NA</td><td>Intergenic</td><td>NA</td><td>NA</td></tr>
<tr><td></td><td></td><td></td><td></td><td></td><td></td><td></td><td></td><td></td><td></td><td></td><td></td></tr>
<tr><td>T2894</td><td>TTCACCGACGTTATCCGACAGGG</td><td>1000</td><td></td><td></td><td></td><td></td><td></td><td></td><td></td><td></td><td></td></tr>
<tr><td>Oligo fwd</td><td>CACCTTCACCGACGTTATCCGACA</td><td></td><td></td><td></td><td></td><td></td><td></td><td></td><td></td><td></td><td></td></tr>
<tr><td>Oligo rev</td><td>AAACTGTCGGATAACGTCGGTGAA</td><td></td><td></td><td></td><td></td><td></td><td></td><td></td><td></td><td></td><td></td></tr>
<tr><td>Chromosome</td><td>start</td><td>end</td><td>strand</td><td>MM</td><td>target_seq</td><td>PAM</td><td>alignment</td><td>distance</td><td>position</td><td>gene name</td><td>gene id</td></tr>
<tr><td>GU305914.1</td><td>45224</td><td>45246</td><td>+</td><td>0</td><td>TTCACCGACGTTATCCGACA</td><td>GGG</td><td>||||||||[||||||||||||]PAM</td><td>NA</td><td>Intergenic</td><td>NA</td><td>NA</td></tr>
<tr><td></td><td></td><td></td><td></td><td></td><td></td><td></td><td></td><td></td><td></td><td></td><td></td></tr>
<tr><td>T2895</td><td>TCACCGACGTTATCCGACAGGGG</td><td>1000</td><td></td><td></td><td></td><td></td><td></td><td></td><td></td><td></td><td></td></tr>
<tr><td>Oligo fwd</td><td>CACCTCACCGACGTTATCCGACAG</td><td></td><td></td><td></td><td></td><td></td><td></td><td></td><td></td><td></td><td></td></tr>
<tr><td>Oligo rev</td><td>AAACCTGTCGGATAACGTCGGTGA</td><td></td><td></td><td></td><td></td><td></td><td></td><td></td><td></td><td></td><td></td></tr>
<tr><td>Chromosome</td><td>start</td><td>end</td><td>strand</td><td>MM</td><td>target_seq</td><td>PAM</td><td>alignment</td><td>distance</td><td>position</td><td>gene name</td><td>gene id</td></tr>
<tr><td>GU305914.1</td><td>45225</td><td>45247</td><td>+</td><td>0</td><td>TCACCGACGTTATCCGACAG</td><td>GGG</td><td>||||||||[||||||||||||]PAM</td><td>NA</td><td>Intergenic</td><td>NA</td><td>NA</td></tr>
<tr><td></td><td></td><td></td><td></td><td></td><td></td><td></td><td></td><td></td><td></td><td></td><td></td></tr>
<tr><td>T2896</td><td>CCTCCTCCGATCGACAGCATCGG</td><td>1000</td><td></td><td></td><td></td><td></td><td></td><td></td><td></td><td></td><td></td></tr>
<tr><td>Oligo fwd</td><td>CACCCCTCCTCCGATCGACAGCAT</td><td></td><td></td><td></td><td></td><td></td><td></td><td></td><td></td><td></td><td></td></tr>
<tr><td>Oligo rev</td><td>AAACATGCTGTCGATCGGAGGAGG</td><td></td><td></td><td></td><td></td><td></td><td></td><td></td><td></td><td></td><td></td></tr>
<tr><td>Chromosome</td><td>start</td><td>end</td><td>strand</td><td>MM</td><td>target_seq</td><td>PAM</td><td>alignment</td><td>distance</td><td>position</td><td>gene name</td><td>gene id</td></tr>
<tr><td>GU305914.1</td><td>45348</td><td>45370</td><td>+</td><td>0</td><td>CCTCCTCCGATCGACAGCAT</td><td>CGG</td><td>||||||||[||||||||||||]PAM</td><td>NA</td><td>Intergenic</td><td>NA</td><td>NA</td></tr>
<tr><td></td><td></td><td></td><td></td><td></td><td></td><td></td><td></td><td></td><td></td><td></td><td></td></tr>
<tr><td>T2897</td><td>CAGCCACGACATGATGACAATGG</td><td>1000</td><td></td><td></td><td></td><td></td><td></td><td></td><td></td><td></td><td></td></tr>
<tr><td>Oligo fwd</td><td>CACCCAGCCACGACATGATGACAA</td><td></td><td></td><td></td><td></td><td></td><td></td><td></td><td></td><td></td><td></td></tr>
<tr><td>Oligo rev</td><td>AAACTTGTCATCATGTCGTGGCTG</td><td></td><td></td><td></td><td></td><td></td><td></td><td></td><td></td><td></td><td></td></tr>
<tr><td>Chromosome</td><td>start</td><td>end</td><td>strand</td><td>MM</td><td>target_seq</td><td>PAM</td><td>alignment</td><td>distance</td><td>position</td><td>gene name</td><td>gene id</td></tr>
<tr><td>GU305914.1</td><td>45371</td><td>45393</td><td>+</td><td>0</td><td>CAGCCACGACATGATGACAA</td><td>TGG</td><td>||||||||[||||||||||||]PAM</td><td>NA</td><td>Intergenic</td><td>NA</td><td>NA</td></tr>
<tr><td></td><td></td><td></td><td></td><td></td><td></td><td></td><td></td><td></td><td></td><td></td><td></td></tr>
<tr><td>T2898</td><td>AGCCACGACATGATGACAATGGG</td><td>1000</td><td></td><td></td><td></td><td></td><td></td><td></td><td></td><td></td><td></td></tr>
<tr><td>Oligo fwd</td><td>CACCAGCCACGACATGATGACAAT</td><td></td><td></td><td></td><td></td><td></td><td></td><td></td><td></td><td></td><td></td></tr>
<tr><td>Oligo rev</td><td>AAACATTGTCATCATGTCGTGGCT</td><td></td><td></td><td></td><td></td><td></td><td></td><td></td><td></td><td></td><td></td></tr>
<tr><td>Chromosome</td><td>start</td><td>end</td><td>strand</td><td>MM</td><td>target_seq</td><td>PAM</td><td>alignment</td><td>distance</td><td>position</td><td>gene name</td><td>gene id</td></tr>
<tr><td>GU305914.1</td><td>45372</td><td>45394</td><td>+</td><td>0</td><td>AGCCACGACATGATGACAAT</td><td>GGG</td><td>||||||||[||||||||||||]PAM</td><td>NA</td><td>Intergenic</td><td>NA</td><td>NA</td></tr>
<tr><td></td><td></td><td></td><td></td><td></td><td></td><td></td><td></td><td></td><td></td><td></td><td></td></tr>
<tr><td>T2899</td><td>GCCACGACATGATGACAATGGGG</td><td>1000</td><td></td><td></td><td></td><td></td><td></td><td></td><td></td><td></td><td></td></tr>
<tr><td>Oligo fwd</td><td>CACCGCCACGACATGATGACAATG</td><td></td><td></td><td></td><td></td><td></td><td></td><td></td><td></td><td></td><td></td></tr>
<tr><td>Oligo rev</td><td>AAACCATTGTCATCATGTCGTGGC</td><td></td><td></td><td></td><td></td><td></td><td></td><td></td><td></td><td></td><td></td></tr>
<tr><td>Chromosome</td><td>start</td><td>end</td><td>strand</td><td>MM</td><td>target_seq</td><td>PAM</td><td>alignment</td><td>distance</td><td>position</td><td>gene name</td><td>gene id</td></tr>
<tr><td>GU305914.1</td><td>45373</td><td>45395</td><td>+</td><td>0</td><td>GCCACGACATGATGACAATG</td><td>GGG</td><td>||||||||[||||||||||||]PAM</td><td>NA</td><td>Intergenic</td><td>NA</td><td>NA</td></tr>
<tr><td></td><td></td><td></td><td></td><td></td><td></td><td></td><td></td><td></td><td></td><td></td><td></td></tr>
<tr><td>T2900</td><td>CCACGACATGATGACAATGGGGG</td><td>1000</td><td></td><td></td><td></td><td></td><td></td><td></td><td></td><td></td><td></td></tr>
<tr><td>Oligo fwd</td><td>CACCCCACGACATGATGACAATGG</td><td></td><td></td><td></td><td></td><td></td><td></td><td></td><td></td><td></td><td></td></tr>
<tr><td>Oligo rev</td><td>AAACCCATTGTCATCATGTCGTGG</td><td></td><td></td><td></td><td></td><td></td><td></td><td></td><td></td><td></td><td></td></tr>
<tr><td>Chromosome</td><td>start</td><td>end</td><td>strand</td><td>MM</td><td>target_seq</td><td>PAM</td><td>alignment</td><td>distance</td><td>position</td><td>gene name</td><td>gene id</td></tr>
<tr><td>GU305914.1</td><td>45374</td><td>45396</td><td>+</td><td>0</td><td>CCACGACATGATGACAATGG</td><td>GGG</td><td>||||||||[||||||||||||]PAM</td><td>NA</td><td>Intergenic</td><td>NA</td><td>NA</td></tr>
<tr><td></td><td></td><td></td><td></td><td></td><td></td><td></td><td></td><td></td><td></td><td></td><td></td></tr>
<tr><td>T2901</td><td>CACGACATGATGACAATGGGGGG</td><td>1000</td><td></td><td></td><td></td><td></td><td></td><td></td><td></td><td></td><td></td></tr>
<tr><td>Oligo fwd</td><td>CACCCACGACATGATGACAATGGG</td><td></td><td></td><td></td><td></td><td></td><td></td><td></td><td></td><td></td><td></td></tr>
<tr><td>Oligo rev</td><td>AAACCCCATTGTCATCATGTCGTG</td><td></td><td></td><td></td><td></td><td></td><td></td><td></td><td></td><td></td><td></td></tr>
<tr><td>Chromosome</td><td>start</td><td>end</td><td>strand</td><td>MM</td><td>target_seq</td><td>PAM</td><td>alignment</td><td>distance</td><td>position</td><td>gene name</td><td>gene id</td></tr>
<tr><td>GU305914.1</td><td>45375</td><td>45397</td><td>+</td><td>0</td><td>CACGACATGATGACAATGGG</td><td>GGG</td><td>||||||||[||||||||||||]PAM</td><td>NA</td><td>Intergenic</td><td>NA</td><td>NA</td></tr>
<tr><td></td><td></td><td></td><td></td><td></td><td></td><td></td><td></td><td></td><td></td><td></td><td></td></tr>
<tr><td>T2902</td><td>GACAATGGGGGGAGAAGAAGAGG</td><td>1000</td><td></td><td></td><td></td><td></td><td></td><td></td><td></td><td></td><td></td></tr>
<tr><td>Oligo fwd</td><td>CACCGACAATGGGGGGAGAAGAAG</td><td></td><td></td><td></td><td></td><td></td><td></td><td></td><td></td><td></td><td></td></tr>
<tr><td>Oligo rev</td><td>AAACCTTCTTCTCCCCCCATTGTC</td><td></td><td></td><td></td><td></td><td></td><td></td><td></td><td></td><td></td><td></td></tr>
<tr><td>Chromosome</td><td>start</td><td>end</td><td>strand</td><td>MM</td><td>target_seq</td><td>PAM</td><td>alignment</td><td>distance</td><td>position</td><td>gene name</td><td>gene id</td></tr>
</table>

<table>
<tr><td>GU305914.1</td><td>45386</td><td>45408</td><td>+</td><td>0</td><td>GACAATGGGGGGAGAAGAAG</td><td>AGG</td><td>||||||||[||||||||||||]PAM</td><td>NA</td><td>Intergenic</td><td>NA</td><td>NA</td></tr>
<tr><td>T2903</td><td>ACAATGGGGGGAGAAGAAGAGGG</td><td>1000</td><td></td><td></td><td></td><td></td><td></td><td></td><td></td><td></td><td></td></tr>
<tr><td>Oligo fwd</td><td>CACCACAATGGGGGGAGAAGAAGA</td><td></td><td></td><td></td><td></td><td></td><td></td><td></td><td></td><td></td><td></td></tr>
<tr><td>Oligo rev</td><td>AAACTCTTCTTCTCCCCCCATTGT</td><td></td><td></td><td></td><td></td><td></td><td></td><td></td><td></td><td></td><td></td></tr>
<tr><td>Chromosome</td><td>start</td><td>end</td><td>strand</td><td>MM</td><td>target_seq</td><td>PAM</td><td>alignment</td><td>distance</td><td>position</td><td>gene name</td><td>gene id</td></tr>
<tr><td>GU305914.1</td><td>45387</td><td>45409</td><td>+</td><td>0</td><td>ACAATGGGGGGAGAAGAAGA</td><td>GGG</td><td>||||||||[||||||||||||]PAM</td><td>NA</td><td>Intergenic</td><td>NA</td><td>NA</td></tr>
<tr><td>T2904</td><td>TCGCCAAGCCCACACGAACTCGG</td><td>1000</td><td></td><td></td><td></td><td></td><td></td><td></td><td></td><td></td><td></td></tr>
<tr><td>Oligo fwd</td><td>CACCTCGCCAAGCCCACACGAACT</td><td></td><td></td><td></td><td></td><td></td><td></td><td></td><td></td><td></td><td></td></tr>
<tr><td>Oligo rev</td><td>AAACAGTTCGTGTGGGCTTGGCGA</td><td></td><td></td><td></td><td></td><td></td><td></td><td></td><td></td><td></td><td></td></tr>
<tr><td>Chromosome</td><td>start</td><td>end</td><td>strand</td><td>MM</td><td>target_seq</td><td>PAM</td><td>alignment</td><td>distance</td><td>position</td><td>gene name</td><td>gene id</td></tr>
<tr><td>GU305914.1</td><td>45486</td><td>45508</td><td>+</td><td>0</td><td>TCGCCAAGCCCACACGAACT</td><td>CGG</td><td>||||||||[||||||||||||]PAM</td><td>NA</td><td>Intergenic</td><td>NA</td><td>NA</td></tr>
<tr><td>T2905</td><td>CGCCAAGCCCACACGAACTCGGG</td><td>1000</td><td></td><td></td><td></td><td></td><td></td><td></td><td></td><td></td><td></td></tr>
<tr><td>Oligo fwd</td><td>CACCCGCCAAGCCCACACGAACTC</td><td></td><td></td><td></td><td></td><td></td><td></td><td></td><td></td><td></td><td></td></tr>
<tr><td>Oligo rev</td><td>AAACGAGTTCGTGTGGGCTTGGCG</td><td></td><td></td><td></td><td></td><td></td><td></td><td></td><td></td><td></td><td></td></tr>
<tr><td>Chromosome</td><td>start</td><td>end</td><td>strand</td><td>MM</td><td>target_seq</td><td>PAM</td><td>alignment</td><td>distance</td><td>position</td><td>gene name</td><td>gene id</td></tr>
<tr><td>GU305914.1</td><td>45487</td><td>45509</td><td>+</td><td>0</td><td>CGCCAAGCCCACACGAACTC</td><td>GGG</td><td>||||||||[||||||||||||]PAM</td><td>NA</td><td>Intergenic</td><td>NA</td><td>NA</td></tr>
<tr><td>T2906</td><td>TTTTCACCGAACGAGATAAACGG</td><td>1000</td><td></td><td></td><td></td><td></td><td></td><td></td><td></td><td></td><td></td></tr>
<tr><td>Oligo fwd</td><td>CACCTTTTCACCGAACGAGATAAA</td><td></td><td></td><td></td><td></td><td></td><td></td><td></td><td></td><td></td><td></td></tr>
<tr><td>Oligo rev</td><td>AAACTTTATCTCGTTCGGTGAAAA</td><td></td><td></td><td></td><td></td><td></td><td></td><td></td><td></td><td></td><td></td></tr>
<tr><td>Chromosome</td><td>start</td><td>end</td><td>strand</td><td>MM</td><td>target_seq</td><td>PAM</td><td>alignment</td><td>distance</td><td>position</td><td>gene name</td><td>gene id</td></tr>
<tr><td>GU305914.1</td><td>45513</td><td>45535</td><td>+</td><td>0</td><td>TTTTCACCGAACGAGATAAA</td><td>CGG</td><td>||||||||[||||||||||||]PAM</td><td>NA</td><td>Intergenic</td><td>NA</td><td>NA</td></tr>
<tr><td>T2907</td><td>AACTTCATCTCCGTCCACGTCGG</td><td>1000</td><td></td><td></td><td></td><td></td><td></td><td></td><td></td><td></td><td></td></tr>
<tr><td>Oligo fwd</td><td>CACCAACTTCATCTCCGTCCACGT</td><td></td><td></td><td></td><td></td><td></td><td></td><td></td><td></td><td></td><td></td></tr>
<tr><td>Oligo rev</td><td>AAACACGTGGACGGAGATGAAGTT</td><td></td><td></td><td></td><td></td><td></td><td></td><td></td><td></td><td></td><td></td></tr>
<tr><td>Chromosome</td><td>start</td><td>end</td><td>strand</td><td>MM</td><td>target_seq</td><td>PAM</td><td>alignment</td><td>distance</td><td>position</td><td>gene name</td><td>gene id</td></tr>
<tr><td>GU305914.1</td><td>45549</td><td>45571</td><td>+</td><td>0</td><td>AACTTCATCTCCGTCCACGT</td><td>CGG</td><td>||||||||[||||||||||||]PAM</td><td>NA</td><td>Intergenic</td><td>NA</td><td>NA</td></tr>
<tr><td>T2908</td><td>TCATCTCCGTCCACGTCGGACGG</td><td>1000</td><td></td><td></td><td></td><td></td><td></td><td></td><td></td><td></td><td></td></tr>
<tr><td>Oligo fwd</td><td>CACCTCATCTCCGTCCACGTCGGA</td><td></td><td></td><td></td><td></td><td></td><td></td><td></td><td></td><td></td><td></td></tr>
<tr><td>Oligo rev</td><td>AAACTCCGACGTGGACGGAGATGA</td><td></td><td></td><td></td><td></td><td></td><td></td><td></td><td></td><td></td><td></td></tr>
<tr><td>Chromosome</td><td>start</td><td>end</td><td>strand</td><td>MM</td><td>target_seq</td><td>PAM</td><td>alignment</td><td>distance</td><td>position</td><td>gene name</td><td>gene id</td></tr>
<tr><td>GU305914.1</td><td>45553</td><td>45575</td><td>+</td><td>0</td><td>TCATCTCCGTCCACGTCGGA</td><td>CGG</td><td>||||||||[||||||||||||]PAM</td><td>NA</td><td>Intergenic</td><td>NA</td><td>NA</td></tr>
<tr><td>T2909</td><td>GTCGGACGGTCTACGCCACCCGG</td><td>1000</td><td></td><td></td><td></td><td></td><td></td><td></td><td></td><td></td><td></td></tr>
<tr><td>Oligo fwd</td><td>CACCGTCGGACGGTCTACGCCACC</td><td></td><td></td><td></td><td></td><td></td><td></td><td></td><td></td><td></td><td></td></tr>
<tr><td>Oligo rev</td><td>AAACGGTGGCGTAGACCGTCCGAC</td><td></td><td></td><td></td><td></td><td></td><td></td><td></td><td></td><td></td><td></td></tr>
<tr><td>Chromosome</td><td>start</td><td>end</td><td>strand</td><td>MM</td><td>target_seq</td><td>PAM</td><td>alignment</td><td>distance</td><td>position</td><td>gene name</td><td>gene id</td></tr>
<tr><td>GU305914.1</td><td>45567</td><td>45589</td><td>+</td><td>0</td><td>GTCGGACGGTCTACGCCACC</td><td>CGG</td><td>||||||||[||||||||||||]PAM</td><td>NA</td><td>Intergenic</td><td>NA</td><td>NA</td></tr>
<tr><td>T2910</td><td>GTCTACGCCACCCGGCGCCGCGG</td><td>1000</td><td></td><td></td><td></td><td></td><td></td><td></td><td></td><td></td><td></td></tr>
<tr><td>Oligo fwd</td><td>CACCGTCTACGCCACCCGGCGCCG</td><td></td><td></td><td></td><td></td><td></td><td></td><td></td><td></td><td></td><td></td></tr>
<tr><td>Oligo rev</td><td>AAACCGGCGCCGGGTGGCGTAGAC</td><td></td><td></td><td></td><td></td><td></td><td></td><td></td><td></td><td></td><td></td></tr>
<tr><td>Chromosome</td><td>start</td><td>end</td><td>strand</td><td>MM</td><td>target_seq</td><td>PAM</td><td>alignment</td><td>distance</td><td>position</td><td>gene name</td><td>gene id</td></tr>
<tr><td>GU305914.1</td><td>45575</td><td>45597</td><td>+</td><td>0</td><td>GTCTACGCCACCCGGCGCCG</td><td>CGG</td><td>||||||||[||||||||||||]PAM</td><td>NA</td><td>Intergenic</td><td>NA</td><td>NA</td></tr>
<tr><td>T2911</td><td>TACGCCACCCGGCGCCGCGGCGG</td><td>1000</td><td></td><td></td><td></td><td></td><td></td><td></td><td></td><td></td><td></td></tr>
<tr><td>Oligo fwd</td><td>CACCTACGCCACCCGGCGCCGCGG</td><td></td><td></td><td></td><td></td><td></td><td></td><td></td><td></td><td></td><td></td></tr>
<tr><td>Oligo rev</td><td>AAACCCGCGGCGCCGGGTGGCGTA</td><td></td><td></td><td></td><td></td><td></td><td></td><td></td><td></td><td></td><td></td></tr>
<tr><td>Chromosome</td><td>start</td><td>end</td><td>strand</td><td>MM</td><td>target_seq</td><td>PAM</td><td>alignment</td><td>distance</td><td>position</td><td>gene name</td><td>gene id</td></tr>
<tr><td>GU305914.1</td><td>45578</td><td>45600</td><td>+</td><td>0</td><td>TACGCCACCCGGCGCCGCGG</td><td>CGG</td><td>||||||||[||||||||||||]PAM</td><td>NA</td><td>Intergenic</td><td>NA</td><td>NA</td></tr>
<tr><td>T2912</td><td>GCCACCCGGCGCCGCGGCGGCGG</td><td>1000</td><td></td><td></td><td></td><td></td><td></td><td></td><td></td><td></td><td></td></tr>
<tr><td>Oligo fwd</td><td>CACCGCCACCCGGCGCCGCGGCGG</td><td></td><td></td><td></td><td></td><td></td><td></td><td></td><td></td><td></td><td></td></tr>
<tr><td>Oligo rev</td><td>AAACCCGCCGCGGCGCCGGGTGGC</td><td></td><td></td><td></td><td></td><td></td><td></td><td></td><td></td><td></td><td></td></tr>
<tr><td>Chromosome</td><td>start</td><td>end</td><td>strand</td><td>MM</td><td>target_seq</td><td>PAM</td><td>alignment</td><td>distance</td><td>position</td><td>gene name</td><td>gene id</td></tr>
<tr><td>GU305914.1</td><td>45581</td><td>45603</td><td>+</td><td>0</td><td>GCCACCCGGCGCCGCGGCGG</td><td>CGG</td><td>||||||||[||||||||||||]PAM</td><td>NA</td><td>Intergenic</td><td>NA</td><td>NA</td></tr>
<tr><td>T2913</td><td>ACCCGGCGCCGCGGCGGCGGCGG</td><td>1000</td><td></td><td></td><td></td><td></td><td></td><td></td><td></td><td></td><td></td></tr>
<tr><td>Oligo fwd</td><td>CACCACCCGGCGCCGCGGCGGCGG</td><td></td><td></td><td></td><td></td><td></td><td></td><td></td><td></td><td></td><td></td></tr>
<tr><td>Oligo rev</td><td>AAACCCGCCGCCGCGGCGCCGGGT</td><td></td><td></td><td></td><td></td><td></td><td></td><td></td><td></td><td></td><td></td></tr>
<tr><td>Chromosome</td><td>start</td><td>end</td><td>strand</td><td>MM</td><td>target_seq</td><td>PAM</td><td>alignment</td><td>distance</td><td>position</td><td>gene name</td><td>gene id</td></tr>
<tr><td>GU305914.1</td><td>45584</td><td>45606</td><td>+</td><td>0</td><td>ACCCGGCGCCGCGGCGGCGG</td><td>CGG</td><td>||||||||[||||||||||||]PAM</td><td>NA</td><td>Intergenic</td><td>NA</td><td>NA</td></tr>
<tr><td>GU305914.1</td><td>56770</td><td>56792</td><td>+</td><td>4</td><td>CTCCGCCGCAGCGGCGGCGG</td><td>CGG</td><td>--|||-||[|-||||||||||]PAM</td><td>NA</td><td>Intergenic</td><td>NA</td><td>NA</td></tr>
<tr><td>T2914</td><td>TGCATCAACATCCTGCTCTAAGG</td><td>1000</td><td></td><td></td><td></td><td></td><td></td><td></td><td></td><td></td><td></td></tr>
<tr><td>Oligo fwd</td><td>CACCTGCATCAACATCCTGCTCTA</td><td></td><td></td><td></td><td></td><td></td><td></td><td></td><td></td><td></td><td></td></tr>
<tr><td>Oligo rev</td><td>AAACTAGAGCAGGATGTTGATGCA</td><td></td><td></td><td></td><td></td><td></td><td></td><td></td><td></td><td></td><td></td></tr>
<tr><td>Chromosome</td><td>start</td><td>end</td><td>strand</td><td>MM</td><td>target_seq</td><td>PAM</td><td>alignment</td><td>distance</td><td>position</td><td>gene name</td><td>gene id</td></tr>
<tr><td>GU305914.1</td><td>45630</td><td>45652</td><td>+</td><td>0</td><td>TGCATCAACATCCTGCTCTA</td><td>AGG</td><td>||||||||[||||||||||||]PAM</td><td>NA</td><td>Intergenic</td><td>NA</td><td>NA</td></tr>
<tr><td>T2915</td><td>GCATCAACATCCTGCTCTAAGGG</td><td>1000</td><td></td><td></td><td></td><td></td><td></td><td></td><td></td><td></td><td></td></tr>
<tr><td>Oligo fwd</td><td>CACCGCATCAACATCCTGCTCTAA</td><td></td><td></td><td></td><td></td><td></td><td></td><td></td><td></td><td></td><td></td></tr>
<tr><td>Oligo rev</td><td>AAACTTAGAGCAGGATGTTGATGC</td><td></td><td></td><td></td><td></td><td></td><td></td><td></td><td></td><td></td><td></td></tr>
<tr><td>Chromosome</td><td>start</td><td>end</td><td>strand</td><td>MM</td><td>target_seq</td><td>PAM</td><td>alignment</td><td>distance</td><td>position</td><td>gene name</td><td>gene id</td></tr>
<tr><td>GU305914.1</td><td>45631</td><td>45653</td><td>+</td><td>0</td><td>GCATCAACATCCTGCTCTAA</td><td>GGG</td><td>||||||||[||||||||||||]PAM</td><td>NA</td><td>Intergenic</td><td>NA</td><td>NA</td></tr>
<tr><td>T2916</td><td>CATCAACATCCTGCTCTAAGGGG</td><td>1000</td><td></td><td></td><td></td><td></td><td></td><td></td><td></td><td></td><td></td></tr>
<tr><td>Oligo fwd</td><td>CACCCATCAACATCCTGCTCTAAG</td><td></td><td></td><td></td><td></td><td></td><td></td><td></td><td></td><td></td><td></td></tr>
<tr><td>Oligo rev</td><td>AAACCTTAGAGCAGGATGTTGATG</td><td></td><td></td><td></td><td></td><td></td><td></td><td></td><td></td><td></td><td></td></tr>
<tr><td>Chromosome</td><td>start</td><td>end</td><td>strand</td><td>MM</td><td>target_seq</td><td>PAM</td><td>alignment</td><td>distance</td><td>position</td><td>gene name</td><td>gene id</td></tr>
<tr><td>GU305914.1</td><td>45632</td><td>45654</td><td>+</td><td>0</td><td>CATCAACATCCTGCTCTAAG</td><td>GGG</td><td>||||||||[||||||||||||]PAM</td><td>NA</td><td>Intergenic</td><td>NA</td><td>NA</td></tr>
<tr><td>T2917</td><td>ATCAACATCCTGCTCTAAGGGGG</td><td>1000</td><td></td><td></td><td></td><td></td><td></td><td></td><td></td><td></td><td></td></tr>
<tr><td>Oligo fwd</td><td>CACCATCAACATCCTGCTCTAAGG</td><td></td><td></td><td></td><td></td><td></td><td></td><td></td><td></td><td></td><td></td></tr>
</table>

<table>
<tr><td>Oligo rev</td><td>AAACCCTTAGAGCAGGATGTTGAT</td><td></td><td></td><td></td><td></td><td></td><td></td><td></td><td></td><td></td><td></td></tr>
<tr><td>Chromosome</td><td>start</td><td>end</td><td>strand</td><td>MM</td><td>target_seq</td><td>PAM</td><td>alignment</td><td>distance</td><td>position</td><td>gene name</td><td>gene id</td></tr>
<tr><td>GU305914.1</td><td>45633</td><td>45655</td><td>+</td><td>0</td><td>ATCAACATCCTGCTCTAAGG</td><td>GGG</td><td>||||||||[||||||||||||]PAM</td><td>NA</td><td>Intergenic</td><td>NA</td><td>NA</td></tr>
<tr><td>T2918</td><td>TCAACATCCTGCTCTAAGGGGGG</td><td>1000</td><td></td><td></td><td></td><td></td><td></td><td></td><td></td><td></td><td></td></tr>
<tr><td>Oligo fwd</td><td>CACCTCAACATCCTGCTCTAAGGG</td><td></td><td></td><td></td><td></td><td></td><td></td><td></td><td></td><td></td><td></td></tr>
<tr><td>Oligo rev</td><td>AAACCCCTTAGAGCAGGATGTTGA</td><td></td><td></td><td></td><td></td><td></td><td></td><td></td><td></td><td></td><td></td></tr>
<tr><td>Chromosome</td><td>start</td><td>end</td><td>strand</td><td>MM</td><td>target_seq</td><td>PAM</td><td>alignment</td><td>distance</td><td>position</td><td>gene name</td><td>gene id</td></tr>
<tr><td>GU305914.1</td><td>45634</td><td>45656</td><td>+</td><td>0</td><td>TCAACATCCTGCTCTAAGGG</td><td>GGG</td><td>||||||||[||||||||||||]PAM</td><td>NA</td><td>Intergenic</td><td>NA</td><td>NA</td></tr>
<tr><td>T2919</td><td>ATCCTGCTCTAAGGGGGGCGCGG</td><td>1000</td><td></td><td></td><td></td><td></td><td></td><td></td><td></td><td></td><td></td></tr>
<tr><td>Oligo fwd</td><td>CACCATCCTGCTCTAAGGGGGGCG</td><td></td><td></td><td></td><td></td><td></td><td></td><td></td><td></td><td></td><td></td></tr>
<tr><td>Oligo rev</td><td>AAACCGCCCCCCTTAGAGCAGGAT</td><td></td><td></td><td></td><td></td><td></td><td></td><td></td><td></td><td></td><td></td></tr>
<tr><td>Chromosome</td><td>start</td><td>end</td><td>strand</td><td>MM</td><td>target_seq</td><td>PAM</td><td>alignment</td><td>distance</td><td>position</td><td>gene name</td><td>gene id</td></tr>
<tr><td>GU305914.1</td><td>45639</td><td>45661</td><td>+</td><td>0</td><td>ATCCTGCTCTAAGGGGGGCG</td><td>CGG</td><td>||||||||[||||||||||||]PAM</td><td>NA</td><td>Intergenic</td><td>NA</td><td>NA</td></tr>
<tr><td>T2920</td><td>TCCTGCTCTAAGGGGGGCGCGGG</td><td>1000</td><td></td><td></td><td></td><td></td><td></td><td></td><td></td><td></td><td></td></tr>
<tr><td>Oligo fwd</td><td>CACCTCCTGCTCTAAGGGGGGCGC</td><td></td><td></td><td></td><td></td><td></td><td></td><td></td><td></td><td></td><td></td></tr>
<tr><td>Oligo rev</td><td>AAACGCGCCCCCCTTAGAGCAGGA</td><td></td><td></td><td></td><td></td><td></td><td></td><td></td><td></td><td></td><td></td></tr>
<tr><td>Chromosome</td><td>start</td><td>end</td><td>strand</td><td>MM</td><td>target_seq</td><td>PAM</td><td>alignment</td><td>distance</td><td>position</td><td>gene name</td><td>gene id</td></tr>
<tr><td>GU305914.1</td><td>45640</td><td>45662</td><td>+</td><td>0</td><td>TCCTGCTCTAAGGGGGGCGC</td><td>GGG</td><td>||||||||[||||||||||||]PAM</td><td>NA</td><td>Intergenic</td><td>NA</td><td>NA</td></tr>
<tr><td>T2921</td><td>TCTAAGGGGGGCGCGGGACGAGG</td><td>1000</td><td></td><td></td><td></td><td></td><td></td><td></td><td></td><td></td><td></td></tr>
<tr><td>Oligo fwd</td><td>CACCTCTAAGGGGGGCGCGGGACG</td><td></td><td></td><td></td><td></td><td></td><td></td><td></td><td></td><td></td><td></td></tr>
<tr><td>Oligo rev</td><td>AAACCGTCCCGCGCCCCCCTTAGA</td><td></td><td></td><td></td><td></td><td></td><td></td><td></td><td></td><td></td><td></td></tr>
<tr><td>Chromosome</td><td>start</td><td>end</td><td>strand</td><td>MM</td><td>target_seq</td><td>PAM</td><td>alignment</td><td>distance</td><td>position</td><td>gene name</td><td>gene id</td></tr>
<tr><td>GU305914.1</td><td>45646</td><td>45668</td><td>+</td><td>0</td><td>TCTAAGGGGGGCGCGGGACG</td><td>AGG</td><td>||||||||[||||||||||||]PAM</td><td>NA</td><td>Intergenic</td><td>NA</td><td>NA</td></tr>
<tr><td>T2922</td><td>CTAAGGGGGGCGCGGGACGAGGG</td><td>1000</td><td></td><td></td><td></td><td></td><td></td><td></td><td></td><td></td><td></td></tr>
<tr><td>Oligo fwd</td><td>CACCCTAAGGGGGGCGCGGGACGA</td><td></td><td></td><td></td><td></td><td></td><td></td><td></td><td></td><td></td><td></td></tr>
<tr><td>Oligo rev</td><td>AAACTCGTCCCGCGCCCCCCTTAG</td><td></td><td></td><td></td><td></td><td></td><td></td><td></td><td></td><td></td><td></td></tr>
<tr><td>Chromosome</td><td>start</td><td>end</td><td>strand</td><td>MM</td><td>target_seq</td><td>PAM</td><td>alignment</td><td>distance</td><td>position</td><td>gene name</td><td>gene id</td></tr>
<tr><td>GU305914.1</td><td>45647</td><td>45669</td><td>+</td><td>0</td><td>CTAAGGGGGGCGCGGGACGA</td><td>GGG</td><td>||||||||[||||||||||||]PAM</td><td>NA</td><td>Intergenic</td><td>NA</td><td>NA</td></tr>
<tr><td>T2923</td><td>TAAGGGGGGCGCGGGACGAGGGG</td><td>1000</td><td></td><td></td><td></td><td></td><td></td><td></td><td></td><td></td><td></td></tr>
<tr><td>Oligo fwd</td><td>CACCTAAGGGGGGCGCGGGACGAG</td><td></td><td></td><td></td><td></td><td></td><td></td><td></td><td></td><td></td><td></td></tr>
<tr><td>Oligo rev</td><td>AAACCTCGTCCCGCGCCCCCCTTA</td><td></td><td></td><td></td><td></td><td></td><td></td><td></td><td></td><td></td><td></td></tr>
<tr><td>Chromosome</td><td>start</td><td>end</td><td>strand</td><td>MM</td><td>target_seq</td><td>PAM</td><td>alignment</td><td>distance</td><td>position</td><td>gene name</td><td>gene id</td></tr>
<tr><td>GU305914.1</td><td>45648</td><td>45670</td><td>+</td><td>0</td><td>TAAGGGGGGCGCGGGACGAG</td><td>GGG</td><td>||||||||[||||||||||||]PAM</td><td>NA</td><td>Intergenic</td><td>NA</td><td>NA</td></tr>
<tr><td>T2924</td><td>GGGGCGCGGGACGAGGGGAACGG</td><td>1000</td><td></td><td></td><td></td><td></td><td></td><td></td><td></td><td></td><td></td></tr>
<tr><td>Oligo fwd</td><td>CACCGGGGCGCGGGACGAGGGGAA</td><td></td><td></td><td></td><td></td><td></td><td></td><td></td><td></td><td></td><td></td></tr>
<tr><td>Oligo rev</td><td>AAACTTCCCCTCGTCCCGCGCCCC</td><td></td><td></td><td></td><td></td><td></td><td></td><td></td><td></td><td></td><td></td></tr>
<tr><td>Chromosome</td><td>start</td><td>end</td><td>strand</td><td>MM</td><td>target_seq</td><td>PAM</td><td>alignment</td><td>distance</td><td>position</td><td>gene name</td><td>gene id</td></tr>
<tr><td>GU305914.1</td><td>45653</td><td>45675</td><td>+</td><td>0</td><td>GGGGCGCGGGACGAGGGGAA</td><td>CGG</td><td>||||||||[||||||||||||]PAM</td><td>NA</td><td>Intergenic</td><td>NA</td><td>NA</td></tr>
<tr><td>T2925</td><td>CGAGGGGAACGGTCGAGACACGG</td><td>1000</td><td></td><td></td><td></td><td></td><td></td><td></td><td></td><td></td><td></td></tr>
<tr><td>Oligo fwd</td><td>CACCCGAGGGGAACGGTCGAGACA</td><td></td><td></td><td></td><td></td><td></td><td></td><td></td><td></td><td></td><td></td></tr>
<tr><td>Oligo rev</td><td>AAACTGTCTCGACCGTTCCCCTCG</td><td></td><td></td><td></td><td></td><td></td><td></td><td></td><td></td><td></td><td></td></tr>
<tr><td>Chromosome</td><td>start</td><td>end</td><td>strand</td><td>MM</td><td>target_seq</td><td>PAM</td><td>alignment</td><td>distance</td><td>position</td><td>gene name</td><td>gene id</td></tr>
<tr><td>GU305914.1</td><td>45664</td><td>45686</td><td>+</td><td>0</td><td>CGAGGGGAACGGTCGAGACA</td><td>CGG</td><td>||||||||[||||||||||||]PAM</td><td>NA</td><td>Intergenic</td><td>NA</td><td>NA</td></tr>
<tr><td>T2926</td><td>ACCGTGTCGTTTTGTAGCATAGG</td><td>1000</td><td></td><td></td><td></td><td></td><td></td><td></td><td></td><td></td><td></td></tr>
<tr><td>Oligo fwd</td><td>CACCACCGTGTCGTTTTGTAGCAT</td><td></td><td></td><td></td><td></td><td></td><td></td><td></td><td></td><td></td><td></td></tr>
<tr><td>Oligo rev</td><td>AAACATGCTACAAAACGACACGGT</td><td></td><td></td><td></td><td></td><td></td><td></td><td></td><td></td><td></td><td></td></tr>
<tr><td>Chromosome</td><td>start</td><td>end</td><td>strand</td><td>MM</td><td>target_seq</td><td>PAM</td><td>alignment</td><td>distance</td><td>position</td><td>gene name</td><td>gene id</td></tr>
<tr><td>GU305914.1</td><td>45715</td><td>45737</td><td>+</td><td>0</td><td>ACCGTGTCGTTTTGTAGCAT</td><td>AGG</td><td>||||||||[||||||||||||]PAM</td><td>NA</td><td>Intergenic</td><td>NA</td><td>NA</td></tr>
<tr><td>T2927</td><td>CCGTGTCGTTTTGTAGCATAGGG</td><td>1000</td><td></td><td></td><td></td><td></td><td></td><td></td><td></td><td></td><td></td></tr>
<tr><td>Oligo fwd</td><td>CACCCCGTGTCGTTTTGTAGCATA</td><td></td><td></td><td></td><td></td><td></td><td></td><td></td><td></td><td></td><td></td></tr>
<tr><td>Oligo rev</td><td>AAACTATGCTACAAAACGACACGG</td><td></td><td></td><td></td><td></td><td></td><td></td><td></td><td></td><td></td><td></td></tr>
<tr><td>Chromosome</td><td>start</td><td>end</td><td>strand</td><td>MM</td><td>target_seq</td><td>PAM</td><td>alignment</td><td>distance</td><td>position</td><td>gene name</td><td>gene id</td></tr>
<tr><td>GU305914.1</td><td>45716</td><td>45738</td><td>+</td><td>0</td><td>CCGTGTCGTTTTGTAGCATA</td><td>GGG</td><td>||||||||[||||||||||||]PAM</td><td>NA</td><td>Intergenic</td><td>NA</td><td>NA</td></tr>
<tr><td>T2928</td><td>CGTGTCGTTTTGTAGCATAGGGG</td><td>1000</td><td></td><td></td><td></td><td></td><td></td><td></td><td></td><td></td><td></td></tr>
<tr><td>Oligo fwd</td><td>CACCCGTGTCGTTTTGTAGCATAG</td><td></td><td></td><td></td><td></td><td></td><td></td><td></td><td></td><td></td><td></td></tr>
<tr><td>Oligo rev</td><td>AAACCTATGCTACAAAACGACACG</td><td></td><td></td><td></td><td></td><td></td><td></td><td></td><td></td><td></td><td></td></tr>
<tr><td>Chromosome</td><td>start</td><td>end</td><td>strand</td><td>MM</td><td>target_seq</td><td>PAM</td><td>alignment</td><td>distance</td><td>position</td><td>gene name</td><td>gene id</td></tr>
<tr><td>GU305914.1</td><td>45717</td><td>45739</td><td>+</td><td>0</td><td>CGTGTCGTTTTGTAGCATAG</td><td>GGG</td><td>||||||||[||||||||||||]PAM</td><td>NA</td><td>Intergenic</td><td>NA</td><td>NA</td></tr>
<tr><td>T2929</td><td>GTTTTGTAGCATAGGGGAAACGG</td><td>1000</td><td></td><td></td><td></td><td></td><td></td><td></td><td></td><td></td><td></td></tr>
<tr><td>Oligo fwd</td><td>CACCGTTTTGTAGCATAGGGGAAA</td><td></td><td></td><td></td><td></td><td></td><td></td><td></td><td></td><td></td><td></td></tr>
<tr><td>Oligo rev</td><td>AAACTTTCCCCTATGCTACAAAAC</td><td></td><td></td><td></td><td></td><td></td><td></td><td></td><td></td><td></td><td></td></tr>
<tr><td>Chromosome</td><td>start</td><td>end</td><td>strand</td><td>MM</td><td>target_seq</td><td>PAM</td><td>alignment</td><td>distance</td><td>position</td><td>gene name</td><td>gene id</td></tr>
<tr><td>GU305914.1</td><td>45723</td><td>45745</td><td>+</td><td>0</td><td>GTTTTGTAGCATAGGGGAAA</td><td>CGG</td><td>||||||||[||||||||||||]PAM</td><td>NA</td><td>Intergenic</td><td>NA</td><td>NA</td></tr>
<tr><td>T2930</td><td>TTTTGTAGCATAGGGGAAACGGG</td><td>1000</td><td></td><td></td><td></td><td></td><td></td><td></td><td></td><td></td><td></td></tr>
<tr><td>Oligo fwd</td><td>CACCTTTTGTAGCATAGGGGAAAC</td><td></td><td></td><td></td><td></td><td></td><td></td><td></td><td></td><td></td><td></td></tr>
<tr><td>Oligo rev</td><td>AAACGTTTCCCCTATGCTACAAAA</td><td></td><td></td><td></td><td></td><td></td><td></td><td></td><td></td><td></td><td></td></tr>
<tr><td>Chromosome</td><td>start</td><td>end</td><td>strand</td><td>MM</td><td>target_seq</td><td>PAM</td><td>alignment</td><td>distance</td><td>position</td><td>gene name</td><td>gene id</td></tr>
<tr><td>GU305914.1</td><td>45724</td><td>45746</td><td>+</td><td>0</td><td>TTTTGTAGCATAGGGGAAAC</td><td>GGG</td><td>||||||||[||||||||||||]PAM</td><td>NA</td><td>Intergenic</td><td>NA</td><td>NA</td></tr>
<tr><td>T2931</td><td>TACATCTAATAAAAACATCTCGG</td><td>1000</td><td></td><td></td><td></td><td></td><td></td><td></td><td></td><td></td><td></td></tr>
<tr><td>Oligo fwd</td><td>CACCTACATCTAATAAAAACATCT</td><td></td><td></td><td></td><td></td><td></td><td></td><td></td><td></td><td></td><td></td></tr>
<tr><td>Oligo rev</td><td>AAACAGATGTTTTTATTAGATGTA</td><td></td><td></td><td></td><td></td><td></td><td></td><td></td><td></td><td></td><td></td></tr>
<tr><td>Chromosome</td><td>start</td><td>end</td><td>strand</td><td>MM</td><td>target_seq</td><td>PAM</td><td>alignment</td><td>distance</td><td>position</td><td>gene name</td><td>gene id</td></tr>
<tr><td>GU305914.1</td><td>45767</td><td>45789</td><td>+</td><td>0</td><td>TACATCTAATAAAAACATCT</td><td>CGG</td><td>||||||||[||||||||||||]PAM</td><td>NA</td><td>Intergenic</td><td>NA</td><td>NA</td></tr>
<tr><td>T2932</td><td>CTAATAAAAACATCTCGGTACGG</td><td>1000</td><td></td><td></td><td></td><td></td><td></td><td></td><td></td><td></td><td></td></tr>
</table>

<table>
<tr><td>Oligo fwd</td><td>CACCCTAATAAAAACATCTCGGTA</td><td></td><td></td><td></td><td></td><td></td><td></td><td></td><td></td><td></td><td></td></tr>
<tr><td>Oligo rev</td><td>AAACTACCGAGATGTTTTTATTAG</td><td></td><td></td><td></td><td></td><td></td><td></td><td></td><td></td><td></td><td></td></tr>
<tr><td>Chromosome</td><td>start</td><td>end</td><td>strand</td><td>MM</td><td>target_seq</td><td>PAM</td><td>alignment</td><td>distance</td><td>position</td><td>gene name</td><td>gene id</td></tr>
<tr><td>GU305914.1</td><td>45772</td><td>45794</td><td>+</td><td>0</td><td>CTAATAAAAACATCTCGGTA</td><td>CGG</td><td>||||||||[||||||||||||]PAM</td><td>NA</td><td>Intergenic</td><td>NA</td><td>NA</td></tr>
<tr><td></td><td></td><td></td><td></td><td></td><td></td><td></td><td></td><td></td><td></td><td></td><td></td></tr>
<tr><td>T2933</td><td>AAACATCTCGGTACGGCCTCTGG</td><td>1000</td><td></td><td></td><td></td><td></td><td></td><td></td><td></td><td></td><td></td></tr>
<tr><td>Oligo fwd</td><td>CACCAAACATCTCGGTACGGCCTC</td><td></td><td></td><td></td><td></td><td></td><td></td><td></td><td></td><td></td><td></td></tr>
<tr><td>Oligo rev</td><td>AAACGAGGCCGTACCGAGATGTTT</td><td></td><td></td><td></td><td></td><td></td><td></td><td></td><td></td><td></td><td></td></tr>
<tr><td>Chromosome</td><td>start</td><td>end</td><td>strand</td><td>MM</td><td>target_seq</td><td>PAM</td><td>alignment</td><td>distance</td><td>position</td><td>gene name</td><td>gene id</td></tr>
<tr><td>GU305914.1</td><td>45779</td><td>45801</td><td>+</td><td>0</td><td>AAACATCTCGGTACGGCCTC</td><td>TGG</td><td>||||||||[||||||||||||]PAM</td><td>NA</td><td>Intergenic</td><td>NA</td><td>NA</td></tr>
<tr><td></td><td></td><td></td><td></td><td></td><td></td><td></td><td></td><td></td><td></td><td></td><td></td></tr>
<tr><td>T2934</td><td>AACATCTCGGTACGGCCTCTGGG</td><td>1000</td><td></td><td></td><td></td><td></td><td></td><td></td><td></td><td></td><td></td></tr>
<tr><td>Oligo fwd</td><td>CACCAACATCTCGGTACGGCCTCT</td><td></td><td></td><td></td><td></td><td></td><td></td><td></td><td></td><td></td><td></td></tr>
<tr><td>Oligo rev</td><td>AAACAGAGGCCGTACCGAGATGTT</td><td></td><td></td><td></td><td></td><td></td><td></td><td></td><td></td><td></td><td></td></tr>
<tr><td>Chromosome</td><td>start</td><td>end</td><td>strand</td><td>MM</td><td>target_seq</td><td>PAM</td><td>alignment</td><td>distance</td><td>position</td><td>gene name</td><td>gene id</td></tr>
<tr><td>GU305914.1</td><td>45780</td><td>45802</td><td>+</td><td>0</td><td>AACATCTCGGTACGGCCTCT</td><td>GGG</td><td>||||||||[||||||||||||]PAM</td><td>NA</td><td>Intergenic</td><td>NA</td><td>NA</td></tr>
<tr><td></td><td></td><td></td><td></td><td></td><td></td><td></td><td></td><td></td><td></td><td></td><td></td></tr>
<tr><td>T2935</td><td>CTCTGGGCCCAAAAGCTAATTGG</td><td>1000</td><td></td><td></td><td></td><td></td><td></td><td></td><td></td><td></td><td></td></tr>
<tr><td>Oligo fwd</td><td>CACCCTCTGGGCCCAAAAGCTAAT</td><td></td><td></td><td></td><td></td><td></td><td></td><td></td><td></td><td></td><td></td></tr>
<tr><td>Oligo rev</td><td>AAACATTAGCTTTTGGGCCCAGAG</td><td></td><td></td><td></td><td></td><td></td><td></td><td></td><td></td><td></td><td></td></tr>
<tr><td>Chromosome</td><td>start</td><td>end</td><td>strand</td><td>MM</td><td>target_seq</td><td>PAM</td><td>alignment</td><td>distance</td><td>position</td><td>gene name</td><td>gene id</td></tr>
<tr><td>GU305914.1</td><td>45796</td><td>45818</td><td>+</td><td>0</td><td>CTCTGGGCCCAAAAGCTAAT</td><td>TGG</td><td>||||||||[||||||||||||]PAM</td><td>NA</td><td>Intergenic</td><td>NA</td><td>NA</td></tr>
<tr><td></td><td></td><td></td><td></td><td></td><td></td><td></td><td></td><td></td><td></td><td></td><td></td></tr>
<tr><td>T2936</td><td>ACATTATTTAACCCATCCATCGG</td><td>1000</td><td></td><td></td><td></td><td></td><td></td><td></td><td></td><td></td><td></td></tr>
<tr><td>Oligo fwd</td><td>CACCACATTATTTAACCCATCCAT</td><td></td><td></td><td></td><td></td><td></td><td></td><td></td><td></td><td></td><td></td></tr>
<tr><td>Oligo rev</td><td>AAACATGGATGGGTTAAATAATGT</td><td></td><td></td><td></td><td></td><td></td><td></td><td></td><td></td><td></td><td></td></tr>
<tr><td>Chromosome</td><td>start</td><td>end</td><td>strand</td><td>MM</td><td>target_seq</td><td>PAM</td><td>alignment</td><td>distance</td><td>position</td><td>gene name</td><td>gene id</td></tr>
<tr><td>GU305914.1</td><td>45829</td><td>45851</td><td>+</td><td>0</td><td>ACATTATTTAACCCATCCAT</td><td>CGG</td><td>||||||||[||||||||||||]PAM</td><td>NA</td><td>Intergenic</td><td>NA</td><td>NA</td></tr>
<tr><td></td><td></td><td></td><td></td><td></td><td></td><td></td><td></td><td></td><td></td><td></td><td></td></tr>
<tr><td>T2937</td><td>CATTATTTAACCCATCCATCGGG</td><td>1000</td><td></td><td></td><td></td><td></td><td></td><td></td><td></td><td></td><td></td></tr>
<tr><td>Oligo fwd</td><td>CACCCATTATTTAACCCATCCATC</td><td></td><td></td><td></td><td></td><td></td><td></td><td></td><td></td><td></td><td></td></tr>
<tr><td>Oligo rev</td><td>AAACGATGGATGGGTTAAATAATG</td><td></td><td></td><td></td><td></td><td></td><td></td><td></td><td></td><td></td><td></td></tr>
<tr><td>Chromosome</td><td>start</td><td>end</td><td>strand</td><td>MM</td><td>target_seq</td><td>PAM</td><td>alignment</td><td>distance</td><td>position</td><td>gene name</td><td>gene id</td></tr>
<tr><td>GU305914.1</td><td>45830</td><td>45852</td><td>+</td><td>0</td><td>CATTATTTAACCCATCCATC</td><td>GGG</td><td>||||||||[||||||||||||]PAM</td><td>NA</td><td>Intergenic</td><td>NA</td><td>NA</td></tr>
<tr><td></td><td></td><td></td><td></td><td></td><td></td><td></td><td></td><td></td><td></td><td></td><td></td></tr>
<tr><td>T2938</td><td>ATTATTTAACCCATCCATCGGGG</td><td>1000</td><td></td><td></td><td></td><td></td><td></td><td></td><td></td><td></td><td></td></tr>
<tr><td>Oligo fwd</td><td>CACCATTATTTAACCCATCCATCG</td><td></td><td></td><td></td><td></td><td></td><td></td><td></td><td></td><td></td><td></td></tr>
<tr><td>Oligo rev</td><td>AAACCGATGGATGGGTTAAATAAT</td><td></td><td></td><td></td><td></td><td></td><td></td><td></td><td></td><td></td><td></td></tr>
<tr><td>Chromosome</td><td>start</td><td>end</td><td>strand</td><td>MM</td><td>target_seq</td><td>PAM</td><td>alignment</td><td>distance</td><td>position</td><td>gene name</td><td>gene id</td></tr>
<tr><td>GU305914.1</td><td>45831</td><td>45853</td><td>+</td><td>0</td><td>ATTATTTAACCCATCCATCG</td><td>GGG</td><td>||||||||[||||||||||||]PAM</td><td>NA</td><td>Intergenic</td><td>NA</td><td>NA</td></tr>
<tr><td></td><td></td><td></td><td></td><td></td><td></td><td></td><td></td><td></td><td></td><td></td><td></td></tr>
<tr><td>T2939</td><td>TTATTTAACCCATCCATCGGGGG</td><td>1000</td><td></td><td></td><td></td><td></td><td></td><td></td><td></td><td></td><td></td></tr>
<tr><td>Oligo fwd</td><td>CACCTTATTTAACCCATCCATCGG</td><td></td><td></td><td></td><td></td><td></td><td></td><td></td><td></td><td></td><td></td></tr>
<tr><td>Oligo rev</td><td>AAACCCGATGGATGGGTTAAATAA</td><td></td><td></td><td></td><td></td><td></td><td></td><td></td><td></td><td></td><td></td></tr>
<tr><td>Chromosome</td><td>start</td><td>end</td><td>strand</td><td>MM</td><td>target_seq</td><td>PAM</td><td>alignment</td><td>distance</td><td>position</td><td>gene name</td><td>gene id</td></tr>
<tr><td>GU305914.1</td><td>45832</td><td>45854</td><td>+</td><td>0</td><td>TTATTTAACCCATCCATCGG</td><td>GGG</td><td>||||||||[||||||||||||]PAM</td><td>NA</td><td>Intergenic</td><td>NA</td><td>NA</td></tr>
<tr><td></td><td></td><td></td><td></td><td></td><td></td><td></td><td></td><td></td><td></td><td></td><td></td></tr>
<tr><td>T2940</td><td>TTAACCCATCCATCGGGGGCTGG</td><td>1000</td><td></td><td></td><td></td><td></td><td></td><td></td><td></td><td></td><td></td></tr>
<tr><td>Oligo fwd</td><td>CACCTTAACCCATCCATCGGGGGC</td><td></td><td></td><td></td><td></td><td></td><td></td><td></td><td></td><td></td><td></td></tr>
<tr><td>Oligo rev</td><td>AAACGCCCCCGATGGATGGGTTAA</td><td></td><td></td><td></td><td></td><td></td><td></td><td></td><td></td><td></td><td></td></tr>
<tr><td>Chromosome</td><td>start</td><td>end</td><td>strand</td><td>MM</td><td>target_seq</td><td>PAM</td><td>alignment</td><td>distance</td><td>position</td><td>gene name</td><td>gene id</td></tr>
<tr><td>GU305914.1</td><td>45836</td><td>45858</td><td>+</td><td>0</td><td>TTAACCCATCCATCGGGGGC</td><td>TGG</td><td>||||||||[||||||||||||]PAM</td><td>NA</td><td>Intergenic</td><td>NA</td><td>NA</td></tr>
<tr><td></td><td></td><td></td><td></td><td></td><td></td><td></td><td></td><td></td><td></td><td></td><td></td></tr>
<tr><td>T2941</td><td>CTGGCGCCACACGCCGCGTGCGG</td><td>1000</td><td></td><td></td><td></td><td></td><td></td><td></td><td></td><td></td><td></td></tr>
<tr><td>Oligo fwd</td><td>CACCCTGGCGCCACACGCCGCGTG</td><td></td><td></td><td></td><td></td><td></td><td></td><td></td><td></td><td></td><td></td></tr>
<tr><td>Oligo rev</td><td>AAACCACGCGGCGTGTGGCGCCAG</td><td></td><td></td><td></td><td></td><td></td><td></td><td></td><td></td><td></td><td></td></tr>
<tr><td>Chromosome</td><td>start</td><td>end</td><td>strand</td><td>MM</td><td>target_seq</td><td>PAM</td><td>alignment</td><td>distance</td><td>position</td><td>gene name</td><td>gene id</td></tr>
<tr><td>GU305914.1</td><td>45855</td><td>45877</td><td>+</td><td>0</td><td>CTGGCGCCACACGCCGCGTG</td><td>CGG</td><td>||||||||[||||||||||||]PAM</td><td>NA</td><td>Intergenic</td><td>NA</td><td>NA</td></tr>
<tr><td></td><td></td><td></td><td></td><td></td><td></td><td></td><td></td><td></td><td></td><td></td><td></td></tr>
<tr><td>T2942</td><td>TGGCGCCACACGCCGCGTGCGGG</td><td>1000</td><td></td><td></td><td></td><td></td><td></td><td></td><td></td><td></td><td></td></tr>
<tr><td>Oligo fwd</td><td>CACCTGGCGCCACACGCCGCGTGC</td><td></td><td></td><td></td><td></td><td></td><td></td><td></td><td></td><td></td><td></td></tr>
<tr><td>Oligo rev</td><td>AAACGCACGCGGCGTGTGGCGCCA</td><td></td><td></td><td></td><td></td><td></td><td></td><td></td><td></td><td></td><td></td></tr>
<tr><td>Chromosome</td><td>start</td><td>end</td><td>strand</td><td>MM</td><td>target_seq</td><td>PAM</td><td>alignment</td><td>distance</td><td>position</td><td>gene name</td><td>gene id</td></tr>
<tr><td>GU305914.1</td><td>45856</td><td>45878</td><td>+</td><td>0</td><td>TGGCGCCACACGCCGCGTGC</td><td>GGG</td><td>||||||||[||||||||||||]PAM</td><td>NA</td><td>Intergenic</td><td>NA</td><td>NA</td></tr>
<tr><td></td><td></td><td></td><td></td><td></td><td></td><td></td><td></td><td></td><td></td><td></td><td></td></tr>
<tr><td>T2943</td><td>CGCCACACGCCGCGTGCGGGTGG</td><td>1000</td><td></td><td></td><td></td><td></td><td></td><td></td><td></td><td></td><td></td></tr>
<tr><td>Oligo fwd</td><td>CACCCGCCACACGCCGCGTGCGGG</td><td></td><td></td><td></td><td></td><td></td><td></td><td></td><td></td><td></td><td></td></tr>
<tr><td>Oligo rev</td><td>AAACCCCGCACGCGGCGTGTGGCG</td><td></td><td></td><td></td><td></td><td></td><td></td><td></td><td></td><td></td><td></td></tr>
<tr><td>Chromosome</td><td>start</td><td>end</td><td>strand</td><td>MM</td><td>target_seq</td><td>PAM</td><td>alignment</td><td>distance</td><td>position</td><td>gene name</td><td>gene id</td></tr>
<tr><td>GU305914.1</td><td>45859</td><td>45881</td><td>+</td><td>0</td><td>CGCCACACGCCGCGTGCGGG</td><td>TGG</td><td>||||||||[||||||||||||]PAM</td><td>NA</td><td>Intergenic</td><td>NA</td><td>NA</td></tr>
<tr><td></td><td></td><td></td><td></td><td></td><td></td><td></td><td></td><td></td><td></td><td></td><td></td></tr>
<tr><td>T2944</td><td>GCCACACGCCGCGTGCGGGTGGG</td><td>1000</td><td></td><td></td><td></td><td></td><td></td><td></td><td></td><td></td><td></td></tr>
<tr><td>Oligo fwd</td><td>CACCGCCACACGCCGCGTGCGGGT</td><td></td><td></td><td></td><td></td><td></td><td></td><td></td><td></td><td></td><td></td></tr>
<tr><td>Oligo rev</td><td>AAACACCCGCACGCGGCGTGTGGC</td><td></td><td></td><td></td><td></td><td></td><td></td><td></td><td></td><td></td><td></td></tr>
<tr><td>Chromosome</td><td>start</td><td>end</td><td>strand</td><td>MM</td><td>target_seq</td><td>PAM</td><td>alignment</td><td>distance</td><td>position</td><td>gene name</td><td>gene id</td></tr>
<tr><td>GU305914.1</td><td>45860</td><td>45882</td><td>+</td><td>0</td><td>GCCACACGCCGCGTGCGGGT</td><td>GGG</td><td>||||||||[||||||||||||]PAM</td><td>NA</td><td>Intergenic</td><td>NA</td><td>NA</td></tr>
<tr><td></td><td></td><td></td><td></td><td></td><td></td><td></td><td></td><td></td><td></td><td></td><td></td></tr>
<tr><td>T2945</td><td>GCGTGCGGGTGGGCATGTATCGG</td><td>1000</td><td></td><td></td><td></td><td></td><td></td><td></td><td></td><td></td><td></td></tr>
<tr><td>Oligo fwd</td><td>CACCGCGTGCGGGTGGGCATGTAT</td><td></td><td></td><td></td><td></td><td></td><td></td><td></td><td></td><td></td><td></td></tr>
<tr><td>Oligo rev</td><td>AAACATACATGCCCACCCGCACGC</td><td></td><td></td><td></td><td></td><td></td><td></td><td></td><td></td><td></td><td></td></tr>
<tr><td>Chromosome</td><td>start</td><td>end</td><td>strand</td><td>MM</td><td>target_seq</td><td>PAM</td><td>alignment</td><td>distance</td><td>position</td><td>gene name</td><td>gene id</td></tr>
<tr><td>GU305914.1</td><td>45870</td><td>45892</td><td>+</td><td>0</td><td>GCGTGCGGGTGGGCATGTAT</td><td>CGG</td><td>||||||||[||||||||||||]PAM</td><td>NA</td><td>Intergenic</td><td>NA</td><td>NA</td></tr>
<tr><td>GU305914.1</td><td>184913</td><td>184935</td><td>-</td><td>4</td><td>GCGTCCTGGGGGGCATCTAT</td><td>AGG</td><td>||||-|-|[|-||||||-|||]PAM</td><td>NA</td><td>Intergenic</td><td>NA</td><td>NA</td></tr>
<tr><td></td><td></td><td></td><td></td><td></td><td></td><td></td><td></td><td></td><td></td><td></td><td></td></tr>
<tr><td>T2946</td><td>CGGGTGGGCATGTATCGGACCGG</td><td>1000</td><td></td><td></td><td></td><td></td><td></td><td></td><td></td><td></td><td></td></tr>
<tr><td>Oligo fwd</td><td>CACCCGGGTGGGCATGTATCGGAC</td><td></td><td></td><td></td><td></td><td></td><td></td><td></td><td></td><td></td><td></td></tr>
<tr><td>Oligo rev</td><td>AAACGTCCGATACATGCCCACCCG</td><td></td><td></td><td></td><td></td><td></td><td></td><td></td><td></td><td></td><td></td></tr>
<tr><td>Chromosome</td><td>start</td><td>end</td><td>strand</td><td>MM</td><td>target_seq</td><td>PAM</td><td>alignment</td><td>distance</td><td>position</td><td>gene name</td><td>gene id</td></tr>
<tr><td>GU305914.1</td><td>45875</td><td>45897</td><td>+</td><td>0</td><td>CGGGTGGGCATGTATCGGAC</td><td>CGG</td><td>||||||||[||||||||||||]PAM</td><td>NA</td><td>Intergenic</td><td>NA</td><td>NA</td></tr>
</table>

<table>
<tr><td>T2947</td><td>CATGTATCGGACCGGCCGCACGG</td><td>1000</td><td></td><td></td><td></td><td></td><td></td><td></td><td></td><td></td><td></td></tr>
<tr><td>Oligo fwd</td><td>CACCCATGTATCGGACCGGCCGCA</td><td></td><td></td><td></td><td></td><td></td><td></td><td></td><td></td><td></td><td></td></tr>
<tr><td>Oligo rev</td><td>AAACTGCGGCCGGTCCGATACATG</td><td></td><td></td><td></td><td></td><td></td><td></td><td></td><td></td><td></td><td></td></tr>
<tr><td>Chromosome</td><td>start</td><td>end</td><td>strand</td><td>MM</td><td>target_seq</td><td>PAM</td><td>alignment</td><td>distance</td><td>position</td><td>gene name</td><td>gene id</td></tr>
<tr><td>GU305914.1</td><td>45883</td><td>45905</td><td>+</td><td>0</td><td>CATGTATCGGACCGGCCGCA</td><td>CGG</td><td>||||||||[||||||||||||]PAM</td><td>NA</td><td>Intergenic</td><td>NA</td><td>NA</td></tr>
<tr><td>T2948</td><td>ATGTATCGGACCGGCCGCACGGG</td><td>1000</td><td></td><td></td><td></td><td></td><td></td><td></td><td></td><td></td><td></td></tr>
<tr><td>Oligo fwd</td><td>CACCATGTATCGGACCGGCCGCAC</td><td></td><td></td><td></td><td></td><td></td><td></td><td></td><td></td><td></td><td></td></tr>
<tr><td>Oligo rev</td><td>AAACGTGCGGCCGGTCCGATACAT</td><td></td><td></td><td></td><td></td><td></td><td></td><td></td><td></td><td></td><td></td></tr>
<tr><td>Chromosome</td><td>start</td><td>end</td><td>strand</td><td>MM</td><td>target_seq</td><td>PAM</td><td>alignment</td><td>distance</td><td>position</td><td>gene name</td><td>gene id</td></tr>
<tr><td>GU305914.1</td><td>45884</td><td>45906</td><td>+</td><td>0</td><td>ATGTATCGGACCGGCCGCAC</td><td>GGG</td><td>||||||||[||||||||||||]PAM</td><td>NA</td><td>Intergenic</td><td>NA</td><td>NA</td></tr>
<tr><td>T2949</td><td>ACCGGCCGCACGGGACGCGATGG</td><td>1000</td><td></td><td></td><td></td><td></td><td></td><td></td><td></td><td></td><td></td></tr>
<tr><td>Oligo fwd</td><td>CACCACCGGCCGCACGGGACGCGA</td><td></td><td></td><td></td><td></td><td></td><td></td><td></td><td></td><td></td><td></td></tr>
<tr><td>Oligo rev</td><td>AAACTCGCGTCCCGTGCGGCCGGT</td><td></td><td></td><td></td><td></td><td></td><td></td><td></td><td></td><td></td><td></td></tr>
<tr><td>Chromosome</td><td>start</td><td>end</td><td>strand</td><td>MM</td><td>target_seq</td><td>PAM</td><td>alignment</td><td>distance</td><td>position</td><td>gene name</td><td>gene id</td></tr>
<tr><td>GU305914.1</td><td>45893</td><td>45915</td><td>+</td><td>0</td><td>ACCGGCCGCACGGGACGCGA</td><td>TGG</td><td>||||||||[||||||||||||]PAM</td><td>NA</td><td>Intergenic</td><td>NA</td><td>NA</td></tr>
<tr><td>T2950</td><td>GGACGCGATGGCCGCTCCGACGG</td><td>1000</td><td></td><td></td><td></td><td></td><td></td><td></td><td></td><td></td><td></td></tr>
<tr><td>Oligo fwd</td><td>CACCGGACGCGATGGCCGCTCCGA</td><td></td><td></td><td></td><td></td><td></td><td></td><td></td><td></td><td></td><td></td></tr>
<tr><td>Oligo rev</td><td>AAACTCGGAGCGGCCATCGCGTCC</td><td></td><td></td><td></td><td></td><td></td><td></td><td></td><td></td><td></td><td></td></tr>
<tr><td>Chromosome</td><td>start</td><td>end</td><td>strand</td><td>MM</td><td>target_seq</td><td>PAM</td><td>alignment</td><td>distance</td><td>position</td><td>gene name</td><td>gene id</td></tr>
<tr><td>GU305914.1</td><td>45905</td><td>45927</td><td>+</td><td>0</td><td>GGACGCGATGGCCGCTCCGA</td><td>CGG</td><td>||||||||[||||||||||||]PAM</td><td>NA</td><td>Intergenic</td><td>NA</td><td>NA</td></tr>
<tr><td>T2951</td><td>CGCGATGGCCGCTCCGACGGAGG</td><td>1000</td><td></td><td></td><td></td><td></td><td></td><td></td><td></td><td></td><td></td></tr>
<tr><td>Oligo fwd</td><td>CACCCGCGATGGCCGCTCCGACGG</td><td></td><td></td><td></td><td></td><td></td><td></td><td></td><td></td><td></td><td></td></tr>
<tr><td>Oligo rev</td><td>AAACCCGTCGGAGCGGCCATCGCG</td><td></td><td></td><td></td><td></td><td></td><td></td><td></td><td></td><td></td><td></td></tr>
<tr><td>Chromosome</td><td>start</td><td>end</td><td>strand</td><td>MM</td><td>target_seq</td><td>PAM</td><td>alignment</td><td>distance</td><td>position</td><td>gene name</td><td>gene id</td></tr>
<tr><td>GU305914.1</td><td>45908</td><td>45930</td><td>+</td><td>0</td><td>CGCGATGGCCGCTCCGACGG</td><td>AGG</td><td>||||||||[||||||||||||]PAM</td><td>NA</td><td>Intergenic</td><td>NA</td><td>NA</td></tr>
<tr><td>GU305914.1</td><td>50346</td><td>50368</td><td>-</td><td>3</td><td>CGCGATCGCCGCGCGGACGG</td><td>CGG</td><td>||||||-|[||||-|-|||||]PAM</td><td>NA</td><td>Intergenic</td><td>NA</td><td>NA</td></tr>
<tr><td>T2952</td><td>TTTCCCCCACAATATGCTGTCGG</td><td>1000</td><td></td><td></td><td></td><td></td><td></td><td></td><td></td><td></td><td></td></tr>
<tr><td>Oligo fwd</td><td>CACCTTTCCCCCACAATATGCTGT</td><td></td><td></td><td></td><td></td><td></td><td></td><td></td><td></td><td></td><td></td></tr>
<tr><td>Oligo rev</td><td>AAACACAGCATATTGTGGGGGAAA</td><td></td><td></td><td></td><td></td><td></td><td></td><td></td><td></td><td></td><td></td></tr>
<tr><td>Chromosome</td><td>start</td><td>end</td><td>strand</td><td>MM</td><td>target_seq</td><td>PAM</td><td>alignment</td><td>distance</td><td>position</td><td>gene name</td><td>gene id</td></tr>
<tr><td>GU305914.1</td><td>45977</td><td>45999</td><td>+</td><td>0</td><td>TTTCCCCCACAATATGCTGT</td><td>CGG</td><td>||||||||[||||||||||||]PAM</td><td>NA</td><td>Intergenic</td><td>NA</td><td>NA</td></tr>
<tr><td>T2953</td><td>CCCCCACAATATGCTGTCGGAGG</td><td>1000</td><td></td><td></td><td></td><td></td><td></td><td></td><td></td><td></td><td></td></tr>
<tr><td>Oligo fwd</td><td>CACCCCCCCACAATATGCTGTCGG</td><td></td><td></td><td></td><td></td><td></td><td></td><td></td><td></td><td></td><td></td></tr>
<tr><td>Oligo rev</td><td>AAACCCGACAGCATATTGTGGGGG</td><td></td><td></td><td></td><td></td><td></td><td></td><td></td><td></td><td></td><td></td></tr>
<tr><td>Chromosome</td><td>start</td><td>end</td><td>strand</td><td>MM</td><td>target_seq</td><td>PAM</td><td>alignment</td><td>distance</td><td>position</td><td>gene name</td><td>gene id</td></tr>
<tr><td>GU305914.1</td><td>45980</td><td>46002</td><td>+</td><td>0</td><td>CCCCCACAATATGCTGTCGG</td><td>AGG</td><td>||||||||[||||||||||||]PAM</td><td>NA</td><td>Intergenic</td><td>NA</td><td>NA</td></tr>
<tr><td>T2954</td><td>GGAAAATATCAATTTCATACAGG</td><td>1000</td><td></td><td></td><td></td><td></td><td></td><td></td><td></td><td></td><td></td></tr>
<tr><td>Oligo fwd</td><td>CACCGGAAAATATCAATTTCATAC</td><td></td><td></td><td></td><td></td><td></td><td></td><td></td><td></td><td></td><td></td></tr>
<tr><td>Oligo rev</td><td>AAACGTATGAAATTGATATTTTCC</td><td></td><td></td><td></td><td></td><td></td><td></td><td></td><td></td><td></td><td></td></tr>
<tr><td>Chromosome</td><td>start</td><td>end</td><td>strand</td><td>MM</td><td>target_seq</td><td>PAM</td><td>alignment</td><td>distance</td><td>position</td><td>gene name</td><td>gene id</td></tr>
<tr><td>GU305914.1</td><td>46001</td><td>46023</td><td>+</td><td>0</td><td>GGAAAATATCAATTTCATAC</td><td>AGG</td><td>||||||||[||||||||||||]PAM</td><td>NA</td><td>Intergenic</td><td>NA</td><td>NA</td></tr>
<tr><td>T2955</td><td>ATACAGGACGCCGTGTGTAACGG</td><td>1000</td><td></td><td></td><td></td><td></td><td></td><td></td><td></td><td></td><td></td></tr>
<tr><td>Oligo fwd</td><td>CACCATACAGGACGCCGTGTGTAA</td><td></td><td></td><td></td><td></td><td></td><td></td><td></td><td></td><td></td><td></td></tr>
<tr><td>Oligo rev</td><td>AAACTTACACACGGCGTCCTGTAT</td><td></td><td></td><td></td><td></td><td></td><td></td><td></td><td></td><td></td><td></td></tr>
<tr><td>Chromosome</td><td>start</td><td>end</td><td>strand</td><td>MM</td><td>target_seq</td><td>PAM</td><td>alignment</td><td>distance</td><td>position</td><td>gene name</td><td>gene id</td></tr>
<tr><td>GU305914.1</td><td>46017</td><td>46039</td><td>+</td><td>0</td><td>ATACAGGACGCCGTGTGTAA</td><td>CGG</td><td>||||||||[||||||||||||]PAM</td><td>NA</td><td>Intergenic</td><td>NA</td><td>NA</td></tr>
<tr><td>T2956</td><td>CGCCGTGTGTAACGGCGACCTGG</td><td>1000</td><td></td><td></td><td></td><td></td><td></td><td></td><td></td><td></td><td></td></tr>
<tr><td>Oligo fwd</td><td>CACCCGCCGTGTGTAACGGCGACC</td><td></td><td></td><td></td><td></td><td></td><td></td><td></td><td></td><td></td><td></td></tr>
<tr><td>Oligo rev</td><td>AAACGGTCGCCGTTACACACGGCG</td><td></td><td></td><td></td><td></td><td></td><td></td><td></td><td></td><td></td><td></td></tr>
<tr><td>Chromosome</td><td>start</td><td>end</td><td>strand</td><td>MM</td><td>target_seq</td><td>PAM</td><td>alignment</td><td>distance</td><td>position</td><td>gene name</td><td>gene id</td></tr>
<tr><td>GU305914.1</td><td>46025</td><td>46047</td><td>+</td><td>0</td><td>CGCCGTGTGTAACGGCGACC</td><td>TGG</td><td>||||||||[||||||||||||]PAM</td><td>NA</td><td>Intergenic</td><td>NA</td><td>NA</td></tr>
<tr><td>T2957</td><td>GCCGTGTGTAACGGCGACCTGGG</td><td>1000</td><td></td><td></td><td></td><td></td><td></td><td></td><td></td><td></td><td></td></tr>
<tr><td>Oligo fwd</td><td>CACCGCCGTGTGTAACGGCGACCT</td><td></td><td></td><td></td><td></td><td></td><td></td><td></td><td></td><td></td><td></td></tr>
<tr><td>Oligo rev</td><td>AAACAGGTCGCCGTTACACACGGC</td><td></td><td></td><td></td><td></td><td></td><td></td><td></td><td></td><td></td><td></td></tr>
<tr><td>Chromosome</td><td>start</td><td>end</td><td>strand</td><td>MM</td><td>target_seq</td><td>PAM</td><td>alignment</td><td>distance</td><td>position</td><td>gene name</td><td>gene id</td></tr>
<tr><td>GU305914.1</td><td>46026</td><td>46048</td><td>+</td><td>0</td><td>GCCGTGTGTAACGGCGACCT</td><td>GGG</td><td>||||||||[||||||||||||]PAM</td><td>NA</td><td>Intergenic</td><td>NA</td><td>NA</td></tr>
<tr><td>T2958</td><td>TAACGGCGACCTGGGCGCCGTGG</td><td>1000</td><td></td><td></td><td></td><td></td><td></td><td></td><td></td><td></td><td></td></tr>
<tr><td>Oligo fwd</td><td>CACCTAACGGCGACCTGGGCGCCG</td><td></td><td></td><td></td><td></td><td></td><td></td><td></td><td></td><td></td><td></td></tr>
<tr><td>Oligo rev</td><td>AAACCGGCGCCCAGGTCGCCGTTA</td><td></td><td></td><td></td><td></td><td></td><td></td><td></td><td></td><td></td><td></td></tr>
<tr><td>Chromosome</td><td>start</td><td>end</td><td>strand</td><td>MM</td><td>target_seq</td><td>PAM</td><td>alignment</td><td>distance</td><td>position</td><td>gene name</td><td>gene id</td></tr>
<tr><td>GU305914.1</td><td>46034</td><td>46056</td><td>+</td><td>0</td><td>TAACGGCGACCTGGGCGCCG</td><td>TGG</td><td>||||||||[||||||||||||]PAM</td><td>NA</td><td>Intergenic</td><td>NA</td><td>NA</td></tr>
<tr><td>T2959</td><td>CGGCGACCTGGGCGCCGTGGCGG</td><td>1000</td><td></td><td></td><td></td><td></td><td></td><td></td><td></td><td></td><td></td></tr>
<tr><td>Oligo fwd</td><td>CACCCGGCGACCTGGGCGCCGTGG</td><td></td><td></td><td></td><td></td><td></td><td></td><td></td><td></td><td></td><td></td></tr>
<tr><td>Oligo rev</td><td>AAACCCACGGCGCCCAGGTCGCCG</td><td></td><td></td><td></td><td></td><td></td><td></td><td></td><td></td><td></td><td></td></tr>
<tr><td>Chromosome</td><td>start</td><td>end</td><td>strand</td><td>MM</td><td>target_seq</td><td>PAM</td><td>alignment</td><td>distance</td><td>position</td><td>gene name</td><td>gene id</td></tr>
<tr><td>GU305914.1</td><td>46037</td><td>46059</td><td>+</td><td>0</td><td>CGGCGACCTGGGCGCCGTGG</td><td>CGG</td><td>||||||||[||||||||||||]PAM</td><td>NA</td><td>Intergenic</td><td>NA</td><td>NA</td></tr>
<tr><td>T2960</td><td>CGCCGTGGCGGCGTTGAACTCGG</td><td>1000</td><td></td><td></td><td></td><td></td><td></td><td></td><td></td><td></td><td></td></tr>
<tr><td>Oligo fwd</td><td>CACCCGCCGTGGCGGCGTTGAACT</td><td></td><td></td><td></td><td></td><td></td><td></td><td></td><td></td><td></td><td></td></tr>
<tr><td>Oligo rev</td><td>AAACAGTTCAACGCCGCCACGGCG</td><td></td><td></td><td></td><td></td><td></td><td></td><td></td><td></td><td></td><td></td></tr>
<tr><td>Chromosome</td><td>start</td><td>end</td><td>strand</td><td>MM</td><td>target_seq</td><td>PAM</td><td>alignment</td><td>distance</td><td>position</td><td>gene name</td><td>gene id</td></tr>
<tr><td>GU305914.1</td><td>46049</td><td>46071</td><td>+</td><td>0</td><td>CGCCGTGGCGGCGTTGAACT</td><td>CGG</td><td>||||||||[||||||||||||]PAM</td><td>NA</td><td>Intergenic</td><td>NA</td><td>NA</td></tr>
<tr><td>T2961</td><td>GCCGTGGCGGCGTTGAACTCGGG</td><td>1000</td><td></td><td></td><td></td><td></td><td></td><td></td><td></td><td></td><td></td></tr>
<tr><td>Oligo fwd</td><td>CACCGCCGTGGCGGCGTTGAACTC</td><td></td><td></td><td></td><td></td><td></td><td></td><td></td><td></td><td></td><td></td></tr>
<tr><td>Oligo rev</td><td>AAACGAGTTCAACGCCGCCACGGC</td><td></td><td></td><td></td><td></td><td></td><td></td><td></td><td></td><td></td><td></td></tr>
</table>

<table>
<tr><td>Chromosome</td><td>start</td><td></td><td>end</td><td>strand</td><td>MM</td><td>target_seq</td><td>PAM</td><td>alignment</td><td>distance</td><td>position</td><td>gene name</td><td>gene id</td></tr>
<tr><td>GU305914.1</td><td></td><td>46050</td><td>46072</td><td>+</td><td>0</td><td>GCCGTGGCGGCGTTGAACTC</td><td>GGG</td><td>||||||||[||||||||||||]PAM</td><td>NA</td><td>Intergenic</td><td>NA</td><td>NA</td></tr>
<tr><td></td></tr>
<tr><td>T2962</td><td>GTTGAACTCGGGTCTGCCGATGG</td><td></td><td>1000</td></tr>
<tr><td>Oligo fwd</td><td>CACCGTTGAACTCGGGTCTGCCGA</td></tr>
<tr><td>Oligo rev</td><td>AAACTCGGCAGACCCGAGTTCAAC</td></tr>
<tr><td>Chromosome</td><td>start</td><td></td><td>end</td><td>strand</td><td>MM</td><td>target_seq</td><td>PAM</td><td>alignment</td><td>distance</td><td>position</td><td>gene name</td><td>gene id</td></tr>
<tr><td>GU305914.1</td><td></td><td>46061</td><td>46083</td><td>+</td><td>0</td><td>GTTGAACTCGGGTCTGCCGA</td><td>TGG</td><td>||||||||[||||||||||||]PAM</td><td>NA</td><td>Intergenic</td><td>NA</td><td>NA</td></tr>
<tr><td></td></tr>
<tr><td>T2963</td><td>GATGGCCCCGTACATGCTAGAGG</td><td></td><td>1000</td></tr>
<tr><td>Oligo fwd</td><td>CACCGATGGCCCCGTACATGCTAG</td></tr>
<tr><td>Oligo rev</td><td>AAACCTAGCATGTACGGGGCCATC</td></tr>
<tr><td>Chromosome</td><td>start</td><td></td><td>end</td><td>strand</td><td>MM</td><td>target_seq</td><td>PAM</td><td>alignment</td><td>distance</td><td>position</td><td>gene name</td><td>gene id</td></tr>
<tr><td>GU305914.1</td><td></td><td>46079</td><td>46101</td><td>+</td><td>0</td><td>GATGGCCCCGTACATGCTAG</td><td>AGG</td><td>||||||||[||||||||||||]PAM</td><td>NA</td><td>Intergenic</td><td>NA</td><td>NA</td></tr>
<tr><td></td></tr>
<tr><td>T2964</td><td>GTACATGCTAGAGGCCCTCCTGG</td><td></td><td>1000</td></tr>
<tr><td>Oligo fwd</td><td>CACCGTACATGCTAGAGGCCCTCC</td></tr>
<tr><td>Oligo rev</td><td>AAACGGAGGGCCTCTAGCATGTAC</td></tr>
<tr><td>Chromosome</td><td>start</td><td></td><td>end</td><td>strand</td><td>MM</td><td>target_seq</td><td>PAM</td><td>alignment</td><td>distance</td><td>position</td><td>gene name</td><td>gene id</td></tr>
<tr><td>GU305914.1</td><td></td><td>46088</td><td>46110</td><td>+</td><td>0</td><td>GTACATGCTAGAGGCCCTCC</td><td>TGG</td><td>||||||||[||||||||||||]PAM</td><td>NA</td><td>Intergenic</td><td>NA</td><td>NA</td></tr>
<tr><td></td></tr>
<tr><td>T2965</td><td>CATGCTAGAGGCCCTCCTGGCGG</td><td></td><td>1000</td></tr>
<tr><td>Oligo fwd</td><td>CACCCATGCTAGAGGCCCTCCTGG</td></tr>
<tr><td>Oligo rev</td><td>AAACCCAGGAGGGCCTCTAGCATG</td></tr>
<tr><td>Chromosome</td><td>start</td><td></td><td>end</td><td>strand</td><td>MM</td><td>target_seq</td><td>PAM</td><td>alignment</td><td>distance</td><td>position</td><td>gene name</td><td>gene id</td></tr>
<tr><td>GU305914.1</td><td></td><td>46091</td><td>46113</td><td>+</td><td>0</td><td>CATGCTAGAGGCCCTCCTGG</td><td>CGG</td><td>||||||||[||||||||||||]PAM</td><td>NA</td><td>Intergenic</td><td>NA</td><td>NA</td></tr>
<tr><td></td></tr>
<tr><td>T2966</td><td>TGGCGGTCCGCGTCAAACACAGG</td><td></td><td>1000</td></tr>
<tr><td>Oligo fwd</td><td>CACCTGGCGGTCCGCGTCAAACAC</td></tr>
<tr><td>Oligo rev</td><td>AAACGTGTTTGACGCGGACCGCCA</td></tr>
<tr><td>Chromosome</td><td>start</td><td></td><td>end</td><td>strand</td><td>MM</td><td>target_seq</td><td>PAM</td><td>alignment</td><td>distance</td><td>position</td><td>gene name</td><td>gene id</td></tr>
<tr><td>GU305914.1</td><td></td><td>46108</td><td>46130</td><td>+</td><td>0</td><td>TGGCGGTCCGCGTCAAACAC</td><td>AGG</td><td>||||||||[||||||||||||]PAM</td><td>NA</td><td>Intergenic</td><td>NA</td><td>NA</td></tr>
<tr><td></td></tr>
<tr><td>T2967</td><td>CGTCAAACACAGGCTGACGAAGG</td><td></td><td>1000</td></tr>
<tr><td>Oligo fwd</td><td>CACCCGTCAAACACAGGCTGACGA</td></tr>
<tr><td>Oligo rev</td><td>AAACTCGTCAGCCTGTGTTTGACG</td></tr>
<tr><td>Chromosome</td><td>start</td><td></td><td>end</td><td>strand</td><td>MM</td><td>target_seq</td><td>PAM</td><td>alignment</td><td>distance</td><td>position</td><td>gene name</td><td>gene id</td></tr>
<tr><td>GU305914.1</td><td></td><td>46118</td><td>46140</td><td>+</td><td>0</td><td>CGTCAAACACAGGCTGACGA</td><td>AGG</td><td>||||||||[||||||||||||]PAM</td><td>NA</td><td>Intergenic</td><td>NA</td><td>NA</td></tr>
<tr><td></td></tr>
<tr><td>T2968</td><td>CATCTGTTACACCATCAGCGTGG</td><td></td><td>1000</td></tr>
<tr><td>Oligo fwd</td><td>CACCCATCTGTTACACCATCAGCG</td></tr>
<tr><td>Oligo rev</td><td>AAACCGCTGATGGTGTAACAGATG</td></tr>
<tr><td>Chromosome</td><td>start</td><td></td><td>end</td><td>strand</td><td>MM</td><td>target_seq</td><td>PAM</td><td>alignment</td><td>distance</td><td>position</td><td>gene name</td><td>gene id</td></tr>
<tr><td>GU305914.1</td><td></td><td>46163</td><td>46185</td><td>+</td><td>0</td><td>CATCTGTTACACCATCAGCG</td><td>TGG</td><td>||||||||[||||||||||||]PAM</td><td>NA</td><td>Intergenic</td><td>NA</td><td>NA</td></tr>
<tr><td></td></tr>
<tr><td>T2969</td><td>AGCGTGGCGCACCTCATCAACGG</td><td></td><td>1000</td></tr>
<tr><td>Oligo fwd</td><td>CACCAGCGTGGCGCACCTCATCAA</td></tr>
<tr><td>Oligo rev</td><td>AAACTTGATGAGGTGCGCCACGCT</td></tr>
<tr><td>Chromosome</td><td>start</td><td></td><td>end</td><td>strand</td><td>MM</td><td>target_seq</td><td>PAM</td><td>alignment</td><td>distance</td><td>position</td><td>gene name</td><td>gene id</td></tr>
<tr><td>GU305914.1</td><td></td><td>46179</td><td>46201</td><td>+</td><td>0</td><td>AGCGTGGCGCACCTCATCAA</td><td>CGG</td><td>||||||||[||||||||||||]PAM</td><td>NA</td><td>Intergenic</td><td>NA</td><td>NA</td></tr>
<tr><td></td></tr>
<tr><td>T2970</td><td>CGGTACGCGCATCTTGCGTTCGG</td><td></td><td>1000</td></tr>
<tr><td>Oligo fwd</td><td>CACCCGGTACGCGCATCTTGCGTT</td></tr>
<tr><td>Oligo rev</td><td>AAACAACGCAAGATGCGCGTACCG</td></tr>
<tr><td>Chromosome</td><td>start</td><td></td><td>end</td><td>strand</td><td>MM</td><td>target_seq</td><td>PAM</td><td>alignment</td><td>distance</td><td>position</td><td>gene name</td><td>gene id</td></tr>
<tr><td>GU305914.1</td><td></td><td>46199</td><td>46221</td><td>+</td><td>0</td><td>CGGTACGCGCATCTTGCGTT</td><td>CGG</td><td>||||||||[||||||||||||]PAM</td><td>NA</td><td>Intergenic</td><td>NA</td><td>NA</td></tr>
<tr><td></td></tr>
<tr><td>T2971</td><td>GCGCATCTTGCGTTCGGCCACGG</td><td></td><td>1000</td></tr>
<tr><td>Oligo fwd</td><td>CACCGCGCATCTTGCGTTCGGCCA</td></tr>
<tr><td>Oligo rev</td><td>AAACTGGCCGAACGCAAGATGCGC</td></tr>
<tr><td>Chromosome</td><td>start</td><td></td><td>end</td><td>strand</td><td>MM</td><td>target_seq</td><td>PAM</td><td>alignment</td><td>distance</td><td>position</td><td>gene name</td><td>gene id</td></tr>
<tr><td>GU305914.1</td><td></td><td>46205</td><td>46227</td><td>+</td><td>0</td><td>GCGCATCTTGCGTTCGGCCA</td><td>CGG</td><td>||||||||[||||||||||||]PAM</td><td>NA</td><td>Intergenic</td><td>NA</td><td>NA</td></tr>
<tr><td></td></tr>
<tr><td>T2972</td><td>GGCCACGGCCAAACACGCGACGG</td><td></td><td>1000</td></tr>
<tr><td>Oligo fwd</td><td>CACCGGCCACGGCCAAACACGCGA</td></tr>
<tr><td>Oligo rev</td><td>AAACTCGCGTGTTTGGCCGTGGCC</td></tr>
<tr><td>Chromosome</td><td>start</td><td></td><td>end</td><td>strand</td><td>MM</td><td>target_seq</td><td>PAM</td><td>alignment</td><td>distance</td><td>position</td><td>gene name</td><td>gene id</td></tr>
<tr><td>GU305914.1</td><td></td><td>46220</td><td>46242</td><td>+</td><td>0</td><td>GGCCACGGCCAAACACGCGA</td><td>CGG</td><td>||||||||[||||||||||||]PAM</td><td>NA</td><td>Intergenic</td><td>NA</td><td>NA</td></tr>
<tr><td></td></tr>
<tr><td>T2973</td><td>CGGCCAAACACGCGACGGCGTGG</td><td></td><td>1000</td></tr>
<tr><td>Oligo fwd</td><td>CACCCGGCCAAACACGCGACGGCG</td></tr>
<tr><td>Oligo rev</td><td>AAACCGCCGTCGCGTGTTTGGCCG</td></tr>
<tr><td>Chromosome</td><td>start</td><td></td><td>end</td><td>strand</td><td>MM</td><td>target_seq</td><td>PAM</td><td>alignment</td><td>distance</td><td>position</td><td>gene name</td><td>gene id</td></tr>
<tr><td>GU305914.1</td><td></td><td>46225</td><td>46247</td><td>+</td><td>0</td><td>CGGCCAAACACGCGACGGCG</td><td>TGG</td><td>||||||||[||||||||||||]PAM</td><td>NA</td><td>Intergenic</td><td>NA</td><td>NA</td></tr>
<tr><td></td></tr>
<tr><td>T2974</td><td>GGCCAAACACGCGACGGCGTGGG</td><td></td><td>1000</td></tr>
<tr><td>Oligo fwd</td><td>CACCGGCCAAACACGCGACGGCGT</td></tr>
<tr><td>Oligo rev</td><td>AAACACGCCGTCGCGTGTTTGGCC</td></tr>
<tr><td>Chromosome</td><td>start</td><td></td><td>end</td><td>strand</td><td>MM</td><td>target_seq</td><td>PAM</td><td>alignment</td><td>distance</td><td>position</td><td>gene name</td><td>gene id</td></tr>
<tr><td>GU305914.1</td><td></td><td>46226</td><td>46248</td><td>+</td><td>0</td><td>GGCCAAACACGCGACGGCGT</td><td>GGG</td><td>||||||||[||||||||||||]PAM</td><td>NA</td><td>Intergenic</td><td>NA</td><td>NA</td></tr>
<tr><td></td></tr>
<tr><td>T2975</td><td>GCCAAACACGCGACGGCGTGGGG</td><td></td><td>1000</td></tr>
<tr><td>Oligo fwd</td><td>CACCGCCAAACACGCGACGGCGTG</td></tr>
<tr><td>Oligo rev</td><td>AAACCACGCCGTCGCGTGTTTGGC</td></tr>
<tr><td>Chromosome</td><td>start</td><td></td><td>end</td><td>strand</td><td>MM</td><td>target_seq</td><td>PAM</td><td>alignment</td><td>distance</td><td>position</td><td>gene name</td><td>gene id</td></tr>
<tr><td>GU305914.1</td><td></td><td>46227</td><td>46249</td><td>+</td><td>0</td><td>GCCAAACACGCGACGGCGTG</td><td>GGG</td><td>||||||||[||||||||||||]PAM</td><td>NA</td><td>Intergenic</td><td>NA</td><td>NA</td></tr>
<tr><td></td></tr>
<tr><td>T2976</td><td>CCAAACACGCGACGGCGTGGGGG</td><td></td><td>1000</td></tr>
<tr><td>Oligo fwd</td><td>CACCCCAAACACGCGACGGCGTGG</td></tr>
</table>

<table>
<tr><td>Oligo rev</td><td>AAACCCACGCCGTCGCGTGTTTGG</td><td></td><td></td><td></td><td></td><td></td><td></td><td></td><td></td><td></td><td></td><td></td></tr>
<tr><td>Chromosome</td><td>start</td><td></td><td>end</td><td>strand</td><td>MM</td><td>target_seq</td><td>PAM</td><td>alignment</td><td>distance</td><td>position</td><td>gene name</td><td>gene id</td></tr>
<tr><td>GU305914.1</td><td></td><td>46228</td><td>46250</td><td>+</td><td>0</td><td>CCAAACACGCGACGGCGTGG</td><td>GGG</td><td>||||||||[||||||||||||]PAM</td><td>NA</td><td>Intergenic</td><td>NA</td><td>NA</td></tr>
<tr><td>T2977</td><td>GCCGAACGACAAACACCGCGCGG</td><td></td><td>1000</td><td></td><td></td><td></td><td></td><td></td><td></td><td></td><td></td><td></td></tr>
<tr><td>Oligo fwd</td><td>CACCGCCGAACGACAAACACCGCG</td><td></td><td></td><td></td><td></td><td></td><td></td><td></td><td></td><td></td><td></td><td></td></tr>
<tr><td>Oligo rev</td><td>AAACCGCGGTGTTTGTCGTTCGGC</td><td></td><td></td><td></td><td></td><td></td><td></td><td></td><td></td><td></td><td></td><td></td></tr>
<tr><td>Chromosome</td><td>start</td><td></td><td>end</td><td>strand</td><td>MM</td><td>target_seq</td><td>PAM</td><td>alignment</td><td>distance</td><td>position</td><td>gene name</td><td>gene id</td></tr>
<tr><td>GU305914.1</td><td></td><td>46250</td><td>46272</td><td>+</td><td>0</td><td>GCCGAACGACAAACACCGCG</td><td>CGG</td><td>||||||||[||||||||||||]PAM</td><td>NA</td><td>Intergenic</td><td>NA</td><td>NA</td></tr>
<tr><td>T2978</td><td>GACAAACACCGCGCGGAGTCCGG</td><td></td><td>1000</td><td></td><td></td><td></td><td></td><td></td><td></td><td></td><td></td><td></td></tr>
<tr><td>Oligo fwd</td><td>CACCGACAAACACCGCGCGGAGTC</td><td></td><td></td><td></td><td></td><td></td><td></td><td></td><td></td><td></td><td></td><td></td></tr>
<tr><td>Oligo rev</td><td>AAACGACTCCGCGCGGTGTTTGTC</td><td></td><td></td><td></td><td></td><td></td><td></td><td></td><td></td><td></td><td></td><td></td></tr>
<tr><td>Chromosome</td><td>start</td><td></td><td>end</td><td>strand</td><td>MM</td><td>target_seq</td><td>PAM</td><td>alignment</td><td>distance</td><td>position</td><td>gene name</td><td>gene id</td></tr>
<tr><td>GU305914.1</td><td></td><td>46257</td><td>46279</td><td>+</td><td>0</td><td>GACAAACACCGCGCGGAGTC</td><td>CGG</td><td>||||||||[||||||||||||]PAM</td><td>NA</td><td>Intergenic</td><td>NA</td><td>NA</td></tr>
<tr><td>T2979</td><td>ACAAACACCGCGCGGAGTCCGGG</td><td></td><td>1000</td><td></td><td></td><td></td><td></td><td></td><td></td><td></td><td></td><td></td></tr>
<tr><td>Oligo fwd</td><td>CACCACAAACACCGCGCGGAGTCC</td><td></td><td></td><td></td><td></td><td></td><td></td><td></td><td></td><td></td><td></td><td></td></tr>
<tr><td>Oligo rev</td><td>AAACGGACTCCGCGCGGTGTTTGT</td><td></td><td></td><td></td><td></td><td></td><td></td><td></td><td></td><td></td><td></td><td></td></tr>
<tr><td>Chromosome</td><td>start</td><td></td><td>end</td><td>strand</td><td>MM</td><td>target_seq</td><td>PAM</td><td>alignment</td><td>distance</td><td>position</td><td>gene name</td><td>gene id</td></tr>
<tr><td>GU305914.1</td><td></td><td>46258</td><td>46280</td><td>+</td><td>0</td><td>ACAAACACCGCGCGGAGTCC</td><td>GGG</td><td>||||||||[||||||||||||]PAM</td><td>NA</td><td>Intergenic</td><td>NA</td><td>NA</td></tr>
<tr><td>T2980</td><td>GCGCGGAGTCCGGGCTCCGAAGG</td><td></td><td>1000</td><td></td><td></td><td></td><td></td><td></td><td></td><td></td><td></td><td></td></tr>
<tr><td>Oligo fwd</td><td>CACCGCGCGGAGTCCGGGCTCCGA</td><td></td><td></td><td></td><td></td><td></td><td></td><td></td><td></td><td></td><td></td><td></td></tr>
<tr><td>Oligo rev</td><td>AAACTCGGAGCCCGGACTCCGCGC</td><td></td><td></td><td></td><td></td><td></td><td></td><td></td><td></td><td></td><td></td><td></td></tr>
<tr><td>Chromosome</td><td>start</td><td></td><td>end</td><td>strand</td><td>MM</td><td>target_seq</td><td>PAM</td><td>alignment</td><td>distance</td><td>position</td><td>gene name</td><td>gene id</td></tr>
<tr><td>GU305914.1</td><td></td><td>46267</td><td>46289</td><td>+</td><td>0</td><td>GCGCGGAGTCCGGGCTCCGA</td><td>AGG</td><td>||||||||[||||||||||||]PAM</td><td>NA</td><td>Intergenic</td><td>NA</td><td>NA</td></tr>
<tr><td>T2981</td><td>GATCTACCGCGCCCTGAACCTGG</td><td></td><td>1000</td><td></td><td></td><td></td><td></td><td></td><td></td><td></td><td></td><td></td></tr>
<tr><td>Oligo fwd</td><td>CACCGATCTACCGCGCCCTGAACC</td><td></td><td></td><td></td><td></td><td></td><td></td><td></td><td></td><td></td><td></td><td></td></tr>
<tr><td>Oligo rev</td><td>AAACGGTTCAGGGCGCGGTAGATC</td><td></td><td></td><td></td><td></td><td></td><td></td><td></td><td></td><td></td><td></td><td></td></tr>
<tr><td>Chromosome</td><td>start</td><td></td><td>end</td><td>strand</td><td>MM</td><td>target_seq</td><td>PAM</td><td>alignment</td><td>distance</td><td>position</td><td>gene name</td><td>gene id</td></tr>
<tr><td>GU305914.1</td><td></td><td>46289</td><td>46311</td><td>+</td><td>0</td><td>GATCTACCGCGCCCTGAACC</td><td>TGG</td><td>||||||||[||||||||||||]PAM</td><td>NA</td><td>Intergenic</td><td>NA</td><td>NA</td></tr>
<tr><td>T2982</td><td>GGAAGATAACCCGTTCGACCTGG</td><td></td><td>1000</td><td></td><td></td><td></td><td></td><td></td><td></td><td></td><td></td><td></td></tr>
<tr><td>Oligo fwd</td><td>CACCGGAAGATAACCCGTTCGACC</td><td></td><td></td><td></td><td></td><td></td><td></td><td></td><td></td><td></td><td></td><td></td></tr>
<tr><td>Oligo rev</td><td>AAACGGTCGAACGGGTTATCTTCC</td><td></td><td></td><td></td><td></td><td></td><td></td><td></td><td></td><td></td><td></td><td></td></tr>
<tr><td>Chromosome</td><td>start</td><td></td><td>end</td><td>strand</td><td>MM</td><td>target_seq</td><td>PAM</td><td>alignment</td><td>distance</td><td>position</td><td>gene name</td><td>gene id</td></tr>
<tr><td>GU305914.1</td><td></td><td>46310</td><td>46332</td><td>+</td><td>0</td><td>GGAAGATAACCCGTTCGACC</td><td>TGG</td><td>||||||||[||||||||||||]PAM</td><td>NA</td><td>Intergenic</td><td>NA</td><td>NA</td></tr>
<tr><td>T2983</td><td>TAACCCGTTCGACCTGGTCGAGG</td><td></td><td>1000</td><td></td><td></td><td></td><td></td><td></td><td></td><td></td><td></td><td></td></tr>
<tr><td>Oligo fwd</td><td>CACCTAACCCGTTCGACCTGGTCG</td><td></td><td></td><td></td><td></td><td></td><td></td><td></td><td></td><td></td><td></td><td></td></tr>
<tr><td>Oligo rev</td><td>AAACCGACCAGGTCGAACGGGTTA</td><td></td><td></td><td></td><td></td><td></td><td></td><td></td><td></td><td></td><td></td><td></td></tr>
<tr><td>Chromosome</td><td>start</td><td></td><td>end</td><td>strand</td><td>MM</td><td>target_seq</td><td>PAM</td><td>alignment</td><td>distance</td><td>position</td><td>gene name</td><td>gene id</td></tr>
<tr><td>GU305914.1</td><td></td><td>46316</td><td>46338</td><td>+</td><td>0</td><td>TAACCCGTTCGACCTGGTCG</td><td>AGG</td><td>||||||||[||||||||||||]PAM</td><td>NA</td><td>Intergenic</td><td>NA</td><td>NA</td></tr>
<tr><td>T2984</td><td>GTTCGACCTGGTCGAGGCCGTGG</td><td></td><td>1000</td><td></td><td></td><td></td><td></td><td></td><td></td><td></td><td></td><td></td></tr>
<tr><td>Oligo fwd</td><td>CACCGTTCGACCTGGTCGAGGCCG</td><td></td><td></td><td></td><td></td><td></td><td></td><td></td><td></td><td></td><td></td><td></td></tr>
<tr><td>Oligo rev</td><td>AAACCGGCCTCGACCAGGTCGAAC</td><td></td><td></td><td></td><td></td><td></td><td></td><td></td><td></td><td></td><td></td><td></td></tr>
<tr><td>Chromosome</td><td>start</td><td></td><td>end</td><td>strand</td><td>MM</td><td>target_seq</td><td>PAM</td><td>alignment</td><td>distance</td><td>position</td><td>gene name</td><td>gene id</td></tr>
<tr><td>GU305914.1</td><td></td><td>46322</td><td>46344</td><td>+</td><td>0</td><td>GTTCGACCTGGTCGAGGCCG</td><td>TGG</td><td>||||||||[||||||||||||]PAM</td><td>NA</td><td>Intergenic</td><td>NA</td><td>NA</td></tr>
<tr><td>T2985</td><td>TTCGACCTGGTCGAGGCCGTGGG</td><td></td><td>1000</td><td></td><td></td><td></td><td></td><td></td><td></td><td></td><td></td><td></td></tr>
<tr><td>Oligo fwd</td><td>CACCTTCGACCTGGTCGAGGCCGT</td><td></td><td></td><td></td><td></td><td></td><td></td><td></td><td></td><td></td><td></td><td></td></tr>
<tr><td>Oligo rev</td><td>AAACACGGCCTCGACCAGGTCGAA</td><td></td><td></td><td></td><td></td><td></td><td></td><td></td><td></td><td></td><td></td><td></td></tr>
<tr><td>Chromosome</td><td>start</td><td></td><td>end</td><td>strand</td><td>MM</td><td>target_seq</td><td>PAM</td><td>alignment</td><td>distance</td><td>position</td><td>gene name</td><td>gene id</td></tr>
<tr><td>GU305914.1</td><td></td><td>46323</td><td>46345</td><td>+</td><td>0</td><td>TTCGACCTGGTCGAGGCCGT</td><td>GGG</td><td>||||||||[||||||||||||]PAM</td><td>NA</td><td>Intergenic</td><td>NA</td><td>NA</td></tr>
<tr><td>T2986</td><td>GGTCGAGGCCGTGGGCGACCTGG</td><td></td><td>1000</td><td></td><td></td><td></td><td></td><td></td><td></td><td></td><td></td><td></td></tr>
<tr><td>Oligo fwd</td><td>CACCGGTCGAGGCCGTGGGCGACC</td><td></td><td></td><td></td><td></td><td></td><td></td><td></td><td></td><td></td><td></td><td></td></tr>
<tr><td>Oligo rev</td><td>AAACGGTCGCCCACGGCCTCGACC</td><td></td><td></td><td></td><td></td><td></td><td></td><td></td><td></td><td></td><td></td><td></td></tr>
<tr><td>Chromosome</td><td>start</td><td></td><td>end</td><td>strand</td><td>MM</td><td>target_seq</td><td>PAM</td><td>alignment</td><td>distance</td><td>position</td><td>gene name</td><td>gene id</td></tr>
<tr><td>GU305914.1</td><td></td><td>46331</td><td>46353</td><td>+</td><td>0</td><td>GGTCGAGGCCGTGGGCGACC</td><td>TGG</td><td>||||||||[||||||||||||]PAM</td><td>NA</td><td>Intergenic</td><td>NA</td><td>NA</td></tr>
<tr><td>GU305914.1</td><td></td><td>148572</td><td>148594</td><td>-</td><td>4</td><td>CGGCGACACCGTGGGCGACC</td><td>TGG</td><td>-|-|||--[||||||||||||]PAM</td><td>NA</td><td>Intergenic</td><td>NA</td><td>NA</td></tr>
<tr><td>T2987</td><td>GGGCGACCTGGACCTGTCTCAGG</td><td></td><td>1000</td><td></td><td></td><td></td><td></td><td></td><td></td><td></td><td></td><td></td></tr>
<tr><td>Oligo fwd</td><td>CACCGGGCGACCTGGACCTGTCTC</td><td></td><td></td><td></td><td></td><td></td><td></td><td></td><td></td><td></td><td></td><td></td></tr>
<tr><td>Oligo rev</td><td>AAACGAGACAGGTCCAGGTCGCCC</td><td></td><td></td><td></td><td></td><td></td><td></td><td></td><td></td><td></td><td></td><td></td></tr>
<tr><td>Chromosome</td><td>start</td><td></td><td>end</td><td>strand</td><td>MM</td><td>target_seq</td><td>PAM</td><td>alignment</td><td>distance</td><td>position</td><td>gene name</td><td>gene id</td></tr>
<tr><td>GU305914.1</td><td></td><td>46343</td><td>46365</td><td>+</td><td>0</td><td>GGGCGACCTGGACCTGTCTC</td><td>AGG</td><td>||||||||[||||||||||||]PAM</td><td>NA</td><td>Intergenic</td><td>NA</td><td>NA</td></tr>
<tr><td>T2988</td><td>GGCGACCTGGACCTGTCTCAGGG</td><td></td><td>1000</td><td></td><td></td><td></td><td></td><td></td><td></td><td></td><td></td><td></td></tr>
<tr><td>Oligo fwd</td><td>CACCGGCGACCTGGACCTGTCTCA</td><td></td><td></td><td></td><td></td><td></td><td></td><td></td><td></td><td></td><td></td><td></td></tr>
<tr><td>Oligo rev</td><td>AAACTGAGACAGGTCCAGGTCGCC</td><td></td><td></td><td></td><td></td><td></td><td></td><td></td><td></td><td></td><td></td><td></td></tr>
<tr><td>Chromosome</td><td>start</td><td></td><td>end</td><td>strand</td><td>MM</td><td>target_seq</td><td>PAM</td><td>alignment</td><td>distance</td><td>position</td><td>gene name</td><td>gene id</td></tr>
<tr><td>GU305914.1</td><td></td><td>46344</td><td>46366</td><td>+</td><td>0</td><td>GGCGACCTGGACCTGTCTCA</td><td>GGG</td><td>||||||||[||||||||||||]PAM</td><td>NA</td><td>Intergenic</td><td>NA</td><td>NA</td></tr>
<tr><td>T2989</td><td>CCTGTCTCAGGGCGCCTACGAGG</td><td></td><td>1000</td><td></td><td></td><td></td><td></td><td></td><td></td><td></td><td></td><td></td></tr>
<tr><td>Oligo fwd</td><td>CACCCCTGTCTCAGGGCGCCTACG</td><td></td><td></td><td></td><td></td><td></td><td></td><td></td><td></td><td></td><td></td><td></td></tr>
<tr><td>Oligo rev</td><td>AAACCGTAGGCGCCCTGAGACAGG</td><td></td><td></td><td></td><td></td><td></td><td></td><td></td><td></td><td></td><td></td><td></td></tr>
<tr><td>Chromosome</td><td>start</td><td></td><td>end</td><td>strand</td><td>MM</td><td>target_seq</td><td>PAM</td><td>alignment</td><td>distance</td><td>position</td><td>gene name</td><td>gene id</td></tr>
<tr><td>GU305914.1</td><td></td><td>46355</td><td>46377</td><td>+</td><td>0</td><td>CCTGTCTCAGGGCGCCTACG</td><td>AGG</td><td>||||||||[||||||||||||]PAM</td><td>NA</td><td>Intergenic</td><td>NA</td><td>NA</td></tr>
<tr><td>T2990</td><td>CTGTCTCAGGGCGCCTACGAGGG</td><td></td><td>1000</td><td></td><td></td><td></td><td></td><td></td><td></td><td></td><td></td><td></td></tr>
<tr><td>Oligo fwd</td><td>CACCCTGTCTCAGGGCGCCTACGA</td><td></td><td></td><td></td><td></td><td></td><td></td><td></td><td></td><td></td><td></td><td></td></tr>
<tr><td>Oligo rev</td><td>AAACTCGTAGGCGCCCTGAGACAG</td><td></td><td></td><td></td><td></td><td></td><td></td><td></td><td></td><td></td><td></td><td></td></tr>
<tr><td>Chromosome</td><td>start</td><td></td><td>end</td><td>strand</td><td>MM</td><td>target_seq</td><td>PAM</td><td>alignment</td><td>distance</td><td>position</td><td>gene name</td><td>gene id</td></tr>
<tr><td>GU305914.1</td><td></td><td>46356</td><td>46378</td><td>+</td><td>0</td><td>CTGTCTCAGGGCGCCTACGA</td><td>GGG</td><td>||||||||[||||||||||||]PAM</td><td>NA</td><td>Intergenic</td><td>NA</td><td>NA</td></tr>
</table>

<table>
<tr><td>T2991</td><td>CTACTCGCTCATGAAGTCCCTGG</td><td>1000</td><td></td><td></td><td></td><td></td><td></td><td></td><td></td><td></td><td></td></tr>
<tr><td>Oligo fwd</td><td>CACCCTACTCGCTCATGAAGTCCC</td><td></td><td></td><td></td><td></td><td></td><td></td><td></td><td></td><td></td><td></td></tr>
<tr><td>Oligo rev</td><td>AAACGGGACTTCATGAGCGAGTAG</td><td></td><td></td><td></td><td></td><td></td><td></td><td></td><td></td><td></td><td></td></tr>
<tr><td>Chromosome</td><td>start</td><td>end</td><td>strand</td><td>MM</td><td>target_seq</td><td>PAM</td><td>alignment</td><td>distance</td><td>position</td><td>gene name</td><td>gene id</td></tr>
<tr><td>GU305914.1</td><td>46394</td><td>46416</td><td>+</td><td>0</td><td>CTACTCGCTCATGAAGTCCC</td><td>TGG</td><td>||||||||[||||||||||||]PAM</td><td>NA</td><td>Intergenic</td><td>NA</td><td>NA</td></tr>
<tr><td>T2992</td><td>TACTCGCTCATGAAGTCCCTGGG</td><td>1000</td><td></td><td></td><td></td><td></td><td></td><td></td><td></td><td></td><td></td></tr>
<tr><td>Oligo fwd</td><td>CACCTACTCGCTCATGAAGTCCCT</td><td></td><td></td><td></td><td></td><td></td><td></td><td></td><td></td><td></td><td></td></tr>
<tr><td>Oligo rev</td><td>AAACAGGGACTTCATGAGCGAGTA</td><td></td><td></td><td></td><td></td><td></td><td></td><td></td><td></td><td></td><td></td></tr>
<tr><td>Chromosome</td><td>start</td><td>end</td><td>strand</td><td>MM</td><td>target_seq</td><td>PAM</td><td>alignment</td><td>distance</td><td>position</td><td>gene name</td><td>gene id</td></tr>
<tr><td>GU305914.1</td><td>46395</td><td>46417</td><td>+</td><td>0</td><td>TACTCGCTCATGAAGTCCCT</td><td>GGG</td><td>||||||||[||||||||||||]PAM</td><td>NA</td><td>Intergenic</td><td>NA</td><td>NA</td></tr>
<tr><td>T2993</td><td>GAAGTCCCTGGGACACGATGTGG</td><td>1000</td><td></td><td></td><td></td><td></td><td></td><td></td><td></td><td></td><td></td></tr>
<tr><td>Oligo fwd</td><td>CACCGAAGTCCCTGGGACACGATG</td><td></td><td></td><td></td><td></td><td></td><td></td><td></td><td></td><td></td><td></td></tr>
<tr><td>Oligo rev</td><td>AAACCATCGTGTCCCAGGGACTTC</td><td></td><td></td><td></td><td></td><td></td><td></td><td></td><td></td><td></td><td></td></tr>
<tr><td>Chromosome</td><td>start</td><td>end</td><td>strand</td><td>MM</td><td>target_seq</td><td>PAM</td><td>alignment</td><td>distance</td><td>position</td><td>gene name</td><td>gene id</td></tr>
<tr><td>GU305914.1</td><td>46406</td><td>46428</td><td>+</td><td>0</td><td>GAAGTCCCTGGGACACGATG</td><td>TGG</td><td>||||||||[||||||||||||]PAM</td><td>NA</td><td>Intergenic</td><td>NA</td><td>NA</td></tr>
<tr><td>T2994</td><td>AAGTCCCTGGGACACGATGTGGG</td><td>1000</td><td></td><td></td><td></td><td></td><td></td><td></td><td></td><td></td><td></td></tr>
<tr><td>Oligo fwd</td><td>CACCAAGTCCCTGGGACACGATGT</td><td></td><td></td><td></td><td></td><td></td><td></td><td></td><td></td><td></td><td></td></tr>
<tr><td>Oligo rev</td><td>AAACACATCGTGTCCCAGGGACTT</td><td></td><td></td><td></td><td></td><td></td><td></td><td></td><td></td><td></td><td></td></tr>
<tr><td>Chromosome</td><td>start</td><td>end</td><td>strand</td><td>MM</td><td>target_seq</td><td>PAM</td><td>alignment</td><td>distance</td><td>position</td><td>gene name</td><td>gene id</td></tr>
<tr><td>GU305914.1</td><td>46407</td><td>46429</td><td>+</td><td>0</td><td>AAGTCCCTGGGACACGATGT</td><td>GGG</td><td>||||||||[||||||||||||]PAM</td><td>NA</td><td>Intergenic</td><td>NA</td><td>NA</td></tr>
<tr><td>T2995</td><td>ACGATGTGGGCCACCTCAGCCGG</td><td>1000</td><td></td><td></td><td></td><td></td><td></td><td></td><td></td><td></td><td></td></tr>
<tr><td>Oligo fwd</td><td>CACCACGATGTGGGCCACCTCAGC</td><td></td><td></td><td></td><td></td><td></td><td></td><td></td><td></td><td></td><td></td></tr>
<tr><td>Oligo rev</td><td>AAACGCTGAGGTGGCCCACATCGT</td><td></td><td></td><td></td><td></td><td></td><td></td><td></td><td></td><td></td><td></td></tr>
<tr><td>Chromosome</td><td>start</td><td>end</td><td>strand</td><td>MM</td><td>target_seq</td><td>PAM</td><td>alignment</td><td>distance</td><td>position</td><td>gene name</td><td>gene id</td></tr>
<tr><td>GU305914.1</td><td>46420</td><td>46442</td><td>+</td><td>0</td><td>ACGATGTGGGCCACCTCAGC</td><td>CGG</td><td>||||||||[||||||||||||]PAM</td><td>NA</td><td>Intergenic</td><td>NA</td><td>NA</td></tr>
<tr><td>T2996</td><td>GGGCCACCTCAGCCGGTCCCTGG</td><td>1000</td><td></td><td></td><td></td><td></td><td></td><td></td><td></td><td></td><td></td></tr>
<tr><td>Oligo fwd</td><td>CACCGGGCCACCTCAGCCGGTCCC</td><td></td><td></td><td></td><td></td><td></td><td></td><td></td><td></td><td></td><td></td></tr>
<tr><td>Oligo rev</td><td>AAACGGGACCGGCTGAGGTGGCCC</td><td></td><td></td><td></td><td></td><td></td><td></td><td></td><td></td><td></td><td></td></tr>
<tr><td>Chromosome</td><td>start</td><td>end</td><td>strand</td><td>MM</td><td>target_seq</td><td>PAM</td><td>alignment</td><td>distance</td><td>position</td><td>gene name</td><td>gene id</td></tr>
<tr><td>GU305914.1</td><td>46427</td><td>46449</td><td>+</td><td>0</td><td>GGGCCACCTCAGCCGGTCCC</td><td>TGG</td><td>||||||||[||||||||||||]PAM</td><td>NA</td><td>Intergenic</td><td>NA</td><td>NA</td></tr>
<tr><td>T2997</td><td>CCTGGACTACTCCACGATGACGG</td><td>1000</td><td></td><td></td><td></td><td></td><td></td><td></td><td></td><td></td><td></td></tr>
<tr><td>Oligo fwd</td><td>CACCCCTGGACTACTCCACGATGA</td><td></td><td></td><td></td><td></td><td></td><td></td><td></td><td></td><td></td><td></td></tr>
<tr><td>Oligo rev</td><td>AAACTCATCGTGGAGTAGTCCAGG</td><td></td><td></td><td></td><td></td><td></td><td></td><td></td><td></td><td></td><td></td></tr>
<tr><td>Chromosome</td><td>start</td><td>end</td><td>strand</td><td>MM</td><td>target_seq</td><td>PAM</td><td>alignment</td><td>distance</td><td>position</td><td>gene name</td><td>gene id</td></tr>
<tr><td>GU305914.1</td><td>46445</td><td>46467</td><td>+</td><td>0</td><td>CCTGGACTACTCCACGATGA</td><td>CGG</td><td>||||||||[||||||||||||]PAM</td><td>NA</td><td>Intergenic</td><td>NA</td><td>NA</td></tr>
<tr><td>T2998</td><td>ATCCCCCCTGTTCACGAGCCAGG</td><td>1000</td><td></td><td></td><td></td><td></td><td></td><td></td><td></td><td></td><td></td></tr>
<tr><td>Oligo fwd</td><td>CACCATCCCCCCTGTTCACGAGCC</td><td></td><td></td><td></td><td></td><td></td><td></td><td></td><td></td><td></td><td></td></tr>
<tr><td>Oligo rev</td><td>AAACGGCTCGTGAACAGGGGGGAT</td><td></td><td></td><td></td><td></td><td></td><td></td><td></td><td></td><td></td><td></td></tr>
<tr><td>Chromosome</td><td>start</td><td>end</td><td>strand</td><td>MM</td><td>target_seq</td><td>PAM</td><td>alignment</td><td>distance</td><td>position</td><td>gene name</td><td>gene id</td></tr>
<tr><td>GU305914.1</td><td>46487</td><td>46509</td><td>+</td><td>0</td><td>ATCCCCCCTGTTCACGAGCC</td><td>AGG</td><td>||||||||[||||||||||||]PAM</td><td>NA</td><td>Intergenic</td><td>NA</td><td>NA</td></tr>
<tr><td>T2999</td><td>CCCCCTGTTCACGAGCCAGGAGG</td><td>1000</td><td></td><td></td><td></td><td></td><td></td><td></td><td></td><td></td><td></td></tr>
<tr><td>Oligo fwd</td><td>CACCCCCCCTGTTCACGAGCCAGG</td><td></td><td></td><td></td><td></td><td></td><td></td><td></td><td></td><td></td><td></td></tr>
<tr><td>Oligo rev</td><td>AAACCCTGGCTCGTGAACAGGGGG</td><td></td><td></td><td></td><td></td><td></td><td></td><td></td><td></td><td></td><td></td></tr>
<tr><td>Chromosome</td><td>start</td><td>end</td><td>strand</td><td>MM</td><td>target_seq</td><td>PAM</td><td>alignment</td><td>distance</td><td>position</td><td>gene name</td><td>gene id</td></tr>
<tr><td>GU305914.1</td><td>46490</td><td>46512</td><td>+</td><td>0</td><td>CCCCCTGTTCACGAGCCAGG</td><td>AGG</td><td>||||||||[||||||||||||]PAM</td><td>NA</td><td>Intergenic</td><td>NA</td><td>NA</td></tr>
<tr><td>T3000</td><td>AGGCCGTGACGATGTACTCCAGG</td><td>1000</td><td></td><td></td><td></td><td></td><td></td><td></td><td></td><td></td><td></td></tr>
<tr><td>Oligo fwd</td><td>CACCAGGCCGTGACGATGTACTCC</td><td></td><td></td><td></td><td></td><td></td><td></td><td></td><td></td><td></td><td></td></tr>
<tr><td>Oligo rev</td><td>AAACGGAGTACATCGTCACGGCCT</td><td></td><td></td><td></td><td></td><td></td><td></td><td></td><td></td><td></td><td></td></tr>
<tr><td>Chromosome</td><td>start</td><td>end</td><td>strand</td><td>MM</td><td>target_seq</td><td>PAM</td><td>alignment</td><td>distance</td><td>position</td><td>gene name</td><td>gene id</td></tr>
<tr><td>GU305914.1</td><td>46510</td><td>46532</td><td>+</td><td>0</td><td>AGGCCGTGACGATGTACTCC</td><td>AGG</td><td>||||||||[||||||||||||]PAM</td><td>NA</td><td>Intergenic</td><td>NA</td><td>NA</td></tr>
<tr><td>T3001</td><td>ATGTACTCCAGGAACCTCGCCGG</td><td>1000</td><td></td><td></td><td></td><td></td><td></td><td></td><td></td><td></td><td></td></tr>
<tr><td>Oligo fwd</td><td>CACCATGTACTCCAGGAACCTCGC</td><td></td><td></td><td></td><td></td><td></td><td></td><td></td><td></td><td></td><td></td></tr>
<tr><td>Oligo rev</td><td>AAACGCGAGGTTCCTGGAGTACAT</td><td></td><td></td><td></td><td></td><td></td><td></td><td></td><td></td><td></td><td></td></tr>
<tr><td>Chromosome</td><td>start</td><td>end</td><td>strand</td><td>MM</td><td>target_seq</td><td>PAM</td><td>alignment</td><td>distance</td><td>position</td><td>gene name</td><td>gene id</td></tr>
<tr><td>GU305914.1</td><td>46521</td><td>46543</td><td>+</td><td>0</td><td>ATGTACTCCAGGAACCTCGC</td><td>CGG</td><td>||||||||[||||||||||||]PAM</td><td>NA</td><td>Intergenic</td><td>NA</td><td>NA</td></tr>
<tr><td>T3002</td><td>CGAGCCGTTCGAGACCCTGTCGG</td><td>1000</td><td></td><td></td><td></td><td></td><td></td><td></td><td></td><td></td><td></td></tr>
<tr><td>Oligo fwd</td><td>CACCCGAGCCGTTCGAGACCCTGT</td><td></td><td></td><td></td><td></td><td></td><td></td><td></td><td></td><td></td><td></td></tr>
<tr><td>Oligo rev</td><td>AAACACAGGGTCTCGAACGGCTCG</td><td></td><td></td><td></td><td></td><td></td><td></td><td></td><td></td><td></td><td></td></tr>
<tr><td>Chromosome</td><td>start</td><td>end</td><td>strand</td><td>MM</td><td>target_seq</td><td>PAM</td><td>alignment</td><td>distance</td><td>position</td><td>gene name</td><td>gene id</td></tr>
<tr><td>GU305914.1</td><td>46562</td><td>46584</td><td>+</td><td>0</td><td>CGAGCCGTTCGAGACCCTGT</td><td>CGG</td><td>||||||||[||||||||||||]PAM</td><td>NA</td><td>Intergenic</td><td>NA</td><td>NA</td></tr>
<tr><td>T3003</td><td>AGACCCTGTCGGTCGTGCACCGG</td><td>1000</td><td></td><td></td><td></td><td></td><td></td><td></td><td></td><td></td><td></td></tr>
<tr><td>Oligo fwd</td><td>CACCAGACCCTGTCGGTCGTGCAC</td><td></td><td></td><td></td><td></td><td></td><td></td><td></td><td></td><td></td><td></td></tr>
<tr><td>Oligo rev</td><td>AAACGTGCACGACCGACAGGGTCT</td><td></td><td></td><td></td><td></td><td></td><td></td><td></td><td></td><td></td><td></td></tr>
<tr><td>Chromosome</td><td>start</td><td>end</td><td>strand</td><td>MM</td><td>target_seq</td><td>PAM</td><td>alignment</td><td>distance</td><td>position</td><td>gene name</td><td>gene id</td></tr>
<tr><td>GU305914.1</td><td>46573</td><td>46595</td><td>+</td><td>0</td><td>AGACCCTGTCGGTCGTGCAC</td><td>CGG</td><td>||||||||[||||||||||||]PAM</td><td>NA</td><td>Intergenic</td><td>NA</td><td>NA</td></tr>
<tr><td>T3004</td><td>CGACATGCTGTTCCTCCTGTCGG</td><td>1000</td><td></td><td></td><td></td><td></td><td></td><td></td><td></td><td></td><td></td></tr>
<tr><td>Oligo fwd</td><td>CACCCGACATGCTGTTCCTCCTGT</td><td></td><td></td><td></td><td></td><td></td><td></td><td></td><td></td><td></td><td></td></tr>
<tr><td>Oligo rev</td><td>AAACACAGGAGGAACAGCATGTCG</td><td></td><td></td><td></td><td></td><td></td><td></td><td></td><td></td><td></td><td></td></tr>
<tr><td>Chromosome</td><td>start</td><td>end</td><td>strand</td><td>MM</td><td>target_seq</td><td>PAM</td><td>alignment</td><td>distance</td><td>position</td><td>gene name</td><td>gene id</td></tr>
<tr><td>GU305914.1</td><td>46622</td><td>46644</td><td>+</td><td>0</td><td>CGACATGCTGTTCCTCCTGT</td><td>CGG</td><td>||||||||[||||||||||||]PAM</td><td>NA</td><td>Intergenic</td><td>NA</td><td>NA</td></tr>
<tr><td>T3005</td><td>ATGCTGTTCCTCCTGTCGGTCGG</td><td>1000</td><td></td><td></td><td></td><td></td><td></td><td></td><td></td><td></td><td></td></tr>
<tr><td>Oligo fwd</td><td>CACCATGCTGTTCCTCCTGTCGGT</td><td></td><td></td><td></td><td></td><td></td><td></td><td></td><td></td><td></td><td></td></tr>
<tr><td>Oligo rev</td><td>AAACACCGACAGGAGGAACAGCAT</td><td></td><td></td><td></td><td></td><td></td><td></td><td></td><td></td><td></td><td></td></tr>
<tr><td>Chromosome</td><td>start</td><td>end</td><td>strand</td><td>MM</td><td>target_seq</td><td>PAM</td><td>alignment</td><td>distance</td><td>position</td><td>gene name</td><td>gene id</td></tr>
<tr><td>GU305914.1</td><td>46626</td><td>46648</td><td>+</td><td>0</td><td>ATGCTGTTCCTCCTGTCGGT</td><td>CGG</td><td>||||||||[||||||||||||]PAM</td><td>NA</td><td>Intergenic</td><td>NA</td><td>NA</td></tr>
</table>

<table>
<tr><td>T3006</td><td>CCGCATGATCGTGCTCCACCAGG</td><td>1000</td><td></td><td></td><td></td><td></td><td></td><td></td><td></td><td></td><td></td></tr>
<tr><td>Oligo fwd</td><td>CACCCCGCATGATCGTGCTCCACC</td><td></td><td></td><td></td><td></td><td></td><td></td><td></td><td></td><td></td><td></td></tr>
<tr><td>Oligo rev</td><td>AAACGGTGGAGCACGATCATGCGG</td><td></td><td></td><td></td><td></td><td></td><td></td><td></td><td></td><td></td><td></td></tr>
<tr><td>Chromosome</td><td>start</td><td>end</td><td>strand</td><td>MM</td><td>target_seq</td><td>PAM</td><td>alignment</td><td>distance</td><td>position</td><td>gene name</td><td>gene id</td></tr>
<tr><td>GU305914.1</td><td>46649</td><td>46671</td><td>+</td><td>0</td><td>CCGCATGATCGTGCTCCACC</td><td>AGG</td><td>||||||||[||||||||||||]PAM</td><td>NA</td><td>Intergenic</td><td>NA</td><td>NA</td></tr>
<tr><td>GU305914.1</td><td>72745</td><td>72767</td><td>+</td><td>4</td><td>CCGGCTGATCGTGCGGCACC</td><td>TGG</td><td>|||--|||[||||||--||||]PAM</td><td>NA</td><td>Intergenic</td><td>NA</td><td>NA</td></tr>
<tr><td>T3007</td><td>GCGCAAGAACCTGATCTTGACGG</td><td>1000</td><td></td><td></td><td></td><td></td><td></td><td></td><td></td><td></td><td></td></tr>
<tr><td>Oligo fwd</td><td>CACCGCGCAAGAACCTGATCTTGA</td><td></td><td></td><td></td><td></td><td></td><td></td><td></td><td></td><td></td><td></td></tr>
<tr><td>Oligo rev</td><td>AAACTCAAGATCAGGTTCTTGCGC</td><td></td><td></td><td></td><td></td><td></td><td></td><td></td><td></td><td></td><td></td></tr>
<tr><td>Chromosome</td><td>start</td><td>end</td><td>strand</td><td>MM</td><td>target_seq</td><td>PAM</td><td>alignment</td><td>distance</td><td>position</td><td>gene name</td><td>gene id</td></tr>
<tr><td>GU305914.1</td><td>46688</td><td>46710</td><td>+</td><td>0</td><td>GCGCAAGAACCTGATCTTGA</td><td>CGG</td><td>||||||||[||||||||||||]PAM</td><td>NA</td><td>Intergenic</td><td>NA</td><td>NA</td></tr>
<tr><td>T3008</td><td>ACTCTACACCGCCTACACTCAGG</td><td>1000</td><td></td><td></td><td></td><td></td><td></td><td></td><td></td><td></td><td></td></tr>
<tr><td>Oligo fwd</td><td>CACCACTCTACACCGCCTACACTC</td><td></td><td></td><td></td><td></td><td></td><td></td><td></td><td></td><td></td><td></td></tr>
<tr><td>Oligo rev</td><td>AAACGAGTGTAGGCGGTGTAGAGT</td><td></td><td></td><td></td><td></td><td></td><td></td><td></td><td></td><td></td><td></td></tr>
<tr><td>Chromosome</td><td>start</td><td>end</td><td>strand</td><td>MM</td><td>target_seq</td><td>PAM</td><td>alignment</td><td>distance</td><td>position</td><td>gene name</td><td>gene id</td></tr>
<tr><td>GU305914.1</td><td>46730</td><td>46752</td><td>+</td><td>0</td><td>ACTCTACACCGCCTACACTC</td><td>AGG</td><td>||||||||[||||||||||||]PAM</td><td>NA</td><td>Intergenic</td><td>NA</td><td>NA</td></tr>
<tr><td>T3009</td><td>CACCGCCTACACTCAGGTACCGG</td><td>1000</td><td></td><td></td><td></td><td></td><td></td><td></td><td></td><td></td><td></td></tr>
<tr><td>Oligo fwd</td><td>CACCCACCGCCTACACTCAGGTAC</td><td></td><td></td><td></td><td></td><td></td><td></td><td></td><td></td><td></td><td></td></tr>
<tr><td>Oligo rev</td><td>AAACGTACCTGAGTGTAGGCGGTG</td><td></td><td></td><td></td><td></td><td></td><td></td><td></td><td></td><td></td><td></td></tr>
<tr><td>Chromosome</td><td>start</td><td>end</td><td>strand</td><td>MM</td><td>target_seq</td><td>PAM</td><td>alignment</td><td>distance</td><td>position</td><td>gene name</td><td>gene id</td></tr>
<tr><td>GU305914.1</td><td>46736</td><td>46758</td><td>+</td><td>0</td><td>CACCGCCTACACTCAGGTAC</td><td>CGG</td><td>||||||||[||||||||||||]PAM</td><td>NA</td><td>Intergenic</td><td>NA</td><td>NA</td></tr>
<tr><td>T3010</td><td>GACGAAATCCCTGTTCCGCGAGG</td><td>1000</td><td></td><td></td><td></td><td></td><td></td><td></td><td></td><td></td><td></td></tr>
<tr><td>Oligo fwd</td><td>CACCGACGAAATCCCTGTTCCGCG</td><td></td><td></td><td></td><td></td><td></td><td></td><td></td><td></td><td></td><td></td></tr>
<tr><td>Oligo rev</td><td>AAACCGCGGAACAGGGATTTCGTC</td><td></td><td></td><td></td><td></td><td></td><td></td><td></td><td></td><td></td><td></td></tr>
<tr><td>Chromosome</td><td>start</td><td>end</td><td>strand</td><td>MM</td><td>target_seq</td><td>PAM</td><td>alignment</td><td>distance</td><td>position</td><td>gene name</td><td>gene id</td></tr>
<tr><td>GU305914.1</td><td>46760</td><td>46782</td><td>+</td><td>0</td><td>GACGAAATCCCTGTTCCGCG</td><td>AGG</td><td>||||||||[||||||||||||]PAM</td><td>NA</td><td>Intergenic</td><td>NA</td><td>NA</td></tr>
<tr><td>T3011</td><td>GTTCCGCGAGGTCGCGCACGAGG</td><td>1000</td><td></td><td></td><td></td><td></td><td></td><td></td><td></td><td></td><td></td></tr>
<tr><td>Oligo fwd</td><td>CACCGTTCCGCGAGGTCGCGCACG</td><td></td><td></td><td></td><td></td><td></td><td></td><td></td><td></td><td></td><td></td></tr>
<tr><td>Oligo rev</td><td>AAACCGTGCGCGACCTCGCGGAAC</td><td></td><td></td><td></td><td></td><td></td><td></td><td></td><td></td><td></td><td></td></tr>
<tr><td>Chromosome</td><td>start</td><td>end</td><td>strand</td><td>MM</td><td>target_seq</td><td>PAM</td><td>alignment</td><td>distance</td><td>position</td><td>gene name</td><td>gene id</td></tr>
<tr><td>GU305914.1</td><td>46772</td><td>46794</td><td>+</td><td>0</td><td>GTTCCGCGAGGTCGCGCACG</td><td>AGG</td><td>||||||||[||||||||||||]PAM</td><td>NA</td><td>Intergenic</td><td>NA</td><td>NA</td></tr>
<tr><td>T3012</td><td>CGCACGCGCTGCTCTCCTCCAGG</td><td>1000</td><td></td><td></td><td></td><td></td><td></td><td></td><td></td><td></td><td></td></tr>
<tr><td>Oligo fwd</td><td>CACCCGCACGCGCTGCTCTCCTCC</td><td></td><td></td><td></td><td></td><td></td><td></td><td></td><td></td><td></td><td></td></tr>
<tr><td>Oligo rev</td><td>AAACGGAGGAGAGCAGCGCGTGCG</td><td></td><td></td><td></td><td></td><td></td><td></td><td></td><td></td><td></td><td></td></tr>
<tr><td>Chromosome</td><td>start</td><td>end</td><td>strand</td><td>MM</td><td>target_seq</td><td>PAM</td><td>alignment</td><td>distance</td><td>position</td><td>gene name</td><td>gene id</td></tr>
<tr><td>GU305914.1</td><td>46795</td><td>46817</td><td>+</td><td>0</td><td>CGCACGCGCTGCTCTCCTCC</td><td>AGG</td><td>||||||||[||||||||||||]PAM</td><td>NA</td><td>Intergenic</td><td>NA</td><td>NA</td></tr>
<tr><td>T3013</td><td>GTTCATCAAGCAGATCATCGCGG</td><td>1000</td><td></td><td></td><td></td><td></td><td></td><td></td><td></td><td></td><td></td></tr>
<tr><td>Oligo fwd</td><td>CACCGTTCATCAAGCAGATCATCG</td><td></td><td></td><td></td><td></td><td></td><td></td><td></td><td></td><td></td><td></td></tr>
<tr><td>Oligo rev</td><td>AAACCGATGATCTGCTTGATGAAC</td><td></td><td></td><td></td><td></td><td></td><td></td><td></td><td></td><td></td><td></td></tr>
<tr><td>Chromosome</td><td>start</td><td>end</td><td>strand</td><td>MM</td><td>target_seq</td><td>PAM</td><td>alignment</td><td>distance</td><td>position</td><td>gene name</td><td>gene id</td></tr>
<tr><td>GU305914.1</td><td>46865</td><td>46887</td><td>+</td><td>0</td><td>GTTCATCAAGCAGATCATCG</td><td>CGG</td><td>||||||||[||||||||||||]PAM</td><td>NA</td><td>Intergenic</td><td>NA</td><td>NA</td></tr>
<tr><td>T3014</td><td>CCGACGTGTACACCTGCCCCAGG</td><td>1000</td><td></td><td></td><td></td><td></td><td></td><td></td><td></td><td></td><td></td></tr>
<tr><td>Oligo fwd</td><td>CACCCCGACGTGTACACCTGCCCC</td><td></td><td></td><td></td><td></td><td></td><td></td><td></td><td></td><td></td><td></td></tr>
<tr><td>Oligo rev</td><td>AAACGGGGCAGGTGTACACGTCGG</td><td></td><td></td><td></td><td></td><td></td><td></td><td></td><td></td><td></td><td></td></tr>
<tr><td>Chromosome</td><td>start</td><td>end</td><td>strand</td><td>MM</td><td>target_seq</td><td>PAM</td><td>alignment</td><td>distance</td><td>position</td><td>gene name</td><td>gene id</td></tr>
<tr><td>GU305914.1</td><td>46888</td><td>46910</td><td>+</td><td>0</td><td>CCGACGTGTACACCTGCCCC</td><td>AGG</td><td>||||||||[||||||||||||]PAM</td><td>NA</td><td>Intergenic</td><td>NA</td><td>NA</td></tr>
<tr><td>T3015</td><td>GTACCTGACCAACCAGATCCTGG</td><td>1000</td><td></td><td></td><td></td><td></td><td></td><td></td><td></td><td></td><td></td></tr>
<tr><td>Oligo fwd</td><td>CACCGTACCTGACCAACCAGATCC</td><td></td><td></td><td></td><td></td><td></td><td></td><td></td><td></td><td></td><td></td></tr>
<tr><td>Oligo rev</td><td>AAACGGATCTGGTTGGTCAGGTAC</td><td></td><td></td><td></td><td></td><td></td><td></td><td></td><td></td><td></td><td></td></tr>
<tr><td>Chromosome</td><td>start</td><td>end</td><td>strand</td><td>MM</td><td>target_seq</td><td>PAM</td><td>alignment</td><td>distance</td><td>position</td><td>gene name</td><td>gene id</td></tr>
<tr><td>GU305914.1</td><td>46910</td><td>46932</td><td>+</td><td>0</td><td>GTACCTGACCAACCAGATCC</td><td>TGG</td><td>||||||||[||||||||||||]PAM</td><td>NA</td><td>Intergenic</td><td>NA</td><td>NA</td></tr>
<tr><td>T3016</td><td>CAACCAGATCCTGGCCGTGACGG</td><td>1000</td><td></td><td></td><td></td><td></td><td></td><td></td><td></td><td></td><td></td></tr>
<tr><td>Oligo fwd</td><td>CACCCAACCAGATCCTGGCCGTGA</td><td></td><td></td><td></td><td></td><td></td><td></td><td></td><td></td><td></td><td></td></tr>
<tr><td>Oligo rev</td><td>AAACTCACGGCCAGGATCTGGTTG</td><td></td><td></td><td></td><td></td><td></td><td></td><td></td><td></td><td></td><td></td></tr>
<tr><td>Chromosome</td><td>start</td><td>end</td><td>strand</td><td>MM</td><td>target_seq</td><td>PAM</td><td>alignment</td><td>distance</td><td>position</td><td>gene name</td><td>gene id</td></tr>
<tr><td>GU305914.1</td><td>46919</td><td>46941</td><td>+</td><td>0</td><td>CAACCAGATCCTGGCCGTGA</td><td>CGG</td><td>||||||||[||||||||||||]PAM</td><td>NA</td><td>Intergenic</td><td>NA</td><td>NA</td></tr>
<tr><td>T3017</td><td>AGATCCTGGCCGTGACGGCGAGG</td><td>1000</td><td></td><td></td><td></td><td></td><td></td><td></td><td></td><td></td><td></td></tr>
<tr><td>Oligo fwd</td><td>CACCAGATCCTGGCCGTGACGGCG</td><td></td><td></td><td></td><td></td><td></td><td></td><td></td><td></td><td></td><td></td></tr>
<tr><td>Oligo rev</td><td>AAACCGCCGTCACGGCCAGGATCT</td><td></td><td></td><td></td><td></td><td></td><td></td><td></td><td></td><td></td><td></td></tr>
<tr><td>Chromosome</td><td>start</td><td>end</td><td>strand</td><td>MM</td><td>target_seq</td><td>PAM</td><td>alignment</td><td>distance</td><td>position</td><td>gene name</td><td>gene id</td></tr>
<tr><td>GU305914.1</td><td>46924</td><td>46946</td><td>+</td><td>0</td><td>AGATCCTGGCCGTGACGGCG</td><td>AGG</td><td>||||||||[||||||||||||]PAM</td><td>NA</td><td>Intergenic</td><td>NA</td><td>NA</td></tr>
<tr><td>T3018</td><td>GCCGTGACGGCGAGGATGCACGG</td><td>1000</td><td></td><td></td><td></td><td></td><td></td><td></td><td></td><td></td><td></td></tr>
<tr><td>Oligo fwd</td><td>CACCGCCGTGACGGCGAGGATGCA</td><td></td><td></td><td></td><td></td><td></td><td></td><td></td><td></td><td></td><td></td></tr>
<tr><td>Oligo rev</td><td>AAACTGCATCCTCGCCGTCACGGC</td><td></td><td></td><td></td><td></td><td></td><td></td><td></td><td></td><td></td><td></td></tr>
<tr><td>Chromosome</td><td>start</td><td>end</td><td>strand</td><td>MM</td><td>target_seq</td><td>PAM</td><td>alignment</td><td>distance</td><td>position</td><td>gene name</td><td>gene id</td></tr>
<tr><td>GU305914.1</td><td>46932</td><td>46954</td><td>+</td><td>0</td><td>GCCGTGACGGCGAGGATGCA</td><td>CGG</td><td>||||||||[||||||||||||]PAM</td><td>NA</td><td>Intergenic</td><td>NA</td><td>NA</td></tr>
<tr><td>T3019</td><td>GACGGCGAGGATGCACGGCCTGG</td><td>1000</td><td></td><td></td><td></td><td></td><td></td><td></td><td></td><td></td><td></td></tr>
<tr><td>Oligo fwd</td><td>CACCGACGGCGAGGATGCACGGCC</td><td></td><td></td><td></td><td></td><td></td><td></td><td></td><td></td><td></td><td></td></tr>
<tr><td>Oligo rev</td><td>AAACGGCCGTGCATCCTCGCCGTC</td><td></td><td></td><td></td><td></td><td></td><td></td><td></td><td></td><td></td><td></td></tr>
<tr><td>Chromosome</td><td>start</td><td>end</td><td>strand</td><td>MM</td><td>target_seq</td><td>PAM</td><td>alignment</td><td>distance</td><td>position</td><td>gene name</td><td>gene id</td></tr>
<tr><td>GU305914.1</td><td>46937</td><td>46959</td><td>+</td><td>0</td><td>GACGGCGAGGATGCACGGCC</td><td>TGG</td><td>||||||||[||||||||||||]PAM</td><td>NA</td><td>Intergenic</td><td>NA</td><td>NA</td></tr>
<tr><td>T3020</td><td>GGCGAGGATGCACGGCCTGGAGG</td><td>1000</td><td></td><td></td><td></td><td></td><td></td><td></td><td></td><td></td><td></td></tr>
<tr><td>Oligo fwd</td><td>CACCGGCGAGGATGCACGGCCTGG</td><td></td><td></td><td></td><td></td><td></td><td></td><td></td><td></td><td></td><td></td></tr>
<tr><td>Oligo rev</td><td>AAACCCAGGCCGTGCATCCTCGCC</td><td></td><td></td><td></td><td></td><td></td><td></td><td></td><td></td><td></td><td></td></tr>
</table>

<table>
<tr><td>Chromosome</td><td>start</td><td>end</td><td>strand</td><td>MM</td><td>target_seq</td><td>PAM</td><td>alignment</td><td>distance</td><td>position</td><td>gene name</td><td>gene id</td></tr>
<tr><td>GU305914.1</td><td>46940</td><td>46962</td><td>+</td><td>0</td><td>GGCGAGGATGCACGGCCTGG</td><td>AGG</td><td>||||||||[||||||||||||]PAM</td><td>NA</td><td>Intergenic</td><td>NA</td><td>NA</td></tr>
<tr><td>T3021</td><td>GCGAGGATGCACGGCCTGGAGGG</td><td>1000</td><td></td><td></td><td></td><td></td><td></td><td></td><td></td><td></td><td></td></tr>
<tr><td>Oligo fwd</td><td>CACCGCGAGGATGCACGGCCTGGA</td><td></td><td></td><td></td><td></td><td></td><td></td><td></td><td></td><td></td><td></td></tr>
<tr><td>Oligo rev</td><td>AAACTCCAGGCCGTGCATCCTCGC</td><td></td><td></td><td></td><td></td><td></td><td></td><td></td><td></td><td></td><td></td></tr>
<tr><td>Chromosome</td><td>start</td><td>end</td><td>strand</td><td>MM</td><td>target_seq</td><td>PAM</td><td>alignment</td><td>distance</td><td>position</td><td>gene name</td><td>gene id</td></tr>
<tr><td>GU305914.1</td><td>46941</td><td>46963</td><td>+</td><td>0</td><td>GCGAGGATGCACGGCCTGGA</td><td>GGG</td><td>||||||||[||||||||||||]PAM</td><td>NA</td><td>Intergenic</td><td>NA</td><td>NA</td></tr>
<tr><td>T3022</td><td>CGAGGATGCACGGCCTGGAGGGG</td><td>1000</td><td></td><td></td><td></td><td></td><td></td><td></td><td></td><td></td><td></td></tr>
<tr><td>Oligo fwd</td><td>CACCCGAGGATGCACGGCCTGGAG</td><td></td><td></td><td></td><td></td><td></td><td></td><td></td><td></td><td></td><td></td></tr>
<tr><td>Oligo rev</td><td>AAACCTCCAGGCCGTGCATCCTCG</td><td></td><td></td><td></td><td></td><td></td><td></td><td></td><td></td><td></td><td></td></tr>
<tr><td>Chromosome</td><td>start</td><td>end</td><td>strand</td><td>MM</td><td>target_seq</td><td>PAM</td><td>alignment</td><td>distance</td><td>position</td><td>gene name</td><td>gene id</td></tr>
<tr><td>GU305914.1</td><td>46942</td><td>46964</td><td>+</td><td>0</td><td>CGAGGATGCACGGCCTGGAG</td><td>GGG</td><td>||||||||[||||||||||||]PAM</td><td>NA</td><td>Intergenic</td><td>NA</td><td>NA</td></tr>
<tr><td>T3023</td><td>GAGGATGCACGGCCTGGAGGGGG</td><td>1000</td><td></td><td></td><td></td><td></td><td></td><td></td><td></td><td></td><td></td></tr>
<tr><td>Oligo fwd</td><td>CACCGAGGATGCACGGCCTGGAGG</td><td></td><td></td><td></td><td></td><td></td><td></td><td></td><td></td><td></td><td></td></tr>
<tr><td>Oligo rev</td><td>AAACCCTCCAGGCCGTGCATCCTC</td><td></td><td></td><td></td><td></td><td></td><td></td><td></td><td></td><td></td><td></td></tr>
<tr><td>Chromosome</td><td>start</td><td>end</td><td>strand</td><td>MM</td><td>target_seq</td><td>PAM</td><td>alignment</td><td>distance</td><td>position</td><td>gene name</td><td>gene id</td></tr>
<tr><td>GU305914.1</td><td>46943</td><td>46965</td><td>+</td><td>0</td><td>GAGGATGCACGGCCTGGAGG</td><td>GGG</td><td>||||||||[||||||||||||]PAM</td><td>NA</td><td>Intergenic</td><td>NA</td><td>NA</td></tr>
<tr><td>T3024</td><td>CGGCCTGGAGGGGGCCGCGATGG</td><td>1000</td><td></td><td></td><td></td><td></td><td></td><td></td><td></td><td></td><td></td></tr>
<tr><td>Oligo fwd</td><td>CACCCGGCCTGGAGGGGGCCGCGA</td><td></td><td></td><td></td><td></td><td></td><td></td><td></td><td></td><td></td><td></td></tr>
<tr><td>Oligo rev</td><td>AAACTCGCGGCCCCCTCCAGGCCG</td><td></td><td></td><td></td><td></td><td></td><td></td><td></td><td></td><td></td><td></td></tr>
<tr><td>Chromosome</td><td>start</td><td>end</td><td>strand</td><td>MM</td><td>target_seq</td><td>PAM</td><td>alignment</td><td>distance</td><td>position</td><td>gene name</td><td>gene id</td></tr>
<tr><td>GU305914.1</td><td>46952</td><td>46974</td><td>+</td><td>0</td><td>CGGCCTGGAGGGGGCCGCGA</td><td>TGG</td><td>||||||||[||||||||||||]PAM</td><td>NA</td><td>Intergenic</td><td>NA</td><td>NA</td></tr>
<tr><td>T3025</td><td>GGCCTGGAGGGGGCCGCGATGGG</td><td>1000</td><td></td><td></td><td></td><td></td><td></td><td></td><td></td><td></td><td></td></tr>
<tr><td>Oligo fwd</td><td>CACCGGCCTGGAGGGGGCCGCGAT</td><td></td><td></td><td></td><td></td><td></td><td></td><td></td><td></td><td></td><td></td></tr>
<tr><td>Oligo rev</td><td>AAACATCGCGGCCCCCTCCAGGCC</td><td></td><td></td><td></td><td></td><td></td><td></td><td></td><td></td><td></td><td></td></tr>
<tr><td>Chromosome</td><td>start</td><td>end</td><td>strand</td><td>MM</td><td>target_seq</td><td>PAM</td><td>alignment</td><td>distance</td><td>position</td><td>gene name</td><td>gene id</td></tr>
<tr><td>GU305914.1</td><td>46953</td><td>46975</td><td>+</td><td>0</td><td>GGCCTGGAGGGGGCCGCGAT</td><td>GGG</td><td>||||||||[||||||||||||]PAM</td><td>NA</td><td>Intergenic</td><td>NA</td><td>NA</td></tr>
<tr><td>T3026</td><td>GATGGGTCACGACGACGACCTGG</td><td>1000</td><td></td><td></td><td></td><td></td><td></td><td></td><td></td><td></td><td></td></tr>
<tr><td>Oligo fwd</td><td>CACCGATGGGTCACGACGACGACC</td><td></td><td></td><td></td><td></td><td></td><td></td><td></td><td></td><td></td><td></td></tr>
<tr><td>Oligo rev</td><td>AAACGGTCGTCGTCGTGACCCATC</td><td></td><td></td><td></td><td></td><td></td><td></td><td></td><td></td><td></td><td></td></tr>
<tr><td>Chromosome</td><td>start</td><td>end</td><td>strand</td><td>MM</td><td>target_seq</td><td>PAM</td><td>alignment</td><td>distance</td><td>position</td><td>gene name</td><td>gene id</td></tr>
<tr><td>GU305914.1</td><td>46970</td><td>46992</td><td>+</td><td>0</td><td>GATGGGTCACGACGACGACC</td><td>TGG</td><td>||||||||[||||||||||||]PAM</td><td>NA</td><td>Intergenic</td><td>NA</td><td>NA</td></tr>
<tr><td>T3027</td><td>CGACGACGACCTGGACATCGAGG</td><td>1000</td><td></td><td></td><td></td><td></td><td></td><td></td><td></td><td></td><td></td></tr>
<tr><td>Oligo fwd</td><td>CACCCGACGACGACCTGGACATCG</td><td></td><td></td><td></td><td></td><td></td><td></td><td></td><td></td><td></td><td></td></tr>
<tr><td>Oligo rev</td><td>AAACCGATGTCCAGGTCGTCGTCG</td><td></td><td></td><td></td><td></td><td></td><td></td><td></td><td></td><td></td><td></td></tr>
<tr><td>Chromosome</td><td>start</td><td>end</td><td>strand</td><td>MM</td><td>target_seq</td><td>PAM</td><td>alignment</td><td>distance</td><td>position</td><td>gene name</td><td>gene id</td></tr>
<tr><td>GU305914.1</td><td>46979</td><td>47001</td><td>+</td><td>0</td><td>CGACGACGACCTGGACATCG</td><td>AGG</td><td>||||||||[||||||||||||]PAM</td><td>NA</td><td>Intergenic</td><td>NA</td><td>NA</td></tr>
<tr><td>T3028</td><td>CGACGACCTGGACATCGAGGTGG</td><td>1000</td><td></td><td></td><td></td><td></td><td></td><td></td><td></td><td></td><td></td></tr>
<tr><td>Oligo fwd</td><td>CACCCGACGACCTGGACATCGAGG</td><td></td><td></td><td></td><td></td><td></td><td></td><td></td><td></td><td></td><td></td></tr>
<tr><td>Oligo rev</td><td>AAACCCTCGATGTCCAGGTCGTCG</td><td></td><td></td><td></td><td></td><td></td><td></td><td></td><td></td><td></td><td></td></tr>
<tr><td>Chromosome</td><td>start</td><td>end</td><td>strand</td><td>MM</td><td>target_seq</td><td>PAM</td><td>alignment</td><td>distance</td><td>position</td><td>gene name</td><td>gene id</td></tr>
<tr><td>GU305914.1</td><td>46982</td><td>47004</td><td>+</td><td>0</td><td>CGACGACCTGGACATCGAGG</td><td>TGG</td><td>||||||||[||||||||||||]PAM</td><td>NA</td><td>Intergenic</td><td>NA</td><td>NA</td></tr>
<tr><td>T3029</td><td>CCTGGACATCGAGGTGGACCCGG</td><td>1000</td><td></td><td></td><td></td><td></td><td></td><td></td><td></td><td></td><td></td></tr>
<tr><td>Oligo fwd</td><td>CACCCCTGGACATCGAGGTGGACC</td><td></td><td></td><td></td><td></td><td></td><td></td><td></td><td></td><td></td><td></td></tr>
<tr><td>Oligo rev</td><td>AAACGGTCCACCTCGATGTCCAGG</td><td></td><td></td><td></td><td></td><td></td><td></td><td></td><td></td><td></td><td></td></tr>
<tr><td>Chromosome</td><td>start</td><td>end</td><td>strand</td><td>MM</td><td>target_seq</td><td>PAM</td><td>alignment</td><td>distance</td><td>position</td><td>gene name</td><td>gene id</td></tr>
<tr><td>GU305914.1</td><td>46988</td><td>47010</td><td>+</td><td>0</td><td>CCTGGACATCGAGGTGGACC</td><td>CGG</td><td>||||||||[||||||||||||]PAM</td><td>NA</td><td>Intergenic</td><td>NA</td><td>NA</td></tr>
<tr><td>GU305914.1</td><td>121337</td><td>121359</td><td>-</td><td>4</td><td>CTCGGACATCTACGTGGACC</td><td>AGG</td><td>|--|||||[||-|-|||||||]PAM</td><td>NA</td><td>Intergenic</td><td>NA</td><td>NA</td></tr>
<tr><td>T3030</td><td>CTGGACATCGAGGTGGACCCGGG</td><td>1000</td><td></td><td></td><td></td><td></td><td></td><td></td><td></td><td></td><td></td></tr>
<tr><td>Oligo fwd</td><td>CACCCTGGACATCGAGGTGGACCC</td><td></td><td></td><td></td><td></td><td></td><td></td><td></td><td></td><td></td><td></td></tr>
<tr><td>Oligo rev</td><td>AAACGGGTCCACCTCGATGTCCAG</td><td></td><td></td><td></td><td></td><td></td><td></td><td></td><td></td><td></td><td></td></tr>
<tr><td>Chromosome</td><td>start</td><td>end</td><td>strand</td><td>MM</td><td>target_seq</td><td>PAM</td><td>alignment</td><td>distance</td><td>position</td><td>gene name</td><td>gene id</td></tr>
<tr><td>GU305914.1</td><td>46989</td><td>47011</td><td>+</td><td>0</td><td>CTGGACATCGAGGTGGACCC</td><td>GGG</td><td>||||||||[||||||||||||]PAM</td><td>NA</td><td>Intergenic</td><td>NA</td><td>NA</td></tr>
<tr><td>T3031</td><td>AGAACATCTTCAAGTGCCCCAGG</td><td>1000</td><td></td><td></td><td></td><td></td><td></td><td></td><td></td><td></td><td></td></tr>
<tr><td>Oligo fwd</td><td>CACCAGAACATCTTCAAGTGCCCC</td><td></td><td></td><td></td><td></td><td></td><td></td><td></td><td></td><td></td><td></td></tr>
<tr><td>Oligo rev</td><td>AAACGGGGCACTTGAAGATGTTCT</td><td></td><td></td><td></td><td></td><td></td><td></td><td></td><td></td><td></td><td></td></tr>
<tr><td>Chromosome</td><td>start</td><td>end</td><td>strand</td><td>MM</td><td>target_seq</td><td>PAM</td><td>alignment</td><td>distance</td><td>position</td><td>gene name</td><td>gene id</td></tr>
<tr><td>GU305914.1</td><td>47059</td><td>47081</td><td>+</td><td>0</td><td>AGAACATCTTCAAGTGCCCC</td><td>AGG</td><td>||||||||[||||||||||||]PAM</td><td>NA</td><td>Intergenic</td><td>NA</td><td>NA</td></tr>
<tr><td>T3032</td><td>CTTCAAGTGCCCCAGGAGCCTGG</td><td>1000</td><td></td><td></td><td></td><td></td><td></td><td></td><td></td><td></td><td></td></tr>
<tr><td>Oligo fwd</td><td>CACCCTTCAAGTGCCCCAGGAGCC</td><td></td><td></td><td></td><td></td><td></td><td></td><td></td><td></td><td></td><td></td></tr>
<tr><td>Oligo rev</td><td>AAACGGCTCCTGGGGCACTTGAAG</td><td></td><td></td><td></td><td></td><td></td><td></td><td></td><td></td><td></td><td></td></tr>
<tr><td>Chromosome</td><td>start</td><td>end</td><td>strand</td><td>MM</td><td>target_seq</td><td>PAM</td><td>alignment</td><td>distance</td><td>position</td><td>gene name</td><td>gene id</td></tr>
<tr><td>GU305914.1</td><td>47066</td><td>47088</td><td>+</td><td>0</td><td>CTTCAAGTGCCCCAGGAGCC</td><td>TGG</td><td>||||||||[||||||||||||]PAM</td><td>NA</td><td>Intergenic</td><td>NA</td><td>NA</td></tr>
<tr><td>T3033</td><td>CAGGAGCCTGGTGCATTACGTGG</td><td>1000</td><td></td><td></td><td></td><td></td><td></td><td></td><td></td><td></td><td></td></tr>
<tr><td>Oligo fwd</td><td>CACCCAGGAGCCTGGTGCATTACG</td><td></td><td></td><td></td><td></td><td></td><td></td><td></td><td></td><td></td><td></td></tr>
<tr><td>Oligo rev</td><td>AAACCGTAATGCACCAGGCTCCTG</td><td></td><td></td><td></td><td></td><td></td><td></td><td></td><td></td><td></td><td></td></tr>
<tr><td>Chromosome</td><td>start</td><td>end</td><td>strand</td><td>MM</td><td>target_seq</td><td>PAM</td><td>alignment</td><td>distance</td><td>position</td><td>gene name</td><td>gene id</td></tr>
<tr><td>GU305914.1</td><td>47078</td><td>47100</td><td>+</td><td>0</td><td>CAGGAGCCTGGTGCATTACG</td><td>TGG</td><td>||||||||[||||||||||||]PAM</td><td>NA</td><td>Intergenic</td><td>NA</td><td>NA</td></tr>
<tr><td>T3034</td><td>AGGAGCCTGGTGCATTACGTGGG</td><td>1000</td><td></td><td></td><td></td><td></td><td></td><td></td><td></td><td></td><td></td></tr>
<tr><td>Oligo fwd</td><td>CACCAGGAGCCTGGTGCATTACGT</td><td></td><td></td><td></td><td></td><td></td><td></td><td></td><td></td><td></td><td></td></tr>
<tr><td>Oligo rev</td><td>AAACACGTAATGCACCAGGCTCCT</td><td></td><td></td><td></td><td></td><td></td><td></td><td></td><td></td><td></td><td></td></tr>
<tr><td>Chromosome</td><td>start</td><td>end</td><td>strand</td><td>MM</td><td>target_seq</td><td>PAM</td><td>alignment</td><td>distance</td><td>position</td><td>gene name</td><td>gene id</td></tr>
<tr><td>GU305914.1</td><td>47079</td><td>47101</td><td>+</td><td>0</td><td>AGGAGCCTGGTGCATTACGT</td><td>GGG</td><td>||||||||[||||||||||||]PAM</td><td>NA</td><td>Intergenic</td><td>NA</td><td>NA</td></tr>
<tr><td>T3035</td><td>GTTCAAGAAGACCATGACCCAGG</td><td>1000</td><td></td><td></td><td></td><td></td><td></td><td></td><td></td><td></td><td></td></tr>
</table>

<table>
<tr><td>Oligo fwd</td><td>CACCGTTCAAGAAGACCATGACCC</td><td></td><td></td><td></td><td></td><td></td><td></td><td></td><td></td><td></td><td></td><td></td></tr>
<tr><td>Oligo rev</td><td>AAACGGGTCATGGTCTTCTTGAAC</td><td></td><td></td><td></td><td></td><td></td><td></td><td></td><td></td><td></td><td></td><td></td></tr>
<tr><td>Chromosome</td><td>start</td><td></td><td>end</td><td>strand</td><td>MM</td><td>target_seq</td><td>PAM</td><td>alignment</td><td>distance</td><td>position</td><td>gene name</td><td>gene id</td></tr>
<tr><td>GU305914.1</td><td></td><td>47111</td><td>47133</td><td>+</td><td>0</td><td>GTTCAAGAAGACCATGACCC</td><td>AGG</td><td>||||||||[||||||||||||]PAM</td><td>NA</td><td>Intergenic</td><td>NA</td><td>NA</td></tr>
<tr><td>T3036</td><td>GGAGCTGCTAGTGAGCTGCACGG</td><td></td><td>1000</td><td></td><td></td><td></td><td></td><td></td><td></td><td></td><td></td><td></td></tr>
<tr><td>Oligo fwd</td><td>CACCGGAGCTGCTAGTGAGCTGCA</td><td></td><td></td><td></td><td></td><td></td><td></td><td></td><td></td><td></td><td></td><td></td></tr>
<tr><td>Oligo rev</td><td>AAACTGCAGCTCACTAGCAGCTCC</td><td></td><td></td><td></td><td></td><td></td><td></td><td></td><td></td><td></td><td></td><td></td></tr>
<tr><td>Chromosome</td><td>start</td><td></td><td>end</td><td>strand</td><td>MM</td><td>target_seq</td><td>PAM</td><td>alignment</td><td>distance</td><td>position</td><td>gene name</td><td>gene id</td></tr>
<tr><td>GU305914.1</td><td></td><td>47132</td><td>47154</td><td>+</td><td>0</td><td>GGAGCTGCTAGTGAGCTGCA</td><td>CGG</td><td>||||||||[||||||||||||]PAM</td><td>NA</td><td>Intergenic</td><td>NA</td><td>NA</td></tr>
<tr><td>T3037</td><td>AGTGAGCTGCACGGACCAAGAGG</td><td></td><td>1000</td><td></td><td></td><td></td><td></td><td></td><td></td><td></td><td></td><td></td></tr>
<tr><td>Oligo fwd</td><td>CACCAGTGAGCTGCACGGACCAAG</td><td></td><td></td><td></td><td></td><td></td><td></td><td></td><td></td><td></td><td></td><td></td></tr>
<tr><td>Oligo rev</td><td>AAACCTTGGTCCGTGCAGCTCACT</td><td></td><td></td><td></td><td></td><td></td><td></td><td></td><td></td><td></td><td></td><td></td></tr>
<tr><td>Chromosome</td><td>start</td><td></td><td>end</td><td>strand</td><td>MM</td><td>target_seq</td><td>PAM</td><td>alignment</td><td>distance</td><td>position</td><td>gene name</td><td>gene id</td></tr>
<tr><td>GU305914.1</td><td></td><td>47141</td><td>47163</td><td>+</td><td>0</td><td>AGTGAGCTGCACGGACCAAG</td><td>AGG</td><td>||||||||[||||||||||||]PAM</td><td>NA</td><td>Intergenic</td><td>NA</td><td>NA</td></tr>
<tr><td>T3038</td><td>CCAGACCTACGAGCTGCCGTCGG</td><td></td><td>1000</td><td></td><td></td><td></td><td></td><td></td><td></td><td></td><td></td><td></td></tr>
<tr><td>Oligo fwd</td><td>CACCCCAGACCTACGAGCTGCCGT</td><td></td><td></td><td></td><td></td><td></td><td></td><td></td><td></td><td></td><td></td><td></td></tr>
<tr><td>Oligo rev</td><td>AAACACGGCAGCTCGTAGGTCTGG</td><td></td><td></td><td></td><td></td><td></td><td></td><td></td><td></td><td></td><td></td><td></td></tr>
<tr><td>Chromosome</td><td>start</td><td></td><td>end</td><td>strand</td><td>MM</td><td>target_seq</td><td>PAM</td><td>alignment</td><td>distance</td><td>position</td><td>gene name</td><td>gene id</td></tr>
<tr><td>GU305914.1</td><td></td><td>47171</td><td>47193</td><td>+</td><td>0</td><td>CCAGACCTACGAGCTGCCGT</td><td>CGG</td><td>||||||||[||||||||||||]PAM</td><td>NA</td><td>Intergenic</td><td>NA</td><td>NA</td></tr>
<tr><td>T3039</td><td>GCCGTCGGTCTCCGAGCTCGTGG</td><td></td><td>1000</td><td></td><td></td><td></td><td></td><td></td><td></td><td></td><td></td><td></td></tr>
<tr><td>Oligo fwd</td><td>CACCGCCGTCGGTCTCCGAGCTCG</td><td></td><td></td><td></td><td></td><td></td><td></td><td></td><td></td><td></td><td></td><td></td></tr>
<tr><td>Oligo rev</td><td>AAACCGAGCTCGGAGACCGACGGC</td><td></td><td></td><td></td><td></td><td></td><td></td><td></td><td></td><td></td><td></td><td></td></tr>
<tr><td>Chromosome</td><td>start</td><td></td><td>end</td><td>strand</td><td>MM</td><td>target_seq</td><td>PAM</td><td>alignment</td><td>distance</td><td>position</td><td>gene name</td><td>gene id</td></tr>
<tr><td>GU305914.1</td><td></td><td>47186</td><td>47208</td><td>+</td><td>0</td><td>GCCGTCGGTCTCCGAGCTCG</td><td>TGG</td><td>||||||||[||||||||||||]PAM</td><td>NA</td><td>Intergenic</td><td>NA</td><td>NA</td></tr>
<tr><td>T3040</td><td>CCGTCGGTCTCCGAGCTCGTGGG</td><td></td><td>1000</td><td></td><td></td><td></td><td></td><td></td><td></td><td></td><td></td><td></td></tr>
<tr><td>Oligo fwd</td><td>CACCCCGTCGGTCTCCGAGCTCGT</td><td></td><td></td><td></td><td></td><td></td><td></td><td></td><td></td><td></td><td></td><td></td></tr>
<tr><td>Oligo rev</td><td>AAACACGAGCTCGGAGACCGACGG</td><td></td><td></td><td></td><td></td><td></td><td></td><td></td><td></td><td></td><td></td><td></td></tr>
<tr><td>Chromosome</td><td>start</td><td></td><td>end</td><td>strand</td><td>MM</td><td>target_seq</td><td>PAM</td><td>alignment</td><td>distance</td><td>position</td><td>gene name</td><td>gene id</td></tr>
<tr><td>GU305914.1</td><td></td><td>47187</td><td>47209</td><td>+</td><td>0</td><td>CCGTCGGTCTCCGAGCTCGT</td><td>GGG</td><td>||||||||[||||||||||||]PAM</td><td>NA</td><td>Intergenic</td><td>NA</td><td>NA</td></tr>
<tr><td>T3041</td><td>GTCTCCGAGCTCGTGGGCGAAGG</td><td></td><td>1000</td><td></td><td></td><td></td><td></td><td></td><td></td><td></td><td></td><td></td></tr>
<tr><td>Oligo fwd</td><td>CACCGTCTCCGAGCTCGTGGGCGA</td><td></td><td></td><td></td><td></td><td></td><td></td><td></td><td></td><td></td><td></td><td></td></tr>
<tr><td>Oligo rev</td><td>AAACTCGCCCACGAGCTCGGAGAC</td><td></td><td></td><td></td><td></td><td></td><td></td><td></td><td></td><td></td><td></td><td></td></tr>
<tr><td>Chromosome</td><td>start</td><td></td><td>end</td><td>strand</td><td>MM</td><td>target_seq</td><td>PAM</td><td>alignment</td><td>distance</td><td>position</td><td>gene name</td><td>gene id</td></tr>
<tr><td>GU305914.1</td><td></td><td>47193</td><td>47215</td><td>+</td><td>0</td><td>GTCTCCGAGCTCGTGGGCGA</td><td>AGG</td><td>||||||||[||||||||||||]PAM</td><td>NA</td><td>Intergenic</td><td>NA</td><td>NA</td></tr>
<tr><td>GU305914.1</td><td></td><td>40508</td><td>40530</td><td>-</td><td>4</td><td>GAGTCCGAGCTCGAGAGCGA</td><td>GGG</td><td>|--|||||[|||||-|-||||]PAM</td><td>NA</td><td>Intergenic</td><td>NA</td><td>NA</td></tr>
<tr><td>T3042</td><td>CGAGCTCGTGGGCGAAGGCGTGG</td><td></td><td>1000</td><td></td><td></td><td></td><td></td><td></td><td></td><td></td><td></td><td></td></tr>
<tr><td>Oligo fwd</td><td>CACCCGAGCTCGTGGGCGAAGGCG</td><td></td><td></td><td></td><td></td><td></td><td></td><td></td><td></td><td></td><td></td><td></td></tr>
<tr><td>Oligo rev</td><td>AAACCGCCTTCGCCCACGAGCTCG</td><td></td><td></td><td></td><td></td><td></td><td></td><td></td><td></td><td></td><td></td><td></td></tr>
<tr><td>Chromosome</td><td>start</td><td></td><td>end</td><td>strand</td><td>MM</td><td>target_seq</td><td>PAM</td><td>alignment</td><td>distance</td><td>position</td><td>gene name</td><td>gene id</td></tr>
<tr><td>GU305914.1</td><td></td><td>47198</td><td>47220</td><td>+</td><td>0</td><td>CGAGCTCGTGGGCGAAGGCG</td><td>TGG</td><td>||||||||[||||||||||||]PAM</td><td>NA</td><td>Intergenic</td><td>NA</td><td>NA</td></tr>
<tr><td>T3043</td><td>TGGGCGAAGGCGTGGCGAAACGG</td><td></td><td>1000</td><td></td><td></td><td></td><td></td><td></td><td></td><td></td><td></td><td></td></tr>
<tr><td>Oligo fwd</td><td>CACCTGGGCGAAGGCGTGGCGAAA</td><td></td><td></td><td></td><td></td><td></td><td></td><td></td><td></td><td></td><td></td><td></td></tr>
<tr><td>Oligo rev</td><td>AAACTTTCGCCACGCCTTCGCCCA</td><td></td><td></td><td></td><td></td><td></td><td></td><td></td><td></td><td></td><td></td><td></td></tr>
<tr><td>Chromosome</td><td>start</td><td></td><td>end</td><td>strand</td><td>MM</td><td>target_seq</td><td>PAM</td><td>alignment</td><td>distance</td><td>position</td><td>gene name</td><td>gene id</td></tr>
<tr><td>GU305914.1</td><td></td><td>47206</td><td>47228</td><td>+</td><td>0</td><td>TGGGCGAAGGCGTGGCGAAA</td><td>CGG</td><td>||||||||[||||||||||||]PAM</td><td>NA</td><td>Intergenic</td><td>NA</td><td>NA</td></tr>
<tr><td>T3044</td><td>GGGCGAAGGCGTGGCGAAACGGG</td><td></td><td>1000</td><td></td><td></td><td></td><td></td><td></td><td></td><td></td><td></td><td></td></tr>
<tr><td>Oligo fwd</td><td>CACCGGGCGAAGGCGTGGCGAAAC</td><td></td><td></td><td></td><td></td><td></td><td></td><td></td><td></td><td></td><td></td><td></td></tr>
<tr><td>Oligo rev</td><td>AAACGTTTCGCCACGCCTTCGCCC</td><td></td><td></td><td></td><td></td><td></td><td></td><td></td><td></td><td></td><td></td><td></td></tr>
<tr><td>Chromosome</td><td>start</td><td></td><td>end</td><td>strand</td><td>MM</td><td>target_seq</td><td>PAM</td><td>alignment</td><td>distance</td><td>position</td><td>gene name</td><td>gene id</td></tr>
<tr><td>GU305914.1</td><td></td><td>47207</td><td>47229</td><td>+</td><td>0</td><td>GGGCGAAGGCGTGGCGAAAC</td><td>GGG</td><td>||||||||[||||||||||||]PAM</td><td>NA</td><td>Intergenic</td><td>NA</td><td>NA</td></tr>
<tr><td>T3045</td><td>GCTGTCGCACTACCTGTCCGTGG</td><td></td><td>1000</td><td></td><td></td><td></td><td></td><td></td><td></td><td></td><td></td><td></td></tr>
<tr><td>Oligo fwd</td><td>CACCGCTGTCGCACTACCTGTCCG</td><td></td><td></td><td></td><td></td><td></td><td></td><td></td><td></td><td></td><td></td><td></td></tr>
<tr><td>Oligo rev</td><td>AAACCGGACAGGTAGTGCGACAGC</td><td></td><td></td><td></td><td></td><td></td><td></td><td></td><td></td><td></td><td></td><td></td></tr>
<tr><td>Chromosome</td><td>start</td><td></td><td>end</td><td>strand</td><td>MM</td><td>target_seq</td><td>PAM</td><td>alignment</td><td>distance</td><td>position</td><td>gene name</td><td>gene id</td></tr>
<tr><td>GU305914.1</td><td></td><td>47249</td><td>47271</td><td>+</td><td>0</td><td>GCTGTCGCACTACCTGTCCG</td><td>TGG</td><td>||||||||[||||||||||||]PAM</td><td>NA</td><td>Intergenic</td><td>NA</td><td>NA</td></tr>
<tr><td>T3046</td><td>CCTGTCCGTGGTAAGCACCCCGG</td><td></td><td>1000</td><td></td><td></td><td></td><td></td><td></td><td></td><td></td><td></td><td></td></tr>
<tr><td>Oligo fwd</td><td>CACCCCTGTCCGTGGTAAGCACCC</td><td></td><td></td><td></td><td></td><td></td><td></td><td></td><td></td><td></td><td></td><td></td></tr>
<tr><td>Oligo rev</td><td>AAACGGGTGCTTACCACGGACAGG</td><td></td><td></td><td></td><td></td><td></td><td></td><td></td><td></td><td></td><td></td><td></td></tr>
<tr><td>Chromosome</td><td>start</td><td></td><td>end</td><td>strand</td><td>MM</td><td>target_seq</td><td>PAM</td><td>alignment</td><td>distance</td><td>position</td><td>gene name</td><td>gene id</td></tr>
<tr><td>GU305914.1</td><td></td><td>47261</td><td>47283</td><td>+</td><td>0</td><td>CCTGTCCGTGGTAAGCACCC</td><td>CGG</td><td>||||||||[||||||||||||]PAM</td><td>NA</td><td>Intergenic</td><td>NA</td><td>NA</td></tr>
<tr><td>T3047</td><td>GTCACAGACGACATCGAAGACGG</td><td></td><td>1000</td><td></td><td></td><td></td><td></td><td></td><td></td><td></td><td></td><td></td></tr>
<tr><td>Oligo fwd</td><td>CACCGTCACAGACGACATCGAAGA</td><td></td><td></td><td></td><td></td><td></td><td></td><td></td><td></td><td></td><td></td><td></td></tr>
<tr><td>Oligo rev</td><td>AAACTCTTCGATGTCGTCTGTGAC</td><td></td><td></td><td></td><td></td><td></td><td></td><td></td><td></td><td></td><td></td><td></td></tr>
<tr><td>Chromosome</td><td>start</td><td></td><td>end</td><td>strand</td><td>MM</td><td>target_seq</td><td>PAM</td><td>alignment</td><td>distance</td><td>position</td><td>gene name</td><td>gene id</td></tr>
<tr><td>GU305914.1</td><td></td><td>47292</td><td>47314</td><td>+</td><td>0</td><td>GTCACAGACGACATCGAAGA</td><td>CGG</td><td>||||||||[||||||||||||]PAM</td><td>NA</td><td>Intergenic</td><td>NA</td><td>NA</td></tr>
<tr><td>T3048</td><td>CGACATCGAAGACGGCGAACAGG</td><td></td><td>1000</td><td></td><td></td><td></td><td></td><td></td><td></td><td></td><td></td><td></td></tr>
<tr><td>Oligo fwd</td><td>CACCCGACATCGAAGACGGCGAAC</td><td></td><td></td><td></td><td></td><td></td><td></td><td></td><td></td><td></td><td></td><td></td></tr>
<tr><td>Oligo rev</td><td>AAACGTTCGCCGTCTTCGATGTCG</td><td></td><td></td><td></td><td></td><td></td><td></td><td></td><td></td><td></td><td></td><td></td></tr>
<tr><td>Chromosome</td><td>start</td><td></td><td>end</td><td>strand</td><td>MM</td><td>target_seq</td><td>PAM</td><td>alignment</td><td>distance</td><td>position</td><td>gene name</td><td>gene id</td></tr>
<tr><td>GU305914.1</td><td></td><td>47300</td><td>47322</td><td>+</td><td>0</td><td>CGACATCGAAGACGGCGAAC</td><td>AGG</td><td>||||||||[||||||||||||]PAM</td><td>NA</td><td>Intergenic</td><td>NA</td><td>NA</td></tr>
<tr><td>T3049</td><td>CATCGAAGACGGCGAACAGGAGG</td><td></td><td>1000</td><td></td><td></td><td></td><td></td><td></td><td></td><td></td><td></td><td></td></tr>
<tr><td>Oligo fwd</td><td>CACCCATCGAAGACGGCGAACAGG</td><td></td><td></td><td></td><td></td><td></td><td></td><td></td><td></td><td></td><td></td><td></td></tr>
<tr><td>Oligo rev</td><td>AAACCCTGTTCGCCGTCTTCGATG</td><td></td><td></td><td></td><td></td><td></td><td></td><td></td><td></td><td></td><td></td><td></td></tr>
<tr><td>Chromosome</td><td>start</td><td></td><td>end</td><td>strand</td><td>MM</td><td>target_seq</td><td>PAM</td><td>alignment</td><td>distance</td><td>position</td><td>gene name</td><td>gene id</td></tr>
<tr><td>GU305914.1</td><td></td><td>47303</td><td>47325</td><td>+</td><td>0</td><td>CATCGAAGACGGCGAACAGG</td><td>AGG</td><td>||||||||[||||||||||||]PAM</td><td>NA</td><td>Intergenic</td><td>NA</td><td>NA</td></tr>
</table>

| | | | | | | | | | | | |
|---|---|---|---|---|---|---|---|---|---|---|---|
| T3050 | CGAAGACGGCGAACAGGAGGAGG | 1000 | | | | | | | | | |
| Oligo fwd | CACCCGAAGACGGCGAACAGGAGG | | | | | | | | | | |
| Oligo rev | AAACCCTCCTGTTCGCCGTCTTCG | | | | | | | | | | |
| Chromosome | start | end | strand | MM | target_seq | PAM | alignment | distance | position | gene name | gene id |
| GU305914.1 | 47306 | 47328 | + | 0 | CGAAGACGGCGAACAGGAGG | AGG | \|\|\|\|\|\|\|\|[\|\|\|\|\|\|\|\|\|\|\|\|]PAM | NA | Intergenic | NA | NA |

| | | | | | | | | | | | |
|---|---|---|---|---|---|---|---|---|---|---|---|
| T3051 | GGATCTGTTCGCCGACGTGTCGG | 1000 | | | | | | | | | |
| Oligo fwd | CACCGGATCTGTTCGCCGACGTGT | | | | | | | | | | |
| Oligo rev | AAACACACGTCGGCGAACAGATCC | | | | | | | | | | |
| Chromosome | start | end | strand | MM | target_seq | PAM | alignment | distance | position | gene name | gene id |
| GU305914.1 | 47327 | 47349 | + | 0 | GGATCTGTTCGCCGACGTGT | CGG | \|\|\|\|\|\|\|\|[\|\|\|\|\|\|\|\|\|\|\|\|]PAM | NA | Intergenic | NA | NA |

| | | | | | | | | | | | |
|---|---|---|---|---|---|---|---|---|---|---|---|
| T3052 | TCTGTTCGCCGACGTGTCGGTGG | 1000 | | | | | | | | | |
| Oligo fwd | CACCTCTGTTCGCCGACGTGTCGG | | | | | | | | | | |
| Oligo rev | AAACCCGACACGTCGGCGAACAGA | | | | | | | | | | |
| Chromosome | start | end | strand | MM | target_seq | PAM | alignment | distance | position | gene name | gene id |
| GU305914.1 | 47330 | 47352 | + | 0 | TCTGTTCGCCGACGTGTCGG | TGG | \|\|\|\|\|\|\|\|[\|\|\|\|\|\|\|\|\|\|\|\|]PAM | NA | Intergenic | NA | NA |

| | | | | | | | | | | | |
|---|---|---|---|---|---|---|---|---|---|---|---|
| T3053 | CGCCGACGTGTCGGTGGCCCCGG | 1000 | | | | | | | | | |
| Oligo fwd | CACCCGCCGACGTGTCGGTGGCCC | | | | | | | | | | |
| Oligo rev | AAACGGGCCACCGACACGTCGGCG | | | | | | | | | | |
| Chromosome | start | end | strand | MM | target_seq | PAM | alignment | distance | position | gene name | gene id |
| GU305914.1 | 47336 | 47358 | + | 0 | CGCCGACGTGTCGGTGGCCC | CGG | \|\|\|\|\|\|\|\|[\|\|\|\|\|\|\|\|\|\|\|\|]PAM | NA | Intergenic | NA | NA |

| | | | | | | | | | | | |
|---|---|---|---|---|---|---|---|---|---|---|---|
| T3054 | TGGCCCCGGCGTCGCCCGTGAGG | 1000 | | | | | | | | | |
| Oligo fwd | CACCTGGCCCCGGCGTCGCCCGTG | | | | | | | | | | |
| Oligo rev | AAACCACGGGCGACGCCGGGGCCA | | | | | | | | | | |
| Chromosome | start | end | strand | MM | target_seq | PAM | alignment | distance | position | gene name | gene id |
| GU305914.1 | 47350 | 47372 | + | 0 | TGGCCCCGGCGTCGCCCGTG | AGG | \|\|\|\|\|\|\|\|[\|\|\|\|\|\|\|\|\|\|\|\|]PAM | NA | Intergenic | NA | NA |

| | | | | | | | | | | | |
|---|---|---|---|---|---|---|---|---|---|---|---|
| T3055 | TCGCCCGTGAGGCCGATCCGCGG | 1000 | | | | | | | | | |
| Oligo fwd | CACCTCGCCCGTGAGGCCGATCCG | | | | | | | | | | |
| Oligo rev | AAACCGGATCGGCCTCACGGGCGA | | | | | | | | | | |
| Chromosome | start | end | strand | MM | target_seq | PAM | alignment | distance | position | gene name | gene id |
| GU305914.1 | 47361 | 47383 | + | 0 | TCGCCCGTGAGGCCGATCCG | CGG | \|\|\|\|\|\|\|\|[\|\|\|\|\|\|\|\|\|\|\|\|]PAM | NA | Intergenic | NA | NA |

| | | | | | | | | | | | |
|---|---|---|---|---|---|---|---|---|---|---|---|
| T3056 | CCCGTGAGGCCGATCCGCGGCGG | 1000 | | | | | | | | | |
| Oligo fwd | CACCCCCGTGAGGCCGATCCGCGG | | | | | | | | | | |
| Oligo rev | AAACCCGCGGATCGGCCTCACGGG | | | | | | | | | | |
| Chromosome | start | end | strand | MM | target_seq | PAM | alignment | distance | position | gene name | gene id |
| GU305914.1 | 47364 | 47386 | + | 0 | CCCGTGAGGCCGATCCGCGG | CGG | \|\|\|\|\|\|\|\|[\|\|\|\|\|\|\|\|\|\|\|\|]PAM | NA | Intergenic | NA | NA |

| | | | | | | | | | | | |
|---|---|---|---|---|---|---|---|---|---|---|---|
| T3057 | CCGTGAGGCCGATCCGCGGCGGG | 1000 | | | | | | | | | |
| Oligo fwd | CACCCCGTGAGGCCGATCCGCGGC | | | | | | | | | | |
| Oligo rev | AAACGCCGCGGATCGGCCTCACGG | | | | | | | | | | |
| Chromosome | start | end | strand | MM | target_seq | PAM | alignment | distance | position | gene name | gene id |
| GU305914.1 | 47365 | 47387 | + | 0 | CCGTGAGGCCGATCCGCGGC | GGG | \|\|\|\|\|\|\|\|[\|\|\|\|\|\|\|\|\|\|\|\|]PAM | NA | Intergenic | NA | NA |

| | | | | | | | | | | | |
|---|---|---|---|---|---|---|---|---|---|---|---|
| T3058 | CGTGAGGCCGATCCGCGGCGGGG | 1000 | | | | | | | | | |
| Oligo fwd | CACCCGTGAGGCCGATCCGCGGCG | | | | | | | | | | |
| Oligo rev | AAACCGCCGCGGATCGGCCTCACG | | | | | | | | | | |
| Chromosome | start | end | strand | MM | target_seq | PAM | alignment | distance | position | gene name | gene id |
| GU305914.1 | 47366 | 47388 | + | 0 | CGTGAGGCCGATCCGCGGCG | GGG | \|\|\|\|\|\|\|\|[\|\|\|\|\|\|\|\|\|\|\|\|]PAM | NA | Intergenic | NA | NA |

| | | | | | | | | | | | |
|---|---|---|---|---|---|---|---|---|---|---|---|
| T3059 | GTGAGGCCGATCCGCGGCGGGGG | 1000 | | | | | | | | | |
| Oligo fwd | CACCGTGAGGCCGATCCGCGGCGG | | | | | | | | | | |
| Oligo rev | AAACCCGCCGCGGATCGGCCTCAC | | | | | | | | | | |
| Chromosome | start | end | strand | MM | target_seq | PAM | alignment | distance | position | gene name | gene id |
| GU305914.1 | 47367 | 47389 | + | 0 | GTGAGGCCGATCCGCGGCGG | GGG | \|\|\|\|\|\|\|\|[\|\|\|\|\|\|\|\|\|\|\|\|]PAM | NA | Intergenic | NA | NA |

| | | | | | | | | | | | |
|---|---|---|---|---|---|---|---|---|---|---|---|
| T3060 | TGAGGCCGATCCGCGGCGGGGGG | 1000 | | | | | | | | | |
| Oligo fwd | CACCTGAGGCCGATCCGCGGCGGG | | | | | | | | | | |
| Oligo rev | AAACCCCGCCGCGGATCGGCCTCA | | | | | | | | | | |
| Chromosome | start | end | strand | MM | target_seq | PAM | alignment | distance | position | gene name | gene id |
| GU305914.1 | 47368 | 47390 | + | 0 | TGAGGCCGATCCGCGGCGGG | GGG | \|\|\|\|\|\|\|\|[\|\|\|\|\|\|\|\|\|\|\|\|]PAM | NA | Intergenic | NA | NA |
| GU305914.1 | 113487 | 113509 | - | 3 | TGAAGACGATCCGCGTCGGG | CGG | \|\|\|-\|-\|\|[\|\|\|\|\|\|\|-\|\|\|\|]PAM | NA | Intergenic | NA | NA |

| | | | | | | | | | | | |
|---|---|---|---|---|---|---|---|---|---|---|---|
| T3061 | CGATCCGCGGCGGGGGGCAACGG | 1000 | | | | | | | | | |
| Oligo fwd | CACCCGATCCGCGGCGGGGGGCAA | | | | | | | | | | |
| Oligo rev | AAACTTGCCCCCCGCCGCGGATCG | | | | | | | | | | |
| Chromosome | start | end | strand | MM | target_seq | PAM | alignment | distance | position | gene name | gene id |
| GU305914.1 | 47374 | 47396 | + | 0 | CGATCCGCGGCGGGGGGCAA | CGG | \|\|\|\|\|\|\|\|[\|\|\|\|\|\|\|\|\|\|\|\|]PAM | NA | Intergenic | NA | NA |

| | | | | | | | | | | | |
|---|---|---|---|---|---|---|---|---|---|---|---|
| T3062 | CCGCGGCGGGGGGCAACGGATGG | 1000 | | | | | | | | | |
| Oligo fwd | CACCCCGCGGCGGGGGGCAACGGA | | | | | | | | | | |
| Oligo rev | AAACTCCGTTGCCCCCCGCCGCGG | | | | | | | | | | |
| Chromosome | start | end | strand | MM | target_seq | PAM | alignment | distance | position | gene name | gene id |
| GU305914.1 | 47378 | 47400 | + | 0 | CCGCGGCGGGGGGCAACGGA | TGG | \|\|\|\|\|\|\|\|[\|\|\|\|\|\|\|\|\|\|\|\|]PAM | NA | Intergenic | NA | NA |

| | | | | | | | | | | | |
|---|---|---|---|---|---|---|---|---|---|---|---|
| T3063 | CAACGGATGGCCAACATGAGAGG | 1000 | | | | | | | | | |
| Oligo fwd | CACCCAACGGATGGCCAACATGAG | | | | | | | | | | |
| Oligo rev | AAACCTCATGTTGGCCATCCGTTG | | | | | | | | | | |
| Chromosome | start | end | strand | MM | target_seq | PAM | alignment | distance | position | gene name | gene id |
| GU305914.1 | 47391 | 47413 | + | 0 | CAACGGATGGCCAACATGAG | AGG | \|\|\|\|\|\|\|\|[\|\|\|\|\|\|\|\|\|\|\|\|]PAM | NA | Intergenic | NA | NA |

| | | |
|---|---|---|
| T3064 | CCGTACTCCACAGTCCAGCGCGG | 1000 |
| Oligo fwd | CACCCCGTACTCCACAGTCCAGCG | |
| Oligo rev | AAACCGCTGGACTGTGGAGTACGG | |

Chromosome start end strand MM target_seq PAM alignment distance position gene name gene id
GU305914.1 47421 47443 + 0 CCGTACTCCACAGTCCAGCG CGG ||||||||[||||||||||||]PAM NA Intergenic NA NA

T3065 CAGTCCAGCGCGGCCGCCGTCGG 1000
Oligo fwd CACCCAGTCCAGCGCGGCCGCCGT
Oligo rev AAACACGGCGGCCGCGCTGGACTG
Chromosome start end strand MM target_seq PAM alignment distance position gene name gene id
GU305914.1 47431 47453 + 0 CAGTCCAGCGCGGCCGCCGT CGG ||||||||[||||||||||||]PAM NA Intergenic NA NA

T3066 CGGCACGAGAGCGAAGTGTGAGG 1000
Oligo fwd CACCCGGCACGAGAGCGAAGTGTG
Oligo rev AAACCACACTTCGCTCTCGTGCCG
Chromosome start end strand MM target_seq PAM alignment distance position gene name gene id
GU305914.1 47451 47473 + 0 CGGCACGAGAGCGAAGTGTG AGG ||||||||[||||||||||||]PAM NA Intergenic NA NA

T3067 GGCACGAGAGCGAAGTGTGAGGG 1000
Oligo fwd CACCGGCACGAGAGCGAAGTGTGA
Oligo rev AAACTCACACTTCGCTCTCGTGCC
Chromosome start end strand MM target_seq PAM alignment distance position gene name gene id
GU305914.1 47452 47474 + 0 GGCACGAGAGCGAAGTGTGA GGG ||||||||[||||||||||||]PAM NA Intergenic NA NA

T3068 CTCTCGATACATCGCCGACACGG 1000
Oligo fwd CACCCTCTCGATACATCGCCGACA
Oligo rev AAACTGTCGGCGATGTATCGAGAG
Chromosome start end strand MM target_seq PAM alignment distance position gene name gene id
GU305914.1 47664 47686 + 0 CTCTCGATACATCGCCGACA CGG ||||||||[||||||||||||]PAM NA Intergenic NA NA

T3069 TCGATACATCGCCGACACGGTGG 1000
Oligo fwd CACCTCGATACATCGCCGACACGG
Oligo rev AAACCCGTGTCGGCGATGTATCGA
Chromosome start end strand MM target_seq PAM alignment distance position gene name gene id
GU305914.1 47667 47689 + 0 TCGATACATCGCCGACACGG TGG ||||||||[||||||||||||]PAM NA Intergenic NA NA

T3070 CGGTGGATCGTTGATCCCGATGG 1000
Oligo fwd CACCCGGTGGATCGTTGATCCCGA
Oligo rev AAACTCGGGATCAACGATCCACCG
Chromosome start end strand MM target_seq PAM alignment distance position gene name gene id
GU305914.1 47684 47706 + 0 CGGTGGATCGTTGATCCCGA TGG ||||||||[||||||||||||]PAM NA Intergenic NA NA

T3071 TGGATCGTTGATCCCGATGGTGG 1000
Oligo fwd CACCTGGATCGTTGATCCCGATGG
Oligo rev AAACCCATCGGGATCAACGATCCA
Chromosome start end strand MM target_seq PAM alignment distance position gene name gene id
GU305914.1 47687 47709 + 0 TGGATCGTTGATCCCGATGG TGG ||||||||[||||||||||||]PAM NA Intergenic NA NA

T3072 CGTTGATCCCGATGGTGGTCAGG 1000
Oligo fwd CACCCGTTGATCCCGATGGTGGTC
Oligo rev AAACGACCACCATCGGGATCAACG
Chromosome start end strand MM target_seq PAM alignment distance position gene name gene id
GU305914.1 47692 47714 + 0 CGTTGATCCCGATGGTGGTC AGG ||||||||[||||||||||||]PAM NA Intergenic NA NA

T3073 GATCCCGATGGTGGTCAGGCTGG 1000
Oligo fwd CACCGATCCCGATGGTGGTCAGGC
Oligo rev AAACGCCTGACCACCATCGGGATC
Chromosome start end strand MM target_seq PAM alignment distance position gene name gene id
GU305914.1 47696 47718 + 0 GATCCCGATGGTGGTCAGGC TGG ||||||||[||||||||||||]PAM NA Intergenic NA NA

T3074 GCTGGTGTCGCAGTCGTCGTCGG 1000
Oligo fwd CACCGCTGGTGTCGCAGTCGTCGT
Oligo rev AAACACGACGACTGCGACACCAGC
Chromosome start end strand MM target_seq PAM alignment distance position gene name gene id
GU305914.1 47714 47736 + 0 GCTGGTGTCGCAGTCGTCGT CGG ||||||||[||||||||||||]PAM NA Intergenic NA NA

T3075 GCTGCGCATCCCCTCGAGCGTGG 1000
Oligo fwd CACCGCTGCGCATCCCCTCGAGCG
Oligo rev AAACCGCTCGAGGGGATGCGCAGC
Chromosome start end strand MM target_seq PAM alignment distance position gene name gene id
GU305914.1 47762 47784 + 0 GCTGCGCATCCCCTCGAGCG TGG ||||||||[||||||||||||]PAM NA Intergenic NA NA

T3076 CTCGAGCGTGGCCTTGATCACGG 1000
Oligo fwd CACCCTCGAGCGTGGCCTTGATCA
Oligo rev AAACTGATCAAGGCCACGCTCGAG
Chromosome start end strand MM target_seq PAM alignment distance position gene name gene id
GU305914.1 47774 47796 + 0 CTCGAGCGTGGCCTTGATCA CGG ||||||||[||||||||||||]PAM NA Intergenic NA NA

T3077 TCGAGCGTGGCCTTGATCACGGG 1000
Oligo fwd CACCTCGAGCGTGGCCTTGATCAC
Oligo rev AAACGTGATCAAGGCCACGCTCGA
Chromosome start end strand MM target_seq PAM alignment distance position gene name gene id
GU305914.1 47775 47797 + 0 TCGAGCGTGGCCTTGATCAC GGG ||||||||[||||||||||||]PAM NA Intergenic NA NA

T3078 CGAGCGTGGCCTTGATCACGGGG 1000
Oligo fwd CACCCGAGCGTGGCCTTGATCACG
Oligo rev AAACCGTGATCAAGGCCACGCTCG
Chromosome start end strand MM target_seq PAM alignment distance position gene name gene id
GU305914.1 47776 47798 + 0 CGAGCGTGGCCTTGATCACG GGG ||||||||[||||||||||||]PAM NA Intergenic NA NA
GU305914.1 135585 135607 - 4 CGCGTCTGGCCTTGATCTCG CGG ||-|--||[|||||||||-||]PAM NA Intergenic NA NA

T3079 GGCCTTGATCACGGGGTCAGAGG 1000

<table>
<tr><td>Oligo fwd</td><td>CACCGGCCTTGATCACGGGGTCAG</td><td></td><td></td><td></td><td></td><td></td><td></td><td></td><td></td><td></td><td></td><td></td></tr>
<tr><td>Oligo rev</td><td>AAACCTGACCCCGTGATCAAGGCC</td><td></td><td></td><td></td><td></td><td></td><td></td><td></td><td></td><td></td><td></td><td></td></tr>
<tr><td>Chromosome</td><td>start</td><td></td><td>end</td><td>strand</td><td>MM</td><td>target_seq</td><td>PAM</td><td>alignment</td><td>distance</td><td>position</td><td>gene name</td><td>gene id</td></tr>
<tr><td>GU305914.1</td><td></td><td>47783</td><td>47805</td><td>+</td><td>0</td><td>GGCCTTGATCACGGGGTCAG</td><td>AGG</td><td>||||||||[||||||||||||]PAM</td><td>NA</td><td>Intergenic</td><td>NA</td><td>NA</td></tr>
<tr><td>T3080</td><td>GATCACGGGGTCAGAGGCCATGG</td><td></td><td>1000</td><td></td><td></td><td></td><td></td><td></td><td></td><td></td><td></td><td></td></tr>
<tr><td>Oligo fwd</td><td>CACCGATCACGGGGTCAGAGGCCA</td><td></td><td></td><td></td><td></td><td></td><td></td><td></td><td></td><td></td><td></td><td></td></tr>
<tr><td>Oligo rev</td><td>AAACTGGCCTCTGACCCCGTGATC</td><td></td><td></td><td></td><td></td><td></td><td></td><td></td><td></td><td></td><td></td><td></td></tr>
<tr><td>Chromosome</td><td>start</td><td></td><td>end</td><td>strand</td><td>MM</td><td>target_seq</td><td>PAM</td><td>alignment</td><td>distance</td><td>position</td><td>gene name</td><td>gene id</td></tr>
<tr><td>GU305914.1</td><td></td><td>47789</td><td>47811</td><td>+</td><td>0</td><td>GATCACGGGGTCAGAGGCCA</td><td>TGG</td><td>||||||||[||||||||||||]PAM</td><td>NA</td><td>Intergenic</td><td>NA</td><td>NA</td></tr>
<tr><td>T3081</td><td>ACGGGGTCAGAGGCCATGGCCGG</td><td></td><td>1000</td><td></td><td></td><td></td><td></td><td></td><td></td><td></td><td></td><td></td></tr>
<tr><td>Oligo fwd</td><td>CACCACGGGGTCAGAGGCCATGGC</td><td></td><td></td><td></td><td></td><td></td><td></td><td></td><td></td><td></td><td></td><td></td></tr>
<tr><td>Oligo rev</td><td>AAACGCCATGGCCTCTGACCCCGT</td><td></td><td></td><td></td><td></td><td></td><td></td><td></td><td></td><td></td><td></td><td></td></tr>
<tr><td>Chromosome</td><td>start</td><td></td><td>end</td><td>strand</td><td>MM</td><td>target_seq</td><td>PAM</td><td>alignment</td><td>distance</td><td>position</td><td>gene name</td><td>gene id</td></tr>
<tr><td>GU305914.1</td><td></td><td>47793</td><td>47815</td><td>+</td><td>0</td><td>ACGGGGTCAGAGGCCATGGC</td><td>CGG</td><td>||||||||[||||||||||||]PAM</td><td>NA</td><td>Intergenic</td><td>NA</td><td>NA</td></tr>
<tr><td>T3082</td><td>CGGGGTCAGAGGCCATGGCCGGG</td><td></td><td>1000</td><td></td><td></td><td></td><td></td><td></td><td></td><td></td><td></td><td></td></tr>
<tr><td>Oligo fwd</td><td>CACCCGGGGTCAGAGGCCATGGCC</td><td></td><td></td><td></td><td></td><td></td><td></td><td></td><td></td><td></td><td></td><td></td></tr>
<tr><td>Oligo rev</td><td>AAACGGCCATGGCCTCTGACCCCG</td><td></td><td></td><td></td><td></td><td></td><td></td><td></td><td></td><td></td><td></td><td></td></tr>
<tr><td>Chromosome</td><td>start</td><td></td><td>end</td><td>strand</td><td>MM</td><td>target_seq</td><td>PAM</td><td>alignment</td><td>distance</td><td>position</td><td>gene name</td><td>gene id</td></tr>
<tr><td>GU305914.1</td><td></td><td>47794</td><td>47816</td><td>+</td><td>0</td><td>CGGGGTCAGAGGCCATGGCC</td><td>GGG</td><td>||||||||[||||||||||||]PAM</td><td>NA</td><td>Intergenic</td><td>NA</td><td>NA</td></tr>
<tr><td>T3083</td><td>GTCAGAGGCCATGGCCGGGCAGG</td><td></td><td>1000</td><td></td><td></td><td></td><td></td><td></td><td></td><td></td><td></td><td></td></tr>
<tr><td>Oligo fwd</td><td>CACCGTCAGAGGCCATGGCCGGGC</td><td></td><td></td><td></td><td></td><td></td><td></td><td></td><td></td><td></td><td></td><td></td></tr>
<tr><td>Oligo rev</td><td>AAACGCCCGGCCATGGCCTCTGAC</td><td></td><td></td><td></td><td></td><td></td><td></td><td></td><td></td><td></td><td></td><td></td></tr>
<tr><td>Chromosome</td><td>start</td><td></td><td>end</td><td>strand</td><td>MM</td><td>target_seq</td><td>PAM</td><td>alignment</td><td>distance</td><td>position</td><td>gene name</td><td>gene id</td></tr>
<tr><td>GU305914.1</td><td></td><td>47798</td><td>47820</td><td>+</td><td>0</td><td>GTCAGAGGCCATGGCCGGGC</td><td>AGG</td><td>||||||||[||||||||||||]PAM</td><td>NA</td><td>Intergenic</td><td>NA</td><td>NA</td></tr>
<tr><td>T3084</td><td>AGAGGCCATGGCCGGGCAGGAGG</td><td></td><td>1000</td><td></td><td></td><td></td><td></td><td></td><td></td><td></td><td></td><td></td></tr>
<tr><td>Oligo fwd</td><td>CACCAGAGGCCATGGCCGGGCAGG</td><td></td><td></td><td></td><td></td><td></td><td></td><td></td><td></td><td></td><td></td><td></td></tr>
<tr><td>Oligo rev</td><td>AAACCCTGCCCGGCCATGGCCTCT</td><td></td><td></td><td></td><td></td><td></td><td></td><td></td><td></td><td></td><td></td><td></td></tr>
<tr><td>Chromosome</td><td>start</td><td></td><td>end</td><td>strand</td><td>MM</td><td>target_seq</td><td>PAM</td><td>alignment</td><td>distance</td><td>position</td><td>gene name</td><td>gene id</td></tr>
<tr><td>GU305914.1</td><td></td><td>47801</td><td>47823</td><td>+</td><td>0</td><td>AGAGGCCATGGCCGGGCAGG</td><td>AGG</td><td>||||||||[||||||||||||]PAM</td><td>NA</td><td>Intergenic</td><td>NA</td><td>NA</td></tr>
<tr><td>T3085</td><td>GAGGCCATGGCCGGGCAGGAGGG</td><td></td><td>1000</td><td></td><td></td><td></td><td></td><td></td><td></td><td></td><td></td><td></td></tr>
<tr><td>Oligo fwd</td><td>CACCGAGGCCATGGCCGGGCAGGA</td><td></td><td></td><td></td><td></td><td></td><td></td><td></td><td></td><td></td><td></td><td></td></tr>
<tr><td>Oligo rev</td><td>AAACTCCTGCCCGGCCATGGCCTC</td><td></td><td></td><td></td><td></td><td></td><td></td><td></td><td></td><td></td><td></td><td></td></tr>
<tr><td>Chromosome</td><td>start</td><td></td><td>end</td><td>strand</td><td>MM</td><td>target_seq</td><td>PAM</td><td>alignment</td><td>distance</td><td>position</td><td>gene name</td><td>gene id</td></tr>
<tr><td>GU305914.1</td><td></td><td>47802</td><td>47824</td><td>+</td><td>0</td><td>GAGGCCATGGCCGGGCAGGA</td><td>GGG</td><td>||||||||[||||||||||||]PAM</td><td>NA</td><td>Intergenic</td><td>NA</td><td>NA</td></tr>
<tr><td>T3086</td><td>AGGCCATGGCCGGGCAGGAGGGG</td><td></td><td>1000</td><td></td><td></td><td></td><td></td><td></td><td></td><td></td><td></td><td></td></tr>
<tr><td>Oligo fwd</td><td>CACCAGGCCATGGCCGGGCAGGAG</td><td></td><td></td><td></td><td></td><td></td><td></td><td></td><td></td><td></td><td></td><td></td></tr>
<tr><td>Oligo rev</td><td>AAACCTCCTGCCCGGCCATGGCCT</td><td></td><td></td><td></td><td></td><td></td><td></td><td></td><td></td><td></td><td></td><td></td></tr>
<tr><td>Chromosome</td><td>start</td><td></td><td>end</td><td>strand</td><td>MM</td><td>target_seq</td><td>PAM</td><td>alignment</td><td>distance</td><td>position</td><td>gene name</td><td>gene id</td></tr>
<tr><td>GU305914.1</td><td></td><td>47803</td><td>47825</td><td>+</td><td>0</td><td>AGGCCATGGCCGGGCAGGAG</td><td>GGG</td><td>||||||||[||||||||||||]PAM</td><td>NA</td><td>Intergenic</td><td>NA</td><td>NA</td></tr>
<tr><td>T3087</td><td>CCATGGCCGGGCAGGAGGGGTGG</td><td></td><td>1000</td><td></td><td></td><td></td><td></td><td></td><td></td><td></td><td></td><td></td></tr>
<tr><td>Oligo fwd</td><td>CACCCCATGGCCGGGCAGGAGGGG</td><td></td><td></td><td></td><td></td><td></td><td></td><td></td><td></td><td></td><td></td><td></td></tr>
<tr><td>Oligo rev</td><td>AAACCCCCTCCTGCCCGGCCATGG</td><td></td><td></td><td></td><td></td><td></td><td></td><td></td><td></td><td></td><td></td><td></td></tr>
<tr><td>Chromosome</td><td>start</td><td></td><td>end</td><td>strand</td><td>MM</td><td>target_seq</td><td>PAM</td><td>alignment</td><td>distance</td><td>position</td><td>gene name</td><td>gene id</td></tr>
<tr><td>GU305914.1</td><td></td><td>47806</td><td>47828</td><td>+</td><td>0</td><td>CCATGGCCGGGCAGGAGGGG</td><td>TGG</td><td>||||||||[||||||||||||]PAM</td><td>NA</td><td>Intergenic</td><td>NA</td><td>NA</td></tr>
<tr><td>T3088</td><td>GGCAGGAGGGGTGGCCGACGAGG</td><td></td><td>1000</td><td></td><td></td><td></td><td></td><td></td><td></td><td></td><td></td><td></td></tr>
<tr><td>Oligo fwd</td><td>CACCGGCAGGAGGGGTGGCCGACG</td><td></td><td></td><td></td><td></td><td></td><td></td><td></td><td></td><td></td><td></td><td></td></tr>
<tr><td>Oligo rev</td><td>AAACCGTCGGCCACCCCTCCTGCC</td><td></td><td></td><td></td><td></td><td></td><td></td><td></td><td></td><td></td><td></td><td></td></tr>
<tr><td>Chromosome</td><td>start</td><td></td><td>end</td><td>strand</td><td>MM</td><td>target_seq</td><td>PAM</td><td>alignment</td><td>distance</td><td>position</td><td>gene name</td><td>gene id</td></tr>
<tr><td>GU305914.1</td><td></td><td>47815</td><td>47837</td><td>+</td><td>0</td><td>GGCAGGAGGGGTGGCCGACG</td><td>AGG</td><td>||||||||[||||||||||||]PAM</td><td>NA</td><td>Intergenic</td><td>NA</td><td>NA</td></tr>
<tr><td>T3089</td><td>GAGGAGCCCCTCGATCTCCTCGG</td><td></td><td>1000</td><td></td><td></td><td></td><td></td><td></td><td></td><td></td><td></td><td></td></tr>
<tr><td>Oligo fwd</td><td>CACCGAGGAGCCCCTCGATCTCCT</td><td></td><td></td><td></td><td></td><td></td><td></td><td></td><td></td><td></td><td></td><td></td></tr>
<tr><td>Oligo rev</td><td>AAACAGGAGATCGAGGGGCTCCTC</td><td></td><td></td><td></td><td></td><td></td><td></td><td></td><td></td><td></td><td></td><td></td></tr>
<tr><td>Chromosome</td><td>start</td><td></td><td>end</td><td>strand</td><td>MM</td><td>target_seq</td><td>PAM</td><td>alignment</td><td>distance</td><td>position</td><td>gene name</td><td>gene id</td></tr>
<tr><td>GU305914.1</td><td></td><td>47834</td><td>47856</td><td>+</td><td>0</td><td>GAGGAGCCCCTCGATCTCCT</td><td>CGG</td><td>||||||||[||||||||||||]PAM</td><td>NA</td><td>Intergenic</td><td>NA</td><td>NA</td></tr>
<tr><td>T3090</td><td>AGGAGCCCCTCGATCTCCTCGGG</td><td></td><td>1000</td><td></td><td></td><td></td><td></td><td></td><td></td><td></td><td></td><td></td></tr>
<tr><td>Oligo fwd</td><td>CACCAGGAGCCCCTCGATCTCCTC</td><td></td><td></td><td></td><td></td><td></td><td></td><td></td><td></td><td></td><td></td><td></td></tr>
<tr><td>Oligo rev</td><td>AAACGAGGAGATCGAGGGGCTCCT</td><td></td><td></td><td></td><td></td><td></td><td></td><td></td><td></td><td></td><td></td><td></td></tr>
<tr><td>Chromosome</td><td>start</td><td></td><td>end</td><td>strand</td><td>MM</td><td>target_seq</td><td>PAM</td><td>alignment</td><td>distance</td><td>position</td><td>gene name</td><td>gene id</td></tr>
<tr><td>GU305914.1</td><td></td><td>47835</td><td>47857</td><td>+</td><td>0</td><td>AGGAGCCCCTCGATCTCCTC</td><td>GGG</td><td>||||||||[||||||||||||]PAM</td><td>NA</td><td>Intergenic</td><td>NA</td><td>NA</td></tr>
<tr><td>T3091</td><td>CCTCGATCTCCTCGGGCCAGAGG</td><td></td><td>1000</td><td></td><td></td><td></td><td></td><td></td><td></td><td></td><td></td><td></td></tr>
<tr><td>Oligo fwd</td><td>CACCCCTCGATCTCCTCGGGCCAG</td><td></td><td></td><td></td><td></td><td></td><td></td><td></td><td></td><td></td><td></td><td></td></tr>
<tr><td>Oligo rev</td><td>AAACCTGGCCCGAGGAGATCGAGG</td><td></td><td></td><td></td><td></td><td></td><td></td><td></td><td></td><td></td><td></td><td></td></tr>
<tr><td>Chromosome</td><td>start</td><td></td><td>end</td><td>strand</td><td>MM</td><td>target_seq</td><td>PAM</td><td>alignment</td><td>distance</td><td>position</td><td>gene name</td><td>gene id</td></tr>
<tr><td>GU305914.1</td><td></td><td>47842</td><td>47864</td><td>+</td><td>0</td><td>CCTCGATCTCCTCGGGCCAG</td><td>AGG</td><td>||||||||[||||||||||||]PAM</td><td>NA</td><td>Intergenic</td><td>NA</td><td>NA</td></tr>
<tr><td>GU305914.1</td><td></td><td>47851</td><td>47873</td><td>-</td><td>4</td><td>CACCGATCTCCTCTGGCCCG</td><td>AGG</td><td>|--|||||[|||||-||||-|]PAM</td><td>NA</td><td>Intergenic</td><td>NA</td><td>NA</td></tr>
<tr><td>T3092</td><td>CTCCTCGGGCCAGAGGAGATCGG</td><td></td><td>1000</td><td></td><td></td><td></td><td></td><td></td><td></td><td></td><td></td><td></td></tr>
<tr><td>Oligo fwd</td><td>CACCCTCCTCGGGCCAGAGGAGAT</td><td></td><td></td><td></td><td></td><td></td><td></td><td></td><td></td><td></td><td></td><td></td></tr>
<tr><td>Oligo rev</td><td>AAACATCTCCTCTGGCCCGAGGAG</td><td></td><td></td><td></td><td></td><td></td><td></td><td></td><td></td><td></td><td></td><td></td></tr>
<tr><td>Chromosome</td><td>start</td><td></td><td>end</td><td>strand</td><td>MM</td><td>target_seq</td><td>PAM</td><td>alignment</td><td>distance</td><td>position</td><td>gene name</td><td>gene id</td></tr>
<tr><td>GU305914.1</td><td></td><td>47849</td><td>47871</td><td>+</td><td>0</td><td>CTCCTCGGGCCAGAGGAGAT</td><td>CGG</td><td>||||||||[||||||||||||]PAM</td><td>NA</td><td>Intergenic</td><td>NA</td><td>NA</td></tr>
<tr><td>T3093</td><td>CTGCTTGAAGCCGAAGTAAGCGG</td><td></td><td>1000</td><td></td><td></td><td></td><td></td><td></td><td></td><td></td><td></td><td></td></tr>
<tr><td>Oligo fwd</td><td>CACCCTGCTTGAAGCCGAAGTAAG</td><td></td><td></td><td></td><td></td><td></td><td></td><td></td><td></td><td></td><td></td><td></td></tr>
<tr><td>Oligo rev</td><td>AAACCTTACTTCGGCTTCAAGCAG</td><td></td><td></td><td></td><td></td><td></td><td></td><td></td><td></td><td></td><td></td><td></td></tr>
<tr><td>Chromosome</td><td>start</td><td></td><td>end</td><td>strand</td><td>MM</td><td>target_seq</td><td>PAM</td><td>alignment</td><td>distance</td><td>position</td><td>gene name</td><td>gene id</td></tr>
<tr><td>GU305914.1</td><td></td><td>47876</td><td>47898</td><td>+</td><td>0</td><td>CTGCTTGAAGCCGAAGTAAG</td><td>CGG</td><td>||||||||[||||||||||||]PAM</td><td>NA</td><td>Intergenic</td><td>NA</td><td>NA</td></tr>
</table>

<table>
<tr><td>T3094</td><td>GTAAGCGGCCCTCATGATGTTGG</td><td>1000</td><td></td><td></td><td></td><td></td><td></td><td></td><td></td><td></td><td></td></tr>
<tr><td>Oligo fwd</td><td>CACCGTAAGCGGCCCTCATGATGT</td><td></td><td></td><td></td><td></td><td></td><td></td><td></td><td></td><td></td><td></td></tr>
<tr><td>Oligo rev</td><td>AAACACATCATGAGGGCCGCTTAC</td><td></td><td></td><td></td><td></td><td></td><td></td><td></td><td></td><td></td><td></td></tr>
<tr><td>Chromosome</td><td>start</td><td>end</td><td>strand</td><td>MM</td><td>target_seq</td><td>PAM</td><td>alignment</td><td>distance</td><td>position</td><td>gene name</td><td>gene id</td></tr>
<tr><td>GU305914.1</td><td>47891</td><td>47913</td><td>+</td><td>0</td><td>GTAAGCGGCCCTCATGATGT</td><td>TGG</td><td>||||||||[||||||||||||]PAM</td><td>NA</td><td>Intergenic</td><td>NA</td><td>NA</td></tr>
<tr><td>T3095</td><td>AGCGGCCCTCATGATGTTGGCGG</td><td>1000</td><td></td><td></td><td></td><td></td><td></td><td></td><td></td><td></td><td></td></tr>
<tr><td>Oligo fwd</td><td>CACCAGCGGCCCTCATGATGTTGG</td><td></td><td></td><td></td><td></td><td></td><td></td><td></td><td></td><td></td><td></td></tr>
<tr><td>Oligo rev</td><td>AAACCCAACATCATGAGGGCCGCT</td><td></td><td></td><td></td><td></td><td></td><td></td><td></td><td></td><td></td><td></td></tr>
<tr><td>Chromosome</td><td>start</td><td>end</td><td>strand</td><td>MM</td><td>target_seq</td><td>PAM</td><td>alignment</td><td>distance</td><td>position</td><td>gene name</td><td>gene id</td></tr>
<tr><td>GU305914.1</td><td>47894</td><td>47916</td><td>+</td><td>0</td><td>AGCGGCCCTCATGATGTTGG</td><td>CGG</td><td>||||||||[||||||||||||]PAM</td><td>NA</td><td>Intergenic</td><td>NA</td><td>NA</td></tr>
<tr><td>T3096</td><td>TGATGTTGGCGGCGTCGTAGTGG</td><td>1000</td><td></td><td></td><td></td><td></td><td></td><td></td><td></td><td></td><td></td></tr>
<tr><td>Oligo fwd</td><td>CACCTGATGTTGGCGGCGTCGTAG</td><td></td><td></td><td></td><td></td><td></td><td></td><td></td><td></td><td></td><td></td></tr>
<tr><td>Oligo rev</td><td>AAACCTACGACGCCGCCAACATCA</td><td></td><td></td><td></td><td></td><td></td><td></td><td></td><td></td><td></td><td></td></tr>
<tr><td>Chromosome</td><td>start</td><td>end</td><td>strand</td><td>MM</td><td>target_seq</td><td>PAM</td><td>alignment</td><td>distance</td><td>position</td><td>gene name</td><td>gene id</td></tr>
<tr><td>GU305914.1</td><td>47905</td><td>47927</td><td>+</td><td>0</td><td>TGATGTTGGCGGCGTCGTAG</td><td>TGG</td><td>||||||||[||||||||||||]PAM</td><td>NA</td><td>Intergenic</td><td>NA</td><td>NA</td></tr>
<tr><td>T3097</td><td>GATGTTGGCGGCGTCGTAGTGGG</td><td>1000</td><td></td><td></td><td></td><td></td><td></td><td></td><td></td><td></td><td></td></tr>
<tr><td>Oligo fwd</td><td>CACCGATGTTGGCGGCGTCGTAGT</td><td></td><td></td><td></td><td></td><td></td><td></td><td></td><td></td><td></td><td></td></tr>
<tr><td>Oligo rev</td><td>AAACACTACGACGCCGCCAACATC</td><td></td><td></td><td></td><td></td><td></td><td></td><td></td><td></td><td></td><td></td></tr>
<tr><td>Chromosome</td><td>start</td><td>end</td><td>strand</td><td>MM</td><td>target_seq</td><td>PAM</td><td>alignment</td><td>distance</td><td>position</td><td>gene name</td><td>gene id</td></tr>
<tr><td>GU305914.1</td><td>47906</td><td>47928</td><td>+</td><td>0</td><td>GATGTTGGCGGCGTCGTAGT</td><td>GGG</td><td>||||||||[||||||||||||]PAM</td><td>NA</td><td>Intergenic</td><td>NA</td><td>NA</td></tr>
<tr><td>T3098</td><td>GTCGTAGTGGGAGAAGACGTCGG</td><td>1000</td><td></td><td></td><td></td><td></td><td></td><td></td><td></td><td></td><td></td></tr>
<tr><td>Oligo fwd</td><td>CACCGTCGTAGTGGGAGAAGACGT</td><td></td><td></td><td></td><td></td><td></td><td></td><td></td><td></td><td></td><td></td></tr>
<tr><td>Oligo rev</td><td>AAACACGTCTTCTCCCACTACGAC</td><td></td><td></td><td></td><td></td><td></td><td></td><td></td><td></td><td></td><td></td></tr>
<tr><td>Chromosome</td><td>start</td><td>end</td><td>strand</td><td>MM</td><td>target_seq</td><td>PAM</td><td>alignment</td><td>distance</td><td>position</td><td>gene name</td><td>gene id</td></tr>
<tr><td>GU305914.1</td><td>47918</td><td>47940</td><td>+</td><td>0</td><td>GTCGTAGTGGGAGAAGACGT</td><td>CGG</td><td>||||||||[||||||||||||]PAM</td><td>NA</td><td>Intergenic</td><td>NA</td><td>NA</td></tr>
<tr><td>T3099</td><td>AGAACTTGTTCTTCCTCCCGAGG</td><td>1000</td><td></td><td></td><td></td><td></td><td></td><td></td><td></td><td></td><td></td></tr>
<tr><td>Oligo fwd</td><td>CACCAGAACTTGTTCTTCCTCCCG</td><td></td><td></td><td></td><td></td><td></td><td></td><td></td><td></td><td></td><td></td></tr>
<tr><td>Oligo rev</td><td>AAACCGGGAGGAAGAACAAGTTCT</td><td></td><td></td><td></td><td></td><td></td><td></td><td></td><td></td><td></td><td></td></tr>
<tr><td>Chromosome</td><td>start</td><td>end</td><td>strand</td><td>MM</td><td>target_seq</td><td>PAM</td><td>alignment</td><td>distance</td><td>position</td><td>gene name</td><td>gene id</td></tr>
<tr><td>GU305914.1</td><td>47989</td><td>48011</td><td>+</td><td>0</td><td>AGAACTTGTTCTTCCTCCCG</td><td>AGG</td><td>||||||||[||||||||||||]PAM</td><td>NA</td><td>Intergenic</td><td>NA</td><td>NA</td></tr>
<tr><td>T3100</td><td>TCCTCCCGAGGAAGAGATTGTGG</td><td>1000</td><td></td><td></td><td></td><td></td><td></td><td></td><td></td><td></td><td></td></tr>
<tr><td>Oligo fwd</td><td>CACCTCCTCCCGAGGAAGAGATTG</td><td></td><td></td><td></td><td></td><td></td><td></td><td></td><td></td><td></td><td></td></tr>
<tr><td>Oligo rev</td><td>AAACCAATCTCTTCCTCGGGAGGA</td><td></td><td></td><td></td><td></td><td></td><td></td><td></td><td></td><td></td><td></td></tr>
<tr><td>Chromosome</td><td>start</td><td>end</td><td>strand</td><td>MM</td><td>target_seq</td><td>PAM</td><td>alignment</td><td>distance</td><td>position</td><td>gene name</td><td>gene id</td></tr>
<tr><td>GU305914.1</td><td>48001</td><td>48023</td><td>+</td><td>0</td><td>TCCTCCCGAGGAAGAGATTG</td><td>TGG</td><td>||||||||[||||||||||||]PAM</td><td>NA</td><td>Intergenic</td><td>NA</td><td>NA</td></tr>
<tr><td>T3101</td><td>GCCGATCTGCTCCCCGAAACTGG</td><td>1000</td><td></td><td></td><td></td><td></td><td></td><td></td><td></td><td></td><td></td></tr>
<tr><td>Oligo fwd</td><td>CACCGCCGATCTGCTCCCCGAAAC</td><td></td><td></td><td></td><td></td><td></td><td></td><td></td><td></td><td></td><td></td></tr>
<tr><td>Oligo rev</td><td>AAACGTTTCGGGGAGCAGATCGGC</td><td></td><td></td><td></td><td></td><td></td><td></td><td></td><td></td><td></td><td></td></tr>
<tr><td>Chromosome</td><td>start</td><td>end</td><td>strand</td><td>MM</td><td>target_seq</td><td>PAM</td><td>alignment</td><td>distance</td><td>position</td><td>gene name</td><td>gene id</td></tr>
<tr><td>GU305914.1</td><td>48023</td><td>48045</td><td>+</td><td>0</td><td>GCCGATCTGCTCCCCGAAAC</td><td>TGG</td><td>||||||||[||||||||||||]PAM</td><td>NA</td><td>Intergenic</td><td>NA</td><td>NA</td></tr>
<tr><td>T3102</td><td>GGTCATCCGCGACATGCAGAAGG</td><td>1000</td><td></td><td></td><td></td><td></td><td></td><td></td><td></td><td></td><td></td></tr>
<tr><td>Oligo fwd</td><td>CACCGGTCATCCGCGACATGCAGA</td><td></td><td></td><td></td><td></td><td></td><td></td><td></td><td></td><td></td><td></td></tr>
<tr><td>Oligo rev</td><td>AAACTCTGCATGTCGCGGATGACC</td><td></td><td></td><td></td><td></td><td></td><td></td><td></td><td></td><td></td><td></td></tr>
<tr><td>Chromosome</td><td>start</td><td>end</td><td>strand</td><td>MM</td><td>target_seq</td><td>PAM</td><td>alignment</td><td>distance</td><td>position</td><td>gene name</td><td>gene id</td></tr>
<tr><td>GU305914.1</td><td>48044</td><td>48066</td><td>+</td><td>0</td><td>GGTCATCCGCGACATGCAGA</td><td>AGG</td><td>||||||||[||||||||||||]PAM</td><td>NA</td><td>Intergenic</td><td>NA</td><td>NA</td></tr>
<tr><td>T3103</td><td>CGACATGCAGAAGGAGCTGTCGG</td><td>1000</td><td></td><td></td><td></td><td></td><td></td><td></td><td></td><td></td><td></td></tr>
<tr><td>Oligo fwd</td><td>CACCCGACATGCAGAAGGAGCTGT</td><td></td><td></td><td></td><td></td><td></td><td></td><td></td><td></td><td></td><td></td></tr>
<tr><td>Oligo rev</td><td>AAACACAGCTCCTTCTGCATGTCG</td><td></td><td></td><td></td><td></td><td></td><td></td><td></td><td></td><td></td><td></td></tr>
<tr><td>Chromosome</td><td>start</td><td>end</td><td>strand</td><td>MM</td><td>target_seq</td><td>PAM</td><td>alignment</td><td>distance</td><td>position</td><td>gene name</td><td>gene id</td></tr>
<tr><td>GU305914.1</td><td>48053</td><td>48075</td><td>+</td><td>0</td><td>CGACATGCAGAAGGAGCTGT</td><td>CGG</td><td>||||||||[||||||||||||]PAM</td><td>NA</td><td>Intergenic</td><td>NA</td><td>NA</td></tr>
<tr><td>T3104</td><td>GACATGCAGAAGGAGCTGTCGGG</td><td>1000</td><td></td><td></td><td></td><td></td><td></td><td></td><td></td><td></td><td></td></tr>
<tr><td>Oligo fwd</td><td>CACCGACATGCAGAAGGAGCTGTC</td><td></td><td></td><td></td><td></td><td></td><td></td><td></td><td></td><td></td><td></td></tr>
<tr><td>Oligo rev</td><td>AAACGACAGCTCCTTCTGCATGTC</td><td></td><td></td><td></td><td></td><td></td><td></td><td></td><td></td><td></td><td></td></tr>
<tr><td>Chromosome</td><td>start</td><td>end</td><td>strand</td><td>MM</td><td>target_seq</td><td>PAM</td><td>alignment</td><td>distance</td><td>position</td><td>gene name</td><td>gene id</td></tr>
<tr><td>GU305914.1</td><td>48054</td><td>48076</td><td>+</td><td>0</td><td>GACATGCAGAAGGAGCTGTC</td><td>GGG</td><td>||||||||[||||||||||||]PAM</td><td>NA</td><td>Intergenic</td><td>NA</td><td>NA</td></tr>
<tr><td>T3105</td><td>ACATGCAGAAGGAGCTGTCGGGG</td><td>1000</td><td></td><td></td><td></td><td></td><td></td><td></td><td></td><td></td><td></td></tr>
<tr><td>Oligo fwd</td><td>CACCACATGCAGAAGGAGCTGTCG</td><td></td><td></td><td></td><td></td><td></td><td></td><td></td><td></td><td></td><td></td></tr>
<tr><td>Oligo rev</td><td>AAACCGACAGCTCCTTCTGCATGT</td><td></td><td></td><td></td><td></td><td></td><td></td><td></td><td></td><td></td><td></td></tr>
<tr><td>Chromosome</td><td>start</td><td>end</td><td>strand</td><td>MM</td><td>target_seq</td><td>PAM</td><td>alignment</td><td>distance</td><td>position</td><td>gene name</td><td>gene id</td></tr>
<tr><td>GU305914.1</td><td>48055</td><td>48077</td><td>+</td><td>0</td><td>ACATGCAGAAGGAGCTGTCG</td><td>GGG</td><td>||||||||[||||||||||||]PAM</td><td>NA</td><td>Intergenic</td><td>NA</td><td>NA</td></tr>
<tr><td>T3106</td><td>GGGTCCCTGACGACGTACTGCGG</td><td>1000</td><td></td><td></td><td></td><td></td><td></td><td></td><td></td><td></td><td></td></tr>
<tr><td>Oligo fwd</td><td>CACCGGGTCCCTGACGACGTACTG</td><td></td><td></td><td></td><td></td><td></td><td></td><td></td><td></td><td></td><td></td></tr>
<tr><td>Oligo rev</td><td>AAACCAGTACGTCGTCAGGGACCC</td><td></td><td></td><td></td><td></td><td></td><td></td><td></td><td></td><td></td><td></td></tr>
<tr><td>Chromosome</td><td>start</td><td>end</td><td>strand</td><td>MM</td><td>target_seq</td><td>PAM</td><td>alignment</td><td>distance</td><td>position</td><td>gene name</td><td>gene id</td></tr>
<tr><td>GU305914.1</td><td>48075</td><td>48097</td><td>+</td><td>0</td><td>GGGTCCCTGACGACGTACTG</td><td>CGG</td><td>||||||||[||||||||||||]PAM</td><td>NA</td><td>Intergenic</td><td>NA</td><td>NA</td></tr>
<tr><td>T3107</td><td>GGTCCCTGACGACGTACTGCGGG</td><td>1000</td><td></td><td></td><td></td><td></td><td></td><td></td><td></td><td></td><td></td></tr>
<tr><td>Oligo fwd</td><td>CACCGGTCCCTGACGACGTACTGC</td><td></td><td></td><td></td><td></td><td></td><td></td><td></td><td></td><td></td><td></td></tr>
<tr><td>Oligo rev</td><td>AAACGCAGTACGTCGTCAGGGACC</td><td></td><td></td><td></td><td></td><td></td><td></td><td></td><td></td><td></td><td></td></tr>
<tr><td>Chromosome</td><td>start</td><td>end</td><td>strand</td><td>MM</td><td>target_seq</td><td>PAM</td><td>alignment</td><td>distance</td><td>position</td><td>gene name</td><td>gene id</td></tr>
<tr><td>GU305914.1</td><td>48076</td><td>48098</td><td>+</td><td>0</td><td>GGTCCCTGACGACGTACTGC</td><td>GGG</td><td>||||||||[||||||||||||]PAM</td><td>NA</td><td>Intergenic</td><td>NA</td><td>NA</td></tr>
<tr><td>T3108</td><td>CTGCGGGTCCTCATCCCCCACGG</td><td>1000</td><td></td><td></td><td></td><td></td><td></td><td></td><td></td><td></td><td></td></tr>
<tr><td>Oligo fwd</td><td>CACCCTGCGGGTCCTCATCCCCCA</td><td></td><td></td><td></td><td></td><td></td><td></td><td></td><td></td><td></td><td></td></tr>
<tr><td>Oligo rev</td><td>AAACTGGGGGATGAGGACCCGCAG</td><td></td><td></td><td></td><td></td><td></td><td></td><td></td><td></td><td></td><td></td></tr>
<tr><td>Chromosome</td><td>start</td><td>end</td><td>strand</td><td>MM</td><td>target_seq</td><td>PAM</td><td>alignment</td><td>distance</td><td>position</td><td>gene name</td><td>gene id</td></tr>
</table>

<table>
<tr><td>GU305914.1</td><td></td><td>48092</td><td>48114</td><td>+</td><td>0</td><td>CTGCGGGTCCTCATCCCCCA</td><td>CGG</td><td>||||||||[||||||||||||]PAM</td><td>NA</td><td>Intergenic</td><td>NA</td><td>NA</td></tr>
<tr><td>T3109</td><td>GAAGAACCACTTCACCCTCGCGG</td><td></td><td>1000</td><td></td><td></td><td></td><td></td><td></td><td></td><td></td><td></td><td></td></tr>
<tr><td>Oligo fwd</td><td>CACCGAAGAACCACTTCACCCTCG</td><td></td><td></td><td></td><td></td><td></td><td></td><td></td><td></td><td></td><td></td><td></td></tr>
<tr><td>Oligo rev</td><td>AAACCGAGGGTGAAGTGGTTCTTC</td><td></td><td></td><td></td><td></td><td></td><td></td><td></td><td></td><td></td><td></td><td></td></tr>
<tr><td>Chromosome</td><td>start</td><td></td><td>end</td><td>strand</td><td>MM</td><td>target_seq</td><td>PAM</td><td>alignment</td><td>distance</td><td>position</td><td>gene name</td><td>gene id</td></tr>
<tr><td>GU305914.1</td><td></td><td>48127</td><td>48149</td><td>+</td><td>0</td><td>GAAGAACCACTTCACCCTCG</td><td>CGG</td><td>||||||||[||||||||||||]PAM</td><td>NA</td><td>Intergenic</td><td>NA</td><td>NA</td></tr>
<tr><td>T3110</td><td>ACTTCACCCTCGCGGACCTGTGG</td><td></td><td>1000</td><td></td><td></td><td></td><td></td><td></td><td></td><td></td><td></td><td></td></tr>
<tr><td>Oligo fwd</td><td>CACCACTTCACCCTCGCGGACCTG</td><td></td><td></td><td></td><td></td><td></td><td></td><td></td><td></td><td></td><td></td><td></td></tr>
<tr><td>Oligo rev</td><td>AAACCAGGTCCGCGAGGGTGAAGT</td><td></td><td></td><td></td><td></td><td></td><td></td><td></td><td></td><td></td><td></td><td></td></tr>
<tr><td>Chromosome</td><td>start</td><td></td><td>end</td><td>strand</td><td>MM</td><td>target_seq</td><td>PAM</td><td>alignment</td><td>distance</td><td>position</td><td>gene name</td><td>gene id</td></tr>
<tr><td>GU305914.1</td><td></td><td>48135</td><td>48157</td><td>+</td><td>0</td><td>ACTTCACCCTCGCGGACCTG</td><td>TGG</td><td>||||||||[||||||||||||]PAM</td><td>NA</td><td>Intergenic</td><td>NA</td><td>NA</td></tr>
<tr><td>T3111</td><td>TCACCCTCGCGGACCTGTGGCGG</td><td></td><td>1000</td><td></td><td></td><td></td><td></td><td></td><td></td><td></td><td></td><td></td></tr>
<tr><td>Oligo fwd</td><td>CACCTCACCCTCGCGGACCTGTGG</td><td></td><td></td><td></td><td></td><td></td><td></td><td></td><td></td><td></td><td></td><td></td></tr>
<tr><td>Oligo rev</td><td>AAACCCACAGGTCCGCGAGGGTGA</td><td></td><td></td><td></td><td></td><td></td><td></td><td></td><td></td><td></td><td></td><td></td></tr>
<tr><td>Chromosome</td><td>start</td><td></td><td>end</td><td>strand</td><td>MM</td><td>target_seq</td><td>PAM</td><td>alignment</td><td>distance</td><td>position</td><td>gene name</td><td>gene id</td></tr>
<tr><td>GU305914.1</td><td></td><td>48138</td><td>48160</td><td>+</td><td>0</td><td>TCACCCTCGCGGACCTGTGG</td><td>CGG</td><td>||||||||[||||||||||||]PAM</td><td>NA</td><td>Intergenic</td><td>NA</td><td>NA</td></tr>
<tr><td>T3112</td><td>CCCTCGCGGACCTGTGGCGGCGG</td><td></td><td>1000</td><td></td><td></td><td></td><td></td><td></td><td></td><td></td><td></td><td></td></tr>
<tr><td>Oligo fwd</td><td>CACCCCCTCGCGGACCTGTGGCGG</td><td></td><td></td><td></td><td></td><td></td><td></td><td></td><td></td><td></td><td></td><td></td></tr>
<tr><td>Oligo rev</td><td>AAACCCGCCACAGGTCCGCGAGGG</td><td></td><td></td><td></td><td></td><td></td><td></td><td></td><td></td><td></td><td></td><td></td></tr>
<tr><td>Chromosome</td><td>start</td><td></td><td>end</td><td>strand</td><td>MM</td><td>target_seq</td><td>PAM</td><td>alignment</td><td>distance</td><td>position</td><td>gene name</td><td>gene id</td></tr>
<tr><td>GU305914.1</td><td></td><td>48141</td><td>48163</td><td>+</td><td>0</td><td>CCCTCGCGGACCTGTGGCGG</td><td>CGG</td><td>||||||||[||||||||||||]PAM</td><td>NA</td><td>Intergenic</td><td>NA</td><td>NA</td></tr>
<tr><td>T3113</td><td>TCGCGGACCTGTGGCGGCGGCGG</td><td></td><td>1000</td><td></td><td></td><td></td><td></td><td></td><td></td><td></td><td></td><td></td></tr>
<tr><td>Oligo fwd</td><td>CACCTCGCGGACCTGTGGCGGCGG</td><td></td><td></td><td></td><td></td><td></td><td></td><td></td><td></td><td></td><td></td><td></td></tr>
<tr><td>Oligo rev</td><td>AAACCCGCCGCCACAGGTCCGCGA</td><td></td><td></td><td></td><td></td><td></td><td></td><td></td><td></td><td></td><td></td><td></td></tr>
<tr><td>Chromosome</td><td>start</td><td></td><td>end</td><td>strand</td><td>MM</td><td>target_seq</td><td>PAM</td><td>alignment</td><td>distance</td><td>position</td><td>gene name</td><td>gene id</td></tr>
<tr><td>GU305914.1</td><td></td><td>48144</td><td>48166</td><td>+</td><td>0</td><td>TCGCGGACCTGTGGCGGCGG</td><td>CGG</td><td>||||||||[||||||||||||]PAM</td><td>NA</td><td>Intergenic</td><td>NA</td><td>NA</td></tr>
<tr><td>GU305914.1</td><td></td><td>67739</td><td>67761</td><td>+</td><td>4</td><td>TCGGGGAGCGGCGGCGGCGG</td><td>CGG</td><td>|||-|||-[|-|-||||||||]PAM</td><td>NA</td><td>Intergenic</td><td>NA</td><td>NA</td></tr>
<tr><td>GU305914.1</td><td></td><td>110035</td><td>110057</td><td>+</td><td>4</td><td>TCGCCGTCCTGCTGCGGCGG</td><td>GGG</td><td>||||-|-|[|||--|||||||]PAM</td><td>NA</td><td>Intergenic</td><td>NA</td><td>NA</td></tr>
<tr><td>T3114</td><td>CGGACCTGTGGCGGCGGCGGTGG</td><td></td><td>1000</td><td></td><td></td><td></td><td></td><td></td><td></td><td></td><td></td><td></td></tr>
<tr><td>Oligo fwd</td><td>CACCCGGACCTGTGGCGGCGGCGG</td><td></td><td></td><td></td><td></td><td></td><td></td><td></td><td></td><td></td><td></td><td></td></tr>
<tr><td>Oligo rev</td><td>AAACCCGCCGCCGCCACAGGTCCG</td><td></td><td></td><td></td><td></td><td></td><td></td><td></td><td></td><td></td><td></td><td></td></tr>
<tr><td>Chromosome</td><td>start</td><td></td><td>end</td><td>strand</td><td>MM</td><td>target_seq</td><td>PAM</td><td>alignment</td><td>distance</td><td>position</td><td>gene name</td><td>gene id</td></tr>
<tr><td>GU305914.1</td><td></td><td>48147</td><td>48169</td><td>+</td><td>0</td><td>CGGACCTGTGGCGGCGGCGG</td><td>TGG</td><td>||||||||[||||||||||||]PAM</td><td>NA</td><td>Intergenic</td><td>NA</td><td>NA</td></tr>
<tr><td>GU305914.1</td><td></td><td>154401</td><td>154423</td><td>+</td><td>4</td><td>CCGGGCGGTGGCGGCGGCGG</td><td>CGG</td><td>|-|--|-|[||||||||||||]PAM</td><td>NA</td><td>Intergenic</td><td>NA</td><td>NA</td></tr>
<tr><td>GU305914.1</td><td></td><td>67742</td><td>67764</td><td>+</td><td>4</td><td>GGGAGCGGCGGCGGCGGCGG</td><td>CGG</td><td>-|||-|-|[-|||||||||||]PAM</td><td>NA</td><td>Intergenic</td><td>NA</td><td>NA</td></tr>
<tr><td>GU305914.1</td><td></td><td>29994</td><td>30016</td><td>+</td><td>4</td><td>GGGACCGGTGCTGGCGGCGG</td><td>CGG</td><td>-|||||-|[||--||||||||]PAM</td><td>NA</td><td>Intergenic</td><td>NA</td><td>NA</td></tr>
<tr><td>T3115</td><td>ACCTGTGGCGGCGGCGGTGGCGG</td><td></td><td>1000</td><td></td><td></td><td></td><td></td><td></td><td></td><td></td><td></td><td></td></tr>
<tr><td>Oligo fwd</td><td>CACCACCTGTGGCGGCGGCGGTGG</td><td></td><td></td><td></td><td></td><td></td><td></td><td></td><td></td><td></td><td></td><td></td></tr>
<tr><td>Oligo rev</td><td>AAACCCACCGCCGCCGCCACAGGT</td><td></td><td></td><td></td><td></td><td></td><td></td><td></td><td></td><td></td><td></td><td></td></tr>
<tr><td>Chromosome</td><td>start</td><td></td><td>end</td><td>strand</td><td>MM</td><td>target_seq</td><td>PAM</td><td>alignment</td><td>distance</td><td>position</td><td>gene name</td><td>gene id</td></tr>
<tr><td>GU305914.1</td><td></td><td>48150</td><td>48172</td><td>+</td><td>0</td><td>ACCTGTGGCGGCGGCGGTGG</td><td>CGG</td><td>||||||||[||||||||||||]PAM</td><td>NA</td><td>Intergenic</td><td>NA</td><td>NA</td></tr>
<tr><td>GU305914.1</td><td></td><td>24624</td><td>24646</td><td>-</td><td>4</td><td>AACACCGGCGGCGGCGGTGG</td><td>CGG</td><td>|-|---||[||||||||||||]PAM</td><td>NA</td><td>Intergenic</td><td>NA</td><td>NA</td></tr>
<tr><td>GU305914.1</td><td></td><td>200150</td><td>200172</td><td>-</td><td>4</td><td>ACGAGTGCCTGCGGCGGTGG</td><td>TGG</td><td>||--|||-[|-||||||||||]PAM</td><td>NA</td><td>Intergenic</td><td>NA</td><td>NA</td></tr>
<tr><td>GU305914.1</td><td></td><td>52719</td><td>52741</td><td>+</td><td>4</td><td>GCCGCTGGCGGTGGCGGTGG</td><td>CGG</td><td>-||--|||[|||-||||||||]PAM</td><td>NA</td><td>Intergenic</td><td>NA</td><td>NA</td></tr>
<tr><td>GU305914.1</td><td></td><td>62437</td><td>62459</td><td>+</td><td>4</td><td>GGCGGTGGCGGCGGGGGTGG</td><td>TGG</td><td>--|-||||[||||||-|||||]PAM</td><td>NA</td><td>Intergenic</td><td>NA</td><td>NA</td></tr>
<tr><td>GU305914.1</td><td></td><td>154404</td><td>154426</td><td>+</td><td>4</td><td>GGCGGTGGCGGCGGCGGCGG</td><td>CGG</td><td>--|-||||[|||||||||-||]PAM</td><td>NA</td><td>Intergenic</td><td>NA</td><td>NA</td></tr>
<tr><td>GU305914.1</td><td></td><td>58126</td><td>58148</td><td>-</td><td>4</td><td>GGCGGTGGCGGCGGCGGCGG</td><td>CGG</td><td>--|-||||[|||||||||-||]PAM</td><td>NA</td><td>Intergenic</td><td>NA</td><td>NA</td></tr>
<tr><td>GU305914.1</td><td></td><td>29997</td><td>30019</td><td>+</td><td>4</td><td>ACCGGTGCTGGCGGCGGCGG</td><td>CGG</td><td>|||-|||-[-||||||||-||]PAM</td><td>NA</td><td>Intergenic</td><td>NA</td><td>NA</td></tr>
<tr><td>GU305914.1</td><td></td><td>152550</td><td>152572</td><td>+</td><td>3</td><td>TCCTGCGGCGGCGGCGGTGC</td><td>CGG</td><td>-||||-||[|||||||||||-]PAM</td><td>NA</td><td>Intergenic</td><td>NA</td><td>NA</td></tr>
<tr><td>GU305914.1</td><td></td><td>35624</td><td>35646</td><td>+</td><td>4</td><td>ATCGGTGGCGTCGGCGGTGT</td><td>CGG</td><td>|-|-||||[||-||||||||-]PAM</td><td>NA</td><td>Intergenic</td><td>NA</td><td>NA</td></tr>
<tr><td>T3116</td><td>TGTGGCGGCGGCGGTGGCGGCGG</td><td></td><td>1000</td><td></td><td></td><td></td><td></td><td></td><td></td><td></td><td></td><td></td></tr>
<tr><td>Oligo fwd</td><td>CACCTGTGGCGGCGGCGGTGGCGG</td><td></td><td></td><td></td><td></td><td></td><td></td><td></td><td></td><td></td><td></td><td></td></tr>
<tr><td>Oligo rev</td><td>AAACCCGCCACCGCCGCCGCCACA</td><td></td><td></td><td></td><td></td><td></td><td></td><td></td><td></td><td></td><td></td><td></td></tr>
<tr><td>Chromosome</td><td>start</td><td></td><td>end</td><td>strand</td><td>MM</td><td>target_seq</td><td>PAM</td><td>alignment</td><td>distance</td><td>position</td><td>gene name</td><td>gene id</td></tr>
<tr><td>GU305914.1</td><td></td><td>48153</td><td>48175</td><td>+</td><td>0</td><td>TGTGGCGGCGGCGGTGGCGG</td><td>CGG</td><td>||||||||[||||||||||||]PAM</td><td>NA</td><td>Intergenic</td><td>NA</td><td>NA</td></tr>
<tr><td>GU305914.1</td><td></td><td>163749</td><td>163771</td><td>+</td><td>4</td><td>GTCGGCGGCGCCGGTGGCGG</td><td>CGG</td><td>---|||||[||-|||||||||]PAM</td><td>NA</td><td>Intergenic</td><td>NA</td><td>NA</td></tr>
<tr><td>GU305914.1</td><td></td><td>154407</td><td>154429</td><td>+</td><td>2</td><td>GGTGGCGGCGGCGGCGGCGG</td><td>CGG</td><td>-|||||||[||||||-|||||]PAM</td><td>NA</td><td>Intergenic</td><td>NA</td><td>NA</td></tr>
<tr><td>GU305914.1</td><td></td><td>58123</td><td>58145</td><td>-</td><td>2</td><td>GGTGGCGGCGGCGGCGGCGG</td><td>AGG</td><td>-|||||||[||||||-|||||]PAM</td><td>NA</td><td>Intergenic</td><td>NA</td><td>NA</td></tr>
<tr><td>GU305914.1</td><td></td><td>62443</td><td>62465</td><td>+</td><td>4</td><td>GGCGGCGGGGGTGGTGGCGG</td><td>CGG</td><td>-|-|||||[-||-||||||||]PAM</td><td>NA</td><td>Intergenic</td><td>NA</td><td>NA</td></tr>
<tr><td>GU305914.1</td><td></td><td>67742</td><td>67764</td><td>+</td><td>4</td><td>GGGAGCGGCGGCGGCGGCGG</td><td>CGG</td><td>-|--||||[||||||-|||||]PAM</td><td>NA</td><td>Intergenic</td><td>NA</td><td>NA</td></tr>
<tr><td>GU305914.1</td><td></td><td>149482</td><td>149504</td><td>-</td><td>4</td><td>GGCGACGGCGGCGGCGGCGG</td><td>CGG</td><td>-|-|-|||[||||||-|||||]PAM</td><td>NA</td><td>Intergenic</td><td>NA</td><td>NA</td></tr>
<tr><td>GU305914.1</td><td></td><td>154404</td><td>154426</td><td>+</td><td>4</td><td>GGCGGTGGCGGCGGCGGCGG</td><td>CGG</td><td>-|-||-||[||||||-|||||]PAM</td><td>NA</td><td>Intergenic</td><td>NA</td><td>NA</td></tr>
<tr><td>GU305914.1</td><td></td><td>58126</td><td>58148</td><td>-</td><td>4</td><td>GGCGGTGGCGGCGGCGGCGG</td><td>CGG</td><td>-|-||-||[||||||-|||||]PAM</td><td>NA</td><td>Intergenic</td><td>NA</td><td>NA</td></tr>
<tr><td>GU305914.1</td><td></td><td>30000</td><td>30022</td><td>+</td><td>4</td><td>GGTGCTGGCGGCGGCGGCGG</td><td>CGG</td><td>-|||--||[||||||-|||||]PAM</td><td>NA</td><td>Intergenic</td><td>NA</td><td>NA</td></tr>
<tr><td>GU305914.1</td><td></td><td>48159</td><td>48181</td><td>+</td><td>4</td><td>GGCGGCGGTGGCGGCGGCGG</td><td>TGG</td><td>-|-|||||[-|||||-|||||]PAM</td><td>NA</td><td>Intergenic</td><td>NA</td><td>NA</td></tr>
<tr><td>GU305914.1</td><td></td><td>152553</td><td>152575</td><td>+</td><td>2</td><td>TGCGGCGGCGGCGGTGCCGG</td><td>CGG</td><td>||-|||||[||||||||-|||]PAM</td><td>NA</td><td>Intergenic</td><td>NA</td><td>NA</td></tr>
<tr><td>GU305914.1</td><td></td><td>168820</td><td>168842</td><td>+</td><td>4</td><td>GGTGGTGGCAGCGGCGGCGG</td><td>CGG</td><td>-||||-||[|-||||-|||||]PAM</td><td>NA</td><td>Intergenic</td><td>NA</td><td>NA</td></tr>
<tr><td>GU305914.1</td><td></td><td>48156</td><td>48178</td><td>+</td><td>4</td><td>GGCGGCGGCGGTGGCGGCGG</td><td>CGG</td><td>-|-|||||[|||-||-|||||]PAM</td><td>NA</td><td>Intergenic</td><td>NA</td><td>NA</td></tr>
<tr><td>GU305914.1</td><td></td><td>62440</td><td>62462</td><td>+</td><td>3</td><td>GGTGGCGGCGGGGGTGGTGG</td><td>CGG</td><td>-|||||||[|||-|||||-||]PAM</td><td>NA</td><td>Intergenic</td><td>NA</td><td>NA</td></tr>
<tr><td>GU305914.1</td><td></td><td>114664</td><td>114686</td><td>-</td><td>4</td><td>TGAGACGGCGGTGGTGGTGG</td><td>CGG</td><td>||-|-|||[|||-|||||-||]PAM</td><td>NA</td><td>Intergenic</td><td>NA</td><td>NA</td></tr>
<tr><td>GU305914.1</td><td></td><td>168811</td><td>168833</td><td>+</td><td>4</td><td>TTTAGCGGCGGTGGTGGCAG</td><td>CGG</td><td>|-|-||||[|||-||||||-|]PAM</td><td>NA</td><td>Intergenic</td><td>NA</td><td>NA</td></tr>
<tr><td>GU305914.1</td><td></td><td>62452</td><td>62474</td><td>+</td><td>4</td><td>GGTGGTGGCGGCGGGGGTGG</td><td>CGG</td><td>-||||-||[||||||-||-||]PAM</td><td>NA</td><td>Intergenic</td><td>NA</td><td>NA</td></tr>
<tr><td>GU305914.1</td><td></td><td>163902</td><td>163924</td><td>+</td><td>4</td><td>AGCGGCGGCGGCGGCGGCAG</td><td>CGG</td><td>-|-|||||[||||||-|||-|]PAM</td><td>NA</td><td>Intergenic</td><td>NA</td><td>NA</td></tr>
<tr><td>GU305914.1</td><td></td><td>30006</td><td>30028</td><td>+</td><td>4</td><td>GGCGGCGGCGGCGGCGGCAG</td><td>TGG</td><td>-|-|||||[||||||-|||-|]PAM</td><td>NA</td><td>Intergenic</td><td>NA</td><td>NA</td></tr>
<tr><td>GU305914.1</td><td></td><td>149476</td><td>149498</td><td>-</td><td>4</td><td>GGCGGCGGCGGCGGCGGCAG</td><td>CGG</td><td>-|-|||||[||||||-|||-|]PAM</td><td>NA</td><td>Intergenic</td><td>NA</td><td>NA</td></tr>
<tr><td>T3117</td><td>GGCGGCGGCGGTGGCGGCGGCGG</td><td></td><td>1000</td><td></td><td></td><td></td><td></td><td></td><td></td><td></td><td></td><td></td></tr>
<tr><td>Oligo fwd</td><td>CACCGGCGGCGGCGGTGGCGGCGG</td><td></td><td></td><td></td><td></td><td></td><td></td><td></td><td></td><td></td><td></td><td></td></tr>
<tr><td>Oligo rev</td><td>AAACCCGCCGCCACCGCCGCCGCC</td><td></td><td></td><td></td><td></td><td></td><td></td><td></td><td></td><td></td><td></td><td></td></tr>
<tr><td>Chromosome</td><td>start</td><td></td><td>end</td><td>strand</td><td>MM</td><td>target_seq</td><td>PAM</td><td>alignment</td><td>distance</td><td>position</td><td>gene name</td><td>gene id</td></tr>
<tr><td>GU305914.1</td><td></td><td>48156</td><td>48178</td><td>+</td><td>0</td><td>GGCGGCGGCGGTGGCGGCGG</td><td>CGG</td><td>||||||||[||||||||||||]PAM</td><td>NA</td><td>Intergenic</td><td>NA</td><td>NA</td></tr>
</table>

<table>
<tr><td>GU305914.1</td><td>62431</td><td>62453</td><td>+</td><td>3</td><td>TTCGACGGCGGTGGCGGCGG</td><td>GGG</td><td>--||-|||[||||||||||||]PAM</td><td>NA</td><td>Intergenic</td><td>NA</td><td>NA</td></tr>
<tr><td>GU305914.1</td><td>62446</td><td>62468</td><td>+</td><td>2</td><td>GGCGGGGGTGGTGGCGGCGG</td><td>GGG</td><td>|||||-||[-|||||||||||]PAM</td><td>NA</td><td>Intergenic</td><td>NA</td><td>NA</td></tr>
<tr><td>GU305914.1</td><td>114658</td><td>114680</td><td>-</td><td>2</td><td>GGCGGTGGTGGTGGCGGCGG</td><td>CGG</td><td>|||||-||[-|||||||||||]PAM</td><td>NA</td><td>Intergenic</td><td>NA</td><td>NA</td></tr>
<tr><td>GU305914.1</td><td>154398</td><td>154420</td><td>+</td><td>4</td><td>GCCCCGGGCGGTGGCGGCGG</td><td>CGG</td><td>|-|---||[||||||||||||]PAM</td><td>NA</td><td>Intergenic</td><td>NA</td><td>NA</td></tr>
<tr><td>GU305914.1</td><td>154407</td><td>154429</td><td>+</td><td>2</td><td>GGTGGCGGCGGCGGCGGCGG</td><td>CGG</td><td>||-|||||[|||-||||||||]PAM</td><td>NA</td><td>Intergenic</td><td>NA</td><td>NA</td></tr>
<tr><td>GU305914.1</td><td>58123</td><td>58145</td><td>-</td><td>2</td><td>GGTGGCGGCGGCGGCGGCGG</td><td>AGG</td><td>||-|||||[|||-||||||||]PAM</td><td>NA</td><td>Intergenic</td><td>NA</td><td>NA</td></tr>
<tr><td>GU305914.1</td><td>149482</td><td>149504</td><td>-</td><td>2</td><td>GGCGACGGCGGCGGCGGCGG</td><td>CGG</td><td>||||-|||[|||-||||||||]PAM</td><td>NA</td><td>Intergenic</td><td>NA</td><td>NA</td></tr>
<tr><td>GU305914.1</td><td>154404</td><td>154426</td><td>+</td><td>2</td><td>GGCGGTGGCGGCGGCGGCGG</td><td>CGG</td><td>|||||-||[|||-||||||||]PAM</td><td>NA</td><td>Intergenic</td><td>NA</td><td>NA</td></tr>
<tr><td>GU305914.1</td><td>58126</td><td>58148</td><td>-</td><td>2</td><td>GGCGGTGGCGGCGGCGGCGG</td><td>CGG</td><td>|||||-||[|||-||||||||]PAM</td><td>NA</td><td>Intergenic</td><td>NA</td><td>NA</td></tr>
<tr><td>GU305914.1</td><td>67742</td><td>67764</td><td>+</td><td>3</td><td>GGGAGCGGCGGCGGCGGCGG</td><td>CGG</td><td>||--||||[|||-||||||||]PAM</td><td>NA</td><td>Intergenic</td><td>NA</td><td>NA</td></tr>
<tr><td>GU305914.1</td><td>48159</td><td>48181</td><td>+</td><td>2</td><td>GGCGGCGGTGGCGGCGGCGG</td><td>TGG</td><td>||||||||[-||-||||||||]PAM</td><td>NA</td><td>Intergenic</td><td>NA</td><td>NA</td></tr>
<tr><td>GU305914.1</td><td>56773</td><td>56795</td><td>+</td><td>4</td><td>CGCCGCAGCGGCGGCGGCGG</td><td>CGG</td><td>-||-||-|[|||-||||||||]PAM</td><td>NA</td><td>Intergenic</td><td>NA</td><td>NA</td></tr>
<tr><td>GU305914.1</td><td>30000</td><td>30022</td><td>+</td><td>4</td><td>GGTGCTGGCGGCGGCGGCGG</td><td>CGG</td><td>||-|--||[|||-||||||||]PAM</td><td>NA</td><td>Intergenic</td><td>NA</td><td>NA</td></tr>
<tr><td>GU305914.1</td><td>149485</td><td>149507</td><td>-</td><td>4</td><td>GCAGGCGACGGCGGCGGCGG</td><td>CGG</td><td>|--||||-[|||-||||||||]PAM</td><td>NA</td><td>Intergenic</td><td>NA</td><td>NA</td></tr>
<tr><td>GU305914.1</td><td>58129</td><td>58151</td><td>-</td><td>4</td><td>GCGGGCGGTGGCGGCGGCGG</td><td>CGG</td><td>|--|||||[-||-||||||||]PAM</td><td>NA</td><td>Intergenic</td><td>NA</td><td>NA</td></tr>
<tr><td>GU305914.1</td><td>161754</td><td>161776</td><td>-</td><td>4</td><td>GGCCGAGGAGGCGGCGGCGG</td><td>AGG</td><td>|||-|-||[-||-||||||||]PAM</td><td>NA</td><td>Intergenic</td><td>NA</td><td>NA</td></tr>
<tr><td>GU305914.1</td><td>161757</td><td>161779</td><td>-</td><td>4</td><td>GGAGGCCGAGGAGGCGGCGG</td><td>CGG</td><td>||-|||-|[-||-||||||||]PAM</td><td>NA</td><td>Intergenic</td><td>NA</td><td>NA</td></tr>
<tr><td>GU305914.1</td><td>168820</td><td>168842</td><td>+</td><td>4</td><td>GGTGGTGGCAGCGGCGGCGG</td><td>CGG</td><td>||-||-||[|-|-||||||||]PAM</td><td>NA</td><td>Intergenic</td><td>NA</td><td>NA</td></tr>
<tr><td>GU305914.1</td><td>62443</td><td>62465</td><td>+</td><td>2</td><td>GGCGGCGGGGGTGGTGGCGG</td><td>CGG</td><td>||||||||[-|||||-|||||]PAM</td><td>NA</td><td>Intergenic</td><td>NA</td><td>NA</td></tr>
<tr><td>GU305914.1</td><td>73792</td><td>73814</td><td>-</td><td>4</td><td>GGCGCCAGCAGCGGCGGCGG</td><td>AGG</td><td>||||-|-|[|-|-||||||||]PAM</td><td>NA</td><td>Intergenic</td><td>NA</td><td>NA</td></tr>
<tr><td>GU305914.1</td><td>114661</td><td>114683</td><td>-</td><td>3</td><td>GACGGCGGTGGTGGTGGCGG</td><td>CGG</td><td>|-||||||[-|||||-|||||]PAM</td><td>NA</td><td>Intergenic</td><td>NA</td><td>NA</td></tr>
<tr><td>GU305914.1</td><td>19936</td><td>19958</td><td>+</td><td>4</td><td>GACGGTAGCGGTGGTGGCGG</td><td>TGG</td><td>|-|||--|[||||||-|||||]PAM</td><td>NA</td><td>Intergenic</td><td>NA</td><td>NA</td></tr>
<tr><td>GU305914.1</td><td>168814</td><td>168836</td><td>+</td><td>3</td><td>AGCGGCGGTGGTGGCAGCGG</td><td>CGG</td><td>-|||||||[-||||||-||||]PAM</td><td>NA</td><td>Intergenic</td><td>NA</td><td>NA</td></tr>
<tr><td>GU305914.1</td><td>48153</td><td>48175</td><td>+</td><td>4</td><td>TGTGGCGGCGGCGGTGGCGG</td><td>CGG</td><td>-|-|||||[|||-||-|||||]PAM</td><td>NA</td><td>Intergenic</td><td>NA</td><td>NA</td></tr>
<tr><td>GU305914.1</td><td>35618</td><td>35640</td><td>+</td><td>4</td><td>GGCGGAATCGGTGGCGTCGG</td><td>CGG</td><td>|||||---[||||||||-|||]PAM</td><td>NA</td><td>Intergenic</td><td>NA</td><td>NA</td></tr>
<tr><td>GU305914.1</td><td>52719</td><td>52741</td><td>+</td><td>4</td><td>GCCGCTGGCGGTGGCGGTGG</td><td>CGG</td><td>|-||--||[|||||||||-||]PAM</td><td>NA</td><td>Intergenic</td><td>NA</td><td>NA</td></tr>
<tr><td>GU305914.1</td><td>74673</td><td>74695</td><td>+</td><td>4</td><td>GGTGCCGTCGGTGGCGGTGG</td><td>CGG</td><td>||-|-||-[|||||||||-||]PAM</td><td>NA</td><td>Intergenic</td><td>NA</td><td>NA</td></tr>
<tr><td>GU305914.1</td><td>18656</td><td>18678</td><td>-</td><td>4</td><td>GACGGCAGAGGTGGCGGAGG</td><td>CGG</td><td>|-||||-|[-||||||||-||]PAM</td><td>NA</td><td>Intergenic</td><td>NA</td><td>NA</td></tr>
<tr><td>GU305914.1</td><td>62458</td><td>62480</td><td>+</td><td>2</td><td>GGCGGCGGGGGTGGCGGCCG</td><td>CGG</td><td>||||||||[-|||||||||-|]PAM</td><td>NA</td><td>Intergenic</td><td>NA</td><td>NA</td></tr>
<tr><td>GU305914.1</td><td>45258</td><td>45280</td><td>-</td><td>4</td><td>AACGGTGGCGGTGGCGGCAG</td><td>CGG</td><td>--|||-||[||||||||||-|]PAM</td><td>NA</td><td>Intergenic</td><td>NA</td><td>NA</td></tr>
<tr><td>GU305914.1</td><td>30006</td><td>30028</td><td>+</td><td>2</td><td>GGCGGCGGCGGCGGCGGCAG</td><td>TGG</td><td>||||||||[|||-||||||-|]PAM</td><td>NA</td><td>Intergenic</td><td>NA</td><td>NA</td></tr>
<tr><td>GU305914.1</td><td>149476</td><td>149498</td><td>-</td><td>2</td><td>GGCGGCGGCGGCGGCGGCAG</td><td>CGG</td><td>||||||||[|||-||||||-|]PAM</td><td>NA</td><td>Intergenic</td><td>NA</td><td>NA</td></tr>
<tr><td>GU305914.1</td><td>163902</td><td>163924</td><td>+</td><td>3</td><td>AGCGGCGGCGGCGGCGGCAG</td><td>CGG</td><td>-|||||||[|||-||||||-|]PAM</td><td>NA</td><td>Intergenic</td><td>NA</td><td>NA</td></tr>
<tr><td>GU305914.1</td><td>113924</td><td>113946</td><td>-</td><td>4</td><td>GTCGGCGTCGGCGGCGGCAG</td><td>CGG</td><td>|-|||||-[|||-||||||-|]PAM</td><td>NA</td><td>Intergenic</td><td>NA</td><td>NA</td></tr>
<tr><td>GU305914.1</td><td>109443</td><td>109465</td><td>+</td><td>3</td><td>GACGGCGGCGGTAGCGGCGT</td><td>CGG</td><td>|-||||||[||||-||||||-]PAM</td><td>NA</td><td>Intergenic</td><td>NA</td><td>NA</td></tr>
</table>

T3118 GGCGGCGGTGGCGGCGGCGGTGG 1000
Oligo fwd CACCGGCGGCGGTGGCGGCGGCGG
Oligo rev AAACCCGCCGCCGCCACCGCCGCC

<table>
<tr><th>Chromosome</th><th>start</th><th>end</th><th>strand</th><th>MM</th><th>target_seq</th><th>PAM</th><th>alignment</th><th>distance</th><th>position</th><th>gene name</th><th>gene id</th></tr>
<tr><td>GU305914.1</td><td>48159</td><td>48181</td><td>+</td><td>0</td><td>GGCGGCGGTGGCGGCGGCGG</td><td>TGG</td><td>||||||||[||||||||||||]PAM</td><td>NA</td><td>Intergenic</td><td>NA</td><td>NA</td></tr>
<tr><td>GU305914.1</td><td>58129</td><td>58151</td><td>-</td><td>2</td><td>GCGGGCGGTGGCGGCGGCGG</td><td>CGG</td><td>|--|||||[||||||||||||]PAM</td><td>NA</td><td>Intergenic</td><td>NA</td><td>NA</td></tr>
<tr><td>GU305914.1</td><td>154401</td><td>154423</td><td>+</td><td>3</td><td>CCGGGCGGTGGCGGCGGCGG</td><td>CGG</td><td>---|||||[||||||||||||]PAM</td><td>NA</td><td>Intergenic</td><td>NA</td><td>NA</td></tr>
<tr><td>GU305914.1</td><td>154407</td><td>154429</td><td>+</td><td>2</td><td>GGTGGCGGCGGCGGCGGCGG</td><td>CGG</td><td>||-|||||[-|||||||||||]PAM</td><td>NA</td><td>Intergenic</td><td>NA</td><td>NA</td></tr>
<tr><td>GU305914.1</td><td>58123</td><td>58145</td><td>-</td><td>2</td><td>GGTGGCGGCGGCGGCGGCGG</td><td>AGG</td><td>||-|||||[-|||||||||||]PAM</td><td>NA</td><td>Intergenic</td><td>NA</td><td>NA</td></tr>
<tr><td>GU305914.1</td><td>149482</td><td>149504</td><td>-</td><td>2</td><td>GGCGACGGCGGCGGCGGCGG</td><td>CGG</td><td>||||-|||[-|||||||||||]PAM</td><td>NA</td><td>Intergenic</td><td>NA</td><td>NA</td></tr>
<tr><td>GU305914.1</td><td>154404</td><td>154426</td><td>+</td><td>2</td><td>GGCGGTGGCGGCGGCGGCGG</td><td>CGG</td><td>|||||-||[-|||||||||||]PAM</td><td>NA</td><td>Intergenic</td><td>NA</td><td>NA</td></tr>
<tr><td>GU305914.1</td><td>58126</td><td>58148</td><td>-</td><td>2</td><td>GGCGGTGGCGGCGGCGGCGG</td><td>CGG</td><td>|||||-||[-|||||||||||]PAM</td><td>NA</td><td>Intergenic</td><td>NA</td><td>NA</td></tr>
<tr><td>GU305914.1</td><td>67742</td><td>67764</td><td>+</td><td>3</td><td>GGGAGCGGCGGCGGCGGCGG</td><td>CGG</td><td>||--||||[-|||||||||||]PAM</td><td>NA</td><td>Intergenic</td><td>NA</td><td>NA</td></tr>
<tr><td>GU305914.1</td><td>29997</td><td>30019</td><td>+</td><td>4</td><td>ACCGGTGCTGGCGGCGGCGG</td><td>CGG</td><td>--|||-|-[||||||||||||]PAM</td><td>NA</td><td>Intergenic</td><td>NA</td><td>NA</td></tr>
<tr><td>GU305914.1</td><td>161754</td><td>161776</td><td>-</td><td>3</td><td>GGCCGAGGAGGCGGCGGCGG</td><td>AGG</td><td>|||-|-||[-|||||||||||]PAM</td><td>NA</td><td>Intergenic</td><td>NA</td><td>NA</td></tr>
<tr><td>GU305914.1</td><td>62446</td><td>62468</td><td>+</td><td>2</td><td>GGCGGGGGTGGTGGCGGCGG</td><td>GGG</td><td>|||||-||[|||-||||||||]PAM</td><td>NA</td><td>Intergenic</td><td>NA</td><td>NA</td></tr>
<tr><td>GU305914.1</td><td>114658</td><td>114680</td><td>-</td><td>2</td><td>GGCGGTGGTGGTGGCGGCGG</td><td>CGG</td><td>|||||-||[|||-||||||||]PAM</td><td>NA</td><td>Intergenic</td><td>NA</td><td>NA</td></tr>
<tr><td>GU305914.1</td><td>56773</td><td>56795</td><td>+</td><td>4</td><td>CGCCGCAGCGGCGGCGGCGG</td><td>CGG</td><td>-||-||-|[-|||||||||||]PAM</td><td>NA</td><td>Intergenic</td><td>NA</td><td>NA</td></tr>
<tr><td>GU305914.1</td><td>30000</td><td>30022</td><td>+</td><td>4</td><td>GGTGCTGGCGGCGGCGGCGG</td><td>CGG</td><td>||-|--||[-|||||||||||]PAM</td><td>NA</td><td>Intergenic</td><td>NA</td><td>NA</td></tr>
<tr><td>GU305914.1</td><td>149485</td><td>149507</td><td>-</td><td>4</td><td>GCAGGCGACGGCGGCGGCGG</td><td>CGG</td><td>|--||||-[-|||||||||||]PAM</td><td>NA</td><td>Intergenic</td><td>NA</td><td>NA</td></tr>
<tr><td>GU305914.1</td><td>168817</td><td>168839</td><td>+</td><td>2</td><td>GGCGGTGGTGGCAGCGGCGG</td><td>CGG</td><td>|||||-||[||||-|||||||]PAM</td><td>NA</td><td>Intergenic</td><td>NA</td><td>NA</td></tr>
<tr><td>GU305914.1</td><td>48156</td><td>48178</td><td>+</td><td>2</td><td>GGCGGCGGCGGTGGCGGCGG</td><td>CGG</td><td>||||||||[-||-||||||||]PAM</td><td>NA</td><td>Intergenic</td><td>NA</td><td>NA</td></tr>
<tr><td>GU305914.1</td><td>168820</td><td>168842</td><td>+</td><td>4</td><td>GGTGGTGGCAGCGGCGGCGG</td><td>CGG</td><td>||-||-||[--||||||||||]PAM</td><td>NA</td><td>Intergenic</td><td>NA</td><td>NA</td></tr>
<tr><td>GU305914.1</td><td>73792</td><td>73814</td><td>-</td><td>4</td><td>GGCGCCAGCAGCGGCGGCGG</td><td>AGG</td><td>||||-|-|[--||||||||||]PAM</td><td>NA</td><td>Intergenic</td><td>NA</td><td>NA</td></tr>
<tr><td>GU305914.1</td><td>161757</td><td>161779</td><td>-</td><td>4</td><td>GGAGGCCGAGGAGGCGGCGG</td><td>CGG</td><td>||-|||-|[-||-||||||||]PAM</td><td>NA</td><td>Intergenic</td><td>NA</td><td>NA</td></tr>
<tr><td>GU305914.1</td><td>26382</td><td>26404</td><td>+</td><td>4</td><td>ATCGGCGCTGGCGACGGCGG</td><td>CGG</td><td>--|||||-[|||||-||||||]PAM</td><td>NA</td><td>Intergenic</td><td>NA</td><td>NA</td></tr>
<tr><td>GU305914.1</td><td>45255</td><td>45277</td><td>-</td><td>2</td><td>GGTGGCGGTGGCGGCAGCGG</td><td>CGG</td><td>||-|||||[|||||||-||||]PAM</td><td>NA</td><td>Intergenic</td><td>NA</td><td>NA</td></tr>
<tr><td>GU305914.1</td><td>18653</td><td>18675</td><td>-</td><td>3</td><td>GGCAGAGGTGGCGGAGGCGG</td><td>GGG</td><td>|||-|-||[||||||-|||||]PAM</td><td>NA</td><td>Intergenic</td><td>NA</td><td>NA</td></tr>
<tr><td>GU305914.1</td><td>74930</td><td>74952</td><td>+</td><td>4</td><td>GGCGCAGGTTGAGGCGGCGG</td><td>TGG</td><td>||||--||[|-|-||||||||]PAM</td><td>NA</td><td>Intergenic</td><td>NA</td><td>NA</td></tr>
<tr><td>GU305914.1</td><td>48153</td><td>48175</td><td>+</td><td>4</td><td>TGTGGCGGCGGCGGTGGCGG</td><td>CGG</td><td>-|-|||||[-|||||-|||||]PAM</td><td>NA</td><td>Intergenic</td><td>NA</td><td>NA</td></tr>
<tr><td>GU305914.1</td><td>52716</td><td>52738</td><td>+</td><td>4</td><td>GCCGCCGCTGGCGGTGGCGG</td><td>TGG</td><td>|-||-||-[||||||-|||||]PAM</td><td>NA</td><td>Intergenic</td><td>NA</td><td>NA</td></tr>
<tr><td>GU305914.1</td><td>114661</td><td>114683</td><td>-</td><td>3</td><td>GACGGCGGTGGTGGTGGCGG</td><td>CGG</td><td>|-||||||[|||-||-|||||]PAM</td><td>NA</td><td>Intergenic</td><td>NA</td><td>NA</td></tr>
<tr><td>GU305914.1</td><td>62434</td><td>62456</td><td>+</td><td>2</td><td>GACGGCGGTGGCGGCGGGGG</td><td>TGG</td><td>|-||||||[|||||||||-||]PAM</td><td>NA</td><td>Intergenic</td><td>NA</td><td>NA</td></tr>
<tr><td>GU305914.1</td><td>168814</td><td>168836</td><td>+</td><td>3</td><td>AGCGGCGGTGGTGGCAGCGG</td><td>CGG</td><td>-|||||||[|||-|||-||||]PAM</td><td>NA</td><td>Intergenic</td><td>NA</td><td>NA</td></tr>
<tr><td>GU305914.1</td><td>62449</td><td>62471</td><td>+</td><td>3</td><td>GGGGGTGGTGGCGGCGGGGG</td><td>TGG</td><td>||-||-||[|||||||||-||]PAM</td><td>NA</td><td>Intergenic</td><td>NA</td><td>NA</td></tr>
<tr><td>GU305914.1</td><td>30006</td><td>30028</td><td>+</td><td>2</td><td>GGCGGCGGCGGCGGCGGCAG</td><td>TGG</td><td>||||||||[-|||||||||-|]PAM</td><td>NA</td><td>Intergenic</td><td>NA</td><td>NA</td></tr>
<tr><td>GU305914.1</td><td>149476</td><td>149498</td><td>-</td><td>2</td><td>GGCGGCGGCGGCGGCGGCAG</td><td>CGG</td><td>||||||||[-|||||||||-|]PAM</td><td>NA</td><td>Intergenic</td><td>NA</td><td>NA</td></tr>
<tr><td>GU305914.1</td><td>163902</td><td>163924</td><td>+</td><td>3</td><td>AGCGGCGGCGGCGGCGGCAG</td><td>CGG</td><td>-|||||||[-|||||||||-|]PAM</td><td>NA</td><td>Intergenic</td><td>NA</td><td>NA</td></tr>
<tr><td>GU305914.1</td><td>109328</td><td>109350</td><td>+</td><td>4</td><td>GACGGAGGTGGCGGGGACGG</td><td>CGG</td><td>|-|||-||[||||||-|-|||]PAM</td><td>NA</td><td>Intergenic</td><td>NA</td><td>NA</td></tr>
<tr><td>GU305914.1</td><td>113924</td><td>113946</td><td>-</td><td>4</td><td>GTCGGCGTCGGCGGCGGCAG</td><td>CGG</td><td>|-|||||-[-|||||||||-|]PAM</td><td>NA</td><td>Intergenic</td><td>NA</td><td>NA</td></tr>
<tr><td>GU305914.1</td><td>56692</td><td>56714</td><td>+</td><td>4</td><td>GGATGCGGTGGCAGCGGCCG</td><td>CGG</td><td>||--||||[||||-|||||-|]PAM</td><td>NA</td><td>Intergenic</td><td>NA</td><td>NA</td></tr>
</table>

T3119 GCGGCGGTGGCGGCGGCGGTGGG 1000
Oligo fwd CACCGCGGCGGTGGCGGCGGCGGT
Oligo rev AAACACCGCCGCCGCCACCGCCGC

<table>
<tr><th>Chromosome</th><th>start</th><th>end</th><th>strand</th><th>MM</th><th>target_seq</th><th>PAM</th><th>alignment</th><th>distance</th><th>position</th><th>gene name</th><th>gene id</th></tr>
<tr><td>GU305914.1</td><td>48160</td><td>48182</td><td>+</td><td>0</td><td>GCGGCGGTGGCGGCGGCGGT</td><td>GGG</td><td>||||||||[||||||||||||]PAM</td><td>NA</td><td>Intergenic</td><td>NA</td><td>NA</td></tr>
<tr><td>GU305914.1</td><td>75852</td><td>75874</td><td>+</td><td>4</td><td>GGGGAGGCGGAGGCGGCGGT</td><td>CGG</td><td>|-||-||-[||-|||||||||]PAM</td><td>NA</td><td>Intergenic</td><td>NA</td><td>NA</td></tr>
<tr><td>GU305914.1</td><td>58122</td><td>58144</td><td>-</td><td>3</td><td>GTGGCGGCGGCGGCGGCGGA</td><td>GGG</td><td>|-|||||-[|||||||||||-]PAM</td><td>NA</td><td>Intergenic</td><td>NA</td><td>NA</td></tr>
<tr><td>GU305914.1</td><td>67743</td><td>67765</td><td>+</td><td>4</td><td>GGAGCGGCGGCGGCGGCGGC</td><td>GGG</td><td>|--||||-[|||||||||||-]PAM</td><td>NA</td><td>Intergenic</td><td>NA</td><td>NA</td></tr>
<tr><td>GU305914.1</td><td>114657</td><td>114679</td><td>-</td><td>3</td><td>GCGGTGGTGGTGGCGGCGGC</td><td>GGG</td><td>||||-|||[||-||||||||-]PAM</td><td>NA</td><td>Intergenic</td><td>NA</td><td>NA</td></tr>
<tr><td>GU305914.1</td><td>221599</td><td>221621</td><td>-</td><td>4</td><td>GAGGCCGTGGTGGCGGCGGC</td><td>AGG</td><td>|-|||-||[||-||||||||-]PAM</td><td>NA</td><td>Intergenic</td><td>NA</td><td>NA</td></tr>
<tr><td>GU305914.1</td><td>26383</td><td>26405</td><td>+</td><td>4</td><td>TCGGCGCTGGCGACGGCGGC</td><td>GGG</td><td>-|||||-|[||||-||||||-]PAM</td><td>NA</td><td>Intergenic</td><td>NA</td><td>NA</td></tr>
</table>

GU305914.1 149475 149497 - 3 GCGGCGGCGGCGGCGGCAGC GGG |||||||-[|||||||||-|-]PAM NA Intergenic NA NA

T3120 TCGCGCACGTTGCCGACGTGCGG 1000
Oligo fwd CACCTCGCGCACGTTGCCGACGTG
Oligo rev AAACCACGTCGGCAACGTGCGCGA
Chromosome start end strand MM target_seq PAM alignment distance position gene name gene id
GU305914.1 48186 48208 + 0 TCGCGCACGTTGCCGACGTG CGG |||||||[|||||||||||||]PAM NA Intergenic NA NA

T3121 GTTGCCGACGTGCGGACAGTAGG 1000
Oligo fwd CACCGTTGCCGACGTGCGGACAGT
Oligo rev AAACACTGTCCGCACGTCGGCAAC
Chromosome start end strand MM target_seq PAM alignment distance position gene name gene id
GU305914.1 48194 48216 + 0 GTTGCCGACGTGCGGACAGT AGG |||||||[|||||||||||||]PAM NA Intergenic NA NA

T3122 TTGCCGACGTGCGGACAGTAGGG 1000
Oligo fwd CACCTTGCCGACGTGCGGACAGTA
Oligo rev AAACTACTGTCCGCACGTCGGCAA
Chromosome start end strand MM target_seq PAM alignment distance position gene name gene id
GU305914.1 48195 48217 + 0 TTGCCGACGTGCGGACAGTA GGG |||||||[|||||||||||||]PAM NA Intergenic NA NA

T3123 GACAGTAGGGAACCTTCTCGAGG 1000
Oligo fwd CACCGACAGTAGGGAACCTTCTCG
Oligo rev AAACCGAGAAGGTTCCCTACTGTC
Chromosome start end strand MM target_seq PAM alignment distance position gene name gene id
GU305914.1 48208 48230 + 0 GACAGTAGGGAACCTTCTCG AGG |||||||[|||||||||||||]PAM NA Intergenic NA NA

T3124 TCTCGAGGCGTCCGATACCGCGG 1000
Oligo fwd CACCTCTCGAGGCGTCCGATACCG
Oligo rev AAACCGGTATCGGACGCCTCGAGA
Chromosome start end strand MM target_seq PAM alignment distance position gene name gene id
GU305914.1 48223 48245 + 0 TCTCGAGGCGTCCGATACCG CGG |||||||[|||||||||||||]PAM NA Intergenic NA NA

T3125 CCTTGTCGTGTCTGAAGTTGAGG 1000
Oligo fwd CACCCCTTGTCGTGTCTGAAGTTG
Oligo rev AAACCAACTTCAGACACGACAAGG
Chromosome start end strand MM target_seq PAM alignment distance position gene name gene id
GU305914.1 48250 48272 + 0 CCTTGTCGTGTCTGAAGTTG AGG |||||||[|||||||||||||]PAM NA Intergenic NA NA

T3126 TGTCGTGTCTGAAGTTGAGGAGG 1000
Oligo fwd CACCTGTCGTGTCTGAAGTTGAGG
Oligo rev AAACCCTCAACTTCAGACACGACA
Chromosome start end strand MM target_seq PAM alignment distance position gene name gene id
GU305914.1 48253 48275 + 0 TGTCGTGTCTGAAGTTGAGG AGG |||||||[|||||||||||||]PAM NA Intergenic NA NA

T3127 TGAAGTTGAGGAGGCTCTTTCGG 1000
Oligo fwd CACCTGAAGTTGAGGAGGCTCTTT
Oligo rev AAACAAAGAGCCTCCTCAACTTCA
Chromosome start end strand MM target_seq PAM alignment distance position gene name gene id
GU305914.1 48262 48284 + 0 TGAAGTTGAGGAGGCTCTTT CGG |||||||[|||||||||||||]PAM NA Intergenic NA NA

T3128 GGCTCTTTCGGCTGCCGATGCGG 1000
Oligo fwd CACCGGCTCTTTCGGCTGCCGATG
Oligo rev AAACCATCGGCAGCCGAAAGAGCC
Chromosome start end strand MM target_seq PAM alignment distance position gene name gene id
GU305914.1 48274 48296 + 0 GGCTCTTTCGGCTGCCGATG CGG |||||||[|||||||||||||]PAM NA Intergenic NA NA

T3129 GATGAACATATTAAAGTTGTCGG 1000
Oligo fwd CACCGATGAACATATTAAAGTTGT
Oligo rev AAACACAACTTTAATATGTTCATC
Chromosome start end strand MM target_seq PAM alignment distance position gene name gene id
GU305914.1 48296 48318 + 0 GATGAACATATTAAAGTTGT CGG |||||||[|||||||||||||]PAM NA Intergenic NA NA

T3130 ACATATTAAAGTTGTCGGCCAGG 1000
Oligo fwd CACCACATATTAAAGTTGTCGGCC
Oligo rev AAACGGCCGACAACTTTAATATGT
Chromosome start end strand MM target_seq PAM alignment distance position gene name gene id
GU305914.1 48301 48323 + 0 ACATATTAAAGTTGTCGGCC AGG |||||||[|||||||||||||]PAM NA Intergenic NA NA

T3131 ACAGTTGTCGTATCCGAAGATGG 1000
Oligo fwd CACCACAGTTGTCGTATCCGAAGA
Oligo rev AAACTCTTCGGATACGACAACTGT
Chromosome start end strand MM target_seq PAM alignment distance position gene name gene id
GU305914.1 48341 48363 + 0 ACAGTTGTCGTATCCGAAGA TGG |||||||[|||||||||||||]PAM NA Intergenic NA NA

T3132 GTATCCGAAGATGGCGCCGAAGG 1000
Oligo fwd CACCGTATCCGAAGATGGCGCCGA
Oligo rev AAACTCGGCGCCATCTTCGGATAC
Chromosome start end strand MM target_seq PAM alignment distance position gene name gene id
GU305914.1 48350 48372 + 0 GTATCCGAAGATGGCGCCGA AGG |||||||[|||||||||||||]PAM NA Intergenic NA NA

T3133 TATCCGAAGATGGCGCCGAAGGG 1000
Oligo fwd CACCTATCCGAAGATGGCGCCGAA
Oligo rev AAACTTCGGCGCCATCTTCGGATA
Chromosome start end strand MM target_seq PAM alignment distance position gene name gene id
GU305914.1 48351 48373 + 0 TATCCGAAGATGGCGCCGAA GGG |||||||[|||||||||||||]PAM NA Intergenic NA NA

T3134 CGCCGAAGGGATCGATGATGAGG 1000
Oligo fwd CACCCGCCGAAGGGATCGATGATG
Oligo rev AAACCATCATCGATCCCTTCGGCG

<table>
<tr><td>Chromosome</td><td>start</td><td>end</td><td>strand</td><td>MM</td><td>target_seq</td><td>PAM</td><td>alignment</td><td>distance</td><td>position</td><td>gene name</td><td>gene id</td></tr>
<tr><td>GU305914.1</td><td>48364</td><td>48386</td><td>+</td><td>0</td><td>CGCCGAAGGGATCGATGATG</td><td>AGG</td><td>||||||||[||||||||||||]PAM</td><td>NA</td><td>Intergenic</td><td>NA</td><td>NA</td></tr>
<tr><td>T3135</td><td>GATCGATGATGAGGATGAGCCGG</td><td>1000</td><td></td><td></td><td></td><td></td><td></td><td></td><td></td><td></td><td></td></tr>
<tr><td>Oligo fwd</td><td>CACCGATCGATGATGAGGATGAGC</td><td></td><td></td><td></td><td></td><td></td><td></td><td></td><td></td><td></td><td></td></tr>
<tr><td>Oligo rev</td><td>AAACGCTCATCCTCATCATCGATC</td><td></td><td></td><td></td><td></td><td></td><td></td><td></td><td></td><td></td><td></td></tr>
<tr><td>Chromosome</td><td>start</td><td>end</td><td>strand</td><td>MM</td><td>target_seq</td><td>PAM</td><td>alignment</td><td>distance</td><td>position</td><td>gene name</td><td>gene id</td></tr>
<tr><td>GU305914.1</td><td>48373</td><td>48395</td><td>+</td><td>0</td><td>GATCGATGATGAGGATGAGC</td><td>CGG</td><td>||||||||[||||||||||||]PAM</td><td>NA</td><td>Intergenic</td><td>NA</td><td>NA</td></tr>
<tr><td>T3136</td><td>ATCGATGATGAGGATGAGCCGGG</td><td>1000</td><td></td><td></td><td></td><td></td><td></td><td></td><td></td><td></td><td></td></tr>
<tr><td>Oligo fwd</td><td>CACCATCGATGATGAGGATGAGCC</td><td></td><td></td><td></td><td></td><td></td><td></td><td></td><td></td><td></td><td></td></tr>
<tr><td>Oligo rev</td><td>AAACGGCTCATCCTCATCATCGAT</td><td></td><td></td><td></td><td></td><td></td><td></td><td></td><td></td><td></td><td></td></tr>
<tr><td>Chromosome</td><td>start</td><td>end</td><td>strand</td><td>MM</td><td>target_seq</td><td>PAM</td><td>alignment</td><td>distance</td><td>position</td><td>gene name</td><td>gene id</td></tr>
<tr><td>GU305914.1</td><td>48374</td><td>48396</td><td>+</td><td>0</td><td>ATCGATGATGAGGATGAGCC</td><td>GGG</td><td>||||||||[||||||||||||]PAM</td><td>NA</td><td>Intergenic</td><td>NA</td><td>NA</td></tr>
<tr><td>GU305914.1</td><td>178105</td><td>178127</td><td>-</td><td>4</td><td>AGCCATCATGAGGATGAGTC</td><td>TGG</td><td>|-|-||-|[||||||||||-|]PAM</td><td>NA</td><td>Intergenic</td><td>NA</td><td>NA</td></tr>
<tr><td>T3137</td><td>TCGATGATGAGGATGAGCCGGGG</td><td>1000</td><td></td><td></td><td></td><td></td><td></td><td></td><td></td><td></td><td></td></tr>
<tr><td>Oligo fwd</td><td>CACCTCGATGATGAGGATGAGCCG</td><td></td><td></td><td></td><td></td><td></td><td></td><td></td><td></td><td></td><td></td></tr>
<tr><td>Oligo rev</td><td>AAACCGGCTCATCCTCATCATCGA</td><td></td><td></td><td></td><td></td><td></td><td></td><td></td><td></td><td></td><td></td></tr>
<tr><td>Chromosome</td><td>start</td><td>end</td><td>strand</td><td>MM</td><td>target_seq</td><td>PAM</td><td>alignment</td><td>distance</td><td>position</td><td>gene name</td><td>gene id</td></tr>
<tr><td>GU305914.1</td><td>48375</td><td>48397</td><td>+</td><td>0</td><td>TCGATGATGAGGATGAGCCG</td><td>GGG</td><td>||||||||[||||||||||||]PAM</td><td>NA</td><td>Intergenic</td><td>NA</td><td>NA</td></tr>
<tr><td>T3138</td><td>TGATGAGGATGAGCCGGGGTCGG</td><td>1000</td><td></td><td></td><td></td><td></td><td></td><td></td><td></td><td></td><td></td></tr>
<tr><td>Oligo fwd</td><td>CACCTGATGAGGATGAGCCGGGGT</td><td></td><td></td><td></td><td></td><td></td><td></td><td></td><td></td><td></td><td></td></tr>
<tr><td>Oligo rev</td><td>AAACACCCCGGCTCATCCTCATCA</td><td></td><td></td><td></td><td></td><td></td><td></td><td></td><td></td><td></td><td></td></tr>
<tr><td>Chromosome</td><td>start</td><td>end</td><td>strand</td><td>MM</td><td>target_seq</td><td>PAM</td><td>alignment</td><td>distance</td><td>position</td><td>gene name</td><td>gene id</td></tr>
<tr><td>GU305914.1</td><td>48379</td><td>48401</td><td>+</td><td>0</td><td>TGATGAGGATGAGCCGGGGT</td><td>CGG</td><td>||||||||[||||||||||||]PAM</td><td>NA</td><td>Intergenic</td><td>NA</td><td>NA</td></tr>
<tr><td>T3139</td><td>GAGCCGGGGTCGGAAGACCGAGG</td><td>1000</td><td></td><td></td><td></td><td></td><td></td><td></td><td></td><td></td><td></td></tr>
<tr><td>Oligo fwd</td><td>CACCGAGCCGGGGTCGGAAGACCG</td><td></td><td></td><td></td><td></td><td></td><td></td><td></td><td></td><td></td><td></td></tr>
<tr><td>Oligo rev</td><td>AAACCGGTCTTCCGACCCCGGCTC</td><td></td><td></td><td></td><td></td><td></td><td></td><td></td><td></td><td></td><td></td></tr>
<tr><td>Chromosome</td><td>start</td><td>end</td><td>strand</td><td>MM</td><td>target_seq</td><td>PAM</td><td>alignment</td><td>distance</td><td>position</td><td>gene name</td><td>gene id</td></tr>
<tr><td>GU305914.1</td><td>48389</td><td>48411</td><td>+</td><td>0</td><td>GAGCCGGGGTCGGAAGACCG</td><td>AGG</td><td>||||||||[||||||||||||]PAM</td><td>NA</td><td>Intergenic</td><td>NA</td><td>NA</td></tr>
<tr><td>T3140</td><td>GGGTCGGAAGACCGAGGTCGCGG</td><td>1000</td><td></td><td></td><td></td><td></td><td></td><td></td><td></td><td></td><td></td></tr>
<tr><td>Oligo fwd</td><td>CACCGGGTCGGAAGACCGAGGTCG</td><td></td><td></td><td></td><td></td><td></td><td></td><td></td><td></td><td></td><td></td></tr>
<tr><td>Oligo rev</td><td>AAACCGACCTCGGTCTTCCGACCC</td><td></td><td></td><td></td><td></td><td></td><td></td><td></td><td></td><td></td><td></td></tr>
<tr><td>Chromosome</td><td>start</td><td>end</td><td>strand</td><td>MM</td><td>target_seq</td><td>PAM</td><td>alignment</td><td>distance</td><td>position</td><td>gene name</td><td>gene id</td></tr>
<tr><td>GU305914.1</td><td>48395</td><td>48417</td><td>+</td><td>0</td><td>GGGTCGGAAGACCGAGGTCG</td><td>CGG</td><td>||||||||[||||||||||||]PAM</td><td>NA</td><td>Intergenic</td><td>NA</td><td>NA</td></tr>
<tr><td>T3141</td><td>CGCGGCCTTGTATCCGACCACGG</td><td>1000</td><td></td><td></td><td></td><td></td><td></td><td></td><td></td><td></td><td></td></tr>
<tr><td>Oligo fwd</td><td>CACCCGCGGCCTTGTATCCGACCA</td><td></td><td></td><td></td><td></td><td></td><td></td><td></td><td></td><td></td><td></td></tr>
<tr><td>Oligo rev</td><td>AAACTGGTCGGATACAAGGCCGCG</td><td></td><td></td><td></td><td></td><td></td><td></td><td></td><td></td><td></td><td></td></tr>
<tr><td>Chromosome</td><td>start</td><td>end</td><td>strand</td><td>MM</td><td>target_seq</td><td>PAM</td><td>alignment</td><td>distance</td><td>position</td><td>gene name</td><td>gene id</td></tr>
<tr><td>GU305914.1</td><td>48413</td><td>48435</td><td>+</td><td>0</td><td>CGCGGCCTTGTATCCGACCA</td><td>CGG</td><td>||||||||[||||||||||||]PAM</td><td>NA</td><td>Intergenic</td><td>NA</td><td>NA</td></tr>
<tr><td>T3142</td><td>GACCACGGCGAAGATGTCAAAGG</td><td>1000</td><td></td><td></td><td></td><td></td><td></td><td></td><td></td><td></td><td></td></tr>
<tr><td>Oligo fwd</td><td>CACCGACCACGGCGAAGATGTCAA</td><td></td><td></td><td></td><td></td><td></td><td></td><td></td><td></td><td></td><td></td></tr>
<tr><td>Oligo rev</td><td>AAACTTGACATCTTCGCCGTGGTC</td><td></td><td></td><td></td><td></td><td></td><td></td><td></td><td></td><td></td><td></td></tr>
<tr><td>Chromosome</td><td>start</td><td>end</td><td>strand</td><td>MM</td><td>target_seq</td><td>PAM</td><td>alignment</td><td>distance</td><td>position</td><td>gene name</td><td>gene id</td></tr>
<tr><td>GU305914.1</td><td>48428</td><td>48450</td><td>+</td><td>0</td><td>GACCACGGCGAAGATGTCAA</td><td>AGG</td><td>||||||||[||||||||||||]PAM</td><td>NA</td><td>Intergenic</td><td>NA</td><td>NA</td></tr>
<tr><td>T3143</td><td>GGCGAAGATGTCAAAGGTGCTGG</td><td>1000</td><td></td><td></td><td></td><td></td><td></td><td></td><td></td><td></td><td></td></tr>
<tr><td>Oligo fwd</td><td>CACCGGCGAAGATGTCAAAGGTGC</td><td></td><td></td><td></td><td></td><td></td><td></td><td></td><td></td><td></td><td></td></tr>
<tr><td>Oligo rev</td><td>AAACGCACCTTTGACATCTTCGCC</td><td></td><td></td><td></td><td></td><td></td><td></td><td></td><td></td><td></td><td></td></tr>
<tr><td>Chromosome</td><td>start</td><td>end</td><td>strand</td><td>MM</td><td>target_seq</td><td>PAM</td><td>alignment</td><td>distance</td><td>position</td><td>gene name</td><td>gene id</td></tr>
<tr><td>GU305914.1</td><td>48434</td><td>48456</td><td>+</td><td>0</td><td>GGCGAAGATGTCAAAGGTGC</td><td>TGG</td><td>||||||||[||||||||||||]PAM</td><td>NA</td><td>Intergenic</td><td>NA</td><td>NA</td></tr>
<tr><td>T3144</td><td>CAAAGGTGCTGGAGAACCTCCGG</td><td>1000</td><td></td><td></td><td></td><td></td><td></td><td></td><td></td><td></td><td></td></tr>
<tr><td>Oligo fwd</td><td>CACCCAAAGGTGCTGGAGAACCTC</td><td></td><td></td><td></td><td></td><td></td><td></td><td></td><td></td><td></td><td></td></tr>
<tr><td>Oligo rev</td><td>AAACGAGGTTCTCCAGCACCTTTG</td><td></td><td></td><td></td><td></td><td></td><td></td><td></td><td></td><td></td><td></td></tr>
<tr><td>Chromosome</td><td>start</td><td>end</td><td>strand</td><td>MM</td><td>target_seq</td><td>PAM</td><td>alignment</td><td>distance</td><td>position</td><td>gene name</td><td>gene id</td></tr>
<tr><td>GU305914.1</td><td>48445</td><td>48467</td><td>+</td><td>0</td><td>CAAAGGTGCTGGAGAACCTC</td><td>CGG</td><td>||||||||[||||||||||||]PAM</td><td>NA</td><td>Intergenic</td><td>NA</td><td>NA</td></tr>
<tr><td>T3145</td><td>GCTGGAGAACCTCCGGACGACGG</td><td>1000</td><td></td><td></td><td></td><td></td><td></td><td></td><td></td><td></td><td></td></tr>
<tr><td>Oligo fwd</td><td>CACCGCTGGAGAACCTCCGGACGA</td><td></td><td></td><td></td><td></td><td></td><td></td><td></td><td></td><td></td><td></td></tr>
<tr><td>Oligo rev</td><td>AAACTCGTCCGGAGGTTCTCCAGC</td><td></td><td></td><td></td><td></td><td></td><td></td><td></td><td></td><td></td><td></td></tr>
<tr><td>Chromosome</td><td>start</td><td>end</td><td>strand</td><td>MM</td><td>target_seq</td><td>PAM</td><td>alignment</td><td>distance</td><td>position</td><td>gene name</td><td>gene id</td></tr>
<tr><td>GU305914.1</td><td>48452</td><td>48474</td><td>+</td><td>0</td><td>GCTGGAGAACCTCCGGACGA</td><td>CGG</td><td>||||||||[||||||||||||]PAM</td><td>NA</td><td>Intergenic</td><td>NA</td><td>NA</td></tr>
<tr><td>T3146</td><td>CCGGACGACGGAGCACATGACGG</td><td>1000</td><td></td><td></td><td></td><td></td><td></td><td></td><td></td><td></td><td></td></tr>
<tr><td>Oligo fwd</td><td>CACCCCGGACGACGGAGCACATGA</td><td></td><td></td><td></td><td></td><td></td><td></td><td></td><td></td><td></td><td></td></tr>
<tr><td>Oligo rev</td><td>AAACTCATGTGCTCCGTCGTCCGG</td><td></td><td></td><td></td><td></td><td></td><td></td><td></td><td></td><td></td><td></td></tr>
<tr><td>Chromosome</td><td>start</td><td>end</td><td>strand</td><td>MM</td><td>target_seq</td><td>PAM</td><td>alignment</td><td>distance</td><td>position</td><td>gene name</td><td>gene id</td></tr>
<tr><td>GU305914.1</td><td>48464</td><td>48486</td><td>+</td><td>0</td><td>CCGGACGACGGAGCACATGA</td><td>CGG</td><td>||||||||[||||||||||||]PAM</td><td>NA</td><td>Intergenic</td><td>NA</td><td>NA</td></tr>
<tr><td>T3147</td><td>GACGACGGAGCACATGACGGAGG</td><td>1000</td><td></td><td></td><td></td><td></td><td></td><td></td><td></td><td></td><td></td></tr>
<tr><td>Oligo fwd</td><td>CACCGACGACGGAGCACATGACGG</td><td></td><td></td><td></td><td></td><td></td><td></td><td></td><td></td><td></td><td></td></tr>
<tr><td>Oligo rev</td><td>AAACCCGTCATGTGCTCCGTCGTC</td><td></td><td></td><td></td><td></td><td></td><td></td><td></td><td></td><td></td><td></td></tr>
<tr><td>Chromosome</td><td>start</td><td>end</td><td>strand</td><td>MM</td><td>target_seq</td><td>PAM</td><td>alignment</td><td>distance</td><td>position</td><td>gene name</td><td>gene id</td></tr>
<tr><td>GU305914.1</td><td>48467</td><td>48489</td><td>+</td><td>0</td><td>GACGACGGAGCACATGACGG</td><td>AGG</td><td>||||||||[||||||||||||]PAM</td><td>NA</td><td>Intergenic</td><td>NA</td><td>NA</td></tr>
<tr><td>T3148</td><td>CGGAGCACATGACGGAGGTGAGG</td><td>1000</td><td></td><td></td><td></td><td></td><td></td><td></td><td></td><td></td><td></td></tr>
<tr><td>Oligo fwd</td><td>CACCCGGAGCACATGACGGAGGTG</td><td></td><td></td><td></td><td></td><td></td><td></td><td></td><td></td><td></td><td></td></tr>
<tr><td>Oligo rev</td><td>AAACCACCTCCGTCATGTGCTCCG</td><td></td><td></td><td></td><td></td><td></td><td></td><td></td><td></td><td></td><td></td></tr>
<tr><td>Chromosome</td><td>start</td><td>end</td><td>strand</td><td>MM</td><td>target_seq</td><td>PAM</td><td>alignment</td><td>distance</td><td>position</td><td>gene name</td><td>gene id</td></tr>
<tr><td>GU305914.1</td><td>48472</td><td>48494</td><td>+</td><td>0</td><td>CGGAGCACATGACGGAGGTG</td><td>AGG</td><td>||||||||[||||||||||||]PAM</td><td>NA</td><td>Intergenic</td><td>NA</td><td>NA</td></tr>
<tr><td>T3149</td><td>TGAGGTTGATGACCGTACCCAGG</td><td>1000</td><td></td><td></td><td></td><td></td><td></td><td></td><td></td><td></td><td></td></tr>
</table>

<table>
<tr><td>Oligo fwd</td><td>CACCTGAGGTTGATGACCGTACCC</td><td></td><td></td><td></td><td></td><td></td><td></td><td></td><td></td><td></td><td></td></tr>
<tr><td>Oligo rev</td><td>AAACGGGTACGGTCATCAACCTCA</td><td></td><td></td><td></td><td></td><td></td><td></td><td></td><td></td><td></td><td></td></tr>
<tr><td>Chromosome</td><td>start</td><td>end</td><td>strand</td><td>MM</td><td>target_seq</td><td>PAM</td><td>alignment</td><td>distance</td><td>position</td><td>gene name</td><td>gene id</td></tr>
<tr><td>GU305914.1</td><td>48490</td><td>48512</td><td>+</td><td>0</td><td>TGAGGTTGATGACCGTACCC</td><td>AGG</td><td>||||||||[||||||||||||]PAM</td><td>NA</td><td>Intergenic</td><td>NA</td><td>NA</td></tr>
<tr><td>T3150</td><td>GATGACCGTACCCAGGCTGCAGG</td><td>1000</td><td></td><td></td><td></td><td></td><td></td><td></td><td></td><td></td><td></td></tr>
<tr><td>Oligo fwd</td><td>CACCGATGACCGTACCCAGGCTGC</td><td></td><td></td><td></td><td></td><td></td><td></td><td></td><td></td><td></td><td></td></tr>
<tr><td>Oligo rev</td><td>AAACGCAGCCTGGGTACGGTCATC</td><td></td><td></td><td></td><td></td><td></td><td></td><td></td><td></td><td></td><td></td></tr>
<tr><td>Chromosome</td><td>start</td><td>end</td><td>strand</td><td>MM</td><td>target_seq</td><td>PAM</td><td>alignment</td><td>distance</td><td>position</td><td>gene name</td><td>gene id</td></tr>
<tr><td>GU305914.1</td><td>48497</td><td>48519</td><td>+</td><td>0</td><td>GATGACCGTACCCAGGCTGC</td><td>AGG</td><td>||||||||[||||||||||||]PAM</td><td>NA</td><td>Intergenic</td><td>NA</td><td>NA</td></tr>
<tr><td>T3151</td><td>GACCGTACCCAGGCTGCAGGAGG</td><td>1000</td><td></td><td></td><td></td><td></td><td></td><td></td><td></td><td></td><td></td></tr>
<tr><td>Oligo fwd</td><td>CACCGACCGTACCCAGGCTGCAGG</td><td></td><td></td><td></td><td></td><td></td><td></td><td></td><td></td><td></td><td></td></tr>
<tr><td>Oligo rev</td><td>AAACCCTGCAGCCTGGGTACGGTC</td><td></td><td></td><td></td><td></td><td></td><td></td><td></td><td></td><td></td><td></td></tr>
<tr><td>Chromosome</td><td>start</td><td>end</td><td>strand</td><td>MM</td><td>target_seq</td><td>PAM</td><td>alignment</td><td>distance</td><td>position</td><td>gene name</td><td>gene id</td></tr>
<tr><td>GU305914.1</td><td>48500</td><td>48522</td><td>+</td><td>0</td><td>GACCGTACCCAGGCTGCAGG</td><td>AGG</td><td>||||||||[||||||||||||]PAM</td><td>NA</td><td>Intergenic</td><td>NA</td><td>NA</td></tr>
<tr><td>T3152</td><td>ACCCAGGCTGCAGGAGGTGACGG</td><td>1000</td><td></td><td></td><td></td><td></td><td></td><td></td><td></td><td></td><td></td></tr>
<tr><td>Oligo fwd</td><td>CACCACCCAGGCTGCAGGAGGTGA</td><td></td><td></td><td></td><td></td><td></td><td></td><td></td><td></td><td></td><td></td></tr>
<tr><td>Oligo rev</td><td>AAACTCACCTCCTGCAGCCTGGGT</td><td></td><td></td><td></td><td></td><td></td><td></td><td></td><td></td><td></td><td></td></tr>
<tr><td>Chromosome</td><td>start</td><td>end</td><td>strand</td><td>MM</td><td>target_seq</td><td>PAM</td><td>alignment</td><td>distance</td><td>position</td><td>gene name</td><td>gene id</td></tr>
<tr><td>GU305914.1</td><td>48506</td><td>48528</td><td>+</td><td>0</td><td>ACCCAGGCTGCAGGAGGTGA</td><td>CGG</td><td>||||||||[||||||||||||]PAM</td><td>NA</td><td>Intergenic</td><td>NA</td><td>NA</td></tr>
<tr><td>T3153</td><td>CGCCGCGATAGAGCATCATCTGG</td><td>1000</td><td></td><td></td><td></td><td></td><td></td><td></td><td></td><td></td><td></td></tr>
<tr><td>Oligo fwd</td><td>CACCCGCCGCGATAGAGCATCATC</td><td></td><td></td><td></td><td></td><td></td><td></td><td></td><td></td><td></td><td></td></tr>
<tr><td>Oligo rev</td><td>AAACGATGATGCTCTATCGCGGCG</td><td></td><td></td><td></td><td></td><td></td><td></td><td></td><td></td><td></td><td></td></tr>
<tr><td>Chromosome</td><td>start</td><td>end</td><td>strand</td><td>MM</td><td>target_seq</td><td>PAM</td><td>alignment</td><td>distance</td><td>position</td><td>gene name</td><td>gene id</td></tr>
<tr><td>GU305914.1</td><td>48538</td><td>48560</td><td>+</td><td>0</td><td>CGCCGCGATAGAGCATCATC</td><td>TGG</td><td>||||||||[||||||||||||]PAM</td><td>NA</td><td>Intergenic</td><td>NA</td><td>NA</td></tr>
<tr><td>T3154</td><td>GCCGCGATAGAGCATCATCTGGG</td><td>1000</td><td></td><td></td><td></td><td></td><td></td><td></td><td></td><td></td><td></td></tr>
<tr><td>Oligo fwd</td><td>CACCGCCGCGATAGAGCATCATCT</td><td></td><td></td><td></td><td></td><td></td><td></td><td></td><td></td><td></td><td></td></tr>
<tr><td>Oligo rev</td><td>AAACAGATGATGCTCTATCGCGGC</td><td></td><td></td><td></td><td></td><td></td><td></td><td></td><td></td><td></td><td></td></tr>
<tr><td>Chromosome</td><td>start</td><td>end</td><td>strand</td><td>MM</td><td>target_seq</td><td>PAM</td><td>alignment</td><td>distance</td><td>position</td><td>gene name</td><td>gene id</td></tr>
<tr><td>GU305914.1</td><td>48539</td><td>48561</td><td>+</td><td>0</td><td>GCCGCGATAGAGCATCATCT</td><td>GGG</td><td>||||||||[||||||||||||]PAM</td><td>NA</td><td>Intergenic</td><td>NA</td><td>NA</td></tr>
<tr><td>T3155</td><td>CATCATCTGGGCGTCGATGCCGG</td><td>1000</td><td></td><td></td><td></td><td></td><td></td><td></td><td></td><td></td><td></td></tr>
<tr><td>Oligo fwd</td><td>CACCCATCATCTGGGCGTCGATGC</td><td></td><td></td><td></td><td></td><td></td><td></td><td></td><td></td><td></td><td></td></tr>
<tr><td>Oligo rev</td><td>AAACGCATCGACGCCCAGATGATG</td><td></td><td></td><td></td><td></td><td></td><td></td><td></td><td></td><td></td><td></td></tr>
<tr><td>Chromosome</td><td>start</td><td>end</td><td>strand</td><td>MM</td><td>target_seq</td><td>PAM</td><td>alignment</td><td>distance</td><td>position</td><td>gene name</td><td>gene id</td></tr>
<tr><td>GU305914.1</td><td>48551</td><td>48573</td><td>+</td><td>0</td><td>CATCATCTGGGCGTCGATGC</td><td>CGG</td><td>||||||||[||||||||||||]PAM</td><td>NA</td><td>Intergenic</td><td>NA</td><td>NA</td></tr>
<tr><td>T3156</td><td>CGGCGATGTTGCCGACCATCTGG</td><td>1000</td><td></td><td></td><td></td><td></td><td></td><td></td><td></td><td></td><td></td></tr>
<tr><td>Oligo fwd</td><td>CACCCGGCGATGTTGCCGACCATC</td><td></td><td></td><td></td><td></td><td></td><td></td><td></td><td></td><td></td><td></td></tr>
<tr><td>Oligo rev</td><td>AAACGATGGTCGGCAACATCGCCG</td><td></td><td></td><td></td><td></td><td></td><td></td><td></td><td></td><td></td><td></td></tr>
<tr><td>Chromosome</td><td>start</td><td>end</td><td>strand</td><td>MM</td><td>target_seq</td><td>PAM</td><td>alignment</td><td>distance</td><td>position</td><td>gene name</td><td>gene id</td></tr>
<tr><td>GU305914.1</td><td>48571</td><td>48593</td><td>+</td><td>0</td><td>CGGCGATGTTGCCGACCATC</td><td>TGG</td><td>||||||||[||||||||||||]PAM</td><td>NA</td><td>Intergenic</td><td>NA</td><td>NA</td></tr>
<tr><td>T3157</td><td>GTTGCCGACCATCTGGTAGACGG</td><td>1000</td><td></td><td></td><td></td><td></td><td></td><td></td><td></td><td></td><td></td></tr>
<tr><td>Oligo fwd</td><td>CACCGTTGCCGACCATCTGGTAGA</td><td></td><td></td><td></td><td></td><td></td><td></td><td></td><td></td><td></td><td></td></tr>
<tr><td>Oligo rev</td><td>AAACTCTACCAGATGGTCGGCAAC</td><td></td><td></td><td></td><td></td><td></td><td></td><td></td><td></td><td></td><td></td></tr>
<tr><td>Chromosome</td><td>start</td><td>end</td><td>strand</td><td>MM</td><td>target_seq</td><td>PAM</td><td>alignment</td><td>distance</td><td>position</td><td>gene name</td><td>gene id</td></tr>
<tr><td>GU305914.1</td><td>48578</td><td>48600</td><td>+</td><td>0</td><td>GTTGCCGACCATCTGGTAGA</td><td>CGG</td><td>||||||||[||||||||||||]PAM</td><td>NA</td><td>Intergenic</td><td>NA</td><td>NA</td></tr>
<tr><td>T3158</td><td>ACCCTGATTCTCCGCGACGATGG</td><td>1000</td><td></td><td></td><td></td><td></td><td></td><td></td><td></td><td></td><td></td></tr>
<tr><td>Oligo fwd</td><td>CACCACCCTGATTCTCCGCGACGA</td><td></td><td></td><td></td><td></td><td></td><td></td><td></td><td></td><td></td><td></td></tr>
<tr><td>Oligo rev</td><td>AAACTCGTCGCGGAGAATCAGGGT</td><td></td><td></td><td></td><td></td><td></td><td></td><td></td><td></td><td></td><td></td></tr>
<tr><td>Chromosome</td><td>start</td><td>end</td><td>strand</td><td>MM</td><td>target_seq</td><td>PAM</td><td>alignment</td><td>distance</td><td>position</td><td>gene name</td><td>gene id</td></tr>
<tr><td>GU305914.1</td><td>48602</td><td>48624</td><td>+</td><td>0</td><td>ACCCTGATTCTCCGCGACGA</td><td>TGG</td><td>||||||||[||||||||||||]PAM</td><td>NA</td><td>Intergenic</td><td>NA</td><td>NA</td></tr>
<tr><td>T3159</td><td>CGCGACGATGGAGATGAAAGCGG</td><td>1000</td><td></td><td></td><td></td><td></td><td></td><td></td><td></td><td></td><td></td></tr>
<tr><td>Oligo fwd</td><td>CACCCGCGACGATGGAGATGAAAG</td><td></td><td></td><td></td><td></td><td></td><td></td><td></td><td></td><td></td><td></td></tr>
<tr><td>Oligo rev</td><td>AAACCTTTCATCTCCATCGTCGCG</td><td></td><td></td><td></td><td></td><td></td><td></td><td></td><td></td><td></td><td></td></tr>
<tr><td>Chromosome</td><td>start</td><td>end</td><td>strand</td><td>MM</td><td>target_seq</td><td>PAM</td><td>alignment</td><td>distance</td><td>position</td><td>gene name</td><td>gene id</td></tr>
<tr><td>GU305914.1</td><td>48614</td><td>48636</td><td>+</td><td>0</td><td>CGCGACGATGGAGATGAAAG</td><td>CGG</td><td>||||||||[||||||||||||]PAM</td><td>NA</td><td>Intergenic</td><td>NA</td><td>NA</td></tr>
<tr><td>T3160</td><td>GCGACGATGGAGATGAAAGCGGG</td><td>1000</td><td></td><td></td><td></td><td></td><td></td><td></td><td></td><td></td><td></td></tr>
<tr><td>Oligo fwd</td><td>CACCGCGACGATGGAGATGAAAGC</td><td></td><td></td><td></td><td></td><td></td><td></td><td></td><td></td><td></td><td></td></tr>
<tr><td>Oligo rev</td><td>AAACGCTTTCATCTCCATCGTCGC</td><td></td><td></td><td></td><td></td><td></td><td></td><td></td><td></td><td></td><td></td></tr>
<tr><td>Chromosome</td><td>start</td><td>end</td><td>strand</td><td>MM</td><td>target_seq</td><td>PAM</td><td>alignment</td><td>distance</td><td>position</td><td>gene name</td><td>gene id</td></tr>
<tr><td>GU305914.1</td><td>48615</td><td>48637</td><td>+</td><td>0</td><td>GCGACGATGGAGATGAAAGC</td><td>GGG</td><td>||||||||[||||||||||||]PAM</td><td>NA</td><td>Intergenic</td><td>NA</td><td>NA</td></tr>
<tr><td>T3161</td><td>GATGGAGATGAAAGCGGGCACGG</td><td>1000</td><td></td><td></td><td></td><td></td><td></td><td></td><td></td><td></td><td></td></tr>
<tr><td>Oligo fwd</td><td>CACCGATGGAGATGAAAGCGGGCA</td><td></td><td></td><td></td><td></td><td></td><td></td><td></td><td></td><td></td><td></td></tr>
<tr><td>Oligo rev</td><td>AAACTGCCCGCTTTCATCTCCATC</td><td></td><td></td><td></td><td></td><td></td><td></td><td></td><td></td><td></td><td></td></tr>
<tr><td>Chromosome</td><td>start</td><td>end</td><td>strand</td><td>MM</td><td>target_seq</td><td>PAM</td><td>alignment</td><td>distance</td><td>position</td><td>gene name</td><td>gene id</td></tr>
<tr><td>GU305914.1</td><td>48620</td><td>48642</td><td>+</td><td>0</td><td>GATGGAGATGAAAGCGGGCA</td><td>CGG</td><td>||||||||[||||||||||||]PAM</td><td>NA</td><td>Intergenic</td><td>NA</td><td>NA</td></tr>
<tr><td>T3162</td><td>GATGAAAGCGGGCACGGAGTCGG</td><td>1000</td><td></td><td></td><td></td><td></td><td></td><td></td><td></td><td></td><td></td></tr>
<tr><td>Oligo fwd</td><td>CACCGATGAAAGCGGGCACGGAGT</td><td></td><td></td><td></td><td></td><td></td><td></td><td></td><td></td><td></td><td></td></tr>
<tr><td>Oligo rev</td><td>AAACACTCCGTGCCCGCTTTCATC</td><td></td><td></td><td></td><td></td><td></td><td></td><td></td><td></td><td></td><td></td></tr>
<tr><td>Chromosome</td><td>start</td><td>end</td><td>strand</td><td>MM</td><td>target_seq</td><td>PAM</td><td>alignment</td><td>distance</td><td>position</td><td>gene name</td><td>gene id</td></tr>
<tr><td>GU305914.1</td><td>48626</td><td>48648</td><td>+</td><td>0</td><td>GATGAAAGCGGGCACGGAGT</td><td>CGG</td><td>||||||||[||||||||||||]PAM</td><td>NA</td><td>Intergenic</td><td>NA</td><td>NA</td></tr>
<tr><td>T3163</td><td>CGGGCACGGAGTCGGCCTCCAGG</td><td>1000</td><td></td><td></td><td></td><td></td><td></td><td></td><td></td><td></td><td></td></tr>
<tr><td>Oligo fwd</td><td>CACCCGGGCACGGAGTCGGCCTCC</td><td></td><td></td><td></td><td></td><td></td><td></td><td></td><td></td><td></td><td></td></tr>
<tr><td>Oligo rev</td><td>AAACGGAGGCCGACTCCGTGCCCG</td><td></td><td></td><td></td><td></td><td></td><td></td><td></td><td></td><td></td><td></td></tr>
<tr><td>Chromosome</td><td>start</td><td>end</td><td>strand</td><td>MM</td><td>target_seq</td><td>PAM</td><td>alignment</td><td>distance</td><td>position</td><td>gene name</td><td>gene id</td></tr>
<tr><td>GU305914.1</td><td>48634</td><td>48656</td><td>+</td><td>0</td><td>CGGGCACGGAGTCGGCCTCC</td><td>AGG</td><td>||||||||[||||||||||||]PAM</td><td>NA</td><td>Intergenic</td><td>NA</td><td>NA</td></tr>
</table>

<table>
<tr><td>T3164</td><td>GCACGGAGTCGGCCTCCAGGAGG</td><td>1000</td><td></td><td></td><td></td><td></td><td></td><td></td><td></td><td></td><td></td></tr>
<tr><td>Oligo fwd</td><td>CACCGCACGGAGTCGGCCTCCAGG</td><td></td><td></td><td></td><td></td><td></td><td></td><td></td><td></td><td></td><td></td></tr>
<tr><td>Oligo rev</td><td>AAACCCTGGAGGCCGACTCCGTGC</td><td></td><td></td><td></td><td></td><td></td><td></td><td></td><td></td><td></td><td></td></tr>
<tr><td>Chromosome</td><td>start</td><td>end</td><td>strand</td><td>MM</td><td>target_seq</td><td>PAM</td><td>alignment</td><td>distance</td><td>position</td><td>gene name</td><td>gene id</td></tr>
<tr><td>GU305914.1</td><td>48637</td><td>48659</td><td>+</td><td>0</td><td>GCACGGAGTCGGCCTCCAGG</td><td>AGG</td><td>||||||||[||||||||||||]PAM</td><td>NA</td><td>Intergenic</td><td>NA</td><td>NA</td></tr>
<tr><td></td><td></td><td></td><td></td><td></td><td></td><td></td><td></td><td></td><td></td><td></td><td></td></tr>
<tr><td>T3165</td><td>GGAGTCGGCCTCCAGGAGGCTGG</td><td>1000</td><td></td><td></td><td></td><td></td><td></td><td></td><td></td><td></td><td></td></tr>
<tr><td>Oligo fwd</td><td>CACCGGAGTCGGCCTCCAGGAGGC</td><td></td><td></td><td></td><td></td><td></td><td></td><td></td><td></td><td></td><td></td></tr>
<tr><td>Oligo rev</td><td>AAACGCCTCCTGGAGGCCGACTCC</td><td></td><td></td><td></td><td></td><td></td><td></td><td></td><td></td><td></td><td></td></tr>
<tr><td>Chromosome</td><td>start</td><td>end</td><td>strand</td><td>MM</td><td>target_seq</td><td>PAM</td><td>alignment</td><td>distance</td><td>position</td><td>gene name</td><td>gene id</td></tr>
<tr><td>GU305914.1</td><td>48641</td><td>48663</td><td>+</td><td>0</td><td>GGAGTCGGCCTCCAGGAGGC</td><td>TGG</td><td>||||||||[||||||||||||]PAM</td><td>NA</td><td>Intergenic</td><td>NA</td><td>NA</td></tr>
<tr><td></td><td></td><td></td><td></td><td></td><td></td><td></td><td></td><td></td><td></td><td></td><td></td></tr>
<tr><td>T3166</td><td>GACACCGTCGACGAGAAGTATGG</td><td>1000</td><td></td><td></td><td></td><td></td><td></td><td></td><td></td><td></td><td></td></tr>
<tr><td>Oligo fwd</td><td>CACCGACACCGTCGACGAGAAGTA</td><td></td><td></td><td></td><td></td><td></td><td></td><td></td><td></td><td></td><td></td></tr>
<tr><td>Oligo rev</td><td>AAACTACTTCTCGTCGACGGTGTC</td><td></td><td></td><td></td><td></td><td></td><td></td><td></td><td></td><td></td><td></td></tr>
<tr><td>Chromosome</td><td>start</td><td>end</td><td>strand</td><td>MM</td><td>target_seq</td><td>PAM</td><td>alignment</td><td>distance</td><td>position</td><td>gene name</td><td>gene id</td></tr>
<tr><td>GU305914.1</td><td>48668</td><td>48690</td><td>+</td><td>0</td><td>GACACCGTCGACGAGAAGTA</td><td>TGG</td><td>||||||||[||||||||||||]PAM</td><td>NA</td><td>Intergenic</td><td>NA</td><td>NA</td></tr>
<tr><td></td><td></td><td></td><td></td><td></td><td></td><td></td><td></td><td></td><td></td><td></td><td></td></tr>
<tr><td>T3167</td><td>ACACCGTCGACGAGAAGTATGGG</td><td>1000</td><td></td><td></td><td></td><td></td><td></td><td></td><td></td><td></td><td></td></tr>
<tr><td>Oligo fwd</td><td>CACCACACCGTCGACGAGAAGTAT</td><td></td><td></td><td></td><td></td><td></td><td></td><td></td><td></td><td></td><td></td></tr>
<tr><td>Oligo rev</td><td>AAACATACTTCTCGTCGACGGTGT</td><td></td><td></td><td></td><td></td><td></td><td></td><td></td><td></td><td></td><td></td></tr>
<tr><td>Chromosome</td><td>start</td><td>end</td><td>strand</td><td>MM</td><td>target_seq</td><td>PAM</td><td>alignment</td><td>distance</td><td>position</td><td>gene name</td><td>gene id</td></tr>
<tr><td>GU305914.1</td><td>48669</td><td>48691</td><td>+</td><td>0</td><td>ACACCGTCGACGAGAAGTAT</td><td>GGG</td><td>||||||||[||||||||||||]PAM</td><td>NA</td><td>Intergenic</td><td>NA</td><td>NA</td></tr>
<tr><td></td><td></td><td></td><td></td><td></td><td></td><td></td><td></td><td></td><td></td><td></td><td></td></tr>
<tr><td>T3168</td><td>TATGGGCGCGACCAGCGACGTGG</td><td>1000</td><td></td><td></td><td></td><td></td><td></td><td></td><td></td><td></td><td></td></tr>
<tr><td>Oligo fwd</td><td>CACCTATGGGCGCGACCAGCGACG</td><td></td><td></td><td></td><td></td><td></td><td></td><td></td><td></td><td></td><td></td></tr>
<tr><td>Oligo rev</td><td>AAACCGTCGCTGGTCGCGCCCATA</td><td></td><td></td><td></td><td></td><td></td><td></td><td></td><td></td><td></td><td></td></tr>
<tr><td>Chromosome</td><td>start</td><td>end</td><td>strand</td><td>MM</td><td>target_seq</td><td>PAM</td><td>alignment</td><td>distance</td><td>position</td><td>gene name</td><td>gene id</td></tr>
<tr><td>GU305914.1</td><td>48686</td><td>48708</td><td>+</td><td>0</td><td>TATGGGCGCGACCAGCGACG</td><td>TGG</td><td>||||||||[||||||||||||]PAM</td><td>NA</td><td>Intergenic</td><td>NA</td><td>NA</td></tr>
<tr><td></td><td></td><td></td><td></td><td></td><td></td><td></td><td></td><td></td><td></td><td></td><td></td></tr>
<tr><td>T3169</td><td>GACCAGCGACGTGGCGAAGATGG</td><td>1000</td><td></td><td></td><td></td><td></td><td></td><td></td><td></td><td></td><td></td></tr>
<tr><td>Oligo fwd</td><td>CACCGACCAGCGACGTGGCGAAGA</td><td></td><td></td><td></td><td></td><td></td><td></td><td></td><td></td><td></td><td></td></tr>
<tr><td>Oligo rev</td><td>AAACTCTTCGCCACGTCGCTGGTC</td><td></td><td></td><td></td><td></td><td></td><td></td><td></td><td></td><td></td><td></td></tr>
<tr><td>Chromosome</td><td>start</td><td>end</td><td>strand</td><td>MM</td><td>target_seq</td><td>PAM</td><td>alignment</td><td>distance</td><td>position</td><td>gene name</td><td>gene id</td></tr>
<tr><td>GU305914.1</td><td>48695</td><td>48717</td><td>+</td><td>0</td><td>GACCAGCGACGTGGCGAAGA</td><td>TGG</td><td>||||||||[||||||||||||]PAM</td><td>NA</td><td>Intergenic</td><td>NA</td><td>NA</td></tr>
<tr><td></td><td></td><td></td><td></td><td></td><td></td><td></td><td></td><td></td><td></td><td></td><td></td></tr>
<tr><td>T3170</td><td>ACCAGCGACGTGGCGAAGATGGG</td><td>1000</td><td></td><td></td><td></td><td></td><td></td><td></td><td></td><td></td><td></td></tr>
<tr><td>Oligo fwd</td><td>CACCACCAGCGACGTGGCGAAGAT</td><td></td><td></td><td></td><td></td><td></td><td></td><td></td><td></td><td></td><td></td></tr>
<tr><td>Oligo rev</td><td>AAACATCTTCGCCACGTCGCTGGT</td><td></td><td></td><td></td><td></td><td></td><td></td><td></td><td></td><td></td><td></td></tr>
<tr><td>Chromosome</td><td>start</td><td>end</td><td>strand</td><td>MM</td><td>target_seq</td><td>PAM</td><td>alignment</td><td>distance</td><td>position</td><td>gene name</td><td>gene id</td></tr>
<tr><td>GU305914.1</td><td>48696</td><td>48718</td><td>+</td><td>0</td><td>ACCAGCGACGTGGCGAAGAT</td><td>GGG</td><td>||||||||[||||||||||||]PAM</td><td>NA</td><td>Intergenic</td><td>NA</td><td>NA</td></tr>
<tr><td></td><td></td><td></td><td></td><td></td><td></td><td></td><td></td><td></td><td></td><td></td><td></td></tr>
<tr><td>T3171</td><td>AGATGGGATCGAGCCGACGCAGG</td><td>1000</td><td></td><td></td><td></td><td></td><td></td><td></td><td></td><td></td><td></td></tr>
<tr><td>Oligo fwd</td><td>CACCAGATGGGATCGAGCCGACGC</td><td></td><td></td><td></td><td></td><td></td><td></td><td></td><td></td><td></td><td></td></tr>
<tr><td>Oligo rev</td><td>AAACGCGTCGGCTCGATCCCATCT</td><td></td><td></td><td></td><td></td><td></td><td></td><td></td><td></td><td></td><td></td></tr>
<tr><td>Chromosome</td><td>start</td><td>end</td><td>strand</td><td>MM</td><td>target_seq</td><td>PAM</td><td>alignment</td><td>distance</td><td>position</td><td>gene name</td><td>gene id</td></tr>
<tr><td>GU305914.1</td><td>48712</td><td>48734</td><td>+</td><td>0</td><td>AGATGGGATCGAGCCGACGC</td><td>AGG</td><td>||||||||[||||||||||||]PAM</td><td>NA</td><td>Intergenic</td><td>NA</td><td>NA</td></tr>
<tr><td>GU305914.1</td><td>65252</td><td>65274</td><td>+</td><td>4</td><td>AGACGTGATCGAGAAGACGC</td><td>GGG</td><td>|||-|-||[|||||--|||||]PAM</td><td>NA</td><td>Intergenic</td><td>NA</td><td>NA</td></tr>
<tr><td></td><td></td><td></td><td></td><td></td><td></td><td></td><td></td><td></td><td></td><td></td><td></td></tr>
<tr><td>T3172</td><td>CGAGCCGACGCAGGCCGTCCTGG</td><td>1000</td><td></td><td></td><td></td><td></td><td></td><td></td><td></td><td></td><td></td></tr>
<tr><td>Oligo fwd</td><td>CACCCGAGCCGACGCAGGCCGTCC</td><td></td><td></td><td></td><td></td><td></td><td></td><td></td><td></td><td></td><td></td></tr>
<tr><td>Oligo rev</td><td>AAACGGACGGCCTGCGTCGGCTCG</td><td></td><td></td><td></td><td></td><td></td><td></td><td></td><td></td><td></td><td></td></tr>
<tr><td>Chromosome</td><td>start</td><td>end</td><td>strand</td><td>MM</td><td>target_seq</td><td>PAM</td><td>alignment</td><td>distance</td><td>position</td><td>gene name</td><td>gene id</td></tr>
<tr><td>GU305914.1</td><td>48721</td><td>48743</td><td>+</td><td>0</td><td>CGAGCCGACGCAGGCCGTCC</td><td>TGG</td><td>||||||||[||||||||||||]PAM</td><td>NA</td><td>Intergenic</td><td>NA</td><td>NA</td></tr>
<tr><td></td><td></td><td></td><td></td><td></td><td></td><td></td><td></td><td></td><td></td><td></td><td></td></tr>
<tr><td>T3173</td><td>ACGCAGGCCGTCCTGGAAGAAGG</td><td>1000</td><td></td><td></td><td></td><td></td><td></td><td></td><td></td><td></td><td></td></tr>
<tr><td>Oligo fwd</td><td>CACCACGCAGGCCGTCCTGGAAGA</td><td></td><td></td><td></td><td></td><td></td><td></td><td></td><td></td><td></td><td></td></tr>
<tr><td>Oligo rev</td><td>AAACTCTTCCAGGACGGCCTGCGT</td><td></td><td></td><td></td><td></td><td></td><td></td><td></td><td></td><td></td><td></td></tr>
<tr><td>Chromosome</td><td>start</td><td>end</td><td>strand</td><td>MM</td><td>target_seq</td><td>PAM</td><td>alignment</td><td>distance</td><td>position</td><td>gene name</td><td>gene id</td></tr>
<tr><td>GU305914.1</td><td>48728</td><td>48750</td><td>+</td><td>0</td><td>ACGCAGGCCGTCCTGGAAGA</td><td>AGG</td><td>||||||||[||||||||||||]PAM</td><td>NA</td><td>Intergenic</td><td>NA</td><td>NA</td></tr>
<tr><td></td><td></td><td></td><td></td><td></td><td></td><td></td><td></td><td></td><td></td><td></td><td></td></tr>
<tr><td>T3174</td><td>ACACTAGACTGTCCCCCAGATGG</td><td>1000</td><td></td><td></td><td></td><td></td><td></td><td></td><td></td><td></td><td></td></tr>
<tr><td>Oligo fwd</td><td>CACCACACTAGACTGTCCCCCAGA</td><td></td><td></td><td></td><td></td><td></td><td></td><td></td><td></td><td></td><td></td></tr>
<tr><td>Oligo rev</td><td>AAACTCTGGGGGACAGTCTAGTGT</td><td></td><td></td><td></td><td></td><td></td><td></td><td></td><td></td><td></td><td></td></tr>
<tr><td>Chromosome</td><td>start</td><td>end</td><td>strand</td><td>MM</td><td>target_seq</td><td>PAM</td><td>alignment</td><td>distance</td><td>position</td><td>gene name</td><td>gene id</td></tr>
<tr><td>GU305914.1</td><td>48751</td><td>48773</td><td>+</td><td>0</td><td>ACACTAGACTGTCCCCCAGA</td><td>TGG</td><td>||||||||[||||||||||||]PAM</td><td>NA</td><td>Intergenic</td><td>NA</td><td>NA</td></tr>
<tr><td></td><td></td><td></td><td></td><td></td><td></td><td></td><td></td><td></td><td></td><td></td><td></td></tr>
<tr><td>T3175</td><td>GTCGTAGCAGTAGATGCCGCCGG</td><td>1000</td><td></td><td></td><td></td><td></td><td></td><td></td><td></td><td></td><td></td></tr>
<tr><td>Oligo fwd</td><td>CACCGTCGTAGCAGTAGATGCCGC</td><td></td><td></td><td></td><td></td><td></td><td></td><td></td><td></td><td></td><td></td></tr>
<tr><td>Oligo rev</td><td>AAACGCGGCATCTACTGCTACGAC</td><td></td><td></td><td></td><td></td><td></td><td></td><td></td><td></td><td></td><td></td></tr>
<tr><td>Chromosome</td><td>start</td><td>end</td><td>strand</td><td>MM</td><td>target_seq</td><td>PAM</td><td>alignment</td><td>distance</td><td>position</td><td>gene name</td><td>gene id</td></tr>
<tr><td>GU305914.1</td><td>48794</td><td>48816</td><td>+</td><td>0</td><td>GTCGTAGCAGTAGATGCCGC</td><td>CGG</td><td>||||||||[||||||||||||]PAM</td><td>NA</td><td>Intergenic</td><td>NA</td><td>NA</td></tr>
<tr><td></td><td></td><td></td><td></td><td></td><td></td><td></td><td></td><td></td><td></td><td></td><td></td></tr>
<tr><td>T3176</td><td>GTAGCAGTAGATGCCGCCGGTGG</td><td>1000</td><td></td><td></td><td></td><td></td><td></td><td></td><td></td><td></td><td></td></tr>
<tr><td>Oligo fwd</td><td>CACCGTAGCAGTAGATGCCGCCGG</td><td></td><td></td><td></td><td></td><td></td><td></td><td></td><td></td><td></td><td></td></tr>
<tr><td>Oligo rev</td><td>AAACCCGGCGGCATCTACTGCTAC</td><td></td><td></td><td></td><td></td><td></td><td></td><td></td><td></td><td></td><td></td></tr>
<tr><td>Chromosome</td><td>start</td><td>end</td><td>strand</td><td>MM</td><td>target_seq</td><td>PAM</td><td>alignment</td><td>distance</td><td>position</td><td>gene name</td><td>gene id</td></tr>
<tr><td>GU305914.1</td><td>48797</td><td>48819</td><td>+</td><td>0</td><td>GTAGCAGTAGATGCCGCCGG</td><td>TGG</td><td>||||||||[||||||||||||]PAM</td><td>NA</td><td>Intergenic</td><td>NA</td><td>NA</td></tr>
<tr><td></td><td></td><td></td><td></td><td></td><td></td><td></td><td></td><td></td><td></td><td></td><td></td></tr>
<tr><td>T3177</td><td>GAAGAGCATCACGAACCTCGTGG</td><td>1000</td><td></td><td></td><td></td><td></td><td></td><td></td><td></td><td></td><td></td></tr>
<tr><td>Oligo fwd</td><td>CACCGAAGAGCATCACGAACCTCG</td><td></td><td></td><td></td><td></td><td></td><td></td><td></td><td></td><td></td><td></td></tr>
<tr><td>Oligo rev</td><td>AAACCGAGGTTCGTGATGCTCTTC</td><td></td><td></td><td></td><td></td><td></td><td></td><td></td><td></td><td></td><td></td></tr>
<tr><td>Chromosome</td><td>start</td><td>end</td><td>strand</td><td>MM</td><td>target_seq</td><td>PAM</td><td>alignment</td><td>distance</td><td>position</td><td>gene name</td><td>gene id</td></tr>
<tr><td>GU305914.1</td><td>48824</td><td>48846</td><td>+</td><td>0</td><td>GAAGAGCATCACGAACCTCG</td><td>TGG</td><td>||||||||[||||||||||||]PAM</td><td>NA</td><td>Intergenic</td><td>NA</td><td>NA</td></tr>
<tr><td></td><td></td><td></td><td></td><td></td><td></td><td></td><td></td><td></td><td></td><td></td><td></td></tr>
<tr><td>T3178</td><td>TCACGAACCTCGTGGTGTCCAGG</td><td>1000</td><td></td><td></td><td></td><td></td><td></td><td></td><td></td><td></td><td></td></tr>
<tr><td>Oligo fwd</td><td>CACCTCACGAACCTCGTGGTGTCC</td><td></td><td></td><td></td><td></td><td></td><td></td><td></td><td></td><td></td><td></td></tr>
<tr><td>Oligo rev</td><td>AAACGGACACCACGAGGTTCGTGA</td><td></td><td></td><td></td><td></td><td></td><td></td><td></td><td></td><td></td><td></td></tr>
<tr><td>Chromosome</td><td>start</td><td>end</td><td>strand</td><td>MM</td><td>target_seq</td><td>PAM</td><td>alignment</td><td>distance</td><td>position</td><td>gene name</td><td>gene id</td></tr>
</table>

<table>
<tr><td>GU305914.1</td><td>48832</td><td>48854</td><td>+</td><td>0</td><td>TCACGAACCTCGTGGTGTCC</td><td>AGG</td><td>||||||||[||||||||||||]PAM</td><td>NA</td><td>Intergenic</td><td>NA</td><td>NA</td></tr>
<tr><td>T3179</td><td>CCAGGTCCCGTCCACGCAGCCGG</td><td>1000</td><td></td><td></td><td></td><td></td><td></td><td></td><td></td><td></td><td></td></tr>
<tr><td>Oligo fwd</td><td>CACCCCAGGTCCCGTCCACGCAGC</td><td></td><td></td><td></td><td></td><td></td><td></td><td></td><td></td><td></td><td></td></tr>
<tr><td>Oligo rev</td><td>AAACGCTGCGTGGACGGGACCTGG</td><td></td><td></td><td></td><td></td><td></td><td></td><td></td><td></td><td></td><td></td></tr>
<tr><td>Chromosome</td><td>start</td><td>end</td><td>strand</td><td>MM</td><td>target_seq</td><td>PAM</td><td>alignment</td><td>distance</td><td>position</td><td>gene name</td><td>gene id</td></tr>
<tr><td>GU305914.1</td><td>48850</td><td>48872</td><td>+</td><td>0</td><td>CCAGGTCCCGTCCACGCAGC</td><td>CGG</td><td>||||||||[||||||||||||]PAM</td><td>NA</td><td>Intergenic</td><td>NA</td><td>NA</td></tr>
<tr><td>T3180</td><td>CGTCCACGCAGCCGGCCGAACGG</td><td>1000</td><td></td><td></td><td></td><td></td><td></td><td></td><td></td><td></td><td></td></tr>
<tr><td>Oligo fwd</td><td>CACCCGTCCACGCAGCCGGCCGAA</td><td></td><td></td><td></td><td></td><td></td><td></td><td></td><td></td><td></td><td></td></tr>
<tr><td>Oligo rev</td><td>AAACTTCGGCCGGCTGCGTGGACG</td><td></td><td></td><td></td><td></td><td></td><td></td><td></td><td></td><td></td><td></td></tr>
<tr><td>Chromosome</td><td>start</td><td>end</td><td>strand</td><td>MM</td><td>target_seq</td><td>PAM</td><td>alignment</td><td>distance</td><td>position</td><td>gene name</td><td>gene id</td></tr>
<tr><td>GU305914.1</td><td>48858</td><td>48880</td><td>+</td><td>0</td><td>CGTCCACGCAGCCGGCCGAA</td><td>CGG</td><td>||||||||[||||||||||||]PAM</td><td>NA</td><td>Intergenic</td><td>NA</td><td>NA</td></tr>
<tr><td>T3181</td><td>AGCAGACGTACTTTCCTTTTAGG</td><td>1000</td><td></td><td></td><td></td><td></td><td></td><td></td><td></td><td></td><td></td></tr>
<tr><td>Oligo fwd</td><td>CACCAGCAGACGTACTTTCCTTTT</td><td></td><td></td><td></td><td></td><td></td><td></td><td></td><td></td><td></td><td></td></tr>
<tr><td>Oligo rev</td><td>AAACAAAAGGAAAGTACGTCTGCT</td><td></td><td></td><td></td><td></td><td></td><td></td><td></td><td></td><td></td><td></td></tr>
<tr><td>Chromosome</td><td>start</td><td>end</td><td>strand</td><td>MM</td><td>target_seq</td><td>PAM</td><td>alignment</td><td>distance</td><td>position</td><td>gene name</td><td>gene id</td></tr>
<tr><td>GU305914.1</td><td>48898</td><td>48920</td><td>+</td><td>0</td><td>AGCAGACGTACTTTCCTTTT</td><td>AGG</td><td>||||||||[||||||||||||]PAM</td><td>NA</td><td>Intergenic</td><td>NA</td><td>NA</td></tr>
<tr><td>T3182</td><td>GCAGACGTACTTTCCTTTTAGGG</td><td>1000</td><td></td><td></td><td></td><td></td><td></td><td></td><td></td><td></td><td></td></tr>
<tr><td>Oligo fwd</td><td>CACCGCAGACGTACTTTCCTTTTA</td><td></td><td></td><td></td><td></td><td></td><td></td><td></td><td></td><td></td><td></td></tr>
<tr><td>Oligo rev</td><td>AAACTAAAAGGAAAGTACGTCTGC</td><td></td><td></td><td></td><td></td><td></td><td></td><td></td><td></td><td></td><td></td></tr>
<tr><td>Chromosome</td><td>start</td><td>end</td><td>strand</td><td>MM</td><td>target_seq</td><td>PAM</td><td>alignment</td><td>distance</td><td>position</td><td>gene name</td><td>gene id</td></tr>
<tr><td>GU305914.1</td><td>48899</td><td>48921</td><td>+</td><td>0</td><td>GCAGACGTACTTTCCTTTTA</td><td>GGG</td><td>||||||||[||||||||||||]PAM</td><td>NA</td><td>Intergenic</td><td>NA</td><td>NA</td></tr>
<tr><td>T3183</td><td>GACGTACTTTCCTTTTAGGGAGG</td><td>1000</td><td></td><td></td><td></td><td></td><td></td><td></td><td></td><td></td><td></td></tr>
<tr><td>Oligo fwd</td><td>CACCGACGTACTTTCCTTTTAGGG</td><td></td><td></td><td></td><td></td><td></td><td></td><td></td><td></td><td></td><td></td></tr>
<tr><td>Oligo rev</td><td>AAACCCCTAAAAGGAAAGTACGTC</td><td></td><td></td><td></td><td></td><td></td><td></td><td></td><td></td><td></td><td></td></tr>
<tr><td>Chromosome</td><td>start</td><td>end</td><td>strand</td><td>MM</td><td>target_seq</td><td>PAM</td><td>alignment</td><td>distance</td><td>position</td><td>gene name</td><td>gene id</td></tr>
<tr><td>GU305914.1</td><td>48902</td><td>48924</td><td>+</td><td>0</td><td>GACGTACTTTCCTTTTAGGG</td><td>AGG</td><td>||||||||[||||||||||||]PAM</td><td>NA</td><td>Intergenic</td><td>NA</td><td>NA</td></tr>
<tr><td>T3184</td><td>TTTAGGGAGGACGTCGTGCGCGG</td><td>1000</td><td></td><td></td><td></td><td></td><td></td><td></td><td></td><td></td><td></td></tr>
<tr><td>Oligo fwd</td><td>CACCTTTAGGGAGGACGTCGTGCG</td><td></td><td></td><td></td><td></td><td></td><td></td><td></td><td></td><td></td><td></td></tr>
<tr><td>Oligo rev</td><td>AAACCGCACGACGTCCTCCCTAAA</td><td></td><td></td><td></td><td></td><td></td><td></td><td></td><td></td><td></td><td></td></tr>
<tr><td>Chromosome</td><td>start</td><td>end</td><td>strand</td><td>MM</td><td>target_seq</td><td>PAM</td><td>alignment</td><td>distance</td><td>position</td><td>gene name</td><td>gene id</td></tr>
<tr><td>GU305914.1</td><td>48915</td><td>48937</td><td>+</td><td>0</td><td>TTTAGGGAGGACGTCGTGCG</td><td>CGG</td><td>||||||||[||||||||||||]PAM</td><td>NA</td><td>Intergenic</td><td>NA</td><td>NA</td></tr>
<tr><td>T3185</td><td>TCCATCCATCCTCCCCTCCGTGG</td><td>1000</td><td></td><td></td><td></td><td></td><td></td><td></td><td></td><td></td><td></td></tr>
<tr><td>Oligo fwd</td><td>CACCTCCATCCATCCTCCCCTCCG</td><td></td><td></td><td></td><td></td><td></td><td></td><td></td><td></td><td></td><td></td></tr>
<tr><td>Oligo rev</td><td>AAACCGGAGGGGAGGATGGATGGA</td><td></td><td></td><td></td><td></td><td></td><td></td><td></td><td></td><td></td><td></td></tr>
<tr><td>Chromosome</td><td>start</td><td>end</td><td>strand</td><td>MM</td><td>target_seq</td><td>PAM</td><td>alignment</td><td>distance</td><td>position</td><td>gene name</td><td>gene id</td></tr>
<tr><td>GU305914.1</td><td>48993</td><td>49015</td><td>+</td><td>0</td><td>TCCATCCATCCTCCCCTCCG</td><td>TGG</td><td>||||||||[||||||||||||]PAM</td><td>NA</td><td>Intergenic</td><td>NA</td><td>NA</td></tr>
<tr><td>T3186</td><td>CCATCCATCCTCCCCTCCGTGGG</td><td>1000</td><td></td><td></td><td></td><td></td><td></td><td></td><td></td><td></td><td></td></tr>
<tr><td>Oligo fwd</td><td>CACCCCATCCATCCTCCCCTCCGT</td><td></td><td></td><td></td><td></td><td></td><td></td><td></td><td></td><td></td><td></td></tr>
<tr><td>Oligo rev</td><td>AAACACGGAGGGGAGGATGGATGG</td><td></td><td></td><td></td><td></td><td></td><td></td><td></td><td></td><td></td><td></td></tr>
<tr><td>Chromosome</td><td>start</td><td>end</td><td>strand</td><td>MM</td><td>target_seq</td><td>PAM</td><td>alignment</td><td>distance</td><td>position</td><td>gene name</td><td>gene id</td></tr>
<tr><td>GU305914.1</td><td>48994</td><td>49016</td><td>+</td><td>0</td><td>CCATCCATCCTCCCCTCCGT</td><td>GGG</td><td>||||||||[||||||||||||]PAM</td><td>NA</td><td>Intergenic</td><td>NA</td><td>NA</td></tr>
<tr><td>T3187</td><td>CTTCAAGAAATACGTATATCTGG</td><td>1000</td><td></td><td></td><td></td><td></td><td></td><td></td><td></td><td></td><td></td></tr>
<tr><td>Oligo fwd</td><td>CACCCTTCAAGAAATACGTATATC</td><td></td><td></td><td></td><td></td><td></td><td></td><td></td><td></td><td></td><td></td></tr>
<tr><td>Oligo rev</td><td>AAACGATATACGTATTTCTTGAAG</td><td></td><td></td><td></td><td></td><td></td><td></td><td></td><td></td><td></td><td></td></tr>
<tr><td>Chromosome</td><td>start</td><td>end</td><td>strand</td><td>MM</td><td>target_seq</td><td>PAM</td><td>alignment</td><td>distance</td><td>position</td><td>gene name</td><td>gene id</td></tr>
<tr><td>GU305914.1</td><td>49064</td><td>49086</td><td>+</td><td>0</td><td>CTTCAAGAAATACGTATATC</td><td>TGG</td><td>||||||||[||||||||||||]PAM</td><td>NA</td><td>Intergenic</td><td>NA</td><td>NA</td></tr>
<tr><td>T3188</td><td>ATCTGGTCATGTCGACGAGCCGG</td><td>1000</td><td></td><td></td><td></td><td></td><td></td><td></td><td></td><td></td><td></td></tr>
<tr><td>Oligo fwd</td><td>CACCATCTGGTCATGTCGACGAGC</td><td></td><td></td><td></td><td></td><td></td><td></td><td></td><td></td><td></td><td></td></tr>
<tr><td>Oligo rev</td><td>AAACGCTCGTCGACATGACCAGAT</td><td></td><td></td><td></td><td></td><td></td><td></td><td></td><td></td><td></td><td></td></tr>
<tr><td>Chromosome</td><td>start</td><td>end</td><td>strand</td><td>MM</td><td>target_seq</td><td>PAM</td><td>alignment</td><td>distance</td><td>position</td><td>gene name</td><td>gene id</td></tr>
<tr><td>GU305914.1</td><td>49081</td><td>49103</td><td>+</td><td>0</td><td>ATCTGGTCATGTCGACGAGC</td><td>CGG</td><td>||||||||[||||||||||||]PAM</td><td>NA</td><td>Intergenic</td><td>NA</td><td>NA</td></tr>
<tr><td>T3189</td><td>AGCCGGACCGAAACCCCGTCCGG</td><td>1000</td><td></td><td></td><td></td><td></td><td></td><td></td><td></td><td></td><td></td></tr>
<tr><td>Oligo fwd</td><td>CACCAGCCGGACCGAAACCCCGTC</td><td></td><td></td><td></td><td></td><td></td><td></td><td></td><td></td><td></td><td></td></tr>
<tr><td>Oligo rev</td><td>AAACGACGGGGTTTCGGTCCGGCT</td><td></td><td></td><td></td><td></td><td></td><td></td><td></td><td></td><td></td><td></td></tr>
<tr><td>Chromosome</td><td>start</td><td>end</td><td>strand</td><td>MM</td><td>target_seq</td><td>PAM</td><td>alignment</td><td>distance</td><td>position</td><td>gene name</td><td>gene id</td></tr>
<tr><td>GU305914.1</td><td>49098</td><td>49120</td><td>+</td><td>0</td><td>AGCCGGACCGAAACCCCGTC</td><td>CGG</td><td>||||||||[||||||||||||]PAM</td><td>NA</td><td>Intergenic</td><td>NA</td><td>NA</td></tr>
<tr><td>T3190</td><td>ACCCCGTCCGGCCAGCTGACCGG</td><td>1000</td><td></td><td></td><td></td><td></td><td></td><td></td><td></td><td></td><td></td></tr>
<tr><td>Oligo fwd</td><td>CACCACCCCGTCCGGCCAGCTGAC</td><td></td><td></td><td></td><td></td><td></td><td></td><td></td><td></td><td></td><td></td></tr>
<tr><td>Oligo rev</td><td>AAACGTCAGCTGGCCGGACGGGGT</td><td></td><td></td><td></td><td></td><td></td><td></td><td></td><td></td><td></td><td></td></tr>
<tr><td>Chromosome</td><td>start</td><td>end</td><td>strand</td><td>MM</td><td>target_seq</td><td>PAM</td><td>alignment</td><td>distance</td><td>position</td><td>gene name</td><td>gene id</td></tr>
<tr><td>GU305914.1</td><td>49110</td><td>49132</td><td>+</td><td>0</td><td>ACCCCGTCCGGCCAGCTGAC</td><td>CGG</td><td>||||||||[||||||||||||]PAM</td><td>NA</td><td>Intergenic</td><td>NA</td><td>NA</td></tr>
<tr><td>T3191</td><td>GTCCGGCCAGCTGACCGGCAAGG</td><td>1000</td><td></td><td></td><td></td><td></td><td></td><td></td><td></td><td></td><td></td></tr>
<tr><td>Oligo fwd</td><td>CACCGTCCGGCCAGCTGACCGGCA</td><td></td><td></td><td></td><td></td><td></td><td></td><td></td><td></td><td></td><td></td></tr>
<tr><td>Oligo rev</td><td>AAACTGCCGGTCAGCTGGCCGGAC</td><td></td><td></td><td></td><td></td><td></td><td></td><td></td><td></td><td></td><td></td></tr>
<tr><td>Chromosome</td><td>start</td><td>end</td><td>strand</td><td>MM</td><td>target_seq</td><td>PAM</td><td>alignment</td><td>distance</td><td>position</td><td>gene name</td><td>gene id</td></tr>
<tr><td>GU305914.1</td><td>49115</td><td>49137</td><td>+</td><td>0</td><td>GTCCGGCCAGCTGACCGGCA</td><td>AGG</td><td>||||||||[||||||||||||]PAM</td><td>NA</td><td>Intergenic</td><td>NA</td><td>NA</td></tr>
<tr><td>T3192</td><td>CGGCCAGCTGACCGGCAAGGAGG</td><td>1000</td><td></td><td></td><td></td><td></td><td></td><td></td><td></td><td></td><td></td></tr>
<tr><td>Oligo fwd</td><td>CACCCGGCCAGCTGACCGGCAAGG</td><td></td><td></td><td></td><td></td><td></td><td></td><td></td><td></td><td></td><td></td></tr>
<tr><td>Oligo rev</td><td>AAACCCTTGCCGGTCAGCTGGCCG</td><td></td><td></td><td></td><td></td><td></td><td></td><td></td><td></td><td></td><td></td></tr>
<tr><td>Chromosome</td><td>start</td><td>end</td><td>strand</td><td>MM</td><td>target_seq</td><td>PAM</td><td>alignment</td><td>distance</td><td>position</td><td>gene name</td><td>gene id</td></tr>
<tr><td>GU305914.1</td><td>49118</td><td>49140</td><td>+</td><td>0</td><td>CGGCCAGCTGACCGGCAAGG</td><td>AGG</td><td>||||||||[||||||||||||]PAM</td><td>NA</td><td>Intergenic</td><td>NA</td><td>NA</td></tr>
<tr><td>T3193</td><td>GCTGACCGGCAAGGAGGTACCGG</td><td>1000</td><td></td><td></td><td></td><td></td><td></td><td></td><td></td><td></td><td></td></tr>
<tr><td>Oligo fwd</td><td>CACCGCTGACCGGCAAGGAGGTAC</td><td></td><td></td><td></td><td></td><td></td><td></td><td></td><td></td><td></td><td></td></tr>
<tr><td>Oligo rev</td><td>AAACGTACCTCCTTGCCGGTCAGC</td><td></td><td></td><td></td><td></td><td></td><td></td><td></td><td></td><td></td><td></td></tr>
</table>

Chromosome start end strand MM target_seq PAM alignment distance position gene name gene id
GU305914.1 49124 49146 + 0 GCTGACCGGCAAGGAGGTAC CGG ||||||||[||||||||||||]PAM NA Intergenic NA NA

T3194 CAAGGAGGTACCGGCGCGCTCGG 1000
Oligo fwd CACCCAAGGAGGTACCGGCGCGCT
Oligo rev AAACAGCGCGCCGGTACCTCCTTG
Chromosome start end strand MM target_seq PAM alignment distance position gene name gene id
GU305914.1 49133 49155 + 0 CAAGGAGGTACCGGCGCGCT CGG ||||||||[||||||||||||]PAM NA Intergenic NA NA

T3195 CTCGGAGACATAGCTATAGATGG 1000
Oligo fwd CACCCTCGGAGACATAGCTATAGA
Oligo rev AAACTCTATAGCTATGTCTCCGAG
Chromosome start end strand MM target_seq PAM alignment distance position gene name gene id
GU305914.1 49151 49173 + 0 CTCGGAGACATAGCTATAGA TGG ||||||||[||||||||||||]PAM NA Intergenic NA NA

T3196 ACATAGCTATAGATGGCCTCCGG 1000
Oligo fwd CACCACATAGCTATAGATGGCCTC
Oligo rev AAACGAGGCCATCTATAGCTATGT
Chromosome start end strand MM target_seq PAM alignment distance position gene name gene id
GU305914.1 49158 49180 + 0 ACATAGCTATAGATGGCCTC CGG ||||||||[||||||||||||]PAM NA Intergenic NA NA

T3197 CTCCGGATTGAAGCCGATGTTGG 1000
Oligo fwd CACCCTCCGGATTGAAGCCGATGT
Oligo rev AAACACATCGGCTTCAATCCGGAG
Chromosome start end strand MM target_seq PAM alignment distance position gene name gene id
GU305914.1 49175 49197 + 0 CTCCGGATTGAAGCCGATGT TGG ||||||||[||||||||||||]PAM NA Intergenic NA NA

T3198 GATAAGCCTCCCCGATGTCCTGG 1000
Oligo fwd CACCGATAAGCCTCCCCGATGTCC
Oligo rev AAACGGACATCGGGGAGGCTTATC
Chromosome start end strand MM target_seq PAM alignment distance position gene name gene id
GU305914.1 49207 49229 + 0 GATAAGCCTCCCCGATGTCC TGG ||||||||[||||||||||||]PAM NA Intergenic NA NA

T3199 GCCTCCCCGATGTCCTGGTAAGG 1000
Oligo fwd CACCGCCTCCCCGATGTCCTGGTA
Oligo rev AAACTACCAGGACATCGGGGAGGC
Chromosome start end strand MM target_seq PAM alignment distance position gene name gene id
GU305914.1 49212 49234 + 0 GCCTCCCCGATGTCCTGGTA AGG ||||||||[||||||||||||]PAM NA Intergenic NA NA

T3200 CCTCCCCGATGTCCTGGTAAGGG 1000
Oligo fwd CACCCCTCCCCGATGTCCTGGTAA
Oligo rev AAACTTACCAGGACATCGGGGAGG
Chromosome start end strand MM target_seq PAM alignment distance position gene name gene id
GU305914.1 49213 49235 + 0 CCTCCCCGATGTCCTGGTAA GGG ||||||||[||||||||||||]PAM NA Intergenic NA NA

T3201 GTCCTGGTAAGGGACGATCTCGG 1000
Oligo fwd CACCGTCCTGGTAAGGGACGATCT
Oligo rev AAACAGATCGTCCCTTACCAGGAC
Chromosome start end strand MM target_seq PAM alignment distance position gene name gene id
GU305914.1 49223 49245 + 0 GTCCTGGTAAGGGACGATCT CGG ||||||||[||||||||||||]PAM NA Intergenic NA NA

T3202 TCCTGGTAAGGGACGATCTCGGG 1000
Oligo fwd CACCTCCTGGTAAGGGACGATCTC
Oligo rev AAACGAGATCGTCCCTTACCAGGA
Chromosome start end strand MM target_seq PAM alignment distance position gene name gene id
GU305914.1 49224 49246 + 0 TCCTGGTAAGGGACGATCTC GGG ||||||||[||||||||||||]PAM NA Intergenic NA NA

T3203 GCAGCAACTTTATTGAGAACTGG 1000
Oligo fwd CACCGCAGCAACTTTATTGAGAAC
Oligo rev AAACGTTCTCAATAAAGTTGCTGC
Chromosome start end strand MM target_seq PAM alignment distance position gene name gene id
GU305914.1 49401 49423 + 0 GCAGCAACTTTATTGAGAAC TGG ||||||||[||||||||||||]PAM NA Intergenic NA NA

T3204 CAGCAACTTTATTGAGAACTGGG 1000
Oligo fwd CACCCAGCAACTTTATTGAGAACT
Oligo rev AAACAGTTCTCAATAAAGTTGCTG
Chromosome start end strand MM target_seq PAM alignment distance position gene name gene id
GU305914.1 49402 49424 + 0 CAGCAACTTTATTGAGAACT GGG ||||||||[||||||||||||]PAM NA Intergenic NA NA

T3205 ATCTATTCACACGTTTTCCGCGG 1000
Oligo fwd CACCATCTATTCACACGTTTTCCG
Oligo rev AAACCGGAAAACGTGTGAATAGAT
Chromosome start end strand MM target_seq PAM alignment distance position gene name gene id
GU305914.1 49438 49460 + 0 ATCTATTCACACGTTTTCCG CGG ||||||||[||||||||||||]PAM NA Intergenic NA NA

T3206 CCGCGGCGAAATCCTCGTTCCGG 1000
Oligo fwd CACCCCGCGGCGAAATCCTCGTTC
Oligo rev AAACGAACGAGGATTTCGCCGCGG
Chromosome start end strand MM target_seq PAM alignment distance position gene name gene id
GU305914.1 49455 49477 + 0 CCGCGGCGAAATCCTCGTTC CGG ||||||||[||||||||||||]PAM NA Intergenic NA NA

T3207 CGAAATCCTCGTTCCGGAGACGG 1000
Oligo fwd CACCCGAAATCCTCGTTCCGGAGA
Oligo rev AAACTCTCCGGAACGAGGATTTCG
Chromosome start end strand MM target_seq PAM alignment distance position gene name gene id
GU305914.1 49461 49483 + 0 CGAAATCCTCGTTCCGGAGA CGG ||||||||[||||||||||||]PAM NA Intergenic NA NA

T3208 AATCCTCGTTCCGGAGACGGAGG 1000
Oligo fwd CACCAATCCTCGTTCCGGAGACGG

Oligo rev AAACCCGTCTCCGGAACGAGGATT
Chromosome start end strand MM target_seq PAM alignment distance position gene name gene id
GU305914.1 49464 49486 + 0 AATCCTCGTTCCGGAGACGG AGG ||||||||[||||||||||||]PAM NA Intergenic NA NA

T3209 ATCCTCGTTCCGGAGACGGAGGG 1000
Oligo fwd CACCATCCTCGTTCCGGAGACGGA
Oligo rev AAACTCCGTCTCCGGAACGAGGAT
Chromosome start end strand MM target_seq PAM alignment distance position gene name gene id
GU305914.1 49465 49487 + 0 ATCCTCGTTCCGGAGACGGA GGG ||||||||[||||||||||||]PAM NA Intergenic NA NA

T3210 TGAACAGCGACTCGCGTTGTCGG 1000
Oligo fwd CACCTGAACAGCGACTCGCGTTGT
Oligo rev AAACACAACGCGAGTCGCTGTTCA
Chromosome start end strand MM target_seq PAM alignment distance position gene name gene id
GU305914.1 49509 49531 + 0 TGAACAGCGACTCGCGTTGT CGG ||||||||[||||||||||||]PAM NA Intergenic NA NA

T3211 CAGCGACTCGCGTTGTCGGATGG 1000
Oligo fwd CACCCAGCGACTCGCGTTGTCGGA
Oligo rev AAACTCCGACAACGCGAGTCGCTG
Chromosome start end strand MM target_seq PAM alignment distance position gene name gene id
GU305914.1 49513 49535 + 0 CAGCGACTCGCGTTGTCGGA TGG ||||||||[||||||||||||]PAM NA Intergenic NA NA

T3212 GGATGGTCATGAAGCAGAAGAGG 1000
Oligo fwd CACCGGATGGTCATGAAGCAGAAG
Oligo rev AAACCTTCTGCTTCATGACCATCC
Chromosome start end strand MM target_seq PAM alignment distance position gene name gene id
GU305914.1 49530 49552 + 0 GGATGGTCATGAAGCAGAAG AGG ||||||||[||||||||||||]PAM NA Intergenic NA NA

T3213 TGAAGCAGAAGAGGATCAACAGG 1000
Oligo fwd CACCTGAAGCAGAAGAGGATCAAC
Oligo rev AAACGTTGATCCTCTTCTGCTTCA
Chromosome start end strand MM target_seq PAM alignment distance position gene name gene id
GU305914.1 49539 49561 + 0 TGAAGCAGAAGAGGATCAAC AGG ||||||||[||||||||||||]PAM NA Intergenic NA NA

T3214 GAAGCAGAAGAGGATCAACAGGG 1000
Oligo fwd CACCGAAGCAGAAGAGGATCAACA
Oligo rev AAACTGTTGATCCTCTTCTGCTTC
Chromosome start end strand MM target_seq PAM alignment distance position gene name gene id
GU305914.1 49540 49562 + 0 GAAGCAGAAGAGGATCAACA GGG ||||||||[||||||||||||]PAM NA Intergenic NA NA

T3215 AGAAGAGGATCAACAGGGCCAGG 1000
Oligo fwd CACCAGAAGAGGATCAACAGGGCC
Oligo rev AAACGGCCCTGTTGATCCTCTTCT
Chromosome start end strand MM target_seq PAM alignment distance position gene name gene id
GU305914.1 49545 49567 + 0 AGAAGAGGATCAACAGGGCC AGG ||||||||[||||||||||||]PAM NA Intergenic NA NA

T3216 GAAGAACATGCCGACCATCTTGG 1000
Oligo fwd CACCGAAGAACATGCCGACCATCT
Oligo rev AAACAGATGGTCGGCATGTTCTTC
Chromosome start end strand MM target_seq PAM alignment distance position gene name gene id
GU305914.1 49582 49604 + 0 GAAGAACATGCCGACCATCT TGG ||||||||[||||||||||||]PAM NA Intergenic NA NA

T3217 AAGAACATGCCGACCATCTTGGG 1000
Oligo fwd CACCAAGAACATGCCGACCATCTT
Oligo rev AAACAAGATGGTCGGCATGTTCTT
Chromosome start end strand MM target_seq PAM alignment distance position gene name gene id
GU305914.1 49583 49605 + 0 AAGAACATGCCGACCATCTT GGG ||||||||[||||||||||||]PAM NA Intergenic NA NA

T3218 ACATGCCGACCATCTTGGGCAGG 1000
Oligo fwd CACCACATGCCGACCATCTTGGGC
Oligo rev AAACGCCCAAGATGGTCGGCATGT
Chromosome start end strand MM target_seq PAM alignment distance position gene name gene id
GU305914.1 49587 49609 + 0 ACATGCCGACCATCTTGGGC AGG ||||||||[||||||||||||]PAM NA Intergenic NA NA

T3219 CATGCCGACCATCTTGGGCAGGG 1000
Oligo fwd CACCCATGCCGACCATCTTGGGCA
Oligo rev AAACTGCCCAAGATGGTCGGCATG
Chromosome start end strand MM target_seq PAM alignment distance position gene name gene id
GU305914.1 49588 49610 + 0 CATGCCGACCATCTTGGGCA GGG ||||||||[||||||||||||]PAM NA Intergenic NA NA

T3220 ATGCCGACCATCTTGGGCAGGGG 1000
Oligo fwd CACCATGCCGACCATCTTGGGCAG
Oligo rev AAACCTGCCCAAGATGGTCGGCAT
Chromosome start end strand MM target_seq PAM alignment distance position gene name gene id
GU305914.1 49589 49611 + 0 ATGCCGACCATCTTGGGCAG GGG ||||||||[||||||||||||]PAM NA Intergenic NA NA

T3221 CATCTTGGGCAGGGGATCGCCGG 1000
Oligo fwd CACCCATCTTGGGCAGGGGATCGC
Oligo rev AAACGCGATCCCCTGCCCAAGATG
Chromosome start end strand MM target_seq PAM alignment distance position gene name gene id
GU305914.1 49597 49619 + 0 CATCTTGGGCAGGGGATCGC CGG ||||||||[||||||||||||]PAM NA Intergenic NA NA

T3222 TGCATCGCGACGAGAAAGCCCGG 1000
Oligo fwd CACCTGCATCGCGACGAGAAAGCC
Oligo rev AAACGGCTTTCTCGTCGCGATGCA
Chromosome start end strand MM target_seq PAM alignment distance position gene name gene id
GU305914.1 49631 49653 + 0 TGCATCGCGACGAGAAAGCC CGG ||||||||[||||||||||||]PAM NA Intergenic NA NA

T3223 GCATCGCGACGAGAAAGCCCGGG 1000

| | | | | | | | | | | | |
|---|---|---|---|---|---|---|---|---|---|---|---|
| Oligo fwd | CACCGCATCGCGACGAGAAAGCCC | | | | | | | | | | |
| Oligo rev | AAACGGGCTTTCTCGTCGCGATGC | | | | | | | | | | |
| Chromosome | start | end | strand | MM | target_seq | PAM | alignment | distance | position | gene name | gene id |
| GU305914.1 | 49632 | 49654 | + | 0 | GCATCGCGACGAGAAAGCCC | GGG | \|\|\|\|\|\|\|\|[\|\|\|\|\|\|\|\|\|\|\|\|]PAM | NA | Intergenic | NA | NA |
| T3224 | CTTCGTCCAGTTGCCCCAGACGG | 1000 | | | | | | | | | |
| Oligo fwd | CACCCTTCGTCCAGTTGCCCCAGA | | | | | | | | | | |
| Oligo rev | AAACTCTGGGGCAACTGGACGAAG | | | | | | | | | | |
| Chromosome | start | end | strand | MM | target_seq | PAM | alignment | distance | position | gene name | gene id |
| GU305914.1 | 49687 | 49709 | + | 0 | CTTCGTCCAGTTGCCCCAGA | CGG | \|\|\|\|\|\|\|\|[\|\|\|\|\|\|\|\|\|\|\|\|]PAM | NA | Intergenic | NA | NA |
| T3225 | CGTCCAGTTGCCCCAGACGGTGG | 1000 | | | | | | | | | |
| Oligo fwd | CACCCGTCCAGTTGCCCCAGACGG | | | | | | | | | | |
| Oligo rev | AAACCCGTCTGGGGCAACTGGACG | | | | | | | | | | |
| Chromosome | start | end | strand | MM | target_seq | PAM | alignment | distance | position | gene name | gene id |
| GU305914.1 | 49690 | 49712 | + | 0 | CGTCCAGTTGCCCCAGACGG | TGG | \|\|\|\|\|\|\|\|[\|\|\|\|\|\|\|\|\|\|\|\|]PAM | NA | Intergenic | NA | NA |
| T3226 | GACGGTGGACCAGTTTCCCGTGG | 1000 | | | | | | | | | |
| Oligo fwd | CACCGACGGTGGACCAGTTTCCCG | | | | | | | | | | |
| Oligo rev | AAACCGGGAAACTGGTCCACCGTC | | | | | | | | | | |
| Chromosome | start | end | strand | MM | target_seq | PAM | alignment | distance | position | gene name | gene id |
| GU305914.1 | 49705 | 49727 | + | 0 | GACGGTGGACCAGTTTCCCG | TGG | \|\|\|\|\|\|\|\|[\|\|\|\|\|\|\|\|\|\|\|\|]PAM | NA | Intergenic | NA | NA |
| T3227 | GGTGGACCAGTTTCCCGTGGCGG | 1000 | | | | | | | | | |
| Oligo fwd | CACCGGTGGACCAGTTTCCCGTGG | | | | | | | | | | |
| Oligo rev | AAACCCACGGGAAACTGGTCCACC | | | | | | | | | | |
| Chromosome | start | end | strand | MM | target_seq | PAM | alignment | distance | position | gene name | gene id |
| GU305914.1 | 49708 | 49730 | + | 0 | GGTGGACCAGTTTCCCGTGG | CGG | \|\|\|\|\|\|\|\|[\|\|\|\|\|\|\|\|\|\|\|\|]PAM | NA | Intergenic | NA | NA |
| T3228 | GTTTCCCGTGGCGGTGCCGCCGG | 1000 | | | | | | | | | |
| Oligo fwd | CACCGTTTCCCGTGGCGGTGCCGC | | | | | | | | | | |
| Oligo rev | AAACGCGGCACCGCCACGGGAAAC | | | | | | | | | | |
| Chromosome | start | end | strand | MM | target_seq | PAM | alignment | distance | position | gene name | gene id |
| GU305914.1 | 49717 | 49739 | + | 0 | GTTTCCCGTGGCGGTGCCGC | CGG | \|\|\|\|\|\|\|\|[\|\|\|\|\|\|\|\|\|\|\|\|]PAM | NA | Intergenic | NA | NA |
| T3229 | CGTGGCGGTGCCGCCGGAGCAGG | 1000 | | | | | | | | | |
| Oligo fwd | CACCCGTGGCGGTGCCGCCGGAGC | | | | | | | | | | |
| Oligo rev | AAACGCTCCGGCGGCACCGCCACG | | | | | | | | | | |
| Chromosome | start | end | strand | MM | target_seq | PAM | alignment | distance | position | gene name | gene id |
| GU305914.1 | 49723 | 49745 | + | 0 | CGTGGCGGTGCCGCCGGAGC | AGG | \|\|\|\|\|\|\|\|[\|\|\|\|\|\|\|\|\|\|\|\|]PAM | NA | Intergenic | NA | NA |
| T3230 | GCAGGCGTCCCTCCTGATGACGG | 1000 | | | | | | | | | |
| Oligo fwd | CACCGCAGGCGTCCCTCCTGATGA | | | | | | | | | | |
| Oligo rev | AAACTCATCAGGAGGGACGCCTGC | | | | | | | | | | |
| Chromosome | start | end | strand | MM | target_seq | PAM | alignment | distance | position | gene name | gene id |
| GU305914.1 | 49741 | 49763 | + | 0 | GCAGGCGTCCCTCCTGATGA | CGG | \|\|\|\|\|\|\|\|[\|\|\|\|\|\|\|\|\|\|\|\|]PAM | NA | Intergenic | NA | NA |
| GU305914.1 | 32556 | 32578 | + | 4 | GACGGCGCCCATCCTGATGA | CGG | \|--\|\|\|\|-[\|\|-\|\|\|\|\|\|\|\|\|]PAM | NA | Intergenic | NA | NA |
| T3231 | GGCGTCCCTCCTGATGACGGCGG | 1000 | | | | | | | | | |
| Oligo fwd | CACCGGCGTCCCTCCTGATGACGG | | | | | | | | | | |
| Oligo rev | AAACCCGTCATCAGGAGGGACGCC | | | | | | | | | | |
| Chromosome | start | end | strand | MM | target_seq | PAM | alignment | distance | position | gene name | gene id |
| GU305914.1 | 49744 | 49766 | + | 0 | GGCGTCCCTCCTGATGACGG | CGG | \|\|\|\|\|\|\|\|[\|\|\|\|\|\|\|\|\|\|\|\|]PAM | NA | Intergenic | NA | NA |
| T3232 | GCGGTCCTCTCCATGATGCCCGG | 1000 | | | | | | | | | |
| Oligo fwd | CACCGCGGTCCTCTCCATGATGCC | | | | | | | | | | |
| Oligo rev | AAACGGCATCATGGAGAGGACCGC | | | | | | | | | | |
| Chromosome | start | end | strand | MM | target_seq | PAM | alignment | distance | position | gene name | gene id |
| GU305914.1 | 49763 | 49785 | + | 0 | GCGGTCCTCTCCATGATGCC | CGG | \|\|\|\|\|\|\|\|[\|\|\|\|\|\|\|\|\|\|\|\|]PAM | NA | Intergenic | NA | NA |
| T3233 | CATGATGCCCGGCGAGAGTTCGG | 1000 | | | | | | | | | |
| Oligo fwd | CACCCATGATGCCCGGCGAGAGTT | | | | | | | | | | |
| Oligo rev | AAACAACTCTCGCCGGGCATCATG | | | | | | | | | | |
| Chromosome | start | end | strand | MM | target_seq | PAM | alignment | distance | position | gene name | gene id |
| GU305914.1 | 49774 | 49796 | + | 0 | CATGATGCCCGGCGAGAGTT | CGG | \|\|\|\|\|\|\|\|[\|\|\|\|\|\|\|\|\|\|\|\|]PAM | NA | Intergenic | NA | NA |
| T3234 | TGCCCGGCGAGAGTTCGGACCGG | 1000 | | | | | | | | | |
| Oligo fwd | CACCTGCCCGGCGAGAGTTCGGAC | | | | | | | | | | |
| Oligo rev | AAACGTCCGAACTCTCGCCGGGCA | | | | | | | | | | |
| Chromosome | start | end | strand | MM | target_seq | PAM | alignment | distance | position | gene name | gene id |
| GU305914.1 | 49779 | 49801 | + | 0 | TGCCCGGCGAGAGTTCGGAC | CGG | \|\|\|\|\|\|\|\|[\|\|\|\|\|\|\|\|\|\|\|\|]PAM | NA | Intergenic | NA | NA |
| T3235 | GGACCGGTGTCCGACACTGACGG | 1000 | | | | | | | | | |
| Oligo fwd | CACCGGACCGGTGTCCGACACTGA | | | | | | | | | | |
| Oligo rev | AAACTCAGTGTCGGACACCGGTCC | | | | | | | | | | |
| Chromosome | start | end | strand | MM | target_seq | PAM | alignment | distance | position | gene name | gene id |
| GU305914.1 | 49795 | 49817 | + | 0 | GGACCGGTGTCCGACACTGA | CGG | \|\|\|\|\|\|\|\|[\|\|\|\|\|\|\|\|\|\|\|\|]PAM | NA | Intergenic | NA | NA |
| T3236 | CGACACTGACGGCGCGCAGCAGG | 1000 | | | | | | | | | |
| Oligo fwd | CACCCGACACTGACGGCGCGCAGC | | | | | | | | | | |
| Oligo rev | AAACGCTGCGCGCCGTCAGTGTCG | | | | | | | | | | |
| Chromosome | start | end | strand | MM | target_seq | PAM | alignment | distance | position | gene name | gene id |
| GU305914.1 | 49806 | 49828 | + | 0 | CGACACTGACGGCGCGCAGC | AGG | \|\|\|\|\|\|\|\|[\|\|\|\|\|\|\|\|\|\|\|\|]PAM | NA | Intergenic | NA | NA |
| T3237 | ACTGACGGCGCGCAGCAGGTCGG | 1000 | | | | | | | | | |
| Oligo fwd | CACCACTGACGGCGCGCAGCAGGT | | | | | | | | | | |
| Oligo rev | AAACACCTGCTGCGCGCCGTCAGT | | | | | | | | | | |
| Chromosome | start | end | strand | MM | target_seq | PAM | alignment | distance | position | gene name | gene id |
| GU305914.1 | 49810 | 49832 | + | 0 | ACTGACGGCGCGCAGCAGGT | CGG | \|\|\|\|\|\|\|\|[\|\|\|\|\|\|\|\|\|\|\|\|]PAM | NA | Intergenic | NA | NA |

<table>
<tr><td>T3238</td><td>CTGACGGCGCGCAGCAGGTCGGG</td><td>1000</td><td></td><td></td><td></td><td></td><td></td><td></td><td></td><td></td><td></td></tr>
<tr><td>Oligo fwd</td><td>CACCCTGACGGCGCGCAGCAGGTC</td><td></td><td></td><td></td><td></td><td></td><td></td><td></td><td></td><td></td><td></td></tr>
<tr><td>Oligo rev</td><td>AAACGACCTGCTGCGCGCCGTCAG</td><td></td><td></td><td></td><td></td><td></td><td></td><td></td><td></td><td></td><td></td></tr>
<tr><td>Chromosome</td><td>start</td><td>end</td><td>strand</td><td>MM</td><td>target_seq</td><td>PAM</td><td>alignment</td><td>distance</td><td>position</td><td>gene name</td><td>gene id</td></tr>
<tr><td>GU305914.1</td><td>49811</td><td>49833</td><td>+</td><td>0</td><td>CTGACGGCGCGCAGCAGGTC</td><td>GGG</td><td>||||||||[||||||||||||]PAM</td><td>NA</td><td>Intergenic</td><td>NA</td><td>NA</td></tr>
<tr><td>T3239</td><td>GGCGCGCAGCAGGTCGGGACAGG</td><td>1000</td><td></td><td></td><td></td><td></td><td></td><td></td><td></td><td></td><td></td></tr>
<tr><td>Oligo fwd</td><td>CACCGGCGCGCAGCAGGTCGGGAC</td><td></td><td></td><td></td><td></td><td></td><td></td><td></td><td></td><td></td><td></td></tr>
<tr><td>Oligo rev</td><td>AAACGTCCCGACCTGCTGCGCGCC</td><td></td><td></td><td></td><td></td><td></td><td></td><td></td><td></td><td></td><td></td></tr>
<tr><td>Chromosome</td><td>start</td><td>end</td><td>strand</td><td>MM</td><td>target_seq</td><td>PAM</td><td>alignment</td><td>distance</td><td>position</td><td>gene name</td><td>gene id</td></tr>
<tr><td>GU305914.1</td><td>49816</td><td>49838</td><td>+</td><td>0</td><td>GGCGCGCAGCAGGTCGGGAC</td><td>AGG</td><td>||||||||[||||||||||||]PAM</td><td>NA</td><td>Intergenic</td><td>NA</td><td>NA</td></tr>
<tr><td>T3240</td><td>GCAGCAGGTCGGGACAGGCACGG</td><td>1000</td><td></td><td></td><td></td><td></td><td></td><td></td><td></td><td></td><td></td></tr>
<tr><td>Oligo fwd</td><td>CACCGCAGCAGGTCGGGACAGGCA</td><td></td><td></td><td></td><td></td><td></td><td></td><td></td><td></td><td></td><td></td></tr>
<tr><td>Oligo rev</td><td>AAACTGCCTGTCCCGACCTGCTGC</td><td></td><td></td><td></td><td></td><td></td><td></td><td></td><td></td><td></td><td></td></tr>
<tr><td>Chromosome</td><td>start</td><td>end</td><td>strand</td><td>MM</td><td>target_seq</td><td>PAM</td><td>alignment</td><td>distance</td><td>position</td><td>gene name</td><td>gene id</td></tr>
<tr><td>GU305914.1</td><td>49821</td><td>49843</td><td>+</td><td>0</td><td>GCAGCAGGTCGGGACAGGCA</td><td>CGG</td><td>||||||||[||||||||||||]PAM</td><td>NA</td><td>Intergenic</td><td>NA</td><td>NA</td></tr>
<tr><td>T3241</td><td>CCAGACTAATGTTTTTCATCAGG</td><td>1000</td><td></td><td></td><td></td><td></td><td></td><td></td><td></td><td></td><td></td></tr>
<tr><td>Oligo fwd</td><td>CACCCCAGACTAATGTTTTTCATC</td><td></td><td></td><td></td><td></td><td></td><td></td><td></td><td></td><td></td><td></td></tr>
<tr><td>Oligo rev</td><td>AAACGATGAAAAACATTAGTCTGG</td><td></td><td></td><td></td><td></td><td></td><td></td><td></td><td></td><td></td><td></td></tr>
<tr><td>Chromosome</td><td>start</td><td>end</td><td>strand</td><td>MM</td><td>target_seq</td><td>PAM</td><td>alignment</td><td>distance</td><td>position</td><td>gene name</td><td>gene id</td></tr>
<tr><td>GU305914.1</td><td>49857</td><td>49879</td><td>+</td><td>0</td><td>CCAGACTAATGTTTTTCATC</td><td>AGG</td><td>||||||||[||||||||||||]PAM</td><td>NA</td><td>Intergenic</td><td>NA</td><td>NA</td></tr>
<tr><td>T3242</td><td>CATCAGGAATTGCACCGTCTCGG</td><td>1000</td><td></td><td></td><td></td><td></td><td></td><td></td><td></td><td></td><td></td></tr>
<tr><td>Oligo fwd</td><td>CACCCATCAGGAATTGCACCGTCT</td><td></td><td></td><td></td><td></td><td></td><td></td><td></td><td></td><td></td><td></td></tr>
<tr><td>Oligo rev</td><td>AAACAGACGGTGCAATTCCTGATG</td><td></td><td></td><td></td><td></td><td></td><td></td><td></td><td></td><td></td><td></td></tr>
<tr><td>Chromosome</td><td>start</td><td>end</td><td>strand</td><td>MM</td><td>target_seq</td><td>PAM</td><td>alignment</td><td>distance</td><td>position</td><td>gene name</td><td>gene id</td></tr>
<tr><td>GU305914.1</td><td>49873</td><td>49895</td><td>+</td><td>0</td><td>CATCAGGAATTGCACCGTCT</td><td>CGG</td><td>||||||||[||||||||||||]PAM</td><td>NA</td><td>Intergenic</td><td>NA</td><td>NA</td></tr>
<tr><td>T3243</td><td>TGCTCGAATCGACCTCCAGACGG</td><td>1000</td><td></td><td></td><td></td><td></td><td></td><td></td><td></td><td></td><td></td></tr>
<tr><td>Oligo fwd</td><td>CACCTGCTCGAATCGACCTCCAGA</td><td></td><td></td><td></td><td></td><td></td><td></td><td></td><td></td><td></td><td></td></tr>
<tr><td>Oligo rev</td><td>AAACTCTGGAGGTCGATTCGAGCA</td><td></td><td></td><td></td><td></td><td></td><td></td><td></td><td></td><td></td><td></td></tr>
<tr><td>Chromosome</td><td>start</td><td>end</td><td>strand</td><td>MM</td><td>target_seq</td><td>PAM</td><td>alignment</td><td>distance</td><td>position</td><td>gene name</td><td>gene id</td></tr>
<tr><td>GU305914.1</td><td>49938</td><td>49960</td><td>+</td><td>0</td><td>TGCTCGAATCGACCTCCAGA</td><td>CGG</td><td>||||||||[||||||||||||]PAM</td><td>NA</td><td>Intergenic</td><td>NA</td><td>NA</td></tr>
<tr><td>T3244</td><td>TCTACCGAACCGCGACCGTCCGG</td><td>1000</td><td></td><td></td><td></td><td></td><td></td><td></td><td></td><td></td><td></td></tr>
<tr><td>Oligo fwd</td><td>CACCTCTACCGAACCGCGACCGTC</td><td></td><td></td><td></td><td></td><td></td><td></td><td></td><td></td><td></td><td></td></tr>
<tr><td>Oligo rev</td><td>AAACGACGGTCGCGGTTCGGTAGA</td><td></td><td></td><td></td><td></td><td></td><td></td><td></td><td></td><td></td><td></td></tr>
<tr><td>Chromosome</td><td>start</td><td>end</td><td>strand</td><td>MM</td><td>target_seq</td><td>PAM</td><td>alignment</td><td>distance</td><td>position</td><td>gene name</td><td>gene id</td></tr>
<tr><td>GU305914.1</td><td>49988</td><td>50010</td><td>+</td><td>0</td><td>TCTACCGAACCGCGACCGTC</td><td>CGG</td><td>||||||||[||||||||||||]PAM</td><td>NA</td><td>Intergenic</td><td>NA</td><td>NA</td></tr>
<tr><td>T3245</td><td>CGGATGACATCTGCCCTTGCCGG</td><td>1000</td><td></td><td></td><td></td><td></td><td></td><td></td><td></td><td></td><td></td></tr>
<tr><td>Oligo fwd</td><td>CACCCGGATGACATCTGCCCTTGC</td><td></td><td></td><td></td><td></td><td></td><td></td><td></td><td></td><td></td><td></td></tr>
<tr><td>Oligo rev</td><td>AAACGCAAGGGCAGATGTCATCCG</td><td></td><td></td><td></td><td></td><td></td><td></td><td></td><td></td><td></td><td></td></tr>
<tr><td>Chromosome</td><td>start</td><td>end</td><td>strand</td><td>MM</td><td>target_seq</td><td>PAM</td><td>alignment</td><td>distance</td><td>position</td><td>gene name</td><td>gene id</td></tr>
<tr><td>GU305914.1</td><td>50008</td><td>50030</td><td>+</td><td>0</td><td>CGGATGACATCTGCCCTTGC</td><td>CGG</td><td>||||||||[||||||||||||]PAM</td><td>NA</td><td>Intergenic</td><td>NA</td><td>NA</td></tr>
<tr><td>T3246</td><td>ACGAAGTCCTTCCATCCGCACGG</td><td>1000</td><td></td><td></td><td></td><td></td><td></td><td></td><td></td><td></td><td></td></tr>
<tr><td>Oligo fwd</td><td>CACCACGAAGTCCTTCCATCCGCA</td><td></td><td></td><td></td><td></td><td></td><td></td><td></td><td></td><td></td><td></td></tr>
<tr><td>Oligo rev</td><td>AAACTGCGGATGGAAGGACTTCGT</td><td></td><td></td><td></td><td></td><td></td><td></td><td></td><td></td><td></td><td></td></tr>
<tr><td>Chromosome</td><td>start</td><td>end</td><td>strand</td><td>MM</td><td>target_seq</td><td>PAM</td><td>alignment</td><td>distance</td><td>position</td><td>gene name</td><td>gene id</td></tr>
<tr><td>GU305914.1</td><td>50037</td><td>50059</td><td>+</td><td>0</td><td>ACGAAGTCCTTCCATCCGCA</td><td>CGG</td><td>||||||||[||||||||||||]PAM</td><td>NA</td><td>Intergenic</td><td>NA</td><td>NA</td></tr>
<tr><td>T3247</td><td>CGCACGGTGCGAGACGAGTCCGG</td><td>1000</td><td></td><td></td><td></td><td></td><td></td><td></td><td></td><td></td><td></td></tr>
<tr><td>Oligo fwd</td><td>CACCCGCACGGTGCGAGACGAGTC</td><td></td><td></td><td></td><td></td><td></td><td></td><td></td><td></td><td></td><td></td></tr>
<tr><td>Oligo rev</td><td>AAACGACTCGTCTCGCACCGTGCG</td><td></td><td></td><td></td><td></td><td></td><td></td><td></td><td></td><td></td><td></td></tr>
<tr><td>Chromosome</td><td>start</td><td>end</td><td>strand</td><td>MM</td><td>target_seq</td><td>PAM</td><td>alignment</td><td>distance</td><td>position</td><td>gene name</td><td>gene id</td></tr>
<tr><td>GU305914.1</td><td>50053</td><td>50075</td><td>+</td><td>0</td><td>CGCACGGTGCGAGACGAGTC</td><td>CGG</td><td>||||||||[||||||||||||]PAM</td><td>NA</td><td>Intergenic</td><td>NA</td><td>NA</td></tr>
<tr><td>T3248</td><td>GCACGGTGCGAGACGAGTCCGGG</td><td>1000</td><td></td><td></td><td></td><td></td><td></td><td></td><td></td><td></td><td></td></tr>
<tr><td>Oligo fwd</td><td>CACCGCACGGTGCGAGACGAGTCC</td><td></td><td></td><td></td><td></td><td></td><td></td><td></td><td></td><td></td><td></td></tr>
<tr><td>Oligo rev</td><td>AAACGGACTCGTCTCGCACCGTGC</td><td></td><td></td><td></td><td></td><td></td><td></td><td></td><td></td><td></td><td></td></tr>
<tr><td>Chromosome</td><td>start</td><td>end</td><td>strand</td><td>MM</td><td>target_seq</td><td>PAM</td><td>alignment</td><td>distance</td><td>position</td><td>gene name</td><td>gene id</td></tr>
<tr><td>GU305914.1</td><td>50054</td><td>50076</td><td>+</td><td>0</td><td>GCACGGTGCGAGACGAGTCC</td><td>GGG</td><td>||||||||[||||||||||||]PAM</td><td>NA</td><td>Intergenic</td><td>NA</td><td>NA</td></tr>
<tr><td>T3249</td><td>CGAGTCCGGGACACGAGCTCAGG</td><td>1000</td><td></td><td></td><td></td><td></td><td></td><td></td><td></td><td></td><td></td></tr>
<tr><td>Oligo fwd</td><td>CACCCGAGTCCGGGACACGAGCTC</td><td></td><td></td><td></td><td></td><td></td><td></td><td></td><td></td><td></td><td></td></tr>
<tr><td>Oligo rev</td><td>AAACGAGCTCGTGTCCCGGACTCG</td><td></td><td></td><td></td><td></td><td></td><td></td><td></td><td></td><td></td><td></td></tr>
<tr><td>Chromosome</td><td>start</td><td>end</td><td>strand</td><td>MM</td><td>target_seq</td><td>PAM</td><td>alignment</td><td>distance</td><td>position</td><td>gene name</td><td>gene id</td></tr>
<tr><td>GU305914.1</td><td>50067</td><td>50089</td><td>+</td><td>0</td><td>CGAGTCCGGGACACGAGCTC</td><td>AGG</td><td>||||||||[||||||||||||]PAM</td><td>NA</td><td>Intergenic</td><td>NA</td><td>NA</td></tr>
<tr><td>T3250</td><td>GAGTCCGGGACACGAGCTCAGGG</td><td>1000</td><td></td><td></td><td></td><td></td><td></td><td></td><td></td><td></td><td></td></tr>
<tr><td>Oligo fwd</td><td>CACCGAGTCCGGGACACGAGCTCA</td><td></td><td></td><td></td><td></td><td></td><td></td><td></td><td></td><td></td><td></td></tr>
<tr><td>Oligo rev</td><td>AAACTGAGCTCGTGTCCCGGACTC</td><td></td><td></td><td></td><td></td><td></td><td></td><td></td><td></td><td></td><td></td></tr>
<tr><td>Chromosome</td><td>start</td><td>end</td><td>strand</td><td>MM</td><td>target_seq</td><td>PAM</td><td>alignment</td><td>distance</td><td>position</td><td>gene name</td><td>gene id</td></tr>
<tr><td>GU305914.1</td><td>50068</td><td>50090</td><td>+</td><td>0</td><td>GAGTCCGGGACACGAGCTCA</td><td>GGG</td><td>||||||||[||||||||||||]PAM</td><td>NA</td><td>Intergenic</td><td>NA</td><td>NA</td></tr>
<tr><td>T3251</td><td>AGTCCGGGACACGAGCTCAGGGG</td><td>1000</td><td></td><td></td><td></td><td></td><td></td><td></td><td></td><td></td><td></td></tr>
<tr><td>Oligo fwd</td><td>CACCAGTCCGGGACACGAGCTCAG</td><td></td><td></td><td></td><td></td><td></td><td></td><td></td><td></td><td></td><td></td></tr>
<tr><td>Oligo rev</td><td>AAACCTGAGCTCGTGTCCCGGACT</td><td></td><td></td><td></td><td></td><td></td><td></td><td></td><td></td><td></td><td></td></tr>
<tr><td>Chromosome</td><td>start</td><td>end</td><td>strand</td><td>MM</td><td>target_seq</td><td>PAM</td><td>alignment</td><td>distance</td><td>position</td><td>gene name</td><td>gene id</td></tr>
<tr><td>GU305914.1</td><td>50069</td><td>50091</td><td>+</td><td>0</td><td>AGTCCGGGACACGAGCTCAG</td><td>GGG</td><td>||||||||[||||||||||||]PAM</td><td>NA</td><td>Intergenic</td><td>NA</td><td>NA</td></tr>
<tr><td>T3252</td><td>GACACGAGCTCAGGGGCCTGAGG</td><td>1000</td><td></td><td></td><td></td><td></td><td></td><td></td><td></td><td></td><td></td></tr>
<tr><td>Oligo fwd</td><td>CACCGACACGAGCTCAGGGGCCTG</td><td></td><td></td><td></td><td></td><td></td><td></td><td></td><td></td><td></td><td></td></tr>
<tr><td>Oligo rev</td><td>AAACCAGGCCCCTGAGCTCGTGTC</td><td></td><td></td><td></td><td></td><td></td><td></td><td></td><td></td><td></td><td></td></tr>
<tr><td>Chromosome</td><td>start</td><td>end</td><td>strand</td><td>MM</td><td>target_seq</td><td>PAM</td><td>alignment</td><td>distance</td><td>position</td><td>gene name</td><td>gene id</td></tr>
</table>

GU305914.1 50076 50098 + 0 GACACGAGCTCAGGGGCCTG AGG ||||||||[||||||||||||]PAM NA Intergenic NA NA

T3253 CAGGGGCCTGAGGATGTAACCGG 1000
Oligo fwd CACCCAGGGGCCTGAGGATGTAAC
Oligo rev AAACGTTACATCCTCAGGCCCCTG
Chromosome start end strand MM target_seq PAM alignment distance position gene name gene id
GU305914.1 50086 50108 + 0 CAGGGGCCTGAGGATGTAAC CGG ||||||||[||||||||||||]PAM NA Intergenic NA NA

T3254 CGATGCCGATGTCAACCAGCTGG 1000
Oligo fwd CACCCGATGCCGATGTCAACCAGC
Oligo rev AAACGCTGGTTGACATCGGCATCG
Chromosome start end strand MM target_seq PAM alignment distance position gene name gene id
GU305914.1 50121 50143 + 0 CGATGCCGATGTCAACCAGC TGG ||||||||[||||||||||||]PAM NA Intergenic NA NA

T3255 TATCCCGAAACGACCAGCGCCGG 1000
Oligo fwd CACCTATCCCGAAACGACCAGCGC
Oligo rev AAACGCGCTGGTCGTTTCGGGATA
Chromosome start end strand MM target_seq PAM alignment distance position gene name gene id
GU305914.1 50145 50167 + 0 TATCCCGAAACGACCAGCGC CGG ||||||||[||||||||||||]PAM NA Intergenic NA NA

T3256 CCGGTTAACCTTGCCGAACCAGG 1000
Oligo fwd CACCCCGGTTAACCTTGCCGAACC
Oligo rev AAACGGTTCGGCAAGGTTAACCGG
Chromosome start end strand MM target_seq PAM alignment distance position gene name gene id
GU305914.1 50164 50186 + 0 CCGGTTAACCTTGCCGAACC AGG ||||||||[||||||||||||]PAM NA Intergenic NA NA

T3257 CCAGGTACTGACCAACGTGAAGG 1000
Oligo fwd CACCCCAGGTACTGACCAACGTGA
Oligo rev AAACTCACGTTGGTCAGTACCTGG
Chromosome start end strand MM target_seq PAM alignment distance position gene name gene id
GU305914.1 50182 50204 + 0 CCAGGTACTGACCAACGTGA AGG ||||||||[||||||||||||]PAM NA Intergenic NA NA

T3258 GAAGGCATCTTGACAGCTGCCGG 1000
Oligo fwd CACCGAAGGCATCTTGACAGCTGC
Oligo rev AAACGCAGCTGTCAAGATGCCTTC
Chromosome start end strand MM target_seq PAM alignment distance position gene name gene id
GU305914.1 50200 50222 + 0 GAAGGCATCTTGACAGCTGC CGG ||||||||[||||||||||||]PAM NA Intergenic NA NA

T3259 GCTGCCGGAGAAGACGAGTTCGG 1000
Oligo fwd CACCGCTGCCGGAGAAGACGAGTT
Oligo rev AAACAACTCGTCTTCTCCGGCAGC
Chromosome start end strand MM target_seq PAM alignment distance position gene name gene id
GU305914.1 50215 50237 + 0 GCTGCCGGAGAAGACGAGTT CGG ||||||||[||||||||||||]PAM NA Intergenic NA NA

T3260 GTCGACGATACAGCCCACACTGG 1000
Oligo fwd CACCGTCGACGATACAGCCCACAC
Oligo rev AAACGTGTGGGCTGTATCGTCGAC
Chromosome start end strand MM target_seq PAM alignment distance position gene name gene id
GU305914.1 50251 50273 + 0 GTCGACGATACAGCCCACAC TGG ||||||||[||||||||||||]PAM NA Intergenic NA NA

T3261 CGATACAGCCCACACTGGTGCGG 1000
Oligo fwd CACCCGATACAGCCCACACTGGTG
Oligo rev AAACCACCAGTGTGGGCTGTATCG
Chromosome start end strand MM target_seq PAM alignment distance position gene name gene id
GU305914.1 50256 50278 + 0 CGATACAGCCCACACTGGTG CGG ||||||||[||||||||||||]PAM NA Intergenic NA NA

T3262 CTGCACGAGACAGACGTCGTAGG 1000
Oligo fwd CACCCTGCACGAGACAGACGTCGT
Oligo rev AAACACGACGTCTGTCTCGTGCAG
Chromosome start end strand MM target_seq PAM alignment distance position gene name gene id
GU305914.1 50290 50312 + 0 CTGCACGAGACAGACGTCGT AGG ||||||||[||||||||||||]PAM NA Intergenic NA NA

T3263 GCACTCGAAGACGCCCGATTCGG 1000
Oligo fwd CACCGCACTCGAAGACGCCCGATT
Oligo rev AAACAATCGGGCGTCTTCGAGTGC
Chromosome start end strand MM target_seq PAM alignment distance position gene name gene id
GU305914.1 50314 50336 + 0 GCACTCGAAGACGCCCGATT CGG ||||||||[||||||||||||]PAM NA Intergenic NA NA

T3264 CACTCGAAGACGCCCGATTCGGG 1000
Oligo fwd CACCCACTCGAAGACGCCCGATTC
Oligo rev AAACGAATCGGGCGTCTTCGAGTG
Chromosome start end strand MM target_seq PAM alignment distance position gene name gene id
GU305914.1 50315 50337 + 0 CACTCGAAGACGCCCGATTC GGG ||||||||[||||||||||||]PAM NA Intergenic NA NA

T3265 CCCCGAGACCGCCGTCCGCGCGG 1000
Oligo fwd CACCCCCCGAGACCGCCGTCCGCG
Oligo rev AAACCGCGGACGGCGGTCTCGGGG
Chromosome start end strand MM target_seq PAM alignment distance position gene name gene id
GU305914.1 50338 50360 + 0 CCCCGAGACCGCCGTCCGCG CGG ||||||||[||||||||||||]PAM NA Intergenic NA NA

T3266 CCGTCCGCGCGGCGATCGCGAGG 1000
Oligo fwd CACCCCGTCCGCGCGGCGATCGCG
Oligo rev AAACCGCGATCGCCGCGCGGACGG
Chromosome start end strand MM target_seq PAM alignment distance position gene name gene id
GU305914.1 50349 50371 + 0 CCGTCCGCGCGGCGATCGCG AGG ||||||||[||||||||||||]PAM NA Intergenic NA NA

T3267 CGCGAGGCAACAACAGAGAGTGG 1000
Oligo fwd CACCCGCGAGGCAACAACAGAGAG
Oligo rev AAACCTCTCTGTTGTTGCCTCGCG

<table>
<tr><td>Chromosome</td><td>start</td><td>end</td><td>strand</td><td>MM</td><td>target_seq</td><td>PAM</td><td>alignment</td><td>distance</td><td>position</td><td>gene name</td><td>gene id</td></tr>
<tr><td>GU305914.1</td><td>50365</td><td>50387</td><td>+</td><td>0</td><td>CGCGAGGCAACAACAGAGAG</td><td>TGG</td><td>||||||||[||||||||||||]PAM</td><td>NA</td><td>Intergenic</td><td>NA</td><td>NA</td></tr>
<tr><td>T3268</td><td>GAGGCAACAACAGAGAGTGGTGG</td><td>1000</td><td></td><td></td><td></td><td></td><td></td><td></td><td></td><td></td><td></td></tr>
<tr><td>Oligo fwd</td><td>CACCGAGGCAACAACAGAGAGTGG</td><td></td><td></td><td></td><td></td><td></td><td></td><td></td><td></td><td></td><td></td></tr>
<tr><td>Oligo rev</td><td>AAACCCACTCTCTGTTGTTGCCTC</td><td></td><td></td><td></td><td></td><td></td><td></td><td></td><td></td><td></td><td></td></tr>
<tr><td>Chromosome</td><td>start</td><td>end</td><td>strand</td><td>MM</td><td>target_seq</td><td>PAM</td><td>alignment</td><td>distance</td><td>position</td><td>gene name</td><td>gene id</td></tr>
<tr><td>GU305914.1</td><td>50368</td><td>50390</td><td>+</td><td>0</td><td>GAGGCAACAACAGAGAGTGG</td><td>TGG</td><td>||||||||[||||||||||||]PAM</td><td>NA</td><td>Intergenic</td><td>NA</td><td>NA</td></tr>
<tr><td>T3269</td><td>ACAGAGAGTGGTGGCACGCGTGG</td><td>1000</td><td></td><td></td><td></td><td></td><td></td><td></td><td></td><td></td><td></td></tr>
<tr><td>Oligo fwd</td><td>CACCACAGAGAGTGGTGGCACGCG</td><td></td><td></td><td></td><td></td><td></td><td></td><td></td><td></td><td></td><td></td></tr>
<tr><td>Oligo rev</td><td>AAACCGCGTGCCACCACTCTCTGT</td><td></td><td></td><td></td><td></td><td></td><td></td><td></td><td></td><td></td><td></td></tr>
<tr><td>Chromosome</td><td>start</td><td>end</td><td>strand</td><td>MM</td><td>target_seq</td><td>PAM</td><td>alignment</td><td>distance</td><td>position</td><td>gene name</td><td>gene id</td></tr>
<tr><td>GU305914.1</td><td>50377</td><td>50399</td><td>+</td><td>0</td><td>ACAGAGAGTGGTGGCACGCG</td><td>TGG</td><td>||||||||[||||||||||||]PAM</td><td>NA</td><td>Intergenic</td><td>NA</td><td>NA</td></tr>
<tr><td>T3270</td><td>GAGAGAATCCGCCCGAGATTCGG</td><td>1000</td><td></td><td></td><td></td><td></td><td></td><td></td><td></td><td></td><td></td></tr>
<tr><td>Oligo fwd</td><td>CACCGAGAGAATCCGCCCGAGATT</td><td></td><td></td><td></td><td></td><td></td><td></td><td></td><td></td><td></td><td></td></tr>
<tr><td>Oligo rev</td><td>AAACAATCTCGGGCGGATTCTCTC</td><td></td><td></td><td></td><td></td><td></td><td></td><td></td><td></td><td></td><td></td></tr>
<tr><td>Chromosome</td><td>start</td><td>end</td><td>strand</td><td>MM</td><td>target_seq</td><td>PAM</td><td>alignment</td><td>distance</td><td>position</td><td>gene name</td><td>gene id</td></tr>
<tr><td>GU305914.1</td><td>50440</td><td>50462</td><td>+</td><td>0</td><td>GAGAGAATCCGCCCGAGATT</td><td>CGG</td><td>||||||||[||||||||||||]PAM</td><td>NA</td><td>Intergenic</td><td>NA</td><td>NA</td></tr>
<tr><td>T3271</td><td>GAGATTCGGACTTATGTGTCAGG</td><td>1000</td><td></td><td></td><td></td><td></td><td></td><td></td><td></td><td></td><td></td></tr>
<tr><td>Oligo fwd</td><td>CACCGAGATTCGGACTTATGTGTC</td><td></td><td></td><td></td><td></td><td></td><td></td><td></td><td></td><td></td><td></td></tr>
<tr><td>Oligo rev</td><td>AAACGACACATAAGTCCGAATCTC</td><td></td><td></td><td></td><td></td><td></td><td></td><td></td><td></td><td></td><td></td></tr>
<tr><td>Chromosome</td><td>start</td><td>end</td><td>strand</td><td>MM</td><td>target_seq</td><td>PAM</td><td>alignment</td><td>distance</td><td>position</td><td>gene name</td><td>gene id</td></tr>
<tr><td>GU305914.1</td><td>50454</td><td>50476</td><td>+</td><td>0</td><td>GAGATTCGGACTTATGTGTC</td><td>AGG</td><td>||||||||[||||||||||||]PAM</td><td>NA</td><td>Intergenic</td><td>NA</td><td>NA</td></tr>
<tr><td>T3272</td><td>TGTGTCAGGACCATCGCTCGAGG</td><td>1000</td><td></td><td></td><td></td><td></td><td></td><td></td><td></td><td></td><td></td></tr>
<tr><td>Oligo fwd</td><td>CACCTGTGTCAGGACCATCGCTCG</td><td></td><td></td><td></td><td></td><td></td><td></td><td></td><td></td><td></td><td></td></tr>
<tr><td>Oligo rev</td><td>AAACCGAGCGATGGTCCTGACACA</td><td></td><td></td><td></td><td></td><td></td><td></td><td></td><td></td><td></td><td></td></tr>
<tr><td>Chromosome</td><td>start</td><td>end</td><td>strand</td><td>MM</td><td>target_seq</td><td>PAM</td><td>alignment</td><td>distance</td><td>position</td><td>gene name</td><td>gene id</td></tr>
<tr><td>GU305914.1</td><td>50468</td><td>50490</td><td>+</td><td>0</td><td>TGTGTCAGGACCATCGCTCG</td><td>AGG</td><td>||||||||[||||||||||||]PAM</td><td>NA</td><td>Intergenic</td><td>NA</td><td>NA</td></tr>
<tr><td>T3273</td><td>CAGGACCATCGCTCGAGGCGCGG</td><td>1000</td><td></td><td></td><td></td><td></td><td></td><td></td><td></td><td></td><td></td></tr>
<tr><td>Oligo fwd</td><td>CACCCAGGACCATCGCTCGAGGCG</td><td></td><td></td><td></td><td></td><td></td><td></td><td></td><td></td><td></td><td></td></tr>
<tr><td>Oligo rev</td><td>AAACCGCCTCGAGCGATGGTCCTG</td><td></td><td></td><td></td><td></td><td></td><td></td><td></td><td></td><td></td><td></td></tr>
<tr><td>Chromosome</td><td>start</td><td>end</td><td>strand</td><td>MM</td><td>target_seq</td><td>PAM</td><td>alignment</td><td>distance</td><td>position</td><td>gene name</td><td>gene id</td></tr>
<tr><td>GU305914.1</td><td>50473</td><td>50495</td><td>+</td><td>0</td><td>CAGGACCATCGCTCGAGGCG</td><td>CGG</td><td>||||||||[||||||||||||]PAM</td><td>NA</td><td>Intergenic</td><td>NA</td><td>NA</td></tr>
<tr><td>T3274</td><td>GAGGCGCGGTCGCACACGCGTGG</td><td>1000</td><td></td><td></td><td></td><td></td><td></td><td></td><td></td><td></td><td></td></tr>
<tr><td>Oligo fwd</td><td>CACCGAGGCGCGGTCGCACACGCG</td><td></td><td></td><td></td><td></td><td></td><td></td><td></td><td></td><td></td><td></td></tr>
<tr><td>Oligo rev</td><td>AAACCGCGTGTGCGACCGCGCCTC</td><td></td><td></td><td></td><td></td><td></td><td></td><td></td><td></td><td></td><td></td></tr>
<tr><td>Chromosome</td><td>start</td><td>end</td><td>strand</td><td>MM</td><td>target_seq</td><td>PAM</td><td>alignment</td><td>distance</td><td>position</td><td>gene name</td><td>gene id</td></tr>
<tr><td>GU305914.1</td><td>50487</td><td>50509</td><td>+</td><td>0</td><td>GAGGCGCGGTCGCACACGCG</td><td>TGG</td><td>||||||||[||||||||||||]PAM</td><td>NA</td><td>Intergenic</td><td>NA</td><td>NA</td></tr>
<tr><td>T3275</td><td>ATGCTCGTCGTCATCACCGCAGG</td><td>1000</td><td></td><td></td><td></td><td></td><td></td><td></td><td></td><td></td><td></td></tr>
<tr><td>Oligo fwd</td><td>CACCATGCTCGTCGTCATCACCGC</td><td></td><td></td><td></td><td></td><td></td><td></td><td></td><td></td><td></td><td></td></tr>
<tr><td>Oligo rev</td><td>AAACGCGGTGATGACGACGAGCAT</td><td></td><td></td><td></td><td></td><td></td><td></td><td></td><td></td><td></td><td></td></tr>
<tr><td>Chromosome</td><td>start</td><td>end</td><td>strand</td><td>MM</td><td>target_seq</td><td>PAM</td><td>alignment</td><td>distance</td><td>position</td><td>gene name</td><td>gene id</td></tr>
<tr><td>GU305914.1</td><td>50532</td><td>50554</td><td>+</td><td>0</td><td>ATGCTCGTCGTCATCACCGC</td><td>AGG</td><td>||||||||[||||||||||||]PAM</td><td>NA</td><td>Intergenic</td><td>NA</td><td>NA</td></tr>
<tr><td>T3276</td><td>TCACCGCAGGTCCCCACAGTTGG</td><td>1000</td><td></td><td></td><td></td><td></td><td></td><td></td><td></td><td></td><td></td></tr>
<tr><td>Oligo fwd</td><td>CACCTCACCGCAGGTCCCCACAGT</td><td></td><td></td><td></td><td></td><td></td><td></td><td></td><td></td><td></td><td></td></tr>
<tr><td>Oligo rev</td><td>AAACACTGTGGGGACCTGCGGTGA</td><td></td><td></td><td></td><td></td><td></td><td></td><td></td><td></td><td></td><td></td></tr>
<tr><td>Chromosome</td><td>start</td><td>end</td><td>strand</td><td>MM</td><td>target_seq</td><td>PAM</td><td>alignment</td><td>distance</td><td>position</td><td>gene name</td><td>gene id</td></tr>
<tr><td>GU305914.1</td><td>50545</td><td>50567</td><td>+</td><td>0</td><td>TCACCGCAGGTCCCCACAGT</td><td>TGG</td><td>||||||||[||||||||||||]PAM</td><td>NA</td><td>Intergenic</td><td>NA</td><td>NA</td></tr>
<tr><td>T3277</td><td>CACCGCAGGTCCCCACAGTTGGG</td><td>1000</td><td></td><td></td><td></td><td></td><td></td><td></td><td></td><td></td><td></td></tr>
<tr><td>Oligo fwd</td><td>CACCCACCGCAGGTCCCCACAGTT</td><td></td><td></td><td></td><td></td><td></td><td></td><td></td><td></td><td></td><td></td></tr>
<tr><td>Oligo rev</td><td>AAACAACTGTGGGGACCTGCGGTG</td><td></td><td></td><td></td><td></td><td></td><td></td><td></td><td></td><td></td><td></td></tr>
<tr><td>Chromosome</td><td>start</td><td>end</td><td>strand</td><td>MM</td><td>target_seq</td><td>PAM</td><td>alignment</td><td>distance</td><td>position</td><td>gene name</td><td>gene id</td></tr>
<tr><td>GU305914.1</td><td>50546</td><td>50568</td><td>+</td><td>0</td><td>CACCGCAGGTCCCCACAGTT</td><td>GGG</td><td>||||||||[||||||||||||]PAM</td><td>NA</td><td>Intergenic</td><td>NA</td><td>NA</td></tr>
<tr><td>T3278</td><td>CTCGAAGCATTCTTCCGTCTTGG</td><td>1000</td><td></td><td></td><td></td><td></td><td></td><td></td><td></td><td></td><td></td></tr>
<tr><td>Oligo fwd</td><td>CACCCTCGAAGCATTCTTCCGTCT</td><td></td><td></td><td></td><td></td><td></td><td></td><td></td><td></td><td></td><td></td></tr>
<tr><td>Oligo rev</td><td>AAACAGACGGAAGAATGCTTCGAG</td><td></td><td></td><td></td><td></td><td></td><td></td><td></td><td></td><td></td><td></td></tr>
<tr><td>Chromosome</td><td>start</td><td>end</td><td>strand</td><td>MM</td><td>target_seq</td><td>PAM</td><td>alignment</td><td>distance</td><td>position</td><td>gene name</td><td>gene id</td></tr>
<tr><td>GU305914.1</td><td>50588</td><td>50610</td><td>+</td><td>0</td><td>CTCGAAGCATTCTTCCGTCT</td><td>TGG</td><td>||||||||[||||||||||||]PAM</td><td>NA</td><td>Intergenic</td><td>NA</td><td>NA</td></tr>
<tr><td>T3279</td><td>TTCTTCCGTCTTGGTGATCTTGG</td><td>1000</td><td></td><td></td><td></td><td></td><td></td><td></td><td></td><td></td><td></td></tr>
<tr><td>Oligo fwd</td><td>CACCTTCTTCCGTCTTGGTGATCT</td><td></td><td></td><td></td><td></td><td></td><td></td><td></td><td></td><td></td><td></td></tr>
<tr><td>Oligo rev</td><td>AAACAGATCACCAAGACGGAAGAA</td><td></td><td></td><td></td><td></td><td></td><td></td><td></td><td></td><td></td><td></td></tr>
<tr><td>Chromosome</td><td>start</td><td>end</td><td>strand</td><td>MM</td><td>target_seq</td><td>PAM</td><td>alignment</td><td>distance</td><td>position</td><td>gene name</td><td>gene id</td></tr>
<tr><td>GU305914.1</td><td>50597</td><td>50619</td><td>+</td><td>0</td><td>TTCTTCCGTCTTGGTGATCT</td><td>TGG</td><td>||||||||[||||||||||||]PAM</td><td>NA</td><td>Intergenic</td><td>NA</td><td>NA</td></tr>
<tr><td>T3280</td><td>CTCGAGCACCCTACTGTCTTCGG</td><td>1000</td><td></td><td></td><td></td><td></td><td></td><td></td><td></td><td></td><td></td></tr>
<tr><td>Oligo fwd</td><td>CACCCTCGAGCACCCTACTGTCTT</td><td></td><td></td><td></td><td></td><td></td><td></td><td></td><td></td><td></td><td></td></tr>
<tr><td>Oligo rev</td><td>AAACAAGACAGTAGGGTGCTCGAG</td><td></td><td></td><td></td><td></td><td></td><td></td><td></td><td></td><td></td><td></td></tr>
<tr><td>Chromosome</td><td>start</td><td>end</td><td>strand</td><td>MM</td><td>target_seq</td><td>PAM</td><td>alignment</td><td>distance</td><td>position</td><td>gene name</td><td>gene id</td></tr>
<tr><td>GU305914.1</td><td>50636</td><td>50658</td><td>+</td><td>0</td><td>CTCGAGCACCCTACTGTCTT</td><td>CGG</td><td>||||||||[||||||||||||]PAM</td><td>NA</td><td>Intergenic</td><td>NA</td><td>NA</td></tr>
<tr><td>T3281</td><td>TCGAGCACCCTACTGTCTTCGGG</td><td>1000</td><td></td><td></td><td></td><td></td><td></td><td></td><td></td><td></td><td></td></tr>
<tr><td>Oligo fwd</td><td>CACCTCGAGCACCCTACTGTCTTC</td><td></td><td></td><td></td><td></td><td></td><td></td><td></td><td></td><td></td><td></td></tr>
<tr><td>Oligo rev</td><td>AAACGAAGACAGTAGGGTGCTCGA</td><td></td><td></td><td></td><td></td><td></td><td></td><td></td><td></td><td></td><td></td></tr>
<tr><td>Chromosome</td><td>start</td><td>end</td><td>strand</td><td>MM</td><td>target_seq</td><td>PAM</td><td>alignment</td><td>distance</td><td>position</td><td>gene name</td><td>gene id</td></tr>
<tr><td>GU305914.1</td><td>50637</td><td>50659</td><td>+</td><td>0</td><td>TCGAGCACCCTACTGTCTTC</td><td>GGG</td><td>||||||||[||||||||||||]PAM</td><td>NA</td><td>Intergenic</td><td>NA</td><td>NA</td></tr>
<tr><td>T3282</td><td>ACCCTACTGTCTTCGGGCGCCGG</td><td>1000</td><td></td><td></td><td></td><td></td><td></td><td></td><td></td><td></td><td></td></tr>
<tr><td>Oligo fwd</td><td>CACCACCCTACTGTCTTCGGGCGC</td><td></td><td></td><td></td><td></td><td></td><td></td><td></td><td></td><td></td><td></td></tr>
</table>

Oligo rev AAACGCGCCCGAAGACAGTAGGGT
Chromosome start end strand MM target_seq PAM alignment distance position gene name gene id
GU305914.1 50643 50665 + 0 ACCCTACTGTCTTCGGGCGC CGG ||||||||[||||||||||||]PAM NA Intergenic NA NA

T3283 TCGGGCGCCGGCCTCTTCCTCGG 1000
Oligo fwd CACCTCGGGCGCCGGCCTCTTCCT
Oligo rev AAACAGGAAGAGGCCGGCGCCCGA
Chromosome start end strand MM target_seq PAM alignment distance position gene name gene id
GU305914.1 50655 50677 + 0 TCGGGCGCCGGCCTCTTCCT CGG ||||||||[||||||||||||]PAM NA Intergenic NA NA

T3284 CGGGCGCCGGCCTCTTCCTCGGG 1000
Oligo fwd CACCCGGGCGCCGGCCTCTTCCTC
Oligo rev AAACGAGGAAGAGGCCGGCGCCCG
Chromosome start end strand MM target_seq PAM alignment distance position gene name gene id
GU305914.1 50656 50678 + 0 CGGGCGCCGGCCTCTTCCTC GGG ||||||||[||||||||||||]PAM NA Intergenic NA NA

T3285 CGCCGGCCTCTTCCTCGGGACGG 1000
Oligo fwd CACCCGCCGGCCTCTTCCTCGGGA
Oligo rev AAACTCCCGAGGAAGAGGCCGGCG
Chromosome start end strand MM target_seq PAM alignment distance position gene name gene id
GU305914.1 50660 50682 + 0 CGCCGGCCTCTTCCTCGGGA CGG ||||||||[||||||||||||]PAM NA Intergenic NA NA

T3286 CGGCCTCTTCCTCGGGACGGAGG 1000
Oligo fwd CACCCGGCCTCTTCCTCGGGACGG
Oligo rev AAACCCGTCCCGAGGAAGAGGCCG
Chromosome start end strand MM target_seq PAM alignment distance position gene name gene id
GU305914.1 50663 50685 + 0 CGGCCTCTTCCTCGGGACGG AGG ||||||||[||||||||||||]PAM NA Intergenic NA NA

T3287 TCCTCGGGACGGAGGCCACCTGG 1000
Oligo fwd CACCTCCTCGGGACGGAGGCCACC
Oligo rev AAACGGTGGCCTCCGTCCCGAGGA
Chromosome start end strand MM target_seq PAM alignment distance position gene name gene id
GU305914.1 50671 50693 + 0 TCCTCGGGACGGAGGCCACC TGG ||||||||[||||||||||||]PAM NA Intergenic NA NA

T3288 GGATACAGCCGTTGTACGACAGG 1000
Oligo fwd CACCGGATACAGCCGTTGTACGAC
Oligo rev AAACGTCGTACAACGGCTGTATCC
Chromosome start end strand MM target_seq PAM alignment distance position gene name gene id
GU305914.1 50692 50714 + 0 GGATACAGCCGTTGTACGAC AGG ||||||||[||||||||||||]PAM NA Intergenic NA NA

T3289 GATACAGCCGTTGTACGACAGGG 1000
Oligo fwd CACCGATACAGCCGTTGTACGACA
Oligo rev AAACTGTCGTACAACGGCTGTATC
Chromosome start end strand MM target_seq PAM alignment distance position gene name gene id
GU305914.1 50693 50715 + 0 GATACAGCCGTTGTACGACA GGG ||||||||[||||||||||||]PAM NA Intergenic NA NA

T3290 CCGTTGTACGACAGGGCCACAGG 1000
Oligo fwd CACCCCGTTGTACGACAGGGCCAC
Oligo rev AAACGTGGCCCTGTCGTACAACGG
Chromosome start end strand MM target_seq PAM alignment distance position gene name gene id
GU305914.1 50700 50722 + 0 CCGTTGTACGACAGGGCCAC AGG ||||||||[||||||||||||]PAM NA Intergenic NA NA

T3291 TTGTACGACAGGGCCACAGGAGG 1000
Oligo fwd CACCTTGTACGACAGGGCCACAGG
Oligo rev AAACCCTGTGGCCCTGTCGTACAA
Chromosome start end strand MM target_seq PAM alignment distance position gene name gene id
GU305914.1 50703 50725 + 0 TTGTACGACAGGGCCACAGG AGG ||||||||[||||||||||||]PAM NA Intergenic NA NA

T3292 CACAGGAGGAGAGCCGTTCACGG 1000
Oligo fwd CACCCACAGGAGGAGAGCCGTTCA
Oligo rev AAACTGAACGGCTCTCCTCCTGTG
Chromosome start end strand MM target_seq PAM alignment distance position gene name gene id
GU305914.1 50717 50739 + 0 CACAGGAGGAGAGCCGTTCA CGG ||||||||[||||||||||||]PAM NA Intergenic NA NA

T3293 GAGGAGAGCCGTTCACGGTGAGG 1000
Oligo fwd CACCGAGGAGAGCCGTTCACGGTG
Oligo rev AAACCACCGTGAACGGCTCTCCTC
Chromosome start end strand MM target_seq PAM alignment distance position gene name gene id
GU305914.1 50722 50744 + 0 GAGGAGAGCCGTTCACGGTG AGG ||||||||[||||||||||||]PAM NA Intergenic NA NA

T3294 GCCGTTCACGGTGAGGTCCATGG 1000
Oligo fwd CACCGCCGTTCACGGTGAGGTCCA
Oligo rev AAACTGGACCTCACCGTGAACGGC
Chromosome start end strand MM target_seq PAM alignment distance position gene name gene id
GU305914.1 50729 50751 + 0 GCCGTTCACGGTGAGGTCCA TGG ||||||||[||||||||||||]PAM NA Intergenic NA NA

T3295 CCGTTCACGGTGAGGTCCATGGG 1000
Oligo fwd CACCCCGTTCACGGTGAGGTCCAT
Oligo rev AAACATGGACCTCACCGTGAACGG
Chromosome start end strand MM target_seq PAM alignment distance position gene name gene id
GU305914.1 50730 50752 + 0 CCGTTCACGGTGAGGTCCAT GGG ||||||||[||||||||||||]PAM NA Intergenic NA NA

T3296 GCTGATCTTCCGAAGAGTCCAGG 1000
Oligo fwd CACCGCTGATCTTCCGAAGAGTCC
Oligo rev AAACGGACTCTTCGGAAGATCAGC
Chromosome start end strand MM target_seq PAM alignment distance position gene name gene id
GU305914.1 50761 50783 + 0 GCTGATCTTCCGAAGAGTCC AGG ||||||||[||||||||||||]PAM NA Intergenic NA NA

T3297 CTGATCTTCCGAAGAGTCCAGGG 1000

<table>
<tr><td>Oligo fwd</td><td>CACCCTGATCTTCCGAAGAGTCCA</td><td></td><td></td><td></td><td></td><td></td><td></td><td></td><td></td><td></td><td></td><td></td></tr>
<tr><td>Oligo rev</td><td>AAACTGGACTCTTCGGAAGATCAG</td><td></td><td></td><td></td><td></td><td></td><td></td><td></td><td></td><td></td><td></td><td></td></tr>
<tr><td>Chromosome</td><td>start</td><td></td><td>end</td><td>strand</td><td>MM</td><td>target_seq</td><td>PAM</td><td>alignment</td><td>distance</td><td>position</td><td>gene name</td><td>gene id</td></tr>
<tr><td>GU305914.1</td><td></td><td>50762</td><td>50784</td><td>+</td><td>0</td><td>CTGATCTTCCGAAGAGTCCA</td><td>GGG</td><td>||||||||[||||||||||||]PAM</td><td>NA</td><td>Intergenic</td><td>NA</td><td>NA</td></tr>
<tr><td></td><td></td><td></td><td></td><td></td><td></td><td></td><td></td><td></td><td></td><td></td><td></td><td></td></tr>
<tr><td>T3298</td><td>CTTCCGAAGAGTCCAGGGTGAGG</td><td></td><td>1000</td><td></td><td></td><td></td><td></td><td></td><td></td><td></td><td></td><td></td></tr>
<tr><td>Oligo fwd</td><td>CACCCTTCCGAAGAGTCCAGGGTG</td><td></td><td></td><td></td><td></td><td></td><td></td><td></td><td></td><td></td><td></td><td></td></tr>
<tr><td>Oligo rev</td><td>AAACCACCCTGGACTCTTCGGAAG</td><td></td><td></td><td></td><td></td><td></td><td></td><td></td><td></td><td></td><td></td><td></td></tr>
<tr><td>Chromosome</td><td>start</td><td></td><td>end</td><td>strand</td><td>MM</td><td>target_seq</td><td>PAM</td><td>alignment</td><td>distance</td><td>position</td><td>gene name</td><td>gene id</td></tr>
<tr><td>GU305914.1</td><td></td><td>50767</td><td>50789</td><td>+</td><td>0</td><td>CTTCCGAAGAGTCCAGGGTG</td><td>AGG</td><td>||||||||[||||||||||||]PAM</td><td>NA</td><td>Intergenic</td><td>NA</td><td>NA</td></tr>
<tr><td></td><td></td><td></td><td></td><td></td><td></td><td></td><td></td><td></td><td></td><td></td><td></td><td></td></tr>
<tr><td>T3299</td><td>TCCAGGGTGAGGTCTATGCACGG</td><td></td><td>1000</td><td></td><td></td><td></td><td></td><td></td><td></td><td></td><td></td><td></td></tr>
<tr><td>Oligo fwd</td><td>CACCTCCAGGGTGAGGTCTATGCA</td><td></td><td></td><td></td><td></td><td></td><td></td><td></td><td></td><td></td><td></td><td></td></tr>
<tr><td>Oligo rev</td><td>AAACTGCATAGACCTCACCCTGGA</td><td></td><td></td><td></td><td></td><td></td><td></td><td></td><td></td><td></td><td></td><td></td></tr>
<tr><td>Chromosome</td><td>start</td><td></td><td>end</td><td>strand</td><td>MM</td><td>target_seq</td><td>PAM</td><td>alignment</td><td>distance</td><td>position</td><td>gene name</td><td>gene id</td></tr>
<tr><td>GU305914.1</td><td></td><td>50778</td><td>50800</td><td>+</td><td>0</td><td>TCCAGGGTGAGGTCTATGCA</td><td>CGG</td><td>||||||||[||||||||||||]PAM</td><td>NA</td><td>Intergenic</td><td>NA</td><td>NA</td></tr>
<tr><td></td><td></td><td></td><td></td><td></td><td></td><td></td><td></td><td></td><td></td><td></td><td></td><td></td></tr>
<tr><td>T3300</td><td>AGGGTGAGGTCTATGCACGGCGG</td><td></td><td>1000</td><td></td><td></td><td></td><td></td><td></td><td></td><td></td><td></td><td></td></tr>
<tr><td>Oligo fwd</td><td>CACCAGGGTGAGGTCTATGCACGG</td><td></td><td></td><td></td><td></td><td></td><td></td><td></td><td></td><td></td><td></td><td></td></tr>
<tr><td>Oligo rev</td><td>AAACCCGTGCATAGACCTCACCCT</td><td></td><td></td><td></td><td></td><td></td><td></td><td></td><td></td><td></td><td></td><td></td></tr>
<tr><td>Chromosome</td><td>start</td><td></td><td>end</td><td>strand</td><td>MM</td><td>target_seq</td><td>PAM</td><td>alignment</td><td>distance</td><td>position</td><td>gene name</td><td>gene id</td></tr>
<tr><td>GU305914.1</td><td></td><td>50781</td><td>50803</td><td>+</td><td>0</td><td>AGGGTGAGGTCTATGCACGG</td><td>CGG</td><td>||||||||[||||||||||||]PAM</td><td>NA</td><td>Intergenic</td><td>NA</td><td>NA</td></tr>
<tr><td></td><td></td><td></td><td></td><td></td><td></td><td></td><td></td><td></td><td></td><td></td><td></td><td></td></tr>
<tr><td>T3301</td><td>CTATGCACGGCGGCTCCGTGTGG</td><td></td><td>1000</td><td></td><td></td><td></td><td></td><td></td><td></td><td></td><td></td><td></td></tr>
<tr><td>Oligo fwd</td><td>CACCCTATGCACGGCGGCTCCGTG</td><td></td><td></td><td></td><td></td><td></td><td></td><td></td><td></td><td></td><td></td><td></td></tr>
<tr><td>Oligo rev</td><td>AAACCACGGAGCCGCCGTGCATAG</td><td></td><td></td><td></td><td></td><td></td><td></td><td></td><td></td><td></td><td></td><td></td></tr>
<tr><td>Chromosome</td><td>start</td><td></td><td>end</td><td>strand</td><td>MM</td><td>target_seq</td><td>PAM</td><td>alignment</td><td>distance</td><td>position</td><td>gene name</td><td>gene id</td></tr>
<tr><td>GU305914.1</td><td></td><td>50791</td><td>50813</td><td>+</td><td>0</td><td>CTATGCACGGCGGCTCCGTG</td><td>TGG</td><td>||||||||[||||||||||||]PAM</td><td>NA</td><td>Intergenic</td><td>NA</td><td>NA</td></tr>
<tr><td></td><td></td><td></td><td></td><td></td><td></td><td></td><td></td><td></td><td></td><td></td><td></td><td></td></tr>
<tr><td>T3302</td><td>GGCGGCTCCGTGTGGTCGCGTGG</td><td></td><td>1000</td><td></td><td></td><td></td><td></td><td></td><td></td><td></td><td></td><td></td></tr>
<tr><td>Oligo fwd</td><td>CACCGGCGGCTCCGTGTGGTCGCG</td><td></td><td></td><td></td><td></td><td></td><td></td><td></td><td></td><td></td><td></td><td></td></tr>
<tr><td>Oligo rev</td><td>AAACCGCGACCACACGGAGCCGCC</td><td></td><td></td><td></td><td></td><td></td><td></td><td></td><td></td><td></td><td></td><td></td></tr>
<tr><td>Chromosome</td><td>start</td><td></td><td>end</td><td>strand</td><td>MM</td><td>target_seq</td><td>PAM</td><td>alignment</td><td>distance</td><td>position</td><td>gene name</td><td>gene id</td></tr>
<tr><td>GU305914.1</td><td></td><td>50799</td><td>50821</td><td>+</td><td>0</td><td>GGCGGCTCCGTGTGGTCGCG</td><td>TGG</td><td>||||||||[||||||||||||]PAM</td><td>NA</td><td>Intergenic</td><td>NA</td><td>NA</td></tr>
<tr><td></td><td></td><td></td><td></td><td></td><td></td><td></td><td></td><td></td><td></td><td></td><td></td><td></td></tr>
<tr><td>T3303</td><td>GGCTCCGTGTGGTCGCGTGGAGG</td><td></td><td>1000</td><td></td><td></td><td></td><td></td><td></td><td></td><td></td><td></td><td></td></tr>
<tr><td>Oligo fwd</td><td>CACCGGCTCCGTGTGGTCGCGTGG</td><td></td><td></td><td></td><td></td><td></td><td></td><td></td><td></td><td></td><td></td><td></td></tr>
<tr><td>Oligo rev</td><td>AAACCCACGCGACCACACGGAGCC</td><td></td><td></td><td></td><td></td><td></td><td></td><td></td><td></td><td></td><td></td><td></td></tr>
<tr><td>Chromosome</td><td>start</td><td></td><td>end</td><td>strand</td><td>MM</td><td>target_seq</td><td>PAM</td><td>alignment</td><td>distance</td><td>position</td><td>gene name</td><td>gene id</td></tr>
<tr><td>GU305914.1</td><td></td><td>50802</td><td>50824</td><td>+</td><td>0</td><td>GGCTCCGTGTGGTCGCGTGG</td><td>AGG</td><td>||||||||[||||||||||||]PAM</td><td>NA</td><td>Intergenic</td><td>NA</td><td>NA</td></tr>
<tr><td></td><td></td><td></td><td></td><td></td><td></td><td></td><td></td><td></td><td></td><td></td><td></td><td></td></tr>
<tr><td>T3304</td><td>GTGGTCGCGTGGAGGAGACACGG</td><td></td><td>1000</td><td></td><td></td><td></td><td></td><td></td><td></td><td></td><td></td><td></td></tr>
<tr><td>Oligo fwd</td><td>CACCGTGGTCGCGTGGAGGAGACA</td><td></td><td></td><td></td><td></td><td></td><td></td><td></td><td></td><td></td><td></td><td></td></tr>
<tr><td>Oligo rev</td><td>AAACTGTCTCCTCCACGCGACCAC</td><td></td><td></td><td></td><td></td><td></td><td></td><td></td><td></td><td></td><td></td><td></td></tr>
<tr><td>Chromosome</td><td>start</td><td></td><td>end</td><td>strand</td><td>MM</td><td>target_seq</td><td>PAM</td><td>alignment</td><td>distance</td><td>position</td><td>gene name</td><td>gene id</td></tr>
<tr><td>GU305914.1</td><td></td><td>50810</td><td>50832</td><td>+</td><td>0</td><td>GTGGTCGCGTGGAGGAGACA</td><td>CGG</td><td>||||||||[||||||||||||]PAM</td><td>NA</td><td>Intergenic</td><td>NA</td><td>NA</td></tr>
<tr><td></td><td></td><td></td><td></td><td></td><td></td><td></td><td></td><td></td><td></td><td></td><td></td><td></td></tr>
<tr><td>T3305</td><td>GGCGAATCGCACCGAGTTGAAGG</td><td></td><td>1000</td><td></td><td></td><td></td><td></td><td></td><td></td><td></td><td></td><td></td></tr>
<tr><td>Oligo fwd</td><td>CACCGGCGAATCGCACCGAGTTGA</td><td></td><td></td><td></td><td></td><td></td><td></td><td></td><td></td><td></td><td></td><td></td></tr>
<tr><td>Oligo rev</td><td>AAACTCAACTCGGTGCGATTCGCC</td><td></td><td></td><td></td><td></td><td></td><td></td><td></td><td></td><td></td><td></td><td></td></tr>
<tr><td>Chromosome</td><td>start</td><td></td><td>end</td><td>strand</td><td>MM</td><td>target_seq</td><td>PAM</td><td>alignment</td><td>distance</td><td>position</td><td>gene name</td><td>gene id</td></tr>
<tr><td>GU305914.1</td><td></td><td>50831</td><td>50853</td><td>+</td><td>0</td><td>GGCGAATCGCACCGAGTTGA</td><td>AGG</td><td>||||||||[||||||||||||]PAM</td><td>NA</td><td>Intergenic</td><td>NA</td><td>NA</td></tr>
<tr><td></td><td></td><td></td><td></td><td></td><td></td><td></td><td></td><td></td><td></td><td></td><td></td><td></td></tr>
<tr><td>T3306</td><td>GTTTTCATCTTCCTCGTCCACGG</td><td></td><td>1000</td><td></td><td></td><td></td><td></td><td></td><td></td><td></td><td></td><td></td></tr>
<tr><td>Oligo fwd</td><td>CACCGTTTTCATCTTCCTCGTCCA</td><td></td><td></td><td></td><td></td><td></td><td></td><td></td><td></td><td></td><td></td><td></td></tr>
<tr><td>Oligo rev</td><td>AAACTGGACGAGGAAGATGAAAAC</td><td></td><td></td><td></td><td></td><td></td><td></td><td></td><td></td><td></td><td></td><td></td></tr>
<tr><td>Chromosome</td><td>start</td><td></td><td>end</td><td>strand</td><td>MM</td><td>target_seq</td><td>PAM</td><td>alignment</td><td>distance</td><td>position</td><td>gene name</td><td>gene id</td></tr>
<tr><td>GU305914.1</td><td></td><td>50864</td><td>50886</td><td>+</td><td>0</td><td>GTTTTCATCTTCCTCGTCCA</td><td>CGG</td><td>||||||||[||||||||||||]PAM</td><td>NA</td><td>Intergenic</td><td>NA</td><td>NA</td></tr>
<tr><td></td><td></td><td></td><td></td><td></td><td></td><td></td><td></td><td></td><td></td><td></td><td></td><td></td></tr>
<tr><td>T3307</td><td>TTTTCATCTTCCTCGTCCACGGG</td><td></td><td>1000</td><td></td><td></td><td></td><td></td><td></td><td></td><td></td><td></td><td></td></tr>
<tr><td>Oligo fwd</td><td>CACCTTTTCATCTTCCTCGTCCAC</td><td></td><td></td><td></td><td></td><td></td><td></td><td></td><td></td><td></td><td></td><td></td></tr>
<tr><td>Oligo rev</td><td>AAACGTGGACGAGGAAGATGAAAA</td><td></td><td></td><td></td><td></td><td></td><td></td><td></td><td></td><td></td><td></td><td></td></tr>
<tr><td>Chromosome</td><td>start</td><td></td><td>end</td><td>strand</td><td>MM</td><td>target_seq</td><td>PAM</td><td>alignment</td><td>distance</td><td>position</td><td>gene name</td><td>gene id</td></tr>
<tr><td>GU305914.1</td><td></td><td>50865</td><td>50887</td><td>+</td><td>0</td><td>TTTTCATCTTCCTCGTCCAC</td><td>GGG</td><td>||||||||[||||||||||||]PAM</td><td>NA</td><td>Intergenic</td><td>NA</td><td>NA</td></tr>
<tr><td></td><td></td><td></td><td></td><td></td><td></td><td></td><td></td><td></td><td></td><td></td><td></td><td></td></tr>
<tr><td>T3308</td><td>CCCGAAGAGAACGTGAACGCCGG</td><td></td><td>1000</td><td></td><td></td><td></td><td></td><td></td><td></td><td></td><td></td><td></td></tr>
<tr><td>Oligo fwd</td><td>CACCCCCGAAGAGAACGTGAACGC</td><td></td><td></td><td></td><td></td><td></td><td></td><td></td><td></td><td></td><td></td><td></td></tr>
<tr><td>Oligo rev</td><td>AAACGCGTTCACGTTCTCTTCGGG</td><td></td><td></td><td></td><td></td><td></td><td></td><td></td><td></td><td></td><td></td><td></td></tr>
<tr><td>Chromosome</td><td>start</td><td></td><td>end</td><td>strand</td><td>MM</td><td>target_seq</td><td>PAM</td><td>alignment</td><td>distance</td><td>position</td><td>gene name</td><td>gene id</td></tr>
<tr><td>GU305914.1</td><td></td><td>50888</td><td>50910</td><td>+</td><td>0</td><td>CCCGAAGAGAACGTGAACGC</td><td>CGG</td><td>||||||||[||||||||||||]PAM</td><td>NA</td><td>Intergenic</td><td>NA</td><td>NA</td></tr>
<tr><td></td><td></td><td></td><td></td><td></td><td></td><td></td><td></td><td></td><td></td><td></td><td></td><td></td></tr>
<tr><td>T3309</td><td>CCGAAGAGAACGTGAACGCCGGG</td><td></td><td>1000</td><td></td><td></td><td></td><td></td><td></td><td></td><td></td><td></td><td></td></tr>
<tr><td>Oligo fwd</td><td>CACCCCGAAGAGAACGTGAACGCC</td><td></td><td></td><td></td><td></td><td></td><td></td><td></td><td></td><td></td><td></td><td></td></tr>
<tr><td>Oligo rev</td><td>AAACGGCGTTCACGTTCTCTTCGG</td><td></td><td></td><td></td><td></td><td></td><td></td><td></td><td></td><td></td><td></td><td></td></tr>
<tr><td>Chromosome</td><td>start</td><td></td><td>end</td><td>strand</td><td>MM</td><td>target_seq</td><td>PAM</td><td>alignment</td><td>distance</td><td>position</td><td>gene name</td><td>gene id</td></tr>
<tr><td>GU305914.1</td><td></td><td>50889</td><td>50911</td><td>+</td><td>0</td><td>CCGAAGAGAACGTGAACGCC</td><td>GGG</td><td>||||||||[||||||||||||]PAM</td><td>NA</td><td>Intergenic</td><td>NA</td><td>NA</td></tr>
<tr><td></td><td></td><td></td><td></td><td></td><td></td><td></td><td></td><td></td><td></td><td></td><td></td><td></td></tr>
<tr><td>T3310</td><td>GAGAACGTGAACGCCGGGCGCGG</td><td></td><td>1000</td><td></td><td></td><td></td><td></td><td></td><td></td><td></td><td></td><td></td></tr>
<tr><td>Oligo fwd</td><td>CACCGAGAACGTGAACGCCGGGCG</td><td></td><td></td><td></td><td></td><td></td><td></td><td></td><td></td><td></td><td></td><td></td></tr>
<tr><td>Oligo rev</td><td>AAACCGCCCGGCGTTCACGTTCTC</td><td></td><td></td><td></td><td></td><td></td><td></td><td></td><td></td><td></td><td></td><td></td></tr>
<tr><td>Chromosome</td><td>start</td><td></td><td>end</td><td>strand</td><td>MM</td><td>target_seq</td><td>PAM</td><td>alignment</td><td>distance</td><td>position</td><td>gene name</td><td>gene id</td></tr>
<tr><td>GU305914.1</td><td></td><td>50894</td><td>50916</td><td>+</td><td>0</td><td>GAGAACGTGAACGCCGGGCG</td><td>CGG</td><td>||||||||[||||||||||||]PAM</td><td>NA</td><td>Intergenic</td><td>NA</td><td>NA</td></tr>
<tr><td></td><td></td><td></td><td></td><td></td><td></td><td></td><td></td><td></td><td></td><td></td><td></td><td></td></tr>
<tr><td>T3311</td><td>AGAACGTGAACGCCGGGCGCGGG</td><td></td><td>1000</td><td></td><td></td><td></td><td></td><td></td><td></td><td></td><td></td><td></td></tr>
<tr><td>Oligo fwd</td><td>CACCAGAACGTGAACGCCGGGCGC</td><td></td><td></td><td></td><td></td><td></td><td></td><td></td><td></td><td></td><td></td><td></td></tr>
<tr><td>Oligo rev</td><td>AAACGCGCCCGGCGTTCACGTTCT</td><td></td><td></td><td></td><td></td><td></td><td></td><td></td><td></td><td></td><td></td><td></td></tr>
<tr><td>Chromosome</td><td>start</td><td></td><td>end</td><td>strand</td><td>MM</td><td>target_seq</td><td>PAM</td><td>alignment</td><td>distance</td><td>position</td><td>gene name</td><td>gene id</td></tr>
<tr><td>GU305914.1</td><td></td><td>50895</td><td>50917</td><td>+</td><td>0</td><td>AGAACGTGAACGCCGGGCGC</td><td>GGG</td><td>||||||||[||||||||||||]PAM</td><td>NA</td><td>Intergenic</td><td>NA</td><td>NA</td></tr>
</table>

T3312 GAACGTGAACGCCGGGCGCGGGG 1000
Oligo fwd CACCGAACGTGAACGCCGGGCGCG
Oligo rev AAACCGCGCCCGGCGTTCACGTTC
Chromosome start end strand MM target_seq PAM alignment distance position gene name gene id
GU305914.1 50896 50918 + 0 GAACGTGAACGCCGGGCGCG GGG ||||||||[||||||||||||]PAM NA Intergenic NA NA

T3313 GCCGGGCGCGGGGCCTCTGACGG 1000
Oligo fwd CACCGCCGGGCGCGGGGCCTCTGA
Oligo rev AAACTCAGAGGCCCCGCGCCCGGC
Chromosome start end strand MM target_seq PAM alignment distance position gene name gene id
GU305914.1 50906 50928 + 0 GCCGGGCGCGGGGCCTCTGA CGG ||||||||[||||||||||||]PAM NA Intergenic NA NA

T3314 GCCTCTGACGGAGAGATCGACGG 1000
Oligo fwd CACCGCCTCTGACGGAGAGATCGA
Oligo rev AAACTCGATCTCTCCGTCAGAGGC
Chromosome start end strand MM target_seq PAM alignment distance position gene name gene id
GU305914.1 50918 50940 + 0 GCCTCTGACGGAGAGATCGA CGG ||||||||[||||||||||||]PAM NA Intergenic NA NA

T3315 CCTCTGACGGAGAGATCGACGGG 1000
Oligo fwd CACCCCTCTGACGGAGAGATCGAC
Oligo rev AAACGTCGATCTCTCCGTCAGAGG
Chromosome start end strand MM target_seq PAM alignment distance position gene name gene id
GU305914.1 50919 50941 + 0 CCTCTGACGGAGAGATCGAC GGG ||||||||[||||||||||||]PAM NA Intergenic NA NA

T3316 GACGGAGAGATCGACGGGCACGG 1000
Oligo fwd CACCGACGGAGAGATCGACGGGCA
Oligo rev AAACTGCCCGTCGATCTCTCCGTC
Chromosome start end strand MM target_seq PAM alignment distance position gene name gene id
GU305914.1 50924 50946 + 0 GACGGAGAGATCGACGGGCA CGG ||||||||[||||||||||||]PAM NA Intergenic NA NA

T3317 ACGGAGAGATCGACGGGCACGGG 1000
Oligo fwd CACCACGGAGAGATCGACGGGCAC
Oligo rev AAACGTGCCCGTCGATCTCTCCGT
Chromosome start end strand MM target_seq PAM alignment distance position gene name gene id
GU305914.1 50925 50947 + 0 ACGGAGAGATCGACGGGCAC GGG ||||||||[||||||||||||]PAM NA Intergenic NA NA

T3318 GAGATCGACGGGCACGGGCGAGG 1000
Oligo fwd CACCGAGATCGACGGGCACGGGCG
Oligo rev AAACCGCCCGTGCCCGTCGATCTC
Chromosome start end strand MM target_seq PAM alignment distance position gene name gene id
GU305914.1 50930 50952 + 0 GAGATCGACGGGCACGGGCG AGG ||||||||[||||||||||||]PAM NA Intergenic NA NA

T3319 TCGACGGGCACGGGCGAGGACGG 1000
Oligo fwd CACCTCGACGGGCACGGGCGAGGA
Oligo rev AAACTCCTCGCCCGTGCCCGTCGA
Chromosome start end strand MM target_seq PAM alignment distance position gene name gene id
GU305914.1 50934 50956 + 0 TCGACGGGCACGGGCGAGGA CGG ||||||||[||||||||||||]PAM NA Intergenic NA NA

T3320 GACGGTACTCCACCGCCACCAGG 1000
Oligo fwd CACCGACGGTACTCCACCGCCACC
Oligo rev AAACGGTGGCGGTGGAGTACCGTC
Chromosome start end strand MM target_seq PAM alignment distance position gene name gene id
GU305914.1 50952 50974 + 0 GACGGTACTCCACCGCCACC AGG ||||||||[||||||||||||]PAM NA Intergenic NA NA

T3321 TACTCCACCGCCACCAGGCACGG 1000
Oligo fwd CACCTACTCCACCGCCACCAGGCA
Oligo rev AAACTGCCTGGTGGCGGTGGAGTA
Chromosome start end strand MM target_seq PAM alignment distance position gene name gene id
GU305914.1 50957 50979 + 0 TACTCCACCGCCACCAGGCA CGG ||||||||[||||||||||||]PAM NA Intergenic NA NA

T3322 ACTCCACCGCCACCAGGCACGGG 1000
Oligo fwd CACCACTCCACCGCCACCAGGCAC
Oligo rev AAACGTGCCTGGTGGCGGTGGAGT
Chromosome start end strand MM target_seq PAM alignment distance position gene name gene id
GU305914.1 50958 50980 + 0 ACTCCACCGCCACCAGGCAC GGG ||||||||[||||||||||||]PAM NA Intergenic NA NA

T3323 CTCCACCGCCACCAGGCACGGGG 1000
Oligo fwd CACCCTCCACCGCCACCAGGCACG
Oligo rev AAACCGTGCCTGGTGGCGGTGGAG
Chromosome start end strand MM target_seq PAM alignment distance position gene name gene id
GU305914.1 50959 50981 + 0 CTCCACCGCCACCAGGCACG GGG ||||||||[||||||||||||]PAM NA Intergenic NA NA

T3324 GCCACCAGGCACGGGGTTCACGG 1000
Oligo fwd CACCGCCACCAGGCACGGGGTTCA
Oligo rev AAACTGAACCCCGTGCCTGGTGGC
Chromosome start end strand MM target_seq PAM alignment distance position gene name gene id
GU305914.1 50966 50988 + 0 GCCACCAGGCACGGGGTTCA CGG ||||||||[||||||||||||]PAM NA Intergenic NA NA

T3325 CCACCAGGCACGGGGTTCACGGG 1000
Oligo fwd CACCCCACCAGGCACGGGGTTCAC
Oligo rev AAACGTGAACCCCGTGCCTGGTGG
Chromosome start end strand MM target_seq PAM alignment distance position gene name gene id
GU305914.1 50967 50989 + 0 CCACCAGGCACGGGGTTCAC GGG ||||||||[||||||||||||]PAM NA Intergenic NA NA

T3326 GTTCACGGGCACGCGAAACATGG 1000
Oligo fwd CACCGTTCACGGGCACGCGAAACA
Oligo rev AAACTGTTTCGCGTGCCCGTGAAC
Chromosome start end strand MM target_seq PAM alignment distance position gene name gene id
GU305914.1 50981 51003 + 0 GTTCACGGGCACGCGAAACA TGG ||||||||[||||||||||||]PAM NA Intergenic NA NA

T3327 TTCACGGGCACGCGAAACATGGG 1000
Oligo fwd CACCTTCACGGGCACGCGAAACAT
Oligo rev AAACATGTTTCGCGTGCCCGTGAA
Chromosome start end strand MM target_seq PAM alignment distance position gene name gene id
GU305914.1 50982 51004 + 0 TTCACGGGCACGCGAAACAT GGG ||||||||[||||||||||||]PAM NA Intergenic NA NA

T3328 ACGGGCACGCGAAACATGGGCGG 1000
Oligo fwd CACCACGGGCACGCGAAACATGGG
Oligo rev AAACCCCATGTTTCGCGTGCCCGT
Chromosome start end strand MM target_seq PAM alignment distance position gene name gene id
GU305914.1 50985 51007 + 0 ACGGGCACGCGAAACATGGG CGG ||||||||[||||||||||||]PAM NA Intergenic NA NA

T3329 CACGCGAAACATGGGCGGCGAGG 1000
Oligo fwd CACCCACGCGAAACATGGGCGGCG
Oligo rev AAACCGCCGCCCATGTTTCGCGTG
Chromosome start end strand MM target_seq PAM alignment distance position gene name gene id
GU305914.1 50990 51012 + 0 CACGCGAAACATGGGCGGCG AGG ||||||||[||||||||||||]PAM NA Intergenic NA NA

T3330 ACATGGGCGGCGAGGAGATGAGG 1000
Oligo fwd CACCACATGGGCGGCGAGGAGATG
Oligo rev AAACCATCTCCTCGCCGCCCATGT
Chromosome start end strand MM target_seq PAM alignment distance position gene name gene id
GU305914.1 50998 51020 + 0 ACATGGGCGGCGAGGAGATG AGG ||||||||[||||||||||||]PAM NA Intergenic NA NA

T3331 CATGGGCGGCGAGGAGATGAGGG 1000
Oligo fwd CACCCATGGGCGGCGAGGAGATGA
Oligo rev AAACTCATCTCCTCGCCGCCCATG
Chromosome start end strand MM target_seq PAM alignment distance position gene name gene id
GU305914.1 50999 51021 + 0 CATGGGCGGCGAGGAGATGA GGG ||||||||[||||||||||||]PAM NA Intergenic NA NA

T3332 CGGCGAGGAGATGAGGGTCACGG 1000
Oligo fwd CACCCGGCGAGGAGATGAGGGTCA
Oligo rev AAACTGACCCTCATCTCCTCGCCG
Chromosome start end strand MM target_seq PAM alignment distance position gene name gene id
GU305914.1 51005 51027 + 0 CGGCGAGGAGATGAGGGTCA CGG ||||||||[||||||||||||]PAM NA Intergenic NA NA
GU305914.1 69052 69074 + 4 GGACGAGGAGATGCGGATCA AGG -|-|||||[|||||-||-|||]PAM NA Intergenic NA NA

T3333 GGAGATGAGGGTCACGGTCATGG 1000
Oligo fwd CACCGGAGATGAGGGTCACGGTCA
Oligo rev AAACTGACCGTGACCCTCATCTCC
Chromosome start end strand MM target_seq PAM alignment distance position gene name gene id
GU305914.1 51011 51033 + 0 GGAGATGAGGGTCACGGTCA TGG ||||||||[||||||||||||]PAM NA Intergenic NA NA

T3334 GAGATGAGGGTCACGGTCATGGG 1000
Oligo fwd CACCGAGATGAGGGTCACGGTCAT
Oligo rev AAACATGACCGTGACCCTCATCTC
Chromosome start end strand MM target_seq PAM alignment distance position gene name gene id
GU305914.1 51012 51034 + 0 GAGATGAGGGTCACGGTCAT GGG ||||||||[||||||||||||]PAM NA Intergenic NA NA

T3335 AGGGTCACGGTCATGGGCATCGG 1000
Oligo fwd CACCAGGGTCACGGTCATGGGCAT
Oligo rev AAACATGCCCATGACCGTGACCCT
Chromosome start end strand MM target_seq PAM alignment distance position gene name gene id
GU305914.1 51018 51040 + 0 AGGGTCACGGTCATGGGCAT CGG ||||||||[||||||||||||]PAM NA Intergenic NA NA

T3336 GTCACGGTCATGGGCATCGGCGG 1000
Oligo fwd CACCGTCACGGTCATGGGCATCGG
Oligo rev AAACCCGATGCCCATGACCGTGAC
Chromosome start end strand MM target_seq PAM alignment distance position gene name gene id
GU305914.1 51021 51043 + 0 GTCACGGTCATGGGCATCGG CGG ||||||||[||||||||||||]PAM NA Intergenic NA NA

T3337 CTCGATAGCCCTCCGAGCGATGG 1000
Oligo fwd CACCCTCGATAGCCCTCCGAGCGA
Oligo rev AAACTCGCTCGGAGGGCTATCGAG
Chromosome start end strand MM target_seq PAM alignment distance position gene name gene id
GU305914.1 51047 51069 + 0 CTCGATAGCCCTCCGAGCGA TGG ||||||||[||||||||||||]PAM NA Intergenic NA NA

T3338 TCGATAGCCCTCCGAGCGATGGG 1000
Oligo fwd CACCTCGATAGCCCTCCGAGCGAT
Oligo rev AAACATCGCTCGGAGGGCTATCGA
Chromosome start end strand MM target_seq PAM alignment distance position gene name gene id
GU305914.1 51048 51070 + 0 TCGATAGCCCTCCGAGCGAT GGG ||||||||[||||||||||||]PAM NA Intergenic NA NA

T3339 ATAGCCCTCCGAGCGATGGGAGG 1000
Oligo fwd CACCATAGCCCTCCGAGCGATGGG
Oligo rev AAACCCCATCGCTCGGAGGGCTAT
Chromosome start end strand MM target_seq PAM alignment distance position gene name gene id
GU305914.1 51051 51073 + 0 ATAGCCCTCCGAGCGATGGG AGG ||||||||[||||||||||||]PAM NA Intergenic NA NA

T3340 AGCGATGGGAGGAGACGCCGCGG 1000
Oligo fwd CACCAGCGATGGGAGGAGACGCCG
Oligo rev AAACCGGCGTCTCCTCCCATCGCT
Chromosome start end strand MM target_seq PAM alignment distance position gene name gene id
GU305914.1 51062 51084 + 0 AGCGATGGGAGGAGACGCCG CGG ||||||||[||||||||||||]PAM NA Intergenic NA NA

T3341 GCCGCGGCGAGATCGATGCACGG 1000
Oligo fwd CACCGCCGCGGCGAGATCGATGCA
Oligo rev AAACTGCATCGATCTCGCCGCGGC

<table>
<tr><td>Chromosome</td><td>start</td><td></td><td>end</td><td>strand</td><td>MM</td><td>target_seq</td><td>PAM</td><td>alignment</td><td>distance</td><td>position</td><td>gene name</td><td>gene id</td></tr>
<tr><td>GU305914.1</td><td></td><td>51078</td><td>51100</td><td>+</td><td>0</td><td>GCCGCGGCGAGATCGATGCA</td><td>CGG</td><td>||||||||[||||||||||||]PAM</td><td>NA</td><td>Intergenic</td><td>NA</td><td>NA</td></tr>
<tr><td>T3342</td><td>CCGCGGCGAGATCGATGCACGGG</td><td></td><td>1000</td><td></td><td></td><td></td><td></td><td></td><td></td><td></td><td></td><td></td></tr>
<tr><td>Oligo fwd</td><td>CACCCCGCGGCGAGATCGATGCAC</td><td></td><td></td><td></td><td></td><td></td><td></td><td></td><td></td><td></td><td></td><td></td></tr>
<tr><td>Oligo rev</td><td>AAACGTGCATCGATCTCGCCGCGG</td><td></td><td></td><td></td><td></td><td></td><td></td><td></td><td></td><td></td><td></td><td></td></tr>
<tr><td>Chromosome</td><td>start</td><td></td><td>end</td><td>strand</td><td>MM</td><td>target_seq</td><td>PAM</td><td>alignment</td><td>distance</td><td>position</td><td>gene name</td><td>gene id</td></tr>
<tr><td>GU305914.1</td><td></td><td>51079</td><td>51101</td><td>+</td><td>0</td><td>CCGCGGCGAGATCGATGCAC</td><td>GGG</td><td>||||||||[||||||||||||]PAM</td><td>NA</td><td>Intergenic</td><td>NA</td><td>NA</td></tr>
<tr><td>T3343</td><td>CGCGGCGAGATCGATGCACGGGG</td><td></td><td>1000</td><td></td><td></td><td></td><td></td><td></td><td></td><td></td><td></td><td></td></tr>
<tr><td>Oligo fwd</td><td>CACCCGCGGCGAGATCGATGCACG</td><td></td><td></td><td></td><td></td><td></td><td></td><td></td><td></td><td></td><td></td><td></td></tr>
<tr><td>Oligo rev</td><td>AAACCGTGCATCGATCTCGCCGCG</td><td></td><td></td><td></td><td></td><td></td><td></td><td></td><td></td><td></td><td></td><td></td></tr>
<tr><td>Chromosome</td><td>start</td><td></td><td>end</td><td>strand</td><td>MM</td><td>target_seq</td><td>PAM</td><td>alignment</td><td>distance</td><td>position</td><td>gene name</td><td>gene id</td></tr>
<tr><td>GU305914.1</td><td></td><td>51080</td><td>51102</td><td>+</td><td>0</td><td>CGCGGCGAGATCGATGCACG</td><td>GGG</td><td>||||||||[||||||||||||]PAM</td><td>NA</td><td>Intergenic</td><td>NA</td><td>NA</td></tr>
<tr><td>T3344</td><td>GATCGATGCACGGGGTGCTCTGG</td><td></td><td>1000</td><td></td><td></td><td></td><td></td><td></td><td></td><td></td><td></td><td></td></tr>
<tr><td>Oligo fwd</td><td>CACCGATCGATGCACGGGGTGCTC</td><td></td><td></td><td></td><td></td><td></td><td></td><td></td><td></td><td></td><td></td><td></td></tr>
<tr><td>Oligo rev</td><td>AAACGAGCACCCCGTGCATCGATC</td><td></td><td></td><td></td><td></td><td></td><td></td><td></td><td></td><td></td><td></td><td></td></tr>
<tr><td>Chromosome</td><td>start</td><td></td><td>end</td><td>strand</td><td>MM</td><td>target_seq</td><td>PAM</td><td>alignment</td><td>distance</td><td>position</td><td>gene name</td><td>gene id</td></tr>
<tr><td>GU305914.1</td><td></td><td>51088</td><td>51110</td><td>+</td><td>0</td><td>GATCGATGCACGGGGTGCTC</td><td>TGG</td><td>||||||||[||||||||||||]PAM</td><td>NA</td><td>Intergenic</td><td>NA</td><td>NA</td></tr>
<tr><td>T3345</td><td>GATGCACGGGGTGCTCTGGATGG</td><td></td><td>1000</td><td></td><td></td><td></td><td></td><td></td><td></td><td></td><td></td><td></td></tr>
<tr><td>Oligo fwd</td><td>CACCGATGCACGGGGTGCTCTGGA</td><td></td><td></td><td></td><td></td><td></td><td></td><td></td><td></td><td></td><td></td><td></td></tr>
<tr><td>Oligo rev</td><td>AAACTCCAGAGCACCCCGTGCATC</td><td></td><td></td><td></td><td></td><td></td><td></td><td></td><td></td><td></td><td></td><td></td></tr>
<tr><td>Chromosome</td><td>start</td><td></td><td>end</td><td>strand</td><td>MM</td><td>target_seq</td><td>PAM</td><td>alignment</td><td>distance</td><td>position</td><td>gene name</td><td>gene id</td></tr>
<tr><td>GU305914.1</td><td></td><td>51092</td><td>51114</td><td>+</td><td>0</td><td>GATGCACGGGGTGCTCTGGA</td><td>TGG</td><td>||||||||[||||||||||||]PAM</td><td>NA</td><td>Intergenic</td><td>NA</td><td>NA</td></tr>
<tr><td>T3346</td><td>TGGATGGACTGCGAACCCCGAGG</td><td></td><td>1000</td><td></td><td></td><td></td><td></td><td></td><td></td><td></td><td></td><td></td></tr>
<tr><td>Oligo fwd</td><td>CACCTGGATGGACTGCGAACCCCG</td><td></td><td></td><td></td><td></td><td></td><td></td><td></td><td></td><td></td><td></td><td></td></tr>
<tr><td>Oligo rev</td><td>AAACCGGGGTTCGCAGTCCATCCA</td><td></td><td></td><td></td><td></td><td></td><td></td><td></td><td></td><td></td><td></td><td></td></tr>
<tr><td>Chromosome</td><td>start</td><td></td><td>end</td><td>strand</td><td>MM</td><td>target_seq</td><td>PAM</td><td>alignment</td><td>distance</td><td>position</td><td>gene name</td><td>gene id</td></tr>
<tr><td>GU305914.1</td><td></td><td>51108</td><td>51130</td><td>+</td><td>0</td><td>TGGATGGACTGCGAACCCCG</td><td>AGG</td><td>||||||||[||||||||||||]PAM</td><td>NA</td><td>Intergenic</td><td>NA</td><td>NA</td></tr>
<tr><td>T3347</td><td>AACCCCGAGGAGTAGATTCGAGG</td><td></td><td>1000</td><td></td><td></td><td></td><td></td><td></td><td></td><td></td><td></td><td></td></tr>
<tr><td>Oligo fwd</td><td>CACCAACCCCGAGGAGTAGATTCG</td><td></td><td></td><td></td><td></td><td></td><td></td><td></td><td></td><td></td><td></td><td></td></tr>
<tr><td>Oligo rev</td><td>AAACCGAATCTACTCCTCGGGGTT</td><td></td><td></td><td></td><td></td><td></td><td></td><td></td><td></td><td></td><td></td><td></td></tr>
<tr><td>Chromosome</td><td>start</td><td></td><td>end</td><td>strand</td><td>MM</td><td>target_seq</td><td>PAM</td><td>alignment</td><td>distance</td><td>position</td><td>gene name</td><td>gene id</td></tr>
<tr><td>GU305914.1</td><td></td><td>51121</td><td>51143</td><td>+</td><td>0</td><td>AACCCCGAGGAGTAGATTCG</td><td>AGG</td><td>||||||||[||||||||||||]PAM</td><td>NA</td><td>Intergenic</td><td>NA</td><td>NA</td></tr>
<tr><td>T3348</td><td>AGGAGTAGATTCGAGGATGATGG</td><td></td><td>1000</td><td></td><td></td><td></td><td></td><td></td><td></td><td></td><td></td><td></td></tr>
<tr><td>Oligo fwd</td><td>CACCAGGAGTAGATTCGAGGATGA</td><td></td><td></td><td></td><td></td><td></td><td></td><td></td><td></td><td></td><td></td><td></td></tr>
<tr><td>Oligo rev</td><td>AAACTCATCCTCGAATCTACTCCT</td><td></td><td></td><td></td><td></td><td></td><td></td><td></td><td></td><td></td><td></td><td></td></tr>
<tr><td>Chromosome</td><td>start</td><td></td><td>end</td><td>strand</td><td>MM</td><td>target_seq</td><td>PAM</td><td>alignment</td><td>distance</td><td>position</td><td>gene name</td><td>gene id</td></tr>
<tr><td>GU305914.1</td><td></td><td>51128</td><td>51150</td><td>+</td><td>0</td><td>AGGAGTAGATTCGAGGATGA</td><td>TGG</td><td>||||||||[||||||||||||]PAM</td><td>NA</td><td>Intergenic</td><td>NA</td><td>NA</td></tr>
<tr><td>T3349</td><td>CCGTTCTGCGCCGATTGCTGCGG</td><td></td><td>1000</td><td></td><td></td><td></td><td></td><td></td><td></td><td></td><td></td><td></td></tr>
<tr><td>Oligo fwd</td><td>CACCCCGTTCTGCGCCGATTGCTG</td><td></td><td></td><td></td><td></td><td></td><td></td><td></td><td></td><td></td><td></td><td></td></tr>
<tr><td>Oligo rev</td><td>AAACCAGCAATCGGCGCAGAACGG</td><td></td><td></td><td></td><td></td><td></td><td></td><td></td><td></td><td></td><td></td><td></td></tr>
<tr><td>Chromosome</td><td>start</td><td></td><td>end</td><td>strand</td><td>MM</td><td>target_seq</td><td>PAM</td><td>alignment</td><td>distance</td><td>position</td><td>gene name</td><td>gene id</td></tr>
<tr><td>GU305914.1</td><td></td><td>51171</td><td>51193</td><td>+</td><td>0</td><td>CCGTTCTGCGCCGATTGCTG</td><td>CGG</td><td>||||||||[||||||||||||]PAM</td><td>NA</td><td>Intergenic</td><td>NA</td><td>NA</td></tr>
<tr><td>T3350</td><td>CCGATTGCTGCGGCACGTAGCGG</td><td></td><td>1000</td><td></td><td></td><td></td><td></td><td></td><td></td><td></td><td></td><td></td></tr>
<tr><td>Oligo fwd</td><td>CACCCCGATTGCTGCGGCACGTAG</td><td></td><td></td><td></td><td></td><td></td><td></td><td></td><td></td><td></td><td></td><td></td></tr>
<tr><td>Oligo rev</td><td>AAACCTACGTGCCGCAGCAATCGG</td><td></td><td></td><td></td><td></td><td></td><td></td><td></td><td></td><td></td><td></td><td></td></tr>
<tr><td>Chromosome</td><td>start</td><td></td><td>end</td><td>strand</td><td>MM</td><td>target_seq</td><td>PAM</td><td>alignment</td><td>distance</td><td>position</td><td>gene name</td><td>gene id</td></tr>
<tr><td>GU305914.1</td><td></td><td>51181</td><td>51203</td><td>+</td><td>0</td><td>CCGATTGCTGCGGCACGTAG</td><td>CGG</td><td>||||||||[||||||||||||]PAM</td><td>NA</td><td>Intergenic</td><td>NA</td><td>NA</td></tr>
<tr><td>T3351</td><td>GCGACTTCCAGTACATCTCGAGG</td><td></td><td>1000</td><td></td><td></td><td></td><td></td><td></td><td></td><td></td><td></td><td></td></tr>
<tr><td>Oligo fwd</td><td>CACCGCGACTTCCAGTACATCTCG</td><td></td><td></td><td></td><td></td><td></td><td></td><td></td><td></td><td></td><td></td><td></td></tr>
<tr><td>Oligo rev</td><td>AAACCGAGATGTACTGGAAGTCGC</td><td></td><td></td><td></td><td></td><td></td><td></td><td></td><td></td><td></td><td></td><td></td></tr>
<tr><td>Chromosome</td><td>start</td><td></td><td>end</td><td>strand</td><td>MM</td><td>target_seq</td><td>PAM</td><td>alignment</td><td>distance</td><td>position</td><td>gene name</td><td>gene id</td></tr>
<tr><td>GU305914.1</td><td></td><td>51205</td><td>51227</td><td>+</td><td>0</td><td>GCGACTTCCAGTACATCTCG</td><td>AGG</td><td>||||||||[||||||||||||]PAM</td><td>NA</td><td>Intergenic</td><td>NA</td><td>NA</td></tr>
<tr><td>T3352</td><td>GGAAGCCCATGTCCACGTTTTGG</td><td></td><td>1000</td><td></td><td></td><td></td><td></td><td></td><td></td><td></td><td></td><td></td></tr>
<tr><td>Oligo fwd</td><td>CACCGGAAGCCCATGTCCACGTTT</td><td></td><td></td><td></td><td></td><td></td><td></td><td></td><td></td><td></td><td></td><td></td></tr>
<tr><td>Oligo rev</td><td>AAACAAACGTGGACATGGGCTTCC</td><td></td><td></td><td></td><td></td><td></td><td></td><td></td><td></td><td></td><td></td><td></td></tr>
<tr><td>Chromosome</td><td>start</td><td></td><td>end</td><td>strand</td><td>MM</td><td>target_seq</td><td>PAM</td><td>alignment</td><td>distance</td><td>position</td><td>gene name</td><td>gene id</td></tr>
<tr><td>GU305914.1</td><td></td><td>51226</td><td>51248</td><td>+</td><td>0</td><td>GGAAGCCCATGTCCACGTTT</td><td>TGG</td><td>||||||||[||||||||||||]PAM</td><td>NA</td><td>Intergenic</td><td>NA</td><td>NA</td></tr>
<tr><td>T3353</td><td>AGCCCATGTCCACGTTTTGGTGG</td><td></td><td>1000</td><td></td><td></td><td></td><td></td><td></td><td></td><td></td><td></td><td></td></tr>
<tr><td>Oligo fwd</td><td>CACCAGCCCATGTCCACGTTTTGG</td><td></td><td></td><td></td><td></td><td></td><td></td><td></td><td></td><td></td><td></td><td></td></tr>
<tr><td>Oligo rev</td><td>AAACCCAAAACGTGGACATGGGCT</td><td></td><td></td><td></td><td></td><td></td><td></td><td></td><td></td><td></td><td></td><td></td></tr>
<tr><td>Chromosome</td><td>start</td><td></td><td>end</td><td>strand</td><td>MM</td><td>target_seq</td><td>PAM</td><td>alignment</td><td>distance</td><td>position</td><td>gene name</td><td>gene id</td></tr>
<tr><td>GU305914.1</td><td></td><td>51229</td><td>51251</td><td>+</td><td>0</td><td>AGCCCATGTCCACGTTTTGG</td><td>TGG</td><td>||||||||[||||||||||||]PAM</td><td>NA</td><td>Intergenic</td><td>NA</td><td>NA</td></tr>
<tr><td>T3354</td><td>CCATGTCCACGTTTTGGTGGAGG</td><td></td><td>1000</td><td></td><td></td><td></td><td></td><td></td><td></td><td></td><td></td><td></td></tr>
<tr><td>Oligo fwd</td><td>CACCCCATGTCCACGTTTTGGTGG</td><td></td><td></td><td></td><td></td><td></td><td></td><td></td><td></td><td></td><td></td><td></td></tr>
<tr><td>Oligo rev</td><td>AAACCCACCAAAACGTGGACATGG</td><td></td><td></td><td></td><td></td><td></td><td></td><td></td><td></td><td></td><td></td><td></td></tr>
<tr><td>Chromosome</td><td>start</td><td></td><td>end</td><td>strand</td><td>MM</td><td>target_seq</td><td>PAM</td><td>alignment</td><td>distance</td><td>position</td><td>gene name</td><td>gene id</td></tr>
<tr><td>GU305914.1</td><td></td><td>51232</td><td>51254</td><td>+</td><td>0</td><td>CCATGTCCACGTTTTGGTGG</td><td>AGG</td><td>||||||||[||||||||||||]PAM</td><td>NA</td><td>Intergenic</td><td>NA</td><td>NA</td></tr>
<tr><td>T3355</td><td>CATGTCCACGTTTTGGTGGAGGG</td><td></td><td>1000</td><td></td><td></td><td></td><td></td><td></td><td></td><td></td><td></td><td></td></tr>
<tr><td>Oligo fwd</td><td>CACCCATGTCCACGTTTTGGTGGA</td><td></td><td></td><td></td><td></td><td></td><td></td><td></td><td></td><td></td><td></td><td></td></tr>
<tr><td>Oligo rev</td><td>AAACTCCACCAAAACGTGGACATG</td><td></td><td></td><td></td><td></td><td></td><td></td><td></td><td></td><td></td><td></td><td></td></tr>
<tr><td>Chromosome</td><td>start</td><td></td><td>end</td><td>strand</td><td>MM</td><td>target_seq</td><td>PAM</td><td>alignment</td><td>distance</td><td>position</td><td>gene name</td><td>gene id</td></tr>
<tr><td>GU305914.1</td><td></td><td>51233</td><td>51255</td><td>+</td><td>0</td><td>CATGTCCACGTTTTGGTGGA</td><td>GGG</td><td>||||||||[||||||||||||]PAM</td><td>NA</td><td>Intergenic</td><td>NA</td><td>NA</td></tr>
<tr><td>T3356</td><td>GTCCACGTTTTGGTGGAGGGTGG</td><td></td><td>1000</td><td></td><td></td><td></td><td></td><td></td><td></td><td></td><td></td><td></td></tr>
<tr><td>Oligo fwd</td><td>CACCGTCCACGTTTTGGTGGAGGG</td><td></td><td></td><td></td><td></td><td></td><td></td><td></td><td></td><td></td><td></td><td></td></tr>
</table>

<table>
<tr><td>Oligo rev</td><td>AAACCCCTCCACCAAAACGTGGAC</td><td></td><td></td><td></td><td></td><td></td><td></td><td></td><td></td><td></td><td></td></tr>
<tr><td>Chromosome</td><td>start</td><td>end</td><td>strand</td><td>MM</td><td>target_seq</td><td>PAM</td><td>alignment</td><td>distance</td><td>position</td><td>gene name</td><td>gene id</td></tr>
<tr><td>GU305914.1</td><td>51236</td><td>51258</td><td>+</td><td>0</td><td>GTCCACGTTTTGGTGGAGGG</td><td>TGG</td><td>||||||||[||||||||||||]PAM</td><td>NA</td><td>Intergenic</td><td>NA</td><td>NA</td></tr>
<tr><td>T3357</td><td>CGTTTTGGTGGAGGGTGGCGCGG</td><td>1000</td><td></td><td></td><td></td><td></td><td></td><td></td><td></td><td></td><td></td></tr>
<tr><td>Oligo fwd</td><td>CACCCGTTTTGGTGGAGGGTGGCG</td><td></td><td></td><td></td><td></td><td></td><td></td><td></td><td></td><td></td><td></td></tr>
<tr><td>Oligo rev</td><td>AAACCGCCACCCTCCACCAAAACG</td><td></td><td></td><td></td><td></td><td></td><td></td><td></td><td></td><td></td><td></td></tr>
<tr><td>Chromosome</td><td>start</td><td>end</td><td>strand</td><td>MM</td><td>target_seq</td><td>PAM</td><td>alignment</td><td>distance</td><td>position</td><td>gene name</td><td>gene id</td></tr>
<tr><td>GU305914.1</td><td>51241</td><td>51263</td><td>+</td><td>0</td><td>CGTTTTGGTGGAGGGTGGCG</td><td>CGG</td><td>||||||||[||||||||||||]PAM</td><td>NA</td><td>Intergenic</td><td>NA</td><td>NA</td></tr>
<tr><td>T3358</td><td>GTGGCGCGGTCAGAGAAGAGCGG</td><td>1000</td><td></td><td></td><td></td><td></td><td></td><td></td><td></td><td></td><td></td></tr>
<tr><td>Oligo fwd</td><td>CACCGTGGCGCGGTCAGAGAAGAG</td><td></td><td></td><td></td><td></td><td></td><td></td><td></td><td></td><td></td><td></td></tr>
<tr><td>Oligo rev</td><td>AAACCTCTTCTCTGACCGCGCCAC</td><td></td><td></td><td></td><td></td><td></td><td></td><td></td><td></td><td></td><td></td></tr>
<tr><td>Chromosome</td><td>start</td><td>end</td><td>strand</td><td>MM</td><td>target_seq</td><td>PAM</td><td>alignment</td><td>distance</td><td>position</td><td>gene name</td><td>gene id</td></tr>
<tr><td>GU305914.1</td><td>51255</td><td>51277</td><td>+</td><td>0</td><td>GTGGCGCGGTCAGAGAAGAG</td><td>CGG</td><td>||||||||[||||||||||||]PAM</td><td>NA</td><td>Intergenic</td><td>NA</td><td>NA</td></tr>
<tr><td>T3359</td><td>GCGCGGTCAGAGAAGAGCGGCGG</td><td>1000</td><td></td><td></td><td></td><td></td><td></td><td></td><td></td><td></td><td></td></tr>
<tr><td>Oligo fwd</td><td>CACCGCGCGGTCAGAGAAGAGCGG</td><td></td><td></td><td></td><td></td><td></td><td></td><td></td><td></td><td></td><td></td></tr>
<tr><td>Oligo rev</td><td>AAACCCGCTCTTCTCTGACCGCGC</td><td></td><td></td><td></td><td></td><td></td><td></td><td></td><td></td><td></td><td></td></tr>
<tr><td>Chromosome</td><td>start</td><td>end</td><td>strand</td><td>MM</td><td>target_seq</td><td>PAM</td><td>alignment</td><td>distance</td><td>position</td><td>gene name</td><td>gene id</td></tr>
<tr><td>GU305914.1</td><td>51258</td><td>51280</td><td>+</td><td>0</td><td>GCGCGGTCAGAGAAGAGCGG</td><td>CGG</td><td>||||||||[||||||||||||]PAM</td><td>NA</td><td>Intergenic</td><td>NA</td><td>NA</td></tr>
<tr><td>T3360</td><td>GTCAGAGAAGAGCGGCGGTATGG</td><td>1000</td><td></td><td></td><td></td><td></td><td></td><td></td><td></td><td></td><td></td></tr>
<tr><td>Oligo fwd</td><td>CACCGTCAGAGAAGAGCGGCGGTA</td><td></td><td></td><td></td><td></td><td></td><td></td><td></td><td></td><td></td><td></td></tr>
<tr><td>Oligo rev</td><td>AAACTACCGCCGCTCTTCTCTGAC</td><td></td><td></td><td></td><td></td><td></td><td></td><td></td><td></td><td></td><td></td></tr>
<tr><td>Chromosome</td><td>start</td><td>end</td><td>strand</td><td>MM</td><td>target_seq</td><td>PAM</td><td>alignment</td><td>distance</td><td>position</td><td>gene name</td><td>gene id</td></tr>
<tr><td>GU305914.1</td><td>51263</td><td>51285</td><td>+</td><td>0</td><td>GTCAGAGAAGAGCGGCGGTA</td><td>TGG</td><td>||||||||[||||||||||||]PAM</td><td>NA</td><td>Intergenic</td><td>NA</td><td>NA</td></tr>
<tr><td>T3361</td><td>GCGGCGGTATGGATCCCTGCTGG</td><td>1000</td><td></td><td></td><td></td><td></td><td></td><td></td><td></td><td></td><td></td></tr>
<tr><td>Oligo fwd</td><td>CACCGCGGCGGTATGGATCCCTGC</td><td></td><td></td><td></td><td></td><td></td><td></td><td></td><td></td><td></td><td></td></tr>
<tr><td>Oligo rev</td><td>AAACGCAGGGATCCATACCGCCGC</td><td></td><td></td><td></td><td></td><td></td><td></td><td></td><td></td><td></td><td></td></tr>
<tr><td>Chromosome</td><td>start</td><td>end</td><td>strand</td><td>MM</td><td>target_seq</td><td>PAM</td><td>alignment</td><td>distance</td><td>position</td><td>gene name</td><td>gene id</td></tr>
<tr><td>GU305914.1</td><td>51274</td><td>51296</td><td>+</td><td>0</td><td>GCGGCGGTATGGATCCCTGC</td><td>TGG</td><td>||||||||[||||||||||||]PAM</td><td>NA</td><td>Intergenic</td><td>NA</td><td>NA</td></tr>
<tr><td>T3362</td><td>CGGTATGGATCCCTGCTGGATGG</td><td>1000</td><td></td><td></td><td></td><td></td><td></td><td></td><td></td><td></td><td></td></tr>
<tr><td>Oligo fwd</td><td>CACCCGGTATGGATCCCTGCTGGA</td><td></td><td></td><td></td><td></td><td></td><td></td><td></td><td></td><td></td><td></td></tr>
<tr><td>Oligo rev</td><td>AAACTCCAGCAGGGATCCATACCG</td><td></td><td></td><td></td><td></td><td></td><td></td><td></td><td></td><td></td><td></td></tr>
<tr><td>Chromosome</td><td>start</td><td>end</td><td>strand</td><td>MM</td><td>target_seq</td><td>PAM</td><td>alignment</td><td>distance</td><td>position</td><td>gene name</td><td>gene id</td></tr>
<tr><td>GU305914.1</td><td>51278</td><td>51300</td><td>+</td><td>0</td><td>CGGTATGGATCCCTGCTGGA</td><td>TGG</td><td>||||||||[||||||||||||]PAM</td><td>NA</td><td>Intergenic</td><td>NA</td><td>NA</td></tr>
<tr><td>T3363</td><td>GGATCCCTGCTGGATGGCGCTGG</td><td>1000</td><td></td><td></td><td></td><td></td><td></td><td></td><td></td><td></td><td></td></tr>
<tr><td>Oligo fwd</td><td>CACCGGATCCCTGCTGGATGGCGC</td><td></td><td></td><td></td><td></td><td></td><td></td><td></td><td></td><td></td><td></td></tr>
<tr><td>Oligo rev</td><td>AAACGCGCCATCCAGCAGGGATCC</td><td></td><td></td><td></td><td></td><td></td><td></td><td></td><td></td><td></td><td></td></tr>
<tr><td>Chromosome</td><td>start</td><td>end</td><td>strand</td><td>MM</td><td>target_seq</td><td>PAM</td><td>alignment</td><td>distance</td><td>position</td><td>gene name</td><td>gene id</td></tr>
<tr><td>GU305914.1</td><td>51284</td><td>51306</td><td>+</td><td>0</td><td>GGATCCCTGCTGGATGGCGC</td><td>TGG</td><td>||||||||[||||||||||||]PAM</td><td>NA</td><td>Intergenic</td><td>NA</td><td>NA</td></tr>
<tr><td>T3364</td><td>CTGCTGGATGGCGCTGGCCGCGG</td><td>1000</td><td></td><td></td><td></td><td></td><td></td><td></td><td></td><td></td><td></td></tr>
<tr><td>Oligo fwd</td><td>CACCCTGCTGGATGGCGCTGGCCG</td><td></td><td></td><td></td><td></td><td></td><td></td><td></td><td></td><td></td><td></td></tr>
<tr><td>Oligo rev</td><td>AAACCGGCCAGCGCCATCCAGCAG</td><td></td><td></td><td></td><td></td><td></td><td></td><td></td><td></td><td></td><td></td></tr>
<tr><td>Chromosome</td><td>start</td><td>end</td><td>strand</td><td>MM</td><td>target_seq</td><td>PAM</td><td>alignment</td><td>distance</td><td>position</td><td>gene name</td><td>gene id</td></tr>
<tr><td>GU305914.1</td><td>51290</td><td>51312</td><td>+</td><td>0</td><td>CTGCTGGATGGCGCTGGCCG</td><td>CGG</td><td>||||||||[||||||||||||]PAM</td><td>NA</td><td>Intergenic</td><td>NA</td><td>NA</td></tr>
<tr><td>T3365</td><td>TGGCGCTGGCCGCGGAAATGAGG</td><td>1000</td><td></td><td></td><td></td><td></td><td></td><td></td><td></td><td></td><td></td></tr>
<tr><td>Oligo fwd</td><td>CACCTGGCGCTGGCCGCGGAAATG</td><td></td><td></td><td></td><td></td><td></td><td></td><td></td><td></td><td></td><td></td></tr>
<tr><td>Oligo rev</td><td>AAACCATTTCCGCGGCCAGCGCCA</td><td></td><td></td><td></td><td></td><td></td><td></td><td></td><td></td><td></td><td></td></tr>
<tr><td>Chromosome</td><td>start</td><td>end</td><td>strand</td><td>MM</td><td>target_seq</td><td>PAM</td><td>alignment</td><td>distance</td><td>position</td><td>gene name</td><td>gene id</td></tr>
<tr><td>GU305914.1</td><td>51298</td><td>51320</td><td>+</td><td>0</td><td>TGGCGCTGGCCGCGGAAATG</td><td>AGG</td><td>||||||||[||||||||||||]PAM</td><td>NA</td><td>Intergenic</td><td>NA</td><td>NA</td></tr>
<tr><td>T3366</td><td>CGGAAATGAGGATGCGCATGAGG</td><td>1000</td><td></td><td></td><td></td><td></td><td></td><td></td><td></td><td></td><td></td></tr>
<tr><td>Oligo fwd</td><td>CACCCGGAAATGAGGATGCGCATG</td><td></td><td></td><td></td><td></td><td></td><td></td><td></td><td></td><td></td><td></td></tr>
<tr><td>Oligo rev</td><td>AAACCATGCGCATCCTCATTTCCG</td><td></td><td></td><td></td><td></td><td></td><td></td><td></td><td></td><td></td><td></td></tr>
<tr><td>Chromosome</td><td>start</td><td>end</td><td>strand</td><td>MM</td><td>target_seq</td><td>PAM</td><td>alignment</td><td>distance</td><td>position</td><td>gene name</td><td>gene id</td></tr>
<tr><td>GU305914.1</td><td>51310</td><td>51332</td><td>+</td><td>0</td><td>CGGAAATGAGGATGCGCATG</td><td>AGG</td><td>||||||||[||||||||||||]PAM</td><td>NA</td><td>Intergenic</td><td>NA</td><td>NA</td></tr>
<tr><td>T3367</td><td>ATGAGGCGATTGCACTTGAGAGG</td><td>1000</td><td></td><td></td><td></td><td></td><td></td><td></td><td></td><td></td><td></td></tr>
<tr><td>Oligo fwd</td><td>CACCATGAGGCGATTGCACTTGAG</td><td></td><td></td><td></td><td></td><td></td><td></td><td></td><td></td><td></td><td></td></tr>
<tr><td>Oligo rev</td><td>AAACCTCAAGTGCAATCGCCTCAT</td><td></td><td></td><td></td><td></td><td></td><td></td><td></td><td></td><td></td><td></td></tr>
<tr><td>Chromosome</td><td>start</td><td>end</td><td>strand</td><td>MM</td><td>target_seq</td><td>PAM</td><td>alignment</td><td>distance</td><td>position</td><td>gene name</td><td>gene id</td></tr>
<tr><td>GU305914.1</td><td>51327</td><td>51349</td><td>+</td><td>0</td><td>ATGAGGCGATTGCACTTGAG</td><td>AGG</td><td>||||||||[||||||||||||]PAM</td><td>NA</td><td>Intergenic</td><td>NA</td><td>NA</td></tr>
<tr><td>T3368</td><td>GGCGATTGCACTTGAGAGGCAGG</td><td>1000</td><td></td><td></td><td></td><td></td><td></td><td></td><td></td><td></td><td></td></tr>
<tr><td>Oligo fwd</td><td>CACCGGCGATTGCACTTGAGAGGC</td><td></td><td></td><td></td><td></td><td></td><td></td><td></td><td></td><td></td><td></td></tr>
<tr><td>Oligo rev</td><td>AAACGCCTCTCAAGTGCAATCGCC</td><td></td><td></td><td></td><td></td><td></td><td></td><td></td><td></td><td></td><td></td></tr>
<tr><td>Chromosome</td><td>start</td><td>end</td><td>strand</td><td>MM</td><td>target_seq</td><td>PAM</td><td>alignment</td><td>distance</td><td>position</td><td>gene name</td><td>gene id</td></tr>
<tr><td>GU305914.1</td><td>51331</td><td>51353</td><td>+</td><td>0</td><td>GGCGATTGCACTTGAGAGGC</td><td>AGG</td><td>||||||||[||||||||||||]PAM</td><td>NA</td><td>Intergenic</td><td>NA</td><td>NA</td></tr>
<tr><td>T3369</td><td>ATTGCACTTGAGAGGCAGGACGG</td><td>1000</td><td></td><td></td><td></td><td></td><td></td><td></td><td></td><td></td><td></td></tr>
<tr><td>Oligo fwd</td><td>CACCATTGCACTTGAGAGGCAGGA</td><td></td><td></td><td></td><td></td><td></td><td></td><td></td><td></td><td></td><td></td></tr>
<tr><td>Oligo rev</td><td>AAACTCCTGCCTCTCAAGTGCAAT</td><td></td><td></td><td></td><td></td><td></td><td></td><td></td><td></td><td></td><td></td></tr>
<tr><td>Chromosome</td><td>start</td><td>end</td><td>strand</td><td>MM</td><td>target_seq</td><td>PAM</td><td>alignment</td><td>distance</td><td>position</td><td>gene name</td><td>gene id</td></tr>
<tr><td>GU305914.1</td><td>51335</td><td>51357</td><td>+</td><td>0</td><td>ATTGCACTTGAGAGGCAGGA</td><td>CGG</td><td>||||||||[||||||||||||]PAM</td><td>NA</td><td>Intergenic</td><td>NA</td><td>NA</td></tr>
<tr><td>T3370</td><td>AGACTTCTGCCTCTTCTCCTTGG</td><td>1000</td><td></td><td></td><td></td><td></td><td></td><td></td><td></td><td></td><td></td></tr>
<tr><td>Oligo fwd</td><td>CACCAGACTTCTGCCTCTTCTCCT</td><td></td><td></td><td></td><td></td><td></td><td></td><td></td><td></td><td></td><td></td></tr>
<tr><td>Oligo rev</td><td>AAACAGGAGAAGAGGCAGAAGTCT</td><td></td><td></td><td></td><td></td><td></td><td></td><td></td><td></td><td></td><td></td></tr>
<tr><td>Chromosome</td><td>start</td><td>end</td><td>strand</td><td>MM</td><td>target_seq</td><td>PAM</td><td>alignment</td><td>distance</td><td>position</td><td>gene name</td><td>gene id</td></tr>
<tr><td>GU305914.1</td><td>51362</td><td>51384</td><td>+</td><td>0</td><td>AGACTTCTGCCTCTTCTCCT</td><td>TGG</td><td>||||||||[||||||||||||]PAM</td><td>NA</td><td>Intergenic</td><td>NA</td><td>NA</td></tr>
<tr><td>T3371</td><td>AGAAGCCCATGAGATCCTTGAGG</td><td>1000</td><td></td><td></td><td></td><td></td><td></td><td></td><td></td><td></td><td></td></tr>
</table>

<table>
<tr><td>Oligo fwd</td><td>CACCAGAAGCCCATGAGATCCTTG</td><td></td><td></td><td></td><td></td><td></td><td></td><td></td><td></td><td></td><td></td></tr>
<tr><td>Oligo rev</td><td>AAACCAAGGATCTCATGGGCTTCT</td><td></td><td></td><td></td><td></td><td></td><td></td><td></td><td></td><td></td><td></td></tr>
<tr><td>Chromosome</td><td>start</td><td>end</td><td>strand</td><td>MM</td><td>target_seq</td><td>PAM</td><td>alignment</td><td>distance</td><td>position</td><td>gene name</td><td>gene id</td></tr>
<tr><td>GU305914.1</td><td>51421</td><td>51443</td><td>+</td><td>0</td><td>AGAAGCCCATGAGATCCTTG</td><td>AGG</td><td>||||||||[||||||||||||]PAM</td><td>NA</td><td>Intergenic</td><td>NA</td><td>NA</td></tr>
<tr><td></td><td></td><td></td><td></td><td></td><td></td><td></td><td></td><td></td><td></td><td></td><td></td></tr>
<tr><td>T3372</td><td>GCTGCTTGACGTACCTCCGCAGG</td><td>1000</td><td></td><td></td><td></td><td></td><td></td><td></td><td></td><td></td><td></td></tr>
<tr><td>Oligo fwd</td><td>CACCGCTGCTTGACGTACCTCCGC</td><td></td><td></td><td></td><td></td><td></td><td></td><td></td><td></td><td></td><td></td></tr>
<tr><td>Oligo rev</td><td>AAACGCGGAGGTACGTCAAGCAGC</td><td></td><td></td><td></td><td></td><td></td><td></td><td></td><td></td><td></td><td></td></tr>
<tr><td>Chromosome</td><td>start</td><td>end</td><td>strand</td><td>MM</td><td>target_seq</td><td>PAM</td><td>alignment</td><td>distance</td><td>position</td><td>gene name</td><td>gene id</td></tr>
<tr><td>GU305914.1</td><td>51454</td><td>51476</td><td>+</td><td>0</td><td>GCTGCTTGACGTACCTCCGC</td><td>AGG</td><td>||||||||[||||||||||||]PAM</td><td>NA</td><td>Intergenic</td><td>NA</td><td>NA</td></tr>
<tr><td></td><td></td><td></td><td></td><td></td><td></td><td></td><td></td><td></td><td></td><td></td><td></td></tr>
<tr><td>T3373</td><td>CTGCTTGACGTACCTCCGCAGGG</td><td>1000</td><td></td><td></td><td></td><td></td><td></td><td></td><td></td><td></td><td></td></tr>
<tr><td>Oligo fwd</td><td>CACCCTGCTTGACGTACCTCCGCA</td><td></td><td></td><td></td><td></td><td></td><td></td><td></td><td></td><td></td><td></td></tr>
<tr><td>Oligo rev</td><td>AAACTGCGGAGGTACGTCAAGCAG</td><td></td><td></td><td></td><td></td><td></td><td></td><td></td><td></td><td></td><td></td></tr>
<tr><td>Chromosome</td><td>start</td><td>end</td><td>strand</td><td>MM</td><td>target_seq</td><td>PAM</td><td>alignment</td><td>distance</td><td>position</td><td>gene name</td><td>gene id</td></tr>
<tr><td>GU305914.1</td><td>51455</td><td>51477</td><td>+</td><td>0</td><td>CTGCTTGACGTACCTCCGCA</td><td>GGG</td><td>||||||||[||||||||||||]PAM</td><td>NA</td><td>Intergenic</td><td>NA</td><td>NA</td></tr>
<tr><td></td><td></td><td></td><td></td><td></td><td></td><td></td><td></td><td></td><td></td><td></td><td></td></tr>
<tr><td>T3374</td><td>TGCTTGACGTACCTCCGCAGGGG</td><td>1000</td><td></td><td></td><td></td><td></td><td></td><td></td><td></td><td></td><td></td></tr>
<tr><td>Oligo fwd</td><td>CACCTGCTTGACGTACCTCCGCAG</td><td></td><td></td><td></td><td></td><td></td><td></td><td></td><td></td><td></td><td></td></tr>
<tr><td>Oligo rev</td><td>AAACCTGCGGAGGTACGTCAAGCA</td><td></td><td></td><td></td><td></td><td></td><td></td><td></td><td></td><td></td><td></td></tr>
<tr><td>Chromosome</td><td>start</td><td>end</td><td>strand</td><td>MM</td><td>target_seq</td><td>PAM</td><td>alignment</td><td>distance</td><td>position</td><td>gene name</td><td>gene id</td></tr>
<tr><td>GU305914.1</td><td>51456</td><td>51478</td><td>+</td><td>0</td><td>TGCTTGACGTACCTCCGCAG</td><td>GGG</td><td>||||||||[||||||||||||]PAM</td><td>NA</td><td>Intergenic</td><td>NA</td><td>NA</td></tr>
<tr><td></td><td></td><td></td><td></td><td></td><td></td><td></td><td></td><td></td><td></td><td></td><td></td></tr>
<tr><td>T3375</td><td>ACGTACCTCCGCAGGGGCGAAGG</td><td>1000</td><td></td><td></td><td></td><td></td><td></td><td></td><td></td><td></td><td></td></tr>
<tr><td>Oligo fwd</td><td>CACCACGTACCTCCGCAGGGGCGA</td><td></td><td></td><td></td><td></td><td></td><td></td><td></td><td></td><td></td><td></td></tr>
<tr><td>Oligo rev</td><td>AAACTCGCCCCTGCGGAGGTACGT</td><td></td><td></td><td></td><td></td><td></td><td></td><td></td><td></td><td></td><td></td></tr>
<tr><td>Chromosome</td><td>start</td><td>end</td><td>strand</td><td>MM</td><td>target_seq</td><td>PAM</td><td>alignment</td><td>distance</td><td>position</td><td>gene name</td><td>gene id</td></tr>
<tr><td>GU305914.1</td><td>51462</td><td>51484</td><td>+</td><td>0</td><td>ACGTACCTCCGCAGGGGCGA</td><td>AGG</td><td>||||||||[||||||||||||]PAM</td><td>NA</td><td>Intergenic</td><td>NA</td><td>NA</td></tr>
<tr><td></td><td></td><td></td><td></td><td></td><td></td><td></td><td></td><td></td><td></td><td></td><td></td></tr>
<tr><td>T3376</td><td>CCGCAGGGGCGAAGGAAAGATGG</td><td>1000</td><td></td><td></td><td></td><td></td><td></td><td></td><td></td><td></td><td></td></tr>
<tr><td>Oligo fwd</td><td>CACCCCGCAGGGGCGAAGGAAAGA</td><td></td><td></td><td></td><td></td><td></td><td></td><td></td><td></td><td></td><td></td></tr>
<tr><td>Oligo rev</td><td>AAACTCTTTCCTTCGCCCCTGCGG</td><td></td><td></td><td></td><td></td><td></td><td></td><td></td><td></td><td></td><td></td></tr>
<tr><td>Chromosome</td><td>start</td><td>end</td><td>strand</td><td>MM</td><td>target_seq</td><td>PAM</td><td>alignment</td><td>distance</td><td>position</td><td>gene name</td><td>gene id</td></tr>
<tr><td>GU305914.1</td><td>51470</td><td>51492</td><td>+</td><td>0</td><td>CCGCAGGGGCGAAGGAAAGA</td><td>TGG</td><td>||||||||[||||||||||||]PAM</td><td>NA</td><td>Intergenic</td><td>NA</td><td>NA</td></tr>
<tr><td></td><td></td><td></td><td></td><td></td><td></td><td></td><td></td><td></td><td></td><td></td><td></td></tr>
<tr><td>T3377</td><td>GTCGTGTTCGAAGACGATGTCGG</td><td>1000</td><td></td><td></td><td></td><td></td><td></td><td></td><td></td><td></td><td></td></tr>
<tr><td>Oligo fwd</td><td>CACCGTCGTGTTCGAAGACGATGT</td><td></td><td></td><td></td><td></td><td></td><td></td><td></td><td></td><td></td><td></td></tr>
<tr><td>Oligo rev</td><td>AAACACATCGTCTTCGAACACGAC</td><td></td><td></td><td></td><td></td><td></td><td></td><td></td><td></td><td></td><td></td></tr>
<tr><td>Chromosome</td><td>start</td><td>end</td><td>strand</td><td>MM</td><td>target_seq</td><td>PAM</td><td>alignment</td><td>distance</td><td>position</td><td>gene name</td><td>gene id</td></tr>
<tr><td>GU305914.1</td><td>51497</td><td>51519</td><td>+</td><td>0</td><td>GTCGTGTTCGAAGACGATGT</td><td>CGG</td><td>||||||||[||||||||||||]PAM</td><td>NA</td><td>Intergenic</td><td>NA</td><td>NA</td></tr>
<tr><td></td><td></td><td></td><td></td><td></td><td></td><td></td><td></td><td></td><td></td><td></td><td></td></tr>
<tr><td>T3378</td><td>CGGCTGTCCAGAAGTCTTGCAGG</td><td>1000</td><td></td><td></td><td></td><td></td><td></td><td></td><td></td><td></td><td></td></tr>
<tr><td>Oligo fwd</td><td>CACCCGGCTGTCCAGAAGTCTTGC</td><td></td><td></td><td></td><td></td><td></td><td></td><td></td><td></td><td></td><td></td></tr>
<tr><td>Oligo rev</td><td>AAACGCAAGACTTCTGGACAGCCG</td><td></td><td></td><td></td><td></td><td></td><td></td><td></td><td></td><td></td><td></td></tr>
<tr><td>Chromosome</td><td>start</td><td>end</td><td>strand</td><td>MM</td><td>target_seq</td><td>PAM</td><td>alignment</td><td>distance</td><td>position</td><td>gene name</td><td>gene id</td></tr>
<tr><td>GU305914.1</td><td>51517</td><td>51539</td><td>+</td><td>0</td><td>CGGCTGTCCAGAAGTCTTGC</td><td>AGG</td><td>||||||||[||||||||||||]PAM</td><td>NA</td><td>Intergenic</td><td>NA</td><td>NA</td></tr>
<tr><td></td><td></td><td></td><td></td><td></td><td></td><td></td><td></td><td></td><td></td><td></td><td></td></tr>
<tr><td>T3379</td><td>TGTCCAGAAGTCTTGCAGGCTGG</td><td>1000</td><td></td><td></td><td></td><td></td><td></td><td></td><td></td><td></td><td></td></tr>
<tr><td>Oligo fwd</td><td>CACCTGTCCAGAAGTCTTGCAGGC</td><td></td><td></td><td></td><td></td><td></td><td></td><td></td><td></td><td></td><td></td></tr>
<tr><td>Oligo rev</td><td>AAACGCCTGCAAGACTTCTGGACA</td><td></td><td></td><td></td><td></td><td></td><td></td><td></td><td></td><td></td><td></td></tr>
<tr><td>Chromosome</td><td>start</td><td>end</td><td>strand</td><td>MM</td><td>target_seq</td><td>PAM</td><td>alignment</td><td>distance</td><td>position</td><td>gene name</td><td>gene id</td></tr>
<tr><td>GU305914.1</td><td>51521</td><td>51543</td><td>+</td><td>0</td><td>TGTCCAGAAGTCTTGCAGGC</td><td>TGG</td><td>||||||||[||||||||||||]PAM</td><td>NA</td><td>Intergenic</td><td>NA</td><td>NA</td></tr>
<tr><td></td><td></td><td></td><td></td><td></td><td></td><td></td><td></td><td></td><td></td><td></td><td></td></tr>
<tr><td>T3380</td><td>GTCCAGAAGTCTTGCAGGCTGGG</td><td>1000</td><td></td><td></td><td></td><td></td><td></td><td></td><td></td><td></td><td></td></tr>
<tr><td>Oligo fwd</td><td>CACCGTCCAGAAGTCTTGCAGGCT</td><td></td><td></td><td></td><td></td><td></td><td></td><td></td><td></td><td></td><td></td></tr>
<tr><td>Oligo rev</td><td>AAACAGCCTGCAAGACTTCTGGAC</td><td></td><td></td><td></td><td></td><td></td><td></td><td></td><td></td><td></td><td></td></tr>
<tr><td>Chromosome</td><td>start</td><td>end</td><td>strand</td><td>MM</td><td>target_seq</td><td>PAM</td><td>alignment</td><td>distance</td><td>position</td><td>gene name</td><td>gene id</td></tr>
<tr><td>GU305914.1</td><td>51522</td><td>51544</td><td>+</td><td>0</td><td>GTCCAGAAGTCTTGCAGGCT</td><td>GGG</td><td>||||||||[||||||||||||]PAM</td><td>NA</td><td>Intergenic</td><td>NA</td><td>NA</td></tr>
<tr><td></td><td></td><td></td><td></td><td></td><td></td><td></td><td></td><td></td><td></td><td></td><td></td></tr>
<tr><td>T3381</td><td>GAACATGACTCCCCTGTCGTAGG</td><td>1000</td><td></td><td></td><td></td><td></td><td></td><td></td><td></td><td></td><td></td></tr>
<tr><td>Oligo fwd</td><td>CACCGAACATGACTCCCCTGTCGT</td><td></td><td></td><td></td><td></td><td></td><td></td><td></td><td></td><td></td><td></td></tr>
<tr><td>Oligo rev</td><td>AAACACGACAGGGGAGTCATGTTC</td><td></td><td></td><td></td><td></td><td></td><td></td><td></td><td></td><td></td><td></td></tr>
<tr><td>Chromosome</td><td>start</td><td>end</td><td>strand</td><td>MM</td><td>target_seq</td><td>PAM</td><td>alignment</td><td>distance</td><td>position</td><td>gene name</td><td>gene id</td></tr>
<tr><td>GU305914.1</td><td>51554</td><td>51576</td><td>+</td><td>0</td><td>GAACATGACTCCCCTGTCGT</td><td>AGG</td><td>||||||||[||||||||||||]PAM</td><td>NA</td><td>Intergenic</td><td>NA</td><td>NA</td></tr>
<tr><td></td><td></td><td></td><td></td><td></td><td></td><td></td><td></td><td></td><td></td><td></td><td></td></tr>
<tr><td>T3382</td><td>GATGAAGATGACCTGTCTCGAGG</td><td>1000</td><td></td><td></td><td></td><td></td><td></td><td></td><td></td><td></td><td></td></tr>
<tr><td>Oligo fwd</td><td>CACCGATGAAGATGACCTGTCTCG</td><td></td><td></td><td></td><td></td><td></td><td></td><td></td><td></td><td></td><td></td></tr>
<tr><td>Oligo rev</td><td>AAACCGAGACAGGTCATCTTCATC</td><td></td><td></td><td></td><td></td><td></td><td></td><td></td><td></td><td></td><td></td></tr>
<tr><td>Chromosome</td><td>start</td><td>end</td><td>strand</td><td>MM</td><td>target_seq</td><td>PAM</td><td>alignment</td><td>distance</td><td>position</td><td>gene name</td><td>gene id</td></tr>
<tr><td>GU305914.1</td><td>51602</td><td>51624</td><td>+</td><td>0</td><td>GATGAAGATGACCTGTCTCG</td><td>AGG</td><td>||||||||[||||||||||||]PAM</td><td>NA</td><td>Intergenic</td><td>NA</td><td>NA</td></tr>
<tr><td></td><td></td><td></td><td></td><td></td><td></td><td></td><td></td><td></td><td></td><td></td><td></td></tr>
<tr><td>T3383</td><td>ATGAAGATGACCTGTCTCGAGGG</td><td>1000</td><td></td><td></td><td></td><td></td><td></td><td></td><td></td><td></td><td></td></tr>
<tr><td>Oligo fwd</td><td>CACCATGAAGATGACCTGTCTCGA</td><td></td><td></td><td></td><td></td><td></td><td></td><td></td><td></td><td></td><td></td></tr>
<tr><td>Oligo rev</td><td>AAACTCGAGACAGGTCATCTTCAT</td><td></td><td></td><td></td><td></td><td></td><td></td><td></td><td></td><td></td><td></td></tr>
<tr><td>Chromosome</td><td>start</td><td>end</td><td>strand</td><td>MM</td><td>target_seq</td><td>PAM</td><td>alignment</td><td>distance</td><td>position</td><td>gene name</td><td>gene id</td></tr>
<tr><td>GU305914.1</td><td>51603</td><td>51625</td><td>+</td><td>0</td><td>ATGAAGATGACCTGTCTCGA</td><td>GGG</td><td>||||||||[||||||||||||]PAM</td><td>NA</td><td>Intergenic</td><td>NA</td><td>NA</td></tr>
<tr><td></td><td></td><td></td><td></td><td></td><td></td><td></td><td></td><td></td><td></td><td></td><td></td></tr>
<tr><td>T3384</td><td>GACCTGTCTCGAGGGTCGATTGG</td><td>1000</td><td></td><td></td><td></td><td></td><td></td><td></td><td></td><td></td><td></td></tr>
<tr><td>Oligo fwd</td><td>CACCGACCTGTCTCGAGGGTCGAT</td><td></td><td></td><td></td><td></td><td></td><td></td><td></td><td></td><td></td><td></td></tr>
<tr><td>Oligo rev</td><td>AAACATCGACCCTCGAGACAGGTC</td><td></td><td></td><td></td><td></td><td></td><td></td><td></td><td></td><td></td><td></td></tr>
<tr><td>Chromosome</td><td>start</td><td>end</td><td>strand</td><td>MM</td><td>target_seq</td><td>PAM</td><td>alignment</td><td>distance</td><td>position</td><td>gene name</td><td>gene id</td></tr>
<tr><td>GU305914.1</td><td>51611</td><td>51633</td><td>+</td><td>0</td><td>GACCTGTCTCGAGGGTCGAT</td><td>TGG</td><td>||||||||[||||||||||||]PAM</td><td>NA</td><td>Intergenic</td><td>NA</td><td>NA</td></tr>
<tr><td>GU305914.1</td><td>145568</td><td>145590</td><td>-</td><td>4</td><td>GCCCTGTGTCGAGTGTCGCT</td><td>CGG</td><td>|-|||||-[|||||-||||-|]PAM</td><td>NA</td><td>Intergenic</td><td>NA</td><td>NA</td></tr>
<tr><td></td><td></td><td></td><td></td><td></td><td></td><td></td><td></td><td></td><td></td><td></td><td></td></tr>
<tr><td>T3385</td><td>CTGTCTCGAGGGTCGATTGGAGG</td><td>1000</td><td></td><td></td><td></td><td></td><td></td><td></td><td></td><td></td><td></td></tr>
<tr><td>Oligo fwd</td><td>CACCCTGTCTCGAGGGTCGATTGG</td><td></td><td></td><td></td><td></td><td></td><td></td><td></td><td></td><td></td><td></td></tr>
<tr><td>Oligo rev</td><td>AAACCCAATCGACCCTCGAGACAG</td><td></td><td></td><td></td><td></td><td></td><td></td><td></td><td></td><td></td><td></td></tr>
<tr><td>Chromosome</td><td>start</td><td>end</td><td>strand</td><td>MM</td><td>target_seq</td><td>PAM</td><td>alignment</td><td>distance</td><td>position</td><td>gene name</td><td>gene id</td></tr>
<tr><td>GU305914.1</td><td>51614</td><td>51636</td><td>+</td><td>0</td><td>CTGTCTCGAGGGTCGATTGG</td><td>AGG</td><td>||||||||[||||||||||||]PAM</td><td>NA</td><td>Intergenic</td><td>NA</td><td>NA</td></tr>
</table>

T3386 GGAGGCGAACCGCCACTCTTCGG 1000
Oligo fwd CACCGGAGGCGAACCGCCACTCTT
Oligo rev AAACAAGAGTGGCGGTTCGCCTCC
Chromosome start end strand MM target_seq PAM alignment distance position gene name gene id
GU305914.1 51632 51654 + 0 GGAGGCGAACCGCCACTCTT CGG ||||||||[||||||||||||]PAM NA Intergenic NA NA

T3387 ACCCCGAAGATGATCATGAGCGG 1000
Oligo fwd CACCACCCCGAAGATGATCATGAG
Oligo rev AAACCTCATGATCATCTTCGGGGT
Chromosome start end strand MM target_seq PAM alignment distance position gene name gene id
GU305914.1 51657 51679 + 0 ACCCCGAAGATGATCATGAG CGG ||||||||[||||||||||||]PAM NA Intergenic NA NA

T3388 GAAGATGATCATGAGCGGTCCGG 1000
Oligo fwd CACCGAAGATGATCATGAGCGGTC
Oligo rev AAACGACCGCTCATGATCATCTTC
Chromosome start end strand MM target_seq PAM alignment distance position gene name gene id
GU305914.1 51662 51684 + 0 GAAGATGATCATGAGCGGTC CGG ||||||||[||||||||||||]PAM NA Intergenic NA NA

T3389 TGATCATGAGCGGTCCGGACAGG 1000
Oligo fwd CACCTGATCATGAGCGGTCCGGAC
Oligo rev AAACGTCCGGACCGCTCATGATCA
Chromosome start end strand MM target_seq PAM alignment distance position gene name gene id
GU305914.1 51667 51689 + 0 TGATCATGAGCGGTCCGGAC AGG ||||||||[||||||||||||]PAM NA Intergenic NA NA

T3390 GATCATGAGCGGTCCGGACAGGG 1000
Oligo fwd CACCGATCATGAGCGGTCCGGACA
Oligo rev AAACTGTCCGGACCGCTCATGATC
Chromosome start end strand MM target_seq PAM alignment distance position gene name gene id
GU305914.1 51668 51690 + 0 GATCATGAGCGGTCCGGACA GGG ||||||||[||||||||||||]PAM NA Intergenic NA NA

T3391 ATCATGAGCGGTCCGGACAGGGG 1000
Oligo fwd CACCATCATGAGCGGTCCGGACAG
Oligo rev AAACCTGTCCGGACCGCTCATGAT
Chromosome start end strand MM target_seq PAM alignment distance position gene name gene id
GU305914.1 51669 51691 + 0 ATCATGAGCGGTCCGGACAG GGG ||||||||[||||||||||||]PAM NA Intergenic NA NA

T3392 TCATGAGCGGTCCGGACAGGGGG 1000
Oligo fwd CACCTCATGAGCGGTCCGGACAGG
Oligo rev AAACCCTGTCCGGACCGCTCATGA
Chromosome start end strand MM target_seq PAM alignment distance position gene name gene id
GU305914.1 51670 51692 + 0 TCATGAGCGGTCCGGACAGG GGG ||||||||[||||||||||||]PAM NA Intergenic NA NA

T3393 ACAGGGGGCAGAAGTCGCGTCGG 1000
Oligo fwd CACCACAGGGGGCAGAAGTCGCGT
Oligo rev AAACACGCGACTTCTGCCCCCTGT
Chromosome start end strand MM target_seq PAM alignment distance position gene name gene id
GU305914.1 51685 51707 + 0 ACAGGGGGCAGAAGTCGCGT CGG ||||||||[||||||||||||]PAM NA Intergenic NA NA

T3394 GCAGAAGTCGCGTCGGATCCAGG 1000
Oligo fwd CACCGCAGAAGTCGCGTCGGATCC
Oligo rev AAACGGATCCGACGCGACTTCTGC
Chromosome start end strand MM target_seq PAM alignment distance position gene name gene id
GU305914.1 51692 51714 + 0 GCAGAAGTCGCGTCGGATCC AGG ||||||||[||||||||||||]PAM NA Intergenic NA NA

T3395 GCGTCGGATCCAGGAGAGCGCGG 1000
Oligo fwd CACCGCGTCGGATCCAGGAGAGCG
Oligo rev AAACCGCTCTCCTGGATCCGACGC
Chromosome start end strand MM target_seq PAM alignment distance position gene name gene id
GU305914.1 51701 51723 + 0 GCGTCGGATCCAGGAGAGCG CGG ||||||||[||||||||||||]PAM NA Intergenic NA NA

T3396 CGTCGGATCCAGGAGAGCGCGGG 1000
Oligo fwd CACCCGTCGGATCCAGGAGAGCGC
Oligo rev AAACGCGCTCTCCTGGATCCGACG
Chromosome start end strand MM target_seq PAM alignment distance position gene name gene id
GU305914.1 51702 51724 + 0 CGTCGGATCCAGGAGAGCGC GGG ||||||||[||||||||||||]PAM NA Intergenic NA NA

T3397 GGAGAGCGCGGGTTCCTTGATGG 1000
Oligo fwd CACCGGAGAGCGCGGGTTCCTTGA
Oligo rev AAACTCAAGGAACCCGCGCTCTCC
Chromosome start end strand MM target_seq PAM alignment distance position gene name gene id
GU305914.1 51713 51735 + 0 GGAGAGCGCGGGTTCCTTGA TGG ||||||||[||||||||||||]PAM NA Intergenic NA NA

T3398 GCGCGGGTTCCTTGATGGTCAGG 1000
Oligo fwd CACCGCGCGGGTTCCTTGATGGTC
Oligo rev AAACGACCATCAAGGAACCCGCGC
Chromosome start end strand MM target_seq PAM alignment distance position gene name gene id
GU305914.1 51718 51740 + 0 GCGCGGGTTCCTTGATGGTC AGG ||||||||[||||||||||||]PAM NA Intergenic NA NA

T3399 CTTGATGGTCAGGTGTTCGTAGG 1000
Oligo fwd CACCCTTGATGGTCAGGTGTTCGT
Oligo rev AAACACGAACACCTGACCATCAAG
Chromosome start end strand MM target_seq PAM alignment distance position gene name gene id
GU305914.1 51728 51750 + 0 CTTGATGGTCAGGTGTTCGT AGG ||||||||[||||||||||||]PAM NA Intergenic NA NA

T3400 TTCGTAGGAGCTATCGCAGACGG 1000
Oligo fwd CACCTTCGTAGGAGCTATCGCAGA
Oligo rev AAACTCTGCGATAGCTCCTACGAA
Chromosome start end strand MM target_seq PAM alignment distance position gene name gene id

GU305914.1 51743 51765 + 0 TTCGTAGGAGCTATCGCAGA CGG ||||||||[||||||||||||]PAM NA Intergenic NA NA

T3401 CAAAGTCCTCACAAACGTATAGG 1000
Oligo fwd CACCCAAAGTCCTCACAAACGTAT
Oligo rev AAACATACGTTTGTGAGGACTTTG
Chromosome start end strand MM target_seq PAM alignment distance position gene name gene id
GU305914.1 51821 51843 + 0 CAAAGTCCTCACAAACGTAT AGG ||||||||[||||||||||||]PAM NA Intergenic NA NA

T3402 CCTGTACCAGTTTCGCCGCCCGG 1000
Oligo fwd CACCCCTGTACCAGTTTCGCCGCC
Oligo rev AAACGGCGGCGAAACTGGTACAGG
Chromosome start end strand MM target_seq PAM alignment distance position gene name gene id
GU305914.1 51859 51881 + 0 CCTGTACCAGTTTCGCCGCC CGG ||||||||[||||||||||||]PAM NA Intergenic NA NA

T3403 TACCAGTTTCGCCGCCCGGATGG 1000
Oligo fwd CACCTACCAGTTTCGCCGCCCGGA
Oligo rev AAACTCCGGGCGGCGAAACTGGTA
Chromosome start end strand MM target_seq PAM alignment distance position gene name gene id
GU305914.1 51863 51885 + 0 TACCAGTTTCGCCGCCCGGA TGG ||||||||[||||||||||||]PAM NA Intergenic NA NA

T3404 CAGTTTCGCCGCCCGGATGGCGG 1000
Oligo fwd CACCCAGTTTCGCCGCCCGGATGG
Oligo rev AAACCCATCCGGGCGGCGAAACTG
Chromosome start end strand MM target_seq PAM alignment distance position gene name gene id
GU305914.1 51866 51888 + 0 CAGTTTCGCCGCCCGGATGG CGG ||||||||[||||||||||||]PAM NA Intergenic NA NA

T3405 GGACTCCCGATCCATCGCGTTGG 1000
Oligo fwd CACCGGACTCCCGATCCATCGCGT
Oligo rev AAACACGCGATGGATCGGGAGTCC
Chromosome start end strand MM target_seq PAM alignment distance position gene name gene id
GU305914.1 51887 51909 + 0 GGACTCCCGATCCATCGCGT TGG ||||||||[||||||||||||]PAM NA Intergenic NA NA

T3406 CGCGTTGGCGAACATCCTCTCGG 1000
Oligo fwd CACCCGCGTTGGCGAACATCCTCT
Oligo rev AAACAGAGGATGTTCGCCAACGCG
Chromosome start end strand MM target_seq PAM alignment distance position gene name gene id
GU305914.1 51902 51924 + 0 CGCGTTGGCGAACATCCTCT CGG ||||||||[||||||||||||]PAM NA Intergenic NA NA

T3407 GCGAACATCCTCTCGGTGTTCGG 1000
Oligo fwd CACCGCGAACATCCTCTCGGTGTT
Oligo rev AAACAACACCGAGAGGATGTTCGC
Chromosome start end strand MM target_seq PAM alignment distance position gene name gene id
GU305914.1 51909 51931 + 0 GCGAACATCCTCTCGGTGTT CGG ||||||||[||||||||||||]PAM NA Intergenic NA NA

T3408 GTTGTCGACACCGCCATCGCCGG 1000
Oligo fwd CACCGTTGTCGACACCGCCATCGC
Oligo rev AAACGCGATGGCGGTGTCGACAAC
Chromosome start end strand MM target_seq PAM alignment distance position gene name gene id
GU305914.1 51942 51964 + 0 GTTGTCGACACCGCCATCGC CGG ||||||||[||||||||||||]PAM NA Intergenic NA NA

T3409 GCTTCCTGACGCTGAGACTGCGG 1000
Oligo fwd CACCGCTTCCTGACGCTGAGACTG
Oligo rev AAACCAGTCTCAGCGTCAGGAAGC
Chromosome start end strand MM target_seq PAM alignment distance position gene name gene id
GU305914.1 51964 51986 + 0 GCTTCCTGACGCTGAGACTG CGG ||||||||[||||||||||||]PAM NA Intergenic NA NA

T3410 TGAGACTGCGGACGAGACTGAGG 1000
Oligo fwd CACCTGAGACTGCGGACGAGACTG
Oligo rev AAACCAGTCTCGTCCGCAGTCTCA
Chromosome start end strand MM target_seq PAM alignment distance position gene name gene id
GU305914.1 51976 51998 + 0 TGAGACTGCGGACGAGACTG AGG ||||||||[||||||||||||]PAM NA Intergenic NA NA

T3411 ACTGCGGACGAGACTGAGGTTGG 1000
Oligo fwd CACCACTGCGGACGAGACTGAGGT
Oligo rev AAACACCTCAGTCTCGTCCGCAGT
Chromosome start end strand MM target_seq PAM alignment distance position gene name gene id
GU305914.1 51980 52002 + 0 ACTGCGGACGAGACTGAGGT TGG ||||||||[||||||||||||]PAM NA Intergenic NA NA

T3412 CTGCGGACGAGACTGAGGTTGGG 1000
Oligo fwd CACCCTGCGGACGAGACTGAGGTT
Oligo rev AAACAACCTCAGTCTCGTCCGCAG
Chromosome start end strand MM target_seq PAM alignment distance position gene name gene id
GU305914.1 51981 52003 + 0 CTGCGGACGAGACTGAGGTT GGG ||||||||[||||||||||||]PAM NA Intergenic NA NA

T3413 TGCGGACGAGACTGAGGTTGGGG 1000
Oligo fwd CACCTGCGGACGAGACTGAGGTTG
Oligo rev AAACCAACCTCAGTCTCGTCCGCA
Chromosome start end strand MM target_seq PAM alignment distance position gene name gene id
GU305914.1 51982 52004 + 0 TGCGGACGAGACTGAGGTTG GGG ||||||||[||||||||||||]PAM NA Intergenic NA NA

T3414 CAAGACGCGTATGAGCTCGTTGG 1000
Oligo fwd CACCCAAGACGCGTATGAGCTCGT
Oligo rev AAACACGAGCTCATACGCGTCTTG
Chromosome start end strand MM target_seq PAM alignment distance position gene name gene id
GU305914.1 52086 52108 + 0 CAAGACGCGTATGAGCTCGT TGG ||||||||[||||||||||||]PAM NA Intergenic NA NA

T3415 GAGCTCGTTGGCAGTCTCCTCGG 1000
Oligo fwd CACCGAGCTCGTTGGCAGTCTCCT
Oligo rev AAACAGGAGACTGCCAACGAGCTC

| | | | | | | | | | | | |
|---|---|---|---|---|---|---|---|---|---|---|---|
| Chromosome | start | end | strand | MM | target_seq | PAM | alignment | distance | position | gene name | gene id |
| GU305914.1 | 52098 | 52120 | + | 0 | GAGCTCGTTGGCAGTCTCCT | CGG | \|\|\|\|\|\|\|\|[\|\|\|\|\|\|\|\|\|\|\|\|]PAM | NA | Intergenic | NA | NA |
| T3416 | TGGCAGTCTCCTCGGCGAAGCGG | 1000 | | | | | | | | | |
| Oligo fwd | CACCTGGCAGTCTCCTCGGCGAAG | | | | | | | | | | |
| Oligo rev | AAACCTTCGCCGAGGAGACTGCCA | | | | | | | | | | |
| Chromosome | start | end | strand | MM | target_seq | PAM | alignment | distance | position | gene name | gene id |
| GU305914.1 | 52106 | 52128 | + | 0 | TGGCAGTCTCCTCGGCGAAG | CGG | \|\|\|\|\|\|\|\|[\|\|\|\|\|\|\|\|\|\|\|\|]PAM | NA | Intergenic | NA | NA |
| T3417 | GCGGCGTTCGCGAGAACCCCAGG | 1000 | | | | | | | | | |
| Oligo fwd | CACCGCGGCGTTCGCGAGAACCCC | | | | | | | | | | |
| Oligo rev | AAACGGGGTTCTCGCGAACGCCGC | | | | | | | | | | |
| Chromosome | start | end | strand | MM | target_seq | PAM | alignment | distance | position | gene name | gene id |
| GU305914.1 | 52125 | 52147 | + | 0 | GCGGCGTTCGCGAGAACCCC | AGG | \|\|\|\|\|\|\|\|[\|\|\|\|\|\|\|\|\|\|\|\|]PAM | NA | Intergenic | NA | NA |
| T3418 | CGCGAGAACCCCAGGAGAAGCGG | 1000 | | | | | | | | | |
| Oligo fwd | CACCCGCGAGAACCCCAGGAGAAG | | | | | | | | | | |
| Oligo rev | AAACCTTCTCCTGGGGTTCTCGCG | | | | | | | | | | |
| Chromosome | start | end | strand | MM | target_seq | PAM | alignment | distance | position | gene name | gene id |
| GU305914.1 | 52133 | 52155 | + | 0 | CGCGAGAACCCCAGGAGAAG | CGG | \|\|\|\|\|\|\|\|[\|\|\|\|\|\|\|\|\|\|\|\|]PAM | NA | Intergenic | NA | NA |
| T3419 | GAGAACCCCAGGAGAAGCGGCGG | 1000 | | | | | | | | | |
| Oligo fwd | CACCGAGAACCCCAGGAGAAGCGG | | | | | | | | | | |
| Oligo rev | AAACCCGCTTCTCCTGGGGTTCTC | | | | | | | | | | |
| Chromosome | start | end | strand | MM | target_seq | PAM | alignment | distance | position | gene name | gene id |
| GU305914.1 | 52136 | 52158 | + | 0 | GAGAACCCCAGGAGAAGCGG | CGG | \|\|\|\|\|\|\|\|[\|\|\|\|\|\|\|\|\|\|\|\|]PAM | NA | Intergenic | NA | NA |
| T3420 | GTCGTCCGTAACGTCTCCAGCGG | 1000 | | | | | | | | | |
| Oligo fwd | CACCGTCGTCCGTAACGTCTCCAG | | | | | | | | | | |
| Oligo rev | AAACCTGGAGACGTTACGGACGAC | | | | | | | | | | |
| Chromosome | start | end | strand | MM | target_seq | PAM | alignment | distance | position | gene name | gene id |
| GU305914.1 | 52188 | 52210 | + | 0 | GTCGTCCGTAACGTCTCCAG | CGG | \|\|\|\|\|\|\|\|[\|\|\|\|\|\|\|\|\|\|\|\|]PAM | NA | Intergenic | NA | NA |
| T3421 | TCGTCCGTAACGTCTCCAGCGGG | 1000 | | | | | | | | | |
| Oligo fwd | CACCTCGTCCGTAACGTCTCCAGC | | | | | | | | | | |
| Oligo rev | AAACGCTGGAGACGTTACGGACGA | | | | | | | | | | |
| Chromosome | start | end | strand | MM | target_seq | PAM | alignment | distance | position | gene name | gene id |
| GU305914.1 | 52189 | 52211 | + | 0 | TCGTCCGTAACGTCTCCAGC | GGG | \|\|\|\|\|\|\|\|[\|\|\|\|\|\|\|\|\|\|\|\|]PAM | NA | Intergenic | NA | NA |
| T3422 | GTTCTGACTGTCCCGCCGCCTGG | 1000 | | | | | | | | | |
| Oligo fwd | CACCGTTCTGACTGTCCCGCCGCC | | | | | | | | | | |
| Oligo rev | AAACGGCGGCGGGACAGTCAGAAC | | | | | | | | | | |
| Chromosome | start | end | strand | MM | target_seq | PAM | alignment | distance | position | gene name | gene id |
| GU305914.1 | 52236 | 52258 | + | 0 | GTTCTGACTGTCCCGCCGCC | TGG | \|\|\|\|\|\|\|\|[\|\|\|\|\|\|\|\|\|\|\|\|]PAM | NA | Intergenic | NA | NA |
| T3423 | CCAATTAGAGAGAATCCAACCGG | 1000 | | | | | | | | | |
| Oligo fwd | CACCCCAATTAGAGAGAATCCAAC | | | | | | | | | | |
| Oligo rev | AAACGTTGGATTCTCTCTAATTGG | | | | | | | | | | |
| Chromosome | start | end | strand | MM | target_seq | PAM | alignment | distance | position | gene name | gene id |
| GU305914.1 | 52263 | 52285 | + | 0 | CCAATTAGAGAGAATCCAAC | CGG | \|\|\|\|\|\|\|\|[\|\|\|\|\|\|\|\|\|\|\|\|]PAM | NA | Intergenic | NA | NA |
| T3424 | TACGAAACCAAGCGCCCCAGAGG | 1000 | | | | | | | | | |
| Oligo fwd | CACCTACGAAACCAAGCGCCCCAG | | | | | | | | | | |
| Oligo rev | AAACCTGGGGCGCTTGGTTTCGTA | | | | | | | | | | |
| Chromosome | start | end | strand | MM | target_seq | PAM | alignment | distance | position | gene name | gene id |
| GU305914.1 | 52287 | 52309 | + | 0 | TACGAAACCAAGCGCCCCAG | AGG | \|\|\|\|\|\|\|\|[\|\|\|\|\|\|\|\|\|\|\|\|]PAM | NA | Intergenic | NA | NA |
| T3425 | CCAGAGGCGAAGAGCAGCGCTGG | 1000 | | | | | | | | | |
| Oligo fwd | CACCCCAGAGGCGAAGAGCAGCGC | | | | | | | | | | |
| Oligo rev | AAACGCGCTGCTCTTCGCCTCTGG | | | | | | | | | | |
| Chromosome | start | end | strand | MM | target_seq | PAM | alignment | distance | position | gene name | gene id |
| GU305914.1 | 52303 | 52325 | + | 0 | CCAGAGGCGAAGAGCAGCGC | TGG | \|\|\|\|\|\|\|\|[\|\|\|\|\|\|\|\|\|\|\|\|]PAM | NA | Intergenic | NA | NA |
| GU305914.1 | 70169 | 70191 | + | 3 | CCCGAGGCGACGAGCAGCAC | CGG | \|\|-\|\|\|\|\|[\|\|-\|\|\|\|\|\|\|-\|]PAM | NA | Intergenic | NA | NA |
| T3426 | TTCGCTTACAAACCCAAAGAAGG | 1000 | | | | | | | | | |
| Oligo fwd | CACCTTCGCTTACAAACCCAAAGA | | | | | | | | | | |
| Oligo rev | AAACTCTTTGGGTTTGTAAGCGAA | | | | | | | | | | |
| Chromosome | start | end | strand | MM | target_seq | PAM | alignment | distance | position | gene name | gene id |
| GU305914.1 | 52329 | 52351 | + | 0 | TTCGCTTACAAACCCAAAGA | AGG | \|\|\|\|\|\|\|\|[\|\|\|\|\|\|\|\|\|\|\|\|]PAM | NA | Intergenic | NA | NA |
| T3427 | TCGCTTACAAACCCAAAGAAGGG | 1000 | | | | | | | | | |
| Oligo fwd | CACCTCGCTTACAAACCCAAAGAA | | | | | | | | | | |
| Oligo rev | AAACTTCTTTGGGTTTGTAAGCGA | | | | | | | | | | |
| Chromosome | start | end | strand | MM | target_seq | PAM | alignment | distance | position | gene name | gene id |
| GU305914.1 | 52330 | 52352 | + | 0 | TCGCTTACAAACCCAAAGAA | GGG | \|\|\|\|\|\|\|\|[\|\|\|\|\|\|\|\|\|\|\|\|]PAM | NA | Intergenic | NA | NA |
| T3428 | TTACAAACCCAAAGAAGGGTCGG | 1000 | | | | | | | | | |
| Oligo fwd | CACCTTACAAACCCAAAGAAGGGT | | | | | | | | | | |
| Oligo rev | AAACACCCTTCTTTGGGTTTGTAA | | | | | | | | | | |
| Chromosome | start | end | strand | MM | target_seq | PAM | alignment | distance | position | gene name | gene id |
| GU305914.1 | 52334 | 52356 | + | 0 | TTACAAACCCAAAGAAGGGT | CGG | \|\|\|\|\|\|\|\|[\|\|\|\|\|\|\|\|\|\|\|\|]PAM | NA | Intergenic | NA | NA |
| T3429 | CCCTCCACAACTACAACCCGCGG | 1000 | | | | | | | | | |
| Oligo fwd | CACCCCCTCCACAACTACAACCCG | | | | | | | | | | |
| Oligo rev | AAACCGGGTTGTAGTTGTGGAGGG | | | | | | | | | | |
| Chromosome | start | end | strand | MM | target_seq | PAM | alignment | distance | position | gene name | gene id |
| GU305914.1 | 52378 | 52400 | + | 0 | CCCTCCACAACTACAACCCG | CGG | \|\|\|\|\|\|\|\|[\|\|\|\|\|\|\|\|\|\|\|\|]PAM | NA | Intergenic | NA | NA |
| T3430 | ATGTGTCCCCCGTCCGTCTCCGG | 1000 | | | | | | | | | |

<table>
<tr><td>Oligo fwd</td><td>CACCATGTGTCCCCCGTCCGTCTC</td><td></td><td></td><td></td><td></td><td></td><td></td><td></td><td></td><td></td><td></td></tr>
<tr><td>Oligo rev</td><td>AAACGAGACGGACGGGGGACACAT</td><td></td><td></td><td></td><td></td><td></td><td></td><td></td><td></td><td></td><td></td></tr>
<tr><td>Chromosome</td><td>start</td><td>end</td><td>strand</td><td>MM</td><td>target_seq</td><td>PAM</td><td>alignment</td><td>distance</td><td>position</td><td>gene name</td><td>gene id</td></tr>
<tr><td>GU305914.1</td><td>52422</td><td>52444</td><td>+</td><td>0</td><td>ATGTGTCCCCCGTCCGTCTC</td><td>CGG</td><td>||||||||[||||||||||||]PAM</td><td>NA</td><td>Intergenic</td><td>NA</td><td>NA</td></tr>
<tr><td></td><td></td><td></td><td></td><td></td><td></td><td></td><td></td><td></td><td></td><td></td><td></td></tr>
<tr><td>T3431</td><td>CCAATCAATCCGAGCCGCCACGG</td><td>1000</td><td></td><td></td><td></td><td></td><td></td><td></td><td></td><td></td><td></td></tr>
<tr><td>Oligo fwd</td><td>CACCCCAATCAATCCGAGCCGCCA</td><td></td><td></td><td></td><td></td><td></td><td></td><td></td><td></td><td></td><td></td></tr>
<tr><td>Oligo rev</td><td>AAACTGGCGGCTCGGATTGATTGG</td><td></td><td></td><td></td><td></td><td></td><td></td><td></td><td></td><td></td><td></td></tr>
<tr><td>Chromosome</td><td>start</td><td>end</td><td>strand</td><td>MM</td><td>target_seq</td><td>PAM</td><td>alignment</td><td>distance</td><td>position</td><td>gene name</td><td>gene id</td></tr>
<tr><td>GU305914.1</td><td>52458</td><td>52480</td><td>+</td><td>0</td><td>CCAATCAATCCGAGCCGCCA</td><td>CGG</td><td>||||||||[||||||||||||]PAM</td><td>NA</td><td>Intergenic</td><td>NA</td><td>NA</td></tr>
<tr><td></td><td></td><td></td><td></td><td></td><td></td><td></td><td></td><td></td><td></td><td></td><td></td></tr>
<tr><td>T3432</td><td>CAATCCGAGCCGCCACGGTCAGG</td><td>1000</td><td></td><td></td><td></td><td></td><td></td><td></td><td></td><td></td><td></td></tr>
<tr><td>Oligo fwd</td><td>CACCCAATCCGAGCCGCCACGGTC</td><td></td><td></td><td></td><td></td><td></td><td></td><td></td><td></td><td></td><td></td></tr>
<tr><td>Oligo rev</td><td>AAACGACCGTGGCGGCTCGGATTG</td><td></td><td></td><td></td><td></td><td></td><td></td><td></td><td></td><td></td><td></td></tr>
<tr><td>Chromosome</td><td>start</td><td>end</td><td>strand</td><td>MM</td><td>target_seq</td><td>PAM</td><td>alignment</td><td>distance</td><td>position</td><td>gene name</td><td>gene id</td></tr>
<tr><td>GU305914.1</td><td>52463</td><td>52485</td><td>+</td><td>0</td><td>CAATCCGAGCCGCCACGGTC</td><td>AGG</td><td>||||||||[||||||||||||]PAM</td><td>NA</td><td>Intergenic</td><td>NA</td><td>NA</td></tr>
<tr><td></td><td></td><td></td><td></td><td></td><td></td><td></td><td></td><td></td><td></td><td></td><td></td></tr>
<tr><td>T3433</td><td>AATCCGAGCCGCCACGGTCAGGG</td><td>1000</td><td></td><td></td><td></td><td></td><td></td><td></td><td></td><td></td><td></td></tr>
<tr><td>Oligo fwd</td><td>CACCAATCCGAGCCGCCACGGTCA</td><td></td><td></td><td></td><td></td><td></td><td></td><td></td><td></td><td></td><td></td></tr>
<tr><td>Oligo rev</td><td>AAACTGACCGTGGCGGCTCGGATT</td><td></td><td></td><td></td><td></td><td></td><td></td><td></td><td></td><td></td><td></td></tr>
<tr><td>Chromosome</td><td>start</td><td>end</td><td>strand</td><td>MM</td><td>target_seq</td><td>PAM</td><td>alignment</td><td>distance</td><td>position</td><td>gene name</td><td>gene id</td></tr>
<tr><td>GU305914.1</td><td>52464</td><td>52486</td><td>+</td><td>0</td><td>AATCCGAGCCGCCACGGTCA</td><td>GGG</td><td>||||||||[||||||||||||]PAM</td><td>NA</td><td>Intergenic</td><td>NA</td><td>NA</td></tr>
<tr><td></td><td></td><td></td><td></td><td></td><td></td><td></td><td></td><td></td><td></td><td></td><td></td></tr>
<tr><td>T3434</td><td>TCAGGGTCAGAAGTTGTGCACGG</td><td>1000</td><td></td><td></td><td></td><td></td><td></td><td></td><td></td><td></td><td></td></tr>
<tr><td>Oligo fwd</td><td>CACCTCAGGGTCAGAAGTTGTGCA</td><td></td><td></td><td></td><td></td><td></td><td></td><td></td><td></td><td></td><td></td></tr>
<tr><td>Oligo rev</td><td>AAACTGCACAACTTCTGACCCTGA</td><td></td><td></td><td></td><td></td><td></td><td></td><td></td><td></td><td></td><td></td></tr>
<tr><td>Chromosome</td><td>start</td><td>end</td><td>strand</td><td>MM</td><td>target_seq</td><td>PAM</td><td>alignment</td><td>distance</td><td>position</td><td>gene name</td><td>gene id</td></tr>
<tr><td>GU305914.1</td><td>52481</td><td>52503</td><td>+</td><td>0</td><td>TCAGGGTCAGAAGTTGTGCA</td><td>CGG</td><td>||||||||[||||||||||||]PAM</td><td>NA</td><td>Intergenic</td><td>NA</td><td>NA</td></tr>
<tr><td></td><td></td><td></td><td></td><td></td><td></td><td></td><td></td><td></td><td></td><td></td><td></td></tr>
<tr><td>T3435</td><td>ATCGTAGTCGTCATCTTCATCGG</td><td>1000</td><td></td><td></td><td></td><td></td><td></td><td></td><td></td><td></td><td></td></tr>
<tr><td>Oligo fwd</td><td>CACCATCGTAGTCGTCATCTTCAT</td><td></td><td></td><td></td><td></td><td></td><td></td><td></td><td></td><td></td><td></td></tr>
<tr><td>Oligo rev</td><td>AAACATGAAGATGACGACTACGAT</td><td></td><td></td><td></td><td></td><td></td><td></td><td></td><td></td><td></td><td></td></tr>
<tr><td>Chromosome</td><td>start</td><td>end</td><td>strand</td><td>MM</td><td>target_seq</td><td>PAM</td><td>alignment</td><td>distance</td><td>position</td><td>gene name</td><td>gene id</td></tr>
<tr><td>GU305914.1</td><td>52535</td><td>52557</td><td>+</td><td>0</td><td>ATCGTAGTCGTCATCTTCAT</td><td>CGG</td><td>||||||||[||||||||||||]PAM</td><td>NA</td><td>Intergenic</td><td>NA</td><td>NA</td></tr>
<tr><td></td><td></td><td></td><td></td><td></td><td></td><td></td><td></td><td></td><td></td><td></td><td></td></tr>
<tr><td>T3436</td><td>GACTGCTGTAGTTGAAACCGAGG</td><td>1000</td><td></td><td></td><td></td><td></td><td></td><td></td><td></td><td></td><td></td></tr>
<tr><td>Oligo fwd</td><td>CACCGACTGCTGTAGTTGAAACCG</td><td></td><td></td><td></td><td></td><td></td><td></td><td></td><td></td><td></td><td></td></tr>
<tr><td>Oligo rev</td><td>AAACCGGTTTCAACTACAGCAGTC</td><td></td><td></td><td></td><td></td><td></td><td></td><td></td><td></td><td></td><td></td></tr>
<tr><td>Chromosome</td><td>start</td><td>end</td><td>strand</td><td>MM</td><td>target_seq</td><td>PAM</td><td>alignment</td><td>distance</td><td>position</td><td>gene name</td><td>gene id</td></tr>
<tr><td>GU305914.1</td><td>52557</td><td>52579</td><td>+</td><td>0</td><td>GACTGCTGTAGTTGAAACCG</td><td>AGG</td><td>||||||||[||||||||||||]PAM</td><td>NA</td><td>Intergenic</td><td>NA</td><td>NA</td></tr>
<tr><td></td><td></td><td></td><td></td><td></td><td></td><td></td><td></td><td></td><td></td><td></td><td></td></tr>
<tr><td>T3437</td><td>CTGTAGTTGAAACCGAGGCAAGG</td><td>1000</td><td></td><td></td><td></td><td></td><td></td><td></td><td></td><td></td><td></td></tr>
<tr><td>Oligo fwd</td><td>CACCCTGTAGTTGAAACCGAGGCA</td><td></td><td></td><td></td><td></td><td></td><td></td><td></td><td></td><td></td><td></td></tr>
<tr><td>Oligo rev</td><td>AAACTGCCTCGGTTTCAACTACAG</td><td></td><td></td><td></td><td></td><td></td><td></td><td></td><td></td><td></td><td></td></tr>
<tr><td>Chromosome</td><td>start</td><td>end</td><td>strand</td><td>MM</td><td>target_seq</td><td>PAM</td><td>alignment</td><td>distance</td><td>position</td><td>gene name</td><td>gene id</td></tr>
<tr><td>GU305914.1</td><td>52562</td><td>52584</td><td>+</td><td>0</td><td>CTGTAGTTGAAACCGAGGCA</td><td>AGG</td><td>||||||||[||||||||||||]PAM</td><td>NA</td><td>Intergenic</td><td>NA</td><td>NA</td></tr>
<tr><td></td><td></td><td></td><td></td><td></td><td></td><td></td><td></td><td></td><td></td><td></td><td></td></tr>
<tr><td>T3438</td><td>TGTAGTTGAAACCGAGGCAAGGG</td><td>1000</td><td></td><td></td><td></td><td></td><td></td><td></td><td></td><td></td><td></td></tr>
<tr><td>Oligo fwd</td><td>CACCTGTAGTTGAAACCGAGGCAA</td><td></td><td></td><td></td><td></td><td></td><td></td><td></td><td></td><td></td><td></td></tr>
<tr><td>Oligo rev</td><td>AAACTTGCCTCGGTTTCAACTACA</td><td></td><td></td><td></td><td></td><td></td><td></td><td></td><td></td><td></td><td></td></tr>
<tr><td>Chromosome</td><td>start</td><td>end</td><td>strand</td><td>MM</td><td>target_seq</td><td>PAM</td><td>alignment</td><td>distance</td><td>position</td><td>gene name</td><td>gene id</td></tr>
<tr><td>GU305914.1</td><td>52563</td><td>52585</td><td>+</td><td>0</td><td>TGTAGTTGAAACCGAGGCAA</td><td>GGG</td><td>||||||||[||||||||||||]PAM</td><td>NA</td><td>Intergenic</td><td>NA</td><td>NA</td></tr>
<tr><td></td><td></td><td></td><td></td><td></td><td></td><td></td><td></td><td></td><td></td><td></td><td></td></tr>
<tr><td>T3439</td><td>AGGCAAGGGATACTGACCCCCGG</td><td>1000</td><td></td><td></td><td></td><td></td><td></td><td></td><td></td><td></td><td></td></tr>
<tr><td>Oligo fwd</td><td>CACCAGGCAAGGGATACTGACCCC</td><td></td><td></td><td></td><td></td><td></td><td></td><td></td><td></td><td></td><td></td></tr>
<tr><td>Oligo rev</td><td>AAACGGGGTCAGTATCCCTTGCCT</td><td></td><td></td><td></td><td></td><td></td><td></td><td></td><td></td><td></td><td></td></tr>
<tr><td>Chromosome</td><td>start</td><td>end</td><td>strand</td><td>MM</td><td>target_seq</td><td>PAM</td><td>alignment</td><td>distance</td><td>position</td><td>gene name</td><td>gene id</td></tr>
<tr><td>GU305914.1</td><td>52577</td><td>52599</td><td>+</td><td>0</td><td>AGGCAAGGGATACTGACCCC</td><td>CGG</td><td>||||||||[||||||||||||]PAM</td><td>NA</td><td>Intergenic</td><td>NA</td><td>NA</td></tr>
<tr><td></td><td></td><td></td><td></td><td></td><td></td><td></td><td></td><td></td><td></td><td></td><td></td></tr>
<tr><td>T3440</td><td>CAAGGGATACTGACCCCCGGCGG</td><td>1000</td><td></td><td></td><td></td><td></td><td></td><td></td><td></td><td></td><td></td></tr>
<tr><td>Oligo fwd</td><td>CACCCAAGGGATACTGACCCCCGG</td><td></td><td></td><td></td><td></td><td></td><td></td><td></td><td></td><td></td><td></td></tr>
<tr><td>Oligo rev</td><td>AAACCCGGGGGTCAGTATCCCTTG</td><td></td><td></td><td></td><td></td><td></td><td></td><td></td><td></td><td></td><td></td></tr>
<tr><td>Chromosome</td><td>start</td><td>end</td><td>strand</td><td>MM</td><td>target_seq</td><td>PAM</td><td>alignment</td><td>distance</td><td>position</td><td>gene name</td><td>gene id</td></tr>
<tr><td>GU305914.1</td><td>52580</td><td>52602</td><td>+</td><td>0</td><td>CAAGGGATACTGACCCCCGG</td><td>CGG</td><td>||||||||[||||||||||||]PAM</td><td>NA</td><td>Intergenic</td><td>NA</td><td>NA</td></tr>
<tr><td></td><td></td><td></td><td></td><td></td><td></td><td></td><td></td><td></td><td></td><td></td><td></td></tr>
<tr><td>T3441</td><td>CATCTGCATCTCCAGAGCATCGG</td><td>1000</td><td></td><td></td><td></td><td></td><td></td><td></td><td></td><td></td><td></td></tr>
<tr><td>Oligo fwd</td><td>CACCCATCTGCATCTCCAGAGCAT</td><td></td><td></td><td></td><td></td><td></td><td></td><td></td><td></td><td></td><td></td></tr>
<tr><td>Oligo rev</td><td>AAACATGCTCTGGAGATGCAGATG</td><td></td><td></td><td></td><td></td><td></td><td></td><td></td><td></td><td></td><td></td></tr>
<tr><td>Chromosome</td><td>start</td><td>end</td><td>strand</td><td>MM</td><td>target_seq</td><td>PAM</td><td>alignment</td><td>distance</td><td>position</td><td>gene name</td><td>gene id</td></tr>
<tr><td>GU305914.1</td><td>52631</td><td>52653</td><td>+</td><td>0</td><td>CATCTGCATCTCCAGAGCAT</td><td>CGG</td><td>||||||||[||||||||||||]PAM</td><td>NA</td><td>Intergenic</td><td>NA</td><td>NA</td></tr>
<tr><td></td><td></td><td></td><td></td><td></td><td></td><td></td><td></td><td></td><td></td><td></td><td></td></tr>
<tr><td>T3442</td><td>CATCTCCAGAGCATCGGCGATGG</td><td>1000</td><td></td><td></td><td></td><td></td><td></td><td></td><td></td><td></td><td></td></tr>
<tr><td>Oligo fwd</td><td>CACCCATCTCCAGAGCATCGGCGA</td><td></td><td></td><td></td><td></td><td></td><td></td><td></td><td></td><td></td><td></td></tr>
<tr><td>Oligo rev</td><td>AAACTCGCCGATGCTCTGGAGATG</td><td></td><td></td><td></td><td></td><td></td><td></td><td></td><td></td><td></td><td></td></tr>
<tr><td>Chromosome</td><td>start</td><td>end</td><td>strand</td><td>MM</td><td>target_seq</td><td>PAM</td><td>alignment</td><td>distance</td><td>position</td><td>gene name</td><td>gene id</td></tr>
<tr><td>GU305914.1</td><td>52637</td><td>52659</td><td>+</td><td>0</td><td>CATCTCCAGAGCATCGGCGA</td><td>TGG</td><td>||||||||[||||||||||||]PAM</td><td>NA</td><td>Intergenic</td><td>NA</td><td>NA</td></tr>
<tr><td>GU305914.1</td><td>71184</td><td>71206</td><td>-</td><td>3</td><td>CTTCTCCAGATCTTCGGCGA</td><td>AGG</td><td>|-||||||[||-|-|||||||]PAM</td><td>NA</td><td>Intergenic</td><td>NA</td><td>NA</td></tr>
<tr><td></td><td></td><td></td><td></td><td></td><td></td><td></td><td></td><td></td><td></td><td></td><td></td></tr>
<tr><td>T3443</td><td>TCGCCTTCGATTTTTTACTCAGG</td><td>1000</td><td></td><td></td><td></td><td></td><td></td><td></td><td></td><td></td><td></td></tr>
<tr><td>Oligo fwd</td><td>CACCTCGCCTTCGATTTTTTACTC</td><td></td><td></td><td></td><td></td><td></td><td></td><td></td><td></td><td></td><td></td></tr>
<tr><td>Oligo rev</td><td>AAACGAGTAAAAAATCGAAGGCGA</td><td></td><td></td><td></td><td></td><td></td><td></td><td></td><td></td><td></td><td></td></tr>
<tr><td>Chromosome</td><td>start</td><td>end</td><td>strand</td><td>MM</td><td>target_seq</td><td>PAM</td><td>alignment</td><td>distance</td><td>position</td><td>gene name</td><td>gene id</td></tr>
<tr><td>GU305914.1</td><td>52672</td><td>52694</td><td>+</td><td>0</td><td>TCGCCTTCGATTTTTTACTC</td><td>AGG</td><td>||||||||[||||||||||||]PAM</td><td>NA</td><td>Intergenic</td><td>NA</td><td>NA</td></tr>
<tr><td></td><td></td><td></td><td></td><td></td><td></td><td></td><td></td><td></td><td></td><td></td><td></td></tr>
<tr><td>T3444</td><td>GTCGCCACCGACGAACACCTTGG</td><td>1000</td><td></td><td></td><td></td><td></td><td></td><td></td><td></td><td></td><td></td></tr>
<tr><td>Oligo fwd</td><td>CACCGTCGCCACCGACGAACACCT</td><td></td><td></td><td></td><td></td><td></td><td></td><td></td><td></td><td></td><td></td></tr>
<tr><td>Oligo rev</td><td>AAACAGGTGTTCGTCGGTGGCGAC</td><td></td><td></td><td></td><td></td><td></td><td></td><td></td><td></td><td></td><td></td></tr>
<tr><td>Chromosome</td><td>start</td><td>end</td><td>strand</td><td>MM</td><td>target_seq</td><td>PAM</td><td>alignment</td><td>distance</td><td>position</td><td>gene name</td><td>gene id</td></tr>
<tr><td>GU305914.1</td><td>52694</td><td>52716</td><td>+</td><td>0</td><td>GTCGCCACCGACGAACACCT</td><td>TGG</td><td>||||||||[||||||||||||]PAM</td><td>NA</td><td>Intergenic</td><td>NA</td><td>NA</td></tr>
</table>

<table>
<tr><td>T3445</td><td>ACGAACACCTTGGCCGCCGCTGG</td><td>1000</td><td></td><td></td><td></td><td></td><td></td><td></td><td></td><td></td><td></td></tr>
<tr><td>Oligo fwd</td><td>CACCACGAACACCTTGGCCGCCGC</td><td></td><td></td><td></td><td></td><td></td><td></td><td></td><td></td><td></td><td></td></tr>
<tr><td>Oligo rev</td><td>AAACGCGGCGGCCAAGGTGTTCGT</td><td></td><td></td><td></td><td></td><td></td><td></td><td></td><td></td><td></td><td></td></tr>
<tr><td>Chromosome</td><td>start</td><td>end</td><td>strand</td><td>MM</td><td>target_seq</td><td>PAM</td><td>alignment</td><td>distance</td><td>position</td><td>gene name</td><td>gene id</td></tr>
<tr><td>GU305914.1</td><td>52704</td><td>52726</td><td>+</td><td>0</td><td>ACGAACACCTTGGCCGCCGC</td><td>TGG</td><td>||||||||[||||||||||||]PAM</td><td>NA</td><td>Intergenic</td><td>NA</td><td>NA</td></tr>
<tr><td></td><td></td><td></td><td></td><td></td><td></td><td></td><td></td><td></td><td></td><td></td><td></td></tr>
<tr><td>T3446</td><td>AACACCTTGGCCGCCGCTGGCGG</td><td>1000</td><td></td><td></td><td></td><td></td><td></td><td></td><td></td><td></td><td></td></tr>
<tr><td>Oligo fwd</td><td>CACCAACACCTTGGCCGCCGCTGG</td><td></td><td></td><td></td><td></td><td></td><td></td><td></td><td></td><td></td><td></td></tr>
<tr><td>Oligo rev</td><td>AAACCCAGCGGCGGCCAAGGTGTT</td><td></td><td></td><td></td><td></td><td></td><td></td><td></td><td></td><td></td><td></td></tr>
<tr><td>Chromosome</td><td>start</td><td>end</td><td>strand</td><td>MM</td><td>target_seq</td><td>PAM</td><td>alignment</td><td>distance</td><td>position</td><td>gene name</td><td>gene id</td></tr>
<tr><td>GU305914.1</td><td>52707</td><td>52729</td><td>+</td><td>0</td><td>AACACCTTGGCCGCCGCTGG</td><td>CGG</td><td>||||||||[||||||||||||]PAM</td><td>NA</td><td>Intergenic</td><td>NA</td><td>NA</td></tr>
<tr><td>GU305914.1</td><td>45160</td><td>45182</td><td>-</td><td>4</td><td>GTCGCCTTCGCCGCCGCTGG</td><td>AGG</td><td>--|-||||[-|||||||||||]PAM</td><td>NA</td><td>Intergenic</td><td>NA</td><td>NA</td></tr>
<tr><td></td><td></td><td></td><td></td><td></td><td></td><td></td><td></td><td></td><td></td><td></td><td></td></tr>
<tr><td>T3447</td><td>ACCTTGGCCGCCGCTGGCGGTGG</td><td>1000</td><td></td><td></td><td></td><td></td><td></td><td></td><td></td><td></td><td></td></tr>
<tr><td>Oligo fwd</td><td>CACCACCTTGGCCGCCGCTGGCGG</td><td></td><td></td><td></td><td></td><td></td><td></td><td></td><td></td><td></td><td></td></tr>
<tr><td>Oligo rev</td><td>AAACCCGCCAGCGGCGGCCAAGGT</td><td></td><td></td><td></td><td></td><td></td><td></td><td></td><td></td><td></td><td></td></tr>
<tr><td>Chromosome</td><td>start</td><td>end</td><td>strand</td><td>MM</td><td>target_seq</td><td>PAM</td><td>alignment</td><td>distance</td><td>position</td><td>gene name</td><td>gene id</td></tr>
<tr><td>GU305914.1</td><td>52710</td><td>52732</td><td>+</td><td>0</td><td>ACCTTGGCCGCCGCTGGCGG</td><td>TGG</td><td>||||||||[||||||||||||]PAM</td><td>NA</td><td>Intergenic</td><td>NA</td><td>NA</td></tr>
<tr><td></td><td></td><td></td><td></td><td></td><td></td><td></td><td></td><td></td><td></td><td></td><td></td></tr>
<tr><td>T3448</td><td>TTGGCCGCCGCTGGCGGTGGCGG</td><td>1000</td><td></td><td></td><td></td><td></td><td></td><td></td><td></td><td></td><td></td></tr>
<tr><td>Oligo fwd</td><td>CACCTTGGCCGCCGCTGGCGGTGG</td><td></td><td></td><td></td><td></td><td></td><td></td><td></td><td></td><td></td><td></td></tr>
<tr><td>Oligo rev</td><td>AAACCCACCGCCAGCGGCGGCCAA</td><td></td><td></td><td></td><td></td><td></td><td></td><td></td><td></td><td></td><td></td></tr>
<tr><td>Chromosome</td><td>start</td><td>end</td><td>strand</td><td>MM</td><td>target_seq</td><td>PAM</td><td>alignment</td><td>distance</td><td>position</td><td>gene name</td><td>gene id</td></tr>
<tr><td>GU305914.1</td><td>52713</td><td>52735</td><td>+</td><td>0</td><td>TTGGCCGCCGCTGGCGGTGG</td><td>CGG</td><td>||||||||[||||||||||||]PAM</td><td>NA</td><td>Intergenic</td><td>NA</td><td>NA</td></tr>
<tr><td></td><td></td><td></td><td></td><td></td><td></td><td></td><td></td><td></td><td></td><td></td><td></td></tr>
<tr><td>T3449</td><td>GCCGCCGCTGGCGGTGGCGGTGG</td><td>1000</td><td></td><td></td><td></td><td></td><td></td><td></td><td></td><td></td><td></td></tr>
<tr><td>Oligo fwd</td><td>CACCGCCGCCGCTGGCGGTGGCGG</td><td></td><td></td><td></td><td></td><td></td><td></td><td></td><td></td><td></td><td></td></tr>
<tr><td>Oligo rev</td><td>AAACCCGCCACCGCCAGCGGCGGC</td><td></td><td></td><td></td><td></td><td></td><td></td><td></td><td></td><td></td><td></td></tr>
<tr><td>Chromosome</td><td>start</td><td>end</td><td>strand</td><td>MM</td><td>target_seq</td><td>PAM</td><td>alignment</td><td>distance</td><td>position</td><td>gene name</td><td>gene id</td></tr>
<tr><td>GU305914.1</td><td>52716</td><td>52738</td><td>+</td><td>0</td><td>GCCGCCGCTGGCGGTGGCGG</td><td>TGG</td><td>||||||||[||||||||||||]PAM</td><td>NA</td><td>Intergenic</td><td>NA</td><td>NA</td></tr>
<tr><td>GU305914.1</td><td>74710</td><td>74732</td><td>-</td><td>4</td><td>GACCCGGGTGGCGGTGGCGG</td><td>CGG</td><td>|-|-|-|-[||||||||||||]PAM</td><td>NA</td><td>Intergenic</td><td>NA</td><td>NA</td></tr>
<tr><td>GU305914.1</td><td>154395</td><td>154417</td><td>+</td><td>4</td><td>ATCGCCCCGGGCGGTGGCGG</td><td>CGG</td><td>--||||-|[-|||||||||||]PAM</td><td>NA</td><td>Intergenic</td><td>NA</td><td>NA</td></tr>
<tr><td>GU305914.1</td><td>114661</td><td>114683</td><td>-</td><td>4</td><td>GACGGCGGTGGTGGTGGCGG</td><td>CGG</td><td>|-||-||-[|||-||||||||]PAM</td><td>NA</td><td>Intergenic</td><td>NA</td><td>NA</td></tr>
<tr><td>GU305914.1</td><td>29997</td><td>30019</td><td>+</td><td>4</td><td>ACCGGTGCTGGCGGCGGCGG</td><td>CGG</td><td>-|||--||[||||||-|||||]PAM</td><td>NA</td><td>Intergenic</td><td>NA</td><td>NA</td></tr>
<tr><td>GU305914.1</td><td>62672</td><td>62694</td><td>-</td><td>3</td><td>GCTGCCGCCGGCGGGGGCGG</td><td>CGG</td><td>||-|||||[-|||||-|||||]PAM</td><td>NA</td><td>Intergenic</td><td>NA</td><td>NA</td></tr>
<tr><td>GU305914.1</td><td>48159</td><td>48181</td><td>+</td><td>4</td><td>GGCGGCGGTGGCGGCGGCGG</td><td>TGG</td><td>|-||-||-[||||||-|||||]PAM</td><td>NA</td><td>Intergenic</td><td>NA</td><td>NA</td></tr>
<tr><td>GU305914.1</td><td>58129</td><td>58151</td><td>-</td><td>4</td><td>GCGGGCGGTGGCGGCGGCGG</td><td>CGG</td><td>||-|-||-[||||||-|||||]PAM</td><td>NA</td><td>Intergenic</td><td>NA</td><td>NA</td></tr>
<tr><td></td><td></td><td></td><td></td><td></td><td></td><td></td><td></td><td></td><td></td><td></td><td></td></tr>
<tr><td>T3450</td><td>GCCGCTGGCGGTGGCGGTGGCGG</td><td>1000</td><td></td><td></td><td></td><td></td><td></td><td></td><td></td><td></td><td></td></tr>
<tr><td>Oligo fwd</td><td>CACCGCCGCTGGCGGTGGCGGTGG</td><td></td><td></td><td></td><td></td><td></td><td></td><td></td><td></td><td></td><td></td></tr>
<tr><td>Oligo rev</td><td>AAACCCACCGCCACCGCCAGCGGC</td><td></td><td></td><td></td><td></td><td></td><td></td><td></td><td></td><td></td><td></td></tr>
<tr><td>Chromosome</td><td>start</td><td>end</td><td>strand</td><td>MM</td><td>target_seq</td><td>PAM</td><td>alignment</td><td>distance</td><td>position</td><td>gene name</td><td>gene id</td></tr>
<tr><td>GU305914.1</td><td>52719</td><td>52741</td><td>+</td><td>0</td><td>GCCGCTGGCGGTGGCGGTGG</td><td>CGG</td><td>||||||||[||||||||||||]PAM</td><td>NA</td><td>Intergenic</td><td>NA</td><td>NA</td></tr>
<tr><td>GU305914.1</td><td>74673</td><td>74695</td><td>+</td><td>4</td><td>GGTGCCGTCGGTGGCGGTGG</td><td>CGG</td><td>|--||-|-[||||||||||||]PAM</td><td>NA</td><td>Intergenic</td><td>NA</td><td>NA</td></tr>
<tr><td>GU305914.1</td><td>48150</td><td>48172</td><td>+</td><td>4</td><td>ACCTGTGGCGGCGGCGGTGG</td><td>CGG</td><td>-||--|||[|||-||||||||]PAM</td><td>NA</td><td>Intergenic</td><td>NA</td><td>NA</td></tr>
<tr><td>GU305914.1</td><td>62437</td><td>62459</td><td>+</td><td>4</td><td>GGCGGTGGCGGCGGGGGTGG</td><td>TGG</td><td>|-||-|||[|||-||-|||||]PAM</td><td>NA</td><td>Intergenic</td><td>NA</td><td>NA</td></tr>
<tr><td>GU305914.1</td><td>225498</td><td>225520</td><td>+</td><td>4</td><td>GCCGAGGGCGGCGGTGGTGG</td><td>TGG</td><td>||||--||[|||-||-|||||]PAM</td><td>NA</td><td>Intergenic</td><td>NA</td><td>NA</td></tr>
<tr><td>GU305914.1</td><td>154398</td><td>154420</td><td>+</td><td>3</td><td>GCCCCGGGCGGTGGCGGCGG</td><td>CGG</td><td>|||-|-||[|||||||||-||]PAM</td><td>NA</td><td>Intergenic</td><td>NA</td><td>NA</td></tr>
<tr><td>GU305914.1</td><td>48156</td><td>48178</td><td>+</td><td>4</td><td>GGCGGCGGCGGTGGCGGCGG</td><td>CGG</td><td>|-||--||[|||||||||-||]PAM</td><td>NA</td><td>Intergenic</td><td>NA</td><td>NA</td></tr>
<tr><td>GU305914.1</td><td>114658</td><td>114680</td><td>-</td><td>4</td><td>GGCGGTGGTGGTGGCGGCGG</td><td>CGG</td><td>|-||-|||[-||||||||-||]PAM</td><td>NA</td><td>Intergenic</td><td>NA</td><td>NA</td></tr>
<tr><td>GU305914.1</td><td>30000</td><td>30022</td><td>+</td><td>4</td><td>GGTGCTGGCGGCGGCGGCGG</td><td>CGG</td><td>|--|||||[|||-|||||-||]PAM</td><td>NA</td><td>Intergenic</td><td>NA</td><td>NA</td></tr>
<tr><td>GU305914.1</td><td>154404</td><td>154426</td><td>+</td><td>4</td><td>GGCGGTGGCGGCGGCGGCGG</td><td>CGG</td><td>|-||-|||[|||-|||||-||]PAM</td><td>NA</td><td>Intergenic</td><td>NA</td><td>NA</td></tr>
<tr><td>GU305914.1</td><td>58126</td><td>58148</td><td>-</td><td>4</td><td>GGCGGTGGCGGCGGCGGCGG</td><td>CGG</td><td>|-||-|||[|||-|||||-||]PAM</td><td>NA</td><td>Intergenic</td><td>NA</td><td>NA</td></tr>
<tr><td>GU305914.1</td><td>36158</td><td>36180</td><td>+</td><td>3</td><td>GCCGCTGCCGGCGGCGGTTG</td><td>TGG</td><td>|||||||-[|||-||||||-|]PAM</td><td>NA</td><td>Intergenic</td><td>NA</td><td>NA</td></tr>
<tr><td>GU305914.1</td><td>37222</td><td>37244</td><td>-</td><td>4</td><td>GCCGCTCTCGGTGACGGCGG</td><td>CGG</td><td>||||||--[|||||-|||-||]PAM</td><td>NA</td><td>Intergenic</td><td>NA</td><td>NA</td></tr>
<tr><td></td><td></td><td></td><td></td><td></td><td></td><td></td><td></td><td></td><td></td><td></td><td></td></tr>
<tr><td>T3451</td><td>GGTGGCGGATGCGACTGTTGAGG</td><td>1000</td><td></td><td></td><td></td><td></td><td></td><td></td><td></td><td></td><td></td></tr>
<tr><td>Oligo fwd</td><td>CACCGGTGGCGGATGCGACTGTTG</td><td></td><td></td><td></td><td></td><td></td><td></td><td></td><td></td><td></td><td></td></tr>
<tr><td>Oligo rev</td><td>AAACCAACAGTCGCATCCGCCACC</td><td></td><td></td><td></td><td></td><td></td><td></td><td></td><td></td><td></td><td></td></tr>
<tr><td>Chromosome</td><td>start</td><td>end</td><td>strand</td><td>MM</td><td>target_seq</td><td>PAM</td><td>alignment</td><td>distance</td><td>position</td><td>gene name</td><td>gene id</td></tr>
<tr><td>GU305914.1</td><td>52734</td><td>52756</td><td>+</td><td>0</td><td>GGTGGCGGATGCGACTGTTG</td><td>AGG</td><td>||||||||[||||||||||||]PAM</td><td>NA</td><td>Intergenic</td><td>NA</td><td>NA</td></tr>
<tr><td></td><td></td><td></td><td></td><td></td><td></td><td></td><td></td><td></td><td></td><td></td><td></td></tr>
<tr><td>T3452</td><td>TGTTGAGGTAGACCCATCTCTGG</td><td>1000</td><td></td><td></td><td></td><td></td><td></td><td></td><td></td><td></td><td></td></tr>
<tr><td>Oligo fwd</td><td>CACCTGTTGAGGTAGACCCATCTC</td><td></td><td></td><td></td><td></td><td></td><td></td><td></td><td></td><td></td><td></td></tr>
<tr><td>Oligo rev</td><td>AAACGAGATGGGTCTACCTCAACA</td><td></td><td></td><td></td><td></td><td></td><td></td><td></td><td></td><td></td><td></td></tr>
<tr><td>Chromosome</td><td>start</td><td>end</td><td>strand</td><td>MM</td><td>target_seq</td><td>PAM</td><td>alignment</td><td>distance</td><td>position</td><td>gene name</td><td>gene id</td></tr>
<tr><td>GU305914.1</td><td>52749</td><td>52771</td><td>+</td><td>0</td><td>TGTTGAGGTAGACCCATCTC</td><td>TGG</td><td>||||||||[||||||||||||]PAM</td><td>NA</td><td>Intergenic</td><td>NA</td><td>NA</td></tr>
<tr><td></td><td></td><td></td><td></td><td></td><td></td><td></td><td></td><td></td><td></td><td></td><td></td></tr>
<tr><td>T3453</td><td>ATCCGTGCAGACCACTTGCACGG</td><td>1000</td><td></td><td></td><td></td><td></td><td></td><td></td><td></td><td></td><td></td></tr>
<tr><td>Oligo fwd</td><td>CACCATCCGTGCAGACCACTTGCA</td><td></td><td></td><td></td><td></td><td></td><td></td><td></td><td></td><td></td><td></td></tr>
<tr><td>Oligo rev</td><td>AAACTGCAAGTGGTCTGCACGGAT</td><td></td><td></td><td></td><td></td><td></td><td></td><td></td><td></td><td></td><td></td></tr>
<tr><td>Chromosome</td><td>start</td><td>end</td><td>strand</td><td>MM</td><td>target_seq</td><td>PAM</td><td>alignment</td><td>distance</td><td>position</td><td>gene name</td><td>gene id</td></tr>
<tr><td>GU305914.1</td><td>52778</td><td>52800</td><td>+</td><td>0</td><td>ATCCGTGCAGACCACTTGCA</td><td>CGG</td><td>||||||||[||||||||||||]PAM</td><td>NA</td><td>Intergenic</td><td>NA</td><td>NA</td></tr>
<tr><td></td><td></td><td></td><td></td><td></td><td></td><td></td><td></td><td></td><td></td><td></td><td></td></tr>
<tr><td>T3454</td><td>CGTGCAGACCACTTGCACGGTGG</td><td>1000</td><td></td><td></td><td></td><td></td><td></td><td></td><td></td><td></td><td></td></tr>
<tr><td>Oligo fwd</td><td>CACCCGTGCAGACCACTTGCACGG</td><td></td><td></td><td></td><td></td><td></td><td></td><td></td><td></td><td></td><td></td></tr>
<tr><td>Oligo rev</td><td>AAACCCGTGCAAGTGGTCTGCACG</td><td></td><td></td><td></td><td></td><td></td><td></td><td></td><td></td><td></td><td></td></tr>
<tr><td>Chromosome</td><td>start</td><td>end</td><td>strand</td><td>MM</td><td>target_seq</td><td>PAM</td><td>alignment</td><td>distance</td><td>position</td><td>gene name</td><td>gene id</td></tr>
<tr><td>GU305914.1</td><td>52781</td><td>52803</td><td>+</td><td>0</td><td>CGTGCAGACCACTTGCACGG</td><td>TGG</td><td>||||||||[||||||||||||]PAM</td><td>NA</td><td>Intergenic</td><td>NA</td><td>NA</td></tr>
<tr><td></td><td></td><td></td><td></td><td></td><td></td><td></td><td></td><td></td><td></td><td></td><td></td></tr>
<tr><td>T3455</td><td>GTGCAGACCACTTGCACGGTGGG</td><td>1000</td><td></td><td></td><td></td><td></td><td></td><td></td><td></td><td></td><td></td></tr>
<tr><td>Oligo fwd</td><td>CACCGTGCAGACCACTTGCACGGT</td><td></td><td></td><td></td><td></td><td></td><td></td><td></td><td></td><td></td><td></td></tr>
<tr><td>Oligo rev</td><td>AAACACCGTGCAAGTGGTCTGCAC</td><td></td><td></td><td></td><td></td><td></td><td></td><td></td><td></td><td></td><td></td></tr>
<tr><td>Chromosome</td><td>start</td><td>end</td><td>strand</td><td>MM</td><td>target_seq</td><td>PAM</td><td>alignment</td><td>distance</td><td>position</td><td>gene name</td><td>gene id</td></tr>
<tr><td>GU305914.1</td><td>52782</td><td>52804</td><td>+</td><td>0</td><td>GTGCAGACCACTTGCACGGT</td><td>GGG</td><td>||||||||[||||||||||||]PAM</td><td>NA</td><td>Intergenic</td><td>NA</td><td>NA</td></tr>
<tr><td></td><td></td><td></td><td></td><td></td><td></td><td></td><td></td><td></td><td></td><td></td><td></td></tr>
<tr><td>T3456</td><td>GTACCATTTCTTCCTGAGAAAGG</td><td>1000</td><td></td><td></td><td></td><td></td><td></td><td></td><td></td><td></td><td></td></tr>
<tr><td>Oligo fwd</td><td>CACCGTACCATTTCTTCCTGAGAA</td><td></td><td></td><td></td><td></td><td></td><td></td><td></td><td></td><td></td><td></td></tr>
</table>

<table>
<tr><td>Oligo rev</td><td>AAACTTCTCAGGAAGAAATGGTAC</td><td></td><td></td><td></td><td></td><td></td><td></td><td></td><td></td><td></td></tr>
<tr><td>Chromosome</td><td>start</td><td>end</td><td>strand</td><td>MM</td><td>target_seq</td><td>PAM</td><td>alignment</td><td>distance</td><td>position</td><td>gene name</td><td>gene id</td></tr>
<tr><td>GU305914.1</td><td>52823</td><td>52845</td><td>+</td><td>0</td><td>GTACCATTTCTTCCTGAGAA</td><td>AGG</td><td>||||||||[||||||||||||]PAM</td><td>NA</td><td>Intergenic</td><td>NA</td><td>NA</td></tr>
<tr><td></td><td></td><td></td><td></td><td></td><td></td><td></td><td></td><td></td><td></td><td></td><td></td></tr>
<tr><td>T3457</td><td>TACCATTTCTTCCTGAGAAAGGG</td><td>1000</td><td></td><td></td><td></td><td></td><td></td><td></td><td></td><td></td><td></td></tr>
<tr><td>Oligo fwd</td><td>CACCTACCATTTCTTCCTGAGAAA</td><td></td><td></td><td></td><td></td><td></td><td></td><td></td><td></td><td></td><td></td></tr>
<tr><td>Oligo rev</td><td>AAACTTTCTCAGGAAGAAATGGTA</td><td></td><td></td><td></td><td></td><td></td><td></td><td></td><td></td><td></td><td></td></tr>
<tr><td>Chromosome</td><td>start</td><td>end</td><td>strand</td><td>MM</td><td>target_seq</td><td>PAM</td><td>alignment</td><td>distance</td><td>position</td><td>gene name</td><td>gene id</td></tr>
<tr><td>GU305914.1</td><td>52824</td><td>52846</td><td>+</td><td>0</td><td>TACCATTTCTTCCTGAGAAA</td><td>GGG</td><td>||||||||[||||||||||||]PAM</td><td>NA</td><td>Intergenic</td><td>NA</td><td>NA</td></tr>
<tr><td></td><td></td><td></td><td></td><td></td><td></td><td></td><td></td><td></td><td></td><td></td><td></td></tr>
<tr><td>T3458</td><td>ACCATTTCTTCCTGAGAAAGGGG</td><td>1000</td><td></td><td></td><td></td><td></td><td></td><td></td><td></td><td></td><td></td></tr>
<tr><td>Oligo fwd</td><td>CACCACCATTTCTTCCTGAGAAAG</td><td></td><td></td><td></td><td></td><td></td><td></td><td></td><td></td><td></td><td></td></tr>
<tr><td>Oligo rev</td><td>AAACCTTTCTCAGGAAGAAATGGT</td><td></td><td></td><td></td><td></td><td></td><td></td><td></td><td></td><td></td><td></td></tr>
<tr><td>Chromosome</td><td>start</td><td>end</td><td>strand</td><td>MM</td><td>target_seq</td><td>PAM</td><td>alignment</td><td>distance</td><td>position</td><td>gene name</td><td>gene id</td></tr>
<tr><td>GU305914.1</td><td>52825</td><td>52847</td><td>+</td><td>0</td><td>ACCATTTCTTCCTGAGAAAG</td><td>GGG</td><td>||||||||[||||||||||||]PAM</td><td>NA</td><td>Intergenic</td><td>NA</td><td>NA</td></tr>
<tr><td></td><td></td><td></td><td></td><td></td><td></td><td></td><td></td><td></td><td></td><td></td><td></td></tr>
<tr><td>T3459</td><td>CCTGAGAAAGGGGAAAGCAGAGG</td><td>1000</td><td></td><td></td><td></td><td></td><td></td><td></td><td></td><td></td><td></td></tr>
<tr><td>Oligo fwd</td><td>CACCCCTGAGAAAGGGGAAAGCAG</td><td></td><td></td><td></td><td></td><td></td><td></td><td></td><td></td><td></td><td></td></tr>
<tr><td>Oligo rev</td><td>AAACCTGCTTTCCCCTTTCTCAGG</td><td></td><td></td><td></td><td></td><td></td><td></td><td></td><td></td><td></td><td></td></tr>
<tr><td>Chromosome</td><td>start</td><td>end</td><td>strand</td><td>MM</td><td>target_seq</td><td>PAM</td><td>alignment</td><td>distance</td><td>position</td><td>gene name</td><td>gene id</td></tr>
<tr><td>GU305914.1</td><td>52835</td><td>52857</td><td>+</td><td>0</td><td>CCTGAGAAAGGGGAAAGCAG</td><td>AGG</td><td>||||||||[||||||||||||]PAM</td><td>NA</td><td>Intergenic</td><td>NA</td><td>NA</td></tr>
<tr><td></td><td></td><td></td><td></td><td></td><td></td><td></td><td></td><td></td><td></td><td></td><td></td></tr>
<tr><td>T3460</td><td>GAGCAGAGCGCGCCCGCCCTCGG</td><td>1000</td><td></td><td></td><td></td><td></td><td></td><td></td><td></td><td></td><td></td></tr>
<tr><td>Oligo fwd</td><td>CACCGAGCAGAGCGCGCCCGCCCT</td><td></td><td></td><td></td><td></td><td></td><td></td><td></td><td></td><td></td><td></td></tr>
<tr><td>Oligo rev</td><td>AAACAGGGCGGGCGCGCTCTGCTC</td><td></td><td></td><td></td><td></td><td></td><td></td><td></td><td></td><td></td><td></td></tr>
<tr><td>Chromosome</td><td>start</td><td>end</td><td>strand</td><td>MM</td><td>target_seq</td><td>PAM</td><td>alignment</td><td>distance</td><td>position</td><td>gene name</td><td>gene id</td></tr>
<tr><td>GU305914.1</td><td>52859</td><td>52881</td><td>+</td><td>0</td><td>GAGCAGAGCGCGCCCGCCCT</td><td>CGG</td><td>||||||||[||||||||||||]PAM</td><td>NA</td><td>Intergenic</td><td>NA</td><td>NA</td></tr>
<tr><td></td><td></td><td></td><td></td><td></td><td></td><td></td><td></td><td></td><td></td><td></td><td></td></tr>
<tr><td>T3461</td><td>CCGCCCTCGGCAAAAGTGACCGG</td><td>1000</td><td></td><td></td><td></td><td></td><td></td><td></td><td></td><td></td><td></td></tr>
<tr><td>Oligo fwd</td><td>CACCCCGCCCTCGGCAAAAGTGAC</td><td></td><td></td><td></td><td></td><td></td><td></td><td></td><td></td><td></td><td></td></tr>
<tr><td>Oligo rev</td><td>AAACGTCACTTTTGCCGAGGGCGG</td><td></td><td></td><td></td><td></td><td></td><td></td><td></td><td></td><td></td><td></td></tr>
<tr><td>Chromosome</td><td>start</td><td>end</td><td>strand</td><td>MM</td><td>target_seq</td><td>PAM</td><td>alignment</td><td>distance</td><td>position</td><td>gene name</td><td>gene id</td></tr>
<tr><td>GU305914.1</td><td>52872</td><td>52894</td><td>+</td><td>0</td><td>CCGCCCTCGGCAAAAGTGAC</td><td>CGG</td><td>||||||||[||||||||||||]PAM</td><td>NA</td><td>Intergenic</td><td>NA</td><td>NA</td></tr>
<tr><td></td><td></td><td></td><td></td><td></td><td></td><td></td><td></td><td></td><td></td><td></td><td></td></tr>
<tr><td>T3462</td><td>CCTCGGCAAAAGTGACCGGTAGG</td><td>1000</td><td></td><td></td><td></td><td></td><td></td><td></td><td></td><td></td><td></td></tr>
<tr><td>Oligo fwd</td><td>CACCCCTCGGCAAAAGTGACCGGT</td><td></td><td></td><td></td><td></td><td></td><td></td><td></td><td></td><td></td><td></td></tr>
<tr><td>Oligo rev</td><td>AAACACCGGTCACTTTTGCCGAGG</td><td></td><td></td><td></td><td></td><td></td><td></td><td></td><td></td><td></td><td></td></tr>
<tr><td>Chromosome</td><td>start</td><td>end</td><td>strand</td><td>MM</td><td>target_seq</td><td>PAM</td><td>alignment</td><td>distance</td><td>position</td><td>gene name</td><td>gene id</td></tr>
<tr><td>GU305914.1</td><td>52876</td><td>52898</td><td>+</td><td>0</td><td>CCTCGGCAAAAGTGACCGGT</td><td>AGG</td><td>||||||||[||||||||||||]PAM</td><td>NA</td><td>Intergenic</td><td>NA</td><td>NA</td></tr>
<tr><td></td><td></td><td></td><td></td><td></td><td></td><td></td><td></td><td></td><td></td><td></td><td></td></tr>
<tr><td>T3463</td><td>CTCGGCAAAAGTGACCGGTAGGG</td><td>1000</td><td></td><td></td><td></td><td></td><td></td><td></td><td></td><td></td><td></td></tr>
<tr><td>Oligo fwd</td><td>CACCCTCGGCAAAAGTGACCGGTA</td><td></td><td></td><td></td><td></td><td></td><td></td><td></td><td></td><td></td><td></td></tr>
<tr><td>Oligo rev</td><td>AAACTACCGGTCACTTTTGCCGAG</td><td></td><td></td><td></td><td></td><td></td><td></td><td></td><td></td><td></td><td></td></tr>
<tr><td>Chromosome</td><td>start</td><td>end</td><td>strand</td><td>MM</td><td>target_seq</td><td>PAM</td><td>alignment</td><td>distance</td><td>position</td><td>gene name</td><td>gene id</td></tr>
<tr><td>GU305914.1</td><td>52877</td><td>52899</td><td>+</td><td>0</td><td>CTCGGCAAAAGTGACCGGTA</td><td>GGG</td><td>||||||||[||||||||||||]PAM</td><td>NA</td><td>Intergenic</td><td>NA</td><td>NA</td></tr>
<tr><td></td><td></td><td></td><td></td><td></td><td></td><td></td><td></td><td></td><td></td><td></td><td></td></tr>
<tr><td>T3464</td><td>TCGGCAAAAGTGACCGGTAGGGG</td><td>1000</td><td></td><td></td><td></td><td></td><td></td><td></td><td></td><td></td><td></td></tr>
<tr><td>Oligo fwd</td><td>CACCTCGGCAAAAGTGACCGGTAG</td><td></td><td></td><td></td><td></td><td></td><td></td><td></td><td></td><td></td><td></td></tr>
<tr><td>Oligo rev</td><td>AAACCTACCGGTCACTTTTGCCGA</td><td></td><td></td><td></td><td></td><td></td><td></td><td></td><td></td><td></td><td></td></tr>
<tr><td>Chromosome</td><td>start</td><td>end</td><td>strand</td><td>MM</td><td>target_seq</td><td>PAM</td><td>alignment</td><td>distance</td><td>position</td><td>gene name</td><td>gene id</td></tr>
<tr><td>GU305914.1</td><td>52878</td><td>52900</td><td>+</td><td>0</td><td>TCGGCAAAAGTGACCGGTAG</td><td>GGG</td><td>||||||||[||||||||||||]PAM</td><td>NA</td><td>Intergenic</td><td>NA</td><td>NA</td></tr>
<tr><td></td><td></td><td></td><td></td><td></td><td></td><td></td><td></td><td></td><td></td><td></td><td></td></tr>
<tr><td>T3465</td><td>GACGATCCTGTCGACGAGAAAGG</td><td>1000</td><td></td><td></td><td></td><td></td><td></td><td></td><td></td><td></td><td></td></tr>
<tr><td>Oligo fwd</td><td>CACCGACGATCCTGTCGACGAGAA</td><td></td><td></td><td></td><td></td><td></td><td></td><td></td><td></td><td></td><td></td></tr>
<tr><td>Oligo rev</td><td>AAACTTCTCGTCGACAGGATCGTC</td><td></td><td></td><td></td><td></td><td></td><td></td><td></td><td></td><td></td><td></td></tr>
<tr><td>Chromosome</td><td>start</td><td>end</td><td>strand</td><td>MM</td><td>target_seq</td><td>PAM</td><td>alignment</td><td>distance</td><td>position</td><td>gene name</td><td>gene id</td></tr>
<tr><td>GU305914.1</td><td>52907</td><td>52929</td><td>+</td><td>0</td><td>GACGATCCTGTCGACGAGAA</td><td>AGG</td><td>||||||||[||||||||||||]PAM</td><td>NA</td><td>Intergenic</td><td>NA</td><td>NA</td></tr>
<tr><td></td><td></td><td></td><td></td><td></td><td></td><td></td><td></td><td></td><td></td><td></td><td></td></tr>
<tr><td>T3466</td><td>AAAGGCGAAGAAGACGCCCGCGG</td><td>1000</td><td></td><td></td><td></td><td></td><td></td><td></td><td></td><td></td><td></td></tr>
<tr><td>Oligo fwd</td><td>CACCAAAGGCGAAGAAGACGCCCG</td><td></td><td></td><td></td><td></td><td></td><td></td><td></td><td></td><td></td><td></td></tr>
<tr><td>Oligo rev</td><td>AAACCGGGCGTCTTCTTCGCCTTT</td><td></td><td></td><td></td><td></td><td></td><td></td><td></td><td></td><td></td><td></td></tr>
<tr><td>Chromosome</td><td>start</td><td>end</td><td>strand</td><td>MM</td><td>target_seq</td><td>PAM</td><td>alignment</td><td>distance</td><td>position</td><td>gene name</td><td>gene id</td></tr>
<tr><td>GU305914.1</td><td>52925</td><td>52947</td><td>+</td><td>0</td><td>AAAGGCGAAGAAGACGCCCG</td><td>CGG</td><td>||||||||[||||||||||||]PAM</td><td>NA</td><td>Intergenic</td><td>NA</td><td>NA</td></tr>
<tr><td>GU305914.1</td><td>147518</td><td>147540</td><td>-</td><td>4</td><td>GAAGGCGCAGAAGACGGCCT</td><td>CGG</td><td>-||||||-[||||||||-||-]PAM</td><td>NA</td><td>Intergenic</td><td>NA</td><td>NA</td></tr>
<tr><td></td><td></td><td></td><td></td><td></td><td></td><td></td><td></td><td></td><td></td><td></td><td></td></tr>
<tr><td>T3467</td><td>ACACACAACACCGCATACTCTGG</td><td>1000</td><td></td><td></td><td></td><td></td><td></td><td></td><td></td><td></td><td></td></tr>
<tr><td>Oligo fwd</td><td>CACCACACACAACACCGCATACTC</td><td></td><td></td><td></td><td></td><td></td><td></td><td></td><td></td><td></td><td></td></tr>
<tr><td>Oligo rev</td><td>AAACGAGTATGCGGTGTTGTGTGT</td><td></td><td></td><td></td><td></td><td></td><td></td><td></td><td></td><td></td><td></td></tr>
<tr><td>Chromosome</td><td>start</td><td>end</td><td>strand</td><td>MM</td><td>target_seq</td><td>PAM</td><td>alignment</td><td>distance</td><td>position</td><td>gene name</td><td>gene id</td></tr>
<tr><td>GU305914.1</td><td>52962</td><td>52984</td><td>+</td><td>0</td><td>ACACACAACACCGCATACTC</td><td>TGG</td><td>||||||||[||||||||||||]PAM</td><td>NA</td><td>Intergenic</td><td>NA</td><td>NA</td></tr>
<tr><td></td><td></td><td></td><td></td><td></td><td></td><td></td><td></td><td></td><td></td><td></td><td></td></tr>
<tr><td>T3468</td><td>CACACAACACCGCATACTCTGGG</td><td>1000</td><td></td><td></td><td></td><td></td><td></td><td></td><td></td><td></td><td></td></tr>
<tr><td>Oligo fwd</td><td>CACCCACACAACACCGCATACTCT</td><td></td><td></td><td></td><td></td><td></td><td></td><td></td><td></td><td></td><td></td></tr>
<tr><td>Oligo rev</td><td>AAACAGAGTATGCGGTGTTGTGTG</td><td></td><td></td><td></td><td></td><td></td><td></td><td></td><td></td><td></td><td></td></tr>
<tr><td>Chromosome</td><td>start</td><td>end</td><td>strand</td><td>MM</td><td>target_seq</td><td>PAM</td><td>alignment</td><td>distance</td><td>position</td><td>gene name</td><td>gene id</td></tr>
<tr><td>GU305914.1</td><td>52963</td><td>52985</td><td>+</td><td>0</td><td>CACACAACACCGCATACTCT</td><td>GGG</td><td>||||||||[||||||||||||]PAM</td><td>NA</td><td>Intergenic</td><td>NA</td><td>NA</td></tr>
<tr><td></td><td></td><td></td><td></td><td></td><td></td><td></td><td></td><td></td><td></td><td></td><td></td></tr>
<tr><td>T3469</td><td>ACACAACACCGCATACTCTGGGG</td><td>1000</td><td></td><td></td><td></td><td></td><td></td><td></td><td></td><td></td><td></td></tr>
<tr><td>Oligo fwd</td><td>CACCACACAACACCGCATACTCTG</td><td></td><td></td><td></td><td></td><td></td><td></td><td></td><td></td><td></td><td></td></tr>
<tr><td>Oligo rev</td><td>AAACCAGAGTATGCGGTGTTGTGT</td><td></td><td></td><td></td><td></td><td></td><td></td><td></td><td></td><td></td><td></td></tr>
<tr><td>Chromosome</td><td>start</td><td>end</td><td>strand</td><td>MM</td><td>target_seq</td><td>PAM</td><td>alignment</td><td>distance</td><td>position</td><td>gene name</td><td>gene id</td></tr>
<tr><td>GU305914.1</td><td>52964</td><td>52986</td><td>+</td><td>0</td><td>ACACAACACCGCATACTCTG</td><td>GGG</td><td>||||||||[||||||||||||]PAM</td><td>NA</td><td>Intergenic</td><td>NA</td><td>NA</td></tr>
<tr><td></td><td></td><td></td><td></td><td></td><td></td><td></td><td></td><td></td><td></td><td></td><td></td></tr>
<tr><td>T3470</td><td>GGCCAATCCTTGTCTCACAATGG</td><td>1000</td><td></td><td></td><td></td><td></td><td></td><td></td><td></td><td></td><td></td></tr>
<tr><td>Oligo fwd</td><td>CACCGGCCAATCCTTGTCTCACAA</td><td></td><td></td><td></td><td></td><td></td><td></td><td></td><td></td><td></td><td></td></tr>
<tr><td>Oligo rev</td><td>AAACTTGTGAGACAAGGATTGGCC</td><td></td><td></td><td></td><td></td><td></td><td></td><td></td><td></td><td></td><td></td></tr>
<tr><td>Chromosome</td><td>start</td><td>end</td><td>strand</td><td>MM</td><td>target_seq</td><td>PAM</td><td>alignment</td><td>distance</td><td>position</td><td>gene name</td><td>gene id</td></tr>
<tr><td>GU305914.1</td><td>52985</td><td>53007</td><td>+</td><td>0</td><td>GGCCAATCCTTGTCTCACAA</td><td>TGG</td><td>||||||||[||||||||||||]PAM</td><td>NA</td><td>Intergenic</td><td>NA</td><td>NA</td></tr>
</table>

| | | | | | | | | | | | | |
|---|---|---|---|---|---|---|---|---|---|---|---|---|
| T3471 | CAATGGTATTGTGTCCGATAAGG | | 1000 | | | | | | | | | |
| Oligo fwd | CACCCAATGGTATTGTGTCCGATA | | | | | | | | | | | |
| Oligo rev | AAACTATCGGACACAATACCATTG | | | | | | | | | | | |
| Chromosome | start | | end | strand | MM | target_seq | PAM | alignment | distance | position | gene name | gene id |
| GU305914.1 | | 53002 | 53024 | + | 0 | CAATGGTATTGTGTCCGATA | AGG | ||||||||[||||||||||||]PAM | NA | Intergenic | NA | NA |
| T3472 | GTCACACGCTCCACAACAGATGG | | 1000 | | | | | | | | | |
| Oligo fwd | CACCGTCACACGCTCCACAACAGA | | | | | | | | | | | |
| Oligo rev | AAACTCTGTTGTGGAGCGTGTGAC | | | | | | | | | | | |
| Chromosome | start | | end | strand | MM | target_seq | PAM | alignment | distance | position | gene name | gene id |
| GU305914.1 | | 53024 | 53046 | + | 0 | GTCACACGCTCCACAACAGA | TGG | ||||||||[||||||||||||]PAM | NA | Intergenic | NA | NA |
| T3473 | CAACAGATGGATGACCCGTTAGG | | 1000 | | | | | | | | | |
| Oligo fwd | CACCCAACAGATGGATGACCCGTT | | | | | | | | | | | |
| Oligo rev | AAACAACGGGTCATCCATCTGTTG | | | | | | | | | | | |
| Chromosome | start | | end | strand | MM | target_seq | PAM | alignment | distance | position | gene name | gene id |
| GU305914.1 | | 53037 | 53059 | + | 0 | CAACAGATGGATGACCCGTT | AGG | ||||||||[||||||||||||]PAM | NA | Intergenic | NA | NA |
| T3474 | AACAGATGGATGACCCGTTAGGG | | 1000 | | | | | | | | | |
| Oligo fwd | CACCAACAGATGGATGACCCGTTA | | | | | | | | | | | |
| Oligo rev | AAACTAACGGGTCATCCATCTGTT | | | | | | | | | | | |
| Chromosome | start | | end | strand | MM | target_seq | PAM | alignment | distance | position | gene name | gene id |
| GU305914.1 | | 53038 | 53060 | + | 0 | AACAGATGGATGACCCGTTA | GGG | ||||||||[||||||||||||]PAM | NA | Intergenic | NA | NA |
| T3475 | GGATGACCCGTTAGGGAAATCGG | | 1000 | | | | | | | | | |
| Oligo fwd | CACCGGATGACCCGTTAGGGAAAT | | | | | | | | | | | |
| Oligo rev | AAACATTTCCCTAACGGGTCATCC | | | | | | | | | | | |
| Chromosome | start | | end | strand | MM | target_seq | PAM | alignment | distance | position | gene name | gene id |
| GU305914.1 | | 53045 | 53067 | + | 0 | GGATGACCCGTTAGGGAAAT | CGG | ||||||||[||||||||||||]PAM | NA | Intergenic | NA | NA |
| T3476 | GATGACCCGTTAGGGAAATCGGG | | 1000 | | | | | | | | | |
| Oligo fwd | CACCGATGACCCGTTAGGGAAATC | | | | | | | | | | | |
| Oligo rev | AAACGATTTCCCTAACGGGTCATC | | | | | | | | | | | |
| Chromosome | start | | end | strand | MM | target_seq | PAM | alignment | distance | position | gene name | gene id |
| GU305914.1 | | 53046 | 53068 | + | 0 | GATGACCCGTTAGGGAAATC | GGG | ||||||||[||||||||||||]PAM | NA | Intergenic | NA | NA |
| T3477 | AAGTTTTGATCACTCGTTCCTGG | | 1000 | | | | | | | | | |
| Oligo fwd | CACCAAGTTTTGATCACTCGTTCC | | | | | | | | | | | |
| Oligo rev | AAACGGAACGAGTGATCAAAACTT | | | | | | | | | | | |
| Chromosome | start | | end | strand | MM | target_seq | PAM | alignment | distance | position | gene name | gene id |
| GU305914.1 | | 53092 | 53114 | + | 0 | AAGTTTTGATCACTCGTTCC | TGG | ||||||||[||||||||||||]PAM | NA | Intergenic | NA | NA |
| T3478 | TGATCACTCGTTCCTGGCCGCGG | | 1000 | | | | | | | | | |
| Oligo fwd | CACCTGATCACTCGTTCCTGGCCG | | | | | | | | | | | |
| Oligo rev | AAACCGGCCAGGAACGAGTGATCA | | | | | | | | | | | |
| Chromosome | start | | end | strand | MM | target_seq | PAM | alignment | distance | position | gene name | gene id |
| GU305914.1 | | 53098 | 53120 | + | 0 | TGATCACTCGTTCCTGGCCG | CGG | ||||||||[||||||||||||]PAM | NA | Intergenic | NA | NA |
| T3479 | GCCGCGGACGAAGATACGAACGG | | 1000 | | | | | | | | | |
| Oligo fwd | CACCGCCGCGGACGAAGATACGAA | | | | | | | | | | | |
| Oligo rev | AAACTTCGTATCTTCGTCCGCGGC | | | | | | | | | | | |
| Chromosome | start | | end | strand | MM | target_seq | PAM | alignment | distance | position | gene name | gene id |
| GU305914.1 | | 53114 | 53136 | + | 0 | GCCGCGGACGAAGATACGAA | CGG | ||||||||[||||||||||||]PAM | NA | Intergenic | NA | NA |
| T3480 | TACGAACGGCGTCTTCGCCGCGG | | 1000 | | | | | | | | | |
| Oligo fwd | CACCTACGAACGGCGTCTTCGCCG | | | | | | | | | | | |
| Oligo rev | AAACCGGCGAAGACGCCGTTCGTA | | | | | | | | | | | |
| Chromosome | start | | end | strand | MM | target_seq | PAM | alignment | distance | position | gene name | gene id |
| GU305914.1 | | 53128 | 53150 | + | 0 | TACGAACGGCGTCTTCGCCG | CGG | ||||||||[||||||||||||]PAM | NA | Intergenic | NA | NA |
| T3481 | CCGACTCTCTTCCCTCAACCTGG | | 1000 | | | | | | | | | |
| Oligo fwd | CACCCCGACTCTCTTCCCTCAACC | | | | | | | | | | | |
| Oligo rev | AAACGGTTGAGGGAAGAGAGTCGG | | | | | | | | | | | |
| Chromosome | start | | end | strand | MM | target_seq | PAM | alignment | distance | position | gene name | gene id |
| GU305914.1 | | 53271 | 53293 | + | 0 | CCGACTCTCTTCCCTCAACC | TGG | ||||||||[||||||||||||]PAM | NA | Intergenic | NA | NA |
| T3482 | CCTGGTCCAACGCGCTCCGCCGG | | 1000 | | | | | | | | | |
| Oligo fwd | CACCCCTGGTCCAACGCGCTCCGC | | | | | | | | | | | |
| Oligo rev | AAACGCGGAGCGCGTTGGACCAGG | | | | | | | | | | | |
| Chromosome | start | | end | strand | MM | target_seq | PAM | alignment | distance | position | gene name | gene id |
| GU305914.1 | | 53289 | 53311 | + | 0 | CCTGGTCCAACGCGCTCCGC | CGG | ||||||||[||||||||||||]PAM | NA | Intergenic | NA | NA |
| T3483 | TCGAACGAATTCACGTCTAACGG | | 1000 | | | | | | | | | |
| Oligo fwd | CACCTCGAACGAATTCACGTCTAA | | | | | | | | | | | |
| Oligo rev | AAACTTAGACGTGAATTCGTTCGA | | | | | | | | | | | |
| Chromosome | start | | end | strand | MM | target_seq | PAM | alignment | distance | position | gene name | gene id |
| GU305914.1 | | 53322 | 53344 | + | 0 | TCGAACGAATTCACGTCTAA | CGG | ||||||||[||||||||||||]PAM | NA | Intergenic | NA | NA |
| T3484 | GTAGATGTTGTACGCGAACCAGG | | 1000 | | | | | | | | | |
| Oligo fwd | CACCGTAGATGTTGTACGCGAACC | | | | | | | | | | | |
| Oligo rev | AAACGGTTCGCGTACAACATCTAC | | | | | | | | | | | |
| Chromosome | start | | end | strand | MM | target_seq | PAM | alignment | distance | position | gene name | gene id |
| GU305914.1 | | 53346 | 53368 | + | 0 | GTAGATGTTGTACGCGAACC | AGG | ||||||||[||||||||||||]PAM | NA | Intergenic | NA | NA |
| T3485 | GGCCCCACACAGCAGCACCACGG | | 1000 | | | | | | | | | |
| Oligo fwd | CACCGGCCCCACACAGCAGCACCA | | | | | | | | | | | |
| Oligo rev | AAACTGGTGCTGCTGTGTGGGGCC | | | | | | | | | | | |
| Chromosome | start | | end | strand | MM | target_seq | PAM | alignment | distance | position | gene name | gene id |
| GU305914.1 | | 53367 | 53389 | + | 0 | GGCCCCACACAGCAGCACCA | CGG | ||||||||[||||||||||||]PAM | NA | Intergenic | NA | NA |

<table>
<tr><td>T3486</td><td>CAGCACCACGGACAAAACCACGG</td><td>1000</td><td></td><td></td><td></td><td></td><td></td><td></td><td></td><td></td><td></td></tr>
<tr><td>Oligo fwd</td><td>CACCCAGCACCACGGACAAAACCA</td><td></td><td></td><td></td><td></td><td></td><td></td><td></td><td></td><td></td><td></td></tr>
<tr><td>Oligo rev</td><td>AAACTGGTTTTGTCCGTGGTGCTG</td><td></td><td></td><td></td><td></td><td></td><td></td><td></td><td></td><td></td><td></td></tr>
<tr><td>Chromosome</td><td>start</td><td>end</td><td>strand</td><td>MM</td><td>target_seq</td><td>PAM</td><td>alignment</td><td>distance</td><td>position</td><td>gene name</td><td>gene id</td></tr>
<tr><td>GU305914.1</td><td>53379</td><td>53401</td><td>+</td><td>0</td><td>CAGCACCACGGACAAAACCA</td><td>CGG</td><td>||||||||[||||||||||||]PAM</td><td>NA</td><td>Intergenic</td><td>NA</td><td>NA</td></tr>
<tr><td>GU305914.1</td><td>152663</td><td>152685</td><td>+</td><td>4</td><td>CGGCGTCACGGCCAAAACCA</td><td>AGG</td><td>|-||--||[|||-||||||||]PAM</td><td>NA</td><td>Intergenic</td><td>NA</td><td>NA</td></tr>
<tr><td>T3487</td><td>CCACGCCAGATGCAAGAGTGAGG</td><td>1000</td><td></td><td></td><td></td><td></td><td></td><td></td><td></td><td></td><td></td></tr>
<tr><td>Oligo fwd</td><td>CACCCCACGCCAGATGCAAGAGTG</td><td></td><td></td><td></td><td></td><td></td><td></td><td></td><td></td><td></td><td></td></tr>
<tr><td>Oligo rev</td><td>AAACCACTCTTGCATCTGGCGTGG</td><td></td><td></td><td></td><td></td><td></td><td></td><td></td><td></td><td></td><td></td></tr>
<tr><td>Chromosome</td><td>start</td><td>end</td><td>strand</td><td>MM</td><td>target_seq</td><td>PAM</td><td>alignment</td><td>distance</td><td>position</td><td>gene name</td><td>gene id</td></tr>
<tr><td>GU305914.1</td><td>53436</td><td>53458</td><td>+</td><td>0</td><td>CCACGCCAGATGCAAGAGTG</td><td>AGG</td><td>||||||||[||||||||||||]PAM</td><td>NA</td><td>Intergenic</td><td>NA</td><td>NA</td></tr>
<tr><td>T3488</td><td>ATGCAAGAGTGAGGCGTCCCCGG</td><td>1000</td><td></td><td></td><td></td><td></td><td></td><td></td><td></td><td></td><td></td></tr>
<tr><td>Oligo fwd</td><td>CACCATGCAAGAGTGAGGCGTCCC</td><td></td><td></td><td></td><td></td><td></td><td></td><td></td><td></td><td></td><td></td></tr>
<tr><td>Oligo rev</td><td>AAACGGGACGCCTCACTCTTGCAT</td><td></td><td></td><td></td><td></td><td></td><td></td><td></td><td></td><td></td><td></td></tr>
<tr><td>Chromosome</td><td>start</td><td>end</td><td>strand</td><td>MM</td><td>target_seq</td><td>PAM</td><td>alignment</td><td>distance</td><td>position</td><td>gene name</td><td>gene id</td></tr>
<tr><td>GU305914.1</td><td>53445</td><td>53467</td><td>+</td><td>0</td><td>ATGCAAGAGTGAGGCGTCCC</td><td>CGG</td><td>||||||||[||||||||||||]PAM</td><td>NA</td><td>Intergenic</td><td>NA</td><td>NA</td></tr>
<tr><td>T3489</td><td>AGGCGTCCCCGGACCCGTCACGG</td><td>1000</td><td></td><td></td><td></td><td></td><td></td><td></td><td></td><td></td><td></td></tr>
<tr><td>Oligo fwd</td><td>CACCAGGCGTCCCCGGACCCGTCA</td><td></td><td></td><td></td><td></td><td></td><td></td><td></td><td></td><td></td><td></td></tr>
<tr><td>Oligo rev</td><td>AAACTGACGGGTCCGGGGACGCCT</td><td></td><td></td><td></td><td></td><td></td><td></td><td></td><td></td><td></td><td></td></tr>
<tr><td>Chromosome</td><td>start</td><td>end</td><td>strand</td><td>MM</td><td>target_seq</td><td>PAM</td><td>alignment</td><td>distance</td><td>position</td><td>gene name</td><td>gene id</td></tr>
<tr><td>GU305914.1</td><td>53456</td><td>53478</td><td>+</td><td>0</td><td>AGGCGTCCCCGGACCCGTCA</td><td>CGG</td><td>||||||||[||||||||||||]PAM</td><td>NA</td><td>Intergenic</td><td>NA</td><td>NA</td></tr>
<tr><td>T3490</td><td>GTCGTTCTCGCCCCGACCGACGG</td><td>1000</td><td></td><td></td><td></td><td></td><td></td><td></td><td></td><td></td><td></td></tr>
<tr><td>Oligo fwd</td><td>CACCGTCGTTCTCGCCCCGACCGA</td><td></td><td></td><td></td><td></td><td></td><td></td><td></td><td></td><td></td><td></td></tr>
<tr><td>Oligo rev</td><td>AAACTCGGTCGGGGCGAGAACGAC</td><td></td><td></td><td></td><td></td><td></td><td></td><td></td><td></td><td></td><td></td></tr>
<tr><td>Chromosome</td><td>start</td><td>end</td><td>strand</td><td>MM</td><td>target_seq</td><td>PAM</td><td>alignment</td><td>distance</td><td>position</td><td>gene name</td><td>gene id</td></tr>
<tr><td>GU305914.1</td><td>53490</td><td>53512</td><td>+</td><td>0</td><td>GTCGTTCTCGCCCCGACCGA</td><td>CGG</td><td>||||||||[||||||||||||]PAM</td><td>NA</td><td>Intergenic</td><td>NA</td><td>NA</td></tr>
<tr><td>GU305914.1</td><td>90624</td><td>90646</td><td>+</td><td>4</td><td>GTACTTCTCGTCCCGCCCGA</td><td>CGG</td><td>||--||||[||-||||-||||]PAM</td><td>NA</td><td>Intergenic</td><td>NA</td><td>NA</td></tr>
<tr><td>T3491</td><td>GTTCTCGCCCCGACCGACGGTGG</td><td>1000</td><td></td><td></td><td></td><td></td><td></td><td></td><td></td><td></td><td></td></tr>
<tr><td>Oligo fwd</td><td>CACCGTTCTCGCCCCGACCGACGG</td><td></td><td></td><td></td><td></td><td></td><td></td><td></td><td></td><td></td><td></td></tr>
<tr><td>Oligo rev</td><td>AAACCCGTCGGTCGGGGCGAGAAC</td><td></td><td></td><td></td><td></td><td></td><td></td><td></td><td></td><td></td><td></td></tr>
<tr><td>Chromosome</td><td>start</td><td>end</td><td>strand</td><td>MM</td><td>target_seq</td><td>PAM</td><td>alignment</td><td>distance</td><td>position</td><td>gene name</td><td>gene id</td></tr>
<tr><td>GU305914.1</td><td>53493</td><td>53515</td><td>+</td><td>0</td><td>GTTCTCGCCCCGACCGACGG</td><td>TGG</td><td>||||||||[||||||||||||]PAM</td><td>NA</td><td>Intergenic</td><td>NA</td><td>NA</td></tr>
<tr><td>T3492</td><td>CGACGGTGGTCGTGCCCGAGAGG</td><td>1000</td><td></td><td></td><td></td><td></td><td></td><td></td><td></td><td></td><td></td></tr>
<tr><td>Oligo fwd</td><td>CACCCGACGGTGGTCGTGCCCGAG</td><td></td><td></td><td></td><td></td><td></td><td></td><td></td><td></td><td></td><td></td></tr>
<tr><td>Oligo rev</td><td>AAACCTCGGGCACGACCACCGTCG</td><td></td><td></td><td></td><td></td><td></td><td></td><td></td><td></td><td></td><td></td></tr>
<tr><td>Chromosome</td><td>start</td><td>end</td><td>strand</td><td>MM</td><td>target_seq</td><td>PAM</td><td>alignment</td><td>distance</td><td>position</td><td>gene name</td><td>gene id</td></tr>
<tr><td>GU305914.1</td><td>53507</td><td>53529</td><td>+</td><td>0</td><td>CGACGGTGGTCGTGCCCGAG</td><td>AGG</td><td>||||||||[||||||||||||]PAM</td><td>NA</td><td>Intergenic</td><td>NA</td><td>NA</td></tr>
<tr><td>T3493</td><td>TTCAAGACCGCCGAGCTCTCCGG</td><td>1000</td><td></td><td></td><td></td><td></td><td></td><td></td><td></td><td></td><td></td></tr>
<tr><td>Oligo fwd</td><td>CACCTTCAAGACCGCCGAGCTCTC</td><td></td><td></td><td></td><td></td><td></td><td></td><td></td><td></td><td></td><td></td></tr>
<tr><td>Oligo rev</td><td>AAACGAGAGCTCGGCGGTCTTGAA</td><td></td><td></td><td></td><td></td><td></td><td></td><td></td><td></td><td></td><td></td></tr>
<tr><td>Chromosome</td><td>start</td><td>end</td><td>strand</td><td>MM</td><td>target_seq</td><td>PAM</td><td>alignment</td><td>distance</td><td>position</td><td>gene name</td><td>gene id</td></tr>
<tr><td>GU305914.1</td><td>53530</td><td>53552</td><td>+</td><td>0</td><td>TTCAAGACCGCCGAGCTCTC</td><td>CGG</td><td>||||||||[||||||||||||]PAM</td><td>NA</td><td>Intergenic</td><td>NA</td><td>NA</td></tr>
<tr><td>T3494</td><td>TCAAGACCGCCGAGCTCTCCGGG</td><td>1000</td><td></td><td></td><td></td><td></td><td></td><td></td><td></td><td></td><td></td></tr>
<tr><td>Oligo fwd</td><td>CACCTCAAGACCGCCGAGCTCTCC</td><td></td><td></td><td></td><td></td><td></td><td></td><td></td><td></td><td></td><td></td></tr>
<tr><td>Oligo rev</td><td>AAACGGAGAGCTCGGCGGTCTTGA</td><td></td><td></td><td></td><td></td><td></td><td></td><td></td><td></td><td></td><td></td></tr>
<tr><td>Chromosome</td><td>start</td><td>end</td><td>strand</td><td>MM</td><td>target_seq</td><td>PAM</td><td>alignment</td><td>distance</td><td>position</td><td>gene name</td><td>gene id</td></tr>
<tr><td>GU305914.1</td><td>53531</td><td>53553</td><td>+</td><td>0</td><td>TCAAGACCGCCGAGCTCTCC</td><td>GGG</td><td>||||||||[||||||||||||]PAM</td><td>NA</td><td>Intergenic</td><td>NA</td><td>NA</td></tr>
<tr><td>T3495</td><td>CGCCGAGCTCTCCGGGAAGATGG</td><td>1000</td><td></td><td></td><td></td><td></td><td></td><td></td><td></td><td></td><td></td></tr>
<tr><td>Oligo fwd</td><td>CACCCGCCGAGCTCTCCGGGAAGA</td><td></td><td></td><td></td><td></td><td></td><td></td><td></td><td></td><td></td><td></td></tr>
<tr><td>Oligo rev</td><td>AAACTCTTCCCGGAGAGCTCGGCG</td><td></td><td></td><td></td><td></td><td></td><td></td><td></td><td></td><td></td><td></td></tr>
<tr><td>Chromosome</td><td>start</td><td>end</td><td>strand</td><td>MM</td><td>target_seq</td><td>PAM</td><td>alignment</td><td>distance</td><td>position</td><td>gene name</td><td>gene id</td></tr>
<tr><td>GU305914.1</td><td>53538</td><td>53560</td><td>+</td><td>0</td><td>CGCCGAGCTCTCCGGGAAGA</td><td>TGG</td><td>||||||||[||||||||||||]PAM</td><td>NA</td><td>Intergenic</td><td>NA</td><td>NA</td></tr>
<tr><td>T3496</td><td>GAAGATGGTGACCTCTTCATCGG</td><td>1000</td><td></td><td></td><td></td><td></td><td></td><td></td><td></td><td></td><td></td></tr>
<tr><td>Oligo fwd</td><td>CACCGAAGATGGTGACCTCTTCAT</td><td></td><td></td><td></td><td></td><td></td><td></td><td></td><td></td><td></td><td></td></tr>
<tr><td>Oligo rev</td><td>AAACATGAAGAGGTCACCATCTTC</td><td></td><td></td><td></td><td></td><td></td><td></td><td></td><td></td><td></td><td></td></tr>
<tr><td>Chromosome</td><td>start</td><td>end</td><td>strand</td><td>MM</td><td>target_seq</td><td>PAM</td><td>alignment</td><td>distance</td><td>position</td><td>gene name</td><td>gene id</td></tr>
<tr><td>GU305914.1</td><td>53553</td><td>53575</td><td>+</td><td>0</td><td>GAAGATGGTGACCTCTTCAT</td><td>CGG</td><td>||||||||[||||||||||||]PAM</td><td>NA</td><td>Intergenic</td><td>NA</td><td>NA</td></tr>
<tr><td>T3497</td><td>TGGTGACCTCTTCATCGGAGAGG</td><td>1000</td><td></td><td></td><td></td><td></td><td></td><td></td><td></td><td></td><td></td></tr>
<tr><td>Oligo fwd</td><td>CACCTGGTGACCTCTTCATCGGAG</td><td></td><td></td><td></td><td></td><td></td><td></td><td></td><td></td><td></td><td></td></tr>
<tr><td>Oligo rev</td><td>AAACCTCCGATGAAGAGGTCACCA</td><td></td><td></td><td></td><td></td><td></td><td></td><td></td><td></td><td></td><td></td></tr>
<tr><td>Chromosome</td><td>start</td><td>end</td><td>strand</td><td>MM</td><td>target_seq</td><td>PAM</td><td>alignment</td><td>distance</td><td>position</td><td>gene name</td><td>gene id</td></tr>
<tr><td>GU305914.1</td><td>53558</td><td>53580</td><td>+</td><td>0</td><td>TGGTGACCTCTTCATCGGAG</td><td>AGG</td><td>||||||||[||||||||||||]PAM</td><td>NA</td><td>Intergenic</td><td>NA</td><td>NA</td></tr>
<tr><td>T3498</td><td>CGGAGAGGATCGTCCGAGACAGG</td><td>1000</td><td></td><td></td><td></td><td></td><td></td><td></td><td></td><td></td><td></td></tr>
<tr><td>Oligo fwd</td><td>CACCCGGAGAGGATCGTCCGAGAC</td><td></td><td></td><td></td><td></td><td></td><td></td><td></td><td></td><td></td><td></td></tr>
<tr><td>Oligo rev</td><td>AAACGTCTCGGACGATCCTCTCCG</td><td></td><td></td><td></td><td></td><td></td><td></td><td></td><td></td><td></td><td></td></tr>
<tr><td>Chromosome</td><td>start</td><td>end</td><td>strand</td><td>MM</td><td>target_seq</td><td>PAM</td><td>alignment</td><td>distance</td><td>position</td><td>gene name</td><td>gene id</td></tr>
<tr><td>GU305914.1</td><td>53573</td><td>53595</td><td>+</td><td>0</td><td>CGGAGAGGATCGTCCGAGAC</td><td>AGG</td><td>||||||||[||||||||||||]PAM</td><td>NA</td><td>Intergenic</td><td>NA</td><td>NA</td></tr>
<tr><td>T3499</td><td>AGACAGGACACCCACCGTCCCGG</td><td>1000</td><td></td><td></td><td></td><td></td><td></td><td></td><td></td><td></td><td></td></tr>
<tr><td>Oligo fwd</td><td>CACCAGACAGGACACCCACCGTCC</td><td></td><td></td><td></td><td></td><td></td><td></td><td></td><td></td><td></td><td></td></tr>
<tr><td>Oligo rev</td><td>AAACGGACGGTGGGTGTCCTGTCT</td><td></td><td></td><td></td><td></td><td></td><td></td><td></td><td></td><td></td><td></td></tr>
<tr><td>Chromosome</td><td>start</td><td>end</td><td>strand</td><td>MM</td><td>target_seq</td><td>PAM</td><td>alignment</td><td>distance</td><td>position</td><td>gene name</td><td>gene id</td></tr>
<tr><td>GU305914.1</td><td>53589</td><td>53611</td><td>+</td><td>0</td><td>AGACAGGACACCCACCGTCC</td><td>CGG</td><td>||||||||[||||||||||||]PAM</td><td>NA</td><td>Intergenic</td><td>NA</td><td>NA</td></tr>
<tr><td>T3500</td><td>GACAGAGAAGACGCGAGTCGAGG</td><td>1000</td><td></td><td></td><td></td><td></td><td></td><td></td><td></td><td></td><td></td></tr>
<tr><td>Oligo fwd</td><td>CACCGACAGAGAAGACGCGAGTCG</td><td></td><td></td><td></td><td></td><td></td><td></td><td></td><td></td><td></td><td></td></tr>
</table>

| | | | | | | | | | | | | |
|---|---|---|---|---|---|---|---|---|---|---|---|---|
| Oligo rev | AAACCGACTCGCGTCTTCTCTGTC | | | | | | | | | | | |
| Chromosome | start | | end | strand | MM | target_seq | PAM | alignment | distance | position | gene name | gene id |
| GU305914.1 | | 53654 | 53676 | + | 0 | GACAGAGAAGACGCGAGTCG | AGG | \|\|\|\|\|\|\|\|[\|\|\|\|\|\|\|\|\|\|\|\|]PAM | NA | Intergenic | NA | NA |
| T3501 | ACAGAGAAGACGCGAGTCGAGGG | | 1000 | | | | | | | | | |
| Oligo fwd | CACCACAGAGAAGACGCGAGTCGA | | | | | | | | | | | |
| Oligo rev | AAACTCGACTCGCGTCTTCTCTGT | | | | | | | | | | | |
| Chromosome | start | | end | strand | MM | target_seq | PAM | alignment | distance | position | gene name | gene id |
| GU305914.1 | | 53655 | 53677 | + | 0 | ACAGAGAAGACGCGAGTCGA | GGG | \|\|\|\|\|\|\|\|[\|\|\|\|\|\|\|\|\|\|\|\|]PAM | NA | Intergenic | NA | NA |
| T3502 | CAGAGAAGACGCGAGTCGAGGGG | | 1000 | | | | | | | | | |
| Oligo fwd | CACCCAGAGAAGACGCGAGTCGAG | | | | | | | | | | | |
| Oligo rev | AAACCTCGACTCGCGTCTTCTCTG | | | | | | | | | | | |
| Chromosome | start | | end | strand | MM | target_seq | PAM | alignment | distance | position | gene name | gene id |
| GU305914.1 | | 53656 | 53678 | + | 0 | CAGAGAAGACGCGAGTCGAG | GGG | \|\|\|\|\|\|\|\|[\|\|\|\|\|\|\|\|\|\|\|\|]PAM | NA | Intergenic | NA | NA |
| T3503 | GAAGACGCGAGTCGAGGGGATGG | | 1000 | | | | | | | | | |
| Oligo fwd | CACCGAAGACGCGAGTCGAGGGGA | | | | | | | | | | | |
| Oligo rev | AAACTCCCCTCGACTCGCGTCTTC | | | | | | | | | | | |
| Chromosome | start | | end | strand | MM | target_seq | PAM | alignment | distance | position | gene name | gene id |
| GU305914.1 | | 53660 | 53682 | + | 0 | GAAGACGCGAGTCGAGGGGA | TGG | \|\|\|\|\|\|\|\|[\|\|\|\|\|\|\|\|\|\|\|\|]PAM | NA | Intergenic | NA | NA |
| T3504 | TGAAGATCAGAGAGATAATTTGG | | 1000 | | | | | | | | | |
| Oligo fwd | CACCTGAAGATCAGAGAGATAATT | | | | | | | | | | | |
| Oligo rev | AAACAATTATCTCTCTGATCTTCA | | | | | | | | | | | |
| Chromosome | start | | end | strand | MM | target_seq | PAM | alignment | distance | position | gene name | gene id |
| GU305914.1 | | 53709 | 53731 | + | 0 | TGAAGATCAGAGAGATAATT | TGG | \|\|\|\|\|\|\|\|[\|\|\|\|\|\|\|\|\|\|\|\|]PAM | NA | Intergenic | NA | NA |
| T3505 | GAAGATCAGAGAGATAATTTGGG | | 1000 | | | | | | | | | |
| Oligo fwd | CACCGAAGATCAGAGAGATAATTT | | | | | | | | | | | |
| Oligo rev | AAACAAATTATCTCTCTGATCTTC | | | | | | | | | | | |
| Chromosome | start | | end | strand | MM | target_seq | PAM | alignment | distance | position | gene name | gene id |
| GU305914.1 | | 53710 | 53732 | + | 0 | GAAGATCAGAGAGATAATTT | GGG | \|\|\|\|\|\|\|\|[\|\|\|\|\|\|\|\|\|\|\|\|]PAM | NA | Intergenic | NA | NA |
| T3506 | TGTCAATGATCACGACGATCAGG | | 1000 | | | | | | | | | |
| Oligo fwd | CACCTGTCAATGATCACGACGATC | | | | | | | | | | | |
| Oligo rev | AAACGATCGTCGTGATCATTGACA | | | | | | | | | | | |
| Chromosome | start | | end | strand | MM | target_seq | PAM | alignment | distance | position | gene name | gene id |
| GU305914.1 | | 53791 | 53813 | + | 0 | TGTCAATGATCACGACGATC | AGG | \|\|\|\|\|\|\|\|[\|\|\|\|\|\|\|\|\|\|\|\|]PAM | NA | Intergenic | NA | NA |
| T3507 | TGACGATGCAGATCACTGCGAGG | | 1000 | | | | | | | | | |
| Oligo fwd | CACCTGACGATGCAGATCACTGCG | | | | | | | | | | | |
| Oligo rev | AAACCGCAGTGATCTGCATCGTCA | | | | | | | | | | | |
| Chromosome | start | | end | strand | MM | target_seq | PAM | alignment | distance | position | gene name | gene id |
| GU305914.1 | | 53815 | 53837 | + | 0 | TGACGATGCAGATCACTGCG | AGG | \|\|\|\|\|\|\|\|[\|\|\|\|\|\|\|\|\|\|\|\|]PAM | NA | Intergenic | NA | NA |
| T3508 | GATGCAGATCACTGCGAGGACGG | | 1000 | | | | | | | | | |
| Oligo fwd | CACCGATGCAGATCACTGCGAGGA | | | | | | | | | | | |
| Oligo rev | AAACTCCTCGCAGTGATCTGCATC | | | | | | | | | | | |
| Chromosome | start | | end | strand | MM | target_seq | PAM | alignment | distance | position | gene name | gene id |
| GU305914.1 | | 53819 | 53841 | + | 0 | GATGCAGATCACTGCGAGGA | CGG | \|\|\|\|\|\|\|\|[\|\|\|\|\|\|\|\|\|\|\|\|]PAM | NA | Intergenic | NA | NA |
| T3509 | GCAGATCACTGCGAGGACGGAGG | | 1000 | | | | | | | | | |
| Oligo fwd | CACCGCAGATCACTGCGAGGACGG | | | | | | | | | | | |
| Oligo rev | AAACCCGTCCTCGCAGTGATCTGC | | | | | | | | | | | |
| Chromosome | start | | end | strand | MM | target_seq | PAM | alignment | distance | position | gene name | gene id |
| GU305914.1 | | 53822 | 53844 | + | 0 | GCAGATCACTGCGAGGACGG | AGG | \|\|\|\|\|\|\|\|[\|\|\|\|\|\|\|\|\|\|\|\|]PAM | NA | Intergenic | NA | NA |
| T3510 | GACGGAGGCGTAAACCACGTCGG | | 1000 | | | | | | | | | |
| Oligo fwd | CACCGACGGAGGCGTAAACCACGT | | | | | | | | | | | |
| Oligo rev | AAACACGTGGTTTACGCCTCCGTC | | | | | | | | | | | |
| Chromosome | start | | end | strand | MM | target_seq | PAM | alignment | distance | position | gene name | gene id |
| GU305914.1 | | 53837 | 53859 | + | 0 | GACGGAGGCGTAAACCACGT | CGG | \|\|\|\|\|\|\|\|[\|\|\|\|\|\|\|\|\|\|\|\|]PAM | NA | Intergenic | NA | NA |
| T3511 | ACGGAGGCGTAAACCACGTCGGG | | 1000 | | | | | | | | | |
| Oligo fwd | CACCACGGAGGCGTAAACCACGTC | | | | | | | | | | | |
| Oligo rev | AAACGACGTGGTTTACGCCTCCGT | | | | | | | | | | | |
| Chromosome | start | | end | strand | MM | target_seq | PAM | alignment | distance | position | gene name | gene id |
| GU305914.1 | | 53838 | 53860 | + | 0 | ACGGAGGCGTAAACCACGTC | GGG | \|\|\|\|\|\|\|\|[\|\|\|\|\|\|\|\|\|\|\|\|]PAM | NA | Intergenic | NA | NA |
| T3512 | ACGTACGCCGTTACAGAGTTTGG | | 1000 | | | | | | | | | |
| Oligo fwd | CACCACGTACGCCGTTACAGAGTT | | | | | | | | | | | |
| Oligo rev | AAACAACTCTGTAACGGCGTACGT | | | | | | | | | | | |
| Chromosome | start | | end | strand | MM | target_seq | PAM | alignment | distance | position | gene name | gene id |
| GU305914.1 | | 53873 | 53895 | + | 0 | ACGTACGCCGTTACAGAGTT | TGG | \|\|\|\|\|\|\|\|[\|\|\|\|\|\|\|\|\|\|\|\|]PAM | NA | Intergenic | NA | NA |
| T3513 | GCGACCGTGCCAACTCTTCGCGG | | 1000 | | | | | | | | | |
| Oligo fwd | CACCGCGACCGTGCCAACTCTTCG | | | | | | | | | | | |
| Oligo rev | AAACCGAAGAGTTGGCACGGTCGC | | | | | | | | | | | |
| Chromosome | start | | end | strand | MM | target_seq | PAM | alignment | distance | position | gene name | gene id |
| GU305914.1 | | 53895 | 53917 | + | 0 | GCGACCGTGCCAACTCTTCG | CGG | \|\|\|\|\|\|\|\|[\|\|\|\|\|\|\|\|\|\|\|\|]PAM | NA | Intergenic | NA | NA |
| T3514 | ACCGTGCCAACTCTTCGCGGCGG | | 1000 | | | | | | | | | |
| Oligo fwd | CACCACCGTGCCAACTCTTCGCGG | | | | | | | | | | | |
| Oligo rev | AAACCCGCGAAGAGTTGGCACGGT | | | | | | | | | | | |
| Chromosome | start | | end | strand | MM | target_seq | PAM | alignment | distance | position | gene name | gene id |
| GU305914.1 | | 53898 | 53920 | + | 0 | ACCGTGCCAACTCTTCGCGG | CGG | \|\|\|\|\|\|\|\|[\|\|\|\|\|\|\|\|\|\|\|\|]PAM | NA | Intergenic | NA | NA |
| T3515 | ACTCTTCGCGGCGGTCGCGCCGG | | 1000 | | | | | | | | | |

Oligo fwd CACCACTCTTCGCGGCGGTCGCGC
Oligo rev AAACGCGCGACCGCCGCGAAGAGT
Chromosome start end strand MM target_seq PAM alignment distance position gene name gene id
GU305914.1 53907 53929 + 0 ACTCTTCGCGGCGGTCGCGC CGG ||||||||[||||||||||||]PAM NA Intergenic NA NA

T3516 CTCTTCGCGGCGGTCGCGCCGGG 1000
Oligo fwd CACCCTCTTCGCGGCGGTCGCGCC
Oligo rev AAACGGCGCGACCGCCGCGAAGAG
Chromosome start end strand MM target_seq PAM alignment distance position gene name gene id
GU305914.1 53908 53930 + 0 CTCTTCGCGGCGGTCGCGCC GGG ||||||||[||||||||||||]PAM NA Intergenic NA NA

T3517 TCGCGGCGGTCGCGCCGGGATGG 1000
Oligo fwd CACCTCGCGGCGGTCGCGCCGGGA
Oligo rev AAACTCCCGGCGCGACCGCCGCGA
Chromosome start end strand MM target_seq PAM alignment distance position gene name gene id
GU305914.1 53912 53934 + 0 TCGCGGCGGTCGCGCCGGGA TGG ||||||||[||||||||||||]PAM NA Intergenic NA NA
GU305914.1 134142 134164 - 3 TCGAGGCGGTCGCTCCGCGA GGG |||-||||[|||||-|||-||]PAM NA Intergenic NA NA

T3518 CGCCGGGATGGCCACTCCGTTGG 1000
Oligo fwd CACCCGCCGGGATGGCCACTCCGT
Oligo rev AAACACGGAGTGGCCATCCCGGCG
Chromosome start end strand MM target_seq PAM alignment distance position gene name gene id
GU305914.1 53924 53946 + 0 CGCCGGGATGGCCACTCCGT TGG ||||||||[||||||||||||]PAM NA Intergenic NA NA

T3519 GGCCACTCCGTTGGCGTCGACGG 1000
Oligo fwd CACCGGCCACTCCGTTGGCGTCGA
Oligo rev AAACTCGACGCCAACGGAGTGGCC
Chromosome start end strand MM target_seq PAM alignment distance position gene name gene id
GU305914.1 53933 53955 + 0 GGCCACTCCGTTGGCGTCGA CGG ||||||||[||||||||||||]PAM NA Intergenic NA NA

T3520 GCCACTCCGTTGGCGTCGACGGG 1000
Oligo fwd CACCGCCACTCCGTTGGCGTCGAC
Oligo rev AAACGTCGACGCCAACGGAGTGGC
Chromosome start end strand MM target_seq PAM alignment distance position gene name gene id
GU305914.1 53934 53956 + 0 GCCACTCCGTTGGCGTCGAC GGG ||||||||[||||||||||||]PAM NA Intergenic NA NA

T3521 CGACGGGAAACAGTTCGAGTCGG 1000
Oligo fwd CACCCGACGGGAAACAGTTCGAGT
Oligo rev AAACACTCGAACTGTTTCCCGTCG
Chromosome start end strand MM target_seq PAM alignment distance position gene name gene id
GU305914.1 53950 53972 + 0 CGACGGGAAACAGTTCGAGT CGG ||||||||[||||||||||||]PAM NA Intergenic NA NA

T3522 GACGGGAAACAGTTCGAGTCGGG 1000
Oligo fwd CACCGACGGGAAACAGTTCGAGTC
Oligo rev AAACGACTCGAACTGTTTCCCGTC
Chromosome start end strand MM target_seq PAM alignment distance position gene name gene id
GU305914.1 53951 53973 + 0 GACGGGAAACAGTTCGAGTC GGG ||||||||[||||||||||||]PAM NA Intergenic NA NA

T3523 GACGAAGATGACGCCTTCCGAGG 1000
Oligo fwd CACCGACGAAGATGACGCCTTCCG
Oligo rev AAACCGGAAGGCGTCATCTTCGTC
Chromosome start end strand MM target_seq PAM alignment distance position gene name gene id
GU305914.1 53984 54006 + 0 GACGAAGATGACGCCTTCCG AGG ||||||||[||||||||||||]PAM NA Intergenic NA NA

T3524 GAAACGTTGTCTCCTAACGACGG 1000
Oligo fwd CACCGAAACGTTGTCTCCTAACGA
Oligo rev AAACTCGTTAGGAGACAACGTTTC
Chromosome start end strand MM target_seq PAM alignment distance position gene name gene id
GU305914.1 54006 54028 + 0 GAAACGTTGTCTCCTAACGA CGG ||||||||[||||||||||||]PAM NA Intergenic NA NA
GU305914.1 163225 163247 - 4 GTAACGTCGTCTCGAAACGA TGG |-|||||-[|||||--|||||]PAM NA Intergenic NA NA

T3525 GCGTAGAGCCCCCGAGCTCACGG 1000
Oligo fwd CACCGCGTAGAGCCCCCGAGCTCA
Oligo rev AAACTGAGCTCGGGGGCTCTACGC
Chromosome start end strand MM target_seq PAM alignment distance position gene name gene id
GU305914.1 54028 54050 + 0 GCGTAGAGCCCCCGAGCTCA CGG ||||||||[||||||||||||]PAM NA Intergenic NA NA

T3526 GAGCCCCCGAGCTCACGGTAAGG 1000
Oligo fwd CACCGAGCCCCCGAGCTCACGGTA
Oligo rev AAACTACCGTGAGCTCGGGGGCTC
Chromosome start end strand MM target_seq PAM alignment distance position gene name gene id
GU305914.1 54033 54055 + 0 GAGCCCCCGAGCTCACGGTA AGG ||||||||[||||||||||||]PAM NA Intergenic NA NA

T3527 CACGGTAAGGCATAGTAAAGAGG 1000
Oligo fwd CACCCACGGTAAGGCATAGTAAAG
Oligo rev AAACCTTTACTATGCCTTACCGTG
Chromosome start end strand MM target_seq PAM alignment distance position gene name gene id
GU305914.1 54046 54068 + 0 CACGGTAAGGCATAGTAAAG AGG ||||||||[||||||||||||]PAM NA Intergenic NA NA

T3528 GCATAGTAAAGAGGCCGCCAAGG 1000
Oligo fwd CACCGCATAGTAAAGAGGCCGCCA
Oligo rev AAACTGGCGGCCTCTTTACTATGC
Chromosome start end strand MM target_seq PAM alignment distance position gene name gene id
GU305914.1 54055 54077 + 0 GCATAGTAAAGAGGCCGCCA AGG ||||||||[||||||||||||]PAM NA Intergenic NA NA

T3529 TAAAGAGGCCGCCAAGGCGACGG 1000
Oligo fwd CACCTAAAGAGGCCGCCAAGGCGA
Oligo rev AAACTCGCCTTGGCGGCCTCTTTA
Chromosome start end strand MM target_seq PAM alignment distance position gene name gene id

<table>
<tr><td>GU305914.1</td><td>54061</td><td>54083</td><td>+</td><td>0</td><td>TAAAGAGGCCGCCAAGGCGA</td><td>CGG</td><td>||||||||[||||||||||||]PAM</td><td>NA</td><td>Intergenic</td><td>NA</td><td>NA</td></tr>
<tr><td></td><td></td><td></td><td></td><td></td><td></td><td></td><td></td><td></td><td></td><td></td><td></td></tr>
<tr><td>T3530</td><td>CCGCCAAGGCGACGGAGATTAGG</td><td>1000</td><td></td><td></td><td></td><td></td><td></td><td></td><td></td><td></td><td></td></tr>
<tr><td>Oligo fwd</td><td>CACCCCGCCAAGGCGACGGAGATT</td><td></td><td></td><td></td><td></td><td></td><td></td><td></td><td></td><td></td><td></td></tr>
<tr><td>Oligo rev</td><td>AAACAATCTCCGTCGCCTTGGCGG</td><td></td><td></td><td></td><td></td><td></td><td></td><td></td><td></td><td></td><td></td></tr>
<tr><td>Chromosome</td><td>start</td><td>end</td><td>strand</td><td>MM</td><td>target_seq</td><td>PAM</td><td>alignment</td><td>distance</td><td>position</td><td>gene name</td><td>gene id</td></tr>
<tr><td>GU305914.1</td><td>54069</td><td>54091</td><td>+</td><td>0</td><td>CCGCCAAGGCGACGGAGATT</td><td>AGG</td><td>||||||||[||||||||||||]PAM</td><td>NA</td><td>Intergenic</td><td>NA</td><td>NA</td></tr>
<tr><td></td><td></td><td></td><td></td><td></td><td></td><td></td><td></td><td></td><td></td><td></td><td></td></tr>
<tr><td>T3531</td><td>CGACGGAGATTAGGTCTTCGCGG</td><td>1000</td><td></td><td></td><td></td><td></td><td></td><td></td><td></td><td></td><td></td></tr>
<tr><td>Oligo fwd</td><td>CACCCGACGGAGATTAGGTCTTCG</td><td></td><td></td><td></td><td></td><td></td><td></td><td></td><td></td><td></td><td></td></tr>
<tr><td>Oligo rev</td><td>AAACCGAAGACCTAATCTCCGTCG</td><td></td><td></td><td></td><td></td><td></td><td></td><td></td><td></td><td></td><td></td></tr>
<tr><td>Chromosome</td><td>start</td><td>end</td><td>strand</td><td>MM</td><td>target_seq</td><td>PAM</td><td>alignment</td><td>distance</td><td>position</td><td>gene name</td><td>gene id</td></tr>
<tr><td>GU305914.1</td><td>54078</td><td>54100</td><td>+</td><td>0</td><td>CGACGGAGATTAGGTCTTCG</td><td>CGG</td><td>||||||||[||||||||||||]PAM</td><td>NA</td><td>Intergenic</td><td>NA</td><td>NA</td></tr>
<tr><td></td><td></td><td></td><td></td><td></td><td></td><td></td><td></td><td></td><td></td><td></td><td></td></tr>
<tr><td>T3532</td><td>CGGAGATTAGGTCTTCGCGGAGG</td><td>1000</td><td></td><td></td><td></td><td></td><td></td><td></td><td></td><td></td><td></td></tr>
<tr><td>Oligo fwd</td><td>CACCCGGAGATTAGGTCTTCGCGG</td><td></td><td></td><td></td><td></td><td></td><td></td><td></td><td></td><td></td><td></td></tr>
<tr><td>Oligo rev</td><td>AAACCCGCGAAGACCTAATCTCCG</td><td></td><td></td><td></td><td></td><td></td><td></td><td></td><td></td><td></td><td></td></tr>
<tr><td>Chromosome</td><td>start</td><td>end</td><td>strand</td><td>MM</td><td>target_seq</td><td>PAM</td><td>alignment</td><td>distance</td><td>position</td><td>gene name</td><td>gene id</td></tr>
<tr><td>GU305914.1</td><td>54081</td><td>54103</td><td>+</td><td>0</td><td>CGGAGATTAGGTCTTCGCGG</td><td>AGG</td><td>||||||||[||||||||||||]PAM</td><td>NA</td><td>Intergenic</td><td>NA</td><td>NA</td></tr>
<tr><td></td><td></td><td></td><td></td><td></td><td></td><td></td><td></td><td></td><td></td><td></td><td></td></tr>
<tr><td>T3533</td><td>GGAGATTAGGTCTTCGCGGAGGG</td><td>1000</td><td></td><td></td><td></td><td></td><td></td><td></td><td></td><td></td><td></td></tr>
<tr><td>Oligo fwd</td><td>CACCGGAGATTAGGTCTTCGCGGA</td><td></td><td></td><td></td><td></td><td></td><td></td><td></td><td></td><td></td><td></td></tr>
<tr><td>Oligo rev</td><td>AAACTCCGCGAAGACCTAATCTCC</td><td></td><td></td><td></td><td></td><td></td><td></td><td></td><td></td><td></td><td></td></tr>
<tr><td>Chromosome</td><td>start</td><td>end</td><td>strand</td><td>MM</td><td>target_seq</td><td>PAM</td><td>alignment</td><td>distance</td><td>position</td><td>gene name</td><td>gene id</td></tr>
<tr><td>GU305914.1</td><td>54082</td><td>54104</td><td>+</td><td>0</td><td>GGAGATTAGGTCTTCGCGGA</td><td>GGG</td><td>||||||||[||||||||||||]PAM</td><td>NA</td><td>Intergenic</td><td>NA</td><td>NA</td></tr>
<tr><td></td><td></td><td></td><td></td><td></td><td></td><td></td><td></td><td></td><td></td><td></td><td></td></tr>
<tr><td>T3534</td><td>TTCGCGGAGGGCGCGACGAAAGG</td><td>1000</td><td></td><td></td><td></td><td></td><td></td><td></td><td></td><td></td><td></td></tr>
<tr><td>Oligo fwd</td><td>CACCTTCGCGGAGGGCGCGACGAA</td><td></td><td></td><td></td><td></td><td></td><td></td><td></td><td></td><td></td><td></td></tr>
<tr><td>Oligo rev</td><td>AAACTTCGTCGCGCCCTCCGCGAA</td><td></td><td></td><td></td><td></td><td></td><td></td><td></td><td></td><td></td><td></td></tr>
<tr><td>Chromosome</td><td>start</td><td>end</td><td>strand</td><td>MM</td><td>target_seq</td><td>PAM</td><td>alignment</td><td>distance</td><td>position</td><td>gene name</td><td>gene id</td></tr>
<tr><td>GU305914.1</td><td>54094</td><td>54116</td><td>+</td><td>0</td><td>TTCGCGGAGGGCGCGACGAA</td><td>AGG</td><td>||||||||[||||||||||||]PAM</td><td>NA</td><td>Intergenic</td><td>NA</td><td>NA</td></tr>
<tr><td></td><td></td><td></td><td></td><td></td><td></td><td></td><td></td><td></td><td></td><td></td><td></td></tr>
<tr><td>T3535</td><td>AGGGCGCGACGAAAGGTTCGCGG</td><td>1000</td><td></td><td></td><td></td><td></td><td></td><td></td><td></td><td></td><td></td></tr>
<tr><td>Oligo fwd</td><td>CACCAGGGCGCGACGAAAGGTTCG</td><td></td><td></td><td></td><td></td><td></td><td></td><td></td><td></td><td></td><td></td></tr>
<tr><td>Oligo rev</td><td>AAACCGAACCTTTCGTCGCGCCCT</td><td></td><td></td><td></td><td></td><td></td><td></td><td></td><td></td><td></td><td></td></tr>
<tr><td>Chromosome</td><td>start</td><td>end</td><td>strand</td><td>MM</td><td>target_seq</td><td>PAM</td><td>alignment</td><td>distance</td><td>position</td><td>gene name</td><td>gene id</td></tr>
<tr><td>GU305914.1</td><td>54101</td><td>54123</td><td>+</td><td>0</td><td>AGGGCGCGACGAAAGGTTCG</td><td>CGG</td><td>||||||||[||||||||||||]PAM</td><td>NA</td><td>Intergenic</td><td>NA</td><td>NA</td></tr>
<tr><td></td><td></td><td></td><td></td><td></td><td></td><td></td><td></td><td></td><td></td><td></td><td></td></tr>
<tr><td>T3536</td><td>GGGCGCGACGAAAGGTTCGCGGG</td><td>1000</td><td></td><td></td><td></td><td></td><td></td><td></td><td></td><td></td><td></td></tr>
<tr><td>Oligo fwd</td><td>CACCGGGCGCGACGAAAGGTTCGC</td><td></td><td></td><td></td><td></td><td></td><td></td><td></td><td></td><td></td><td></td></tr>
<tr><td>Oligo rev</td><td>AAACGCGAACCTTTCGTCGCGCCC</td><td></td><td></td><td></td><td></td><td></td><td></td><td></td><td></td><td></td><td></td></tr>
<tr><td>Chromosome</td><td>start</td><td>end</td><td>strand</td><td>MM</td><td>target_seq</td><td>PAM</td><td>alignment</td><td>distance</td><td>position</td><td>gene name</td><td>gene id</td></tr>
<tr><td>GU305914.1</td><td>54102</td><td>54124</td><td>+</td><td>0</td><td>GGGCGCGACGAAAGGTTCGC</td><td>GGG</td><td>||||||||[||||||||||||]PAM</td><td>NA</td><td>Intergenic</td><td>NA</td><td>NA</td></tr>
<tr><td></td><td></td><td></td><td></td><td></td><td></td><td></td><td></td><td></td><td></td><td></td><td></td></tr>
<tr><td>T3537</td><td>CGACGAAAGGTTCGCGGGAGCGG</td><td>1000</td><td></td><td></td><td></td><td></td><td></td><td></td><td></td><td></td><td></td></tr>
<tr><td>Oligo fwd</td><td>CACCCGACGAAAGGTTCGCGGGAG</td><td></td><td></td><td></td><td></td><td></td><td></td><td></td><td></td><td></td><td></td></tr>
<tr><td>Oligo rev</td><td>AAACCTCCCGCGAACCTTTCGTCG</td><td></td><td></td><td></td><td></td><td></td><td></td><td></td><td></td><td></td><td></td></tr>
<tr><td>Chromosome</td><td>start</td><td>end</td><td>strand</td><td>MM</td><td>target_seq</td><td>PAM</td><td>alignment</td><td>distance</td><td>position</td><td>gene name</td><td>gene id</td></tr>
<tr><td>GU305914.1</td><td>54107</td><td>54129</td><td>+</td><td>0</td><td>CGACGAAAGGTTCGCGGGAG</td><td>CGG</td><td>||||||||[||||||||||||]PAM</td><td>NA</td><td>Intergenic</td><td>NA</td><td>NA</td></tr>
<tr><td></td><td></td><td></td><td></td><td></td><td></td><td></td><td></td><td></td><td></td><td></td><td></td></tr>
<tr><td>T3538</td><td>AGCGGCAATCTTCTCACCGTCGG</td><td>1000</td><td></td><td></td><td></td><td></td><td></td><td></td><td></td><td></td><td></td></tr>
<tr><td>Oligo fwd</td><td>CACCAGCGGCAATCTTCTCACCGT</td><td></td><td></td><td></td><td></td><td></td><td></td><td></td><td></td><td></td><td></td></tr>
<tr><td>Oligo rev</td><td>AAACACGGTGAGAAGATTGCCGCT</td><td></td><td></td><td></td><td></td><td></td><td></td><td></td><td></td><td></td><td></td></tr>
<tr><td>Chromosome</td><td>start</td><td>end</td><td>strand</td><td>MM</td><td>target_seq</td><td>PAM</td><td>alignment</td><td>distance</td><td>position</td><td>gene name</td><td>gene id</td></tr>
<tr><td>GU305914.1</td><td>54125</td><td>54147</td><td>+</td><td>0</td><td>AGCGGCAATCTTCTCACCGT</td><td>CGG</td><td>||||||||[||||||||||||]PAM</td><td>NA</td><td>Intergenic</td><td>NA</td><td>NA</td></tr>
<tr><td></td><td></td><td></td><td></td><td></td><td></td><td></td><td></td><td></td><td></td><td></td><td></td></tr>
<tr><td>T3539</td><td>CGATCATCGTCTCCCATCAAAGG</td><td>1000</td><td></td><td></td><td></td><td></td><td></td><td></td><td></td><td></td><td></td></tr>
<tr><td>Oligo fwd</td><td>CACCCGATCATCGTCTCCCATCAA</td><td></td><td></td><td></td><td></td><td></td><td></td><td></td><td></td><td></td><td></td></tr>
<tr><td>Oligo rev</td><td>AAACTTGATGGGAGACGATGATCG</td><td></td><td></td><td></td><td></td><td></td><td></td><td></td><td></td><td></td><td></td></tr>
<tr><td>Chromosome</td><td>start</td><td>end</td><td>strand</td><td>MM</td><td>target_seq</td><td>PAM</td><td>alignment</td><td>distance</td><td>position</td><td>gene name</td><td>gene id</td></tr>
<tr><td>GU305914.1</td><td>54186</td><td>54208</td><td>+</td><td>0</td><td>CGATCATCGTCTCCCATCAA</td><td>AGG</td><td>||||||||[||||||||||||]PAM</td><td>NA</td><td>Intergenic</td><td>NA</td><td>NA</td></tr>
<tr><td></td><td></td><td></td><td></td><td></td><td></td><td></td><td></td><td></td><td></td><td></td><td></td></tr>
<tr><td>T3540</td><td>ATCGTCTCCCATCAAAGGTTCGG</td><td>1000</td><td></td><td></td><td></td><td></td><td></td><td></td><td></td><td></td><td></td></tr>
<tr><td>Oligo fwd</td><td>CACCATCGTCTCCCATCAAAGGTT</td><td></td><td></td><td></td><td></td><td></td><td></td><td></td><td></td><td></td><td></td></tr>
<tr><td>Oligo rev</td><td>AAACAACCTTTGATGGGAGACGAT</td><td></td><td></td><td></td><td></td><td></td><td></td><td></td><td></td><td></td><td></td></tr>
<tr><td>Chromosome</td><td>start</td><td>end</td><td>strand</td><td>MM</td><td>target_seq</td><td>PAM</td><td>alignment</td><td>distance</td><td>position</td><td>gene name</td><td>gene id</td></tr>
<tr><td>GU305914.1</td><td>54191</td><td>54213</td><td>+</td><td>0</td><td>ATCGTCTCCCATCAAAGGTT</td><td>CGG</td><td>||||||||[||||||||||||]PAM</td><td>NA</td><td>Intergenic</td><td>NA</td><td>NA</td></tr>
<tr><td></td><td></td><td></td><td></td><td></td><td></td><td></td><td></td><td></td><td></td><td></td><td></td></tr>
<tr><td>T3541</td><td>GTCTCCCATCAAAGGTTCGGAGG</td><td>1000</td><td></td><td></td><td></td><td></td><td></td><td></td><td></td><td></td><td></td></tr>
<tr><td>Oligo fwd</td><td>CACCGTCTCCCATCAAAGGTTCGG</td><td></td><td></td><td></td><td></td><td></td><td></td><td></td><td></td><td></td><td></td></tr>
<tr><td>Oligo rev</td><td>AAACCCGAACCTTTGATGGGAGAC</td><td></td><td></td><td></td><td></td><td></td><td></td><td></td><td></td><td></td><td></td></tr>
<tr><td>Chromosome</td><td>start</td><td>end</td><td>strand</td><td>MM</td><td>target_seq</td><td>PAM</td><td>alignment</td><td>distance</td><td>position</td><td>gene name</td><td>gene id</td></tr>
<tr><td>GU305914.1</td><td>54194</td><td>54216</td><td>+</td><td>0</td><td>GTCTCCCATCAAAGGTTCGG</td><td>AGG</td><td>||||||||[||||||||||||]PAM</td><td>NA</td><td>Intergenic</td><td>NA</td><td>NA</td></tr>
<tr><td></td><td></td><td></td><td></td><td></td><td></td><td></td><td></td><td></td><td></td><td></td><td></td></tr>
<tr><td>T3542</td><td>GAGGAACCTGCGCACGACAACGG</td><td>1000</td><td></td><td></td><td></td><td></td><td></td><td></td><td></td><td></td><td></td></tr>
<tr><td>Oligo fwd</td><td>CACCGAGGAACCTGCGCACGACAA</td><td></td><td></td><td></td><td></td><td></td><td></td><td></td><td></td><td></td><td></td></tr>
<tr><td>Oligo rev</td><td>AAACTTGTCGTGCGCAGGTTCCTC</td><td></td><td></td><td></td><td></td><td></td><td></td><td></td><td></td><td></td><td></td></tr>
<tr><td>Chromosome</td><td>start</td><td>end</td><td>strand</td><td>MM</td><td>target_seq</td><td>PAM</td><td>alignment</td><td>distance</td><td>position</td><td>gene name</td><td>gene id</td></tr>
<tr><td>GU305914.1</td><td>54213</td><td>54235</td><td>+</td><td>0</td><td>GAGGAACCTGCGCACGACAA</td><td>CGG</td><td>||||||||[||||||||||||]PAM</td><td>NA</td><td>Intergenic</td><td>NA</td><td>NA</td></tr>
<tr><td></td><td></td><td></td><td></td><td></td><td></td><td></td><td></td><td></td><td></td><td></td><td></td></tr>
<tr><td>T3543</td><td>GCGCACGACAACGGAATGTGAGG</td><td>1000</td><td></td><td></td><td></td><td></td><td></td><td></td><td></td><td></td><td></td></tr>
<tr><td>Oligo fwd</td><td>CACCGCGCACGACAACGGAATGTG</td><td></td><td></td><td></td><td></td><td></td><td></td><td></td><td></td><td></td><td></td></tr>
<tr><td>Oligo rev</td><td>AAACCACATTCCGTTGTCGTGCGC</td><td></td><td></td><td></td><td></td><td></td><td></td><td></td><td></td><td></td><td></td></tr>
<tr><td>Chromosome</td><td>start</td><td>end</td><td>strand</td><td>MM</td><td>target_seq</td><td>PAM</td><td>alignment</td><td>distance</td><td>position</td><td>gene name</td><td>gene id</td></tr>
<tr><td>GU305914.1</td><td>54222</td><td>54244</td><td>+</td><td>0</td><td>GCGCACGACAACGGAATGTG</td><td>AGG</td><td>||||||||[||||||||||||]PAM</td><td>NA</td><td>Intergenic</td><td>NA</td><td>NA</td></tr>
<tr><td></td><td></td><td></td><td></td><td></td><td></td><td></td><td></td><td></td><td></td><td></td><td></td></tr>
<tr><td>T3544</td><td>GACAACGGAATGTGAGGAAACGG</td><td>1000</td><td></td><td></td><td></td><td></td><td></td><td></td><td></td><td></td><td></td></tr>
<tr><td>Oligo fwd</td><td>CACCGACAACGGAATGTGAGGAAA</td><td></td><td></td><td></td><td></td><td></td><td></td><td></td><td></td><td></td><td></td></tr>
<tr><td>Oligo rev</td><td>AAACTTTCCTCACATTCCGTTGTC</td><td></td><td></td><td></td><td></td><td></td><td></td><td></td><td></td><td></td><td></td></tr>
</table>

<table>
<tr><td>Chromosome</td><td>start</td><td></td><td>end</td><td>strand</td><td>MM</td><td>target_seq</td><td>PAM</td><td>alignment</td><td>distance</td><td>position</td><td>gene name</td><td>gene id</td></tr>
<tr><td>GU305914.1</td><td></td><td>54228</td><td>54250</td><td>+</td><td>0</td><td>GACAACGGAATGTGAGGAAA</td><td>CGG</td><td>||||||||[||||||||||||]PAM</td><td>NA</td><td>Intergenic</td><td>NA</td><td>NA</td></tr>
<tr><td colspan="13"></td></tr>
<tr><td>T3545</td><td>GCGTTTAAAATAGTTTTGTAAGG</td><td></td><td>1000</td><td colspan="9"></td></tr>
<tr><td>Oligo fwd</td><td>CACCGCGTTTAAAATAGTTTTGTA</td><td colspan="11"></td></tr>
<tr><td>Oligo rev</td><td>AAACTACAAAACTATTTTAAACGC</td><td colspan="11"></td></tr>
<tr><td>Chromosome</td><td>start</td><td></td><td>end</td><td>strand</td><td>MM</td><td>target_seq</td><td>PAM</td><td>alignment</td><td>distance</td><td>position</td><td>gene name</td><td>gene id</td></tr>
<tr><td>GU305914.1</td><td></td><td>54250</td><td>54272</td><td>+</td><td>0</td><td>GCGTTTAAAATAGTTTTGTA</td><td>AGG</td><td>||||||||[||||||||||||]PAM</td><td>NA</td><td>Intergenic</td><td>NA</td><td>NA</td></tr>
<tr><td colspan="13"></td></tr>
<tr><td>T3546</td><td>ACACATCGTCATCTCCCGTCCGG</td><td></td><td>1000</td><td colspan="9"></td></tr>
<tr><td>Oligo fwd</td><td>CACCACACATCGTCATCTCCCGTC</td><td colspan="11"></td></tr>
<tr><td>Oligo rev</td><td>AAACGACGGGAGATGACGATGTGT</td><td colspan="11"></td></tr>
<tr><td>Chromosome</td><td>start</td><td></td><td>end</td><td>strand</td><td>MM</td><td>target_seq</td><td>PAM</td><td>alignment</td><td>distance</td><td>position</td><td>gene name</td><td>gene id</td></tr>
<tr><td>GU305914.1</td><td></td><td>54309</td><td>54331</td><td>+</td><td>0</td><td>ACACATCGTCATCTCCCGTC</td><td>CGG</td><td>||||||||[||||||||||||]PAM</td><td>NA</td><td>Intergenic</td><td>NA</td><td>NA</td></tr>
<tr><td colspan="13"></td></tr>
<tr><td>T3547</td><td>GTCATCTCCCGTCCGGCGACCGG</td><td></td><td>1000</td><td colspan="9"></td></tr>
<tr><td>Oligo fwd</td><td>CACCGTCATCTCCCGTCCGGCGAC</td><td colspan="11"></td></tr>
<tr><td>Oligo rev</td><td>AAACGTCGCCGGACGGGAGATGAC</td><td colspan="11"></td></tr>
<tr><td>Chromosome</td><td>start</td><td></td><td>end</td><td>strand</td><td>MM</td><td>target_seq</td><td>PAM</td><td>alignment</td><td>distance</td><td>position</td><td>gene name</td><td>gene id</td></tr>
<tr><td>GU305914.1</td><td></td><td>54316</td><td>54338</td><td>+</td><td>0</td><td>GTCATCTCCCGTCCGGCGAC</td><td>CGG</td><td>||||||||[||||||||||||]PAM</td><td>NA</td><td>Intergenic</td><td>NA</td><td>NA</td></tr>
<tr><td colspan="13"></td></tr>
<tr><td>T3548</td><td>ACCTCCCCGACCAGTGACCACGG</td><td></td><td>1000</td><td colspan="9"></td></tr>
<tr><td>Oligo fwd</td><td>CACCACCTCCCCGACCAGTGACCA</td><td colspan="11"></td></tr>
<tr><td>Oligo rev</td><td>AAACTGGTCACTGGTCGGGGAGGT</td><td colspan="11"></td></tr>
<tr><td>Chromosome</td><td>start</td><td></td><td>end</td><td>strand</td><td>MM</td><td>target_seq</td><td>PAM</td><td>alignment</td><td>distance</td><td>position</td><td>gene name</td><td>gene id</td></tr>
<tr><td>GU305914.1</td><td></td><td>54370</td><td>54392</td><td>+</td><td>0</td><td>ACCTCCCCGACCAGTGACCA</td><td>CGG</td><td>||||||||[||||||||||||]PAM</td><td>NA</td><td>Intergenic</td><td>NA</td><td>NA</td></tr>
<tr><td colspan="13"></td></tr>
<tr><td>T3549</td><td>CGAGAACGAGTATGACCAAGAGG</td><td></td><td>1000</td><td colspan="9"></td></tr>
<tr><td>Oligo fwd</td><td>CACCCGAGAACGAGTATGACCAAG</td><td colspan="11"></td></tr>
<tr><td>Oligo rev</td><td>AAACCTTGGTCATACTCGTTCTCG</td><td colspan="11"></td></tr>
<tr><td>Chromosome</td><td>start</td><td></td><td>end</td><td>strand</td><td>MM</td><td>target_seq</td><td>PAM</td><td>alignment</td><td>distance</td><td>position</td><td>gene name</td><td>gene id</td></tr>
<tr><td>GU305914.1</td><td></td><td>54402</td><td>54424</td><td>+</td><td>0</td><td>CGAGAACGAGTATGACCAAG</td><td>AGG</td><td>||||||||[||||||||||||]PAM</td><td>NA</td><td>Intergenic</td><td>NA</td><td>NA</td></tr>
<tr><td colspan="13"></td></tr>
<tr><td>T3550</td><td>GAGAACGAGTATGACCAAGAGGG</td><td></td><td>1000</td><td colspan="9"></td></tr>
<tr><td>Oligo fwd</td><td>CACCGAGAACGAGTATGACCAAGA</td><td colspan="11"></td></tr>
<tr><td>Oligo rev</td><td>AAACTCTTGGTCATACTCGTTCTC</td><td colspan="11"></td></tr>
<tr><td>Chromosome</td><td>start</td><td></td><td>end</td><td>strand</td><td>MM</td><td>target_seq</td><td>PAM</td><td>alignment</td><td>distance</td><td>position</td><td>gene name</td><td>gene id</td></tr>
<tr><td>GU305914.1</td><td></td><td>54403</td><td>54425</td><td>+</td><td>0</td><td>GAGAACGAGTATGACCAAGA</td><td>GGG</td><td>||||||||[||||||||||||]PAM</td><td>NA</td><td>Intergenic</td><td>NA</td><td>NA</td></tr>
<tr><td colspan="13"></td></tr>
<tr><td>T3551</td><td>CACACAGACTCCGACCGTGATGG</td><td></td><td>1000</td><td colspan="9"></td></tr>
<tr><td>Oligo fwd</td><td>CACCCACACAGACTCCGACCGTGA</td><td colspan="11"></td></tr>
<tr><td>Oligo rev</td><td>AAACTCACGGTCGGAGTCTGTGTG</td><td colspan="11"></td></tr>
<tr><td>Chromosome</td><td>start</td><td></td><td>end</td><td>strand</td><td>MM</td><td>target_seq</td><td>PAM</td><td>alignment</td><td>distance</td><td>position</td><td>gene name</td><td>gene id</td></tr>
<tr><td>GU305914.1</td><td></td><td>54427</td><td>54449</td><td>+</td><td>0</td><td>CACACAGACTCCGACCGTGA</td><td>TGG</td><td>||||||||[||||||||||||]PAM</td><td>NA</td><td>Intergenic</td><td>NA</td><td>NA</td></tr>
<tr><td colspan="13"></td></tr>
<tr><td>T3552</td><td>ACAGACTCCGACCGTGATGGCGG</td><td></td><td>1000</td><td colspan="9"></td></tr>
<tr><td>Oligo fwd</td><td>CACCACAGACTCCGACCGTGATGG</td><td colspan="11"></td></tr>
<tr><td>Oligo rev</td><td>AAACCCATCACGGTCGGAGTCTGT</td><td colspan="11"></td></tr>
<tr><td>Chromosome</td><td>start</td><td></td><td>end</td><td>strand</td><td>MM</td><td>target_seq</td><td>PAM</td><td>alignment</td><td>distance</td><td>position</td><td>gene name</td><td>gene id</td></tr>
<tr><td>GU305914.1</td><td></td><td>54430</td><td>54452</td><td>+</td><td>0</td><td>ACAGACTCCGACCGTGATGG</td><td>CGG</td><td>||||||||[||||||||||||]PAM</td><td>NA</td><td>Intergenic</td><td>NA</td><td>NA</td></tr>
<tr><td colspan="13"></td></tr>
<tr><td>T3553</td><td>TATCCATGCGTTCGAGAGACAGG</td><td></td><td>1000</td><td colspan="9"></td></tr>
<tr><td>Oligo fwd</td><td>CACCTATCCATGCGTTCGAGAGAC</td><td colspan="11"></td></tr>
<tr><td>Oligo rev</td><td>AAACGTCTCTCGAACGCATGGATA</td><td colspan="11"></td></tr>
<tr><td>Chromosome</td><td>start</td><td></td><td>end</td><td>strand</td><td>MM</td><td>target_seq</td><td>PAM</td><td>alignment</td><td>distance</td><td>position</td><td>gene name</td><td>gene id</td></tr>
<tr><td>GU305914.1</td><td></td><td>54468</td><td>54490</td><td>+</td><td>0</td><td>TATCCATGCGTTCGAGAGAC</td><td>AGG</td><td>||||||||[||||||||||||]PAM</td><td>NA</td><td>Intergenic</td><td>NA</td><td>NA</td></tr>
<tr><td colspan="13"></td></tr>
<tr><td>T3554</td><td>ATCCATGCGTTCGAGAGACAGGG</td><td></td><td>1000</td><td colspan="9"></td></tr>
<tr><td>Oligo fwd</td><td>CACCATCCATGCGTTCGAGAGACA</td><td colspan="11"></td></tr>
<tr><td>Oligo rev</td><td>AAACTGTCTCTCGAACGCATGGAT</td><td colspan="11"></td></tr>
<tr><td>Chromosome</td><td>start</td><td></td><td>end</td><td>strand</td><td>MM</td><td>target_seq</td><td>PAM</td><td>alignment</td><td>distance</td><td>position</td><td>gene name</td><td>gene id</td></tr>
<tr><td>GU305914.1</td><td></td><td>54469</td><td>54491</td><td>+</td><td>0</td><td>ATCCATGCGTTCGAGAGACA</td><td>GGG</td><td>||||||||[||||||||||||]PAM</td><td>NA</td><td>Intergenic</td><td>NA</td><td>NA</td></tr>
<tr><td colspan="13"></td></tr>
<tr><td>T3555</td><td>TCTCGCTCGCTGTTCGCGCGCGG</td><td></td><td>1000</td><td colspan="9"></td></tr>
<tr><td>Oligo fwd</td><td>CACCTCTCGCTCGCTGTTCGCGCG</td><td colspan="11"></td></tr>
<tr><td>Oligo rev</td><td>AAACCGCGCGAACAGCGAGCGAGA</td><td colspan="11"></td></tr>
<tr><td>Chromosome</td><td>start</td><td></td><td>end</td><td>strand</td><td>MM</td><td>target_seq</td><td>PAM</td><td>alignment</td><td>distance</td><td>position</td><td>gene name</td><td>gene id</td></tr>
<tr><td>GU305914.1</td><td></td><td>54504</td><td>54526</td><td>+</td><td>0</td><td>TCTCGCTCGCTGTTCGCGCG</td><td>CGG</td><td>||||||||[||||||||||||]PAM</td><td>NA</td><td>Intergenic</td><td>NA</td><td>NA</td></tr>
<tr><td colspan="13"></td></tr>
<tr><td>T3556</td><td>ACGCGCAGCCACCTCCGTCGAGG</td><td></td><td>1000</td><td colspan="9"></td></tr>
<tr><td>Oligo fwd</td><td>CACCACGCGCAGCCACCTCCGTCG</td><td colspan="11"></td></tr>
<tr><td>Oligo rev</td><td>AAACCGACGGAGGTGGCTGCGCGT</td><td colspan="11"></td></tr>
<tr><td>Chromosome</td><td>start</td><td></td><td>end</td><td>strand</td><td>MM</td><td>target_seq</td><td>PAM</td><td>alignment</td><td>distance</td><td>position</td><td>gene name</td><td>gene id</td></tr>
<tr><td>GU305914.1</td><td></td><td>54541</td><td>54563</td><td>+</td><td>0</td><td>ACGCGCAGCCACCTCCGTCG</td><td>AGG</td><td>||||||||[||||||||||||]PAM</td><td>NA</td><td>Intergenic</td><td>NA</td><td>NA</td></tr>
<tr><td colspan="13"></td></tr>
<tr><td>T3557</td><td>AGCCACCTCCGTCGAGGACGAGG</td><td></td><td>1000</td><td colspan="9"></td></tr>
<tr><td>Oligo fwd</td><td>CACCAGCCACCTCCGTCGAGGACG</td><td colspan="11"></td></tr>
<tr><td>Oligo rev</td><td>AAACCGTCCTCGACGGAGGTGGCT</td><td colspan="11"></td></tr>
<tr><td>Chromosome</td><td>start</td><td></td><td>end</td><td>strand</td><td>MM</td><td>target_seq</td><td>PAM</td><td>alignment</td><td>distance</td><td>position</td><td>gene name</td><td>gene id</td></tr>
<tr><td>GU305914.1</td><td></td><td>54547</td><td>54569</td><td>+</td><td>0</td><td>AGCCACCTCCGTCGAGGACG</td><td>AGG</td><td>||||||||[||||||||||||]PAM</td><td>NA</td><td>Intergenic</td><td>NA</td><td>NA</td></tr>
<tr><td colspan="13"></td></tr>
<tr><td>T3558</td><td>CACCTCCGTCGAGGACGAGGTGG</td><td></td><td>1000</td><td colspan="9"></td></tr>
<tr><td>Oligo fwd</td><td>CACCCACCTCCGTCGAGGACGAGG</td><td colspan="11"></td></tr>
<tr><td>Oligo rev</td><td>AAACCCTCGTCCTCGACGGAGGTG</td><td colspan="11"></td></tr>
<tr><td>Chromosome</td><td>start</td><td></td><td>end</td><td>strand</td><td>MM</td><td>target_seq</td><td>PAM</td><td>alignment</td><td>distance</td><td>position</td><td>gene name</td><td>gene id</td></tr>
<tr><td>GU305914.1</td><td></td><td>54550</td><td>54572</td><td>+</td><td>0</td><td>CACCTCCGTCGAGGACGAGG</td><td>TGG</td><td>||||||||[||||||||||||]PAM</td><td>NA</td><td>Intergenic</td><td>NA</td><td>NA</td></tr>
<tr><td colspan="13"></td></tr>
<tr><td>T3559</td><td>CTCCGTCGAGGACGAGGTGGTGG</td><td></td><td>1000</td><td colspan="9"></td></tr>
<tr><td>Oligo fwd</td><td>CACCCTCCGTCGAGGACGAGGTGG</td><td colspan="11"></td></tr>
</table>

<table>
<tr><td>Oligo rev</td><td>AAACCCACCTCGTCCTCGACGGAG</td><td></td><td></td><td></td><td></td><td></td><td></td><td></td><td></td><td></td><td></td></tr>
<tr><td>Chromosome</td><td>start</td><td>end</td><td>strand</td><td>MM</td><td>target_seq</td><td>PAM</td><td>alignment</td><td>distance</td><td>position</td><td>gene name</td><td>gene id</td></tr>
<tr><td>GU305914.1</td><td>54553</td><td>54575</td><td>+</td><td>0</td><td>CTCCGTCGAGGACGAGGTGG</td><td>TGG</td><td>||||||||[||||||||||||]PAM</td><td>NA</td><td>Intergenic</td><td>NA</td><td>NA</td></tr>
<tr><td>T3560</td><td>CGTCGAGGACGAGGTGGTGGAGG</td><td>1000</td><td></td><td></td><td></td><td></td><td></td><td></td><td></td><td></td><td></td></tr>
<tr><td>Oligo fwd</td><td>CACCCGTCGAGGACGAGGTGGTGG</td><td></td><td></td><td></td><td></td><td></td><td></td><td></td><td></td><td></td><td></td></tr>
<tr><td>Oligo rev</td><td>AAACCCACCACCTCGTCCTCGACG</td><td></td><td></td><td></td><td></td><td></td><td></td><td></td><td></td><td></td><td></td></tr>
<tr><td>Chromosome</td><td>start</td><td>end</td><td>strand</td><td>MM</td><td>target_seq</td><td>PAM</td><td>alignment</td><td>distance</td><td>position</td><td>gene name</td><td>gene id</td></tr>
<tr><td>GU305914.1</td><td>54556</td><td>54578</td><td>+</td><td>0</td><td>CGTCGAGGACGAGGTGGTGG</td><td>AGG</td><td>||||||||[||||||||||||]PAM</td><td>NA</td><td>Intergenic</td><td>NA</td><td>NA</td></tr>
<tr><td>GU305914.1</td><td>196806</td><td>196828</td><td>+</td><td>4</td><td>TGTCGATGACGAGCAGGTGG</td><td>TGG</td><td>-|||||-|[|||||--|||||]PAM</td><td>NA</td><td>Intergenic</td><td>NA</td><td>NA</td></tr>
<tr><td>T3561</td><td>CGAGGACGAGGTGGTGGAGGCGG</td><td>1000</td><td></td><td></td><td></td><td></td><td></td><td></td><td></td><td></td><td></td></tr>
<tr><td>Oligo fwd</td><td>CACCCGAGGACGAGGTGGTGGAGG</td><td></td><td></td><td></td><td></td><td></td><td></td><td></td><td></td><td></td><td></td></tr>
<tr><td>Oligo rev</td><td>AAACCCTCCACCACCTCGTCCTCG</td><td></td><td></td><td></td><td></td><td></td><td></td><td></td><td></td><td></td><td></td></tr>
<tr><td>Chromosome</td><td>start</td><td>end</td><td>strand</td><td>MM</td><td>target_seq</td><td>PAM</td><td>alignment</td><td>distance</td><td>position</td><td>gene name</td><td>gene id</td></tr>
<tr><td>GU305914.1</td><td>54559</td><td>54581</td><td>+</td><td>0</td><td>CGAGGACGAGGTGGTGGAGG</td><td>CGG</td><td>||||||||[||||||||||||]PAM</td><td>NA</td><td>Intergenic</td><td>NA</td><td>NA</td></tr>
<tr><td>T3562</td><td>GGTGGTGGAGGCGGTTGACGAGG</td><td>1000</td><td></td><td></td><td></td><td></td><td></td><td></td><td></td><td></td><td></td></tr>
<tr><td>Oligo fwd</td><td>CACCGGTGGTGGAGGCGGTTGACG</td><td></td><td></td><td></td><td></td><td></td><td></td><td></td><td></td><td></td><td></td></tr>
<tr><td>Oligo rev</td><td>AAACCGTCAACCGCCTCCACCACC</td><td></td><td></td><td></td><td></td><td></td><td></td><td></td><td></td><td></td><td></td></tr>
<tr><td>Chromosome</td><td>start</td><td>end</td><td>strand</td><td>MM</td><td>target_seq</td><td>PAM</td><td>alignment</td><td>distance</td><td>position</td><td>gene name</td><td>gene id</td></tr>
<tr><td>GU305914.1</td><td>54568</td><td>54590</td><td>+</td><td>0</td><td>GGTGGTGGAGGCGGTTGACG</td><td>AGG</td><td>||||||||[||||||||||||]PAM</td><td>NA</td><td>Intergenic</td><td>NA</td><td>NA</td></tr>
<tr><td>T3563</td><td>GGAGGCGGTTGACGAGGATACGG</td><td>1000</td><td></td><td></td><td></td><td></td><td></td><td></td><td></td><td></td><td></td></tr>
<tr><td>Oligo fwd</td><td>CACCGGAGGCGGTTGACGAGGATA</td><td></td><td></td><td></td><td></td><td></td><td></td><td></td><td></td><td></td><td></td></tr>
<tr><td>Oligo rev</td><td>AAACTATCCTCGTCAACCGCCTCC</td><td></td><td></td><td></td><td></td><td></td><td></td><td></td><td></td><td></td><td></td></tr>
<tr><td>Chromosome</td><td>start</td><td>end</td><td>strand</td><td>MM</td><td>target_seq</td><td>PAM</td><td>alignment</td><td>distance</td><td>position</td><td>gene name</td><td>gene id</td></tr>
<tr><td>GU305914.1</td><td>54574</td><td>54596</td><td>+</td><td>0</td><td>GGAGGCGGTTGACGAGGATA</td><td>CGG</td><td>||||||||[||||||||||||]PAM</td><td>NA</td><td>Intergenic</td><td>NA</td><td>NA</td></tr>
<tr><td>GU305914.1</td><td>77043</td><td>77065</td><td>-</td><td>3</td><td>GGAGTCGGTCGACGAGGATG</td><td>AGG</td><td>||||-|||[|-|||||||||-]PAM</td><td>NA</td><td>Intergenic</td><td>NA</td><td>NA</td></tr>
<tr><td>T3564</td><td>GGTTGACGAGGATACGGAGATGG</td><td>1000</td><td></td><td></td><td></td><td></td><td></td><td></td><td></td><td></td><td></td></tr>
<tr><td>Oligo fwd</td><td>CACCGGTTGACGAGGATACGGAGA</td><td></td><td></td><td></td><td></td><td></td><td></td><td></td><td></td><td></td><td></td></tr>
<tr><td>Oligo rev</td><td>AAACTCTCCGTATCCTCGTCAACC</td><td></td><td></td><td></td><td></td><td></td><td></td><td></td><td></td><td></td><td></td></tr>
<tr><td>Chromosome</td><td>start</td><td>end</td><td>strand</td><td>MM</td><td>target_seq</td><td>PAM</td><td>alignment</td><td>distance</td><td>position</td><td>gene name</td><td>gene id</td></tr>
<tr><td>GU305914.1</td><td>54580</td><td>54602</td><td>+</td><td>0</td><td>GGTTGACGAGGATACGGAGA</td><td>TGG</td><td>||||||||[||||||||||||]PAM</td><td>NA</td><td>Intergenic</td><td>NA</td><td>NA</td></tr>
<tr><td>T3565</td><td>CGAGGATACGGAGATGGAGATGG</td><td>1000</td><td></td><td></td><td></td><td></td><td></td><td></td><td></td><td></td><td></td></tr>
<tr><td>Oligo fwd</td><td>CACCCGAGGATACGGAGATGGAGA</td><td></td><td></td><td></td><td></td><td></td><td></td><td></td><td></td><td></td><td></td></tr>
<tr><td>Oligo rev</td><td>AAACTCTCCATCTCCGTATCCTCG</td><td></td><td></td><td></td><td></td><td></td><td></td><td></td><td></td><td></td><td></td></tr>
<tr><td>Chromosome</td><td>start</td><td>end</td><td>strand</td><td>MM</td><td>target_seq</td><td>PAM</td><td>alignment</td><td>distance</td><td>position</td><td>gene name</td><td>gene id</td></tr>
<tr><td>GU305914.1</td><td>54586</td><td>54608</td><td>+</td><td>0</td><td>CGAGGATACGGAGATGGAGA</td><td>TGG</td><td>||||||||[||||||||||||]PAM</td><td>NA</td><td>Intergenic</td><td>NA</td><td>NA</td></tr>
<tr><td>T3566</td><td>GGAGTTCTCTCCGCGTAACGCGG</td><td>1000</td><td></td><td></td><td></td><td></td><td></td><td></td><td></td><td></td><td></td></tr>
<tr><td>Oligo fwd</td><td>CACCGGAGTTCTCTCCGCGTAACG</td><td></td><td></td><td></td><td></td><td></td><td></td><td></td><td></td><td></td><td></td></tr>
<tr><td>Oligo rev</td><td>AAACCGTTACGCGGAGAGAACTCC</td><td></td><td></td><td></td><td></td><td></td><td></td><td></td><td></td><td></td><td></td></tr>
<tr><td>Chromosome</td><td>start</td><td>end</td><td>strand</td><td>MM</td><td>target_seq</td><td>PAM</td><td>alignment</td><td>distance</td><td>position</td><td>gene name</td><td>gene id</td></tr>
<tr><td>GU305914.1</td><td>54607</td><td>54629</td><td>+</td><td>0</td><td>GGAGTTCTCTCCGCGTAACG</td><td>CGG</td><td>||||||||[||||||||||||]PAM</td><td>NA</td><td>Intergenic</td><td>NA</td><td>NA</td></tr>
<tr><td>T3567</td><td>TCTCTCCGCGTAACGCGGCGCGG</td><td>1000</td><td></td><td></td><td></td><td></td><td></td><td></td><td></td><td></td><td></td></tr>
<tr><td>Oligo fwd</td><td>CACCTCTCTCCGCGTAACGCGGCG</td><td></td><td></td><td></td><td></td><td></td><td></td><td></td><td></td><td></td><td></td></tr>
<tr><td>Oligo rev</td><td>AAACCGCCGCGTTACGCGGAGAGA</td><td></td><td></td><td></td><td></td><td></td><td></td><td></td><td></td><td></td><td></td></tr>
<tr><td>Chromosome</td><td>start</td><td>end</td><td>strand</td><td>MM</td><td>target_seq</td><td>PAM</td><td>alignment</td><td>distance</td><td>position</td><td>gene name</td><td>gene id</td></tr>
<tr><td>GU305914.1</td><td>54612</td><td>54634</td><td>+</td><td>0</td><td>TCTCTCCGCGTAACGCGGCG</td><td>CGG</td><td>||||||||[||||||||||||]PAM</td><td>NA</td><td>Intergenic</td><td>NA</td><td>NA</td></tr>
<tr><td>T3568</td><td>GTTACCAAAGAGACTCTCGTAGG</td><td>1000</td><td></td><td></td><td></td><td></td><td></td><td></td><td></td><td></td><td></td></tr>
<tr><td>Oligo fwd</td><td>CACCGTTACCAAAGAGACTCTCGT</td><td></td><td></td><td></td><td></td><td></td><td></td><td></td><td></td><td></td><td></td></tr>
<tr><td>Oligo rev</td><td>AAACACGAGAGTCTCTTTGGTAAC</td><td></td><td></td><td></td><td></td><td></td><td></td><td></td><td></td><td></td><td></td></tr>
<tr><td>Chromosome</td><td>start</td><td>end</td><td>strand</td><td>MM</td><td>target_seq</td><td>PAM</td><td>alignment</td><td>distance</td><td>position</td><td>gene name</td><td>gene id</td></tr>
<tr><td>GU305914.1</td><td>54670</td><td>54692</td><td>+</td><td>0</td><td>GTTACCAAAGAGACTCTCGT</td><td>AGG</td><td>||||||||[||||||||||||]PAM</td><td>NA</td><td>Intergenic</td><td>NA</td><td>NA</td></tr>
<tr><td>T3569</td><td>CCAAAGAGACTCTCGTAGGACGG</td><td>1000</td><td></td><td></td><td></td><td></td><td></td><td></td><td></td><td></td><td></td></tr>
<tr><td>Oligo fwd</td><td>CACCCCAAAGAGACTCTCGTAGGA</td><td></td><td></td><td></td><td></td><td></td><td></td><td></td><td></td><td></td><td></td></tr>
<tr><td>Oligo rev</td><td>AAACTCCTACGAGAGTCTCTTTGG</td><td></td><td></td><td></td><td></td><td></td><td></td><td></td><td></td><td></td><td></td></tr>
<tr><td>Chromosome</td><td>start</td><td>end</td><td>strand</td><td>MM</td><td>target_seq</td><td>PAM</td><td>alignment</td><td>distance</td><td>position</td><td>gene name</td><td>gene id</td></tr>
<tr><td>GU305914.1</td><td>54674</td><td>54696</td><td>+</td><td>0</td><td>CCAAAGAGACTCTCGTAGGA</td><td>CGG</td><td>||||||||[||||||||||||]PAM</td><td>NA</td><td>Intergenic</td><td>NA</td><td>NA</td></tr>
<tr><td>T3570</td><td>AAGAGACTCTCGTAGGACGGTGG</td><td>1000</td><td></td><td></td><td></td><td></td><td></td><td></td><td></td><td></td><td></td></tr>
<tr><td>Oligo fwd</td><td>CACCAAGAGACTCTCGTAGGACGG</td><td></td><td></td><td></td><td></td><td></td><td></td><td></td><td></td><td></td><td></td></tr>
<tr><td>Oligo rev</td><td>AAACCCGTCCTACGAGAGTCTCTT</td><td></td><td></td><td></td><td></td><td></td><td></td><td></td><td></td><td></td><td></td></tr>
<tr><td>Chromosome</td><td>start</td><td>end</td><td>strand</td><td>MM</td><td>target_seq</td><td>PAM</td><td>alignment</td><td>distance</td><td>position</td><td>gene name</td><td>gene id</td></tr>
<tr><td>GU305914.1</td><td>54677</td><td>54699</td><td>+</td><td>0</td><td>AAGAGACTCTCGTAGGACGG</td><td>TGG</td><td>||||||||[||||||||||||]PAM</td><td>NA</td><td>Intergenic</td><td>NA</td><td>NA</td></tr>
<tr><td>T3571</td><td>AGAGACTCTCGTAGGACGGTGGG</td><td>1000</td><td></td><td></td><td></td><td></td><td></td><td></td><td></td><td></td><td></td></tr>
<tr><td>Oligo fwd</td><td>CACCAGAGACTCTCGTAGGACGGT</td><td></td><td></td><td></td><td></td><td></td><td></td><td></td><td></td><td></td><td></td></tr>
<tr><td>Oligo rev</td><td>AAACACCGTCCTACGAGAGTCTCT</td><td></td><td></td><td></td><td></td><td></td><td></td><td></td><td></td><td></td><td></td></tr>
<tr><td>Chromosome</td><td>start</td><td>end</td><td>strand</td><td>MM</td><td>target_seq</td><td>PAM</td><td>alignment</td><td>distance</td><td>position</td><td>gene name</td><td>gene id</td></tr>
<tr><td>GU305914.1</td><td>54678</td><td>54700</td><td>+</td><td>0</td><td>AGAGACTCTCGTAGGACGGT</td><td>GGG</td><td>||||||||[||||||||||||]PAM</td><td>NA</td><td>Intergenic</td><td>NA</td><td>NA</td></tr>
<tr><td>T3572</td><td>GAGACTCTCGTAGGACGGTGGGG</td><td>1000</td><td></td><td></td><td></td><td></td><td></td><td></td><td></td><td></td><td></td></tr>
<tr><td>Oligo fwd</td><td>CACCGAGACTCTCGTAGGACGGTG</td><td></td><td></td><td></td><td></td><td></td><td></td><td></td><td></td><td></td><td></td></tr>
<tr><td>Oligo rev</td><td>AAACCACCGTCCTACGAGAGTCTC</td><td></td><td></td><td></td><td></td><td></td><td></td><td></td><td></td><td></td><td></td></tr>
<tr><td>Chromosome</td><td>start</td><td>end</td><td>strand</td><td>MM</td><td>target_seq</td><td>PAM</td><td>alignment</td><td>distance</td><td>position</td><td>gene name</td><td>gene id</td></tr>
<tr><td>GU305914.1</td><td>54679</td><td>54701</td><td>+</td><td>0</td><td>GAGACTCTCGTAGGACGGTG</td><td>GGG</td><td>||||||||[||||||||||||]PAM</td><td>NA</td><td>Intergenic</td><td>NA</td><td>NA</td></tr>
<tr><td>T3573</td><td>AGACTCTCGTAGGACGGTGGGGG</td><td>1000</td><td></td><td></td><td></td><td></td><td></td><td></td><td></td><td></td><td></td></tr>
<tr><td>Oligo fwd</td><td>CACCAGACTCTCGTAGGACGGTGG</td><td></td><td></td><td></td><td></td><td></td><td></td><td></td><td></td><td></td><td></td></tr>
<tr><td>Oligo rev</td><td>AAACCCACCGTCCTACGAGAGTCT</td><td></td><td></td><td></td><td></td><td></td><td></td><td></td><td></td><td></td><td></td></tr>
<tr><td>Chromosome</td><td>start</td><td>end</td><td>strand</td><td>MM</td><td>target_seq</td><td>PAM</td><td>alignment</td><td>distance</td><td>position</td><td>gene name</td><td>gene id</td></tr>
<tr><td>GU305914.1</td><td>54680</td><td>54702</td><td>+</td><td>0</td><td>AGACTCTCGTAGGACGGTGG</td><td>GGG</td><td>||||||||[||||||||||||]PAM</td><td>NA</td><td>Intergenic</td><td>NA</td><td>NA</td></tr>
</table>

| | | | | | | | | | | | |
|---|---|---|---|---|---|---|---|---|---|---|---|
| T3574 | ATTCGTGACCACCAGTCGCGTGG | 1000 | | | | | | | | | |
| Oligo fwd | CACCATTCGTGACCACCAGTCGCG | | | | | | | | | | |
| Oligo rev | AAACCGCGACTGGTGGTCACGAAT | | | | | | | | | | |
| Chromosome | start | end | strand | MM | target_seq | PAM | alignment | distance | position | gene name | gene id |
| GU305914.1 | 54727 | 54749 | + | 0 | ATTCGTGACCACCAGTCGCG | TGG | \|\|\|\|\|\|\|\|[\|\|\|\|\|\|\|\|\|\|\|\|]PAM | NA | Intergenic | NA | NA |
| T3575 | CGTGACCACCAGTCGCGTGGTGG | 1000 | | | | | | | | | |
| Oligo fwd | CACCCGTGACCACCAGTCGCGTGG | | | | | | | | | | |
| Oligo rev | AAACCCACGCGACTGGTGGTCACG | | | | | | | | | | |
| Chromosome | start | end | strand | MM | target_seq | PAM | alignment | distance | position | gene name | gene id |
| GU305914.1 | 54730 | 54752 | + | 0 | CGTGACCACCAGTCGCGTGG | TGG | \|\|\|\|\|\|\|\|[\|\|\|\|\|\|\|\|\|\|\|\|]PAM | NA | Intergenic | NA | NA |
| T3576 | CCGACCGCTTCCAGATAAGTAGG | 1000 | | | | | | | | | |
| Oligo fwd | CACCCCGACCGCTTCCAGATAAGT | | | | | | | | | | |
| Oligo rev | AAACACTTATCTGGAAGCGGTCGG | | | | | | | | | | |
| Chromosome | start | end | strand | MM | target_seq | PAM | alignment | distance | position | gene name | gene id |
| GU305914.1 | 54794 | 54816 | + | 0 | CCGACCGCTTCCAGATAAGT | AGG | \|\|\|\|\|\|\|\|[\|\|\|\|\|\|\|\|\|\|\|\|]PAM | NA | Intergenic | NA | NA |
| T3577 | CTTCCAGATAAGTAGGCAAACGG | 1000 | | | | | | | | | |
| Oligo fwd | CACCCTTCCAGATAAGTAGGCAAA | | | | | | | | | | |
| Oligo rev | AAACTTTGCCTACTTATCTGGAAG | | | | | | | | | | |
| Chromosome | start | end | strand | MM | target_seq | PAM | alignment | distance | position | gene name | gene id |
| GU305914.1 | 54801 | 54823 | + | 0 | CTTCCAGATAAGTAGGCAAA | CGG | \|\|\|\|\|\|\|\|[\|\|\|\|\|\|\|\|\|\|\|\|]PAM | NA | Intergenic | NA | NA |
| T3578 | TAAGTAGGCAAACGGTGCTCCGG | 1000 | | | | | | | | | |
| Oligo fwd | CACCTAAGTAGGCAAACGGTGCTC | | | | | | | | | | |
| Oligo rev | AAACGAGCACCGTTTGCCTACTTA | | | | | | | | | | |
| Chromosome | start | end | strand | MM | target_seq | PAM | alignment | distance | position | gene name | gene id |
| GU305914.1 | 54809 | 54831 | + | 0 | TAAGTAGGCAAACGGTGCTC | CGG | \|\|\|\|\|\|\|\|[\|\|\|\|\|\|\|\|\|\|\|\|]PAM | NA | Intergenic | NA | NA |
| T3579 | AGGCAAACGGTGCTCCGGTTCGG | 1000 | | | | | | | | | |
| Oligo fwd | CACCAGGCAAACGGTGCTCCGGTT | | | | | | | | | | |
| Oligo rev | AAACAACCGGAGCACCGTTTGCCT | | | | | | | | | | |
| Chromosome | start | end | strand | MM | target_seq | PAM | alignment | distance | position | gene name | gene id |
| GU305914.1 | 54814 | 54836 | + | 0 | AGGCAAACGGTGCTCCGGTT | CGG | \|\|\|\|\|\|\|\|[\|\|\|\|\|\|\|\|\|\|\|\|]PAM | NA | Intergenic | NA | NA |
| T3580 | TATCAGCATATATAGTATCAAGG | 1000 | | | | | | | | | |
| Oligo fwd | CACCTATCAGCATATATAGTATCA | | | | | | | | | | |
| Oligo rev | AAACTGATACTATATATGCTGATA | | | | | | | | | | |
| Chromosome | start | end | strand | MM | target_seq | PAM | alignment | distance | position | gene name | gene id |
| GU305914.1 | 54894 | 54916 | + | 0 | TATCAGCATATATAGTATCA | AGG | \|\|\|\|\|\|\|\|[\|\|\|\|\|\|\|\|\|\|\|\|]PAM | NA | Intergenic | NA | NA |
| T3581 | CAATGAGAGCCACGATCTTCAGG | 1000 | | | | | | | | | |
| Oligo fwd | CACCCAATGAGAGCCACGATCTTC | | | | | | | | | | |
| Oligo rev | AAACGAAGATCGTGGCTCTCATTG | | | | | | | | | | |
| Chromosome | start | end | strand | MM | target_seq | PAM | alignment | distance | position | gene name | gene id |
| GU305914.1 | 54926 | 54948 | + | 0 | CAATGAGAGCCACGATCTTC | AGG | \|\|\|\|\|\|\|\|[\|\|\|\|\|\|\|\|\|\|\|\|]PAM | NA | Intergenic | NA | NA |
| T3582 | ATCTTCAGGAGATCGTACCACGG | 1000 | | | | | | | | | |
| Oligo fwd | CACCATCTTCAGGAGATCGTACCA | | | | | | | | | | |
| Oligo rev | AAACTGGTACGATCTCCTGAAGAT | | | | | | | | | | |
| Chromosome | start | end | strand | MM | target_seq | PAM | alignment | distance | position | gene name | gene id |
| GU305914.1 | 54940 | 54962 | + | 0 | ATCTTCAGGAGATCGTACCA | CGG | \|\|\|\|\|\|\|\|[\|\|\|\|\|\|\|\|\|\|\|\|]PAM | NA | Intergenic | NA | NA |
| T3583 | TCTTCAGGAGATCGTACCACGGG | 1000 | | | | | | | | | |
| Oligo fwd | CACCTCTTCAGGAGATCGTACCAC | | | | | | | | | | |
| Oligo rev | AAACGTGGTACGATCTCCTGAAGA | | | | | | | | | | |
| Chromosome | start | end | strand | MM | target_seq | PAM | alignment | distance | position | gene name | gene id |
| GU305914.1 | 54941 | 54963 | + | 0 | TCTTCAGGAGATCGTACCAC | GGG | \|\|\|\|\|\|\|\|[\|\|\|\|\|\|\|\|\|\|\|\|]PAM | NA | Intergenic | NA | NA |
| T3584 | GGAGATCGTACCACGGGTTGTGG | 1000 | | | | | | | | | |
| Oligo fwd | CACCGGAGATCGTACCACGGGTTG | | | | | | | | | | |
| Oligo rev | AAACCAACCCGTGGTACGATCTCC | | | | | | | | | | |
| Chromosome | start | end | strand | MM | target_seq | PAM | alignment | distance | position | gene name | gene id |
| GU305914.1 | 54947 | 54969 | + | 0 | GGAGATCGTACCACGGGTTG | TGG | \|\|\|\|\|\|\|\|[\|\|\|\|\|\|\|\|\|\|\|\|]PAM | NA | Intergenic | NA | NA |
| T3585 | TGGCCCTGCCTGACCCGCCTAGG | 1000 | | | | | | | | | |
| Oligo fwd | CACCTGGCCCTGCCTGACCCGCCT | | | | | | | | | | |
| Oligo rev | AAACAGGCGGGTCAGGCAGGGCCA | | | | | | | | | | |
| Chromosome | start | end | strand | MM | target_seq | PAM | alignment | distance | position | gene name | gene id |
| GU305914.1 | 54967 | 54989 | + | 0 | TGGCCCTGCCTGACCCGCCT | AGG | \|\|\|\|\|\|\|\|[\|\|\|\|\|\|\|\|\|\|\|\|]PAM | NA | Intergenic | NA | NA |
| T3586 | CATACTCCGTCTATCCCGAAAGG | 1000 | | | | | | | | | |
| Oligo fwd | CACCCATACTCCGTCTATCCCGAA | | | | | | | | | | |
| Oligo rev | AAACTTCGGGATAGACGGAGTATG | | | | | | | | | | |
| Chromosome | start | end | strand | MM | target_seq | PAM | alignment | distance | position | gene name | gene id |
| GU305914.1 | 54990 | 55012 | + | 0 | CATACTCCGTCTATCCCGAA | AGG | \|\|\|\|\|\|\|\|[\|\|\|\|\|\|\|\|\|\|\|\|]PAM | NA | Intergenic | NA | NA |
| T3587 | CGTCTATCCCGAAAGGCAATCGG | 1000 | | | | | | | | | |
| Oligo fwd | CACCCGTCTATCCCGAAAGGCAAT | | | | | | | | | | |
| Oligo rev | AAACATTGCCTTTCGGGATAGACG | | | | | | | | | | |
| Chromosome | start | end | strand | MM | target_seq | PAM | alignment | distance | position | gene name | gene id |
| GU305914.1 | 54997 | 55019 | + | 0 | CGTCTATCCCGAAAGGCAAT | CGG | \|\|\|\|\|\|\|\|[\|\|\|\|\|\|\|\|\|\|\|\|]PAM | NA | Intergenic | NA | NA |
| T3588 | ACCTTCCGAAGCGTTCAGAGAGG | 1000 | | | | | | | | | |
| Oligo fwd | CACCACCTTCCGAAGCGTTCAGAG | | | | | | | | | | |
| Oligo rev | AAACCTCTGAACGCTTCGGAAGGT | | | | | | | | | | |
| Chromosome | start | end | strand | MM | target_seq | PAM | alignment | distance | position | gene name | gene id |

GU305914.1 55067 55089 + 0 ACCTTCCGAAGCGTTCAGAG AGG ||||||||[||||||||||||]PAM NA Intergenic NA NA

T3589 ACGAGCAAACACCAGAGTCTCGG 1000
Oligo fwd CACCACGAGCAAACACCAGAGTCT
Oligo rev AAACAGACTCTGGTGTTTGCTCGT
Chromosome start end strand MM target_seq PAM alignment distance position gene name gene id
GU305914.1 55096 55118 + 0 ACGAGCAAACACCAGAGTCT CGG ||||||||[||||||||||||]PAM NA Intergenic NA NA

T3590 GTCTCGGTTCCCGTACCTATCGG 1000
Oligo fwd CACCGTCTCGGTTCCCGTACCTAT
Oligo rev AAACATAGGTACGGGAACCGAGAC
Chromosome start end strand MM target_seq PAM alignment distance position gene name gene id
GU305914.1 55112 55134 + 0 GTCTCGGTTCCCGTACCTAT CGG ||||||||[||||||||||||]PAM NA Intergenic NA NA

T3591 ACAGCGTTAAATAATAATCCTGG 1000
Oligo fwd CACCACAGCGTTAAATAATAATCC
Oligo rev AAACGGATTATTATTTAACGCTGT
Chromosome start end strand MM target_seq PAM alignment distance position gene name gene id
GU305914.1 55149 55171 + 0 ACAGCGTTAAATAATAATCC TGG ||||||||[||||||||||||]PAM NA Intergenic NA NA

T3592 AATAATAATCCTGGCGCCCGCGG 1000
Oligo fwd CACCAATAATAATCCTGGCGCCCG
Oligo rev AAACCGGGCGCCAGGATTATTATT
Chromosome start end strand MM target_seq PAM alignment distance position gene name gene id
GU305914.1 55158 55180 + 0 AATAATAATCCTGGCGCCCG CGG ||||||||[||||||||||||]PAM NA Intergenic NA NA

T3593 AATAATCCTGGCGCCCGCGGCGG 1000
Oligo fwd CACCAATAATCCTGGCGCCCGCGG
Oligo rev AAACCCGCGGGCGCCAGGATTATT
Chromosome start end strand MM target_seq PAM alignment distance position gene name gene id
GU305914.1 55161 55183 + 0 AATAATCCTGGCGCCCGCGG CGG ||||||||[||||||||||||]PAM NA Intergenic NA NA

T3594 GGCGCCGAGACAGTTATCTACGG 1000
Oligo fwd CACCGGCGCCGAGACAGTTATCTA
Oligo rev AAACTAGATAACTGTCTCGGCGCC
Chromosome start end strand MM target_seq PAM alignment distance position gene name gene id
GU305914.1 55182 55204 + 0 GGCGCCGAGACAGTTATCTA CGG ||||||||[||||||||||||]PAM NA Intergenic NA NA

T3595 CCGAGACAGTTATCTACGGAAGG 1000
Oligo fwd CACCCCGAGACAGTTATCTACGGA
Oligo rev AAACTCCGTAGATAACTGTCTCGG
Chromosome start end strand MM target_seq PAM alignment distance position gene name gene id
GU305914.1 55186 55208 + 0 CCGAGACAGTTATCTACGGA AGG ||||||||[||||||||||||]PAM NA Intergenic NA NA

T3596 CGAGACAGTTATCTACGGAAGGG 1000
Oligo fwd CACCCGAGACAGTTATCTACGGAA
Oligo rev AAACTTCCGTAGATAACTGTCTCG
Chromosome start end strand MM target_seq PAM alignment distance position gene name gene id
GU305914.1 55187 55209 + 0 CGAGACAGTTATCTACGGAA GGG ||||||||[||||||||||||]PAM NA Intergenic NA NA

T3597 CGGAAGGGAACCGCCGTTCCCGG 1000
Oligo fwd CACCCGGAAGGGAACCGCCGTTCC
Oligo rev AAACGGAACGGCGGTTCCCTTCCG
Chromosome start end strand MM target_seq PAM alignment distance position gene name gene id
GU305914.1 55202 55224 + 0 CGGAAGGGAACCGCCGTTCC CGG ||||||||[||||||||||||]PAM NA Intergenic NA NA

T3598 GTTCCCGGCCCTCTTCTCCCCGG 1000
Oligo fwd CACCGTTCCCGGCCCTCTTCTCCC
Oligo rev AAACGGGAGAAGAGGGCCGGGAAC
Chromosome start end strand MM target_seq PAM alignment distance position gene name gene id
GU305914.1 55217 55239 + 0 GTTCCCGGCCCTCTTCTCCC CGG ||||||||[||||||||||||]PAM NA Intergenic NA NA

T3599 TTCCCGGCCCTCTTCTCCCCGGG 1000
Oligo fwd CACCTTCCCGGCCCTCTTCTCCCC
Oligo rev AAACGGGGAGAAGAGGGCCGGGAA
Chromosome start end strand MM target_seq PAM alignment distance position gene name gene id
GU305914.1 55218 55240 + 0 TTCCCGGCCCTCTTCTCCCC GGG ||||||||[||||||||||||]PAM NA Intergenic NA NA

T3600 CCCTCTTCTCCCCGGGAGAGAGG 1000
Oligo fwd CACCCCCTCTTCTCCCCGGGAGAG
Oligo rev AAACCTCTCCCGGGGAGAAGAGGG
Chromosome start end strand MM target_seq PAM alignment distance position gene name gene id
GU305914.1 55225 55247 + 0 CCCTCTTCTCCCCGGGAGAG AGG ||||||||[||||||||||||]PAM NA Intergenic NA NA

T3601 CTTCTCCCCGGGAGAGAGGAAGG 1000
Oligo fwd CACCCTTCTCCCCGGGAGAGAGGA
Oligo rev AAACTCCTCTCTCCCGGGGAGAAG
Chromosome start end strand MM target_seq PAM alignment distance position gene name gene id
GU305914.1 55229 55251 + 0 CTTCTCCCCGGGAGAGAGGA AGG ||||||||[||||||||||||]PAM NA Intergenic NA NA
GU305914.1 57789 57811 + 4 ATTCACCGCGGGCGAGAGGA GGG -|||-||-[||||-|||||||]PAM NA Intergenic NA NA

T3602 CGTCCCGTTTTTATTGATGTCGG 1000
Oligo fwd CACCCGTCCCGTTTTTATTGATGT
Oligo rev AAACACATCAATAAAAACGGGACG
Chromosome start end strand MM target_seq PAM alignment distance position gene name gene id
GU305914.1 55344 55366 + 0 CGTCCCGTTTTTATTGATGT CGG ||||||||[||||||||||||]PAM NA Intergenic NA NA

T3603 TCGAGACCCTTCTCTTGCGTCGG 1000
Oligo fwd CACCTCGAGACCCTTCTCTTGCGT

Oligo rev AAACACGCAAGAGAAGGGTCTCGA
Chromosome start end strand MM target_seq PAM alignment distance position gene name gene id
GU305914.1 55425 55447 + 0 TCGAGACCCTTCTCTTGCGT CGG ||||||||[||||||||||||]PAM NA Intergenic NA NA

T3604 CGTCGGCACCTCTGCCCGAGCGG 1000
Oligo fwd CACCCGTCGGCACCTCTGCCCGAG
Oligo rev AAACCTCGGGCAGAGGTGCCGACG
Chromosome start end strand MM target_seq PAM alignment distance position gene name gene id
GU305914.1 55442 55464 + 0 CGTCGGCACCTCTGCCCGAG CGG ||||||||[||||||||||||]PAM NA Intergenic NA NA

T3605 TGCCCGAGCGGACAGAAGACCGG 1000
Oligo fwd CACCTGCCCGAGCGGACAGAAGAC
Oligo rev AAACGTCTTCTGTCCGCTCGGGCA
Chromosome start end strand MM target_seq PAM alignment distance position gene name gene id
GU305914.1 55454 55476 + 0 TGCCCGAGCGGACAGAAGAC CGG ||||||||[||||||||||||]PAM NA Intergenic NA NA

T3606 CGGACAGAAGACCGGCCTGATGG 1000
Oligo fwd CACCCGGACAGAAGACCGGCCTGA
Oligo rev AAACTCAGGCCGGTCTTCTGTCCG
Chromosome start end strand MM target_seq PAM alignment distance position gene name gene id
GU305914.1 55462 55484 + 0 CGGACAGAAGACCGGCCTGA TGG ||||||||[||||||||||||]PAM NA Intergenic NA NA

T3607 GGACAGAAGACCGGCCTGATGGG 1000
Oligo fwd CACCGGACAGAAGACCGGCCTGAT
Oligo rev AAACATCAGGCCGGTCTTCTGTCC
Chromosome start end strand MM target_seq PAM alignment distance position gene name gene id
GU305914.1 55463 55485 + 0 GGACAGAAGACCGGCCTGAT GGG ||||||||[||||||||||||]PAM NA Intergenic NA NA

T3608 GACCGGCCTGATGGGCTGTCCGG 1000
Oligo fwd CACCGACCGGCCTGATGGGCTGTC
Oligo rev AAACGACAGCCCATCAGGCCGGTC
Chromosome start end strand MM target_seq PAM alignment distance position gene name gene id
GU305914.1 55471 55493 + 0 GACCGGCCTGATGGGCTGTC CGG ||||||||[||||||||||||]PAM NA Intergenic NA NA

T3609 ACCGGCCTGATGGGCTGTCCGGG 1000
Oligo fwd CACCACCGGCCTGATGGGCTGTCC
Oligo rev AAACGGACAGCCCATCAGGCCGGT
Chromosome start end strand MM target_seq PAM alignment distance position gene name gene id
GU305914.1 55472 55494 + 0 ACCGGCCTGATGGGCTGTCC GGG ||||||||[||||||||||||]PAM NA Intergenic NA NA

T3610 CCGGCCTGATGGGCTGTCCGGGG 1000
Oligo fwd CACCCCGGCCTGATGGGCTGTCCG
Oligo rev AAACCGGACAGCCCATCAGGCCGG
Chromosome start end strand MM target_seq PAM alignment distance position gene name gene id
GU305914.1 55473 55495 + 0 CCGGCCTGATGGGCTGTCCG GGG ||||||||[||||||||||||]PAM NA Intergenic NA NA

T3611 CTGATGGGCTGTCCGGGGCGCGG 1000
Oligo fwd CACCCTGATGGGCTGTCCGGGGCG
Oligo rev AAACCGCCCCGGACAGCCCATCAG
Chromosome start end strand MM target_seq PAM alignment distance position gene name gene id
GU305914.1 55478 55500 + 0 CTGATGGGCTGTCCGGGGCG CGG ||||||||[||||||||||||]PAM NA Intergenic NA NA

T3612 TGATGGGCTGTCCGGGGCGCGGG 1000
Oligo fwd CACCTGATGGGCTGTCCGGGGCGC
Oligo rev AAACGCGCCCCGGACAGCCCATCA
Chromosome start end strand MM target_seq PAM alignment distance position gene name gene id
GU305914.1 55479 55501 + 0 TGATGGGCTGTCCGGGGCGC GGG ||||||||[||||||||||||]PAM NA Intergenic NA NA

T3613 CGGGGCGCGGGTCTTCGAGTCGG 1000
Oligo fwd CACCCGGGGCGCGGGTCTTCGAGT
Oligo rev AAACACTCGAAGACCCGCGCCCCG
Chromosome start end strand MM target_seq PAM alignment distance position gene name gene id
GU305914.1 55491 55513 + 0 CGGGGCGCGGGTCTTCGAGT CGG ||||||||[||||||||||||]PAM NA Intergenic NA NA

T3614 GCGGGTCTTCGAGTCGGCGCTGG 1000
Oligo fwd CACCGCGGGTCTTCGAGTCGGCGC
Oligo rev AAACGCGCCGACTCGAAGACCCGC
Chromosome start end strand MM target_seq PAM alignment distance position gene name gene id
GU305914.1 55497 55519 + 0 GCGGGTCTTCGAGTCGGCGC TGG ||||||||[||||||||||||]PAM NA Intergenic NA NA

T3615 CGGGTCTTCGAGTCGGCGCTGGG 1000
Oligo fwd CACCCGGGTCTTCGAGTCGGCGCT
Oligo rev AAACAGCGCCGACTCGAAGACCCG
Chromosome start end strand MM target_seq PAM alignment distance position gene name gene id
GU305914.1 55498 55520 + 0 CGGGTCTTCGAGTCGGCGCT GGG ||||||||[||||||||||||]PAM NA Intergenic NA NA

T3616 GGGTCTTCGAGTCGGCGCTGGGG 1000
Oligo fwd CACCGGGTCTTCGAGTCGGCGCTG
Oligo rev AAACCAGCGCCGACTCGAAGACCC
Chromosome start end strand MM target_seq PAM alignment distance position gene name gene id
GU305914.1 55499 55521 + 0 GGGTCTTCGAGTCGGCGCTG GGG ||||||||[||||||||||||]PAM NA Intergenic NA NA

T3617 GGTCTTCGAGTCGGCGCTGGGGG 1000
Oligo fwd CACCGGTCTTCGAGTCGGCGCTGG
Oligo rev AAACCCAGCGCCGACTCGAAGACC
Chromosome start end strand MM target_seq PAM alignment distance position gene name gene id
GU305914.1 55500 55522 + 0 GGTCTTCGAGTCGGCGCTGG GGG ||||||||[||||||||||||]PAM NA Intergenic NA NA

T3618 TTCGAGTCGGCGCTGGGGGAAGG 1000

<table>
<tr><td>Oligo fwd</td><td>CACCTTCGAGTCGGCGCTGGGGGA</td><td></td><td></td><td></td><td></td><td></td><td></td><td></td><td></td><td></td><td></td></tr>
<tr><td>Oligo rev</td><td>AAACTCCCCCAGCGCCGACTCGAA</td><td></td><td></td><td></td><td></td><td></td><td></td><td></td><td></td><td></td><td></td></tr>
<tr><td>Chromosome</td><td>start</td><td>end</td><td>strand</td><td>MM</td><td>target_seq</td><td>PAM</td><td>alignment</td><td>distance</td><td>position</td><td>gene name</td><td>gene id</td></tr>
<tr><td>GU305914.1</td><td>55504</td><td>55526</td><td>+</td><td>0</td><td>TTCGAGTCGGCGCTGGGGGA</td><td>AGG</td><td>||||||||[||||||||||||]PAM</td><td>NA</td><td>Intergenic</td><td>NA</td><td>NA</td></tr>
<tr><td></td><td></td><td></td><td></td><td></td><td></td><td></td><td></td><td></td><td></td><td></td><td></td></tr>
<tr><td>T3619</td><td>TCGGCGCTGGGGGAAGGTGTTGG</td><td>1000</td><td></td><td></td><td></td><td></td><td></td><td></td><td></td><td></td><td></td></tr>
<tr><td>Oligo fwd</td><td>CACCTCGGCGCTGGGGGAAGGTGT</td><td></td><td></td><td></td><td></td><td></td><td></td><td></td><td></td><td></td><td></td></tr>
<tr><td>Oligo rev</td><td>AAACACACCTTCCCCCAGCGCCGA</td><td></td><td></td><td></td><td></td><td></td><td></td><td></td><td></td><td></td><td></td></tr>
<tr><td>Chromosome</td><td>start</td><td>end</td><td>strand</td><td>MM</td><td>target_seq</td><td>PAM</td><td>alignment</td><td>distance</td><td>position</td><td>gene name</td><td>gene id</td></tr>
<tr><td>GU305914.1</td><td>55510</td><td>55532</td><td>+</td><td>0</td><td>TCGGCGCTGGGGGAAGGTGT</td><td>TGG</td><td>||||||||[||||||||||||]PAM</td><td>NA</td><td>Intergenic</td><td>NA</td><td>NA</td></tr>
<tr><td></td><td></td><td></td><td></td><td></td><td></td><td></td><td></td><td></td><td></td><td></td><td></td></tr>
<tr><td>T3620</td><td>CGGCGCTGGGGGAAGGTGTTGGG</td><td>1000</td><td></td><td></td><td></td><td></td><td></td><td></td><td></td><td></td><td></td></tr>
<tr><td>Oligo fwd</td><td>CACCCGGCGCTGGGGGAAGGTGTT</td><td></td><td></td><td></td><td></td><td></td><td></td><td></td><td></td><td></td><td></td></tr>
<tr><td>Oligo rev</td><td>AAACAACACCTTCCCCCAGCGCCG</td><td></td><td></td><td></td><td></td><td></td><td></td><td></td><td></td><td></td><td></td></tr>
<tr><td>Chromosome</td><td>start</td><td>end</td><td>strand</td><td>MM</td><td>target_seq</td><td>PAM</td><td>alignment</td><td>distance</td><td>position</td><td>gene name</td><td>gene id</td></tr>
<tr><td>GU305914.1</td><td>55511</td><td>55533</td><td>+</td><td>0</td><td>CGGCGCTGGGGGAAGGTGTT</td><td>GGG</td><td>||||||||[||||||||||||]PAM</td><td>NA</td><td>Intergenic</td><td>NA</td><td>NA</td></tr>
<tr><td></td><td></td><td></td><td></td><td></td><td></td><td></td><td></td><td></td><td></td><td></td><td></td></tr>
<tr><td>T3621</td><td>GGCGCTGGGGGAAGGTGTTGGGG</td><td>1000</td><td></td><td></td><td></td><td></td><td></td><td></td><td></td><td></td><td></td></tr>
<tr><td>Oligo fwd</td><td>CACCGGCGCTGGGGGAAGGTGTTG</td><td></td><td></td><td></td><td></td><td></td><td></td><td></td><td></td><td></td><td></td></tr>
<tr><td>Oligo rev</td><td>AAACCAACACCTTCCCCCAGCGCC</td><td></td><td></td><td></td><td></td><td></td><td></td><td></td><td></td><td></td><td></td></tr>
<tr><td>Chromosome</td><td>start</td><td>end</td><td>strand</td><td>MM</td><td>target_seq</td><td>PAM</td><td>alignment</td><td>distance</td><td>position</td><td>gene name</td><td>gene id</td></tr>
<tr><td>GU305914.1</td><td>55512</td><td>55534</td><td>+</td><td>0</td><td>GGCGCTGGGGGAAGGTGTTG</td><td>GGG</td><td>||||||||[||||||||||||]PAM</td><td>NA</td><td>Intergenic</td><td>NA</td><td>NA</td></tr>
<tr><td></td><td></td><td></td><td></td><td></td><td></td><td></td><td></td><td></td><td></td><td></td><td></td></tr>
<tr><td>T3622</td><td>GGAGAAAGCAGTTGCTGTTGCGG</td><td>1000</td><td></td><td></td><td></td><td></td><td></td><td></td><td></td><td></td><td></td></tr>
<tr><td>Oligo fwd</td><td>CACCGGAGAAAGCAGTTGCTGTTG</td><td></td><td></td><td></td><td></td><td></td><td></td><td></td><td></td><td></td><td></td></tr>
<tr><td>Oligo rev</td><td>AAACCAACAGCAACTGCTTTCTCC</td><td></td><td></td><td></td><td></td><td></td><td></td><td></td><td></td><td></td><td></td></tr>
<tr><td>Chromosome</td><td>start</td><td>end</td><td>strand</td><td>MM</td><td>target_seq</td><td>PAM</td><td>alignment</td><td>distance</td><td>position</td><td>gene name</td><td>gene id</td></tr>
<tr><td>GU305914.1</td><td>55533</td><td>55555</td><td>+</td><td>0</td><td>GGAGAAAGCAGTTGCTGTTG</td><td>CGG</td><td>||||||||[||||||||||||]PAM</td><td>NA</td><td>Intergenic</td><td>NA</td><td>NA</td></tr>
<tr><td></td><td></td><td></td><td></td><td></td><td></td><td></td><td></td><td></td><td></td><td></td><td></td></tr>
<tr><td>T3623</td><td>ATCGCATGACGCCGAGCACGAGG</td><td>1000</td><td></td><td></td><td></td><td></td><td></td><td></td><td></td><td></td><td></td></tr>
<tr><td>Oligo fwd</td><td>CACCATCGCATGACGCCGAGCACG</td><td></td><td></td><td></td><td></td><td></td><td></td><td></td><td></td><td></td><td></td></tr>
<tr><td>Oligo rev</td><td>AAACCGTGCTCGGCGTCATGCGAT</td><td></td><td></td><td></td><td></td><td></td><td></td><td></td><td></td><td></td><td></td></tr>
<tr><td>Chromosome</td><td>start</td><td>end</td><td>strand</td><td>MM</td><td>target_seq</td><td>PAM</td><td>alignment</td><td>distance</td><td>position</td><td>gene name</td><td>gene id</td></tr>
<tr><td>GU305914.1</td><td>55557</td><td>55579</td><td>+</td><td>0</td><td>ATCGCATGACGCCGAGCACG</td><td>AGG</td><td>||||||||[||||||||||||]PAM</td><td>NA</td><td>Intergenic</td><td>NA</td><td>NA</td></tr>
<tr><td></td><td></td><td></td><td></td><td></td><td></td><td></td><td></td><td></td><td></td><td></td><td></td></tr>
<tr><td>T3624</td><td>CATGACGCCGAGCACGAGGAAGG</td><td>1000</td><td></td><td></td><td></td><td></td><td></td><td></td><td></td><td></td><td></td></tr>
<tr><td>Oligo fwd</td><td>CACCCATGACGCCGAGCACGAGGA</td><td></td><td></td><td></td><td></td><td></td><td></td><td></td><td></td><td></td><td></td></tr>
<tr><td>Oligo rev</td><td>AAACTCCTCGTGCTCGGCGTCATG</td><td></td><td></td><td></td><td></td><td></td><td></td><td></td><td></td><td></td><td></td></tr>
<tr><td>Chromosome</td><td>start</td><td>end</td><td>strand</td><td>MM</td><td>target_seq</td><td>PAM</td><td>alignment</td><td>distance</td><td>position</td><td>gene name</td><td>gene id</td></tr>
<tr><td>GU305914.1</td><td>55561</td><td>55583</td><td>+</td><td>0</td><td>CATGACGCCGAGCACGAGGA</td><td>AGG</td><td>||||||||[||||||||||||]PAM</td><td>NA</td><td>Intergenic</td><td>NA</td><td>NA</td></tr>
<tr><td></td><td></td><td></td><td></td><td></td><td></td><td></td><td></td><td></td><td></td><td></td><td></td></tr>
<tr><td>T3625</td><td>CGAGCACGAGGAAGGTGAAGAGG</td><td>1000</td><td></td><td></td><td></td><td></td><td></td><td></td><td></td><td></td><td></td></tr>
<tr><td>Oligo fwd</td><td>CACCCGAGCACGAGGAAGGTGAAG</td><td></td><td></td><td></td><td></td><td></td><td></td><td></td><td></td><td></td><td></td></tr>
<tr><td>Oligo rev</td><td>AAACCTTCACCTTCCTCGTGCTCG</td><td></td><td></td><td></td><td></td><td></td><td></td><td></td><td></td><td></td><td></td></tr>
<tr><td>Chromosome</td><td>start</td><td>end</td><td>strand</td><td>MM</td><td>target_seq</td><td>PAM</td><td>alignment</td><td>distance</td><td>position</td><td>gene name</td><td>gene id</td></tr>
<tr><td>GU305914.1</td><td>55569</td><td>55591</td><td>+</td><td>0</td><td>CGAGCACGAGGAAGGTGAAG</td><td>AGG</td><td>||||||||[||||||||||||]PAM</td><td>NA</td><td>Intergenic</td><td>NA</td><td>NA</td></tr>
<tr><td></td><td></td><td></td><td></td><td></td><td></td><td></td><td></td><td></td><td></td><td></td><td></td></tr>
<tr><td>T3626</td><td>GAGGAAGGTGAAGAGGTCTGCGG</td><td>1000</td><td></td><td></td><td></td><td></td><td></td><td></td><td></td><td></td><td></td></tr>
<tr><td>Oligo fwd</td><td>CACCGAGGAAGGTGAAGAGGTCTG</td><td></td><td></td><td></td><td></td><td></td><td></td><td></td><td></td><td></td><td></td></tr>
<tr><td>Oligo rev</td><td>AAACCAGACCTCTTCACCTTCCTC</td><td></td><td></td><td></td><td></td><td></td><td></td><td></td><td></td><td></td><td></td></tr>
<tr><td>Chromosome</td><td>start</td><td>end</td><td>strand</td><td>MM</td><td>target_seq</td><td>PAM</td><td>alignment</td><td>distance</td><td>position</td><td>gene name</td><td>gene id</td></tr>
<tr><td>GU305914.1</td><td>55576</td><td>55598</td><td>+</td><td>0</td><td>GAGGAAGGTGAAGAGGTCTG</td><td>CGG</td><td>||||||||[||||||||||||]PAM</td><td>NA</td><td>Intergenic</td><td>NA</td><td>NA</td></tr>
<tr><td></td><td></td><td></td><td></td><td></td><td></td><td></td><td></td><td></td><td></td><td></td><td></td></tr>
<tr><td>T3627</td><td>AGGTGAAGAGGTCTGCGGCGAGG</td><td>1000</td><td></td><td></td><td></td><td></td><td></td><td></td><td></td><td></td><td></td></tr>
<tr><td>Oligo fwd</td><td>CACCAGGTGAAGAGGTCTGCGGCG</td><td></td><td></td><td></td><td></td><td></td><td></td><td></td><td></td><td></td><td></td></tr>
<tr><td>Oligo rev</td><td>AAACCGCCGCAGACCTCTTCACCT</td><td></td><td></td><td></td><td></td><td></td><td></td><td></td><td></td><td></td><td></td></tr>
<tr><td>Chromosome</td><td>start</td><td>end</td><td>strand</td><td>MM</td><td>target_seq</td><td>PAM</td><td>alignment</td><td>distance</td><td>position</td><td>gene name</td><td>gene id</td></tr>
<tr><td>GU305914.1</td><td>55581</td><td>55603</td><td>+</td><td>0</td><td>AGGTGAAGAGGTCTGCGGCG</td><td>AGG</td><td>||||||||[||||||||||||]PAM</td><td>NA</td><td>Intergenic</td><td>NA</td><td>NA</td></tr>
<tr><td></td><td></td><td></td><td></td><td></td><td></td><td></td><td></td><td></td><td></td><td></td><td></td></tr>
<tr><td>T3628</td><td>AACTTGATGTTACCGTGCCCCGG</td><td>1000</td><td></td><td></td><td></td><td></td><td></td><td></td><td></td><td></td><td></td></tr>
<tr><td>Oligo fwd</td><td>CACCAACTTGATGTTACCGTGCCC</td><td></td><td></td><td></td><td></td><td></td><td></td><td></td><td></td><td></td><td></td></tr>
<tr><td>Oligo rev</td><td>AAACGGGCACGGTAACATCAAGTT</td><td></td><td></td><td></td><td></td><td></td><td></td><td></td><td></td><td></td><td></td></tr>
<tr><td>Chromosome</td><td>start</td><td>end</td><td>strand</td><td>MM</td><td>target_seq</td><td>PAM</td><td>alignment</td><td>distance</td><td>position</td><td>gene name</td><td>gene id</td></tr>
<tr><td>GU305914.1</td><td>55604</td><td>55626</td><td>+</td><td>0</td><td>AACTTGATGTTACCGTGCCC</td><td>CGG</td><td>||||||||[||||||||||||]PAM</td><td>NA</td><td>Intergenic</td><td>NA</td><td>NA</td></tr>
<tr><td></td><td></td><td></td><td></td><td></td><td></td><td></td><td></td><td></td><td></td><td></td><td></td></tr>
<tr><td>T3629</td><td>ACTTGATGTTACCGTGCCCCGGG</td><td>1000</td><td></td><td></td><td></td><td></td><td></td><td></td><td></td><td></td><td></td></tr>
<tr><td>Oligo fwd</td><td>CACCACTTGATGTTACCGTGCCCC</td><td></td><td></td><td></td><td></td><td></td><td></td><td></td><td></td><td></td><td></td></tr>
<tr><td>Oligo rev</td><td>AAACGGGGCACGGTAACATCAAGT</td><td></td><td></td><td></td><td></td><td></td><td></td><td></td><td></td><td></td><td></td></tr>
<tr><td>Chromosome</td><td>start</td><td>end</td><td>strand</td><td>MM</td><td>target_seq</td><td>PAM</td><td>alignment</td><td>distance</td><td>position</td><td>gene name</td><td>gene id</td></tr>
<tr><td>GU305914.1</td><td>55605</td><td>55627</td><td>+</td><td>0</td><td>ACTTGATGTTACCGTGCCCC</td><td>GGG</td><td>||||||||[||||||||||||]PAM</td><td>NA</td><td>Intergenic</td><td>NA</td><td>NA</td></tr>
<tr><td></td><td></td><td></td><td></td><td></td><td></td><td></td><td></td><td></td><td></td><td></td><td></td></tr>
<tr><td>T3630</td><td>CCCCGGGTCGACGTAGTAGACGG</td><td>1000</td><td></td><td></td><td></td><td></td><td></td><td></td><td></td><td></td><td></td></tr>
<tr><td>Oligo fwd</td><td>CACCCCCCGGGTCGACGTAGTAGA</td><td></td><td></td><td></td><td></td><td></td><td></td><td></td><td></td><td></td><td></td></tr>
<tr><td>Oligo rev</td><td>AAACTCTACTACGTCGACCCGGGG</td><td></td><td></td><td></td><td></td><td></td><td></td><td></td><td></td><td></td><td></td></tr>
<tr><td>Chromosome</td><td>start</td><td>end</td><td>strand</td><td>MM</td><td>target_seq</td><td>PAM</td><td>alignment</td><td>distance</td><td>position</td><td>gene name</td><td>gene id</td></tr>
<tr><td>GU305914.1</td><td>55621</td><td>55643</td><td>+</td><td>0</td><td>CCCCGGGTCGACGTAGTAGA</td><td>CGG</td><td>||||||||[||||||||||||]PAM</td><td>NA</td><td>Intergenic</td><td>NA</td><td>NA</td></tr>
<tr><td></td><td></td><td></td><td></td><td></td><td></td><td></td><td></td><td></td><td></td><td></td><td></td></tr>
<tr><td>T3631</td><td>GTAGTAGACGGTCCCGCCGTCGG</td><td>1000</td><td></td><td></td><td></td><td></td><td></td><td></td><td></td><td></td><td></td></tr>
<tr><td>Oligo fwd</td><td>CACCGTAGTAGACGGTCCCGCCGT</td><td></td><td></td><td></td><td></td><td></td><td></td><td></td><td></td><td></td><td></td></tr>
<tr><td>Oligo rev</td><td>AAACACGGCGGGACCGTCTACTAC</td><td></td><td></td><td></td><td></td><td></td><td></td><td></td><td></td><td></td><td></td></tr>
<tr><td>Chromosome</td><td>start</td><td>end</td><td>strand</td><td>MM</td><td>target_seq</td><td>PAM</td><td>alignment</td><td>distance</td><td>position</td><td>gene name</td><td>gene id</td></tr>
<tr><td>GU305914.1</td><td>55633</td><td>55655</td><td>+</td><td>0</td><td>GTAGTAGACGGTCCCGCCGT</td><td>CGG</td><td>||||||||[||||||||||||]PAM</td><td>NA</td><td>Intergenic</td><td>NA</td><td>NA</td></tr>
<tr><td></td><td></td><td></td><td></td><td></td><td></td><td></td><td></td><td></td><td></td><td></td><td></td></tr>
<tr><td>T3632</td><td>CGGTCCCGCCGTCGGACACGAGG</td><td>1000</td><td></td><td></td><td></td><td></td><td></td><td></td><td></td><td></td><td></td></tr>
<tr><td>Oligo fwd</td><td>CACCCGGTCCCGCCGTCGGACACG</td><td></td><td></td><td></td><td></td><td></td><td></td><td></td><td></td><td></td><td></td></tr>
<tr><td>Oligo rev</td><td>AAACCGTGTCCGACGGCGGGACCG</td><td></td><td></td><td></td><td></td><td></td><td></td><td></td><td></td><td></td><td></td></tr>
<tr><td>Chromosome</td><td>start</td><td>end</td><td>strand</td><td>MM</td><td>target_seq</td><td>PAM</td><td>alignment</td><td>distance</td><td>position</td><td>gene name</td><td>gene id</td></tr>
<tr><td>GU305914.1</td><td>55641</td><td>55663</td><td>+</td><td>0</td><td>CGGTCCCGCCGTCGGACACG</td><td>AGG</td><td>||||||||[||||||||||||]PAM</td><td>NA</td><td>Intergenic</td><td>NA</td><td>NA</td></tr>
</table>

<table>
<tr><td>T3633</td><td>CCCGCCGTCGGACACGAGGATGG</td><td></td><td>1000</td></tr>
<tr><td>Oligo fwd</td><td>CACCCCCGCCGTCGGACACGAGGA</td></tr>
<tr><td>Oligo rev</td><td>AAACTCCTCGTGTCCGACGGCGGG</td></tr>
<tr><td>Chromosome</td><td>start</td><td></td><td>end</td><td>strand</td><td>MM</td><td>target_seq</td><td>PAM</td><td>alignment</td><td>distance</td><td>position</td><td>gene name</td><td>gene id</td></tr>
<tr><td>GU305914.1</td><td></td><td>55645</td><td>55667</td><td>+</td><td>0</td><td>CCCGCCGTCGGACACGAGGA</td><td>TGG</td><td>||||||||[||||||||||||]PAM</td><td>NA</td><td>Intergenic</td><td>NA</td><td>NA</td></tr>
<tr><td>T3634</td><td>CCGCCGTCGGACACGAGGATGGG</td><td></td><td>1000</td></tr>
<tr><td>Oligo fwd</td><td>CACCCCGCCGTCGGACACGAGGAT</td></tr>
<tr><td>Oligo rev</td><td>AAACATCCTCGTGTCCGACGGCGG</td></tr>
<tr><td>Chromosome</td><td>start</td><td></td><td>end</td><td>strand</td><td>MM</td><td>target_seq</td><td>PAM</td><td>alignment</td><td>distance</td><td>position</td><td>gene name</td><td>gene id</td></tr>
<tr><td>GU305914.1</td><td></td><td>55646</td><td>55668</td><td>+</td><td>0</td><td>CCGCCGTCGGACACGAGGAT</td><td>GGG</td><td>||||||||[||||||||||||]PAM</td><td>NA</td><td>Intergenic</td><td>NA</td><td>NA</td></tr>
<tr><td>T3635</td><td>GGGCGCGTTCACCCAGACCCCGG</td><td></td><td>1000</td></tr>
<tr><td>Oligo fwd</td><td>CACCGGGCGCGTTCACCCAGACCC</td></tr>
<tr><td>Oligo rev</td><td>AAACGGGTCTGGGTGAACGCGCCC</td></tr>
<tr><td>Chromosome</td><td>start</td><td></td><td>end</td><td>strand</td><td>MM</td><td>target_seq</td><td>PAM</td><td>alignment</td><td>distance</td><td>position</td><td>gene name</td><td>gene id</td></tr>
<tr><td>GU305914.1</td><td></td><td>55666</td><td>55688</td><td>+</td><td>0</td><td>GGGCGCGTTCACCCAGACCC</td><td>CGG</td><td>||||||||[||||||||||||]PAM</td><td>NA</td><td>Intergenic</td><td>NA</td><td>NA</td></tr>
<tr><td>T3636</td><td>GCACGCCTTGACCATGAAGACGG</td><td></td><td>1000</td></tr>
<tr><td>Oligo fwd</td><td>CACCGCACGCCTTGACCATGAAGA</td></tr>
<tr><td>Oligo rev</td><td>AAACTCTTCATGGTCAAGGCGTGC</td></tr>
<tr><td>Chromosome</td><td>start</td><td></td><td>end</td><td>strand</td><td>MM</td><td>target_seq</td><td>PAM</td><td>alignment</td><td>distance</td><td>position</td><td>gene name</td><td>gene id</td></tr>
<tr><td>GU305914.1</td><td></td><td>55693</td><td>55715</td><td>+</td><td>0</td><td>GCACGCCTTGACCATGAAGA</td><td>CGG</td><td>||||||||[||||||||||||]PAM</td><td>NA</td><td>Intergenic</td><td>NA</td><td>NA</td></tr>
<tr><td>T3637</td><td>CACGCCTTGACCATGAAGACGGG</td><td></td><td>1000</td></tr>
<tr><td>Oligo fwd</td><td>CACCCACGCCTTGACCATGAAGAC</td></tr>
<tr><td>Oligo rev</td><td>AAACGTCTTCATGGTCAAGGCGTG</td></tr>
<tr><td>Chromosome</td><td>start</td><td></td><td>end</td><td>strand</td><td>MM</td><td>target_seq</td><td>PAM</td><td>alignment</td><td>distance</td><td>position</td><td>gene name</td><td>gene id</td></tr>
<tr><td>GU305914.1</td><td></td><td>55694</td><td>55716</td><td>+</td><td>0</td><td>CACGCCTTGACCATGAAGAC</td><td>GGG</td><td>||||||||[||||||||||||]PAM</td><td>NA</td><td>Intergenic</td><td>NA</td><td>NA</td></tr>
<tr><td>T3638</td><td>CCATGAAGACGGGCTCGAGTCGG</td><td></td><td>1000</td></tr>
<tr><td>Oligo fwd</td><td>CACCCCATGAAGACGGGCTCGAGT</td></tr>
<tr><td>Oligo rev</td><td>AAACACTCGAGCCCGTCTTCATGG</td></tr>
<tr><td>Chromosome</td><td>start</td><td></td><td>end</td><td>strand</td><td>MM</td><td>target_seq</td><td>PAM</td><td>alignment</td><td>distance</td><td>position</td><td>gene name</td><td>gene id</td></tr>
<tr><td>GU305914.1</td><td></td><td>55704</td><td>55726</td><td>+</td><td>0</td><td>CCATGAAGACGGGCTCGAGT</td><td>CGG</td><td>||||||||[||||||||||||]PAM</td><td>NA</td><td>Intergenic</td><td>NA</td><td>NA</td></tr>
<tr><td>GU305914.1</td><td></td><td>75134</td><td>75156</td><td>+</td><td>4</td><td>TCTTGAAGACGTGCTCGAGG</td><td>CGG</td><td>-|-|||||[|||-|||||||-]PAM</td><td>NA</td><td>Intergenic</td><td>NA</td><td>NA</td></tr>
<tr><td>T3639</td><td>AGACGGGCTCGAGTCGGCACTGG</td><td></td><td>1000</td></tr>
<tr><td>Oligo fwd</td><td>CACCAGACGGGCTCGAGTCGGCAC</td></tr>
<tr><td>Oligo rev</td><td>AAACGTGCCGACTCGAGCCCGTCT</td></tr>
<tr><td>Chromosome</td><td>start</td><td></td><td>end</td><td>strand</td><td>MM</td><td>target_seq</td><td>PAM</td><td>alignment</td><td>distance</td><td>position</td><td>gene name</td><td>gene id</td></tr>
<tr><td>GU305914.1</td><td></td><td>55710</td><td>55732</td><td>+</td><td>0</td><td>AGACGGGCTCGAGTCGGCAC</td><td>TGG</td><td>||||||||[||||||||||||]PAM</td><td>NA</td><td>Intergenic</td><td>NA</td><td>NA</td></tr>
<tr><td>T3640</td><td>GGGCTCGAGTCGGCACTGGCCGG</td><td></td><td>1000</td></tr>
<tr><td>Oligo fwd</td><td>CACCGGGCTCGAGTCGGCACTGGC</td></tr>
<tr><td>Oligo rev</td><td>AAACGCCAGTGCCGACTCGAGCCC</td></tr>
<tr><td>Chromosome</td><td>start</td><td></td><td>end</td><td>strand</td><td>MM</td><td>target_seq</td><td>PAM</td><td>alignment</td><td>distance</td><td>position</td><td>gene name</td><td>gene id</td></tr>
<tr><td>GU305914.1</td><td></td><td>55714</td><td>55736</td><td>+</td><td>0</td><td>GGGCTCGAGTCGGCACTGGC</td><td>CGG</td><td>||||||||[||||||||||||]PAM</td><td>NA</td><td>Intergenic</td><td>NA</td><td>NA</td></tr>
<tr><td>T3641</td><td>GTAGTCGAAGCTGTCGTAGCCGG</td><td></td><td>1000</td></tr>
<tr><td>Oligo fwd</td><td>CACCGTAGTCGAAGCTGTCGTAGC</td></tr>
<tr><td>Oligo rev</td><td>AAACGCTACGACAGCTTCGACTAC</td></tr>
<tr><td>Chromosome</td><td>start</td><td></td><td>end</td><td>strand</td><td>MM</td><td>target_seq</td><td>PAM</td><td>alignment</td><td>distance</td><td>position</td><td>gene name</td><td>gene id</td></tr>
<tr><td>GU305914.1</td><td></td><td>55756</td><td>55778</td><td>+</td><td>0</td><td>GTAGTCGAAGCTGTCGTAGC</td><td>CGG</td><td>||||||||[||||||||||||]PAM</td><td>NA</td><td>Intergenic</td><td>NA</td><td>NA</td></tr>
<tr><td>T3642</td><td>TGTCGTAGCCGGTGTCGCACAGG</td><td></td><td>1000</td></tr>
<tr><td>Oligo fwd</td><td>CACCTGTCGTAGCCGGTGTCGCAC</td></tr>
<tr><td>Oligo rev</td><td>AAACGTGCGACACCGGCTACGACA</td></tr>
<tr><td>Chromosome</td><td>start</td><td></td><td>end</td><td>strand</td><td>MM</td><td>target_seq</td><td>PAM</td><td>alignment</td><td>distance</td><td>position</td><td>gene name</td><td>gene id</td></tr>
<tr><td>GU305914.1</td><td></td><td>55767</td><td>55789</td><td>+</td><td>0</td><td>TGTCGTAGCCGGTGTCGCAC</td><td>AGG</td><td>||||||||[||||||||||||]PAM</td><td>NA</td><td>Intergenic</td><td>NA</td><td>NA</td></tr>
<tr><td>T3643</td><td>GGTGTCGCACAGGTTGCTGACGG</td><td></td><td>1000</td></tr>
<tr><td>Oligo fwd</td><td>CACCGGTGTCGCACAGGTTGCTGA</td></tr>
<tr><td>Oligo rev</td><td>AAACTCAGCAACCTGTGCGACACC</td></tr>
<tr><td>Chromosome</td><td>start</td><td></td><td>end</td><td>strand</td><td>MM</td><td>target_seq</td><td>PAM</td><td>alignment</td><td>distance</td><td>position</td><td>gene name</td><td>gene id</td></tr>
<tr><td>GU305914.1</td><td></td><td>55777</td><td>55799</td><td>+</td><td>0</td><td>GGTGTCGCACAGGTTGCTGA</td><td>CGG</td><td>||||||||[||||||||||||]PAM</td><td>NA</td><td>Intergenic</td><td>NA</td><td>NA</td></tr>
<tr><td>T3644</td><td>ACAGGTTGCTGACGGTGAGCAGG</td><td></td><td>1000</td></tr>
<tr><td>Oligo fwd</td><td>CACCACAGGTTGCTGACGGTGAGC</td></tr>
<tr><td>Oligo rev</td><td>AAACGCTCACCGTCAGCAACCTGT</td></tr>
<tr><td>Chromosome</td><td>start</td><td></td><td>end</td><td>strand</td><td>MM</td><td>target_seq</td><td>PAM</td><td>alignment</td><td>distance</td><td>position</td><td>gene name</td><td>gene id</td></tr>
<tr><td>GU305914.1</td><td></td><td>55785</td><td>55807</td><td>+</td><td>0</td><td>ACAGGTTGCTGACGGTGAGC</td><td>AGG</td><td>||||||||[||||||||||||]PAM</td><td>NA</td><td>Intergenic</td><td>NA</td><td>NA</td></tr>
<tr><td>T3645</td><td>CAGGTTGCTGACGGTGAGCAGGG</td><td></td><td>1000</td></tr>
<tr><td>Oligo fwd</td><td>CACCCAGGTTGCTGACGGTGAGCA</td></tr>
<tr><td>Oligo rev</td><td>AAACTGCTCACCGTCAGCAACCTG</td></tr>
<tr><td>Chromosome</td><td>start</td><td></td><td>end</td><td>strand</td><td>MM</td><td>target_seq</td><td>PAM</td><td>alignment</td><td>distance</td><td>position</td><td>gene name</td><td>gene id</td></tr>
<tr><td>GU305914.1</td><td></td><td>55786</td><td>55808</td><td>+</td><td>0</td><td>CAGGTTGCTGACGGTGAGCA</td><td>GGG</td><td>||||||||[||||||||||||]PAM</td><td>NA</td><td>Intergenic</td><td>NA</td><td>NA</td></tr>
<tr><td>T3646</td><td>GGGTCCACATGCTATCCGAGTGG</td><td></td><td>1000</td></tr>
<tr><td>Oligo fwd</td><td>CACCGGGTCCACATGCTATCCGAG</td></tr>
<tr><td>Oligo rev</td><td>AAACCTCGGATAGCATGTGGACCC</td></tr>
<tr><td>Chromosome</td><td>start</td><td></td><td>end</td><td>strand</td><td>MM</td><td>target_seq</td><td>PAM</td><td>alignment</td><td>distance</td><td>position</td><td>gene name</td><td>gene id</td></tr>
<tr><td>GU305914.1</td><td></td><td>55806</td><td>55828</td><td>+</td><td>0</td><td>GGGTCCACATGCTATCCGAG</td><td>TGG</td><td>||||||||[||||||||||||]PAM</td><td>NA</td><td>Intergenic</td><td>NA</td><td>NA</td></tr>
<tr><td>T3647</td><td>CATGCTATCCGAGTGGAAGATGG</td><td></td><td>1000</td></tr>
<tr><td>Oligo fwd</td><td>CACCCATGCTATCCGAGTGGAAGA</td></tr>
<tr><td>Oligo rev</td><td>AAACTCTTCCACTCGGATAGCATG</td></tr>
<tr><td>Chromosome</td><td>start</td><td></td><td>end</td><td>strand</td><td>MM</td><td>target_seq</td><td>PAM</td><td>alignment</td><td>distance</td><td>position</td><td>gene name</td><td>gene id</td></tr>
</table>

GU305914.1 55813 55835 + 0 CATGCTATCCGAGTGGAAGA TGG ||||||||[||||||||||||]PAM NA Intergenic NA NA

T3648 GCTATCCGAGTGGAAGATGGCGG 1000
Oligo fwd CACCGCTATCCGAGTGGAAGATGG
Oligo rev AAACCCATCTTCCACTCGGATAGC
Chromosome start end strand MM target_seq PAM alignment distance position gene name gene id
GU305914.1 55816 55838 + 0 GCTATCCGAGTGGAAGATGG CGG ||||||||[||||||||||||]PAM NA Intergenic NA NA

T3649 CGAGTGGAAGATGGCGGTCTCGG 1000
Oligo fwd CACCCGAGTGGAAGATGGCGGTCT
Oligo rev AAACAGACCGCCATCTTCCACTCG
Chromosome start end strand MM target_seq PAM alignment distance position gene name gene id
GU305914.1 55822 55844 + 0 CGAGTGGAAGATGGCGGTCT CGG ||||||||[||||||||||||]PAM NA Intergenic NA NA

T3650 TGGCGGTCTCGGCGCCGAGCAGG 1000
Oligo fwd CACCTGGCGGTCTCGGCGCCGAGC
Oligo rev AAACGCTCGGCGCCGAGACCGCCA
Chromosome start end strand MM target_seq PAM alignment distance position gene name gene id
GU305914.1 55833 55855 + 0 TGGCGGTCTCGGCGCCGAGC AGG ||||||||[||||||||||||]PAM NA Intergenic NA NA
GU305914.1 124338 124360 - 4 TGTCGATCGCGGCGCTGAGC CGG ||-||-||[-||||||-||||]PAM NA Intergenic NA NA

T3651 CGGCGCCGAGCAGGCGATCGCGG 1000
Oligo fwd CACCCGGCGCCGAGCAGGCGATCG
Oligo rev AAACCGATCGCCTGCTCGGCGCCG
Chromosome start end strand MM target_seq PAM alignment distance position gene name gene id
GU305914.1 55842 55864 + 0 CGGCGCCGAGCAGGCGATCG CGG ||||||||[||||||||||||]PAM NA Intergenic NA NA

T3652 GGCGATCGCGGCTGACGACCAGG 1000
Oligo fwd CACCGGCGATCGCGGCTGACGACC
Oligo rev AAACGGTCGTCAGCCGCGATCGCC
Chromosome start end strand MM target_seq PAM alignment distance position gene name gene id
GU305914.1 55854 55876 + 0 GGCGATCGCGGCTGACGACC AGG ||||||||[||||||||||||]PAM NA Intergenic NA NA
GU305914.1 110116 110138 - 4 GGCGGCCGCGGCGGACGAGC CGG ||||--||[||||-|||||-|]PAM NA Intergenic NA NA

T3653 CAGGTCGTCTATCGAGCTCCTGG 1000
Oligo fwd CACCCAGGTCGTCTATCGAGCTCC
Oligo rev AAACGGAGCTCGATAGACGACCTG
Chromosome start end strand MM target_seq PAM alignment distance position gene name gene id
GU305914.1 55873 55895 + 0 CAGGTCGTCTATCGAGCTCC TGG ||||||||[||||||||||||]PAM NA Intergenic NA NA

T3654 CTATCGAGCTCCTGGATTCGAGG 1000
Oligo fwd CACCCTATCGAGCTCCTGGATTCG
Oligo rev AAACCGAATCCAGGAGCTCGATAG
Chromosome start end strand MM target_seq PAM alignment distance position gene name gene id
GU305914.1 55881 55903 + 0 CTATCGAGCTCCTGGATTCG AGG ||||||||[||||||||||||]PAM NA Intergenic NA NA

T3655 CGAGCTCCTGGATTCGAGGATGG 1000
Oligo fwd CACCCGAGCTCCTGGATTCGAGGA
Oligo rev AAACTCCTCGAATCCAGGAGCTCG
Chromosome start end strand MM target_seq PAM alignment distance position gene name gene id
GU305914.1 55885 55907 + 0 CGAGCTCCTGGATTCGAGGA TGG ||||||||[||||||||||||]PAM NA Intergenic NA NA

T3656 GATTCGAGGATGGACTTGAGCGG 1000
Oligo fwd CACCGATTCGAGGATGGACTTGAG
Oligo rev AAACCTCAAGTCCATCCTCGAATC
Chromosome start end strand MM target_seq PAM alignment distance position gene name gene id
GU305914.1 55895 55917 + 0 GATTCGAGGATGGACTTGAG CGG ||||||||[||||||||||||]PAM NA Intergenic NA NA

T3657 GGATCCTCCGTATAGTCGATCGG 1000
Oligo fwd CACCGGATCCTCCGTATAGTCGAT
Oligo rev AAACATCGACTATACGGAGGATCC
Chromosome start end strand MM target_seq PAM alignment distance position gene name gene id
GU305914.1 55916 55938 + 0 GGATCCTCCGTATAGTCGAT CGG ||||||||[||||||||||||]PAM NA Intergenic NA NA

T3658 ATAGTCGATCGGCGTCGATCCGG 1000
Oligo fwd CACCATAGTCGATCGGCGTCGATC
Oligo rev AAACGATCGACGCCGATCGACTAT
Chromosome start end strand MM target_seq PAM alignment distance position gene name gene id
GU305914.1 55927 55949 + 0 ATAGTCGATCGGCGTCGATC CGG ||||||||[||||||||||||]PAM NA Intergenic NA NA
GU305914.1 163206 163228 - 3 ATGGGCGATCCGCGTCGATC TGG ||-|-|||[||-|||||||||]PAM NA Intergenic NA NA

T3659 TCGGCGTCGATCCGGCGTCCAGG 1000
Oligo fwd CACCTCGGCGTCGATCCGGCGTCC
Oligo rev AAACGGACGCCGGATCGACGCCGA
Chromosome start end strand MM target_seq PAM alignment distance position gene name gene id
GU305914.1 55935 55957 + 0 TCGGCGTCGATCCGGCGTCC AGG ||||||||[||||||||||||]PAM NA Intergenic NA NA

T3660 GGCCGATGCCGCACCTCTCACGG 1000
Oligo fwd CACCGGCCGATGCCGCACCTCTCA
Oligo rev AAACTGAGAGGTGCGGCATCGGCC
Chromosome start end strand MM target_seq PAM alignment distance position gene name gene id
GU305914.1 55956 55978 + 0 GGCCGATGCCGCACCTCTCA CGG ||||||||[||||||||||||]PAM NA Intergenic NA NA

T3661 ACCTCTCACGGAGTCCGCGCCGG 1000
Oligo fwd CACCACCTCTCACGGAGTCCGCGC
Oligo rev AAACGCGCGGACTCCGTGAGAGGT
Chromosome start end strand MM target_seq PAM alignment distance position gene name gene id
GU305914.1 55968 55990 + 0 ACCTCTCACGGAGTCCGCGC CGG ||||||||[||||||||||||]PAM NA Intergenic NA NA

| | | | | | | | | | | | |
|---|---|---|---|---|---|---|---|---|---|---|---|
| T3662 | GAGTCCGCGCCGGCAGAAGTCGG | 1000 | | | | | | | | | |
| Oligo fwd | CACCGAGTCCGCGCCGGCAGAAGT | | | | | | | | | | |
| Oligo rev | AAACACTTCTGCCGGCGCGGACTC | | | | | | | | | | |
| Chromosome | start | end | strand | MM | target_seq | PAM | alignment | distance | position | gene name | gene id |
| GU305914.1 | 55978 | 56000 | + | 0 | GAGTCCGCGCCGGCAGAAGT | CGG | \|\|\|\|\|\|\|\|[\|\|\|\|\|\|\|\|\|\|\|\|]PAM | NA | Intergenic | NA | NA |
| T3663 | GAAGTCGGAGAAGCCGCACTTGG | 1000 | | | | | | | | | |
| Oligo fwd | CACCGAAGTCGGAGAAGCCGCACT | | | | | | | | | | |
| Oligo rev | AAACAGTGCGGCTTCTCCGACTTC | | | | | | | | | | |
| Chromosome | start | end | strand | MM | target_seq | PAM | alignment | distance | position | gene name | gene id |
| GU305914.1 | 55993 | 56015 | + | 0 | GAAGTCGGAGAAGCCGCACT | TGG | \|\|\|\|\|\|\|\|[\|\|\|\|\|\|\|\|\|\|\|\|]PAM | NA | Intergenic | NA | NA |
| T3664 | AGAAGCCGCACTTGGTGAGCAGG | 1000 | | | | | | | | | |
| Oligo fwd | CACCAGAAGCCGCACTTGGTGAGC | | | | | | | | | | |
| Oligo rev | AAACGCTCACCAAGTGCGGCTTCT | | | | | | | | | | |
| Chromosome | start | end | strand | MM | target_seq | PAM | alignment | distance | position | gene name | gene id |
| GU305914.1 | 56001 | 56023 | + | 0 | AGAAGCCGCACTTGGTGAGC | AGG | \|\|\|\|\|\|\|\|[\|\|\|\|\|\|\|\|\|\|\|\|]PAM | NA | Intergenic | NA | NA |
| T3665 | CTTGGTGAGCAGGTGCAGCACGG | 1000 | | | | | | | | | |
| Oligo fwd | CACCCTTGGTGAGCAGGTGCAGCA | | | | | | | | | | |
| Oligo rev | AAACTGCTGCACCTGCTCACCAAG | | | | | | | | | | |
| Chromosome | start | end | strand | MM | target_seq | PAM | alignment | distance | position | gene name | gene id |
| GU305914.1 | 56011 | 56033 | + | 0 | CTTGGTGAGCAGGTGCAGCA | CGG | \|\|\|\|\|\|\|\|[\|\|\|\|\|\|\|\|\|\|\|\|]PAM | NA | Intergenic | NA | NA |
| T3666 | TTGGTGAGCAGGTGCAGCACGGG | 1000 | | | | | | | | | |
| Oligo fwd | CACCTTGGTGAGCAGGTGCAGCAC | | | | | | | | | | |
| Oligo rev | AAACGTGCTGCACCTGCTCACCAA | | | | | | | | | | |
| Chromosome | start | end | strand | MM | target_seq | PAM | alignment | distance | position | gene name | gene id |
| GU305914.1 | 56012 | 56034 | + | 0 | TTGGTGAGCAGGTGCAGCAC | GGG | \|\|\|\|\|\|\|\|[\|\|\|\|\|\|\|\|\|\|\|\|]PAM | NA | Intergenic | NA | NA |
| T3667 | TGGTGAGCAGGTGCAGCACGGGG | 1000 | | | | | | | | | |
| Oligo fwd | CACCTGGTGAGCAGGTGCAGCACG | | | | | | | | | | |
| Oligo rev | AAACCGTGCTGCACCTGCTCACCA | | | | | | | | | | |
| Chromosome | start | end | strand | MM | target_seq | PAM | alignment | distance | position | gene name | gene id |
| GU305914.1 | 56013 | 56035 | + | 0 | TGGTGAGCAGGTGCAGCACG | GGG | \|\|\|\|\|\|\|\|[\|\|\|\|\|\|\|\|\|\|\|\|]PAM | NA | Intergenic | NA | NA |
| T3668 | ACATGTACACGAAGCCATCGAGG | 1000 | | | | | | | | | |
| Oligo fwd | CACCACATGTACACGAAGCCATCG | | | | | | | | | | |
| Oligo rev | AAACCGATGGCTTCGTGTACATGT | | | | | | | | | | |
| Chromosome | start | end | strand | MM | target_seq | PAM | alignment | distance | position | gene name | gene id |
| GU305914.1 | 56055 | 56077 | + | 0 | ACATGTACACGAAGCCATCG | AGG | \|\|\|\|\|\|\|\|[\|\|\|\|\|\|\|\|\|\|\|\|]PAM | NA | Intergenic | NA | NA |
| T3669 | CGAAGCCATCGAGGTTGACGAGG | 1000 | | | | | | | | | |
| Oligo fwd | CACCCGAAGCCATCGAGGTTGACG | | | | | | | | | | |
| Oligo rev | AAACCGTCAACCTCGATGGCTTCG | | | | | | | | | | |
| Chromosome | start | end | strand | MM | target_seq | PAM | alignment | distance | position | gene name | gene id |
| GU305914.1 | 56064 | 56086 | + | 0 | CGAAGCCATCGAGGTTGACG | AGG | \|\|\|\|\|\|\|\|[\|\|\|\|\|\|\|\|\|\|\|\|]PAM | NA | Intergenic | NA | NA |
| T3670 | TCGAGGTTGACGAGGATGACCGG | 1000 | | | | | | | | | |
| Oligo fwd | CACCTCGAGGTTGACGAGGATGAC | | | | | | | | | | |
| Oligo rev | AAACGTCATCCTCGTCAACCTCGA | | | | | | | | | | |
| Chromosome | start | end | strand | MM | target_seq | PAM | alignment | distance | position | gene name | gene id |
| GU305914.1 | 56072 | 56094 | + | 0 | TCGAGGTTGACGAGGATGAC | CGG | \|\|\|\|\|\|\|\|[\|\|\|\|\|\|\|\|\|\|\|\|]PAM | NA | Intergenic | NA | NA |
| T3671 | CGAGGTTGACGAGGATGACCGGG | 1000 | | | | | | | | | |
| Oligo fwd | CACCCGAGGTTGACGAGGATGACC | | | | | | | | | | |
| Oligo rev | AAACGGTCATCCTCGTCAACCTCG | | | | | | | | | | |
| Chromosome | start | end | strand | MM | target_seq | PAM | alignment | distance | position | gene name | gene id |
| GU305914.1 | 56073 | 56095 | + | 0 | CGAGGTTGACGAGGATGACC | GGG | \|\|\|\|\|\|\|\|[\|\|\|\|\|\|\|\|\|\|\|\|]PAM | NA | Intergenic | NA | NA |
| T3672 | GACGAGGATGACCGGGTACTCGG | 1000 | | | | | | | | | |
| Oligo fwd | CACCGACGAGGATGACCGGGTACT | | | | | | | | | | |
| Oligo rev | AAACAGTACCCGGTCATCCTCGTC | | | | | | | | | | |
| Chromosome | start | end | strand | MM | target_seq | PAM | alignment | distance | position | gene name | gene id |
| GU305914.1 | 56080 | 56102 | + | 0 | GACGAGGATGACCGGGTACT | CGG | \|\|\|\|\|\|\|\|[\|\|\|\|\|\|\|\|\|\|\|\|]PAM | NA | Intergenic | NA | NA |
| T3673 | GATGACCGGGTACTCGGCCTCGG | 1000 | | | | | | | | | |
| Oligo fwd | CACCGATGACCGGGTACTCGGCCT | | | | | | | | | | |
| Oligo rev | AAACAGGCCGAGTACCCGGTCATC | | | | | | | | | | |
| Chromosome | start | end | strand | MM | target_seq | PAM | alignment | distance | position | gene name | gene id |
| GU305914.1 | 56086 | 56108 | + | 0 | GATGACCGGGTACTCGGCCT | CGG | \|\|\|\|\|\|\|\|[\|\|\|\|\|\|\|\|\|\|\|\|]PAM | NA | Intergenic | NA | NA |
| T3674 | GACCGGGTACTCGGCCTCGGAGG | 1000 | | | | | | | | | |
| Oligo fwd | CACCGACCGGGTACTCGGCCTCGG | | | | | | | | | | |
| Oligo rev | AAACCCGAGGCCGAGTACCCGGTC | | | | | | | | | | |
| Chromosome | start | end | strand | MM | target_seq | PAM | alignment | distance | position | gene name | gene id |
| GU305914.1 | 56089 | 56111 | + | 0 | GACCGGGTACTCGGCCTCGG | AGG | \|\|\|\|\|\|\|\|[\|\|\|\|\|\|\|\|\|\|\|\|]PAM | NA | Intergenic | NA | NA |
| T3675 | ACCGGGTACTCGGCCTCGGAGGG | 1000 | | | | | | | | | |
| Oligo fwd | CACCACCGGGTACTCGGCCTCGGA | | | | | | | | | | |
| Oligo rev | AAACTCCGAGGCCGAGTACCCGGT | | | | | | | | | | |
| Chromosome | start | end | strand | MM | target_seq | PAM | alignment | distance | position | gene name | gene id |
| GU305914.1 | 56090 | 56112 | + | 0 | ACCGGGTACTCGGCCTCGGA | GGG | \|\|\|\|\|\|\|\|[\|\|\|\|\|\|\|\|\|\|\|\|]PAM | NA | Intergenic | NA | NA |
| T3676 | CCTCGGAGGGTTCGCGCCCCAGG | 1000 | | | | | | | | | |
| Oligo fwd | CACCCCTCGGAGGGTTCGCGCCCC | | | | | | | | | | |
| Oligo rev | AAACGGGGCGCGAACCCTCCGAGG | | | | | | | | | | |
| Chromosome | start | end | strand | MM | target_seq | PAM | alignment | distance | position | gene name | gene id |
| GU305914.1 | 56103 | 56125 | + | 0 | CCTCGGAGGGTTCGCGCCCC | AGG | \|\|\|\|\|\|\|\|[\|\|\|\|\|\|\|\|\|\|\|\|]PAM | NA | Intergenic | NA | NA |

T3677 AGGGTTCGCGCCCCAGGTCGTGG 1000
Oligo fwd CACCAGGGTTCGCGCCCCAGGTCG
Oligo rev AAACCGACCTGGGGCGCGAACCCT
Chromosome start end strand MM target_seq PAM alignment distance position gene name gene id
GU305914.1 56109 56131 + 0 AGGGTTCGCGCCCCAGGTCG TGG ||||||||[||||||||||||]PAM NA Intergenic NA NA

T3678 CAGGTCGTGGACCACCCCGATGG 1000
Oligo fwd CACCCAGGTCGTGGACCACCCCGA
Oligo rev AAACTCGGGGTGGTCCACGACCTG
Chromosome start end strand MM target_seq PAM alignment distance position gene name gene id
GU305914.1 56122 56144 + 0 CAGGTCGTGGACCACCCCGA TGG ||||||||[||||||||||||]PAM NA Intergenic NA NA

T3679 GGACCACCCCGATGGCGATGAGG 1000
Oligo fwd CACCGGACCACCCCGATGGCGATG
Oligo rev AAACCATCGCCATCGGGGTGGTCC
Chromosome start end strand MM target_seq PAM alignment distance position gene name gene id
GU305914.1 56130 56152 + 0 GGACCACCCCGATGGCGATG AGG ||||||||[||||||||||||]PAM NA Intergenic NA NA

T3680 CGCAGCAGACGTGACCGTCCCGG 1000
Oligo fwd CACCCGCAGCAGACGTGACCGTCC
Oligo rev AAACGGACGGTCACGTCTGCTGCG
Chromosome start end strand MM target_seq PAM alignment distance position gene name gene id
GU305914.1 56160 56182 + 0 CGCAGCAGACGTGACCGTCC CGG ||||||||[||||||||||||]PAM NA Intergenic NA NA

T3681 GACGTGACCGTCCCGGACGACGG 1000
Oligo fwd CACCGACGTGACCGTCCCGGACGA
Oligo rev AAACTCGTCCGGGACGGTCACGTC
Chromosome start end strand MM target_seq PAM alignment distance position gene name gene id
GU305914.1 56167 56189 + 0 GACGTGACCGTCCCGGACGA CGG ||||||||[||||||||||||]PAM NA Intergenic NA NA

T3682 GTGACCGTCCCGGACGACGGCGG 1000
Oligo fwd CACCGTGACCGTCCCGGACGACGG
Oligo rev AAACCCGTCGTCCGGGACGGTCAC
Chromosome start end strand MM target_seq PAM alignment distance position gene name gene id
GU305914.1 56170 56192 + 0 GTGACCGTCCCGGACGACGG CGG ||||||||[||||||||||||]PAM NA Intergenic NA NA

T3683 GCGCTCGCCTTCTGCAGCCTCGG 1000
Oligo fwd CACCGCGCTCGCCTTCTGCAGCCT
Oligo rev AAACAGGCTGCAGAAGGCGAGCGC
Chromosome start end strand MM target_seq PAM alignment distance position gene name gene id
GU305914.1 56192 56214 + 0 GCGCTCGCCTTCTGCAGCCT CGG ||||||||[||||||||||||]PAM NA Intergenic NA NA

T3684 CGAGCGAGTCGCATCGATCACGG 1000
Oligo fwd CACCCGAGCGAGTCGCATCGATCA
Oligo rev AAACTGATCGATGCGACTCGCTCG
Chromosome start end strand MM target_seq PAM alignment distance position gene name gene id
GU305914.1 56215 56237 + 0 CGAGCGAGTCGCATCGATCA CGG ||||||||[||||||||||||]PAM NA Intergenic NA NA

T3685 GATCACGGTGAAGACCCAGCCGG 1000
Oligo fwd CACCGATCACGGTGAAGACCCAGC
Oligo rev AAACGCTGGGTCTTCACCGTGATC
Chromosome start end strand MM target_seq PAM alignment distance position gene name gene id
GU305914.1 56230 56252 + 0 GATCACGGTGAAGACCCAGC CGG ||||||||[||||||||||||]PAM NA Intergenic NA NA

T3686 ATCACGGTGAAGACCCAGCCGGG 1000
Oligo fwd CACCATCACGGTGAAGACCCAGCC
Oligo rev AAACGGCTGGGTCTTCACCGTGAT
Chromosome start end strand MM target_seq PAM alignment distance position gene name gene id
GU305914.1 56231 56253 + 0 ATCACGGTGAAGACCCAGCC GGG ||||||||[||||||||||||]PAM NA Intergenic NA NA

T3687 TCACGGTGAAGACCCAGCCGGGG 1000
Oligo fwd CACCTCACGGTGAAGACCCAGCCG
Oligo rev AAACCGGCTGGGTCTTCACCGTGA
Chromosome start end strand MM target_seq PAM alignment distance position gene name gene id
GU305914.1 56232 56254 + 0 TCACGGTGAAGACCCAGCCG GGG ||||||||[||||||||||||]PAM NA Intergenic NA NA

T3688 CACGGTGAAGACCCAGCCGGGGG 1000
Oligo fwd CACCCACGGTGAAGACCCAGCCGG
Oligo rev AAACCCGGCTGGGTCTTCACCGTG
Chromosome start end strand MM target_seq PAM alignment distance position gene name gene id
GU305914.1 56233 56255 + 0 CACGGTGAAGACCCAGCCGG GGG ||||||||[||||||||||||]PAM NA Intergenic NA NA

T3689 ACGGTGAAGACCCAGCCGGGGGG 1000
Oligo fwd CACCACGGTGAAGACCCAGCCGGG
Oligo rev AAACCCCGGCTGGGTCTTCACCGT
Chromosome start end strand MM target_seq PAM alignment distance position gene name gene id
GU305914.1 56234 56256 + 0 ACGGTGAAGACCCAGCCGGG GGG ||||||||[||||||||||||]PAM NA Intergenic NA NA

T3690 GAAGACCCAGCCGGGGGGCCAGG 1000
Oligo fwd CACCGAAGACCCAGCCGGGGGGCC
Oligo rev AAACGGCCCCCCGGCTGGGTCTTC
Chromosome start end strand MM target_seq PAM alignment distance position gene name gene id
GU305914.1 56239 56261 + 0 GAAGACCCAGCCGGGGGGCC AGG ||||||||[||||||||||||]PAM NA Intergenic NA NA

T3691 GCCAGGTCAACAGCACGTGATGG 1000
Oligo fwd CACCGCCAGGTCAACAGCACGTGA
Oligo rev AAACTCACGTGCTGTTGACCTGGC
Chromosome start end strand MM target_seq PAM alignment distance position gene name gene id

| | | | | | | | | | | | |
|---|---|---|---|---|---|---|---|---|---|---|---|
| GU305914.1 | 56256 | 56278 | + | 0 | GCCAGGTCAACAGCACGTGA | TGG | \|\|\|\|\|\|\|\|[\|\|\|\|\|\|\|\|\|\|\|\|]PAM | NA | Intergenic | NA | NA |
| T3692 | GGAGATGAAACCTGTAGATGAGG | 1000 | | | | | | | | | |
| Oligo fwd | CACCGGAGATGAAACCTGTAGATG | | | | | | | | | | |
| Oligo rev | AAACCATCTACAGGTTTCATCTCC | | | | | | | | | | |
| Chromosome | start | end | strand | MM | target_seq | PAM | alignment | distance | position | gene name | gene id |
| GU305914.1 | 56277 | 56299 | + | 0 | GGAGATGAAACCTGTAGATG | AGG | \|\|\|\|\|\|\|\|[\|\|\|\|\|\|\|\|\|\|\|\|]PAM | NA | Intergenic | NA | NA |
| T3693 | AGATGAGGCAGTTGAGTCGCCGG | 1000 | | | | | | | | | |
| Oligo fwd | CACCAGATGAGGCAGTTGAGTCGC | | | | | | | | | | |
| Oligo rev | AAACGCGACTCAACTGCCTCATCT | | | | | | | | | | |
| Chromosome | start | end | strand | MM | target_seq | PAM | alignment | distance | position | gene name | gene id |
| GU305914.1 | 56292 | 56314 | + | 0 | AGATGAGGCAGTTGAGTCGC | CGG | \|\|\|\|\|\|\|\|[\|\|\|\|\|\|\|\|\|\|\|\|]PAM | NA | Intergenic | NA | NA |
| T3694 | GTTGAGTCGCCGGATGTCCCTGG | 1000 | | | | | | | | | |
| Oligo fwd | CACCGTTGAGTCGCCGGATGTCCC | | | | | | | | | | |
| Oligo rev | AAACGGGACATCCGGCGACTCAAC | | | | | | | | | | |
| Chromosome | start | end | strand | MM | target_seq | PAM | alignment | distance | position | gene name | gene id |
| GU305914.1 | 56302 | 56324 | + | 0 | GTTGAGTCGCCGGATGTCCC | TGG | \|\|\|\|\|\|\|\|[\|\|\|\|\|\|\|\|\|\|\|\|]PAM | NA | Intergenic | NA | NA |
| T3695 | GAGTCGCCGGATGTCCCTGGCGG | 1000 | | | | | | | | | |
| Oligo fwd | CACCGAGTCGCCGGATGTCCCTGG | | | | | | | | | | |
| Oligo rev | AAACCCAGGGACATCCGGCGACTC | | | | | | | | | | |
| Chromosome | start | end | strand | MM | target_seq | PAM | alignment | distance | position | gene name | gene id |
| GU305914.1 | 56305 | 56327 | + | 0 | GAGTCGCCGGATGTCCCTGG | CGG | \|\|\|\|\|\|\|\|[\|\|\|\|\|\|\|\|\|\|\|\|]PAM | NA | Intergenic | NA | NA |
| T3696 | CCCTGGCGGCGATCAATTCGTGG | 1000 | | | | | | | | | |
| Oligo fwd | CACCCCCTGGCGGCGATCAATTCG | | | | | | | | | | |
| Oligo rev | AAACCGAATTGATCGCCGCCAGGG | | | | | | | | | | |
| Chromosome | start | end | strand | MM | target_seq | PAM | alignment | distance | position | gene name | gene id |
| GU305914.1 | 56319 | 56341 | + | 0 | CCCTGGCGGCGATCAATTCG | TGG | \|\|\|\|\|\|\|\|[\|\|\|\|\|\|\|\|\|\|\|\|]PAM | NA | Intergenic | NA | NA |
| T3697 | AGAGTCGACCTGAGAGAGTCTGG | 1000 | | | | | | | | | |
| Oligo fwd | CACCAGAGTCGACCTGAGAGAGTC | | | | | | | | | | |
| Oligo rev | AAACGACTCTCTCAGGTCGACTCT | | | | | | | | | | |
| Chromosome | start | end | strand | MM | target_seq | PAM | alignment | distance | position | gene name | gene id |
| GU305914.1 | 56377 | 56399 | + | 0 | AGAGTCGACCTGAGAGAGTC | TGG | \|\|\|\|\|\|\|\|[\|\|\|\|\|\|\|\|\|\|\|\|]PAM | NA | Intergenic | NA | NA |
| T3698 | CGACGAAATCTTTACTGAGCTGG | 1000 | | | | | | | | | |
| Oligo fwd | CACCCGACGAAATCTTTACTGAGC | | | | | | | | | | |
| Oligo rev | AAACGCTCAGTAAAGATTTCGTCG | | | | | | | | | | |
| Chromosome | start | end | strand | MM | target_seq | PAM | alignment | distance | position | gene name | gene id |
| GU305914.1 | 56412 | 56434 | + | 0 | CGACGAAATCTTTACTGAGC | TGG | \|\|\|\|\|\|\|\|[\|\|\|\|\|\|\|\|\|\|\|\|]PAM | NA | Intergenic | NA | NA |
| T3699 | GAAATCTTTACTGAGCTGGACGG | 1000 | | | | | | | | | |
| Oligo fwd | CACCGAAATCTTTACTGAGCTGGA | | | | | | | | | | |
| Oligo rev | AAACTCCAGCTCAGTAAAGATTTC | | | | | | | | | | |
| Chromosome | start | end | strand | MM | target_seq | PAM | alignment | distance | position | gene name | gene id |
| GU305914.1 | 56416 | 56438 | + | 0 | GAAATCTTTACTGAGCTGGA | CGG | \|\|\|\|\|\|\|\|[\|\|\|\|\|\|\|\|\|\|\|\|]PAM | NA | Intergenic | NA | NA |
| T3700 | AAATCTTTACTGAGCTGGACGGG | 1000 | | | | | | | | | |
| Oligo fwd | CACCAAATCTTTACTGAGCTGGAC | | | | | | | | | | |
| Oligo rev | AAACGTCCAGCTCAGTAAAGATTT | | | | | | | | | | |
| Chromosome | start | end | strand | MM | target_seq | PAM | alignment | distance | position | gene name | gene id |
| GU305914.1 | 56417 | 56439 | + | 0 | AAATCTTTACTGAGCTGGAC | GGG | \|\|\|\|\|\|\|\|[\|\|\|\|\|\|\|\|\|\|\|\|]PAM | NA | Intergenic | NA | NA |
| T3701 | AATCTTTACTGAGCTGGACGGGG | 1000 | | | | | | | | | |
| Oligo fwd | CACCAATCTTTACTGAGCTGGACG | | | | | | | | | | |
| Oligo rev | AAACCGTCCAGCTCAGTAAAGATT | | | | | | | | | | |
| Chromosome | start | end | strand | MM | target_seq | PAM | alignment | distance | position | gene name | gene id |
| GU305914.1 | 56418 | 56440 | + | 0 | AATCTTTACTGAGCTGGACG | GGG | \|\|\|\|\|\|\|\|[\|\|\|\|\|\|\|\|\|\|\|\|]PAM | NA | Intergenic | NA | NA |
| T3702 | TTACTGAGCTGGACGGGGTCCGG | 1000 | | | | | | | | | |
| Oligo fwd | CACCTTACTGAGCTGGACGGGGTC | | | | | | | | | | |
| Oligo rev | AAACGACCCCGTCCAGCTCAGTAA | | | | | | | | | | |
| Chromosome | start | end | strand | MM | target_seq | PAM | alignment | distance | position | gene name | gene id |
| GU305914.1 | 56423 | 56445 | + | 0 | TTACTGAGCTGGACGGGGTC | CGG | \|\|\|\|\|\|\|\|[\|\|\|\|\|\|\|\|\|\|\|\|]PAM | NA | Intergenic | NA | NA |
| T3703 | TGAGCTGGACGGGGTCCGGCAGG | 1000 | | | | | | | | | |
| Oligo fwd | CACCTGAGCTGGACGGGGTCCGGC | | | | | | | | | | |
| Oligo rev | AAACGCCGGACCCCGTCCAGCTCA | | | | | | | | | | |
| Chromosome | start | end | strand | MM | target_seq | PAM | alignment | distance | position | gene name | gene id |
| GU305914.1 | 56427 | 56449 | + | 0 | TGAGCTGGACGGGGTCCGGC | AGG | \|\|\|\|\|\|\|\|[\|\|\|\|\|\|\|\|\|\|\|\|]PAM | NA | Intergenic | NA | NA |
| T3704 | GAGCTGGACGGGGTCCGGCAGGG | 1000 | | | | | | | | | |
| Oligo fwd | CACCGAGCTGGACGGGGTCCGGCA | | | | | | | | | | |
| Oligo rev | AAACTGCCGGACCCCGTCCAGCTC | | | | | | | | | | |
| Chromosome | start | end | strand | MM | target_seq | PAM | alignment | distance | position | gene name | gene id |
| GU305914.1 | 56428 | 56450 | + | 0 | GAGCTGGACGGGGTCCGGCA | GGG | \|\|\|\|\|\|\|\|[\|\|\|\|\|\|\|\|\|\|\|\|]PAM | NA | Intergenic | NA | NA |
| T3705 | TGGACGGGGTCCGGCAGGGTCGG | 1000 | | | | | | | | | |
| Oligo fwd | CACCTGGACGGGGTCCGGCAGGGT | | | | | | | | | | |
| Oligo rev | AAACACCCTGCCGGACCCCGTCCA | | | | | | | | | | |
| Chromosome | start | end | strand | MM | target_seq | PAM | alignment | distance | position | gene name | gene id |
| GU305914.1 | 56432 | 56454 | + | 0 | TGGACGGGGTCCGGCAGGGT | CGG | \|\|\|\|\|\|\|\|[\|\|\|\|\|\|\|\|\|\|\|\|]PAM | NA | Intergenic | NA | NA |
| T3706 | ACGGGGTCCGGCAGGGTCGGAGG | 1000 | | | | | | | | | |
| Oligo fwd | CACCACGGGGTCCGGCAGGGTCGG | | | | | | | | | | |
| Oligo rev | AAACCCGACCCTGCCGGACCCCGT | | | | | | | | | | |

<table>
<tr><td>Chromosome</td><td>start</td><td>end</td><td>strand</td><td>MM</td><td>target_seq</td><td>PAM</td><td>alignment</td><td>distance</td><td>position</td><td>gene name</td><td>gene id</td></tr>
<tr><td>GU305914.1</td><td>56435</td><td>56457</td><td>+</td><td>0</td><td>ACGGGGTCCGGCAGGGTCGG</td><td>AGG</td><td>||||||||[||||||||||||]PAM</td><td>NA</td><td>Intergenic</td><td>NA</td><td>NA</td></tr>
<tr><td>T3707</td><td>GAGGCGCTCCGATCAGCGAGTGG</td><td>1000</td></tr>
<tr><td>Oligo fwd</td><td>CACCGAGGCGCTCCGATCAGCGAG</td></tr>
<tr><td>Oligo rev</td><td>AAACCTCGCTGATCGGAGCGCCTC</td></tr>
<tr><td>Chromosome</td><td>start</td><td>end</td><td>strand</td><td>MM</td><td>target_seq</td><td>PAM</td><td>alignment</td><td>distance</td><td>position</td><td>gene name</td><td>gene id</td></tr>
<tr><td>GU305914.1</td><td>56454</td><td>56476</td><td>+</td><td>0</td><td>GAGGCGCTCCGATCAGCGAG</td><td>TGG</td><td>||||||||[||||||||||||]PAM</td><td>NA</td><td>Intergenic</td><td>NA</td><td>NA</td></tr>
<tr><td>T3708</td><td>CGAGTGGCGATCGAGAGTAAAGG</td><td>1000</td></tr>
<tr><td>Oligo fwd</td><td>CACCCGAGTGGCGATCGAGAGTAA</td></tr>
<tr><td>Oligo rev</td><td>AAACTTACTCTCGATCGCCACTCG</td></tr>
<tr><td>Chromosome</td><td>start</td><td>end</td><td>strand</td><td>MM</td><td>target_seq</td><td>PAM</td><td>alignment</td><td>distance</td><td>position</td><td>gene name</td><td>gene id</td></tr>
<tr><td>GU305914.1</td><td>56470</td><td>56492</td><td>+</td><td>0</td><td>CGAGTGGCGATCGAGAGTAA</td><td>AGG</td><td>||||||||[||||||||||||]PAM</td><td>NA</td><td>Intergenic</td><td>NA</td><td>NA</td></tr>
<tr><td>T3709</td><td>GGCGATCGAGAGTAAAGGTCAGG</td><td>1000</td></tr>
<tr><td>Oligo fwd</td><td>CACCGGCGATCGAGAGTAAAGGTC</td></tr>
<tr><td>Oligo rev</td><td>AAACGACCTTTACTCTCGATCGCC</td></tr>
<tr><td>Chromosome</td><td>start</td><td>end</td><td>strand</td><td>MM</td><td>target_seq</td><td>PAM</td><td>alignment</td><td>distance</td><td>position</td><td>gene name</td><td>gene id</td></tr>
<tr><td>GU305914.1</td><td>56475</td><td>56497</td><td>+</td><td>0</td><td>GGCGATCGAGAGTAAAGGTC</td><td>AGG</td><td>||||||||[||||||||||||]PAM</td><td>NA</td><td>Intergenic</td><td>NA</td><td>NA</td></tr>
<tr><td>T3710</td><td>GATCGAGAGTAAAGGTCAGGCGG</td><td>1000</td></tr>
<tr><td>Oligo fwd</td><td>CACCGATCGAGAGTAAAGGTCAGG</td></tr>
<tr><td>Oligo rev</td><td>AAACCCTGACCTTTACTCTCGATC</td></tr>
<tr><td>Chromosome</td><td>start</td><td>end</td><td>strand</td><td>MM</td><td>target_seq</td><td>PAM</td><td>alignment</td><td>distance</td><td>position</td><td>gene name</td><td>gene id</td></tr>
<tr><td>GU305914.1</td><td>56478</td><td>56500</td><td>+</td><td>0</td><td>GATCGAGAGTAAAGGTCAGG</td><td>CGG</td><td>||||||||[||||||||||||]PAM</td><td>NA</td><td>Intergenic</td><td>NA</td><td>NA</td></tr>
<tr><td>T3711</td><td>TGTCGTTCCTAGTACTGCTGCGG</td><td>1000</td></tr>
<tr><td>Oligo fwd</td><td>CACCTGTCGTTCCTAGTACTGCTG</td></tr>
<tr><td>Oligo rev</td><td>AAACCAGCAGTACTAGGAACGACA</td></tr>
<tr><td>Chromosome</td><td>start</td><td>end</td><td>strand</td><td>MM</td><td>target_seq</td><td>PAM</td><td>alignment</td><td>distance</td><td>position</td><td>gene name</td><td>gene id</td></tr>
<tr><td>GU305914.1</td><td>56553</td><td>56575</td><td>+</td><td>0</td><td>TGTCGTTCCTAGTACTGCTG</td><td>CGG</td><td>||||||||[||||||||||||]PAM</td><td>NA</td><td>Intergenic</td><td>NA</td><td>NA</td></tr>
<tr><td>T3712</td><td>TCCTAGTACTGCTGCGGCGACGG</td><td>1000</td></tr>
<tr><td>Oligo fwd</td><td>CACCTCCTAGTACTGCTGCGGCGA</td></tr>
<tr><td>Oligo rev</td><td>AAACTCGCCGCAGCAGTACTAGGA</td></tr>
<tr><td>Chromosome</td><td>start</td><td>end</td><td>strand</td><td>MM</td><td>target_seq</td><td>PAM</td><td>alignment</td><td>distance</td><td>position</td><td>gene name</td><td>gene id</td></tr>
<tr><td>GU305914.1</td><td>56559</td><td>56581</td><td>+</td><td>0</td><td>TCCTAGTACTGCTGCGGCGA</td><td>CGG</td><td>||||||||[||||||||||||]PAM</td><td>NA</td><td>Intergenic</td><td>NA</td><td>NA</td></tr>
<tr><td>T3713</td><td>GACGCGAGATTTCTTCAGAGTGG</td><td>1000</td></tr>
<tr><td>Oligo fwd</td><td>CACCGACGCGAGATTTCTTCAGAG</td></tr>
<tr><td>Oligo rev</td><td>AAACCTCTGAAGAAATCTCGCGTC</td></tr>
<tr><td>Chromosome</td><td>start</td><td>end</td><td>strand</td><td>MM</td><td>target_seq</td><td>PAM</td><td>alignment</td><td>distance</td><td>position</td><td>gene name</td><td>gene id</td></tr>
<tr><td>GU305914.1</td><td>56581</td><td>56603</td><td>+</td><td>0</td><td>GACGCGAGATTTCTTCAGAG</td><td>TGG</td><td>||||||||[||||||||||||]PAM</td><td>NA</td><td>Intergenic</td><td>NA</td><td>NA</td></tr>
<tr><td>T3714</td><td>GCGAGATTTCTTCAGAGTGGCGG</td><td>1000</td></tr>
<tr><td>Oligo fwd</td><td>CACCGCGAGATTTCTTCAGAGTGG</td></tr>
<tr><td>Oligo rev</td><td>AAACCCACTCTGAAGAAATCTCGC</td></tr>
<tr><td>Chromosome</td><td>start</td><td>end</td><td>strand</td><td>MM</td><td>target_seq</td><td>PAM</td><td>alignment</td><td>distance</td><td>position</td><td>gene name</td><td>gene id</td></tr>
<tr><td>GU305914.1</td><td>56584</td><td>56606</td><td>+</td><td>0</td><td>GCGAGATTTCTTCAGAGTGG</td><td>CGG</td><td>||||||||[||||||||||||]PAM</td><td>NA</td><td>Intergenic</td><td>NA</td><td>NA</td></tr>
<tr><td>T3715</td><td>CAGAGTGGCGGCGACACCGCAGG</td><td>1000</td></tr>
<tr><td>Oligo fwd</td><td>CACCCAGAGTGGCGGCGACACCGC</td></tr>
<tr><td>Oligo rev</td><td>AAACGCGGTGTCGCCGCCACTCTG</td></tr>
<tr><td>Chromosome</td><td>start</td><td>end</td><td>strand</td><td>MM</td><td>target_seq</td><td>PAM</td><td>alignment</td><td>distance</td><td>position</td><td>gene name</td><td>gene id</td></tr>
<tr><td>GU305914.1</td><td>56596</td><td>56618</td><td>+</td><td>0</td><td>CAGAGTGGCGGCGACACCGC</td><td>AGG</td><td>||||||||[||||||||||||]PAM</td><td>NA</td><td>Intergenic</td><td>NA</td><td>NA</td></tr>
<tr><td>GU305914.1</td><td>195416</td><td>195438</td><td>-</td><td>4</td><td>CTGTGCTGCGGCGACACCGC</td><td>CGG</td><td>|-|-|--|[||||||||||||]PAM</td><td>NA</td><td>Intergenic</td><td>NA</td><td>NA</td></tr>
<tr><td>T3716</td><td>TCTGCCGTCGTCGAGCGAGCAGG</td><td>1000</td></tr>
<tr><td>Oligo fwd</td><td>CACCTCTGCCGTCGTCGAGCGAGC</td></tr>
<tr><td>Oligo rev</td><td>AAACGCTCGCTCGACGACGGCAGA</td></tr>
<tr><td>Chromosome</td><td>start</td><td>end</td><td>strand</td><td>MM</td><td>target_seq</td><td>PAM</td><td>alignment</td><td>distance</td><td>position</td><td>gene name</td><td>gene id</td></tr>
<tr><td>GU305914.1</td><td>56656</td><td>56678</td><td>+</td><td>0</td><td>TCTGCCGTCGTCGAGCGAGC</td><td>AGG</td><td>||||||||[||||||||||||]PAM</td><td>NA</td><td>Intergenic</td><td>NA</td><td>NA</td></tr>
<tr><td>T3717</td><td>GCCGTCGTCGAGCGAGCAGGCGG</td><td>1000</td></tr>
<tr><td>Oligo fwd</td><td>CACCGCCGTCGTCGAGCGAGCAGG</td></tr>
<tr><td>Oligo rev</td><td>AAACCCTGCTCGCTCGACGACGGC</td></tr>
<tr><td>Chromosome</td><td>start</td><td>end</td><td>strand</td><td>MM</td><td>target_seq</td><td>PAM</td><td>alignment</td><td>distance</td><td>position</td><td>gene name</td><td>gene id</td></tr>
<tr><td>GU305914.1</td><td>56659</td><td>56681</td><td>+</td><td>0</td><td>GCCGTCGTCGAGCGAGCAGG</td><td>CGG</td><td>||||||||[||||||||||||]PAM</td><td>NA</td><td>Intergenic</td><td>NA</td><td>NA</td></tr>
<tr><td>T3718</td><td>GTCGAGCGAGCAGGCGGCCGCGG</td><td>1000</td></tr>
<tr><td>Oligo fwd</td><td>CACCGTCGAGCGAGCAGGCGGCCG</td></tr>
<tr><td>Oligo rev</td><td>AAACCGGCCGCCTGCTCGCTCGAC</td></tr>
<tr><td>Chromosome</td><td>start</td><td>end</td><td>strand</td><td>MM</td><td>target_seq</td><td>PAM</td><td>alignment</td><td>distance</td><td>position</td><td>gene name</td><td>gene id</td></tr>
<tr><td>GU305914.1</td><td>56665</td><td>56687</td><td>+</td><td>0</td><td>GTCGAGCGAGCAGGCGGCCG</td><td>CGG</td><td>||||||||[||||||||||||]PAM</td><td>NA</td><td>Intergenic</td><td>NA</td><td>NA</td></tr>
<tr><td>T3719</td><td>CGAGCAGGCGGCCGCGGATGTGG</td><td>1000</td></tr>
<tr><td>Oligo fwd</td><td>CACCCGAGCAGGCGGCCGCGGATG</td></tr>
<tr><td>Oligo rev</td><td>AAACCATCCGCGGCCGCCTGCTCG</td></tr>
<tr><td>Chromosome</td><td>start</td><td>end</td><td>strand</td><td>MM</td><td>target_seq</td><td>PAM</td><td>alignment</td><td>distance</td><td>position</td><td>gene name</td><td>gene id</td></tr>
<tr><td>GU305914.1</td><td>56671</td><td>56693</td><td>+</td><td>0</td><td>CGAGCAGGCGGCCGCGGATG</td><td>TGG</td><td>||||||||[||||||||||||]PAM</td><td>NA</td><td>Intergenic</td><td>NA</td><td>NA</td></tr>
<tr><td>T3720</td><td>GGCGGCCGCGGATGTGGATGCGG</td><td>1000</td></tr>
<tr><td>Oligo fwd</td><td>CACCGGCGGCCGCGGATGTGGATG</td></tr>
<tr><td>Oligo rev</td><td>AAACCATCCACATCCGCGGCCGCC</td></tr>
<tr><td>Chromosome</td><td>start</td><td>end</td><td>strand</td><td>MM</td><td>target_seq</td><td>PAM</td><td>alignment</td><td>distance</td><td>position</td><td>gene name</td><td>gene id</td></tr>
<tr><td>GU305914.1</td><td>56677</td><td>56699</td><td>+</td><td>0</td><td>GGCGGCCGCGGATGTGGATG</td><td>CGG</td><td>||||||||[||||||||||||]PAM</td><td>NA</td><td>Intergenic</td><td>NA</td><td>NA</td></tr>
<tr><td>T3721</td><td>GGCCGCGGATGTGGATGCGGTGG</td><td>1000</td></tr>
</table>

<table>
<tr><td>Oligo fwd</td><td>CACCGGCCGCGGATGTGGATGCGG</td><td></td><td></td><td></td><td></td><td></td><td></td><td></td><td></td><td></td><td></td></tr>
<tr><td>Oligo rev</td><td>AAACCCGCATCCACATCCGCGGCC</td><td></td><td></td><td></td><td></td><td></td><td></td><td></td><td></td><td></td><td></td></tr>
<tr><td>Chromosome</td><td>start</td><td>end</td><td>strand</td><td>MM</td><td>target_seq</td><td>PAM</td><td>alignment</td><td>distance</td><td>position</td><td>gene name</td><td>gene id</td></tr>
<tr><td>GU305914.1</td><td>56680</td><td>56702</td><td>+</td><td>0</td><td>GGCCGCGGATGTGGATGCGG</td><td>TGG</td><td>|||||||[|||||||||||||]PAM</td><td>NA</td><td>Intergenic</td><td>NA</td><td>NA</td></tr>
<tr><td></td><td></td><td></td><td></td><td></td><td></td><td></td><td></td><td></td><td></td><td></td><td></td></tr>
<tr><td>T3722</td><td>GGATGTGGATGCGGTGGCAGCGG</td><td>1000</td><td></td><td></td><td></td><td></td><td></td><td></td><td></td><td></td><td></td></tr>
<tr><td>Oligo fwd</td><td>CACCGGATGTGGATGCGGTGGCAG</td><td></td><td></td><td></td><td></td><td></td><td></td><td></td><td></td><td></td><td></td></tr>
<tr><td>Oligo rev</td><td>AAACCTGCCACCGCATCCACATCC</td><td></td><td></td><td></td><td></td><td></td><td></td><td></td><td></td><td></td><td></td></tr>
<tr><td>Chromosome</td><td>start</td><td>end</td><td>strand</td><td>MM</td><td>target_seq</td><td>PAM</td><td>alignment</td><td>distance</td><td>position</td><td>gene name</td><td>gene id</td></tr>
<tr><td>GU305914.1</td><td>56686</td><td>56708</td><td>+</td><td>0</td><td>GGATGTGGATGCGGTGGCAG</td><td>CGG</td><td>|||||||[|||||||||||||]PAM</td><td>NA</td><td>Intergenic</td><td>NA</td><td>NA</td></tr>
<tr><td></td><td></td><td></td><td></td><td></td><td></td><td></td><td></td><td></td><td></td><td></td><td></td></tr>
<tr><td>T3723</td><td>GGATGCGGTGGCAGCGGCCGCGG</td><td>1000</td><td></td><td></td><td></td><td></td><td></td><td></td><td></td><td></td><td></td></tr>
<tr><td>Oligo fwd</td><td>CACCGGATGCGGTGGCAGCGGCCG</td><td></td><td></td><td></td><td></td><td></td><td></td><td></td><td></td><td></td><td></td></tr>
<tr><td>Oligo rev</td><td>AAACCGGCCGCTGCCACCGCATCC</td><td></td><td></td><td></td><td></td><td></td><td></td><td></td><td></td><td></td><td></td></tr>
<tr><td>Chromosome</td><td>start</td><td>end</td><td>strand</td><td>MM</td><td>target_seq</td><td>PAM</td><td>alignment</td><td>distance</td><td>position</td><td>gene name</td><td>gene id</td></tr>
<tr><td>GU305914.1</td><td>56692</td><td>56714</td><td>+</td><td>0</td><td>GGATGCGGTGGCAGCGGCCG</td><td>CGG</td><td>|||||||[|||||||||||||]PAM</td><td>NA</td><td>Intergenic</td><td>NA</td><td>NA</td></tr>
<tr><td>GU305914.1</td><td>168817</td><td>168839</td><td>+</td><td>4</td><td>GGCGGTGGTGGCAGCGGCGG</td><td>CGG</td><td>||--|-||[|||||||||-|]PAM</td><td>NA</td><td>Intergenic</td><td>NA</td><td>NA</td></tr>
<tr><td>GU305914.1</td><td>163908</td><td>163930</td><td>+</td><td>4</td><td>GGCGGCGGCGGCAGCGGCAG</td><td>CGG</td><td>||--||||[-|||||||||-|]PAM</td><td>NA</td><td>Intergenic</td><td>NA</td><td>NA</td></tr>
<tr><td>GU305914.1</td><td>48159</td><td>48181</td><td>+</td><td>4</td><td>GGCGGCGGTGGCGGCGGCGG</td><td>TGG</td><td>||--||||[||||-|||||-|]PAM</td><td>NA</td><td>Intergenic</td><td>NA</td><td>NA</td></tr>
<tr><td></td><td></td><td></td><td></td><td></td><td></td><td></td><td></td><td></td><td></td><td></td><td></td></tr>
<tr><td>T3724</td><td>GCCGCGGCACCCTTCACCGTCGG</td><td>1000</td><td></td><td></td><td></td><td></td><td></td><td></td><td></td><td></td><td></td></tr>
<tr><td>Oligo fwd</td><td>CACCGCCGCGGCACCCTTCACCGT</td><td></td><td></td><td></td><td></td><td></td><td></td><td></td><td></td><td></td><td></td></tr>
<tr><td>Oligo rev</td><td>AAACACGGTGAAGGGTGCCGCGGC</td><td></td><td></td><td></td><td></td><td></td><td></td><td></td><td></td><td></td><td></td></tr>
<tr><td>Chromosome</td><td>start</td><td>end</td><td>strand</td><td>MM</td><td>target_seq</td><td>PAM</td><td>alignment</td><td>distance</td><td>position</td><td>gene name</td><td>gene id</td></tr>
<tr><td>GU305914.1</td><td>56708</td><td>56730</td><td>+</td><td>0</td><td>GCCGCGGCACCCTTCACCGT</td><td>CGG</td><td>|||||||[|||||||||||||]PAM</td><td>NA</td><td>Intergenic</td><td>NA</td><td>NA</td></tr>
<tr><td></td><td></td><td></td><td></td><td></td><td></td><td></td><td></td><td></td><td></td><td></td><td></td></tr>
<tr><td>T3725</td><td>ACGACTCTTCTCCGCCGCAGCGG</td><td>1000</td><td></td><td></td><td></td><td></td><td></td><td></td><td></td><td></td><td></td></tr>
<tr><td>Oligo fwd</td><td>CACCACGACTCTTCTCCGCCGCAG</td><td></td><td></td><td></td><td></td><td></td><td></td><td></td><td></td><td></td><td></td></tr>
<tr><td>Oligo rev</td><td>AAACCTGCGGCGGAGAAGAGTCGT</td><td></td><td></td><td></td><td></td><td></td><td></td><td></td><td></td><td></td><td></td></tr>
<tr><td>Chromosome</td><td>start</td><td>end</td><td>strand</td><td>MM</td><td>target_seq</td><td>PAM</td><td>alignment</td><td>distance</td><td>position</td><td>gene name</td><td>gene id</td></tr>
<tr><td>GU305914.1</td><td>56761</td><td>56783</td><td>+</td><td>0</td><td>ACGACTCTTCTCCGCCGCAG</td><td>CGG</td><td>|||||||[|||||||||||||]PAM</td><td>NA</td><td>Intergenic</td><td>NA</td><td>NA</td></tr>
<tr><td></td><td></td><td></td><td></td><td></td><td></td><td></td><td></td><td></td><td></td><td></td><td></td></tr>
<tr><td>T3726</td><td>ACTCTTCTCCGCCGCAGCGGCGG</td><td>1000</td><td></td><td></td><td></td><td></td><td></td><td></td><td></td><td></td><td></td></tr>
<tr><td>Oligo fwd</td><td>CACCACTCTTCTCCGCCGCAGCGG</td><td></td><td></td><td></td><td></td><td></td><td></td><td></td><td></td><td></td><td></td></tr>
<tr><td>Oligo rev</td><td>AAACCCGCTGCGGCGGAGAAGAGT</td><td></td><td></td><td></td><td></td><td></td><td></td><td></td><td></td><td></td><td></td></tr>
<tr><td>Chromosome</td><td>start</td><td>end</td><td>strand</td><td>MM</td><td>target_seq</td><td>PAM</td><td>alignment</td><td>distance</td><td>position</td><td>gene name</td><td>gene id</td></tr>
<tr><td>GU305914.1</td><td>56764</td><td>56786</td><td>+</td><td>0</td><td>ACTCTTCTCCGCCGCAGCGG</td><td>CGG</td><td>|||||||[|||||||||||||]PAM</td><td>NA</td><td>Intergenic</td><td>NA</td><td>NA</td></tr>
<tr><td></td><td></td><td></td><td></td><td></td><td></td><td></td><td></td><td></td><td></td><td></td><td></td></tr>
<tr><td>T3727</td><td>CTTCTCCGCCGCAGCGGCGGCGG</td><td>1000</td><td></td><td></td><td></td><td></td><td></td><td></td><td></td><td></td><td></td></tr>
<tr><td>Oligo fwd</td><td>CACCCTTCTCCGCCGCAGCGGCGG</td><td></td><td></td><td></td><td></td><td></td><td></td><td></td><td></td><td></td><td></td></tr>
<tr><td>Oligo rev</td><td>AAACCCGCCGCTGCGGCGGAGAAG</td><td></td><td></td><td></td><td></td><td></td><td></td><td></td><td></td><td></td><td></td></tr>
<tr><td>Chromosome</td><td>start</td><td>end</td><td>strand</td><td>MM</td><td>target_seq</td><td>PAM</td><td>alignment</td><td>distance</td><td>position</td><td>gene name</td><td>gene id</td></tr>
<tr><td>GU305914.1</td><td>56767</td><td>56789</td><td>+</td><td>0</td><td>CTTCTCCGCCGCAGCGGCGG</td><td>CGG</td><td>|||||||[|||||||||||||]PAM</td><td>NA</td><td>Intergenic</td><td>NA</td><td>NA</td></tr>
<tr><td>GU305914.1</td><td>56770</td><td>56792</td><td>+</td><td>4</td><td>CTCCGCCGCAGCGGCGGCGG</td><td>CGG</td><td>||-|-|||[|-||-|||||||]PAM</td><td>NA</td><td>Intergenic</td><td>NA</td><td>NA</td></tr>
<tr><td>GU305914.1</td><td>26436</td><td>26458</td><td>+</td><td>4</td><td>CAACTCCGCCACGGCGGCGG</td><td>CGG</td><td>|--|||||[||-|-|||||||]PAM</td><td>NA</td><td>Intergenic</td><td>NA</td><td>NA</td></tr>
<tr><td>GU305914.1</td><td>71744</td><td>71766</td><td>+</td><td>4</td><td>CTCCTCAGCAGCAGCAGCGG</td><td>CGG</td><td>||-|||-|[|-|||||-||||]PAM</td><td>NA</td><td>Intergenic</td><td>NA</td><td>NA</td></tr>
<tr><td></td><td></td><td></td><td></td><td></td><td></td><td></td><td></td><td></td><td></td><td></td><td></td></tr>
<tr><td>T3728</td><td>CTCCGCCGCAGCGGCGGCGGCGG</td><td>1000</td><td></td><td></td><td></td><td></td><td></td><td></td><td></td><td></td><td></td></tr>
<tr><td>Oligo fwd</td><td>CACCCTCCGCCGCAGCGGCGGCGG</td><td></td><td></td><td></td><td></td><td></td><td></td><td></td><td></td><td></td><td></td></tr>
<tr><td>Oligo rev</td><td>AAACCCGCCGCCGCTGCGGCGGAG</td><td></td><td></td><td></td><td></td><td></td><td></td><td></td><td></td><td></td><td></td></tr>
<tr><td>Chromosome</td><td>start</td><td>end</td><td>strand</td><td>MM</td><td>target_seq</td><td>PAM</td><td>alignment</td><td>distance</td><td>position</td><td>gene name</td><td>gene id</td></tr>
<tr><td>GU305914.1</td><td>56770</td><td>56792</td><td>+</td><td>0</td><td>CTCCGCCGCAGCGGCGGCGG</td><td>CGG</td><td>|||||||[|||||||||||||]PAM</td><td>NA</td><td>Intergenic</td><td>NA</td><td>NA</td></tr>
<tr><td>GU305914.1</td><td>56773</td><td>56795</td><td>+</td><td>3</td><td>CGCCGCAGCGGCGGCGGCGG</td><td>CGG</td><td>|-||||-|[|-||||||||||]PAM</td><td>NA</td><td>Intergenic</td><td>NA</td><td>NA</td></tr>
<tr><td>GU305914.1</td><td>45584</td><td>45606</td><td>+</td><td>4</td><td>ACCCGGCGCCGCGGCGGCGG</td><td>CGG</td><td>--|||-||[|-||||||||||]PAM</td><td>NA</td><td>Intergenic</td><td>NA</td><td>NA</td></tr>
<tr><td>GU305914.1</td><td>35653</td><td>35675</td><td>+</td><td>4</td><td>ATCCACGGCGGCGGCGGCGG</td><td>CGG</td><td>-|||-|-|[|-||||||||||]PAM</td><td>NA</td><td>Intergenic</td><td>NA</td><td>NA</td></tr>
<tr><td>GU305914.1</td><td>71747</td><td>71769</td><td>+</td><td>3</td><td>CTCAGCAGCAGCAGCGGCGG</td><td>CGG</td><td>|||-||-|[||||-|||||||]PAM</td><td>NA</td><td>Intergenic</td><td>NA</td><td>NA</td></tr>
<tr><td>GU305914.1</td><td>56767</td><td>56789</td><td>+</td><td>4</td><td>CTTCTCCGCCGCAGCGGCGG</td><td>CGG</td><td>||-|-|||[|-||-|||||||]PAM</td><td>NA</td><td>Intergenic</td><td>NA</td><td>NA</td></tr>
<tr><td>GU305914.1</td><td>71744</td><td>71766</td><td>+</td><td>4</td><td>CTCCTCAGCAGCAGCAGCGG</td><td>CGG</td><td>||||-|-|[||||-||-||||]PAM</td><td>NA</td><td>Intergenic</td><td>NA</td><td>NA</td></tr>
<tr><td>GU305914.1</td><td>66846</td><td>66868</td><td>-</td><td>4</td><td>CGTCGGCGCAGCGGCGGCGC</td><td>CGG</td><td>|--||-||[||||||||||||-]PAM</td><td>NA</td><td>Intergenic</td><td>NA</td><td>NA</td></tr>
<tr><td></td><td></td><td></td><td></td><td></td><td></td><td></td><td></td><td></td><td></td><td></td><td></td></tr>
<tr><td>T3729</td><td>CGCCGCAGCGGCGGCGGCGGCGG</td><td>1000</td><td></td><td></td><td></td><td></td><td></td><td></td><td></td><td></td><td></td></tr>
<tr><td>Oligo fwd</td><td>CACCCGCCGCAGCGGCGGCGGCGG</td><td></td><td></td><td></td><td></td><td></td><td></td><td></td><td></td><td></td><td></td></tr>
<tr><td>Oligo rev</td><td>AAACCCGCCGCCGCCGCTGCGGCG</td><td></td><td></td><td></td><td></td><td></td><td></td><td></td><td></td><td></td><td></td></tr>
<tr><td>Chromosome</td><td>start</td><td>end</td><td>strand</td><td>MM</td><td>target_seq</td><td>PAM</td><td>alignment</td><td>distance</td><td>position</td><td>gene name</td><td>gene id</td></tr>
<tr><td>GU305914.1</td><td>56773</td><td>56795</td><td>+</td><td>0</td><td>CGCCGCAGCGGCGGCGGCGG</td><td>CGG</td><td>|||||||[|||||||||||||]PAM</td><td>NA</td><td>Intergenic</td><td>NA</td><td>NA</td></tr>
<tr><td>GU305914.1</td><td>67742</td><td>67764</td><td>+</td><td>4</td><td>GGGAGCGGCGGCGGCGGCGG</td><td>CGG</td><td>-|--||-|[||||||||||||]PAM</td><td>NA</td><td>Intergenic</td><td>NA</td><td>NA</td></tr>
<tr><td>GU305914.1</td><td>154407</td><td>154429</td><td>+</td><td>4</td><td>GGTGGCGGCGGCGGCGGCGG</td><td>CGG</td><td>-|--||-|[||||||||||||]PAM</td><td>NA</td><td>Intergenic</td><td>NA</td><td>NA</td></tr>
<tr><td>GU305914.1</td><td>58123</td><td>58145</td><td>-</td><td>4</td><td>GGTGGCGGCGGCGGCGGCGG</td><td>AGG</td><td>-|--||-|[||||||||||||]PAM</td><td>NA</td><td>Intergenic</td><td>NA</td><td>NA</td></tr>
<tr><td>GU305914.1</td><td>35653</td><td>35675</td><td>+</td><td>4</td><td>ATCCACGGCGGCGGCGGCGG</td><td>CGG</td><td>--||-|-|[||||||||||||]PAM</td><td>NA</td><td>Intergenic</td><td>NA</td><td>NA</td></tr>
<tr><td>GU305914.1</td><td>24627</td><td>24649</td><td>-</td><td>3</td><td>CGCAACACCGGCGGCGGCGG</td><td>TGG</td><td>|||--||-[||||||||||||]PAM</td><td>NA</td><td>Intergenic</td><td>NA</td><td>NA</td></tr>
<tr><td>GU305914.1</td><td>149482</td><td>149504</td><td>-</td><td>4</td><td>GGCGACGGCGGCGGCGGCGG</td><td>CGG</td><td>-||--|-|[||||||||||||]PAM</td><td>NA</td><td>Intergenic</td><td>NA</td><td>NA</td></tr>
<tr><td>GU305914.1</td><td>154404</td><td>154426</td><td>+</td><td>4</td><td>GGCGGTGGCGGCGGCGGCGG</td><td>CGG</td><td>-||-|--|[||||||||||||]PAM</td><td>NA</td><td>Intergenic</td><td>NA</td><td>NA</td></tr>
<tr><td>GU305914.1</td><td>58126</td><td>58148</td><td>-</td><td>4</td><td>GGCGGTGGCGGCGGCGGCGG</td><td>CGG</td><td>-||-|--|[||||||||||||]PAM</td><td>NA</td><td>Intergenic</td><td>NA</td><td>NA</td></tr>
<tr><td>GU305914.1</td><td>56770</td><td>56792</td><td>+</td><td>3</td><td>CTCCGCCGCAGCGGCGGCGG</td><td>CGG</td><td>|-||||-|[|-||||||||||]PAM</td><td>NA</td><td>Intergenic</td><td>NA</td><td>NA</td></tr>
<tr><td>GU305914.1</td><td>73792</td><td>73814</td><td>-</td><td>4</td><td>GGCGCCAGCAGCGGCGGCGG</td><td>AGG</td><td>-||--|||[|-||||||||||]PAM</td><td>NA</td><td>Intergenic</td><td>NA</td><td>NA</td></tr>
<tr><td>GU305914.1</td><td>48159</td><td>48181</td><td>+</td><td>4</td><td>GGCGGCGGTGGCGGCGGCGG</td><td>TGG</td><td>-||-||-|[-||||||||||]PAM</td><td>NA</td><td>Intergenic</td><td>NA</td><td>NA</td></tr>
<tr><td>GU305914.1</td><td>161754</td><td>161776</td><td>-</td><td>4</td><td>GGCCGAGGAGGCGGCGGCGG</td><td>AGG</td><td>-||||--|[-||||||||||]PAM</td><td>NA</td><td>Intergenic</td><td>NA</td><td>NA</td></tr>
<tr><td>GU305914.1</td><td>48156</td><td>48178</td><td>+</td><td>4</td><td>GGCGGCGGCGGTGGCGGCGG</td><td>CGG</td><td>-||-||-|[|||-||||||||]PAM</td><td>NA</td><td>Intergenic</td><td>NA</td><td>NA</td></tr>
<tr><td>GU305914.1</td><td>71747</td><td>71769</td><td>+</td><td>4</td><td>CTCAGCAGCAGCAGCGGCGG</td><td>CGG</td><td>|-|-||||[|-||-|||||||]PAM</td><td>NA</td><td>Intergenic</td><td>NA</td><td>NA</td></tr>
<tr><td>GU305914.1</td><td>73789</td><td>73811</td><td>-</td><td>4</td><td>GCCAGCAGCGGCGGCGGAGG</td><td>AGG</td><td>--|-||||[|||||||||-||]PAM</td><td>NA</td><td>Intergenic</td><td>NA</td><td>NA</td></tr>
<tr><td>GU305914.1</td><td>110090</td><td>110112</td><td>+</td><td>4</td><td>CGTCGCCGCGGCCGCGTCGG</td><td>GGG</td><td>||-|||-|[||||-|||-|||]PAM</td><td>NA</td><td>Intergenic</td><td>NA</td><td>NA</td></tr>
<tr><td>GU305914.1</td><td>163902</td><td>163924</td><td>+</td><td>4</td><td>AGCGGCGGCGGCGGCGGCAG</td><td>CGG</td><td>-||-||-|[||||||||||-|]PAM</td><td>NA</td><td>Intergenic</td><td>NA</td><td>NA</td></tr>
<tr><td>GU305914.1</td><td>30006</td><td>30028</td><td>+</td><td>4</td><td>GGCGGCGGCGGCGGCGGCAG</td><td>TGG</td><td>-||-||-|[||||||||||-|]PAM</td><td>NA</td><td>Intergenic</td><td>NA</td><td>NA</td></tr>
<tr><td>GU305914.1</td><td>149476</td><td>149498</td><td>-</td><td>4</td><td>GGCGGCGGCGGCGGCGGCAG</td><td>CGG</td><td>-||-||-|[||||||||||-|]PAM</td><td>NA</td><td>Intergenic</td><td>NA</td><td>NA</td></tr>
<tr><td>GU305914.1</td><td>110083</td><td>110105</td><td>-</td><td>4</td><td>CGCGGCCGCGGCGACGCCGG</td><td>TGG</td><td>|||-||-|[|||||-||-|||]PAM</td><td>NA</td><td>Intergenic</td><td>NA</td><td>NA</td></tr>
<tr><td>GU305914.1</td><td>113912</td><td>113934</td><td>-</td><td>4</td><td>GGCGGCAGCGGCGACGGCGT</td><td>CGG</td><td>-||-||||[|||||-|||||-]PAM</td><td>NA</td><td>Intergenic</td><td>NA</td><td>NA</td></tr>
<tr><td>GU305914.1</td><td>71753</td><td>71775</td><td>+</td><td>4</td><td>AGCAGCAGCGGCGGCGGCAA</td><td>CGG</td><td>-||-||||[||||||||||--]PAM</td><td>NA</td><td>Intergenic</td><td>NA</td><td>NA</td></tr>
<tr><td></td><td></td><td></td><td></td><td></td><td></td><td></td><td></td><td></td><td></td><td></td><td></td></tr>
<tr><td>T3730</td><td>TTTCGAAAAGATCTCTCTCTCGG</td><td>1000</td><td></td><td></td><td></td><td></td><td></td><td></td><td></td><td></td><td></td></tr>
</table>

<table>
<tr><td>Oligo fwd</td><td>CACCTTTCGAAAAGATCTCTCTCT</td><td></td><td></td><td></td><td></td><td></td><td></td><td></td><td></td><td></td><td></td></tr>
<tr><td>Oligo rev</td><td>AAACAGAGAGAGATCTTTTCGAAA</td><td></td><td></td><td></td><td></td><td></td><td></td><td></td><td></td><td></td><td></td></tr>
<tr><td>Chromosome</td><td>start</td><td>end</td><td>strand</td><td>MM</td><td>target_seq</td><td>PAM</td><td>alignment</td><td>distance</td><td>position</td><td>gene name</td><td>gene id</td></tr>
<tr><td>GU305914.1</td><td>56825</td><td>56847</td><td>+</td><td>0</td><td>TTTCGAAAAGATCTCTCTCT</td><td>CGG</td><td>||||||||[||||||||||||]PAM</td><td>NA</td><td>Intergenic</td><td>NA</td><td>NA</td></tr>
<tr><td>T3731</td><td>GATGAGCGCCGTGAGATCGTCGG</td><td>1000</td><td></td><td></td><td></td><td></td><td></td><td></td><td></td><td></td><td></td></tr>
<tr><td>Oligo fwd</td><td>CACCGATGAGCGCCGTGAGATCGT</td><td></td><td></td><td></td><td></td><td></td><td></td><td></td><td></td><td></td><td></td></tr>
<tr><td>Oligo rev</td><td>AAACACGATCTCACGGCGCTCATC</td><td></td><td></td><td></td><td></td><td></td><td></td><td></td><td></td><td></td><td></td></tr>
<tr><td>Chromosome</td><td>start</td><td>end</td><td>strand</td><td>MM</td><td>target_seq</td><td>PAM</td><td>alignment</td><td>distance</td><td>position</td><td>gene name</td><td>gene id</td></tr>
<tr><td>GU305914.1</td><td>56860</td><td>56882</td><td>+</td><td>0</td><td>GATGAGCGCCGTGAGATCGT</td><td>CGG</td><td>||||||||[||||||||||||]PAM</td><td>NA</td><td>Intergenic</td><td>NA</td><td>NA</td></tr>
<tr><td>T3732</td><td>AGCGCCGTGAGATCGTCGGACGG</td><td>1000</td><td></td><td></td><td></td><td></td><td></td><td></td><td></td><td></td><td></td></tr>
<tr><td>Oligo fwd</td><td>CACCAGCGCCGTGAGATCGTCGGA</td><td></td><td></td><td></td><td></td><td></td><td></td><td></td><td></td><td></td><td></td></tr>
<tr><td>Oligo rev</td><td>AAACTCCGACGATCTCACGGCGCT</td><td></td><td></td><td></td><td></td><td></td><td></td><td></td><td></td><td></td><td></td></tr>
<tr><td>Chromosome</td><td>start</td><td>end</td><td>strand</td><td>MM</td><td>target_seq</td><td>PAM</td><td>alignment</td><td>distance</td><td>position</td><td>gene name</td><td>gene id</td></tr>
<tr><td>GU305914.1</td><td>56864</td><td>56886</td><td>+</td><td>0</td><td>AGCGCCGTGAGATCGTCGGA</td><td>CGG</td><td>||||||||[||||||||||||]PAM</td><td>NA</td><td>Intergenic</td><td>NA</td><td>NA</td></tr>
<tr><td>T3733</td><td>GGAGACCAATCTCGTCCACGAGG</td><td>1000</td><td></td><td></td><td></td><td></td><td></td><td></td><td></td><td></td><td></td></tr>
<tr><td>Oligo fwd</td><td>CACCGGAGACCAATCTCGTCCACG</td><td></td><td></td><td></td><td></td><td></td><td></td><td></td><td></td><td></td><td></td></tr>
<tr><td>Oligo rev</td><td>AAACCGTGGACGAGATTGGTCTCC</td><td></td><td></td><td></td><td></td><td></td><td></td><td></td><td></td><td></td><td></td></tr>
<tr><td>Chromosome</td><td>start</td><td>end</td><td>strand</td><td>MM</td><td>target_seq</td><td>PAM</td><td>alignment</td><td>distance</td><td>position</td><td>gene name</td><td>gene id</td></tr>
<tr><td>GU305914.1</td><td>56885</td><td>56907</td><td>+</td><td>0</td><td>GGAGACCAATCTCGTCCACG</td><td>AGG</td><td>||||||||[||||||||||||]PAM</td><td>NA</td><td>Intergenic</td><td>NA</td><td>NA</td></tr>
<tr><td>T3734</td><td>AGGTGCCGAAAATCGCACGACGG</td><td>1000</td><td></td><td></td><td></td><td></td><td></td><td></td><td></td><td></td><td></td></tr>
<tr><td>Oligo fwd</td><td>CACCAGGTGCCGAAAATCGCACGA</td><td></td><td></td><td></td><td></td><td></td><td></td><td></td><td></td><td></td><td></td></tr>
<tr><td>Oligo rev</td><td>AAACTCGTGCGATTTTCGGCACCT</td><td></td><td></td><td></td><td></td><td></td><td></td><td></td><td></td><td></td><td></td></tr>
<tr><td>Chromosome</td><td>start</td><td>end</td><td>strand</td><td>MM</td><td>target_seq</td><td>PAM</td><td>alignment</td><td>distance</td><td>position</td><td>gene name</td><td>gene id</td></tr>
<tr><td>GU305914.1</td><td>56905</td><td>56927</td><td>+</td><td>0</td><td>AGGTGCCGAAAATCGCACGA</td><td>CGG</td><td>||||||||[||||||||||||]PAM</td><td>NA</td><td>Intergenic</td><td>NA</td><td>NA</td></tr>
<tr><td>T3735</td><td>ATCGCACGACGGCCCTCCCACGG</td><td>1000</td><td></td><td></td><td></td><td></td><td></td><td></td><td></td><td></td><td></td></tr>
<tr><td>Oligo fwd</td><td>CACCATCGCACGACGGCCCTCCCA</td><td></td><td></td><td></td><td></td><td></td><td></td><td></td><td></td><td></td><td></td></tr>
<tr><td>Oligo rev</td><td>AAACTGGGAGGGCCGTCGTGCGAT</td><td></td><td></td><td></td><td></td><td></td><td></td><td></td><td></td><td></td><td></td></tr>
<tr><td>Chromosome</td><td>start</td><td>end</td><td>strand</td><td>MM</td><td>target_seq</td><td>PAM</td><td>alignment</td><td>distance</td><td>position</td><td>gene name</td><td>gene id</td></tr>
<tr><td>GU305914.1</td><td>56916</td><td>56938</td><td>+</td><td>0</td><td>ATCGCACGACGGCCCTCCCA</td><td>CGG</td><td>||||||||[||||||||||||]PAM</td><td>NA</td><td>Intergenic</td><td>NA</td><td>NA</td></tr>
<tr><td>T3736</td><td>TCGCACGACGGCCCTCCCACGGG</td><td>1000</td><td></td><td></td><td></td><td></td><td></td><td></td><td></td><td></td><td></td></tr>
<tr><td>Oligo fwd</td><td>CACCTCGCACGACGGCCCTCCCAC</td><td></td><td></td><td></td><td></td><td></td><td></td><td></td><td></td><td></td><td></td></tr>
<tr><td>Oligo rev</td><td>AAACGTGGGAGGGCCGTCGTGCGA</td><td></td><td></td><td></td><td></td><td></td><td></td><td></td><td></td><td></td><td></td></tr>
<tr><td>Chromosome</td><td>start</td><td>end</td><td>strand</td><td>MM</td><td>target_seq</td><td>PAM</td><td>alignment</td><td>distance</td><td>position</td><td>gene name</td><td>gene id</td></tr>
<tr><td>GU305914.1</td><td>56917</td><td>56939</td><td>+</td><td>0</td><td>TCGCACGACGGCCCTCCCAC</td><td>GGG</td><td>||||||||[||||||||||||]PAM</td><td>NA</td><td>Intergenic</td><td>NA</td><td>NA</td></tr>
<tr><td>T3737</td><td>CCTCCCACGGGCCTCCGACCTGG</td><td>1000</td><td></td><td></td><td></td><td></td><td></td><td></td><td></td><td></td><td></td></tr>
<tr><td>Oligo fwd</td><td>CACCCCTCCCACGGGCCTCCGACC</td><td></td><td></td><td></td><td></td><td></td><td></td><td></td><td></td><td></td><td></td></tr>
<tr><td>Oligo rev</td><td>AAACGGTCGGAGGCCCGTGGGAGG</td><td></td><td></td><td></td><td></td><td></td><td></td><td></td><td></td><td></td><td></td></tr>
<tr><td>Chromosome</td><td>start</td><td>end</td><td>strand</td><td>MM</td><td>target_seq</td><td>PAM</td><td>alignment</td><td>distance</td><td>position</td><td>gene name</td><td>gene id</td></tr>
<tr><td>GU305914.1</td><td>56929</td><td>56951</td><td>+</td><td>0</td><td>CCTCCCACGGGCCTCCGACC</td><td>TGG</td><td>||||||||[||||||||||||]PAM</td><td>NA</td><td>Intergenic</td><td>NA</td><td>NA</td></tr>
<tr><td>GU305914.1</td><td>64214</td><td>64236</td><td>+</td><td>3</td><td>CCTCCCGCTGGCCGCCGACC</td><td>TGG</td><td>||||||-|[-||||-||||||]PAM</td><td>NA</td><td>Intergenic</td><td>NA</td><td>NA</td></tr>
<tr><td>T3738</td><td>CCGACCTGGTATCCAGCGTGCGG</td><td>1000</td><td></td><td></td><td></td><td></td><td></td><td></td><td></td><td></td><td></td></tr>
<tr><td>Oligo fwd</td><td>CACCCCGACCTGGTATCCAGCGTG</td><td></td><td></td><td></td><td></td><td></td><td></td><td></td><td></td><td></td><td></td></tr>
<tr><td>Oligo rev</td><td>AAACCACGCTGGATACCAGGTCGG</td><td></td><td></td><td></td><td></td><td></td><td></td><td></td><td></td><td></td><td></td></tr>
<tr><td>Chromosome</td><td>start</td><td>end</td><td>strand</td><td>MM</td><td>target_seq</td><td>PAM</td><td>alignment</td><td>distance</td><td>position</td><td>gene name</td><td>gene id</td></tr>
<tr><td>GU305914.1</td><td>56943</td><td>56965</td><td>+</td><td>0</td><td>CCGACCTGGTATCCAGCGTG</td><td>CGG</td><td>||||||||[||||||||||||]PAM</td><td>NA</td><td>Intergenic</td><td>NA</td><td>NA</td></tr>
<tr><td>T3739</td><td>TCCAGCGTGCGGATGATCCTCGG</td><td>1000</td><td></td><td></td><td></td><td></td><td></td><td></td><td></td><td></td><td></td></tr>
<tr><td>Oligo fwd</td><td>CACCTCCAGCGTGCGGATGATCCT</td><td></td><td></td><td></td><td></td><td></td><td></td><td></td><td></td><td></td><td></td></tr>
<tr><td>Oligo rev</td><td>AAACAGGATCATCCGCACGCTGGA</td><td></td><td></td><td></td><td></td><td></td><td></td><td></td><td></td><td></td><td></td></tr>
<tr><td>Chromosome</td><td>start</td><td>end</td><td>strand</td><td>MM</td><td>target_seq</td><td>PAM</td><td>alignment</td><td>distance</td><td>position</td><td>gene name</td><td>gene id</td></tr>
<tr><td>GU305914.1</td><td>56954</td><td>56976</td><td>+</td><td>0</td><td>TCCAGCGTGCGGATGATCCT</td><td>CGG</td><td>||||||||[||||||||||||]PAM</td><td>NA</td><td>Intergenic</td><td>NA</td><td>NA</td></tr>
<tr><td>T3740</td><td>GTGCGGATGATCCTCGGCTGCGG</td><td>1000</td><td></td><td></td><td></td><td></td><td></td><td></td><td></td><td></td><td></td></tr>
<tr><td>Oligo fwd</td><td>CACCGTGCGGATGATCCTCGGCTG</td><td></td><td></td><td></td><td></td><td></td><td></td><td></td><td></td><td></td><td></td></tr>
<tr><td>Oligo rev</td><td>AAACCAGCCGAGGATCATCCGCAC</td><td></td><td></td><td></td><td></td><td></td><td></td><td></td><td></td><td></td><td></td></tr>
<tr><td>Chromosome</td><td>start</td><td>end</td><td>strand</td><td>MM</td><td>target_seq</td><td>PAM</td><td>alignment</td><td>distance</td><td>position</td><td>gene name</td><td>gene id</td></tr>
<tr><td>GU305914.1</td><td>56960</td><td>56982</td><td>+</td><td>0</td><td>GTGCGGATGATCCTCGGCTG</td><td>CGG</td><td>||||||||[||||||||||||]PAM</td><td>NA</td><td>Intergenic</td><td>NA</td><td>NA</td></tr>
<tr><td>T3741</td><td>GGATGATCCTCGGCTGCGGTCGG</td><td>1000</td><td></td><td></td><td></td><td></td><td></td><td></td><td></td><td></td><td></td></tr>
<tr><td>Oligo fwd</td><td>CACCGGATGATCCTCGGCTGCGGT</td><td></td><td></td><td></td><td></td><td></td><td></td><td></td><td></td><td></td><td></td></tr>
<tr><td>Oligo rev</td><td>AAACACCGCAGCCGAGGATCATCC</td><td></td><td></td><td></td><td></td><td></td><td></td><td></td><td></td><td></td><td></td></tr>
<tr><td>Chromosome</td><td>start</td><td>end</td><td>strand</td><td>MM</td><td>target_seq</td><td>PAM</td><td>alignment</td><td>distance</td><td>position</td><td>gene name</td><td>gene id</td></tr>
<tr><td>GU305914.1</td><td>56964</td><td>56986</td><td>+</td><td>0</td><td>GGATGATCCTCGGCTGCGGT</td><td>CGG</td><td>||||||||[||||||||||||]PAM</td><td>NA</td><td>Intergenic</td><td>NA</td><td>NA</td></tr>
<tr><td>T3742</td><td>CTGCGGTCGGAACCGCGCGTCGG</td><td>1000</td><td></td><td></td><td></td><td></td><td></td><td></td><td></td><td></td><td></td></tr>
<tr><td>Oligo fwd</td><td>CACCCTGCGGTCGGAACCGCGCGT</td><td></td><td></td><td></td><td></td><td></td><td></td><td></td><td></td><td></td><td></td></tr>
<tr><td>Oligo rev</td><td>AAACACGCGCGGTTCCGACCGCAG</td><td></td><td></td><td></td><td></td><td></td><td></td><td></td><td></td><td></td><td></td></tr>
<tr><td>Chromosome</td><td>start</td><td>end</td><td>strand</td><td>MM</td><td>target_seq</td><td>PAM</td><td>alignment</td><td>distance</td><td>position</td><td>gene name</td><td>gene id</td></tr>
<tr><td>GU305914.1</td><td>56977</td><td>56999</td><td>+</td><td>0</td><td>CTGCGGTCGGAACCGCGCGT</td><td>CGG</td><td>||||||||[||||||||||||]PAM</td><td>NA</td><td>Intergenic</td><td>NA</td><td>NA</td></tr>
<tr><td>T3743</td><td>CCGCGCGTCGGCCAGTCGTTCGG</td><td>1000</td><td></td><td></td><td></td><td></td><td></td><td></td><td></td><td></td><td></td></tr>
<tr><td>Oligo fwd</td><td>CACCCCGCGCGTCGGCCAGTCGTT</td><td></td><td></td><td></td><td></td><td></td><td></td><td></td><td></td><td></td><td></td></tr>
<tr><td>Oligo rev</td><td>AAACAACGACTGGCCGACGCGCGG</td><td></td><td></td><td></td><td></td><td></td><td></td><td></td><td></td><td></td><td></td></tr>
<tr><td>Chromosome</td><td>start</td><td>end</td><td>strand</td><td>MM</td><td>target_seq</td><td>PAM</td><td>alignment</td><td>distance</td><td>position</td><td>gene name</td><td>gene id</td></tr>
<tr><td>GU305914.1</td><td>56989</td><td>57011</td><td>+</td><td>0</td><td>CCGCGCGTCGGCCAGTCGTT</td><td>CGG</td><td>||||||||[||||||||||||]PAM</td><td>NA</td><td>Intergenic</td><td>NA</td><td>NA</td></tr>
<tr><td>T3744</td><td>CGCGCGTCGGCCAGTCGTTCGGG</td><td>1000</td><td></td><td></td><td></td><td></td><td></td><td></td><td></td><td></td><td></td></tr>
<tr><td>Oligo fwd</td><td>CACCCGCGCGTCGGCCAGTCGTTC</td><td></td><td></td><td></td><td></td><td></td><td></td><td></td><td></td><td></td><td></td></tr>
<tr><td>Oligo rev</td><td>AAACGAACGACTGGCCGACGCGCG</td><td></td><td></td><td></td><td></td><td></td><td></td><td></td><td></td><td></td><td></td></tr>
<tr><td>Chromosome</td><td>start</td><td>end</td><td>strand</td><td>MM</td><td>target_seq</td><td>PAM</td><td>alignment</td><td>distance</td><td>position</td><td>gene name</td><td>gene id</td></tr>
<tr><td>GU305914.1</td><td>56990</td><td>57012</td><td>+</td><td>0</td><td>CGCGCGTCGGCCAGTCGTTC</td><td>GGG</td><td>||||||||[||||||||||||]PAM</td><td>NA</td><td>Intergenic</td><td>NA</td><td>NA</td></tr>
</table>

<table>
<tr><td>T3745</td><td>CGTTCGGGATACATAGAGAACGG</td><td>1000</td><td></td><td></td><td></td><td></td><td></td><td></td><td></td><td></td><td></td></tr>
<tr><td>Oligo fwd</td><td>CACCCGTTCGGGATACATAGAGAA</td><td></td><td></td><td></td><td></td><td></td><td></td><td></td><td></td><td></td><td></td></tr>
<tr><td>Oligo rev</td><td>AAACTTCTCTATGTATCCCGAACG</td><td></td><td></td><td></td><td></td><td></td><td></td><td></td><td></td><td></td><td></td></tr>
<tr><td>Chromosome</td><td>start</td><td>end</td><td>strand</td><td>MM</td><td>target_seq</td><td>PAM</td><td>alignment</td><td>distance</td><td>position</td><td>gene name</td><td>gene id</td></tr>
<tr><td>GU305914.1</td><td>57005</td><td>57027</td><td>+</td><td>0</td><td>CGTTCGGGATACATAGAGAA</td><td>CGG</td><td>||||||||[||||||||||||]PAM</td><td>NA</td><td>Intergenic</td><td>NA</td><td>NA</td></tr>
<tr><td>T3746</td><td>GGGATACATAGAGAACGGCGCGG</td><td>1000</td><td></td><td></td><td></td><td></td><td></td><td></td><td></td><td></td><td></td></tr>
<tr><td>Oligo fwd</td><td>CACCGGGATACATAGAGAACGGCG</td><td></td><td></td><td></td><td></td><td></td><td></td><td></td><td></td><td></td><td></td></tr>
<tr><td>Oligo rev</td><td>AAACCGCCGTTCTCTATGTATCCC</td><td></td><td></td><td></td><td></td><td></td><td></td><td></td><td></td><td></td><td></td></tr>
<tr><td>Chromosome</td><td>start</td><td>end</td><td>strand</td><td>MM</td><td>target_seq</td><td>PAM</td><td>alignment</td><td>distance</td><td>position</td><td>gene name</td><td>gene id</td></tr>
<tr><td>GU305914.1</td><td>57010</td><td>57032</td><td>+</td><td>0</td><td>GGGATACATAGAGAACGGCG</td><td>CGG</td><td>||||||||[||||||||||||]PAM</td><td>NA</td><td>Intergenic</td><td>NA</td><td>NA</td></tr>
<tr><td>T3747</td><td>GGATACATAGAGAACGGCGCGGG</td><td>1000</td><td></td><td></td><td></td><td></td><td></td><td></td><td></td><td></td><td></td></tr>
<tr><td>Oligo fwd</td><td>CACCGGATACATAGAGAACGGCGC</td><td></td><td></td><td></td><td></td><td></td><td></td><td></td><td></td><td></td><td></td></tr>
<tr><td>Oligo rev</td><td>AAACGCGCCGTTCTCTATGTATCC</td><td></td><td></td><td></td><td></td><td></td><td></td><td></td><td></td><td></td><td></td></tr>
<tr><td>Chromosome</td><td>start</td><td>end</td><td>strand</td><td>MM</td><td>target_seq</td><td>PAM</td><td>alignment</td><td>distance</td><td>position</td><td>gene name</td><td>gene id</td></tr>
<tr><td>GU305914.1</td><td>57011</td><td>57033</td><td>+</td><td>0</td><td>GGATACATAGAGAACGGCGC</td><td>GGG</td><td>||||||||[||||||||||||]PAM</td><td>NA</td><td>Intergenic</td><td>NA</td><td>NA</td></tr>
<tr><td>T3748</td><td>GATACATAGAGAACGGCGCGGGG</td><td>1000</td><td></td><td></td><td></td><td></td><td></td><td></td><td></td><td></td><td></td></tr>
<tr><td>Oligo fwd</td><td>CACCGATACATAGAGAACGGCGCG</td><td></td><td></td><td></td><td></td><td></td><td></td><td></td><td></td><td></td><td></td></tr>
<tr><td>Oligo rev</td><td>AAACCGCGCCGTTCTCTATGTATC</td><td></td><td></td><td></td><td></td><td></td><td></td><td></td><td></td><td></td><td></td></tr>
<tr><td>Chromosome</td><td>start</td><td>end</td><td>strand</td><td>MM</td><td>target_seq</td><td>PAM</td><td>alignment</td><td>distance</td><td>position</td><td>gene name</td><td>gene id</td></tr>
<tr><td>GU305914.1</td><td>57012</td><td>57034</td><td>+</td><td>0</td><td>GATACATAGAGAACGGCGCG</td><td>GGG</td><td>||||||||[||||||||||||]PAM</td><td>NA</td><td>Intergenic</td><td>NA</td><td>NA</td></tr>
<tr><td>T3749</td><td>AGAGAACGGCGCGGGGACGCTGG</td><td>1000</td><td></td><td></td><td></td><td></td><td></td><td></td><td></td><td></td><td></td></tr>
<tr><td>Oligo fwd</td><td>CACCAGAGAACGGCGCGGGGACGC</td><td></td><td></td><td></td><td></td><td></td><td></td><td></td><td></td><td></td><td></td></tr>
<tr><td>Oligo rev</td><td>AAACGCGTCCCCGCGCCGTTCTCT</td><td></td><td></td><td></td><td></td><td></td><td></td><td></td><td></td><td></td><td></td></tr>
<tr><td>Chromosome</td><td>start</td><td>end</td><td>strand</td><td>MM</td><td>target_seq</td><td>PAM</td><td>alignment</td><td>distance</td><td>position</td><td>gene name</td><td>gene id</td></tr>
<tr><td>GU305914.1</td><td>57019</td><td>57041</td><td>+</td><td>0</td><td>AGAGAACGGCGCGGGGACGC</td><td>TGG</td><td>||||||||[||||||||||||]PAM</td><td>NA</td><td>Intergenic</td><td>NA</td><td>NA</td></tr>
<tr><td>T3750</td><td>GATGTTCCGTCGCCACCTAGCGG</td><td>1000</td><td></td><td></td><td></td><td></td><td></td><td></td><td></td><td></td><td></td></tr>
<tr><td>Oligo fwd</td><td>CACCGATGTTCCGTCGCCACCTAG</td><td></td><td></td><td></td><td></td><td></td><td></td><td></td><td></td><td></td><td></td></tr>
<tr><td>Oligo rev</td><td>AAACCTAGGTGGCGACGGAACATC</td><td></td><td></td><td></td><td></td><td></td><td></td><td></td><td></td><td></td><td></td></tr>
<tr><td>Chromosome</td><td>start</td><td>end</td><td>strand</td><td>MM</td><td>target_seq</td><td>PAM</td><td>alignment</td><td>distance</td><td>position</td><td>gene name</td><td>gene id</td></tr>
<tr><td>GU305914.1</td><td>57043</td><td>57065</td><td>+</td><td>0</td><td>GATGTTCCGTCGCCACCTAG</td><td>CGG</td><td>||||||||[||||||||||||]PAM</td><td>NA</td><td>Intergenic</td><td>NA</td><td>NA</td></tr>
<tr><td>T3751</td><td>GTACCCGAGCTCTCGAGCCATGG</td><td>1000</td><td></td><td></td><td></td><td></td><td></td><td></td><td></td><td></td><td></td></tr>
<tr><td>Oligo fwd</td><td>CACCGTACCCGAGCTCTCGAGCCA</td><td></td><td></td><td></td><td></td><td></td><td></td><td></td><td></td><td></td><td></td></tr>
<tr><td>Oligo rev</td><td>AAACTGGCTCGAGAGCTCGGGTAC</td><td></td><td></td><td></td><td></td><td></td><td></td><td></td><td></td><td></td><td></td></tr>
<tr><td>Chromosome</td><td>start</td><td>end</td><td>strand</td><td>MM</td><td>target_seq</td><td>PAM</td><td>alignment</td><td>distance</td><td>position</td><td>gene name</td><td>gene id</td></tr>
<tr><td>GU305914.1</td><td>57079</td><td>57101</td><td>+</td><td>0</td><td>GTACCCGAGCTCTCGAGCCA</td><td>TGG</td><td>||||||||[||||||||||||]PAM</td><td>NA</td><td>Intergenic</td><td>NA</td><td>NA</td></tr>
<tr><td>T3752</td><td>CTCGAGCCATGGCTCGTCGTCGG</td><td>1000</td><td></td><td></td><td></td><td></td><td></td><td></td><td></td><td></td><td></td></tr>
<tr><td>Oligo fwd</td><td>CACCCTCGAGCCATGGCTCGTCGT</td><td></td><td></td><td></td><td></td><td></td><td></td><td></td><td></td><td></td><td></td></tr>
<tr><td>Oligo rev</td><td>AAACACGACGAGCCATGGCTCGAG</td><td></td><td></td><td></td><td></td><td></td><td></td><td></td><td></td><td></td><td></td></tr>
<tr><td>Chromosome</td><td>start</td><td>end</td><td>strand</td><td>MM</td><td>target_seq</td><td>PAM</td><td>alignment</td><td>distance</td><td>position</td><td>gene name</td><td>gene id</td></tr>
<tr><td>GU305914.1</td><td>57090</td><td>57112</td><td>+</td><td>0</td><td>CTCGAGCCATGGCTCGTCGT</td><td>CGG</td><td>||||||||[||||||||||||]PAM</td><td>NA</td><td>Intergenic</td><td>NA</td><td>NA</td></tr>
<tr><td>T3753</td><td>ATGGCTCGTCGTCGGCACGACGG</td><td>1000</td><td></td><td></td><td></td><td></td><td></td><td></td><td></td><td></td><td></td></tr>
<tr><td>Oligo fwd</td><td>CACCATGGCTCGTCGTCGGCACGA</td><td></td><td></td><td></td><td></td><td></td><td></td><td></td><td></td><td></td><td></td></tr>
<tr><td>Oligo rev</td><td>AAACTCGTGCCGACGACGAGCCAT</td><td></td><td></td><td></td><td></td><td></td><td></td><td></td><td></td><td></td><td></td></tr>
<tr><td>Chromosome</td><td>start</td><td>end</td><td>strand</td><td>MM</td><td>target_seq</td><td>PAM</td><td>alignment</td><td>distance</td><td>position</td><td>gene name</td><td>gene id</td></tr>
<tr><td>GU305914.1</td><td>57098</td><td>57120</td><td>+</td><td>0</td><td>ATGGCTCGTCGTCGGCACGA</td><td>CGG</td><td>||||||||[||||||||||||]PAM</td><td>NA</td><td>Intergenic</td><td>NA</td><td>NA</td></tr>
<tr><td>T3754</td><td>TGGCTCGTCGTCGGCACGACGGG</td><td>1000</td><td></td><td></td><td></td><td></td><td></td><td></td><td></td><td></td><td></td></tr>
<tr><td>Oligo fwd</td><td>CACCTGGCTCGTCGTCGGCACGAC</td><td></td><td></td><td></td><td></td><td></td><td></td><td></td><td></td><td></td><td></td></tr>
<tr><td>Oligo rev</td><td>AAACGTCGTGCCGACGACGAGCCA</td><td></td><td></td><td></td><td></td><td></td><td></td><td></td><td></td><td></td><td></td></tr>
<tr><td>Chromosome</td><td>start</td><td>end</td><td>strand</td><td>MM</td><td>target_seq</td><td>PAM</td><td>alignment</td><td>distance</td><td>position</td><td>gene name</td><td>gene id</td></tr>
<tr><td>GU305914.1</td><td>57099</td><td>57121</td><td>+</td><td>0</td><td>TGGCTCGTCGTCGGCACGAC</td><td>GGG</td><td>||||||||[||||||||||||]PAM</td><td>NA</td><td>Intergenic</td><td>NA</td><td>NA</td></tr>
<tr><td>T3755</td><td>CGTCGTCGGCACGACGGGCAAGG</td><td>1000</td><td></td><td></td><td></td><td></td><td></td><td></td><td></td><td></td><td></td></tr>
<tr><td>Oligo fwd</td><td>CACCCGTCGTCGGCACGACGGGCA</td><td></td><td></td><td></td><td></td><td></td><td></td><td></td><td></td><td></td><td></td></tr>
<tr><td>Oligo rev</td><td>AAACTGCCCGTCGTGCCGACGACG</td><td></td><td></td><td></td><td></td><td></td><td></td><td></td><td></td><td></td><td></td></tr>
<tr><td>Chromosome</td><td>start</td><td>end</td><td>strand</td><td>MM</td><td>target_seq</td><td>PAM</td><td>alignment</td><td>distance</td><td>position</td><td>gene name</td><td>gene id</td></tr>
<tr><td>GU305914.1</td><td>57104</td><td>57126</td><td>+</td><td>0</td><td>CGTCGTCGGCACGACGGGCA</td><td>AGG</td><td>||||||||[||||||||||||]PAM</td><td>NA</td><td>Intergenic</td><td>NA</td><td>NA</td></tr>
<tr><td>T3756</td><td>TTGTCATAGACCAACGAACGTGG</td><td>1000</td><td></td><td></td><td></td><td></td><td></td><td></td><td></td><td></td><td></td></tr>
<tr><td>Oligo fwd</td><td>CACCTTGTCATAGACCAACGAACG</td><td></td><td></td><td></td><td></td><td></td><td></td><td></td><td></td><td></td><td></td></tr>
<tr><td>Oligo rev</td><td>AAACCGTTCGTTGGTCTATGACAA</td><td></td><td></td><td></td><td></td><td></td><td></td><td></td><td></td><td></td><td></td></tr>
<tr><td>Chromosome</td><td>start</td><td>end</td><td>strand</td><td>MM</td><td>target_seq</td><td>PAM</td><td>alignment</td><td>distance</td><td>position</td><td>gene name</td><td>gene id</td></tr>
<tr><td>GU305914.1</td><td>57141</td><td>57163</td><td>+</td><td>0</td><td>TTGTCATAGACCAACGAACG</td><td>TGG</td><td>||||||||[||||||||||||]PAM</td><td>NA</td><td>Intergenic</td><td>NA</td><td>NA</td></tr>
<tr><td>T3757</td><td>ATAGACCAACGAACGTGGCGAGG</td><td>1000</td><td></td><td></td><td></td><td></td><td></td><td></td><td></td><td></td><td></td></tr>
<tr><td>Oligo fwd</td><td>CACCATAGACCAACGAACGTGGCG</td><td></td><td></td><td></td><td></td><td></td><td></td><td></td><td></td><td></td><td></td></tr>
<tr><td>Oligo rev</td><td>AAACCGCCACGTTCGTTGGTCTAT</td><td></td><td></td><td></td><td></td><td></td><td></td><td></td><td></td><td></td><td></td></tr>
<tr><td>Chromosome</td><td>start</td><td>end</td><td>strand</td><td>MM</td><td>target_seq</td><td>PAM</td><td>alignment</td><td>distance</td><td>position</td><td>gene name</td><td>gene id</td></tr>
<tr><td>GU305914.1</td><td>57146</td><td>57168</td><td>+</td><td>0</td><td>ATAGACCAACGAACGTGGCG</td><td>AGG</td><td>||||||||[||||||||||||]PAM</td><td>NA</td><td>Intergenic</td><td>NA</td><td>NA</td></tr>
<tr><td>T3758</td><td>ACGAACGTGGCGAGGTCTACTGG</td><td>1000</td><td></td><td></td><td></td><td></td><td></td><td></td><td></td><td></td><td></td></tr>
<tr><td>Oligo fwd</td><td>CACCACGAACGTGGCGAGGTCTAC</td><td></td><td></td><td></td><td></td><td></td><td></td><td></td><td></td><td></td><td></td></tr>
<tr><td>Oligo rev</td><td>AAACGTAGACCTCGCCACGTTCGT</td><td></td><td></td><td></td><td></td><td></td><td></td><td></td><td></td><td></td><td></td></tr>
<tr><td>Chromosome</td><td>start</td><td>end</td><td>strand</td><td>MM</td><td>target_seq</td><td>PAM</td><td>alignment</td><td>distance</td><td>position</td><td>gene name</td><td>gene id</td></tr>
<tr><td>GU305914.1</td><td>57154</td><td>57176</td><td>+</td><td>0</td><td>ACGAACGTGGCGAGGTCTAC</td><td>TGG</td><td>||||||||[||||||||||||]PAM</td><td>NA</td><td>Intergenic</td><td>NA</td><td>NA</td></tr>
<tr><td>T3759</td><td>CACAGAGTAGTAGTGTAAAACGG</td><td>1000</td><td></td><td></td><td></td><td></td><td></td><td></td><td></td><td></td><td></td></tr>
<tr><td>Oligo fwd</td><td>CACCCACAGAGTAGTAGTGTAAAA</td><td></td><td></td><td></td><td></td><td></td><td></td><td></td><td></td><td></td><td></td></tr>
<tr><td>Oligo rev</td><td>AAACTTTTACACTACTACTCTGTG</td><td></td><td></td><td></td><td></td><td></td><td></td><td></td><td></td><td></td><td></td></tr>
<tr><td>Chromosome</td><td>start</td><td>end</td><td>strand</td><td>MM</td><td>target_seq</td><td>PAM</td><td>alignment</td><td>distance</td><td>position</td><td>gene name</td><td>gene id</td></tr>
</table>

<table>
<tr><td>GU305914.1</td><td>57191</td><td>57213</td><td>+</td><td>0</td><td>CACAGAGTAGTAGTGTAAAA</td><td>CGG</td><td>||||||||[||||||||||||]PAM</td><td>NA</td><td>Intergenic</td><td>NA</td><td>NA</td></tr>
<tr><td></td><td></td><td></td><td></td><td></td><td></td><td></td><td></td><td></td><td></td><td></td><td></td></tr>
<tr><td>T3760</td><td>ACAGAGTAGTAGTGTAAAACGGG</td><td>1000</td><td></td><td></td><td></td><td></td><td></td><td></td><td></td><td></td><td></td></tr>
<tr><td>Oligo fwd</td><td>CACCACAGAGTAGTAGTGTAAAAC</td><td></td><td></td><td></td><td></td><td></td><td></td><td></td><td></td><td></td><td></td></tr>
<tr><td>Oligo rev</td><td>AAACGTTTTACACTACTACTCTGT</td><td></td><td></td><td></td><td></td><td></td><td></td><td></td><td></td><td></td><td></td></tr>
<tr><td>Chromosome</td><td>start</td><td>end</td><td>strand</td><td>MM</td><td>target_seq</td><td>PAM</td><td>alignment</td><td>distance</td><td>position</td><td>gene name</td><td>gene id</td></tr>
<tr><td>GU305914.1</td><td>57192</td><td>57214</td><td>+</td><td>0</td><td>ACAGAGTAGTAGTGTAAAAC</td><td>GGG</td><td>||||||||[||||||||||||]PAM</td><td>NA</td><td>Intergenic</td><td>NA</td><td>NA</td></tr>
<tr><td></td><td></td><td></td><td></td><td></td><td></td><td></td><td></td><td></td><td></td><td></td><td></td></tr>
<tr><td>T3761</td><td>AAAACGGGACGCCGTCACGCCGG</td><td>1000</td><td></td><td></td><td></td><td></td><td></td><td></td><td></td><td></td><td></td></tr>
<tr><td>Oligo fwd</td><td>CACCAAAACGGGACGCCGTCACGC</td><td></td><td></td><td></td><td></td><td></td><td></td><td></td><td></td><td></td><td></td></tr>
<tr><td>Oligo rev</td><td>AAACGCGTGACGGCGTCCCGTTTT</td><td></td><td></td><td></td><td></td><td></td><td></td><td></td><td></td><td></td><td></td></tr>
<tr><td>Chromosome</td><td>start</td><td>end</td><td>strand</td><td>MM</td><td>target_seq</td><td>PAM</td><td>alignment</td><td>distance</td><td>position</td><td>gene name</td><td>gene id</td></tr>
<tr><td>GU305914.1</td><td>57207</td><td>57229</td><td>+</td><td>0</td><td>AAAACGGGACGCCGTCACGC</td><td>CGG</td><td>||||||||[||||||||||||]PAM</td><td>NA</td><td>Intergenic</td><td>NA</td><td>NA</td></tr>
<tr><td></td><td></td><td></td><td></td><td></td><td></td><td></td><td></td><td></td><td></td><td></td><td></td></tr>
<tr><td>T3762</td><td>CGAGCGTAGACGACAATTATAGG</td><td>1000</td><td></td><td></td><td></td><td></td><td></td><td></td><td></td><td></td><td></td></tr>
<tr><td>Oligo fwd</td><td>CACCCGAGCGTAGACGACAATTAT</td><td></td><td></td><td></td><td></td><td></td><td></td><td></td><td></td><td></td><td></td></tr>
<tr><td>Oligo rev</td><td>AAACATAATTGTCGTCTACGCTCG</td><td></td><td></td><td></td><td></td><td></td><td></td><td></td><td></td><td></td><td></td></tr>
<tr><td>Chromosome</td><td>start</td><td>end</td><td>strand</td><td>MM</td><td>target_seq</td><td>PAM</td><td>alignment</td><td>distance</td><td>position</td><td>gene name</td><td>gene id</td></tr>
<tr><td>GU305914.1</td><td>57233</td><td>57255</td><td>+</td><td>0</td><td>CGAGCGTAGACGACAATTAT</td><td>AGG</td><td>||||||||[||||||||||||]PAM</td><td>NA</td><td>Intergenic</td><td>NA</td><td>NA</td></tr>
<tr><td></td><td></td><td></td><td></td><td></td><td></td><td></td><td></td><td></td><td></td><td></td><td></td></tr>
<tr><td>T3763</td><td>CGTAGACGACAATTATAGGTAGG</td><td>1000</td><td></td><td></td><td></td><td></td><td></td><td></td><td></td><td></td><td></td></tr>
<tr><td>Oligo fwd</td><td>CACCCGTAGACGACAATTATAGGT</td><td></td><td></td><td></td><td></td><td></td><td></td><td></td><td></td><td></td><td></td></tr>
<tr><td>Oligo rev</td><td>AAACACCTATAATTGTCGTCTACG</td><td></td><td></td><td></td><td></td><td></td><td></td><td></td><td></td><td></td><td></td></tr>
<tr><td>Chromosome</td><td>start</td><td>end</td><td>strand</td><td>MM</td><td>target_seq</td><td>PAM</td><td>alignment</td><td>distance</td><td>position</td><td>gene name</td><td>gene id</td></tr>
<tr><td>GU305914.1</td><td>57237</td><td>57259</td><td>+</td><td>0</td><td>CGTAGACGACAATTATAGGT</td><td>AGG</td><td>||||||||[||||||||||||]PAM</td><td>NA</td><td>Intergenic</td><td>NA</td><td>NA</td></tr>
<tr><td></td><td></td><td></td><td></td><td></td><td></td><td></td><td></td><td></td><td></td><td></td><td></td></tr>
<tr><td>T3764</td><td>GAAAACCCCCTCGAAGACGCCGG</td><td>1000</td><td></td><td></td><td></td><td></td><td></td><td></td><td></td><td></td><td></td></tr>
<tr><td>Oligo fwd</td><td>CACCGAAAACCCCCTCGAAGACGC</td><td></td><td></td><td></td><td></td><td></td><td></td><td></td><td></td><td></td><td></td></tr>
<tr><td>Oligo rev</td><td>AAACGCGTCTTCGAGGGGGTTTTC</td><td></td><td></td><td></td><td></td><td></td><td></td><td></td><td></td><td></td><td></td></tr>
<tr><td>Chromosome</td><td>start</td><td>end</td><td>strand</td><td>MM</td><td>target_seq</td><td>PAM</td><td>alignment</td><td>distance</td><td>position</td><td>gene name</td><td>gene id</td></tr>
<tr><td>GU305914.1</td><td>57302</td><td>57324</td><td>+</td><td>0</td><td>GAAAACCCCCTCGAAGACGC</td><td>CGG</td><td>||||||||[||||||||||||]PAM</td><td>NA</td><td>Intergenic</td><td>NA</td><td>NA</td></tr>
<tr><td></td><td></td><td></td><td></td><td></td><td></td><td></td><td></td><td></td><td></td><td></td><td></td></tr>
<tr><td>T3765</td><td>AGACGCCGGTCTCTAGCTAACGG</td><td>1000</td><td></td><td></td><td></td><td></td><td></td><td></td><td></td><td></td><td></td></tr>
<tr><td>Oligo fwd</td><td>CACCAGACGCCGGTCTCTAGCTAA</td><td></td><td></td><td></td><td></td><td></td><td></td><td></td><td></td><td></td><td></td></tr>
<tr><td>Oligo rev</td><td>AAACTTAGCTAGAGACCGGCGTCT</td><td></td><td></td><td></td><td></td><td></td><td></td><td></td><td></td><td></td><td></td></tr>
<tr><td>Chromosome</td><td>start</td><td>end</td><td>strand</td><td>MM</td><td>target_seq</td><td>PAM</td><td>alignment</td><td>distance</td><td>position</td><td>gene name</td><td>gene id</td></tr>
<tr><td>GU305914.1</td><td>57316</td><td>57338</td><td>+</td><td>0</td><td>AGACGCCGGTCTCTAGCTAA</td><td>CGG</td><td>||||||||[||||||||||||]PAM</td><td>NA</td><td>Intergenic</td><td>NA</td><td>NA</td></tr>
<tr><td></td><td></td><td></td><td></td><td></td><td></td><td></td><td></td><td></td><td></td><td></td><td></td></tr>
<tr><td>T3766</td><td>CCGGTCTCTAGCTAACGGATCGG</td><td>1000</td><td></td><td></td><td></td><td></td><td></td><td></td><td></td><td></td><td></td></tr>
<tr><td>Oligo fwd</td><td>CACCCCGGTCTCTAGCTAACGGAT</td><td></td><td></td><td></td><td></td><td></td><td></td><td></td><td></td><td></td><td></td></tr>
<tr><td>Oligo rev</td><td>AAACATCCGTTAGCTAGAGACCGG</td><td></td><td></td><td></td><td></td><td></td><td></td><td></td><td></td><td></td><td></td></tr>
<tr><td>Chromosome</td><td>start</td><td>end</td><td>strand</td><td>MM</td><td>target_seq</td><td>PAM</td><td>alignment</td><td>distance</td><td>position</td><td>gene name</td><td>gene id</td></tr>
<tr><td>GU305914.1</td><td>57321</td><td>57343</td><td>+</td><td>0</td><td>CCGGTCTCTAGCTAACGGAT</td><td>CGG</td><td>||||||||[||||||||||||]PAM</td><td>NA</td><td>Intergenic</td><td>NA</td><td>NA</td></tr>
<tr><td></td><td></td><td></td><td></td><td></td><td></td><td></td><td></td><td></td><td></td><td></td><td></td></tr>
<tr><td>T3767</td><td>CGGATCGGACGCCGTGAAGACGG</td><td>1000</td><td></td><td></td><td></td><td></td><td></td><td></td><td></td><td></td><td></td></tr>
<tr><td>Oligo fwd</td><td>CACCCGGATCGGACGCCGTGAAGA</td><td></td><td></td><td></td><td></td><td></td><td></td><td></td><td></td><td></td><td></td></tr>
<tr><td>Oligo rev</td><td>AAACTCTTCACGGCGTCCGATCCG</td><td></td><td></td><td></td><td></td><td></td><td></td><td></td><td></td><td></td><td></td></tr>
<tr><td>Chromosome</td><td>start</td><td>end</td><td>strand</td><td>MM</td><td>target_seq</td><td>PAM</td><td>alignment</td><td>distance</td><td>position</td><td>gene name</td><td>gene id</td></tr>
<tr><td>GU305914.1</td><td>57336</td><td>57358</td><td>+</td><td>0</td><td>CGGATCGGACGCCGTGAAGA</td><td>CGG</td><td>||||||||[||||||||||||]PAM</td><td>NA</td><td>Intergenic</td><td>NA</td><td>NA</td></tr>
<tr><td></td><td></td><td></td><td></td><td></td><td></td><td></td><td></td><td></td><td></td><td></td><td></td></tr>
<tr><td>T3768</td><td>TATCTGTTGTATACGATTGCCGG</td><td>1000</td><td></td><td></td><td></td><td></td><td></td><td></td><td></td><td></td><td></td></tr>
<tr><td>Oligo fwd</td><td>CACCTATCTGTTGTATACGATTGC</td><td></td><td></td><td></td><td></td><td></td><td></td><td></td><td></td><td></td><td></td></tr>
<tr><td>Oligo rev</td><td>AAACGCAATCGTATACAACAGATA</td><td></td><td></td><td></td><td></td><td></td><td></td><td></td><td></td><td></td><td></td></tr>
<tr><td>Chromosome</td><td>start</td><td>end</td><td>strand</td><td>MM</td><td>target_seq</td><td>PAM</td><td>alignment</td><td>distance</td><td>position</td><td>gene name</td><td>gene id</td></tr>
<tr><td>GU305914.1</td><td>57377</td><td>57399</td><td>+</td><td>0</td><td>TATCTGTTGTATACGATTGC</td><td>CGG</td><td>||||||||[||||||||||||]PAM</td><td>NA</td><td>Intergenic</td><td>NA</td><td>NA</td></tr>
<tr><td></td><td></td><td></td><td></td><td></td><td></td><td></td><td></td><td></td><td></td><td></td><td></td></tr>
<tr><td>T3769</td><td>ACGATTGCCGGCGACGACCGCGG</td><td>1000</td><td></td><td></td><td></td><td></td><td></td><td></td><td></td><td></td><td></td></tr>
<tr><td>Oligo fwd</td><td>CACCACGATTGCCGGCGACGACCG</td><td></td><td></td><td></td><td></td><td></td><td></td><td></td><td></td><td></td><td></td></tr>
<tr><td>Oligo rev</td><td>AAACCGGTCGTCGCCGGCAATCGT</td><td></td><td></td><td></td><td></td><td></td><td></td><td></td><td></td><td></td><td></td></tr>
<tr><td>Chromosome</td><td>start</td><td>end</td><td>strand</td><td>MM</td><td>target_seq</td><td>PAM</td><td>alignment</td><td>distance</td><td>position</td><td>gene name</td><td>gene id</td></tr>
<tr><td>GU305914.1</td><td>57389</td><td>57411</td><td>+</td><td>0</td><td>ACGATTGCCGGCGACGACCG</td><td>CGG</td><td>||||||||[||||||||||||]PAM</td><td>NA</td><td>Intergenic</td><td>NA</td><td>NA</td></tr>
<tr><td></td><td></td><td></td><td></td><td></td><td></td><td></td><td></td><td></td><td></td><td></td><td></td></tr>
<tr><td>T3770</td><td>GCCGGCGACGACCGCGGCTCTGG</td><td>1000</td><td></td><td></td><td></td><td></td><td></td><td></td><td></td><td></td><td></td></tr>
<tr><td>Oligo fwd</td><td>CACCGCCGGCGACGACCGCGGCTC</td><td></td><td></td><td></td><td></td><td></td><td></td><td></td><td></td><td></td><td></td></tr>
<tr><td>Oligo rev</td><td>AAACGAGCCGCGGTCGTCGCCGGC</td><td></td><td></td><td></td><td></td><td></td><td></td><td></td><td></td><td></td><td></td></tr>
<tr><td>Chromosome</td><td>start</td><td>end</td><td>strand</td><td>MM</td><td>target_seq</td><td>PAM</td><td>alignment</td><td>distance</td><td>position</td><td>gene name</td><td>gene id</td></tr>
<tr><td>GU305914.1</td><td>57395</td><td>57417</td><td>+</td><td>0</td><td>GCCGGCGACGACCGCGGCTC</td><td>TGG</td><td>||||||||[||||||||||||]PAM</td><td>NA</td><td>Intergenic</td><td>NA</td><td>NA</td></tr>
<tr><td></td><td></td><td></td><td></td><td></td><td></td><td></td><td></td><td></td><td></td><td></td><td></td></tr>
<tr><td>T3771</td><td>GCTCTGGAAAAAGATACGATCGG</td><td>1000</td><td></td><td></td><td></td><td></td><td></td><td></td><td></td><td></td><td></td></tr>
<tr><td>Oligo fwd</td><td>CACCGCTCTGGAAAAAGATACGAT</td><td></td><td></td><td></td><td></td><td></td><td></td><td></td><td></td><td></td><td></td></tr>
<tr><td>Oligo rev</td><td>AAACATCGTATCTTTTTCCAGAGC</td><td></td><td></td><td></td><td></td><td></td><td></td><td></td><td></td><td></td><td></td></tr>
<tr><td>Chromosome</td><td>start</td><td>end</td><td>strand</td><td>MM</td><td>target_seq</td><td>PAM</td><td>alignment</td><td>distance</td><td>position</td><td>gene name</td><td>gene id</td></tr>
<tr><td>GU305914.1</td><td>57411</td><td>57433</td><td>+</td><td>0</td><td>GCTCTGGAAAAAGATACGAT</td><td>CGG</td><td>||||||||[||||||||||||]PAM</td><td>NA</td><td>Intergenic</td><td>NA</td><td>NA</td></tr>
<tr><td></td><td></td><td></td><td></td><td></td><td></td><td></td><td></td><td></td><td></td><td></td><td></td></tr>
<tr><td>T3772</td><td>GGAGCGACAGTTTCCTTCTCCGG</td><td>1000</td><td></td><td></td><td></td><td></td><td></td><td></td><td></td><td></td><td></td></tr>
<tr><td>Oligo fwd</td><td>CACCGGAGCGACAGTTTCCTTCTC</td><td></td><td></td><td></td><td></td><td></td><td></td><td></td><td></td><td></td><td></td></tr>
<tr><td>Oligo rev</td><td>AAACGAGAAGGAAACTGTCGCTCC</td><td></td><td></td><td></td><td></td><td></td><td></td><td></td><td></td><td></td><td></td></tr>
<tr><td>Chromosome</td><td>start</td><td>end</td><td>strand</td><td>MM</td><td>target_seq</td><td>PAM</td><td>alignment</td><td>distance</td><td>position</td><td>gene name</td><td>gene id</td></tr>
<tr><td>GU305914.1</td><td>57432</td><td>57454</td><td>+</td><td>0</td><td>GGAGCGACAGTTTCCTTCTC</td><td>CGG</td><td>||||||||[||||||||||||]PAM</td><td>NA</td><td>Intergenic</td><td>NA</td><td>NA</td></tr>
<tr><td></td><td></td><td></td><td></td><td></td><td></td><td></td><td></td><td></td><td></td><td></td><td></td></tr>
<tr><td>T3773</td><td>GTTTCCTTCTCCGGAGCTCGCGG</td><td>1000</td><td></td><td></td><td></td><td></td><td></td><td></td><td></td><td></td><td></td></tr>
<tr><td>Oligo fwd</td><td>CACCGTTTCCTTCTCCGGAGCTCG</td><td></td><td></td><td></td><td></td><td></td><td></td><td></td><td></td><td></td><td></td></tr>
<tr><td>Oligo rev</td><td>AAACCGAGCTCCGGAGAAGGAAAC</td><td></td><td></td><td></td><td></td><td></td><td></td><td></td><td></td><td></td><td></td></tr>
<tr><td>Chromosome</td><td>start</td><td>end</td><td>strand</td><td>MM</td><td>target_seq</td><td>PAM</td><td>alignment</td><td>distance</td><td>position</td><td>gene name</td><td>gene id</td></tr>
<tr><td>GU305914.1</td><td>57441</td><td>57463</td><td>+</td><td>0</td><td>GTTTCCTTCTCCGGAGCTCG</td><td>CGG</td><td>||||||||[||||||||||||]PAM</td><td>NA</td><td>Intergenic</td><td>NA</td><td>NA</td></tr>
<tr><td></td><td></td><td></td><td></td><td></td><td></td><td></td><td></td><td></td><td></td><td></td><td></td></tr>
<tr><td>T3774</td><td>TCCGGAGCTCGCGGTCGTCACGG</td><td>1000</td><td></td><td></td><td></td><td></td><td></td><td></td><td></td><td></td><td></td></tr>
<tr><td>Oligo fwd</td><td>CACCTCCGGAGCTCGCGGTCGTCA</td><td></td><td></td><td></td><td></td><td></td><td></td><td></td><td></td><td></td><td></td></tr>
<tr><td>Oligo rev</td><td>AAACTGACGACCGCGAGCTCCGGA</td><td></td><td></td><td></td><td></td><td></td><td></td><td></td><td></td><td></td><td></td></tr>
</table>

<table>
<tr><td>Chromosome</td><td>start</td><td>end</td><td>strand</td><td>MM</td><td>target_seq</td><td>PAM</td><td>alignment</td><td>distance</td><td>position</td><td>gene name</td><td>gene id</td></tr>
<tr><td>GU305914.1</td><td>57450</td><td>57472</td><td>+</td><td>0</td><td>TCCGGAGCTCGCGGTCGTCA</td><td>CGG</td><td>||||||||[||||||||||||]PAM</td><td>NA</td><td>Intergenic</td><td>NA</td><td>NA</td></tr>
<tr><td></td><td></td><td></td><td></td><td></td><td></td><td></td><td></td><td></td><td></td><td></td><td></td></tr>
<tr><td>T3775</td><td>GGAGCTCGCGGTCGTCACGGCGG</td><td>1000</td><td></td><td></td><td></td><td></td><td></td><td></td><td></td><td></td><td></td></tr>
<tr><td>Oligo fwd</td><td>CACCGGAGCTCGCGGTCGTCACGG</td><td></td><td></td><td></td><td></td><td></td><td></td><td></td><td></td><td></td><td></td></tr>
<tr><td>Oligo rev</td><td>AAACCCGTGACGACCGCGAGCTCC</td><td></td><td></td><td></td><td></td><td></td><td></td><td></td><td></td><td></td><td></td></tr>
<tr><td>Chromosome</td><td>start</td><td>end</td><td>strand</td><td>MM</td><td>target_seq</td><td>PAM</td><td>alignment</td><td>distance</td><td>position</td><td>gene name</td><td>gene id</td></tr>
<tr><td>GU305914.1</td><td>57453</td><td>57475</td><td>+</td><td>0</td><td>GGAGCTCGCGGTCGTCACGG</td><td>CGG</td><td>||||||||[||||||||||||]PAM</td><td>NA</td><td>Intergenic</td><td>NA</td><td>NA</td></tr>
<tr><td></td><td></td><td></td><td></td><td></td><td></td><td></td><td></td><td></td><td></td><td></td><td></td></tr>
<tr><td>T3776</td><td>TAACACAGACGTACGTCGATCGG</td><td>1000</td><td></td><td></td><td></td><td></td><td></td><td></td><td></td><td></td><td></td></tr>
<tr><td>Oligo fwd</td><td>CACCTAACACAGACGTACGTCGAT</td><td></td><td></td><td></td><td></td><td></td><td></td><td></td><td></td><td></td><td></td></tr>
<tr><td>Oligo rev</td><td>AAACATCGACGTACGTCTGTGTTA</td><td></td><td></td><td></td><td></td><td></td><td></td><td></td><td></td><td></td><td></td></tr>
<tr><td>Chromosome</td><td>start</td><td>end</td><td>strand</td><td>MM</td><td>target_seq</td><td>PAM</td><td>alignment</td><td>distance</td><td>position</td><td>gene name</td><td>gene id</td></tr>
<tr><td>GU305914.1</td><td>57480</td><td>57502</td><td>+</td><td>0</td><td>TAACACAGACGTACGTCGAT</td><td>CGG</td><td>||||||||[||||||||||||]PAM</td><td>NA</td><td>Intergenic</td><td>NA</td><td>NA</td></tr>
<tr><td></td><td></td><td></td><td></td><td></td><td></td><td></td><td></td><td></td><td></td><td></td><td></td></tr>
<tr><td>T3777</td><td>TCGCTTCTTCGACGCCGTCGTGG</td><td>1000</td><td></td><td></td><td></td><td></td><td></td><td></td><td></td><td></td><td></td></tr>
<tr><td>Oligo fwd</td><td>CACCTCGCTTCTTCGACGCCGTCG</td><td></td><td></td><td></td><td></td><td></td><td></td><td></td><td></td><td></td><td></td></tr>
<tr><td>Oligo rev</td><td>AAACCGACGGCGTCGAAGAAGCGA</td><td></td><td></td><td></td><td></td><td></td><td></td><td></td><td></td><td></td><td></td></tr>
<tr><td>Chromosome</td><td>start</td><td>end</td><td>strand</td><td>MM</td><td>target_seq</td><td>PAM</td><td>alignment</td><td>distance</td><td>position</td><td>gene name</td><td>gene id</td></tr>
<tr><td>GU305914.1</td><td>57508</td><td>57530</td><td>+</td><td>0</td><td>TCGCTTCTTCGACGCCGTCG</td><td>TGG</td><td>||||||||[||||||||||||]PAM</td><td>NA</td><td>Intergenic</td><td>NA</td><td>NA</td></tr>
<tr><td></td><td></td><td></td><td></td><td></td><td></td><td></td><td></td><td></td><td></td><td></td><td></td></tr>
<tr><td>T3778</td><td>CTTCGACGCCGTCGTGGCGATGG</td><td>1000</td><td></td><td></td><td></td><td></td><td></td><td></td><td></td><td></td><td></td></tr>
<tr><td>Oligo fwd</td><td>CACCCTTCGACGCCGTCGTGGCGA</td><td></td><td></td><td></td><td></td><td></td><td></td><td></td><td></td><td></td><td></td></tr>
<tr><td>Oligo rev</td><td>AAACTCGCCACGACGGCGTCGAAG</td><td></td><td></td><td></td><td></td><td></td><td></td><td></td><td></td><td></td><td></td></tr>
<tr><td>Chromosome</td><td>start</td><td>end</td><td>strand</td><td>MM</td><td>target_seq</td><td>PAM</td><td>alignment</td><td>distance</td><td>position</td><td>gene name</td><td>gene id</td></tr>
<tr><td>GU305914.1</td><td>57514</td><td>57536</td><td>+</td><td>0</td><td>CTTCGACGCCGTCGTGGCGA</td><td>TGG</td><td>||||||||[||||||||||||]PAM</td><td>NA</td><td>Intergenic</td><td>NA</td><td>NA</td></tr>
<tr><td></td><td></td><td></td><td></td><td></td><td></td><td></td><td></td><td></td><td></td><td></td><td></td></tr>
<tr><td>T3779</td><td>ACGCCGTCGTGGCGATGGCGCGG</td><td>1000</td><td></td><td></td><td></td><td></td><td></td><td></td><td></td><td></td><td></td></tr>
<tr><td>Oligo fwd</td><td>CACCACGCCGTCGTGGCGATGGCG</td><td></td><td></td><td></td><td></td><td></td><td></td><td></td><td></td><td></td><td></td></tr>
<tr><td>Oligo rev</td><td>AAACCGCCATCGCCACGACGGCGT</td><td></td><td></td><td></td><td></td><td></td><td></td><td></td><td></td><td></td><td></td></tr>
<tr><td>Chromosome</td><td>start</td><td>end</td><td>strand</td><td>MM</td><td>target_seq</td><td>PAM</td><td>alignment</td><td>distance</td><td>position</td><td>gene name</td><td>gene id</td></tr>
<tr><td>GU305914.1</td><td>57519</td><td>57541</td><td>+</td><td>0</td><td>ACGCCGTCGTGGCGATGGCG</td><td>CGG</td><td>||||||||[||||||||||||]PAM</td><td>NA</td><td>Intergenic</td><td>NA</td><td>NA</td></tr>
<tr><td></td><td></td><td></td><td></td><td></td><td></td><td></td><td></td><td></td><td></td><td></td><td></td></tr>
<tr><td>T3780</td><td>CCGTCGTGGCGATGGCGCGGCGG</td><td>1000</td><td></td><td></td><td></td><td></td><td></td><td></td><td></td><td></td><td></td></tr>
<tr><td>Oligo fwd</td><td>CACCCCGTCGTGGCGATGGCGCGG</td><td></td><td></td><td></td><td></td><td></td><td></td><td></td><td></td><td></td><td></td></tr>
<tr><td>Oligo rev</td><td>AAACCCGCGCCATCGCCACGACGG</td><td></td><td></td><td></td><td></td><td></td><td></td><td></td><td></td><td></td><td></td></tr>
<tr><td>Chromosome</td><td>start</td><td>end</td><td>strand</td><td>MM</td><td>target_seq</td><td>PAM</td><td>alignment</td><td>distance</td><td>position</td><td>gene name</td><td>gene id</td></tr>
<tr><td>GU305914.1</td><td>57522</td><td>57544</td><td>+</td><td>0</td><td>CCGTCGTGGCGATGGCGCGG</td><td>CGG</td><td>||||||||[||||||||||||]PAM</td><td>NA</td><td>Intergenic</td><td>NA</td><td>NA</td></tr>
<tr><td></td><td></td><td></td><td></td><td></td><td></td><td></td><td></td><td></td><td></td><td></td><td></td></tr>
<tr><td>T3781</td><td>TCGTGGCGATGGCGCGGCGGTGG</td><td>1000</td><td></td><td></td><td></td><td></td><td></td><td></td><td></td><td></td><td></td></tr>
<tr><td>Oligo fwd</td><td>CACCTCGTGGCGATGGCGCGGCGG</td><td></td><td></td><td></td><td></td><td></td><td></td><td></td><td></td><td></td><td></td></tr>
<tr><td>Oligo rev</td><td>AAACCCGCCGCGCCATCGCCACGA</td><td></td><td></td><td></td><td></td><td></td><td></td><td></td><td></td><td></td><td></td></tr>
<tr><td>Chromosome</td><td>start</td><td>end</td><td>strand</td><td>MM</td><td>target_seq</td><td>PAM</td><td>alignment</td><td>distance</td><td>position</td><td>gene name</td><td>gene id</td></tr>
<tr><td>GU305914.1</td><td>57525</td><td>57547</td><td>+</td><td>0</td><td>TCGTGGCGATGGCGCGGCGG</td><td>TGG</td><td>||||||||[||||||||||||]PAM</td><td>NA</td><td>Intergenic</td><td>NA</td><td>NA</td></tr>
<tr><td></td><td></td><td></td><td></td><td></td><td></td><td></td><td></td><td></td><td></td><td></td><td></td></tr>
<tr><td>T3782</td><td>TGGCGATGGCGCGGCGGTGGTGG</td><td>1000</td><td></td><td></td><td></td><td></td><td></td><td></td><td></td><td></td><td></td></tr>
<tr><td>Oligo fwd</td><td>CACCTGGCGATGGCGCGGCGGTGG</td><td></td><td></td><td></td><td></td><td></td><td></td><td></td><td></td><td></td><td></td></tr>
<tr><td>Oligo rev</td><td>AAACCCACCGCCGCGCCATCGCCA</td><td></td><td></td><td></td><td></td><td></td><td></td><td></td><td></td><td></td><td></td></tr>
<tr><td>Chromosome</td><td>start</td><td>end</td><td>strand</td><td>MM</td><td>target_seq</td><td>PAM</td><td>alignment</td><td>distance</td><td>position</td><td>gene name</td><td>gene id</td></tr>
<tr><td>GU305914.1</td><td>57528</td><td>57550</td><td>+</td><td>0</td><td>TGGCGATGGCGCGGCGGTGG</td><td>TGG</td><td>||||||||[||||||||||||]PAM</td><td>NA</td><td>Intergenic</td><td>NA</td><td>NA</td></tr>
<tr><td>GU305914.1</td><td>125354</td><td>125376</td><td>-</td><td>4</td><td>GGGCGACGGCGCGGTGGTGC</td><td>CGG</td><td>-|||||-|[||||||-||||-]PAM</td><td>NA</td><td>Intergenic</td><td>NA</td><td>NA</td></tr>
<tr><td></td><td></td><td></td><td></td><td></td><td></td><td></td><td></td><td></td><td></td><td></td><td></td></tr>
<tr><td>T3783</td><td>GGTGGTGAGCTTTTCTCAAACGG</td><td>1000</td><td></td><td></td><td></td><td></td><td></td><td></td><td></td><td></td><td></td></tr>
<tr><td>Oligo fwd</td><td>CACCGGTGGTGAGCTTTTCTCAAA</td><td></td><td></td><td></td><td></td><td></td><td></td><td></td><td></td><td></td><td></td></tr>
<tr><td>Oligo rev</td><td>AAACTTTGAGAAAAGCTCACCACC</td><td></td><td></td><td></td><td></td><td></td><td></td><td></td><td></td><td></td><td></td></tr>
<tr><td>Chromosome</td><td>start</td><td>end</td><td>strand</td><td>MM</td><td>target_seq</td><td>PAM</td><td>alignment</td><td>distance</td><td>position</td><td>gene name</td><td>gene id</td></tr>
<tr><td>GU305914.1</td><td>57546</td><td>57568</td><td>+</td><td>0</td><td>GGTGGTGAGCTTTTCTCAAA</td><td>CGG</td><td>||||||||[||||||||||||]PAM</td><td>NA</td><td>Intergenic</td><td>NA</td><td>NA</td></tr>
<tr><td></td><td></td><td></td><td></td><td></td><td></td><td></td><td></td><td></td><td></td><td></td><td></td></tr>
<tr><td>T3784</td><td>TTCTCAAACGGTGATGATACAGG</td><td>1000</td><td></td><td></td><td></td><td></td><td></td><td></td><td></td><td></td><td></td></tr>
<tr><td>Oligo fwd</td><td>CACCTTCTCAAACGGTGATGATAC</td><td></td><td></td><td></td><td></td><td></td><td></td><td></td><td></td><td></td><td></td></tr>
<tr><td>Oligo rev</td><td>AAACGTATCATCACCGTTTGAGAA</td><td></td><td></td><td></td><td></td><td></td><td></td><td></td><td></td><td></td><td></td></tr>
<tr><td>Chromosome</td><td>start</td><td>end</td><td>strand</td><td>MM</td><td>target_seq</td><td>PAM</td><td>alignment</td><td>distance</td><td>position</td><td>gene name</td><td>gene id</td></tr>
<tr><td>GU305914.1</td><td>57558</td><td>57580</td><td>+</td><td>0</td><td>TTCTCAAACGGTGATGATAC</td><td>AGG</td><td>||||||||[||||||||||||]PAM</td><td>NA</td><td>Intergenic</td><td>NA</td><td>NA</td></tr>
<tr><td></td><td></td><td></td><td></td><td></td><td></td><td></td><td></td><td></td><td></td><td></td><td></td></tr>
<tr><td>T3785</td><td>CAAACGGTGATGATACAGGTAGG</td><td>1000</td><td></td><td></td><td></td><td></td><td></td><td></td><td></td><td></td><td></td></tr>
<tr><td>Oligo fwd</td><td>CACCCAAACGGTGATGATACAGGT</td><td></td><td></td><td></td><td></td><td></td><td></td><td></td><td></td><td></td><td></td></tr>
<tr><td>Oligo rev</td><td>AAACACCTGTATCATCACCGTTTG</td><td></td><td></td><td></td><td></td><td></td><td></td><td></td><td></td><td></td><td></td></tr>
<tr><td>Chromosome</td><td>start</td><td>end</td><td>strand</td><td>MM</td><td>target_seq</td><td>PAM</td><td>alignment</td><td>distance</td><td>position</td><td>gene name</td><td>gene id</td></tr>
<tr><td>GU305914.1</td><td>57562</td><td>57584</td><td>+</td><td>0</td><td>CAAACGGTGATGATACAGGT</td><td>AGG</td><td>||||||||[||||||||||||]PAM</td><td>NA</td><td>Intergenic</td><td>NA</td><td>NA</td></tr>
<tr><td></td><td></td><td></td><td></td><td></td><td></td><td></td><td></td><td></td><td></td><td></td><td></td></tr>
<tr><td>T3786</td><td>ATAAAATGATCTAACTATTATGG</td><td>1000</td><td></td><td></td><td></td><td></td><td></td><td></td><td></td><td></td><td></td></tr>
<tr><td>Oligo fwd</td><td>CACCATAAAATGATCTAACTATTA</td><td></td><td></td><td></td><td></td><td></td><td></td><td></td><td></td><td></td><td></td></tr>
<tr><td>Oligo rev</td><td>AAACTAATAGTTAGATCATTTTAT</td><td></td><td></td><td></td><td></td><td></td><td></td><td></td><td></td><td></td><td></td></tr>
<tr><td>Chromosome</td><td>start</td><td>end</td><td>strand</td><td>MM</td><td>target_seq</td><td>PAM</td><td>alignment</td><td>distance</td><td>position</td><td>gene name</td><td>gene id</td></tr>
<tr><td>GU305914.1</td><td>57596</td><td>57618</td><td>+</td><td>0</td><td>ATAAAATGATCTAACTATTA</td><td>TGG</td><td>||||||||[||||||||||||]PAM</td><td>NA</td><td>Intergenic</td><td>NA</td><td>NA</td></tr>
<tr><td></td><td></td><td></td><td></td><td></td><td></td><td></td><td></td><td></td><td></td><td></td><td></td></tr>
<tr><td>T3787</td><td>CTATTATGGATTCTAAATCACGG</td><td>1000</td><td></td><td></td><td></td><td></td><td></td><td></td><td></td><td></td><td></td></tr>
<tr><td>Oligo fwd</td><td>CACCCTATTATGGATTCTAAATCA</td><td></td><td></td><td></td><td></td><td></td><td></td><td></td><td></td><td></td><td></td></tr>
<tr><td>Oligo rev</td><td>AAACTGATTTAGAATCCATAATAG</td><td></td><td></td><td></td><td></td><td></td><td></td><td></td><td></td><td></td><td></td></tr>
<tr><td>Chromosome</td><td>start</td><td>end</td><td>strand</td><td>MM</td><td>target_seq</td><td>PAM</td><td>alignment</td><td>distance</td><td>position</td><td>gene name</td><td>gene id</td></tr>
<tr><td>GU305914.1</td><td>57610</td><td>57632</td><td>+</td><td>0</td><td>CTATTATGGATTCTAAATCA</td><td>CGG</td><td>||||||||[||||||||||||]PAM</td><td>NA</td><td>Intergenic</td><td>NA</td><td>NA</td></tr>
<tr><td></td><td></td><td></td><td></td><td></td><td></td><td></td><td></td><td></td><td></td><td></td><td></td></tr>
<tr><td>T3788</td><td>ATGGATTCTAAATCACGGAGTGG</td><td>1000</td><td></td><td></td><td></td><td></td><td></td><td></td><td></td><td></td><td></td></tr>
<tr><td>Oligo fwd</td><td>CACCATGGATTCTAAATCACGGAG</td><td></td><td></td><td></td><td></td><td></td><td></td><td></td><td></td><td></td><td></td></tr>
<tr><td>Oligo rev</td><td>AAACCTCCGTGATTTAGAATCCAT</td><td></td><td></td><td></td><td></td><td></td><td></td><td></td><td></td><td></td><td></td></tr>
<tr><td>Chromosome</td><td>start</td><td>end</td><td>strand</td><td>MM</td><td>target_seq</td><td>PAM</td><td>alignment</td><td>distance</td><td>position</td><td>gene name</td><td>gene id</td></tr>
<tr><td>GU305914.1</td><td>57615</td><td>57637</td><td>+</td><td>0</td><td>ATGGATTCTAAATCACGGAG</td><td>TGG</td><td>||||||||[||||||||||||]PAM</td><td>NA</td><td>Intergenic</td><td>NA</td><td>NA</td></tr>
<tr><td></td><td></td><td></td><td></td><td></td><td></td><td></td><td></td><td></td><td></td><td></td><td></td></tr>
<tr><td>T3789</td><td>GGAGTGGTTGTGATTCTTTTCGG</td><td>1000</td><td></td><td></td><td></td><td></td><td></td><td></td><td></td><td></td><td></td></tr>
</table>

<table>
<tr><td>Oligo fwd</td><td>CACCGGAGTGGTTGTGATTCTTTT</td><td></td><td></td><td></td><td></td><td></td><td></td><td></td><td></td><td></td><td></td></tr>
<tr><td>Oligo rev</td><td>AAACAAAAGAATCACAACCACTCC</td><td></td><td></td><td></td><td></td><td></td><td></td><td></td><td></td><td></td><td></td></tr>
<tr><td>Chromosome</td><td>start</td><td>end</td><td>strand</td><td>MM</td><td>target_seq</td><td>PAM</td><td>alignment</td><td>distance</td><td>position</td><td>gene name</td><td>gene id</td></tr>
<tr><td>GU305914.1</td><td>57631</td><td>57653</td><td>+</td><td>0</td><td>GGAGTGGTTGTGATTCTTTT</td><td>CGG</td><td>||||||||[||||||||||||]PAM</td><td>NA</td><td>Intergenic</td><td>NA</td><td>NA</td></tr>
<tr><td></td><td></td><td></td><td></td><td></td><td></td><td></td><td></td><td></td><td></td><td></td><td></td></tr>
<tr><td>T3790</td><td>GAGTGGTTGTGATTCTTTTCGGG</td><td>1000</td><td></td><td></td><td></td><td></td><td></td><td></td><td></td><td></td><td></td></tr>
<tr><td>Oligo fwd</td><td>CACCGAGTGGTTGTGATTCTTTTC</td><td></td><td></td><td></td><td></td><td></td><td></td><td></td><td></td><td></td><td></td></tr>
<tr><td>Oligo rev</td><td>AAACGAAAAGAATCACAACCACTC</td><td></td><td></td><td></td><td></td><td></td><td></td><td></td><td></td><td></td><td></td></tr>
<tr><td>Chromosome</td><td>start</td><td>end</td><td>strand</td><td>MM</td><td>target_seq</td><td>PAM</td><td>alignment</td><td>distance</td><td>position</td><td>gene name</td><td>gene id</td></tr>
<tr><td>GU305914.1</td><td>57632</td><td>57654</td><td>+</td><td>0</td><td>GAGTGGTTGTGATTCTTTTC</td><td>GGG</td><td>||||||||[||||||||||||]PAM</td><td>NA</td><td>Intergenic</td><td>NA</td><td>NA</td></tr>
<tr><td></td><td></td><td></td><td></td><td></td><td></td><td></td><td></td><td></td><td></td><td></td><td></td></tr>
<tr><td>T3791</td><td>AGTGGTTGTGATTCTTTTCGGGG</td><td>1000</td><td></td><td></td><td></td><td></td><td></td><td></td><td></td><td></td><td></td></tr>
<tr><td>Oligo fwd</td><td>CACCAGTGGTTGTGATTCTTTTCG</td><td></td><td></td><td></td><td></td><td></td><td></td><td></td><td></td><td></td><td></td></tr>
<tr><td>Oligo rev</td><td>AAACCGAAAAGAATCACAACCACT</td><td></td><td></td><td></td><td></td><td></td><td></td><td></td><td></td><td></td><td></td></tr>
<tr><td>Chromosome</td><td>start</td><td>end</td><td>strand</td><td>MM</td><td>target_seq</td><td>PAM</td><td>alignment</td><td>distance</td><td>position</td><td>gene name</td><td>gene id</td></tr>
<tr><td>GU305914.1</td><td>57633</td><td>57655</td><td>+</td><td>0</td><td>AGTGGTTGTGATTCTTTTCG</td><td>GGG</td><td>||||||||[||||||||||||]PAM</td><td>NA</td><td>Intergenic</td><td>NA</td><td>NA</td></tr>
<tr><td></td><td></td><td></td><td></td><td></td><td></td><td></td><td></td><td></td><td></td><td></td><td></td></tr>
<tr><td>T3792</td><td>TGTGATTCTTTTCGGGGATCCGG</td><td>1000</td><td></td><td></td><td></td><td></td><td></td><td></td><td></td><td></td><td></td></tr>
<tr><td>Oligo fwd</td><td>CACCTGTGATTCTTTTCGGGGATC</td><td></td><td></td><td></td><td></td><td></td><td></td><td></td><td></td><td></td><td></td></tr>
<tr><td>Oligo rev</td><td>AAACGATCCCCGAAAAGAATCACA</td><td></td><td></td><td></td><td></td><td></td><td></td><td></td><td></td><td></td><td></td></tr>
<tr><td>Chromosome</td><td>start</td><td>end</td><td>strand</td><td>MM</td><td>target_seq</td><td>PAM</td><td>alignment</td><td>distance</td><td>position</td><td>gene name</td><td>gene id</td></tr>
<tr><td>GU305914.1</td><td>57639</td><td>57661</td><td>+</td><td>0</td><td>TGTGATTCTTTTCGGGGATC</td><td>CGG</td><td>||||||||[||||||||||||]PAM</td><td>NA</td><td>Intergenic</td><td>NA</td><td>NA</td></tr>
<tr><td></td><td></td><td></td><td></td><td></td><td></td><td></td><td></td><td></td><td></td><td></td><td></td></tr>
<tr><td>T3793</td><td>TTTCGGGGATCCGGCACCTGAGG</td><td>1000</td><td></td><td></td><td></td><td></td><td></td><td></td><td></td><td></td><td></td></tr>
<tr><td>Oligo fwd</td><td>CACCTTTCGGGGATCCGGCACCTG</td><td></td><td></td><td></td><td></td><td></td><td></td><td></td><td></td><td></td><td></td></tr>
<tr><td>Oligo rev</td><td>AAACCAGGTGCCGGATCCCCGAAA</td><td></td><td></td><td></td><td></td><td></td><td></td><td></td><td></td><td></td><td></td></tr>
<tr><td>Chromosome</td><td>start</td><td>end</td><td>strand</td><td>MM</td><td>target_seq</td><td>PAM</td><td>alignment</td><td>distance</td><td>position</td><td>gene name</td><td>gene id</td></tr>
<tr><td>GU305914.1</td><td>57648</td><td>57670</td><td>+</td><td>0</td><td>TTTCGGGGATCCGGCACCTG</td><td>AGG</td><td>||||||||[||||||||||||]PAM</td><td>NA</td><td>Intergenic</td><td>NA</td><td>NA</td></tr>
<tr><td></td><td></td><td></td><td></td><td></td><td></td><td></td><td></td><td></td><td></td><td></td><td></td></tr>
<tr><td>T3794</td><td>TTCGGGGATCCGGCACCTGAGGG</td><td>1000</td><td></td><td></td><td></td><td></td><td></td><td></td><td></td><td></td><td></td></tr>
<tr><td>Oligo fwd</td><td>CACCTTCGGGGATCCGGCACCTGA</td><td></td><td></td><td></td><td></td><td></td><td></td><td></td><td></td><td></td><td></td></tr>
<tr><td>Oligo rev</td><td>AAACTCAGGTGCCGGATCCCCGAA</td><td></td><td></td><td></td><td></td><td></td><td></td><td></td><td></td><td></td><td></td></tr>
<tr><td>Chromosome</td><td>start</td><td>end</td><td>strand</td><td>MM</td><td>target_seq</td><td>PAM</td><td>alignment</td><td>distance</td><td>position</td><td>gene name</td><td>gene id</td></tr>
<tr><td>GU305914.1</td><td>57649</td><td>57671</td><td>+</td><td>0</td><td>TTCGGGGATCCGGCACCTGA</td><td>GGG</td><td>||||||||[||||||||||||]PAM</td><td>NA</td><td>Intergenic</td><td>NA</td><td>NA</td></tr>
<tr><td></td><td></td><td></td><td></td><td></td><td></td><td></td><td></td><td></td><td></td><td></td><td></td></tr>
<tr><td>T3795</td><td>TCGGGGATCCGGCACCTGAGGGG</td><td>1000</td><td></td><td></td><td></td><td></td><td></td><td></td><td></td><td></td><td></td></tr>
<tr><td>Oligo fwd</td><td>CACCTCGGGGATCCGGCACCTGAG</td><td></td><td></td><td></td><td></td><td></td><td></td><td></td><td></td><td></td><td></td></tr>
<tr><td>Oligo rev</td><td>AAACCTCAGGTGCCGGATCCCCGA</td><td></td><td></td><td></td><td></td><td></td><td></td><td></td><td></td><td></td><td></td></tr>
<tr><td>Chromosome</td><td>start</td><td>end</td><td>strand</td><td>MM</td><td>target_seq</td><td>PAM</td><td>alignment</td><td>distance</td><td>position</td><td>gene name</td><td>gene id</td></tr>
<tr><td>GU305914.1</td><td>57650</td><td>57672</td><td>+</td><td>0</td><td>TCGGGGATCCGGCACCTGAG</td><td>GGG</td><td>||||||||[||||||||||||]PAM</td><td>NA</td><td>Intergenic</td><td>NA</td><td>NA</td></tr>
<tr><td></td><td></td><td></td><td></td><td></td><td></td><td></td><td></td><td></td><td></td><td></td><td></td></tr>
<tr><td>T3796</td><td>CGGGGATCCGGCACCTGAGGGGG</td><td>1000</td><td></td><td></td><td></td><td></td><td></td><td></td><td></td><td></td><td></td></tr>
<tr><td>Oligo fwd</td><td>CACCCGGGGATCCGGCACCTGAGG</td><td></td><td></td><td></td><td></td><td></td><td></td><td></td><td></td><td></td><td></td></tr>
<tr><td>Oligo rev</td><td>AAACCCTCAGGTGCCGGATCCCCG</td><td></td><td></td><td></td><td></td><td></td><td></td><td></td><td></td><td></td><td></td></tr>
<tr><td>Chromosome</td><td>start</td><td>end</td><td>strand</td><td>MM</td><td>target_seq</td><td>PAM</td><td>alignment</td><td>distance</td><td>position</td><td>gene name</td><td>gene id</td></tr>
<tr><td>GU305914.1</td><td>57651</td><td>57673</td><td>+</td><td>0</td><td>CGGGGATCCGGCACCTGAGG</td><td>GGG</td><td>||||||||[||||||||||||]PAM</td><td>NA</td><td>Intergenic</td><td>NA</td><td>NA</td></tr>
<tr><td>GU305914.1</td><td>62476</td><td>62498</td><td>+</td><td>3</td><td>CGCGGATCCAGCACCAGAGG</td><td>TGG</td><td>||-|||||[|-|||||-||||]PAM</td><td>NA</td><td>Intergenic</td><td>NA</td><td>NA</td></tr>
<tr><td></td><td></td><td></td><td></td><td></td><td></td><td></td><td></td><td></td><td></td><td></td><td></td></tr>
<tr><td>T3797</td><td>GGGGATCCGGCACCTGAGGGGGG</td><td>1000</td><td></td><td></td><td></td><td></td><td></td><td></td><td></td><td></td><td></td></tr>
<tr><td>Oligo fwd</td><td>CACCGGGGATCCGGCACCTGAGGG</td><td></td><td></td><td></td><td></td><td></td><td></td><td></td><td></td><td></td><td></td></tr>
<tr><td>Oligo rev</td><td>AAACCCCTCAGGTGCCGGATCCCC</td><td></td><td></td><td></td><td></td><td></td><td></td><td></td><td></td><td></td><td></td></tr>
<tr><td>Chromosome</td><td>start</td><td>end</td><td>strand</td><td>MM</td><td>target_seq</td><td>PAM</td><td>alignment</td><td>distance</td><td>position</td><td>gene name</td><td>gene id</td></tr>
<tr><td>GU305914.1</td><td>57652</td><td>57674</td><td>+</td><td>0</td><td>GGGGATCCGGCACCTGAGGG</td><td>GGG</td><td>||||||||[||||||||||||]PAM</td><td>NA</td><td>Intergenic</td><td>NA</td><td>NA</td></tr>
<tr><td></td><td></td><td></td><td></td><td></td><td></td><td></td><td></td><td></td><td></td><td></td><td></td></tr>
<tr><td>T3798</td><td>AGGGGGGTTCAGATCCTCCTCGG</td><td>1000</td><td></td><td></td><td></td><td></td><td></td><td></td><td></td><td></td><td></td></tr>
<tr><td>Oligo fwd</td><td>CACCAGGGGGGTTCAGATCCTCCT</td><td></td><td></td><td></td><td></td><td></td><td></td><td></td><td></td><td></td><td></td></tr>
<tr><td>Oligo rev</td><td>AAACAGGAGGATCTGAACCCCCCT</td><td></td><td></td><td></td><td></td><td></td><td></td><td></td><td></td><td></td><td></td></tr>
<tr><td>Chromosome</td><td>start</td><td>end</td><td>strand</td><td>MM</td><td>target_seq</td><td>PAM</td><td>alignment</td><td>distance</td><td>position</td><td>gene name</td><td>gene id</td></tr>
<tr><td>GU305914.1</td><td>57668</td><td>57690</td><td>+</td><td>0</td><td>AGGGGGGTTCAGATCCTCCT</td><td>CGG</td><td>||||||||[||||||||||||]PAM</td><td>NA</td><td>Intergenic</td><td>NA</td><td>NA</td></tr>
<tr><td></td><td></td><td></td><td></td><td></td><td></td><td></td><td></td><td></td><td></td><td></td><td></td></tr>
<tr><td>T3799</td><td>TGAAAGAAAGACCGCACGCTCGG</td><td>1000</td><td></td><td></td><td></td><td></td><td></td><td></td><td></td><td></td><td></td></tr>
<tr><td>Oligo fwd</td><td>CACCTGAAAGAAAGACCGCACGCT</td><td></td><td></td><td></td><td></td><td></td><td></td><td></td><td></td><td></td><td></td></tr>
<tr><td>Oligo rev</td><td>AAACAGCGTGCGGTCTTTCTTTCA</td><td></td><td></td><td></td><td></td><td></td><td></td><td></td><td></td><td></td><td></td></tr>
<tr><td>Chromosome</td><td>start</td><td>end</td><td>strand</td><td>MM</td><td>target_seq</td><td>PAM</td><td>alignment</td><td>distance</td><td>position</td><td>gene name</td><td>gene id</td></tr>
<tr><td>GU305914.1</td><td>57693</td><td>57715</td><td>+</td><td>0</td><td>TGAAAGAAAGACCGCACGCT</td><td>CGG</td><td>||||||||[||||||||||||]PAM</td><td>NA</td><td>Intergenic</td><td>NA</td><td>NA</td></tr>
<tr><td></td><td></td><td></td><td></td><td></td><td></td><td></td><td></td><td></td><td></td><td></td><td></td></tr>
<tr><td>T3800</td><td>GAAAGAAAGACCGCACGCTCGGG</td><td>1000</td><td></td><td></td><td></td><td></td><td></td><td></td><td></td><td></td><td></td></tr>
<tr><td>Oligo fwd</td><td>CACCGAAAGAAAGACCGCACGCTC</td><td></td><td></td><td></td><td></td><td></td><td></td><td></td><td></td><td></td><td></td></tr>
<tr><td>Oligo rev</td><td>AAACGAGCGTGCGGTCTTTCTTTC</td><td></td><td></td><td></td><td></td><td></td><td></td><td></td><td></td><td></td><td></td></tr>
<tr><td>Chromosome</td><td>start</td><td>end</td><td>strand</td><td>MM</td><td>target_seq</td><td>PAM</td><td>alignment</td><td>distance</td><td>position</td><td>gene name</td><td>gene id</td></tr>
<tr><td>GU305914.1</td><td>57694</td><td>57716</td><td>+</td><td>0</td><td>GAAAGAAAGACCGCACGCTC</td><td>GGG</td><td>||||||||[||||||||||||]PAM</td><td>NA</td><td>Intergenic</td><td>NA</td><td>NA</td></tr>
<tr><td></td><td></td><td></td><td></td><td></td><td></td><td></td><td></td><td></td><td></td><td></td><td></td></tr>
<tr><td>T3801</td><td>AAAGAAAGACCGCACGCTCGGGG</td><td>1000</td><td></td><td></td><td></td><td></td><td></td><td></td><td></td><td></td><td></td></tr>
<tr><td>Oligo fwd</td><td>CACCAAAGAAAGACCGCACGCTCG</td><td></td><td></td><td></td><td></td><td></td><td></td><td></td><td></td><td></td><td></td></tr>
<tr><td>Oligo rev</td><td>AAACCGAGCGTGCGGTCTTTCTTT</td><td></td><td></td><td></td><td></td><td></td><td></td><td></td><td></td><td></td><td></td></tr>
<tr><td>Chromosome</td><td>start</td><td>end</td><td>strand</td><td>MM</td><td>target_seq</td><td>PAM</td><td>alignment</td><td>distance</td><td>position</td><td>gene name</td><td>gene id</td></tr>
<tr><td>GU305914.1</td><td>57695</td><td>57717</td><td>+</td><td>0</td><td>AAAGAAAGACCGCACGCTCG</td><td>GGG</td><td>||||||||[||||||||||||]PAM</td><td>NA</td><td>Intergenic</td><td>NA</td><td>NA</td></tr>
<tr><td></td><td></td><td></td><td></td><td></td><td></td><td></td><td></td><td></td><td></td><td></td><td></td></tr>
<tr><td>T3802</td><td>TCGGGGAATTTATCTCGATTAGG</td><td>1000</td><td></td><td></td><td></td><td></td><td></td><td></td><td></td><td></td><td></td></tr>
<tr><td>Oligo fwd</td><td>CACCTCGGGGAATTTATCTCGATT</td><td></td><td></td><td></td><td></td><td></td><td></td><td></td><td></td><td></td><td></td></tr>
<tr><td>Oligo rev</td><td>AAACAATCGAGATAAATTCCCCGA</td><td></td><td></td><td></td><td></td><td></td><td></td><td></td><td></td><td></td><td></td></tr>
<tr><td>Chromosome</td><td>start</td><td>end</td><td>strand</td><td>MM</td><td>target_seq</td><td>PAM</td><td>alignment</td><td>distance</td><td>position</td><td>gene name</td><td>gene id</td></tr>
<tr><td>GU305914.1</td><td>57712</td><td>57734</td><td>+</td><td>0</td><td>TCGGGGAATTTATCTCGATT</td><td>AGG</td><td>||||||||[||||||||||||]PAM</td><td>NA</td><td>Intergenic</td><td>NA</td><td>NA</td></tr>
<tr><td></td><td></td><td></td><td></td><td></td><td></td><td></td><td></td><td></td><td></td><td></td><td></td></tr>
<tr><td>T3803</td><td>TTCGTTCGTGCGATTCACCGCGG</td><td>1000</td><td></td><td></td><td></td><td></td><td></td><td></td><td></td><td></td><td></td></tr>
<tr><td>Oligo fwd</td><td>CACCTTCGTTCGTGCGATTCACCG</td><td></td><td></td><td></td><td></td><td></td><td></td><td></td><td></td><td></td><td></td></tr>
<tr><td>Oligo rev</td><td>AAACCGGTGAATCGCACGAACGAA</td><td></td><td></td><td></td><td></td><td></td><td></td><td></td><td></td><td></td><td></td></tr>
<tr><td>Chromosome</td><td>start</td><td>end</td><td>strand</td><td>MM</td><td>target_seq</td><td>PAM</td><td>alignment</td><td>distance</td><td>position</td><td>gene name</td><td>gene id</td></tr>
<tr><td>GU305914.1</td><td>57777</td><td>57799</td><td>+</td><td>0</td><td>TTCGTTCGTGCGATTCACCG</td><td>CGG</td><td>||||||||[||||||||||||]PAM</td><td>NA</td><td>Intergenic</td><td>NA</td><td>NA</td></tr>
</table>

| | | | | | | | | | | | |
|---|---|---|---|---|---|---|---|---|---|---|---|
| T3804 | TCGTTCGTGCGATTCACCGCGGG | 1000 | | | | | | | | | |
| Oligo fwd | CACCTCGTTCGTGCGATTCACCGC | | | | | | | | | | |
| Oligo rev | AAACGCGGTGAATCGCACGAACGA | | | | | | | | | | |
| Chromosome | start | end | strand | MM | target_seq | PAM | alignment | distance | position | gene name | gene id |
| GU305914.1 | 57778 | 57800 | + | 0 | TCGTTCGTGCGATTCACCGC | GGG | ||||||||[||||||||||||]PAM | NA | Intergenic | NA | NA |
| T3805 | TGCGATTCACCGCGGGCGAGAGG | 1000 | | | | | | | | | |
| Oligo fwd | CACCTGCGATTCACCGCGGGCGAG | | | | | | | | | | |
| Oligo rev | AAACCTCGCCCGCGGTGAATCGCA | | | | | | | | | | |
| Chromosome | start | end | strand | MM | target_seq | PAM | alignment | distance | position | gene name | gene id |
| GU305914.1 | 57785 | 57807 | + | 0 | TGCGATTCACCGCGGGCGAG | AGG | ||||||||[||||||||||||]PAM | NA | Intergenic | NA | NA |
| T3806 | GATTCACCGCGGGCGAGAGGAGG | 1000 | | | | | | | | | |
| Oligo fwd | CACCGATTCACCGCGGGCGAGAGG | | | | | | | | | | |
| Oligo rev | AAACCCTCTCGCCCGCGGTGAATC | | | | | | | | | | |
| Chromosome | start | end | strand | MM | target_seq | PAM | alignment | distance | position | gene name | gene id |
| GU305914.1 | 57788 | 57810 | + | 0 | GATTCACCGCGGGCGAGAGG | AGG | ||||||||[||||||||||||]PAM | NA | Intergenic | NA | NA |
| T3807 | ATTCACCGCGGGCGAGAGGAGGG | 1000 | | | | | | | | | |
| Oligo fwd | CACCATTCACCGCGGGCGAGAGGA | | | | | | | | | | |
| Oligo rev | AAACTCCTCTCGCCCGCGGTGAAT | | | | | | | | | | |
| Chromosome | start | end | strand | MM | target_seq | PAM | alignment | distance | position | gene name | gene id |
| GU305914.1 | 57789 | 57811 | + | 0 | ATTCACCGCGGGCGAGAGGA | GGG | ||||||||[||||||||||||]PAM | NA | Intergenic | NA | NA |
| GU305914.1 | 55229 | 55251 | + | 4 | CTTCTCCCCGGGAGAGAGGA | AGG | -|||-||-[||||-|||||||]PAM | NA | Intergenic | NA | NA |
| T3808 | TTCACCGCGGGCGAGAGGAGGGG | 1000 | | | | | | | | | |
| Oligo fwd | CACCTTCACCGCGGGCGAGAGGAG | | | | | | | | | | |
| Oligo rev | AAACCTCCTCTCGCCCGCGGTGAA | | | | | | | | | | |
| Chromosome | start | end | strand | MM | target_seq | PAM | alignment | distance | position | gene name | gene id |
| GU305914.1 | 57790 | 57812 | + | 0 | TTCACCGCGGGCGAGAGGAG | GGG | ||||||||[||||||||||||]PAM | NA | Intergenic | NA | NA |
| T3809 | ACCGCGGGCGAGAGGAGGGGAGG | 1000 | | | | | | | | | |
| Oligo fwd | CACCACCGCGGGCGAGAGGAGGGG | | | | | | | | | | |
| Oligo rev | AAACCCCCTCCTCTCGCCCGCGGT | | | | | | | | | | |
| Chromosome | start | end | strand | MM | target_seq | PAM | alignment | distance | position | gene name | gene id |
| GU305914.1 | 57793 | 57815 | + | 0 | ACCGCGGGCGAGAGGAGGGG | AGG | ||||||||[||||||||||||]PAM | NA | Intergenic | NA | NA |
| T3810 | GTCGCCGCCGCCGTCGCTCTCGG | 1000 | | | | | | | | | |
| Oligo fwd | CACCGTCGCCGCCGCCGTCGCTCT | | | | | | | | | | |
| Oligo rev | AAACAGAGCGACGGCGGCGGCGAC | | | | | | | | | | |
| Chromosome | start | end | strand | MM | target_seq | PAM | alignment | distance | position | gene name | gene id |
| GU305914.1 | 57828 | 57850 | + | 0 | GTCGCCGCCGCCGTCGCTCT | CGG | ||||||||[||||||||||||]PAM | NA | Intergenic | NA | NA |
| T3811 | GCCGCCGCCGTCGCTCTCGGTGG | 1000 | | | | | | | | | |
| Oligo fwd | CACCGCCGCCGCCGTCGCTCTCGG | | | | | | | | | | |
| Oligo rev | AAACCCGAGAGCGACGGCGGCGGC | | | | | | | | | | |
| Chromosome | start | end | strand | MM | target_seq | PAM | alignment | distance | position | gene name | gene id |
| GU305914.1 | 57831 | 57853 | + | 0 | GCCGCCGCCGTCGCTCTCGG | TGG | ||||||||[||||||||||||]PAM | NA | Intergenic | NA | NA |
| GU305914.1 | 24134 | 24156 | + | 4 | GGCGCCGACGTCGGCCTCGG | CGG | |-|||||-[|||||--|||||]PAM | NA | Intergenic | NA | NA |
| T3812 | CGTCGCTCTCGGTGGTCAGTCGG | 1000 | | | | | | | | | |
| Oligo fwd | CACCCGTCGCTCTCGGTGGTCAGT | | | | | | | | | | |
| Oligo rev | AAACACTGACCACCGAGAGCGACG | | | | | | | | | | |
| Chromosome | start | end | strand | MM | target_seq | PAM | alignment | distance | position | gene name | gene id |
| GU305914.1 | 57839 | 57861 | + | 0 | CGTCGCTCTCGGTGGTCAGT | CGG | ||||||||[||||||||||||]PAM | NA | Intergenic | NA | NA |
| T3813 | GTGGTCAGTCGGTCGATCCCCGG | 1000 | | | | | | | | | |
| Oligo fwd | CACCGTGGTCAGTCGGTCGATCCC | | | | | | | | | | |
| Oligo rev | AAACGGGATCGACCGACTGACCAC | | | | | | | | | | |
| Chromosome | start | end | strand | MM | target_seq | PAM | alignment | distance | position | gene name | gene id |
| GU305914.1 | 57850 | 57872 | + | 0 | GTGGTCAGTCGGTCGATCCC | CGG | ||||||||[||||||||||||]PAM | NA | Intergenic | NA | NA |
| T3814 | TGGTCAGTCGGTCGATCCCCGGG | 1000 | | | | | | | | | |
| Oligo fwd | CACCTGGTCAGTCGGTCGATCCCC | | | | | | | | | | |
| Oligo rev | AAACGGGGATCGACCGACTGACCA | | | | | | | | | | |
| Chromosome | start | end | strand | MM | target_seq | PAM | alignment | distance | position | gene name | gene id |
| GU305914.1 | 57851 | 57873 | + | 0 | TGGTCAGTCGGTCGATCCCC | GGG | ||||||||[||||||||||||]PAM | NA | Intergenic | NA | NA |
| T3815 | TCGATCCCCGGGCCAGAATAAGG | 1000 | | | | | | | | | |
| Oligo fwd | CACCTCGATCCCCGGGCCAGAATA | | | | | | | | | | |
| Oligo rev | AAACTATTCTGGCCCGGGGATCGA | | | | | | | | | | |
| Chromosome | start | end | strand | MM | target_seq | PAM | alignment | distance | position | gene name | gene id |
| GU305914.1 | 57862 | 57884 | + | 0 | TCGATCCCCGGGCCAGAATA | AGG | ||||||||[||||||||||||]PAM | NA | Intergenic | NA | NA |
| T3816 | CGGGCCAGAATAAGGCATTCAGG | 1000 | | | | | | | | | |
| Oligo fwd | CACCCGGGCCAGAATAAGGCATTC | | | | | | | | | | |
| Oligo rev | AAACGAATGCCTTATTCTGGCCCG | | | | | | | | | | |
| Chromosome | start | end | strand | MM | target_seq | PAM | alignment | distance | position | gene name | gene id |
| GU305914.1 | 57870 | 57892 | + | 0 | CGGGCCAGAATAAGGCATTC | AGG | ||||||||[||||||||||||]PAM | NA | Intergenic | NA | NA |
| T3817 | GCACTTTTGTTTTTTAGTGTTGG | 1000 | | | | | | | | | |
| Oligo fwd | CACCGCACTTTTGTTTTTTAGTGT | | | | | | | | | | |
| Oligo rev | AAACACACTAAAAAACAAAAGTGC | | | | | | | | | | |
| Chromosome | start | end | strand | MM | target_seq | PAM | alignment | distance | position | gene name | gene id |
| GU305914.1 | 57897 | 57919 | + | 0 | GCACTTTTGTTTTTTAGTGT | TGG | ||||||||[||||||||||||]PAM | NA | Intergenic | NA | NA |
| T3818 | CACTTTTGTTTTTTAGTGTTGGG | 1000 | | | | | | | | | |
| Oligo fwd | CACCCACTTTTGTTTTTTAGTGTT | | | | | | | | | | |

<table>
<tr><td>Oligo rev</td><td>AAACAACACTAAAAAACAAAAGTG</td><td></td><td></td><td></td><td></td><td></td><td></td><td></td><td></td><td></td><td></td><td></td></tr>
<tr><td>Chromosome</td><td>start</td><td></td><td>end</td><td>strand</td><td>MM</td><td>target_seq</td><td>PAM</td><td>alignment</td><td>distance</td><td>position</td><td>gene name</td><td>gene id</td></tr>
<tr><td>GU305914.1</td><td></td><td>57898</td><td>57920</td><td>+</td><td>0</td><td>CACTTTTGTTTTTTAGTGTT</td><td>GGG</td><td>||||||||[||||||||||||]PAM</td><td>NA</td><td>Intergenic</td><td>NA</td><td>NA</td></tr>
<tr><td>T3819</td><td>AGTGTTGGGAGTGTATTCGAAGG</td><td></td><td>1000</td><td></td><td></td><td></td><td></td><td></td><td></td><td></td><td></td><td></td></tr>
<tr><td>Oligo fwd</td><td>CACCAGTGTTGGGAGTGTATTCGA</td><td></td><td></td><td></td><td></td><td></td><td></td><td></td><td></td><td></td><td></td><td></td></tr>
<tr><td>Oligo rev</td><td>AAACTCGAATACACTCCCAACACT</td><td></td><td></td><td></td><td></td><td></td><td></td><td></td><td></td><td></td><td></td><td></td></tr>
<tr><td>Chromosome</td><td>start</td><td></td><td>end</td><td>strand</td><td>MM</td><td>target_seq</td><td>PAM</td><td>alignment</td><td>distance</td><td>position</td><td>gene name</td><td>gene id</td></tr>
<tr><td>GU305914.1</td><td></td><td>57912</td><td>57934</td><td>+</td><td>0</td><td>AGTGTTGGGAGTGTATTCGA</td><td>AGG</td><td>||||||||[||||||||||||]PAM</td><td>NA</td><td>Intergenic</td><td>NA</td><td>NA</td></tr>
<tr><td>T3820</td><td>TTCACTGTCGCTCTCTTCCTTGG</td><td></td><td>1000</td><td></td><td></td><td></td><td></td><td></td><td></td><td></td><td></td><td></td></tr>
<tr><td>Oligo fwd</td><td>CACCTTCACTGTCGCTCTCTTCCT</td><td></td><td></td><td></td><td></td><td></td><td></td><td></td><td></td><td></td><td></td><td></td></tr>
<tr><td>Oligo rev</td><td>AAACAGGAAGAGAGCGACAGTGAA</td><td></td><td></td><td></td><td></td><td></td><td></td><td></td><td></td><td></td><td></td><td></td></tr>
<tr><td>Chromosome</td><td>start</td><td></td><td>end</td><td>strand</td><td>MM</td><td>target_seq</td><td>PAM</td><td>alignment</td><td>distance</td><td>position</td><td>gene name</td><td>gene id</td></tr>
<tr><td>GU305914.1</td><td></td><td>57945</td><td>57967</td><td>+</td><td>0</td><td>TTCACTGTCGCTCTCTTCCT</td><td>TGG</td><td>||||||||[||||||||||||]PAM</td><td>NA</td><td>Intergenic</td><td>NA</td><td>NA</td></tr>
<tr><td>T3821</td><td>TGTCGTTGAAGTAGCCACCGCGG</td><td></td><td>1000</td><td></td><td></td><td></td><td></td><td></td><td></td><td></td><td></td><td></td></tr>
<tr><td>Oligo fwd</td><td>CACCTGTCGTTGAAGTAGCCACCG</td><td></td><td></td><td></td><td></td><td></td><td></td><td></td><td></td><td></td><td></td><td></td></tr>
<tr><td>Oligo rev</td><td>AAACCGGTGGCTACTTCAACGACA</td><td></td><td></td><td></td><td></td><td></td><td></td><td></td><td></td><td></td><td></td><td></td></tr>
<tr><td>Chromosome</td><td>start</td><td></td><td>end</td><td>strand</td><td>MM</td><td>target_seq</td><td>PAM</td><td>alignment</td><td>distance</td><td>position</td><td>gene name</td><td>gene id</td></tr>
<tr><td>GU305914.1</td><td></td><td>57968</td><td>57990</td><td>+</td><td>0</td><td>TGTCGTTGAAGTAGCCACCG</td><td>CGG</td><td>||||||||[||||||||||||]PAM</td><td>NA</td><td>Intergenic</td><td>NA</td><td>NA</td></tr>
<tr><td>T3822</td><td>GCGGTCAGAGCCGCCTCCCCCGG</td><td></td><td>1000</td><td></td><td></td><td></td><td></td><td></td><td></td><td></td><td></td><td></td></tr>
<tr><td>Oligo fwd</td><td>CACCGCGGTCAGAGCCGCCTCCCC</td><td></td><td></td><td></td><td></td><td></td><td></td><td></td><td></td><td></td><td></td><td></td></tr>
<tr><td>Oligo rev</td><td>AAACGGGGAGGCGGCTCTGACCGC</td><td></td><td></td><td></td><td></td><td></td><td></td><td></td><td></td><td></td><td></td><td></td></tr>
<tr><td>Chromosome</td><td>start</td><td></td><td>end</td><td>strand</td><td>MM</td><td>target_seq</td><td>PAM</td><td>alignment</td><td>distance</td><td>position</td><td>gene name</td><td>gene id</td></tr>
<tr><td>GU305914.1</td><td></td><td>57987</td><td>58009</td><td>+</td><td>0</td><td>GCGGTCAGAGCCGCCTCCCC</td><td>CGG</td><td>||||||||[||||||||||||]PAM</td><td>NA</td><td>Intergenic</td><td>NA</td><td>NA</td></tr>
<tr><td>T3823</td><td>CCCTCCACCCACAGCCCCCTTGG</td><td></td><td>1000</td><td></td><td></td><td></td><td></td><td></td><td></td><td></td><td></td><td></td></tr>
<tr><td>Oligo fwd</td><td>CACCCCCTCCACCCACAGCCCCCT</td><td></td><td></td><td></td><td></td><td></td><td></td><td></td><td></td><td></td><td></td><td></td></tr>
<tr><td>Oligo rev</td><td>AAACAGGGGGCTGTGGGTGGAGGG</td><td></td><td></td><td></td><td></td><td></td><td></td><td></td><td></td><td></td><td></td><td></td></tr>
<tr><td>Chromosome</td><td>start</td><td></td><td>end</td><td>strand</td><td>MM</td><td>target_seq</td><td>PAM</td><td>alignment</td><td>distance</td><td>position</td><td>gene name</td><td>gene id</td></tr>
<tr><td>GU305914.1</td><td></td><td>58011</td><td>58033</td><td>+</td><td>0</td><td>CCCTCCACCCACAGCCCCCT</td><td>TGG</td><td>||||||||[||||||||||||]PAM</td><td>NA</td><td>Intergenic</td><td>NA</td><td>NA</td></tr>
<tr><td>T3824</td><td>CCCCTTGGAGACCATGAAACTGG</td><td></td><td>1000</td><td></td><td></td><td></td><td></td><td></td><td></td><td></td><td></td><td></td></tr>
<tr><td>Oligo fwd</td><td>CACCCCCCTTGGAGACCATGAAAC</td><td></td><td></td><td></td><td></td><td></td><td></td><td></td><td></td><td></td><td></td><td></td></tr>
<tr><td>Oligo rev</td><td>AAACGTTTCATGGTCTCCAAGGGG</td><td></td><td></td><td></td><td></td><td></td><td></td><td></td><td></td><td></td><td></td><td></td></tr>
<tr><td>Chromosome</td><td>start</td><td></td><td>end</td><td>strand</td><td>MM</td><td>target_seq</td><td>PAM</td><td>alignment</td><td>distance</td><td>position</td><td>gene name</td><td>gene id</td></tr>
<tr><td>GU305914.1</td><td></td><td>58026</td><td>58048</td><td>+</td><td>0</td><td>CCCCTTGGAGACCATGAAAC</td><td>TGG</td><td>||||||||[||||||||||||]PAM</td><td>NA</td><td>Intergenic</td><td>NA</td><td>NA</td></tr>
<tr><td>T3825</td><td>CGCTTCCGCCGCCACCTTGATGG</td><td></td><td>1000</td><td></td><td></td><td></td><td></td><td></td><td></td><td></td><td></td><td></td></tr>
<tr><td>Oligo fwd</td><td>CACCCGCTTCCGCCGCCACCTTGA</td><td></td><td></td><td></td><td></td><td></td><td></td><td></td><td></td><td></td><td></td><td></td></tr>
<tr><td>Oligo rev</td><td>AAACTCAAGGTGGCGGCGGAAGCG</td><td></td><td></td><td></td><td></td><td></td><td></td><td></td><td></td><td></td><td></td><td></td></tr>
<tr><td>Chromosome</td><td>start</td><td></td><td>end</td><td>strand</td><td>MM</td><td>target_seq</td><td>PAM</td><td>alignment</td><td>distance</td><td>position</td><td>gene name</td><td>gene id</td></tr>
<tr><td>GU305914.1</td><td></td><td>58082</td><td>58104</td><td>+</td><td>0</td><td>CGCTTCCGCCGCCACCTTGA</td><td>TGG</td><td>||||||||[||||||||||||]PAM</td><td>NA</td><td>Intergenic</td><td>NA</td><td>NA</td></tr>
<tr><td>T3826</td><td>CGAGACCCTCTCGTGTTTCTTGG</td><td></td><td>1000</td><td></td><td></td><td></td><td></td><td></td><td></td><td></td><td></td><td></td></tr>
<tr><td>Oligo fwd</td><td>CACCCGAGACCCTCTCGTGTTTCT</td><td></td><td></td><td></td><td></td><td></td><td></td><td></td><td></td><td></td><td></td><td></td></tr>
<tr><td>Oligo rev</td><td>AAACAGAAACACGAGAGGGTCTCG</td><td></td><td></td><td></td><td></td><td></td><td></td><td></td><td></td><td></td><td></td><td></td></tr>
<tr><td>Chromosome</td><td>start</td><td></td><td>end</td><td>strand</td><td>MM</td><td>target_seq</td><td>PAM</td><td>alignment</td><td>distance</td><td>position</td><td>gene name</td><td>gene id</td></tr>
<tr><td>GU305914.1</td><td></td><td>58158</td><td>58180</td><td>+</td><td>0</td><td>CGAGACCCTCTCGTGTTTCT</td><td>TGG</td><td>||||||||[||||||||||||]PAM</td><td>NA</td><td>Intergenic</td><td>NA</td><td>NA</td></tr>
<tr><td>T3827</td><td>GTCGATGACTTGATCGACGCAGG</td><td></td><td>1000</td><td></td><td></td><td></td><td></td><td></td><td></td><td></td><td></td><td></td></tr>
<tr><td>Oligo fwd</td><td>CACCGTCGATGACTTGATCGACGC</td><td></td><td></td><td></td><td></td><td></td><td></td><td></td><td></td><td></td><td></td><td></td></tr>
<tr><td>Oligo rev</td><td>AAACGCGTCGATCAAGTCATCGAC</td><td></td><td></td><td></td><td></td><td></td><td></td><td></td><td></td><td></td><td></td><td></td></tr>
<tr><td>Chromosome</td><td>start</td><td></td><td>end</td><td>strand</td><td>MM</td><td>target_seq</td><td>PAM</td><td>alignment</td><td>distance</td><td>position</td><td>gene name</td><td>gene id</td></tr>
<tr><td>GU305914.1</td><td></td><td>58188</td><td>58210</td><td>+</td><td>0</td><td>GTCGATGACTTGATCGACGC</td><td>AGG</td><td>||||||||[||||||||||||]PAM</td><td>NA</td><td>Intergenic</td><td>NA</td><td>NA</td></tr>
<tr><td>T3828</td><td>GATGACTTGATCGACGCAGGCGG</td><td></td><td>1000</td><td></td><td></td><td></td><td></td><td></td><td></td><td></td><td></td><td></td></tr>
<tr><td>Oligo fwd</td><td>CACCGATGACTTGATCGACGCAGG</td><td></td><td></td><td></td><td></td><td></td><td></td><td></td><td></td><td></td><td></td><td></td></tr>
<tr><td>Oligo rev</td><td>AAACCCTGCGTCGATCAAGTCATC</td><td></td><td></td><td></td><td></td><td></td><td></td><td></td><td></td><td></td><td></td><td></td></tr>
<tr><td>Chromosome</td><td>start</td><td></td><td>end</td><td>strand</td><td>MM</td><td>target_seq</td><td>PAM</td><td>alignment</td><td>distance</td><td>position</td><td>gene name</td><td>gene id</td></tr>
<tr><td>GU305914.1</td><td></td><td>58191</td><td>58213</td><td>+</td><td>0</td><td>GATGACTTGATCGACGCAGG</td><td>CGG</td><td>||||||||[||||||||||||]PAM</td><td>NA</td><td>Intergenic</td><td>NA</td><td>NA</td></tr>
<tr><td>T3829</td><td>GCAGGCGGCGAAATCGTCGCCGG</td><td></td><td>1000</td><td></td><td></td><td></td><td></td><td></td><td></td><td></td><td></td><td></td></tr>
<tr><td>Oligo fwd</td><td>CACCGCAGGCGGCGAAATCGTCGC</td><td></td><td></td><td></td><td></td><td></td><td></td><td></td><td></td><td></td><td></td><td></td></tr>
<tr><td>Oligo rev</td><td>AAACGCGACGATTTCGCCGCCTGC</td><td></td><td></td><td></td><td></td><td></td><td></td><td></td><td></td><td></td><td></td><td></td></tr>
<tr><td>Chromosome</td><td>start</td><td></td><td>end</td><td>strand</td><td>MM</td><td>target_seq</td><td>PAM</td><td>alignment</td><td>distance</td><td>position</td><td>gene name</td><td>gene id</td></tr>
<tr><td>GU305914.1</td><td></td><td>58206</td><td>58228</td><td>+</td><td>0</td><td>GCAGGCGGCGAAATCGTCGC</td><td>CGG</td><td>||||||||[||||||||||||]PAM</td><td>NA</td><td>Intergenic</td><td>NA</td><td>NA</td></tr>
<tr><td>T3830</td><td>GGCGGCGAAATCGTCGCCGGCGG</td><td></td><td>1000</td><td></td><td></td><td></td><td></td><td></td><td></td><td></td><td></td><td></td></tr>
<tr><td>Oligo fwd</td><td>CACCGGCGGCGAAATCGTCGCCGG</td><td></td><td></td><td></td><td></td><td></td><td></td><td></td><td></td><td></td><td></td><td></td></tr>
<tr><td>Oligo rev</td><td>AAACCCGGCGACGATTTCGCCGCC</td><td></td><td></td><td></td><td></td><td></td><td></td><td></td><td></td><td></td><td></td><td></td></tr>
<tr><td>Chromosome</td><td>start</td><td></td><td>end</td><td>strand</td><td>MM</td><td>target_seq</td><td>PAM</td><td>alignment</td><td>distance</td><td>position</td><td>gene name</td><td>gene id</td></tr>
<tr><td>GU305914.1</td><td></td><td>58209</td><td>58231</td><td>+</td><td>0</td><td>GGCGGCGAAATCGTCGCCGG</td><td>CGG</td><td>||||||||[||||||||||||]PAM</td><td>NA</td><td>Intergenic</td><td>NA</td><td>NA</td></tr>
<tr><td>T3831</td><td>ATCGTCGCCGGCGGAGCTGCTGG</td><td></td><td>1000</td><td></td><td></td><td></td><td></td><td></td><td></td><td></td><td></td><td></td></tr>
<tr><td>Oligo fwd</td><td>CACCATCGTCGCCGGCGGAGCTGC</td><td></td><td></td><td></td><td></td><td></td><td></td><td></td><td></td><td></td><td></td><td></td></tr>
<tr><td>Oligo rev</td><td>AAACGCAGCTCCGCCGGCGACGAT</td><td></td><td></td><td></td><td></td><td></td><td></td><td></td><td></td><td></td><td></td><td></td></tr>
<tr><td>Chromosome</td><td>start</td><td></td><td>end</td><td>strand</td><td>MM</td><td>target_seq</td><td>PAM</td><td>alignment</td><td>distance</td><td>position</td><td>gene name</td><td>gene id</td></tr>
<tr><td>GU305914.1</td><td></td><td>58218</td><td>58240</td><td>+</td><td>0</td><td>ATCGTCGCCGGCGGAGCTGC</td><td>TGG</td><td>||||||||[||||||||||||]PAM</td><td>NA</td><td>Intergenic</td><td>NA</td><td>NA</td></tr>
<tr><td>T3832</td><td>CGGCGGAGCTGCTGGAGTTCTGG</td><td></td><td>1000</td><td></td><td></td><td></td><td></td><td></td><td></td><td></td><td></td><td></td></tr>
<tr><td>Oligo fwd</td><td>CACCCGGCGGAGCTGCTGGAGTTC</td><td></td><td></td><td></td><td></td><td></td><td></td><td></td><td></td><td></td><td></td><td></td></tr>
<tr><td>Oligo rev</td><td>AAACGAACTCCAGCAGCTCCGCCG</td><td></td><td></td><td></td><td></td><td></td><td></td><td></td><td></td><td></td><td></td><td></td></tr>
<tr><td>Chromosome</td><td>start</td><td></td><td>end</td><td>strand</td><td>MM</td><td>target_seq</td><td>PAM</td><td>alignment</td><td>distance</td><td>position</td><td>gene name</td><td>gene id</td></tr>
<tr><td>GU305914.1</td><td></td><td>58226</td><td>58248</td><td>+</td><td>0</td><td>CGGCGGAGCTGCTGGAGTTC</td><td>TGG</td><td>||||||||[||||||||||||]PAM</td><td>NA</td><td>Intergenic</td><td>NA</td><td>NA</td></tr>
<tr><td>T3833</td><td>GTTCTGGTAGTTGTTGCTGTTGG</td><td></td><td>1000</td><td></td><td></td><td></td><td></td><td></td><td></td><td></td><td></td><td></td></tr>
</table>

<table>
<tr><td>Oligo fwd</td><td>CACCGTTCTGGTAGTTGTTGCTGT</td><td></td><td></td><td></td><td></td><td></td><td></td><td></td><td></td><td></td><td></td></tr>
<tr><td>Oligo rev</td><td>AAACACAGCAACAACTACCAGAAC</td><td></td><td></td><td></td><td></td><td></td><td></td><td></td><td></td><td></td><td></td></tr>
<tr><td>Chromosome</td><td>start</td><td>end</td><td>strand</td><td>MM</td><td>target_seq</td><td>PAM</td><td>alignment</td><td>distance</td><td>position</td><td>gene name</td><td>gene id</td></tr>
<tr><td>GU305914.1</td><td>58242</td><td>58264</td><td>+</td><td>0</td><td>GTTCTGGTAGTTGTTGCTGT</td><td>TGG</td><td>||||||||[||||||||||||]PAM</td><td>NA</td><td>Intergenic</td><td>NA</td><td>NA</td></tr>
<tr><td></td><td></td><td></td><td></td><td></td><td></td><td></td><td></td><td></td><td></td><td></td><td></td></tr>
<tr><td>T3834</td><td>GTAGTTGTTGCTGTTGGCGACGG</td><td>1000</td><td></td><td></td><td></td><td></td><td></td><td></td><td></td><td></td><td></td></tr>
<tr><td>Oligo fwd</td><td>CACCGTAGTTGTTGCTGTTGGCGA</td><td></td><td></td><td></td><td></td><td></td><td></td><td></td><td></td><td></td><td></td></tr>
<tr><td>Oligo rev</td><td>AAACTCGCCAACAGCAACAACTAC</td><td></td><td></td><td></td><td></td><td></td><td></td><td></td><td></td><td></td><td></td></tr>
<tr><td>Chromosome</td><td>start</td><td>end</td><td>strand</td><td>MM</td><td>target_seq</td><td>PAM</td><td>alignment</td><td>distance</td><td>position</td><td>gene name</td><td>gene id</td></tr>
<tr><td>GU305914.1</td><td>58248</td><td>58270</td><td>+</td><td>0</td><td>GTAGTTGTTGCTGTTGGCGA</td><td>CGG</td><td>||||||||[||||||||||||]PAM</td><td>NA</td><td>Intergenic</td><td>NA</td><td>NA</td></tr>
<tr><td></td><td></td><td></td><td></td><td></td><td></td><td></td><td></td><td></td><td></td><td></td><td></td></tr>
<tr><td>T3835</td><td>TAGTTGTTGCTGTTGGCGACGGG</td><td>1000</td><td></td><td></td><td></td><td></td><td></td><td></td><td></td><td></td><td></td></tr>
<tr><td>Oligo fwd</td><td>CACCTAGTTGTTGCTGTTGGCGAC</td><td></td><td></td><td></td><td></td><td></td><td></td><td></td><td></td><td></td><td></td></tr>
<tr><td>Oligo rev</td><td>AAACGTCGCCAACAGCAACAACTA</td><td></td><td></td><td></td><td></td><td></td><td></td><td></td><td></td><td></td><td></td></tr>
<tr><td>Chromosome</td><td>start</td><td>end</td><td>strand</td><td>MM</td><td>target_seq</td><td>PAM</td><td>alignment</td><td>distance</td><td>position</td><td>gene name</td><td>gene id</td></tr>
<tr><td>GU305914.1</td><td>58249</td><td>58271</td><td>+</td><td>0</td><td>TAGTTGTTGCTGTTGGCGAC</td><td>GGG</td><td>||||||||[||||||||||||]PAM</td><td>NA</td><td>Intergenic</td><td>NA</td><td>NA</td></tr>
<tr><td></td><td></td><td></td><td></td><td></td><td></td><td></td><td></td><td></td><td></td><td></td><td></td></tr>
<tr><td>T3836</td><td>TGCTGTTGGCGACGGGCATGCGG</td><td>1000</td><td></td><td></td><td></td><td></td><td></td><td></td><td></td><td></td><td></td></tr>
<tr><td>Oligo fwd</td><td>CACCTGCTGTTGGCGACGGGCATG</td><td></td><td></td><td></td><td></td><td></td><td></td><td></td><td></td><td></td><td></td></tr>
<tr><td>Oligo rev</td><td>AAACCATGCCCGTCGCCAACAGCA</td><td></td><td></td><td></td><td></td><td></td><td></td><td></td><td></td><td></td><td></td></tr>
<tr><td>Chromosome</td><td>start</td><td>end</td><td>strand</td><td>MM</td><td>target_seq</td><td>PAM</td><td>alignment</td><td>distance</td><td>position</td><td>gene name</td><td>gene id</td></tr>
<tr><td>GU305914.1</td><td>58256</td><td>58278</td><td>+</td><td>0</td><td>TGCTGTTGGCGACGGGCATG</td><td>CGG</td><td>||||||||[||||||||||||]PAM</td><td>NA</td><td>Intergenic</td><td>NA</td><td>NA</td></tr>
<tr><td></td><td></td><td></td><td></td><td></td><td></td><td></td><td></td><td></td><td></td><td></td><td></td></tr>
<tr><td>T3837</td><td>GCCCACGTCGCCGCGCACGAAGG</td><td>1000</td><td></td><td></td><td></td><td></td><td></td><td></td><td></td><td></td><td></td></tr>
<tr><td>Oligo fwd</td><td>CACCGCCCACGTCGCCGCGCACGA</td><td></td><td></td><td></td><td></td><td></td><td></td><td></td><td></td><td></td><td></td></tr>
<tr><td>Oligo rev</td><td>AAACTCGTGCGCGGCGACGTGGGC</td><td></td><td></td><td></td><td></td><td></td><td></td><td></td><td></td><td></td><td></td></tr>
<tr><td>Chromosome</td><td>start</td><td>end</td><td>strand</td><td>MM</td><td>target_seq</td><td>PAM</td><td>alignment</td><td>distance</td><td>position</td><td>gene name</td><td>gene id</td></tr>
<tr><td>GU305914.1</td><td>58284</td><td>58306</td><td>+</td><td>0</td><td>GCCCACGTCGCCGCGCACGA</td><td>AGG</td><td>||||||||[||||||||||||]PAM</td><td>NA</td><td>Intergenic</td><td>NA</td><td>NA</td></tr>
<tr><td></td><td></td><td></td><td></td><td></td><td></td><td></td><td></td><td></td><td></td><td></td><td></td></tr>
<tr><td>T3838</td><td>CCCACGTCGCCGCGCACGAAGGG</td><td>1000</td><td></td><td></td><td></td><td></td><td></td><td></td><td></td><td></td><td></td></tr>
<tr><td>Oligo fwd</td><td>CACCCCCACGTCGCCGCGCACGAA</td><td></td><td></td><td></td><td></td><td></td><td></td><td></td><td></td><td></td><td></td></tr>
<tr><td>Oligo rev</td><td>AAACTTCGTGCGCGGCGACGTGGG</td><td></td><td></td><td></td><td></td><td></td><td></td><td></td><td></td><td></td><td></td></tr>
<tr><td>Chromosome</td><td>start</td><td>end</td><td>strand</td><td>MM</td><td>target_seq</td><td>PAM</td><td>alignment</td><td>distance</td><td>position</td><td>gene name</td><td>gene id</td></tr>
<tr><td>GU305914.1</td><td>58285</td><td>58307</td><td>+</td><td>0</td><td>CCCACGTCGCCGCGCACGAA</td><td>GGG</td><td>||||||||[||||||||||||]PAM</td><td>NA</td><td>Intergenic</td><td>NA</td><td>NA</td></tr>
<tr><td>GU305914.1</td><td>147085</td><td>147107</td><td>-</td><td>4</td><td>GCGACGTCGCTGAGCACGAA</td><td>GGG</td><td>-|-|||||[||-|-|||||||]PAM</td><td>NA</td><td>Intergenic</td><td>NA</td><td>NA</td></tr>
<tr><td></td><td></td><td></td><td></td><td></td><td></td><td></td><td></td><td></td><td></td><td></td><td></td></tr>
<tr><td>T3839</td><td>GCACGAAGGGCTCCTCGCTGAGG</td><td>1000</td><td></td><td></td><td></td><td></td><td></td><td></td><td></td><td></td><td></td></tr>
<tr><td>Oligo fwd</td><td>CACCGCACGAAGGGCTCCTCGCTG</td><td></td><td></td><td></td><td></td><td></td><td></td><td></td><td></td><td></td><td></td></tr>
<tr><td>Oligo rev</td><td>AAACCAGCGAGGAGCCCTTCGTGC</td><td></td><td></td><td></td><td></td><td></td><td></td><td></td><td></td><td></td><td></td></tr>
<tr><td>Chromosome</td><td>start</td><td>end</td><td>strand</td><td>MM</td><td>target_seq</td><td>PAM</td><td>alignment</td><td>distance</td><td>position</td><td>gene name</td><td>gene id</td></tr>
<tr><td>GU305914.1</td><td>58298</td><td>58320</td><td>+</td><td>0</td><td>GCACGAAGGGCTCCTCGCTG</td><td>AGG</td><td>||||||||[||||||||||||]PAM</td><td>NA</td><td>Intergenic</td><td>NA</td><td>NA</td></tr>
<tr><td></td><td></td><td></td><td></td><td></td><td></td><td></td><td></td><td></td><td></td><td></td><td></td></tr>
<tr><td>T3840</td><td>CTCGCTGAGGAAGTTCTCGATGG</td><td>1000</td><td></td><td></td><td></td><td></td><td></td><td></td><td></td><td></td><td></td></tr>
<tr><td>Oligo fwd</td><td>CACCCTCGCTGAGGAAGTTCTCGA</td><td></td><td></td><td></td><td></td><td></td><td></td><td></td><td></td><td></td><td></td></tr>
<tr><td>Oligo rev</td><td>AAACTCGAGAACTTCCTCAGCGAG</td><td></td><td></td><td></td><td></td><td></td><td></td><td></td><td></td><td></td><td></td></tr>
<tr><td>Chromosome</td><td>start</td><td>end</td><td>strand</td><td>MM</td><td>target_seq</td><td>PAM</td><td>alignment</td><td>distance</td><td>position</td><td>gene name</td><td>gene id</td></tr>
<tr><td>GU305914.1</td><td>58311</td><td>58333</td><td>+</td><td>0</td><td>CTCGCTGAGGAAGTTCTCGA</td><td>TGG</td><td>||||||||[||||||||||||]PAM</td><td>NA</td><td>Intergenic</td><td>NA</td><td>NA</td></tr>
<tr><td></td><td></td><td></td><td></td><td></td><td></td><td></td><td></td><td></td><td></td><td></td><td></td></tr>
<tr><td>T3841</td><td>GGTGAAGTTAGCGTTCTTGCTGG</td><td>1000</td><td></td><td></td><td></td><td></td><td></td><td></td><td></td><td></td><td></td></tr>
<tr><td>Oligo fwd</td><td>CACCGGTGAAGTTAGCGTTCTTGC</td><td></td><td></td><td></td><td></td><td></td><td></td><td></td><td></td><td></td><td></td></tr>
<tr><td>Oligo rev</td><td>AAACGCAAGAACGCTAACTTCACC</td><td></td><td></td><td></td><td></td><td></td><td></td><td></td><td></td><td></td><td></td></tr>
<tr><td>Chromosome</td><td>start</td><td>end</td><td>strand</td><td>MM</td><td>target_seq</td><td>PAM</td><td>alignment</td><td>distance</td><td>position</td><td>gene name</td><td>gene id</td></tr>
<tr><td>GU305914.1</td><td>58332</td><td>58354</td><td>+</td><td>0</td><td>GGTGAAGTTAGCGTTCTTGC</td><td>TGG</td><td>||||||||[||||||||||||]PAM</td><td>NA</td><td>Intergenic</td><td>NA</td><td>NA</td></tr>
<tr><td></td><td></td><td></td><td></td><td></td><td></td><td></td><td></td><td></td><td></td><td></td><td></td></tr>
<tr><td>T3842</td><td>CTTGCTGGCGACGTAGAGCATGG</td><td>1000</td><td></td><td></td><td></td><td></td><td></td><td></td><td></td><td></td><td></td></tr>
<tr><td>Oligo fwd</td><td>CACCCTTGCTGGCGACGTAGAGCA</td><td></td><td></td><td></td><td></td><td></td><td></td><td></td><td></td><td></td><td></td></tr>
<tr><td>Oligo rev</td><td>AAACTGCTCTACGTCGCCAGCAAG</td><td></td><td></td><td></td><td></td><td></td><td></td><td></td><td></td><td></td><td></td></tr>
<tr><td>Chromosome</td><td>start</td><td>end</td><td>strand</td><td>MM</td><td>target_seq</td><td>PAM</td><td>alignment</td><td>distance</td><td>position</td><td>gene name</td><td>gene id</td></tr>
<tr><td>GU305914.1</td><td>58347</td><td>58369</td><td>+</td><td>0</td><td>CTTGCTGGCGACGTAGAGCA</td><td>TGG</td><td>||||||||[||||||||||||]PAM</td><td>NA</td><td>Intergenic</td><td>NA</td><td>NA</td></tr>
<tr><td></td><td></td><td></td><td></td><td></td><td></td><td></td><td></td><td></td><td></td><td></td><td></td></tr>
<tr><td>T3843</td><td>CGACGTAGAGCATGGTCTCGTGG</td><td>1000</td><td></td><td></td><td></td><td></td><td></td><td></td><td></td><td></td><td></td></tr>
<tr><td>Oligo fwd</td><td>CACCCGACGTAGAGCATGGTCTCG</td><td></td><td></td><td></td><td></td><td></td><td></td><td></td><td></td><td></td><td></td></tr>
<tr><td>Oligo rev</td><td>AAACCGAGACCATGCTCTACGTCG</td><td></td><td></td><td></td><td></td><td></td><td></td><td></td><td></td><td></td><td></td></tr>
<tr><td>Chromosome</td><td>start</td><td>end</td><td>strand</td><td>MM</td><td>target_seq</td><td>PAM</td><td>alignment</td><td>distance</td><td>position</td><td>gene name</td><td>gene id</td></tr>
<tr><td>GU305914.1</td><td>58355</td><td>58377</td><td>+</td><td>0</td><td>CGACGTAGAGCATGGTCTCG</td><td>TGG</td><td>||||||||[||||||||||||]PAM</td><td>NA</td><td>Intergenic</td><td>NA</td><td>NA</td></tr>
<tr><td></td><td></td><td></td><td></td><td></td><td></td><td></td><td></td><td></td><td></td><td></td><td></td></tr>
<tr><td>T3844</td><td>CGTCATGACGCGCAGCGTGCAGG</td><td>1000</td><td></td><td></td><td></td><td></td><td></td><td></td><td></td><td></td><td></td></tr>
<tr><td>Oligo fwd</td><td>CACCCGTCATGACGCGCAGCGTGC</td><td></td><td></td><td></td><td></td><td></td><td></td><td></td><td></td><td></td><td></td></tr>
<tr><td>Oligo rev</td><td>AAACGCACGCTGCGCGTCATGACG</td><td></td><td></td><td></td><td></td><td></td><td></td><td></td><td></td><td></td><td></td></tr>
<tr><td>Chromosome</td><td>start</td><td>end</td><td>strand</td><td>MM</td><td>target_seq</td><td>PAM</td><td>alignment</td><td>distance</td><td>position</td><td>gene name</td><td>gene id</td></tr>
<tr><td>GU305914.1</td><td>58380</td><td>58402</td><td>+</td><td>0</td><td>CGTCATGACGCGCAGCGTGC</td><td>AGG</td><td>||||||||[||||||||||||]PAM</td><td>NA</td><td>Intergenic</td><td>NA</td><td>NA</td></tr>
<tr><td></td><td></td><td></td><td></td><td></td><td></td><td></td><td></td><td></td><td></td><td></td><td></td></tr>
<tr><td>T3845</td><td>CCACACAGTTACAGAGCGCCTGG</td><td>1000</td><td></td><td></td><td></td><td></td><td></td><td></td><td></td><td></td><td></td></tr>
<tr><td>Oligo fwd</td><td>CACCCCACACAGTTACAGAGCGCC</td><td></td><td></td><td></td><td></td><td></td><td></td><td></td><td></td><td></td><td></td></tr>
<tr><td>Oligo rev</td><td>AAACGGCGCTCTGTAACTGTGTGG</td><td></td><td></td><td></td><td></td><td></td><td></td><td></td><td></td><td></td><td></td></tr>
<tr><td>Chromosome</td><td>start</td><td>end</td><td>strand</td><td>MM</td><td>target_seq</td><td>PAM</td><td>alignment</td><td>distance</td><td>position</td><td>gene name</td><td>gene id</td></tr>
<tr><td>GU305914.1</td><td>58415</td><td>58437</td><td>+</td><td>0</td><td>CCACACAGTTACAGAGCGCC</td><td>TGG</td><td>||||||||[||||||||||||]PAM</td><td>NA</td><td>Intergenic</td><td>NA</td><td>NA</td></tr>
<tr><td></td><td></td><td></td><td></td><td></td><td></td><td></td><td></td><td></td><td></td><td></td><td></td></tr>
<tr><td>T3846</td><td>AGTTACAGAGCGCCTGGTGCAGG</td><td>1000</td><td></td><td></td><td></td><td></td><td></td><td></td><td></td><td></td><td></td></tr>
<tr><td>Oligo fwd</td><td>CACCAGTTACAGAGCGCCTGGTGC</td><td></td><td></td><td></td><td></td><td></td><td></td><td></td><td></td><td></td><td></td></tr>
<tr><td>Oligo rev</td><td>AAACGCACCAGGCGCTCTGTAACT</td><td></td><td></td><td></td><td></td><td></td><td></td><td></td><td></td><td></td><td></td></tr>
<tr><td>Chromosome</td><td>start</td><td>end</td><td>strand</td><td>MM</td><td>target_seq</td><td>PAM</td><td>alignment</td><td>distance</td><td>position</td><td>gene name</td><td>gene id</td></tr>
<tr><td>GU305914.1</td><td>58421</td><td>58443</td><td>+</td><td>0</td><td>AGTTACAGAGCGCCTGGTGC</td><td>AGG</td><td>||||||||[||||||||||||]PAM</td><td>NA</td><td>Intergenic</td><td>NA</td><td>NA</td></tr>
<tr><td></td><td></td><td></td><td></td><td></td><td></td><td></td><td></td><td></td><td></td><td></td><td></td></tr>
<tr><td>T3847</td><td>GAGCGCCTGGTGCAGGTTCTTGG</td><td>1000</td><td></td><td></td><td></td><td></td><td></td><td></td><td></td><td></td><td></td></tr>
<tr><td>Oligo fwd</td><td>CACCGAGCGCCTGGTGCAGGTTCT</td><td></td><td></td><td></td><td></td><td></td><td></td><td></td><td></td><td></td><td></td></tr>
<tr><td>Oligo rev</td><td>AAACAGAACCTGCACCAGGCGCTC</td><td></td><td></td><td></td><td></td><td></td><td></td><td></td><td></td><td></td><td></td></tr>
<tr><td>Chromosome</td><td>start</td><td>end</td><td>strand</td><td>MM</td><td>target_seq</td><td>PAM</td><td>alignment</td><td>distance</td><td>position</td><td>gene name</td><td>gene id</td></tr>
<tr><td>GU305914.1</td><td>58428</td><td>58450</td><td>+</td><td>0</td><td>GAGCGCCTGGTGCAGGTTCT</td><td>TGG</td><td>||||||||[||||||||||||]PAM</td><td>NA</td><td>Intergenic</td><td>NA</td><td>NA</td></tr>
</table>

<table>
<tr><td>T3848</td><td>TGGCCTGCACCGTGATGCGCAGG</td><td>1000</td><td></td><td></td><td></td><td></td><td></td><td></td><td></td><td></td><td></td></tr>
<tr><td>Oligo fwd</td><td>CACCTGGCCTGCACCGTGATGCGC</td><td></td><td></td><td></td><td></td><td></td><td></td><td></td><td></td><td></td><td></td></tr>
<tr><td>Oligo rev</td><td>AAACGCGCATCACGGTGCAGGCCA</td><td></td><td></td><td></td><td></td><td></td><td></td><td></td><td></td><td></td><td></td></tr>
<tr><td>Chromosome</td><td>start</td><td>end</td><td>strand</td><td>MM</td><td>target_seq</td><td>PAM</td><td>alignment</td><td>distance</td><td>position</td><td>gene name</td><td>gene id</td></tr>
<tr><td>GU305914.1</td><td>58448</td><td>58470</td><td>+</td><td>0</td><td>TGGCCTGCACCGTGATGCGC</td><td>AGG</td><td>||||||||[||||||||||||]PAM</td><td>NA</td><td>Intergenic</td><td>NA</td><td>NA</td></tr>
<tr><td></td><td></td><td></td><td></td><td></td><td></td><td></td><td></td><td></td><td></td><td></td><td></td></tr>
<tr><td>T3849</td><td>CAGGTTCTTGACCTCGTGAAAGG</td><td>1000</td><td></td><td></td><td></td><td></td><td></td><td></td><td></td><td></td><td></td></tr>
<tr><td>Oligo fwd</td><td>CACCCAGGTTCTTGACCTCGTGAA</td><td></td><td></td><td></td><td></td><td></td><td></td><td></td><td></td><td></td><td></td></tr>
<tr><td>Oligo rev</td><td>AAACTTCACGAGGTCAAGAACCTG</td><td></td><td></td><td></td><td></td><td></td><td></td><td></td><td></td><td></td><td></td></tr>
<tr><td>Chromosome</td><td>start</td><td>end</td><td>strand</td><td>MM</td><td>target_seq</td><td>PAM</td><td>alignment</td><td>distance</td><td>position</td><td>gene name</td><td>gene id</td></tr>
<tr><td>GU305914.1</td><td>58467</td><td>58489</td><td>+</td><td>0</td><td>CAGGTTCTTGACCTCGTGAA</td><td>AGG</td><td>||||||||[||||||||||||]PAM</td><td>NA</td><td>Intergenic</td><td>NA</td><td>NA</td></tr>
<tr><td></td><td></td><td></td><td></td><td></td><td></td><td></td><td></td><td></td><td></td><td></td><td></td></tr>
<tr><td>T3850</td><td>TGACCTCGTGAAAGGCGATGCGG</td><td>1000</td><td></td><td></td><td></td><td></td><td></td><td></td><td></td><td></td><td></td></tr>
<tr><td>Oligo fwd</td><td>CACCTGACCTCGTGAAAGGCGATG</td><td></td><td></td><td></td><td></td><td></td><td></td><td></td><td></td><td></td><td></td></tr>
<tr><td>Oligo rev</td><td>AAACCATCGCCTTTCACGAGGTCA</td><td></td><td></td><td></td><td></td><td></td><td></td><td></td><td></td><td></td><td></td></tr>
<tr><td>Chromosome</td><td>start</td><td>end</td><td>strand</td><td>MM</td><td>target_seq</td><td>PAM</td><td>alignment</td><td>distance</td><td>position</td><td>gene name</td><td>gene id</td></tr>
<tr><td>GU305914.1</td><td>58475</td><td>58497</td><td>+</td><td>0</td><td>TGACCTCGTGAAAGGCGATG</td><td>CGG</td><td>||||||||[||||||||||||]PAM</td><td>NA</td><td>Intergenic</td><td>NA</td><td>NA</td></tr>
<tr><td></td><td></td><td></td><td></td><td></td><td></td><td></td><td></td><td></td><td></td><td></td><td></td></tr>
<tr><td>T3851</td><td>GTGAAAGGCGATGCGGTTGCTGG</td><td>1000</td><td></td><td></td><td></td><td></td><td></td><td></td><td></td><td></td><td></td></tr>
<tr><td>Oligo fwd</td><td>CACCGTGAAAGGCGATGCGGTTGC</td><td></td><td></td><td></td><td></td><td></td><td></td><td></td><td></td><td></td><td></td></tr>
<tr><td>Oligo rev</td><td>AAACGCAACCGCATCGCCTTTCAC</td><td></td><td></td><td></td><td></td><td></td><td></td><td></td><td></td><td></td><td></td></tr>
<tr><td>Chromosome</td><td>start</td><td>end</td><td>strand</td><td>MM</td><td>target_seq</td><td>PAM</td><td>alignment</td><td>distance</td><td>position</td><td>gene name</td><td>gene id</td></tr>
<tr><td>GU305914.1</td><td>58482</td><td>58504</td><td>+</td><td>0</td><td>GTGAAAGGCGATGCGGTTGC</td><td>TGG</td><td>||||||||[||||||||||||]PAM</td><td>NA</td><td>Intergenic</td><td>NA</td><td>NA</td></tr>
<tr><td></td><td></td><td></td><td></td><td></td><td></td><td></td><td></td><td></td><td></td><td></td><td></td></tr>
<tr><td>T3852</td><td>AAAGGCGATGCGGTTGCTGGCGG</td><td>1000</td><td></td><td></td><td></td><td></td><td></td><td></td><td></td><td></td><td></td></tr>
<tr><td>Oligo fwd</td><td>CACCAAAGGCGATGCGGTTGCTGG</td><td></td><td></td><td></td><td></td><td></td><td></td><td></td><td></td><td></td><td></td></tr>
<tr><td>Oligo rev</td><td>AAACCCAGCAACCGCATCGCCTTT</td><td></td><td></td><td></td><td></td><td></td><td></td><td></td><td></td><td></td><td></td></tr>
<tr><td>Chromosome</td><td>start</td><td>end</td><td>strand</td><td>MM</td><td>target_seq</td><td>PAM</td><td>alignment</td><td>distance</td><td>position</td><td>gene name</td><td>gene id</td></tr>
<tr><td>GU305914.1</td><td>58485</td><td>58507</td><td>+</td><td>0</td><td>AAAGGCGATGCGGTTGCTGG</td><td>CGG</td><td>||||||||[||||||||||||]PAM</td><td>NA</td><td>Intergenic</td><td>NA</td><td>NA</td></tr>
<tr><td></td><td></td><td></td><td></td><td></td><td></td><td></td><td></td><td></td><td></td><td></td><td></td></tr>
<tr><td>T3853</td><td>ACCGAGCGAGAACTTGACCGTGG</td><td>1000</td><td></td><td></td><td></td><td></td><td></td><td></td><td></td><td></td><td></td></tr>
<tr><td>Oligo fwd</td><td>CACCACCGAGCGAGAACTTGACCG</td><td></td><td></td><td></td><td></td><td></td><td></td><td></td><td></td><td></td><td></td></tr>
<tr><td>Oligo rev</td><td>AAACCGGTCAAGTTCTCGCTCGGT</td><td></td><td></td><td></td><td></td><td></td><td></td><td></td><td></td><td></td><td></td></tr>
<tr><td>Chromosome</td><td>start</td><td>end</td><td>strand</td><td>MM</td><td>target_seq</td><td>PAM</td><td>alignment</td><td>distance</td><td>position</td><td>gene name</td><td>gene id</td></tr>
<tr><td>GU305914.1</td><td>58530</td><td>58552</td><td>+</td><td>0</td><td>ACCGAGCGAGAACTTGACCG</td><td>TGG</td><td>||||||||[||||||||||||]PAM</td><td>NA</td><td>Intergenic</td><td>NA</td><td>NA</td></tr>
<tr><td></td><td></td><td></td><td></td><td></td><td></td><td></td><td></td><td></td><td></td><td></td><td></td></tr>
<tr><td>T3854</td><td>CCGAGCGAGAACTTGACCGTGGG</td><td>1000</td><td></td><td></td><td></td><td></td><td></td><td></td><td></td><td></td><td></td></tr>
<tr><td>Oligo fwd</td><td>CACCCCGAGCGAGAACTTGACCGT</td><td></td><td></td><td></td><td></td><td></td><td></td><td></td><td></td><td></td><td></td></tr>
<tr><td>Oligo rev</td><td>AAACACGGTCAAGTTCTCGCTCGG</td><td></td><td></td><td></td><td></td><td></td><td></td><td></td><td></td><td></td><td></td></tr>
<tr><td>Chromosome</td><td>start</td><td>end</td><td>strand</td><td>MM</td><td>target_seq</td><td>PAM</td><td>alignment</td><td>distance</td><td>position</td><td>gene name</td><td>gene id</td></tr>
<tr><td>GU305914.1</td><td>58531</td><td>58553</td><td>+</td><td>0</td><td>CCGAGCGAGAACTTGACCGT</td><td>GGG</td><td>||||||||[||||||||||||]PAM</td><td>NA</td><td>Intergenic</td><td>NA</td><td>NA</td></tr>
<tr><td></td><td></td><td></td><td></td><td></td><td></td><td></td><td></td><td></td><td></td><td></td><td></td></tr>
<tr><td>T3855</td><td>AGCGAGAACTTGACCGTGGGCGG</td><td>1000</td><td></td><td></td><td></td><td></td><td></td><td></td><td></td><td></td><td></td></tr>
<tr><td>Oligo fwd</td><td>CACCAGCGAGAACTTGACCGTGGG</td><td></td><td></td><td></td><td></td><td></td><td></td><td></td><td></td><td></td><td></td></tr>
<tr><td>Oligo rev</td><td>AAACCCCACGGTCAAGTTCTCGCT</td><td></td><td></td><td></td><td></td><td></td><td></td><td></td><td></td><td></td><td></td></tr>
<tr><td>Chromosome</td><td>start</td><td>end</td><td>strand</td><td>MM</td><td>target_seq</td><td>PAM</td><td>alignment</td><td>distance</td><td>position</td><td>gene name</td><td>gene id</td></tr>
<tr><td>GU305914.1</td><td>58534</td><td>58556</td><td>+</td><td>0</td><td>AGCGAGAACTTGACCGTGGG</td><td>CGG</td><td>||||||||[||||||||||||]PAM</td><td>NA</td><td>Intergenic</td><td>NA</td><td>NA</td></tr>
<tr><td></td><td></td><td></td><td></td><td></td><td></td><td></td><td></td><td></td><td></td><td></td><td></td></tr>
<tr><td>T3856</td><td>CGTGATCTGTACCGTGCTGCTGG</td><td>1000</td><td></td><td></td><td></td><td></td><td></td><td></td><td></td><td></td><td></td></tr>
<tr><td>Oligo fwd</td><td>CACCCGTGATCTGTACCGTGCTGC</td><td></td><td></td><td></td><td></td><td></td><td></td><td></td><td></td><td></td><td></td></tr>
<tr><td>Oligo rev</td><td>AAACGCAGCACGGTACAGATCACG</td><td></td><td></td><td></td><td></td><td></td><td></td><td></td><td></td><td></td><td></td></tr>
<tr><td>Chromosome</td><td>start</td><td>end</td><td>strand</td><td>MM</td><td>target_seq</td><td>PAM</td><td>alignment</td><td>distance</td><td>position</td><td>gene name</td><td>gene id</td></tr>
<tr><td>GU305914.1</td><td>58569</td><td>58591</td><td>+</td><td>0</td><td>CGTGATCTGTACCGTGCTGC</td><td>TGG</td><td>||||||||[||||||||||||]PAM</td><td>NA</td><td>Intergenic</td><td>NA</td><td>NA</td></tr>
<tr><td></td><td></td><td></td><td></td><td></td><td></td><td></td><td></td><td></td><td></td><td></td><td></td></tr>
<tr><td>T3857</td><td>GATCTGTACCGTGCTGCTGGAGG</td><td>1000</td><td></td><td></td><td></td><td></td><td></td><td></td><td></td><td></td><td></td></tr>
<tr><td>Oligo fwd</td><td>CACCGATCTGTACCGTGCTGCTGG</td><td></td><td></td><td></td><td></td><td></td><td></td><td></td><td></td><td></td><td></td></tr>
<tr><td>Oligo rev</td><td>AAACCCAGCAGCACGGTACAGATC</td><td></td><td></td><td></td><td></td><td></td><td></td><td></td><td></td><td></td><td></td></tr>
<tr><td>Chromosome</td><td>start</td><td>end</td><td>strand</td><td>MM</td><td>target_seq</td><td>PAM</td><td>alignment</td><td>distance</td><td>position</td><td>gene name</td><td>gene id</td></tr>
<tr><td>GU305914.1</td><td>58572</td><td>58594</td><td>+</td><td>0</td><td>GATCTGTACCGTGCTGCTGG</td><td>AGG</td><td>||||||||[||||||||||||]PAM</td><td>NA</td><td>Intergenic</td><td>NA</td><td>NA</td></tr>
<tr><td></td><td></td><td></td><td></td><td></td><td></td><td></td><td></td><td></td><td></td><td></td><td></td></tr>
<tr><td>T3858</td><td>CTGTACCGTGCTGCTGGAGGTGG</td><td>1000</td><td></td><td></td><td></td><td></td><td></td><td></td><td></td><td></td><td></td></tr>
<tr><td>Oligo fwd</td><td>CACCCTGTACCGTGCTGCTGGAGG</td><td></td><td></td><td></td><td></td><td></td><td></td><td></td><td></td><td></td><td></td></tr>
<tr><td>Oligo rev</td><td>AAACCCTCCAGCAGCACGGTACAG</td><td></td><td></td><td></td><td></td><td></td><td></td><td></td><td></td><td></td><td></td></tr>
<tr><td>Chromosome</td><td>start</td><td>end</td><td>strand</td><td>MM</td><td>target_seq</td><td>PAM</td><td>alignment</td><td>distance</td><td>position</td><td>gene name</td><td>gene id</td></tr>
<tr><td>GU305914.1</td><td>58575</td><td>58597</td><td>+</td><td>0</td><td>CTGTACCGTGCTGCTGGAGG</td><td>TGG</td><td>||||||||[||||||||||||]PAM</td><td>NA</td><td>Intergenic</td><td>NA</td><td>NA</td></tr>
<tr><td>GU305914.1</td><td>102094</td><td>102116</td><td>+</td><td>4</td><td>CTCCTCCGAGCTGCTGGAGG</td><td>AGG</td><td>||---|||[-|||||||||||]PAM</td><td>NA</td><td>Intergenic</td><td>NA</td><td>NA</td></tr>
<tr><td></td><td></td><td></td><td></td><td></td><td></td><td></td><td></td><td></td><td></td><td></td><td></td></tr>
<tr><td>T3859</td><td>GCTGGAGGTGGATTGCTTCTTGG</td><td>1000</td><td></td><td></td><td></td><td></td><td></td><td></td><td></td><td></td><td></td></tr>
<tr><td>Oligo fwd</td><td>CACCGCTGGAGGTGGATTGCTTCT</td><td></td><td></td><td></td><td></td><td></td><td></td><td></td><td></td><td></td><td></td></tr>
<tr><td>Oligo rev</td><td>AAACAGAAGCAATCCACCTCCAGC</td><td></td><td></td><td></td><td></td><td></td><td></td><td></td><td></td><td></td><td></td></tr>
<tr><td>Chromosome</td><td>start</td><td>end</td><td>strand</td><td>MM</td><td>target_seq</td><td>PAM</td><td>alignment</td><td>distance</td><td>position</td><td>gene name</td><td>gene id</td></tr>
<tr><td>GU305914.1</td><td>58587</td><td>58609</td><td>+</td><td>0</td><td>GCTGGAGGTGGATTGCTTCT</td><td>TGG</td><td>||||||||[||||||||||||]PAM</td><td>NA</td><td>Intergenic</td><td>NA</td><td>NA</td></tr>
<tr><td></td><td></td><td></td><td></td><td></td><td></td><td></td><td></td><td></td><td></td><td></td><td></td></tr>
<tr><td>T3860</td><td>TTGCTTCTTGGAGTTTCGCTTGG</td><td>1000</td><td></td><td></td><td></td><td></td><td></td><td></td><td></td><td></td><td></td></tr>
<tr><td>Oligo fwd</td><td>CACCTTGCTTCTTGGAGTTTCGCT</td><td></td><td></td><td></td><td></td><td></td><td></td><td></td><td></td><td></td><td></td></tr>
<tr><td>Oligo rev</td><td>AAACAGCGAAACTCCAAGAAGCAA</td><td></td><td></td><td></td><td></td><td></td><td></td><td></td><td></td><td></td><td></td></tr>
<tr><td>Chromosome</td><td>start</td><td>end</td><td>strand</td><td>MM</td><td>target_seq</td><td>PAM</td><td>alignment</td><td>distance</td><td>position</td><td>gene name</td><td>gene id</td></tr>
<tr><td>GU305914.1</td><td>58599</td><td>58621</td><td>+</td><td>0</td><td>TTGCTTCTTGGAGTTTCGCT</td><td>TGG</td><td>||||||||[||||||||||||]PAM</td><td>NA</td><td>Intergenic</td><td>NA</td><td>NA</td></tr>
<tr><td></td><td></td><td></td><td></td><td></td><td></td><td></td><td></td><td></td><td></td><td></td><td></td></tr>
<tr><td>T3861</td><td>TGCTTCTTGGAGTTTCGCTTGGG</td><td>1000</td><td></td><td></td><td></td><td></td><td></td><td></td><td></td><td></td><td></td></tr>
<tr><td>Oligo fwd</td><td>CACCTGCTTCTTGGAGTTTCGCTT</td><td></td><td></td><td></td><td></td><td></td><td></td><td></td><td></td><td></td><td></td></tr>
<tr><td>Oligo rev</td><td>AAACAAGCGAAACTCCAAGAAGCA</td><td></td><td></td><td></td><td></td><td></td><td></td><td></td><td></td><td></td><td></td></tr>
<tr><td>Chromosome</td><td>start</td><td>end</td><td>strand</td><td>MM</td><td>target_seq</td><td>PAM</td><td>alignment</td><td>distance</td><td>position</td><td>gene name</td><td>gene id</td></tr>
<tr><td>GU305914.1</td><td>58600</td><td>58622</td><td>+</td><td>0</td><td>TGCTTCTTGGAGTTTCGCTT</td><td>GGG</td><td>||||||||[||||||||||||]PAM</td><td>NA</td><td>Intergenic</td><td>NA</td><td>NA</td></tr>
<tr><td></td><td></td><td></td><td></td><td></td><td></td><td></td><td></td><td></td><td></td><td></td><td></td></tr>
<tr><td>T3862</td><td>GCTTCTTGGAGTTTCGCTTGGGG</td><td>1000</td><td></td><td></td><td></td><td></td><td></td><td></td><td></td><td></td><td></td></tr>
<tr><td>Oligo fwd</td><td>CACCGCTTCTTGGAGTTTCGCTTG</td><td></td><td></td><td></td><td></td><td></td><td></td><td></td><td></td><td></td><td></td></tr>
<tr><td>Oligo rev</td><td>AAACCAAGCGAAACTCCAAGAAGC</td><td></td><td></td><td></td><td></td><td></td><td></td><td></td><td></td><td></td><td></td></tr>
</table>

| | | | | | | | | | | | |
|---|---|---|---|---|---|---|---|---|---|---|---|
| Chromosome | start | end | strand | MM | target_seq | PAM | alignment | distance | position | gene name | gene id |
| GU305914.1 | 58601 | 58623 | + | 0 | GCTTCTTGGAGTTTCGCTTG | GGG | \|\|\|\|\|\|\|\|[\|\|\|\|\|\|\|\|\|\|\|\|]PAM | NA | Intergenic | NA | NA |
| T3863 | TCTTGGAGTTTCGCTTGGGGCGG | 1000 | | | | | | | | | |
| Oligo fwd | CACCTCTTGGAGTTTCGCTTGGGG | | | | | | | | | | |
| Oligo rev | AAACCCCCAAGCGAAACTCCAAGA | | | | | | | | | | |
| Chromosome | start | end | strand | MM | target_seq | PAM | alignment | distance | position | gene name | gene id |
| GU305914.1 | 58604 | 58626 | + | 0 | TCTTGGAGTTTCGCTTGGGG | CGG | \|\|\|\|\|\|\|\|[\|\|\|\|\|\|\|\|\|\|\|\|]PAM | NA | Intergenic | NA | NA |
| T3864 | TTTCGCTTGGGGCGGATGTTCGG | 1000 | | | | | | | | | |
| Oligo fwd | CACCTTTCGCTTGGGGCGGATGTT | | | | | | | | | | |
| Oligo rev | AAACAACATCCGCCCCAAGCGAAA | | | | | | | | | | |
| Chromosome | start | end | strand | MM | target_seq | PAM | alignment | distance | position | gene name | gene id |
| GU305914.1 | 58612 | 58634 | + | 0 | TTTCGCTTGGGGCGGATGTT | CGG | \|\|\|\|\|\|\|\|[\|\|\|\|\|\|\|\|\|\|\|\|]PAM | NA | Intergenic | NA | NA |
| T3865 | GGATGTTCGGCGCGATCCAGCGG | 1000 | | | | | | | | | |
| Oligo fwd | CACCGGATGTTCGGCGCGATCCAG | | | | | | | | | | |
| Oligo rev | AAACCTGGATCGCGCCGAACATCC | | | | | | | | | | |
| Chromosome | start | end | strand | MM | target_seq | PAM | alignment | distance | position | gene name | gene id |
| GU305914.1 | 58625 | 58647 | + | 0 | GGATGTTCGGCGCGATCCAG | CGG | \|\|\|\|\|\|\|\|[\|\|\|\|\|\|\|\|\|\|\|\|]PAM | NA | Intergenic | NA | NA |
| T3866 | GCGGATGAGCTCGCCGACGACGG | 1000 | | | | | | | | | |
| Oligo fwd | CACCGCGGATGAGCTCGCCGACGA | | | | | | | | | | |
| Oligo rev | AAACTCGTCGGCGAGCTCATCCGC | | | | | | | | | | |
| Chromosome | start | end | strand | MM | target_seq | PAM | alignment | distance | position | gene name | gene id |
| GU305914.1 | 58644 | 58666 | + | 0 | GCGGATGAGCTCGCCGACGA | CGG | \|\|\|\|\|\|\|\|[\|\|\|\|\|\|\|\|\|\|\|\|]PAM | NA | Intergenic | NA | NA |
| GU305914.1 | 151350 | 151372 | + | 4 | GCGCCCGAGCTCGGCGACGA | CGG | \|\|\|---\|\|[\|\|\|\|\|-\|\|\|\|\|\|]PAM | NA | Intergenic | NA | NA |
| T3867 | GATGAGCTCGCCGACGACGGCGG | 1000 | | | | | | | | | |
| Oligo fwd | CACCGATGAGCTCGCCGACGACGG | | | | | | | | | | |
| Oligo rev | AAACCCGTCGTCGGCGAGCTCATC | | | | | | | | | | |
| Chromosome | start | end | strand | MM | target_seq | PAM | alignment | distance | position | gene name | gene id |
| GU305914.1 | 58647 | 58669 | + | 0 | GATGAGCTCGCCGACGACGG | CGG | \|\|\|\|\|\|\|\|[\|\|\|\|\|\|\|\|\|\|\|\|]PAM | NA | Intergenic | NA | NA |
| GU305914.1 | 154117 | 154139 | + | 4 | GCTGATCGCGACGACGACGG | TGG | \|-\|\|\|-\|-[\|\|-\|\|\|\|\|\|\|\|\|]PAM | NA | Intergenic | NA | NA |
| T3868 | GGAGTCGAGATCGACGCGCGCGG | 1000 | | | | | | | | | |
| Oligo fwd | CACCGGAGTCGAGATCGACGCGCG | | | | | | | | | | |
| Oligo rev | AAACCGCGCGTCGATCTCGACTCC | | | | | | | | | | |
| Chromosome | start | end | strand | MM | target_seq | PAM | alignment | distance | position | gene name | gene id |
| GU305914.1 | 58668 | 58690 | + | 0 | GGAGTCGAGATCGACGCGCG | CGG | \|\|\|\|\|\|\|\|[\|\|\|\|\|\|\|\|\|\|\|\|]PAM | NA | Intergenic | NA | NA |
| T3869 | GTCGAGATCGACGCGCGCGGCGG | 1000 | | | | | | | | | |
| Oligo fwd | CACCGTCGAGATCGACGCGCGCGG | | | | | | | | | | |
| Oligo rev | AAACCCGCGCGCGTCGATCTCGAC | | | | | | | | | | |
| Chromosome | start | end | strand | MM | target_seq | PAM | alignment | distance | position | gene name | gene id |
| GU305914.1 | 58671 | 58693 | + | 0 | GTCGAGATCGACGCGCGCGG | CGG | \|\|\|\|\|\|\|\|[\|\|\|\|\|\|\|\|\|\|\|\|]PAM | NA | Intergenic | NA | NA |
| T3870 | GACGCGCGCGGCGGAGTCGCCGG | 1000 | | | | | | | | | |
| Oligo fwd | CACCGACGCGCGCGGCGGAGTCGC | | | | | | | | | | |
| Oligo rev | AAACGCGACTCCGCCGCGCGCGTC | | | | | | | | | | |
| Chromosome | start | end | strand | MM | target_seq | PAM | alignment | distance | position | gene name | gene id |
| GU305914.1 | 58680 | 58702 | + | 0 | GACGCGCGCGGCGGAGTCGC | CGG | \|\|\|\|\|\|\|\|[\|\|\|\|\|\|\|\|\|\|\|\|]PAM | NA | Intergenic | NA | NA |
| GU305914.1 | 225704 | 225726 | + | 3 | GACGCCCGCGGCGGGGGCGC | TGG | \|\|\|\|\|-\|\|[\|\|\|\|\|\|-\|-\|\|\|]PAM | NA | Intergenic | NA | NA |
| T3871 | CGGCGGAGTCGCCGGTCTCGCGG | 1000 | | | | | | | | | |
| Oligo fwd | CACCCGGCGGAGTCGCCGGTCTCG | | | | | | | | | | |
| Oligo rev | AAACCGAGACCGGCGACTCCGCCG | | | | | | | | | | |
| Chromosome | start | end | strand | MM | target_seq | PAM | alignment | distance | position | gene name | gene id |
| GU305914.1 | 58688 | 58710 | + | 0 | CGGCGGAGTCGCCGGTCTCG | CGG | \|\|\|\|\|\|\|\|[\|\|\|\|\|\|\|\|\|\|\|\|]PAM | NA | Intergenic | NA | NA |
| T3872 | CGGTCTCGCGGATGATGTCCTGG | 1000 | | | | | | | | | |
| Oligo fwd | CACCCGGTCTCGCGGATGATGTCC | | | | | | | | | | |
| Oligo rev | AAACGGACATCATCCGCGAGACCG | | | | | | | | | | |
| Chromosome | start | end | strand | MM | target_seq | PAM | alignment | distance | position | gene name | gene id |
| GU305914.1 | 58700 | 58722 | + | 0 | CGGTCTCGCGGATGATGTCC | TGG | \|\|\|\|\|\|\|\|[\|\|\|\|\|\|\|\|\|\|\|\|]PAM | NA | Intergenic | NA | NA |
| T3873 | GATGTCCTGGTTGTGCACGCAGG | 1000 | | | | | | | | | |
| Oligo fwd | CACCGATGTCCTGGTTGTGCACGC | | | | | | | | | | |
| Oligo rev | AAACGCGTGCACAACCAGGACATC | | | | | | | | | | |
| Chromosome | start | end | strand | MM | target_seq | PAM | alignment | distance | position | gene name | gene id |
| GU305914.1 | 58713 | 58735 | + | 0 | GATGTCCTGGTTGTGCACGC | AGG | \|\|\|\|\|\|\|\|[\|\|\|\|\|\|\|\|\|\|\|\|]PAM | NA | Intergenic | NA | NA |
| T3874 | GTCCTGGTTGTGCACGCAGGCGG | 1000 | | | | | | | | | |
| Oligo fwd | CACCGTCCTGGTTGTGCACGCAGG | | | | | | | | | | |
| Oligo rev | AAACCCTGCGTGCACAACCAGGAC | | | | | | | | | | |
| Chromosome | start | end | strand | MM | target_seq | PAM | alignment | distance | position | gene name | gene id |
| GU305914.1 | 58716 | 58738 | + | 0 | GTCCTGGTTGTGCACGCAGG | CGG | \|\|\|\|\|\|\|\|[\|\|\|\|\|\|\|\|\|\|\|\|]PAM | NA | Intergenic | NA | NA |
| T3875 | GCTGAACTCCATGTTGTAGTCGG | 1000 | | | | | | | | | |
| Oligo fwd | CACCGCTGAACTCCATGTTGTAGT | | | | | | | | | | |
| Oligo rev | AAACACTACAACATGGAGTTCAGC | | | | | | | | | | |
| Chromosome | start | end | strand | MM | target_seq | PAM | alignment | distance | position | gene name | gene id |
| GU305914.1 | 58740 | 58762 | + | 0 | GCTGAACTCCATGTTGTAGT | CGG | \|\|\|\|\|\|\|\|[\|\|\|\|\|\|\|\|\|\|\|\|]PAM | NA | Intergenic | NA | NA |
| T3876 | CTGAACTCCATGTTGTAGTCGGG | 1000 | | | | | | | | | |
| Oligo fwd | CACCCTGAACTCCATGTTGTAGTC | | | | | | | | | | |
| Oligo rev | AAACGACTACAACATGGAGTTCAG | | | | | | | | | | |
| Chromosome | start | end | strand | MM | target_seq | PAM | alignment | distance | position | gene name | gene id |
| GU305914.1 | 58741 | 58763 | + | 0 | CTGAACTCCATGTTGTAGTC | GGG | \|\|\|\|\|\|\|\|[\|\|\|\|\|\|\|\|\|\|\|\|]PAM | NA | Intergenic | NA | NA |

<table>
<tr><td>T3877</td><td>GGGCGCCGACATGCAGACCTTGG</td><td>1000</td><td></td><td></td><td></td><td></td><td></td><td></td><td></td><td></td><td></td></tr>
<tr><td>Oligo fwd</td><td>CACCGGGCGCCGACATGCAGACCT</td><td></td><td></td><td></td><td></td><td></td><td></td><td></td><td></td><td></td><td></td></tr>
<tr><td>Oligo rev</td><td>AAACAGGTCTGCATGTCGGCGCCC</td><td></td><td></td><td></td><td></td><td></td><td></td><td></td><td></td><td></td><td></td></tr>
<tr><td>Chromosome</td><td>start</td><td>end</td><td>strand</td><td>MM</td><td>target_seq</td><td>PAM</td><td>alignment</td><td>distance</td><td>position</td><td>gene name</td><td>gene id</td></tr>
<tr><td>GU305914.1</td><td>58761</td><td>58783</td><td>+</td><td>0</td><td>GGGCGCCGACATGCAGACCT</td><td>TGG</td><td>||||||||[||||||||||||]PAM</td><td>NA</td><td>Intergenic</td><td>NA</td><td>NA</td></tr>
<tr><td>T3878</td><td>ACATGCAGACCTTGGCGCTGAGG</td><td>1000</td><td></td><td></td><td></td><td></td><td></td><td></td><td></td><td></td><td></td></tr>
<tr><td>Oligo fwd</td><td>CACCACATGCAGACCTTGGCGCTG</td><td></td><td></td><td></td><td></td><td></td><td></td><td></td><td></td><td></td><td></td></tr>
<tr><td>Oligo rev</td><td>AAACCAGCGCCAAGGTCTGCATGT</td><td></td><td></td><td></td><td></td><td></td><td></td><td></td><td></td><td></td><td></td></tr>
<tr><td>Chromosome</td><td>start</td><td>end</td><td>strand</td><td>MM</td><td>target_seq</td><td>PAM</td><td>alignment</td><td>distance</td><td>position</td><td>gene name</td><td>gene id</td></tr>
<tr><td>GU305914.1</td><td>58769</td><td>58791</td><td>+</td><td>0</td><td>ACATGCAGACCTTGGCGCTG</td><td>AGG</td><td>||||||||[||||||||||||]PAM</td><td>NA</td><td>Intergenic</td><td>NA</td><td>NA</td></tr>
<tr><td>T3879</td><td>GACCTTGGCGCTGAGGTCCGAGG</td><td>1000</td><td></td><td></td><td></td><td></td><td></td><td></td><td></td><td></td><td></td></tr>
<tr><td>Oligo fwd</td><td>CACCGACCTTGGCGCTGAGGTCCG</td><td></td><td></td><td></td><td></td><td></td><td></td><td></td><td></td><td></td><td></td></tr>
<tr><td>Oligo rev</td><td>AAACCGGACCTCAGCGCCAAGGTC</td><td></td><td></td><td></td><td></td><td></td><td></td><td></td><td></td><td></td><td></td></tr>
<tr><td>Chromosome</td><td>start</td><td>end</td><td>strand</td><td>MM</td><td>target_seq</td><td>PAM</td><td>alignment</td><td>distance</td><td>position</td><td>gene name</td><td>gene id</td></tr>
<tr><td>GU305914.1</td><td>58776</td><td>58798</td><td>+</td><td>0</td><td>GACCTTGGCGCTGAGGTCCG</td><td>AGG</td><td>||||||||[||||||||||||]PAM</td><td>NA</td><td>Intergenic</td><td>NA</td><td>NA</td></tr>
<tr><td>T3880</td><td>GAGGTCCGAGGTGTCCTGCACGG</td><td>1000</td><td></td><td></td><td></td><td></td><td></td><td></td><td></td><td></td><td></td></tr>
<tr><td>Oligo fwd</td><td>CACCGAGGTCCGAGGTGTCCTGCA</td><td></td><td></td><td></td><td></td><td></td><td></td><td></td><td></td><td></td><td></td></tr>
<tr><td>Oligo rev</td><td>AAACTGCAGGACACCTCGGACCTC</td><td></td><td></td><td></td><td></td><td></td><td></td><td></td><td></td><td></td><td></td></tr>
<tr><td>Chromosome</td><td>start</td><td>end</td><td>strand</td><td>MM</td><td>target_seq</td><td>PAM</td><td>alignment</td><td>distance</td><td>position</td><td>gene name</td><td>gene id</td></tr>
<tr><td>GU305914.1</td><td>58788</td><td>58810</td><td>+</td><td>0</td><td>GAGGTCCGAGGTGTCCTGCA</td><td>CGG</td><td>||||||||[||||||||||||]PAM</td><td>NA</td><td>Intergenic</td><td>NA</td><td>NA</td></tr>
<tr><td>T3881</td><td>GGTGTCCTGCACGGTGAATTTGG</td><td>1000</td><td></td><td></td><td></td><td></td><td></td><td></td><td></td><td></td><td></td></tr>
<tr><td>Oligo fwd</td><td>CACCGGTGTCCTGCACGGTGAATT</td><td></td><td></td><td></td><td></td><td></td><td></td><td></td><td></td><td></td><td></td></tr>
<tr><td>Oligo rev</td><td>AAACAATTCACCGTGCAGGACACC</td><td></td><td></td><td></td><td></td><td></td><td></td><td></td><td></td><td></td><td></td></tr>
<tr><td>Chromosome</td><td>start</td><td>end</td><td>strand</td><td>MM</td><td>target_seq</td><td>PAM</td><td>alignment</td><td>distance</td><td>position</td><td>gene name</td><td>gene id</td></tr>
<tr><td>GU305914.1</td><td>58797</td><td>58819</td><td>+</td><td>0</td><td>GGTGTCCTGCACGGTGAATT</td><td>TGG</td><td>||||||||[||||||||||||]PAM</td><td>NA</td><td>Intergenic</td><td>NA</td><td>NA</td></tr>
<tr><td>T3882</td><td>CGGTGAATTTGGTCAGATCGCGG</td><td>1000</td><td></td><td></td><td></td><td></td><td></td><td></td><td></td><td></td><td></td></tr>
<tr><td>Oligo fwd</td><td>CACCCGGTGAATTTGGTCAGATCG</td><td></td><td></td><td></td><td></td><td></td><td></td><td></td><td></td><td></td><td></td></tr>
<tr><td>Oligo rev</td><td>AAACCGATCTGACCAAATTCACCG</td><td></td><td></td><td></td><td></td><td></td><td></td><td></td><td></td><td></td><td></td></tr>
<tr><td>Chromosome</td><td>start</td><td>end</td><td>strand</td><td>MM</td><td>target_seq</td><td>PAM</td><td>alignment</td><td>distance</td><td>position</td><td>gene name</td><td>gene id</td></tr>
<tr><td>GU305914.1</td><td>58808</td><td>58830</td><td>+</td><td>0</td><td>CGGTGAATTTGGTCAGATCG</td><td>CGG</td><td>||||||||[||||||||||||]PAM</td><td>NA</td><td>Intergenic</td><td>NA</td><td>NA</td></tr>
<tr><td>T3883</td><td>TTTGGTCAGATCGCGGCTCGAGG</td><td>1000</td><td></td><td></td><td></td><td></td><td></td><td></td><td></td><td></td><td></td></tr>
<tr><td>Oligo fwd</td><td>CACCTTTGGTCAGATCGCGGCTCG</td><td></td><td></td><td></td><td></td><td></td><td></td><td></td><td></td><td></td><td></td></tr>
<tr><td>Oligo rev</td><td>AAACCGAGCCGCGATCTGACCAAA</td><td></td><td></td><td></td><td></td><td></td><td></td><td></td><td></td><td></td><td></td></tr>
<tr><td>Chromosome</td><td>start</td><td>end</td><td>strand</td><td>MM</td><td>target_seq</td><td>PAM</td><td>alignment</td><td>distance</td><td>position</td><td>gene name</td><td>gene id</td></tr>
<tr><td>GU305914.1</td><td>58815</td><td>58837</td><td>+</td><td>0</td><td>TTTGGTCAGATCGCGGCTCG</td><td>AGG</td><td>||||||||[||||||||||||]PAM</td><td>NA</td><td>Intergenic</td><td>NA</td><td>NA</td></tr>
<tr><td>T3884</td><td>GGCTCGAGGTCATGTACATGAGG</td><td>1000</td><td></td><td></td><td></td><td></td><td></td><td></td><td></td><td></td><td></td></tr>
<tr><td>Oligo fwd</td><td>CACCGGCTCGAGGTCATGTACATG</td><td></td><td></td><td></td><td></td><td></td><td></td><td></td><td></td><td></td><td></td></tr>
<tr><td>Oligo rev</td><td>AAACCATGTACATGACCTCGAGCC</td><td></td><td></td><td></td><td></td><td></td><td></td><td></td><td></td><td></td><td></td></tr>
<tr><td>Chromosome</td><td>start</td><td>end</td><td>strand</td><td>MM</td><td>target_seq</td><td>PAM</td><td>alignment</td><td>distance</td><td>position</td><td>gene name</td><td>gene id</td></tr>
<tr><td>GU305914.1</td><td>58829</td><td>58851</td><td>+</td><td>0</td><td>GGCTCGAGGTCATGTACATG</td><td>AGG</td><td>||||||||[||||||||||||]PAM</td><td>NA</td><td>Intergenic</td><td>NA</td><td>NA</td></tr>
<tr><td>T3885</td><td>TGTACATGAGGTTTCCCAGCAGG</td><td>1000</td><td></td><td></td><td></td><td></td><td></td><td></td><td></td><td></td><td></td></tr>
<tr><td>Oligo fwd</td><td>CACCTGTACATGAGGTTTCCCAGC</td><td></td><td></td><td></td><td></td><td></td><td></td><td></td><td></td><td></td><td></td></tr>
<tr><td>Oligo rev</td><td>AAACGCTGGGAAACCTCATGTACA</td><td></td><td></td><td></td><td></td><td></td><td></td><td></td><td></td><td></td><td></td></tr>
<tr><td>Chromosome</td><td>start</td><td>end</td><td>strand</td><td>MM</td><td>target_seq</td><td>PAM</td><td>alignment</td><td>distance</td><td>position</td><td>gene name</td><td>gene id</td></tr>
<tr><td>GU305914.1</td><td>58841</td><td>58863</td><td>+</td><td>0</td><td>TGTACATGAGGTTTCCCAGC</td><td>AGG</td><td>||||||||[||||||||||||]PAM</td><td>NA</td><td>Intergenic</td><td>NA</td><td>NA</td></tr>
<tr><td>T3886</td><td>GTACATGAGGTTTCCCAGCAGGG</td><td>1000</td><td></td><td></td><td></td><td></td><td></td><td></td><td></td><td></td><td></td></tr>
<tr><td>Oligo fwd</td><td>CACCGTACATGAGGTTTCCCAGCA</td><td></td><td></td><td></td><td></td><td></td><td></td><td></td><td></td><td></td><td></td></tr>
<tr><td>Oligo rev</td><td>AAACTGCTGGGAAACCTCATGTAC</td><td></td><td></td><td></td><td></td><td></td><td></td><td></td><td></td><td></td><td></td></tr>
<tr><td>Chromosome</td><td>start</td><td>end</td><td>strand</td><td>MM</td><td>target_seq</td><td>PAM</td><td>alignment</td><td>distance</td><td>position</td><td>gene name</td><td>gene id</td></tr>
<tr><td>GU305914.1</td><td>58842</td><td>58864</td><td>+</td><td>0</td><td>GTACATGAGGTTTCCCAGCA</td><td>GGG</td><td>||||||||[||||||||||||]PAM</td><td>NA</td><td>Intergenic</td><td>NA</td><td>NA</td></tr>
<tr><td>T3887</td><td>TACATGAGGTTTCCCAGCAGGGG</td><td>1000</td><td></td><td></td><td></td><td></td><td></td><td></td><td></td><td></td><td></td></tr>
<tr><td>Oligo fwd</td><td>CACCTACATGAGGTTTCCCAGCAG</td><td></td><td></td><td></td><td></td><td></td><td></td><td></td><td></td><td></td><td></td></tr>
<tr><td>Oligo rev</td><td>AAACCTGCTGGGAAACCTCATGTA</td><td></td><td></td><td></td><td></td><td></td><td></td><td></td><td></td><td></td><td></td></tr>
<tr><td>Chromosome</td><td>start</td><td>end</td><td>strand</td><td>MM</td><td>target_seq</td><td>PAM</td><td>alignment</td><td>distance</td><td>position</td><td>gene name</td><td>gene id</td></tr>
<tr><td>GU305914.1</td><td>58843</td><td>58865</td><td>+</td><td>0</td><td>TACATGAGGTTTCCCAGCAG</td><td>GGG</td><td>||||||||[||||||||||||]PAM</td><td>NA</td><td>Intergenic</td><td>NA</td><td>NA</td></tr>
<tr><td>T3888</td><td>ACATGAGGTTTCCCAGCAGGGGG</td><td>1000</td><td></td><td></td><td></td><td></td><td></td><td></td><td></td><td></td><td></td></tr>
<tr><td>Oligo fwd</td><td>CACCACATGAGGTTTCCCAGCAGG</td><td></td><td></td><td></td><td></td><td></td><td></td><td></td><td></td><td></td><td></td></tr>
<tr><td>Oligo rev</td><td>AAACCCTGCTGGGAAACCTCATGT</td><td></td><td></td><td></td><td></td><td></td><td></td><td></td><td></td><td></td><td></td></tr>
<tr><td>Chromosome</td><td>start</td><td>end</td><td>strand</td><td>MM</td><td>target_seq</td><td>PAM</td><td>alignment</td><td>distance</td><td>position</td><td>gene name</td><td>gene id</td></tr>
<tr><td>GU305914.1</td><td>58844</td><td>58866</td><td>+</td><td>0</td><td>ACATGAGGTTTCCCAGCAGG</td><td>GGG</td><td>||||||||[||||||||||||]PAM</td><td>NA</td><td>Intergenic</td><td>NA</td><td>NA</td></tr>
<tr><td>T3889</td><td>GAGGTTTCCCAGCAGGGGGATGG</td><td>1000</td><td></td><td></td><td></td><td></td><td></td><td></td><td></td><td></td><td></td></tr>
<tr><td>Oligo fwd</td><td>CACCGAGGTTTCCCAGCAGGGGGA</td><td></td><td></td><td></td><td></td><td></td><td></td><td></td><td></td><td></td><td></td></tr>
<tr><td>Oligo rev</td><td>AAACTCCCCCTGCTGGGAAACCTC</td><td></td><td></td><td></td><td></td><td></td><td></td><td></td><td></td><td></td><td></td></tr>
<tr><td>Chromosome</td><td>start</td><td>end</td><td>strand</td><td>MM</td><td>target_seq</td><td>PAM</td><td>alignment</td><td>distance</td><td>position</td><td>gene name</td><td>gene id</td></tr>
<tr><td>GU305914.1</td><td>58848</td><td>58870</td><td>+</td><td>0</td><td>GAGGTTTCCCAGCAGGGGGA</td><td>TGG</td><td>||||||||[||||||||||||]PAM</td><td>NA</td><td>Intergenic</td><td>NA</td><td>NA</td></tr>
<tr><td>T3890</td><td>CAGGGGGATGGAGTTGTTGATGG</td><td>1000</td><td></td><td></td><td></td><td></td><td></td><td></td><td></td><td></td><td></td></tr>
<tr><td>Oligo fwd</td><td>CACCCAGGGGGATGGAGTTGTTGA</td><td></td><td></td><td></td><td></td><td></td><td></td><td></td><td></td><td></td><td></td></tr>
<tr><td>Oligo rev</td><td>AAACTCAACAACTCCATCCCCCTG</td><td></td><td></td><td></td><td></td><td></td><td></td><td></td><td></td><td></td><td></td></tr>
<tr><td>Chromosome</td><td>start</td><td>end</td><td>strand</td><td>MM</td><td>target_seq</td><td>PAM</td><td>alignment</td><td>distance</td><td>position</td><td>gene name</td><td>gene id</td></tr>
<tr><td>GU305914.1</td><td>58860</td><td>58882</td><td>+</td><td>0</td><td>CAGGGGGATGGAGTTGTTGA</td><td>TGG</td><td>||||||||[||||||||||||]PAM</td><td>NA</td><td>Intergenic</td><td>NA</td><td>NA</td></tr>
<tr><td>T3891</td><td>GATGGAGTTGTTGATGGTCTTGG</td><td>1000</td><td></td><td></td><td></td><td></td><td></td><td></td><td></td><td></td><td></td></tr>
<tr><td>Oligo fwd</td><td>CACCGATGGAGTTGTTGATGGTCT</td><td></td><td></td><td></td><td></td><td></td><td></td><td></td><td></td><td></td><td></td></tr>
<tr><td>Oligo rev</td><td>AAACAGACCATCAACAACTCCATC</td><td></td><td></td><td></td><td></td><td></td><td></td><td></td><td></td><td></td><td></td></tr>
<tr><td>Chromosome</td><td>start</td><td>end</td><td>strand</td><td>MM</td><td>target_seq</td><td>PAM</td><td>alignment</td><td>distance</td><td>position</td><td>gene name</td><td>gene id</td></tr>
</table>

<table>
<tr><td>GU305914.1</td><td>58866</td><td>58888</td><td>+</td><td>0</td><td>GATGGAGTTGTTGATGGTCT</td><td>TGG</td><td>||||||||[||||||||||||]PAM</td><td>NA</td><td>Intergenic</td><td>NA</td><td>NA</td></tr>
<tr><td>GU305914.1</td><td>173685</td><td>173707</td><td>+</td><td>4</td><td>GATGATGTTGATGATGGGCT</td><td>GGG</td><td>||||--||[||-||||||-||]PAM</td><td>NA</td><td>Intergenic</td><td>NA</td><td>NA</td></tr>
<tr><td>T3892</td><td>CTTGGCCGAGAAAGACTTGTCGG</td><td>1000</td><td></td><td></td><td></td><td></td><td></td><td></td><td></td><td></td><td></td></tr>
<tr><td>Oligo fwd</td><td>CACCCTTGGCCGAGAAAGACTTGT</td><td></td><td></td><td></td><td></td><td></td><td></td><td></td><td></td><td></td><td></td></tr>
<tr><td>Oligo rev</td><td>AAACACAAGTCTTTCTCGGCCAAG</td><td></td><td></td><td></td><td></td><td></td><td></td><td></td><td></td><td></td><td></td></tr>
<tr><td>Chromosome</td><td>start</td><td>end</td><td>strand</td><td>MM</td><td>target_seq</td><td>PAM</td><td>alignment</td><td>distance</td><td>position</td><td>gene name</td><td>gene id</td></tr>
<tr><td>GU305914.1</td><td>58884</td><td>58906</td><td>+</td><td>0</td><td>CTTGGCCGAGAAAGACTTGT</td><td>CGG</td><td>||||||||[||||||||||||]PAM</td><td>NA</td><td>Intergenic</td><td>NA</td><td>NA</td></tr>
<tr><td>T3893</td><td>CTTGTCGGTGATGTACAGACAGG</td><td>1000</td><td></td><td></td><td></td><td></td><td></td><td></td><td></td><td></td><td></td></tr>
<tr><td>Oligo fwd</td><td>CACCCTTGTCGGTGATGTACAGAC</td><td></td><td></td><td></td><td></td><td></td><td></td><td></td><td></td><td></td><td></td></tr>
<tr><td>Oligo rev</td><td>AAACGTCTGTACATCACCGACAAG</td><td></td><td></td><td></td><td></td><td></td><td></td><td></td><td></td><td></td><td></td></tr>
<tr><td>Chromosome</td><td>start</td><td>end</td><td>strand</td><td>MM</td><td>target_seq</td><td>PAM</td><td>alignment</td><td>distance</td><td>position</td><td>gene name</td><td>gene id</td></tr>
<tr><td>GU305914.1</td><td>58899</td><td>58921</td><td>+</td><td>0</td><td>CTTGTCGGTGATGTACAGAC</td><td>AGG</td><td>||||||||[||||||||||||]PAM</td><td>NA</td><td>Intergenic</td><td>NA</td><td>NA</td></tr>
<tr><td>T3894</td><td>GGAGCTGTTGAAGACGATCTTGG</td><td>1000</td><td></td><td></td><td></td><td></td><td></td><td></td><td></td><td></td><td></td></tr>
<tr><td>Oligo fwd</td><td>CACCGGAGCTGTTGAAGACGATCT</td><td></td><td></td><td></td><td></td><td></td><td></td><td></td><td></td><td></td><td></td></tr>
<tr><td>Oligo rev</td><td>AAACAGATCGTCTTCAACAGCTCC</td><td></td><td></td><td></td><td></td><td></td><td></td><td></td><td></td><td></td><td></td></tr>
<tr><td>Chromosome</td><td>start</td><td>end</td><td>strand</td><td>MM</td><td>target_seq</td><td>PAM</td><td>alignment</td><td>distance</td><td>position</td><td>gene name</td><td>gene id</td></tr>
<tr><td>GU305914.1</td><td>58920</td><td>58942</td><td>+</td><td>0</td><td>GGAGCTGTTGAAGACGATCT</td><td>TGG</td><td>||||||||[||||||||||||]PAM</td><td>NA</td><td>Intergenic</td><td>NA</td><td>NA</td></tr>
<tr><td>GU305914.1</td><td>69640</td><td>69662</td><td>+</td><td>3</td><td>GAAGCTGCTGAAGACGAGCT</td><td>GGG</td><td>|-|||||-[|||||||||-||]PAM</td><td>NA</td><td>Intergenic</td><td>NA</td><td>NA</td></tr>
<tr><td>T3895</td><td>AGACGATCTTGGCGATGAACTGG</td><td>1000</td><td></td><td></td><td></td><td></td><td></td><td></td><td></td><td></td><td></td></tr>
<tr><td>Oligo fwd</td><td>CACCAGACGATCTTGGCGATGAAC</td><td></td><td></td><td></td><td></td><td></td><td></td><td></td><td></td><td></td><td></td></tr>
<tr><td>Oligo rev</td><td>AAACGTTCATCGCCAAGATCGTCT</td><td></td><td></td><td></td><td></td><td></td><td></td><td></td><td></td><td></td><td></td></tr>
<tr><td>Chromosome</td><td>start</td><td>end</td><td>strand</td><td>MM</td><td>target_seq</td><td>PAM</td><td>alignment</td><td>distance</td><td>position</td><td>gene name</td><td>gene id</td></tr>
<tr><td>GU305914.1</td><td>58931</td><td>58953</td><td>+</td><td>0</td><td>AGACGATCTTGGCGATGAAC</td><td>TGG</td><td>||||||||[||||||||||||]PAM</td><td>NA</td><td>Intergenic</td><td>NA</td><td>NA</td></tr>
<tr><td>T3896</td><td>GGCGATGAACTGGTTCTTGACGG</td><td>1000</td><td></td><td></td><td></td><td></td><td></td><td></td><td></td><td></td><td></td></tr>
<tr><td>Oligo fwd</td><td>CACCGGCGATGAACTGGTTCTTGA</td><td></td><td></td><td></td><td></td><td></td><td></td><td></td><td></td><td></td><td></td></tr>
<tr><td>Oligo rev</td><td>AAACTCAAGAACCAGTTCATCGCC</td><td></td><td></td><td></td><td></td><td></td><td></td><td></td><td></td><td></td><td></td></tr>
<tr><td>Chromosome</td><td>start</td><td>end</td><td>strand</td><td>MM</td><td>target_seq</td><td>PAM</td><td>alignment</td><td>distance</td><td>position</td><td>gene name</td><td>gene id</td></tr>
<tr><td>GU305914.1</td><td>58941</td><td>58963</td><td>+</td><td>0</td><td>GGCGATGAACTGGTTCTTGA</td><td>CGG</td><td>||||||||[||||||||||||]PAM</td><td>NA</td><td>Intergenic</td><td>NA</td><td>NA</td></tr>
<tr><td>T3897</td><td>TGAACTGGTTCTTGACGGTCTGG</td><td>1000</td><td></td><td></td><td></td><td></td><td></td><td></td><td></td><td></td><td></td></tr>
<tr><td>Oligo fwd</td><td>CACCTGAACTGGTTCTTGACGGTC</td><td></td><td></td><td></td><td></td><td></td><td></td><td></td><td></td><td></td><td></td></tr>
<tr><td>Oligo rev</td><td>AAACGACCGTCAAGAACCAGTTCA</td><td></td><td></td><td></td><td></td><td></td><td></td><td></td><td></td><td></td><td></td></tr>
<tr><td>Chromosome</td><td>start</td><td>end</td><td>strand</td><td>MM</td><td>target_seq</td><td>PAM</td><td>alignment</td><td>distance</td><td>position</td><td>gene name</td><td>gene id</td></tr>
<tr><td>GU305914.1</td><td>58946</td><td>58968</td><td>+</td><td>0</td><td>TGAACTGGTTCTTGACGGTC</td><td>TGG</td><td>||||||||[||||||||||||]PAM</td><td>NA</td><td>Intergenic</td><td>NA</td><td>NA</td></tr>
<tr><td>GU305914.1</td><td>73030</td><td>73052</td><td>-</td><td>4</td><td>AGAACGGGTTCTTGACCGTT</td><td>AGG</td><td>-||||-||[||||||||-||-]PAM</td><td>NA</td><td>Intergenic</td><td>NA</td><td>NA</td></tr>
<tr><td>T3898</td><td>GACGGTCTGGACGATGAGCGCGG</td><td>1000</td><td></td><td></td><td></td><td></td><td></td><td></td><td></td><td></td><td></td></tr>
<tr><td>Oligo fwd</td><td>CACCGACGGTCTGGACGATGAGCG</td><td></td><td></td><td></td><td></td><td></td><td></td><td></td><td></td><td></td><td></td></tr>
<tr><td>Oligo rev</td><td>AAACCGCTCATCGTCCAGACCGTC</td><td></td><td></td><td></td><td></td><td></td><td></td><td></td><td></td><td></td><td></td></tr>
<tr><td>Chromosome</td><td>start</td><td>end</td><td>strand</td><td>MM</td><td>target_seq</td><td>PAM</td><td>alignment</td><td>distance</td><td>position</td><td>gene name</td><td>gene id</td></tr>
<tr><td>GU305914.1</td><td>58959</td><td>58981</td><td>+</td><td>0</td><td>GACGGTCTGGACGATGAGCG</td><td>CGG</td><td>||||||||[||||||||||||]PAM</td><td>NA</td><td>Intergenic</td><td>NA</td><td>NA</td></tr>
<tr><td>T3899</td><td>ACGGTCTGGACGATGAGCGCGGG</td><td>1000</td><td></td><td></td><td></td><td></td><td></td><td></td><td></td><td></td><td></td></tr>
<tr><td>Oligo fwd</td><td>CACCACGGTCTGGACGATGAGCGC</td><td></td><td></td><td></td><td></td><td></td><td></td><td></td><td></td><td></td><td></td></tr>
<tr><td>Oligo rev</td><td>AAACGCGCTCATCGTCCAGACCGT</td><td></td><td></td><td></td><td></td><td></td><td></td><td></td><td></td><td></td><td></td></tr>
<tr><td>Chromosome</td><td>start</td><td>end</td><td>strand</td><td>MM</td><td>target_seq</td><td>PAM</td><td>alignment</td><td>distance</td><td>position</td><td>gene name</td><td>gene id</td></tr>
<tr><td>GU305914.1</td><td>58960</td><td>58982</td><td>+</td><td>0</td><td>ACGGTCTGGACGATGAGCGC</td><td>GGG</td><td>||||||||[||||||||||||]PAM</td><td>NA</td><td>Intergenic</td><td>NA</td><td>NA</td></tr>
<tr><td>T3900</td><td>CGGTCTGGACGATGAGCGCGGGG</td><td>1000</td><td></td><td></td><td></td><td></td><td></td><td></td><td></td><td></td><td></td></tr>
<tr><td>Oligo fwd</td><td>CACCCGGTCTGGACGATGAGCGCG</td><td></td><td></td><td></td><td></td><td></td><td></td><td></td><td></td><td></td><td></td></tr>
<tr><td>Oligo rev</td><td>AAACCGCGCTCATCGTCCAGACCG</td><td></td><td></td><td></td><td></td><td></td><td></td><td></td><td></td><td></td><td></td></tr>
<tr><td>Chromosome</td><td>start</td><td>end</td><td>strand</td><td>MM</td><td>target_seq</td><td>PAM</td><td>alignment</td><td>distance</td><td>position</td><td>gene name</td><td>gene id</td></tr>
<tr><td>GU305914.1</td><td>58961</td><td>58983</td><td>+</td><td>0</td><td>CGGTCTGGACGATGAGCGCG</td><td>GGG</td><td>||||||||[||||||||||||]PAM</td><td>NA</td><td>Intergenic</td><td>NA</td><td>NA</td></tr>
<tr><td>T3901</td><td>GGTCTGGACGATGAGCGCGGGGG</td><td>1000</td><td></td><td></td><td></td><td></td><td></td><td></td><td></td><td></td><td></td></tr>
<tr><td>Oligo fwd</td><td>CACCGGTCTGGACGATGAGCGCGG</td><td></td><td></td><td></td><td></td><td></td><td></td><td></td><td></td><td></td><td></td></tr>
<tr><td>Oligo rev</td><td>AAACCCGCGCTCATCGTCCAGACC</td><td></td><td></td><td></td><td></td><td></td><td></td><td></td><td></td><td></td><td></td></tr>
<tr><td>Chromosome</td><td>start</td><td>end</td><td>strand</td><td>MM</td><td>target_seq</td><td>PAM</td><td>alignment</td><td>distance</td><td>position</td><td>gene name</td><td>gene id</td></tr>
<tr><td>GU305914.1</td><td>58962</td><td>58984</td><td>+</td><td>0</td><td>GGTCTGGACGATGAGCGCGG</td><td>GGG</td><td>||||||||[||||||||||||]PAM</td><td>NA</td><td>Intergenic</td><td>NA</td><td>NA</td></tr>
<tr><td>T3902</td><td>CTGGACGATGAGCGCGGGGGTGG</td><td>1000</td><td></td><td></td><td></td><td></td><td></td><td></td><td></td><td></td><td></td></tr>
<tr><td>Oligo fwd</td><td>CACCCTGGACGATGAGCGCGGGGG</td><td></td><td></td><td></td><td></td><td></td><td></td><td></td><td></td><td></td><td></td></tr>
<tr><td>Oligo rev</td><td>AAACCCCCCGCGCTCATCGTCCAG</td><td></td><td></td><td></td><td></td><td></td><td></td><td></td><td></td><td></td><td></td></tr>
<tr><td>Chromosome</td><td>start</td><td>end</td><td>strand</td><td>MM</td><td>target_seq</td><td>PAM</td><td>alignment</td><td>distance</td><td>position</td><td>gene name</td><td>gene id</td></tr>
<tr><td>GU305914.1</td><td>58965</td><td>58987</td><td>+</td><td>0</td><td>CTGGACGATGAGCGCGGGGG</td><td>TGG</td><td>||||||||[||||||||||||]PAM</td><td>NA</td><td>Intergenic</td><td>NA</td><td>NA</td></tr>
<tr><td>GU305914.1</td><td>41545</td><td>41567</td><td>+</td><td>4</td><td>CGGGACGGGGAGCGCGCGGG</td><td>CGG</td><td>|-|||||-[-|||||||-|||]PAM</td><td>NA</td><td>Intergenic</td><td>NA</td><td>NA</td></tr>
<tr><td>T3903</td><td>TGGACGATGAGCGCGGGGGTGGG</td><td>1000</td><td></td><td></td><td></td><td></td><td></td><td></td><td></td><td></td><td></td></tr>
<tr><td>Oligo fwd</td><td>CACCTGGACGATGAGCGCGGGGGT</td><td></td><td></td><td></td><td></td><td></td><td></td><td></td><td></td><td></td><td></td></tr>
<tr><td>Oligo rev</td><td>AAACACCCCCGCGCTCATCGTCCA</td><td></td><td></td><td></td><td></td><td></td><td></td><td></td><td></td><td></td><td></td></tr>
<tr><td>Chromosome</td><td>start</td><td>end</td><td>strand</td><td>MM</td><td>target_seq</td><td>PAM</td><td>alignment</td><td>distance</td><td>position</td><td>gene name</td><td>gene id</td></tr>
<tr><td>GU305914.1</td><td>58966</td><td>58988</td><td>+</td><td>0</td><td>TGGACGATGAGCGCGGGGGT</td><td>GGG</td><td>||||||||[||||||||||||]PAM</td><td>NA</td><td>Intergenic</td><td>NA</td><td>NA</td></tr>
<tr><td>T3904</td><td>CGATGAGCGCGGGGGTGGGCAGG</td><td>1000</td><td></td><td></td><td></td><td></td><td></td><td></td><td></td><td></td><td></td></tr>
<tr><td>Oligo fwd</td><td>CACCCGATGAGCGCGGGGGTGGGC</td><td></td><td></td><td></td><td></td><td></td><td></td><td></td><td></td><td></td><td></td></tr>
<tr><td>Oligo rev</td><td>AAACGCCCACCCCCGCGCTCATCG</td><td></td><td></td><td></td><td></td><td></td><td></td><td></td><td></td><td></td><td></td></tr>
<tr><td>Chromosome</td><td>start</td><td>end</td><td>strand</td><td>MM</td><td>target_seq</td><td>PAM</td><td>alignment</td><td>distance</td><td>position</td><td>gene name</td><td>gene id</td></tr>
<tr><td>GU305914.1</td><td>58970</td><td>58992</td><td>+</td><td>0</td><td>CGATGAGCGCGGGGGTGGGC</td><td>AGG</td><td>||||||||[||||||||||||]PAM</td><td>NA</td><td>Intergenic</td><td>NA</td><td>NA</td></tr>
<tr><td>T3905</td><td>GGTGGGCAGGAAGCTGACCGTGG</td><td>1000</td><td></td><td></td><td></td><td></td><td></td><td></td><td></td><td></td><td></td></tr>
<tr><td>Oligo fwd</td><td>CACCGGTGGGCAGGAAGCTGACCG</td><td></td><td></td><td></td><td></td><td></td><td></td><td></td><td></td><td></td><td></td></tr>
<tr><td>Oligo rev</td><td>AAACCGGTCAGCTTCCTGCCCACC</td><td></td><td></td><td></td><td></td><td></td><td></td><td></td><td></td><td></td><td></td></tr>
<tr><td>Chromosome</td><td>start</td><td>end</td><td>strand</td><td>MM</td><td>target_seq</td><td>PAM</td><td>alignment</td><td>distance</td><td>position</td><td>gene name</td><td>gene id</td></tr>
<tr><td>GU305914.1</td><td>58983</td><td>59005</td><td>+</td><td>0</td><td>GGTGGGCAGGAAGCTGACCG</td><td>TGG</td><td>||||||||[||||||||||||]PAM</td><td>NA</td><td>Intergenic</td><td>NA</td><td>NA</td></tr>
</table>

| | | | | | | | | | | | |
|---|---|---|---|---|---|---|---|---|---|---|---|
| T3906 | GTACGACGCAACGCAACTGCTGG | 1000 | | | | | | | | | |
| Oligo fwd | CACCGTACGACGCAACGCAACTGC | | | | | | | | | | |
| Oligo rev | AAACGCAGTTGCGTTGCGTCGTAC | | | | | | | | | | |
| Chromosome | start | end | strand | MM | target_seq | PAM | alignment | distance | position | gene name | gene id |
| GU305914.1 | 59024 | 59046 | + | 0 | GTACGACGCAACGCAACTGC | TGG | \|\|\|\|\|\|\|\|[\|\|\|\|\|\|\|\|\|\|\|\|]PAM | NA | Intergenic | NA | NA |
| T3907 | GACGCAACGCAACTGCTGGATGG | 1000 | | | | | | | | | |
| Oligo fwd | CACCGACGCAACGCAACTGCTGGA | | | | | | | | | | |
| Oligo rev | AAACTCCAGCAGTTGCGTTGCGTC | | | | | | | | | | |
| Chromosome | start | end | strand | MM | target_seq | PAM | alignment | distance | position | gene name | gene id |
| GU305914.1 | 59028 | 59050 | + | 0 | GACGCAACGCAACTGCTGGA | TGG | \|\|\|\|\|\|\|\|[\|\|\|\|\|\|\|\|\|\|\|\|]PAM | NA | Intergenic | NA | NA |
| T3908 | GCAACGCAACTGCTGGATGGCGG | 1000 | | | | | | | | | |
| Oligo fwd | CACCGCAACGCAACTGCTGGATGG | | | | | | | | | | |
| Oligo rev | AAACCCATCCAGCAGTTGCGTTGC | | | | | | | | | | |
| Chromosome | start | end | strand | MM | target_seq | PAM | alignment | distance | position | gene name | gene id |
| GU305914.1 | 59031 | 59053 | + | 0 | GCAACGCAACTGCTGGATGG | CGG | \|\|\|\|\|\|\|\|[\|\|\|\|\|\|\|\|\|\|\|\|]PAM | NA | Intergenic | NA | NA |
| T3909 | TGGCGGTCTTGTACGACTTGAGG | 1000 | | | | | | | | | |
| Oligo fwd | CACCTGGCGGTCTTGTACGACTTG | | | | | | | | | | |
| Oligo rev | AAACCAAGTCGTACAAGACCGCCA | | | | | | | | | | |
| Chromosome | start | end | strand | MM | target_seq | PAM | alignment | distance | position | gene name | gene id |
| GU305914.1 | 59048 | 59070 | + | 0 | TGGCGGTCTTGTACGACTTG | AGG | \|\|\|\|\|\|\|\|[\|\|\|\|\|\|\|\|\|\|\|\|]PAM | NA | Intergenic | NA | NA |
| T3910 | CTTGTACGACTTGAGGCGAAAGG | 1000 | | | | | | | | | |
| Oligo fwd | CACCCTTGTACGACTTGAGGCGAA | | | | | | | | | | |
| Oligo rev | AAACTTCGCCTCAAGTCGTACAAG | | | | | | | | | | |
| Chromosome | start | end | strand | MM | target_seq | PAM | alignment | distance | position | gene name | gene id |
| GU305914.1 | 59055 | 59077 | + | 0 | CTTGTACGACTTGAGGCGAA | AGG | \|\|\|\|\|\|\|\|[\|\|\|\|\|\|\|\|\|\|\|\|]PAM | NA | Intergenic | NA | NA |
| T3911 | ACGACTTGAGGCGAAAGGCCAGG | 1000 | | | | | | | | | |
| Oligo fwd | CACCACGACTTGAGGCGAAAGGCC | | | | | | | | | | |
| Oligo rev | AAACGGCCTTTCGCCTCAAGTCGT | | | | | | | | | | |
| Chromosome | start | end | strand | MM | target_seq | PAM | alignment | distance | position | gene name | gene id |
| GU305914.1 | 59060 | 59082 | + | 0 | ACGACTTGAGGCGAAAGGCC | AGG | \|\|\|\|\|\|\|\|[\|\|\|\|\|\|\|\|\|\|\|\|]PAM | NA | Intergenic | NA | NA |
| T3912 | CGACTTGAGGCGAAAGGCCAGGG | 1000 | | | | | | | | | |
| Oligo fwd | CACCCGACTTGAGGCGAAAGGCCA | | | | | | | | | | |
| Oligo rev | AAACTGGCCTTTCGCCTCAAGTCG | | | | | | | | | | |
| Chromosome | start | end | strand | MM | target_seq | PAM | alignment | distance | position | gene name | gene id |
| GU305914.1 | 59061 | 59083 | + | 0 | CGACTTGAGGCGAAAGGCCA | GGG | \|\|\|\|\|\|\|\|[\|\|\|\|\|\|\|\|\|\|\|\|]PAM | NA | Intergenic | NA | NA |
| T3913 | TTGAGGCGAAAGGCCAGGGTCGG | 1000 | | | | | | | | | |
| Oligo fwd | CACCTTGAGGCGAAAGGCCAGGGT | | | | | | | | | | |
| Oligo rev | AAACACCCTGGCCTTTCGCCTCAA | | | | | | | | | | |
| Chromosome | start | end | strand | MM | target_seq | PAM | alignment | distance | position | gene name | gene id |
| GU305914.1 | 59065 | 59087 | + | 0 | TTGAGGCGAAAGGCCAGGGT | CGG | \|\|\|\|\|\|\|\|[\|\|\|\|\|\|\|\|\|\|\|\|]PAM | NA | Intergenic | NA | NA |
| T3914 | AGGCGAAAGGCCAGGGTCGGCGG | 1000 | | | | | | | | | |
| Oligo fwd | CACCAGGCGAAAGGCCAGGGTCGG | | | | | | | | | | |
| Oligo rev | AAACCCGACCCTGGCCTTTCGCCT | | | | | | | | | | |
| Chromosome | start | end | strand | MM | target_seq | PAM | alignment | distance | position | gene name | gene id |
| GU305914.1 | 59068 | 59090 | + | 0 | AGGCGAAAGGCCAGGGTCGG | CGG | \|\|\|\|\|\|\|\|[\|\|\|\|\|\|\|\|\|\|\|\|]PAM | NA | Intergenic | NA | NA |
| T3915 | CCTACCACCCTCCATTCCCGCGG | 1000 | | | | | | | | | |
| Oligo fwd | CACCCCTACCACCCTCCATTCCCG | | | | | | | | | | |
| Oligo rev | AAACCGGGAATGGAGGGTGGTAGG | | | | | | | | | | |
| Chromosome | start | end | strand | MM | target_seq | PAM | alignment | distance | position | gene name | gene id |
| GU305914.1 | 59109 | 59131 | + | 0 | CCTACCACCCTCCATTCCCG | CGG | \|\|\|\|\|\|\|\|[\|\|\|\|\|\|\|\|\|\|\|\|]PAM | NA | Intergenic | NA | NA |
| T3916 | GATGTATGTGAGCGAGATCGCGG | 1000 | | | | | | | | | |
| Oligo fwd | CACCGATGTATGTGAGCGAGATCG | | | | | | | | | | |
| Oligo rev | AAACCGATCTCGCTCACATACATC | | | | | | | | | | |
| Chromosome | start | end | strand | MM | target_seq | PAM | alignment | distance | position | gene name | gene id |
| GU305914.1 | 59133 | 59155 | + | 0 | GATGTATGTGAGCGAGATCG | CGG | \|\|\|\|\|\|\|\|[\|\|\|\|\|\|\|\|\|\|\|\|]PAM | NA | Intergenic | NA | NA |
| T3917 | TGAGCGAGATCGCGGCGCGCCGG | 1000 | | | | | | | | | |
| Oligo fwd | CACCTGAGCGAGATCGCGGCGCGC | | | | | | | | | | |
| Oligo rev | AAACGCGCGCCGCGATCTCGCTCA | | | | | | | | | | |
| Chromosome | start | end | strand | MM | target_seq | PAM | alignment | distance | position | gene name | gene id |
| GU305914.1 | 59141 | 59163 | + | 0 | TGAGCGAGATCGCGGCGCGC | CGG | \|\|\|\|\|\|\|\|[\|\|\|\|\|\|\|\|\|\|\|\|]PAM | NA | Intergenic | NA | NA |
| T3918 | CGCGGCGCGCCGGCGAAAGAAGG | 1000 | | | | | | | | | |
| Oligo fwd | CACCCGCGGCGCGCCGGCGAAAGA | | | | | | | | | | |
| Oligo rev | AAACTCTTTCGCCGGCGCGCCGCG | | | | | | | | | | |
| Chromosome | start | end | strand | MM | target_seq | PAM | alignment | distance | position | gene name | gene id |
| GU305914.1 | 59151 | 59173 | + | 0 | CGCGGCGCGCCGGCGAAAGA | AGG | \|\|\|\|\|\|\|\|[\|\|\|\|\|\|\|\|\|\|\|\|]PAM | NA | Intergenic | NA | NA |
| T3919 | CGACCGAGCGAGCGAGCGCGAGG | 1000 | | | | | | | | | |
| Oligo fwd | CACCCGACCGAGCGAGCGAGCGCG | | | | | | | | | | |
| Oligo rev | AAACCGCGCTCGCTCGCTCGGTCG | | | | | | | | | | |
| Chromosome | start | end | strand | MM | target_seq | PAM | alignment | distance | position | gene name | gene id |
| GU305914.1 | 59191 | 59213 | + | 0 | CGACCGAGCGAGCGAGCGCG | AGG | \|\|\|\|\|\|\|\|[\|\|\|\|\|\|\|\|\|\|\|\|]PAM | NA | Intergenic | NA | NA |
| T3920 | GAGCGAGCGCGAGGCGACGACGG | 1000 | | | | | | | | | |
| Oligo fwd | CACCGAGCGAGCGCGAGGCGACGA | | | | | | | | | | |
| Oligo rev | AAACTCGTCGCCTCGCGCTCGCTC | | | | | | | | | | |
| Chromosome | start | end | strand | MM | target_seq | PAM | alignment | distance | position | gene name | gene id |

GU305914.1 59200 59222 + 0 GAGCGAGCGCGAGGCGACGA CGG ||||||||[||||||||||||]PAM NA Intergenic NA NA

T3921 GACGACGGCGAGAGTCCGATCGG 1000
Oligo fwd CACCGACGACGGCGAGAGTCCGAT
Oligo rev AAACATCGGACTCTCGCCGTCGTC
Chromosome start end strand MM target_seq PAM alignment distance position gene name gene id
GU305914.1 59215 59237 + 0 GACGACGGCGAGAGTCCGAT CGG ||||||||[||||||||||||]PAM NA Intergenic NA NA

T3922 ACGACGGCGAGAGTCCGATCGGG 1000
Oligo fwd CACCACGACGGCGAGAGTCCGATC
Oligo rev AAACGATCGGACTCTCGCCGTCGT
Chromosome start end strand MM target_seq PAM alignment distance position gene name gene id
GU305914.1 59216 59238 + 0 ACGACGGCGAGAGTCCGATC GGG ||||||||[||||||||||||]PAM NA Intergenic NA NA

T3923 CGAGAGTCCGATCGGGCGCGG 1000
Oligo fwd CACCCGAGAGTCCGATCGGGCGCG
Oligo rev AAACCGCGCCCGATCGGACTCTCG
Chromosome start end strand MM target_seq PAM alignment distance position gene name gene id
GU305914.1 59223 59245 + 0 CGAGAGTCCGATCGGGCGCG CGG ||||||||[||||||||||||]PAM NA Intergenic NA NA

T3924 AACAGAGCGAAAACGAACCACGG 1000
Oligo fwd CACCAACAGAGCGAAAACGAACCA
Oligo rev AAACTGGTTCGTTTTCGCTCTGTT
Chromosome start end strand MM target_seq PAM alignment distance position gene name gene id
GU305914.1 59248 59270 + 0 AACAGAGCGAAAACGAACCA CGG ||||||||[||||||||||||]PAM NA Intergenic NA NA

T3925 ACAGAGCGAAAACGAACCACGGG 1000
Oligo fwd CACCACAGAGCGAAAACGAACCAC
Oligo rev AAACGTGGTTCGTTTTCGCTCTGT
Chromosome start end strand MM target_seq PAM alignment distance position gene name gene id
GU305914.1 59249 59271 + 0 ACAGAGCGAAAACGAACCAC GGG ||||||||[||||||||||||]PAM NA Intergenic NA NA

T3926 GATGCGTCAGAAAAAAAAGTCGG 1000
Oligo fwd CACCGATGCGTCAGAAAAAAAAGT
Oligo rev AAACACTTTTTTTTCTGACGCATC
Chromosome start end strand MM target_seq PAM alignment distance position gene name gene id
GU305914.1 59271 59293 + 0 GATGCGTCAGAAAAAAAAGT CGG ||||||||[||||||||||||]PAM NA Intergenic NA NA

T3927 AGCAGAGAGACCCTGCACCGCGG 1000
Oligo fwd CACCAGCAGAGAGACCCTGCACCG
Oligo rev AAACCGGTGCAGGGTCTCTCTGCT
Chromosome start end strand MM target_seq PAM alignment distance position gene name gene id
GU305914.1 59312 59334 + 0 AGCAGAGAGACCCTGCACCG CGG ||||||||[||||||||||||]PAM NA Intergenic NA NA

T3928 TGCACCGCGGATCTCCTCGAAGG 1000
Oligo fwd CACCTGCACCGCGGATCTCCTCGA
Oligo rev AAACTCGAGGAGATCCGCGGTGCA
Chromosome start end strand MM target_seq PAM alignment distance position gene name gene id
GU305914.1 59325 59347 + 0 TGCACCGCGGATCTCCTCGA AGG ||||||||[||||||||||||]PAM NA Intergenic NA NA

T3929 GCACCGCGGATCTCCTCGAAGGG 1000
Oligo fwd CACCGCACCGCGGATCTCCTCGAA
Oligo rev AAACTTCGAGGAGATCCGCGGTGC
Chromosome start end strand MM target_seq PAM alignment distance position gene name gene id
GU305914.1 59326 59348 + 0 GCACCGCGGATCTCCTCGAA GGG ||||||||[||||||||||||]PAM NA Intergenic NA NA

T3930 CACCGCGGATCTCCTCGAAGGGG 1000
Oligo fwd CACCCACCGCGGATCTCCTCGAAG
Oligo rev AAACCTTCGAGGAGATCCGCGGTG
Chromosome start end strand MM target_seq PAM alignment distance position gene name gene id
GU305914.1 59327 59349 + 0 CACCGCGGATCTCCTCGAAG GGG ||||||||[||||||||||||]PAM NA Intergenic NA NA

T3931 TGTCACGAAAACCAATCGTCTGG 1000
Oligo fwd CACCTGTCACGAAAACCAATCGTC
Oligo rev AAACGACGATTGGTTTTCGTGACA
Chromosome start end strand MM target_seq PAM alignment distance position gene name gene id
GU305914.1 59398 59420 + 0 TGTCACGAAAACCAATCGTC TGG ||||||||[||||||||||||]PAM NA Intergenic NA NA

T3932 GAGCAATAGAACTCGTTTTTTGG 1000
Oligo fwd CACCGAGCAATAGAACTCGTTTTT
Oligo rev AAACAAAAACGAGTTCTATTGCTC
Chromosome start end strand MM target_seq PAM alignment distance position gene name gene id
GU305914.1 59481 59503 + 0 GAGCAATAGAACTCGTTTTT TGG ||||||||[||||||||||||]PAM NA Intergenic NA NA

T3933 TCGCCGTCAGCGATAATTCACGG 1000
Oligo fwd CACCTCGCCGTCAGCGATAATTCA
Oligo rev AAACTGAATTATCGCTGACGGCGA
Chromosome start end strand MM target_seq PAM alignment distance position gene name gene id
GU305914.1 59512 59534 + 0 TCGCCGTCAGCGATAATTCA CGG ||||||||[||||||||||||]PAM NA Intergenic NA NA

T3934 CGTCAGCGATAATTCACGGAAGG 1000
Oligo fwd CACCCGTCAGCGATAATTCACGGA
Oligo rev AAACTCCGTGAATTATCGCTGACG
Chromosome start end strand MM target_seq PAM alignment distance position gene name gene id
GU305914.1 59516 59538 + 0 CGTCAGCGATAATTCACGGA AGG ||||||||[||||||||||||]PAM NA Intergenic NA NA

T3935 GTCAGCGATAATTCACGGAAGGG 1000
Oligo fwd CACCGTCAGCGATAATTCACGGAA
Oligo rev AAACTTCCGTGAATTATCGCTGAC

<table>
<tr><td>Chromosome</td><td>start</td><td>end</td><td>strand</td><td>MM</td><td>target_seq</td><td>PAM</td><td>alignment</td><td>distance</td><td>position</td><td>gene name</td><td>gene id</td></tr>
<tr><td>GU305914.1</td><td>59517</td><td>59539</td><td>+</td><td>0</td><td>GTCAGCGATAATTCACGGAA</td><td>GGG</td><td>||||||||[||||||||||||]PAM</td><td>NA</td><td>Intergenic</td><td>NA</td><td>NA</td></tr>
<tr><td></td><td></td><td></td><td></td><td></td><td></td><td></td><td></td><td></td><td></td><td></td><td></td></tr>
<tr><td>T3936</td><td>TCAGCGATAATTCACGGAAGGGG</td><td>1000</td><td></td><td></td><td></td><td></td><td></td><td></td><td></td><td></td><td></td></tr>
<tr><td>Oligo fwd</td><td>CACCTCAGCGATAATTCACGGAAG</td><td></td><td></td><td></td><td></td><td></td><td></td><td></td><td></td><td></td><td></td></tr>
<tr><td>Oligo rev</td><td>AAACCTTCCGTGAATTATCGCTGA</td><td></td><td></td><td></td><td></td><td></td><td></td><td></td><td></td><td></td><td></td></tr>
<tr><td>Chromosome</td><td>start</td><td>end</td><td>strand</td><td>MM</td><td>target_seq</td><td>PAM</td><td>alignment</td><td>distance</td><td>position</td><td>gene name</td><td>gene id</td></tr>
<tr><td>GU305914.1</td><td>59518</td><td>59540</td><td>+</td><td>0</td><td>TCAGCGATAATTCACGGAAG</td><td>GGG</td><td>||||||||[||||||||||||]PAM</td><td>NA</td><td>Intergenic</td><td>NA</td><td>NA</td></tr>
<tr><td></td><td></td><td></td><td></td><td></td><td></td><td></td><td></td><td></td><td></td><td></td><td></td></tr>
<tr><td>T3937</td><td>CAGCGATAATTCACGGAAGGGGG</td><td>1000</td><td></td><td></td><td></td><td></td><td></td><td></td><td></td><td></td><td></td></tr>
<tr><td>Oligo fwd</td><td>CACCCAGCGATAATTCACGGAAGG</td><td></td><td></td><td></td><td></td><td></td><td></td><td></td><td></td><td></td><td></td></tr>
<tr><td>Oligo rev</td><td>AAACCCTTCCGTGAATTATCGCTG</td><td></td><td></td><td></td><td></td><td></td><td></td><td></td><td></td><td></td><td></td></tr>
<tr><td>Chromosome</td><td>start</td><td>end</td><td>strand</td><td>MM</td><td>target_seq</td><td>PAM</td><td>alignment</td><td>distance</td><td>position</td><td>gene name</td><td>gene id</td></tr>
<tr><td>GU305914.1</td><td>59519</td><td>59541</td><td>+</td><td>0</td><td>CAGCGATAATTCACGGAAGG</td><td>GGG</td><td>||||||||[||||||||||||]PAM</td><td>NA</td><td>Intergenic</td><td>NA</td><td>NA</td></tr>
<tr><td></td><td></td><td></td><td></td><td></td><td></td><td></td><td></td><td></td><td></td><td></td><td></td></tr>
<tr><td>T3938</td><td>ACTACATTTATATACTCCGACGG</td><td>1000</td><td></td><td></td><td></td><td></td><td></td><td></td><td></td><td></td><td></td></tr>
<tr><td>Oligo fwd</td><td>CACCACTACATTTATATACTCCGA</td><td></td><td></td><td></td><td></td><td></td><td></td><td></td><td></td><td></td><td></td></tr>
<tr><td>Oligo rev</td><td>AAACTCGGAGTATATAAATGTAGT</td><td></td><td></td><td></td><td></td><td></td><td></td><td></td><td></td><td></td><td></td></tr>
<tr><td>Chromosome</td><td>start</td><td>end</td><td>strand</td><td>MM</td><td>target_seq</td><td>PAM</td><td>alignment</td><td>distance</td><td>position</td><td>gene name</td><td>gene id</td></tr>
<tr><td>GU305914.1</td><td>59545</td><td>59567</td><td>+</td><td>0</td><td>ACTACATTTATATACTCCGA</td><td>CGG</td><td>||||||||[||||||||||||]PAM</td><td>NA</td><td>Intergenic</td><td>NA</td><td>NA</td></tr>
<tr><td></td><td></td><td></td><td></td><td></td><td></td><td></td><td></td><td></td><td></td><td></td><td></td></tr>
<tr><td>T3939</td><td>CGAAGACCCCCCGAAGCCGATGG</td><td>1000</td><td></td><td></td><td></td><td></td><td></td><td></td><td></td><td></td><td></td></tr>
<tr><td>Oligo fwd</td><td>CACCCGAAGACCCCCCGAAGCCGA</td><td></td><td></td><td></td><td></td><td></td><td></td><td></td><td></td><td></td><td></td></tr>
<tr><td>Oligo rev</td><td>AAACTCGGCTTCGGGGGGTCTTCG</td><td></td><td></td><td></td><td></td><td></td><td></td><td></td><td></td><td></td><td></td></tr>
<tr><td>Chromosome</td><td>start</td><td>end</td><td>strand</td><td>MM</td><td>target_seq</td><td>PAM</td><td>alignment</td><td>distance</td><td>position</td><td>gene name</td><td>gene id</td></tr>
<tr><td>GU305914.1</td><td>59614</td><td>59636</td><td>+</td><td>0</td><td>CGAAGACCCCCCGAAGCCGA</td><td>TGG</td><td>||||||||[||||||||||||]PAM</td><td>NA</td><td>Intergenic</td><td>NA</td><td>NA</td></tr>
<tr><td></td><td></td><td></td><td></td><td></td><td></td><td></td><td></td><td></td><td></td><td></td><td></td></tr>
<tr><td>T3940</td><td>CCCCCCGAAGCCGATGGCGACGG</td><td>1000</td><td></td><td></td><td></td><td></td><td></td><td></td><td></td><td></td><td></td></tr>
<tr><td>Oligo fwd</td><td>CACCCCCCCCGAAGCCGATGGCGA</td><td></td><td></td><td></td><td></td><td></td><td></td><td></td><td></td><td></td><td></td></tr>
<tr><td>Oligo rev</td><td>AAACTCGCCATCGGCTTCGGGGGG</td><td></td><td></td><td></td><td></td><td></td><td></td><td></td><td></td><td></td><td></td></tr>
<tr><td>Chromosome</td><td>start</td><td>end</td><td>strand</td><td>MM</td><td>target_seq</td><td>PAM</td><td>alignment</td><td>distance</td><td>position</td><td>gene name</td><td>gene id</td></tr>
<tr><td>GU305914.1</td><td>59620</td><td>59642</td><td>+</td><td>0</td><td>CCCCCCGAAGCCGATGGCGA</td><td>CGG</td><td>||||||||[||||||||||||]PAM</td><td>NA</td><td>Intergenic</td><td>NA</td><td>NA</td></tr>
<tr><td></td><td></td><td></td><td></td><td></td><td></td><td></td><td></td><td></td><td></td><td></td><td></td></tr>
<tr><td>T3941</td><td>CCCCCGAAGCCGATGGCGACGGG</td><td>1000</td><td></td><td></td><td></td><td></td><td></td><td></td><td></td><td></td><td></td></tr>
<tr><td>Oligo fwd</td><td>CACCCCCCCGAAGCCGATGGCGAC</td><td></td><td></td><td></td><td></td><td></td><td></td><td></td><td></td><td></td><td></td></tr>
<tr><td>Oligo rev</td><td>AAACGTCGCCATCGGCTTCGGGGG</td><td></td><td></td><td></td><td></td><td></td><td></td><td></td><td></td><td></td><td></td></tr>
<tr><td>Chromosome</td><td>start</td><td>end</td><td>strand</td><td>MM</td><td>target_seq</td><td>PAM</td><td>alignment</td><td>distance</td><td>position</td><td>gene name</td><td>gene id</td></tr>
<tr><td>GU305914.1</td><td>59621</td><td>59643</td><td>+</td><td>0</td><td>CCCCCGAAGCCGATGGCGAC</td><td>GGG</td><td>||||||||[||||||||||||]PAM</td><td>NA</td><td>Intergenic</td><td>NA</td><td>NA</td></tr>
<tr><td></td><td></td><td></td><td></td><td></td><td></td><td></td><td></td><td></td><td></td><td></td><td></td></tr>
<tr><td>T3942</td><td>CGACGGGCAGCGACTGCGCATGG</td><td>1000</td><td></td><td></td><td></td><td></td><td></td><td></td><td></td><td></td><td></td></tr>
<tr><td>Oligo fwd</td><td>CACCCGACGGGCAGCGACTGCGCA</td><td></td><td></td><td></td><td></td><td></td><td></td><td></td><td></td><td></td><td></td></tr>
<tr><td>Oligo rev</td><td>AAACTGCGCAGTCGCTGCCCGTCG</td><td></td><td></td><td></td><td></td><td></td><td></td><td></td><td></td><td></td><td></td></tr>
<tr><td>Chromosome</td><td>start</td><td>end</td><td>strand</td><td>MM</td><td>target_seq</td><td>PAM</td><td>alignment</td><td>distance</td><td>position</td><td>gene name</td><td>gene id</td></tr>
<tr><td>GU305914.1</td><td>59637</td><td>59659</td><td>+</td><td>0</td><td>CGACGGGCAGCGACTGCGCA</td><td>TGG</td><td>||||||||[||||||||||||]PAM</td><td>NA</td><td>Intergenic</td><td>NA</td><td>NA</td></tr>
<tr><td></td><td></td><td></td><td></td><td></td><td></td><td></td><td></td><td></td><td></td><td></td><td></td></tr>
<tr><td>T3943</td><td>GACTGCGCATGGTCGACGAACGG</td><td>1000</td><td></td><td></td><td></td><td></td><td></td><td></td><td></td><td></td><td></td></tr>
<tr><td>Oligo fwd</td><td>CACCGACTGCGCATGGTCGACGAA</td><td></td><td></td><td></td><td></td><td></td><td></td><td></td><td></td><td></td><td></td></tr>
<tr><td>Oligo rev</td><td>AAACTTCGTCGACCATGCGCAGTC</td><td></td><td></td><td></td><td></td><td></td><td></td><td></td><td></td><td></td><td></td></tr>
<tr><td>Chromosome</td><td>start</td><td>end</td><td>strand</td><td>MM</td><td>target_seq</td><td>PAM</td><td>alignment</td><td>distance</td><td>position</td><td>gene name</td><td>gene id</td></tr>
<tr><td>GU305914.1</td><td>59648</td><td>59670</td><td>+</td><td>0</td><td>GACTGCGCATGGTCGACGAA</td><td>CGG</td><td>||||||||[||||||||||||]PAM</td><td>NA</td><td>Intergenic</td><td>NA</td><td>NA</td></tr>
<tr><td></td><td></td><td></td><td></td><td></td><td></td><td></td><td></td><td></td><td></td><td></td><td></td></tr>
<tr><td>T3944</td><td>GCACATCTTGATGACCGCGTCGG</td><td>1000</td><td></td><td></td><td></td><td></td><td></td><td></td><td></td><td></td><td></td></tr>
<tr><td>Oligo fwd</td><td>CACCGCACATCTTGATGACCGCGT</td><td></td><td></td><td></td><td></td><td></td><td></td><td></td><td></td><td></td><td></td></tr>
<tr><td>Oligo rev</td><td>AAACACGCGGTCATCAAGATGTGC</td><td></td><td></td><td></td><td></td><td></td><td></td><td></td><td></td><td></td><td></td></tr>
<tr><td>Chromosome</td><td>start</td><td>end</td><td>strand</td><td>MM</td><td>target_seq</td><td>PAM</td><td>alignment</td><td>distance</td><td>position</td><td>gene name</td><td>gene id</td></tr>
<tr><td>GU305914.1</td><td>59683</td><td>59705</td><td>+</td><td>0</td><td>GCACATCTTGATGACCGCGT</td><td>CGG</td><td>||||||||[||||||||||||]PAM</td><td>NA</td><td>Intergenic</td><td>NA</td><td>NA</td></tr>
<tr><td></td><td></td><td></td><td></td><td></td><td></td><td></td><td></td><td></td><td></td><td></td><td></td></tr>
<tr><td>T3945</td><td>GATGACCGCGTCGGTCGACACGG</td><td>1000</td><td></td><td></td><td></td><td></td><td></td><td></td><td></td><td></td><td></td></tr>
<tr><td>Oligo fwd</td><td>CACCGATGACCGCGTCGGTCGACA</td><td></td><td></td><td></td><td></td><td></td><td></td><td></td><td></td><td></td><td></td></tr>
<tr><td>Oligo rev</td><td>AAACTGTCGACCGACGCGGTCATC</td><td></td><td></td><td></td><td></td><td></td><td></td><td></td><td></td><td></td><td></td></tr>
<tr><td>Chromosome</td><td>start</td><td>end</td><td>strand</td><td>MM</td><td>target_seq</td><td>PAM</td><td>alignment</td><td>distance</td><td>position</td><td>gene name</td><td>gene id</td></tr>
<tr><td>GU305914.1</td><td>59692</td><td>59714</td><td>+</td><td>0</td><td>GATGACCGCGTCGGTCGACA</td><td>CGG</td><td>||||||||[||||||||||||]PAM</td><td>NA</td><td>Intergenic</td><td>NA</td><td>NA</td></tr>
<tr><td></td><td></td><td></td><td></td><td></td><td></td><td></td><td></td><td></td><td></td><td></td><td></td></tr>
<tr><td>T3946</td><td>GTCGGTCGACACGGTCCAGCCGG</td><td>1000</td><td></td><td></td><td></td><td></td><td></td><td></td><td></td><td></td><td></td></tr>
<tr><td>Oligo fwd</td><td>CACCGTCGGTCGACACGGTCCAGC</td><td></td><td></td><td></td><td></td><td></td><td></td><td></td><td></td><td></td><td></td></tr>
<tr><td>Oligo rev</td><td>AAACGCTGGACCGTGTCGACCGAC</td><td></td><td></td><td></td><td></td><td></td><td></td><td></td><td></td><td></td><td></td></tr>
<tr><td>Chromosome</td><td>start</td><td>end</td><td>strand</td><td>MM</td><td>target_seq</td><td>PAM</td><td>alignment</td><td>distance</td><td>position</td><td>gene name</td><td>gene id</td></tr>
<tr><td>GU305914.1</td><td>59701</td><td>59723</td><td>+</td><td>0</td><td>GTCGGTCGACACGGTCCAGC</td><td>CGG</td><td>||||||||[||||||||||||]PAM</td><td>NA</td><td>Intergenic</td><td>NA</td><td>NA</td></tr>
<tr><td></td><td></td><td></td><td></td><td></td><td></td><td></td><td></td><td></td><td></td><td></td><td></td></tr>
<tr><td>T3947</td><td>CGACACGGTCCAGCCGGTCTCGG</td><td>1000</td><td></td><td></td><td></td><td></td><td></td><td></td><td></td><td></td><td></td></tr>
<tr><td>Oligo fwd</td><td>CACCCGACACGGTCCAGCCGGTCT</td><td></td><td></td><td></td><td></td><td></td><td></td><td></td><td></td><td></td><td></td></tr>
<tr><td>Oligo rev</td><td>AAACAGACCGGCTGGACCGTGTCG</td><td></td><td></td><td></td><td></td><td></td><td></td><td></td><td></td><td></td><td></td></tr>
<tr><td>Chromosome</td><td>start</td><td>end</td><td>strand</td><td>MM</td><td>target_seq</td><td>PAM</td><td>alignment</td><td>distance</td><td>position</td><td>gene name</td><td>gene id</td></tr>
<tr><td>GU305914.1</td><td>59707</td><td>59729</td><td>+</td><td>0</td><td>CGACACGGTCCAGCCGGTCT</td><td>CGG</td><td>||||||||[||||||||||||]PAM</td><td>NA</td><td>Intergenic</td><td>NA</td><td>NA</td></tr>
<tr><td></td><td></td><td></td><td></td><td></td><td></td><td></td><td></td><td></td><td></td><td></td><td></td></tr>
<tr><td>T3948</td><td>GTCGTCCGAGTCGTCGTAGAAGG</td><td>1000</td><td></td><td></td><td></td><td></td><td></td><td></td><td></td><td></td><td></td></tr>
<tr><td>Oligo fwd</td><td>CACCGTCGTCCGAGTCGTCGTAGA</td><td></td><td></td><td></td><td></td><td></td><td></td><td></td><td></td><td></td><td></td></tr>
<tr><td>Oligo rev</td><td>AAACTCTACGACGACTCGGACGAC</td><td></td><td></td><td></td><td></td><td></td><td></td><td></td><td></td><td></td><td></td></tr>
<tr><td>Chromosome</td><td>start</td><td>end</td><td>strand</td><td>MM</td><td>target_seq</td><td>PAM</td><td>alignment</td><td>distance</td><td>position</td><td>gene name</td><td>gene id</td></tr>
<tr><td>GU305914.1</td><td>59749</td><td>59771</td><td>+</td><td>0</td><td>GTCGTCCGAGTCGTCGTAGA</td><td>AGG</td><td>||||||||[||||||||||||]PAM</td><td>NA</td><td>Intergenic</td><td>NA</td><td>NA</td></tr>
<tr><td></td><td></td><td></td><td></td><td></td><td></td><td></td><td></td><td></td><td></td><td></td><td></td></tr>
<tr><td>T3949</td><td>TCGTCCGAGTCGTCGTAGAAGGG</td><td>1000</td><td></td><td></td><td></td><td></td><td></td><td></td><td></td><td></td><td></td></tr>
<tr><td>Oligo fwd</td><td>CACCTCGTCCGAGTCGTCGTAGAA</td><td></td><td></td><td></td><td></td><td></td><td></td><td></td><td></td><td></td><td></td></tr>
<tr><td>Oligo rev</td><td>AAACTTCTACGACGACTCGGACGA</td><td></td><td></td><td></td><td></td><td></td><td></td><td></td><td></td><td></td><td></td></tr>
<tr><td>Chromosome</td><td>start</td><td>end</td><td>strand</td><td>MM</td><td>target_seq</td><td>PAM</td><td>alignment</td><td>distance</td><td>position</td><td>gene name</td><td>gene id</td></tr>
<tr><td>GU305914.1</td><td>59750</td><td>59772</td><td>+</td><td>0</td><td>TCGTCCGAGTCGTCGTAGAA</td><td>GGG</td><td>||||||||[||||||||||||]PAM</td><td>NA</td><td>Intergenic</td><td>NA</td><td>NA</td></tr>
<tr><td></td><td></td><td></td><td></td><td></td><td></td><td></td><td></td><td></td><td></td><td></td><td></td></tr>
<tr><td>T3950</td><td>CGTCGTAGAAGGGCGCCGAGCGG</td><td>1000</td><td></td><td></td><td></td><td></td><td></td><td></td><td></td><td></td><td></td></tr>
<tr><td>Oligo fwd</td><td>CACCCGTCGTAGAAGGGCGCCGAG</td><td></td><td></td><td></td><td></td><td></td><td></td><td></td><td></td><td></td><td></td></tr>
</table>

Oligo rev AAACCTCGGCGCCCTTCTACGACG
Chromosome start end strand MM target_seq PAM alignment distance position gene name gene id
GU305914.1 59760 59782 + 0 CGTCGTAGAAGGGCGCCGAG CGG ||||||||[||||||||||||]PAM NA Intergenic NA NA

T3951 AGAAGGGCGCCGAGCGGCCGCGG 1000
Oligo fwd CACCAGAAGGGCGCCGAGCGGCCG
Oligo rev AAACCGGCCGCTCGGCGCCCTTCT
Chromosome start end strand MM target_seq PAM alignment distance position gene name gene id
GU305914.1 59766 59788 + 0 AGAAGGGCGCCGAGCGGCCG CGG ||||||||[||||||||||||]PAM NA Intergenic NA NA

T3952 CGCCGAGCGGCCGCGGCGCACGG 1000
Oligo fwd CACCCGCCGAGCGGCCGCGGCGCA
Oligo rev AAACTGCGCCGCGGCCGCTCGGCG
Chromosome start end strand MM target_seq PAM alignment distance position gene name gene id
GU305914.1 59773 59795 + 0 CGCCGAGCGGCCGCGGCGCA CGG ||||||||[||||||||||||]PAM NA Intergenic NA NA

T3953 GCGGCGCACGGCGACCGTCGCGG 1000
Oligo fwd CACCGCGGCGCACGGCGACCGTCG
Oligo rev AAACCGACGGTCGCCGTGCGCCGC
Chromosome start end strand MM target_seq PAM alignment distance position gene name gene id
GU305914.1 59785 59807 + 0 GCGGCGCACGGCGACCGTCG CGG ||||||||[||||||||||||]PAM NA Intergenic NA NA

T3954 ACTCTTTCGAGATCTGCCTGAGG 1000
Oligo fwd CACCACTCTTTCGAGATCTGCCTG
Oligo rev AAACCAGGCAGATCTCGAAAGAGT
Chromosome start end strand MM target_seq PAM alignment distance position gene name gene id
GU305914.1 59808 59830 + 0 ACTCTTTCGAGATCTGCCTG AGG ||||||||[||||||||||||]PAM NA Intergenic NA NA

T3955 CTTTCGAGATCTGCCTGAGGCGG 1000
Oligo fwd CACCCTTTCGAGATCTGCCTGAGG
Oligo rev AAACCCTCAGGCAGATCTCGAAAG
Chromosome start end strand MM target_seq PAM alignment distance position gene name gene id
GU305914.1 59811 59833 + 0 CTTTCGAGATCTGCCTGAGG CGG ||||||||[||||||||||||]PAM NA Intergenic NA NA

T3956 CCTGAGGCGGCTGTTGAGCACGG 1000
Oligo fwd CACCCCTGAGGCGGCTGTTGAGCA
Oligo rev AAACTGCTCAACAGCCGCCTCAGG
Chromosome start end strand MM target_seq PAM alignment distance position gene name gene id
GU305914.1 59824 59846 + 0 CCTGAGGCGGCTGTTGAGCA CGG ||||||||[||||||||||||]PAM NA Intergenic NA NA

T3957 CTGAGGCGGCTGTTGAGCACGGG 1000
Oligo fwd CACCCTGAGGCGGCTGTTGAGCAC
Oligo rev AAACGTGCTCAACAGCCGCCTCAG
Chromosome start end strand MM target_seq PAM alignment distance position gene name gene id
GU305914.1 59825 59847 + 0 CTGAGGCGGCTGTTGAGCAC GGG ||||||||[||||||||||||]PAM NA Intergenic NA NA

T3958 GTTGAGCACGGGCAGCTCGACGG 1000
Oligo fwd CACCGTTGAGCACGGGCAGCTCGA
Oligo rev AAACTCGAGCTGCCCGTGCTCAAC
Chromosome start end strand MM target_seq PAM alignment distance position gene name gene id
GU305914.1 59836 59858 + 0 GTTGAGCACGGGCAGCTCGA CGG ||||||||[||||||||||||]PAM NA Intergenic NA NA

T3959 TTGAGCACGGGCAGCTCGACGGG 1000
Oligo fwd CACCTTGAGCACGGGCAGCTCGAC
Oligo rev AAACGTCGAGCTGCCCGTGCTCAA
Chromosome start end strand MM target_seq PAM alignment distance position gene name gene id
GU305914.1 59837 59859 + 0 TTGAGCACGGGCAGCTCGAC GGG ||||||||[||||||||||||]PAM NA Intergenic NA NA

T3960 GCACGGGCAGCTCGACGGGCCGG 1000
Oligo fwd CACCGCACGGGCAGCTCGACGGGC
Oligo rev AAACGCCCGTCGAGCTGCCCGTGC
Chromosome start end strand MM target_seq PAM alignment distance position gene name gene id
GU305914.1 59841 59863 + 0 GCACGGGCAGCTCGACGGGC CGG ||||||||[||||||||||||]PAM NA Intergenic NA NA

T3961 GGGCCGGTTCTCGCCGAGACAGG 1000
Oligo fwd CACCGGGCCGGTTCTCGCCGAGAC
Oligo rev AAACGTCTCGGCGAGAACCGGCCC
Chromosome start end strand MM target_seq PAM alignment distance position gene name gene id
GU305914.1 59857 59879 + 0 GGGCCGGTTCTCGCCGAGAC AGG ||||||||[||||||||||||]PAM NA Intergenic NA NA

T3962 AGGAGAGATCCTCCTCGTGCAGG 1000
Oligo fwd CACCAGGAGAGATCCTCCTCGTGC
Oligo rev AAACGCACGAGGAGGATCTCTCCT
Chromosome start end strand MM target_seq PAM alignment distance position gene name gene id
GU305914.1 59877 59899 + 0 AGGAGAGATCCTCCTCGTGC AGG ||||||||[||||||||||||]PAM NA Intergenic NA NA

T3963 GATCCTCCTCGTGCAGGCGATGG 1000
Oligo fwd CACCGATCCTCCTCGTGCAGGCGA
Oligo rev AAACTCGCCTGCACGAGGAGGATC
Chromosome start end strand MM target_seq PAM alignment distance position gene name gene id
GU305914.1 59883 59905 + 0 GATCCTCCTCGTGCAGGCGA TGG ||||||||[||||||||||||]PAM NA Intergenic NA NA

T3964 ATCCTCCTCGTGCAGGCGATGGG 1000
Oligo fwd CACCATCCTCCTCGTGCAGGCGAT
Oligo rev AAACATCGCCTGCACGAGGAGGAT
Chromosome start end strand MM target_seq PAM alignment distance position gene name gene id
GU305914.1 59884 59906 + 0 ATCCTCCTCGTGCAGGCGAT GGG ||||||||[||||||||||||]PAM NA Intergenic NA NA

T3965 TCCTCCTCGTGCAGGCGATGGGG 1000

<table>
<tr><td>Oligo fwd</td><td>CACCTCCTCCTCGTGCAGGCGATG</td><td></td><td></td><td></td><td></td><td></td><td></td><td></td><td></td><td></td><td></td></tr>
<tr><td>Oligo rev</td><td>AAACCATCGCCTGCACGAGGAGGA</td><td></td><td></td><td></td><td></td><td></td><td></td><td></td><td></td><td></td><td></td></tr>
<tr><td>Chromosome</td><td>start</td><td>end</td><td>strand</td><td>MM</td><td>target_seq</td><td>PAM</td><td>alignment</td><td>distance</td><td>position</td><td>gene name</td><td>gene id</td></tr>
<tr><td>GU305914.1</td><td>59885</td><td>59907</td><td>+</td><td>0</td><td>TCCTCCTCGTGCAGGCGATG</td><td>GGG</td><td>||||||||[||||||||||||]PAM</td><td>NA</td><td>Intergenic</td><td>NA</td><td>NA</td></tr>
<tr><td>T3966</td><td>GCGATGGGGCTCGATCTTGACGG</td><td>1000</td><td></td><td></td><td></td><td></td><td></td><td></td><td></td><td></td><td></td></tr>
<tr><td>Oligo fwd</td><td>CACCGCGATGGGGCTCGATCTTGA</td><td></td><td></td><td></td><td></td><td></td><td></td><td></td><td></td><td></td><td></td></tr>
<tr><td>Oligo rev</td><td>AAACTCAAGATCGAGCCCCATCGC</td><td></td><td></td><td></td><td></td><td></td><td></td><td></td><td></td><td></td><td></td></tr>
<tr><td>Chromosome</td><td>start</td><td>end</td><td>strand</td><td>MM</td><td>target_seq</td><td>PAM</td><td>alignment</td><td>distance</td><td>position</td><td>gene name</td><td>gene id</td></tr>
<tr><td>GU305914.1</td><td>59899</td><td>59921</td><td>+</td><td>0</td><td>GCGATGGGGCTCGATCTTGA</td><td>CGG</td><td>||||||||[||||||||||||]PAM</td><td>NA</td><td>Intergenic</td><td>NA</td><td>NA</td></tr>
<tr><td>T3967</td><td>TCGAGCATCTGATCCTTCGTCGG</td><td>1000</td><td></td><td></td><td></td><td></td><td></td><td></td><td></td><td></td><td></td></tr>
<tr><td>Oligo fwd</td><td>CACCTCGAGCATCTGATCCTTCGT</td><td></td><td></td><td></td><td></td><td></td><td></td><td></td><td></td><td></td><td></td></tr>
<tr><td>Oligo rev</td><td>AAACACGAAGGATCAGATGCTCGA</td><td></td><td></td><td></td><td></td><td></td><td></td><td></td><td></td><td></td><td></td></tr>
<tr><td>Chromosome</td><td>start</td><td>end</td><td>strand</td><td>MM</td><td>target_seq</td><td>PAM</td><td>alignment</td><td>distance</td><td>position</td><td>gene name</td><td>gene id</td></tr>
<tr><td>GU305914.1</td><td>59936</td><td>59958</td><td>+</td><td>0</td><td>TCGAGCATCTGATCCTTCGT</td><td>CGG</td><td>||||||||[||||||||||||]PAM</td><td>NA</td><td>Intergenic</td><td>NA</td><td>NA</td></tr>
<tr><td>T3968</td><td>CGAGCATCTGATCCTTCGTCGGG</td><td>1000</td><td></td><td></td><td></td><td></td><td></td><td></td><td></td><td></td><td></td></tr>
<tr><td>Oligo fwd</td><td>CACCCGAGCATCTGATCCTTCGTC</td><td></td><td></td><td></td><td></td><td></td><td></td><td></td><td></td><td></td><td></td></tr>
<tr><td>Oligo rev</td><td>AAACGACGAAGGATCAGATGCTCG</td><td></td><td></td><td></td><td></td><td></td><td></td><td></td><td></td><td></td><td></td></tr>
<tr><td>Chromosome</td><td>start</td><td>end</td><td>strand</td><td>MM</td><td>target_seq</td><td>PAM</td><td>alignment</td><td>distance</td><td>position</td><td>gene name</td><td>gene id</td></tr>
<tr><td>GU305914.1</td><td>59937</td><td>59959</td><td>+</td><td>0</td><td>CGAGCATCTGATCCTTCGTC</td><td>GGG</td><td>||||||||[||||||||||||]PAM</td><td>NA</td><td>Intergenic</td><td>NA</td><td>NA</td></tr>
<tr><td>T3969</td><td>GAGCATCTGATCCTTCGTCGGGG</td><td>1000</td><td></td><td></td><td></td><td></td><td></td><td></td><td></td><td></td><td></td></tr>
<tr><td>Oligo fwd</td><td>CACCGAGCATCTGATCCTTCGTCG</td><td></td><td></td><td></td><td></td><td></td><td></td><td></td><td></td><td></td><td></td></tr>
<tr><td>Oligo rev</td><td>AAACCGACGAAGGATCAGATGCTC</td><td></td><td></td><td></td><td></td><td></td><td></td><td></td><td></td><td></td><td></td></tr>
<tr><td>Chromosome</td><td>start</td><td>end</td><td>strand</td><td>MM</td><td>target_seq</td><td>PAM</td><td>alignment</td><td>distance</td><td>position</td><td>gene name</td><td>gene id</td></tr>
<tr><td>GU305914.1</td><td>59938</td><td>59960</td><td>+</td><td>0</td><td>GAGCATCTGATCCTTCGTCG</td><td>GGG</td><td>||||||||[||||||||||||]PAM</td><td>NA</td><td>Intergenic</td><td>NA</td><td>NA</td></tr>
<tr><td>T3970</td><td>TTCGTCGGGGTCATCATGAGCGG</td><td>1000</td><td></td><td></td><td></td><td></td><td></td><td></td><td></td><td></td><td></td></tr>
<tr><td>Oligo fwd</td><td>CACCTTCGTCGGGGTCATCATGAG</td><td></td><td></td><td></td><td></td><td></td><td></td><td></td><td></td><td></td><td></td></tr>
<tr><td>Oligo rev</td><td>AAACCTCATGATGACCCCGACGAA</td><td></td><td></td><td></td><td></td><td></td><td></td><td></td><td></td><td></td><td></td></tr>
<tr><td>Chromosome</td><td>start</td><td>end</td><td>strand</td><td>MM</td><td>target_seq</td><td>PAM</td><td>alignment</td><td>distance</td><td>position</td><td>gene name</td><td>gene id</td></tr>
<tr><td>GU305914.1</td><td>59951</td><td>59973</td><td>+</td><td>0</td><td>TTCGTCGGGGTCATCATGAG</td><td>CGG</td><td>||||||||[||||||||||||]PAM</td><td>NA</td><td>Intergenic</td><td>NA</td><td>NA</td></tr>
<tr><td>T3971</td><td>GCTCGAGAAAGACGTTGTCGCGG</td><td>1000</td><td></td><td></td><td></td><td></td><td></td><td></td><td></td><td></td><td></td></tr>
<tr><td>Oligo fwd</td><td>CACCGCTCGAGAAAGACGTTGTCG</td><td></td><td></td><td></td><td></td><td></td><td></td><td></td><td></td><td></td><td></td></tr>
<tr><td>Oligo rev</td><td>AAACCGACAACGTCTTTCTCGAGC</td><td></td><td></td><td></td><td></td><td></td><td></td><td></td><td></td><td></td><td></td></tr>
<tr><td>Chromosome</td><td>start</td><td>end</td><td>strand</td><td>MM</td><td>target_seq</td><td>PAM</td><td>alignment</td><td>distance</td><td>position</td><td>gene name</td><td>gene id</td></tr>
<tr><td>GU305914.1</td><td>59979</td><td>60001</td><td>+</td><td>0</td><td>GCTCGAGAAAGACGTTGTCG</td><td>CGG</td><td>||||||||[||||||||||||]PAM</td><td>NA</td><td>Intergenic</td><td>NA</td><td>NA</td></tr>
<tr><td>GU305914.1</td><td>77878</td><td>77900</td><td>-</td><td>4</td><td>GCTGCAGGAACACGTTGTCG</td><td>CGG</td><td>|||--||-[||-|||||||||]PAM</td><td>NA</td><td>Intergenic</td><td>NA</td><td>NA</td></tr>
<tr><td>T3972</td><td>AGAAAGACGTTGTCGCGGCGCGG</td><td>1000</td><td></td><td></td><td></td><td></td><td></td><td></td><td></td><td></td><td></td></tr>
<tr><td>Oligo fwd</td><td>CACCAGAAAGACGTTGTCGCGGCG</td><td></td><td></td><td></td><td></td><td></td><td></td><td></td><td></td><td></td><td></td></tr>
<tr><td>Oligo rev</td><td>AAACCGCCGCGACAACGTCTTTCT</td><td></td><td></td><td></td><td></td><td></td><td></td><td></td><td></td><td></td><td></td></tr>
<tr><td>Chromosome</td><td>start</td><td>end</td><td>strand</td><td>MM</td><td>target_seq</td><td>PAM</td><td>alignment</td><td>distance</td><td>position</td><td>gene name</td><td>gene id</td></tr>
<tr><td>GU305914.1</td><td>59984</td><td>60006</td><td>+</td><td>0</td><td>AGAAAGACGTTGTCGCGGCG</td><td>CGG</td><td>||||||||[||||||||||||]PAM</td><td>NA</td><td>Intergenic</td><td>NA</td><td>NA</td></tr>
<tr><td>T3973</td><td>TCGCGGCGCGGCCAGAAGCCCGG</td><td>1000</td><td></td><td></td><td></td><td></td><td></td><td></td><td></td><td></td><td></td></tr>
<tr><td>Oligo fwd</td><td>CACCTCGCGGCGCGGCCAGAAGCC</td><td></td><td></td><td></td><td></td><td></td><td></td><td></td><td></td><td></td><td></td></tr>
<tr><td>Oligo rev</td><td>AAACGGCTTCTGGCCGCGCCGCGA</td><td></td><td></td><td></td><td></td><td></td><td></td><td></td><td></td><td></td><td></td></tr>
<tr><td>Chromosome</td><td>start</td><td>end</td><td>strand</td><td>MM</td><td>target_seq</td><td>PAM</td><td>alignment</td><td>distance</td><td>position</td><td>gene name</td><td>gene id</td></tr>
<tr><td>GU305914.1</td><td>59996</td><td>60018</td><td>+</td><td>0</td><td>TCGCGGCGCGGCCAGAAGCC</td><td>CGG</td><td>||||||||[||||||||||||]PAM</td><td>NA</td><td>Intergenic</td><td>NA</td><td>NA</td></tr>
<tr><td>T3974</td><td>GCGCGGCCAGAAGCCCGGAGTGG</td><td>1000</td><td></td><td></td><td></td><td></td><td></td><td></td><td></td><td></td><td></td></tr>
<tr><td>Oligo fwd</td><td>CACCGCGCGGCCAGAAGCCCGGAG</td><td></td><td></td><td></td><td></td><td></td><td></td><td></td><td></td><td></td><td></td></tr>
<tr><td>Oligo rev</td><td>AAACCTCCGGGCTTCTGGCCGCGC</td><td></td><td></td><td></td><td></td><td></td><td></td><td></td><td></td><td></td><td></td></tr>
<tr><td>Chromosome</td><td>start</td><td>end</td><td>strand</td><td>MM</td><td>target_seq</td><td>PAM</td><td>alignment</td><td>distance</td><td>position</td><td>gene name</td><td>gene id</td></tr>
<tr><td>GU305914.1</td><td>60001</td><td>60023</td><td>+</td><td>0</td><td>GCGCGGCCAGAAGCCCGGAG</td><td>TGG</td><td>||||||||[||||||||||||]PAM</td><td>NA</td><td>Intergenic</td><td>NA</td><td>NA</td></tr>
<tr><td>T3975</td><td>AGCCCGGAGTGGTGCCCGCCAGG</td><td>1000</td><td></td><td></td><td></td><td></td><td></td><td></td><td></td><td></td><td></td></tr>
<tr><td>Oligo fwd</td><td>CACCAGCCCGGAGTGGTGCCCGCC</td><td></td><td></td><td></td><td></td><td></td><td></td><td></td><td></td><td></td><td></td></tr>
<tr><td>Oligo rev</td><td>AAACGGCGGGCACCACTCCGGGCT</td><td></td><td></td><td></td><td></td><td></td><td></td><td></td><td></td><td></td><td></td></tr>
<tr><td>Chromosome</td><td>start</td><td>end</td><td>strand</td><td>MM</td><td>target_seq</td><td>PAM</td><td>alignment</td><td>distance</td><td>position</td><td>gene name</td><td>gene id</td></tr>
<tr><td>GU305914.1</td><td>60012</td><td>60034</td><td>+</td><td>0</td><td>AGCCCGGAGTGGTGCCCGCC</td><td>AGG</td><td>||||||||[||||||||||||]PAM</td><td>NA</td><td>Intergenic</td><td>NA</td><td>NA</td></tr>
<tr><td>T3976</td><td>CGGAGTGGTGCCCGCCAGGTTGG</td><td>1000</td><td></td><td></td><td></td><td></td><td></td><td></td><td></td><td></td><td></td></tr>
<tr><td>Oligo fwd</td><td>CACCCGGAGTGGTGCCCGCCAGGT</td><td></td><td></td><td></td><td></td><td></td><td></td><td></td><td></td><td></td><td></td></tr>
<tr><td>Oligo rev</td><td>AAACACCTGGCGGGCACCACTCCG</td><td></td><td></td><td></td><td></td><td></td><td></td><td></td><td></td><td></td><td></td></tr>
<tr><td>Chromosome</td><td>start</td><td>end</td><td>strand</td><td>MM</td><td>target_seq</td><td>PAM</td><td>alignment</td><td>distance</td><td>position</td><td>gene name</td><td>gene id</td></tr>
<tr><td>GU305914.1</td><td>60016</td><td>60038</td><td>+</td><td>0</td><td>CGGAGTGGTGCCCGCCAGGT</td><td>TGG</td><td>||||||||[||||||||||||]PAM</td><td>NA</td><td>Intergenic</td><td>NA</td><td>NA</td></tr>
<tr><td>T3977</td><td>AGTGGTGCCCGCCAGGTTGGCGG</td><td>1000</td><td></td><td></td><td></td><td></td><td></td><td></td><td></td><td></td><td></td></tr>
<tr><td>Oligo fwd</td><td>CACCAGTGGTGCCCGCCAGGTTGG</td><td></td><td></td><td></td><td></td><td></td><td></td><td></td><td></td><td></td><td></td></tr>
<tr><td>Oligo rev</td><td>AAACCCAACCTGGCGGGCACCACT</td><td></td><td></td><td></td><td></td><td></td><td></td><td></td><td></td><td></td><td></td></tr>
<tr><td>Chromosome</td><td>start</td><td>end</td><td>strand</td><td>MM</td><td>target_seq</td><td>PAM</td><td>alignment</td><td>distance</td><td>position</td><td>gene name</td><td>gene id</td></tr>
<tr><td>GU305914.1</td><td>60019</td><td>60041</td><td>+</td><td>0</td><td>AGTGGTGCCCGCCAGGTTGG</td><td>CGG</td><td>||||||||[||||||||||||]PAM</td><td>NA</td><td>Intergenic</td><td>NA</td><td>NA</td></tr>
<tr><td>T3978</td><td>CGCCAGGTTGGCGGCTTCGTCGG</td><td>1000</td><td></td><td></td><td></td><td></td><td></td><td></td><td></td><td></td><td></td></tr>
<tr><td>Oligo fwd</td><td>CACCCGCCAGGTTGGCGGCTTCGT</td><td></td><td></td><td></td><td></td><td></td><td></td><td></td><td></td><td></td><td></td></tr>
<tr><td>Oligo rev</td><td>AAACACGAAGCCGCCAACCTGGCG</td><td></td><td></td><td></td><td></td><td></td><td></td><td></td><td></td><td></td><td></td></tr>
<tr><td>Chromosome</td><td>start</td><td>end</td><td>strand</td><td>MM</td><td>target_seq</td><td>PAM</td><td>alignment</td><td>distance</td><td>position</td><td>gene name</td><td>gene id</td></tr>
<tr><td>GU305914.1</td><td>60028</td><td>60050</td><td>+</td><td>0</td><td>CGCCAGGTTGGCGGCTTCGT</td><td>CGG</td><td>||||||||[||||||||||||]PAM</td><td>NA</td><td>Intergenic</td><td>NA</td><td>NA</td></tr>
<tr><td>T3979</td><td>GGCTTCGTCGGCCGAGCAGCCGG</td><td>1000</td><td></td><td></td><td></td><td></td><td></td><td></td><td></td><td></td><td></td></tr>
<tr><td>Oligo fwd</td><td>CACCGGCTTCGTCGGCCGAGCAGC</td><td></td><td></td><td></td><td></td><td></td><td></td><td></td><td></td><td></td><td></td></tr>
<tr><td>Oligo rev</td><td>AAACGCTGCTCGGCCGACGAAGCC</td><td></td><td></td><td></td><td></td><td></td><td></td><td></td><td></td><td></td><td></td></tr>
<tr><td>Chromosome</td><td>start</td><td>end</td><td>strand</td><td>MM</td><td>target_seq</td><td>PAM</td><td>alignment</td><td>distance</td><td>position</td><td>gene name</td><td>gene id</td></tr>
<tr><td>GU305914.1</td><td>60040</td><td>60062</td><td>+</td><td>0</td><td>GGCTTCGTCGGCCGAGCAGC</td><td>CGG</td><td>||||||||[||||||||||||]PAM</td><td>NA</td><td>Intergenic</td><td>NA</td><td>NA</td></tr>
</table>

| | | | | | | | | | | | |
|---|---|---|---|---|---|---|---|---|---|---|---|
| T3980 | GGCCGAGCAGCCGGTGATGAAGG | 1000 | | | | | | | | | |
| Oligo fwd | CACCGGCCGAGCAGCCGGTGATGA | | | | | | | | | | |
| Oligo rev | AAACTCATCACCGGCTGCTCGGCC | | | | | | | | | | |
| Chromosome | start | end | strand | MM | target_seq | PAM | alignment | distance | position | gene name | gene id |
| GU305914.1 | 60049 | 60071 | + | 0 | GGCCGAGCAGCCGGTGATGA | AGG | \|\|\|\|\|\|\|\|[\|\|\|\|\|\|\|\|\|\|\|\|]PAM | NA | Intergenic | NA | NA |
| GU305914.1 | 77455 | 77477 | + | 4 | CGACGAGCACCCGGTGATGG | TGG | -\|-\|\|\|\|\|[\|-\|\|\|\|\|\|\|\|\|-]PAM | NA | Intergenic | NA | NA |
| T3981 | CGAGCAGCCGGTGATGAAGGAGG | 1000 | | | | | | | | | |
| Oligo fwd | CACCCGAGCAGCCGGTGATGAAGG | | | | | | | | | | |
| Oligo rev | AAACCCTTCATCACCGGCTGCTCG | | | | | | | | | | |
| Chromosome | start | end | strand | MM | target_seq | PAM | alignment | distance | position | gene name | gene id |
| GU305914.1 | 60052 | 60074 | + | 0 | CGAGCAGCCGGTGATGAAGG | AGG | \|\|\|\|\|\|\|\|[\|\|\|\|\|\|\|\|\|\|\|\|]PAM | NA | Intergenic | NA | NA |
| T3982 | CGTCTTCACGTAGTTTCTCGAGG | 1000 | | | | | | | | | |
| Oligo fwd | CACCCGTCTTCACGTAGTTTCTCG | | | | | | | | | | |
| Oligo rev | AAACCGAGAAACTACGTGAAGACG | | | | | | | | | | |
| Chromosome | start | end | strand | MM | target_seq | PAM | alignment | distance | position | gene name | gene id |
| GU305914.1 | 60099 | 60121 | + | 0 | CGTCTTCACGTAGTTTCTCG | AGG | \|\|\|\|\|\|\|\|[\|\|\|\|\|\|\|\|\|\|\|\|]PAM | NA | Intergenic | NA | NA |
| T3983 | TAGTTTCTCGAGGAACTCCTTGG | 1000 | | | | | | | | | |
| Oligo fwd | CACCTAGTTTCTCGAGGAACTCCT | | | | | | | | | | |
| Oligo rev | AAACAGGAGTTCCTCGAGAAACTA | | | | | | | | | | |
| Chromosome | start | end | strand | MM | target_seq | PAM | alignment | distance | position | gene name | gene id |
| GU305914.1 | 60109 | 60131 | + | 0 | TAGTTTCTCGAGGAACTCCT | TGG | \|\|\|\|\|\|\|\|[\|\|\|\|\|\|\|\|\|\|\|\|]PAM | NA | Intergenic | NA | NA |
| T3984 | AACTCCTTGGTCATGCGACGCGG | 1000 | | | | | | | | | |
| Oligo fwd | CACCAACTCCTTGGTCATGCGACG | | | | | | | | | | |
| Oligo rev | AAACCGTCGCATGACCAAGGAGTT | | | | | | | | | | |
| Chromosome | start | end | strand | MM | target_seq | PAM | alignment | distance | position | gene name | gene id |
| GU305914.1 | 60122 | 60144 | + | 0 | AACTCCTTGGTCATGCGACG | CGG | \|\|\|\|\|\|\|\|[\|\|\|\|\|\|\|\|\|\|\|\|]PAM | NA | Intergenic | NA | NA |
| T3985 | ACTCCTTGGTCATGCGACGCGGG | 1000 | | | | | | | | | |
| Oligo fwd | CACCACTCCTTGGTCATGCGACGC | | | | | | | | | | |
| Oligo rev | AAACGCGTCGCATGACCAAGGAGT | | | | | | | | | | |
| Chromosome | start | end | strand | MM | target_seq | PAM | alignment | distance | position | gene name | gene id |
| GU305914.1 | 60123 | 60145 | + | 0 | ACTCCTTGGTCATGCGACGC | GGG | \|\|\|\|\|\|\|\|[\|\|\|\|\|\|\|\|\|\|\|\|]PAM | NA | Intergenic | NA | NA |
| T3986 | CGCGCTCGTCGTCGTCGAAGAGG | 1000 | | | | | | | | | |
| Oligo fwd | CACCCGCGCTCGTCGTCGTCGAAG | | | | | | | | | | |
| Oligo rev | AAACCTTCGACGACGACGAGCGCG | | | | | | | | | | |
| Chromosome | start | end | strand | MM | target_seq | PAM | alignment | distance | position | gene name | gene id |
| GU305914.1 | 60147 | 60169 | + | 0 | CGCGCTCGTCGTCGTCGAAG | AGG | \|\|\|\|\|\|\|\|[\|\|\|\|\|\|\|\|\|\|\|\|]PAM | NA | Intergenic | NA | NA |
| T3987 | CTCGTCGTCGTCGAAGAGGTCGG | 1000 | | | | | | | | | |
| Oligo fwd | CACCCTCGTCGTCGTCGAAGAGGT | | | | | | | | | | |
| Oligo rev | AAACACCTCTTCGACGACGACGAG | | | | | | | | | | |
| Chromosome | start | end | strand | MM | target_seq | PAM | alignment | distance | position | gene name | gene id |
| GU305914.1 | 60151 | 60173 | + | 0 | CTCGTCGTCGTCGAAGAGGT | CGG | \|\|\|\|\|\|\|\|[\|\|\|\|\|\|\|\|\|\|\|\|]PAM | NA | Intergenic | NA | NA |
| T3988 | TCGTCGTCGTCGAAGAGGTCGGG | 1000 | | | | | | | | | |
| Oligo fwd | CACCTCGTCGTCGTCGAAGAGGTC | | | | | | | | | | |
| Oligo rev | AAACGACCTCTTCGACGACGACGA | | | | | | | | | | |
| Chromosome | start | end | strand | MM | target_seq | PAM | alignment | distance | position | gene name | gene id |
| GU305914.1 | 60152 | 60174 | + | 0 | TCGTCGTCGTCGAAGAGGTC | GGG | \|\|\|\|\|\|\|\|[\|\|\|\|\|\|\|\|\|\|\|\|]PAM | NA | Intergenic | NA | NA |
| GU305914.1 | 18033 | 18055 | + | 4 | GTGTCGTAGTCGTAGAGGTC | TGG | --\|\|\|\|\|-[\|\|\|\|-\|\|\|\|\|\|\|]PAM | NA | Intergenic | NA | NA |
| T3989 | CGTCGTCGTCGAAGAGGTCGGGG | 1000 | | | | | | | | | |
| Oligo fwd | CACCCGTCGTCGTCGAAGAGGTCG | | | | | | | | | | |
| Oligo rev | AAACCGACCTCTTCGACGACGACG | | | | | | | | | | |
| Chromosome | start | end | strand | MM | target_seq | PAM | alignment | distance | position | gene name | gene id |
| GU305914.1 | 60153 | 60175 | + | 0 | CGTCGTCGTCGAAGAGGTCG | GGG | \|\|\|\|\|\|\|\|[\|\|\|\|\|\|\|\|\|\|\|\|]PAM | NA | Intergenic | NA | NA |
| T3990 | CGAAGAGGTCGGGGTAGAAGCGG | 1000 | | | | | | | | | |
| Oligo fwd | CACCCGAAGAGGTCGGGGTAGAAG | | | | | | | | | | |
| Oligo rev | AAACCTTCTACCCCGACCTCTTCG | | | | | | | | | | |
| Chromosome | start | end | strand | MM | target_seq | PAM | alignment | distance | position | gene name | gene id |
| GU305914.1 | 60162 | 60184 | + | 0 | CGAAGAGGTCGGGGTAGAAG | CGG | \|\|\|\|\|\|\|\|[\|\|\|\|\|\|\|\|\|\|\|\|]PAM | NA | Intergenic | NA | NA |
| T3991 | GAGGTCGGGGTAGAAGCGGCCGG | 1000 | | | | | | | | | |
| Oligo fwd | CACCGAGGTCGGGGTAGAAGCGGC | | | | | | | | | | |
| Oligo rev | AAACGCCGCTTCTACCCCGACCTC | | | | | | | | | | |
| Chromosome | start | end | strand | MM | target_seq | PAM | alignment | distance | position | gene name | gene id |
| GU305914.1 | 60166 | 60188 | + | 0 | GAGGTCGGGGTAGAAGCGGC | CGG | \|\|\|\|\|\|\|\|[\|\|\|\|\|\|\|\|\|\|\|\|]PAM | NA | Intergenic | NA | NA |
| T3992 | AGAAGCGGCCGGCCGCGTACAGG | 1000 | | | | | | | | | |
| Oligo fwd | CACCAGAAGCGGCCGGCCGCGTAC | | | | | | | | | | |
| Oligo rev | AAACGTACGCGGCCGGCCGCTTCT | | | | | | | | | | |
| Chromosome | start | end | strand | MM | target_seq | PAM | alignment | distance | position | gene name | gene id |
| GU305914.1 | 60177 | 60199 | + | 0 | AGAAGCGGCCGGCCGCGTAC | AGG | \|\|\|\|\|\|\|\|[\|\|\|\|\|\|\|\|\|\|\|\|]PAM | NA | Intergenic | NA | NA |
| T3993 | CCGGCCGCGTACAGGCTCTTAGG | 1000 | | | | | | | | | |
| Oligo fwd | CACCCCGGCCGCGTACAGGCTCTT | | | | | | | | | | |
| Oligo rev | AAACAAGAGCCTGTACGCGGCCGG | | | | | | | | | | |
| Chromosome | start | end | strand | MM | target_seq | PAM | alignment | distance | position | gene name | gene id |
| GU305914.1 | 60185 | 60207 | + | 0 | CCGGCCGCGTACAGGCTCTT | AGG | \|\|\|\|\|\|\|\|[\|\|\|\|\|\|\|\|\|\|\|\|]PAM | NA | Intergenic | NA | NA |
| T3994 | CGGCCGCGTACAGGCTCTTAGGG | 1000 | | | | | | | | | |
| Oligo fwd | CACCCGGCCGCGTACAGGCTCTTA | | | | | | | | | | |

| | | | | | | | | | | | |
|---|---|---|---|---|---|---|---|---|---|---|---|
| Oligo rev | AAACTAAGAGCCTGTACGCGGCCG | | | | | | | | | | |
| Chromosome | start | end | strand | MM | target_seq | PAM | alignment | distance | position | gene name | gene id |
| GU305914.1 | 60186 | 60208 | + | 0 | CGGCCGCGTACAGGCTCTTA | GGG | \|\|\|\|\|\|\|\|[\|\|\|\|\|\|\|\|\|\|\|\|]PAM | NA | Intergenic | NA | NA |
| T3995 | GTACAGGCTCTTAGGGAACTTGG | 1000 | | | | | | | | | |
| Oligo fwd | CACCGTACAGGCTCTTAGGGAACT | | | | | | | | | | |
| Oligo rev | AAACAGTTCCCTAAGAGCCTGTAC | | | | | | | | | | |
| Chromosome | start | end | strand | MM | target_seq | PAM | alignment | distance | position | gene name | gene id |
| GU305914.1 | 60193 | 60215 | + | 0 | GTACAGGCTCTTAGGGAACT | TGG | \|\|\|\|\|\|\|\|[\|\|\|\|\|\|\|\|\|\|\|\|]PAM | NA | Intergenic | NA | NA |
| T3996 | TACAGGCTCTTAGGGAACTTGGG | 1000 | | | | | | | | | |
| Oligo fwd | CACCTACAGGCTCTTAGGGAACTT | | | | | | | | | | |
| Oligo rev | AAACAAGTTCCCTAAGAGCCTGTA | | | | | | | | | | |
| Chromosome | start | end | strand | MM | target_seq | PAM | alignment | distance | position | gene name | gene id |
| GU305914.1 | 60194 | 60216 | + | 0 | TACAGGCTCTTAGGGAACTT | GGG | \|\|\|\|\|\|\|\|[\|\|\|\|\|\|\|\|\|\|\|\|]PAM | NA | Intergenic | NA | NA |
| T3997 | ACAGGCTCTTAGGGAACTTGGGG | 1000 | | | | | | | | | |
| Oligo fwd | CACCACAGGCTCTTAGGGAACTTG | | | | | | | | | | |
| Oligo rev | AAACCAAGTTCCCTAAGAGCCTGT | | | | | | | | | | |
| Chromosome | start | end | strand | MM | target_seq | PAM | alignment | distance | position | gene name | gene id |
| GU305914.1 | 60195 | 60217 | + | 0 | ACAGGCTCTTAGGGAACTTG | GGG | \|\|\|\|\|\|\|\|[\|\|\|\|\|\|\|\|\|\|\|\|]PAM | NA | Intergenic | NA | NA |
| T3998 | CTCTTAGGGAACTTGGGGAACGG | 1000 | | | | | | | | | |
| Oligo fwd | CACCCTCTTAGGGAACTTGGGGAA | | | | | | | | | | |
| Oligo rev | AAACTTCCCCAAGTTCCCTAAGAG | | | | | | | | | | |
| Chromosome | start | end | strand | MM | target_seq | PAM | alignment | distance | position | gene name | gene id |
| GU305914.1 | 60200 | 60222 | + | 0 | CTCTTAGGGAACTTGGGGAA | CGG | \|\|\|\|\|\|\|\|[\|\|\|\|\|\|\|\|\|\|\|\|]PAM | NA | Intergenic | NA | NA |
| T3999 | AGGGAACTTGGGGAACGGATCGG | 1000 | | | | | | | | | |
| Oligo fwd | CACCAGGGAACTTGGGGAACGGAT | | | | | | | | | | |
| Oligo rev | AAACATCCGTTCCCCAAGTTCCCT | | | | | | | | | | |
| Chromosome | start | end | strand | MM | target_seq | PAM | alignment | distance | position | gene name | gene id |
| GU305914.1 | 60205 | 60227 | + | 0 | AGGGAACTTGGGGAACGGAT | CGG | \|\|\|\|\|\|\|\|[\|\|\|\|\|\|\|\|\|\|\|\|]PAM | NA | Intergenic | NA | NA |
| T4000 | GATCGGCGCCCGCCATGCACAGG | 1000 | | | | | | | | | |
| Oligo fwd | CACCGATCGGCGCCCGCCATGCAC | | | | | | | | | | |
| Oligo rev | AAACGTGCATGGCGGGCGCCGATC | | | | | | | | | | |
| Chromosome | start | end | strand | MM | target_seq | PAM | alignment | distance | position | gene name | gene id |
| GU305914.1 | 60222 | 60244 | + | 0 | GATCGGCGCCCGCCATGCAC | AGG | \|\|\|\|\|\|\|\|[\|\|\|\|\|\|\|\|\|\|\|\|]PAM | NA | Intergenic | NA | NA |
| T4001 | CGCCATGCACAGGTCGACGCTGG | 1000 | | | | | | | | | |
| Oligo fwd | CACCCGCCATGCACAGGTCGACGC | | | | | | | | | | |
| Oligo rev | AAACGCGTCGACCTGTGCATGGCG | | | | | | | | | | |
| Chromosome | start | end | strand | MM | target_seq | PAM | alignment | distance | position | gene name | gene id |
| GU305914.1 | 60232 | 60254 | + | 0 | CGCCATGCACAGGTCGACGC | TGG | \|\|\|\|\|\|\|\|[\|\|\|\|\|\|\|\|\|\|\|\|]PAM | NA | Intergenic | NA | NA |
| T4002 | CAGGTCGACGCTGGCCCGCACGG | 1000 | | | | | | | | | |
| Oligo fwd | CACCCAGGTCGACGCTGGCCCGCA | | | | | | | | | | |
| Oligo rev | AAACTGCGGGCCAGCGTCGACCTG | | | | | | | | | | |
| Chromosome | start | end | strand | MM | target_seq | PAM | alignment | distance | position | gene name | gene id |
| GU305914.1 | 60241 | 60263 | + | 0 | CAGGTCGACGCTGGCCCGCA | CGG | \|\|\|\|\|\|\|\|[\|\|\|\|\|\|\|\|\|\|\|\|]PAM | NA | Intergenic | NA | NA |
| GU305914.1 | 141904 | 141926 | - | 4 | CATGTCGTCGCTGCCCAGCA | AGG | \|\|-\|\|\|\|-[\|\|\|\|\|-\|\|-\|\|\|]PAM | NA | Intergenic | NA | NA |
| T4003 | GTCGACGCTGGCCCGCACGGCGG | 1000 | | | | | | | | | |
| Oligo fwd | CACCGTCGACGCTGGCCCGCACGG | | | | | | | | | | |
| Oligo rev | AAACCCGTGCGGGCCAGCGTCGAC | | | | | | | | | | |
| Chromosome | start | end | strand | MM | target_seq | PAM | alignment | distance | position | gene name | gene id |
| GU305914.1 | 60244 | 60266 | + | 0 | GTCGACGCTGGCCCGCACGG | CGG | \|\|\|\|\|\|\|\|[\|\|\|\|\|\|\|\|\|\|\|\|]PAM | NA | Intergenic | NA | NA |
| T4004 | AGATAAACTCCGCGATACGCTGG | 1000 | | | | | | | | | |
| Oligo fwd | CACCAGATAAACTCCGCGATACGC | | | | | | | | | | |
| Oligo rev | AAACGCGTATCGCGGAGTTTATCT | | | | | | | | | | |
| Chromosome | start | end | strand | MM | target_seq | PAM | alignment | distance | position | gene name | gene id |
| GU305914.1 | 60273 | 60295 | + | 0 | AGATAAACTCCGCGATACGC | TGG | \|\|\|\|\|\|\|\|[\|\|\|\|\|\|\|\|\|\|\|\|]PAM | NA | Intergenic | NA | NA |
| T4005 | ACTCCGCGATACGCTGGTTGAGG | 1000 | | | | | | | | | |
| Oligo fwd | CACCACTCCGCGATACGCTGGTTG | | | | | | | | | | |
| Oligo rev | AAACCAACCAGCGTATCGCGGAGT | | | | | | | | | | |
| Chromosome | start | end | strand | MM | target_seq | PAM | alignment | distance | position | gene name | gene id |
| GU305914.1 | 60279 | 60301 | + | 0 | ACTCCGCGATACGCTGGTTG | AGG | \|\|\|\|\|\|\|\|[\|\|\|\|\|\|\|\|\|\|\|\|]PAM | NA | Intergenic | NA | NA |
| T4006 | GAGGTCGTAAGACTCCATGTCGG | 1000 | | | | | | | | | |
| Oligo fwd | CACCGAGGTCGTAAGACTCCATGT | | | | | | | | | | |
| Oligo rev | AAACACATGGAGTCTTACGACCTC | | | | | | | | | | |
| Chromosome | start | end | strand | MM | target_seq | PAM | alignment | distance | position | gene name | gene id |
| GU305914.1 | 60298 | 60320 | + | 0 | GAGGTCGTAAGACTCCATGT | CGG | \|\|\|\|\|\|\|\|[\|\|\|\|\|\|\|\|\|\|\|\|]PAM | NA | Intergenic | NA | NA |
| T4007 | TGTCGGCGAACTTCTGCCCGAGG | 1000 | | | | | | | | | |
| Oligo fwd | CACCTGTCGGCGAACTTCTGCCCG | | | | | | | | | | |
| Oligo rev | AAACCGGGCAGAAGTTCGCCGACA | | | | | | | | | | |
| Chromosome | start | end | strand | MM | target_seq | PAM | alignment | distance | position | gene name | gene id |
| GU305914.1 | 60315 | 60337 | + | 0 | TGTCGGCGAACTTCTGCCCG | AGG | \|\|\|\|\|\|\|\|[\|\|\|\|\|\|\|\|\|\|\|\|]PAM | NA | Intergenic | NA | NA |
| T4008 | CGAGGCGATTCATCACAGCGTGG | 1000 | | | | | | | | | |
| Oligo fwd | CACCCGAGGCGATTCATCACAGCG | | | | | | | | | | |
| Oligo rev | AAACCGCTGTGATGAATCGCCTCG | | | | | | | | | | |
| Chromosome | start | end | strand | MM | target_seq | PAM | alignment | distance | position | gene name | gene id |
| GU305914.1 | 60333 | 60355 | + | 0 | CGAGGCGATTCATCACAGCG | TGG | \|\|\|\|\|\|\|\|[\|\|\|\|\|\|\|\|\|\|\|\|]PAM | NA | Intergenic | NA | NA |

<table>
<tr><td>T4009</td><td>AGCCGACGACCCCGATGTGCAGG</td><td>1000</td><td></td><td></td><td></td><td></td><td></td><td></td><td></td><td></td><td></td></tr>
<tr><td>Oligo fwd</td><td>CACCAGCCGACGACCCCGATGTGC</td><td></td><td></td><td></td><td></td><td></td><td></td><td></td><td></td><td></td><td></td></tr>
<tr><td>Oligo rev</td><td>AAACGCACATCGGGGTCGTCGGCT</td><td></td><td></td><td></td><td></td><td></td><td></td><td></td><td></td><td></td><td></td></tr>
<tr><td>Chromosome</td><td>start</td><td>end</td><td>strand</td><td>MM</td><td>target_seq</td><td>PAM</td><td>alignment</td><td>distance</td><td>position</td><td>gene name</td><td>gene id</td></tr>
<tr><td>GU305914.1</td><td>60357</td><td>60379</td><td>+</td><td>0</td><td>AGCCGACGACCCCGATGTGC</td><td>AGG</td><td>||||||||[||||||||||||]PAM</td><td>NA</td><td>Intergenic</td><td>NA</td><td>NA</td></tr>
<tr><td>T4010</td><td>GCAGGTCGCGATAGAGCGTCAGG</td><td>1000</td><td></td><td></td><td></td><td></td><td></td><td></td><td></td><td></td><td></td></tr>
<tr><td>Oligo fwd</td><td>CACCGCAGGTCGCGATAGAGCGTC</td><td></td><td></td><td></td><td></td><td></td><td></td><td></td><td></td><td></td><td></td></tr>
<tr><td>Oligo rev</td><td>AAACGACGCTCTATCGCGACCTGC</td><td></td><td></td><td></td><td></td><td></td><td></td><td></td><td></td><td></td><td></td></tr>
<tr><td>Chromosome</td><td>start</td><td>end</td><td>strand</td><td>MM</td><td>target_seq</td><td>PAM</td><td>alignment</td><td>distance</td><td>position</td><td>gene name</td><td>gene id</td></tr>
<tr><td>GU305914.1</td><td>60375</td><td>60397</td><td>+</td><td>0</td><td>GCAGGTCGCGATAGAGCGTC</td><td>AGG</td><td>||||||||[||||||||||||]PAM</td><td>NA</td><td>Intergenic</td><td>NA</td><td>NA</td></tr>
<tr><td>T4011</td><td>ATAGAGCGTCAGGAGCTTGCCGG</td><td>1000</td><td></td><td></td><td></td><td></td><td></td><td></td><td></td><td></td><td></td></tr>
<tr><td>Oligo fwd</td><td>CACCATAGAGCGTCAGGAGCTTGC</td><td></td><td></td><td></td><td></td><td></td><td></td><td></td><td></td><td></td><td></td></tr>
<tr><td>Oligo rev</td><td>AAACGCAAGCTCCTGACGCTCTAT</td><td></td><td></td><td></td><td></td><td></td><td></td><td></td><td></td><td></td><td></td></tr>
<tr><td>Chromosome</td><td>start</td><td>end</td><td>strand</td><td>MM</td><td>target_seq</td><td>PAM</td><td>alignment</td><td>distance</td><td>position</td><td>gene name</td><td>gene id</td></tr>
<tr><td>GU305914.1</td><td>60385</td><td>60407</td><td>+</td><td>0</td><td>ATAGAGCGTCAGGAGCTTGC</td><td>CGG</td><td>||||||||[||||||||||||]PAM</td><td>NA</td><td>Intergenic</td><td>NA</td><td>NA</td></tr>
<tr><td>T4012</td><td>CTTGCCGGCGCCGTACGCGTTGG</td><td>1000</td><td></td><td></td><td></td><td></td><td></td><td></td><td></td><td></td><td></td></tr>
<tr><td>Oligo fwd</td><td>CACCCTTGCCGGCGCCGTACGCGT</td><td></td><td></td><td></td><td></td><td></td><td></td><td></td><td></td><td></td><td></td></tr>
<tr><td>Oligo rev</td><td>AAACACGCGTACGGCGCCGGCAAG</td><td></td><td></td><td></td><td></td><td></td><td></td><td></td><td></td><td></td><td></td></tr>
<tr><td>Chromosome</td><td>start</td><td>end</td><td>strand</td><td>MM</td><td>target_seq</td><td>PAM</td><td>alignment</td><td>distance</td><td>position</td><td>gene name</td><td>gene id</td></tr>
<tr><td>GU305914.1</td><td>60400</td><td>60422</td><td>+</td><td>0</td><td>CTTGCCGGCGCCGTACGCGT</td><td>TGG</td><td>||||||||[||||||||||||]PAM</td><td>NA</td><td>Intergenic</td><td>NA</td><td>NA</td></tr>
<tr><td>T4013</td><td>GCCGTACGCGTTGGTGCGCACGG</td><td>1000</td><td></td><td></td><td></td><td></td><td></td><td></td><td></td><td></td><td></td></tr>
<tr><td>Oligo fwd</td><td>CACCGCCGTACGCGTTGGTGCGCA</td><td></td><td></td><td></td><td></td><td></td><td></td><td></td><td></td><td></td><td></td></tr>
<tr><td>Oligo rev</td><td>AAACTGCGCACCAACGCGTACGGC</td><td></td><td></td><td></td><td></td><td></td><td></td><td></td><td></td><td></td><td></td></tr>
<tr><td>Chromosome</td><td>start</td><td>end</td><td>strand</td><td>MM</td><td>target_seq</td><td>PAM</td><td>alignment</td><td>distance</td><td>position</td><td>gene name</td><td>gene id</td></tr>
<tr><td>GU305914.1</td><td>60409</td><td>60431</td><td>+</td><td>0</td><td>GCCGTACGCGTTGGTGCGCA</td><td>CGG</td><td>||||||||[||||||||||||]PAM</td><td>NA</td><td>Intergenic</td><td>NA</td><td>NA</td></tr>
<tr><td>T4014</td><td>GCACGGTCATGTCGATGAGCAGG</td><td>1000</td><td></td><td></td><td></td><td></td><td></td><td></td><td></td><td></td><td></td></tr>
<tr><td>Oligo fwd</td><td>CACCGCACGGTCATGTCGATGAGC</td><td></td><td></td><td></td><td></td><td></td><td></td><td></td><td></td><td></td><td></td></tr>
<tr><td>Oligo rev</td><td>AAACGCTCATCGACATGACCGTGC</td><td></td><td></td><td></td><td></td><td></td><td></td><td></td><td></td><td></td><td></td></tr>
<tr><td>Chromosome</td><td>start</td><td>end</td><td>strand</td><td>MM</td><td>target_seq</td><td>PAM</td><td>alignment</td><td>distance</td><td>position</td><td>gene name</td><td>gene id</td></tr>
<tr><td>GU305914.1</td><td>60426</td><td>60448</td><td>+</td><td>0</td><td>GCACGGTCATGTCGATGAGC</td><td>AGG</td><td>||||||||[||||||||||||]PAM</td><td>NA</td><td>Intergenic</td><td>NA</td><td>NA</td></tr>
<tr><td>T4015</td><td>TACCCAGCACCACCGCGTCGCGG</td><td>1000</td><td></td><td></td><td></td><td></td><td></td><td></td><td></td><td></td><td></td></tr>
<tr><td>Oligo fwd</td><td>CACCTACCCAGCACCACCGCGTCG</td><td></td><td></td><td></td><td></td><td></td><td></td><td></td><td></td><td></td><td></td></tr>
<tr><td>Oligo rev</td><td>AAACCGACGCGGTGGTGCTGGGTA</td><td></td><td></td><td></td><td></td><td></td><td></td><td></td><td></td><td></td><td></td></tr>
<tr><td>Chromosome</td><td>start</td><td>end</td><td>strand</td><td>MM</td><td>target_seq</td><td>PAM</td><td>alignment</td><td>distance</td><td>position</td><td>gene name</td><td>gene id</td></tr>
<tr><td>GU305914.1</td><td>60450</td><td>60472</td><td>+</td><td>0</td><td>TACCCAGCACCACCGCGTCG</td><td>CGG</td><td>||||||||[||||||||||||]PAM</td><td>NA</td><td>Intergenic</td><td>NA</td><td>NA</td></tr>
<tr><td>T4016</td><td>GCACCACCGCGTCGCGGACCAGG</td><td>1000</td><td></td><td></td><td></td><td></td><td></td><td></td><td></td><td></td><td></td></tr>
<tr><td>Oligo fwd</td><td>CACCGCACCACCGCGTCGCGGACC</td><td></td><td></td><td></td><td></td><td></td><td></td><td></td><td></td><td></td><td></td></tr>
<tr><td>Oligo rev</td><td>AAACGGTCCGCGACGCGGTGGTGC</td><td></td><td></td><td></td><td></td><td></td><td></td><td></td><td></td><td></td><td></td></tr>
<tr><td>Chromosome</td><td>start</td><td>end</td><td>strand</td><td>MM</td><td>target_seq</td><td>PAM</td><td>alignment</td><td>distance</td><td>position</td><td>gene name</td><td>gene id</td></tr>
<tr><td>GU305914.1</td><td>60456</td><td>60478</td><td>+</td><td>0</td><td>GCACCACCGCGTCGCGGACC</td><td>AGG</td><td>||||||||[||||||||||||]PAM</td><td>NA</td><td>Intergenic</td><td>NA</td><td>NA</td></tr>
<tr><td>T4017</td><td>CACCACCGCGTCGCGGACCAGGG</td><td>1000</td><td></td><td></td><td></td><td></td><td></td><td></td><td></td><td></td><td></td></tr>
<tr><td>Oligo fwd</td><td>CACCCACCACCGCGTCGCGGACCA</td><td></td><td></td><td></td><td></td><td></td><td></td><td></td><td></td><td></td><td></td></tr>
<tr><td>Oligo rev</td><td>AAACTGGTCCGCGACGCGGTGGTG</td><td></td><td></td><td></td><td></td><td></td><td></td><td></td><td></td><td></td><td></td></tr>
<tr><td>Chromosome</td><td>start</td><td>end</td><td>strand</td><td>MM</td><td>target_seq</td><td>PAM</td><td>alignment</td><td>distance</td><td>position</td><td>gene name</td><td>gene id</td></tr>
<tr><td>GU305914.1</td><td>60457</td><td>60479</td><td>+</td><td>0</td><td>CACCACCGCGTCGCGGACCA</td><td>GGG</td><td>||||||||[||||||||||||]PAM</td><td>NA</td><td>Intergenic</td><td>NA</td><td>NA</td></tr>
<tr><td>T4018</td><td>CGTCGCGGACCAGGGACTCCAGG</td><td>1000</td><td></td><td></td><td></td><td></td><td></td><td></td><td></td><td></td><td></td></tr>
<tr><td>Oligo fwd</td><td>CACCCGTCGCGGACCAGGGACTCC</td><td></td><td></td><td></td><td></td><td></td><td></td><td></td><td></td><td></td><td></td></tr>
<tr><td>Oligo rev</td><td>AAACGGAGTCCCTGGTCCGCGACG</td><td></td><td></td><td></td><td></td><td></td><td></td><td></td><td></td><td></td><td></td></tr>
<tr><td>Chromosome</td><td>start</td><td>end</td><td>strand</td><td>MM</td><td>target_seq</td><td>PAM</td><td>alignment</td><td>distance</td><td>position</td><td>gene name</td><td>gene id</td></tr>
<tr><td>GU305914.1</td><td>60465</td><td>60487</td><td>+</td><td>0</td><td>CGTCGCGGACCAGGGACTCC</td><td>AGG</td><td>||||||||[||||||||||||]PAM</td><td>NA</td><td>Intergenic</td><td>NA</td><td>NA</td></tr>
<tr><td>T4019</td><td>GTCGCGGACCAGGGACTCCAGGG</td><td>1000</td><td></td><td></td><td></td><td></td><td></td><td></td><td></td><td></td><td></td></tr>
<tr><td>Oligo fwd</td><td>CACCGTCGCGGACCAGGGACTCCA</td><td></td><td></td><td></td><td></td><td></td><td></td><td></td><td></td><td></td><td></td></tr>
<tr><td>Oligo rev</td><td>AAACTGGAGTCCCTGGTCCGCGAC</td><td></td><td></td><td></td><td></td><td></td><td></td><td></td><td></td><td></td><td></td></tr>
<tr><td>Chromosome</td><td>start</td><td>end</td><td>strand</td><td>MM</td><td>target_seq</td><td>PAM</td><td>alignment</td><td>distance</td><td>position</td><td>gene name</td><td>gene id</td></tr>
<tr><td>GU305914.1</td><td>60466</td><td>60488</td><td>+</td><td>0</td><td>GTCGCGGACCAGGGACTCCA</td><td>GGG</td><td>||||||||[||||||||||||]PAM</td><td>NA</td><td>Intergenic</td><td>NA</td><td>NA</td></tr>
<tr><td>T4020</td><td>CCAGGGCCTTGAAGCTGAAGCGG</td><td>1000</td><td></td><td></td><td></td><td></td><td></td><td></td><td></td><td></td><td></td></tr>
<tr><td>Oligo fwd</td><td>CACCCCAGGGCCTTGAAGCTGAAG</td><td></td><td></td><td></td><td></td><td></td><td></td><td></td><td></td><td></td><td></td></tr>
<tr><td>Oligo rev</td><td>AAACCTTCAGCTTCAAGGCCCTGG</td><td></td><td></td><td></td><td></td><td></td><td></td><td></td><td></td><td></td><td></td></tr>
<tr><td>Chromosome</td><td>start</td><td>end</td><td>strand</td><td>MM</td><td>target_seq</td><td>PAM</td><td>alignment</td><td>distance</td><td>position</td><td>gene name</td><td>gene id</td></tr>
<tr><td>GU305914.1</td><td>60483</td><td>60505</td><td>+</td><td>0</td><td>CCAGGGCCTTGAAGCTGAAG</td><td>CGG</td><td>||||||||[||||||||||||]PAM</td><td>NA</td><td>Intergenic</td><td>NA</td><td>NA</td></tr>
<tr><td>T4021</td><td>TGAAGCGGCAACCGAGACCGAGG</td><td>1000</td><td></td><td></td><td></td><td></td><td></td><td></td><td></td><td></td><td></td></tr>
<tr><td>Oligo fwd</td><td>CACCTGAAGCGGCAACCGAGACCG</td><td></td><td></td><td></td><td></td><td></td><td></td><td></td><td></td><td></td><td></td></tr>
<tr><td>Oligo rev</td><td>AAACCGGTCTCGGTTGCCGCTTCA</td><td></td><td></td><td></td><td></td><td></td><td></td><td></td><td></td><td></td><td></td></tr>
<tr><td>Chromosome</td><td>start</td><td>end</td><td>strand</td><td>MM</td><td>target_seq</td><td>PAM</td><td>alignment</td><td>distance</td><td>position</td><td>gene name</td><td>gene id</td></tr>
<tr><td>GU305914.1</td><td>60498</td><td>60520</td><td>+</td><td>0</td><td>TGAAGCGGCAACCGAGACCG</td><td>AGG</td><td>||||||||[||||||||||||]PAM</td><td>NA</td><td>Intergenic</td><td>NA</td><td>NA</td></tr>
<tr><td>T4022</td><td>GAGACCGAGGACCACGTCGACGG</td><td>1000</td><td></td><td></td><td></td><td></td><td></td><td></td><td></td><td></td><td></td></tr>
<tr><td>Oligo fwd</td><td>CACCGAGACCGAGGACCACGTCGA</td><td></td><td></td><td></td><td></td><td></td><td></td><td></td><td></td><td></td><td></td></tr>
<tr><td>Oligo rev</td><td>AAACTCGACGTGGTCCTCGGTCTC</td><td></td><td></td><td></td><td></td><td></td><td></td><td></td><td></td><td></td><td></td></tr>
<tr><td>Chromosome</td><td>start</td><td>end</td><td>strand</td><td>MM</td><td>target_seq</td><td>PAM</td><td>alignment</td><td>distance</td><td>position</td><td>gene name</td><td>gene id</td></tr>
<tr><td>GU305914.1</td><td>60511</td><td>60533</td><td>+</td><td>0</td><td>GAGACCGAGGACCACGTCGA</td><td>CGG</td><td>||||||||[||||||||||||]PAM</td><td>NA</td><td>Intergenic</td><td>NA</td><td>NA</td></tr>
<tr><td>T4023</td><td>TCTCGACTGCCGCGCCCTCCAGG</td><td>1000</td><td></td><td></td><td></td><td></td><td></td><td></td><td></td><td></td><td></td></tr>
<tr><td>Oligo fwd</td><td>CACCTCTCGACTGCCGCGCCCTCC</td><td></td><td></td><td></td><td></td><td></td><td></td><td></td><td></td><td></td><td></td></tr>
<tr><td>Oligo rev</td><td>AAACGGAGGGCGCGGCAGTCGAGA</td><td></td><td></td><td></td><td></td><td></td><td></td><td></td><td></td><td></td><td></td></tr>
<tr><td>Chromosome</td><td>start</td><td>end</td><td>strand</td><td>MM</td><td>target_seq</td><td>PAM</td><td>alignment</td><td>distance</td><td>position</td><td>gene name</td><td>gene id</td></tr>
<tr><td>GU305914.1</td><td>60537</td><td>60559</td><td>+</td><td>0</td><td>TCTCGACTGCCGCGCCCTCC</td><td>AGG</td><td>||||||||[||||||||||||]PAM</td><td>NA</td><td>Intergenic</td><td>NA</td><td>NA</td></tr>
</table>

<table>
<tr><td>T4024</td><td>CCTCCAGGACGCAGTTCTCCAGG</td><td>1000</td><td></td><td></td><td></td><td></td><td></td><td></td><td></td><td></td><td></td></tr>
<tr><td>Oligo fwd</td><td>CACCCCTCCAGGACGCAGTTCTCC</td><td></td><td></td><td></td><td></td><td></td><td></td><td></td><td></td><td></td><td></td></tr>
<tr><td>Oligo rev</td><td>AAACGGAGAACTGCGTCCTGGAGG</td><td></td><td></td><td></td><td></td><td></td><td></td><td></td><td></td><td></td><td></td></tr>
<tr><td>Chromosome</td><td>start</td><td>end</td><td>strand</td><td>MM</td><td>target_seq</td><td>PAM</td><td>alignment</td><td>distance</td><td>position</td><td>gene name</td><td>gene id</td></tr>
<tr><td>GU305914.1</td><td>60552</td><td>60574</td><td>+</td><td>0</td><td>CCTCCAGGACGCAGTTCTCC</td><td>AGG</td><td>||||||||[||||||||||||]PAM</td><td>NA</td><td>Intergenic</td><td>NA</td><td>NA</td></tr>
<tr><td>T4025</td><td>CCAGGTTGATGCACAGCTCGTGG</td><td>1000</td><td></td><td></td><td></td><td></td><td></td><td></td><td></td><td></td><td></td></tr>
<tr><td>Oligo fwd</td><td>CACCCCAGGTTGATGCACAGCTCG</td><td></td><td></td><td></td><td></td><td></td><td></td><td></td><td></td><td></td><td></td></tr>
<tr><td>Oligo rev</td><td>AAACCGAGCTGTGCATCAACCTGG</td><td></td><td></td><td></td><td></td><td></td><td></td><td></td><td></td><td></td><td></td></tr>
<tr><td>Chromosome</td><td>start</td><td>end</td><td>strand</td><td>MM</td><td>target_seq</td><td>PAM</td><td>alignment</td><td>distance</td><td>position</td><td>gene name</td><td>gene id</td></tr>
<tr><td>GU305914.1</td><td>60570</td><td>60592</td><td>+</td><td>0</td><td>CCAGGTTGATGCACAGCTCG</td><td>TGG</td><td>||||||||[||||||||||||]PAM</td><td>NA</td><td>Intergenic</td><td>NA</td><td>NA</td></tr>
<tr><td>T4026</td><td>GTTGATGCACAGCTCGTGGATGG</td><td>1000</td><td></td><td></td><td></td><td></td><td></td><td></td><td></td><td></td><td></td></tr>
<tr><td>Oligo fwd</td><td>CACCGTTGATGCACAGCTCGTGGA</td><td></td><td></td><td></td><td></td><td></td><td></td><td></td><td></td><td></td><td></td></tr>
<tr><td>Oligo rev</td><td>AAACTCCACGAGCTGTGCATCAAC</td><td></td><td></td><td></td><td></td><td></td><td></td><td></td><td></td><td></td><td></td></tr>
<tr><td>Chromosome</td><td>start</td><td>end</td><td>strand</td><td>MM</td><td>target_seq</td><td>PAM</td><td>alignment</td><td>distance</td><td>position</td><td>gene name</td><td>gene id</td></tr>
<tr><td>GU305914.1</td><td>60574</td><td>60596</td><td>+</td><td>0</td><td>GTTGATGCACAGCTCGTGGA</td><td>TGG</td><td>||||||||[||||||||||||]PAM</td><td>NA</td><td>Intergenic</td><td>NA</td><td>NA</td></tr>
<tr><td>T4027</td><td>TTGATGCACAGCTCGTGGATGGG</td><td>1000</td><td></td><td></td><td></td><td></td><td></td><td></td><td></td><td></td><td></td></tr>
<tr><td>Oligo fwd</td><td>CACCTTGATGCACAGCTCGTGGAT</td><td></td><td></td><td></td><td></td><td></td><td></td><td></td><td></td><td></td><td></td></tr>
<tr><td>Oligo rev</td><td>AAACATCCACGAGCTGTGCATCAA</td><td></td><td></td><td></td><td></td><td></td><td></td><td></td><td></td><td></td><td></td></tr>
<tr><td>Chromosome</td><td>start</td><td>end</td><td>strand</td><td>MM</td><td>target_seq</td><td>PAM</td><td>alignment</td><td>distance</td><td>position</td><td>gene name</td><td>gene id</td></tr>
<tr><td>GU305914.1</td><td>60575</td><td>60597</td><td>+</td><td>0</td><td>TTGATGCACAGCTCGTGGAT</td><td>GGG</td><td>||||||||[||||||||||||]PAM</td><td>NA</td><td>Intergenic</td><td>NA</td><td>NA</td></tr>
<tr><td>T4028</td><td>TGATGCACAGCTCGTGGATGGGG</td><td>1000</td><td></td><td></td><td></td><td></td><td></td><td></td><td></td><td></td><td></td></tr>
<tr><td>Oligo fwd</td><td>CACCTGATGCACAGCTCGTGGATG</td><td></td><td></td><td></td><td></td><td></td><td></td><td></td><td></td><td></td><td></td></tr>
<tr><td>Oligo rev</td><td>AAACCATCCACGAGCTGTGCATCA</td><td></td><td></td><td></td><td></td><td></td><td></td><td></td><td></td><td></td><td></td></tr>
<tr><td>Chromosome</td><td>start</td><td>end</td><td>strand</td><td>MM</td><td>target_seq</td><td>PAM</td><td>alignment</td><td>distance</td><td>position</td><td>gene name</td><td>gene id</td></tr>
<tr><td>GU305914.1</td><td>60576</td><td>60598</td><td>+</td><td>0</td><td>TGATGCACAGCTCGTGGATG</td><td>GGG</td><td>||||||||[||||||||||||]PAM</td><td>NA</td><td>Intergenic</td><td>NA</td><td>NA</td></tr>
<tr><td>T4029</td><td>GATGCACAGCTCGTGGATGGGGG</td><td>1000</td><td></td><td></td><td></td><td></td><td></td><td></td><td></td><td></td><td></td></tr>
<tr><td>Oligo fwd</td><td>CACCGATGCACAGCTCGTGGATGG</td><td></td><td></td><td></td><td></td><td></td><td></td><td></td><td></td><td></td><td></td></tr>
<tr><td>Oligo rev</td><td>AAACCCATCCACGAGCTGTGCATC</td><td></td><td></td><td></td><td></td><td></td><td></td><td></td><td></td><td></td><td></td></tr>
<tr><td>Chromosome</td><td>start</td><td>end</td><td>strand</td><td>MM</td><td>target_seq</td><td>PAM</td><td>alignment</td><td>distance</td><td>position</td><td>gene name</td><td>gene id</td></tr>
<tr><td>GU305914.1</td><td>60577</td><td>60599</td><td>+</td><td>0</td><td>GATGCACAGCTCGTGGATGG</td><td>GGG</td><td>||||||||[||||||||||||]PAM</td><td>NA</td><td>Intergenic</td><td>NA</td><td>NA</td></tr>
<tr><td>T4030</td><td>GGTACAGCCGACGAAGCCGCCGG</td><td>1000</td><td></td><td></td><td></td><td></td><td></td><td></td><td></td><td></td><td></td></tr>
<tr><td>Oligo fwd</td><td>CACCGGTACAGCCGACGAAGCCGC</td><td></td><td></td><td></td><td></td><td></td><td></td><td></td><td></td><td></td><td></td></tr>
<tr><td>Oligo rev</td><td>AAACGCGGCTTCGTCGGCTGTACC</td><td></td><td></td><td></td><td></td><td></td><td></td><td></td><td></td><td></td><td></td></tr>
<tr><td>Chromosome</td><td>start</td><td>end</td><td>strand</td><td>MM</td><td>target_seq</td><td>PAM</td><td>alignment</td><td>distance</td><td>position</td><td>gene name</td><td>gene id</td></tr>
<tr><td>GU305914.1</td><td>60598</td><td>60620</td><td>+</td><td>0</td><td>GGTACAGCCGACGAAGCCGC</td><td>CGG</td><td>||||||||[||||||||||||]PAM</td><td>NA</td><td>Intergenic</td><td>NA</td><td>NA</td></tr>
<tr><td>GU305914.1</td><td>137037</td><td>137059</td><td>-</td><td>4</td><td>GACGCAGCCGACGAAGCGGC</td><td>CGG</td><td>|---||||[|||||||||-||]PAM</td><td>NA</td><td>Intergenic</td><td>NA</td><td>NA</td></tr>
<tr><td>T4031</td><td>GACGAAGCCGCCGGAGACGACGG</td><td>1000</td><td></td><td></td><td></td><td></td><td></td><td></td><td></td><td></td><td></td></tr>
<tr><td>Oligo fwd</td><td>CACCGACGAAGCCGCCGGAGACGA</td><td></td><td></td><td></td><td></td><td></td><td></td><td></td><td></td><td></td><td></td></tr>
<tr><td>Oligo rev</td><td>AAACTCGTCTCCGGCGGCTTCGTC</td><td></td><td></td><td></td><td></td><td></td><td></td><td></td><td></td><td></td><td></td></tr>
<tr><td>Chromosome</td><td>start</td><td>end</td><td>strand</td><td>MM</td><td>target_seq</td><td>PAM</td><td>alignment</td><td>distance</td><td>position</td><td>gene name</td><td>gene id</td></tr>
<tr><td>GU305914.1</td><td>60607</td><td>60629</td><td>+</td><td>0</td><td>GACGAAGCCGCCGGAGACGA</td><td>CGG</td><td>||||||||[||||||||||||]PAM</td><td>NA</td><td>Intergenic</td><td>NA</td><td>NA</td></tr>
<tr><td>GU305914.1</td><td>192825</td><td>192847</td><td>+</td><td>4</td><td>GATGGAGTCGTCGGAGACGA</td><td>AGG</td><td>||-|-||-[||-|||||||||]PAM</td><td>NA</td><td>Intergenic</td><td>NA</td><td>NA</td></tr>
<tr><td>T4032</td><td>GCCGGAGACGACGGAGACGCCGG</td><td>1000</td><td></td><td></td><td></td><td></td><td></td><td></td><td></td><td></td><td></td></tr>
<tr><td>Oligo fwd</td><td>CACCGCCGGAGACGACGGAGACGC</td><td></td><td></td><td></td><td></td><td></td><td></td><td></td><td></td><td></td><td></td></tr>
<tr><td>Oligo rev</td><td>AAACGCGTCTCCGTCGTCTCCGGC</td><td></td><td></td><td></td><td></td><td></td><td></td><td></td><td></td><td></td><td></td></tr>
<tr><td>Chromosome</td><td>start</td><td>end</td><td>strand</td><td>MM</td><td>target_seq</td><td>PAM</td><td>alignment</td><td>distance</td><td>position</td><td>gene name</td><td>gene id</td></tr>
<tr><td>GU305914.1</td><td>60616</td><td>60638</td><td>+</td><td>0</td><td>GCCGGAGACGACGGAGACGC</td><td>CGG</td><td>||||||||[||||||||||||]PAM</td><td>NA</td><td>Intergenic</td><td>NA</td><td>NA</td></tr>
<tr><td>T4033</td><td>CCGGAGACGACGGAGACGCCGGG</td><td>1000</td><td></td><td></td><td></td><td></td><td></td><td></td><td></td><td></td><td></td></tr>
<tr><td>Oligo fwd</td><td>CACCCCGGAGACGACGGAGACGCC</td><td></td><td></td><td></td><td></td><td></td><td></td><td></td><td></td><td></td><td></td></tr>
<tr><td>Oligo rev</td><td>AAACGGCGTCTCCGTCGTCTCCGG</td><td></td><td></td><td></td><td></td><td></td><td></td><td></td><td></td><td></td><td></td></tr>
<tr><td>Chromosome</td><td>start</td><td>end</td><td>strand</td><td>MM</td><td>target_seq</td><td>PAM</td><td>alignment</td><td>distance</td><td>position</td><td>gene name</td><td>gene id</td></tr>
<tr><td>GU305914.1</td><td>60617</td><td>60639</td><td>+</td><td>0</td><td>CCGGAGACGACGGAGACGCC</td><td>GGG</td><td>||||||||[||||||||||||]PAM</td><td>NA</td><td>Intergenic</td><td>NA</td><td>NA</td></tr>
<tr><td>T4034</td><td>GGAGACGCCGGGTCCGCACACGG</td><td>1000</td><td></td><td></td><td></td><td></td><td></td><td></td><td></td><td></td><td></td></tr>
<tr><td>Oligo fwd</td><td>CACCGGAGACGCCGGGTCCGCACA</td><td></td><td></td><td></td><td></td><td></td><td></td><td></td><td></td><td></td><td></td></tr>
<tr><td>Oligo rev</td><td>AAACTGTGCGGACCCGGCGTCTCC</td><td></td><td></td><td></td><td></td><td></td><td></td><td></td><td></td><td></td><td></td></tr>
<tr><td>Chromosome</td><td>start</td><td>end</td><td>strand</td><td>MM</td><td>target_seq</td><td>PAM</td><td>alignment</td><td>distance</td><td>position</td><td>gene name</td><td>gene id</td></tr>
<tr><td>GU305914.1</td><td>60628</td><td>60650</td><td>+</td><td>0</td><td>GGAGACGCCGGGTCCGCACA</td><td>CGG</td><td>||||||||[||||||||||||]PAM</td><td>NA</td><td>Intergenic</td><td>NA</td><td>NA</td></tr>
<tr><td>T4035</td><td>GACGCCGGGTCCGCACACGGCGG</td><td>1000</td><td></td><td></td><td></td><td></td><td></td><td></td><td></td><td></td><td></td></tr>
<tr><td>Oligo fwd</td><td>CACCGACGCCGGGTCCGCACACGG</td><td></td><td></td><td></td><td></td><td></td><td></td><td></td><td></td><td></td><td></td></tr>
<tr><td>Oligo rev</td><td>AAACCCGTGTGCGGACCCGGCGTC</td><td></td><td></td><td></td><td></td><td></td><td></td><td></td><td></td><td></td><td></td></tr>
<tr><td>Chromosome</td><td>start</td><td>end</td><td>strand</td><td>MM</td><td>target_seq</td><td>PAM</td><td>alignment</td><td>distance</td><td>position</td><td>gene name</td><td>gene id</td></tr>
<tr><td>GU305914.1</td><td>60631</td><td>60653</td><td>+</td><td>0</td><td>GACGCCGGGTCCGCACACGG</td><td>CGG</td><td>||||||||[||||||||||||]PAM</td><td>NA</td><td>Intergenic</td><td>NA</td><td>NA</td></tr>
<tr><td>T4036</td><td>ACGCCGGGTCCGCACACGGCGGG</td><td>1000</td><td></td><td></td><td></td><td></td><td></td><td></td><td></td><td></td><td></td></tr>
<tr><td>Oligo fwd</td><td>CACCACGCCGGGTCCGCACACGGC</td><td></td><td></td><td></td><td></td><td></td><td></td><td></td><td></td><td></td><td></td></tr>
<tr><td>Oligo rev</td><td>AAACGCCGTGTGCGGACCCGGCGT</td><td></td><td></td><td></td><td></td><td></td><td></td><td></td><td></td><td></td><td></td></tr>
<tr><td>Chromosome</td><td>start</td><td>end</td><td>strand</td><td>MM</td><td>target_seq</td><td>PAM</td><td>alignment</td><td>distance</td><td>position</td><td>gene name</td><td>gene id</td></tr>
<tr><td>GU305914.1</td><td>60632</td><td>60654</td><td>+</td><td>0</td><td>ACGCCGGGTCCGCACACGGC</td><td>GGG</td><td>||||||||[||||||||||||]PAM</td><td>NA</td><td>Intergenic</td><td>NA</td><td>NA</td></tr>
<tr><td>T4037</td><td>CGCCGGGTCCGCACACGGCGGGG</td><td>1000</td><td></td><td></td><td></td><td></td><td></td><td></td><td></td><td></td><td></td></tr>
<tr><td>Oligo fwd</td><td>CACCCGCCGGGTCCGCACACGGCG</td><td></td><td></td><td></td><td></td><td></td><td></td><td></td><td></td><td></td><td></td></tr>
<tr><td>Oligo rev</td><td>AAACCGCCGTGTGCGGACCCGGCG</td><td></td><td></td><td></td><td></td><td></td><td></td><td></td><td></td><td></td><td></td></tr>
<tr><td>Chromosome</td><td>start</td><td>end</td><td>strand</td><td>MM</td><td>target_seq</td><td>PAM</td><td>alignment</td><td>distance</td><td>position</td><td>gene name</td><td>gene id</td></tr>
<tr><td>GU305914.1</td><td>60633</td><td>60655</td><td>+</td><td>0</td><td>CGCCGGGTCCGCACACGGCG</td><td>GGG</td><td>||||||||[||||||||||||]PAM</td><td>NA</td><td>Intergenic</td><td>NA</td><td>NA</td></tr>
<tr><td>T4038</td><td>GGCGGGGCGCCCGATGTTCTCGG</td><td>1000</td><td></td><td></td><td></td><td></td><td></td><td></td><td></td><td></td><td></td></tr>
<tr><td>Oligo fwd</td><td>CACCGGCGGGGCGCCCGATGTTCT</td><td></td><td></td><td></td><td></td><td></td><td></td><td></td><td></td><td></td><td></td></tr>
</table>

Oligo rev AAACAGAACATCGGGCGCCCCGCC
Chromosome start end strand MM target_seq PAM alignment distance position gene name gene id
GU305914.1 60649 60671 + 0 GGCGGGGCGCCCGATGTTCT CGG ||||||||[||||||||||||]PAM NA Intergenic NA NA

T4039 GCGGGGCGCCCGATGTTCTCGGG 1000
Oligo fwd CACCGCGGGGCGCCCGATGTTCTC
Oligo rev AAACGAGAACATCGGGCGCCCCGC
Chromosome start end strand MM target_seq PAM alignment distance position gene name gene id
GU305914.1 60650 60672 + 0 GCGGGGCGCCCGATGTTCTC GGG ||||||||[||||||||||||]PAM NA Intergenic NA NA

T4040 CGGGGCGCCCGATGTTCTCGGGG 1000
Oligo fwd CACCCGGGGCGCCCGATGTTCTCG
Oligo rev AAACCGAGAACATCGGGCGCCCCG
Chromosome start end strand MM target_seq PAM alignment distance position gene name gene id
GU305914.1 60651 60673 + 0 CGGGGCGCCCGATGTTCTCG GGG ||||||||[||||||||||||]PAM NA Intergenic NA NA
GU305914.1 197031 197053 + 4 CGTCGCGCCCGCTGTTCTCC TGG ||--||||[|||-|||||||-]PAM NA Intergenic NA NA

T4041 ATCATACTCTTGACGTTGTGAGG 1000
Oligo fwd CACCATCATACTCTTGACGTTGTG
Oligo rev AAACCACAACGTCAAGAGTATGAT
Chromosome start end strand MM target_seq PAM alignment distance position gene name gene id
GU305914.1 60674 60696 + 0 ATCATACTCTTGACGTTGTG AGG ||||||||[||||||||||||]PAM NA Intergenic NA NA

T4042 TCATACTCTTGACGTTGTGAGGG 1000
Oligo fwd CACCTCATACTCTTGACGTTGTGA
Oligo rev AAACTCACAACGTCAAGAGTATGA
Chromosome start end strand MM target_seq PAM alignment distance position gene name gene id
GU305914.1 60675 60697 + 0 TCATACTCTTGACGTTGTGA GGG ||||||||[||||||||||||]PAM NA Intergenic NA NA

T4043 GACGTTGTGAGGGAAGACCACGG 1000
Oligo fwd CACCGACGTTGTGAGGGAAGACCA
Oligo rev AAACTGGTCTTCCCTCACAACGTC
Chromosome start end strand MM target_seq PAM alignment distance position gene name gene id
GU305914.1 60685 60707 + 0 GACGTTGTGAGGGAAGACCA CGG ||||||||[||||||||||||]PAM NA Intergenic NA NA

T4044 GCACGCGTCGAGCTGACAGACGG 1000
Oligo fwd CACCGCACGCGTCGAGCTGACAGA
Oligo rev AAACTCTGTCAGCTCGACGCGTGC
Chromosome start end strand MM target_seq PAM alignment distance position gene name gene id
GU305914.1 60727 60749 + 0 GCACGCGTCGAGCTGACAGA CGG ||||||||[||||||||||||]PAM NA Intergenic NA NA

T4045 GAGCTGACAGACGGCCCACCAGG 1000
Oligo fwd CACCGAGCTGACAGACGGCCCACC
Oligo rev AAACGGTGGGCCGTCTGTCAGCTC
Chromosome start end strand MM target_seq PAM alignment distance position gene name gene id
GU305914.1 60736 60758 + 0 GAGCTGACAGACGGCCCACC AGG ||||||||[||||||||||||]PAM NA Intergenic NA NA

T4046 AGCTGACAGACGGCCCACCAGGG 1000
Oligo fwd CACCAGCTGACAGACGGCCCACCA
Oligo rev AAACTGGTGGGCCGTCTGTCAGCT
Chromosome start end strand MM target_seq PAM alignment distance position gene name gene id
GU305914.1 60737 60759 + 0 AGCTGACAGACGGCCCACCA GGG ||||||||[||||||||||||]PAM NA Intergenic NA NA

T4047 GACGGCCCACCAGGGCGAGACGG 1000
Oligo fwd CACCGACGGCCCACCAGGGCGAGA
Oligo rev AAACTCTCGCCCTGGTGGGCCGTC
Chromosome start end strand MM target_seq PAM alignment distance position gene name gene id
GU305914.1 60745 60767 + 0 GACGGCCCACCAGGGCGAGA CGG ||||||||[||||||||||||]PAM NA Intergenic NA NA

T4048 CCACCAGGGCGAGACGGTGATGG 1000
Oligo fwd CACCCCACCAGGGCGAGACGGTGA
Oligo rev AAACTCACCGTCTCGCCCTGGTGG
Chromosome start end strand MM target_seq PAM alignment distance position gene name gene id
GU305914.1 60751 60773 + 0 CCACCAGGGCGAGACGGTGA TGG ||||||||[||||||||||||]PAM NA Intergenic NA NA

T4049 CCATCTTCTCGTAGACGTTACGG 1000
Oligo fwd CACCCCATCTTCTCGTAGACGTTA
Oligo rev AAACTAACGTCTACGAGAAGATGG
Chromosome start end strand MM target_seq PAM alignment distance position gene name gene id
GU305914.1 60792 60814 + 0 CCATCTTCTCGTAGACGTTA CGG ||||||||[||||||||||||]PAM NA Intergenic NA NA

T4050 CTTCTCGTAGACGTTACGGAAGG 1000
Oligo fwd CACCCTTCTCGTAGACGTTACGGA
Oligo rev AAACTCCGTAACGTCTACGAGAAG
Chromosome start end strand MM target_seq PAM alignment distance position gene name gene id
GU305914.1 60796 60818 + 0 CTTCTCGTAGACGTTACGGA AGG ||||||||[||||||||||||]PAM NA Intergenic NA NA

T4051 AGACGTTACGGAAGGACTTGAGG 1000
Oligo fwd CACCAGACGTTACGGAAGGACTTG
Oligo rev AAACCAAGTCCTTCCGTAACGTCT
Chromosome start end strand MM target_seq PAM alignment distance position gene name gene id
GU305914.1 60804 60826 + 0 AGACGTTACGGAAGGACTTG AGG ||||||||[||||||||||||]PAM NA Intergenic NA NA

T4052 TACGGAAGGACTTGAGGTCGAGG 1000
Oligo fwd CACCTACGGAAGGACTTGAGGTCG
Oligo rev AAACCGACCTCAAGTCCTTCCGTA
Chromosome start end strand MM target_seq PAM alignment distance position gene name gene id
GU305914.1 60810 60832 + 0 TACGGAAGGACTTGAGGTCG AGG ||||||||[||||||||||||]PAM NA Intergenic NA NA

<table>
<tr><td>T4053</td><td>ACTTGAGGTCGAGGCCGCACAGG</td><td></td><td>1000</td><td></td><td></td><td></td><td></td><td></td><td></td><td></td><td></td><td></td></tr>
<tr><td>Oligo fwd</td><td>CACCACTTGAGGTCGAGGCCGCAC</td><td></td><td></td><td></td><td></td><td></td><td></td><td></td><td></td><td></td><td></td><td></td></tr>
<tr><td>Oligo rev</td><td>AAACGTGCGGCCTCGACCTCAAGT</td><td></td><td></td><td></td><td></td><td></td><td></td><td></td><td></td><td></td><td></td><td></td></tr>
<tr><td>Chromosome</td><td>start</td><td></td><td>end</td><td>strand</td><td>MM</td><td>target_seq</td><td>PAM</td><td>alignment</td><td>distance</td><td>position</td><td>gene name</td><td>gene id</td></tr>
<tr><td>GU305914.1</td><td></td><td>60819</td><td>60841</td><td>+</td><td>0</td><td>ACTTGAGGTCGAGGCCGCAC</td><td>AGG</td><td>||||||||[||||||||||||]PAM</td><td>NA</td><td>Intergenic</td><td>NA</td><td>NA</td></tr>
<tr><td>T4054</td><td>CTTGAGGTCGAGGCCGCACAGGG</td><td></td><td>1000</td><td></td><td></td><td></td><td></td><td></td><td></td><td></td><td></td><td></td></tr>
<tr><td>Oligo fwd</td><td>CACCCTTGAGGTCGAGGCCGCACA</td><td></td><td></td><td></td><td></td><td></td><td></td><td></td><td></td><td></td><td></td><td></td></tr>
<tr><td>Oligo rev</td><td>AAACTGTGCGGCCTCGACCTCAAG</td><td></td><td></td><td></td><td></td><td></td><td></td><td></td><td></td><td></td><td></td><td></td></tr>
<tr><td>Chromosome</td><td>start</td><td></td><td>end</td><td>strand</td><td>MM</td><td>target_seq</td><td>PAM</td><td>alignment</td><td>distance</td><td>position</td><td>gene name</td><td>gene id</td></tr>
<tr><td>GU305914.1</td><td></td><td>60820</td><td>60842</td><td>+</td><td>0</td><td>CTTGAGGTCGAGGCCGCACA</td><td>GGG</td><td>||||||||[||||||||||||]PAM</td><td>NA</td><td>Intergenic</td><td>NA</td><td>NA</td></tr>
<tr><td>T4055</td><td>GTCGAGGCCGCACAGGGCCGCGG</td><td></td><td>1000</td><td></td><td></td><td></td><td></td><td></td><td></td><td></td><td></td><td></td></tr>
<tr><td>Oligo fwd</td><td>CACCGTCGAGGCCGCACAGGGCCG</td><td></td><td></td><td></td><td></td><td></td><td></td><td></td><td></td><td></td><td></td><td></td></tr>
<tr><td>Oligo rev</td><td>AAACCGGCCCTGTGCGGCCTCGAC</td><td></td><td></td><td></td><td></td><td></td><td></td><td></td><td></td><td></td><td></td><td></td></tr>
<tr><td>Chromosome</td><td>start</td><td></td><td>end</td><td>strand</td><td>MM</td><td>target_seq</td><td>PAM</td><td>alignment</td><td>distance</td><td>position</td><td>gene name</td><td>gene id</td></tr>
<tr><td>GU305914.1</td><td></td><td>60826</td><td>60848</td><td>+</td><td>0</td><td>GTCGAGGCCGCACAGGGCCG</td><td>CGG</td><td>||||||||[||||||||||||]PAM</td><td>NA</td><td>Intergenic</td><td>NA</td><td>NA</td></tr>
<tr><td>T4056</td><td>CATGAGCCACATCTTGTCGCCGG</td><td></td><td>1000</td><td></td><td></td><td></td><td></td><td></td><td></td><td></td><td></td><td></td></tr>
<tr><td>Oligo fwd</td><td>CACCCATGAGCCACATCTTGTCGC</td><td></td><td></td><td></td><td></td><td></td><td></td><td></td><td></td><td></td><td></td><td></td></tr>
<tr><td>Oligo rev</td><td>AAACGCGACAAGATGTGGCTCATG</td><td></td><td></td><td></td><td></td><td></td><td></td><td></td><td></td><td></td><td></td><td></td></tr>
<tr><td>Chromosome</td><td>start</td><td></td><td>end</td><td>strand</td><td>MM</td><td>target_seq</td><td>PAM</td><td>alignment</td><td>distance</td><td>position</td><td>gene name</td><td>gene id</td></tr>
<tr><td>GU305914.1</td><td></td><td>60865</td><td>60887</td><td>+</td><td>0</td><td>CATGAGCCACATCTTGTCGC</td><td>CGG</td><td>||||||||[||||||||||||]PAM</td><td>NA</td><td>Intergenic</td><td>NA</td><td>NA</td></tr>
<tr><td>T4057</td><td>TCTTGTCGCCGGAGCTGTCTAGG</td><td></td><td>1000</td><td></td><td></td><td></td><td></td><td></td><td></td><td></td><td></td><td></td></tr>
<tr><td>Oligo fwd</td><td>CACCTCTTGTCGCCGGAGCTGTCT</td><td></td><td></td><td></td><td></td><td></td><td></td><td></td><td></td><td></td><td></td><td></td></tr>
<tr><td>Oligo rev</td><td>AAACAGACAGCTCCGGCGACAAGA</td><td></td><td></td><td></td><td></td><td></td><td></td><td></td><td></td><td></td><td></td><td></td></tr>
<tr><td>Chromosome</td><td>start</td><td></td><td>end</td><td>strand</td><td>MM</td><td>target_seq</td><td>PAM</td><td>alignment</td><td>distance</td><td>position</td><td>gene name</td><td>gene id</td></tr>
<tr><td>GU305914.1</td><td></td><td>60876</td><td>60898</td><td>+</td><td>0</td><td>TCTTGTCGCCGGAGCTGTCT</td><td>AGG</td><td>||||||||[||||||||||||]PAM</td><td>NA</td><td>Intergenic</td><td>NA</td><td>NA</td></tr>
<tr><td>T4058</td><td>TAGCGATCCCAGAAGATCCTCGG</td><td></td><td>1000</td><td></td><td></td><td></td><td></td><td></td><td></td><td></td><td></td><td></td></tr>
<tr><td>Oligo fwd</td><td>CACCTAGCGATCCCAGAAGATCCT</td><td></td><td></td><td></td><td></td><td></td><td></td><td></td><td></td><td></td><td></td><td></td></tr>
<tr><td>Oligo rev</td><td>AAACAGGATCTTCTGGGATCGCTA</td><td></td><td></td><td></td><td></td><td></td><td></td><td></td><td></td><td></td><td></td><td></td></tr>
<tr><td>Chromosome</td><td>start</td><td></td><td>end</td><td>strand</td><td>MM</td><td>target_seq</td><td>PAM</td><td>alignment</td><td>distance</td><td>position</td><td>gene name</td><td>gene id</td></tr>
<tr><td>GU305914.1</td><td></td><td>60899</td><td>60921</td><td>+</td><td>0</td><td>TAGCGATCCCAGAAGATCCT</td><td>CGG</td><td>||||||||[||||||||||||]PAM</td><td>NA</td><td>Intergenic</td><td>NA</td><td>NA</td></tr>
<tr><td>T4059</td><td>CCTCGGCGTGTTGACCGCGAAGG</td><td></td><td>1000</td><td></td><td></td><td></td><td></td><td></td><td></td><td></td><td></td><td></td></tr>
<tr><td>Oligo fwd</td><td>CACCCCTCGGCGTGTTGACCGCGA</td><td></td><td></td><td></td><td></td><td></td><td></td><td></td><td></td><td></td><td></td><td></td></tr>
<tr><td>Oligo rev</td><td>AAACTCGCGGTCAACACGCCGAGG</td><td></td><td></td><td></td><td></td><td></td><td></td><td></td><td></td><td></td><td></td><td></td></tr>
<tr><td>Chromosome</td><td>start</td><td></td><td>end</td><td>strand</td><td>MM</td><td>target_seq</td><td>PAM</td><td>alignment</td><td>distance</td><td>position</td><td>gene name</td><td>gene id</td></tr>
<tr><td>GU305914.1</td><td></td><td>60916</td><td>60938</td><td>+</td><td>0</td><td>CCTCGGCGTGTTGACCGCGA</td><td>AGG</td><td>||||||||[||||||||||||]PAM</td><td>NA</td><td>Intergenic</td><td>NA</td><td>NA</td></tr>
<tr><td>T4060</td><td>TGTTGACCGCGAAGGCCATCAGG</td><td></td><td>1000</td><td></td><td></td><td></td><td></td><td></td><td></td><td></td><td></td><td></td></tr>
<tr><td>Oligo fwd</td><td>CACCTGTTGACCGCGAAGGCCATC</td><td></td><td></td><td></td><td></td><td></td><td></td><td></td><td></td><td></td><td></td><td></td></tr>
<tr><td>Oligo rev</td><td>AAACGATGGCCTTCGCGGTCAACA</td><td></td><td></td><td></td><td></td><td></td><td></td><td></td><td></td><td></td><td></td><td></td></tr>
<tr><td>Chromosome</td><td>start</td><td></td><td>end</td><td>strand</td><td>MM</td><td>target_seq</td><td>PAM</td><td>alignment</td><td>distance</td><td>position</td><td>gene name</td><td>gene id</td></tr>
<tr><td>GU305914.1</td><td></td><td>60924</td><td>60946</td><td>+</td><td>0</td><td>TGTTGACCGCGAAGGCCATC</td><td>AGG</td><td>||||||||[||||||||||||]PAM</td><td>NA</td><td>Intergenic</td><td>NA</td><td>NA</td></tr>
<tr><td>T4061</td><td>CGTATTCCCGCCCGACGTCGAGG</td><td></td><td>1000</td><td></td><td></td><td></td><td></td><td></td><td></td><td></td><td></td><td></td></tr>
<tr><td>Oligo fwd</td><td>CACCCGTATTCCCGCCCGACGTCG</td><td></td><td></td><td></td><td></td><td></td><td></td><td></td><td></td><td></td><td></td><td></td></tr>
<tr><td>Oligo rev</td><td>AAACCGACGTCGGGCGGGAATACG</td><td></td><td></td><td></td><td></td><td></td><td></td><td></td><td></td><td></td><td></td><td></td></tr>
<tr><td>Chromosome</td><td>start</td><td></td><td>end</td><td>strand</td><td>MM</td><td>target_seq</td><td>PAM</td><td>alignment</td><td>distance</td><td>position</td><td>gene name</td><td>gene id</td></tr>
<tr><td>GU305914.1</td><td></td><td>60951</td><td>60973</td><td>+</td><td>0</td><td>CGTATTCCCGCCCGACGTCG</td><td>AGG</td><td>||||||||[||||||||||||]PAM</td><td>NA</td><td>Intergenic</td><td>NA</td><td>NA</td></tr>
<tr><td>T4062</td><td>GAAGATCTTGAGAGCGTCGACGG</td><td></td><td>1000</td><td></td><td></td><td></td><td></td><td></td><td></td><td></td><td></td><td></td></tr>
<tr><td>Oligo fwd</td><td>CACCGAAGATCTTGAGAGCGTCGA</td><td></td><td></td><td></td><td></td><td></td><td></td><td></td><td></td><td></td><td></td><td></td></tr>
<tr><td>Oligo rev</td><td>AAACTCGACGCTCTCAAGATCTTC</td><td></td><td></td><td></td><td></td><td></td><td></td><td></td><td></td><td></td><td></td><td></td></tr>
<tr><td>Chromosome</td><td>start</td><td></td><td>end</td><td>strand</td><td>MM</td><td>target_seq</td><td>PAM</td><td>alignment</td><td>distance</td><td>position</td><td>gene name</td><td>gene id</td></tr>
<tr><td>GU305914.1</td><td></td><td>60982</td><td>61004</td><td>+</td><td>0</td><td>GAAGATCTTGAGAGCGTCGA</td><td>CGG</td><td>||||||||[||||||||||||]PAM</td><td>NA</td><td>Intergenic</td><td>NA</td><td>NA</td></tr>
<tr><td>T4063</td><td>GTCGAAGTAGATCCTCAAGACGG</td><td></td><td>1000</td><td></td><td></td><td></td><td></td><td></td><td></td><td></td><td></td><td></td></tr>
<tr><td>Oligo fwd</td><td>CACCGTCGAAGTAGATCCTCAAGA</td><td></td><td></td><td></td><td></td><td></td><td></td><td></td><td></td><td></td><td></td><td></td></tr>
<tr><td>Oligo rev</td><td>AAACTCTTGAGGATCTACTTCGAC</td><td></td><td></td><td></td><td></td><td></td><td></td><td></td><td></td><td></td><td></td><td></td></tr>
<tr><td>Chromosome</td><td>start</td><td></td><td>end</td><td>strand</td><td>MM</td><td>target_seq</td><td>PAM</td><td>alignment</td><td>distance</td><td>position</td><td>gene name</td><td>gene id</td></tr>
<tr><td>GU305914.1</td><td></td><td>61012</td><td>61034</td><td>+</td><td>0</td><td>GTCGAAGTAGATCCTCAAGA</td><td>CGG</td><td>||||||||[||||||||||||]PAM</td><td>NA</td><td>Intergenic</td><td>NA</td><td>NA</td></tr>
<tr><td>T4064</td><td>ACGGTCTTGAATCGCGCGTACGG</td><td></td><td>1000</td><td></td><td></td><td></td><td></td><td></td><td></td><td></td><td></td><td></td></tr>
<tr><td>Oligo fwd</td><td>CACCACGGTCTTGAATCGCGCGTA</td><td></td><td></td><td></td><td></td><td></td><td></td><td></td><td></td><td></td><td></td><td></td></tr>
<tr><td>Oligo rev</td><td>AAACTACGCGCGATTCAAGACCGT</td><td></td><td></td><td></td><td></td><td></td><td></td><td></td><td></td><td></td><td></td><td></td></tr>
<tr><td>Chromosome</td><td>start</td><td></td><td>end</td><td>strand</td><td>MM</td><td>target_seq</td><td>PAM</td><td>alignment</td><td>distance</td><td>position</td><td>gene name</td><td>gene id</td></tr>
<tr><td>GU305914.1</td><td></td><td>61031</td><td>61053</td><td>+</td><td>0</td><td>ACGGTCTTGAATCGCGCGTA</td><td>CGG</td><td>||||||||[||||||||||||]PAM</td><td>NA</td><td>Intergenic</td><td>NA</td><td>NA</td></tr>
<tr><td>T4065</td><td>GCGCCTTGCACTGCTGTCCGAGG</td><td></td><td>1000</td><td></td><td></td><td></td><td></td><td></td><td></td><td></td><td></td><td></td></tr>
<tr><td>Oligo fwd</td><td>CACCGCGCCTTGCACTGCTGTCCG</td><td></td><td></td><td></td><td></td><td></td><td></td><td></td><td></td><td></td><td></td><td></td></tr>
<tr><td>Oligo rev</td><td>AAACCGGACAGCAGTGCAAGGCGC</td><td></td><td></td><td></td><td></td><td></td><td></td><td></td><td></td><td></td><td></td><td></td></tr>
<tr><td>Chromosome</td><td>start</td><td></td><td>end</td><td>strand</td><td>MM</td><td>target_seq</td><td>PAM</td><td>alignment</td><td>distance</td><td>position</td><td>gene name</td><td>gene id</td></tr>
<tr><td>GU305914.1</td><td></td><td>61059</td><td>61081</td><td>+</td><td>0</td><td>GCGCCTTGCACTGCTGTCCG</td><td>AGG</td><td>||||||||[||||||||||||]PAM</td><td>NA</td><td>Intergenic</td><td>NA</td><td>NA</td></tr>
<tr><td>T4066</td><td>CTGTCCGAGGATGTGCAGAGAGG</td><td></td><td>1000</td><td></td><td></td><td></td><td></td><td></td><td></td><td></td><td></td><td></td></tr>
<tr><td>Oligo fwd</td><td>CACCCTGTCCGAGGATGTGCAGAG</td><td></td><td></td><td></td><td></td><td></td><td></td><td></td><td></td><td></td><td></td><td></td></tr>
<tr><td>Oligo rev</td><td>AAACCTCTGCACATCCTCGGACAG</td><td></td><td></td><td></td><td></td><td></td><td></td><td></td><td></td><td></td><td></td><td></td></tr>
<tr><td>Chromosome</td><td>start</td><td></td><td>end</td><td>strand</td><td>MM</td><td>target_seq</td><td>PAM</td><td>alignment</td><td>distance</td><td>position</td><td>gene name</td><td>gene id</td></tr>
<tr><td>GU305914.1</td><td></td><td>61072</td><td>61094</td><td>+</td><td>0</td><td>CTGTCCGAGGATGTGCAGAG</td><td>AGG</td><td>||||||||[||||||||||||]PAM</td><td>NA</td><td>Intergenic</td><td>NA</td><td>NA</td></tr>
<tr><td>T4067</td><td>AGGCCACGATGTCGCCTCCGAGG</td><td></td><td>1000</td><td></td><td></td><td></td><td></td><td></td><td></td><td></td><td></td><td></td></tr>
<tr><td>Oligo fwd</td><td>CACCAGGCCACGATGTCGCCTCCG</td><td></td><td></td><td></td><td></td><td></td><td></td><td></td><td></td><td></td><td></td><td></td></tr>
<tr><td>Oligo rev</td><td>AAACCGGAGGCGACATCGTGGCCT</td><td></td><td></td><td></td><td></td><td></td><td></td><td></td><td></td><td></td><td></td><td></td></tr>
<tr><td>Chromosome</td><td>start</td><td></td><td>end</td><td>strand</td><td>MM</td><td>target_seq</td><td>PAM</td><td>alignment</td><td>distance</td><td>position</td><td>gene name</td><td>gene id</td></tr>
<tr><td>GU305914.1</td><td></td><td>61092</td><td>61114</td><td>+</td><td>0</td><td>AGGCCACGATGTCGCCTCCG</td><td>AGG</td><td>||||||||[||||||||||||]PAM</td><td>NA</td><td>Intergenic</td><td>NA</td><td>NA</td></tr>
</table>

<table>
<tr><td>T4068</td><td>TCCGAGGTCCGCCGCGCTGACGG</td><td>1000</td><td></td><td></td><td></td><td></td><td></td><td></td><td></td><td></td><td></td></tr>
<tr><td>Oligo fwd</td><td>CACCTCCGAGGTCCGCCGCGCTGA</td><td></td><td></td><td></td><td></td><td></td><td></td><td></td><td></td><td></td><td></td></tr>
<tr><td>Oligo rev</td><td>AAACTCAGCGCGGCGGACCTCGGA</td><td></td><td></td><td></td><td></td><td></td><td></td><td></td><td></td><td></td><td></td></tr>
<tr><td>Chromosome</td><td>start</td><td>end</td><td>strand</td><td>MM</td><td>target_seq</td><td>PAM</td><td>alignment</td><td>distance</td><td>position</td><td>gene name</td><td>gene id</td></tr>
<tr><td>GU305914.1</td><td>61108</td><td>61130</td><td>+</td><td>0</td><td>TCCGAGGTCCGCCGCGCTGA</td><td>CGG</td><td>||||||||[||||||||||||]PAM</td><td>NA</td><td>Intergenic</td><td>NA</td><td>NA</td></tr>
<tr><td>T4069</td><td>CCGCGCTGACGGACACCGAGAGG</td><td>1000</td><td></td><td></td><td></td><td></td><td></td><td></td><td></td><td></td><td></td></tr>
<tr><td>Oligo fwd</td><td>CACCCCGCGCTGACGGACACCGAG</td><td></td><td></td><td></td><td></td><td></td><td></td><td></td><td></td><td></td><td></td></tr>
<tr><td>Oligo rev</td><td>AAACCTCGGTGTCCGTCAGCGCGG</td><td></td><td></td><td></td><td></td><td></td><td></td><td></td><td></td><td></td><td></td></tr>
<tr><td>Chromosome</td><td>start</td><td>end</td><td>strand</td><td>MM</td><td>target_seq</td><td>PAM</td><td>alignment</td><td>distance</td><td>position</td><td>gene name</td><td>gene id</td></tr>
<tr><td>GU305914.1</td><td>61119</td><td>61141</td><td>+</td><td>0</td><td>CCGCGCTGACGGACACCGAG</td><td>AGG</td><td>||||||||[||||||||||||]PAM</td><td>NA</td><td>Intergenic</td><td>NA</td><td>NA</td></tr>
<tr><td>T4070</td><td>GACGGACACCGAGAGGCCGTTGG</td><td>1000</td><td></td><td></td><td></td><td></td><td></td><td></td><td></td><td></td><td></td></tr>
<tr><td>Oligo fwd</td><td>CACCGACGGACACCGAGAGGCCGT</td><td></td><td></td><td></td><td></td><td></td><td></td><td></td><td></td><td></td><td></td></tr>
<tr><td>Oligo rev</td><td>AAACACGGCCTCTCGGTGTCCGTC</td><td></td><td></td><td></td><td></td><td></td><td></td><td></td><td></td><td></td><td></td></tr>
<tr><td>Chromosome</td><td>start</td><td>end</td><td>strand</td><td>MM</td><td>target_seq</td><td>PAM</td><td>alignment</td><td>distance</td><td>position</td><td>gene name</td><td>gene id</td></tr>
<tr><td>GU305914.1</td><td>61126</td><td>61148</td><td>+</td><td>0</td><td>GACGGACACCGAGAGGCCGT</td><td>TGG</td><td>||||||||[||||||||||||]PAM</td><td>NA</td><td>Intergenic</td><td>NA</td><td>NA</td></tr>
<tr><td>T4071</td><td>CCATGTCGCAGATGTCGTGCAGG</td><td>1000</td><td></td><td></td><td></td><td></td><td></td><td></td><td></td><td></td><td></td></tr>
<tr><td>Oligo fwd</td><td>CACCCCATGTCGCAGATGTCGTGC</td><td></td><td></td><td></td><td></td><td></td><td></td><td></td><td></td><td></td><td></td></tr>
<tr><td>Oligo rev</td><td>AAACGCACGACATCTGCGACATGG</td><td></td><td></td><td></td><td></td><td></td><td></td><td></td><td></td><td></td><td></td></tr>
<tr><td>Chromosome</td><td>start</td><td>end</td><td>strand</td><td>MM</td><td>target_seq</td><td>PAM</td><td>alignment</td><td>distance</td><td>position</td><td>gene name</td><td>gene id</td></tr>
<tr><td>GU305914.1</td><td>61152</td><td>61174</td><td>+</td><td>0</td><td>CCATGTCGCAGATGTCGTGC</td><td>AGG</td><td>||||||||[||||||||||||]PAM</td><td>NA</td><td>Intergenic</td><td>NA</td><td>NA</td></tr>
<tr><td>GU305914.1</td><td>69531</td><td>69553</td><td>+</td><td>3</td><td>TCGTGTCGCAGATGTCGCGC</td><td>CGG</td><td>-|-|||||[|||||||||-||]PAM</td><td>NA</td><td>Intergenic</td><td>NA</td><td>NA</td></tr>
<tr><td>T4072</td><td>CATGTCGCAGATGTCGTGCAGGG</td><td>1000</td><td></td><td></td><td></td><td></td><td></td><td></td><td></td><td></td><td></td></tr>
<tr><td>Oligo fwd</td><td>CACCCATGTCGCAGATGTCGTGCA</td><td></td><td></td><td></td><td></td><td></td><td></td><td></td><td></td><td></td><td></td></tr>
<tr><td>Oligo rev</td><td>AAACTGCACGACATCTGCGACATG</td><td></td><td></td><td></td><td></td><td></td><td></td><td></td><td></td><td></td><td></td></tr>
<tr><td>Chromosome</td><td>start</td><td>end</td><td>strand</td><td>MM</td><td>target_seq</td><td>PAM</td><td>alignment</td><td>distance</td><td>position</td><td>gene name</td><td>gene id</td></tr>
<tr><td>GU305914.1</td><td>61153</td><td>61175</td><td>+</td><td>0</td><td>CATGTCGCAGATGTCGTGCA</td><td>GGG</td><td>||||||||[||||||||||||]PAM</td><td>NA</td><td>Intergenic</td><td>NA</td><td>NA</td></tr>
<tr><td>T4073</td><td>GGTGCCGTTGATCGTGCTCTCGG</td><td>1000</td><td></td><td></td><td></td><td></td><td></td><td></td><td></td><td></td><td></td></tr>
<tr><td>Oligo fwd</td><td>CACCGGTGCCGTTGATCGTGCTCT</td><td></td><td></td><td></td><td></td><td></td><td></td><td></td><td></td><td></td><td></td></tr>
<tr><td>Oligo rev</td><td>AAACAGAGCACGATCAACGGCACC</td><td></td><td></td><td></td><td></td><td></td><td></td><td></td><td></td><td></td><td></td></tr>
<tr><td>Chromosome</td><td>start</td><td>end</td><td>strand</td><td>MM</td><td>target_seq</td><td>PAM</td><td>alignment</td><td>distance</td><td>position</td><td>gene name</td><td>gene id</td></tr>
<tr><td>GU305914.1</td><td>61174</td><td>61196</td><td>+</td><td>0</td><td>GGTGCCGTTGATCGTGCTCT</td><td>CGG</td><td>||||||||[||||||||||||]PAM</td><td>NA</td><td>Intergenic</td><td>NA</td><td>NA</td></tr>
<tr><td>GU305914.1</td><td>122665</td><td>122687</td><td>+</td><td>4</td><td>CGTGTCGTCGATCGTGATCT</td><td>CGG</td><td>-|||-|||[-|||||||-|||]PAM</td><td>NA</td><td>Intergenic</td><td>NA</td><td>NA</td></tr>
<tr><td>T4074</td><td>CGGCCGAGTAGAACTGAGCGCGG</td><td>1000</td><td></td><td></td><td></td><td></td><td></td><td></td><td></td><td></td><td></td></tr>
<tr><td>Oligo fwd</td><td>CACCCGGCCGAGTAGAACTGAGCG</td><td></td><td></td><td></td><td></td><td></td><td></td><td></td><td></td><td></td><td></td></tr>
<tr><td>Oligo rev</td><td>AAACCGCTCAGTTCTACTCGGCCG</td><td></td><td></td><td></td><td></td><td></td><td></td><td></td><td></td><td></td><td></td></tr>
<tr><td>Chromosome</td><td>start</td><td>end</td><td>strand</td><td>MM</td><td>target_seq</td><td>PAM</td><td>alignment</td><td>distance</td><td>position</td><td>gene name</td><td>gene id</td></tr>
<tr><td>GU305914.1</td><td>61194</td><td>61216</td><td>+</td><td>0</td><td>CGGCCGAGTAGAACTGAGCG</td><td>CGG</td><td>||||||||[||||||||||||]PAM</td><td>NA</td><td>Intergenic</td><td>NA</td><td>NA</td></tr>
<tr><td>T4075</td><td>GAACTGAGCGCGGTAAGAAACGG</td><td>1000</td><td></td><td></td><td></td><td></td><td></td><td></td><td></td><td></td><td></td></tr>
<tr><td>Oligo fwd</td><td>CACCGAACTGAGCGCGGTAAGAAA</td><td></td><td></td><td></td><td></td><td></td><td></td><td></td><td></td><td></td><td></td></tr>
<tr><td>Oligo rev</td><td>AAACTTTCTTACCGCGCTCAGTTC</td><td></td><td></td><td></td><td></td><td></td><td></td><td></td><td></td><td></td><td></td></tr>
<tr><td>Chromosome</td><td>start</td><td>end</td><td>strand</td><td>MM</td><td>target_seq</td><td>PAM</td><td>alignment</td><td>distance</td><td>position</td><td>gene name</td><td>gene id</td></tr>
<tr><td>GU305914.1</td><td>61204</td><td>61226</td><td>+</td><td>0</td><td>GAACTGAGCGCGGTAAGAAA</td><td>CGG</td><td>||||||||[||||||||||||]PAM</td><td>NA</td><td>Intergenic</td><td>NA</td><td>NA</td></tr>
<tr><td>T4076</td><td>TAAGAAACGGCGTCGAAGATCGG</td><td>1000</td><td></td><td></td><td></td><td></td><td></td><td></td><td></td><td></td><td></td></tr>
<tr><td>Oligo fwd</td><td>CACCTAAGAAACGGCGTCGAAGAT</td><td></td><td></td><td></td><td></td><td></td><td></td><td></td><td></td><td></td><td></td></tr>
<tr><td>Oligo rev</td><td>AAACATCTTCGACGCCGTTTCTTA</td><td></td><td></td><td></td><td></td><td></td><td></td><td></td><td></td><td></td><td></td></tr>
<tr><td>Chromosome</td><td>start</td><td>end</td><td>strand</td><td>MM</td><td>target_seq</td><td>PAM</td><td>alignment</td><td>distance</td><td>position</td><td>gene name</td><td>gene id</td></tr>
<tr><td>GU305914.1</td><td>61217</td><td>61239</td><td>+</td><td>0</td><td>TAAGAAACGGCGTCGAAGAT</td><td>CGG</td><td>||||||||[||||||||||||]PAM</td><td>NA</td><td>Intergenic</td><td>NA</td><td>NA</td></tr>
<tr><td>T4077</td><td>AAGAAACGGCGTCGAAGATCGGG</td><td>1000</td><td></td><td></td><td></td><td></td><td></td><td></td><td></td><td></td><td></td></tr>
<tr><td>Oligo fwd</td><td>CACCAAGAAACGGCGTCGAAGATC</td><td></td><td></td><td></td><td></td><td></td><td></td><td></td><td></td><td></td><td></td></tr>
<tr><td>Oligo rev</td><td>AAACGATCTTCGACGCCGTTTCTT</td><td></td><td></td><td></td><td></td><td></td><td></td><td></td><td></td><td></td><td></td></tr>
<tr><td>Chromosome</td><td>start</td><td>end</td><td>strand</td><td>MM</td><td>target_seq</td><td>PAM</td><td>alignment</td><td>distance</td><td>position</td><td>gene name</td><td>gene id</td></tr>
<tr><td>GU305914.1</td><td>61218</td><td>61240</td><td>+</td><td>0</td><td>AAGAAACGGCGTCGAAGATC</td><td>GGG</td><td>||||||||[||||||||||||]PAM</td><td>NA</td><td>Intergenic</td><td>NA</td><td>NA</td></tr>
<tr><td>T4078</td><td>AACGGCGTCGAAGATCGGGTTGG</td><td>1000</td><td></td><td></td><td></td><td></td><td></td><td></td><td></td><td></td><td></td></tr>
<tr><td>Oligo fwd</td><td>CACCAACGGCGTCGAAGATCGGGT</td><td></td><td></td><td></td><td></td><td></td><td></td><td></td><td></td><td></td><td></td></tr>
<tr><td>Oligo rev</td><td>AAACACCCGATCTTCGACGCCGTT</td><td></td><td></td><td></td><td></td><td></td><td></td><td></td><td></td><td></td><td></td></tr>
<tr><td>Chromosome</td><td>start</td><td>end</td><td>strand</td><td>MM</td><td>target_seq</td><td>PAM</td><td>alignment</td><td>distance</td><td>position</td><td>gene name</td><td>gene id</td></tr>
<tr><td>GU305914.1</td><td>61222</td><td>61244</td><td>+</td><td>0</td><td>AACGGCGTCGAAGATCGGGT</td><td>TGG</td><td>||||||||[||||||||||||]PAM</td><td>NA</td><td>Intergenic</td><td>NA</td><td>NA</td></tr>
<tr><td>T4079</td><td>CGGGTTGGAACCGCCGAACCCGG</td><td>1000</td><td></td><td></td><td></td><td></td><td></td><td></td><td></td><td></td><td></td></tr>
<tr><td>Oligo fwd</td><td>CACCCGGGTTGGAACCGCCGAACC</td><td></td><td></td><td></td><td></td><td></td><td></td><td></td><td></td><td></td><td></td></tr>
<tr><td>Oligo rev</td><td>AAACGGTTCGGCGGTTCCAACCCG</td><td></td><td></td><td></td><td></td><td></td><td></td><td></td><td></td><td></td><td></td></tr>
<tr><td>Chromosome</td><td>start</td><td>end</td><td>strand</td><td>MM</td><td>target_seq</td><td>PAM</td><td>alignment</td><td>distance</td><td>position</td><td>gene name</td><td>gene id</td></tr>
<tr><td>GU305914.1</td><td>61237</td><td>61259</td><td>+</td><td>0</td><td>CGGGTTGGAACCGCCGAACC</td><td>CGG</td><td>||||||||[||||||||||||]PAM</td><td>NA</td><td>Intergenic</td><td>NA</td><td>NA</td></tr>
<tr><td>T4080</td><td>GGAACCGCCGAACCCGGCGTAGG</td><td>1000</td><td></td><td></td><td></td><td></td><td></td><td></td><td></td><td></td><td></td></tr>
<tr><td>Oligo fwd</td><td>CACCGGAACCGCCGAACCCGGCGT</td><td></td><td></td><td></td><td></td><td></td><td></td><td></td><td></td><td></td><td></td></tr>
<tr><td>Oligo rev</td><td>AAACACGCCGGGTTCGGCGGTTCC</td><td></td><td></td><td></td><td></td><td></td><td></td><td></td><td></td><td></td><td></td></tr>
<tr><td>Chromosome</td><td>start</td><td>end</td><td>strand</td><td>MM</td><td>target_seq</td><td>PAM</td><td>alignment</td><td>distance</td><td>position</td><td>gene name</td><td>gene id</td></tr>
<tr><td>GU305914.1</td><td>61243</td><td>61265</td><td>+</td><td>0</td><td>GGAACCGCCGAACCCGGCGT</td><td>AGG</td><td>||||||||[||||||||||||]PAM</td><td>NA</td><td>Intergenic</td><td>NA</td><td>NA</td></tr>
<tr><td>T4081</td><td>CGCCGAACCCGGCGTAGGCCAGG</td><td>1000</td><td></td><td></td><td></td><td></td><td></td><td></td><td></td><td></td><td></td></tr>
<tr><td>Oligo fwd</td><td>CACCCGCCGAACCCGGCGTAGGCC</td><td></td><td></td><td></td><td></td><td></td><td></td><td></td><td></td><td></td><td></td></tr>
<tr><td>Oligo rev</td><td>AAACGGCCTACGCCGGGTTCGGCG</td><td></td><td></td><td></td><td></td><td></td><td></td><td></td><td></td><td></td><td></td></tr>
<tr><td>Chromosome</td><td>start</td><td>end</td><td>strand</td><td>MM</td><td>target_seq</td><td>PAM</td><td>alignment</td><td>distance</td><td>position</td><td>gene name</td><td>gene id</td></tr>
<tr><td>GU305914.1</td><td>61248</td><td>61270</td><td>+</td><td>0</td><td>CGCCGAACCCGGCGTAGGCC</td><td>AGG</td><td>||||||||[||||||||||||]PAM</td><td>NA</td><td>Intergenic</td><td>NA</td><td>NA</td></tr>
<tr><td>T4082</td><td>CCGGCGTAGGCCAGGACCCGCGG</td><td>1000</td><td></td><td></td><td></td><td></td><td></td><td></td><td></td><td></td><td></td></tr>
<tr><td>Oligo fwd</td><td>CACCCCGGCGTAGGCCAGGACCCG</td><td></td><td></td><td></td><td></td><td></td><td></td><td></td><td></td><td></td><td></td></tr>
</table>

<table>
<tr><td>Oligo rev</td><td>AAACCGGGTCCTGGCCTACGCCGG</td><td></td><td></td><td></td><td></td><td></td><td></td><td></td><td></td><td></td><td></td><td></td></tr>
<tr><td>Chromosome</td><td>start</td><td></td><td>end</td><td>strand</td><td>MM</td><td>target_seq</td><td>PAM</td><td>alignment</td><td>distance</td><td>position</td><td>gene name</td><td>gene id</td></tr>
<tr><td>GU305914.1</td><td></td><td>61256</td><td>61278</td><td>+</td><td>0</td><td>CCGGCGTAGGCCAGGACCCG</td><td>CGG</td><td>||||||||[||||||||||||]PAM</td><td>NA</td><td>Intergenic</td><td>NA</td><td>NA</td></tr>
<tr><td>T4083</td><td>GTAGGCCAGGACCCGCGGCGAGG</td><td></td><td>1000</td><td></td><td></td><td></td><td></td><td></td><td></td><td></td><td></td><td></td></tr>
<tr><td>Oligo fwd</td><td>CACCGTAGGCCAGGACCCGCGGCG</td><td></td><td></td><td></td><td></td><td></td><td></td><td></td><td></td><td></td><td></td><td></td></tr>
<tr><td>Oligo rev</td><td>AAACCGCCGCGGGTCCTGGCCTAC</td><td></td><td></td><td></td><td></td><td></td><td></td><td></td><td></td><td></td><td></td><td></td></tr>
<tr><td>Chromosome</td><td>start</td><td></td><td>end</td><td>strand</td><td>MM</td><td>target_seq</td><td>PAM</td><td>alignment</td><td>distance</td><td>position</td><td>gene name</td><td>gene id</td></tr>
<tr><td>GU305914.1</td><td></td><td>61261</td><td>61283</td><td>+</td><td>0</td><td>GTAGGCCAGGACCCGCGGCG</td><td>AGG</td><td>||||||||[||||||||||||]PAM</td><td>NA</td><td>Intergenic</td><td>NA</td><td>NA</td></tr>
<tr><td>T4084</td><td>TAGGCCAGGACCCGCGGCGAGGG</td><td></td><td>1000</td><td></td><td></td><td></td><td></td><td></td><td></td><td></td><td></td><td></td></tr>
<tr><td>Oligo fwd</td><td>CACCTAGGCCAGGACCCGCGGCGA</td><td></td><td></td><td></td><td></td><td></td><td></td><td></td><td></td><td></td><td></td><td></td></tr>
<tr><td>Oligo rev</td><td>AAACTCGCCGCGGGTCCTGGCCTA</td><td></td><td></td><td></td><td></td><td></td><td></td><td></td><td></td><td></td><td></td><td></td></tr>
<tr><td>Chromosome</td><td>start</td><td></td><td>end</td><td>strand</td><td>MM</td><td>target_seq</td><td>PAM</td><td>alignment</td><td>distance</td><td>position</td><td>gene name</td><td>gene id</td></tr>
<tr><td>GU305914.1</td><td></td><td>61262</td><td>61284</td><td>+</td><td>0</td><td>TAGGCCAGGACCCGCGGCGA</td><td>GGG</td><td>||||||||[||||||||||||]PAM</td><td>NA</td><td>Intergenic</td><td>NA</td><td>NA</td></tr>
<tr><td>T4085</td><td>AGGCCAGGACCCGCGGCGAGGGG</td><td></td><td>1000</td><td></td><td></td><td></td><td></td><td></td><td></td><td></td><td></td><td></td></tr>
<tr><td>Oligo fwd</td><td>CACCAGGCCAGGACCCGCGGCGAG</td><td></td><td></td><td></td><td></td><td></td><td></td><td></td><td></td><td></td><td></td><td></td></tr>
<tr><td>Oligo rev</td><td>AAACCTCGCCGCGGGTCCTGGCCT</td><td></td><td></td><td></td><td></td><td></td><td></td><td></td><td></td><td></td><td></td><td></td></tr>
<tr><td>Chromosome</td><td>start</td><td></td><td>end</td><td>strand</td><td>MM</td><td>target_seq</td><td>PAM</td><td>alignment</td><td>distance</td><td>position</td><td>gene name</td><td>gene id</td></tr>
<tr><td>GU305914.1</td><td></td><td>61263</td><td>61285</td><td>+</td><td>0</td><td>AGGCCAGGACCCGCGGCGAG</td><td>GGG</td><td>||||||||[||||||||||||]PAM</td><td>NA</td><td>Intergenic</td><td>NA</td><td>NA</td></tr>
<tr><td>T4086</td><td>GACCCGCGGCGAGGGGATGATGG</td><td></td><td>1000</td><td></td><td></td><td></td><td></td><td></td><td></td><td></td><td></td><td></td></tr>
<tr><td>Oligo fwd</td><td>CACCGACCCGCGGCGAGGGGATGA</td><td></td><td></td><td></td><td></td><td></td><td></td><td></td><td></td><td></td><td></td><td></td></tr>
<tr><td>Oligo rev</td><td>AAACTCATCCCCTCGCCGCGGGTC</td><td></td><td></td><td></td><td></td><td></td><td></td><td></td><td></td><td></td><td></td><td></td></tr>
<tr><td>Chromosome</td><td>start</td><td></td><td>end</td><td>strand</td><td>MM</td><td>target_seq</td><td>PAM</td><td>alignment</td><td>distance</td><td>position</td><td>gene name</td><td>gene id</td></tr>
<tr><td>GU305914.1</td><td></td><td>61270</td><td>61292</td><td>+</td><td>0</td><td>GACCCGCGGCGAGGGGATGA</td><td>TGG</td><td>||||||||[||||||||||||]PAM</td><td>NA</td><td>Intergenic</td><td>NA</td><td>NA</td></tr>
<tr><td>T4087</td><td>CGGCGAGGGGATGATGGCGAAGG</td><td></td><td>1000</td><td></td><td></td><td></td><td></td><td></td><td></td><td></td><td></td><td></td></tr>
<tr><td>Oligo fwd</td><td>CACCCGGCGAGGGGATGATGGCGA</td><td></td><td></td><td></td><td></td><td></td><td></td><td></td><td></td><td></td><td></td><td></td></tr>
<tr><td>Oligo rev</td><td>AAACTCGCCATCATCCCCTCGCCG</td><td></td><td></td><td></td><td></td><td></td><td></td><td></td><td></td><td></td><td></td><td></td></tr>
<tr><td>Chromosome</td><td>start</td><td></td><td>end</td><td>strand</td><td>MM</td><td>target_seq</td><td>PAM</td><td>alignment</td><td>distance</td><td>position</td><td>gene name</td><td>gene id</td></tr>
<tr><td>GU305914.1</td><td></td><td>61276</td><td>61298</td><td>+</td><td>0</td><td>CGGCGAGGGGATGATGGCGA</td><td>AGG</td><td>||||||||[||||||||||||]PAM</td><td>NA</td><td>Intergenic</td><td>NA</td><td>NA</td></tr>
<tr><td>T4088</td><td>GGATGATGGCGAAGGAGCAGAGG</td><td></td><td>1000</td><td></td><td></td><td></td><td></td><td></td><td></td><td></td><td></td><td></td></tr>
<tr><td>Oligo fwd</td><td>CACCGGATGATGGCGAAGGAGCAG</td><td></td><td></td><td></td><td></td><td></td><td></td><td></td><td></td><td></td><td></td><td></td></tr>
<tr><td>Oligo rev</td><td>AAACCTGCTCCTTCGCCATCATCC</td><td></td><td></td><td></td><td></td><td></td><td></td><td></td><td></td><td></td><td></td><td></td></tr>
<tr><td>Chromosome</td><td>start</td><td></td><td>end</td><td>strand</td><td>MM</td><td>target_seq</td><td>PAM</td><td>alignment</td><td>distance</td><td>position</td><td>gene name</td><td>gene id</td></tr>
<tr><td>GU305914.1</td><td></td><td>61284</td><td>61306</td><td>+</td><td>0</td><td>GGATGATGGCGAAGGAGCAG</td><td>AGG</td><td>||||||||[||||||||||||]PAM</td><td>NA</td><td>Intergenic</td><td>NA</td><td>NA</td></tr>
<tr><td>T4089</td><td>GATGATGGCGAAGGAGCAGAGGG</td><td></td><td>1000</td><td></td><td></td><td></td><td></td><td></td><td></td><td></td><td></td><td></td></tr>
<tr><td>Oligo fwd</td><td>CACCGATGATGGCGAAGGAGCAGA</td><td></td><td></td><td></td><td></td><td></td><td></td><td></td><td></td><td></td><td></td><td></td></tr>
<tr><td>Oligo rev</td><td>AAACTCTGCTCCTTCGCCATCATC</td><td></td><td></td><td></td><td></td><td></td><td></td><td></td><td></td><td></td><td></td><td></td></tr>
<tr><td>Chromosome</td><td>start</td><td></td><td>end</td><td>strand</td><td>MM</td><td>target_seq</td><td>PAM</td><td>alignment</td><td>distance</td><td>position</td><td>gene name</td><td>gene id</td></tr>
<tr><td>GU305914.1</td><td></td><td>61285</td><td>61307</td><td>+</td><td>0</td><td>GATGATGGCGAAGGAGCAGA</td><td>GGG</td><td>||||||||[||||||||||||]PAM</td><td>NA</td><td>Intergenic</td><td>NA</td><td>NA</td></tr>
<tr><td>T4090</td><td>GATGGCGAAGGAGCAGAGGGCGG</td><td></td><td>1000</td><td></td><td></td><td></td><td></td><td></td><td></td><td></td><td></td><td></td></tr>
<tr><td>Oligo fwd</td><td>CACCGATGGCGAAGGAGCAGAGGG</td><td></td><td></td><td></td><td></td><td></td><td></td><td></td><td></td><td></td><td></td><td></td></tr>
<tr><td>Oligo rev</td><td>AAACCCCTCTGCTCCTTCGCCATC</td><td></td><td></td><td></td><td></td><td></td><td></td><td></td><td></td><td></td><td></td><td></td></tr>
<tr><td>Chromosome</td><td>start</td><td></td><td>end</td><td>strand</td><td>MM</td><td>target_seq</td><td>PAM</td><td>alignment</td><td>distance</td><td>position</td><td>gene name</td><td>gene id</td></tr>
<tr><td>GU305914.1</td><td></td><td>61288</td><td>61310</td><td>+</td><td>0</td><td>GATGGCGAAGGAGCAGAGGG</td><td>CGG</td><td>||||||||[||||||||||||]PAM</td><td>NA</td><td>Intergenic</td><td>NA</td><td>NA</td></tr>
<tr><td>T4091</td><td>CGAAGGAGCAGAGGGCGGAGAGG</td><td></td><td>1000</td><td></td><td></td><td></td><td></td><td></td><td></td><td></td><td></td><td></td></tr>
<tr><td>Oligo fwd</td><td>CACCCGAAGGAGCAGAGGGCGGAG</td><td></td><td></td><td></td><td></td><td></td><td></td><td></td><td></td><td></td><td></td><td></td></tr>
<tr><td>Oligo rev</td><td>AAACCTCCGCCCTCTGCTCCTTCG</td><td></td><td></td><td></td><td></td><td></td><td></td><td></td><td></td><td></td><td></td><td></td></tr>
<tr><td>Chromosome</td><td>start</td><td></td><td>end</td><td>strand</td><td>MM</td><td>target_seq</td><td>PAM</td><td>alignment</td><td>distance</td><td>position</td><td>gene name</td><td>gene id</td></tr>
<tr><td>GU305914.1</td><td></td><td>61293</td><td>61315</td><td>+</td><td>0</td><td>CGAAGGAGCAGAGGGCGGAG</td><td>AGG</td><td>||||||||[||||||||||||]PAM</td><td>NA</td><td>Intergenic</td><td>NA</td><td>NA</td></tr>
<tr><td>T4092</td><td>GGAGCAGAGGGCGGAGAGGAAGG</td><td></td><td>1000</td><td></td><td></td><td></td><td></td><td></td><td></td><td></td><td></td><td></td></tr>
<tr><td>Oligo fwd</td><td>CACCGGAGCAGAGGGCGGAGAGGA</td><td></td><td></td><td></td><td></td><td></td><td></td><td></td><td></td><td></td><td></td><td></td></tr>
<tr><td>Oligo rev</td><td>AAACTCCTCTCCGCCCTCTGCTCC</td><td></td><td></td><td></td><td></td><td></td><td></td><td></td><td></td><td></td><td></td><td></td></tr>
<tr><td>Chromosome</td><td>start</td><td></td><td>end</td><td>strand</td><td>MM</td><td>target_seq</td><td>PAM</td><td>alignment</td><td>distance</td><td>position</td><td>gene name</td><td>gene id</td></tr>
<tr><td>GU305914.1</td><td></td><td>61297</td><td>61319</td><td>+</td><td>0</td><td>GGAGCAGAGGGCGGAGAGGA</td><td>AGG</td><td>||||||||[||||||||||||]PAM</td><td>NA</td><td>Intergenic</td><td>NA</td><td>NA</td></tr>
<tr><td>T4093</td><td>GGCGGAGAGGAAGGCGTCGAAGG</td><td></td><td>1000</td><td></td><td></td><td></td><td></td><td></td><td></td><td></td><td></td><td></td></tr>
<tr><td>Oligo fwd</td><td>CACCGGCGGAGAGGAAGGCGTCGA</td><td></td><td></td><td></td><td></td><td></td><td></td><td></td><td></td><td></td><td></td><td></td></tr>
<tr><td>Oligo rev</td><td>AAACTCGACGCCTTCCTCTCCGCC</td><td></td><td></td><td></td><td></td><td></td><td></td><td></td><td></td><td></td><td></td><td></td></tr>
<tr><td>Chromosome</td><td>start</td><td></td><td>end</td><td>strand</td><td>MM</td><td>target_seq</td><td>PAM</td><td>alignment</td><td>distance</td><td>position</td><td>gene name</td><td>gene id</td></tr>
<tr><td>GU305914.1</td><td></td><td>61306</td><td>61328</td><td>+</td><td>0</td><td>GGCGGAGAGGAAGGCGTCGA</td><td>AGG</td><td>||||||||[||||||||||||]PAM</td><td>NA</td><td>Intergenic</td><td>NA</td><td>NA</td></tr>
<tr><td>GU305914.1</td><td></td><td>74988</td><td>75010</td><td>+</td><td>4</td><td>GGCGCGGAGGAAGGCGACGC</td><td>AGG</td><td>||||--||[||||||||-||-]PAM</td><td>NA</td><td>Intergenic</td><td>NA</td><td>NA</td></tr>
<tr><td>T4094</td><td>GAAGGCGTCGAAGGCCTGCGCGG</td><td></td><td>1000</td><td></td><td></td><td></td><td></td><td></td><td></td><td></td><td></td><td></td></tr>
<tr><td>Oligo fwd</td><td>CACCGAAGGCGTCGAAGGCCTGCG</td><td></td><td></td><td></td><td></td><td></td><td></td><td></td><td></td><td></td><td></td><td></td></tr>
<tr><td>Oligo rev</td><td>AAACCGCAGGCCTTCGACGCCTTC</td><td></td><td></td><td></td><td></td><td></td><td></td><td></td><td></td><td></td><td></td><td></td></tr>
<tr><td>Chromosome</td><td>start</td><td></td><td>end</td><td>strand</td><td>MM</td><td>target_seq</td><td>PAM</td><td>alignment</td><td>distance</td><td>position</td><td>gene name</td><td>gene id</td></tr>
<tr><td>GU305914.1</td><td></td><td>61315</td><td>61337</td><td>+</td><td>0</td><td>GAAGGCGTCGAAGGCCTGCG</td><td>CGG</td><td>||||||||[||||||||||||]PAM</td><td>NA</td><td>Intergenic</td><td>NA</td><td>NA</td></tr>
<tr><td>T4095</td><td>GAAGGCCTGCGCGGCGTCGACGG</td><td></td><td>1000</td><td></td><td></td><td></td><td></td><td></td><td></td><td></td><td></td><td></td></tr>
<tr><td>Oligo fwd</td><td>CACCGAAGGCCTGCGCGGCGTCGA</td><td></td><td></td><td></td><td></td><td></td><td></td><td></td><td></td><td></td><td></td><td></td></tr>
<tr><td>Oligo rev</td><td>AAACTCGACGCCGCGCAGGCCTTC</td><td></td><td></td><td></td><td></td><td></td><td></td><td></td><td></td><td></td><td></td><td></td></tr>
<tr><td>Chromosome</td><td>start</td><td></td><td>end</td><td>strand</td><td>MM</td><td>target_seq</td><td>PAM</td><td>alignment</td><td>distance</td><td>position</td><td>gene name</td><td>gene id</td></tr>
<tr><td>GU305914.1</td><td></td><td>61324</td><td>61346</td><td>+</td><td>0</td><td>GAAGGCCTGCGCGGCGTCGA</td><td>CGG</td><td>||||||||[||||||||||||]PAM</td><td>NA</td><td>Intergenic</td><td>NA</td><td>NA</td></tr>
<tr><td>GU305914.1</td><td></td><td>128755</td><td>128777</td><td>+</td><td>4</td><td>CGATGCCTCCGCGGCGTCGA</td><td>CGG</td><td>--|-||||[-|||||||||||]PAM</td><td>NA</td><td>Intergenic</td><td>NA</td><td>NA</td></tr>
<tr><td>T4096</td><td>AAGGCCTGCGCGGCGTCGACGGG</td><td></td><td>1000</td><td></td><td></td><td></td><td></td><td></td><td></td><td></td><td></td><td></td></tr>
<tr><td>Oligo fwd</td><td>CACCAAGGCCTGCGCGGCGTCGAC</td><td></td><td></td><td></td><td></td><td></td><td></td><td></td><td></td><td></td><td></td><td></td></tr>
<tr><td>Oligo rev</td><td>AAACGTCGACGCCGCGCAGGCCTT</td><td></td><td></td><td></td><td></td><td></td><td></td><td></td><td></td><td></td><td></td><td></td></tr>
<tr><td>Chromosome</td><td>start</td><td></td><td>end</td><td>strand</td><td>MM</td><td>target_seq</td><td>PAM</td><td>alignment</td><td>distance</td><td>position</td><td>gene name</td><td>gene id</td></tr>
<tr><td>GU305914.1</td><td></td><td>61325</td><td>61347</td><td>+</td><td>0</td><td>AAGGCCTGCGCGGCGTCGAC</td><td>GGG</td><td>||||||||[||||||||||||]PAM</td><td>NA</td><td>Intergenic</td><td>NA</td><td>NA</td></tr>
</table>

<table>
<tr><td>T4097</td><td>CTGCGCGGCGTCGACGGGCTTGG</td><td>1000</td><td></td><td></td><td></td><td></td><td></td><td></td><td></td><td></td><td></td></tr>
<tr><td>Oligo fwd</td><td>CACCCTGCGCGGCGTCGACGGGCT</td><td></td><td></td><td></td><td></td><td></td><td></td><td></td><td></td><td></td><td></td></tr>
<tr><td>Oligo rev</td><td>AAACAGCCCGTCGACGCCGCGCAG</td><td></td><td></td><td></td><td></td><td></td><td></td><td></td><td></td><td></td><td></td></tr>
<tr><td>Chromosome</td><td>start</td><td>end</td><td>strand</td><td>MM</td><td>target_seq</td><td>PAM</td><td>alignment</td><td>distance</td><td>position</td><td>gene name</td><td>gene id</td></tr>
<tr><td>GU305914.1</td><td>61330</td><td>61352</td><td>+</td><td>0</td><td>CTGCGCGGCGTCGACGGGCT</td><td>TGG</td><td>||||||||[||||||||||||]PAM</td><td>NA</td><td>Intergenic</td><td>NA</td><td>NA</td></tr>
<tr><td>T4098</td><td>CGTCGACGGGCTTGGAGCTGAGG</td><td>1000</td><td></td><td></td><td></td><td></td><td></td><td></td><td></td><td></td><td></td></tr>
<tr><td>Oligo fwd</td><td>CACCCGTCGACGGGCTTGGAGCTG</td><td></td><td></td><td></td><td></td><td></td><td></td><td></td><td></td><td></td><td></td></tr>
<tr><td>Oligo rev</td><td>AAACCAGCTCCAAGCCCGTCGACG</td><td></td><td></td><td></td><td></td><td></td><td></td><td></td><td></td><td></td><td></td></tr>
<tr><td>Chromosome</td><td>start</td><td>end</td><td>strand</td><td>MM</td><td>target_seq</td><td>PAM</td><td>alignment</td><td>distance</td><td>position</td><td>gene name</td><td>gene id</td></tr>
<tr><td>GU305914.1</td><td>61338</td><td>61360</td><td>+</td><td>0</td><td>CGTCGACGGGCTTGGAGCTG</td><td>AGG</td><td>||||||||[||||||||||||]PAM</td><td>NA</td><td>Intergenic</td><td>NA</td><td>NA</td></tr>
<tr><td>T4099</td><td>CCCGTCGACCTCGCACTCGACGG</td><td>1000</td><td></td><td></td><td></td><td></td><td></td><td></td><td></td><td></td><td></td></tr>
<tr><td>Oligo fwd</td><td>CACCCCCGTCGACCTCGCACTCGA</td><td></td><td></td><td></td><td></td><td></td><td></td><td></td><td></td><td></td><td></td></tr>
<tr><td>Oligo rev</td><td>AAACTCGAGTGCGAGGTCGACGGG</td><td></td><td></td><td></td><td></td><td></td><td></td><td></td><td></td><td></td><td></td></tr>
<tr><td>Chromosome</td><td>start</td><td>end</td><td>strand</td><td>MM</td><td>target_seq</td><td>PAM</td><td>alignment</td><td>distance</td><td>position</td><td>gene name</td><td>gene id</td></tr>
<tr><td>GU305914.1</td><td>61378</td><td>61400</td><td>+</td><td>0</td><td>CCCGTCGACCTCGCACTCGA</td><td>CGG</td><td>||||||||[||||||||||||]PAM</td><td>NA</td><td>Intergenic</td><td>NA</td><td>NA</td></tr>
<tr><td>T4100</td><td>GCACTCGACGGCGAGAGCCGCGG</td><td>1000</td><td></td><td></td><td></td><td></td><td></td><td></td><td></td><td></td><td></td></tr>
<tr><td>Oligo fwd</td><td>CACCGCACTCGACGGCGAGAGCCG</td><td></td><td></td><td></td><td></td><td></td><td></td><td></td><td></td><td></td><td></td></tr>
<tr><td>Oligo rev</td><td>AAACCGGCTCTCGCCGTCGAGTGC</td><td></td><td></td><td></td><td></td><td></td><td></td><td></td><td></td><td></td><td></td></tr>
<tr><td>Chromosome</td><td>start</td><td>end</td><td>strand</td><td>MM</td><td>target_seq</td><td>PAM</td><td>alignment</td><td>distance</td><td>position</td><td>gene name</td><td>gene id</td></tr>
<tr><td>GU305914.1</td><td>61390</td><td>61412</td><td>+</td><td>0</td><td>GCACTCGACGGCGAGAGCCG</td><td>CGG</td><td>||||||||[||||||||||||]PAM</td><td>NA</td><td>Intergenic</td><td>NA</td><td>NA</td></tr>
<tr><td>T4101</td><td>CGGCGAGAGCCGCGGCGACCCGG</td><td>1000</td><td></td><td></td><td></td><td></td><td></td><td></td><td></td><td></td><td></td></tr>
<tr><td>Oligo fwd</td><td>CACCCGGCGAGAGCCGCGGCGACC</td><td></td><td></td><td></td><td></td><td></td><td></td><td></td><td></td><td></td><td></td></tr>
<tr><td>Oligo rev</td><td>AAACGGTCGCCGCGGCTCTCGCCG</td><td></td><td></td><td></td><td></td><td></td><td></td><td></td><td></td><td></td><td></td></tr>
<tr><td>Chromosome</td><td>start</td><td>end</td><td>strand</td><td>MM</td><td>target_seq</td><td>PAM</td><td>alignment</td><td>distance</td><td>position</td><td>gene name</td><td>gene id</td></tr>
<tr><td>GU305914.1</td><td>61398</td><td>61420</td><td>+</td><td>0</td><td>CGGCGAGAGCCGCGGCGACC</td><td>CGG</td><td>||||||||[||||||||||||]PAM</td><td>NA</td><td>Intergenic</td><td>NA</td><td>NA</td></tr>
<tr><td>T4102</td><td>GGCGAGAGCCGCGGCGACCCGGG</td><td>1000</td><td></td><td></td><td></td><td></td><td></td><td></td><td></td><td></td><td></td></tr>
<tr><td>Oligo fwd</td><td>CACCGGCGAGAGCCGCGGCGACCC</td><td></td><td></td><td></td><td></td><td></td><td></td><td></td><td></td><td></td><td></td></tr>
<tr><td>Oligo rev</td><td>AAACGGGTCGCCGCGGCTCTCGCC</td><td></td><td></td><td></td><td></td><td></td><td></td><td></td><td></td><td></td><td></td></tr>
<tr><td>Chromosome</td><td>start</td><td>end</td><td>strand</td><td>MM</td><td>target_seq</td><td>PAM</td><td>alignment</td><td>distance</td><td>position</td><td>gene name</td><td>gene id</td></tr>
<tr><td>GU305914.1</td><td>61399</td><td>61421</td><td>+</td><td>0</td><td>GGCGAGAGCCGCGGCGACCC</td><td>GGG</td><td>||||||||[||||||||||||]PAM</td><td>NA</td><td>Intergenic</td><td>NA</td><td>NA</td></tr>
<tr><td>T4103</td><td>CGCGGCGACCCGGGCGAAGACGG</td><td>1000</td><td></td><td></td><td></td><td></td><td></td><td></td><td></td><td></td><td></td></tr>
<tr><td>Oligo fwd</td><td>CACCCGCGGCGACCCGGGCGAAGA</td><td></td><td></td><td></td><td></td><td></td><td></td><td></td><td></td><td></td><td></td></tr>
<tr><td>Oligo rev</td><td>AAACTCTTCGCCCGGGTCGCCGCG</td><td></td><td></td><td></td><td></td><td></td><td></td><td></td><td></td><td></td><td></td></tr>
<tr><td>Chromosome</td><td>start</td><td>end</td><td>strand</td><td>MM</td><td>target_seq</td><td>PAM</td><td>alignment</td><td>distance</td><td>position</td><td>gene name</td><td>gene id</td></tr>
<tr><td>GU305914.1</td><td>61408</td><td>61430</td><td>+</td><td>0</td><td>CGCGGCGACCCGGGCGAAGA</td><td>CGG</td><td>||||||||[||||||||||||]PAM</td><td>NA</td><td>Intergenic</td><td>NA</td><td>NA</td></tr>
<tr><td>T4104</td><td>GACCCGGGCGAAGACGGAGATGG</td><td>1000</td><td></td><td></td><td></td><td></td><td></td><td></td><td></td><td></td><td></td></tr>
<tr><td>Oligo fwd</td><td>CACCGACCCGGGCGAAGACGGAGA</td><td></td><td></td><td></td><td></td><td></td><td></td><td></td><td></td><td></td><td></td></tr>
<tr><td>Oligo rev</td><td>AAACTCTCCGTCTTCGCCCGGGTC</td><td></td><td></td><td></td><td></td><td></td><td></td><td></td><td></td><td></td><td></td></tr>
<tr><td>Chromosome</td><td>start</td><td>end</td><td>strand</td><td>MM</td><td>target_seq</td><td>PAM</td><td>alignment</td><td>distance</td><td>position</td><td>gene name</td><td>gene id</td></tr>
<tr><td>GU305914.1</td><td>61414</td><td>61436</td><td>+</td><td>0</td><td>GACCCGGGCGAAGACGGAGA</td><td>TGG</td><td>||||||||[||||||||||||]PAM</td><td>NA</td><td>Intergenic</td><td>NA</td><td>NA</td></tr>
<tr><td>T4105</td><td>ACCCGGGCGAAGACGGAGATGGG</td><td>1000</td><td></td><td></td><td></td><td></td><td></td><td></td><td></td><td></td><td></td></tr>
<tr><td>Oligo fwd</td><td>CACCACCCGGGCGAAGACGGAGAT</td><td></td><td></td><td></td><td></td><td></td><td></td><td></td><td></td><td></td><td></td></tr>
<tr><td>Oligo rev</td><td>AAACATCTCCGTCTTCGCCCGGGT</td><td></td><td></td><td></td><td></td><td></td><td></td><td></td><td></td><td></td><td></td></tr>
<tr><td>Chromosome</td><td>start</td><td>end</td><td>strand</td><td>MM</td><td>target_seq</td><td>PAM</td><td>alignment</td><td>distance</td><td>position</td><td>gene name</td><td>gene id</td></tr>
<tr><td>GU305914.1</td><td>61415</td><td>61437</td><td>+</td><td>0</td><td>ACCCGGGCGAAGACGGAGAT</td><td>GGG</td><td>||||||||[||||||||||||]PAM</td><td>NA</td><td>Intergenic</td><td>NA</td><td>NA</td></tr>
<tr><td>T4106</td><td>CCCGGGCGAAGACGGAGATGGGG</td><td>1000</td><td></td><td></td><td></td><td></td><td></td><td></td><td></td><td></td><td></td></tr>
<tr><td>Oligo fwd</td><td>CACCCCCGGGCGAAGACGGAGATG</td><td></td><td></td><td></td><td></td><td></td><td></td><td></td><td></td><td></td><td></td></tr>
<tr><td>Oligo rev</td><td>AAACCATCTCCGTCTTCGCCCGGG</td><td></td><td></td><td></td><td></td><td></td><td></td><td></td><td></td><td></td><td></td></tr>
<tr><td>Chromosome</td><td>start</td><td>end</td><td>strand</td><td>MM</td><td>target_seq</td><td>PAM</td><td>alignment</td><td>distance</td><td>position</td><td>gene name</td><td>gene id</td></tr>
<tr><td>GU305914.1</td><td>61416</td><td>61438</td><td>+</td><td>0</td><td>CCCGGGCGAAGACGGAGATG</td><td>GGG</td><td>||||||||[||||||||||||]PAM</td><td>NA</td><td>Intergenic</td><td>NA</td><td>NA</td></tr>
<tr><td>T4107</td><td>CCGGGCGAAGACGGAGATGGGGG</td><td>1000</td><td></td><td></td><td></td><td></td><td></td><td></td><td></td><td></td><td></td></tr>
<tr><td>Oligo fwd</td><td>CACCCCGGGCGAAGACGGAGATGG</td><td></td><td></td><td></td><td></td><td></td><td></td><td></td><td></td><td></td><td></td></tr>
<tr><td>Oligo rev</td><td>AAACCCATCTCCGTCTTCGCCCGG</td><td></td><td></td><td></td><td></td><td></td><td></td><td></td><td></td><td></td><td></td></tr>
<tr><td>Chromosome</td><td>start</td><td>end</td><td>strand</td><td>MM</td><td>target_seq</td><td>PAM</td><td>alignment</td><td>distance</td><td>position</td><td>gene name</td><td>gene id</td></tr>
<tr><td>GU305914.1</td><td>61417</td><td>61439</td><td>+</td><td>0</td><td>CCGGGCGAAGACGGAGATGG</td><td>GGG</td><td>||||||||[||||||||||||]PAM</td><td>NA</td><td>Intergenic</td><td>NA</td><td>NA</td></tr>
<tr><td>T4108</td><td>TGGGGGTCTCCTCGCGTCCCTGG</td><td>1000</td><td></td><td></td><td></td><td></td><td></td><td></td><td></td><td></td><td></td></tr>
<tr><td>Oligo fwd</td><td>CACCTGGGGGTCTCCTCGCGTCCC</td><td></td><td></td><td></td><td></td><td></td><td></td><td></td><td></td><td></td><td></td></tr>
<tr><td>Oligo rev</td><td>AAACGGGACGCGAGGAGACCCCCA</td><td></td><td></td><td></td><td></td><td></td><td></td><td></td><td></td><td></td><td></td></tr>
<tr><td>Chromosome</td><td>start</td><td>end</td><td>strand</td><td>MM</td><td>target_seq</td><td>PAM</td><td>alignment</td><td>distance</td><td>position</td><td>gene name</td><td>gene id</td></tr>
<tr><td>GU305914.1</td><td>61434</td><td>61456</td><td>+</td><td>0</td><td>TGGGGGTCTCCTCGCGTCCC</td><td>TGG</td><td>||||||||[||||||||||||]PAM</td><td>NA</td><td>Intergenic</td><td>NA</td><td>NA</td></tr>
<tr><td>T4109</td><td>GCAGCGATCCATCATGACGACGG</td><td>1000</td><td></td><td></td><td></td><td></td><td></td><td></td><td></td><td></td><td></td></tr>
<tr><td>Oligo fwd</td><td>CACCGCAGCGATCCATCATGACGA</td><td></td><td></td><td></td><td></td><td></td><td></td><td></td><td></td><td></td><td></td></tr>
<tr><td>Oligo rev</td><td>AAACTCGTCATGATGGATCGCTGC</td><td></td><td></td><td></td><td></td><td></td><td></td><td></td><td></td><td></td><td></td></tr>
<tr><td>Chromosome</td><td>start</td><td>end</td><td>strand</td><td>MM</td><td>target_seq</td><td>PAM</td><td>alignment</td><td>distance</td><td>position</td><td>gene name</td><td>gene id</td></tr>
<tr><td>GU305914.1</td><td>61468</td><td>61490</td><td>+</td><td>0</td><td>GCAGCGATCCATCATGACGA</td><td>CGG</td><td>||||||||[||||||||||||]PAM</td><td>NA</td><td>Intergenic</td><td>NA</td><td>NA</td></tr>
<tr><td>T4110</td><td>TCATGACGACGGTCGAGTAGAGG</td><td>1000</td><td></td><td></td><td></td><td></td><td></td><td></td><td></td><td></td><td></td></tr>
<tr><td>Oligo fwd</td><td>CACCTCATGACGACGGTCGAGTAG</td><td></td><td></td><td></td><td></td><td></td><td></td><td></td><td></td><td></td><td></td></tr>
<tr><td>Oligo rev</td><td>AAACCTACTCGACCGTCGTCATGA</td><td></td><td></td><td></td><td></td><td></td><td></td><td></td><td></td><td></td><td></td></tr>
<tr><td>Chromosome</td><td>start</td><td>end</td><td>strand</td><td>MM</td><td>target_seq</td><td>PAM</td><td>alignment</td><td>distance</td><td>position</td><td>gene name</td><td>gene id</td></tr>
<tr><td>GU305914.1</td><td>61479</td><td>61501</td><td>+</td><td>0</td><td>TCATGACGACGGTCGAGTAG</td><td>AGG</td><td>||||||||[||||||||||||]PAM</td><td>NA</td><td>Intergenic</td><td>NA</td><td>NA</td></tr>
<tr><td>T4111</td><td>TCGAGTAGAGGCCCCTACCCAGG</td><td>1000</td><td></td><td></td><td></td><td></td><td></td><td></td><td></td><td></td><td></td></tr>
<tr><td>Oligo fwd</td><td>CACCTCGAGTAGAGGCCCCTACCC</td><td></td><td></td><td></td><td></td><td></td><td></td><td></td><td></td><td></td><td></td></tr>
<tr><td>Oligo rev</td><td>AAACGGGTAGGGGCCTCTACTCGA</td><td></td><td></td><td></td><td></td><td></td><td></td><td></td><td></td><td></td><td></td></tr>
<tr><td>Chromosome</td><td>start</td><td>end</td><td>strand</td><td>MM</td><td>target_seq</td><td>PAM</td><td>alignment</td><td>distance</td><td>position</td><td>gene name</td><td>gene id</td></tr>
</table>

<table>
<tr><td>GU305914.1</td><td>61491</td><td>61513</td><td>+</td><td>0</td><td>TCGAGTAGAGGCCCCTACCC</td><td>AGG</td><td>||||||||[||||||||||||]PAM</td><td>NA</td><td>Intergenic</td><td>NA</td><td>NA</td></tr>
<tr><td></td><td></td><td></td><td></td><td></td><td></td><td></td><td></td><td></td><td></td><td></td><td></td></tr>
<tr><td>T4112</td><td>CGAGTAGAGGCCCCTACCCAGGG</td><td>1000</td><td></td><td></td><td></td><td></td><td></td><td></td><td></td><td></td><td></td></tr>
<tr><td>Oligo fwd</td><td>CACCCGAGTAGAGGCCCCTACCCA</td><td></td><td></td><td></td><td></td><td></td><td></td><td></td><td></td><td></td><td></td></tr>
<tr><td>Oligo rev</td><td>AAACTGGGTAGGGGCCTCTACTCG</td><td></td><td></td><td></td><td></td><td></td><td></td><td></td><td></td><td></td><td></td></tr>
<tr><td>Chromosome</td><td>start</td><td>end</td><td>strand</td><td>MM</td><td>target_seq</td><td>PAM</td><td>alignment</td><td>distance</td><td>position</td><td>gene name</td><td>gene id</td></tr>
<tr><td>GU305914.1</td><td>61492</td><td>61514</td><td>+</td><td>0</td><td>CGAGTAGAGGCCCCTACCCA</td><td>GGG</td><td>||||||||[||||||||||||]PAM</td><td>NA</td><td>Intergenic</td><td>NA</td><td>NA</td></tr>
<tr><td></td><td></td><td></td><td></td><td></td><td></td><td></td><td></td><td></td><td></td><td></td><td></td></tr>
<tr><td>T4113</td><td>TAGAGGCCCCTACCCAGGGCCGG</td><td>1000</td><td></td><td></td><td></td><td></td><td></td><td></td><td></td><td></td><td></td></tr>
<tr><td>Oligo fwd</td><td>CACCTAGAGGCCCCTACCCAGGGC</td><td></td><td></td><td></td><td></td><td></td><td></td><td></td><td></td><td></td><td></td></tr>
<tr><td>Oligo rev</td><td>AAACGCCCTGGGTAGGGGCCTCTA</td><td></td><td></td><td></td><td></td><td></td><td></td><td></td><td></td><td></td><td></td></tr>
<tr><td>Chromosome</td><td>start</td><td>end</td><td>strand</td><td>MM</td><td>target_seq</td><td>PAM</td><td>alignment</td><td>distance</td><td>position</td><td>gene name</td><td>gene id</td></tr>
<tr><td>GU305914.1</td><td>61496</td><td>61518</td><td>+</td><td>0</td><td>TAGAGGCCCCTACCCAGGGC</td><td>CGG</td><td>||||||||[||||||||||||]PAM</td><td>NA</td><td>Intergenic</td><td>NA</td><td>NA</td></tr>
<tr><td></td><td></td><td></td><td></td><td></td><td></td><td></td><td></td><td></td><td></td><td></td><td></td></tr>
<tr><td>T4114</td><td>CTCCGTGATGCGCTGCGCCATGG</td><td>1000</td><td></td><td></td><td></td><td></td><td></td><td></td><td></td><td></td><td></td></tr>
<tr><td>Oligo fwd</td><td>CACCCTCCGTGATGCGCTGCGCCA</td><td></td><td></td><td></td><td></td><td></td><td></td><td></td><td></td><td></td><td></td></tr>
<tr><td>Oligo rev</td><td>AAACTGGCGCAGCGCATCACGGAG</td><td></td><td></td><td></td><td></td><td></td><td></td><td></td><td></td><td></td><td></td></tr>
<tr><td>Chromosome</td><td>start</td><td>end</td><td>strand</td><td>MM</td><td>target_seq</td><td>PAM</td><td>alignment</td><td>distance</td><td>position</td><td>gene name</td><td>gene id</td></tr>
<tr><td>GU305914.1</td><td>61528</td><td>61550</td><td>+</td><td>0</td><td>CTCCGTGATGCGCTGCGCCA</td><td>TGG</td><td>||||||||[||||||||||||]PAM</td><td>NA</td><td>Intergenic</td><td>NA</td><td>NA</td></tr>
<tr><td></td><td></td><td></td><td></td><td></td><td></td><td></td><td></td><td></td><td></td><td></td><td></td></tr>
<tr><td>T4115</td><td>GGCGTCCACCACCATGCCGTAGG</td><td>1000</td><td></td><td></td><td></td><td></td><td></td><td></td><td></td><td></td><td></td></tr>
<tr><td>Oligo fwd</td><td>CACCGGCGTCCACCACCATGCCGT</td><td></td><td></td><td></td><td></td><td></td><td></td><td></td><td></td><td></td><td></td></tr>
<tr><td>Oligo rev</td><td>AAACACGGCATGGTGGTGGACGCC</td><td></td><td></td><td></td><td></td><td></td><td></td><td></td><td></td><td></td><td></td></tr>
<tr><td>Chromosome</td><td>start</td><td>end</td><td>strand</td><td>MM</td><td>target_seq</td><td>PAM</td><td>alignment</td><td>distance</td><td>position</td><td>gene name</td><td>gene id</td></tr>
<tr><td>GU305914.1</td><td>61549</td><td>61571</td><td>+</td><td>0</td><td>GGCGTCCACCACCATGCCGT</td><td>AGG</td><td>||||||||[||||||||||||]PAM</td><td>NA</td><td>Intergenic</td><td>NA</td><td>NA</td></tr>
<tr><td>GU305914.1</td><td>148010</td><td>148032</td><td>-</td><td>4</td><td>GCGTTCCACCACCATGCCGA</td><td>CGG</td><td>|---||||[|||||||||||-]PAM</td><td>NA</td><td>Intergenic</td><td>NA</td><td>NA</td></tr>
<tr><td></td><td></td><td></td><td></td><td></td><td></td><td></td><td></td><td></td><td></td><td></td><td></td></tr>
<tr><td>T4116</td><td>GCCGTAGGTCTTGAAGAACGCGG</td><td>1000</td><td></td><td></td><td></td><td></td><td></td><td></td><td></td><td></td><td></td></tr>
<tr><td>Oligo fwd</td><td>CACCGCCGTAGGTCTTGAAGAACG</td><td></td><td></td><td></td><td></td><td></td><td></td><td></td><td></td><td></td><td></td></tr>
<tr><td>Oligo rev</td><td>AAACCGTTCTTCAAGACCTACGGC</td><td></td><td></td><td></td><td></td><td></td><td></td><td></td><td></td><td></td><td></td></tr>
<tr><td>Chromosome</td><td>start</td><td>end</td><td>strand</td><td>MM</td><td>target_seq</td><td>PAM</td><td>alignment</td><td>distance</td><td>position</td><td>gene name</td><td>gene id</td></tr>
<tr><td>GU305914.1</td><td>61564</td><td>61586</td><td>+</td><td>0</td><td>GCCGTAGGTCTTGAAGAACG</td><td>CGG</td><td>||||||||[||||||||||||]PAM</td><td>NA</td><td>Intergenic</td><td>NA</td><td>NA</td></tr>
<tr><td></td><td></td><td></td><td></td><td></td><td></td><td></td><td></td><td></td><td></td><td></td><td></td></tr>
<tr><td>T4117</td><td>ACGCGGCGACAGCATCCCGCAGG</td><td>1000</td><td></td><td></td><td></td><td></td><td></td><td></td><td></td><td></td><td></td></tr>
<tr><td>Oligo fwd</td><td>CACCACGCGGCGACAGCATCCCGC</td><td></td><td></td><td></td><td></td><td></td><td></td><td></td><td></td><td></td><td></td></tr>
<tr><td>Oligo rev</td><td>AAACGCGGGATGCTGTCGCCGCGT</td><td></td><td></td><td></td><td></td><td></td><td></td><td></td><td></td><td></td><td></td></tr>
<tr><td>Chromosome</td><td>start</td><td>end</td><td>strand</td><td>MM</td><td>target_seq</td><td>PAM</td><td>alignment</td><td>distance</td><td>position</td><td>gene name</td><td>gene id</td></tr>
<tr><td>GU305914.1</td><td>61581</td><td>61603</td><td>+</td><td>0</td><td>ACGCGGCGACAGCATCCCGC</td><td>AGG</td><td>||||||||[||||||||||||]PAM</td><td>NA</td><td>Intergenic</td><td>NA</td><td>NA</td></tr>
<tr><td></td><td></td><td></td><td></td><td></td><td></td><td></td><td></td><td></td><td></td><td></td><td></td></tr>
<tr><td>T4118</td><td>CCCGCAGGCGATCGCAGTCGCGG</td><td>1000</td><td></td><td></td><td></td><td></td><td></td><td></td><td></td><td></td><td></td></tr>
<tr><td>Oligo fwd</td><td>CACCCCCGCAGGCGATCGCAGTCG</td><td></td><td></td><td></td><td></td><td></td><td></td><td></td><td></td><td></td><td></td></tr>
<tr><td>Oligo rev</td><td>AAACCGACTGCGATCGCCTGCGGG</td><td></td><td></td><td></td><td></td><td></td><td></td><td></td><td></td><td></td><td></td></tr>
<tr><td>Chromosome</td><td>start</td><td>end</td><td>strand</td><td>MM</td><td>target_seq</td><td>PAM</td><td>alignment</td><td>distance</td><td>position</td><td>gene name</td><td>gene id</td></tr>
<tr><td>GU305914.1</td><td>61596</td><td>61618</td><td>+</td><td>0</td><td>CCCGCAGGCGATCGCAGTCG</td><td>CGG</td><td>||||||||[||||||||||||]PAM</td><td>NA</td><td>Intergenic</td><td>NA</td><td>NA</td></tr>
<tr><td></td><td></td><td></td><td></td><td></td><td></td><td></td><td></td><td></td><td></td><td></td><td></td></tr>
<tr><td>T4119</td><td>GCGATCGCAGTCGCGGTAGAAGG</td><td>1000</td><td></td><td></td><td></td><td></td><td></td><td></td><td></td><td></td><td></td></tr>
<tr><td>Oligo fwd</td><td>CACCGCGATCGCAGTCGCGGTAGA</td><td></td><td></td><td></td><td></td><td></td><td></td><td></td><td></td><td></td><td></td></tr>
<tr><td>Oligo rev</td><td>AAACTCTACCGCGACTGCGATCGC</td><td></td><td></td><td></td><td></td><td></td><td></td><td></td><td></td><td></td><td></td></tr>
<tr><td>Chromosome</td><td>start</td><td>end</td><td>strand</td><td>MM</td><td>target_seq</td><td>PAM</td><td>alignment</td><td>distance</td><td>position</td><td>gene name</td><td>gene id</td></tr>
<tr><td>GU305914.1</td><td>61603</td><td>61625</td><td>+</td><td>0</td><td>GCGATCGCAGTCGCGGTAGA</td><td>AGG</td><td>||||||||[||||||||||||]PAM</td><td>NA</td><td>Intergenic</td><td>NA</td><td>NA</td></tr>
<tr><td></td><td></td><td></td><td></td><td></td><td></td><td></td><td></td><td></td><td></td><td></td><td></td></tr>
<tr><td>T4120</td><td>AGTCGCGGTAGAAGGCCGAGAGG</td><td>1000</td><td></td><td></td><td></td><td></td><td></td><td></td><td></td><td></td><td></td></tr>
<tr><td>Oligo fwd</td><td>CACCAGTCGCGGTAGAAGGCCGAG</td><td></td><td></td><td></td><td></td><td></td><td></td><td></td><td></td><td></td><td></td></tr>
<tr><td>Oligo rev</td><td>AAACCTCGGCCTTCTACCGCGACT</td><td></td><td></td><td></td><td></td><td></td><td></td><td></td><td></td><td></td><td></td></tr>
<tr><td>Chromosome</td><td>start</td><td>end</td><td>strand</td><td>MM</td><td>target_seq</td><td>PAM</td><td>alignment</td><td>distance</td><td>position</td><td>gene name</td><td>gene id</td></tr>
<tr><td>GU305914.1</td><td>61611</td><td>61633</td><td>+</td><td>0</td><td>AGTCGCGGTAGAAGGCCGAG</td><td>AGG</td><td>||||||||[||||||||||||]PAM</td><td>NA</td><td>Intergenic</td><td>NA</td><td>NA</td></tr>
<tr><td></td><td></td><td></td><td></td><td></td><td></td><td></td><td></td><td></td><td></td><td></td><td></td></tr>
<tr><td>T4121</td><td>GTCGCGGTAGAAGGCCGAGAGGG</td><td>1000</td><td></td><td></td><td></td><td></td><td></td><td></td><td></td><td></td><td></td></tr>
<tr><td>Oligo fwd</td><td>CACCGTCGCGGTAGAAGGCCGAGA</td><td></td><td></td><td></td><td></td><td></td><td></td><td></td><td></td><td></td><td></td></tr>
<tr><td>Oligo rev</td><td>AAACTCTCGGCCTTCTACCGCGAC</td><td></td><td></td><td></td><td></td><td></td><td></td><td></td><td></td><td></td><td></td></tr>
<tr><td>Chromosome</td><td>start</td><td>end</td><td>strand</td><td>MM</td><td>target_seq</td><td>PAM</td><td>alignment</td><td>distance</td><td>position</td><td>gene name</td><td>gene id</td></tr>
<tr><td>GU305914.1</td><td>61612</td><td>61634</td><td>+</td><td>0</td><td>GTCGCGGTAGAAGGCCGAGA</td><td>GGG</td><td>||||||||[||||||||||||]PAM</td><td>NA</td><td>Intergenic</td><td>NA</td><td>NA</td></tr>
<tr><td></td><td></td><td></td><td></td><td></td><td></td><td></td><td></td><td></td><td></td><td></td><td></td></tr>
<tr><td>T4122</td><td>TCGCGGTAGAAGGCCGAGAGGGG</td><td>1000</td><td></td><td></td><td></td><td></td><td></td><td></td><td></td><td></td><td></td></tr>
<tr><td>Oligo fwd</td><td>CACCTCGCGGTAGAAGGCCGAGAG</td><td></td><td></td><td></td><td></td><td></td><td></td><td></td><td></td><td></td><td></td></tr>
<tr><td>Oligo rev</td><td>AAACCTCTCGGCCTTCTACCGCGA</td><td></td><td></td><td></td><td></td><td></td><td></td><td></td><td></td><td></td><td></td></tr>
<tr><td>Chromosome</td><td>start</td><td>end</td><td>strand</td><td>MM</td><td>target_seq</td><td>PAM</td><td>alignment</td><td>distance</td><td>position</td><td>gene name</td><td>gene id</td></tr>
<tr><td>GU305914.1</td><td>61613</td><td>61635</td><td>+</td><td>0</td><td>TCGCGGTAGAAGGCCGAGAG</td><td>GGG</td><td>||||||||[||||||||||||]PAM</td><td>NA</td><td>Intergenic</td><td>NA</td><td>NA</td></tr>
<tr><td></td><td></td><td></td><td></td><td></td><td></td><td></td><td></td><td></td><td></td><td></td><td></td></tr>
<tr><td>T4123</td><td>CGCGGTAGAAGGCCGAGAGGGGG</td><td>1000</td><td></td><td></td><td></td><td></td><td></td><td></td><td></td><td></td><td></td></tr>
<tr><td>Oligo fwd</td><td>CACCCGCGGTAGAAGGCCGAGAGG</td><td></td><td></td><td></td><td></td><td></td><td></td><td></td><td></td><td></td><td></td></tr>
<tr><td>Oligo rev</td><td>AAACCCTCTCGGCCTTCTACCGCG</td><td></td><td></td><td></td><td></td><td></td><td></td><td></td><td></td><td></td><td></td></tr>
<tr><td>Chromosome</td><td>start</td><td>end</td><td>strand</td><td>MM</td><td>target_seq</td><td>PAM</td><td>alignment</td><td>distance</td><td>position</td><td>gene name</td><td>gene id</td></tr>
<tr><td>GU305914.1</td><td>61614</td><td>61636</td><td>+</td><td>0</td><td>CGCGGTAGAAGGCCGAGAGG</td><td>GGG</td><td>||||||||[||||||||||||]PAM</td><td>NA</td><td>Intergenic</td><td>NA</td><td>NA</td></tr>
<tr><td></td><td></td><td></td><td></td><td></td><td></td><td></td><td></td><td></td><td></td><td></td><td></td></tr>
<tr><td>T4124</td><td>GTAGAAGGCCGAGAGGGGGACGG</td><td>1000</td><td></td><td></td><td></td><td></td><td></td><td></td><td></td><td></td><td></td></tr>
<tr><td>Oligo fwd</td><td>CACCGTAGAAGGCCGAGAGGGGGA</td><td></td><td></td><td></td><td></td><td></td><td></td><td></td><td></td><td></td><td></td></tr>
<tr><td>Oligo rev</td><td>AAACTCCCCCTCTCGGCCTTCTAC</td><td></td><td></td><td></td><td></td><td></td><td></td><td></td><td></td><td></td><td></td></tr>
<tr><td>Chromosome</td><td>start</td><td>end</td><td>strand</td><td>MM</td><td>target_seq</td><td>PAM</td><td>alignment</td><td>distance</td><td>position</td><td>gene name</td><td>gene id</td></tr>
<tr><td>GU305914.1</td><td>61618</td><td>61640</td><td>+</td><td>0</td><td>GTAGAAGGCCGAGAGGGGGA</td><td>CGG</td><td>||||||||[||||||||||||]PAM</td><td>NA</td><td>Intergenic</td><td>NA</td><td>NA</td></tr>
<tr><td></td><td></td><td></td><td></td><td></td><td></td><td></td><td></td><td></td><td></td><td></td><td></td></tr>
<tr><td>T4125</td><td>GAGGGGGACGGCGTTCCTCTTGG</td><td>1000</td><td></td><td></td><td></td><td></td><td></td><td></td><td></td><td></td><td></td></tr>
<tr><td>Oligo fwd</td><td>CACCGAGGGGGACGGCGTTCCTCT</td><td></td><td></td><td></td><td></td><td></td><td></td><td></td><td></td><td></td><td></td></tr>
<tr><td>Oligo rev</td><td>AAACAGAGGAACGCCGTCCCCCTC</td><td></td><td></td><td></td><td></td><td></td><td></td><td></td><td></td><td></td><td></td></tr>
<tr><td>Chromosome</td><td>start</td><td>end</td><td>strand</td><td>MM</td><td>target_seq</td><td>PAM</td><td>alignment</td><td>distance</td><td>position</td><td>gene name</td><td>gene id</td></tr>
<tr><td>GU305914.1</td><td>61630</td><td>61652</td><td>+</td><td>0</td><td>GAGGGGGACGGCGTTCCTCT</td><td>TGG</td><td>||||||||[||||||||||||]PAM</td><td>NA</td><td>Intergenic</td><td>NA</td><td>NA</td></tr>
<tr><td></td><td></td><td></td><td></td><td></td><td></td><td></td><td></td><td></td><td></td><td></td><td></td></tr>
<tr><td>T4126</td><td>CGCCGTCTCGACGTAACATTCGG</td><td>1000</td><td></td><td></td><td></td><td></td><td></td><td></td><td></td><td></td><td></td></tr>
<tr><td>Oligo fwd</td><td>CACCCGCCGTCTCGACGTAACATT</td><td></td><td></td><td></td><td></td><td></td><td></td><td></td><td></td><td></td><td></td></tr>
</table>

<table>
<tr><td>Oligo rev</td><td>AAACAATGTTACGTCGAGACGGCG</td><td></td><td></td><td></td><td></td><td></td><td></td><td></td><td></td><td></td><td></td></tr>
<tr><td>Chromosome</td><td>start</td><td>end</td><td>strand</td><td>MM</td><td>target_seq</td><td>PAM</td><td>alignment</td><td>distance</td><td>position</td><td>gene name</td><td>gene id</td></tr>
<tr><td>GU305914.1</td><td>61654</td><td>61676</td><td>+</td><td>0</td><td>CGCCGTCTCGACGTAACATT</td><td>CGG</td><td>||||||||[||||||||||||]PAM</td><td>NA</td><td>Intergenic</td><td>NA</td><td>NA</td></tr>
<tr><td>T4127</td><td>CGTAACATTCGGCGAGATCCAGG</td><td>1000</td><td></td><td></td><td></td><td></td><td></td><td></td><td></td><td></td><td></td></tr>
<tr><td>Oligo fwd</td><td>CACCCGTAACATTCGGCGAGATCC</td><td></td><td></td><td></td><td></td><td></td><td></td><td></td><td></td><td></td><td></td></tr>
<tr><td>Oligo rev</td><td>AAACGGATCTCGCCGAATGTTACG</td><td></td><td></td><td></td><td></td><td></td><td></td><td></td><td></td><td></td><td></td></tr>
<tr><td>Chromosome</td><td>start</td><td>end</td><td>strand</td><td>MM</td><td>target_seq</td><td>PAM</td><td>alignment</td><td>distance</td><td>position</td><td>gene name</td><td>gene id</td></tr>
<tr><td>GU305914.1</td><td>61665</td><td>61687</td><td>+</td><td>0</td><td>CGTAACATTCGGCGAGATCC</td><td>AGG</td><td>||||||||[||||||||||||]PAM</td><td>NA</td><td>Intergenic</td><td>NA</td><td>NA</td></tr>
<tr><td>T4128</td><td>GCGAGATCCAGGTCGCACGACGG</td><td>1000</td><td></td><td></td><td></td><td></td><td></td><td></td><td></td><td></td><td></td></tr>
<tr><td>Oligo fwd</td><td>CACCGCGAGATCCAGGTCGCACGA</td><td></td><td></td><td></td><td></td><td></td><td></td><td></td><td></td><td></td><td></td></tr>
<tr><td>Oligo rev</td><td>AAACTCGTGCGACCTGGATCTCGC</td><td></td><td></td><td></td><td></td><td></td><td></td><td></td><td></td><td></td><td></td></tr>
<tr><td>Chromosome</td><td>start</td><td>end</td><td>strand</td><td>MM</td><td>target_seq</td><td>PAM</td><td>alignment</td><td>distance</td><td>position</td><td>gene name</td><td>gene id</td></tr>
<tr><td>GU305914.1</td><td>61676</td><td>61698</td><td>+</td><td>0</td><td>GCGAGATCCAGGTCGCACGA</td><td>CGG</td><td>||||||||[||||||||||||]PAM</td><td>NA</td><td>Intergenic</td><td>NA</td><td>NA</td></tr>
<tr><td>T4129</td><td>CGAGATCCAGGTCGCACGACGGG</td><td>1000</td><td></td><td></td><td></td><td></td><td></td><td></td><td></td><td></td><td></td></tr>
<tr><td>Oligo fwd</td><td>CACCCGAGATCCAGGTCGCACGAC</td><td></td><td></td><td></td><td></td><td></td><td></td><td></td><td></td><td></td><td></td></tr>
<tr><td>Oligo rev</td><td>AAACGTCGTGCGACCTGGATCTCG</td><td></td><td></td><td></td><td></td><td></td><td></td><td></td><td></td><td></td><td></td></tr>
<tr><td>Chromosome</td><td>start</td><td>end</td><td>strand</td><td>MM</td><td>target_seq</td><td>PAM</td><td>alignment</td><td>distance</td><td>position</td><td>gene name</td><td>gene id</td></tr>
<tr><td>GU305914.1</td><td>61677</td><td>61699</td><td>+</td><td>0</td><td>CGAGATCCAGGTCGCACGAC</td><td>GGG</td><td>||||||||[||||||||||||]PAM</td><td>NA</td><td>Intergenic</td><td>NA</td><td>NA</td></tr>
<tr><td>T4130</td><td>ATCCAGGTCGCACGACGGGTCGG</td><td>1000</td><td></td><td></td><td></td><td></td><td></td><td></td><td></td><td></td><td></td></tr>
<tr><td>Oligo fwd</td><td>CACCATCCAGGTCGCACGACGGGT</td><td></td><td></td><td></td><td></td><td></td><td></td><td></td><td></td><td></td><td></td></tr>
<tr><td>Oligo rev</td><td>AAACACCCGTCGTGCGACCTGGAT</td><td></td><td></td><td></td><td></td><td></td><td></td><td></td><td></td><td></td><td></td></tr>
<tr><td>Chromosome</td><td>start</td><td>end</td><td>strand</td><td>MM</td><td>target_seq</td><td>PAM</td><td>alignment</td><td>distance</td><td>position</td><td>gene name</td><td>gene id</td></tr>
<tr><td>GU305914.1</td><td>61681</td><td>61703</td><td>+</td><td>0</td><td>ATCCAGGTCGCACGACGGGT</td><td>CGG</td><td>||||||||[||||||||||||]PAM</td><td>NA</td><td>Intergenic</td><td>NA</td><td>NA</td></tr>
<tr><td>T4131</td><td>GGTCGCACGACGGGTCGGTCTGG</td><td>1000</td><td></td><td></td><td></td><td></td><td></td><td></td><td></td><td></td><td></td></tr>
<tr><td>Oligo fwd</td><td>CACCGGTCGCACGACGGGTCGGTC</td><td></td><td></td><td></td><td></td><td></td><td></td><td></td><td></td><td></td><td></td></tr>
<tr><td>Oligo rev</td><td>AAACGACCGACCCGTCGTGCGACC</td><td></td><td></td><td></td><td></td><td></td><td></td><td></td><td></td><td></td><td></td></tr>
<tr><td>Chromosome</td><td>start</td><td>end</td><td>strand</td><td>MM</td><td>target_seq</td><td>PAM</td><td>alignment</td><td>distance</td><td>position</td><td>gene name</td><td>gene id</td></tr>
<tr><td>GU305914.1</td><td>61686</td><td>61708</td><td>+</td><td>0</td><td>GGTCGCACGACGGGTCGGTC</td><td>TGG</td><td>||||||||[||||||||||||]PAM</td><td>NA</td><td>Intergenic</td><td>NA</td><td>NA</td></tr>
<tr><td>T4132</td><td>ACGGGTCGGTCTGGATGAGACGG</td><td>1000</td><td></td><td></td><td></td><td></td><td></td><td></td><td></td><td></td><td></td></tr>
<tr><td>Oligo fwd</td><td>CACCACGGGTCGGTCTGGATGAGA</td><td></td><td></td><td></td><td></td><td></td><td></td><td></td><td></td><td></td><td></td></tr>
<tr><td>Oligo rev</td><td>AAACTCTCATCCAGACCGACCCGT</td><td></td><td></td><td></td><td></td><td></td><td></td><td></td><td></td><td></td><td></td></tr>
<tr><td>Chromosome</td><td>start</td><td>end</td><td>strand</td><td>MM</td><td>target_seq</td><td>PAM</td><td>alignment</td><td>distance</td><td>position</td><td>gene name</td><td>gene id</td></tr>
<tr><td>GU305914.1</td><td>61695</td><td>61717</td><td>+</td><td>0</td><td>ACGGGTCGGTCTGGATGAGA</td><td>CGG</td><td>||||||||[||||||||||||]PAM</td><td>NA</td><td>Intergenic</td><td>NA</td><td>NA</td></tr>
<tr><td>T4133</td><td>GTCGGTCTGGATGAGACGGTTGG</td><td>1000</td><td></td><td></td><td></td><td></td><td></td><td></td><td></td><td></td><td></td></tr>
<tr><td>Oligo fwd</td><td>CACCGTCGGTCTGGATGAGACGGT</td><td></td><td></td><td></td><td></td><td></td><td></td><td></td><td></td><td></td><td></td></tr>
<tr><td>Oligo rev</td><td>AAACACCGTCTCATCCAGACCGAC</td><td></td><td></td><td></td><td></td><td></td><td></td><td></td><td></td><td></td><td></td></tr>
<tr><td>Chromosome</td><td>start</td><td>end</td><td>strand</td><td>MM</td><td>target_seq</td><td>PAM</td><td>alignment</td><td>distance</td><td>position</td><td>gene name</td><td>gene id</td></tr>
<tr><td>GU305914.1</td><td>61699</td><td>61721</td><td>+</td><td>0</td><td>GTCGGTCTGGATGAGACGGT</td><td>TGG</td><td>||||||||[||||||||||||]PAM</td><td>NA</td><td>Intergenic</td><td>NA</td><td>NA</td></tr>
<tr><td>T4134</td><td>CTGGATGAGACGGTTGGCGCTGG</td><td>1000</td><td></td><td></td><td></td><td></td><td></td><td></td><td></td><td></td><td></td></tr>
<tr><td>Oligo fwd</td><td>CACCCTGGATGAGACGGTTGGCGC</td><td></td><td></td><td></td><td></td><td></td><td></td><td></td><td></td><td></td><td></td></tr>
<tr><td>Oligo rev</td><td>AAACGCGCCAACCGTCTCATCCAG</td><td></td><td></td><td></td><td></td><td></td><td></td><td></td><td></td><td></td><td></td></tr>
<tr><td>Chromosome</td><td>start</td><td>end</td><td>strand</td><td>MM</td><td>target_seq</td><td>PAM</td><td>alignment</td><td>distance</td><td>position</td><td>gene name</td><td>gene id</td></tr>
<tr><td>GU305914.1</td><td>61705</td><td>61727</td><td>+</td><td>0</td><td>CTGGATGAGACGGTTGGCGC</td><td>TGG</td><td>||||||||[||||||||||||]PAM</td><td>NA</td><td>Intergenic</td><td>NA</td><td>NA</td></tr>
<tr><td>T4135</td><td>ACGGTTGGCGCTGGAGACGATGG</td><td>1000</td><td></td><td></td><td></td><td></td><td></td><td></td><td></td><td></td><td></td></tr>
<tr><td>Oligo fwd</td><td>CACCACGGTTGGCGCTGGAGACGA</td><td></td><td></td><td></td><td></td><td></td><td></td><td></td><td></td><td></td><td></td></tr>
<tr><td>Oligo rev</td><td>AAACTCGTCTCCAGCGCCAACCGT</td><td></td><td></td><td></td><td></td><td></td><td></td><td></td><td></td><td></td><td></td></tr>
<tr><td>Chromosome</td><td>start</td><td>end</td><td>strand</td><td>MM</td><td>target_seq</td><td>PAM</td><td>alignment</td><td>distance</td><td>position</td><td>gene name</td><td>gene id</td></tr>
<tr><td>GU305914.1</td><td>61714</td><td>61736</td><td>+</td><td>0</td><td>ACGGTTGGCGCTGGAGACGA</td><td>TGG</td><td>||||||||[||||||||||||]PAM</td><td>NA</td><td>Intergenic</td><td>NA</td><td>NA</td></tr>
<tr><td>T4136</td><td>GTTGGCGCTGGAGACGATGGCGG</td><td>1000</td><td></td><td></td><td></td><td></td><td></td><td></td><td></td><td></td><td></td></tr>
<tr><td>Oligo fwd</td><td>CACCGTTGGCGCTGGAGACGATGG</td><td></td><td></td><td></td><td></td><td></td><td></td><td></td><td></td><td></td><td></td></tr>
<tr><td>Oligo rev</td><td>AAACCCATCGTCTCCAGCGCCAAC</td><td></td><td></td><td></td><td></td><td></td><td></td><td></td><td></td><td></td><td></td></tr>
<tr><td>Chromosome</td><td>start</td><td>end</td><td>strand</td><td>MM</td><td>target_seq</td><td>PAM</td><td>alignment</td><td>distance</td><td>position</td><td>gene name</td><td>gene id</td></tr>
<tr><td>GU305914.1</td><td>61717</td><td>61739</td><td>+</td><td>0</td><td>GTTGGCGCTGGAGACGATGG</td><td>CGG</td><td>||||||||[||||||||||||]PAM</td><td>NA</td><td>Intergenic</td><td>NA</td><td>NA</td></tr>
<tr><td>T4137</td><td>CGATGGCGGTGTAGAGCGCGAGG</td><td>1000</td><td></td><td></td><td></td><td></td><td></td><td></td><td></td><td></td><td></td></tr>
<tr><td>Oligo fwd</td><td>CACCCGATGGCGGTGTAGAGCGCG</td><td></td><td></td><td></td><td></td><td></td><td></td><td></td><td></td><td></td><td></td></tr>
<tr><td>Oligo rev</td><td>AAACCGCGCTCTACACCGCCATCG</td><td></td><td></td><td></td><td></td><td></td><td></td><td></td><td></td><td></td><td></td></tr>
<tr><td>Chromosome</td><td>start</td><td>end</td><td>strand</td><td>MM</td><td>target_seq</td><td>PAM</td><td>alignment</td><td>distance</td><td>position</td><td>gene name</td><td>gene id</td></tr>
<tr><td>GU305914.1</td><td>61731</td><td>61753</td><td>+</td><td>0</td><td>CGATGGCGGTGTAGAGCGCG</td><td>AGG</td><td>||||||||[||||||||||||]PAM</td><td>NA</td><td>Intergenic</td><td>NA</td><td>NA</td></tr>
<tr><td>T4138</td><td>GATGGCGGTGTAGAGCGCGAGGG</td><td>1000</td><td></td><td></td><td></td><td></td><td></td><td></td><td></td><td></td><td></td></tr>
<tr><td>Oligo fwd</td><td>CACCGATGGCGGTGTAGAGCGCGA</td><td></td><td></td><td></td><td></td><td></td><td></td><td></td><td></td><td></td><td></td></tr>
<tr><td>Oligo rev</td><td>AAACTCGCGCTCTACACCGCCATC</td><td></td><td></td><td></td><td></td><td></td><td></td><td></td><td></td><td></td><td></td></tr>
<tr><td>Chromosome</td><td>start</td><td>end</td><td>strand</td><td>MM</td><td>target_seq</td><td>PAM</td><td>alignment</td><td>distance</td><td>position</td><td>gene name</td><td>gene id</td></tr>
<tr><td>GU305914.1</td><td>61732</td><td>61754</td><td>+</td><td>0</td><td>GATGGCGGTGTAGAGCGCGA</td><td>GGG</td><td>||||||||[||||||||||||]PAM</td><td>NA</td><td>Intergenic</td><td>NA</td><td>NA</td></tr>
<tr><td>T4139</td><td>ATGGCGGTGTAGAGCGCGAGGGG</td><td>1000</td><td></td><td></td><td></td><td></td><td></td><td></td><td></td><td></td><td></td></tr>
<tr><td>Oligo fwd</td><td>CACCATGGCGGTGTAGAGCGCGAG</td><td></td><td></td><td></td><td></td><td></td><td></td><td></td><td></td><td></td><td></td></tr>
<tr><td>Oligo rev</td><td>AAACCTCGCGCTCTACACCGCCAT</td><td></td><td></td><td></td><td></td><td></td><td></td><td></td><td></td><td></td><td></td></tr>
<tr><td>Chromosome</td><td>start</td><td>end</td><td>strand</td><td>MM</td><td>target_seq</td><td>PAM</td><td>alignment</td><td>distance</td><td>position</td><td>gene name</td><td>gene id</td></tr>
<tr><td>GU305914.1</td><td>61733</td><td>61755</td><td>+</td><td>0</td><td>ATGGCGGTGTAGAGCGCGAG</td><td>GGG</td><td>||||||||[||||||||||||]PAM</td><td>NA</td><td>Intergenic</td><td>NA</td><td>NA</td></tr>
<tr><td>T4140</td><td>TCCGCAGCGCGTCACGTCCGCGG</td><td>1000</td><td></td><td></td><td></td><td></td><td></td><td></td><td></td><td></td><td></td></tr>
<tr><td>Oligo fwd</td><td>CACCTCCGCAGCGCGTCACGTCCG</td><td></td><td></td><td></td><td></td><td></td><td></td><td></td><td></td><td></td><td></td></tr>
<tr><td>Oligo rev</td><td>AAACCGGACGTGACGCGCTGCGGA</td><td></td><td></td><td></td><td></td><td></td><td></td><td></td><td></td><td></td><td></td></tr>
<tr><td>Chromosome</td><td>start</td><td>end</td><td>strand</td><td>MM</td><td>target_seq</td><td>PAM</td><td>alignment</td><td>distance</td><td>position</td><td>gene name</td><td>gene id</td></tr>
<tr><td>GU305914.1</td><td>61756</td><td>61778</td><td>+</td><td>0</td><td>TCCGCAGCGCGTCACGTCCG</td><td>CGG</td><td>||||||||[||||||||||||]PAM</td><td>NA</td><td>Intergenic</td><td>NA</td><td>NA</td></tr>
<tr><td>T4141</td><td>GCGTCACGTCCGCGGCGTAGCGG</td><td>1000</td><td></td><td></td><td></td><td></td><td></td><td></td><td></td><td></td><td></td></tr>
</table>

<table>
<tr><td>Oligo fwd</td><td>CACCGCGTCACGTCCGCGGCGTAG</td><td></td><td></td><td></td><td></td><td></td><td></td><td></td><td></td><td></td><td></td></tr>
<tr><td>Oligo rev</td><td>AAACCTACGCCGCGGACGTGACGC</td><td></td><td></td><td></td><td></td><td></td><td></td><td></td><td></td><td></td><td></td></tr>
<tr><td>Chromosome</td><td>start</td><td>end</td><td>strand</td><td>MM</td><td>target_seq</td><td>PAM</td><td>alignment</td><td>distance</td><td>position</td><td>gene name</td><td>gene id</td></tr>
<tr><td>GU305914.1</td><td>61764</td><td>61786</td><td>+</td><td>0</td><td>GCGTCACGTCCGCGGCGTAG</td><td>CGG</td><td>||||||||[||||||||||||]PAM</td><td>NA</td><td>Intergenic</td><td>NA</td><td>NA</td></tr>
<tr><td></td><td></td><td></td><td></td><td></td><td></td><td></td><td></td><td></td><td></td><td></td><td></td></tr>
<tr><td>T4142</td><td>TCACGTCCGCGGCGTAGCGGCGG</td><td>1000</td><td></td><td></td><td></td><td></td><td></td><td></td><td></td><td></td><td></td></tr>
<tr><td>Oligo fwd</td><td>CACCTCACGTCCGCGGCGTAGCGG</td><td></td><td></td><td></td><td></td><td></td><td></td><td></td><td></td><td></td><td></td></tr>
<tr><td>Oligo rev</td><td>AAACCCGCTACGCCGCGGACGTGA</td><td></td><td></td><td></td><td></td><td></td><td></td><td></td><td></td><td></td><td></td></tr>
<tr><td>Chromosome</td><td>start</td><td>end</td><td>strand</td><td>MM</td><td>target_seq</td><td>PAM</td><td>alignment</td><td>distance</td><td>position</td><td>gene name</td><td>gene id</td></tr>
<tr><td>GU305914.1</td><td>61767</td><td>61789</td><td>+</td><td>0</td><td>TCACGTCCGCGGCGTAGCGG</td><td>CGG</td><td>||||||||[||||||||||||]PAM</td><td>NA</td><td>Intergenic</td><td>NA</td><td>NA</td></tr>
<tr><td></td><td></td><td></td><td></td><td></td><td></td><td></td><td></td><td></td><td></td><td></td><td></td></tr>
<tr><td>T4143</td><td>GATGCGAGCCATGCGCTCCTTGG</td><td>1000</td><td></td><td></td><td></td><td></td><td></td><td></td><td></td><td></td><td></td></tr>
<tr><td>Oligo fwd</td><td>CACCGATGCGAGCCATGCGCTCCT</td><td></td><td></td><td></td><td></td><td></td><td></td><td></td><td></td><td></td><td></td></tr>
<tr><td>Oligo rev</td><td>AAACAGGAGCGCATGGCTCGCATC</td><td></td><td></td><td></td><td></td><td></td><td></td><td></td><td></td><td></td><td></td></tr>
<tr><td>Chromosome</td><td>start</td><td>end</td><td>strand</td><td>MM</td><td>target_seq</td><td>PAM</td><td>alignment</td><td>distance</td><td>position</td><td>gene name</td><td>gene id</td></tr>
<tr><td>GU305914.1</td><td>61789</td><td>61811</td><td>+</td><td>0</td><td>GATGCGAGCCATGCGCTCCT</td><td>TGG</td><td>||||||||[||||||||||||]PAM</td><td>NA</td><td>Intergenic</td><td>NA</td><td>NA</td></tr>
<tr><td></td><td></td><td></td><td></td><td></td><td></td><td></td><td></td><td></td><td></td><td></td><td></td></tr>
<tr><td>T4144</td><td>GCGAGCCATGCGCTCCTTGGCGG</td><td>1000</td><td></td><td></td><td></td><td></td><td></td><td></td><td></td><td></td><td></td></tr>
<tr><td>Oligo fwd</td><td>CACCGCGAGCCATGCGCTCCTTGG</td><td></td><td></td><td></td><td></td><td></td><td></td><td></td><td></td><td></td><td></td></tr>
<tr><td>Oligo rev</td><td>AAACCCAAGGAGCGCATGGCTCGC</td><td></td><td></td><td></td><td></td><td></td><td></td><td></td><td></td><td></td><td></td></tr>
<tr><td>Chromosome</td><td>start</td><td>end</td><td>strand</td><td>MM</td><td>target_seq</td><td>PAM</td><td>alignment</td><td>distance</td><td>position</td><td>gene name</td><td>gene id</td></tr>
<tr><td>GU305914.1</td><td>61792</td><td>61814</td><td>+</td><td>0</td><td>GCGAGCCATGCGCTCCTTGG</td><td>CGG</td><td>||||||||[||||||||||||]PAM</td><td>NA</td><td>Intergenic</td><td>NA</td><td>NA</td></tr>
<tr><td></td><td></td><td></td><td></td><td></td><td></td><td></td><td></td><td></td><td></td><td></td><td></td></tr>
<tr><td>T4145</td><td>CGAGCCATGCGCTCCTTGGCGGG</td><td>1000</td><td></td><td></td><td></td><td></td><td></td><td></td><td></td><td></td><td></td></tr>
<tr><td>Oligo fwd</td><td>CACCCGAGCCATGCGCTCCTTGGC</td><td></td><td></td><td></td><td></td><td></td><td></td><td></td><td></td><td></td><td></td></tr>
<tr><td>Oligo rev</td><td>AAACGCCAAGGAGCGCATGGCTCG</td><td></td><td></td><td></td><td></td><td></td><td></td><td></td><td></td><td></td><td></td></tr>
<tr><td>Chromosome</td><td>start</td><td>end</td><td>strand</td><td>MM</td><td>target_seq</td><td>PAM</td><td>alignment</td><td>distance</td><td>position</td><td>gene name</td><td>gene id</td></tr>
<tr><td>GU305914.1</td><td>61793</td><td>61815</td><td>+</td><td>0</td><td>CGAGCCATGCGCTCCTTGGC</td><td>GGG</td><td>||||||||[||||||||||||]PAM</td><td>NA</td><td>Intergenic</td><td>NA</td><td>NA</td></tr>
<tr><td></td><td></td><td></td><td></td><td></td><td></td><td></td><td></td><td></td><td></td><td></td><td></td></tr>
<tr><td>T4146</td><td>CATGCGCTCCTTGGCGGGTTCGG</td><td>1000</td><td></td><td></td><td></td><td></td><td></td><td></td><td></td><td></td><td></td></tr>
<tr><td>Oligo fwd</td><td>CACCCATGCGCTCCTTGGCGGGTT</td><td></td><td></td><td></td><td></td><td></td><td></td><td></td><td></td><td></td><td></td></tr>
<tr><td>Oligo rev</td><td>AAACAACCCGCCAAGGAGCGCATG</td><td></td><td></td><td></td><td></td><td></td><td></td><td></td><td></td><td></td><td></td></tr>
<tr><td>Chromosome</td><td>start</td><td>end</td><td>strand</td><td>MM</td><td>target_seq</td><td>PAM</td><td>alignment</td><td>distance</td><td>position</td><td>gene name</td><td>gene id</td></tr>
<tr><td>GU305914.1</td><td>61798</td><td>61820</td><td>+</td><td>0</td><td>CATGCGCTCCTTGGCGGGTT</td><td>CGG</td><td>||||||||[||||||||||||]PAM</td><td>NA</td><td>Intergenic</td><td>NA</td><td>NA</td></tr>
<tr><td></td><td></td><td></td><td></td><td></td><td></td><td></td><td></td><td></td><td></td><td></td><td></td></tr>
<tr><td>T4147</td><td>CGGAGCACTTCTCGTTGACGAGG</td><td>1000</td><td></td><td></td><td></td><td></td><td></td><td></td><td></td><td></td><td></td></tr>
<tr><td>Oligo fwd</td><td>CACCCGGAGCACTTCTCGTTGACG</td><td></td><td></td><td></td><td></td><td></td><td></td><td></td><td></td><td></td><td></td></tr>
<tr><td>Oligo rev</td><td>AAACCGTCAACGAGAAGTGCTCCG</td><td></td><td></td><td></td><td></td><td></td><td></td><td></td><td></td><td></td><td></td></tr>
<tr><td>Chromosome</td><td>start</td><td>end</td><td>strand</td><td>MM</td><td>target_seq</td><td>PAM</td><td>alignment</td><td>distance</td><td>position</td><td>gene name</td><td>gene id</td></tr>
<tr><td>GU305914.1</td><td>61818</td><td>61840</td><td>+</td><td>0</td><td>CGGAGCACTTCTCGTTGACG</td><td>AGG</td><td>||||||||[||||||||||||]PAM</td><td>NA</td><td>Intergenic</td><td>NA</td><td>NA</td></tr>
<tr><td>GU305914.1</td><td>220892</td><td>220914</td><td>-</td><td>4</td><td>CGAAACACGTCTCGTTGGCG</td><td>AGG</td><td>||-|-|||[-||||||||-||]PAM</td><td>NA</td><td>Intergenic</td><td>NA</td><td>NA</td></tr>
<tr><td></td><td></td><td></td><td></td><td></td><td></td><td></td><td></td><td></td><td></td><td></td><td></td></tr>
<tr><td>T4148</td><td>GGAGCACTTCTCGTTGACGAGGG</td><td>1000</td><td></td><td></td><td></td><td></td><td></td><td></td><td></td><td></td><td></td></tr>
<tr><td>Oligo fwd</td><td>CACCGGAGCACTTCTCGTTGACGA</td><td></td><td></td><td></td><td></td><td></td><td></td><td></td><td></td><td></td><td></td></tr>
<tr><td>Oligo rev</td><td>AAACTCGTCAACGAGAAGTGCTCC</td><td></td><td></td><td></td><td></td><td></td><td></td><td></td><td></td><td></td><td></td></tr>
<tr><td>Chromosome</td><td>start</td><td>end</td><td>strand</td><td>MM</td><td>target_seq</td><td>PAM</td><td>alignment</td><td>distance</td><td>position</td><td>gene name</td><td>gene id</td></tr>
<tr><td>GU305914.1</td><td>61819</td><td>61841</td><td>+</td><td>0</td><td>GGAGCACTTCTCGTTGACGA</td><td>GGG</td><td>||||||||[||||||||||||]PAM</td><td>NA</td><td>Intergenic</td><td>NA</td><td>NA</td></tr>
<tr><td></td><td></td><td></td><td></td><td></td><td></td><td></td><td></td><td></td><td></td><td></td><td></td></tr>
<tr><td>T4149</td><td>CGTTGACGAGGGCGCAGAATCGG</td><td>1000</td><td></td><td></td><td></td><td></td><td></td><td></td><td></td><td></td><td></td></tr>
<tr><td>Oligo fwd</td><td>CACCCGTTGACGAGGGCGCAGAAT</td><td></td><td></td><td></td><td></td><td></td><td></td><td></td><td></td><td></td><td></td></tr>
<tr><td>Oligo rev</td><td>AAACATTCTGCGCCCTCGTCAACG</td><td></td><td></td><td></td><td></td><td></td><td></td><td></td><td></td><td></td><td></td></tr>
<tr><td>Chromosome</td><td>start</td><td>end</td><td>strand</td><td>MM</td><td>target_seq</td><td>PAM</td><td>alignment</td><td>distance</td><td>position</td><td>gene name</td><td>gene id</td></tr>
<tr><td>GU305914.1</td><td>61830</td><td>61852</td><td>+</td><td>0</td><td>CGTTGACGAGGGCGCAGAAT</td><td>CGG</td><td>||||||||[||||||||||||]PAM</td><td>NA</td><td>Intergenic</td><td>NA</td><td>NA</td></tr>
<tr><td></td><td></td><td></td><td></td><td></td><td></td><td></td><td></td><td></td><td></td><td></td><td></td></tr>
<tr><td>T4150</td><td>CGCAGAATCGGCTGTTGCCGAGG</td><td>1000</td><td></td><td></td><td></td><td></td><td></td><td></td><td></td><td></td><td></td></tr>
<tr><td>Oligo fwd</td><td>CACCCGCAGAATCGGCTGTTGCCG</td><td></td><td></td><td></td><td></td><td></td><td></td><td></td><td></td><td></td><td></td></tr>
<tr><td>Oligo rev</td><td>AAACCGGCAACAGCCGATTCTGCG</td><td></td><td></td><td></td><td></td><td></td><td></td><td></td><td></td><td></td><td></td></tr>
<tr><td>Chromosome</td><td>start</td><td>end</td><td>strand</td><td>MM</td><td>target_seq</td><td>PAM</td><td>alignment</td><td>distance</td><td>position</td><td>gene name</td><td>gene id</td></tr>
<tr><td>GU305914.1</td><td>61842</td><td>61864</td><td>+</td><td>0</td><td>CGCAGAATCGGCTGTTGCCG</td><td>AGG</td><td>||||||||[||||||||||||]PAM</td><td>NA</td><td>Intergenic</td><td>NA</td><td>NA</td></tr>
<tr><td></td><td></td><td></td><td></td><td></td><td></td><td></td><td></td><td></td><td></td><td></td><td></td></tr>
<tr><td>T4151</td><td>GGCTGTTGCCGAGGATGCCGTGG</td><td>1000</td><td></td><td></td><td></td><td></td><td></td><td></td><td></td><td></td><td></td></tr>
<tr><td>Oligo fwd</td><td>CACCGGCTGTTGCCGAGGATGCCG</td><td></td><td></td><td></td><td></td><td></td><td></td><td></td><td></td><td></td><td></td></tr>
<tr><td>Oligo rev</td><td>AAACCGGCATCCTCGGCAACAGCC</td><td></td><td></td><td></td><td></td><td></td><td></td><td></td><td></td><td></td><td></td></tr>
<tr><td>Chromosome</td><td>start</td><td>end</td><td>strand</td><td>MM</td><td>target_seq</td><td>PAM</td><td>alignment</td><td>distance</td><td>position</td><td>gene name</td><td>gene id</td></tr>
<tr><td>GU305914.1</td><td>61851</td><td>61873</td><td>+</td><td>0</td><td>GGCTGTTGCCGAGGATGCCG</td><td>TGG</td><td>||||||||[||||||||||||]PAM</td><td>NA</td><td>Intergenic</td><td>NA</td><td>NA</td></tr>
<tr><td></td><td></td><td></td><td></td><td></td><td></td><td></td><td></td><td></td><td></td><td></td><td></td></tr>
<tr><td>T4152</td><td>GCCGTGGCAGTAGCAGCGCACGG</td><td>1000</td><td></td><td></td><td></td><td></td><td></td><td></td><td></td><td></td><td></td></tr>
<tr><td>Oligo fwd</td><td>CACCGCCGTGGCAGTAGCAGCGCA</td><td></td><td></td><td></td><td></td><td></td><td></td><td></td><td></td><td></td><td></td></tr>
<tr><td>Oligo rev</td><td>AAACTGCGCTGCTACTGCCACGGC</td><td></td><td></td><td></td><td></td><td></td><td></td><td></td><td></td><td></td><td></td></tr>
<tr><td>Chromosome</td><td>start</td><td>end</td><td>strand</td><td>MM</td><td>target_seq</td><td>PAM</td><td>alignment</td><td>distance</td><td>position</td><td>gene name</td><td>gene id</td></tr>
<tr><td>GU305914.1</td><td>61867</td><td>61889</td><td>+</td><td>0</td><td>GCCGTGGCAGTAGCAGCGCA</td><td>CGG</td><td>||||||||[||||||||||||]PAM</td><td>NA</td><td>Intergenic</td><td>NA</td><td>NA</td></tr>
<tr><td>GU305914.1</td><td>127779</td><td>127801</td><td>-</td><td>4</td><td>CTCGTCGCAGTAGCAGGGCA</td><td>CGG</td><td>--|||-||[||||||||-|||]PAM</td><td>NA</td><td>Intergenic</td><td>NA</td><td>NA</td></tr>
<tr><td></td><td></td><td></td><td></td><td></td><td></td><td></td><td></td><td></td><td></td><td></td><td></td></tr>
<tr><td>T4153</td><td>GTAGCAGCGCACGGACTCGACGG</td><td>1000</td><td></td><td></td><td></td><td></td><td></td><td></td><td></td><td></td><td></td></tr>
<tr><td>Oligo fwd</td><td>CACCGTAGCAGCGCACGGACTCGA</td><td></td><td></td><td></td><td></td><td></td><td></td><td></td><td></td><td></td><td></td></tr>
<tr><td>Oligo rev</td><td>AAACTCGAGTCCGTGCGCTGCTAC</td><td></td><td></td><td></td><td></td><td></td><td></td><td></td><td></td><td></td><td></td></tr>
<tr><td>Chromosome</td><td>start</td><td>end</td><td>strand</td><td>MM</td><td>target_seq</td><td>PAM</td><td>alignment</td><td>distance</td><td>position</td><td>gene name</td><td>gene id</td></tr>
<tr><td>GU305914.1</td><td>61876</td><td>61898</td><td>+</td><td>0</td><td>GTAGCAGCGCACGGACTCGA</td><td>CGG</td><td>||||||||[||||||||||||]PAM</td><td>NA</td><td>Intergenic</td><td>NA</td><td>NA</td></tr>
<tr><td></td><td></td><td></td><td></td><td></td><td></td><td></td><td></td><td></td><td></td><td></td><td></td></tr>
<tr><td>T4154</td><td>CTCGACGGCGTCGTCAAACAAGG</td><td>1000</td><td></td><td></td><td></td><td></td><td></td><td></td><td></td><td></td><td></td></tr>
<tr><td>Oligo fwd</td><td>CACCCTCGACGGCGTCGTCAAACA</td><td></td><td></td><td></td><td></td><td></td><td></td><td></td><td></td><td></td><td></td></tr>
<tr><td>Oligo rev</td><td>AAACTGTTTGACGACGCCGTCGAG</td><td></td><td></td><td></td><td></td><td></td><td></td><td></td><td></td><td></td><td></td></tr>
<tr><td>Chromosome</td><td>start</td><td>end</td><td>strand</td><td>MM</td><td>target_seq</td><td>PAM</td><td>alignment</td><td>distance</td><td>position</td><td>gene name</td><td>gene id</td></tr>
<tr><td>GU305914.1</td><td>61891</td><td>61913</td><td>+</td><td>0</td><td>CTCGACGGCGTCGTCAAACA</td><td>AGG</td><td>||||||||[||||||||||||]PAM</td><td>NA</td><td>Intergenic</td><td>NA</td><td>NA</td></tr>
<tr><td></td><td></td><td></td><td></td><td></td><td></td><td></td><td></td><td></td><td></td><td></td><td></td></tr>
<tr><td>T4155</td><td>GCCCGCTGTCGCGCCCCGCGCGG</td><td>1000</td><td></td><td></td><td></td><td></td><td></td><td></td><td></td><td></td><td></td></tr>
<tr><td>Oligo fwd</td><td>CACCGCCCGCTGTCGCGCCCCGCG</td><td></td><td></td><td></td><td></td><td></td><td></td><td></td><td></td><td></td><td></td></tr>
<tr><td>Oligo rev</td><td>AAACCGCGGGGCGCGACAGCGGGC</td><td></td><td></td><td></td><td></td><td></td><td></td><td></td><td></td><td></td><td></td></tr>
<tr><td>Chromosome</td><td>start</td><td>end</td><td>strand</td><td>MM</td><td>target_seq</td><td>PAM</td><td>alignment</td><td>distance</td><td>position</td><td>gene name</td><td>gene id</td></tr>
</table>

GU305914.1 61920 61942 + 0 GCCCGCTGTCGCGCCCCGCG CGG ||||||||[||||||||||||]PAM NA Intergenic NA NA
GU305914.1 110132 110154 + 4 GGCCGCCGTCGCGTCCCGGG AGG |-||||-|[|||||-||||-|]PAM NA Intergenic NA NA

T4156 GCTGTCGCGCCCCGCGCGGAAGG 1000
Oligo fwd CACCGCTGTCGCGCCCCGCGCGGA
Oligo rev AAACTCCGCGCGGGGCGCGACAGC
Chromosome start end strand MM target_seq PAM alignment distance position gene name gene id
GU305914.1 61924 61946 + 0 GCTGTCGCGCCCCGCGCGGA AGG ||||||||[||||||||||||]PAM NA Intergenic NA NA

T4157 CTGTCGCGCCCCGCGCGGAAGGG 1000
Oligo fwd CACCCTGTCGCGCCCCGCGCGGAA
Oligo rev AAACTTCCGCGCGGGGCGCGACAG
Chromosome start end strand MM target_seq PAM alignment distance position gene name gene id
GU305914.1 61925 61947 + 0 CTGTCGCGCCCCGCGCGGAA GGG ||||||||[||||||||||||]PAM NA Intergenic NA NA

T4158 CGCGCGGAAGGGCCTGCTGACGG 1000
Oligo fwd CACCCGCGCGGAAGGGCCTGCTGA
Oligo rev AAACTCAGCAGGCCCTTCCGCGCG
Chromosome start end strand MM target_seq PAM alignment distance position gene name gene id
GU305914.1 61936 61958 + 0 CGCGCGGAAGGGCCTGCTGA CGG ||||||||[||||||||||||]PAM NA Intergenic NA NA

T4159 GCGCGGAAGGGCCTGCTGACGGG 1000
Oligo fwd CACCGCGCGGAAGGGCCTGCTGAC
Oligo rev AAACGTCAGCAGGCCCTTCCGCGC
Chromosome start end strand MM target_seq PAM alignment distance position gene name gene id
GU305914.1 61937 61959 + 0 GCGCGGAAGGGCCTGCTGAC GGG ||||||||[||||||||||||]PAM NA Intergenic NA NA

T4160 ACGGGAATGACCGTCATCGTCGG 1000
Oligo fwd CACCACGGGAATGACCGTCATCGT
Oligo rev AAACACGATGACGGTCATTCCCGT
Chromosome start end strand MM target_seq PAM alignment distance position gene name gene id
GU305914.1 61955 61977 + 0 ACGGGAATGACCGTCATCGT CGG ||||||||[||||||||||||]PAM NA Intergenic NA NA

T4161 CGGGAATGACCGTCATCGTCGGG 1000
Oligo fwd CACCCGGGAATGACCGTCATCGTC
Oligo rev AAACGACGATGACGGTCATTCCCG
Chromosome start end strand MM target_seq PAM alignment distance position gene name gene id
GU305914.1 61956 61978 + 0 CGGGAATGACCGTCATCGTC GGG ||||||||[||||||||||||]PAM NA Intergenic NA NA

T4162 GGGAATGACCGTCATCGTCGGGG 1000
Oligo fwd CACCGGGAATGACCGTCATCGTCG
Oligo rev AAACCGACGATGACGGTCATTCCC
Chromosome start end strand MM target_seq PAM alignment distance position gene name gene id
GU305914.1 61957 61979 + 0 GGGAATGACCGTCATCGTCG GGG ||||||||[||||||||||||]PAM NA Intergenic NA NA

T4163 ATGACCGTCATCGTCGGGGTCGG 1000
Oligo fwd CACCATGACCGTCATCGTCGGGGT
Oligo rev AAACACCCCGACGATGACGGTCAT
Chromosome start end strand MM target_seq PAM alignment distance position gene name gene id
GU305914.1 61961 61983 + 0 ATGACCGTCATCGTCGGGGT CGG ||||||||[||||||||||||]PAM NA Intergenic NA NA

T4164 TGACCGTCATCGTCGGGGTCGGG 1000
Oligo fwd CACCTGACCGTCATCGTCGGGGTC
Oligo rev AAACGACCCCGACGATGACGGTCA
Chromosome start end strand MM target_seq PAM alignment distance position gene name gene id
GU305914.1 61962 61984 + 0 TGACCGTCATCGTCGGGGTC GGG ||||||||[||||||||||||]PAM NA Intergenic NA NA

T4165 GACCGTCATCGTCGGGGTCGGGG 1000
Oligo fwd CACCGACCGTCATCGTCGGGGTCG
Oligo rev AAACCGACCCCGACGATGACGGTC
Chromosome start end strand MM target_seq PAM alignment distance position gene name gene id
GU305914.1 61963 61985 + 0 GACCGTCATCGTCGGGGTCG GGG ||||||||[||||||||||||]PAM NA Intergenic NA NA
GU305914.1 146796 146818 + 3 GCCCGACATCGTCGGCGTCG CGG |-|||-||[|||||||-||||]PAM NA Intergenic NA NA

T4166 ACCGTCATCGTCGGGGTCGGGGG 1000
Oligo fwd CACCACCGTCATCGTCGGGGTCGG
Oligo rev AAACCCGACCCCGACGATGACGGT
Chromosome start end strand MM target_seq PAM alignment distance position gene name gene id
GU305914.1 61964 61986 + 0 ACCGTCATCGTCGGGGTCGG GGG ||||||||[||||||||||||]PAM NA Intergenic NA NA
GU305914.1 146797 146819 + 4 CCCGACATCGTCGGCGTCGC GGG -|||-|||[||||||-||||-]PAM NA Intergenic NA NA

T4167 CCGTCATCGTCGGGGTCGGGGGG 1000
Oligo fwd CACCCCGTCATCGTCGGGGTCGGG
Oligo rev AAACCCCGACCCCGACGATGACGG
Chromosome start end strand MM target_seq PAM alignment distance position gene name gene id
GU305914.1 61965 61987 + 0 CCGTCATCGTCGGGGTCGGG GGG ||||||||[||||||||||||]PAM NA Intergenic NA NA

T4168 GGGGCGCATCTCGTCCCTGACGG 1000
Oligo fwd CACCGGGGCGCATCTCGTCCCTGA
Oligo rev AAACTCAGGGACGAGATGCGCCCC
Chromosome start end strand MM target_seq PAM alignment distance position gene name gene id
GU305914.1 61984 62006 + 0 GGGGCGCATCTCGTCCCTGA CGG ||||||||[||||||||||||]PAM NA Intergenic NA NA

T4169 GGGCGCATCTCGTCCCTGACGGG 1000
Oligo fwd CACCGGGCGCATCTCGTCCCTGAC
Oligo rev AAACGTCAGGGACGAGATGCGCCC
Chromosome start end strand MM target_seq PAM alignment distance position gene name gene id
GU305914.1 61985 62007 + 0 GGGCGCATCTCGTCCCTGAC GGG ||||||||[||||||||||||]PAM NA Intergenic NA NA

<table>
<tr><td>T4170</td><td>GGCGCATCTCGTCCCTGACGGGG</td><td>1000</td><td></td><td></td><td></td><td></td><td></td><td></td><td></td><td></td><td></td></tr>
<tr><td>Oligo fwd</td><td>CACCGGCGCATCTCGTCCCTGACG</td><td></td><td></td><td></td><td></td><td></td><td></td><td></td><td></td><td></td><td></td></tr>
<tr><td>Oligo rev</td><td>AAACCGTCAGGGACGAGATGCGCC</td><td></td><td></td><td></td><td></td><td></td><td></td><td></td><td></td><td></td><td></td></tr>
<tr><td>Chromosome</td><td>start</td><td>end</td><td>strand</td><td>MM</td><td>target_seq</td><td>PAM</td><td>alignment</td><td>distance</td><td>position</td><td>gene name</td><td>gene id</td></tr>
<tr><td>GU305914.1</td><td>61986</td><td>62008</td><td>+</td><td>0</td><td>GGCGCATCTCGTCCCTGACG</td><td>GGG</td><td>||||||||[||||||||||||]PAM</td><td>NA</td><td>Intergenic</td><td>NA</td><td>NA</td></tr>
<tr><td>GU305914.1</td><td>80211</td><td>80233</td><td>+</td><td>4</td><td>GGTGCAGGTCGTCCCTGGCG</td><td>CGG</td><td>||-|||--[|||||||||-||]PAM</td><td>NA</td><td>Intergenic</td><td>NA</td><td>NA</td></tr>
<tr><td>T4171</td><td>CTCGTCCCTGACGGGGAGCCCGG</td><td>1000</td><td></td><td></td><td></td><td></td><td></td><td></td><td></td><td></td><td></td></tr>
<tr><td>Oligo fwd</td><td>CACCCTCGTCCCTGACGGGGAGCC</td><td></td><td></td><td></td><td></td><td></td><td></td><td></td><td></td><td></td><td></td></tr>
<tr><td>Oligo rev</td><td>AAACGGCTCCCCGTCAGGGACGAG</td><td></td><td></td><td></td><td></td><td></td><td></td><td></td><td></td><td></td><td></td></tr>
<tr><td>Chromosome</td><td>start</td><td>end</td><td>strand</td><td>MM</td><td>target_seq</td><td>PAM</td><td>alignment</td><td>distance</td><td>position</td><td>gene name</td><td>gene id</td></tr>
<tr><td>GU305914.1</td><td>61993</td><td>62015</td><td>+</td><td>0</td><td>CTCGTCCCTGACGGGGAGCC</td><td>CGG</td><td>||||||||[||||||||||||]PAM</td><td>NA</td><td>Intergenic</td><td>NA</td><td>NA</td></tr>
<tr><td>T4172</td><td>GCAGATCTTCGAATTCTCCGAGG</td><td>1000</td><td></td><td></td><td></td><td></td><td></td><td></td><td></td><td></td><td></td></tr>
<tr><td>Oligo fwd</td><td>CACCGCAGATCTTCGAATTCTCCG</td><td></td><td></td><td></td><td></td><td></td><td></td><td></td><td></td><td></td><td></td></tr>
<tr><td>Oligo rev</td><td>AAACCGGAGAATTCGAAGATCTGC</td><td></td><td></td><td></td><td></td><td></td><td></td><td></td><td></td><td></td><td></td></tr>
<tr><td>Chromosome</td><td>start</td><td>end</td><td>strand</td><td>MM</td><td>target_seq</td><td>PAM</td><td>alignment</td><td>distance</td><td>position</td><td>gene name</td><td>gene id</td></tr>
<tr><td>GU305914.1</td><td>62022</td><td>62044</td><td>+</td><td>0</td><td>GCAGATCTTCGAATTCTCCG</td><td>AGG</td><td>||||||||[||||||||||||]PAM</td><td>NA</td><td>Intergenic</td><td>NA</td><td>NA</td></tr>
<tr><td>T4173</td><td>CTGATCTTCGTCGTCTCCCCCGG</td><td>1000</td><td></td><td></td><td></td><td></td><td></td><td></td><td></td><td></td><td></td></tr>
<tr><td>Oligo fwd</td><td>CACCCTGATCTTCGTCGTCTCCCC</td><td></td><td></td><td></td><td></td><td></td><td></td><td></td><td></td><td></td><td></td></tr>
<tr><td>Oligo rev</td><td>AAACGGGGAGACGACGAAGATCAG</td><td></td><td></td><td></td><td></td><td></td><td></td><td></td><td></td><td></td><td></td></tr>
<tr><td>Chromosome</td><td>start</td><td>end</td><td>strand</td><td>MM</td><td>target_seq</td><td>PAM</td><td>alignment</td><td>distance</td><td>position</td><td>gene name</td><td>gene id</td></tr>
<tr><td>GU305914.1</td><td>62074</td><td>62096</td><td>+</td><td>0</td><td>CTGATCTTCGTCGTCTCCCC</td><td>CGG</td><td>||||||||[||||||||||||]PAM</td><td>NA</td><td>Intergenic</td><td>NA</td><td>NA</td></tr>
<tr><td>T4174</td><td>GTCGCCATCGCTGCTGCTGCTGG</td><td>1000</td><td></td><td></td><td></td><td></td><td></td><td></td><td></td><td></td><td></td></tr>
<tr><td>Oligo fwd</td><td>CACCGTCGCCATCGCTGCTGCTGC</td><td></td><td></td><td></td><td></td><td></td><td></td><td></td><td></td><td></td><td></td></tr>
<tr><td>Oligo rev</td><td>AAACGCAGCAGCAGCGATGGCGAC</td><td></td><td></td><td></td><td></td><td></td><td></td><td></td><td></td><td></td><td></td></tr>
<tr><td>Chromosome</td><td>start</td><td>end</td><td>strand</td><td>MM</td><td>target_seq</td><td>PAM</td><td>alignment</td><td>distance</td><td>position</td><td>gene name</td><td>gene id</td></tr>
<tr><td>GU305914.1</td><td>62096</td><td>62118</td><td>+</td><td>0</td><td>GTCGCCATCGCTGCTGCTGC</td><td>TGG</td><td>||||||||[||||||||||||]PAM</td><td>NA</td><td>Intergenic</td><td>NA</td><td>NA</td></tr>
<tr><td>GU305914.1</td><td>128801</td><td>128823</td><td>-</td><td>4</td><td>GCCGACGTCGCTGCCGCTGC</td><td>GGG</td><td>|-||-|-|[||||||-|||||]PAM</td><td>NA</td><td>Intergenic</td><td>NA</td><td>NA</td></tr>
<tr><td>T4175</td><td>GCCATCGCTGCTGCTGCTGGTGG</td><td>1000</td><td></td><td></td><td></td><td></td><td></td><td></td><td></td><td></td><td></td></tr>
<tr><td>Oligo fwd</td><td>CACCGCCATCGCTGCTGCTGCTGG</td><td></td><td></td><td></td><td></td><td></td><td></td><td></td><td></td><td></td><td></td></tr>
<tr><td>Oligo rev</td><td>AAACCCAGCAGCAGCAGCGATGGC</td><td></td><td></td><td></td><td></td><td></td><td></td><td></td><td></td><td></td><td></td></tr>
<tr><td>Chromosome</td><td>start</td><td>end</td><td>strand</td><td>MM</td><td>target_seq</td><td>PAM</td><td>alignment</td><td>distance</td><td>position</td><td>gene name</td><td>gene id</td></tr>
<tr><td>GU305914.1</td><td>62099</td><td>62121</td><td>+</td><td>0</td><td>GCCATCGCTGCTGCTGCTGG</td><td>TGG</td><td>||||||||[||||||||||||]PAM</td><td>NA</td><td>Intergenic</td><td>NA</td><td>NA</td></tr>
<tr><td>T4176</td><td>TGCTGCTGGTGGTGCCGAAGCGG</td><td>1000</td><td></td><td></td><td></td><td></td><td></td><td></td><td></td><td></td><td></td></tr>
<tr><td>Oligo fwd</td><td>CACCTGCTGCTGGTGGTGCCGAAG</td><td></td><td></td><td></td><td></td><td></td><td></td><td></td><td></td><td></td><td></td></tr>
<tr><td>Oligo rev</td><td>AAACCTTCGGCACCACCAGCAGCA</td><td></td><td></td><td></td><td></td><td></td><td></td><td></td><td></td><td></td><td></td></tr>
<tr><td>Chromosome</td><td>start</td><td>end</td><td>strand</td><td>MM</td><td>target_seq</td><td>PAM</td><td>alignment</td><td>distance</td><td>position</td><td>gene name</td><td>gene id</td></tr>
<tr><td>GU305914.1</td><td>62110</td><td>62132</td><td>+</td><td>0</td><td>TGCTGCTGGTGGTGCCGAAG</td><td>CGG</td><td>||||||||[||||||||||||]PAM</td><td>NA</td><td>Intergenic</td><td>NA</td><td>NA</td></tr>
<tr><td>T4177</td><td>GCTGGTGGTGCCGAAGCGGCAGG</td><td>1000</td><td></td><td></td><td></td><td></td><td></td><td></td><td></td><td></td><td></td></tr>
<tr><td>Oligo fwd</td><td>CACCGCTGGTGGTGCCGAAGCGGC</td><td></td><td></td><td></td><td></td><td></td><td></td><td></td><td></td><td></td><td></td></tr>
<tr><td>Oligo rev</td><td>AAACGCCGCTTCGGCACCACCAGC</td><td></td><td></td><td></td><td></td><td></td><td></td><td></td><td></td><td></td><td></td></tr>
<tr><td>Chromosome</td><td>start</td><td>end</td><td>strand</td><td>MM</td><td>target_seq</td><td>PAM</td><td>alignment</td><td>distance</td><td>position</td><td>gene name</td><td>gene id</td></tr>
<tr><td>GU305914.1</td><td>62114</td><td>62136</td><td>+</td><td>0</td><td>GCTGGTGGTGCCGAAGCGGC</td><td>AGG</td><td>||||||||[||||||||||||]PAM</td><td>NA</td><td>Intergenic</td><td>NA</td><td>NA</td></tr>
<tr><td>T4178</td><td>TGGTGCCGAAGCGGCAGGTGAGG</td><td>1000</td><td></td><td></td><td></td><td></td><td></td><td></td><td></td><td></td><td></td></tr>
<tr><td>Oligo fwd</td><td>CACCTGGTGCCGAAGCGGCAGGTG</td><td></td><td></td><td></td><td></td><td></td><td></td><td></td><td></td><td></td><td></td></tr>
<tr><td>Oligo rev</td><td>AAACCACCTGCCGCTTCGGCACCA</td><td></td><td></td><td></td><td></td><td></td><td></td><td></td><td></td><td></td><td></td></tr>
<tr><td>Chromosome</td><td>start</td><td>end</td><td>strand</td><td>MM</td><td>target_seq</td><td>PAM</td><td>alignment</td><td>distance</td><td>position</td><td>gene name</td><td>gene id</td></tr>
<tr><td>GU305914.1</td><td>62119</td><td>62141</td><td>+</td><td>0</td><td>TGGTGCCGAAGCGGCAGGTG</td><td>AGG</td><td>||||||||[||||||||||||]PAM</td><td>NA</td><td>Intergenic</td><td>NA</td><td>NA</td></tr>
<tr><td>T4179</td><td>CGAAGCGGCAGGTGAGGATTCGG</td><td>1000</td><td></td><td></td><td></td><td></td><td></td><td></td><td></td><td></td><td></td></tr>
<tr><td>Oligo fwd</td><td>CACCCGAAGCGGCAGGTGAGGATT</td><td></td><td></td><td></td><td></td><td></td><td></td><td></td><td></td><td></td><td></td></tr>
<tr><td>Oligo rev</td><td>AAACAATCCTCACCTGCCGCTTCG</td><td></td><td></td><td></td><td></td><td></td><td></td><td></td><td></td><td></td><td></td></tr>
<tr><td>Chromosome</td><td>start</td><td>end</td><td>strand</td><td>MM</td><td>target_seq</td><td>PAM</td><td>alignment</td><td>distance</td><td>position</td><td>gene name</td><td>gene id</td></tr>
<tr><td>GU305914.1</td><td>62125</td><td>62147</td><td>+</td><td>0</td><td>CGAAGCGGCAGGTGAGGATT</td><td>CGG</td><td>||||||||[||||||||||||]PAM</td><td>NA</td><td>Intergenic</td><td>NA</td><td>NA</td></tr>
<tr><td>T4180</td><td>GCAGGTGAGGATTCGGATTCAGG</td><td>1000</td><td></td><td></td><td></td><td></td><td></td><td></td><td></td><td></td><td></td></tr>
<tr><td>Oligo fwd</td><td>CACCGCAGGTGAGGATTCGGATTC</td><td></td><td></td><td></td><td></td><td></td><td></td><td></td><td></td><td></td><td></td></tr>
<tr><td>Oligo rev</td><td>AAACGAATCCGAATCCTCACCTGC</td><td></td><td></td><td></td><td></td><td></td><td></td><td></td><td></td><td></td><td></td></tr>
<tr><td>Chromosome</td><td>start</td><td>end</td><td>strand</td><td>MM</td><td>target_seq</td><td>PAM</td><td>alignment</td><td>distance</td><td>position</td><td>gene name</td><td>gene id</td></tr>
<tr><td>GU305914.1</td><td>62132</td><td>62154</td><td>+</td><td>0</td><td>GCAGGTGAGGATTCGGATTC</td><td>AGG</td><td>||||||||[||||||||||||]PAM</td><td>NA</td><td>Intergenic</td><td>NA</td><td>NA</td></tr>
<tr><td>T4181</td><td>CAGGTGAGGATTCGGATTCAGGG</td><td>1000</td><td></td><td></td><td></td><td></td><td></td><td></td><td></td><td></td><td></td></tr>
<tr><td>Oligo fwd</td><td>CACCCAGGTGAGGATTCGGATTCA</td><td></td><td></td><td></td><td></td><td></td><td></td><td></td><td></td><td></td><td></td></tr>
<tr><td>Oligo rev</td><td>AAACTGAATCCGAATCCTCACCTG</td><td></td><td></td><td></td><td></td><td></td><td></td><td></td><td></td><td></td><td></td></tr>
<tr><td>Chromosome</td><td>start</td><td>end</td><td>strand</td><td>MM</td><td>target_seq</td><td>PAM</td><td>alignment</td><td>distance</td><td>position</td><td>gene name</td><td>gene id</td></tr>
<tr><td>GU305914.1</td><td>62133</td><td>62155</td><td>+</td><td>0</td><td>CAGGTGAGGATTCGGATTCA</td><td>GGG</td><td>||||||||[||||||||||||]PAM</td><td>NA</td><td>Intergenic</td><td>NA</td><td>NA</td></tr>
<tr><td>T4182</td><td>AGGTGAGGATTCGGATTCAGGGG</td><td>1000</td><td></td><td></td><td></td><td></td><td></td><td></td><td></td><td></td><td></td></tr>
<tr><td>Oligo fwd</td><td>CACCAGGTGAGGATTCGGATTCAG</td><td></td><td></td><td></td><td></td><td></td><td></td><td></td><td></td><td></td><td></td></tr>
<tr><td>Oligo rev</td><td>AAACCTGAATCCGAATCCTCACCT</td><td></td><td></td><td></td><td></td><td></td><td></td><td></td><td></td><td></td><td></td></tr>
<tr><td>Chromosome</td><td>start</td><td>end</td><td>strand</td><td>MM</td><td>target_seq</td><td>PAM</td><td>alignment</td><td>distance</td><td>position</td><td>gene name</td><td>gene id</td></tr>
<tr><td>GU305914.1</td><td>62134</td><td>62156</td><td>+</td><td>0</td><td>AGGTGAGGATTCGGATTCAG</td><td>GGG</td><td>||||||||[||||||||||||]PAM</td><td>NA</td><td>Intergenic</td><td>NA</td><td>NA</td></tr>
<tr><td>T4183</td><td>GGATTCGGATTCAGGGGCGACGG</td><td>1000</td><td></td><td></td><td></td><td></td><td></td><td></td><td></td><td></td><td></td></tr>
<tr><td>Oligo fwd</td><td>CACCGGATTCGGATTCAGGGGCGA</td><td></td><td></td><td></td><td></td><td></td><td></td><td></td><td></td><td></td><td></td></tr>
<tr><td>Oligo rev</td><td>AAACTCGCCCCTGAATCCGAATCC</td><td></td><td></td><td></td><td></td><td></td><td></td><td></td><td></td><td></td><td></td></tr>
<tr><td>Chromosome</td><td>start</td><td>end</td><td>strand</td><td>MM</td><td>target_seq</td><td>PAM</td><td>alignment</td><td>distance</td><td>position</td><td>gene name</td><td>gene id</td></tr>
<tr><td>GU305914.1</td><td>62140</td><td>62162</td><td>+</td><td>0</td><td>GGATTCGGATTCAGGGGCGA</td><td>CGG</td><td>||||||||[||||||||||||]PAM</td><td>NA</td><td>Intergenic</td><td>NA</td><td>NA</td></tr>
<tr><td>T4184</td><td>GGATTCAGGGGCGACGGCAGCGG</td><td>1000</td><td></td><td></td><td></td><td></td><td></td><td></td><td></td><td></td><td></td></tr>
<tr><td>Oligo fwd</td><td>CACCGGATTCAGGGGCGACGGCAG</td><td></td><td></td><td></td><td></td><td></td><td></td><td></td><td></td><td></td><td></td></tr>
<tr><td>Oligo rev</td><td>AAACCTGCCGTCGCCCCTGAATCC</td><td></td><td></td><td></td><td></td><td></td><td></td><td></td><td></td><td></td><td></td></tr>
</table>

| | | | | | | | | | | | |
|---|---|---|---|---|---|---|---|---|---|---|---|
| Chromosome | start | end | strand | MM | target_seq | PAM | alignment | distance | position | gene name | gene id |
| GU305914.1 | 62146 | 62168 | + | 0 | GGATTCAGGGGCGACGGCAG | CGG | \|\|\|\|\|\|\|\|[\|\|\|\|\|\|\|\|\|\|\|\|]PAM | NA | Intergenic | NA | NA |
| T4185 | AGGGGCGACGGCAGCGGCAGCGG | 1000 | | | | | | | | | |
| Oligo fwd | CACCAGGGGCGACGGCAGCGGCAG | | | | | | | | | | |
| Oligo rev | AAACCTGCCGCTGCCGTCGCCCCT | | | | | | | | | | |
| Chromosome | start | end | strand | MM | target_seq | PAM | alignment | distance | position | gene name | gene id |
| GU305914.1 | 62152 | 62174 | + | 0 | AGGGGCGACGGCAGCGGCAG | CGG | \|\|\|\|\|\|\|\|[\|\|\|\|\|\|\|\|\|\|\|\|]PAM | NA | Intergenic | NA | NA |
| GU305914.1 | 163908 | 163930 | + | 3 | GGCGGCGGCGGCAGCGGCAG | CGG | -\|-\|\|\|\|-[\|\|\|\|\|\|\|\|\|\|\|\|]PAM | NA | Intergenic | NA | NA |
| GU305914.1 | 163902 | 163924 | + | 3 | AGCGGCGGCGGCGGCGGCAG | CGG | \|\|-\|\|\|\|-[\|\|\|\|-\|\|\|\|\|\|\|]PAM | NA | Intergenic | NA | NA |
| GU305914.1 | 30006 | 30028 | + | 4 | GGCGGCGGCGGCGGCGGCAG | TGG | -\|-\|\|\|\|-[\|\|\|\|-\|\|\|\|\|\|\|]PAM | NA | Intergenic | NA | NA |
| GU305914.1 | 149476 | 149498 | - | 4 | GGCGGCGGCGGCGGCGGCAG | CGG | -\|-\|\|\|\|-[\|\|\|\|-\|\|\|\|\|\|\|]PAM | NA | Intergenic | NA | NA |
| GU305914.1 | 168829 | 168851 | + | 4 | AGCGGCGGCGGCGGCGACAG | CGG | \|\|-\|\|\|\|-[\|\|\|\|-\|\|\|-\|\|\|]PAM | NA | Intergenic | NA | NA |
| GU305914.1 | 109437 | 109459 | + | 4 | ACGGGAGACGGCGGCGGTAG | CGG | \|-\|\|\|-\|\|[\|\|\|\|-\|\|\|\|-\|\|]PAM | NA | Intergenic | NA | NA |
| GU305914.1 | 73795 | 73817 | - | 4 | AGTGGCGCCAGCAGCGGCGG | CGG | \|\|-\|\|\|\|-[\|-\|\|\|\|\|\|\|\|-\|]PAM | NA | Intergenic | NA | NA |
| GU305914.1 | 71759 | 71781 | + | 4 | AGCGGCGGCGGCAACGGGAG | CGG | \|\|-\|\|\|\|-[\|\|\|\|\|-\|\|\|-\|\|]PAM | NA | Intergenic | NA | NA |
| T4186 | GGCGACGGCAGCGGCAGCGGCGG | 1000 | | | | | | | | | |
| Oligo fwd | CACCGGCGACGGCAGCGGCAGCGG | | | | | | | | | | |
| Oligo rev | AAACCCGCTGCCGCTGCCGTCGCC | | | | | | | | | | |
| Chromosome | start | end | strand | MM | target_seq | PAM | alignment | distance | position | gene name | gene id |
| GU305914.1 | 62155 | 62177 | + | 0 | GGCGACGGCAGCGGCAGCGG | CGG | \|\|\|\|\|\|\|\|[\|\|\|\|\|\|\|\|\|\|\|\|]PAM | NA | Intergenic | NA | NA |
| GU305914.1 | 163911 | 163933 | + | 1 | GGCGGCGGCAGCGGCAGCGG | AGG | \|\|\|\|-\|\|\|[\|\|\|\|\|\|\|\|\|\|\|\|]PAM | NA | Intergenic | NA | NA |
| GU305914.1 | 45255 | 45277 | - | 4 | GGTGGCGGTGGCGGCAGCGG | CGG | \|\|-\|-\|\|\|[--\|\|\|\|\|\|\|\|\|\|]PAM | NA | Intergenic | NA | NA |
| GU305914.1 | 73792 | 73814 | - | 3 | GGCGCCAGCAGCGGCGGCGG | AGG | \|\|\|\|-\|-\|[\|\|\|\|\|\|\|-\|\|\|\|]PAM | NA | Intergenic | NA | NA |
| GU305914.1 | 149482 | 149504 | - | 2 | GGCGACGGCGGCGGCGGCGG | CGG | \|\|\|\|\|\|\|\|[\|-\|\|\|\|\|-\|\|\|\|]PAM | NA | Intergenic | NA | NA |
| GU305914.1 | 168820 | 168842 | + | 4 | GGTGGTGGCAGCGGCGGCGG | CGG | \|\|-\|--\|\|[\|\|\|\|\|\|\|-\|\|\|\|]PAM | NA | Intergenic | NA | NA |
| GU305914.1 | 154407 | 154429 | + | 4 | GGTGGCGGCGGCGGCGGCGG | CGG | \|\|-\|-\|\|\|[\|-\|\|\|\|\|-\|\|\|\|]PAM | NA | Intergenic | NA | NA |
| GU305914.1 | 58123 | 58145 | - | 4 | GGTGGCGGCGGCGGCGGCGG | AGG | \|\|-\|-\|\|\|[\|-\|\|\|\|\|-\|\|\|\|]PAM | NA | Intergenic | NA | NA |
| GU305914.1 | 154404 | 154426 | + | 4 | GGCGGTGGCGGCGGCGGCGG | CGG | \|\|\|\|--\|\|[\|-\|\|\|\|\|-\|\|\|\|]PAM | NA | Intergenic | NA | NA |
| GU305914.1 | 58126 | 58148 | - | 4 | GGCGGTGGCGGCGGCGGCGG | CGG | \|\|\|\|--\|\|[\|-\|\|\|\|\|-\|\|\|\|]PAM | NA | Intergenic | NA | NA |
| T4187 | AGCGGCGGTCGTTACGCTCGCGG | 1000 | | | | | | | | | |
| Oligo fwd | CACCAGCGGCGGTCGTTACGCTCG | | | | | | | | | | |
| Oligo rev | AAACCGAGCGTAACGACCGCCGCT | | | | | | | | | | |
| Chromosome | start | end | strand | MM | target_seq | PAM | alignment | distance | position | gene name | gene id |
| GU305914.1 | 62170 | 62192 | + | 0 | AGCGGCGGTCGTTACGCTCG | CGG | \|\|\|\|\|\|\|\|[\|\|\|\|\|\|\|\|\|\|\|\|]PAM | NA | Intergenic | NA | NA |
| T4188 | GCGGCGGTCGTTACGCTCGCGGG | 1000 | | | | | | | | | |
| Oligo fwd | CACCGCGGCGGTCGTTACGCTCGC | | | | | | | | | | |
| Oligo rev | AAACGCGAGCGTAACGACCGCCGC | | | | | | | | | | |
| Chromosome | start | end | strand | MM | target_seq | PAM | alignment | distance | position | gene name | gene id |
| GU305914.1 | 62171 | 62193 | + | 0 | GCGGCGGTCGTTACGCTCGC | GGG | \|\|\|\|\|\|\|\|[\|\|\|\|\|\|\|\|\|\|\|\|]PAM | NA | Intergenic | NA | NA |
| GU305914.1 | 123247 | 123269 | - | 4 | CCGTCGGACGTCACGCTCGC | CGG | -\|\|-\|\|\|-[\|\|\|-\|\|\|\|\|\|\|\|]PAM | NA | Intergenic | NA | NA |
| T4189 | GCGGTCGTTACGCTCGCGGGAGG | 1000 | | | | | | | | | |
| Oligo fwd | CACCGCGGTCGTTACGCTCGCGGG | | | | | | | | | | |
| Oligo rev | AAACCCCGCGAGCGTAACGACCGC | | | | | | | | | | |
| Chromosome | start | end | strand | MM | target_seq | PAM | alignment | distance | position | gene name | gene id |
| GU305914.1 | 62174 | 62196 | + | 0 | GCGGTCGTTACGCTCGCGGG | AGG | \|\|\|\|\|\|\|\|[\|\|\|\|\|\|\|\|\|\|\|\|]PAM | NA | Intergenic | NA | NA |
| T4190 | GTCGTTACGCTCGCGGGAGGCGG | 1000 | | | | | | | | | |
| Oligo fwd | CACCGTCGTTACGCTCGCGGGAGG | | | | | | | | | | |
| Oligo rev | AAACCCTCCCGCGAGCGTAACGAC | | | | | | | | | | |
| Chromosome | start | end | strand | MM | target_seq | PAM | alignment | distance | position | gene name | gene id |
| GU305914.1 | 62177 | 62199 | + | 0 | GTCGTTACGCTCGCGGGAGG | CGG | \|\|\|\|\|\|\|\|[\|\|\|\|\|\|\|\|\|\|\|\|]PAM | NA | Intergenic | NA | NA |
| T4191 | GCGGCATCGTCGCCGCGTACAGG | 1000 | | | | | | | | | |
| Oligo fwd | CACCGCGGCATCGTCGCCGCGTAC | | | | | | | | | | |
| Oligo rev | AAACGTACGCGGCGACGATGCCGC | | | | | | | | | | |
| Chromosome | start | end | strand | MM | target_seq | PAM | alignment | distance | position | gene name | gene id |
| GU305914.1 | 62196 | 62218 | + | 0 | GCGGCATCGTCGCCGCGTAC | AGG | \|\|\|\|\|\|\|\|[\|\|\|\|\|\|\|\|\|\|\|\|]PAM | NA | Intergenic | NA | NA |
| T4192 | GCATCGTCGCCGCGTACAGGAGG | 1000 | | | | | | | | | |
| Oligo fwd | CACCGCATCGTCGCCGCGTACAGG | | | | | | | | | | |
| Oligo rev | AAACCCTGTACGCGGCGACGATGC | | | | | | | | | | |
| Chromosome | start | end | strand | MM | target_seq | PAM | alignment | distance | position | gene name | gene id |
| GU305914.1 | 62199 | 62221 | + | 0 | GCATCGTCGCCGCGTACAGG | AGG | \|\|\|\|\|\|\|\|[\|\|\|\|\|\|\|\|\|\|\|\|]PAM | NA | Intergenic | NA | NA |
| T4193 | CATCGTCGCCGCGTACAGGAGGG | 1000 | | | | | | | | | |
| Oligo fwd | CACCCATCGTCGCCGCGTACAGGA | | | | | | | | | | |
| Oligo rev | AAACTCCTGTACGCGGCGACGATG | | | | | | | | | | |
| Chromosome | start | end | strand | MM | target_seq | PAM | alignment | distance | position | gene name | gene id |
| GU305914.1 | 62200 | 62222 | + | 0 | CATCGTCGCCGCGTACAGGA | GGG | \|\|\|\|\|\|\|\|[\|\|\|\|\|\|\|\|\|\|\|\|]PAM | NA | Intergenic | NA | NA |
| T4194 | ATCGTCGCCGCGTACAGGAGGGG | 1000 | | | | | | | | | |
| Oligo fwd | CACCATCGTCGCCGCGTACAGGAG | | | | | | | | | | |
| Oligo rev | AAACCTCCTGTACGCGGCGACGAT | | | | | | | | | | |
| Chromosome | start | end | strand | MM | target_seq | PAM | alignment | distance | position | gene name | gene id |
| GU305914.1 | 62201 | 62223 | + | 0 | ATCGTCGCCGCGTACAGGAG | GGG | \|\|\|\|\|\|\|\|[\|\|\|\|\|\|\|\|\|\|\|\|]PAM | NA | Intergenic | NA | NA |
| T4195 | CAGGAGGGGATCAGCTTTAGCGG | 1000 | | | | | | | | | |
| Oligo fwd | CACCCAGGAGGGGATCAGCTTTAG | | | | | | | | | | |
| Oligo rev | AAACCTAAAGCTGATCCCCTCCTG | | | | | | | | | | |
| Chromosome | start | end | strand | MM | target_seq | PAM | alignment | distance | position | gene name | gene id |
| GU305914.1 | 62215 | 62237 | + | 0 | CAGGAGGGGATCAGCTTTAG | CGG | \|\|\|\|\|\|\|\|[\|\|\|\|\|\|\|\|\|\|\|\|]PAM | NA | Intergenic | NA | NA |
| T4196 | AGGAGGGGATCAGCTTTAGCGGG | 1000 | | | | | | | | | |
| Oligo fwd | CACCAGGAGGGGATCAGCTTTAGC | | | | | | | | | | |

<table>
<tr><td>Oligo rev</td><td>AAACGCTAAAGCTGATCCCCTCCT</td><td></td><td></td><td></td><td></td><td></td><td></td><td></td><td></td><td></td><td></td></tr>
<tr><td>Chromosome</td><td>start</td><td>end</td><td>strand</td><td>MM</td><td>target_seq</td><td>PAM</td><td>alignment</td><td>distance</td><td>position</td><td>gene name</td><td>gene id</td></tr>
<tr><td>GU305914.1</td><td>62216</td><td>62238</td><td>+</td><td>0</td><td>AGGAGGGGATCAGCTTTAGC</td><td>GGG</td><td>||||||||[||||||||||||]PAM</td><td>NA</td><td>Intergenic</td><td>NA</td><td>NA</td></tr>
<tr><td></td><td></td><td></td><td></td><td></td><td></td><td></td><td></td><td></td><td></td><td></td><td></td></tr>
<tr><td>T4197</td><td>TTTAGCGGGCTTAAGCTTCTTGG</td><td>1000</td><td></td><td></td><td></td><td></td><td></td><td></td><td></td><td></td><td></td></tr>
<tr><td>Oligo fwd</td><td>CACCTTTAGCGGGCTTAAGCTTCT</td><td></td><td></td><td></td><td></td><td></td><td></td><td></td><td></td><td></td><td></td></tr>
<tr><td>Oligo rev</td><td>AAACAGAAGCTTAAGCCCGCTAAA</td><td></td><td></td><td></td><td></td><td></td><td></td><td></td><td></td><td></td><td></td></tr>
<tr><td>Chromosome</td><td>start</td><td>end</td><td>strand</td><td>MM</td><td>target_seq</td><td>PAM</td><td>alignment</td><td>distance</td><td>position</td><td>gene name</td><td>gene id</td></tr>
<tr><td>GU305914.1</td><td>62230</td><td>62252</td><td>+</td><td>0</td><td>TTTAGCGGGCTTAAGCTTCT</td><td>TGG</td><td>||||||||[||||||||||||]PAM</td><td>NA</td><td>Intergenic</td><td>NA</td><td>NA</td></tr>
<tr><td></td><td></td><td></td><td></td><td></td><td></td><td></td><td></td><td></td><td></td><td></td><td></td></tr>
<tr><td>T4198</td><td>GGCTTAAGCTTCTTGGCTTGAGG</td><td>1000</td><td></td><td></td><td></td><td></td><td></td><td></td><td></td><td></td><td></td></tr>
<tr><td>Oligo fwd</td><td>CACCGGCTTAAGCTTCTTGGCTTG</td><td></td><td></td><td></td><td></td><td></td><td></td><td></td><td></td><td></td><td></td></tr>
<tr><td>Oligo rev</td><td>AAACCAAGCCAAGAAGCTTAAGCC</td><td></td><td></td><td></td><td></td><td></td><td></td><td></td><td></td><td></td><td></td></tr>
<tr><td>Chromosome</td><td>start</td><td>end</td><td>strand</td><td>MM</td><td>target_seq</td><td>PAM</td><td>alignment</td><td>distance</td><td>position</td><td>gene name</td><td>gene id</td></tr>
<tr><td>GU305914.1</td><td>62237</td><td>62259</td><td>+</td><td>0</td><td>GGCTTAAGCTTCTTGGCTTG</td><td>AGG</td><td>||||||||[||||||||||||]PAM</td><td>NA</td><td>Intergenic</td><td>NA</td><td>NA</td></tr>
<tr><td></td><td></td><td></td><td></td><td></td><td></td><td></td><td></td><td></td><td></td><td></td><td></td></tr>
<tr><td>T4199</td><td>CTTGAGGTTGCTGTTGTTGTTGG</td><td>1000</td><td></td><td></td><td></td><td></td><td></td><td></td><td></td><td></td><td></td></tr>
<tr><td>Oligo fwd</td><td>CACCCTTGAGGTTGCTGTTGTTGT</td><td></td><td></td><td></td><td></td><td></td><td></td><td></td><td></td><td></td><td></td></tr>
<tr><td>Oligo rev</td><td>AAACACAACAACAGCAACCTCAAG</td><td></td><td></td><td></td><td></td><td></td><td></td><td></td><td></td><td></td><td></td></tr>
<tr><td>Chromosome</td><td>start</td><td>end</td><td>strand</td><td>MM</td><td>target_seq</td><td>PAM</td><td>alignment</td><td>distance</td><td>position</td><td>gene name</td><td>gene id</td></tr>
<tr><td>GU305914.1</td><td>62253</td><td>62275</td><td>+</td><td>0</td><td>CTTGAGGTTGCTGTTGTTGT</td><td>TGG</td><td>||||||||[||||||||||||]PAM</td><td>NA</td><td>Intergenic</td><td>NA</td><td>NA</td></tr>
<tr><td></td><td></td><td></td><td></td><td></td><td></td><td></td><td></td><td></td><td></td><td></td><td></td></tr>
<tr><td>T4200</td><td>GTTGTTGGCCTGTGCTGCACCGG</td><td>1000</td><td></td><td></td><td></td><td></td><td></td><td></td><td></td><td></td><td></td></tr>
<tr><td>Oligo fwd</td><td>CACCGTTGTTGGCCTGTGCTGCAC</td><td></td><td></td><td></td><td></td><td></td><td></td><td></td><td></td><td></td><td></td></tr>
<tr><td>Oligo rev</td><td>AAACGTGCAGCACAGGCCAACAAC</td><td></td><td></td><td></td><td></td><td></td><td></td><td></td><td></td><td></td><td></td></tr>
<tr><td>Chromosome</td><td>start</td><td>end</td><td>strand</td><td>MM</td><td>target_seq</td><td>PAM</td><td>alignment</td><td>distance</td><td>position</td><td>gene name</td><td>gene id</td></tr>
<tr><td>GU305914.1</td><td>62268</td><td>62290</td><td>+</td><td>0</td><td>GTTGTTGGCCTGTGCTGCAC</td><td>CGG</td><td>||||||||[||||||||||||]PAM</td><td>NA</td><td>Intergenic</td><td>NA</td><td>NA</td></tr>
<tr><td></td><td></td><td></td><td></td><td></td><td></td><td></td><td></td><td></td><td></td><td></td><td></td></tr>
<tr><td>T4201</td><td>TTGTTGGCCTGTGCTGCACCGGG</td><td>1000</td><td></td><td></td><td></td><td></td><td></td><td></td><td></td><td></td><td></td></tr>
<tr><td>Oligo fwd</td><td>CACCTTGTTGGCCTGTGCTGCACC</td><td></td><td></td><td></td><td></td><td></td><td></td><td></td><td></td><td></td><td></td></tr>
<tr><td>Oligo rev</td><td>AAACGGTGCAGCACAGGCCAACAA</td><td></td><td></td><td></td><td></td><td></td><td></td><td></td><td></td><td></td><td></td></tr>
<tr><td>Chromosome</td><td>start</td><td>end</td><td>strand</td><td>MM</td><td>target_seq</td><td>PAM</td><td>alignment</td><td>distance</td><td>position</td><td>gene name</td><td>gene id</td></tr>
<tr><td>GU305914.1</td><td>62269</td><td>62291</td><td>+</td><td>0</td><td>TTGTTGGCCTGTGCTGCACC</td><td>GGG</td><td>||||||||[||||||||||||]PAM</td><td>NA</td><td>Intergenic</td><td>NA</td><td>NA</td></tr>
<tr><td></td><td></td><td></td><td></td><td></td><td></td><td></td><td></td><td></td><td></td><td></td><td></td></tr>
<tr><td>T4202</td><td>TGTTGGCCTGTGCTGCACCGGGG</td><td>1000</td><td></td><td></td><td></td><td></td><td></td><td></td><td></td><td></td><td></td></tr>
<tr><td>Oligo fwd</td><td>CACCTGTTGGCCTGTGCTGCACCG</td><td></td><td></td><td></td><td></td><td></td><td></td><td></td><td></td><td></td><td></td></tr>
<tr><td>Oligo rev</td><td>AAACCGGTGCAGCACAGGCCAACA</td><td></td><td></td><td></td><td></td><td></td><td></td><td></td><td></td><td></td><td></td></tr>
<tr><td>Chromosome</td><td>start</td><td>end</td><td>strand</td><td>MM</td><td>target_seq</td><td>PAM</td><td>alignment</td><td>distance</td><td>position</td><td>gene name</td><td>gene id</td></tr>
<tr><td>GU305914.1</td><td>62270</td><td>62292</td><td>+</td><td>0</td><td>TGTTGGCCTGTGCTGCACCG</td><td>GGG</td><td>||||||||[||||||||||||]PAM</td><td>NA</td><td>Intergenic</td><td>NA</td><td>NA</td></tr>
<tr><td></td><td></td><td></td><td></td><td></td><td></td><td></td><td></td><td></td><td></td><td></td><td></td></tr>
<tr><td>T4203</td><td>GCCTGTGCTGCACCGGGGAGCGG</td><td>1000</td><td></td><td></td><td></td><td></td><td></td><td></td><td></td><td></td><td></td></tr>
<tr><td>Oligo fwd</td><td>CACCGCCTGTGCTGCACCGGGGAG</td><td></td><td></td><td></td><td></td><td></td><td></td><td></td><td></td><td></td><td></td></tr>
<tr><td>Oligo rev</td><td>AAACCTCCCCGGTGCAGCACAGGC</td><td></td><td></td><td></td><td></td><td></td><td></td><td></td><td></td><td></td><td></td></tr>
<tr><td>Chromosome</td><td>start</td><td>end</td><td>strand</td><td>MM</td><td>target_seq</td><td>PAM</td><td>alignment</td><td>distance</td><td>position</td><td>gene name</td><td>gene id</td></tr>
<tr><td>GU305914.1</td><td>62275</td><td>62297</td><td>+</td><td>0</td><td>GCCTGTGCTGCACCGGGGAG</td><td>CGG</td><td>||||||||[||||||||||||]PAM</td><td>NA</td><td>Intergenic</td><td>NA</td><td>NA</td></tr>
<tr><td></td><td></td><td></td><td></td><td></td><td></td><td></td><td></td><td></td><td></td><td></td><td></td></tr>
<tr><td>T4204</td><td>CCTGTGCTGCACCGGGGAGCGGG</td><td>1000</td><td></td><td></td><td></td><td></td><td></td><td></td><td></td><td></td><td></td></tr>
<tr><td>Oligo fwd</td><td>CACCCCTGTGCTGCACCGGGGAGC</td><td></td><td></td><td></td><td></td><td></td><td></td><td></td><td></td><td></td><td></td></tr>
<tr><td>Oligo rev</td><td>AAACGCTCCCCGGTGCAGCACAGG</td><td></td><td></td><td></td><td></td><td></td><td></td><td></td><td></td><td></td><td></td></tr>
<tr><td>Chromosome</td><td>start</td><td>end</td><td>strand</td><td>MM</td><td>target_seq</td><td>PAM</td><td>alignment</td><td>distance</td><td>position</td><td>gene name</td><td>gene id</td></tr>
<tr><td>GU305914.1</td><td>62276</td><td>62298</td><td>+</td><td>0</td><td>CCTGTGCTGCACCGGGGAGC</td><td>GGG</td><td>||||||||[||||||||||||]PAM</td><td>NA</td><td>Intergenic</td><td>NA</td><td>NA</td></tr>
<tr><td></td><td></td><td></td><td></td><td></td><td></td><td></td><td></td><td></td><td></td><td></td><td></td></tr>
<tr><td>T4205</td><td>CTGCACCGGGGAGCGGGAATCGG</td><td>1000</td><td></td><td></td><td></td><td></td><td></td><td></td><td></td><td></td><td></td></tr>
<tr><td>Oligo fwd</td><td>CACCCTGCACCGGGGAGCGGGAAT</td><td></td><td></td><td></td><td></td><td></td><td></td><td></td><td></td><td></td><td></td></tr>
<tr><td>Oligo rev</td><td>AAACATTCCCGCTCCCCGGTGCAG</td><td></td><td></td><td></td><td></td><td></td><td></td><td></td><td></td><td></td><td></td></tr>
<tr><td>Chromosome</td><td>start</td><td>end</td><td>strand</td><td>MM</td><td>target_seq</td><td>PAM</td><td>alignment</td><td>distance</td><td>position</td><td>gene name</td><td>gene id</td></tr>
<tr><td>GU305914.1</td><td>62282</td><td>62304</td><td>+</td><td>0</td><td>CTGCACCGGGGAGCGGGAAT</td><td>CGG</td><td>||||||||[||||||||||||]PAM</td><td>NA</td><td>Intergenic</td><td>NA</td><td>NA</td></tr>
<tr><td></td><td></td><td></td><td></td><td></td><td></td><td></td><td></td><td></td><td></td><td></td><td></td></tr>
<tr><td>T4206</td><td>TGCACCGGGGAGCGGGAATCGGG</td><td>1000</td><td></td><td></td><td></td><td></td><td></td><td></td><td></td><td></td><td></td></tr>
<tr><td>Oligo fwd</td><td>CACCTGCACCGGGGAGCGGGAATC</td><td></td><td></td><td></td><td></td><td></td><td></td><td></td><td></td><td></td><td></td></tr>
<tr><td>Oligo rev</td><td>AAACGATTCCCGCTCCCCGGTGCA</td><td></td><td></td><td></td><td></td><td></td><td></td><td></td><td></td><td></td><td></td></tr>
<tr><td>Chromosome</td><td>start</td><td>end</td><td>strand</td><td>MM</td><td>target_seq</td><td>PAM</td><td>alignment</td><td>distance</td><td>position</td><td>gene name</td><td>gene id</td></tr>
<tr><td>GU305914.1</td><td>62283</td><td>62305</td><td>+</td><td>0</td><td>TGCACCGGGGAGCGGGAATC</td><td>GGG</td><td>||||||||[||||||||||||]PAM</td><td>NA</td><td>Intergenic</td><td>NA</td><td>NA</td></tr>
<tr><td></td><td></td><td></td><td></td><td></td><td></td><td></td><td></td><td></td><td></td><td></td><td></td></tr>
<tr><td>T4207</td><td>CGGGGAGCGGGAATCGGGATAGG</td><td>1000</td><td></td><td></td><td></td><td></td><td></td><td></td><td></td><td></td><td></td></tr>
<tr><td>Oligo fwd</td><td>CACCCGGGGAGCGGGAATCGGGAT</td><td></td><td></td><td></td><td></td><td></td><td></td><td></td><td></td><td></td><td></td></tr>
<tr><td>Oligo rev</td><td>AAACATCCCGATTCCCGCTCCCCG</td><td></td><td></td><td></td><td></td><td></td><td></td><td></td><td></td><td></td><td></td></tr>
<tr><td>Chromosome</td><td>start</td><td>end</td><td>strand</td><td>MM</td><td>target_seq</td><td>PAM</td><td>alignment</td><td>distance</td><td>position</td><td>gene name</td><td>gene id</td></tr>
<tr><td>GU305914.1</td><td>62288</td><td>62310</td><td>+</td><td>0</td><td>CGGGGAGCGGGAATCGGGAT</td><td>AGG</td><td>||||||||[||||||||||||]PAM</td><td>NA</td><td>Intergenic</td><td>NA</td><td>NA</td></tr>
<tr><td>GU305914.1</td><td>170397</td><td>170419</td><td>+</td><td>4</td><td>CCGGGATCGGGTATCGAGAT</td><td>GGG</td><td>|-||||-|[|||-||||-|||]PAM</td><td>NA</td><td>Intergenic</td><td>NA</td><td>NA</td></tr>
<tr><td></td><td></td><td></td><td></td><td></td><td></td><td></td><td></td><td></td><td></td><td></td><td></td></tr>
<tr><td>T4208</td><td>GGAATCGGGATAGGAGTCGACGG</td><td>1000</td><td></td><td></td><td></td><td></td><td></td><td></td><td></td><td></td><td></td></tr>
<tr><td>Oligo fwd</td><td>CACCGGAATCGGGATAGGAGTCGA</td><td></td><td></td><td></td><td></td><td></td><td></td><td></td><td></td><td></td><td></td></tr>
<tr><td>Oligo rev</td><td>AAACTCGACTCCTATCCCGATTCC</td><td></td><td></td><td></td><td></td><td></td><td></td><td></td><td></td><td></td><td></td></tr>
<tr><td>Chromosome</td><td>start</td><td>end</td><td>strand</td><td>MM</td><td>target_seq</td><td>PAM</td><td>alignment</td><td>distance</td><td>position</td><td>gene name</td><td>gene id</td></tr>
<tr><td>GU305914.1</td><td>62297</td><td>62319</td><td>+</td><td>0</td><td>GGAATCGGGATAGGAGTCGA</td><td>CGG</td><td>||||||||[||||||||||||]PAM</td><td>NA</td><td>Intergenic</td><td>NA</td><td>NA</td></tr>
<tr><td></td><td></td><td></td><td></td><td></td><td></td><td></td><td></td><td></td><td></td><td></td><td></td></tr>
<tr><td>T4209</td><td>GTGCCGACACTAACGTGACTAGG</td><td>1000</td><td></td><td></td><td></td><td></td><td></td><td></td><td></td><td></td><td></td></tr>
<tr><td>Oligo fwd</td><td>CACCGTGCCGACACTAACGTGACT</td><td></td><td></td><td></td><td></td><td></td><td></td><td></td><td></td><td></td><td></td></tr>
<tr><td>Oligo rev</td><td>AAACAGTCACGTTAGTGTCGGCAC</td><td></td><td></td><td></td><td></td><td></td><td></td><td></td><td></td><td></td><td></td></tr>
<tr><td>Chromosome</td><td>start</td><td>end</td><td>strand</td><td>MM</td><td>target_seq</td><td>PAM</td><td>alignment</td><td>distance</td><td>position</td><td>gene name</td><td>gene id</td></tr>
<tr><td>GU305914.1</td><td>62330</td><td>62352</td><td>+</td><td>0</td><td>GTGCCGACACTAACGTGACT</td><td>AGG</td><td>||||||||[||||||||||||]PAM</td><td>NA</td><td>Intergenic</td><td>NA</td><td>NA</td></tr>
<tr><td></td><td></td><td></td><td></td><td></td><td></td><td></td><td></td><td></td><td></td><td></td><td></td></tr>
<tr><td>T4210</td><td>CACTAACGTGACTAGGACGATGG</td><td>1000</td><td></td><td></td><td></td><td></td><td></td><td></td><td></td><td></td><td></td></tr>
<tr><td>Oligo fwd</td><td>CACCCACTAACGTGACTAGGACGA</td><td></td><td></td><td></td><td></td><td></td><td></td><td></td><td></td><td></td><td></td></tr>
<tr><td>Oligo rev</td><td>AAACTCGTCCTAGTCACGTTAGTG</td><td></td><td></td><td></td><td></td><td></td><td></td><td></td><td></td><td></td><td></td></tr>
<tr><td>Chromosome</td><td>start</td><td>end</td><td>strand</td><td>MM</td><td>target_seq</td><td>PAM</td><td>alignment</td><td>distance</td><td>position</td><td>gene name</td><td>gene id</td></tr>
<tr><td>GU305914.1</td><td>62337</td><td>62359</td><td>+</td><td>0</td><td>CACTAACGTGACTAGGACGA</td><td>TGG</td><td>||||||||[||||||||||||]PAM</td><td>NA</td><td>Intergenic</td><td>NA</td><td>NA</td></tr>
</table>

<table>
<tr><td>T4211</td><td>TAACGTGACTAGGACGATGGTGG</td><td>1000</td><td></td><td></td><td></td><td></td><td></td><td></td><td></td><td></td><td></td></tr>
<tr><td>Oligo fwd</td><td>CACCTAACGTGACTAGGACGATGG</td><td></td><td></td><td></td><td></td><td></td><td></td><td></td><td></td><td></td><td></td></tr>
<tr><td>Oligo rev</td><td>AAACCCATCGTCCTAGTCACGTTA</td><td></td><td></td><td></td><td></td><td></td><td></td><td></td><td></td><td></td><td></td></tr>
<tr><td>Chromosome</td><td>start</td><td>end</td><td>strand</td><td>MM</td><td>target_seq</td><td>PAM</td><td>alignment</td><td>distance</td><td>position</td><td>gene name</td><td>gene id</td></tr>
<tr><td>GU305914.1</td><td>62340</td><td>62362</td><td>+</td><td>0</td><td>TAACGTGACTAGGACGATGG</td><td>TGG</td><td>||||||||[||||||||||||]PAM</td><td>NA</td><td>Intergenic</td><td>NA</td><td>NA</td></tr>
<tr><td>T4212</td><td>TGACTAGGACGATGGTGGCGAGG</td><td>1000</td><td></td><td></td><td></td><td></td><td></td><td></td><td></td><td></td><td></td></tr>
<tr><td>Oligo fwd</td><td>CACCTGACTAGGACGATGGTGGCG</td><td></td><td></td><td></td><td></td><td></td><td></td><td></td><td></td><td></td><td></td></tr>
<tr><td>Oligo rev</td><td>AAACCGCCACCATCGTCCTAGTCA</td><td></td><td></td><td></td><td></td><td></td><td></td><td></td><td></td><td></td><td></td></tr>
<tr><td>Chromosome</td><td>start</td><td>end</td><td>strand</td><td>MM</td><td>target_seq</td><td>PAM</td><td>alignment</td><td>distance</td><td>position</td><td>gene name</td><td>gene id</td></tr>
<tr><td>GU305914.1</td><td>62345</td><td>62367</td><td>+</td><td>0</td><td>TGACTAGGACGATGGTGGCG</td><td>AGG</td><td>||||||||[||||||||||||]PAM</td><td>NA</td><td>Intergenic</td><td>NA</td><td>NA</td></tr>
<tr><td>T4213</td><td>AGGAGAAGATCCCTCGTCACAGG</td><td>1000</td><td></td><td></td><td></td><td></td><td></td><td></td><td></td><td></td><td></td></tr>
<tr><td>Oligo fwd</td><td>CACCAGGAGAAGATCCCTCGTCAC</td><td></td><td></td><td></td><td></td><td></td><td></td><td></td><td></td><td></td><td></td></tr>
<tr><td>Oligo rev</td><td>AAACGTGACGAGGGATCTTCTCCT</td><td></td><td></td><td></td><td></td><td></td><td></td><td></td><td></td><td></td><td></td></tr>
<tr><td>Chromosome</td><td>start</td><td>end</td><td>strand</td><td>MM</td><td>target_seq</td><td>PAM</td><td>alignment</td><td>distance</td><td>position</td><td>gene name</td><td>gene id</td></tr>
<tr><td>GU305914.1</td><td>62365</td><td>62387</td><td>+</td><td>0</td><td>AGGAGAAGATCCCTCGTCAC</td><td>AGG</td><td>||||||||[||||||||||||]PAM</td><td>NA</td><td>Intergenic</td><td>NA</td><td>NA</td></tr>
<tr><td>T4214</td><td>GGAGAAGATCCCTCGTCACAGGG</td><td>1000</td><td></td><td></td><td></td><td></td><td></td><td></td><td></td><td></td><td></td></tr>
<tr><td>Oligo fwd</td><td>CACCGGAGAAGATCCCTCGTCACA</td><td></td><td></td><td></td><td></td><td></td><td></td><td></td><td></td><td></td><td></td></tr>
<tr><td>Oligo rev</td><td>AAACTGTGACGAGGGATCTTCTCC</td><td></td><td></td><td></td><td></td><td></td><td></td><td></td><td></td><td></td><td></td></tr>
<tr><td>Chromosome</td><td>start</td><td>end</td><td>strand</td><td>MM</td><td>target_seq</td><td>PAM</td><td>alignment</td><td>distance</td><td>position</td><td>gene name</td><td>gene id</td></tr>
<tr><td>GU305914.1</td><td>62366</td><td>62388</td><td>+</td><td>0</td><td>GGAGAAGATCCCTCGTCACA</td><td>GGG</td><td>||||||||[||||||||||||]PAM</td><td>NA</td><td>Intergenic</td><td>NA</td><td>NA</td></tr>
<tr><td>T4215</td><td>CTCGTCACAGGGCGCAGTCGAGG</td><td>1000</td><td></td><td></td><td></td><td></td><td></td><td></td><td></td><td></td><td></td></tr>
<tr><td>Oligo fwd</td><td>CACCCTCGTCACAGGGCGCAGTCG</td><td></td><td></td><td></td><td></td><td></td><td></td><td></td><td></td><td></td><td></td></tr>
<tr><td>Oligo rev</td><td>AAACCGACTGCGCCCTGTGACGAG</td><td></td><td></td><td></td><td></td><td></td><td></td><td></td><td></td><td></td><td></td></tr>
<tr><td>Chromosome</td><td>start</td><td>end</td><td>strand</td><td>MM</td><td>target_seq</td><td>PAM</td><td>alignment</td><td>distance</td><td>position</td><td>gene name</td><td>gene id</td></tr>
<tr><td>GU305914.1</td><td>62377</td><td>62399</td><td>+</td><td>0</td><td>CTCGTCACAGGGCGCAGTCG</td><td>AGG</td><td>||||||||[||||||||||||]PAM</td><td>NA</td><td>Intergenic</td><td>NA</td><td>NA</td></tr>
<tr><td>T4216</td><td>GGGCGCAGTCGAGGCTGTAGAGG</td><td>1000</td><td></td><td></td><td></td><td></td><td></td><td></td><td></td><td></td><td></td></tr>
<tr><td>Oligo fwd</td><td>CACCGGGCGCAGTCGAGGCTGTAG</td><td></td><td></td><td></td><td></td><td></td><td></td><td></td><td></td><td></td><td></td></tr>
<tr><td>Oligo rev</td><td>AAACCTACAGCCTCGACTGCGCCC</td><td></td><td></td><td></td><td></td><td></td><td></td><td></td><td></td><td></td><td></td></tr>
<tr><td>Chromosome</td><td>start</td><td>end</td><td>strand</td><td>MM</td><td>target_seq</td><td>PAM</td><td>alignment</td><td>distance</td><td>position</td><td>gene name</td><td>gene id</td></tr>
<tr><td>GU305914.1</td><td>62386</td><td>62408</td><td>+</td><td>0</td><td>GGGCGCAGTCGAGGCTGTAG</td><td>AGG</td><td>||||||||[||||||||||||]PAM</td><td>NA</td><td>Intergenic</td><td>NA</td><td>NA</td></tr>
<tr><td>T4217</td><td>GGCGCAGTCGAGGCTGTAGAGGG</td><td>1000</td><td></td><td></td><td></td><td></td><td></td><td></td><td></td><td></td><td></td></tr>
<tr><td>Oligo fwd</td><td>CACCGGCGCAGTCGAGGCTGTAGA</td><td></td><td></td><td></td><td></td><td></td><td></td><td></td><td></td><td></td><td></td></tr>
<tr><td>Oligo rev</td><td>AAACTCTACAGCCTCGACTGCGCC</td><td></td><td></td><td></td><td></td><td></td><td></td><td></td><td></td><td></td><td></td></tr>
<tr><td>Chromosome</td><td>start</td><td>end</td><td>strand</td><td>MM</td><td>target_seq</td><td>PAM</td><td>alignment</td><td>distance</td><td>position</td><td>gene name</td><td>gene id</td></tr>
<tr><td>GU305914.1</td><td>62387</td><td>62409</td><td>+</td><td>0</td><td>GGCGCAGTCGAGGCTGTAGA</td><td>GGG</td><td>||||||||[||||||||||||]PAM</td><td>NA</td><td>Intergenic</td><td>NA</td><td>NA</td></tr>
<tr><td>T4218</td><td>GCGCAGTCGAGGCTGTAGAGGGG</td><td>1000</td><td></td><td></td><td></td><td></td><td></td><td></td><td></td><td></td><td></td></tr>
<tr><td>Oligo fwd</td><td>CACCGCGCAGTCGAGGCTGTAGAG</td><td></td><td></td><td></td><td></td><td></td><td></td><td></td><td></td><td></td><td></td></tr>
<tr><td>Oligo rev</td><td>AAACCTCTACAGCCTCGACTGCGC</td><td></td><td></td><td></td><td></td><td></td><td></td><td></td><td></td><td></td><td></td></tr>
<tr><td>Chromosome</td><td>start</td><td>end</td><td>strand</td><td>MM</td><td>target_seq</td><td>PAM</td><td>alignment</td><td>distance</td><td>position</td><td>gene name</td><td>gene id</td></tr>
<tr><td>GU305914.1</td><td>62388</td><td>62410</td><td>+</td><td>0</td><td>GCGCAGTCGAGGCTGTAGAG</td><td>GGG</td><td>||||||||[||||||||||||]PAM</td><td>NA</td><td>Intergenic</td><td>NA</td><td>NA</td></tr>
<tr><td>T4219</td><td>GTCGAGGCTGTAGAGGGGCTCGG</td><td>1000</td><td></td><td></td><td></td><td></td><td></td><td></td><td></td><td></td><td></td></tr>
<tr><td>Oligo fwd</td><td>CACCGTCGAGGCTGTAGAGGGGCT</td><td></td><td></td><td></td><td></td><td></td><td></td><td></td><td></td><td></td><td></td></tr>
<tr><td>Oligo rev</td><td>AAACAGCCCCTCTACAGCCTCGAC</td><td></td><td></td><td></td><td></td><td></td><td></td><td></td><td></td><td></td><td></td></tr>
<tr><td>Chromosome</td><td>start</td><td>end</td><td>strand</td><td>MM</td><td>target_seq</td><td>PAM</td><td>alignment</td><td>distance</td><td>position</td><td>gene name</td><td>gene id</td></tr>
<tr><td>GU305914.1</td><td>62393</td><td>62415</td><td>+</td><td>0</td><td>GTCGAGGCTGTAGAGGGGCT</td><td>CGG</td><td>||||||||[||||||||||||]PAM</td><td>NA</td><td>Intergenic</td><td>NA</td><td>NA</td></tr>
<tr><td>T4220</td><td>TCGAGGCTGTAGAGGGGCTCGGG</td><td>1000</td><td></td><td></td><td></td><td></td><td></td><td></td><td></td><td></td><td></td></tr>
<tr><td>Oligo fwd</td><td>CACCTCGAGGCTGTAGAGGGGCTC</td><td></td><td></td><td></td><td></td><td></td><td></td><td></td><td></td><td></td><td></td></tr>
<tr><td>Oligo rev</td><td>AAACGAGCCCCTCTACAGCCTCGA</td><td></td><td></td><td></td><td></td><td></td><td></td><td></td><td></td><td></td><td></td></tr>
<tr><td>Chromosome</td><td>start</td><td>end</td><td>strand</td><td>MM</td><td>target_seq</td><td>PAM</td><td>alignment</td><td>distance</td><td>position</td><td>gene name</td><td>gene id</td></tr>
<tr><td>GU305914.1</td><td>62394</td><td>62416</td><td>+</td><td>0</td><td>TCGAGGCTGTAGAGGGGCTC</td><td>GGG</td><td>||||||||[||||||||||||]PAM</td><td>NA</td><td>Intergenic</td><td>NA</td><td>NA</td></tr>
<tr><td>GU305914.1</td><td>120752</td><td>120774</td><td>+</td><td>4</td><td>TCGACGATGTCGAGGGTCTC</td><td>CGG</td><td>||||-|-|[||-|||||-|||]PAM</td><td>NA</td><td>Intergenic</td><td>NA</td><td>NA</td></tr>
<tr><td>T4221</td><td>GGCTGTAGAGGGGCTCGGGCTGG</td><td>1000</td><td></td><td></td><td></td><td></td><td></td><td></td><td></td><td></td><td></td></tr>
<tr><td>Oligo fwd</td><td>CACCGGCTGTAGAGGGGCTCGGGC</td><td></td><td></td><td></td><td></td><td></td><td></td><td></td><td></td><td></td><td></td></tr>
<tr><td>Oligo rev</td><td>AAACGCCCGAGCCCCTCTACAGCC</td><td></td><td></td><td></td><td></td><td></td><td></td><td></td><td></td><td></td><td></td></tr>
<tr><td>Chromosome</td><td>start</td><td>end</td><td>strand</td><td>MM</td><td>target_seq</td><td>PAM</td><td>alignment</td><td>distance</td><td>position</td><td>gene name</td><td>gene id</td></tr>
<tr><td>GU305914.1</td><td>62398</td><td>62420</td><td>+</td><td>0</td><td>GGCTGTAGAGGGGCTCGGGC</td><td>TGG</td><td>||||||||[||||||||||||]PAM</td><td>NA</td><td>Intergenic</td><td>NA</td><td>NA</td></tr>
<tr><td>T4222</td><td>GCTGTAGAGGGGCTCGGGCTGGG</td><td>1000</td><td></td><td></td><td></td><td></td><td></td><td></td><td></td><td></td><td></td></tr>
<tr><td>Oligo fwd</td><td>CACCGCTGTAGAGGGGCTCGGGCT</td><td></td><td></td><td></td><td></td><td></td><td></td><td></td><td></td><td></td><td></td></tr>
<tr><td>Oligo rev</td><td>AAACAGCCCGAGCCCCTCTACAGC</td><td></td><td></td><td></td><td></td><td></td><td></td><td></td><td></td><td></td><td></td></tr>
<tr><td>Chromosome</td><td>start</td><td>end</td><td>strand</td><td>MM</td><td>target_seq</td><td>PAM</td><td>alignment</td><td>distance</td><td>position</td><td>gene name</td><td>gene id</td></tr>
<tr><td>GU305914.1</td><td>62399</td><td>62421</td><td>+</td><td>0</td><td>GCTGTAGAGGGGCTCGGGCT</td><td>GGG</td><td>||||||||[||||||||||||]PAM</td><td>NA</td><td>Intergenic</td><td>NA</td><td>NA</td></tr>
<tr><td>T4223</td><td>AGGGGCTCGGGCTGGGTCGTCGG</td><td>1000</td><td></td><td></td><td></td><td></td><td></td><td></td><td></td><td></td><td></td></tr>
<tr><td>Oligo fwd</td><td>CACCAGGGGCTCGGGCTGGGTCGT</td><td></td><td></td><td></td><td></td><td></td><td></td><td></td><td></td><td></td><td></td></tr>
<tr><td>Oligo rev</td><td>AAACACGACCCAGCCCGAGCCCCT</td><td></td><td></td><td></td><td></td><td></td><td></td><td></td><td></td><td></td><td></td></tr>
<tr><td>Chromosome</td><td>start</td><td>end</td><td>strand</td><td>MM</td><td>target_seq</td><td>PAM</td><td>alignment</td><td>distance</td><td>position</td><td>gene name</td><td>gene id</td></tr>
<tr><td>GU305914.1</td><td>62406</td><td>62428</td><td>+</td><td>0</td><td>AGGGGCTCGGGCTGGGTCGT</td><td>CGG</td><td>||||||||[||||||||||||]PAM</td><td>NA</td><td>Intergenic</td><td>NA</td><td>NA</td></tr>
<tr><td>T4224</td><td>GCTGGGTCGTCGGTTTTCGACGG</td><td>1000</td><td></td><td></td><td></td><td></td><td></td><td></td><td></td><td></td><td></td></tr>
<tr><td>Oligo fwd</td><td>CACCGCTGGGTCGTCGGTTTTCGA</td><td></td><td></td><td></td><td></td><td></td><td></td><td></td><td></td><td></td><td></td></tr>
<tr><td>Oligo rev</td><td>AAACTCGAAAACCGACGACCCAGC</td><td></td><td></td><td></td><td></td><td></td><td></td><td></td><td></td><td></td><td></td></tr>
<tr><td>Chromosome</td><td>start</td><td>end</td><td>strand</td><td>MM</td><td>target_seq</td><td>PAM</td><td>alignment</td><td>distance</td><td>position</td><td>gene name</td><td>gene id</td></tr>
<tr><td>GU305914.1</td><td>62416</td><td>62438</td><td>+</td><td>0</td><td>GCTGGGTCGTCGGTTTTCGA</td><td>CGG</td><td>||||||||[||||||||||||]PAM</td><td>NA</td><td>Intergenic</td><td>NA</td><td>NA</td></tr>
<tr><td>T4225</td><td>GGGTCGTCGGTTTTCGACGGCGG</td><td>1000</td><td></td><td></td><td></td><td></td><td></td><td></td><td></td><td></td><td></td></tr>
<tr><td>Oligo fwd</td><td>CACCGGGTCGTCGGTTTTCGACGG</td><td></td><td></td><td></td><td></td><td></td><td></td><td></td><td></td><td></td><td></td></tr>
<tr><td>Oligo rev</td><td>AAACCCGTCGAAAACCGACGACCC</td><td></td><td></td><td></td><td></td><td></td><td></td><td></td><td></td><td></td><td></td></tr>
<tr><td>Chromosome</td><td>start</td><td>end</td><td>strand</td><td>MM</td><td>target_seq</td><td>PAM</td><td>alignment</td><td>distance</td><td>position</td><td>gene name</td><td>gene id</td></tr>
</table>

<table>
<tr><td>GU305914.1</td><td>62419</td><td>62441</td><td>+</td><td>0</td><td>GGGTCGTCGGTTTTCGACGG</td><td>CGG</td><td>||||||||[||||||||||||]PAM</td><td>NA</td><td>Intergenic</td><td>NA</td><td>NA</td></tr>
<tr><td></td><td></td><td></td><td></td><td></td><td></td><td></td><td></td><td></td><td></td><td></td><td></td></tr>
<tr><td>T4226</td><td>TCGTCGGTTTTCGACGGCGGTGG</td><td>1000</td><td></td><td></td><td></td><td></td><td></td><td></td><td></td><td></td><td></td></tr>
<tr><td>Oligo fwd</td><td>CACCTCGTCGGTTTTCGACGGCGG</td><td></td><td></td><td></td><td></td><td></td><td></td><td></td><td></td><td></td><td></td></tr>
<tr><td>Oligo rev</td><td>AAACCCGCCGTCGAAAACCGACGA</td><td></td><td></td><td></td><td></td><td></td><td></td><td></td><td></td><td></td><td></td></tr>
<tr><td>Chromosome</td><td>start</td><td>end</td><td>strand</td><td>MM</td><td>target_seq</td><td>PAM</td><td>alignment</td><td>distance</td><td>position</td><td>gene name</td><td>gene id</td></tr>
<tr><td>GU305914.1</td><td>62422</td><td>62444</td><td>+</td><td>0</td><td>TCGTCGGTTTTCGACGGCGG</td><td>TGG</td><td>||||||||[||||||||||||]PAM</td><td>NA</td><td>Intergenic</td><td>NA</td><td>NA</td></tr>
<tr><td></td><td></td><td></td><td></td><td></td><td></td><td></td><td></td><td></td><td></td><td></td><td></td></tr>
<tr><td>T4227</td><td>TCGGTTTTCGACGGCGGTGGCGG</td><td>1000</td><td></td><td></td><td></td><td></td><td></td><td></td><td></td><td></td><td></td></tr>
<tr><td>Oligo fwd</td><td>CACCTCGGTTTTCGACGGCGGTGG</td><td></td><td></td><td></td><td></td><td></td><td></td><td></td><td></td><td></td><td></td></tr>
<tr><td>Oligo rev</td><td>AAACCCACCGCCGTCGAAAACCGA</td><td></td><td></td><td></td><td></td><td></td><td></td><td></td><td></td><td></td><td></td></tr>
<tr><td>Chromosome</td><td>start</td><td>end</td><td>strand</td><td>MM</td><td>target_seq</td><td>PAM</td><td>alignment</td><td>distance</td><td>position</td><td>gene name</td><td>gene id</td></tr>
<tr><td>GU305914.1</td><td>62425</td><td>62447</td><td>+</td><td>0</td><td>TCGGTTTTCGACGGCGGTGG</td><td>CGG</td><td>||||||||[||||||||||||]PAM</td><td>NA</td><td>Intergenic</td><td>NA</td><td>NA</td></tr>
<tr><td></td><td></td><td></td><td></td><td></td><td></td><td></td><td></td><td></td><td></td><td></td><td></td></tr>
<tr><td>T4228</td><td>GTTTTCGACGGCGGTGGCGGCGG</td><td>1000</td><td></td><td></td><td></td><td></td><td></td><td></td><td></td><td></td><td></td></tr>
<tr><td>Oligo fwd</td><td>CACCGTTTTCGACGGCGGTGGCGG</td><td></td><td></td><td></td><td></td><td></td><td></td><td></td><td></td><td></td><td></td></tr>
<tr><td>Oligo rev</td><td>AAACCCGCCACCGCCGTCGAAAAC</td><td></td><td></td><td></td><td></td><td></td><td></td><td></td><td></td><td></td><td></td></tr>
<tr><td>Chromosome</td><td>start</td><td>end</td><td>strand</td><td>MM</td><td>target_seq</td><td>PAM</td><td>alignment</td><td>distance</td><td>position</td><td>gene name</td><td>gene id</td></tr>
<tr><td>GU305914.1</td><td>62428</td><td>62450</td><td>+</td><td>0</td><td>GTTTTCGACGGCGGTGGCGG</td><td>CGG</td><td>||||||||[||||||||||||]PAM</td><td>NA</td><td>Intergenic</td><td>NA</td><td>NA</td></tr>
<tr><td></td><td></td><td></td><td></td><td></td><td></td><td></td><td></td><td></td><td></td><td></td><td></td></tr>
<tr><td>T4229</td><td>TTTTCGACGGCGGTGGCGGCGGG</td><td>1000</td><td></td><td></td><td></td><td></td><td></td><td></td><td></td><td></td><td></td></tr>
<tr><td>Oligo fwd</td><td>CACCTTTTCGACGGCGGTGGCGGC</td><td></td><td></td><td></td><td></td><td></td><td></td><td></td><td></td><td></td><td></td></tr>
<tr><td>Oligo rev</td><td>AAACGCCGCCACCGCCGTCGAAAA</td><td></td><td></td><td></td><td></td><td></td><td></td><td></td><td></td><td></td><td></td></tr>
<tr><td>Chromosome</td><td>start</td><td>end</td><td>strand</td><td>MM</td><td>target_seq</td><td>PAM</td><td>alignment</td><td>distance</td><td>position</td><td>gene name</td><td>gene id</td></tr>
<tr><td>GU305914.1</td><td>62429</td><td>62451</td><td>+</td><td>0</td><td>TTTTCGACGGCGGTGGCGGC</td><td>GGG</td><td>||||||||[||||||||||||]PAM</td><td>NA</td><td>Intergenic</td><td>NA</td><td>NA</td></tr>
<tr><td></td><td></td><td></td><td></td><td></td><td></td><td></td><td></td><td></td><td></td><td></td><td></td></tr>
<tr><td>T4230</td><td>TTTCGACGGCGGTGGCGGCGGGG</td><td>1000</td><td></td><td></td><td></td><td></td><td></td><td></td><td></td><td></td><td></td></tr>
<tr><td>Oligo fwd</td><td>CACCTTTCGACGGCGGTGGCGGCG</td><td></td><td></td><td></td><td></td><td></td><td></td><td></td><td></td><td></td><td></td></tr>
<tr><td>Oligo rev</td><td>AAACCGCCGCCACCGCCGTCGAAA</td><td></td><td></td><td></td><td></td><td></td><td></td><td></td><td></td><td></td><td></td></tr>
<tr><td>Chromosome</td><td>start</td><td>end</td><td>strand</td><td>MM</td><td>target_seq</td><td>PAM</td><td>alignment</td><td>distance</td><td>position</td><td>gene name</td><td>gene id</td></tr>
<tr><td>GU305914.1</td><td>62430</td><td>62452</td><td>+</td><td>0</td><td>TTTCGACGGCGGTGGCGGCG</td><td>GGG</td><td>||||||||[||||||||||||]PAM</td><td>NA</td><td>Intergenic</td><td>NA</td><td>NA</td></tr>
<tr><td>GU305914.1</td><td>159954</td><td>159976</td><td>-</td><td>4</td><td>TATCGGCGGCGACGGCGGCG</td><td>AGG</td><td>|-|||-||[|||--|||||||]PAM</td><td>NA</td><td>Intergenic</td><td>NA</td><td>NA</td></tr>
<tr><td></td><td></td><td></td><td></td><td></td><td></td><td></td><td></td><td></td><td></td><td></td><td></td></tr>
<tr><td>T4231</td><td>TTCGACGGCGGTGGCGGCGGGGG</td><td>1000</td><td></td><td></td><td></td><td></td><td></td><td></td><td></td><td></td><td></td></tr>
<tr><td>Oligo fwd</td><td>CACCTTCGACGGCGGTGGCGGCGG</td><td></td><td></td><td></td><td></td><td></td><td></td><td></td><td></td><td></td><td></td></tr>
<tr><td>Oligo rev</td><td>AAACCCGCCGCCACCGCCGTCGAA</td><td></td><td></td><td></td><td></td><td></td><td></td><td></td><td></td><td></td><td></td></tr>
<tr><td>Chromosome</td><td>start</td><td>end</td><td>strand</td><td>MM</td><td>target_seq</td><td>PAM</td><td>alignment</td><td>distance</td><td>position</td><td>gene name</td><td>gene id</td></tr>
<tr><td>GU305914.1</td><td>62431</td><td>62453</td><td>+</td><td>0</td><td>TTCGACGGCGGTGGCGGCGG</td><td>GGG</td><td>||||||||[||||||||||||]PAM</td><td>NA</td><td>Intergenic</td><td>NA</td><td>NA</td></tr>
<tr><td>GU305914.1</td><td>48156</td><td>48178</td><td>+</td><td>3</td><td>GGCGGCGGCGGTGGCGGCGG</td><td>CGG</td><td>--||-|||[||||||||||||]PAM</td><td>NA</td><td>Intergenic</td><td>NA</td><td>NA</td></tr>
<tr><td>GU305914.1</td><td>149482</td><td>149504</td><td>-</td><td>3</td><td>GGCGACGGCGGCGGCGGCGG</td><td>CGG</td><td>--||||||[|||-||||||||]PAM</td><td>NA</td><td>Intergenic</td><td>NA</td><td>NA</td></tr>
<tr><td>GU305914.1</td><td>35653</td><td>35675</td><td>+</td><td>3</td><td>ATCCACGGCGGCGGCGGCGG</td><td>CGG</td><td>-||-||||[|||-||||||||]PAM</td><td>NA</td><td>Intergenic</td><td>NA</td><td>NA</td></tr>
<tr><td>GU305914.1</td><td>73189</td><td>73211</td><td>+</td><td>4</td><td>TTCCTCGTCGGTGCCGGCGG</td><td>AGG</td><td>|||--||-[|||||-||||||]PAM</td><td>NA</td><td>Intergenic</td><td>NA</td><td>NA</td></tr>
<tr><td>GU305914.1</td><td>114664</td><td>114686</td><td>-</td><td>4</td><td>TGAGACGGCGGTGGTGGTGG</td><td>CGG</td><td>|--|||||[||||||-||-||]PAM</td><td>NA</td><td>Intergenic</td><td>NA</td><td>NA</td></tr>
<tr><td></td><td></td><td></td><td></td><td></td><td></td><td></td><td></td><td></td><td></td><td></td><td></td></tr>
<tr><td>T4232</td><td>GACGGCGGTGGCGGCGGGGGTGG</td><td>1000</td><td></td><td></td><td></td><td></td><td></td><td></td><td></td><td></td><td></td></tr>
<tr><td>Oligo fwd</td><td>CACCGACGGCGGTGGCGGCGGGGG</td><td></td><td></td><td></td><td></td><td></td><td></td><td></td><td></td><td></td><td></td></tr>
<tr><td>Oligo rev</td><td>AAACCCCCCGCCGCCACCGCCGTC</td><td></td><td></td><td></td><td></td><td></td><td></td><td></td><td></td><td></td><td></td></tr>
<tr><td>Chromosome</td><td>start</td><td>end</td><td>strand</td><td>MM</td><td>target_seq</td><td>PAM</td><td>alignment</td><td>distance</td><td>position</td><td>gene name</td><td>gene id</td></tr>
<tr><td>GU305914.1</td><td>62434</td><td>62456</td><td>+</td><td>0</td><td>GACGGCGGTGGCGGCGGGGG</td><td>TGG</td><td>||||||||[||||||||||||]PAM</td><td>NA</td><td>Intergenic</td><td>NA</td><td>NA</td></tr>
<tr><td>GU305914.1</td><td>62449</td><td>62471</td><td>+</td><td>3</td><td>GGGGGTGGTGGCGGCGGGGG</td><td>TGG</td><td>|--||-||[||||||||||||]PAM</td><td>NA</td><td>Intergenic</td><td>NA</td><td>NA</td></tr>
<tr><td>GU305914.1</td><td>143022</td><td>143044</td><td>+</td><td>4</td><td>GCTGGTGGTGGCGCCGGGGG</td><td>CGG</td><td>|--||-||[|||||-||||||]PAM</td><td>NA</td><td>Intergenic</td><td>NA</td><td>NA</td></tr>
<tr><td>GU305914.1</td><td>48159</td><td>48181</td><td>+</td><td>2</td><td>GGCGGCGGTGGCGGCGGCGG</td><td>TGG</td><td>|-||||||[|||||||||-||]PAM</td><td>NA</td><td>Intergenic</td><td>NA</td><td>NA</td></tr>
<tr><td>GU305914.1</td><td>58129</td><td>58151</td><td>-</td><td>3</td><td>GCGGGCGGTGGCGGCGGCGG</td><td>CGG</td><td>|--|||||[|||||||||-||]PAM</td><td>NA</td><td>Intergenic</td><td>NA</td><td>NA</td></tr>
<tr><td>GU305914.1</td><td>154401</td><td>154423</td><td>+</td><td>4</td><td>CCGGGCGGTGGCGGCGGCGG</td><td>CGG</td><td>---|||||[|||||||||-||]PAM</td><td>NA</td><td>Intergenic</td><td>NA</td><td>NA</td></tr>
<tr><td>GU305914.1</td><td>154407</td><td>154429</td><td>+</td><td>4</td><td>GGTGGCGGCGGCGGCGGCGG</td><td>CGG</td><td>|--|||||[-||||||||-||]PAM</td><td>NA</td><td>Intergenic</td><td>NA</td><td>NA</td></tr>
<tr><td>GU305914.1</td><td>58123</td><td>58145</td><td>-</td><td>4</td><td>GGTGGCGGCGGCGGCGGCGG</td><td>AGG</td><td>|--|||||[-||||||||-||]PAM</td><td>NA</td><td>Intergenic</td><td>NA</td><td>NA</td></tr>
<tr><td>GU305914.1</td><td>149482</td><td>149504</td><td>-</td><td>4</td><td>GGCGACGGCGGCGGCGGCGG</td><td>CGG</td><td>|-||-|||[-||||||||-||]PAM</td><td>NA</td><td>Intergenic</td><td>NA</td><td>NA</td></tr>
<tr><td>GU305914.1</td><td>154404</td><td>154426</td><td>+</td><td>4</td><td>GGCGGTGGCGGCGGCGGCGG</td><td>CGG</td><td>|-|||-||[-||||||||-||]PAM</td><td>NA</td><td>Intergenic</td><td>NA</td><td>NA</td></tr>
<tr><td>GU305914.1</td><td>58126</td><td>58148</td><td>-</td><td>4</td><td>GGCGGTGGCGGCGGCGGCGG</td><td>CGG</td><td>|-|||-||[-||||||||-||]PAM</td><td>NA</td><td>Intergenic</td><td>NA</td><td>NA</td></tr>
<tr><td>GU305914.1</td><td>62446</td><td>62468</td><td>+</td><td>4</td><td>GGCGGGGGTGGTGGCGGCGG</td><td>GGG</td><td>|-|||-||[|||-|||||-||]PAM</td><td>NA</td><td>Intergenic</td><td>NA</td><td>NA</td></tr>
<tr><td>GU305914.1</td><td>114658</td><td>114680</td><td>-</td><td>4</td><td>GGCGGTGGTGGTGGCGGCGG</td><td>CGG</td><td>|-|||-||[|||-|||||-||]PAM</td><td>NA</td><td>Intergenic</td><td>NA</td><td>NA</td></tr>
<tr><td>GU305914.1</td><td>168817</td><td>168839</td><td>+</td><td>4</td><td>GGCGGTGGTGGCAGCGGCGG</td><td>CGG</td><td>|-|||-||[||||-||||-||]PAM</td><td>NA</td><td>Intergenic</td><td>NA</td><td>NA</td></tr>
<tr><td>GU305914.1</td><td>45255</td><td>45277</td><td>-</td><td>4</td><td>GGTGGCGGTGGCGGCAGCGG</td><td>CGG</td><td>|--|||||[|||||||-|-||]PAM</td><td>NA</td><td>Intergenic</td><td>NA</td><td>NA</td></tr>
<tr><td></td><td></td><td></td><td></td><td></td><td></td><td></td><td></td><td></td><td></td><td></td><td></td></tr>
<tr><td>T4233</td><td>GGCGGTGGCGGCGGGGGTGGTGG</td><td>1000</td><td></td><td></td><td></td><td></td><td></td><td></td><td></td><td></td><td></td></tr>
<tr><td>Oligo fwd</td><td>CACCGGCGGTGGCGGCGGGGGTGG</td><td></td><td></td><td></td><td></td><td></td><td></td><td></td><td></td><td></td><td></td></tr>
<tr><td>Oligo rev</td><td>AAACCCACCCCCGCCGCCACCGCC</td><td></td><td></td><td></td><td></td><td></td><td></td><td></td><td></td><td></td><td></td></tr>
<tr><td>Chromosome</td><td>start</td><td>end</td><td>strand</td><td>MM</td><td>target_seq</td><td>PAM</td><td>alignment</td><td>distance</td><td>position</td><td>gene name</td><td>gene id</td></tr>
<tr><td>GU305914.1</td><td>62437</td><td>62459</td><td>+</td><td>0</td><td>GGCGGTGGCGGCGGGGGTGG</td><td>TGG</td><td>||||||||[||||||||||||]PAM</td><td>NA</td><td>Intergenic</td><td>NA</td><td>NA</td></tr>
<tr><td>GU305914.1</td><td>62452</td><td>62474</td><td>+</td><td>1</td><td>GGTGGTGGCGGCGGGGGTGG</td><td>CGG</td><td>||-|||||[||||||||||||]PAM</td><td>NA</td><td>Intergenic</td><td>NA</td><td>NA</td></tr>
<tr><td>GU305914.1</td><td>48150</td><td>48172</td><td>+</td><td>4</td><td>ACCTGTGGCGGCGGCGGTGG</td><td>CGG</td><td>--|-||||[||||||-|||||]PAM</td><td>NA</td><td>Intergenic</td><td>NA</td><td>NA</td></tr>
<tr><td>GU305914.1</td><td>225498</td><td>225520</td><td>+</td><td>4</td><td>GCCGAGGGCGGCGGTGGTGG</td><td>TGG</td><td>|-||--||[||||||-|||||]PAM</td><td>NA</td><td>Intergenic</td><td>NA</td><td>NA</td></tr>
<tr><td>GU305914.1</td><td>52719</td><td>52741</td><td>+</td><td>4</td><td>GCCGCTGGCGGTGGCGGTGG</td><td>CGG</td><td>|-||-|||[|||-||-|||||]PAM</td><td>NA</td><td>Intergenic</td><td>NA</td><td>NA</td></tr>
<tr><td>GU305914.1</td><td>62440</td><td>62462</td><td>+</td><td>4</td><td>GGTGGCGGCGGGGGTGGTGG</td><td>CGG</td><td>||-||-||[|||-||-|||||]PAM</td><td>NA</td><td>Intergenic</td><td>NA</td><td>NA</td></tr>
<tr><td>GU305914.1</td><td>114470</td><td>114492</td><td>-</td><td>4</td><td>AGCGGAGGCGGCAGGGTTGG</td><td>CGG</td><td>-||||-||[||||-|||-|||]PAM</td><td>NA</td><td>Intergenic</td><td>NA</td><td>NA</td></tr>
<tr><td>GU305914.1</td><td>154404</td><td>154426</td><td>+</td><td>2</td><td>GGCGGTGGCGGCGGCGGCGG</td><td>CGG</td><td>||||||||[||||||-||-||]PAM</td><td>NA</td><td>Intergenic</td><td>NA</td><td>NA</td></tr>
<tr><td>GU305914.1</td><td>58126</td><td>58148</td><td>-</td><td>2</td><td>GGCGGTGGCGGCGGCGGCGG</td><td>CGG</td><td>||||||||[||||||-||-||]PAM</td><td>NA</td><td>Intergenic</td><td>NA</td><td>NA</td></tr>
<tr><td>GU305914.1</td><td>30000</td><td>30022</td><td>+</td><td>4</td><td>GGTGCTGGCGGCGGCGGCGG</td><td>CGG</td><td>||-|-|||[||||||-||-||]PAM</td><td>NA</td><td>Intergenic</td><td>NA</td><td>NA</td></tr>
<tr><td>GU305914.1</td><td>154407</td><td>154429</td><td>+</td><td>4</td><td>GGTGGCGGCGGCGGCGGCGG</td><td>CGG</td><td>||-||-||[||||||-||-||]PAM</td><td>NA</td><td>Intergenic</td><td>NA</td><td>NA</td></tr>
<tr><td>GU305914.1</td><td>58123</td><td>58145</td><td>-</td><td>4</td><td>GGTGGCGGCGGCGGCGGCGG</td><td>AGG</td><td>||-||-||[||||||-||-||]PAM</td><td>NA</td><td>Intergenic</td><td>NA</td><td>NA</td></tr>
<tr><td>GU305914.1</td><td>149482</td><td>149504</td><td>-</td><td>4</td><td>GGCGACGGCGGCGGCGGCGG</td><td>CGG</td><td>||||--||[||||||-||-||]PAM</td><td>NA</td><td>Intergenic</td><td>NA</td><td>NA</td></tr>
<tr><td>GU305914.1</td><td>161748</td><td>161770</td><td>-</td><td>4</td><td>GGAGGCGGCGGCGGAGGTGA</td><td>CGG</td><td>||-||-||[||||||-||||-]PAM</td><td>NA</td><td>Intergenic</td><td>NA</td><td>NA</td></tr>
<tr><td>GU305914.1</td><td>81240</td><td>81262</td><td>-</td><td>4</td><td>GGAGGCGGCGGCGGGGGGTG</td><td>AGG</td><td>||-||-||[|||||||||--|]PAM</td><td>NA</td><td>Intergenic</td><td>NA</td><td>NA</td></tr>
<tr><td></td><td></td><td></td><td></td><td></td><td></td><td></td><td></td><td></td><td></td><td></td><td></td></tr>
<tr><td>T4234</td><td>GGTGGCGGCGGGGGTGGTGGCGG</td><td>1000</td><td></td><td></td><td></td><td></td><td></td><td></td><td></td><td></td><td></td></tr>
<tr><td>Oligo fwd</td><td>CACCGGTGGCGGCGGGGGTGGTGG</td><td></td><td></td><td></td><td></td><td></td><td></td><td></td><td></td><td></td><td></td></tr>
<tr><td>Oligo rev</td><td>AAACCCACCACCCCCGCCGCCACC</td><td></td><td></td><td></td><td></td><td></td><td></td><td></td><td></td><td></td><td></td></tr>
<tr><td>Chromosome</td><td>start</td><td>end</td><td>strand</td><td>MM</td><td>target_seq</td><td>PAM</td><td>alignment</td><td>distance</td><td>position</td><td>gene name</td><td>gene id</td></tr>
<tr><td>GU305914.1</td><td>62440</td><td>62462</td><td>+</td><td>0</td><td>GGTGGCGGCGGGGGTGGTGG</td><td>CGG</td><td>||||||||[||||||||||||]PAM</td><td>NA</td><td>Intergenic</td><td>NA</td><td>NA</td></tr>
</table>

<table>
<tr><td>GU305914.1</td><td>114664</td><td>114686</td><td>-</td><td>4</td><td>TGAGACGGCGGTGGTGGTGG</td><td>CGG</td><td>-|-|-|||[|||-||||||||]PAM</td><td>NA</td><td>Intergenic</td><td>NA</td><td>NA</td></tr>
<tr><td>GU305914.1</td><td>62452</td><td>62474</td><td>+</td><td>3</td><td>GGTGGTGGCGGCGGGGGTGG</td><td>CGG</td><td>|||||-||[|||-||-|||||]PAM</td><td>NA</td><td>Intergenic</td><td>NA</td><td>NA</td></tr>
<tr><td>GU305914.1</td><td>62437</td><td>62459</td><td>+</td><td>4</td><td>GGCGGTGGCGGCGGGGGTGG</td><td>TGG</td><td>||-||-||[|||-||-|||||]PAM</td><td>NA</td><td>Intergenic</td><td>NA</td><td>NA</td></tr>
<tr><td>GU305914.1</td><td>74673</td><td>74695</td><td>+</td><td>4</td><td>GGTGCCGTCGGTGGCGGTGG</td><td>CGG</td><td>||||-||-[|||-||-|||||]PAM</td><td>NA</td><td>Intergenic</td><td>NA</td><td>NA</td></tr>
<tr><td>GU305914.1</td><td>36161</td><td>36183</td><td>+</td><td>4</td><td>GCTGCCGGCGGCGGTTGTGG</td><td>TGG</td><td>|-||-|||[|||-|||-||||]PAM</td><td>NA</td><td>Intergenic</td><td>NA</td><td>NA</td></tr>
<tr><td>GU305914.1</td><td>48153</td><td>48175</td><td>+</td><td>3</td><td>TGTGGCGGCGGCGGTGGCGG</td><td>CGG</td><td>-|||||||[|||-|||||-||]PAM</td><td>NA</td><td>Intergenic</td><td>NA</td><td>NA</td></tr>
<tr><td></td><td></td><td></td><td></td><td></td><td></td><td></td><td></td><td></td><td></td><td></td><td></td></tr>
<tr><td>T4235</td><td>GGCGGCGGGGGTGGTGGCGGCGG</td><td>1000</td><td></td><td></td><td></td><td></td><td></td><td></td><td></td><td></td><td></td></tr>
<tr><td>Oligo fwd</td><td>CACCGGCGGCGGGGGTGGTGGCGG</td><td></td><td></td><td></td><td></td><td></td><td></td><td></td><td></td><td></td><td></td></tr>
<tr><td>Oligo rev</td><td>AAACCCGCCACCACCCCCGCCGCC</td><td></td><td></td><td></td><td></td><td></td><td></td><td></td><td></td><td></td><td></td></tr>
<tr><td>Chromosome</td><td>start</td><td>end</td><td>strand</td><td>MM</td><td>target_seq</td><td>PAM</td><td>alignment</td><td>distance</td><td>position</td><td>gene name</td><td>gene id</td></tr>
<tr><td>GU305914.1</td><td>62443</td><td>62465</td><td>+</td><td>0</td><td>GGCGGCGGGGGTGGTGGCGG</td><td>CGG</td><td>||||||||[||||||||||||]PAM</td><td>NA</td><td>Intergenic</td><td>NA</td><td>NA</td></tr>
<tr><td>GU305914.1</td><td>114661</td><td>114683</td><td>-</td><td>2</td><td>GACGGCGGTGGTGGTGGCGG</td><td>CGG</td><td>|-||||||[-|||||||||||]PAM</td><td>NA</td><td>Intergenic</td><td>NA</td><td>NA</td></tr>
<tr><td>GU305914.1</td><td>19936</td><td>19958</td><td>+</td><td>4</td><td>GACGGTAGCGGTGGTGGCGG</td><td>TGG</td><td>|-|||--|[-|||||||||||]PAM</td><td>NA</td><td>Intergenic</td><td>NA</td><td>NA</td></tr>
<tr><td>GU305914.1</td><td>48153</td><td>48175</td><td>+</td><td>4</td><td>TGTGGCGGCGGCGGTGGCGG</td><td>CGG</td><td>-|-|||||[-||-||||||||]PAM</td><td>NA</td><td>Intergenic</td><td>NA</td><td>NA</td></tr>
<tr><td>GU305914.1</td><td>48156</td><td>48178</td><td>+</td><td>2</td><td>GGCGGCGGCGGTGGCGGCGG</td><td>CGG</td><td>||||||||[-|||||-|||||]PAM</td><td>NA</td><td>Intergenic</td><td>NA</td><td>NA</td></tr>
<tr><td>GU305914.1</td><td>62446</td><td>62468</td><td>+</td><td>3</td><td>GGCGGGGGTGGTGGCGGCGG</td><td>GGG</td><td>|||||-||[-|||||-|||||]PAM</td><td>NA</td><td>Intergenic</td><td>NA</td><td>NA</td></tr>
<tr><td>GU305914.1</td><td>114658</td><td>114680</td><td>-</td><td>3</td><td>GGCGGTGGTGGTGGCGGCGG</td><td>CGG</td><td>|||||-||[-|||||-|||||]PAM</td><td>NA</td><td>Intergenic</td><td>NA</td><td>NA</td></tr>
<tr><td>GU305914.1</td><td>62458</td><td>62480</td><td>+</td><td>2</td><td>GGCGGCGGGGGTGGCGGCCG</td><td>CGG</td><td>||||||||[||||||-|||-|]PAM</td><td>NA</td><td>Intergenic</td><td>NA</td><td>NA</td></tr>
<tr><td></td><td></td><td></td><td></td><td></td><td></td><td></td><td></td><td></td><td></td><td></td><td></td></tr>
<tr><td>T4236</td><td>GCGGCGGGGGTGGTGGCGGCGGG</td><td>1000</td><td></td><td></td><td></td><td></td><td></td><td></td><td></td><td></td><td></td></tr>
<tr><td>Oligo fwd</td><td>CACCGCGGCGGGGGTGGTGGCGGC</td><td></td><td></td><td></td><td></td><td></td><td></td><td></td><td></td><td></td><td></td></tr>
<tr><td>Oligo rev</td><td>AAACGCCGCCACCACCCCCGCCGC</td><td></td><td></td><td></td><td></td><td></td><td></td><td></td><td></td><td></td><td></td></tr>
<tr><td>Chromosome</td><td>start</td><td>end</td><td>strand</td><td>MM</td><td>target_seq</td><td>PAM</td><td>alignment</td><td>distance</td><td>position</td><td>gene name</td><td>gene id</td></tr>
<tr><td>GU305914.1</td><td>62444</td><td>62466</td><td>+</td><td>0</td><td>GCGGCGGGGGTGGTGGCGGC</td><td>GGG</td><td>||||||||[||||||||||||]PAM</td><td>NA</td><td>Intergenic</td><td>NA</td><td>NA</td></tr>
<tr><td>GU305914.1</td><td>114657</td><td>114679</td><td>-</td><td>3</td><td>GCGGTGGTGGTGGCGGCGGC</td><td>GGG</td><td>||||-||-[|||||-||||||]PAM</td><td>NA</td><td>Intergenic</td><td>NA</td><td>NA</td></tr>
<tr><td>GU305914.1</td><td>221599</td><td>221621</td><td>-</td><td>4</td><td>GAGGCCGTGGTGGCGGCGGC</td><td>AGG</td><td>|-|||-|-[|||||-||||||]PAM</td><td>NA</td><td>Intergenic</td><td>NA</td><td>NA</td></tr>
<tr><td>GU305914.1</td><td>152554</td><td>152576</td><td>+</td><td>3</td><td>GCGGCGGCGGCGGTGCCGGC</td><td>GGG</td><td>|||||||-[||-||||-||||]PAM</td><td>NA</td><td>Intergenic</td><td>NA</td><td>NA</td></tr>
<tr><td></td><td></td><td></td><td></td><td></td><td></td><td></td><td></td><td></td><td></td><td></td><td></td></tr>
<tr><td>T4237</td><td>CGGCGGGGGTGGTGGCGGCGGGG</td><td>1000</td><td></td><td></td><td></td><td></td><td></td><td></td><td></td><td></td><td></td></tr>
<tr><td>Oligo fwd</td><td>CACCCGGCGGGGGTGGTGGCGGCG</td><td></td><td></td><td></td><td></td><td></td><td></td><td></td><td></td><td></td><td></td></tr>
<tr><td>Oligo rev</td><td>AAACCGCCGCCACCACCCCCGCCG</td><td></td><td></td><td></td><td></td><td></td><td></td><td></td><td></td><td></td><td></td></tr>
<tr><td>Chromosome</td><td>start</td><td>end</td><td>strand</td><td>MM</td><td>target_seq</td><td>PAM</td><td>alignment</td><td>distance</td><td>position</td><td>gene name</td><td>gene id</td></tr>
<tr><td>GU305914.1</td><td>62445</td><td>62467</td><td>+</td><td>0</td><td>CGGCGGGGGTGGTGGCGGCG</td><td>GGG</td><td>||||||||[||||||||||||]PAM</td><td>NA</td><td>Intergenic</td><td>NA</td><td>NA</td></tr>
<tr><td>GU305914.1</td><td>152555</td><td>152577</td><td>+</td><td>3</td><td>CGGCGGCGGCGGTGCCGGCG</td><td>GGG</td><td>||||||-|[|-||||-|||||]PAM</td><td>NA</td><td>Intergenic</td><td>NA</td><td>NA</td></tr>
<tr><td>GU305914.1</td><td>18654</td><td>18676</td><td>-</td><td>4</td><td>CGGCAGAGGTGGCGGAGGCG</td><td>GGG</td><td>||||-|-|[||||-||-||||]PAM</td><td>NA</td><td>Intergenic</td><td>NA</td><td>NA</td></tr>
<tr><td></td><td></td><td></td><td></td><td></td><td></td><td></td><td></td><td></td><td></td><td></td><td></td></tr>
<tr><td>T4238</td><td>GGCGGGGGTGGTGGCGGCGGGGG</td><td>1000</td><td></td><td></td><td></td><td></td><td></td><td></td><td></td><td></td><td></td></tr>
<tr><td>Oligo fwd</td><td>CACCGGCGGGGGTGGTGGCGGCGG</td><td></td><td></td><td></td><td></td><td></td><td></td><td></td><td></td><td></td><td></td></tr>
<tr><td>Oligo rev</td><td>AAACCCGCCGCCACCACCCCCGCC</td><td></td><td></td><td></td><td></td><td></td><td></td><td></td><td></td><td></td><td></td></tr>
<tr><td>Chromosome</td><td>start</td><td>end</td><td>strand</td><td>MM</td><td>target_seq</td><td>PAM</td><td>alignment</td><td>distance</td><td>position</td><td>gene name</td><td>gene id</td></tr>
<tr><td>GU305914.1</td><td>62446</td><td>62468</td><td>+</td><td>0</td><td>GGCGGGGGTGGTGGCGGCGG</td><td>GGG</td><td>||||||||[||||||||||||]PAM</td><td>NA</td><td>Intergenic</td><td>NA</td><td>NA</td></tr>
<tr><td>GU305914.1</td><td>114658</td><td>114680</td><td>-</td><td>1</td><td>GGCGGTGGTGGTGGCGGCGG</td><td>CGG</td><td>|||||-||[||||||||||||]PAM</td><td>NA</td><td>Intergenic</td><td>NA</td><td>NA</td></tr>
<tr><td>GU305914.1</td><td>48156</td><td>48178</td><td>+</td><td>2</td><td>GGCGGCGGCGGTGGCGGCGG</td><td>CGG</td><td>|||||-||[-|||||||||||]PAM</td><td>NA</td><td>Intergenic</td><td>NA</td><td>NA</td></tr>
<tr><td>GU305914.1</td><td>154398</td><td>154420</td><td>+</td><td>4</td><td>GCCCCGGGCGGTGGCGGCGG</td><td>CGG</td><td>|-|--|||[-|||||||||||]PAM</td><td>NA</td><td>Intergenic</td><td>NA</td><td>NA</td></tr>
<tr><td>GU305914.1</td><td>48159</td><td>48181</td><td>+</td><td>2</td><td>GGCGGCGGTGGCGGCGGCGG</td><td>TGG</td><td>|||||-||[|||-||||||||]PAM</td><td>NA</td><td>Intergenic</td><td>NA</td><td>NA</td></tr>
<tr><td>GU305914.1</td><td>58129</td><td>58151</td><td>-</td><td>4</td><td>GCGGGCGGTGGCGGCGGCGG</td><td>CGG</td><td>|--||-||[|||-||||||||]PAM</td><td>NA</td><td>Intergenic</td><td>NA</td><td>NA</td></tr>
<tr><td>GU305914.1</td><td>154404</td><td>154426</td><td>+</td><td>3</td><td>GGCGGTGGCGGCGGCGGCGG</td><td>CGG</td><td>|||||-||[-||-||||||||]PAM</td><td>NA</td><td>Intergenic</td><td>NA</td><td>NA</td></tr>
<tr><td>GU305914.1</td><td>58126</td><td>58148</td><td>-</td><td>3</td><td>GGCGGTGGCGGCGGCGGCGG</td><td>CGG</td><td>|||||-||[-||-||||||||]PAM</td><td>NA</td><td>Intergenic</td><td>NA</td><td>NA</td></tr>
<tr><td>GU305914.1</td><td>154407</td><td>154429</td><td>+</td><td>4</td><td>GGTGGCGGCGGCGGCGGCGG</td><td>CGG</td><td>||-||-||[-||-||||||||]PAM</td><td>NA</td><td>Intergenic</td><td>NA</td><td>NA</td></tr>
<tr><td>GU305914.1</td><td>58123</td><td>58145</td><td>-</td><td>4</td><td>GGTGGCGGCGGCGGCGGCGG</td><td>AGG</td><td>||-||-||[-||-||||||||]PAM</td><td>NA</td><td>Intergenic</td><td>NA</td><td>NA</td></tr>
<tr><td>GU305914.1</td><td>161754</td><td>161776</td><td>-</td><td>4</td><td>GGCCGAGGAGGCGGCGGCGG</td><td>AGG</td><td>|||-|-||[-||-||||||||]PAM</td><td>NA</td><td>Intergenic</td><td>NA</td><td>NA</td></tr>
<tr><td>GU305914.1</td><td>149482</td><td>149504</td><td>-</td><td>4</td><td>GGCGACGGCGGCGGCGGCGG</td><td>CGG</td><td>||||--||[-||-||||||||]PAM</td><td>NA</td><td>Intergenic</td><td>NA</td><td>NA</td></tr>
<tr><td>GU305914.1</td><td>114661</td><td>114683</td><td>-</td><td>3</td><td>GACGGCGGTGGTGGTGGCGG</td><td>CGG</td><td>|-|||-||[||||||-|||||]PAM</td><td>NA</td><td>Intergenic</td><td>NA</td><td>NA</td></tr>
<tr><td>GU305914.1</td><td>74930</td><td>74952</td><td>+</td><td>4</td><td>GGCGCAGGTTGAGGCGGCGG</td><td>TGG</td><td>||||--||[|-|-||||||||]PAM</td><td>NA</td><td>Intergenic</td><td>NA</td><td>NA</td></tr>
<tr><td>GU305914.1</td><td>168817</td><td>168839</td><td>+</td><td>3</td><td>GGCGGTGGTGGCAGCGGCGG</td><td>CGG</td><td>|||||-||[|||--|||||||]PAM</td><td>NA</td><td>Intergenic</td><td>NA</td><td>NA</td></tr>
<tr><td>GU305914.1</td><td>168814</td><td>168836</td><td>+</td><td>3</td><td>AGCGGCGGTGGTGGCAGCGG</td><td>CGG</td><td>-||||-||[|||||||-||||]PAM</td><td>NA</td><td>Intergenic</td><td>NA</td><td>NA</td></tr>
<tr><td>GU305914.1</td><td>62443</td><td>62465</td><td>+</td><td>3</td><td>GGCGGCGGGGGTGGTGGCGG</td><td>CGG</td><td>|||||-||[-|||||-|||||]PAM</td><td>NA</td><td>Intergenic</td><td>NA</td><td>NA</td></tr>
<tr><td>GU305914.1</td><td>18653</td><td>18675</td><td>-</td><td>4</td><td>GGCAGAGGTGGCGGAGGCGG</td><td>GGG</td><td>|||-|-||[|||-||-|||||]PAM</td><td>NA</td><td>Intergenic</td><td>NA</td><td>NA</td></tr>
<tr><td>GU305914.1</td><td>45255</td><td>45277</td><td>-</td><td>4</td><td>GGTGGCGGTGGCGGCAGCGG</td><td>CGG</td><td>||-||-||[|||-|||-||||]PAM</td><td>NA</td><td>Intergenic</td><td>NA</td><td>NA</td></tr>
<tr><td>GU305914.1</td><td>62434</td><td>62456</td><td>+</td><td>4</td><td>GACGGCGGTGGCGGCGGGGG</td><td>TGG</td><td>|-|||-||[|||-|||||-||]PAM</td><td>NA</td><td>Intergenic</td><td>NA</td><td>NA</td></tr>
<tr><td>GU305914.1</td><td>62458</td><td>62480</td><td>+</td><td>3</td><td>GGCGGCGGGGGTGGCGGCCG</td><td>CGG</td><td>|||||-||[-|||||||||-|]PAM</td><td>NA</td><td>Intergenic</td><td>NA</td><td>NA</td></tr>
<tr><td>GU305914.1</td><td>62449</td><td>62471</td><td>+</td><td>4</td><td>GGGGGTGGTGGCGGCGGGGG</td><td>TGG</td><td>||-||-||[|||-|||||-||]PAM</td><td>NA</td><td>Intergenic</td><td>NA</td><td>NA</td></tr>
<tr><td></td><td></td><td></td><td></td><td></td><td></td><td></td><td></td><td></td><td></td><td></td><td></td></tr>
<tr><td>T4239</td><td>GGGGGTGGTGGCGGCGGGGGTGG</td><td>1000</td><td></td><td></td><td></td><td></td><td></td><td></td><td></td><td></td><td></td></tr>
<tr><td>Oligo fwd</td><td>CACCGGGGGTGGTGGCGGCGGGGG</td><td></td><td></td><td></td><td></td><td></td><td></td><td></td><td></td><td></td><td></td></tr>
<tr><td>Oligo rev</td><td>AAACCCCCCGCCGCCACCACCCCC</td><td></td><td></td><td></td><td></td><td></td><td></td><td></td><td></td><td></td><td></td></tr>
<tr><td>Chromosome</td><td>start</td><td>end</td><td>strand</td><td>MM</td><td>target_seq</td><td>PAM</td><td>alignment</td><td>distance</td><td>position</td><td>gene name</td><td>gene id</td></tr>
<tr><td>GU305914.1</td><td>62449</td><td>62471</td><td>+</td><td>0</td><td>GGGGGTGGTGGCGGCGGGGG</td><td>TGG</td><td>||||||||[||||||||||||]PAM</td><td>NA</td><td>Intergenic</td><td>NA</td><td>NA</td></tr>
<tr><td>GU305914.1</td><td>62434</td><td>62456</td><td>+</td><td>3</td><td>GACGGCGGTGGCGGCGGGGG</td><td>TGG</td><td>|--||-||[||||||||||||]PAM</td><td>NA</td><td>Intergenic</td><td>NA</td><td>NA</td></tr>
<tr><td>GU305914.1</td><td>143022</td><td>143044</td><td>+</td><td>3</td><td>GCTGGTGGTGGCGCCGGGGG</td><td>CGG</td><td>|--|||||[|||||-||||||]PAM</td><td>NA</td><td>Intergenic</td><td>NA</td><td>NA</td></tr>
<tr><td>GU305914.1</td><td>58129</td><td>58151</td><td>-</td><td>3</td><td>GCGGGCGGTGGCGGCGGCGG</td><td>CGG</td><td>|-|||-||[|||||||||-||]PAM</td><td>NA</td><td>Intergenic</td><td>NA</td><td>NA</td></tr>
<tr><td>GU305914.1</td><td>48159</td><td>48181</td><td>+</td><td>3</td><td>GGCGGCGGTGGCGGCGGCGG</td><td>TGG</td><td>||-||-||[|||||||||-||]PAM</td><td>NA</td><td>Intergenic</td><td>NA</td><td>NA</td></tr>
<tr><td>GU305914.1</td><td>154401</td><td>154423</td><td>+</td><td>4</td><td>CCGGGCGGTGGCGGCGGCGG</td><td>CGG</td><td>--|||-||[|||||||||-||]PAM</td><td>NA</td><td>Intergenic</td><td>NA</td><td>NA</td></tr>
<tr><td>GU305914.1</td><td>154404</td><td>154426</td><td>+</td><td>3</td><td>GGCGGTGGCGGCGGCGGCGG</td><td>CGG</td><td>||-|||||[-||||||||-||]PAM</td><td>NA</td><td>Intergenic</td><td>NA</td><td>NA</td></tr>
<tr><td>GU305914.1</td><td>58126</td><td>58148</td><td>-</td><td>3</td><td>GGCGGTGGCGGCGGCGGCGG</td><td>CGG</td><td>||-|||||[-||||||||-||]PAM</td><td>NA</td><td>Intergenic</td><td>NA</td><td>NA</td></tr>
<tr><td>GU305914.1</td><td>30000</td><td>30022</td><td>+</td><td>4</td><td>GGTGCTGGCGGCGGCGGCGG</td><td>CGG</td><td>||-|-|||[-||||||||-||]PAM</td><td>NA</td><td>Intergenic</td><td>NA</td><td>NA</td></tr>
<tr><td>GU305914.1</td><td>154407</td><td>154429</td><td>+</td><td>4</td><td>GGTGGCGGCGGCGGCGGCGG</td><td>CGG</td><td>||-||-||[-||||||||-||]PAM</td><td>NA</td><td>Intergenic</td><td>NA</td><td>NA</td></tr>
<tr><td>GU305914.1</td><td>58123</td><td>58145</td><td>-</td><td>4</td><td>GGTGGCGGCGGCGGCGGCGG</td><td>AGG</td><td>||-||-||[-||||||||-||]PAM</td><td>NA</td><td>Intergenic</td><td>NA</td><td>NA</td></tr>
<tr><td>GU305914.1</td><td>67742</td><td>67764</td><td>+</td><td>4</td><td>GGGAGCGGCGGCGGCGGCGG</td><td>CGG</td><td>|||-|-||[-||||||||-||]PAM</td><td>NA</td><td>Intergenic</td><td>NA</td><td>NA</td></tr>
<tr><td>GU305914.1</td><td>114658</td><td>114680</td><td>-</td><td>3</td><td>GGCGGTGGTGGTGGCGGCGG</td><td>CGG</td><td>||-|||||[|||-|||||-||]PAM</td><td>NA</td><td>Intergenic</td><td>NA</td><td>NA</td></tr>
<tr><td>GU305914.1</td><td>168817</td><td>168839</td><td>+</td><td>3</td><td>GGCGGTGGTGGCAGCGGCGG</td><td>CGG</td><td>||-|||||[||||-||||-||]PAM</td><td>NA</td><td>Intergenic</td><td>NA</td><td>NA</td></tr>
<tr><td>GU305914.1</td><td>62446</td><td>62468</td><td>+</td><td>4</td><td>GGCGGGGGTGGTGGCGGCGG</td><td>GGG</td><td>||-||-||[|||-|||||-||]PAM</td><td>NA</td><td>Intergenic</td><td>NA</td><td>NA</td></tr>
<tr><td>GU305914.1</td><td>45255</td><td>45277</td><td>-</td><td>4</td><td>GGTGGCGGTGGCGGCAGCGG</td><td>CGG</td><td>||-||-||[|||||||-|-||]PAM</td><td>NA</td><td>Intergenic</td><td>NA</td><td>NA</td></tr>
<tr><td>GU305914.1</td><td>98053</td><td>98075</td><td>-</td><td>4</td><td>GGACGTGGTGGCGGCGATGG</td><td>CGG</td><td>||--||||[||||||||--||]PAM</td><td>NA</td><td>Intergenic</td><td>NA</td><td>NA</td></tr>
<tr><td></td><td></td><td></td><td></td><td></td><td></td><td></td><td></td><td></td><td></td><td></td><td></td></tr>
<tr><td>T4240</td><td>GGTGGTGGCGGCGGGGGTGGCGG</td><td>1000</td><td></td><td></td><td></td><td></td><td></td><td></td><td></td><td></td><td></td></tr>
<tr><td>Oligo fwd</td><td>CACCGGTGGTGGCGGCGGGGGTGG</td><td></td><td></td><td></td><td></td><td></td><td></td><td></td><td></td><td></td><td></td></tr>
<tr><td>Oligo rev</td><td>AAACCCACCCCCGCCGCCACCACC</td><td></td><td></td><td></td><td></td><td></td><td></td><td></td><td></td><td></td><td></td></tr>
</table>

<table>
<tr><td>Chromosome</td><td>start</td><td>end</td><td>strand</td><td>MM</td><td>target_seq</td><td>PAM</td><td>alignment</td><td>distance</td><td>position</td><td>gene name</td><td>gene id</td></tr>
<tr><td>GU305914.1</td><td>62452</td><td>62474</td><td>+</td><td>0</td><td>GGTGGTGGCGGCGGGGGTGG</td><td>CGG</td><td>||||||||[||||||||||||]PAM</td><td>NA</td><td>Intergenic</td><td>NA</td><td>NA</td></tr>
<tr><td>GU305914.1</td><td>62437</td><td>62459</td><td>+</td><td>1</td><td>GGCGGTGGCGGCGGGGGTGG</td><td>TGG</td><td>||-|||||[||||||||||||]PAM</td><td>NA</td><td>Intergenic</td><td>NA</td><td>NA</td></tr>
<tr><td>GU305914.1</td><td>62440</td><td>62462</td><td>+</td><td>3</td><td>GGTGGCGGCGGGGGTGGTGG</td><td>CGG</td><td>|||||-||[|||-||-|||||]PAM</td><td>NA</td><td>Intergenic</td><td>NA</td><td>NA</td></tr>
<tr><td>GU305914.1</td><td>154404</td><td>154426</td><td>+</td><td>3</td><td>GGCGGTGGCGGCGGCGGCGG</td><td>CGG</td><td>||-|||||[||||||-||-||]PAM</td><td>NA</td><td>Intergenic</td><td>NA</td><td>NA</td></tr>
<tr><td>GU305914.1</td><td>58126</td><td>58148</td><td>-</td><td>3</td><td>GGCGGTGGCGGCGGCGGCGG</td><td>CGG</td><td>||-|||||[||||||-||-||]PAM</td><td>NA</td><td>Intergenic</td><td>NA</td><td>NA</td></tr>
<tr><td>GU305914.1</td><td>30000</td><td>30022</td><td>+</td><td>3</td><td>GGTGCTGGCGGCGGCGGCGG</td><td>CGG</td><td>||||-|||[||||||-||-||]PAM</td><td>NA</td><td>Intergenic</td><td>NA</td><td>NA</td></tr>
<tr><td>GU305914.1</td><td>154407</td><td>154429</td><td>+</td><td>3</td><td>GGTGGCGGCGGCGGCGGCGG</td><td>CGG</td><td>|||||-||[||||||-||-||]PAM</td><td>NA</td><td>Intergenic</td><td>NA</td><td>NA</td></tr>
<tr><td>GU305914.1</td><td>58123</td><td>58145</td><td>-</td><td>3</td><td>GGTGGCGGCGGCGGCGGCGG</td><td>AGG</td><td>|||||-||[||||||-||-||]PAM</td><td>NA</td><td>Intergenic</td><td>NA</td><td>NA</td></tr>
<tr><td>GU305914.1</td><td>48153</td><td>48175</td><td>+</td><td>4</td><td>TGTGGCGGCGGCGGTGGCGG</td><td>CGG</td><td>-||||-||[||||||-||-||]PAM</td><td>NA</td><td>Intergenic</td><td>NA</td><td>NA</td></tr>
<tr><td>GU305914.1</td><td>161748</td><td>161770</td><td>-</td><td>4</td><td>GGAGGCGGCGGCGGAGGTGA</td><td>CGG</td><td>||-||-||[||||||-||||-]PAM</td><td>NA</td><td>Intergenic</td><td>NA</td><td>NA</td></tr>
<tr><td>GU305914.1</td><td>81240</td><td>81262</td><td>-</td><td>4</td><td>GGAGGCGGCGGCGGGGGGTG</td><td>AGG</td><td>||-||-||[|||||||||--|]PAM</td><td>NA</td><td>Intergenic</td><td>NA</td><td>NA</td></tr>
<tr><td>T4241</td><td>GGCGGCGGGGGTGGCGGCCGCGG</td><td>1000</td><td></td><td></td><td></td><td></td><td></td><td></td><td></td><td></td><td></td></tr>
<tr><td>Oligo fwd</td><td>CACCGGCGGCGGGGGTGGCGGCCG</td><td></td><td></td><td></td><td></td><td></td><td></td><td></td><td></td><td></td><td></td></tr>
<tr><td>Oligo rev</td><td>AAACCGGCCGCCACCCCCGCCGCC</td><td></td><td></td><td></td><td></td><td></td><td></td><td></td><td></td><td></td><td></td></tr>
<tr><td>Chromosome</td><td>start</td><td>end</td><td>strand</td><td>MM</td><td>target_seq</td><td>PAM</td><td>alignment</td><td>distance</td><td>position</td><td>gene name</td><td>gene id</td></tr>
<tr><td>GU305914.1</td><td>62458</td><td>62480</td><td>+</td><td>0</td><td>GGCGGCGGGGGTGGCGGCCG</td><td>CGG</td><td>||||||||[||||||||||||]PAM</td><td>NA</td><td>Intergenic</td><td>NA</td><td>NA</td></tr>
<tr><td>GU305914.1</td><td>48156</td><td>48178</td><td>+</td><td>2</td><td>GGCGGCGGCGGTGGCGGCGG</td><td>CGG</td><td>||||||||[-|||||||||-|]PAM</td><td>NA</td><td>Intergenic</td><td>NA</td><td>NA</td></tr>
<tr><td>GU305914.1</td><td>100225</td><td>100247</td><td>-</td><td>4</td><td>CGCCGCGGGGGCGGCGGACG</td><td>AGG</td><td>-||-||||[|||-|||||-||]PAM</td><td>NA</td><td>Intergenic</td><td>NA</td><td>NA</td></tr>
<tr><td>GU305914.1</td><td>62446</td><td>62468</td><td>+</td><td>3</td><td>GGCGGGGGTGGTGGCGGCGG</td><td>GGG</td><td>|||||-||[-|||||||||-|]PAM</td><td>NA</td><td>Intergenic</td><td>NA</td><td>NA</td></tr>
<tr><td>GU305914.1</td><td>114658</td><td>114680</td><td>-</td><td>3</td><td>GGCGGTGGTGGTGGCGGCGG</td><td>CGG</td><td>|||||-||[-|||||||||-|]PAM</td><td>NA</td><td>Intergenic</td><td>NA</td><td>NA</td></tr>
<tr><td>GU305914.1</td><td>62443</td><td>62465</td><td>+</td><td>2</td><td>GGCGGCGGGGGTGGTGGCGG</td><td>CGG</td><td>||||||||[||||||-|||-|]PAM</td><td>NA</td><td>Intergenic</td><td>NA</td><td>NA</td></tr>
<tr><td>T4242</td><td>GGCCGCGGATCCAGCACCAGAGG</td><td>1000</td><td></td><td></td><td></td><td></td><td></td><td></td><td></td><td></td><td></td></tr>
<tr><td>Oligo fwd</td><td>CACCGGCCGCGGATCCAGCACCAG</td><td></td><td></td><td></td><td></td><td></td><td></td><td></td><td></td><td></td><td></td></tr>
<tr><td>Oligo rev</td><td>AAACCTGGTGCTGGATCCGCGGCC</td><td></td><td></td><td></td><td></td><td></td><td></td><td></td><td></td><td></td><td></td></tr>
<tr><td>Chromosome</td><td>start</td><td>end</td><td>strand</td><td>MM</td><td>target_seq</td><td>PAM</td><td>alignment</td><td>distance</td><td>position</td><td>gene name</td><td>gene id</td></tr>
<tr><td>GU305914.1</td><td>62473</td><td>62495</td><td>+</td><td>0</td><td>GGCCGCGGATCCAGCACCAG</td><td>AGG</td><td>||||||||[||||||||||||]PAM</td><td>NA</td><td>Intergenic</td><td>NA</td><td>NA</td></tr>
<tr><td>T4243</td><td>CGCGGATCCAGCACCAGAGGTGG</td><td>1000</td><td></td><td></td><td></td><td></td><td></td><td></td><td></td><td></td><td></td></tr>
<tr><td>Oligo fwd</td><td>CACCCGCGGATCCAGCACCAGAGG</td><td></td><td></td><td></td><td></td><td></td><td></td><td></td><td></td><td></td><td></td></tr>
<tr><td>Oligo rev</td><td>AAACCCTCTGGTGCTGGATCCGCG</td><td></td><td></td><td></td><td></td><td></td><td></td><td></td><td></td><td></td><td></td></tr>
<tr><td>Chromosome</td><td>start</td><td>end</td><td>strand</td><td>MM</td><td>target_seq</td><td>PAM</td><td>alignment</td><td>distance</td><td>position</td><td>gene name</td><td>gene id</td></tr>
<tr><td>GU305914.1</td><td>62476</td><td>62498</td><td>+</td><td>0</td><td>CGCGGATCCAGCACCAGAGG</td><td>TGG</td><td>||||||||[||||||||||||]PAM</td><td>NA</td><td>Intergenic</td><td>NA</td><td>NA</td></tr>
<tr><td>GU305914.1</td><td>57651</td><td>57673</td><td>+</td><td>3</td><td>CGGGGATCCGGCACCTGAGG</td><td>GGG</td><td>||-|||||[|-|||||-||||]PAM</td><td>NA</td><td>Intergenic</td><td>NA</td><td>NA</td></tr>
<tr><td>T4244</td><td>AGCGTGATTGTTGTCACTCTTGG</td><td>1000</td><td></td><td></td><td></td><td></td><td></td><td></td><td></td><td></td><td></td></tr>
<tr><td>Oligo fwd</td><td>CACCAGCGTGATTGTTGTCACTCT</td><td></td><td></td><td></td><td></td><td></td><td></td><td></td><td></td><td></td><td></td></tr>
<tr><td>Oligo rev</td><td>AAACAGAGTGACAACAATCACGCT</td><td></td><td></td><td></td><td></td><td></td><td></td><td></td><td></td><td></td><td></td></tr>
<tr><td>Chromosome</td><td>start</td><td>end</td><td>strand</td><td>MM</td><td>target_seq</td><td>PAM</td><td>alignment</td><td>distance</td><td>position</td><td>gene name</td><td>gene id</td></tr>
<tr><td>GU305914.1</td><td>62503</td><td>62525</td><td>+</td><td>0</td><td>AGCGTGATTGTTGTCACTCT</td><td>TGG</td><td>||||||||[||||||||||||]PAM</td><td>NA</td><td>Intergenic</td><td>NA</td><td>NA</td></tr>
<tr><td>T4245</td><td>ATTGTTGTCACTCTTGGCGTTGG</td><td>1000</td><td></td><td></td><td></td><td></td><td></td><td></td><td></td><td></td><td></td></tr>
<tr><td>Oligo fwd</td><td>CACCATTGTTGTCACTCTTGGCGT</td><td></td><td></td><td></td><td></td><td></td><td></td><td></td><td></td><td></td><td></td></tr>
<tr><td>Oligo rev</td><td>AAACACGCCAAGAGTGACAACAAT</td><td></td><td></td><td></td><td></td><td></td><td></td><td></td><td></td><td></td><td></td></tr>
<tr><td>Chromosome</td><td>start</td><td>end</td><td>strand</td><td>MM</td><td>target_seq</td><td>PAM</td><td>alignment</td><td>distance</td><td>position</td><td>gene name</td><td>gene id</td></tr>
<tr><td>GU305914.1</td><td>62509</td><td>62531</td><td>+</td><td>0</td><td>ATTGTTGTCACTCTTGGCGT</td><td>TGG</td><td>||||||||[||||||||||||]PAM</td><td>NA</td><td>Intergenic</td><td>NA</td><td>NA</td></tr>
<tr><td>T4246</td><td>ACTCTTGGCGTTGGAAGCACCGG</td><td>1000</td><td></td><td></td><td></td><td></td><td></td><td></td><td></td><td></td><td></td></tr>
<tr><td>Oligo fwd</td><td>CACCACTCTTGGCGTTGGAAGCAC</td><td></td><td></td><td></td><td></td><td></td><td></td><td></td><td></td><td></td><td></td></tr>
<tr><td>Oligo rev</td><td>AAACGTGCTTCCAACGCCAAGAGT</td><td></td><td></td><td></td><td></td><td></td><td></td><td></td><td></td><td></td><td></td></tr>
<tr><td>Chromosome</td><td>start</td><td>end</td><td>strand</td><td>MM</td><td>target_seq</td><td>PAM</td><td>alignment</td><td>distance</td><td>position</td><td>gene name</td><td>gene id</td></tr>
<tr><td>GU305914.1</td><td>62518</td><td>62540</td><td>+</td><td>0</td><td>ACTCTTGGCGTTGGAAGCAC</td><td>CGG</td><td>||||||||[||||||||||||]PAM</td><td>NA</td><td>Intergenic</td><td>NA</td><td>NA</td></tr>
<tr><td>T4247</td><td>GGCGTTGGAAGCACCGGCGACGG</td><td>1000</td><td></td><td></td><td></td><td></td><td></td><td></td><td></td><td></td><td></td></tr>
<tr><td>Oligo fwd</td><td>CACCGGCGTTGGAAGCACCGGCGA</td><td></td><td></td><td></td><td></td><td></td><td></td><td></td><td></td><td></td><td></td></tr>
<tr><td>Oligo rev</td><td>AAACTCGCCGGTGCTTCCAACGCC</td><td></td><td></td><td></td><td></td><td></td><td></td><td></td><td></td><td></td><td></td></tr>
<tr><td>Chromosome</td><td>start</td><td>end</td><td>strand</td><td>MM</td><td>target_seq</td><td>PAM</td><td>alignment</td><td>distance</td><td>position</td><td>gene name</td><td>gene id</td></tr>
<tr><td>GU305914.1</td><td>62524</td><td>62546</td><td>+</td><td>0</td><td>GGCGTTGGAAGCACCGGCGA</td><td>CGG</td><td>||||||||[||||||||||||]PAM</td><td>NA</td><td>Intergenic</td><td>NA</td><td>NA</td></tr>
<tr><td>T4248</td><td>TTGGAAGCACCGGCGACGGCAGG</td><td>1000</td><td></td><td></td><td></td><td></td><td></td><td></td><td></td><td></td><td></td></tr>
<tr><td>Oligo fwd</td><td>CACCTTGGAAGCACCGGCGACGGC</td><td></td><td></td><td></td><td></td><td></td><td></td><td></td><td></td><td></td><td></td></tr>
<tr><td>Oligo rev</td><td>AAACGCCGTCGCCGGTGCTTCCAA</td><td></td><td></td><td></td><td></td><td></td><td></td><td></td><td></td><td></td><td></td></tr>
<tr><td>Chromosome</td><td>start</td><td>end</td><td>strand</td><td>MM</td><td>target_seq</td><td>PAM</td><td>alignment</td><td>distance</td><td>position</td><td>gene name</td><td>gene id</td></tr>
<tr><td>GU305914.1</td><td>62528</td><td>62550</td><td>+</td><td>0</td><td>TTGGAAGCACCGGCGACGGC</td><td>AGG</td><td>||||||||[||||||||||||]PAM</td><td>NA</td><td>Intergenic</td><td>NA</td><td>NA</td></tr>
<tr><td>T4249</td><td>AGCACCGGCGACGGCAGGAGCGG</td><td>1000</td><td></td><td></td><td></td><td></td><td></td><td></td><td></td><td></td><td></td></tr>
<tr><td>Oligo fwd</td><td>CACCAGCACCGGCGACGGCAGGAG</td><td></td><td></td><td></td><td></td><td></td><td></td><td></td><td></td><td></td><td></td></tr>
<tr><td>Oligo rev</td><td>AAACCTCCTGCCGTCGCCGGTGCT</td><td></td><td></td><td></td><td></td><td></td><td></td><td></td><td></td><td></td><td></td></tr>
<tr><td>Chromosome</td><td>start</td><td>end</td><td>strand</td><td>MM</td><td>target_seq</td><td>PAM</td><td>alignment</td><td>distance</td><td>position</td><td>gene name</td><td>gene id</td></tr>
<tr><td>GU305914.1</td><td>62533</td><td>62555</td><td>+</td><td>0</td><td>AGCACCGGCGACGGCAGGAG</td><td>CGG</td><td>||||||||[||||||||||||]PAM</td><td>NA</td><td>Intergenic</td><td>NA</td><td>NA</td></tr>
<tr><td>T4250</td><td>GGCGACGGCAGGAGCGGCCGCGG</td><td>1000</td><td></td><td></td><td></td><td></td><td></td><td></td><td></td><td></td><td></td></tr>
<tr><td>Oligo fwd</td><td>CACCGGCGACGGCAGGAGCGGCCG</td><td></td><td></td><td></td><td></td><td></td><td></td><td></td><td></td><td></td><td></td></tr>
<tr><td>Oligo rev</td><td>AAACCGGCCGCTCCTGCCGTCGCC</td><td></td><td></td><td></td><td></td><td></td><td></td><td></td><td></td><td></td><td></td></tr>
<tr><td>Chromosome</td><td>start</td><td>end</td><td>strand</td><td>MM</td><td>target_seq</td><td>PAM</td><td>alignment</td><td>distance</td><td>position</td><td>gene name</td><td>gene id</td></tr>
<tr><td>GU305914.1</td><td>62539</td><td>62561</td><td>+</td><td>0</td><td>GGCGACGGCAGGAGCGGCCG</td><td>CGG</td><td>||||||||[||||||||||||]PAM</td><td>NA</td><td>Intergenic</td><td>NA</td><td>NA</td></tr>
<tr><td>T4251</td><td>GACGGCAGGAGCGGCCGCGGCGG</td><td>1000</td><td></td><td></td><td></td><td></td><td></td><td></td><td></td><td></td><td></td></tr>
<tr><td>Oligo fwd</td><td>CACCGACGGCAGGAGCGGCCGCGG</td><td></td><td></td><td></td><td></td><td></td><td></td><td></td><td></td><td></td><td></td></tr>
<tr><td>Oligo rev</td><td>AAACCCGCGGCCGCTCCTGCCGTC</td><td></td><td></td><td></td><td></td><td></td><td></td><td></td><td></td><td></td><td></td></tr>
<tr><td>Chromosome</td><td>start</td><td>end</td><td>strand</td><td>MM</td><td>target_seq</td><td>PAM</td><td>alignment</td><td>distance</td><td>position</td><td>gene name</td><td>gene id</td></tr>
<tr><td>GU305914.1</td><td>62542</td><td>62564</td><td>+</td><td>0</td><td>GACGGCAGGAGCGGCCGCGG</td><td>CGG</td><td>||||||||[||||||||||||]PAM</td><td>NA</td><td>Intergenic</td><td>NA</td><td>NA</td></tr>
<tr><td>GU305914.1</td><td>73792</td><td>73814</td><td>-</td><td>4</td><td>GGCGCCAGCAGCGGCGGCGG</td><td>AGG</td><td>|-||-|||[-||||||-||||]PAM</td><td>NA</td><td>Intergenic</td><td>NA</td><td>NA</td></tr>
<tr><td>GU305914.1</td><td>163911</td><td>163933</td><td>+</td><td>4</td><td>GGCGGCGGCAGCGGCAGCGG</td><td>AGG</td><td>|-||||-|[-||||||-||||]PAM</td><td>NA</td><td>Intergenic</td><td>NA</td><td>NA</td></tr>
<tr><td>T4252</td><td>AGCGGCCGCGGCGGTAGCGTTGG</td><td>1000</td><td></td><td></td><td></td><td></td><td></td><td></td><td></td><td></td><td></td></tr>
<tr><td>Oligo fwd</td><td>CACCAGCGGCCGCGGCGGTAGCGT</td><td></td><td></td><td></td><td></td><td></td><td></td><td></td><td></td><td></td><td></td></tr>
</table>

<table>
<tr><td>Oligo rev</td><td>AAACACGCTACCGCCGCGGCCGCT</td></tr>
<tr><td>Chromosome</td><td>start</td><td>end</td><td>strand</td><td>MM</td><td>target_seq</td><td>PAM</td><td>alignment</td><td>distance</td><td>position</td><td>gene name</td><td>gene id</td></tr>
<tr><td>GU305914.1</td><td>62551</td><td>62573</td><td>+</td><td>0</td><td>AGCGGCCGCGGCGGTAGCGT</td><td>TGG</td><td>||||||||[||||||||||||]PAM</td><td>NA</td><td>Intergenic</td><td>NA</td><td>NA</td></tr>
<tr><td>GU305914.1</td><td>19589</td><td>19611</td><td>+</td><td>4</td><td>GTCGGTGGCGGCGGTAGCGT</td><td>CGG</td><td>--|||--|[||||||||||||]PAM</td><td>NA</td><td>Intergenic</td><td>NA</td><td>NA</td></tr>
<tr><td>T4253</td><td>GCGGCGGTAGCGTTGGTCTTCGG</td><td>1000</td></tr>
<tr><td>Oligo fwd</td><td>CACCGCGGCGGTAGCGTTGGTCTT</td></tr>
<tr><td>Oligo rev</td><td>AAACAAGACCAACGCTACCGCCGC</td></tr>
<tr><td>Chromosome</td><td>start</td><td>end</td><td>strand</td><td>MM</td><td>target_seq</td><td>PAM</td><td>alignment</td><td>distance</td><td>position</td><td>gene name</td><td>gene id</td></tr>
<tr><td>GU305914.1</td><td>62558</td><td>62580</td><td>+</td><td>0</td><td>GCGGCGGTAGCGTTGGTCTT</td><td>CGG</td><td>||||||||[||||||||||||]PAM</td><td>NA</td><td>Intergenic</td><td>NA</td><td>NA</td></tr>
<tr><td>T4254</td><td>TAGCGTTGGTCTTCGGCATTCGG</td><td>1000</td></tr>
<tr><td>Oligo fwd</td><td>CACCTAGCGTTGGTCTTCGGCATT</td></tr>
<tr><td>Oligo rev</td><td>AAACAATGCCGAAGACCAACGCTA</td></tr>
<tr><td>Chromosome</td><td>start</td><td>end</td><td>strand</td><td>MM</td><td>target_seq</td><td>PAM</td><td>alignment</td><td>distance</td><td>position</td><td>gene name</td><td>gene id</td></tr>
<tr><td>GU305914.1</td><td>62565</td><td>62587</td><td>+</td><td>0</td><td>TAGCGTTGGTCTTCGGCATT</td><td>CGG</td><td>||||||||[||||||||||||]PAM</td><td>NA</td><td>Intergenic</td><td>NA</td><td>NA</td></tr>
<tr><td>T4255</td><td>AGCGTTGGTCTTCGGCATTCGGG</td><td>1000</td></tr>
<tr><td>Oligo fwd</td><td>CACCAGCGTTGGTCTTCGGCATTC</td></tr>
<tr><td>Oligo rev</td><td>AAACGAATGCCGAAGACCAACGCT</td></tr>
<tr><td>Chromosome</td><td>start</td><td>end</td><td>strand</td><td>MM</td><td>target_seq</td><td>PAM</td><td>alignment</td><td>distance</td><td>position</td><td>gene name</td><td>gene id</td></tr>
<tr><td>GU305914.1</td><td>62566</td><td>62588</td><td>+</td><td>0</td><td>AGCGTTGGTCTTCGGCATTC</td><td>GGG</td><td>||||||||[||||||||||||]PAM</td><td>NA</td><td>Intergenic</td><td>NA</td><td>NA</td></tr>
<tr><td>T4256</td><td>GTTGGTCTTCGGCATTCGGGTGG</td><td>1000</td></tr>
<tr><td>Oligo fwd</td><td>CACCGTTGGTCTTCGGCATTCGGG</td></tr>
<tr><td>Oligo rev</td><td>AAACCCCGAATGCCGAAGACCAAC</td></tr>
<tr><td>Chromosome</td><td>start</td><td>end</td><td>strand</td><td>MM</td><td>target_seq</td><td>PAM</td><td>alignment</td><td>distance</td><td>position</td><td>gene name</td><td>gene id</td></tr>
<tr><td>GU305914.1</td><td>62569</td><td>62591</td><td>+</td><td>0</td><td>GTTGGTCTTCGGCATTCGGG</td><td>TGG</td><td>||||||||[||||||||||||]PAM</td><td>NA</td><td>Intergenic</td><td>NA</td><td>NA</td></tr>
<tr><td>T4257</td><td>TTGGTCTTCGGCATTCGGGTGGG</td><td>1000</td></tr>
<tr><td>Oligo fwd</td><td>CACCTTGGTCTTCGGCATTCGGGT</td></tr>
<tr><td>Oligo rev</td><td>AAACACCCGAATGCCGAAGACCAA</td></tr>
<tr><td>Chromosome</td><td>start</td><td>end</td><td>strand</td><td>MM</td><td>target_seq</td><td>PAM</td><td>alignment</td><td>distance</td><td>position</td><td>gene name</td><td>gene id</td></tr>
<tr><td>GU305914.1</td><td>62570</td><td>62592</td><td>+</td><td>0</td><td>TTGGTCTTCGGCATTCGGGT</td><td>GGG</td><td>||||||||[||||||||||||]PAM</td><td>NA</td><td>Intergenic</td><td>NA</td><td>NA</td></tr>
<tr><td>T4258</td><td>GTCTTCGGCATTCGGGTGGGAGG</td><td>1000</td></tr>
<tr><td>Oligo fwd</td><td>CACCGTCTTCGGCATTCGGGTGGG</td></tr>
<tr><td>Oligo rev</td><td>AAACCCCACCCGAATGCCGAAGAC</td></tr>
<tr><td>Chromosome</td><td>start</td><td>end</td><td>strand</td><td>MM</td><td>target_seq</td><td>PAM</td><td>alignment</td><td>distance</td><td>position</td><td>gene name</td><td>gene id</td></tr>
<tr><td>GU305914.1</td><td>62573</td><td>62595</td><td>+</td><td>0</td><td>GTCTTCGGCATTCGGGTGGG</td><td>AGG</td><td>||||||||[||||||||||||]PAM</td><td>NA</td><td>Intergenic</td><td>NA</td><td>NA</td></tr>
<tr><td>T4259</td><td>TCTTCGGCATTCGGGTGGGAGGG</td><td>1000</td></tr>
<tr><td>Oligo fwd</td><td>CACCTCTTCGGCATTCGGGTGGGA</td></tr>
<tr><td>Oligo rev</td><td>AAACTCCCACCCGAATGCCGAAGA</td></tr>
<tr><td>Chromosome</td><td>start</td><td>end</td><td>strand</td><td>MM</td><td>target_seq</td><td>PAM</td><td>alignment</td><td>distance</td><td>position</td><td>gene name</td><td>gene id</td></tr>
<tr><td>GU305914.1</td><td>62574</td><td>62596</td><td>+</td><td>0</td><td>TCTTCGGCATTCGGGTGGGA</td><td>GGG</td><td>||||||||[||||||||||||]PAM</td><td>NA</td><td>Intergenic</td><td>NA</td><td>NA</td></tr>
<tr><td>T4260</td><td>TCGGCATTCGGGTGGGAGGGAGG</td><td>1000</td></tr>
<tr><td>Oligo fwd</td><td>CACCTCGGCATTCGGGTGGGAGGG</td></tr>
<tr><td>Oligo rev</td><td>AAACCCCTCCCACCCGAATGCCGA</td></tr>
<tr><td>Chromosome</td><td>start</td><td>end</td><td>strand</td><td>MM</td><td>target_seq</td><td>PAM</td><td>alignment</td><td>distance</td><td>position</td><td>gene name</td><td>gene id</td></tr>
<tr><td>GU305914.1</td><td>62577</td><td>62599</td><td>+</td><td>0</td><td>TCGGCATTCGGGTGGGAGGG</td><td>AGG</td><td>||||||||[||||||||||||]PAM</td><td>NA</td><td>Intergenic</td><td>NA</td><td>NA</td></tr>
<tr><td>T4261</td><td>CGGCATTCGGGTGGGAGGGAGGG</td><td>1000</td></tr>
<tr><td>Oligo fwd</td><td>CACCCGGCATTCGGGTGGGAGGGA</td></tr>
<tr><td>Oligo rev</td><td>AAACTCCCTCCCACCCGAATGCCG</td></tr>
<tr><td>Chromosome</td><td>start</td><td>end</td><td>strand</td><td>MM</td><td>target_seq</td><td>PAM</td><td>alignment</td><td>distance</td><td>position</td><td>gene name</td><td>gene id</td></tr>
<tr><td>GU305914.1</td><td>62578</td><td>62600</td><td>+</td><td>0</td><td>CGGCATTCGGGTGGGAGGGA</td><td>GGG</td><td>||||||||[||||||||||||]PAM</td><td>NA</td><td>Intergenic</td><td>NA</td><td>NA</td></tr>
<tr><td>T4262</td><td>AAGTGACAGCAGCAGCCAGTAGG</td><td>1000</td></tr>
<tr><td>Oligo fwd</td><td>CACCAAGTGACAGCAGCAGCCAGT</td></tr>
<tr><td>Oligo rev</td><td>AAACACTGGCTGCTGCTGTCACTT</td></tr>
<tr><td>Chromosome</td><td>start</td><td>end</td><td>strand</td><td>MM</td><td>target_seq</td><td>PAM</td><td>alignment</td><td>distance</td><td>position</td><td>gene name</td><td>gene id</td></tr>
<tr><td>GU305914.1</td><td>62643</td><td>62665</td><td>+</td><td>0</td><td>AAGTGACAGCAGCAGCCAGT</td><td>AGG</td><td>||||||||[||||||||||||]PAM</td><td>NA</td><td>Intergenic</td><td>NA</td><td>NA</td></tr>
<tr><td>T4263</td><td>GTTAGCGCCGCCGCCCCCGCCGG</td><td>1000</td></tr>
<tr><td>Oligo fwd</td><td>CACCGTTAGCGCCGCCGCCCCCGC</td></tr>
<tr><td>Oligo rev</td><td>AAACGCGGGGGCGGCGGCGCTAAC</td></tr>
<tr><td>Chromosome</td><td>start</td><td>end</td><td>strand</td><td>MM</td><td>target_seq</td><td>PAM</td><td>alignment</td><td>distance</td><td>position</td><td>gene name</td><td>gene id</td></tr>
<tr><td>GU305914.1</td><td>62665</td><td>62687</td><td>+</td><td>0</td><td>GTTAGCGCCGCCGCCCCCGC</td><td>CGG</td><td>||||||||[||||||||||||]PAM</td><td>NA</td><td>Intergenic</td><td>NA</td><td>NA</td></tr>
<tr><td>T4264</td><td>AGCGCCGCCGCCCCCGCCGGCGG</td><td>1000</td></tr>
<tr><td>Oligo fwd</td><td>CACCAGCGCCGCCGCCCCCGCCGG</td></tr>
<tr><td>Oligo rev</td><td>AAACCCGGCGGGGGCGGCGGCGCT</td></tr>
<tr><td>Chromosome</td><td>start</td><td>end</td><td>strand</td><td>MM</td><td>target_seq</td><td>PAM</td><td>alignment</td><td>distance</td><td>position</td><td>gene name</td><td>gene id</td></tr>
<tr><td>GU305914.1</td><td>62668</td><td>62690</td><td>+</td><td>0</td><td>AGCGCCGCCGCCCCCGCCGG</td><td>CGG</td><td>||||||||[||||||||||||]PAM</td><td>NA</td><td>Intergenic</td><td>NA</td><td>NA</td></tr>
<tr><td>GU305914.1</td><td>154399</td><td>154421</td><td>-</td><td>4</td><td>GCCGCCGCCGCCACCGCCCG</td><td>GGG</td><td>--||||||[||||-|||||-|]PAM</td><td>NA</td><td>Intergenic</td><td>NA</td><td>NA</td></tr>
<tr><td>GU305914.1</td><td>225708</td><td>225730</td><td>-</td><td>3</td><td>AGCGCCAGCGCCCCCGCCGC</td><td>GGG</td><td>||||||--[|||||||||||-]PAM</td><td>NA</td><td>Intergenic</td><td>NA</td><td>NA</td></tr>
<tr><td>T4265</td><td>CCCGCCGGCGGCAGCAGCCGCGG</td><td>1000</td></tr>
<tr><td>Oligo fwd</td><td>CACCCCCGCCGGCGGCAGCAGCCG</td></tr>
<tr><td>Oligo rev</td><td>AAACCGGCTGCTGCCGCCGGCGGG</td></tr>
<tr><td>Chromosome</td><td>start</td><td>end</td><td>strand</td><td>MM</td><td>target_seq</td><td>PAM</td><td>alignment</td><td>distance</td><td>position</td><td>gene name</td><td>gene id</td></tr>
<tr><td>GU305914.1</td><td>62680</td><td>62702</td><td>+</td><td>0</td><td>CCCGCCGGCGGCAGCAGCCG</td><td>CGG</td><td>||||||||[||||||||||||]PAM</td><td>NA</td><td>Intergenic</td><td>NA</td><td>NA</td></tr>
<tr><td>GU305914.1</td><td>164959</td><td>164981</td><td>-</td><td>4</td><td>GGCGGCGGCGGCAGCAGCTG</td><td>CGG</td><td>--||-|||[||||||||||-|]PAM</td><td>NA</td><td>Intergenic</td><td>NA</td><td>NA</td></tr>
<tr><td>T4266</td><td>AGCAGCATTGACAGCAGCATTGG</td><td>1000</td></tr>
<tr><td>Oligo fwd</td><td>CACCAGCAGCATTGACAGCAGCAT</td></tr>
<tr><td>Oligo rev</td><td>AAACATGCTGCTGTCAATGCTGCT</td></tr>
</table>

| | | | | | | | | | | | |
|---|---|---|---|---|---|---|---|---|---|---|---|
| Chromosome | start | end | strand | MM | target_seq | PAM | alignment | distance | position | gene name | gene id |
| GU305914.1 | 62704 | 62726 | + | 0 | AGCAGCATTGACAGCAGCAT | TGG | \|\|\|\|\|\|\|\|[\|\|\|\|\|\|\|\|\|\|\|\|]PAM | NA | Intergenic | NA | NA |
| T4267 | AGCATTGACAGCAGCATTGGCGG | 1000 | | | | | | | | | |
| Oligo fwd | CACCAGCATTGACAGCAGCATTGG | | | | | | | | | | |
| Oligo rev | AAACCCAATGCTGCTGTCAATGCT | | | | | | | | | | |
| Chromosome | start | end | strand | MM | target_seq | PAM | alignment | distance | position | gene name | gene id |
| GU305914.1 | 62707 | 62729 | + | 0 | AGCATTGACAGCAGCATTGG | CGG | \|\|\|\|\|\|\|\|[\|\|\|\|\|\|\|\|\|\|\|\|]PAM | NA | Intergenic | NA | NA |
| T4268 | ATTGACAGCAGCATTGGCGGCGG | 1000 | | | | | | | | | |
| Oligo fwd | CACCATTGACAGCAGCATTGGCGG | | | | | | | | | | |
| Oligo rev | AAACCCGCCAATGCTGCTGTCAAT | | | | | | | | | | |
| Chromosome | start | end | strand | MM | target_seq | PAM | alignment | distance | position | gene name | gene id |
| GU305914.1 | 62710 | 62732 | + | 0 | ATTGACAGCAGCATTGGCGG | CGG | \|\|\|\|\|\|\|\|[\|\|\|\|\|\|\|\|\|\|\|\|]PAM | NA | Intergenic | NA | NA |
| T4269 | AGCAGCATTGGCGGCGGAGACGG | 1000 | | | | | | | | | |
| Oligo fwd | CACCAGCAGCATTGGCGGCGGAGA | | | | | | | | | | |
| Oligo rev | AAACTCTCCGCCGCCAATGCTGCT | | | | | | | | | | |
| Chromosome | start | end | strand | MM | target_seq | PAM | alignment | distance | position | gene name | gene id |
| GU305914.1 | 62716 | 62738 | + | 0 | AGCAGCATTGGCGGCGGAGA | CGG | \|\|\|\|\|\|\|\|[\|\|\|\|\|\|\|\|\|\|\|\|]PAM | NA | Intergenic | NA | NA |
| T4270 | ATTGGCGGCGGAGACGGCAGCGG | 1000 | | | | | | | | | |
| Oligo fwd | CACCATTGGCGGCGGAGACGGCAG | | | | | | | | | | |
| Oligo rev | AAACCTGCCGTCTCCGCCGCCAAT | | | | | | | | | | |
| Chromosome | start | end | strand | MM | target_seq | PAM | alignment | distance | position | gene name | gene id |
| GU305914.1 | 62722 | 62744 | + | 0 | ATTGGCGGCGGAGACGGCAG | CGG | \|\|\|\|\|\|\|\|[\|\|\|\|\|\|\|\|\|\|\|\|]PAM | NA | Intergenic | NA | NA |
| GU305914.1 | 163902 | 163924 | + | 4 | AGCGGCGGCGGCGGCGGCAG | CGG | \|--\|\|\|\|\|[\|\|\|-\|-\|\|\|\|\|\|]PAM | NA | Intergenic | NA | NA |
| T4271 | GACCCTGTCTGCCGCCGCGTTGG | 1000 | | | | | | | | | |
| Oligo fwd | CACCGACCCTGTCTGCCGCCGCGT | | | | | | | | | | |
| Oligo rev | AAACACGCGGCGGCAGACAGGGTC | | | | | | | | | | |
| Chromosome | start | end | strand | MM | target_seq | PAM | alignment | distance | position | gene name | gene id |
| GU305914.1 | 62746 | 62768 | + | 0 | GACCCTGTCTGCCGCCGCGT | TGG | \|\|\|\|\|\|\|\|[\|\|\|\|\|\|\|\|\|\|\|\|]PAM | NA | Intergenic | NA | NA |
| T4272 | GTCTGCCGCCGCGTTGGTCGCGG | 1000 | | | | | | | | | |
| Oligo fwd | CACCGTCTGCCGCCGCGTTGGTCG | | | | | | | | | | |
| Oligo rev | AAACCGACCAACGCGGCGGCAGAC | | | | | | | | | | |
| Chromosome | start | end | strand | MM | target_seq | PAM | alignment | distance | position | gene name | gene id |
| GU305914.1 | 62752 | 62774 | + | 0 | GTCTGCCGCCGCGTTGGTCG | CGG | \|\|\|\|\|\|\|\|[\|\|\|\|\|\|\|\|\|\|\|\|]PAM | NA | Intergenic | NA | NA |
| T4273 | CGCCGCGTTGGTCGCGGTCTTGG | 1000 | | | | | | | | | |
| Oligo fwd | CACCCGCCGCGTTGGTCGCGGTCT | | | | | | | | | | |
| Oligo rev | AAACAGACCGCGACCAACGCGGCG | | | | | | | | | | |
| Chromosome | start | end | strand | MM | target_seq | PAM | alignment | distance | position | gene name | gene id |
| GU305914.1 | 62758 | 62780 | + | 0 | CGCCGCGTTGGTCGCGGTCT | TGG | \|\|\|\|\|\|\|\|[\|\|\|\|\|\|\|\|\|\|\|\|]PAM | NA | Intergenic | NA | NA |
| T4274 | GCCGCGTTGGTCGCGGTCTTGGG | 1000 | | | | | | | | | |
| Oligo fwd | CACCGCCGCGTTGGTCGCGGTCTT | | | | | | | | | | |
| Oligo rev | AAACAAGACCGCGACCAACGCGGC | | | | | | | | | | |
| Chromosome | start | end | strand | MM | target_seq | PAM | alignment | distance | position | gene name | gene id |
| GU305914.1 | 62759 | 62781 | + | 0 | GCCGCGTTGGTCGCGGTCTT | GGG | \|\|\|\|\|\|\|\|[\|\|\|\|\|\|\|\|\|\|\|\|]PAM | NA | Intergenic | NA | NA |
| T4275 | CCGCGTTGGTCGCGGTCTTGGGG | 1000 | | | | | | | | | |
| Oligo fwd | CACCCCGCGTTGGTCGCGGTCTTG | | | | | | | | | | |
| Oligo rev | AAACCAAGACCGCGACCAACGCGG | | | | | | | | | | |
| Chromosome | start | end | strand | MM | target_seq | PAM | alignment | distance | position | gene name | gene id |
| GU305914.1 | 62760 | 62782 | + | 0 | CCGCGTTGGTCGCGGTCTTG | GGG | \|\|\|\|\|\|\|\|[\|\|\|\|\|\|\|\|\|\|\|\|]PAM | NA | Intergenic | NA | NA |
| T4276 | CGGTCTTGGGGTAGCTCCTGCGG | 1000 | | | | | | | | | |
| Oligo fwd | CACCCGGTCTTGGGGTAGCTCCTG | | | | | | | | | | |
| Oligo rev | AAACCAGGAGCTACCCCAAGACCG | | | | | | | | | | |
| Chromosome | start | end | strand | MM | target_seq | PAM | alignment | distance | position | gene name | gene id |
| GU305914.1 | 62772 | 62794 | + | 0 | CGGTCTTGGGGTAGCTCCTG | CGG | \|\|\|\|\|\|\|\|[\|\|\|\|\|\|\|\|\|\|\|\|]PAM | NA | Intergenic | NA | NA |
| T4277 | TGGGGTAGCTCCTGCGGCGTCGG | 1000 | | | | | | | | | |
| Oligo fwd | CACCTGGGGTAGCTCCTGCGGCGT | | | | | | | | | | |
| Oligo rev | AAACACGCCGCAGGAGCTACCCCA | | | | | | | | | | |
| Chromosome | start | end | strand | MM | target_seq | PAM | alignment | distance | position | gene name | gene id |
| GU305914.1 | 62778 | 62800 | + | 0 | TGGGGTAGCTCCTGCGGCGT | CGG | \|\|\|\|\|\|\|\|[\|\|\|\|\|\|\|\|\|\|\|\|]PAM | NA | Intergenic | NA | NA |
| T4278 | CCTGCGGCGTCGGTCAGTGTAGG | 1000 | | | | | | | | | |
| Oligo fwd | CACCCCTGCGGCGTCGGTCAGTGT | | | | | | | | | | |
| Oligo rev | AAACACACTGACCGACGCCGCAGG | | | | | | | | | | |
| Chromosome | start | end | strand | MM | target_seq | PAM | alignment | distance | position | gene name | gene id |
| GU305914.1 | 62788 | 62810 | + | 0 | CCTGCGGCGTCGGTCAGTGT | AGG | \|\|\|\|\|\|\|\|[\|\|\|\|\|\|\|\|\|\|\|\|]PAM | NA | Intergenic | NA | NA |
| T4279 | GCGGCGTCGGTCAGTGTAGGAGG | 1000 | | | | | | | | | |
| Oligo fwd | CACCGCGGCGTCGGTCAGTGTAGG | | | | | | | | | | |
| Oligo rev | AAACCCTACACTGACCGACGCCGC | | | | | | | | | | |
| Chromosome | start | end | strand | MM | target_seq | PAM | alignment | distance | position | gene name | gene id |
| GU305914.1 | 62791 | 62813 | + | 0 | GCGGCGTCGGTCAGTGTAGG | AGG | \|\|\|\|\|\|\|\|[\|\|\|\|\|\|\|\|\|\|\|\|]PAM | NA | Intergenic | NA | NA |
| T4280 | CGGCGTCGGTCAGTGTAGGAGGG | 1000 | | | | | | | | | |
| Oligo fwd | CACCCGGCGTCGGTCAGTGTAGGA | | | | | | | | | | |
| Oligo rev | AAACTCCTACACTGACCGACGCCG | | | | | | | | | | |
| Chromosome | start | end | strand | MM | target_seq | PAM | alignment | distance | position | gene name | gene id |
| GU305914.1 | 62792 | 62814 | + | 0 | CGGCGTCGGTCAGTGTAGGA | GGG | \|\|\|\|\|\|\|\|[\|\|\|\|\|\|\|\|\|\|\|\|]PAM | NA | Intergenic | NA | NA |
| T4281 | GTAGGAGGGACTCCAGCAAGAGG | 1000 | | | | | | | | | |

<table>
<tr><td>Oligo fwd</td><td>CACCGTAGGAGGGACTCCAGCAAG</td><td></td><td></td><td></td><td></td><td></td><td></td><td></td><td></td><td></td><td></td></tr>
<tr><td>Oligo rev</td><td>AAACCTTGCTGGAGTCCCTCCTAC</td><td></td><td></td><td></td><td></td><td></td><td></td><td></td><td></td><td></td><td></td></tr>
<tr><td>Chromosome</td><td>start</td><td>end</td><td>strand</td><td>MM</td><td>target_seq</td><td>PAM</td><td>alignment</td><td>distance</td><td>position</td><td>gene name</td><td>gene id</td></tr>
<tr><td>GU305914.1</td><td>62806</td><td>62828</td><td>+</td><td>0</td><td>GTAGGAGGGACTCCAGCAAG</td><td>AGG</td><td>||||||||[||||||||||||]PAM</td><td>NA</td><td>Intergenic</td><td>NA</td><td>NA</td></tr>
<tr><td>T4282</td><td>AGGCGATGCTCATAGCATTATGG</td><td>1000</td><td></td><td></td><td></td><td></td><td></td><td></td><td></td><td></td><td></td></tr>
<tr><td>Oligo fwd</td><td>CACCAGGCGATGCTCATAGCATTA</td><td></td><td></td><td></td><td></td><td></td><td></td><td></td><td></td><td></td><td></td></tr>
<tr><td>Oligo rev</td><td>AAACTAATGCTATGAGCATCGCCT</td><td></td><td></td><td></td><td></td><td></td><td></td><td></td><td></td><td></td><td></td></tr>
<tr><td>Chromosome</td><td>start</td><td>end</td><td>strand</td><td>MM</td><td>target_seq</td><td>PAM</td><td>alignment</td><td>distance</td><td>position</td><td>gene name</td><td>gene id</td></tr>
<tr><td>GU305914.1</td><td>62826</td><td>62848</td><td>+</td><td>0</td><td>AGGCGATGCTCATAGCATTA</td><td>TGG</td><td>||||||||[||||||||||||]PAM</td><td>NA</td><td>Intergenic</td><td>NA</td><td>NA</td></tr>
<tr><td>T4283</td><td>TCCATTCATGATCCTGACGTAGG</td><td>1000</td><td></td><td></td><td></td><td></td><td></td><td></td><td></td><td></td><td></td></tr>
<tr><td>Oligo fwd</td><td>CACCTCCATTCATGATCCTGACGT</td><td></td><td></td><td></td><td></td><td></td><td></td><td></td><td></td><td></td><td></td></tr>
<tr><td>Oligo rev</td><td>AAACACGTCAGGATCATGAATGGA</td><td></td><td></td><td></td><td></td><td></td><td></td><td></td><td></td><td></td><td></td></tr>
<tr><td>Chromosome</td><td>start</td><td>end</td><td>strand</td><td>MM</td><td>target_seq</td><td>PAM</td><td>alignment</td><td>distance</td><td>position</td><td>gene name</td><td>gene id</td></tr>
<tr><td>GU305914.1</td><td>62872</td><td>62894</td><td>+</td><td>0</td><td>TCCATTCATGATCCTGACGT</td><td>AGG</td><td>||||||||[||||||||||||]PAM</td><td>NA</td><td>Intergenic</td><td>NA</td><td>NA</td></tr>
<tr><td>T4284</td><td>CCATTCATGATCCTGACGTAGGG</td><td>1000</td><td></td><td></td><td></td><td></td><td></td><td></td><td></td><td></td><td></td></tr>
<tr><td>Oligo fwd</td><td>CACCCCATTCATGATCCTGACGTA</td><td></td><td></td><td></td><td></td><td></td><td></td><td></td><td></td><td></td><td></td></tr>
<tr><td>Oligo rev</td><td>AAACTACGTCAGGATCATGAATGG</td><td></td><td></td><td></td><td></td><td></td><td></td><td></td><td></td><td></td><td></td></tr>
<tr><td>Chromosome</td><td>start</td><td>end</td><td>strand</td><td>MM</td><td>target_seq</td><td>PAM</td><td>alignment</td><td>distance</td><td>position</td><td>gene name</td><td>gene id</td></tr>
<tr><td>GU305914.1</td><td>62873</td><td>62895</td><td>+</td><td>0</td><td>CCATTCATGATCCTGACGTA</td><td>GGG</td><td>||||||||[||||||||||||]PAM</td><td>NA</td><td>Intergenic</td><td>NA</td><td>NA</td></tr>
<tr><td>T4285</td><td>GTGCCCCCCACCCACCCCGAAGG</td><td>1000</td><td></td><td></td><td></td><td></td><td></td><td></td><td></td><td></td><td></td></tr>
<tr><td>Oligo fwd</td><td>CACCGTGCCCCCCACCCACCCCGA</td><td></td><td></td><td></td><td></td><td></td><td></td><td></td><td></td><td></td><td></td></tr>
<tr><td>Oligo rev</td><td>AAACTCGGGGTGGGTGGGGGGCAC</td><td></td><td></td><td></td><td></td><td></td><td></td><td></td><td></td><td></td><td></td></tr>
<tr><td>Chromosome</td><td>start</td><td>end</td><td>strand</td><td>MM</td><td>target_seq</td><td>PAM</td><td>alignment</td><td>distance</td><td>position</td><td>gene name</td><td>gene id</td></tr>
<tr><td>GU305914.1</td><td>62902</td><td>62924</td><td>+</td><td>0</td><td>GTGCCCCCCACCCACCCCGA</td><td>AGG</td><td>||||||||[||||||||||||]PAM</td><td>NA</td><td>Intergenic</td><td>NA</td><td>NA</td></tr>
<tr><td>T4286</td><td>ACCCACCCCGAAGGCTCCCCCGG</td><td>1000</td><td></td><td></td><td></td><td></td><td></td><td></td><td></td><td></td><td></td></tr>
<tr><td>Oligo fwd</td><td>CACCACCCACCCCGAAGGCTCCCC</td><td></td><td></td><td></td><td></td><td></td><td></td><td></td><td></td><td></td><td></td></tr>
<tr><td>Oligo rev</td><td>AAACGGGGAGCCTTCGGGGTGGGT</td><td></td><td></td><td></td><td></td><td></td><td></td><td></td><td></td><td></td><td></td></tr>
<tr><td>Chromosome</td><td>start</td><td>end</td><td>strand</td><td>MM</td><td>target_seq</td><td>PAM</td><td>alignment</td><td>distance</td><td>position</td><td>gene name</td><td>gene id</td></tr>
<tr><td>GU305914.1</td><td>62911</td><td>62933</td><td>+</td><td>0</td><td>ACCCACCCCGAAGGCTCCCC</td><td>CGG</td><td>||||||||[||||||||||||]PAM</td><td>NA</td><td>Intergenic</td><td>NA</td><td>NA</td></tr>
<tr><td>T4287</td><td>CCCGAAGGCTCCCCCGGCAGCGG</td><td>1000</td><td></td><td></td><td></td><td></td><td></td><td></td><td></td><td></td><td></td></tr>
<tr><td>Oligo fwd</td><td>CACCCCCGAAGGCTCCCCCGGCAG</td><td></td><td></td><td></td><td></td><td></td><td></td><td></td><td></td><td></td><td></td></tr>
<tr><td>Oligo rev</td><td>AAACCTGCCGGGGGAGCCTTCGGG</td><td></td><td></td><td></td><td></td><td></td><td></td><td></td><td></td><td></td><td></td></tr>
<tr><td>Chromosome</td><td>start</td><td>end</td><td>strand</td><td>MM</td><td>target_seq</td><td>PAM</td><td>alignment</td><td>distance</td><td>position</td><td>gene name</td><td>gene id</td></tr>
<tr><td>GU305914.1</td><td>62917</td><td>62939</td><td>+</td><td>0</td><td>CCCGAAGGCTCCCCCGGCAG</td><td>CGG</td><td>||||||||[||||||||||||]PAM</td><td>NA</td><td>Intergenic</td><td>NA</td><td>NA</td></tr>
<tr><td>T4288</td><td>GAAGGCTCCCCCGGCAGCGGTGG</td><td>1000</td><td></td><td></td><td></td><td></td><td></td><td></td><td></td><td></td><td></td></tr>
<tr><td>Oligo fwd</td><td>CACCGAAGGCTCCCCCGGCAGCGG</td><td></td><td></td><td></td><td></td><td></td><td></td><td></td><td></td><td></td><td></td></tr>
<tr><td>Oligo rev</td><td>AAACCCGCTGCCGGGGGAGCCTTC</td><td></td><td></td><td></td><td></td><td></td><td></td><td></td><td></td><td></td><td></td></tr>
<tr><td>Chromosome</td><td>start</td><td>end</td><td>strand</td><td>MM</td><td>target_seq</td><td>PAM</td><td>alignment</td><td>distance</td><td>position</td><td>gene name</td><td>gene id</td></tr>
<tr><td>GU305914.1</td><td>62920</td><td>62942</td><td>+</td><td>0</td><td>GAAGGCTCCCCCGGCAGCGG</td><td>TGG</td><td>||||||||[||||||||||||]PAM</td><td>NA</td><td>Intergenic</td><td>NA</td><td>NA</td></tr>
<tr><td>T4289</td><td>GGCTCCCCCGGCAGCGGTGGTGG</td><td>1000</td><td></td><td></td><td></td><td></td><td></td><td></td><td></td><td></td><td></td></tr>
<tr><td>Oligo fwd</td><td>CACCGGCTCCCCCGGCAGCGGTGG</td><td></td><td></td><td></td><td></td><td></td><td></td><td></td><td></td><td></td><td></td></tr>
<tr><td>Oligo rev</td><td>AAACCCACCGCTGCCGGGGGAGCC</td><td></td><td></td><td></td><td></td><td></td><td></td><td></td><td></td><td></td><td></td></tr>
<tr><td>Chromosome</td><td>start</td><td>end</td><td>strand</td><td>MM</td><td>target_seq</td><td>PAM</td><td>alignment</td><td>distance</td><td>position</td><td>gene name</td><td>gene id</td></tr>
<tr><td>GU305914.1</td><td>62923</td><td>62945</td><td>+</td><td>0</td><td>GGCTCCCCCGGCAGCGGTGG</td><td>TGG</td><td>||||||||[||||||||||||]PAM</td><td>NA</td><td>Intergenic</td><td>NA</td><td>NA</td></tr>
<tr><td>T4290</td><td>TCCCCCGGCAGCGGTGGTGGCGG</td><td>1000</td><td></td><td></td><td></td><td></td><td></td><td></td><td></td><td></td><td></td></tr>
<tr><td>Oligo fwd</td><td>CACCTCCCCCGGCAGCGGTGGTGG</td><td></td><td></td><td></td><td></td><td></td><td></td><td></td><td></td><td></td><td></td></tr>
<tr><td>Oligo rev</td><td>AAACCCACCACCGCTGCCGGGGGA</td><td></td><td></td><td></td><td></td><td></td><td></td><td></td><td></td><td></td><td></td></tr>
<tr><td>Chromosome</td><td>start</td><td>end</td><td>strand</td><td>MM</td><td>target_seq</td><td>PAM</td><td>alignment</td><td>distance</td><td>position</td><td>gene name</td><td>gene id</td></tr>
<tr><td>GU305914.1</td><td>62926</td><td>62948</td><td>+</td><td>0</td><td>TCCCCCGGCAGCGGTGGTGG</td><td>CGG</td><td>||||||||[||||||||||||]PAM</td><td>NA</td><td>Intergenic</td><td>NA</td><td>NA</td></tr>
<tr><td>T4291</td><td>CAGCGGTGGTGGCGGTCTGCTGG</td><td>1000</td><td></td><td></td><td></td><td></td><td></td><td></td><td></td><td></td><td></td></tr>
<tr><td>Oligo fwd</td><td>CACCCAGCGGTGGTGGCGGTCTGC</td><td></td><td></td><td></td><td></td><td></td><td></td><td></td><td></td><td></td><td></td></tr>
<tr><td>Oligo rev</td><td>AAACGCAGACCGCCACCACCGCTG</td><td></td><td></td><td></td><td></td><td></td><td></td><td></td><td></td><td></td><td></td></tr>
<tr><td>Chromosome</td><td>start</td><td>end</td><td>strand</td><td>MM</td><td>target_seq</td><td>PAM</td><td>alignment</td><td>distance</td><td>position</td><td>gene name</td><td>gene id</td></tr>
<tr><td>GU305914.1</td><td>62934</td><td>62956</td><td>+</td><td>0</td><td>CAGCGGTGGTGGCGGTCTGC</td><td>TGG</td><td>||||||||[||||||||||||]PAM</td><td>NA</td><td>Intergenic</td><td>NA</td><td>NA</td></tr>
<tr><td>T4292</td><td>TGGCCTCCCTGCCCGTCGTCAGG</td><td>1000</td><td></td><td></td><td></td><td></td><td></td><td></td><td></td><td></td><td></td></tr>
<tr><td>Oligo fwd</td><td>CACCTGGCCTCCCTGCCCGTCGTC</td><td></td><td></td><td></td><td></td><td></td><td></td><td></td><td></td><td></td><td></td></tr>
<tr><td>Oligo rev</td><td>AAACGACGACGGGCAGGGAGGCCA</td><td></td><td></td><td></td><td></td><td></td><td></td><td></td><td></td><td></td><td></td></tr>
<tr><td>Chromosome</td><td>start</td><td>end</td><td>strand</td><td>MM</td><td>target_seq</td><td>PAM</td><td>alignment</td><td>distance</td><td>position</td><td>gene name</td><td>gene id</td></tr>
<tr><td>GU305914.1</td><td>62954</td><td>62976</td><td>+</td><td>0</td><td>TGGCCTCCCTGCCCGTCGTC</td><td>AGG</td><td>||||||||[||||||||||||]PAM</td><td>NA</td><td>Intergenic</td><td>NA</td><td>NA</td></tr>
<tr><td>T4293</td><td>TCCCTGCCCGTCGTCAGGCATGG</td><td>1000</td><td></td><td></td><td></td><td></td><td></td><td></td><td></td><td></td><td></td></tr>
<tr><td>Oligo fwd</td><td>CACCTCCCTGCCCGTCGTCAGGCA</td><td></td><td></td><td></td><td></td><td></td><td></td><td></td><td></td><td></td><td></td></tr>
<tr><td>Oligo rev</td><td>AAACTGCCTGACGACGGGCAGGGA</td><td></td><td></td><td></td><td></td><td></td><td></td><td></td><td></td><td></td><td></td></tr>
<tr><td>Chromosome</td><td>start</td><td>end</td><td>strand</td><td>MM</td><td>target_seq</td><td>PAM</td><td>alignment</td><td>distance</td><td>position</td><td>gene name</td><td>gene id</td></tr>
<tr><td>GU305914.1</td><td>62959</td><td>62981</td><td>+</td><td>0</td><td>TCCCTGCCCGTCGTCAGGCA</td><td>TGG</td><td>||||||||[||||||||||||]PAM</td><td>NA</td><td>Intergenic</td><td>NA</td><td>NA</td></tr>
<tr><td>T4294</td><td>CCCTGCCCGTCGTCAGGCATGGG</td><td>1000</td><td></td><td></td><td></td><td></td><td></td><td></td><td></td><td></td><td></td></tr>
<tr><td>Oligo fwd</td><td>CACCCCCTGCCCGTCGTCAGGCAT</td><td></td><td></td><td></td><td></td><td></td><td></td><td></td><td></td><td></td><td></td></tr>
<tr><td>Oligo rev</td><td>AAACATGCCTGACGACGGGCAGGG</td><td></td><td></td><td></td><td></td><td></td><td></td><td></td><td></td><td></td><td></td></tr>
<tr><td>Chromosome</td><td>start</td><td>end</td><td>strand</td><td>MM</td><td>target_seq</td><td>PAM</td><td>alignment</td><td>distance</td><td>position</td><td>gene name</td><td>gene id</td></tr>
<tr><td>GU305914.1</td><td>62960</td><td>62982</td><td>+</td><td>0</td><td>CCCTGCCCGTCGTCAGGCAT</td><td>GGG</td><td>||||||||[||||||||||||]PAM</td><td>NA</td><td>Intergenic</td><td>NA</td><td>NA</td></tr>
<tr><td>T4295</td><td>CCTGCCCGTCGTCAGGCATGGGG</td><td>1000</td><td></td><td></td><td></td><td></td><td></td><td></td><td></td><td></td><td></td></tr>
<tr><td>Oligo fwd</td><td>CACCCCTGCCCGTCGTCAGGCATG</td><td></td><td></td><td></td><td></td><td></td><td></td><td></td><td></td><td></td><td></td></tr>
<tr><td>Oligo rev</td><td>AAACCATGCCTGACGACGGGCAGG</td><td></td><td></td><td></td><td></td><td></td><td></td><td></td><td></td><td></td><td></td></tr>
<tr><td>Chromosome</td><td>start</td><td>end</td><td>strand</td><td>MM</td><td>target_seq</td><td>PAM</td><td>alignment</td><td>distance</td><td>position</td><td>gene name</td><td>gene id</td></tr>
<tr><td>GU305914.1</td><td>62961</td><td>62983</td><td>+</td><td>0</td><td>CCTGCCCGTCGTCAGGCATG</td><td>GGG</td><td>||||||||[||||||||||||]PAM</td><td>NA</td><td>Intergenic</td><td>NA</td><td>NA</td></tr>
</table>

T4296 AAAAAGAAGAATCCGTTGTCCGG 1000
Oligo fwd CACCAAAAAGAAGAATCCGTTGTC
Oligo rev AAACGACAACGGATTCTTCTTTTT
Chromosome start end strand MM target_seq PAM alignment distance position gene name gene id
GU305914.1 62990 63012 + 0 AAAAAGAAGAATCCGTTGTC CGG ||||||||[||||||||||||]PAM NA Intergenic NA NA

T4297 AAAAGAAGAATCCGTTGTCCGGG 1000
Oligo fwd CACCAAAAGAAGAATCCGTTGTCC
Oligo rev AAACGGACAACGGATTCTTCTTTT
Chromosome start end strand MM target_seq PAM alignment distance position gene name gene id
GU305914.1 62991 63013 + 0 AAAAGAAGAATCCGTTGTCC GGG ||||||||[||||||||||||]PAM NA Intergenic NA NA

T4298 GAAGAATCCGTTGTCCGGGTCGG 1000
Oligo fwd CACCGAAGAATCCGTTGTCCGGGT
Oligo rev AAACACCCGGACAACGGATTCTTC
Chromosome start end strand MM target_seq PAM alignment distance position gene name gene id
GU305914.1 62995 63017 + 0 GAAGAATCCGTTGTCCGGGT CGG ||||||||[||||||||||||]PAM NA Intergenic NA NA

T4299 GTCCGGGTCGGAGTAGACTTTGG 1000
Oligo fwd CACCGTCCGGGTCGGAGTAGACTT
Oligo rev AAACAAGTCTACTCCGACCCGGAC
Chromosome start end strand MM target_seq PAM alignment distance position gene name gene id
GU305914.1 63007 63029 + 0 GTCCGGGTCGGAGTAGACTT TGG ||||||||[||||||||||||]PAM NA Intergenic NA NA

T4300 TCCGGGTCGGAGTAGACTTTGGG 1000
Oligo fwd CACCTCCGGGTCGGAGTAGACTTT
Oligo rev AAACAAAGTCTACTCCGACCCGGA
Chromosome start end strand MM target_seq PAM alignment distance position gene name gene id
GU305914.1 63008 63030 + 0 TCCGGGTCGGAGTAGACTTT GGG ||||||||[||||||||||||]PAM NA Intergenic NA NA

T4301 GGTCGGAGTAGACTTTGGGCTGG 1000
Oligo fwd CACCGGTCGGAGTAGACTTTGGGC
Oligo rev AAACGCCCAAAGTCTACTCCGACC
Chromosome start end strand MM target_seq PAM alignment distance position gene name gene id
GU305914.1 63012 63034 + 0 GGTCGGAGTAGACTTTGGGC TGG ||||||||[||||||||||||]PAM NA Intergenic NA NA

T4302 CATGTCACGCGAACTCGACGAGG 1000
Oligo fwd CACCCATGTCACGCGAACTCGACG
Oligo rev AAACCGTCGAGTTCGCGTGACATG
Chromosome start end strand MM target_seq PAM alignment distance position gene name gene id
GU305914.1 63040 63062 + 0 CATGTCACGCGAACTCGACG AGG ||||||||[||||||||||||]PAM NA Intergenic NA NA

T4303 ATGTCACGCGAACTCGACGAGGG 1000
Oligo fwd CACCATGTCACGCGAACTCGACGA
Oligo rev AAACTCGTCGAGTTCGCGTGACAT
Chromosome start end strand MM target_seq PAM alignment distance position gene name gene id
GU305914.1 63041 63063 + 0 ATGTCACGCGAACTCGACGA GGG ||||||||[||||||||||||]PAM NA Intergenic NA NA

T4304 TGTCACGCGAACTCGACGAGGGG 1000
Oligo fwd CACCTGTCACGCGAACTCGACGAG
Oligo rev AAACCTCGTCGAGTTCGCGTGACA
Chromosome start end strand MM target_seq PAM alignment distance position gene name gene id
GU305914.1 63042 63064 + 0 TGTCACGCGAACTCGACGAG GGG ||||||||[||||||||||||]PAM NA Intergenic NA NA

T4305 GAACTTCTCTAGCTTAACCACGG 1000
Oligo fwd CACCGAACTTCTCTAGCTTAACCA
Oligo rev AAACTGGTTAAGCTAGAGAAGTTC
Chromosome start end strand MM target_seq PAM alignment distance position gene name gene id
GU305914.1 63080 63102 + 0 GAACTTCTCTAGCTTAACCA CGG ||||||||[||||||||||||]PAM NA Intergenic NA NA

T4306 CTTAACCACGGAGAGCTGTCCGG 1000
Oligo fwd CACCCTTAACCACGGAGAGCTGTC
Oligo rev AAACGACAGCTCTCCGTGGTTAAG
Chromosome start end strand MM target_seq PAM alignment distance position gene name gene id
GU305914.1 63092 63114 + 0 CTTAACCACGGAGAGCTGTC CGG ||||||||[||||||||||||]PAM NA Intergenic NA NA

T4307 GGAGAGCTGTCCGGCGCGCGAGG 1000
Oligo fwd CACCGGAGAGCTGTCCGGCGCGCG
Oligo rev AAACCGCGCGCCGGACAGCTCTCC
Chromosome start end strand MM target_seq PAM alignment distance position gene name gene id
GU305914.1 63101 63123 + 0 GGAGAGCTGTCCGGCGCGCG AGG ||||||||[||||||||||||]PAM NA Intergenic NA NA

T4308 GGCGCGCGAGGCGACGATCCCGG 1000
Oligo fwd CACCGGCGCGCGAGGCGACGATCC
Oligo rev AAACGGATCGTCGCCTCGCGCGCC
Chromosome start end strand MM target_seq PAM alignment distance position gene name gene id
GU305914.1 63113 63135 + 0 GGCGCGCGAGGCGACGATCC CGG ||||||||[||||||||||||]PAM NA Intergenic NA NA

T4309 CTGTAGCGTGCCGAGACAGCCGG 1000
Oligo fwd CACCCTGTAGCGTGCCGAGACAGC
Oligo rev AAACGCTGTCTCGGCACGCTACAG
Chromosome start end strand MM target_seq PAM alignment distance position gene name gene id
GU305914.1 63170 63192 + 0 CTGTAGCGTGCCGAGACAGC CGG ||||||||[||||||||||||]PAM NA Intergenic NA NA

T4310 TAGCGTGCCGAGACAGCCGGTGG 1000
Oligo fwd CACCTAGCGTGCCGAGACAGCCGG
Oligo rev AAACCCGGCTGTCTCGGCACGCTA
Chromosome start end strand MM target_seq PAM alignment distance position gene name gene id
GU305914.1 63173 63195 + 0 TAGCGTGCCGAGACAGCCGG TGG ||||||||[||||||||||||]PAM NA Intergenic NA NA

T4311 GTGCCGAGACAGCCGGTGGACGG 1000
Oligo fwd CACCGTGCCGAGACAGCCGGTGGA
Oligo rev AAACTCCACCGGCTGTCTCGGCAC
Chromosome start end strand MM target_seq PAM alignment distance position gene name gene id
GU305914.1 63177 63199 + 0 GTGCCGAGACAGCCGGTGGA CGG ||||||||[||||||||||||]PAM NA Intergenic NA NA

T4312 GTTATTTAAGAAGTGAAATCTGG 1000
Oligo fwd CACCGTTATTTAAGAAGTGAAATC
Oligo rev AAACGATTTCACTTCTTAAATAAC
Chromosome start end strand MM target_seq PAM alignment distance position gene name gene id
GU305914.1 63218 63240 + 0 GTTATTTAAGAAGTGAAATC TGG ||||||||[||||||||||||]PAM NA Intergenic NA NA

T4313 AGACCCTGAACCTATAGCGCAGG 1000
Oligo fwd CACCAGACCCTGAACCTATAGCGC
Oligo rev AAACGCGCTATAGGTTCAGGGTCT
Chromosome start end strand MM target_seq PAM alignment distance position gene name gene id
GU305914.1 63241 63263 + 0 AGACCCTGAACCTATAGCGC AGG ||||||||[||||||||||||]PAM NA Intergenic NA NA

T4314 CGTATGCCTGTGTGTGATCCTGG 1000
Oligo fwd CACCCGTATGCCTGTGTGTGATCC
Oligo rev AAACGGATCACACACAGGCATACG
Chromosome start end strand MM target_seq PAM alignment distance position gene name gene id
GU305914.1 63275 63297 + 0 CGTATGCCTGTGTGTGATCC TGG ||||||||[||||||||||||]PAM NA Intergenic NA NA

T4315 GATCCTGGAGACCACCACAAAGG 1000
Oligo fwd CACCGATCCTGGAGACCACCACAA
Oligo rev AAACTTGTGGTGGTCTCCAGGATC
Chromosome start end strand MM target_seq PAM alignment distance position gene name gene id
GU305914.1 63290 63312 + 0 GATCCTGGAGACCACCACAA AGG ||||||||[||||||||||||]PAM NA Intergenic NA NA

T4316 CCTGGAGACCACCACAAAGGCGG 1000
Oligo fwd CACCCCTGGAGACCACCACAAAGG
Oligo rev AAACCCTTTGTGGTGGTCTCCAGG
Chromosome start end strand MM target_seq PAM alignment distance position gene name gene id
GU305914.1 63293 63315 + 0 CCTGGAGACCACCACAAAGG CGG ||||||||[||||||||||||]PAM NA Intergenic NA NA

T4317 GAGACCACCACAAAGGCGGACGG 1000
Oligo fwd CACCGAGACCACCACAAAGGCGGA
Oligo rev AAACTCCGCCTTTGTGGTGGTCTC
Chromosome start end strand MM target_seq PAM alignment distance position gene name gene id
GU305914.1 63297 63319 + 0 GAGACCACCACAAAGGCGGA CGG ||||||||[||||||||||||]PAM NA Intergenic NA NA

T4318 ATAAACGAATCTGATCTCATCGG 1000
Oligo fwd CACCATAAACGAATCTGATCTCAT
Oligo rev AAACATGAGATCAGATTCGTTTAT
Chromosome start end strand MM target_seq PAM alignment distance position gene name gene id
GU305914.1 63362 63384 + 0 ATAAACGAATCTGATCTCAT CGG ||||||||[||||||||||||]PAM NA Intergenic NA NA

T4319 CATCGGTGCTTTCCCCATCGAGG 1000
Oligo fwd CACCCATCGGTGCTTTCCCCATCG
Oligo rev AAACCGATGGGGAAAGCACCGATG
Chromosome start end strand MM target_seq PAM alignment distance position gene name gene id
GU305914.1 63379 63401 + 0 CATCGGTGCTTTCCCCATCG AGG ||||||||[||||||||||||]PAM NA Intergenic NA NA

T4320 TGCTTTCCCCATCGAGGTCGAGG 1000
Oligo fwd CACCTGCTTTCCCCATCGAGGTCG
Oligo rev AAACCGACCTCGATGGGGAAAGCA
Chromosome start end strand MM target_seq PAM alignment distance position gene name gene id
GU305914.1 63385 63407 + 0 TGCTTTCCCCATCGAGGTCG AGG ||||||||[||||||||||||]PAM NA Intergenic NA NA

T4321 GAGGTCGAGGTAGATCTCCTTGG 1000
Oligo fwd CACCGAGGTCGAGGTAGATCTCCT
Oligo rev AAACAGGAGATCTACCTCGACCTC
Chromosome start end strand MM target_seq PAM alignment distance position gene name gene id
GU305914.1 63398 63420 + 0 GAGGTCGAGGTAGATCTCCT TGG ||||||||[||||||||||||]PAM NA Intergenic NA NA

T4322 AGGTCGAGGTAGATCTCCTTGGG 1000
Oligo fwd CACCAGGTCGAGGTAGATCTCCTT
Oligo rev AAACAAGGAGATCTACCTCGACCT
Chromosome start end strand MM target_seq PAM alignment distance position gene name gene id
GU305914.1 63399 63421 + 0 AGGTCGAGGTAGATCTCCTT GGG ||||||||[||||||||||||]PAM NA Intergenic NA NA

T4323 AGATCTCCTTGGGAACGAGCAGG 1000
Oligo fwd CACCAGATCTCCTTGGGAACGAGC
Oligo rev AAACGCTCGTTCCCAAGGAGATCT
Chromosome start end strand MM target_seq PAM alignment distance position gene name gene id
GU305914.1 63409 63431 + 0 AGATCTCCTTGGGAACGAGC AGG ||||||||[||||||||||||]PAM NA Intergenic NA NA

T4324 GATCTCCTTGGGAACGAGCAGGG 1000
Oligo fwd CACCGATCTCCTTGGGAACGAGCA
Oligo rev AAACTGCTCGTTCCCAAGGAGATC
Chromosome start end strand MM target_seq PAM alignment distance position gene name gene id
GU305914.1 63410 63432 + 0 GATCTCCTTGGGAACGAGCA GGG ||||||||[||||||||||||]PAM NA Intergenic NA NA

T4325 ATCTCCTTGGGAACGAGCAGGGG 1000
Oligo fwd CACCATCTCCTTGGGAACGAGCAG
Oligo rev AAACCTGCTCGTTCCCAAGGAGAT
Chromosome start end strand MM target_seq PAM alignment distance position gene name gene id

<table>
<tr><td>GU305914.1</td><td>63411</td><td>63433</td><td>+</td><td>0</td><td>ATCTCCTTGGGAACGAGCAG</td><td>GGG</td><td>||||||||[||||||||||||]PAM</td><td>NA</td><td>Intergenic</td><td>NA</td><td>NA</td></tr>
<tr><td>T4326</td><td>AACGAGCAGGGGATAGATCTCGG</td><td>1000</td><td></td><td></td><td></td><td></td><td></td><td></td><td></td><td></td><td></td></tr>
<tr><td>Oligo fwd</td><td>CACCAACGAGCAGGGGATAGATCT</td><td></td><td></td><td></td><td></td><td></td><td></td><td></td><td></td><td></td><td></td></tr>
<tr><td>Oligo rev</td><td>AAACAGATCTATCCCCTGCTCGTT</td><td></td><td></td><td></td><td></td><td></td><td></td><td></td><td></td><td></td><td></td></tr>
<tr><td>Chromosome</td><td>start</td><td>end</td><td>strand</td><td>MM</td><td>target_seq</td><td>PAM</td><td>alignment</td><td>distance</td><td>position</td><td>gene name</td><td>gene id</td></tr>
<tr><td>GU305914.1</td><td>63422</td><td>63444</td><td>+</td><td>0</td><td>AACGAGCAGGGGATAGATCT</td><td>CGG</td><td>||||||||[||||||||||||]PAM</td><td>NA</td><td>Intergenic</td><td>NA</td><td>NA</td></tr>
<tr><td>T4327</td><td>CGAGTGTCCGCGCATCGAGCAGG</td><td>1000</td><td></td><td></td><td></td><td></td><td></td><td></td><td></td><td></td><td></td></tr>
<tr><td>Oligo fwd</td><td>CACCCGAGTGTCCGCGCATCGAGC</td><td></td><td></td><td></td><td></td><td></td><td></td><td></td><td></td><td></td><td></td></tr>
<tr><td>Oligo rev</td><td>AAACGCTCGATGCGCGGACACTCG</td><td></td><td></td><td></td><td></td><td></td><td></td><td></td><td></td><td></td><td></td></tr>
<tr><td>Chromosome</td><td>start</td><td>end</td><td>strand</td><td>MM</td><td>target_seq</td><td>PAM</td><td>alignment</td><td>distance</td><td>position</td><td>gene name</td><td>gene id</td></tr>
<tr><td>GU305914.1</td><td>63461</td><td>63483</td><td>+</td><td>0</td><td>CGAGTGTCCGCGCATCGAGC</td><td>AGG</td><td>||||||||[||||||||||||]PAM</td><td>NA</td><td>Intergenic</td><td>NA</td><td>NA</td></tr>
<tr><td>T4328</td><td>CGCGCATCGAGCAGGTGATCAGG</td><td>1000</td><td></td><td></td><td></td><td></td><td></td><td></td><td></td><td></td><td></td></tr>
<tr><td>Oligo fwd</td><td>CACCCGCGCATCGAGCAGGTGATC</td><td></td><td></td><td></td><td></td><td></td><td></td><td></td><td></td><td></td><td></td></tr>
<tr><td>Oligo rev</td><td>AAACGATCACCTGCTCGATGCGCG</td><td></td><td></td><td></td><td></td><td></td><td></td><td></td><td></td><td></td><td></td></tr>
<tr><td>Chromosome</td><td>start</td><td>end</td><td>strand</td><td>MM</td><td>target_seq</td><td>PAM</td><td>alignment</td><td>distance</td><td>position</td><td>gene name</td><td>gene id</td></tr>
<tr><td>GU305914.1</td><td>63469</td><td>63491</td><td>+</td><td>0</td><td>CGCGCATCGAGCAGGTGATC</td><td>AGG</td><td>||||||||[||||||||||||]PAM</td><td>NA</td><td>Intergenic</td><td>NA</td><td>NA</td></tr>
<tr><td>T4329</td><td>GGTGATCAGGCACGTCGCTCCGG</td><td>1000</td><td></td><td></td><td></td><td></td><td></td><td></td><td></td><td></td><td></td></tr>
<tr><td>Oligo fwd</td><td>CACCGGTGATCAGGCACGTCGCTC</td><td></td><td></td><td></td><td></td><td></td><td></td><td></td><td></td><td></td><td></td></tr>
<tr><td>Oligo rev</td><td>AAACGAGCGACGTGCCTGATCACC</td><td></td><td></td><td></td><td></td><td></td><td></td><td></td><td></td><td></td><td></td></tr>
<tr><td>Chromosome</td><td>start</td><td>end</td><td>strand</td><td>MM</td><td>target_seq</td><td>PAM</td><td>alignment</td><td>distance</td><td>position</td><td>gene name</td><td>gene id</td></tr>
<tr><td>GU305914.1</td><td>63482</td><td>63504</td><td>+</td><td>0</td><td>GGTGATCAGGCACGTCGCTC</td><td>CGG</td><td>||||||||[||||||||||||]PAM</td><td>NA</td><td>Intergenic</td><td>NA</td><td>NA</td></tr>
<tr><td>T4330</td><td>GTGATCAGGCACGTCGCTCCGGG</td><td>1000</td><td></td><td></td><td></td><td></td><td></td><td></td><td></td><td></td><td></td></tr>
<tr><td>Oligo fwd</td><td>CACCGTGATCAGGCACGTCGCTCC</td><td></td><td></td><td></td><td></td><td></td><td></td><td></td><td></td><td></td><td></td></tr>
<tr><td>Oligo rev</td><td>AAACGGAGCGACGTGCCTGATCAC</td><td></td><td></td><td></td><td></td><td></td><td></td><td></td><td></td><td></td><td></td></tr>
<tr><td>Chromosome</td><td>start</td><td>end</td><td>strand</td><td>MM</td><td>target_seq</td><td>PAM</td><td>alignment</td><td>distance</td><td>position</td><td>gene name</td><td>gene id</td></tr>
<tr><td>GU305914.1</td><td>63483</td><td>63505</td><td>+</td><td>0</td><td>GTGATCAGGCACGTCGCTCC</td><td>GGG</td><td>||||||||[||||||||||||]PAM</td><td>NA</td><td>Intergenic</td><td>NA</td><td>NA</td></tr>
<tr><td>T4331</td><td>CGCTCCGGGCGTGATCGAGTTGG</td><td>1000</td><td></td><td></td><td></td><td></td><td></td><td></td><td></td><td></td><td></td></tr>
<tr><td>Oligo fwd</td><td>CACCCGCTCCGGGCGTGATCGAGT</td><td></td><td></td><td></td><td></td><td></td><td></td><td></td><td></td><td></td><td></td></tr>
<tr><td>Oligo rev</td><td>AAACACTCGATCACGCCCGGAGCG</td><td></td><td></td><td></td><td></td><td></td><td></td><td></td><td></td><td></td><td></td></tr>
<tr><td>Chromosome</td><td>start</td><td>end</td><td>strand</td><td>MM</td><td>target_seq</td><td>PAM</td><td>alignment</td><td>distance</td><td>position</td><td>gene name</td><td>gene id</td></tr>
<tr><td>GU305914.1</td><td>63497</td><td>63519</td><td>+</td><td>0</td><td>CGCTCCGGGCGTGATCGAGT</td><td>TGG</td><td>||||||||[||||||||||||]PAM</td><td>NA</td><td>Intergenic</td><td>NA</td><td>NA</td></tr>
<tr><td>T4332</td><td>ATCGAGTTGGTGCATCCGAGCGG</td><td>1000</td><td></td><td></td><td></td><td></td><td></td><td></td><td></td><td></td><td></td></tr>
<tr><td>Oligo fwd</td><td>CACCATCGAGTTGGTGCATCCGAG</td><td></td><td></td><td></td><td></td><td></td><td></td><td></td><td></td><td></td><td></td></tr>
<tr><td>Oligo rev</td><td>AAACCTCGGATGCACCAACTCGAT</td><td></td><td></td><td></td><td></td><td></td><td></td><td></td><td></td><td></td><td></td></tr>
<tr><td>Chromosome</td><td>start</td><td>end</td><td>strand</td><td>MM</td><td>target_seq</td><td>PAM</td><td>alignment</td><td>distance</td><td>position</td><td>gene name</td><td>gene id</td></tr>
<tr><td>GU305914.1</td><td>63510</td><td>63532</td><td>+</td><td>0</td><td>ATCGAGTTGGTGCATCCGAG</td><td>CGG</td><td>||||||||[||||||||||||]PAM</td><td>NA</td><td>Intergenic</td><td>NA</td><td>NA</td></tr>
<tr><td>T4333</td><td>TCGAGTTGGTGCATCCGAGCGGG</td><td>1000</td><td></td><td></td><td></td><td></td><td></td><td></td><td></td><td></td><td></td></tr>
<tr><td>Oligo fwd</td><td>CACCTCGAGTTGGTGCATCCGAGC</td><td></td><td></td><td></td><td></td><td></td><td></td><td></td><td></td><td></td><td></td></tr>
<tr><td>Oligo rev</td><td>AAACGCTCGGATGCACCAACTCGA</td><td></td><td></td><td></td><td></td><td></td><td></td><td></td><td></td><td></td><td></td></tr>
<tr><td>Chromosome</td><td>start</td><td>end</td><td>strand</td><td>MM</td><td>target_seq</td><td>PAM</td><td>alignment</td><td>distance</td><td>position</td><td>gene name</td><td>gene id</td></tr>
<tr><td>GU305914.1</td><td>63511</td><td>63533</td><td>+</td><td>0</td><td>TCGAGTTGGTGCATCCGAGC</td><td>GGG</td><td>||||||||[||||||||||||]PAM</td><td>NA</td><td>Intergenic</td><td>NA</td><td>NA</td></tr>
<tr><td>T4334</td><td>AGTTGGTGCATCCGAGCGGGTGG</td><td>1000</td><td></td><td></td><td></td><td></td><td></td><td></td><td></td><td></td><td></td></tr>
<tr><td>Oligo fwd</td><td>CACCAGTTGGTGCATCCGAGCGGG</td><td></td><td></td><td></td><td></td><td></td><td></td><td></td><td></td><td></td><td></td></tr>
<tr><td>Oligo rev</td><td>AAACCCCGCTCGGATGCACCAACT</td><td></td><td></td><td></td><td></td><td></td><td></td><td></td><td></td><td></td><td></td></tr>
<tr><td>Chromosome</td><td>start</td><td>end</td><td>strand</td><td>MM</td><td>target_seq</td><td>PAM</td><td>alignment</td><td>distance</td><td>position</td><td>gene name</td><td>gene id</td></tr>
<tr><td>GU305914.1</td><td>63514</td><td>63536</td><td>+</td><td>0</td><td>AGTTGGTGCATCCGAGCGGG</td><td>TGG</td><td>||||||||[||||||||||||]PAM</td><td>NA</td><td>Intergenic</td><td>NA</td><td>NA</td></tr>
<tr><td>T4335</td><td>ATCCGAGCGGGTGGACGACCCGG</td><td>1000</td><td></td><td></td><td></td><td></td><td></td><td></td><td></td><td></td><td></td></tr>
<tr><td>Oligo fwd</td><td>CACCATCCGAGCGGGTGGACGACC</td><td></td><td></td><td></td><td></td><td></td><td></td><td></td><td></td><td></td><td></td></tr>
<tr><td>Oligo rev</td><td>AAACGGTCGTCCACCCGCTCGGAT</td><td></td><td></td><td></td><td></td><td></td><td></td><td></td><td></td><td></td><td></td></tr>
<tr><td>Chromosome</td><td>start</td><td>end</td><td>strand</td><td>MM</td><td>target_seq</td><td>PAM</td><td>alignment</td><td>distance</td><td>position</td><td>gene name</td><td>gene id</td></tr>
<tr><td>GU305914.1</td><td>63523</td><td>63545</td><td>+</td><td>0</td><td>ATCCGAGCGGGTGGACGACC</td><td>CGG</td><td>||||||||[||||||||||||]PAM</td><td>NA</td><td>Intergenic</td><td>NA</td><td>NA</td></tr>
<tr><td>T4336</td><td>GACGACCCGGTGCAGCCGCCAGG</td><td>1000</td><td></td><td></td><td></td><td></td><td></td><td></td><td></td><td></td><td></td></tr>
<tr><td>Oligo fwd</td><td>CACCGACGACCCGGTGCAGCCGCC</td><td></td><td></td><td></td><td></td><td></td><td></td><td></td><td></td><td></td><td></td></tr>
<tr><td>Oligo rev</td><td>AAACGGCGGCTGCACCGGGTCGTC</td><td></td><td></td><td></td><td></td><td></td><td></td><td></td><td></td><td></td><td></td></tr>
<tr><td>Chromosome</td><td>start</td><td>end</td><td>strand</td><td>MM</td><td>target_seq</td><td>PAM</td><td>alignment</td><td>distance</td><td>position</td><td>gene name</td><td>gene id</td></tr>
<tr><td>GU305914.1</td><td>63536</td><td>63558</td><td>+</td><td>0</td><td>GACGACCCGGTGCAGCCGCC</td><td>AGG</td><td>||||||||[||||||||||||]PAM</td><td>NA</td><td>Intergenic</td><td>NA</td><td>NA</td></tr>
<tr><td>T4337</td><td>CCAGGCGCCGTCGCGCAGCCAGG</td><td>1000</td><td></td><td></td><td></td><td></td><td></td><td></td><td></td><td></td><td></td></tr>
<tr><td>Oligo fwd</td><td>CACCCCAGGCGCCGTCGCGCAGCC</td><td></td><td></td><td></td><td></td><td></td><td></td><td></td><td></td><td></td><td></td></tr>
<tr><td>Oligo rev</td><td>AAACGGCTGCGCGACGGCGCCTGG</td><td></td><td></td><td></td><td></td><td></td><td></td><td></td><td></td><td></td><td></td></tr>
<tr><td>Chromosome</td><td>start</td><td>end</td><td>strand</td><td>MM</td><td>target_seq</td><td>PAM</td><td>alignment</td><td>distance</td><td>position</td><td>gene name</td><td>gene id</td></tr>
<tr><td>GU305914.1</td><td>63554</td><td>63576</td><td>+</td><td>0</td><td>CCAGGCGCCGTCGCGCAGCC</td><td>AGG</td><td>||||||||[||||||||||||]PAM</td><td>NA</td><td>Intergenic</td><td>NA</td><td>NA</td></tr>
<tr><td>T4338</td><td>CGCCGTCGCGCAGCCAGGTGCGG</td><td>1000</td><td></td><td></td><td></td><td></td><td></td><td></td><td></td><td></td><td></td></tr>
<tr><td>Oligo fwd</td><td>CACCCGCCGTCGCGCAGCCAGGTG</td><td></td><td></td><td></td><td></td><td></td><td></td><td></td><td></td><td></td><td></td></tr>
<tr><td>Oligo rev</td><td>AAACCACCTGGCTGCGCGACGGCG</td><td></td><td></td><td></td><td></td><td></td><td></td><td></td><td></td><td></td><td></td></tr>
<tr><td>Chromosome</td><td>start</td><td>end</td><td>strand</td><td>MM</td><td>target_seq</td><td>PAM</td><td>alignment</td><td>distance</td><td>position</td><td>gene name</td><td>gene id</td></tr>
<tr><td>GU305914.1</td><td>63559</td><td>63581</td><td>+</td><td>0</td><td>CGCCGTCGCGCAGCCAGGTG</td><td>CGG</td><td>||||||||[||||||||||||]PAM</td><td>NA</td><td>Intergenic</td><td>NA</td><td>NA</td></tr>
<tr><td>T4339</td><td>GTCGCGCAGCCAGGTGCGGATGG</td><td>1000</td><td></td><td></td><td></td><td></td><td></td><td></td><td></td><td></td><td></td></tr>
<tr><td>Oligo fwd</td><td>CACCGTCGCGCAGCCAGGTGCGGA</td><td></td><td></td><td></td><td></td><td></td><td></td><td></td><td></td><td></td><td></td></tr>
<tr><td>Oligo rev</td><td>AAACTCCGCACCTGGCTGCGCGAC</td><td></td><td></td><td></td><td></td><td></td><td></td><td></td><td></td><td></td><td></td></tr>
<tr><td>Chromosome</td><td>start</td><td>end</td><td>strand</td><td>MM</td><td>target_seq</td><td>PAM</td><td>alignment</td><td>distance</td><td>position</td><td>gene name</td><td>gene id</td></tr>
<tr><td>GU305914.1</td><td>63563</td><td>63585</td><td>+</td><td>0</td><td>GTCGCGCAGCCAGGTGCGGA</td><td>TGG</td><td>||||||||[||||||||||||]PAM</td><td>NA</td><td>Intergenic</td><td>NA</td><td>NA</td></tr>
<tr><td>T4340</td><td>TGCGGATGGCCTCGCCCTCGCGG</td><td>1000</td><td></td><td></td><td></td><td></td><td></td><td></td><td></td><td></td><td></td></tr>
<tr><td>Oligo fwd</td><td>CACCTGCGGATGGCCTCGCCCTCG</td><td></td><td></td><td></td><td></td><td></td><td></td><td></td><td></td><td></td><td></td></tr>
<tr><td>Oligo rev</td><td>AAACCGAGGGCGAGGCCATCCGCA</td><td></td><td></td><td></td><td></td><td></td><td></td><td></td><td></td><td></td><td></td></tr>
</table>

<table>
<tr><td>Chromosome</td><td>start</td><td>end</td><td>strand</td><td>MM</td><td>target_seq</td><td>PAM</td><td>alignment</td><td>distance</td><td>position</td><td>gene name</td><td>gene id</td></tr>
<tr><td>GU305914.1</td><td>63577</td><td>63599</td><td>+</td><td>0</td><td>TGCGGATGGCCTCGCCCTCG</td><td>CGG</td><td>||||||||[||||||||||||]PAM</td><td>NA</td><td>Intergenic</td><td>NA</td><td>NA</td></tr>
<tr><td>GU305914.1</td><td>200933</td><td>200955</td><td>+</td><td>4</td><td>TACCGATGGCGTCGCGCTCG</td><td>AGG</td><td>|-|-||||[||-||||-||||]PAM</td><td>NA</td><td>Intergenic</td><td>NA</td><td>NA</td></tr>
<tr><td></td><td></td><td></td><td></td><td></td><td></td><td></td><td></td><td></td><td></td><td></td><td></td></tr>
<tr><td>T4341</td><td>GATCGCCGCGAGCGTGTTCTTGG</td><td>1000</td><td></td><td></td><td></td><td></td><td></td><td></td><td></td><td></td><td></td></tr>
<tr><td>Oligo fwd</td><td>CACCGATCGCCGCGAGCGTGTTCT</td><td></td><td></td><td></td><td></td><td></td><td></td><td></td><td></td><td></td><td></td></tr>
<tr><td>Oligo rev</td><td>AAACAGAACACGCTCGCGGCGATC</td><td></td><td></td><td></td><td></td><td></td><td></td><td></td><td></td><td></td><td></td></tr>
<tr><td>Chromosome</td><td>start</td><td>end</td><td>strand</td><td>MM</td><td>target_seq</td><td>PAM</td><td>alignment</td><td>distance</td><td>position</td><td>gene name</td><td>gene id</td></tr>
<tr><td>GU305914.1</td><td>63605</td><td>63627</td><td>+</td><td>0</td><td>GATCGCCGCGAGCGTGTTCT</td><td>TGG</td><td>||||||||[||||||||||||]PAM</td><td>NA</td><td>Intergenic</td><td>NA</td><td>NA</td></tr>
<tr><td></td><td></td><td></td><td></td><td></td><td></td><td></td><td></td><td></td><td></td><td></td><td></td></tr>
<tr><td>T4342</td><td>CGCCGCGAGCGTGTTCTTGGTGG</td><td>1000</td><td></td><td></td><td></td><td></td><td></td><td></td><td></td><td></td><td></td></tr>
<tr><td>Oligo fwd</td><td>CACCCGCCGCGAGCGTGTTCTTGG</td><td></td><td></td><td></td><td></td><td></td><td></td><td></td><td></td><td></td><td></td></tr>
<tr><td>Oligo rev</td><td>AAACCCAAGAACACGCTCGCGGCG</td><td></td><td></td><td></td><td></td><td></td><td></td><td></td><td></td><td></td><td></td></tr>
<tr><td>Chromosome</td><td>start</td><td>end</td><td>strand</td><td>MM</td><td>target_seq</td><td>PAM</td><td>alignment</td><td>distance</td><td>position</td><td>gene name</td><td>gene id</td></tr>
<tr><td>GU305914.1</td><td>63608</td><td>63630</td><td>+</td><td>0</td><td>CGCCGCGAGCGTGTTCTTGG</td><td>TGG</td><td>||||||||[||||||||||||]PAM</td><td>NA</td><td>Intergenic</td><td>NA</td><td>NA</td></tr>
<tr><td></td><td></td><td></td><td></td><td></td><td></td><td></td><td></td><td></td><td></td><td></td><td></td></tr>
<tr><td>T4343</td><td>CGCGAGCGTGTTCTTGGTGGCGG</td><td>1000</td><td></td><td></td><td></td><td></td><td></td><td></td><td></td><td></td><td></td></tr>
<tr><td>Oligo fwd</td><td>CACCCGCGAGCGTGTTCTTGGTGG</td><td></td><td></td><td></td><td></td><td></td><td></td><td></td><td></td><td></td><td></td></tr>
<tr><td>Oligo rev</td><td>AAACCCACCAAGAACACGCTCGCG</td><td></td><td></td><td></td><td></td><td></td><td></td><td></td><td></td><td></td><td></td></tr>
<tr><td>Chromosome</td><td>start</td><td>end</td><td>strand</td><td>MM</td><td>target_seq</td><td>PAM</td><td>alignment</td><td>distance</td><td>position</td><td>gene name</td><td>gene id</td></tr>
<tr><td>GU305914.1</td><td>63611</td><td>63633</td><td>+</td><td>0</td><td>CGCGAGCGTGTTCTTGGTGG</td><td>CGG</td><td>||||||||[||||||||||||]PAM</td><td>NA</td><td>Intergenic</td><td>NA</td><td>NA</td></tr>
<tr><td></td><td></td><td></td><td></td><td></td><td></td><td></td><td></td><td></td><td></td><td></td><td></td></tr>
<tr><td>T4344</td><td>GAGCGTGTTCTTGGTGGCGGCGG</td><td>1000</td><td></td><td></td><td></td><td></td><td></td><td></td><td></td><td></td><td></td></tr>
<tr><td>Oligo fwd</td><td>CACCGAGCGTGTTCTTGGTGGCGG</td><td></td><td></td><td></td><td></td><td></td><td></td><td></td><td></td><td></td><td></td></tr>
<tr><td>Oligo rev</td><td>AAACCCGCCACCAAGAACACGCTC</td><td></td><td></td><td></td><td></td><td></td><td></td><td></td><td></td><td></td><td></td></tr>
<tr><td>Chromosome</td><td>start</td><td>end</td><td>strand</td><td>MM</td><td>target_seq</td><td>PAM</td><td>alignment</td><td>distance</td><td>position</td><td>gene name</td><td>gene id</td></tr>
<tr><td>GU305914.1</td><td>63614</td><td>63636</td><td>+</td><td>0</td><td>GAGCGTGTTCTTGGTGGCGG</td><td>CGG</td><td>||||||||[||||||||||||]PAM</td><td>NA</td><td>Intergenic</td><td>NA</td><td>NA</td></tr>
<tr><td></td><td></td><td></td><td></td><td></td><td></td><td></td><td></td><td></td><td></td><td></td><td></td></tr>
<tr><td>T4345</td><td>GGTGGCGGCGGAGACGTTCTCGG</td><td>1000</td><td></td><td></td><td></td><td></td><td></td><td></td><td></td><td></td><td></td></tr>
<tr><td>Oligo fwd</td><td>CACCGGTGGCGGCGGAGACGTTCT</td><td></td><td></td><td></td><td></td><td></td><td></td><td></td><td></td><td></td><td></td></tr>
<tr><td>Oligo rev</td><td>AAACAGAACGTCTCCGCCGCCACC</td><td></td><td></td><td></td><td></td><td></td><td></td><td></td><td></td><td></td><td></td></tr>
<tr><td>Chromosome</td><td>start</td><td>end</td><td>strand</td><td>MM</td><td>target_seq</td><td>PAM</td><td>alignment</td><td>distance</td><td>position</td><td>gene name</td><td>gene id</td></tr>
<tr><td>GU305914.1</td><td>63626</td><td>63648</td><td>+</td><td>0</td><td>GGTGGCGGCGGAGACGTTCT</td><td>CGG</td><td>||||||||[||||||||||||]PAM</td><td>NA</td><td>Intergenic</td><td>NA</td><td>NA</td></tr>
<tr><td></td><td></td><td></td><td></td><td></td><td></td><td></td><td></td><td></td><td></td><td></td><td></td></tr>
<tr><td>T4346</td><td>CGTTCTCGGCCACCATCGAGAGG</td><td>1000</td><td></td><td></td><td></td><td></td><td></td><td></td><td></td><td></td><td></td></tr>
<tr><td>Oligo fwd</td><td>CACCCGTTCTCGGCCACCATCGAG</td><td></td><td></td><td></td><td></td><td></td><td></td><td></td><td></td><td></td><td></td></tr>
<tr><td>Oligo rev</td><td>AAACCTCGATGGTGGCCGAGAACG</td><td></td><td></td><td></td><td></td><td></td><td></td><td></td><td></td><td></td><td></td></tr>
<tr><td>Chromosome</td><td>start</td><td>end</td><td>strand</td><td>MM</td><td>target_seq</td><td>PAM</td><td>alignment</td><td>distance</td><td>position</td><td>gene name</td><td>gene id</td></tr>
<tr><td>GU305914.1</td><td>63640</td><td>63662</td><td>+</td><td>0</td><td>CGTTCTCGGCCACCATCGAG</td><td>AGG</td><td>||||||||[||||||||||||]PAM</td><td>NA</td><td>Intergenic</td><td>NA</td><td>NA</td></tr>
<tr><td></td><td></td><td></td><td></td><td></td><td></td><td></td><td></td><td></td><td></td><td></td><td></td></tr>
<tr><td>T4347</td><td>CCACCATCGAGAGGAAGAAGAGG</td><td>1000</td><td></td><td></td><td></td><td></td><td></td><td></td><td></td><td></td><td></td></tr>
<tr><td>Oligo fwd</td><td>CACCCCACCATCGAGAGGAAGAAG</td><td></td><td></td><td></td><td></td><td></td><td></td><td></td><td></td><td></td><td></td></tr>
<tr><td>Oligo rev</td><td>AAACCTTCTTCCTCTCGATGGTGG</td><td></td><td></td><td></td><td></td><td></td><td></td><td></td><td></td><td></td><td></td></tr>
<tr><td>Chromosome</td><td>start</td><td>end</td><td>strand</td><td>MM</td><td>target_seq</td><td>PAM</td><td>alignment</td><td>distance</td><td>position</td><td>gene name</td><td>gene id</td></tr>
<tr><td>GU305914.1</td><td>63649</td><td>63671</td><td>+</td><td>0</td><td>CCACCATCGAGAGGAAGAAG</td><td>AGG</td><td>||||||||[||||||||||||]PAM</td><td>NA</td><td>Intergenic</td><td>NA</td><td>NA</td></tr>
<tr><td></td><td></td><td></td><td></td><td></td><td></td><td></td><td></td><td></td><td></td><td></td><td></td></tr>
<tr><td>T4348</td><td>GAAGAAGAGGCTGACGAGCACGG</td><td>1000</td><td></td><td></td><td></td><td></td><td></td><td></td><td></td><td></td><td></td></tr>
<tr><td>Oligo fwd</td><td>CACCGAAGAAGAGGCTGACGAGCA</td><td></td><td></td><td></td><td></td><td></td><td></td><td></td><td></td><td></td><td></td></tr>
<tr><td>Oligo rev</td><td>AAACTGCTCGTCAGCCTCTTCTTC</td><td></td><td></td><td></td><td></td><td></td><td></td><td></td><td></td><td></td><td></td></tr>
<tr><td>Chromosome</td><td>start</td><td>end</td><td>strand</td><td>MM</td><td>target_seq</td><td>PAM</td><td>alignment</td><td>distance</td><td>position</td><td>gene name</td><td>gene id</td></tr>
<tr><td>GU305914.1</td><td>63662</td><td>63684</td><td>+</td><td>0</td><td>GAAGAAGAGGCTGACGAGCA</td><td>CGG</td><td>||||||||[||||||||||||]PAM</td><td>NA</td><td>Intergenic</td><td>NA</td><td>NA</td></tr>
<tr><td></td><td></td><td></td><td></td><td></td><td></td><td></td><td></td><td></td><td></td><td></td><td></td></tr>
<tr><td>T4349</td><td>GAGGCTGACGAGCACGGTGCTGG</td><td>1000</td><td></td><td></td><td></td><td></td><td></td><td></td><td></td><td></td><td></td></tr>
<tr><td>Oligo fwd</td><td>CACCGAGGCTGACGAGCACGGTGC</td><td></td><td></td><td></td><td></td><td></td><td></td><td></td><td></td><td></td><td></td></tr>
<tr><td>Oligo rev</td><td>AAACGCACCGTGCTCGTCAGCCTC</td><td></td><td></td><td></td><td></td><td></td><td></td><td></td><td></td><td></td><td></td></tr>
<tr><td>Chromosome</td><td>start</td><td>end</td><td>strand</td><td>MM</td><td>target_seq</td><td>PAM</td><td>alignment</td><td>distance</td><td>position</td><td>gene name</td><td>gene id</td></tr>
<tr><td>GU305914.1</td><td>63668</td><td>63690</td><td>+</td><td>0</td><td>GAGGCTGACGAGCACGGTGC</td><td>TGG</td><td>||||||||[||||||||||||]PAM</td><td>NA</td><td>Intergenic</td><td>NA</td><td>NA</td></tr>
<tr><td></td><td></td><td></td><td></td><td></td><td></td><td></td><td></td><td></td><td></td><td></td><td></td></tr>
<tr><td>T4350</td><td>AGGCTGACGAGCACGGTGCTGGG</td><td>1000</td><td></td><td></td><td></td><td></td><td></td><td></td><td></td><td></td><td></td></tr>
<tr><td>Oligo fwd</td><td>CACCAGGCTGACGAGCACGGTGCT</td><td></td><td></td><td></td><td></td><td></td><td></td><td></td><td></td><td></td><td></td></tr>
<tr><td>Oligo rev</td><td>AAACAGCACCGTGCTCGTCAGCCT</td><td></td><td></td><td></td><td></td><td></td><td></td><td></td><td></td><td></td><td></td></tr>
<tr><td>Chromosome</td><td>start</td><td>end</td><td>strand</td><td>MM</td><td>target_seq</td><td>PAM</td><td>alignment</td><td>distance</td><td>position</td><td>gene name</td><td>gene id</td></tr>
<tr><td>GU305914.1</td><td>63669</td><td>63691</td><td>+</td><td>0</td><td>AGGCTGACGAGCACGGTGCT</td><td>GGG</td><td>||||||||[||||||||||||]PAM</td><td>NA</td><td>Intergenic</td><td>NA</td><td>NA</td></tr>
<tr><td></td><td></td><td></td><td></td><td></td><td></td><td></td><td></td><td></td><td></td><td></td><td></td></tr>
<tr><td>T4351</td><td>CCGTGTCGAGCCTGAACATCAGG</td><td>1000</td><td></td><td></td><td></td><td></td><td></td><td></td><td></td><td></td><td></td></tr>
<tr><td>Oligo fwd</td><td>CACCCCGTGTCGAGCCTGAACATC</td><td></td><td></td><td></td><td></td><td></td><td></td><td></td><td></td><td></td><td></td></tr>
<tr><td>Oligo rev</td><td>AAACGATGTTCAGGCTCGACACGG</td><td></td><td></td><td></td><td></td><td></td><td></td><td></td><td></td><td></td><td></td></tr>
<tr><td>Chromosome</td><td>start</td><td>end</td><td>strand</td><td>MM</td><td>target_seq</td><td>PAM</td><td>alignment</td><td>distance</td><td>position</td><td>gene name</td><td>gene id</td></tr>
<tr><td>GU305914.1</td><td>63703</td><td>63725</td><td>+</td><td>0</td><td>CCGTGTCGAGCCTGAACATC</td><td>AGG</td><td>||||||||[||||||||||||]PAM</td><td>NA</td><td>Intergenic</td><td>NA</td><td>NA</td></tr>
<tr><td></td><td></td><td></td><td></td><td></td><td></td><td></td><td></td><td></td><td></td><td></td><td></td></tr>
<tr><td>T4352</td><td>GCCTGAACATCAGGACCCCCAGG</td><td>1000</td><td></td><td></td><td></td><td></td><td></td><td></td><td></td><td></td><td></td></tr>
<tr><td>Oligo fwd</td><td>CACCGCCTGAACATCAGGACCCCC</td><td></td><td></td><td></td><td></td><td></td><td></td><td></td><td></td><td></td><td></td></tr>
<tr><td>Oligo rev</td><td>AAACGGGGGTCCTGATGTTCAGGC</td><td></td><td></td><td></td><td></td><td></td><td></td><td></td><td></td><td></td><td></td></tr>
<tr><td>Chromosome</td><td>start</td><td>end</td><td>strand</td><td>MM</td><td>target_seq</td><td>PAM</td><td>alignment</td><td>distance</td><td>position</td><td>gene name</td><td>gene id</td></tr>
<tr><td>GU305914.1</td><td>63712</td><td>63734</td><td>+</td><td>0</td><td>GCCTGAACATCAGGACCCCC</td><td>AGG</td><td>||||||||[||||||||||||]PAM</td><td>NA</td><td>Intergenic</td><td>NA</td><td>NA</td></tr>
<tr><td></td><td></td><td></td><td></td><td></td><td></td><td></td><td></td><td></td><td></td><td></td><td></td></tr>
<tr><td>T4353</td><td>GACTCTCCCGCACGAAGAAGCGG</td><td>1000</td><td></td><td></td><td></td><td></td><td></td><td></td><td></td><td></td><td></td></tr>
<tr><td>Oligo fwd</td><td>CACCGACTCTCCCGCACGAAGAAG</td><td></td><td></td><td></td><td></td><td></td><td></td><td></td><td></td><td></td><td></td></tr>
<tr><td>Oligo rev</td><td>AAACCTTCTTCGTGCGGGAGAGTC</td><td></td><td></td><td></td><td></td><td></td><td></td><td></td><td></td><td></td><td></td></tr>
<tr><td>Chromosome</td><td>start</td><td>end</td><td>strand</td><td>MM</td><td>target_seq</td><td>PAM</td><td>alignment</td><td>distance</td><td>position</td><td>gene name</td><td>gene id</td></tr>
<tr><td>GU305914.1</td><td>63757</td><td>63779</td><td>+</td><td>0</td><td>GACTCTCCCGCACGAAGAAG</td><td>CGG</td><td>||||||||[||||||||||||]PAM</td><td>NA</td><td>Intergenic</td><td>NA</td><td>NA</td></tr>
<tr><td></td><td></td><td></td><td></td><td></td><td></td><td></td><td></td><td></td><td></td><td></td><td></td></tr>
<tr><td>T4354</td><td>ACTCTCCCGCACGAAGAAGCGGG</td><td>1000</td><td></td><td></td><td></td><td></td><td></td><td></td><td></td><td></td><td></td></tr>
<tr><td>Oligo fwd</td><td>CACCACTCTCCCGCACGAAGAAGC</td><td></td><td></td><td></td><td></td><td></td><td></td><td></td><td></td><td></td><td></td></tr>
<tr><td>Oligo rev</td><td>AAACGCTTCTTCGTGCGGGAGAGT</td><td></td><td></td><td></td><td></td><td></td><td></td><td></td><td></td><td></td><td></td></tr>
<tr><td>Chromosome</td><td>start</td><td>end</td><td>strand</td><td>MM</td><td>target_seq</td><td>PAM</td><td>alignment</td><td>distance</td><td>position</td><td>gene name</td><td>gene id</td></tr>
<tr><td>GU305914.1</td><td>63758</td><td>63780</td><td>+</td><td>0</td><td>ACTCTCCCGCACGAAGAAGC</td><td>GGG</td><td>||||||||[||||||||||||]PAM</td><td>NA</td><td>Intergenic</td><td>NA</td><td>NA</td></tr>
<tr><td></td><td></td><td></td><td></td><td></td><td></td><td></td><td></td><td></td><td></td><td></td><td></td></tr>
<tr><td>T4355</td><td>AGAAGCGGGCCGTGATCGCCTGG</td><td>1000</td><td></td><td></td><td></td><td></td><td></td><td></td><td></td><td></td><td></td></tr>
</table>

Oligo fwd CACCAGAAGCGGGCCGTGATCGCC
Oligo rev AAACGGCGATCACGGCCCGCTTCT
Chromosome start end strand MM target_seq PAM alignment distance position gene name gene id
GU305914.1 63772 63794 + 0 AGAAGCGGGCCGTGATCGCC TGG ||||||||[||||||||||||]PAM NA Intergenic NA NA

T4356 AGCGGGCCGTGATCGCCTGGTGG 1000
Oligo fwd CACCAGCGGGCCGTGATCGCCTGG
Oligo rev AAACCCAGGCGATCACGGCCCGCT
Chromosome start end strand MM target_seq PAM alignment distance position gene name gene id
GU305914.1 63775 63797 + 0 AGCGGGCCGTGATCGCCTGG TGG ||||||||[||||||||||||]PAM NA Intergenic NA NA

T4357 GGCCGTGATCGCCTGGTGGTCGG 1000
Oligo fwd CACCGGCCGTGATCGCCTGGTGGT
Oligo rev AAACACCACCAGGCGATCACGGCC
Chromosome start end strand MM target_seq PAM alignment distance position gene name gene id
GU305914.1 63779 63801 + 0 GGCCGTGATCGCCTGGTGGT CGG ||||||||[||||||||||||]PAM NA Intergenic NA NA

T4358 GGCGCCGTAGACATAGCCCTTGG 1000
Oligo fwd CACCGGCGCCGTAGACATAGCCCT
Oligo rev AAACAGGGCTATGTCTACGGCGCC
Chromosome start end strand MM target_seq PAM alignment distance position gene name gene id
GU305914.1 63800 63822 + 0 GGCGCCGTAGACATAGCCCT TGG ||||||||[||||||||||||]PAM NA Intergenic NA NA

T4359 CCTCATCTTAGACACCTTGTAGG 1000
Oligo fwd CACCCCTCATCTTAGACACCTTGT
Oligo rev AAACACAAGGTGTCTAAGATGAGG
Chromosome start end strand MM target_seq PAM alignment distance position gene name gene id
GU305914.1 63842 63864 + 0 CCTCATCTTAGACACCTTGT AGG ||||||||[||||||||||||]PAM NA Intergenic NA NA

T4360 CTCATCTTAGACACCTTGTAGGG 1000
Oligo fwd CACCCTCATCTTAGACACCTTGTA
Oligo rev AAACTACAAGGTGTCTAAGATGAG
Chromosome start end strand MM target_seq PAM alignment distance position gene name gene id
GU305914.1 63843 63865 + 0 CTCATCTTAGACACCTTGTA GGG ||||||||[||||||||||||]PAM NA Intergenic NA NA

T4361 TCATCTTAGACACCTTGTAGGGG 1000
Oligo fwd CACCTCATCTTAGACACCTTGTAG
Oligo rev AAACCTACAAGGTGTCTAAGATGA
Chromosome start end strand MM target_seq PAM alignment distance position gene name gene id
GU305914.1 63844 63866 + 0 TCATCTTAGACACCTTGTAG GGG ||||||||[||||||||||||]PAM NA Intergenic NA NA

T4362 AGACACCTTGTAGGGGTCCTCGG 1000
Oligo fwd CACCAGACACCTTGTAGGGGTCCT
Oligo rev AAACAGGACCCCTACAAGGTGTCT
Chromosome start end strand MM target_seq PAM alignment distance position gene name gene id
GU305914.1 63851 63873 + 0 AGACACCTTGTAGGGGTCCT CGG ||||||||[||||||||||||]PAM NA Intergenic NA NA

T4363 GACACCTTGTAGGGGTCCTCGGG 1000
Oligo fwd CACCGACACCTTGTAGGGGTCCTC
Oligo rev AAACGAGGACCCCTACAAGGTGTC
Chromosome start end strand MM target_seq PAM alignment distance position gene name gene id
GU305914.1 63852 63874 + 0 GACACCTTGTAGGGGTCCTC GGG ||||||||[||||||||||||]PAM NA Intergenic NA NA

T4364 ACACCTTGTAGGGGTCCTCGGGG 1000
Oligo fwd CACCACACCTTGTAGGGGTCCTCG
Oligo rev AAACCGAGGACCCCTACAAGGTGT
Chromosome start end strand MM target_seq PAM alignment distance position gene name gene id
GU305914.1 63853 63875 + 0 ACACCTTGTAGGGGTCCTCG GGG ||||||||[||||||||||||]PAM NA Intergenic NA NA

T4365 TCCATCACCTCCCGCTTCATCGG 1000
Oligo fwd CACCTCCATCACCTCCCGCTTCAT
Oligo rev AAACATGAAGCGGGAGGTGATGGA
Chromosome start end strand MM target_seq PAM alignment distance position gene name gene id
GU305914.1 63882 63904 + 0 TCCATCACCTCCCGCTTCAT CGG ||||||||[||||||||||||]PAM NA Intergenic NA NA

T4366 CCATCACCTCCCGCTTCATCGGG 1000
Oligo fwd CACCCCATCACCTCCCGCTTCATC
Oligo rev AAACGATGAAGCGGGAGGTGATGG
Chromosome start end strand MM target_seq PAM alignment distance position gene name gene id
GU305914.1 63883 63905 + 0 CCATCACCTCCCGCTTCATC GGG ||||||||[||||||||||||]PAM NA Intergenic NA NA

T4367 GAAGCTGCTGACGTTCGCCATGG 1000
Oligo fwd CACCGAAGCTGCTGACGTTCGCCA
Oligo rev AAACTGGCGAACGTCAGCAGCTTC
Chromosome start end strand MM target_seq PAM alignment distance position gene name gene id
GU305914.1 63908 63930 + 0 GAAGCTGCTGACGTTCGCCA TGG ||||||||[||||||||||||]PAM NA Intergenic NA NA

T4368 CGCCATGGCCGCGTTCGCGAAGG 1000
Oligo fwd CACCCGCCATGGCCGCGTTCGCGA
Oligo rev AAACTCGCGAACGCGGCCATGGCG
Chromosome start end strand MM target_seq PAM alignment distance position gene name gene id
GU305914.1 63923 63945 + 0 CGCCATGGCCGCGTTCGCGA AGG ||||||||[||||||||||||]PAM NA Intergenic NA NA

T4369 CGTCACCCGAGCCCTCAACCTGG 1000
Oligo fwd CACCCGTCACCCGAGCCCTCAACC
Oligo rev AAACGGTTGAGGGCTCGGGTGACG
Chromosome start end strand MM target_seq PAM alignment distance position gene name gene id
GU305914.1 63959 63981 + 0 CGTCACCCGAGCCCTCAACC TGG ||||||||[||||||||||||]PAM NA Intergenic NA NA

| | | | | | | | | | | | |
|---|---|---|---|---|---|---|---|---|---|---|---|
| T4370 | CCTCAACCTGGAACAGACGCTGG | 1000 | | | | | | | | | |
| Oligo fwd | CACCCCTCAACCTGGAACAGACGC | | | | | | | | | | |
| Oligo rev | AAACGCGTCTGTTCCAGGTTGAGG | | | | | | | | | | |
| Chromosome | start | end | strand | MM | target_seq | PAM | alignment | distance | position | gene name | gene id |
| GU305914.1 | 63971 | 63993 | + | 0 | CCTCAACCTGGAACAGACGC | TGG | \|\|\|\|\|\|\|\|[\|\|\|\|\|\|\|\|\|\|\|\|]PAM | NA | Intergenic | NA | NA |

| | | | | | | | | | | | |
|---|---|---|---|---|---|---|---|---|---|---|---|
| T4371 | AACAGACGCTGGAAGATCTGCGG | 1000 | | | | | | | | | |
| Oligo fwd | CACCAACAGACGCTGGAAGATCTG | | | | | | | | | | |
| Oligo rev | AAACCAGATCTTCCAGCGTCTGTT | | | | | | | | | | |
| Chromosome | start | end | strand | MM | target_seq | PAM | alignment | distance | position | gene name | gene id |
| GU305914.1 | 63982 | 64004 | + | 0 | AACAGACGCTGGAAGATCTG | CGG | \|\|\|\|\|\|\|\|[\|\|\|\|\|\|\|\|\|\|\|\|]PAM | NA | Intergenic | NA | NA |

| | | | | | | | | | | | |
|---|---|---|---|---|---|---|---|---|---|---|---|
| T4372 | GGAAGATCTGCGGATGAAAACGG | 1000 | | | | | | | | | |
| Oligo fwd | CACCGGAAGATCTGCGGATGAAAA | | | | | | | | | | |
| Oligo rev | AAACTTTTCATCCGCAGATCTTCC | | | | | | | | | | |
| Chromosome | start | end | strand | MM | target_seq | PAM | alignment | distance | position | gene name | gene id |
| GU305914.1 | 63992 | 64014 | + | 0 | GGAAGATCTGCGGATGAAAA | CGG | \|\|\|\|\|\|\|\|[\|\|\|\|\|\|\|\|\|\|\|\|]PAM | NA | Intergenic | NA | NA |

| | | | | | | | | | | | |
|---|---|---|---|---|---|---|---|---|---|---|---|
| T4373 | GATGAAAACGGCGTGTGACGAGG | 1000 | | | | | | | | | |
| Oligo fwd | CACCGATGAAAACGGCGTGTGACG | | | | | | | | | | |
| Oligo rev | AAACCGTCACACGCCGTTTTCATC | | | | | | | | | | |
| Chromosome | start | end | strand | MM | target_seq | PAM | alignment | distance | position | gene name | gene id |
| GU305914.1 | 64004 | 64026 | + | 0 | GATGAAAACGGCGTGTGACG | AGG | \|\|\|\|\|\|\|\|[\|\|\|\|\|\|\|\|\|\|\|\|]PAM | NA | Intergenic | NA | NA |

| | | | | | | | | | | | |
|---|---|---|---|---|---|---|---|---|---|---|---|
| T4374 | CGAGGATCTACTTAATATTCTGG | 1000 | | | | | | | | | |
| Oligo fwd | CACCCGAGGATCTACTTAATATTC | | | | | | | | | | |
| Oligo rev | AAACGAATATTAAGTAGATCCTCG | | | | | | | | | | |
| Chromosome | start | end | strand | MM | target_seq | PAM | alignment | distance | position | gene name | gene id |
| GU305914.1 | 64022 | 64044 | + | 0 | CGAGGATCTACTTAATATTC | TGG | \|\|\|\|\|\|\|\|[\|\|\|\|\|\|\|\|\|\|\|\|]PAM | NA | Intergenic | NA | NA |

| | | | | | | | | | | | |
|---|---|---|---|---|---|---|---|---|---|---|---|
| T4375 | CTCCGCCGTGCAACTGCAGACGG | 1000 | | | | | | | | | |
| Oligo fwd | CACCCTCCGCCGTGCAACTGCAGA | | | | | | | | | | |
| Oligo rev | AAACTCTGCAGTTGCACGGCGGAG | | | | | | | | | | |
| Chromosome | start | end | strand | MM | target_seq | PAM | alignment | distance | position | gene name | gene id |
| GU305914.1 | 64058 | 64080 | + | 0 | CTCCGCCGTGCAACTGCAGA | CGG | \|\|\|\|\|\|\|\|[\|\|\|\|\|\|\|\|\|\|\|\|]PAM | NA | Intergenic | NA | NA |

| | | | | | | | | | | | |
|---|---|---|---|---|---|---|---|---|---|---|---|
| T4376 | CGTGCAACTGCAGACGGTGACGG | 1000 | | | | | | | | | |
| Oligo fwd | CACCCGTGCAACTGCAGACGGTGA | | | | | | | | | | |
| Oligo rev | AAACTCACCGTCTGCAGTTGCACG | | | | | | | | | | |
| Chromosome | start | end | strand | MM | target_seq | PAM | alignment | distance | position | gene name | gene id |
| GU305914.1 | 64064 | 64086 | + | 0 | CGTGCAACTGCAGACGGTGA | CGG | \|\|\|\|\|\|\|\|[\|\|\|\|\|\|\|\|\|\|\|\|]PAM | NA | Intergenic | NA | NA |

| | | | | | | | | | | | |
|---|---|---|---|---|---|---|---|---|---|---|---|
| T4377 | CCTGCAGCACGTCCCCGAGACGG | 1000 | | | | | | | | | |
| Oligo fwd | CACCCCTGCAGCACGTCCCCGAGA | | | | | | | | | | |
| Oligo rev | AAACTCTCGGGGACGTGCTGCAGG | | | | | | | | | | |
| Chromosome | start | end | strand | MM | target_seq | PAM | alignment | distance | position | gene name | gene id |
| GU305914.1 | 64106 | 64128 | + | 0 | CCTGCAGCACGTCCCCGAGA | CGG | \|\|\|\|\|\|\|\|[\|\|\|\|\|\|\|\|\|\|\|\|]PAM | NA | Intergenic | NA | NA |

| | | | | | | | | | | | |
|---|---|---|---|---|---|---|---|---|---|---|---|
| T4378 | CTGCAGCACGTCCCCGAGACGGG | 1000 | | | | | | | | | |
| Oligo fwd | CACCCTGCAGCACGTCCCCGAGAC | | | | | | | | | | |
| Oligo rev | AAACGTCTCGGGGACGTGCTGCAG | | | | | | | | | | |
| Chromosome | start | end | strand | MM | target_seq | PAM | alignment | distance | position | gene name | gene id |
| GU305914.1 | 64107 | 64129 | + | 0 | CTGCAGCACGTCCCCGAGAC | GGG | \|\|\|\|\|\|\|\|[\|\|\|\|\|\|\|\|\|\|\|\|]PAM | NA | Intergenic | NA | NA |

| | | | | | | | | | | | |
|---|---|---|---|---|---|---|---|---|---|---|---|
| T4379 | CGAGTTCATCCGCAGACACCCGG | 1000 | | | | | | | | | |
| Oligo fwd | CACCCGAGTTCATCCGCAGACACC | | | | | | | | | | |
| Oligo rev | AAACGGTGTCTGCGGATGAACTCG | | | | | | | | | | |
| Chromosome | start | end | strand | MM | target_seq | PAM | alignment | distance | position | gene name | gene id |
| GU305914.1 | 64139 | 64161 | + | 0 | CGAGTTCATCCGCAGACACC | CGG | \|\|\|\|\|\|\|\|[\|\|\|\|\|\|\|\|\|\|\|\|]PAM | NA | Intergenic | NA | NA |

| | | | | | | | | | | | |
|---|---|---|---|---|---|---|---|---|---|---|---|
| T4380 | GTGTTCTACTTTCTGAACCACGG | 1000 | | | | | | | | | |
| Oligo fwd | CACCGTGTTCTACTTTCTGAACCA | | | | | | | | | | |
| Oligo rev | AAACTGGTTCAGAAAGTAGAACAC | | | | | | | | | | |
| Chromosome | start | end | strand | MM | target_seq | PAM | alignment | distance | position | gene name | gene id |
| GU305914.1 | 64161 | 64183 | + | 0 | GTGTTCTACTTTCTGAACCA | CGG | \|\|\|\|\|\|\|\|[\|\|\|\|\|\|\|\|\|\|\|\|]PAM | NA | Intergenic | NA | NA |

| | | | | | | | | | | | |
|---|---|---|---|---|---|---|---|---|---|---|---|
| T4381 | TTTCTGAACCACGGCACCTTCGG | 1000 | | | | | | | | | |
| Oligo fwd | CACCTTTCTGAACCACGGCACCTT | | | | | | | | | | |
| Oligo rev | AAACAAGGTGCCGTGGTTCAGAAA | | | | | | | | | | |
| Chromosome | start | end | strand | MM | target_seq | PAM | alignment | distance | position | gene name | gene id |
| GU305914.1 | 64170 | 64192 | + | 0 | TTTCTGAACCACGGCACCTT | CGG | \|\|\|\|\|\|\|\|[\|\|\|\|\|\|\|\|\|\|\|\|]PAM | NA | Intergenic | NA | NA |

| | | | | | | | | | | | |
|---|---|---|---|---|---|---|---|---|---|---|---|
| T4382 | TTCTGAACCACGGCACCTTCGGG | 1000 | | | | | | | | | |
| Oligo fwd | CACCTTCTGAACCACGGCACCTTC | | | | | | | | | | |
| Oligo rev | AAACGAAGGTGCCGTGGTTCAGAA | | | | | | | | | | |
| Chromosome | start | end | strand | MM | target_seq | PAM | alignment | distance | position | gene name | gene id |
| GU305914.1 | 64171 | 64193 | + | 0 | TTCTGAACCACGGCACCTTC | GGG | \|\|\|\|\|\|\|\|[\|\|\|\|\|\|\|\|\|\|\|\|]PAM | NA | Intergenic | NA | NA |

| | | | | | | | | | | | |
|---|---|---|---|---|---|---|---|---|---|---|---|
| T4383 | GAACCACGGCACCTTCGGGCCGG | 1000 | | | | | | | | | |
| Oligo fwd | CACCGAACCACGGCACCTTCGGGC | | | | | | | | | | |
| Oligo rev | AAACGCCCGAAGGTGCCGTGGTTC | | | | | | | | | | |
| Chromosome | start | end | strand | MM | target_seq | PAM | alignment | distance | position | gene name | gene id |
| GU305914.1 | 64175 | 64197 | + | 0 | GAACCACGGCACCTTCGGGC | CGG | \|\|\|\|\|\|\|\|[\|\|\|\|\|\|\|\|\|\|\|\|]PAM | NA | Intergenic | NA | NA |

| | | | | | | | | | | | |
|---|---|---|---|---|---|---|---|---|---|---|---|
| T4384 | ACCTTCGGGCCGGCGCAGAAAGG | 1000 | | | | | | | | | |
| Oligo fwd | CACCACCTTCGGGCCGGCGCAGAA | | | | | | | | | | |
| Oligo rev | AAACTTCTGCGCCGGCCCGAAGGT | | | | | | | | | | |
| Chromosome | start | end | strand | MM | target_seq | PAM | alignment | distance | position | gene name | gene id |
| GU305914.1 | 64185 | 64207 | + | 0 | ACCTTCGGGCCGGCGCAGAA | AGG | \|\|\|\|\|\|\|\|[\|\|\|\|\|\|\|\|\|\|\|\|]PAM | NA | Intergenic | NA | NA |

| | | | | | | | | | | | |
|---|---|---|---|---|---|---|---|---|---|---|---|
| T4385 | CCTTCGGGCCGGCGCAGAAAGGG | 1000 | | | | | | | | | |
| Oligo fwd | CACCCCTTCGGGCCGGCGCAGAAA | | | | | | | | | | |
| Oligo rev | AAACTTTCTGCGCCGGCCCGAAGG | | | | | | | | | | |
| Chromosome | start | end | strand | MM | target_seq | PAM | alignment | distance | position | gene name | gene id |
| GU305914.1 | 64186 | 64208 | + | 0 | CCTTCGGGCCGGCGCAGAAA | GGG | \|\|\|\|\|\|\|\|[\|\|\|\|\|\|\|\|\|\|\|\|]PAM | NA | Intergenic | NA | NA |

| | | | | | | | | | | | |
|---|---|---|---|---|---|---|---|---|---|---|---|
| T4386 | GAAAGGGCTTCTCCTCCCGCTGG | 1000 | | | | | | | | | |
| Oligo fwd | CACCGAAAGGGCTTCTCCTCCCGC | | | | | | | | | | |
| Oligo rev | AAACGCGGGAGGAGAAGCCCTTTC | | | | | | | | | | |
| Chromosome | start | end | strand | MM | target_seq | PAM | alignment | distance | position | gene name | gene id |
| GU305914.1 | 64202 | 64224 | + | 0 | GAAAGGGCTTCTCCTCCCGC | TGG | \|\|\|\|\|\|\|\|[\|\|\|\|\|\|\|\|\|\|\|\|]PAM | NA | Intergenic | NA | NA |

| | | | | | | | | | | | |
|---|---|---|---|---|---|---|---|---|---|---|---|
| T4387 | CCTCCCGCTGGCCGCCGACCTGG | 1000 | | | | | | | | | |
| Oligo fwd | CACCCCTCCCGCTGGCCGCCGACC | | | | | | | | | | |
| Oligo rev | AAACGGTCGGCGGCCAGCGGGAGG | | | | | | | | | | |
| Chromosome | start | end | strand | MM | target_seq | PAM | alignment | distance | position | gene name | gene id |
| GU305914.1 | 64214 | 64236 | + | 0 | CCTCCCGCTGGCCGCCGACC | TGG | \|\|\|\|\|\|\|\|[\|\|\|\|\|\|\|\|\|\|\|\|]PAM | NA | Intergenic | NA | NA |
| GU305914.1 | 56929 | 56951 | + | 3 | CCTCCCACGGGCCTCCGACC | TGG | \|\|\|\|\|\|-\|[-\|\|\|\|-\|\|\|\|\|\|]PAM | NA | Intergenic | NA | NA |

| | | | | | | | | | | | |
|---|---|---|---|---|---|---|---|---|---|---|---|
| T4388 | GCTGGCCGCCGACCTGGTCTCGG | 1000 | | | | | | | | | |
| Oligo fwd | CACCGCTGGCCGCCGACCTGGTCT | | | | | | | | | | |
| Oligo rev | AAACAGACCAGGTCGGCGGCCAGC | | | | | | | | | | |
| Chromosome | start | end | strand | MM | target_seq | PAM | alignment | distance | position | gene name | gene id |
| GU305914.1 | 64220 | 64242 | + | 0 | GCTGGCCGCCGACCTGGTCT | CGG | \|\|\|\|\|\|\|\|[\|\|\|\|\|\|\|\|\|\|\|\|]PAM | NA | Intergenic | NA | NA |

| | | | | | | | | | | | |
|---|---|---|---|---|---|---|---|---|---|---|---|
| T4389 | CAAGTATACCGAGAAGTCGCAGG | 1000 | | | | | | | | | |
| Oligo fwd | CACCCAAGTATACCGAGAAGTCGC | | | | | | | | | | |
| Oligo rev | AAACGCGACTTCTCGGTATACTTG | | | | | | | | | | |
| Chromosome | start | end | strand | MM | target_seq | PAM | alignment | distance | position | gene name | gene id |
| GU305914.1 | 64250 | 64272 | + | 0 | CAAGTATACCGAGAAGTCGC | AGG | \|\|\|\|\|\|\|\|[\|\|\|\|\|\|\|\|\|\|\|\|]PAM | NA | Intergenic | NA | NA |

| | | | | | | | | | | | |
|---|---|---|---|---|---|---|---|---|---|---|---|
| T4390 | GTATACCGAGAAGTCGCAGGCGG | 1000 | | | | | | | | | |
| Oligo fwd | CACCGTATACCGAGAAGTCGCAGG | | | | | | | | | | |
| Oligo rev | AAACCCTGCGACTTCTCGGTATAC | | | | | | | | | | |
| Chromosome | start | end | strand | MM | target_seq | PAM | alignment | distance | position | gene name | gene id |
| GU305914.1 | 64253 | 64275 | + | 0 | GTATACCGAGAAGTCGCAGG | CGG | \|\|\|\|\|\|\|\|[\|\|\|\|\|\|\|\|\|\|\|\|]PAM | NA | Intergenic | NA | NA |

| | | | | | | | | | | | |
|---|---|---|---|---|---|---|---|---|---|---|---|
| T4391 | TACCGAGAAGTCGCAGGCGGCGG | 1000 | | | | | | | | | |
| Oligo fwd | CACCTACCGAGAAGTCGCAGGCGG | | | | | | | | | | |
| Oligo rev | AAACCCGCCTGCGACTTCTCGGTA | | | | | | | | | | |
| Chromosome | start | end | strand | MM | target_seq | PAM | alignment | distance | position | gene name | gene id |
| GU305914.1 | 64256 | 64278 | + | 0 | TACCGAGAAGTCGCAGGCGG | CGG | \|\|\|\|\|\|\|\|[\|\|\|\|\|\|\|\|\|\|\|\|]PAM | NA | Intergenic | NA | NA |

| | | | | | | | | | | | |
|---|---|---|---|---|---|---|---|---|---|---|---|
| T4392 | ACCGAGAAGTCGCAGGCGGCGGG | 1000 | | | | | | | | | |
| Oligo fwd | CACCACCGAGAAGTCGCAGGCGGC | | | | | | | | | | |
| Oligo rev | AAACGCCGCCTGCGACTTCTCGGT | | | | | | | | | | |
| Chromosome | start | end | strand | MM | target_seq | PAM | alignment | distance | position | gene name | gene id |
| GU305914.1 | 64257 | 64279 | + | 0 | ACCGAGAAGTCGCAGGCGGC | GGG | \|\|\|\|\|\|\|\|[\|\|\|\|\|\|\|\|\|\|\|\|]PAM | NA | Intergenic | NA | NA |

| | | | | | | | | | | | |
|---|---|---|---|---|---|---|---|---|---|---|---|
| T4393 | GAGAAGTCGCAGGCGGCGGGCGG | 1000 | | | | | | | | | |
| Oligo fwd | CACCGAGAAGTCGCAGGCGGCGGG | | | | | | | | | | |
| Oligo rev | AAACCCCGCCGCCTGCGACTTCTC | | | | | | | | | | |
| Chromosome | start | end | strand | MM | target_seq | PAM | alignment | distance | position | gene name | gene id |
| GU305914.1 | 64260 | 64282 | + | 0 | GAGAAGTCGCAGGCGGCGGG | CGG | \|\|\|\|\|\|\|\|[\|\|\|\|\|\|\|\|\|\|\|\|]PAM | NA | Intergenic | NA | NA |

| | | | | | | | | | | | |
|---|---|---|---|---|---|---|---|---|---|---|---|
| T4394 | AAGTCGCAGGCGGCGGGCGGAGG | 1000 | | | | | | | | | |
| Oligo fwd | CACCAAGTCGCAGGCGGCGGGCGG | | | | | | | | | | |
| Oligo rev | AAACCCGCCCGCCGCCTGCGACTT | | | | | | | | | | |
| Chromosome | start | end | strand | MM | target_seq | PAM | alignment | distance | position | gene name | gene id |
| GU305914.1 | 64263 | 64285 | + | 0 | AAGTCGCAGGCGGCGGGCGG | AGG | \|\|\|\|\|\|\|\|[\|\|\|\|\|\|\|\|\|\|\|\|]PAM | NA | Intergenic | NA | NA |

| | | | | | | | | | | | |
|---|---|---|---|---|---|---|---|---|---|---|---|
| T4395 | GCGGCGGGCGGAGGCAGTGTCGG | 1000 | | | | | | | | | |
| Oligo fwd | CACCGCGGCGGGCGGAGGCAGTGT | | | | | | | | | | |
| Oligo rev | AAACACACTGCCTCCGCCCGCCGC | | | | | | | | | | |
| Chromosome | start | end | strand | MM | target_seq | PAM | alignment | distance | position | gene name | gene id |
| GU305914.1 | 64272 | 64294 | + | 0 | GCGGCGGGCGGAGGCAGTGT | CGG | \|\|\|\|\|\|\|\|[\|\|\|\|\|\|\|\|\|\|\|\|]PAM | NA | Intergenic | NA | NA |

| | | | | | | | | | | | |
|---|---|---|---|---|---|---|---|---|---|---|---|
| T4396 | GCGGGCGGAGGCAGTGTCGGCGG | 1000 | | | | | | | | | |
| Oligo fwd | CACCGCGGGCGGAGGCAGTGTCGG | | | | | | | | | | |
| Oligo rev | AAACCCGACACTGCCTCCGCCCGC | | | | | | | | | | |
| Chromosome | start | end | strand | MM | target_seq | PAM | alignment | distance | position | gene name | gene id |
| GU305914.1 | 64275 | 64297 | + | 0 | GCGGGCGGAGGCAGTGTCGG | CGG | \|\|\|\|\|\|\|\|[\|\|\|\|\|\|\|\|\|\|\|\|]PAM | NA | Intergenic | NA | NA |

| | | | | | | | | | | | |
|---|---|---|---|---|---|---|---|---|---|---|---|
| T4397 | CGGAGGCAGTGTCGGCGGCGAGG | 1000 | | | | | | | | | |
| Oligo fwd | CACCCGGAGGCAGTGTCGGCGGCG | | | | | | | | | | |
| Oligo rev | AAACCGCCGCCGACACTGCCTCCG | | | | | | | | | | |
| Chromosome | start | end | strand | MM | target_seq | PAM | alignment | distance | position | gene name | gene id |
| GU305914.1 | 64280 | 64302 | + | 0 | CGGAGGCAGTGTCGGCGGCG | AGG | \|\|\|\|\|\|\|\|[\|\|\|\|\|\|\|\|\|\|\|\|]PAM | NA | Intergenic | NA | NA |

| | | | | | | | | | | | |
|---|---|---|---|---|---|---|---|---|---|---|---|
| T4398 | CCTCGAAACTCTCCAACGAGCGG | 1000 | | | | | | | | | |
| Oligo fwd | CACCCCTCGAAACTCTCCAACGAG | | | | | | | | | | |
| Oligo rev | AAACCTCGTTGGAGAGTTTCGAGG | | | | | | | | | | |
| Chromosome | start | end | strand | MM | target_seq | PAM | alignment | distance | position | gene name | gene id |
| GU305914.1 | 64309 | 64331 | + | 0 | CCTCGAAACTCTCCAACGAG | CGG | \|\|\|\|\|\|\|\|[\|\|\|\|\|\|\|\|\|\|\|\|]PAM | NA | Intergenic | NA | NA |

| | | |
|---|---|---|
| T4399 | CTCGAAACTCTCCAACGAGCGGG | 1000 |
| Oligo fwd | CACCCTCGAAACTCTCCAACGAGC | |
| Oligo rev | AAACGCTCGTTGGAGAGTTTCGAG | |

| Chromosome | start | end | strand | MM | target_seq | PAM | alignment | distance | position | gene name | gene id |
|---|---|---|---|---|---|---|---|---|---|---|---|
| GU305914.1 | 64310 | 64332 | + | 0 | CTCGAAACTCTCCAACGAGC | GGG | \|\|\|\|\|\|\|\|[\|\|\|\|\|\|\|\|\|\|\|\|]PAM | NA | Intergenic | NA | NA |
| T4400 | ACAGACGCTGTCCGAGTTCCTGG | 1000 | | | | | | | | | |
| Oligo fwd | CACCACAGACGCTGTCCGAGTTCC | | | | | | | | | | |
| Oligo rev | AAACGGAACTCGGACAGCGTCTGT | | | | | | | | | | |
| Chromosome | start | end | strand | MM | target_seq | PAM | alignment | distance | position | gene name | gene id |
| GU305914.1 | 64337 | 64359 | + | 0 | ACAGACGCTGTCCGAGTTCC | TGG | \|\|\|\|\|\|\|\|[\|\|\|\|\|\|\|\|\|\|\|\|]PAM | NA | Intergenic | NA | NA |
| T4401 | ACGCTGTCCGAGTTCCTGGAAGG | 1000 | | | | | | | | | |
| Oligo fwd | CACCACGCTGTCCGAGTTCCTGGA | | | | | | | | | | |
| Oligo rev | AAACTCCAGGAACTCGGACAGCGT | | | | | | | | | | |
| Chromosome | start | end | strand | MM | target_seq | PAM | alignment | distance | position | gene name | gene id |
| GU305914.1 | 64341 | 64363 | + | 0 | ACGCTGTCCGAGTTCCTGGA | AGG | \|\|\|\|\|\|\|\|[\|\|\|\|\|\|\|\|\|\|\|\|]PAM | NA | Intergenic | NA | NA |
| T4402 | GTCCGAGTTCCTGGAAGGCGCGG | 1000 | | | | | | | | | |
| Oligo fwd | CACCGTCCGAGTTCCTGGAAGGCG | | | | | | | | | | |
| Oligo rev | AAACCGCCTTCCAGGAACTCGGAC | | | | | | | | | | |
| Chromosome | start | end | strand | MM | target_seq | PAM | alignment | distance | position | gene name | gene id |
| GU305914.1 | 64346 | 64368 | + | 0 | GTCCGAGTTCCTGGAAGGCG | CGG | \|\|\|\|\|\|\|\|[\|\|\|\|\|\|\|\|\|\|\|\|]PAM | NA | Intergenic | NA | NA |
| T4403 | GGAAGGCGCGGCGCGCGTCTCGG | 1000 | | | | | | | | | |
| Oligo fwd | CACCGGAAGGCGCGGCGCGCGTCT | | | | | | | | | | |
| Oligo rev | AAACAGACGCGCGCCGCGCCTTCC | | | | | | | | | | |
| Chromosome | start | end | strand | MM | target_seq | PAM | alignment | distance | position | gene name | gene id |
| GU305914.1 | 64358 | 64380 | + | 0 | GGAAGGCGCGGCGCGCGTCT | CGG | \|\|\|\|\|\|\|\|[\|\|\|\|\|\|\|\|\|\|\|\|]PAM | NA | Intergenic | NA | NA |
| T4404 | GCGCGTCTCGGACATCCAACAGG | 1000 | | | | | | | | | |
| Oligo fwd | CACCGCGCGTCTCGGACATCCAAC | | | | | | | | | | |
| Oligo rev | AAACGTTGGATGTCCGAGACGCGC | | | | | | | | | | |
| Chromosome | start | end | strand | MM | target_seq | PAM | alignment | distance | position | gene name | gene id |
| GU305914.1 | 64370 | 64392 | + | 0 | GCGCGTCTCGGACATCCAAC | AGG | \|\|\|\|\|\|\|\|[\|\|\|\|\|\|\|\|\|\|\|\|]PAM | NA | Intergenic | NA | NA |
| T4405 | CGCGTCTCGGACATCCAACAGGG | 1000 | | | | | | | | | |
| Oligo fwd | CACCCGCGTCTCGGACATCCAACA | | | | | | | | | | |
| Oligo rev | AAACTGTTGGATGTCCGAGACGCG | | | | | | | | | | |
| Chromosome | start | end | strand | MM | target_seq | PAM | alignment | distance | position | gene name | gene id |
| GU305914.1 | 64371 | 64393 | + | 0 | CGCGTCTCGGACATCCAACA | GGG | \|\|\|\|\|\|\|\|[\|\|\|\|\|\|\|\|\|\|\|\|]PAM | NA | Intergenic | NA | NA |
| T4406 | GGACATCCAACAGGGTCTGCTGG | 1000 | | | | | | | | | |
| Oligo fwd | CACCGGACATCCAACAGGGTCTGC | | | | | | | | | | |
| Oligo rev | AAACGCAGACCCTGTTGGATGTCC | | | | | | | | | | |
| Chromosome | start | end | strand | MM | target_seq | PAM | alignment | distance | position | gene name | gene id |
| GU305914.1 | 64379 | 64401 | + | 0 | GGACATCCAACAGGGTCTGC | TGG | \|\|\|\|\|\|\|\|[\|\|\|\|\|\|\|\|\|\|\|\|]PAM | NA | Intergenic | NA | NA |
| T4407 | CCAACAGGGTCTGCTGGACTCGG | 1000 | | | | | | | | | |
| Oligo fwd | CACCCCAACAGGGTCTGCTGGACT | | | | | | | | | | |
| Oligo rev | AAACAGTCCAGCAGACCCTGTTGG | | | | | | | | | | |
| Chromosome | start | end | strand | MM | target_seq | PAM | alignment | distance | position | gene name | gene id |
| GU305914.1 | 64385 | 64407 | + | 0 | CCAACAGGGTCTGCTGGACT | CGG | \|\|\|\|\|\|\|\|[\|\|\|\|\|\|\|\|\|\|\|\|]PAM | NA | Intergenic | NA | NA |
| T4408 | CGAAGAAACCGACCGTCCTCAGG | 1000 | | | | | | | | | |
| Oligo fwd | CACCCGAAGAAACCGACCGTCCTC | | | | | | | | | | |
| Oligo rev | AAACGAGGACGGTCGGTTTCTTCG | | | | | | | | | | |
| Chromosome | start | end | strand | MM | target_seq | PAM | alignment | distance | position | gene name | gene id |
| GU305914.1 | 64424 | 64446 | + | 0 | CGAAGAAACCGACCGTCCTC | AGG | \|\|\|\|\|\|\|\|[\|\|\|\|\|\|\|\|\|\|\|\|]PAM | NA | Intergenic | NA | NA |
| T4409 | GAAGAAACCGACCGTCCTCAGGG | 1000 | | | | | | | | | |
| Oligo fwd | CACCGAAGAAACCGACCGTCCTCA | | | | | | | | | | |
| Oligo rev | AAACTGAGGACGGTCGGTTTCTTC | | | | | | | | | | |
| Chromosome | start | end | strand | MM | target_seq | PAM | alignment | distance | position | gene name | gene id |
| GU305914.1 | 64425 | 64447 | + | 0 | GAAGAAACCGACCGTCCTCA | GGG | \|\|\|\|\|\|\|\|[\|\|\|\|\|\|\|\|\|\|\|\|]PAM | NA | Intergenic | NA | NA |
| T4410 | ACCGACCGTCCTCAGGGCCGCGG | 1000 | | | | | | | | | |
| Oligo fwd | CACCACCGACCGTCCTCAGGGCCG | | | | | | | | | | |
| Oligo rev | AAACCGGCCCTGAGGACGGTCGGT | | | | | | | | | | |
| Chromosome | start | end | strand | MM | target_seq | PAM | alignment | distance | position | gene name | gene id |
| GU305914.1 | 64431 | 64453 | + | 0 | ACCGACCGTCCTCAGGGCCG | CGG | \|\|\|\|\|\|\|\|[\|\|\|\|\|\|\|\|\|\|\|\|]PAM | NA | Intergenic | NA | NA |
| T4411 | ACCGTCCTCAGGGCCGCGGCCGG | 1000 | | | | | | | | | |
| Oligo fwd | CACCACCGTCCTCAGGGCCGCGGC | | | | | | | | | | |
| Oligo rev | AAACGCCGCGGCCCTGAGGACGGT | | | | | | | | | | |
| Chromosome | start | end | strand | MM | target_seq | PAM | alignment | distance | position | gene name | gene id |
| GU305914.1 | 64435 | 64457 | + | 0 | ACCGTCCTCAGGGCCGCGGC | CGG | \|\|\|\|\|\|\|\|[\|\|\|\|\|\|\|\|\|\|\|\|]PAM | NA | Intergenic | NA | NA |
| T4412 | GTCCTCAGGGCCGCGGCCGGAGG | 1000 | | | | | | | | | |
| Oligo fwd | CACCGTCCTCAGGGCCGCGGCCGG | | | | | | | | | | |
| Oligo rev | AAACCCGGCCGCGGCCCTGAGGAC | | | | | | | | | | |
| Chromosome | start | end | strand | MM | target_seq | PAM | alignment | distance | position | gene name | gene id |
| GU305914.1 | 64438 | 64460 | + | 0 | GTCCTCAGGGCCGCGGCCGG | AGG | \|\|\|\|\|\|\|\|[\|\|\|\|\|\|\|\|\|\|\|\|]PAM | NA | Intergenic | NA | NA |
| GU305914.1 | 31058 | 31080 | - | 4 | CTCGGCAGGGCCGCGGCCGA | GGG | -\|\|--\|\|\|[\|\|\|\|\|\|\|\|\|\|\|-]PAM | NA | Intergenic | NA | NA |
| T4413 | TCCTCAGGGCCGCGGCCGGAGGG | 1000 | | | | | | | | | |
| Oligo fwd | CACCTCCTCAGGGCCGCGGCCGGA | | | | | | | | | | |
| Oligo rev | AAACTCCGGCCGCGGCCCTGAGGA | | | | | | | | | | |
| Chromosome | start | end | strand | MM | target_seq | PAM | alignment | distance | position | gene name | gene id |
| GU305914.1 | 64439 | 64461 | + | 0 | TCCTCAGGGCCGCGGCCGGA | GGG | \|\|\|\|\|\|\|\|[\|\|\|\|\|\|\|\|\|\|\|\|]PAM | NA | Intergenic | NA | NA |
| T4414 | CACCGACGCCGACCTAAAGACGG | 1000 | | | | | | | | | |

<table>
<tr><td>Oligo fwd</td><td>CACCCACCGACGCCGACCTAAAGA</td><td></td><td></td><td></td><td></td><td></td><td></td><td></td><td></td><td></td><td></td><td></td></tr>
<tr><td>Oligo rev</td><td>AAACTCTTTAGGTCGGCGTCGGTG</td><td></td><td></td><td></td><td></td><td></td><td></td><td></td><td></td><td></td><td></td><td></td></tr>
<tr><td>Chromosome</td><td>start</td><td></td><td>end</td><td>strand</td><td>MM</td><td>target_seq</td><td>PAM</td><td>alignment</td><td>distance</td><td>position</td><td>gene name</td><td>gene id</td></tr>
<tr><td>GU305914.1</td><td></td><td>64466</td><td>64488</td><td>+</td><td>0</td><td>CACCGACGCCGACCTAAAGA</td><td>CGG</td><td>||||||||[||||||||||||]PAM</td><td>NA</td><td>Intergenic</td><td>NA</td><td>NA</td></tr>
<tr><td></td><td></td><td></td><td></td><td></td><td></td><td></td><td></td><td></td><td></td><td></td><td></td><td></td></tr>
<tr><td>T4415</td><td>CCTAAAGACGGTCGAGAAGCTGG</td><td></td><td>1000</td><td></td><td></td><td></td><td></td><td></td><td></td><td></td><td></td><td></td></tr>
<tr><td>Oligo fwd</td><td>CACCCCTAAAGACGGTCGAGAAGC</td><td></td><td></td><td></td><td></td><td></td><td></td><td></td><td></td><td></td><td></td><td></td></tr>
<tr><td>Oligo rev</td><td>AAACGCTTCTCGACCGTCTTTAGG</td><td></td><td></td><td></td><td></td><td></td><td></td><td></td><td></td><td></td><td></td><td></td></tr>
<tr><td>Chromosome</td><td>start</td><td></td><td>end</td><td>strand</td><td>MM</td><td>target_seq</td><td>PAM</td><td>alignment</td><td>distance</td><td>position</td><td>gene name</td><td>gene id</td></tr>
<tr><td>GU305914.1</td><td></td><td>64478</td><td>64500</td><td>+</td><td>0</td><td>CCTAAAGACGGTCGAGAAGC</td><td>TGG</td><td>||||||||[||||||||||||]PAM</td><td>NA</td><td>Intergenic</td><td>NA</td><td>NA</td></tr>
<tr><td>GU305914.1</td><td></td><td>131546</td><td>131568</td><td>+</td><td>4</td><td>CGTCAAGGTGGTCGAGAAGC</td><td>TGG</td><td>|-|-|||-[-|||||||||||]PAM</td><td>NA</td><td>Intergenic</td><td>NA</td><td>NA</td></tr>
<tr><td></td><td></td><td></td><td></td><td></td><td></td><td></td><td></td><td></td><td></td><td></td><td></td><td></td></tr>
<tr><td>T4416</td><td>GACGGTCGAGAAGCTGGTCGCGG</td><td></td><td>1000</td><td></td><td></td><td></td><td></td><td></td><td></td><td></td><td></td><td></td></tr>
<tr><td>Oligo fwd</td><td>CACCGACGGTCGAGAAGCTGGTCG</td><td></td><td></td><td></td><td></td><td></td><td></td><td></td><td></td><td></td><td></td><td></td></tr>
<tr><td>Oligo rev</td><td>AAACCGACCAGCTTCTCGACCGTC</td><td></td><td></td><td></td><td></td><td></td><td></td><td></td><td></td><td></td><td></td><td></td></tr>
<tr><td>Chromosome</td><td>start</td><td></td><td>end</td><td>strand</td><td>MM</td><td>target_seq</td><td>PAM</td><td>alignment</td><td>distance</td><td>position</td><td>gene name</td><td>gene id</td></tr>
<tr><td>GU305914.1</td><td></td><td>64484</td><td>64506</td><td>+</td><td>0</td><td>GACGGTCGAGAAGCTGGTCG</td><td>CGG</td><td>||||||||[||||||||||||]PAM</td><td>NA</td><td>Intergenic</td><td>NA</td><td>NA</td></tr>
<tr><td></td><td></td><td></td><td></td><td></td><td></td><td></td><td></td><td></td><td></td><td></td><td></td><td></td></tr>
<tr><td>T4417</td><td>TCGCGGCCACGCAGACGCTGAGG</td><td></td><td>1000</td><td></td><td></td><td></td><td></td><td></td><td></td><td></td><td></td><td></td></tr>
<tr><td>Oligo fwd</td><td>CACCTCGCGGCCACGCAGACGCTG</td><td></td><td></td><td></td><td></td><td></td><td></td><td></td><td></td><td></td><td></td><td></td></tr>
<tr><td>Oligo rev</td><td>AAACCAGCGTCTGCGTGGCCGCGA</td><td></td><td></td><td></td><td></td><td></td><td></td><td></td><td></td><td></td><td></td><td></td></tr>
<tr><td>Chromosome</td><td>start</td><td></td><td>end</td><td>strand</td><td>MM</td><td>target_seq</td><td>PAM</td><td>alignment</td><td>distance</td><td>position</td><td>gene name</td><td>gene id</td></tr>
<tr><td>GU305914.1</td><td></td><td>64501</td><td>64523</td><td>+</td><td>0</td><td>TCGCGGCCACGCAGACGCTG</td><td>AGG</td><td>||||||||[||||||||||||]PAM</td><td>NA</td><td>Intergenic</td><td>NA</td><td>NA</td></tr>
<tr><td></td><td></td><td></td><td></td><td></td><td></td><td></td><td></td><td></td><td></td><td></td><td></td><td></td></tr>
<tr><td>T4418</td><td>GCTGAGGAACTGTCGCGCCGTGG</td><td></td><td>1000</td><td></td><td></td><td></td><td></td><td></td><td></td><td></td><td></td><td></td></tr>
<tr><td>Oligo fwd</td><td>CACCGCTGAGGAACTGTCGCGCCG</td><td></td><td></td><td></td><td></td><td></td><td></td><td></td><td></td><td></td><td></td><td></td></tr>
<tr><td>Oligo rev</td><td>AAACCGGCGCGACAGTTCCTCAGC</td><td></td><td></td><td></td><td></td><td></td><td></td><td></td><td></td><td></td><td></td><td></td></tr>
<tr><td>Chromosome</td><td>start</td><td></td><td>end</td><td>strand</td><td>MM</td><td>target_seq</td><td>PAM</td><td>alignment</td><td>distance</td><td>position</td><td>gene name</td><td>gene id</td></tr>
<tr><td>GU305914.1</td><td></td><td>64517</td><td>64539</td><td>+</td><td>0</td><td>GCTGAGGAACTGTCGCGCCG</td><td>TGG</td><td>||||||||[||||||||||||]PAM</td><td>NA</td><td>Intergenic</td><td>NA</td><td>NA</td></tr>
<tr><td></td><td></td><td></td><td></td><td></td><td></td><td></td><td></td><td></td><td></td><td></td><td></td><td></td></tr>
<tr><td>T4419</td><td>TCGCGCCGTGGCCGAGCTGCTGG</td><td></td><td>1000</td><td></td><td></td><td></td><td></td><td></td><td></td><td></td><td></td><td></td></tr>
<tr><td>Oligo fwd</td><td>CACCTCGCGCCGTGGCCGAGCTGC</td><td></td><td></td><td></td><td></td><td></td><td></td><td></td><td></td><td></td><td></td><td></td></tr>
<tr><td>Oligo rev</td><td>AAACGCAGCTCGGCCACGGCGCGA</td><td></td><td></td><td></td><td></td><td></td><td></td><td></td><td></td><td></td><td></td><td></td></tr>
<tr><td>Chromosome</td><td>start</td><td></td><td>end</td><td>strand</td><td>MM</td><td>target_seq</td><td>PAM</td><td>alignment</td><td>distance</td><td>position</td><td>gene name</td><td>gene id</td></tr>
<tr><td>GU305914.1</td><td></td><td>64529</td><td>64551</td><td>+</td><td>0</td><td>TCGCGCCGTGGCCGAGCTGC</td><td>TGG</td><td>||||||||[||||||||||||]PAM</td><td>NA</td><td>Intergenic</td><td>NA</td><td>NA</td></tr>
<tr><td></td><td></td><td></td><td></td><td></td><td></td><td></td><td></td><td></td><td></td><td></td><td></td><td></td></tr>
<tr><td>T4420</td><td>CGCCGTGGCCGAGCTGCTGGAGG</td><td></td><td>1000</td><td></td><td></td><td></td><td></td><td></td><td></td><td></td><td></td><td></td></tr>
<tr><td>Oligo fwd</td><td>CACCCGCCGTGGCCGAGCTGCTGG</td><td></td><td></td><td></td><td></td><td></td><td></td><td></td><td></td><td></td><td></td><td></td></tr>
<tr><td>Oligo rev</td><td>AAACCCAGCAGCTCGGCCACGGCG</td><td></td><td></td><td></td><td></td><td></td><td></td><td></td><td></td><td></td><td></td><td></td></tr>
<tr><td>Chromosome</td><td>start</td><td></td><td>end</td><td>strand</td><td>MM</td><td>target_seq</td><td>PAM</td><td>alignment</td><td>distance</td><td>position</td><td>gene name</td><td>gene id</td></tr>
<tr><td>GU305914.1</td><td></td><td>64532</td><td>64554</td><td>+</td><td>0</td><td>CGCCGTGGCCGAGCTGCTGG</td><td>AGG</td><td>||||||||[||||||||||||]PAM</td><td>NA</td><td>Intergenic</td><td>NA</td><td>NA</td></tr>
<tr><td>GU305914.1</td><td></td><td>26061</td><td>26083</td><td>-</td><td>4</td><td>CGCCGCGTGCGAGCCGCTGG</td><td>TGG</td><td>|||||-|-[-|||||-|||||]PAM</td><td>NA</td><td>Intergenic</td><td>NA</td><td>NA</td></tr>
<tr><td></td><td></td><td></td><td></td><td></td><td></td><td></td><td></td><td></td><td></td><td></td><td></td><td></td></tr>
<tr><td>T4421</td><td>CGCTCACCACGCAGCAGCTGCGG</td><td></td><td>1000</td><td></td><td></td><td></td><td></td><td></td><td></td><td></td><td></td><td></td></tr>
<tr><td>Oligo fwd</td><td>CACCCGCTCACCACGCAGCAGCTG</td><td></td><td></td><td></td><td></td><td></td><td></td><td></td><td></td><td></td><td></td><td></td></tr>
<tr><td>Oligo rev</td><td>AAACCAGCTGCTGCGTGGTGAGCG</td><td></td><td></td><td></td><td></td><td></td><td></td><td></td><td></td><td></td><td></td><td></td></tr>
<tr><td>Chromosome</td><td>start</td><td></td><td>end</td><td>strand</td><td>MM</td><td>target_seq</td><td>PAM</td><td>alignment</td><td>distance</td><td>position</td><td>gene name</td><td>gene id</td></tr>
<tr><td>GU305914.1</td><td></td><td>64591</td><td>64613</td><td>+</td><td>0</td><td>CGCTCACCACGCAGCAGCTG</td><td>CGG</td><td>||||||||[||||||||||||]PAM</td><td>NA</td><td>Intergenic</td><td>NA</td><td>NA</td></tr>
<tr><td></td><td></td><td></td><td></td><td></td><td></td><td></td><td></td><td></td><td></td><td></td><td></td><td></td></tr>
<tr><td>T4422</td><td>ACGCAGCAGCTGCGGCGCGACGG</td><td></td><td>1000</td><td></td><td></td><td></td><td></td><td></td><td></td><td></td><td></td><td></td></tr>
<tr><td>Oligo fwd</td><td>CACCACGCAGCAGCTGCGGCGCGA</td><td></td><td></td><td></td><td></td><td></td><td></td><td></td><td></td><td></td><td></td><td></td></tr>
<tr><td>Oligo rev</td><td>AAACTCGCGCCGCAGCTGCTGCGT</td><td></td><td></td><td></td><td></td><td></td><td></td><td></td><td></td><td></td><td></td><td></td></tr>
<tr><td>Chromosome</td><td>start</td><td></td><td>end</td><td>strand</td><td>MM</td><td>target_seq</td><td>PAM</td><td>alignment</td><td>distance</td><td>position</td><td>gene name</td><td>gene id</td></tr>
<tr><td>GU305914.1</td><td></td><td>64599</td><td>64621</td><td>+</td><td>0</td><td>ACGCAGCAGCTGCGGCGCGA</td><td>CGG</td><td>||||||||[||||||||||||]PAM</td><td>NA</td><td>Intergenic</td><td>NA</td><td>NA</td></tr>
<tr><td></td><td></td><td></td><td></td><td></td><td></td><td></td><td></td><td></td><td></td><td></td><td></td><td></td></tr>
<tr><td>T4423</td><td>GCGCGACGGCGACACACAGCTGG</td><td></td><td>1000</td><td></td><td></td><td></td><td></td><td></td><td></td><td></td><td></td><td></td></tr>
<tr><td>Oligo fwd</td><td>CACCGCGCGACGGCGACACACAGC</td><td></td><td></td><td></td><td></td><td></td><td></td><td></td><td></td><td></td><td></td><td></td></tr>
<tr><td>Oligo rev</td><td>AAACGCTGTGTGTCGCCGTCGCGC</td><td></td><td></td><td></td><td></td><td></td><td></td><td></td><td></td><td></td><td></td><td></td></tr>
<tr><td>Chromosome</td><td>start</td><td></td><td>end</td><td>strand</td><td>MM</td><td>target_seq</td><td>PAM</td><td>alignment</td><td>distance</td><td>position</td><td>gene name</td><td>gene id</td></tr>
<tr><td>GU305914.1</td><td></td><td>64613</td><td>64635</td><td>+</td><td>0</td><td>GCGCGACGGCGACACACAGC</td><td>TGG</td><td>||||||||[||||||||||||]PAM</td><td>NA</td><td>Intergenic</td><td>NA</td><td>NA</td></tr>
<tr><td></td><td></td><td></td><td></td><td></td><td></td><td></td><td></td><td></td><td></td><td></td><td></td><td></td></tr>
<tr><td>T4424</td><td>CGGCGACACACAGCTGGACGCGG</td><td></td><td>1000</td><td></td><td></td><td></td><td></td><td></td><td></td><td></td><td></td><td></td></tr>
<tr><td>Oligo fwd</td><td>CACCCGGCGACACACAGCTGGACG</td><td></td><td></td><td></td><td></td><td></td><td></td><td></td><td></td><td></td><td></td><td></td></tr>
<tr><td>Oligo rev</td><td>AAACCGTCCAGCTGTGTGTCGCCG</td><td></td><td></td><td></td><td></td><td></td><td></td><td></td><td></td><td></td><td></td><td></td></tr>
<tr><td>Chromosome</td><td>start</td><td></td><td>end</td><td>strand</td><td>MM</td><td>target_seq</td><td>PAM</td><td>alignment</td><td>distance</td><td>position</td><td>gene name</td><td>gene id</td></tr>
<tr><td>GU305914.1</td><td></td><td>64619</td><td>64641</td><td>+</td><td>0</td><td>CGGCGACACACAGCTGGACG</td><td>CGG</td><td>||||||||[||||||||||||]PAM</td><td>NA</td><td>Intergenic</td><td>NA</td><td>NA</td></tr>
<tr><td></td><td></td><td></td><td></td><td></td><td></td><td></td><td></td><td></td><td></td><td></td><td></td><td></td></tr>
<tr><td>T4425</td><td>CCACTGCCACGATAACCACGCGG</td><td></td><td>1000</td><td></td><td></td><td></td><td></td><td></td><td></td><td></td><td></td><td></td></tr>
<tr><td>Oligo fwd</td><td>CACCCCACTGCCACGATAACCACG</td><td></td><td></td><td></td><td></td><td></td><td></td><td></td><td></td><td></td><td></td><td></td></tr>
<tr><td>Oligo rev</td><td>AAACCGTGGTTATCGTGGCAGTGG</td><td></td><td></td><td></td><td></td><td></td><td></td><td></td><td></td><td></td><td></td><td></td></tr>
<tr><td>Chromosome</td><td>start</td><td></td><td>end</td><td>strand</td><td>MM</td><td>target_seq</td><td>PAM</td><td>alignment</td><td>distance</td><td>position</td><td>gene name</td><td>gene id</td></tr>
<tr><td>GU305914.1</td><td></td><td>64655</td><td>64677</td><td>+</td><td>0</td><td>CCACTGCCACGATAACCACG</td><td>CGG</td><td>||||||||[||||||||||||]PAM</td><td>NA</td><td>Intergenic</td><td>NA</td><td>NA</td></tr>
<tr><td></td><td></td><td></td><td></td><td></td><td></td><td></td><td></td><td></td><td></td><td></td><td></td><td></td></tr>
<tr><td>T4426</td><td>GGCGACGCCCGCCCTAGAGACGG</td><td></td><td>1000</td><td></td><td></td><td></td><td></td><td></td><td></td><td></td><td></td><td></td></tr>
<tr><td>Oligo fwd</td><td>CACCGGCGACGCCCGCCCTAGAGA</td><td></td><td></td><td></td><td></td><td></td><td></td><td></td><td></td><td></td><td></td><td></td></tr>
<tr><td>Oligo rev</td><td>AAACTCTCTAGGGCGGGCGTCGCC</td><td></td><td></td><td></td><td></td><td></td><td></td><td></td><td></td><td></td><td></td><td></td></tr>
<tr><td>Chromosome</td><td>start</td><td></td><td>end</td><td>strand</td><td>MM</td><td>target_seq</td><td>PAM</td><td>alignment</td><td>distance</td><td>position</td><td>gene name</td><td>gene id</td></tr>
<tr><td>GU305914.1</td><td></td><td>64676</td><td>64698</td><td>+</td><td>0</td><td>GGCGACGCCCGCCCTAGAGA</td><td>CGG</td><td>||||||||[||||||||||||]PAM</td><td>NA</td><td>Intergenic</td><td>NA</td><td>NA</td></tr>
<tr><td></td><td></td><td></td><td></td><td></td><td></td><td></td><td></td><td></td><td></td><td></td><td></td><td></td></tr>
<tr><td>T4427</td><td>CCTAGAGACGGACTTCCAGAAGG</td><td></td><td>1000</td><td></td><td></td><td></td><td></td><td></td><td></td><td></td><td></td><td></td></tr>
<tr><td>Oligo fwd</td><td>CACCCCTAGAGACGGACTTCCAGA</td><td></td><td></td><td></td><td></td><td></td><td></td><td></td><td></td><td></td><td></td><td></td></tr>
<tr><td>Oligo rev</td><td>AAACTCTGGAAGTCCGTCTCTAGG</td><td></td><td></td><td></td><td></td><td></td><td></td><td></td><td></td><td></td><td></td><td></td></tr>
<tr><td>Chromosome</td><td>start</td><td></td><td>end</td><td>strand</td><td>MM</td><td>target_seq</td><td>PAM</td><td>alignment</td><td>distance</td><td>position</td><td>gene name</td><td>gene id</td></tr>
<tr><td>GU305914.1</td><td></td><td>64688</td><td>64710</td><td>+</td><td>0</td><td>CCTAGAGACGGACTTCCAGA</td><td>AGG</td><td>||||||||[||||||||||||]PAM</td><td>NA</td><td>Intergenic</td><td>NA</td><td>NA</td></tr>
<tr><td></td><td></td><td></td><td></td><td></td><td></td><td></td><td></td><td></td><td></td><td></td><td></td><td></td></tr>
<tr><td>T4428</td><td>CGACCGCGCGTCCGCCGTGCTGG</td><td></td><td>1000</td><td></td><td></td><td></td><td></td><td></td><td></td><td></td><td></td><td></td></tr>
<tr><td>Oligo fwd</td><td>CACCCGACCGCGCGTCCGCCGTGC</td><td></td><td></td><td></td><td></td><td></td><td></td><td></td><td></td><td></td><td></td><td></td></tr>
<tr><td>Oligo rev</td><td>AAACGCACGGCGGACGCGCGGTCG</td><td></td><td></td><td></td><td></td><td></td><td></td><td></td><td></td><td></td><td></td><td></td></tr>
<tr><td>Chromosome</td><td>start</td><td></td><td>end</td><td>strand</td><td>MM</td><td>target_seq</td><td>PAM</td><td>alignment</td><td>distance</td><td>position</td><td>gene name</td><td>gene id</td></tr>
</table>

<table>
<tr><td>GU305914.1</td><td>64715</td><td>64737</td><td>+</td><td>0</td><td>CGACCGCGCGTCCGCCGTGC</td><td>TGG</td><td>||||||||[||||||||||||]PAM</td><td>NA</td><td>Intergenic</td><td>NA</td><td>NA</td></tr>
<tr><td></td><td></td><td></td><td></td><td></td><td></td><td></td><td></td><td></td><td></td><td></td><td></td></tr>
<tr><td>T4429</td><td>ATCCACCAGATCCAGCAGCTCGG</td><td>1000</td><td></td><td></td><td></td><td></td><td></td><td></td><td></td><td></td><td></td></tr>
<tr><td>Oligo fwd</td><td>CACCATCCACCAGATCCAGCAGCT</td><td></td><td></td><td></td><td></td><td></td><td></td><td></td><td></td><td></td><td></td></tr>
<tr><td>Oligo rev</td><td>AAACAGCTGCTGGATCTGGTGGAT</td><td></td><td></td><td></td><td></td><td></td><td></td><td></td><td></td><td></td><td></td></tr>
<tr><td>Chromosome</td><td>start</td><td>end</td><td>strand</td><td>MM</td><td>target_seq</td><td>PAM</td><td>alignment</td><td>distance</td><td>position</td><td>gene name</td><td>gene id</td></tr>
<tr><td>GU305914.1</td><td>64758</td><td>64780</td><td>+</td><td>0</td><td>ATCCACCAGATCCAGCAGCT</td><td>CGG</td><td>||||||||[||||||||||||]PAM</td><td>NA</td><td>Intergenic</td><td>NA</td><td>NA</td></tr>
<tr><td></td><td></td><td></td><td></td><td></td><td></td><td></td><td></td><td></td><td></td><td></td><td></td></tr>
<tr><td>T4430</td><td>CCAGATCCAGCAGCTCGGCTCGG</td><td>1000</td><td></td><td></td><td></td><td></td><td></td><td></td><td></td><td></td><td></td></tr>
<tr><td>Oligo fwd</td><td>CACCCCAGATCCAGCAGCTCGGCT</td><td></td><td></td><td></td><td></td><td></td><td></td><td></td><td></td><td></td><td></td></tr>
<tr><td>Oligo rev</td><td>AAACAGCCGAGCTGCTGGATCTGG</td><td></td><td></td><td></td><td></td><td></td><td></td><td></td><td></td><td></td><td></td></tr>
<tr><td>Chromosome</td><td>start</td><td>end</td><td>strand</td><td>MM</td><td>target_seq</td><td>PAM</td><td>alignment</td><td>distance</td><td>position</td><td>gene name</td><td>gene id</td></tr>
<tr><td>GU305914.1</td><td>64763</td><td>64785</td><td>+</td><td>0</td><td>CCAGATCCAGCAGCTCGGCT</td><td>CGG</td><td>||||||||[||||||||||||]PAM</td><td>NA</td><td>Intergenic</td><td>NA</td><td>NA</td></tr>
<tr><td></td><td></td><td></td><td></td><td></td><td></td><td></td><td></td><td></td><td></td><td></td><td></td></tr>
<tr><td>T4431</td><td>GATCTCCTCGCGCTCGCTGTCGG</td><td>1000</td><td></td><td></td><td></td><td></td><td></td><td></td><td></td><td></td><td></td></tr>
<tr><td>Oligo fwd</td><td>CACCGATCTCCTCGCGCTCGCTGT</td><td></td><td></td><td></td><td></td><td></td><td></td><td></td><td></td><td></td><td></td></tr>
<tr><td>Oligo rev</td><td>AAACACAGCGAGCGCGAGGAGATC</td><td></td><td></td><td></td><td></td><td></td><td></td><td></td><td></td><td></td><td></td></tr>
<tr><td>Chromosome</td><td>start</td><td>end</td><td>strand</td><td>MM</td><td>target_seq</td><td>PAM</td><td>alignment</td><td>distance</td><td>position</td><td>gene name</td><td>gene id</td></tr>
<tr><td>GU305914.1</td><td>64811</td><td>64833</td><td>+</td><td>0</td><td>GATCTCCTCGCGCTCGCTGT</td><td>CGG</td><td>||||||||[||||||||||||]PAM</td><td>NA</td><td>Intergenic</td><td>NA</td><td>NA</td></tr>
<tr><td></td><td></td><td></td><td></td><td></td><td></td><td></td><td></td><td></td><td></td><td></td><td></td></tr>
<tr><td>T4432</td><td>CTCGCGCTCGCTGTCGGCCAAGG</td><td>1000</td><td></td><td></td><td></td><td></td><td></td><td></td><td></td><td></td><td></td></tr>
<tr><td>Oligo fwd</td><td>CACCCTCGCGCTCGCTGTCGGCCA</td><td></td><td></td><td></td><td></td><td></td><td></td><td></td><td></td><td></td><td></td></tr>
<tr><td>Oligo rev</td><td>AAACTGGCCGACAGCGAGCGCGAG</td><td></td><td></td><td></td><td></td><td></td><td></td><td></td><td></td><td></td><td></td></tr>
<tr><td>Chromosome</td><td>start</td><td>end</td><td>strand</td><td>MM</td><td>target_seq</td><td>PAM</td><td>alignment</td><td>distance</td><td>position</td><td>gene name</td><td>gene id</td></tr>
<tr><td>GU305914.1</td><td>64817</td><td>64839</td><td>+</td><td>0</td><td>CTCGCGCTCGCTGTCGGCCA</td><td>AGG</td><td>||||||||[||||||||||||]PAM</td><td>NA</td><td>Intergenic</td><td>NA</td><td>NA</td></tr>
<tr><td></td><td></td><td></td><td></td><td></td><td></td><td></td><td></td><td></td><td></td><td></td><td></td></tr>
<tr><td>T4433</td><td>GCCAACGTGCTCTTCTACCCCGG</td><td>1000</td><td></td><td></td><td></td><td></td><td></td><td></td><td></td><td></td><td></td></tr>
<tr><td>Oligo fwd</td><td>CACCGCCAACGTGCTCTTCTACCC</td><td></td><td></td><td></td><td></td><td></td><td></td><td></td><td></td><td></td><td></td></tr>
<tr><td>Oligo rev</td><td>AAACGGGTAGAAGAGCACGTTGGC</td><td></td><td></td><td></td><td></td><td></td><td></td><td></td><td></td><td></td><td></td></tr>
<tr><td>Chromosome</td><td>start</td><td>end</td><td>strand</td><td>MM</td><td>target_seq</td><td>PAM</td><td>alignment</td><td>distance</td><td>position</td><td>gene name</td><td>gene id</td></tr>
<tr><td>GU305914.1</td><td>64905</td><td>64927</td><td>+</td><td>0</td><td>GCCAACGTGCTCTTCTACCC</td><td>CGG</td><td>||||||||[||||||||||||]PAM</td><td>NA</td><td>Intergenic</td><td>NA</td><td>NA</td></tr>
<tr><td></td><td></td><td></td><td></td><td></td><td></td><td></td><td></td><td></td><td></td><td></td><td></td></tr>
<tr><td>T4434</td><td>CGTGCTCTTCTACCCCGGCGTGG</td><td>1000</td><td></td><td></td><td></td><td></td><td></td><td></td><td></td><td></td><td></td></tr>
<tr><td>Oligo fwd</td><td>CACCCGTGCTCTTCTACCCCGGCG</td><td></td><td></td><td></td><td></td><td></td><td></td><td></td><td></td><td></td><td></td></tr>
<tr><td>Oligo rev</td><td>AAACCGCCGGGGTAGAAGAGCACG</td><td></td><td></td><td></td><td></td><td></td><td></td><td></td><td></td><td></td><td></td></tr>
<tr><td>Chromosome</td><td>start</td><td>end</td><td>strand</td><td>MM</td><td>target_seq</td><td>PAM</td><td>alignment</td><td>distance</td><td>position</td><td>gene name</td><td>gene id</td></tr>
<tr><td>GU305914.1</td><td>64910</td><td>64932</td><td>+</td><td>0</td><td>CGTGCTCTTCTACCCCGGCG</td><td>TGG</td><td>||||||||[||||||||||||]PAM</td><td>NA</td><td>Intergenic</td><td>NA</td><td>NA</td></tr>
<tr><td></td><td></td><td></td><td></td><td></td><td></td><td></td><td></td><td></td><td></td><td></td><td></td></tr>
<tr><td>T4435</td><td>GCGTGGTGTTCGCCATCCTGCGG</td><td>1000</td><td></td><td></td><td></td><td></td><td></td><td></td><td></td><td></td><td></td></tr>
<tr><td>Oligo fwd</td><td>CACCGCGTGGTGTTCGCCATCCTG</td><td></td><td></td><td></td><td></td><td></td><td></td><td></td><td></td><td></td><td></td></tr>
<tr><td>Oligo rev</td><td>AAACCAGGATGGCGAACACCACGC</td><td></td><td></td><td></td><td></td><td></td><td></td><td></td><td></td><td></td><td></td></tr>
<tr><td>Chromosome</td><td>start</td><td>end</td><td>strand</td><td>MM</td><td>target_seq</td><td>PAM</td><td>alignment</td><td>distance</td><td>position</td><td>gene name</td><td>gene id</td></tr>
<tr><td>GU305914.1</td><td>64927</td><td>64949</td><td>+</td><td>0</td><td>GCGTGGTGTTCGCCATCCTG</td><td>CGG</td><td>||||||||[||||||||||||]PAM</td><td>NA</td><td>Intergenic</td><td>NA</td><td>NA</td></tr>
<tr><td></td><td></td><td></td><td></td><td></td><td></td><td></td><td></td><td></td><td></td><td></td><td></td></tr>
<tr><td>T4436</td><td>GTTCGCCATCCTGCGGACCGCGG</td><td>1000</td><td></td><td></td><td></td><td></td><td></td><td></td><td></td><td></td><td></td></tr>
<tr><td>Oligo fwd</td><td>CACCGTTCGCCATCCTGCGGACCG</td><td></td><td></td><td></td><td></td><td></td><td></td><td></td><td></td><td></td><td></td></tr>
<tr><td>Oligo rev</td><td>AAACCGGTCCGCAGGATGGCGAAC</td><td></td><td></td><td></td><td></td><td></td><td></td><td></td><td></td><td></td><td></td></tr>
<tr><td>Chromosome</td><td>start</td><td>end</td><td>strand</td><td>MM</td><td>target_seq</td><td>PAM</td><td>alignment</td><td>distance</td><td>position</td><td>gene name</td><td>gene id</td></tr>
<tr><td>GU305914.1</td><td>64934</td><td>64956</td><td>+</td><td>0</td><td>GTTCGCCATCCTGCGGACCG</td><td>CGG</td><td>||||||||[||||||||||||]PAM</td><td>NA</td><td>Intergenic</td><td>NA</td><td>NA</td></tr>
<tr><td></td><td></td><td></td><td></td><td></td><td></td><td></td><td></td><td></td><td></td><td></td><td></td></tr>
<tr><td>T4437</td><td>CTGCGGACCGCGGCGCGCGTCGG</td><td>1000</td><td></td><td></td><td></td><td></td><td></td><td></td><td></td><td></td><td></td></tr>
<tr><td>Oligo fwd</td><td>CACCCTGCGGACCGCGGCGCGCGT</td><td></td><td></td><td></td><td></td><td></td><td></td><td></td><td></td><td></td><td></td></tr>
<tr><td>Oligo rev</td><td>AAACACGCGCGCCGCGGTCCGCAG</td><td></td><td></td><td></td><td></td><td></td><td></td><td></td><td></td><td></td><td></td></tr>
<tr><td>Chromosome</td><td>start</td><td>end</td><td>strand</td><td>MM</td><td>target_seq</td><td>PAM</td><td>alignment</td><td>distance</td><td>position</td><td>gene name</td><td>gene id</td></tr>
<tr><td>GU305914.1</td><td>64944</td><td>64966</td><td>+</td><td>0</td><td>CTGCGGACCGCGGCGCGCGT</td><td>CGG</td><td>||||||||[||||||||||||]PAM</td><td>NA</td><td>Intergenic</td><td>NA</td><td>NA</td></tr>
<tr><td></td><td></td><td></td><td></td><td></td><td></td><td></td><td></td><td></td><td></td><td></td><td></td></tr>
<tr><td>T4438</td><td>CGGCGCGCGTCGGAGAAGAGCGG</td><td>1000</td><td></td><td></td><td></td><td></td><td></td><td></td><td></td><td></td><td></td></tr>
<tr><td>Oligo fwd</td><td>CACCCGGCGCGCGTCGGAGAAGAG</td><td></td><td></td><td></td><td></td><td></td><td></td><td></td><td></td><td></td><td></td></tr>
<tr><td>Oligo rev</td><td>AAACCTCTTCTCCGACGCGCGCCG</td><td></td><td></td><td></td><td></td><td></td><td></td><td></td><td></td><td></td><td></td></tr>
<tr><td>Chromosome</td><td>start</td><td>end</td><td>strand</td><td>MM</td><td>target_seq</td><td>PAM</td><td>alignment</td><td>distance</td><td>position</td><td>gene name</td><td>gene id</td></tr>
<tr><td>GU305914.1</td><td>64954</td><td>64976</td><td>+</td><td>0</td><td>CGGCGCGCGTCGGAGAAGAG</td><td>CGG</td><td>||||||||[||||||||||||]PAM</td><td>NA</td><td>Intergenic</td><td>NA</td><td>NA</td></tr>
<tr><td></td><td></td><td></td><td></td><td></td><td></td><td></td><td></td><td></td><td></td><td></td><td></td></tr>
<tr><td>T4439</td><td>AGAAGAGCGGCGCCAGTCGCCGG</td><td>1000</td><td></td><td></td><td></td><td></td><td></td><td></td><td></td><td></td><td></td></tr>
<tr><td>Oligo fwd</td><td>CACCAGAAGAGCGGCGCCAGTCGC</td><td></td><td></td><td></td><td></td><td></td><td></td><td></td><td></td><td></td><td></td></tr>
<tr><td>Oligo rev</td><td>AAACGCGACTGGCGCCGCTCTTCT</td><td></td><td></td><td></td><td></td><td></td><td></td><td></td><td></td><td></td><td></td></tr>
<tr><td>Chromosome</td><td>start</td><td>end</td><td>strand</td><td>MM</td><td>target_seq</td><td>PAM</td><td>alignment</td><td>distance</td><td>position</td><td>gene name</td><td>gene id</td></tr>
<tr><td>GU305914.1</td><td>64967</td><td>64989</td><td>+</td><td>0</td><td>AGAAGAGCGGCGCCAGTCGC</td><td>CGG</td><td>||||||||[||||||||||||]PAM</td><td>NA</td><td>Intergenic</td><td>NA</td><td>NA</td></tr>
<tr><td></td><td></td><td></td><td></td><td></td><td></td><td></td><td></td><td></td><td></td><td></td><td></td></tr>
<tr><td>T4440</td><td>GAAGAGCGGCGCCAGTCGCCGGG</td><td>1000</td><td></td><td></td><td></td><td></td><td></td><td></td><td></td><td></td><td></td></tr>
<tr><td>Oligo fwd</td><td>CACCGAAGAGCGGCGCCAGTCGCC</td><td></td><td></td><td></td><td></td><td></td><td></td><td></td><td></td><td></td><td></td></tr>
<tr><td>Oligo rev</td><td>AAACGGCGACTGGCGCCGCTCTTC</td><td></td><td></td><td></td><td></td><td></td><td></td><td></td><td></td><td></td><td></td></tr>
<tr><td>Chromosome</td><td>start</td><td>end</td><td>strand</td><td>MM</td><td>target_seq</td><td>PAM</td><td>alignment</td><td>distance</td><td>position</td><td>gene name</td><td>gene id</td></tr>
<tr><td>GU305914.1</td><td>64968</td><td>64990</td><td>+</td><td>0</td><td>GAAGAGCGGCGCCAGTCGCC</td><td>GGG</td><td>||||||||[||||||||||||]PAM</td><td>NA</td><td>Intergenic</td><td>NA</td><td>NA</td></tr>
<tr><td></td><td></td><td></td><td></td><td></td><td></td><td></td><td></td><td></td><td></td><td></td><td></td></tr>
<tr><td>T4441</td><td>GAGCGGCGCCAGTCGCCGGGCGG</td><td>1000</td><td></td><td></td><td></td><td></td><td></td><td></td><td></td><td></td><td></td></tr>
<tr><td>Oligo fwd</td><td>CACCGAGCGGCGCCAGTCGCCGGG</td><td></td><td></td><td></td><td></td><td></td><td></td><td></td><td></td><td></td><td></td></tr>
<tr><td>Oligo rev</td><td>AAACCCCGGCGACTGGCGCCGCTC</td><td></td><td></td><td></td><td></td><td></td><td></td><td></td><td></td><td></td><td></td></tr>
<tr><td>Chromosome</td><td>start</td><td>end</td><td>strand</td><td>MM</td><td>target_seq</td><td>PAM</td><td>alignment</td><td>distance</td><td>position</td><td>gene name</td><td>gene id</td></tr>
<tr><td>GU305914.1</td><td>64971</td><td>64993</td><td>+</td><td>0</td><td>GAGCGGCGCCAGTCGCCGGG</td><td>CGG</td><td>||||||||[||||||||||||]PAM</td><td>NA</td><td>Intergenic</td><td>NA</td><td>NA</td></tr>
<tr><td></td><td></td><td></td><td></td><td></td><td></td><td></td><td></td><td></td><td></td><td></td><td></td></tr>
<tr><td>T4442</td><td>AGCGGCGCCAGTCGCCGGGCGGG</td><td>1000</td><td></td><td></td><td></td><td></td><td></td><td></td><td></td><td></td><td></td></tr>
<tr><td>Oligo fwd</td><td>CACCAGCGGCGCCAGTCGCCGGGC</td><td></td><td></td><td></td><td></td><td></td><td></td><td></td><td></td><td></td><td></td></tr>
<tr><td>Oligo rev</td><td>AAACGCCCGGCGACTGGCGCCGCT</td><td></td><td></td><td></td><td></td><td></td><td></td><td></td><td></td><td></td><td></td></tr>
<tr><td>Chromosome</td><td>start</td><td>end</td><td>strand</td><td>MM</td><td>target_seq</td><td>PAM</td><td>alignment</td><td>distance</td><td>position</td><td>gene name</td><td>gene id</td></tr>
<tr><td>GU305914.1</td><td>64972</td><td>64994</td><td>+</td><td>0</td><td>AGCGGCGCCAGTCGCCGGGC</td><td>GGG</td><td>||||||||[||||||||||||]PAM</td><td>NA</td><td>Intergenic</td><td>NA</td><td>NA</td></tr>
<tr><td></td><td></td><td></td><td></td><td></td><td></td><td></td><td></td><td></td><td></td><td></td><td></td></tr>
<tr><td>T4443</td><td>GGACGCCGCCTCAGAGACGACGG</td><td>1000</td><td></td><td></td><td></td><td></td><td></td><td></td><td></td><td></td><td></td></tr>
<tr><td>Oligo fwd</td><td>CACCGGACGCCGCCTCAGAGACGA</td><td></td><td></td><td></td><td></td><td></td><td></td><td></td><td></td><td></td><td></td></tr>
<tr><td>Oligo rev</td><td>AAACTCGTCTCTGAGGCGGCGTCC</td><td></td><td></td><td></td><td></td><td></td><td></td><td></td><td></td><td></td><td></td></tr>
</table>

| Chromosome | start | end | strand | MM | target_seq | PAM | alignment | distance | position | gene name | gene id |
|---|---|---|---|---|---|---|---|---|---|---|---|
| GU305914.1 | 64993 | 65015 | + | 0 | GGACGCCGCCTCAGAGACGA | CGG | \|\|\|\|\|\|\|\|[\|\|\|\|\|\|\|\|\|\|\|\|]PAM | NA | Intergenic | NA | NA |
| T4444 | GATAGCAGAGAACGACGAAAAGG | 1000 | | | | | | | | | |
| Oligo fwd | CACCGATAGCAGAGAACGACGAAA | | | | | | | | | | |
| Oligo rev | AAACTTTCGTCGTTCTCTGCTATC | | | | | | | | | | |
| Chromosome | start | end | strand | MM | target_seq | PAM | alignment | distance | position | gene name | gene id |
| GU305914.1 | 65075 | 65097 | + | 0 | GATAGCAGAGAACGACGAAA | AGG | \|\|\|\|\|\|\|\|[\|\|\|\|\|\|\|\|\|\|\|\|]PAM | NA | Intergenic | NA | NA |
| T4445 | AGCAGAGAACGACGAAAAGGAGG | 1000 | | | | | | | | | |
| Oligo fwd | CACCAGCAGAGAACGACGAAAAGG | | | | | | | | | | |
| Oligo rev | AAACCCTTTTCGTCGTTCTCTGCT | | | | | | | | | | |
| Chromosome | start | end | strand | MM | target_seq | PAM | alignment | distance | position | gene name | gene id |
| GU305914.1 | 65078 | 65100 | + | 0 | AGCAGAGAACGACGAAAAGG | AGG | \|\|\|\|\|\|\|\|[\|\|\|\|\|\|\|\|\|\|\|\|]PAM | NA | Intergenic | NA | NA |
| T4446 | CGACGAAAAGGAGGATGAAGAGG | 1000 | | | | | | | | | |
| Oligo fwd | CACCCGACGAAAAGGAGGATGAAG | | | | | | | | | | |
| Oligo rev | AAACCTTCATCCTCCTTTTCGTCG | | | | | | | | | | |
| Chromosome | start | end | strand | MM | target_seq | PAM | alignment | distance | position | gene name | gene id |
| GU305914.1 | 65087 | 65109 | + | 0 | CGACGAAAAGGAGGATGAAG | AGG | \|\|\|\|\|\|\|\|[\|\|\|\|\|\|\|\|\|\|\|\|]PAM | NA | Intergenic | NA | NA |
| T4447 | CGAAAAGGAGGATGAAGAGGAGG | 1000 | | | | | | | | | |
| Oligo fwd | CACCCGAAAAGGAGGATGAAGAGG | | | | | | | | | | |
| Oligo rev | AAACCCTCTTCATCCTCCTTTTCG | | | | | | | | | | |
| Chromosome | start | end | strand | MM | target_seq | PAM | alignment | distance | position | gene name | gene id |
| GU305914.1 | 65090 | 65112 | + | 0 | CGAAAAGGAGGATGAAGAGG | AGG | \|\|\|\|\|\|\|\|[\|\|\|\|\|\|\|\|\|\|\|\|]PAM | NA | Intergenic | NA | NA |
| GU305914.1 | 27345 | 27367 | + | 4 | CGAAGATGATGATGAAGAAG | AGG | \|\|\|\|-\|-\|[\|-\|\|\|\|\|\|\|\|-\|]PAM | NA | Intergenic | NA | NA |
| T4448 | GGAGGAAGAAGACGACGACGAGG | 1000 | | | | | | | | | |
| Oligo fwd | CACCGGAGGAAGAAGACGACGACG | | | | | | | | | | |
| Oligo rev | AAACCGTCGTCGTCTTCTTCCTCC | | | | | | | | | | |
| Chromosome | start | end | strand | MM | target_seq | PAM | alignment | distance | position | gene name | gene id |
| GU305914.1 | 65108 | 65130 | + | 0 | GGAGGAAGAAGACGACGACG | AGG | \|\|\|\|\|\|\|\|[\|\|\|\|\|\|\|\|\|\|\|\|]PAM | NA | Intergenic | NA | NA |
| GU305914.1 | 118494 | 118516 | - | 2 | GGAGGACGAGGACGACGACG | AGG | \|\|\|\|\|\|-\|[\|-\|\|\|\|\|\|\|\|\|\|]PAM | NA | Intergenic | NA | NA |
| GU305914.1 | 118842 | 118864 | - | 4 | AGAAGACGAAGAGGACGACG | AGG | -\|\|-\|\|-\|[\|\|\|\|-\|\|\|\|\|\|\|]PAM | NA | Intergenic | NA | NA |
| GU305914.1 | 65111 | 65133 | + | 3 | GGAAGAAGACGACGACGAGG | AGG | \|\|\|-\|\|\|\|[\|-\|\|\|\|\|\|\|\|-\|]PAM | NA | Intergenic | NA | NA |
| GU305914.1 | 118851 | 118873 | - | 3 | GGAGGAGGAAGAAGACGAAG | AGG | \|\|\|\|\|\|-\|[\|\|\|\|-\|\|\|\|\|-\|]PAM | NA | Intergenic | NA | NA |
| T4449 | GGAAGAAGACGACGACGAGGAGG | 1000 | | | | | | | | | |
| Oligo fwd | CACCGGAAGAAGACGACGACGAGG | | | | | | | | | | |
| Oligo rev | AAACCCTCGTCGTCGTCTTCTTCC | | | | | | | | | | |
| Chromosome | start | end | strand | MM | target_seq | PAM | alignment | distance | position | gene name | gene id |
| GU305914.1 | 65111 | 65133 | + | 0 | GGAAGAAGACGACGACGAGG | AGG | \|\|\|\|\|\|\|\|[\|\|\|\|\|\|\|\|\|\|\|\|]PAM | NA | Intergenic | NA | NA |
| GU305914.1 | 118809 | 118831 | - | 4 | CGACGATGATGACGACGAGG | AGG | -\|\|-\|\|-\|[\|-\|\|\|\|\|\|\|\|\|\|]PAM | NA | Intergenic | NA | NA |
| GU305914.1 | 65114 | 65136 | + | 3 | AGAAGACGACGACGAGGAGG | AGG | -\|\|\|\|\|-\|[\|\|\|\|\|\|\|-\|\|\|\|]PAM | NA | Intergenic | NA | NA |
| GU305914.1 | 65108 | 65130 | + | 3 | GGAGGAAGAAGACGACGACG | AGG | \|\|\|-\|\|\|\|[\|-\|\|\|\|\|\|\|\|-\|]PAM | NA | Intergenic | NA | NA |
| GU305914.1 | 118494 | 118516 | - | 4 | GGAGGACGAGGACGACGACG | AGG | \|\|\|-\|\|-\|[\|-\|\|\|\|\|\|\|\|-\|]PAM | NA | Intergenic | NA | NA |
| GU305914.1 | 52531 | 52553 | - | 4 | TGAAGATGACGACTACGATG | GGG | -\|\|\|\|\|-\|[\|\|\|\|\|-\|\|\|\|-\|]PAM | NA | Intergenic | NA | NA |
| T4450 | AGAAGACGACGACGAGGAGGAGG | 1000 | | | | | | | | | |
| Oligo fwd | CACCAGAAGACGACGACGAGGAGG | | | | | | | | | | |
| Oligo rev | AAACCCTCCTCGTCGTCGTCTTCT | | | | | | | | | | |
| Chromosome | start | end | strand | MM | target_seq | PAM | alignment | distance | position | gene name | gene id |
| GU305914.1 | 65114 | 65136 | + | 0 | AGAAGACGACGACGAGGAGG | AGG | \|\|\|\|\|\|\|\|[\|\|\|\|\|\|\|\|\|\|\|\|]PAM | NA | Intergenic | NA | NA |
| GU305914.1 | 65111 | 65133 | + | 3 | GGAAGAAGACGACGACGAGG | AGG | -\|\|\|\|\|-\|[\|\|\|\|\|\|\|-\|\|\|\|]PAM | NA | Intergenic | NA | NA |
| GU305914.1 | 140717 | 140739 | + | 4 | AGATGAGGACGAGGAGGATG | AGG | \|\|\|-\|\|-\|[\|\|\|\|-\|\|\|\|\|-\|]PAM | NA | Intergenic | NA | NA |
| T4451 | GAAGACGACGACGAGGAGGAGGG | 1000 | | | | | | | | | |
| Oligo fwd | CACCGAAGACGACGACGAGGAGGA | | | | | | | | | | |
| Oligo rev | AAACTCCTCCTCGTCGTCGTCTTC | | | | | | | | | | |
| Chromosome | start | end | strand | MM | target_seq | PAM | alignment | distance | position | gene name | gene id |
| GU305914.1 | 65115 | 65137 | + | 0 | GAAGACGACGACGAGGAGGA | GGG | \|\|\|\|\|\|\|\|[\|\|\|\|\|\|\|\|\|\|\|\|]PAM | NA | Intergenic | NA | NA |
| GU305914.1 | 118805 | 118827 | - | 2 | GATGATGACGACGAGGAGGA | CGG | \|\|-\|\|-\|\|[\|\|\|\|\|\|\|\|\|\|\|\|]PAM | NA | Intergenic | NA | NA |
| GU305914.1 | 77033 | 77055 | - | 4 | GACGAGGATGAGGAGGAGGA | CGG | \|\|-\|\|-\|\|[-\|\|-\|\|\|\|\|\|\|\|]PAM | NA | Intergenic | NA | NA |
| GU305914.1 | 65118 | 65140 | + | 3 | GACGACGACGAGGAGGAGGG | CGG | \|\|-\|\|\|\|\|[\|\|\|-\|\|\|\|\|\|\|-]PAM | NA | Intergenic | NA | NA |
| GU305914.1 | 25768 | 25790 | + | 3 | GAAGACGTCGACGAGGACCA | GGG | \|\|\|\|\|\|\|-[\|\|\|\|\|\|\|\|\|--\|]PAM | NA | Intergenic | NA | NA |
| T4452 | GACGACGACGAGGAGGAGGGCGG | 1000 | | | | | | | | | |
| Oligo fwd | CACCGACGACGACGAGGAGGAGGG | | | | | | | | | | |
| Oligo rev | AAACCCCTCCTCCTCGTCGTCGTC | | | | | | | | | | |
| Chromosome | start | end | strand | MM | target_seq | PAM | alignment | distance | position | gene name | gene id |
| GU305914.1 | 65118 | 65140 | + | 0 | GACGACGACGAGGAGGAGGG | CGG | \|\|\|\|\|\|\|\|[\|\|\|\|\|\|\|\|\|\|\|\|]PAM | NA | Intergenic | NA | NA |
| GU305914.1 | 77033 | 77055 | - | 3 | GACGAGGATGAGGAGGAGGA | CGG | \|\|\|\|\|-\|\|[-\|\|\|\|\|\|\|\|\|\|-]PAM | NA | Intergenic | NA | NA |
| GU305914.1 | 65115 | 65137 | + | 3 | GAAGACGACGACGAGGAGGA | GGG | \|\|-\|\|\|\|\|[\|\|\|-\|\|\|\|\|\|\|-]PAM | NA | Intergenic | NA | NA |
| GU305914.1 | 118805 | 118827 | - | 4 | GATGATGACGACGAGGAGGA | CGG | \|\|-\|\|-\|\|[\|\|\|-\|\|\|\|\|\|\|-]PAM | NA | Intergenic | NA | NA |
| T4453 | CGACGAGGAGGAGGGCGGCGAGG | 1000 | | | | | | | | | |
| Oligo fwd | CACCCGACGAGGAGGAGGGCGGCG | | | | | | | | | | |
| Oligo rev | AAACCGCCGCCCTCCTCCTCGTCG | | | | | | | | | | |
| Chromosome | start | end | strand | MM | target_seq | PAM | alignment | distance | position | gene name | gene id |
| GU305914.1 | 65123 | 65145 | + | 0 | CGACGAGGAGGAGGGCGGCG | AGG | \|\|\|\|\|\|\|\|[\|\|\|\|\|\|\|\|\|\|\|\|]PAM | NA | Intergenic | NA | NA |
| GU305914.1 | 150931 | 150953 | - | 3 | CGCGGAGGACGAGGGCGGCG | CGG | \|\|--\|\|\|\|[\|-\|\|\|\|\|\|\|\|\|\|]PAM | NA | Intergenic | NA | NA |
| T4454 | GACGAGGAGGAGGGCGGCGAGGG | 1000 | | | | | | | | | |
| Oligo fwd | CACCGACGAGGAGGAGGGCGGCGA | | | | | | | | | | |
| Oligo rev | AAACTCGCCGCCCTCCTCCTCGTC | | | | | | | | | | |
| Chromosome | start | end | strand | MM | target_seq | PAM | alignment | distance | position | gene name | gene id |
| GU305914.1 | 65124 | 65146 | + | 0 | GACGAGGAGGAGGGCGGCGA | GGG | \|\|\|\|\|\|\|\|[\|\|\|\|\|\|\|\|\|\|\|\|]PAM | NA | Intergenic | NA | NA |

| | | | | | | | | | | | |
|---|---|---|---|---|---|---|---|---|---|---|---|
| T4455 | ACGAGGAGGAGGGCGGCGAGGGG | 1000 | | | | | | | | | |
| Oligo fwd | CACCACGAGGAGGAGGGCGGCGAG | | | | | | | | | | |
| Oligo rev | AAACCTCGCCGCCCTCCTCCTCGT | | | | | | | | | | |
| Chromosome | start | end | strand | MM | target_seq | PAM | alignment | distance | position | gene name | gene id |
| GU305914.1 | 65125 | 65147 | + | 0 | ACGAGGAGGAGGGCGGCGAG | GGG | \|\|\|\|\|\|\|\|[\|\|\|\|\|\|\|\|\|\|\|\|]PAM | NA | Intergenic | NA | NA |
| T4456 | CGAGGAGGAGGGCGGCGAGGGGG | 1000 | | | | | | | | | |
| Oligo fwd | CACCCGAGGAGGAGGGCGGCGAGG | | | | | | | | | | |
| Oligo rev | AAACCCTCGCCGCCCTCCTCCTCG | | | | | | | | | | |
| Chromosome | start | end | strand | MM | target_seq | PAM | alignment | distance | position | gene name | gene id |
| GU305914.1 | 65126 | 65148 | + | 0 | CGAGGAGGAGGGCGGCGAGG | GGG | \|\|\|\|\|\|\|\|[\|\|\|\|\|\|\|\|\|\|\|\|]PAM | NA | Intergenic | NA | NA |
| GU305914.1 | 75736 | 75758 | + | 4 | GGAAGTGGAGGTCGGCGAGG | CGG | -\|\|-\|-\|\|[\|\|\|-\|\|\|\|\|\|\|\|]PAM | NA | Intergenic | NA | NA |
| GU305914.1 | 75860 | 75882 | + | 4 | GGAGGCGGCGGTCGGCGAGG | GGG | -\|\|\|\|-\|\|[-\|\|-\|\|\|\|\|\|\|\|]PAM | NA | Intergenic | NA | NA |
| T4457 | GAGGAGGAGGGCGGCGAGGGGGG | 1000 | | | | | | | | | |
| Oligo fwd | CACCGAGGAGGAGGGCGGCGAGGG | | | | | | | | | | |
| Oligo rev | AAACCCCTCGCCGCCCTCCTCCTC | | | | | | | | | | |
| Chromosome | start | end | strand | MM | target_seq | PAM | alignment | distance | position | gene name | gene id |
| GU305914.1 | 65127 | 65149 | + | 0 | GAGGAGGAGGGCGGCGAGGG | GGG | \|\|\|\|\|\|\|\|[\|\|\|\|\|\|\|\|\|\|\|\|]PAM | NA | Intergenic | NA | NA |
| T4458 | CGACTTCTGTGTGACCATCGCGG | 1000 | | | | | | | | | |
| Oligo fwd | CACCCGACTTCTGTGTGACCATCG | | | | | | | | | | |
| Oligo rev | AAACCGATGGTCACACAGAAGTCG | | | | | | | | | | |
| Chromosome | start | end | strand | MM | target_seq | PAM | alignment | distance | position | gene name | gene id |
| GU305914.1 | 65174 | 65196 | + | 0 | CGACTTCTGTGTGACCATCG | CGG | \|\|\|\|\|\|\|\|[\|\|\|\|\|\|\|\|\|\|\|\|]PAM | NA | Intergenic | NA | NA |
| T4459 | CATTTTATTCACCCAGACCCAGG | 1000 | | | | | | | | | |
| Oligo fwd | CACCCATTTTATTCACCCAGACCC | | | | | | | | | | |
| Oligo rev | AAACGGGTCTGGGTGAATAAAATG | | | | | | | | | | |
| Chromosome | start | end | strand | MM | target_seq | PAM | alignment | distance | position | gene name | gene id |
| GU305914.1 | 65198 | 65220 | + | 0 | CATTTTATTCACCCAGACCC | AGG | \|\|\|\|\|\|\|\|[\|\|\|\|\|\|\|\|\|\|\|\|]PAM | NA | Intergenic | NA | NA |
| T4460 | CCCAGGAATCGTCCCGCTCGCGG | 1000 | | | | | | | | | |
| Oligo fwd | CACCCCCAGGAATCGTCCCGCTCG | | | | | | | | | | |
| Oligo rev | AAACCGAGCGGGACGATTCCTGGG | | | | | | | | | | |
| Chromosome | start | end | strand | MM | target_seq | PAM | alignment | distance | position | gene name | gene id |
| GU305914.1 | 65215 | 65237 | + | 0 | CCCAGGAATCGTCCCGCTCG | CGG | \|\|\|\|\|\|\|\|[\|\|\|\|\|\|\|\|\|\|\|\|]PAM | NA | Intergenic | NA | NA |
| T4461 | AAGACGTGATCGAGAAGACGCGG | 1000 | | | | | | | | | |
| Oligo fwd | CACCAAGACGTGATCGAGAAGACG | | | | | | | | | | |
| Oligo rev | AAACCGTCTTCTCGATCACGTCTT | | | | | | | | | | |
| Chromosome | start | end | strand | MM | target_seq | PAM | alignment | distance | position | gene name | gene id |
| GU305914.1 | 65251 | 65273 | + | 0 | AAGACGTGATCGAGAAGACG | CGG | \|\|\|\|\|\|\|\|[\|\|\|\|\|\|\|\|\|\|\|\|]PAM | NA | Intergenic | NA | NA |
| T4462 | AGACGTGATCGAGAAGACGCGGG | 1000 | | | | | | | | | |
| Oligo fwd | CACCAGACGTGATCGAGAAGACGC | | | | | | | | | | |
| Oligo rev | AAACGCGTCTTCTCGATCACGTCT | | | | | | | | | | |
| Chromosome | start | end | strand | MM | target_seq | PAM | alignment | distance | position | gene name | gene id |
| GU305914.1 | 65252 | 65274 | + | 0 | AGACGTGATCGAGAAGACGC | GGG | \|\|\|\|\|\|\|\|[\|\|\|\|\|\|\|\|\|\|\|\|]PAM | NA | Intergenic | NA | NA |
| GU305914.1 | 48712 | 48734 | + | 4 | AGATGGGATCGAGCCGACGC | AGG | \|\|\|-\|-\|\|[\|\|\|\|\|--\|\|\|\|\|]PAM | NA | Intergenic | NA | NA |
| T4463 | ACGCGGGCGTTCTACCGCCTCGG | 1000 | | | | | | | | | |
| Oligo fwd | CACCACGCGGGCGTTCTACCGCCT | | | | | | | | | | |
| Oligo rev | AAACAGGCGGTAGAACGCCCGCGT | | | | | | | | | | |
| Chromosome | start | end | strand | MM | target_seq | PAM | alignment | distance | position | gene name | gene id |
| GU305914.1 | 65268 | 65290 | + | 0 | ACGCGGGCGTTCTACCGCCT | CGG | \|\|\|\|\|\|\|\|[\|\|\|\|\|\|\|\|\|\|\|\|]PAM | NA | Intergenic | NA | NA |
| T4464 | CGCGGGCGTTCTACCGCCTCGGG | 1000 | | | | | | | | | |
| Oligo fwd | CACCCGCGGGCGTTCTACCGCCTC | | | | | | | | | | |
| Oligo rev | AAACGAGGCGGTAGAACGCCCGCG | | | | | | | | | | |
| Chromosome | start | end | strand | MM | target_seq | PAM | alignment | distance | position | gene name | gene id |
| GU305914.1 | 65269 | 65291 | + | 0 | CGCGGGCGTTCTACCGCCTC | GGG | \|\|\|\|\|\|\|\|[\|\|\|\|\|\|\|\|\|\|\|\|]PAM | NA | Intergenic | NA | NA |
| T4465 | CGGGTTGACCCCGAAGAGCACGG | 1000 | | | | | | | | | |
| Oligo fwd | CACCCGGGTTGACCCCGAAGAGCA | | | | | | | | | | |
| Oligo rev | AAACTGCTCTTCGGGGTCAACCCG | | | | | | | | | | |
| Chromosome | start | end | strand | MM | target_seq | PAM | alignment | distance | position | gene name | gene id |
| GU305914.1 | 65288 | 65310 | + | 0 | CGGGTTGACCCCGAAGAGCA | CGG | \|\|\|\|\|\|\|\|[\|\|\|\|\|\|\|\|\|\|\|\|]PAM | NA | Intergenic | NA | NA |
| T4466 | CCTGCACGCCACGTACCGCACGG | 1000 | | | | | | | | | |
| Oligo fwd | CACCCCTGCACGCCACGTACCGCA | | | | | | | | | | |
| Oligo rev | AAACTGCGGTACGTGGCGTGCAGG | | | | | | | | | | |
| Chromosome | start | end | strand | MM | target_seq | PAM | alignment | distance | position | gene name | gene id |
| GU305914.1 | 65333 | 65355 | + | 0 | CCTGCACGCCACGTACCGCA | CGG | \|\|\|\|\|\|\|\|[\|\|\|\|\|\|\|\|\|\|\|\|]PAM | NA | Intergenic | NA | NA |
| T4467 | CGCCACGTACCGCACGGCCGCGG | 1000 | | | | | | | | | |
| Oligo fwd | CACCCGCCACGTACCGCACGGCCG | | | | | | | | | | |
| Oligo rev | AAACCGGCCGTGCGGTACGTGGCG | | | | | | | | | | |
| Chromosome | start | end | strand | MM | target_seq | PAM | alignment | distance | position | gene name | gene id |
| GU305914.1 | 65339 | 65361 | + | 0 | CGCCACGTACCGCACGGCCG | CGG | \|\|\|\|\|\|\|\|[\|\|\|\|\|\|\|\|\|\|\|\|]PAM | NA | Intergenic | NA | NA |
| GU305914.1 | 195888 | 195910 | - | 3 | CTCCACGGATCGCACGGCCG | GGG | \|-\|\|\|\|\|-[\|-\|\|\|\|\|\|\|\|\|\|]PAM | NA | Intergenic | NA | NA |
| T4468 | AAGCATCGACGAGTTCAGCGAGG | 1000 | | | | | | | | | |
| Oligo fwd | CACCAAGCATCGACGAGTTCAGCG | | | | | | | | | | |
| Oligo rev | AAACCGCTGAACTCGTCGATGCTT | | | | | | | | | | |
| Chromosome | start | end | strand | MM | target_seq | PAM | alignment | distance | position | gene name | gene id |
| GU305914.1 | 65369 | 65391 | + | 0 | AAGCATCGACGAGTTCAGCG | AGG | \|\|\|\|\|\|\|\|[\|\|\|\|\|\|\|\|\|\|\|\|]PAM | NA | Intergenic | NA | NA |
| T4469 | CATCGACGAGTTCAGCGAGGCGG | 1000 | | | | | | | | | |

<table>
<tr><td>Oligo fwd</td><td>CACCCATCGACGAGTTCAGCGAGG</td><td></td><td></td><td></td><td></td><td></td><td></td><td></td><td></td><td></td><td></td></tr>
<tr><td>Oligo rev</td><td>AAACCCTCGCTGAACTCGTCGATG</td><td></td><td></td><td></td><td></td><td></td><td></td><td></td><td></td><td></td><td></td></tr>
<tr><td>Chromosome</td><td>start</td><td>end</td><td>strand</td><td>MM</td><td>target_seq</td><td>PAM</td><td>alignment</td><td>distance</td><td>position</td><td>gene name</td><td>gene id</td></tr>
<tr><td>GU305914.1</td><td>65372</td><td>65394</td><td>+</td><td>0</td><td>CATCGACGAGTTCAGCGAGG</td><td>CGG</td><td>||||||||[||||||||||||]PAM</td><td>NA</td><td>Intergenic</td><td>NA</td><td>NA</td></tr>
<tr><td></td><td></td><td></td><td></td><td></td><td></td><td></td><td></td><td></td><td></td><td></td><td></td></tr>
<tr><td>T4470</td><td>CGACGAGTTCAGCGAGGCGGCGG</td><td>1000</td><td></td><td></td><td></td><td></td><td></td><td></td><td></td><td></td><td></td></tr>
<tr><td>Oligo fwd</td><td>CACCCGACGAGTTCAGCGAGGCGG</td><td></td><td></td><td></td><td></td><td></td><td></td><td></td><td></td><td></td><td></td></tr>
<tr><td>Oligo rev</td><td>AAACCCGCCTCGCTGAACTCGTCG</td><td></td><td></td><td></td><td></td><td></td><td></td><td></td><td></td><td></td><td></td></tr>
<tr><td>Chromosome</td><td>start</td><td>end</td><td>strand</td><td>MM</td><td>target_seq</td><td>PAM</td><td>alignment</td><td>distance</td><td>position</td><td>gene name</td><td>gene id</td></tr>
<tr><td>GU305914.1</td><td>65375</td><td>65397</td><td>+</td><td>0</td><td>CGACGAGTTCAGCGAGGCGG</td><td>CGG</td><td>||||||||[||||||||||||]PAM</td><td>NA</td><td>Intergenic</td><td>NA</td><td>NA</td></tr>
<tr><td></td><td></td><td></td><td></td><td></td><td></td><td></td><td></td><td></td><td></td><td></td><td></td></tr>
<tr><td>T4471</td><td>GGCGGACCACATCACCTTTCTGG</td><td>1000</td><td></td><td></td><td></td><td></td><td></td><td></td><td></td><td></td><td></td></tr>
<tr><td>Oligo fwd</td><td>CACCGGCGGACCACATCACCTTTC</td><td></td><td></td><td></td><td></td><td></td><td></td><td></td><td></td><td></td><td></td></tr>
<tr><td>Oligo rev</td><td>AAACGAAAGGTGATGTGGTCCGCC</td><td></td><td></td><td></td><td></td><td></td><td></td><td></td><td></td><td></td><td></td></tr>
<tr><td>Chromosome</td><td>start</td><td>end</td><td>strand</td><td>MM</td><td>target_seq</td><td>PAM</td><td>alignment</td><td>distance</td><td>position</td><td>gene name</td><td>gene id</td></tr>
<tr><td>GU305914.1</td><td>65393</td><td>65415</td><td>+</td><td>0</td><td>GGCGGACCACATCACCTTTC</td><td>TGG</td><td>||||||||[||||||||||||]PAM</td><td>NA</td><td>Intergenic</td><td>NA</td><td>NA</td></tr>
<tr><td></td><td></td><td></td><td></td><td></td><td></td><td></td><td></td><td></td><td></td><td></td><td></td></tr>
<tr><td>T4472</td><td>TCCTGTTTTACCACGCGAAGCGG</td><td>1000</td><td></td><td></td><td></td><td></td><td></td><td></td><td></td><td></td><td></td></tr>
<tr><td>Oligo fwd</td><td>CACCTCCTGTTTTACCACGCGAAG</td><td></td><td></td><td></td><td></td><td></td><td></td><td></td><td></td><td></td><td></td></tr>
<tr><td>Oligo rev</td><td>AAACCTTCGCGTGGTAAAACAGGA</td><td></td><td></td><td></td><td></td><td></td><td></td><td></td><td></td><td></td><td></td></tr>
<tr><td>Chromosome</td><td>start</td><td>end</td><td>strand</td><td>MM</td><td>target_seq</td><td>PAM</td><td>alignment</td><td>distance</td><td>position</td><td>gene name</td><td>gene id</td></tr>
<tr><td>GU305914.1</td><td>65467</td><td>65489</td><td>+</td><td>0</td><td>TCCTGTTTTACCACGCGAAG</td><td>CGG</td><td>||||||||[||||||||||||]PAM</td><td>NA</td><td>Intergenic</td><td>NA</td><td>NA</td></tr>
<tr><td></td><td></td><td></td><td></td><td></td><td></td><td></td><td></td><td></td><td></td><td></td><td></td></tr>
<tr><td>T4473</td><td>CCACGCGAAGCGGCTGATCCTGG</td><td>1000</td><td></td><td></td><td></td><td></td><td></td><td></td><td></td><td></td><td></td></tr>
<tr><td>Oligo fwd</td><td>CACCCCACGCGAAGCGGCTGATCC</td><td></td><td></td><td></td><td></td><td></td><td></td><td></td><td></td><td></td><td></td></tr>
<tr><td>Oligo rev</td><td>AAACGGATCAGCCGCTTCGCGTGG</td><td></td><td></td><td></td><td></td><td></td><td></td><td></td><td></td><td></td><td></td></tr>
<tr><td>Chromosome</td><td>start</td><td>end</td><td>strand</td><td>MM</td><td>target_seq</td><td>PAM</td><td>alignment</td><td>distance</td><td>position</td><td>gene name</td><td>gene id</td></tr>
<tr><td>GU305914.1</td><td>65477</td><td>65499</td><td>+</td><td>0</td><td>CCACGCGAAGCGGCTGATCC</td><td>TGG</td><td>||||||||[||||||||||||]PAM</td><td>NA</td><td>Intergenic</td><td>NA</td><td>NA</td></tr>
<tr><td></td><td></td><td></td><td></td><td></td><td></td><td></td><td></td><td></td><td></td><td></td><td></td></tr>
<tr><td>T4474</td><td>ACAACAGCGCTCCCTGTTGACGG</td><td>1000</td><td></td><td></td><td></td><td></td><td></td><td></td><td></td><td></td><td></td></tr>
<tr><td>Oligo fwd</td><td>CACCACAACAGCGCTCCCTGTTGA</td><td></td><td></td><td></td><td></td><td></td><td></td><td></td><td></td><td></td><td></td></tr>
<tr><td>Oligo rev</td><td>AAACTCAACAGGGAGCGCTGTTGT</td><td></td><td></td><td></td><td></td><td></td><td></td><td></td><td></td><td></td><td></td></tr>
<tr><td>Chromosome</td><td>start</td><td>end</td><td>strand</td><td>MM</td><td>target_seq</td><td>PAM</td><td>alignment</td><td>distance</td><td>position</td><td>gene name</td><td>gene id</td></tr>
<tr><td>GU305914.1</td><td>65501</td><td>65523</td><td>+</td><td>0</td><td>ACAACAGCGCTCCCTGTTGA</td><td>CGG</td><td>||||||||[||||||||||||]PAM</td><td>NA</td><td>Intergenic</td><td>NA</td><td>NA</td></tr>
<tr><td></td><td></td><td></td><td></td><td></td><td></td><td></td><td></td><td></td><td></td><td></td><td></td></tr>
<tr><td>T4475</td><td>CAACAGCGCTCCCTGTTGACGGG</td><td>1000</td><td></td><td></td><td></td><td></td><td></td><td></td><td></td><td></td><td></td></tr>
<tr><td>Oligo fwd</td><td>CACCCAACAGCGCTCCCTGTTGAC</td><td></td><td></td><td></td><td></td><td></td><td></td><td></td><td></td><td></td><td></td></tr>
<tr><td>Oligo rev</td><td>AAACGTCAACAGGGAGCGCTGTTG</td><td></td><td></td><td></td><td></td><td></td><td></td><td></td><td></td><td></td><td></td></tr>
<tr><td>Chromosome</td><td>start</td><td>end</td><td>strand</td><td>MM</td><td>target_seq</td><td>PAM</td><td>alignment</td><td>distance</td><td>position</td><td>gene name</td><td>gene id</td></tr>
<tr><td>GU305914.1</td><td>65502</td><td>65524</td><td>+</td><td>0</td><td>CAACAGCGCTCCCTGTTGAC</td><td>GGG</td><td>||||||||[||||||||||||]PAM</td><td>NA</td><td>Intergenic</td><td>NA</td><td>NA</td></tr>
<tr><td></td><td></td><td></td><td></td><td></td><td></td><td></td><td></td><td></td><td></td><td></td><td></td></tr>
<tr><td>T4476</td><td>GCTCCCTGTTGACGGGCAACCGG</td><td>1000</td><td></td><td></td><td></td><td></td><td></td><td></td><td></td><td></td><td></td></tr>
<tr><td>Oligo fwd</td><td>CACCGCTCCCTGTTGACGGGCAAC</td><td></td><td></td><td></td><td></td><td></td><td></td><td></td><td></td><td></td><td></td></tr>
<tr><td>Oligo rev</td><td>AAACGTTGCCCGTCAACAGGGAGC</td><td></td><td></td><td></td><td></td><td></td><td></td><td></td><td></td><td></td><td></td></tr>
<tr><td>Chromosome</td><td>start</td><td>end</td><td>strand</td><td>MM</td><td>target_seq</td><td>PAM</td><td>alignment</td><td>distance</td><td>position</td><td>gene name</td><td>gene id</td></tr>
<tr><td>GU305914.1</td><td>65509</td><td>65531</td><td>+</td><td>0</td><td>GCTCCCTGTTGACGGGCAAC</td><td>CGG</td><td>||||||||[||||||||||||]PAM</td><td>NA</td><td>Intergenic</td><td>NA</td><td>NA</td></tr>
<tr><td></td><td></td><td></td><td></td><td></td><td></td><td></td><td></td><td></td><td></td><td></td><td></td></tr>
<tr><td>T4477</td><td>GACGGGCAACCGGTCCCTGCAGG</td><td>1000</td><td></td><td></td><td></td><td></td><td></td><td></td><td></td><td></td><td></td></tr>
<tr><td>Oligo fwd</td><td>CACCGACGGGCAACCGGTCCCTGC</td><td></td><td></td><td></td><td></td><td></td><td></td><td></td><td></td><td></td><td></td></tr>
<tr><td>Oligo rev</td><td>AAACGCAGGGACCGGTTGCCCGTC</td><td></td><td></td><td></td><td></td><td></td><td></td><td></td><td></td><td></td><td></td></tr>
<tr><td>Chromosome</td><td>start</td><td>end</td><td>strand</td><td>MM</td><td>target_seq</td><td>PAM</td><td>alignment</td><td>distance</td><td>position</td><td>gene name</td><td>gene id</td></tr>
<tr><td>GU305914.1</td><td>65519</td><td>65541</td><td>+</td><td>0</td><td>GACGGGCAACCGGTCCCTGC</td><td>AGG</td><td>||||||||[||||||||||||]PAM</td><td>NA</td><td>Intergenic</td><td>NA</td><td>NA</td></tr>
<tr><td></td><td></td><td></td><td></td><td></td><td></td><td></td><td></td><td></td><td></td><td></td><td></td></tr>
<tr><td>T4478</td><td>ACCGGTCCCTGCAGGACGTGTGG</td><td>1000</td><td></td><td></td><td></td><td></td><td></td><td></td><td></td><td></td><td></td></tr>
<tr><td>Oligo fwd</td><td>CACCACCGGTCCCTGCAGGACGTG</td><td></td><td></td><td></td><td></td><td></td><td></td><td></td><td></td><td></td><td></td></tr>
<tr><td>Oligo rev</td><td>AAACCACGTCCTGCAGGGACCGGT</td><td></td><td></td><td></td><td></td><td></td><td></td><td></td><td></td><td></td><td></td></tr>
<tr><td>Chromosome</td><td>start</td><td>end</td><td>strand</td><td>MM</td><td>target_seq</td><td>PAM</td><td>alignment</td><td>distance</td><td>position</td><td>gene name</td><td>gene id</td></tr>
<tr><td>GU305914.1</td><td>65527</td><td>65549</td><td>+</td><td>0</td><td>ACCGGTCCCTGCAGGACGTG</td><td>TGG</td><td>||||||||[||||||||||||]PAM</td><td>NA</td><td>Intergenic</td><td>NA</td><td>NA</td></tr>
<tr><td></td><td></td><td></td><td></td><td></td><td></td><td></td><td></td><td></td><td></td><td></td><td></td></tr>
<tr><td>T4479</td><td>CCGGTCCCTGCAGGACGTGTGGG</td><td>1000</td><td></td><td></td><td></td><td></td><td></td><td></td><td></td><td></td><td></td></tr>
<tr><td>Oligo fwd</td><td>CACCCCGGTCCCTGCAGGACGTGT</td><td></td><td></td><td></td><td></td><td></td><td></td><td></td><td></td><td></td><td></td></tr>
<tr><td>Oligo rev</td><td>AAACACACGTCCTGCAGGGACCGG</td><td></td><td></td><td></td><td></td><td></td><td></td><td></td><td></td><td></td><td></td></tr>
<tr><td>Chromosome</td><td>start</td><td>end</td><td>strand</td><td>MM</td><td>target_seq</td><td>PAM</td><td>alignment</td><td>distance</td><td>position</td><td>gene name</td><td>gene id</td></tr>
<tr><td>GU305914.1</td><td>65528</td><td>65550</td><td>+</td><td>0</td><td>CCGGTCCCTGCAGGACGTGT</td><td>GGG</td><td>||||||||[||||||||||||]PAM</td><td>NA</td><td>Intergenic</td><td>NA</td><td>NA</td></tr>
<tr><td></td><td></td><td></td><td></td><td></td><td></td><td></td><td></td><td></td><td></td><td></td><td></td></tr>
<tr><td>T4480</td><td>GCAGGACGTGTGGGCCAACGTGG</td><td>1000</td><td></td><td></td><td></td><td></td><td></td><td></td><td></td><td></td><td></td></tr>
<tr><td>Oligo fwd</td><td>CACCGCAGGACGTGTGGGCCAACG</td><td></td><td></td><td></td><td></td><td></td><td></td><td></td><td></td><td></td><td></td></tr>
<tr><td>Oligo rev</td><td>AAACCGTTGGCCCACACGTCCTGC</td><td></td><td></td><td></td><td></td><td></td><td></td><td></td><td></td><td></td><td></td></tr>
<tr><td>Chromosome</td><td>start</td><td>end</td><td>strand</td><td>MM</td><td>target_seq</td><td>PAM</td><td>alignment</td><td>distance</td><td>position</td><td>gene name</td><td>gene id</td></tr>
<tr><td>GU305914.1</td><td>65537</td><td>65559</td><td>+</td><td>0</td><td>GCAGGACGTGTGGGCCAACG</td><td>TGG</td><td>||||||||[||||||||||||]PAM</td><td>NA</td><td>Intergenic</td><td>NA</td><td>NA</td></tr>
<tr><td></td><td></td><td></td><td></td><td></td><td></td><td></td><td></td><td></td><td></td><td></td><td></td></tr>
<tr><td>T4481</td><td>ACGTGGCGTTCAACATCAACCGG</td><td>1000</td><td></td><td></td><td></td><td></td><td></td><td></td><td></td><td></td><td></td></tr>
<tr><td>Oligo fwd</td><td>CACCACGTGGCGTTCAACATCAAC</td><td></td><td></td><td></td><td></td><td></td><td></td><td></td><td></td><td></td><td></td></tr>
<tr><td>Oligo rev</td><td>AAACGTTGATGTTGAACGCCACGT</td><td></td><td></td><td></td><td></td><td></td><td></td><td></td><td></td><td></td><td></td></tr>
<tr><td>Chromosome</td><td>start</td><td>end</td><td>strand</td><td>MM</td><td>target_seq</td><td>PAM</td><td>alignment</td><td>distance</td><td>position</td><td>gene name</td><td>gene id</td></tr>
<tr><td>GU305914.1</td><td>65554</td><td>65576</td><td>+</td><td>0</td><td>ACGTGGCGTTCAACATCAAC</td><td>CGG</td><td>||||||||[||||||||||||]PAM</td><td>NA</td><td>Intergenic</td><td>NA</td><td>NA</td></tr>
<tr><td></td><td></td><td></td><td></td><td></td><td></td><td></td><td></td><td></td><td></td><td></td><td></td></tr>
<tr><td>T4482</td><td>AGAGTTCACGCAGCTGACGCTGG</td><td>1000</td><td></td><td></td><td></td><td></td><td></td><td></td><td></td><td></td><td></td></tr>
<tr><td>Oligo fwd</td><td>CACCAGAGTTCACGCAGCTGACGC</td><td></td><td></td><td></td><td></td><td></td><td></td><td></td><td></td><td></td><td></td></tr>
<tr><td>Oligo rev</td><td>AAACGCGTCAGCTGCGTGAACTCT</td><td></td><td></td><td></td><td></td><td></td><td></td><td></td><td></td><td></td><td></td></tr>
<tr><td>Chromosome</td><td>start</td><td>end</td><td>strand</td><td>MM</td><td>target_seq</td><td>PAM</td><td>alignment</td><td>distance</td><td>position</td><td>gene name</td><td>gene id</td></tr>
<tr><td>GU305914.1</td><td>65603</td><td>65625</td><td>+</td><td>0</td><td>AGAGTTCACGCAGCTGACGC</td><td>TGG</td><td>||||||||[||||||||||||]PAM</td><td>NA</td><td>Intergenic</td><td>NA</td><td>NA</td></tr>
<tr><td></td><td></td><td></td><td></td><td></td><td></td><td></td><td></td><td></td><td></td><td></td><td></td></tr>
<tr><td>T4483</td><td>GAGTTCACGCAGCTGACGCTGGG</td><td>1000</td><td></td><td></td><td></td><td></td><td></td><td></td><td></td><td></td><td></td></tr>
<tr><td>Oligo fwd</td><td>CACCGAGTTCACGCAGCTGACGCT</td><td></td><td></td><td></td><td></td><td></td><td></td><td></td><td></td><td></td><td></td></tr>
<tr><td>Oligo rev</td><td>AAACAGCGTCAGCTGCGTGAACTC</td><td></td><td></td><td></td><td></td><td></td><td></td><td></td><td></td><td></td><td></td></tr>
<tr><td>Chromosome</td><td>start</td><td>end</td><td>strand</td><td>MM</td><td>target_seq</td><td>PAM</td><td>alignment</td><td>distance</td><td>position</td><td>gene name</td><td>gene id</td></tr>
<tr><td>GU305914.1</td><td>65604</td><td>65626</td><td>+</td><td>0</td><td>GAGTTCACGCAGCTGACGCT</td><td>GGG</td><td>||||||||[||||||||||||]PAM</td><td>NA</td><td>Intergenic</td><td>NA</td><td>NA</td></tr>
</table>

<table>
<tr><td>T4484</td><td>GCTGACGCTGGGCGCGTCGCCGG</td><td>1000</td></tr>
<tr><td>Oligo fwd</td><td>CACCGCTGACGCTGGGCGCGTCGC</td></tr>
<tr><td>Oligo rev</td><td>AAACGCGACGCGCCCAGCGTCAGC</td></tr>
<tr><td>Chromosome</td><td>start</td><td>end</td><td>strand</td><td>MM</td><td>target_seq</td><td>PAM</td><td>alignment</td><td>distance</td><td>position</td><td>gene name</td><td>gene id</td></tr>
<tr><td>GU305914.1</td><td>65615</td><td>65637</td><td>+</td><td>0</td><td>GCTGACGCTGGGCGCGTCGC</td><td>CGG</td><td>||||||||[||||||||||||]PAM</td><td>NA</td><td>Intergenic</td><td>NA</td><td>NA</td></tr>
<tr><td></td></tr>
<tr><td>T4485</td><td>GACGCTGGGCGCGTCGCCGGCGG</td><td>1000</td></tr>
<tr><td>Oligo fwd</td><td>CACCGACGCTGGGCGCGTCGCCGG</td></tr>
<tr><td>Oligo rev</td><td>AAACCCGGCGACGCGCCCAGCGTC</td></tr>
<tr><td>Chromosome</td><td>start</td><td>end</td><td>strand</td><td>MM</td><td>target_seq</td><td>PAM</td><td>alignment</td><td>distance</td><td>position</td><td>gene name</td><td>gene id</td></tr>
<tr><td>GU305914.1</td><td>65618</td><td>65640</td><td>+</td><td>0</td><td>GACGCTGGGCGCGTCGCCGG</td><td>CGG</td><td>||||||||[||||||||||||]PAM</td><td>NA</td><td>Intergenic</td><td>NA</td><td>NA</td></tr>
<tr><td>GU305914.1</td><td>27320</td><td>27342</td><td>-</td><td>4</td><td>GACGATAGACGCGTCGTCGG</td><td>CGG</td><td>||||-|-|[-|||||||-|||]PAM</td><td>NA</td><td>Intergenic</td><td>NA</td><td>NA</td></tr>
<tr><td></td></tr>
<tr><td>T4486</td><td>ACGCTGGGCGCGTCGCCGGCGGG</td><td>1000</td></tr>
<tr><td>Oligo fwd</td><td>CACCACGCTGGGCGCGTCGCCGGC</td></tr>
<tr><td>Oligo rev</td><td>AAACGCCGGCGACGCGCCCAGCGT</td></tr>
<tr><td>Chromosome</td><td>start</td><td>end</td><td>strand</td><td>MM</td><td>target_seq</td><td>PAM</td><td>alignment</td><td>distance</td><td>position</td><td>gene name</td><td>gene id</td></tr>
<tr><td>GU305914.1</td><td>65619</td><td>65641</td><td>+</td><td>0</td><td>ACGCTGGGCGCGTCGCCGGC</td><td>GGG</td><td>||||||||[||||||||||||]PAM</td><td>NA</td><td>Intergenic</td><td>NA</td><td>NA</td></tr>
<tr><td></td></tr>
<tr><td>T4487</td><td>CGAGTACCTGTACCGCGACGCGG</td><td>1000</td></tr>
<tr><td>Oligo fwd</td><td>CACCCGAGTACCTGTACCGCGACG</td></tr>
<tr><td>Oligo rev</td><td>AAACCGTCGCGGTACAGGTACTCG</td></tr>
<tr><td>Chromosome</td><td>start</td><td>end</td><td>strand</td><td>MM</td><td>target_seq</td><td>PAM</td><td>alignment</td><td>distance</td><td>position</td><td>gene name</td><td>gene id</td></tr>
<tr><td>GU305914.1</td><td>65645</td><td>65667</td><td>+</td><td>0</td><td>CGAGTACCTGTACCGCGACG</td><td>CGG</td><td>||||||||[||||||||||||]PAM</td><td>NA</td><td>Intergenic</td><td>NA</td><td>NA</td></tr>
<tr><td>GU305914.1</td><td>60462</td><td>60484</td><td>-</td><td>4</td><td>GGAGTCCCTGGTCCGCGACG</td><td>CGG</td><td>-||||-||[||--||||||||]PAM</td><td>NA</td><td>Intergenic</td><td>NA</td><td>NA</td></tr>
<tr><td></td></tr>
<tr><td>T4488</td><td>ACCGCGACGCGGCGAACAAGTGG</td><td>1000</td></tr>
<tr><td>Oligo fwd</td><td>CACCACCGCGACGCGGCGAACAAG</td></tr>
<tr><td>Oligo rev</td><td>AAACCTTGTTCGCCGCGTCGCGGT</td></tr>
<tr><td>Chromosome</td><td>start</td><td>end</td><td>strand</td><td>MM</td><td>target_seq</td><td>PAM</td><td>alignment</td><td>distance</td><td>position</td><td>gene name</td><td>gene id</td></tr>
<tr><td>GU305914.1</td><td>65656</td><td>65678</td><td>+</td><td>0</td><td>ACCGCGACGCGGCGAACAAG</td><td>TGG</td><td>||||||||[||||||||||||]PAM</td><td>NA</td><td>Intergenic</td><td>NA</td><td>NA</td></tr>
<tr><td></td></tr>
<tr><td>T4489</td><td>CCGCGACGCGGCGAACAAGTGGG</td><td>1000</td></tr>
<tr><td>Oligo fwd</td><td>CACCCCGCGACGCGGCGAACAAGT</td></tr>
<tr><td>Oligo rev</td><td>AAACACTTGTTCGCCGCGTCGCGG</td></tr>
<tr><td>Chromosome</td><td>start</td><td>end</td><td>strand</td><td>MM</td><td>target_seq</td><td>PAM</td><td>alignment</td><td>distance</td><td>position</td><td>gene name</td><td>gene id</td></tr>
<tr><td>GU305914.1</td><td>65657</td><td>65679</td><td>+</td><td>0</td><td>CCGCGACGCGGCGAACAAGT</td><td>GGG</td><td>||||||||[||||||||||||]PAM</td><td>NA</td><td>Intergenic</td><td>NA</td><td>NA</td></tr>
<tr><td></td></tr>
<tr><td>T4490</td><td>GGACGACATCGCGTTCACGCTGG</td><td>1000</td></tr>
<tr><td>Oligo fwd</td><td>CACCGGACGACATCGCGTTCACGC</td></tr>
<tr><td>Oligo rev</td><td>AAACGCGTGAACGCGATGTCGTCC</td></tr>
<tr><td>Chromosome</td><td>start</td><td>end</td><td>strand</td><td>MM</td><td>target_seq</td><td>PAM</td><td>alignment</td><td>distance</td><td>position</td><td>gene name</td><td>gene id</td></tr>
<tr><td>GU305914.1</td><td>65678</td><td>65700</td><td>+</td><td>0</td><td>GGACGACATCGCGTTCACGC</td><td>TGG</td><td>||||||||[||||||||||||]PAM</td><td>NA</td><td>Intergenic</td><td>NA</td><td>NA</td></tr>
<tr><td>GU305914.1</td><td>192902</td><td>192924</td><td>-</td><td>4</td><td>GTCCGACGGCGCGTTCACGC</td><td>TGG</td><td>|--||||-[-|||||||||||]PAM</td><td>NA</td><td>Intergenic</td><td>NA</td><td>NA</td></tr>
<tr><td></td></tr>
<tr><td>T4491</td><td>CATCGCGTTCACGCTGGACCAGG</td><td>1000</td></tr>
<tr><td>Oligo fwd</td><td>CACCCATCGCGTTCACGCTGGACC</td></tr>
<tr><td>Oligo rev</td><td>AAACGGTCCAGCGTGAACGCGATG</td></tr>
<tr><td>Chromosome</td><td>start</td><td>end</td><td>strand</td><td>MM</td><td>target_seq</td><td>PAM</td><td>alignment</td><td>distance</td><td>position</td><td>gene name</td><td>gene id</td></tr>
<tr><td>GU305914.1</td><td>65684</td><td>65706</td><td>+</td><td>0</td><td>CATCGCGTTCACGCTGGACC</td><td>AGG</td><td>||||||||[||||||||||||]PAM</td><td>NA</td><td>Intergenic</td><td>NA</td><td>NA</td></tr>
<tr><td></td></tr>
<tr><td>T4492</td><td>CGCGTTCACGCTGGACCAGGAGG</td><td>1000</td></tr>
<tr><td>Oligo fwd</td><td>CACCCGCGTTCACGCTGGACCAGG</td></tr>
<tr><td>Oligo rev</td><td>AAACCCTGGTCCAGCGTGAACGCG</td></tr>
<tr><td>Chromosome</td><td>start</td><td>end</td><td>strand</td><td>MM</td><td>target_seq</td><td>PAM</td><td>alignment</td><td>distance</td><td>position</td><td>gene name</td><td>gene id</td></tr>
<tr><td>GU305914.1</td><td>65687</td><td>65709</td><td>+</td><td>0</td><td>CGCGTTCACGCTGGACCAGG</td><td>AGG</td><td>||||||||[||||||||||||]PAM</td><td>NA</td><td>Intergenic</td><td>NA</td><td>NA</td></tr>
<tr><td></td></tr>
<tr><td>T4493</td><td>GCTGGACCAGGAGGACCTGACGG</td><td>1000</td></tr>
<tr><td>Oligo fwd</td><td>CACCGCTGGACCAGGAGGACCTGA</td></tr>
<tr><td>Oligo rev</td><td>AAACTCAGGTCCTCCTGGTCCAGC</td></tr>
<tr><td>Chromosome</td><td>start</td><td>end</td><td>strand</td><td>MM</td><td>target_seq</td><td>PAM</td><td>alignment</td><td>distance</td><td>position</td><td>gene name</td><td>gene id</td></tr>
<tr><td>GU305914.1</td><td>65696</td><td>65718</td><td>+</td><td>0</td><td>GCTGGACCAGGAGGACCTGA</td><td>CGG</td><td>||||||||[||||||||||||]PAM</td><td>NA</td><td>Intergenic</td><td>NA</td><td>NA</td></tr>
<tr><td>GU305914.1</td><td>124607</td><td>124629</td><td>-</td><td>4</td><td>GCTGCTGCAGGAGTACCTGA</td><td>CGG</td><td>||||---|[|||||-||||||]PAM</td><td>NA</td><td>Intergenic</td><td>NA</td><td>NA</td></tr>
<tr><td></td></tr>
<tr><td>T4494</td><td>CCCGCTGCCGCTGCCGCCGCAGG</td><td>1000</td></tr>
<tr><td>Oligo fwd</td><td>CACCCCCGCTGCCGCTGCCGCCGC</td></tr>
<tr><td>Oligo rev</td><td>AAACGCGGCGGCAGCGGCAGCGGG</td></tr>
<tr><td>Chromosome</td><td>start</td><td>end</td><td>strand</td><td>MM</td><td>target_seq</td><td>PAM</td><td>alignment</td><td>distance</td><td>position</td><td>gene name</td><td>gene id</td></tr>
<tr><td>GU305914.1</td><td>65720</td><td>65742</td><td>+</td><td>0</td><td>CCCGCTGCCGCTGCCGCCGC</td><td>AGG</td><td>||||||||[||||||||||||]PAM</td><td>NA</td><td>Intergenic</td><td>NA</td><td>NA</td></tr>
<tr><td>GU305914.1</td><td>164964</td><td>164986</td><td>+</td><td>3</td><td>GCTGCTGCCGCCGCCGCCGC</td><td>CGG</td><td>-|-|||||[|||-||||||||]PAM</td><td>NA</td><td>Intergenic</td><td>NA</td><td>NA</td></tr>
<tr><td>GU305914.1</td><td>103496</td><td>103518</td><td>-</td><td>4</td><td>ACCGCTGAGCCTGCCGCCGC</td><td>GGG</td><td>-||||||-[--||||||||||]PAM</td><td>NA</td><td>Intergenic</td><td>NA</td><td>NA</td></tr>
<tr><td>GU305914.1</td><td>62681</td><td>62703</td><td>-</td><td>4</td><td>GCCGCGGCTGCTGCCGCCGG</td><td>CGG</td><td>-||||-||[-||||||||||-]PAM</td><td>NA</td><td>Intergenic</td><td>NA</td><td>NA</td></tr>
<tr><td></td></tr>
<tr><td>T4495</td><td>GCAGGAAGATCCGACGCCCGAGG</td><td>1000</td></tr>
<tr><td>Oligo fwd</td><td>CACCGCAGGAAGATCCGACGCCCG</td></tr>
<tr><td>Oligo rev</td><td>AAACCGGGCGTCGGATCTTCCTGC</td></tr>
<tr><td>Chromosome</td><td>start</td><td>end</td><td>strand</td><td>MM</td><td>target_seq</td><td>PAM</td><td>alignment</td><td>distance</td><td>position</td><td>gene name</td><td>gene id</td></tr>
<tr><td>GU305914.1</td><td>65738</td><td>65760</td><td>+</td><td>0</td><td>GCAGGAAGATCCGACGCCCG</td><td>AGG</td><td>||||||||[||||||||||||]PAM</td><td>NA</td><td>Intergenic</td><td>NA</td><td>NA</td></tr>
<tr><td></td></tr>
<tr><td>T4496</td><td>CGACGCCCGAGGACATCGCGCGG</td><td>1000</td></tr>
<tr><td>Oligo fwd</td><td>CACCCGACGCCCGAGGACATCGCG</td></tr>
<tr><td>Oligo rev</td><td>AAACCGCGATGTCCTCGGGCGTCG</td></tr>
<tr><td>Chromosome</td><td>start</td><td>end</td><td>strand</td><td>MM</td><td>target_seq</td><td>PAM</td><td>alignment</td><td>distance</td><td>position</td><td>gene name</td><td>gene id</td></tr>
<tr><td>GU305914.1</td><td>65749</td><td>65771</td><td>+</td><td>0</td><td>CGACGCCCGAGGACATCGCG</td><td>CGG</td><td>||||||||[||||||||||||]PAM</td><td>NA</td><td>Intergenic</td><td>NA</td><td>NA</td></tr>
<tr><td></td></tr>
<tr><td>T4497</td><td>GACGCCCGAGGACATCGCGCGGG</td><td>1000</td></tr>
<tr><td>Oligo fwd</td><td>CACCGACGCCCGAGGACATCGCGC</td></tr>
<tr><td>Oligo rev</td><td>AAACGCGCGATGTCCTCGGGCGTC</td></tr>
<tr><td>Chromosome</td><td>start</td><td>end</td><td>strand</td><td>MM</td><td>target_seq</td><td>PAM</td><td>alignment</td><td>distance</td><td>position</td><td>gene name</td><td>gene id</td></tr>
</table>

GU305914.1 65750 65772 + 0 GACGCCCGAGGACATCGCGC GGG ||||||||[||||||||||||]PAM NA Intergenic NA NA
GU305914.1 29944 29966 - 4 CGAGCCCGAGGACTTCGCGC GGG ---|||||[|||||-||||||]PAM NA Intergenic NA NA

T4498 CATCGCGCGGGCGTGCGAGCTGG 1000
Oligo fwd CACCCATCGCGCGGGCGTGCGAGC
Oligo rev AAACGCTCGCACGCCCGCGCGATG
Chromosome start end strand MM target_seq PAM alignment distance position gene name gene id
GU305914.1 65762 65784 + 0 CATCGCGCGGGCGTGCGAGC TGG ||||||||[||||||||||||]PAM NA Intergenic NA NA

T4499 CGCGCGGGCGTGCGAGCTGGTGG 1000
Oligo fwd CACCCGCGCGGGCGTGCGAGCTGG
Oligo rev AAACCCAGCTCGCACGCCCGCGCG
Chromosome start end strand MM target_seq PAM alignment distance position gene name gene id
GU305914.1 65765 65787 + 0 CGCGCGGGCGTGCGAGCTGG TGG ||||||||[||||||||||||]PAM NA Intergenic NA NA

T4500 GTGCGAGCTGGTGGACGCGCAGG 1000
Oligo fwd CACCGTGCGAGCTGGTGGACGCGC
Oligo rev AAACGCGCGTCCACCAGCTCGCAC
Chromosome start end strand MM target_seq PAM alignment distance position gene name gene id
GU305914.1 65774 65796 + 0 GTGCGAGCTGGTGGACGCGC AGG ||||||||[||||||||||||]PAM NA Intergenic NA NA

T4501 CGACTTCGACAAGACGCTGACGG 1000
Oligo fwd CACCCGACTTCGACAAGACGCTGA
Oligo rev AAACTCAGCGTCTTGTCGAAGTCG
Chromosome start end strand MM target_seq PAM alignment distance position gene name gene id
GU305914.1 65840 65862 + 0 CGACTTCGACAAGACGCTGA CGG ||||||||[||||||||||||]PAM NA Intergenic NA NA

T4502 CGGAGAAGATCATCATCCCGCGG 1000
Oligo fwd CACCCGGAGAAGATCATCATCCCG
Oligo rev AAACCGGGATGATGATCTTCTCCG
Chromosome start end strand MM target_seq PAM alignment distance position gene name gene id
GU305914.1 65860 65882 + 0 CGGAGAAGATCATCATCCCG CGG ||||||||[||||||||||||]PAM NA Intergenic NA NA

T4503 GATCATCATCCCGCGGATCGAGG 1000
Oligo fwd CACCGATCATCATCCCGCGGATCG
Oligo rev AAACCGATCCGCGGGATGATGATC
Chromosome start end strand MM target_seq PAM alignment distance position gene name gene id
GU305914.1 65867 65889 + 0 GATCATCATCCCGCGGATCG AGG ||||||||[||||||||||||]PAM NA Intergenic NA NA

T4504 GGCGATCGTGAACGCGATGCCGG 1000
Oligo fwd CACCGGCGATCGTGAACGCGATGC
Oligo rev AAACGCATCGCGTTCACGATCGCC
Chromosome start end strand MM target_seq PAM alignment distance position gene name gene id
GU305914.1 65888 65910 + 0 GGCGATCGTGAACGCGATGC CGG ||||||||[||||||||||||]PAM NA Intergenic NA NA

T4505 GATCGTGAACGCGATGCCGGCGG 1000
Oligo fwd CACCGATCGTGAACGCGATGCCGG
Oligo rev AAACCCGGCATCGCGTTCACGATC
Chromosome start end strand MM target_seq PAM alignment distance position gene name gene id
GU305914.1 65891 65913 + 0 GATCGTGAACGCGATGCCGG CGG ||||||||[||||||||||||]PAM NA Intergenic NA NA

T4506 TCCGCGCGCACGACGACATGCGG 1000
Oligo fwd CACCTCCGCGCGCACGACGACATG
Oligo rev AAACCATGTCGTCGTGCGCGCGGA
Chromosome start end strand MM target_seq PAM alignment distance position gene name gene id
GU305914.1 65917 65939 + 0 TCCGCGCGCACGACGACATG CGG ||||||||[||||||||||||]PAM NA Intergenic NA NA

T4507 TCTGCCGCCTGCTGATGCCGCGG 1000
Oligo fwd CACCTCTGCCGCCTGCTGATGCCG
Oligo rev AAACCGGCATCAGCAGGCGGCAGA
Chromosome start end strand MM target_seq PAM alignment distance position gene name gene id
GU305914.1 65959 65981 + 0 TCTGCCGCCTGCTGATGCCG CGG ||||||||[||||||||||||]PAM NA Intergenic NA NA

T4508 CCTGCTGATGCCGCGGCGCCTGG 1000
Oligo fwd CACCCCTGCTGATGCCGCGGCGCC
Oligo rev AAACGGCGCCGCGGCATCAGCAGG
Chromosome start end strand MM target_seq PAM alignment distance position gene name gene id
GU305914.1 65966 65988 + 0 CCTGCTGATGCCGCGGCGCC TGG ||||||||[||||||||||||]PAM NA Intergenic NA NA

T4509 CCTGGAACTGTACCGCAACCTGG 1000
Oligo fwd CACCCCTGGAACTGTACCGCAACC
Oligo rev AAACGGTTGCGGTACAGTTCCAGG
Chromosome start end strand MM target_seq PAM alignment distance position gene name gene id
GU305914.1 65984 66006 + 0 CCTGGAACTGTACCGCAACC TGG ||||||||[||||||||||||]PAM NA Intergenic NA NA

T4510 GCTGCACTACGTGAGCCAGACGG 1000
Oligo fwd CACCGCTGCACTACGTGAGCCAGA
Oligo rev AAACTCTGGCTCACGTAGTGCAGC
Chromosome start end strand MM target_seq PAM alignment distance position gene name gene id
GU305914.1 66023 66045 + 0 GCTGCACTACGTGAGCCAGA CGG ||||||||[||||||||||||]PAM NA Intergenic NA NA
GU305914.1 22753 22775 - 4 GGAGCAATTCGTGAGCCAGA CGG |--|||-|[-|||||||||||]PAM NA Intergenic NA NA

T4511 TACGTGAGCCAGACGGACCTCGG 1000
Oligo fwd CACCTACGTGAGCCAGACGGACCT
Oligo rev AAACAGGTCCGTCTGGCTCACGTA
Chromosome start end strand MM target_seq PAM alignment distance position gene name gene id
GU305914.1 66030 66052 + 0 TACGTGAGCCAGACGGACCT CGG ||||||||[||||||||||||]PAM NA Intergenic NA NA

T4512 ACGTGAGCCAGACGGACCTCGGG 1000

Oligo fwd CACCACGTGAGCCAGACGGACCTC
Oligo rev AAACGAGGTCCGTCTGGCTCACGT
Chromosome start end strand MM target_seq PAM alignment distance position gene name gene id
GU305914.1 66031 66053 + 0 ACGTGAGCCAGACGGACCTC GGG ||||||||[||||||||||||]PAM NA Intergenic NA NA

T4513 GAGCCAGACGGACCTCGGGCTGG 1000
Oligo fwd CACCGAGCCAGACGGACCTCGGGC
Oligo rev AAACGCCCGAGGTCCGTCTGGCTC
Chromosome start end strand MM target_seq PAM alignment distance position gene name gene id
GU305914.1 66035 66057 + 0 GAGCCAGACGGACCTCGGGC TGG ||||||||[||||||||||||]PAM NA Intergenic NA NA

T4514 GACGGACCTCGGGCTGGTGAAGG 1000
Oligo fwd CACCGACGGACCTCGGGCTGGTGA
Oligo rev AAACTCACCAGCCCGAGGTCCGTC
Chromosome start end strand MM target_seq PAM alignment distance position gene name gene id
GU305914.1 66041 66063 + 0 GACGGACCTCGGGCTGGTGA AGG ||||||||[||||||||||||]PAM NA Intergenic NA NA

T4515 GGGCTGGTGAAGGTGCTGTACGG 1000
Oligo fwd CACCGGGCTGGTGAAGGTGCTGTA
Oligo rev AAACTACAGCACCTTCACCAGCCC
Chromosome start end strand MM target_seq PAM alignment distance position gene name gene id
GU305914.1 66051 66073 + 0 GGGCTGGTGAAGGTGCTGTA CGG ||||||||[||||||||||||]PAM NA Intergenic NA NA

T4516 GTACGGCGTGATCCGCAACGTGG 1000
Oligo fwd CACCGTACGGCGTGATCCGCAACG
Oligo rev AAACCGTTGCGGATCACGCCGTAC
Chromosome start end strand MM target_seq PAM alignment distance position gene name gene id
GU305914.1 66068 66090 + 0 GTACGGCGTGATCCGCAACG TGG ||||||||[||||||||||||]PAM NA Intergenic NA NA

T4517 CGCAACGTGGTCCTGCACCTCGG 1000
Oligo fwd CACCCGCAACGTGGTCCTGCACCT
Oligo rev AAACAGGTGCAGGACCACGTTGCG
Chromosome start end strand MM target_seq PAM alignment distance position gene name gene id
GU305914.1 66081 66103 + 0 CGCAACGTGGTCCTGCACCT CGG ||||||||[||||||||||||]PAM NA Intergenic NA NA

T4518 CTGCACCTCGGCGACATCACCGG 1000
Oligo fwd CACCCTGCACCTCGGCGACATCAC
Oligo rev AAACGTGATGTCGCCGAGGTGCAG
Chromosome start end strand MM target_seq PAM alignment distance position gene name gene id
GU305914.1 66093 66115 + 0 CTGCACCTCGGCGACATCAC CGG ||||||||[||||||||||||]PAM NA Intergenic NA NA

T4519 TGCACCTCGGCGACATCACCGGG 1000
Oligo fwd CACCTGCACCTCGGCGACATCACC
Oligo rev AAACGGTGATGTCGCCGAGGTGCA
Chromosome start end strand MM target_seq PAM alignment distance position gene name gene id
GU305914.1 66094 66116 + 0 TGCACCTCGGCGACATCACC GGG ||||||||[||||||||||||]PAM NA Intergenic NA NA

T4520 GCACCTCGGCGACATCACCGGGG 1000
Oligo fwd CACCGCACCTCGGCGACATCACCG
Oligo rev AAACCGGTGATGTCGCCGAGGTGC
Chromosome start end strand MM target_seq PAM alignment distance position gene name gene id
GU305914.1 66095 66117 + 0 GCACCTCGGCGACATCACCG GGG ||||||||[||||||||||||]PAM NA Intergenic NA NA

T4521 GGACTCGTACTCGTTCAACTCGG 1000
Oligo fwd CACCGGACTCGTACTCGTTCAACT
Oligo rev AAACAGTTGAACGAGTACGAGTCC
Chromosome start end strand MM target_seq PAM alignment distance position gene name gene id
GU305914.1 66116 66138 + 0 GGACTCGTACTCGTTCAACT CGG ||||||||[||||||||||||]PAM NA Intergenic NA NA

T4522 GTTCAACTCGGAGCTGCTCGAGG 1000
Oligo fwd CACCGTTCAACTCGGAGCTGCTCG
Oligo rev AAACCGAGCAGCTCCGAGTTGAAC
Chromosome start end strand MM target_seq PAM alignment distance position gene name gene id
GU305914.1 66128 66150 + 0 GTTCAACTCGGAGCTGCTCG AGG ||||||||[||||||||||||]PAM NA Intergenic NA NA

T4523 CTCGGAGCTGCTCGAGGACCTGG 1000
Oligo fwd CACCCTCGGAGCTGCTCGAGGACC
Oligo rev AAACGGTCCTCGAGCAGCTCCGAG
Chromosome start end strand MM target_seq PAM alignment distance position gene name gene id
GU305914.1 66134 66156 + 0 CTCGGAGCTGCTCGAGGACC TGG ||||||||[||||||||||||]PAM NA Intergenic NA NA

T4524 CGAGTCGTTCCTGCACGCGCTGG 1000
Oligo fwd CACCCGAGTCGTTCCTGCACGCGC
Oligo rev AAACGCGCGTGCAGGAACGACTCG
Chromosome start end strand MM target_seq PAM alignment distance position gene name gene id
GU305914.1 66161 66183 + 0 CGAGTCGTTCCTGCACGCGC TGG ||||||||[||||||||||||]PAM NA Intergenic NA NA

T4525 GAGTCGTTCCTGCACGCGCTGGG 1000
Oligo fwd CACCGAGTCGTTCCTGCACGCGCT
Oligo rev AAACAGCGCGTGCAGGAACGACTC
Chromosome start end strand MM target_seq PAM alignment distance position gene name gene id
GU305914.1 66162 66184 + 0 GAGTCGTTCCTGCACGCGCT GGG ||||||||[||||||||||||]PAM NA Intergenic NA NA

T4526 AGTCGTTCCTGCACGCGCTGGGG 1000
Oligo fwd CACCAGTCGTTCCTGCACGCGCTG
Oligo rev AAACCAGCGCGTGCAGGAACGACT
Chromosome start end strand MM target_seq PAM alignment distance position gene name gene id
GU305914.1 66163 66185 + 0 AGTCGTTCCTGCACGCGCTG GGG ||||||||[||||||||||||]PAM NA Intergenic NA NA

```
T4527        GTCGTTCCTGCACGCGCTGGGGG    1000
Oligo fwd    CACCGTCGTTCCTGCACGCGCTGG
Oligo rev    AAACCCAGCGCGTGCAGGAACGAC
Chromosome   start    end      strand  MM  target_seq            PAM  alignment                    distance  position    gene name  gene id
GU305914.1   66164    66186    +       0   GTCGTTCCTGCACGCGCTGG  GGG  ||||||||[||||||||||||]PAM    NA        Intergenic  NA         NA

T4528        TCGTTCCTGCACGCGCTGGGGGG    1000
Oligo fwd    CACCTCGTTCCTGCACGCGCTGGG
Oligo rev    AAACCCCAGCGCGTGCAGGAACGA
Chromosome   start    end      strand  MM  target_seq            PAM  alignment                    distance  position    gene name  gene id
GU305914.1   66165    66187    +       0   TCGTTCCTGCACGCGCTGGG  GGG  ||||||||[||||||||||||]PAM    NA        Intergenic  NA         NA
GU305914.1   126496   126518   -       4   GCGCTGCTGAACGCGCTGGG  CGG  -||-|-||[|-||||||||||]PAM    NA        Intergenic  NA         NA

T4529        GCTGGGGGGCGACATCACCGAGG    1000
Oligo fwd    CACCGCTGGGGGGCGACATCACCG
Oligo rev    AAACCGGTGATGTCGCCCCCCAGC
Chromosome   start    end      strand  MM  target_seq            PAM  alignment                    distance  position    gene name  gene id
GU305914.1   66179    66201    +       0   GCTGGGGGGCGACATCACCG  AGG  ||||||||[||||||||||||]PAM    NA        Intergenic  NA         NA

T4530        GGGCGACATCACCGAGGCGCTGG    1000
Oligo fwd    CACCGGGCGACATCACCGAGGCGC
Oligo rev    AAACGCGCCTCGGTGATGTCGCCC
Chromosome   start    end      strand  MM  target_seq            PAM  alignment                    distance  position    gene name  gene id
GU305914.1   66185    66207    +       0   GGGCGACATCACCGAGGCGC  TGG  ||||||||[||||||||||||]PAM    NA        Intergenic  NA         NA

T4531        TCACCGAGGCGCTGGCGCGCTGG    1000
Oligo fwd    CACCTCACCGAGGCGCTGGCGCGC
Oligo rev    AAACGCGCGCCAGCGCCTCGGTGA
Chromosome   start    end      strand  MM  target_seq            PAM  alignment                    distance  position    gene name  gene id
GU305914.1   66193    66215    +       0   TCACCGAGGCGCTGGCGCGC  TGG  ||||||||[||||||||||||]PAM    NA        Intergenic  NA         NA

T4532        CACCGAGGCGCTGGCGCGCTGGG    1000
Oligo fwd    CACCCACCGAGGCGCTGGCGCGCT
Oligo rev    AAACAGCGCGCCAGCGCCTCGGTG
Chromosome   start    end      strand  MM  target_seq            PAM  alignment                    distance  position    gene name  gene id
GU305914.1   66194    66216    +       0   CACCGAGGCGCTGGCGCGCT  GGG  ||||||||[||||||||||||]PAM    NA        Intergenic  NA         NA

T4533        GGCGCTGGCGCGCTGGGCCGCGG    1000
Oligo fwd    CACCGGCGCTGGCGCGCTGGGCCG
Oligo rev    AAACCGGCCCAGCGCGCCAGCGCC
Chromosome   start    end      strand  MM  target_seq            PAM  alignment                    distance  position    gene name  gene id
GU305914.1   66200    66222    +       0   GGCGCTGGCGCGCTGGGCCG  CGG  ||||||||[||||||||||||]PAM    NA        Intergenic  NA         NA

T4534        TGGCGCGCTGGGCCGCGGACCGG    1000
Oligo fwd    CACCTGGCGCGCTGGGCCGCGGAC
Oligo rev    AAACGTCCGCGGCCCAGCGCGCCA
Chromosome   start    end      strand  MM  target_seq            PAM  alignment                    distance  position    gene name  gene id
GU305914.1   66205    66227    +       0   TGGCGCGCTGGGCCGCGGAC  CGG  ||||||||[||||||||||||]PAM    NA        Intergenic  NA         NA

T4535        GGCCGCGGACCGGCGCCAGCTGG    1000
Oligo fwd    CACCGGCCGCGGACCGGCGCCAGC
Oligo rev    AAACGCTGGCGCCGGTCCGCGGCC
Chromosome   start    end      strand  MM  target_seq            PAM  alignment                    distance  position    gene name  gene id
GU305914.1   66215    66237    +       0   GGCCGCGGACCGGCGCCAGC  TGG  ||||||||[||||||||||||]PAM    NA        Intergenic  NA         NA

T4536        ACCGGCGCCAGCTGGCCGAGCGG    1000
Oligo fwd    CACCACCGGCGCCAGCTGGCCGAG
Oligo rev    AAACCTCGGCCAGCTGGCGCCGGT
Chromosome   start    end      strand  MM  target_seq            PAM  alignment                    distance  position    gene name  gene id
GU305914.1   66223    66245    +       0   ACCGGCGCCAGCTGGCCGAG  CGG  ||||||||[||||||||||||]PAM    NA        Intergenic  NA         NA

T4537        CCAGCTGGCCGAGCGGTTCGCGG    1000
Oligo fwd    CACCCCAGCTGGCCGAGCGGTTCG
Oligo rev    AAACCGAACCGCTCGGCCAGCTGG
Chromosome   start    end      strand  MM  target_seq            PAM  alignment                    distance  position    gene name  gene id
GU305914.1   66230    66252    +       0   CCAGCTGGCCGAGCGGTTCG  CGG  ||||||||[||||||||||||]PAM    NA        Intergenic  NA         NA
GU305914.1   101308   101330   -       4   CAAGCAGACCGCGCGGTTCG  CGG  |-|||-|-[|||-||||||||]PAM    NA        Intergenic  NA         NA

T4538        CTACTCGCTGACGAAGAGCGTGG    1000
Oligo fwd    CACCCTACTCGCTGACGAAGAGCG
Oligo rev    AAACCGCTCTTCGTCAGCGAGTAG
Chromosome   start    end      strand  MM  target_seq            PAM  alignment                    distance  position    gene name  gene id
GU305914.1   66269    66291    +       0   CTACTCGCTGACGAAGAGCG  TGG  ||||||||[||||||||||||]PAM    NA        Intergenic  NA         NA

T4539        GCTGTACACGAACACGCACTCGG    1000
Oligo fwd    CACCGCTGTACACGAACACGCACT
Oligo rev    AAACAGTGCGTGTTCGTGTACAGC
Chromosome   start    end      strand  MM  target_seq            PAM  alignment                    distance  position    gene name  gene id
GU305914.1   66296    66318    +       0   GCTGTACACGAACACGCACT  CGG  ||||||||[||||||||||||]PAM    NA        Intergenic  NA         NA

T4540        GTACACGAACACGCACTCGGTGG    1000
Oligo fwd    CACCGTACACGAACACGCACTCGG
Oligo rev    AAACCCGAGTGCGTGTTCGTGTAC
Chromosome   start    end      strand  MM  target_seq            PAM  alignment                    distance  position    gene name  gene id
GU305914.1   66299    66321    +       0   GTACACGAACACGCACTCGG  TGG  ||||||||[||||||||||||]PAM    NA        Intergenic  NA         NA

T4541        CACGCACTCGGTGGTGCTGACGG    1000
Oligo fwd    CACCCACGCACTCGGTGGTGCTGA
Oligo rev    AAACTCAGCACCACCGAGTGCGTG
```

| | | | | | | | | | | | |
|---|---|---|---|---|---|---|---|---|---|---|---|
| Chromosome | start | end | strand | MM | target_seq | PAM | alignment | distance | position | gene name | gene id |
| GU305914.1 | 66308 | 66330 | + | 0 | CACGCACTCGGTGGTGCTGA | CGG | \|\|\|\|\|\|\|\|[\|\|\|\|\|\|\|\|\|\|\|\|]PAM | NA | Intergenic | NA | NA |
| T4542 | ACGCACTCGGTGGTGCTGACGGG | 1000 | | | | | | | | | |
| Oligo fwd | CACCACGCACTCGGTGGTGCTGAC | | | | | | | | | | |
| Oligo rev | AAACGTCAGCACCACCGAGTGCGT | | | | | | | | | | |
| Chromosome | start | end | strand | MM | target_seq | PAM | alignment | distance | position | gene name | gene id |
| GU305914.1 | 66309 | 66331 | + | 0 | ACGCACTCGGTGGTGCTGAC | GGG | \|\|\|\|\|\|\|\|[\|\|\|\|\|\|\|\|\|\|\|\|]PAM | NA | Intergenic | NA | NA |
| T4543 | CAAGAACGAGATCGTGACGATGG | 1000 | | | | | | | | | |
| Oligo fwd | CACCCAAGAACGAGATCGTGACGA | | | | | | | | | | |
| Oligo rev | AAACTCGTCACGATCTCGTTCTTG | | | | | | | | | | |
| Chromosome | start | end | strand | MM | target_seq | PAM | alignment | distance | position | gene name | gene id |
| GU305914.1 | 66332 | 66354 | + | 0 | CAAGAACGAGATCGTGACGA | TGG | \|\|\|\|\|\|\|\|[\|\|\|\|\|\|\|\|\|\|\|\|]PAM | NA | Intergenic | NA | NA |
| T4544 | GAACGAGATCGTGACGATGGTGG | 1000 | | | | | | | | | |
| Oligo fwd | CACCGAACGAGATCGTGACGATGG | | | | | | | | | | |
| Oligo rev | AAACCCATCGTCACGATCTCGTTC | | | | | | | | | | |
| Chromosome | start | end | strand | MM | target_seq | PAM | alignment | distance | position | gene name | gene id |
| GU305914.1 | 66335 | 66357 | + | 0 | GAACGAGATCGTGACGATGG | TGG | \|\|\|\|\|\|\|\|[\|\|\|\|\|\|\|\|\|\|\|\|]PAM | NA | Intergenic | NA | NA |
| T4545 | GGTGGCCCTGCCCGAGTTCACGG | 1000 | | | | | | | | | |
| Oligo fwd | CACCGGTGGCCCTGCCCGAGTTCA | | | | | | | | | | |
| Oligo rev | AAACTGAACTCGGGCAGGGCCACC | | | | | | | | | | |
| Chromosome | start | end | strand | MM | target_seq | PAM | alignment | distance | position | gene name | gene id |
| GU305914.1 | 66353 | 66375 | + | 0 | GGTGGCCCTGCCCGAGTTCA | CGG | \|\|\|\|\|\|\|\|[\|\|\|\|\|\|\|\|\|\|\|\|]PAM | NA | Intergenic | NA | NA |
| T4546 | CCGAGTTCACGGACACGCTGCGG | 1000 | | | | | | | | | |
| Oligo fwd | CACCCCGAGTTCACGGACACGCTG | | | | | | | | | | |
| Oligo rev | AAACCAGCGTGTCCGTGAACTCGG | | | | | | | | | | |
| Chromosome | start | end | strand | MM | target_seq | PAM | alignment | distance | position | gene name | gene id |
| GU305914.1 | 66364 | 66386 | + | 0 | CCGAGTTCACGGACACGCTG | CGG | \|\|\|\|\|\|\|\|[\|\|\|\|\|\|\|\|\|\|\|\|]PAM | NA | Intergenic | NA | NA |
| T4547 | GGACACGCTGCGGAAGATCGCGG | 1000 | | | | | | | | | |
| Oligo fwd | CACCGGACACGCTGCGGAAGATCG | | | | | | | | | | |
| Oligo rev | AAACCGATCTTCCGCAGCGTGTCC | | | | | | | | | | |
| Chromosome | start | end | strand | MM | target_seq | PAM | alignment | distance | position | gene name | gene id |
| GU305914.1 | 66374 | 66396 | + | 0 | GGACACGCTGCGGAAGATCG | CGG | \|\|\|\|\|\|\|\|[\|\|\|\|\|\|\|\|\|\|\|\|]PAM | NA | Intergenic | NA | NA |
| T4548 | CACGCTGCGGAAGATCGCGGCGG | 1000 | | | | | | | | | |
| Oligo fwd | CACCCACGCTGCGGAAGATCGCGG | | | | | | | | | | |
| Oligo rev | AAACCCGCGATCTTCCGCAGCGTG | | | | | | | | | | |
| Chromosome | start | end | strand | MM | target_seq | PAM | alignment | distance | position | gene name | gene id |
| GU305914.1 | 66377 | 66399 | + | 0 | CACGCTGCGGAAGATCGCGG | CGG | \|\|\|\|\|\|\|\|[\|\|\|\|\|\|\|\|\|\|\|\|]PAM | NA | Intergenic | NA | NA |
| T4549 | GAAGATCGCGGCGGACAACGCGG | 1000 | | | | | | | | | |
| Oligo fwd | CACCGAAGATCGCGGCGGACAACG | | | | | | | | | | |
| Oligo rev | AAACCGTTGTCCGCCGCGATCTTC | | | | | | | | | | |
| Chromosome | start | end | strand | MM | target_seq | PAM | alignment | distance | position | gene name | gene id |
| GU305914.1 | 66386 | 66408 | + | 0 | GAAGATCGCGGCGGACAACG | CGG | \|\|\|\|\|\|\|\|[\|\|\|\|\|\|\|\|\|\|\|\|]PAM | NA | Intergenic | NA | NA |
| T4550 | CGCGGCGGACAACGCGGAGCTGG | 1000 | | | | | | | | | |
| Oligo fwd | CACCCGCGGCGGACAACGCGGAGC | | | | | | | | | | |
| Oligo rev | AAACGCTCCGCGTTGTCCGCCGCG | | | | | | | | | | |
| Chromosome | start | end | strand | MM | target_seq | PAM | alignment | distance | position | gene name | gene id |
| GU305914.1 | 66392 | 66414 | + | 0 | CGCGGCGGACAACGCGGAGC | TGG | \|\|\|\|\|\|\|\|[\|\|\|\|\|\|\|\|\|\|\|\|]PAM | NA | Intergenic | NA | NA |
| GU305914.1 | 121214 | 121236 | - | 4 | CGAGGAGGACGACGCGCAGC | TGG | \|\|-\|\|-\|\|[\|\|-\|\|\|\|\|-\|\|\|]PAM | NA | Intergenic | NA | NA |
| T4551 | GGCGGACAACGCGGAGCTGGAGG | 1000 | | | | | | | | | |
| Oligo fwd | CACCGGCGGACAACGCGGAGCTGG | | | | | | | | | | |
| Oligo rev | AAACCCAGCTCCGCGTTGTCCGCC | | | | | | | | | | |
| Chromosome | start | end | strand | MM | target_seq | PAM | alignment | distance | position | gene name | gene id |
| GU305914.1 | 66395 | 66417 | + | 0 | GGCGGACAACGCGGAGCTGG | AGG | \|\|\|\|\|\|\|\|[\|\|\|\|\|\|\|\|\|\|\|\|]PAM | NA | Intergenic | NA | NA |
| T4552 | ACAACGCGGAGCTGGAGGAGCGG | 1000 | | | | | | | | | |
| Oligo fwd | CACCACAACGCGGAGCTGGAGGAG | | | | | | | | | | |
| Oligo rev | AAACCTCCTCCAGCTCCGCGTTGT | | | | | | | | | | |
| Chromosome | start | end | strand | MM | target_seq | PAM | alignment | distance | position | gene name | gene id |
| GU305914.1 | 66400 | 66422 | + | 0 | ACAACGCGGAGCTGGAGGAG | CGG | \|\|\|\|\|\|\|\|[\|\|\|\|\|\|\|\|\|\|\|\|]PAM | NA | Intergenic | NA | NA |
| T4553 | CTGCGTTTCCTGAGCAACACCGG | 1000 | | | | | | | | | |
| Oligo fwd | CACCCTGCGTTTCCTGAGCAACAC | | | | | | | | | | |
| Oligo rev | AAACGTGTTGCTCAGGAAACGCAG | | | | | | | | | | |
| Chromosome | start | end | strand | MM | target_seq | PAM | alignment | distance | position | gene name | gene id |
| GU305914.1 | 66423 | 66445 | + | 0 | CTGCGTTTCCTGAGCAACAC | CGG | \|\|\|\|\|\|\|\|[\|\|\|\|\|\|\|\|\|\|\|\|]PAM | NA | Intergenic | NA | NA |
| T4554 | CCGGCGCGCTGATCGTCGAGCGG | 1000 | | | | | | | | | |
| Oligo fwd | CACCCCGGCGCGCTGATCGTCGAG | | | | | | | | | | |
| Oligo rev | AAACCTCGACGATCAGCGCGCCGG | | | | | | | | | | |
| Chromosome | start | end | strand | MM | target_seq | PAM | alignment | distance | position | gene name | gene id |
| GU305914.1 | 66442 | 66464 | + | 0 | CCGGCGCGCTGATCGTCGAG | CGG | \|\|\|\|\|\|\|\|[\|\|\|\|\|\|\|\|\|\|\|\|]PAM | NA | Intergenic | NA | NA |
| T4555 | GCGGCTGCGCTCGCTGACCGAGG | 1000 | | | | | | | | | |
| Oligo fwd | CACCGCGGCTGCGCTCGCTGACCG | | | | | | | | | | |
| Oligo rev | AAACCGGTCAGCGAGCGCAGCCGC | | | | | | | | | | |
| Chromosome | start | end | strand | MM | target_seq | PAM | alignment | distance | position | gene name | gene id |
| GU305914.1 | 66461 | 66483 | + | 0 | GCGGCTGCGCTCGCTGACCG | AGG | \|\|\|\|\|\|\|\|[\|\|\|\|\|\|\|\|\|\|\|\|]PAM | NA | Intergenic | NA | NA |
| T4556 | CCGACCTGACTTTCTCGACGCGG | 1000 | | | | | | | | | |

<table>
<tr><td>Oligo fwd</td><td>CACCCCGACCTGACTTTCTCGACG</td><td></td><td></td><td></td><td></td><td></td><td></td><td></td><td></td><td></td><td></td></tr>
<tr><td>Oligo rev</td><td>AAACCGTCGAGAAAGTCAGGTCGG</td><td></td><td></td><td></td><td></td><td></td><td></td><td></td><td></td><td></td><td></td></tr>
<tr><td>Chromosome</td><td>start</td><td>end</td><td>strand</td><td>MM</td><td>target_seq</td><td>PAM</td><td>alignment</td><td>distance</td><td>position</td><td>gene name</td><td>gene id</td></tr>
<tr><td>GU305914.1</td><td>66508</td><td>66530</td><td>+</td><td>0</td><td>CCGACCTGACTTTCTCGACG</td><td>CGG</td><td>||||||||[||||||||||||]PAM</td><td>NA</td><td>Intergenic</td><td>NA</td><td>NA</td></tr>
<tr><td>T4557</td><td>CGATCACGAAGCTGTACGCGCGG</td><td>1000</td><td></td><td></td><td></td><td></td><td></td><td></td><td></td><td></td><td></td></tr>
<tr><td>Oligo fwd</td><td>CACCCGATCACGAAGCTGTACGCG</td><td></td><td></td><td></td><td></td><td></td><td></td><td></td><td></td><td></td><td></td></tr>
<tr><td>Oligo rev</td><td>AAACCGCGTACAGCTTCGTGATCG</td><td></td><td></td><td></td><td></td><td></td><td></td><td></td><td></td><td></td><td></td></tr>
<tr><td>Chromosome</td><td>start</td><td>end</td><td>strand</td><td>MM</td><td>target_seq</td><td>PAM</td><td>alignment</td><td>distance</td><td>position</td><td>gene name</td><td>gene id</td></tr>
<tr><td>GU305914.1</td><td>66532</td><td>66554</td><td>+</td><td>0</td><td>CGATCACGAAGCTGTACGCG</td><td>CGG</td><td>||||||||[||||||||||||]PAM</td><td>NA</td><td>Intergenic</td><td>NA</td><td>NA</td></tr>
<tr><td>T4558</td><td>GAAGCTGTACGCGCGGCTGTCGG</td><td>1000</td><td></td><td></td><td></td><td></td><td></td><td></td><td></td><td></td><td></td></tr>
<tr><td>Oligo fwd</td><td>CACCGAAGCTGTACGCGCGGCTGT</td><td></td><td></td><td></td><td></td><td></td><td></td><td></td><td></td><td></td><td></td></tr>
<tr><td>Oligo rev</td><td>AAACACAGCCGCGCGTACAGCTTC</td><td></td><td></td><td></td><td></td><td></td><td></td><td></td><td></td><td></td><td></td></tr>
<tr><td>Chromosome</td><td>start</td><td>end</td><td>strand</td><td>MM</td><td>target_seq</td><td>PAM</td><td>alignment</td><td>distance</td><td>position</td><td>gene name</td><td>gene id</td></tr>
<tr><td>GU305914.1</td><td>66539</td><td>66561</td><td>+</td><td>0</td><td>GAAGCTGTACGCGCGGCTGT</td><td>CGG</td><td>||||||||[||||||||||||]PAM</td><td>NA</td><td>Intergenic</td><td>NA</td><td>NA</td></tr>
<tr><td>T4559</td><td>GGAGACGCTGCAGACGAACCAGG</td><td>1000</td><td></td><td></td><td></td><td></td><td></td><td></td><td></td><td></td><td></td></tr>
<tr><td>Oligo fwd</td><td>CACCGGAGACGCTGCAGACGAACC</td><td></td><td></td><td></td><td></td><td></td><td></td><td></td><td></td><td></td><td></td></tr>
<tr><td>Oligo rev</td><td>AAACGGTTCGTCTGCAGCGTCTCC</td><td></td><td></td><td></td><td></td><td></td><td></td><td></td><td></td><td></td><td></td></tr>
<tr><td>Chromosome</td><td>start</td><td>end</td><td>strand</td><td>MM</td><td>target_seq</td><td>PAM</td><td>alignment</td><td>distance</td><td>position</td><td>gene name</td><td>gene id</td></tr>
<tr><td>GU305914.1</td><td>66560</td><td>66582</td><td>+</td><td>0</td><td>GGAGACGCTGCAGACGAACC</td><td>AGG</td><td>||||||||[||||||||||||]PAM</td><td>NA</td><td>Intergenic</td><td>NA</td><td>NA</td></tr>
<tr><td>T4560</td><td>GCAGACGAACCAGGCGACGCTGG</td><td>1000</td><td></td><td></td><td></td><td></td><td></td><td></td><td></td><td></td><td></td></tr>
<tr><td>Oligo fwd</td><td>CACCGCAGACGAACCAGGCGACGC</td><td></td><td></td><td></td><td></td><td></td><td></td><td></td><td></td><td></td><td></td></tr>
<tr><td>Oligo rev</td><td>AAACGCGTCGCCTGGTTCGTCTGC</td><td></td><td></td><td></td><td></td><td></td><td></td><td></td><td></td><td></td><td></td></tr>
<tr><td>Chromosome</td><td>start</td><td>end</td><td>strand</td><td>MM</td><td>target_seq</td><td>PAM</td><td>alignment</td><td>distance</td><td>position</td><td>gene name</td><td>gene id</td></tr>
<tr><td>GU305914.1</td><td>66569</td><td>66591</td><td>+</td><td>0</td><td>GCAGACGAACCAGGCGACGC</td><td>TGG</td><td>||||||||[||||||||||||]PAM</td><td>NA</td><td>Intergenic</td><td>NA</td><td>NA</td></tr>
<tr><td>T4561</td><td>GACGCTGGAATCGCTGTGCGTGG</td><td>1000</td><td></td><td></td><td></td><td></td><td></td><td></td><td></td><td></td><td></td></tr>
<tr><td>Oligo fwd</td><td>CACCGACGCTGGAATCGCTGTGCG</td><td></td><td></td><td></td><td></td><td></td><td></td><td></td><td></td><td></td><td></td></tr>
<tr><td>Oligo rev</td><td>AAACCGCACAGCGATTCCAGCGTC</td><td></td><td></td><td></td><td></td><td></td><td></td><td></td><td></td><td></td><td></td></tr>
<tr><td>Chromosome</td><td>start</td><td>end</td><td>strand</td><td>MM</td><td>target_seq</td><td>PAM</td><td>alignment</td><td>distance</td><td>position</td><td>gene name</td><td>gene id</td></tr>
<tr><td>GU305914.1</td><td>66584</td><td>66606</td><td>+</td><td>0</td><td>GACGCTGGAATCGCTGTGCG</td><td>TGG</td><td>||||||||[||||||||||||]PAM</td><td>NA</td><td>Intergenic</td><td>NA</td><td>NA</td></tr>
<tr><td>T4562</td><td>GCGCCTGCGCTTCAACCGCGCGG</td><td>1000</td><td></td><td></td><td></td><td></td><td></td><td></td><td></td><td></td><td></td></tr>
<tr><td>Oligo fwd</td><td>CACCGCGCCTGCGCTTCAACCGCG</td><td></td><td></td><td></td><td></td><td></td><td></td><td></td><td></td><td></td><td></td></tr>
<tr><td>Oligo rev</td><td>AAACCGCGGTTGAAGCGCAGGCGC</td><td></td><td></td><td></td><td></td><td></td><td></td><td></td><td></td><td></td><td></td></tr>
<tr><td>Chromosome</td><td>start</td><td>end</td><td>strand</td><td>MM</td><td>target_seq</td><td>PAM</td><td>alignment</td><td>distance</td><td>position</td><td>gene name</td><td>gene id</td></tr>
<tr><td>GU305914.1</td><td>66608</td><td>66630</td><td>+</td><td>0</td><td>GCGCCTGCGCTTCAACCGCG</td><td>CGG</td><td>||||||||[||||||||||||]PAM</td><td>NA</td><td>Intergenic</td><td>NA</td><td>NA</td></tr>
<tr><td>T4563</td><td>GTGCGAGTCGTTCTCGTCGCTGG</td><td>1000</td><td></td><td></td><td></td><td></td><td></td><td></td><td></td><td></td><td></td></tr>
<tr><td>Oligo fwd</td><td>CACCGTGCGAGTCGTTCTCGTCGC</td><td></td><td></td><td></td><td></td><td></td><td></td><td></td><td></td><td></td><td></td></tr>
<tr><td>Oligo rev</td><td>AAACGCGACGAGAACGACTCGCAC</td><td></td><td></td><td></td><td></td><td></td><td></td><td></td><td></td><td></td><td></td></tr>
<tr><td>Chromosome</td><td>start</td><td>end</td><td>strand</td><td>MM</td><td>target_seq</td><td>PAM</td><td>alignment</td><td>distance</td><td>position</td><td>gene name</td><td>gene id</td></tr>
<tr><td>GU305914.1</td><td>66632</td><td>66654</td><td>+</td><td>0</td><td>GTGCGAGTCGTTCTCGTCGC</td><td>TGG</td><td>||||||||[||||||||||||]PAM</td><td>NA</td><td>Intergenic</td><td>NA</td><td>NA</td></tr>
<tr><td>T4564</td><td>GGCGAGCGCCTGCCGCACGATGG</td><td>1000</td><td></td><td></td><td></td><td></td><td></td><td></td><td></td><td></td><td></td></tr>
<tr><td>Oligo fwd</td><td>CACCGGCGAGCGCCTGCCGCACGA</td><td></td><td></td><td></td><td></td><td></td><td></td><td></td><td></td><td></td><td></td></tr>
<tr><td>Oligo rev</td><td>AAACTCGTGCGGCAGGCGCTCGCC</td><td></td><td></td><td></td><td></td><td></td><td></td><td></td><td></td><td></td><td></td></tr>
<tr><td>Chromosome</td><td>start</td><td>end</td><td>strand</td><td>MM</td><td>target_seq</td><td>PAM</td><td>alignment</td><td>distance</td><td>position</td><td>gene name</td><td>gene id</td></tr>
<tr><td>GU305914.1</td><td>66653</td><td>66675</td><td>+</td><td>0</td><td>GGCGAGCGCCTGCCGCACGA</td><td>TGG</td><td>||||||||[||||||||||||]PAM</td><td>NA</td><td>Intergenic</td><td>NA</td><td>NA</td></tr>
<tr><td>T4565</td><td>CGCCTGCCGCACGATGGACGCGG</td><td>1000</td><td></td><td></td><td></td><td></td><td></td><td></td><td></td><td></td><td></td></tr>
<tr><td>Oligo fwd</td><td>CACCCGCCTGCCGCACGATGGACG</td><td></td><td></td><td></td><td></td><td></td><td></td><td></td><td></td><td></td><td></td></tr>
<tr><td>Oligo rev</td><td>AAACCGTCCATCGTGCGGCAGGCG</td><td></td><td></td><td></td><td></td><td></td><td></td><td></td><td></td><td></td><td></td></tr>
<tr><td>Chromosome</td><td>start</td><td>end</td><td>strand</td><td>MM</td><td>target_seq</td><td>PAM</td><td>alignment</td><td>distance</td><td>position</td><td>gene name</td><td>gene id</td></tr>
<tr><td>GU305914.1</td><td>66659</td><td>66681</td><td>+</td><td>0</td><td>CGCCTGCCGCACGATGGACG</td><td>CGG</td><td>||||||||[||||||||||||]PAM</td><td>NA</td><td>Intergenic</td><td>NA</td><td>NA</td></tr>
<tr><td>T4566</td><td>GCCGCACGATGGACGCGGACAGG</td><td>1000</td><td></td><td></td><td></td><td></td><td></td><td></td><td></td><td></td><td></td></tr>
<tr><td>Oligo fwd</td><td>CACCGCCGCACGATGGACGCGGAC</td><td></td><td></td><td></td><td></td><td></td><td></td><td></td><td></td><td></td><td></td></tr>
<tr><td>Oligo rev</td><td>AAACGTCCGCGTCCATCGTGCGGC</td><td></td><td></td><td></td><td></td><td></td><td></td><td></td><td></td><td></td><td></td></tr>
<tr><td>Chromosome</td><td>start</td><td>end</td><td>strand</td><td>MM</td><td>target_seq</td><td>PAM</td><td>alignment</td><td>distance</td><td>position</td><td>gene name</td><td>gene id</td></tr>
<tr><td>GU305914.1</td><td>66664</td><td>66686</td><td>+</td><td>0</td><td>GCCGCACGATGGACGCGGAC</td><td>AGG</td><td>||||||||[||||||||||||]PAM</td><td>NA</td><td>Intergenic</td><td>NA</td><td>NA</td></tr>
<tr><td>T4567</td><td>CGATGGACGCGGACAGGATCCGG</td><td>1000</td><td></td><td></td><td></td><td></td><td></td><td></td><td></td><td></td><td></td></tr>
<tr><td>Oligo fwd</td><td>CACCCGATGGACGCGGACAGGATC</td><td></td><td></td><td></td><td></td><td></td><td></td><td></td><td></td><td></td><td></td></tr>
<tr><td>Oligo rev</td><td>AAACGATCCTGTCCGCGTCCATCG</td><td></td><td></td><td></td><td></td><td></td><td></td><td></td><td></td><td></td><td></td></tr>
<tr><td>Chromosome</td><td>start</td><td>end</td><td>strand</td><td>MM</td><td>target_seq</td><td>PAM</td><td>alignment</td><td>distance</td><td>position</td><td>gene name</td><td>gene id</td></tr>
<tr><td>GU305914.1</td><td>66670</td><td>66692</td><td>+</td><td>0</td><td>CGATGGACGCGGACAGGATC</td><td>CGG</td><td>||||||||[||||||||||||]PAM</td><td>NA</td><td>Intergenic</td><td>NA</td><td>NA</td></tr>
<tr><td>GU305914.1</td><td>75564</td><td>75586</td><td>+</td><td>4</td><td>CGATGGCGGCGGCGAGGATC</td><td>GGG</td><td>||||||--[||||--||||||]PAM</td><td>NA</td><td>Intergenic</td><td>NA</td><td>NA</td></tr>
<tr><td>T4568</td><td>GCGGACAGGATCCGGAGCCGCGG</td><td>1000</td><td></td><td></td><td></td><td></td><td></td><td></td><td></td><td></td><td></td></tr>
<tr><td>Oligo fwd</td><td>CACCGCGGACAGGATCCGGAGCCG</td><td></td><td></td><td></td><td></td><td></td><td></td><td></td><td></td><td></td><td></td></tr>
<tr><td>Oligo rev</td><td>AAACCGGCTCCGGATCCTGTCCGC</td><td></td><td></td><td></td><td></td><td></td><td></td><td></td><td></td><td></td><td></td></tr>
<tr><td>Chromosome</td><td>start</td><td>end</td><td>strand</td><td>MM</td><td>target_seq</td><td>PAM</td><td>alignment</td><td>distance</td><td>position</td><td>gene name</td><td>gene id</td></tr>
<tr><td>GU305914.1</td><td>66678</td><td>66700</td><td>+</td><td>0</td><td>GCGGACAGGATCCGGAGCCG</td><td>CGG</td><td>||||||||[||||||||||||]PAM</td><td>NA</td><td>Intergenic</td><td>NA</td><td>NA</td></tr>
<tr><td>T4569</td><td>CGGACAGGATCCGGAGCCGCGGG</td><td>1000</td><td></td><td></td><td></td><td></td><td></td><td></td><td></td><td></td><td></td></tr>
<tr><td>Oligo fwd</td><td>CACCCGGACAGGATCCGGAGCCGC</td><td></td><td></td><td></td><td></td><td></td><td></td><td></td><td></td><td></td><td></td></tr>
<tr><td>Oligo rev</td><td>AAACGCGGCTCCGGATCCTGTCCG</td><td></td><td></td><td></td><td></td><td></td><td></td><td></td><td></td><td></td><td></td></tr>
<tr><td>Chromosome</td><td>start</td><td>end</td><td>strand</td><td>MM</td><td>target_seq</td><td>PAM</td><td>alignment</td><td>distance</td><td>position</td><td>gene name</td><td>gene id</td></tr>
<tr><td>GU305914.1</td><td>66679</td><td>66701</td><td>+</td><td>0</td><td>CGGACAGGATCCGGAGCCGC</td><td>GGG</td><td>||||||||[||||||||||||]PAM</td><td>NA</td><td>Intergenic</td><td>NA</td><td>NA</td></tr>
<tr><td>T4570</td><td>GGATCCGGAGCCGCGGGCTGCGG</td><td>1000</td><td></td><td></td><td></td><td></td><td></td><td></td><td></td><td></td><td></td></tr>
<tr><td>Oligo fwd</td><td>CACCGGATCCGGAGCCGCGGGCTG</td><td></td><td></td><td></td><td></td><td></td><td></td><td></td><td></td><td></td><td></td></tr>
<tr><td>Oligo rev</td><td>AAACCAGCCCGCGGCTCCGGATCC</td><td></td><td></td><td></td><td></td><td></td><td></td><td></td><td></td><td></td><td></td></tr>
<tr><td>Chromosome</td><td>start</td><td>end</td><td>strand</td><td>MM</td><td>target_seq</td><td>PAM</td><td>alignment</td><td>distance</td><td>position</td><td>gene name</td><td>gene id</td></tr>
<tr><td>GU305914.1</td><td>66685</td><td>66707</td><td>+</td><td>0</td><td>GGATCCGGAGCCGCGGGCTG</td><td>CGG</td><td>||||||||[||||||||||||]PAM</td><td>NA</td><td>Intergenic</td><td>NA</td><td>NA</td></tr>
</table>

<table>
<tr><td>GU305914.1</td><td></td><td>213767</td><td>213789</td><td>+</td><td>4</td><td>GGCTCCGTAGTCTCGGGCTG</td><td>AGG</td><td>||-||||-[||-|-|||||||]PAM</td><td>NA</td><td>Intergenic</td><td>NA</td><td>NA</td></tr>
<tr><td></td></tr>
<tr><td>T4571</td><td>CCGCGGGCTGCGGCGCTGCGTGG</td><td></td><td>1000</td></tr>
<tr><td>Oligo fwd</td><td>CACCCCGCGGGCTGCGGCGCTGCG</td></tr>
<tr><td>Oligo rev</td><td>AAACCGCAGCGCCGCAGCCCGCGG</td></tr>
<tr><td>Chromosome</td><td>start</td><td></td><td>end</td><td>strand</td><td>MM</td><td>target_seq</td><td>PAM</td><td>alignment</td><td>distance</td><td>position</td><td>gene name</td><td>gene id</td></tr>
<tr><td>GU305914.1</td><td></td><td>66695</td><td>66717</td><td>+</td><td>0</td><td>CCGCGGGCTGCGGCGCTGCG</td><td>TGG</td><td>||||||||[||||||||||||]PAM</td><td>NA</td><td>Intergenic</td><td>NA</td><td>NA</td></tr>
<tr><td></td></tr>
<tr><td>T4572</td><td>CGGGCTGCGGCGCTGCGTGGCGG</td><td></td><td>1000</td></tr>
<tr><td>Oligo fwd</td><td>CACCCGGGCTGCGGCGCTGCGTGG</td></tr>
<tr><td>Oligo rev</td><td>AAACCCACGCAGCGCCGCAGCCCG</td></tr>
<tr><td>Chromosome</td><td>start</td><td></td><td>end</td><td>strand</td><td>MM</td><td>target_seq</td><td>PAM</td><td>alignment</td><td>distance</td><td>position</td><td>gene name</td><td>gene id</td></tr>
<tr><td>GU305914.1</td><td></td><td>66698</td><td>66720</td><td>+</td><td>0</td><td>CGGGCTGCGGCGCTGCGTGG</td><td>CGG</td><td>||||||||[||||||||||||]PAM</td><td>NA</td><td>Intergenic</td><td>NA</td><td>NA</td></tr>
<tr><td></td></tr>
<tr><td>T4573</td><td>GGCGGACGCGCTGACGCTGATGG</td><td></td><td>1000</td></tr>
<tr><td>Oligo fwd</td><td>CACCGGCGGACGCGCTGACGCTGA</td></tr>
<tr><td>Oligo rev</td><td>AAACTCAGCGTCAGCGCGTCCGCC</td></tr>
<tr><td>Chromosome</td><td>start</td><td></td><td>end</td><td>strand</td><td>MM</td><td>target_seq</td><td>PAM</td><td>alignment</td><td>distance</td><td>position</td><td>gene name</td><td>gene id</td></tr>
<tr><td>GU305914.1</td><td></td><td>66716</td><td>66738</td><td>+</td><td>0</td><td>GGCGGACGCGCTGACGCTGA</td><td>TGG</td><td>||||||||[||||||||||||]PAM</td><td>NA</td><td>Intergenic</td><td>NA</td><td>NA</td></tr>
<tr><td>GU305914.1</td><td></td><td>225713</td><td>225735</td><td>+</td><td>3</td><td>GGCGGGGGCGCTGGCGCTGA</td><td>TGG</td><td>|||||--|[|||||-||||||]PAM</td><td>NA</td><td>Intergenic</td><td>NA</td><td>NA</td></tr>
<tr><td></td></tr>
<tr><td>T4574</td><td>TGACGCTGATGGAGAGCCACCGG</td><td></td><td>1000</td></tr>
<tr><td>Oligo fwd</td><td>CACCTGACGCTGATGGAGAGCCAC</td></tr>
<tr><td>Oligo rev</td><td>AAACGTGGCTCTCCATCAGCGTCA</td></tr>
<tr><td>Chromosome</td><td>start</td><td></td><td>end</td><td>strand</td><td>MM</td><td>target_seq</td><td>PAM</td><td>alignment</td><td>distance</td><td>position</td><td>gene name</td><td>gene id</td></tr>
<tr><td>GU305914.1</td><td></td><td>66727</td><td>66749</td><td>+</td><td>0</td><td>TGACGCTGATGGAGAGCCAC</td><td>CGG</td><td>||||||||[||||||||||||]PAM</td><td>NA</td><td>Intergenic</td><td>NA</td><td>NA</td></tr>
<tr><td></td></tr>
<tr><td>T4575</td><td>GACGCTGATGGAGAGCCACCGGG</td><td></td><td>1000</td></tr>
<tr><td>Oligo fwd</td><td>CACCGACGCTGATGGAGAGCCACC</td></tr>
<tr><td>Oligo rev</td><td>AAACGGTGGCTCTCCATCAGCGTC</td></tr>
<tr><td>Chromosome</td><td>start</td><td></td><td>end</td><td>strand</td><td>MM</td><td>target_seq</td><td>PAM</td><td>alignment</td><td>distance</td><td>position</td><td>gene name</td><td>gene id</td></tr>
<tr><td>GU305914.1</td><td></td><td>66728</td><td>66750</td><td>+</td><td>0</td><td>GACGCTGATGGAGAGCCACC</td><td>GGG</td><td>||||||||[||||||||||||]PAM</td><td>NA</td><td>Intergenic</td><td>NA</td><td>NA</td></tr>
<tr><td></td></tr>
<tr><td>T4576</td><td>CCACAGCCCCCCCAAGAGACAGG</td><td></td><td>1000</td></tr>
<tr><td>Oligo fwd</td><td>CACCCCACAGCCCCCCCAAGAGAC</td></tr>
<tr><td>Oligo rev</td><td>AAACGTCTCTTGGGGGGGCTGTGG</td></tr>
<tr><td>Chromosome</td><td>start</td><td></td><td>end</td><td>strand</td><td>MM</td><td>target_seq</td><td>PAM</td><td>alignment</td><td>distance</td><td>position</td><td>gene name</td><td>gene id</td></tr>
<tr><td>GU305914.1</td><td></td><td>66755</td><td>66777</td><td>+</td><td>0</td><td>CCACAGCCCCCCCAAGAGAC</td><td>AGG</td><td>||||||||[||||||||||||]PAM</td><td>NA</td><td>Intergenic</td><td>NA</td><td>NA</td></tr>
<tr><td></td></tr>
<tr><td>T4577</td><td>CCAAGAGACAGGAGCTCGACCGG</td><td></td><td>1000</td></tr>
<tr><td>Oligo fwd</td><td>CACCCCAAGAGACAGGAGCTCGAC</td></tr>
<tr><td>Oligo rev</td><td>AAACGTCGAGCTCCTGTCTCTTGG</td></tr>
<tr><td>Chromosome</td><td>start</td><td></td><td>end</td><td>strand</td><td>MM</td><td>target_seq</td><td>PAM</td><td>alignment</td><td>distance</td><td>position</td><td>gene name</td><td>gene id</td></tr>
<tr><td>GU305914.1</td><td></td><td>66766</td><td>66788</td><td>+</td><td>0</td><td>CCAAGAGACAGGAGCTCGAC</td><td>CGG</td><td>||||||||[||||||||||||]PAM</td><td>NA</td><td>Intergenic</td><td>NA</td><td>NA</td></tr>
<tr><td></td></tr>
<tr><td>T4578</td><td>CAAGAGACAGGAGCTCGACCGGG</td><td></td><td>1000</td></tr>
<tr><td>Oligo fwd</td><td>CACCCAAGAGACAGGAGCTCGACC</td></tr>
<tr><td>Oligo rev</td><td>AAACGGTCGAGCTCCTGTCTCTTG</td></tr>
<tr><td>Chromosome</td><td>start</td><td></td><td>end</td><td>strand</td><td>MM</td><td>target_seq</td><td>PAM</td><td>alignment</td><td>distance</td><td>position</td><td>gene name</td><td>gene id</td></tr>
<tr><td>GU305914.1</td><td></td><td>66767</td><td>66789</td><td>+</td><td>0</td><td>CAAGAGACAGGAGCTCGACC</td><td>GGG</td><td>||||||||[||||||||||||]PAM</td><td>NA</td><td>Intergenic</td><td>NA</td><td>NA</td></tr>
<tr><td></td></tr>
<tr><td>T4579</td><td>GAGCTCGACCGGGAATCGATAGG</td><td></td><td>1000</td></tr>
<tr><td>Oligo fwd</td><td>CACCGAGCTCGACCGGGAATCGAT</td></tr>
<tr><td>Oligo rev</td><td>AAACATCGATTCCCGGTCGAGCTC</td></tr>
<tr><td>Chromosome</td><td>start</td><td></td><td>end</td><td>strand</td><td>MM</td><td>target_seq</td><td>PAM</td><td>alignment</td><td>distance</td><td>position</td><td>gene name</td><td>gene id</td></tr>
<tr><td>GU305914.1</td><td></td><td>66777</td><td>66799</td><td>+</td><td>0</td><td>GAGCTCGACCGGGAATCGAT</td><td>AGG</td><td>||||||||[||||||||||||]PAM</td><td>NA</td><td>Intergenic</td><td>NA</td><td>NA</td></tr>
<tr><td></td></tr>
<tr><td>T4580</td><td>AGAAGCTCTTCGATCCGTACCGG</td><td></td><td>1000</td></tr>
<tr><td>Oligo fwd</td><td>CACCAGAAGCTCTTCGATCCGTAC</td></tr>
<tr><td>Oligo rev</td><td>AAACGTACGGATCGAAGAGCTTCT</td></tr>
<tr><td>Chromosome</td><td>start</td><td></td><td>end</td><td>strand</td><td>MM</td><td>target_seq</td><td>PAM</td><td>alignment</td><td>distance</td><td>position</td><td>gene name</td><td>gene id</td></tr>
<tr><td>GU305914.1</td><td></td><td>66808</td><td>66830</td><td>+</td><td>0</td><td>AGAAGCTCTTCGATCCGTAC</td><td>CGG</td><td>||||||||[||||||||||||]PAM</td><td>NA</td><td>Intergenic</td><td>NA</td><td>NA</td></tr>
<tr><td></td></tr>
<tr><td>T4581</td><td>CGATCCGTACCGGAATCGCGAGG</td><td></td><td>1000</td></tr>
<tr><td>Oligo fwd</td><td>CACCCGATCCGTACCGGAATCGCG</td></tr>
<tr><td>Oligo rev</td><td>AAACCGCGATTCCGGTACGGATCG</td></tr>
<tr><td>Chromosome</td><td>start</td><td></td><td>end</td><td>strand</td><td>MM</td><td>target_seq</td><td>PAM</td><td>alignment</td><td>distance</td><td>position</td><td>gene name</td><td>gene id</td></tr>
<tr><td>GU305914.1</td><td></td><td>66818</td><td>66840</td><td>+</td><td>0</td><td>CGATCCGTACCGGAATCGCG</td><td>AGG</td><td>||||||||[||||||||||||]PAM</td><td>NA</td><td>Intergenic</td><td>NA</td><td>NA</td></tr>
<tr><td></td></tr>
<tr><td>T4582</td><td>TCCGTACCGGAATCGCGAGGAGG</td><td></td><td>1000</td></tr>
<tr><td>Oligo fwd</td><td>CACCTCCGTACCGGAATCGCGAGG</td></tr>
<tr><td>Oligo rev</td><td>AAACCCTCGCGATTCCGGTACGGA</td></tr>
<tr><td>Chromosome</td><td>start</td><td></td><td>end</td><td>strand</td><td>MM</td><td>target_seq</td><td>PAM</td><td>alignment</td><td>distance</td><td>position</td><td>gene name</td><td>gene id</td></tr>
<tr><td>GU305914.1</td><td></td><td>66821</td><td>66843</td><td>+</td><td>0</td><td>TCCGTACCGGAATCGCGAGG</td><td>AGG</td><td>||||||||[||||||||||||]PAM</td><td>NA</td><td>Intergenic</td><td>NA</td><td>NA</td></tr>
<tr><td></td></tr>
<tr><td>T4583</td><td>CCGGAATCGCGAGGAGGCGCCGG</td><td></td><td>1000</td></tr>
<tr><td>Oligo fwd</td><td>CACCCCGGAATCGCGAGGAGGCGC</td></tr>
<tr><td>Oligo rev</td><td>AAACGCGCCTCCTCGCGATTCCGG</td></tr>
<tr><td>Chromosome</td><td>start</td><td></td><td>end</td><td>strand</td><td>MM</td><td>target_seq</td><td>PAM</td><td>alignment</td><td>distance</td><td>position</td><td>gene name</td><td>gene id</td></tr>
<tr><td>GU305914.1</td><td></td><td>66827</td><td>66849</td><td>+</td><td>0</td><td>CCGGAATCGCGAGGAGGCGC</td><td>CGG</td><td>||||||||[||||||||||||]PAM</td><td>NA</td><td>Intergenic</td><td>NA</td><td>NA</td></tr>
<tr><td></td></tr>
<tr><td>T4584</td><td>CTGCGCCGACGAAACATGAGCGG</td><td></td><td>1000</td></tr>
<tr><td>Oligo fwd</td><td>CACCCTGCGCCGACGAAACATGAG</td></tr>
<tr><td>Oligo rev</td><td>AAACCTCATGTTTCGTCGGCGCAG</td></tr>
<tr><td>Chromosome</td><td>start</td><td></td><td>end</td><td>strand</td><td>MM</td><td>target_seq</td><td>PAM</td><td>alignment</td><td>distance</td><td>position</td><td>gene name</td><td>gene id</td></tr>
<tr><td>GU305914.1</td><td></td><td>66858</td><td>66880</td><td>+</td><td>0</td><td>CTGCGCCGACGAAACATGAG</td><td>CGG</td><td>||||||||[||||||||||||]PAM</td><td>NA</td><td>Intergenic</td><td>NA</td><td>NA</td></tr>
<tr><td></td></tr>
<tr><td>T4585</td><td>GAAACATGAGCGGACGTCCGAGG</td><td></td><td>1000</td></tr>
<tr><td>Oligo fwd</td><td>CACCGAAACATGAGCGGACGTCCG</td></tr>
</table>

| | | | | | | | | | | | |
|---|---|---|---|---|---|---|---|---|---|---|---|
| Oligo rev | AAACCGGACGTCCGCTCATGTTTC | | | | | | | | | | |
| Chromosome | start | end | strand | MM | target_seq | PAM | alignment | distance | position | gene name | gene id |
| GU305914.1 | 66868 | 66890 | + | 0 | GAAACATGAGCGGACGTCCG | AGG | \|\|\|\|\|\|\|\|[\|\|\|\|\|\|\|\|\|\|\|\|]PAM | NA | Intergenic | NA | NA |
| T4586 | AAACATGAGCGGACGTCCGAGGG | 1000 | | | | | | | | | |
| Oligo fwd | CACCAAACATGAGCGGACGTCCGA | | | | | | | | | | |
| Oligo rev | AAACTCGGACGTCCGCTCATGTTT | | | | | | | | | | |
| Chromosome | start | end | strand | MM | target_seq | PAM | alignment | distance | position | gene name | gene id |
| GU305914.1 | 66869 | 66891 | + | 0 | AAACATGAGCGGACGTCCGA | GGG | \|\|\|\|\|\|\|\|[\|\|\|\|\|\|\|\|\|\|\|\|]PAM | NA | Intergenic | NA | NA |
| T4587 | GGGACGCGTCCGTTGATTTCCGG | 1000 | | | | | | | | | |
| Oligo fwd | CACCGGGACGCGTCCGTTGATTTC | | | | | | | | | | |
| Oligo rev | AAACGAAATCAACGGACGCGTCCC | | | | | | | | | | |
| Chromosome | start | end | strand | MM | target_seq | PAM | alignment | distance | position | gene name | gene id |
| GU305914.1 | 66889 | 66911 | + | 0 | GGGACGCGTCCGTTGATTTC | CGG | \|\|\|\|\|\|\|\|[\|\|\|\|\|\|\|\|\|\|\|\|]PAM | NA | Intergenic | NA | NA |
| T4588 | GTCCGTTGATTTCCGGTACGAGG | 1000 | | | | | | | | | |
| Oligo fwd | CACCGTCCGTTGATTTCCGGTACG | | | | | | | | | | |
| Oligo rev | AAACCGTACCGGAAATCAACGGAC | | | | | | | | | | |
| Chromosome | start | end | strand | MM | target_seq | PAM | alignment | distance | position | gene name | gene id |
| GU305914.1 | 66896 | 66918 | + | 0 | GTCCGTTGATTTCCGGTACG | AGG | \|\|\|\|\|\|\|\|[\|\|\|\|\|\|\|\|\|\|\|\|]PAM | NA | Intergenic | NA | NA |
| T4589 | CGTTGATTTCCGGTACGAGGCGG | 1000 | | | | | | | | | |
| Oligo fwd | CACCCGTTGATTTCCGGTACGAGG | | | | | | | | | | |
| Oligo rev | AAACCCTCGTACCGGAAATCAACG | | | | | | | | | | |
| Chromosome | start | end | strand | MM | target_seq | PAM | alignment | distance | position | gene name | gene id |
| GU305914.1 | 66899 | 66921 | + | 0 | CGTTGATTTCCGGTACGAGG | CGG | \|\|\|\|\|\|\|\|[\|\|\|\|\|\|\|\|\|\|\|\|]PAM | NA | Intergenic | NA | NA |
| T4590 | ATTTCCGGTACGAGGCGGAGAGG | 1000 | | | | | | | | | |
| Oligo fwd | CACCATTTCCGGTACGAGGCGGAG | | | | | | | | | | |
| Oligo rev | AAACCTCCGCCTCGTACCGGAAAT | | | | | | | | | | |
| Chromosome | start | end | strand | MM | target_seq | PAM | alignment | distance | position | gene name | gene id |
| GU305914.1 | 66904 | 66926 | + | 0 | ATTTCCGGTACGAGGCGGAG | AGG | \|\|\|\|\|\|\|\|[\|\|\|\|\|\|\|\|\|\|\|\|]PAM | NA | Intergenic | NA | NA |
| T4591 | CGAGGCGGAGAGGTACGTGCCGG | 1000 | | | | | | | | | |
| Oligo fwd | CACCCGAGGCGGAGAGGTACGTGC | | | | | | | | | | |
| Oligo rev | AAACGCACGTACCTCTCCGCCTCG | | | | | | | | | | |
| Chromosome | start | end | strand | MM | target_seq | PAM | alignment | distance | position | gene name | gene id |
| GU305914.1 | 66914 | 66936 | + | 0 | CGAGGCGGAGAGGTACGTGC | CGG | \|\|\|\|\|\|\|\|[\|\|\|\|\|\|\|\|\|\|\|\|]PAM | NA | Intergenic | NA | NA |
| T4592 | CGCCGAACGACCTGCGCGAGTGG | 1000 | | | | | | | | | |
| Oligo fwd | CACCCGCCGAACGACCTGCGCGAG | | | | | | | | | | |
| Oligo rev | AAACCTCGCGCAGGTCGTTCGGCG | | | | | | | | | | |
| Chromosome | start | end | strand | MM | target_seq | PAM | alignment | distance | position | gene name | gene id |
| GU305914.1 | 66943 | 66965 | + | 0 | CGCCGAACGACCTGCGCGAG | TGG | \|\|\|\|\|\|\|\|[\|\|\|\|\|\|\|\|\|\|\|\|]PAM | NA | Intergenic | NA | NA |
| GU305914.1 | 110585 | 110607 | - | 4 | GTTCGAACGGCCTGCGCGAG | AGG | ---\|\|\|\|\|[\|-\|\|\|\|\|\|\|\|\|\|]PAM | NA | Intergenic | NA | NA |
| T4593 | CGACCTGCGCGAGTGGTACGTGG | 1000 | | | | | | | | | |
| Oligo fwd | CACCCGACCTGCGCGAGTGGTACG | | | | | | | | | | |
| Oligo rev | AAACCGTACCACTCGCGCAGGTCG | | | | | | | | | | |
| Chromosome | start | end | strand | MM | target_seq | PAM | alignment | distance | position | gene name | gene id |
| GU305914.1 | 66950 | 66972 | + | 0 | CGACCTGCGCGAGTGGTACG | TGG | \|\|\|\|\|\|\|\|[\|\|\|\|\|\|\|\|\|\|\|\|]PAM | NA | Intergenic | NA | NA |
| T4594 | CGAGTGGTACGTGGAGACGAAGG | 1000 | | | | | | | | | |
| Oligo fwd | CACCCGAGTGGTACGTGGAGACGA | | | | | | | | | | |
| Oligo rev | AAACTCGTCTCCACGTACCACTCG | | | | | | | | | | |
| Chromosome | start | end | strand | MM | target_seq | PAM | alignment | distance | position | gene name | gene id |
| GU305914.1 | 66959 | 66981 | + | 0 | CGAGTGGTACGTGGAGACGA | AGG | \|\|\|\|\|\|\|\|[\|\|\|\|\|\|\|\|\|\|\|\|]PAM | NA | Intergenic | NA | NA |
| GU305914.1 | 27478 | 27500 | - | 4 | CGCGTGATGCGCGGAGACGA | CGG | \|\|-\|\|\|-\|[-\|\|-\|\|\|\|\|\|\|\|]PAM | NA | Intergenic | NA | NA |
| T4595 | GTACGTGGAGACGAAGGAGAAGG | 1000 | | | | | | | | | |
| Oligo fwd | CACCGTACGTGGAGACGAAGGAGA | | | | | | | | | | |
| Oligo rev | AAACTCTCCTTCGTCTCCACGTAC | | | | | | | | | | |
| Chromosome | start | end | strand | MM | target_seq | PAM | alignment | distance | position | gene name | gene id |
| GU305914.1 | 66965 | 66987 | + | 0 | GTACGTGGAGACGAAGGAGA | AGG | \|\|\|\|\|\|\|\|[\|\|\|\|\|\|\|\|\|\|\|\|]PAM | NA | Intergenic | NA | NA |
| GU305914.1 | 111427 | 111449 | - | 4 | GTACAGGTAGAAGAAGGAGA | CGG | \|\|\|\|--\|-[\|\|\|-\|\|\|\|\|\|\|\|]PAM | NA | Intergenic | NA | NA |
| T4596 | CCTGACGAACCCGCTACGCCTGG | 1000 | | | | | | | | | |
| Oligo fwd | CACCCCTGACGAACCCGCTACGCC | | | | | | | | | | |
| Oligo rev | AAACGGCGTAGCGGGTTCGTCAGG | | | | | | | | | | |
| Chromosome | start | end | strand | MM | target_seq | PAM | alignment | distance | position | gene name | gene id |
| GU305914.1 | 66998 | 67020 | + | 0 | CCTGACGAACCCGCTACGCC | TGG | \|\|\|\|\|\|\|\|[\|\|\|\|\|\|\|\|\|\|\|\|]PAM | NA | Intergenic | NA | NA |
| T4597 | CCTGGCCAGAAACGTACAGAAGG | 1000 | | | | | | | | | |
| Oligo fwd | CACCCCTGGCCAGAAACGTACAGA | | | | | | | | | | |
| Oligo rev | AAACTCTGTACGTTTCTGGCCAGG | | | | | | | | | | |
| Chromosome | start | end | strand | MM | target_seq | PAM | alignment | distance | position | gene name | gene id |
| GU305914.1 | 67016 | 67038 | + | 0 | CCTGGCCAGAAACGTACAGA | AGG | \|\|\|\|\|\|\|\|[\|\|\|\|\|\|\|\|\|\|\|\|]PAM | NA | Intergenic | NA | NA |
| T4598 | CCGCGCAAGCCGTGACCAGTCGG | 1000 | | | | | | | | | |
| Oligo fwd | CACCCCGCGCAAGCCGTGACCAGT | | | | | | | | | | |
| Oligo rev | AAACACTGGTCACGGCTTGCGCGG | | | | | | | | | | |
| Chromosome | start | end | strand | MM | target_seq | PAM | alignment | distance | position | gene name | gene id |
| GU305914.1 | 67057 | 67079 | + | 0 | CCGCGCAAGCCGTGACCAGT | CGG | \|\|\|\|\|\|\|\|[\|\|\|\|\|\|\|\|\|\|\|\|]PAM | NA | Intergenic | NA | NA |
| T4599 | GACCAGTCGGCGCCCGTGTACGG | 1000 | | | | | | | | | |
| Oligo fwd | CACCGACCAGTCGGCGCCCGTGTA | | | | | | | | | | |
| Oligo rev | AAACTACACGGGCGCCGACTGGTC | | | | | | | | | | |
| Chromosome | start | end | strand | MM | target_seq | PAM | alignment | distance | position | gene name | gene id |

GU305914.1	67070	67092	+	0	GACCAGTCGGCGCCCGTGTA	CGG	||||||||[||||||||||||]PAM	NA	Intergenic	NA	NA

T4600	ACCAGTCGGCGCCCGTGTACGGG	1000
Oligo fwd	CACCACCAGTCGGCGCCCGTGTAC
Oligo rev	AAACGTACACGGGCGCCGACTGGT
Chromosome	start	end	strand	MM	target_seq	PAM	alignment	distance	position	gene name	gene id
GU305914.1	67071	67093	+	0	ACCAGTCGGCGCCCGTGTAC	GGG	||||||||[||||||||||||]PAM	NA	Intergenic	NA	NA

T4601	GCCCGTGTACGGGCCCCGCGCGG	1000
Oligo fwd	CACCGCCCGTGTACGGGCCCCGCG
Oligo rev	AAACCGCGGGGCCCGTACACGGGC
Chromosome	start	end	strand	MM	target_seq	PAM	alignment	distance	position	gene name	gene id
GU305914.1	67081	67103	+	0	GCCCGTGTACGGGCCCCGCG	CGG	||||||||[||||||||||||]PAM	NA	Intergenic	NA	NA

T4602	CCCGTGTACGGGCCCCGCGCGGG	1000
Oligo fwd	CACCCCCGTGTACGGGCCCCGCGC
Oligo rev	AAACGCGCGGGGCCCGTACACGGG
Chromosome	start	end	strand	MM	target_seq	PAM	alignment	distance	position	gene name	gene id
GU305914.1	67082	67104	+	0	CCCGTGTACGGGCCCCGCGC	GGG	||||||||[||||||||||||]PAM	NA	Intergenic	NA	NA

T4603	CCTACACACCTGTTACCTGATGG	1000
Oligo fwd	CACCCCTACACACCTGTTACCTGA
Oligo rev	AAACTCAGGTAACAGGTGTGTAGG
Chromosome	start	end	strand	MM	target_seq	PAM	alignment	distance	position	gene name	gene id
GU305914.1	67135	67157	+	0	CCTACACACCTGTTACCTGA	TGG	||||||||[||||||||||||]PAM	NA	Intergenic	NA	NA

T4604	CTACACACCTGTTACCTGATGGG	1000
Oligo fwd	CACCCTACACACCTGTTACCTGAT
Oligo rev	AAACATCAGGTAACAGGTGTGTAG
Chromosome	start	end	strand	MM	target_seq	PAM	alignment	distance	position	gene name	gene id
GU305914.1	67136	67158	+	0	CTACACACCTGTTACCTGAT	GGG	||||||||[||||||||||||]PAM	NA	Intergenic	NA	NA
GU305914.1	186983	187005	+	4	GGACACACCTGGTACCTGAA	CGG	--||||||[|||-|||||||-]PAM	NA	Intergenic	NA	NA

T4605	TTACCTGATGGGCATCGATCCGG	1000
Oligo fwd	CACCTTACCTGATGGGCATCGATC
Oligo rev	AAACGATCGATGCCCATCAGGTAA
Chromosome	start	end	strand	MM	target_seq	PAM	alignment	distance	position	gene name	gene id
GU305914.1	67147	67169	+	0	TTACCTGATGGGCATCGATC	CGG	||||||||[||||||||||||]PAM	NA	Intergenic	NA	NA

T4606	GATGGGCATCGATCCGGTGCTGG	1000
Oligo fwd	CACCGATGGGCATCGATCCGGTGC
Oligo rev	AAACGCACCGGATCGATGCCCATC
Chromosome	start	end	strand	MM	target_seq	PAM	alignment	distance	position	gene name	gene id
GU305914.1	67153	67175	+	0	GATGGGCATCGATCCGGTGC	TGG	||||||||[||||||||||||]PAM	NA	Intergenic	NA	NA

T4607	GGTGCTGGACACGACGTCGCTGG	1000
Oligo fwd	CACCGGTGCTGGACACGACGTCGC
Oligo rev	AAACGCGACGTCGTGTCCAGCACC
Chromosome	start	end	strand	MM	target_seq	PAM	alignment	distance	position	gene name	gene id
GU305914.1	67168	67190	+	0	GGTGCTGGACACGACGTCGC	TGG	||||||||[||||||||||||]PAM	NA	Intergenic	NA	NA

T4608	GACGTCGCTGGACGCCGTGCTGG	1000
Oligo fwd	CACCGACGTCGCTGGACGCCGTGC
Oligo rev	AAACGCACGGCGTCCAGCGACGTC
Chromosome	start	end	strand	MM	target_seq	PAM	alignment	distance	position	gene name	gene id
GU305914.1	67180	67202	+	0	GACGTCGCTGGACGCCGTGC	TGG	||||||||[||||||||||||]PAM	NA	Intergenic	NA	NA

T4609	CTGGACGCCGTGCTGGACAGCGG	1000
Oligo fwd	CACCCTGGACGCCGTGCTGGACAG
Oligo rev	AAACCTGTCCAGCACGGCGTCCAG
Chromosome	start	end	strand	MM	target_seq	PAM	alignment	distance	position	gene name	gene id
GU305914.1	67187	67209	+	0	CTGGACGCCGTGCTGGACAG	CGG	||||||||[||||||||||||]PAM	NA	Intergenic	NA	NA

T4610	GCTGGACAGCGGCGCGCGCCTGG	1000
Oligo fwd	CACCGCTGGACAGCGGCGCGCGCC
Oligo rev	AAACGGCGCGCGCCGCTGTCCAGC
Chromosome	start	end	strand	MM	target_seq	PAM	alignment	distance	position	gene name	gene id
GU305914.1	67198	67220	+	0	GCTGGACAGCGGCGCGCGCC	TGG	||||||||[||||||||||||]PAM	NA	Intergenic	NA	NA

T4611	CGCGCGCCTGGACGCGATCGCGG	1000
Oligo fwd	CACCCGCGCGCCTGGACGCGATCG
Oligo rev	AAACCGATCGCGTCCAGGCGCGCG
Chromosome	start	end	strand	MM	target_seq	PAM	alignment	distance	position	gene name	gene id
GU305914.1	67210	67232	+	0	CGCGCGCCTGGACGCGATCG	CGG	||||||||[||||||||||||]PAM	NA	Intergenic	NA	NA

T4612	GGACGCGATCGCGGACGAGAAGG	1000
Oligo fwd	CACCGGACGCGATCGCGGACGAGA
Oligo rev	AAACTCTCGTCCGCGATCGCGTCC
Chromosome	start	end	strand	MM	target_seq	PAM	alignment	distance	position	gene name	gene id
GU305914.1	67219	67241	+	0	GGACGCGATCGCGGACGAGA	AGG	||||||||[||||||||||||]PAM	NA	Intergenic	NA	NA
GU305914.1	229967	229989	-	4	GGCCGGGATGGCGCACGAGA	CGG	||-||-||[|-|||-||||||]PAM	NA	Intergenic	NA	NA

T4613	TCGCGGACGAGAAGGTGAAGAGG	1000
Oligo fwd	CACCTCGCGGACGAGAAGGTGAAG
Oligo rev	AAACCTTCACCTTCTCGTCCGCGA
Chromosome	start	end	strand	MM	target_seq	PAM	alignment	distance	position	gene name	gene id
GU305914.1	67227	67249	+	0	TCGCGGACGAGAAGGTGAAG	AGG	||||||||[||||||||||||]PAM	NA	Intergenic	NA	NA

T4614	GGACGAGAAGGTGAAGAGGCAGG	1000

Oligo fwd CACCGGACGAGAAGGTGAAGAGGC
Oligo rev AAACGCCTCTTCACCTTCTCGTCC
Chromosome start end strand MM target_seq PAM alignment distance position gene name gene id
GU305914.1 67231 67253 + 0 GGACGAGAAGGTGAAGAGGC AGG ||||||||[||||||||||||]PAM NA Intergenic NA NA

T4615 GGTGAAGAGGCAGGCGCTGACGG 1000
Oligo fwd CACCGGTGAAGAGGCAGGCGCTGA
Oligo rev AAACTCAGCGCCTGCCTCTTCACC
Chromosome start end strand MM target_seq PAM alignment distance position gene name gene id
GU305914.1 67240 67262 + 0 GGTGAAGAGGCAGGCGCTGA CGG ||||||||[||||||||||||]PAM NA Intergenic NA NA

T4616 GACGGACCACCCGTACCGTCTGG 1000
Oligo fwd CACCGACGGACCACCCGTACCGTC
Oligo rev AAACGACGGTACGGGTGGTCCGTC
Chromosome start end strand MM target_seq PAM alignment distance position gene name gene id
GU305914.1 67258 67280 + 0 GACGGACCACCCGTACCGTC TGG ||||||||[||||||||||||]PAM NA Intergenic NA NA

T4617 ACGGACCACCCGTACCGTCTGGG 1000
Oligo fwd CACCACGGACCACCCGTACCGTCT
Oligo rev AAACAGACGGTACGGGTGGTCCGT
Chromosome start end strand MM target_seq PAM alignment distance position gene name gene id
GU305914.1 67259 67281 + 0 ACGGACCACCCGTACCGTCT GGG ||||||||[||||||||||||]PAM NA Intergenic NA NA

T4618 CGGACCACCCGTACCGTCTGGGG 1000
Oligo fwd CACCCGGACCACCCGTACCGTCTG
Oligo rev AAACCAGACGGTACGGGTGGTCCG
Chromosome start end strand MM target_seq PAM alignment distance position gene name gene id
GU305914.1 67260 67282 + 0 CGGACCACCCGTACCGTCTG GGG ||||||||[||||||||||||]PAM NA Intergenic NA NA

T4619 CCACCCGTACCGTCTGGGGACGG 1000
Oligo fwd CACCCCACCCGTACCGTCTGGGGA
Oligo rev AAACTCCCCAGACGGTACGGGTGG
Chromosome start end strand MM target_seq PAM alignment distance position gene name gene id
GU305914.1 67264 67286 + 0 CCACCCGTACCGTCTGGGGA CGG ||||||||[||||||||||||]PAM NA Intergenic NA NA

T4620 TCTGGGGACGGAGATCCCCACGG 1000
Oligo fwd CACCTCTGGGGACGGAGATCCCCA
Oligo rev AAACTGGGGATCTCCGTCCCCAGA
Chromosome start end strand MM target_seq PAM alignment distance position gene name gene id
GU305914.1 67276 67298 + 0 TCTGGGGACGGAGATCCCCA CGG ||||||||[||||||||||||]PAM NA Intergenic NA NA

T4621 CACGGTGATCGAGACGCCCGCGG 1000
Oligo fwd CACCCACGGTGATCGAGACGCCCG
Oligo rev AAACCGGGCGTCTCGATCACCGTG
Chromosome start end strand MM target_seq PAM alignment distance position gene name gene id
GU305914.1 67294 67316 + 0 CACGGTGATCGAGACGCCCG CGG ||||||||[||||||||||||]PAM NA Intergenic NA NA

T4622 ACGGTGATCGAGACGCCCGCGGG 1000
Oligo fwd CACCACGGTGATCGAGACGCCCGC
Oligo rev AAACGCGGGCGTCTCGATCACCGT
Chromosome start end strand MM target_seq PAM alignment distance position gene name gene id
GU305914.1 67295 67317 + 0 ACGGTGATCGAGACGCCCGC GGG ||||||||[||||||||||||]PAM NA Intergenic NA NA

T4623 CGAGACGCCCGCGGGTATCACGG 1000
Oligo fwd CACCCGAGACGCCCGCGGGTATCA
Oligo rev AAACTGATACCCGCGGGCGTCTCG
Chromosome start end strand MM target_seq PAM alignment distance position gene name gene id
GU305914.1 67303 67325 + 0 CGAGACGCCCGCGGGTATCA CGG ||||||||[||||||||||||]PAM NA Intergenic NA NA

T4624 GAGACGCCCGCGGGTATCACGGG 1000
Oligo fwd CACCGAGACGCCCGCGGGTATCAC
Oligo rev AAACGTGATACCCGCGGGCGTCTC
Chromosome start end strand MM target_seq PAM alignment distance position gene name gene id
GU305914.1 67304 67326 + 0 GAGACGCCCGCGGGTATCAC GGG ||||||||[||||||||||||]PAM NA Intergenic NA NA

T4625 AGACGCCCGCGGGTATCACGGGG 1000
Oligo fwd CACCAGACGCCCGCGGGTATCACG
Oligo rev AAACCGTGATACCCGCGGGCGTCT
Chromosome start end strand MM target_seq PAM alignment distance position gene name gene id
GU305914.1 67305 67327 + 0 AGACGCCCGCGGGTATCACG GGG ||||||||[||||||||||||]PAM NA Intergenic NA NA

T4626 TCACGGGGCACGCGTTATCGCGG 1000
Oligo fwd CACCTCACGGGGCACGCGTTATCG
Oligo rev AAACCGATAACGCGTGCCCCGTGA
Chromosome start end strand MM target_seq PAM alignment distance position gene name gene id
GU305914.1 67320 67342 + 0 TCACGGGGCACGCGTTATCG CGG ||||||||[||||||||||||]PAM NA Intergenic NA NA

T4627 GCGTTATCGCGGCCGTTCAACGG 1000
Oligo fwd CACCGCGTTATCGCGGCCGTTCAA
Oligo rev AAACTTGAACGGCCGCGATAACGC
Chromosome start end strand MM target_seq PAM alignment distance position gene name gene id
GU305914.1 67331 67353 + 0 GCGTTATCGCGGCCGTTCAA CGG ||||||||[||||||||||||]PAM NA Intergenic NA NA

T4628 ATCGCGGCCGTTCAACGGCACGG 1000
Oligo fwd CACCATCGCGGCCGTTCAACGGCA
Oligo rev AAACTGCCGTTGAACGGCCGCGAT
Chromosome start end strand MM target_seq PAM alignment distance position gene name gene id
GU305914.1 67336 67358 + 0 ATCGCGGCCGTTCAACGGCA CGG ||||||||[||||||||||||]PAM NA Intergenic NA NA

| | | | | | | | | | | | |
|---|---|---|---|---|---|---|---|---|---|---|---|
| T4629 | GCGGCCGTTCAACGGCACGGCGG | 1000 | | | | | | | | | |
| Oligo fwd | CACCGCGGCCGTTCAACGGCACGG | | | | | | | | | | |
| Oligo rev | AAACCCGTGCCGTTGAACGGCCGC | | | | | | | | | | |
| Chromosome | start | end | strand | MM | target_seq | PAM | alignment | distance | position | gene name | gene id |
| GU305914.1 | 67339 | 67361 | + | 0 | GCGGCCGTTCAACGGCACGG | CGG | \|\|\|\|\|\|\|\|[\|\|\|\|\|\|\|\|\|\|\|\|]PAM | NA | Intergenic | NA | NA |
| T4630 | CAACGGCACGGCGGAGACGCAGG | 1000 | | | | | | | | | |
| Oligo fwd | CACCCAACGGCACGGCGGAGACGC | | | | | | | | | | |
| Oligo rev | AAACGCGTCTCCGCCGTGCCGTTG | | | | | | | | | | |
| Chromosome | start | end | strand | MM | target_seq | PAM | alignment | distance | position | gene name | gene id |
| GU305914.1 | 67348 | 67370 | + | 0 | CAACGGCACGGCGGAGACGC | AGG | \|\|\|\|\|\|\|\|[\|\|\|\|\|\|\|\|\|\|\|\|]PAM | NA | Intergenic | NA | NA |
| T4631 | CACGGCGGAGACGCAGGACCTGG | 1000 | | | | | | | | | |
| Oligo fwd | CACCCACGGCGGAGACGCAGGACC | | | | | | | | | | |
| Oligo rev | AAACGGTCCTGCGTCTCCGCCGTG | | | | | | | | | | |
| Chromosome | start | end | strand | MM | target_seq | PAM | alignment | distance | position | gene name | gene id |
| GU305914.1 | 67354 | 67376 | + | 0 | CACGGCGGAGACGCAGGACC | TGG | \|\|\|\|\|\|\|\|[\|\|\|\|\|\|\|\|\|\|\|\|]PAM | NA | Intergenic | NA | NA |
| T4632 | ACGGCGGAGACGCAGGACCTGGG | 1000 | | | | | | | | | |
| Oligo fwd | CACCACGGCGGAGACGCAGGACCT | | | | | | | | | | |
| Oligo rev | AAACAGGTCCTGCGTCTCCGCCGT | | | | | | | | | | |
| Chromosome | start | end | strand | MM | target_seq | PAM | alignment | distance | position | gene name | gene id |
| GU305914.1 | 67355 | 67377 | + | 0 | ACGGCGGAGACGCAGGACCT | GGG | \|\|\|\|\|\|\|\|[\|\|\|\|\|\|\|\|\|\|\|\|]PAM | NA | Intergenic | NA | NA |
| T4633 | CGGCGGAGACGCAGGACCTGGGG | 1000 | | | | | | | | | |
| Oligo fwd | CACCCGGCGGAGACGCAGGACCTG | | | | | | | | | | |
| Oligo rev | AAACCAGGTCCTGCGTCTCCGCCG | | | | | | | | | | |
| Chromosome | start | end | strand | MM | target_seq | PAM | alignment | distance | position | gene name | gene id |
| GU305914.1 | 67356 | 67378 | + | 0 | CGGCGGAGACGCAGGACCTG | GGG | \|\|\|\|\|\|\|\|[\|\|\|\|\|\|\|\|\|\|\|\|]PAM | NA | Intergenic | NA | NA |
| T4634 | GGCGGAGACGCAGGACCTGGGGG | 1000 | | | | | | | | | |
| Oligo fwd | CACCGGCGGAGACGCAGGACCTGG | | | | | | | | | | |
| Oligo rev | AAACCCAGGTCCTGCGTCTCCGCC | | | | | | | | | | |
| Chromosome | start | end | strand | MM | target_seq | PAM | alignment | distance | position | gene name | gene id |
| GU305914.1 | 67357 | 67379 | + | 0 | GGCGGAGACGCAGGACCTGG | GGG | \|\|\|\|\|\|\|\|[\|\|\|\|\|\|\|\|\|\|\|\|]PAM | NA | Intergenic | NA | NA |
| GU305914.1 | 72061 | 72083 | + | 4 | GGAGGCGACGCATGCCCTGG | AGG | \|\|-\|\|-\|\|[\|\|\|\|-\|-\|\|\|\|\|]PAM | NA | Intergenic | NA | NA |
| T4635 | GCGGAGACGCAGGACCTGGGGGG | 1000 | | | | | | | | | |
| Oligo fwd | CACCGCGGAGACGCAGGACCTGGG | | | | | | | | | | |
| Oligo rev | AAACCCCAGGTCCTGCGTCTCCGC | | | | | | | | | | |
| Chromosome | start | end | strand | MM | target_seq | PAM | alignment | distance | position | gene name | gene id |
| GU305914.1 | 67358 | 67380 | + | 0 | GCGGAGACGCAGGACCTGGG | GGG | \|\|\|\|\|\|\|\|[\|\|\|\|\|\|\|\|\|\|\|\|]PAM | NA | Intergenic | NA | NA |
| T4636 | CCTGGGGGGCTACAAGTGCCTGG | 1000 | | | | | | | | | |
| Oligo fwd | CACCCCTGGGGGGCTACAAGTGCC | | | | | | | | | | |
| Oligo rev | AAACGGCACTTGTAGCCCCCCAGG | | | | | | | | | | |
| Chromosome | start | end | strand | MM | target_seq | PAM | alignment | distance | position | gene name | gene id |
| GU305914.1 | 67372 | 67394 | + | 0 | CCTGGGGGGCTACAAGTGCC | TGG | \|\|\|\|\|\|\|\|[\|\|\|\|\|\|\|\|\|\|\|\|]PAM | NA | Intergenic | NA | NA |
| T4637 | CTGGGGGGCTACAAGTGCCTGGG | 1000 | | | | | | | | | |
| Oligo fwd | CACCCTGGGGGGCTACAAGTGCCT | | | | | | | | | | |
| Oligo rev | AAACAGGCACTTGTAGCCCCCCAG | | | | | | | | | | |
| Chromosome | start | end | strand | MM | target_seq | PAM | alignment | distance | position | gene name | gene id |
| GU305914.1 | 67373 | 67395 | + | 0 | CTGGGGGGCTACAAGTGCCT | GGG | \|\|\|\|\|\|\|\|[\|\|\|\|\|\|\|\|\|\|\|\|]PAM | NA | Intergenic | NA | NA |
| T4638 | CTACAAGTGCCTGGGCATCCTGG | 1000 | | | | | | | | | |
| Oligo fwd | CACCCTACAAGTGCCTGGGCATCC | | | | | | | | | | |
| Oligo rev | AAACGGATGCCCAGGCACTTGTAG | | | | | | | | | | |
| Chromosome | start | end | strand | MM | target_seq | PAM | alignment | distance | position | gene name | gene id |
| GU305914.1 | 67381 | 67403 | + | 0 | CTACAAGTGCCTGGGCATCC | TGG | \|\|\|\|\|\|\|\|[\|\|\|\|\|\|\|\|\|\|\|\|]PAM | NA | Intergenic | NA | NA |
| T4639 | GACTTCCTGACGTACGCGAGAGG | 1000 | | | | | | | | | |
| Oligo fwd | CACCGACTTCCTGACGTACGCGAG | | | | | | | | | | |
| Oligo rev | AAACCTCGCGTACGTCAGGAAGTC | | | | | | | | | | |
| Chromosome | start | end | strand | MM | target_seq | PAM | alignment | distance | position | gene name | gene id |
| GU305914.1 | 67403 | 67425 | + | 0 | GACTTCCTGACGTACGCGAG | AGG | \|\|\|\|\|\|\|\|[\|\|\|\|\|\|\|\|\|\|\|\|]PAM | NA | Intergenic | NA | NA |
| T4640 | ACTTCCTGACGTACGCGAGAGGG | 1000 | | | | | | | | | |
| Oligo fwd | CACCACTTCCTGACGTACGCGAGA | | | | | | | | | | |
| Oligo rev | AAACTCTCGCGTACGTCAGGAAGT | | | | | | | | | | |
| Chromosome | start | end | strand | MM | target_seq | PAM | alignment | distance | position | gene name | gene id |
| GU305914.1 | 67404 | 67426 | + | 0 | ACTTCCTGACGTACGCGAGA | GGG | \|\|\|\|\|\|\|\|[\|\|\|\|\|\|\|\|\|\|\|\|]PAM | NA | Intergenic | NA | NA |
| T4641 | CGCGAGAGGGAAGCCGCTGCCGG | 1000 | | | | | | | | | |
| Oligo fwd | CACCCGCGAGAGGGAAGCCGCTGC | | | | | | | | | | |
| Oligo rev | AAACGCAGCGGCTTCCCTCTCGCG | | | | | | | | | | |
| Chromosome | start | end | strand | MM | target_seq | PAM | alignment | distance | position | gene name | gene id |
| GU305914.1 | 67417 | 67439 | + | 0 | CGCGAGAGGGAAGCCGCTGC | CGG | \|\|\|\|\|\|\|\|[\|\|\|\|\|\|\|\|\|\|\|\|]PAM | NA | Intergenic | NA | NA |
| T4642 | GCCGGTGTACATCATCGTGACGG | 1000 | | | | | | | | | |
| Oligo fwd | CACCGCCGGTGTACATCATCGTGA | | | | | | | | | | |
| Oligo rev | AAACTCACGATGATGTACACCGGC | | | | | | | | | | |
| Chromosome | start | end | strand | MM | target_seq | PAM | alignment | distance | position | gene name | gene id |
| GU305914.1 | 67435 | 67457 | + | 0 | GCCGGTGTACATCATCGTGA | CGG | \|\|\|\|\|\|\|\|[\|\|\|\|\|\|\|\|\|\|\|\|]PAM | NA | Intergenic | NA | NA |
| T4643 | GTGTACATCATCGTGACGGTCGG | 1000 | | | | | | | | | |
| Oligo fwd | CACCGTGTACATCATCGTGACGGT | | | | | | | | | | |
| Oligo rev | AAACACCGTCACGATGATGTACAC | | | | | | | | | | |
| Chromosome | start | end | strand | MM | target_seq | PAM | alignment | distance | position | gene name | gene id |

| | | | | | | | | | | | |
|---|---|---|---|---|---|---|---|---|---|---|---|
| GU305914.1 | 67439 | 67461 | + | 0 | GTGTACATCATCGTGACGGT | CGG | \|\|\|\|\|\|\|\|[\|\|\|\|\|\|\|\|\|\|\|\|]PAM | NA | Intergenic | NA | NA |
| T4644 | ACGGTCGGCGTGCACACGAGAGG | 1000 | | | | | | | | | |
| Oligo fwd | CACCACGGTCGGCGTGCACACGAG | | | | | | | | | | |
| Oligo rev | AAACCTCGTGTGCACGCCGACCGT | | | | | | | | | | |
| Chromosome | start | end | strand | MM | target_seq | PAM | alignment | distance | position | gene name | gene id |
| GU305914.1 | 67454 | 67476 | + | 0 | ACGGTCGGCGTGCACACGAG | AGG | \|\|\|\|\|\|\|\|[\|\|\|\|\|\|\|\|\|\|\|\|]PAM | NA | Intergenic | NA | NA |
| T4645 | CGGTCGGCGTGCACACGAGAGGG | 1000 | | | | | | | | | |
| Oligo fwd | CACCCGGTCGGCGTGCACACGAGA | | | | | | | | | | |
| Oligo rev | AAACTCTCGTGTGCACGCCGACCG | | | | | | | | | | |
| Chromosome | start | end | strand | MM | target_seq | PAM | alignment | distance | position | gene name | gene id |
| GU305914.1 | 67455 | 67477 | + | 0 | CGGTCGGCGTGCACACGAGA | GGG | \|\|\|\|\|\|\|\|[\|\|\|\|\|\|\|\|\|\|\|\|]PAM | NA | Intergenic | NA | NA |
| T4646 | GGTCGGCGTGCACACGAGAGGGG | 1000 | | | | | | | | | |
| Oligo fwd | CACCGGTCGGCGTGCACACGAGAG | | | | | | | | | | |
| Oligo rev | AAACCTCTCGTGTGCACGCCGACC | | | | | | | | | | |
| Chromosome | start | end | strand | MM | target_seq | PAM | alignment | distance | position | gene name | gene id |
| GU305914.1 | 67456 | 67478 | + | 0 | GGTCGGCGTGCACACGAGAG | GGG | \|\|\|\|\|\|\|\|[\|\|\|\|\|\|\|\|\|\|\|\|]PAM | NA | Intergenic | NA | NA |
| T4647 | GCACACGAGAGGGGTGATCGTGG | 1000 | | | | | | | | | |
| Oligo fwd | CACCGCACACGAGAGGGGTGATCG | | | | | | | | | | |
| Oligo rev | AAACCGATCACCCCTCTCGTGTGC | | | | | | | | | | |
| Chromosome | start | end | strand | MM | target_seq | PAM | alignment | distance | position | gene name | gene id |
| GU305914.1 | 67465 | 67487 | + | 0 | GCACACGAGAGGGGTGATCG | TGG | \|\|\|\|\|\|\|\|[\|\|\|\|\|\|\|\|\|\|\|\|]PAM | NA | Intergenic | NA | NA |
| T4648 | CGAGAGGGGTGATCGTGGCGCGG | 1000 | | | | | | | | | |
| Oligo fwd | CACCCGAGAGGGGTGATCGTGGCG | | | | | | | | | | |
| Oligo rev | AAACCGCCACGATCACCCCTCTCG | | | | | | | | | | |
| Chromosome | start | end | strand | MM | target_seq | PAM | alignment | distance | position | gene name | gene id |
| GU305914.1 | 67470 | 67492 | + | 0 | CGAGAGGGGTGATCGTGGCG | CGG | \|\|\|\|\|\|\|\|[\|\|\|\|\|\|\|\|\|\|\|\|]PAM | NA | Intergenic | NA | NA |
| T4649 | GAGAGGGGTGATCGTGGCGCGGG | 1000 | | | | | | | | | |
| Oligo fwd | CACCGAGAGGGGTGATCGTGGCGC | | | | | | | | | | |
| Oligo rev | AAACGCGCCACGATCACCCCTCTC | | | | | | | | | | |
| Chromosome | start | end | strand | MM | target_seq | PAM | alignment | distance | position | gene name | gene id |
| GU305914.1 | 67471 | 67493 | + | 0 | GAGAGGGGTGATCGTGGCGC | GGG | \|\|\|\|\|\|\|\|[\|\|\|\|\|\|\|\|\|\|\|\|]PAM | NA | Intergenic | NA | NA |
| T4650 | AGAGGGGTGATCGTGGCGCGGGG | 1000 | | | | | | | | | |
| Oligo fwd | CACCAGAGGGGTGATCGTGGCGCG | | | | | | | | | | |
| Oligo rev | AAACCGCGCCACGATCACCCCTCT | | | | | | | | | | |
| Chromosome | start | end | strand | MM | target_seq | PAM | alignment | distance | position | gene name | gene id |
| GU305914.1 | 67472 | 67494 | + | 0 | AGAGGGGTGATCGTGGCGCG | GGG | \|\|\|\|\|\|\|\|[\|\|\|\|\|\|\|\|\|\|\|\|]PAM | NA | Intergenic | NA | NA |
| T4651 | GAGGGGTGATCGTGGCGCGGGGG | 1000 | | | | | | | | | |
| Oligo fwd | CACCGAGGGGTGATCGTGGCGCGG | | | | | | | | | | |
| Oligo rev | AAACCCGCGCCACGATCACCCCTC | | | | | | | | | | |
| Chromosome | start | end | strand | MM | target_seq | PAM | alignment | distance | position | gene name | gene id |
| GU305914.1 | 67473 | 67495 | + | 0 | GAGGGGTGATCGTGGCGCGG | GGG | \|\|\|\|\|\|\|\|[\|\|\|\|\|\|\|\|\|\|\|\|]PAM | NA | Intergenic | NA | NA |
| T4652 | AGGGGTGATCGTGGCGCGGGGGG | 1000 | | | | | | | | | |
| Oligo fwd | CACCAGGGGTGATCGTGGCGCGGG | | | | | | | | | | |
| Oligo rev | AAACCCCGCGCCACGATCACCCCT | | | | | | | | | | |
| Chromosome | start | end | strand | MM | target_seq | PAM | alignment | distance | position | gene name | gene id |
| GU305914.1 | 67474 | 67496 | + | 0 | AGGGGTGATCGTGGCGCGGG | GGG | \|\|\|\|\|\|\|\|[\|\|\|\|\|\|\|\|\|\|\|\|]PAM | NA | Intergenic | NA | NA |
| T4653 | CGTGTTCGACCCGCACACGACGG | 1000 | | | | | | | | | |
| Oligo fwd | CACCCGTGTTCGACCCGCACACGA | | | | | | | | | | |
| Oligo rev | AAACTCGTGTGCGGGTCGAACACG | | | | | | | | | | |
| Chromosome | start | end | strand | MM | target_seq | PAM | alignment | distance | position | gene name | gene id |
| GU305914.1 | 67504 | 67526 | + | 0 | CGTGTTCGACCCGCACACGA | CGG | \|\|\|\|\|\|\|\|[\|\|\|\|\|\|\|\|\|\|\|\|]PAM | NA | Intergenic | NA | NA |
| T4654 | CCCGCACACGACGGACCTGTCGG | 1000 | | | | | | | | | |
| Oligo fwd | CACCCCCGCACACGACGGACCTGT | | | | | | | | | | |
| Oligo rev | AAACACAGGTCCGTCGTGTGCGGG | | | | | | | | | | |
| Chromosome | start | end | strand | MM | target_seq | PAM | alignment | distance | position | gene name | gene id |
| GU305914.1 | 67513 | 67535 | + | 0 | CCCGCACACGACGGACCTGT | CGG | \|\|\|\|\|\|\|\|[\|\|\|\|\|\|\|\|\|\|\|\|]PAM | NA | Intergenic | NA | NA |
| T4655 | CACGACGGACCTGTCGGCCGAGG | 1000 | | | | | | | | | |
| Oligo fwd | CACCCACGACGGACCTGTCGGCCG | | | | | | | | | | |
| Oligo rev | AAACCGGCCGACAGGTCCGTCGTG | | | | | | | | | | |
| Chromosome | start | end | strand | MM | target_seq | PAM | alignment | distance | position | gene name | gene id |
| GU305914.1 | 67519 | 67541 | + | 0 | CACGACGGACCTGTCGGCCG | AGG | \|\|\|\|\|\|\|\|[\|\|\|\|\|\|\|\|\|\|\|\|]PAM | NA | Intergenic | NA | NA |
| T4656 | GACGGACCTGTCGGCCGAGGCGG | 1000 | | | | | | | | | |
| Oligo fwd | CACCGACGGACCTGTCGGCCGAGG | | | | | | | | | | |
| Oligo rev | AAACCCTCGGCCGACAGGTCCGTC | | | | | | | | | | |
| Chromosome | start | end | strand | MM | target_seq | PAM | alignment | distance | position | gene name | gene id |
| GU305914.1 | 67522 | 67544 | + | 0 | GACGGACCTGTCGGCCGAGG | CGG | \|\|\|\|\|\|\|\|[\|\|\|\|\|\|\|\|\|\|\|\|]PAM | NA | Intergenic | NA | NA |
| T4657 | GGACCTGTCGGCCGAGGCGGCGG | 1000 | | | | | | | | | |
| Oligo fwd | CACCGGACCTGTCGGCCGAGGCGG | | | | | | | | | | |
| Oligo rev | AAACCCGCCTCGGCCGACAGGTCC | | | | | | | | | | |
| Chromosome | start | end | strand | MM | target_seq | PAM | alignment | distance | position | gene name | gene id |
| GU305914.1 | 67525 | 67547 | + | 0 | GGACCTGTCGGCCGAGGCGG | CGG | \|\|\|\|\|\|\|\|[\|\|\|\|\|\|\|\|\|\|\|\|]PAM | NA | Intergenic | NA | NA |
| T4658 | CGTGTGCGACGACTTCACCGAGG | 1000 | | | | | | | | | |
| Oligo fwd | CACCCGTGTGCGACGACTTCACCG | | | | | | | | | | |
| Oligo rev | AAACCGGTGAAGTCGTCGCACACG | | | | | | | | | | |

Chromosome start end strand MM target_seq PAM alignment distance position gene name gene id
GU305914.1 67552 67574 + 0 CGTGTGCGACGACTTCACCG AGG ||||||||[||||||||||||]PAM NA Intergenic NA NA

T4659 CGACTTCACCGAGGCGATCTCGG 1000
Oligo fwd CACCCGACTTCACCGAGGCGATCT
Oligo rev AAACAGATCGCCTCGGTGAAGTCG
Chromosome start end strand MM target_seq PAM alignment distance position gene name gene id
GU305914.1 67561 67583 + 0 CGACTTCACCGAGGCGATCT CGG ||||||||[||||||||||||]PAM NA Intergenic NA NA

T4660 ATCTCGGCGCTGTCGTTCTTCGG 1000
Oligo fwd CACCATCTCGGCGCTGTCGTTCTT
Oligo rev AAACAAGAACGACAGCGCCGAGAT
Chromosome start end strand MM target_seq PAM alignment distance position gene name gene id
GU305914.1 67577 67599 + 0 ATCTCGGCGCTGTCGTTCTT CGG ||||||||[||||||||||||]PAM NA Intergenic NA NA

T4661 TCTCGGCGCTGTCGTTCTTCGGG 1000
Oligo fwd CACCTCTCGGCGCTGTCGTTCTTC
Oligo rev AAACGAAGAACGACAGCGCCGAGA
Chromosome start end strand MM target_seq PAM alignment distance position gene name gene id
GU305914.1 67578 67600 + 0 TCTCGGCGCTGTCGTTCTTC GGG ||||||||[||||||||||||]PAM NA Intergenic NA NA

T4662 CTCGGCGCTGTCGTTCTTCGGGG 1000
Oligo fwd CACCCTCGGCGCTGTCGTTCTTCG
Oligo rev AAACCGAAGAACGACAGCGCCGAG
Chromosome start end strand MM target_seq PAM alignment distance position gene name gene id
GU305914.1 67579 67601 + 0 CTCGGCGCTGTCGTTCTTCG GGG ||||||||[||||||||||||]PAM NA Intergenic NA NA

T4663 TCGTTCTTCGGGGCCATGATCGG 1000
Oligo fwd CACCTCGTTCTTCGGGGCCATGAT
Oligo rev AAACATCATGGCCCCGAAGAACGA
Chromosome start end strand MM target_seq PAM alignment distance position gene name gene id
GU305914.1 67589 67611 + 0 TCGTTCTTCGGGGCCATGAT CGG ||||||||[||||||||||||]PAM NA Intergenic NA NA

T4664 CGGAGACTTCTACTACGACGCGG 1000
Oligo fwd CACCCGGAGACTTCTACTACGACG
Oligo rev AAACCGTCGTAGTAGAAGTCTCCG
Chromosome start end strand MM target_seq PAM alignment distance position gene name gene id
GU305914.1 67609 67631 + 0 CGGAGACTTCTACTACGACG CGG ||||||||[||||||||||||]PAM NA Intergenic NA NA

T4665 CTTCTACTACGACGCGGTCCTGG 1000
Oligo fwd CACCCTTCTACTACGACGCGGTCC
Oligo rev AAACGGACCGCGTCGTAGTAGAAG
Chromosome start end strand MM target_seq PAM alignment distance position gene name gene id
GU305914.1 67615 67637 + 0 CTTCTACTACGACGCGGTCC TGG ||||||||[||||||||||||]PAM NA Intergenic NA NA

T4666 CACGACGCTGATCTCGCCGTCGG 1000
Oligo fwd CACCCACGACGCTGATCTCGCCGT
Oligo rev AAACACGGCGAGATCAGCGTCGTG
Chromosome start end strand MM target_seq PAM alignment distance position gene name gene id
GU305914.1 67657 67679 + 0 CACGACGCTGATCTCGCCGT CGG ||||||||[||||||||||||]PAM NA Intergenic NA NA

T4667 GATCTCGCCGTCGGAGCTGCTGG 1000
Oligo fwd CACCGATCTCGCCGTCGGAGCTGC
Oligo rev AAACGCAGCTCCGACGGCGAGATC
Chromosome start end strand MM target_seq PAM alignment distance position gene name gene id
GU305914.1 67666 67688 + 0 GATCTCGCCGTCGGAGCTGC TGG ||||||||[||||||||||||]PAM NA Intergenic NA NA

T4668 GGAGCTGCTGGTGCAGATCATGG 1000
Oligo fwd CACCGGAGCTGCTGGTGCAGATCA
Oligo rev AAACTGATCTGCACCAGCAGCTCC
Chromosome start end strand MM target_seq PAM alignment distance position gene name gene id
GU305914.1 67678 67700 + 0 GGAGCTGCTGGTGCAGATCA TGG ||||||||[||||||||||||]PAM NA Intergenic NA NA

T4669 GCAGATCATGGACCAGTACAAGG 1000
Oligo fwd CACCGCAGATCATGGACCAGTACA
Oligo rev AAACTGTACTGGTCCATGATCTGC
Chromosome start end strand MM target_seq PAM alignment distance position gene name gene id
GU305914.1 67690 67712 + 0 GCAGATCATGGACCAGTACA AGG ||||||||[||||||||||||]PAM NA Intergenic NA NA

T4670 CATGGACCAGTACAAGGACCCGG 1000
Oligo fwd CACCCATGGACCAGTACAAGGACC
Oligo rev AAACGGTCCTTGTACTGGTCCATG
Chromosome start end strand MM target_seq PAM alignment distance position gene name gene id
GU305914.1 67696 67718 + 0 CATGGACCAGTACAAGGACC CGG ||||||||[||||||||||||]PAM NA Intergenic NA NA

T4671 GGACCCGGACATCGACGCGTCGG 1000
Oligo fwd CACCGGACCCGGACATCGACGCGT
Oligo rev AAACACGCGTCGATGTCCGGGTCC
Chromosome start end strand MM target_seq PAM alignment distance position gene name gene id
GU305914.1 67711 67733 + 0 GGACCCGGACATCGACGCGT CGG ||||||||[||||||||||||]PAM NA Intergenic NA NA

T4672 CATCGACGCGTCGGTGATGTCGG 1000
Oligo fwd CACCCATCGACGCGTCGGTGATGT
Oligo rev AAACACATCACCGACGCGTCGATG
Chromosome start end strand MM target_seq PAM alignment distance position gene name gene id
GU305914.1 67720 67742 + 0 CATCGACGCGTCGGTGATGT CGG ||||||||[||||||||||||]PAM NA Intergenic NA NA

T4673 ATCGACGCGTCGGTGATGTCGGG 1000
Oligo fwd CACCATCGACGCGTCGGTGATGTC

<table>
<tr><td>Oligo rev</td><td>AAACGACATCACCGACGCGTCGAT</td><td></td><td></td><td></td><td></td><td></td><td></td><td></td><td></td><td></td><td></td></tr>
<tr><td>Chromosome</td><td>start</td><td>end</td><td>strand</td><td>MM</td><td>target_seq</td><td>PAM</td><td>alignment</td><td>distance</td><td>position</td><td>gene name</td><td>gene id</td></tr>
<tr><td>GU305914.1</td><td>67721</td><td>67743</td><td>+</td><td>0</td><td>ATCGACGCGTCGGTGATGTC</td><td>GGG</td><td>||||||||[||||||||||||]PAM</td><td>NA</td><td>Intergenic</td><td>NA</td><td>NA</td></tr>
<tr><td></td><td></td><td></td><td></td><td></td><td></td><td></td><td></td><td></td><td></td><td></td><td></td></tr>
<tr><td>T4674</td><td>TCGACGCGTCGGTGATGTCGGGG</td><td>1000</td><td></td><td></td><td></td><td></td><td></td><td></td><td></td><td></td><td></td></tr>
<tr><td>Oligo fwd</td><td>CACCTCGACGCGTCGGTGATGTCG</td><td></td><td></td><td></td><td></td><td></td><td></td><td></td><td></td><td></td><td></td></tr>
<tr><td>Oligo rev</td><td>AAACCGACATCACCGACGCGTCGA</td><td></td><td></td><td></td><td></td><td></td><td></td><td></td><td></td><td></td><td></td></tr>
<tr><td>Chromosome</td><td>start</td><td>end</td><td>strand</td><td>MM</td><td>target_seq</td><td>PAM</td><td>alignment</td><td>distance</td><td>position</td><td>gene name</td><td>gene id</td></tr>
<tr><td>GU305914.1</td><td>67722</td><td>67744</td><td>+</td><td>0</td><td>TCGACGCGTCGGTGATGTCG</td><td>GGG</td><td>||||||||[||||||||||||]PAM</td><td>NA</td><td>Intergenic</td><td>NA</td><td>NA</td></tr>
<tr><td></td><td></td><td></td><td></td><td></td><td></td><td></td><td></td><td></td><td></td><td></td><td></td></tr>
<tr><td>T4675</td><td>GCGTCGGTGATGTCGGGGAGCGG</td><td>1000</td><td></td><td></td><td></td><td></td><td></td><td></td><td></td><td></td><td></td></tr>
<tr><td>Oligo fwd</td><td>CACCGCGTCGGTGATGTCGGGGAG</td><td></td><td></td><td></td><td></td><td></td><td></td><td></td><td></td><td></td><td></td></tr>
<tr><td>Oligo rev</td><td>AAACCTCCCCGACATCACCGACGC</td><td></td><td></td><td></td><td></td><td></td><td></td><td></td><td></td><td></td><td></td></tr>
<tr><td>Chromosome</td><td>start</td><td>end</td><td>strand</td><td>MM</td><td>target_seq</td><td>PAM</td><td>alignment</td><td>distance</td><td>position</td><td>gene name</td><td>gene id</td></tr>
<tr><td>GU305914.1</td><td>67727</td><td>67749</td><td>+</td><td>0</td><td>GCGTCGGTGATGTCGGGGAG</td><td>CGG</td><td>||||||||[||||||||||||]PAM</td><td>NA</td><td>Intergenic</td><td>NA</td><td>NA</td></tr>
<tr><td></td><td></td><td></td><td></td><td></td><td></td><td></td><td></td><td></td><td></td><td></td><td></td></tr>
<tr><td>T4676</td><td>TCGGTGATGTCGGGGAGCGGCGG</td><td>1000</td><td></td><td></td><td></td><td></td><td></td><td></td><td></td><td></td><td></td></tr>
<tr><td>Oligo fwd</td><td>CACCTCGGTGATGTCGGGGAGCGG</td><td></td><td></td><td></td><td></td><td></td><td></td><td></td><td></td><td></td><td></td></tr>
<tr><td>Oligo rev</td><td>AAACCCGCTCCCCGACATCACCGA</td><td></td><td></td><td></td><td></td><td></td><td></td><td></td><td></td><td></td><td></td></tr>
<tr><td>Chromosome</td><td>start</td><td>end</td><td>strand</td><td>MM</td><td>target_seq</td><td>PAM</td><td>alignment</td><td>distance</td><td>position</td><td>gene name</td><td>gene id</td></tr>
<tr><td>GU305914.1</td><td>67730</td><td>67752</td><td>+</td><td>0</td><td>TCGGTGATGTCGGGGAGCGG</td><td>CGG</td><td>||||||||[||||||||||||]PAM</td><td>NA</td><td>Intergenic</td><td>NA</td><td>NA</td></tr>
<tr><td>GU305914.1</td><td>225229</td><td>225251</td><td>+</td><td>4</td><td>ACGGTGTCGTCGGTGAGCGG</td><td>AGG</td><td>-|||||--[|||||-||||||]PAM</td><td>NA</td><td>Intergenic</td><td>NA</td><td>NA</td></tr>
<tr><td></td><td></td><td></td><td></td><td></td><td></td><td></td><td></td><td></td><td></td><td></td><td></td></tr>
<tr><td>T4677</td><td>GTGATGTCGGGGAGCGGCGGCGG</td><td>1000</td><td></td><td></td><td></td><td></td><td></td><td></td><td></td><td></td><td></td></tr>
<tr><td>Oligo fwd</td><td>CACCGTGATGTCGGGGAGCGGCGG</td><td></td><td></td><td></td><td></td><td></td><td></td><td></td><td></td><td></td><td></td></tr>
<tr><td>Oligo rev</td><td>AAACCCGCCGCTCCCCGACATCAC</td><td></td><td></td><td></td><td></td><td></td><td></td><td></td><td></td><td></td><td></td></tr>
<tr><td>Chromosome</td><td>start</td><td>end</td><td>strand</td><td>MM</td><td>target_seq</td><td>PAM</td><td>alignment</td><td>distance</td><td>position</td><td>gene name</td><td>gene id</td></tr>
<tr><td>GU305914.1</td><td>67733</td><td>67755</td><td>+</td><td>0</td><td>GTGATGTCGGGGAGCGGCGG</td><td>CGG</td><td>||||||||[||||||||||||]PAM</td><td>NA</td><td>Intergenic</td><td>NA</td><td>NA</td></tr>
<tr><td>GU305914.1</td><td>53614</td><td>53636</td><td>-</td><td>3</td><td>GGGATGTCGGGCAGAGGCGG</td><td>AGG</td><td>|-||||||[|||-||-|||||]PAM</td><td>NA</td><td>Intergenic</td><td>NA</td><td>NA</td></tr>
<tr><td></td><td></td><td></td><td></td><td></td><td></td><td></td><td></td><td></td><td></td><td></td><td></td></tr>
<tr><td>T4678</td><td>ATGTCGGGGAGCGGCGGCGGCGG</td><td>1000</td><td></td><td></td><td></td><td></td><td></td><td></td><td></td><td></td><td></td></tr>
<tr><td>Oligo fwd</td><td>CACCATGTCGGGGAGCGGCGGCGG</td><td></td><td></td><td></td><td></td><td></td><td></td><td></td><td></td><td></td><td></td></tr>
<tr><td>Oligo rev</td><td>AAACCCGCCGCCGCTCCCCGACAT</td><td></td><td></td><td></td><td></td><td></td><td></td><td></td><td></td><td></td><td></td></tr>
<tr><td>Chromosome</td><td>start</td><td>end</td><td>strand</td><td>MM</td><td>target_seq</td><td>PAM</td><td>alignment</td><td>distance</td><td>position</td><td>gene name</td><td>gene id</td></tr>
<tr><td>GU305914.1</td><td>67736</td><td>67758</td><td>+</td><td>0</td><td>ATGTCGGGGAGCGGCGGCGG</td><td>CGG</td><td>||||||||[||||||||||||]PAM</td><td>NA</td><td>Intergenic</td><td>NA</td><td>NA</td></tr>
<tr><td></td><td></td><td></td><td></td><td></td><td></td><td></td><td></td><td></td><td></td><td></td><td></td></tr>
<tr><td>T4679</td><td>TCGGGGAGCGGCGGCGGCGGCGG</td><td>1000</td><td></td><td></td><td></td><td></td><td></td><td></td><td></td><td></td><td></td></tr>
<tr><td>Oligo fwd</td><td>CACCTCGGGGAGCGGCGGCGGCGG</td><td></td><td></td><td></td><td></td><td></td><td></td><td></td><td></td><td></td><td></td></tr>
<tr><td>Oligo rev</td><td>AAACCCGCCGCCGCCGCTCCCCGA</td><td></td><td></td><td></td><td></td><td></td><td></td><td></td><td></td><td></td><td></td></tr>
<tr><td>Chromosome</td><td>start</td><td>end</td><td>strand</td><td>MM</td><td>target_seq</td><td>PAM</td><td>alignment</td><td>distance</td><td>position</td><td>gene name</td><td>gene id</td></tr>
<tr><td>GU305914.1</td><td>67739</td><td>67761</td><td>+</td><td>0</td><td>TCGGGGAGCGGCGGCGGCGG</td><td>CGG</td><td>||||||||[||||||||||||]PAM</td><td>NA</td><td>Intergenic</td><td>NA</td><td>NA</td></tr>
<tr><td>GU305914.1</td><td>164971</td><td>164993</td><td>-</td><td>4</td><td>GGTGGGACCGGCGGCGGCGG</td><td>CGG</td><td>---||||-[||||||||||||]PAM</td><td>NA</td><td>Intergenic</td><td>NA</td><td>NA</td></tr>
<tr><td>GU305914.1</td><td>154401</td><td>154423</td><td>+</td><td>4</td><td>CCGGGCGGTGGCGGCGGCGG</td><td>CGG</td><td>-||||--|[-|||||||||||]PAM</td><td>NA</td><td>Intergenic</td><td>NA</td><td>NA</td></tr>
<tr><td>GU305914.1</td><td>58129</td><td>58151</td><td>-</td><td>4</td><td>GCGGGCGGTGGCGGCGGCGG</td><td>CGG</td><td>-||||--|[-|||||||||||]PAM</td><td>NA</td><td>Intergenic</td><td>NA</td><td>NA</td></tr>
<tr><td>GU305914.1</td><td>48144</td><td>48166</td><td>+</td><td>4</td><td>TCGCGGACCTGTGGCGGCGG</td><td>CGG</td><td>|||-|||-[|-|-||||||||]PAM</td><td>NA</td><td>Intergenic</td><td>NA</td><td>NA</td></tr>
<tr><td></td><td></td><td></td><td></td><td></td><td></td><td></td><td></td><td></td><td></td><td></td><td></td></tr>
<tr><td>T4680</td><td>GGGAGCGGCGGCGGCGGCGGCGG</td><td>1000</td><td></td><td></td><td></td><td></td><td></td><td></td><td></td><td></td><td></td></tr>
<tr><td>Oligo fwd</td><td>CACCGGGAGCGGCGGCGGCGGCGG</td><td></td><td></td><td></td><td></td><td></td><td></td><td></td><td></td><td></td><td></td></tr>
<tr><td>Oligo rev</td><td>AAACCCGCCGCCGCCGCCGCTCCC</td><td></td><td></td><td></td><td></td><td></td><td></td><td></td><td></td><td></td><td></td></tr>
<tr><td>Chromosome</td><td>start</td><td>end</td><td>strand</td><td>MM</td><td>target_seq</td><td>PAM</td><td>alignment</td><td>distance</td><td>position</td><td>gene name</td><td>gene id</td></tr>
<tr><td>GU305914.1</td><td>67742</td><td>67764</td><td>+</td><td>0</td><td>GGGAGCGGCGGCGGCGGCGG</td><td>CGG</td><td>||||||||[||||||||||||]PAM</td><td>NA</td><td>Intergenic</td><td>NA</td><td>NA</td></tr>
<tr><td>GU305914.1</td><td>154407</td><td>154429</td><td>+</td><td>2</td><td>GGTGGCGGCGGCGGCGGCGG</td><td>CGG</td><td>||--||||[||||||||||||]PAM</td><td>NA</td><td>Intergenic</td><td>NA</td><td>NA</td></tr>
<tr><td>GU305914.1</td><td>58123</td><td>58145</td><td>-</td><td>2</td><td>GGTGGCGGCGGCGGCGGCGG</td><td>AGG</td><td>||--||||[||||||||||||]PAM</td><td>NA</td><td>Intergenic</td><td>NA</td><td>NA</td></tr>
<tr><td>GU305914.1</td><td>149482</td><td>149504</td><td>-</td><td>3</td><td>GGCGACGGCGGCGGCGGCGG</td><td>CGG</td><td>||---|||[||||||||||||]PAM</td><td>NA</td><td>Intergenic</td><td>NA</td><td>NA</td></tr>
<tr><td>GU305914.1</td><td>154404</td><td>154426</td><td>+</td><td>3</td><td>GGCGGTGGCGGCGGCGGCGG</td><td>CGG</td><td>||--|-||[||||||||||||]PAM</td><td>NA</td><td>Intergenic</td><td>NA</td><td>NA</td></tr>
<tr><td>GU305914.1</td><td>58126</td><td>58148</td><td>-</td><td>3</td><td>GGCGGTGGCGGCGGCGGCGG</td><td>CGG</td><td>||--|-||[||||||||||||]PAM</td><td>NA</td><td>Intergenic</td><td>NA</td><td>NA</td></tr>
<tr><td>GU305914.1</td><td>56773</td><td>56795</td><td>+</td><td>4</td><td>CGCCGCAGCGGCGGCGGCGG</td><td>CGG</td><td>-|--||-|[||||||||||||]PAM</td><td>NA</td><td>Intergenic</td><td>NA</td><td>NA</td></tr>
<tr><td>GU305914.1</td><td>58129</td><td>58151</td><td>-</td><td>3</td><td>GCGGGCGGTGGCGGCGGCGG</td><td>CGG</td><td>|-|-||||[-|||||||||||]PAM</td><td>NA</td><td>Intergenic</td><td>NA</td><td>NA</td></tr>
<tr><td>GU305914.1</td><td>48159</td><td>48181</td><td>+</td><td>3</td><td>GGCGGCGGTGGCGGCGGCGG</td><td>TGG</td><td>||--||||[-|||||||||||]PAM</td><td>NA</td><td>Intergenic</td><td>NA</td><td>NA</td></tr>
<tr><td>GU305914.1</td><td>30000</td><td>30022</td><td>+</td><td>4</td><td>GGTGCTGGCGGCGGCGGCGG</td><td>CGG</td><td>||----||[||||||||||||]PAM</td><td>NA</td><td>Intergenic</td><td>NA</td><td>NA</td></tr>
<tr><td>GU305914.1</td><td>149485</td><td>149507</td><td>-</td><td>4</td><td>GCAGGCGACGGCGGCGGCGG</td><td>CGG</td><td>|---|||-[||||||||||||]PAM</td><td>NA</td><td>Intergenic</td><td>NA</td><td>NA</td></tr>
<tr><td>GU305914.1</td><td>154401</td><td>154423</td><td>+</td><td>4</td><td>CCGGGCGGTGGCGGCGGCGG</td><td>CGG</td><td>--|-||||[-|||||||||||]PAM</td><td>NA</td><td>Intergenic</td><td>NA</td><td>NA</td></tr>
<tr><td>GU305914.1</td><td>35650</td><td>35672</td><td>+</td><td>3</td><td>GGGATCCACGGCGGCGGCGG</td><td>CGG</td><td>||||-|--[||||||||||||]PAM</td><td>NA</td><td>Intergenic</td><td>NA</td><td>NA</td></tr>
<tr><td>GU305914.1</td><td>161754</td><td>161776</td><td>-</td><td>4</td><td>GGCCGAGGAGGCGGCGGCGG</td><td>AGG</td><td>||--|-||[-|||||||||||]PAM</td><td>NA</td><td>Intergenic</td><td>NA</td><td>NA</td></tr>
<tr><td>GU305914.1</td><td>48147</td><td>48169</td><td>+</td><td>4</td><td>CGGACCTGTGGCGGCGGCGG</td><td>TGG</td><td>-|||-|-|[-|||||||||||]PAM</td><td>NA</td><td>Intergenic</td><td>NA</td><td>NA</td></tr>
<tr><td>GU305914.1</td><td>48156</td><td>48178</td><td>+</td><td>3</td><td>GGCGGCGGCGGTGGCGGCGG</td><td>CGG</td><td>||--||||[|||-||||||||]PAM</td><td>NA</td><td>Intergenic</td><td>NA</td><td>NA</td></tr>
<tr><td>GU305914.1</td><td>168820</td><td>168842</td><td>+</td><td>4</td><td>GGTGGTGGCAGCGGCGGCGG</td><td>CGG</td><td>||--|-||[|-||||||||||]PAM</td><td>NA</td><td>Intergenic</td><td>NA</td><td>NA</td></tr>
<tr><td>GU305914.1</td><td>48153</td><td>48175</td><td>+</td><td>4</td><td>TGTGGCGGCGGCGGTGGCGG</td><td>CGG</td><td>-|--||||[||||||-|||||]PAM</td><td>NA</td><td>Intergenic</td><td>NA</td><td>NA</td></tr>
<tr><td>GU305914.1</td><td>90259</td><td>90281</td><td>-</td><td>4</td><td>GCGGGCGACGGCGGAGGCGG</td><td>CGG</td><td>|-|-|||-[||||||-|||||]PAM</td><td>NA</td><td>Intergenic</td><td>NA</td><td>NA</td></tr>
<tr><td>GU305914.1</td><td>86784</td><td>86806</td><td>-</td><td>4</td><td>GGAAGAGGGGGCGGGGGCGG</td><td>CGG</td><td>||-||-||[-|||||-|||||]PAM</td><td>NA</td><td>Intergenic</td><td>NA</td><td>NA</td></tr>
<tr><td>GU305914.1</td><td>18653</td><td>18675</td><td>-</td><td>4</td><td>GGCAGAGGTGGCGGAGGCGG</td><td>GGG</td><td>||-||-||[-|||||-|||||]PAM</td><td>NA</td><td>Intergenic</td><td>NA</td><td>NA</td></tr>
<tr><td>GU305914.1</td><td>110125</td><td>110147</td><td>-</td><td>4</td><td>GGACGCGACGGCGGCCGCGG</td><td>CGG</td><td>||--|||-[|||||||-||||]PAM</td><td>NA</td><td>Intergenic</td><td>NA</td><td>NA</td></tr>
<tr><td>GU305914.1</td><td>45255</td><td>45277</td><td>-</td><td>4</td><td>GGTGGCGGTGGCGGCAGCGG</td><td>CGG</td><td>||--||||[-||||||-||||]PAM</td><td>NA</td><td>Intergenic</td><td>NA</td><td>NA</td></tr>
<tr><td>GU305914.1</td><td>163911</td><td>163933</td><td>+</td><td>4</td><td>GGCGGCGGCAGCGGCAGCGG</td><td>AGG</td><td>||--||||[|-|||||-||||]PAM</td><td>NA</td><td>Intergenic</td><td>NA</td><td>NA</td></tr>
<tr><td>GU305914.1</td><td>73789</td><td>73811</td><td>-</td><td>4</td><td>GCCAGCAGCGGCGGCGGAGG</td><td>AGG</td><td>|--|||-|[|||||||||-||]PAM</td><td>NA</td><td>Intergenic</td><td>NA</td><td>NA</td></tr>
<tr><td>GU305914.1</td><td>30006</td><td>30028</td><td>+</td><td>3</td><td>GGCGGCGGCGGCGGCGGCAG</td><td>TGG</td><td>||--||||[||||||||||-|]PAM</td><td>NA</td><td>Intergenic</td><td>NA</td><td>NA</td></tr>
<tr><td>GU305914.1</td><td>149476</td><td>149498</td><td>-</td><td>3</td><td>GGCGGCGGCGGCGGCGGCAG</td><td>CGG</td><td>||--||||[||||||||||-|]PAM</td><td>NA</td><td>Intergenic</td><td>NA</td><td>NA</td></tr>
<tr><td>GU305914.1</td><td>62449</td><td>62471</td><td>+</td><td>4</td><td>GGGGGTGGTGGCGGCGGGGG</td><td>TGG</td><td>|||-|-||[-||||||||-||]PAM</td><td>NA</td><td>Intergenic</td><td>NA</td><td>NA</td></tr>
<tr><td>GU305914.1</td><td>163902</td><td>163924</td><td>+</td><td>4</td><td>AGCGGCGGCGGCGGCGGCAG</td><td>CGG</td><td>-|--||||[||||||||||-|]PAM</td><td>NA</td><td>Intergenic</td><td>NA</td><td>NA</td></tr>
<tr><td>GU305914.1</td><td>73786</td><td>73808</td><td>-</td><td>4</td><td>AGCAGCGGCGGCGGAGGAGG</td><td>AGG</td><td>-|-|||||[||||||-||-||]PAM</td><td>NA</td><td>Intergenic</td><td>NA</td><td>NA</td></tr>
<tr><td>GU305914.1</td><td>163908</td><td>163930</td><td>+</td><td>4</td><td>GGCGGCGGCGGCAGCGGCAG</td><td>CGG</td><td>||--||||[||||-|||||-|]PAM</td><td>NA</td><td>Intergenic</td><td>NA</td><td>NA</td></tr>
<tr><td>GU305914.1</td><td>128809</td><td>128831</td><td>+</td><td>4</td><td>GGCAGCGACGTCGGCGGCGA</td><td>AGG</td><td>||-||||-[||-||||||||-]PAM</td><td>NA</td><td>Intergenic</td><td>NA</td><td>NA</td></tr>
<tr><td>GU305914.1</td><td>73813</td><td>73835</td><td>-</td><td>4</td><td>GGGAGTAGCGGCGGCAGCAG</td><td>TGG</td><td>|||||--|[|||||||-||-|]PAM</td><td>NA</td><td>Intergenic</td><td>NA</td><td>NA</td></tr>
<tr><td></td><td></td><td></td><td></td><td></td><td></td><td></td><td></td><td></td><td></td><td></td><td></td></tr>
<tr><td>T4681</td><td>GGAGCGGCGGCGGCGGCGGCGGG</td><td>1000</td><td></td><td></td><td></td><td></td><td></td><td></td><td></td><td></td><td></td></tr>
<tr><td>Oligo fwd</td><td>CACCGGAGCGGCGGCGGCGGCGGC</td><td></td><td></td><td></td><td></td><td></td><td></td><td></td><td></td><td></td><td></td></tr>
<tr><td>Oligo rev</td><td>AAACGCCGCCGCCGCCGCCGCTCC</td><td></td><td></td><td></td><td></td><td></td><td></td><td></td><td></td><td></td><td></td></tr>
<tr><td>Chromosome</td><td>start</td><td>end</td><td>strand</td><td>MM</td><td>target_seq</td><td>PAM</td><td>alignment</td><td>distance</td><td>position</td><td>gene name</td><td>gene id</td></tr>
<tr><td>GU305914.1</td><td>67743</td><td>67765</td><td>+</td><td>0</td><td>GGAGCGGCGGCGGCGGCGGC</td><td>GGG</td><td>||||||||[||||||||||||]PAM</td><td>NA</td><td>Intergenic</td><td>NA</td><td>NA</td></tr>
</table>

| | | | | | | | | | | | |
|---|---|---|---|---|---|---|---|---|---|---|---|
| GU305914.1 | 35654 | 35676 | + | 4 | TCCACGGCGGCGGCGGCGGC | GGG | ----\|\|\|\|[\|\|\|\|\|\|\|\|\|\|\|\|]PAM | NA | Intergenic | NA | NA |
| GU305914.1 | 149296 | 149318 | + | 4 | CGGGCGACGACGGCGGCGGC | AGG | -\|-\|\|\|-\|[\|-\|\|\|\|\|\|\|\|\|\|]PAM | NA | Intergenic | NA | NA |
| GU305914.1 | 86780 | 86802 | - | 4 | GAGGGGGCGGGGGCGGCGGC | AGG | \|--\|-\|\|\|[\|\|-\|\|\|\|\|\|\|\|\|]PAM | NA | Intergenic | NA | NA |
| GU305914.1 | 90246 | 90268 | - | 4 | GAGGCGGCGGAGCCGGCGGC | AGG | \|--\|\|\|\|\|[\|\|-\|-\|\|\|\|\|\|\|]PAM | NA | Intergenic | NA | NA |
| GU305914.1 | 149475 | 149497 | - | 3 | GCGGCGGCGGCGGCGGCAGC | GGG | \|--\|\|\|\|\|[\|\|\|\|\|\|\|\|\|-\|\|]PAM | NA | Intergenic | NA | NA |
| GU305914.1 | 152554 | 152576 | + | 4 | GCGGCGGCGGCGGTGCCGGC | GGG | \|--\|\|\|\|\|[\|\|\|\|\|-\|-\|\|\|\|]PAM | NA | Intergenic | NA | NA |
| GU305914.1 | 58122 | 58144 | - | 3 | GTGGCGGCGGCGGCGGCGGA | GGG | \|--\|\|\|\|\|[\|\|\|\|\|\|\|\|\|\|\|-]PAM | NA | Intergenic | NA | NA |
| GU305914.1 | 48160 | 48182 | + | 4 | GCGGCGGTGGCGGCGGCGGT | GGG | \|--\|\|\|\|-[\|\|\|\|\|\|\|\|\|\|\|-]PAM | NA | Intergenic | NA | NA |
| GU305914.1 | 168830 | 168852 | + | 4 | GCGGCGGCGGCGGCGACAGC | GGG | \|--\|\|\|\|\|[\|\|\|\|\|\|\|-\|-\|\|]PAM | NA | Intergenic | NA | NA |
| GU305914.1 | 75852 | 75874 | + | 4 | GGGGAGGCGGAGGCGGCGGT | CGG | \|\|-\|-\|\|\|[\|\|-\|\|\|\|\|\|\|\|-]PAM | NA | Intergenic | NA | NA |
| GU305914.1 | 71754 | 71776 | + | 4 | GCAGCAGCGGCGGCGGCAAC | GGG | \|-\|\|\|-\|\|[\|\|\|\|\|\|\|\|\|--\|]PAM | NA | Intergenic | NA | NA |
| T4682 | TCTGTTTCTCCTTTGCCGAGCGG | 1000 | | | | | | | | | |
| Oligo fwd | CACCTCTGTTTCTCCTTTGCCGAG | | | | | | | | | | |
| Oligo rev | AAACCTCGGCAAAGGAGAAACAGA | | | | | | | | | | |
| Chromosome | start | end | strand | MM | target_seq | PAM | alignment | distance | position | gene name | gene id |
| GU305914.1 | 67802 | 67824 | + | 0 | TCTGTTTCTCCTTTGCCGAG | CGG | \|\|\|\|\|\|\|\|[\|\|\|\|\|\|\|\|\|\|\|\|]PAM | NA | Intergenic | NA | NA |
| T4683 | CCGAGCGGCGCCGCGTCCGACGG | 1000 | | | | | | | | | |
| Oligo fwd | CACCCCGAGCGGCGCCGCGTCCGA | | | | | | | | | | |
| Oligo rev | AAACTCGGACGCGGCGCCGCTCGG | | | | | | | | | | |
| Chromosome | start | end | strand | MM | target_seq | PAM | alignment | distance | position | gene name | gene id |
| GU305914.1 | 67817 | 67839 | + | 0 | CCGAGCGGCGCCGCGTCCGA | CGG | \|\|\|\|\|\|\|\|[\|\|\|\|\|\|\|\|\|\|\|\|]PAM | NA | Intergenic | NA | NA |
| T4684 | GCGTCCGACGGCGTGACGAACGG | 1000 | | | | | | | | | |
| Oligo fwd | CACCGCGTCCGACGGCGTGACGAA | | | | | | | | | | |
| Oligo rev | AAACTTCGTCACGCCGTCGGACGC | | | | | | | | | | |
| Chromosome | start | end | strand | MM | target_seq | PAM | alignment | distance | position | gene name | gene id |
| GU305914.1 | 67829 | 67851 | + | 0 | GCGTCCGACGGCGTGACGAA | CGG | \|\|\|\|\|\|\|\|[\|\|\|\|\|\|\|\|\|\|\|\|]PAM | NA | Intergenic | NA | NA |
| T4685 | CGACGGCGTGACGAACGGCGCGG | 1000 | | | | | | | | | |
| Oligo fwd | CACCCGACGGCGTGACGAACGGCG | | | | | | | | | | |
| Oligo rev | AAACCGCCGTTCGTCACGCCGTCG | | | | | | | | | | |
| Chromosome | start | end | strand | MM | target_seq | PAM | alignment | distance | position | gene name | gene id |
| GU305914.1 | 67834 | 67856 | + | 0 | CGACGGCGTGACGAACGGCG | CGG | \|\|\|\|\|\|\|\|[\|\|\|\|\|\|\|\|\|\|\|\|]PAM | NA | Intergenic | NA | NA |
| T4686 | GACGGCGTGACGAACGGCGCGGG | 1000 | | | | | | | | | |
| Oligo fwd | CACCGACGGCGTGACGAACGGCGC | | | | | | | | | | |
| Oligo rev | AAACGCGCCGTTCGTCACGCCGTC | | | | | | | | | | |
| Chromosome | start | end | strand | MM | target_seq | PAM | alignment | distance | position | gene name | gene id |
| GU305914.1 | 67835 | 67857 | + | 0 | GACGGCGTGACGAACGGCGC | GGG | \|\|\|\|\|\|\|\|[\|\|\|\|\|\|\|\|\|\|\|\|]PAM | NA | Intergenic | NA | NA |
| T4687 | CGCCCAGACTTCGTCCAAGTCGG | 1000 | | | | | | | | | |
| Oligo fwd | CACCCGCCCAGACTTCGTCCAAGT | | | | | | | | | | |
| Oligo rev | AAACACTTGGACGAAGTCTGGGCG | | | | | | | | | | |
| Chromosome | start | end | strand | MM | target_seq | PAM | alignment | distance | position | gene name | gene id |
| GU305914.1 | 67858 | 67880 | + | 0 | CGCCCAGACTTCGTCCAAGT | CGG | \|\|\|\|\|\|\|\|[\|\|\|\|\|\|\|\|\|\|\|\|]PAM | NA | Intergenic | NA | NA |
| T4688 | GCCCAGACTTCGTCCAAGTCGGG | 1000 | | | | | | | | | |
| Oligo fwd | CACCGCCCAGACTTCGTCCAAGTC | | | | | | | | | | |
| Oligo rev | AAACGACTTGGACGAAGTCTGGGC | | | | | | | | | | |
| Chromosome | start | end | strand | MM | target_seq | PAM | alignment | distance | position | gene name | gene id |
| GU305914.1 | 67859 | 67881 | + | 0 | GCCCAGACTTCGTCCAAGTC | GGG | \|\|\|\|\|\|\|\|[\|\|\|\|\|\|\|\|\|\|\|\|]PAM | NA | Intergenic | NA | NA |
| T4689 | CCAAGTCGGGCAAGAAGAGACGG | 1000 | | | | | | | | | |
| Oligo fwd | CACCCCAAGTCGGGCAAGAAGAGA | | | | | | | | | | |
| Oligo rev | AAACTCTCTTCTTGCCCGACTTGG | | | | | | | | | | |
| Chromosome | start | end | strand | MM | target_seq | PAM | alignment | distance | position | gene name | gene id |
| GU305914.1 | 67872 | 67894 | + | 0 | CCAAGTCGGGCAAGAAGAGA | CGG | \|\|\|\|\|\|\|\|[\|\|\|\|\|\|\|\|\|\|\|\|]PAM | NA | Intergenic | NA | NA |
| T4690 | CAAGTCGGGCAAGAAGAGACGGG | 1000 | | | | | | | | | |
| Oligo fwd | CACCCAAGTCGGGCAAGAAGAGAC | | | | | | | | | | |
| Oligo rev | AAACGTCTCTTCTTGCCCGACTTG | | | | | | | | | | |
| Chromosome | start | end | strand | MM | target_seq | PAM | alignment | distance | position | gene name | gene id |
| GU305914.1 | 67873 | 67895 | + | 0 | CAAGTCGGGCAAGAAGAGAC | GGG | \|\|\|\|\|\|\|\|[\|\|\|\|\|\|\|\|\|\|\|\|]PAM | NA | Intergenic | NA | NA |
| T4691 | GGGCAAGAAGAGACGGGCGCCGG | 1000 | | | | | | | | | |
| Oligo fwd | CACCGGGCAAGAAGAGACGGGCGC | | | | | | | | | | |
| Oligo rev | AAACGCGCCCGTCTCTTCTTGCCC | | | | | | | | | | |
| Chromosome | start | end | strand | MM | target_seq | PAM | alignment | distance | position | gene name | gene id |
| GU305914.1 | 67879 | 67901 | + | 0 | GGGCAAGAAGAGACGGGCGC | CGG | \|\|\|\|\|\|\|\|[\|\|\|\|\|\|\|\|\|\|\|\|]PAM | NA | Intergenic | NA | NA |
| T4692 | GGACGTGCCGCCGCAGCAGAAGG | 1000 | | | | | | | | | |
| Oligo fwd | CACCGGACGTGCCGCCGCAGCAGA | | | | | | | | | | |
| Oligo rev | AAACTCTGCTGCGGCGGCACGTCC | | | | | | | | | | |
| Chromosome | start | end | strand | MM | target_seq | PAM | alignment | distance | position | gene name | gene id |
| GU305914.1 | 67900 | 67922 | + | 0 | GGACGTGCCGCCGCAGCAGA | AGG | \|\|\|\|\|\|\|\|[\|\|\|\|\|\|\|\|\|\|\|\|]PAM | NA | Intergenic | NA | NA |
| T4693 | GACGTGCCGCCGCAGCAGAAGGG | 1000 | | | | | | | | | |
| Oligo fwd | CACCGACGTGCCGCCGCAGCAGAA | | | | | | | | | | |
| Oligo rev | AAACTTCTGCTGCGGCGGCACGTC | | | | | | | | | | |
| Chromosome | start | end | strand | MM | target_seq | PAM | alignment | distance | position | gene name | gene id |
| GU305914.1 | 67901 | 67923 | + | 0 | GACGTGCCGCCGCAGCAGAA | GGG | \|\|\|\|\|\|\|\|[\|\|\|\|\|\|\|\|\|\|\|\|]PAM | NA | Intergenic | NA | NA |
| T4694 | GCCGCCGCAGCAGAAGGGCACGG | 1000 | | | | | | | | | |
| Oligo fwd | CACCGCCGCCGCAGCAGAAGGGCA | | | | | | | | | | |
| Oligo rev | AAACTGCCCTTCTGCTGCGGCGGC | | | | | | | | | | |
| Chromosome | start | end | strand | MM | target_seq | PAM | alignment | distance | position | gene name | gene id |
| GU305914.1 | 67906 | 67928 | + | 0 | GCCGCCGCAGCAGAAGGGCA | CGG | \|\|\|\|\|\|\|\|[\|\|\|\|\|\|\|\|\|\|\|\|]PAM | NA | Intergenic | NA | NA |

| | | | | | | | | | | | |
|---|---|---|---|---|---|---|---|---|---|---|---|
| T4695 | CGAAGACGCTGCGCCGCAGCCGG | 1000 | | | | | | | | | |
| Oligo fwd | CACCCGAAGACGCTGCGCCGCAGC | | | | | | | | | | |
| Oligo rev | AAACGCTGCGGCGCAGCGTCTTCG | | | | | | | | | | |
| Chromosome | start | end | strand | MM | target_seq | PAM | alignment | distance | position | gene name | gene id |
| GU305914.1 | 67929 | 67951 | + | 0 | CGAAGACGCTGCGCCGCAGC | CGG | ||||||||[||||||||||||]PAM | NA | Intergenic | NA | NA |

| | | | | | | | | | | | |
|---|---|---|---|---|---|---|---|---|---|---|---|
| T4696 | AGACGCTGCGCCGCAGCCGGCGG | 1000 | | | | | | | | | |
| Oligo fwd | CACCAGACGCTGCGCCGCAGCCGG | | | | | | | | | | |
| Oligo rev | AAACCCGGCTGCGGCGCAGCGTCT | | | | | | | | | | |
| Chromosome | start | end | strand | MM | target_seq | PAM | alignment | distance | position | gene name | gene id |
| GU305914.1 | 67932 | 67954 | + | 0 | AGACGCTGCGCCGCAGCCGG | CGG | ||||||||[||||||||||||]PAM | NA | Intergenic | NA | NA |
| GU305914.1 | 66695 | 66717 | - | 4 | CCACGCAGCGCCGCAGCCCG | CGG | --||||-|[||||||||||-|]PAM | NA | Intergenic | NA | NA |

| | | | | | | | | | | | |
|---|---|---|---|---|---|---|---|---|---|---|---|
| T4697 | GCCGCAGCCGGCGGCCGATCAGG | 1000 | | | | | | | | | |
| Oligo fwd | CACCGCCGCAGCCGGCGGCCGATC | | | | | | | | | | |
| Oligo rev | AAACGATCGGCCGCCGGCTGCGGC | | | | | | | | | | |
| Chromosome | start | end | strand | MM | target_seq | PAM | alignment | distance | position | gene name | gene id |
| GU305914.1 | 67941 | 67963 | + | 0 | GCCGCAGCCGGCGGCCGATC | AGG | ||||||||[||||||||||||]PAM | NA | Intergenic | NA | NA |

| | | | | | | | | | | | |
|---|---|---|---|---|---|---|---|---|---|---|---|
| T4698 | GTTGCCCGAGCGTCTGTCCGAGG | 1000 | | | | | | | | | |
| Oligo fwd | CACCGTTGCCCGAGCGTCTGTCCG | | | | | | | | | | |
| Oligo rev | AAACCGGACAGACGCTCGGGCAAC | | | | | | | | | | |
| Chromosome | start | end | strand | MM | target_seq | PAM | alignment | distance | position | gene name | gene id |
| GU305914.1 | 67963 | 67985 | + | 0 | GTTGCCCGAGCGTCTGTCCG | AGG | ||||||||[||||||||||||]PAM | NA | Intergenic | NA | NA |

| | | | | | | | | | | | |
|---|---|---|---|---|---|---|---|---|---|---|---|
| T4699 | GCGTCTGTCCGAGGTGATCGTGG | 1000 | | | | | | | | | |
| Oligo fwd | CACCGCGTCTGTCCGAGGTGATCG | | | | | | | | | | |
| Oligo rev | AAACCGATCACCTCGGACAGACGC | | | | | | | | | | |
| Chromosome | start | end | strand | MM | target_seq | PAM | alignment | distance | position | gene name | gene id |
| GU305914.1 | 67972 | 67994 | + | 0 | GCGTCTGTCCGAGGTGATCG | TGG | ||||||||[||||||||||||]PAM | NA | Intergenic | NA | NA |

| | | | | | | | | | | | |
|---|---|---|---|---|---|---|---|---|---|---|---|
| T4700 | CACCGCCACGCAGTCGCTGCCGG | 1000 | | | | | | | | | |
| Oligo fwd | CACCCACCGCCACGCAGTCGCTGC | | | | | | | | | | |
| Oligo rev | AAACGCAGCGACTGCGTGGCGGTG | | | | | | | | | | |
| Chromosome | start | end | strand | MM | target_seq | PAM | alignment | distance | position | gene name | gene id |
| GU305914.1 | 68023 | 68045 | + | 0 | CACCGCCACGCAGTCGCTGC | CGG | ||||||||[||||||||||||]PAM | NA | Intergenic | NA | NA |

| | | | | | | | | | | | |
|---|---|---|---|---|---|---|---|---|---|---|---|
| T4701 | GCTGCCGGTGAAACCCCCGCAGG | 1000 | | | | | | | | | |
| Oligo fwd | CACCGCTGCCGGTGAAACCCCCGC | | | | | | | | | | |
| Oligo rev | AAACGCGGGGGTTTCACCGGCAGC | | | | | | | | | | |
| Chromosome | start | end | strand | MM | target_seq | PAM | alignment | distance | position | gene name | gene id |
| GU305914.1 | 68038 | 68060 | + | 0 | GCTGCCGGTGAAACCCCCGC | AGG | ||||||||[||||||||||||]PAM | NA | Intergenic | NA | NA |

| | | | | | | | | | | | |
|---|---|---|---|---|---|---|---|---|---|---|---|
| T4702 | CGGTGAAACCCCCGCAGGAATGG | 1000 | | | | | | | | | |
| Oligo fwd | CACCCGGTGAAACCCCCGCAGGAA | | | | | | | | | | |
| Oligo rev | AAACTTCCTGCGGGGGTTTCACCG | | | | | | | | | | |
| Chromosome | start | end | strand | MM | target_seq | PAM | alignment | distance | position | gene name | gene id |
| GU305914.1 | 68043 | 68065 | + | 0 | CGGTGAAACCCCCGCAGGAA | TGG | ||||||||[||||||||||||]PAM | NA | Intergenic | NA | NA |

| | | | | | | | | | | | |
|---|---|---|---|---|---|---|---|---|---|---|---|
| T4703 | CAGGAATGGACCATGTACTGCGG | 1000 | | | | | | | | | |
| Oligo fwd | CACCCAGGAATGGACCATGTACTG | | | | | | | | | | |
| Oligo rev | AAACCAGTACATGGTCCATTCCTG | | | | | | | | | | |
| Chromosome | start | end | strand | MM | target_seq | PAM | alignment | distance | position | gene name | gene id |
| GU305914.1 | 68057 | 68079 | + | 0 | CAGGAATGGACCATGTACTG | CGG | ||||||||[||||||||||||]PAM | NA | Intergenic | NA | NA |

| | | | | | | | | | | | |
|---|---|---|---|---|---|---|---|---|---|---|---|
| T4704 | AGGAATGGACCATGTACTGCGGG | 1000 | | | | | | | | | |
| Oligo fwd | CACCAGGAATGGACCATGTACTGC | | | | | | | | | | |
| Oligo rev | AAACGCAGTACATGGTCCATTCCT | | | | | | | | | | |
| Chromosome | start | end | strand | MM | target_seq | PAM | alignment | distance | position | gene name | gene id |
| GU305914.1 | 68058 | 68080 | + | 0 | AGGAATGGACCATGTACTGC | GGG | ||||||||[||||||||||||]PAM | NA | Intergenic | NA | NA |

| | | | | | | | | | | | |
|---|---|---|---|---|---|---|---|---|---|---|---|
| T4705 | GGAATGGACCATGTACTGCGGGG | 1000 | | | | | | | | | |
| Oligo fwd | CACCGGAATGGACCATGTACTGCG | | | | | | | | | | |
| Oligo rev | AAACCGCAGTACATGGTCCATTCC | | | | | | | | | | |
| Chromosome | start | end | strand | MM | target_seq | PAM | alignment | distance | position | gene name | gene id |
| GU305914.1 | 68059 | 68081 | + | 0 | GGAATGGACCATGTACTGCG | GGG | ||||||||[||||||||||||]PAM | NA | Intergenic | NA | NA |

| | | | | | | | | | | | |
|---|---|---|---|---|---|---|---|---|---|---|---|
| T4706 | ATGGACCATGTACTGCGGGGAGG | 1000 | | | | | | | | | |
| Oligo fwd | CACCATGGACCATGTACTGCGGGG | | | | | | | | | | |
| Oligo rev | AAACCCCCGCAGTACATGGTCCAT | | | | | | | | | | |
| Chromosome | start | end | strand | MM | target_seq | PAM | alignment | distance | position | gene name | gene id |
| GU305914.1 | 68062 | 68084 | + | 0 | ATGGACCATGTACTGCGGGG | AGG | ||||||||[||||||||||||]PAM | NA | Intergenic | NA | NA |

| | | | | | | | | | | | |
|---|---|---|---|---|---|---|---|---|---|---|---|
| T4707 | GCGGGGAGGAGCCGTTCTACCGG | 1000 | | | | | | | | | |
| Oligo fwd | CACCGCGGGGAGGAGCCGTTCTAC | | | | | | | | | | |
| Oligo rev | AAACGTAGAACGGCTCCTCCCCGC | | | | | | | | | | |
| Chromosome | start | end | strand | MM | target_seq | PAM | alignment | distance | position | gene name | gene id |
| GU305914.1 | 68076 | 68098 | + | 0 | GCGGGGAGGAGCCGTTCTAC | CGG | ||||||||[||||||||||||]PAM | NA | Intergenic | NA | NA |
| GU305914.1 | 71544 | 71566 | + | 4 | CAGGGCAGGAGCCGTTCGAC | TGG | --|||-||[|||||||||-||]PAM | NA | Intergenic | NA | NA |

| | | | | | | | | | | | |
|---|---|---|---|---|---|---|---|---|---|---|---|
| T4708 | GTTCTACCGGCAGTACTTCGTGG | 1000 | | | | | | | | | |
| Oligo fwd | CACCGTTCTACCGGCAGTACTTCG | | | | | | | | | | |
| Oligo rev | AAACCGAAGTACTGCCGGTAGAAC | | | | | | | | | | |
| Chromosome | start | end | strand | MM | target_seq | PAM | alignment | distance | position | gene name | gene id |
| GU305914.1 | 68089 | 68111 | + | 0 | GTTCTACCGGCAGTACTTCG | TGG | ||||||||[||||||||||||]PAM | NA | Intergenic | NA | NA |

| | | |
|---|---|---|
| T4709 | CCGGCAGTACTTCGTGGACGTGG | 1000 |
| Oligo fwd | CACCCCGGCAGTACTTCGTGGACG | |

<table>
<tr><td>Oligo rev</td><td>AAACCGTCCACGAAGTACTGCCGG</td><td></td><td></td><td></td><td></td><td></td><td></td><td></td><td></td><td></td><td></td></tr>
<tr><td>Chromosome</td><td>start</td><td>end</td><td>strand</td><td>MM</td><td>target_seq</td><td>PAM</td><td>alignment</td><td>distance</td><td>position</td><td>gene name</td><td>gene id</td></tr>
<tr><td>GU305914.1</td><td>68095</td><td>68117</td><td>+</td><td>0</td><td>CCGGCAGTACTTCGTGGACG</td><td>TGG</td><td>||||||||[||||||||||||]PAM</td><td>NA</td><td>Intergenic</td><td>NA</td><td>NA</td></tr>
<tr><td>T4710</td><td>CGGCAGTACTTCGTGGACGTGGG</td><td>1000</td><td></td><td></td><td></td><td></td><td></td><td></td><td></td><td></td><td></td></tr>
<tr><td>Oligo fwd</td><td>CACCCGGCAGTACTTCGTGGACGT</td><td></td><td></td><td></td><td></td><td></td><td></td><td></td><td></td><td></td><td></td></tr>
<tr><td>Oligo rev</td><td>AAACACGTCCACGAAGTACTGCCG</td><td></td><td></td><td></td><td></td><td></td><td></td><td></td><td></td><td></td><td></td></tr>
<tr><td>Chromosome</td><td>start</td><td>end</td><td>strand</td><td>MM</td><td>target_seq</td><td>PAM</td><td>alignment</td><td>distance</td><td>position</td><td>gene name</td><td>gene id</td></tr>
<tr><td>GU305914.1</td><td>68096</td><td>68118</td><td>+</td><td>0</td><td>CGGCAGTACTTCGTGGACGT</td><td>GGG</td><td>||||||||[||||||||||||]PAM</td><td>NA</td><td>Intergenic</td><td>NA</td><td>NA</td></tr>
<tr><td>T4711</td><td>GGACGTGGGCCGCCAGCTGCTGG</td><td>1000</td><td></td><td></td><td></td><td></td><td></td><td></td><td></td><td></td><td></td></tr>
<tr><td>Oligo fwd</td><td>CACCGGACGTGGGCCGCCAGCTGC</td><td></td><td></td><td></td><td></td><td></td><td></td><td></td><td></td><td></td><td></td></tr>
<tr><td>Oligo rev</td><td>AAACGCAGCTGGCGGCCCACGTCC</td><td></td><td></td><td></td><td></td><td></td><td></td><td></td><td></td><td></td><td></td></tr>
<tr><td>Chromosome</td><td>start</td><td>end</td><td>strand</td><td>MM</td><td>target_seq</td><td>PAM</td><td>alignment</td><td>distance</td><td>position</td><td>gene name</td><td>gene id</td></tr>
<tr><td>GU305914.1</td><td>68110</td><td>68132</td><td>+</td><td>0</td><td>GGACGTGGGCCGCCAGCTGC</td><td>TGG</td><td>||||||||[||||||||||||]PAM</td><td>NA</td><td>Intergenic</td><td>NA</td><td>NA</td></tr>
<tr><td>T4712</td><td>GATCGATACTTACGTGAGTCTGG</td><td>1000</td><td></td><td></td><td></td><td></td><td></td><td></td><td></td><td></td><td></td></tr>
<tr><td>Oligo fwd</td><td>CACCGATCGATACTTACGTGAGTC</td><td></td><td></td><td></td><td></td><td></td><td></td><td></td><td></td><td></td><td></td></tr>
<tr><td>Oligo rev</td><td>AAACGACTCACGTAAGTATCGATC</td><td></td><td></td><td></td><td></td><td></td><td></td><td></td><td></td><td></td><td></td></tr>
<tr><td>Chromosome</td><td>start</td><td>end</td><td>strand</td><td>MM</td><td>target_seq</td><td>PAM</td><td>alignment</td><td>distance</td><td>position</td><td>gene name</td><td>gene id</td></tr>
<tr><td>GU305914.1</td><td>68140</td><td>68162</td><td>+</td><td>0</td><td>GATCGATACTTACGTGAGTC</td><td>TGG</td><td>||||||||[||||||||||||]PAM</td><td>NA</td><td>Intergenic</td><td>NA</td><td>NA</td></tr>
<tr><td>T4713</td><td>CGTGAGTCTGGACAAGACGACGG</td><td>1000</td><td></td><td></td><td></td><td></td><td></td><td></td><td></td><td></td><td></td></tr>
<tr><td>Oligo fwd</td><td>CACCCGTGAGTCTGGACAAGACGA</td><td></td><td></td><td></td><td></td><td></td><td></td><td></td><td></td><td></td><td></td></tr>
<tr><td>Oligo rev</td><td>AAACTCGTCTTGTCCAGACTCACG</td><td></td><td></td><td></td><td></td><td></td><td></td><td></td><td></td><td></td><td></td></tr>
<tr><td>Chromosome</td><td>start</td><td>end</td><td>strand</td><td>MM</td><td>target_seq</td><td>PAM</td><td>alignment</td><td>distance</td><td>position</td><td>gene name</td><td>gene id</td></tr>
<tr><td>GU305914.1</td><td>68152</td><td>68174</td><td>+</td><td>0</td><td>CGTGAGTCTGGACAAGACGA</td><td>CGG</td><td>||||||||[||||||||||||]PAM</td><td>NA</td><td>Intergenic</td><td>NA</td><td>NA</td></tr>
<tr><td>T4714</td><td>GGACAAGACGACGGACGAGTCGG</td><td>1000</td><td></td><td></td><td></td><td></td><td></td><td></td><td></td><td></td><td></td></tr>
<tr><td>Oligo fwd</td><td>CACCGGACAAGACGACGGACGAGT</td><td></td><td></td><td></td><td></td><td></td><td></td><td></td><td></td><td></td><td></td></tr>
<tr><td>Oligo rev</td><td>AAACACTCGTCCGTCGTCTTGTCC</td><td></td><td></td><td></td><td></td><td></td><td></td><td></td><td></td><td></td><td></td></tr>
<tr><td>Chromosome</td><td>start</td><td>end</td><td>strand</td><td>MM</td><td>target_seq</td><td>PAM</td><td>alignment</td><td>distance</td><td>position</td><td>gene name</td><td>gene id</td></tr>
<tr><td>GU305914.1</td><td>68161</td><td>68183</td><td>+</td><td>0</td><td>GGACAAGACGACGGACGAGT</td><td>CGG</td><td>||||||||[||||||||||||]PAM</td><td>NA</td><td>Intergenic</td><td>NA</td><td>NA</td></tr>
<tr><td>T4715</td><td>CAAGACGACGGACGAGTCGGCGG</td><td>1000</td><td></td><td></td><td></td><td></td><td></td><td></td><td></td><td></td><td></td></tr>
<tr><td>Oligo fwd</td><td>CACCCAAGACGACGGACGAGTCGG</td><td></td><td></td><td></td><td></td><td></td><td></td><td></td><td></td><td></td><td></td></tr>
<tr><td>Oligo rev</td><td>AAACCCGACTCGTCCGTCGTCTTG</td><td></td><td></td><td></td><td></td><td></td><td></td><td></td><td></td><td></td><td></td></tr>
<tr><td>Chromosome</td><td>start</td><td>end</td><td>strand</td><td>MM</td><td>target_seq</td><td>PAM</td><td>alignment</td><td>distance</td><td>position</td><td>gene name</td><td>gene id</td></tr>
<tr><td>GU305914.1</td><td>68164</td><td>68186</td><td>+</td><td>0</td><td>CAAGACGACGGACGAGTCGG</td><td>CGG</td><td>||||||||[||||||||||||]PAM</td><td>NA</td><td>Intergenic</td><td>NA</td><td>NA</td></tr>
<tr><td>T4716</td><td>CGGACGAGTCGGCGGTGCAGCGG</td><td>1000</td><td></td><td></td><td></td><td></td><td></td><td></td><td></td><td></td><td></td></tr>
<tr><td>Oligo fwd</td><td>CACCCGGACGAGTCGGCGGTGCAG</td><td></td><td></td><td></td><td></td><td></td><td></td><td></td><td></td><td></td><td></td></tr>
<tr><td>Oligo rev</td><td>AAACCTGCACCGCCGACTCGTCCG</td><td></td><td></td><td></td><td></td><td></td><td></td><td></td><td></td><td></td><td></td></tr>
<tr><td>Chromosome</td><td>start</td><td>end</td><td>strand</td><td>MM</td><td>target_seq</td><td>PAM</td><td>alignment</td><td>distance</td><td>position</td><td>gene name</td><td>gene id</td></tr>
<tr><td>GU305914.1</td><td>68172</td><td>68194</td><td>+</td><td>0</td><td>CGGACGAGTCGGCGGTGCAG</td><td>CGG</td><td>||||||||[||||||||||||]PAM</td><td>NA</td><td>Intergenic</td><td>NA</td><td>NA</td></tr>
<tr><td>T4717</td><td>GCAGCGGTTCGAGAGCCTGATGG</td><td>1000</td><td></td><td></td><td></td><td></td><td></td><td></td><td></td><td></td><td></td></tr>
<tr><td>Oligo fwd</td><td>CACCGCAGCGGTTCGAGAGCCTGA</td><td></td><td></td><td></td><td></td><td></td><td></td><td></td><td></td><td></td><td></td></tr>
<tr><td>Oligo rev</td><td>AAACTCAGGCTCTCGAACCGCTGC</td><td></td><td></td><td></td><td></td><td></td><td></td><td></td><td></td><td></td><td></td></tr>
<tr><td>Chromosome</td><td>start</td><td>end</td><td>strand</td><td>MM</td><td>target_seq</td><td>PAM</td><td>alignment</td><td>distance</td><td>position</td><td>gene name</td><td>gene id</td></tr>
<tr><td>GU305914.1</td><td>68188</td><td>68210</td><td>+</td><td>0</td><td>GCAGCGGTTCGAGAGCCTGA</td><td>TGG</td><td>||||||||[||||||||||||]PAM</td><td>NA</td><td>Intergenic</td><td>NA</td><td>NA</td></tr>
<tr><td>T4718</td><td>CAGCGGTTCGAGAGCCTGATGGG</td><td>1000</td><td></td><td></td><td></td><td></td><td></td><td></td><td></td><td></td><td></td></tr>
<tr><td>Oligo fwd</td><td>CACCCAGCGGTTCGAGAGCCTGAT</td><td></td><td></td><td></td><td></td><td></td><td></td><td></td><td></td><td></td><td></td></tr>
<tr><td>Oligo rev</td><td>AAACATCAGGCTCTCGAACCGCTG</td><td></td><td></td><td></td><td></td><td></td><td></td><td></td><td></td><td></td><td></td></tr>
<tr><td>Chromosome</td><td>start</td><td>end</td><td>strand</td><td>MM</td><td>target_seq</td><td>PAM</td><td>alignment</td><td>distance</td><td>position</td><td>gene name</td><td>gene id</td></tr>
<tr><td>GU305914.1</td><td>68189</td><td>68211</td><td>+</td><td>0</td><td>CAGCGGTTCGAGAGCCTGAT</td><td>GGG</td><td>||||||||[||||||||||||]PAM</td><td>NA</td><td>Intergenic</td><td>NA</td><td>NA</td></tr>
<tr><td>T4719</td><td>CGAGAGCCTGATGGGTCTGTCGG</td><td>1000</td><td></td><td></td><td></td><td></td><td></td><td></td><td></td><td></td><td></td></tr>
<tr><td>Oligo fwd</td><td>CACCCGAGAGCCTGATGGGTCTGT</td><td></td><td></td><td></td><td></td><td></td><td></td><td></td><td></td><td></td><td></td></tr>
<tr><td>Oligo rev</td><td>AAACACAGACCCATCAGGCTCTCG</td><td></td><td></td><td></td><td></td><td></td><td></td><td></td><td></td><td></td><td></td></tr>
<tr><td>Chromosome</td><td>start</td><td>end</td><td>strand</td><td>MM</td><td>target_seq</td><td>PAM</td><td>alignment</td><td>distance</td><td>position</td><td>gene name</td><td>gene id</td></tr>
<tr><td>GU305914.1</td><td>68197</td><td>68219</td><td>+</td><td>0</td><td>CGAGAGCCTGATGGGTCTGT</td><td>CGG</td><td>||||||||[||||||||||||]PAM</td><td>NA</td><td>Intergenic</td><td>NA</td><td>NA</td></tr>
<tr><td>T4720</td><td>GATGGGTCTGTCGGACGAGCTGG</td><td>1000</td><td></td><td></td><td></td><td></td><td></td><td></td><td></td><td></td><td></td></tr>
<tr><td>Oligo fwd</td><td>CACCGATGGGTCTGTCGGACGAGC</td><td></td><td></td><td></td><td></td><td></td><td></td><td></td><td></td><td></td><td></td></tr>
<tr><td>Oligo rev</td><td>AAACGCTCGTCCGACAGACCCATC</td><td></td><td></td><td></td><td></td><td></td><td></td><td></td><td></td><td></td><td></td></tr>
<tr><td>Chromosome</td><td>start</td><td>end</td><td>strand</td><td>MM</td><td>target_seq</td><td>PAM</td><td>alignment</td><td>distance</td><td>position</td><td>gene name</td><td>gene id</td></tr>
<tr><td>GU305914.1</td><td>68206</td><td>68228</td><td>+</td><td>0</td><td>GATGGGTCTGTCGGACGAGC</td><td>TGG</td><td>||||||||[||||||||||||]PAM</td><td>NA</td><td>Intergenic</td><td>NA</td><td>NA</td></tr>
<tr><td>T4721</td><td>TCTGTCGGACGAGCTGGACGCGG</td><td>1000</td><td></td><td></td><td></td><td></td><td></td><td></td><td></td><td></td><td></td></tr>
<tr><td>Oligo fwd</td><td>CACCTCTGTCGGACGAGCTGGACG</td><td></td><td></td><td></td><td></td><td></td><td></td><td></td><td></td><td></td><td></td></tr>
<tr><td>Oligo rev</td><td>AAACCGTCCAGCTCGTCCGACAGA</td><td></td><td></td><td></td><td></td><td></td><td></td><td></td><td></td><td></td><td></td></tr>
<tr><td>Chromosome</td><td>start</td><td>end</td><td>strand</td><td>MM</td><td>target_seq</td><td>PAM</td><td>alignment</td><td>distance</td><td>position</td><td>gene name</td><td>gene id</td></tr>
<tr><td>GU305914.1</td><td>68212</td><td>68234</td><td>+</td><td>0</td><td>TCTGTCGGACGAGCTGGACG</td><td>CGG</td><td>||||||||[||||||||||||]PAM</td><td>NA</td><td>Intergenic</td><td>NA</td><td>NA</td></tr>
<tr><td>T4722</td><td>CGAGCTGGACGCGGTGATCGCGG</td><td>1000</td><td></td><td></td><td></td><td></td><td></td><td></td><td></td><td></td><td></td></tr>
<tr><td>Oligo fwd</td><td>CACCCGAGCTGGACGCGGTGATCG</td><td></td><td></td><td></td><td></td><td></td><td></td><td></td><td></td><td></td><td></td></tr>
<tr><td>Oligo rev</td><td>AAACCGATCACCGCGTCCAGCTCG</td><td></td><td></td><td></td><td></td><td></td><td></td><td></td><td></td><td></td><td></td></tr>
<tr><td>Chromosome</td><td>start</td><td>end</td><td>strand</td><td>MM</td><td>target_seq</td><td>PAM</td><td>alignment</td><td>distance</td><td>position</td><td>gene name</td><td>gene id</td></tr>
<tr><td>GU305914.1</td><td>68221</td><td>68243</td><td>+</td><td>0</td><td>CGAGCTGGACGCGGTGATCG</td><td>CGG</td><td>||||||||[||||||||||||]PAM</td><td>NA</td><td>Intergenic</td><td>NA</td><td>NA</td></tr>
<tr><td>T4723</td><td>GCTGGACGCGGTGATCGCGGCGG</td><td>1000</td><td></td><td></td><td></td><td></td><td></td><td></td><td></td><td></td><td></td></tr>
<tr><td>Oligo fwd</td><td>CACCGCTGGACGCGGTGATCGCGG</td><td></td><td></td><td></td><td></td><td></td><td></td><td></td><td></td><td></td><td></td></tr>
<tr><td>Oligo rev</td><td>AAACCCGCGATCACCGCGTCCAGC</td><td></td><td></td><td></td><td></td><td></td><td></td><td></td><td></td><td></td><td></td></tr>
<tr><td>Chromosome</td><td>start</td><td>end</td><td>strand</td><td>MM</td><td>target_seq</td><td>PAM</td><td>alignment</td><td>distance</td><td>position</td><td>gene name</td><td>gene id</td></tr>
<tr><td>GU305914.1</td><td>68224</td><td>68246</td><td>+</td><td>0</td><td>GCTGGACGCGGTGATCGCGG</td><td>CGG</td><td>||||||||[||||||||||||]PAM</td><td>NA</td><td>Intergenic</td><td>NA</td><td>NA</td></tr>
<tr><td>GU305914.1</td><td>68824</td><td>68846</td><td>+</td><td>4</td><td>GCCGGACTCGGACATCGCGG</td><td>CGG</td><td>||-||||-[|||--|||||||]PAM</td><td>NA</td><td>Intergenic</td><td>NA</td><td>NA</td></tr>
</table>

| | | | | | | | | | | | |
|---|---|---|---|---|---|---|---|---|---|---|---|
| T4724 | GATCGCGGCGGCGCGCGACGG | 1000 | | | | | | | | | |
| Oligo fwd | CACCGATCGCGGCGGCGCGCGA | | | | | | | | | | |
| Oligo rev | AAACTCGCGCGCGCCGCCGCGATC | | | | | | | | | | |
| Chromosome | start | end | strand | MM | target_seq | PAM | alignment | distance | position | gene name | gene id |
| GU305914.1 | 68236 | 68258 | + | 0 | GATCGCGGCGGCGCGCGCGA | CGG | \|\|\|\|\|\|\|\|[\|\|\|\|\|\|\|\|\|\|\|\|]PAM | NA | Intergenic | NA | NA |
| T4725 | GGCGCGCGCGACGGAGCTGTCGG | 1000 | | | | | | | | | |
| Oligo fwd | CACCGGCGCGCGCGACGGAGCTGT | | | | | | | | | | |
| Oligo rev | AAACACAGCTCCGTCGCGCGCGCC | | | | | | | | | | |
| Chromosome | start | end | strand | MM | target_seq | PAM | alignment | distance | position | gene name | gene id |
| GU305914.1 | 68245 | 68267 | + | 0 | GGCGCGCGCGACGGAGCTGT | CGG | \|\|\|\|\|\|\|\|[\|\|\|\|\|\|\|\|\|\|\|\|]PAM | NA | Intergenic | NA | NA |
| T4726 | ACAAGACTTACCTGTCTCAGCGG | 1000 | | | | | | | | | |
| Oligo fwd | CACCACAAGACTTACCTGTCTCAG | | | | | | | | | | |
| Oligo rev | AAACCTGAGACAGGTAAGTCTTGT | | | | | | | | | | |
| Chromosome | start | end | strand | MM | target_seq | PAM | alignment | distance | position | gene name | gene id |
| GU305914.1 | 68280 | 68302 | + | 0 | ACAAGACTTACCTGTCTCAG | CGG | \|\|\|\|\|\|\|\|[\|\|\|\|\|\|\|\|\|\|\|\|]PAM | NA | Intergenic | NA | NA |
| T4727 | CAAGACTTACCTGTCTCAGCGGG | 1000 | | | | | | | | | |
| Oligo fwd | CACCCAAGACTTACCTGTCTCAGC | | | | | | | | | | |
| Oligo rev | AAACGCTGAGACAGGTAAGTCTTG | | | | | | | | | | |
| Chromosome | start | end | strand | MM | target_seq | PAM | alignment | distance | position | gene name | gene id |
| GU305914.1 | 68281 | 68303 | + | 0 | CAAGACTTACCTGTCTCAGC | GGG | \|\|\|\|\|\|\|\|[\|\|\|\|\|\|\|\|\|\|\|\|]PAM | NA | Intergenic | NA | NA |
| T4728 | GTCTCAGCGGGTGAGCGCGTCGG | 1000 | | | | | | | | | |
| Oligo fwd | CACCGTCTCAGCGGGTGAGCGCGT | | | | | | | | | | |
| Oligo rev | AAACACGCGCTCACCCGCTGAGAC | | | | | | | | | | |
| Chromosome | start | end | strand | MM | target_seq | PAM | alignment | distance | position | gene name | gene id |
| GU305914.1 | 68293 | 68315 | + | 0 | GTCTCAGCGGGTGAGCGCGT | CGG | \|\|\|\|\|\|\|\|[\|\|\|\|\|\|\|\|\|\|\|\|]PAM | NA | Intergenic | NA | NA |
| T4729 | TCAGCGGGTGAGCGCGTCGGCGG | 1000 | | | | | | | | | |
| Oligo fwd | CACCTCAGCGGGTGAGCGCGTCGG | | | | | | | | | | |
| Oligo rev | AAACCCGACGCGCTCACCCGCTGA | | | | | | | | | | |
| Chromosome | start | end | strand | MM | target_seq | PAM | alignment | distance | position | gene name | gene id |
| GU305914.1 | 68296 | 68318 | + | 0 | TCAGCGGGTGAGCGCGTCGG | CGG | \|\|\|\|\|\|\|\|[\|\|\|\|\|\|\|\|\|\|\|\|]PAM | NA | Intergenic | NA | NA |
| T4730 | TGAGCGCGTCGGCGGTGACGAGG | 1000 | | | | | | | | | |
| Oligo fwd | CACCTGAGCGCGTCGGCGGTGACG | | | | | | | | | | |
| Oligo rev | AAACCGTCACCGCCGACGCGCTCA | | | | | | | | | | |
| Chromosome | start | end | strand | MM | target_seq | PAM | alignment | distance | position | gene name | gene id |
| GU305914.1 | 68304 | 68326 | + | 0 | TGAGCGCGTCGGCGGTGACG | AGG | \|\|\|\|\|\|\|\|[\|\|\|\|\|\|\|\|\|\|\|\|]PAM | NA | Intergenic | NA | NA |
| T4731 | CGCGTCGGCGGTGACGAGGACGG | 1000 | | | | | | | | | |
| Oligo fwd | CACCCGCGTCGGCGGTGACGAGGA | | | | | | | | | | |
| Oligo rev | AAACTCCTCGTCACCGCCGACGCG | | | | | | | | | | |
| Chromosome | start | end | strand | MM | target_seq | PAM | alignment | distance | position | gene name | gene id |
| GU305914.1 | 68308 | 68330 | + | 0 | CGCGTCGGCGGTGACGAGGA | CGG | \|\|\|\|\|\|\|\|[\|\|\|\|\|\|\|\|\|\|\|\|]PAM | NA | Intergenic | NA | NA |
| T4732 | CGGCGGTGACGAGGACGGACCGG | 1000 | | | | | | | | | |
| Oligo fwd | CACCCGGCGGTGACGAGGACGGAC | | | | | | | | | | |
| Oligo rev | AAACGTCCGTCCTCGTCACCGCCG | | | | | | | | | | |
| Chromosome | start | end | strand | MM | target_seq | PAM | alignment | distance | position | gene name | gene id |
| GU305914.1 | 68313 | 68335 | + | 0 | CGGCGGTGACGAGGACGGAC | CGG | \|\|\|\|\|\|\|\|[\|\|\|\|\|\|\|\|\|\|\|\|]PAM | NA | Intergenic | NA | NA |
| GU305914.1 | 174261 | 174283 | + | 4 | CGACGGCGACGATGACGAAC | AGG | \|\|-\|\|\|-\|[\|\|\|\|-\|\|\|\|-\|\|]PAM | NA | Intergenic | NA | NA |
| GU305914.1 | 32951 | 32973 | + | 4 | CGGAGGCGACGAAGACGGGC | CGG | \|\|\|-\|\|-\|[\|\|\|\|-\|\|\|\|\|-\|]PAM | NA | Intergenic | NA | NA |
| GU305914.1 | 96811 | 96833 | - | 4 | CGCCGGCGACGAGGACGAAG | GGG | \|\|-\|\|\|-\|[\|\|\|\|\|\|\|\|\|-\|-]PAM | NA | Intergenic | NA | NA |
| T4733 | GACGAGGACGGACCGGCTGCTGG | 1000 | | | | | | | | | |
| Oligo fwd | CACCGACGAGGACGGACCGGCTGC | | | | | | | | | | |
| Oligo rev | AAACGCAGCCGGTCCGTCCTCGTC | | | | | | | | | | |
| Chromosome | start | end | strand | MM | target_seq | PAM | alignment | distance | position | gene name | gene id |
| GU305914.1 | 68320 | 68342 | + | 0 | GACGAGGACGGACCGGCTGC | TGG | \|\|\|\|\|\|\|\|[\|\|\|\|\|\|\|\|\|\|\|\|]PAM | NA | Intergenic | NA | NA |
| T4734 | GCTGCTGGCGTCGAAGATCCTGG | 1000 | | | | | | | | | |
| Oligo fwd | CACCGCTGCTGGCGTCGAAGATCC | | | | | | | | | | |
| Oligo rev | AAACGGATCTTCGACGCCAGCAGC | | | | | | | | | | |
| Chromosome | start | end | strand | MM | target_seq | PAM | alignment | distance | position | gene name | gene id |
| GU305914.1 | 68335 | 68357 | + | 0 | GCTGCTGGCGTCGAAGATCC | TGG | \|\|\|\|\|\|\|\|[\|\|\|\|\|\|\|\|\|\|\|\|]PAM | NA | Intergenic | NA | NA |
| T4735 | GAAGATCCTGGCGCTGTTCGAGG | 1000 | | | | | | | | | |
| Oligo fwd | CACCGAAGATCCTGGCGCTGTTCG | | | | | | | | | | |
| Oligo rev | AAACCGAACAGCGCCAGGATCTTC | | | | | | | | | | |
| Chromosome | start | end | strand | MM | target_seq | PAM | alignment | distance | position | gene name | gene id |
| GU305914.1 | 68347 | 68369 | + | 0 | GAAGATCCTGGCGCTGTTCG | AGG | \|\|\|\|\|\|\|\|[\|\|\|\|\|\|\|\|\|\|\|\|]PAM | NA | Intergenic | NA | NA |
| T4736 | GATCCTGGCGCTGTTCGAGGAGG | 1000 | | | | | | | | | |
| Oligo fwd | CACCGATCCTGGCGCTGTTCGAGG | | | | | | | | | | |
| Oligo rev | AAACCCTCGAACAGCGCCAGGATC | | | | | | | | | | |
| Chromosome | start | end | strand | MM | target_seq | PAM | alignment | distance | position | gene name | gene id |
| GU305914.1 | 68350 | 68372 | + | 0 | GATCCTGGCGCTGTTCGAGG | AGG | \|\|\|\|\|\|\|\|[\|\|\|\|\|\|\|\|\|\|\|\|]PAM | NA | Intergenic | NA | NA |
| T4737 | ATCCTGGCGCTGTTCGAGGAGGG | 1000 | | | | | | | | | |
| Oligo fwd | CACCATCCTGGCGCTGTTCGAGGA | | | | | | | | | | |
| Oligo rev | AAACTCCTCGAACAGCGCCAGGAT | | | | | | | | | | |
| Chromosome | start | end | strand | MM | target_seq | PAM | alignment | distance | position | gene name | gene id |
| GU305914.1 | 68351 | 68373 | + | 0 | ATCCTGGCGCTGTTCGAGGA | GGG | \|\|\|\|\|\|\|\|[\|\|\|\|\|\|\|\|\|\|\|\|]PAM | NA | Intergenic | NA | NA |
| T4738 | GGCGCTGTTCGAGGAGGGCTCGG | 1000 | | | | | | | | | |
| Oligo fwd | CACCGGCGCTGTTCGAGGAGGGCT | | | | | | | | | | |

| | | | | | | | | | | | |
|---|---|---|---|---|---|---|---|---|---|---|---|
| Oligo rev | AAACAGCCCTCCTCGAACAGCGCC | | | | | | | | | | |
| Chromosome | start | end | strand | MM | target_seq | PAM | alignment | distance | position | gene name | gene id |
| GU305914.1 | 68356 | 68378 | + | 0 | GGCGCTGTTCGAGGAGGGCT | CGG | ||||||||[||||||||||||]PAM | NA | Intergenic | NA | NA |
| T4739 | GTTCGAGGAGGGCTCGGCGACGG | 1000 | | | | | | | | | |
| Oligo fwd | CACCGTTCGAGGAGGGCTCGGCGA | | | | | | | | | | |
| Oligo rev | AAACTCGCCGAGCCCTCCTCGAAC | | | | | | | | | | |
| Chromosome | start | end | strand | MM | target_seq | PAM | alignment | distance | position | gene name | gene id |
| GU305914.1 | 68362 | 68384 | + | 0 | GTTCGAGGAGGGCTCGGCGA | CGG | ||||||||[||||||||||||]PAM | NA | Intergenic | NA | NA |
| T4740 | CTCGGCGACGGACTTCGAGACGG | 1000 | | | | | | | | | |
| Oligo fwd | CACCCTCGGCGACGGACTTCGAGA | | | | | | | | | | |
| Oligo rev | AAACTCTCGAAGTCCGTCGCCGAG | | | | | | | | | | |
| Chromosome | start | end | strand | MM | target_seq | PAM | alignment | distance | position | gene name | gene id |
| GU305914.1 | 68374 | 68396 | + | 0 | CTCGGCGACGGACTTCGAGA | CGG | ||||||||[||||||||||||]PAM | NA | Intergenic | NA | NA |
| T4741 | ACTTCGAGACGGTGTCGTCGTGG | 1000 | | | | | | | | | |
| Oligo fwd | CACCACTTCGAGACGGTGTCGTCG | | | | | | | | | | |
| Oligo rev | AAACCGACGACACCGTCTCGAAGT | | | | | | | | | | |
| Chromosome | start | end | strand | MM | target_seq | PAM | alignment | distance | position | gene name | gene id |
| GU305914.1 | 68385 | 68407 | + | 0 | ACTTCGAGACGGTGTCGTCG | TGG | ||||||||[||||||||||||]PAM | NA | Intergenic | NA | NA |
| T4742 | AGACGGTGTCGTCGTGGCTGCGG | 1000 | | | | | | | | | |
| Oligo fwd | CACCAGACGGTGTCGTCGTGGCTG | | | | | | | | | | |
| Oligo rev | AAACCAGCCACGACGACACCGTCT | | | | | | | | | | |
| Chromosome | start | end | strand | MM | target_seq | PAM | alignment | distance | position | gene name | gene id |
| GU305914.1 | 68391 | 68413 | + | 0 | AGACGGTGTCGTCGTGGCTG | CGG | ||||||||[||||||||||||]PAM | NA | Intergenic | NA | NA |
| T4743 | GACGGTGTCGTCGTGGCTGCGGG | 1000 | | | | | | | | | |
| Oligo fwd | CACCGACGGTGTCGTCGTGGCTGC | | | | | | | | | | |
| Oligo rev | AAACGCAGCCACGACGACACCGTC | | | | | | | | | | |
| Chromosome | start | end | strand | MM | target_seq | PAM | alignment | distance | position | gene name | gene id |
| GU305914.1 | 68392 | 68414 | + | 0 | GACGGTGTCGTCGTGGCTGC | GGG | ||||||||[||||||||||||]PAM | NA | Intergenic | NA | NA |
| T4744 | GCTGCTGCAGCAGCTGCCCGTGG | 1000 | | | | | | | | | |
| Oligo fwd | CACCGCTGCTGCAGCAGCTGCCCG | | | | | | | | | | |
| Oligo rev | AAACCGGGCAGCTGCTGCAGCAGC | | | | | | | | | | |
| Chromosome | start | end | strand | MM | target_seq | PAM | alignment | distance | position | gene name | gene id |
| GU305914.1 | 68416 | 68438 | + | 0 | GCTGCTGCAGCAGCTGCCCG | TGG | ||||||||[||||||||||||]PAM | NA | Intergenic | NA | NA |
| GU305914.1 | 164955 | 164977 | - | 4 | GCGGCGGCAGCAGCTGCGGG | TGG | ||-||-||[|||||||||--|]PAM | NA | Intergenic | NA | NA |
| T4745 | GGAGAACACGACGACGACCGAGG | 1000 | | | | | | | | | |
| Oligo fwd | CACCGGAGAACACGACGACGACCG | | | | | | | | | | |
| Oligo rev | AAACCGGTCGTCGTCGTGTTCTCC | | | | | | | | | | |
| Chromosome | start | end | strand | MM | target_seq | PAM | alignment | distance | position | gene name | gene id |
| GU305914.1 | 68437 | 68459 | + | 0 | GGAGAACACGACGACGACCG | AGG | ||||||||[||||||||||||]PAM | NA | Intergenic | NA | NA |
| GU305914.1 | 212416 | 212438 | - | 4 | GGAATACACGACGAAGAACG | AGG | |||--|||[||||||-||-||]PAM | NA | Intergenic | NA | NA |
| GU305914.1 | 40445 | 40467 | - | 4 | GAAGAAGACGACGATGACGG | CGG | |-||||-|[||||||-|||-|]PAM | NA | Intergenic | NA | NA |
| T4746 | GGCGAAGATCGCCGCGTTCGTGG | 1000 | | | | | | | | | |
| Oligo fwd | CACCGGCGAAGATCGCCGCGTTCG | | | | | | | | | | |
| Oligo rev | AAACCGAACGCGGCGATCTTCGCC | | | | | | | | | | |
| Chromosome | start | end | strand | MM | target_seq | PAM | alignment | distance | position | gene name | gene id |
| GU305914.1 | 68458 | 68480 | + | 0 | GGCGAAGATCGCCGCGTTCG | TGG | ||||||||[||||||||||||]PAM | NA | Intergenic | NA | NA |
| T4747 | GTGGCGCAGCACCCCATCCCCGG | 1000 | | | | | | | | | |
| Oligo fwd | CACCGTGGCGCAGCACCCCATCCC | | | | | | | | | | |
| Oligo rev | AAACGGGATGGGGTGCTGCGCCAC | | | | | | | | | | |
| Chromosome | start | end | strand | MM | target_seq | PAM | alignment | distance | position | gene name | gene id |
| GU305914.1 | 68477 | 68499 | + | 0 | GTGGCGCAGCACCCCATCCC | CGG | ||||||||[||||||||||||]PAM | NA | Intergenic | NA | NA |
| T4748 | TGGCGCAGCACCCCATCCCCGGG | 1000 | | | | | | | | | |
| Oligo fwd | CACCTGGCGCAGCACCCCATCCCC | | | | | | | | | | |
| Oligo rev | AAACGGGGATGGGGTGCTGCGCCA | | | | | | | | | | |
| Chromosome | start | end | strand | MM | target_seq | PAM | alignment | distance | position | gene name | gene id |
| GU305914.1 | 68478 | 68500 | + | 0 | TGGCGCAGCACCCCATCCCC | GGG | ||||||||[||||||||||||]PAM | NA | Intergenic | NA | NA |
| T4749 | GGCGCAGCACCCCATCCCCGGGG | 1000 | | | | | | | | | |
| Oligo fwd | CACCGGCGCAGCACCCCATCCCCG | | | | | | | | | | |
| Oligo rev | AAACCGGGGATGGGGTGCTGCGCC | | | | | | | | | | |
| Chromosome | start | end | strand | MM | target_seq | PAM | alignment | distance | position | gene name | gene id |
| GU305914.1 | 68479 | 68501 | + | 0 | GGCGCAGCACCCCATCCCCG | GGG | ||||||||[||||||||||||]PAM | NA | Intergenic | NA | NA |
| GU305914.1 | 720 | 742 | + | 4 | GGCGGGGCACCCCCCCCCCG | CGG | ||||--||[|||||--|||||]PAM | NA | Intergenic | NA | NA |
| T4750 | CGTGTGCCTGCGCAACTCGCAGG | 1000 | | | | | | | | | |
| Oligo fwd | CACCCGTGTGCCTGCGCAACTCGC | | | | | | | | | | |
| Oligo rev | AAACGCGAGTTGCGCAGGCACACG | | | | | | | | | | |
| Chromosome | start | end | strand | MM | target_seq | PAM | alignment | distance | position | gene name | gene id |
| GU305914.1 | 68512 | 68534 | + | 0 | CGTGTGCCTGCGCAACTCGC | AGG | ||||||||[||||||||||||]PAM | NA | Intergenic | NA | NA |
| T4751 | CAACTCGCAGGTGAAGAGCCTGG | 1000 | | | | | | | | | |
| Oligo fwd | CACCCAACTCGCAGGTGAAGAGCC | | | | | | | | | | |
| Oligo rev | AAACGGCTCTTCACCTGCGAGTTG | | | | | | | | | | |
| Chromosome | start | end | strand | MM | target_seq | PAM | alignment | distance | position | gene name | gene id |
| GU305914.1 | 68524 | 68546 | + | 0 | CAACTCGCAGGTGAAGAGCC | TGG | ||||||||[||||||||||||]PAM | NA | Intergenic | NA | NA |
| T4752 | AACTCGCAGGTGAAGAGCCTGGG | 1000 | | | | | | | | | |
| Oligo fwd | CACCAACTCGCAGGTGAAGAGCCT | | | | | | | | | | |
| Oligo rev | AAACAGGCTCTTCACCTGCGAGTT | | | | | | | | | | |

<table>
<tr><td>Chromosome</td><td>start</td><td>end</td><td>strand</td><td>MM</td><td>target_seq</td><td>PAM</td><td>alignment</td><td>distance</td><td>position</td><td>gene name</td><td>gene id</td></tr>
<tr><td>GU305914.1</td><td>68525</td><td>68547</td><td>+</td><td>0</td><td>AACTCGCAGGTGAAGAGCCT</td><td>GGG</td><td>||||||||[||||||||||||]PAM</td><td>NA</td><td>Intergenic</td><td>NA</td><td>NA</td></tr>
<tr><td>T4753</td><td>GAGCCTGGGCGAGCTGCTGTCGG</td><td>1000</td><td></td><td></td><td></td><td></td><td></td><td></td><td></td><td></td><td></td></tr>
<tr><td>Oligo fwd</td><td>CACCGAGCCTGGGCGAGCTGCTGT</td><td></td><td></td><td></td><td></td><td></td><td></td><td></td><td></td><td></td><td></td></tr>
<tr><td>Oligo rev</td><td>AAACACAGCAGCTCGCCCAGGCTC</td><td></td><td></td><td></td><td></td><td></td><td></td><td></td><td></td><td></td><td></td></tr>
<tr><td>Chromosome</td><td>start</td><td>end</td><td>strand</td><td>MM</td><td>target_seq</td><td>PAM</td><td>alignment</td><td>distance</td><td>position</td><td>gene name</td><td>gene id</td></tr>
<tr><td>GU305914.1</td><td>68539</td><td>68561</td><td>+</td><td>0</td><td>GAGCCTGGGCGAGCTGCTGT</td><td>CGG</td><td>||||||||[||||||||||||]PAM</td><td>NA</td><td>Intergenic</td><td>NA</td><td>NA</td></tr>
<tr><td>T4754</td><td>GCGAGCTGCTGTCGGTGAAGAGG</td><td>1000</td><td></td><td></td><td></td><td></td><td></td><td></td><td></td><td></td><td></td></tr>
<tr><td>Oligo fwd</td><td>CACCGCGAGCTGCTGTCGGTGAAG</td><td></td><td></td><td></td><td></td><td></td><td></td><td></td><td></td><td></td><td></td></tr>
<tr><td>Oligo rev</td><td>AAACCTTCACCGACAGCAGCTCGC</td><td></td><td></td><td></td><td></td><td></td><td></td><td></td><td></td><td></td><td></td></tr>
<tr><td>Chromosome</td><td>start</td><td>end</td><td>strand</td><td>MM</td><td>target_seq</td><td>PAM</td><td>alignment</td><td>distance</td><td>position</td><td>gene name</td><td>gene id</td></tr>
<tr><td>GU305914.1</td><td>68547</td><td>68569</td><td>+</td><td>0</td><td>GCGAGCTGCTGTCGGTGAAG</td><td>AGG</td><td>||||||||[||||||||||||]PAM</td><td>NA</td><td>Intergenic</td><td>NA</td><td>NA</td></tr>
<tr><td>T4755</td><td>CGAGCTGCTGTCGGTGAAGAGGG</td><td>1000</td><td></td><td></td><td></td><td></td><td></td><td></td><td></td><td></td><td></td></tr>
<tr><td>Oligo fwd</td><td>CACCCGAGCTGCTGTCGGTGAAGA</td><td></td><td></td><td></td><td></td><td></td><td></td><td></td><td></td><td></td><td></td></tr>
<tr><td>Oligo rev</td><td>AAACTCTTCACCGACAGCAGCTCG</td><td></td><td></td><td></td><td></td><td></td><td></td><td></td><td></td><td></td><td></td></tr>
<tr><td>Chromosome</td><td>start</td><td>end</td><td>strand</td><td>MM</td><td>target_seq</td><td>PAM</td><td>alignment</td><td>distance</td><td>position</td><td>gene name</td><td>gene id</td></tr>
<tr><td>GU305914.1</td><td>68548</td><td>68570</td><td>+</td><td>0</td><td>CGAGCTGCTGTCGGTGAAGA</td><td>GGG</td><td>||||||||[||||||||||||]PAM</td><td>NA</td><td>Intergenic</td><td>NA</td><td>NA</td></tr>
<tr><td>T4756</td><td>GGTGAAGAGGGAGACCCTGAAGG</td><td>1000</td><td></td><td></td><td></td><td></td><td></td><td></td><td></td><td></td><td></td></tr>
<tr><td>Oligo fwd</td><td>CACCGGTGAAGAGGGAGACCCTGA</td><td></td><td></td><td></td><td></td><td></td><td></td><td></td><td></td><td></td><td></td></tr>
<tr><td>Oligo rev</td><td>AAACTCAGGGTCTCCCTCTTCACC</td><td></td><td></td><td></td><td></td><td></td><td></td><td></td><td></td><td></td><td></td></tr>
<tr><td>Chromosome</td><td>start</td><td>end</td><td>strand</td><td>MM</td><td>target_seq</td><td>PAM</td><td>alignment</td><td>distance</td><td>position</td><td>gene name</td><td>gene id</td></tr>
<tr><td>GU305914.1</td><td>68560</td><td>68582</td><td>+</td><td>0</td><td>GGTGAAGAGGGAGACCCTGA</td><td>AGG</td><td>||||||||[||||||||||||]PAM</td><td>NA</td><td>Intergenic</td><td>NA</td><td>NA</td></tr>
<tr><td>T4757</td><td>GACCCTGAAGGTGAAATACATGG</td><td>1000</td><td></td><td></td><td></td><td></td><td></td><td></td><td></td><td></td><td></td></tr>
<tr><td>Oligo fwd</td><td>CACCGACCCTGAAGGTGAAATACA</td><td></td><td></td><td></td><td></td><td></td><td></td><td></td><td></td><td></td><td></td></tr>
<tr><td>Oligo rev</td><td>AAACTGTATTTCACCTTCAGGGTC</td><td></td><td></td><td></td><td></td><td></td><td></td><td></td><td></td><td></td><td></td></tr>
<tr><td>Chromosome</td><td>start</td><td>end</td><td>strand</td><td>MM</td><td>target_seq</td><td>PAM</td><td>alignment</td><td>distance</td><td>position</td><td>gene name</td><td>gene id</td></tr>
<tr><td>GU305914.1</td><td>68572</td><td>68594</td><td>+</td><td>0</td><td>GACCCTGAAGGTGAAATACA</td><td>TGG</td><td>||||||||[||||||||||||]PAM</td><td>NA</td><td>Intergenic</td><td>NA</td><td>NA</td></tr>
<tr><td>T4758</td><td>CGCGACTTACCGCAAGATCCTGG</td><td>1000</td><td></td><td></td><td></td><td></td><td></td><td></td><td></td><td></td><td></td></tr>
<tr><td>Oligo fwd</td><td>CACCCGCGACTTACCGCAAGATCC</td><td></td><td></td><td></td><td></td><td></td><td></td><td></td><td></td><td></td><td></td></tr>
<tr><td>Oligo rev</td><td>AAACGGATCTTGCGGTAAGTCGCG</td><td></td><td></td><td></td><td></td><td></td><td></td><td></td><td></td><td></td><td></td></tr>
<tr><td>Chromosome</td><td>start</td><td>end</td><td>strand</td><td>MM</td><td>target_seq</td><td>PAM</td><td>alignment</td><td>distance</td><td>position</td><td>gene name</td><td>gene id</td></tr>
<tr><td>GU305914.1</td><td>68602</td><td>68624</td><td>+</td><td>0</td><td>CGCGACTTACCGCAAGATCC</td><td>TGG</td><td>||||||||[||||||||||||]PAM</td><td>NA</td><td>Intergenic</td><td>NA</td><td>NA</td></tr>
<tr><td>T4759</td><td>CCTGGACGCGATCTCGAAGCTGG</td><td>1000</td><td></td><td></td><td></td><td></td><td></td><td></td><td></td><td></td><td></td></tr>
<tr><td>Oligo fwd</td><td>CACCCCTGGACGCGATCTCGAAGC</td><td></td><td></td><td></td><td></td><td></td><td></td><td></td><td></td><td></td><td></td></tr>
<tr><td>Oligo rev</td><td>AAACGCTTCGAGATCGCGTCCAGG</td><td></td><td></td><td></td><td></td><td></td><td></td><td></td><td></td><td></td><td></td></tr>
<tr><td>Chromosome</td><td>start</td><td>end</td><td>strand</td><td>MM</td><td>target_seq</td><td>PAM</td><td>alignment</td><td>distance</td><td>position</td><td>gene name</td><td>gene id</td></tr>
<tr><td>GU305914.1</td><td>68620</td><td>68642</td><td>+</td><td>0</td><td>CCTGGACGCGATCTCGAAGC</td><td>TGG</td><td>||||||||[||||||||||||]PAM</td><td>NA</td><td>Intergenic</td><td>NA</td><td>NA</td></tr>
<tr><td>T4760</td><td>CTGGACGCGATCTCGAAGCTGGG</td><td>1000</td><td></td><td></td><td></td><td></td><td></td><td></td><td></td><td></td><td></td></tr>
<tr><td>Oligo fwd</td><td>CACCCTGGACGCGATCTCGAAGCT</td><td></td><td></td><td></td><td></td><td></td><td></td><td></td><td></td><td></td><td></td></tr>
<tr><td>Oligo rev</td><td>AAACAGCTTCGAGATCGCGTCCAG</td><td></td><td></td><td></td><td></td><td></td><td></td><td></td><td></td><td></td><td></td></tr>
<tr><td>Chromosome</td><td>start</td><td>end</td><td>strand</td><td>MM</td><td>target_seq</td><td>PAM</td><td>alignment</td><td>distance</td><td>position</td><td>gene name</td><td>gene id</td></tr>
<tr><td>GU305914.1</td><td>68621</td><td>68643</td><td>+</td><td>0</td><td>CTGGACGCGATCTCGAAGCT</td><td>GGG</td><td>||||||||[||||||||||||]PAM</td><td>NA</td><td>Intergenic</td><td>NA</td><td>NA</td></tr>
<tr><td>T4761</td><td>CGCGATCTCGAAGCTGGGCCTGG</td><td>1000</td><td></td><td></td><td></td><td></td><td></td><td></td><td></td><td></td><td></td></tr>
<tr><td>Oligo fwd</td><td>CACCCGCGATCTCGAAGCTGGGCC</td><td></td><td></td><td></td><td></td><td></td><td></td><td></td><td></td><td></td><td></td></tr>
<tr><td>Oligo rev</td><td>AAACGGCCCAGCTTCGAGATCGCG</td><td></td><td></td><td></td><td></td><td></td><td></td><td></td><td></td><td></td><td></td></tr>
<tr><td>Chromosome</td><td>start</td><td>end</td><td>strand</td><td>MM</td><td>target_seq</td><td>PAM</td><td>alignment</td><td>distance</td><td>position</td><td>gene name</td><td>gene id</td></tr>
<tr><td>GU305914.1</td><td>68626</td><td>68648</td><td>+</td><td>0</td><td>CGCGATCTCGAAGCTGGGCC</td><td>TGG</td><td>||||||||[||||||||||||]PAM</td><td>NA</td><td>Intergenic</td><td>NA</td><td>NA</td></tr>
<tr><td>T4762</td><td>GCTGGGCCTGGCGAGCAACGCGG</td><td>1000</td><td></td><td></td><td></td><td></td><td></td><td></td><td></td><td></td><td></td></tr>
<tr><td>Oligo fwd</td><td>CACCGCTGGGCCTGGCGAGCAACG</td><td></td><td></td><td></td><td></td><td></td><td></td><td></td><td></td><td></td><td></td></tr>
<tr><td>Oligo rev</td><td>AAACCGTTGCTCGCCAGGCCCAGC</td><td></td><td></td><td></td><td></td><td></td><td></td><td></td><td></td><td></td><td></td></tr>
<tr><td>Chromosome</td><td>start</td><td>end</td><td>strand</td><td>MM</td><td>target_seq</td><td>PAM</td><td>alignment</td><td>distance</td><td>position</td><td>gene name</td><td>gene id</td></tr>
<tr><td>GU305914.1</td><td>68638</td><td>68660</td><td>+</td><td>0</td><td>GCTGGGCCTGGCGAGCAACG</td><td>CGG</td><td>||||||||[||||||||||||]PAM</td><td>NA</td><td>Intergenic</td><td>NA</td><td>NA</td></tr>
<tr><td>T4763</td><td>GGGCCTGGCGAGCAACGCGGCGG</td><td>1000</td><td></td><td></td><td></td><td></td><td></td><td></td><td></td><td></td><td></td></tr>
<tr><td>Oligo fwd</td><td>CACCGGGCCTGGCGAGCAACGCGG</td><td></td><td></td><td></td><td></td><td></td><td></td><td></td><td></td><td></td><td></td></tr>
<tr><td>Oligo rev</td><td>AAACCCGCGTTGCTCGCCAGGCCC</td><td></td><td></td><td></td><td></td><td></td><td></td><td></td><td></td><td></td><td></td></tr>
<tr><td>Chromosome</td><td>start</td><td>end</td><td>strand</td><td>MM</td><td>target_seq</td><td>PAM</td><td>alignment</td><td>distance</td><td>position</td><td>gene name</td><td>gene id</td></tr>
<tr><td>GU305914.1</td><td>68641</td><td>68663</td><td>+</td><td>0</td><td>GGGCCTGGCGAGCAACGCGG</td><td>CGG</td><td>||||||||[||||||||||||]PAM</td><td>NA</td><td>Intergenic</td><td>NA</td><td>NA</td></tr>
<tr><td>T4764</td><td>CCTGGCGAGCAACGCGGCGGCGG</td><td>1000</td><td></td><td></td><td></td><td></td><td></td><td></td><td></td><td></td><td></td></tr>
<tr><td>Oligo fwd</td><td>CACCCCTGGCGAGCAACGCGGCGG</td><td></td><td></td><td></td><td></td><td></td><td></td><td></td><td></td><td></td><td></td></tr>
<tr><td>Oligo rev</td><td>AAACCCGCCGCGTTGCTCGCCAGG</td><td></td><td></td><td></td><td></td><td></td><td></td><td></td><td></td><td></td><td></td></tr>
<tr><td>Chromosome</td><td>start</td><td>end</td><td>strand</td><td>MM</td><td>target_seq</td><td>PAM</td><td>alignment</td><td>distance</td><td>position</td><td>gene name</td><td>gene id</td></tr>
<tr><td>GU305914.1</td><td>68644</td><td>68666</td><td>+</td><td>0</td><td>CCTGGCGAGCAACGCGGCGG</td><td>CGG</td><td>||||||||[||||||||||||]PAM</td><td>NA</td><td>Intergenic</td><td>NA</td><td>NA</td></tr>
<tr><td>T4765</td><td>GAGCGCCGACACGTCGCACCTGG</td><td>1000</td><td></td><td></td><td></td><td></td><td></td><td></td><td></td><td></td><td></td></tr>
<tr><td>Oligo fwd</td><td>CACCGAGCGCCGACACGTCGCACC</td><td></td><td></td><td></td><td></td><td></td><td></td><td></td><td></td><td></td><td></td></tr>
<tr><td>Oligo rev</td><td>AAACGGTGCGACGTGTCGGCGCTC</td><td></td><td></td><td></td><td></td><td></td><td></td><td></td><td></td><td></td><td></td></tr>
<tr><td>Chromosome</td><td>start</td><td>end</td><td>strand</td><td>MM</td><td>target_seq</td><td>PAM</td><td>alignment</td><td>distance</td><td>position</td><td>gene name</td><td>gene id</td></tr>
<tr><td>GU305914.1</td><td>68674</td><td>68696</td><td>+</td><td>0</td><td>GAGCGCCGACACGTCGCACC</td><td>TGG</td><td>||||||||[||||||||||||]PAM</td><td>NA</td><td>Intergenic</td><td>NA</td><td>NA</td></tr>
<tr><td>T4766</td><td>GCACCTGGACAGCGAGCAGCTGG</td><td>1000</td><td></td><td></td><td></td><td></td><td></td><td></td><td></td><td></td><td></td></tr>
<tr><td>Oligo fwd</td><td>CACCGCACCTGGACAGCGAGCAGC</td><td></td><td></td><td></td><td></td><td></td><td></td><td></td><td></td><td></td><td></td></tr>
<tr><td>Oligo rev</td><td>AAACGCTGCTCGCTGTCCAGGTGC</td><td></td><td></td><td></td><td></td><td></td><td></td><td></td><td></td><td></td><td></td></tr>
<tr><td>Chromosome</td><td>start</td><td>end</td><td>strand</td><td>MM</td><td>target_seq</td><td>PAM</td><td>alignment</td><td>distance</td><td>position</td><td>gene name</td><td>gene id</td></tr>
<tr><td>GU305914.1</td><td>68689</td><td>68711</td><td>+</td><td>0</td><td>GCACCTGGACAGCGAGCAGC</td><td>TGG</td><td>||||||||[||||||||||||]PAM</td><td>NA</td><td>Intergenic</td><td>NA</td><td>NA</td></tr>
<tr><td>T4767</td><td>CAGCGAGCAGCTGGCGTCGCTGG</td><td>1000</td><td></td><td></td><td></td><td></td><td></td><td></td><td></td><td></td><td></td></tr>
<tr><td>Oligo fwd</td><td>CACCCAGCGAGCAGCTGGCGTCGC</td><td></td><td></td><td></td><td></td><td></td><td></td><td></td><td></td><td></td><td></td></tr>
</table>

<table>
<tr><td>Oligo rev</td><td>AAACGCGACGCCAGCTGCTCGCTG</td><td></td><td></td><td></td><td></td><td></td><td></td><td></td><td></td><td></td><td></td></tr>
<tr><td>Chromosome</td><td>start</td><td>end</td><td>strand</td><td>MM</td><td>target_seq</td><td>PAM</td><td>alignment</td><td>distance</td><td>position</td><td>gene name</td><td>gene id</td></tr>
<tr><td>GU305914.1</td><td>68698</td><td>68720</td><td>+</td><td>0</td><td>CAGCGAGCAGCTGGCGTCGC</td><td>TGG</td><td>||||||||[||||||||||||]PAM</td><td>NA</td><td>Intergenic</td><td>NA</td><td>NA</td></tr>
<tr><td>GU305914.1</td><td>197469</td><td>197491</td><td>-</td><td>4</td><td>CCGCGGGGAGCTGGCGTCCC</td><td>TGG</td><td>|-|||-|-[||||||||||-|]PAM</td><td>NA</td><td>Intergenic</td><td>NA</td><td>NA</td></tr>
<tr><td></td><td></td><td></td><td></td><td></td><td></td><td></td><td></td><td></td><td></td><td></td><td></td></tr>
<tr><td>T4768</td><td>GCAGCTGGCGTCGCTGGCACAGG</td><td>1000</td><td></td><td></td><td></td><td></td><td></td><td></td><td></td><td></td><td></td></tr>
<tr><td>Oligo fwd</td><td>CACCGCAGCTGGCGTCGCTGGCAC</td><td></td><td></td><td></td><td></td><td></td><td></td><td></td><td></td><td></td><td></td></tr>
<tr><td>Oligo rev</td><td>AAACGTGCCAGCGACGCCAGCTGC</td><td></td><td></td><td></td><td></td><td></td><td></td><td></td><td></td><td></td><td></td></tr>
<tr><td>Chromosome</td><td>start</td><td>end</td><td>strand</td><td>MM</td><td>target_seq</td><td>PAM</td><td>alignment</td><td>distance</td><td>position</td><td>gene name</td><td>gene id</td></tr>
<tr><td>GU305914.1</td><td>68704</td><td>68726</td><td>+</td><td>0</td><td>GCAGCTGGCGTCGCTGGCAC</td><td>AGG</td><td>||||||||[||||||||||||]PAM</td><td>NA</td><td>Intergenic</td><td>NA</td><td>NA</td></tr>
<tr><td></td><td></td><td></td><td></td><td></td><td></td><td></td><td></td><td></td><td></td><td></td><td></td></tr>
<tr><td>T4769</td><td>GCTGGCGTCGCTGGCACAGGCGG</td><td>1000</td><td></td><td></td><td></td><td></td><td></td><td></td><td></td><td></td><td></td></tr>
<tr><td>Oligo fwd</td><td>CACCGCTGGCGTCGCTGGCACAGG</td><td></td><td></td><td></td><td></td><td></td><td></td><td></td><td></td><td></td><td></td></tr>
<tr><td>Oligo rev</td><td>AAACCCTGTGCCAGCGACGCCAGC</td><td></td><td></td><td></td><td></td><td></td><td></td><td></td><td></td><td></td><td></td></tr>
<tr><td>Chromosome</td><td>start</td><td>end</td><td>strand</td><td>MM</td><td>target_seq</td><td>PAM</td><td>alignment</td><td>distance</td><td>position</td><td>gene name</td><td>gene id</td></tr>
<tr><td>GU305914.1</td><td>68707</td><td>68729</td><td>+</td><td>0</td><td>GCTGGCGTCGCTGGCACAGG</td><td>CGG</td><td>||||||||[||||||||||||]PAM</td><td>NA</td><td>Intergenic</td><td>NA</td><td>NA</td></tr>
<tr><td></td><td></td><td></td><td></td><td></td><td></td><td></td><td></td><td></td><td></td><td></td><td></td></tr>
<tr><td>T4770</td><td>GTCGCTGGCACAGGCGGCCGAGG</td><td>1000</td><td></td><td></td><td></td><td></td><td></td><td></td><td></td><td></td><td></td></tr>
<tr><td>Oligo fwd</td><td>CACCGTCGCTGGCACAGGCGGCCG</td><td></td><td></td><td></td><td></td><td></td><td></td><td></td><td></td><td></td><td></td></tr>
<tr><td>Oligo rev</td><td>AAACCGGCCGCCTGTGCCAGCGAC</td><td></td><td></td><td></td><td></td><td></td><td></td><td></td><td></td><td></td><td></td></tr>
<tr><td>Chromosome</td><td>start</td><td>end</td><td>strand</td><td>MM</td><td>target_seq</td><td>PAM</td><td>alignment</td><td>distance</td><td>position</td><td>gene name</td><td>gene id</td></tr>
<tr><td>GU305914.1</td><td>68713</td><td>68735</td><td>+</td><td>0</td><td>GTCGCTGGCACAGGCGGCCG</td><td>AGG</td><td>||||||||[||||||||||||]PAM</td><td>NA</td><td>Intergenic</td><td>NA</td><td>NA</td></tr>
<tr><td></td><td></td><td></td><td></td><td></td><td></td><td></td><td></td><td></td><td></td><td></td><td></td></tr>
<tr><td>T4771</td><td>GGCCGAGGCGTACGCGCAGTCGG</td><td>1000</td><td></td><td></td><td></td><td></td><td></td><td></td><td></td><td></td><td></td></tr>
<tr><td>Oligo fwd</td><td>CACCGGCCGAGGCGTACGCGCAGT</td><td></td><td></td><td></td><td></td><td></td><td></td><td></td><td></td><td></td><td></td></tr>
<tr><td>Oligo rev</td><td>AAACACTGCGCGTACGCCTCGGCC</td><td></td><td></td><td></td><td></td><td></td><td></td><td></td><td></td><td></td><td></td></tr>
<tr><td>Chromosome</td><td>start</td><td>end</td><td>strand</td><td>MM</td><td>target_seq</td><td>PAM</td><td>alignment</td><td>distance</td><td>position</td><td>gene name</td><td>gene id</td></tr>
<tr><td>GU305914.1</td><td>68728</td><td>68750</td><td>+</td><td>0</td><td>GGCCGAGGCGTACGCGCAGT</td><td>CGG</td><td>||||||||[||||||||||||]PAM</td><td>NA</td><td>Intergenic</td><td>NA</td><td>NA</td></tr>
<tr><td></td><td></td><td></td><td></td><td></td><td></td><td></td><td></td><td></td><td></td><td></td><td></td></tr>
<tr><td>T4772</td><td>GTCGGCGTACGAGTCGTGCCAGG</td><td>1000</td><td></td><td></td><td></td><td></td><td></td><td></td><td></td><td></td><td></td></tr>
<tr><td>Oligo fwd</td><td>CACCGTCGGCGTACGAGTCGTGCC</td><td></td><td></td><td></td><td></td><td></td><td></td><td></td><td></td><td></td><td></td></tr>
<tr><td>Oligo rev</td><td>AAACGGCACGACTCGTACGCCGAC</td><td></td><td></td><td></td><td></td><td></td><td></td><td></td><td></td><td></td><td></td></tr>
<tr><td>Chromosome</td><td>start</td><td>end</td><td>strand</td><td>MM</td><td>target_seq</td><td>PAM</td><td>alignment</td><td>distance</td><td>position</td><td>gene name</td><td>gene id</td></tr>
<tr><td>GU305914.1</td><td>68746</td><td>68768</td><td>+</td><td>0</td><td>GTCGGCGTACGAGTCGTGCC</td><td>AGG</td><td>||||||||[||||||||||||]PAM</td><td>NA</td><td>Intergenic</td><td>NA</td><td>NA</td></tr>
<tr><td></td><td></td><td></td><td></td><td></td><td></td><td></td><td></td><td></td><td></td><td></td><td></td></tr>
<tr><td>T4773</td><td>GTCGTGCCAGGCGAAGATGACGG</td><td>1000</td><td></td><td></td><td></td><td></td><td></td><td></td><td></td><td></td><td></td></tr>
<tr><td>Oligo fwd</td><td>CACCGTCGTGCCAGGCGAAGATGA</td><td></td><td></td><td></td><td></td><td></td><td></td><td></td><td></td><td></td><td></td></tr>
<tr><td>Oligo rev</td><td>AAACTCATCTTCGCCTGGCACGAC</td><td></td><td></td><td></td><td></td><td></td><td></td><td></td><td></td><td></td><td></td></tr>
<tr><td>Chromosome</td><td>start</td><td>end</td><td>strand</td><td>MM</td><td>target_seq</td><td>PAM</td><td>alignment</td><td>distance</td><td>position</td><td>gene name</td><td>gene id</td></tr>
<tr><td>GU305914.1</td><td>68758</td><td>68780</td><td>+</td><td>0</td><td>GTCGTGCCAGGCGAAGATGA</td><td>CGG</td><td>||||||||[||||||||||||]PAM</td><td>NA</td><td>Intergenic</td><td>NA</td><td>NA</td></tr>
<tr><td></td><td></td><td></td><td></td><td></td><td></td><td></td><td></td><td></td><td></td><td></td><td></td></tr>
<tr><td>T4774</td><td>GAACCACAACCGCATCCTGACGG</td><td>1000</td><td></td><td></td><td></td><td></td><td></td><td></td><td></td><td></td><td></td></tr>
<tr><td>Oligo fwd</td><td>CACCGAACCACAACCGCATCCTGA</td><td></td><td></td><td></td><td></td><td></td><td></td><td></td><td></td><td></td><td></td></tr>
<tr><td>Oligo rev</td><td>AAACTCAGGATGCGGTTGTGGTTC</td><td></td><td></td><td></td><td></td><td></td><td></td><td></td><td></td><td></td><td></td></tr>
<tr><td>Chromosome</td><td>start</td><td>end</td><td>strand</td><td>MM</td><td>target_seq</td><td>PAM</td><td>alignment</td><td>distance</td><td>position</td><td>gene name</td><td>gene id</td></tr>
<tr><td>GU305914.1</td><td>68794</td><td>68816</td><td>+</td><td>0</td><td>GAACCACAACCGCATCCTGA</td><td>CGG</td><td>||||||||[||||||||||||]PAM</td><td>NA</td><td>Intergenic</td><td>NA</td><td>NA</td></tr>
<tr><td></td><td></td><td></td><td></td><td></td><td></td><td></td><td></td><td></td><td></td><td></td><td></td></tr>
<tr><td>T4775</td><td>AACCACAACCGCATCCTGACGGG</td><td>1000</td><td></td><td></td><td></td><td></td><td></td><td></td><td></td><td></td><td></td></tr>
<tr><td>Oligo fwd</td><td>CACCAACCACAACCGCATCCTGAC</td><td></td><td></td><td></td><td></td><td></td><td></td><td></td><td></td><td></td><td></td></tr>
<tr><td>Oligo rev</td><td>AAACGTCAGGATGCGGTTGTGGTT</td><td></td><td></td><td></td><td></td><td></td><td></td><td></td><td></td><td></td><td></td></tr>
<tr><td>Chromosome</td><td>start</td><td>end</td><td>strand</td><td>MM</td><td>target_seq</td><td>PAM</td><td>alignment</td><td>distance</td><td>position</td><td>gene name</td><td>gene id</td></tr>
<tr><td>GU305914.1</td><td>68795</td><td>68817</td><td>+</td><td>0</td><td>AACCACAACCGCATCCTGAC</td><td>GGG</td><td>||||||||[||||||||||||]PAM</td><td>NA</td><td>Intergenic</td><td>NA</td><td>NA</td></tr>
<tr><td></td><td></td><td></td><td></td><td></td><td></td><td></td><td></td><td></td><td></td><td></td><td></td></tr>
<tr><td>T4776</td><td>ACCACAACCGCATCCTGACGGGG</td><td>1000</td><td></td><td></td><td></td><td></td><td></td><td></td><td></td><td></td><td></td></tr>
<tr><td>Oligo fwd</td><td>CACCACCACAACCGCATCCTGACG</td><td></td><td></td><td></td><td></td><td></td><td></td><td></td><td></td><td></td><td></td></tr>
<tr><td>Oligo rev</td><td>AAACCGTCAGGATGCGGTTGTGGT</td><td></td><td></td><td></td><td></td><td></td><td></td><td></td><td></td><td></td><td></td></tr>
<tr><td>Chromosome</td><td>start</td><td>end</td><td>strand</td><td>MM</td><td>target_seq</td><td>PAM</td><td>alignment</td><td>distance</td><td>position</td><td>gene name</td><td>gene id</td></tr>
<tr><td>GU305914.1</td><td>68796</td><td>68818</td><td>+</td><td>0</td><td>ACCACAACCGCATCCTGACG</td><td>GGG</td><td>||||||||[||||||||||||]PAM</td><td>NA</td><td>Intergenic</td><td>NA</td><td>NA</td></tr>
<tr><td></td><td></td><td></td><td></td><td></td><td></td><td></td><td></td><td></td><td></td><td></td><td></td></tr>
<tr><td>T4777</td><td>CATCCTGACGGGGTCGCTGCCGG</td><td>1000</td><td></td><td></td><td></td><td></td><td></td><td></td><td></td><td></td><td></td></tr>
<tr><td>Oligo fwd</td><td>CACCCATCCTGACGGGGTCGCTGC</td><td></td><td></td><td></td><td></td><td></td><td></td><td></td><td></td><td></td><td></td></tr>
<tr><td>Oligo rev</td><td>AAACGCAGCGACCCCGTCAGGATG</td><td></td><td></td><td></td><td></td><td></td><td></td><td></td><td></td><td></td><td></td></tr>
<tr><td>Chromosome</td><td>start</td><td>end</td><td>strand</td><td>MM</td><td>target_seq</td><td>PAM</td><td>alignment</td><td>distance</td><td>position</td><td>gene name</td><td>gene id</td></tr>
<tr><td>GU305914.1</td><td>68806</td><td>68828</td><td>+</td><td>0</td><td>CATCCTGACGGGGTCGCTGC</td><td>CGG</td><td>||||||||[||||||||||||]PAM</td><td>NA</td><td>Intergenic</td><td>NA</td><td>NA</td></tr>
<tr><td></td><td></td><td></td><td></td><td></td><td></td><td></td><td></td><td></td><td></td><td></td><td></td></tr>
<tr><td>T4778</td><td>GACGGGGTCGCTGCCGGACTCGG</td><td>1000</td><td></td><td></td><td></td><td></td><td></td><td></td><td></td><td></td><td></td></tr>
<tr><td>Oligo fwd</td><td>CACCGACGGGGTCGCTGCCGGACT</td><td></td><td></td><td></td><td></td><td></td><td></td><td></td><td></td><td></td><td></td></tr>
<tr><td>Oligo rev</td><td>AAACAGTCCGGCAGCGACCCCGTC</td><td></td><td></td><td></td><td></td><td></td><td></td><td></td><td></td><td></td><td></td></tr>
<tr><td>Chromosome</td><td>start</td><td>end</td><td>strand</td><td>MM</td><td>target_seq</td><td>PAM</td><td>alignment</td><td>distance</td><td>position</td><td>gene name</td><td>gene id</td></tr>
<tr><td>GU305914.1</td><td>68812</td><td>68834</td><td>+</td><td>0</td><td>GACGGGGTCGCTGCCGGACT</td><td>CGG</td><td>||||||||[||||||||||||]PAM</td><td>NA</td><td>Intergenic</td><td>NA</td><td>NA</td></tr>
<tr><td></td><td></td><td></td><td></td><td></td><td></td><td></td><td></td><td></td><td></td><td></td><td></td></tr>
<tr><td>T4779</td><td>GCTGCCGGACTCGGACATCGCGG</td><td>1000</td><td></td><td></td><td></td><td></td><td></td><td></td><td></td><td></td><td></td></tr>
<tr><td>Oligo fwd</td><td>CACCGCTGCCGGACTCGGACATCG</td><td></td><td></td><td></td><td></td><td></td><td></td><td></td><td></td><td></td><td></td></tr>
<tr><td>Oligo rev</td><td>AAACCGATGTCCGAGTCCGGCAGC</td><td></td><td></td><td></td><td></td><td></td><td></td><td></td><td></td><td></td><td></td></tr>
<tr><td>Chromosome</td><td>start</td><td>end</td><td>strand</td><td>MM</td><td>target_seq</td><td>PAM</td><td>alignment</td><td>distance</td><td>position</td><td>gene name</td><td>gene id</td></tr>
<tr><td>GU305914.1</td><td>68821</td><td>68843</td><td>+</td><td>0</td><td>GCTGCCGGACTCGGACATCG</td><td>CGG</td><td>||||||||[||||||||||||]PAM</td><td>NA</td><td>Intergenic</td><td>NA</td><td>NA</td></tr>
<tr><td></td><td></td><td></td><td></td><td></td><td></td><td></td><td></td><td></td><td></td><td></td><td></td></tr>
<tr><td>T4780</td><td>GCCGGACTCGGACATCGCGGCGG</td><td>1000</td><td></td><td></td><td></td><td></td><td></td><td></td><td></td><td></td><td></td></tr>
<tr><td>Oligo fwd</td><td>CACCGCCGGACTCGGACATCGCGG</td><td></td><td></td><td></td><td></td><td></td><td></td><td></td><td></td><td></td><td></td></tr>
<tr><td>Oligo rev</td><td>AAACCCGCGATGTCCGAGTCCGGC</td><td></td><td></td><td></td><td></td><td></td><td></td><td></td><td></td><td></td><td></td></tr>
<tr><td>Chromosome</td><td>start</td><td>end</td><td>strand</td><td>MM</td><td>target_seq</td><td>PAM</td><td>alignment</td><td>distance</td><td>position</td><td>gene name</td><td>gene id</td></tr>
<tr><td>GU305914.1</td><td>68824</td><td>68846</td><td>+</td><td>0</td><td>GCCGGACTCGGACATCGCGG</td><td>CGG</td><td>||||||||[||||||||||||]PAM</td><td>NA</td><td>Intergenic</td><td>NA</td><td>NA</td></tr>
<tr><td>GU305914.1</td><td>68224</td><td>68246</td><td>+</td><td>4</td><td>GCTGGACGCGGTGATCGCGG</td><td>CGG</td><td>||-||||-[|||--|||||||]PAM</td><td>NA</td><td>Intergenic</td><td>NA</td><td>NA</td></tr>
<tr><td></td><td></td><td></td><td></td><td></td><td></td><td></td><td></td><td></td><td></td><td></td><td></td></tr>
<tr><td>T4781</td><td>CTCGGACATCGCGGCGGTGCTGG</td><td>1000</td><td></td><td></td><td></td><td></td><td></td><td></td><td></td><td></td><td></td></tr>
<tr><td>Oligo fwd</td><td>CACCCTCGGACATCGCGGCGGTGC</td><td></td><td></td><td></td><td></td><td></td><td></td><td></td><td></td><td></td><td></td></tr>
<tr><td>Oligo rev</td><td>AAACGCACCGCCGCGATGTCCGAG</td><td></td><td></td><td></td><td></td><td></td><td></td><td></td><td></td><td></td><td></td></tr>
<tr><td>Chromosome</td><td>start</td><td>end</td><td>strand</td><td>MM</td><td>target_seq</td><td>PAM</td><td>alignment</td><td>distance</td><td>position</td><td>gene name</td><td>gene id</td></tr>
<tr><td>GU305914.1</td><td>68830</td><td>68852</td><td>+</td><td>0</td><td>CTCGGACATCGCGGCGGTGC</td><td>TGG</td><td>||||||||[||||||||||||]PAM</td><td>NA</td><td>Intergenic</td><td>NA</td><td>NA</td></tr>
</table>

T4782 GGACATCGCGGCGGTGCTGGCGG 1000
Oligo fwd CACCGGACATCGCGGCGGTGCTGG
Oligo rev AAACCCAGCACCGCCGCGATGTCC
Chromosome start end strand MM target_seq PAM alignment distance position gene name gene id
GU305914.1 68833 68855 + 0 GGACATCGCGGCGGTGCTGG CGG ||||||||[||||||||||||||]PAM NA Intergenic NA NA

T4783 GGCGGTGCTGGCGGCCTTCAAGG 1000
Oligo fwd CACCGGCGGTGCTGGCGGCCTTCA
Oligo rev AAACTGAAGGCCGCCAGCACCGCC
Chromosome start end strand MM target_seq PAM alignment distance position gene name gene id
GU305914.1 68842 68864 + 0 GGCGGTGCTGGCGGCCTTCA AGG ||||||||[||||||||||||||]PAM NA Intergenic NA NA

T4784 GGCGGCCTTCAAGGCCGTGTCGG 1000
Oligo fwd CACCGGCGGCCTTCAAGGCCGTGT
Oligo rev AAACACACGGCCTTGAAGGCCGCC
Chromosome start end strand MM target_seq PAM alignment distance position gene name gene id
GU305914.1 68851 68873 + 0 GGCGGCCTTCAAGGCCGTGT CGG ||||||||[||||||||||||||]PAM NA Intergenic NA NA

T4785 AGGCCGTGTCGGAGAACGTGCGG 1000
Oligo fwd CACCAGGCCGTGTCGGAGAACGTG
Oligo rev AAACCACGTTCTCCGACACGGCCT
Chromosome start end strand MM target_seq PAM alignment distance position gene name gene id
GU305914.1 68862 68884 + 0 AGGCCGTGTCGGAGAACGTG CGG ||||||||[||||||||||||||]PAM NA Intergenic NA NA

T4786 GGCCGTGTCGGAGAACGTGCGGG 1000
Oligo fwd CACCGGCCGTGTCGGAGAACGTGC
Oligo rev AAACGCACGTTCTCCGACACGGCC
Chromosome start end strand MM target_seq PAM alignment distance position gene name gene id
GU305914.1 68863 68885 + 0 GGCCGTGTCGGAGAACGTGC GGG ||||||||[||||||||||||||]PAM NA Intergenic NA NA

T4787 GAACGTGCGGGCGCTGCGCGAGG 1000
Oligo fwd CACCGAACGTGCGGGCGCTGCGCG
Oligo rev AAACCGCGCAGCGCCCGCACGTTC
Chromosome start end strand MM target_seq PAM alignment distance position gene name gene id
GU305914.1 68875 68897 + 0 GAACGTGCGGGCGCTGCGCG AGG ||||||||[||||||||||||||]PAM NA Intergenic NA NA

T4788 GGTCGAGCTGCTGAAGTCGCAGG 1000
Oligo fwd CACCGGTCGAGCTGCTGAAGTCGC
Oligo rev AAACGCGACTTCAGCAGCTCGACC
Chromosome start end strand MM target_seq PAM alignment distance position gene name gene id
GU305914.1 68896 68918 + 0 GGTCGAGCTGCTGAAGTCGC AGG ||||||||[||||||||||||||]PAM NA Intergenic NA NA
GU305914.1 108389 108411 + 4 GGTCGGGGTGCGGACGTCGC AGG |||||-|-[|||-||-|||||]PAM NA Intergenic NA NA

T4789 GCAACTCGCTCAGCTGACCGAGG 1000
Oligo fwd CACCGCAACTCGCTCAGCTGACCG
Oligo rev AAACCGGTCAGCTGAGCGAGTTGC
Chromosome start end strand MM target_seq PAM alignment distance position gene name gene id
GU305914.1 68926 68948 + 0 GCAACTCGCTCAGCTGACCG AGG ||||||||[||||||||||||||]PAM NA Intergenic NA NA

T4790 GGACCTGCTGTACCTGCAGACGG 1000
Oligo fwd CACCGGACCTGCTGTACCTGCAGA
Oligo rev AAACTCTGCAGGTACAGCAGGTCC
Chromosome start end strand MM target_seq PAM alignment distance position gene name gene id
GU305914.1 68947 68969 + 0 GGACCTGCTGTACCTGCAGA CGG ||||||||[||||||||||||||]PAM NA Intergenic NA NA

T4791 GCTGTACCTGCAGACGGCCGAGG 1000
Oligo fwd CACCGCTGTACCTGCAGACGGCCG
Oligo rev AAACCGGCCGTCTGCAGGTACAGC
Chromosome start end strand MM target_seq PAM alignment distance position gene name gene id
GU305914.1 68953 68975 + 0 GCTGTACCTGCAGACGGCCG AGG ||||||||[||||||||||||||]PAM NA Intergenic NA NA

T4792 GCAGACGGCCGAGGTGAAGCTGG 1000
Oligo fwd CACCGCAGACGGCCGAGGTGAAGC
Oligo rev AAACGCTTCACCTCGGCCGTCTGC
Chromosome start end strand MM target_seq PAM alignment distance position gene name gene id
GU305914.1 68962 68984 + 0 GCAGACGGCCGAGGTGAAGC TGG ||||||||[||||||||||||||]PAM NA Intergenic NA NA

T4793 GACGGCCGAGGTGAAGCTGGAGG 1000
Oligo fwd CACCGACGGCCGAGGTGAAGCTGG
Oligo rev AAACCCAGCTTCACCTCGGCCGTC
Chromosome start end strand MM target_seq PAM alignment distance position gene name gene id
GU305914.1 68965 68987 + 0 GACGGCCGAGGTGAAGCTGG AGG ||||||||[||||||||||||||]PAM NA Intergenic NA NA

T4794 GCTGGAGGTGACGCCGAGCGCGG 1000
Oligo fwd CACCGCTGGAGGTGACGCCGAGCG
Oligo rev AAACCGCTCGGCGTCACCTCCAGC
Chromosome start end strand MM target_seq PAM alignment distance position gene name gene id
GU305914.1 68980 69002 + 0 GCTGGAGGTGACGCCGAGCG CGG ||||||||[||||||||||||||]PAM NA Intergenic NA NA

T4795 GGAGATCAAGAAGTTGCGCGCGG 1000
Oligo fwd CACCGGAGATCAAGAAGTTGCGCG
Oligo rev AAACCGCGCAACTTCTTGATCTCC
Chromosome start end strand MM target_seq PAM alignment distance position gene name gene id
GU305914.1 69001 69023 + 0 GGAGATCAAGAAGTTGCGCG CGG ||||||||[||||||||||||||]PAM NA Intergenic NA NA

T4796 GTTGCGCGCGGAGTACGAGACGG 1000
Oligo fwd CACCGTTGCGCGCGGAGTACGAGA
Oligo rev AAACTCTCGTACTCCGCGCGCAAC

<table>
<tr><td>Chromosome</td><td>start</td><td>end</td><td>strand</td><td>MM</td><td>target_seq</td><td>PAM</td><td>alignment</td><td>distance</td><td>position</td><td>gene name</td><td>gene id</td></tr>
<tr><td>GU305914.1</td><td>69013</td><td>69035</td><td>+</td><td>0</td><td>GTTGCGCGCGGAGTACGAGA</td><td>CGG</td><td>||||||||[||||||||||||]PAM</td><td>NA</td><td>Intergenic</td><td>NA</td><td>NA</td></tr>
<tr><td>T4797</td><td>GACGGCGCGCAAGCAGATCTCGG</td><td>1000</td><td></td><td></td><td></td><td></td><td></td><td></td><td></td><td></td><td></td></tr>
<tr><td>Oligo fwd</td><td>CACCGACGGCGCGCAAGCAGATCT</td><td></td><td></td><td></td><td></td><td></td><td></td><td></td><td></td><td></td><td></td></tr>
<tr><td>Oligo rev</td><td>AAACAGATCTGCTTGCGCGCCGTC</td><td></td><td></td><td></td><td></td><td></td><td></td><td></td><td></td><td></td><td></td></tr>
<tr><td>Chromosome</td><td>start</td><td>end</td><td>strand</td><td>MM</td><td>target_seq</td><td>PAM</td><td>alignment</td><td>distance</td><td>position</td><td>gene name</td><td>gene id</td></tr>
<tr><td>GU305914.1</td><td>69031</td><td>69053</td><td>+</td><td>0</td><td>GACGGCGCGCAAGCAGATCT</td><td>CGG</td><td>||||||||[||||||||||||]PAM</td><td>NA</td><td>Intergenic</td><td>NA</td><td>NA</td></tr>
<tr><td>T4798</td><td>GCGCAAGCAGATCTCGGACGAGG</td><td>1000</td><td></td><td></td><td></td><td></td><td></td><td></td><td></td><td></td><td></td></tr>
<tr><td>Oligo fwd</td><td>CACCGCGCAAGCAGATCTCGGACG</td><td></td><td></td><td></td><td></td><td></td><td></td><td></td><td></td><td></td><td></td></tr>
<tr><td>Oligo rev</td><td>AAACCGTCCGAGATCTGCTTGCGC</td><td></td><td></td><td></td><td></td><td></td><td></td><td></td><td></td><td></td><td></td></tr>
<tr><td>Chromosome</td><td>start</td><td>end</td><td>strand</td><td>MM</td><td>target_seq</td><td>PAM</td><td>alignment</td><td>distance</td><td>position</td><td>gene name</td><td>gene id</td></tr>
<tr><td>GU305914.1</td><td>69037</td><td>69059</td><td>+</td><td>0</td><td>GCGCAAGCAGATCTCGGACG</td><td>AGG</td><td>||||||||[||||||||||||]PAM</td><td>NA</td><td>Intergenic</td><td>NA</td><td>NA</td></tr>
<tr><td>GU305914.1</td><td>106735</td><td>106757</td><td>-</td><td>3</td><td>GCGCTCGCAGATGTCGGACG</td><td>AGG</td><td>||||--||[||||-|||||||]PAM</td><td>NA</td><td>Intergenic</td><td>NA</td><td>NA</td></tr>
<tr><td>T4799</td><td>AGATCTCGGACGAGGAGATGCGG</td><td>1000</td><td></td><td></td><td></td><td></td><td></td><td></td><td></td><td></td><td></td></tr>
<tr><td>Oligo fwd</td><td>CACCAGATCTCGGACGAGGAGATG</td><td></td><td></td><td></td><td></td><td></td><td></td><td></td><td></td><td></td><td></td></tr>
<tr><td>Oligo rev</td><td>AAACCATCTCCTCGTCCGAGATCT</td><td></td><td></td><td></td><td></td><td></td><td></td><td></td><td></td><td></td><td></td></tr>
<tr><td>Chromosome</td><td>start</td><td>end</td><td>strand</td><td>MM</td><td>target_seq</td><td>PAM</td><td>alignment</td><td>distance</td><td>position</td><td>gene name</td><td>gene id</td></tr>
<tr><td>GU305914.1</td><td>69045</td><td>69067</td><td>+</td><td>0</td><td>AGATCTCGGACGAGGAGATG</td><td>CGG</td><td>||||||||[||||||||||||]PAM</td><td>NA</td><td>Intergenic</td><td>NA</td><td>NA</td></tr>
<tr><td>T4800</td><td>GGACGAGGAGATGCGGATCAAGG</td><td>1000</td><td></td><td></td><td></td><td></td><td></td><td></td><td></td><td></td><td></td></tr>
<tr><td>Oligo fwd</td><td>CACCGGACGAGGAGATGCGGATCA</td><td></td><td></td><td></td><td></td><td></td><td></td><td></td><td></td><td></td><td></td></tr>
<tr><td>Oligo rev</td><td>AAACTGATCCGCATCTCCTCGTCC</td><td></td><td></td><td></td><td></td><td></td><td></td><td></td><td></td><td></td><td></td></tr>
<tr><td>Chromosome</td><td>start</td><td>end</td><td>strand</td><td>MM</td><td>target_seq</td><td>PAM</td><td>alignment</td><td>distance</td><td>position</td><td>gene name</td><td>gene id</td></tr>
<tr><td>GU305914.1</td><td>69052</td><td>69074</td><td>+</td><td>0</td><td>GGACGAGGAGATGCGGATCA</td><td>AGG</td><td>||||||||[||||||||||||]PAM</td><td>NA</td><td>Intergenic</td><td>NA</td><td>NA</td></tr>
<tr><td>GU305914.1</td><td>51005</td><td>51027</td><td>+</td><td>4</td><td>CGGCGAGGAGATGAGGGTCA</td><td>CGG</td><td>-|-|||||[|||||-||-|||]PAM</td><td>NA</td><td>Intergenic</td><td>NA</td><td>NA</td></tr>
<tr><td>T4801</td><td>CAAGGAACTGATCGAGAACATGG</td><td>1000</td><td></td><td></td><td></td><td></td><td></td><td></td><td></td><td></td><td></td></tr>
<tr><td>Oligo fwd</td><td>CACCCAAGGAACTGATCGAGAACA</td><td></td><td></td><td></td><td></td><td></td><td></td><td></td><td></td><td></td><td></td></tr>
<tr><td>Oligo rev</td><td>AAACTGTTCTCGATCAGTTCCTTG</td><td></td><td></td><td></td><td></td><td></td><td></td><td></td><td></td><td></td><td></td></tr>
<tr><td>Chromosome</td><td>start</td><td>end</td><td>strand</td><td>MM</td><td>target_seq</td><td>PAM</td><td>alignment</td><td>distance</td><td>position</td><td>gene name</td><td>gene id</td></tr>
<tr><td>GU305914.1</td><td>69070</td><td>69092</td><td>+</td><td>0</td><td>CAAGGAACTGATCGAGAACA</td><td>TGG</td><td>||||||||[||||||||||||]PAM</td><td>NA</td><td>Intergenic</td><td>NA</td><td>NA</td></tr>
<tr><td>T4802</td><td>GGAACTGATCGAGAACATGGAGG</td><td>1000</td><td></td><td></td><td></td><td></td><td></td><td></td><td></td><td></td><td></td></tr>
<tr><td>Oligo fwd</td><td>CACCGGAACTGATCGAGAACATGG</td><td></td><td></td><td></td><td></td><td></td><td></td><td></td><td></td><td></td><td></td></tr>
<tr><td>Oligo rev</td><td>AAACCCATGTTCTCGATCAGTTCC</td><td></td><td></td><td></td><td></td><td></td><td></td><td></td><td></td><td></td><td></td></tr>
<tr><td>Chromosome</td><td>start</td><td>end</td><td>strand</td><td>MM</td><td>target_seq</td><td>PAM</td><td>alignment</td><td>distance</td><td>position</td><td>gene name</td><td>gene id</td></tr>
<tr><td>GU305914.1</td><td>69073</td><td>69095</td><td>+</td><td>0</td><td>GGAACTGATCGAGAACATGG</td><td>AGG</td><td>||||||||[||||||||||||]PAM</td><td>NA</td><td>Intergenic</td><td>NA</td><td>NA</td></tr>
<tr><td>T4803</td><td>GAACATGGAGGACATGATCACGG</td><td>1000</td><td></td><td></td><td></td><td></td><td></td><td></td><td></td><td></td><td></td></tr>
<tr><td>Oligo fwd</td><td>CACCGAACATGGAGGACATGATCA</td><td></td><td></td><td></td><td></td><td></td><td></td><td></td><td></td><td></td><td></td></tr>
<tr><td>Oligo rev</td><td>AAACTGATCATGTCCTCCATGTTC</td><td></td><td></td><td></td><td></td><td></td><td></td><td></td><td></td><td></td><td></td></tr>
<tr><td>Chromosome</td><td>start</td><td>end</td><td>strand</td><td>MM</td><td>target_seq</td><td>PAM</td><td>alignment</td><td>distance</td><td>position</td><td>gene name</td><td>gene id</td></tr>
<tr><td>GU305914.1</td><td>69085</td><td>69107</td><td>+</td><td>0</td><td>GAACATGGAGGACATGATCA</td><td>CGG</td><td>||||||||[||||||||||||]PAM</td><td>NA</td><td>Intergenic</td><td>NA</td><td>NA</td></tr>
<tr><td>T4804</td><td>CAGCAGCAGCCCGCCGCCGCCGG</td><td>1000</td><td></td><td></td><td></td><td></td><td></td><td></td><td></td><td></td><td></td></tr>
<tr><td>Oligo fwd</td><td>CACCCAGCAGCAGCCCGCCGCCGC</td><td></td><td></td><td></td><td></td><td></td><td></td><td></td><td></td><td></td><td></td></tr>
<tr><td>Oligo rev</td><td>AAACGCGGCGGCGGGCTGCTGCTG</td><td></td><td></td><td></td><td></td><td></td><td></td><td></td><td></td><td></td><td></td></tr>
<tr><td>Chromosome</td><td>start</td><td>end</td><td>strand</td><td>MM</td><td>target_seq</td><td>PAM</td><td>alignment</td><td>distance</td><td>position</td><td>gene name</td><td>gene id</td></tr>
<tr><td>GU305914.1</td><td>69112</td><td>69134</td><td>+</td><td>0</td><td>CAGCAGCAGCCCGCCGCCGC</td><td>CGG</td><td>||||||||[||||||||||||]PAM</td><td>NA</td><td>Intergenic</td><td>NA</td><td>NA</td></tr>
<tr><td>T4805</td><td>CCCGCCGCCGCCGGAGATGCTGG</td><td>1000</td><td></td><td></td><td></td><td></td><td></td><td></td><td></td><td></td><td></td></tr>
<tr><td>Oligo fwd</td><td>CACCCCCGCCGCCGCCGGAGATGC</td><td></td><td></td><td></td><td></td><td></td><td></td><td></td><td></td><td></td><td></td></tr>
<tr><td>Oligo rev</td><td>AAACGCATCTCCGGCGGCGGCGGG</td><td></td><td></td><td></td><td></td><td></td><td></td><td></td><td></td><td></td><td></td></tr>
<tr><td>Chromosome</td><td>start</td><td>end</td><td>strand</td><td>MM</td><td>target_seq</td><td>PAM</td><td>alignment</td><td>distance</td><td>position</td><td>gene name</td><td>gene id</td></tr>
<tr><td>GU305914.1</td><td>69121</td><td>69143</td><td>+</td><td>0</td><td>CCCGCCGCCGCCGGAGATGC</td><td>TGG</td><td>||||||||[||||||||||||]PAM</td><td>NA</td><td>Intergenic</td><td>NA</td><td>NA</td></tr>
<tr><td>GU305914.1</td><td>70188</td><td>70210</td><td>-</td><td>4</td><td>GGCGCCGCCGCCGGCGCTGC</td><td>CGG</td><td>--||||||[||||||-|-|||]PAM</td><td>NA</td><td>Intergenic</td><td>NA</td><td>NA</td></tr>
<tr><td>T4806</td><td>TATGTTACGCACGCAGATCGAGG</td><td>1000</td><td></td><td></td><td></td><td></td><td></td><td></td><td></td><td></td><td></td></tr>
<tr><td>Oligo fwd</td><td>CACCTATGTTACGCACGCAGATCG</td><td></td><td></td><td></td><td></td><td></td><td></td><td></td><td></td><td></td><td></td></tr>
<tr><td>Oligo rev</td><td>AAACCGATCTGCGTGCGTAACATA</td><td></td><td></td><td></td><td></td><td></td><td></td><td></td><td></td><td></td><td></td></tr>
<tr><td>Chromosome</td><td>start</td><td>end</td><td>strand</td><td>MM</td><td>target_seq</td><td>PAM</td><td>alignment</td><td>distance</td><td>position</td><td>gene name</td><td>gene id</td></tr>
<tr><td>GU305914.1</td><td>69145</td><td>69167</td><td>+</td><td>0</td><td>TATGTTACGCACGCAGATCG</td><td>AGG</td><td>||||||||[||||||||||||]PAM</td><td>NA</td><td>Intergenic</td><td>NA</td><td>NA</td></tr>
<tr><td>T4807</td><td>GTTACGCACGCAGATCGAGGAGG</td><td>1000</td><td></td><td></td><td></td><td></td><td></td><td></td><td></td><td></td><td></td></tr>
<tr><td>Oligo fwd</td><td>CACCGTTACGCACGCAGATCGAGG</td><td></td><td></td><td></td><td></td><td></td><td></td><td></td><td></td><td></td><td></td></tr>
<tr><td>Oligo rev</td><td>AAACCCTCGATCTGCGTGCGTAAC</td><td></td><td></td><td></td><td></td><td></td><td></td><td></td><td></td><td></td><td></td></tr>
<tr><td>Chromosome</td><td>start</td><td>end</td><td>strand</td><td>MM</td><td>target_seq</td><td>PAM</td><td>alignment</td><td>distance</td><td>position</td><td>gene name</td><td>gene id</td></tr>
<tr><td>GU305914.1</td><td>69148</td><td>69170</td><td>+</td><td>0</td><td>GTTACGCACGCAGATCGAGG</td><td>AGG</td><td>||||||||[||||||||||||]PAM</td><td>NA</td><td>Intergenic</td><td>NA</td><td>NA</td></tr>
<tr><td>T4808</td><td>ACGCACGCAGATCGAGGAGGTGG</td><td>1000</td><td></td><td></td><td></td><td></td><td></td><td></td><td></td><td></td><td></td></tr>
<tr><td>Oligo fwd</td><td>CACCACGCACGCAGATCGAGGAGG</td><td></td><td></td><td></td><td></td><td></td><td></td><td></td><td></td><td></td><td></td></tr>
<tr><td>Oligo rev</td><td>AAACCCTCCTCGATCTGCGTGCGT</td><td></td><td></td><td></td><td></td><td></td><td></td><td></td><td></td><td></td><td></td></tr>
<tr><td>Chromosome</td><td>start</td><td>end</td><td>strand</td><td>MM</td><td>target_seq</td><td>PAM</td><td>alignment</td><td>distance</td><td>position</td><td>gene name</td><td>gene id</td></tr>
<tr><td>GU305914.1</td><td>69151</td><td>69173</td><td>+</td><td>0</td><td>ACGCACGCAGATCGAGGAGG</td><td>TGG</td><td>||||||||[||||||||||||]PAM</td><td>NA</td><td>Intergenic</td><td>NA</td><td>NA</td></tr>
<tr><td>T4809</td><td>CGAGGAGGTGGAGAGCATGACGG</td><td>1000</td><td></td><td></td><td></td><td></td><td></td><td></td><td></td><td></td><td></td></tr>
<tr><td>Oligo fwd</td><td>CACCCGAGGAGGTGGAGAGCATGA</td><td></td><td></td><td></td><td></td><td></td><td></td><td></td><td></td><td></td><td></td></tr>
<tr><td>Oligo rev</td><td>AAACTCATGCTCTCCACCTCCTCG</td><td></td><td></td><td></td><td></td><td></td><td></td><td></td><td></td><td></td><td></td></tr>
<tr><td>Chromosome</td><td>start</td><td>end</td><td>strand</td><td>MM</td><td>target_seq</td><td>PAM</td><td>alignment</td><td>distance</td><td>position</td><td>gene name</td><td>gene id</td></tr>
<tr><td>GU305914.1</td><td>69163</td><td>69185</td><td>+</td><td>0</td><td>CGAGGAGGTGGAGAGCATGA</td><td>CGG</td><td>||||||||[||||||||||||]PAM</td><td>NA</td><td>Intergenic</td><td>NA</td><td>NA</td></tr>
<tr><td>GU305914.1</td><td>28747</td><td>28769</td><td>-</td><td>4</td><td>GGGGGAGGAGGAGAGGATGA</td><td>TGG</td><td>-|-|||||[-||||||-||||]PAM</td><td>NA</td><td>Intergenic</td><td>NA</td><td>NA</td></tr>
<tr><td>T4810</td><td>GGAGGTGGAGAGCATGACGGTGG</td><td>1000</td><td></td><td></td><td></td><td></td><td></td><td></td><td></td><td></td><td></td></tr>
<tr><td>Oligo fwd</td><td>CACCGGAGGTGGAGAGCATGACGG</td><td></td><td></td><td></td><td></td><td></td><td></td><td></td><td></td><td></td><td></td></tr>
<tr><td>Oligo rev</td><td>AAACCCGTCATGCTCTCCACCTCC</td><td></td><td></td><td></td><td></td><td></td><td></td><td></td><td></td><td></td><td></td></tr>
<tr><td>Chromosome</td><td>start</td><td>end</td><td>strand</td><td>MM</td><td>target_seq</td><td>PAM</td><td>alignment</td><td>distance</td><td>position</td><td>gene name</td><td>gene id</td></tr>
</table>

| | | | | | | | | | | | |
|---|---|---|---|---|---|---|---|---|---|---|---|
| GU305914.1 | 69166 | 69188 | + | 0 | GGAGGTGGAGAGCATGACGG | TGG | \|\|\|\|\|\|\|\|[\|\|\|\|\|\|\|\|\|\|\|\|]PAM | NA | Intergenic | NA | NA |
| GU305914.1 | 28744 | 28766 | - | 3 | GGAGGAGGAGAGGATGATGG | TGG | \|\|\|\|\|-\|\|[\|\|\|\|-\|\|\|\|-\|\|]PAM | NA | Intergenic | NA | NA |
| T4811 | GAGCATGACGGTGGACGAGCAGG | 1000 | | | | | | | | | |
| Oligo fwd | CACCGAGCATGACGGTGGACGAGC | | | | | | | | | | |
| Oligo rev | AAACGCTCGTCCACCGTCATGCTC | | | | | | | | | | |
| Chromosome | start | end | strand | MM | target_seq | PAM | alignment | distance | position | gene name | gene id |
| GU305914.1 | 69175 | 69197 | + | 0 | GAGCATGACGGTGGACGAGC | AGG | \|\|\|\|\|\|\|\|[\|\|\|\|\|\|\|\|\|\|\|\|]PAM | NA | Intergenic | NA | NA |
| T4812 | CGGTGGACGAGCAGGACGCGCGG | 1000 | | | | | | | | | |
| Oligo fwd | CACCCGGTGGACGAGCAGGACGCG | | | | | | | | | | |
| Oligo rev | AAACCGCGTCCTGCTCGTCCACCG | | | | | | | | | | |
| Chromosome | start | end | strand | MM | target_seq | PAM | alignment | distance | position | gene name | gene id |
| GU305914.1 | 69183 | 69205 | + | 0 | CGGTGGACGAGCAGGACGCG | CGG | \|\|\|\|\|\|\|\|[\|\|\|\|\|\|\|\|\|\|\|\|]PAM | NA | Intergenic | NA | NA |
| T4813 | GGACGCGCGGCGCGCCGACAAGG | 1000 | | | | | | | | | |
| Oligo fwd | CACCGGACGCGCGGCGCGCCGACA | | | | | | | | | | |
| Oligo rev | AAACTGTCGGCGCGCCGCGCGTCC | | | | | | | | | | |
| Chromosome | start | end | strand | MM | target_seq | PAM | alignment | distance | position | gene name | gene id |
| GU305914.1 | 69196 | 69218 | + | 0 | GGACGCGCGGCGCGCCGACA | AGG | \|\|\|\|\|\|\|\|[\|\|\|\|\|\|\|\|\|\|\|\|]PAM | NA | Intergenic | NA | NA |
| GU305914.1 | 201365 | 201387 | - | 4 | GGAGGCGGGGCGCTACGACA | CGG | \|\|\|-\|\|\|-[\|\|\|\|\|--\|\|\|\|\|]PAM | NA | Intergenic | NA | NA |
| T4814 | GCGGCGCGCCGACAAGGTGCTGG | 1000 | | | | | | | | | |
| Oligo fwd | CACCGCGGCGCGCCGACAAGGTGC | | | | | | | | | | |
| Oligo rev | AAACGCACCTTGTCGGCGCGCCGC | | | | | | | | | | |
| Chromosome | start | end | strand | MM | target_seq | PAM | alignment | distance | position | gene name | gene id |
| GU305914.1 | 69202 | 69224 | + | 0 | GCGGCGCGCCGACAAGGTGC | TGG | \|\|\|\|\|\|\|\|[\|\|\|\|\|\|\|\|\|\|\|\|]PAM | NA | Intergenic | NA | NA |
| GU305914.1 | 197552 | 197574 | + | 4 | GTGGCGTGCCGATAGGGTGC | CGG | \|-\|\|\|\|-\|[\|\|\|\|-\|-\|\|\|\|\|]PAM | NA | Intergenic | NA | NA |
| T4815 | CGGCGCGCCGACAAGGTGCTGGG | 1000 | | | | | | | | | |
| Oligo fwd | CACCCGGCGCGCCGACAAGGTGCT | | | | | | | | | | |
| Oligo rev | AAACAGCACCTTGTCGGCGCGCCG | | | | | | | | | | |
| Chromosome | start | end | strand | MM | target_seq | PAM | alignment | distance | position | gene name | gene id |
| GU305914.1 | 69203 | 69225 | + | 0 | CGGCGCGCCGACAAGGTGCT | GGG | \|\|\|\|\|\|\|\|[\|\|\|\|\|\|\|\|\|\|\|\|]PAM | NA | Intergenic | NA | NA |
| T4816 | GGCGCGCCGACAAGGTGCTGGGG | 1000 | | | | | | | | | |
| Oligo fwd | CACCGGCGCGCCGACAAGGTGCTG | | | | | | | | | | |
| Oligo rev | AAACCAGCACCTTGTCGGCGCGCC | | | | | | | | | | |
| Chromosome | start | end | strand | MM | target_seq | PAM | alignment | distance | position | gene name | gene id |
| GU305914.1 | 69204 | 69226 | + | 0 | GGCGCGCCGACAAGGTGCTG | GGG | \|\|\|\|\|\|\|\|[\|\|\|\|\|\|\|\|\|\|\|\|]PAM | NA | Intergenic | NA | NA |
| T4817 | GCTGGGGAAGCTGTCGACGCTGG | 1000 | | | | | | | | | |
| Oligo fwd | CACCGCTGGGGAAGCTGTCGACGC | | | | | | | | | | |
| Oligo rev | AAACGCGTCGACAGCTTCCCCAGC | | | | | | | | | | |
| Chromosome | start | end | strand | MM | target_seq | PAM | alignment | distance | position | gene name | gene id |
| GU305914.1 | 69220 | 69242 | + | 0 | GCTGGGGAAGCTGTCGACGC | TGG | \|\|\|\|\|\|\|\|[\|\|\|\|\|\|\|\|\|\|\|\|]PAM | NA | Intergenic | NA | NA |
| T4818 | GAAGCTGTCGACGCTGGACGAGG | 1000 | | | | | | | | | |
| Oligo fwd | CACCGAAGCTGTCGACGCTGGACG | | | | | | | | | | |
| Oligo rev | AAACCGTCCAGCGTCGACAGCTTC | | | | | | | | | | |
| Chromosome | start | end | strand | MM | target_seq | PAM | alignment | distance | position | gene name | gene id |
| GU305914.1 | 69226 | 69248 | + | 0 | GAAGCTGTCGACGCTGGACG | AGG | \|\|\|\|\|\|\|\|[\|\|\|\|\|\|\|\|\|\|\|\|]PAM | NA | Intergenic | NA | NA |
| T4819 | GCTGTCGACGCTGGACGAGGCGG | 1000 | | | | | | | | | |
| Oligo fwd | CACCGCTGTCGACGCTGGACGAGG | | | | | | | | | | |
| Oligo rev | AAACCCTCGTCCAGCGTCGACAGC | | | | | | | | | | |
| Chromosome | start | end | strand | MM | target_seq | PAM | alignment | distance | position | gene name | gene id |
| GU305914.1 | 69229 | 69251 | + | 0 | GCTGTCGACGCTGGACGAGG | CGG | \|\|\|\|\|\|\|\|[\|\|\|\|\|\|\|\|\|\|\|\|]PAM | NA | Intergenic | NA | NA |
| GU305914.1 | 87687 | 87709 | - | 4 | CCTGGCGACGCTGAACGACG | TGG | -\|\|\|-\|\|\|[\|\|\|\|\|-\|\|\|\|-\|]PAM | NA | Intergenic | NA | NA |
| T4820 | GTCGACGCTGGACGAGGCGGAGG | 1000 | | | | | | | | | |
| Oligo fwd | CACCGTCGACGCTGGACGAGGCGG | | | | | | | | | | |
| Oligo rev | AAACCCGCCTCGTCCAGCGTCGAC | | | | | | | | | | |
| Chromosome | start | end | strand | MM | target_seq | PAM | alignment | distance | position | gene name | gene id |
| GU305914.1 | 69232 | 69254 | + | 0 | GTCGACGCTGGACGAGGCGG | AGG | \|\|\|\|\|\|\|\|[\|\|\|\|\|\|\|\|\|\|\|\|]PAM | NA | Intergenic | NA | NA |
| GU305914.1 | 77074 | 77096 | + | 4 | GACGATGCTGGAGGAGGAGG | AGG | \|-\|\|\|-\|\|[\|\|\|\|-\|\|\|\|-\|\|]PAM | NA | Intergenic | NA | NA |
| T4821 | GCTGGACGAGGCGGAGGCGAAGG | 1000 | | | | | | | | | |
| Oligo fwd | CACCGCTGGACGAGGCGGAGGCGA | | | | | | | | | | |
| Oligo rev | AAACTCGCCTCCGCCTCGTCCAGC | | | | | | | | | | |
| Chromosome | start | end | strand | MM | target_seq | PAM | alignment | distance | position | gene name | gene id |
| GU305914.1 | 69238 | 69260 | + | 0 | GCTGGACGAGGCGGAGGCGA | AGG | \|\|\|\|\|\|\|\|[\|\|\|\|\|\|\|\|\|\|\|\|]PAM | NA | Intergenic | NA | NA |
| T4822 | CGAGGCGGAGGCGAAGGCGACGG | 1000 | | | | | | | | | |
| Oligo fwd | CACCCGAGGCGGAGGCGAAGGCGA | | | | | | | | | | |
| Oligo rev | AAACTCGCCTTCGCCTCCGCCTCG | | | | | | | | | | |
| Chromosome | start | end | strand | MM | target_seq | PAM | alignment | distance | position | gene name | gene id |
| GU305914.1 | 69244 | 69266 | + | 0 | CGAGGCGGAGGCGAAGGCGA | CGG | \|\|\|\|\|\|\|\|[\|\|\|\|\|\|\|\|\|\|\|\|]PAM | NA | Intergenic | NA | NA |
| GU305914.1 | 111940 | 111962 | - | 4 | CACGTCGCAGGCGAAGGCGA | CGG | \|--\|-\|\|-[\|\|\|\|\|\|\|\|\|\|\|\|]PAM | NA | Intergenic | NA | NA |
| T4823 | GTTCGCACAGTCGCTGTCGACGG | 1000 | | | | | | | | | |
| Oligo fwd | CACCGTTCGCACAGTCGCTGTCGA | | | | | | | | | | |
| Oligo rev | AAACTCGACAGCGACTGTGCGAAC | | | | | | | | | | |
| Chromosome | start | end | strand | MM | target_seq | PAM | alignment | distance | position | gene name | gene id |
| GU305914.1 | 69268 | 69290 | + | 0 | GTTCGCACAGTCGCTGTCGA | CGG | \|\|\|\|\|\|\|\|[\|\|\|\|\|\|\|\|\|\|\|\|]PAM | NA | Intergenic | NA | NA |
| T4824 | CCCGTCGCTGAACGAGATCAAGG | 1000 | | | | | | | | | |
| Oligo fwd | CACCCCCGTCGCTGAACGAGATCA | | | | | | | | | | |
| Oligo rev | AAACTGATCTCGTTCAGCGACGGG | | | | | | | | | | |

<table>
<tr><td>Chromosome</td><td>start</td><td>end</td><td>strand</td><td>MM</td><td>target_seq</td><td>PAM</td><td>alignment</td><td>distance</td><td>position</td><td>gene name</td><td>gene id</td></tr>
<tr><td>GU305914.1</td><td>69298</td><td>69320</td><td>+</td><td>0</td><td>CCCGTCGCTGAACGAGATCA</td><td>AGG</td><td>||||||||[||||||||||||]PAM</td><td>NA</td><td>Intergenic</td><td>NA</td><td>NA</td></tr>
<tr><td>T4825</td><td>CCGTCGCTGAACGAGATCAAGGG</td><td>1000</td><td></td><td></td><td></td><td></td><td></td><td></td><td></td><td></td><td></td></tr>
<tr><td>Oligo fwd</td><td>CACCCCGTCGCTGAACGAGATCAA</td><td></td><td></td><td></td><td></td><td></td><td></td><td></td><td></td><td></td><td></td></tr>
<tr><td>Oligo rev</td><td>AAACTTGATCTCGTTCAGCGACGG</td><td></td><td></td><td></td><td></td><td></td><td></td><td></td><td></td><td></td><td></td></tr>
<tr><td>Chromosome</td><td>start</td><td>end</td><td>strand</td><td>MM</td><td>target_seq</td><td>PAM</td><td>alignment</td><td>distance</td><td>position</td><td>gene name</td><td>gene id</td></tr>
<tr><td>GU305914.1</td><td>69299</td><td>69321</td><td>+</td><td>0</td><td>CCGTCGCTGAACGAGATCAA</td><td>GGG</td><td>||||||||[||||||||||||]PAM</td><td>NA</td><td>Intergenic</td><td>NA</td><td>NA</td></tr>
<tr><td>T4826</td><td>GAAGATGCTGAAGAGCGTGCTGG</td><td>1000</td><td></td><td></td><td></td><td></td><td></td><td></td><td></td><td></td><td></td></tr>
<tr><td>Oligo fwd</td><td>CACCGAAGATGCTGAAGAGCGTGC</td><td></td><td></td><td></td><td></td><td></td><td></td><td></td><td></td><td></td><td></td></tr>
<tr><td>Oligo rev</td><td>AAACGCACGCTCTTCAGCATCTTC</td><td></td><td></td><td></td><td></td><td></td><td></td><td></td><td></td><td></td><td></td></tr>
<tr><td>Chromosome</td><td>start</td><td>end</td><td>strand</td><td>MM</td><td>target_seq</td><td>PAM</td><td>alignment</td><td>distance</td><td>position</td><td>gene name</td><td>gene id</td></tr>
<tr><td>GU305914.1</td><td>69325</td><td>69347</td><td>+</td><td>0</td><td>GAAGATGCTGAAGAGCGTGC</td><td>TGG</td><td>||||||||[||||||||||||]PAM</td><td>NA</td><td>Intergenic</td><td>NA</td><td>NA</td></tr>
<tr><td>T4827</td><td>GAGCGTGCTGGAGACGAACGCGG</td><td>1000</td><td></td><td></td><td></td><td></td><td></td><td></td><td></td><td></td><td></td></tr>
<tr><td>Oligo fwd</td><td>CACCGAGCGTGCTGGAGACGAACG</td><td></td><td></td><td></td><td></td><td></td><td></td><td></td><td></td><td></td><td></td></tr>
<tr><td>Oligo rev</td><td>AAACCGTTCGTCTCCAGCACGCTC</td><td></td><td></td><td></td><td></td><td></td><td></td><td></td><td></td><td></td><td></td></tr>
<tr><td>Chromosome</td><td>start</td><td>end</td><td>strand</td><td>MM</td><td>target_seq</td><td>PAM</td><td>alignment</td><td>distance</td><td>position</td><td>gene name</td><td>gene id</td></tr>
<tr><td>GU305914.1</td><td>69337</td><td>69359</td><td>+</td><td>0</td><td>GAGCGTGCTGGAGACGAACG</td><td>CGG</td><td>||||||||[||||||||||||]PAM</td><td>NA</td><td>Intergenic</td><td>NA</td><td>NA</td></tr>
<tr><td>T4828</td><td>TGGAGACGAACGCGGATCTGAGG</td><td>1000</td><td></td><td></td><td></td><td></td><td></td><td></td><td></td><td></td><td></td></tr>
<tr><td>Oligo fwd</td><td>CACCTGGAGACGAACGCGGATCTG</td><td></td><td></td><td></td><td></td><td></td><td></td><td></td><td></td><td></td><td></td></tr>
<tr><td>Oligo rev</td><td>AAACCAGATCCGCGTTCGTCTCCA</td><td></td><td></td><td></td><td></td><td></td><td></td><td></td><td></td><td></td><td></td></tr>
<tr><td>Chromosome</td><td>start</td><td>end</td><td>strand</td><td>MM</td><td>target_seq</td><td>PAM</td><td>alignment</td><td>distance</td><td>position</td><td>gene name</td><td>gene id</td></tr>
<tr><td>GU305914.1</td><td>69345</td><td>69367</td><td>+</td><td>0</td><td>TGGAGACGAACGCGGATCTG</td><td>AGG</td><td>||||||||[||||||||||||]PAM</td><td>NA</td><td>Intergenic</td><td>NA</td><td>NA</td></tr>
<tr><td>T4829</td><td>TCTGAGGCGCGCGTACGTGCAGG</td><td>1000</td><td></td><td></td><td></td><td></td><td></td><td></td><td></td><td></td><td></td></tr>
<tr><td>Oligo fwd</td><td>CACCTCTGAGGCGCGCGTACGTGC</td><td></td><td></td><td></td><td></td><td></td><td></td><td></td><td></td><td></td><td></td></tr>
<tr><td>Oligo rev</td><td>AAACGCACGTACGCGCGCCTCAGA</td><td></td><td></td><td></td><td></td><td></td><td></td><td></td><td></td><td></td><td></td></tr>
<tr><td>Chromosome</td><td>start</td><td>end</td><td>strand</td><td>MM</td><td>target_seq</td><td>PAM</td><td>alignment</td><td>distance</td><td>position</td><td>gene name</td><td>gene id</td></tr>
<tr><td>GU305914.1</td><td>69361</td><td>69383</td><td>+</td><td>0</td><td>TCTGAGGCGCGCGTACGTGC</td><td>AGG</td><td>||||||||[||||||||||||]PAM</td><td>NA</td><td>Intergenic</td><td>NA</td><td>NA</td></tr>
<tr><td>T4830</td><td>GAGGCGCGCGTACGTGCAGGCGG</td><td>1000</td><td></td><td></td><td></td><td></td><td></td><td></td><td></td><td></td><td></td></tr>
<tr><td>Oligo fwd</td><td>CACCGAGGCGCGCGTACGTGCAGG</td><td></td><td></td><td></td><td></td><td></td><td></td><td></td><td></td><td></td><td></td></tr>
<tr><td>Oligo rev</td><td>AAACCCTGCACGTACGCGCGCCTC</td><td></td><td></td><td></td><td></td><td></td><td></td><td></td><td></td><td></td><td></td></tr>
<tr><td>Chromosome</td><td>start</td><td>end</td><td>strand</td><td>MM</td><td>target_seq</td><td>PAM</td><td>alignment</td><td>distance</td><td>position</td><td>gene name</td><td>gene id</td></tr>
<tr><td>GU305914.1</td><td>69364</td><td>69386</td><td>+</td><td>0</td><td>GAGGCGCGCGTACGTGCAGG</td><td>CGG</td><td>||||||||[||||||||||||]PAM</td><td>NA</td><td>Intergenic</td><td>NA</td><td>NA</td></tr>
<tr><td>T4831</td><td>CGCGTACGTGCAGGCGGTGACGG</td><td>1000</td><td></td><td></td><td></td><td></td><td></td><td></td><td></td><td></td><td></td></tr>
<tr><td>Oligo fwd</td><td>CACCCGCGTACGTGCAGGCGGTGA</td><td></td><td></td><td></td><td></td><td></td><td></td><td></td><td></td><td></td><td></td></tr>
<tr><td>Oligo rev</td><td>AAACTCACCGCCTGCACGTACGCG</td><td></td><td></td><td></td><td></td><td></td><td></td><td></td><td></td><td></td><td></td></tr>
<tr><td>Chromosome</td><td>start</td><td>end</td><td>strand</td><td>MM</td><td>target_seq</td><td>PAM</td><td>alignment</td><td>distance</td><td>position</td><td>gene name</td><td>gene id</td></tr>
<tr><td>GU305914.1</td><td>69370</td><td>69392</td><td>+</td><td>0</td><td>CGCGTACGTGCAGGCGGTGA</td><td>CGG</td><td>||||||||[||||||||||||]PAM</td><td>NA</td><td>Intergenic</td><td>NA</td><td>NA</td></tr>
<tr><td>GU305914.1</td><td>215365</td><td>215387</td><td>-</td><td>4</td><td>CACGTACCTGCAGACCGTGA</td><td>GGG</td><td>|-|||||-[|||||-|-||||]PAM</td><td>NA</td><td>Intergenic</td><td>NA</td><td>NA</td></tr>
<tr><td>T4832</td><td>GCGTACGTGCAGGCGGTGACGGG</td><td>1000</td><td></td><td></td><td></td><td></td><td></td><td></td><td></td><td></td><td></td></tr>
<tr><td>Oligo fwd</td><td>CACCGCGTACGTGCAGGCGGTGAC</td><td></td><td></td><td></td><td></td><td></td><td></td><td></td><td></td><td></td><td></td></tr>
<tr><td>Oligo rev</td><td>AAACGTCACCGCCTGCACGTACGC</td><td></td><td></td><td></td><td></td><td></td><td></td><td></td><td></td><td></td><td></td></tr>
<tr><td>Chromosome</td><td>start</td><td>end</td><td>strand</td><td>MM</td><td>target_seq</td><td>PAM</td><td>alignment</td><td>distance</td><td>position</td><td>gene name</td><td>gene id</td></tr>
<tr><td>GU305914.1</td><td>69371</td><td>69393</td><td>+</td><td>0</td><td>GCGTACGTGCAGGCGGTGAC</td><td>GGG</td><td>||||||||[||||||||||||]PAM</td><td>NA</td><td>Intergenic</td><td>NA</td><td>NA</td></tr>
<tr><td>T4833</td><td>GCAGGCGGTGACGGGCATGCTGG</td><td>1000</td><td></td><td></td><td></td><td></td><td></td><td></td><td></td><td></td><td></td></tr>
<tr><td>Oligo fwd</td><td>CACCGCAGGCGGTGACGGGCATGC</td><td></td><td></td><td></td><td></td><td></td><td></td><td></td><td></td><td></td><td></td></tr>
<tr><td>Oligo rev</td><td>AAACGCATGCCCGTCACCGCCTGC</td><td></td><td></td><td></td><td></td><td></td><td></td><td></td><td></td><td></td><td></td></tr>
<tr><td>Chromosome</td><td>start</td><td>end</td><td>strand</td><td>MM</td><td>target_seq</td><td>PAM</td><td>alignment</td><td>distance</td><td>position</td><td>gene name</td><td>gene id</td></tr>
<tr><td>GU305914.1</td><td>69379</td><td>69401</td><td>+</td><td>0</td><td>GCAGGCGGTGACGGGCATGC</td><td>TGG</td><td>||||||||[||||||||||||]PAM</td><td>NA</td><td>Intergenic</td><td>NA</td><td>NA</td></tr>
<tr><td>T4834</td><td>GGAGAACGCGCTGAAGCAGCTGG</td><td>1000</td><td></td><td></td><td></td><td></td><td></td><td></td><td></td><td></td><td></td></tr>
<tr><td>Oligo fwd</td><td>CACCGGAGAACGCGCTGAAGCAGC</td><td></td><td></td><td></td><td></td><td></td><td></td><td></td><td></td><td></td><td></td></tr>
<tr><td>Oligo rev</td><td>AAACGCTGCTTCAGCGCGTTCTCC</td><td></td><td></td><td></td><td></td><td></td><td></td><td></td><td></td><td></td><td></td></tr>
<tr><td>Chromosome</td><td>start</td><td>end</td><td>strand</td><td>MM</td><td>target_seq</td><td>PAM</td><td>alignment</td><td>distance</td><td>position</td><td>gene name</td><td>gene id</td></tr>
<tr><td>GU305914.1</td><td>69400</td><td>69422</td><td>+</td><td>0</td><td>GGAGAACGCGCTGAAGCAGC</td><td>TGG</td><td>||||||||[||||||||||||]PAM</td><td>NA</td><td>Intergenic</td><td>NA</td><td>NA</td></tr>
<tr><td>T4835</td><td>GAACGCGCTGAAGCAGCTGGCGG</td><td>1000</td><td></td><td></td><td></td><td></td><td></td><td></td><td></td><td></td><td></td></tr>
<tr><td>Oligo fwd</td><td>CACCGAACGCGCTGAAGCAGCTGG</td><td></td><td></td><td></td><td></td><td></td><td></td><td></td><td></td><td></td><td></td></tr>
<tr><td>Oligo rev</td><td>AAACCCAGCTGCTTCAGCGCGTTC</td><td></td><td></td><td></td><td></td><td></td><td></td><td></td><td></td><td></td><td></td></tr>
<tr><td>Chromosome</td><td>start</td><td>end</td><td>strand</td><td>MM</td><td>target_seq</td><td>PAM</td><td>alignment</td><td>distance</td><td>position</td><td>gene name</td><td>gene id</td></tr>
<tr><td>GU305914.1</td><td>69403</td><td>69425</td><td>+</td><td>0</td><td>GAACGCGCTGAAGCAGCTGG</td><td>CGG</td><td>||||||||[||||||||||||]PAM</td><td>NA</td><td>Intergenic</td><td>NA</td><td>NA</td></tr>
<tr><td>T4836</td><td>CGCGCTGAAGCAGCTGGCGGAGG</td><td>1000</td><td></td><td></td><td></td><td></td><td></td><td></td><td></td><td></td><td></td></tr>
<tr><td>Oligo fwd</td><td>CACCCGCGCTGAAGCAGCTGGCGG</td><td></td><td></td><td></td><td></td><td></td><td></td><td></td><td></td><td></td><td></td></tr>
<tr><td>Oligo rev</td><td>AAACCCGCCAGCTGCTTCAGCGCG</td><td></td><td></td><td></td><td></td><td></td><td></td><td></td><td></td><td></td><td></td></tr>
<tr><td>Chromosome</td><td>start</td><td>end</td><td>strand</td><td>MM</td><td>target_seq</td><td>PAM</td><td>alignment</td><td>distance</td><td>position</td><td>gene name</td><td>gene id</td></tr>
<tr><td>GU305914.1</td><td>69406</td><td>69428</td><td>+</td><td>0</td><td>CGCGCTGAAGCAGCTGGCGG</td><td>AGG</td><td>||||||||[||||||||||||]PAM</td><td>NA</td><td>Intergenic</td><td>NA</td><td>NA</td></tr>
<tr><td>T4837</td><td>GCGCTGAAGCAGCTGGCGGAGGG</td><td>1000</td><td></td><td></td><td></td><td></td><td></td><td></td><td></td><td></td><td></td></tr>
<tr><td>Oligo fwd</td><td>CACCGCGCTGAAGCAGCTGGCGGA</td><td></td><td></td><td></td><td></td><td></td><td></td><td></td><td></td><td></td><td></td></tr>
<tr><td>Oligo rev</td><td>AAACTCCGCCAGCTGCTTCAGCGC</td><td></td><td></td><td></td><td></td><td></td><td></td><td></td><td></td><td></td><td></td></tr>
<tr><td>Chromosome</td><td>start</td><td>end</td><td>strand</td><td>MM</td><td>target_seq</td><td>PAM</td><td>alignment</td><td>distance</td><td>position</td><td>gene name</td><td>gene id</td></tr>
<tr><td>GU305914.1</td><td>69407</td><td>69429</td><td>+</td><td>0</td><td>GCGCTGAAGCAGCTGGCGGA</td><td>GGG</td><td>||||||||[||||||||||||]PAM</td><td>NA</td><td>Intergenic</td><td>NA</td><td>NA</td></tr>
<tr><td>T4838</td><td>GGCGGAGGGCAACCTGCCCTCGG</td><td>1000</td><td></td><td></td><td></td><td></td><td></td><td></td><td></td><td></td><td></td></tr>
<tr><td>Oligo fwd</td><td>CACCGGCGGAGGGCAACCTGCCCT</td><td></td><td></td><td></td><td></td><td></td><td></td><td></td><td></td><td></td><td></td></tr>
<tr><td>Oligo rev</td><td>AAACAGGGCAGGTTGCCCTCCGCC</td><td></td><td></td><td></td><td></td><td></td><td></td><td></td><td></td><td></td><td></td></tr>
<tr><td>Chromosome</td><td>start</td><td>end</td><td>strand</td><td>MM</td><td>target_seq</td><td>PAM</td><td>alignment</td><td>distance</td><td>position</td><td>gene name</td><td>gene id</td></tr>
<tr><td>GU305914.1</td><td>69421</td><td>69443</td><td>+</td><td>0</td><td>GGCGGAGGGCAACCTGCCCT</td><td>CGG</td><td>||||||||[||||||||||||]PAM</td><td>NA</td><td>Intergenic</td><td>NA</td><td>NA</td></tr>
<tr><td>T4839</td><td>CCTCGGACGACATGATGTCGAGG</td><td>1000</td><td></td><td></td><td></td><td></td><td></td><td></td><td></td><td></td><td></td></tr>
</table>

<table>
<tr><td>Oligo fwd</td><td>CACCCCTCGGACGACATGATGTCG</td><td></td><td></td><td></td><td></td><td></td><td></td><td></td><td></td><td></td><td></td></tr>
<tr><td>Oligo rev</td><td>AAACCGACATCATGTCGTCCGAGG</td><td></td><td></td><td></td><td></td><td></td><td></td><td></td><td></td><td></td><td></td></tr>
<tr><td>Chromosome</td><td>start</td><td>end</td><td>strand</td><td>MM</td><td>target_seq</td><td>PAM</td><td>alignment</td><td>distance</td><td>position</td><td>gene name</td><td>gene id</td></tr>
<tr><td>GU305914.1</td><td>69438</td><td>69460</td><td>+</td><td>0</td><td>CCTCGGACGACATGATGTCG</td><td>AGG</td><td>||||||||[||||||||||||]PAM</td><td>NA</td><td>Intergenic</td><td>NA</td><td>NA</td></tr>
<tr><td></td><td></td><td></td><td></td><td></td><td></td><td></td><td></td><td></td><td></td><td></td><td></td></tr>
<tr><td>T4840</td><td>CGACATGATGTCGAGGATCACGG</td><td>1000</td><td></td><td></td><td></td><td></td><td></td><td></td><td></td><td></td><td></td></tr>
<tr><td>Oligo fwd</td><td>CACCCGACATGATGTCGAGGATCA</td><td></td><td></td><td></td><td></td><td></td><td></td><td></td><td></td><td></td><td></td></tr>
<tr><td>Oligo rev</td><td>AAACTGATCCTCGACATCATGTCG</td><td></td><td></td><td></td><td></td><td></td><td></td><td></td><td></td><td></td><td></td></tr>
<tr><td>Chromosome</td><td>start</td><td>end</td><td>strand</td><td>MM</td><td>target_seq</td><td>PAM</td><td>alignment</td><td>distance</td><td>position</td><td>gene name</td><td>gene id</td></tr>
<tr><td>GU305914.1</td><td>69445</td><td>69467</td><td>+</td><td>0</td><td>CGACATGATGTCGAGGATCA</td><td>CGG</td><td>||||||||[||||||||||||]PAM</td><td>NA</td><td>Intergenic</td><td>NA</td><td>NA</td></tr>
<tr><td></td><td></td><td></td><td></td><td></td><td></td><td></td><td></td><td></td><td></td><td></td><td></td></tr>
<tr><td>T4841</td><td>GATGTCGAGGATCACGGCGCTGG</td><td>1000</td><td></td><td></td><td></td><td></td><td></td><td></td><td></td><td></td><td></td></tr>
<tr><td>Oligo fwd</td><td>CACCGATGTCGAGGATCACGGCGC</td><td></td><td></td><td></td><td></td><td></td><td></td><td></td><td></td><td></td><td></td></tr>
<tr><td>Oligo rev</td><td>AAACGCGCCGTGATCCTCGACATC</td><td></td><td></td><td></td><td></td><td></td><td></td><td></td><td></td><td></td><td></td></tr>
<tr><td>Chromosome</td><td>start</td><td>end</td><td>strand</td><td>MM</td><td>target_seq</td><td>PAM</td><td>alignment</td><td>distance</td><td>position</td><td>gene name</td><td>gene id</td></tr>
<tr><td>GU305914.1</td><td>69451</td><td>69473</td><td>+</td><td>0</td><td>GATGTCGAGGATCACGGCGC</td><td>TGG</td><td>||||||||[||||||||||||]PAM</td><td>NA</td><td>Intergenic</td><td>NA</td><td>NA</td></tr>
<tr><td></td><td></td><td></td><td></td><td></td><td></td><td></td><td></td><td></td><td></td><td></td><td></td></tr>
<tr><td>T4842</td><td>GTCGAGGATCACGGCGCTGGCGG</td><td>1000</td><td></td><td></td><td></td><td></td><td></td><td></td><td></td><td></td><td></td></tr>
<tr><td>Oligo fwd</td><td>CACCGTCGAGGATCACGGCGCTGG</td><td></td><td></td><td></td><td></td><td></td><td></td><td></td><td></td><td></td><td></td></tr>
<tr><td>Oligo rev</td><td>AAACCCAGCGCCGTGATCCTCGAC</td><td></td><td></td><td></td><td></td><td></td><td></td><td></td><td></td><td></td><td></td></tr>
<tr><td>Chromosome</td><td>start</td><td>end</td><td>strand</td><td>MM</td><td>target_seq</td><td>PAM</td><td>alignment</td><td>distance</td><td>position</td><td>gene name</td><td>gene id</td></tr>
<tr><td>GU305914.1</td><td>69454</td><td>69476</td><td>+</td><td>0</td><td>GTCGAGGATCACGGCGCTGG</td><td>CGG</td><td>||||||||[||||||||||||]PAM</td><td>NA</td><td>Intergenic</td><td>NA</td><td>NA</td></tr>
<tr><td></td><td></td><td></td><td></td><td></td><td></td><td></td><td></td><td></td><td></td><td></td><td></td></tr>
<tr><td>T4843</td><td>GCTGGCGGAACAGCTGCCGACGG</td><td>1000</td><td></td><td></td><td></td><td></td><td></td><td></td><td></td><td></td><td></td></tr>
<tr><td>Oligo fwd</td><td>CACCGCTGGCGGAACAGCTGCCGA</td><td></td><td></td><td></td><td></td><td></td><td></td><td></td><td></td><td></td><td></td></tr>
<tr><td>Oligo rev</td><td>AAACTCGGCAGCTGTTCCGCCAGC</td><td></td><td></td><td></td><td></td><td></td><td></td><td></td><td></td><td></td><td></td></tr>
<tr><td>Chromosome</td><td>start</td><td>end</td><td>strand</td><td>MM</td><td>target_seq</td><td>PAM</td><td>alignment</td><td>distance</td><td>position</td><td>gene name</td><td>gene id</td></tr>
<tr><td>GU305914.1</td><td>69469</td><td>69491</td><td>+</td><td>0</td><td>GCTGGCGGAACAGCTGCCGA</td><td>CGG</td><td>||||||||[||||||||||||]PAM</td><td>NA</td><td>Intergenic</td><td>NA</td><td>NA</td></tr>
<tr><td></td><td></td><td></td><td></td><td></td><td></td><td></td><td></td><td></td><td></td><td></td><td></td></tr>
<tr><td>T4844</td><td>CTGGCGGAACAGCTGCCGACGGG</td><td>1000</td><td></td><td></td><td></td><td></td><td></td><td></td><td></td><td></td><td></td></tr>
<tr><td>Oligo fwd</td><td>CACCCTGGCGGAACAGCTGCCGAC</td><td></td><td></td><td></td><td></td><td></td><td></td><td></td><td></td><td></td><td></td></tr>
<tr><td>Oligo rev</td><td>AAACGTCGGCAGCTGTTCCGCCAG</td><td></td><td></td><td></td><td></td><td></td><td></td><td></td><td></td><td></td><td></td></tr>
<tr><td>Chromosome</td><td>start</td><td>end</td><td>strand</td><td>MM</td><td>target_seq</td><td>PAM</td><td>alignment</td><td>distance</td><td>position</td><td>gene name</td><td>gene id</td></tr>
<tr><td>GU305914.1</td><td>69470</td><td>69492</td><td>+</td><td>0</td><td>CTGGCGGAACAGCTGCCGAC</td><td>GGG</td><td>||||||||[||||||||||||]PAM</td><td>NA</td><td>Intergenic</td><td>NA</td><td>NA</td></tr>
<tr><td>GU305914.1</td><td>19529</td><td>19551</td><td>+</td><td>4</td><td>ACGGCGGAACCGCTGCCGAA</td><td>CGG</td><td>--||||||[||-||||||||-]PAM</td><td>NA</td><td>Intergenic</td><td>NA</td><td>NA</td></tr>
<tr><td>GU305914.1</td><td>109272</td><td>109294</td><td>+</td><td>3</td><td>CAGGCGGAACAGCTGCTGTC</td><td>CGG</td><td>|-||||||[||||||||-|-|]PAM</td><td>NA</td><td>Intergenic</td><td>NA</td><td>NA</td></tr>
<tr><td></td><td></td><td></td><td></td><td></td><td></td><td></td><td></td><td></td><td></td><td></td><td></td></tr>
<tr><td>T4845</td><td>TGGCGGAACAGCTGCCGACGGGG</td><td>1000</td><td></td><td></td><td></td><td></td><td></td><td></td><td></td><td></td><td></td></tr>
<tr><td>Oligo fwd</td><td>CACCTGGCGGAACAGCTGCCGACG</td><td></td><td></td><td></td><td></td><td></td><td></td><td></td><td></td><td></td><td></td></tr>
<tr><td>Oligo rev</td><td>AAACCGTCGGCAGCTGTTCCGCCA</td><td></td><td></td><td></td><td></td><td></td><td></td><td></td><td></td><td></td><td></td></tr>
<tr><td>Chromosome</td><td>start</td><td>end</td><td>strand</td><td>MM</td><td>target_seq</td><td>PAM</td><td>alignment</td><td>distance</td><td>position</td><td>gene name</td><td>gene id</td></tr>
<tr><td>GU305914.1</td><td>69471</td><td>69493</td><td>+</td><td>0</td><td>TGGCGGAACAGCTGCCGACG</td><td>GGG</td><td>||||||||[||||||||||||]PAM</td><td>NA</td><td>Intergenic</td><td>NA</td><td>NA</td></tr>
<tr><td></td><td></td><td></td><td></td><td></td><td></td><td></td><td></td><td></td><td></td><td></td><td></td></tr>
<tr><td>T4846</td><td>ACAGCTGCCGACGGGGCGCCAGG</td><td>1000</td><td></td><td></td><td></td><td></td><td></td><td></td><td></td><td></td><td></td></tr>
<tr><td>Oligo fwd</td><td>CACCACAGCTGCCGACGGGGCGCC</td><td></td><td></td><td></td><td></td><td></td><td></td><td></td><td></td><td></td><td></td></tr>
<tr><td>Oligo rev</td><td>AAACGGCGCCCCGTCGGCAGCTGT</td><td></td><td></td><td></td><td></td><td></td><td></td><td></td><td></td><td></td><td></td></tr>
<tr><td>Chromosome</td><td>start</td><td>end</td><td>strand</td><td>MM</td><td>target_seq</td><td>PAM</td><td>alignment</td><td>distance</td><td>position</td><td>gene name</td><td>gene id</td></tr>
<tr><td>GU305914.1</td><td>69478</td><td>69500</td><td>+</td><td>0</td><td>ACAGCTGCCGACGGGGCGCC</td><td>AGG</td><td>||||||||[||||||||||||]PAM</td><td>NA</td><td>Intergenic</td><td>NA</td><td>NA</td></tr>
<tr><td></td><td></td><td></td><td></td><td></td><td></td><td></td><td></td><td></td><td></td><td></td><td></td></tr>
<tr><td>T4847</td><td>CCAGGTGCGCGCCGACCTGCTGG</td><td>1000</td><td></td><td></td><td></td><td></td><td></td><td></td><td></td><td></td><td></td></tr>
<tr><td>Oligo fwd</td><td>CACCCCAGGTGCGCGCCGACCTGC</td><td></td><td></td><td></td><td></td><td></td><td></td><td></td><td></td><td></td><td></td></tr>
<tr><td>Oligo rev</td><td>AAACGCAGGTCGGCGCGCACCTGG</td><td></td><td></td><td></td><td></td><td></td><td></td><td></td><td></td><td></td><td></td></tr>
<tr><td>Chromosome</td><td>start</td><td>end</td><td>strand</td><td>MM</td><td>target_seq</td><td>PAM</td><td>alignment</td><td>distance</td><td>position</td><td>gene name</td><td>gene id</td></tr>
<tr><td>GU305914.1</td><td>69496</td><td>69518</td><td>+</td><td>0</td><td>CCAGGTGCGCGCCGACCTGC</td><td>TGG</td><td>||||||||[||||||||||||]PAM</td><td>NA</td><td>Intergenic</td><td>NA</td><td>NA</td></tr>
<tr><td>GU305914.1</td><td>100384</td><td>100406</td><td>-</td><td>4</td><td>CAAGCTGCGCGAGGACCTGC</td><td>AGG</td><td>|-||-|||[|||--|||||||]PAM</td><td>NA</td><td>Intergenic</td><td>NA</td><td>NA</td></tr>
<tr><td></td><td></td><td></td><td></td><td></td><td></td><td></td><td></td><td></td><td></td><td></td><td></td></tr>
<tr><td>T4848</td><td>CGCCGACCTGCTGGACTCGACGG</td><td>1000</td><td></td><td></td><td></td><td></td><td></td><td></td><td></td><td></td><td></td></tr>
<tr><td>Oligo fwd</td><td>CACCCGCCGACCTGCTGGACTCGA</td><td></td><td></td><td></td><td></td><td></td><td></td><td></td><td></td><td></td><td></td></tr>
<tr><td>Oligo rev</td><td>AAACTCGAGTCCAGCAGGTCGGCG</td><td></td><td></td><td></td><td></td><td></td><td></td><td></td><td></td><td></td><td></td></tr>
<tr><td>Chromosome</td><td>start</td><td>end</td><td>strand</td><td>MM</td><td>target_seq</td><td>PAM</td><td>alignment</td><td>distance</td><td>position</td><td>gene name</td><td>gene id</td></tr>
<tr><td>GU305914.1</td><td>69505</td><td>69527</td><td>+</td><td>0</td><td>CGCCGACCTGCTGGACTCGA</td><td>CGG</td><td>||||||||[||||||||||||]PAM</td><td>NA</td><td>Intergenic</td><td>NA</td><td>NA</td></tr>
<tr><td>GU305914.1</td><td>123974</td><td>123996</td><td>-</td><td>4</td><td>CACCGAGGTGCTGGACTCGG</td><td>AGG</td><td>|-||||--[|||||||||||-]PAM</td><td>NA</td><td>Intergenic</td><td>NA</td><td>NA</td></tr>
<tr><td></td><td></td><td></td><td></td><td></td><td></td><td></td><td></td><td></td><td></td><td></td><td></td></tr>
<tr><td>T4849</td><td>TCGTGTCGCAGATGTCGCGCCGG</td><td>1000</td><td></td><td></td><td></td><td></td><td></td><td></td><td></td><td></td><td></td></tr>
<tr><td>Oligo fwd</td><td>CACCTCGTGTCGCAGATGTCGCGC</td><td></td><td></td><td></td><td></td><td></td><td></td><td></td><td></td><td></td><td></td></tr>
<tr><td>Oligo rev</td><td>AAACGCGCGACATCTGCGACACGA</td><td></td><td></td><td></td><td></td><td></td><td></td><td></td><td></td><td></td><td></td></tr>
<tr><td>Chromosome</td><td>start</td><td>end</td><td>strand</td><td>MM</td><td>target_seq</td><td>PAM</td><td>alignment</td><td>distance</td><td>position</td><td>gene name</td><td>gene id</td></tr>
<tr><td>GU305914.1</td><td>69531</td><td>69553</td><td>+</td><td>0</td><td>TCGTGTCGCAGATGTCGCGC</td><td>CGG</td><td>||||||||[||||||||||||]PAM</td><td>NA</td><td>Intergenic</td><td>NA</td><td>NA</td></tr>
<tr><td>GU305914.1</td><td>61152</td><td>61174</td><td>+</td><td>3</td><td>CCATGTCGCAGATGTCGTGC</td><td>AGG</td><td>-|-|||||[|||||||||-||]PAM</td><td>NA</td><td>Intergenic</td><td>NA</td><td>NA</td></tr>
<tr><td></td><td></td><td></td><td></td><td></td><td></td><td></td><td></td><td></td><td></td><td></td><td></td></tr>
<tr><td>T4850</td><td>CGCAGATGTCGCGCCGGCTGCGG</td><td>1000</td><td></td><td></td><td></td><td></td><td></td><td></td><td></td><td></td><td></td></tr>
<tr><td>Oligo fwd</td><td>CACCCGCAGATGTCGCGCCGGCTG</td><td></td><td></td><td></td><td></td><td></td><td></td><td></td><td></td><td></td><td></td></tr>
<tr><td>Oligo rev</td><td>AAACCAGCCGGCGCGACATCTGCG</td><td></td><td></td><td></td><td></td><td></td><td></td><td></td><td></td><td></td><td></td></tr>
<tr><td>Chromosome</td><td>start</td><td>end</td><td>strand</td><td>MM</td><td>target_seq</td><td>PAM</td><td>alignment</td><td>distance</td><td>position</td><td>gene name</td><td>gene id</td></tr>
<tr><td>GU305914.1</td><td>69537</td><td>69559</td><td>+</td><td>0</td><td>CGCAGATGTCGCGCCGGCTG</td><td>CGG</td><td>||||||||[||||||||||||]PAM</td><td>NA</td><td>Intergenic</td><td>NA</td><td>NA</td></tr>
<tr><td></td><td></td><td></td><td></td><td></td><td></td><td></td><td></td><td></td><td></td><td></td><td></td></tr>
<tr><td>T4851</td><td>GTCGCGCCGGCTGCGGTACGCGG</td><td>1000</td><td></td><td></td><td></td><td></td><td></td><td></td><td></td><td></td><td></td></tr>
<tr><td>Oligo fwd</td><td>CACCGTCGCGCCGGCTGCGGTACG</td><td></td><td></td><td></td><td></td><td></td><td></td><td></td><td></td><td></td><td></td></tr>
<tr><td>Oligo rev</td><td>AAACCGTACCGCAGCCGGCGCGAC</td><td></td><td></td><td></td><td></td><td></td><td></td><td></td><td></td><td></td><td></td></tr>
<tr><td>Chromosome</td><td>start</td><td>end</td><td>strand</td><td>MM</td><td>target_seq</td><td>PAM</td><td>alignment</td><td>distance</td><td>position</td><td>gene name</td><td>gene id</td></tr>
<tr><td>GU305914.1</td><td>69544</td><td>69566</td><td>+</td><td>0</td><td>GTCGCGCCGGCTGCGGTACG</td><td>CGG</td><td>||||||||[||||||||||||]PAM</td><td>NA</td><td>Intergenic</td><td>NA</td><td>NA</td></tr>
<tr><td>GU305914.1</td><td>42353</td><td>42375</td><td>-</td><td>4</td><td>GTGGCTGCGCCTGCGGTACG</td><td>AGG</td><td>||-||--|[|-||||||||||]PAM</td><td>NA</td><td>Intergenic</td><td>NA</td><td>NA</td></tr>
<tr><td></td><td></td><td></td><td></td><td></td><td></td><td></td><td></td><td></td><td></td><td></td><td></td></tr>
<tr><td>T4852</td><td>GAAGAACTCGACGCAGTCGCTGG</td><td>1000</td><td></td><td></td><td></td><td></td><td></td><td></td><td></td><td></td><td></td></tr>
<tr><td>Oligo fwd</td><td>CACCGAAGAACTCGACGCAGTCGC</td><td></td><td></td><td></td><td></td><td></td><td></td><td></td><td></td><td></td><td></td></tr>
<tr><td>Oligo rev</td><td>AAACGCGACTGCGTCGAGTTCTTC</td><td></td><td></td><td></td><td></td><td></td><td></td><td></td><td></td><td></td><td></td></tr>
<tr><td>Chromosome</td><td>start</td><td>end</td><td>strand</td><td>MM</td><td>target_seq</td><td>PAM</td><td>alignment</td><td>distance</td><td>position</td><td>gene name</td><td>gene id</td></tr>
<tr><td>GU305914.1</td><td>69574</td><td>69596</td><td>+</td><td>0</td><td>GAAGAACTCGACGCAGTCGC</td><td>TGG</td><td>||||||||[||||||||||||]PAM</td><td>NA</td><td>Intergenic</td><td>NA</td><td>NA</td></tr>
<tr><td>GU305914.1</td><td>110440</td><td>110462</td><td>-</td><td>4</td><td>GAAGTTCTTGACGCAGCCGC</td><td>CGG</td><td>||||--||[-|||||||-|||]PAM</td><td>NA</td><td>Intergenic</td><td>NA</td><td>NA</td></tr>
</table>

| | | | | | | | | | | | |
|---|---|---|---|---|---|---|---|---|---|---|---|
| GU305914.1 | | 141520 | 141542 | - | 4 | GAAGACCACGACGCAGTGTC | CGG | \|\|\|\|\|-\|-[\|\|\|\|\|\|\|\|\|--\|]PAM | NA | Intergenic | NA | NA |
| T4853 | GAACTCGACGCAGTCGCTGGAGG | | 1000 | | | | | | | | |
| Oligo fwd | CACCGAACTCGACGCAGTCGCTGG | | | | | | | | | | |
| Oligo rev | AAACCCAGCGACTGCGTCGAGTTC | | | | | | | | | | |
| Chromosome | start | | end | strand | MM | target_seq | PAM | alignment | distance | position | gene name | gene id |
| GU305914.1 | | 69577 | 69599 | + | 0 | GAACTCGACGCAGTCGCTGG | AGG | \|\|\|\|\|\|\|\|[\|\|\|\|\|\|\|\|\|\|\|\|]PAM | NA | Intergenic | NA | NA |
| T4854 | CCGAGAACGACGACATGATCAGG | | 1000 | | | | | | | | |
| Oligo fwd | CACCCCGAGAACGACGACATGATC | | | | | | | | | | |
| Oligo rev | AAACGATCATGTCGTCGTTCTCGG | | | | | | | | | | |
| Chromosome | start | | end | strand | MM | target_seq | PAM | alignment | distance | position | gene name | gene id |
| GU305914.1 | | 69618 | 69640 | + | 0 | CCGAGAACGACGACATGATC | AGG | \|\|\|\|\|\|\|\|[\|\|\|\|\|\|\|\|\|\|\|\|]PAM | NA | Intergenic | NA | NA |
| T4855 | GGAAGCTGCTGAAGACGAGCTGG | | 1000 | | | | | | | | |
| Oligo fwd | CACCGGAAGCTGCTGAAGACGAGC | | | | | | | | | | |
| Oligo rev | AAACGCTCGTCTTCAGCAGCTTCC | | | | | | | | | | |
| Chromosome | start | | end | strand | MM | target_seq | PAM | alignment | distance | position | gene name | gene id |
| GU305914.1 | | 69639 | 69661 | + | 0 | GGAAGCTGCTGAAGACGAGC | TGG | \|\|\|\|\|\|\|\|[\|\|\|\|\|\|\|\|\|\|\|\|]PAM | NA | Intergenic | NA | NA |
| T4856 | GAAGCTGCTGAAGACGAGCTGGG | | 1000 | | | | | | | | |
| Oligo fwd | CACCGAAGCTGCTGAAGACGAGCT | | | | | | | | | | |
| Oligo rev | AAACAGCTCGTCTTCAGCAGCTTC | | | | | | | | | | |
| Chromosome | start | | end | strand | MM | target_seq | PAM | alignment | distance | position | gene name | gene id |
| GU305914.1 | | 69640 | 69662 | + | 0 | GAAGCTGCTGAAGACGAGCT | GGG | \|\|\|\|\|\|\|\|[\|\|\|\|\|\|\|\|\|\|\|\|]PAM | NA | Intergenic | NA | NA |
| GU305914.1 | | 58920 | 58942 | + | 3 | GGAGCTGTTGAAGACGATCT | TGG | \|-\|\|\|\|\|-[\|\|\|\|\|\|\|\|\|-\|\|]PAM | NA | Intergenic | NA | NA |
| T4857 | AAGCTGCTGAAGACGAGCTGGGG | | 1000 | | | | | | | | |
| Oligo fwd | CACCAAGCTGCTGAAGACGAGCTG | | | | | | | | | | |
| Oligo rev | AAACCAGCTCGTCTTCAGCAGCTT | | | | | | | | | | |
| Chromosome | start | | end | strand | MM | target_seq | PAM | alignment | distance | position | gene name | gene id |
| GU305914.1 | | 69641 | 69663 | + | 0 | AAGCTGCTGAAGACGAGCTG | GGG | \|\|\|\|\|\|\|\|[\|\|\|\|\|\|\|\|\|\|\|\|]PAM | NA | Intergenic | NA | NA |
| T4858 | GACGAGCTGGGGTAAGCCCGTGG | | 1000 | | | | | | | | |
| Oligo fwd | CACCGACGAGCTGGGGTAAGCCCG | | | | | | | | | | |
| Oligo rev | AAACCGGGCTTACCCCAGCTCGTC | | | | | | | | | | |
| Chromosome | start | | end | strand | MM | target_seq | PAM | alignment | distance | position | gene name | gene id |
| GU305914.1 | | 69652 | 69674 | + | 0 | GACGAGCTGGGGTAAGCCCG | TGG | \|\|\|\|\|\|\|\|[\|\|\|\|\|\|\|\|\|\|\|\|]PAM | NA | Intergenic | NA | NA |
| T4859 | CTGGGGTAAGCCCGTGGCGACGG | | 1000 | | | | | | | | |
| Oligo fwd | CACCCTGGGGTAAGCCCGTGGCGA | | | | | | | | | | |
| Oligo rev | AAACTCGCCACGGGCTTACCCCAG | | | | | | | | | | |
| Chromosome | start | | end | strand | MM | target_seq | PAM | alignment | distance | position | gene name | gene id |
| GU305914.1 | | 69658 | 69680 | + | 0 | CTGGGGTAAGCCCGTGGCGA | CGG | \|\|\|\|\|\|\|\|[\|\|\|\|\|\|\|\|\|\|\|\|]PAM | NA | Intergenic | NA | NA |
| T4860 | CCGTGGCGACGGTGTACCGCCGG | | 1000 | | | | | | | | |
| Oligo fwd | CACCCCGTGGCGACGGTGTACCGC | | | | | | | | | | |
| Oligo rev | AAACGCGGTACACCGTCGCCACGG | | | | | | | | | | |
| Chromosome | start | | end | strand | MM | target_seq | PAM | alignment | distance | position | gene name | gene id |
| GU305914.1 | | 69669 | 69691 | + | 0 | CCGTGGCGACGGTGTACCGC | CGG | \|\|\|\|\|\|\|\|[\|\|\|\|\|\|\|\|\|\|\|\|]PAM | NA | Intergenic | NA | NA |
| T4861 | CGTGGCGACGGTGTACCGCCGGG | | 1000 | | | | | | | | |
| Oligo fwd | CACCCGTGGCGACGGTGTACCGCC | | | | | | | | | | |
| Oligo rev | AAACGGCGGTACACCGTCGCCACG | | | | | | | | | | |
| Chromosome | start | | end | strand | MM | target_seq | PAM | alignment | distance | position | gene name | gene id |
| GU305914.1 | | 69670 | 69692 | + | 0 | CGTGGCGACGGTGTACCGCC | GGG | \|\|\|\|\|\|\|\|[\|\|\|\|\|\|\|\|\|\|\|\|]PAM | NA | Intergenic | NA | NA |
| T4862 | GACGGTGTACCGCCGGGTGCAGG | | 1000 | | | | | | | | |
| Oligo fwd | CACCGACGGTGTACCGCCGGGTGC | | | | | | | | | | |
| Oligo rev | AAACGCACCCGGCGGTACACCGTC | | | | | | | | | | |
| Chromosome | start | | end | strand | MM | target_seq | PAM | alignment | distance | position | gene name | gene id |
| GU305914.1 | | 69676 | 69698 | + | 0 | GACGGTGTACCGCCGGGTGC | AGG | \|\|\|\|\|\|\|\|[\|\|\|\|\|\|\|\|\|\|\|\|]PAM | NA | Intergenic | NA | NA |
| T4863 | GGTGTACCGCCGGGTGCAGGTGG | | 1000 | | | | | | | | |
| Oligo fwd | CACCGGTGTACCGCCGGGTGCAGG | | | | | | | | | | |
| Oligo rev | AAACCCTGCACCCGGCGGTACACC | | | | | | | | | | |
| Chromosome | start | | end | strand | MM | target_seq | PAM | alignment | distance | position | gene name | gene id |
| GU305914.1 | | 69679 | 69701 | + | 0 | GGTGTACCGCCGGGTGCAGG | TGG | \|\|\|\|\|\|\|\|[\|\|\|\|\|\|\|\|\|\|\|\|]PAM | NA | Intergenic | NA | NA |
| T4864 | GGTGGAATACGACCGCAAGATGG | | 1000 | | | | | | | | |
| Oligo fwd | CACCGGTGGAATACGACCGCAAGA | | | | | | | | | | |
| Oligo rev | AAACTCTTGCGGTCGTATTCCACC | | | | | | | | | | |
| Chromosome | start | | end | strand | MM | target_seq | PAM | alignment | distance | position | gene name | gene id |
| GU305914.1 | | 69697 | 69719 | + | 0 | GGTGGAATACGACCGCAAGA | TGG | \|\|\|\|\|\|\|\|[\|\|\|\|\|\|\|\|\|\|\|\|]PAM | NA | Intergenic | NA | NA |
| T4865 | GGAATACGACCGCAAGATGGAGG | | 1000 | | | | | | | | |
| Oligo fwd | CACCGGAATACGACCGCAAGATGG | | | | | | | | | | |
| Oligo rev | AAACCCATCTTGCGGTCGTATTCC | | | | | | | | | | |
| Chromosome | start | | end | strand | MM | target_seq | PAM | alignment | distance | position | gene name | gene id |
| GU305914.1 | | 69700 | 69722 | + | 0 | GGAATACGACCGCAAGATGG | AGG | \|\|\|\|\|\|\|\|[\|\|\|\|\|\|\|\|\|\|\|\|]PAM | NA | Intergenic | NA | NA |
| T4866 | AGGAGCTGCGCGAATCCGAGTGG | | 1000 | | | | | | | | |
| Oligo fwd | CACCAGGAGCTGCGCGAATCCGAG | | | | | | | | | | |
| Oligo rev | AAACCTCGGATTCGCGCAGCTCCT | | | | | | | | | | |
| Chromosome | start | | end | strand | MM | target_seq | PAM | alignment | distance | position | gene name | gene id |
| GU305914.1 | | 69720 | 69742 | + | 0 | AGGAGCTGCGCGAATCCGAG | TGG | \|\|\|\|\|\|\|\|[\|\|\|\|\|\|\|\|\|\|\|\|]PAM | NA | Intergenic | NA | NA |
| T4867 | CGAGTGGCTGAAGCGCGTGAAGG | | 1000 | | | | | | | | |
| Oligo fwd | CACCCGAGTGGCTGAAGCGCGTGA | | | | | | | | | | |

<table>
<tr><td>Oligo rev</td><td>AAACTCACGCGCTTCAGCCACTCG</td><td></td><td></td><td></td><td></td><td></td><td></td><td></td><td></td><td></td><td></td><td></td></tr>
<tr><td>Chromosome</td><td>start</td><td></td><td>end</td><td>strand</td><td>MM</td><td>target_seq</td><td>PAM</td><td>alignment</td><td>distance</td><td>position</td><td>gene name</td><td>gene id</td></tr>
<tr><td>GU305914.1</td><td></td><td>69736</td><td>69758</td><td>+</td><td>0</td><td>CGAGTGGCTGAAGCGCGTGA</td><td>AGG</td><td>||||||||[||||||||||||]PAM</td><td>NA</td><td>Intergenic</td><td>NA</td><td>NA</td></tr>
<tr><td>T4868</td><td>GCTGAAGCGCGTGAAGGAGACGG</td><td></td><td>1000</td><td></td><td></td><td></td><td></td><td></td><td></td><td></td><td></td><td></td></tr>
<tr><td>Oligo fwd</td><td>CACCGCTGAAGCGCGTGAAGGAGA</td><td></td><td></td><td></td><td></td><td></td><td></td><td></td><td></td><td></td><td></td><td></td></tr>
<tr><td>Oligo rev</td><td>AAACTCTCCTTCACGCGCTTCAGC</td><td></td><td></td><td></td><td></td><td></td><td></td><td></td><td></td><td></td><td></td><td></td></tr>
<tr><td>Chromosome</td><td>start</td><td></td><td>end</td><td>strand</td><td>MM</td><td>target_seq</td><td>PAM</td><td>alignment</td><td>distance</td><td>position</td><td>gene name</td><td>gene id</td></tr>
<tr><td>GU305914.1</td><td></td><td>69742</td><td>69764</td><td>+</td><td>0</td><td>GCTGAAGCGCGTGAAGGAGA</td><td>CGG</td><td>||||||||[||||||||||||]PAM</td><td>NA</td><td>Intergenic</td><td>NA</td><td>NA</td></tr>
<tr><td>T4869</td><td>CATCGACTCCCCGCAGACGCTGG</td><td></td><td>1000</td><td></td><td></td><td></td><td></td><td></td><td></td><td></td><td></td><td></td></tr>
<tr><td>Oligo fwd</td><td>CACCCATCGACTCCCCGCAGACGC</td><td></td><td></td><td></td><td></td><td></td><td></td><td></td><td></td><td></td><td></td><td></td></tr>
<tr><td>Oligo rev</td><td>AAACGCGTCTGCGGGGAGTCGATG</td><td></td><td></td><td></td><td></td><td></td><td></td><td></td><td></td><td></td><td></td><td></td></tr>
<tr><td>Chromosome</td><td>start</td><td></td><td>end</td><td>strand</td><td>MM</td><td>target_seq</td><td>PAM</td><td>alignment</td><td>distance</td><td>position</td><td>gene name</td><td>gene id</td></tr>
<tr><td>GU305914.1</td><td></td><td>69766</td><td>69788</td><td>+</td><td>0</td><td>CATCGACTCCCCGCAGACGC</td><td>TGG</td><td>||||||||[||||||||||||]PAM</td><td>NA</td><td>Intergenic</td><td>NA</td><td>NA</td></tr>
<tr><td>T4870</td><td>ACTCCCCGCAGACGCTGGAGAGG</td><td></td><td>1000</td><td></td><td></td><td></td><td></td><td></td><td></td><td></td><td></td><td></td></tr>
<tr><td>Oligo fwd</td><td>CACCACTCCCCGCAGACGCTGGAG</td><td></td><td></td><td></td><td></td><td></td><td></td><td></td><td></td><td></td><td></td><td></td></tr>
<tr><td>Oligo rev</td><td>AAACCTCCAGCGTCTGCGGGGAGT</td><td></td><td></td><td></td><td></td><td></td><td></td><td></td><td></td><td></td><td></td><td></td></tr>
<tr><td>Chromosome</td><td>start</td><td></td><td>end</td><td>strand</td><td>MM</td><td>target_seq</td><td>PAM</td><td>alignment</td><td>distance</td><td>position</td><td>gene name</td><td>gene id</td></tr>
<tr><td>GU305914.1</td><td></td><td>69771</td><td>69793</td><td>+</td><td>0</td><td>ACTCCCCGCAGACGCTGGAG</td><td>AGG</td><td>||||||||[||||||||||||]PAM</td><td>NA</td><td>Intergenic</td><td>NA</td><td>NA</td></tr>
<tr><td>T4871</td><td>GCTGGAGAGGCTGCTGAAGACGG</td><td></td><td>1000</td><td></td><td></td><td></td><td></td><td></td><td></td><td></td><td></td><td></td></tr>
<tr><td>Oligo fwd</td><td>CACCGCTGGAGAGGCTGCTGAAGA</td><td></td><td></td><td></td><td></td><td></td><td></td><td></td><td></td><td></td><td></td><td></td></tr>
<tr><td>Oligo rev</td><td>AAACTCTTCAGCAGCCTCTCCAGC</td><td></td><td></td><td></td><td></td><td></td><td></td><td></td><td></td><td></td><td></td><td></td></tr>
<tr><td>Chromosome</td><td>start</td><td></td><td>end</td><td>strand</td><td>MM</td><td>target_seq</td><td>PAM</td><td>alignment</td><td>distance</td><td>position</td><td>gene name</td><td>gene id</td></tr>
<tr><td>GU305914.1</td><td></td><td>69784</td><td>69806</td><td>+</td><td>0</td><td>GCTGGAGAGGCTGCTGAAGA</td><td>CGG</td><td>||||||||[||||||||||||]PAM</td><td>NA</td><td>Intergenic</td><td>NA</td><td>NA</td></tr>
<tr><td>GU305914.1</td><td></td><td>80230</td><td>80252</td><td>+</td><td>4</td><td>GCGGCAGAGGCGGCTGACGA</td><td>CGG</td><td>||-|-|||[|||-|||||-||]PAM</td><td>NA</td><td>Intergenic</td><td>NA</td><td>NA</td></tr>
<tr><td>T4872</td><td>GGCGCCGAACGAAACGATCCTGG</td><td></td><td>1000</td><td></td><td></td><td></td><td></td><td></td><td></td><td></td><td></td><td></td></tr>
<tr><td>Oligo fwd</td><td>CACCGGCGCCGAACGAAACGATCC</td><td></td><td></td><td></td><td></td><td></td><td></td><td></td><td></td><td></td><td></td><td></td></tr>
<tr><td>Oligo rev</td><td>AAACGGATCGTTTCGTTCGGCGCC</td><td></td><td></td><td></td><td></td><td></td><td></td><td></td><td></td><td></td><td></td><td></td></tr>
<tr><td>Chromosome</td><td>start</td><td></td><td>end</td><td>strand</td><td>MM</td><td>target_seq</td><td>PAM</td><td>alignment</td><td>distance</td><td>position</td><td>gene name</td><td>gene id</td></tr>
<tr><td>GU305914.1</td><td></td><td>69805</td><td>69827</td><td>+</td><td>0</td><td>GGCGCCGAACGAAACGATCC</td><td>TGG</td><td>||||||||[||||||||||||]PAM</td><td>NA</td><td>Intergenic</td><td>NA</td><td>NA</td></tr>
<tr><td>T4873</td><td>GATCCTGGCGAAGCACGCGCCGG</td><td></td><td>1000</td><td></td><td></td><td></td><td></td><td></td><td></td><td></td><td></td><td></td></tr>
<tr><td>Oligo fwd</td><td>CACCGATCCTGGCGAAGCACGCGC</td><td></td><td></td><td></td><td></td><td></td><td></td><td></td><td></td><td></td><td></td><td></td></tr>
<tr><td>Oligo rev</td><td>AAACGCGCGTGCTTCGCCAGGATC</td><td></td><td></td><td></td><td></td><td></td><td></td><td></td><td></td><td></td><td></td><td></td></tr>
<tr><td>Chromosome</td><td>start</td><td></td><td>end</td><td>strand</td><td>MM</td><td>target_seq</td><td>PAM</td><td>alignment</td><td>distance</td><td>position</td><td>gene name</td><td>gene id</td></tr>
<tr><td>GU305914.1</td><td></td><td>69820</td><td>69842</td><td>+</td><td>0</td><td>GATCCTGGCGAAGCACGCGC</td><td>CGG</td><td>||||||||[||||||||||||]PAM</td><td>NA</td><td>Intergenic</td><td>NA</td><td>NA</td></tr>
<tr><td>T4874</td><td>TGCACGCGCGACTGAAGAAGAGG</td><td></td><td>1000</td><td></td><td></td><td></td><td></td><td></td><td></td><td></td><td></td><td></td></tr>
<tr><td>Oligo fwd</td><td>CACCTGCACGCGCGACTGAAGAAG</td><td></td><td></td><td></td><td></td><td></td><td></td><td></td><td></td><td></td><td></td><td></td></tr>
<tr><td>Oligo rev</td><td>AAACCTTCTTCAGTCGCGCGTGCA</td><td></td><td></td><td></td><td></td><td></td><td></td><td></td><td></td><td></td><td></td><td></td></tr>
<tr><td>Chromosome</td><td>start</td><td></td><td>end</td><td>strand</td><td>MM</td><td>target_seq</td><td>PAM</td><td>alignment</td><td>distance</td><td>position</td><td>gene name</td><td>gene id</td></tr>
<tr><td>GU305914.1</td><td></td><td>69846</td><td>69868</td><td>+</td><td>0</td><td>TGCACGCGCGACTGAAGAAG</td><td>AGG</td><td>||||||||[||||||||||||]PAM</td><td>NA</td><td>Intergenic</td><td>NA</td><td>NA</td></tr>
<tr><td>T4875</td><td>GAAGAAGAGGATGCAGAGCGAGG</td><td></td><td>1000</td><td></td><td></td><td></td><td></td><td></td><td></td><td></td><td></td><td></td></tr>
<tr><td>Oligo fwd</td><td>CACCGAAGAAGAGGATGCAGAGCG</td><td></td><td></td><td></td><td></td><td></td><td></td><td></td><td></td><td></td><td></td><td></td></tr>
<tr><td>Oligo rev</td><td>AAACCGCTCTGCATCCTCTTCTTC</td><td></td><td></td><td></td><td></td><td></td><td></td><td></td><td></td><td></td><td></td><td></td></tr>
<tr><td>Chromosome</td><td>start</td><td></td><td>end</td><td>strand</td><td>MM</td><td>target_seq</td><td>PAM</td><td>alignment</td><td>distance</td><td>position</td><td>gene name</td><td>gene id</td></tr>
<tr><td>GU305914.1</td><td></td><td>69859</td><td>69881</td><td>+</td><td>0</td><td>GAAGAAGAGGATGCAGAGCG</td><td>AGG</td><td>||||||||[||||||||||||]PAM</td><td>NA</td><td>Intergenic</td><td>NA</td><td>NA</td></tr>
<tr><td>T4876</td><td>GAAGAGGATGCAGAGCGAGGCGG</td><td></td><td>1000</td><td></td><td></td><td></td><td></td><td></td><td></td><td></td><td></td><td></td></tr>
<tr><td>Oligo fwd</td><td>CACCGAAGAGGATGCAGAGCGAGG</td><td></td><td></td><td></td><td></td><td></td><td></td><td></td><td></td><td></td><td></td><td></td></tr>
<tr><td>Oligo rev</td><td>AAACCCTCGCTCTGCATCCTCTTC</td><td></td><td></td><td></td><td></td><td></td><td></td><td></td><td></td><td></td><td></td><td></td></tr>
<tr><td>Chromosome</td><td>start</td><td></td><td>end</td><td>strand</td><td>MM</td><td>target_seq</td><td>PAM</td><td>alignment</td><td>distance</td><td>position</td><td>gene name</td><td>gene id</td></tr>
<tr><td>GU305914.1</td><td></td><td>69862</td><td>69884</td><td>+</td><td>0</td><td>GAAGAGGATGCAGAGCGAGG</td><td>CGG</td><td>||||||||[||||||||||||]PAM</td><td>NA</td><td>Intergenic</td><td>NA</td><td>NA</td></tr>
<tr><td>T4877</td><td>TGCAGAGCGAGGCGGAGAAGAGG</td><td></td><td>1000</td><td></td><td></td><td></td><td></td><td></td><td></td><td></td><td></td><td></td></tr>
<tr><td>Oligo fwd</td><td>CACCTGCAGAGCGAGGCGGAGAAG</td><td></td><td></td><td></td><td></td><td></td><td></td><td></td><td></td><td></td><td></td><td></td></tr>
<tr><td>Oligo rev</td><td>AAACCTTCTCCGCCTCGCTCTGCA</td><td></td><td></td><td></td><td></td><td></td><td></td><td></td><td></td><td></td><td></td><td></td></tr>
<tr><td>Chromosome</td><td>start</td><td></td><td>end</td><td>strand</td><td>MM</td><td>target_seq</td><td>PAM</td><td>alignment</td><td>distance</td><td>position</td><td>gene name</td><td>gene id</td></tr>
<tr><td>GU305914.1</td><td></td><td>69870</td><td>69892</td><td>+</td><td>0</td><td>TGCAGAGCGAGGCGGAGAAG</td><td>AGG</td><td>||||||||[||||||||||||]PAM</td><td>NA</td><td>Intergenic</td><td>NA</td><td>NA</td></tr>
<tr><td>T4878</td><td>GAGCGAGGCGGAGAAGAGGACGG</td><td></td><td>1000</td><td></td><td></td><td></td><td></td><td></td><td></td><td></td><td></td><td></td></tr>
<tr><td>Oligo fwd</td><td>CACCGAGCGAGGCGGAGAAGAGGA</td><td></td><td></td><td></td><td></td><td></td><td></td><td></td><td></td><td></td><td></td><td></td></tr>
<tr><td>Oligo rev</td><td>AAACTCCTCTTCTCCGCCTCGCTC</td><td></td><td></td><td></td><td></td><td></td><td></td><td></td><td></td><td></td><td></td><td></td></tr>
<tr><td>Chromosome</td><td>start</td><td></td><td>end</td><td>strand</td><td>MM</td><td>target_seq</td><td>PAM</td><td>alignment</td><td>distance</td><td>position</td><td>gene name</td><td>gene id</td></tr>
<tr><td>GU305914.1</td><td></td><td>69874</td><td>69896</td><td>+</td><td>0</td><td>GAGCGAGGCGGAGAAGAGGA</td><td>CGG</td><td>||||||||[||||||||||||]PAM</td><td>NA</td><td>Intergenic</td><td>NA</td><td>NA</td></tr>
<tr><td>T4879</td><td>CGAGGCGGAGAAGAGGACGGCGG</td><td></td><td>1000</td><td></td><td></td><td></td><td></td><td></td><td></td><td></td><td></td><td></td></tr>
<tr><td>Oligo fwd</td><td>CACCCGAGGCGGAGAAGAGGACGG</td><td></td><td></td><td></td><td></td><td></td><td></td><td></td><td></td><td></td><td></td><td></td></tr>
<tr><td>Oligo rev</td><td>AAACCCGTCCTCTTCTCCGCCTCG</td><td></td><td></td><td></td><td></td><td></td><td></td><td></td><td></td><td></td><td></td><td></td></tr>
<tr><td>Chromosome</td><td>start</td><td></td><td>end</td><td>strand</td><td>MM</td><td>target_seq</td><td>PAM</td><td>alignment</td><td>distance</td><td>position</td><td>gene name</td><td>gene id</td></tr>
<tr><td>GU305914.1</td><td></td><td>69877</td><td>69899</td><td>+</td><td>0</td><td>CGAGGCGGAGAAGAGGACGG</td><td>CGG</td><td>||||||||[||||||||||||]PAM</td><td>NA</td><td>Intergenic</td><td>NA</td><td>NA</td></tr>
<tr><td>T4880</td><td>GGACATGAAGCGACTGTACGAGG</td><td></td><td>1000</td><td></td><td></td><td></td><td></td><td></td><td></td><td></td><td></td><td></td></tr>
<tr><td>Oligo fwd</td><td>CACCGGACATGAAGCGACTGTACG</td><td></td><td></td><td></td><td></td><td></td><td></td><td></td><td></td><td></td><td></td><td></td></tr>
<tr><td>Oligo rev</td><td>AAACCGTACAGTCGCTTCATGTCC</td><td></td><td></td><td></td><td></td><td></td><td></td><td></td><td></td><td></td><td></td><td></td></tr>
<tr><td>Chromosome</td><td>start</td><td></td><td>end</td><td>strand</td><td>MM</td><td>target_seq</td><td>PAM</td><td>alignment</td><td>distance</td><td>position</td><td>gene name</td><td>gene id</td></tr>
<tr><td>GU305914.1</td><td></td><td>69898</td><td>69920</td><td>+</td><td>0</td><td>GGACATGAAGCGACTGTACG</td><td>AGG</td><td>||||||||[||||||||||||]PAM</td><td>NA</td><td>Intergenic</td><td>NA</td><td>NA</td></tr>
<tr><td>T4881</td><td>GTACGAGGAGATGCGCAAGAAGG</td><td></td><td>1000</td><td></td><td></td><td></td><td></td><td></td><td></td><td></td><td></td><td></td></tr>
<tr><td>Oligo fwd</td><td>CACCGTACGAGGAGATGCGCAAGA</td><td></td><td></td><td></td><td></td><td></td><td></td><td></td><td></td><td></td><td></td><td></td></tr>
<tr><td>Oligo rev</td><td>AAACTCTTGCGCATCTCCTCGTAC</td><td></td><td></td><td></td><td></td><td></td><td></td><td></td><td></td><td></td><td></td><td></td></tr>
<tr><td>Chromosome</td><td>start</td><td></td><td>end</td><td>strand</td><td>MM</td><td>target_seq</td><td>PAM</td><td>alignment</td><td>distance</td><td>position</td><td>gene name</td><td>gene id</td></tr>
<tr><td>GU305914.1</td><td></td><td>69913</td><td>69935</td><td>+</td><td>0</td><td>GTACGAGGAGATGCGCAAGA</td><td>AGG</td><td>||||||||[||||||||||||]PAM</td><td>NA</td><td>Intergenic</td><td>NA</td><td>NA</td></tr>
</table>

<table>
<tr><td>T4882</td><td>CGAGGAGATGCGCAAGAAGGTGG</td><td>1000</td><td></td><td></td><td></td><td></td><td></td><td></td><td></td><td></td><td></td></tr>
<tr><td>Oligo fwd</td><td>CACCCGAGGAGATGCGCAAGAAGG</td><td></td><td></td><td></td><td></td><td></td><td></td><td></td><td></td><td></td><td></td></tr>
<tr><td>Oligo rev</td><td>AAACCCTTCTTGCGCATCTCCTCG</td><td></td><td></td><td></td><td></td><td></td><td></td><td></td><td></td><td></td><td></td></tr>
<tr><td>Chromosome</td><td>start</td><td>end</td><td>strand</td><td>MM</td><td>target_seq</td><td>PAM</td><td>alignment</td><td>distance</td><td>position</td><td>gene name</td><td>gene id</td></tr>
<tr><td>GU305914.1</td><td>69916</td><td>69938</td><td>+</td><td>0</td><td>CGAGGAGATGCGCAAGAAGG</td><td>TGG</td><td>||||||||[||||||||||||]PAM</td><td>NA</td><td>Intergenic</td><td>NA</td><td>NA</td></tr>
<tr><td></td><td></td><td></td><td></td><td></td><td></td><td></td><td></td><td></td><td></td><td></td><td></td></tr>
<tr><td>T4883</td><td>GAAGGTGGACACCGACCTGAAGG</td><td>1000</td><td></td><td></td><td></td><td></td><td></td><td></td><td></td><td></td><td></td></tr>
<tr><td>Oligo fwd</td><td>CACCGAAGGTGGACACCGACCTGA</td><td></td><td></td><td></td><td></td><td></td><td></td><td></td><td></td><td></td><td></td></tr>
<tr><td>Oligo rev</td><td>AAACTCAGGTCGGTGTCCACCTTC</td><td></td><td></td><td></td><td></td><td></td><td></td><td></td><td></td><td></td><td></td></tr>
<tr><td>Chromosome</td><td>start</td><td>end</td><td>strand</td><td>MM</td><td>target_seq</td><td>PAM</td><td>alignment</td><td>distance</td><td>position</td><td>gene name</td><td>gene id</td></tr>
<tr><td>GU305914.1</td><td>69931</td><td>69953</td><td>+</td><td>0</td><td>GAAGGTGGACACCGACCTGA</td><td>AGG</td><td>||||||||[||||||||||||]PAM</td><td>NA</td><td>Intergenic</td><td>NA</td><td>NA</td></tr>
<tr><td></td><td></td><td></td><td></td><td></td><td></td><td></td><td></td><td></td><td></td><td></td><td></td></tr>
<tr><td>T4884</td><td>GGTGGACACCGACCTGAAGGTGG</td><td>1000</td><td></td><td></td><td></td><td></td><td></td><td></td><td></td><td></td><td></td></tr>
<tr><td>Oligo fwd</td><td>CACCGGTGGACACCGACCTGAAGG</td><td></td><td></td><td></td><td></td><td></td><td></td><td></td><td></td><td></td><td></td></tr>
<tr><td>Oligo rev</td><td>AAACCCTTCAGGTCGGTGTCCACC</td><td></td><td></td><td></td><td></td><td></td><td></td><td></td><td></td><td></td><td></td></tr>
<tr><td>Chromosome</td><td>start</td><td>end</td><td>strand</td><td>MM</td><td>target_seq</td><td>PAM</td><td>alignment</td><td>distance</td><td>position</td><td>gene name</td><td>gene id</td></tr>
<tr><td>GU305914.1</td><td>69934</td><td>69956</td><td>+</td><td>0</td><td>GGTGGACACCGACCTGAAGG</td><td>TGG</td><td>||||||||[||||||||||||]PAM</td><td>NA</td><td>Intergenic</td><td>NA</td><td>NA</td></tr>
<tr><td></td><td></td><td></td><td></td><td></td><td></td><td></td><td></td><td></td><td></td><td></td><td></td></tr>
<tr><td>T4885</td><td>CACCGACCTGAAGGTGGTGTCGG</td><td>1000</td><td></td><td></td><td></td><td></td><td></td><td></td><td></td><td></td><td></td></tr>
<tr><td>Oligo fwd</td><td>CACCCACCGACCTGAAGGTGGTGT</td><td></td><td></td><td></td><td></td><td></td><td></td><td></td><td></td><td></td><td></td></tr>
<tr><td>Oligo rev</td><td>AAACACACCACCTTCAGGTCGGTG</td><td></td><td></td><td></td><td></td><td></td><td></td><td></td><td></td><td></td><td></td></tr>
<tr><td>Chromosome</td><td>start</td><td>end</td><td>strand</td><td>MM</td><td>target_seq</td><td>PAM</td><td>alignment</td><td>distance</td><td>position</td><td>gene name</td><td>gene id</td></tr>
<tr><td>GU305914.1</td><td>69940</td><td>69962</td><td>+</td><td>0</td><td>CACCGACCTGAAGGTGGTGT</td><td>CGG</td><td>||||||||[||||||||||||]PAM</td><td>NA</td><td>Intergenic</td><td>NA</td><td>NA</td></tr>
<tr><td></td><td></td><td></td><td></td><td></td><td></td><td></td><td></td><td></td><td></td><td></td><td></td></tr>
<tr><td>T4886</td><td>TCGATCGACCTGAAAAGCTGCGG</td><td>1000</td><td></td><td></td><td></td><td></td><td></td><td></td><td></td><td></td><td></td></tr>
<tr><td>Oligo fwd</td><td>CACCTCGATCGACCTGAAAAGCTG</td><td></td><td></td><td></td><td></td><td></td><td></td><td></td><td></td><td></td><td></td></tr>
<tr><td>Oligo rev</td><td>AAACCAGCTTTTCAGGTCGATCGA</td><td></td><td></td><td></td><td></td><td></td><td></td><td></td><td></td><td></td><td></td></tr>
<tr><td>Chromosome</td><td>start</td><td>end</td><td>strand</td><td>MM</td><td>target_seq</td><td>PAM</td><td>alignment</td><td>distance</td><td>position</td><td>gene name</td><td>gene id</td></tr>
<tr><td>GU305914.1</td><td>69998</td><td>70020</td><td>+</td><td>0</td><td>TCGATCGACCTGAAAAGCTG</td><td>CGG</td><td>||||||||[||||||||||||]PAM</td><td>NA</td><td>Intergenic</td><td>NA</td><td>NA</td></tr>
<tr><td></td><td></td><td></td><td></td><td></td><td></td><td></td><td></td><td></td><td></td><td></td><td></td></tr>
<tr><td>T4887</td><td>GCTGCGGCACCTCGCTCGTGAGG</td><td>1000</td><td></td><td></td><td></td><td></td><td></td><td></td><td></td><td></td><td></td></tr>
<tr><td>Oligo fwd</td><td>CACCGCTGCGGCACCTCGCTCGTG</td><td></td><td></td><td></td><td></td><td></td><td></td><td></td><td></td><td></td><td></td></tr>
<tr><td>Oligo rev</td><td>AAACCACGAGCGAGGTGCCGCAGC</td><td></td><td></td><td></td><td></td><td></td><td></td><td></td><td></td><td></td><td></td></tr>
<tr><td>Chromosome</td><td>start</td><td>end</td><td>strand</td><td>MM</td><td>target_seq</td><td>PAM</td><td>alignment</td><td>distance</td><td>position</td><td>gene name</td><td>gene id</td></tr>
<tr><td>GU305914.1</td><td>70014</td><td>70036</td><td>+</td><td>0</td><td>GCTGCGGCACCTCGCTCGTG</td><td>AGG</td><td>||||||||[||||||||||||]PAM</td><td>NA</td><td>Intergenic</td><td>NA</td><td>NA</td></tr>
<tr><td></td><td></td><td></td><td></td><td></td><td></td><td></td><td></td><td></td><td></td><td></td><td></td></tr>
<tr><td>T4888</td><td>CTCGCTCGTGAGGCTGACCAAGG</td><td>1000</td><td></td><td></td><td></td><td></td><td></td><td></td><td></td><td></td><td></td></tr>
<tr><td>Oligo fwd</td><td>CACCCTCGCTCGTGAGGCTGACCA</td><td></td><td></td><td></td><td></td><td></td><td></td><td></td><td></td><td></td><td></td></tr>
<tr><td>Oligo rev</td><td>AAACTGGTCAGCCTCACGAGCGAG</td><td></td><td></td><td></td><td></td><td></td><td></td><td></td><td></td><td></td><td></td></tr>
<tr><td>Chromosome</td><td>start</td><td>end</td><td>strand</td><td>MM</td><td>target_seq</td><td>PAM</td><td>alignment</td><td>distance</td><td>position</td><td>gene name</td><td>gene id</td></tr>
<tr><td>GU305914.1</td><td>70024</td><td>70046</td><td>+</td><td>0</td><td>CTCGCTCGTGAGGCTGACCA</td><td>AGG</td><td>||||||||[||||||||||||]PAM</td><td>NA</td><td>Intergenic</td><td>NA</td><td>NA</td></tr>
<tr><td></td><td></td><td></td><td></td><td></td><td></td><td></td><td></td><td></td><td></td><td></td><td></td></tr>
<tr><td>T4889</td><td>GCTCGTGAGGCTGACCAAGGAGG</td><td>1000</td><td></td><td></td><td></td><td></td><td></td><td></td><td></td><td></td><td></td></tr>
<tr><td>Oligo fwd</td><td>CACCGCTCGTGAGGCTGACCAAGG</td><td></td><td></td><td></td><td></td><td></td><td></td><td></td><td></td><td></td><td></td></tr>
<tr><td>Oligo rev</td><td>AAACCCTTGGTCAGCCTCACGAGC</td><td></td><td></td><td></td><td></td><td></td><td></td><td></td><td></td><td></td><td></td></tr>
<tr><td>Chromosome</td><td>start</td><td>end</td><td>strand</td><td>MM</td><td>target_seq</td><td>PAM</td><td>alignment</td><td>distance</td><td>position</td><td>gene name</td><td>gene id</td></tr>
<tr><td>GU305914.1</td><td>70027</td><td>70049</td><td>+</td><td>0</td><td>GCTCGTGAGGCTGACCAAGG</td><td>AGG</td><td>||||||||[||||||||||||]PAM</td><td>NA</td><td>Intergenic</td><td>NA</td><td>NA</td></tr>
<tr><td></td><td></td><td></td><td></td><td></td><td></td><td></td><td></td><td></td><td></td><td></td><td></td></tr>
<tr><td>T4890</td><td>TGAGGCTGACCAAGGAGGACAGG</td><td>1000</td><td></td><td></td><td></td><td></td><td></td><td></td><td></td><td></td><td></td></tr>
<tr><td>Oligo fwd</td><td>CACCTGAGGCTGACCAAGGAGGAC</td><td></td><td></td><td></td><td></td><td></td><td></td><td></td><td></td><td></td><td></td></tr>
<tr><td>Oligo rev</td><td>AAACGTCCTCCTTGGTCAGCCTCA</td><td></td><td></td><td></td><td></td><td></td><td></td><td></td><td></td><td></td><td></td></tr>
<tr><td>Chromosome</td><td>start</td><td>end</td><td>strand</td><td>MM</td><td>target_seq</td><td>PAM</td><td>alignment</td><td>distance</td><td>position</td><td>gene name</td><td>gene id</td></tr>
<tr><td>GU305914.1</td><td>70032</td><td>70054</td><td>+</td><td>0</td><td>TGAGGCTGACCAAGGAGGAC</td><td>AGG</td><td>||||||||[||||||||||||]PAM</td><td>NA</td><td>Intergenic</td><td>NA</td><td>NA</td></tr>
<tr><td>GU305914.1</td><td>124206</td><td>124228</td><td>-</td><td>4</td><td>CGATCGTGACCAAGGAGGAC</td><td>CGG</td><td>-||---||[||||||||||||]PAM</td><td>NA</td><td>Intergenic</td><td>NA</td><td>NA</td></tr>
<tr><td></td><td></td><td></td><td></td><td></td><td></td><td></td><td></td><td></td><td></td><td></td><td></td></tr>
<tr><td>T4891</td><td>CAAATCCCTGAACTCCCTGCTGG</td><td>1000</td><td></td><td></td><td></td><td></td><td></td><td></td><td></td><td></td><td></td></tr>
<tr><td>Oligo fwd</td><td>CACCCAAATCCCTGAACTCCCTGC</td><td></td><td></td><td></td><td></td><td></td><td></td><td></td><td></td><td></td><td></td></tr>
<tr><td>Oligo rev</td><td>AAACGCAGGGAGTTCAGGGATTTG</td><td></td><td></td><td></td><td></td><td></td><td></td><td></td><td></td><td></td><td></td></tr>
<tr><td>Chromosome</td><td>start</td><td>end</td><td>strand</td><td>MM</td><td>target_seq</td><td>PAM</td><td>alignment</td><td>distance</td><td>position</td><td>gene name</td><td>gene id</td></tr>
<tr><td>GU305914.1</td><td>70090</td><td>70112</td><td>+</td><td>0</td><td>CAAATCCCTGAACTCCCTGC</td><td>TGG</td><td>||||||||[||||||||||||]PAM</td><td>NA</td><td>Intergenic</td><td>NA</td><td>NA</td></tr>
<tr><td></td><td></td><td></td><td></td><td></td><td></td><td></td><td></td><td></td><td></td><td></td><td></td></tr>
<tr><td>T4892</td><td>CCTGCTGGCGTCGCTCGTCACGG</td><td>1000</td><td></td><td></td><td></td><td></td><td></td><td></td><td></td><td></td><td></td></tr>
<tr><td>Oligo fwd</td><td>CACCCCTGCTGGCGTCGCTCGTCA</td><td></td><td></td><td></td><td></td><td></td><td></td><td></td><td></td><td></td><td></td></tr>
<tr><td>Oligo rev</td><td>AAACTGACGAGCGACGCCAGCAGG</td><td></td><td></td><td></td><td></td><td></td><td></td><td></td><td></td><td></td><td></td></tr>
<tr><td>Chromosome</td><td>start</td><td>end</td><td>strand</td><td>MM</td><td>target_seq</td><td>PAM</td><td>alignment</td><td>distance</td><td>position</td><td>gene name</td><td>gene id</td></tr>
<tr><td>GU305914.1</td><td>70105</td><td>70127</td><td>+</td><td>0</td><td>CCTGCTGGCGTCGCTCGTCA</td><td>CGG</td><td>||||||||[||||||||||||]PAM</td><td>NA</td><td>Intergenic</td><td>NA</td><td>NA</td></tr>
<tr><td></td><td></td><td></td><td></td><td></td><td></td><td></td><td></td><td></td><td></td><td></td><td></td></tr>
<tr><td>T4893</td><td>GCTGGCGTCGCTCGTCACGGTGG</td><td>1000</td><td></td><td></td><td></td><td></td><td></td><td></td><td></td><td></td><td></td></tr>
<tr><td>Oligo fwd</td><td>CACCGCTGGCGTCGCTCGTCACGG</td><td></td><td></td><td></td><td></td><td></td><td></td><td></td><td></td><td></td><td></td></tr>
<tr><td>Oligo rev</td><td>AAACCCGTGACGAGCGACGCCAGC</td><td></td><td></td><td></td><td></td><td></td><td></td><td></td><td></td><td></td><td></td></tr>
<tr><td>Chromosome</td><td>start</td><td>end</td><td>strand</td><td>MM</td><td>target_seq</td><td>PAM</td><td>alignment</td><td>distance</td><td>position</td><td>gene name</td><td>gene id</td></tr>
<tr><td>GU305914.1</td><td>70108</td><td>70130</td><td>+</td><td>0</td><td>GCTGGCGTCGCTCGTCACGG</td><td>TGG</td><td>||||||||[||||||||||||]PAM</td><td>NA</td><td>Intergenic</td><td>NA</td><td>NA</td></tr>
<tr><td></td><td></td><td></td><td></td><td></td><td></td><td></td><td></td><td></td><td></td><td></td><td></td></tr>
<tr><td>T4894</td><td>GTCGCTCGTCACGGTGGAGACGG</td><td>1000</td><td></td><td></td><td></td><td></td><td></td><td></td><td></td><td></td><td></td></tr>
<tr><td>Oligo fwd</td><td>CACCGTCGCTCGTCACGGTGGAGA</td><td></td><td></td><td></td><td></td><td></td><td></td><td></td><td></td><td></td><td></td></tr>
<tr><td>Oligo rev</td><td>AAACTCTCCACCGTGACGAGCGAC</td><td></td><td></td><td></td><td></td><td></td><td></td><td></td><td></td><td></td><td></td></tr>
<tr><td>Chromosome</td><td>start</td><td>end</td><td>strand</td><td>MM</td><td>target_seq</td><td>PAM</td><td>alignment</td><td>distance</td><td>position</td><td>gene name</td><td>gene id</td></tr>
<tr><td>GU305914.1</td><td>70114</td><td>70136</td><td>+</td><td>0</td><td>GTCGCTCGTCACGGTGGAGA</td><td>CGG</td><td>||||||||[||||||||||||]PAM</td><td>NA</td><td>Intergenic</td><td>NA</td><td>NA</td></tr>
<tr><td></td><td></td><td></td><td></td><td></td><td></td><td></td><td></td><td></td><td></td><td></td><td></td></tr>
<tr><td>T4895</td><td>CACGATCGCCGCGATCCTCAAGG</td><td>1000</td><td></td><td></td><td></td><td></td><td></td><td></td><td></td><td></td><td></td></tr>
<tr><td>Oligo fwd</td><td>CACCCACGATCGCCGCGATCCTCA</td><td></td><td></td><td></td><td></td><td></td><td></td><td></td><td></td><td></td><td></td></tr>
<tr><td>Oligo rev</td><td>AAACTGAGGATCGCGGCGATCGTG</td><td></td><td></td><td></td><td></td><td></td><td></td><td></td><td></td><td></td><td></td></tr>
<tr><td>Chromosome</td><td>start</td><td>end</td><td>strand</td><td>MM</td><td>target_seq</td><td>PAM</td><td>alignment</td><td>distance</td><td>position</td><td>gene name</td><td>gene id</td></tr>
<tr><td>GU305914.1</td><td>70138</td><td>70160</td><td>+</td><td>0</td><td>CACGATCGCCGCGATCCTCA</td><td>AGG</td><td>||||||||[||||||||||||]PAM</td><td>NA</td><td>Intergenic</td><td>NA</td><td>NA</td></tr>
<tr><td></td><td></td><td></td><td></td><td></td><td></td><td></td><td></td><td></td><td></td><td></td><td></td></tr>
<tr><td>T4896</td><td>ACGATCGCCGCGATCCTCAAGGG</td><td>1000</td><td></td><td></td><td></td><td></td><td></td><td></td><td></td><td></td><td></td></tr>
<tr><td>Oligo fwd</td><td>CACCACGATCGCCGCGATCCTCAA</td><td></td><td></td><td></td><td></td><td></td><td></td><td></td><td></td><td></td><td></td></tr>
<tr><td>Oligo rev</td><td>AAACTTGAGGATCGCGGCGATCGT</td><td></td><td></td><td></td><td></td><td></td><td></td><td></td><td></td><td></td><td></td></tr>
<tr><td>Chromosome</td><td>start</td><td>end</td><td>strand</td><td>MM</td><td>target_seq</td><td>PAM</td><td>alignment</td><td>distance</td><td>position</td><td>gene name</td><td>gene id</td></tr>
</table>

<table>
<tr><td>GU305914.1</td><td></td><td>70139</td><td>70161</td><td>+</td><td>0</td><td>ACGATCGCCGCGATCCTCAA</td><td>GGG</td><td>||||||||[||||||||||||]PAM</td><td>NA</td><td>Intergenic</td><td>NA</td><td>NA</td></tr>
<tr><td>T4897</td><td>CGCCGCGATCCTCAAGGGCGCGG</td><td></td><td>1000</td><td></td><td></td><td></td><td></td><td></td><td></td><td></td><td></td><td></td></tr>
<tr><td>Oligo fwd</td><td>CACCCGCCGCGATCCTCAAGGGCG</td><td></td><td></td><td></td><td></td><td></td><td></td><td></td><td></td><td></td><td></td><td></td></tr>
<tr><td>Oligo rev</td><td>AAACCGCCCTTGAGGATCGCGGCG</td><td></td><td></td><td></td><td></td><td></td><td></td><td></td><td></td><td></td><td></td><td></td></tr>
<tr><td>Chromosome</td><td>start</td><td></td><td>end</td><td>strand</td><td>MM</td><td>target_seq</td><td>PAM</td><td>alignment</td><td>distance</td><td>position</td><td>gene name</td><td>gene id</td></tr>
<tr><td>GU305914.1</td><td></td><td>70144</td><td>70166</td><td>+</td><td>0</td><td>CGCCGCGATCCTCAAGGGCG</td><td>CGG</td><td>||||||||[||||||||||||]PAM</td><td>NA</td><td>Intergenic</td><td>NA</td><td>NA</td></tr>
<tr><td>T4898</td><td>CCTCAAGGGCGCGGACCCCGAGG</td><td></td><td>1000</td><td></td><td></td><td></td><td></td><td></td><td></td><td></td><td></td><td></td></tr>
<tr><td>Oligo fwd</td><td>CACCCCTCAAGGGCGCGGACCCCG</td><td></td><td></td><td></td><td></td><td></td><td></td><td></td><td></td><td></td><td></td><td></td></tr>
<tr><td>Oligo rev</td><td>AAACCGGGGTCCGCGCCCTTGAGG</td><td></td><td></td><td></td><td></td><td></td><td></td><td></td><td></td><td></td><td></td><td></td></tr>
<tr><td>Chromosome</td><td>start</td><td></td><td>end</td><td>strand</td><td>MM</td><td>target_seq</td><td>PAM</td><td>alignment</td><td>distance</td><td>position</td><td>gene name</td><td>gene id</td></tr>
<tr><td>GU305914.1</td><td></td><td>70153</td><td>70175</td><td>+</td><td>0</td><td>CCTCAAGGGCGCGGACCCCG</td><td>AGG</td><td>||||||||[||||||||||||]PAM</td><td>NA</td><td>Intergenic</td><td>NA</td><td>NA</td></tr>
<tr><td>T4899</td><td>CCCGAGGCGACGAGCAGCACCGG</td><td></td><td>1000</td><td></td><td></td><td></td><td></td><td></td><td></td><td></td><td></td><td></td></tr>
<tr><td>Oligo fwd</td><td>CACCCCCGAGGCGACGAGCAGCAC</td><td></td><td></td><td></td><td></td><td></td><td></td><td></td><td></td><td></td><td></td><td></td></tr>
<tr><td>Oligo rev</td><td>AAACGTGCTGCTCGTCGCCTCGGG</td><td></td><td></td><td></td><td></td><td></td><td></td><td></td><td></td><td></td><td></td><td></td></tr>
<tr><td>Chromosome</td><td>start</td><td></td><td>end</td><td>strand</td><td>MM</td><td>target_seq</td><td>PAM</td><td>alignment</td><td>distance</td><td>position</td><td>gene name</td><td>gene id</td></tr>
<tr><td>GU305914.1</td><td></td><td>70169</td><td>70191</td><td>+</td><td>0</td><td>CCCGAGGCGACGAGCAGCAC</td><td>CGG</td><td>||||||||[||||||||||||]PAM</td><td>NA</td><td>Intergenic</td><td>NA</td><td>NA</td></tr>
<tr><td>GU305914.1</td><td></td><td>52303</td><td>52325</td><td>+</td><td>3</td><td>CCAGAGGCGAAGAGCAGCGC</td><td>TGG</td><td>||-|||||[||-|||||||-|]PAM</td><td>NA</td><td>Intergenic</td><td>NA</td><td>NA</td></tr>
<tr><td>T4900</td><td>ACGAGCAGCACCGGCAGCGCCGG</td><td></td><td>1000</td><td></td><td></td><td></td><td></td><td></td><td></td><td></td><td></td><td></td></tr>
<tr><td>Oligo fwd</td><td>CACCACGAGCAGCACCGGCAGCGC</td><td></td><td></td><td></td><td></td><td></td><td></td><td></td><td></td><td></td><td></td><td></td></tr>
<tr><td>Oligo rev</td><td>AAACGCGCTGCCGGTGCTGCTCGT</td><td></td><td></td><td></td><td></td><td></td><td></td><td></td><td></td><td></td><td></td><td></td></tr>
<tr><td>Chromosome</td><td>start</td><td></td><td>end</td><td>strand</td><td>MM</td><td>target_seq</td><td>PAM</td><td>alignment</td><td>distance</td><td>position</td><td>gene name</td><td>gene id</td></tr>
<tr><td>GU305914.1</td><td></td><td>70178</td><td>70200</td><td>+</td><td>0</td><td>ACGAGCAGCACCGGCAGCGC</td><td>CGG</td><td>||||||||[||||||||||||]PAM</td><td>NA</td><td>Intergenic</td><td>NA</td><td>NA</td></tr>
<tr><td>T4901</td><td>AGCAGCACCGGCAGCGCCGGCGG</td><td></td><td>1000</td><td></td><td></td><td></td><td></td><td></td><td></td><td></td><td></td><td></td></tr>
<tr><td>Oligo fwd</td><td>CACCAGCAGCACCGGCAGCGCCGG</td><td></td><td></td><td></td><td></td><td></td><td></td><td></td><td></td><td></td><td></td><td></td></tr>
<tr><td>Oligo rev</td><td>AAACCCGGCGCTGCCGGTGCTGCT</td><td></td><td></td><td></td><td></td><td></td><td></td><td></td><td></td><td></td><td></td><td></td></tr>
<tr><td>Chromosome</td><td>start</td><td></td><td>end</td><td>strand</td><td>MM</td><td>target_seq</td><td>PAM</td><td>alignment</td><td>distance</td><td>position</td><td>gene name</td><td>gene id</td></tr>
<tr><td>GU305914.1</td><td></td><td>70181</td><td>70203</td><td>+</td><td>0</td><td>AGCAGCACCGGCAGCGCCGG</td><td>CGG</td><td>||||||||[||||||||||||]PAM</td><td>NA</td><td>Intergenic</td><td>NA</td><td>NA</td></tr>
<tr><td>GU305914.1</td><td></td><td>24627</td><td>24649</td><td>-</td><td>4</td><td>CGCAACACCGGCGGCGGCGG</td><td>TGG</td><td>-|||-|||[||||-|||-|||]PAM</td><td>NA</td><td>Intergenic</td><td>NA</td><td>NA</td></tr>
<tr><td>GU305914.1</td><td></td><td>117379</td><td>117401</td><td>+</td><td>4</td><td>AACGGCACCGACAGCGCCGA</td><td>CGG</td><td>|-|-||||[||-||||||||-]PAM</td><td>NA</td><td>Intergenic</td><td>NA</td><td>NA</td></tr>
<tr><td>T4902</td><td>AGCACCGGCAGCGCCGGCGGCGG</td><td></td><td>1000</td><td></td><td></td><td></td><td></td><td></td><td></td><td></td><td></td><td></td></tr>
<tr><td>Oligo fwd</td><td>CACCAGCACCGGCAGCGCCGGCGG</td><td></td><td></td><td></td><td></td><td></td><td></td><td></td><td></td><td></td><td></td><td></td></tr>
<tr><td>Oligo rev</td><td>AAACCCGCCGGCGCTGCCGGTGCT</td><td></td><td></td><td></td><td></td><td></td><td></td><td></td><td></td><td></td><td></td><td></td></tr>
<tr><td>Chromosome</td><td>start</td><td></td><td>end</td><td>strand</td><td>MM</td><td>target_seq</td><td>PAM</td><td>alignment</td><td>distance</td><td>position</td><td>gene name</td><td>gene id</td></tr>
<tr><td>GU305914.1</td><td></td><td>70184</td><td>70206</td><td>+</td><td>0</td><td>AGCACCGGCAGCGCCGGCGG</td><td>CGG</td><td>||||||||[||||||||||||]PAM</td><td>NA</td><td>Intergenic</td><td>NA</td><td>NA</td></tr>
<tr><td>GU305914.1</td><td></td><td>73792</td><td>73814</td><td>-</td><td>4</td><td>GGCGCCAGCAGCGGCGGCGG</td><td>AGG</td><td>-||-||-|[|||||-||||||]PAM</td><td>NA</td><td>Intergenic</td><td>NA</td><td>NA</td></tr>
<tr><td>GU305914.1</td><td></td><td>57832</td><td>57854</td><td>-</td><td>4</td><td>ACCACCGAGAGCGACGGCGG</td><td>CGG</td><td>|-|||||-[-||||-||||||]PAM</td><td>NA</td><td>Intergenic</td><td>NA</td><td>NA</td></tr>
<tr><td>GU305914.1</td><td></td><td>128803</td><td>128825</td><td>+</td><td>4</td><td>CGCAGCGGCAGCGACGTCGG</td><td>CGG</td><td>-|||-|||[|||||-||-|||]PAM</td><td>NA</td><td>Intergenic</td><td>NA</td><td>NA</td></tr>
<tr><td>T4903</td><td>GAACAACATCTCGACGTTGCTGG</td><td></td><td>1000</td><td></td><td></td><td></td><td></td><td></td><td></td><td></td><td></td><td></td></tr>
<tr><td>Oligo fwd</td><td>CACCGAACAACATCTCGACGTTGC</td><td></td><td></td><td></td><td></td><td></td><td></td><td></td><td></td><td></td><td></td><td></td></tr>
<tr><td>Oligo rev</td><td>AAACGCAACGTCGAGATGTTGTTC</td><td></td><td></td><td></td><td></td><td></td><td></td><td></td><td></td><td></td><td></td><td></td></tr>
<tr><td>Chromosome</td><td>start</td><td></td><td>end</td><td>strand</td><td>MM</td><td>target_seq</td><td>PAM</td><td>alignment</td><td>distance</td><td>position</td><td>gene name</td><td>gene id</td></tr>
<tr><td>GU305914.1</td><td></td><td>70240</td><td>70262</td><td>+</td><td>0</td><td>GAACAACATCTCGACGTTGC</td><td>TGG</td><td>||||||||[||||||||||||]PAM</td><td>NA</td><td>Intergenic</td><td>NA</td><td>NA</td></tr>
<tr><td>T4904</td><td>AACAACATCTCGACGTTGCTGGG</td><td></td><td>1000</td><td></td><td></td><td></td><td></td><td></td><td></td><td></td><td></td><td></td></tr>
<tr><td>Oligo fwd</td><td>CACCAACAACATCTCGACGTTGCT</td><td></td><td></td><td></td><td></td><td></td><td></td><td></td><td></td><td></td><td></td><td></td></tr>
<tr><td>Oligo rev</td><td>AAACAGCAACGTCGAGATGTTGTT</td><td></td><td></td><td></td><td></td><td></td><td></td><td></td><td></td><td></td><td></td><td></td></tr>
<tr><td>Chromosome</td><td>start</td><td></td><td>end</td><td>strand</td><td>MM</td><td>target_seq</td><td>PAM</td><td>alignment</td><td>distance</td><td>position</td><td>gene name</td><td>gene id</td></tr>
<tr><td>GU305914.1</td><td></td><td>70241</td><td>70263</td><td>+</td><td>0</td><td>AACAACATCTCGACGTTGCT</td><td>GGG</td><td>||||||||[||||||||||||]PAM</td><td>NA</td><td>Intergenic</td><td>NA</td><td>NA</td></tr>
<tr><td>T4905</td><td>ACATCTCGACGTTGCTGGGCTGG</td><td></td><td>1000</td><td></td><td></td><td></td><td></td><td></td><td></td><td></td><td></td><td></td></tr>
<tr><td>Oligo fwd</td><td>CACCACATCTCGACGTTGCTGGGC</td><td></td><td></td><td></td><td></td><td></td><td></td><td></td><td></td><td></td><td></td><td></td></tr>
<tr><td>Oligo rev</td><td>AAACGCCCAGCAACGTCGAGATGT</td><td></td><td></td><td></td><td></td><td></td><td></td><td></td><td></td><td></td><td></td><td></td></tr>
<tr><td>Chromosome</td><td>start</td><td></td><td>end</td><td>strand</td><td>MM</td><td>target_seq</td><td>PAM</td><td>alignment</td><td>distance</td><td>position</td><td>gene name</td><td>gene id</td></tr>
<tr><td>GU305914.1</td><td></td><td>70245</td><td>70267</td><td>+</td><td>0</td><td>ACATCTCGACGTTGCTGGGC</td><td>TGG</td><td>||||||||[||||||||||||]PAM</td><td>NA</td><td>Intergenic</td><td>NA</td><td>NA</td></tr>
<tr><td>T4906</td><td>GTTGCTGGGCTGGCAGCAGAAGG</td><td></td><td>1000</td><td></td><td></td><td></td><td></td><td></td><td></td><td></td><td></td><td></td></tr>
<tr><td>Oligo fwd</td><td>CACCGTTGCTGGGCTGGCAGCAGA</td><td></td><td></td><td></td><td></td><td></td><td></td><td></td><td></td><td></td><td></td><td></td></tr>
<tr><td>Oligo rev</td><td>AAACTCTGCTGCCAGCCCAGCAAC</td><td></td><td></td><td></td><td></td><td></td><td></td><td></td><td></td><td></td><td></td><td></td></tr>
<tr><td>Chromosome</td><td>start</td><td></td><td>end</td><td>strand</td><td>MM</td><td>target_seq</td><td>PAM</td><td>alignment</td><td>distance</td><td>position</td><td>gene name</td><td>gene id</td></tr>
<tr><td>GU305914.1</td><td></td><td>70255</td><td>70277</td><td>+</td><td>0</td><td>GTTGCTGGGCTGGCAGCAGA</td><td>AGG</td><td>||||||||[||||||||||||]PAM</td><td>NA</td><td>Intergenic</td><td>NA</td><td>NA</td></tr>
<tr><td>T4907</td><td>GCAGCAGAAGGTGCTGCTGCCGG</td><td></td><td>1000</td><td></td><td></td><td></td><td></td><td></td><td></td><td></td><td></td><td></td></tr>
<tr><td>Oligo fwd</td><td>CACCGCAGCAGAAGGTGCTGCTGC</td><td></td><td></td><td></td><td></td><td></td><td></td><td></td><td></td><td></td><td></td><td></td></tr>
<tr><td>Oligo rev</td><td>AAACGCAGCAGCACCTTCTGCTGC</td><td></td><td></td><td></td><td></td><td></td><td></td><td></td><td></td><td></td><td></td><td></td></tr>
<tr><td>Chromosome</td><td>start</td><td></td><td>end</td><td>strand</td><td>MM</td><td>target_seq</td><td>PAM</td><td>alignment</td><td>distance</td><td>position</td><td>gene name</td><td>gene id</td></tr>
<tr><td>GU305914.1</td><td></td><td>70267</td><td>70289</td><td>+</td><td>0</td><td>GCAGCAGAAGGTGCTGCTGC</td><td>CGG</td><td>||||||||[||||||||||||]PAM</td><td>NA</td><td>Intergenic</td><td>NA</td><td>NA</td></tr>
<tr><td>T4908</td><td>GAAGGTGCTGCTGCCGGAGACGG</td><td></td><td>1000</td><td></td><td></td><td></td><td></td><td></td><td></td><td></td><td></td><td></td></tr>
<tr><td>Oligo fwd</td><td>CACCGAAGGTGCTGCTGCCGGAGA</td><td></td><td></td><td></td><td></td><td></td><td></td><td></td><td></td><td></td><td></td><td></td></tr>
<tr><td>Oligo rev</td><td>AAACTCTCCGGCAGCAGCACCTTC</td><td></td><td></td><td></td><td></td><td></td><td></td><td></td><td></td><td></td><td></td><td></td></tr>
<tr><td>Chromosome</td><td>start</td><td></td><td>end</td><td>strand</td><td>MM</td><td>target_seq</td><td>PAM</td><td>alignment</td><td>distance</td><td>position</td><td>gene name</td><td>gene id</td></tr>
<tr><td>GU305914.1</td><td></td><td>70273</td><td>70295</td><td>+</td><td>0</td><td>GAAGGTGCTGCTGCCGGAGA</td><td>CGG</td><td>||||||||[||||||||||||]PAM</td><td>NA</td><td>Intergenic</td><td>NA</td><td>NA</td></tr>
<tr><td>T4909</td><td>TGACGCACATGCGAAAGAACTGG</td><td></td><td>1000</td><td></td><td></td><td></td><td></td><td></td><td></td><td></td><td></td><td></td></tr>
<tr><td>Oligo fwd</td><td>CACCTGACGCACATGCGAAAGAAC</td><td></td><td></td><td></td><td></td><td></td><td></td><td></td><td></td><td></td><td></td><td></td></tr>
<tr><td>Oligo rev</td><td>AAACGTTCTTTCGCATGTGCGTCA</td><td></td><td></td><td></td><td></td><td></td><td></td><td></td><td></td><td></td><td></td><td></td></tr>
<tr><td>Chromosome</td><td>start</td><td></td><td>end</td><td>strand</td><td>MM</td><td>target_seq</td><td>PAM</td><td>alignment</td><td>distance</td><td>position</td><td>gene name</td><td>gene id</td></tr>
<tr><td>GU305914.1</td><td></td><td>70335</td><td>70357</td><td>+</td><td>0</td><td>TGACGCACATGCGAAAGAAC</td><td>TGG</td><td>||||||||[||||||||||||]PAM</td><td>NA</td><td>Intergenic</td><td>NA</td><td>NA</td></tr>
<tr><td>T4910</td><td>AAAGAACTGGCGCAACCCCGCGG</td><td></td><td>1000</td><td></td><td></td><td></td><td></td><td></td><td></td><td></td><td></td><td></td></tr>
<tr><td>Oligo fwd</td><td>CACCAAAGAACTGGCGCAACCCCG</td><td></td><td></td><td></td><td></td><td></td><td></td><td></td><td></td><td></td><td></td><td></td></tr>
<tr><td>Oligo rev</td><td>AAACCGGGGTTGCGCCAGTTCTTT</td><td></td><td></td><td></td><td></td><td></td><td></td><td></td><td></td><td></td><td></td><td></td></tr>
</table>

<table>
<tr><td>Chromosome</td><td>start</td><td>end</td><td>strand</td><td>MM</td><td>target_seq</td><td>PAM</td><td>alignment</td><td>distance</td><td>position</td><td>gene name</td><td>gene id</td></tr>
<tr><td>GU305914.1</td><td>70348</td><td>70370</td><td>+</td><td>0</td><td>AAAGAACTGGCGCAACCCCG</td><td>CGG</td><td>||||||||[||||||||||||]PAM</td><td>NA</td><td>Intergenic</td><td>NA</td><td>NA</td></tr>
<tr><td></td><td></td><td></td><td></td><td></td><td></td><td></td><td></td><td></td><td></td><td></td><td></td></tr>
<tr><td>T4911</td><td>GAACTGGCGCAACCCCGCGGCGG</td><td>1000</td><td></td><td></td><td></td><td></td><td></td><td></td><td></td><td></td><td></td></tr>
<tr><td>Oligo fwd</td><td>CACCGAACTGGCGCAACCCCGCGG</td><td></td><td></td><td></td><td></td><td></td><td></td><td></td><td></td><td></td><td></td></tr>
<tr><td>Oligo rev</td><td>AAACCCGCGGGGTTGCGCCAGTTC</td><td></td><td></td><td></td><td></td><td></td><td></td><td></td><td></td><td></td><td></td></tr>
<tr><td>Chromosome</td><td>start</td><td>end</td><td>strand</td><td>MM</td><td>target_seq</td><td>PAM</td><td>alignment</td><td>distance</td><td>position</td><td>gene name</td><td>gene id</td></tr>
<tr><td>GU305914.1</td><td>70351</td><td>70373</td><td>+</td><td>0</td><td>GAACTGGCGCAACCCCGCGG</td><td>CGG</td><td>||||||||[||||||||||||]PAM</td><td>NA</td><td>Intergenic</td><td>NA</td><td>NA</td></tr>
<tr><td></td><td></td><td></td><td></td><td></td><td></td><td></td><td></td><td></td><td></td><td></td><td></td></tr>
<tr><td>T4912</td><td>CTTCAGCAGCACCCCCCACACGG</td><td>1000</td><td></td><td></td><td></td><td></td><td></td><td></td><td></td><td></td><td></td></tr>
<tr><td>Oligo fwd</td><td>CACCCTTCAGCAGCACCCCCCACA</td><td></td><td></td><td></td><td></td><td></td><td></td><td></td><td></td><td></td><td></td></tr>
<tr><td>Oligo rev</td><td>AAACTGTGGGGGGTGCTGCTGAAG</td><td></td><td></td><td></td><td></td><td></td><td></td><td></td><td></td><td></td><td></td></tr>
<tr><td>Chromosome</td><td>start</td><td>end</td><td>strand</td><td>MM</td><td>target_seq</td><td>PAM</td><td>alignment</td><td>distance</td><td>position</td><td>gene name</td><td>gene id</td></tr>
<tr><td>GU305914.1</td><td>70375</td><td>70397</td><td>+</td><td>0</td><td>CTTCAGCAGCACCCCCCACA</td><td>CGG</td><td>||||||||[||||||||||||]PAM</td><td>NA</td><td>Intergenic</td><td>NA</td><td>NA</td></tr>
<tr><td></td><td></td><td></td><td></td><td></td><td></td><td></td><td></td><td></td><td></td><td></td><td></td></tr>
<tr><td>T4913</td><td>CCCCCCACACGGCCGCGTACAGG</td><td>1000</td><td></td><td></td><td></td><td></td><td></td><td></td><td></td><td></td><td></td></tr>
<tr><td>Oligo fwd</td><td>CACCCCCCCCACACGGCCGCGTAC</td><td></td><td></td><td></td><td></td><td></td><td></td><td></td><td></td><td></td><td></td></tr>
<tr><td>Oligo rev</td><td>AAACGTACGCGGCCGTGTGGGGGG</td><td></td><td></td><td></td><td></td><td></td><td></td><td></td><td></td><td></td><td></td></tr>
<tr><td>Chromosome</td><td>start</td><td>end</td><td>strand</td><td>MM</td><td>target_seq</td><td>PAM</td><td>alignment</td><td>distance</td><td>position</td><td>gene name</td><td>gene id</td></tr>
<tr><td>GU305914.1</td><td>70386</td><td>70408</td><td>+</td><td>0</td><td>CCCCCCACACGGCCGCGTAC</td><td>AGG</td><td>||||||||[||||||||||||]PAM</td><td>NA</td><td>Intergenic</td><td>NA</td><td>NA</td></tr>
<tr><td></td><td></td><td></td><td></td><td></td><td></td><td></td><td></td><td></td><td></td><td></td><td></td></tr>
<tr><td>T4914</td><td>GTACAGGAACTTCGCCGAACTGG</td><td>1000</td><td></td><td></td><td></td><td></td><td></td><td></td><td></td><td></td><td></td></tr>
<tr><td>Oligo fwd</td><td>CACCGTACAGGAACTTCGCCGAAC</td><td></td><td></td><td></td><td></td><td></td><td></td><td></td><td></td><td></td><td></td></tr>
<tr><td>Oligo rev</td><td>AAACGTTCGGCGAAGTTCCTGTAC</td><td></td><td></td><td></td><td></td><td></td><td></td><td></td><td></td><td></td><td></td></tr>
<tr><td>Chromosome</td><td>start</td><td>end</td><td>strand</td><td>MM</td><td>target_seq</td><td>PAM</td><td>alignment</td><td>distance</td><td>position</td><td>gene name</td><td>gene id</td></tr>
<tr><td>GU305914.1</td><td>70402</td><td>70424</td><td>+</td><td>0</td><td>GTACAGGAACTTCGCCGAAC</td><td>TGG</td><td>||||||||[||||||||||||]PAM</td><td>NA</td><td>Intergenic</td><td>NA</td><td>NA</td></tr>
<tr><td></td><td></td><td></td><td></td><td></td><td></td><td></td><td></td><td></td><td></td><td></td><td></td></tr>
<tr><td>T4915</td><td>CGAACGAGATCGAGAGCCGCAGG</td><td>1000</td><td></td><td></td><td></td><td></td><td></td><td></td><td></td><td></td><td></td></tr>
<tr><td>Oligo fwd</td><td>CACCCGAACGAGATCGAGAGCCGC</td><td></td><td></td><td></td><td></td><td></td><td></td><td></td><td></td><td></td><td></td></tr>
<tr><td>Oligo rev</td><td>AAACGCGGCTCTCGATCTCGTTCG</td><td></td><td></td><td></td><td></td><td></td><td></td><td></td><td></td><td></td><td></td></tr>
<tr><td>Chromosome</td><td>start</td><td>end</td><td>strand</td><td>MM</td><td>target_seq</td><td>PAM</td><td>alignment</td><td>distance</td><td>position</td><td>gene name</td><td>gene id</td></tr>
<tr><td>GU305914.1</td><td>70425</td><td>70447</td><td>+</td><td>0</td><td>CGAACGAGATCGAGAGCCGC</td><td>AGG</td><td>||||||||[||||||||||||]PAM</td><td>NA</td><td>Intergenic</td><td>NA</td><td>NA</td></tr>
<tr><td></td><td></td><td></td><td></td><td></td><td></td><td></td><td></td><td></td><td></td><td></td><td></td></tr>
<tr><td>T4916</td><td>CGAGATCGAGAGCCGCAGGACGG</td><td>1000</td><td></td><td></td><td></td><td></td><td></td><td></td><td></td><td></td><td></td></tr>
<tr><td>Oligo fwd</td><td>CACCCGAGATCGAGAGCCGCAGGA</td><td></td><td></td><td></td><td></td><td></td><td></td><td></td><td></td><td></td><td></td></tr>
<tr><td>Oligo rev</td><td>AAACTCCTGCGGCTCTCGATCTCG</td><td></td><td></td><td></td><td></td><td></td><td></td><td></td><td></td><td></td><td></td></tr>
<tr><td>Chromosome</td><td>start</td><td>end</td><td>strand</td><td>MM</td><td>target_seq</td><td>PAM</td><td>alignment</td><td>distance</td><td>position</td><td>gene name</td><td>gene id</td></tr>
<tr><td>GU305914.1</td><td>70429</td><td>70451</td><td>+</td><td>0</td><td>CGAGATCGAGAGCCGCAGGA</td><td>CGG</td><td>||||||||[||||||||||||]PAM</td><td>NA</td><td>Intergenic</td><td>NA</td><td>NA</td></tr>
<tr><td></td><td></td><td></td><td></td><td></td><td></td><td></td><td></td><td></td><td></td><td></td><td></td></tr>
<tr><td>T4917</td><td>GAGCCGCAGGACGGAGACGCTGG</td><td>1000</td><td></td><td></td><td></td><td></td><td></td><td></td><td></td><td></td><td></td></tr>
<tr><td>Oligo fwd</td><td>CACCGAGCCGCAGGACGGAGACGC</td><td></td><td></td><td></td><td></td><td></td><td></td><td></td><td></td><td></td><td></td></tr>
<tr><td>Oligo rev</td><td>AAACGCGTCTCCGTCCTGCGGCTC</td><td></td><td></td><td></td><td></td><td></td><td></td><td></td><td></td><td></td><td></td></tr>
<tr><td>Chromosome</td><td>start</td><td>end</td><td>strand</td><td>MM</td><td>target_seq</td><td>PAM</td><td>alignment</td><td>distance</td><td>position</td><td>gene name</td><td>gene id</td></tr>
<tr><td>GU305914.1</td><td>70438</td><td>70460</td><td>+</td><td>0</td><td>GAGCCGCAGGACGGAGACGC</td><td>TGG</td><td>||||||||[||||||||||||]PAM</td><td>NA</td><td>Intergenic</td><td>NA</td><td>NA</td></tr>
<tr><td></td><td></td><td></td><td></td><td></td><td></td><td></td><td></td><td></td><td></td><td></td><td></td></tr>
<tr><td>T4918</td><td>GCGATACAAGAGCAAGTACGCGG</td><td>1000</td><td></td><td></td><td></td><td></td><td></td><td></td><td></td><td></td><td></td></tr>
<tr><td>Oligo fwd</td><td>CACCGCGATACAAGAGCAAGTACG</td><td></td><td></td><td></td><td></td><td></td><td></td><td></td><td></td><td></td><td></td></tr>
<tr><td>Oligo rev</td><td>AAACCGTACTTGCTCTTGTATCGC</td><td></td><td></td><td></td><td></td><td></td><td></td><td></td><td></td><td></td><td></td></tr>
<tr><td>Chromosome</td><td>start</td><td>end</td><td>strand</td><td>MM</td><td>target_seq</td><td>PAM</td><td>alignment</td><td>distance</td><td>position</td><td>gene name</td><td>gene id</td></tr>
<tr><td>GU305914.1</td><td>70462</td><td>70484</td><td>+</td><td>0</td><td>GCGATACAAGAGCAAGTACG</td><td>CGG</td><td>||||||||[||||||||||||]PAM</td><td>NA</td><td>Intergenic</td><td>NA</td><td>NA</td></tr>
<tr><td></td><td></td><td></td><td></td><td></td><td></td><td></td><td></td><td></td><td></td><td></td><td></td></tr>
<tr><td>T4919</td><td>CGACGTGACGATCAAACCCGAGG</td><td>1000</td><td></td><td></td><td></td><td></td><td></td><td></td><td></td><td></td><td></td></tr>
<tr><td>Oligo fwd</td><td>CACCCGACGTGACGATCAAACCCG</td><td></td><td></td><td></td><td></td><td></td><td></td><td></td><td></td><td></td><td></td></tr>
<tr><td>Oligo rev</td><td>AAACCGGGTTTGATCGTCACGTCG</td><td></td><td></td><td></td><td></td><td></td><td></td><td></td><td></td><td></td><td></td></tr>
<tr><td>Chromosome</td><td>start</td><td>end</td><td>strand</td><td>MM</td><td>target_seq</td><td>PAM</td><td>alignment</td><td>distance</td><td>position</td><td>gene name</td><td>gene id</td></tr>
<tr><td>GU305914.1</td><td>70522</td><td>70544</td><td>+</td><td>0</td><td>CGACGTGACGATCAAACCCG</td><td>AGG</td><td>||||||||[||||||||||||]PAM</td><td>NA</td><td>Intergenic</td><td>NA</td><td>NA</td></tr>
<tr><td></td><td></td><td></td><td></td><td></td><td></td><td></td><td></td><td></td><td></td><td></td><td></td></tr>
<tr><td>T4920</td><td>CAAACCCGAGGACACCTTCACGG</td><td>1000</td><td></td><td></td><td></td><td></td><td></td><td></td><td></td><td></td><td></td></tr>
<tr><td>Oligo fwd</td><td>CACCCAAACCCGAGGACACCTTCA</td><td></td><td></td><td></td><td></td><td></td><td></td><td></td><td></td><td></td><td></td></tr>
<tr><td>Oligo rev</td><td>AAACTGAAGGTGTCCTCGGGTTTG</td><td></td><td></td><td></td><td></td><td></td><td></td><td></td><td></td><td></td><td></td></tr>
<tr><td>Chromosome</td><td>start</td><td>end</td><td>strand</td><td>MM</td><td>target_seq</td><td>PAM</td><td>alignment</td><td>distance</td><td>position</td><td>gene name</td><td>gene id</td></tr>
<tr><td>GU305914.1</td><td>70534</td><td>70556</td><td>+</td><td>0</td><td>CAAACCCGAGGACACCTTCA</td><td>CGG</td><td>||||||||[||||||||||||]PAM</td><td>NA</td><td>Intergenic</td><td>NA</td><td>NA</td></tr>
<tr><td>GU305914.1</td><td>63856</td><td>63878</td><td>-</td><td>4</td><td>CGACCCCGAGGACCCCTACA</td><td>AGG</td><td>|-|-||||[|||||-|||-||]PAM</td><td>NA</td><td>Intergenic</td><td>NA</td><td>NA</td></tr>
<tr><td></td><td></td><td></td><td></td><td></td><td></td><td></td><td></td><td></td><td></td><td></td><td></td></tr>
<tr><td>T4921</td><td>ACCCGAGGACACCTTCACGGCGG</td><td>1000</td><td></td><td></td><td></td><td></td><td></td><td></td><td></td><td></td><td></td></tr>
<tr><td>Oligo fwd</td><td>CACCACCCGAGGACACCTTCACGG</td><td></td><td></td><td></td><td></td><td></td><td></td><td></td><td></td><td></td><td></td></tr>
<tr><td>Oligo rev</td><td>AAACCCGTGAAGGTGTCCTCGGGT</td><td></td><td></td><td></td><td></td><td></td><td></td><td></td><td></td><td></td><td></td></tr>
<tr><td>Chromosome</td><td>start</td><td>end</td><td>strand</td><td>MM</td><td>target_seq</td><td>PAM</td><td>alignment</td><td>distance</td><td>position</td><td>gene name</td><td>gene id</td></tr>
<tr><td>GU305914.1</td><td>70537</td><td>70559</td><td>+</td><td>0</td><td>ACCCGAGGACACCTTCACGG</td><td>CGG</td><td>||||||||[||||||||||||]PAM</td><td>NA</td><td>Intergenic</td><td>NA</td><td>NA</td></tr>
<tr><td></td><td></td><td></td><td></td><td></td><td></td><td></td><td></td><td></td><td></td><td></td><td></td></tr>
<tr><td>T4922</td><td>ACACCTTCACGGCGGCGTTCAGG</td><td>1000</td><td></td><td></td><td></td><td></td><td></td><td></td><td></td><td></td><td></td></tr>
<tr><td>Oligo fwd</td><td>CACCACACCTTCACGGCGGCGTTC</td><td></td><td></td><td></td><td></td><td></td><td></td><td></td><td></td><td></td><td></td></tr>
<tr><td>Oligo rev</td><td>AAACGAACGCCGCCGTGAAGGTGT</td><td></td><td></td><td></td><td></td><td></td><td></td><td></td><td></td><td></td><td></td></tr>
<tr><td>Chromosome</td><td>start</td><td>end</td><td>strand</td><td>MM</td><td>target_seq</td><td>PAM</td><td>alignment</td><td>distance</td><td>position</td><td>gene name</td><td>gene id</td></tr>
<tr><td>GU305914.1</td><td>70545</td><td>70567</td><td>+</td><td>0</td><td>ACACCTTCACGGCGGCGTTC</td><td>AGG</td><td>||||||||[||||||||||||]PAM</td><td>NA</td><td>Intergenic</td><td>NA</td><td>NA</td></tr>
<tr><td></td><td></td><td></td><td></td><td></td><td></td><td></td><td></td><td></td><td></td><td></td><td></td></tr>
<tr><td>T4923</td><td>CACCTTCACGGCGGCGTTCAGGG</td><td>1000</td><td></td><td></td><td></td><td></td><td></td><td></td><td></td><td></td><td></td></tr>
<tr><td>Oligo fwd</td><td>CACCCACCTTCACGGCGGCGTTCA</td><td></td><td></td><td></td><td></td><td></td><td></td><td></td><td></td><td></td><td></td></tr>
<tr><td>Oligo rev</td><td>AAACTGAACGCCGCCGTGAAGGTG</td><td></td><td></td><td></td><td></td><td></td><td></td><td></td><td></td><td></td><td></td></tr>
<tr><td>Chromosome</td><td>start</td><td>end</td><td>strand</td><td>MM</td><td>target_seq</td><td>PAM</td><td>alignment</td><td>distance</td><td>position</td><td>gene name</td><td>gene id</td></tr>
<tr><td>GU305914.1</td><td>70546</td><td>70568</td><td>+</td><td>0</td><td>CACCTTCACGGCGGCGTTCA</td><td>GGG</td><td>||||||||[||||||||||||]PAM</td><td>NA</td><td>Intergenic</td><td>NA</td><td>NA</td></tr>
<tr><td></td><td></td><td></td><td></td><td></td><td></td><td></td><td></td><td></td><td></td><td></td><td></td></tr>
<tr><td>T4924</td><td>GGCGTTCAGGGACACCGTGAAGG</td><td>1000</td><td></td><td></td><td></td><td></td><td></td><td></td><td></td><td></td><td></td></tr>
<tr><td>Oligo fwd</td><td>CACCGGCGTTCAGGGACACCGTGA</td><td></td><td></td><td></td><td></td><td></td><td></td><td></td><td></td><td></td><td></td></tr>
<tr><td>Oligo rev</td><td>AAACTCACGGTGTCCCTGAACGCC</td><td></td><td></td><td></td><td></td><td></td><td></td><td></td><td></td><td></td><td></td></tr>
<tr><td>Chromosome</td><td>start</td><td>end</td><td>strand</td><td>MM</td><td>target_seq</td><td>PAM</td><td>alignment</td><td>distance</td><td>position</td><td>gene name</td><td>gene id</td></tr>
<tr><td>GU305914.1</td><td>70558</td><td>70580</td><td>+</td><td>0</td><td>GGCGTTCAGGGACACCGTGA</td><td>AGG</td><td>||||||||[||||||||||||]PAM</td><td>NA</td><td>Intergenic</td><td>NA</td><td>NA</td></tr>
<tr><td>GU305914.1</td><td>195667</td><td>195689</td><td>+</td><td>3</td><td>GGCCTTCCTGGACACCGTGA</td><td>CGG</td><td>|||-|||-[-|||||||||||]PAM</td><td>NA</td><td>Intergenic</td><td>NA</td><td>NA</td></tr>
</table>

<table>
<tr><td>T4925</td><td>CAGGGACACCGTGAAGGCGATGG</td><td>1000</td></tr>
<tr><td>Oligo fwd</td><td>CACCCAGGGACACCGTGAAGGCGA</td></tr>
<tr><td>Oligo rev</td><td>AAACTCGCCTTCACGGTGTCCCTG</td></tr>
<tr><td>Chromosome</td><td>start</td><td>end</td><td>strand</td><td>MM</td><td>target_seq</td><td>PAM</td><td>alignment</td><td>distance</td><td>position</td><td>gene name</td><td>gene id</td></tr>
<tr><td>GU305914.1</td><td>70564</td><td>70586</td><td>+</td><td>0</td><td>CAGGGACACCGTGAAGGCGA</td><td>TGG</td><td>||||||||[||||||||||||]PAM</td><td>NA</td><td>Intergenic</td><td>NA</td><td>NA</td></tr>
<tr><td>T4926</td><td>GGACACCGTGAAGGCGATGGCGG</td><td>1000</td></tr>
<tr><td>Oligo fwd</td><td>CACCGGACACCGTGAAGGCGATGG</td></tr>
<tr><td>Oligo rev</td><td>AAACCCATCGCCTTCACGGTGTCC</td></tr>
<tr><td>Chromosome</td><td>start</td><td>end</td><td>strand</td><td>MM</td><td>target_seq</td><td>PAM</td><td>alignment</td><td>distance</td><td>position</td><td>gene name</td><td>gene id</td></tr>
<tr><td>GU305914.1</td><td>70567</td><td>70589</td><td>+</td><td>0</td><td>GGACACCGTGAAGGCGATGG</td><td>CGG</td><td>||||||||[||||||||||||]PAM</td><td>NA</td><td>Intergenic</td><td>NA</td><td>NA</td></tr>
<tr><td>T4927</td><td>GATGGCGGCGCCCTTCTCGACGG</td><td>1000</td></tr>
<tr><td>Oligo fwd</td><td>CACCGATGGCGGCGCCCTTCTCGA</td></tr>
<tr><td>Oligo rev</td><td>AAACTCGAGAAGGGCGCCGCCATC</td></tr>
<tr><td>Chromosome</td><td>start</td><td>end</td><td>strand</td><td>MM</td><td>target_seq</td><td>PAM</td><td>alignment</td><td>distance</td><td>position</td><td>gene name</td><td>gene id</td></tr>
<tr><td>GU305914.1</td><td>70582</td><td>70604</td><td>+</td><td>0</td><td>GATGGCGGCGCCCTTCTCGA</td><td>CGG</td><td>||||||||[||||||||||||]PAM</td><td>NA</td><td>Intergenic</td><td>NA</td><td>NA</td></tr>
<tr><td>T4928</td><td>GCCCTTCTCGACGGAGCTGCAGG</td><td>1000</td></tr>
<tr><td>Oligo fwd</td><td>CACCGCCCTTCTCGACGGAGCTGC</td></tr>
<tr><td>Oligo rev</td><td>AAACGCAGCTCCGTCGAGAAGGGC</td></tr>
<tr><td>Chromosome</td><td>start</td><td>end</td><td>strand</td><td>MM</td><td>target_seq</td><td>PAM</td><td>alignment</td><td>distance</td><td>position</td><td>gene name</td><td>gene id</td></tr>
<tr><td>GU305914.1</td><td>70591</td><td>70613</td><td>+</td><td>0</td><td>GCCCTTCTCGACGGAGCTGC</td><td>AGG</td><td>||||||||[||||||||||||]PAM</td><td>NA</td><td>Intergenic</td><td>NA</td><td>NA</td></tr>
<tr><td>T4929</td><td>GGAGCTGCAGGCGCGCCTGCAGG</td><td>1000</td></tr>
<tr><td>Oligo fwd</td><td>CACCGGAGCTGCAGGCGCGCCTGC</td></tr>
<tr><td>Oligo rev</td><td>AAACGCAGGCGCGCCTGCAGCTCC</td></tr>
<tr><td>Chromosome</td><td>start</td><td>end</td><td>strand</td><td>MM</td><td>target_seq</td><td>PAM</td><td>alignment</td><td>distance</td><td>position</td><td>gene name</td><td>gene id</td></tr>
<tr><td>GU305914.1</td><td>70603</td><td>70625</td><td>+</td><td>0</td><td>GGAGCTGCAGGCGCGCCTGC</td><td>AGG</td><td>||||||||[||||||||||||]PAM</td><td>NA</td><td>Intergenic</td><td>NA</td><td>NA</td></tr>
<tr><td>T4930</td><td>GGCGCGAGAGAACGAGCTCAAGG</td><td>1000</td></tr>
<tr><td>Oligo fwd</td><td>CACCGGCGCGAGAGAACGAGCTCA</td></tr>
<tr><td>Oligo rev</td><td>AAACTGAGCTCGTTCTCTCGCGCC</td></tr>
<tr><td>Chromosome</td><td>start</td><td>end</td><td>strand</td><td>MM</td><td>target_seq</td><td>PAM</td><td>alignment</td><td>distance</td><td>position</td><td>gene name</td><td>gene id</td></tr>
<tr><td>GU305914.1</td><td>70624</td><td>70646</td><td>+</td><td>0</td><td>GGCGCGAGAGAACGAGCTCA</td><td>AGG</td><td>||||||||[||||||||||||]PAM</td><td>NA</td><td>Intergenic</td><td>NA</td><td>NA</td></tr>
<tr><td>T4931</td><td>CGAGCTCAAGGACGAGCTGTCGG</td><td>1000</td></tr>
<tr><td>Oligo fwd</td><td>CACCCGAGCTCAAGGACGAGCTGT</td></tr>
<tr><td>Oligo rev</td><td>AAACACAGCTCGTCCTTGAGCTCG</td></tr>
<tr><td>Chromosome</td><td>start</td><td>end</td><td>strand</td><td>MM</td><td>target_seq</td><td>PAM</td><td>alignment</td><td>distance</td><td>position</td><td>gene name</td><td>gene id</td></tr>
<tr><td>GU305914.1</td><td>70636</td><td>70658</td><td>+</td><td>0</td><td>CGAGCTCAAGGACGAGCTGT</td><td>CGG</td><td>||||||||[||||||||||||]PAM</td><td>NA</td><td>Intergenic</td><td>NA</td><td>NA</td></tr>
<tr><td>T4932</td><td>CGTGAAGCTCAAGACGAAGCTGG</td><td>1000</td></tr>
<tr><td>Oligo fwd</td><td>CACCCGTGAAGCTCAAGACGAAGC</td></tr>
<tr><td>Oligo rev</td><td>AAACGCTTCGTCTTGAGCTTCACG</td></tr>
<tr><td>Chromosome</td><td>start</td><td>end</td><td>strand</td><td>MM</td><td>target_seq</td><td>PAM</td><td>alignment</td><td>distance</td><td>position</td><td>gene name</td><td>gene id</td></tr>
<tr><td>GU305914.1</td><td>70666</td><td>70688</td><td>+</td><td>0</td><td>CGTGAAGCTCAAGACGAAGC</td><td>TGG</td><td>||||||||[||||||||||||]PAM</td><td>NA</td><td>Intergenic</td><td>NA</td><td>NA</td></tr>
<tr><td>T4933</td><td>GACGAAGCTGGAGCGCTACCTGG</td><td>1000</td></tr>
<tr><td>Oligo fwd</td><td>CACCGACGAAGCTGGAGCGCTACC</td></tr>
<tr><td>Oligo rev</td><td>AAACGGTAGCGCTCCAGCTTCGTC</td></tr>
<tr><td>Chromosome</td><td>start</td><td>end</td><td>strand</td><td>MM</td><td>target_seq</td><td>PAM</td><td>alignment</td><td>distance</td><td>position</td><td>gene name</td><td>gene id</td></tr>
<tr><td>GU305914.1</td><td>70678</td><td>70700</td><td>+</td><td>0</td><td>GACGAAGCTGGAGCGCTACC</td><td>TGG</td><td>||||||||[||||||||||||]PAM</td><td>NA</td><td>Intergenic</td><td>NA</td><td>NA</td></tr>
<tr><td>T4934</td><td>CCTGGCTAAGCGCAGCTCTCAGG</td><td>1000</td></tr>
<tr><td>Oligo fwd</td><td>CACCCCTGGCTAAGCGCAGCTCTC</td></tr>
<tr><td>Oligo rev</td><td>AAACGAGAGCTGCGCTTAGCCAGG</td></tr>
<tr><td>Chromosome</td><td>start</td><td>end</td><td>strand</td><td>MM</td><td>target_seq</td><td>PAM</td><td>alignment</td><td>distance</td><td>position</td><td>gene name</td><td>gene id</td></tr>
<tr><td>GU305914.1</td><td>70696</td><td>70718</td><td>+</td><td>0</td><td>CCTGGCTAAGCGCAGCTCTC</td><td>AGG</td><td>||||||||[||||||||||||]PAM</td><td>NA</td><td>Intergenic</td><td>NA</td><td>NA</td></tr>
<tr><td>T4935</td><td>AGCGCAGCTCTCAGGACGCTAGG</td><td>1000</td></tr>
<tr><td>Oligo fwd</td><td>CACCAGCGCAGCTCTCAGGACGCT</td></tr>
<tr><td>Oligo rev</td><td>AAACAGCGTCCTGAGAGCTGCGCT</td></tr>
<tr><td>Chromosome</td><td>start</td><td>end</td><td>strand</td><td>MM</td><td>target_seq</td><td>PAM</td><td>alignment</td><td>distance</td><td>position</td><td>gene name</td><td>gene id</td></tr>
<tr><td>GU305914.1</td><td>70704</td><td>70726</td><td>+</td><td>0</td><td>AGCGCAGCTCTCAGGACGCT</td><td>AGG</td><td>||||||||[||||||||||||]PAM</td><td>NA</td><td>Intergenic</td><td>NA</td><td>NA</td></tr>
<tr><td>T4936</td><td>GCAGCTCTCAGGACGCTAGGTGG</td><td>1000</td></tr>
<tr><td>Oligo fwd</td><td>CACCGCAGCTCTCAGGACGCTAGG</td></tr>
<tr><td>Oligo rev</td><td>AAACCCTAGCGTCCTGAGAGCTGC</td></tr>
<tr><td>Chromosome</td><td>start</td><td>end</td><td>strand</td><td>MM</td><td>target_seq</td><td>PAM</td><td>alignment</td><td>distance</td><td>position</td><td>gene name</td><td>gene id</td></tr>
<tr><td>GU305914.1</td><td>70707</td><td>70729</td><td>+</td><td>0</td><td>GCAGCTCTCAGGACGCTAGG</td><td>TGG</td><td>||||||||[||||||||||||]PAM</td><td>NA</td><td>Intergenic</td><td>NA</td><td>NA</td></tr>
<tr><td>T4937</td><td>TGATCACCCAGCACCGCATCAGG</td><td>1000</td></tr>
<tr><td>Oligo fwd</td><td>CACCTGATCACCCAGCACCGCATC</td></tr>
<tr><td>Oligo rev</td><td>AAACGATGCGGTGCTGGGTGATCA</td></tr>
<tr><td>Chromosome</td><td>start</td><td>end</td><td>strand</td><td>MM</td><td>target_seq</td><td>PAM</td><td>alignment</td><td>distance</td><td>position</td><td>gene name</td><td>gene id</td></tr>
<tr><td>GU305914.1</td><td>70737</td><td>70759</td><td>+</td><td>0</td><td>TGATCACCCAGCACCGCATC</td><td>AGG</td><td>||||||||[||||||||||||]PAM</td><td>NA</td><td>Intergenic</td><td>NA</td><td>NA</td></tr>
<tr><td>T4938</td><td>GCACCGCATCAGGCTGCCCGAGG</td><td>1000</td></tr>
<tr><td>Oligo fwd</td><td>CACCGCACCGCATCAGGCTGCCCG</td></tr>
<tr><td>Oligo rev</td><td>AAACCGGGCAGCCTGATGCGGTGC</td></tr>
<tr><td>Chromosome</td><td>start</td><td>end</td><td>strand</td><td>MM</td><td>target_seq</td><td>PAM</td><td>alignment</td><td>distance</td><td>position</td><td>gene name</td><td>gene id</td></tr>
<tr><td>GU305914.1</td><td>70747</td><td>70769</td><td>+</td><td>0</td><td>GCACCGCATCAGGCTGCCCG</td><td>AGG</td><td>||||||||[||||||||||||]PAM</td><td>NA</td><td>Intergenic</td><td>NA</td><td>NA</td></tr>
<tr><td>T4939</td><td>CACCGCATCAGGCTGCCCGAGGG</td><td>1000</td></tr>
<tr><td>Oligo fwd</td><td>CACCCACCGCATCAGGCTGCCCGA</td></tr>
<tr><td>Oligo rev</td><td>AAACTCGGGCAGCCTGATGCGGTG</td></tr>
<tr><td>Chromosome</td><td>start</td><td>end</td><td>strand</td><td>MM</td><td>target_seq</td><td>PAM</td><td>alignment</td><td>distance</td><td>position</td><td>gene name</td><td>gene id</td></tr>
<tr><td>GU305914.1</td><td>70748</td><td>70770</td><td>+</td><td>0</td><td>CACCGCATCAGGCTGCCCGA</td><td>GGG</td><td>||||||||[||||||||||||]PAM</td><td>NA</td><td>Intergenic</td><td>NA</td><td>NA</td></tr>
</table>

| | | | | | | | | | | | | |
|---|---|---|---|---|---|---|---|---|---|---|---|---|
| T4940 | CATCAGGCTGCCCGAGGGCCTGG | | 1000 | | | | | | | | | |
| Oligo fwd | CACCCATCAGGCTGCCCGAGGGCC | | | | | | | | | | | |
| Oligo rev | AAACGGCCCTCGGGCAGCCTGATG | | | | | | | | | | | |
| Chromosome | start | | end | strand | MM | target_seq | PAM | alignment | distance | position | gene name | gene id |
| GU305914.1 | | 70753 | 70775 | + | 0 | CATCAGGCTGCCCGAGGGCC | TGG | ||||||||[||||||||||||]PAM | NA | Intergenic | NA | NA |
| T4941 | GCTGCCCGAGGGCCTGGACGTGG | | 1000 | | | | | | | | | |
| Oligo fwd | CACCGCTGCCCGAGGGCCTGGACG | | | | | | | | | | | |
| Oligo rev | AAACCGTCCAGGCCCTCGGGCAGC | | | | | | | | | | | |
| Chromosome | start | | end | strand | MM | target_seq | PAM | alignment | distance | position | gene name | gene id |
| GU305914.1 | | 70759 | 70781 | + | 0 | GCTGCCCGAGGGCCTGGACG | TGG | ||||||||[||||||||||||]PAM | NA | Intergenic | NA | NA |
| T4942 | GGACACCAACCGCCTGCGCACGG | | 1000 | | | | | | | | | |
| Oligo fwd | CACCGGACACCAACCGCCTGCGCA | | | | | | | | | | | |
| Oligo rev | AAACTGCGCAGGCGGTTGGTGTCC | | | | | | | | | | | |
| Chromosome | start | | end | strand | MM | target_seq | PAM | alignment | distance | position | gene name | gene id |
| GU305914.1 | | 70780 | 70802 | + | 0 | GGACACCAACCGCCTGCGCA | CGG | ||||||||[||||||||||||]PAM | NA | Intergenic | NA | NA |
| T4943 | CAACCGCCTGCGCACGGACACGG | | 1000 | | | | | | | | | |
| Oligo fwd | CACCCAACCGCCTGCGCACGGACA | | | | | | | | | | | |
| Oligo rev | AAACTGTCCGTGCGCAGGCGGTTG | | | | | | | | | | | |
| Chromosome | start | | end | strand | MM | target_seq | PAM | alignment | distance | position | gene name | gene id |
| GU305914.1 | | 70786 | 70808 | + | 0 | CAACCGCCTGCGCACGGACA | CGG | ||||||||[||||||||||||]PAM | NA | Intergenic | NA | NA |
| T4944 | CCGCCTGCGCACGGACACGGTGG | | 1000 | | | | | | | | | |
| Oligo fwd | CACCCCGCCTGCGCACGGACACGG | | | | | | | | | | | |
| Oligo rev | AAACCCGTGTCCGTGCGCAGGCGG | | | | | | | | | | | |
| Chromosome | start | | end | strand | MM | target_seq | PAM | alignment | distance | position | gene name | gene id |
| GU305914.1 | | 70789 | 70811 | + | 0 | CCGCCTGCGCACGGACACGG | TGG | ||||||||[||||||||||||]PAM | NA | Intergenic | NA | NA |
| T4945 | GACACGGTGGTGACGCTGAACGG | | 1000 | | | | | | | | | |
| Oligo fwd | CACCGACACGGTGGTGACGCTGAA | | | | | | | | | | | |
| Oligo rev | AAACTTCAGCGTCACCACCGTGTC | | | | | | | | | | | |
| Chromosome | start | | end | strand | MM | target_seq | PAM | alignment | distance | position | gene name | gene id |
| GU305914.1 | | 70802 | 70824 | + | 0 | GACACGGTGGTGACGCTGAA | CGG | ||||||||[||||||||||||]PAM | NA | Intergenic | NA | NA |
| T4946 | GCTGAACGGCATCATCCGCTCGG | | 1000 | | | | | | | | | |
| Oligo fwd | CACCGCTGAACGGCATCATCCGCT | | | | | | | | | | | |
| Oligo rev | AAACAGCGGATGATGCCGTTCAGC | | | | | | | | | | | |
| Chromosome | start | | end | strand | MM | target_seq | PAM | alignment | distance | position | gene name | gene id |
| GU305914.1 | | 70816 | 70838 | + | 0 | GCTGAACGGCATCATCCGCT | CGG | ||||||||[||||||||||||]PAM | NA | Intergenic | NA | NA |
| T4947 | GAACGGCATCATCCGCTCGGCGG | | 1000 | | | | | | | | | |
| Oligo fwd | CACCGAACGGCATCATCCGCTCGG | | | | | | | | | | | |
| Oligo rev | AAACCCGAGCGGATGATGCCGTTC | | | | | | | | | | | |
| Chromosome | start | | end | strand | MM | target_seq | PAM | alignment | distance | position | gene name | gene id |
| GU305914.1 | | 70819 | 70841 | + | 0 | GAACGGCATCATCCGCTCGG | CGG | ||||||||[||||||||||||]PAM | NA | Intergenic | NA | NA |
| T4948 | CACCACCCTGCCCTACATCACGG | | 1000 | | | | | | | | | |
| Oligo fwd | CACCCACCACCCTGCCCTACATCA | | | | | | | | | | | |
| Oligo rev | AAACTGATGTAGGGCAGGGTGGTG | | | | | | | | | | | |
| Chromosome | start | | end | strand | MM | target_seq | PAM | alignment | distance | position | gene name | gene id |
| GU305914.1 | | 70843 | 70865 | + | 0 | CACCACCCTGCCCTACATCA | CGG | ||||||||[||||||||||||]PAM | NA | Intergenic | NA | NA |
| T4949 | TGCCCTACATCACGGCCAAGAGG | | 1000 | | | | | | | | | |
| Oligo fwd | CACCTGCCCTACATCACGGCCAAG | | | | | | | | | | | |
| Oligo rev | AAACCTTGGCCGTGATGTAGGGCA | | | | | | | | | | | |
| Chromosome | start | | end | strand | MM | target_seq | PAM | alignment | distance | position | gene name | gene id |
| GU305914.1 | | 70851 | 70873 | + | 0 | TGCCCTACATCACGGCCAAG | AGG | ||||||||[||||||||||||]PAM | NA | Intergenic | NA | NA |
| T4950 | GCCCTACATCACGGCCAAGAGGG | | 1000 | | | | | | | | | |
| Oligo fwd | CACCGCCCTACATCACGGCCAAGA | | | | | | | | | | | |
| Oligo rev | AAACTCTTGGCCGTGATGTAGGGC | | | | | | | | | | | |
| Chromosome | start | | end | strand | MM | target_seq | PAM | alignment | distance | position | gene name | gene id |
| GU305914.1 | | 70852 | 70874 | + | 0 | GCCCTACATCACGGCCAAGA | GGG | ||||||||[||||||||||||]PAM | NA | Intergenic | NA | NA |
| T4951 | CCCTACATCACGGCCAAGAGGGG | | 1000 | | | | | | | | | |
| Oligo fwd | CACCCCCTACATCACGGCCAAGAG | | | | | | | | | | | |
| Oligo rev | AAACCTCTTGGCCGTGATGTAGGG | | | | | | | | | | | |
| Chromosome | start | | end | strand | MM | target_seq | PAM | alignment | distance | position | gene name | gene id |
| GU305914.1 | | 70853 | 70875 | + | 0 | CCCTACATCACGGCCAAGAG | GGG | ||||||||[||||||||||||]PAM | NA | Intergenic | NA | NA |
| T4952 | CATCACGGCCAAGAGGGGCCTGG | | 1000 | | | | | | | | | |
| Oligo fwd | CACCCATCACGGCCAAGAGGGGCC | | | | | | | | | | | |
| Oligo rev | AAACGGCCCCTCTTGGCCGTGATG | | | | | | | | | | | |
| Chromosome | start | | end | strand | MM | target_seq | PAM | alignment | distance | position | gene name | gene id |
| GU305914.1 | | 70858 | 70880 | + | 0 | CATCACGGCCAAGAGGGGCC | TGG | ||||||||[||||||||||||]PAM | NA | Intergenic | NA | NA |
| T4953 | CGGCCAAGAGGGGCCTGGAGTGG | | 1000 | | | | | | | | | |
| Oligo fwd | CACCCGGCCAAGAGGGGCCTGGAG | | | | | | | | | | | |
| Oligo rev | AAACCTCCAGGCCCCTCTTGGCCG | | | | | | | | | | | |
| Chromosome | start | | end | strand | MM | target_seq | PAM | alignment | distance | position | gene name | gene id |
| GU305914.1 | | 70863 | 70885 | + | 0 | CGGCCAAGAGGGGCCTGGAG | TGG | ||||||||[||||||||||||]PAM | NA | Intergenic | NA | NA |
| T4954 | GAGGGGCCTGGAGTGGACGACGG | | 1000 | | | | | | | | | |
| Oligo fwd | CACCGAGGGGCCTGGAGTGGACGA | | | | | | | | | | | |
| Oligo rev | AAACTCGTCCACTCCAGGCCCCTC | | | | | | | | | | | |
| Chromosome | start | | end | strand | MM | target_seq | PAM | alignment | distance | position | gene name | gene id |

<table>
<tr><td>GU305914.1</td><td>70870</td><td>70892</td><td>+</td><td>0</td><td>GAGGGGCCTGGAGTGGACGA</td><td>CGG</td><td>||||||||[||||||||||||]PAM</td><td>NA</td><td>Intergenic</td><td>NA</td><td>NA</td></tr>
<tr><td></td><td></td><td></td><td></td><td></td><td></td><td></td><td></td><td></td><td></td><td></td><td></td></tr>
<tr><td>T4955</td><td>GTGGACGACGGAATTCATCCTGG</td><td>1000</td><td></td><td></td><td></td><td></td><td></td><td></td><td></td><td></td><td></td></tr>
<tr><td>Oligo fwd</td><td>CACCGTGGACGACGGAATTCATCC</td><td></td><td></td><td></td><td></td><td></td><td></td><td></td><td></td><td></td><td></td></tr>
<tr><td>Oligo rev</td><td>AAACGGATGAATTCCGTCGTCCAC</td><td></td><td></td><td></td><td></td><td></td><td></td><td></td><td></td><td></td><td></td></tr>
<tr><td>Chromosome</td><td>start</td><td>end</td><td>strand</td><td>MM</td><td>target_seq</td><td>PAM</td><td>alignment</td><td>distance</td><td>position</td><td>gene name</td><td>gene id</td></tr>
<tr><td>GU305914.1</td><td>70882</td><td>70904</td><td>+</td><td>0</td><td>GTGGACGACGGAATTCATCC</td><td>TGG</td><td>||||||||[||||||||||||]PAM</td><td>NA</td><td>Intergenic</td><td>NA</td><td>NA</td></tr>
<tr><td></td><td></td><td></td><td></td><td></td><td></td><td></td><td></td><td></td><td></td><td></td><td></td></tr>
<tr><td>T4956</td><td>GACGACGGAATTCATCCTGGCGG</td><td>1000</td><td></td><td></td><td></td><td></td><td></td><td></td><td></td><td></td><td></td></tr>
<tr><td>Oligo fwd</td><td>CACCGACGACGGAATTCATCCTGG</td><td></td><td></td><td></td><td></td><td></td><td></td><td></td><td></td><td></td><td></td></tr>
<tr><td>Oligo rev</td><td>AAACCCAGGATGAATTCCGTCGTC</td><td></td><td></td><td></td><td></td><td></td><td></td><td></td><td></td><td></td><td></td></tr>
<tr><td>Chromosome</td><td>start</td><td>end</td><td>strand</td><td>MM</td><td>target_seq</td><td>PAM</td><td>alignment</td><td>distance</td><td>position</td><td>gene name</td><td>gene id</td></tr>
<tr><td>GU305914.1</td><td>70885</td><td>70907</td><td>+</td><td>0</td><td>GACGACGGAATTCATCCTGG</td><td>CGG</td><td>||||||||[||||||||||||]PAM</td><td>NA</td><td>Intergenic</td><td>NA</td><td>NA</td></tr>
<tr><td></td><td></td><td></td><td></td><td></td><td></td><td></td><td></td><td></td><td></td><td></td><td></td></tr>
<tr><td>T4957</td><td>GATCGAAGAGAAGAAAGACGCGG</td><td>1000</td><td></td><td></td><td></td><td></td><td></td><td></td><td></td><td></td><td></td></tr>
<tr><td>Oligo fwd</td><td>CACCGATCGAAGAGAAGAAAGACG</td><td></td><td></td><td></td><td></td><td></td><td></td><td></td><td></td><td></td><td></td></tr>
<tr><td>Oligo rev</td><td>AAACCGTCTTTCTTCTCTTCGATC</td><td></td><td></td><td></td><td></td><td></td><td></td><td></td><td></td><td></td><td></td></tr>
<tr><td>Chromosome</td><td>start</td><td>end</td><td>strand</td><td>MM</td><td>target_seq</td><td>PAM</td><td>alignment</td><td>distance</td><td>position</td><td>gene name</td><td>gene id</td></tr>
<tr><td>GU305914.1</td><td>70909</td><td>70931</td><td>+</td><td>0</td><td>GATCGAAGAGAAGAAAGACG</td><td>CGG</td><td>||||||||[||||||||||||]PAM</td><td>NA</td><td>Intergenic</td><td>NA</td><td>NA</td></tr>
<tr><td></td><td></td><td></td><td></td><td></td><td></td><td></td><td></td><td></td><td></td><td></td><td></td></tr>
<tr><td>T4958</td><td>GAGAAGAAAGACGCGGACCCCGG</td><td>1000</td><td></td><td></td><td></td><td></td><td></td><td></td><td></td><td></td><td></td></tr>
<tr><td>Oligo fwd</td><td>CACCGAGAAGAAAGACGCGGACCC</td><td></td><td></td><td></td><td></td><td></td><td></td><td></td><td></td><td></td><td></td></tr>
<tr><td>Oligo rev</td><td>AAACGGGTCCGCGTCTTTCTTCTC</td><td></td><td></td><td></td><td></td><td></td><td></td><td></td><td></td><td></td><td></td></tr>
<tr><td>Chromosome</td><td>start</td><td>end</td><td>strand</td><td>MM</td><td>target_seq</td><td>PAM</td><td>alignment</td><td>distance</td><td>position</td><td>gene name</td><td>gene id</td></tr>
<tr><td>GU305914.1</td><td>70916</td><td>70938</td><td>+</td><td>0</td><td>GAGAAGAAAGACGCGGACCC</td><td>CGG</td><td>||||||||[||||||||||||]PAM</td><td>NA</td><td>Intergenic</td><td>NA</td><td>NA</td></tr>
<tr><td></td><td></td><td></td><td></td><td></td><td></td><td></td><td></td><td></td><td></td><td></td><td></td></tr>
<tr><td>T4959</td><td>AGAAGAAAGACGCGGACCCCGGG</td><td>1000</td><td></td><td></td><td></td><td></td><td></td><td></td><td></td><td></td><td></td></tr>
<tr><td>Oligo fwd</td><td>CACCAGAAGAAAGACGCGGACCCC</td><td></td><td></td><td></td><td></td><td></td><td></td><td></td><td></td><td></td><td></td></tr>
<tr><td>Oligo rev</td><td>AAACGGGGTCCGCGTCTTTCTTCT</td><td></td><td></td><td></td><td></td><td></td><td></td><td></td><td></td><td></td><td></td></tr>
<tr><td>Chromosome</td><td>start</td><td>end</td><td>strand</td><td>MM</td><td>target_seq</td><td>PAM</td><td>alignment</td><td>distance</td><td>position</td><td>gene name</td><td>gene id</td></tr>
<tr><td>GU305914.1</td><td>70917</td><td>70939</td><td>+</td><td>0</td><td>AGAAGAAAGACGCGGACCCC</td><td>GGG</td><td>||||||||[||||||||||||]PAM</td><td>NA</td><td>Intergenic</td><td>NA</td><td>NA</td></tr>
<tr><td></td><td></td><td></td><td></td><td></td><td></td><td></td><td></td><td></td><td></td><td></td><td></td></tr>
<tr><td>T4960</td><td>CCCCGGGATGTTCGCCCAGCTGG</td><td>1000</td><td></td><td></td><td></td><td></td><td></td><td></td><td></td><td></td><td></td></tr>
<tr><td>Oligo fwd</td><td>CACCCCCCGGGATGTTCGCCCAGC</td><td></td><td></td><td></td><td></td><td></td><td></td><td></td><td></td><td></td><td></td></tr>
<tr><td>Oligo rev</td><td>AAACGCTGGGCGAACATCCCGGGG</td><td></td><td></td><td></td><td></td><td></td><td></td><td></td><td></td><td></td><td></td></tr>
<tr><td>Chromosome</td><td>start</td><td>end</td><td>strand</td><td>MM</td><td>target_seq</td><td>PAM</td><td>alignment</td><td>distance</td><td>position</td><td>gene name</td><td>gene id</td></tr>
<tr><td>GU305914.1</td><td>70933</td><td>70955</td><td>+</td><td>0</td><td>CCCCGGGATGTTCGCCCAGC</td><td>TGG</td><td>||||||||[||||||||||||]PAM</td><td>NA</td><td>Intergenic</td><td>NA</td><td>NA</td></tr>
<tr><td></td><td></td><td></td><td></td><td></td><td></td><td></td><td></td><td></td><td></td><td></td><td></td></tr>
<tr><td>T4961</td><td>CGCCCAGCTGGCGACCCTGCTGG</td><td>1000</td><td></td><td></td><td></td><td></td><td></td><td></td><td></td><td></td><td></td></tr>
<tr><td>Oligo fwd</td><td>CACCCGCCCAGCTGGCGACCCTGC</td><td></td><td></td><td></td><td></td><td></td><td></td><td></td><td></td><td></td><td></td></tr>
<tr><td>Oligo rev</td><td>AAACGCAGGGTCGCCAGCTGGGCG</td><td></td><td></td><td></td><td></td><td></td><td></td><td></td><td></td><td></td><td></td></tr>
<tr><td>Chromosome</td><td>start</td><td>end</td><td>strand</td><td>MM</td><td>target_seq</td><td>PAM</td><td>alignment</td><td>distance</td><td>position</td><td>gene name</td><td>gene id</td></tr>
<tr><td>GU305914.1</td><td>70945</td><td>70967</td><td>+</td><td>0</td><td>CGCCCAGCTGGCGACCCTGC</td><td>TGG</td><td>||||||||[||||||||||||]PAM</td><td>NA</td><td>Intergenic</td><td>NA</td><td>NA</td></tr>
<tr><td></td><td></td><td></td><td></td><td></td><td></td><td></td><td></td><td></td><td></td><td></td><td></td></tr>
<tr><td>T4962</td><td>GACCCTGCTGGACAACTCGCAGG</td><td>1000</td><td></td><td></td><td></td><td></td><td></td><td></td><td></td><td></td><td></td></tr>
<tr><td>Oligo fwd</td><td>CACCGACCCTGCTGGACAACTCGC</td><td></td><td></td><td></td><td></td><td></td><td></td><td></td><td></td><td></td><td></td></tr>
<tr><td>Oligo rev</td><td>AAACGCGAGTTGTCCAGCAGGGTC</td><td></td><td></td><td></td><td></td><td></td><td></td><td></td><td></td><td></td><td></td></tr>
<tr><td>Chromosome</td><td>start</td><td>end</td><td>strand</td><td>MM</td><td>target_seq</td><td>PAM</td><td>alignment</td><td>distance</td><td>position</td><td>gene name</td><td>gene id</td></tr>
<tr><td>GU305914.1</td><td>70957</td><td>70979</td><td>+</td><td>0</td><td>GACCCTGCTGGACAACTCGC</td><td>AGG</td><td>||||||||[||||||||||||]PAM</td><td>NA</td><td>Intergenic</td><td>NA</td><td>NA</td></tr>
<tr><td></td><td></td><td></td><td></td><td></td><td></td><td></td><td></td><td></td><td></td><td></td><td></td></tr>
<tr><td>T4963</td><td>ACAACTCGCAGGCGATCGCCAGG</td><td>1000</td><td></td><td></td><td></td><td></td><td></td><td></td><td></td><td></td><td></td></tr>
<tr><td>Oligo fwd</td><td>CACCACAACTCGCAGGCGATCGCC</td><td></td><td></td><td></td><td></td><td></td><td></td><td></td><td></td><td></td><td></td></tr>
<tr><td>Oligo rev</td><td>AAACGGCGATCGCCTGCGAGTTGT</td><td></td><td></td><td></td><td></td><td></td><td></td><td></td><td></td><td></td><td></td></tr>
<tr><td>Chromosome</td><td>start</td><td>end</td><td>strand</td><td>MM</td><td>target_seq</td><td>PAM</td><td>alignment</td><td>distance</td><td>position</td><td>gene name</td><td>gene id</td></tr>
<tr><td>GU305914.1</td><td>70968</td><td>70990</td><td>+</td><td>0</td><td>ACAACTCGCAGGCGATCGCC</td><td>AGG</td><td>||||||||[||||||||||||]PAM</td><td>NA</td><td>Intergenic</td><td>NA</td><td>NA</td></tr>
<tr><td></td><td></td><td></td><td></td><td></td><td></td><td></td><td></td><td></td><td></td><td></td><td></td></tr>
<tr><td>T4964</td><td>GGCGATCGCCAGGAAGATCGAGG</td><td>1000</td><td></td><td></td><td></td><td></td><td></td><td></td><td></td><td></td><td></td></tr>
<tr><td>Oligo fwd</td><td>CACCGGCGATCGCCAGGAAGATCG</td><td></td><td></td><td></td><td></td><td></td><td></td><td></td><td></td><td></td><td></td></tr>
<tr><td>Oligo rev</td><td>AAACCGATCTTCCTGGCGATCGCC</td><td></td><td></td><td></td><td></td><td></td><td></td><td></td><td></td><td></td><td></td></tr>
<tr><td>Chromosome</td><td>start</td><td>end</td><td>strand</td><td>MM</td><td>target_seq</td><td>PAM</td><td>alignment</td><td>distance</td><td>position</td><td>gene name</td><td>gene id</td></tr>
<tr><td>GU305914.1</td><td>70978</td><td>71000</td><td>+</td><td>0</td><td>GGCGATCGCCAGGAAGATCG</td><td>AGG</td><td>||||||||[||||||||||||]PAM</td><td>NA</td><td>Intergenic</td><td>NA</td><td>NA</td></tr>
<tr><td>GU305914.1</td><td>31494</td><td>31516</td><td>+</td><td>4</td><td>GGCGGTAGCCAGGGAGACCG</td><td>CGG</td><td>||||-|-|[|||||-|||-||]PAM</td><td>NA</td><td>Intergenic</td><td>NA</td><td>NA</td></tr>
<tr><td>GU305914.1</td><td>83427</td><td>83449</td><td>+</td><td>4</td><td>GCCGATCACCAGGAGGATGG</td><td>TGG</td><td>|-|||||-[||||||-|||-|]PAM</td><td>NA</td><td>Intergenic</td><td>NA</td><td>NA</td></tr>
<tr><td></td><td></td><td></td><td></td><td></td><td></td><td></td><td></td><td></td><td></td><td></td><td></td></tr>
<tr><td>T4965</td><td>GAAGATCGTGTACAACACGAAGG</td><td>1000</td><td></td><td></td><td></td><td></td><td></td><td></td><td></td><td></td><td></td></tr>
<tr><td>Oligo fwd</td><td>CACCGAAGATCGTGTACAACACGA</td><td></td><td></td><td></td><td></td><td></td><td></td><td></td><td></td><td></td><td></td></tr>
<tr><td>Oligo rev</td><td>AAACTCGTGTTGTACACGATCTTC</td><td></td><td></td><td></td><td></td><td></td><td></td><td></td><td></td><td></td><td></td></tr>
<tr><td>Chromosome</td><td>start</td><td>end</td><td>strand</td><td>MM</td><td>target_seq</td><td>PAM</td><td>alignment</td><td>distance</td><td>position</td><td>gene name</td><td>gene id</td></tr>
<tr><td>GU305914.1</td><td>71002</td><td>71024</td><td>+</td><td>0</td><td>GAAGATCGTGTACAACACGA</td><td>AGG</td><td>||||||||[||||||||||||]PAM</td><td>NA</td><td>Intergenic</td><td>NA</td><td>NA</td></tr>
<tr><td></td><td></td><td></td><td></td><td></td><td></td><td></td><td></td><td></td><td></td><td></td><td></td></tr>
<tr><td>T4966</td><td>GATCGTGTACAACACGAAGGTGG</td><td>1000</td><td></td><td></td><td></td><td></td><td></td><td></td><td></td><td></td><td></td></tr>
<tr><td>Oligo fwd</td><td>CACCGATCGTGTACAACACGAAGG</td><td></td><td></td><td></td><td></td><td></td><td></td><td></td><td></td><td></td><td></td></tr>
<tr><td>Oligo rev</td><td>AAACCCTTCGTGTTGTACACGATC</td><td></td><td></td><td></td><td></td><td></td><td></td><td></td><td></td><td></td><td></td></tr>
<tr><td>Chromosome</td><td>start</td><td>end</td><td>strand</td><td>MM</td><td>target_seq</td><td>PAM</td><td>alignment</td><td>distance</td><td>position</td><td>gene name</td><td>gene id</td></tr>
<tr><td>GU305914.1</td><td>71005</td><td>71027</td><td>+</td><td>0</td><td>GATCGTGTACAACACGAAGG</td><td>TGG</td><td>||||||||[||||||||||||]PAM</td><td>NA</td><td>Intergenic</td><td>NA</td><td>NA</td></tr>
<tr><td></td><td></td><td></td><td></td><td></td><td></td><td></td><td></td><td></td><td></td><td></td><td></td></tr>
<tr><td>T4967</td><td>CGTGTACAACACGAAGGTGGAGG</td><td>1000</td><td></td><td></td><td></td><td></td><td></td><td></td><td></td><td></td><td></td></tr>
<tr><td>Oligo fwd</td><td>CACCCGTGTACAACACGAAGGTGG</td><td></td><td></td><td></td><td></td><td></td><td></td><td></td><td></td><td></td><td></td></tr>
<tr><td>Oligo rev</td><td>AAACCCACCTTCGTGTTGTACACG</td><td></td><td></td><td></td><td></td><td></td><td></td><td></td><td></td><td></td><td></td></tr>
<tr><td>Chromosome</td><td>start</td><td>end</td><td>strand</td><td>MM</td><td>target_seq</td><td>PAM</td><td>alignment</td><td>distance</td><td>position</td><td>gene name</td><td>gene id</td></tr>
<tr><td>GU305914.1</td><td>71008</td><td>71030</td><td>+</td><td>0</td><td>CGTGTACAACACGAAGGTGG</td><td>AGG</td><td>||||||||[||||||||||||]PAM</td><td>NA</td><td>Intergenic</td><td>NA</td><td>NA</td></tr>
<tr><td></td><td></td><td></td><td></td><td></td><td></td><td></td><td></td><td></td><td></td><td></td><td></td></tr>
<tr><td>T4968</td><td>GTACAACACGAAGGTGGAGGCGG</td><td>1000</td><td></td><td></td><td></td><td></td><td></td><td></td><td></td><td></td><td></td></tr>
<tr><td>Oligo fwd</td><td>CACCGTACAACACGAAGGTGGAGG</td><td></td><td></td><td></td><td></td><td></td><td></td><td></td><td></td><td></td><td></td></tr>
<tr><td>Oligo rev</td><td>AAACCCTCCACCTTCGTGTTGTAC</td><td></td><td></td><td></td><td></td><td></td><td></td><td></td><td></td><td></td><td></td></tr>
<tr><td>Chromosome</td><td>start</td><td>end</td><td>strand</td><td>MM</td><td>target_seq</td><td>PAM</td><td>alignment</td><td>distance</td><td>position</td><td>gene name</td><td>gene id</td></tr>
<tr><td>GU305914.1</td><td>71011</td><td>71033</td><td>+</td><td>0</td><td>GTACAACACGAAGGTGGAGG</td><td>CGG</td><td>||||||||[||||||||||||]PAM</td><td>NA</td><td>Intergenic</td><td>NA</td><td>NA</td></tr>
<tr><td></td><td></td><td></td><td></td><td></td><td></td><td></td><td></td><td></td><td></td><td></td><td></td></tr>
<tr><td>T4969</td><td>GAAGGTGGAGGCGGAGATGTTGG</td><td>1000</td><td></td><td></td><td></td><td></td><td></td><td></td><td></td><td></td><td></td></tr>
</table>

<table>
<tr><td>Oligo fwd</td><td>CACCGAAGGTGGAGGCGGAGATGT</td><td></td><td></td><td></td><td></td><td></td><td></td><td></td><td></td><td></td><td></td></tr>
<tr><td>Oligo rev</td><td>AAACACATCTCCGCCTCCACCTTC</td><td></td><td></td><td></td><td></td><td></td><td></td><td></td><td></td><td></td><td></td></tr>
<tr><td>Chromosome</td><td>start</td><td>end</td><td>strand</td><td>MM</td><td>target_seq</td><td>PAM</td><td>alignment</td><td>distance</td><td>position</td><td>gene name</td><td>gene id</td></tr>
<tr><td>GU305914.1</td><td>71020</td><td>71042</td><td>+</td><td>0</td><td>GAAGGTGGAGGCGGAGATGT</td><td>TGG</td><td>||||||||[||||||||||||]PAM</td><td>NA</td><td>Intergenic</td><td>NA</td><td>NA</td></tr>
<tr><td>T4970</td><td>GGAGGCGGAGATGTTGGAGACGG</td><td>1000</td><td></td><td></td><td></td><td></td><td></td><td></td><td></td><td></td><td></td></tr>
<tr><td>Oligo fwd</td><td>CACCGGAGGCGGAGATGTTGGAGA</td><td></td><td></td><td></td><td></td><td></td><td></td><td></td><td></td><td></td><td></td></tr>
<tr><td>Oligo rev</td><td>AAACTCTCCAACATCTCCGCCTCC</td><td></td><td></td><td></td><td></td><td></td><td></td><td></td><td></td><td></td><td></td></tr>
<tr><td>Chromosome</td><td>start</td><td>end</td><td>strand</td><td>MM</td><td>target_seq</td><td>PAM</td><td>alignment</td><td>distance</td><td>position</td><td>gene name</td><td>gene id</td></tr>
<tr><td>GU305914.1</td><td>71026</td><td>71048</td><td>+</td><td>0</td><td>GGAGGCGGAGATGTTGGAGA</td><td>CGG</td><td>||||||||[||||||||||||]PAM</td><td>NA</td><td>Intergenic</td><td>NA</td><td>NA</td></tr>
<tr><td>GU305914.1</td><td>74637</td><td>74659</td><td>+</td><td>4</td><td>GAAGGTGGAGAGCTTGGAGA</td><td>GGG</td><td>|-|||-||[|||--|||||||]PAM</td><td>NA</td><td>Intergenic</td><td>NA</td><td>NA</td></tr>
<tr><td>GU305914.1</td><td>111499</td><td>111521</td><td>-</td><td>4</td><td>GCCGGCGGAGGTGTTGAAGA</td><td>TGG</td><td>|--|||||[||-|||||-|||]PAM</td><td>NA</td><td>Intergenic</td><td>NA</td><td>NA</td></tr>
<tr><td>T4971</td><td>GGAGATGTTGGAGACGGCGCCGG</td><td>1000</td><td></td><td></td><td></td><td></td><td></td><td></td><td></td><td></td><td></td></tr>
<tr><td>Oligo fwd</td><td>CACCGGAGATGTTGGAGACGGCGC</td><td></td><td></td><td></td><td></td><td></td><td></td><td></td><td></td><td></td><td></td></tr>
<tr><td>Oligo rev</td><td>AAACGCGCCGTCTCCAACATCTCC</td><td></td><td></td><td></td><td></td><td></td><td></td><td></td><td></td><td></td><td></td></tr>
<tr><td>Chromosome</td><td>start</td><td>end</td><td>strand</td><td>MM</td><td>target_seq</td><td>PAM</td><td>alignment</td><td>distance</td><td>position</td><td>gene name</td><td>gene id</td></tr>
<tr><td>GU305914.1</td><td>71032</td><td>71054</td><td>+</td><td>0</td><td>GGAGATGTTGGAGACGGCGC</td><td>CGG</td><td>||||||||[||||||||||||]PAM</td><td>NA</td><td>Intergenic</td><td>NA</td><td>NA</td></tr>
<tr><td>T4972</td><td>AGACGGCGCCGGAGCACACGAGG</td><td>1000</td><td></td><td></td><td></td><td></td><td></td><td></td><td></td><td></td><td></td></tr>
<tr><td>Oligo fwd</td><td>CACCAGACGGCGCCGGAGCACACG</td><td></td><td></td><td></td><td></td><td></td><td></td><td></td><td></td><td></td><td></td></tr>
<tr><td>Oligo rev</td><td>AAACCGTGTGCTCCGGCGCCGTCT</td><td></td><td></td><td></td><td></td><td></td><td></td><td></td><td></td><td></td><td></td></tr>
<tr><td>Chromosome</td><td>start</td><td>end</td><td>strand</td><td>MM</td><td>target_seq</td><td>PAM</td><td>alignment</td><td>distance</td><td>position</td><td>gene name</td><td>gene id</td></tr>
<tr><td>GU305914.1</td><td>71043</td><td>71065</td><td>+</td><td>0</td><td>AGACGGCGCCGGAGCACACG</td><td>AGG</td><td>||||||||[||||||||||||]PAM</td><td>NA</td><td>Intergenic</td><td>NA</td><td>NA</td></tr>
<tr><td>T4973</td><td>GACGGCGCCGGAGCACACGAGGG</td><td>1000</td><td></td><td></td><td></td><td></td><td></td><td></td><td></td><td></td><td></td></tr>
<tr><td>Oligo fwd</td><td>CACCGACGGCGCCGGAGCACACGA</td><td></td><td></td><td></td><td></td><td></td><td></td><td></td><td></td><td></td><td></td></tr>
<tr><td>Oligo rev</td><td>AAACTCGTGTGCTCCGGCGCCGTC</td><td></td><td></td><td></td><td></td><td></td><td></td><td></td><td></td><td></td><td></td></tr>
<tr><td>Chromosome</td><td>start</td><td>end</td><td>strand</td><td>MM</td><td>target_seq</td><td>PAM</td><td>alignment</td><td>distance</td><td>position</td><td>gene name</td><td>gene id</td></tr>
<tr><td>GU305914.1</td><td>71044</td><td>71066</td><td>+</td><td>0</td><td>GACGGCGCCGGAGCACACGA</td><td>GGG</td><td>||||||||[||||||||||||]PAM</td><td>NA</td><td>Intergenic</td><td>NA</td><td>NA</td></tr>
<tr><td>T4974</td><td>GTCGAACCTGCAGCTGATGCTGG</td><td>1000</td><td></td><td></td><td></td><td></td><td></td><td></td><td></td><td></td><td></td></tr>
<tr><td>Oligo fwd</td><td>CACCGTCGAACCTGCAGCTGATGC</td><td></td><td></td><td></td><td></td><td></td><td></td><td></td><td></td><td></td><td></td></tr>
<tr><td>Oligo rev</td><td>AAACGCATCAGCTGCAGGTTCGAC</td><td></td><td></td><td></td><td></td><td></td><td></td><td></td><td></td><td></td><td></td></tr>
<tr><td>Chromosome</td><td>start</td><td>end</td><td>strand</td><td>MM</td><td>target_seq</td><td>PAM</td><td>alignment</td><td>distance</td><td>position</td><td>gene name</td><td>gene id</td></tr>
<tr><td>GU305914.1</td><td>71074</td><td>71096</td><td>+</td><td>0</td><td>GTCGAACCTGCAGCTGATGC</td><td>TGG</td><td>||||||||[||||||||||||]PAM</td><td>NA</td><td>Intergenic</td><td>NA</td><td>NA</td></tr>
<tr><td>T4975</td><td>TCGAACCTGCAGCTGATGCTGGG</td><td>1000</td><td></td><td></td><td></td><td></td><td></td><td></td><td></td><td></td><td></td></tr>
<tr><td>Oligo fwd</td><td>CACCTCGAACCTGCAGCTGATGCT</td><td></td><td></td><td></td><td></td><td></td><td></td><td></td><td></td><td></td><td></td></tr>
<tr><td>Oligo rev</td><td>AAACAGCATCAGCTGCAGGTTCGA</td><td></td><td></td><td></td><td></td><td></td><td></td><td></td><td></td><td></td><td></td></tr>
<tr><td>Chromosome</td><td>start</td><td>end</td><td>strand</td><td>MM</td><td>target_seq</td><td>PAM</td><td>alignment</td><td>distance</td><td>position</td><td>gene name</td><td>gene id</td></tr>
<tr><td>GU305914.1</td><td>71075</td><td>71097</td><td>+</td><td>0</td><td>TCGAACCTGCAGCTGATGCT</td><td>GGG</td><td>||||||||[||||||||||||]PAM</td><td>NA</td><td>Intergenic</td><td>NA</td><td>NA</td></tr>
<tr><td>T4976</td><td>GCAGCTGATGCTGGGTCAGCTGG</td><td>1000</td><td></td><td></td><td></td><td></td><td></td><td></td><td></td><td></td><td></td></tr>
<tr><td>Oligo fwd</td><td>CACCGCAGCTGATGCTGGGTCAGC</td><td></td><td></td><td></td><td></td><td></td><td></td><td></td><td></td><td></td><td></td></tr>
<tr><td>Oligo rev</td><td>AAACGCTGACCCAGCATCAGCTGC</td><td></td><td></td><td></td><td></td><td></td><td></td><td></td><td></td><td></td><td></td></tr>
<tr><td>Chromosome</td><td>start</td><td>end</td><td>strand</td><td>MM</td><td>target_seq</td><td>PAM</td><td>alignment</td><td>distance</td><td>position</td><td>gene name</td><td>gene id</td></tr>
<tr><td>GU305914.1</td><td>71083</td><td>71105</td><td>+</td><td>0</td><td>GCAGCTGATGCTGGGTCAGC</td><td>TGG</td><td>||||||||[||||||||||||]PAM</td><td>NA</td><td>Intergenic</td><td>NA</td><td>NA</td></tr>
<tr><td>T4977</td><td>GCTGATGCTGGGTCAGCTGGAGG</td><td>1000</td><td></td><td></td><td></td><td></td><td></td><td></td><td></td><td></td><td></td></tr>
<tr><td>Oligo fwd</td><td>CACCGCTGATGCTGGGTCAGCTGG</td><td></td><td></td><td></td><td></td><td></td><td></td><td></td><td></td><td></td><td></td></tr>
<tr><td>Oligo rev</td><td>AAACCCAGCTGACCCAGCATCAGC</td><td></td><td></td><td></td><td></td><td></td><td></td><td></td><td></td><td></td><td></td></tr>
<tr><td>Chromosome</td><td>start</td><td>end</td><td>strand</td><td>MM</td><td>target_seq</td><td>PAM</td><td>alignment</td><td>distance</td><td>position</td><td>gene name</td><td>gene id</td></tr>
<tr><td>GU305914.1</td><td>71086</td><td>71108</td><td>+</td><td>0</td><td>GCTGATGCTGGGTCAGCTGG</td><td>AGG</td><td>||||||||[||||||||||||]PAM</td><td>NA</td><td>Intergenic</td><td>NA</td><td>NA</td></tr>
<tr><td>T4978</td><td>TCAGCTGGAGGCCAAGAGAGTGG</td><td>1000</td><td></td><td></td><td></td><td></td><td></td><td></td><td></td><td></td><td></td></tr>
<tr><td>Oligo fwd</td><td>CACCTCAGCTGGAGGCCAAGAGAG</td><td></td><td></td><td></td><td></td><td></td><td></td><td></td><td></td><td></td><td></td></tr>
<tr><td>Oligo rev</td><td>AAACCTCTCTTGGCCTCCAGCTGA</td><td></td><td></td><td></td><td></td><td></td><td></td><td></td><td></td><td></td><td></td></tr>
<tr><td>Chromosome</td><td>start</td><td>end</td><td>strand</td><td>MM</td><td>target_seq</td><td>PAM</td><td>alignment</td><td>distance</td><td>position</td><td>gene name</td><td>gene id</td></tr>
<tr><td>GU305914.1</td><td>71098</td><td>71120</td><td>+</td><td>0</td><td>TCAGCTGGAGGCCAAGAGAG</td><td>TGG</td><td>||||||||[||||||||||||]PAM</td><td>NA</td><td>Intergenic</td><td>NA</td><td>NA</td></tr>
<tr><td>T4979</td><td>CTGGAGGCCAAGAGAGTGGTCGG</td><td>1000</td><td></td><td></td><td></td><td></td><td></td><td></td><td></td><td></td><td></td></tr>
<tr><td>Oligo fwd</td><td>CACCCTGGAGGCCAAGAGAGTGGT</td><td></td><td></td><td></td><td></td><td></td><td></td><td></td><td></td><td></td><td></td></tr>
<tr><td>Oligo rev</td><td>AAACACCACTCTCTTGGCCTCCAG</td><td></td><td></td><td></td><td></td><td></td><td></td><td></td><td></td><td></td><td></td></tr>
<tr><td>Chromosome</td><td>start</td><td>end</td><td>strand</td><td>MM</td><td>target_seq</td><td>PAM</td><td>alignment</td><td>distance</td><td>position</td><td>gene name</td><td>gene id</td></tr>
<tr><td>GU305914.1</td><td>71102</td><td>71124</td><td>+</td><td>0</td><td>CTGGAGGCCAAGAGAGTGGT</td><td>CGG</td><td>||||||||[||||||||||||]PAM</td><td>NA</td><td>Intergenic</td><td>NA</td><td>NA</td></tr>
<tr><td>T4980</td><td>GAGGCCAAGAGAGTGGTCGGCGG</td><td>1000</td><td></td><td></td><td></td><td></td><td></td><td></td><td></td><td></td><td></td></tr>
<tr><td>Oligo fwd</td><td>CACCGAGGCCAAGAGAGTGGTCGG</td><td></td><td></td><td></td><td></td><td></td><td></td><td></td><td></td><td></td><td></td></tr>
<tr><td>Oligo rev</td><td>AAACCCGACCACTCTCTTGGCCTC</td><td></td><td></td><td></td><td></td><td></td><td></td><td></td><td></td><td></td><td></td></tr>
<tr><td>Chromosome</td><td>start</td><td>end</td><td>strand</td><td>MM</td><td>target_seq</td><td>PAM</td><td>alignment</td><td>distance</td><td>position</td><td>gene name</td><td>gene id</td></tr>
<tr><td>GU305914.1</td><td>71105</td><td>71127</td><td>+</td><td>0</td><td>GAGGCCAAGAGAGTGGTCGG</td><td>CGG</td><td>||||||||[||||||||||||]PAM</td><td>NA</td><td>Intergenic</td><td>NA</td><td>NA</td></tr>
<tr><td>T4981</td><td>AGGCCAAGAGAGTGGTCGGCGGG</td><td>1000</td><td></td><td></td><td></td><td></td><td></td><td></td><td></td><td></td><td></td></tr>
<tr><td>Oligo fwd</td><td>CACCAGGCCAAGAGAGTGGTCGGC</td><td></td><td></td><td></td><td></td><td></td><td></td><td></td><td></td><td></td><td></td></tr>
<tr><td>Oligo rev</td><td>AAACGCCGACCACTCTCTTGGCCT</td><td></td><td></td><td></td><td></td><td></td><td></td><td></td><td></td><td></td><td></td></tr>
<tr><td>Chromosome</td><td>start</td><td>end</td><td>strand</td><td>MM</td><td>target_seq</td><td>PAM</td><td>alignment</td><td>distance</td><td>position</td><td>gene name</td><td>gene id</td></tr>
<tr><td>GU305914.1</td><td>71106</td><td>71128</td><td>+</td><td>0</td><td>AGGCCAAGAGAGTGGTCGGC</td><td>GGG</td><td>||||||||[||||||||||||]PAM</td><td>NA</td><td>Intergenic</td><td>NA</td><td>NA</td></tr>
<tr><td>T4982</td><td>GGCCAAGAGAGTGGTCGGCGGGG</td><td>1000</td><td></td><td></td><td></td><td></td><td></td><td></td><td></td><td></td><td></td></tr>
<tr><td>Oligo fwd</td><td>CACCGGCCAAGAGAGTGGTCGGCG</td><td></td><td></td><td></td><td></td><td></td><td></td><td></td><td></td><td></td><td></td></tr>
<tr><td>Oligo rev</td><td>AAACCGCCGACCACTCTCTTGGCC</td><td></td><td></td><td></td><td></td><td></td><td></td><td></td><td></td><td></td><td></td></tr>
<tr><td>Chromosome</td><td>start</td><td>end</td><td>strand</td><td>MM</td><td>target_seq</td><td>PAM</td><td>alignment</td><td>distance</td><td>position</td><td>gene name</td><td>gene id</td></tr>
<tr><td>GU305914.1</td><td>71107</td><td>71129</td><td>+</td><td>0</td><td>GGCCAAGAGAGTGGTCGGCG</td><td>GGG</td><td>||||||||[||||||||||||]PAM</td><td>NA</td><td>Intergenic</td><td>NA</td><td>NA</td></tr>
<tr><td>T4983</td><td>CAAGAGAGTGGTCGGCGGGGAGG</td><td>1000</td><td></td><td></td><td></td><td></td><td></td><td></td><td></td><td></td><td></td></tr>
<tr><td>Oligo fwd</td><td>CACCCAAGAGAGTGGTCGGCGGGG</td><td></td><td></td><td></td><td></td><td></td><td></td><td></td><td></td><td></td><td></td></tr>
<tr><td>Oligo rev</td><td>AAACCCCCGCCGACCACTCTCTTG</td><td></td><td></td><td></td><td></td><td></td><td></td><td></td><td></td><td></td><td></td></tr>
<tr><td>Chromosome</td><td>start</td><td>end</td><td>strand</td><td>MM</td><td>target_seq</td><td>PAM</td><td>alignment</td><td>distance</td><td>position</td><td>gene name</td><td>gene id</td></tr>
</table>

<table>
<tr><td>GU305914.1</td><td>71110</td><td>71132</td><td>+</td><td>0</td><td>CAAGAGAGTGGTCGGCGGGG</td><td>AGG</td><td>||||||||[||||||||||||]PAM</td><td>NA</td><td>Intergenic</td><td>NA</td><td>NA</td></tr>
<tr><td colspan="12"></td></tr>
<tr><td>T4984</td><td>CCTGACCAAACAGAACAAACTGG</td><td>1000</td><td colspan="9"></td></tr>
<tr><td>Oligo fwd</td><td>CACCCCTGACCAAACAGAACAAAC</td><td colspan="10"></td></tr>
<tr><td>Oligo rev</td><td>AAACGTTTGTTCTGTTTGGTCAGG</td><td colspan="10"></td></tr>
<tr><td>Chromosome</td><td>start</td><td>end</td><td>strand</td><td>MM</td><td>target_seq</td><td>PAM</td><td>alignment</td><td>distance</td><td>position</td><td>gene name</td><td>gene id</td></tr>
<tr><td>GU305914.1</td><td>71161</td><td>71183</td><td>+</td><td>0</td><td>CCTGACCAAACAGAACAAAC</td><td>TGG</td><td>||||||||[||||||||||||]PAM</td><td>NA</td><td>Intergenic</td><td>NA</td><td>NA</td></tr>
<tr><td colspan="12"></td></tr>
<tr><td>T4985</td><td>ACTGGCCTTCGCCGAAGATCTGG</td><td>1000</td><td colspan="9"></td></tr>
<tr><td>Oligo fwd</td><td>CACCACTGGCCTTCGCCGAAGATC</td><td colspan="10"></td></tr>
<tr><td>Oligo rev</td><td>AAACGATCTTCGGCGAAGGCCAGT</td><td colspan="10"></td></tr>
<tr><td>Chromosome</td><td>start</td><td>end</td><td>strand</td><td>MM</td><td>target_seq</td><td>PAM</td><td>alignment</td><td>distance</td><td>position</td><td>gene name</td><td>gene id</td></tr>
<tr><td>GU305914.1</td><td>71179</td><td>71201</td><td>+</td><td>0</td><td>ACTGGCCTTCGCCGAAGATC</td><td>TGG</td><td>||||||||[||||||||||||]PAM</td><td>NA</td><td>Intergenic</td><td>NA</td><td>NA</td></tr>
<tr><td colspan="12"></td></tr>
<tr><td>T4986</td><td>CGAAGATCTGGAGAAGCTGTCGG</td><td>1000</td><td colspan="9"></td></tr>
<tr><td>Oligo fwd</td><td>CACCCGAAGATCTGGAGAAGCTGT</td><td colspan="10"></td></tr>
<tr><td>Oligo rev</td><td>AAACACAGCTTCTCCAGATCTTCG</td><td colspan="10"></td></tr>
<tr><td>Chromosome</td><td>start</td><td>end</td><td>strand</td><td>MM</td><td>target_seq</td><td>PAM</td><td>alignment</td><td>distance</td><td>position</td><td>gene name</td><td>gene id</td></tr>
<tr><td>GU305914.1</td><td>71191</td><td>71213</td><td>+</td><td>0</td><td>CGAAGATCTGGAGAAGCTGT</td><td>CGG</td><td>||||||||[||||||||||||]PAM</td><td>NA</td><td>Intergenic</td><td>NA</td><td>NA</td></tr>
<tr><td colspan="12"></td></tr>
<tr><td>T4987</td><td>ATCTGGAGAAGCTGTCGGCGCGG</td><td>1000</td><td colspan="9"></td></tr>
<tr><td>Oligo fwd</td><td>CACCATCTGGAGAAGCTGTCGGCG</td><td colspan="10"></td></tr>
<tr><td>Oligo rev</td><td>AAACCGCCGACAGCTTCTCCAGAT</td><td colspan="10"></td></tr>
<tr><td>Chromosome</td><td>start</td><td>end</td><td>strand</td><td>MM</td><td>target_seq</td><td>PAM</td><td>alignment</td><td>distance</td><td>position</td><td>gene name</td><td>gene id</td></tr>
<tr><td>GU305914.1</td><td>71196</td><td>71218</td><td>+</td><td>0</td><td>ATCTGGAGAAGCTGTCGGCG</td><td>CGG</td><td>||||||||[||||||||||||]PAM</td><td>NA</td><td>Intergenic</td><td>NA</td><td>NA</td></tr>
<tr><td>GU305914.1</td><td>130607</td><td>130629</td><td>-</td><td>4</td><td>AGATGGAGAAGCCGGCGGCG</td><td>CGG</td><td>|--|||||[||||-|-|||||]PAM</td><td>NA</td><td>Intergenic</td><td>NA</td><td>NA</td></tr>
<tr><td colspan="12"></td></tr>
<tr><td>T4988</td><td>GTCGGCGCGGTACTTCGAACTGG</td><td>1000</td><td colspan="9"></td></tr>
<tr><td>Oligo fwd</td><td>CACCGTCGGCGCGGTACTTCGAAC</td><td colspan="10"></td></tr>
<tr><td>Oligo rev</td><td>AAACGTTCGAAGTACCGCGCCGAC</td><td colspan="10"></td></tr>
<tr><td>Chromosome</td><td>start</td><td>end</td><td>strand</td><td>MM</td><td>target_seq</td><td>PAM</td><td>alignment</td><td>distance</td><td>position</td><td>gene name</td><td>gene id</td></tr>
<tr><td>GU305914.1</td><td>71209</td><td>71231</td><td>+</td><td>0</td><td>GTCGGCGCGGTACTTCGAAC</td><td>TGG</td><td>||||||||[||||||||||||]PAM</td><td>NA</td><td>Intergenic</td><td>NA</td><td>NA</td></tr>
<tr><td colspan="12"></td></tr>
<tr><td>T4989</td><td>GACATCCGCAGCTCGAAGTACGG</td><td>1000</td><td colspan="9"></td></tr>
<tr><td>Oligo fwd</td><td>CACCGACATCCGCAGCTCGAAGTA</td><td colspan="10"></td></tr>
<tr><td>Oligo rev</td><td>AAACTACTTCGAGCTGCGGATGTC</td><td colspan="10"></td></tr>
<tr><td>Chromosome</td><td>start</td><td>end</td><td>strand</td><td>MM</td><td>target_seq</td><td>PAM</td><td>alignment</td><td>distance</td><td>position</td><td>gene name</td><td>gene id</td></tr>
<tr><td>GU305914.1</td><td>71237</td><td>71259</td><td>+</td><td>0</td><td>GACATCCGCAGCTCGAAGTA</td><td>CGG</td><td>||||||||[||||||||||||]PAM</td><td>NA</td><td>Intergenic</td><td>NA</td><td>NA</td></tr>
<tr><td colspan="12"></td></tr>
<tr><td>T4990</td><td>ACATCCGCAGCTCGAAGTACGGG</td><td>1000</td><td colspan="9"></td></tr>
<tr><td>Oligo fwd</td><td>CACCACATCCGCAGCTCGAAGTAC</td><td colspan="10"></td></tr>
<tr><td>Oligo rev</td><td>AAACGTACTTCGAGCTGCGGATGT</td><td colspan="10"></td></tr>
<tr><td>Chromosome</td><td>start</td><td>end</td><td>strand</td><td>MM</td><td>target_seq</td><td>PAM</td><td>alignment</td><td>distance</td><td>position</td><td>gene name</td><td>gene id</td></tr>
<tr><td>GU305914.1</td><td>71238</td><td>71260</td><td>+</td><td>0</td><td>ACATCCGCAGCTCGAAGTAC</td><td>GGG</td><td>||||||||[||||||||||||]PAM</td><td>NA</td><td>Intergenic</td><td>NA</td><td>NA</td></tr>
<tr><td colspan="12"></td></tr>
<tr><td>T4991</td><td>CCGCAGCTCGAAGTACGGGCTGG</td><td>1000</td><td colspan="9"></td></tr>
<tr><td>Oligo fwd</td><td>CACCCCGCAGCTCGAAGTACGGGC</td><td colspan="10"></td></tr>
<tr><td>Oligo rev</td><td>AAACGCCCGTACTTCGAGCTGCGG</td><td colspan="10"></td></tr>
<tr><td>Chromosome</td><td>start</td><td>end</td><td>strand</td><td>MM</td><td>target_seq</td><td>PAM</td><td>alignment</td><td>distance</td><td>position</td><td>gene name</td><td>gene id</td></tr>
<tr><td>GU305914.1</td><td>71242</td><td>71264</td><td>+</td><td>0</td><td>CCGCAGCTCGAAGTACGGGC</td><td>TGG</td><td>||||||||[||||||||||||]PAM</td><td>NA</td><td>Intergenic</td><td>NA</td><td>NA</td></tr>
<tr><td colspan="12"></td></tr>
<tr><td>T4992</td><td>GAAGACGAGCCTCCTGAAGTCGG</td><td>1000</td><td colspan="9"></td></tr>
<tr><td>Oligo fwd</td><td>CACCGAAGACGAGCCTCCTGAAGT</td><td colspan="10"></td></tr>
<tr><td>Oligo rev</td><td>AAACACTTCAGGAGGCTCGTCTTC</td><td colspan="10"></td></tr>
<tr><td>Chromosome</td><td>start</td><td>end</td><td>strand</td><td>MM</td><td>target_seq</td><td>PAM</td><td>alignment</td><td>distance</td><td>position</td><td>gene name</td><td>gene id</td></tr>
<tr><td>GU305914.1</td><td>71284</td><td>71306</td><td>+</td><td>0</td><td>GAAGACGAGCCTCCTGAAGT</td><td>CGG</td><td>||||||||[||||||||||||]PAM</td><td>NA</td><td>Intergenic</td><td>NA</td><td>NA</td></tr>
<tr><td colspan="12"></td></tr>
<tr><td>T4993</td><td>AGCCTCCTGAAGTCGGAGATCGG</td><td>1000</td><td colspan="9"></td></tr>
<tr><td>Oligo fwd</td><td>CACCAGCCTCCTGAAGTCGGAGAT</td><td colspan="10"></td></tr>
<tr><td>Oligo rev</td><td>AAACATCTCCGACTTCAGGAGGCT</td><td colspan="10"></td></tr>
<tr><td>Chromosome</td><td>start</td><td>end</td><td>strand</td><td>MM</td><td>target_seq</td><td>PAM</td><td>alignment</td><td>distance</td><td>position</td><td>gene name</td><td>gene id</td></tr>
<tr><td>GU305914.1</td><td>71291</td><td>71313</td><td>+</td><td>0</td><td>AGCCTCCTGAAGTCGGAGAT</td><td>CGG</td><td>||||||||[||||||||||||]PAM</td><td>NA</td><td>Intergenic</td><td>NA</td><td>NA</td></tr>
<tr><td colspan="12"></td></tr>
<tr><td>T4994</td><td>GCCTCCTGAAGTCGGAGATCGGG</td><td>1000</td><td colspan="9"></td></tr>
<tr><td>Oligo fwd</td><td>CACCGCCTCCTGAAGTCGGAGATC</td><td colspan="10"></td></tr>
<tr><td>Oligo rev</td><td>AAACGATCTCCGACTTCAGGAGGC</td><td colspan="10"></td></tr>
<tr><td>Chromosome</td><td>start</td><td>end</td><td>strand</td><td>MM</td><td>target_seq</td><td>PAM</td><td>alignment</td><td>distance</td><td>position</td><td>gene name</td><td>gene id</td></tr>
<tr><td>GU305914.1</td><td>71292</td><td>71314</td><td>+</td><td>0</td><td>GCCTCCTGAAGTCGGAGATC</td><td>GGG</td><td>||||||||[||||||||||||]PAM</td><td>NA</td><td>Intergenic</td><td>NA</td><td>NA</td></tr>
<tr><td colspan="12"></td></tr>
<tr><td>T4995</td><td>CAAGAAAGATCCCCCCTCGACGG</td><td>1000</td><td colspan="9"></td></tr>
<tr><td>Oligo fwd</td><td>CACCCAAGAAAGATCCCCCCTCGA</td><td colspan="10"></td></tr>
<tr><td>Oligo rev</td><td>AAACTCGAGGGGGGATCTTTCTTG</td><td colspan="10"></td></tr>
<tr><td>Chromosome</td><td>start</td><td>end</td><td>strand</td><td>MM</td><td>target_seq</td><td>PAM</td><td>alignment</td><td>distance</td><td>position</td><td>gene name</td><td>gene id</td></tr>
<tr><td>GU305914.1</td><td>71320</td><td>71342</td><td>+</td><td>0</td><td>CAAGAAAGATCCCCCCTCGA</td><td>CGG</td><td>||||||||[||||||||||||]PAM</td><td>NA</td><td>Intergenic</td><td>NA</td><td>NA</td></tr>
<tr><td colspan="12"></td></tr>
<tr><td>T4996</td><td>AAGAAAGATCCCCCCTCGACGGG</td><td>1000</td><td colspan="9"></td></tr>
<tr><td>Oligo fwd</td><td>CACCAAGAAAGATCCCCCCTCGAC</td><td colspan="10"></td></tr>
<tr><td>Oligo rev</td><td>AAACGTCGAGGGGGGATCTTTCTT</td><td colspan="10"></td></tr>
<tr><td>Chromosome</td><td>start</td><td>end</td><td>strand</td><td>MM</td><td>target_seq</td><td>PAM</td><td>alignment</td><td>distance</td><td>position</td><td>gene name</td><td>gene id</td></tr>
<tr><td>GU305914.1</td><td>71321</td><td>71343</td><td>+</td><td>0</td><td>AAGAAAGATCCCCCCTCGAC</td><td>GGG</td><td>||||||||[||||||||||||]PAM</td><td>NA</td><td>Intergenic</td><td>NA</td><td>NA</td></tr>
<tr><td colspan="12"></td></tr>
<tr><td>T4997</td><td>AGAAAGATCCCCCCTCGACGGGG</td><td>1000</td><td colspan="9"></td></tr>
<tr><td>Oligo fwd</td><td>CACCAGAAAGATCCCCCCTCGACG</td><td colspan="10"></td></tr>
<tr><td>Oligo rev</td><td>AAACCGTCGAGGGGGGATCTTTCT</td><td colspan="10"></td></tr>
<tr><td>Chromosome</td><td>start</td><td>end</td><td>strand</td><td>MM</td><td>target_seq</td><td>PAM</td><td>alignment</td><td>distance</td><td>position</td><td>gene name</td><td>gene id</td></tr>
<tr><td>GU305914.1</td><td>71322</td><td>71344</td><td>+</td><td>0</td><td>AGAAAGATCCCCCCTCGACG</td><td>GGG</td><td>||||||||[||||||||||||]PAM</td><td>NA</td><td>Intergenic</td><td>NA</td><td>NA</td></tr>
<tr><td colspan="12"></td></tr>
<tr><td>T4998</td><td>GAAAGATCCCCCCTCGACGGGGG</td><td>1000</td><td colspan="9"></td></tr>
<tr><td>Oligo fwd</td><td>CACCGAAAGATCCCCCCTCGACGG</td><td colspan="10"></td></tr>
</table>

<table>
<tr><td>Oligo rev</td><td>AAACCCGTCGAGGGGGGATCTTTC</td><td></td><td></td><td></td><td></td><td></td><td></td><td></td><td></td><td></td><td></td></tr>
<tr><td>Chromosome</td><td>start</td><td>end</td><td>strand</td><td>MM</td><td>target_seq</td><td>PAM</td><td>alignment</td><td>distance</td><td>position</td><td>gene name</td><td>gene id</td></tr>
<tr><td>GU305914.1</td><td>71323</td><td>71345</td><td>+</td><td>0</td><td>GAAAGATCCCCCCTCGACGG</td><td>GGG</td><td>||||||||[||||||||||||]PAM</td><td>NA</td><td>Intergenic</td><td>NA</td><td>NA</td></tr>
<tr><td>T4999</td><td>AGATCCCCCCTCGACGGGGGAGG</td><td>1000</td><td></td><td></td><td></td><td></td><td></td><td></td><td></td><td></td><td></td></tr>
<tr><td>Oligo fwd</td><td>CACCAGATCCCCCCTCGACGGGGG</td><td></td><td></td><td></td><td></td><td></td><td></td><td></td><td></td><td></td><td></td></tr>
<tr><td>Oligo rev</td><td>AAACCCCCCGTCGAGGGGGGATCT</td><td></td><td></td><td></td><td></td><td></td><td></td><td></td><td></td><td></td><td></td></tr>
<tr><td>Chromosome</td><td>start</td><td>end</td><td>strand</td><td>MM</td><td>target_seq</td><td>PAM</td><td>alignment</td><td>distance</td><td>position</td><td>gene name</td><td>gene id</td></tr>
<tr><td>GU305914.1</td><td>71326</td><td>71348</td><td>+</td><td>0</td><td>AGATCCCCCCTCGACGGGGG</td><td>AGG</td><td>||||||||[||||||||||||]PAM</td><td>NA</td><td>Intergenic</td><td>NA</td><td>NA</td></tr>
<tr><td>T5000</td><td>TCCCCCCTCGACGGGGGAGGAGG</td><td>1000</td><td></td><td></td><td></td><td></td><td></td><td></td><td></td><td></td><td></td></tr>
<tr><td>Oligo fwd</td><td>CACCTCCCCCCTCGACGGGGGAGG</td><td></td><td></td><td></td><td></td><td></td><td></td><td></td><td></td><td></td><td></td></tr>
<tr><td>Oligo rev</td><td>AAACCCTCCCCCGTCGAGGGGGGA</td><td></td><td></td><td></td><td></td><td></td><td></td><td></td><td></td><td></td><td></td></tr>
<tr><td>Chromosome</td><td>start</td><td>end</td><td>strand</td><td>MM</td><td>target_seq</td><td>PAM</td><td>alignment</td><td>distance</td><td>position</td><td>gene name</td><td>gene id</td></tr>
<tr><td>GU305914.1</td><td>71329</td><td>71351</td><td>+</td><td>0</td><td>TCCCCCCTCGACGGGGGAGG</td><td>AGG</td><td>||||||||[||||||||||||]PAM</td><td>NA</td><td>Intergenic</td><td>NA</td><td>NA</td></tr>
<tr><td>T5001</td><td>CCCCTCGACGGGGGAGGAGGTGG</td><td>1000</td><td></td><td></td><td></td><td></td><td></td><td></td><td></td><td></td><td></td></tr>
<tr><td>Oligo fwd</td><td>CACCCCCCTCGACGGGGGAGGAGG</td><td></td><td></td><td></td><td></td><td></td><td></td><td></td><td></td><td></td><td></td></tr>
<tr><td>Oligo rev</td><td>AAACCCTCCTCCCCCGTCGAGGGG</td><td></td><td></td><td></td><td></td><td></td><td></td><td></td><td></td><td></td><td></td></tr>
<tr><td>Chromosome</td><td>start</td><td>end</td><td>strand</td><td>MM</td><td>target_seq</td><td>PAM</td><td>alignment</td><td>distance</td><td>position</td><td>gene name</td><td>gene id</td></tr>
<tr><td>GU305914.1</td><td>71332</td><td>71354</td><td>+</td><td>0</td><td>CCCCTCGACGGGGGAGGAGG</td><td>TGG</td><td>||||||||[||||||||||||]PAM</td><td>NA</td><td>Intergenic</td><td>NA</td><td>NA</td></tr>
<tr><td>GU305914.1</td><td>99537</td><td>99559</td><td>-</td><td>4</td><td>ACGCTCGTCGGGCGAGGAGG</td><td>CGG</td><td>-|-||||-[||||-|||||||]PAM</td><td>NA</td><td>Intergenic</td><td>NA</td><td>NA</td></tr>
<tr><td>T5002</td><td>CCCTCGACGGGGGAGGAGGTGGG</td><td>1000</td><td></td><td></td><td></td><td></td><td></td><td></td><td></td><td></td><td></td></tr>
<tr><td>Oligo fwd</td><td>CACCCCCTCGACGGGGGAGGAGGT</td><td></td><td></td><td></td><td></td><td></td><td></td><td></td><td></td><td></td><td></td></tr>
<tr><td>Oligo rev</td><td>AAACACCTCCTCCCCCGTCGAGGG</td><td></td><td></td><td></td><td></td><td></td><td></td><td></td><td></td><td></td><td></td></tr>
<tr><td>Chromosome</td><td>start</td><td>end</td><td>strand</td><td>MM</td><td>target_seq</td><td>PAM</td><td>alignment</td><td>distance</td><td>position</td><td>gene name</td><td>gene id</td></tr>
<tr><td>GU305914.1</td><td>71333</td><td>71355</td><td>+</td><td>0</td><td>CCCTCGACGGGGGAGGAGGT</td><td>GGG</td><td>||||||||[||||||||||||]PAM</td><td>NA</td><td>Intergenic</td><td>NA</td><td>NA</td></tr>
<tr><td>T5003</td><td>CTTACCTGAAGAGAGCACGACGG</td><td>1000</td><td></td><td></td><td></td><td></td><td></td><td></td><td></td><td></td><td></td></tr>
<tr><td>Oligo fwd</td><td>CACCCTTACCTGAAGAGAGCACGA</td><td></td><td></td><td></td><td></td><td></td><td></td><td></td><td></td><td></td><td></td></tr>
<tr><td>Oligo rev</td><td>AAACTCGTGCTCTCTTCAGGTAAG</td><td></td><td></td><td></td><td></td><td></td><td></td><td></td><td></td><td></td><td></td></tr>
<tr><td>Chromosome</td><td>start</td><td>end</td><td>strand</td><td>MM</td><td>target_seq</td><td>PAM</td><td>alignment</td><td>distance</td><td>position</td><td>gene name</td><td>gene id</td></tr>
<tr><td>GU305914.1</td><td>71356</td><td>71378</td><td>+</td><td>0</td><td>CTTACCTGAAGAGAGCACGA</td><td>CGG</td><td>||||||||[||||||||||||]PAM</td><td>NA</td><td>Intergenic</td><td>NA</td><td>NA</td></tr>
<tr><td>T5004</td><td>ACCTGAAGAGAGCACGACGGCGG</td><td>1000</td><td></td><td></td><td></td><td></td><td></td><td></td><td></td><td></td><td></td></tr>
<tr><td>Oligo fwd</td><td>CACCACCTGAAGAGAGCACGACGG</td><td></td><td></td><td></td><td></td><td></td><td></td><td></td><td></td><td></td><td></td></tr>
<tr><td>Oligo rev</td><td>AAACCCGTCGTGCTCTCTTCAGGT</td><td></td><td></td><td></td><td></td><td></td><td></td><td></td><td></td><td></td><td></td></tr>
<tr><td>Chromosome</td><td>start</td><td>end</td><td>strand</td><td>MM</td><td>target_seq</td><td>PAM</td><td>alignment</td><td>distance</td><td>position</td><td>gene name</td><td>gene id</td></tr>
<tr><td>GU305914.1</td><td>71359</td><td>71381</td><td>+</td><td>0</td><td>ACCTGAAGAGAGCACGACGG</td><td>CGG</td><td>||||||||[||||||||||||]PAM</td><td>NA</td><td>Intergenic</td><td>NA</td><td>NA</td></tr>
<tr><td>T5005</td><td>CTCGATCTCCTCGCTCCTGTTGG</td><td>1000</td><td></td><td></td><td></td><td></td><td></td><td></td><td></td><td></td><td></td></tr>
<tr><td>Oligo fwd</td><td>CACCCTCGATCTCCTCGCTCCTGT</td><td></td><td></td><td></td><td></td><td></td><td></td><td></td><td></td><td></td><td></td></tr>
<tr><td>Oligo rev</td><td>AAACACAGGAGCGAGGAGATCGAG</td><td></td><td></td><td></td><td></td><td></td><td></td><td></td><td></td><td></td><td></td></tr>
<tr><td>Chromosome</td><td>start</td><td>end</td><td>strand</td><td>MM</td><td>target_seq</td><td>PAM</td><td>alignment</td><td>distance</td><td>position</td><td>gene name</td><td>gene id</td></tr>
<tr><td>GU305914.1</td><td>71389</td><td>71411</td><td>+</td><td>0</td><td>CTCGATCTCCTCGCTCCTGT</td><td>TGG</td><td>||||||||[||||||||||||]PAM</td><td>NA</td><td>Intergenic</td><td>NA</td><td>NA</td></tr>
<tr><td>T5006</td><td>TCGATCTCCTCGCTCCTGTTGGG</td><td>1000</td><td></td><td></td><td></td><td></td><td></td><td></td><td></td><td></td><td></td></tr>
<tr><td>Oligo fwd</td><td>CACCTCGATCTCCTCGCTCCTGTT</td><td></td><td></td><td></td><td></td><td></td><td></td><td></td><td></td><td></td><td></td></tr>
<tr><td>Oligo rev</td><td>AAACAACAGGAGCGAGGAGATCGA</td><td></td><td></td><td></td><td></td><td></td><td></td><td></td><td></td><td></td><td></td></tr>
<tr><td>Chromosome</td><td>start</td><td>end</td><td>strand</td><td>MM</td><td>target_seq</td><td>PAM</td><td>alignment</td><td>distance</td><td>position</td><td>gene name</td><td>gene id</td></tr>
<tr><td>GU305914.1</td><td>71390</td><td>71412</td><td>+</td><td>0</td><td>TCGATCTCCTCGCTCCTGTT</td><td>GGG</td><td>||||||||[||||||||||||]PAM</td><td>NA</td><td>Intergenic</td><td>NA</td><td>NA</td></tr>
<tr><td>T5007</td><td>CTCGCTCCTGTTGGGCATCGCGG</td><td>1000</td><td></td><td></td><td></td><td></td><td></td><td></td><td></td><td></td><td></td></tr>
<tr><td>Oligo fwd</td><td>CACCCTCGCTCCTGTTGGGCATCG</td><td></td><td></td><td></td><td></td><td></td><td></td><td></td><td></td><td></td><td></td></tr>
<tr><td>Oligo rev</td><td>AAACCGATGCCCAACAGGAGCGAG</td><td></td><td></td><td></td><td></td><td></td><td></td><td></td><td></td><td></td><td></td></tr>
<tr><td>Chromosome</td><td>start</td><td>end</td><td>strand</td><td>MM</td><td>target_seq</td><td>PAM</td><td>alignment</td><td>distance</td><td>position</td><td>gene name</td><td>gene id</td></tr>
<tr><td>GU305914.1</td><td>71398</td><td>71420</td><td>+</td><td>0</td><td>CTCGCTCCTGTTGGGCATCG</td><td>CGG</td><td>||||||||[||||||||||||]PAM</td><td>NA</td><td>Intergenic</td><td>NA</td><td>NA</td></tr>
<tr><td>T5008</td><td>CCTGTTGGGCATCGCGGCGCTGG</td><td>1000</td><td></td><td></td><td></td><td></td><td></td><td></td><td></td><td></td><td></td></tr>
<tr><td>Oligo fwd</td><td>CACCCCTGTTGGGCATCGCGGCGC</td><td></td><td></td><td></td><td></td><td></td><td></td><td></td><td></td><td></td><td></td></tr>
<tr><td>Oligo rev</td><td>AAACGCGCCGCGATGCCCAACAGG</td><td></td><td></td><td></td><td></td><td></td><td></td><td></td><td></td><td></td><td></td></tr>
<tr><td>Chromosome</td><td>start</td><td>end</td><td>strand</td><td>MM</td><td>target_seq</td><td>PAM</td><td>alignment</td><td>distance</td><td>position</td><td>gene name</td><td>gene id</td></tr>
<tr><td>GU305914.1</td><td>71404</td><td>71426</td><td>+</td><td>0</td><td>CCTGTTGGGCATCGCGGCGC</td><td>TGG</td><td>||||||||[||||||||||||]PAM</td><td>NA</td><td>Intergenic</td><td>NA</td><td>NA</td></tr>
<tr><td>T5009</td><td>GGCGCTGGAGAAGTACATCGTGG</td><td>1000</td><td></td><td></td><td></td><td></td><td></td><td></td><td></td><td></td><td></td></tr>
<tr><td>Oligo fwd</td><td>CACCGGCGCTGGAGAAGTACATCG</td><td></td><td></td><td></td><td></td><td></td><td></td><td></td><td></td><td></td><td></td></tr>
<tr><td>Oligo rev</td><td>AAACCGATGTACTTCTCCAGCGCC</td><td></td><td></td><td></td><td></td><td></td><td></td><td></td><td></td><td></td><td></td></tr>
<tr><td>Chromosome</td><td>start</td><td>end</td><td>strand</td><td>MM</td><td>target_seq</td><td>PAM</td><td>alignment</td><td>distance</td><td>position</td><td>gene name</td><td>gene id</td></tr>
<tr><td>GU305914.1</td><td>71419</td><td>71441</td><td>+</td><td>0</td><td>GGCGCTGGAGAAGTACATCG</td><td>TGG</td><td>||||||||[||||||||||||]PAM</td><td>NA</td><td>Intergenic</td><td>NA</td><td>NA</td></tr>
<tr><td>T5010</td><td>CGTGGCGCACCGCACCCTGCTGG</td><td>1000</td><td></td><td></td><td></td><td></td><td></td><td></td><td></td><td></td><td></td></tr>
<tr><td>Oligo fwd</td><td>CACCCGTGGCGCACCGCACCCTGC</td><td></td><td></td><td></td><td></td><td></td><td></td><td></td><td></td><td></td><td></td></tr>
<tr><td>Oligo rev</td><td>AAACGCAGGGTGCGGTGCGCCACG</td><td></td><td></td><td></td><td></td><td></td><td></td><td></td><td></td><td></td><td></td></tr>
<tr><td>Chromosome</td><td>start</td><td>end</td><td>strand</td><td>MM</td><td>target_seq</td><td>PAM</td><td>alignment</td><td>distance</td><td>position</td><td>gene name</td><td>gene id</td></tr>
<tr><td>GU305914.1</td><td>71437</td><td>71459</td><td>+</td><td>0</td><td>CGTGGCGCACCGCACCCTGC</td><td>TGG</td><td>||||||||[||||||||||||]PAM</td><td>NA</td><td>Intergenic</td><td>NA</td><td>NA</td></tr>
<tr><td>T5011</td><td>CTCGCAGCCGTTGATCAGTCAGG</td><td>1000</td><td></td><td></td><td></td><td></td><td></td><td></td><td></td><td></td><td></td></tr>
<tr><td>Oligo fwd</td><td>CACCCTCGCAGCCGTTGATCAGTC</td><td></td><td></td><td></td><td></td><td></td><td></td><td></td><td></td><td></td><td></td></tr>
<tr><td>Oligo rev</td><td>AAACGACTGATCAACGGCTGCGAG</td><td></td><td></td><td></td><td></td><td></td><td></td><td></td><td></td><td></td><td></td></tr>
<tr><td>Chromosome</td><td>start</td><td>end</td><td>strand</td><td>MM</td><td>target_seq</td><td>PAM</td><td>alignment</td><td>distance</td><td>position</td><td>gene name</td><td>gene id</td></tr>
<tr><td>GU305914.1</td><td>71473</td><td>71495</td><td>+</td><td>0</td><td>CTCGCAGCCGTTGATCAGTC</td><td>AGG</td><td>||||||||[||||||||||||]PAM</td><td>NA</td><td>Intergenic</td><td>NA</td><td>NA</td></tr>
<tr><td>T5012</td><td>GCAGCCGTTGATCAGTCAGGCGG</td><td>1000</td><td></td><td></td><td></td><td></td><td></td><td></td><td></td><td></td><td></td></tr>
<tr><td>Oligo fwd</td><td>CACCGCAGCCGTTGATCAGTCAGG</td><td></td><td></td><td></td><td></td><td></td><td></td><td></td><td></td><td></td><td></td></tr>
<tr><td>Oligo rev</td><td>AAACCCTGACTGATCAACGGCTGC</td><td></td><td></td><td></td><td></td><td></td><td></td><td></td><td></td><td></td><td></td></tr>
<tr><td>Chromosome</td><td>start</td><td>end</td><td>strand</td><td>MM</td><td>target_seq</td><td>PAM</td><td>alignment</td><td>distance</td><td>position</td><td>gene name</td><td>gene id</td></tr>
<tr><td>GU305914.1</td><td>71476</td><td>71498</td><td>+</td><td>0</td><td>GCAGCCGTTGATCAGTCAGG</td><td>CGG</td><td>||||||||[||||||||||||]PAM</td><td>NA</td><td>Intergenic</td><td>NA</td><td>NA</td></tr>
</table>

T5013 CAGTCAGGCGGAAAACATCCCGG 1000
Oligo fwd CACCCAGTCAGGCGGAAAACATCC
Oligo rev AAACGGATGTTTTCCGCCTGACTG
Chromosome start end strand MM target_seq PAM alignment distance position gene name gene id
GU305914.1 71488 71510 + 0 CAGTCAGGCGGAAAACATCC CGG ||||||||[||||||||||||]PAM NA Intergenic NA NA

T5014 GGCGGAAAACATCCCGGCTCTGG 1000
Oligo fwd CACCGGCGGAAAACATCCCGGCTC
Oligo rev AAACGAGCCGGGATGTTTTCCGCC
Chromosome start end strand MM target_seq PAM alignment distance position gene name gene id
GU305914.1 71494 71516 + 0 GGCGGAAAACATCCCGGCTC TGG ||||||||[||||||||||||]PAM NA Intergenic NA NA

T5015 GGAAAACATCCCGGCTCTGGTGG 1000
Oligo fwd CACCGGAAAACATCCCGGCTCTGG
Oligo rev AAACCCAGAGCCGGGATGTTTTCC
Chromosome start end strand MM target_seq PAM alignment distance position gene name gene id
GU305914.1 71497 71519 + 0 GGAAAACATCCCGGCTCTGG TGG ||||||||[||||||||||||]PAM NA Intergenic NA NA

T5016 GAAAACATCCCGGCTCTGGTGGG 1000
Oligo fwd CACCGAAAACATCCCGGCTCTGGT
Oligo rev AAACACCAGAGCCGGGATGTTTTC
Chromosome start end strand MM target_seq PAM alignment distance position gene name gene id
GU305914.1 71498 71520 + 0 GAAAACATCCCGGCTCTGGT GGG ||||||||[||||||||||||]PAM NA Intergenic NA NA

T5017 AACATCCCGGCTCTGGTGGGTGG 1000
Oligo fwd CACCAACATCCCGGCTCTGGTGGG
Oligo rev AAACCCCACCAGAGCCGGGATGTT
Chromosome start end strand MM target_seq PAM alignment distance position gene name gene id
GU305914.1 71501 71523 + 0 AACATCCCGGCTCTGGTGGG TGG ||||||||[||||||||||||]PAM NA Intergenic NA NA

T5018 CCGGCTCTGGTGGGTGGCCCCGG 1000
Oligo fwd CACCCCGGCTCTGGTGGGTGGCCC
Oligo rev AAACGGGCCACCCACCAGAGCCGG
Chromosome start end strand MM target_seq PAM alignment distance position gene name gene id
GU305914.1 71507 71529 + 0 CCGGCTCTGGTGGGTGGCCC CGG ||||||||[||||||||||||]PAM NA Intergenic NA NA
GU305914.1 150962 150984 + 4 CCGCCGCCGGTGGGAGGCCC AGG |||-|-|-[||||||-|||||]PAM NA Intergenic NA NA

T5019 TCTGGTGGGTGGCCCCGGTGAGG 1000
Oligo fwd CACCTCTGGTGGGTGGCCCCGGTG
Oligo rev AAACCACCGGGGCCACCCACCAGA
Chromosome start end strand MM target_seq PAM alignment distance position gene name gene id
GU305914.1 71512 71534 + 0 TCTGGTGGGTGGCCCCGGTG AGG ||||||||[||||||||||||]PAM NA Intergenic NA NA

T5020 CCCGGTGAGGACGATAAGCCAGG 1000
Oligo fwd CACCCCCGGTGAGGACGATAAGCC
Oligo rev AAACGGCTTATCGTCCTCACCGGG
Chromosome start end strand MM target_seq PAM alignment distance position gene name gene id
GU305914.1 71525 71547 + 0 CCCGGTGAGGACGATAAGCC AGG ||||||||[||||||||||||]PAM NA Intergenic NA NA

T5021 CCGGTGAGGACGATAAGCCAGGG 1000
Oligo fwd CACCCCGGTGAGGACGATAAGCCA
Oligo rev AAACTGGCTTATCGTCCTCACCGG
Chromosome start end strand MM target_seq PAM alignment distance position gene name gene id
GU305914.1 71526 71548 + 0 CCGGTGAGGACGATAAGCCA GGG ||||||||[||||||||||||]PAM NA Intergenic NA NA

T5022 TGAGGACGATAAGCCAGGGCAGG 1000
Oligo fwd CACCTGAGGACGATAAGCCAGGGC
Oligo rev AAACGCCCTGGCTTATCGTCCTCA
Chromosome start end strand MM target_seq PAM alignment distance position gene name gene id
GU305914.1 71530 71552 + 0 TGAGGACGATAAGCCAGGGC AGG ||||||||[||||||||||||]PAM NA Intergenic NA NA

T5023 CAGGGCAGGAGCCGTTCGACTGG 1000
Oligo fwd CACCCAGGGCAGGAGCCGTTCGAC
Oligo rev AAACGTCGAACGGCTCCTGCCCTG
Chromosome start end strand MM target_seq PAM alignment distance position gene name gene id
GU305914.1 71544 71566 + 0 CAGGGCAGGAGCCGTTCGAC TGG ||||||||[||||||||||||]PAM NA Intergenic NA NA
GU305914.1 68076 68098 + 4 GCGGGGAGGAGCCGTTCTAC CGG --|||-||[|||||||||-||]PAM NA Intergenic NA NA

T5024 GTCAGTACCGCCTTATATCGCGG 1000
Oligo fwd CACCGTCAGTACCGCCTTATATCG
Oligo rev AAACCGATATAAGGCGGTACTGAC
Chromosome start end strand MM target_seq PAM alignment distance position gene name gene id
GU305914.1 71603 71625 + 0 GTCAGTACCGCCTTATATCG CGG ||||||||[||||||||||||]PAM NA Intergenic NA NA

T5025 TCAGTACCGCCTTATATCGCGGG 1000
Oligo fwd CACCTCAGTACCGCCTTATATCGC
Oligo rev AAACGCGATATAAGGCGGTACTGA
Chromosome start end strand MM target_seq PAM alignment distance position gene name gene id
GU305914.1 71604 71626 + 0 TCAGTACCGCCTTATATCGC GGG ||||||||[||||||||||||]PAM NA Intergenic NA NA

T5026 TACCGCCTTATATCGCGGGACGG 1000
Oligo fwd CACCTACCGCCTTATATCGCGGGA
Oligo rev AAACTCCCGCGATATAAGGCGGTA
Chromosome start end strand MM target_seq PAM alignment distance position gene name gene id
GU305914.1 71608 71630 + 0 TACCGCCTTATATCGCGGGA CGG ||||||||[||||||||||||]PAM NA Intergenic NA NA

T5027 TATCGCGGGACGGACGTGTTCGG 1000
Oligo fwd CACCTATCGCGGGACGGACGTGTT
Oligo rev AAACAACACGTCCGTCCCGCGATA

<table>
<tr><td>Chromosome</td><td>start</td><td>end</td><td>strand</td><td>MM</td><td>target_seq</td><td>PAM</td><td>alignment</td><td>distance</td><td>position</td><td>gene name</td><td>gene id</td></tr>
<tr><td>GU305914.1</td><td>71618</td><td>71640</td><td>+</td><td>0</td><td>TATCGCGGGACGGACGTGTT</td><td>CGG</td><td>||||||||[||||||||||||]PAM</td><td>NA</td><td>Intergenic</td><td>NA</td><td>NA</td></tr>
<tr><td></td><td></td><td></td><td></td><td></td><td></td><td></td><td></td><td></td><td></td><td></td><td></td></tr>
<tr><td>T5028</td><td>ATCGCGGGACGGACGTGTTCGGG</td><td>1000</td><td></td><td></td><td></td><td></td><td></td><td></td><td></td><td></td><td></td></tr>
<tr><td>Oligo fwd</td><td>CACCATCGCGGGACGGACGTGTTC</td><td></td><td></td><td></td><td></td><td></td><td></td><td></td><td></td><td></td><td></td></tr>
<tr><td>Oligo rev</td><td>AAACGAACACGTCCGTCCCGCGAT</td><td></td><td></td><td></td><td></td><td></td><td></td><td></td><td></td><td></td><td></td></tr>
<tr><td>Chromosome</td><td>start</td><td>end</td><td>strand</td><td>MM</td><td>target_seq</td><td>PAM</td><td>alignment</td><td>distance</td><td>position</td><td>gene name</td><td>gene id</td></tr>
<tr><td>GU305914.1</td><td>71619</td><td>71641</td><td>+</td><td>0</td><td>ATCGCGGGACGGACGTGTTC</td><td>GGG</td><td>||||||||[||||||||||||]PAM</td><td>NA</td><td>Intergenic</td><td>NA</td><td>NA</td></tr>
<tr><td></td><td></td><td></td><td></td><td></td><td></td><td></td><td></td><td></td><td></td><td></td><td></td></tr>
<tr><td>T5029</td><td>TCGCGGGACGGACGTGTTCGGGG</td><td>1000</td><td></td><td></td><td></td><td></td><td></td><td></td><td></td><td></td><td></td></tr>
<tr><td>Oligo fwd</td><td>CACCTCGCGGGACGGACGTGTTCG</td><td></td><td></td><td></td><td></td><td></td><td></td><td></td><td></td><td></td><td></td></tr>
<tr><td>Oligo rev</td><td>AAACCGAACACGTCCGTCCCGCGA</td><td></td><td></td><td></td><td></td><td></td><td></td><td></td><td></td><td></td><td></td></tr>
<tr><td>Chromosome</td><td>start</td><td>end</td><td>strand</td><td>MM</td><td>target_seq</td><td>PAM</td><td>alignment</td><td>distance</td><td>position</td><td>gene name</td><td>gene id</td></tr>
<tr><td>GU305914.1</td><td>71620</td><td>71642</td><td>+</td><td>0</td><td>TCGCGGGACGGACGTGTTCG</td><td>GGG</td><td>||||||||[||||||||||||]PAM</td><td>NA</td><td>Intergenic</td><td>NA</td><td>NA</td></tr>
<tr><td>GU305914.1</td><td>139922</td><td>139944</td><td>+</td><td>4</td><td>TCGCGACGCGGACGAGTTCG</td><td>CGG</td><td>|||||---[||||||-|||||]PAM</td><td>NA</td><td>Intergenic</td><td>NA</td><td>NA</td></tr>
<tr><td></td><td></td><td></td><td></td><td></td><td></td><td></td><td></td><td></td><td></td><td></td><td></td></tr>
<tr><td>T5030</td><td>GGACGGACGTGTTCGGGGAGAGG</td><td>1000</td><td></td><td></td><td></td><td></td><td></td><td></td><td></td><td></td><td></td></tr>
<tr><td>Oligo fwd</td><td>CACCGGACGGACGTGTTCGGGGAG</td><td></td><td></td><td></td><td></td><td></td><td></td><td></td><td></td><td></td><td></td></tr>
<tr><td>Oligo rev</td><td>AAACCTCCCCGAACACGTCCGTCC</td><td></td><td></td><td></td><td></td><td></td><td></td><td></td><td></td><td></td><td></td></tr>
<tr><td>Chromosome</td><td>start</td><td>end</td><td>strand</td><td>MM</td><td>target_seq</td><td>PAM</td><td>alignment</td><td>distance</td><td>position</td><td>gene name</td><td>gene id</td></tr>
<tr><td>GU305914.1</td><td>71625</td><td>71647</td><td>+</td><td>0</td><td>GGACGGACGTGTTCGGGGAG</td><td>AGG</td><td>||||||||[||||||||||||]PAM</td><td>NA</td><td>Intergenic</td><td>NA</td><td>NA</td></tr>
<tr><td></td><td></td><td></td><td></td><td></td><td></td><td></td><td></td><td></td><td></td><td></td><td></td></tr>
<tr><td>T5031</td><td>CGGGGAGAGGCGCGTGATGACGG</td><td>1000</td><td></td><td></td><td></td><td></td><td></td><td></td><td></td><td></td><td></td></tr>
<tr><td>Oligo fwd</td><td>CACCCGGGGAGAGGCGCGTGATGA</td><td></td><td></td><td></td><td></td><td></td><td></td><td></td><td></td><td></td><td></td></tr>
<tr><td>Oligo rev</td><td>AAACTCATCACGCGCCTCTCCCCG</td><td></td><td></td><td></td><td></td><td></td><td></td><td></td><td></td><td></td><td></td></tr>
<tr><td>Chromosome</td><td>start</td><td>end</td><td>strand</td><td>MM</td><td>target_seq</td><td>PAM</td><td>alignment</td><td>distance</td><td>position</td><td>gene name</td><td>gene id</td></tr>
<tr><td>GU305914.1</td><td>71638</td><td>71660</td><td>+</td><td>0</td><td>CGGGGAGAGGCGCGTGATGA</td><td>CGG</td><td>||||||||[||||||||||||]PAM</td><td>NA</td><td>Intergenic</td><td>NA</td><td>NA</td></tr>
<tr><td></td><td></td><td></td><td></td><td></td><td></td><td></td><td></td><td></td><td></td><td></td><td></td></tr>
<tr><td>T5032</td><td>AGAGGCGCGTGATGACGGCACGG</td><td>1000</td><td></td><td></td><td></td><td></td><td></td><td></td><td></td><td></td><td></td></tr>
<tr><td>Oligo fwd</td><td>CACCAGAGGCGCGTGATGACGGCA</td><td></td><td></td><td></td><td></td><td></td><td></td><td></td><td></td><td></td><td></td></tr>
<tr><td>Oligo rev</td><td>AAACTGCCGTCATCACGCGCCTCT</td><td></td><td></td><td></td><td></td><td></td><td></td><td></td><td></td><td></td><td></td></tr>
<tr><td>Chromosome</td><td>start</td><td>end</td><td>strand</td><td>MM</td><td>target_seq</td><td>PAM</td><td>alignment</td><td>distance</td><td>position</td><td>gene name</td><td>gene id</td></tr>
<tr><td>GU305914.1</td><td>71643</td><td>71665</td><td>+</td><td>0</td><td>AGAGGCGCGTGATGACGGCA</td><td>CGG</td><td>||||||||[||||||||||||]PAM</td><td>NA</td><td>Intergenic</td><td>NA</td><td>NA</td></tr>
<tr><td></td><td></td><td></td><td></td><td></td><td></td><td></td><td></td><td></td><td></td><td></td><td></td></tr>
<tr><td>T5033</td><td>GAGGCGCGTGATGACGGCACGGG</td><td>1000</td><td></td><td></td><td></td><td></td><td></td><td></td><td></td><td></td><td></td></tr>
<tr><td>Oligo fwd</td><td>CACCGAGGCGCGTGATGACGGCAC</td><td></td><td></td><td></td><td></td><td></td><td></td><td></td><td></td><td></td><td></td></tr>
<tr><td>Oligo rev</td><td>AAACGTGCCGTCATCACGCGCCTC</td><td></td><td></td><td></td><td></td><td></td><td></td><td></td><td></td><td></td><td></td></tr>
<tr><td>Chromosome</td><td>start</td><td>end</td><td>strand</td><td>MM</td><td>target_seq</td><td>PAM</td><td>alignment</td><td>distance</td><td>position</td><td>gene name</td><td>gene id</td></tr>
<tr><td>GU305914.1</td><td>71644</td><td>71666</td><td>+</td><td>0</td><td>GAGGCGCGTGATGACGGCAC</td><td>GGG</td><td>||||||||[||||||||||||]PAM</td><td>NA</td><td>Intergenic</td><td>NA</td><td>NA</td></tr>
<tr><td></td><td></td><td></td><td></td><td></td><td></td><td></td><td></td><td></td><td></td><td></td><td></td></tr>
<tr><td>T5034</td><td>AGGCGCGTGATGACGGCACGGGG</td><td>1000</td><td></td><td></td><td></td><td></td><td></td><td></td><td></td><td></td><td></td></tr>
<tr><td>Oligo fwd</td><td>CACCAGGCGCGTGATGACGGCACG</td><td></td><td></td><td></td><td></td><td></td><td></td><td></td><td></td><td></td><td></td></tr>
<tr><td>Oligo rev</td><td>AAACCGTGCCGTCATCACGCGCCT</td><td></td><td></td><td></td><td></td><td></td><td></td><td></td><td></td><td></td><td></td></tr>
<tr><td>Chromosome</td><td>start</td><td>end</td><td>strand</td><td>MM</td><td>target_seq</td><td>PAM</td><td>alignment</td><td>distance</td><td>position</td><td>gene name</td><td>gene id</td></tr>
<tr><td>GU305914.1</td><td>71645</td><td>71667</td><td>+</td><td>0</td><td>AGGCGCGTGATGACGGCACG</td><td>GGG</td><td>||||||||[||||||||||||]PAM</td><td>NA</td><td>Intergenic</td><td>NA</td><td>NA</td></tr>
<tr><td></td><td></td><td></td><td></td><td></td><td></td><td></td><td></td><td></td><td></td><td></td><td></td></tr>
<tr><td>T5035</td><td>TTGTACCTGTACGCGACGCACGG</td><td>1000</td><td></td><td></td><td></td><td></td><td></td><td></td><td></td><td></td><td></td></tr>
<tr><td>Oligo fwd</td><td>CACCTTGTACCTGTACGCGACGCA</td><td></td><td></td><td></td><td></td><td></td><td></td><td></td><td></td><td></td><td></td></tr>
<tr><td>Oligo rev</td><td>AAACTGCGTCGCGTACAGGTACAA</td><td></td><td></td><td></td><td></td><td></td><td></td><td></td><td></td><td></td><td></td></tr>
<tr><td>Chromosome</td><td>start</td><td>end</td><td>strand</td><td>MM</td><td>target_seq</td><td>PAM</td><td>alignment</td><td>distance</td><td>position</td><td>gene name</td><td>gene id</td></tr>
<tr><td>GU305914.1</td><td>71675</td><td>71697</td><td>+</td><td>0</td><td>TTGTACCTGTACGCGACGCA</td><td>CGG</td><td>||||||||[||||||||||||]PAM</td><td>NA</td><td>Intergenic</td><td>NA</td><td>NA</td></tr>
<tr><td></td><td></td><td></td><td></td><td></td><td></td><td></td><td></td><td></td><td></td><td></td><td></td></tr>
<tr><td>T5036</td><td>TGTACCTGTACGCGACGCACGGG</td><td>1000</td><td></td><td></td><td></td><td></td><td></td><td></td><td></td><td></td><td></td></tr>
<tr><td>Oligo fwd</td><td>CACCTGTACCTGTACGCGACGCAC</td><td></td><td></td><td></td><td></td><td></td><td></td><td></td><td></td><td></td><td></td></tr>
<tr><td>Oligo rev</td><td>AAACGTGCGTCGCGTACAGGTACA</td><td></td><td></td><td></td><td></td><td></td><td></td><td></td><td></td><td></td><td></td></tr>
<tr><td>Chromosome</td><td>start</td><td>end</td><td>strand</td><td>MM</td><td>target_seq</td><td>PAM</td><td>alignment</td><td>distance</td><td>position</td><td>gene name</td><td>gene id</td></tr>
<tr><td>GU305914.1</td><td>71676</td><td>71698</td><td>+</td><td>0</td><td>TGTACCTGTACGCGACGCAC</td><td>GGG</td><td>||||||||[||||||||||||]PAM</td><td>NA</td><td>Intergenic</td><td>NA</td><td>NA</td></tr>
<tr><td></td><td></td><td></td><td></td><td></td><td></td><td></td><td></td><td></td><td></td><td></td><td></td></tr>
<tr><td>T5037</td><td>GCACGGGAACTTCGTGTTCGAGG</td><td>1000</td><td></td><td></td><td></td><td></td><td></td><td></td><td></td><td></td><td></td></tr>
<tr><td>Oligo fwd</td><td>CACCGCACGGGAACTTCGTGTTCG</td><td></td><td></td><td></td><td></td><td></td><td></td><td></td><td></td><td></td><td></td></tr>
<tr><td>Oligo rev</td><td>AAACCGAACACGAAGTTCCCGTGC</td><td></td><td></td><td></td><td></td><td></td><td></td><td></td><td></td><td></td><td></td></tr>
<tr><td>Chromosome</td><td>start</td><td>end</td><td>strand</td><td>MM</td><td>target_seq</td><td>PAM</td><td>alignment</td><td>distance</td><td>position</td><td>gene name</td><td>gene id</td></tr>
<tr><td>GU305914.1</td><td>71692</td><td>71714</td><td>+</td><td>0</td><td>GCACGGGAACTTCGTGTTCG</td><td>AGG</td><td>||||||||[||||||||||||]PAM</td><td>NA</td><td>Intergenic</td><td>NA</td><td>NA</td></tr>
<tr><td></td><td></td><td></td><td></td><td></td><td></td><td></td><td></td><td></td><td></td><td></td><td></td></tr>
<tr><td>T5038</td><td>GAGGCGTTCTCACACGTCCGCGG</td><td>1000</td><td></td><td></td><td></td><td></td><td></td><td></td><td></td><td></td><td></td></tr>
<tr><td>Oligo fwd</td><td>CACCGAGGCGTTCTCACACGTCCG</td><td></td><td></td><td></td><td></td><td></td><td></td><td></td><td></td><td></td><td></td></tr>
<tr><td>Oligo rev</td><td>AAACCGGACGTGTGAGAACGCCTC</td><td></td><td></td><td></td><td></td><td></td><td></td><td></td><td></td><td></td><td></td></tr>
<tr><td>Chromosome</td><td>start</td><td>end</td><td>strand</td><td>MM</td><td>target_seq</td><td>PAM</td><td>alignment</td><td>distance</td><td>position</td><td>gene name</td><td>gene id</td></tr>
<tr><td>GU305914.1</td><td>71711</td><td>71733</td><td>+</td><td>0</td><td>GAGGCGTTCTCACACGTCCG</td><td>CGG</td><td>||||||||[||||||||||||]PAM</td><td>NA</td><td>Intergenic</td><td>NA</td><td>NA</td></tr>
<tr><td></td><td></td><td></td><td></td><td></td><td></td><td></td><td></td><td></td><td></td><td></td><td></td></tr>
<tr><td>T5039</td><td>AGGCGTTCTCACACGTCCGCGGG</td><td>1000</td><td></td><td></td><td></td><td></td><td></td><td></td><td></td><td></td><td></td></tr>
<tr><td>Oligo fwd</td><td>CACCAGGCGTTCTCACACGTCCGC</td><td></td><td></td><td></td><td></td><td></td><td></td><td></td><td></td><td></td><td></td></tr>
<tr><td>Oligo rev</td><td>AAACGCGGACGTGTGAGAACGCCT</td><td></td><td></td><td></td><td></td><td></td><td></td><td></td><td></td><td></td><td></td></tr>
<tr><td>Chromosome</td><td>start</td><td>end</td><td>strand</td><td>MM</td><td>target_seq</td><td>PAM</td><td>alignment</td><td>distance</td><td>position</td><td>gene name</td><td>gene id</td></tr>
<tr><td>GU305914.1</td><td>71712</td><td>71734</td><td>+</td><td>0</td><td>AGGCGTTCTCACACGTCCGC</td><td>GGG</td><td>||||||||[||||||||||||]PAM</td><td>NA</td><td>Intergenic</td><td>NA</td><td>NA</td></tr>
<tr><td></td><td></td><td></td><td></td><td></td><td></td><td></td><td></td><td></td><td></td><td></td><td></td></tr>
<tr><td>T5040</td><td>CTCACACGTCCGCGGGACGAAGG</td><td>1000</td><td></td><td></td><td></td><td></td><td></td><td></td><td></td><td></td><td></td></tr>
<tr><td>Oligo fwd</td><td>CACCCTCACACGTCCGCGGGACGA</td><td></td><td></td><td></td><td></td><td></td><td></td><td></td><td></td><td></td><td></td></tr>
<tr><td>Oligo rev</td><td>AAACTCGTCCCGCGGACGTGTGAG</td><td></td><td></td><td></td><td></td><td></td><td></td><td></td><td></td><td></td><td></td></tr>
<tr><td>Chromosome</td><td>start</td><td>end</td><td>strand</td><td>MM</td><td>target_seq</td><td>PAM</td><td>alignment</td><td>distance</td><td>position</td><td>gene name</td><td>gene id</td></tr>
<tr><td>GU305914.1</td><td>71719</td><td>71741</td><td>+</td><td>0</td><td>CTCACACGTCCGCGGGACGA</td><td>AGG</td><td>||||||||[||||||||||||]PAM</td><td>NA</td><td>Intergenic</td><td>NA</td><td>NA</td></tr>
<tr><td></td><td></td><td></td><td></td><td></td><td></td><td></td><td></td><td></td><td></td><td></td><td></td></tr>
<tr><td>T5041</td><td>TCACACGTCCGCGGGACGAAGGG</td><td>1000</td><td></td><td></td><td></td><td></td><td></td><td></td><td></td><td></td><td></td></tr>
<tr><td>Oligo fwd</td><td>CACCTCACACGTCCGCGGGACGAA</td><td></td><td></td><td></td><td></td><td></td><td></td><td></td><td></td><td></td><td></td></tr>
<tr><td>Oligo rev</td><td>AAACTTCGTCCCGCGGACGTGTGA</td><td></td><td></td><td></td><td></td><td></td><td></td><td></td><td></td><td></td><td></td></tr>
<tr><td>Chromosome</td><td>start</td><td>end</td><td>strand</td><td>MM</td><td>target_seq</td><td>PAM</td><td>alignment</td><td>distance</td><td>position</td><td>gene name</td><td>gene id</td></tr>
<tr><td>GU305914.1</td><td>71720</td><td>71742</td><td>+</td><td>0</td><td>TCACACGTCCGCGGGACGAA</td><td>GGG</td><td>||||||||[||||||||||||]PAM</td><td>NA</td><td>Intergenic</td><td>NA</td><td>NA</td></tr>
<tr><td></td><td></td><td></td><td></td><td></td><td></td><td></td><td></td><td></td><td></td><td></td><td></td></tr>
<tr><td>T5042</td><td>GGCCTCCTCAGCAGCAGCAGCGG</td><td>1000</td><td></td><td></td><td></td><td></td><td></td><td></td><td></td><td></td><td></td></tr>
</table>

| | | | | | | | | | | | |
|---|---|---|---|---|---|---|---|---|---|---|---|
| Oligo fwd | CACCGGCCTCCTCAGCAGCAGCAG | | | | | | | | | | |
| Oligo rev | AAACCTGCTGCTGCTGAGGAGGCC | | | | | | | | | | |
| Chromosome | start | end | strand | MM | target_seq | PAM | alignment | distance | position | gene name | gene id |
| GU305914.1 | 71741 | 71763 | + | 0 | GGCCTCCTCAGCAGCAGCAG | CGG | \|\|\|\|\|\|\|\|[\|\|\|\|\|\|\|\|\|\|\|\|]PAM | NA | Intergenic | NA | NA |
| GU305914.1 | 27408 | 27430 | + | 4 | GGCAGCAGCAGCAGCAGCAG | CGG | \|\|\|--\|--[\|\|\|\|\|\|\|\|\|\|\|\|]PAM | NA | Intergenic | NA | NA |

| | | | | | | | | | | | |
|---|---|---|---|---|---|---|---|---|---|---|---|
| T5043 | CTCCTCAGCAGCAGCAGCGGCGG | 1000 | | | | | | | | | |
| Oligo fwd | CACCCTCCTCAGCAGCAGCAGCGG | | | | | | | | | | |
| Oligo rev | AAACCCGCTGCTGCTGCTGAGGAG | | | | | | | | | | |
| Chromosome | start | end | strand | MM | target_seq | PAM | alignment | distance | position | gene name | gene id |
| GU305914.1 | 71744 | 71766 | + | 0 | CTCCTCAGCAGCAGCAGCGG | CGG | \|\|\|\|\|\|\|\|[\|\|\|\|\|\|\|\|\|\|\|\|]PAM | NA | Intergenic | NA | NA |
| GU305914.1 | 27411 | 27433 | + | 4 | AGCAGCAGCAGCAGCAGCGG | CGG | --\|--\|\|\|[\|\|\|\|\|\|\|\|\|\|\|\|]PAM | NA | Intergenic | NA | NA |
| GU305914.1 | 71747 | 71769 | + | 3 | CTCAGCAGCAGCAGCGGCGG | CGG | \|\|\|--\|\|\|[\|\|\|\|\|\|\|-\|\|\|\|]PAM | NA | Intergenic | NA | NA |
| GU305914.1 | 56767 | 56789 | + | 4 | CTTCTCCGCCGCAGCGGCGG | CGG | \|\|-\|\|\|-\|[\|-\|\|\|\|\|-\|\|\|\|]PAM | NA | Intergenic | NA | NA |
| GU305914.1 | 56770 | 56792 | + | 4 | CTCCGCCGCAGCGGCGGCGG | CGG | \|\|\|\|-\|-\|[\|\|\|\|-\|\|-\|\|\|\|]PAM | NA | Intergenic | NA | NA |

| | | | | | | | | | | | |
|---|---|---|---|---|---|---|---|---|---|---|---|
| T5044 | CTCAGCAGCAGCAGCGGCGGCGG | 1000 | | | | | | | | | |
| Oligo fwd | CACCCTCAGCAGCAGCAGCGGCGG | | | | | | | | | | |
| Oligo rev | AAACCCGCCGCTGCTGCTGCTGAG | | | | | | | | | | |
| Chromosome | start | end | strand | MM | target_seq | PAM | alignment | distance | position | gene name | gene id |
| GU305914.1 | 71747 | 71769 | + | 0 | CTCAGCAGCAGCAGCGGCGG | CGG | \|\|\|\|\|\|\|\|[\|\|\|\|\|\|\|\|\|\|\|\|]PAM | NA | Intergenic | NA | NA |
| GU305914.1 | 56770 | 56792 | + | 3 | CTCCGCCGCAGCGGCGGCGG | CGG | \|\|\|-\|\|-\|[\|\|\|\|-\|\|\|\|\|\|\|]PAM | NA | Intergenic | NA | NA |
| GU305914.1 | 56773 | 56795 | + | 4 | CGCCGCAGCGGCGGCGGCGG | CGG | \|-\|-\|\|\|\|[\|-\|\|-\|\|\|\|\|\|\|]PAM | NA | Intergenic | NA | NA |
| GU305914.1 | 27411 | 27433 | + | 3 | AGCAGCAGCAGCAGCAGCGG | CGG | --\|\|\|\|\|\|[\|\|\|\|\|\|\|-\|\|\|\|]PAM | NA | Intergenic | NA | NA |
| GU305914.1 | 71744 | 71766 | + | 3 | CTCCTCAGCAGCAGCAGCGG | CGG | \|\|\|--\|\|\|[\|\|\|\|\|\|\|-\|\|\|\|]PAM | NA | Intergenic | NA | NA |
| GU305914.1 | 128721 | 128743 | - | 4 | GGCGGCAGCAGCAGCGGCGC | TGG | --\|-\|\|\|\|[\|\|\|\|\|\|\|\|\|\|\|-]PAM | NA | Intergenic | NA | NA |
| GU305914.1 | 27408 | 27430 | + | 4 | GGCAGCAGCAGCAGCAGCAG | CGG | --\|\|\|\|\|\|[\|\|\|\|\|\|\|-\|\|-\|]PAM | NA | Intergenic | NA | NA |

| | | | | | | | | | | | |
|---|---|---|---|---|---|---|---|---|---|---|---|
| T5045 | AGCAGCAGCGGCGGCGGCAACGG | 1000 | | | | | | | | | |
| Oligo fwd | CACCAGCAGCAGCGGCGGCGGCAA | | | | | | | | | | |
| Oligo rev | AAACTTGCCGCCGCCGCTGCTGCT | | | | | | | | | | |
| Chromosome | start | end | strand | MM | target_seq | PAM | alignment | distance | position | gene name | gene id |
| GU305914.1 | 71753 | 71775 | + | 0 | AGCAGCAGCGGCGGCGGCAA | CGG | \|\|\|\|\|\|\|\|[\|\|\|\|\|\|\|\|\|\|\|\|]PAM | NA | Intergenic | NA | NA |
| GU305914.1 | 163902 | 163924 | + | 3 | AGCGGCGGCGGCGGCGGCAG | CGG | \|\|\|-\|\|-\|[\|\|\|\|\|\|\|\|\|\|\|-]PAM | NA | Intergenic | NA | NA |
| GU305914.1 | 30006 | 30028 | + | 4 | GGCGGCGGCGGCGGCGGCAG | TGG | -\|\|-\|\|-\|[\|\|\|\|\|\|\|\|\|\|\|-]PAM | NA | Intergenic | NA | NA |
| GU305914.1 | 149476 | 149498 | - | 4 | GGCGGCGGCGGCGGCGGCAG | CGG | -\|\|-\|\|-\|[\|\|\|\|\|\|\|\|\|\|\|-]PAM | NA | Intergenic | NA | NA |
| GU305914.1 | 168829 | 168851 | + | 4 | AGCGGCGGCGGCGGCGACAG | CGG | \|\|\|-\|\|-\|[\|\|\|\|\|\|\|\|-\|\|-]PAM | NA | Intergenic | NA | NA |
| GU305914.1 | 56773 | 56795 | + | 4 | CGCCGCAGCGGCGGCGGCGG | CGG | -\|\|-\|\|\|\|[\|\|\|\|\|\|\|\|\|\|--]PAM | NA | Intergenic | NA | NA |

| | | | | | | | | | | | |
|---|---|---|---|---|---|---|---|---|---|---|---|
| T5046 | GCAGCAGCGGCGGCGGCAACGGG | 1000 | | | | | | | | | |
| Oligo fwd | CACCGCAGCAGCGGCGGCGGCAAC | | | | | | | | | | |
| Oligo rev | AAACGTTGCCGCCGCCGCTGCTGC | | | | | | | | | | |
| Chromosome | start | end | strand | MM | target_seq | PAM | alignment | distance | position | gene name | gene id |
| GU305914.1 | 71754 | 71776 | + | 0 | GCAGCAGCGGCGGCGGCAAC | GGG | \|\|\|\|\|\|\|\|[\|\|\|\|\|\|\|\|\|\|\|\|]PAM | NA | Intergenic | NA | NA |
| GU305914.1 | 149475 | 149497 | - | 3 | GCGGCGGCGGCGGCGGCAGC | GGG | \|\|-\|\|-\|\|[\|\|\|\|\|\|\|\|\|\|-\|]PAM | NA | Intergenic | NA | NA |
| GU305914.1 | 168830 | 168852 | + | 4 | GCGGCGGCGGCGGCGACAGC | GGG | \|\|-\|\|-\|\|[\|\|\|\|\|\|\|-\|\|-\|]PAM | NA | Intergenic | NA | NA |
| GU305914.1 | 67743 | 67765 | + | 4 | GGAGCGGCGGCGGCGGCGGC | GGG | \|-\|\|\|-\|\|[\|\|\|\|\|\|\|\|\|--\|]PAM | NA | Intergenic | NA | NA |

| | | | | | | | | | | | |
|---|---|---|---|---|---|---|---|---|---|---|---|
| T5047 | AGCGGCGGCGGCAACGGGAGCGG | 1000 | | | | | | | | | |
| Oligo fwd | CACCAGCGGCGGCGGCAACGGGAG | | | | | | | | | | |
| Oligo rev | AAACCTCCCGTTGCCGCCGCCGCT | | | | | | | | | | |
| Chromosome | start | end | strand | MM | target_seq | PAM | alignment | distance | position | gene name | gene id |
| GU305914.1 | 71759 | 71781 | + | 0 | AGCGGCGGCGGCAACGGGAG | CGG | \|\|\|\|\|\|\|\|[\|\|\|\|\|\|\|\|\|\|\|\|]PAM | NA | Intergenic | NA | NA |
| GU305914.1 | 163908 | 163930 | + | 3 | GGCGGCGGCGGCAGCGGCAG | CGG | -\|\|\|\|\|\|\|[\|\|\|\|\|-\|\|\|-\|\|]PAM | NA | Intergenic | NA | NA |
| GU305914.1 | 62152 | 62174 | + | 4 | AGGGGCGACGGCAGCGGCAG | CGG | \|\|-\|\|\|\|-[\|\|\|\|\|-\|\|\|-\|\|]PAM | NA | Intergenic | NA | NA |

| | | | | | | | | | | | |
|---|---|---|---|---|---|---|---|---|---|---|---|
| T5048 | GAAAGACCAGCAGAATGCGACGG | 1000 | | | | | | | | | |
| Oligo fwd | CACCGAAAGACCAGCAGAATGCGA | | | | | | | | | | |
| Oligo rev | AAACTCGCATTCTGCTGGTCTTTC | | | | | | | | | | |
| Chromosome | start | end | strand | MM | target_seq | PAM | alignment | distance | position | gene name | gene id |
| GU305914.1 | 71797 | 71819 | + | 0 | GAAAGACCAGCAGAATGCGA | CGG | \|\|\|\|\|\|\|\|[\|\|\|\|\|\|\|\|\|\|\|\|]PAM | NA | Intergenic | NA | NA |

| | | | | | | | | | | | |
|---|---|---|---|---|---|---|---|---|---|---|---|
| T5049 | AGCAGAATGCGACGGTGACGCGG | 1000 | | | | | | | | | |
| Oligo fwd | CACCAGCAGAATGCGACGGTGACG | | | | | | | | | | |
| Oligo rev | AAACCGTCACCGTCGCATTCTGCT | | | | | | | | | | |
| Chromosome | start | end | strand | MM | target_seq | PAM | alignment | distance | position | gene name | gene id |
| GU305914.1 | 71805 | 71827 | + | 0 | AGCAGAATGCGACGGTGACG | CGG | \|\|\|\|\|\|\|\|[\|\|\|\|\|\|\|\|\|\|\|\|]PAM | NA | Intergenic | NA | NA |

| | | | | | | | | | | | |
|---|---|---|---|---|---|---|---|---|---|---|---|
| T5050 | CGACGGTGACGCGGCGCTACCGG | 1000 | | | | | | | | | |
| Oligo fwd | CACCCGACGGTGACGCGGCGCTAC | | | | | | | | | | |
| Oligo rev | AAACGTAGCGCCGCGTCACCGTCG | | | | | | | | | | |
| Chromosome | start | end | strand | MM | target_seq | PAM | alignment | distance | position | gene name | gene id |
| GU305914.1 | 71814 | 71836 | + | 0 | CGACGGTGACGCGGCGCTAC | CGG | \|\|\|\|\|\|\|\|[\|\|\|\|\|\|\|\|\|\|\|\|]PAM | NA | Intergenic | NA | NA |

| | | | | | | | | | | | |
|---|---|---|---|---|---|---|---|---|---|---|---|
| T5051 | GGTGACGCGGCGCTACCGGTCGG | 1000 | | | | | | | | | |
| Oligo fwd | CACCGGTGACGCGGCGCTACCGGT | | | | | | | | | | |
| Oligo rev | AAACACCGGTAGCGCCGCGTCACC | | | | | | | | | | |
| Chromosome | start | end | strand | MM | target_seq | PAM | alignment | distance | position | gene name | gene id |
| GU305914.1 | 71818 | 71840 | + | 0 | GGTGACGCGGCGCTACCGGT | CGG | \|\|\|\|\|\|\|\|[\|\|\|\|\|\|\|\|\|\|\|\|]PAM | NA | Intergenic | NA | NA |

| | | | | | | | | | | | |
|---|---|---|---|---|---|---|---|---|---|---|---|
| T5052 | CTACCGGTCGGTGACCGTGTTGG | 1000 | | | | | | | | | |
| Oligo fwd | CACCCTACCGGTCGGTGACCGTGT | | | | | | | | | | |
| Oligo rev | AAACACACGGTCACCGACCGGTAG | | | | | | | | | | |
| Chromosome | start | end | strand | MM | target_seq | PAM | alignment | distance | position | gene name | gene id |
| GU305914.1 | 71830 | 71852 | + | 0 | CTACCGGTCGGTGACCGTGT | TGG | \|\|\|\|\|\|\|\|[\|\|\|\|\|\|\|\|\|\|\|\|]PAM | NA | Intergenic | NA | NA |

| | | |
|---|---|---|
| T5053 | CCGCCACTCTGCAGACGTTCTGG | 1000 |
| Oligo fwd | CACCCCGCCACTCTGCAGACGTTC | |
| Oligo rev | AAACGAACGTCTGCAGAGTGGCGG | |

| | | | | | | | | | | | |
|---|---|---|---|---|---|---|---|---|---|---|---|
| Chromosome | start | end | strand | MM | target_seq | PAM | alignment | distance | position | gene name | gene id |
| GU305914.1 | 71862 | 71884 | + | 0 | CCGCCACTCTGCAGACGTTC | TGG | \|\|\|\|\|\|\|\|[\|\|\|\|\|\|\|\|\|\|\|\|]PAM | NA | Intergenic | NA | NA |
| T5054 | CACTCTGCAGACGTTCTGGTCGG | 1000 | | | | | | | | | |
| Oligo fwd | CACCCACTCTGCAGACGTTCTGGT | | | | | | | | | | |
| Oligo rev | AAACACCAGAACGTCTGCAGAGTG | | | | | | | | | | |
| Chromosome | start | end | strand | MM | target_seq | PAM | alignment | distance | position | gene name | gene id |
| GU305914.1 | 71866 | 71888 | + | 0 | CACTCTGCAGACGTTCTGGT | CGG | \|\|\|\|\|\|\|\|[\|\|\|\|\|\|\|\|\|\|\|\|]PAM | NA | Intergenic | NA | NA |
| T5055 | CGTTCTGGTCGGAGATCTCGAGG | 1000 | | | | | | | | | |
| Oligo fwd | CACCCGTTCTGGTCGGAGATCTCG | | | | | | | | | | |
| Oligo rev | AAACCGAGATCTCCGACCAGAACG | | | | | | | | | | |
| Chromosome | start | end | strand | MM | target_seq | PAM | alignment | distance | position | gene name | gene id |
| GU305914.1 | 71877 | 71899 | + | 0 | CGTTCTGGTCGGAGATCTCG | AGG | \|\|\|\|\|\|\|\|[\|\|\|\|\|\|\|\|\|\|\|\|]PAM | NA | Intergenic | NA | NA |
| T5056 | GTACGACATACGCGAGCTGCTGG | 1000 | | | | | | | | | |
| Oligo fwd | CACCGTACGACATACGCGAGCTGC | | | | | | | | | | |
| Oligo rev | AAACGCAGCTCGCGTATGTCGTAC | | | | | | | | | | |
| Chromosome | start | end | strand | MM | target_seq | PAM | alignment | distance | position | gene name | gene id |
| GU305914.1 | 71899 | 71921 | + | 0 | GTACGACATACGCGAGCTGC | TGG | \|\|\|\|\|\|\|\|[\|\|\|\|\|\|\|\|\|\|\|\|]PAM | NA | Intergenic | NA | NA |
| T5057 | CGACATACGCGAGCTGCTGGTGG | 1000 | | | | | | | | | |
| Oligo fwd | CACCCGACATACGCGAGCTGCTGG | | | | | | | | | | |
| Oligo rev | AAACCCAGCAGCTCGCGTATGTCG | | | | | | | | | | |
| Chromosome | start | end | strand | MM | target_seq | PAM | alignment | distance | position | gene name | gene id |
| GU305914.1 | 71902 | 71924 | + | 0 | CGACATACGCGAGCTGCTGG | TGG | \|\|\|\|\|\|\|\|[\|\|\|\|\|\|\|\|\|\|\|\|]PAM | NA | Intergenic | NA | NA |
| T5058 | CGCGAGCTGCTGGTGGAGAGCGG | 1000 | | | | | | | | | |
| Oligo fwd | CACCCGCGAGCTGCTGGTGGAGAG | | | | | | | | | | |
| Oligo rev | AAACCTCTCCACCAGCAGCTCGCG | | | | | | | | | | |
| Chromosome | start | end | strand | MM | target_seq | PAM | alignment | distance | position | gene name | gene id |
| GU305914.1 | 71909 | 71931 | + | 0 | CGCGAGCTGCTGGTGGAGAG | CGG | \|\|\|\|\|\|\|\|[\|\|\|\|\|\|\|\|\|\|\|\|]PAM | NA | Intergenic | NA | NA |
| T5059 | AGCTGCTGGTGGAGAGCGGCAGG | 1000 | | | | | | | | | |
| Oligo fwd | CACCAGCTGCTGGTGGAGAGCGGC | | | | | | | | | | |
| Oligo rev | AAACGCCGCTCTCCACCAGCAGCT | | | | | | | | | | |
| Chromosome | start | end | strand | MM | target_seq | PAM | alignment | distance | position | gene name | gene id |
| GU305914.1 | 71913 | 71935 | + | 0 | AGCTGCTGGTGGAGAGCGGC | AGG | \|\|\|\|\|\|\|\|[\|\|\|\|\|\|\|\|\|\|\|\|]PAM | NA | Intergenic | NA | NA |
| T5060 | GCTGCTGGTGGAGAGCGGCAGGG | 1000 | | | | | | | | | |
| Oligo fwd | CACCGCTGCTGGTGGAGAGCGGCA | | | | | | | | | | |
| Oligo rev | AAACTGCCGCTCTCCACCAGCAGC | | | | | | | | | | |
| Chromosome | start | end | strand | MM | target_seq | PAM | alignment | distance | position | gene name | gene id |
| GU305914.1 | 71914 | 71936 | + | 0 | GCTGCTGGTGGAGAGCGGCA | GGG | \|\|\|\|\|\|\|\|[\|\|\|\|\|\|\|\|\|\|\|\|]PAM | NA | Intergenic | NA | NA |
| T5061 | GGTGGAGAGCGGCAGGGATCCGG | 1000 | | | | | | | | | |
| Oligo fwd | CACCGGTGGAGAGCGGCAGGGATC | | | | | | | | | | |
| Oligo rev | AAACGATCCCTGCCGCTCTCCACC | | | | | | | | | | |
| Chromosome | start | end | strand | MM | target_seq | PAM | alignment | distance | position | gene name | gene id |
| GU305914.1 | 71920 | 71942 | + | 0 | GGTGGAGAGCGGCAGGGATC | CGG | \|\|\|\|\|\|\|\|[\|\|\|\|\|\|\|\|\|\|\|\|]PAM | NA | Intergenic | NA | NA |
| T5062 | CGGCAGGGATCCGGAAAGCGCGG | 1000 | | | | | | | | | |
| Oligo fwd | CACCCGGCAGGGATCCGGAAAGCG | | | | | | | | | | |
| Oligo rev | AAACCGCTTTCCGGATCCCTGCCG | | | | | | | | | | |
| Chromosome | start | end | strand | MM | target_seq | PAM | alignment | distance | position | gene name | gene id |
| GU305914.1 | 71929 | 71951 | + | 0 | CGGCAGGGATCCGGAAAGCG | CGG | \|\|\|\|\|\|\|\|[\|\|\|\|\|\|\|\|\|\|\|\|]PAM | NA | Intergenic | NA | NA |
| T5063 | CGGCCGACCAGCGCGACAGACGG | 1000 | | | | | | | | | |
| Oligo fwd | CACCCGGCCGACCAGCGCGACAGA | | | | | | | | | | |
| Oligo rev | AAACTCTGTCGCGCTGGTCGGCCG | | | | | | | | | | |
| Chromosome | start | end | strand | MM | target_seq | PAM | alignment | distance | position | gene name | gene id |
| GU305914.1 | 71949 | 71971 | + | 0 | CGGCCGACCAGCGCGACAGA | CGG | \|\|\|\|\|\|\|\|[\|\|\|\|\|\|\|\|\|\|\|\|]PAM | NA | Intergenic | NA | NA |
| T5064 | GCGCGACAGACGGCTGAACACGG | 1000 | | | | | | | | | |
| Oligo fwd | CACCGCGCGACAGACGGCTGAACA | | | | | | | | | | |
| Oligo rev | AAACTGTTCAGCCGTCTGTCGCGC | | | | | | | | | | |
| Chromosome | start | end | strand | MM | target_seq | PAM | alignment | distance | position | gene name | gene id |
| GU305914.1 | 71959 | 71981 | + | 0 | GCGCGACAGACGGCTGAACA | CGG | \|\|\|\|\|\|\|\|[\|\|\|\|\|\|\|\|\|\|\|\|]PAM | NA | Intergenic | NA | NA |
| T5065 | ACTGTTCGTGTATGTGATGACGG | 1000 | | | | | | | | | |
| Oligo fwd | CACCACTGTTCGTGTATGTGATGA | | | | | | | | | | |
| Oligo rev | AAACTCATCACATACACGAACAGT | | | | | | | | | | |
| Chromosome | start | end | strand | MM | target_seq | PAM | alignment | distance | position | gene name | gene id |
| GU305914.1 | 71995 | 72017 | + | 0 | ACTGTTCGTGTATGTGATGA | CGG | \|\|\|\|\|\|\|\|[\|\|\|\|\|\|\|\|\|\|\|\|]PAM | NA | Intergenic | NA | NA |
| T5066 | GTTCGTGTATGTGATGACGGTGG | 1000 | | | | | | | | | |
| Oligo fwd | CACCGTTCGTGTATGTGATGACGG | | | | | | | | | | |
| Oligo rev | AAACCCGTCATCACATACACGAAC | | | | | | | | | | |
| Chromosome | start | end | strand | MM | target_seq | PAM | alignment | distance | position | gene name | gene id |
| GU305914.1 | 71998 | 72020 | + | 0 | GTTCGTGTATGTGATGACGG | TGG | \|\|\|\|\|\|\|\|[\|\|\|\|\|\|\|\|\|\|\|\|]PAM | NA | Intergenic | NA | NA |
| T5067 | TGTATGTGATGACGGTGGCCTGG | 1000 | | | | | | | | | |
| Oligo fwd | CACCTGTATGTGATGACGGTGGCC | | | | | | | | | | |
| Oligo rev | AAACGGCCACCGTCATCACATACA | | | | | | | | | | |
| Chromosome | start | end | strand | MM | target_seq | PAM | alignment | distance | position | gene name | gene id |
| GU305914.1 | 72003 | 72025 | + | 0 | TGTATGTGATGACGGTGGCC | TGG | \|\|\|\|\|\|\|\|[\|\|\|\|\|\|\|\|\|\|\|\|]PAM | NA | Intergenic | NA | NA |
| T5068 | TGTGATGACGGTGGCCTGGTCGG | 1000 | | | | | | | | | |
| Oligo fwd | CACCTGTGATGACGGTGGCCTGGT | | | | | | | | | | |

<table>
<tr><td>Oligo rev</td><td>AAACACCAGGCCACCGTCATCACA</td><td></td><td></td><td></td><td></td><td></td><td></td><td></td><td></td><td></td><td></td><td></td></tr>
<tr><td>Chromosome</td><td>start</td><td></td><td>end</td><td>strand</td><td>MM</td><td>target_seq</td><td>PAM</td><td>alignment</td><td>distance</td><td>position</td><td>gene name</td><td>gene id</td></tr>
<tr><td>GU305914.1</td><td></td><td>72007</td><td>72029</td><td>+</td><td>0</td><td>TGTGATGACGGTGGCCTGGT</td><td>CGG</td><td>||||||||[||||||||||||]PAM</td><td>NA</td><td>Intergenic</td><td>NA</td><td>NA</td></tr>
<tr><td>T5069</td><td>GATGACGGTGGCCTGGTCGGAGG</td><td></td><td>1000</td><td></td><td></td><td></td><td></td><td></td><td></td><td></td><td></td><td></td></tr>
<tr><td>Oligo fwd</td><td>CACCGATGACGGTGGCCTGGTCGG</td><td></td><td></td><td></td><td></td><td></td><td></td><td></td><td></td><td></td><td></td><td></td></tr>
<tr><td>Oligo rev</td><td>AAACCCGACCAGGCCACCGTCATC</td><td></td><td></td><td></td><td></td><td></td><td></td><td></td><td></td><td></td><td></td><td></td></tr>
<tr><td>Chromosome</td><td>start</td><td></td><td>end</td><td>strand</td><td>MM</td><td>target_seq</td><td>PAM</td><td>alignment</td><td>distance</td><td>position</td><td>gene name</td><td>gene id</td></tr>
<tr><td>GU305914.1</td><td></td><td>72010</td><td>72032</td><td>+</td><td>0</td><td>GATGACGGTGGCCTGGTCGG</td><td>AGG</td><td>||||||||[||||||||||||]PAM</td><td>NA</td><td>Intergenic</td><td>NA</td><td>NA</td></tr>
<tr><td>T5070</td><td>CTGGTCGGAGGCGACGCCGCCGG</td><td></td><td>1000</td><td></td><td></td><td></td><td></td><td></td><td></td><td></td><td></td><td></td></tr>
<tr><td>Oligo fwd</td><td>CACCCTGGTCGGAGGCGACGCCGC</td><td></td><td></td><td></td><td></td><td></td><td></td><td></td><td></td><td></td><td></td><td></td></tr>
<tr><td>Oligo rev</td><td>AAACGCGGCGTCGCCTCCGACCAG</td><td></td><td></td><td></td><td></td><td></td><td></td><td></td><td></td><td></td><td></td><td></td></tr>
<tr><td>Chromosome</td><td>start</td><td></td><td>end</td><td>strand</td><td>MM</td><td>target_seq</td><td>PAM</td><td>alignment</td><td>distance</td><td>position</td><td>gene name</td><td>gene id</td></tr>
<tr><td>GU305914.1</td><td></td><td>72022</td><td>72044</td><td>+</td><td>0</td><td>CTGGTCGGAGGCGACGCCGC</td><td>CGG</td><td>||||||||[||||||||||||]PAM</td><td>NA</td><td>Intergenic</td><td>NA</td><td>NA</td></tr>
<tr><td>GU305914.1</td><td></td><td>163541</td><td>163563</td><td>+</td><td>4</td><td>CTCTTCGGTCGCGACGCCGC</td><td>CGG</td><td>||--||||[--||||||||||]PAM</td><td>NA</td><td>Intergenic</td><td>NA</td><td>NA</td></tr>
<tr><td>T5071</td><td>GTCGGAGGCGACGCCGCCGGTGG</td><td></td><td>1000</td><td></td><td></td><td></td><td></td><td></td><td></td><td></td><td></td><td></td></tr>
<tr><td>Oligo fwd</td><td>CACCGTCGGAGGCGACGCCGCCGG</td><td></td><td></td><td></td><td></td><td></td><td></td><td></td><td></td><td></td><td></td><td></td></tr>
<tr><td>Oligo rev</td><td>AAACCCGGCGGCGTCGCCTCCGAC</td><td></td><td></td><td></td><td></td><td></td><td></td><td></td><td></td><td></td><td></td><td></td></tr>
<tr><td>Chromosome</td><td>start</td><td></td><td>end</td><td>strand</td><td>MM</td><td>target_seq</td><td>PAM</td><td>alignment</td><td>distance</td><td>position</td><td>gene name</td><td>gene id</td></tr>
<tr><td>GU305914.1</td><td></td><td>72025</td><td>72047</td><td>+</td><td>0</td><td>GTCGGAGGCGACGCCGCCGG</td><td>TGG</td><td>||||||||[||||||||||||]PAM</td><td>NA</td><td>Intergenic</td><td>NA</td><td>NA</td></tr>
<tr><td>T5072</td><td>GGCGACGCCGCCGGTGGAGCCGG</td><td></td><td>1000</td><td></td><td></td><td></td><td></td><td></td><td></td><td></td><td></td><td></td></tr>
<tr><td>Oligo fwd</td><td>CACCGGCGACGCCGCCGGTGGAGC</td><td></td><td></td><td></td><td></td><td></td><td></td><td></td><td></td><td></td><td></td><td></td></tr>
<tr><td>Oligo rev</td><td>AAACGCTCCACCGGCGGCGTCGCC</td><td></td><td></td><td></td><td></td><td></td><td></td><td></td><td></td><td></td><td></td><td></td></tr>
<tr><td>Chromosome</td><td>start</td><td></td><td>end</td><td>strand</td><td>MM</td><td>target_seq</td><td>PAM</td><td>alignment</td><td>distance</td><td>position</td><td>gene name</td><td>gene id</td></tr>
<tr><td>GU305914.1</td><td></td><td>72031</td><td>72053</td><td>+</td><td>0</td><td>GGCGACGCCGCCGGTGGAGC</td><td>CGG</td><td>||||||||[||||||||||||]PAM</td><td>NA</td><td>Intergenic</td><td>NA</td><td>NA</td></tr>
<tr><td>T5073</td><td>GCGACGCCGCCGGTGGAGCCGGG</td><td></td><td>1000</td><td></td><td></td><td></td><td></td><td></td><td></td><td></td><td></td><td></td></tr>
<tr><td>Oligo fwd</td><td>CACCGCGACGCCGCCGGTGGAGCC</td><td></td><td></td><td></td><td></td><td></td><td></td><td></td><td></td><td></td><td></td><td></td></tr>
<tr><td>Oligo rev</td><td>AAACGGCTCCACCGGCGGCGTCGC</td><td></td><td></td><td></td><td></td><td></td><td></td><td></td><td></td><td></td><td></td><td></td></tr>
<tr><td>Chromosome</td><td>start</td><td></td><td>end</td><td>strand</td><td>MM</td><td>target_seq</td><td>PAM</td><td>alignment</td><td>distance</td><td>position</td><td>gene name</td><td>gene id</td></tr>
<tr><td>GU305914.1</td><td></td><td>72032</td><td>72054</td><td>+</td><td>0</td><td>GCGACGCCGCCGGTGGAGCC</td><td>GGG</td><td>||||||||[||||||||||||]PAM</td><td>NA</td><td>Intergenic</td><td>NA</td><td>NA</td></tr>
<tr><td>T5074</td><td>GCCGGTGGAGCCGGGCAGCCCGG</td><td></td><td>1000</td><td></td><td></td><td></td><td></td><td></td><td></td><td></td><td></td><td></td></tr>
<tr><td>Oligo fwd</td><td>CACCGCCGGTGGAGCCGGGCAGCC</td><td></td><td></td><td></td><td></td><td></td><td></td><td></td><td></td><td></td><td></td><td></td></tr>
<tr><td>Oligo rev</td><td>AAACGGCTGCCCGGCTCCACCGGC</td><td></td><td></td><td></td><td></td><td></td><td></td><td></td><td></td><td></td><td></td><td></td></tr>
<tr><td>Chromosome</td><td>start</td><td></td><td>end</td><td>strand</td><td>MM</td><td>target_seq</td><td>PAM</td><td>alignment</td><td>distance</td><td>position</td><td>gene name</td><td>gene id</td></tr>
<tr><td>GU305914.1</td><td></td><td>72040</td><td>72062</td><td>+</td><td>0</td><td>GCCGGTGGAGCCGGGCAGCC</td><td>CGG</td><td>||||||||[||||||||||||]PAM</td><td>NA</td><td>Intergenic</td><td>NA</td><td>NA</td></tr>
<tr><td>T5075</td><td>GGTGGAGCCGGGCAGCCCGGAGG</td><td></td><td>1000</td><td></td><td></td><td></td><td></td><td></td><td></td><td></td><td></td><td></td></tr>
<tr><td>Oligo fwd</td><td>CACCGGTGGAGCCGGGCAGCCCGG</td><td></td><td></td><td></td><td></td><td></td><td></td><td></td><td></td><td></td><td></td><td></td></tr>
<tr><td>Oligo rev</td><td>AAACCCGGGCTGCCCGGCTCCACC</td><td></td><td></td><td></td><td></td><td></td><td></td><td></td><td></td><td></td><td></td><td></td></tr>
<tr><td>Chromosome</td><td>start</td><td></td><td>end</td><td>strand</td><td>MM</td><td>target_seq</td><td>PAM</td><td>alignment</td><td>distance</td><td>position</td><td>gene name</td><td>gene id</td></tr>
<tr><td>GU305914.1</td><td></td><td>72043</td><td>72065</td><td>+</td><td>0</td><td>GGTGGAGCCGGGCAGCCCGG</td><td>AGG</td><td>||||||||[||||||||||||]PAM</td><td>NA</td><td>Intergenic</td><td>NA</td><td>NA</td></tr>
<tr><td>T5076</td><td>CCCGGAGGCGACGCATGCCCTGG</td><td></td><td>1000</td><td></td><td></td><td></td><td></td><td></td><td></td><td></td><td></td><td></td></tr>
<tr><td>Oligo fwd</td><td>CACCCCCGGAGGCGACGCATGCCC</td><td></td><td></td><td></td><td></td><td></td><td></td><td></td><td></td><td></td><td></td><td></td></tr>
<tr><td>Oligo rev</td><td>AAACGGGCATGCGTCGCCTCCGGG</td><td></td><td></td><td></td><td></td><td></td><td></td><td></td><td></td><td></td><td></td><td></td></tr>
<tr><td>Chromosome</td><td>start</td><td></td><td>end</td><td>strand</td><td>MM</td><td>target_seq</td><td>PAM</td><td>alignment</td><td>distance</td><td>position</td><td>gene name</td><td>gene id</td></tr>
<tr><td>GU305914.1</td><td></td><td>72058</td><td>72080</td><td>+</td><td>0</td><td>CCCGGAGGCGACGCATGCCC</td><td>TGG</td><td>||||||||[||||||||||||]PAM</td><td>NA</td><td>Intergenic</td><td>NA</td><td>NA</td></tr>
<tr><td>T5077</td><td>GGAGGCGACGCATGCCCTGGAGG</td><td></td><td>1000</td><td></td><td></td><td></td><td></td><td></td><td></td><td></td><td></td><td></td></tr>
<tr><td>Oligo fwd</td><td>CACCGGAGGCGACGCATGCCCTGG</td><td></td><td></td><td></td><td></td><td></td><td></td><td></td><td></td><td></td><td></td><td></td></tr>
<tr><td>Oligo rev</td><td>AAACCCAGGGCATGCGTCGCCTCC</td><td></td><td></td><td></td><td></td><td></td><td></td><td></td><td></td><td></td><td></td><td></td></tr>
<tr><td>Chromosome</td><td>start</td><td></td><td>end</td><td>strand</td><td>MM</td><td>target_seq</td><td>PAM</td><td>alignment</td><td>distance</td><td>position</td><td>gene name</td><td>gene id</td></tr>
<tr><td>GU305914.1</td><td></td><td>72061</td><td>72083</td><td>+</td><td>0</td><td>GGAGGCGACGCATGCCCTGG</td><td>AGG</td><td>||||||||[||||||||||||]PAM</td><td>NA</td><td>Intergenic</td><td>NA</td><td>NA</td></tr>
<tr><td>GU305914.1</td><td></td><td>67357</td><td>67379</td><td>+</td><td>4</td><td>GGCGGAGACGCAGGACCTGG</td><td>GGG</td><td>||-||-||[||||-|-|||||]PAM</td><td>NA</td><td>Intergenic</td><td>NA</td><td>NA</td></tr>
<tr><td>T5078</td><td>TGCCCTGGAGGTGAGCCTGCTGG</td><td></td><td>1000</td><td></td><td></td><td></td><td></td><td></td><td></td><td></td><td></td><td></td></tr>
<tr><td>Oligo fwd</td><td>CACCTGCCCTGGAGGTGAGCCTGC</td><td></td><td></td><td></td><td></td><td></td><td></td><td></td><td></td><td></td><td></td><td></td></tr>
<tr><td>Oligo rev</td><td>AAACGCAGGCTCACCTCCAGGGCA</td><td></td><td></td><td></td><td></td><td></td><td></td><td></td><td></td><td></td><td></td><td></td></tr>
<tr><td>Chromosome</td><td>start</td><td></td><td>end</td><td>strand</td><td>MM</td><td>target_seq</td><td>PAM</td><td>alignment</td><td>distance</td><td>position</td><td>gene name</td><td>gene id</td></tr>
<tr><td>GU305914.1</td><td></td><td>72073</td><td>72095</td><td>+</td><td>0</td><td>TGCCCTGGAGGTGAGCCTGC</td><td>TGG</td><td>||||||||[||||||||||||]PAM</td><td>NA</td><td>Intergenic</td><td>NA</td><td>NA</td></tr>
<tr><td>T5079</td><td>GCTGGACTTCTCGACGCTGATGG</td><td></td><td>1000</td><td></td><td></td><td></td><td></td><td></td><td></td><td></td><td></td><td></td></tr>
<tr><td>Oligo fwd</td><td>CACCGCTGGACTTCTCGACGCTGA</td><td></td><td></td><td></td><td></td><td></td><td></td><td></td><td></td><td></td><td></td><td></td></tr>
<tr><td>Oligo rev</td><td>AAACTCAGCGTCGAGAAGTCCAGC</td><td></td><td></td><td></td><td></td><td></td><td></td><td></td><td></td><td></td><td></td><td></td></tr>
<tr><td>Chromosome</td><td>start</td><td></td><td>end</td><td>strand</td><td>MM</td><td>target_seq</td><td>PAM</td><td>alignment</td><td>distance</td><td>position</td><td>gene name</td><td>gene id</td></tr>
<tr><td>GU305914.1</td><td></td><td>72091</td><td>72113</td><td>+</td><td>0</td><td>GCTGGACTTCTCGACGCTGA</td><td>TGG</td><td>||||||||[||||||||||||]PAM</td><td>NA</td><td>Intergenic</td><td>NA</td><td>NA</td></tr>
<tr><td>T5080</td><td>GGACTTCTCGACGCTGATGGCGG</td><td></td><td>1000</td><td></td><td></td><td></td><td></td><td></td><td></td><td></td><td></td><td></td></tr>
<tr><td>Oligo fwd</td><td>CACCGGACTTCTCGACGCTGATGG</td><td></td><td></td><td></td><td></td><td></td><td></td><td></td><td></td><td></td><td></td><td></td></tr>
<tr><td>Oligo rev</td><td>AAACCCATCAGCGTCGAGAAGTCC</td><td></td><td></td><td></td><td></td><td></td><td></td><td></td><td></td><td></td><td></td><td></td></tr>
<tr><td>Chromosome</td><td>start</td><td></td><td>end</td><td>strand</td><td>MM</td><td>target_seq</td><td>PAM</td><td>alignment</td><td>distance</td><td>position</td><td>gene name</td><td>gene id</td></tr>
<tr><td>GU305914.1</td><td></td><td>72094</td><td>72116</td><td>+</td><td>0</td><td>GGACTTCTCGACGCTGATGG</td><td>CGG</td><td>||||||||[||||||||||||]PAM</td><td>NA</td><td>Intergenic</td><td>NA</td><td>NA</td></tr>
<tr><td>T5081</td><td>GCTGATGGCGGCGCTGCACCCGG</td><td></td><td>1000</td><td></td><td></td><td></td><td></td><td></td><td></td><td></td><td></td><td></td></tr>
<tr><td>Oligo fwd</td><td>CACCGCTGATGGCGGCGCTGCACC</td><td></td><td></td><td></td><td></td><td></td><td></td><td></td><td></td><td></td><td></td><td></td></tr>
<tr><td>Oligo rev</td><td>AAACGGTGCAGCGCCGCCATCAGC</td><td></td><td></td><td></td><td></td><td></td><td></td><td></td><td></td><td></td><td></td><td></td></tr>
<tr><td>Chromosome</td><td>start</td><td></td><td>end</td><td>strand</td><td>MM</td><td>target_seq</td><td>PAM</td><td>alignment</td><td>distance</td><td>position</td><td>gene name</td><td>gene id</td></tr>
<tr><td>GU305914.1</td><td></td><td>72106</td><td>72128</td><td>+</td><td>0</td><td>GCTGATGGCGGCGCTGCACC</td><td>CGG</td><td>||||||||[||||||||||||]PAM</td><td>NA</td><td>Intergenic</td><td>NA</td><td>NA</td></tr>
<tr><td>T5082</td><td>GTACGCCATCACGACCCAGCCGG</td><td></td><td>1000</td><td></td><td></td><td></td><td></td><td></td><td></td><td></td><td></td><td></td></tr>
<tr><td>Oligo fwd</td><td>CACCGTACGCCATCACGACCCAGC</td><td></td><td></td><td></td><td></td><td></td><td></td><td></td><td></td><td></td><td></td><td></td></tr>
<tr><td>Oligo rev</td><td>AAACGCTGGGTCGTGATGGCGTAC</td><td></td><td></td><td></td><td></td><td></td><td></td><td></td><td></td><td></td><td></td><td></td></tr>
<tr><td>Chromosome</td><td>start</td><td></td><td>end</td><td>strand</td><td>MM</td><td>target_seq</td><td>PAM</td><td>alignment</td><td>distance</td><td>position</td><td>gene name</td><td>gene id</td></tr>
<tr><td>GU305914.1</td><td></td><td>72136</td><td>72158</td><td>+</td><td>0</td><td>GTACGCCATCACGACCCAGC</td><td>CGG</td><td>||||||||[||||||||||||]PAM</td><td>NA</td><td>Intergenic</td><td>NA</td><td>NA</td></tr>
</table>

<table>
<tr><td>T5083</td><td>CGCCATCACGACCCAGCCGGTGG</td><td>1000</td><td></td><td></td><td></td><td></td><td></td><td></td><td></td><td></td><td></td></tr>
<tr><td>Oligo fwd</td><td>CACCCGCCATCACGACCCAGCCGG</td><td></td><td></td><td></td><td></td><td></td><td></td><td></td><td></td><td></td><td></td></tr>
<tr><td>Oligo rev</td><td>AAACCCGGCTGGGTCGTGATGGCG</td><td></td><td></td><td></td><td></td><td></td><td></td><td></td><td></td><td></td><td></td></tr>
<tr><td>Chromosome</td><td>start</td><td>end</td><td>strand</td><td>MM</td><td>target_seq</td><td>PAM</td><td>alignment</td><td>distance</td><td>position</td><td>gene name</td><td>gene id</td></tr>
<tr><td>GU305914.1</td><td>72139</td><td>72161</td><td>+</td><td>0</td><td>CGCCATCACGACCCAGCCGG</td><td>TGG</td><td>||||||||[||||||||||||]PAM</td><td>NA</td><td>Intergenic</td><td>NA</td><td>NA</td></tr>
<tr><td>T5084</td><td>CCGGTGGACGCCACCCTGCGCGG</td><td>1000</td><td></td><td></td><td></td><td></td><td></td><td></td><td></td><td></td><td></td></tr>
<tr><td>Oligo fwd</td><td>CACCCCGGTGGACGCCACCCTGCG</td><td></td><td></td><td></td><td></td><td></td><td></td><td></td><td></td><td></td><td></td></tr>
<tr><td>Oligo rev</td><td>AAACCGCAGGGTGGCGTCCACCGG</td><td></td><td></td><td></td><td></td><td></td><td></td><td></td><td></td><td></td><td></td></tr>
<tr><td>Chromosome</td><td>start</td><td>end</td><td>strand</td><td>MM</td><td>target_seq</td><td>PAM</td><td>alignment</td><td>distance</td><td>position</td><td>gene name</td><td>gene id</td></tr>
<tr><td>GU305914.1</td><td>72155</td><td>72177</td><td>+</td><td>0</td><td>CCGGTGGACGCCACCCTGCG</td><td>CGG</td><td>||||||||[||||||||||||]PAM</td><td>NA</td><td>Intergenic</td><td>NA</td><td>NA</td></tr>
<tr><td>T5085</td><td>GCGCGGCTTGATCGCCAGACTGG</td><td>1000</td><td></td><td></td><td></td><td></td><td></td><td></td><td></td><td></td><td></td></tr>
<tr><td>Oligo fwd</td><td>CACCGCGCGGCTTGATCGCCAGAC</td><td></td><td></td><td></td><td></td><td></td><td></td><td></td><td></td><td></td><td></td></tr>
<tr><td>Oligo rev</td><td>AAACGTCTGGCGATCAAGCCGCGC</td><td></td><td></td><td></td><td></td><td></td><td></td><td></td><td></td><td></td><td></td></tr>
<tr><td>Chromosome</td><td>start</td><td>end</td><td>strand</td><td>MM</td><td>target_seq</td><td>PAM</td><td>alignment</td><td>distance</td><td>position</td><td>gene name</td><td>gene id</td></tr>
<tr><td>GU305914.1</td><td>72172</td><td>72194</td><td>+</td><td>0</td><td>GCGCGGCTTGATCGCCAGAC</td><td>TGG</td><td>||||||||[||||||||||||]PAM</td><td>NA</td><td>Intergenic</td><td>NA</td><td>NA</td></tr>
<tr><td>T5086</td><td>GCTTGATCGCCAGACTGGACAGG</td><td>1000</td><td></td><td></td><td></td><td></td><td></td><td></td><td></td><td></td><td></td></tr>
<tr><td>Oligo fwd</td><td>CACCGCTTGATCGCCAGACTGGAC</td><td></td><td></td><td></td><td></td><td></td><td></td><td></td><td></td><td></td><td></td></tr>
<tr><td>Oligo rev</td><td>AAACGTCCAGTCTGGCGATCAAGC</td><td></td><td></td><td></td><td></td><td></td><td></td><td></td><td></td><td></td><td></td></tr>
<tr><td>Chromosome</td><td>start</td><td>end</td><td>strand</td><td>MM</td><td>target_seq</td><td>PAM</td><td>alignment</td><td>distance</td><td>position</td><td>gene name</td><td>gene id</td></tr>
<tr><td>GU305914.1</td><td>72177</td><td>72199</td><td>+</td><td>0</td><td>GCTTGATCGCCAGACTGGAC</td><td>AGG</td><td>||||||||[||||||||||||]PAM</td><td>NA</td><td>Intergenic</td><td>NA</td><td>NA</td></tr>
<tr><td>T5087</td><td>TGATCGCCAGACTGGACAGGAGG</td><td>1000</td><td></td><td></td><td></td><td></td><td></td><td></td><td></td><td></td><td></td></tr>
<tr><td>Oligo fwd</td><td>CACCTGATCGCCAGACTGGACAGG</td><td></td><td></td><td></td><td></td><td></td><td></td><td></td><td></td><td></td><td></td></tr>
<tr><td>Oligo rev</td><td>AAACCCTGTCCAGTCTGGCGATCA</td><td></td><td></td><td></td><td></td><td></td><td></td><td></td><td></td><td></td><td></td></tr>
<tr><td>Chromosome</td><td>start</td><td>end</td><td>strand</td><td>MM</td><td>target_seq</td><td>PAM</td><td>alignment</td><td>distance</td><td>position</td><td>gene name</td><td>gene id</td></tr>
<tr><td>GU305914.1</td><td>72180</td><td>72202</td><td>+</td><td>0</td><td>TGATCGCCAGACTGGACAGG</td><td>AGG</td><td>||||||||[||||||||||||]PAM</td><td>NA</td><td>Intergenic</td><td>NA</td><td>NA</td></tr>
<tr><td>T5088</td><td>CGCCAGACTGGACAGGAGGACGG</td><td>1000</td><td></td><td></td><td></td><td></td><td></td><td></td><td></td><td></td><td></td></tr>
<tr><td>Oligo fwd</td><td>CACCCGCCAGACTGGACAGGAGGA</td><td></td><td></td><td></td><td></td><td></td><td></td><td></td><td></td><td></td><td></td></tr>
<tr><td>Oligo rev</td><td>AAACTCCTCCTGTCCAGTCTGGCG</td><td></td><td></td><td></td><td></td><td></td><td></td><td></td><td></td><td></td><td></td></tr>
<tr><td>Chromosome</td><td>start</td><td>end</td><td>strand</td><td>MM</td><td>target_seq</td><td>PAM</td><td>alignment</td><td>distance</td><td>position</td><td>gene name</td><td>gene id</td></tr>
<tr><td>GU305914.1</td><td>72184</td><td>72206</td><td>+</td><td>0</td><td>CGCCAGACTGGACAGGAGGA</td><td>CGG</td><td>||||||||[||||||||||||]PAM</td><td>NA</td><td>Intergenic</td><td>NA</td><td>NA</td></tr>
<tr><td>T5089</td><td>CAGACTGGACAGGAGGACGGTGG</td><td>1000</td><td></td><td></td><td></td><td></td><td></td><td></td><td></td><td></td><td></td></tr>
<tr><td>Oligo fwd</td><td>CACCCAGACTGGACAGGAGGACGG</td><td></td><td></td><td></td><td></td><td></td><td></td><td></td><td></td><td></td><td></td></tr>
<tr><td>Oligo rev</td><td>AAACCCGTCCTCCTGTCCAGTCTG</td><td></td><td></td><td></td><td></td><td></td><td></td><td></td><td></td><td></td><td></td></tr>
<tr><td>Chromosome</td><td>start</td><td>end</td><td>strand</td><td>MM</td><td>target_seq</td><td>PAM</td><td>alignment</td><td>distance</td><td>position</td><td>gene name</td><td>gene id</td></tr>
<tr><td>GU305914.1</td><td>72187</td><td>72209</td><td>+</td><td>0</td><td>CAGACTGGACAGGAGGACGG</td><td>TGG</td><td>||||||||[||||||||||||]PAM</td><td>NA</td><td>Intergenic</td><td>NA</td><td>NA</td></tr>
<tr><td>T5090</td><td>GGACAGGAGGACGGTGGACGCGG</td><td>1000</td><td></td><td></td><td></td><td></td><td></td><td></td><td></td><td></td><td></td></tr>
<tr><td>Oligo fwd</td><td>CACCGGACAGGAGGACGGTGGACG</td><td></td><td></td><td></td><td></td><td></td><td></td><td></td><td></td><td></td><td></td></tr>
<tr><td>Oligo rev</td><td>AAACCGTCCACCGTCCTCCTGTCC</td><td></td><td></td><td></td><td></td><td></td><td></td><td></td><td></td><td></td><td></td></tr>
<tr><td>Chromosome</td><td>start</td><td>end</td><td>strand</td><td>MM</td><td>target_seq</td><td>PAM</td><td>alignment</td><td>distance</td><td>position</td><td>gene name</td><td>gene id</td></tr>
<tr><td>GU305914.1</td><td>72193</td><td>72215</td><td>+</td><td>0</td><td>GGACAGGAGGACGGTGGACG</td><td>CGG</td><td>||||||||[||||||||||||]PAM</td><td>NA</td><td>Intergenic</td><td>NA</td><td>NA</td></tr>
<tr><td>T5091</td><td>GGACGCGGCGATGAACACGCAGG</td><td>1000</td><td></td><td></td><td></td><td></td><td></td><td></td><td></td><td></td><td></td></tr>
<tr><td>Oligo fwd</td><td>CACCGGACGCGGCGATGAACACGC</td><td></td><td></td><td></td><td></td><td></td><td></td><td></td><td></td><td></td><td></td></tr>
<tr><td>Oligo rev</td><td>AAACGCGTGTTCATCGCCGCGTCC</td><td></td><td></td><td></td><td></td><td></td><td></td><td></td><td></td><td></td><td></td></tr>
<tr><td>Chromosome</td><td>start</td><td>end</td><td>strand</td><td>MM</td><td>target_seq</td><td>PAM</td><td>alignment</td><td>distance</td><td>position</td><td>gene name</td><td>gene id</td></tr>
<tr><td>GU305914.1</td><td>72208</td><td>72230</td><td>+</td><td>0</td><td>GGACGCGGCGATGAACACGC</td><td>AGG</td><td>||||||||[||||||||||||]PAM</td><td>NA</td><td>Intergenic</td><td>NA</td><td>NA</td></tr>
<tr><td>T5092</td><td>CGAATTGAAAGCGTTCTGCTTGG</td><td>1000</td><td></td><td></td><td></td><td></td><td></td><td></td><td></td><td></td><td></td></tr>
<tr><td>Oligo fwd</td><td>CACCCGAATTGAAAGCGTTCTGCT</td><td></td><td></td><td></td><td></td><td></td><td></td><td></td><td></td><td></td><td></td></tr>
<tr><td>Oligo rev</td><td>AAACAGCAGAACGCTTTCAATTCG</td><td></td><td></td><td></td><td></td><td></td><td></td><td></td><td></td><td></td><td></td></tr>
<tr><td>Chromosome</td><td>start</td><td>end</td><td>strand</td><td>MM</td><td>target_seq</td><td>PAM</td><td>alignment</td><td>distance</td><td>position</td><td>gene name</td><td>gene id</td></tr>
<tr><td>GU305914.1</td><td>72256</td><td>72278</td><td>+</td><td>0</td><td>CGAATTGAAAGCGTTCTGCT</td><td>TGG</td><td>||||||||[||||||||||||]PAM</td><td>NA</td><td>Intergenic</td><td>NA</td><td>NA</td></tr>
<tr><td>T5093</td><td>TCTGCTTGGATACGAAGCAGTGG</td><td>1000</td><td></td><td></td><td></td><td></td><td></td><td></td><td></td><td></td><td></td></tr>
<tr><td>Oligo fwd</td><td>CACCTCTGCTTGGATACGAAGCAG</td><td></td><td></td><td></td><td></td><td></td><td></td><td></td><td></td><td></td><td></td></tr>
<tr><td>Oligo rev</td><td>AAACCTGCTTCGTATCCAAGCAGA</td><td></td><td></td><td></td><td></td><td></td><td></td><td></td><td></td><td></td><td></td></tr>
<tr><td>Chromosome</td><td>start</td><td>end</td><td>strand</td><td>MM</td><td>target_seq</td><td>PAM</td><td>alignment</td><td>distance</td><td>position</td><td>gene name</td><td>gene id</td></tr>
<tr><td>GU305914.1</td><td>72270</td><td>72292</td><td>+</td><td>0</td><td>TCTGCTTGGATACGAAGCAG</td><td>TGG</td><td>||||||||[||||||||||||]PAM</td><td>NA</td><td>Intergenic</td><td>NA</td><td>NA</td></tr>
<tr><td>T5094</td><td>GGATACGAAGCAGTGGACGCAGG</td><td>1000</td><td></td><td></td><td></td><td></td><td></td><td></td><td></td><td></td><td></td></tr>
<tr><td>Oligo fwd</td><td>CACCGGATACGAAGCAGTGGACGC</td><td></td><td></td><td></td><td></td><td></td><td></td><td></td><td></td><td></td><td></td></tr>
<tr><td>Oligo rev</td><td>AAACGCGTCCACTGCTTCGTATCC</td><td></td><td></td><td></td><td></td><td></td><td></td><td></td><td></td><td></td><td></td></tr>
<tr><td>Chromosome</td><td>start</td><td>end</td><td>strand</td><td>MM</td><td>target_seq</td><td>PAM</td><td>alignment</td><td>distance</td><td>position</td><td>gene name</td><td>gene id</td></tr>
<tr><td>GU305914.1</td><td>72277</td><td>72299</td><td>+</td><td>0</td><td>GGATACGAAGCAGTGGACGC</td><td>AGG</td><td>||||||||[||||||||||||]PAM</td><td>NA</td><td>Intergenic</td><td>NA</td><td>NA</td></tr>
<tr><td>T5095</td><td>TACGAAGCAGTGGACGCAGGAGG</td><td>1000</td><td></td><td></td><td></td><td></td><td></td><td></td><td></td><td></td><td></td></tr>
<tr><td>Oligo fwd</td><td>CACCTACGAAGCAGTGGACGCAGG</td><td></td><td></td><td></td><td></td><td></td><td></td><td></td><td></td><td></td><td></td></tr>
<tr><td>Oligo rev</td><td>AAACCCTGCGTCCACTGCTTCGTA</td><td></td><td></td><td></td><td></td><td></td><td></td><td></td><td></td><td></td><td></td></tr>
<tr><td>Chromosome</td><td>start</td><td>end</td><td>strand</td><td>MM</td><td>target_seq</td><td>PAM</td><td>alignment</td><td>distance</td><td>position</td><td>gene name</td><td>gene id</td></tr>
<tr><td>GU305914.1</td><td>72280</td><td>72302</td><td>+</td><td>0</td><td>TACGAAGCAGTGGACGCAGG</td><td>AGG</td><td>||||||||[||||||||||||]PAM</td><td>NA</td><td>Intergenic</td><td>NA</td><td>NA</td></tr>
<tr><td>T5096</td><td>AGGACATCCGCCCGCAGATGTGG</td><td>1000</td><td></td><td></td><td></td><td></td><td></td><td></td><td></td><td></td><td></td></tr>
<tr><td>Oligo fwd</td><td>CACCAGGACATCCGCCCGCAGATG</td><td></td><td></td><td></td><td></td><td></td><td></td><td></td><td></td><td></td><td></td></tr>
<tr><td>Oligo rev</td><td>AAACCATCTGCGGGCGGATGTCCT</td><td></td><td></td><td></td><td></td><td></td><td></td><td></td><td></td><td></td><td></td></tr>
<tr><td>Chromosome</td><td>start</td><td>end</td><td>strand</td><td>MM</td><td>target_seq</td><td>PAM</td><td>alignment</td><td>distance</td><td>position</td><td>gene name</td><td>gene id</td></tr>
<tr><td>GU305914.1</td><td>72300</td><td>72322</td><td>+</td><td>0</td><td>AGGACATCCGCCCGCAGATG</td><td>TGG</td><td>||||||||[||||||||||||]PAM</td><td>NA</td><td>Intergenic</td><td>NA</td><td>NA</td></tr>
<tr><td>T5097</td><td>CCGCCCGCAGATGTGGATGTCGG</td><td>1000</td><td></td><td></td><td></td><td></td><td></td><td></td><td></td><td></td><td></td></tr>
<tr><td>Oligo fwd</td><td>CACCCCGCCCGCAGATGTGGATGT</td><td></td><td></td><td></td><td></td><td></td><td></td><td></td><td></td><td></td><td></td></tr>
<tr><td>Oligo rev</td><td>AAACACATCCACATCTGCGGGCGG</td><td></td><td></td><td></td><td></td><td></td><td></td><td></td><td></td><td></td><td></td></tr>
<tr><td>Chromosome</td><td>start</td><td>end</td><td>strand</td><td>MM</td><td>target_seq</td><td>PAM</td><td>alignment</td><td>distance</td><td>position</td><td>gene name</td><td>gene id</td></tr>
</table>

<table>
<tr><td>GU305914.1</td><td>72307</td><td>72329</td><td>+</td><td>0</td><td>CCGCCCGCAGATGTGGATGT</td><td>CGG</td><td>||||||||[||||||||||||]PAM</td><td>NA</td><td>Intergenic</td><td>NA</td><td>NA</td></tr>
<tr><td>T5098</td><td>GCAGATGTGGATGTCGGACCTGG</td><td>1000</td><td></td><td></td><td></td><td></td><td></td><td></td><td></td><td></td><td></td></tr>
<tr><td>Oligo fwd</td><td>CACCGCAGATGTGGATGTCGGACC</td><td></td><td></td><td></td><td></td><td></td><td></td><td></td><td></td><td></td><td></td></tr>
<tr><td>Oligo rev</td><td>AAACGGTCCGACATCCACATCTGC</td><td></td><td></td><td></td><td></td><td></td><td></td><td></td><td></td><td></td><td></td></tr>
<tr><td>Chromosome</td><td>start</td><td>end</td><td>strand</td><td>MM</td><td>target_seq</td><td>PAM</td><td>alignment</td><td>distance</td><td>position</td><td>gene name</td><td>gene id</td></tr>
<tr><td>GU305914.1</td><td>72313</td><td>72335</td><td>+</td><td>0</td><td>GCAGATGTGGATGTCGGACC</td><td>TGG</td><td>||||||||[||||||||||||]PAM</td><td>NA</td><td>Intergenic</td><td>NA</td><td>NA</td></tr>
<tr><td>T5099</td><td>TCTATACATGTTAGCCACAAAGG</td><td>1000</td><td></td><td></td><td></td><td></td><td></td><td></td><td></td><td></td><td></td></tr>
<tr><td>Oligo fwd</td><td>CACCTCTATACATGTTAGCCACAA</td><td></td><td></td><td></td><td></td><td></td><td></td><td></td><td></td><td></td><td></td></tr>
<tr><td>Oligo rev</td><td>AAACTTGTGGCTAACATGTATAGA</td><td></td><td></td><td></td><td></td><td></td><td></td><td></td><td></td><td></td><td></td></tr>
<tr><td>Chromosome</td><td>start</td><td>end</td><td>strand</td><td>MM</td><td>target_seq</td><td>PAM</td><td>alignment</td><td>distance</td><td>position</td><td>gene name</td><td>gene id</td></tr>
<tr><td>GU305914.1</td><td>72397</td><td>72419</td><td>+</td><td>0</td><td>TCTATACATGTTAGCCACAA</td><td>AGG</td><td>||||||||[||||||||||||]PAM</td><td>NA</td><td>Intergenic</td><td>NA</td><td>NA</td></tr>
<tr><td>T5100</td><td>TAGCCACAAAGGTGCTGCCCAGG</td><td>1000</td><td></td><td></td><td></td><td></td><td></td><td></td><td></td><td></td><td></td></tr>
<tr><td>Oligo fwd</td><td>CACCTAGCCACAAAGGTGCTGCCC</td><td></td><td></td><td></td><td></td><td></td><td></td><td></td><td></td><td></td><td></td></tr>
<tr><td>Oligo rev</td><td>AAACGGGCAGCACCTTTGTGGCTA</td><td></td><td></td><td></td><td></td><td></td><td></td><td></td><td></td><td></td><td></td></tr>
<tr><td>Chromosome</td><td>start</td><td>end</td><td>strand</td><td>MM</td><td>target_seq</td><td>PAM</td><td>alignment</td><td>distance</td><td>position</td><td>gene name</td><td>gene id</td></tr>
<tr><td>GU305914.1</td><td>72408</td><td>72430</td><td>+</td><td>0</td><td>TAGCCACAAAGGTGCTGCCC</td><td>AGG</td><td>||||||||[||||||||||||]PAM</td><td>NA</td><td>Intergenic</td><td>NA</td><td>NA</td></tr>
<tr><td>T5101</td><td>AGCCACAAAGGTGCTGCCCAGGG</td><td>1000</td><td></td><td></td><td></td><td></td><td></td><td></td><td></td><td></td><td></td></tr>
<tr><td>Oligo fwd</td><td>CACCAGCCACAAAGGTGCTGCCCA</td><td></td><td></td><td></td><td></td><td></td><td></td><td></td><td></td><td></td><td></td></tr>
<tr><td>Oligo rev</td><td>AAACTGGGCAGCACCTTTGTGGCT</td><td></td><td></td><td></td><td></td><td></td><td></td><td></td><td></td><td></td><td></td></tr>
<tr><td>Chromosome</td><td>start</td><td>end</td><td>strand</td><td>MM</td><td>target_seq</td><td>PAM</td><td>alignment</td><td>distance</td><td>position</td><td>gene name</td><td>gene id</td></tr>
<tr><td>GU305914.1</td><td>72409</td><td>72431</td><td>+</td><td>0</td><td>AGCCACAAAGGTGCTGCCCA</td><td>GGG</td><td>||||||||[||||||||||||]PAM</td><td>NA</td><td>Intergenic</td><td>NA</td><td>NA</td></tr>
<tr><td>T5102</td><td>TGCTGCCCAGGGACATCCTTCGG</td><td>1000</td><td></td><td></td><td></td><td></td><td></td><td></td><td></td><td></td><td></td></tr>
<tr><td>Oligo fwd</td><td>CACCTGCTGCCCAGGGACATCCTT</td><td></td><td></td><td></td><td></td><td></td><td></td><td></td><td></td><td></td><td></td></tr>
<tr><td>Oligo rev</td><td>AAACAAGGATGTCCCTGGGCAGCA</td><td></td><td></td><td></td><td></td><td></td><td></td><td></td><td></td><td></td><td></td></tr>
<tr><td>Chromosome</td><td>start</td><td>end</td><td>strand</td><td>MM</td><td>target_seq</td><td>PAM</td><td>alignment</td><td>distance</td><td>position</td><td>gene name</td><td>gene id</td></tr>
<tr><td>GU305914.1</td><td>72420</td><td>72442</td><td>+</td><td>0</td><td>TGCTGCCCAGGGACATCCTT</td><td>CGG</td><td>||||||||[||||||||||||]PAM</td><td>NA</td><td>Intergenic</td><td>NA</td><td>NA</td></tr>
<tr><td>T5103</td><td>GGGACATCCTTCGGTGTCTCTGG</td><td>1000</td><td></td><td></td><td></td><td></td><td></td><td></td><td></td><td></td><td></td></tr>
<tr><td>Oligo fwd</td><td>CACCGGGACATCCTTCGGTGTCTC</td><td></td><td></td><td></td><td></td><td></td><td></td><td></td><td></td><td></td><td></td></tr>
<tr><td>Oligo rev</td><td>AAACGAGACACCGAAGGATGTCCC</td><td></td><td></td><td></td><td></td><td></td><td></td><td></td><td></td><td></td><td></td></tr>
<tr><td>Chromosome</td><td>start</td><td>end</td><td>strand</td><td>MM</td><td>target_seq</td><td>PAM</td><td>alignment</td><td>distance</td><td>position</td><td>gene name</td><td>gene id</td></tr>
<tr><td>GU305914.1</td><td>72429</td><td>72451</td><td>+</td><td>0</td><td>GGGACATCCTTCGGTGTCTC</td><td>TGG</td><td>||||||||[||||||||||||]PAM</td><td>NA</td><td>Intergenic</td><td>NA</td><td>NA</td></tr>
<tr><td>T5104</td><td>GGACATCCTTCGGTGTCTCTGGG</td><td>1000</td><td></td><td></td><td></td><td></td><td></td><td></td><td></td><td></td><td></td></tr>
<tr><td>Oligo fwd</td><td>CACCGGACATCCTTCGGTGTCTCT</td><td></td><td></td><td></td><td></td><td></td><td></td><td></td><td></td><td></td><td></td></tr>
<tr><td>Oligo rev</td><td>AAACAGAGACACCGAAGGATGTCC</td><td></td><td></td><td></td><td></td><td></td><td></td><td></td><td></td><td></td><td></td></tr>
<tr><td>Chromosome</td><td>start</td><td>end</td><td>strand</td><td>MM</td><td>target_seq</td><td>PAM</td><td>alignment</td><td>distance</td><td>position</td><td>gene name</td><td>gene id</td></tr>
<tr><td>GU305914.1</td><td>72430</td><td>72452</td><td>+</td><td>0</td><td>GGACATCCTTCGGTGTCTCT</td><td>GGG</td><td>||||||||[||||||||||||]PAM</td><td>NA</td><td>Intergenic</td><td>NA</td><td>NA</td></tr>
<tr><td>T5105</td><td>GGTGTCTCTGGGTGCAGTTCAGG</td><td>1000</td><td></td><td></td><td></td><td></td><td></td><td></td><td></td><td></td><td></td></tr>
<tr><td>Oligo fwd</td><td>CACCGGTGTCTCTGGGTGCAGTTC</td><td></td><td></td><td></td><td></td><td></td><td></td><td></td><td></td><td></td><td></td></tr>
<tr><td>Oligo rev</td><td>AAACGAACTGCACCCAGAGACACC</td><td></td><td></td><td></td><td></td><td></td><td></td><td></td><td></td><td></td><td></td></tr>
<tr><td>Chromosome</td><td>start</td><td>end</td><td>strand</td><td>MM</td><td>target_seq</td><td>PAM</td><td>alignment</td><td>distance</td><td>position</td><td>gene name</td><td>gene id</td></tr>
<tr><td>GU305914.1</td><td>72441</td><td>72463</td><td>+</td><td>0</td><td>GGTGTCTCTGGGTGCAGTTC</td><td>AGG</td><td>||||||||[||||||||||||]PAM</td><td>NA</td><td>Intergenic</td><td>NA</td><td>NA</td></tr>
<tr><td>T5106</td><td>TCTCTGGGTGCAGTTCAGGCCGG</td><td>1000</td><td></td><td></td><td></td><td></td><td></td><td></td><td></td><td></td><td></td></tr>
<tr><td>Oligo fwd</td><td>CACCTCTCTGGGTGCAGTTCAGGC</td><td></td><td></td><td></td><td></td><td></td><td></td><td></td><td></td><td></td><td></td></tr>
<tr><td>Oligo rev</td><td>AAACGCCTGAACTGCACCCAGAGA</td><td></td><td></td><td></td><td></td><td></td><td></td><td></td><td></td><td></td><td></td></tr>
<tr><td>Chromosome</td><td>start</td><td>end</td><td>strand</td><td>MM</td><td>target_seq</td><td>PAM</td><td>alignment</td><td>distance</td><td>position</td><td>gene name</td><td>gene id</td></tr>
<tr><td>GU305914.1</td><td>72445</td><td>72467</td><td>+</td><td>0</td><td>TCTCTGGGTGCAGTTCAGGC</td><td>CGG</td><td>||||||||[||||||||||||]PAM</td><td>NA</td><td>Intergenic</td><td>NA</td><td>NA</td></tr>
<tr><td>T5107</td><td>GATAGCGTCCTTAGAAGAGCTGG</td><td>1000</td><td></td><td></td><td></td><td></td><td></td><td></td><td></td><td></td><td></td></tr>
<tr><td>Oligo fwd</td><td>CACCGATAGCGTCCTTAGAAGAGC</td><td></td><td></td><td></td><td></td><td></td><td></td><td></td><td></td><td></td><td></td></tr>
<tr><td>Oligo rev</td><td>AAACGCTCTTCTAAGGACGCTATC</td><td></td><td></td><td></td><td></td><td></td><td></td><td></td><td></td><td></td><td></td></tr>
<tr><td>Chromosome</td><td>start</td><td>end</td><td>strand</td><td>MM</td><td>target_seq</td><td>PAM</td><td>alignment</td><td>distance</td><td>position</td><td>gene name</td><td>gene id</td></tr>
<tr><td>GU305914.1</td><td>72481</td><td>72503</td><td>+</td><td>0</td><td>GATAGCGTCCTTAGAAGAGC</td><td>TGG</td><td>||||||||[||||||||||||]PAM</td><td>NA</td><td>Intergenic</td><td>NA</td><td>NA</td></tr>
<tr><td>T5108</td><td>GACTCCTTCTTCAAGATCTACGG</td><td>1000</td><td></td><td></td><td></td><td></td><td></td><td></td><td></td><td></td><td></td></tr>
<tr><td>Oligo fwd</td><td>CACCGACTCCTTCTTCAAGATCTA</td><td></td><td></td><td></td><td></td><td></td><td></td><td></td><td></td><td></td><td></td></tr>
<tr><td>Oligo rev</td><td>AAACTAGATCTTGAAGAAGGAGTC</td><td></td><td></td><td></td><td></td><td></td><td></td><td></td><td></td><td></td><td></td></tr>
<tr><td>Chromosome</td><td>start</td><td>end</td><td>strand</td><td>MM</td><td>target_seq</td><td>PAM</td><td>alignment</td><td>distance</td><td>position</td><td>gene name</td><td>gene id</td></tr>
<tr><td>GU305914.1</td><td>72518</td><td>72540</td><td>+</td><td>0</td><td>GACTCCTTCTTCAAGATCTA</td><td>CGG</td><td>||||||||[||||||||||||]PAM</td><td>NA</td><td>Intergenic</td><td>NA</td><td>NA</td></tr>
<tr><td>GU305914.1</td><td>61565</td><td>61587</td><td>-</td><td>4</td><td>GCCGCGTTCTTCAAGACCTA</td><td>CGG</td><td>|-|-|-||[||||||||-|||]PAM</td><td>NA</td><td>Intergenic</td><td>NA</td><td>NA</td></tr>
<tr><td>T5109</td><td>CTACGGTACGACGAGCGAGACGG</td><td>1000</td><td></td><td></td><td></td><td></td><td></td><td></td><td></td><td></td><td></td></tr>
<tr><td>Oligo fwd</td><td>CACCCTACGGTACGACGAGCGAGA</td><td></td><td></td><td></td><td></td><td></td><td></td><td></td><td></td><td></td><td></td></tr>
<tr><td>Oligo rev</td><td>AAACTCTCGCTCGTCGTACCGTAG</td><td></td><td></td><td></td><td></td><td></td><td></td><td></td><td></td><td></td><td></td></tr>
<tr><td>Chromosome</td><td>start</td><td>end</td><td>strand</td><td>MM</td><td>target_seq</td><td>PAM</td><td>alignment</td><td>distance</td><td>position</td><td>gene name</td><td>gene id</td></tr>
<tr><td>GU305914.1</td><td>72535</td><td>72557</td><td>+</td><td>0</td><td>CTACGGTACGACGAGCGAGA</td><td>CGG</td><td>||||||||[||||||||||||]PAM</td><td>NA</td><td>Intergenic</td><td>NA</td><td>NA</td></tr>
<tr><td>T5110</td><td>TACGACGAGCGAGACGGTGTCGG</td><td>1000</td><td></td><td></td><td></td><td></td><td></td><td></td><td></td><td></td><td></td></tr>
<tr><td>Oligo fwd</td><td>CACCTACGACGAGCGAGACGGTGT</td><td></td><td></td><td></td><td></td><td></td><td></td><td></td><td></td><td></td><td></td></tr>
<tr><td>Oligo rev</td><td>AAACACACCGTCTCGCTCGTCGTA</td><td></td><td></td><td></td><td></td><td></td><td></td><td></td><td></td><td></td><td></td></tr>
<tr><td>Chromosome</td><td>start</td><td>end</td><td>strand</td><td>MM</td><td>target_seq</td><td>PAM</td><td>alignment</td><td>distance</td><td>position</td><td>gene name</td><td>gene id</td></tr>
<tr><td>GU305914.1</td><td>72541</td><td>72563</td><td>+</td><td>0</td><td>TACGACGAGCGAGACGGTGT</td><td>CGG</td><td>||||||||[||||||||||||]PAM</td><td>NA</td><td>Intergenic</td><td>NA</td><td>NA</td></tr>
<tr><td>T5111</td><td>CGAGCGAGACGGTGTCGGCGCGG</td><td>1000</td><td></td><td></td><td></td><td></td><td></td><td></td><td></td><td></td><td></td></tr>
<tr><td>Oligo fwd</td><td>CACCCGAGCGAGACGGTGTCGGCG</td><td></td><td></td><td></td><td></td><td></td><td></td><td></td><td></td><td></td><td></td></tr>
<tr><td>Oligo rev</td><td>AAACCGCCGACACCGTCTCGCTCG</td><td></td><td></td><td></td><td></td><td></td><td></td><td></td><td></td><td></td><td></td></tr>
<tr><td>Chromosome</td><td>start</td><td>end</td><td>strand</td><td>MM</td><td>target_seq</td><td>PAM</td><td>alignment</td><td>distance</td><td>position</td><td>gene name</td><td>gene id</td></tr>
<tr><td>GU305914.1</td><td>72546</td><td>72568</td><td>+</td><td>0</td><td>CGAGCGAGACGGTGTCGGCG</td><td>CGG</td><td>||||||||[||||||||||||]PAM</td><td>NA</td><td>Intergenic</td><td>NA</td><td>NA</td></tr>
<tr><td>T5112</td><td>GTGTCGGCGCGGCTGAAGACCGG</td><td>1000</td><td></td><td></td><td></td><td></td><td></td><td></td><td></td><td></td><td></td></tr>
<tr><td>Oligo fwd</td><td>CACCGTGTCGGCGCGGCTGAAGAC</td><td></td><td></td><td></td><td></td><td></td><td></td><td></td><td></td><td></td><td></td></tr>
</table>

<table>
<tr><td>Oligo rev</td><td>AAACGTCTTCAGCCGCGCCGACAC</td><td></td><td></td><td></td><td></td><td></td><td></td><td></td><td></td><td></td><td></td><td></td></tr>
<tr><td>Chromosome</td><td>start</td><td></td><td>end</td><td>strand</td><td>MM</td><td>target_seq</td><td>PAM</td><td>alignment</td><td>distance</td><td>position</td><td>gene name</td><td>gene id</td></tr>
<tr><td>GU305914.1</td><td></td><td>72557</td><td>72579</td><td>+</td><td>0</td><td>GTGTCGGCGCGGCTGAAGAC</td><td>CGG</td><td>||||||||[||||||||||||]PAM</td><td>NA</td><td>Intergenic</td><td>NA</td><td>NA</td></tr>
<tr><td>T5113</td><td>TGTCGGCGCGGCTGAAGACCGGG</td><td></td><td>1000</td><td></td><td></td><td></td><td></td><td></td><td></td><td></td><td></td><td></td></tr>
<tr><td>Oligo fwd</td><td>CACCTGTCGGCGCGGCTGAAGACC</td><td></td><td></td><td></td><td></td><td></td><td></td><td></td><td></td><td></td><td></td><td></td></tr>
<tr><td>Oligo rev</td><td>AAACGGTCTTCAGCCGCGCCGACA</td><td></td><td></td><td></td><td></td><td></td><td></td><td></td><td></td><td></td><td></td><td></td></tr>
<tr><td>Chromosome</td><td>start</td><td></td><td>end</td><td>strand</td><td>MM</td><td>target_seq</td><td>PAM</td><td>alignment</td><td>distance</td><td>position</td><td>gene name</td><td>gene id</td></tr>
<tr><td>GU305914.1</td><td></td><td>72558</td><td>72580</td><td>+</td><td>0</td><td>TGTCGGCGCGGCTGAAGACC</td><td>GGG</td><td>||||||||[||||||||||||]PAM</td><td>NA</td><td>Intergenic</td><td>NA</td><td>NA</td></tr>
<tr><td>T5114</td><td>GTCGGCGCGGCTGAAGACCGGGG</td><td></td><td>1000</td><td></td><td></td><td></td><td></td><td></td><td></td><td></td><td></td><td></td></tr>
<tr><td>Oligo fwd</td><td>CACCGTCGGCGCGGCTGAAGACCG</td><td></td><td></td><td></td><td></td><td></td><td></td><td></td><td></td><td></td><td></td><td></td></tr>
<tr><td>Oligo rev</td><td>AAACCGGTCTTCAGCCGCGCCGAC</td><td></td><td></td><td></td><td></td><td></td><td></td><td></td><td></td><td></td><td></td><td></td></tr>
<tr><td>Chromosome</td><td>start</td><td></td><td>end</td><td>strand</td><td>MM</td><td>target_seq</td><td>PAM</td><td>alignment</td><td>distance</td><td>position</td><td>gene name</td><td>gene id</td></tr>
<tr><td>GU305914.1</td><td></td><td>72559</td><td>72581</td><td>+</td><td>0</td><td>GTCGGCGCGGCTGAAGACCG</td><td>GGG</td><td>||||||||[||||||||||||]PAM</td><td>NA</td><td>Intergenic</td><td>NA</td><td>NA</td></tr>
<tr><td>T5115</td><td>GCGGCTGAAGACCGGGGAGAAGG</td><td></td><td>1000</td><td></td><td></td><td></td><td></td><td></td><td></td><td></td><td></td><td></td></tr>
<tr><td>Oligo fwd</td><td>CACCGCGGCTGAAGACCGGGGAGA</td><td></td><td></td><td></td><td></td><td></td><td></td><td></td><td></td><td></td><td></td><td></td></tr>
<tr><td>Oligo rev</td><td>AAACTCTCCCCGGTCTTCAGCCGC</td><td></td><td></td><td></td><td></td><td></td><td></td><td></td><td></td><td></td><td></td><td></td></tr>
<tr><td>Chromosome</td><td>start</td><td></td><td>end</td><td>strand</td><td>MM</td><td>target_seq</td><td>PAM</td><td>alignment</td><td>distance</td><td>position</td><td>gene name</td><td>gene id</td></tr>
<tr><td>GU305914.1</td><td></td><td>72565</td><td>72587</td><td>+</td><td>0</td><td>GCGGCTGAAGACCGGGGAGA</td><td>AGG</td><td>||||||||[||||||||||||]PAM</td><td>NA</td><td>Intergenic</td><td>NA</td><td>NA</td></tr>
<tr><td>T5116</td><td>GCTGAAGACCGGGGAGAAGGTGG</td><td></td><td>1000</td><td></td><td></td><td></td><td></td><td></td><td></td><td></td><td></td><td></td></tr>
<tr><td>Oligo fwd</td><td>CACCGCTGAAGACCGGGGAGAAGG</td><td></td><td></td><td></td><td></td><td></td><td></td><td></td><td></td><td></td><td></td><td></td></tr>
<tr><td>Oligo rev</td><td>AAACCCTTCTCCCCGGTCTTCAGC</td><td></td><td></td><td></td><td></td><td></td><td></td><td></td><td></td><td></td><td></td><td></td></tr>
<tr><td>Chromosome</td><td>start</td><td></td><td>end</td><td>strand</td><td>MM</td><td>target_seq</td><td>PAM</td><td>alignment</td><td>distance</td><td>position</td><td>gene name</td><td>gene id</td></tr>
<tr><td>GU305914.1</td><td></td><td>72568</td><td>72590</td><td>+</td><td>0</td><td>GCTGAAGACCGGGGAGAAGG</td><td>TGG</td><td>||||||||[||||||||||||]PAM</td><td>NA</td><td>Intergenic</td><td>NA</td><td>NA</td></tr>
<tr><td>T5117</td><td>AGACCGGGGAGAAGGTGGAACGG</td><td></td><td>1000</td><td></td><td></td><td></td><td></td><td></td><td></td><td></td><td></td><td></td></tr>
<tr><td>Oligo fwd</td><td>CACCAGACCGGGGAGAAGGTGGAA</td><td></td><td></td><td></td><td></td><td></td><td></td><td></td><td></td><td></td><td></td><td></td></tr>
<tr><td>Oligo rev</td><td>AAACTTCCACCTTCTCCCCGGTCT</td><td></td><td></td><td></td><td></td><td></td><td></td><td></td><td></td><td></td><td></td><td></td></tr>
<tr><td>Chromosome</td><td>start</td><td></td><td>end</td><td>strand</td><td>MM</td><td>target_seq</td><td>PAM</td><td>alignment</td><td>distance</td><td>position</td><td>gene name</td><td>gene id</td></tr>
<tr><td>GU305914.1</td><td></td><td>72573</td><td>72595</td><td>+</td><td>0</td><td>AGACCGGGGAGAAGGTGGAA</td><td>CGG</td><td>||||||||[||||||||||||]PAM</td><td>NA</td><td>Intergenic</td><td>NA</td><td>NA</td></tr>
<tr><td>T5118</td><td>CGGGGAGAAGGTGGAACGGCAGG</td><td></td><td>1000</td><td></td><td></td><td></td><td></td><td></td><td></td><td></td><td></td><td></td></tr>
<tr><td>Oligo fwd</td><td>CACCCGGGGAGAAGGTGGAACGGC</td><td></td><td></td><td></td><td></td><td></td><td></td><td></td><td></td><td></td><td></td><td></td></tr>
<tr><td>Oligo rev</td><td>AAACGCCGTTCCACCTTCTCCCCG</td><td></td><td></td><td></td><td></td><td></td><td></td><td></td><td></td><td></td><td></td><td></td></tr>
<tr><td>Chromosome</td><td>start</td><td></td><td>end</td><td>strand</td><td>MM</td><td>target_seq</td><td>PAM</td><td>alignment</td><td>distance</td><td>position</td><td>gene name</td><td>gene id</td></tr>
<tr><td>GU305914.1</td><td></td><td>72577</td><td>72599</td><td>+</td><td>0</td><td>CGGGGAGAAGGTGGAACGGC</td><td>AGG</td><td>||||||||[||||||||||||]PAM</td><td>NA</td><td>Intergenic</td><td>NA</td><td>NA</td></tr>
<tr><td>T5119</td><td>GGAGAAGGTGGAACGGCAGGTGG</td><td></td><td>1000</td><td></td><td></td><td></td><td></td><td></td><td></td><td></td><td></td><td></td></tr>
<tr><td>Oligo fwd</td><td>CACCGGAGAAGGTGGAACGGCAGG</td><td></td><td></td><td></td><td></td><td></td><td></td><td></td><td></td><td></td><td></td><td></td></tr>
<tr><td>Oligo rev</td><td>AAACCCTGCCGTTCCACCTTCTCC</td><td></td><td></td><td></td><td></td><td></td><td></td><td></td><td></td><td></td><td></td><td></td></tr>
<tr><td>Chromosome</td><td>start</td><td></td><td>end</td><td>strand</td><td>MM</td><td>target_seq</td><td>PAM</td><td>alignment</td><td>distance</td><td>position</td><td>gene name</td><td>gene id</td></tr>
<tr><td>GU305914.1</td><td></td><td>72580</td><td>72602</td><td>+</td><td>0</td><td>GGAGAAGGTGGAACGGCAGG</td><td>TGG</td><td>||||||||[||||||||||||]PAM</td><td>NA</td><td>Intergenic</td><td>NA</td><td>NA</td></tr>
<tr><td>GU305914.1</td><td></td><td>150313</td><td>150335</td><td>-</td><td>3</td><td>GGAGAAGCGGGACCGGCAGG</td><td>TGG</td><td>|||||||-[-|||-|||||||]PAM</td><td>NA</td><td>Intergenic</td><td>NA</td><td>NA</td></tr>
<tr><td>T5120</td><td>CAAGCCGACGATGTCGCTGCTGG</td><td></td><td>1000</td><td></td><td></td><td></td><td></td><td></td><td></td><td></td><td></td><td></td></tr>
<tr><td>Oligo fwd</td><td>CACCCAAGCCGACGATGTCGCTGC</td><td></td><td></td><td></td><td></td><td></td><td></td><td></td><td></td><td></td><td></td><td></td></tr>
<tr><td>Oligo rev</td><td>AAACGCAGCGACATCGTCGGCTTG</td><td></td><td></td><td></td><td></td><td></td><td></td><td></td><td></td><td></td><td></td><td></td></tr>
<tr><td>Chromosome</td><td>start</td><td></td><td>end</td><td>strand</td><td>MM</td><td>target_seq</td><td>PAM</td><td>alignment</td><td>distance</td><td>position</td><td>gene name</td><td>gene id</td></tr>
<tr><td>GU305914.1</td><td></td><td>72613</td><td>72635</td><td>+</td><td>0</td><td>CAAGCCGACGATGTCGCTGC</td><td>TGG</td><td>||||||||[||||||||||||]PAM</td><td>NA</td><td>Intergenic</td><td>NA</td><td>NA</td></tr>
<tr><td>T5121</td><td>CGAGTTCTCGCAGCAAGAAGCGG</td><td></td><td>1000</td><td></td><td></td><td></td><td></td><td></td><td></td><td></td><td></td><td></td></tr>
<tr><td>Oligo fwd</td><td>CACCCGAGTTCTCGCAGCAAGAAG</td><td></td><td></td><td></td><td></td><td></td><td></td><td></td><td></td><td></td><td></td><td></td></tr>
<tr><td>Oligo rev</td><td>AAACCTTCTTGCTGCGAGAACTCG</td><td></td><td></td><td></td><td></td><td></td><td></td><td></td><td></td><td></td><td></td><td></td></tr>
<tr><td>Chromosome</td><td>start</td><td></td><td>end</td><td>strand</td><td>MM</td><td>target_seq</td><td>PAM</td><td>alignment</td><td>distance</td><td>position</td><td>gene name</td><td>gene id</td></tr>
<tr><td>GU305914.1</td><td></td><td>72637</td><td>72659</td><td>+</td><td>0</td><td>CGAGTTCTCGCAGCAAGAAG</td><td>CGG</td><td>||||||||[||||||||||||]PAM</td><td>NA</td><td>Intergenic</td><td>NA</td><td>NA</td></tr>
<tr><td>T5122</td><td>CTCGCAGCAAGAAGCGGTGCTGG</td><td></td><td>1000</td><td></td><td></td><td></td><td></td><td></td><td></td><td></td><td></td><td></td></tr>
<tr><td>Oligo fwd</td><td>CACCCTCGCAGCAAGAAGCGGTGC</td><td></td><td></td><td></td><td></td><td></td><td></td><td></td><td></td><td></td><td></td><td></td></tr>
<tr><td>Oligo rev</td><td>AAACGCACCGCTTCTTGCTGCGAG</td><td></td><td></td><td></td><td></td><td></td><td></td><td></td><td></td><td></td><td></td><td></td></tr>
<tr><td>Chromosome</td><td>start</td><td></td><td>end</td><td>strand</td><td>MM</td><td>target_seq</td><td>PAM</td><td>alignment</td><td>distance</td><td>position</td><td>gene name</td><td>gene id</td></tr>
<tr><td>GU305914.1</td><td></td><td>72643</td><td>72665</td><td>+</td><td>0</td><td>CTCGCAGCAAGAAGCGGTGC</td><td>TGG</td><td>||||||||[||||||||||||]PAM</td><td>NA</td><td>Intergenic</td><td>NA</td><td>NA</td></tr>
<tr><td>T5123</td><td>AGCGGTGCTGGACTACATCCTGG</td><td></td><td>1000</td><td></td><td></td><td></td><td></td><td></td><td></td><td></td><td></td><td></td></tr>
<tr><td>Oligo fwd</td><td>CACCAGCGGTGCTGGACTACATCC</td><td></td><td></td><td></td><td></td><td></td><td></td><td></td><td></td><td></td><td></td><td></td></tr>
<tr><td>Oligo rev</td><td>AAACGGATGTAGTCCAGCACCGCT</td><td></td><td></td><td></td><td></td><td></td><td></td><td></td><td></td><td></td><td></td><td></td></tr>
<tr><td>Chromosome</td><td>start</td><td></td><td>end</td><td>strand</td><td>MM</td><td>target_seq</td><td>PAM</td><td>alignment</td><td>distance</td><td>position</td><td>gene name</td><td>gene id</td></tr>
<tr><td>GU305914.1</td><td></td><td>72655</td><td>72677</td><td>+</td><td>0</td><td>AGCGGTGCTGGACTACATCC</td><td>TGG</td><td>||||||||[||||||||||||]PAM</td><td>NA</td><td>Intergenic</td><td>NA</td><td>NA</td></tr>
<tr><td>T5124</td><td>GCGGTGCTGGACTACATCCTGGG</td><td></td><td>1000</td><td></td><td></td><td></td><td></td><td></td><td></td><td></td><td></td><td></td></tr>
<tr><td>Oligo fwd</td><td>CACCGCGGTGCTGGACTACATCCT</td><td></td><td></td><td></td><td></td><td></td><td></td><td></td><td></td><td></td><td></td><td></td></tr>
<tr><td>Oligo rev</td><td>AAACAGGATGTAGTCCAGCACCGC</td><td></td><td></td><td></td><td></td><td></td><td></td><td></td><td></td><td></td><td></td><td></td></tr>
<tr><td>Chromosome</td><td>start</td><td></td><td>end</td><td>strand</td><td>MM</td><td>target_seq</td><td>PAM</td><td>alignment</td><td>distance</td><td>position</td><td>gene name</td><td>gene id</td></tr>
<tr><td>GU305914.1</td><td></td><td>72656</td><td>72678</td><td>+</td><td>0</td><td>GCGGTGCTGGACTACATCCT</td><td>GGG</td><td>||||||||[||||||||||||]PAM</td><td>NA</td><td>Intergenic</td><td>NA</td><td>NA</td></tr>
<tr><td>T5125</td><td>CGGTGCTGGACTACATCCTGGGG</td><td></td><td>1000</td><td></td><td></td><td></td><td></td><td></td><td></td><td></td><td></td><td></td></tr>
<tr><td>Oligo fwd</td><td>CACCCGGTGCTGGACTACATCCTG</td><td></td><td></td><td></td><td></td><td></td><td></td><td></td><td></td><td></td><td></td><td></td></tr>
<tr><td>Oligo rev</td><td>AAACCAGGATGTAGTCCAGCACCG</td><td></td><td></td><td></td><td></td><td></td><td></td><td></td><td></td><td></td><td></td><td></td></tr>
<tr><td>Chromosome</td><td>start</td><td></td><td>end</td><td>strand</td><td>MM</td><td>target_seq</td><td>PAM</td><td>alignment</td><td>distance</td><td>position</td><td>gene name</td><td>gene id</td></tr>
<tr><td>GU305914.1</td><td></td><td>72657</td><td>72679</td><td>+</td><td>0</td><td>CGGTGCTGGACTACATCCTG</td><td>GGG</td><td>||||||||[||||||||||||]PAM</td><td>NA</td><td>Intergenic</td><td>NA</td><td>NA</td></tr>
<tr><td>T5126</td><td>CGTGTTCGCGATCCCGATGACGG</td><td></td><td>1000</td><td></td><td></td><td></td><td></td><td></td><td></td><td></td><td></td><td></td></tr>
<tr><td>Oligo fwd</td><td>CACCCGTGTTCGCGATCCCGATGA</td><td></td><td></td><td></td><td></td><td></td><td></td><td></td><td></td><td></td><td></td><td></td></tr>
<tr><td>Oligo rev</td><td>AAACTCATCGGGATCGCGAACACG</td><td></td><td></td><td></td><td></td><td></td><td></td><td></td><td></td><td></td><td></td><td></td></tr>
<tr><td>Chromosome</td><td>start</td><td></td><td>end</td><td>strand</td><td>MM</td><td>target_seq</td><td>PAM</td><td>alignment</td><td>distance</td><td>position</td><td>gene name</td><td>gene id</td></tr>
<tr><td>GU305914.1</td><td></td><td>72685</td><td>72707</td><td>+</td><td>0</td><td>CGTGTTCGCGATCCCGATGA</td><td>CGG</td><td>||||||||[||||||||||||]PAM</td><td>NA</td><td>Intergenic</td><td>NA</td><td>NA</td></tr>
</table>

<table>
<tr><td>T5127</td><td>GTTCGCGATCCCGATGACGGTGG</td><td></td><td>1000</td><td></td><td></td><td></td><td></td><td></td><td></td><td></td><td></td><td></td></tr>
<tr><td>Oligo fwd</td><td>CACCGTTCGCGATCCCGATGACGG</td><td></td><td></td><td></td><td></td><td></td><td></td><td></td><td></td><td></td><td></td><td></td></tr>
<tr><td>Oligo rev</td><td>AAACCCGTCATCGGGATCGCGAAC</td><td></td><td></td><td></td><td></td><td></td><td></td><td></td><td></td><td></td><td></td><td></td></tr>
<tr><td>Chromosome</td><td>start</td><td></td><td>end</td><td>strand</td><td>MM</td><td>target_seq</td><td>PAM</td><td>alignment</td><td>distance</td><td>position</td><td>gene name</td><td>gene id</td></tr>
<tr><td>GU305914.1</td><td></td><td>72688</td><td>72710</td><td>+</td><td>0</td><td>GTTCGCGATCCCGATGACGG</td><td>TGG</td><td>||||||||[||||||||||||]PAM</td><td>NA</td><td>Intergenic</td><td>NA</td><td>NA</td></tr>
<tr><td>T5128</td><td>TTCGCGATCCCGATGACGGTGGG</td><td></td><td>1000</td><td></td><td></td><td></td><td></td><td></td><td></td><td></td><td></td><td></td></tr>
<tr><td>Oligo fwd</td><td>CACCTTCGCGATCCCGATGACGGT</td><td></td><td></td><td></td><td></td><td></td><td></td><td></td><td></td><td></td><td></td><td></td></tr>
<tr><td>Oligo rev</td><td>AAACACCGTCATCGGGATCGCGAA</td><td></td><td></td><td></td><td></td><td></td><td></td><td></td><td></td><td></td><td></td><td></td></tr>
<tr><td>Chromosome</td><td>start</td><td></td><td>end</td><td>strand</td><td>MM</td><td>target_seq</td><td>PAM</td><td>alignment</td><td>distance</td><td>position</td><td>gene name</td><td>gene id</td></tr>
<tr><td>GU305914.1</td><td></td><td>72689</td><td>72711</td><td>+</td><td>0</td><td>TTCGCGATCCCGATGACGGT</td><td>GGG</td><td>||||||||[||||||||||||]PAM</td><td>NA</td><td>Intergenic</td><td>NA</td><td>NA</td></tr>
<tr><td>T5129</td><td>CATGTCACGAACATCGTAAACGG</td><td></td><td>1000</td><td></td><td></td><td></td><td></td><td></td><td></td><td></td><td></td><td></td></tr>
<tr><td>Oligo fwd</td><td>CACCCATGTCACGAACATCGTAAA</td><td></td><td></td><td></td><td></td><td></td><td></td><td></td><td></td><td></td><td></td><td></td></tr>
<tr><td>Oligo rev</td><td>AAACTTTACGATGTTCGTGACATG</td><td></td><td></td><td></td><td></td><td></td><td></td><td></td><td></td><td></td><td></td><td></td></tr>
<tr><td>Chromosome</td><td>start</td><td></td><td>end</td><td>strand</td><td>MM</td><td>target_seq</td><td>PAM</td><td>alignment</td><td>distance</td><td>position</td><td>gene name</td><td>gene id</td></tr>
<tr><td>GU305914.1</td><td></td><td>72716</td><td>72738</td><td>+</td><td>0</td><td>CATGTCACGAACATCGTAAA</td><td>CGG</td><td>||||||||[||||||||||||]PAM</td><td>NA</td><td>Intergenic</td><td>NA</td><td>NA</td></tr>
<tr><td>T5130</td><td>ACATCGTAAACGGCCGATACCGG</td><td></td><td>1000</td><td></td><td></td><td></td><td></td><td></td><td></td><td></td><td></td><td></td></tr>
<tr><td>Oligo fwd</td><td>CACCACATCGTAAACGGCCGATAC</td><td></td><td></td><td></td><td></td><td></td><td></td><td></td><td></td><td></td><td></td><td></td></tr>
<tr><td>Oligo rev</td><td>AAACGTATCGGCCGTTTACGATGT</td><td></td><td></td><td></td><td></td><td></td><td></td><td></td><td></td><td></td><td></td><td></td></tr>
<tr><td>Chromosome</td><td>start</td><td></td><td>end</td><td>strand</td><td>MM</td><td>target_seq</td><td>PAM</td><td>alignment</td><td>distance</td><td>position</td><td>gene name</td><td>gene id</td></tr>
<tr><td>GU305914.1</td><td></td><td>72726</td><td>72748</td><td>+</td><td>0</td><td>ACATCGTAAACGGCCGATAC</td><td>CGG</td><td>||||||||[||||||||||||]PAM</td><td>NA</td><td>Intergenic</td><td>NA</td><td>NA</td></tr>
<tr><td>T5131</td><td>GCCGATACCGGCTGATCGTGCGG</td><td></td><td>1000</td><td></td><td></td><td></td><td></td><td></td><td></td><td></td><td></td><td></td></tr>
<tr><td>Oligo fwd</td><td>CACCGCCGATACCGGCTGATCGTG</td><td></td><td></td><td></td><td></td><td></td><td></td><td></td><td></td><td></td><td></td><td></td></tr>
<tr><td>Oligo rev</td><td>AAACCACGATCAGCCGGTATCGGC</td><td></td><td></td><td></td><td></td><td></td><td></td><td></td><td></td><td></td><td></td><td></td></tr>
<tr><td>Chromosome</td><td>start</td><td></td><td>end</td><td>strand</td><td>MM</td><td>target_seq</td><td>PAM</td><td>alignment</td><td>distance</td><td>position</td><td>gene name</td><td>gene id</td></tr>
<tr><td>GU305914.1</td><td></td><td>72738</td><td>72760</td><td>+</td><td>0</td><td>GCCGATACCGGCTGATCGTG</td><td>CGG</td><td>||||||||[||||||||||||]PAM</td><td>NA</td><td>Intergenic</td><td>NA</td><td>NA</td></tr>
<tr><td>T5132</td><td>CCGGCTGATCGTGCGGCACCTGG</td><td></td><td>1000</td><td></td><td></td><td></td><td></td><td></td><td></td><td></td><td></td><td></td></tr>
<tr><td>Oligo fwd</td><td>CACCCCGGCTGATCGTGCGGCACC</td><td></td><td></td><td></td><td></td><td></td><td></td><td></td><td></td><td></td><td></td><td></td></tr>
<tr><td>Oligo rev</td><td>AAACGGTGCCGCACGATCAGCCGG</td><td></td><td></td><td></td><td></td><td></td><td></td><td></td><td></td><td></td><td></td><td></td></tr>
<tr><td>Chromosome</td><td>start</td><td></td><td>end</td><td>strand</td><td>MM</td><td>target_seq</td><td>PAM</td><td>alignment</td><td>distance</td><td>position</td><td>gene name</td><td>gene id</td></tr>
<tr><td>GU305914.1</td><td></td><td>72745</td><td>72767</td><td>+</td><td>0</td><td>CCGGCTGATCGTGCGGCACC</td><td>TGG</td><td>||||||||[||||||||||||]PAM</td><td>NA</td><td>Intergenic</td><td>NA</td><td>NA</td></tr>
<tr><td>GU305914.1</td><td></td><td>46649</td><td>46671</td><td>+</td><td>4</td><td>CCGCATGATCGTGCTCCACC</td><td>AGG</td><td>|||--|||[||||||--||||]PAM</td><td>NA</td><td>Intergenic</td><td>NA</td><td>NA</td></tr>
<tr><td>T5133</td><td>GGAGAACCTGCCGAGCGACCCGG</td><td></td><td>1000</td><td></td><td></td><td></td><td></td><td></td><td></td><td></td><td></td><td></td></tr>
<tr><td>Oligo fwd</td><td>CACCGGAGAACCTGCCGAGCGACC</td><td></td><td></td><td></td><td></td><td></td><td></td><td></td><td></td><td></td><td></td><td></td></tr>
<tr><td>Oligo rev</td><td>AAACGGTCGCTCGGCAGGTTCTCC</td><td></td><td></td><td></td><td></td><td></td><td></td><td></td><td></td><td></td><td></td><td></td></tr>
<tr><td>Chromosome</td><td>start</td><td></td><td>end</td><td>strand</td><td>MM</td><td>target_seq</td><td>PAM</td><td>alignment</td><td>distance</td><td>position</td><td>gene name</td><td>gene id</td></tr>
<tr><td>GU305914.1</td><td></td><td>72766</td><td>72788</td><td>+</td><td>0</td><td>GGAGAACCTGCCGAGCGACC</td><td>CGG</td><td>||||||||[||||||||||||]PAM</td><td>NA</td><td>Intergenic</td><td>NA</td><td>NA</td></tr>
<tr><td>T5134</td><td>GAGCGACCCGGACTTCGTCAAGG</td><td></td><td>1000</td><td></td><td></td><td></td><td></td><td></td><td></td><td></td><td></td><td></td></tr>
<tr><td>Oligo fwd</td><td>CACCGAGCGACCCGGACTTCGTCA</td><td></td><td></td><td></td><td></td><td></td><td></td><td></td><td></td><td></td><td></td><td></td></tr>
<tr><td>Oligo rev</td><td>AAACTGACGAAGTCCGGGTCGCTC</td><td></td><td></td><td></td><td></td><td></td><td></td><td></td><td></td><td></td><td></td><td></td></tr>
<tr><td>Chromosome</td><td>start</td><td></td><td>end</td><td>strand</td><td>MM</td><td>target_seq</td><td>PAM</td><td>alignment</td><td>distance</td><td>position</td><td>gene name</td><td>gene id</td></tr>
<tr><td>GU305914.1</td><td></td><td>72778</td><td>72800</td><td>+</td><td>0</td><td>GAGCGACCCGGACTTCGTCA</td><td>AGG</td><td>||||||||[||||||||||||]PAM</td><td>NA</td><td>Intergenic</td><td>NA</td><td>NA</td></tr>
<tr><td>GU305914.1</td><td></td><td>195765</td><td>195787</td><td>-</td><td>3</td><td>GAGCGACTCAGACTTCGACA</td><td>TGG</td><td>|||||||-[|-|||||||-||]PAM</td><td>NA</td><td>Intergenic</td><td>NA</td><td>NA</td></tr>
<tr><td>T5135</td><td>CGACCCGGACTTCGTCAAGGTGG</td><td></td><td>1000</td><td></td><td></td><td></td><td></td><td></td><td></td><td></td><td></td><td></td></tr>
<tr><td>Oligo fwd</td><td>CACCCGACCCGGACTTCGTCAAGG</td><td></td><td></td><td></td><td></td><td></td><td></td><td></td><td></td><td></td><td></td><td></td></tr>
<tr><td>Oligo rev</td><td>AAACCCTTGACGAAGTCCGGGTCG</td><td></td><td></td><td></td><td></td><td></td><td></td><td></td><td></td><td></td><td></td><td></td></tr>
<tr><td>Chromosome</td><td>start</td><td></td><td>end</td><td>strand</td><td>MM</td><td>target_seq</td><td>PAM</td><td>alignment</td><td>distance</td><td>position</td><td>gene name</td><td>gene id</td></tr>
<tr><td>GU305914.1</td><td></td><td>72781</td><td>72803</td><td>+</td><td>0</td><td>CGACCCGGACTTCGTCAAGG</td><td>TGG</td><td>||||||||[||||||||||||]PAM</td><td>NA</td><td>Intergenic</td><td>NA</td><td>NA</td></tr>
<tr><td>T5136</td><td>GACCTGAGCTTCGACCGCTTCGG</td><td></td><td>1000</td><td></td><td></td><td></td><td></td><td></td><td></td><td></td><td></td><td></td></tr>
<tr><td>Oligo fwd</td><td>CACCGACCTGAGCTTCGACCGCTT</td><td></td><td></td><td></td><td></td><td></td><td></td><td></td><td></td><td></td><td></td><td></td></tr>
<tr><td>Oligo rev</td><td>AAACAAGCGGTCGAAGCTCAGGTC</td><td></td><td></td><td></td><td></td><td></td><td></td><td></td><td></td><td></td><td></td><td></td></tr>
<tr><td>Chromosome</td><td>start</td><td></td><td>end</td><td>strand</td><td>MM</td><td>target_seq</td><td>PAM</td><td>alignment</td><td>distance</td><td>position</td><td>gene name</td><td>gene id</td></tr>
<tr><td>GU305914.1</td><td></td><td>72815</td><td>72837</td><td>+</td><td>0</td><td>GACCTGAGCTTCGACCGCTT</td><td>CGG</td><td>||||||||[||||||||||||]PAM</td><td>NA</td><td>Intergenic</td><td>NA</td><td>NA</td></tr>
<tr><td>T5137</td><td>TGAGCTTCGACCGCTTCGGATGG</td><td></td><td>1000</td><td></td><td></td><td></td><td></td><td></td><td></td><td></td><td></td><td></td></tr>
<tr><td>Oligo fwd</td><td>CACCTGAGCTTCGACCGCTTCGGA</td><td></td><td></td><td></td><td></td><td></td><td></td><td></td><td></td><td></td><td></td><td></td></tr>
<tr><td>Oligo rev</td><td>AAACTCCGAAGCGGTCGAAGCTCA</td><td></td><td></td><td></td><td></td><td></td><td></td><td></td><td></td><td></td><td></td><td></td></tr>
<tr><td>Chromosome</td><td>start</td><td></td><td>end</td><td>strand</td><td>MM</td><td>target_seq</td><td>PAM</td><td>alignment</td><td>distance</td><td>position</td><td>gene name</td><td>gene id</td></tr>
<tr><td>GU305914.1</td><td></td><td>72819</td><td>72841</td><td>+</td><td>0</td><td>TGAGCTTCGACCGCTTCGGA</td><td>TGG</td><td>||||||||[||||||||||||]PAM</td><td>NA</td><td>Intergenic</td><td>NA</td><td>NA</td></tr>
<tr><td>T5138</td><td>AAAACCCCGTCGAACGCTCCTGG</td><td></td><td>1000</td><td></td><td></td><td></td><td></td><td></td><td></td><td></td><td></td><td></td></tr>
<tr><td>Oligo fwd</td><td>CACCAAAACCCCGTCGAACGCTCC</td><td></td><td></td><td></td><td></td><td></td><td></td><td></td><td></td><td></td><td></td><td></td></tr>
<tr><td>Oligo rev</td><td>AAACGGAGCGTTCGACGGGGTTTT</td><td></td><td></td><td></td><td></td><td></td><td></td><td></td><td></td><td></td><td></td><td></td></tr>
<tr><td>Chromosome</td><td>start</td><td></td><td>end</td><td>strand</td><td>MM</td><td>target_seq</td><td>PAM</td><td>alignment</td><td>distance</td><td>position</td><td>gene name</td><td>gene id</td></tr>
<tr><td>GU305914.1</td><td></td><td>72855</td><td>72877</td><td>+</td><td>0</td><td>AAAACCCCGTCGAACGCTCC</td><td>TGG</td><td>||||||||[||||||||||||]PAM</td><td>NA</td><td>Intergenic</td><td>NA</td><td>NA</td></tr>
<tr><td>T5139</td><td>CTCCTGGTTCTCGCTGCAAGAGG</td><td></td><td>1000</td><td></td><td></td><td></td><td></td><td></td><td></td><td></td><td></td><td></td></tr>
<tr><td>Oligo fwd</td><td>CACCCTCCTGGTTCTCGCTGCAAG</td><td></td><td></td><td></td><td></td><td></td><td></td><td></td><td></td><td></td><td></td><td></td></tr>
<tr><td>Oligo rev</td><td>AAACCTTGCAGCGAGAACCAGGAG</td><td></td><td></td><td></td><td></td><td></td><td></td><td></td><td></td><td></td><td></td><td></td></tr>
<tr><td>Chromosome</td><td>start</td><td></td><td>end</td><td>strand</td><td>MM</td><td>target_seq</td><td>PAM</td><td>alignment</td><td>distance</td><td>position</td><td>gene name</td><td>gene id</td></tr>
<tr><td>GU305914.1</td><td></td><td>72871</td><td>72893</td><td>+</td><td>0</td><td>CTCCTGGTTCTCGCTGCAAG</td><td>AGG</td><td>||||||||[||||||||||||]PAM</td><td>NA</td><td>Intergenic</td><td>NA</td><td>NA</td></tr>
<tr><td>T5140</td><td>GGTTCTCGCTGCAAGAGGATCGG</td><td></td><td>1000</td><td></td><td></td><td></td><td></td><td></td><td></td><td></td><td></td><td></td></tr>
<tr><td>Oligo fwd</td><td>CACCGGTTCTCGCTGCAAGAGGAT</td><td></td><td></td><td></td><td></td><td></td><td></td><td></td><td></td><td></td><td></td><td></td></tr>
<tr><td>Oligo rev</td><td>AAACATCCTCTTGCAGCGAGAACC</td><td></td><td></td><td></td><td></td><td></td><td></td><td></td><td></td><td></td><td></td><td></td></tr>
<tr><td>Chromosome</td><td>start</td><td></td><td>end</td><td>strand</td><td>MM</td><td>target_seq</td><td>PAM</td><td>alignment</td><td>distance</td><td>position</td><td>gene name</td><td>gene id</td></tr>
<tr><td>GU305914.1</td><td></td><td>72876</td><td>72898</td><td>+</td><td>0</td><td>GGTTCTCGCTGCAAGAGGAT</td><td>CGG</td><td>||||||||[||||||||||||]PAM</td><td>NA</td><td>Intergenic</td><td>NA</td><td>NA</td></tr>
<tr><td>T5141</td><td>GCTGACTAACCCCCCACAGCAGG</td><td></td><td>1000</td><td></td><td></td><td></td><td></td><td></td><td></td><td></td><td></td><td></td></tr>
<tr><td>Oligo fwd</td><td>CACCGCTGACTAACCCCCCACAGC</td><td></td><td></td><td></td><td></td><td></td><td></td><td></td><td></td><td></td><td></td><td></td></tr>
<tr><td>Oligo rev</td><td>AAACGCTGTGGGGGGTTAGTCAGC</td><td></td><td></td><td></td><td></td><td></td><td></td><td></td><td></td><td></td><td></td><td></td></tr>
</table>

<table>
<tr><td>Chromosome</td><td>start</td><td>end</td><td>strand</td><td>MM</td><td>target_seq</td><td>PAM</td><td>alignment</td><td>distance</td><td>position</td><td>gene name</td><td>gene id</td></tr>
<tr><td>GU305914.1</td><td>72910</td><td>72932</td><td>+</td><td>0</td><td>GCTGACTAACCCCCCACAGC</td><td>AGG</td><td>||||||||[||||||||||||]PAM</td><td>NA</td><td>Intergenic</td><td>NA</td><td>NA</td></tr>
<tr><td></td><td></td><td></td><td></td><td></td><td></td><td></td><td></td><td></td><td></td><td></td><td></td></tr>
<tr><td>T5142</td><td>ACAGCAGGACAGAACGCCGTTGG</td><td>1000</td><td></td><td></td><td></td><td></td><td></td><td></td><td></td><td></td><td></td></tr>
<tr><td>Oligo fwd</td><td>CACCACAGCAGGACAGAACGCCGT</td><td></td><td></td><td></td><td></td><td></td><td></td><td></td><td></td><td></td><td></td></tr>
<tr><td>Oligo rev</td><td>AAACACGGCGTTCTGTCCTGCTGT</td><td></td><td></td><td></td><td></td><td></td><td></td><td></td><td></td><td></td><td></td></tr>
<tr><td>Chromosome</td><td>start</td><td>end</td><td>strand</td><td>MM</td><td>target_seq</td><td>PAM</td><td>alignment</td><td>distance</td><td>position</td><td>gene name</td><td>gene id</td></tr>
<tr><td>GU305914.1</td><td>72925</td><td>72947</td><td>+</td><td>0</td><td>ACAGCAGGACAGAACGCCGT</td><td>TGG</td><td>||||||||[||||||||||||]PAM</td><td>NA</td><td>Intergenic</td><td>NA</td><td>NA</td></tr>
<tr><td></td><td></td><td></td><td></td><td></td><td></td><td></td><td></td><td></td><td></td><td></td><td></td></tr>
<tr><td>T5143</td><td>GCAGGACAGAACGCCGTTGGTGG</td><td>1000</td><td></td><td></td><td></td><td></td><td></td><td></td><td></td><td></td><td></td></tr>
<tr><td>Oligo fwd</td><td>CACCGCAGGACAGAACGCCGTTGG</td><td></td><td></td><td></td><td></td><td></td><td></td><td></td><td></td><td></td><td></td></tr>
<tr><td>Oligo rev</td><td>AAACCCAACGGCGTTCTGTCCTGC</td><td></td><td></td><td></td><td></td><td></td><td></td><td></td><td></td><td></td><td></td></tr>
<tr><td>Chromosome</td><td>start</td><td>end</td><td>strand</td><td>MM</td><td>target_seq</td><td>PAM</td><td>alignment</td><td>distance</td><td>position</td><td>gene name</td><td>gene id</td></tr>
<tr><td>GU305914.1</td><td>72928</td><td>72950</td><td>+</td><td>0</td><td>GCAGGACAGAACGCCGTTGG</td><td>TGG</td><td>||||||||[||||||||||||]PAM</td><td>NA</td><td>Intergenic</td><td>NA</td><td>NA</td></tr>
<tr><td></td><td></td><td></td><td></td><td></td><td></td><td></td><td></td><td></td><td></td><td></td><td></td></tr>
<tr><td>T5144</td><td>CGACTCCAACACCAACTACGCGG</td><td>1000</td><td></td><td></td><td></td><td></td><td></td><td></td><td></td><td></td><td></td></tr>
<tr><td>Oligo fwd</td><td>CACCCGACTCCAACACCAACTACG</td><td></td><td></td><td></td><td></td><td></td><td></td><td></td><td></td><td></td><td></td></tr>
<tr><td>Oligo rev</td><td>AAACCGTAGTTGGTGTTGGAGTCG</td><td></td><td></td><td></td><td></td><td></td><td></td><td></td><td></td><td></td><td></td></tr>
<tr><td>Chromosome</td><td>start</td><td>end</td><td>strand</td><td>MM</td><td>target_seq</td><td>PAM</td><td>alignment</td><td>distance</td><td>position</td><td>gene name</td><td>gene id</td></tr>
<tr><td>GU305914.1</td><td>72955</td><td>72977</td><td>+</td><td>0</td><td>CGACTCCAACACCAACTACG</td><td>CGG</td><td>||||||||[||||||||||||]PAM</td><td>NA</td><td>Intergenic</td><td>NA</td><td>NA</td></tr>
<tr><td></td><td></td><td></td><td></td><td></td><td></td><td></td><td></td><td></td><td></td><td></td><td></td></tr>
<tr><td>T5145</td><td>CGCGATGATGCCCCCGCTGAAGG</td><td>1000</td><td></td><td></td><td></td><td></td><td></td><td></td><td></td><td></td><td></td></tr>
<tr><td>Oligo fwd</td><td>CACCCGCGATGATGCCCCCGCTGA</td><td></td><td></td><td></td><td></td><td></td><td></td><td></td><td></td><td></td><td></td></tr>
<tr><td>Oligo rev</td><td>AAACTCAGCGGGGGCATCATCGCG</td><td></td><td></td><td></td><td></td><td></td><td></td><td></td><td></td><td></td><td></td></tr>
<tr><td>Chromosome</td><td>start</td><td>end</td><td>strand</td><td>MM</td><td>target_seq</td><td>PAM</td><td>alignment</td><td>distance</td><td>position</td><td>gene name</td><td>gene id</td></tr>
<tr><td>GU305914.1</td><td>72982</td><td>73004</td><td>+</td><td>0</td><td>CGCGATGATGCCCCCGCTGA</td><td>AGG</td><td>||||||||[||||||||||||]PAM</td><td>NA</td><td>Intergenic</td><td>NA</td><td>NA</td></tr>
<tr><td></td><td></td><td></td><td></td><td></td><td></td><td></td><td></td><td></td><td></td><td></td><td></td></tr>
<tr><td>T5146</td><td>GCCCCCGCTGAAGGCGCCCCCGG</td><td>1000</td><td></td><td></td><td></td><td></td><td></td><td></td><td></td><td></td><td></td></tr>
<tr><td>Oligo fwd</td><td>CACCGCCCCCGCTGAAGGCGCCCC</td><td></td><td></td><td></td><td></td><td></td><td></td><td></td><td></td><td></td><td></td></tr>
<tr><td>Oligo rev</td><td>AAACGGGGCGCCTTCAGCGGGGGC</td><td></td><td></td><td></td><td></td><td></td><td></td><td></td><td></td><td></td><td></td></tr>
<tr><td>Chromosome</td><td>start</td><td>end</td><td>strand</td><td>MM</td><td>target_seq</td><td>PAM</td><td>alignment</td><td>distance</td><td>position</td><td>gene name</td><td>gene id</td></tr>
<tr><td>GU305914.1</td><td>72991</td><td>73013</td><td>+</td><td>0</td><td>GCCCCCGCTGAAGGCGCCCC</td><td>CGG</td><td>||||||||[||||||||||||]PAM</td><td>NA</td><td>Intergenic</td><td>NA</td><td>NA</td></tr>
<tr><td></td><td></td><td></td><td></td><td></td><td></td><td></td><td></td><td></td><td></td><td></td><td></td></tr>
<tr><td>T5147</td><td>AGGCGCCCCCGGCGACGAGTCGG</td><td>1000</td><td></td><td></td><td></td><td></td><td></td><td></td><td></td><td></td><td></td></tr>
<tr><td>Oligo fwd</td><td>CACCAGGCGCCCCCGGCGACGAGT</td><td></td><td></td><td></td><td></td><td></td><td></td><td></td><td></td><td></td><td></td></tr>
<tr><td>Oligo rev</td><td>AAACACTCGTCGCCGGGGGCGCCT</td><td></td><td></td><td></td><td></td><td></td><td></td><td></td><td></td><td></td><td></td></tr>
<tr><td>Chromosome</td><td>start</td><td>end</td><td>strand</td><td>MM</td><td>target_seq</td><td>PAM</td><td>alignment</td><td>distance</td><td>position</td><td>gene name</td><td>gene id</td></tr>
<tr><td>GU305914.1</td><td>73002</td><td>73024</td><td>+</td><td>0</td><td>AGGCGCCCCCGGCGACGAGT</td><td>CGG</td><td>||||||||[||||||||||||]PAM</td><td>NA</td><td>Intergenic</td><td>NA</td><td>NA</td></tr>
<tr><td></td><td></td><td></td><td></td><td></td><td></td><td></td><td></td><td></td><td></td><td></td><td></td></tr>
<tr><td>T5148</td><td>GGCGCCCCCGGCGACGAGTCGGG</td><td>1000</td><td></td><td></td><td></td><td></td><td></td><td></td><td></td><td></td><td></td></tr>
<tr><td>Oligo fwd</td><td>CACCGGCGCCCCCGGCGACGAGTC</td><td></td><td></td><td></td><td></td><td></td><td></td><td></td><td></td><td></td><td></td></tr>
<tr><td>Oligo rev</td><td>AAACGACTCGTCGCCGGGGGCGCC</td><td></td><td></td><td></td><td></td><td></td><td></td><td></td><td></td><td></td><td></td></tr>
<tr><td>Chromosome</td><td>start</td><td>end</td><td>strand</td><td>MM</td><td>target_seq</td><td>PAM</td><td>alignment</td><td>distance</td><td>position</td><td>gene name</td><td>gene id</td></tr>
<tr><td>GU305914.1</td><td>73003</td><td>73025</td><td>+</td><td>0</td><td>GGCGCCCCCGGCGACGAGTC</td><td>GGG</td><td>||||||||[||||||||||||]PAM</td><td>NA</td><td>Intergenic</td><td>NA</td><td>NA</td></tr>
<tr><td>GU305914.1</td><td>203821</td><td>203843</td><td>+</td><td>4</td><td>GGCTTCCCCGTCGACGTGTC</td><td>CGG</td><td>|||--|||[||-|||||-|||]PAM</td><td>NA</td><td>Intergenic</td><td>NA</td><td>NA</td></tr>
<tr><td></td><td></td><td></td><td></td><td></td><td></td><td></td><td></td><td></td><td></td><td></td><td></td></tr>
<tr><td>T5149</td><td>GACGAGTCGGGTCCACCTAACGG</td><td>1000</td><td></td><td></td><td></td><td></td><td></td><td></td><td></td><td></td><td></td></tr>
<tr><td>Oligo fwd</td><td>CACCGACGAGTCGGGTCCACCTAA</td><td></td><td></td><td></td><td></td><td></td><td></td><td></td><td></td><td></td><td></td></tr>
<tr><td>Oligo rev</td><td>AAACTTAGGTGGACCCGACTCGTC</td><td></td><td></td><td></td><td></td><td></td><td></td><td></td><td></td><td></td><td></td></tr>
<tr><td>Chromosome</td><td>start</td><td>end</td><td>strand</td><td>MM</td><td>target_seq</td><td>PAM</td><td>alignment</td><td>distance</td><td>position</td><td>gene name</td><td>gene id</td></tr>
<tr><td>GU305914.1</td><td>73015</td><td>73037</td><td>+</td><td>0</td><td>GACGAGTCGGGTCCACCTAA</td><td>CGG</td><td>||||||||[||||||||||||]PAM</td><td>NA</td><td>Intergenic</td><td>NA</td><td>NA</td></tr>
<tr><td></td><td></td><td></td><td></td><td></td><td></td><td></td><td></td><td></td><td></td><td></td><td></td></tr>
<tr><td>T5150</td><td>AGAACCCGTTCTCGACGATGCGG</td><td>1000</td><td></td><td></td><td></td><td></td><td></td><td></td><td></td><td></td><td></td></tr>
<tr><td>Oligo fwd</td><td>CACCAGAACCCGTTCTCGACGATG</td><td></td><td></td><td></td><td></td><td></td><td></td><td></td><td></td><td></td><td></td></tr>
<tr><td>Oligo rev</td><td>AAACCATCGTCGAGAACGGGTTCT</td><td></td><td></td><td></td><td></td><td></td><td></td><td></td><td></td><td></td><td></td></tr>
<tr><td>Chromosome</td><td>start</td><td>end</td><td>strand</td><td>MM</td><td>target_seq</td><td>PAM</td><td>alignment</td><td>distance</td><td>position</td><td>gene name</td><td>gene id</td></tr>
<tr><td>GU305914.1</td><td>73041</td><td>73063</td><td>+</td><td>0</td><td>AGAACCCGTTCTCGACGATG</td><td>CGG</td><td>||||||||[||||||||||||]PAM</td><td>NA</td><td>Intergenic</td><td>NA</td><td>NA</td></tr>
<tr><td></td><td></td><td></td><td></td><td></td><td></td><td></td><td></td><td></td><td></td><td></td><td></td></tr>
<tr><td>T5151</td><td>GATGCGGATGATCCCCCACGAGG</td><td>1000</td><td></td><td></td><td></td><td></td><td></td><td></td><td></td><td></td><td></td></tr>
<tr><td>Oligo fwd</td><td>CACCGATGCGGATGATCCCCCACG</td><td></td><td></td><td></td><td></td><td></td><td></td><td></td><td></td><td></td><td></td></tr>
<tr><td>Oligo rev</td><td>AAACCGTGGGGGATCATCCGCATC</td><td></td><td></td><td></td><td></td><td></td><td></td><td></td><td></td><td></td><td></td></tr>
<tr><td>Chromosome</td><td>start</td><td>end</td><td>strand</td><td>MM</td><td>target_seq</td><td>PAM</td><td>alignment</td><td>distance</td><td>position</td><td>gene name</td><td>gene id</td></tr>
<tr><td>GU305914.1</td><td>73057</td><td>73079</td><td>+</td><td>0</td><td>GATGCGGATGATCCCCCACG</td><td>AGG</td><td>||||||||[||||||||||||]PAM</td><td>NA</td><td>Intergenic</td><td>NA</td><td>NA</td></tr>
<tr><td></td><td></td><td></td><td></td><td></td><td></td><td></td><td></td><td></td><td></td><td></td><td></td></tr>
<tr><td>T5152</td><td>GATGATCCCCCACGAGGACGAGG</td><td>1000</td><td></td><td></td><td></td><td></td><td></td><td></td><td></td><td></td><td></td></tr>
<tr><td>Oligo fwd</td><td>CACCGATGATCCCCCACGAGGACG</td><td></td><td></td><td></td><td></td><td></td><td></td><td></td><td></td><td></td><td></td></tr>
<tr><td>Oligo rev</td><td>AAACCGTCCTCGTGGGGGATCATC</td><td></td><td></td><td></td><td></td><td></td><td></td><td></td><td></td><td></td><td></td></tr>
<tr><td>Chromosome</td><td>start</td><td>end</td><td>strand</td><td>MM</td><td>target_seq</td><td>PAM</td><td>alignment</td><td>distance</td><td>position</td><td>gene name</td><td>gene id</td></tr>
<tr><td>GU305914.1</td><td>73063</td><td>73085</td><td>+</td><td>0</td><td>GATGATCCCCCACGAGGACG</td><td>AGG</td><td>||||||||[||||||||||||]PAM</td><td>NA</td><td>Intergenic</td><td>NA</td><td>NA</td></tr>
<tr><td></td><td></td><td></td><td></td><td></td><td></td><td></td><td></td><td></td><td></td><td></td><td></td></tr>
<tr><td>T5153</td><td>ATGATCCCCCACGAGGACGAGGG</td><td>1000</td><td></td><td></td><td></td><td></td><td></td><td></td><td></td><td></td><td></td></tr>
<tr><td>Oligo fwd</td><td>CACCATGATCCCCCACGAGGACGA</td><td></td><td></td><td></td><td></td><td></td><td></td><td></td><td></td><td></td><td></td></tr>
<tr><td>Oligo rev</td><td>AAACTCGTCCTCGTGGGGGATCAT</td><td></td><td></td><td></td><td></td><td></td><td></td><td></td><td></td><td></td><td></td></tr>
<tr><td>Chromosome</td><td>start</td><td>end</td><td>strand</td><td>MM</td><td>target_seq</td><td>PAM</td><td>alignment</td><td>distance</td><td>position</td><td>gene name</td><td>gene id</td></tr>
<tr><td>GU305914.1</td><td>73064</td><td>73086</td><td>+</td><td>0</td><td>ATGATCCCCCACGAGGACGA</td><td>GGG</td><td>||||||||[||||||||||||]PAM</td><td>NA</td><td>Intergenic</td><td>NA</td><td>NA</td></tr>
<tr><td></td><td></td><td></td><td></td><td></td><td></td><td></td><td></td><td></td><td></td><td></td><td></td></tr>
<tr><td>T5154</td><td>TCCCCGACTGAAGAGAGCGTCGG</td><td>1000</td><td></td><td></td><td></td><td></td><td></td><td></td><td></td><td></td><td></td></tr>
<tr><td>Oligo fwd</td><td>CACCTCCCCGACTGAAGAGAGCGT</td><td></td><td></td><td></td><td></td><td></td><td></td><td></td><td></td><td></td><td></td></tr>
<tr><td>Oligo rev</td><td>AAACACGCTCTCTTCAGTCGGGGA</td><td></td><td></td><td></td><td></td><td></td><td></td><td></td><td></td><td></td><td></td></tr>
<tr><td>Chromosome</td><td>start</td><td>end</td><td>strand</td><td>MM</td><td>target_seq</td><td>PAM</td><td>alignment</td><td>distance</td><td>position</td><td>gene name</td><td>gene id</td></tr>
<tr><td>GU305914.1</td><td>73159</td><td>73181</td><td>+</td><td>0</td><td>TCCCCGACTGAAGAGAGCGT</td><td>CGG</td><td>||||||||[||||||||||||]PAM</td><td>NA</td><td>Intergenic</td><td>NA</td><td>NA</td></tr>
<tr><td></td><td></td><td></td><td></td><td></td><td></td><td></td><td></td><td></td><td></td><td></td><td></td></tr>
<tr><td>T5155</td><td>CCCCGACTGAAGAGAGCGTCGGG</td><td>1000</td><td></td><td></td><td></td><td></td><td></td><td></td><td></td><td></td><td></td></tr>
<tr><td>Oligo fwd</td><td>CACCCCCCGACTGAAGAGAGCGTC</td><td></td><td></td><td></td><td></td><td></td><td></td><td></td><td></td><td></td><td></td></tr>
<tr><td>Oligo rev</td><td>AAACGACGCTCTCTTCAGTCGGGG</td><td></td><td></td><td></td><td></td><td></td><td></td><td></td><td></td><td></td><td></td></tr>
<tr><td>Chromosome</td><td>start</td><td>end</td><td>strand</td><td>MM</td><td>target_seq</td><td>PAM</td><td>alignment</td><td>distance</td><td>position</td><td>gene name</td><td>gene id</td></tr>
<tr><td>GU305914.1</td><td>73160</td><td>73182</td><td>+</td><td>0</td><td>CCCCGACTGAAGAGAGCGTC</td><td>GGG</td><td>||||||||[||||||||||||]PAM</td><td>NA</td><td>Intergenic</td><td>NA</td><td>NA</td></tr>
<tr><td></td><td></td><td></td><td></td><td></td><td></td><td></td><td></td><td></td><td></td><td></td><td></td></tr>
<tr><td>T5156</td><td>GTCGGGAGACTCTTCCTCGTCGG</td><td>1000</td><td></td><td></td><td></td><td></td><td></td><td></td><td></td><td></td><td></td></tr>
</table>

<table>
<tr><td>Oligo fwd</td><td>CACCGTCGGGAGACTCTTCCTCGT</td><td></td><td></td><td></td><td></td><td></td><td></td><td></td><td></td><td></td><td></td></tr>
<tr><td>Oligo rev</td><td>AAACACGAGGAAGAGTCTCCCGAC</td><td></td><td></td><td></td><td></td><td></td><td></td><td></td><td></td><td></td><td></td></tr>
<tr><td>Chromosome</td><td>start</td><td>end</td><td>strand</td><td>MM</td><td>target_seq</td><td>PAM</td><td>alignment</td><td>distance</td><td>position</td><td>gene name</td><td>gene id</td></tr>
<tr><td>GU305914.1</td><td>73177</td><td>73199</td><td>+</td><td>0</td><td>GTCGGGAGACTCTTCCTCGT</td><td>CGG</td><td>||||||||[||||||||||||]PAM</td><td>NA</td><td>Intergenic</td><td>NA</td><td>NA</td></tr>
<tr><td>T5157</td><td>AGACTCTTCCTCGTCGGTGCCGG</td><td>1000</td><td></td><td></td><td></td><td></td><td></td><td></td><td></td><td></td><td></td></tr>
<tr><td>Oligo fwd</td><td>CACCAGACTCTTCCTCGTCGGTGC</td><td></td><td></td><td></td><td></td><td></td><td></td><td></td><td></td><td></td><td></td></tr>
<tr><td>Oligo rev</td><td>AAACGCACCGACGAGGAAGAGTCT</td><td></td><td></td><td></td><td></td><td></td><td></td><td></td><td></td><td></td><td></td></tr>
<tr><td>Chromosome</td><td>start</td><td>end</td><td>strand</td><td>MM</td><td>target_seq</td><td>PAM</td><td>alignment</td><td>distance</td><td>position</td><td>gene name</td><td>gene id</td></tr>
<tr><td>GU305914.1</td><td>73183</td><td>73205</td><td>+</td><td>0</td><td>AGACTCTTCCTCGTCGGTGC</td><td>CGG</td><td>||||||||[||||||||||||]PAM</td><td>NA</td><td>Intergenic</td><td>NA</td><td>NA</td></tr>
<tr><td>T5158</td><td>CTCTTCCTCGTCGGTGCCGGCGG</td><td>1000</td><td></td><td></td><td></td><td></td><td></td><td></td><td></td><td></td><td></td></tr>
<tr><td>Oligo fwd</td><td>CACCCTCTTCCTCGTCGGTGCCGG</td><td></td><td></td><td></td><td></td><td></td><td></td><td></td><td></td><td></td><td></td></tr>
<tr><td>Oligo rev</td><td>AAACCCGGCACCGACGAGGAAGAG</td><td></td><td></td><td></td><td></td><td></td><td></td><td></td><td></td><td></td><td></td></tr>
<tr><td>Chromosome</td><td>start</td><td>end</td><td>strand</td><td>MM</td><td>target_seq</td><td>PAM</td><td>alignment</td><td>distance</td><td>position</td><td>gene name</td><td>gene id</td></tr>
<tr><td>GU305914.1</td><td>73186</td><td>73208</td><td>+</td><td>0</td><td>CTCTTCCTCGTCGGTGCCGG</td><td>CGG</td><td>||||||||[||||||||||||]PAM</td><td>NA</td><td>Intergenic</td><td>NA</td><td>NA</td></tr>
<tr><td>GU305914.1</td><td>127716</td><td>127738</td><td>-</td><td>4</td><td>CTTGTCCTCGTCGGTGGCGC</td><td>CGG</td><td>||--||||[||||||||-||-]PAM</td><td>NA</td><td>Intergenic</td><td>NA</td><td>NA</td></tr>
<tr><td>T5159</td><td>TTCCTCGTCGGTGCCGGCGGAGG</td><td>1000</td><td></td><td></td><td></td><td></td><td></td><td></td><td></td><td></td><td></td></tr>
<tr><td>Oligo fwd</td><td>CACCTTCCTCGTCGGTGCCGGCGG</td><td></td><td></td><td></td><td></td><td></td><td></td><td></td><td></td><td></td><td></td></tr>
<tr><td>Oligo rev</td><td>AAACCCGCCGGCACCGACGAGGAA</td><td></td><td></td><td></td><td></td><td></td><td></td><td></td><td></td><td></td><td></td></tr>
<tr><td>Chromosome</td><td>start</td><td>end</td><td>strand</td><td>MM</td><td>target_seq</td><td>PAM</td><td>alignment</td><td>distance</td><td>position</td><td>gene name</td><td>gene id</td></tr>
<tr><td>GU305914.1</td><td>73189</td><td>73211</td><td>+</td><td>0</td><td>TTCCTCGTCGGTGCCGGCGG</td><td>AGG</td><td>||||||||[||||||||||||]PAM</td><td>NA</td><td>Intergenic</td><td>NA</td><td>NA</td></tr>
<tr><td>GU305914.1</td><td>62431</td><td>62453</td><td>+</td><td>4</td><td>TTCGACGGCGGTGGCGGCGG</td><td>GGG</td><td>|||--||-[|||||-||||||]PAM</td><td>NA</td><td>Intergenic</td><td>NA</td><td>NA</td></tr>
<tr><td>T5160</td><td>CGTCGGTGCCGGCGGAGGATCGG</td><td>1000</td><td></td><td></td><td></td><td></td><td></td><td></td><td></td><td></td><td></td></tr>
<tr><td>Oligo fwd</td><td>CACCCGTCGGTGCCGGCGGAGGAT</td><td></td><td></td><td></td><td></td><td></td><td></td><td></td><td></td><td></td><td></td></tr>
<tr><td>Oligo rev</td><td>AAACATCCTCCGCCGGCACCGACG</td><td></td><td></td><td></td><td></td><td></td><td></td><td></td><td></td><td></td><td></td></tr>
<tr><td>Chromosome</td><td>start</td><td>end</td><td>strand</td><td>MM</td><td>target_seq</td><td>PAM</td><td>alignment</td><td>distance</td><td>position</td><td>gene name</td><td>gene id</td></tr>
<tr><td>GU305914.1</td><td>73194</td><td>73216</td><td>+</td><td>0</td><td>CGTCGGTGCCGGCGGAGGAT</td><td>CGG</td><td>||||||||[||||||||||||]PAM</td><td>NA</td><td>Intergenic</td><td>NA</td><td>NA</td></tr>
<tr><td>T5161</td><td>GTCGGTGCCGGCGGAGGATCGGG</td><td>1000</td><td></td><td></td><td></td><td></td><td></td><td></td><td></td><td></td><td></td></tr>
<tr><td>Oligo fwd</td><td>CACCGTCGGTGCCGGCGGAGGATC</td><td></td><td></td><td></td><td></td><td></td><td></td><td></td><td></td><td></td><td></td></tr>
<tr><td>Oligo rev</td><td>AAACGATCCTCCGCCGGCACCGAC</td><td></td><td></td><td></td><td></td><td></td><td></td><td></td><td></td><td></td><td></td></tr>
<tr><td>Chromosome</td><td>start</td><td>end</td><td>strand</td><td>MM</td><td>target_seq</td><td>PAM</td><td>alignment</td><td>distance</td><td>position</td><td>gene name</td><td>gene id</td></tr>
<tr><td>GU305914.1</td><td>73195</td><td>73217</td><td>+</td><td>0</td><td>GTCGGTGCCGGCGGAGGATC</td><td>GGG</td><td>||||||||[||||||||||||]PAM</td><td>NA</td><td>Intergenic</td><td>NA</td><td>NA</td></tr>
<tr><td>GU305914.1</td><td>40436</td><td>40458</td><td>-</td><td>4</td><td>GACGATGACGGCGGAGGACC</td><td>CGG</td><td>|-||-||-[||||||||||-|]PAM</td><td>NA</td><td>Intergenic</td><td>NA</td><td>NA</td></tr>
<tr><td>T5162</td><td>GGACCCCGACGCGCGACCCCAGG</td><td>1000</td><td></td><td></td><td></td><td></td><td></td><td></td><td></td><td></td><td></td></tr>
<tr><td>Oligo fwd</td><td>CACCGGACCCCGACGCGCGACCCC</td><td></td><td></td><td></td><td></td><td></td><td></td><td></td><td></td><td></td><td></td></tr>
<tr><td>Oligo rev</td><td>AAACGGGGTCGCGCGTCGGGGTCC</td><td></td><td></td><td></td><td></td><td></td><td></td><td></td><td></td><td></td><td></td></tr>
<tr><td>Chromosome</td><td>start</td><td>end</td><td>strand</td><td>MM</td><td>target_seq</td><td>PAM</td><td>alignment</td><td>distance</td><td>position</td><td>gene name</td><td>gene id</td></tr>
<tr><td>GU305914.1</td><td>73216</td><td>73238</td><td>+</td><td>0</td><td>GGACCCCGACGCGCGACCCC</td><td>AGG</td><td>||||||||[||||||||||||]PAM</td><td>NA</td><td>Intergenic</td><td>NA</td><td>NA</td></tr>
<tr><td>T5163</td><td>CGACGCGCGACCCCAGGATTCGG</td><td>1000</td><td></td><td></td><td></td><td></td><td></td><td></td><td></td><td></td><td></td></tr>
<tr><td>Oligo fwd</td><td>CACCCGACGCGCGACCCCAGGATT</td><td></td><td></td><td></td><td></td><td></td><td></td><td></td><td></td><td></td><td></td></tr>
<tr><td>Oligo rev</td><td>AAACAATCCTGGGGTCGCGCGTCG</td><td></td><td></td><td></td><td></td><td></td><td></td><td></td><td></td><td></td><td></td></tr>
<tr><td>Chromosome</td><td>start</td><td>end</td><td>strand</td><td>MM</td><td>target_seq</td><td>PAM</td><td>alignment</td><td>distance</td><td>position</td><td>gene name</td><td>gene id</td></tr>
<tr><td>GU305914.1</td><td>73222</td><td>73244</td><td>+</td><td>0</td><td>CGACGCGCGACCCCAGGATT</td><td>CGG</td><td>||||||||[||||||||||||]PAM</td><td>NA</td><td>Intergenic</td><td>NA</td><td>NA</td></tr>
<tr><td>T5164</td><td>ACCCCTGTTCTCGCAGACCAAGG</td><td>1000</td><td></td><td></td><td></td><td></td><td></td><td></td><td></td><td></td><td></td></tr>
<tr><td>Oligo fwd</td><td>CACCACCCCTGTTCTCGCAGACCA</td><td></td><td></td><td></td><td></td><td></td><td></td><td></td><td></td><td></td><td></td></tr>
<tr><td>Oligo rev</td><td>AAACTGGTCTGCGAGAACAGGGGT</td><td></td><td></td><td></td><td></td><td></td><td></td><td></td><td></td><td></td><td></td></tr>
<tr><td>Chromosome</td><td>start</td><td>end</td><td>strand</td><td>MM</td><td>target_seq</td><td>PAM</td><td>alignment</td><td>distance</td><td>position</td><td>gene name</td><td>gene id</td></tr>
<tr><td>GU305914.1</td><td>73267</td><td>73289</td><td>+</td><td>0</td><td>ACCCCTGTTCTCGCAGACCA</td><td>AGG</td><td>||||||||[||||||||||||]PAM</td><td>NA</td><td>Intergenic</td><td>NA</td><td>NA</td></tr>
<tr><td>T5165</td><td>GGTGTCCAGCATCGTGCCGAAGG</td><td>1000</td><td></td><td></td><td></td><td></td><td></td><td></td><td></td><td></td><td></td></tr>
<tr><td>Oligo fwd</td><td>CACCGGTGTCCAGCATCGTGCCGA</td><td></td><td></td><td></td><td></td><td></td><td></td><td></td><td></td><td></td><td></td></tr>
<tr><td>Oligo rev</td><td>AAACTCGGCACGATGCTGGACACC</td><td></td><td></td><td></td><td></td><td></td><td></td><td></td><td></td><td></td><td></td></tr>
<tr><td>Chromosome</td><td>start</td><td>end</td><td>strand</td><td>MM</td><td>target_seq</td><td>PAM</td><td>alignment</td><td>distance</td><td>position</td><td>gene name</td><td>gene id</td></tr>
<tr><td>GU305914.1</td><td>73288</td><td>73310</td><td>+</td><td>0</td><td>GGTGTCCAGCATCGTGCCGA</td><td>AGG</td><td>||||||||[||||||||||||]PAM</td><td>NA</td><td>Intergenic</td><td>NA</td><td>NA</td></tr>
<tr><td>T5166</td><td>CAGCATCGTGCCGAAGGACGCGG</td><td>1000</td><td></td><td></td><td></td><td></td><td></td><td></td><td></td><td></td><td></td></tr>
<tr><td>Oligo fwd</td><td>CACCCAGCATCGTGCCGAAGGACG</td><td></td><td></td><td></td><td></td><td></td><td></td><td></td><td></td><td></td><td></td></tr>
<tr><td>Oligo rev</td><td>AAACCGTCCTTCGGCACGATGCTG</td><td></td><td></td><td></td><td></td><td></td><td></td><td></td><td></td><td></td><td></td></tr>
<tr><td>Chromosome</td><td>start</td><td>end</td><td>strand</td><td>MM</td><td>target_seq</td><td>PAM</td><td>alignment</td><td>distance</td><td>position</td><td>gene name</td><td>gene id</td></tr>
<tr><td>GU305914.1</td><td>73294</td><td>73316</td><td>+</td><td>0</td><td>CAGCATCGTGCCGAAGGACG</td><td>CGG</td><td>||||||||[||||||||||||]PAM</td><td>NA</td><td>Intergenic</td><td>NA</td><td>NA</td></tr>
<tr><td>T5167</td><td>AGCCCTGTCTAGCGCGATCATGG</td><td>1000</td><td></td><td></td><td></td><td></td><td></td><td></td><td></td><td></td><td></td></tr>
<tr><td>Oligo fwd</td><td>CACCAGCCCTGTCTAGCGCGATCA</td><td></td><td></td><td></td><td></td><td></td><td></td><td></td><td></td><td></td><td></td></tr>
<tr><td>Oligo rev</td><td>AAACTGATCGCGCTAGACAGGGCT</td><td></td><td></td><td></td><td></td><td></td><td></td><td></td><td></td><td></td><td></td></tr>
<tr><td>Chromosome</td><td>start</td><td>end</td><td>strand</td><td>MM</td><td>target_seq</td><td>PAM</td><td>alignment</td><td>distance</td><td>position</td><td>gene name</td><td>gene id</td></tr>
<tr><td>GU305914.1</td><td>73372</td><td>73394</td><td>+</td><td>0</td><td>AGCCCTGTCTAGCGCGATCA</td><td>TGG</td><td>||||||||[||||||||||||]PAM</td><td>NA</td><td>Intergenic</td><td>NA</td><td>NA</td></tr>
<tr><td>T5168</td><td>CCTGTCTAGCGCGATCATGGCGG</td><td>1000</td><td></td><td></td><td></td><td></td><td></td><td></td><td></td><td></td><td></td></tr>
<tr><td>Oligo fwd</td><td>CACCCCTGTCTAGCGCGATCATGG</td><td></td><td></td><td></td><td></td><td></td><td></td><td></td><td></td><td></td><td></td></tr>
<tr><td>Oligo rev</td><td>AAACCCATGATCGCGCTAGACAGG</td><td></td><td></td><td></td><td></td><td></td><td></td><td></td><td></td><td></td><td></td></tr>
<tr><td>Chromosome</td><td>start</td><td>end</td><td>strand</td><td>MM</td><td>target_seq</td><td>PAM</td><td>alignment</td><td>distance</td><td>position</td><td>gene name</td><td>gene id</td></tr>
<tr><td>GU305914.1</td><td>73375</td><td>73397</td><td>+</td><td>0</td><td>CCTGTCTAGCGCGATCATGG</td><td>CGG</td><td>||||||||[||||||||||||]PAM</td><td>NA</td><td>Intergenic</td><td>NA</td><td>NA</td></tr>
<tr><td>T5169</td><td>GCAAGAGACCCGACTGCAGCTGG</td><td>1000</td><td></td><td></td><td></td><td></td><td></td><td></td><td></td><td></td><td></td></tr>
<tr><td>Oligo fwd</td><td>CACCGCAAGAGACCCGACTGCAGC</td><td></td><td></td><td></td><td></td><td></td><td></td><td></td><td></td><td></td><td></td></tr>
<tr><td>Oligo rev</td><td>AAACGCTGCAGTCGGGTCTCTTGC</td><td></td><td></td><td></td><td></td><td></td><td></td><td></td><td></td><td></td><td></td></tr>
<tr><td>Chromosome</td><td>start</td><td>end</td><td>strand</td><td>MM</td><td>target_seq</td><td>PAM</td><td>alignment</td><td>distance</td><td>position</td><td>gene name</td><td>gene id</td></tr>
<tr><td>GU305914.1</td><td>73411</td><td>73433</td><td>+</td><td>0</td><td>GCAAGAGACCCGACTGCAGC</td><td>TGG</td><td>||||||||[||||||||||||]PAM</td><td>NA</td><td>Intergenic</td><td>NA</td><td>NA</td></tr>
<tr><td>T5170</td><td>TTTCGAGAGCGACATGTGCGAGG</td><td>1000</td><td></td><td></td><td></td><td></td><td></td><td></td><td></td><td></td><td></td></tr>
<tr><td>Oligo fwd</td><td>CACCTTTCGAGAGCGACATGTGCG</td><td></td><td></td><td></td><td></td><td></td><td></td><td></td><td></td><td></td><td></td></tr>
<tr><td>Oligo rev</td><td>AAACCGCACATGTCGCTCTCGAAA</td><td></td><td></td><td></td><td></td><td></td><td></td><td></td><td></td><td></td><td></td></tr>
</table>

<table>
<tr><td>Chromosome</td><td>start</td><td>end</td><td>strand</td><td>MM</td><td>target_seq</td><td>PAM</td><td>alignment</td><td>distance</td><td>position</td><td>gene name</td><td>gene id</td></tr>
<tr><td>GU305914.1</td><td>73438</td><td>73460</td><td>+</td><td>0</td><td>TTTCGAGAGCGACATGTGCG</td><td>AGG</td><td>||||||||[||||||||||||]PAM</td><td>NA</td><td>Intergenic</td><td>NA</td><td>NA</td></tr>
<tr><td>T5171</td><td>GCGACATGTGCGAGGCCATCAGG</td><td>1000</td><td></td><td></td><td></td><td></td><td></td><td></td><td></td><td></td><td></td></tr>
<tr><td>Oligo fwd</td><td>CACCGCGACATGTGCGAGGCCATC</td><td></td><td></td><td></td><td></td><td></td><td></td><td></td><td></td><td></td><td></td></tr>
<tr><td>Oligo rev</td><td>AAACGATGGCCTCGCACATGTCGC</td><td></td><td></td><td></td><td></td><td></td><td></td><td></td><td></td><td></td><td></td></tr>
<tr><td>Chromosome</td><td>start</td><td>end</td><td>strand</td><td>MM</td><td>target_seq</td><td>PAM</td><td>alignment</td><td>distance</td><td>position</td><td>gene name</td><td>gene id</td></tr>
<tr><td>GU305914.1</td><td>73446</td><td>73468</td><td>+</td><td>0</td><td>GCGACATGTGCGAGGCCATC</td><td>AGG</td><td>||||||||[||||||||||||]PAM</td><td>NA</td><td>Intergenic</td><td>NA</td><td>NA</td></tr>
<tr><td>T5172</td><td>ACATGTGCGAGGCCATCAGGAGG</td><td>1000</td><td></td><td></td><td></td><td></td><td></td><td></td><td></td><td></td><td></td></tr>
<tr><td>Oligo fwd</td><td>CACCACATGTGCGAGGCCATCAGG</td><td></td><td></td><td></td><td></td><td></td><td></td><td></td><td></td><td></td><td></td></tr>
<tr><td>Oligo rev</td><td>AAACCCTGATGGCCTCGCACATGT</td><td></td><td></td><td></td><td></td><td></td><td></td><td></td><td></td><td></td><td></td></tr>
<tr><td>Chromosome</td><td>start</td><td>end</td><td>strand</td><td>MM</td><td>target_seq</td><td>PAM</td><td>alignment</td><td>distance</td><td>position</td><td>gene name</td><td>gene id</td></tr>
<tr><td>GU305914.1</td><td>73449</td><td>73471</td><td>+</td><td>0</td><td>ACATGTGCGAGGCCATCAGG</td><td>AGG</td><td>||||||||[||||||||||||]PAM</td><td>NA</td><td>Intergenic</td><td>NA</td><td>NA</td></tr>
<tr><td>T5173</td><td>AAAACCCACACCACACACAGAGG</td><td>1000</td><td></td><td></td><td></td><td></td><td></td><td></td><td></td><td></td><td></td></tr>
<tr><td>Oligo fwd</td><td>CACCAAAACCCACACCACACACAG</td><td></td><td></td><td></td><td></td><td></td><td></td><td></td><td></td><td></td><td></td></tr>
<tr><td>Oligo rev</td><td>AAACCTGTGTGTGGTGTGGGTTTT</td><td></td><td></td><td></td><td></td><td></td><td></td><td></td><td></td><td></td><td></td></tr>
<tr><td>Chromosome</td><td>start</td><td>end</td><td>strand</td><td>MM</td><td>target_seq</td><td>PAM</td><td>alignment</td><td>distance</td><td>position</td><td>gene name</td><td>gene id</td></tr>
<tr><td>GU305914.1</td><td>73507</td><td>73529</td><td>+</td><td>0</td><td>AAAACCCACACCACACACAG</td><td>AGG</td><td>||||||||[||||||||||||]PAM</td><td>NA</td><td>Intergenic</td><td>NA</td><td>NA</td></tr>
<tr><td>T5174</td><td>ATGTGTGTCTTATTGTATGACGG</td><td>1000</td><td></td><td></td><td></td><td></td><td></td><td></td><td></td><td></td><td></td></tr>
<tr><td>Oligo fwd</td><td>CACCATGTGTGTCTTATTGTATGA</td><td></td><td></td><td></td><td></td><td></td><td></td><td></td><td></td><td></td><td></td></tr>
<tr><td>Oligo rev</td><td>AAACTCATACAATAAGACACACAT</td><td></td><td></td><td></td><td></td><td></td><td></td><td></td><td></td><td></td><td></td></tr>
<tr><td>Chromosome</td><td>start</td><td>end</td><td>strand</td><td>MM</td><td>target_seq</td><td>PAM</td><td>alignment</td><td>distance</td><td>position</td><td>gene name</td><td>gene id</td></tr>
<tr><td>GU305914.1</td><td>73566</td><td>73588</td><td>+</td><td>0</td><td>ATGTGTGTCTTATTGTATGA</td><td>CGG</td><td>||||||||[||||||||||||]PAM</td><td>NA</td><td>Intergenic</td><td>NA</td><td>NA</td></tr>
<tr><td>T5175</td><td>TGTGTCTTATTGTATGACGGTGG</td><td>1000</td><td></td><td></td><td></td><td></td><td></td><td></td><td></td><td></td><td></td></tr>
<tr><td>Oligo fwd</td><td>CACCTGTGTCTTATTGTATGACGG</td><td></td><td></td><td></td><td></td><td></td><td></td><td></td><td></td><td></td><td></td></tr>
<tr><td>Oligo rev</td><td>AAACCCGTCATACAATAAGACACA</td><td></td><td></td><td></td><td></td><td></td><td></td><td></td><td></td><td></td><td></td></tr>
<tr><td>Chromosome</td><td>start</td><td>end</td><td>strand</td><td>MM</td><td>target_seq</td><td>PAM</td><td>alignment</td><td>distance</td><td>position</td><td>gene name</td><td>gene id</td></tr>
<tr><td>GU305914.1</td><td>73569</td><td>73591</td><td>+</td><td>0</td><td>TGTGTCTTATTGTATGACGG</td><td>TGG</td><td>||||||||[||||||||||||]PAM</td><td>NA</td><td>Intergenic</td><td>NA</td><td>NA</td></tr>
<tr><td>T5176</td><td>CGATCGAGAGCATCCGCAGAAGG</td><td>1000</td><td></td><td></td><td></td><td></td><td></td><td></td><td></td><td></td><td></td></tr>
<tr><td>Oligo fwd</td><td>CACCCGATCGAGAGCATCCGCAGA</td><td></td><td></td><td></td><td></td><td></td><td></td><td></td><td></td><td></td><td></td></tr>
<tr><td>Oligo rev</td><td>AAACTCTGCGGATGCTCTCGATCG</td><td></td><td></td><td></td><td></td><td></td><td></td><td></td><td></td><td></td><td></td></tr>
<tr><td>Chromosome</td><td>start</td><td>end</td><td>strand</td><td>MM</td><td>target_seq</td><td>PAM</td><td>alignment</td><td>distance</td><td>position</td><td>gene name</td><td>gene id</td></tr>
<tr><td>GU305914.1</td><td>73601</td><td>73623</td><td>+</td><td>0</td><td>CGATCGAGAGCATCCGCAGA</td><td>AGG</td><td>||||||||[||||||||||||]PAM</td><td>NA</td><td>Intergenic</td><td>NA</td><td>NA</td></tr>
<tr><td>T5177</td><td>GCATCCGCAGAAGGTCCAGCTGG</td><td>1000</td><td></td><td></td><td></td><td></td><td></td><td></td><td></td><td></td><td></td></tr>
<tr><td>Oligo fwd</td><td>CACCGCATCCGCAGAAGGTCCAGC</td><td></td><td></td><td></td><td></td><td></td><td></td><td></td><td></td><td></td><td></td></tr>
<tr><td>Oligo rev</td><td>AAACGCTGGACCTTCTGCGGATGC</td><td></td><td></td><td></td><td></td><td></td><td></td><td></td><td></td><td></td><td></td></tr>
<tr><td>Chromosome</td><td>start</td><td>end</td><td>strand</td><td>MM</td><td>target_seq</td><td>PAM</td><td>alignment</td><td>distance</td><td>position</td><td>gene name</td><td>gene id</td></tr>
<tr><td>GU305914.1</td><td>73610</td><td>73632</td><td>+</td><td>0</td><td>GCATCCGCAGAAGGTCCAGC</td><td>TGG</td><td>||||||||[||||||||||||]PAM</td><td>NA</td><td>Intergenic</td><td>NA</td><td>NA</td></tr>
<tr><td>T5178</td><td>CCGCAGAAGGTCCAGCTGGTAGG</td><td>1000</td><td></td><td></td><td></td><td></td><td></td><td></td><td></td><td></td><td></td></tr>
<tr><td>Oligo fwd</td><td>CACCCCGCAGAAGGTCCAGCTGGT</td><td></td><td></td><td></td><td></td><td></td><td></td><td></td><td></td><td></td><td></td></tr>
<tr><td>Oligo rev</td><td>AAACACCAGCTGGACCTTCTGCGG</td><td></td><td></td><td></td><td></td><td></td><td></td><td></td><td></td><td></td><td></td></tr>
<tr><td>Chromosome</td><td>start</td><td>end</td><td>strand</td><td>MM</td><td>target_seq</td><td>PAM</td><td>alignment</td><td>distance</td><td>position</td><td>gene name</td><td>gene id</td></tr>
<tr><td>GU305914.1</td><td>73614</td><td>73636</td><td>+</td><td>0</td><td>CCGCAGAAGGTCCAGCTGGT</td><td>AGG</td><td>||||||||[||||||||||||]PAM</td><td>NA</td><td>Intergenic</td><td>NA</td><td>NA</td></tr>
<tr><td>T5179</td><td>CCTCCCATCTTCATATATTTCGG</td><td>1000</td><td></td><td></td><td></td><td></td><td></td><td></td><td></td><td></td><td></td></tr>
<tr><td>Oligo fwd</td><td>CACCCCTCCCATCTTCATATATTT</td><td></td><td></td><td></td><td></td><td></td><td></td><td></td><td></td><td></td><td></td></tr>
<tr><td>Oligo rev</td><td>AAACAAATATATGAAGATGGGAGG</td><td></td><td></td><td></td><td></td><td></td><td></td><td></td><td></td><td></td><td></td></tr>
<tr><td>Chromosome</td><td>start</td><td>end</td><td>strand</td><td>MM</td><td>target_seq</td><td>PAM</td><td>alignment</td><td>distance</td><td>position</td><td>gene name</td><td>gene id</td></tr>
<tr><td>GU305914.1</td><td>73654</td><td>73676</td><td>+</td><td>0</td><td>CCTCCCATCTTCATATATTT</td><td>CGG</td><td>||||||||[||||||||||||]PAM</td><td>NA</td><td>Intergenic</td><td>NA</td><td>NA</td></tr>
<tr><td>T5180</td><td>CTTCATATATTTCGGAATCATGG</td><td>1000</td><td></td><td></td><td></td><td></td><td></td><td></td><td></td><td></td><td></td></tr>
<tr><td>Oligo fwd</td><td>CACCCTTCATATATTTCGGAATCA</td><td></td><td></td><td></td><td></td><td></td><td></td><td></td><td></td><td></td><td></td></tr>
<tr><td>Oligo rev</td><td>AAACTGATTCCGAAATATATGAAG</td><td></td><td></td><td></td><td></td><td></td><td></td><td></td><td></td><td></td><td></td></tr>
<tr><td>Chromosome</td><td>start</td><td>end</td><td>strand</td><td>MM</td><td>target_seq</td><td>PAM</td><td>alignment</td><td>distance</td><td>position</td><td>gene name</td><td>gene id</td></tr>
<tr><td>GU305914.1</td><td>73662</td><td>73684</td><td>+</td><td>0</td><td>CTTCATATATTTCGGAATCA</td><td>TGG</td><td>||||||||[||||||||||||]PAM</td><td>NA</td><td>Intergenic</td><td>NA</td><td>NA</td></tr>
<tr><td>T5181</td><td>CGGAATCATGGTCGAGATGACGG</td><td>1000</td><td></td><td></td><td></td><td></td><td></td><td></td><td></td><td></td><td></td></tr>
<tr><td>Oligo fwd</td><td>CACCCGGAATCATGGTCGAGATGA</td><td></td><td></td><td></td><td></td><td></td><td></td><td></td><td></td><td></td><td></td></tr>
<tr><td>Oligo rev</td><td>AAACTCATCTCGACCATGATTCCG</td><td></td><td></td><td></td><td></td><td></td><td></td><td></td><td></td><td></td><td></td></tr>
<tr><td>Chromosome</td><td>start</td><td>end</td><td>strand</td><td>MM</td><td>target_seq</td><td>PAM</td><td>alignment</td><td>distance</td><td>position</td><td>gene name</td><td>gene id</td></tr>
<tr><td>GU305914.1</td><td>73674</td><td>73696</td><td>+</td><td>0</td><td>CGGAATCATGGTCGAGATGA</td><td>CGG</td><td>||||||||[||||||||||||]PAM</td><td>NA</td><td>Intergenic</td><td>NA</td><td>NA</td></tr>
<tr><td>T5182</td><td>GGAATCATGGTCGAGATGACGGG</td><td>1000</td><td></td><td></td><td></td><td></td><td></td><td></td><td></td><td></td><td></td></tr>
<tr><td>Oligo fwd</td><td>CACCGGAATCATGGTCGAGATGAC</td><td></td><td></td><td></td><td></td><td></td><td></td><td></td><td></td><td></td><td></td></tr>
<tr><td>Oligo rev</td><td>AAACGTCATCTCGACCATGATTCC</td><td></td><td></td><td></td><td></td><td></td><td></td><td></td><td></td><td></td><td></td></tr>
<tr><td>Chromosome</td><td>start</td><td>end</td><td>strand</td><td>MM</td><td>target_seq</td><td>PAM</td><td>alignment</td><td>distance</td><td>position</td><td>gene name</td><td>gene id</td></tr>
<tr><td>GU305914.1</td><td>73675</td><td>73697</td><td>+</td><td>0</td><td>GGAATCATGGTCGAGATGAC</td><td>GGG</td><td>||||||||[||||||||||||]PAM</td><td>NA</td><td>Intergenic</td><td>NA</td><td>NA</td></tr>
<tr><td>T5183</td><td>ACGGGTTGTGCAACCATCGCAGG</td><td>1000</td><td></td><td></td><td></td><td></td><td></td><td></td><td></td><td></td><td></td></tr>
<tr><td>Oligo fwd</td><td>CACCACGGGTTGTGCAACCATCGC</td><td></td><td></td><td></td><td></td><td></td><td></td><td></td><td></td><td></td><td></td></tr>
<tr><td>Oligo rev</td><td>AAACGCGATGGTTGCACAACCCGT</td><td></td><td></td><td></td><td></td><td></td><td></td><td></td><td></td><td></td><td></td></tr>
<tr><td>Chromosome</td><td>start</td><td>end</td><td>strand</td><td>MM</td><td>target_seq</td><td>PAM</td><td>alignment</td><td>distance</td><td>position</td><td>gene name</td><td>gene id</td></tr>
<tr><td>GU305914.1</td><td>73693</td><td>73715</td><td>+</td><td>0</td><td>ACGGGTTGTGCAACCATCGC</td><td>AGG</td><td>||||||||[||||||||||||]PAM</td><td>NA</td><td>Intergenic</td><td>NA</td><td>NA</td></tr>
<tr><td>T5184</td><td>GTGCAACCATCGCAGGCGCCAGG</td><td>1000</td><td></td><td></td><td></td><td></td><td></td><td></td><td></td><td></td><td></td></tr>
<tr><td>Oligo fwd</td><td>CACCGTGCAACCATCGCAGGCGCC</td><td></td><td></td><td></td><td></td><td></td><td></td><td></td><td></td><td></td><td></td></tr>
<tr><td>Oligo rev</td><td>AAACGGCGCCTGCGATGGTTGCAC</td><td></td><td></td><td></td><td></td><td></td><td></td><td></td><td></td><td></td><td></td></tr>
<tr><td>Chromosome</td><td>start</td><td>end</td><td>strand</td><td>MM</td><td>target_seq</td><td>PAM</td><td>alignment</td><td>distance</td><td>position</td><td>gene name</td><td>gene id</td></tr>
<tr><td>GU305914.1</td><td>73700</td><td>73722</td><td>+</td><td>0</td><td>GTGCAACCATCGCAGGCGCC</td><td>AGG</td><td>||||||||[||||||||||||]PAM</td><td>NA</td><td>Intergenic</td><td>NA</td><td>NA</td></tr>
<tr><td>T5185</td><td>AGGCGCCAGGTTGAGTACGTTGG</td><td>1000</td><td></td><td></td><td></td><td></td><td></td><td></td><td></td><td></td><td></td></tr>
<tr><td>Oligo fwd</td><td>CACCAGGCGCCAGGTTGAGTACGT</td><td></td><td></td><td></td><td></td><td></td><td></td><td></td><td></td><td></td><td></td></tr>
</table>

<table>
<tr><td>Oligo rev</td><td>AAACACGTACTCAACCTGGCGCCT</td><td></td><td></td><td></td><td></td><td></td><td></td><td></td><td></td><td></td><td></td></tr>
<tr><td>Chromosome</td><td>start</td><td>end</td><td>strand</td><td>MM</td><td>target_seq</td><td>PAM</td><td>alignment</td><td>distance</td><td>position</td><td>gene name</td><td>gene id</td></tr>
<tr><td>GU305914.1</td><td>73713</td><td>73735</td><td>+</td><td>0</td><td>AGGCGCCAGGTTGAGTACGT</td><td>TGG</td><td>||||||||[||||||||||||]PAM</td><td>NA</td><td>Intergenic</td><td>NA</td><td>NA</td></tr>
<tr><td>T5186</td><td>TCCTCCTCCGCCGCCGCTGCTGG</td><td>1000</td><td></td><td></td><td></td><td></td><td></td><td></td><td></td><td></td><td></td></tr>
<tr><td>Oligo fwd</td><td>CACCTCCTCCTCCGCCGCCGCTGC</td><td></td><td></td><td></td><td></td><td></td><td></td><td></td><td></td><td></td><td></td></tr>
<tr><td>Oligo rev</td><td>AAACGCAGCGGCGGCGGAGGAGGA</td><td></td><td></td><td></td><td></td><td></td><td></td><td></td><td></td><td></td><td></td></tr>
<tr><td>Chromosome</td><td>start</td><td>end</td><td>strand</td><td>MM</td><td>target_seq</td><td>PAM</td><td>alignment</td><td>distance</td><td>position</td><td>gene name</td><td>gene id</td></tr>
<tr><td>GU305914.1</td><td>73788</td><td>73810</td><td>+</td><td>0</td><td>TCCTCCTCCGCCGCCGCTGC</td><td>TGG</td><td>||||||||[||||||||||||]PAM</td><td>NA</td><td>Intergenic</td><td>NA</td><td>NA</td></tr>
<tr><td>T5187</td><td>GCCGCTACTCCCGCCTCCCGAGG</td><td>1000</td><td></td><td></td><td></td><td></td><td></td><td></td><td></td><td></td><td></td></tr>
<tr><td>Oligo fwd</td><td>CACCGCCGCTACTCCCGCCTCCCG</td><td></td><td></td><td></td><td></td><td></td><td></td><td></td><td></td><td></td><td></td></tr>
<tr><td>Oligo rev</td><td>AAACCGGGAGGCGGGAGTAGCGGC</td><td></td><td></td><td></td><td></td><td></td><td></td><td></td><td></td><td></td><td></td></tr>
<tr><td>Chromosome</td><td>start</td><td>end</td><td>strand</td><td>MM</td><td>target_seq</td><td>PAM</td><td>alignment</td><td>distance</td><td>position</td><td>gene name</td><td>gene id</td></tr>
<tr><td>GU305914.1</td><td>73824</td><td>73846</td><td>+</td><td>0</td><td>GCCGCTACTCCCGCCTCCCG</td><td>AGG</td><td>||||||||[||||||||||||]PAM</td><td>NA</td><td>Intergenic</td><td>NA</td><td>NA</td></tr>
<tr><td>T5188</td><td>CGAGGCCGCGCTACTAACGTTGG</td><td>1000</td><td></td><td></td><td></td><td></td><td></td><td></td><td></td><td></td><td></td></tr>
<tr><td>Oligo fwd</td><td>CACCCGAGGCCGCGCTACTAACGT</td><td></td><td></td><td></td><td></td><td></td><td></td><td></td><td></td><td></td><td></td></tr>
<tr><td>Oligo rev</td><td>AAACACGTTAGTAGCGCGGCCTCG</td><td></td><td></td><td></td><td></td><td></td><td></td><td></td><td></td><td></td><td></td></tr>
<tr><td>Chromosome</td><td>start</td><td>end</td><td>strand</td><td>MM</td><td>target_seq</td><td>PAM</td><td>alignment</td><td>distance</td><td>position</td><td>gene name</td><td>gene id</td></tr>
<tr><td>GU305914.1</td><td>73842</td><td>73864</td><td>+</td><td>0</td><td>CGAGGCCGCGCTACTAACGT</td><td>TGG</td><td>||||||||[||||||||||||]PAM</td><td>NA</td><td>Intergenic</td><td>NA</td><td>NA</td></tr>
<tr><td>T5189</td><td>TCAGTGATAATAGTGTTGATCGG</td><td>1000</td><td></td><td></td><td></td><td></td><td></td><td></td><td></td><td></td><td></td></tr>
<tr><td>Oligo fwd</td><td>CACCTCAGTGATAATAGTGTTGAT</td><td></td><td></td><td></td><td></td><td></td><td></td><td></td><td></td><td></td><td></td></tr>
<tr><td>Oligo rev</td><td>AAACATCAACACTATTATCACTGA</td><td></td><td></td><td></td><td></td><td></td><td></td><td></td><td></td><td></td><td></td></tr>
<tr><td>Chromosome</td><td>start</td><td>end</td><td>strand</td><td>MM</td><td>target_seq</td><td>PAM</td><td>alignment</td><td>distance</td><td>position</td><td>gene name</td><td>gene id</td></tr>
<tr><td>GU305914.1</td><td>73933</td><td>73955</td><td>+</td><td>0</td><td>TCAGTGATAATAGTGTTGAT</td><td>CGG</td><td>||||||||[||||||||||||]PAM</td><td>NA</td><td>Intergenic</td><td>NA</td><td>NA</td></tr>
<tr><td>T5190</td><td>GATAATAGTGTTGATCGGACCGG</td><td>1000</td><td></td><td></td><td></td><td></td><td></td><td></td><td></td><td></td><td></td></tr>
<tr><td>Oligo fwd</td><td>CACCGATAATAGTGTTGATCGGAC</td><td></td><td></td><td></td><td></td><td></td><td></td><td></td><td></td><td></td><td></td></tr>
<tr><td>Oligo rev</td><td>AAACGTCCGATCAACACTATTATC</td><td></td><td></td><td></td><td></td><td></td><td></td><td></td><td></td><td></td><td></td></tr>
<tr><td>Chromosome</td><td>start</td><td>end</td><td>strand</td><td>MM</td><td>target_seq</td><td>PAM</td><td>alignment</td><td>distance</td><td>position</td><td>gene name</td><td>gene id</td></tr>
<tr><td>GU305914.1</td><td>73938</td><td>73960</td><td>+</td><td>0</td><td>GATAATAGTGTTGATCGGAC</td><td>CGG</td><td>||||||||[||||||||||||]PAM</td><td>NA</td><td>Intergenic</td><td>NA</td><td>NA</td></tr>
<tr><td>T5191</td><td>GTTGATCGGACCGGCAGCGCCGG</td><td>1000</td><td></td><td></td><td></td><td></td><td></td><td></td><td></td><td></td><td></td></tr>
<tr><td>Oligo fwd</td><td>CACCGTTGATCGGACCGGCAGCGC</td><td></td><td></td><td></td><td></td><td></td><td></td><td></td><td></td><td></td><td></td></tr>
<tr><td>Oligo rev</td><td>AAACGCGCTGCCGGTCCGATCAAC</td><td></td><td></td><td></td><td></td><td></td><td></td><td></td><td></td><td></td><td></td></tr>
<tr><td>Chromosome</td><td>start</td><td>end</td><td>strand</td><td>MM</td><td>target_seq</td><td>PAM</td><td>alignment</td><td>distance</td><td>position</td><td>gene name</td><td>gene id</td></tr>
<tr><td>GU305914.1</td><td>73947</td><td>73969</td><td>+</td><td>0</td><td>GTTGATCGGACCGGCAGCGC</td><td>CGG</td><td>||||||||[||||||||||||]PAM</td><td>NA</td><td>Intergenic</td><td>NA</td><td>NA</td></tr>
<tr><td>T5192</td><td>CCTACGCCTGATCTCCAGACAGG</td><td>1000</td><td></td><td></td><td></td><td></td><td></td><td></td><td></td><td></td><td></td></tr>
<tr><td>Oligo fwd</td><td>CACCCCTACGCCTGATCTCCAGAC</td><td></td><td></td><td></td><td></td><td></td><td></td><td></td><td></td><td></td><td></td></tr>
<tr><td>Oligo rev</td><td>AAACGTCTGGAGATCAGGCGTAGG</td><td></td><td></td><td></td><td></td><td></td><td></td><td></td><td></td><td></td><td></td></tr>
<tr><td>Chromosome</td><td>start</td><td>end</td><td>strand</td><td>MM</td><td>target_seq</td><td>PAM</td><td>alignment</td><td>distance</td><td>position</td><td>gene name</td><td>gene id</td></tr>
<tr><td>GU305914.1</td><td>73989</td><td>74011</td><td>+</td><td>0</td><td>CCTACGCCTGATCTCCAGAC</td><td>AGG</td><td>||||||||[||||||||||||]PAM</td><td>NA</td><td>Intergenic</td><td>NA</td><td>NA</td></tr>
<tr><td>T5193</td><td>CCAGACAGGAACAGCGAGAGTGG</td><td>1000</td><td></td><td></td><td></td><td></td><td></td><td></td><td></td><td></td><td></td></tr>
<tr><td>Oligo fwd</td><td>CACCCCAGACAGGAACAGCGAGAG</td><td></td><td></td><td></td><td></td><td></td><td></td><td></td><td></td><td></td><td></td></tr>
<tr><td>Oligo rev</td><td>AAACCTCTCGCTGTTCCTGTCTGG</td><td></td><td></td><td></td><td></td><td></td><td></td><td></td><td></td><td></td><td></td></tr>
<tr><td>Chromosome</td><td>start</td><td>end</td><td>strand</td><td>MM</td><td>target_seq</td><td>PAM</td><td>alignment</td><td>distance</td><td>position</td><td>gene name</td><td>gene id</td></tr>
<tr><td>GU305914.1</td><td>74003</td><td>74025</td><td>+</td><td>0</td><td>CCAGACAGGAACAGCGAGAG</td><td>TGG</td><td>||||||||[||||||||||||]PAM</td><td>NA</td><td>Intergenic</td><td>NA</td><td>NA</td></tr>
<tr><td>T5194</td><td>AGGAACAGCGAGAGTGGATCTGG</td><td>1000</td><td></td><td></td><td></td><td></td><td></td><td></td><td></td><td></td><td></td></tr>
<tr><td>Oligo fwd</td><td>CACCAGGAACAGCGAGAGTGGATC</td><td></td><td></td><td></td><td></td><td></td><td></td><td></td><td></td><td></td><td></td></tr>
<tr><td>Oligo rev</td><td>AAACGATCCACTCTCGCTGTTCCT</td><td></td><td></td><td></td><td></td><td></td><td></td><td></td><td></td><td></td><td></td></tr>
<tr><td>Chromosome</td><td>start</td><td>end</td><td>strand</td><td>MM</td><td>target_seq</td><td>PAM</td><td>alignment</td><td>distance</td><td>position</td><td>gene name</td><td>gene id</td></tr>
<tr><td>GU305914.1</td><td>74009</td><td>74031</td><td>+</td><td>0</td><td>AGGAACAGCGAGAGTGGATC</td><td>TGG</td><td>||||||||[||||||||||||]PAM</td><td>NA</td><td>Intergenic</td><td>NA</td><td>NA</td></tr>
<tr><td>T5195</td><td>GTGGATCTGGCAGCCGAGACAGG</td><td>1000</td><td></td><td></td><td></td><td></td><td></td><td></td><td></td><td></td><td></td></tr>
<tr><td>Oligo fwd</td><td>CACCGTGGATCTGGCAGCCGAGAC</td><td></td><td></td><td></td><td></td><td></td><td></td><td></td><td></td><td></td><td></td></tr>
<tr><td>Oligo rev</td><td>AAACGTCTCGGCTGCCAGATCCAC</td><td></td><td></td><td></td><td></td><td></td><td></td><td></td><td></td><td></td><td></td></tr>
<tr><td>Chromosome</td><td>start</td><td>end</td><td>strand</td><td>MM</td><td>target_seq</td><td>PAM</td><td>alignment</td><td>distance</td><td>position</td><td>gene name</td><td>gene id</td></tr>
<tr><td>GU305914.1</td><td>74022</td><td>74044</td><td>+</td><td>0</td><td>GTGGATCTGGCAGCCGAGAC</td><td>AGG</td><td>||||||||[||||||||||||]PAM</td><td>NA</td><td>Intergenic</td><td>NA</td><td>NA</td></tr>
<tr><td>T5196</td><td>CTGGCAGCCGAGACAGGCGTCGG</td><td>1000</td><td></td><td></td><td></td><td></td><td></td><td></td><td></td><td></td><td></td></tr>
<tr><td>Oligo fwd</td><td>CACCCTGGCAGCCGAGACAGGCGT</td><td></td><td></td><td></td><td></td><td></td><td></td><td></td><td></td><td></td><td></td></tr>
<tr><td>Oligo rev</td><td>AAACACGCCTGTCTCGGCTGCCAG</td><td></td><td></td><td></td><td></td><td></td><td></td><td></td><td></td><td></td><td></td></tr>
<tr><td>Chromosome</td><td>start</td><td>end</td><td>strand</td><td>MM</td><td>target_seq</td><td>PAM</td><td>alignment</td><td>distance</td><td>position</td><td>gene name</td><td>gene id</td></tr>
<tr><td>GU305914.1</td><td>74028</td><td>74050</td><td>+</td><td>0</td><td>CTGGCAGCCGAGACAGGCGT</td><td>CGG</td><td>||||||||[||||||||||||]PAM</td><td>NA</td><td>Intergenic</td><td>NA</td><td>NA</td></tr>
<tr><td>GU305914.1</td><td>128969</td><td>128991</td><td>+</td><td>3</td><td>CTGGCCGCCGAGGCAGCCGT</td><td>CGG</td><td>|||||-||[||||-|||-|||]PAM</td><td>NA</td><td>Intergenic</td><td>NA</td><td>NA</td></tr>
<tr><td>T5197</td><td>AGCCGAGACAGGCGTCGGCGAGG</td><td>1000</td><td></td><td></td><td></td><td></td><td></td><td></td><td></td><td></td><td></td></tr>
<tr><td>Oligo fwd</td><td>CACCAGCCGAGACAGGCGTCGGCG</td><td></td><td></td><td></td><td></td><td></td><td></td><td></td><td></td><td></td><td></td></tr>
<tr><td>Oligo rev</td><td>AAACCGCCGACGCCTGTCTCGGCT</td><td></td><td></td><td></td><td></td><td></td><td></td><td></td><td></td><td></td><td></td></tr>
<tr><td>Chromosome</td><td>start</td><td>end</td><td>strand</td><td>MM</td><td>target_seq</td><td>PAM</td><td>alignment</td><td>distance</td><td>position</td><td>gene name</td><td>gene id</td></tr>
<tr><td>GU305914.1</td><td>74033</td><td>74055</td><td>+</td><td>0</td><td>AGCCGAGACAGGCGTCGGCG</td><td>AGG</td><td>||||||||[||||||||||||]PAM</td><td>NA</td><td>Intergenic</td><td>NA</td><td>NA</td></tr>
<tr><td>T5198</td><td>GACAGGCGTCGGCGAGGTCGAGG</td><td>1000</td><td></td><td></td><td></td><td></td><td></td><td></td><td></td><td></td><td></td></tr>
<tr><td>Oligo fwd</td><td>CACCGACAGGCGTCGGCGAGGTCG</td><td></td><td></td><td></td><td></td><td></td><td></td><td></td><td></td><td></td><td></td></tr>
<tr><td>Oligo rev</td><td>AAACCGACCTCGCCGACGCCTGTC</td><td></td><td></td><td></td><td></td><td></td><td></td><td></td><td></td><td></td><td></td></tr>
<tr><td>Chromosome</td><td>start</td><td>end</td><td>strand</td><td>MM</td><td>target_seq</td><td>PAM</td><td>alignment</td><td>distance</td><td>position</td><td>gene name</td><td>gene id</td></tr>
<tr><td>GU305914.1</td><td>74039</td><td>74061</td><td>+</td><td>0</td><td>GACAGGCGTCGGCGAGGTCG</td><td>AGG</td><td>||||||||[||||||||||||]PAM</td><td>NA</td><td>Intergenic</td><td>NA</td><td>NA</td></tr>
<tr><td>T5199</td><td>CGGCGAGGTCGAGGTCGTCGCGG</td><td>1000</td><td></td><td></td><td></td><td></td><td></td><td></td><td></td><td></td><td></td></tr>
<tr><td>Oligo fwd</td><td>CACCCGGCGAGGTCGAGGTCGTCG</td><td></td><td></td><td></td><td></td><td></td><td></td><td></td><td></td><td></td><td></td></tr>
<tr><td>Oligo rev</td><td>AAACCGACGACCTCGACCTCGCCG</td><td></td><td></td><td></td><td></td><td></td><td></td><td></td><td></td><td></td><td></td></tr>
<tr><td>Chromosome</td><td>start</td><td>end</td><td>strand</td><td>MM</td><td>target_seq</td><td>PAM</td><td>alignment</td><td>distance</td><td>position</td><td>gene name</td><td>gene id</td></tr>
<tr><td>GU305914.1</td><td>74048</td><td>74070</td><td>+</td><td>0</td><td>CGGCGAGGTCGAGGTCGTCG</td><td>CGG</td><td>||||||||[||||||||||||]PAM</td><td>NA</td><td>Intergenic</td><td>NA</td><td>NA</td></tr>
<tr><td>GU305914.1</td><td>61377</td><td>61399</td><td>-</td><td>4</td><td>CGTCGAGTGCGAGGTCGACG</td><td>GGG</td><td>||-||||-[-||||||||-||]PAM</td><td>NA</td><td>Intergenic</td><td>NA</td><td>NA</td></tr>
</table>

<table>
<tr><td>T5200</td><td>GGTCGAGGTCGTCGCGGTCCAGG</td><td>1000</td><td></td><td></td><td></td><td></td><td></td><td></td><td></td><td></td><td></td></tr>
<tr><td>Oligo fwd</td><td>CACCGGTCGAGGTCGTCGCGGTCC</td><td></td><td></td><td></td><td></td><td></td><td></td><td></td><td></td><td></td><td></td></tr>
<tr><td>Oligo rev</td><td>AAACGGACCGCGACGACCTCGACC</td><td></td><td></td><td></td><td></td><td></td><td></td><td></td><td></td><td></td><td></td></tr>
<tr><td>Chromosome</td><td>start</td><td>end</td><td>strand</td><td>MM</td><td>target_seq</td><td>PAM</td><td>alignment</td><td>distance</td><td>position</td><td>gene name</td><td>gene id</td></tr>
<tr><td>GU305914.1</td><td>74054</td><td>74076</td><td>+</td><td>0</td><td>GGTCGAGGTCGTCGCGGTCC</td><td>AGG</td><td>||||||||[||||||||||||]PAM</td><td>NA</td><td>Intergenic</td><td>NA</td><td>NA</td></tr>
<tr><td>T5201</td><td>GAGGTCGTCGCGGTCCAGGCAGG</td><td>1000</td><td></td><td></td><td></td><td></td><td></td><td></td><td></td><td></td><td></td></tr>
<tr><td>Oligo fwd</td><td>CACCGAGGTCGTCGCGGTCCAGGC</td><td></td><td></td><td></td><td></td><td></td><td></td><td></td><td></td><td></td><td></td></tr>
<tr><td>Oligo rev</td><td>AAACGCCTGGACCGCGACGACCTC</td><td></td><td></td><td></td><td></td><td></td><td></td><td></td><td></td><td></td><td></td></tr>
<tr><td>Chromosome</td><td>start</td><td>end</td><td>strand</td><td>MM</td><td>target_seq</td><td>PAM</td><td>alignment</td><td>distance</td><td>position</td><td>gene name</td><td>gene id</td></tr>
<tr><td>GU305914.1</td><td>74058</td><td>74080</td><td>+</td><td>0</td><td>GAGGTCGTCGCGGTCCAGGC</td><td>AGG</td><td>||||||||[||||||||||||]PAM</td><td>NA</td><td>Intergenic</td><td>NA</td><td>NA</td></tr>
<tr><td>T5202</td><td>CTCCCTGTAGACGTTCTGACAGG</td><td>1000</td><td></td><td></td><td></td><td></td><td></td><td></td><td></td><td></td><td></td></tr>
<tr><td>Oligo fwd</td><td>CACCCTCCCTGTAGACGTTCTGAC</td><td></td><td></td><td></td><td></td><td></td><td></td><td></td><td></td><td></td><td></td></tr>
<tr><td>Oligo rev</td><td>AAACGTCAGAACGTCTACAGGGAG</td><td></td><td></td><td></td><td></td><td></td><td></td><td></td><td></td><td></td><td></td></tr>
<tr><td>Chromosome</td><td>start</td><td>end</td><td>strand</td><td>MM</td><td>target_seq</td><td>PAM</td><td>alignment</td><td>distance</td><td>position</td><td>gene name</td><td>gene id</td></tr>
<tr><td>GU305914.1</td><td>74082</td><td>74104</td><td>+</td><td>0</td><td>CTCCCTGTAGACGTTCTGAC</td><td>AGG</td><td>||||||||[||||||||||||]PAM</td><td>NA</td><td>Intergenic</td><td>NA</td><td>NA</td></tr>
<tr><td>T5203</td><td>TGTAGACGTTCTGACAGGACTGG</td><td>1000</td><td></td><td></td><td></td><td></td><td></td><td></td><td></td><td></td><td></td></tr>
<tr><td>Oligo fwd</td><td>CACCTGTAGACGTTCTGACAGGAC</td><td></td><td></td><td></td><td></td><td></td><td></td><td></td><td></td><td></td><td></td></tr>
<tr><td>Oligo rev</td><td>AAACGTCCTGTCAGAACGTCTACA</td><td></td><td></td><td></td><td></td><td></td><td></td><td></td><td></td><td></td><td></td></tr>
<tr><td>Chromosome</td><td>start</td><td>end</td><td>strand</td><td>MM</td><td>target_seq</td><td>PAM</td><td>alignment</td><td>distance</td><td>position</td><td>gene name</td><td>gene id</td></tr>
<tr><td>GU305914.1</td><td>74087</td><td>74109</td><td>+</td><td>0</td><td>TGTAGACGTTCTGACAGGAC</td><td>TGG</td><td>||||||||[||||||||||||]PAM</td><td>NA</td><td>Intergenic</td><td>NA</td><td>NA</td></tr>
<tr><td>T5204</td><td>GGACTGGCAGAAGAACTCGCCGG</td><td>1000</td><td></td><td></td><td></td><td></td><td></td><td></td><td></td><td></td><td></td></tr>
<tr><td>Oligo fwd</td><td>CACCGGACTGGCAGAAGAACTCGC</td><td></td><td></td><td></td><td></td><td></td><td></td><td></td><td></td><td></td><td></td></tr>
<tr><td>Oligo rev</td><td>AAACGCGAGTTCTTCTGCCAGTCC</td><td></td><td></td><td></td><td></td><td></td><td></td><td></td><td></td><td></td><td></td></tr>
<tr><td>Chromosome</td><td>start</td><td>end</td><td>strand</td><td>MM</td><td>target_seq</td><td>PAM</td><td>alignment</td><td>distance</td><td>position</td><td>gene name</td><td>gene id</td></tr>
<tr><td>GU305914.1</td><td>74103</td><td>74125</td><td>+</td><td>0</td><td>GGACTGGCAGAAGAACTCGC</td><td>CGG</td><td>||||||||[||||||||||||]PAM</td><td>NA</td><td>Intergenic</td><td>NA</td><td>NA</td></tr>
<tr><td>T5205</td><td>ACTCGCCGGAGTGCGAGACGAGG</td><td>1000</td><td></td><td></td><td></td><td></td><td></td><td></td><td></td><td></td><td></td></tr>
<tr><td>Oligo fwd</td><td>CACCACTCGCCGGAGTGCGAGACG</td><td></td><td></td><td></td><td></td><td></td><td></td><td></td><td></td><td></td><td></td></tr>
<tr><td>Oligo rev</td><td>AAACCGTCTCGCACTCCGGCGAGT</td><td></td><td></td><td></td><td></td><td></td><td></td><td></td><td></td><td></td><td></td></tr>
<tr><td>Chromosome</td><td>start</td><td>end</td><td>strand</td><td>MM</td><td>target_seq</td><td>PAM</td><td>alignment</td><td>distance</td><td>position</td><td>gene name</td><td>gene id</td></tr>
<tr><td>GU305914.1</td><td>74117</td><td>74139</td><td>+</td><td>0</td><td>ACTCGCCGGAGTGCGAGACG</td><td>AGG</td><td>||||||||[||||||||||||]PAM</td><td>NA</td><td>Intergenic</td><td>NA</td><td>NA</td></tr>
<tr><td>T5206</td><td>CGCCGGAGTGCGAGACGAGGCGG</td><td>1000</td><td></td><td></td><td></td><td></td><td></td><td></td><td></td><td></td><td></td></tr>
<tr><td>Oligo fwd</td><td>CACCCGCCGGAGTGCGAGACGAGG</td><td></td><td></td><td></td><td></td><td></td><td></td><td></td><td></td><td></td><td></td></tr>
<tr><td>Oligo rev</td><td>AAACCCTCGTCTCGCACTCCGGCG</td><td></td><td></td><td></td><td></td><td></td><td></td><td></td><td></td><td></td><td></td></tr>
<tr><td>Chromosome</td><td>start</td><td>end</td><td>strand</td><td>MM</td><td>target_seq</td><td>PAM</td><td>alignment</td><td>distance</td><td>position</td><td>gene name</td><td>gene id</td></tr>
<tr><td>GU305914.1</td><td>74120</td><td>74142</td><td>+</td><td>0</td><td>CGCCGGAGTGCGAGACGAGG</td><td>CGG</td><td>||||||||[||||||||||||]PAM</td><td>NA</td><td>Intergenic</td><td>NA</td><td>NA</td></tr>
<tr><td>T5207</td><td>CGGAGTGCGAGACGAGGCGGAGG</td><td>1000</td><td></td><td></td><td></td><td></td><td></td><td></td><td></td><td></td><td></td></tr>
<tr><td>Oligo fwd</td><td>CACCCGGAGTGCGAGACGAGGCGG</td><td></td><td></td><td></td><td></td><td></td><td></td><td></td><td></td><td></td><td></td></tr>
<tr><td>Oligo rev</td><td>AAACCCGCCTCGTCTCGCACTCCG</td><td></td><td></td><td></td><td></td><td></td><td></td><td></td><td></td><td></td><td></td></tr>
<tr><td>Chromosome</td><td>start</td><td>end</td><td>strand</td><td>MM</td><td>target_seq</td><td>PAM</td><td>alignment</td><td>distance</td><td>position</td><td>gene name</td><td>gene id</td></tr>
<tr><td>GU305914.1</td><td>74123</td><td>74145</td><td>+</td><td>0</td><td>CGGAGTGCGAGACGAGGCGG</td><td>AGG</td><td>||||||||[||||||||||||]PAM</td><td>NA</td><td>Intergenic</td><td>NA</td><td>NA</td></tr>
<tr><td>T5208</td><td>GAGGCGGAGGAGCTTCTCGACGG</td><td>1000</td><td></td><td></td><td></td><td></td><td></td><td></td><td></td><td></td><td></td></tr>
<tr><td>Oligo fwd</td><td>CACCGAGGCGGAGGAGCTTCTCGA</td><td></td><td></td><td></td><td></td><td></td><td></td><td></td><td></td><td></td><td></td></tr>
<tr><td>Oligo rev</td><td>AAACTCGAGAAGCTCCTCCGCCTC</td><td></td><td></td><td></td><td></td><td></td><td></td><td></td><td></td><td></td><td></td></tr>
<tr><td>Chromosome</td><td>start</td><td>end</td><td>strand</td><td>MM</td><td>target_seq</td><td>PAM</td><td>alignment</td><td>distance</td><td>position</td><td>gene name</td><td>gene id</td></tr>
<tr><td>GU305914.1</td><td>74136</td><td>74158</td><td>+</td><td>0</td><td>GAGGCGGAGGAGCTTCTCGA</td><td>CGG</td><td>||||||||[||||||||||||]PAM</td><td>NA</td><td>Intergenic</td><td>NA</td><td>NA</td></tr>
<tr><td>T5209</td><td>GGATGACTCGCGCAGCACGACGG</td><td>1000</td><td></td><td></td><td></td><td></td><td></td><td></td><td></td><td></td><td></td></tr>
<tr><td>Oligo fwd</td><td>CACCGGATGACTCGCGCAGCACGA</td><td></td><td></td><td></td><td></td><td></td><td></td><td></td><td></td><td></td><td></td></tr>
<tr><td>Oligo rev</td><td>AAACTCGTGCTGCGCGAGTCATCC</td><td></td><td></td><td></td><td></td><td></td><td></td><td></td><td></td><td></td><td></td></tr>
<tr><td>Chromosome</td><td>start</td><td>end</td><td>strand</td><td>MM</td><td>target_seq</td><td>PAM</td><td>alignment</td><td>distance</td><td>position</td><td>gene name</td><td>gene id</td></tr>
<tr><td>GU305914.1</td><td>74157</td><td>74179</td><td>+</td><td>0</td><td>GGATGACTCGCGCAGCACGA</td><td>CGG</td><td>||||||||[||||||||||||]PAM</td><td>NA</td><td>Intergenic</td><td>NA</td><td>NA</td></tr>
<tr><td>T5210</td><td>CAGCACGACGGTGCCGAAACAGG</td><td>1000</td><td></td><td></td><td></td><td></td><td></td><td></td><td></td><td></td><td></td></tr>
<tr><td>Oligo fwd</td><td>CACCCAGCACGACGGTGCCGAAAC</td><td></td><td></td><td></td><td></td><td></td><td></td><td></td><td></td><td></td><td></td></tr>
<tr><td>Oligo rev</td><td>AAACGTTTCGGCACCGTCGTGCTG</td><td></td><td></td><td></td><td></td><td></td><td></td><td></td><td></td><td></td><td></td></tr>
<tr><td>Chromosome</td><td>start</td><td>end</td><td>strand</td><td>MM</td><td>target_seq</td><td>PAM</td><td>alignment</td><td>distance</td><td>position</td><td>gene name</td><td>gene id</td></tr>
<tr><td>GU305914.1</td><td>74169</td><td>74191</td><td>+</td><td>0</td><td>CAGCACGACGGTGCCGAAAC</td><td>AGG</td><td>||||||||[||||||||||||]PAM</td><td>NA</td><td>Intergenic</td><td>NA</td><td>NA</td></tr>
<tr><td>T5211</td><td>GGTGCCGAAACAGGTCCTAGAGG</td><td>1000</td><td></td><td></td><td></td><td></td><td></td><td></td><td></td><td></td><td></td></tr>
<tr><td>Oligo fwd</td><td>CACCGGTGCCGAAACAGGTCCTAG</td><td></td><td></td><td></td><td></td><td></td><td></td><td></td><td></td><td></td><td></td></tr>
<tr><td>Oligo rev</td><td>AAACCTAGGACCTGTTTCGGCACC</td><td></td><td></td><td></td><td></td><td></td><td></td><td></td><td></td><td></td><td></td></tr>
<tr><td>Chromosome</td><td>start</td><td>end</td><td>strand</td><td>MM</td><td>target_seq</td><td>PAM</td><td>alignment</td><td>distance</td><td>position</td><td>gene name</td><td>gene id</td></tr>
<tr><td>GU305914.1</td><td>74178</td><td>74200</td><td>+</td><td>0</td><td>GGTGCCGAAACAGGTCCTAG</td><td>AGG</td><td>||||||||[||||||||||||]PAM</td><td>NA</td><td>Intergenic</td><td>NA</td><td>NA</td></tr>
<tr><td>T5212</td><td>GAAACAGGTCCTAGAGGAGCAGG</td><td>1000</td><td></td><td></td><td></td><td></td><td></td><td></td><td></td><td></td><td></td></tr>
<tr><td>Oligo fwd</td><td>CACCGAAACAGGTCCTAGAGGAGC</td><td></td><td></td><td></td><td></td><td></td><td></td><td></td><td></td><td></td><td></td></tr>
<tr><td>Oligo rev</td><td>AAACGCTCCTCTAGGACCTGTTTC</td><td></td><td></td><td></td><td></td><td></td><td></td><td></td><td></td><td></td><td></td></tr>
<tr><td>Chromosome</td><td>start</td><td>end</td><td>strand</td><td>MM</td><td>target_seq</td><td>PAM</td><td>alignment</td><td>distance</td><td>position</td><td>gene name</td><td>gene id</td></tr>
<tr><td>GU305914.1</td><td>74184</td><td>74206</td><td>+</td><td>0</td><td>GAAACAGGTCCTAGAGGAGC</td><td>AGG</td><td>||||||||[||||||||||||]PAM</td><td>NA</td><td>Intergenic</td><td>NA</td><td>NA</td></tr>
<tr><td>T5213</td><td>AAACAGGTCCTAGAGGAGCAGGG</td><td>1000</td><td></td><td></td><td></td><td></td><td></td><td></td><td></td><td></td><td></td></tr>
<tr><td>Oligo fwd</td><td>CACCAAACAGGTCCTAGAGGAGCA</td><td></td><td></td><td></td><td></td><td></td><td></td><td></td><td></td><td></td><td></td></tr>
<tr><td>Oligo rev</td><td>AAACTGCTCCTCTAGGACCTGTTT</td><td></td><td></td><td></td><td></td><td></td><td></td><td></td><td></td><td></td><td></td></tr>
<tr><td>Chromosome</td><td>start</td><td>end</td><td>strand</td><td>MM</td><td>target_seq</td><td>PAM</td><td>alignment</td><td>distance</td><td>position</td><td>gene name</td><td>gene id</td></tr>
<tr><td>GU305914.1</td><td>74185</td><td>74207</td><td>+</td><td>0</td><td>AAACAGGTCCTAGAGGAGCA</td><td>GGG</td><td>||||||||[||||||||||||]PAM</td><td>NA</td><td>Intergenic</td><td>NA</td><td>NA</td></tr>
<tr><td>T5214</td><td>TCCTAGAGGAGCAGGGCACGAGG</td><td>1000</td><td></td><td></td><td></td><td></td><td></td><td></td><td></td><td></td><td></td></tr>
<tr><td>Oligo fwd</td><td>CACCTCCTAGAGGAGCAGGGCACG</td><td></td><td></td><td></td><td></td><td></td><td></td><td></td><td></td><td></td><td></td></tr>
<tr><td>Oligo rev</td><td>AAACCGTGCCCTGCTCCTCTAGGA</td><td></td><td></td><td></td><td></td><td></td><td></td><td></td><td></td><td></td><td></td></tr>
<tr><td>Chromosome</td><td>start</td><td>end</td><td>strand</td><td>MM</td><td>target_seq</td><td>PAM</td><td>alignment</td><td>distance</td><td>position</td><td>gene name</td><td>gene id</td></tr>
</table>

GU305914.1 74192 74214 + 0 TCCTAGAGGAGCAGGGCACG AGG ||||||||[||||||||||||]PAM NA Intergenic NA NA

T5215 AGCAGGGCACGAGGACGTCGAGG 1000
Oligo fwd CACCAGCAGGGCACGAGGACGTCG
Oligo rev AAACCGACGTCCTCGTGCCCTGCT
Chromosome start end strand MM target_seq PAM alignment distance position gene name gene id
GU305914.1 74201 74223 + 0 AGCAGGGCACGAGGACGTCG AGG ||||||||[||||||||||||]PAM NA Intergenic NA NA

T5216 AGGGCACGAGGACGTCGAGGCGG 1000
Oligo fwd CACCAGGGCACGAGGACGTCGAGG
Oligo rev AAACCCTCGACGTCCTCGTGCCCT
Chromosome start end strand MM target_seq PAM alignment distance position gene name gene id
GU305914.1 74204 74226 + 0 AGGGCACGAGGACGTCGAGG CGG ||||||||[||||||||||||]PAM NA Intergenic NA NA

T5217 GGGCACGAGGACGTCGAGGCGGG 1000
Oligo fwd CACCGGGCACGAGGACGTCGAGGC
Oligo rev AAACGCCTCGACGTCCTCGTGCCC
Chromosome start end strand MM target_seq PAM alignment distance position gene name gene id
GU305914.1 74205 74227 + 0 GGGCACGAGGACGTCGAGGC GGG ||||||||[||||||||||||]PAM NA Intergenic NA NA

T5218 AGGACGTCGAGGCGGGTGCCCGG 1000
Oligo fwd CACCAGGACGTCGAGGCGGGTGCC
Oligo rev AAACGGCACCCGCCTCGACGTCCT
Chromosome start end strand MM target_seq PAM alignment distance position gene name gene id
GU305914.1 74212 74234 + 0 AGGACGTCGAGGCGGGTGCC CGG ||||||||[||||||||||||]PAM NA Intergenic NA NA

T5219 GGACGTCGAGGCGGGTGCCCGGG 1000
Oligo fwd CACCGGACGTCGAGGCGGGTGCCC
Oligo rev AAACGGGCACCCGCCTCGACGTCC
Chromosome start end strand MM target_seq PAM alignment distance position gene name gene id
GU305914.1 74213 74235 + 0 GGACGTCGAGGCGGGTGCCC GGG ||||||||[||||||||||||]PAM NA Intergenic NA NA

T5220 GCCCGGGCCGAGAACCCCGCAGG 1000
Oligo fwd CACCGCCCGGGCCGAGAACCCCGC
Oligo rev AAACGCGGGGTTCTCGGCCCGGGC
Chromosome start end strand MM target_seq PAM alignment distance position gene name gene id
GU305914.1 74229 74251 + 0 GCCCGGGCCGAGAACCCCGC AGG ||||||||[||||||||||||]PAM NA Intergenic NA NA

T5221 CGGGCCGAGAACCCCGCAGGCGG 1000
Oligo fwd CACCCGGGCCGAGAACCCCGCAGG
Oligo rev AAACCCTGCGGGGTTCTCGGCCCG
Chromosome start end strand MM target_seq PAM alignment distance position gene name gene id
GU305914.1 74232 74254 + 0 CGGGCCGAGAACCCCGCAGG CGG ||||||||[||||||||||||]PAM NA Intergenic NA NA

T5222 TCGCGAACCTGAGCCCGTGATGG 1000
Oligo fwd CACCTCGCGAACCTGAGCCCGTGA
Oligo rev AAACTCACGGGCTCAGGTTCGCGA
Chromosome start end strand MM target_seq PAM alignment distance position gene name gene id
GU305914.1 74288 74310 + 0 TCGCGAACCTGAGCCCGTGA TGG ||||||||[||||||||||||]PAM NA Intergenic NA NA

T5223 CCTGAGCCCGTGATGGTAATCGG 1000
Oligo fwd CACCCCTGAGCCCGTGATGGTAAT
Oligo rev AAACATTACCATCACGGGCTCAGG
Chromosome start end strand MM target_seq PAM alignment distance position gene name gene id
GU305914.1 74295 74317 + 0 CCTGAGCCCGTGATGGTAAT CGG ||||||||[||||||||||||]PAM NA Intergenic NA NA

T5224 TGACCTCGTAGATCCGCTCTCGG 1000
Oligo fwd CACCTGACCTCGTAGATCCGCTCT
Oligo rev AAACAGAGCGGATCTACGAGGTCA
Chromosome start end strand MM target_seq PAM alignment distance position gene name gene id
GU305914.1 74324 74346 + 0 TGACCTCGTAGATCCGCTCT CGG ||||||||[||||||||||||]PAM NA Intergenic NA NA

T5225 TCCGCTCTCGGCTTAAATACTGG 1000
Oligo fwd CACCTCCGCTCTCGGCTTAAATAC
Oligo rev AAACGTATTTAAGCCGAGAGCGGA
Chromosome start end strand MM target_seq PAM alignment distance position gene name gene id
GU305914.1 74336 74358 + 0 TCCGCTCTCGGCTTAAATAC TGG ||||||||[||||||||||||]PAM NA Intergenic NA NA

T5226 TTAAATACTGGTTCCCACACAGG 1000
Oligo fwd CACCTTAAATACTGGTTCCCACAC
Oligo rev AAACGTGTGGGAACCAGTATTTAA
Chromosome start end strand MM target_seq PAM alignment distance position gene name gene id
GU305914.1 74348 74370 + 0 TTAAATACTGGTTCCCACAC AGG ||||||||[||||||||||||]PAM NA Intergenic NA NA

T5227 TAAATACTGGTTCCCACACAGGG 1000
Oligo fwd CACCTAAATACTGGTTCCCACACA
Oligo rev AAACTGTGTGGGAACCAGTATTTA
Chromosome start end strand MM target_seq PAM alignment distance position gene name gene id
GU305914.1 74349 74371 + 0 TAAATACTGGTTCCCACACA GGG ||||||||[||||||||||||]PAM NA Intergenic NA NA

T5228 CCCACACAGGGAACAGTGAGCGG 1000
Oligo fwd CACCCCCACACAGGGAACAGTGAG
Oligo rev AAACCTCACTGTTCCCTGTGTGGG
Chromosome start end strand MM target_seq PAM alignment distance position gene name gene id
GU305914.1 74361 74383 + 0 CCCACACAGGGAACAGTGAG CGG ||||||||[||||||||||||]PAM NA Intergenic NA NA

T5229 AACAGTGAGCGGTGTCGTTGTGG 1000
Oligo fwd CACCAACAGTGAGCGGTGTCGTTG
Oligo rev AAACCAACGACACCGCTCACTGTT

<table>
<tr><td>Chromosome</td><td>start</td><td>end</td><td>strand</td><td>MM</td><td>target_seq</td><td>PAM</td><td>alignment</td><td>distance</td><td>position</td><td>gene name</td><td>gene id</td></tr>
<tr><td>GU305914.1</td><td>74372</td><td>74394</td><td>+</td><td>0</td><td>AACAGTGAGCGGTGTCGTTG</td><td>TGG</td><td>||||||||[||||||||||||]PAM</td><td>NA</td><td>Intergenic</td><td>NA</td><td>NA</td></tr>
<tr><td></td><td></td><td></td><td></td><td></td><td></td><td></td><td></td><td></td><td></td><td></td><td></td></tr>
<tr><td>T5230</td><td>ACAGTGAGCGGTGTCGTTGTGGG</td><td>1000</td><td></td><td></td><td></td><td></td><td></td><td></td><td></td><td></td><td></td></tr>
<tr><td>Oligo fwd</td><td>CACCACAGTGAGCGGTGTCGTTGT</td><td></td><td></td><td></td><td></td><td></td><td></td><td></td><td></td><td></td><td></td></tr>
<tr><td>Oligo rev</td><td>AAACACAACGACACCGCTCACTGT</td><td></td><td></td><td></td><td></td><td></td><td></td><td></td><td></td><td></td><td></td></tr>
<tr><td>Chromosome</td><td>start</td><td>end</td><td>strand</td><td>MM</td><td>target_seq</td><td>PAM</td><td>alignment</td><td>distance</td><td>position</td><td>gene name</td><td>gene id</td></tr>
<tr><td>GU305914.1</td><td>74373</td><td>74395</td><td>+</td><td>0</td><td>ACAGTGAGCGGTGTCGTTGT</td><td>GGG</td><td>||||||||[||||||||||||]PAM</td><td>NA</td><td>Intergenic</td><td>NA</td><td>NA</td></tr>
<tr><td></td><td></td><td></td><td></td><td></td><td></td><td></td><td></td><td></td><td></td><td></td><td></td></tr>
<tr><td>T5231</td><td>TGTAGATGACACTCTTCTCCTGG</td><td>1000</td><td></td><td></td><td></td><td></td><td></td><td></td><td></td><td></td><td></td></tr>
<tr><td>Oligo fwd</td><td>CACCTGTAGATGACACTCTTCTCC</td><td></td><td></td><td></td><td></td><td></td><td></td><td></td><td></td><td></td><td></td></tr>
<tr><td>Oligo rev</td><td>AAACGGAGAAGAGTGTCATCTACA</td><td></td><td></td><td></td><td></td><td></td><td></td><td></td><td></td><td></td><td></td></tr>
<tr><td>Chromosome</td><td>start</td><td>end</td><td>strand</td><td>MM</td><td>target_seq</td><td>PAM</td><td>alignment</td><td>distance</td><td>position</td><td>gene name</td><td>gene id</td></tr>
<tr><td>GU305914.1</td><td>74399</td><td>74421</td><td>+</td><td>0</td><td>TGTAGATGACACTCTTCTCC</td><td>TGG</td><td>||||||||[||||||||||||]PAM</td><td>NA</td><td>Intergenic</td><td>NA</td><td>NA</td></tr>
<tr><td>GU305914.1</td><td>10666</td><td>10688</td><td>+</td><td>4</td><td>GGTAGATCACACTCTGCTCA</td><td>AGG</td><td>-||||||-[|||||||-|||-]PAM</td><td>NA</td><td>Intergenic</td><td>NA</td><td>NA</td></tr>
<tr><td></td><td></td><td></td><td></td><td></td><td></td><td></td><td></td><td></td><td></td><td></td><td></td></tr>
<tr><td>T5232</td><td>AGATGACACTCTTCTCCTGGCGG</td><td>1000</td><td></td><td></td><td></td><td></td><td></td><td></td><td></td><td></td><td></td></tr>
<tr><td>Oligo fwd</td><td>CACCAGATGACACTCTTCTCCTGG</td><td></td><td></td><td></td><td></td><td></td><td></td><td></td><td></td><td></td><td></td></tr>
<tr><td>Oligo rev</td><td>AAACCCAGGAGAAGAGTGTCATCT</td><td></td><td></td><td></td><td></td><td></td><td></td><td></td><td></td><td></td><td></td></tr>
<tr><td>Chromosome</td><td>start</td><td>end</td><td>strand</td><td>MM</td><td>target_seq</td><td>PAM</td><td>alignment</td><td>distance</td><td>position</td><td>gene name</td><td>gene id</td></tr>
<tr><td>GU305914.1</td><td>74402</td><td>74424</td><td>+</td><td>0</td><td>AGATGACACTCTTCTCCTGG</td><td>CGG</td><td>||||||||[||||||||||||]PAM</td><td>NA</td><td>Intergenic</td><td>NA</td><td>NA</td></tr>
<tr><td></td><td></td><td></td><td></td><td></td><td></td><td></td><td></td><td></td><td></td><td></td><td></td></tr>
<tr><td>T5233</td><td>AGTGTCCGCAGTTGTCACAGAGG</td><td>1000</td><td></td><td></td><td></td><td></td><td></td><td></td><td></td><td></td><td></td></tr>
<tr><td>Oligo fwd</td><td>CACCAGTGTCCGCAGTTGTCACAG</td><td></td><td></td><td></td><td></td><td></td><td></td><td></td><td></td><td></td><td></td></tr>
<tr><td>Oligo rev</td><td>AAACCTGTGACAACTGCGGACACT</td><td></td><td></td><td></td><td></td><td></td><td></td><td></td><td></td><td></td><td></td></tr>
<tr><td>Chromosome</td><td>start</td><td>end</td><td>strand</td><td>MM</td><td>target_seq</td><td>PAM</td><td>alignment</td><td>distance</td><td>position</td><td>gene name</td><td>gene id</td></tr>
<tr><td>GU305914.1</td><td>74498</td><td>74520</td><td>+</td><td>0</td><td>AGTGTCCGCAGTTGTCACAG</td><td>AGG</td><td>||||||||[||||||||||||]PAM</td><td>NA</td><td>Intergenic</td><td>NA</td><td>NA</td></tr>
<tr><td></td><td></td><td></td><td></td><td></td><td></td><td></td><td></td><td></td><td></td><td></td><td></td></tr>
<tr><td>T5234</td><td>TCCGCAGTTGTCACAGAGGATGG</td><td>1000</td><td></td><td></td><td></td><td></td><td></td><td></td><td></td><td></td><td></td></tr>
<tr><td>Oligo fwd</td><td>CACCTCCGCAGTTGTCACAGAGGA</td><td></td><td></td><td></td><td></td><td></td><td></td><td></td><td></td><td></td><td></td></tr>
<tr><td>Oligo rev</td><td>AAACTCCTCTGTGACAACTGCGGA</td><td></td><td></td><td></td><td></td><td></td><td></td><td></td><td></td><td></td><td></td></tr>
<tr><td>Chromosome</td><td>start</td><td>end</td><td>strand</td><td>MM</td><td>target_seq</td><td>PAM</td><td>alignment</td><td>distance</td><td>position</td><td>gene name</td><td>gene id</td></tr>
<tr><td>GU305914.1</td><td>74502</td><td>74524</td><td>+</td><td>0</td><td>TCCGCAGTTGTCACAGAGGA</td><td>TGG</td><td>||||||||[||||||||||||]PAM</td><td>NA</td><td>Intergenic</td><td>NA</td><td>NA</td></tr>
<tr><td></td><td></td><td></td><td></td><td></td><td></td><td></td><td></td><td></td><td></td><td></td><td></td></tr>
<tr><td>T5235</td><td>GGCCTCGAGCACGCCGTCCTCGG</td><td>1000</td><td></td><td></td><td></td><td></td><td></td><td></td><td></td><td></td><td></td></tr>
<tr><td>Oligo fwd</td><td>CACCGGCCTCGAGCACGCCGTCCT</td><td></td><td></td><td></td><td></td><td></td><td></td><td></td><td></td><td></td><td></td></tr>
<tr><td>Oligo rev</td><td>AAACAGGACGGCGTGCTCGAGGCC</td><td></td><td></td><td></td><td></td><td></td><td></td><td></td><td></td><td></td><td></td></tr>
<tr><td>Chromosome</td><td>start</td><td>end</td><td>strand</td><td>MM</td><td>target_seq</td><td>PAM</td><td>alignment</td><td>distance</td><td>position</td><td>gene name</td><td>gene id</td></tr>
<tr><td>GU305914.1</td><td>74523</td><td>74545</td><td>+</td><td>0</td><td>GGCCTCGAGCACGCCGTCCT</td><td>CGG</td><td>||||||||[||||||||||||]PAM</td><td>NA</td><td>Intergenic</td><td>NA</td><td>NA</td></tr>
<tr><td></td><td></td><td></td><td></td><td></td><td></td><td></td><td></td><td></td><td></td><td></td><td></td></tr>
<tr><td>T5236</td><td>CTCGGCGTAGCGCTGACACTTGG</td><td>1000</td><td></td><td></td><td></td><td></td><td></td><td></td><td></td><td></td><td></td></tr>
<tr><td>Oligo fwd</td><td>CACCCTCGGCGTAGCGCTGACACT</td><td></td><td></td><td></td><td></td><td></td><td></td><td></td><td></td><td></td><td></td></tr>
<tr><td>Oligo rev</td><td>AAACAGTGTCAGCGCTACGCCGAG</td><td></td><td></td><td></td><td></td><td></td><td></td><td></td><td></td><td></td><td></td></tr>
<tr><td>Chromosome</td><td>start</td><td>end</td><td>strand</td><td>MM</td><td>target_seq</td><td>PAM</td><td>alignment</td><td>distance</td><td>position</td><td>gene name</td><td>gene id</td></tr>
<tr><td>GU305914.1</td><td>74541</td><td>74563</td><td>+</td><td>0</td><td>CTCGGCGTAGCGCTGACACT</td><td>TGG</td><td>||||||||[||||||||||||]PAM</td><td>NA</td><td>Intergenic</td><td>NA</td><td>NA</td></tr>
<tr><td></td><td></td><td></td><td></td><td></td><td></td><td></td><td></td><td></td><td></td><td></td><td></td></tr>
<tr><td>T5237</td><td>GGTCTTGCAGAAGCACCACACGG</td><td>1000</td><td></td><td></td><td></td><td></td><td></td><td></td><td></td><td></td><td></td></tr>
<tr><td>Oligo fwd</td><td>CACCGGTCTTGCAGAAGCACCACA</td><td></td><td></td><td></td><td></td><td></td><td></td><td></td><td></td><td></td><td></td></tr>
<tr><td>Oligo rev</td><td>AAACTGTGGTGCTTCTGCAAGACC</td><td></td><td></td><td></td><td></td><td></td><td></td><td></td><td></td><td></td><td></td></tr>
<tr><td>Chromosome</td><td>start</td><td>end</td><td>strand</td><td>MM</td><td>target_seq</td><td>PAM</td><td>alignment</td><td>distance</td><td>position</td><td>gene name</td><td>gene id</td></tr>
<tr><td>GU305914.1</td><td>74562</td><td>74584</td><td>+</td><td>0</td><td>GGTCTTGCAGAAGCACCACA</td><td>CGG</td><td>||||||||[||||||||||||]PAM</td><td>NA</td><td>Intergenic</td><td>NA</td><td>NA</td></tr>
<tr><td>GU305914.1</td><td>153359</td><td>153381</td><td>+</td><td>4</td><td>GATCATGCAGAAGCACACCA</td><td>AGG</td><td>|-||-|||[||||||||--||]PAM</td><td>NA</td><td>Intergenic</td><td>NA</td><td>NA</td></tr>
<tr><td></td><td></td><td></td><td></td><td></td><td></td><td></td><td></td><td></td><td></td><td></td><td></td></tr>
<tr><td>T5238</td><td>GTCTTGCAGAAGCACCACACGGG</td><td>1000</td><td></td><td></td><td></td><td></td><td></td><td></td><td></td><td></td><td></td></tr>
<tr><td>Oligo fwd</td><td>CACCGTCTTGCAGAAGCACCACAC</td><td></td><td></td><td></td><td></td><td></td><td></td><td></td><td></td><td></td><td></td></tr>
<tr><td>Oligo rev</td><td>AAACGTGTGGTGCTTCTGCAAGAC</td><td></td><td></td><td></td><td></td><td></td><td></td><td></td><td></td><td></td><td></td></tr>
<tr><td>Chromosome</td><td>start</td><td>end</td><td>strand</td><td>MM</td><td>target_seq</td><td>PAM</td><td>alignment</td><td>distance</td><td>position</td><td>gene name</td><td>gene id</td></tr>
<tr><td>GU305914.1</td><td>74563</td><td>74585</td><td>+</td><td>0</td><td>GTCTTGCAGAAGCACCACAC</td><td>GGG</td><td>||||||||[||||||||||||]PAM</td><td>NA</td><td>Intergenic</td><td>NA</td><td>NA</td></tr>
<tr><td></td><td></td><td></td><td></td><td></td><td></td><td></td><td></td><td></td><td></td><td></td><td></td></tr>
<tr><td>T5239</td><td>TCTTGCAGAAGCACCACACGGGG</td><td>1000</td><td></td><td></td><td></td><td></td><td></td><td></td><td></td><td></td><td></td></tr>
<tr><td>Oligo fwd</td><td>CACCTCTTGCAGAAGCACCACACG</td><td></td><td></td><td></td><td></td><td></td><td></td><td></td><td></td><td></td><td></td></tr>
<tr><td>Oligo rev</td><td>AAACCGTGTGGTGCTTCTGCAAGA</td><td></td><td></td><td></td><td></td><td></td><td></td><td></td><td></td><td></td><td></td></tr>
<tr><td>Chromosome</td><td>start</td><td>end</td><td>strand</td><td>MM</td><td>target_seq</td><td>PAM</td><td>alignment</td><td>distance</td><td>position</td><td>gene name</td><td>gene id</td></tr>
<tr><td>GU305914.1</td><td>74564</td><td>74586</td><td>+</td><td>0</td><td>TCTTGCAGAAGCACCACACG</td><td>GGG</td><td>||||||||[||||||||||||]PAM</td><td>NA</td><td>Intergenic</td><td>NA</td><td>NA</td></tr>
<tr><td></td><td></td><td></td><td></td><td></td><td></td><td></td><td></td><td></td><td></td><td></td><td></td></tr>
<tr><td>T5240</td><td>TGCAGAAGCACCACACGGGGAGG</td><td>1000</td><td></td><td></td><td></td><td></td><td></td><td></td><td></td><td></td><td></td></tr>
<tr><td>Oligo fwd</td><td>CACCTGCAGAAGCACCACACGGGG</td><td></td><td></td><td></td><td></td><td></td><td></td><td></td><td></td><td></td><td></td></tr>
<tr><td>Oligo rev</td><td>AAACCCCCGTGTGGTGCTTCTGCA</td><td></td><td></td><td></td><td></td><td></td><td></td><td></td><td></td><td></td><td></td></tr>
<tr><td>Chromosome</td><td>start</td><td>end</td><td>strand</td><td>MM</td><td>target_seq</td><td>PAM</td><td>alignment</td><td>distance</td><td>position</td><td>gene name</td><td>gene id</td></tr>
<tr><td>GU305914.1</td><td>74567</td><td>74589</td><td>+</td><td>0</td><td>TGCAGAAGCACCACACGGGG</td><td>AGG</td><td>||||||||[||||||||||||]PAM</td><td>NA</td><td>Intergenic</td><td>NA</td><td>NA</td></tr>
<tr><td></td><td></td><td></td><td></td><td></td><td></td><td></td><td></td><td></td><td></td><td></td><td></td></tr>
<tr><td>T5241</td><td>ACCACACGGGGAGGTCGATGAGG</td><td>1000</td><td></td><td></td><td></td><td></td><td></td><td></td><td></td><td></td><td></td></tr>
<tr><td>Oligo fwd</td><td>CACCACCACACGGGGAGGTCGATG</td><td></td><td></td><td></td><td></td><td></td><td></td><td></td><td></td><td></td><td></td></tr>
<tr><td>Oligo rev</td><td>AAACCATCGACCTCCCCGTGTGGT</td><td></td><td></td><td></td><td></td><td></td><td></td><td></td><td></td><td></td><td></td></tr>
<tr><td>Chromosome</td><td>start</td><td>end</td><td>strand</td><td>MM</td><td>target_seq</td><td>PAM</td><td>alignment</td><td>distance</td><td>position</td><td>gene name</td><td>gene id</td></tr>
<tr><td>GU305914.1</td><td>74576</td><td>74598</td><td>+</td><td>0</td><td>ACCACACGGGGAGGTCGATG</td><td>AGG</td><td>||||||||[||||||||||||]PAM</td><td>NA</td><td>Intergenic</td><td>NA</td><td>NA</td></tr>
<tr><td></td><td></td><td></td><td></td><td></td><td></td><td></td><td></td><td></td><td></td><td></td><td></td></tr>
<tr><td>T5242</td><td>CACGGGGAGGTCGATGAGGCTGG</td><td>1000</td><td></td><td></td><td></td><td></td><td></td><td></td><td></td><td></td><td></td></tr>
<tr><td>Oligo fwd</td><td>CACCCACGGGGAGGTCGATGAGGC</td><td></td><td></td><td></td><td></td><td></td><td></td><td></td><td></td><td></td><td></td></tr>
<tr><td>Oligo rev</td><td>AAACGCCTCATCGACCTCCCCGTG</td><td></td><td></td><td></td><td></td><td></td><td></td><td></td><td></td><td></td><td></td></tr>
<tr><td>Chromosome</td><td>start</td><td>end</td><td>strand</td><td>MM</td><td>target_seq</td><td>PAM</td><td>alignment</td><td>distance</td><td>position</td><td>gene name</td><td>gene id</td></tr>
<tr><td>GU305914.1</td><td>74580</td><td>74602</td><td>+</td><td>0</td><td>CACGGGGAGGTCGATGAGGC</td><td>TGG</td><td>||||||||[||||||||||||]PAM</td><td>NA</td><td>Intergenic</td><td>NA</td><td>NA</td></tr>
<tr><td></td><td></td><td></td><td></td><td></td><td></td><td></td><td></td><td></td><td></td><td></td><td></td></tr>
<tr><td>T5243</td><td>GGTCGATGAGGCTGGAGATGAGG</td><td>1000</td><td></td><td></td><td></td><td></td><td></td><td></td><td></td><td></td><td></td></tr>
<tr><td>Oligo fwd</td><td>CACCGGTCGATGAGGCTGGAGATG</td><td></td><td></td><td></td><td></td><td></td><td></td><td></td><td></td><td></td><td></td></tr>
<tr><td>Oligo rev</td><td>AAACCATCTCCAGCCTCATCGACC</td><td></td><td></td><td></td><td></td><td></td><td></td><td></td><td></td><td></td><td></td></tr>
<tr><td>Chromosome</td><td>start</td><td>end</td><td>strand</td><td>MM</td><td>target_seq</td><td>PAM</td><td>alignment</td><td>distance</td><td>position</td><td>gene name</td><td>gene id</td></tr>
<tr><td>GU305914.1</td><td>74588</td><td>74610</td><td>+</td><td>0</td><td>GGTCGATGAGGCTGGAGATG</td><td>AGG</td><td>||||||||[||||||||||||]PAM</td><td>NA</td><td>Intergenic</td><td>NA</td><td>NA</td></tr>
</table>

<table>
<tr><td>T5244</td><td>GCCGTTTCTGAGCACGACCGAGG</td><td></td><td>1000</td><td></td><td></td><td></td><td></td><td></td><td></td><td></td><td></td><td></td></tr>
<tr><td>Oligo fwd</td><td>CACCGCCGTTTCTGAGCACGACCG</td><td></td><td></td><td></td><td></td><td></td><td></td><td></td><td></td><td></td><td></td><td></td></tr>
<tr><td>Oligo rev</td><td>AAACCGGTCGTGCTCAGAAACGGC</td><td></td><td></td><td></td><td></td><td></td><td></td><td></td><td></td><td></td><td></td><td></td></tr>
<tr><td>Chromosome</td><td>start</td><td></td><td>end</td><td>strand</td><td>MM</td><td>target_seq</td><td>PAM</td><td>alignment</td><td>distance</td><td>position</td><td>gene name</td><td>gene id</td></tr>
<tr><td>GU305914.1</td><td></td><td>74610</td><td>74632</td><td>+</td><td>0</td><td>GCCGTTTCTGAGCACGACCG</td><td>AGG</td><td>||||||||[||||||||||||]PAM</td><td>NA</td><td>Intergenic</td><td>NA</td><td>NA</td></tr>
<tr><td>T5245</td><td>GTTTCTGAGCACGACCGAGGTGG</td><td></td><td>1000</td><td></td><td></td><td></td><td></td><td></td><td></td><td></td><td></td><td></td></tr>
<tr><td>Oligo fwd</td><td>CACCGTTTCTGAGCACGACCGAGG</td><td></td><td></td><td></td><td></td><td></td><td></td><td></td><td></td><td></td><td></td><td></td></tr>
<tr><td>Oligo rev</td><td>AAACCCTCGGTCGTGCTCAGAAAC</td><td></td><td></td><td></td><td></td><td></td><td></td><td></td><td></td><td></td><td></td><td></td></tr>
<tr><td>Chromosome</td><td>start</td><td></td><td>end</td><td>strand</td><td>MM</td><td>target_seq</td><td>PAM</td><td>alignment</td><td>distance</td><td>position</td><td>gene name</td><td>gene id</td></tr>
<tr><td>GU305914.1</td><td></td><td>74613</td><td>74635</td><td>+</td><td>0</td><td>GTTTCTGAGCACGACCGAGG</td><td>TGG</td><td>||||||||[||||||||||||]PAM</td><td>NA</td><td>Intergenic</td><td>NA</td><td>NA</td></tr>
<tr><td>T5246</td><td>GAGCACGACCGAGGTGGCGAAGG</td><td></td><td>1000</td><td></td><td></td><td></td><td></td><td></td><td></td><td></td><td></td><td></td></tr>
<tr><td>Oligo fwd</td><td>CACCGAGCACGACCGAGGTGGCGA</td><td></td><td></td><td></td><td></td><td></td><td></td><td></td><td></td><td></td><td></td><td></td></tr>
<tr><td>Oligo rev</td><td>AAACTCGCCACCTCGGTCGTGCTC</td><td></td><td></td><td></td><td></td><td></td><td></td><td></td><td></td><td></td><td></td><td></td></tr>
<tr><td>Chromosome</td><td>start</td><td></td><td>end</td><td>strand</td><td>MM</td><td>target_seq</td><td>PAM</td><td>alignment</td><td>distance</td><td>position</td><td>gene name</td><td>gene id</td></tr>
<tr><td>GU305914.1</td><td></td><td>74619</td><td>74641</td><td>+</td><td>0</td><td>GAGCACGACCGAGGTGGCGA</td><td>AGG</td><td>||||||||[||||||||||||]PAM</td><td>NA</td><td>Intergenic</td><td>NA</td><td>NA</td></tr>
<tr><td>T5247</td><td>CACGACCGAGGTGGCGAAGGTGG</td><td></td><td>1000</td><td></td><td></td><td></td><td></td><td></td><td></td><td></td><td></td><td></td></tr>
<tr><td>Oligo fwd</td><td>CACCCACGACCGAGGTGGCGAAGG</td><td></td><td></td><td></td><td></td><td></td><td></td><td></td><td></td><td></td><td></td><td></td></tr>
<tr><td>Oligo rev</td><td>AAACCCTTCGCCACCTCGGTCGTG</td><td></td><td></td><td></td><td></td><td></td><td></td><td></td><td></td><td></td><td></td><td></td></tr>
<tr><td>Chromosome</td><td>start</td><td></td><td>end</td><td>strand</td><td>MM</td><td>target_seq</td><td>PAM</td><td>alignment</td><td>distance</td><td>position</td><td>gene name</td><td>gene id</td></tr>
<tr><td>GU305914.1</td><td></td><td>74622</td><td>74644</td><td>+</td><td>0</td><td>CACGACCGAGGTGGCGAAGG</td><td>TGG</td><td>||||||||[||||||||||||]PAM</td><td>NA</td><td>Intergenic</td><td>NA</td><td>NA</td></tr>
<tr><td>GU305914.1</td><td></td><td>18656</td><td>18678</td><td>-</td><td>4</td><td>GACGGCAGAGGTGGCGGAGG</td><td>CGG</td><td>-|||-|-|[||||||||-|||]PAM</td><td>NA</td><td>Intergenic</td><td>NA</td><td>NA</td></tr>
<tr><td>T5248</td><td>GGTGGCGAAGGTGGAGAGCTTGG</td><td></td><td>1000</td><td></td><td></td><td></td><td></td><td></td><td></td><td></td><td></td><td></td></tr>
<tr><td>Oligo fwd</td><td>CACCGGTGGCGAAGGTGGAGAGCT</td><td></td><td></td><td></td><td></td><td></td><td></td><td></td><td></td><td></td><td></td><td></td></tr>
<tr><td>Oligo rev</td><td>AAACAGCTCTCCACCTTCGCCACC</td><td></td><td></td><td></td><td></td><td></td><td></td><td></td><td></td><td></td><td></td><td></td></tr>
<tr><td>Chromosome</td><td>start</td><td></td><td>end</td><td>strand</td><td>MM</td><td>target_seq</td><td>PAM</td><td>alignment</td><td>distance</td><td>position</td><td>gene name</td><td>gene id</td></tr>
<tr><td>GU305914.1</td><td></td><td>74631</td><td>74653</td><td>+</td><td>0</td><td>GGTGGCGAAGGTGGAGAGCT</td><td>TGG</td><td>||||||||[||||||||||||]PAM</td><td>NA</td><td>Intergenic</td><td>NA</td><td>NA</td></tr>
<tr><td>T5249</td><td>CGAAGGTGGAGAGCTTGGAGAGG</td><td></td><td>1000</td><td></td><td></td><td></td><td></td><td></td><td></td><td></td><td></td><td></td></tr>
<tr><td>Oligo fwd</td><td>CACCCGAAGGTGGAGAGCTTGGAG</td><td></td><td></td><td></td><td></td><td></td><td></td><td></td><td></td><td></td><td></td><td></td></tr>
<tr><td>Oligo rev</td><td>AAACCTCCAAGCTCTCCACCTTCG</td><td></td><td></td><td></td><td></td><td></td><td></td><td></td><td></td><td></td><td></td><td></td></tr>
<tr><td>Chromosome</td><td>start</td><td></td><td>end</td><td>strand</td><td>MM</td><td>target_seq</td><td>PAM</td><td>alignment</td><td>distance</td><td>position</td><td>gene name</td><td>gene id</td></tr>
<tr><td>GU305914.1</td><td></td><td>74636</td><td>74658</td><td>+</td><td>0</td><td>CGAAGGTGGAGAGCTTGGAG</td><td>AGG</td><td>||||||||[||||||||||||]PAM</td><td>NA</td><td>Intergenic</td><td>NA</td><td>NA</td></tr>
<tr><td>T5250</td><td>GAAGGTGGAGAGCTTGGAGAGGG</td><td></td><td>1000</td><td></td><td></td><td></td><td></td><td></td><td></td><td></td><td></td><td></td></tr>
<tr><td>Oligo fwd</td><td>CACCGAAGGTGGAGAGCTTGGAGA</td><td></td><td></td><td></td><td></td><td></td><td></td><td></td><td></td><td></td><td></td><td></td></tr>
<tr><td>Oligo rev</td><td>AAACTCTCCAAGCTCTCCACCTTC</td><td></td><td></td><td></td><td></td><td></td><td></td><td></td><td></td><td></td><td></td><td></td></tr>
<tr><td>Chromosome</td><td>start</td><td></td><td>end</td><td>strand</td><td>MM</td><td>target_seq</td><td>PAM</td><td>alignment</td><td>distance</td><td>position</td><td>gene name</td><td>gene id</td></tr>
<tr><td>GU305914.1</td><td></td><td>74637</td><td>74659</td><td>+</td><td>0</td><td>GAAGGTGGAGAGCTTGGAGA</td><td>GGG</td><td>||||||||[||||||||||||]PAM</td><td>NA</td><td>Intergenic</td><td>NA</td><td>NA</td></tr>
<tr><td>GU305914.1</td><td></td><td>71026</td><td>71048</td><td>+</td><td>4</td><td>GGAGGCGGAGATGTTGGAGA</td><td>CGG</td><td>|-|||-||[|||--|||||||]PAM</td><td>NA</td><td>Intergenic</td><td>NA</td><td>NA</td></tr>
<tr><td>T5251</td><td>AAGGTGGAGAGCTTGGAGAGGGG</td><td></td><td>1000</td><td></td><td></td><td></td><td></td><td></td><td></td><td></td><td></td><td></td></tr>
<tr><td>Oligo fwd</td><td>CACCAAGGTGGAGAGCTTGGAGAG</td><td></td><td></td><td></td><td></td><td></td><td></td><td></td><td></td><td></td><td></td><td></td></tr>
<tr><td>Oligo rev</td><td>AAACCTCTCCAAGCTCTCCACCTT</td><td></td><td></td><td></td><td></td><td></td><td></td><td></td><td></td><td></td><td></td><td></td></tr>
<tr><td>Chromosome</td><td>start</td><td></td><td>end</td><td>strand</td><td>MM</td><td>target_seq</td><td>PAM</td><td>alignment</td><td>distance</td><td>position</td><td>gene name</td><td>gene id</td></tr>
<tr><td>GU305914.1</td><td></td><td>74638</td><td>74660</td><td>+</td><td>0</td><td>AAGGTGGAGAGCTTGGAGAG</td><td>GGG</td><td>||||||||[||||||||||||]PAM</td><td>NA</td><td>Intergenic</td><td>NA</td><td>NA</td></tr>
<tr><td>T5252</td><td>GGAGAGCTTGGAGAGGGGCCAGG</td><td></td><td>1000</td><td></td><td></td><td></td><td></td><td></td><td></td><td></td><td></td><td></td></tr>
<tr><td>Oligo fwd</td><td>CACCGGAGAGCTTGGAGAGGGGCC</td><td></td><td></td><td></td><td></td><td></td><td></td><td></td><td></td><td></td><td></td><td></td></tr>
<tr><td>Oligo rev</td><td>AAACGGCCCCTCTCCAAGCTCTCC</td><td></td><td></td><td></td><td></td><td></td><td></td><td></td><td></td><td></td><td></td><td></td></tr>
<tr><td>Chromosome</td><td>start</td><td></td><td>end</td><td>strand</td><td>MM</td><td>target_seq</td><td>PAM</td><td>alignment</td><td>distance</td><td>position</td><td>gene name</td><td>gene id</td></tr>
<tr><td>GU305914.1</td><td></td><td>74643</td><td>74665</td><td>+</td><td>0</td><td>GGAGAGCTTGGAGAGGGGCC</td><td>AGG</td><td>||||||||[||||||||||||]PAM</td><td>NA</td><td>Intergenic</td><td>NA</td><td>NA</td></tr>
<tr><td>T5253</td><td>GAGAGCTTGGAGAGGGGCCAGGG</td><td></td><td>1000</td><td></td><td></td><td></td><td></td><td></td><td></td><td></td><td></td><td></td></tr>
<tr><td>Oligo fwd</td><td>CACCGAGAGCTTGGAGAGGGGCCA</td><td></td><td></td><td></td><td></td><td></td><td></td><td></td><td></td><td></td><td></td><td></td></tr>
<tr><td>Oligo rev</td><td>AAACTGGCCCCTCTCCAAGCTCTC</td><td></td><td></td><td></td><td></td><td></td><td></td><td></td><td></td><td></td><td></td><td></td></tr>
<tr><td>Chromosome</td><td>start</td><td></td><td>end</td><td>strand</td><td>MM</td><td>target_seq</td><td>PAM</td><td>alignment</td><td>distance</td><td>position</td><td>gene name</td><td>gene id</td></tr>
<tr><td>GU305914.1</td><td></td><td>74644</td><td>74666</td><td>+</td><td>0</td><td>GAGAGCTTGGAGAGGGGCCA</td><td>GGG</td><td>||||||||[||||||||||||]PAM</td><td>NA</td><td>Intergenic</td><td>NA</td><td>NA</td></tr>
<tr><td>T5254</td><td>GGAGAGGGGCCAGGGACAGTTGG</td><td></td><td>1000</td><td></td><td></td><td></td><td></td><td></td><td></td><td></td><td></td><td></td></tr>
<tr><td>Oligo fwd</td><td>CACCGGAGAGGGGCCAGGGACAGT</td><td></td><td></td><td></td><td></td><td></td><td></td><td></td><td></td><td></td><td></td><td></td></tr>
<tr><td>Oligo rev</td><td>AAACACTGTCCCTGGCCCCTCTCC</td><td></td><td></td><td></td><td></td><td></td><td></td><td></td><td></td><td></td><td></td><td></td></tr>
<tr><td>Chromosome</td><td>start</td><td></td><td>end</td><td>strand</td><td>MM</td><td>target_seq</td><td>PAM</td><td>alignment</td><td>distance</td><td>position</td><td>gene name</td><td>gene id</td></tr>
<tr><td>GU305914.1</td><td></td><td>74652</td><td>74674</td><td>+</td><td>0</td><td>GGAGAGGGGCCAGGGACAGT</td><td>TGG</td><td>||||||||[||||||||||||]PAM</td><td>NA</td><td>Intergenic</td><td>NA</td><td>NA</td></tr>
<tr><td>T5255</td><td>CCAGGGACAGTTGGTGCCGTCGG</td><td></td><td>1000</td><td></td><td></td><td></td><td></td><td></td><td></td><td></td><td></td><td></td></tr>
<tr><td>Oligo fwd</td><td>CACCCCAGGGACAGTTGGTGCCGT</td><td></td><td></td><td></td><td></td><td></td><td></td><td></td><td></td><td></td><td></td><td></td></tr>
<tr><td>Oligo rev</td><td>AAACACGGCACCAACTGTCCCTGG</td><td></td><td></td><td></td><td></td><td></td><td></td><td></td><td></td><td></td><td></td><td></td></tr>
<tr><td>Chromosome</td><td>start</td><td></td><td>end</td><td>strand</td><td>MM</td><td>target_seq</td><td>PAM</td><td>alignment</td><td>distance</td><td>position</td><td>gene name</td><td>gene id</td></tr>
<tr><td>GU305914.1</td><td></td><td>74661</td><td>74683</td><td>+</td><td>0</td><td>CCAGGGACAGTTGGTGCCGT</td><td>CGG</td><td>||||||||[||||||||||||]PAM</td><td>NA</td><td>Intergenic</td><td>NA</td><td>NA</td></tr>
<tr><td>T5256</td><td>GGGACAGTTGGTGCCGTCGGTGG</td><td></td><td>1000</td><td></td><td></td><td></td><td></td><td></td><td></td><td></td><td></td><td></td></tr>
<tr><td>Oligo fwd</td><td>CACCGGGACAGTTGGTGCCGTCGG</td><td></td><td></td><td></td><td></td><td></td><td></td><td></td><td></td><td></td><td></td><td></td></tr>
<tr><td>Oligo rev</td><td>AAACCCGACGGCACCAACTGTCCC</td><td></td><td></td><td></td><td></td><td></td><td></td><td></td><td></td><td></td><td></td><td></td></tr>
<tr><td>Chromosome</td><td>start</td><td></td><td>end</td><td>strand</td><td>MM</td><td>target_seq</td><td>PAM</td><td>alignment</td><td>distance</td><td>position</td><td>gene name</td><td>gene id</td></tr>
<tr><td>GU305914.1</td><td></td><td>74664</td><td>74686</td><td>+</td><td>0</td><td>GGGACAGTTGGTGCCGTCGG</td><td>TGG</td><td>||||||||[||||||||||||]PAM</td><td>NA</td><td>Intergenic</td><td>NA</td><td>NA</td></tr>
<tr><td>T5257</td><td>ACAGTTGGTGCCGTCGGTGGCGG</td><td></td><td>1000</td><td></td><td></td><td></td><td></td><td></td><td></td><td></td><td></td><td></td></tr>
<tr><td>Oligo fwd</td><td>CACCACAGTTGGTGCCGTCGGTGG</td><td></td><td></td><td></td><td></td><td></td><td></td><td></td><td></td><td></td><td></td><td></td></tr>
<tr><td>Oligo rev</td><td>AAACCCACCGACGGCACCAACTGT</td><td></td><td></td><td></td><td></td><td></td><td></td><td></td><td></td><td></td><td></td><td></td></tr>
<tr><td>Chromosome</td><td>start</td><td></td><td>end</td><td>strand</td><td>MM</td><td>target_seq</td><td>PAM</td><td>alignment</td><td>distance</td><td>position</td><td>gene name</td><td>gene id</td></tr>
<tr><td>GU305914.1</td><td></td><td>74667</td><td>74689</td><td>+</td><td>0</td><td>ACAGTTGGTGCCGTCGGTGG</td><td>CGG</td><td>||||||||[||||||||||||]PAM</td><td>NA</td><td>Intergenic</td><td>NA</td><td>NA</td></tr>
<tr><td>T5258</td><td>GTTGGTGCCGTCGGTGGCGGTGG</td><td></td><td>1000</td><td></td><td></td><td></td><td></td><td></td><td></td><td></td><td></td><td></td></tr>
<tr><td>Oligo fwd</td><td>CACCGTTGGTGCCGTCGGTGGCGG</td><td></td><td></td><td></td><td></td><td></td><td></td><td></td><td></td><td></td><td></td><td></td></tr>
<tr><td>Oligo rev</td><td>AAACCCGCCACCGACGGCACCAAC</td><td></td><td></td><td></td><td></td><td></td><td></td><td></td><td></td><td></td><td></td><td></td></tr>
</table>

| | | | | | | | | | | | |
|---|---|---|---|---|---|---|---|---|---|---|---|
| Chromosome | start | end | strand | MM | target_seq | PAM | alignment | distance | position | gene name | gene id |
| GU305914.1 | 74670 | 74692 | + | 0 | GTTGGTGCCGTCGGTGGCGG | TGG | \|\|\|\|\|\|\|\|[\|\|\|\|\|\|\|\|\|\|\|\|]PAM | NA | Intergenic | NA | NA |
| GU305914.1 | 163749 | 163771 | + | 4 | GTCGGCGGCGCCGGTGGCGG | CGG | \|\|-\|\|-\|-[\|\|-\|\|\|\|\|\|\|\|\|]PAM | NA | Intergenic | NA | NA |
| T5259 | GGTGCCGTCGGTGGCGGTGGCGG | 1000 | | | | | | | | | |
| Oligo fwd | CACCGGTGCCGTCGGTGGCGGTGG | | | | | | | | | | |
| Oligo rev | AAACCCACCGCCACCGACGGCACC | | | | | | | | | | |
| Chromosome | start | end | strand | MM | target_seq | PAM | alignment | distance | position | gene name | gene id |
| GU305914.1 | 74673 | 74695 | + | 0 | GGTGCCGTCGGTGGCGGTGG | CGG | \|\|\|\|\|\|\|\|[\|\|\|\|\|\|\|\|\|\|\|\|]PAM | NA | Intergenic | NA | NA |
| GU305914.1 | 52719 | 52741 | + | 4 | GCCGCTGGCGGTGGCGGTGG | CGG | \|--\|\|-\|-[\|\|\|\|\|\|\|\|\|\|\|\|]PAM | NA | Intergenic | NA | NA |
| GU305914.1 | 62440 | 62462 | + | 4 | GGTGGCGGCGGGGGTGGTGG | CGG | \|\|\|\|-\|\|-[\|\|\|-\|\|-\|\|\|\|\|]PAM | NA | Intergenic | NA | NA |
| GU305914.1 | 48156 | 48178 | + | 4 | GGCGGCGGCGGTGGCGGCGG | CGG | \|\|-\|-\|\|-[\|\|\|\|\|\|\|\|\|-\|\|]PAM | NA | Intergenic | NA | NA |
| GU305914.1 | 154407 | 154429 | + | 4 | GGTGGCGGCGGCGGCGGCGG | CGG | \|\|\|\|-\|\|-[\|\|\|-\|\|\|\|\|-\|\|]PAM | NA | Intergenic | NA | NA |
| GU305914.1 | 58123 | 58145 | - | 4 | GGTGGCGGCGGCGGCGGCGG | AGG | \|\|\|\|-\|\|-[\|\|\|-\|\|\|\|\|-\|\|]PAM | NA | Intergenic | NA | NA |
| GU305914.1 | 30000 | 30022 | + | 4 | GGTGCTGGCGGCGGCGGCGG | CGG | \|\|\|\|\|-\|-[\|\|\|-\|\|\|\|\|-\|\|]PAM | NA | Intergenic | NA | NA |
| T5260 | GGCGGTGGCGGCCGAGTTCCCGG | 1000 | | | | | | | | | |
| Oligo fwd | CACCGGCGGTGGCGGCCGAGTTCC | | | | | | | | | | |
| Oligo rev | AAACGGAACTCGGCCGCCACCGCC | | | | | | | | | | |
| Chromosome | start | end | strand | MM | target_seq | PAM | alignment | distance | position | gene name | gene id |
| GU305914.1 | 74685 | 74707 | + | 0 | GGCGGTGGCGGCCGAGTTCC | CGG | \|\|\|\|\|\|\|\|[\|\|\|\|\|\|\|\|\|\|\|\|]PAM | NA | Intergenic | NA | NA |
| T5261 | GAACCGCCGCCACCGCCACCCGG | 1000 | | | | | | | | | |
| Oligo fwd | CACCGAACCGCCGCCACCGCCACC | | | | | | | | | | |
| Oligo rev | AAACGGTGGCGGTGGCGGCGGTTC | | | | | | | | | | |
| Chromosome | start | end | strand | MM | target_seq | PAM | alignment | distance | position | gene name | gene id |
| GU305914.1 | 74707 | 74729 | + | 0 | GAACCGCCGCCACCGCCACC | CGG | \|\|\|\|\|\|\|\|[\|\|\|\|\|\|\|\|\|\|\|\|]PAM | NA | Intergenic | NA | NA |
| T5262 | AACCGCCGCCACCGCCACCCGGG | 1000 | | | | | | | | | |
| Oligo fwd | CACCAACCGCCGCCACCGCCACCC | | | | | | | | | | |
| Oligo rev | AAACGGGTGGCGGTGGCGGCGGTT | | | | | | | | | | |
| Chromosome | start | end | strand | MM | target_seq | PAM | alignment | distance | position | gene name | gene id |
| GU305914.1 | 74708 | 74730 | + | 0 | AACCGCCGCCACCGCCACCC | GGG | \|\|\|\|\|\|\|\|[\|\|\|\|\|\|\|\|\|\|\|\|]PAM | NA | Intergenic | NA | NA |
| GU305914.1 | 98350 | 98372 | + | 4 | CGCCGCCGCCACCGCCGCCT | AGG | --\|\|\|\|\|\|[\|\|\|\|\|\|\|\|-\|\|-]PAM | NA | Intergenic | NA | NA |
| T5263 | GCCACCGCCACCCGGGTCTGAGG | 1000 | | | | | | | | | |
| Oligo fwd | CACCGCCACCGCCACCCGGGTCTG | | | | | | | | | | |
| Oligo rev | AAACCAGACCCGGGTGGCGGTGGC | | | | | | | | | | |
| Chromosome | start | end | strand | MM | target_seq | PAM | alignment | distance | position | gene name | gene id |
| GU305914.1 | 74715 | 74737 | + | 0 | GCCACCGCCACCCGGGTCTG | AGG | \|\|\|\|\|\|\|\|[\|\|\|\|\|\|\|\|\|\|\|\|]PAM | NA | Intergenic | NA | NA |
| T5264 | CGCCACCCGGGTCTGAGGTCCGG | 1000 | | | | | | | | | |
| Oligo fwd | CACCCGCCACCCGGGTCTGAGGTC | | | | | | | | | | |
| Oligo rev | AAACGACCTCAGACCCGGGTGGCG | | | | | | | | | | |
| Chromosome | start | end | strand | MM | target_seq | PAM | alignment | distance | position | gene name | gene id |
| GU305914.1 | 74720 | 74742 | + | 0 | CGCCACCCGGGTCTGAGGTC | CGG | \|\|\|\|\|\|\|\|[\|\|\|\|\|\|\|\|\|\|\|\|]PAM | NA | Intergenic | NA | NA |
| T5265 | CGGGTCTGAGGTCCGGACCGCGG | 1000 | | | | | | | | | |
| Oligo fwd | CACCCGGGTCTGAGGTCCGGACCG | | | | | | | | | | |
| Oligo rev | AAACCGGTCCGGACCTCAGACCCG | | | | | | | | | | |
| Chromosome | start | end | strand | MM | target_seq | PAM | alignment | distance | position | gene name | gene id |
| GU305914.1 | 74727 | 74749 | + | 0 | CGGGTCTGAGGTCCGGACCG | CGG | \|\|\|\|\|\|\|\|[\|\|\|\|\|\|\|\|\|\|\|\|]PAM | NA | Intergenic | NA | NA |
| T5266 | CGGCGCGTCCGCCGTAGCGCAGG | 1000 | | | | | | | | | |
| Oligo fwd | CACCCGGCGCGTCCGCCGTAGCGC | | | | | | | | | | |
| Oligo rev | AAACGCGCTACGGCGGACGCGCCG | | | | | | | | | | |
| Chromosome | start | end | strand | MM | target_seq | PAM | alignment | distance | position | gene name | gene id |
| GU305914.1 | 74747 | 74769 | + | 0 | CGGCGCGTCCGCCGTAGCGC | AGG | \|\|\|\|\|\|\|\|[\|\|\|\|\|\|\|\|\|\|\|\|]PAM | NA | Intergenic | NA | NA |
| T5267 | GGCGCGTCCGCCGTAGCGCAGGG | 1000 | | | | | | | | | |
| Oligo fwd | CACCGGCGCGTCCGCCGTAGCGCA | | | | | | | | | | |
| Oligo rev | AAACTGCGCTACGGCGGACGCGCC | | | | | | | | | | |
| Chromosome | start | end | strand | MM | target_seq | PAM | alignment | distance | position | gene name | gene id |
| GU305914.1 | 74748 | 74770 | + | 0 | GGCGCGTCCGCCGTAGCGCA | GGG | \|\|\|\|\|\|\|\|[\|\|\|\|\|\|\|\|\|\|\|\|]PAM | NA | Intergenic | NA | NA |
| T5268 | GCGCAGGGTCCGATTGCCGATGG | 1000 | | | | | | | | | |
| Oligo fwd | CACCGCGCAGGGTCCGATTGCCGA | | | | | | | | | | |
| Oligo rev | AAACTCGGCAATCGGACCCTGCGC | | | | | | | | | | |
| Chromosome | start | end | strand | MM | target_seq | PAM | alignment | distance | position | gene name | gene id |
| GU305914.1 | 74763 | 74785 | + | 0 | GCGCAGGGTCCGATTGCCGA | TGG | \|\|\|\|\|\|\|\|[\|\|\|\|\|\|\|\|\|\|\|\|]PAM | NA | Intergenic | NA | NA |
| T5269 | CGCAGGGTCCGATTGCCGATGGG | 1000 | | | | | | | | | |
| Oligo fwd | CACCCGCAGGGTCCGATTGCCGAT | | | | | | | | | | |
| Oligo rev | AAACATCGGCAATCGGACCCTGCG | | | | | | | | | | |
| Chromosome | start | end | strand | MM | target_seq | PAM | alignment | distance | position | gene name | gene id |
| GU305914.1 | 74764 | 74786 | + | 0 | CGCAGGGTCCGATTGCCGAT | GGG | \|\|\|\|\|\|\|\|[\|\|\|\|\|\|\|\|\|\|\|\|]PAM | NA | Intergenic | NA | NA |
| T5270 | GCAGGGTCCGATTGCCGATGGGG | 1000 | | | | | | | | | |
| Oligo fwd | CACCGCAGGGTCCGATTGCCGATG | | | | | | | | | | |
| Oligo rev | AAACCATCGGCAATCGGACCCTGC | | | | | | | | | | |
| Chromosome | start | end | strand | MM | target_seq | PAM | alignment | distance | position | gene name | gene id |
| GU305914.1 | 74765 | 74787 | + | 0 | GCAGGGTCCGATTGCCGATG | GGG | \|\|\|\|\|\|\|\|[\|\|\|\|\|\|\|\|\|\|\|\|]PAM | NA | Intergenic | NA | NA |
| T5271 | TCCGATTGCCGATGGGGATGAGG | 1000 | | | | | | | | | |
| Oligo fwd | CACCTCCGATTGCCGATGGGGATG | | | | | | | | | | |
| Oligo rev | AAACCATCCCCATCGGCAATCGGA | | | | | | | | | | |
| Chromosome | start | end | strand | MM | target_seq | PAM | alignment | distance | position | gene name | gene id |
| GU305914.1 | 74771 | 74793 | + | 0 | TCCGATTGCCGATGGGGATG | AGG | \|\|\|\|\|\|\|\|[\|\|\|\|\|\|\|\|\|\|\|\|]PAM | NA | Intergenic | NA | NA |

| | | | | | | | | | | | |
|---|---|---|---|---|---|---|---|---|---|---|---|
| T5272 | CCGATGGGGATGAGGAAGCGCGG | 1000 | | | | | | | | | |
| Oligo fwd | CACCCCGATGGGGATGAGGAAGCG | | | | | | | | | | |
| Oligo rev | AAACCGCTTCCTCATCCCCATCGG | | | | | | | | | | |
| Chromosome | start | end | strand | MM | target_seq | PAM | alignment | distance | position | gene name | gene id |
| GU305914.1 | 74779 | 74801 | + | 0 | CCGATGGGGATGAGGAAGCG | CGG | \|\|\|\|\|\|\|\|[\|\|\|\|\|\|\|\|\|\|\|\|]PAM | NA | Intergenic | NA | NA |

| | | | | | | | | | | | |
|---|---|---|---|---|---|---|---|---|---|---|---|
| T5273 | CGATGGGGATGAGGAAGCGCGGG | 1000 | | | | | | | | | |
| Oligo fwd | CACCCGATGGGGATGAGGAAGCGC | | | | | | | | | | |
| Oligo rev | AAACGCGCTTCCTCATCCCCATCG | | | | | | | | | | |
| Chromosome | start | end | strand | MM | target_seq | PAM | alignment | distance | position | gene name | gene id |
| GU305914.1 | 74780 | 74802 | + | 0 | CGATGGGGATGAGGAAGCGC | GGG | \|\|\|\|\|\|\|\|[\|\|\|\|\|\|\|\|\|\|\|\|]PAM | NA | Intergenic | NA | NA |

| | | | | | | | | | | | |
|---|---|---|---|---|---|---|---|---|---|---|---|
| T5274 | AACTTTAGCCATGACGTAGAAGG | 1000 | | | | | | | | | |
| Oligo fwd | CACCAACTTTAGCCATGACGTAGA | | | | | | | | | | |
| Oligo rev | AAACTCTACGTCATGGCTAAAGTT | | | | | | | | | | |
| Chromosome | start | end | strand | MM | target_seq | PAM | alignment | distance | position | gene name | gene id |
| GU305914.1 | 74829 | 74851 | + | 0 | AACTTTAGCCATGACGTAGA | AGG | \|\|\|\|\|\|\|\|[\|\|\|\|\|\|\|\|\|\|\|\|]PAM | NA | Intergenic | NA | NA |

| | | | | | | | | | | | |
|---|---|---|---|---|---|---|---|---|---|---|---|
| T5275 | ACTTTAGCCATGACGTAGAAGGG | 1000 | | | | | | | | | |
| Oligo fwd | CACCACTTTAGCCATGACGTAGAA | | | | | | | | | | |
| Oligo rev | AAACTTCTACGTCATGGCTAAAGT | | | | | | | | | | |
| Chromosome | start | end | strand | MM | target_seq | PAM | alignment | distance | position | gene name | gene id |
| GU305914.1 | 74830 | 74852 | + | 0 | ACTTTAGCCATGACGTAGAA | GGG | \|\|\|\|\|\|\|\|[\|\|\|\|\|\|\|\|\|\|\|\|]PAM | NA | Intergenic | NA | NA |

| | | | | | | | | | | | |
|---|---|---|---|---|---|---|---|---|---|---|---|
| T5276 | CTTTAGCCATGACGTAGAAGGGG | 1000 | | | | | | | | | |
| Oligo fwd | CACCCTTTAGCCATGACGTAGAAG | | | | | | | | | | |
| Oligo rev | AAACCTTCTACGTCATGGCTAAAG | | | | | | | | | | |
| Chromosome | start | end | strand | MM | target_seq | PAM | alignment | distance | position | gene name | gene id |
| GU305914.1 | 74831 | 74853 | + | 0 | CTTTAGCCATGACGTAGAAG | GGG | \|\|\|\|\|\|\|\|[\|\|\|\|\|\|\|\|\|\|\|\|]PAM | NA | Intergenic | NA | NA |

| | | | | | | | | | | | |
|---|---|---|---|---|---|---|---|---|---|---|---|
| T5277 | GACGTAGAAGGGGTTTCCGCAGG | 1000 | | | | | | | | | |
| Oligo fwd | CACCGACGTAGAAGGGGTTTCCGC | | | | | | | | | | |
| Oligo rev | AAACGCGGAAACCCCTTCTACGTC | | | | | | | | | | |
| Chromosome | start | end | strand | MM | target_seq | PAM | alignment | distance | position | gene name | gene id |
| GU305914.1 | 74841 | 74863 | + | 0 | GACGTAGAAGGGGTTTCCGC | AGG | \|\|\|\|\|\|\|\|[\|\|\|\|\|\|\|\|\|\|\|\|]PAM | NA | Intergenic | NA | NA |

| | | | | | | | | | | | |
|---|---|---|---|---|---|---|---|---|---|---|---|
| T5278 | ACGTAGAAGGGGTTTCCGCAGGG | 1000 | | | | | | | | | |
| Oligo fwd | CACCACGTAGAAGGGGTTTCCGCA | | | | | | | | | | |
| Oligo rev | AAACTGCGGAAACCCCTTCTACGT | | | | | | | | | | |
| Chromosome | start | end | strand | MM | target_seq | PAM | alignment | distance | position | gene name | gene id |
| GU305914.1 | 74842 | 74864 | + | 0 | ACGTAGAAGGGGTTTCCGCA | GGG | \|\|\|\|\|\|\|\|[\|\|\|\|\|\|\|\|\|\|\|\|]PAM | NA | Intergenic | NA | NA |

| | | | | | | | | | | | |
|---|---|---|---|---|---|---|---|---|---|---|---|
| T5279 | GGTTTCCGCAGGGAGTCTCGCGG | 1000 | | | | | | | | | |
| Oligo fwd | CACCGGTTTCCGCAGGGAGTCTCG | | | | | | | | | | |
| Oligo rev | AAACCGAGACTCCCTGCGGAAACC | | | | | | | | | | |
| Chromosome | start | end | strand | MM | target_seq | PAM | alignment | distance | position | gene name | gene id |
| GU305914.1 | 74852 | 74874 | + | 0 | GGTTTCCGCAGGGAGTCTCG | CGG | \|\|\|\|\|\|\|\|[\|\|\|\|\|\|\|\|\|\|\|\|]PAM | NA | Intergenic | NA | NA |

| | | | | | | | | | | | |
|---|---|---|---|---|---|---|---|---|---|---|---|
| T5280 | GCGCAAGATGCTGACGAGCGTGG | 1000 | | | | | | | | | |
| Oligo fwd | CACCGCGCAAGATGCTGACGAGCG | | | | | | | | | | |
| Oligo rev | AAACCGCTCGTCAGCATCTTGCGC | | | | | | | | | | |
| Chromosome | start | end | strand | MM | target_seq | PAM | alignment | distance | position | gene name | gene id |
| GU305914.1 | 74883 | 74905 | + | 0 | GCGCAAGATGCTGACGAGCG | TGG | \|\|\|\|\|\|\|\|[\|\|\|\|\|\|\|\|\|\|\|\|]PAM | NA | Intergenic | NA | NA |

| | | | | | | | | | | | |
|---|---|---|---|---|---|---|---|---|---|---|---|
| T5281 | CTGACGAGCGTGGAGACGCCCGG | 1000 | | | | | | | | | |
| Oligo fwd | CACCCTGACGAGCGTGGAGACGCC | | | | | | | | | | |
| Oligo rev | AAACGGCGTCTCCACGCTCGTCAG | | | | | | | | | | |
| Chromosome | start | end | strand | MM | target_seq | PAM | alignment | distance | position | gene name | gene id |
| GU305914.1 | 74893 | 74915 | + | 0 | CTGACGAGCGTGGAGACGCC | CGG | \|\|\|\|\|\|\|\|[\|\|\|\|\|\|\|\|\|\|\|\|]PAM | NA | Intergenic | NA | NA |

| | | | | | | | | | | | |
|---|---|---|---|---|---|---|---|---|---|---|---|
| T5282 | TGACGAGCGTGGAGACGCCCGGG | 1000 | | | | | | | | | |
| Oligo fwd | CACCTGACGAGCGTGGAGACGCCC | | | | | | | | | | |
| Oligo rev | AAACGGGCGTCTCCACGCTCGTCA | | | | | | | | | | |
| Chromosome | start | end | strand | MM | target_seq | PAM | alignment | distance | position | gene name | gene id |
| GU305914.1 | 74894 | 74916 | + | 0 | TGACGAGCGTGGAGACGCCC | GGG | \|\|\|\|\|\|\|\|[\|\|\|\|\|\|\|\|\|\|\|\|]PAM | NA | Intergenic | NA | NA |
| GU305914.1 | 129276 | 129298 | + | 4 | CGACCATCGTGCAGACGCCC | AGG | -\|\|\|-\|-\|[\|\|\|-\|\|\|\|\|\|\|\|]PAM | NA | Intergenic | NA | NA |

| | | | | | | | | | | | |
|---|---|---|---|---|---|---|---|---|---|---|---|
| T5283 | GAGCGTGGAGACGCCCGGGTCGG | 1000 | | | | | | | | | |
| Oligo fwd | CACCGAGCGTGGAGACGCCCGGGT | | | | | | | | | | |
| Oligo rev | AAACACCCGGGCGTCTCCACGCTC | | | | | | | | | | |
| Chromosome | start | end | strand | MM | target_seq | PAM | alignment | distance | position | gene name | gene id |
| GU305914.1 | 74898 | 74920 | + | 0 | GAGCGTGGAGACGCCCGGGT | CGG | \|\|\|\|\|\|\|\|[\|\|\|\|\|\|\|\|\|\|\|\|]PAM | NA | Intergenic | NA | NA |

| | | | | | | | | | | | |
|---|---|---|---|---|---|---|---|---|---|---|---|
| T5284 | AGCGTGGAGACGCCCGGGTCGGG | 1000 | | | | | | | | | |
| Oligo fwd | CACCAGCGTGGAGACGCCCGGGTC | | | | | | | | | | |
| Oligo rev | AAACGACCCGGGCGTCTCCACGCT | | | | | | | | | | |
| Chromosome | start | end | strand | MM | target_seq | PAM | alignment | distance | position | gene name | gene id |
| GU305914.1 | 74899 | 74921 | + | 0 | AGCGTGGAGACGCCCGGGTC | GGG | \|\|\|\|\|\|\|\|[\|\|\|\|\|\|\|\|\|\|\|\|]PAM | NA | Intergenic | NA | NA |

| | | | | | | | | | | | |
|---|---|---|---|---|---|---|---|---|---|---|---|
| T5285 | AGACGCCCGGGTCGGGACCGAGG | 1000 | | | | | | | | | |
| Oligo fwd | CACCAGACGCCCGGGTCGGGACCG | | | | | | | | | | |
| Oligo rev | AAACCGGTCCCGACCCGGGCGTCT | | | | | | | | | | |
| Chromosome | start | end | strand | MM | target_seq | PAM | alignment | distance | position | gene name | gene id |
| GU305914.1 | 74906 | 74928 | + | 0 | AGACGCCCGGGTCGGGACCG | AGG | \|\|\|\|\|\|\|\|[\|\|\|\|\|\|\|\|\|\|\|\|]PAM | NA | Intergenic | NA | NA |

| | | | | | | | | | | | |
|---|---|---|---|---|---|---|---|---|---|---|---|
| T5286 | CGCCCGGGTCGGGACCGAGGCGG | 1000 | | | | | | | | | |
| Oligo fwd | CACCCGCCCGGGTCGGGACCGAGG | | | | | | | | | | |
| Oligo rev | AAACCCTCGGTCCCGACCCGGGCG | | | | | | | | | | |
| Chromosome | start | end | strand | MM | target_seq | PAM | alignment | distance | position | gene name | gene id |

| | | | | | | | | | | | | |
|---|---|---|---|---|---|---|---|---|---|---|---|---|
| GU305914.1 | | 74909 | 74931 | + | 0 | CGCCCGGGTCGGGACCGAGG | CGG | \|\|\|\|\|\|\|\|[\|\|\|\|\|\|\|\|\|\|\|\|]PAM | NA | Intergenic | NA | NA |
| T5287 | GGTCGGGACCGAGGCGGCGCAGG | | 1000 | | | | | | | | | |
| Oligo fwd | CACCGGTCGGGACCGAGGCGGCGC | | | | | | | | | | | |
| Oligo rev | AAACGCGCCGCCTCGGTCCCGACC | | | | | | | | | | | |
| Chromosome | start | | end | strand | MM | target_seq | PAM | alignment | distance | position | gene name | gene id |
| GU305914.1 | | 74915 | 74937 | + | 0 | GGTCGGGACCGAGGCGGCGC | AGG | \|\|\|\|\|\|\|\|[\|\|\|\|\|\|\|\|\|\|\|\|]PAM | NA | Intergenic | NA | NA |
| T5288 | GACCGAGGCGGCGCAGGTTGAGG | | 1000 | | | | | | | | | |
| Oligo fwd | CACCGACCGAGGCGGCGCAGGTTG | | | | | | | | | | | |
| Oligo rev | AAACCAACCTGCGCCGCCTCGGTC | | | | | | | | | | | |
| Chromosome | start | | end | strand | MM | target_seq | PAM | alignment | distance | position | gene name | gene id |
| GU305914.1 | | 74921 | 74943 | + | 0 | GACCGAGGCGGCGCAGGTTG | AGG | \|\|\|\|\|\|\|\|[\|\|\|\|\|\|\|\|\|\|\|\|]PAM | NA | Intergenic | NA | NA |
| T5289 | CGAGGCGGCGCAGGTTGAGGCGG | | 1000 | | | | | | | | | |
| Oligo fwd | CACCCGAGGCGGCGCAGGTTGAGG | | | | | | | | | | | |
| Oligo rev | AAACCCTCAACCTGCGCCGCCTCG | | | | | | | | | | | |
| Chromosome | start | | end | strand | MM | target_seq | PAM | alignment | distance | position | gene name | gene id |
| GU305914.1 | | 74924 | 74946 | + | 0 | CGAGGCGGCGCAGGTTGAGG | CGG | \|\|\|\|\|\|\|\|[\|\|\|\|\|\|\|\|\|\|\|\|]PAM | NA | Intergenic | NA | NA |
| T5290 | GGCGGCGCAGGTTGAGGCGGCGG | | 1000 | | | | | | | | | |
| Oligo fwd | CACCGGCGGCGCAGGTTGAGGCGG | | | | | | | | | | | |
| Oligo rev | AAACCCGCCTCAACCTGCGCCGCC | | | | | | | | | | | |
| Chromosome | start | | end | strand | MM | target_seq | PAM | alignment | distance | position | gene name | gene id |
| GU305914.1 | | 74927 | 74949 | + | 0 | GGCGGCGCAGGTTGAGGCGG | CGG | \|\|\|\|\|\|\|\|[\|\|\|\|\|\|\|\|\|\|\|\|]PAM | NA | Intergenic | NA | NA |
| T5291 | GGCGCAGGTTGAGGCGGCGGTGG | | 1000 | | | | | | | | | |
| Oligo fwd | CACCGGCGCAGGTTGAGGCGGCGG | | | | | | | | | | | |
| Oligo rev | AAACCCGCCGCCTCAACCTGCGCC | | | | | | | | | | | |
| Chromosome | start | | end | strand | MM | target_seq | PAM | alignment | distance | position | gene name | gene id |
| GU305914.1 | | 74930 | 74952 | + | 0 | GGCGCAGGTTGAGGCGGCGG | TGG | \|\|\|\|\|\|\|\|[\|\|\|\|\|\|\|\|\|\|\|\|]PAM | NA | Intergenic | NA | NA |
| GU305914.1 | | 48159 | 48181 | + | 4 | GGCGGCGGTGGCGGCGGCGG | TGG | \|\|\|\|--\|\|[\|-\|-\|\|\|\|\|\|\|\|]PAM | NA | Intergenic | NA | NA |
| GU305914.1 | | 62446 | 62468 | + | 4 | GGCGGGGGTGGTGGCGGCGG | GGG | \|\|\|\|--\|\|[\|-\|-\|\|\|\|\|\|\|\|]PAM | NA | Intergenic | NA | NA |
| GU305914.1 | | 114658 | 114680 | - | 4 | GGCGGTGGTGGTGGCGGCGG | CGG | \|\|\|\|--\|\|[\|-\|-\|\|\|\|\|\|\|\|]PAM | NA | Intergenic | NA | NA |
| GU305914.1 | | 153117 | 153139 | - | 4 | GGTGGAGGTTGAAGCTGCGG | TGG | \|\|-\|-\|\|\|[\|\|\|\|-\|\|-\|\|\|\|]PAM | NA | Intergenic | NA | NA |
| T5292 | GTTGAGGCGGCGGTGGTAGACGG | | 1000 | | | | | | | | | |
| Oligo fwd | CACCGTTGAGGCGGCGGTGGTAGA | | | | | | | | | | | |
| Oligo rev | AAACTCTACCACCGCCGCCTCAAC | | | | | | | | | | | |
| Chromosome | start | | end | strand | MM | target_seq | PAM | alignment | distance | position | gene name | gene id |
| GU305914.1 | | 74937 | 74959 | + | 0 | GTTGAGGCGGCGGTGGTAGA | CGG | \|\|\|\|\|\|\|\|[\|\|\|\|\|\|\|\|\|\|\|\|]PAM | NA | Intergenic | NA | NA |
| T5293 | TTGAGGCGGCGGTGGTAGACGGG | | 1000 | | | | | | | | | |
| Oligo fwd | CACCTTGAGGCGGCGGTGGTAGAC | | | | | | | | | | | |
| Oligo rev | AAACGTCTACCACCGCCGCCTCAA | | | | | | | | | | | |
| Chromosome | start | | end | strand | MM | target_seq | PAM | alignment | distance | position | gene name | gene id |
| GU305914.1 | | 74938 | 74960 | + | 0 | TTGAGGCGGCGGTGGTAGAC | GGG | \|\|\|\|\|\|\|\|[\|\|\|\|\|\|\|\|\|\|\|\|]PAM | NA | Intergenic | NA | NA |
| T5294 | GGCGGCGGTGGTAGACGGGCAGG | | 1000 | | | | | | | | | |
| Oligo fwd | CACCGGCGGCGGTGGTAGACGGGC | | | | | | | | | | | |
| Oligo rev | AAACGCCCGTCTACCACCGCCGCC | | | | | | | | | | | |
| Chromosome | start | | end | strand | MM | target_seq | PAM | alignment | distance | position | gene name | gene id |
| GU305914.1 | | 74942 | 74964 | + | 0 | GGCGGCGGTGGTAGACGGGC | AGG | \|\|\|\|\|\|\|\|[\|\|\|\|\|\|\|\|\|\|\|\|]PAM | NA | Intergenic | NA | NA |
| T5295 | CGCAGCCGACCTCGTACTCGCGG | | 1000 | | | | | | | | | |
| Oligo fwd | CACCCGCAGCCGACCTCGTACTCG | | | | | | | | | | | |
| Oligo rev | AAACCGAGTACGAGGTCGGCTGCG | | | | | | | | | | | |
| Chromosome | start | | end | strand | MM | target_seq | PAM | alignment | distance | position | gene name | gene id |
| GU305914.1 | | 74966 | 74988 | + | 0 | CGCAGCCGACCTCGTACTCG | CGG | \|\|\|\|\|\|\|\|[\|\|\|\|\|\|\|\|\|\|\|\|]PAM | NA | Intergenic | NA | NA |
| T5296 | GCAGCCGACCTCGTACTCGCGGG | | 1000 | | | | | | | | | |
| Oligo fwd | CACCGCAGCCGACCTCGTACTCGC | | | | | | | | | | | |
| Oligo rev | AAACGCGAGTACGAGGTCGGCTGC | | | | | | | | | | | |
| Chromosome | start | | end | strand | MM | target_seq | PAM | alignment | distance | position | gene name | gene id |
| GU305914.1 | | 74967 | 74989 | + | 0 | GCAGCCGACCTCGTACTCGC | GGG | \|\|\|\|\|\|\|\|[\|\|\|\|\|\|\|\|\|\|\|\|]PAM | NA | Intergenic | NA | NA |
| T5297 | CGACCTCGTACTCGCGGGCGCGG | | 1000 | | | | | | | | | |
| Oligo fwd | CACCCGACCTCGTACTCGCGGGCG | | | | | | | | | | | |
| Oligo rev | AAACCGCCCGCGAGTACGAGGTCG | | | | | | | | | | | |
| Chromosome | start | | end | strand | MM | target_seq | PAM | alignment | distance | position | gene name | gene id |
| GU305914.1 | | 74972 | 74994 | + | 0 | CGACCTCGTACTCGCGGGCG | CGG | \|\|\|\|\|\|\|\|[\|\|\|\|\|\|\|\|\|\|\|\|]PAM | NA | Intergenic | NA | NA |
| T5298 | CCTCGTACTCGCGGGCGCGGAGG | | 1000 | | | | | | | | | |
| Oligo fwd | CACCCCTCGTACTCGCGGGCGCGG | | | | | | | | | | | |
| Oligo rev | AAACCCGCGCCCGCGAGTACGAGG | | | | | | | | | | | |
| Chromosome | start | | end | strand | MM | target_seq | PAM | alignment | distance | position | gene name | gene id |
| GU305914.1 | | 74975 | 74997 | + | 0 | CCTCGTACTCGCGGGCGCGG | AGG | \|\|\|\|\|\|\|\|[\|\|\|\|\|\|\|\|\|\|\|\|]PAM | NA | Intergenic | NA | NA |
| T5299 | GTACTCGCGGGCGCGGAGGAAGG | | 1000 | | | | | | | | | |
| Oligo fwd | CACCGTACTCGCGGGCGCGGAGGA | | | | | | | | | | | |
| Oligo rev | AAACTCCTCCGCGCCCGCGAGTAC | | | | | | | | | | | |
| Chromosome | start | | end | strand | MM | target_seq | PAM | alignment | distance | position | gene name | gene id |
| GU305914.1 | | 74979 | 75001 | + | 0 | GTACTCGCGGGCGCGGAGGA | AGG | \|\|\|\|\|\|\|\|[\|\|\|\|\|\|\|\|\|\|\|\|]PAM | NA | Intergenic | NA | NA |
| T5300 | GGCGCGGAGGAAGGCGACGCAGG | | 1000 | | | | | | | | | |
| Oligo fwd | CACCGGCGCGGAGGAAGGCGACGC | | | | | | | | | | | |
| Oligo rev | AAACGCGTCGCCTTCCTCCGCGCC | | | | | | | | | | | |
| Chromosome | start | | end | strand | MM | target_seq | PAM | alignment | distance | position | gene name | gene id |
| GU305914.1 | | 74988 | 75010 | + | 0 | GGCGCGGAGGAAGGCGACGC | AGG | \|\|\|\|\|\|\|\|[\|\|\|\|\|\|\|\|\|\|\|\|]PAM | NA | Intergenic | NA | NA |

<table>
<tr><td>GU305914.1</td><td>126734</td><td>126756</td><td>-</td><td>4</td><td>CGCGCGGCGCATGGCGACGC</td><td>GGG</td><td>-||||||-[|-|-||||||||]PAM</td><td>NA</td><td>Intergenic</td><td>NA</td><td>NA</td></tr>
<tr><td>GU305914.1</td><td>61306</td><td>61328</td><td>+</td><td>4</td><td>GGCGGAGAGGAAGGCGTCGA</td><td>AGG</td><td>||||--||[||||||||-||-]PAM</td><td>NA</td><td>Intergenic</td><td>NA</td><td>NA</td></tr>
<tr><td>T5301</td><td>GCGCGGAGGAAGGCGACGCAGGG</td><td>1000</td><td></td><td></td><td></td><td></td><td></td><td></td><td></td><td></td><td></td></tr>
<tr><td>Oligo fwd</td><td>CACCGCGCGGAGGAAGGCGACGCA</td><td></td><td></td><td></td><td></td><td></td><td></td><td></td><td></td><td></td><td></td></tr>
<tr><td>Oligo rev</td><td>AAACTGCGTCGCCTTCCTCCGCGC</td><td></td><td></td><td></td><td></td><td></td><td></td><td></td><td></td><td></td><td></td></tr>
<tr><td>Chromosome</td><td>start</td><td>end</td><td>strand</td><td>MM</td><td>target_seq</td><td>PAM</td><td>alignment</td><td>distance</td><td>position</td><td>gene name</td><td>gene id</td></tr>
<tr><td>GU305914.1</td><td>74989</td><td>75011</td><td>+</td><td>0</td><td>GCGCGGAGGAAGGCGACGCA</td><td>GGG</td><td>||||||||[||||||||||||]PAM</td><td>NA</td><td>Intergenic</td><td>NA</td><td>NA</td></tr>
<tr><td>T5302</td><td>CGCGGAGGAAGGCGACGCAGGGG</td><td>1000</td><td></td><td></td><td></td><td></td><td></td><td></td><td></td><td></td><td></td></tr>
<tr><td>Oligo fwd</td><td>CACCCGCGGAGGAAGGCGACGCAG</td><td></td><td></td><td></td><td></td><td></td><td></td><td></td><td></td><td></td><td></td></tr>
<tr><td>Oligo rev</td><td>AAACCTGCGTCGCCTTCCTCCGCG</td><td></td><td></td><td></td><td></td><td></td><td></td><td></td><td></td><td></td><td></td></tr>
<tr><td>Chromosome</td><td>start</td><td>end</td><td>strand</td><td>MM</td><td>target_seq</td><td>PAM</td><td>alignment</td><td>distance</td><td>position</td><td>gene name</td><td>gene id</td></tr>
<tr><td>GU305914.1</td><td>74990</td><td>75012</td><td>+</td><td>0</td><td>CGCGGAGGAAGGCGACGCAG</td><td>GGG</td><td>||||||||[||||||||||||]PAM</td><td>NA</td><td>Intergenic</td><td>NA</td><td>NA</td></tr>
<tr><td>T5303</td><td>GCGGAGGAAGGCGACGCAGGGGG</td><td>1000</td><td></td><td></td><td></td><td></td><td></td><td></td><td></td><td></td><td></td></tr>
<tr><td>Oligo fwd</td><td>CACCGCGGAGGAAGGCGACGCAGG</td><td></td><td></td><td></td><td></td><td></td><td></td><td></td><td></td><td></td><td></td></tr>
<tr><td>Oligo rev</td><td>AAACCCTGCGTCGCCTTCCTCCGC</td><td></td><td></td><td></td><td></td><td></td><td></td><td></td><td></td><td></td><td></td></tr>
<tr><td>Chromosome</td><td>start</td><td>end</td><td>strand</td><td>MM</td><td>target_seq</td><td>PAM</td><td>alignment</td><td>distance</td><td>position</td><td>gene name</td><td>gene id</td></tr>
<tr><td>GU305914.1</td><td>74991</td><td>75013</td><td>+</td><td>0</td><td>GCGGAGGAAGGCGACGCAGG</td><td>GGG</td><td>||||||||[||||||||||||]PAM</td><td>NA</td><td>Intergenic</td><td>NA</td><td>NA</td></tr>
<tr><td>T5304</td><td>GAAGGCGACGCAGGGGGAGCTGG</td><td>1000</td><td></td><td></td><td></td><td></td><td></td><td></td><td></td><td></td><td></td></tr>
<tr><td>Oligo fwd</td><td>CACCGAAGGCGACGCAGGGGGAGC</td><td></td><td></td><td></td><td></td><td></td><td></td><td></td><td></td><td></td><td></td></tr>
<tr><td>Oligo rev</td><td>AAACGCTCCCCCTGCGTCGCCTTC</td><td></td><td></td><td></td><td></td><td></td><td></td><td></td><td></td><td></td><td></td></tr>
<tr><td>Chromosome</td><td>start</td><td>end</td><td>strand</td><td>MM</td><td>target_seq</td><td>PAM</td><td>alignment</td><td>distance</td><td>position</td><td>gene name</td><td>gene id</td></tr>
<tr><td>GU305914.1</td><td>74997</td><td>75019</td><td>+</td><td>0</td><td>GAAGGCGACGCAGGGGGAGC</td><td>TGG</td><td>||||||||[||||||||||||]PAM</td><td>NA</td><td>Intergenic</td><td>NA</td><td>NA</td></tr>
<tr><td>T5305</td><td>CGACGCAGGGGGAGCTGGCCTGG</td><td>1000</td><td></td><td></td><td></td><td></td><td></td><td></td><td></td><td></td><td></td></tr>
<tr><td>Oligo fwd</td><td>CACCCGACGCAGGGGGAGCTGGCC</td><td></td><td></td><td></td><td></td><td></td><td></td><td></td><td></td><td></td><td></td></tr>
<tr><td>Oligo rev</td><td>AAACGGCCAGCTCCCCCTGCGTCG</td><td></td><td></td><td></td><td></td><td></td><td></td><td></td><td></td><td></td><td></td></tr>
<tr><td>Chromosome</td><td>start</td><td>end</td><td>strand</td><td>MM</td><td>target_seq</td><td>PAM</td><td>alignment</td><td>distance</td><td>position</td><td>gene name</td><td>gene id</td></tr>
<tr><td>GU305914.1</td><td>75002</td><td>75024</td><td>+</td><td>0</td><td>CGACGCAGGGGGAGCTGGCC</td><td>TGG</td><td>||||||||[||||||||||||]PAM</td><td>NA</td><td>Intergenic</td><td>NA</td><td>NA</td></tr>
<tr><td>T5306</td><td>GCAGGGGGAGCTGGCCTGGTTGG</td><td>1000</td><td></td><td></td><td></td><td></td><td></td><td></td><td></td><td></td><td></td></tr>
<tr><td>Oligo fwd</td><td>CACCGCAGGGGGAGCTGGCCTGGT</td><td></td><td></td><td></td><td></td><td></td><td></td><td></td><td></td><td></td><td></td></tr>
<tr><td>Oligo rev</td><td>AAACACCAGGCCAGCTCCCCCTGC</td><td></td><td></td><td></td><td></td><td></td><td></td><td></td><td></td><td></td><td></td></tr>
<tr><td>Chromosome</td><td>start</td><td>end</td><td>strand</td><td>MM</td><td>target_seq</td><td>PAM</td><td>alignment</td><td>distance</td><td>position</td><td>gene name</td><td>gene id</td></tr>
<tr><td>GU305914.1</td><td>75006</td><td>75028</td><td>+</td><td>0</td><td>GCAGGGGGAGCTGGCCTGGT</td><td>TGG</td><td>||||||||[||||||||||||]PAM</td><td>NA</td><td>Intergenic</td><td>NA</td><td>NA</td></tr>
<tr><td>T5307</td><td>CAGGGGGAGCTGGCCTGGTTGGG</td><td>1000</td><td></td><td></td><td></td><td></td><td></td><td></td><td></td><td></td><td></td></tr>
<tr><td>Oligo fwd</td><td>CACCCAGGGGGAGCTGGCCTGGTT</td><td></td><td></td><td></td><td></td><td></td><td></td><td></td><td></td><td></td><td></td></tr>
<tr><td>Oligo rev</td><td>AAACAACCAGGCCAGCTCCCCCTG</td><td></td><td></td><td></td><td></td><td></td><td></td><td></td><td></td><td></td><td></td></tr>
<tr><td>Chromosome</td><td>start</td><td>end</td><td>strand</td><td>MM</td><td>target_seq</td><td>PAM</td><td>alignment</td><td>distance</td><td>position</td><td>gene name</td><td>gene id</td></tr>
<tr><td>GU305914.1</td><td>75007</td><td>75029</td><td>+</td><td>0</td><td>CAGGGGGAGCTGGCCTGGTT</td><td>GGG</td><td>||||||||[||||||||||||]PAM</td><td>NA</td><td>Intergenic</td><td>NA</td><td>NA</td></tr>
<tr><td>T5308</td><td>GGAGCTGGCCTGGTTGGGAATGG</td><td>1000</td><td></td><td></td><td></td><td></td><td></td><td></td><td></td><td></td><td></td></tr>
<tr><td>Oligo fwd</td><td>CACCGGAGCTGGCCTGGTTGGGAA</td><td></td><td></td><td></td><td></td><td></td><td></td><td></td><td></td><td></td><td></td></tr>
<tr><td>Oligo rev</td><td>AAACTTCCCAACCAGGCCAGCTCC</td><td></td><td></td><td></td><td></td><td></td><td></td><td></td><td></td><td></td><td></td></tr>
<tr><td>Chromosome</td><td>start</td><td>end</td><td>strand</td><td>MM</td><td>target_seq</td><td>PAM</td><td>alignment</td><td>distance</td><td>position</td><td>gene name</td><td>gene id</td></tr>
<tr><td>GU305914.1</td><td>75012</td><td>75034</td><td>+</td><td>0</td><td>GGAGCTGGCCTGGTTGGGAA</td><td>TGG</td><td>||||||||[||||||||||||]PAM</td><td>NA</td><td>Intergenic</td><td>NA</td><td>NA</td></tr>
<tr><td>T5309</td><td>GAGCTGGCCTGGTTGGGAATGGG</td><td>1000</td><td></td><td></td><td></td><td></td><td></td><td></td><td></td><td></td><td></td></tr>
<tr><td>Oligo fwd</td><td>CACCGAGCTGGCCTGGTTGGGAAT</td><td></td><td></td><td></td><td></td><td></td><td></td><td></td><td></td><td></td><td></td></tr>
<tr><td>Oligo rev</td><td>AAACATTCCCAACCAGGCCAGCTC</td><td></td><td></td><td></td><td></td><td></td><td></td><td></td><td></td><td></td><td></td></tr>
<tr><td>Chromosome</td><td>start</td><td>end</td><td>strand</td><td>MM</td><td>target_seq</td><td>PAM</td><td>alignment</td><td>distance</td><td>position</td><td>gene name</td><td>gene id</td></tr>
<tr><td>GU305914.1</td><td>75013</td><td>75035</td><td>+</td><td>0</td><td>GAGCTGGCCTGGTTGGGAAT</td><td>GGG</td><td>||||||||[||||||||||||]PAM</td><td>NA</td><td>Intergenic</td><td>NA</td><td>NA</td></tr>
<tr><td>T5310</td><td>AGCTGGCCTGGTTGGGAATGGGG</td><td>1000</td><td></td><td></td><td></td><td></td><td></td><td></td><td></td><td></td><td></td></tr>
<tr><td>Oligo fwd</td><td>CACCAGCTGGCCTGGTTGGGAATG</td><td></td><td></td><td></td><td></td><td></td><td></td><td></td><td></td><td></td><td></td></tr>
<tr><td>Oligo rev</td><td>AAACCATTCCCAACCAGGCCAGCT</td><td></td><td></td><td></td><td></td><td></td><td></td><td></td><td></td><td></td><td></td></tr>
<tr><td>Chromosome</td><td>start</td><td>end</td><td>strand</td><td>MM</td><td>target_seq</td><td>PAM</td><td>alignment</td><td>distance</td><td>position</td><td>gene name</td><td>gene id</td></tr>
<tr><td>GU305914.1</td><td>75014</td><td>75036</td><td>+</td><td>0</td><td>AGCTGGCCTGGTTGGGAATG</td><td>GGG</td><td>||||||||[||||||||||||]PAM</td><td>NA</td><td>Intergenic</td><td>NA</td><td>NA</td></tr>
<tr><td>T5311</td><td>GTTGGGAATGGGGATGCCGAAGG</td><td>1000</td><td></td><td></td><td></td><td></td><td></td><td></td><td></td><td></td><td></td></tr>
<tr><td>Oligo fwd</td><td>CACCGTTGGGAATGGGGATGCCGA</td><td></td><td></td><td></td><td></td><td></td><td></td><td></td><td></td><td></td><td></td></tr>
<tr><td>Oligo rev</td><td>AAACTCGGCATCCCCATTCCCAAC</td><td></td><td></td><td></td><td></td><td></td><td></td><td></td><td></td><td></td><td></td></tr>
<tr><td>Chromosome</td><td>start</td><td>end</td><td>strand</td><td>MM</td><td>target_seq</td><td>PAM</td><td>alignment</td><td>distance</td><td>position</td><td>gene name</td><td>gene id</td></tr>
<tr><td>GU305914.1</td><td>75024</td><td>75046</td><td>+</td><td>0</td><td>GTTGGGAATGGGGATGCCGA</td><td>AGG</td><td>||||||||[||||||||||||]PAM</td><td>NA</td><td>Intergenic</td><td>NA</td><td>NA</td></tr>
<tr><td>T5312</td><td>TTGGGAATGGGGATGCCGAAGGG</td><td>1000</td><td></td><td></td><td></td><td></td><td></td><td></td><td></td><td></td><td></td></tr>
<tr><td>Oligo fwd</td><td>CACCTTGGGAATGGGGATGCCGAA</td><td></td><td></td><td></td><td></td><td></td><td></td><td></td><td></td><td></td><td></td></tr>
<tr><td>Oligo rev</td><td>AAACTTCGGCATCCCCATTCCCAA</td><td></td><td></td><td></td><td></td><td></td><td></td><td></td><td></td><td></td><td></td></tr>
<tr><td>Chromosome</td><td>start</td><td>end</td><td>strand</td><td>MM</td><td>target_seq</td><td>PAM</td><td>alignment</td><td>distance</td><td>position</td><td>gene name</td><td>gene id</td></tr>
<tr><td>GU305914.1</td><td>75025</td><td>75047</td><td>+</td><td>0</td><td>TTGGGAATGGGGATGCCGAA</td><td>GGG</td><td>||||||||[||||||||||||]PAM</td><td>NA</td><td>Intergenic</td><td>NA</td><td>NA</td></tr>
<tr><td>T5313</td><td>TGGGAATGGGGATGCCGAAGGGG</td><td>1000</td><td></td><td></td><td></td><td></td><td></td><td></td><td></td><td></td><td></td></tr>
<tr><td>Oligo fwd</td><td>CACCTGGGAATGGGGATGCCGAAG</td><td></td><td></td><td></td><td></td><td></td><td></td><td></td><td></td><td></td><td></td></tr>
<tr><td>Oligo rev</td><td>AAACCTTCGGCATCCCCATTCCCA</td><td></td><td></td><td></td><td></td><td></td><td></td><td></td><td></td><td></td><td></td></tr>
<tr><td>Chromosome</td><td>start</td><td>end</td><td>strand</td><td>MM</td><td>target_seq</td><td>PAM</td><td>alignment</td><td>distance</td><td>position</td><td>gene name</td><td>gene id</td></tr>
<tr><td>GU305914.1</td><td>75026</td><td>75048</td><td>+</td><td>0</td><td>TGGGAATGGGGATGCCGAAG</td><td>GGG</td><td>||||||||[||||||||||||]PAM</td><td>NA</td><td>Intergenic</td><td>NA</td><td>NA</td></tr>
<tr><td>T5314</td><td>TGGGGATGCCGAAGGGGAAGTGG</td><td>1000</td><td></td><td></td><td></td><td></td><td></td><td></td><td></td><td></td><td></td></tr>
<tr><td>Oligo fwd</td><td>CACCTGGGGATGCCGAAGGGGAAG</td><td></td><td></td><td></td><td></td><td></td><td></td><td></td><td></td><td></td><td></td></tr>
<tr><td>Oligo rev</td><td>AAACCTTCCCCTTCGGCATCCCCA</td><td></td><td></td><td></td><td></td><td></td><td></td><td></td><td></td><td></td><td></td></tr>
<tr><td>Chromosome</td><td>start</td><td>end</td><td>strand</td><td>MM</td><td>target_seq</td><td>PAM</td><td>alignment</td><td>distance</td><td>position</td><td>gene name</td><td>gene id</td></tr>
<tr><td>GU305914.1</td><td>75032</td><td>75054</td><td>+</td><td>0</td><td>TGGGGATGCCGAAGGGGAAG</td><td>TGG</td><td>||||||||[||||||||||||]PAM</td><td>NA</td><td>Intergenic</td><td>NA</td><td>NA</td></tr>
<tr><td>T5315</td><td>AGGGGAAGTGGAAGACGAACTGG</td><td>1000</td><td></td><td></td><td></td><td></td><td></td><td></td><td></td><td></td><td></td></tr>
<tr><td>Oligo fwd</td><td>CACCAGGGGAAGTGGAAGACGAAC</td><td></td><td></td><td></td><td></td><td></td><td></td><td></td><td></td><td></td><td></td></tr>
</table>

| | | | | | | | | | | | |
|---|---|---|---|---|---|---|---|---|---|---|---|
| Oligo rev | AAACGTTCGTCTTCCACTTCCCCT | | | | | | | | | | |
| Chromosome | start | end | strand | MM | target_seq | PAM | alignment | distance | position | gene name | gene id |
| GU305914.1 | 75044 | 75066 | + | 0 | AGGGGAAGTGGAAGACGAAC | TGG | \|\|\|\|\|\|\|\|[\|\|\|\|\|\|\|\|\|\|\|\|]PAM | NA | Intergenic | NA | NA |
| T5316 | AGTGGAAGACGAACTGGTGCAGG | 1000 | | | | | | | | | |
| Oligo fwd | CACCAGTGGAAGACGAACTGGTGC | | | | | | | | | | |
| Oligo rev | AAACGCACCAGTTCGTCTTCCACT | | | | | | | | | | |
| Chromosome | start | end | strand | MM | target_seq | PAM | alignment | distance | position | gene name | gene id |
| GU305914.1 | 75050 | 75072 | + | 0 | AGTGGAAGACGAACTGGTGC | AGG | \|\|\|\|\|\|\|\|[\|\|\|\|\|\|\|\|\|\|\|\|]PAM | NA | Intergenic | NA | NA |
| T5317 | GAAGACGAACTGGTGCAGGTCGG | 1000 | | | | | | | | | |
| Oligo fwd | CACCGAAGACGAACTGGTGCAGGT | | | | | | | | | | |
| Oligo rev | AAACACCTGCACCAGTTCGTCTTC | | | | | | | | | | |
| Chromosome | start | end | strand | MM | target_seq | PAM | alignment | distance | position | gene name | gene id |
| GU305914.1 | 75054 | 75076 | + | 0 | GAAGACGAACTGGTGCAGGT | CGG | \|\|\|\|\|\|\|\|[\|\|\|\|\|\|\|\|\|\|\|\|]PAM | NA | Intergenic | NA | NA |
| T5318 | CGAACTGGTGCAGGTCGGTGAGG | 1000 | | | | | | | | | |
| Oligo fwd | CACCCGAACTGGTGCAGGTCGGTG | | | | | | | | | | |
| Oligo rev | AAACCACCGACCTGCACCAGTTCG | | | | | | | | | | |
| Chromosome | start | end | strand | MM | target_seq | PAM | alignment | distance | position | gene name | gene id |
| GU305914.1 | 75059 | 75081 | + | 0 | CGAACTGGTGCAGGTCGGTG | AGG | \|\|\|\|\|\|\|\|[\|\|\|\|\|\|\|\|\|\|\|\|]PAM | NA | Intergenic | NA | NA |
| T5319 | GGTGCAGGTCGGTGAGGCCCTGG | 1000 | | | | | | | | | |
| Oligo fwd | CACCGGTGCAGGTCGGTGAGGCCC | | | | | | | | | | |
| Oligo rev | AAACGGGCCTCACCGACCTGCACC | | | | | | | | | | |
| Chromosome | start | end | strand | MM | target_seq | PAM | alignment | distance | position | gene name | gene id |
| GU305914.1 | 75065 | 75087 | + | 0 | GGTGCAGGTCGGTGAGGCCC | TGG | \|\|\|\|\|\|\|\|[\|\|\|\|\|\|\|\|\|\|\|\|]PAM | NA | Intergenic | NA | NA |
| T5320 | GTCGGTGAGGCCCTGGTAGCTGG | 1000 | | | | | | | | | |
| Oligo fwd | CACCGTCGGTGAGGCCCTGGTAGC | | | | | | | | | | |
| Oligo rev | AAACGCTACCAGGGCCTCACCGAC | | | | | | | | | | |
| Chromosome | start | end | strand | MM | target_seq | PAM | alignment | distance | position | gene name | gene id |
| GU305914.1 | 75072 | 75094 | + | 0 | GTCGGTGAGGCCCTGGTAGC | TGG | \|\|\|\|\|\|\|\|[\|\|\|\|\|\|\|\|\|\|\|\|]PAM | NA | Intergenic | NA | NA |
| T5321 | GGTAGCTGGCGTTGTCGACCAGG | 1000 | | | | | | | | | |
| Oligo fwd | CACCGGTAGCTGGCGTTGTCGACC | | | | | | | | | | |
| Oligo rev | AAACGGTCGACAACGCCAGCTACC | | | | | | | | | | |
| Chromosome | start | end | strand | MM | target_seq | PAM | alignment | distance | position | gene name | gene id |
| GU305914.1 | 75086 | 75108 | + | 0 | GGTAGCTGGCGTTGTCGACC | AGG | \|\|\|\|\|\|\|\|[\|\|\|\|\|\|\|\|\|\|\|\|]PAM | NA | Intergenic | NA | NA |
| T5322 | GTAGCTGGCGTTGTCGACCAGGG | 1000 | | | | | | | | | |
| Oligo fwd | CACCGTAGCTGGCGTTGTCGACCA | | | | | | | | | | |
| Oligo rev | AAACTGGTCGACAACGCCAGCTAC | | | | | | | | | | |
| Chromosome | start | end | strand | MM | target_seq | PAM | alignment | distance | position | gene name | gene id |
| GU305914.1 | 75087 | 75109 | + | 0 | GTAGCTGGCGTTGTCGACCA | GGG | \|\|\|\|\|\|\|\|[\|\|\|\|\|\|\|\|\|\|\|\|]PAM | NA | Intergenic | NA | NA |
| T5323 | GTCGACCAGGGCGTCGACCTTGG | 1000 | | | | | | | | | |
| Oligo fwd | CACCGTCGACCAGGGCGTCGACCT | | | | | | | | | | |
| Oligo rev | AAACAGGTCGACGCCCTGGTCGAC | | | | | | | | | | |
| Chromosome | start | end | strand | MM | target_seq | PAM | alignment | distance | position | gene name | gene id |
| GU305914.1 | 75099 | 75121 | + | 0 | GTCGACCAGGGCGTCGACCT | TGG | \|\|\|\|\|\|\|\|[\|\|\|\|\|\|\|\|\|\|\|\|]PAM | NA | Intergenic | NA | NA |
| T5324 | CCAGGGCGTCGACCTTGGAGCGG | 1000 | | | | | | | | | |
| Oligo fwd | CACCCCAGGGCGTCGACCTTGGAG | | | | | | | | | | |
| Oligo rev | AAACCTCCAAGGTCGACGCCCTGG | | | | | | | | | | |
| Chromosome | start | end | strand | MM | target_seq | PAM | alignment | distance | position | gene name | gene id |
| GU305914.1 | 75104 | 75126 | + | 0 | CCAGGGCGTCGACCTTGGAG | CGG | \|\|\|\|\|\|\|\|[\|\|\|\|\|\|\|\|\|\|\|\|]PAM | NA | Intergenic | NA | NA |
| T5325 | ACCTCTTGAAGACGTGCTCGAGG | 1000 | | | | | | | | | |
| Oligo fwd | CACCACCTCTTGAAGACGTGCTCG | | | | | | | | | | |
| Oligo rev | AAACCGAGCACGTCTTCAAGAGGT | | | | | | | | | | |
| Chromosome | start | end | strand | MM | target_seq | PAM | alignment | distance | position | gene name | gene id |
| GU305914.1 | 75131 | 75153 | + | 0 | ACCTCTTGAAGACGTGCTCG | AGG | \|\|\|\|\|\|\|\|[\|\|\|\|\|\|\|\|\|\|\|\|]PAM | NA | Intergenic | NA | NA |
| T5326 | TCTTGAAGACGTGCTCGAGGCGG | 1000 | | | | | | | | | |
| Oligo fwd | CACCTCTTGAAGACGTGCTCGAGG | | | | | | | | | | |
| Oligo rev | AAACCCTCGAGCACGTCTTCAAGA | | | | | | | | | | |
| Chromosome | start | end | strand | MM | target_seq | PAM | alignment | distance | position | gene name | gene id |
| GU305914.1 | 75134 | 75156 | + | 0 | TCTTGAAGACGTGCTCGAGG | CGG | \|\|\|\|\|\|\|\|[\|\|\|\|\|\|\|\|\|\|\|\|]PAM | NA | Intergenic | NA | NA |
| GU305914.1 | 55704 | 55726 | + | 4 | CCATGAAGACGGGCTCGAGT | CGG | -\|-\|\|\|\|\|[\|\|\|-\|\|\|\|\|\|\|-]PAM | NA | Intergenic | NA | NA |
| T5327 | GGCTGCTCATGCCGCTCCTGAGG | 1000 | | | | | | | | | |
| Oligo fwd | CACCGGCTGCTCATGCCGCTCCTG | | | | | | | | | | |
| Oligo rev | AAACCAGGAGCGGCATGAGCAGCC | | | | | | | | | | |
| Chromosome | start | end | strand | MM | target_seq | PAM | alignment | distance | position | gene name | gene id |
| GU305914.1 | 75155 | 75177 | + | 0 | GGCTGCTCATGCCGCTCCTG | AGG | \|\|\|\|\|\|\|\|[\|\|\|\|\|\|\|\|\|\|\|\|]PAM | NA | Intergenic | NA | NA |
| T5328 | TCATGCCGCTCCTGAGGTAGAGG | 1000 | | | | | | | | | |
| Oligo fwd | CACCTCATGCCGCTCCTGAGGTAG | | | | | | | | | | |
| Oligo rev | AAACCTACCTCAGGAGCGGCATGA | | | | | | | | | | |
| Chromosome | start | end | strand | MM | target_seq | PAM | alignment | distance | position | gene name | gene id |
| GU305914.1 | 75161 | 75183 | + | 0 | TCATGCCGCTCCTGAGGTAG | AGG | \|\|\|\|\|\|\|\|[\|\|\|\|\|\|\|\|\|\|\|\|]PAM | NA | Intergenic | NA | NA |
| T5329 | GCCGCTCCTGAGGTAGAGGCAGG | 1000 | | | | | | | | | |
| Oligo fwd | CACCGCCGCTCCTGAGGTAGAGGC | | | | | | | | | | |
| Oligo rev | AAACGCCTCTACCTCAGGAGCGGC | | | | | | | | | | |
| Chromosome | start | end | strand | MM | target_seq | PAM | alignment | distance | position | gene name | gene id |
| GU305914.1 | 75165 | 75187 | + | 0 | GCCGCTCCTGAGGTAGAGGC | AGG | \|\|\|\|\|\|\|\|[\|\|\|\|\|\|\|\|\|\|\|\|]PAM | NA | Intergenic | NA | NA |

<table>
<tr><td>T5330</td><td>GACGCGCTCCATGCAGCGCACGG</td><td>1000</td><td></td><td></td><td></td><td></td><td></td><td></td><td></td><td></td><td></td></tr>
<tr><td>Oligo fwd</td><td>CACCGACGCGCTCCATGCAGCGCA</td><td></td><td></td><td></td><td></td><td></td><td></td><td></td><td></td><td></td><td></td></tr>
<tr><td>Oligo rev</td><td>AAACTGCGCTGCATGGAGCGCGTC</td><td></td><td></td><td></td><td></td><td></td><td></td><td></td><td></td><td></td><td></td></tr>
<tr><td>Chromosome</td><td>start</td><td>end</td><td>strand</td><td>MM</td><td>target_seq</td><td>PAM</td><td>alignment</td><td>distance</td><td>position</td><td>gene name</td><td>gene id</td></tr>
<tr><td>GU305914.1</td><td>75201</td><td>75223</td><td>+</td><td>0</td><td>GACGCGCTCCATGCAGCGCA</td><td>CGG</td><td>||||||||[||||||||||||]PAM</td><td>NA</td><td>Intergenic</td><td>NA</td><td>NA</td></tr>
<tr><td>T5331</td><td>GCAGCGCACGGCGTAAAAGTAGG</td><td>1000</td><td></td><td></td><td></td><td></td><td></td><td></td><td></td><td></td><td></td></tr>
<tr><td>Oligo fwd</td><td>CACCGCAGCGCACGGCGTAAAAGT</td><td></td><td></td><td></td><td></td><td></td><td></td><td></td><td></td><td></td><td></td></tr>
<tr><td>Oligo rev</td><td>AAACACTTTTACGCCGTGCGCTGC</td><td></td><td></td><td></td><td></td><td></td><td></td><td></td><td></td><td></td><td></td></tr>
<tr><td>Chromosome</td><td>start</td><td>end</td><td>strand</td><td>MM</td><td>target_seq</td><td>PAM</td><td>alignment</td><td>distance</td><td>position</td><td>gene name</td><td>gene id</td></tr>
<tr><td>GU305914.1</td><td>75213</td><td>75235</td><td>+</td><td>0</td><td>GCAGCGCACGGCGTAAAAGT</td><td>AGG</td><td>||||||||[||||||||||||]PAM</td><td>NA</td><td>Intergenic</td><td>NA</td><td>NA</td></tr>
<tr><td>T5332</td><td>CGGCGTAAAAGTAGGCGAGCAGG</td><td>1000</td><td></td><td></td><td></td><td></td><td></td><td></td><td></td><td></td><td></td></tr>
<tr><td>Oligo fwd</td><td>CACCCGGCGTAAAAGTAGGCGAGC</td><td></td><td></td><td></td><td></td><td></td><td></td><td></td><td></td><td></td><td></td></tr>
<tr><td>Oligo rev</td><td>AAACGCTCGCCTACTTTTACGCCG</td><td></td><td></td><td></td><td></td><td></td><td></td><td></td><td></td><td></td><td></td></tr>
<tr><td>Chromosome</td><td>start</td><td>end</td><td>strand</td><td>MM</td><td>target_seq</td><td>PAM</td><td>alignment</td><td>distance</td><td>position</td><td>gene name</td><td>gene id</td></tr>
<tr><td>GU305914.1</td><td>75221</td><td>75243</td><td>+</td><td>0</td><td>CGGCGTAAAAGTAGGCGAGC</td><td>AGG</td><td>||||||||[||||||||||||]PAM</td><td>NA</td><td>Intergenic</td><td>NA</td><td>NA</td></tr>
<tr><td>T5333</td><td>CCTCTTTCTGCGAGATCTCGCGG</td><td>1000</td><td></td><td></td><td></td><td></td><td></td><td></td><td></td><td></td><td></td></tr>
<tr><td>Oligo fwd</td><td>CACCCCTCTTTCTGCGAGATCTCG</td><td></td><td></td><td></td><td></td><td></td><td></td><td></td><td></td><td></td><td></td></tr>
<tr><td>Oligo rev</td><td>AAACCGAGATCTCGCAGAAAGAGG</td><td></td><td></td><td></td><td></td><td></td><td></td><td></td><td></td><td></td><td></td></tr>
<tr><td>Chromosome</td><td>start</td><td>end</td><td>strand</td><td>MM</td><td>target_seq</td><td>PAM</td><td>alignment</td><td>distance</td><td>position</td><td>gene name</td><td>gene id</td></tr>
<tr><td>GU305914.1</td><td>75266</td><td>75288</td><td>+</td><td>0</td><td>CCTCTTTCTGCGAGATCTCG</td><td>CGG</td><td>||||||||[||||||||||||]PAM</td><td>NA</td><td>Intergenic</td><td>NA</td><td>NA</td></tr>
<tr><td>T5334</td><td>CTGCGAGATCTCGCGGACGATGG</td><td>1000</td><td></td><td></td><td></td><td></td><td></td><td></td><td></td><td></td><td></td></tr>
<tr><td>Oligo fwd</td><td>CACCCTGCGAGATCTCGCGGACGA</td><td></td><td></td><td></td><td></td><td></td><td></td><td></td><td></td><td></td><td></td></tr>
<tr><td>Oligo rev</td><td>AAACTCGTCCGCGAGATCTCGCAG</td><td></td><td></td><td></td><td></td><td></td><td></td><td></td><td></td><td></td><td></td></tr>
<tr><td>Chromosome</td><td>start</td><td>end</td><td>strand</td><td>MM</td><td>target_seq</td><td>PAM</td><td>alignment</td><td>distance</td><td>position</td><td>gene name</td><td>gene id</td></tr>
<tr><td>GU305914.1</td><td>75273</td><td>75295</td><td>+</td><td>0</td><td>CTGCGAGATCTCGCGGACGA</td><td>TGG</td><td>||||||||[||||||||||||]PAM</td><td>NA</td><td>Intergenic</td><td>NA</td><td>NA</td></tr>
<tr><td>T5335</td><td>CGCGGACGATGGACTTGACGAGG</td><td>1000</td><td></td><td></td><td></td><td></td><td></td><td></td><td></td><td></td><td></td></tr>
<tr><td>Oligo fwd</td><td>CACCCGCGGACGATGGACTTGACG</td><td></td><td></td><td></td><td></td><td></td><td></td><td></td><td></td><td></td><td></td></tr>
<tr><td>Oligo rev</td><td>AAACCGTCAAGTCCATCGTCCGCG</td><td></td><td></td><td></td><td></td><td></td><td></td><td></td><td></td><td></td><td></td></tr>
<tr><td>Chromosome</td><td>start</td><td>end</td><td>strand</td><td>MM</td><td>target_seq</td><td>PAM</td><td>alignment</td><td>distance</td><td>position</td><td>gene name</td><td>gene id</td></tr>
<tr><td>GU305914.1</td><td>75284</td><td>75306</td><td>+</td><td>0</td><td>CGCGGACGATGGACTTGACG</td><td>AGG</td><td>||||||||[||||||||||||]PAM</td><td>NA</td><td>Intergenic</td><td>NA</td><td>NA</td></tr>
<tr><td>T5336</td><td>GCGGACGATGGACTTGACGAGGG</td><td>1000</td><td></td><td></td><td></td><td></td><td></td><td></td><td></td><td></td><td></td></tr>
<tr><td>Oligo fwd</td><td>CACCGCGGACGATGGACTTGACGA</td><td></td><td></td><td></td><td></td><td></td><td></td><td></td><td></td><td></td><td></td></tr>
<tr><td>Oligo rev</td><td>AAACTCGTCAAGTCCATCGTCCGC</td><td></td><td></td><td></td><td></td><td></td><td></td><td></td><td></td><td></td><td></td></tr>
<tr><td>Chromosome</td><td>start</td><td>end</td><td>strand</td><td>MM</td><td>target_seq</td><td>PAM</td><td>alignment</td><td>distance</td><td>position</td><td>gene name</td><td>gene id</td></tr>
<tr><td>GU305914.1</td><td>75285</td><td>75307</td><td>+</td><td>0</td><td>GCGGACGATGGACTTGACGA</td><td>GGG</td><td>||||||||[||||||||||||]PAM</td><td>NA</td><td>Intergenic</td><td>NA</td><td>NA</td></tr>
<tr><td>T5337</td><td>GGACTTGACGAGGGCCGAGCAGG</td><td>1000</td><td></td><td></td><td></td><td></td><td></td><td></td><td></td><td></td><td></td></tr>
<tr><td>Oligo fwd</td><td>CACCGGACTTGACGAGGGCCGAGC</td><td></td><td></td><td></td><td></td><td></td><td></td><td></td><td></td><td></td><td></td></tr>
<tr><td>Oligo rev</td><td>AAACGCTCGGCCCTCGTCAAGTCC</td><td></td><td></td><td></td><td></td><td></td><td></td><td></td><td></td><td></td><td></td></tr>
<tr><td>Chromosome</td><td>start</td><td>end</td><td>strand</td><td>MM</td><td>target_seq</td><td>PAM</td><td>alignment</td><td>distance</td><td>position</td><td>gene name</td><td>gene id</td></tr>
<tr><td>GU305914.1</td><td>75294</td><td>75316</td><td>+</td><td>0</td><td>GGACTTGACGAGGGCCGAGC</td><td>AGG</td><td>||||||||[||||||||||||]PAM</td><td>NA</td><td>Intergenic</td><td>NA</td><td>NA</td></tr>
<tr><td>T5338</td><td>CTTGACGAGGGCCGAGCAGGAGG</td><td>1000</td><td></td><td></td><td></td><td></td><td></td><td></td><td></td><td></td><td></td></tr>
<tr><td>Oligo fwd</td><td>CACCCTTGACGAGGGCCGAGCAGG</td><td></td><td></td><td></td><td></td><td></td><td></td><td></td><td></td><td></td><td></td></tr>
<tr><td>Oligo rev</td><td>AAACCCTGCTCGGCCCTCGTCAAG</td><td></td><td></td><td></td><td></td><td></td><td></td><td></td><td></td><td></td><td></td></tr>
<tr><td>Chromosome</td><td>start</td><td>end</td><td>strand</td><td>MM</td><td>target_seq</td><td>PAM</td><td>alignment</td><td>distance</td><td>position</td><td>gene name</td><td>gene id</td></tr>
<tr><td>GU305914.1</td><td>75297</td><td>75319</td><td>+</td><td>0</td><td>CTTGACGAGGGCCGAGCAGG</td><td>AGG</td><td>||||||||[||||||||||||]PAM</td><td>NA</td><td>Intergenic</td><td>NA</td><td>NA</td></tr>
<tr><td>T5339</td><td>AGCAGGAGGACCAGCGTTCGAGG</td><td>1000</td><td></td><td></td><td></td><td></td><td></td><td></td><td></td><td></td><td></td></tr>
<tr><td>Oligo fwd</td><td>CACCAGCAGGAGGACCAGCGTTCG</td><td></td><td></td><td></td><td></td><td></td><td></td><td></td><td></td><td></td><td></td></tr>
<tr><td>Oligo rev</td><td>AAACCGAACGCTGGTCCTCCTGCT</td><td></td><td></td><td></td><td></td><td></td><td></td><td></td><td></td><td></td><td></td></tr>
<tr><td>Chromosome</td><td>start</td><td>end</td><td>strand</td><td>MM</td><td>target_seq</td><td>PAM</td><td>alignment</td><td>distance</td><td>position</td><td>gene name</td><td>gene id</td></tr>
<tr><td>GU305914.1</td><td>75311</td><td>75333</td><td>+</td><td>0</td><td>AGCAGGAGGACCAGCGTTCG</td><td>AGG</td><td>||||||||[||||||||||||]PAM</td><td>NA</td><td>Intergenic</td><td>NA</td><td>NA</td></tr>
<tr><td>T5340</td><td>TGCCGTTCATGAACATGTTGAGG</td><td>1000</td><td></td><td></td><td></td><td></td><td></td><td></td><td></td><td></td><td></td></tr>
<tr><td>Oligo fwd</td><td>CACCTGCCGTTCATGAACATGTTG</td><td></td><td></td><td></td><td></td><td></td><td></td><td></td><td></td><td></td><td></td></tr>
<tr><td>Oligo rev</td><td>AAACCAACATGTTCATGAACGGCA</td><td></td><td></td><td></td><td></td><td></td><td></td><td></td><td></td><td></td><td></td></tr>
<tr><td>Chromosome</td><td>start</td><td>end</td><td>strand</td><td>MM</td><td>target_seq</td><td>PAM</td><td>alignment</td><td>distance</td><td>position</td><td>gene name</td><td>gene id</td></tr>
<tr><td>GU305914.1</td><td>75380</td><td>75402</td><td>+</td><td>0</td><td>TGCCGTTCATGAACATGTTG</td><td>AGG</td><td>||||||||[||||||||||||]PAM</td><td>NA</td><td>Intergenic</td><td>NA</td><td>NA</td></tr>
<tr><td>T5341</td><td>TGTTGAGGACGCAGTAGACGCGG</td><td>1000</td><td></td><td></td><td></td><td></td><td></td><td></td><td></td><td></td><td></td></tr>
<tr><td>Oligo fwd</td><td>CACCTGTTGAGGACGCAGTAGACG</td><td></td><td></td><td></td><td></td><td></td><td></td><td></td><td></td><td></td><td></td></tr>
<tr><td>Oligo rev</td><td>AAACCGTCTACTGCGTCCTCAACA</td><td></td><td></td><td></td><td></td><td></td><td></td><td></td><td></td><td></td><td></td></tr>
<tr><td>Chromosome</td><td>start</td><td>end</td><td>strand</td><td>MM</td><td>target_seq</td><td>PAM</td><td>alignment</td><td>distance</td><td>position</td><td>gene name</td><td>gene id</td></tr>
<tr><td>GU305914.1</td><td>75395</td><td>75417</td><td>+</td><td>0</td><td>TGTTGAGGACGCAGTAGACG</td><td>CGG</td><td>||||||||[||||||||||||]PAM</td><td>NA</td><td>Intergenic</td><td>NA</td><td>NA</td></tr>
<tr><td>T5342</td><td>TCCGTCGCGTTCGTCCTCGACGG</td><td>1000</td><td></td><td></td><td></td><td></td><td></td><td></td><td></td><td></td><td></td></tr>
<tr><td>Oligo fwd</td><td>CACCTCCGTCGCGTTCGTCCTCGA</td><td></td><td></td><td></td><td></td><td></td><td></td><td></td><td></td><td></td><td></td></tr>
<tr><td>Oligo rev</td><td>AAACTCGAGGACGAACGCGACGGA</td><td></td><td></td><td></td><td></td><td></td><td></td><td></td><td></td><td></td><td></td></tr>
<tr><td>Chromosome</td><td>start</td><td>end</td><td>strand</td><td>MM</td><td>target_seq</td><td>PAM</td><td>alignment</td><td>distance</td><td>position</td><td>gene name</td><td>gene id</td></tr>
<tr><td>GU305914.1</td><td>75420</td><td>75442</td><td>+</td><td>0</td><td>TCCGTCGCGTTCGTCCTCGA</td><td>CGG</td><td>||||||||[||||||||||||]PAM</td><td>NA</td><td>Intergenic</td><td>NA</td><td>NA</td></tr>
<tr><td>T5343</td><td>CGTCCTCGACGGTGCCCGAGAGG</td><td>1000</td><td></td><td></td><td></td><td></td><td></td><td></td><td></td><td></td><td></td></tr>
<tr><td>Oligo fwd</td><td>CACCCGTCCTCGACGGTGCCCGAG</td><td></td><td></td><td></td><td></td><td></td><td></td><td></td><td></td><td></td><td></td></tr>
<tr><td>Oligo rev</td><td>AAACCTCGGGCACCGTCGAGGACG</td><td></td><td></td><td></td><td></td><td></td><td></td><td></td><td></td><td></td><td></td></tr>
<tr><td>Chromosome</td><td>start</td><td>end</td><td>strand</td><td>MM</td><td>target_seq</td><td>PAM</td><td>alignment</td><td>distance</td><td>position</td><td>gene name</td><td>gene id</td></tr>
<tr><td>GU305914.1</td><td>75431</td><td>75453</td><td>+</td><td>0</td><td>CGTCCTCGACGGTGCCCGAG</td><td>AGG</td><td>||||||||[||||||||||||]PAM</td><td>NA</td><td>Intergenic</td><td>NA</td><td>NA</td></tr>
<tr><td>T5344</td><td>CGGTGCCCGAGAGGAGCAGCTGG</td><td>1000</td><td></td><td></td><td></td><td></td><td></td><td></td><td></td><td></td><td></td></tr>
<tr><td>Oligo fwd</td><td>CACCCGGTGCCCGAGAGGAGCAGC</td><td></td><td></td><td></td><td></td><td></td><td></td><td></td><td></td><td></td><td></td></tr>
<tr><td>Oligo rev</td><td>AAACGCTGCTCCTCTCGGGCACCG</td><td></td><td></td><td></td><td></td><td></td><td></td><td></td><td></td><td></td><td></td></tr>
<tr><td>Chromosome</td><td>start</td><td>end</td><td>strand</td><td>MM</td><td>target_seq</td><td>PAM</td><td>alignment</td><td>distance</td><td>position</td><td>gene name</td><td>gene id</td></tr>
<tr><td>GU305914.1</td><td>75440</td><td>75462</td><td>+</td><td>0</td><td>CGGTGCCCGAGAGGAGCAGC</td><td>TGG</td><td>||||||||[||||||||||||]PAM</td><td>NA</td><td>Intergenic</td><td>NA</td><td>NA</td></tr>
</table>

T5345 TGCCCGAGAGGAGCAGCTGGAGG 1000
Oligo fwd CACCTGCCCGAGAGGAGCAGCTGG
Oligo rev AAACCCAGCTGCTCCTCTCGGGCA
Chromosome start end strand MM target_seq PAM alignment distance position gene name gene id
GU305914.1 75443 75465 + 0 TGCCCGAGAGGAGCAGCTGG AGG ||||||||[||||||||||||]PAM NA Intergenic NA NA

T5346 GCTGGAGGTGATACTTGTCGCGG 1000
Oligo fwd CACCGCTGGAGGTGATACTTGTCG
Oligo rev AAACCGACAAGTATCACCTCCAGC
Chromosome start end strand MM target_seq PAM alignment distance position gene name gene id
GU305914.1 75458 75480 + 0 GCTGGAGGTGATACTTGTCG CGG ||||||||[||||||||||||]PAM NA Intergenic NA NA

T5347 GGTGATACTTGTCGCGGTGCTGG 1000
Oligo fwd CACCGGTGATACTTGTCGCGGTGC
Oligo rev AAACGCACCGCGACAAGTATCACC
Chromosome start end strand MM target_seq PAM alignment distance position gene name gene id
GU305914.1 75464 75486 + 0 GGTGATACTTGTCGCGGTGC TGG ||||||||[||||||||||||]PAM NA Intergenic NA NA

T5348 AGACGCCCCCGAAGAGCTCACGG 1000
Oligo fwd CACCAGACGCCCCCGAAGAGCTCA
Oligo rev AAACTGAGCTCTTCGGGGGCGTCT
Chromosome start end strand MM target_seq PAM alignment distance position gene name gene id
GU305914.1 75488 75510 + 0 AGACGCCCCCGAAGAGCTCA CGG ||||||||[||||||||||||]PAM NA Intergenic NA NA

T5349 CTCACGGATGACCCGCCGAACGG 1000
Oligo fwd CACCCTCACGGATGACCCGCCGAA
Oligo rev AAACTTCGGCGGGTCATCCGTGAG
Chromosome start end strand MM target_seq PAM alignment distance position gene name gene id
GU305914.1 75504 75526 + 0 CTCACGGATGACCCGCCGAA CGG ||||||||[||||||||||||]PAM NA Intergenic NA NA

T5350 TCACGGATGACCCGCCGAACGGG 1000
Oligo fwd CACCTCACGGATGACCCGCCGAAC
Oligo rev AAACGTTCGGCGGGTCATCCGTGA
Chromosome start end strand MM target_seq PAM alignment distance position gene name gene id
GU305914.1 75505 75527 + 0 TCACGGATGACCCGCCGAAC GGG ||||||||[||||||||||||]PAM NA Intergenic NA NA

T5351 AACGGGCCGCATGCCAGTAGAGG 1000
Oligo fwd CACCAACGGGCCGCATGCCAGTAG
Oligo rev AAACCTACTGGCATGCGGCCCGTT
Chromosome start end strand MM target_seq PAM alignment distance position gene name gene id
GU305914.1 75522 75544 + 0 AACGGGCCGCATGCCAGTAG AGG ||||||||[||||||||||||]PAM NA Intergenic NA NA

T5352 GCCAGTAGAGGAGCCAGAGCAGG 1000
Oligo fwd CACCGCCAGTAGAGGAGCCAGAGC
Oligo rev AAACGCTCTGGCTCCTCTACTGGC
Chromosome start end strand MM target_seq PAM alignment distance position gene name gene id
GU305914.1 75534 75556 + 0 GCCAGTAGAGGAGCCAGAGC AGG ||||||||[||||||||||||]PAM NA Intergenic NA NA

T5353 AGAGGAGCCAGAGCAGGACGAGG 1000
Oligo fwd CACCAGAGGAGCCAGAGCAGGACG
Oligo rev AAACCGTCCTGCTCTGGCTCCTCT
Chromosome start end strand MM target_seq PAM alignment distance position gene name gene id
GU305914.1 75540 75562 + 0 AGAGGAGCCAGAGCAGGACG AGG ||||||||[||||||||||||]PAM NA Intergenic NA NA

T5354 GAGGAGCCAGAGCAGGACGAGGG 1000
Oligo fwd CACCGAGGAGCCAGAGCAGGACGA
Oligo rev AAACTCGTCCTGCTCTGGCTCCTC
Chromosome start end strand MM target_seq PAM alignment distance position gene name gene id
GU305914.1 75541 75563 + 0 GAGGAGCCAGAGCAGGACGA GGG ||||||||[||||||||||||]PAM NA Intergenic NA NA

T5355 CCAGAGCAGGACGAGGGCGATGG 1000
Oligo fwd CACCCCAGAGCAGGACGAGGGCGA
Oligo rev AAACTCGCCCTCGTCCTGCTCTGG
Chromosome start end strand MM target_seq PAM alignment distance position gene name gene id
GU305914.1 75547 75569 + 0 CCAGAGCAGGACGAGGGCGA TGG ||||||||[||||||||||||]PAM NA Intergenic NA NA

T5356 GAGCAGGACGAGGGCGATGGCGG 1000
Oligo fwd CACCGAGCAGGACGAGGGCGATGG
Oligo rev AAACCCATCGCCCTCGTCCTGCTC
Chromosome start end strand MM target_seq PAM alignment distance position gene name gene id
GU305914.1 75550 75572 + 0 GAGCAGGACGAGGGCGATGG CGG ||||||||[||||||||||||]PAM NA Intergenic NA NA

T5357 CAGGACGAGGGCGATGGCGGCGG 1000
Oligo fwd CACCCAGGACGAGGGCGATGGCGG
Oligo rev AAACCCGCCATCGCCCTCGTCCTG
Chromosome start end strand MM target_seq PAM alignment distance position gene name gene id
GU305914.1 75553 75575 + 0 CAGGACGAGGGCGATGGCGG CGG ||||||||[||||||||||||]PAM NA Intergenic NA NA
GU305914.1 90265 90287 - 4 CGGGAGGCGGGCGACGGCGG AGG |-|||-|-[||||||-|||||]PAM NA Intergenic NA NA

T5358 CGAGGGCGATGGCGGCGGCGAGG 1000
Oligo fwd CACCCGAGGGCGATGGCGGCGGCG
Oligo rev AAACCGCCGCCGCCATCGCCCTCG
Chromosome start end strand MM target_seq PAM alignment distance position gene name gene id
GU305914.1 75558 75580 + 0 CGAGGGCGATGGCGGCGGCG AGG ||||||||[||||||||||||]PAM NA Intergenic NA NA
GU305914.1 35655 35677 + 4 CCACGGCGGCGGCGGCGGCG GGG |-|-||||[--||||||||||]PAM NA Intergenic NA NA

T5359 GCGATGGCGGCGGCGAGGATCGG 1000
Oligo fwd CACCGCGATGGCGGCGGCGAGGAT

<table>
<tr><td>Oligo rev</td><td>AAACATCCTCGCCGCCGCCATCGC</td><td></td><td></td><td></td><td></td><td></td><td></td><td></td><td></td><td></td><td></td><td></td></tr>
<tr><td>Chromosome</td><td>start</td><td></td><td>end</td><td>strand</td><td>MM</td><td>target_seq</td><td>PAM</td><td>alignment</td><td>distance</td><td>position</td><td>gene name</td><td>gene id</td></tr>
<tr><td>GU305914.1</td><td></td><td>75563</td><td>75585</td><td>+</td><td>0</td><td>GCGATGGCGGCGGCGAGGAT</td><td>CGG</td><td>||||||||[||||||||||||]PAM</td><td>NA</td><td>Intergenic</td><td>NA</td><td>NA</td></tr>
<tr><td></td><td></td><td></td><td></td><td></td><td></td><td></td><td></td><td></td><td></td><td></td><td></td><td></td></tr>
<tr><td>T5360</td><td>CGATGGCGGCGGCGAGGATCGGG</td><td></td><td>1000</td><td></td><td></td><td></td><td></td><td></td><td></td><td></td><td></td><td></td></tr>
<tr><td>Oligo fwd</td><td>CACCCGATGGCGGCGGCGAGGATC</td><td></td><td></td><td></td><td></td><td></td><td></td><td></td><td></td><td></td><td></td><td></td></tr>
<tr><td>Oligo rev</td><td>AAACGATCCTCGCCGCCGCCATCG</td><td></td><td></td><td></td><td></td><td></td><td></td><td></td><td></td><td></td><td></td><td></td></tr>
<tr><td>Chromosome</td><td>start</td><td></td><td>end</td><td>strand</td><td>MM</td><td>target_seq</td><td>PAM</td><td>alignment</td><td>distance</td><td>position</td><td>gene name</td><td>gene id</td></tr>
<tr><td>GU305914.1</td><td></td><td>75564</td><td>75586</td><td>+</td><td>0</td><td>CGATGGCGGCGGCGAGGATC</td><td>GGG</td><td>||||||||[||||||||||||]PAM</td><td>NA</td><td>Intergenic</td><td>NA</td><td>NA</td></tr>
<tr><td>GU305914.1</td><td></td><td>66670</td><td>66692</td><td>+</td><td>4</td><td>CGATGGACGCGGACAGGATC</td><td>CGG</td><td>||||||--[||||--||||||]PAM</td><td>NA</td><td>Intergenic</td><td>NA</td><td>NA</td></tr>
<tr><td></td><td></td><td></td><td></td><td></td><td></td><td></td><td></td><td></td><td></td><td></td><td></td><td></td></tr>
<tr><td>T5361</td><td>GATGGCGGCGGCGAGGATCGGGG</td><td></td><td>1000</td><td></td><td></td><td></td><td></td><td></td><td></td><td></td><td></td><td></td></tr>
<tr><td>Oligo fwd</td><td>CACCGATGGCGGCGGCGAGGATCG</td><td></td><td></td><td></td><td></td><td></td><td></td><td></td><td></td><td></td><td></td><td></td></tr>
<tr><td>Oligo rev</td><td>AAACCGATCCTCGCCGCCGCCATC</td><td></td><td></td><td></td><td></td><td></td><td></td><td></td><td></td><td></td><td></td><td></td></tr>
<tr><td>Chromosome</td><td>start</td><td></td><td>end</td><td>strand</td><td>MM</td><td>target_seq</td><td>PAM</td><td>alignment</td><td>distance</td><td>position</td><td>gene name</td><td>gene id</td></tr>
<tr><td>GU305914.1</td><td></td><td>75565</td><td>75587</td><td>+</td><td>0</td><td>GATGGCGGCGGCGAGGATCG</td><td>GGG</td><td>||||||||[||||||||||||]PAM</td><td>NA</td><td>Intergenic</td><td>NA</td><td>NA</td></tr>
<tr><td></td><td></td><td></td><td></td><td></td><td></td><td></td><td></td><td></td><td></td><td></td><td></td><td></td></tr>
<tr><td>T5362</td><td>CGGGGCGCAGTGAATAGCGTTGG</td><td></td><td>1000</td><td></td><td></td><td></td><td></td><td></td><td></td><td></td><td></td><td></td></tr>
<tr><td>Oligo fwd</td><td>CACCCGGGGCGCAGTGAATAGCGT</td><td></td><td></td><td></td><td></td><td></td><td></td><td></td><td></td><td></td><td></td><td></td></tr>
<tr><td>Oligo rev</td><td>AAACACGCTATTCACTGCGCCCCG</td><td></td><td></td><td></td><td></td><td></td><td></td><td></td><td></td><td></td><td></td><td></td></tr>
<tr><td>Chromosome</td><td>start</td><td></td><td>end</td><td>strand</td><td>MM</td><td>target_seq</td><td>PAM</td><td>alignment</td><td>distance</td><td>position</td><td>gene name</td><td>gene id</td></tr>
<tr><td>GU305914.1</td><td></td><td>75583</td><td>75605</td><td>+</td><td>0</td><td>CGGGGCGCAGTGAATAGCGT</td><td>TGG</td><td>||||||||[||||||||||||]PAM</td><td>NA</td><td>Intergenic</td><td>NA</td><td>NA</td></tr>
<tr><td></td><td></td><td></td><td></td><td></td><td></td><td></td><td></td><td></td><td></td><td></td><td></td><td></td></tr>
<tr><td>T5363</td><td>GTGAATAGCGTTGGCGTGCCTGG</td><td></td><td>1000</td><td></td><td></td><td></td><td></td><td></td><td></td><td></td><td></td><td></td></tr>
<tr><td>Oligo fwd</td><td>CACCGTGAATAGCGTTGGCGTGCC</td><td></td><td></td><td></td><td></td><td></td><td></td><td></td><td></td><td></td><td></td><td></td></tr>
<tr><td>Oligo rev</td><td>AAACGGCACGCCAACGCTATTCAC</td><td></td><td></td><td></td><td></td><td></td><td></td><td></td><td></td><td></td><td></td><td></td></tr>
<tr><td>Chromosome</td><td>start</td><td></td><td>end</td><td>strand</td><td>MM</td><td>target_seq</td><td>PAM</td><td>alignment</td><td>distance</td><td>position</td><td>gene name</td><td>gene id</td></tr>
<tr><td>GU305914.1</td><td></td><td>75592</td><td>75614</td><td>+</td><td>0</td><td>GTGAATAGCGTTGGCGTGCC</td><td>TGG</td><td>||||||||[||||||||||||]PAM</td><td>NA</td><td>Intergenic</td><td>NA</td><td>NA</td></tr>
<tr><td></td><td></td><td></td><td></td><td></td><td></td><td></td><td></td><td></td><td></td><td></td><td></td><td></td></tr>
<tr><td>T5364</td><td>GTTGGCGTGCCTGGTCAAAAAGG</td><td></td><td>1000</td><td></td><td></td><td></td><td></td><td></td><td></td><td></td><td></td><td></td></tr>
<tr><td>Oligo fwd</td><td>CACCGTTGGCGTGCCTGGTCAAAA</td><td></td><td></td><td></td><td></td><td></td><td></td><td></td><td></td><td></td><td></td><td></td></tr>
<tr><td>Oligo rev</td><td>AAACTTTTGACCAGGCACGCCAAC</td><td></td><td></td><td></td><td></td><td></td><td></td><td></td><td></td><td></td><td></td><td></td></tr>
<tr><td>Chromosome</td><td>start</td><td></td><td>end</td><td>strand</td><td>MM</td><td>target_seq</td><td>PAM</td><td>alignment</td><td>distance</td><td>position</td><td>gene name</td><td>gene id</td></tr>
<tr><td>GU305914.1</td><td></td><td>75601</td><td>75623</td><td>+</td><td>0</td><td>GTTGGCGTGCCTGGTCAAAA</td><td>AGG</td><td>||||||||[||||||||||||]PAM</td><td>NA</td><td>Intergenic</td><td>NA</td><td>NA</td></tr>
<tr><td>GU305914.1</td><td></td><td>117909</td><td>117931</td><td>-</td><td>4</td><td>GTCGGTGTGCCTGGACGAAA</td><td>CGG</td><td>||-||-||[||||||-|-|||]PAM</td><td>NA</td><td>Intergenic</td><td>NA</td><td>NA</td></tr>
<tr><td></td><td></td><td></td><td></td><td></td><td></td><td></td><td></td><td></td><td></td><td></td><td></td><td></td></tr>
<tr><td>T5365</td><td>CAAAAAGGAAATAGTTCTTTCGG</td><td></td><td>1000</td><td></td><td></td><td></td><td></td><td></td><td></td><td></td><td></td><td></td></tr>
<tr><td>Oligo fwd</td><td>CACCCAAAAAGGAAATAGTTCTTT</td><td></td><td></td><td></td><td></td><td></td><td></td><td></td><td></td><td></td><td></td><td></td></tr>
<tr><td>Oligo rev</td><td>AAACAAAGAACTATTTCCTTTTTG</td><td></td><td></td><td></td><td></td><td></td><td></td><td></td><td></td><td></td><td></td><td></td></tr>
<tr><td>Chromosome</td><td>start</td><td></td><td>end</td><td>strand</td><td>MM</td><td>target_seq</td><td>PAM</td><td>alignment</td><td>distance</td><td>position</td><td>gene name</td><td>gene id</td></tr>
<tr><td>GU305914.1</td><td></td><td>75616</td><td>75638</td><td>+</td><td>0</td><td>CAAAAAGGAAATAGTTCTTT</td><td>CGG</td><td>||||||||[||||||||||||]PAM</td><td>NA</td><td>Intergenic</td><td>NA</td><td>NA</td></tr>
<tr><td></td><td></td><td></td><td></td><td></td><td></td><td></td><td></td><td></td><td></td><td></td><td></td><td></td></tr>
<tr><td>T5366</td><td>AATAGTTCTTTCGGAGTACGTGG</td><td></td><td>1000</td><td></td><td></td><td></td><td></td><td></td><td></td><td></td><td></td><td></td></tr>
<tr><td>Oligo fwd</td><td>CACCAATAGTTCTTTCGGAGTACG</td><td></td><td></td><td></td><td></td><td></td><td></td><td></td><td></td><td></td><td></td><td></td></tr>
<tr><td>Oligo rev</td><td>AAACCGTACTCCGAAAGAACTATT</td><td></td><td></td><td></td><td></td><td></td><td></td><td></td><td></td><td></td><td></td><td></td></tr>
<tr><td>Chromosome</td><td>start</td><td></td><td>end</td><td>strand</td><td>MM</td><td>target_seq</td><td>PAM</td><td>alignment</td><td>distance</td><td>position</td><td>gene name</td><td>gene id</td></tr>
<tr><td>GU305914.1</td><td></td><td>75625</td><td>75647</td><td>+</td><td>0</td><td>AATAGTTCTTTCGGAGTACG</td><td>TGG</td><td>||||||||[||||||||||||]PAM</td><td>NA</td><td>Intergenic</td><td>NA</td><td>NA</td></tr>
<tr><td></td><td></td><td></td><td></td><td></td><td></td><td></td><td></td><td></td><td></td><td></td><td></td><td></td></tr>
<tr><td>T5367</td><td>TCTTTCGGAGTACGTGGTCTCGG</td><td></td><td>1000</td><td></td><td></td><td></td><td></td><td></td><td></td><td></td><td></td><td></td></tr>
<tr><td>Oligo fwd</td><td>CACCTCTTTCGGAGTACGTGGTCT</td><td></td><td></td><td></td><td></td><td></td><td></td><td></td><td></td><td></td><td></td><td></td></tr>
<tr><td>Oligo rev</td><td>AAACAGACCACGTACTCCGAAAGA</td><td></td><td></td><td></td><td></td><td></td><td></td><td></td><td></td><td></td><td></td><td></td></tr>
<tr><td>Chromosome</td><td>start</td><td></td><td>end</td><td>strand</td><td>MM</td><td>target_seq</td><td>PAM</td><td>alignment</td><td>distance</td><td>position</td><td>gene name</td><td>gene id</td></tr>
<tr><td>GU305914.1</td><td></td><td>75631</td><td>75653</td><td>+</td><td>0</td><td>TCTTTCGGAGTACGTGGTCT</td><td>CGG</td><td>||||||||[||||||||||||]PAM</td><td>NA</td><td>Intergenic</td><td>NA</td><td>NA</td></tr>
<tr><td></td><td></td><td></td><td></td><td></td><td></td><td></td><td></td><td></td><td></td><td></td><td></td><td></td></tr>
<tr><td>T5368</td><td>TTCGGAGTACGTGGTCTCGGCGG</td><td></td><td>1000</td><td></td><td></td><td></td><td></td><td></td><td></td><td></td><td></td><td></td></tr>
<tr><td>Oligo fwd</td><td>CACCTTCGGAGTACGTGGTCTCGG</td><td></td><td></td><td></td><td></td><td></td><td></td><td></td><td></td><td></td><td></td><td></td></tr>
<tr><td>Oligo rev</td><td>AAACCCGAGACCACGTACTCCGAA</td><td></td><td></td><td></td><td></td><td></td><td></td><td></td><td></td><td></td><td></td><td></td></tr>
<tr><td>Chromosome</td><td>start</td><td></td><td>end</td><td>strand</td><td>MM</td><td>target_seq</td><td>PAM</td><td>alignment</td><td>distance</td><td>position</td><td>gene name</td><td>gene id</td></tr>
<tr><td>GU305914.1</td><td></td><td>75634</td><td>75656</td><td>+</td><td>0</td><td>TTCGGAGTACGTGGTCTCGG</td><td>CGG</td><td>||||||||[||||||||||||]PAM</td><td>NA</td><td>Intergenic</td><td>NA</td><td>NA</td></tr>
<tr><td></td><td></td><td></td><td></td><td></td><td></td><td></td><td></td><td></td><td></td><td></td><td></td><td></td></tr>
<tr><td>T5369</td><td>GGCCGTGACGTTCGTGTCGCCGG</td><td></td><td>1000</td><td></td><td></td><td></td><td></td><td></td><td></td><td></td><td></td><td></td></tr>
<tr><td>Oligo fwd</td><td>CACCGGCCGTGACGTTCGTGTCGC</td><td></td><td></td><td></td><td></td><td></td><td></td><td></td><td></td><td></td><td></td><td></td></tr>
<tr><td>Oligo rev</td><td>AAACGCGACACGAACGTCACGGCC</td><td></td><td></td><td></td><td></td><td></td><td></td><td></td><td></td><td></td><td></td><td></td></tr>
<tr><td>Chromosome</td><td>start</td><td></td><td>end</td><td>strand</td><td>MM</td><td>target_seq</td><td>PAM</td><td>alignment</td><td>distance</td><td>position</td><td>gene name</td><td>gene id</td></tr>
<tr><td>GU305914.1</td><td></td><td>75655</td><td>75677</td><td>+</td><td>0</td><td>GGCCGTGACGTTCGTGTCGC</td><td>CGG</td><td>||||||||[||||||||||||]PAM</td><td>NA</td><td>Intergenic</td><td>NA</td><td>NA</td></tr>
<tr><td></td><td></td><td></td><td></td><td></td><td></td><td></td><td></td><td></td><td></td><td></td><td></td><td></td></tr>
<tr><td>T5370</td><td>CGTGACGTTCGTGTCGCCGGCGG</td><td></td><td>1000</td><td></td><td></td><td></td><td></td><td></td><td></td><td></td><td></td><td></td></tr>
<tr><td>Oligo fwd</td><td>CACCCGTGACGTTCGTGTCGCCGG</td><td></td><td></td><td></td><td></td><td></td><td></td><td></td><td></td><td></td><td></td><td></td></tr>
<tr><td>Oligo rev</td><td>AAACCCGGCGACACGAACGTCACG</td><td></td><td></td><td></td><td></td><td></td><td></td><td></td><td></td><td></td><td></td><td></td></tr>
<tr><td>Chromosome</td><td>start</td><td></td><td>end</td><td>strand</td><td>MM</td><td>target_seq</td><td>PAM</td><td>alignment</td><td>distance</td><td>position</td><td>gene name</td><td>gene id</td></tr>
<tr><td>GU305914.1</td><td></td><td>75658</td><td>75680</td><td>+</td><td>0</td><td>CGTGACGTTCGTGTCGCCGG</td><td>CGG</td><td>||||||||[||||||||||||]PAM</td><td>NA</td><td>Intergenic</td><td>NA</td><td>NA</td></tr>
<tr><td></td><td></td><td></td><td></td><td></td><td></td><td></td><td></td><td></td><td></td><td></td><td></td><td></td></tr>
<tr><td>T5371</td><td>GTGACGTTCGTGTCGCCGGCGGG</td><td></td><td>1000</td><td></td><td></td><td></td><td></td><td></td><td></td><td></td><td></td><td></td></tr>
<tr><td>Oligo fwd</td><td>CACCGTGACGTTCGTGTCGCCGGC</td><td></td><td></td><td></td><td></td><td></td><td></td><td></td><td></td><td></td><td></td><td></td></tr>
<tr><td>Oligo rev</td><td>AAACGCCGGCGACACGAACGTCAC</td><td></td><td></td><td></td><td></td><td></td><td></td><td></td><td></td><td></td><td></td><td></td></tr>
<tr><td>Chromosome</td><td>start</td><td></td><td>end</td><td>strand</td><td>MM</td><td>target_seq</td><td>PAM</td><td>alignment</td><td>distance</td><td>position</td><td>gene name</td><td>gene id</td></tr>
<tr><td>GU305914.1</td><td></td><td>75659</td><td>75681</td><td>+</td><td>0</td><td>GTGACGTTCGTGTCGCCGGC</td><td>GGG</td><td>||||||||[||||||||||||]PAM</td><td>NA</td><td>Intergenic</td><td>NA</td><td>NA</td></tr>
<tr><td></td><td></td><td></td><td></td><td></td><td></td><td></td><td></td><td></td><td></td><td></td><td></td><td></td></tr>
<tr><td>T5372</td><td>TGACGTTCGTGTCGCCGGCGGGG</td><td></td><td>1000</td><td></td><td></td><td></td><td></td><td></td><td></td><td></td><td></td><td></td></tr>
<tr><td>Oligo fwd</td><td>CACCTGACGTTCGTGTCGCCGGCG</td><td></td><td></td><td></td><td></td><td></td><td></td><td></td><td></td><td></td><td></td><td></td></tr>
<tr><td>Oligo rev</td><td>AAACCGCCGGCGACACGAACGTCA</td><td></td><td></td><td></td><td></td><td></td><td></td><td></td><td></td><td></td><td></td><td></td></tr>
<tr><td>Chromosome</td><td>start</td><td></td><td>end</td><td>strand</td><td>MM</td><td>target_seq</td><td>PAM</td><td>alignment</td><td>distance</td><td>position</td><td>gene name</td><td>gene id</td></tr>
<tr><td>GU305914.1</td><td></td><td>75660</td><td>75682</td><td>+</td><td>0</td><td>TGACGTTCGTGTCGCCGGCG</td><td>GGG</td><td>||||||||[||||||||||||]PAM</td><td>NA</td><td>Intergenic</td><td>NA</td><td>NA</td></tr>
<tr><td></td><td></td><td></td><td></td><td></td><td></td><td></td><td></td><td></td><td></td><td></td><td></td><td></td></tr>
<tr><td>T5373</td><td>GACGTTCGTGTCGCCGGCGGGGG</td><td></td><td>1000</td><td></td><td></td><td></td><td></td><td></td><td></td><td></td><td></td><td></td></tr>
<tr><td>Oligo fwd</td><td>CACCGACGTTCGTGTCGCCGGCGG</td><td></td><td></td><td></td><td></td><td></td><td></td><td></td><td></td><td></td><td></td><td></td></tr>
<tr><td>Oligo rev</td><td>AAACCCGCCGGCGACACGAACGTC</td><td></td><td></td><td></td><td></td><td></td><td></td><td></td><td></td><td></td><td></td><td></td></tr>
<tr><td>Chromosome</td><td>start</td><td></td><td>end</td><td>strand</td><td>MM</td><td>target_seq</td><td>PAM</td><td>alignment</td><td>distance</td><td>position</td><td>gene name</td><td>gene id</td></tr>
<tr><td>GU305914.1</td><td></td><td>75661</td><td>75683</td><td>+</td><td>0</td><td>GACGTTCGTGTCGCCGGCGG</td><td>GGG</td><td>||||||||[||||||||||||]PAM</td><td>NA</td><td>Intergenic</td><td>NA</td><td>NA</td></tr>
</table>

<table>
<tr><td>GU305914.1</td><td>110632</td><td>110654</td><td>-</td><td>4</td><td>GACGTGCAGGACGCCGGCGG</td><td>CGG</td><td>|||||-|-[-|-|||||||||]PAM</td><td>NA</td><td>Intergenic</td><td>NA</td><td>NA</td></tr>
<tr><td>T5374</td><td>ACGTTCGTGTCGCCGGCGGGGGG</td><td>1000</td><td></td><td></td><td></td><td></td><td></td><td></td><td></td><td></td><td></td></tr>
<tr><td>Oligo fwd</td><td>CACCACGTTCGTGTCGCCGGCGGG</td><td></td><td></td><td></td><td></td><td></td><td></td><td></td><td></td><td></td><td></td></tr>
<tr><td>Oligo rev</td><td>AAACCCCGCCGGCGACACGAACGT</td><td></td><td></td><td></td><td></td><td></td><td></td><td></td><td></td><td></td><td></td></tr>
<tr><td>Chromosome</td><td>start</td><td>end</td><td>strand</td><td>MM</td><td>target_seq</td><td>PAM</td><td>alignment</td><td>distance</td><td>position</td><td>gene name</td><td>gene id</td></tr>
<tr><td>GU305914.1</td><td>75662</td><td>75684</td><td>+</td><td>0</td><td>ACGTTCGTGTCGCCGGCGGG</td><td>GGG</td><td>||||||||[||||||||||||]PAM</td><td>NA</td><td>Intergenic</td><td>NA</td><td>NA</td></tr>
<tr><td>T5375</td><td>TTCGTGTCGCCGGCGGGGGGAGG</td><td>1000</td><td></td><td></td><td></td><td></td><td></td><td></td><td></td><td></td><td></td></tr>
<tr><td>Oligo fwd</td><td>CACCTTCGTGTCGCCGGCGGGGGG</td><td></td><td></td><td></td><td></td><td></td><td></td><td></td><td></td><td></td><td></td></tr>
<tr><td>Oligo rev</td><td>AAACCCCCCCGCCGGCGACACGAA</td><td></td><td></td><td></td><td></td><td></td><td></td><td></td><td></td><td></td><td></td></tr>
<tr><td>Chromosome</td><td>start</td><td>end</td><td>strand</td><td>MM</td><td>target_seq</td><td>PAM</td><td>alignment</td><td>distance</td><td>position</td><td>gene name</td><td>gene id</td></tr>
<tr><td>GU305914.1</td><td>75665</td><td>75687</td><td>+</td><td>0</td><td>TTCGTGTCGCCGGCGGGGGG</td><td>AGG</td><td>||||||||[||||||||||||]PAM</td><td>NA</td><td>Intergenic</td><td>NA</td><td>NA</td></tr>
<tr><td>T5376</td><td>CGCCGGCGGGGGGAGGCCGCTGG</td><td>1000</td><td></td><td></td><td></td><td></td><td></td><td></td><td></td><td></td><td></td></tr>
<tr><td>Oligo fwd</td><td>CACCCGCCGGCGGGGGGAGGCCGC</td><td></td><td></td><td></td><td></td><td></td><td></td><td></td><td></td><td></td><td></td></tr>
<tr><td>Oligo rev</td><td>AAACGCGGCCTCCCCCCGCCGGCG</td><td></td><td></td><td></td><td></td><td></td><td></td><td></td><td></td><td></td><td></td></tr>
<tr><td>Chromosome</td><td>start</td><td>end</td><td>strand</td><td>MM</td><td>target_seq</td><td>PAM</td><td>alignment</td><td>distance</td><td>position</td><td>gene name</td><td>gene id</td></tr>
<tr><td>GU305914.1</td><td>75672</td><td>75694</td><td>+</td><td>0</td><td>CGCCGGCGGGGGGAGGCCGC</td><td>TGG</td><td>||||||||[||||||||||||]PAM</td><td>NA</td><td>Intergenic</td><td>NA</td><td>NA</td></tr>
<tr><td>T5377</td><td>GCCGGCGGGGGGAGGCCGCTGGG</td><td>1000</td><td></td><td></td><td></td><td></td><td></td><td></td><td></td><td></td><td></td></tr>
<tr><td>Oligo fwd</td><td>CACCGCCGGCGGGGGGAGGCCGCT</td><td></td><td></td><td></td><td></td><td></td><td></td><td></td><td></td><td></td><td></td></tr>
<tr><td>Oligo rev</td><td>AAACAGCGGCCTCCCCCCGCCGGC</td><td></td><td></td><td></td><td></td><td></td><td></td><td></td><td></td><td></td><td></td></tr>
<tr><td>Chromosome</td><td>start</td><td>end</td><td>strand</td><td>MM</td><td>target_seq</td><td>PAM</td><td>alignment</td><td>distance</td><td>position</td><td>gene name</td><td>gene id</td></tr>
<tr><td>GU305914.1</td><td>75673</td><td>75695</td><td>+</td><td>0</td><td>GCCGGCGGGGGGAGGCCGCT</td><td>GGG</td><td>||||||||[||||||||||||]PAM</td><td>NA</td><td>Intergenic</td><td>NA</td><td>NA</td></tr>
<tr><td>T5378</td><td>CGGGGGGAGGCCGCTGGGTCTGG</td><td>1000</td><td></td><td></td><td></td><td></td><td></td><td></td><td></td><td></td><td></td></tr>
<tr><td>Oligo fwd</td><td>CACCCGGGGGGAGGCCGCTGGGTC</td><td></td><td></td><td></td><td></td><td></td><td></td><td></td><td></td><td></td><td></td></tr>
<tr><td>Oligo rev</td><td>AAACGACCCAGCGGCCTCCCCCCG</td><td></td><td></td><td></td><td></td><td></td><td></td><td></td><td></td><td></td><td></td></tr>
<tr><td>Chromosome</td><td>start</td><td>end</td><td>strand</td><td>MM</td><td>target_seq</td><td>PAM</td><td>alignment</td><td>distance</td><td>position</td><td>gene name</td><td>gene id</td></tr>
<tr><td>GU305914.1</td><td>75678</td><td>75700</td><td>+</td><td>0</td><td>CGGGGGGAGGCCGCTGGGTC</td><td>TGG</td><td>||||||||[||||||||||||]PAM</td><td>NA</td><td>Intergenic</td><td>NA</td><td>NA</td></tr>
<tr><td>T5379</td><td>GGTCTGGTCCTGATCCCGCGAGG</td><td>1000</td><td></td><td></td><td></td><td></td><td></td><td></td><td></td><td></td><td></td></tr>
<tr><td>Oligo fwd</td><td>CACCGGTCTGGTCCTGATCCCGCG</td><td></td><td></td><td></td><td></td><td></td><td></td><td></td><td></td><td></td><td></td></tr>
<tr><td>Oligo rev</td><td>AAACCGCGGGATCAGGACCAGACC</td><td></td><td></td><td></td><td></td><td></td><td></td><td></td><td></td><td></td><td></td></tr>
<tr><td>Chromosome</td><td>start</td><td>end</td><td>strand</td><td>MM</td><td>target_seq</td><td>PAM</td><td>alignment</td><td>distance</td><td>position</td><td>gene name</td><td>gene id</td></tr>
<tr><td>GU305914.1</td><td>75694</td><td>75716</td><td>+</td><td>0</td><td>GGTCTGGTCCTGATCCCGCG</td><td>AGG</td><td>||||||||[||||||||||||]PAM</td><td>NA</td><td>Intergenic</td><td>NA</td><td>NA</td></tr>
<tr><td>T5380</td><td>GTCTGGTCCTGATCCCGCGAGGG</td><td>1000</td><td></td><td></td><td></td><td></td><td></td><td></td><td></td><td></td><td></td></tr>
<tr><td>Oligo fwd</td><td>CACCGTCTGGTCCTGATCCCGCGA</td><td></td><td></td><td></td><td></td><td></td><td></td><td></td><td></td><td></td><td></td></tr>
<tr><td>Oligo rev</td><td>AAACTCGCGGGATCAGGACCAGAC</td><td></td><td></td><td></td><td></td><td></td><td></td><td></td><td></td><td></td><td></td></tr>
<tr><td>Chromosome</td><td>start</td><td>end</td><td>strand</td><td>MM</td><td>target_seq</td><td>PAM</td><td>alignment</td><td>distance</td><td>position</td><td>gene name</td><td>gene id</td></tr>
<tr><td>GU305914.1</td><td>75695</td><td>75717</td><td>+</td><td>0</td><td>GTCTGGTCCTGATCCCGCGA</td><td>GGG</td><td>||||||||[||||||||||||]PAM</td><td>NA</td><td>Intergenic</td><td>NA</td><td>NA</td></tr>
<tr><td>T5381</td><td>TGATCCCGCGAGGGAGCCGCCGG</td><td>1000</td><td></td><td></td><td></td><td></td><td></td><td></td><td></td><td></td><td></td></tr>
<tr><td>Oligo fwd</td><td>CACCTGATCCCGCGAGGGAGCCGC</td><td></td><td></td><td></td><td></td><td></td><td></td><td></td><td></td><td></td><td></td></tr>
<tr><td>Oligo rev</td><td>AAACGCGGCTCCCTCGCGGGATCA</td><td></td><td></td><td></td><td></td><td></td><td></td><td></td><td></td><td></td><td></td></tr>
<tr><td>Chromosome</td><td>start</td><td>end</td><td>strand</td><td>MM</td><td>target_seq</td><td>PAM</td><td>alignment</td><td>distance</td><td>position</td><td>gene name</td><td>gene id</td></tr>
<tr><td>GU305914.1</td><td>75704</td><td>75726</td><td>+</td><td>0</td><td>TGATCCCGCGAGGGAGCCGC</td><td>CGG</td><td>||||||||[||||||||||||]PAM</td><td>NA</td><td>Intergenic</td><td>NA</td><td>NA</td></tr>
<tr><td>GU305914.1</td><td>113607</td><td>113629</td><td>-</td><td>4</td><td>CGACGTCGCGAGGGAGCCGC</td><td>AGG</td><td>-||---||[||||||||||||]PAM</td><td>NA</td><td>Intergenic</td><td>NA</td><td>NA</td></tr>
<tr><td>T5382</td><td>GATCCCGCGAGGGAGCCGCCGGG</td><td>1000</td><td></td><td></td><td></td><td></td><td></td><td></td><td></td><td></td><td></td></tr>
<tr><td>Oligo fwd</td><td>CACCGATCCCGCGAGGGAGCCGCC</td><td></td><td></td><td></td><td></td><td></td><td></td><td></td><td></td><td></td><td></td></tr>
<tr><td>Oligo rev</td><td>AAACGGCGGCTCCCTCGCGGGATC</td><td></td><td></td><td></td><td></td><td></td><td></td><td></td><td></td><td></td><td></td></tr>
<tr><td>Chromosome</td><td>start</td><td>end</td><td>strand</td><td>MM</td><td>target_seq</td><td>PAM</td><td>alignment</td><td>distance</td><td>position</td><td>gene name</td><td>gene id</td></tr>
<tr><td>GU305914.1</td><td>75705</td><td>75727</td><td>+</td><td>0</td><td>GATCCCGCGAGGGAGCCGCC</td><td>GGG</td><td>||||||||[||||||||||||]PAM</td><td>NA</td><td>Intergenic</td><td>NA</td><td>NA</td></tr>
<tr><td>T5383</td><td>ATCCCGCGAGGGAGCCGCCGGGG</td><td>1000</td><td></td><td></td><td></td><td></td><td></td><td></td><td></td><td></td><td></td></tr>
<tr><td>Oligo fwd</td><td>CACCATCCCGCGAGGGAGCCGCCG</td><td></td><td></td><td></td><td></td><td></td><td></td><td></td><td></td><td></td><td></td></tr>
<tr><td>Oligo rev</td><td>AAACCGGCGGCTCCCTCGCGGGAT</td><td></td><td></td><td></td><td></td><td></td><td></td><td></td><td></td><td></td><td></td></tr>
<tr><td>Chromosome</td><td>start</td><td>end</td><td>strand</td><td>MM</td><td>target_seq</td><td>PAM</td><td>alignment</td><td>distance</td><td>position</td><td>gene name</td><td>gene id</td></tr>
<tr><td>GU305914.1</td><td>75706</td><td>75728</td><td>+</td><td>0</td><td>ATCCCGCGAGGGAGCCGCCG</td><td>GGG</td><td>||||||||[||||||||||||]PAM</td><td>NA</td><td>Intergenic</td><td>NA</td><td>NA</td></tr>
<tr><td>T5384</td><td>CCGCGAGGGAGCCGCCGGGGAGG</td><td>1000</td><td></td><td></td><td></td><td></td><td></td><td></td><td></td><td></td><td></td></tr>
<tr><td>Oligo fwd</td><td>CACCCCGCGAGGGAGCCGCCGGGG</td><td></td><td></td><td></td><td></td><td></td><td></td><td></td><td></td><td></td><td></td></tr>
<tr><td>Oligo rev</td><td>AAACCCCCGGCGGCTCCCTCGCGG</td><td></td><td></td><td></td><td></td><td></td><td></td><td></td><td></td><td></td><td></td></tr>
<tr><td>Chromosome</td><td>start</td><td>end</td><td>strand</td><td>MM</td><td>target_seq</td><td>PAM</td><td>alignment</td><td>distance</td><td>position</td><td>gene name</td><td>gene id</td></tr>
<tr><td>GU305914.1</td><td>75709</td><td>75731</td><td>+</td><td>0</td><td>CCGCGAGGGAGCCGCCGGGG</td><td>AGG</td><td>||||||||[||||||||||||]PAM</td><td>NA</td><td>Intergenic</td><td>NA</td><td>NA</td></tr>
<tr><td>T5385</td><td>CGAGGGAGCCGCCGGGGAGGAGG</td><td>1000</td><td></td><td></td><td></td><td></td><td></td><td></td><td></td><td></td><td></td></tr>
<tr><td>Oligo fwd</td><td>CACCCGAGGGAGCCGCCGGGGAGG</td><td></td><td></td><td></td><td></td><td></td><td></td><td></td><td></td><td></td><td></td></tr>
<tr><td>Oligo rev</td><td>AAACCCTCCCCGGCGGCTCCCTCG</td><td></td><td></td><td></td><td></td><td></td><td></td><td></td><td></td><td></td><td></td></tr>
<tr><td>Chromosome</td><td>start</td><td>end</td><td>strand</td><td>MM</td><td>target_seq</td><td>PAM</td><td>alignment</td><td>distance</td><td>position</td><td>gene name</td><td>gene id</td></tr>
<tr><td>GU305914.1</td><td>75712</td><td>75734</td><td>+</td><td>0</td><td>CGAGGGAGCCGCCGGGGAGG</td><td>AGG</td><td>||||||||[||||||||||||]PAM</td><td>NA</td><td>Intergenic</td><td>NA</td><td>NA</td></tr>
<tr><td>T5386</td><td>GGGAGCCGCCGGGGAGGAGGTGG</td><td>1000</td><td></td><td></td><td></td><td></td><td></td><td></td><td></td><td></td><td></td></tr>
<tr><td>Oligo fwd</td><td>CACCGGGAGCCGCCGGGGAGGAGG</td><td></td><td></td><td></td><td></td><td></td><td></td><td></td><td></td><td></td><td></td></tr>
<tr><td>Oligo rev</td><td>AAACCCTCCTCCCCGGCGGCTCCC</td><td></td><td></td><td></td><td></td><td></td><td></td><td></td><td></td><td></td><td></td></tr>
<tr><td>Chromosome</td><td>start</td><td>end</td><td>strand</td><td>MM</td><td>target_seq</td><td>PAM</td><td>alignment</td><td>distance</td><td>position</td><td>gene name</td><td>gene id</td></tr>
<tr><td>GU305914.1</td><td>75715</td><td>75737</td><td>+</td><td>0</td><td>GGGAGCCGCCGGGGAGGAGG</td><td>TGG</td><td>||||||||[||||||||||||]PAM</td><td>NA</td><td>Intergenic</td><td>NA</td><td>NA</td></tr>
<tr><td>T5387</td><td>CGCCGGGGAGGAGGTGGAAGTGG</td><td>1000</td><td></td><td></td><td></td><td></td><td></td><td></td><td></td><td></td><td></td></tr>
<tr><td>Oligo fwd</td><td>CACCCGCCGGGGAGGAGGTGGAAG</td><td></td><td></td><td></td><td></td><td></td><td></td><td></td><td></td><td></td><td></td></tr>
<tr><td>Oligo rev</td><td>AAACCTTCCACCTCCTCCCCGGCG</td><td></td><td></td><td></td><td></td><td></td><td></td><td></td><td></td><td></td><td></td></tr>
<tr><td>Chromosome</td><td>start</td><td>end</td><td>strand</td><td>MM</td><td>target_seq</td><td>PAM</td><td>alignment</td><td>distance</td><td>position</td><td>gene name</td><td>gene id</td></tr>
<tr><td>GU305914.1</td><td>75721</td><td>75743</td><td>+</td><td>0</td><td>CGCCGGGGAGGAGGTGGAAG</td><td>TGG</td><td>||||||||[||||||||||||]PAM</td><td>NA</td><td>Intergenic</td><td>NA</td><td>NA</td></tr>
<tr><td>GU305914.1</td><td>73780</td><td>73802</td><td>-</td><td>4</td><td>GGCGGCGGAGGAGGAGGAAG</td><td>CGG</td><td>-||-|-||[||||||-|||||]PAM</td><td>NA</td><td>Intergenic</td><td>NA</td><td>NA</td></tr>
<tr><td>T5388</td><td>CGGGGAGGAGGTGGAAGTGGAGG</td><td>1000</td><td></td><td></td><td></td><td></td><td></td><td></td><td></td><td></td><td></td></tr>
</table>

<table>
<tr><td>Oligo fwd</td><td>CACCCGGGGAGGAGGTGGAAGTGG</td><td></td><td></td><td></td><td></td><td></td><td></td><td></td><td></td><td></td><td></td></tr>
<tr><td>Oligo rev</td><td>AAACCCACTTCCACCTCCTCCCCG</td><td></td><td></td><td></td><td></td><td></td><td></td><td></td><td></td><td></td><td></td></tr>
<tr><td>Chromosome</td><td>start</td><td>end</td><td>strand</td><td>MM</td><td>target_seq</td><td>PAM</td><td>alignment</td><td>distance</td><td>position</td><td>gene name</td><td>gene id</td></tr>
<tr><td>GU305914.1</td><td>75724</td><td>75746</td><td>+</td><td>0</td><td>CGGGGAGGAGGTGGAAGTGG</td><td>AGG</td><td>||||||||[||||||||||||]PAM</td><td>NA</td><td>Intergenic</td><td>NA</td><td>NA</td></tr>
<tr><td>GU305914.1</td><td>73777</td><td>73799</td><td>-</td><td>4</td><td>GGCGGAGGAGGAGGAAGCGG</td><td>CGG</td><td>-|-|||||[|||-|||||-||]PAM</td><td>NA</td><td>Intergenic</td><td>NA</td><td>NA</td></tr>
<tr><td></td><td></td><td></td><td></td><td></td><td></td><td></td><td></td><td></td><td></td><td></td><td></td></tr>
<tr><td>T5389</td><td>GAGGAGGTGGAAGTGGAGGTCGG</td><td>1000</td><td></td><td></td><td></td><td></td><td></td><td></td><td></td><td></td><td></td></tr>
<tr><td>Oligo fwd</td><td>CACCGAGGAGGTGGAAGTGGAGGT</td><td></td><td></td><td></td><td></td><td></td><td></td><td></td><td></td><td></td><td></td></tr>
<tr><td>Oligo rev</td><td>AAACACCTCCACTTCCACCTCCTC</td><td></td><td></td><td></td><td></td><td></td><td></td><td></td><td></td><td></td><td></td></tr>
<tr><td>Chromosome</td><td>start</td><td>end</td><td>strand</td><td>MM</td><td>target_seq</td><td>PAM</td><td>alignment</td><td>distance</td><td>position</td><td>gene name</td><td>gene id</td></tr>
<tr><td>GU305914.1</td><td>75728</td><td>75750</td><td>+</td><td>0</td><td>GAGGAGGTGGAAGTGGAGGT</td><td>CGG</td><td>||||||||[||||||||||||]PAM</td><td>NA</td><td>Intergenic</td><td>NA</td><td>NA</td></tr>
<tr><td></td><td></td><td></td><td></td><td></td><td></td><td></td><td></td><td></td><td></td><td></td><td></td></tr>
<tr><td>T5390</td><td>GGTGGAAGTGGAGGTCGGCGAGG</td><td>1000</td><td></td><td></td><td></td><td></td><td></td><td></td><td></td><td></td><td></td></tr>
<tr><td>Oligo fwd</td><td>CACCGGTGGAAGTGGAGGTCGGCG</td><td></td><td></td><td></td><td></td><td></td><td></td><td></td><td></td><td></td><td></td></tr>
<tr><td>Oligo rev</td><td>AAACCGCCGACCTCCACTTCCACC</td><td></td><td></td><td></td><td></td><td></td><td></td><td></td><td></td><td></td><td></td></tr>
<tr><td>Chromosome</td><td>start</td><td>end</td><td>strand</td><td>MM</td><td>target_seq</td><td>PAM</td><td>alignment</td><td>distance</td><td>position</td><td>gene name</td><td>gene id</td></tr>
<tr><td>GU305914.1</td><td>75733</td><td>75755</td><td>+</td><td>0</td><td>GGTGGAAGTGGAGGTCGGCG</td><td>AGG</td><td>||||||||[||||||||||||]PAM</td><td>NA</td><td>Intergenic</td><td>NA</td><td>NA</td></tr>
<tr><td>GU305914.1</td><td>91358</td><td>91380</td><td>-</td><td>4</td><td>CGTGAAGGTGCAGGTCGGCG</td><td>CGG</td><td>-|||-|-|[||-|||||||||]PAM</td><td>NA</td><td>Intergenic</td><td>NA</td><td>NA</td></tr>
<tr><td>GU305914.1</td><td>75857</td><td>75879</td><td>+</td><td>4</td><td>GGCGGAGGCGGCGGTCGGCG</td><td>AGG</td><td>||-|||-|[-||-||||||||]PAM</td><td>NA</td><td>Intergenic</td><td>NA</td><td>NA</td></tr>
<tr><td></td><td></td><td></td><td></td><td></td><td></td><td></td><td></td><td></td><td></td><td></td><td></td></tr>
<tr><td>T5391</td><td>GGAAGTGGAGGTCGGCGAGGCGG</td><td>1000</td><td></td><td></td><td></td><td></td><td></td><td></td><td></td><td></td><td></td></tr>
<tr><td>Oligo fwd</td><td>CACCGGAAGTGGAGGTCGGCGAGG</td><td></td><td></td><td></td><td></td><td></td><td></td><td></td><td></td><td></td><td></td></tr>
<tr><td>Oligo rev</td><td>AAACCCTCGCCGACCTCCACTTCC</td><td></td><td></td><td></td><td></td><td></td><td></td><td></td><td></td><td></td><td></td></tr>
<tr><td>Chromosome</td><td>start</td><td>end</td><td>strand</td><td>MM</td><td>target_seq</td><td>PAM</td><td>alignment</td><td>distance</td><td>position</td><td>gene name</td><td>gene id</td></tr>
<tr><td>GU305914.1</td><td>75736</td><td>75758</td><td>+</td><td>0</td><td>GGAAGTGGAGGTCGGCGAGG</td><td>CGG</td><td>||||||||[||||||||||||]PAM</td><td>NA</td><td>Intergenic</td><td>NA</td><td>NA</td></tr>
<tr><td>GU305914.1</td><td>75860</td><td>75882</td><td>+</td><td>3</td><td>GGAGGCGGCGGTCGGCGAGG</td><td>GGG</td><td>|||-|-||[-|||||||||||]PAM</td><td>NA</td><td>Intergenic</td><td>NA</td><td>NA</td></tr>
<tr><td>GU305914.1</td><td>65126</td><td>65148</td><td>+</td><td>4</td><td>CGAGGAGGAGGGCGGCGAGG</td><td>GGG</td><td>-||-|-||[|||-||||||||]PAM</td><td>NA</td><td>Intergenic</td><td>NA</td><td>NA</td></tr>
<tr><td>GU305914.1</td><td>213268</td><td>213290</td><td>+</td><td>3</td><td>GGAGGTGGCGGTCGGAGAGG</td><td>CGG</td><td>|||-||||[-||||||-||||]PAM</td><td>NA</td><td>Intergenic</td><td>NA</td><td>NA</td></tr>
<tr><td></td><td></td><td></td><td></td><td></td><td></td><td></td><td></td><td></td><td></td><td></td><td></td></tr>
<tr><td>T5392</td><td>GGAGGTCGGCGAGGCGGCCGCGG</td><td>1000</td><td></td><td></td><td></td><td></td><td></td><td></td><td></td><td></td><td></td></tr>
<tr><td>Oligo fwd</td><td>CACCGGAGGTCGGCGAGGCGGCCG</td><td></td><td></td><td></td><td></td><td></td><td></td><td></td><td></td><td></td><td></td></tr>
<tr><td>Oligo rev</td><td>AAACCGGCCGCCTCGCCGACCTCC</td><td></td><td></td><td></td><td></td><td></td><td></td><td></td><td></td><td></td><td></td></tr>
<tr><td>Chromosome</td><td>start</td><td>end</td><td>strand</td><td>MM</td><td>target_seq</td><td>PAM</td><td>alignment</td><td>distance</td><td>position</td><td>gene name</td><td>gene id</td></tr>
<tr><td>GU305914.1</td><td>75742</td><td>75764</td><td>+</td><td>0</td><td>GGAGGTCGGCGAGGCGGCCG</td><td>CGG</td><td>||||||||[||||||||||||]PAM</td><td>NA</td><td>Intergenic</td><td>NA</td><td>NA</td></tr>
<tr><td></td><td></td><td></td><td></td><td></td><td></td><td></td><td></td><td></td><td></td><td></td><td></td></tr>
<tr><td>T5393</td><td>GAGGTCGGCGAGGCGGCCGCGGG</td><td>1000</td><td></td><td></td><td></td><td></td><td></td><td></td><td></td><td></td><td></td></tr>
<tr><td>Oligo fwd</td><td>CACCGAGGTCGGCGAGGCGGCCGC</td><td></td><td></td><td></td><td></td><td></td><td></td><td></td><td></td><td></td><td></td></tr>
<tr><td>Oligo rev</td><td>AAACGCGGCCGCCTCGCCGACCTC</td><td></td><td></td><td></td><td></td><td></td><td></td><td></td><td></td><td></td><td></td></tr>
<tr><td>Chromosome</td><td>start</td><td>end</td><td>strand</td><td>MM</td><td>target_seq</td><td>PAM</td><td>alignment</td><td>distance</td><td>position</td><td>gene name</td><td>gene id</td></tr>
<tr><td>GU305914.1</td><td>75743</td><td>75765</td><td>+</td><td>0</td><td>GAGGTCGGCGAGGCGGCCGC</td><td>GGG</td><td>||||||||[||||||||||||]PAM</td><td>NA</td><td>Intergenic</td><td>NA</td><td>NA</td></tr>
<tr><td></td><td></td><td></td><td></td><td></td><td></td><td></td><td></td><td></td><td></td><td></td><td></td></tr>
<tr><td>T5394</td><td>GGCGGCCGCGGGCTGATTCTTGG</td><td>1000</td><td></td><td></td><td></td><td></td><td></td><td></td><td></td><td></td><td></td></tr>
<tr><td>Oligo fwd</td><td>CACCGGCGGCCGCGGGCTGATTCT</td><td></td><td></td><td></td><td></td><td></td><td></td><td></td><td></td><td></td><td></td></tr>
<tr><td>Oligo rev</td><td>AAACAGAATCAGCCCGCGGCCGCC</td><td></td><td></td><td></td><td></td><td></td><td></td><td></td><td></td><td></td><td></td></tr>
<tr><td>Chromosome</td><td>start</td><td>end</td><td>strand</td><td>MM</td><td>target_seq</td><td>PAM</td><td>alignment</td><td>distance</td><td>position</td><td>gene name</td><td>gene id</td></tr>
<tr><td>GU305914.1</td><td>75754</td><td>75776</td><td>+</td><td>0</td><td>GGCGGCCGCGGGCTGATTCT</td><td>TGG</td><td>||||||||[||||||||||||]PAM</td><td>NA</td><td>Intergenic</td><td>NA</td><td>NA</td></tr>
<tr><td></td><td></td><td></td><td></td><td></td><td></td><td></td><td></td><td></td><td></td><td></td><td></td></tr>
<tr><td>T5395</td><td>GCGTATCTTCGCGTGAGAACCGG</td><td>1000</td><td></td><td></td><td></td><td></td><td></td><td></td><td></td><td></td><td></td></tr>
<tr><td>Oligo fwd</td><td>CACCGCGTATCTTCGCGTGAGAAC</td><td></td><td></td><td></td><td></td><td></td><td></td><td></td><td></td><td></td><td></td></tr>
<tr><td>Oligo rev</td><td>AAACGTTCTCACGCGAAGATACGC</td><td></td><td></td><td></td><td></td><td></td><td></td><td></td><td></td><td></td><td></td></tr>
<tr><td>Chromosome</td><td>start</td><td>end</td><td>strand</td><td>MM</td><td>target_seq</td><td>PAM</td><td>alignment</td><td>distance</td><td>position</td><td>gene name</td><td>gene id</td></tr>
<tr><td>GU305914.1</td><td>75794</td><td>75816</td><td>+</td><td>0</td><td>GCGTATCTTCGCGTGAGAAC</td><td>CGG</td><td>||||||||[||||||||||||]PAM</td><td>NA</td><td>Intergenic</td><td>NA</td><td>NA</td></tr>
<tr><td></td><td></td><td></td><td></td><td></td><td></td><td></td><td></td><td></td><td></td><td></td><td></td></tr>
<tr><td>T5396</td><td>CGTATCTTCGCGTGAGAACCGGG</td><td>1000</td><td></td><td></td><td></td><td></td><td></td><td></td><td></td><td></td><td></td></tr>
<tr><td>Oligo fwd</td><td>CACCCGTATCTTCGCGTGAGAACC</td><td></td><td></td><td></td><td></td><td></td><td></td><td></td><td></td><td></td><td></td></tr>
<tr><td>Oligo rev</td><td>AAACGGTTCTCACGCGAAGATACG</td><td></td><td></td><td></td><td></td><td></td><td></td><td></td><td></td><td></td><td></td></tr>
<tr><td>Chromosome</td><td>start</td><td>end</td><td>strand</td><td>MM</td><td>target_seq</td><td>PAM</td><td>alignment</td><td>distance</td><td>position</td><td>gene name</td><td>gene id</td></tr>
<tr><td>GU305914.1</td><td>75795</td><td>75817</td><td>+</td><td>0</td><td>CGTATCTTCGCGTGAGAACC</td><td>GGG</td><td>||||||||[||||||||||||]PAM</td><td>NA</td><td>Intergenic</td><td>NA</td><td>NA</td></tr>
<tr><td></td><td></td><td></td><td></td><td></td><td></td><td></td><td></td><td></td><td></td><td></td><td></td></tr>
<tr><td>T5397</td><td>GAACCGGGCCGTGTTTGCGCAGG</td><td>1000</td><td></td><td></td><td></td><td></td><td></td><td></td><td></td><td></td><td></td></tr>
<tr><td>Oligo fwd</td><td>CACCGAACCGGGCCGTGTTTGCGC</td><td></td><td></td><td></td><td></td><td></td><td></td><td></td><td></td><td></td><td></td></tr>
<tr><td>Oligo rev</td><td>AAACGCGCAAACACGGCCCGGTTC</td><td></td><td></td><td></td><td></td><td></td><td></td><td></td><td></td><td></td><td></td></tr>
<tr><td>Chromosome</td><td>start</td><td>end</td><td>strand</td><td>MM</td><td>target_seq</td><td>PAM</td><td>alignment</td><td>distance</td><td>position</td><td>gene name</td><td>gene id</td></tr>
<tr><td>GU305914.1</td><td>75810</td><td>75832</td><td>+</td><td>0</td><td>GAACCGGGCCGTGTTTGCGC</td><td>AGG</td><td>||||||||[||||||||||||]PAM</td><td>NA</td><td>Intergenic</td><td>NA</td><td>NA</td></tr>
<tr><td></td><td></td><td></td><td></td><td></td><td></td><td></td><td></td><td></td><td></td><td></td><td></td></tr>
<tr><td>T5398</td><td>AGGCTGCCCCAGCGCTGACCCGG</td><td>1000</td><td></td><td></td><td></td><td></td><td></td><td></td><td></td><td></td><td></td></tr>
<tr><td>Oligo fwd</td><td>CACCAGGCTGCCCCAGCGCTGACC</td><td></td><td></td><td></td><td></td><td></td><td></td><td></td><td></td><td></td><td></td></tr>
<tr><td>Oligo rev</td><td>AAACGGTCAGCGCTGGGGCAGCCT</td><td></td><td></td><td></td><td></td><td></td><td></td><td></td><td></td><td></td><td></td></tr>
<tr><td>Chromosome</td><td>start</td><td>end</td><td>strand</td><td>MM</td><td>target_seq</td><td>PAM</td><td>alignment</td><td>distance</td><td>position</td><td>gene name</td><td>gene id</td></tr>
<tr><td>GU305914.1</td><td>75830</td><td>75852</td><td>+</td><td>0</td><td>AGGCTGCCCCAGCGCTGACC</td><td>CGG</td><td>||||||||[||||||||||||]PAM</td><td>NA</td><td>Intergenic</td><td>NA</td><td>NA</td></tr>
<tr><td></td><td></td><td></td><td></td><td></td><td></td><td></td><td></td><td></td><td></td><td></td><td></td></tr>
<tr><td>T5399</td><td>GGCTGCCCCAGCGCTGACCCGGG</td><td>1000</td><td></td><td></td><td></td><td></td><td></td><td></td><td></td><td></td><td></td></tr>
<tr><td>Oligo fwd</td><td>CACCGGCTGCCCCAGCGCTGACCC</td><td></td><td></td><td></td><td></td><td></td><td></td><td></td><td></td><td></td><td></td></tr>
<tr><td>Oligo rev</td><td>AAACGGGTCAGCGCTGGGGCAGCC</td><td></td><td></td><td></td><td></td><td></td><td></td><td></td><td></td><td></td><td></td></tr>
<tr><td>Chromosome</td><td>start</td><td>end</td><td>strand</td><td>MM</td><td>target_seq</td><td>PAM</td><td>alignment</td><td>distance</td><td>position</td><td>gene name</td><td>gene id</td></tr>
<tr><td>GU305914.1</td><td>75831</td><td>75853</td><td>+</td><td>0</td><td>GGCTGCCCCAGCGCTGACCC</td><td>GGG</td><td>||||||||[||||||||||||]PAM</td><td>NA</td><td>Intergenic</td><td>NA</td><td>NA</td></tr>
<tr><td></td><td></td><td></td><td></td><td></td><td></td><td></td><td></td><td></td><td></td><td></td><td></td></tr>
<tr><td>T5400</td><td>GCTGCCCCAGCGCTGACCCGGGG</td><td>1000</td><td></td><td></td><td></td><td></td><td></td><td></td><td></td><td></td><td></td></tr>
<tr><td>Oligo fwd</td><td>CACCGCTGCCCCAGCGCTGACCCG</td><td></td><td></td><td></td><td></td><td></td><td></td><td></td><td></td><td></td><td></td></tr>
<tr><td>Oligo rev</td><td>AAACCGGGTCAGCGCTGGGGCAGC</td><td></td><td></td><td></td><td></td><td></td><td></td><td></td><td></td><td></td><td></td></tr>
<tr><td>Chromosome</td><td>start</td><td>end</td><td>strand</td><td>MM</td><td>target_seq</td><td>PAM</td><td>alignment</td><td>distance</td><td>position</td><td>gene name</td><td>gene id</td></tr>
<tr><td>GU305914.1</td><td>75832</td><td>75854</td><td>+</td><td>0</td><td>GCTGCCCCAGCGCTGACCCG</td><td>GGG</td><td>||||||||[||||||||||||]PAM</td><td>NA</td><td>Intergenic</td><td>NA</td><td>NA</td></tr>
<tr><td></td><td></td><td></td><td></td><td></td><td></td><td></td><td></td><td></td><td></td><td></td><td></td></tr>
<tr><td>T5401</td><td>CTGCCCCAGCGCTGACCCGGGGG</td><td>1000</td><td></td><td></td><td></td><td></td><td></td><td></td><td></td><td></td><td></td></tr>
<tr><td>Oligo fwd</td><td>CACCCTGCCCCAGCGCTGACCCGG</td><td></td><td></td><td></td><td></td><td></td><td></td><td></td><td></td><td></td><td></td></tr>
<tr><td>Oligo rev</td><td>AAACCCGGGTCAGCGCTGGGGCAG</td><td></td><td></td><td></td><td></td><td></td><td></td><td></td><td></td><td></td><td></td></tr>
<tr><td>Chromosome</td><td>start</td><td>end</td><td>strand</td><td>MM</td><td>target_seq</td><td>PAM</td><td>alignment</td><td>distance</td><td>position</td><td>gene name</td><td>gene id</td></tr>
<tr><td>GU305914.1</td><td>75833</td><td>75855</td><td>+</td><td>0</td><td>CTGCCCCAGCGCTGACCCGG</td><td>GGG</td><td>||||||||[||||||||||||]PAM</td><td>NA</td><td>Intergenic</td><td>NA</td><td>NA</td></tr>
</table>

<table>
<tr><td>T5402</td><td>CCCCAGCGCTGACCCGGGGGAGG</td><td></td><td>1000</td><td></td><td></td><td></td><td></td><td></td><td></td><td></td><td></td><td></td></tr>
<tr><td>Oligo fwd</td><td>CACCCCCCAGCGCTGACCCGGGGG</td><td></td><td></td><td></td><td></td><td></td><td></td><td></td><td></td><td></td><td></td><td></td></tr>
<tr><td>Oligo rev</td><td>AAACCCCCCGGGTCAGCGCTGGGG</td><td></td><td></td><td></td><td></td><td></td><td></td><td></td><td></td><td></td><td></td><td></td></tr>
<tr><td>Chromosome</td><td>start</td><td></td><td>end</td><td>strand</td><td>MM</td><td>target_seq</td><td>PAM</td><td>alignment</td><td>distance</td><td>position</td><td>gene name</td><td>gene id</td></tr>
<tr><td>GU305914.1</td><td></td><td>75836</td><td>75858</td><td>+</td><td>0</td><td>CCCCAGCGCTGACCCGGGGG</td><td>AGG</td><td>||||||||[||||||||||||]PAM</td><td>NA</td><td>Intergenic</td><td>NA</td><td>NA</td></tr>
<tr><td></td><td></td><td></td><td></td><td></td><td></td><td></td><td></td><td></td><td></td><td></td><td></td><td></td></tr>
<tr><td>T5403</td><td>CAGCGCTGACCCGGGGGAGGCGG</td><td></td><td>1000</td><td></td><td></td><td></td><td></td><td></td><td></td><td></td><td></td><td></td></tr>
<tr><td>Oligo fwd</td><td>CACCCAGCGCTGACCCGGGGGAGG</td><td></td><td></td><td></td><td></td><td></td><td></td><td></td><td></td><td></td><td></td><td></td></tr>
<tr><td>Oligo rev</td><td>AAACCCTCCCCCGGGTCAGCGCTG</td><td></td><td></td><td></td><td></td><td></td><td></td><td></td><td></td><td></td><td></td><td></td></tr>
<tr><td>Chromosome</td><td>start</td><td></td><td>end</td><td>strand</td><td>MM</td><td>target_seq</td><td>PAM</td><td>alignment</td><td>distance</td><td>position</td><td>gene name</td><td>gene id</td></tr>
<tr><td>GU305914.1</td><td></td><td>75839</td><td>75861</td><td>+</td><td>0</td><td>CAGCGCTGACCCGGGGGAGG</td><td>CGG</td><td>||||||||[||||||||||||]PAM</td><td>NA</td><td>Intergenic</td><td>NA</td><td>NA</td></tr>
<tr><td></td><td></td><td></td><td></td><td></td><td></td><td></td><td></td><td></td><td></td><td></td><td></td><td></td></tr>
<tr><td>T5404</td><td>CGCTGACCCGGGGGAGGCGGAGG</td><td></td><td>1000</td><td></td><td></td><td></td><td></td><td></td><td></td><td></td><td></td><td></td></tr>
<tr><td>Oligo fwd</td><td>CACCCGCTGACCCGGGGGAGGCGG</td><td></td><td></td><td></td><td></td><td></td><td></td><td></td><td></td><td></td><td></td><td></td></tr>
<tr><td>Oligo rev</td><td>AAACCCGCCTCCCCCGGGTCAGCG</td><td></td><td></td><td></td><td></td><td></td><td></td><td></td><td></td><td></td><td></td><td></td></tr>
<tr><td>Chromosome</td><td>start</td><td></td><td>end</td><td>strand</td><td>MM</td><td>target_seq</td><td>PAM</td><td>alignment</td><td>distance</td><td>position</td><td>gene name</td><td>gene id</td></tr>
<tr><td>GU305914.1</td><td></td><td>75842</td><td>75864</td><td>+</td><td>0</td><td>CGCTGACCCGGGGGAGGCGG</td><td>AGG</td><td>||||||||[||||||||||||]PAM</td><td>NA</td><td>Intergenic</td><td>NA</td><td>NA</td></tr>
<tr><td></td><td></td><td></td><td></td><td></td><td></td><td></td><td></td><td></td><td></td><td></td><td></td><td></td></tr>
<tr><td>T5405</td><td>TGACCCGGGGGAGGCGGAGGCGG</td><td></td><td>1000</td><td></td><td></td><td></td><td></td><td></td><td></td><td></td><td></td><td></td></tr>
<tr><td>Oligo fwd</td><td>CACCTGACCCGGGGGAGGCGGAGG</td><td></td><td></td><td></td><td></td><td></td><td></td><td></td><td></td><td></td><td></td><td></td></tr>
<tr><td>Oligo rev</td><td>AAACCCTCCGCCTCCCCCGGGTCA</td><td></td><td></td><td></td><td></td><td></td><td></td><td></td><td></td><td></td><td></td><td></td></tr>
<tr><td>Chromosome</td><td>start</td><td></td><td>end</td><td>strand</td><td>MM</td><td>target_seq</td><td>PAM</td><td>alignment</td><td>distance</td><td>position</td><td>gene name</td><td>gene id</td></tr>
<tr><td>GU305914.1</td><td></td><td>75845</td><td>75867</td><td>+</td><td>0</td><td>TGACCCGGGGGAGGCGGAGG</td><td>CGG</td><td>||||||||[||||||||||||]PAM</td><td>NA</td><td>Intergenic</td><td>NA</td><td>NA</td></tr>
<tr><td></td><td></td><td></td><td></td><td></td><td></td><td></td><td></td><td></td><td></td><td></td><td></td><td></td></tr>
<tr><td>T5406</td><td>CCCGGGGGAGGCGGAGGCGGCGG</td><td></td><td>1000</td><td></td><td></td><td></td><td></td><td></td><td></td><td></td><td></td><td></td></tr>
<tr><td>Oligo fwd</td><td>CACCCCCGGGGGAGGCGGAGGCGG</td><td></td><td></td><td></td><td></td><td></td><td></td><td></td><td></td><td></td><td></td><td></td></tr>
<tr><td>Oligo rev</td><td>AAACCCGCCTCCGCCTCCCCCGGG</td><td></td><td></td><td></td><td></td><td></td><td></td><td></td><td></td><td></td><td></td><td></td></tr>
<tr><td>Chromosome</td><td>start</td><td></td><td>end</td><td>strand</td><td>MM</td><td>target_seq</td><td>PAM</td><td>alignment</td><td>distance</td><td>position</td><td>gene name</td><td>gene id</td></tr>
<tr><td>GU305914.1</td><td></td><td>75848</td><td>75870</td><td>+</td><td>0</td><td>CCCGGGGGAGGCGGAGGCGG</td><td>CGG</td><td>||||||||[||||||||||||]PAM</td><td>NA</td><td>Intergenic</td><td>NA</td><td>NA</td></tr>
<tr><td>GU305914.1</td><td></td><td>154401</td><td>154423</td><td>+</td><td>4</td><td>CCGGGCGGTGGCGGCGGCGG</td><td>CGG</td><td>||-||-||[-|||||-|||||]PAM</td><td>NA</td><td>Intergenic</td><td>NA</td><td>NA</td></tr>
<tr><td></td><td></td><td></td><td></td><td></td><td></td><td></td><td></td><td></td><td></td><td></td><td></td><td></td></tr>
<tr><td>T5407</td><td>GGGGAGGCGGAGGCGGCGGTCGG</td><td></td><td>1000</td><td></td><td></td><td></td><td></td><td></td><td></td><td></td><td></td><td></td></tr>
<tr><td>Oligo fwd</td><td>CACCGGGGAGGCGGAGGCGGCGGT</td><td></td><td></td><td></td><td></td><td></td><td></td><td></td><td></td><td></td><td></td><td></td></tr>
<tr><td>Oligo rev</td><td>AAACACCGCCGCCTCCGCCTCCCC</td><td></td><td></td><td></td><td></td><td></td><td></td><td></td><td></td><td></td><td></td><td></td></tr>
<tr><td>Chromosome</td><td>start</td><td></td><td>end</td><td>strand</td><td>MM</td><td>target_seq</td><td>PAM</td><td>alignment</td><td>distance</td><td>position</td><td>gene name</td><td>gene id</td></tr>
<tr><td>GU305914.1</td><td></td><td>75852</td><td>75874</td><td>+</td><td>0</td><td>GGGGAGGCGGAGGCGGCGGT</td><td>CGG</td><td>||||||||[||||||||||||]PAM</td><td>NA</td><td>Intergenic</td><td>NA</td><td>NA</td></tr>
<tr><td>GU305914.1</td><td></td><td>48160</td><td>48182</td><td>+</td><td>4</td><td>GCGGCGGTGGCGGCGGCGGT</td><td>GGG</td><td>|-||-||-[||-|||||||||]PAM</td><td>NA</td><td>Intergenic</td><td>NA</td><td>NA</td></tr>
<tr><td>GU305914.1</td><td></td><td>58122</td><td>58144</td><td>-</td><td>4</td><td>GTGGCGGCGGCGGCGGCGGA</td><td>GGG</td><td>|-||-|||[||-||||||||-]PAM</td><td>NA</td><td>Intergenic</td><td>NA</td><td>NA</td></tr>
<tr><td>GU305914.1</td><td></td><td>86780</td><td>86802</td><td>-</td><td>4</td><td>GAGGGGGCGGGGGCGGCGGC</td><td>AGG</td><td>|-||-|||[||-||||||||-]PAM</td><td>NA</td><td>Intergenic</td><td>NA</td><td>NA</td></tr>
<tr><td>GU305914.1</td><td></td><td>67743</td><td>67765</td><td>+</td><td>4</td><td>GGAGCGGCGGCGGCGGCGGC</td><td>GGG</td><td>||-|-|||[||-||||||||-]PAM</td><td>NA</td><td>Intergenic</td><td>NA</td><td>NA</td></tr>
<tr><td>GU305914.1</td><td></td><td>90246</td><td>90268</td><td>-</td><td>4</td><td>GAGGCGGCGGAGCCGGCGGC</td><td>AGG</td><td>|-||-|||[||||-||||||-]PAM</td><td>NA</td><td>Intergenic</td><td>NA</td><td>NA</td></tr>
<tr><td></td><td></td><td></td><td></td><td></td><td></td><td></td><td></td><td></td><td></td><td></td><td></td><td></td></tr>
<tr><td>T5408</td><td>GGCGGAGGCGGCGGTCGGCGAGG</td><td></td><td>1000</td><td></td><td></td><td></td><td></td><td></td><td></td><td></td><td></td><td></td></tr>
<tr><td>Oligo fwd</td><td>CACCGGCGGAGGCGGCGGTCGGCG</td><td></td><td></td><td></td><td></td><td></td><td></td><td></td><td></td><td></td><td></td><td></td></tr>
<tr><td>Oligo rev</td><td>AAACCGCCGACCGCCGCCTCCGCC</td><td></td><td></td><td></td><td></td><td></td><td></td><td></td><td></td><td></td><td></td><td></td></tr>
<tr><td>Chromosome</td><td>start</td><td></td><td>end</td><td>strand</td><td>MM</td><td>target_seq</td><td>PAM</td><td>alignment</td><td>distance</td><td>position</td><td>gene name</td><td>gene id</td></tr>
<tr><td>GU305914.1</td><td></td><td>75857</td><td>75879</td><td>+</td><td>0</td><td>GGCGGAGGCGGCGGTCGGCG</td><td>AGG</td><td>||||||||[||||||||||||]PAM</td><td>NA</td><td>Intergenic</td><td>NA</td><td>NA</td></tr>
<tr><td>GU305914.1</td><td></td><td>75733</td><td>75755</td><td>+</td><td>4</td><td>GGTGGAAGTGGAGGTCGGCG</td><td>AGG</td><td>||-|||-|[-||-||||||||]PAM</td><td>NA</td><td>Intergenic</td><td>NA</td><td>NA</td></tr>
<tr><td>GU305914.1</td><td></td><td>213265</td><td>213287</td><td>+</td><td>3</td><td>GTCGGAGGTGGCGGTCGGAG</td><td>AGG</td><td>|-||||||[-|||||||||-|]PAM</td><td>NA</td><td>Intergenic</td><td>NA</td><td>NA</td></tr>
<tr><td></td><td></td><td></td><td></td><td></td><td></td><td></td><td></td><td></td><td></td><td></td><td></td><td></td></tr>
<tr><td>T5409</td><td>GCGGAGGCGGCGGTCGGCGAGGG</td><td></td><td>1000</td><td></td><td></td><td></td><td></td><td></td><td></td><td></td><td></td><td></td></tr>
<tr><td>Oligo fwd</td><td>CACCGCGGAGGCGGCGGTCGGCGA</td><td></td><td></td><td></td><td></td><td></td><td></td><td></td><td></td><td></td><td></td><td></td></tr>
<tr><td>Oligo rev</td><td>AAACTCGCCGACCGCCGCCTCCGC</td><td></td><td></td><td></td><td></td><td></td><td></td><td></td><td></td><td></td><td></td><td></td></tr>
<tr><td>Chromosome</td><td>start</td><td></td><td>end</td><td>strand</td><td>MM</td><td>target_seq</td><td>PAM</td><td>alignment</td><td>distance</td><td>position</td><td>gene name</td><td>gene id</td></tr>
<tr><td>GU305914.1</td><td></td><td>75858</td><td>75880</td><td>+</td><td>0</td><td>GCGGAGGCGGCGGTCGGCGA</td><td>GGG</td><td>||||||||[||||||||||||]PAM</td><td>NA</td><td>Intergenic</td><td>NA</td><td>NA</td></tr>
<tr><td></td><td></td><td></td><td></td><td></td><td></td><td></td><td></td><td></td><td></td><td></td><td></td><td></td></tr>
<tr><td>T5410</td><td>CGGAGGCGGCGGTCGGCGAGGGG</td><td></td><td>1000</td><td></td><td></td><td></td><td></td><td></td><td></td><td></td><td></td><td></td></tr>
<tr><td>Oligo fwd</td><td>CACCCGGAGGCGGCGGTCGGCGAG</td><td></td><td></td><td></td><td></td><td></td><td></td><td></td><td></td><td></td><td></td><td></td></tr>
<tr><td>Oligo rev</td><td>AAACCTCGCCGACCGCCGCCTCCG</td><td></td><td></td><td></td><td></td><td></td><td></td><td></td><td></td><td></td><td></td><td></td></tr>
<tr><td>Chromosome</td><td>start</td><td></td><td>end</td><td>strand</td><td>MM</td><td>target_seq</td><td>PAM</td><td>alignment</td><td>distance</td><td>position</td><td>gene name</td><td>gene id</td></tr>
<tr><td>GU305914.1</td><td></td><td>75859</td><td>75881</td><td>+</td><td>0</td><td>CGGAGGCGGCGGTCGGCGAG</td><td>GGG</td><td>||||||||[||||||||||||]PAM</td><td>NA</td><td>Intergenic</td><td>NA</td><td>NA</td></tr>
<tr><td></td><td></td><td></td><td></td><td></td><td></td><td></td><td></td><td></td><td></td><td></td><td></td><td></td></tr>
<tr><td>T5411</td><td>GGAGGCGGCGGTCGGCGAGGGGG</td><td></td><td>1000</td><td></td><td></td><td></td><td></td><td></td><td></td><td></td><td></td><td></td></tr>
<tr><td>Oligo fwd</td><td>CACCGGAGGCGGCGGTCGGCGAGG</td><td></td><td></td><td></td><td></td><td></td><td></td><td></td><td></td><td></td><td></td><td></td></tr>
<tr><td>Oligo rev</td><td>AAACCCTCGCCGACCGCCGCCTCC</td><td></td><td></td><td></td><td></td><td></td><td></td><td></td><td></td><td></td><td></td><td></td></tr>
<tr><td>Chromosome</td><td>start</td><td></td><td>end</td><td>strand</td><td>MM</td><td>target_seq</td><td>PAM</td><td>alignment</td><td>distance</td><td>position</td><td>gene name</td><td>gene id</td></tr>
<tr><td>GU305914.1</td><td></td><td>75860</td><td>75882</td><td>+</td><td>0</td><td>GGAGGCGGCGGTCGGCGAGG</td><td>GGG</td><td>||||||||[||||||||||||]PAM</td><td>NA</td><td>Intergenic</td><td>NA</td><td>NA</td></tr>
<tr><td>GU305914.1</td><td></td><td>75736</td><td>75758</td><td>+</td><td>3</td><td>GGAAGTGGAGGTCGGCGAGG</td><td>CGG</td><td>|||-|-||[-|||||||||||]PAM</td><td>NA</td><td>Intergenic</td><td>NA</td><td>NA</td></tr>
<tr><td>GU305914.1</td><td></td><td>65126</td><td>65148</td><td>+</td><td>4</td><td>CGAGGAGGAGGGCGGCGAGG</td><td>GGG</td><td>-||||-||[-||-||||||||]PAM</td><td>NA</td><td>Intergenic</td><td>NA</td><td>NA</td></tr>
<tr><td>GU305914.1</td><td></td><td>213268</td><td>213290</td><td>+</td><td>2</td><td>GGAGGTGGCGGTCGGAGAGG</td><td>CGG</td><td>|||||-||[|||||||-||||]PAM</td><td>NA</td><td>Intergenic</td><td>NA</td><td>NA</td></tr>
<tr><td>GU305914.1</td><td></td><td>173582</td><td>173604</td><td>+</td><td>4</td><td>CGAGGTGTCGGTCGGCGACG</td><td>AGG</td><td>-||||-|-[||||||||||-|]PAM</td><td>NA</td><td>Intergenic</td><td>NA</td><td>NA</td></tr>
<tr><td></td><td></td><td></td><td></td><td></td><td></td><td></td><td></td><td></td><td></td><td></td><td></td><td></td></tr>
<tr><td>T5412</td><td>CGGCGGTCGGCGAGGGGGAGCGG</td><td></td><td>1000</td><td></td><td></td><td></td><td></td><td></td><td></td><td></td><td></td><td></td></tr>
<tr><td>Oligo fwd</td><td>CACCCGGCGGTCGGCGAGGGGGAG</td><td></td><td></td><td></td><td></td><td></td><td></td><td></td><td></td><td></td><td></td><td></td></tr>
<tr><td>Oligo rev</td><td>AAACCTCCCCCTCGCCGACCGCCG</td><td></td><td></td><td></td><td></td><td></td><td></td><td></td><td></td><td></td><td></td><td></td></tr>
<tr><td>Chromosome</td><td>start</td><td></td><td>end</td><td>strand</td><td>MM</td><td>target_seq</td><td>PAM</td><td>alignment</td><td>distance</td><td>position</td><td>gene name</td><td>gene id</td></tr>
<tr><td>GU305914.1</td><td></td><td>75865</td><td>75887</td><td>+</td><td>0</td><td>CGGCGGTCGGCGAGGGGGAG</td><td>CGG</td><td>||||||||[||||||||||||]PAM</td><td>NA</td><td>Intergenic</td><td>NA</td><td>NA</td></tr>
<tr><td>GU305914.1</td><td></td><td>224570</td><td>224592</td><td>+</td><td>4</td><td>CTGCGCCCGGCGAGCGGGAG</td><td>GGG</td><td>|-|||--|[||||||-|||||]PAM</td><td>NA</td><td>Intergenic</td><td>NA</td><td>NA</td></tr>
<tr><td></td><td></td><td></td><td></td><td></td><td></td><td></td><td></td><td></td><td></td><td></td><td></td><td></td></tr>
<tr><td>T5413</td><td>CGGTCGGCGAGGGGGAGCGGCGG</td><td></td><td>1000</td><td></td><td></td><td></td><td></td><td></td><td></td><td></td><td></td><td></td></tr>
<tr><td>Oligo fwd</td><td>CACCCGGTCGGCGAGGGGGAGCGG</td><td></td><td></td><td></td><td></td><td></td><td></td><td></td><td></td><td></td><td></td><td></td></tr>
<tr><td>Oligo rev</td><td>AAACCCGCTCCCCCTCGCCGACCG</td><td></td><td></td><td></td><td></td><td></td><td></td><td></td><td></td><td></td><td></td><td></td></tr>
<tr><td>Chromosome</td><td>start</td><td></td><td>end</td><td>strand</td><td>MM</td><td>target_seq</td><td>PAM</td><td>alignment</td><td>distance</td><td>position</td><td>gene name</td><td>gene id</td></tr>
<tr><td>GU305914.1</td><td></td><td>75868</td><td>75890</td><td>+</td><td>0</td><td>CGGTCGGCGAGGGGGAGCGG</td><td>CGG</td><td>||||||||[||||||||||||]PAM</td><td>NA</td><td>Intergenic</td><td>NA</td><td>NA</td></tr>
<tr><td></td><td></td><td></td><td></td><td></td><td></td><td></td><td></td><td></td><td></td><td></td><td></td><td></td></tr>
<tr><td>T5414</td><td>GGCGAGGGGGAGCGGCGGCGAGG</td><td></td><td>1000</td><td></td><td></td><td></td><td></td><td></td><td></td><td></td><td></td><td></td></tr>
<tr><td>Oligo fwd</td><td>CACCGGCGAGGGGGAGCGGCGGCG</td><td></td><td></td><td></td><td></td><td></td><td></td><td></td><td></td><td></td><td></td><td></td></tr>
<tr><td>Oligo rev</td><td>AAACCGCCGCCGCTCCCCCTCGCC</td><td></td><td></td><td></td><td></td><td></td><td></td><td></td><td></td><td></td><td></td><td></td></tr>
<tr><td>Chromosome</td><td>start</td><td></td><td>end</td><td>strand</td><td>MM</td><td>target_seq</td><td>PAM</td><td>alignment</td><td>distance</td><td>position</td><td>gene name</td><td>gene id</td></tr>
</table>

<table>
<tr><td>GU305914.1</td><td>75873</td><td>75895</td><td>+</td><td>0</td><td>GGCGAGGGGGAGCGGCGGCG</td><td>AGG</td><td>||||||||[||||||||||||]PAM</td><td>NA</td><td>Intergenic</td><td>NA</td><td>NA</td></tr>
<tr><td>T5415</td><td>GGGAGCGGCGGCGAGGCACCTGG</td><td>1000</td><td></td><td></td><td></td><td></td><td></td><td></td><td></td><td></td><td></td></tr>
<tr><td>Oligo fwd</td><td>CACCGGGAGCGGCGGCGAGGCACC</td><td></td><td></td><td></td><td></td><td></td><td></td><td></td><td></td><td></td><td></td></tr>
<tr><td>Oligo rev</td><td>AAACGGTGCCTCGCCGCCGCTCCC</td><td></td><td></td><td></td><td></td><td></td><td></td><td></td><td></td><td></td><td></td></tr>
<tr><td>Chromosome</td><td>start</td><td>end</td><td>strand</td><td>MM</td><td>target_seq</td><td>PAM</td><td>alignment</td><td>distance</td><td>position</td><td>gene name</td><td>gene id</td></tr>
<tr><td>GU305914.1</td><td>75880</td><td>75902</td><td>+</td><td>0</td><td>GGGAGCGGCGGCGAGGCACC</td><td>TGG</td><td>||||||||[||||||||||||]PAM</td><td>NA</td><td>Intergenic</td><td>NA</td><td>NA</td></tr>
<tr><td>T5416</td><td>CGGCGGCGAGGCACCTGGCACGG</td><td>1000</td><td></td><td></td><td></td><td></td><td></td><td></td><td></td><td></td><td></td></tr>
<tr><td>Oligo fwd</td><td>CACCCGGCGGCGAGGCACCTGGCA</td><td></td><td></td><td></td><td></td><td></td><td></td><td></td><td></td><td></td><td></td></tr>
<tr><td>Oligo rev</td><td>AAACTGCCAGGTGCCTCGCCGCCG</td><td></td><td></td><td></td><td></td><td></td><td></td><td></td><td></td><td></td><td></td></tr>
<tr><td>Chromosome</td><td>start</td><td>end</td><td>strand</td><td>MM</td><td>target_seq</td><td>PAM</td><td>alignment</td><td>distance</td><td>position</td><td>gene name</td><td>gene id</td></tr>
<tr><td>GU305914.1</td><td>75885</td><td>75907</td><td>+</td><td>0</td><td>CGGCGGCGAGGCACCTGGCA</td><td>CGG</td><td>||||||||[||||||||||||]PAM</td><td>NA</td><td>Intergenic</td><td>NA</td><td>NA</td></tr>
<tr><td>T5417</td><td>ACCTGGCACGGCGACAGATGAGG</td><td>1000</td><td></td><td></td><td></td><td></td><td></td><td></td><td></td><td></td><td></td></tr>
<tr><td>Oligo fwd</td><td>CACCACCTGGCACGGCGACAGATG</td><td></td><td></td><td></td><td></td><td></td><td></td><td></td><td></td><td></td><td></td></tr>
<tr><td>Oligo rev</td><td>AAACCATCTGTCGCCGTGCCAGGT</td><td></td><td></td><td></td><td></td><td></td><td></td><td></td><td></td><td></td><td></td></tr>
<tr><td>Chromosome</td><td>start</td><td>end</td><td>strand</td><td>MM</td><td>target_seq</td><td>PAM</td><td>alignment</td><td>distance</td><td>position</td><td>gene name</td><td>gene id</td></tr>
<tr><td>GU305914.1</td><td>75897</td><td>75919</td><td>+</td><td>0</td><td>ACCTGGCACGGCGACAGATG</td><td>AGG</td><td>||||||||[||||||||||||]PAM</td><td>NA</td><td>Intergenic</td><td>NA</td><td>NA</td></tr>
<tr><td>T5418</td><td>CCTGGCACGGCGACAGATGAGGG</td><td>1000</td><td></td><td></td><td></td><td></td><td></td><td></td><td></td><td></td><td></td></tr>
<tr><td>Oligo fwd</td><td>CACCCCTGGCACGGCGACAGATGA</td><td></td><td></td><td></td><td></td><td></td><td></td><td></td><td></td><td></td><td></td></tr>
<tr><td>Oligo rev</td><td>AAACTCATCTGTCGCCGTGCCAGG</td><td></td><td></td><td></td><td></td><td></td><td></td><td></td><td></td><td></td><td></td></tr>
<tr><td>Chromosome</td><td>start</td><td>end</td><td>strand</td><td>MM</td><td>target_seq</td><td>PAM</td><td>alignment</td><td>distance</td><td>position</td><td>gene name</td><td>gene id</td></tr>
<tr><td>GU305914.1</td><td>75898</td><td>75920</td><td>+</td><td>0</td><td>CCTGGCACGGCGACAGATGA</td><td>GGG</td><td>||||||||[||||||||||||]PAM</td><td>NA</td><td>Intergenic</td><td>NA</td><td>NA</td></tr>
<tr><td>GU305914.1</td><td>24311</td><td>24333</td><td>+</td><td>4</td><td>CGCGGCCCCGCGACAGATGA</td><td>TGG</td><td>|--|||-|[-|||||||||||]PAM</td><td>NA</td><td>Intergenic</td><td>NA</td><td>NA</td></tr>
<tr><td>T5419</td><td>ACAGATGAGGGACCCGCAGTCGG</td><td>1000</td><td></td><td></td><td></td><td></td><td></td><td></td><td></td><td></td><td></td></tr>
<tr><td>Oligo fwd</td><td>CACCACAGATGAGGGACCCGCAGT</td><td></td><td></td><td></td><td></td><td></td><td></td><td></td><td></td><td></td><td></td></tr>
<tr><td>Oligo rev</td><td>AAACACTGCGGGTCCCTCATCTGT</td><td></td><td></td><td></td><td></td><td></td><td></td><td></td><td></td><td></td><td></td></tr>
<tr><td>Chromosome</td><td>start</td><td>end</td><td>strand</td><td>MM</td><td>target_seq</td><td>PAM</td><td>alignment</td><td>distance</td><td>position</td><td>gene name</td><td>gene id</td></tr>
<tr><td>GU305914.1</td><td>75910</td><td>75932</td><td>+</td><td>0</td><td>ACAGATGAGGGACCCGCAGT</td><td>CGG</td><td>||||||||[||||||||||||]PAM</td><td>NA</td><td>Intergenic</td><td>NA</td><td>NA</td></tr>
<tr><td>T5420</td><td>GATGAGGGACCCGCAGTCGGTGG</td><td>1000</td><td></td><td></td><td></td><td></td><td></td><td></td><td></td><td></td><td></td></tr>
<tr><td>Oligo fwd</td><td>CACCGATGAGGGACCCGCAGTCGG</td><td></td><td></td><td></td><td></td><td></td><td></td><td></td><td></td><td></td><td></td></tr>
<tr><td>Oligo rev</td><td>AAACCCGACTGCGGGTCCCTCATC</td><td></td><td></td><td></td><td></td><td></td><td></td><td></td><td></td><td></td><td></td></tr>
<tr><td>Chromosome</td><td>start</td><td>end</td><td>strand</td><td>MM</td><td>target_seq</td><td>PAM</td><td>alignment</td><td>distance</td><td>position</td><td>gene name</td><td>gene id</td></tr>
<tr><td>GU305914.1</td><td>75913</td><td>75935</td><td>+</td><td>0</td><td>GATGAGGGACCCGCAGTCGG</td><td>TGG</td><td>||||||||[||||||||||||]PAM</td><td>NA</td><td>Intergenic</td><td>NA</td><td>NA</td></tr>
<tr><td>T5421</td><td>ATGAGGGACCCGCAGTCGGTGGG</td><td>1000</td><td></td><td></td><td></td><td></td><td></td><td></td><td></td><td></td><td></td></tr>
<tr><td>Oligo fwd</td><td>CACCATGAGGGACCCGCAGTCGGT</td><td></td><td></td><td></td><td></td><td></td><td></td><td></td><td></td><td></td><td></td></tr>
<tr><td>Oligo rev</td><td>AAACACCGACTGCGGGTCCCTCAT</td><td></td><td></td><td></td><td></td><td></td><td></td><td></td><td></td><td></td><td></td></tr>
<tr><td>Chromosome</td><td>start</td><td>end</td><td>strand</td><td>MM</td><td>target_seq</td><td>PAM</td><td>alignment</td><td>distance</td><td>position</td><td>gene name</td><td>gene id</td></tr>
<tr><td>GU305914.1</td><td>75914</td><td>75936</td><td>+</td><td>0</td><td>ATGAGGGACCCGCAGTCGGT</td><td>GGG</td><td>||||||||[||||||||||||]PAM</td><td>NA</td><td>Intergenic</td><td>NA</td><td>NA</td></tr>
<tr><td>T5422</td><td>GGACCCGCAGTCGGTGGGAACGG</td><td>1000</td><td></td><td></td><td></td><td></td><td></td><td></td><td></td><td></td><td></td></tr>
<tr><td>Oligo fwd</td><td>CACCGGACCCGCAGTCGGTGGGAA</td><td></td><td></td><td></td><td></td><td></td><td></td><td></td><td></td><td></td><td></td></tr>
<tr><td>Oligo rev</td><td>AAACTTCCCACCGACTGCGGGTCC</td><td></td><td></td><td></td><td></td><td></td><td></td><td></td><td></td><td></td><td></td></tr>
<tr><td>Chromosome</td><td>start</td><td>end</td><td>strand</td><td>MM</td><td>target_seq</td><td>PAM</td><td>alignment</td><td>distance</td><td>position</td><td>gene name</td><td>gene id</td></tr>
<tr><td>GU305914.1</td><td>75919</td><td>75941</td><td>+</td><td>0</td><td>GGACCCGCAGTCGGTGGGAA</td><td>CGG</td><td>||||||||[||||||||||||]PAM</td><td>NA</td><td>Intergenic</td><td>NA</td><td>NA</td></tr>
<tr><td>T5423</td><td>TCCGCGCGTAGATGTCGCGCAGG</td><td>1000</td><td></td><td></td><td></td><td></td><td></td><td></td><td></td><td></td><td></td></tr>
<tr><td>Oligo fwd</td><td>CACCTCCGCGCGTAGATGTCGCGC</td><td></td><td></td><td></td><td></td><td></td><td></td><td></td><td></td><td></td><td></td></tr>
<tr><td>Oligo rev</td><td>AAACGCGCGACATCTACGCGCGGA</td><td></td><td></td><td></td><td></td><td></td><td></td><td></td><td></td><td></td><td></td></tr>
<tr><td>Chromosome</td><td>start</td><td>end</td><td>strand</td><td>MM</td><td>target_seq</td><td>PAM</td><td>alignment</td><td>distance</td><td>position</td><td>gene name</td><td>gene id</td></tr>
<tr><td>GU305914.1</td><td>75951</td><td>75973</td><td>+</td><td>0</td><td>TCCGCGCGTAGATGTCGCGC</td><td>AGG</td><td>||||||||[||||||||||||]PAM</td><td>NA</td><td>Intergenic</td><td>NA</td><td>NA</td></tr>
<tr><td>T5424</td><td>GCGCGTAGATGTCGCGCAGGAGG</td><td>1000</td><td></td><td></td><td></td><td></td><td></td><td></td><td></td><td></td><td></td></tr>
<tr><td>Oligo fwd</td><td>CACCGCGCGTAGATGTCGCGCAGG</td><td></td><td></td><td></td><td></td><td></td><td></td><td></td><td></td><td></td><td></td></tr>
<tr><td>Oligo rev</td><td>AAACCCTGCGCGACATCTACGCGC</td><td></td><td></td><td></td><td></td><td></td><td></td><td></td><td></td><td></td><td></td></tr>
<tr><td>Chromosome</td><td>start</td><td>end</td><td>strand</td><td>MM</td><td>target_seq</td><td>PAM</td><td>alignment</td><td>distance</td><td>position</td><td>gene name</td><td>gene id</td></tr>
<tr><td>GU305914.1</td><td>75954</td><td>75976</td><td>+</td><td>0</td><td>GCGCGTAGATGTCGCGCAGG</td><td>AGG</td><td>||||||||[||||||||||||]PAM</td><td>NA</td><td>Intergenic</td><td>NA</td><td>NA</td></tr>
<tr><td>T5425</td><td>CGCGTAGATGTCGCGCAGGAGGG</td><td>1000</td><td></td><td></td><td></td><td></td><td></td><td></td><td></td><td></td><td></td></tr>
<tr><td>Oligo fwd</td><td>CACCCGCGTAGATGTCGCGCAGGA</td><td></td><td></td><td></td><td></td><td></td><td></td><td></td><td></td><td></td><td></td></tr>
<tr><td>Oligo rev</td><td>AAACTCCTGCGCGACATCTACGCG</td><td></td><td></td><td></td><td></td><td></td><td></td><td></td><td></td><td></td><td></td></tr>
<tr><td>Chromosome</td><td>start</td><td>end</td><td>strand</td><td>MM</td><td>target_seq</td><td>PAM</td><td>alignment</td><td>distance</td><td>position</td><td>gene name</td><td>gene id</td></tr>
<tr><td>GU305914.1</td><td>75955</td><td>75977</td><td>+</td><td>0</td><td>CGCGTAGATGTCGCGCAGGA</td><td>GGG</td><td>||||||||[||||||||||||]PAM</td><td>NA</td><td>Intergenic</td><td>NA</td><td>NA</td></tr>
<tr><td>T5426</td><td>AGATGTCGCGCAGGAGGGCCTGG</td><td>1000</td><td></td><td></td><td></td><td></td><td></td><td></td><td></td><td></td><td></td></tr>
<tr><td>Oligo fwd</td><td>CACCAGATGTCGCGCAGGAGGGCC</td><td></td><td></td><td></td><td></td><td></td><td></td><td></td><td></td><td></td><td></td></tr>
<tr><td>Oligo rev</td><td>AAACGGCCCTCCTGCGCGACATCT</td><td></td><td></td><td></td><td></td><td></td><td></td><td></td><td></td><td></td><td></td></tr>
<tr><td>Chromosome</td><td>start</td><td>end</td><td>strand</td><td>MM</td><td>target_seq</td><td>PAM</td><td>alignment</td><td>distance</td><td>position</td><td>gene name</td><td>gene id</td></tr>
<tr><td>GU305914.1</td><td>75960</td><td>75982</td><td>+</td><td>0</td><td>AGATGTCGCGCAGGAGGGCC</td><td>TGG</td><td>||||||||[||||||||||||]PAM</td><td>NA</td><td>Intergenic</td><td>NA</td><td>NA</td></tr>
<tr><td>T5427</td><td>GCGCAGGAGGGCCTGGTACTCGG</td><td>1000</td><td></td><td></td><td></td><td></td><td></td><td></td><td></td><td></td><td></td></tr>
<tr><td>Oligo fwd</td><td>CACCGCGCAGGAGGGCCTGGTACT</td><td></td><td></td><td></td><td></td><td></td><td></td><td></td><td></td><td></td><td></td></tr>
<tr><td>Oligo rev</td><td>AAACAGTACCAGGCCCTCCTGCGC</td><td></td><td></td><td></td><td></td><td></td><td></td><td></td><td></td><td></td><td></td></tr>
<tr><td>Chromosome</td><td>start</td><td>end</td><td>strand</td><td>MM</td><td>target_seq</td><td>PAM</td><td>alignment</td><td>distance</td><td>position</td><td>gene name</td><td>gene id</td></tr>
<tr><td>GU305914.1</td><td>75967</td><td>75989</td><td>+</td><td>0</td><td>GCGCAGGAGGGCCTGGTACT</td><td>CGG</td><td>||||||||[||||||||||||]PAM</td><td>NA</td><td>Intergenic</td><td>NA</td><td>NA</td></tr>
<tr><td>T5428</td><td>CTGGTACTCGGCCTCGTTGTCGG</td><td>1000</td><td></td><td></td><td></td><td></td><td></td><td></td><td></td><td></td><td></td></tr>
<tr><td>Oligo fwd</td><td>CACCCTGGTACTCGGCCTCGTTGT</td><td></td><td></td><td></td><td></td><td></td><td></td><td></td><td></td><td></td><td></td></tr>
<tr><td>Oligo rev</td><td>AAACACAACGAGGCCGAGTACCAG</td><td></td><td></td><td></td><td></td><td></td><td></td><td></td><td></td><td></td><td></td></tr>
<tr><td>Chromosome</td><td>start</td><td>end</td><td>strand</td><td>MM</td><td>target_seq</td><td>PAM</td><td>alignment</td><td>distance</td><td>position</td><td>gene name</td><td>gene id</td></tr>
<tr><td>GU305914.1</td><td>75979</td><td>76001</td><td>+</td><td>0</td><td>CTGGTACTCGGCCTCGTTGT</td><td>CGG</td><td>||||||||[||||||||||||]PAM</td><td>NA</td><td>Intergenic</td><td>NA</td><td>NA</td></tr>
<tr><td>T5429</td><td>TGGTACTCGGCCTCGTTGTCGGG</td><td>1000</td><td></td><td></td><td></td><td></td><td></td><td></td><td></td><td></td><td></td></tr>
<tr><td>Oligo fwd</td><td>CACCTGGTACTCGGCCTCGTTGTC</td><td></td><td></td><td></td><td></td><td></td><td></td><td></td><td></td><td></td><td></td></tr>
</table>

<table>
<tr><td>Oligo rev</td><td>AAACGACAACGAGGCCGAGTACCA</td></tr>
<tr><td>Chromosome</td><td>start</td><td></td><td>end</td><td>strand</td><td>MM</td><td>target_seq</td><td>PAM</td><td>alignment</td><td>distance</td><td>position</td><td>gene name</td><td>gene id</td></tr>
<tr><td>GU305914.1</td><td></td><td>75980</td><td>76002</td><td>+</td><td>0</td><td>TGGTACTCGGCCTCGTTGTC</td><td>GGG</td><td>||||||||[||||||||||||]PAM</td><td>NA</td><td>Intergenic</td><td>NA</td><td>NA</td></tr>
<tr><td>T5430</td><td>GGCCTCGTTGTCGGGTCCGAAGG</td><td></td><td>1000</td></tr>
<tr><td>Oligo fwd</td><td>CACCGGCCTCGTTGTCGGGTCCGA</td></tr>
<tr><td>Oligo rev</td><td>AAACTCGGACCCGACAACGAGGCC</td></tr>
<tr><td>Chromosome</td><td>start</td><td></td><td>end</td><td>strand</td><td>MM</td><td>target_seq</td><td>PAM</td><td>alignment</td><td>distance</td><td>position</td><td>gene name</td><td>gene id</td></tr>
<tr><td>GU305914.1</td><td></td><td>75988</td><td>76010</td><td>+</td><td>0</td><td>GGCCTCGTTGTCGGGTCCGA</td><td>AGG</td><td>||||||||[||||||||||||]PAM</td><td>NA</td><td>Intergenic</td><td>NA</td><td>NA</td></tr>
<tr><td>T5431</td><td>GGCGACGACCTCGATGTTGTAGG</td><td></td><td>1000</td></tr>
<tr><td>Oligo fwd</td><td>CACCGGCGACGACCTCGATGTTGT</td></tr>
<tr><td>Oligo rev</td><td>AAACACAACATCGAGGTCGTCGCC</td></tr>
<tr><td>Chromosome</td><td>start</td><td></td><td>end</td><td>strand</td><td>MM</td><td>target_seq</td><td>PAM</td><td>alignment</td><td>distance</td><td>position</td><td>gene name</td><td>gene id</td></tr>
<tr><td>GU305914.1</td><td></td><td>76009</td><td>76031</td><td>+</td><td>0</td><td>GGCGACGACCTCGATGTTGT</td><td>AGG</td><td>||||||||[||||||||||||]PAM</td><td>NA</td><td>Intergenic</td><td>NA</td><td>NA</td></tr>
<tr><td>T5432</td><td>GTTGTAGGACATGCCGCTGTTGG</td><td></td><td>1000</td></tr>
<tr><td>Oligo fwd</td><td>CACCGTTGTAGGACATGCCGCTGT</td></tr>
<tr><td>Oligo rev</td><td>AAACACAGCGGCATGTCCTACAAC</td></tr>
<tr><td>Chromosome</td><td>start</td><td></td><td>end</td><td>strand</td><td>MM</td><td>target_seq</td><td>PAM</td><td>alignment</td><td>distance</td><td>position</td><td>gene name</td><td>gene id</td></tr>
<tr><td>GU305914.1</td><td></td><td>76024</td><td>76046</td><td>+</td><td>0</td><td>GTTGTAGGACATGCCGCTGT</td><td>TGG</td><td>||||||||[||||||||||||]PAM</td><td>NA</td><td>Intergenic</td><td>NA</td><td>NA</td></tr>
<tr><td>T5433</td><td>GGACTTCCTGACGACGCAGTTGG</td><td></td><td>1000</td></tr>
<tr><td>Oligo fwd</td><td>CACCGGACTTCCTGACGACGCAGT</td></tr>
<tr><td>Oligo rev</td><td>AAACACTGCGTCGTCAGGAAGTCC</td></tr>
<tr><td>Chromosome</td><td>start</td><td></td><td>end</td><td>strand</td><td>MM</td><td>target_seq</td><td>PAM</td><td>alignment</td><td>distance</td><td>position</td><td>gene name</td><td>gene id</td></tr>
<tr><td>GU305914.1</td><td></td><td>76045</td><td>76067</td><td>+</td><td>0</td><td>GGACTTCCTGACGACGCAGT</td><td>TGG</td><td>||||||||[||||||||||||]PAM</td><td>NA</td><td>Intergenic</td><td>NA</td><td>NA</td></tr>
<tr><td>T5434</td><td>GGTGAGCACCCCCTTGTCGCAGG</td><td></td><td>1000</td></tr>
<tr><td>Oligo fwd</td><td>CACCGGTGAGCACCCCCTTGTCGC</td></tr>
<tr><td>Oligo rev</td><td>AAACGCGACAAGGGGGTGCTCACC</td></tr>
<tr><td>Chromosome</td><td>start</td><td></td><td>end</td><td>strand</td><td>MM</td><td>target_seq</td><td>PAM</td><td>alignment</td><td>distance</td><td>position</td><td>gene name</td><td>gene id</td></tr>
<tr><td>GU305914.1</td><td></td><td>76066</td><td>76088</td><td>+</td><td>0</td><td>GGTGAGCACCCCCTTGTCGC</td><td>AGG</td><td>||||||||[||||||||||||]PAM</td><td>NA</td><td>Intergenic</td><td>NA</td><td>NA</td></tr>
<tr><td>T5435</td><td>GTGAGCACCCCCTTGTCGCAGGG</td><td></td><td>1000</td></tr>
<tr><td>Oligo fwd</td><td>CACCGTGAGCACCCCCTTGTCGCA</td></tr>
<tr><td>Oligo rev</td><td>AAACTGCGACAAGGGGGTGCTCAC</td></tr>
<tr><td>Chromosome</td><td>start</td><td></td><td>end</td><td>strand</td><td>MM</td><td>target_seq</td><td>PAM</td><td>alignment</td><td>distance</td><td>position</td><td>gene name</td><td>gene id</td></tr>
<tr><td>GU305914.1</td><td></td><td>76067</td><td>76089</td><td>+</td><td>0</td><td>GTGAGCACCCCCTTGTCGCA</td><td>GGG</td><td>||||||||[||||||||||||]PAM</td><td>NA</td><td>Intergenic</td><td>NA</td><td>NA</td></tr>
<tr><td>T5436</td><td>GGGCTTGATCCTCTCGATGTCGG</td><td></td><td>1000</td></tr>
<tr><td>Oligo fwd</td><td>CACCGGGCTTGATCCTCTCGATGT</td></tr>
<tr><td>Oligo rev</td><td>AAACACATCGAGAGGATCAAGCCC</td></tr>
<tr><td>Chromosome</td><td>start</td><td></td><td>end</td><td>strand</td><td>MM</td><td>target_seq</td><td>PAM</td><td>alignment</td><td>distance</td><td>position</td><td>gene name</td><td>gene id</td></tr>
<tr><td>GU305914.1</td><td></td><td>76087</td><td>76109</td><td>+</td><td>0</td><td>GGGCTTGATCCTCTCGATGT</td><td>CGG</td><td>||||||||[||||||||||||]PAM</td><td>NA</td><td>Intergenic</td><td>NA</td><td>NA</td></tr>
<tr><td>T5437</td><td>TGTCGGCCAGCGTGACCACGAGG</td><td></td><td>1000</td></tr>
<tr><td>Oligo fwd</td><td>CACCTGTCGGCCAGCGTGACCACG</td></tr>
<tr><td>Oligo rev</td><td>AAACCGTGGTCACGCTGGCCGACA</td></tr>
<tr><td>Chromosome</td><td>start</td><td></td><td>end</td><td>strand</td><td>MM</td><td>target_seq</td><td>PAM</td><td>alignment</td><td>distance</td><td>position</td><td>gene name</td><td>gene id</td></tr>
<tr><td>GU305914.1</td><td></td><td>76104</td><td>76126</td><td>+</td><td>0</td><td>TGTCGGCCAGCGTGACCACG</td><td>AGG</td><td>||||||||[||||||||||||]PAM</td><td>NA</td><td>Intergenic</td><td>NA</td><td>NA</td></tr>
<tr><td>T5438</td><td>GTCGGCCAGCGTGACCACGAGGG</td><td></td><td>1000</td></tr>
<tr><td>Oligo fwd</td><td>CACCGTCGGCCAGCGTGACCACGA</td></tr>
<tr><td>Oligo rev</td><td>AAACTCGTGGTCACGCTGGCCGAC</td></tr>
<tr><td>Chromosome</td><td>start</td><td></td><td>end</td><td>strand</td><td>MM</td><td>target_seq</td><td>PAM</td><td>alignment</td><td>distance</td><td>position</td><td>gene name</td><td>gene id</td></tr>
<tr><td>GU305914.1</td><td></td><td>76105</td><td>76127</td><td>+</td><td>0</td><td>GTCGGCCAGCGTGACCACGA</td><td>GGG</td><td>||||||||[||||||||||||]PAM</td><td>NA</td><td>Intergenic</td><td>NA</td><td>NA</td></tr>
<tr><td>T5439</td><td>CGAGGGCTTCGTTGACGCTCAGG</td><td></td><td>1000</td></tr>
<tr><td>Oligo fwd</td><td>CACCCGAGGGCTTCGTTGACGCTC</td></tr>
<tr><td>Oligo rev</td><td>AAACGAGCGTCAACGAAGCCCTCG</td></tr>
<tr><td>Chromosome</td><td>start</td><td></td><td>end</td><td>strand</td><td>MM</td><td>target_seq</td><td>PAM</td><td>alignment</td><td>distance</td><td>position</td><td>gene name</td><td>gene id</td></tr>
<tr><td>GU305914.1</td><td></td><td>76122</td><td>76144</td><td>+</td><td>0</td><td>CGAGGGCTTCGTTGACGCTC</td><td>AGG</td><td>||||||||[||||||||||||]PAM</td><td>NA</td><td>Intergenic</td><td>NA</td><td>NA</td></tr>
<tr><td>T5440</td><td>TAAAATCGCAGACCCTCTTGAGG</td><td></td><td>1000</td></tr>
<tr><td>Oligo fwd</td><td>CACCTAAAATCGCAGACCCTCTTG</td></tr>
<tr><td>Oligo rev</td><td>AAACCAAGAGGGTCTGCGATTTTA</td></tr>
<tr><td>Chromosome</td><td>start</td><td></td><td>end</td><td>strand</td><td>MM</td><td>target_seq</td><td>PAM</td><td>alignment</td><td>distance</td><td>position</td><td>gene name</td><td>gene id</td></tr>
<tr><td>GU305914.1</td><td></td><td>76149</td><td>76171</td><td>+</td><td>0</td><td>TAAAATCGCAGACCCTCTTG</td><td>AGG</td><td>||||||||[||||||||||||]PAM</td><td>NA</td><td>Intergenic</td><td>NA</td><td>NA</td></tr>
<tr><td>T5441</td><td>AGACCCTCTTGAGGTCCTTGAGG</td><td></td><td>1000</td></tr>
<tr><td>Oligo fwd</td><td>CACCAGACCCTCTTGAGGTCCTTG</td></tr>
<tr><td>Oligo rev</td><td>AAACCAAGGACCTCAAGAGGGTCT</td></tr>
<tr><td>Chromosome</td><td>start</td><td></td><td>end</td><td>strand</td><td>MM</td><td>target_seq</td><td>PAM</td><td>alignment</td><td>distance</td><td>position</td><td>gene name</td><td>gene id</td></tr>
<tr><td>GU305914.1</td><td></td><td>76158</td><td>76180</td><td>+</td><td>0</td><td>AGACCCTCTTGAGGTCCTTG</td><td>AGG</td><td>||||||||[||||||||||||]PAM</td><td>NA</td><td>Intergenic</td><td>NA</td><td>NA</td></tr>
<tr><td>T5442</td><td>CTTGAGGTCCTTGAGGCGCTCGG</td><td></td><td>1000</td></tr>
<tr><td>Oligo fwd</td><td>CACCCTTGAGGTCCTTGAGGCGCT</td></tr>
<tr><td>Oligo rev</td><td>AAACAGCGCCTCAAGGACCTCAAG</td></tr>
<tr><td>Chromosome</td><td>start</td><td></td><td>end</td><td>strand</td><td>MM</td><td>target_seq</td><td>PAM</td><td>alignment</td><td>distance</td><td>position</td><td>gene name</td><td>gene id</td></tr>
<tr><td>GU305914.1</td><td></td><td>76165</td><td>76187</td><td>+</td><td>0</td><td>CTTGAGGTCCTTGAGGCGCT</td><td>CGG</td><td>||||||||[||||||||||||]PAM</td><td>NA</td><td>Intergenic</td><td>NA</td><td>NA</td></tr>
<tr><td>T5443</td><td>TTGAGGTCCTTGAGGCGCTCGGG</td><td></td><td>1000</td></tr>
<tr><td>Oligo fwd</td><td>CACCTTGAGGTCCTTGAGGCGCTC</td></tr>
<tr><td>Oligo rev</td><td>AAACGAGCGCCTCAAGGACCTCAA</td></tr>
<tr><td>Chromosome</td><td>start</td><td></td><td>end</td><td>strand</td><td>MM</td><td>target_seq</td><td>PAM</td><td>alignment</td><td>distance</td><td>position</td><td>gene name</td><td>gene id</td></tr>
<tr><td>GU305914.1</td><td></td><td>76166</td><td>76188</td><td>+</td><td>0</td><td>TTGAGGTCCTTGAGGCGCTC</td><td>GGG</td><td>||||||||[||||||||||||]PAM</td><td>NA</td><td>Intergenic</td><td>NA</td><td>NA</td></tr>
<tr><td>T5444</td><td>CTCGGGAAAGCCTATATAGCAGG</td><td></td><td>1000</td></tr>
</table>

<table>
<tr><td>Oligo fwd</td><td>CACCCTCGGGAAAGCCTATATAGC</td><td></td><td></td><td></td><td></td><td></td><td></td><td></td><td></td><td></td><td></td></tr>
<tr><td>Oligo rev</td><td>AAACGCTATATAGGCTTTCCCGAG</td><td></td><td></td><td></td><td></td><td></td><td></td><td></td><td></td><td></td><td></td></tr>
<tr><td>Chromosome</td><td>start</td><td>end</td><td>strand</td><td>MM</td><td>target_seq</td><td>PAM</td><td>alignment</td><td>distance</td><td>position</td><td>gene name</td><td>gene id</td></tr>
<tr><td>GU305914.1</td><td>76183</td><td>76205</td><td>+</td><td>0</td><td>CTCGGGAAAGCCTATATAGC</td><td>AGG</td><td>||||||||[||||||||||||]PAM</td><td>NA</td><td>Intergenic</td><td>NA</td><td>NA</td></tr>
<tr><td>T5445</td><td>GAAAGCCTATATAGCAGGAGAGG</td><td>1000</td><td></td><td></td><td></td><td></td><td></td><td></td><td></td><td></td><td></td></tr>
<tr><td>Oligo fwd</td><td>CACCGAAAGCCTATATAGCAGGAG</td><td></td><td></td><td></td><td></td><td></td><td></td><td></td><td></td><td></td><td></td></tr>
<tr><td>Oligo rev</td><td>AAACCTCCTGCTATATAGGCTTTC</td><td></td><td></td><td></td><td></td><td></td><td></td><td></td><td></td><td></td><td></td></tr>
<tr><td>Chromosome</td><td>start</td><td>end</td><td>strand</td><td>MM</td><td>target_seq</td><td>PAM</td><td>alignment</td><td>distance</td><td>position</td><td>gene name</td><td>gene id</td></tr>
<tr><td>GU305914.1</td><td>76188</td><td>76210</td><td>+</td><td>0</td><td>GAAAGCCTATATAGCAGGAG</td><td>AGG</td><td>||||||||[||||||||||||]PAM</td><td>NA</td><td>Intergenic</td><td>NA</td><td>NA</td></tr>
<tr><td>T5446</td><td>AAAGCCTATATAGCAGGAGAGGG</td><td>1000</td><td></td><td></td><td></td><td></td><td></td><td></td><td></td><td></td><td></td></tr>
<tr><td>Oligo fwd</td><td>CACCAAAGCCTATATAGCAGGAGA</td><td></td><td></td><td></td><td></td><td></td><td></td><td></td><td></td><td></td><td></td></tr>
<tr><td>Oligo rev</td><td>AAACTCTCCTGCTATATAGGCTTT</td><td></td><td></td><td></td><td></td><td></td><td></td><td></td><td></td><td></td><td></td></tr>
<tr><td>Chromosome</td><td>start</td><td>end</td><td>strand</td><td>MM</td><td>target_seq</td><td>PAM</td><td>alignment</td><td>distance</td><td>position</td><td>gene name</td><td>gene id</td></tr>
<tr><td>GU305914.1</td><td>76189</td><td>76211</td><td>+</td><td>0</td><td>AAAGCCTATATAGCAGGAGA</td><td>GGG</td><td>||||||||[||||||||||||]PAM</td><td>NA</td><td>Intergenic</td><td>NA</td><td>NA</td></tr>
<tr><td>T5447</td><td>GGAGAGGGAGACCCGACAGCCGG</td><td>1000</td><td></td><td></td><td></td><td></td><td></td><td></td><td></td><td></td><td></td></tr>
<tr><td>Oligo fwd</td><td>CACCGGAGAGGGAGACCCGACAGC</td><td></td><td></td><td></td><td></td><td></td><td></td><td></td><td></td><td></td><td></td></tr>
<tr><td>Oligo rev</td><td>AAACGCTGTCGGGTCTCCCTCTCC</td><td></td><td></td><td></td><td></td><td></td><td></td><td></td><td></td><td></td><td></td></tr>
<tr><td>Chromosome</td><td>start</td><td>end</td><td>strand</td><td>MM</td><td>target_seq</td><td>PAM</td><td>alignment</td><td>distance</td><td>position</td><td>gene name</td><td>gene id</td></tr>
<tr><td>GU305914.1</td><td>76204</td><td>76226</td><td>+</td><td>0</td><td>GGAGAGGGAGACCCGACAGC</td><td>CGG</td><td>||||||||[||||||||||||]PAM</td><td>NA</td><td>Intergenic</td><td>NA</td><td>NA</td></tr>
<tr><td>T5448</td><td>GTTTTTGAACACGAAGCAAGCGG</td><td>1000</td><td></td><td></td><td></td><td></td><td></td><td></td><td></td><td></td><td></td></tr>
<tr><td>Oligo fwd</td><td>CACCGTTTTTGAACACGAAGCAAG</td><td></td><td></td><td></td><td></td><td></td><td></td><td></td><td></td><td></td><td></td></tr>
<tr><td>Oligo rev</td><td>AAACCTTGCTTCGTGTTCAAAAAC</td><td></td><td></td><td></td><td></td><td></td><td></td><td></td><td></td><td></td><td></td></tr>
<tr><td>Chromosome</td><td>start</td><td>end</td><td>strand</td><td>MM</td><td>target_seq</td><td>PAM</td><td>alignment</td><td>distance</td><td>position</td><td>gene name</td><td>gene id</td></tr>
<tr><td>GU305914.1</td><td>76228</td><td>76250</td><td>+</td><td>0</td><td>GTTTTTGAACACGAAGCAAG</td><td>CGG</td><td>||||||||[||||||||||||]PAM</td><td>NA</td><td>Intergenic</td><td>NA</td><td>NA</td></tr>
<tr><td>T5449</td><td>TCCTACACTCCCAATAGCTGAGG</td><td>1000</td><td></td><td></td><td></td><td></td><td></td><td></td><td></td><td></td><td></td></tr>
<tr><td>Oligo fwd</td><td>CACCTCCTACACTCCCAATAGCTG</td><td></td><td></td><td></td><td></td><td></td><td></td><td></td><td></td><td></td><td></td></tr>
<tr><td>Oligo rev</td><td>AAACCAGCTATTGGGAGTGTAGGA</td><td></td><td></td><td></td><td></td><td></td><td></td><td></td><td></td><td></td><td></td></tr>
<tr><td>Chromosome</td><td>start</td><td>end</td><td>strand</td><td>MM</td><td>target_seq</td><td>PAM</td><td>alignment</td><td>distance</td><td>position</td><td>gene name</td><td>gene id</td></tr>
<tr><td>GU305914.1</td><td>76260</td><td>76282</td><td>+</td><td>0</td><td>TCCTACACTCCCAATAGCTG</td><td>AGG</td><td>||||||||[||||||||||||]PAM</td><td>NA</td><td>Intergenic</td><td>NA</td><td>NA</td></tr>
<tr><td>T5450</td><td>TAGCTGAGGAGATACTCGACCGG</td><td>1000</td><td></td><td></td><td></td><td></td><td></td><td></td><td></td><td></td><td></td></tr>
<tr><td>Oligo fwd</td><td>CACCTAGCTGAGGAGATACTCGAC</td><td></td><td></td><td></td><td></td><td></td><td></td><td></td><td></td><td></td><td></td></tr>
<tr><td>Oligo rev</td><td>AAACGTCGAGTATCTCCTCAGCTA</td><td></td><td></td><td></td><td></td><td></td><td></td><td></td><td></td><td></td><td></td></tr>
<tr><td>Chromosome</td><td>start</td><td>end</td><td>strand</td><td>MM</td><td>target_seq</td><td>PAM</td><td>alignment</td><td>distance</td><td>position</td><td>gene name</td><td>gene id</td></tr>
<tr><td>GU305914.1</td><td>76274</td><td>76296</td><td>+</td><td>0</td><td>TAGCTGAGGAGATACTCGAC</td><td>CGG</td><td>||||||||[||||||||||||]PAM</td><td>NA</td><td>Intergenic</td><td>NA</td><td>NA</td></tr>
<tr><td>T5451</td><td>CTCGACCGGATAGACGATATCGG</td><td>1000</td><td></td><td></td><td></td><td></td><td></td><td></td><td></td><td></td><td></td></tr>
<tr><td>Oligo fwd</td><td>CACCCTCGACCGGATAGACGATAT</td><td></td><td></td><td></td><td></td><td></td><td></td><td></td><td></td><td></td><td></td></tr>
<tr><td>Oligo rev</td><td>AAACATATCGTCTATCCGGTCGAG</td><td></td><td></td><td></td><td></td><td></td><td></td><td></td><td></td><td></td><td></td></tr>
<tr><td>Chromosome</td><td>start</td><td>end</td><td>strand</td><td>MM</td><td>target_seq</td><td>PAM</td><td>alignment</td><td>distance</td><td>position</td><td>gene name</td><td>gene id</td></tr>
<tr><td>GU305914.1</td><td>76288</td><td>76310</td><td>+</td><td>0</td><td>CTCGACCGGATAGACGATAT</td><td>CGG</td><td>||||||||[||||||||||||]PAM</td><td>NA</td><td>Intergenic</td><td>NA</td><td>NA</td></tr>
<tr><td>T5452</td><td>GACCGGATAGACGATATCGGTGG</td><td>1000</td><td></td><td></td><td></td><td></td><td></td><td></td><td></td><td></td><td></td></tr>
<tr><td>Oligo fwd</td><td>CACCGACCGGATAGACGATATCGG</td><td></td><td></td><td></td><td></td><td></td><td></td><td></td><td></td><td></td><td></td></tr>
<tr><td>Oligo rev</td><td>AAACCCGATATCGTCTATCCGGTC</td><td></td><td></td><td></td><td></td><td></td><td></td><td></td><td></td><td></td><td></td></tr>
<tr><td>Chromosome</td><td>start</td><td>end</td><td>strand</td><td>MM</td><td>target_seq</td><td>PAM</td><td>alignment</td><td>distance</td><td>position</td><td>gene name</td><td>gene id</td></tr>
<tr><td>GU305914.1</td><td>76291</td><td>76313</td><td>+</td><td>0</td><td>GACCGGATAGACGATATCGG</td><td>TGG</td><td>||||||||[||||||||||||]PAM</td><td>NA</td><td>Intergenic</td><td>NA</td><td>NA</td></tr>
<tr><td>T5453</td><td>CGCGTCGCAGAGCGAATAGTTGG</td><td>1000</td><td></td><td></td><td></td><td></td><td></td><td></td><td></td><td></td><td></td></tr>
<tr><td>Oligo fwd</td><td>CACCCGCGTCGCAGAGCGAATAGT</td><td></td><td></td><td></td><td></td><td></td><td></td><td></td><td></td><td></td><td></td></tr>
<tr><td>Oligo rev</td><td>AAACACTATTCGCTCTGCGACGCG</td><td></td><td></td><td></td><td></td><td></td><td></td><td></td><td></td><td></td><td></td></tr>
<tr><td>Chromosome</td><td>start</td><td>end</td><td>strand</td><td>MM</td><td>target_seq</td><td>PAM</td><td>alignment</td><td>distance</td><td>position</td><td>gene name</td><td>gene id</td></tr>
<tr><td>GU305914.1</td><td>76321</td><td>76343</td><td>+</td><td>0</td><td>CGCGTCGCAGAGCGAATAGT</td><td>TGG</td><td>||||||||[||||||||||||]PAM</td><td>NA</td><td>Intergenic</td><td>NA</td><td>NA</td></tr>
<tr><td>T5454</td><td>GCGTCGCAGAGCGAATAGTTGGG</td><td>1000</td><td></td><td></td><td></td><td></td><td></td><td></td><td></td><td></td><td></td></tr>
<tr><td>Oligo fwd</td><td>CACCGCGTCGCAGAGCGAATAGTT</td><td></td><td></td><td></td><td></td><td></td><td></td><td></td><td></td><td></td><td></td></tr>
<tr><td>Oligo rev</td><td>AAACAACTATTCGCTCTGCGACGC</td><td></td><td></td><td></td><td></td><td></td><td></td><td></td><td></td><td></td><td></td></tr>
<tr><td>Chromosome</td><td>start</td><td>end</td><td>strand</td><td>MM</td><td>target_seq</td><td>PAM</td><td>alignment</td><td>distance</td><td>position</td><td>gene name</td><td>gene id</td></tr>
<tr><td>GU305914.1</td><td>76322</td><td>76344</td><td>+</td><td>0</td><td>GCGTCGCAGAGCGAATAGTT</td><td>GGG</td><td>||||||||[||||||||||||]PAM</td><td>NA</td><td>Intergenic</td><td>NA</td><td>NA</td></tr>
<tr><td>T5455</td><td>CGTCGCAGAGCGAATAGTTGGGG</td><td>1000</td><td></td><td></td><td></td><td></td><td></td><td></td><td></td><td></td><td></td></tr>
<tr><td>Oligo fwd</td><td>CACCCGTCGCAGAGCGAATAGTTG</td><td></td><td></td><td></td><td></td><td></td><td></td><td></td><td></td><td></td><td></td></tr>
<tr><td>Oligo rev</td><td>AAACCAACTATTCGCTCTGCGACG</td><td></td><td></td><td></td><td></td><td></td><td></td><td></td><td></td><td></td><td></td></tr>
<tr><td>Chromosome</td><td>start</td><td>end</td><td>strand</td><td>MM</td><td>target_seq</td><td>PAM</td><td>alignment</td><td>distance</td><td>position</td><td>gene name</td><td>gene id</td></tr>
<tr><td>GU305914.1</td><td>76323</td><td>76345</td><td>+</td><td>0</td><td>CGTCGCAGAGCGAATAGTTG</td><td>GGG</td><td>||||||||[||||||||||||]PAM</td><td>NA</td><td>Intergenic</td><td>NA</td><td>NA</td></tr>
<tr><td>T5456</td><td>GGGGTTTTTGCATATTAAAGCGG</td><td>1000</td><td></td><td></td><td></td><td></td><td></td><td></td><td></td><td></td><td></td></tr>
<tr><td>Oligo fwd</td><td>CACCGGGGTTTTTGCATATTAAAG</td><td></td><td></td><td></td><td></td><td></td><td></td><td></td><td></td><td></td><td></td></tr>
<tr><td>Oligo rev</td><td>AAACCTTTAATATGCAAAAACCCC</td><td></td><td></td><td></td><td></td><td></td><td></td><td></td><td></td><td></td><td></td></tr>
<tr><td>Chromosome</td><td>start</td><td>end</td><td>strand</td><td>MM</td><td>target_seq</td><td>PAM</td><td>alignment</td><td>distance</td><td>position</td><td>gene name</td><td>gene id</td></tr>
<tr><td>GU305914.1</td><td>76342</td><td>76364</td><td>+</td><td>0</td><td>GGGGTTTTTGCATATTAAAG</td><td>CGG</td><td>||||||||[||||||||||||]PAM</td><td>NA</td><td>Intergenic</td><td>NA</td><td>NA</td></tr>
<tr><td>T5457</td><td>TTTGCATATTAAAGCGGTGTCGG</td><td>1000</td><td></td><td></td><td></td><td></td><td></td><td></td><td></td><td></td><td></td></tr>
<tr><td>Oligo fwd</td><td>CACCTTTGCATATTAAAGCGGTGT</td><td></td><td></td><td></td><td></td><td></td><td></td><td></td><td></td><td></td><td></td></tr>
<tr><td>Oligo rev</td><td>AAACACACCGCTTTAATATGCAAA</td><td></td><td></td><td></td><td></td><td></td><td></td><td></td><td></td><td></td><td></td></tr>
<tr><td>Chromosome</td><td>start</td><td>end</td><td>strand</td><td>MM</td><td>target_seq</td><td>PAM</td><td>alignment</td><td>distance</td><td>position</td><td>gene name</td><td>gene id</td></tr>
<tr><td>GU305914.1</td><td>76348</td><td>76370</td><td>+</td><td>0</td><td>TTTGCATATTAAAGCGGTGT</td><td>CGG</td><td>||||||||[||||||||||||]PAM</td><td>NA</td><td>Intergenic</td><td>NA</td><td>NA</td></tr>
<tr><td>T5458</td><td>TGAGCTCGTTCTCGTCGAGCCGG</td><td>1000</td><td></td><td></td><td></td><td></td><td></td><td></td><td></td><td></td><td></td></tr>
<tr><td>Oligo fwd</td><td>CACCTGAGCTCGTTCTCGTCGAGC</td><td></td><td></td><td></td><td></td><td></td><td></td><td></td><td></td><td></td><td></td></tr>
<tr><td>Oligo rev</td><td>AAACGCTCGACGAGAACGAGCTCA</td><td></td><td></td><td></td><td></td><td></td><td></td><td></td><td></td><td></td><td></td></tr>
<tr><td>Chromosome</td><td>start</td><td>end</td><td>strand</td><td>MM</td><td>target_seq</td><td>PAM</td><td>alignment</td><td>distance</td><td>position</td><td>gene name</td><td>gene id</td></tr>
<tr><td>GU305914.1</td><td>76377</td><td>76399</td><td>+</td><td>0</td><td>TGAGCTCGTTCTCGTCGAGC</td><td>CGG</td><td>||||||||[||||||||||||]PAM</td><td>NA</td><td>Intergenic</td><td>NA</td><td>NA</td></tr>
</table>

<table>
<tr><td>T5459</td><td>GCTCGTTCTCGTCGAGCCGGAGG</td><td>1000</td><td></td><td></td><td></td><td></td><td></td><td></td><td></td><td></td><td></td></tr>
<tr><td>Oligo fwd</td><td>CACCGCTCGTTCTCGTCGAGCCGG</td><td></td><td></td><td></td><td></td><td></td><td></td><td></td><td></td><td></td><td></td></tr>
<tr><td>Oligo rev</td><td>AAACCCGGCTCGACGAGAACGAGC</td><td></td><td></td><td></td><td></td><td></td><td></td><td></td><td></td><td></td><td></td></tr>
<tr><td>Chromosome</td><td>start</td><td>end</td><td>strand</td><td>MM</td><td>target_seq</td><td>PAM</td><td>alignment</td><td>distance</td><td>position</td><td>gene name</td><td>gene id</td></tr>
<tr><td>GU305914.1</td><td>76380</td><td>76402</td><td>+</td><td>0</td><td>GCTCGTTCTCGTCGAGCCGG</td><td>AGG</td><td>||||||||[||||||||||||]PAM</td><td>NA</td><td>Intergenic</td><td>NA</td><td>NA</td></tr>
<tr><td>T5460</td><td>CCGGAGGATCCGCCGCGTGTTGG</td><td>1000</td><td></td><td></td><td></td><td></td><td></td><td></td><td></td><td></td><td></td></tr>
<tr><td>Oligo fwd</td><td>CACCCCGGAGGATCCGCCGCGTGT</td><td></td><td></td><td></td><td></td><td></td><td></td><td></td><td></td><td></td><td></td></tr>
<tr><td>Oligo rev</td><td>AAACACACGCGGCGGATCCTCCGG</td><td></td><td></td><td></td><td></td><td></td><td></td><td></td><td></td><td></td><td></td></tr>
<tr><td>Chromosome</td><td>start</td><td>end</td><td>strand</td><td>MM</td><td>target_seq</td><td>PAM</td><td>alignment</td><td>distance</td><td>position</td><td>gene name</td><td>gene id</td></tr>
<tr><td>GU305914.1</td><td>76396</td><td>76418</td><td>+</td><td>0</td><td>CCGGAGGATCCGCCGCGTGT</td><td>TGG</td><td>||||||||[||||||||||||]PAM</td><td>NA</td><td>Intergenic</td><td>NA</td><td>NA</td></tr>
<tr><td>T5461</td><td>TCCGCCGCGTGTTGGAAATCAGG</td><td>1000</td><td></td><td></td><td></td><td></td><td></td><td></td><td></td><td></td><td></td></tr>
<tr><td>Oligo fwd</td><td>CACCTCCGCCGCGTGTTGGAAATC</td><td></td><td></td><td></td><td></td><td></td><td></td><td></td><td></td><td></td><td></td></tr>
<tr><td>Oligo rev</td><td>AAACGATTTCCAACACGCGGCGGA</td><td></td><td></td><td></td><td></td><td></td><td></td><td></td><td></td><td></td><td></td></tr>
<tr><td>Chromosome</td><td>start</td><td>end</td><td>strand</td><td>MM</td><td>target_seq</td><td>PAM</td><td>alignment</td><td>distance</td><td>position</td><td>gene name</td><td>gene id</td></tr>
<tr><td>GU305914.1</td><td>76404</td><td>76426</td><td>+</td><td>0</td><td>TCCGCCGCGTGTTGGAAATC</td><td>AGG</td><td>||||||||[||||||||||||]PAM</td><td>NA</td><td>Intergenic</td><td>NA</td><td>NA</td></tr>
<tr><td>T5462</td><td>CCGCGTGTTGGAAATCAGGTCGG</td><td>1000</td><td></td><td></td><td></td><td></td><td></td><td></td><td></td><td></td><td></td></tr>
<tr><td>Oligo fwd</td><td>CACCCCGCGTGTTGGAAATCAGGT</td><td></td><td></td><td></td><td></td><td></td><td></td><td></td><td></td><td></td><td></td></tr>
<tr><td>Oligo rev</td><td>AAACACCTGATTTCCAACACGCGG</td><td></td><td></td><td></td><td></td><td></td><td></td><td></td><td></td><td></td><td></td></tr>
<tr><td>Chromosome</td><td>start</td><td>end</td><td>strand</td><td>MM</td><td>target_seq</td><td>PAM</td><td>alignment</td><td>distance</td><td>position</td><td>gene name</td><td>gene id</td></tr>
<tr><td>GU305914.1</td><td>76408</td><td>76430</td><td>+</td><td>0</td><td>CCGCGTGTTGGAAATCAGGT</td><td>CGG</td><td>||||||||[||||||||||||]PAM</td><td>NA</td><td>Intergenic</td><td>NA</td><td>NA</td></tr>
<tr><td>T5463</td><td>CTTGTCGATCTCCATCGCGCCGG</td><td>1000</td><td></td><td></td><td></td><td></td><td></td><td></td><td></td><td></td><td></td></tr>
<tr><td>Oligo fwd</td><td>CACCCTTGTCGATCTCCATCGCGC</td><td></td><td></td><td></td><td></td><td></td><td></td><td></td><td></td><td></td><td></td></tr>
<tr><td>Oligo rev</td><td>AAACGCGCGATGGAGATCGACAAG</td><td></td><td></td><td></td><td></td><td></td><td></td><td></td><td></td><td></td><td></td></tr>
<tr><td>Chromosome</td><td>start</td><td>end</td><td>strand</td><td>MM</td><td>target_seq</td><td>PAM</td><td>alignment</td><td>distance</td><td>position</td><td>gene name</td><td>gene id</td></tr>
<tr><td>GU305914.1</td><td>76441</td><td>76463</td><td>+</td><td>0</td><td>CTTGTCGATCTCCATCGCGC</td><td>CGG</td><td>||||||||[||||||||||||]PAM</td><td>NA</td><td>Intergenic</td><td>NA</td><td>NA</td></tr>
<tr><td>GU305914.1</td><td>32891</td><td>32913</td><td>-</td><td>4</td><td>CCTGTCCATCGCCAACGCGC</td><td>TGG</td><td>|-||||-|[||-|||-|||||]PAM</td><td>NA</td><td>Intergenic</td><td>NA</td><td>NA</td></tr>
<tr><td>T5464</td><td>CCATCGCGCCGGCGCCGCCGCGG</td><td>1000</td><td></td><td></td><td></td><td></td><td></td><td></td><td></td><td></td><td></td></tr>
<tr><td>Oligo fwd</td><td>CACCCCATCGCGCCGGCGCCGCCG</td><td></td><td></td><td></td><td></td><td></td><td></td><td></td><td></td><td></td><td></td></tr>
<tr><td>Oligo rev</td><td>AAACCGGCGGCGCCGGCGCGATGG</td><td></td><td></td><td></td><td></td><td></td><td></td><td></td><td></td><td></td><td></td></tr>
<tr><td>Chromosome</td><td>start</td><td>end</td><td>strand</td><td>MM</td><td>target_seq</td><td>PAM</td><td>alignment</td><td>distance</td><td>position</td><td>gene name</td><td>gene id</td></tr>
<tr><td>GU305914.1</td><td>76452</td><td>76474</td><td>+</td><td>0</td><td>CCATCGCGCCGGCGCCGCCG</td><td>CGG</td><td>||||||||[||||||||||||]PAM</td><td>NA</td><td>Intergenic</td><td>NA</td><td>NA</td></tr>
<tr><td>T5465</td><td>TGAGCTCGCCGCCGTCCGAGTGG</td><td>1000</td><td></td><td></td><td></td><td></td><td></td><td></td><td></td><td></td><td></td></tr>
<tr><td>Oligo fwd</td><td>CACCTGAGCTCGCCGCCGTCCGAG</td><td></td><td></td><td></td><td></td><td></td><td></td><td></td><td></td><td></td><td></td></tr>
<tr><td>Oligo rev</td><td>AAACCTCGGACGGCGGCGAGCTCA</td><td></td><td></td><td></td><td></td><td></td><td></td><td></td><td></td><td></td><td></td></tr>
<tr><td>Chromosome</td><td>start</td><td>end</td><td>strand</td><td>MM</td><td>target_seq</td><td>PAM</td><td>alignment</td><td>distance</td><td>position</td><td>gene name</td><td>gene id</td></tr>
<tr><td>GU305914.1</td><td>76476</td><td>76498</td><td>+</td><td>0</td><td>TGAGCTCGCCGCCGTCCGAG</td><td>TGG</td><td>||||||||[||||||||||||]PAM</td><td>NA</td><td>Intergenic</td><td>NA</td><td>NA</td></tr>
<tr><td>GU305914.1</td><td>114393</td><td>114415</td><td>+</td><td>4</td><td>GGAGCGCGCCGCCGGCCGCG</td><td>AGG</td><td>-||||-||[||||||-|||-|]PAM</td><td>NA</td><td>Intergenic</td><td>NA</td><td>NA</td></tr>
<tr><td>T5466</td><td>CCCGAGCCGACCCGATCAGTCGG</td><td>1000</td><td></td><td></td><td></td><td></td><td></td><td></td><td></td><td></td><td></td></tr>
<tr><td>Oligo fwd</td><td>CACCCCCGAGCCGACCCGATCAGT</td><td></td><td></td><td></td><td></td><td></td><td></td><td></td><td></td><td></td><td></td></tr>
<tr><td>Oligo rev</td><td>AAACACTGATCGGGTCGGCTCGGG</td><td></td><td></td><td></td><td></td><td></td><td></td><td></td><td></td><td></td><td></td></tr>
<tr><td>Chromosome</td><td>start</td><td>end</td><td>strand</td><td>MM</td><td>target_seq</td><td>PAM</td><td>alignment</td><td>distance</td><td>position</td><td>gene name</td><td>gene id</td></tr>
<tr><td>GU305914.1</td><td>76504</td><td>76526</td><td>+</td><td>0</td><td>CCCGAGCCGACCCGATCAGT</td><td>CGG</td><td>||||||||[||||||||||||]PAM</td><td>NA</td><td>Intergenic</td><td>NA</td><td>NA</td></tr>
<tr><td>T5467</td><td>GTCTAATAATTTAGTGATCACGG</td><td>1000</td><td></td><td></td><td></td><td></td><td></td><td></td><td></td><td></td><td></td></tr>
<tr><td>Oligo fwd</td><td>CACCGTCTAATAATTTAGTGATCA</td><td></td><td></td><td></td><td></td><td></td><td></td><td></td><td></td><td></td><td></td></tr>
<tr><td>Oligo rev</td><td>AAACTGATCACTAAATTATTAGAC</td><td></td><td></td><td></td><td></td><td></td><td></td><td></td><td></td><td></td><td></td></tr>
<tr><td>Chromosome</td><td>start</td><td>end</td><td>strand</td><td>MM</td><td>target_seq</td><td>PAM</td><td>alignment</td><td>distance</td><td>position</td><td>gene name</td><td>gene id</td></tr>
<tr><td>GU305914.1</td><td>76558</td><td>76580</td><td>+</td><td>0</td><td>GTCTAATAATTTAGTGATCA</td><td>CGG</td><td>||||||||[||||||||||||]PAM</td><td>NA</td><td>Intergenic</td><td>NA</td><td>NA</td></tr>
<tr><td>T5468</td><td>AGAGCTTCATCTTCTCGTTGAGG</td><td>1000</td><td></td><td></td><td></td><td></td><td></td><td></td><td></td><td></td><td></td></tr>
<tr><td>Oligo fwd</td><td>CACCAGAGCTTCATCTTCTCGTTG</td><td></td><td></td><td></td><td></td><td></td><td></td><td></td><td></td><td></td><td></td></tr>
<tr><td>Oligo rev</td><td>AAACCAACGAGAAGATGAAGCTCT</td><td></td><td></td><td></td><td></td><td></td><td></td><td></td><td></td><td></td><td></td></tr>
<tr><td>Chromosome</td><td>start</td><td>end</td><td>strand</td><td>MM</td><td>target_seq</td><td>PAM</td><td>alignment</td><td>distance</td><td>position</td><td>gene name</td><td>gene id</td></tr>
<tr><td>GU305914.1</td><td>76587</td><td>76609</td><td>+</td><td>0</td><td>AGAGCTTCATCTTCTCGTTG</td><td>AGG</td><td>||||||||[||||||||||||]PAM</td><td>NA</td><td>Intergenic</td><td>NA</td><td>NA</td></tr>
<tr><td>GU305914.1</td><td>111223</td><td>111245</td><td>+</td><td>4</td><td>GGAGCCTCATCCTCTCGATG</td><td>TGG</td><td>-||||-||[|||-|||||-||]PAM</td><td>NA</td><td>Intergenic</td><td>NA</td><td>NA</td></tr>
<tr><td>T5469</td><td>CTTCATCTTCTCGTTGAGGACGG</td><td>1000</td><td></td><td></td><td></td><td></td><td></td><td></td><td></td><td></td><td></td></tr>
<tr><td>Oligo fwd</td><td>CACCCTTCATCTTCTCGTTGAGGA</td><td></td><td></td><td></td><td></td><td></td><td></td><td></td><td></td><td></td><td></td></tr>
<tr><td>Oligo rev</td><td>AAACTCCTCAACGAGAAGATGAAG</td><td></td><td></td><td></td><td></td><td></td><td></td><td></td><td></td><td></td><td></td></tr>
<tr><td>Chromosome</td><td>start</td><td>end</td><td>strand</td><td>MM</td><td>target_seq</td><td>PAM</td><td>alignment</td><td>distance</td><td>position</td><td>gene name</td><td>gene id</td></tr>
<tr><td>GU305914.1</td><td>76591</td><td>76613</td><td>+</td><td>0</td><td>CTTCATCTTCTCGTTGAGGA</td><td>CGG</td><td>||||||||[||||||||||||]PAM</td><td>NA</td><td>Intergenic</td><td>NA</td><td>NA</td></tr>
<tr><td>T5470</td><td>CATCTTCTCGTTGAGGACGGCGG</td><td>1000</td><td></td><td></td><td></td><td></td><td></td><td></td><td></td><td></td><td></td></tr>
<tr><td>Oligo fwd</td><td>CACCCATCTTCTCGTTGAGGACGG</td><td></td><td></td><td></td><td></td><td></td><td></td><td></td><td></td><td></td><td></td></tr>
<tr><td>Oligo rev</td><td>AAACCCGTCCTCAACGAGAAGATG</td><td></td><td></td><td></td><td></td><td></td><td></td><td></td><td></td><td></td><td></td></tr>
<tr><td>Chromosome</td><td>start</td><td>end</td><td>strand</td><td>MM</td><td>target_seq</td><td>PAM</td><td>alignment</td><td>distance</td><td>position</td><td>gene name</td><td>gene id</td></tr>
<tr><td>GU305914.1</td><td>76594</td><td>76616</td><td>+</td><td>0</td><td>CATCTTCTCGTTGAGGACGG</td><td>CGG</td><td>||||||||[||||||||||||]PAM</td><td>NA</td><td>Intergenic</td><td>NA</td><td>NA</td></tr>
<tr><td>GU305914.1</td><td>19513</td><td>19535</td><td>+</td><td>4</td><td>CATCGTCGCGATGGGGACGG</td><td>CGG</td><td>||||-||-[||-||-||||||]PAM</td><td>NA</td><td>Intergenic</td><td>NA</td><td>NA</td></tr>
<tr><td>T5471</td><td>GTACACATGCCGCAGAAAAGCGG</td><td>1000</td><td></td><td></td><td></td><td></td><td></td><td></td><td></td><td></td><td></td></tr>
<tr><td>Oligo fwd</td><td>CACCGTACACATGCCGCAGAAAAG</td><td></td><td></td><td></td><td></td><td></td><td></td><td></td><td></td><td></td><td></td></tr>
<tr><td>Oligo rev</td><td>AAACCTTTTCTGCGGCATGTGTAC</td><td></td><td></td><td></td><td></td><td></td><td></td><td></td><td></td><td></td><td></td></tr>
<tr><td>Chromosome</td><td>start</td><td>end</td><td>strand</td><td>MM</td><td>target_seq</td><td>PAM</td><td>alignment</td><td>distance</td><td>position</td><td>gene name</td><td>gene id</td></tr>
<tr><td>GU305914.1</td><td>76642</td><td>76664</td><td>+</td><td>0</td><td>GTACACATGCCGCAGAAAAG</td><td>CGG</td><td>||||||||[||||||||||||]PAM</td><td>NA</td><td>Intergenic</td><td>NA</td><td>NA</td></tr>
<tr><td>T5472</td><td>TTTAACCTCGAGTCGATAGACGG</td><td>1000</td><td></td><td></td><td></td><td></td><td></td><td></td><td></td><td></td><td></td></tr>
<tr><td>Oligo fwd</td><td>CACCTTTAACCTCGAGTCGATAGA</td><td></td><td></td><td></td><td></td><td></td><td></td><td></td><td></td><td></td><td></td></tr>
<tr><td>Oligo rev</td><td>AAACTCTATCGACTCGAGGTTAAA</td><td></td><td></td><td></td><td></td><td></td><td></td><td></td><td></td><td></td><td></td></tr>
<tr><td>Chromosome</td><td>start</td><td>end</td><td>strand</td><td>MM</td><td>target_seq</td><td>PAM</td><td>alignment</td><td>distance</td><td>position</td><td>gene name</td><td>gene id</td></tr>
<tr><td>GU305914.1</td><td>76691</td><td>76713</td><td>+</td><td>0</td><td>TTTAACCTCGAGTCGATAGA</td><td>CGG</td><td>||||||||[||||||||||||]PAM</td><td>NA</td><td>Intergenic</td><td>NA</td><td>NA</td></tr>
<tr><td>T5473</td><td>AACGTCAGTTCGTAGACCCTAGG</td><td>1000</td><td></td><td></td><td></td><td></td><td></td><td></td><td></td><td></td><td></td></tr>
</table>

| | | | | | | | | | | | |
|---|---|---|---|---|---|---|---|---|---|---|---|
| Oligo fwd | CACCAACGTCAGTTCGTAGACCCT | | | | | | | | | | |
| Oligo rev | AAACAGGGTCTACGAACTGACGTT | | | | | | | | | | |
| Chromosome | start | end | strand | MM | target_seq | PAM | alignment | distance | position | gene name | gene id |
| GU305914.1 | 76715 | 76737 | + | 0 | AACGTCAGTTCGTAGACCCT | AGG | \|\|\|\|\|\|\|\|[\|\|\|\|\|\|\|\|\|\|\|\|]PAM | NA | Intergenic | NA | NA |
| T5474 | TCGTAGACCCTAGGCAGCATCGG | 1000 | | | | | | | | | |
| Oligo fwd | CACCTCGTAGACCCTAGGCAGCAT | | | | | | | | | | |
| Oligo rev | AAACATGCTGCCTAGGGTCTACGA | | | | | | | | | | |
| Chromosome | start | end | strand | MM | target_seq | PAM | alignment | distance | position | gene name | gene id |
| GU305914.1 | 76724 | 76746 | + | 0 | TCGTAGACCCTAGGCAGCAT | CGG | \|\|\|\|\|\|\|\|[\|\|\|\|\|\|\|\|\|\|\|\|]PAM | NA | Intergenic | NA | NA |
| T5475 | TAGGCAGCATCGGCTCGAAACGG | 1000 | | | | | | | | | |
| Oligo fwd | CACCTAGGCAGCATCGGCTCGAAA | | | | | | | | | | |
| Oligo rev | AAACTTTCGAGCCGATGCTGCCTA | | | | | | | | | | |
| Chromosome | start | end | strand | MM | target_seq | PAM | alignment | distance | position | gene name | gene id |
| GU305914.1 | 76734 | 76756 | + | 0 | TAGGCAGCATCGGCTCGAAA | CGG | \|\|\|\|\|\|\|\|[\|\|\|\|\|\|\|\|\|\|\|\|]PAM | NA | Intergenic | NA | NA |
| T5476 | CGGCTCGAAACGGATCGCGTCGG | 1000 | | | | | | | | | |
| Oligo fwd | CACCCGGCTCGAAACGGATCGCGT | | | | | | | | | | |
| Oligo rev | AAACACGCGATCCGTTTCGAGCCG | | | | | | | | | | |
| Chromosome | start | end | strand | MM | target_seq | PAM | alignment | distance | position | gene name | gene id |
| GU305914.1 | 76744 | 76766 | + | 0 | CGGCTCGAAACGGATCGCGT | CGG | \|\|\|\|\|\|\|\|[\|\|\|\|\|\|\|\|\|\|\|\|]PAM | NA | Intergenic | NA | NA |
| T5477 | AGCGTTCGAAGAGCGCGTCGAGG | 1000 | | | | | | | | | |
| Oligo fwd | CACCAGCGTTCGAAGAGCGCGTCG | | | | | | | | | | |
| Oligo rev | AAACCGACGCGCTCTTCGAACGCT | | | | | | | | | | |
| Chromosome | start | end | strand | MM | target_seq | PAM | alignment | distance | position | gene name | gene id |
| GU305914.1 | 76785 | 76807 | + | 0 | AGCGTTCGAAGAGCGCGTCG | AGG | \|\|\|\|\|\|\|\|[\|\|\|\|\|\|\|\|\|\|\|\|]PAM | NA | Intergenic | NA | NA |
| T5478 | CGCGTCGAGGTTGCACTCCTCGG | 1000 | | | | | | | | | |
| Oligo fwd | CACCCGCGTCGAGGTTGCACTCCT | | | | | | | | | | |
| Oligo rev | AAACAGGAGTGCAACCTCGACGCG | | | | | | | | | | |
| Chromosome | start | end | strand | MM | target_seq | PAM | alignment | distance | position | gene name | gene id |
| GU305914.1 | 76798 | 76820 | + | 0 | CGCGTCGAGGTTGCACTCCT | CGG | \|\|\|\|\|\|\|\|[\|\|\|\|\|\|\|\|\|\|\|\|]PAM | NA | Intergenic | NA | NA |
| T5479 | GTCGAGGTTGCACTCCTCGGCGG | 1000 | | | | | | | | | |
| Oligo fwd | CACCGTCGAGGTTGCACTCCTCGG | | | | | | | | | | |
| Oligo rev | AAACCCGAGGAGTGCAACCTCGAC | | | | | | | | | | |
| Chromosome | start | end | strand | MM | target_seq | PAM | alignment | distance | position | gene name | gene id |
| GU305914.1 | 76801 | 76823 | + | 0 | GTCGAGGTTGCACTCCTCGG | CGG | \|\|\|\|\|\|\|\|[\|\|\|\|\|\|\|\|\|\|\|\|]PAM | NA | Intergenic | NA | NA |
| T5480 | TCGAGGTTGCACTCCTCGGCGGG | 1000 | | | | | | | | | |
| Oligo fwd | CACCTCGAGGTTGCACTCCTCGGC | | | | | | | | | | |
| Oligo rev | AAACGCCGAGGAGTGCAACCTCGA | | | | | | | | | | |
| Chromosome | start | end | strand | MM | target_seq | PAM | alignment | distance | position | gene name | gene id |
| GU305914.1 | 76802 | 76824 | + | 0 | TCGAGGTTGCACTCCTCGGC | GGG | \|\|\|\|\|\|\|\|[\|\|\|\|\|\|\|\|\|\|\|\|]PAM | NA | Intergenic | NA | NA |
| T5481 | CGAGGTTGCACTCCTCGGCGGGG | 1000 | | | | | | | | | |
| Oligo fwd | CACCCGAGGTTGCACTCCTCGGCG | | | | | | | | | | |
| Oligo rev | AAACCGCCGAGGAGTGCAACCTCG | | | | | | | | | | |
| Chromosome | start | end | strand | MM | target_seq | PAM | alignment | distance | position | gene name | gene id |
| GU305914.1 | 76803 | 76825 | + | 0 | CGAGGTTGCACTCCTCGGCG | GGG | \|\|\|\|\|\|\|\|[\|\|\|\|\|\|\|\|\|\|\|\|]PAM | NA | Intergenic | NA | NA |
| GU305914.1 | 201680 | 201702 | - | 4 | CGACGTGGCACCCCCCGGCG | AGG | \|\|\|-\|\|-\|[\|\|\|-\|\|-\|\|\|\|\|]PAM | NA | Intergenic | NA | NA |
| T5482 | GTTGCACTCCTCGGCGGGGACGG | 1000 | | | | | | | | | |
| Oligo fwd | CACCGTTGCACTCCTCGGCGGGGA | | | | | | | | | | |
| Oligo rev | AAACTCCCCGCCGAGGAGTGCAAC | | | | | | | | | | |
| Chromosome | start | end | strand | MM | target_seq | PAM | alignment | distance | position | gene name | gene id |
| GU305914.1 | 76807 | 76829 | + | 0 | GTTGCACTCCTCGGCGGGGA | CGG | \|\|\|\|\|\|\|\|[\|\|\|\|\|\|\|\|\|\|\|\|]PAM | NA | Intergenic | NA | NA |
| T5483 | TTGCACTCCTCGGCGGGGACGGG | 1000 | | | | | | | | | |
| Oligo fwd | CACCTTGCACTCCTCGGCGGGGAC | | | | | | | | | | |
| Oligo rev | AAACGTCCCCGCCGAGGAGTGCAA | | | | | | | | | | |
| Chromosome | start | end | strand | MM | target_seq | PAM | alignment | distance | position | gene name | gene id |
| GU305914.1 | 76808 | 76830 | + | 0 | TTGCACTCCTCGGCGGGGAC | GGG | \|\|\|\|\|\|\|\|[\|\|\|\|\|\|\|\|\|\|\|\|]PAM | NA | Intergenic | NA | NA |
| GU305914.1 | 19913 | 19935 | - | 4 | TTCGATTCCTCGGCGGCGAC | GGG | \|\|--\|-\|\|[\|\|\|\|\|\|\|\|-\|\|\|]PAM | NA | Intergenic | NA | NA |
| T5484 | CTCCTCGGCGGGGACGGGCGCGG | 1000 | | | | | | | | | |
| Oligo fwd | CACCCTCCTCGGCGGGGACGGGCG | | | | | | | | | | |
| Oligo rev | AAACCGCCCGTCCCCGCCGAGGAG | | | | | | | | | | |
| Chromosome | start | end | strand | MM | target_seq | PAM | alignment | distance | position | gene name | gene id |
| GU305914.1 | 76813 | 76835 | + | 0 | CTCCTCGGCGGGGACGGGCG | CGG | \|\|\|\|\|\|\|\|[\|\|\|\|\|\|\|\|\|\|\|\|]PAM | NA | Intergenic | NA | NA |
| T5485 | GGCGGGGACGGGCGCGGCGCCGG | 1000 | | | | | | | | | |
| Oligo fwd | CACCGGCGGGGACGGGCGCGGCGC | | | | | | | | | | |
| Oligo rev | AAACGCGCCGCGCCCGTCCCCGCC | | | | | | | | | | |
| Chromosome | start | end | strand | MM | target_seq | PAM | alignment | distance | position | gene name | gene id |
| GU305914.1 | 76819 | 76841 | + | 0 | GGCGGGGACGGGCGCGGCGC | CGG | \|\|\|\|\|\|\|\|[\|\|\|\|\|\|\|\|\|\|\|\|]PAM | NA | Intergenic | NA | NA |
| T5486 | GGGGACGGGCGCGGCGCCGGCGG | 1000 | | | | | | | | | |
| Oligo fwd | CACCGGGGACGGGCGCGGCGCCGG | | | | | | | | | | |
| Oligo rev | AAACCCGGCGCCGCGCCCGTCCCC | | | | | | | | | | |
| Chromosome | start | end | strand | MM | target_seq | PAM | alignment | distance | position | gene name | gene id |
| GU305914.1 | 76822 | 76844 | + | 0 | GGGGACGGGCGCGGCGCCGG | CGG | \|\|\|\|\|\|\|\|[\|\|\|\|\|\|\|\|\|\|\|\|]PAM | NA | Intergenic | NA | NA |
| T5487 | CCGGCGGCCTCGTCGTCGTCCGG | 1000 | | | | | | | | | |
| Oligo fwd | CACCCCGGCGGCCTCGTCGTCGTC | | | | | | | | | | |
| Oligo rev | AAACGACGACGACGAGGCCGCCGG | | | | | | | | | | |
| Chromosome | start | end | strand | MM | target_seq | PAM | alignment | distance | position | gene name | gene id |

GU305914.1 76838 76860 + 0 CCGGCGGCCTCGTCGTCGTC CGG ||||||||[||||||||||||]PAM NA Intergenic NA NA

T5488 CCGGCGCATCGCCGCCGCGACGG 1000
Oligo fwd CACCCCGGCGCATCGCCGCCGCGA
Oligo rev AAACTCGCGGCGGCGATGCGCCGG
Chromosome start end strand MM target_seq PAM alignment distance position gene name gene id
GU305914.1 76857 76879 + 0 CCGGCGCATCGCCGCCGCGA CGG ||||||||[||||||||||||]PAM NA Intergenic NA NA

T5489 ATCACCGTTCATCGCCGTCGCGG 1000
Oligo fwd CACCATCACCGTTCATCGCCGTCG
Oligo rev AAACCGACGGCGATGAACGGTGAT
Chromosome start end strand MM target_seq PAM alignment distance position gene name gene id
GU305914.1 76888 76910 + 0 ATCACCGTTCATCGCCGTCG CGG ||||||||[||||||||||||]PAM NA Intergenic NA NA

T5490 TCGCGGCGCTCGCGATGTACAGG 1000
Oligo fwd CACCTCGCGGCGCTCGCGATGTAC
Oligo rev AAACGTACATCGCGAGCGCCGCGA
Chromosome start end strand MM target_seq PAM alignment distance position gene name gene id
GU305914.1 76905 76927 + 0 TCGCGGCGCTCGCGATGTAC AGG ||||||||[||||||||||||]PAM NA Intergenic NA NA

T5491 CGCGGCGCTCGCGATGTACAGGG 1000
Oligo fwd CACCCGCGGCGCTCGCGATGTACA
Oligo rev AAACTGTACATCGCGAGCGCCGCG
Chromosome start end strand MM target_seq PAM alignment distance position gene name gene id
GU305914.1 76906 76928 + 0 CGCGGCGCTCGCGATGTACA GGG ||||||||[||||||||||||]PAM NA Intergenic NA NA

T5492 CGCTCGCGATGTACAGGGCCTGG 1000
Oligo fwd CACCCGCTCGCGATGTACAGGGCC
Oligo rev AAACGGCCCTGTACATCGCGAGCG
Chromosome start end strand MM target_seq PAM alignment distance position gene name gene id
GU305914.1 76911 76933 + 0 CGCTCGCGATGTACAGGGCC TGG ||||||||[||||||||||||]PAM NA Intergenic NA NA

T5493 GCTCGCGATGTACAGGGCCTGGG 1000
Oligo fwd CACCGCTCGCGATGTACAGGGCCT
Oligo rev AAACAGGCCCTGTACATCGCGAGC
Chromosome start end strand MM target_seq PAM alignment distance position gene name gene id
GU305914.1 76912 76934 + 0 GCTCGCGATGTACAGGGCCT GGG ||||||||[||||||||||||]PAM NA Intergenic NA NA

T5494 CAAAACCCCTCCAGAAGTGACGG 1000
Oligo fwd CACCCAAAACCCCTCCAGAAGTGA
Oligo rev AAACTCACTTCTGGAGGGGTTTTG
Chromosome start end strand MM target_seq PAM alignment distance position gene name gene id
GU305914.1 76999 77021 + 0 CAAAACCCCTCCAGAAGTGA CGG ||||||||[||||||||||||]PAM NA Intergenic NA NA

T5495 AACCCCTCCAGAAGTGACGGAGG 1000
Oligo fwd CACCAACCCCTCCAGAAGTGACGG
Oligo rev AAACCCGTCACTTCTGGAGGGGTT
Chromosome start end strand MM target_seq PAM alignment distance position gene name gene id
GU305914.1 77002 77024 + 0 AACCCCTCCAGAAGTGACGG AGG ||||||||[||||||||||||]PAM NA Intergenic NA NA

T5496 ACCCCTCCAGAAGTGACGGAGGG 1000
Oligo fwd CACCACCCCTCCAGAAGTGACGGA
Oligo rev AAACTCCGTCACTTCTGGAGGGGT
Chromosome start end strand MM target_seq PAM alignment distance position gene name gene id
GU305914.1 77003 77025 + 0 ACCCCTCCAGAAGTGACGGA GGG ||||||||[||||||||||||]PAM NA Intergenic NA NA

T5497 TCATCCTCGTCGACCGACTCCGG 1000
Oligo fwd CACCTCATCCTCGTCGACCGACTC
Oligo rev AAACGAGTCGGTCGACGAGGATGA
Chromosome start end strand MM target_seq PAM alignment distance position gene name gene id
GU305914.1 77045 77067 + 0 TCATCCTCGTCGACCGACTC CGG ||||||||[||||||||||||]PAM NA Intergenic NA NA
GU305914.1 114964 114986 - 3 TCATCGTCGTCGTCCGAATC GGG |||||-||[||||-||||-||]PAM NA Intergenic NA NA

T5498 CATCCTCGTCGACCGACTCCGGG 1000
Oligo fwd CACCCATCCTCGTCGACCGACTCC
Oligo rev AAACGGAGTCGGTCGACGAGGATG
Chromosome start end strand MM target_seq PAM alignment distance position gene name gene id
GU305914.1 77046 77068 + 0 CATCCTCGTCGACCGACTCC GGG ||||||||[||||||||||||]PAM NA Intergenic NA NA

T5499 CCTCGTCGACCGACTCCGGGCGG 1000
Oligo fwd CACCCCTCGTCGACCGACTCCGGG
Oligo rev AAACCCCGGAGTCGGTCGACGAGG
Chromosome start end strand MM target_seq PAM alignment distance position gene name gene id
GU305914.1 77049 77071 + 0 CCTCGTCGACCGACTCCGGG CGG ||||||||[||||||||||||]PAM NA Intergenic NA NA

T5500 CTCGTCGACCGACTCCGGGCGGG 1000
Oligo fwd CACCCTCGTCGACCGACTCCGGGC
Oligo rev AAACGCCCGGAGTCGGTCGACGAG
Chromosome start end strand MM target_seq PAM alignment distance position gene name gene id
GU305914.1 77050 77072 + 0 CTCGTCGACCGACTCCGGGC GGG ||||||||[||||||||||||]PAM NA Intergenic NA NA

T5501 CTCCGGGCGGGAGACGATGCTGG 1000
Oligo fwd CACCCTCCGGGCGGGAGACGATGC
Oligo rev AAACGCATCGTCTCCCGCCCGGAG
Chromosome start end strand MM target_seq PAM alignment distance position gene name gene id
GU305914.1 77062 77084 + 0 CTCCGGGCGGGAGACGATGC TGG ||||||||[||||||||||||]PAM NA Intergenic NA NA
GU305914.1 18264 18286 - 4 CTCCAGTCGGGAGACGACTC CGG ||||-|-|[|||||||||--|]PAM NA Intergenic NA NA

T5502 CGGGCGGGAGACGATGCTGGAGG 1000

Oligo fwd CACCCGGGCGGGAGACGATGCTGG
Oligo rev AAACCCAGCATCGTCTCCCGCCCG
Chromosome start end strand MM target_seq PAM alignment distance position gene name gene id
GU305914.1 77065 77087 + 0 CGGGCGGGAGACGATGCTGG AGG ||||||||[||||||||||||]PAM NA Intergenic NA NA

T5503 GCGGGAGACGATGCTGGAGGAGG 1000
Oligo fwd CACCGCGGGAGACGATGCTGGAGG
Oligo rev AAACCCTCCAGCATCGTCTCCCGC
Chromosome start end strand MM target_seq PAM alignment distance position gene name gene id
GU305914.1 77068 77090 + 0 GCGGGAGACGATGCTGGAGG AGG ||||||||[||||||||||||]PAM NA Intergenic NA NA

T5504 GGAGACGATGCTGGAGGAGGAGG 1000
Oligo fwd CACCGGAGACGATGCTGGAGGAGG
Oligo rev AAACCCTCCTCCAGCATCGTCTCC
Chromosome start end strand MM target_seq PAM alignment distance position gene name gene id
GU305914.1 77071 77093 + 0 GGAGACGATGCTGGAGGAGG AGG ||||||||[||||||||||||]PAM NA Intergenic NA NA
GU305914.1 145963 145985 + 4 GGAGAGGTCGGTGGAGGAGG CGG |||||-|-[-|-|||||||||]PAM NA Intergenic NA NA

T5505 GACGATGCTGGAGGAGGAGGAGG 1000
Oligo fwd CACCGACGATGCTGGAGGAGGAGG
Oligo rev AAACCCTCCTCCTCCAGCATCGTC
Chromosome start end strand MM target_seq PAM alignment distance position gene name gene id
GU305914.1 77074 77096 + 0 GACGATGCTGGAGGAGGAGG AGG ||||||||[||||||||||||]PAM NA Intergenic NA NA
GU305914.1 77077 77099 + 4 GATGCTGGAGGAGGAGGAGG CGG ||-|-||-[-|||||||||||]PAM NA Intergenic NA NA
GU305914.1 69232 69254 + 4 GTCGACGCTGGACGAGGCGG AGG |-|||-||[||||-||||-||]PAM NA Intergenic NA NA

T5506 GATGCTGGAGGAGGAGGAGGCGG 1000
Oligo fwd CACCGATGCTGGAGGAGGAGGAGG
Oligo rev AAACCCTCCTCCTCCTCCAGCATC
Chromosome start end strand MM target_seq PAM alignment distance position gene name gene id
GU305914.1 77077 77099 + 0 GATGCTGGAGGAGGAGGAGG CGG ||||||||[||||||||||||]PAM NA Intergenic NA NA
GU305914.1 77074 77096 + 4 GACGATGCTGGAGGAGGAGG AGG ||-|-||-[-|||||||||||]PAM NA Intergenic NA NA

T5507 GGCGTGCTGCGACCTCGACTCGG 1000
Oligo fwd CACCGGCGTGCTGCGACCTCGACT
Oligo rev AAACAGTCGAGGTCGCAGCACGCC
Chromosome start end strand MM target_seq PAM alignment distance position gene name gene id
GU305914.1 77098 77120 + 0 GGCGTGCTGCGACCTCGACT CGG ||||||||[||||||||||||]PAM NA Intergenic NA NA

T5508 CTGCGACCTCGACTCGGAGCTGG 1000
Oligo fwd CACCCTGCGACCTCGACTCGGAGC
Oligo rev AAACGCTCCGAGTCGAGGTCGCAG
Chromosome start end strand MM target_seq PAM alignment distance position gene name gene id
GU305914.1 77104 77126 + 0 CTGCGACCTCGACTCGGAGC TGG ||||||||[||||||||||||]PAM NA Intergenic NA NA

T5509 ACCTCGACTCGGAGCTGGCGCGG 1000
Oligo fwd CACCACCTCGACTCGGAGCTGGCG
Oligo rev AAACCGCCAGCTCCGAGTCGAGGT
Chromosome start end strand MM target_seq PAM alignment distance position gene name gene id
GU305914.1 77109 77131 + 0 ACCTCGACTCGGAGCTGGCG CGG ||||||||[||||||||||||]PAM NA Intergenic NA NA

T5510 GACTCGGAGCTGGCGCGGATCGG 1000
Oligo fwd CACCGACTCGGAGCTGGCGCGGAT
Oligo rev AAACATCCGCGCCAGCTCCGAGTC
Chromosome start end strand MM target_seq PAM alignment distance position gene name gene id
GU305914.1 77114 77136 + 0 GACTCGGAGCTGGCGCGGAT CGG ||||||||[||||||||||||]PAM NA Intergenic NA NA

T5511 GCGGATCGGCGACGAGAACACGG 1000
Oligo fwd CACCGCGGATCGGCGACGAGAACA
Oligo rev AAACTGTTCTCGTCGCCGATCCGC
Chromosome start end strand MM target_seq PAM alignment distance position gene name gene id
GU305914.1 77128 77150 + 0 GCGGATCGGCGACGAGAACA CGG ||||||||[||||||||||||]PAM NA Intergenic NA NA

T5512 CCGCGAGCTGTGCCTGCCGCTGG 1000
Oligo fwd CACCCCGCGAGCTGTGCCTGCCGC
Oligo rev AAACGCGGCAGGCACAGCTCGCGG
Chromosome start end strand MM target_seq PAM alignment distance position gene name gene id
GU305914.1 77158 77180 + 0 CCGCGAGCTGTGCCTGCCGC TGG ||||||||[||||||||||||]PAM NA Intergenic NA NA

T5513 CGATCGTCAGCATCTGCCTGCGG 1000
Oligo fwd CACCCGATCGTCAGCATCTGCCTG
Oligo rev AAACCAGGCAGATGCTGACGATCG
Chromosome start end strand MM target_seq PAM alignment distance position gene name gene id
GU305914.1 77208 77230 + 0 CGATCGTCAGCATCTGCCTG CGG ||||||||[||||||||||||]PAM NA Intergenic NA NA

T5514 GGCGCGACCCGCAGCAGAAATGG 1000
Oligo fwd CACCGGCGCGACCCGCAGCAGAAA
Oligo rev AAACTTTCTGCTGCGGGTCGCGCC
Chromosome start end strand MM target_seq PAM alignment distance position gene name gene id
GU305914.1 77229 77251 + 0 GGCGCGACCCGCAGCAGAAA TGG ||||||||[||||||||||||]PAM NA Intergenic NA NA

T5515 CCCGCAGCAGAAATGGCTGCTGG 1000
Oligo fwd CACCCCCGCAGCAGAAATGGCTGC
Oligo rev AAACGCAGCCATTTCTGCTGCGGG
Chromosome start end strand MM target_seq PAM alignment distance position gene name gene id
GU305914.1 77236 77258 + 0 CCCGCAGCAGAAATGGCTGC TGG ||||||||[||||||||||||]PAM NA Intergenic NA NA

T5516 CTTCCTGTGCCACAAGTGCGCGG 1000
Oligo fwd CACCCTTCCTGTGCCACAAGTGCG

<table>
<tr><td>Oligo rev</td><td>AAACCGCACTTGTGGCACAGGAAG</td><td></td><td></td><td></td><td></td><td></td><td></td><td></td><td></td><td></td><td></td></tr>
<tr><td>Chromosome</td><td>start</td><td>end</td><td>strand</td><td>MM</td><td>target_seq</td><td>PAM</td><td>alignment</td><td>distance</td><td>position</td><td>gene name</td><td>gene id</td></tr>
<tr><td>GU305914.1</td><td>77266</td><td>77288</td><td>+</td><td>0</td><td>CTTCCTGTGCCACAAGTGCG</td><td>CGG</td><td>||||||||[||||||||||||]PAM</td><td>NA</td><td>Intergenic</td><td>NA</td><td>NA</td></tr>
<tr><td>T5517</td><td>CCTGTGCCACAAGTGCGCGGCGG</td><td>1000</td><td></td><td></td><td></td><td></td><td></td><td></td><td></td><td></td><td></td></tr>
<tr><td>Oligo fwd</td><td>CACCCCTGTGCCACAAGTGCGCGG</td><td></td><td></td><td></td><td></td><td></td><td></td><td></td><td></td><td></td><td></td></tr>
<tr><td>Oligo rev</td><td>AAACCCGCGCACTTGTGGCACAGG</td><td></td><td></td><td></td><td></td><td></td><td></td><td></td><td></td><td></td><td></td></tr>
<tr><td>Chromosome</td><td>start</td><td>end</td><td>strand</td><td>MM</td><td>target_seq</td><td>PAM</td><td>alignment</td><td>distance</td><td>position</td><td>gene name</td><td>gene id</td></tr>
<tr><td>GU305914.1</td><td>77269</td><td>77291</td><td>+</td><td>0</td><td>CCTGTGCCACAAGTGCGCGG</td><td>CGG</td><td>||||||||[||||||||||||]PAM</td><td>NA</td><td>Intergenic</td><td>NA</td><td>NA</td></tr>
<tr><td>T5518</td><td>GTGCGCGGCGGCGCCCCGCACGG</td><td>1000</td><td></td><td></td><td></td><td></td><td></td><td></td><td></td><td></td><td></td></tr>
<tr><td>Oligo fwd</td><td>CACCGTGCGCGGCGGCGCCCCGCA</td><td></td><td></td><td></td><td></td><td></td><td></td><td></td><td></td><td></td><td></td></tr>
<tr><td>Oligo rev</td><td>AAACTGCGGGGCGCCGCCGCGCAC</td><td></td><td></td><td></td><td></td><td></td><td></td><td></td><td></td><td></td><td></td></tr>
<tr><td>Chromosome</td><td>start</td><td>end</td><td>strand</td><td>MM</td><td>target_seq</td><td>PAM</td><td>alignment</td><td>distance</td><td>position</td><td>gene name</td><td>gene id</td></tr>
<tr><td>GU305914.1</td><td>77281</td><td>77303</td><td>+</td><td>0</td><td>GTGCGCGGCGGCGCCCCGCA</td><td>CGG</td><td>||||||||[||||||||||||]PAM</td><td>NA</td><td>Intergenic</td><td>NA</td><td>NA</td></tr>
<tr><td>T5519</td><td>GGCGGCGCCCCGCACGGCGATGG</td><td>1000</td><td></td><td></td><td></td><td></td><td></td><td></td><td></td><td></td><td></td></tr>
<tr><td>Oligo fwd</td><td>CACCGGCGGCGCCCCGCACGGCGA</td><td></td><td></td><td></td><td></td><td></td><td></td><td></td><td></td><td></td><td></td></tr>
<tr><td>Oligo rev</td><td>AAACTCGCCGTGCGGGGCGCCGCC</td><td></td><td></td><td></td><td></td><td></td><td></td><td></td><td></td><td></td><td></td></tr>
<tr><td>Chromosome</td><td>start</td><td>end</td><td>strand</td><td>MM</td><td>target_seq</td><td>PAM</td><td>alignment</td><td>distance</td><td>position</td><td>gene name</td><td>gene id</td></tr>
<tr><td>GU305914.1</td><td>77287</td><td>77309</td><td>+</td><td>0</td><td>GGCGGCGCCCCGCACGGCGA</td><td>TGG</td><td>||||||||[||||||||||||]PAM</td><td>NA</td><td>Intergenic</td><td>NA</td><td>NA</td></tr>
<tr><td>T5520</td><td>GGCGATGGCCACGCTGATCGTGG</td><td>1000</td><td></td><td></td><td></td><td></td><td></td><td></td><td></td><td></td><td></td></tr>
<tr><td>Oligo fwd</td><td>CACCGGCGATGGCCACGCTGATCG</td><td></td><td></td><td></td><td></td><td></td><td></td><td></td><td></td><td></td><td></td></tr>
<tr><td>Oligo rev</td><td>AAACCGATCAGCGTGGCCATCGCC</td><td></td><td></td><td></td><td></td><td></td><td></td><td></td><td></td><td></td><td></td></tr>
<tr><td>Chromosome</td><td>start</td><td>end</td><td>strand</td><td>MM</td><td>target_seq</td><td>PAM</td><td>alignment</td><td>distance</td><td>position</td><td>gene name</td><td>gene id</td></tr>
<tr><td>GU305914.1</td><td>77302</td><td>77324</td><td>+</td><td>0</td><td>GGCGATGGCCACGCTGATCG</td><td>TGG</td><td>||||||||[||||||||||||]PAM</td><td>NA</td><td>Intergenic</td><td>NA</td><td>NA</td></tr>
<tr><td>T5521</td><td>CGCCTTCGACAACATCTTCAAGG</td><td>1000</td><td></td><td></td><td></td><td></td><td></td><td></td><td></td><td></td><td></td></tr>
<tr><td>Oligo fwd</td><td>CACCCGCCTTCGACAACATCTTCA</td><td></td><td></td><td></td><td></td><td></td><td></td><td></td><td></td><td></td><td></td></tr>
<tr><td>Oligo rev</td><td>AAACTGAAGATGTTGTCGAAGGCG</td><td></td><td></td><td></td><td></td><td></td><td></td><td></td><td></td><td></td><td></td></tr>
<tr><td>Chromosome</td><td>start</td><td>end</td><td>strand</td><td>MM</td><td>target_seq</td><td>PAM</td><td>alignment</td><td>distance</td><td>position</td><td>gene name</td><td>gene id</td></tr>
<tr><td>GU305914.1</td><td>77368</td><td>77390</td><td>+</td><td>0</td><td>CGCCTTCGACAACATCTTCA</td><td>AGG</td><td>||||||||[||||||||||||]PAM</td><td>NA</td><td>Intergenic</td><td>NA</td><td>NA</td></tr>
<tr><td>GU305914.1</td><td>147312</td><td>147334</td><td>+</td><td>4</td><td>CAACTTCGACACCATCGTCA</td><td>GGG</td><td>|--|||||[|||-||||-|||]PAM</td><td>NA</td><td>Intergenic</td><td>NA</td><td>NA</td></tr>
<tr><td>T5522</td><td>TCGACAACATCTTCAAGGAGCGG</td><td>1000</td><td></td><td></td><td></td><td></td><td></td><td></td><td></td><td></td><td></td></tr>
<tr><td>Oligo fwd</td><td>CACCTCGACAACATCTTCAAGGAG</td><td></td><td></td><td></td><td></td><td></td><td></td><td></td><td></td><td></td><td></td></tr>
<tr><td>Oligo rev</td><td>AAACCTCCTTGAAGATGTTGTCGA</td><td></td><td></td><td></td><td></td><td></td><td></td><td></td><td></td><td></td><td></td></tr>
<tr><td>Chromosome</td><td>start</td><td>end</td><td>strand</td><td>MM</td><td>target_seq</td><td>PAM</td><td>alignment</td><td>distance</td><td>position</td><td>gene name</td><td>gene id</td></tr>
<tr><td>GU305914.1</td><td>77373</td><td>77395</td><td>+</td><td>0</td><td>TCGACAACATCTTCAAGGAG</td><td>CGG</td><td>||||||||[||||||||||||]PAM</td><td>NA</td><td>Intergenic</td><td>NA</td><td>NA</td></tr>
<tr><td>T5523</td><td>ACGCGCACTTCTTCATCAACCGG</td><td>1000</td><td></td><td></td><td></td><td></td><td></td><td></td><td></td><td></td><td></td></tr>
<tr><td>Oligo fwd</td><td>CACCACGCGCACTTCTTCATCAAC</td><td></td><td></td><td></td><td></td><td></td><td></td><td></td><td></td><td></td><td></td></tr>
<tr><td>Oligo rev</td><td>AAACGTTGATGAAGAAGTGCGCGT</td><td></td><td></td><td></td><td></td><td></td><td></td><td></td><td></td><td></td><td></td></tr>
<tr><td>Chromosome</td><td>start</td><td>end</td><td>strand</td><td>MM</td><td>target_seq</td><td>PAM</td><td>alignment</td><td>distance</td><td>position</td><td>gene name</td><td>gene id</td></tr>
<tr><td>GU305914.1</td><td>77418</td><td>77440</td><td>+</td><td>0</td><td>ACGCGCACTTCTTCATCAAC</td><td>CGG</td><td>||||||||[||||||||||||]PAM</td><td>NA</td><td>Intergenic</td><td>NA</td><td>NA</td></tr>
<tr><td>T5524</td><td>CACGCAGCGCGACGAGCACCCGG</td><td>1000</td><td></td><td></td><td></td><td></td><td></td><td></td><td></td><td></td><td></td></tr>
<tr><td>Oligo fwd</td><td>CACCCACGCAGCGCGACGAGCACC</td><td></td><td></td><td></td><td></td><td></td><td></td><td></td><td></td><td></td><td></td></tr>
<tr><td>Oligo rev</td><td>AAACGGTGCTCGTCGCGCTGCGTG</td><td></td><td></td><td></td><td></td><td></td><td></td><td></td><td></td><td></td><td></td></tr>
<tr><td>Chromosome</td><td>start</td><td>end</td><td>strand</td><td>MM</td><td>target_seq</td><td>PAM</td><td>alignment</td><td>distance</td><td>position</td><td>gene name</td><td>gene id</td></tr>
<tr><td>GU305914.1</td><td>77446</td><td>77468</td><td>+</td><td>0</td><td>CACGCAGCGCGACGAGCACC</td><td>CGG</td><td>||||||||[||||||||||||]PAM</td><td>NA</td><td>Intergenic</td><td>NA</td><td>NA</td></tr>
<tr><td>T5525</td><td>GCGCGACGAGCACCCGGTGATGG</td><td>1000</td><td></td><td></td><td></td><td></td><td></td><td></td><td></td><td></td><td></td></tr>
<tr><td>Oligo fwd</td><td>CACCGCGCGACGAGCACCCGGTGA</td><td></td><td></td><td></td><td></td><td></td><td></td><td></td><td></td><td></td><td></td></tr>
<tr><td>Oligo rev</td><td>AAACTCACCGGGTGCTCGTCGCGC</td><td></td><td></td><td></td><td></td><td></td><td></td><td></td><td></td><td></td><td></td></tr>
<tr><td>Chromosome</td><td>start</td><td>end</td><td>strand</td><td>MM</td><td>target_seq</td><td>PAM</td><td>alignment</td><td>distance</td><td>position</td><td>gene name</td><td>gene id</td></tr>
<tr><td>GU305914.1</td><td>77452</td><td>77474</td><td>+</td><td>0</td><td>GCGCGACGAGCACCCGGTGA</td><td>TGG</td><td>||||||||[||||||||||||]PAM</td><td>NA</td><td>Intergenic</td><td>NA</td><td>NA</td></tr>
<tr><td>T5526</td><td>CGACGAGCACCCGGTGATGGTGG</td><td>1000</td><td></td><td></td><td></td><td></td><td></td><td></td><td></td><td></td><td></td></tr>
<tr><td>Oligo fwd</td><td>CACCCGACGAGCACCCGGTGATGG</td><td></td><td></td><td></td><td></td><td></td><td></td><td></td><td></td><td></td><td></td></tr>
<tr><td>Oligo rev</td><td>AAACCCATCACCGGGTGCTCGTCG</td><td></td><td></td><td></td><td></td><td></td><td></td><td></td><td></td><td></td><td></td></tr>
<tr><td>Chromosome</td><td>start</td><td>end</td><td>strand</td><td>MM</td><td>target_seq</td><td>PAM</td><td>alignment</td><td>distance</td><td>position</td><td>gene name</td><td>gene id</td></tr>
<tr><td>GU305914.1</td><td>77455</td><td>77477</td><td>+</td><td>0</td><td>CGACGAGCACCCGGTGATGG</td><td>TGG</td><td>||||||||[||||||||||||]PAM</td><td>NA</td><td>Intergenic</td><td>NA</td><td>NA</td></tr>
<tr><td>GU305914.1</td><td>60049</td><td>60071</td><td>+</td><td>4</td><td>GGCCGAGCAGCCGGTGATGA</td><td>AGG</td><td>-|-|||||[|-|||||||||-]PAM</td><td>NA</td><td>Intergenic</td><td>NA</td><td>NA</td></tr>
<tr><td>T5527</td><td>GATGGTGGAGAACATCACGCTGG</td><td>1000</td><td></td><td></td><td></td><td></td><td></td><td></td><td></td><td></td><td></td></tr>
<tr><td>Oligo fwd</td><td>CACCGATGGTGGAGAACATCACGC</td><td></td><td></td><td></td><td></td><td></td><td></td><td></td><td></td><td></td><td></td></tr>
<tr><td>Oligo rev</td><td>AAACGCGTGATGTTCTCCACCATC</td><td></td><td></td><td></td><td></td><td></td><td></td><td></td><td></td><td></td><td></td></tr>
<tr><td>Chromosome</td><td>start</td><td>end</td><td>strand</td><td>MM</td><td>target_seq</td><td>PAM</td><td>alignment</td><td>distance</td><td>position</td><td>gene name</td><td>gene id</td></tr>
<tr><td>GU305914.1</td><td>77470</td><td>77492</td><td>+</td><td>0</td><td>GATGGTGGAGAACATCACGC</td><td>TGG</td><td>||||||||[||||||||||||]PAM</td><td>NA</td><td>Intergenic</td><td>NA</td><td>NA</td></tr>
<tr><td>GU305914.1</td><td>201539</td><td>201561</td><td>-</td><td>4</td><td>GACGGAGGAGAACAGCTCGC</td><td>GGG</td><td>||-||-||[||||||-|-|||]PAM</td><td>NA</td><td>Intergenic</td><td>NA</td><td>NA</td></tr>
<tr><td>T5528</td><td>GAACATCACGCTGGCGCACATGG</td><td>1000</td><td></td><td></td><td></td><td></td><td></td><td></td><td></td><td></td><td></td></tr>
<tr><td>Oligo fwd</td><td>CACCGAACATCACGCTGGCGCACA</td><td></td><td></td><td></td><td></td><td></td><td></td><td></td><td></td><td></td><td></td></tr>
<tr><td>Oligo rev</td><td>AAACTGTGCGCCAGCGTGATGTTC</td><td></td><td></td><td></td><td></td><td></td><td></td><td></td><td></td><td></td><td></td></tr>
<tr><td>Chromosome</td><td>start</td><td>end</td><td>strand</td><td>MM</td><td>target_seq</td><td>PAM</td><td>alignment</td><td>distance</td><td>position</td><td>gene name</td><td>gene id</td></tr>
<tr><td>GU305914.1</td><td>77479</td><td>77501</td><td>+</td><td>0</td><td>GAACATCACGCTGGCGCACA</td><td>TGG</td><td>||||||||[||||||||||||]PAM</td><td>NA</td><td>Intergenic</td><td>NA</td><td>NA</td></tr>
<tr><td>T5529</td><td>CATCACGCTGGCGCACATGGCGG</td><td>1000</td><td></td><td></td><td></td><td></td><td></td><td></td><td></td><td></td><td></td></tr>
<tr><td>Oligo fwd</td><td>CACCCATCACGCTGGCGCACATGG</td><td></td><td></td><td></td><td></td><td></td><td></td><td></td><td></td><td></td><td></td></tr>
<tr><td>Oligo rev</td><td>AAACCCATGTGCGCCAGCGTGATG</td><td></td><td></td><td></td><td></td><td></td><td></td><td></td><td></td><td></td><td></td></tr>
<tr><td>Chromosome</td><td>start</td><td>end</td><td>strand</td><td>MM</td><td>target_seq</td><td>PAM</td><td>alignment</td><td>distance</td><td>position</td><td>gene name</td><td>gene id</td></tr>
<tr><td>GU305914.1</td><td>77482</td><td>77504</td><td>+</td><td>0</td><td>CATCACGCTGGCGCACATGG</td><td>CGG</td><td>||||||||[||||||||||||]PAM</td><td>NA</td><td>Intergenic</td><td>NA</td><td>NA</td></tr>
<tr><td>T5530</td><td>TGGCGCACATGGCGGTGACCCGG</td><td>1000</td><td></td><td></td><td></td><td></td><td></td><td></td><td></td><td></td><td></td></tr>
<tr><td>Oligo fwd</td><td>CACCTGGCGCACATGGCGGTGACC</td><td></td><td></td><td></td><td></td><td></td><td></td><td></td><td></td><td></td><td></td></tr>
<tr><td>Oligo rev</td><td>AAACGGTCACCGCCATGTGCGCCA</td><td></td><td></td><td></td><td></td><td></td><td></td><td></td><td></td><td></td><td></td></tr>
<tr><td>Chromosome</td><td>start</td><td>end</td><td>strand</td><td>MM</td><td>target_seq</td><td>PAM</td><td>alignment</td><td>distance</td><td>position</td><td>gene name</td><td>gene id</td></tr>
</table>

| | | | | | | | | | | | |
|---|---|---|---|---|---|---|---|---|---|---|---|
| GU305914.1 | | 77490 | 77512 + | 0 | TGGCGCACATGGCGGTGACC | CGG | \|\|\|\|\|\|\|\|[\|\|\|\|\|\|\|\|\|\|\|\|]PAM | NA | Intergenic | NA | NA |
| T5531 | GGCGCACATGGCGGTGACCCGGG | 1000 | | | | | | | | | |
| Oligo fwd | CACCGGCGCACATGGCGGTGACCC | | | | | | | | | | |
| Oligo rev | AAACGGGTCACCGCCATGTGCGCC | | | | | | | | | | |
| Chromosome | start | end | strand | MM | target_seq | PAM | alignment | distance | position | gene name | gene id |
| GU305914.1 | 77491 | 77513 | + | 0 | GGCGCACATGGCGGTGACCC | GGG | \|\|\|\|\|\|\|\|[\|\|\|\|\|\|\|\|\|\|\|\|]PAM | NA | Intergenic | NA | NA |
| T5532 | GGTGACCCGGGCGCTGCTGACGG | 1000 | | | | | | | | | |
| Oligo fwd | CACCGGTGACCCGGGCGCTGCTGA | | | | | | | | | | |
| Oligo rev | AAACTCAGCAGCGCCCGGGTCACC | | | | | | | | | | |
| Chromosome | start | end | strand | MM | target_seq | PAM | alignment | distance | position | gene name | gene id |
| GU305914.1 | 77503 | 77525 | + | 0 | GGTGACCCGGGCGCTGCTGA | CGG | \|\|\|\|\|\|\|\|[\|\|\|\|\|\|\|\|\|\|\|\|]PAM | NA | Intergenic | NA | NA |
| T5533 | GCTGCTGACGGACGACGACGCGG | 1000 | | | | | | | | | |
| Oligo fwd | CACCGCTGCTGACGGACGACGACG | | | | | | | | | | |
| Oligo rev | AAACCGTCGTCGTCCGTCAGCAGC | | | | | | | | | | |
| Chromosome | start | end | strand | MM | target_seq | PAM | alignment | distance | position | gene name | gene id |
| GU305914.1 | 77515 | 77537 | + | 0 | GCTGCTGACGGACGACGACG | CGG | \|\|\|\|\|\|\|\|[\|\|\|\|\|\|\|\|\|\|\|\|]PAM | NA | Intergenic | NA | NA |
| GU305914.1 | 26049 | 26071 | - | 4 | GCCGCTGGTGGACGACGCCG | AGG | \|\|-\|\|\|\|-[-\|\|\|\|\|\|\|\|-\|\|]PAM | NA | Intergenic | NA | NA |
| T5534 | ACGCGGTGCCGTACACCAAGCGG | 1000 | | | | | | | | | |
| Oligo fwd | CACCACGCGGTGCCGTACACCAAG | | | | | | | | | | |
| Oligo rev | AAACCTTGGTGTACGGCACCGCGT | | | | | | | | | | |
| Chromosome | start | end | strand | MM | target_seq | PAM | alignment | distance | position | gene name | gene id |
| GU305914.1 | 77532 | 77554 | + | 0 | ACGCGGTGCCGTACACCAAG | CGG | \|\|\|\|\|\|\|\|[\|\|\|\|\|\|\|\|\|\|\|\|]PAM | NA | Intergenic | NA | NA |
| T5535 | GTACAAGCTGCCGAAGAAGCCGG | 1000 | | | | | | | | | |
| Oligo fwd | CACCGTACAAGCTGCCGAAGAAGC | | | | | | | | | | |
| Oligo rev | AAACGCTTCTTCGGCAGCTTGTAC | | | | | | | | | | |
| Chromosome | start | end | strand | MM | target_seq | PAM | alignment | distance | position | gene name | gene id |
| GU305914.1 | 77566 | 77588 | + | 0 | GTACAAGCTGCCGAAGAAGC | CGG | \|\|\|\|\|\|\|\|[\|\|\|\|\|\|\|\|\|\|\|\|]PAM | NA | Intergenic | NA | NA |
| T5536 | GCTGCCGAAGAAGCCGGCGCCGG | 1000 | | | | | | | | | |
| Oligo fwd | CACCGCTGCCGAAGAAGCCGGCGC | | | | | | | | | | |
| Oligo rev | AAACGCGCCGGCTTCTTCGGCAGC | | | | | | | | | | |
| Chromosome | start | end | strand | MM | target_seq | PAM | alignment | distance | position | gene name | gene id |
| GU305914.1 | 77572 | 77594 | + | 0 | GCTGCCGAAGAAGCCGGCGC | CGG | \|\|\|\|\|\|\|\|[\|\|\|\|\|\|\|\|\|\|\|\|]PAM | NA | Intergenic | NA | NA |
| T5537 | GCCGAAGAAGCCGGCGCCGGCGG | 1000 | | | | | | | | | |
| Oligo fwd | CACCGCCGAAGAAGCCGGCGCCGG | | | | | | | | | | |
| Oligo rev | AAACCCGGCGCCGGCTTCTTCGGC | | | | | | | | | | |
| Chromosome | start | end | strand | MM | target_seq | PAM | alignment | distance | position | gene name | gene id |
| GU305914.1 | 77575 | 77597 | + | 0 | GCCGAAGAAGCCGGCGCCGG | CGG | \|\|\|\|\|\|\|\|[\|\|\|\|\|\|\|\|\|\|\|\|]PAM | NA | Intergenic | NA | NA |
| T5538 | GCCGGCGCCGGCGGAGCCCGAGG | 1000 | | | | | | | | | |
| Oligo fwd | CACCGCCGGCGCCGGCGGAGCCCG | | | | | | | | | | |
| Oligo rev | AAACCGGGCTCCGCCGGCGCCGGC | | | | | | | | | | |
| Chromosome | start | end | strand | MM | target_seq | PAM | alignment | distance | position | gene name | gene id |
| GU305914.1 | 77584 | 77606 | + | 0 | GCCGGCGCCGGCGGAGCCCG | AGG | \|\|\|\|\|\|\|\|[\|\|\|\|\|\|\|\|\|\|\|\|]PAM | NA | Intergenic | NA | NA |
| GU305914.1 | 90250 | 90272 | - | 4 | GGCGGAGGCGGCGGAGCCGG | CGG | \|-\|\|\|-\|-[\|\|\|\|\|\|\|\|\|\|-\|]PAM | NA | Intergenic | NA | NA |
| GU305914.1 | 90259 | 90281 | - | 4 | GCGGGCGACGGCGGAGGCGG | CGG | \|\|-\|\|\|\|-[\|\|\|\|\|\|\|\|-\|-\|]PAM | NA | Intergenic | NA | NA |
| T5539 | CGGCGGAGCCCGAGGAGCTGCGG | 1000 | | | | | | | | | |
| Oligo fwd | CACCCGGCGGAGCCCGAGGAGCTG | | | | | | | | | | |
| Oligo rev | AAACCAGCTCCTCGGGCTCCGCCG | | | | | | | | | | |
| Chromosome | start | end | strand | MM | target_seq | PAM | alignment | distance | position | gene name | gene id |
| GU305914.1 | 77592 | 77614 | + | 0 | CGGCGGAGCCCGAGGAGCTG | CGG | \|\|\|\|\|\|\|\|[\|\|\|\|\|\|\|\|\|\|\|\|]PAM | NA | Intergenic | NA | NA |
| T5540 | GCCCGAGGAGCTGCGGCTGCTGG | 1000 | | | | | | | | | |
| Oligo fwd | CACCGCCCGAGGAGCTGCGGCTGC | | | | | | | | | | |
| Oligo rev | AAACGCAGCCGCAGCTCCTCGGGC | | | | | | | | | | |
| Chromosome | start | end | strand | MM | target_seq | PAM | alignment | distance | position | gene name | gene id |
| GU305914.1 | 77599 | 77621 | + | 0 | GCCCGAGGAGCTGCGGCTGC | TGG | \|\|\|\|\|\|\|\|[\|\|\|\|\|\|\|\|\|\|\|\|]PAM | NA | Intergenic | NA | NA |
| T5541 | TGCGGCTGCTGGACAAGTACCGG | 1000 | | | | | | | | | |
| Oligo fwd | CACCTGCGGCTGCTGGACAAGTAC | | | | | | | | | | |
| Oligo rev | AAACGTACTTGTCCAGCAGCCGCA | | | | | | | | | | |
| Chromosome | start | end | strand | MM | target_seq | PAM | alignment | distance | position | gene name | gene id |
| GU305914.1 | 77610 | 77632 | + | 0 | TGCGGCTGCTGGACAAGTAC | CGG | \|\|\|\|\|\|\|\|[\|\|\|\|\|\|\|\|\|\|\|\|]PAM | NA | Intergenic | NA | NA |
| T5542 | CAAGTACCGGCGCGCGACCGAGG | 1000 | | | | | | | | | |
| Oligo fwd | CACCCAAGTACCGGCGCGCGACCG | | | | | | | | | | |
| Oligo rev | AAACCGGTCGCGCGCCGGTACTTG | | | | | | | | | | |
| Chromosome | start | end | strand | MM | target_seq | PAM | alignment | distance | position | gene name | gene id |
| GU305914.1 | 77623 | 77645 | + | 0 | CAAGTACCGGCGCGCGACCG | AGG | \|\|\|\|\|\|\|\|[\|\|\|\|\|\|\|\|\|\|\|\|]PAM | NA | Intergenic | NA | NA |
| T5543 | AAGTACCGGCGCGCGACCGAGGG | 1000 | | | | | | | | | |
| Oligo fwd | CACCAAGTACCGGCGCGCGACCGA | | | | | | | | | | |
| Oligo rev | AAACTCGGTCGCGCGCCGGTACTT | | | | | | | | | | |
| Chromosome | start | end | strand | MM | target_seq | PAM | alignment | distance | position | gene name | gene id |
| GU305914.1 | 77624 | 77646 | + | 0 | AAGTACCGGCGCGCGACCGA | GGG | \|\|\|\|\|\|\|\|[\|\|\|\|\|\|\|\|\|\|\|\|]PAM | NA | Intergenic | NA | NA |
| T5544 | CGACCGAGGGCTCGTTCGCGCGG | 1000 | | | | | | | | | |
| Oligo fwd | CACCCGACCGAGGGCTCGTTCGCG | | | | | | | | | | |
| Oligo rev | AAACCGCGAACGAGCCCTCGGTCG | | | | | | | | | | |
| Chromosome | start | end | strand | MM | target_seq | PAM | alignment | distance | position | gene name | gene id |
| GU305914.1 | 77637 | 77659 | + | 0 | CGACCGAGGGCTCGTTCGCG | CGG | \|\|\|\|\|\|\|\|[\|\|\|\|\|\|\|\|\|\|\|\|]PAM | NA | Intergenic | NA | NA |
| GU305914.1 | 170839 | 170861 | + | 3 | CGAACGTGGTCTCGTTCGCG | CGG | \|\|\|-\|\|-\|[\|-\|\|\|\|\|\|\|\|\|\|]PAM | NA | Intergenic | NA | NA |

| | | | | | | | | | | | |
|---|---|---|---|---|---|---|---|---|---|---|---|
| T5545 | GACCGAGGGCTCGTTCGCGCGGG | 1000 | | | | | | | | | |
| Oligo fwd | CACCGACCGAGGGCTCGTTCGCGC | | | | | | | | | | |
| Oligo rev | AAACGCGCGAACGAGCCCTCGGTC | | | | | | | | | | |
| Chromosome | start | end | strand | MM | target_seq | PAM | alignment | distance | position | gene name | gene id |
| GU305914.1 | 77638 | 77660 | + | 0 | GACCGAGGGCTCGTTCGCGC | GGG | ||||||||[||||||||||||]PAM | NA | Intergenic | NA | NA |
| GU305914.1 | 192902 | 192924 | - | 4 | GTCCGACGGCGCGTTCACGC | TGG | |-||||-|[||-|||||-|||]PAM | NA | Intergenic | NA | NA |
| T5546 | CGCGGGTGCTCTTCTACATCTGG | 1000 | | | | | | | | | |
| Oligo fwd | CACCCGCGGGTGCTCTTCTACATC | | | | | | | | | | |
| Oligo rev | AAACGATGTAGAAGAGCACCCGCG | | | | | | | | | | |
| Chromosome | start | end | strand | MM | target_seq | PAM | alignment | distance | position | gene name | gene id |
| GU305914.1 | 77655 | 77677 | + | 0 | CGCGGGTGCTCTTCTACATC | TGG | ||||||||[||||||||||||]PAM | NA | Intergenic | NA | NA |
| T5547 | GGTGCTCTTCTACATCTGGTCGG | 1000 | | | | | | | | | |
| Oligo fwd | CACCGGTGCTCTTCTACATCTGGT | | | | | | | | | | |
| Oligo rev | AAACACCAGATGTAGAAGAGCACC | | | | | | | | | | |
| Chromosome | start | end | strand | MM | target_seq | PAM | alignment | distance | position | gene name | gene id |
| GU305914.1 | 77659 | 77681 | + | 0 | GGTGCTCTTCTACATCTGGT | CGG | ||||||||[||||||||||||]PAM | NA | Intergenic | NA | NA |
| T5548 | GTGCTCTTCTACATCTGGTCGGG | 1000 | | | | | | | | | |
| Oligo fwd | CACCGTGCTCTTCTACATCTGGTC | | | | | | | | | | |
| Oligo rev | AAACGACCAGATGTAGAAGAGCAC | | | | | | | | | | |
| Chromosome | start | end | strand | MM | target_seq | PAM | alignment | distance | position | gene name | gene id |
| GU305914.1 | 77660 | 77682 | + | 0 | GTGCTCTTCTACATCTGGTC | GGG | ||||||||[||||||||||||]PAM | NA | Intergenic | NA | NA |
| T5549 | GATGTTCAACACGACGCTGACGG | 1000 | | | | | | | | | |
| Oligo fwd | CACCGATGTTCAACACGACGCTGA | | | | | | | | | | |
| Oligo rev | AAACTCAGCGTCGTGTTGAACATC | | | | | | | | | | |
| Chromosome | start | end | strand | MM | target_seq | PAM | alignment | distance | position | gene name | gene id |
| GU305914.1 | 77692 | 77714 | + | 0 | GATGTTCAACACGACGCTGA | CGG | ||||||||[||||||||||||]PAM | NA | Intergenic | NA | NA |
| GU305914.1 | 135143 | 135165 | - | 4 | GAGGTGCAGCACCACGCTGA | AGG | ||-||-||[-|||-||||||||]PAM | NA | Intergenic | NA | NA |
| T5550 | CAACACGACGCTGACGGACCTGG | 1000 | | | | | | | | | |
| Oligo fwd | CACCCAACACGACGCTGACGGACC | | | | | | | | | | |
| Oligo rev | AAACGGTCCGTCAGCGTCGTGTTG | | | | | | | | | | |
| Chromosome | start | end | strand | MM | target_seq | PAM | alignment | distance | position | gene name | gene id |
| GU305914.1 | 77698 | 77720 | + | 0 | CAACACGACGCTGACGGACC | TGG | ||||||||[||||||||||||]PAM | NA | Intergenic | NA | NA |
| GU305914.1 | 213222 | 213244 | - | 4 | CCTCACGACGCTGATCGACC | GGG | |--|||||[||||||--||||]PAM | NA | Intergenic | NA | NA |
| T5551 | CCTGGCGATCAAGAAGAGCAAGG | 1000 | | | | | | | | | |
| Oligo fwd | CACCCCTGGCGATCAAGAAGAGCA | | | | | | | | | | |
| Oligo rev | AAACTGCTCTTCTTGATCGCCAGG | | | | | | | | | | |
| Chromosome | start | end | strand | MM | target_seq | PAM | alignment | distance | position | gene name | gene id |
| GU305914.1 | 77716 | 77738 | + | 0 | CCTGGCGATCAAGAAGAGCA | AGG | ||||||||[||||||||||||]PAM | NA | Intergenic | NA | NA |
| T5552 | CAAGAAGAGCAAGGCGCTGAAGG | 1000 | | | | | | | | | |
| Oligo fwd | CACCCAAGAAGAGCAAGGCGCTGA | | | | | | | | | | |
| Oligo rev | AAACTCAGCGCCTTGCTCTTCTTG | | | | | | | | | | |
| Chromosome | start | end | strand | MM | target_seq | PAM | alignment | distance | position | gene name | gene id |
| GU305914.1 | 77725 | 77747 | + | 0 | CAAGAAGAGCAAGGCGCTGA | AGG | ||||||||[||||||||||||]PAM | NA | Intergenic | NA | NA |
| T5553 | GGCGCTCAAGACGCGACAGTCGG | 1000 | | | | | | | | | |
| Oligo fwd | CACCGGCGCTCAAGACGCGACAGT | | | | | | | | | | |
| Oligo rev | AAACACTGTCGCGTCTTGAGCGCC | | | | | | | | | | |
| Chromosome | start | end | strand | MM | target_seq | PAM | alignment | distance | position | gene name | gene id |
| GU305914.1 | 77746 | 77768 | + | 0 | GGCGCTCAAGACGCGACAGT | CGG | ||||||||[||||||||||||]PAM | NA | Intergenic | NA | NA |
| GU305914.1 | 99574 | 99596 | - | 4 | GTCGCTCGAGTCGGGACAGT | CGG | |-|||||-[||-||-||||||]PAM | NA | Intergenic | NA | NA |
| T5554 | GTCGGAGATCGAGCCGAGCGTGG | 1000 | | | | | | | | | |
| Oligo fwd | CACCGTCGGAGATCGAGCCGAGCG | | | | | | | | | | |
| Oligo rev | AAACCGCTCGGCTCGATCTCCGAC | | | | | | | | | | |
| Chromosome | start | end | strand | MM | target_seq | PAM | alignment | distance | position | gene name | gene id |
| GU305914.1 | 77764 | 77786 | + | 0 | GTCGGAGATCGAGCCGAGCG | TGG | ||||||||[||||||||||||]PAM | NA | Intergenic | NA | NA |
| T5555 | TCGGAGATCGAGCCGAGCGTGGG | 1000 | | | | | | | | | |
| Oligo fwd | CACCTCGGAGATCGAGCCGAGCGT | | | | | | | | | | |
| Oligo rev | AAACACGCTCGGCTCGATCTCCGA | | | | | | | | | | |
| Chromosome | start | end | strand | MM | target_seq | PAM | alignment | distance | position | gene name | gene id |
| GU305914.1 | 77765 | 77787 | + | 0 | TCGGAGATCGAGCCGAGCGT | GGG | ||||||||[||||||||||||]PAM | NA | Intergenic | NA | NA |
| GU305914.1 | 197989 | 198011 | - | 4 | TCGGACGTCGAGTCGAGCGG | CGG | |||||--|[||||-||||||-]PAM | NA | Intergenic | NA | NA |
| T5556 | CCTGCGCAACGCCACGACCACGG | 1000 | | | | | | | | | |
| Oligo fwd | CACCCCTGCGCAACGCCACGACCA | | | | | | | | | | |
| Oligo rev | AAACTGGTCGTGGCGTTGCGCAGG | | | | | | | | | | |
| Chromosome | start | end | strand | MM | target_seq | PAM | alignment | distance | position | gene name | gene id |
| GU305914.1 | 77821 | 77843 | + | 0 | CCTGCGCAACGCCACGACCA | CGG | ||||||||[||||||||||||]PAM | NA | Intergenic | NA | NA |
| T5557 | GTGCCTGCTGTGCGAGCTGATGG | 1000 | | | | | | | | | |
| Oligo fwd | CACCGTGCCTGCTGTGCGAGCTGA | | | | | | | | | | |
| Oligo rev | AAACTCAGCTCGCACAGCAGGCAC | | | | | | | | | | |
| Chromosome | start | end | strand | MM | target_seq | PAM | alignment | distance | position | gene name | gene id |
| GU305914.1 | 77845 | 77867 | + | 0 | GTGCCTGCTGTGCGAGCTGA | TGG | ||||||||[||||||||||||]PAM | NA | Intergenic | NA | NA |
| T5558 | ACGTGTTCCTGCAGCAGCTGCGG | 1000 | | | | | | | | | |
| Oligo fwd | CACCACGTGTTCCTGCAGCAGCTG | | | | | | | | | | |
| Oligo rev | AAACCAGCTGCTGCAGGAACACGT | | | | | | | | | | |
| Chromosome | start | end | strand | MM | target_seq | PAM | alignment | distance | position | gene name | gene id |
| GU305914.1 | 77886 | 77908 | + | 0 | ACGTGTTCCTGCAGCAGCTG | CGG | ||||||||[||||||||||||]PAM | NA | Intergenic | NA | NA |

GU305914.1 124612 124634 + 4 AGGTACTCCTGCAGCAGCTT CGG |-||--||[|||||||||||-]PAM NA Intergenic NA NA

T5559 CGTGTTCCTGCAGCAGCTGCGGG 1000
Oligo fwd CACCCGTGTTCCTGCAGCAGCTGC
Oligo rev AAACGCAGCTGCTGCAGGAACACG
Chromosome start end strand MM target_seq PAM alignment distance position gene name gene id
GU305914.1 77887 77909 + 0 CGTGTTCCTGCAGCAGCTGC GGG ||||||||[||||||||||||]PAM NA Intergenic NA NA

T5560 ACAACCTGAAGATCATCGACCGG 1000
Oligo fwd CACCACAACCTGAAGATCATCGAC
Oligo rev AAACGTCGATGATCTTCAGGTTGT
Chromosome start end strand MM target_seq PAM alignment distance position gene name gene id
GU305914.1 77934 77956 + 0 ACAACCTGAAGATCATCGAC CGG ||||||||[||||||||||||]PAM NA Intergenic NA NA

T5561 CGACCGGACGCAGCTGACCATGG 1000
Oligo fwd CACCCGACCGGACGCAGCTGACCA
Oligo rev AAACTGGTCAGCTGCGTCCGGTCG
Chromosome start end strand MM target_seq PAM alignment distance position gene name gene id
GU305914.1 77950 77972 + 0 CGACCGGACGCAGCTGACCA TGG ||||||||[||||||||||||]PAM NA Intergenic NA NA

T5562 CCGGACGCAGCTGACCATGGCGG 1000
Oligo fwd CACCCCGGACGCAGCTGACCATGG
Oligo rev AAACCCATGGTCAGCTGCGTCCGG
Chromosome start end strand MM target_seq PAM alignment distance position gene name gene id
GU305914.1 77953 77975 + 0 CCGGACGCAGCTGACCATGG CGG ||||||||[||||||||||||]PAM NA Intergenic NA NA

T5563 ATGGCGGAGATCCTGTCGCACGG 1000
Oligo fwd CACCATGGCGGAGATCCTGTCGCA
Oligo rev AAACTGCGACAGGATCTCCGCCAT
Chromosome start end strand MM target_seq PAM alignment distance position gene name gene id
GU305914.1 77969 77991 + 0 ATGGCGGAGATCCTGTCGCA CGG ||||||||[||||||||||||]PAM NA Intergenic NA NA

T5564 CCTGTCGCACGGACGCAACTCGG 1000
Oligo fwd CACCCCTGTCGCACGGACGCAACT
Oligo rev AAACAGTTGCGTCCGTGCGACAGG
Chromosome start end strand MM target_seq PAM alignment distance position gene name gene id
GU305914.1 77980 78002 + 0 CCTGTCGCACGGACGCAACT CGG ||||||||[||||||||||||]PAM NA Intergenic NA NA

T5565 CCCGCAGCAGCTCAAGAACAAGG 1000
Oligo fwd CACCCCCGCAGCAGCTCAAGAACA
Oligo rev AAACTGTTCTTGAGCTGCTGCGGG
Chromosome start end strand MM target_seq PAM alignment distance position gene name gene id
GU305914.1 78007 78029 + 0 CCCGCAGCAGCTCAAGAACA AGG ||||||||[||||||||||||]PAM NA Intergenic NA NA

T5566 GACGTCGAGCCCCGAGACGCTGG 1000
Oligo fwd CACCGACGTCGAGCCCCGAGACGC
Oligo rev AAACGCGTCTCGGGGCTCGACGTC
Chromosome start end strand MM target_seq PAM alignment distance position gene name gene id
GU305914.1 78046 78068 + 0 GACGTCGAGCCCCGAGACGC TGG ||||||||[||||||||||||]PAM NA Intergenic NA NA

T5567 GAGCCCCGAGACGCTGGCCGCGG 1000
Oligo fwd CACCGAGCCCCGAGACGCTGGCCG
Oligo rev AAACCGGCCAGCGTCTCGGGGCTC
Chromosome start end strand MM target_seq PAM alignment distance position gene name gene id
GU305914.1 78052 78074 + 0 GAGCCCCGAGACGCTGGCCG CGG ||||||||[||||||||||||]PAM NA Intergenic NA NA

T5568 CTGGCCGCGGCGAGCCGCCCCGG 1000
Oligo fwd CACCCTGGCCGCGGCGAGCCGCCC
Oligo rev AAACGGGCGGCTCGCCGCGGCCAG
Chromosome start end strand MM target_seq PAM alignment distance position gene name gene id
GU305914.1 78065 78087 + 0 CTGGCCGCGGCGAGCCGCCC CGG ||||||||[||||||||||||]PAM NA Intergenic NA NA

T5569 TGGCCGCGGCGAGCCGCCCCGGG 1000
Oligo fwd CACCTGGCCGCGGCGAGCCGCCCC
Oligo rev AAACGGGGCGGCTCGCCGCGGCCA
Chromosome start end strand MM target_seq PAM alignment distance position gene name gene id
GU305914.1 78066 78088 + 0 TGGCCGCGGCGAGCCGCCCC GGG ||||||||[||||||||||||]PAM NA Intergenic NA NA

T5570 CGCGGCGAGCCGCCCCGGGACGG 1000
Oligo fwd CACCCGCGGCGAGCCGCCCCGGGA
Oligo rev AAACTCCCGGGGCGGCTCGCCGCG
Chromosome start end strand MM target_seq PAM alignment distance position gene name gene id
GU305914.1 78070 78092 + 0 CGCGGCGAGCCGCCCCGGGA CGG ||||||||[||||||||||||]PAM NA Intergenic NA NA

T5571 GGCGAGCCGCCCCGGGACGGAGG 1000
Oligo fwd CACCGGCGAGCCGCCCCGGGACGG
Oligo rev AAACCCGTCCCGGGGCGGCTCGCC
Chromosome start end strand MM target_seq PAM alignment distance position gene name gene id
GU305914.1 78073 78095 + 0 GGCGAGCCGCCCCGGGACGG AGG ||||||||[||||||||||||]PAM NA Intergenic NA NA

T5572 GCTGAGCGCGCTGACCTACCTGG 1000
Oligo fwd CACCGCTGAGCGCGCTGACCTACC
Oligo rev AAACGGTAGGTCAGCGCGCTCAGC
Chromosome start end strand MM target_seq PAM alignment distance position gene name gene id
GU305914.1 78106 78128 + 0 GCTGAGCGCGCTGACCTACC TGG ||||||||[||||||||||||]PAM NA Intergenic NA NA

T5573 GACCTACCTGGTCCTGCGTCAGG 1000
Oligo fwd CACCGACCTACCTGGTCCTGCGTC
Oligo rev AAACGACGCAGGACCAGGTAGGTC

```
Chromosome	start	end	strand	MM	target_seq	PAM	alignment	distance	position	gene name	gene id
GU305914.1	78118	78140	+	0	GACCTACCTGGTCCTGCGTC	AGG	||||||||[||||||||||||]PAM	NA	Intergenic	NA	NA

T5574	TACCTGGTCCTGCGTCAGGTCGG	1000
Oligo fwd	CACCTACCTGGTCCTGCGTCAGGT
Oligo rev	AAACACCTGACGCAGGACCAGGTA
Chromosome	start	end	strand	MM	target_seq	PAM	alignment	distance	position	gene name	gene id
GU305914.1	78122	78144	+	0	TACCTGGTCCTGCGTCAGGT	CGG	||||||||[||||||||||||]PAM	NA	Intergenic	NA	NA

T5575	CTGCGTCAGGTCGGCGTGATCGG	1000
Oligo fwd	CACCCTGCGTCAGGTCGGCGTGAT
Oligo rev	AAACATCACGCCGACCTGACGCAG
Chromosome	start	end	strand	MM	target_seq	PAM	alignment	distance	position	gene name	gene id
GU305914.1	78131	78153	+	0	CTGCGTCAGGTCGGCGTGAT	CGG	||||||||[||||||||||||]PAM	NA	Intergenic	NA	NA

T5576	TGCGTCAGGTCGGCGTGATCGGG	1000
Oligo fwd	CACCTGCGTCAGGTCGGCGTGATC
Oligo rev	AAACGATCACGCCGACCTGACGCA
Chromosome	start	end	strand	MM	target_seq	PAM	alignment	distance	position	gene name	gene id
GU305914.1	78132	78154	+	0	TGCGTCAGGTCGGCGTGATC	GGG	||||||||[||||||||||||]PAM	NA	Intergenic	NA	NA

T5577	GCGTCAGGTCGGCGTGATCGGGG	1000
Oligo fwd	CACCGCGTCAGGTCGGCGTGATCG
Oligo rev	AAACCGATCACGCCGACCTGACGC
Chromosome	start	end	strand	MM	target_seq	PAM	alignment	distance	position	gene name	gene id
GU305914.1	78133	78155	+	0	GCGTCAGGTCGGCGTGATCG	GGG	||||||||[||||||||||||]PAM	NA	Intergenic	NA	NA

T5578	GGTGTACAAACACCTGTTCGCGG	1000
Oligo fwd	CACCGGTGTACAAACACCTGTTCG
Oligo rev	AAACCGAACAGGTGTTTGTACACC
Chromosome	start	end	strand	MM	target_seq	PAM	alignment	distance	position	gene name	gene id
GU305914.1	78154	78176	+	0	GGTGTACAAACACCTGTTCG	CGG	||||||||[||||||||||||]PAM	NA	Intergenic	NA	NA

T5579	GTTCGCGGACCCGCTGTGCGCGG	1000
Oligo fwd	CACCGTTCGCGGACCCGCTGTGCG
Oligo rev	AAACCGCACAGCGGGTCCGCGAAC
Chromosome	start	end	strand	MM	target_seq	PAM	alignment	distance	position	gene name	gene id
GU305914.1	78169	78191	+	0	GTTCGCGGACCCGCTGTGCG	CGG	||||||||[||||||||||||]PAM	NA	Intergenic	NA	NA

T5580	CGCGGCTAACATGCGATCGACGG	1000
Oligo fwd	CACCCGCGGCTAACATGCGATCGA
Oligo rev	AAACTCGATCGCATGTTAGCCGCG
Chromosome	start	end	strand	MM	target_seq	PAM	alignment	distance	position	gene name	gene id
GU305914.1	78187	78209	+	0	CGCGGCTAACATGCGATCGA	CGG	||||||||[||||||||||||]PAM	NA	Intergenic	NA	NA

T5581	GTATTTAAACGAAGCGAAGCTGG	1000
Oligo fwd	CACCGTATTTAAACGAAGCGAAGC
Oligo rev	AAACGCTTCGCTTCGTTTAAATAC
Chromosome	start	end	strand	MM	target_seq	PAM	alignment	distance	position	gene name	gene id
GU305914.1	78244	78266	+	0	GTATTTAAACGAAGCGAAGC	TGG	||||||||[||||||||||||]PAM	NA	Intergenic	NA	NA

T5582	CGCGCGTCGAGCGCGATTTCTGG	1000
Oligo fwd	CACCCGCGCGTCGAGCGCGATTTC
Oligo rev	AAACGAAATCGCGCTCGACGCGCG
Chromosome	start	end	strand	MM	target_seq	PAM	alignment	distance	position	gene name	gene id
GU305914.1	78294	78316	+	0	CGCGCGTCGAGCGCGATTTC	TGG	||||||||[||||||||||||]PAM	NA	Intergenic	NA	NA

T5583	CGCAACTCAGCGACTTTCTGCGG	1000
Oligo fwd	CACCCGCAACTCAGCGACTTTCTG
Oligo rev	AAACCAGAAAGTCGCTGAGTTGCG
Chromosome	start	end	strand	MM	target_seq	PAM	alignment	distance	position	gene name	gene id
GU305914.1	78375	78397	+	0	CGCAACTCAGCGACTTTCTG	CGG	||||||||[||||||||||||]PAM	NA	Intergenic	NA	NA

T5584	GCAACTCAGCGACTTTCTGCGGG	1000
Oligo fwd	CACCGCAACTCAGCGACTTTCTGC
Oligo rev	AAACGCAGAAAGTCGCTGAGTTGC
Chromosome	start	end	strand	MM	target_seq	PAM	alignment	distance	position	gene name	gene id
GU305914.1	78376	78398	+	0	GCAACTCAGCGACTTTCTGC	GGG	||||||||[||||||||||||]PAM	NA	Intergenic	NA	NA

T5585	CTTTCTGCGGGACTTCTCGCAGG	1000
Oligo fwd	CACCCTTTCTGCGGGACTTCTCGC
Oligo rev	AAACGCGAGAAGTCCCGCAGAAAG
Chromosome	start	end	strand	MM	target_seq	PAM	alignment	distance	position	gene name	gene id
GU305914.1	78388	78410	+	0	CTTTCTGCGGGACTTCTCGC	AGG	||||||||[||||||||||||]PAM	NA	Intergenic	NA	NA

T5586	GCGGGACTTCTCGCAGGTGCTGG	1000
Oligo fwd	CACCGCGGGACTTCTCGCAGGTGC
Oligo rev	AAACGCACCTGCGAGAAGTCCCGC
Chromosome	start	end	strand	MM	target_seq	PAM	alignment	distance	position	gene name	gene id
GU305914.1	78394	78416	+	0	GCGGGACTTCTCGCAGGTGC	TGG	||||||||[||||||||||||]PAM	NA	Intergenic	NA	NA
GU305914.1	110054	110076	+	4	GGGGGCCATCGCGCAGGTGC	CGG	|-|||-|-[||-|||||||||]PAM	NA	Intergenic	NA	NA

T5587	ATCGCACGATTTCAGCCTCGTGG	1000
Oligo fwd	CACCATCGCACGATTTCAGCCTCG
Oligo rev	AAACCGAGGCTGAAATCGTGCGAT
Chromosome	start	end	strand	MM	target_seq	PAM	alignment	distance	position	gene name	gene id
GU305914.1	78418	78440	+	0	ATCGCACGATTTCAGCCTCG	TGG	||||||||[||||||||||||]PAM	NA	Intergenic	NA	NA

T5588	CACCGTCGAGAAGTATGTTTAGG	1000
```

<table>
<tr><td>Oligo fwd</td><td>CACCCACCGTCGAGAAGTATGTTT</td><td></td><td></td><td></td><td></td><td></td><td></td><td></td><td></td><td></td><td></td><td></td></tr>
<tr><td>Oligo rev</td><td>AAACAAACATACTTCTCGACGGTG</td><td></td><td></td><td></td><td></td><td></td><td></td><td></td><td></td><td></td><td></td><td></td></tr>
<tr><td>Chromosome</td><td>start</td><td></td><td>end</td><td>strand</td><td>MM</td><td>target_seq</td><td>PAM</td><td>alignment</td><td>distance</td><td>position</td><td>gene name</td><td>gene id</td></tr>
<tr><td>GU305914.1</td><td></td><td>78451</td><td>78473</td><td>+</td><td>0</td><td>CACCGTCGAGAAGTATGTTT</td><td>AGG</td><td>||||||||[||||||||||||]PAM</td><td>NA</td><td>Intergenic</td><td>NA</td><td>NA</td></tr>
<tr><td>T5589</td><td>GAGAAGTATGTTTAGGAGCCCGG</td><td></td><td>1000</td><td></td><td></td><td></td><td></td><td></td><td></td><td></td><td></td><td></td></tr>
<tr><td>Oligo fwd</td><td>CACCGAGAAGTATGTTTAGGAGCC</td><td></td><td></td><td></td><td></td><td></td><td></td><td></td><td></td><td></td><td></td><td></td></tr>
<tr><td>Oligo rev</td><td>AAACGGCTCCTAAACATACTTCTC</td><td></td><td></td><td></td><td></td><td></td><td></td><td></td><td></td><td></td><td></td><td></td></tr>
<tr><td>Chromosome</td><td>start</td><td></td><td>end</td><td>strand</td><td>MM</td><td>target_seq</td><td>PAM</td><td>alignment</td><td>distance</td><td>position</td><td>gene name</td><td>gene id</td></tr>
<tr><td>GU305914.1</td><td></td><td>78458</td><td>78480</td><td>+</td><td>0</td><td>GAGAAGTATGTTTAGGAGCC</td><td>CGG</td><td>||||||||[||||||||||||]PAM</td><td>NA</td><td>Intergenic</td><td>NA</td><td>NA</td></tr>
<tr><td>T5590</td><td>AAGTATGTTTAGGAGCCCGGAGG</td><td></td><td>1000</td><td></td><td></td><td></td><td></td><td></td><td></td><td></td><td></td><td></td></tr>
<tr><td>Oligo fwd</td><td>CACCAAGTATGTTTAGGAGCCCGG</td><td></td><td></td><td></td><td></td><td></td><td></td><td></td><td></td><td></td><td></td><td></td></tr>
<tr><td>Oligo rev</td><td>AAACCCGGGCTCCTAAACATACTT</td><td></td><td></td><td></td><td></td><td></td><td></td><td></td><td></td><td></td><td></td><td></td></tr>
<tr><td>Chromosome</td><td>start</td><td></td><td>end</td><td>strand</td><td>MM</td><td>target_seq</td><td>PAM</td><td>alignment</td><td>distance</td><td>position</td><td>gene name</td><td>gene id</td></tr>
<tr><td>GU305914.1</td><td></td><td>78461</td><td>78483</td><td>+</td><td>0</td><td>AAGTATGTTTAGGAGCCCGG</td><td>AGG</td><td>||||||||[||||||||||||]PAM</td><td>NA</td><td>Intergenic</td><td>NA</td><td>NA</td></tr>
<tr><td>T5591</td><td>AGTATGTTTAGGAGCCCGGAGGG</td><td></td><td>1000</td><td></td><td></td><td></td><td></td><td></td><td></td><td></td><td></td><td></td></tr>
<tr><td>Oligo fwd</td><td>CACCAGTATGTTTAGGAGCCCGGA</td><td></td><td></td><td></td><td></td><td></td><td></td><td></td><td></td><td></td><td></td><td></td></tr>
<tr><td>Oligo rev</td><td>AAACTCCGGGCTCCTAAACATACT</td><td></td><td></td><td></td><td></td><td></td><td></td><td></td><td></td><td></td><td></td><td></td></tr>
<tr><td>Chromosome</td><td>start</td><td></td><td>end</td><td>strand</td><td>MM</td><td>target_seq</td><td>PAM</td><td>alignment</td><td>distance</td><td>position</td><td>gene name</td><td>gene id</td></tr>
<tr><td>GU305914.1</td><td></td><td>78462</td><td>78484</td><td>+</td><td>0</td><td>AGTATGTTTAGGAGCCCGGA</td><td>GGG</td><td>||||||||[||||||||||||]PAM</td><td>NA</td><td>Intergenic</td><td>NA</td><td>NA</td></tr>
<tr><td>T5592</td><td>GTTTAGGAGCCCGGAGGGAGAGG</td><td></td><td>1000</td><td></td><td></td><td></td><td></td><td></td><td></td><td></td><td></td><td></td></tr>
<tr><td>Oligo fwd</td><td>CACCGTTTAGGAGCCCGGAGGGAG</td><td></td><td></td><td></td><td></td><td></td><td></td><td></td><td></td><td></td><td></td><td></td></tr>
<tr><td>Oligo rev</td><td>AAACCTCCCTCCGGGCTCCTAAAC</td><td></td><td></td><td></td><td></td><td></td><td></td><td></td><td></td><td></td><td></td><td></td></tr>
<tr><td>Chromosome</td><td>start</td><td></td><td>end</td><td>strand</td><td>MM</td><td>target_seq</td><td>PAM</td><td>alignment</td><td>distance</td><td>position</td><td>gene name</td><td>gene id</td></tr>
<tr><td>GU305914.1</td><td></td><td>78467</td><td>78489</td><td>+</td><td>0</td><td>GTTTAGGAGCCCGGAGGGAG</td><td>AGG</td><td>||||||||[||||||||||||]PAM</td><td>NA</td><td>Intergenic</td><td>NA</td><td>NA</td></tr>
<tr><td>T5593</td><td>GGAGCCCGGAGGGAGAGGAACGG</td><td></td><td>1000</td><td></td><td></td><td></td><td></td><td></td><td></td><td></td><td></td><td></td></tr>
<tr><td>Oligo fwd</td><td>CACCGGAGCCCGGAGGGAGAGGAA</td><td></td><td></td><td></td><td></td><td></td><td></td><td></td><td></td><td></td><td></td><td></td></tr>
<tr><td>Oligo rev</td><td>AAACTTCCTCTCCCTCCGGGCTCC</td><td></td><td></td><td></td><td></td><td></td><td></td><td></td><td></td><td></td><td></td><td></td></tr>
<tr><td>Chromosome</td><td>start</td><td></td><td>end</td><td>strand</td><td>MM</td><td>target_seq</td><td>PAM</td><td>alignment</td><td>distance</td><td>position</td><td>gene name</td><td>gene id</td></tr>
<tr><td>GU305914.1</td><td></td><td>78472</td><td>78494</td><td>+</td><td>0</td><td>GGAGCCCGGAGGGAGAGGAA</td><td>CGG</td><td>||||||||[||||||||||||]PAM</td><td>NA</td><td>Intergenic</td><td>NA</td><td>NA</td></tr>
<tr><td>T5594</td><td>GAGCCCGGAGGGAGAGGAACGGG</td><td></td><td>1000</td><td></td><td></td><td></td><td></td><td></td><td></td><td></td><td></td><td></td></tr>
<tr><td>Oligo fwd</td><td>CACCGAGCCCGGAGGGAGAGGAAC</td><td></td><td></td><td></td><td></td><td></td><td></td><td></td><td></td><td></td><td></td><td></td></tr>
<tr><td>Oligo rev</td><td>AAACGTTCCTCTCCCTCCGGGCTC</td><td></td><td></td><td></td><td></td><td></td><td></td><td></td><td></td><td></td><td></td><td></td></tr>
<tr><td>Chromosome</td><td>start</td><td></td><td>end</td><td>strand</td><td>MM</td><td>target_seq</td><td>PAM</td><td>alignment</td><td>distance</td><td>position</td><td>gene name</td><td>gene id</td></tr>
<tr><td>GU305914.1</td><td></td><td>78473</td><td>78495</td><td>+</td><td>0</td><td>GAGCCCGGAGGGAGAGGAAC</td><td>GGG</td><td>||||||||[||||||||||||]PAM</td><td>NA</td><td>Intergenic</td><td>NA</td><td>NA</td></tr>
<tr><td>T5595</td><td>GGACGCCGCCGACCGAGAAGAGG</td><td></td><td>1000</td><td></td><td></td><td></td><td></td><td></td><td></td><td></td><td></td><td></td></tr>
<tr><td>Oligo fwd</td><td>CACCGGACGCCGCCGACCGAGAAG</td><td></td><td></td><td></td><td></td><td></td><td></td><td></td><td></td><td></td><td></td><td></td></tr>
<tr><td>Oligo rev</td><td>AAACCTTCTCGGTCGGCGGCGTCC</td><td></td><td></td><td></td><td></td><td></td><td></td><td></td><td></td><td></td><td></td><td></td></tr>
<tr><td>Chromosome</td><td>start</td><td></td><td>end</td><td>strand</td><td>MM</td><td>target_seq</td><td>PAM</td><td>alignment</td><td>distance</td><td>position</td><td>gene name</td><td>gene id</td></tr>
<tr><td>GU305914.1</td><td></td><td>78494</td><td>78516</td><td>+</td><td>0</td><td>GGACGCCGCCGACCGAGAAG</td><td>AGG</td><td>||||||||[||||||||||||]PAM</td><td>NA</td><td>Intergenic</td><td>NA</td><td>NA</td></tr>
<tr><td>T5596</td><td>CGCCGCCGACCGAGAAGAGGAGG</td><td></td><td>1000</td><td></td><td></td><td></td><td></td><td></td><td></td><td></td><td></td><td></td></tr>
<tr><td>Oligo fwd</td><td>CACCCGCCGCCGACCGAGAAGAGG</td><td></td><td></td><td></td><td></td><td></td><td></td><td></td><td></td><td></td><td></td><td></td></tr>
<tr><td>Oligo rev</td><td>AAACCCTCTTCTCGGTCGGCGGCG</td><td></td><td></td><td></td><td></td><td></td><td></td><td></td><td></td><td></td><td></td><td></td></tr>
<tr><td>Chromosome</td><td>start</td><td></td><td>end</td><td>strand</td><td>MM</td><td>target_seq</td><td>PAM</td><td>alignment</td><td>distance</td><td>position</td><td>gene name</td><td>gene id</td></tr>
<tr><td>GU305914.1</td><td></td><td>78497</td><td>78519</td><td>+</td><td>0</td><td>CGCCGCCGACCGAGAAGAGG</td><td>AGG</td><td>||||||||[||||||||||||]PAM</td><td>NA</td><td>Intergenic</td><td>NA</td><td>NA</td></tr>
<tr><td>T5597</td><td>GCCGACCGAGAAGAGGAGGAAGG</td><td></td><td>1000</td><td></td><td></td><td></td><td></td><td></td><td></td><td></td><td></td><td></td></tr>
<tr><td>Oligo fwd</td><td>CACCGCCGACCGAGAAGAGGAGGA</td><td></td><td></td><td></td><td></td><td></td><td></td><td></td><td></td><td></td><td></td><td></td></tr>
<tr><td>Oligo rev</td><td>AAACTCCTCCTCTTCTCGGTCGGC</td><td></td><td></td><td></td><td></td><td></td><td></td><td></td><td></td><td></td><td></td><td></td></tr>
<tr><td>Chromosome</td><td>start</td><td></td><td>end</td><td>strand</td><td>MM</td><td>target_seq</td><td>PAM</td><td>alignment</td><td>distance</td><td>position</td><td>gene name</td><td>gene id</td></tr>
<tr><td>GU305914.1</td><td></td><td>78501</td><td>78523</td><td>+</td><td>0</td><td>GCCGACCGAGAAGAGGAGGA</td><td>AGG</td><td>||||||||[||||||||||||]PAM</td><td>NA</td><td>Intergenic</td><td>NA</td><td>NA</td></tr>
<tr><td>GU305914.1</td><td></td><td>39672</td><td>39694</td><td>+</td><td>4</td><td>GCCCTCCGAGGGGAGGAGGA</td><td>AGG</td><td>|||--|||[||--||||||||]PAM</td><td>NA</td><td>Intergenic</td><td>NA</td><td>NA</td></tr>
<tr><td>T5598</td><td>GACCGAGAAGAGGAGGAAGGAGG</td><td></td><td>1000</td><td></td><td></td><td></td><td></td><td></td><td></td><td></td><td></td><td></td></tr>
<tr><td>Oligo fwd</td><td>CACCGACCGAGAAGAGGAGGAAGG</td><td></td><td></td><td></td><td></td><td></td><td></td><td></td><td></td><td></td><td></td><td></td></tr>
<tr><td>Oligo rev</td><td>AAACCCTTCCTCCTCTTCTCGGTC</td><td></td><td></td><td></td><td></td><td></td><td></td><td></td><td></td><td></td><td></td><td></td></tr>
<tr><td>Chromosome</td><td>start</td><td></td><td>end</td><td>strand</td><td>MM</td><td>target_seq</td><td>PAM</td><td>alignment</td><td>distance</td><td>position</td><td>gene name</td><td>gene id</td></tr>
<tr><td>GU305914.1</td><td></td><td>78504</td><td>78526</td><td>+</td><td>0</td><td>GACCGAGAAGAGGAGGAAGG</td><td>AGG</td><td>||||||||[||||||||||||]PAM</td><td>NA</td><td>Intergenic</td><td>NA</td><td>NA</td></tr>
<tr><td>T5599</td><td>AGAAGAGGAGGAAGGAGGAGAGG</td><td></td><td>1000</td><td></td><td></td><td></td><td></td><td></td><td></td><td></td><td></td><td></td></tr>
<tr><td>Oligo fwd</td><td>CACCAGAAGAGGAGGAAGGAGGAG</td><td></td><td></td><td></td><td></td><td></td><td></td><td></td><td></td><td></td><td></td><td></td></tr>
<tr><td>Oligo rev</td><td>AAACCTCCTCCTTCCTCCTCTTCT</td><td></td><td></td><td></td><td></td><td></td><td></td><td></td><td></td><td></td><td></td><td></td></tr>
<tr><td>Chromosome</td><td>start</td><td></td><td>end</td><td>strand</td><td>MM</td><td>target_seq</td><td>PAM</td><td>alignment</td><td>distance</td><td>position</td><td>gene name</td><td>gene id</td></tr>
<tr><td>GU305914.1</td><td></td><td>78509</td><td>78531</td><td>+</td><td>0</td><td>AGAAGAGGAGGAAGGAGGAG</td><td>AGG</td><td>||||||||[||||||||||||]PAM</td><td>NA</td><td>Intergenic</td><td>NA</td><td>NA</td></tr>
<tr><td>T5600</td><td>AAGGAGGAGAGGCTCGACGACGG</td><td></td><td>1000</td><td></td><td></td><td></td><td></td><td></td><td></td><td></td><td></td><td></td></tr>
<tr><td>Oligo fwd</td><td>CACCAAGGAGGAGAGGCTCGACGA</td><td></td><td></td><td></td><td></td><td></td><td></td><td></td><td></td><td></td><td></td><td></td></tr>
<tr><td>Oligo rev</td><td>AAACTCGTCGAGCCTCTCCTCCTT</td><td></td><td></td><td></td><td></td><td></td><td></td><td></td><td></td><td></td><td></td><td></td></tr>
<tr><td>Chromosome</td><td>start</td><td></td><td>end</td><td>strand</td><td>MM</td><td>target_seq</td><td>PAM</td><td>alignment</td><td>distance</td><td>position</td><td>gene name</td><td>gene id</td></tr>
<tr><td>GU305914.1</td><td></td><td>78520</td><td>78542</td><td>+</td><td>0</td><td>AAGGAGGAGAGGCTCGACGA</td><td>CGG</td><td>||||||||[||||||||||||]PAM</td><td>NA</td><td>Intergenic</td><td>NA</td><td>NA</td></tr>
<tr><td>T5601</td><td>GAGAGGCTCGACGACGGTCGAGG</td><td></td><td>1000</td><td></td><td></td><td></td><td></td><td></td><td></td><td></td><td></td><td></td></tr>
<tr><td>Oligo fwd</td><td>CACCGAGAGGCTCGACGACGGTCG</td><td></td><td></td><td></td><td></td><td></td><td></td><td></td><td></td><td></td><td></td><td></td></tr>
<tr><td>Oligo rev</td><td>AAACCGACCGTCGTCGAGCCTCTC</td><td></td><td></td><td></td><td></td><td></td><td></td><td></td><td></td><td></td><td></td><td></td></tr>
<tr><td>Chromosome</td><td>start</td><td></td><td>end</td><td>strand</td><td>MM</td><td>target_seq</td><td>PAM</td><td>alignment</td><td>distance</td><td>position</td><td>gene name</td><td>gene id</td></tr>
<tr><td>GU305914.1</td><td></td><td>78526</td><td>78548</td><td>+</td><td>0</td><td>GAGAGGCTCGACGACGGTCG</td><td>AGG</td><td>||||||||[||||||||||||]PAM</td><td>NA</td><td>Intergenic</td><td>NA</td><td>NA</td></tr>
<tr><td>T5602</td><td>CAAGCGCGCCCGACACCGCCCGG</td><td></td><td>1000</td><td></td><td></td><td></td><td></td><td></td><td></td><td></td><td></td><td></td></tr>
<tr><td>Oligo fwd</td><td>CACCCAAGCGCGCCCGACACCGCC</td><td></td><td></td><td></td><td></td><td></td><td></td><td></td><td></td><td></td><td></td><td></td></tr>
<tr><td>Oligo rev</td><td>AAACGGCGGTGTCGGGCGCGCTTG</td><td></td><td></td><td></td><td></td><td></td><td></td><td></td><td></td><td></td><td></td><td></td></tr>
<tr><td>Chromosome</td><td>start</td><td></td><td>end</td><td>strand</td><td>MM</td><td>target_seq</td><td>PAM</td><td>alignment</td><td>distance</td><td>position</td><td>gene name</td><td>gene id</td></tr>
<tr><td>GU305914.1</td><td></td><td>78572</td><td>78594</td><td>+</td><td>0</td><td>CAAGCGCGCCCGACACCGCC</td><td>CGG</td><td>||||||||[||||||||||||]PAM</td><td>NA</td><td>Intergenic</td><td>NA</td><td>NA</td></tr>
</table>

| | | | | | | | | | | | | |
|---|---|---|---|---|---|---|---|---|---|---|---|---|
| T5603 | CGCGCCCGACACCGCCCGGCCGG | 1000 | | | | | | | | | | |
| Oligo fwd | CACCCGCGCCCGACACCGCCCGGC | | | | | | | | | | | |
| Oligo rev | AAACGCCGGGCGGTGTCGGGCGCG | | | | | | | | | | | |
| Chromosome | start | | end | strand | MM | target_seq | PAM | alignment | distance | position | gene name | gene id |
| GU305914.1 | | 78576 | 78598 | + | 0 | CGCGCCCGACACCGCCCGGC | CGG | \|\|\|\|\|\|\|\|[\|\|\|\|\|\|\|\|\|\|\|\|]PAM | NA | Intergenic | NA | NA |
| T5604 | CGACACCGCCCGGCCGGCAGCGG | 1000 | | | | | | | | | | |
| Oligo fwd | CACCCGACACCGCCCGGCCGGCAG | | | | | | | | | | | |
| Oligo rev | AAACCTGCCGGCCGGGCGGTGTCG | | | | | | | | | | | |
| Chromosome | start | | end | strand | MM | target_seq | PAM | alignment | distance | position | gene name | gene id |
| GU305914.1 | | 78582 | 78604 | + | 0 | CGACACCGCCCGGCCGGCAG | CGG | \|\|\|\|\|\|\|\|[\|\|\|\|\|\|\|\|\|\|\|\|]PAM | NA | Intergenic | NA | NA |
| T5605 | GCGGCCTCAGAACACCGCTGAGG | 1000 | | | | | | | | | | |
| Oligo fwd | CACCGCGGCCTCAGAACACCGCTG | | | | | | | | | | | |
| Oligo rev | AAACCAGCGGTGTTCTGAGGCCGC | | | | | | | | | | | |
| Chromosome | start | | end | strand | MM | target_seq | PAM | alignment | distance | position | gene name | gene id |
| GU305914.1 | | 78601 | 78623 | + | 0 | GCGGCCTCAGAACACCGCTG | AGG | \|\|\|\|\|\|\|\|[\|\|\|\|\|\|\|\|\|\|\|\|]PAM | NA | Intergenic | NA | NA |
| T5606 | CCGCTGCTCCTCTCCCTCTCCGG | 1000 | | | | | | | | | | |
| Oligo fwd | CACCCCGCTGCTCCTCTCCCTCTC | | | | | | | | | | | |
| Oligo rev | AAACGAGAGGGAGAGGAGCAGCGG | | | | | | | | | | | |
| Chromosome | start | | end | strand | MM | target_seq | PAM | alignment | distance | position | gene name | gene id |
| GU305914.1 | | 78638 | 78660 | + | 0 | CCGCTGCTCCTCTCCCTCTC | CGG | \|\|\|\|\|\|\|\|[\|\|\|\|\|\|\|\|\|\|\|\|]PAM | NA | Intergenic | NA | NA |
| T5607 | CTCCCTCTCCGGAGCGACAGTGG | 1000 | | | | | | | | | | |
| Oligo fwd | CACCCTCCCTCTCCGGAGCGACAG | | | | | | | | | | | |
| Oligo rev | AAACCTGTCGCTCCGGAGAGGGAG | | | | | | | | | | | |
| Chromosome | start | | end | strand | MM | target_seq | PAM | alignment | distance | position | gene name | gene id |
| GU305914.1 | | 78649 | 78671 | + | 0 | CTCCCTCTCCGGAGCGACAG | TGG | \|\|\|\|\|\|\|\|[\|\|\|\|\|\|\|\|\|\|\|\|]PAM | NA | Intergenic | NA | NA |
| T5608 | AGCGACAGTGGCAACAGCGACGG | 1000 | | | | | | | | | | |
| Oligo fwd | CACCAGCGACAGTGGCAACAGCGA | | | | | | | | | | | |
| Oligo rev | AAACTCGCTGTTGCCACTGTCGCT | | | | | | | | | | | |
| Chromosome | start | | end | strand | MM | target_seq | PAM | alignment | distance | position | gene name | gene id |
| GU305914.1 | | 78661 | 78683 | + | 0 | AGCGACAGTGGCAACAGCGA | CGG | \|\|\|\|\|\|\|\|[\|\|\|\|\|\|\|\|\|\|\|\|]PAM | NA | Intergenic | NA | NA |
| T5609 | GACAGTGGCAACAGCGACGGCGG | 1000 | | | | | | | | | | |
| Oligo fwd | CACCGACAGTGGCAACAGCGACGG | | | | | | | | | | | |
| Oligo rev | AAACCCGTCGCTGTTGCCACTGTC | | | | | | | | | | | |
| Chromosome | start | | end | strand | MM | target_seq | PAM | alignment | distance | position | gene name | gene id |
| GU305914.1 | | 78664 | 78686 | + | 0 | GACAGTGGCAACAGCGACGG | CGG | \|\|\|\|\|\|\|\|[\|\|\|\|\|\|\|\|\|\|\|\|]PAM | NA | Intergenic | NA | NA |
| T5610 | ACAGTGGCAACAGCGACGGCGGG | 1000 | | | | | | | | | | |
| Oligo fwd | CACCACAGTGGCAACAGCGACGGC | | | | | | | | | | | |
| Oligo rev | AAACGCCGTCGCTGTTGCCACTGT | | | | | | | | | | | |
| Chromosome | start | | end | strand | MM | target_seq | PAM | alignment | distance | position | gene name | gene id |
| GU305914.1 | | 78665 | 78687 | + | 0 | ACAGTGGCAACAGCGACGGC | GGG | \|\|\|\|\|\|\|\|[\|\|\|\|\|\|\|\|\|\|\|\|]PAM | NA | Intergenic | NA | NA |
| T5611 | AGCGACGGCGGGCAGAAAAGAGG | 1000 | | | | | | | | | | |
| Oligo fwd | CACCAGCGACGGCGGGCAGAAAAG | | | | | | | | | | | |
| Oligo rev | AAACCTTTTCTGCCCGCCGTCGCT | | | | | | | | | | | |
| Chromosome | start | | end | strand | MM | target_seq | PAM | alignment | distance | position | gene name | gene id |
| GU305914.1 | | 78676 | 78698 | + | 0 | AGCGACGGCGGGCAGAAAAG | AGG | \|\|\|\|\|\|\|\|[\|\|\|\|\|\|\|\|\|\|\|\|]PAM | NA | Intergenic | NA | NA |
| T5612 | GCCTCCGCCACCCAAGAGATCGG | 1000 | | | | | | | | | | |
| Oligo fwd | CACCGCCTCCGCCACCCAAGAGAT | | | | | | | | | | | |
| Oligo rev | AAACATCTCTTGGGTGGCGGAGGC | | | | | | | | | | | |
| Chromosome | start | | end | strand | MM | target_seq | PAM | alignment | distance | position | gene name | gene id |
| GU305914.1 | | 78719 | 78741 | + | 0 | GCCTCCGCCACCCAAGAGAT | CGG | \|\|\|\|\|\|\|\|[\|\|\|\|\|\|\|\|\|\|\|\|]PAM | NA | Intergenic | NA | NA |
| T5613 | GAGATCGGCCGCCAGCGCAGCGG | 1000 | | | | | | | | | | |
| Oligo fwd | CACCGAGATCGGCCGCCAGCGCAG | | | | | | | | | | | |
| Oligo rev | AAACCTGCGCTGGCGGCCGATCTC | | | | | | | | | | | |
| Chromosome | start | | end | strand | MM | target_seq | PAM | alignment | distance | position | gene name | gene id |
| GU305914.1 | | 78734 | 78756 | + | 0 | GAGATCGGCCGCCAGCGCAG | CGG | \|\|\|\|\|\|\|\|[\|\|\|\|\|\|\|\|\|\|\|\|]PAM | NA | Intergenic | NA | NA |
| T5614 | TCGGCCGCCAGCGCAGCGGCCGG | 1000 | | | | | | | | | | |
| Oligo fwd | CACCTCGGCCGCCAGCGCAGCGGC | | | | | | | | | | | |
| Oligo rev | AAACGCCGCTGCGCTGGCGGCCGA | | | | | | | | | | | |
| Chromosome | start | | end | strand | MM | target_seq | PAM | alignment | distance | position | gene name | gene id |
| GU305914.1 | | 78738 | 78760 | + | 0 | TCGGCCGCCAGCGCAGCGGC | CGG | \|\|\|\|\|\|\|\|[\|\|\|\|\|\|\|\|\|\|\|\|]PAM | NA | Intergenic | NA | NA |
| T5615 | CGCCAGCGCAGCGGCCGGCGCGG | 1000 | | | | | | | | | | |
| Oligo fwd | CACCCGCCAGCGCAGCGGCCGGCG | | | | | | | | | | | |
| Oligo rev | AAACCGCCGGCCGCTGCGCTGGCG | | | | | | | | | | | |
| Chromosome | start | | end | strand | MM | target_seq | PAM | alignment | distance | position | gene name | gene id |
| GU305914.1 | | 78743 | 78765 | + | 0 | CGCCAGCGCAGCGGCCGGCG | CGG | \|\|\|\|\|\|\|\|[\|\|\|\|\|\|\|\|\|\|\|\|]PAM | NA | Intergenic | NA | NA |
| T5616 | GATCATGAAGCTACCGATCACGG | 1000 | | | | | | | | | | |
| Oligo fwd | CACCGATCATGAAGCTACCGATCA | | | | | | | | | | | |
| Oligo rev | AAACTGATCGGTAGCTTCATGATC | | | | | | | | | | | |
| Chromosome | start | | end | strand | MM | target_seq | PAM | alignment | distance | position | gene name | gene id |
| GU305914.1 | | 78854 | 78876 | + | 0 | GATCATGAAGCTACCGATCA | CGG | \|\|\|\|\|\|\|\|[\|\|\|\|\|\|\|\|\|\|\|\|]PAM | NA | Intergenic | NA | NA |
| T5617 | ATCATGAAGCTACCGATCACGGG | 1000 | | | | | | | | | | |
| Oligo fwd | CACCATCATGAAGCTACCGATCAC | | | | | | | | | | | |
| Oligo rev | AAACGTGATCGGTAGCTTCATGAT | | | | | | | | | | | |
| Chromosome | start | | end | strand | MM | target_seq | PAM | alignment | distance | position | gene name | gene id |

<table>
<tr><td>GU305914.1</td><td>78855</td><td>78877</td><td>+</td><td>0</td><td>ATCATGAAGCTACCGATCAC</td><td>GGG</td><td>||||||||[||||||||||||]PAM</td><td>NA</td><td>Intergenic</td><td>NA</td><td>NA</td></tr>
<tr><td></td><td></td><td></td><td></td><td></td><td></td><td></td><td></td><td></td><td></td><td></td><td></td></tr>
<tr><td>T5618</td><td>GAAGCTACCGATCACGGGCAAGG</td><td>1000</td><td></td><td></td><td></td><td></td><td></td><td></td><td></td><td></td><td></td></tr>
<tr><td>Oligo fwd</td><td>CACCGAAGCTACCGATCACGGGCA</td><td></td><td></td><td></td><td></td><td></td><td></td><td></td><td></td><td></td><td></td></tr>
<tr><td>Oligo rev</td><td>AAACTGCCCGTGATCGGTAGCTTC</td><td></td><td></td><td></td><td></td><td></td><td></td><td></td><td></td><td></td><td></td></tr>
<tr><td>Chromosome</td><td>start</td><td>end</td><td>strand</td><td>MM</td><td>target_seq</td><td>PAM</td><td>alignment</td><td>distance</td><td>position</td><td>gene name</td><td>gene id</td></tr>
<tr><td>GU305914.1</td><td>78860</td><td>78882</td><td>+</td><td>0</td><td>GAAGCTACCGATCACGGGCA</td><td>AGG</td><td>||||||||[||||||||||||]PAM</td><td>NA</td><td>Intergenic</td><td>NA</td><td>NA</td></tr>
<tr><td></td><td></td><td></td><td></td><td></td><td></td><td></td><td></td><td></td><td></td><td></td><td></td></tr>
<tr><td>T5619</td><td>CGTTCGACTTCAAGTCGCACCGG</td><td>1000</td><td></td><td></td><td></td><td></td><td></td><td></td><td></td><td></td><td></td></tr>
<tr><td>Oligo fwd</td><td>CACCCGTTCGACTTCAAGTCGCAC</td><td></td><td></td><td></td><td></td><td></td><td></td><td></td><td></td><td></td><td></td></tr>
<tr><td>Oligo rev</td><td>AAACGTGCGACTTGAAGTCGAACG</td><td></td><td></td><td></td><td></td><td></td><td></td><td></td><td></td><td></td><td></td></tr>
<tr><td>Chromosome</td><td>start</td><td>end</td><td>strand</td><td>MM</td><td>target_seq</td><td>PAM</td><td>alignment</td><td>distance</td><td>position</td><td>gene name</td><td>gene id</td></tr>
<tr><td>GU305914.1</td><td>78898</td><td>78920</td><td>+</td><td>0</td><td>CGTTCGACTTCAAGTCGCAC</td><td>CGG</td><td>||||||||[||||||||||||]PAM</td><td>NA</td><td>Intergenic</td><td>NA</td><td>NA</td></tr>
<tr><td>GU305914.1</td><td>78896</td><td>78918</td><td>-</td><td>4</td><td>GGTGCGACTTGAAGTCGAAC</td><td>GGG</td><td>-||-||||[||-||||||-||]PAM</td><td>NA</td><td>Intergenic</td><td>NA</td><td>NA</td></tr>
<tr><td></td><td></td><td></td><td></td><td></td><td></td><td></td><td></td><td></td><td></td><td></td><td></td></tr>
<tr><td>T5620</td><td>GCACCGGCAGCACACGTGTCTGG</td><td>1000</td><td></td><td></td><td></td><td></td><td></td><td></td><td></td><td></td><td></td></tr>
<tr><td>Oligo fwd</td><td>CACCGCACCGGCAGCACACGTGTC</td><td></td><td></td><td></td><td></td><td></td><td></td><td></td><td></td><td></td><td></td></tr>
<tr><td>Oligo rev</td><td>AAACGACACGTGTGCTGCCGGTGC</td><td></td><td></td><td></td><td></td><td></td><td></td><td></td><td></td><td></td><td></td></tr>
<tr><td>Chromosome</td><td>start</td><td>end</td><td>strand</td><td>MM</td><td>target_seq</td><td>PAM</td><td>alignment</td><td>distance</td><td>position</td><td>gene name</td><td>gene id</td></tr>
<tr><td>GU305914.1</td><td>78914</td><td>78936</td><td>+</td><td>0</td><td>GCACCGGCAGCACACGTGTC</td><td>TGG</td><td>||||||||[||||||||||||]PAM</td><td>NA</td><td>Intergenic</td><td>NA</td><td>NA</td></tr>
<tr><td></td><td></td><td></td><td></td><td></td><td></td><td></td><td></td><td></td><td></td><td></td><td></td></tr>
<tr><td>T5621</td><td>TGTCTGGACCTCTCCCCCTACGG</td><td>1000</td><td></td><td></td><td></td><td></td><td></td><td></td><td></td><td></td><td></td></tr>
<tr><td>Oligo fwd</td><td>CACCTGTCTGGACCTCTCCCCCTA</td><td></td><td></td><td></td><td></td><td></td><td></td><td></td><td></td><td></td><td></td></tr>
<tr><td>Oligo rev</td><td>AAACTAGGGGGAGAGGTCCAGACA</td><td></td><td></td><td></td><td></td><td></td><td></td><td></td><td></td><td></td><td></td></tr>
<tr><td>Chromosome</td><td>start</td><td>end</td><td>strand</td><td>MM</td><td>target_seq</td><td>PAM</td><td>alignment</td><td>distance</td><td>position</td><td>gene name</td><td>gene id</td></tr>
<tr><td>GU305914.1</td><td>78930</td><td>78952</td><td>+</td><td>0</td><td>TGTCTGGACCTCTCCCCCTA</td><td>CGG</td><td>||||||||[||||||||||||]PAM</td><td>NA</td><td>Intergenic</td><td>NA</td><td>NA</td></tr>
<tr><td></td><td></td><td></td><td></td><td></td><td></td><td></td><td></td><td></td><td></td><td></td><td></td></tr>
<tr><td>T5622</td><td>GTCTGGACCTCTCCCCCTACGGG</td><td>1000</td><td></td><td></td><td></td><td></td><td></td><td></td><td></td><td></td><td></td></tr>
<tr><td>Oligo fwd</td><td>CACCGTCTGGACCTCTCCCCCTAC</td><td></td><td></td><td></td><td></td><td></td><td></td><td></td><td></td><td></td><td></td></tr>
<tr><td>Oligo rev</td><td>AAACGTAGGGGGAGAGGTCCAGAC</td><td></td><td></td><td></td><td></td><td></td><td></td><td></td><td></td><td></td><td></td></tr>
<tr><td>Chromosome</td><td>start</td><td>end</td><td>strand</td><td>MM</td><td>target_seq</td><td>PAM</td><td>alignment</td><td>distance</td><td>position</td><td>gene name</td><td>gene id</td></tr>
<tr><td>GU305914.1</td><td>78931</td><td>78953</td><td>+</td><td>0</td><td>GTCTGGACCTCTCCCCCTAC</td><td>GGG</td><td>||||||||[||||||||||||]PAM</td><td>NA</td><td>Intergenic</td><td>NA</td><td>NA</td></tr>
<tr><td></td><td></td><td></td><td></td><td></td><td></td><td></td><td></td><td></td><td></td><td></td><td></td></tr>
<tr><td>T5623</td><td>CTCCCCCTACGGGAACGACCAGG</td><td>1000</td><td></td><td></td><td></td><td></td><td></td><td></td><td></td><td></td><td></td></tr>
<tr><td>Oligo fwd</td><td>CACCCTCCCCCTACGGGAACGACC</td><td></td><td></td><td></td><td></td><td></td><td></td><td></td><td></td><td></td><td></td></tr>
<tr><td>Oligo rev</td><td>AAACGGTCGTTCCCGTAGGGGGAG</td><td></td><td></td><td></td><td></td><td></td><td></td><td></td><td></td><td></td><td></td></tr>
<tr><td>Chromosome</td><td>start</td><td>end</td><td>strand</td><td>MM</td><td>target_seq</td><td>PAM</td><td>alignment</td><td>distance</td><td>position</td><td>gene name</td><td>gene id</td></tr>
<tr><td>GU305914.1</td><td>78941</td><td>78963</td><td>+</td><td>0</td><td>CTCCCCCTACGGGAACGACC</td><td>AGG</td><td>||||||||[||||||||||||]PAM</td><td>NA</td><td>Intergenic</td><td>NA</td><td>NA</td></tr>
<tr><td></td><td></td><td></td><td></td><td></td><td></td><td></td><td></td><td></td><td></td><td></td><td></td></tr>
<tr><td>T5624</td><td>ACGGGAACGACCAGGTGTCGCGG</td><td>1000</td><td></td><td></td><td></td><td></td><td></td><td></td><td></td><td></td><td></td></tr>
<tr><td>Oligo fwd</td><td>CACCACGGGAACGACCAGGTGTCG</td><td></td><td></td><td></td><td></td><td></td><td></td><td></td><td></td><td></td><td></td></tr>
<tr><td>Oligo rev</td><td>AAACCGACACCTGGTCGTTCCCGT</td><td></td><td></td><td></td><td></td><td></td><td></td><td></td><td></td><td></td><td></td></tr>
<tr><td>Chromosome</td><td>start</td><td>end</td><td>strand</td><td>MM</td><td>target_seq</td><td>PAM</td><td>alignment</td><td>distance</td><td>position</td><td>gene name</td><td>gene id</td></tr>
<tr><td>GU305914.1</td><td>78949</td><td>78971</td><td>+</td><td>0</td><td>ACGGGAACGACCAGGTGTCG</td><td>CGG</td><td>||||||||[||||||||||||]PAM</td><td>NA</td><td>Intergenic</td><td>NA</td><td>NA</td></tr>
<tr><td></td><td></td><td></td><td></td><td></td><td></td><td></td><td></td><td></td><td></td><td></td><td></td></tr>
<tr><td>T5625</td><td>GAGCGCGTGCACCACCTGCAAGG</td><td>1000</td><td></td><td></td><td></td><td></td><td></td><td></td><td></td><td></td><td></td></tr>
<tr><td>Oligo fwd</td><td>CACCGAGCGCGTGCACCACCTGCA</td><td></td><td></td><td></td><td></td><td></td><td></td><td></td><td></td><td></td><td></td></tr>
<tr><td>Oligo rev</td><td>AAACTGCAGGTGGTGCACGCGCTC</td><td></td><td></td><td></td><td></td><td></td><td></td><td></td><td></td><td></td><td></td></tr>
<tr><td>Chromosome</td><td>start</td><td>end</td><td>strand</td><td>MM</td><td>target_seq</td><td>PAM</td><td>alignment</td><td>distance</td><td>position</td><td>gene name</td><td>gene id</td></tr>
<tr><td>GU305914.1</td><td>78971</td><td>78993</td><td>+</td><td>0</td><td>GAGCGCGTGCACCACCTGCA</td><td>AGG</td><td>||||||||[||||||||||||]PAM</td><td>NA</td><td>Intergenic</td><td>NA</td><td>NA</td></tr>
<tr><td></td><td></td><td></td><td></td><td></td><td></td><td></td><td></td><td></td><td></td><td></td><td></td></tr>
<tr><td>T5626</td><td>GGAGACGACGAGACTGCCGACGG</td><td>1000</td><td></td><td></td><td></td><td></td><td></td><td></td><td></td><td></td><td></td></tr>
<tr><td>Oligo fwd</td><td>CACCGGAGACGACGAGACTGCCGA</td><td></td><td></td><td></td><td></td><td></td><td></td><td></td><td></td><td></td><td></td></tr>
<tr><td>Oligo rev</td><td>AAACTCGGCAGTCTCGTCGTCTCC</td><td></td><td></td><td></td><td></td><td></td><td></td><td></td><td></td><td></td><td></td></tr>
<tr><td>Chromosome</td><td>start</td><td>end</td><td>strand</td><td>MM</td><td>target_seq</td><td>PAM</td><td>alignment</td><td>distance</td><td>position</td><td>gene name</td><td>gene id</td></tr>
<tr><td>GU305914.1</td><td>78992</td><td>79014</td><td>+</td><td>0</td><td>GGAGACGACGAGACTGCCGA</td><td>CGG</td><td>||||||||[||||||||||||]PAM</td><td>NA</td><td>Intergenic</td><td>NA</td><td>NA</td></tr>
<tr><td>GU305914.1</td><td>83084</td><td>83106</td><td>-</td><td>4</td><td>GAAAGCGACTAGACTGCCGA</td><td>CGG</td><td>|-|--|||[|-||||||||||]PAM</td><td>NA</td><td>Intergenic</td><td>NA</td><td>NA</td></tr>
<tr><td></td><td></td><td></td><td></td><td></td><td></td><td></td><td></td><td></td><td></td><td></td><td></td></tr>
<tr><td>T5627</td><td>GACGAGACTGCCGACGGCGTCGG</td><td>1000</td><td></td><td></td><td></td><td></td><td></td><td></td><td></td><td></td><td></td></tr>
<tr><td>Oligo fwd</td><td>CACCGACGAGACTGCCGACGGCGT</td><td></td><td></td><td></td><td></td><td></td><td></td><td></td><td></td><td></td><td></td></tr>
<tr><td>Oligo rev</td><td>AAACACGCCGTCGGCAGTCTCGTC</td><td></td><td></td><td></td><td></td><td></td><td></td><td></td><td></td><td></td><td></td></tr>
<tr><td>Chromosome</td><td>start</td><td>end</td><td>strand</td><td>MM</td><td>target_seq</td><td>PAM</td><td>alignment</td><td>distance</td><td>position</td><td>gene name</td><td>gene id</td></tr>
<tr><td>GU305914.1</td><td>78998</td><td>79020</td><td>+</td><td>0</td><td>GACGAGACTGCCGACGGCGT</td><td>CGG</td><td>||||||||[||||||||||||]PAM</td><td>NA</td><td>Intergenic</td><td>NA</td><td>NA</td></tr>
<tr><td></td><td></td><td></td><td></td><td></td><td></td><td></td><td></td><td></td><td></td><td></td><td></td></tr>
<tr><td>T5628</td><td>GCCGACGGCGTCGGACAGCATGG</td><td>1000</td><td></td><td></td><td></td><td></td><td></td><td></td><td></td><td></td><td></td></tr>
<tr><td>Oligo fwd</td><td>CACCGCCGACGGCGTCGGACAGCA</td><td></td><td></td><td></td><td></td><td></td><td></td><td></td><td></td><td></td><td></td></tr>
<tr><td>Oligo rev</td><td>AAACTGCTGTCCGACGCCGTCGGC</td><td></td><td></td><td></td><td></td><td></td><td></td><td></td><td></td><td></td><td></td></tr>
<tr><td>Chromosome</td><td>start</td><td>end</td><td>strand</td><td>MM</td><td>target_seq</td><td>PAM</td><td>alignment</td><td>distance</td><td>position</td><td>gene name</td><td>gene id</td></tr>
<tr><td>GU305914.1</td><td>79007</td><td>79029</td><td>+</td><td>0</td><td>GCCGACGGCGTCGGACAGCA</td><td>TGG</td><td>||||||||[||||||||||||]PAM</td><td>NA</td><td>Intergenic</td><td>NA</td><td>NA</td></tr>
<tr><td></td><td></td><td></td><td></td><td></td><td></td><td></td><td></td><td></td><td></td><td></td><td></td></tr>
<tr><td>T5629</td><td>GACGGCGTCGGACAGCATGGTGG</td><td>1000</td><td></td><td></td><td></td><td></td><td></td><td></td><td></td><td></td><td></td></tr>
<tr><td>Oligo fwd</td><td>CACCGACGGCGTCGGACAGCATGG</td><td></td><td></td><td></td><td></td><td></td><td></td><td></td><td></td><td></td><td></td></tr>
<tr><td>Oligo rev</td><td>AAACCCATGCTGTCCGACGCCGTC</td><td></td><td></td><td></td><td></td><td></td><td></td><td></td><td></td><td></td><td></td></tr>
<tr><td>Chromosome</td><td>start</td><td>end</td><td>strand</td><td>MM</td><td>target_seq</td><td>PAM</td><td>alignment</td><td>distance</td><td>position</td><td>gene name</td><td>gene id</td></tr>
<tr><td>GU305914.1</td><td>79010</td><td>79032</td><td>+</td><td>0</td><td>GACGGCGTCGGACAGCATGG</td><td>TGG</td><td>||||||||[||||||||||||]PAM</td><td>NA</td><td>Intergenic</td><td>NA</td><td>NA</td></tr>
<tr><td></td><td></td><td></td><td></td><td></td><td></td><td></td><td></td><td></td><td></td><td></td><td></td></tr>
<tr><td>T5630</td><td>AAGCACCGCAAGTTCTACTTCGG</td><td>1000</td><td></td><td></td><td></td><td></td><td></td><td></td><td></td><td></td><td></td></tr>
<tr><td>Oligo fwd</td><td>CACCAAGCACCGCAAGTTCTACTT</td><td></td><td></td><td></td><td></td><td></td><td></td><td></td><td></td><td></td><td></td></tr>
<tr><td>Oligo rev</td><td>AAACAAGTAGAACTTGCGGTGCTT</td><td></td><td></td><td></td><td></td><td></td><td></td><td></td><td></td><td></td><td></td></tr>
<tr><td>Chromosome</td><td>start</td><td>end</td><td>strand</td><td>MM</td><td>target_seq</td><td>PAM</td><td>alignment</td><td>distance</td><td>position</td><td>gene name</td><td>gene id</td></tr>
<tr><td>GU305914.1</td><td>79062</td><td>79084</td><td>+</td><td>0</td><td>AAGCACCGCAAGTTCTACTT</td><td>CGG</td><td>||||||||[||||||||||||]PAM</td><td>NA</td><td>Intergenic</td><td>NA</td><td>NA</td></tr>
<tr><td></td><td></td><td></td><td></td><td></td><td></td><td></td><td></td><td></td><td></td><td></td><td></td></tr>
<tr><td>T5631</td><td>AGCACCGCAAGTTCTACTTCGGG</td><td>1000</td><td></td><td></td><td></td><td></td><td></td><td></td><td></td><td></td><td></td></tr>
<tr><td>Oligo fwd</td><td>CACCAGCACCGCAAGTTCTACTTC</td><td></td><td></td><td></td><td></td><td></td><td></td><td></td><td></td><td></td><td></td></tr>
<tr><td>Oligo rev</td><td>AAACGAAGTAGAACTTGCGGTGCT</td><td></td><td></td><td></td><td></td><td></td><td></td><td></td><td></td><td></td><td></td></tr>
<tr><td>Chromosome</td><td>start</td><td>end</td><td>strand</td><td>MM</td><td>target_seq</td><td>PAM</td><td>alignment</td><td>distance</td><td>position</td><td>gene name</td><td>gene id</td></tr>
<tr><td>GU305914.1</td><td>79063</td><td>79085</td><td>+</td><td>0</td><td>AGCACCGCAAGTTCTACTTC</td><td>GGG</td><td>||||||||[||||||||||||]PAM</td><td>NA</td><td>Intergenic</td><td>NA</td><td>NA</td></tr>
<tr><td></td><td></td><td></td><td></td><td></td><td></td><td></td><td></td><td></td><td></td><td></td><td></td></tr>
<tr><td>T5632</td><td>CTTCGGGTTCCGTAAGAACATGG</td><td>1000</td><td></td><td></td><td></td><td></td><td></td><td></td><td></td><td></td><td></td></tr>
</table>

| | | | | | | | | | | | |
|---|---|---|---|---|---|---|---|---|---|---|---|
| Oligo fwd | CACCCTTCGGGTTCCGTAAGAACA | | | | | | | | | | |
| Oligo rev | AAACTGTTCTTACGGAACCCGAAG | | | | | | | | | | |
| Chromosome | start | end | strand | MM | target_seq | PAM | alignment | distance | position | gene name | gene id |
| GU305914.1 | 79079 | 79101 | + | 0 | CTTCGGGTTCCGTAAGAACA | TGG | \|\|\|\|\|\|\|\|[\|\|\|\|\|\|\|\|\|\|\|\|]PAM | NA | Intergenic | NA | NA |
| T5633 | GAACATGGAACTGCTGAAGATGG | 1000 | | | | | | | | | |
| Oligo fwd | CACCGAACATGGAACTGCTGAAGA | | | | | | | | | | |
| Oligo rev | AAACTCTTCAGCAGTTCCATGTTC | | | | | | | | | | |
| Chromosome | start | end | strand | MM | target_seq | PAM | alignment | distance | position | gene name | gene id |
| GU305914.1 | 79094 | 79116 | + | 0 | GAACATGGAACTGCTGAAGA | TGG | \|\|\|\|\|\|\|\|[\|\|\|\|\|\|\|\|\|\|\|\|]PAM | NA | Intergenic | NA | NA |
| T5634 | GATCTACTACGACCGAGAGATGG | 1000 | | | | | | | | | |
| Oligo fwd | CACCGATCTACTACGACCGAGAGA | | | | | | | | | | |
| Oligo rev | AAACTCTCTCGGTCGTAGTAGATC | | | | | | | | | | |
| Chromosome | start | end | strand | MM | target_seq | PAM | alignment | distance | position | gene name | gene id |
| GU305914.1 | 79187 | 79209 | + | 0 | GATCTACTACGACCGAGAGA | TGG | \|\|\|\|\|\|\|\|[\|\|\|\|\|\|\|\|\|\|\|\|]PAM | NA | Intergenic | NA | NA |
| T5635 | CATGCAACTGATTTTCGAGAAGG | 1000 | | | | | | | | | |
| Oligo fwd | CACCCATGCAACTGATTTTCGAGA | | | | | | | | | | |
| Oligo rev | AAACTCTCGAAAATCAGTTGCATG | | | | | | | | | | |
| Chromosome | start | end | strand | MM | target_seq | PAM | alignment | distance | position | gene name | gene id |
| GU305914.1 | 79214 | 79236 | + | 0 | CATGCAACTGATTTTCGAGA | AGG | \|\|\|\|\|\|\|\|[\|\|\|\|\|\|\|\|\|\|\|\|]PAM | NA | Intergenic | NA | NA |
| T5636 | ACTGATTTTCGAGAAGGAGACGG | 1000 | | | | | | | | | |
| Oligo fwd | CACCACTGATTTTCGAGAAGGAGA | | | | | | | | | | |
| Oligo rev | AAACTCTCCTTCTCGAAAATCAGT | | | | | | | | | | |
| Chromosome | start | end | strand | MM | target_seq | PAM | alignment | distance | position | gene name | gene id |
| GU305914.1 | 79220 | 79242 | + | 0 | ACTGATTTTCGAGAAGGAGA | CGG | \|\|\|\|\|\|\|\|[\|\|\|\|\|\|\|\|\|\|\|\|]PAM | NA | Intergenic | NA | NA |
| T5637 | GTGCATCGAGCAGATCCTAACGG | 1000 | | | | | | | | | |
| Oligo fwd | CACCGTGCATCGAGCAGATCCTAA | | | | | | | | | | |
| Oligo rev | AAACTTAGGATCTGCTCGATGCAC | | | | | | | | | | |
| Chromosome | start | end | strand | MM | target_seq | PAM | alignment | distance | position | gene name | gene id |
| GU305914.1 | 79259 | 79281 | + | 0 | GTGCATCGAGCAGATCCTAA | CGG | \|\|\|\|\|\|\|\|[\|\|\|\|\|\|\|\|\|\|\|\|]PAM | NA | Intergenic | NA | NA |
| T5638 | CATCGAGCAGATCCTAACGGTGG | 1000 | | | | | | | | | |
| Oligo fwd | CACCCATCGAGCAGATCCTAACGG | | | | | | | | | | |
| Oligo rev | AAACCCGTTAGGATCTGCTCGATG | | | | | | | | | | |
| Chromosome | start | end | strand | MM | target_seq | PAM | alignment | distance | position | gene name | gene id |
| GU305914.1 | 79262 | 79284 | + | 0 | CATCGAGCAGATCCTAACGG | TGG | \|\|\|\|\|\|\|\|[\|\|\|\|\|\|\|\|\|\|\|\|]PAM | NA | Intergenic | NA | NA |
| T5639 | GCAGATCCTAACGGTGGCGAAGG | 1000 | | | | | | | | | |
| Oligo fwd | CACCGCAGATCCTAACGGTGGCGA | | | | | | | | | | |
| Oligo rev | AAACTCGCCACCGTTAGGATCTGC | | | | | | | | | | |
| Chromosome | start | end | strand | MM | target_seq | PAM | alignment | distance | position | gene name | gene id |
| GU305914.1 | 79268 | 79290 | + | 0 | GCAGATCCTAACGGTGGCGA | AGG | \|\|\|\|\|\|\|\|[\|\|\|\|\|\|\|\|\|\|\|\|]PAM | NA | Intergenic | NA | NA |
| T5640 | ACGGTGGCGAAGGACGCGTACGG | 1000 | | | | | | | | | |
| Oligo fwd | CACCACGGTGGCGAAGGACGCGTA | | | | | | | | | | |
| Oligo rev | AAACTACGCGTCCTTCGCCACCGT | | | | | | | | | | |
| Chromosome | start | end | strand | MM | target_seq | PAM | alignment | distance | position | gene name | gene id |
| GU305914.1 | 79278 | 79300 | + | 0 | ACGGTGGCGAAGGACGCGTA | CGG | \|\|\|\|\|\|\|\|[\|\|\|\|\|\|\|\|\|\|\|\|]PAM | NA | Intergenic | NA | NA |
| T5641 | CGGTGGCGAAGGACGCGTACGGG | 1000 | | | | | | | | | |
| Oligo fwd | CACCCGGTGGCGAAGGACGCGTAC | | | | | | | | | | |
| Oligo rev | AAACGTACGCGTCCTTCGCCACCG | | | | | | | | | | |
| Chromosome | start | end | strand | MM | target_seq | PAM | alignment | distance | position | gene name | gene id |
| GU305914.1 | 79279 | 79301 | + | 0 | CGGTGGCGAAGGACGCGTAC | GGG | \|\|\|\|\|\|\|\|[\|\|\|\|\|\|\|\|\|\|\|\|]PAM | NA | Intergenic | NA | NA |
| T5642 | GGTGGCGAAGGACGCGTACGGGG | 1000 | | | | | | | | | |
| Oligo fwd | CACCGGTGGCGAAGGACGCGTACG | | | | | | | | | | |
| Oligo rev | AAACCGTACGCGTCCTTCGCCACC | | | | | | | | | | |
| Chromosome | start | end | strand | MM | target_seq | PAM | alignment | distance | position | gene name | gene id |
| GU305914.1 | 79280 | 79302 | + | 0 | GGTGGCGAAGGACGCGTACG | GGG | \|\|\|\|\|\|\|\|[\|\|\|\|\|\|\|\|\|\|\|\|]PAM | NA | Intergenic | NA | NA |
| T5643 | GGACGCGTACGGGGTGAGCCTGG | 1000 | | | | | | | | | |
| Oligo fwd | CACCGGACGCGTACGGGGTGAGCC | | | | | | | | | | |
| Oligo rev | AAACGGCTCACCCCGTACGCGTCC | | | | | | | | | | |
| Chromosome | start | end | strand | MM | target_seq | PAM | alignment | distance | position | gene name | gene id |
| GU305914.1 | 79289 | 79311 | + | 0 | GGACGCGTACGGGGTGAGCC | TGG | \|\|\|\|\|\|\|\|[\|\|\|\|\|\|\|\|\|\|\|\|]PAM | NA | Intergenic | NA | NA |
| T5644 | GCCTGGACATCGCGCACCAGCGG | 1000 | | | | | | | | | |
| Oligo fwd | CACCGCCTGGACATCGCGCACCAG | | | | | | | | | | |
| Oligo rev | AAACCTGGTGCGCGATGTCCAGGC | | | | | | | | | | |
| Chromosome | start | end | strand | MM | target_seq | PAM | alignment | distance | position | gene name | gene id |
| GU305914.1 | 79306 | 79328 | + | 0 | GCCTGGACATCGCGCACCAG | CGG | \|\|\|\|\|\|\|\|[\|\|\|\|\|\|\|\|\|\|\|\|]PAM | NA | Intergenic | NA | NA |
| T5645 | GCACCAGCGGATCACGCTGACGG | 1000 | | | | | | | | | |
| Oligo fwd | CACCGCACCAGCGGATCACGCTGA | | | | | | | | | | |
| Oligo rev | AAACTCAGCGTGATCCGCTGGTGC | | | | | | | | | | |
| Chromosome | start | end | strand | MM | target_seq | PAM | alignment | distance | position | gene name | gene id |
| GU305914.1 | 79319 | 79341 | + | 0 | GCACCAGCGGATCACGCTGA | CGG | \|\|\|\|\|\|\|\|[\|\|\|\|\|\|\|\|\|\|\|\|]PAM | NA | Intergenic | NA | NA |
| T5646 | CGCTGACGGCGCGATGTCTGCGG | 1000 | | | | | | | | | |
| Oligo fwd | CACCCGCTGACGGCGCGATGTCTG | | | | | | | | | | |
| Oligo rev | AAACCAGACATCGCGCCGTCAGCG | | | | | | | | | | |
| Chromosome | start | end | strand | MM | target_seq | PAM | alignment | distance | position | gene name | gene id |
| GU305914.1 | 79333 | 79355 | + | 0 | CGCTGACGGCGCGATGTCTG | CGG | \|\|\|\|\|\|\|\|[\|\|\|\|\|\|\|\|\|\|\|\|]PAM | NA | Intergenic | NA | NA |

<table>
<tr><td>T5647</td><td>TCGACGTGCTGATGCTGCAGAGG</td><td>1000</td><td></td><td></td><td></td><td></td><td></td><td></td><td></td><td></td><td></td></tr>
<tr><td>Oligo fwd</td><td>CACCTCGACGTGCTGATGCTGCAG</td><td></td><td></td><td></td><td></td><td></td><td></td><td></td><td></td><td></td><td></td></tr>
<tr><td>Oligo rev</td><td>AAACCTGCAGCATCAGCACGTCGA</td><td></td><td></td><td></td><td></td><td></td><td></td><td></td><td></td><td></td><td></td></tr>
<tr><td>Chromosome</td><td>start</td><td>end</td><td>strand</td><td>MM</td><td>target_seq</td><td>PAM</td><td>alignment</td><td>distance</td><td>position</td><td>gene name</td><td>gene id</td></tr>
<tr><td>GU305914.1</td><td>79378</td><td>79400</td><td>+</td><td>0</td><td>TCGACGTGCTGATGCTGCAG</td><td>AGG</td><td>||||||||[||||||||||||]PAM</td><td>NA</td><td>Intergenic</td><td>NA</td><td>NA</td></tr>
<tr><td>T5648</td><td>CGTGCTGATGCTGCAGAGGAAGG</td><td>1000</td><td></td><td></td><td></td><td></td><td></td><td></td><td></td><td></td><td></td></tr>
<tr><td>Oligo fwd</td><td>CACCCGTGCTGATGCTGCAGAGGA</td><td></td><td></td><td></td><td></td><td></td><td></td><td></td><td></td><td></td><td></td></tr>
<tr><td>Oligo rev</td><td>AAACTCCTCTGCAGCATCAGCACG</td><td></td><td></td><td></td><td></td><td></td><td></td><td></td><td></td><td></td><td></td></tr>
<tr><td>Chromosome</td><td>start</td><td>end</td><td>strand</td><td>MM</td><td>target_seq</td><td>PAM</td><td>alignment</td><td>distance</td><td>position</td><td>gene name</td><td>gene id</td></tr>
<tr><td>GU305914.1</td><td>79382</td><td>79404</td><td>+</td><td>0</td><td>CGTGCTGATGCTGCAGAGGA</td><td>AGG</td><td>||||||||[||||||||||||]PAM</td><td>NA</td><td>Intergenic</td><td>NA</td><td>NA</td></tr>
<tr><td>GU305914.1</td><td>133230</td><td>133252</td><td>+</td><td>3</td><td>CGTGCCGATGCAGAAGAGGA</td><td>AGG</td><td>|||||-||[|||-|-||||||]PAM</td><td>NA</td><td>Intergenic</td><td>NA</td><td>NA</td></tr>
<tr><td>T5649</td><td>GCTGATGCTGCAGAGGAAGGTGG</td><td>1000</td><td></td><td></td><td></td><td></td><td></td><td></td><td></td><td></td><td></td></tr>
<tr><td>Oligo fwd</td><td>CACCGCTGATGCTGCAGAGGAAGG</td><td></td><td></td><td></td><td></td><td></td><td></td><td></td><td></td><td></td><td></td></tr>
<tr><td>Oligo rev</td><td>AAACCCTTCCTCTGCAGCATCAGC</td><td></td><td></td><td></td><td></td><td></td><td></td><td></td><td></td><td></td><td></td></tr>
<tr><td>Chromosome</td><td>start</td><td>end</td><td>strand</td><td>MM</td><td>target_seq</td><td>PAM</td><td>alignment</td><td>distance</td><td>position</td><td>gene name</td><td>gene id</td></tr>
<tr><td>GU305914.1</td><td>79385</td><td>79407</td><td>+</td><td>0</td><td>GCTGATGCTGCAGAGGAAGG</td><td>TGG</td><td>||||||||[||||||||||||]PAM</td><td>NA</td><td>Intergenic</td><td>NA</td><td>NA</td></tr>
<tr><td>T5650</td><td>GAGAGTTACTCGTTGTGACGAGG</td><td>1000</td><td></td><td></td><td></td><td></td><td></td><td></td><td></td><td></td><td></td></tr>
<tr><td>Oligo fwd</td><td>CACCGAGAGTTACTCGTTGTGACG</td><td></td><td></td><td></td><td></td><td></td><td></td><td></td><td></td><td></td><td></td></tr>
<tr><td>Oligo rev</td><td>AAACCGTCACAACGAGTAACTCTC</td><td></td><td></td><td></td><td></td><td></td><td></td><td></td><td></td><td></td><td></td></tr>
<tr><td>Chromosome</td><td>start</td><td>end</td><td>strand</td><td>MM</td><td>target_seq</td><td>PAM</td><td>alignment</td><td>distance</td><td>position</td><td>gene name</td><td>gene id</td></tr>
<tr><td>GU305914.1</td><td>79446</td><td>79468</td><td>+</td><td>0</td><td>GAGAGTTACTCGTTGTGACG</td><td>AGG</td><td>||||||||[||||||||||||]PAM</td><td>NA</td><td>Intergenic</td><td>NA</td><td>NA</td></tr>
<tr><td>T5651</td><td>AATTACCGCGACATTGTTGTCGG</td><td>1000</td><td></td><td></td><td></td><td></td><td></td><td></td><td></td><td></td><td></td></tr>
<tr><td>Oligo fwd</td><td>CACCAATTACCGCGACATTGTTGT</td><td></td><td></td><td></td><td></td><td></td><td></td><td></td><td></td><td></td><td></td></tr>
<tr><td>Oligo rev</td><td>AAACACAACAATGTCGCGGTAATT</td><td></td><td></td><td></td><td></td><td></td><td></td><td></td><td></td><td></td><td></td></tr>
<tr><td>Chromosome</td><td>start</td><td>end</td><td>strand</td><td>MM</td><td>target_seq</td><td>PAM</td><td>alignment</td><td>distance</td><td>position</td><td>gene name</td><td>gene id</td></tr>
<tr><td>GU305914.1</td><td>79570</td><td>79592</td><td>+</td><td>0</td><td>AATTACCGCGACATTGTTGT</td><td>CGG</td><td>||||||||[||||||||||||]PAM</td><td>NA</td><td>Intergenic</td><td>NA</td><td>NA</td></tr>
<tr><td>T5652</td><td>ATTGTTGTCGGTCACACAATAGG</td><td>1000</td><td></td><td></td><td></td><td></td><td></td><td></td><td></td><td></td><td></td></tr>
<tr><td>Oligo fwd</td><td>CACCATTGTTGTCGGTCACACAAT</td><td></td><td></td><td></td><td></td><td></td><td></td><td></td><td></td><td></td><td></td></tr>
<tr><td>Oligo rev</td><td>AAACATTGTGTGACCGACAACAAT</td><td></td><td></td><td></td><td></td><td></td><td></td><td></td><td></td><td></td><td></td></tr>
<tr><td>Chromosome</td><td>start</td><td>end</td><td>strand</td><td>MM</td><td>target_seq</td><td>PAM</td><td>alignment</td><td>distance</td><td>position</td><td>gene name</td><td>gene id</td></tr>
<tr><td>GU305914.1</td><td>79582</td><td>79604</td><td>+</td><td>0</td><td>ATTGTTGTCGGTCACACAAT</td><td>AGG</td><td>||||||||[||||||||||||]PAM</td><td>NA</td><td>Intergenic</td><td>NA</td><td>NA</td></tr>
<tr><td>T5653</td><td>TTGTTGTCGGTCACACAATAGGG</td><td>1000</td><td></td><td></td><td></td><td></td><td></td><td></td><td></td><td></td><td></td></tr>
<tr><td>Oligo fwd</td><td>CACCTTGTTGTCGGTCACACAATA</td><td></td><td></td><td></td><td></td><td></td><td></td><td></td><td></td><td></td><td></td></tr>
<tr><td>Oligo rev</td><td>AAACTATTGTGTGACCGACAACAA</td><td></td><td></td><td></td><td></td><td></td><td></td><td></td><td></td><td></td><td></td></tr>
<tr><td>Chromosome</td><td>start</td><td>end</td><td>strand</td><td>MM</td><td>target_seq</td><td>PAM</td><td>alignment</td><td>distance</td><td>position</td><td>gene name</td><td>gene id</td></tr>
<tr><td>GU305914.1</td><td>79583</td><td>79605</td><td>+</td><td>0</td><td>TTGTTGTCGGTCACACAATA</td><td>GGG</td><td>||||||||[||||||||||||]PAM</td><td>NA</td><td>Intergenic</td><td>NA</td><td>NA</td></tr>
<tr><td>T5654</td><td>TGTTGTCGGTCACACAATAGGGG</td><td>1000</td><td></td><td></td><td></td><td></td><td></td><td></td><td></td><td></td><td></td></tr>
<tr><td>Oligo fwd</td><td>CACCTGTTGTCGGTCACACAATAG</td><td></td><td></td><td></td><td></td><td></td><td></td><td></td><td></td><td></td><td></td></tr>
<tr><td>Oligo rev</td><td>AAACCTATTGTGTGACCGACAACA</td><td></td><td></td><td></td><td></td><td></td><td></td><td></td><td></td><td></td><td></td></tr>
<tr><td>Chromosome</td><td>start</td><td>end</td><td>strand</td><td>MM</td><td>target_seq</td><td>PAM</td><td>alignment</td><td>distance</td><td>position</td><td>gene name</td><td>gene id</td></tr>
<tr><td>GU305914.1</td><td>79584</td><td>79606</td><td>+</td><td>0</td><td>TGTTGTCGGTCACACAATAG</td><td>GGG</td><td>||||||||[||||||||||||]PAM</td><td>NA</td><td>Intergenic</td><td>NA</td><td>NA</td></tr>
<tr><td>T5655</td><td>ACACAATAGGGGAGAATAATAGG</td><td>1000</td><td></td><td></td><td></td><td></td><td></td><td></td><td></td><td></td><td></td></tr>
<tr><td>Oligo fwd</td><td>CACCACACAATAGGGGAGAATAAT</td><td></td><td></td><td></td><td></td><td></td><td></td><td></td><td></td><td></td><td></td></tr>
<tr><td>Oligo rev</td><td>AAACATTATTCTCCCCTATTGTGT</td><td></td><td></td><td></td><td></td><td></td><td></td><td></td><td></td><td></td><td></td></tr>
<tr><td>Chromosome</td><td>start</td><td>end</td><td>strand</td><td>MM</td><td>target_seq</td><td>PAM</td><td>alignment</td><td>distance</td><td>position</td><td>gene name</td><td>gene id</td></tr>
<tr><td>GU305914.1</td><td>79595</td><td>79617</td><td>+</td><td>0</td><td>ACACAATAGGGGAGAATAAT</td><td>AGG</td><td>||||||||[||||||||||||]PAM</td><td>NA</td><td>Intergenic</td><td>NA</td><td>NA</td></tr>
<tr><td>T5656</td><td>GGGGAGAATAATAGGATCAAAGG</td><td>1000</td><td></td><td></td><td></td><td></td><td></td><td></td><td></td><td></td><td></td></tr>
<tr><td>Oligo fwd</td><td>CACCGGGGAGAATAATAGGATCAA</td><td></td><td></td><td></td><td></td><td></td><td></td><td></td><td></td><td></td><td></td></tr>
<tr><td>Oligo rev</td><td>AAACTTGATCCTATTATTCTCCCC</td><td></td><td></td><td></td><td></td><td></td><td></td><td></td><td></td><td></td><td></td></tr>
<tr><td>Chromosome</td><td>start</td><td>end</td><td>strand</td><td>MM</td><td>target_seq</td><td>PAM</td><td>alignment</td><td>distance</td><td>position</td><td>gene name</td><td>gene id</td></tr>
<tr><td>GU305914.1</td><td>79603</td><td>79625</td><td>+</td><td>0</td><td>GGGGAGAATAATAGGATCAA</td><td>AGG</td><td>||||||||[||||||||||||]PAM</td><td>NA</td><td>Intergenic</td><td>NA</td><td>NA</td></tr>
<tr><td>T5657</td><td>GGGAGAATAATAGGATCAAAGGG</td><td>1000</td><td></td><td></td><td></td><td></td><td></td><td></td><td></td><td></td><td></td></tr>
<tr><td>Oligo fwd</td><td>CACCGGGAGAATAATAGGATCAAA</td><td></td><td></td><td></td><td></td><td></td><td></td><td></td><td></td><td></td><td></td></tr>
<tr><td>Oligo rev</td><td>AAACTTTGATCCTATTATTCTCCC</td><td></td><td></td><td></td><td></td><td></td><td></td><td></td><td></td><td></td><td></td></tr>
<tr><td>Chromosome</td><td>start</td><td>end</td><td>strand</td><td>MM</td><td>target_seq</td><td>PAM</td><td>alignment</td><td>distance</td><td>position</td><td>gene name</td><td>gene id</td></tr>
<tr><td>GU305914.1</td><td>79604</td><td>79626</td><td>+</td><td>0</td><td>GGGAGAATAATAGGATCAAA</td><td>GGG</td><td>||||||||[||||||||||||]PAM</td><td>NA</td><td>Intergenic</td><td>NA</td><td>NA</td></tr>
<tr><td>T5658</td><td>ATAATAGGATCAAAGGGAATCGG</td><td>1000</td><td></td><td></td><td></td><td></td><td></td><td></td><td></td><td></td><td></td></tr>
<tr><td>Oligo fwd</td><td>CACCATAATAGGATCAAAGGGAAT</td><td></td><td></td><td></td><td></td><td></td><td></td><td></td><td></td><td></td><td></td></tr>
<tr><td>Oligo rev</td><td>AAACATTCCCTTTGATCCTATTAT</td><td></td><td></td><td></td><td></td><td></td><td></td><td></td><td></td><td></td><td></td></tr>
<tr><td>Chromosome</td><td>start</td><td>end</td><td>strand</td><td>MM</td><td>target_seq</td><td>PAM</td><td>alignment</td><td>distance</td><td>position</td><td>gene name</td><td>gene id</td></tr>
<tr><td>GU305914.1</td><td>79610</td><td>79632</td><td>+</td><td>0</td><td>ATAATAGGATCAAAGGGAAT</td><td>CGG</td><td>||||||||[||||||||||||]PAM</td><td>NA</td><td>Intergenic</td><td>NA</td><td>NA</td></tr>
<tr><td>T5659</td><td>TAATAGGATCAAAGGGAATCGGG</td><td>1000</td><td></td><td></td><td></td><td></td><td></td><td></td><td></td><td></td><td></td></tr>
<tr><td>Oligo fwd</td><td>CACCTAATAGGATCAAAGGGAATC</td><td></td><td></td><td></td><td></td><td></td><td></td><td></td><td></td><td></td><td></td></tr>
<tr><td>Oligo rev</td><td>AAACGATTCCCTTTGATCCTATTA</td><td></td><td></td><td></td><td></td><td></td><td></td><td></td><td></td><td></td><td></td></tr>
<tr><td>Chromosome</td><td>start</td><td>end</td><td>strand</td><td>MM</td><td>target_seq</td><td>PAM</td><td>alignment</td><td>distance</td><td>position</td><td>gene name</td><td>gene id</td></tr>
<tr><td>GU305914.1</td><td>79611</td><td>79633</td><td>+</td><td>0</td><td>TAATAGGATCAAAGGGAATC</td><td>GGG</td><td>||||||||[||||||||||||]PAM</td><td>NA</td><td>Intergenic</td><td>NA</td><td>NA</td></tr>
<tr><td>T5660</td><td>ACAATATTTATTATTGTTATCGG</td><td>1000</td><td></td><td></td><td></td><td></td><td></td><td></td><td></td><td></td><td></td></tr>
<tr><td>Oligo fwd</td><td>CACCACAATATTTATTATTGTTAT</td><td></td><td></td><td></td><td></td><td></td><td></td><td></td><td></td><td></td><td></td></tr>
<tr><td>Oligo rev</td><td>AAACATAACAATAATAAATATTGT</td><td></td><td></td><td></td><td></td><td></td><td></td><td></td><td></td><td></td><td></td></tr>
<tr><td>Chromosome</td><td>start</td><td>end</td><td>strand</td><td>MM</td><td>target_seq</td><td>PAM</td><td>alignment</td><td>distance</td><td>position</td><td>gene name</td><td>gene id</td></tr>
<tr><td>GU305914.1</td><td>79636</td><td>79658</td><td>+</td><td>0</td><td>ACAATATTTATTATTGTTAT</td><td>CGG</td><td>||||||||[||||||||||||]PAM</td><td>NA</td><td>Intergenic</td><td>NA</td><td>NA</td></tr>
<tr><td>T5661</td><td>ACACGATATCAATCGAGATAAGG</td><td>1000</td><td></td><td></td><td></td><td></td><td></td><td></td><td></td><td></td><td></td></tr>
<tr><td>Oligo fwd</td><td>CACCACACGATATCAATCGAGATA</td><td></td><td></td><td></td><td></td><td></td><td></td><td></td><td></td><td></td><td></td></tr>
<tr><td>Oligo rev</td><td>AAACTATCTCGATTGATATCGTGT</td><td></td><td></td><td></td><td></td><td></td><td></td><td></td><td></td><td></td><td></td></tr>
<tr><td>Chromosome</td><td>start</td><td>end</td><td>strand</td><td>MM</td><td>target_seq</td><td>PAM</td><td>alignment</td><td>distance</td><td>position</td><td>gene name</td><td>gene id</td></tr>
</table>

<table>
<tr><td>GU305914.1</td><td>79663</td><td>79685</td><td>+</td><td>0</td><td>ACACGATATCAATCGAGATA</td><td>AGG</td><td>||||||||[||||||||||||]PAM</td><td>NA</td><td>Intergenic</td><td>NA</td><td>NA</td></tr>
<tr><td></td><td></td><td></td><td></td><td></td><td></td><td></td><td></td><td></td><td></td><td></td><td></td></tr>
<tr><td>T5662</td><td>CACGATATCAATCGAGATAAGGG</td><td>1000</td><td></td><td></td><td></td><td></td><td></td><td></td><td></td><td></td><td></td></tr>
<tr><td>Oligo fwd</td><td>CACCCACGATATCAATCGAGATAA</td><td></td><td></td><td></td><td></td><td></td><td></td><td></td><td></td><td></td><td></td></tr>
<tr><td>Oligo rev</td><td>AAACTTATCTCGATTGATATCGTG</td><td></td><td></td><td></td><td></td><td></td><td></td><td></td><td></td><td></td><td></td></tr>
<tr><td>Chromosome</td><td>start</td><td>end</td><td>strand</td><td>MM</td><td>target_seq</td><td>PAM</td><td>alignment</td><td>distance</td><td>position</td><td>gene name</td><td>gene id</td></tr>
<tr><td>GU305914.1</td><td>79664</td><td>79686</td><td>+</td><td>0</td><td>CACGATATCAATCGAGATAA</td><td>GGG</td><td>||||||||[||||||||||||]PAM</td><td>NA</td><td>Intergenic</td><td>NA</td><td>NA</td></tr>
<tr><td></td><td></td><td></td><td></td><td></td><td></td><td></td><td></td><td></td><td></td><td></td><td></td></tr>
<tr><td>T5663</td><td>GATATCAATCGAGATAAGGGAGG</td><td>1000</td><td></td><td></td><td></td><td></td><td></td><td></td><td></td><td></td><td></td></tr>
<tr><td>Oligo fwd</td><td>CACCGATATCAATCGAGATAAGGG</td><td></td><td></td><td></td><td></td><td></td><td></td><td></td><td></td><td></td><td></td></tr>
<tr><td>Oligo rev</td><td>AAACCCCTTATCTCGATTGATATC</td><td></td><td></td><td></td><td></td><td></td><td></td><td></td><td></td><td></td><td></td></tr>
<tr><td>Chromosome</td><td>start</td><td>end</td><td>strand</td><td>MM</td><td>target_seq</td><td>PAM</td><td>alignment</td><td>distance</td><td>position</td><td>gene name</td><td>gene id</td></tr>
<tr><td>GU305914.1</td><td>79667</td><td>79689</td><td>+</td><td>0</td><td>GATATCAATCGAGATAAGGG</td><td>AGG</td><td>||||||||[||||||||||||]PAM</td><td>NA</td><td>Intergenic</td><td>NA</td><td>NA</td></tr>
<tr><td></td><td></td><td></td><td></td><td></td><td></td><td></td><td></td><td></td><td></td><td></td><td></td></tr>
<tr><td>T5664</td><td>ATATCAATCGAGATAAGGGAGGG</td><td>1000</td><td></td><td></td><td></td><td></td><td></td><td></td><td></td><td></td><td></td></tr>
<tr><td>Oligo fwd</td><td>CACCATATCAATCGAGATAAGGGA</td><td></td><td></td><td></td><td></td><td></td><td></td><td></td><td></td><td></td><td></td></tr>
<tr><td>Oligo rev</td><td>AAACTCCCTTATCTCGATTGATAT</td><td></td><td></td><td></td><td></td><td></td><td></td><td></td><td></td><td></td><td></td></tr>
<tr><td>Chromosome</td><td>start</td><td>end</td><td>strand</td><td>MM</td><td>target_seq</td><td>PAM</td><td>alignment</td><td>distance</td><td>position</td><td>gene name</td><td>gene id</td></tr>
<tr><td>GU305914.1</td><td>79668</td><td>79690</td><td>+</td><td>0</td><td>ATATCAATCGAGATAAGGGA</td><td>GGG</td><td>||||||||[||||||||||||]PAM</td><td>NA</td><td>Intergenic</td><td>NA</td><td>NA</td></tr>
<tr><td></td><td></td><td></td><td></td><td></td><td></td><td></td><td></td><td></td><td></td><td></td><td></td></tr>
<tr><td>T5665</td><td>GAGATAAGGGAGGGCGCCCCCGG</td><td>1000</td><td></td><td></td><td></td><td></td><td></td><td></td><td></td><td></td><td></td></tr>
<tr><td>Oligo fwd</td><td>CACCGAGATAAGGGAGGGCGCCCC</td><td></td><td></td><td></td><td></td><td></td><td></td><td></td><td></td><td></td><td></td></tr>
<tr><td>Oligo rev</td><td>AAACGGGGCGCCCTCCCTTATCTC</td><td></td><td></td><td></td><td></td><td></td><td></td><td></td><td></td><td></td><td></td></tr>
<tr><td>Chromosome</td><td>start</td><td>end</td><td>strand</td><td>MM</td><td>target_seq</td><td>PAM</td><td>alignment</td><td>distance</td><td>position</td><td>gene name</td><td>gene id</td></tr>
<tr><td>GU305914.1</td><td>79677</td><td>79699</td><td>+</td><td>0</td><td>GAGATAAGGGAGGGCGCCCC</td><td>CGG</td><td>||||||||[||||||||||||]PAM</td><td>NA</td><td>Intergenic</td><td>NA</td><td>NA</td></tr>
<tr><td></td><td></td><td></td><td></td><td></td><td></td><td></td><td></td><td></td><td></td><td></td><td></td></tr>
<tr><td>T5666</td><td>AGATAAGGGAGGGCGCCCCCGGG</td><td>1000</td><td></td><td></td><td></td><td></td><td></td><td></td><td></td><td></td><td></td></tr>
<tr><td>Oligo fwd</td><td>CACCAGATAAGGGAGGGCGCCCCC</td><td></td><td></td><td></td><td></td><td></td><td></td><td></td><td></td><td></td><td></td></tr>
<tr><td>Oligo rev</td><td>AAACGGGGGCGCCCTCCCTTATCT</td><td></td><td></td><td></td><td></td><td></td><td></td><td></td><td></td><td></td><td></td></tr>
<tr><td>Chromosome</td><td>start</td><td>end</td><td>strand</td><td>MM</td><td>target_seq</td><td>PAM</td><td>alignment</td><td>distance</td><td>position</td><td>gene name</td><td>gene id</td></tr>
<tr><td>GU305914.1</td><td>79678</td><td>79700</td><td>+</td><td>0</td><td>AGATAAGGGAGGGCGCCCCC</td><td>GGG</td><td>||||||||[||||||||||||]PAM</td><td>NA</td><td>Intergenic</td><td>NA</td><td>NA</td></tr>
<tr><td></td><td></td><td></td><td></td><td></td><td></td><td></td><td></td><td></td><td></td><td></td><td></td></tr>
<tr><td>T5667</td><td>CGTCTCCATCGCCTCGTGCGAGG</td><td>1000</td><td></td><td></td><td></td><td></td><td></td><td></td><td></td><td></td><td></td></tr>
<tr><td>Oligo fwd</td><td>CACCCGTCTCCATCGCCTCGTGCG</td><td></td><td></td><td></td><td></td><td></td><td></td><td></td><td></td><td></td><td></td></tr>
<tr><td>Oligo rev</td><td>AAACCGCACGAGGCGATGGAGACG</td><td></td><td></td><td></td><td></td><td></td><td></td><td></td><td></td><td></td><td></td></tr>
<tr><td>Chromosome</td><td>start</td><td>end</td><td>strand</td><td>MM</td><td>target_seq</td><td>PAM</td><td>alignment</td><td>distance</td><td>position</td><td>gene name</td><td>gene id</td></tr>
<tr><td>GU305914.1</td><td>79705</td><td>79727</td><td>+</td><td>0</td><td>CGTCTCCATCGCCTCGTGCG</td><td>AGG</td><td>||||||||[||||||||||||]PAM</td><td>NA</td><td>Intergenic</td><td>NA</td><td>NA</td></tr>
<tr><td></td><td></td><td></td><td></td><td></td><td></td><td></td><td></td><td></td><td></td><td></td><td></td></tr>
<tr><td>T5668</td><td>GTTCGTTCACGTTTCCACAGAGG</td><td>1000</td><td></td><td></td><td></td><td></td><td></td><td></td><td></td><td></td><td></td></tr>
<tr><td>Oligo fwd</td><td>CACCGTTCGTTCACGTTTCCACAG</td><td></td><td></td><td></td><td></td><td></td><td></td><td></td><td></td><td></td><td></td></tr>
<tr><td>Oligo rev</td><td>AAACCTGTGGAAACGTGAACGAAC</td><td></td><td></td><td></td><td></td><td></td><td></td><td></td><td></td><td></td><td></td></tr>
<tr><td>Chromosome</td><td>start</td><td>end</td><td>strand</td><td>MM</td><td>target_seq</td><td>PAM</td><td>alignment</td><td>distance</td><td>position</td><td>gene name</td><td>gene id</td></tr>
<tr><td>GU305914.1</td><td>79797</td><td>79819</td><td>+</td><td>0</td><td>GTTCGTTCACGTTTCCACAG</td><td>AGG</td><td>||||||||[||||||||||||]PAM</td><td>NA</td><td>Intergenic</td><td>NA</td><td>NA</td></tr>
<tr><td></td><td></td><td></td><td></td><td></td><td></td><td></td><td></td><td></td><td></td><td></td><td></td></tr>
<tr><td>T5669</td><td>TCACGTTTCCACAGAGGTGTAGG</td><td>1000</td><td></td><td></td><td></td><td></td><td></td><td></td><td></td><td></td><td></td></tr>
<tr><td>Oligo fwd</td><td>CACCTCACGTTTCCACAGAGGTGT</td><td></td><td></td><td></td><td></td><td></td><td></td><td></td><td></td><td></td><td></td></tr>
<tr><td>Oligo rev</td><td>AAACACACCTCTGTGGAAACGTGA</td><td></td><td></td><td></td><td></td><td></td><td></td><td></td><td></td><td></td><td></td></tr>
<tr><td>Chromosome</td><td>start</td><td>end</td><td>strand</td><td>MM</td><td>target_seq</td><td>PAM</td><td>alignment</td><td>distance</td><td>position</td><td>gene name</td><td>gene id</td></tr>
<tr><td>GU305914.1</td><td>79803</td><td>79825</td><td>+</td><td>0</td><td>TCACGTTTCCACAGAGGTGT</td><td>AGG</td><td>||||||||[||||||||||||]PAM</td><td>NA</td><td>Intergenic</td><td>NA</td><td>NA</td></tr>
<tr><td></td><td></td><td></td><td></td><td></td><td></td><td></td><td></td><td></td><td></td><td></td><td></td></tr>
<tr><td>T5670</td><td>TTCCACAGAGGTGTAGGAACTGG</td><td>1000</td><td></td><td></td><td></td><td></td><td></td><td></td><td></td><td></td><td></td></tr>
<tr><td>Oligo fwd</td><td>CACCTTCCACAGAGGTGTAGGAAC</td><td></td><td></td><td></td><td></td><td></td><td></td><td></td><td></td><td></td><td></td></tr>
<tr><td>Oligo rev</td><td>AAACGTTCCTACACCTCTGTGGAA</td><td></td><td></td><td></td><td></td><td></td><td></td><td></td><td></td><td></td><td></td></tr>
<tr><td>Chromosome</td><td>start</td><td>end</td><td>strand</td><td>MM</td><td>target_seq</td><td>PAM</td><td>alignment</td><td>distance</td><td>position</td><td>gene name</td><td>gene id</td></tr>
<tr><td>GU305914.1</td><td>79809</td><td>79831</td><td>+</td><td>0</td><td>TTCCACAGAGGTGTAGGAAC</td><td>TGG</td><td>||||||||[||||||||||||]PAM</td><td>NA</td><td>Intergenic</td><td>NA</td><td>NA</td></tr>
<tr><td></td><td></td><td></td><td></td><td></td><td></td><td></td><td></td><td></td><td></td><td></td><td></td></tr>
<tr><td>T5671</td><td>ACAGAGGTGTAGGAACTGGTCGG</td><td>1000</td><td></td><td></td><td></td><td></td><td></td><td></td><td></td><td></td><td></td></tr>
<tr><td>Oligo fwd</td><td>CACCACAGAGGTGTAGGAACTGGT</td><td></td><td></td><td></td><td></td><td></td><td></td><td></td><td></td><td></td><td></td></tr>
<tr><td>Oligo rev</td><td>AAACACCAGTTCCTACACCTCTGT</td><td></td><td></td><td></td><td></td><td></td><td></td><td></td><td></td><td></td><td></td></tr>
<tr><td>Chromosome</td><td>start</td><td>end</td><td>strand</td><td>MM</td><td>target_seq</td><td>PAM</td><td>alignment</td><td>distance</td><td>position</td><td>gene name</td><td>gene id</td></tr>
<tr><td>GU305914.1</td><td>79813</td><td>79835</td><td>+</td><td>0</td><td>ACAGAGGTGTAGGAACTGGT</td><td>CGG</td><td>||||||||[||||||||||||]PAM</td><td>NA</td><td>Intergenic</td><td>NA</td><td>NA</td></tr>
<tr><td></td><td></td><td></td><td></td><td></td><td></td><td></td><td></td><td></td><td></td><td></td><td></td></tr>
<tr><td>T5672</td><td>CAGAGGTGTAGGAACTGGTCGGG</td><td>1000</td><td></td><td></td><td></td><td></td><td></td><td></td><td></td><td></td><td></td></tr>
<tr><td>Oligo fwd</td><td>CACCCAGAGGTGTAGGAACTGGTC</td><td></td><td></td><td></td><td></td><td></td><td></td><td></td><td></td><td></td><td></td></tr>
<tr><td>Oligo rev</td><td>AAACGACCAGTTCCTACACCTCTG</td><td></td><td></td><td></td><td></td><td></td><td></td><td></td><td></td><td></td><td></td></tr>
<tr><td>Chromosome</td><td>start</td><td>end</td><td>strand</td><td>MM</td><td>target_seq</td><td>PAM</td><td>alignment</td><td>distance</td><td>position</td><td>gene name</td><td>gene id</td></tr>
<tr><td>GU305914.1</td><td>79814</td><td>79836</td><td>+</td><td>0</td><td>CAGAGGTGTAGGAACTGGTC</td><td>GGG</td><td>||||||||[||||||||||||]PAM</td><td>NA</td><td>Intergenic</td><td>NA</td><td>NA</td></tr>
<tr><td></td><td></td><td></td><td></td><td></td><td></td><td></td><td></td><td></td><td></td><td></td><td></td></tr>
<tr><td>T5673</td><td>AGAGGTGTAGGAACTGGTCGGGG</td><td>1000</td><td></td><td></td><td></td><td></td><td></td><td></td><td></td><td></td><td></td></tr>
<tr><td>Oligo fwd</td><td>CACCAGAGGTGTAGGAACTGGTCG</td><td></td><td></td><td></td><td></td><td></td><td></td><td></td><td></td><td></td><td></td></tr>
<tr><td>Oligo rev</td><td>AAACCGACCAGTTCCTACACCTCT</td><td></td><td></td><td></td><td></td><td></td><td></td><td></td><td></td><td></td><td></td></tr>
<tr><td>Chromosome</td><td>start</td><td>end</td><td>strand</td><td>MM</td><td>target_seq</td><td>PAM</td><td>alignment</td><td>distance</td><td>position</td><td>gene name</td><td>gene id</td></tr>
<tr><td>GU305914.1</td><td>79815</td><td>79837</td><td>+</td><td>0</td><td>AGAGGTGTAGGAACTGGTCG</td><td>GGG</td><td>||||||||[||||||||||||]PAM</td><td>NA</td><td>Intergenic</td><td>NA</td><td>NA</td></tr>
<tr><td></td><td></td><td></td><td></td><td></td><td></td><td></td><td></td><td></td><td></td><td></td><td></td></tr>
<tr><td>T5674</td><td>GGTCGGGGATGTAGATCCTCTGG</td><td>1000</td><td></td><td></td><td></td><td></td><td></td><td></td><td></td><td></td><td></td></tr>
<tr><td>Oligo fwd</td><td>CACCGGTCGGGGATGTAGATCCTC</td><td></td><td></td><td></td><td></td><td></td><td></td><td></td><td></td><td></td><td></td></tr>
<tr><td>Oligo rev</td><td>AAACGAGGATCTACATCCCCGACC</td><td></td><td></td><td></td><td></td><td></td><td></td><td></td><td></td><td></td><td></td></tr>
<tr><td>Chromosome</td><td>start</td><td>end</td><td>strand</td><td>MM</td><td>target_seq</td><td>PAM</td><td>alignment</td><td>distance</td><td>position</td><td>gene name</td><td>gene id</td></tr>
<tr><td>GU305914.1</td><td>79830</td><td>79852</td><td>+</td><td>0</td><td>GGTCGGGGATGTAGATCCTC</td><td>TGG</td><td>||||||||[||||||||||||]PAM</td><td>NA</td><td>Intergenic</td><td>NA</td><td>NA</td></tr>
<tr><td></td><td></td><td></td><td></td><td></td><td></td><td></td><td></td><td></td><td></td><td></td><td></td></tr>
<tr><td>T5675</td><td>GTCGGGGATGTAGATCCTCTGGG</td><td>1000</td><td></td><td></td><td></td><td></td><td></td><td></td><td></td><td></td><td></td></tr>
<tr><td>Oligo fwd</td><td>CACCGTCGGGGATGTAGATCCTCT</td><td></td><td></td><td></td><td></td><td></td><td></td><td></td><td></td><td></td><td></td></tr>
<tr><td>Oligo rev</td><td>AAACAGAGGATCTACATCCCCGAC</td><td></td><td></td><td></td><td></td><td></td><td></td><td></td><td></td><td></td><td></td></tr>
<tr><td>Chromosome</td><td>start</td><td>end</td><td>strand</td><td>MM</td><td>target_seq</td><td>PAM</td><td>alignment</td><td>distance</td><td>position</td><td>gene name</td><td>gene id</td></tr>
<tr><td>GU305914.1</td><td>79831</td><td>79853</td><td>+</td><td>0</td><td>GTCGGGGATGTAGATCCTCT</td><td>GGG</td><td>||||||||[||||||||||||]PAM</td><td>NA</td><td>Intergenic</td><td>NA</td><td>NA</td></tr>
<tr><td></td><td></td><td></td><td></td><td></td><td></td><td></td><td></td><td></td><td></td><td></td><td></td></tr>
<tr><td>T5676</td><td>TCGGGGATGTAGATCCTCTGGGG</td><td>1000</td><td></td><td></td><td></td><td></td><td></td><td></td><td></td><td></td><td></td></tr>
<tr><td>Oligo fwd</td><td>CACCTCGGGGATGTAGATCCTCTG</td><td></td><td></td><td></td><td></td><td></td><td></td><td></td><td></td><td></td><td></td></tr>
<tr><td>Oligo rev</td><td>AAACCAGAGGATCTACATCCCCGA</td><td></td><td></td><td></td><td></td><td></td><td></td><td></td><td></td><td></td><td></td></tr>
</table>

Chromosome start end strand MM target_seq PAM alignment distance position gene name gene id

GU305914.1 79832 79854 + 0 TCGGGGATGTAGATCCTCTG GGG ||||||||[||||||||||||]PAM NA Intergenic NA NA

T5677 AGATCCTCTGGGGCACCACGAGG 1000

Oligo fwd CACCAGATCCTCTGGGGCACCACG

Oligo rev AAACCGTGGTGCCCCAGAGGATCT

Chromosome start end strand MM target_seq PAM alignment distance position gene name gene id

GU305914.1 79842 79864 + 0 AGATCCTCTGGGGCACCACG AGG ||||||||[||||||||||||]PAM NA Intergenic NA NA

T5678 GATCCTCTGGGGCACCACGAGGG 1000

Oligo fwd CACCGATCCTCTGGGGCACCACGA

Oligo rev AAACTCGTGGTGCCCCAGAGGATC

Chromosome start end strand MM target_seq PAM alignment distance position gene name gene id

GU305914.1 79843 79865 + 0 GATCCTCTGGGGCACCACGA GGG ||||||||[||||||||||||]PAM NA Intergenic NA NA

T5679 GGAGAAGAACTTCTCCCTCTTGG 1000

Oligo fwd CACCGGAGAAGAACTTCTCCCTCT

Oligo rev AAACAGAGGGAGAAGTTCTTCTCC

Chromosome start end strand MM target_seq PAM alignment distance position gene name gene id

GU305914.1 79864 79886 + 0 GGAGAAGAACTTCTCCCTCT TGG ||||||||[||||||||||||]PAM NA Intergenic NA NA

GU305914.1 79857 79879 - 3 GGAGAAGTTCTTCTCCCTCG TGG |||||||-[-||||||||||-]PAM NA Intergenic NA NA

T5680 GAGAAGAACTTCTCCCTCTTGGG 1000

Oligo fwd CACCGAGAAGAACTTCTCCCTCTT

Oligo rev AAACAAGAGGGAGAAGTTCTTCTC

Chromosome start end strand MM target_seq PAM alignment distance position gene name gene id

GU305914.1 79865 79887 + 0 GAGAAGAACTTCTCCCTCTT GGG ||||||||[||||||||||||]PAM NA Intergenic NA NA

T5681 TCTCCCTCTTGGGCATGTCCCGG 1000

Oligo fwd CACCTCTCCCTCTTGGGCATGTCC

Oligo rev AAACGGACATGCCCAAGAGGGAGA

Chromosome start end strand MM target_seq PAM alignment distance position gene name gene id

GU305914.1 79875 79897 + 0 TCTCCCTCTTGGGCATGTCC CGG ||||||||[||||||||||||]PAM NA Intergenic NA NA

T5682 CTCCCTCTTGGGCATGTCCCGGG 1000

Oligo fwd CACCCTCCCTCTTGGGCATGTCCC

Oligo rev AAACGGGACATGCCCAAGAGGGAG

Chromosome start end strand MM target_seq PAM alignment distance position gene name gene id

GU305914.1 79876 79898 + 0 CTCCCTCTTGGGCATGTCCC GGG ||||||||[||||||||||||]PAM NA Intergenic NA NA

T5683 TCCCTCTTGGGCATGTCCCGGGG 1000

Oligo fwd CACCTCCCTCTTGGGCATGTCCCG

Oligo rev AAACCGGGACATGCCCAAGAGGGA

Chromosome start end strand MM target_seq PAM alignment distance position gene name gene id

GU305914.1 79877 79899 + 0 TCCCTCTTGGGCATGTCCCG GGG ||||||||[||||||||||||]PAM NA Intergenic NA NA

T5684 CCCTCTTGGGCATGTCCCGGGGG 1000

Oligo fwd CACCCCCTCTTGGGCATGTCCCGG

Oligo rev AAACCCGGGACATGCCCAAGAGGG

Chromosome start end strand MM target_seq PAM alignment distance position gene name gene id

GU305914.1 79878 79900 + 0 CCCTCTTGGGCATGTCCCGG GGG ||||||||[||||||||||||]PAM NA Intergenic NA NA

T5685 GGGCATGTCCCGGGGGAAGACGG 1000

Oligo fwd CACCGGGCATGTCCCGGGGGAAGA

Oligo rev AAACTCTTCCCCCGGGACATGCCC

Chromosome start end strand MM target_seq PAM alignment distance position gene name gene id

GU305914.1 79885 79907 + 0 GGGCATGTCCCGGGGGAAGA CGG ||||||||[||||||||||||]PAM NA Intergenic NA NA

T5686 GGCATGTCCCGGGGGAAGACGGG 1000

Oligo fwd CACCGGCATGTCCCGGGGGAAGAC

Oligo rev AAACGTCTTCCCCCGGGACATGCC

Chromosome start end strand MM target_seq PAM alignment distance position gene name gene id

GU305914.1 79886 79908 + 0 GGCATGTCCCGGGGGAAGAC GGG ||||||||[||||||||||||]PAM NA Intergenic NA NA

T5687 GAAGACGGGCATGAGCGTGTTGG 1000

Oligo fwd CACCGAAGACGGGCATGAGCGTGT

Oligo rev AAACACACGCTCATGCCCGTCTTC

Chromosome start end strand MM target_seq PAM alignment distance position gene name gene id

GU305914.1 79900 79922 + 0 GAAGACGGGCATGAGCGTGT TGG ||||||||[||||||||||||]PAM NA Intergenic NA NA

T5688 GGGCATGAGCGTGTTGGTGACGG 1000

Oligo fwd CACCGGGCATGAGCGTGTTGGTGA

Oligo rev AAACTCACCAACACGCTCATGCCC

Chromosome start end strand MM target_seq PAM alignment distance position gene name gene id

GU305914.1 79906 79928 + 0 GGGCATGAGCGTGTTGGTGA CGG ||||||||[||||||||||||]PAM NA Intergenic NA NA

GU305914.1 19474 19496 + 4 GGCGATGAGCGTGGTGGGGA CGG ||--||||[|||||-|||-||]PAM NA Intergenic NA NA

T5689 GACCTGCTCGTAGTACTTTTCGG 1000

Oligo fwd CACCGACCTGCTCGTAGTACTTTT

Oligo rev AAACAAAAGTACTACGAGCAGGTC

Chromosome start end strand MM target_seq PAM alignment distance position gene name gene id

GU305914.1 79936 79958 + 0 GACCTGCTCGTAGTACTTTT CGG ||||||||[||||||||||||]PAM NA Intergenic NA NA

T5690 TTCGGCGTTGATACTGAGTTTGG 1000

Oligo fwd CACCTTCGGCGTTGATACTGAGTT

Oligo rev AAACAACTCAGTATCAACGCCGAA

Chromosome start end strand MM target_seq PAM alignment distance position gene name gene id

GU305914.1 79954 79976 + 0 TTCGGCGTTGATACTGAGTT TGG ||||||||[||||||||||||]PAM NA Intergenic NA NA

<table>
<tr><td>T5691</td><td>TTGGCCTCGACGACGTGTCGCGG</td><td></td><td>1000</td><td></td><td></td><td></td><td></td><td></td><td></td><td></td><td></td><td></td></tr>
<tr><td>Oligo fwd</td><td>CACCTTGGCCTCGACGACGTGTCG</td><td></td><td></td><td></td><td></td><td></td><td></td><td></td><td></td><td></td><td></td><td></td></tr>
<tr><td>Oligo rev</td><td>AAACCGACACGTCGTCGAGGCCAA</td><td></td><td></td><td></td><td></td><td></td><td></td><td></td><td></td><td></td><td></td><td></td></tr>
<tr><td>Chromosome</td><td>start</td><td></td><td>end</td><td>strand</td><td>MM</td><td>target_seq</td><td>PAM</td><td>alignment</td><td>distance</td><td>position</td><td>gene name</td><td>gene id</td></tr>
<tr><td>GU305914.1</td><td></td><td>79973</td><td>79995</td><td>+</td><td>0</td><td>TTGGCCTCGACGACGTGTCG</td><td>CGG</td><td>||||||||[||||||||||||]PAM</td><td>NA</td><td>Intergenic</td><td>NA</td><td>NA</td></tr>
<tr><td>T5692</td><td>GACGACGTGTCGCGGATCCTCGG</td><td></td><td>1000</td><td></td><td></td><td></td><td></td><td></td><td></td><td></td><td></td><td></td></tr>
<tr><td>Oligo fwd</td><td>CACCGACGACGTGTCGCGGATCCT</td><td></td><td></td><td></td><td></td><td></td><td></td><td></td><td></td><td></td><td></td><td></td></tr>
<tr><td>Oligo rev</td><td>AAACAGGATCCGCGACACGTCGTC</td><td></td><td></td><td></td><td></td><td></td><td></td><td></td><td></td><td></td><td></td><td></td></tr>
<tr><td>Chromosome</td><td>start</td><td></td><td>end</td><td>strand</td><td>MM</td><td>target_seq</td><td>PAM</td><td>alignment</td><td>distance</td><td>position</td><td>gene name</td><td>gene id</td></tr>
<tr><td>GU305914.1</td><td></td><td>79981</td><td>80003</td><td>+</td><td>0</td><td>GACGACGTGTCGCGGATCCT</td><td>CGG</td><td>||||||||[||||||||||||]PAM</td><td>NA</td><td>Intergenic</td><td>NA</td><td>NA</td></tr>
<tr><td>T5693</td><td>ATCCTCGGCGATCTCGTAGTTGG</td><td></td><td>1000</td><td></td><td></td><td></td><td></td><td></td><td></td><td></td><td></td><td></td></tr>
<tr><td>Oligo fwd</td><td>CACCATCCTCGGCGATCTCGTAGT</td><td></td><td></td><td></td><td></td><td></td><td></td><td></td><td></td><td></td><td></td><td></td></tr>
<tr><td>Oligo rev</td><td>AAACACTACGAGATCGCCGAGGAT</td><td></td><td></td><td></td><td></td><td></td><td></td><td></td><td></td><td></td><td></td><td></td></tr>
<tr><td>Chromosome</td><td>start</td><td></td><td>end</td><td>strand</td><td>MM</td><td>target_seq</td><td>PAM</td><td>alignment</td><td>distance</td><td>position</td><td>gene name</td><td>gene id</td></tr>
<tr><td>GU305914.1</td><td></td><td>79996</td><td>80018</td><td>+</td><td>0</td><td>ATCCTCGGCGATCTCGTAGT</td><td>TGG</td><td>||||||||[||||||||||||]PAM</td><td>NA</td><td>Intergenic</td><td>NA</td><td>NA</td></tr>
<tr><td>T5694</td><td>TCCTCGGCGATCTCGTAGTTGGG</td><td></td><td>1000</td><td></td><td></td><td></td><td></td><td></td><td></td><td></td><td></td><td></td></tr>
<tr><td>Oligo fwd</td><td>CACCTCCTCGGCGATCTCGTAGTT</td><td></td><td></td><td></td><td></td><td></td><td></td><td></td><td></td><td></td><td></td><td></td></tr>
<tr><td>Oligo rev</td><td>AAACAACTACGAGATCGCCGAGGA</td><td></td><td></td><td></td><td></td><td></td><td></td><td></td><td></td><td></td><td></td><td></td></tr>
<tr><td>Chromosome</td><td>start</td><td></td><td>end</td><td>strand</td><td>MM</td><td>target_seq</td><td>PAM</td><td>alignment</td><td>distance</td><td>position</td><td>gene name</td><td>gene id</td></tr>
<tr><td>GU305914.1</td><td></td><td>79997</td><td>80019</td><td>+</td><td>0</td><td>TCCTCGGCGATCTCGTAGTT</td><td>GGG</td><td>||||||||[||||||||||||]PAM</td><td>NA</td><td>Intergenic</td><td>NA</td><td>NA</td></tr>
<tr><td>T5695</td><td>AACGTTCTTTTTGCAGCCGTCGG</td><td></td><td>1000</td><td></td><td></td><td></td><td></td><td></td><td></td><td></td><td></td><td></td></tr>
<tr><td>Oligo fwd</td><td>CACCAACGTTCTTTTTGCAGCCGT</td><td></td><td></td><td></td><td></td><td></td><td></td><td></td><td></td><td></td><td></td><td></td></tr>
<tr><td>Oligo rev</td><td>AAACACGGCTGCAAAAAGAACGTT</td><td></td><td></td><td></td><td></td><td></td><td></td><td></td><td></td><td></td><td></td><td></td></tr>
<tr><td>Chromosome</td><td>start</td><td></td><td>end</td><td>strand</td><td>MM</td><td>target_seq</td><td>PAM</td><td>alignment</td><td>distance</td><td>position</td><td>gene name</td><td>gene id</td></tr>
<tr><td>GU305914.1</td><td></td><td>80020</td><td>80042</td><td>+</td><td>0</td><td>AACGTTCTTTTTGCAGCCGT</td><td>CGG</td><td>||||||||[||||||||||||]PAM</td><td>NA</td><td>Intergenic</td><td>NA</td><td>NA</td></tr>
<tr><td>T5696</td><td>ACGTTCTTTTTGCAGCCGTCGGG</td><td></td><td>1000</td><td></td><td></td><td></td><td></td><td></td><td></td><td></td><td></td><td></td></tr>
<tr><td>Oligo fwd</td><td>CACCACGTTCTTTTTGCAGCCGTC</td><td></td><td></td><td></td><td></td><td></td><td></td><td></td><td></td><td></td><td></td><td></td></tr>
<tr><td>Oligo rev</td><td>AAACGACGGCTGCAAAAAGAACGT</td><td></td><td></td><td></td><td></td><td></td><td></td><td></td><td></td><td></td><td></td><td></td></tr>
<tr><td>Chromosome</td><td>start</td><td></td><td>end</td><td>strand</td><td>MM</td><td>target_seq</td><td>PAM</td><td>alignment</td><td>distance</td><td>position</td><td>gene name</td><td>gene id</td></tr>
<tr><td>GU305914.1</td><td></td><td>80021</td><td>80043</td><td>+</td><td>0</td><td>ACGTTCTTTTTGCAGCCGTC</td><td>GGG</td><td>||||||||[||||||||||||]PAM</td><td>NA</td><td>Intergenic</td><td>NA</td><td>NA</td></tr>
<tr><td>T5697</td><td>TCTTTTTGCAGCCGTCGGGCAGG</td><td></td><td>1000</td><td></td><td></td><td></td><td></td><td></td><td></td><td></td><td></td><td></td></tr>
<tr><td>Oligo fwd</td><td>CACCTCTTTTTGCAGCCGTCGGGC</td><td></td><td></td><td></td><td></td><td></td><td></td><td></td><td></td><td></td><td></td><td></td></tr>
<tr><td>Oligo rev</td><td>AAACGCCCGACGGCTGCAAAAAGA</td><td></td><td></td><td></td><td></td><td></td><td></td><td></td><td></td><td></td><td></td><td></td></tr>
<tr><td>Chromosome</td><td>start</td><td></td><td>end</td><td>strand</td><td>MM</td><td>target_seq</td><td>PAM</td><td>alignment</td><td>distance</td><td>position</td><td>gene name</td><td>gene id</td></tr>
<tr><td>GU305914.1</td><td></td><td>80025</td><td>80047</td><td>+</td><td>0</td><td>TCTTTTTGCAGCCGTCGGGC</td><td>AGG</td><td>||||||||[||||||||||||]PAM</td><td>NA</td><td>Intergenic</td><td>NA</td><td>NA</td></tr>
<tr><td>T5698</td><td>CTTTTTGCAGCCGTCGGGCAGGG</td><td></td><td>1000</td><td></td><td></td><td></td><td></td><td></td><td></td><td></td><td></td><td></td></tr>
<tr><td>Oligo fwd</td><td>CACCCTTTTTGCAGCCGTCGGGCA</td><td></td><td></td><td></td><td></td><td></td><td></td><td></td><td></td><td></td><td></td><td></td></tr>
<tr><td>Oligo rev</td><td>AAACTGCCCGACGGCTGCAAAAAG</td><td></td><td></td><td></td><td></td><td></td><td></td><td></td><td></td><td></td><td></td><td></td></tr>
<tr><td>Chromosome</td><td>start</td><td></td><td>end</td><td>strand</td><td>MM</td><td>target_seq</td><td>PAM</td><td>alignment</td><td>distance</td><td>position</td><td>gene name</td><td>gene id</td></tr>
<tr><td>GU305914.1</td><td></td><td>80026</td><td>80048</td><td>+</td><td>0</td><td>CTTTTTGCAGCCGTCGGGCA</td><td>GGG</td><td>||||||||[||||||||||||]PAM</td><td>NA</td><td>Intergenic</td><td>NA</td><td>NA</td></tr>
<tr><td>T5699</td><td>TGCAGCCGTCGGGCAGGGTGAGG</td><td></td><td>1000</td><td></td><td></td><td></td><td></td><td></td><td></td><td></td><td></td><td></td></tr>
<tr><td>Oligo fwd</td><td>CACCTGCAGCCGTCGGGCAGGGTG</td><td></td><td></td><td></td><td></td><td></td><td></td><td></td><td></td><td></td><td></td><td></td></tr>
<tr><td>Oligo rev</td><td>AAACCACCCTGCCCGACGGCTGCA</td><td></td><td></td><td></td><td></td><td></td><td></td><td></td><td></td><td></td><td></td><td></td></tr>
<tr><td>Chromosome</td><td>start</td><td></td><td>end</td><td>strand</td><td>MM</td><td>target_seq</td><td>PAM</td><td>alignment</td><td>distance</td><td>position</td><td>gene name</td><td>gene id</td></tr>
<tr><td>GU305914.1</td><td></td><td>80031</td><td>80053</td><td>+</td><td>0</td><td>TGCAGCCGTCGGGCAGGGTG</td><td>AGG</td><td>||||||||[||||||||||||]PAM</td><td>NA</td><td>Intergenic</td><td>NA</td><td>NA</td></tr>
<tr><td>T5700</td><td>GGGTGAGGACGTACTCGACACGG</td><td></td><td>1000</td><td></td><td></td><td></td><td></td><td></td><td></td><td></td><td></td><td></td></tr>
<tr><td>Oligo fwd</td><td>CACCGGGTGAGGACGTACTCGACA</td><td></td><td></td><td></td><td></td><td></td><td></td><td></td><td></td><td></td><td></td><td></td></tr>
<tr><td>Oligo rev</td><td>AAACTGTCGAGTACGTCCTCACCC</td><td></td><td></td><td></td><td></td><td></td><td></td><td></td><td></td><td></td><td></td><td></td></tr>
<tr><td>Chromosome</td><td>start</td><td></td><td>end</td><td>strand</td><td>MM</td><td>target_seq</td><td>PAM</td><td>alignment</td><td>distance</td><td>position</td><td>gene name</td><td>gene id</td></tr>
<tr><td>GU305914.1</td><td></td><td>80046</td><td>80068</td><td>+</td><td>0</td><td>GGGTGAGGACGTACTCGACA</td><td>CGG</td><td>||||||||[||||||||||||]PAM</td><td>NA</td><td>Intergenic</td><td>NA</td><td>NA</td></tr>
<tr><td>T5701</td><td>CTCGACACGGTCGCCGACCGCGG</td><td></td><td>1000</td><td></td><td></td><td></td><td></td><td></td><td></td><td></td><td></td><td></td></tr>
<tr><td>Oligo fwd</td><td>CACCCTCGACACGGTCGCCGACCG</td><td></td><td></td><td></td><td></td><td></td><td></td><td></td><td></td><td></td><td></td><td></td></tr>
<tr><td>Oligo rev</td><td>AAACCGGTCGGCGACCGTGTCGAG</td><td></td><td></td><td></td><td></td><td></td><td></td><td></td><td></td><td></td><td></td><td></td></tr>
<tr><td>Chromosome</td><td>start</td><td></td><td>end</td><td>strand</td><td>MM</td><td>target_seq</td><td>PAM</td><td>alignment</td><td>distance</td><td>position</td><td>gene name</td><td>gene id</td></tr>
<tr><td>GU305914.1</td><td></td><td>80059</td><td>80081</td><td>+</td><td>0</td><td>CTCGACACGGTCGCCGACCG</td><td>CGG</td><td>||||||||[||||||||||||]PAM</td><td>NA</td><td>Intergenic</td><td>NA</td><td>NA</td></tr>
<tr><td>T5702</td><td>TCGACACGGTCGCCGACCGCGGG</td><td></td><td>1000</td><td></td><td></td><td></td><td></td><td></td><td></td><td></td><td></td><td></td></tr>
<tr><td>Oligo fwd</td><td>CACCTCGACACGGTCGCCGACCGC</td><td></td><td></td><td></td><td></td><td></td><td></td><td></td><td></td><td></td><td></td><td></td></tr>
<tr><td>Oligo rev</td><td>AAACGCGGTCGGCGACCGTGTCGA</td><td></td><td></td><td></td><td></td><td></td><td></td><td></td><td></td><td></td><td></td><td></td></tr>
<tr><td>Chromosome</td><td>start</td><td></td><td>end</td><td>strand</td><td>MM</td><td>target_seq</td><td>PAM</td><td>alignment</td><td>distance</td><td>position</td><td>gene name</td><td>gene id</td></tr>
<tr><td>GU305914.1</td><td></td><td>80060</td><td>80082</td><td>+</td><td>0</td><td>TCGACACGGTCGCCGACCGC</td><td>GGG</td><td>||||||||[||||||||||||]PAM</td><td>NA</td><td>Intergenic</td><td>NA</td><td>NA</td></tr>
<tr><td>T5703</td><td>CCTTCCGAGCCGCCAGACGTCGG</td><td></td><td>1000</td><td></td><td></td><td></td><td></td><td></td><td></td><td></td><td></td><td></td></tr>
<tr><td>Oligo fwd</td><td>CACCCCTTCCGAGCCGCCAGACGT</td><td></td><td></td><td></td><td></td><td></td><td></td><td></td><td></td><td></td><td></td><td></td></tr>
<tr><td>Oligo rev</td><td>AAACACGTCTGGCGGCTCGGAAGG</td><td></td><td></td><td></td><td></td><td></td><td></td><td></td><td></td><td></td><td></td><td></td></tr>
<tr><td>Chromosome</td><td>start</td><td></td><td>end</td><td>strand</td><td>MM</td><td>target_seq</td><td>PAM</td><td>alignment</td><td>distance</td><td>position</td><td>gene name</td><td>gene id</td></tr>
<tr><td>GU305914.1</td><td></td><td>80091</td><td>80113</td><td>+</td><td>0</td><td>CCTTCCGAGCCGCCAGACGT</td><td>CGG</td><td>||||||||[||||||||||||]PAM</td><td>NA</td><td>Intergenic</td><td>NA</td><td>NA</td></tr>
<tr><td>T5704</td><td>AGCCGCCAGACGTCGGATGACGG</td><td></td><td>1000</td><td></td><td></td><td></td><td></td><td></td><td></td><td></td><td></td><td></td></tr>
<tr><td>Oligo fwd</td><td>CACCAGCCGCCAGACGTCGGATGA</td><td></td><td></td><td></td><td></td><td></td><td></td><td></td><td></td><td></td><td></td><td></td></tr>
<tr><td>Oligo rev</td><td>AAACTCATCCGACGTCTGGCGGCT</td><td></td><td></td><td></td><td></td><td></td><td></td><td></td><td></td><td></td><td></td><td></td></tr>
<tr><td>Chromosome</td><td>start</td><td></td><td>end</td><td>strand</td><td>MM</td><td>target_seq</td><td>PAM</td><td>alignment</td><td>distance</td><td>position</td><td>gene name</td><td>gene id</td></tr>
<tr><td>GU305914.1</td><td></td><td>80098</td><td>80120</td><td>+</td><td>0</td><td>AGCCGCCAGACGTCGGATGA</td><td>CGG</td><td>||||||||[||||||||||||]PAM</td><td>NA</td><td>Intergenic</td><td>NA</td><td>NA</td></tr>
<tr><td>T5705</td><td>CCAGACGTCGGATGACGGCAAGG</td><td></td><td>1000</td><td></td><td></td><td></td><td></td><td></td><td></td><td></td><td></td><td></td></tr>
<tr><td>Oligo fwd</td><td>CACCCCAGACGTCGGATGACGGCA</td><td></td><td></td><td></td><td></td><td></td><td></td><td></td><td></td><td></td><td></td><td></td></tr>
<tr><td>Oligo rev</td><td>AAACTGCCGTCATCCGACGTCTGG</td><td></td><td></td><td></td><td></td><td></td><td></td><td></td><td></td><td></td><td></td><td></td></tr>
<tr><td>Chromosome</td><td>start</td><td></td><td>end</td><td>strand</td><td>MM</td><td>target_seq</td><td>PAM</td><td>alignment</td><td>distance</td><td>position</td><td>gene name</td><td>gene id</td></tr>
<tr><td>GU305914.1</td><td></td><td>80103</td><td>80125</td><td>+</td><td>0</td><td>CCAGACGTCGGATGACGGCA</td><td>AGG</td><td>||||||||[||||||||||||]PAM</td><td>NA</td><td>Intergenic</td><td>NA</td><td>NA</td></tr>
</table>

<table>
<tr><td>T5706</td><td>CGTCGGATGACGGCAAGGTGAGG</td><td>1000</td><td></td><td></td><td></td><td></td><td></td><td></td><td></td><td></td><td></td></tr>
<tr><td>Oligo fwd</td><td>CACCCGTCGGATGACGGCAAGGTG</td><td></td><td></td><td></td><td></td><td></td><td></td><td></td><td></td><td></td><td></td></tr>
<tr><td>Oligo rev</td><td>AAACCACCTTGCCGTCATCCGACG</td><td></td><td></td><td></td><td></td><td></td><td></td><td></td><td></td><td></td><td></td></tr>
<tr><td>Chromosome</td><td>start</td><td>end</td><td>strand</td><td>MM</td><td>target_seq</td><td>PAM</td><td>alignment</td><td>distance</td><td>position</td><td>gene name</td><td>gene id</td></tr>
<tr><td>GU305914.1</td><td>80108</td><td>80130</td><td>+</td><td>0</td><td>CGTCGGATGACGGCAAGGTG</td><td>AGG</td><td>||||||||[||||||||||||]PAM</td><td>NA</td><td>Intergenic</td><td>NA</td><td>NA</td></tr>
<tr><td>T5707</td><td>GGATGACGGCAAGGTGAGGCAGG</td><td>1000</td><td></td><td></td><td></td><td></td><td></td><td></td><td></td><td></td><td></td></tr>
<tr><td>Oligo fwd</td><td>CACCGGATGACGGCAAGGTGAGGC</td><td></td><td></td><td></td><td></td><td></td><td></td><td></td><td></td><td></td><td></td></tr>
<tr><td>Oligo rev</td><td>AAACGCCTCACCTTGCCGTCATCC</td><td></td><td></td><td></td><td></td><td></td><td></td><td></td><td></td><td></td><td></td></tr>
<tr><td>Chromosome</td><td>start</td><td>end</td><td>strand</td><td>MM</td><td>target_seq</td><td>PAM</td><td>alignment</td><td>distance</td><td>position</td><td>gene name</td><td>gene id</td></tr>
<tr><td>GU305914.1</td><td>80112</td><td>80134</td><td>+</td><td>0</td><td>GGATGACGGCAAGGTGAGGC</td><td>AGG</td><td>||||||||[||||||||||||]PAM</td><td>NA</td><td>Intergenic</td><td>NA</td><td>NA</td></tr>
<tr><td>T5708</td><td>CAGGTTTTTCTGTTTATAGCAGG</td><td>1000</td><td></td><td></td><td></td><td></td><td></td><td></td><td></td><td></td><td></td></tr>
<tr><td>Oligo fwd</td><td>CACCCAGGTTTTTCTGTTTATAGC</td><td></td><td></td><td></td><td></td><td></td><td></td><td></td><td></td><td></td><td></td></tr>
<tr><td>Oligo rev</td><td>AAACGCTATAAACAGAAAAACCTG</td><td></td><td></td><td></td><td></td><td></td><td></td><td></td><td></td><td></td><td></td></tr>
<tr><td>Chromosome</td><td>start</td><td>end</td><td>strand</td><td>MM</td><td>target_seq</td><td>PAM</td><td>alignment</td><td>distance</td><td>position</td><td>gene name</td><td>gene id</td></tr>
<tr><td>GU305914.1</td><td>80131</td><td>80153</td><td>+</td><td>0</td><td>CAGGTTTTTCTGTTTATAGC</td><td>AGG</td><td>||||||||[||||||||||||]PAM</td><td>NA</td><td>Intergenic</td><td>NA</td><td>NA</td></tr>
<tr><td>T5709</td><td>GTTTATAGCAGGAGAGCTCTTGG</td><td>1000</td><td></td><td></td><td></td><td></td><td></td><td></td><td></td><td></td><td></td></tr>
<tr><td>Oligo fwd</td><td>CACCGTTTATAGCAGGAGAGCTCT</td><td></td><td></td><td></td><td></td><td></td><td></td><td></td><td></td><td></td><td></td></tr>
<tr><td>Oligo rev</td><td>AAACAGAGCTCTCCTGCTATAAAC</td><td></td><td></td><td></td><td></td><td></td><td></td><td></td><td></td><td></td><td></td></tr>
<tr><td>Chromosome</td><td>start</td><td>end</td><td>strand</td><td>MM</td><td>target_seq</td><td>PAM</td><td>alignment</td><td>distance</td><td>position</td><td>gene name</td><td>gene id</td></tr>
<tr><td>GU305914.1</td><td>80142</td><td>80164</td><td>+</td><td>0</td><td>GTTTATAGCAGGAGAGCTCT</td><td>TGG</td><td>||||||||[||||||||||||]PAM</td><td>NA</td><td>Intergenic</td><td>NA</td><td>NA</td></tr>
<tr><td>T5710</td><td>TTTATAGCAGGAGAGCTCTTGGG</td><td>1000</td><td></td><td></td><td></td><td></td><td></td><td></td><td></td><td></td><td></td></tr>
<tr><td>Oligo fwd</td><td>CACCTTTATAGCAGGAGAGCTCTT</td><td></td><td></td><td></td><td></td><td></td><td></td><td></td><td></td><td></td><td></td></tr>
<tr><td>Oligo rev</td><td>AAACAAGAGCTCTCCTGCTATAAA</td><td></td><td></td><td></td><td></td><td></td><td></td><td></td><td></td><td></td><td></td></tr>
<tr><td>Chromosome</td><td>start</td><td>end</td><td>strand</td><td>MM</td><td>target_seq</td><td>PAM</td><td>alignment</td><td>distance</td><td>position</td><td>gene name</td><td>gene id</td></tr>
<tr><td>GU305914.1</td><td>80143</td><td>80165</td><td>+</td><td>0</td><td>TTTATAGCAGGAGAGCTCTT</td><td>GGG</td><td>||||||||[||||||||||||]PAM</td><td>NA</td><td>Intergenic</td><td>NA</td><td>NA</td></tr>
<tr><td>T5711</td><td>AGCAGGAGAGCTCTTGGGAGAGG</td><td>1000</td><td></td><td></td><td></td><td></td><td></td><td></td><td></td><td></td><td></td></tr>
<tr><td>Oligo fwd</td><td>CACCAGCAGGAGAGCTCTTGGGAG</td><td></td><td></td><td></td><td></td><td></td><td></td><td></td><td></td><td></td><td></td></tr>
<tr><td>Oligo rev</td><td>AAACCTCCCAAGAGCTCTCCTGCT</td><td></td><td></td><td></td><td></td><td></td><td></td><td></td><td></td><td></td><td></td></tr>
<tr><td>Chromosome</td><td>start</td><td>end</td><td>strand</td><td>MM</td><td>target_seq</td><td>PAM</td><td>alignment</td><td>distance</td><td>position</td><td>gene name</td><td>gene id</td></tr>
<tr><td>GU305914.1</td><td>80148</td><td>80170</td><td>+</td><td>0</td><td>AGCAGGAGAGCTCTTGGGAG</td><td>AGG</td><td>||||||||[||||||||||||]PAM</td><td>NA</td><td>Intergenic</td><td>NA</td><td>NA</td></tr>
<tr><td>T5712</td><td>GGAGAGCTCTTGGGAGAGGACGG</td><td>1000</td><td></td><td></td><td></td><td></td><td></td><td></td><td></td><td></td><td></td></tr>
<tr><td>Oligo fwd</td><td>CACCGGAGAGCTCTTGGGAGAGGA</td><td></td><td></td><td></td><td></td><td></td><td></td><td></td><td></td><td></td><td></td></tr>
<tr><td>Oligo rev</td><td>AAACTCCTCTCCCAAGAGCTCTCC</td><td></td><td></td><td></td><td></td><td></td><td></td><td></td><td></td><td></td><td></td></tr>
<tr><td>Chromosome</td><td>start</td><td>end</td><td>strand</td><td>MM</td><td>target_seq</td><td>PAM</td><td>alignment</td><td>distance</td><td>position</td><td>gene name</td><td>gene id</td></tr>
<tr><td>GU305914.1</td><td>80152</td><td>80174</td><td>+</td><td>0</td><td>GGAGAGCTCTTGGGAGAGGA</td><td>CGG</td><td>||||||||[||||||||||||]PAM</td><td>NA</td><td>Intergenic</td><td>NA</td><td>NA</td></tr>
<tr><td>T5713</td><td>GAGCTCTTGGGAGAGGACGGAGG</td><td>1000</td><td></td><td></td><td></td><td></td><td></td><td></td><td></td><td></td><td></td></tr>
<tr><td>Oligo fwd</td><td>CACCGAGCTCTTGGGAGAGGACGG</td><td></td><td></td><td></td><td></td><td></td><td></td><td></td><td></td><td></td><td></td></tr>
<tr><td>Oligo rev</td><td>AAACCCGTCCTCTCCCAAGAGCTC</td><td></td><td></td><td></td><td></td><td></td><td></td><td></td><td></td><td></td><td></td></tr>
<tr><td>Chromosome</td><td>start</td><td>end</td><td>strand</td><td>MM</td><td>target_seq</td><td>PAM</td><td>alignment</td><td>distance</td><td>position</td><td>gene name</td><td>gene id</td></tr>
<tr><td>GU305914.1</td><td>80155</td><td>80177</td><td>+</td><td>0</td><td>GAGCTCTTGGGAGAGGACGG</td><td>AGG</td><td>||||||||[||||||||||||]PAM</td><td>NA</td><td>Intergenic</td><td>NA</td><td>NA</td></tr>
<tr><td>T5714</td><td>CTTGGGAGAGGACGGAGGAGAGG</td><td>1000</td><td></td><td></td><td></td><td></td><td></td><td></td><td></td><td></td><td></td></tr>
<tr><td>Oligo fwd</td><td>CACCCTTGGGAGAGGACGGAGGAG</td><td></td><td></td><td></td><td></td><td></td><td></td><td></td><td></td><td></td><td></td></tr>
<tr><td>Oligo rev</td><td>AAACCTCCTCCGTCCTCTCCCAAG</td><td></td><td></td><td></td><td></td><td></td><td></td><td></td><td></td><td></td><td></td></tr>
<tr><td>Chromosome</td><td>start</td><td>end</td><td>strand</td><td>MM</td><td>target_seq</td><td>PAM</td><td>alignment</td><td>distance</td><td>position</td><td>gene name</td><td>gene id</td></tr>
<tr><td>GU305914.1</td><td>80160</td><td>80182</td><td>+</td><td>0</td><td>CTTGGGAGAGGACGGAGGAG</td><td>AGG</td><td>||||||||[||||||||||||]PAM</td><td>NA</td><td>Intergenic</td><td>NA</td><td>NA</td></tr>
<tr><td>T5715</td><td>TTGGGAGAGGACGGAGGAGAGGG</td><td>1000</td><td></td><td></td><td></td><td></td><td></td><td></td><td></td><td></td><td></td></tr>
<tr><td>Oligo fwd</td><td>CACCTTGGGAGAGGACGGAGGAGA</td><td></td><td></td><td></td><td></td><td></td><td></td><td></td><td></td><td></td><td></td></tr>
<tr><td>Oligo rev</td><td>AAACTCTCCTCCGTCCTCTCCCAA</td><td></td><td></td><td></td><td></td><td></td><td></td><td></td><td></td><td></td><td></td></tr>
<tr><td>Chromosome</td><td>start</td><td>end</td><td>strand</td><td>MM</td><td>target_seq</td><td>PAM</td><td>alignment</td><td>distance</td><td>position</td><td>gene name</td><td>gene id</td></tr>
<tr><td>GU305914.1</td><td>80161</td><td>80183</td><td>+</td><td>0</td><td>TTGGGAGAGGACGGAGGAGA</td><td>GGG</td><td>||||||||[||||||||||||]PAM</td><td>NA</td><td>Intergenic</td><td>NA</td><td>NA</td></tr>
<tr><td>T5716</td><td>GACGGAGGAGAGGGTGAGTTCGG</td><td>1000</td><td></td><td></td><td></td><td></td><td></td><td></td><td></td><td></td><td></td></tr>
<tr><td>Oligo fwd</td><td>CACCGACGGAGGAGAGGGTGAGTT</td><td></td><td></td><td></td><td></td><td></td><td></td><td></td><td></td><td></td><td></td></tr>
<tr><td>Oligo rev</td><td>AAACAACTCACCCTCTCCTCCGTC</td><td></td><td></td><td></td><td></td><td></td><td></td><td></td><td></td><td></td><td></td></tr>
<tr><td>Chromosome</td><td>start</td><td>end</td><td>strand</td><td>MM</td><td>target_seq</td><td>PAM</td><td>alignment</td><td>distance</td><td>position</td><td>gene name</td><td>gene id</td></tr>
<tr><td>GU305914.1</td><td>80170</td><td>80192</td><td>+</td><td>0</td><td>GACGGAGGAGAGGGTGAGTT</td><td>CGG</td><td>||||||||[||||||||||||]PAM</td><td>NA</td><td>Intergenic</td><td>NA</td><td>NA</td></tr>
<tr><td>T5717</td><td>ACGGAGGAGAGGGTGAGTTCGGG</td><td>1000</td><td></td><td></td><td></td><td></td><td></td><td></td><td></td><td></td><td></td></tr>
<tr><td>Oligo fwd</td><td>CACCACGGAGGAGAGGGTGAGTTC</td><td></td><td></td><td></td><td></td><td></td><td></td><td></td><td></td><td></td><td></td></tr>
<tr><td>Oligo rev</td><td>AAACGAACTCACCCTCTCCTCCGT</td><td></td><td></td><td></td><td></td><td></td><td></td><td></td><td></td><td></td><td></td></tr>
<tr><td>Chromosome</td><td>start</td><td>end</td><td>strand</td><td>MM</td><td>target_seq</td><td>PAM</td><td>alignment</td><td>distance</td><td>position</td><td>gene name</td><td>gene id</td></tr>
<tr><td>GU305914.1</td><td>80171</td><td>80193</td><td>+</td><td>0</td><td>ACGGAGGAGAGGGTGAGTTC</td><td>GGG</td><td>||||||||[||||||||||||]PAM</td><td>NA</td><td>Intergenic</td><td>NA</td><td>NA</td></tr>
<tr><td>T5718</td><td>GGAGAGGGTGAGTTCGGGCACGG</td><td>1000</td><td></td><td></td><td></td><td></td><td></td><td></td><td></td><td></td><td></td></tr>
<tr><td>Oligo fwd</td><td>CACCGGAGAGGGTGAGTTCGGGCA</td><td></td><td></td><td></td><td></td><td></td><td></td><td></td><td></td><td></td><td></td></tr>
<tr><td>Oligo rev</td><td>AAACTGCCCGAACTCACCCTCTCC</td><td></td><td></td><td></td><td></td><td></td><td></td><td></td><td></td><td></td><td></td></tr>
<tr><td>Chromosome</td><td>start</td><td>end</td><td>strand</td><td>MM</td><td>target_seq</td><td>PAM</td><td>alignment</td><td>distance</td><td>position</td><td>gene name</td><td>gene id</td></tr>
<tr><td>GU305914.1</td><td>80176</td><td>80198</td><td>+</td><td>0</td><td>GGAGAGGGTGAGTTCGGGCA</td><td>CGG</td><td>||||||||[||||||||||||]PAM</td><td>NA</td><td>Intergenic</td><td>NA</td><td>NA</td></tr>
<tr><td>T5719</td><td>GAGAGGGTGAGTTCGGGCACGGG</td><td>1000</td><td></td><td></td><td></td><td></td><td></td><td></td><td></td><td></td><td></td></tr>
<tr><td>Oligo fwd</td><td>CACCGAGAGGGTGAGTTCGGGCAC</td><td></td><td></td><td></td><td></td><td></td><td></td><td></td><td></td><td></td><td></td></tr>
<tr><td>Oligo rev</td><td>AAACGTGCCCGAACTCACCCTCTC</td><td></td><td></td><td></td><td></td><td></td><td></td><td></td><td></td><td></td><td></td></tr>
<tr><td>Chromosome</td><td>start</td><td>end</td><td>strand</td><td>MM</td><td>target_seq</td><td>PAM</td><td>alignment</td><td>distance</td><td>position</td><td>gene name</td><td>gene id</td></tr>
<tr><td>GU305914.1</td><td>80177</td><td>80199</td><td>+</td><td>0</td><td>GAGAGGGTGAGTTCGGGCAC</td><td>GGG</td><td>||||||||[||||||||||||]PAM</td><td>NA</td><td>Intergenic</td><td>NA</td><td>NA</td></tr>
<tr><td>T5720</td><td>CGGGCACGGGCACCTTCTTCAGG</td><td>1000</td><td></td><td></td><td></td><td></td><td></td><td></td><td></td><td></td><td></td></tr>
<tr><td>Oligo fwd</td><td>CACCCGGGCACGGGCACCTTCTTC</td><td></td><td></td><td></td><td></td><td></td><td></td><td></td><td></td><td></td><td></td></tr>
<tr><td>Oligo rev</td><td>AAACGAAGAAGGTGCCCGTGCCCG</td><td></td><td></td><td></td><td></td><td></td><td></td><td></td><td></td><td></td><td></td></tr>
<tr><td>Chromosome</td><td>start</td><td>end</td><td>strand</td><td>MM</td><td>target_seq</td><td>PAM</td><td>alignment</td><td>distance</td><td>position</td><td>gene name</td><td>gene id</td></tr>
</table>

| | | | | | | | | | | | |
|---|---|---|---|---|---|---|---|---|---|---|---|
| GU305914.1 | 80190 | 80212 | + | 0 | CGGGCACGGGCACCTTCTTC | AGG | \|\|\|\|\|\|\|\|[\|\|\|\|\|\|\|\|\|\|\|\|]PAM | NA | Intergenic | NA | NA |
| T5721 | CGGGCACCTTCTTCAGGTGCAGG | 1000 | | | | | | | | | |
| Oligo fwd | CACCCGGGCACCTTCTTCAGGTGC | | | | | | | | | | |
| Oligo rev | AAACGCACCTGAAGAAGGTGCCCG | | | | | | | | | | |
| Chromosome | start | end | strand | MM | target_seq | PAM | alignment | distance | position | gene name | gene id |
| GU305914.1 | 80196 | 80218 | + | 0 | CGGGCACCTTCTTCAGGTGC | AGG | \|\|\|\|\|\|\|\|[\|\|\|\|\|\|\|\|\|\|\|\|]PAM | NA | Intergenic | NA | NA |
| GU305914.1 | 77335 | 77357 | - | 4 | AGTGCAGCTTCATCAGGTGC | AGG | -\|-\|\|\|-\|[\|\|\|-\|\|\|\|\|\|\|\|]PAM | NA | Intergenic | NA | NA |
| T5722 | CTTCAGGTGCAGGTCGTCCCTGG | 1000 | | | | | | | | | |
| Oligo fwd | CACCCTTCAGGTGCAGGTCGTCCC | | | | | | | | | | |
| Oligo rev | AAACGGGACGACCTGCACCTGAAG | | | | | | | | | | |
| Chromosome | start | end | strand | MM | target_seq | PAM | alignment | distance | position | gene name | gene id |
| GU305914.1 | 80206 | 80228 | + | 0 | CTTCAGGTGCAGGTCGTCCC | TGG | \|\|\|\|\|\|\|\|[\|\|\|\|\|\|\|\|\|\|\|\|]PAM | NA | Intergenic | NA | NA |
| T5723 | GGTGCAGGTCGTCCCTGGCGCGG | 1000 | | | | | | | | | |
| Oligo fwd | CACCGGTGCAGGTCGTCCCTGGCG | | | | | | | | | | |
| Oligo rev | AAACCGCCAGGGACGACCTGCACC | | | | | | | | | | |
| Chromosome | start | end | strand | MM | target_seq | PAM | alignment | distance | position | gene name | gene id |
| GU305914.1 | 80211 | 80233 | + | 0 | GGTGCAGGTCGTCCCTGGCG | CGG | \|\|\|\|\|\|\|\|[\|\|\|\|\|\|\|\|\|\|\|\|]PAM | NA | Intergenic | NA | NA |
| GU305914.1 | 61986 | 62008 | + | 4 | GGCGCATCTCGTCCCTGACG | GGG | \|\|-\|\|\|--[\|\|\|\|\|\|\|\|\|-\|\|]PAM | NA | Intergenic | NA | NA |
| T5724 | GGTCGTCCCTGGCGCGGCAGAGG | 1000 | | | | | | | | | |
| Oligo fwd | CACCGGTCGTCCCTGGCGCGGCAG | | | | | | | | | | |
| Oligo rev | AAACCTGCCGCGCCAGGGACGACC | | | | | | | | | | |
| Chromosome | start | end | strand | MM | target_seq | PAM | alignment | distance | position | gene name | gene id |
| GU305914.1 | 80217 | 80239 | + | 0 | GGTCGTCCCTGGCGCGGCAG | AGG | \|\|\|\|\|\|\|\|[\|\|\|\|\|\|\|\|\|\|\|\|]PAM | NA | Intergenic | NA | NA |
| T5725 | CGTCCCTGGCGCGGCAGAGGCGG | 1000 | | | | | | | | | |
| Oligo fwd | CACCCGTCCCTGGCGCGGCAGAGG | | | | | | | | | | |
| Oligo rev | AAACCCTCTGCCGCGCCAGGGACG | | | | | | | | | | |
| Chromosome | start | end | strand | MM | target_seq | PAM | alignment | distance | position | gene name | gene id |
| GU305914.1 | 80220 | 80242 | + | 0 | CGTCCCTGGCGCGGCAGAGG | CGG | \|\|\|\|\|\|\|\|[\|\|\|\|\|\|\|\|\|\|\|\|]PAM | NA | Intergenic | NA | NA |
| GU305914.1 | 128089 | 128111 | - | 4 | GGTACCTGGCGAGGCAGACG | AGG | -\|\|-\|\|\|\|[\|\|\|-\|\|\|\|\|\|-\|]PAM | NA | Intergenic | NA | NA |
| T5726 | GCGGCAGAGGCGGCTGACGACGG | 1000 | | | | | | | | | |
| Oligo fwd | CACCGCGGCAGAGGCGGCTGACGA | | | | | | | | | | |
| Oligo rev | AAACTCGTCAGCCGCCTCTGCCGC | | | | | | | | | | |
| Chromosome | start | end | strand | MM | target_seq | PAM | alignment | distance | position | gene name | gene id |
| GU305914.1 | 80230 | 80252 | + | 0 | GCGGCAGAGGCGGCTGACGA | CGG | \|\|\|\|\|\|\|\|[\|\|\|\|\|\|\|\|\|\|\|\|]PAM | NA | Intergenic | NA | NA |
| GU305914.1 | 69784 | 69806 | + | 4 | GCTGGAGAGGCTGCTGAAGA | CGG | \|\|-\|-\|\|\|[\|\|\|-\|\|\|\|\|-\|\|]PAM | NA | Intergenic | NA | NA |
| T5727 | CGGCAGAGGCGGCTGACGACGGG | 1000 | | | | | | | | | |
| Oligo fwd | CACCCGGCAGAGGCGGCTGACGAC | | | | | | | | | | |
| Oligo rev | AAACGTCGTCAGCCGCCTCTGCCG | | | | | | | | | | |
| Chromosome | start | end | strand | MM | target_seq | PAM | alignment | distance | position | gene name | gene id |
| GU305914.1 | 80231 | 80253 | + | 0 | CGGCAGAGGCGGCTGACGAC | GGG | \|\|\|\|\|\|\|\|[\|\|\|\|\|\|\|\|\|\|\|\|]PAM | NA | Intergenic | NA | NA |
| T5728 | ACGACGGGCCCGAACTCACATGG | 1000 | | | | | | | | | |
| Oligo fwd | CACCACGACGGGCCCGAACTCACA | | | | | | | | | | |
| Oligo rev | AAACTGTGAGTTCGGGCCCGTCGT | | | | | | | | | | |
| Chromosome | start | end | strand | MM | target_seq | PAM | alignment | distance | position | gene name | gene id |
| GU305914.1 | 80246 | 80268 | + | 0 | ACGACGGGCCCGAACTCACA | TGG | \|\|\|\|\|\|\|\|[\|\|\|\|\|\|\|\|\|\|\|\|]PAM | NA | Intergenic | NA | NA |
| T5729 | CGACGGGCCCGAACTCACATGGG | 1000 | | | | | | | | | |
| Oligo fwd | CACCCGACGGGCCCGAACTCACAT | | | | | | | | | | |
| Oligo rev | AAACATGTGAGTTCGGGCCCGTCG | | | | | | | | | | |
| Chromosome | start | end | strand | MM | target_seq | PAM | alignment | distance | position | gene name | gene id |
| GU305914.1 | 80247 | 80269 | + | 0 | CGACGGGCCCGAACTCACAT | GGG | \|\|\|\|\|\|\|\|[\|\|\|\|\|\|\|\|\|\|\|\|]PAM | NA | Intergenic | NA | NA |
| T5730 | ATGGGATACCGTTCTTCTTGAGG | 1000 | | | | | | | | | |
| Oligo fwd | CACCATGGGATACCGTTCTTCTTG | | | | | | | | | | |
| Oligo rev | AAACCAAGAAGAACGGTATCCCAT | | | | | | | | | | |
| Chromosome | start | end | strand | MM | target_seq | PAM | alignment | distance | position | gene name | gene id |
| GU305914.1 | 80265 | 80287 | + | 0 | ATGGGATACCGTTCTTCTTG | AGG | \|\|\|\|\|\|\|\|[\|\|\|\|\|\|\|\|\|\|\|\|]PAM | NA | Intergenic | NA | NA |
| T5731 | TGAGGTCGTCGAAAGTCATCCGG | 1000 | | | | | | | | | |
| Oligo fwd | CACCTGAGGTCGTCGAAAGTCATC | | | | | | | | | | |
| Oligo rev | AAACGATGACTTTCGACGACCTCA | | | | | | | | | | |
| Chromosome | start | end | strand | MM | target_seq | PAM | alignment | distance | position | gene name | gene id |
| GU305914.1 | 80283 | 80305 | + | 0 | TGAGGTCGTCGAAAGTCATC | CGG | \|\|\|\|\|\|\|\|[\|\|\|\|\|\|\|\|\|\|\|\|]PAM | NA | Intergenic | NA | NA |
| T5732 | GAGGTCGTCGAAAGTCATCCGGG | 1000 | | | | | | | | | |
| Oligo fwd | CACCGAGGTCGTCGAAAGTCATCC | | | | | | | | | | |
| Oligo rev | AAACGGATGACTTTCGACGACCTC | | | | | | | | | | |
| Chromosome | start | end | strand | MM | target_seq | PAM | alignment | distance | position | gene name | gene id |
| GU305914.1 | 80284 | 80306 | + | 0 | GAGGTCGTCGAAAGTCATCC | GGG | \|\|\|\|\|\|\|\|[\|\|\|\|\|\|\|\|\|\|\|\|]PAM | NA | Intergenic | NA | NA |
| T5733 | CTTCATAGTCCCCTCCGAAACGG | 1000 | | | | | | | | | |
| Oligo fwd | CACCCTTCATAGTCCCCTCCGAAA | | | | | | | | | | |
| Oligo rev | AAACTTTCGGAGGGGACTATGAAG | | | | | | | | | | |
| Chromosome | start | end | strand | MM | target_seq | PAM | alignment | distance | position | gene name | gene id |
| GU305914.1 | 80311 | 80333 | + | 0 | CTTCATAGTCCCCTCCGAAA | CGG | \|\|\|\|\|\|\|\|[\|\|\|\|\|\|\|\|\|\|\|\|]PAM | NA | Intergenic | NA | NA |
| T5734 | TCTCGTCGTTGAAAACCATATGG | 1000 | | | | | | | | | |
| Oligo fwd | CACCTCTCGTCGTTGAAAACCATA | | | | | | | | | | |
| Oligo rev | AAACTATGGTTTTCAACGACGAGA | | | | | | | | | | |
| Chromosome | start | end | strand | MM | target_seq | PAM | alignment | distance | position | gene name | gene id |
| GU305914.1 | 80334 | 80356 | + | 0 | TCTCGTCGTTGAAAACCATA | TGG | \|\|\|\|\|\|\|\|[\|\|\|\|\|\|\|\|\|\|\|\|]PAM | NA | Intergenic | NA | NA |

T5735 CGTCGTTGAAAACCATATGGAGG 1000
Oligo fwd CACCCGTCGTTGAAAACCATATGG
Oligo rev AAACCCATATGGTTTTCAACGACG
Chromosome start end strand MM target_seq PAM alignment distance position gene name gene id
GU305914.1 80337 80359 + 0 CGTCGTTGAAAACCATATGG AGG ||||||||[||||||||||||]PAM NA Intergenic NA NA

T5736 CGCCTTGACGAAACCGCATGCGG 1000
Oligo fwd CACCCGCCTTGACGAAACCGCATG
Oligo rev AAACCATGCGGTTTCGTCAAGGCG
Chromosome start end strand MM target_seq PAM alignment distance position gene name gene id
GU305914.1 80374 80396 + 0 CGCCTTGACGAAACCGCATG CGG ||||||||[||||||||||||]PAM NA Intergenic NA NA

T5737 TGACGAAACCGCATGCGGTGCGG 1000
Oligo fwd CACCTGACGAAACCGCATGCGGTG
Oligo rev AAACCACCGCATGCGGTTTCGTCA
Chromosome start end strand MM target_seq PAM alignment distance position gene name gene id
GU305914.1 80379 80401 + 0 TGACGAAACCGCATGCGGTG CGG ||||||||[||||||||||||]PAM NA Intergenic NA NA

T5738 GATCGACGCCTTTCATACTGAGG 1000
Oligo fwd CACCGATCGACGCCTTTCATACTG
Oligo rev AAACCAGTATGAAAGGCGTCGATC
Chromosome start end strand MM target_seq PAM alignment distance position gene name gene id
GU305914.1 80409 80431 + 0 GATCGACGCCTTTCATACTG AGG ||||||||[||||||||||||]PAM NA Intergenic NA NA

T5739 CGCCTTTCATACTGAGGTTCTGG 1000
Oligo fwd CACCCGCCTTTCATACTGAGGTTC
Oligo rev AAACGAACCTCAGTATGAAAGGCG
Chromosome start end strand MM target_seq PAM alignment distance position gene name gene id
GU305914.1 80415 80437 + 0 CGCCTTTCATACTGAGGTTC TGG ||||||||[||||||||||||]PAM NA Intergenic NA NA

T5740 GCCTTTCATACTGAGGTTCTGGG 1000
Oligo fwd CACCGCCTTTCATACTGAGGTTCT
Oligo rev AAACAGAACCTCAGTATGAAAGGC
Chromosome start end strand MM target_seq PAM alignment distance position gene name gene id
GU305914.1 80416 80438 + 0 GCCTTTCATACTGAGGTTCT GGG ||||||||[||||||||||||]PAM NA Intergenic NA NA

T5741 TACTGAGGTTCTGGGAACCGTGG 1000
Oligo fwd CACCTACTGAGGTTCTGGGAACCG
Oligo rev AAACCGGTTCCCAGAACCTCAGTA
Chromosome start end strand MM target_seq PAM alignment distance position gene name gene id
GU305914.1 80424 80446 + 0 TACTGAGGTTCTGGGAACCG TGG ||||||||[||||||||||||]PAM NA Intergenic NA NA

T5742 TCTTCTTGCATATCATCATAAGG 1000
Oligo fwd CACCTCTTCTTGCATATCATCATA
Oligo rev AAACTATGATGATATGCAAGAAGA
Chromosome start end strand MM target_seq PAM alignment distance position gene name gene id
GU305914.1 80463 80485 + 0 TCTTCTTGCATATCATCATA AGG ||||||||[||||||||||||]PAM NA Intergenic NA NA

T5743 CTTCTTGCATATCATCATAAGGG 1000
Oligo fwd CACCCTTCTTGCATATCATCATAA
Oligo rev AAACTTATGATGATATGCAAGAAG
Chromosome start end strand MM target_seq PAM alignment distance position gene name gene id
GU305914.1 80464 80486 + 0 CTTCTTGCATATCATCATAA GGG ||||||||[||||||||||||]PAM NA Intergenic NA NA

T5744 CGAACATCTTCTCGAACTCGAGG 1000
Oligo fwd CACCCGAACATCTTCTCGAACTCG
Oligo rev AAACCGAGTTCGAGAAGATGTTCG
Chromosome start end strand MM target_seq PAM alignment distance position gene name gene id
GU305914.1 80490 80512 + 0 CGAACATCTTCTCGAACTCG AGG ||||||||[||||||||||||]PAM NA Intergenic NA NA

T5745 CTTCTCGAACTCGAGGCGAACGG 1000
Oligo fwd CACCCTTCTCGAACTCGAGGCGAA
Oligo rev AAACTTCGCCTCGAGTTCGAGAAG
Chromosome start end strand MM target_seq PAM alignment distance position gene name gene id
GU305914.1 80497 80519 + 0 CTTCTCGAACTCGAGGCGAA CGG ||||||||[||||||||||||]PAM NA Intergenic NA NA

T5746 TTCTCGAACTCGAGGCGAACGGG 1000
Oligo fwd CACCTTCTCGAACTCGAGGCGAAC
Oligo rev AAACGTTCGCCTCGAGTTCGAGAA
Chromosome start end strand MM target_seq PAM alignment distance position gene name gene id
GU305914.1 80498 80520 + 0 TTCTCGAACTCGAGGCGAAC GGG ||||||||[||||||||||||]PAM NA Intergenic NA NA

T5747 GGCGAACGGGCTCTTTGAAAAGG 1000
Oligo fwd CACCGGCGAACGGGCTCTTTGAAA
Oligo rev AAACTTTCAAAGAGCCCGTTCGCC
Chromosome start end strand MM target_seq PAM alignment distance position gene name gene id
GU305914.1 80511 80533 + 0 GGCGAACGGGCTCTTTGAAA AGG ||||||||[||||||||||||]PAM NA Intergenic NA NA

T5748 GCGAACGGGCTCTTTGAAAAGGG 1000
Oligo fwd CACCGCGAACGGGCTCTTTGAAAA
Oligo rev AAACTTTTCAAAGAGCCCGTTCGC
Chromosome start end strand MM target_seq PAM alignment distance position gene name gene id
GU305914.1 80512 80534 + 0 GCGAACGGGCTCTTTGAAAA GGG ||||||||[||||||||||||]PAM NA Intergenic NA NA

T5749 AAGGGCGTTCGTAACGTTCGCGG 1000
Oligo fwd CACCAAGGGCGTTCGTAACGTTCG
Oligo rev AAACCGAACGTTACGAACGCCCTT
Chromosome start end strand MM target_seq PAM alignment distance position gene name gene id

<table>
<tr><td>GU305914.1</td><td>80530</td><td>80552</td><td>+</td><td>0</td><td>AAGGGCGTTCGTAACGTTCG</td><td>CGG</td><td>||||||||[||||||||||||]PAM</td><td>NA</td><td>Intergenic</td><td>NA</td><td>NA</td></tr>
<tr><td>T5750</td><td>CGTTCGTAACGTTCGCGGCGAGG</td><td>1000</td><td></td><td></td><td></td><td></td><td></td><td></td><td></td><td></td><td></td></tr>
<tr><td>Oligo fwd</td><td>CACCGTTCGTAACGTTCGCGGCG</td><td></td><td></td><td></td><td></td><td></td><td></td><td></td><td></td><td></td><td></td></tr>
<tr><td>Oligo rev</td><td>AAACCGCCGCGAACGTTACGAACG</td><td></td><td></td><td></td><td></td><td></td><td></td><td></td><td></td><td></td><td></td></tr>
<tr><td>Chromosome</td><td>start</td><td>end</td><td>strand</td><td>MM</td><td>target_seq</td><td>PAM</td><td>alignment</td><td>distance</td><td>position</td><td>gene name</td><td>gene id</td></tr>
<tr><td>GU305914.1</td><td>80535</td><td>80557</td><td>+</td><td>0</td><td>CGTTCGTAACGTTCGCGGCG</td><td>AGG</td><td>||||||||[||||||||||||]PAM</td><td>NA</td><td>Intergenic</td><td>NA</td><td>NA</td></tr>
<tr><td>T5751</td><td>GTTCGTAACGTTCGCGGCGAGGG</td><td>1000</td><td></td><td></td><td></td><td></td><td></td><td></td><td></td><td></td><td></td></tr>
<tr><td>Oligo fwd</td><td>CACCGTTCGTAACGTTCGCGGCGA</td><td></td><td></td><td></td><td></td><td></td><td></td><td></td><td></td><td></td><td></td></tr>
<tr><td>Oligo rev</td><td>AAACTCGCCGCGAACGTTACGAAC</td><td></td><td></td><td></td><td></td><td></td><td></td><td></td><td></td><td></td><td></td></tr>
<tr><td>Chromosome</td><td>start</td><td>end</td><td>strand</td><td>MM</td><td>target_seq</td><td>PAM</td><td>alignment</td><td>distance</td><td>position</td><td>gene name</td><td>gene id</td></tr>
<tr><td>GU305914.1</td><td>80536</td><td>80558</td><td>+</td><td>0</td><td>GTTCGTAACGTTCGCGGCGA</td><td>GGG</td><td>||||||||[||||||||||||]PAM</td><td>NA</td><td>Intergenic</td><td>NA</td><td>NA</td></tr>
<tr><td>T5752</td><td>GTTCGCGGCGAGGGCTCCAACGG</td><td>1000</td><td></td><td></td><td></td><td></td><td></td><td></td><td></td><td></td><td></td></tr>
<tr><td>Oligo fwd</td><td>CACCGTTCGCGGCGAGGGCTCCAA</td><td></td><td></td><td></td><td></td><td></td><td></td><td></td><td></td><td></td><td></td></tr>
<tr><td>Oligo rev</td><td>AAACTTGGAGCCCTCGCCGCGAAC</td><td></td><td></td><td></td><td></td><td></td><td></td><td></td><td></td><td></td><td></td></tr>
<tr><td>Chromosome</td><td>start</td><td>end</td><td>strand</td><td>MM</td><td>target_seq</td><td>PAM</td><td>alignment</td><td>distance</td><td>position</td><td>gene name</td><td>gene id</td></tr>
<tr><td>GU305914.1</td><td>80545</td><td>80567</td><td>+</td><td>0</td><td>GTTCGCGGCGAGGGCTCCAA</td><td>CGG</td><td>||||||||[||||||||||||]PAM</td><td>NA</td><td>Intergenic</td><td>NA</td><td>NA</td></tr>
<tr><td>T5753</td><td>GGGCTCCAACGGCCTTCAAAAGG</td><td>1000</td><td></td><td></td><td></td><td></td><td></td><td></td><td></td><td></td><td></td></tr>
<tr><td>Oligo fwd</td><td>CACCGGGCTCCAACGGCCTTCAAA</td><td></td><td></td><td></td><td></td><td></td><td></td><td></td><td></td><td></td><td></td></tr>
<tr><td>Oligo rev</td><td>AAACTTTGAAGGCCGTTGGAGCCC</td><td></td><td></td><td></td><td></td><td></td><td></td><td></td><td></td><td></td><td></td></tr>
<tr><td>Chromosome</td><td>start</td><td>end</td><td>strand</td><td>MM</td><td>target_seq</td><td>PAM</td><td>alignment</td><td>distance</td><td>position</td><td>gene name</td><td>gene id</td></tr>
<tr><td>GU305914.1</td><td>80556</td><td>80578</td><td>+</td><td>0</td><td>GGGCTCCAACGGCCTTCAAA</td><td>AGG</td><td>||||||||[||||||||||||]PAM</td><td>NA</td><td>Intergenic</td><td>NA</td><td>NA</td></tr>
<tr><td>T5754</td><td>GGCTCCAACGGCCTTCAAAAGGG</td><td>1000</td><td></td><td></td><td></td><td></td><td></td><td></td><td></td><td></td><td></td></tr>
<tr><td>Oligo fwd</td><td>CACCGGCTCCAACGGCCTTCAAAA</td><td></td><td></td><td></td><td></td><td></td><td></td><td></td><td></td><td></td><td></td></tr>
<tr><td>Oligo rev</td><td>AAACTTTTGAAGGCCGTTGGAGCC</td><td></td><td></td><td></td><td></td><td></td><td></td><td></td><td></td><td></td><td></td></tr>
<tr><td>Chromosome</td><td>start</td><td>end</td><td>strand</td><td>MM</td><td>target_seq</td><td>PAM</td><td>alignment</td><td>distance</td><td>position</td><td>gene name</td><td>gene id</td></tr>
<tr><td>GU305914.1</td><td>80557</td><td>80579</td><td>+</td><td>0</td><td>GGCTCCAACGGCCTTCAAAA</td><td>GGG</td><td>||||||||[||||||||||||]PAM</td><td>NA</td><td>Intergenic</td><td>NA</td><td>NA</td></tr>
<tr><td>T5755</td><td>GAAAGCCGCAAACACACTGTCGG</td><td>1000</td><td></td><td></td><td></td><td></td><td></td><td></td><td></td><td></td><td></td></tr>
<tr><td>Oligo fwd</td><td>CACCGAAAGCCGCAAACACACTGT</td><td></td><td></td><td></td><td></td><td></td><td></td><td></td><td></td><td></td><td></td></tr>
<tr><td>Oligo rev</td><td>AAACACAGTGTGTTTGCGGCTTTC</td><td></td><td></td><td></td><td></td><td></td><td></td><td></td><td></td><td></td><td></td></tr>
<tr><td>Chromosome</td><td>start</td><td>end</td><td>strand</td><td>MM</td><td>target_seq</td><td>PAM</td><td>alignment</td><td>distance</td><td>position</td><td>gene name</td><td>gene id</td></tr>
<tr><td>GU305914.1</td><td>80599</td><td>80621</td><td>+</td><td>0</td><td>GAAAGCCGCAAACACACTGT</td><td>CGG</td><td>||||||||[||||||||||||]PAM</td><td>NA</td><td>Intergenic</td><td>NA</td><td>NA</td></tr>
<tr><td>T5756</td><td>CGCCGTAGATGATCTTGACCCGG</td><td>1000</td><td></td><td></td><td></td><td></td><td></td><td></td><td></td><td></td><td></td></tr>
<tr><td>Oligo fwd</td><td>CACCCGCCGTAGATGATCTTGACC</td><td></td><td></td><td></td><td></td><td></td><td></td><td></td><td></td><td></td><td></td></tr>
<tr><td>Oligo rev</td><td>AAACGGTCAAGATCATCTACGGCG</td><td></td><td></td><td></td><td></td><td></td><td></td><td></td><td></td><td></td><td></td></tr>
<tr><td>Chromosome</td><td>start</td><td>end</td><td>strand</td><td>MM</td><td>target_seq</td><td>PAM</td><td>alignment</td><td>distance</td><td>position</td><td>gene name</td><td>gene id</td></tr>
<tr><td>GU305914.1</td><td>80625</td><td>80647</td><td>+</td><td>0</td><td>CGCCGTAGATGATCTTGACC</td><td>CGG</td><td>||||||||[||||||||||||]PAM</td><td>NA</td><td>Intergenic</td><td>NA</td><td>NA</td></tr>
<tr><td>T5757</td><td>CGTAGATGATCTTGACCCGGAGG</td><td>1000</td><td></td><td></td><td></td><td></td><td></td><td></td><td></td><td></td><td></td></tr>
<tr><td>Oligo fwd</td><td>CACCCGTAGATGATCTTGACCCGG</td><td></td><td></td><td></td><td></td><td></td><td></td><td></td><td></td><td></td><td></td></tr>
<tr><td>Oligo rev</td><td>AAACCCGGGTCAAGATCATCTACG</td><td></td><td></td><td></td><td></td><td></td><td></td><td></td><td></td><td></td><td></td></tr>
<tr><td>Chromosome</td><td>start</td><td>end</td><td>strand</td><td>MM</td><td>target_seq</td><td>PAM</td><td>alignment</td><td>distance</td><td>position</td><td>gene name</td><td>gene id</td></tr>
<tr><td>GU305914.1</td><td>80628</td><td>80650</td><td>+</td><td>0</td><td>CGTAGATGATCTTGACCCGG</td><td>AGG</td><td>||||||||[||||||||||||]PAM</td><td>NA</td><td>Intergenic</td><td>NA</td><td>NA</td></tr>
<tr><td>T5758</td><td>GTAGATGATCTTGACCCGGAGGG</td><td>1000</td><td></td><td></td><td></td><td></td><td></td><td></td><td></td><td></td><td></td></tr>
<tr><td>Oligo fwd</td><td>CACCGTAGATGATCTTGACCCGGA</td><td></td><td></td><td></td><td></td><td></td><td></td><td></td><td></td><td></td><td></td></tr>
<tr><td>Oligo rev</td><td>AAACTCCGGGTCAAGATCATCTAC</td><td></td><td></td><td></td><td></td><td></td><td></td><td></td><td></td><td></td><td></td></tr>
<tr><td>Chromosome</td><td>start</td><td>end</td><td>strand</td><td>MM</td><td>target_seq</td><td>PAM</td><td>alignment</td><td>distance</td><td>position</td><td>gene name</td><td>gene id</td></tr>
<tr><td>GU305914.1</td><td>80629</td><td>80651</td><td>+</td><td>0</td><td>GTAGATGATCTTGACCCGGA</td><td>GGG</td><td>||||||||[||||||||||||]PAM</td><td>NA</td><td>Intergenic</td><td>NA</td><td>NA</td></tr>
<tr><td>T5759</td><td>GATCTTGACCCGGAGGGCTTCGG</td><td>1000</td><td></td><td></td><td></td><td></td><td></td><td></td><td></td><td></td><td></td></tr>
<tr><td>Oligo fwd</td><td>CACCGATCTTGACCCGGAGGGCTT</td><td></td><td></td><td></td><td></td><td></td><td></td><td></td><td></td><td></td><td></td></tr>
<tr><td>Oligo rev</td><td>AAACAAGCCCTCCGGGTCAAGATC</td><td></td><td></td><td></td><td></td><td></td><td></td><td></td><td></td><td></td><td></td></tr>
<tr><td>Chromosome</td><td>start</td><td>end</td><td>strand</td><td>MM</td><td>target_seq</td><td>PAM</td><td>alignment</td><td>distance</td><td>position</td><td>gene name</td><td>gene id</td></tr>
<tr><td>GU305914.1</td><td>80635</td><td>80657</td><td>+</td><td>0</td><td>GATCTTGACCCGGAGGGCTT</td><td>CGG</td><td>||||||||[||||||||||||]PAM</td><td>NA</td><td>Intergenic</td><td>NA</td><td>NA</td></tr>
<tr><td>T5760</td><td>ATCTTGACCCGGAGGGCTTCGGG</td><td>1000</td><td></td><td></td><td></td><td></td><td></td><td></td><td></td><td></td><td></td></tr>
<tr><td>Oligo fwd</td><td>CACCATCTTGACCCGGAGGGCTTC</td><td></td><td></td><td></td><td></td><td></td><td></td><td></td><td></td><td></td><td></td></tr>
<tr><td>Oligo rev</td><td>AAACGAAGCCCTCCGGGTCAAGAT</td><td></td><td></td><td></td><td></td><td></td><td></td><td></td><td></td><td></td><td></td></tr>
<tr><td>Chromosome</td><td>start</td><td>end</td><td>strand</td><td>MM</td><td>target_seq</td><td>PAM</td><td>alignment</td><td>distance</td><td>position</td><td>gene name</td><td>gene id</td></tr>
<tr><td>GU305914.1</td><td>80636</td><td>80658</td><td>+</td><td>0</td><td>ATCTTGACCCGGAGGGCTTC</td><td>GGG</td><td>||||||||[||||||||||||]PAM</td><td>NA</td><td>Intergenic</td><td>NA</td><td>NA</td></tr>
<tr><td>T5761</td><td>TCTTGACCCGGAGGGCTTCGGGG</td><td>1000</td><td></td><td></td><td></td><td></td><td></td><td></td><td></td><td></td><td></td></tr>
<tr><td>Oligo fwd</td><td>CACCTCTTGACCCGGAGGGCTTCG</td><td></td><td></td><td></td><td></td><td></td><td></td><td></td><td></td><td></td><td></td></tr>
<tr><td>Oligo rev</td><td>AAACCGAAGCCCTCCGGGTCAAGA</td><td></td><td></td><td></td><td></td><td></td><td></td><td></td><td></td><td></td><td></td></tr>
<tr><td>Chromosome</td><td>start</td><td>end</td><td>strand</td><td>MM</td><td>target_seq</td><td>PAM</td><td>alignment</td><td>distance</td><td>position</td><td>gene name</td><td>gene id</td></tr>
<tr><td>GU305914.1</td><td>80637</td><td>80659</td><td>+</td><td>0</td><td>TCTTGACCCGGAGGGCTTCG</td><td>GGG</td><td>||||||||[||||||||||||]PAM</td><td>NA</td><td>Intergenic</td><td>NA</td><td>NA</td></tr>
<tr><td>T5762</td><td>GGCTTCGGGGTCGATAGCCTCGG</td><td>1000</td><td></td><td></td><td></td><td></td><td></td><td></td><td></td><td></td><td></td></tr>
<tr><td>Oligo fwd</td><td>CACCGGCTTCGGGGTCGATAGCCT</td><td></td><td></td><td></td><td></td><td></td><td></td><td></td><td></td><td></td><td></td></tr>
<tr><td>Oligo rev</td><td>AAACAGGCTATCGACCCCGAAGCC</td><td></td><td></td><td></td><td></td><td></td><td></td><td></td><td></td><td></td><td></td></tr>
<tr><td>Chromosome</td><td>start</td><td>end</td><td>strand</td><td>MM</td><td>target_seq</td><td>PAM</td><td>alignment</td><td>distance</td><td>position</td><td>gene name</td><td>gene id</td></tr>
<tr><td>GU305914.1</td><td>80650</td><td>80672</td><td>+</td><td>0</td><td>GGCTTCGGGGTCGATAGCCT</td><td>CGG</td><td>||||||||[||||||||||||]PAM</td><td>NA</td><td>Intergenic</td><td>NA</td><td>NA</td></tr>
<tr><td>T5763</td><td>GCTTCGGGGTCGATAGCCTCGGG</td><td>1000</td><td></td><td></td><td></td><td></td><td></td><td></td><td></td><td></td><td></td></tr>
<tr><td>Oligo fwd</td><td>CACCGCTTCGGGGTCGATAGCCTC</td><td></td><td></td><td></td><td></td><td></td><td></td><td></td><td></td><td></td><td></td></tr>
<tr><td>Oligo rev</td><td>AAACGAGGCTATCGACCCCGAAGC</td><td></td><td></td><td></td><td></td><td></td><td></td><td></td><td></td><td></td><td></td></tr>
<tr><td>Chromosome</td><td>start</td><td>end</td><td>strand</td><td>MM</td><td>target_seq</td><td>PAM</td><td>alignment</td><td>distance</td><td>position</td><td>gene name</td><td>gene id</td></tr>
<tr><td>GU305914.1</td><td>80651</td><td>80673</td><td>+</td><td>0</td><td>GCTTCGGGGTCGATAGCCTC</td><td>GGG</td><td>||||||||[||||||||||||]PAM</td><td>NA</td><td>Intergenic</td><td>NA</td><td>NA</td></tr>
<tr><td>T5764</td><td>CTTCGGGGTCGATAGCCTCGGGG</td><td>1000</td><td></td><td></td><td></td><td></td><td></td><td></td><td></td><td></td><td></td></tr>
<tr><td>Oligo fwd</td><td>CACCCTTCGGGGTCGATAGCCTCG</td><td></td><td></td><td></td><td></td><td></td><td></td><td></td><td></td><td></td><td></td></tr>
<tr><td>Oligo rev</td><td>AAACCGAGGCTATCGACCCCGAAG</td><td></td><td></td><td></td><td></td><td></td><td></td><td></td><td></td><td></td><td></td></tr>
</table>

<table>
<tr><td>Chromosome</td><td>start</td><td>end</td><td>strand</td><td>MM</td><td>target_seq</td><td>PAM</td><td>alignment</td><td>distance</td><td>position</td><td>gene name</td><td>gene id</td></tr>
<tr><td>GU305914.1</td><td>80652</td><td>80674</td><td>+</td><td>0</td><td>CTTCGGGGTCGATAGCCTCG</td><td>GGG</td><td>||||||||[||||||||||||]PAM</td><td>NA</td><td>Intergenic</td><td>NA</td><td>NA</td></tr>
<tr><td>T5765</td><td>TTCGGGGTCGATAGCCTCGGGGG</td><td>1000</td><td></td><td></td><td></td><td></td><td></td><td></td><td></td><td></td><td></td></tr>
<tr><td>Oligo fwd</td><td>CACCTTCGGGGTCGATAGCCTCGG</td><td></td><td></td><td></td><td></td><td></td><td></td><td></td><td></td><td></td><td></td></tr>
<tr><td>Oligo rev</td><td>AAACCCGAGGCTATCGACCCCGAA</td><td></td><td></td><td></td><td></td><td></td><td></td><td></td><td></td><td></td><td></td></tr>
<tr><td>Chromosome</td><td>start</td><td>end</td><td>strand</td><td>MM</td><td>target_seq</td><td>PAM</td><td>alignment</td><td>distance</td><td>position</td><td>gene name</td><td>gene id</td></tr>
<tr><td>GU305914.1</td><td>80653</td><td>80675</td><td>+</td><td>0</td><td>TTCGGGGTCGATAGCCTCGG</td><td>GGG</td><td>||||||||[||||||||||||]PAM</td><td>NA</td><td>Intergenic</td><td>NA</td><td>NA</td></tr>
<tr><td>T5766</td><td>TCGGGGTCGATAGCCTCGGGGGG</td><td>1000</td><td></td><td></td><td></td><td></td><td></td><td></td><td></td><td></td><td></td></tr>
<tr><td>Oligo fwd</td><td>CACCTCGGGGTCGATAGCCTCGGG</td><td></td><td></td><td></td><td></td><td></td><td></td><td></td><td></td><td></td><td></td></tr>
<tr><td>Oligo rev</td><td>AAACCCCGAGGCTATCGACCCCGA</td><td></td><td></td><td></td><td></td><td></td><td></td><td></td><td></td><td></td><td></td></tr>
<tr><td>Chromosome</td><td>start</td><td>end</td><td>strand</td><td>MM</td><td>target_seq</td><td>PAM</td><td>alignment</td><td>distance</td><td>position</td><td>gene name</td><td>gene id</td></tr>
<tr><td>GU305914.1</td><td>80654</td><td>80676</td><td>+</td><td>0</td><td>TCGGGGTCGATAGCCTCGGG</td><td>GGG</td><td>||||||||[||||||||||||]PAM</td><td>NA</td><td>Intergenic</td><td>NA</td><td>NA</td></tr>
<tr><td>T5767</td><td>CGGGGTCGATAGCCTCGGGGGGG</td><td>1000</td><td></td><td></td><td></td><td></td><td></td><td></td><td></td><td></td><td></td></tr>
<tr><td>Oligo fwd</td><td>CACCCGGGGTCGATAGCCTCGGGG</td><td></td><td></td><td></td><td></td><td></td><td></td><td></td><td></td><td></td><td></td></tr>
<tr><td>Oligo rev</td><td>AAACCCCCGAGGCTATCGACCCCG</td><td></td><td></td><td></td><td></td><td></td><td></td><td></td><td></td><td></td><td></td></tr>
<tr><td>Chromosome</td><td>start</td><td>end</td><td>strand</td><td>MM</td><td>target_seq</td><td>PAM</td><td>alignment</td><td>distance</td><td>position</td><td>gene name</td><td>gene id</td></tr>
<tr><td>GU305914.1</td><td>80655</td><td>80677</td><td>+</td><td>0</td><td>CGGGGTCGATAGCCTCGGGG</td><td>GGG</td><td>||||||||[||||||||||||]PAM</td><td>NA</td><td>Intergenic</td><td>NA</td><td>NA</td></tr>
<tr><td>T5768</td><td>TAGCCTCGGGGGGGAGTCCGAGG</td><td>1000</td><td></td><td></td><td></td><td></td><td></td><td></td><td></td><td></td><td></td></tr>
<tr><td>Oligo fwd</td><td>CACCTAGCCTCGGGGGGGAGTCCG</td><td></td><td></td><td></td><td></td><td></td><td></td><td></td><td></td><td></td><td></td></tr>
<tr><td>Oligo rev</td><td>AAACCGGACTCCCCCCCGAGGCTA</td><td></td><td></td><td></td><td></td><td></td><td></td><td></td><td></td><td></td><td></td></tr>
<tr><td>Chromosome</td><td>start</td><td>end</td><td>strand</td><td>MM</td><td>target_seq</td><td>PAM</td><td>alignment</td><td>distance</td><td>position</td><td>gene name</td><td>gene id</td></tr>
<tr><td>GU305914.1</td><td>80664</td><td>80686</td><td>+</td><td>0</td><td>TAGCCTCGGGGGGGAGTCCG</td><td>AGG</td><td>||||||||[||||||||||||]PAM</td><td>NA</td><td>Intergenic</td><td>NA</td><td>NA</td></tr>
<tr><td>T5769</td><td>CTCGGGGGGGAGTCCGAGGACGG</td><td>1000</td><td></td><td></td><td></td><td></td><td></td><td></td><td></td><td></td><td></td></tr>
<tr><td>Oligo fwd</td><td>CACCCTCGGGGGGGAGTCCGAGGA</td><td></td><td></td><td></td><td></td><td></td><td></td><td></td><td></td><td></td><td></td></tr>
<tr><td>Oligo rev</td><td>AAACTCCTCGGACTCCCCCCCGAG</td><td></td><td></td><td></td><td></td><td></td><td></td><td></td><td></td><td></td><td></td></tr>
<tr><td>Chromosome</td><td>start</td><td>end</td><td>strand</td><td>MM</td><td>target_seq</td><td>PAM</td><td>alignment</td><td>distance</td><td>position</td><td>gene name</td><td>gene id</td></tr>
<tr><td>GU305914.1</td><td>80668</td><td>80690</td><td>+</td><td>0</td><td>CTCGGGGGGGAGTCCGAGGA</td><td>CGG</td><td>||||||||[||||||||||||]PAM</td><td>NA</td><td>Intergenic</td><td>NA</td><td>NA</td></tr>
<tr><td>T5770</td><td>GGGGGAGTCCGAGGACGGTACGG</td><td>1000</td><td></td><td></td><td></td><td></td><td></td><td></td><td></td><td></td><td></td></tr>
<tr><td>Oligo fwd</td><td>CACCGGGGGAGTCCGAGGACGGTA</td><td></td><td></td><td></td><td></td><td></td><td></td><td></td><td></td><td></td><td></td></tr>
<tr><td>Oligo rev</td><td>AAACTACCGTCCTCGGACTCCCCC</td><td></td><td></td><td></td><td></td><td></td><td></td><td></td><td></td><td></td><td></td></tr>
<tr><td>Chromosome</td><td>start</td><td>end</td><td>strand</td><td>MM</td><td>target_seq</td><td>PAM</td><td>alignment</td><td>distance</td><td>position</td><td>gene name</td><td>gene id</td></tr>
<tr><td>GU305914.1</td><td>80673</td><td>80695</td><td>+</td><td>0</td><td>GGGGGAGTCCGAGGACGGTA</td><td>CGG</td><td>||||||||[||||||||||||]PAM</td><td>NA</td><td>Intergenic</td><td>NA</td><td>NA</td></tr>
<tr><td>T5771</td><td>GAGGACGGTACGGAGAAAATCGG</td><td>1000</td><td></td><td></td><td></td><td></td><td></td><td></td><td></td><td></td><td></td></tr>
<tr><td>Oligo fwd</td><td>CACCGAGGACGGTACGGAGAAAAT</td><td></td><td></td><td></td><td></td><td></td><td></td><td></td><td></td><td></td><td></td></tr>
<tr><td>Oligo rev</td><td>AAACATTTTCTCCGTACCGTCCTC</td><td></td><td></td><td></td><td></td><td></td><td></td><td></td><td></td><td></td><td></td></tr>
<tr><td>Chromosome</td><td>start</td><td>end</td><td>strand</td><td>MM</td><td>target_seq</td><td>PAM</td><td>alignment</td><td>distance</td><td>position</td><td>gene name</td><td>gene id</td></tr>
<tr><td>GU305914.1</td><td>80683</td><td>80705</td><td>+</td><td>0</td><td>GAGGACGGTACGGAGAAAAT</td><td>CGG</td><td>||||||||[||||||||||||]PAM</td><td>NA</td><td>Intergenic</td><td>NA</td><td>NA</td></tr>
<tr><td>T5772</td><td>AGGACGGTACGGAGAAAATCGGG</td><td>1000</td><td></td><td></td><td></td><td></td><td></td><td></td><td></td><td></td><td></td></tr>
<tr><td>Oligo fwd</td><td>CACCAGGACGGTACGGAGAAAATC</td><td></td><td></td><td></td><td></td><td></td><td></td><td></td><td></td><td></td><td></td></tr>
<tr><td>Oligo rev</td><td>AAACGATTTTCTCCGTACCGTCCT</td><td></td><td></td><td></td><td></td><td></td><td></td><td></td><td></td><td></td><td></td></tr>
<tr><td>Chromosome</td><td>start</td><td>end</td><td>strand</td><td>MM</td><td>target_seq</td><td>PAM</td><td>alignment</td><td>distance</td><td>position</td><td>gene name</td><td>gene id</td></tr>
<tr><td>GU305914.1</td><td>80684</td><td>80706</td><td>+</td><td>0</td><td>AGGACGGTACGGAGAAAATC</td><td>GGG</td><td>||||||||[||||||||||||]PAM</td><td>NA</td><td>Intergenic</td><td>NA</td><td>NA</td></tr>
<tr><td>T5773</td><td>GGACGGTACGGAGAAAATCGGGG</td><td>1000</td><td></td><td></td><td></td><td></td><td></td><td></td><td></td><td></td><td></td></tr>
<tr><td>Oligo fwd</td><td>CACCGGACGGTACGGAGAAAATCG</td><td></td><td></td><td></td><td></td><td></td><td></td><td></td><td></td><td></td><td></td></tr>
<tr><td>Oligo rev</td><td>AAACCGATTTTCTCCGTACCGTCC</td><td></td><td></td><td></td><td></td><td></td><td></td><td></td><td></td><td></td><td></td></tr>
<tr><td>Chromosome</td><td>start</td><td>end</td><td>strand</td><td>MM</td><td>target_seq</td><td>PAM</td><td>alignment</td><td>distance</td><td>position</td><td>gene name</td><td>gene id</td></tr>
<tr><td>GU305914.1</td><td>80685</td><td>80707</td><td>+</td><td>0</td><td>GGACGGTACGGAGAAAATCG</td><td>GGG</td><td>||||||||[||||||||||||]PAM</td><td>NA</td><td>Intergenic</td><td>NA</td><td>NA</td></tr>
<tr><td>T5774</td><td>CGGTACGGAGAAAATCGGGGCGG</td><td>1000</td><td></td><td></td><td></td><td></td><td></td><td></td><td></td><td></td><td></td></tr>
<tr><td>Oligo fwd</td><td>CACCCGGTACGGAGAAAATCGGGG</td><td></td><td></td><td></td><td></td><td></td><td></td><td></td><td></td><td></td><td></td></tr>
<tr><td>Oligo rev</td><td>AAACCCCCGATTTTCTCCGTACCG</td><td></td><td></td><td></td><td></td><td></td><td></td><td></td><td></td><td></td><td></td></tr>
<tr><td>Chromosome</td><td>start</td><td>end</td><td>strand</td><td>MM</td><td>target_seq</td><td>PAM</td><td>alignment</td><td>distance</td><td>position</td><td>gene name</td><td>gene id</td></tr>
<tr><td>GU305914.1</td><td>80688</td><td>80710</td><td>+</td><td>0</td><td>CGGTACGGAGAAAATCGGGG</td><td>CGG</td><td>||||||||[||||||||||||]PAM</td><td>NA</td><td>Intergenic</td><td>NA</td><td>NA</td></tr>
<tr><td>T5775</td><td>GAAAATCGGGGCGGTTGCACCGG</td><td>1000</td><td></td><td></td><td></td><td></td><td></td><td></td><td></td><td></td><td></td></tr>
<tr><td>Oligo fwd</td><td>CACCGAAAATCGGGGCGGTTGCAC</td><td></td><td></td><td></td><td></td><td></td><td></td><td></td><td></td><td></td><td></td></tr>
<tr><td>Oligo rev</td><td>AAACGTGCAACCGCCCCGATTTTC</td><td></td><td></td><td></td><td></td><td></td><td></td><td></td><td></td><td></td><td></td></tr>
<tr><td>Chromosome</td><td>start</td><td>end</td><td>strand</td><td>MM</td><td>target_seq</td><td>PAM</td><td>alignment</td><td>distance</td><td>position</td><td>gene name</td><td>gene id</td></tr>
<tr><td>GU305914.1</td><td>80697</td><td>80719</td><td>+</td><td>0</td><td>GAAAATCGGGGCGGTTGCAC</td><td>CGG</td><td>||||||||[||||||||||||]PAM</td><td>NA</td><td>Intergenic</td><td>NA</td><td>NA</td></tr>
<tr><td>T5776</td><td>GGTTGCACCGGTCCTCGATATGG</td><td>1000</td><td></td><td></td><td></td><td></td><td></td><td></td><td></td><td></td><td></td></tr>
<tr><td>Oligo fwd</td><td>CACCGGTTGCACCGGTCCTCGATA</td><td></td><td></td><td></td><td></td><td></td><td></td><td></td><td></td><td></td><td></td></tr>
<tr><td>Oligo rev</td><td>AAACTATCGAGGACCGGTGCAACC</td><td></td><td></td><td></td><td></td><td></td><td></td><td></td><td></td><td></td><td></td></tr>
<tr><td>Chromosome</td><td>start</td><td>end</td><td>strand</td><td>MM</td><td>target_seq</td><td>PAM</td><td>alignment</td><td>distance</td><td>position</td><td>gene name</td><td>gene id</td></tr>
<tr><td>GU305914.1</td><td>80709</td><td>80731</td><td>+</td><td>0</td><td>GGTTGCACCGGTCCTCGATA</td><td>TGG</td><td>||||||||[||||||||||||]PAM</td><td>NA</td><td>Intergenic</td><td>NA</td><td>NA</td></tr>
<tr><td>T5777</td><td>GCACCGGTCCTCGATATGGCCGG</td><td>1000</td><td></td><td></td><td></td><td></td><td></td><td></td><td></td><td></td><td></td></tr>
<tr><td>Oligo fwd</td><td>CACCGCACCGGTCCTCGATATGGC</td><td></td><td></td><td></td><td></td><td></td><td></td><td></td><td></td><td></td><td></td></tr>
<tr><td>Oligo rev</td><td>AAACGCCATATCGAGGACCGGTGC</td><td></td><td></td><td></td><td></td><td></td><td></td><td></td><td></td><td></td><td></td></tr>
<tr><td>Chromosome</td><td>start</td><td>end</td><td>strand</td><td>MM</td><td>target_seq</td><td>PAM</td><td>alignment</td><td>distance</td><td>position</td><td>gene name</td><td>gene id</td></tr>
<tr><td>GU305914.1</td><td>80713</td><td>80735</td><td>+</td><td>0</td><td>GCACCGGTCCTCGATATGGC</td><td>CGG</td><td>||||||||[||||||||||||]PAM</td><td>NA</td><td>Intergenic</td><td>NA</td><td>NA</td></tr>
<tr><td>T5778</td><td>CCGGTCCTCGATATGGCCGGCGG</td><td>1000</td><td></td><td></td><td></td><td></td><td></td><td></td><td></td><td></td><td></td></tr>
<tr><td>Oligo fwd</td><td>CACCCCGGTCCTCGATATGGCCGG</td><td></td><td></td><td></td><td></td><td></td><td></td><td></td><td></td><td></td><td></td></tr>
<tr><td>Oligo rev</td><td>AAACCCGGCCATATCGAGGACCGG</td><td></td><td></td><td></td><td></td><td></td><td></td><td></td><td></td><td></td><td></td></tr>
<tr><td>Chromosome</td><td>start</td><td>end</td><td>strand</td><td>MM</td><td>target_seq</td><td>PAM</td><td>alignment</td><td>distance</td><td>position</td><td>gene name</td><td>gene id</td></tr>
<tr><td>GU305914.1</td><td>80716</td><td>80738</td><td>+</td><td>0</td><td>CCGGTCCTCGATATGGCCGG</td><td>CGG</td><td>||||||||[||||||||||||]PAM</td><td>NA</td><td>Intergenic</td><td>NA</td><td>NA</td></tr>
<tr><td>T5779</td><td>CAACATATCCCTGCCGATCTTGG</td><td>1000</td><td></td><td></td><td></td><td></td><td></td><td></td><td></td><td></td><td></td></tr>
<tr><td>Oligo fwd</td><td>CACCCAACATATCCCTGCCGATCT</td><td></td><td></td><td></td><td></td><td></td><td></td><td></td><td></td><td></td><td></td></tr>
</table>

<table>
<tr><td>Oligo rev</td><td>AAACAGATCGGCAGGGATATGTTG</td><td></td><td></td><td></td><td></td><td></td><td></td><td></td><td></td><td></td><td></td><td></td></tr>
<tr><td>Chromosome</td><td>start</td><td></td><td>end</td><td>strand</td><td>MM</td><td>target_seq</td><td>PAM</td><td>alignment</td><td>distance</td><td>position</td><td>gene name</td><td>gene id</td></tr>
<tr><td>GU305914.1</td><td></td><td>80746</td><td>80768</td><td>+</td><td>0</td><td>CAACATATCCCTGCCGATCT</td><td>TGG</td><td>||||||||[||||||||||||]PAM</td><td>NA</td><td>Intergenic</td><td>NA</td><td>NA</td></tr>
<tr><td>T5780</td><td>GCCGATCTTGGTGATAGAAGCGG</td><td></td><td>1000</td><td></td><td></td><td></td><td></td><td></td><td></td><td></td><td></td><td></td></tr>
<tr><td>Oligo fwd</td><td>CACCGCCGATCTTGGTGATAGAAG</td><td></td><td></td><td></td><td></td><td></td><td></td><td></td><td></td><td></td><td></td><td></td></tr>
<tr><td>Oligo rev</td><td>AAACCTTCTATCACCAAGATCGGC</td><td></td><td></td><td></td><td></td><td></td><td></td><td></td><td></td><td></td><td></td><td></td></tr>
<tr><td>Chromosome</td><td>start</td><td></td><td>end</td><td>strand</td><td>MM</td><td>target_seq</td><td>PAM</td><td>alignment</td><td>distance</td><td>position</td><td>gene name</td><td>gene id</td></tr>
<tr><td>GU305914.1</td><td></td><td>80758</td><td>80780</td><td>+</td><td>0</td><td>GCCGATCTTGGTGATAGAAG</td><td>CGG</td><td>||||||||[||||||||||||]PAM</td><td>NA</td><td>Intergenic</td><td>NA</td><td>NA</td></tr>
<tr><td>T5781</td><td>CTTGGTGATAGAAGCGGCGATGG</td><td></td><td>1000</td><td></td><td></td><td></td><td></td><td></td><td></td><td></td><td></td><td></td></tr>
<tr><td>Oligo fwd</td><td>CACCCTTGGTGATAGAAGCGGCGA</td><td></td><td></td><td></td><td></td><td></td><td></td><td></td><td></td><td></td><td></td><td></td></tr>
<tr><td>Oligo rev</td><td>AAACTCGCCGCTTCTATCACCAAG</td><td></td><td></td><td></td><td></td><td></td><td></td><td></td><td></td><td></td><td></td><td></td></tr>
<tr><td>Chromosome</td><td>start</td><td></td><td>end</td><td>strand</td><td>MM</td><td>target_seq</td><td>PAM</td><td>alignment</td><td>distance</td><td>position</td><td>gene name</td><td>gene id</td></tr>
<tr><td>GU305914.1</td><td></td><td>80764</td><td>80786</td><td>+</td><td>0</td><td>CTTGGTGATAGAAGCGGCGA</td><td>TGG</td><td>||||||||[||||||||||||]PAM</td><td>NA</td><td>Intergenic</td><td>NA</td><td>NA</td></tr>
<tr><td>T5782</td><td>TTGGTGATAGAAGCGGCGATGGG</td><td></td><td>1000</td><td></td><td></td><td></td><td></td><td></td><td></td><td></td><td></td><td></td></tr>
<tr><td>Oligo fwd</td><td>CACCTTGGTGATAGAAGCGGCGAT</td><td></td><td></td><td></td><td></td><td></td><td></td><td></td><td></td><td></td><td></td><td></td></tr>
<tr><td>Oligo rev</td><td>AAACATCGCCGCTTCTATCACCAA</td><td></td><td></td><td></td><td></td><td></td><td></td><td></td><td></td><td></td><td></td><td></td></tr>
<tr><td>Chromosome</td><td>start</td><td></td><td>end</td><td>strand</td><td>MM</td><td>target_seq</td><td>PAM</td><td>alignment</td><td>distance</td><td>position</td><td>gene name</td><td>gene id</td></tr>
<tr><td>GU305914.1</td><td></td><td>80765</td><td>80787</td><td>+</td><td>0</td><td>TTGGTGATAGAAGCGGCGAT</td><td>GGG</td><td>||||||||[||||||||||||]PAM</td><td>NA</td><td>Intergenic</td><td>NA</td><td>NA</td></tr>
<tr><td>T5783</td><td>TGGTGATAGAAGCGGCGATGGGG</td><td></td><td>1000</td><td></td><td></td><td></td><td></td><td></td><td></td><td></td><td></td><td></td></tr>
<tr><td>Oligo fwd</td><td>CACCTGGTGATAGAAGCGGCGATG</td><td></td><td></td><td></td><td></td><td></td><td></td><td></td><td></td><td></td><td></td><td></td></tr>
<tr><td>Oligo rev</td><td>AAACCATCGCCGCTTCTATCACCA</td><td></td><td></td><td></td><td></td><td></td><td></td><td></td><td></td><td></td><td></td><td></td></tr>
<tr><td>Chromosome</td><td>start</td><td></td><td>end</td><td>strand</td><td>MM</td><td>target_seq</td><td>PAM</td><td>alignment</td><td>distance</td><td>position</td><td>gene name</td><td>gene id</td></tr>
<tr><td>GU305914.1</td><td></td><td>80766</td><td>80788</td><td>+</td><td>0</td><td>TGGTGATAGAAGCGGCGATG</td><td>GGG</td><td>||||||||[||||||||||||]PAM</td><td>NA</td><td>Intergenic</td><td>NA</td><td>NA</td></tr>
<tr><td>T5784</td><td>GAAGCGGCGATGGGGAGACACGG</td><td></td><td>1000</td><td></td><td></td><td></td><td></td><td></td><td></td><td></td><td></td><td></td></tr>
<tr><td>Oligo fwd</td><td>CACCGAAGCGGCGATGGGGAGACA</td><td></td><td></td><td></td><td></td><td></td><td></td><td></td><td></td><td></td><td></td><td></td></tr>
<tr><td>Oligo rev</td><td>AAACTGTCTCCCCATCGCCGCTTC</td><td></td><td></td><td></td><td></td><td></td><td></td><td></td><td></td><td></td><td></td><td></td></tr>
<tr><td>Chromosome</td><td>start</td><td></td><td>end</td><td>strand</td><td>MM</td><td>target_seq</td><td>PAM</td><td>alignment</td><td>distance</td><td>position</td><td>gene name</td><td>gene id</td></tr>
<tr><td>GU305914.1</td><td></td><td>80774</td><td>80796</td><td>+</td><td>0</td><td>GAAGCGGCGATGGGGAGACA</td><td>CGG</td><td>||||||||[||||||||||||]PAM</td><td>NA</td><td>Intergenic</td><td>NA</td><td>NA</td></tr>
<tr><td>T5785</td><td>AAGCGGCGATGGGGAGACACGGG</td><td></td><td>1000</td><td></td><td></td><td></td><td></td><td></td><td></td><td></td><td></td><td></td></tr>
<tr><td>Oligo fwd</td><td>CACCAAGCGGCGATGGGGAGACAC</td><td></td><td></td><td></td><td></td><td></td><td></td><td></td><td></td><td></td><td></td><td></td></tr>
<tr><td>Oligo rev</td><td>AAACGTGTCTCCCCATCGCCGCTT</td><td></td><td></td><td></td><td></td><td></td><td></td><td></td><td></td><td></td><td></td><td></td></tr>
<tr><td>Chromosome</td><td>start</td><td></td><td>end</td><td>strand</td><td>MM</td><td>target_seq</td><td>PAM</td><td>alignment</td><td>distance</td><td>position</td><td>gene name</td><td>gene id</td></tr>
<tr><td>GU305914.1</td><td></td><td>80775</td><td>80797</td><td>+</td><td>0</td><td>AAGCGGCGATGGGGAGACAC</td><td>GGG</td><td>||||||||[||||||||||||]PAM</td><td>NA</td><td>Intergenic</td><td>NA</td><td>NA</td></tr>
<tr><td>T5786</td><td>GGGGAGACACGGGAGCATCCCGG</td><td></td><td>1000</td><td></td><td></td><td></td><td></td><td></td><td></td><td></td><td></td><td></td></tr>
<tr><td>Oligo fwd</td><td>CACCGGGGAGACACGGGAGCATCC</td><td></td><td></td><td></td><td></td><td></td><td></td><td></td><td></td><td></td><td></td><td></td></tr>
<tr><td>Oligo rev</td><td>AAACGGATGCTCCCGTGTCTCCCC</td><td></td><td></td><td></td><td></td><td></td><td></td><td></td><td></td><td></td><td></td><td></td></tr>
<tr><td>Chromosome</td><td>start</td><td></td><td>end</td><td>strand</td><td>MM</td><td>target_seq</td><td>PAM</td><td>alignment</td><td>distance</td><td>position</td><td>gene name</td><td>gene id</td></tr>
<tr><td>GU305914.1</td><td></td><td>80785</td><td>80807</td><td>+</td><td>0</td><td>GGGGAGACACGGGAGCATCC</td><td>CGG</td><td>||||||||[||||||||||||]PAM</td><td>NA</td><td>Intergenic</td><td>NA</td><td>NA</td></tr>
<tr><td>T5787</td><td>AAGCGTTGCACGTTACTTTGAGG</td><td></td><td>1000</td><td></td><td></td><td></td><td></td><td></td><td></td><td></td><td></td><td></td></tr>
<tr><td>Oligo fwd</td><td>CACCAAGCGTTGCACGTTACTTTG</td><td></td><td></td><td></td><td></td><td></td><td></td><td></td><td></td><td></td><td></td><td></td></tr>
<tr><td>Oligo rev</td><td>AAACCAAAGTAACGTGCAACGCTT</td><td></td><td></td><td></td><td></td><td></td><td></td><td></td><td></td><td></td><td></td><td></td></tr>
<tr><td>Chromosome</td><td>start</td><td></td><td>end</td><td>strand</td><td>MM</td><td>target_seq</td><td>PAM</td><td>alignment</td><td>distance</td><td>position</td><td>gene name</td><td>gene id</td></tr>
<tr><td>GU305914.1</td><td></td><td>80832</td><td>80854</td><td>+</td><td>0</td><td>AAGCGTTGCACGTTACTTTG</td><td>AGG</td><td>||||||||[||||||||||||]PAM</td><td>NA</td><td>Intergenic</td><td>NA</td><td>NA</td></tr>
<tr><td>T5788</td><td>AGCGTTGCACGTTACTTTGAGGG</td><td></td><td>1000</td><td></td><td></td><td></td><td></td><td></td><td></td><td></td><td></td><td></td></tr>
<tr><td>Oligo fwd</td><td>CACCAGCGTTGCACGTTACTTTGA</td><td></td><td></td><td></td><td></td><td></td><td></td><td></td><td></td><td></td><td></td><td></td></tr>
<tr><td>Oligo rev</td><td>AAACTCAAAGTAACGTGCAACGCT</td><td></td><td></td><td></td><td></td><td></td><td></td><td></td><td></td><td></td><td></td><td></td></tr>
<tr><td>Chromosome</td><td>start</td><td></td><td>end</td><td>strand</td><td>MM</td><td>target_seq</td><td>PAM</td><td>alignment</td><td>distance</td><td>position</td><td>gene name</td><td>gene id</td></tr>
<tr><td>GU305914.1</td><td></td><td>80833</td><td>80855</td><td>+</td><td>0</td><td>AGCGTTGCACGTTACTTTGA</td><td>GGG</td><td>||||||||[||||||||||||]PAM</td><td>NA</td><td>Intergenic</td><td>NA</td><td>NA</td></tr>
<tr><td>T5789</td><td>GATCATCCGTTGCATCTCGTTGG</td><td></td><td>1000</td><td></td><td></td><td></td><td></td><td></td><td></td><td></td><td></td><td></td></tr>
<tr><td>Oligo fwd</td><td>CACCGATCATCCGTTGCATCTCGT</td><td></td><td></td><td></td><td></td><td></td><td></td><td></td><td></td><td></td><td></td><td></td></tr>
<tr><td>Oligo rev</td><td>AAACACGAGATGCAACGGATGATC</td><td></td><td></td><td></td><td></td><td></td><td></td><td></td><td></td><td></td><td></td><td></td></tr>
<tr><td>Chromosome</td><td>start</td><td></td><td>end</td><td>strand</td><td>MM</td><td>target_seq</td><td>PAM</td><td>alignment</td><td>distance</td><td>position</td><td>gene name</td><td>gene id</td></tr>
<tr><td>GU305914.1</td><td></td><td>80875</td><td>80897</td><td>+</td><td>0</td><td>GATCATCCGTTGCATCTCGT</td><td>TGG</td><td>||||||||[||||||||||||]PAM</td><td>NA</td><td>Intergenic</td><td>NA</td><td>NA</td></tr>
<tr><td>T5790</td><td>TGCACTGCTTCATCGCCTCCCGG</td><td></td><td>1000</td><td></td><td></td><td></td><td></td><td></td><td></td><td></td><td></td><td></td></tr>
<tr><td>Oligo fwd</td><td>CACCTGCACTGCTTCATCGCCTCC</td><td></td><td></td><td></td><td></td><td></td><td></td><td></td><td></td><td></td><td></td><td></td></tr>
<tr><td>Oligo rev</td><td>AAACGGAGGCGATGAAGCAGTGCA</td><td></td><td></td><td></td><td></td><td></td><td></td><td></td><td></td><td></td><td></td><td></td></tr>
<tr><td>Chromosome</td><td>start</td><td></td><td>end</td><td>strand</td><td>MM</td><td>target_seq</td><td>PAM</td><td>alignment</td><td>distance</td><td>position</td><td>gene name</td><td>gene id</td></tr>
<tr><td>GU305914.1</td><td></td><td>80898</td><td>80920</td><td>+</td><td>0</td><td>TGCACTGCTTCATCGCCTCC</td><td>CGG</td><td>||||||||[||||||||||||]PAM</td><td>NA</td><td>Intergenic</td><td>NA</td><td>NA</td></tr>
<tr><td>T5791</td><td>CCTCCCGGACCAGCTTGCGTTGG</td><td></td><td>1000</td><td></td><td></td><td></td><td></td><td></td><td></td><td></td><td></td><td></td></tr>
<tr><td>Oligo fwd</td><td>CACCCCTCCCGGACCAGCTTGCGT</td><td></td><td></td><td></td><td></td><td></td><td></td><td></td><td></td><td></td><td></td><td></td></tr>
<tr><td>Oligo rev</td><td>AAACACGCAAGCTGGTCCGGGAGG</td><td></td><td></td><td></td><td></td><td></td><td></td><td></td><td></td><td></td><td></td><td></td></tr>
<tr><td>Chromosome</td><td>start</td><td></td><td>end</td><td>strand</td><td>MM</td><td>target_seq</td><td>PAM</td><td>alignment</td><td>distance</td><td>position</td><td>gene name</td><td>gene id</td></tr>
<tr><td>GU305914.1</td><td></td><td>80913</td><td>80935</td><td>+</td><td>0</td><td>CCTCCCGGACCAGCTTGCGT</td><td>TGG</td><td>||||||||[||||||||||||]PAM</td><td>NA</td><td>Intergenic</td><td>NA</td><td>NA</td></tr>
<tr><td>T5792</td><td>CTCCCGGACCAGCTTGCGTTGGG</td><td></td><td>1000</td><td></td><td></td><td></td><td></td><td></td><td></td><td></td><td></td><td></td></tr>
<tr><td>Oligo fwd</td><td>CACCCTCCCGGACCAGCTTGCGTT</td><td></td><td></td><td></td><td></td><td></td><td></td><td></td><td></td><td></td><td></td><td></td></tr>
<tr><td>Oligo rev</td><td>AAACAACGCAAGCTGGTCCGGGAG</td><td></td><td></td><td></td><td></td><td></td><td></td><td></td><td></td><td></td><td></td><td></td></tr>
<tr><td>Chromosome</td><td>start</td><td></td><td>end</td><td>strand</td><td>MM</td><td>target_seq</td><td>PAM</td><td>alignment</td><td>distance</td><td>position</td><td>gene name</td><td>gene id</td></tr>
<tr><td>GU305914.1</td><td></td><td>80914</td><td>80936</td><td>+</td><td>0</td><td>CTCCCGGACCAGCTTGCGTT</td><td>GGG</td><td>||||||||[||||||||||||]PAM</td><td>NA</td><td>Intergenic</td><td>NA</td><td>NA</td></tr>
<tr><td>T5793</td><td>TAGCCATCGTACCAGCAGCTCGG</td><td></td><td>1000</td><td></td><td></td><td></td><td></td><td></td><td></td><td></td><td></td><td></td></tr>
<tr><td>Oligo fwd</td><td>CACCTAGCCATCGTACCAGCAGCT</td><td></td><td></td><td></td><td></td><td></td><td></td><td></td><td></td><td></td><td></td><td></td></tr>
<tr><td>Oligo rev</td><td>AAACAGCTGCTGGTACGATGGCTA</td><td></td><td></td><td></td><td></td><td></td><td></td><td></td><td></td><td></td><td></td><td></td></tr>
<tr><td>Chromosome</td><td>start</td><td></td><td>end</td><td>strand</td><td>MM</td><td>target_seq</td><td>PAM</td><td>alignment</td><td>distance</td><td>position</td><td>gene name</td><td>gene id</td></tr>
<tr><td>GU305914.1</td><td></td><td>80938</td><td>80960</td><td>+</td><td>0</td><td>TAGCCATCGTACCAGCAGCT</td><td>CGG</td><td>||||||||[||||||||||||]PAM</td><td>NA</td><td>Intergenic</td><td>NA</td><td>NA</td></tr>
<tr><td>T5794</td><td>GTACGTTCTCCCGCACAAACCGG</td><td></td><td>1000</td><td></td><td></td><td></td><td></td><td></td><td></td><td></td><td></td><td></td></tr>
</table>

<table>
<tr><td>Oligo fwd</td><td>CACCGTACGTTCTCCCGCACAAAC</td><td></td><td></td><td></td><td></td><td></td><td></td><td></td><td></td><td></td><td></td><td></td></tr>
<tr><td>Oligo rev</td><td>AAACGTTTGTGCGGGAGAACGTAC</td><td></td><td></td><td></td><td></td><td></td><td></td><td></td><td></td><td></td><td></td><td></td></tr>
<tr><td>Chromosome</td><td>start</td><td></td><td>end</td><td>strand</td><td>MM</td><td>target_seq</td><td>PAM</td><td>alignment</td><td>distance</td><td>position</td><td>gene name</td><td>gene id</td></tr>
<tr><td>GU305914.1</td><td></td><td>80976</td><td>80998</td><td>+</td><td>0</td><td>GTACGTTCTCCCGCACAAAC</td><td>CGG</td><td>||||||||[||||||||||||]PAM</td><td>NA</td><td>Intergenic</td><td>NA</td><td>NA</td></tr>
<tr><td>T5795</td><td>GATCACCGATTTCTACCCTCAGG</td><td></td><td>1000</td><td></td><td></td><td></td><td></td><td></td><td></td><td></td><td></td><td></td></tr>
<tr><td>Oligo fwd</td><td>CACCGATCACCGATTTCTACCCTC</td><td></td><td></td><td></td><td></td><td></td><td></td><td></td><td></td><td></td><td></td><td></td></tr>
<tr><td>Oligo rev</td><td>AAACGAGGGTAGAAATCGGTGATC</td><td></td><td></td><td></td><td></td><td></td><td></td><td></td><td></td><td></td><td></td><td></td></tr>
<tr><td>Chromosome</td><td>start</td><td></td><td>end</td><td>strand</td><td>MM</td><td>target_seq</td><td>PAM</td><td>alignment</td><td>distance</td><td>position</td><td>gene name</td><td>gene id</td></tr>
<tr><td>GU305914.1</td><td></td><td>81006</td><td>81028</td><td>+</td><td>0</td><td>GATCACCGATTTCTACCCTC</td><td>AGG</td><td>||||||||[||||||||||||]PAM</td><td>NA</td><td>Intergenic</td><td>NA</td><td>NA</td></tr>
<tr><td>T5796</td><td>ACCCTCAGGACGTCCTTTTCAGG</td><td></td><td>1000</td><td></td><td></td><td></td><td></td><td></td><td></td><td></td><td></td><td></td></tr>
<tr><td>Oligo fwd</td><td>CACCACCCTCAGGACGTCCTTTTC</td><td></td><td></td><td></td><td></td><td></td><td></td><td></td><td></td><td></td><td></td><td></td></tr>
<tr><td>Oligo rev</td><td>AAACGAAAAGGACGTCCTGAGGGT</td><td></td><td></td><td></td><td></td><td></td><td></td><td></td><td></td><td></td><td></td><td></td></tr>
<tr><td>Chromosome</td><td>start</td><td></td><td>end</td><td>strand</td><td>MM</td><td>target_seq</td><td>PAM</td><td>alignment</td><td>distance</td><td>position</td><td>gene name</td><td>gene id</td></tr>
<tr><td>GU305914.1</td><td></td><td>81020</td><td>81042</td><td>+</td><td>0</td><td>ACCCTCAGGACGTCCTTTTC</td><td>AGG</td><td>||||||||[||||||||||||]PAM</td><td>NA</td><td>Intergenic</td><td>NA</td><td>NA</td></tr>
<tr><td>T5797</td><td>GACGTCCTTTTCAGGCACCTCGG</td><td></td><td>1000</td><td></td><td></td><td></td><td></td><td></td><td></td><td></td><td></td><td></td></tr>
<tr><td>Oligo fwd</td><td>CACCGACGTCCTTTTCAGGCACCT</td><td></td><td></td><td></td><td></td><td></td><td></td><td></td><td></td><td></td><td></td><td></td></tr>
<tr><td>Oligo rev</td><td>AAACAGGTGCCTGAAAAGGACGTC</td><td></td><td></td><td></td><td></td><td></td><td></td><td></td><td></td><td></td><td></td><td></td></tr>
<tr><td>Chromosome</td><td>start</td><td></td><td>end</td><td>strand</td><td>MM</td><td>target_seq</td><td>PAM</td><td>alignment</td><td>distance</td><td>position</td><td>gene name</td><td>gene id</td></tr>
<tr><td>GU305914.1</td><td></td><td>81028</td><td>81050</td><td>+</td><td>0</td><td>GACGTCCTTTTCAGGCACCT</td><td>CGG</td><td>||||||||[||||||||||||]PAM</td><td>NA</td><td>Intergenic</td><td>NA</td><td>NA</td></tr>
<tr><td>T5798</td><td>ACGTCCTTTTCAGGCACCTCGGG</td><td></td><td>1000</td><td></td><td></td><td></td><td></td><td></td><td></td><td></td><td></td><td></td></tr>
<tr><td>Oligo fwd</td><td>CACCACGTCCTTTTCAGGCACCTC</td><td></td><td></td><td></td><td></td><td></td><td></td><td></td><td></td><td></td><td></td><td></td></tr>
<tr><td>Oligo rev</td><td>AAACGAGGTGCCTGAAAAGGACGT</td><td></td><td></td><td></td><td></td><td></td><td></td><td></td><td></td><td></td><td></td><td></td></tr>
<tr><td>Chromosome</td><td>start</td><td></td><td>end</td><td>strand</td><td>MM</td><td>target_seq</td><td>PAM</td><td>alignment</td><td>distance</td><td>position</td><td>gene name</td><td>gene id</td></tr>
<tr><td>GU305914.1</td><td></td><td>81029</td><td>81051</td><td>+</td><td>0</td><td>ACGTCCTTTTCAGGCACCTC</td><td>GGG</td><td>||||||||[||||||||||||]PAM</td><td>NA</td><td>Intergenic</td><td>NA</td><td>NA</td></tr>
<tr><td>T5799</td><td>CGTCCTTTTCAGGCACCTCGGGG</td><td></td><td>1000</td><td></td><td></td><td></td><td></td><td></td><td></td><td></td><td></td><td></td></tr>
<tr><td>Oligo fwd</td><td>CACCCGTCCTTTTCAGGCACCTCG</td><td></td><td></td><td></td><td></td><td></td><td></td><td></td><td></td><td></td><td></td><td></td></tr>
<tr><td>Oligo rev</td><td>AAACCGAGGTGCCTGAAAAGGACG</td><td></td><td></td><td></td><td></td><td></td><td></td><td></td><td></td><td></td><td></td><td></td></tr>
<tr><td>Chromosome</td><td>start</td><td></td><td>end</td><td>strand</td><td>MM</td><td>target_seq</td><td>PAM</td><td>alignment</td><td>distance</td><td>position</td><td>gene name</td><td>gene id</td></tr>
<tr><td>GU305914.1</td><td></td><td>81030</td><td>81052</td><td>+</td><td>0</td><td>CGTCCTTTTCAGGCACCTCG</td><td>GGG</td><td>||||||||[||||||||||||]PAM</td><td>NA</td><td>Intergenic</td><td>NA</td><td>NA</td></tr>
<tr><td>T5800</td><td>GTCCTTTTCAGGCACCTCGGGGG</td><td></td><td>1000</td><td></td><td></td><td></td><td></td><td></td><td></td><td></td><td></td><td></td></tr>
<tr><td>Oligo fwd</td><td>CACCGTCCTTTTCAGGCACCTCGG</td><td></td><td></td><td></td><td></td><td></td><td></td><td></td><td></td><td></td><td></td><td></td></tr>
<tr><td>Oligo rev</td><td>AAACCCGAGGTGCCTGAAAAGGAC</td><td></td><td></td><td></td><td></td><td></td><td></td><td></td><td></td><td></td><td></td><td></td></tr>
<tr><td>Chromosome</td><td>start</td><td></td><td>end</td><td>strand</td><td>MM</td><td>target_seq</td><td>PAM</td><td>alignment</td><td>distance</td><td>position</td><td>gene name</td><td>gene id</td></tr>
<tr><td>GU305914.1</td><td></td><td>81031</td><td>81053</td><td>+</td><td>0</td><td>GTCCTTTTCAGGCACCTCGG</td><td>GGG</td><td>||||||||[||||||||||||]PAM</td><td>NA</td><td>Intergenic</td><td>NA</td><td>NA</td></tr>
<tr><td>T5801</td><td>CGGGGGAGCCCTCCACCAAGAGG</td><td></td><td>1000</td><td></td><td></td><td></td><td></td><td></td><td></td><td></td><td></td><td></td></tr>
<tr><td>Oligo fwd</td><td>CACCCGGGGGAGCCCTCCACCAAG</td><td></td><td></td><td></td><td></td><td></td><td></td><td></td><td></td><td></td><td></td><td></td></tr>
<tr><td>Oligo rev</td><td>AAACCTTGGTGGAGGGCTCCCCCG</td><td></td><td></td><td></td><td></td><td></td><td></td><td></td><td></td><td></td><td></td><td></td></tr>
<tr><td>Chromosome</td><td>start</td><td></td><td>end</td><td>strand</td><td>MM</td><td>target_seq</td><td>PAM</td><td>alignment</td><td>distance</td><td>position</td><td>gene name</td><td>gene id</td></tr>
<tr><td>GU305914.1</td><td></td><td>81048</td><td>81070</td><td>+</td><td>0</td><td>CGGGGGAGCCCTCCACCAAG</td><td>AGG</td><td>||||||||[||||||||||||]PAM</td><td>NA</td><td>Intergenic</td><td>NA</td><td>NA</td></tr>
<tr><td>T5802</td><td>GGGGGAGCCCTCCACCAAGAGGG</td><td></td><td>1000</td><td></td><td></td><td></td><td></td><td></td><td></td><td></td><td></td><td></td></tr>
<tr><td>Oligo fwd</td><td>CACCGGGGGAGCCCTCCACCAAGA</td><td></td><td></td><td></td><td></td><td></td><td></td><td></td><td></td><td></td><td></td><td></td></tr>
<tr><td>Oligo rev</td><td>AAACTCTTGGTGGAGGGCTCCCCC</td><td></td><td></td><td></td><td></td><td></td><td></td><td></td><td></td><td></td><td></td><td></td></tr>
<tr><td>Chromosome</td><td>start</td><td></td><td>end</td><td>strand</td><td>MM</td><td>target_seq</td><td>PAM</td><td>alignment</td><td>distance</td><td>position</td><td>gene name</td><td>gene id</td></tr>
<tr><td>GU305914.1</td><td></td><td>81049</td><td>81071</td><td>+</td><td>0</td><td>GGGGGAGCCCTCCACCAAGA</td><td>GGG</td><td>||||||||[||||||||||||]PAM</td><td>NA</td><td>Intergenic</td><td>NA</td><td>NA</td></tr>
<tr><td>T5803</td><td>GGAGCCCTCCACCAAGAGGGTGG</td><td></td><td>1000</td><td></td><td></td><td></td><td></td><td></td><td></td><td></td><td></td><td></td></tr>
<tr><td>Oligo fwd</td><td>CACCGGAGCCCTCCACCAAGAGGG</td><td></td><td></td><td></td><td></td><td></td><td></td><td></td><td></td><td></td><td></td><td></td></tr>
<tr><td>Oligo rev</td><td>AAACCCCTCTTGGTGGAGGGCTCC</td><td></td><td></td><td></td><td></td><td></td><td></td><td></td><td></td><td></td><td></td><td></td></tr>
<tr><td>Chromosome</td><td>start</td><td></td><td>end</td><td>strand</td><td>MM</td><td>target_seq</td><td>PAM</td><td>alignment</td><td>distance</td><td>position</td><td>gene name</td><td>gene id</td></tr>
<tr><td>GU305914.1</td><td></td><td>81052</td><td>81074</td><td>+</td><td>0</td><td>GGAGCCCTCCACCAAGAGGG</td><td>TGG</td><td>||||||||[||||||||||||]PAM</td><td>NA</td><td>Intergenic</td><td>NA</td><td>NA</td></tr>
<tr><td>T5804</td><td>GGTGGAGTAGCACAGATTGTTGG</td><td></td><td>1000</td><td></td><td></td><td></td><td></td><td></td><td></td><td></td><td></td><td></td></tr>
<tr><td>Oligo fwd</td><td>CACCGGTGGAGTAGCACAGATTGT</td><td></td><td></td><td></td><td></td><td></td><td></td><td></td><td></td><td></td><td></td><td></td></tr>
<tr><td>Oligo rev</td><td>AAACACAATCTGTGCTACTCCACC</td><td></td><td></td><td></td><td></td><td></td><td></td><td></td><td></td><td></td><td></td><td></td></tr>
<tr><td>Chromosome</td><td>start</td><td></td><td>end</td><td>strand</td><td>MM</td><td>target_seq</td><td>PAM</td><td>alignment</td><td>distance</td><td>position</td><td>gene name</td><td>gene id</td></tr>
<tr><td>GU305914.1</td><td></td><td>81070</td><td>81092</td><td>+</td><td>0</td><td>GGTGGAGTAGCACAGATTGT</td><td>TGG</td><td>||||||||[||||||||||||]PAM</td><td>NA</td><td>Intergenic</td><td>NA</td><td>NA</td></tr>
<tr><td>T5805</td><td>CAGATTGTTGGACATGATGATGG</td><td></td><td>1000</td><td></td><td></td><td></td><td></td><td></td><td></td><td></td><td></td><td></td></tr>
<tr><td>Oligo fwd</td><td>CACCCAGATTGTTGGACATGATGA</td><td></td><td></td><td></td><td></td><td></td><td></td><td></td><td></td><td></td><td></td><td></td></tr>
<tr><td>Oligo rev</td><td>AAACTCATCATGTCCAACAATCTG</td><td></td><td></td><td></td><td></td><td></td><td></td><td></td><td></td><td></td><td></td><td></td></tr>
<tr><td>Chromosome</td><td>start</td><td></td><td>end</td><td>strand</td><td>MM</td><td>target_seq</td><td>PAM</td><td>alignment</td><td>distance</td><td>position</td><td>gene name</td><td>gene id</td></tr>
<tr><td>GU305914.1</td><td></td><td>81082</td><td>81104</td><td>+</td><td>0</td><td>CAGATTGTTGGACATGATGA</td><td>TGG</td><td>||||||||[||||||||||||]PAM</td><td>NA</td><td>Intergenic</td><td>NA</td><td>NA</td></tr>
<tr><td>T5806</td><td>TTGTTGGACATGATGATGGACGG</td><td></td><td>1000</td><td></td><td></td><td></td><td></td><td></td><td></td><td></td><td></td><td></td></tr>
<tr><td>Oligo fwd</td><td>CACCTTGTTGGACATGATGATGGA</td><td></td><td></td><td></td><td></td><td></td><td></td><td></td><td></td><td></td><td></td><td></td></tr>
<tr><td>Oligo rev</td><td>AAACTCCATCATCATGTCCAACAA</td><td></td><td></td><td></td><td></td><td></td><td></td><td></td><td></td><td></td><td></td><td></td></tr>
<tr><td>Chromosome</td><td>start</td><td></td><td>end</td><td>strand</td><td>MM</td><td>target_seq</td><td>PAM</td><td>alignment</td><td>distance</td><td>position</td><td>gene name</td><td>gene id</td></tr>
<tr><td>GU305914.1</td><td></td><td>81086</td><td>81108</td><td>+</td><td>0</td><td>TTGTTGGACATGATGATGGA</td><td>CGG</td><td>||||||||[||||||||||||]PAM</td><td>NA</td><td>Intergenic</td><td>NA</td><td>NA</td></tr>
<tr><td>T5807</td><td>GATGATGGACGGATACAGACTGG</td><td></td><td>1000</td><td></td><td></td><td></td><td></td><td></td><td></td><td></td><td></td><td></td></tr>
<tr><td>Oligo fwd</td><td>CACCGATGATGGACGGATACAGAC</td><td></td><td></td><td></td><td></td><td></td><td></td><td></td><td></td><td></td><td></td><td></td></tr>
<tr><td>Oligo rev</td><td>AAACGTCTGTATCCGTCCATCATC</td><td></td><td></td><td></td><td></td><td></td><td></td><td></td><td></td><td></td><td></td><td></td></tr>
<tr><td>Chromosome</td><td>start</td><td></td><td>end</td><td>strand</td><td>MM</td><td>target_seq</td><td>PAM</td><td>alignment</td><td>distance</td><td>position</td><td>gene name</td><td>gene id</td></tr>
<tr><td>GU305914.1</td><td></td><td>81097</td><td>81119</td><td>+</td><td>0</td><td>GATGATGGACGGATACAGAC</td><td>TGG</td><td>||||||||[||||||||||||]PAM</td><td>NA</td><td>Intergenic</td><td>NA</td><td>NA</td></tr>
<tr><td>T5808</td><td>GGCGAAATCGAACACGCAGACGG</td><td></td><td>1000</td><td></td><td></td><td></td><td></td><td></td><td></td><td></td><td></td><td></td></tr>
<tr><td>Oligo fwd</td><td>CACCGGCGAAATCGAACACGCAGA</td><td></td><td></td><td></td><td></td><td></td><td></td><td></td><td></td><td></td><td></td><td></td></tr>
<tr><td>Oligo rev</td><td>AAACTCTGCGTGTTCGATTTCGCC</td><td></td><td></td><td></td><td></td><td></td><td></td><td></td><td></td><td></td><td></td><td></td></tr>
<tr><td>Chromosome</td><td>start</td><td></td><td>end</td><td>strand</td><td>MM</td><td>target_seq</td><td>PAM</td><td>alignment</td><td>distance</td><td>position</td><td>gene name</td><td>gene id</td></tr>
<tr><td>GU305914.1</td><td></td><td>81118</td><td>81140</td><td>+</td><td>0</td><td>GGCGAAATCGAACACGCAGA</td><td>CGG</td><td>||||||||[||||||||||||]PAM</td><td>NA</td><td>Intergenic</td><td>NA</td><td>NA</td></tr>
</table>

T5809 GCGAAATCGAACACGCAGACGGG 1000
Oligo fwd CACCGCGAAATCGAACACGCAGAC
Oligo rev AAACGTCTGCGTGTTCGATTTCGC

| Chromosome | start | end | strand | MM | target_seq | PAM | alignment | distance | position | gene name | gene id |
|---|---|---|---|---|---|---|---|---|---|---|---|
| GU305914.1 | 81119 | 81141 | + | 0 | GCGAAATCGAACACGCAGAC | GGG | ||||||||[||||||||||||]PAM | NA | Intergenic | NA | NA |

T5810 CGAACACGCAGACGGGAACGTGG 1000
Oligo fwd CACCCGAACACGCAGACGGGAACG
Oligo rev AAACCGTTCCCGTCTGCGTGTTCG

| Chromosome | start | end | strand | MM | target_seq | PAM | alignment | distance | position | gene name | gene id |
|---|---|---|---|---|---|---|---|---|---|---|---|
| GU305914.1 | 81126 | 81148 | + | 0 | CGAACACGCAGACGGGAACG | TGG | ||||||||[||||||||||||]PAM | NA | Intergenic | NA | NA |

T5811 ACACGCAGACGGGAACGTGGTGG 1000
Oligo fwd CACCACACGCAGACGGGAACGTGG
Oligo rev AAACCCACGTTCCCGTCTGCGTGT

| Chromosome | start | end | strand | MM | target_seq | PAM | alignment | distance | position | gene name | gene id |
|---|---|---|---|---|---|---|---|---|---|---|---|
| GU305914.1 | 81129 | 81151 | + | 0 | ACACGCAGACGGGAACGTGG | TGG | ||||||||[||||||||||||]PAM | NA | Intergenic | NA | NA |

T5812 ACGTGGTGGAACCCGCACTCCGG 1000
Oligo fwd CACCACGTGGTGGAACCCGCACTC
Oligo rev AAACGAGTGCGGGTTCCACCACGT

| Chromosome | start | end | strand | MM | target_seq | PAM | alignment | distance | position | gene name | gene id |
|---|---|---|---|---|---|---|---|---|---|---|---|
| GU305914.1 | 81143 | 81165 | + | 0 | ACGTGGTGGAACCCGCACTC | CGG | ||||||||[||||||||||||]PAM | NA | Intergenic | NA | NA |

T5813 GTTCGAGCACCGTAGCTCCCTGG 1000
Oligo fwd CACCGTTCGAGCACCGTAGCTCCC
Oligo rev AAACGGGAGCTACGGTGCTCGAAC

| Chromosome | start | end | strand | MM | target_seq | PAM | alignment | distance | position | gene name | gene id |
|---|---|---|---|---|---|---|---|---|---|---|---|
| GU305914.1 | 81165 | 81187 | + | 0 | GTTCGAGCACCGTAGCTCCC | TGG | ||||||||[||||||||||||]PAM | NA | Intergenic | NA | NA |

T5814 GTTCGAGTTCTCGCCCTCCGAGG 1000
Oligo fwd CACCGTTCGAGTTCTCGCCCTCCG
Oligo rev AAACCGGAGGGCGAGAACTCGAAC

| Chromosome | start | end | strand | MM | target_seq | PAM | alignment | distance | position | gene name | gene id |
|---|---|---|---|---|---|---|---|---|---|---|---|
| GU305914.1 | 81208 | 81230 | + | 0 | GTTCGAGTTCTCGCCCTCCG | AGG | ||||||||[||||||||||||]PAM | NA | Intergenic | NA | NA |
| GU305914.1 | 226825 | 226847 | + | 4 | TTTCGAGGTCTCGCTCTCCC | AGG | -||||||-[||||||-||||-]PAM | NA | Intergenic | NA | NA |

T5815 CGCCGCCGCCTCCTCGATCGCGG 1000
Oligo fwd CACCCGCCGCCGCCTCCTCGATCG
Oligo rev AAACCGATCGAGGAGGCGGCGGCG

| Chromosome | start | end | strand | MM | target_seq | PAM | alignment | distance | position | gene name | gene id |
|---|---|---|---|---|---|---|---|---|---|---|---|
| GU305914.1 | 81250 | 81272 | + | 0 | CGCCGCCGCCTCCTCGATCG | CGG | ||||||||[||||||||||||]PAM | NA | Intergenic | NA | NA |

T5816 CGCCGCCTCCTCGATCGCGGCGG 1000
Oligo fwd CACCCGCCGCCTCCTCGATCGCGG
Oligo rev AAACCCGCGATCGAGGAGGCGGCG

| Chromosome | start | end | strand | MM | target_seq | PAM | alignment | distance | position | gene name | gene id |
|---|---|---|---|---|---|---|---|---|---|---|---|
| GU305914.1 | 81253 | 81275 | + | 0 | CGCCGCCTCCTCGATCGCGG | CGG | ||||||||[||||||||||||]PAM | NA | Intergenic | NA | NA |

T5817 CGATCGCGGCGGCCGCCTCGTGG 1000
Oligo fwd CACCCGATCGCGGCGGCCGCCTCG
Oligo rev AAACCGAGGCGGCCGCCGCGATCG

| Chromosome | start | end | strand | MM | target_seq | PAM | alignment | distance | position | gene name | gene id |
|---|---|---|---|---|---|---|---|---|---|---|---|
| GU305914.1 | 81264 | 81286 | + | 0 | CGATCGCGGCGGCCGCCTCG | TGG | ||||||||[||||||||||||]PAM | NA | Intergenic | NA | NA |
| GU305914.1 | 199811 | 199833 | - | 4 | CGCACGTGGCGGCCGCCGCG | TGG | ||--||-|[|||||||||-||]PAM | NA | Intergenic | NA | NA |

T5818 GGCCGCCTCGTGGCCGAGCGAGG 1000
Oligo fwd CACCGGCCGCCTCGTGGCCGAGCG
Oligo rev AAACCGCTCGGCCACGAGGCGGCC

| Chromosome | start | end | strand | MM | target_seq | PAM | alignment | distance | position | gene name | gene id |
|---|---|---|---|---|---|---|---|---|---|---|---|
| GU305914.1 | 81274 | 81296 | + | 0 | GGCCGCCTCGTGGCCGAGCG | AGG | ||||||||[||||||||||||]PAM | NA | Intergenic | NA | NA |

T5819 GCCGCCTCGTGGCCGAGCGAGGG 1000
Oligo fwd CACCGCCGCCTCGTGGCCGAGCGA
Oligo rev AAACTCGCTCGGCCACGAGGCGGC

| Chromosome | start | end | strand | MM | target_seq | PAM | alignment | distance | position | gene name | gene id |
|---|---|---|---|---|---|---|---|---|---|---|---|
| GU305914.1 | 81275 | 81297 | + | 0 | GCCGCCTCGTGGCCGAGCGA | GGG | ||||||||[||||||||||||]PAM | NA | Intergenic | NA | NA |

T5820 GTGGCCGAGCGAGGGCATGTTGG 1000
Oligo fwd CACCGTGGCCGAGCGAGGGCATGT
Oligo rev AAACACATGCCCTCGCTCGGCCAC

| Chromosome | start | end | strand | MM | target_seq | PAM | alignment | distance | position | gene name | gene id |
|---|---|---|---|---|---|---|---|---|---|---|---|
| GU305914.1 | 81283 | 81305 | + | 0 | GTGGCCGAGCGAGGGCATGT | TGG | ||||||||[||||||||||||]PAM | NA | Intergenic | NA | NA |

T5821 TGGCCGAGCGAGGGCATGTTGGG 1000
Oligo fwd CACCTGGCCGAGCGAGGGCATGTT
Oligo rev AAACAACATGCCCTCGCTCGGCCA

| Chromosome | start | end | strand | MM | target_seq | PAM | alignment | distance | position | gene name | gene id |
|---|---|---|---|---|---|---|---|---|---|---|---|
| GU305914.1 | 81284 | 81306 | + | 0 | TGGCCGAGCGAGGGCATGTT | GGG | ||||||||[||||||||||||]PAM | NA | Intergenic | NA | NA |

T5822 CGAGCGAGGGCATGTTGGGCAGG 1000
Oligo fwd CACCCGAGCGAGGGCATGTTGGGC
Oligo rev AAACGCCCAACATGCCCTCGCTCG

| Chromosome | start | end | strand | MM | target_seq | PAM | alignment | distance | position | gene name | gene id |
|---|---|---|---|---|---|---|---|---|---|---|---|
| GU305914.1 | 81288 | 81310 | + | 0 | CGAGCGAGGGCATGTTGGGC | AGG | ||||||||[||||||||||||]PAM | NA | Intergenic | NA | NA |

T5823 CGAACACTCCTCGAGCAGACAGG 1000
Oligo fwd CACCCGAACACTCCTCGAGCAGAC
Oligo rev AAACGTCTGCTCGAGGAGTGTTCG

| | | | | | | | | | | | |
|---|---|---|---|---|---|---|---|---|---|---|---|
| Chromosome | start | end | strand | MM | target_seq | PAM | alignment | distance | position | gene name | gene id |
| GU305914.1 | 81325 | 81347 | + | 0 | CGAACACTCCTCGAGCAGAC | AGG | \|\|\|\|\|\|\|\|[\|\|\|\|\|\|\|\|\|\|\|\|]PAM | NA | Intergenic | NA | NA |
| T5824 | CGAGCAGACAGGTGTAGATCCGG | 1000 | | | | | | | | | |
| Oligo fwd | CACCCGAGCAGACAGGTGTAGATC | | | | | | | | | | |
| Oligo rev | AAACGATCTACACCTGTCTGCTCG | | | | | | | | | | |
| Chromosome | start | end | strand | MM | target_seq | PAM | alignment | distance | position | gene name | gene id |
| GU305914.1 | 81336 | 81358 | + | 0 | CGAGCAGACAGGTGTAGATC | CGG | \|\|\|\|\|\|\|\|[\|\|\|\|\|\|\|\|\|\|\|\|]PAM | NA | Intergenic | NA | NA |
| GU305914.1 | 126818 | 126840 | + | 4 | CGCGCAGGAAGGTGTCGATC | AGG | \|\|-\|\|\|\|-[-\|\|\|\|\|\|-\|\|\|\|]PAM | NA | Intergenic | NA | NA |
| T5825 | CGTCGAAGATGACCTTCCTGAGG | 1000 | | | | | | | | | |
| Oligo fwd | CACCCGTCGAAGATGACCTTCCTG | | | | | | | | | | |
| Oligo rev | AAACCAGGAAGGTCATCTTCGACG | | | | | | | | | | |
| Chromosome | start | end | strand | MM | target_seq | PAM | alignment | distance | position | gene name | gene id |
| GU305914.1 | 81369 | 81391 | + | 0 | CGTCGAAGATGACCTTCCTG | AGG | \|\|\|\|\|\|\|\|[\|\|\|\|\|\|\|\|\|\|\|\|]PAM | NA | Intergenic | NA | NA |
| T5826 | GTCGAAGATGACCTTCCTGAGGG | 1000 | | | | | | | | | |
| Oligo fwd | CACCGTCGAAGATGACCTTCCTGA | | | | | | | | | | |
| Oligo rev | AAACTCAGGAAGGTCATCTTCGAC | | | | | | | | | | |
| Chromosome | start | end | strand | MM | target_seq | PAM | alignment | distance | position | gene name | gene id |
| GU305914.1 | 81370 | 81392 | + | 0 | GTCGAAGATGACCTTCCTGA | GGG | \|\|\|\|\|\|\|\|[\|\|\|\|\|\|\|\|\|\|\|\|]PAM | NA | Intergenic | NA | NA |
| GU305914.1 | 147821 | 147843 | - | 4 | GACGATGATGACCTTCTCGA | GGG | \|-\|\|\|-\|\|[\|\|\|\|\|\|\|\|--\|\|]PAM | NA | Intergenic | NA | NA |
| T5827 | TCGAAGATGACCTTCCTGAGGGG | 1000 | | | | | | | | | |
| Oligo fwd | CACCTCGAAGATGACCTTCCTGAG | | | | | | | | | | |
| Oligo rev | AAACCTCAGGAAGGTCATCTTCGA | | | | | | | | | | |
| Chromosome | start | end | strand | MM | target_seq | PAM | alignment | distance | position | gene name | gene id |
| GU305914.1 | 81371 | 81393 | + | 0 | TCGAAGATGACCTTCCTGAG | GGG | \|\|\|\|\|\|\|\|[\|\|\|\|\|\|\|\|\|\|\|\|]PAM | NA | Intergenic | NA | NA |
| T5828 | CGAAGATGACCTTCCTGAGGGGG | 1000 | | | | | | | | | |
| Oligo fwd | CACCCGAAGATGACCTTCCTGAGG | | | | | | | | | | |
| Oligo rev | AAACCCTCAGGAAGGTCATCTTCG | | | | | | | | | | |
| Chromosome | start | end | strand | MM | target_seq | PAM | alignment | distance | position | gene name | gene id |
| GU305914.1 | 81372 | 81394 | + | 0 | CGAAGATGACCTTCCTGAGG | GGG | \|\|\|\|\|\|\|\|[\|\|\|\|\|\|\|\|\|\|\|\|]PAM | NA | Intergenic | NA | NA |
| T5829 | GAGGGGGATCCTCGCCAACCTGG | 1000 | | | | | | | | | |
| Oligo fwd | CACCGAGGGGGATCCTCGCCAACC | | | | | | | | | | |
| Oligo rev | AAACGGTTGGCGAGGATCCCCCTC | | | | | | | | | | |
| Chromosome | start | end | strand | MM | target_seq | PAM | alignment | distance | position | gene name | gene id |
| GU305914.1 | 81388 | 81410 | + | 0 | GAGGGGGATCCTCGCCAACC | TGG | \|\|\|\|\|\|\|\|[\|\|\|\|\|\|\|\|\|\|\|\|]PAM | NA | Intergenic | NA | NA |
| T5830 | GATCCTCGCCAACCTGGCGATGG | 1000 | | | | | | | | | |
| Oligo fwd | CACCGATCCTCGCCAACCTGGCGA | | | | | | | | | | |
| Oligo rev | AAACTCGCCAGGTTGGCGAGGATC | | | | | | | | | | |
| Chromosome | start | end | strand | MM | target_seq | PAM | alignment | distance | position | gene name | gene id |
| GU305914.1 | 81394 | 81416 | + | 0 | GATCCTCGCCAACCTGGCGA | TGG | \|\|\|\|\|\|\|\|[\|\|\|\|\|\|\|\|\|\|\|\|]PAM | NA | Intergenic | NA | NA |
| T5831 | CGCCAACCTGGCGATGGCCGCGG | 1000 | | | | | | | | | |
| Oligo fwd | CACCCGCCAACCTGGCGATGGCCG | | | | | | | | | | |
| Oligo rev | AAACCGGCCATCGCCAGGTTGGCG | | | | | | | | | | |
| Chromosome | start | end | strand | MM | target_seq | PAM | alignment | distance | position | gene name | gene id |
| GU305914.1 | 81400 | 81422 | + | 0 | CGCCAACCTGGCGATGGCCG | CGG | \|\|\|\|\|\|\|\|[\|\|\|\|\|\|\|\|\|\|\|\|]PAM | NA | Intergenic | NA | NA |
| T5832 | GAAGAGTTCGCGCACGAGCACGG | 1000 | | | | | | | | | |
| Oligo fwd | CACCGAAGAGTTCGCGCACGAGCA | | | | | | | | | | |
| Oligo rev | AAACTGCTCGTGCGCGAACTCTTC | | | | | | | | | | |
| Chromosome | start | end | strand | MM | target_seq | PAM | alignment | distance | position | gene name | gene id |
| GU305914.1 | 81448 | 81470 | + | 0 | GAAGAGTTCGCGCACGAGCA | CGG | \|\|\|\|\|\|\|\|[\|\|\|\|\|\|\|\|\|\|\|\|]PAM | NA | Intergenic | NA | NA |
| T5833 | GCACGAGCACGGCGTCCTGCAGG | 1000 | | | | | | | | | |
| Oligo fwd | CACCGCACGAGCACGGCGTCCTGC | | | | | | | | | | |
| Oligo rev | AAACGCAGGACGCCGTGCTCGTGC | | | | | | | | | | |
| Chromosome | start | end | strand | MM | target_seq | PAM | alignment | distance | position | gene name | gene id |
| GU305914.1 | 81459 | 81481 | + | 0 | GCACGAGCACGGCGTCCTGC | AGG | \|\|\|\|\|\|\|\|[\|\|\|\|\|\|\|\|\|\|\|\|]PAM | NA | Intergenic | NA | NA |
| T5834 | GCAGGCAGTACTTGCCCACCCGG | 1000 | | | | | | | | | |
| Oligo fwd | CACCGCAGGCAGTACTTGCCCACC | | | | | | | | | | |
| Oligo rev | AAACGGTGGGCAAGTACTGCCTGC | | | | | | | | | | |
| Chromosome | start | end | strand | MM | target_seq | PAM | alignment | distance | position | gene name | gene id |
| GU305914.1 | 81477 | 81499 | + | 0 | GCAGGCAGTACTTGCCCACC | CGG | \|\|\|\|\|\|\|\|[\|\|\|\|\|\|\|\|\|\|\|\|]PAM | NA | Intergenic | NA | NA |
| T5835 | CAGGCAGTACTTGCCCACCCGGG | 1000 | | | | | | | | | |
| Oligo fwd | CACCCAGGCAGTACTTGCCCACCC | | | | | | | | | | |
| Oligo rev | AAACGGGTGGGCAAGTACTGCCTG | | | | | | | | | | |
| Chromosome | start | end | strand | MM | target_seq | PAM | alignment | distance | position | gene name | gene id |
| GU305914.1 | 81478 | 81500 | + | 0 | CAGGCAGTACTTGCCCACCC | GGG | \|\|\|\|\|\|\|\|[\|\|\|\|\|\|\|\|\|\|\|\|]PAM | NA | Intergenic | NA | NA |
| T5836 | CCGGGCCCGTCCCTCGTCCCCGG | 1000 | | | | | | | | | |
| Oligo fwd | CACCCCGGGCCCGTCCCTCGTCCC | | | | | | | | | | |
| Oligo rev | AAACGGGACGAGGGACGGGCCCGG | | | | | | | | | | |
| Chromosome | start | end | strand | MM | target_seq | PAM | alignment | distance | position | gene name | gene id |
| GU305914.1 | 81496 | 81518 | + | 0 | CCGGGCCCGTCCCTCGTCCC | CGG | \|\|\|\|\|\|\|\|[\|\|\|\|\|\|\|\|\|\|\|\|]PAM | NA | Intergenic | NA | NA |
| T5837 | GGCCCGTCCCTCGTCCCCGGCGG | 1000 | | | | | | | | | |
| Oligo fwd | CACCGGCCCGTCCCTCGTCCCCGG | | | | | | | | | | |
| Oligo rev | AAACCCGGGGACGAGGGACGGGCC | | | | | | | | | | |
| Chromosome | start | end | strand | MM | target_seq | PAM | alignment | distance | position | gene name | gene id |
| GU305914.1 | 81499 | 81521 | + | 0 | GGCCCGTCCCTCGTCCCCGG | CGG | \|\|\|\|\|\|\|\|[\|\|\|\|\|\|\|\|\|\|\|\|]PAM | NA | Intergenic | NA | NA |

<table>
<tr><td>T5838</td><td>CCTCGTCCCCGGCGGAGAACTGG</td><td>1000</td><td></td><td></td><td></td><td></td><td></td><td></td><td></td><td></td><td></td></tr>
<tr><td>Oligo fwd</td><td>CACCCCTCGTCCCCGGCGGAGAAC</td><td></td><td></td><td></td><td></td><td></td><td></td><td></td><td></td><td></td><td></td></tr>
<tr><td>Oligo rev</td><td>AAACGTTCTCCGCCGGGGACGAGG</td><td></td><td></td><td></td><td></td><td></td><td></td><td></td><td></td><td></td><td></td></tr>
<tr><td>Chromosome</td><td>start</td><td>end</td><td>strand</td><td>MM</td><td>target_seq</td><td>PAM</td><td>alignment</td><td>distance</td><td>position</td><td>gene name</td><td>gene id</td></tr>
<tr><td>GU305914.1</td><td>81507</td><td>81529</td><td>+</td><td>0</td><td>CCTCGTCCCCGGCGGAGAAC</td><td>TGG</td><td>||||||||[||||||||||||]PAM</td><td>NA</td><td>Intergenic</td><td>NA</td><td>NA</td></tr>
<tr><td>T5839</td><td>GTCCCCGGCGGAGAACTGGACGG</td><td>1000</td><td></td><td></td><td></td><td></td><td></td><td></td><td></td><td></td><td></td></tr>
<tr><td>Oligo fwd</td><td>CACCGTCCCCGGCGGAGAACTGGA</td><td></td><td></td><td></td><td></td><td></td><td></td><td></td><td></td><td></td><td></td></tr>
<tr><td>Oligo rev</td><td>AAACTCCAGTTCTCCGCCGGGGAC</td><td></td><td></td><td></td><td></td><td></td><td></td><td></td><td></td><td></td><td></td></tr>
<tr><td>Chromosome</td><td>start</td><td>end</td><td>strand</td><td>MM</td><td>target_seq</td><td>PAM</td><td>alignment</td><td>distance</td><td>position</td><td>gene name</td><td>gene id</td></tr>
<tr><td>GU305914.1</td><td>81511</td><td>81533</td><td>+</td><td>0</td><td>GTCCCCGGCGGAGAACTGGA</td><td>CGG</td><td>||||||||[||||||||||||]PAM</td><td>NA</td><td>Intergenic</td><td>NA</td><td>NA</td></tr>
<tr><td>T5840</td><td>TCCCCGGCGGAGAACTGGACGGG</td><td>1000</td><td></td><td></td><td></td><td></td><td></td><td></td><td></td><td></td><td></td></tr>
<tr><td>Oligo fwd</td><td>CACCTCCCCGGCGGAGAACTGGAC</td><td></td><td></td><td></td><td></td><td></td><td></td><td></td><td></td><td></td><td></td></tr>
<tr><td>Oligo rev</td><td>AAACGTCCAGTTCTCCGCCGGGGA</td><td></td><td></td><td></td><td></td><td></td><td></td><td></td><td></td><td></td><td></td></tr>
<tr><td>Chromosome</td><td>start</td><td>end</td><td>strand</td><td>MM</td><td>target_seq</td><td>PAM</td><td>alignment</td><td>distance</td><td>position</td><td>gene name</td><td>gene id</td></tr>
<tr><td>GU305914.1</td><td>81512</td><td>81534</td><td>+</td><td>0</td><td>TCCCCGGCGGAGAACTGGAC</td><td>GGG</td><td>||||||||[||||||||||||]PAM</td><td>NA</td><td>Intergenic</td><td>NA</td><td>NA</td></tr>
<tr><td>T5841</td><td>CCCCGGCGGAGAACTGGACGGGG</td><td>1000</td><td></td><td></td><td></td><td></td><td></td><td></td><td></td><td></td><td></td></tr>
<tr><td>Oligo fwd</td><td>CACCCCCCGGCGGAGAACTGGACG</td><td></td><td></td><td></td><td></td><td></td><td></td><td></td><td></td><td></td><td></td></tr>
<tr><td>Oligo rev</td><td>AAACCGTCCAGTTCTCCGCCGGGG</td><td></td><td></td><td></td><td></td><td></td><td></td><td></td><td></td><td></td><td></td></tr>
<tr><td>Chromosome</td><td>start</td><td>end</td><td>strand</td><td>MM</td><td>target_seq</td><td>PAM</td><td>alignment</td><td>distance</td><td>position</td><td>gene name</td><td>gene id</td></tr>
<tr><td>GU305914.1</td><td>81513</td><td>81535</td><td>+</td><td>0</td><td>CCCCGGCGGAGAACTGGACG</td><td>GGG</td><td>||||||||[||||||||||||]PAM</td><td>NA</td><td>Intergenic</td><td>NA</td><td>NA</td></tr>
<tr><td>T5842</td><td>CGGGGATCTCTTTGTAGCTCAGG</td><td>1000</td><td></td><td></td><td></td><td></td><td></td><td></td><td></td><td></td><td></td></tr>
<tr><td>Oligo fwd</td><td>CACCCGGGGATCTCTTTGTAGCTC</td><td></td><td></td><td></td><td></td><td></td><td></td><td></td><td></td><td></td><td></td></tr>
<tr><td>Oligo rev</td><td>AAACGAGCTACAAAGAGATCCCCG</td><td></td><td></td><td></td><td></td><td></td><td></td><td></td><td></td><td></td><td></td></tr>
<tr><td>Chromosome</td><td>start</td><td>end</td><td>strand</td><td>MM</td><td>target_seq</td><td>PAM</td><td>alignment</td><td>distance</td><td>position</td><td>gene name</td><td>gene id</td></tr>
<tr><td>GU305914.1</td><td>81531</td><td>81553</td><td>+</td><td>0</td><td>CGGGGATCTCTTTGTAGCTC</td><td>AGG</td><td>||||||||[||||||||||||]PAM</td><td>NA</td><td>Intergenic</td><td>NA</td><td>NA</td></tr>
<tr><td>T5843</td><td>GGTCGTCTTTCTTCTTCTTGAGG</td><td>1000</td><td></td><td></td><td></td><td></td><td></td><td></td><td></td><td></td><td></td></tr>
<tr><td>Oligo fwd</td><td>CACCGGTCGTCTTTCTTCTTCTTG</td><td></td><td></td><td></td><td></td><td></td><td></td><td></td><td></td><td></td><td></td></tr>
<tr><td>Oligo rev</td><td>AAACCAAGAAGAAGAAAGACGACC</td><td></td><td></td><td></td><td></td><td></td><td></td><td></td><td></td><td></td><td></td></tr>
<tr><td>Chromosome</td><td>start</td><td>end</td><td>strand</td><td>MM</td><td>target_seq</td><td>PAM</td><td>alignment</td><td>distance</td><td>position</td><td>gene name</td><td>gene id</td></tr>
<tr><td>GU305914.1</td><td>81552</td><td>81574</td><td>+</td><td>0</td><td>GGTCGTCTTTCTTCTTCTTG</td><td>AGG</td><td>||||||||[||||||||||||]PAM</td><td>NA</td><td>Intergenic</td><td>NA</td><td>NA</td></tr>
<tr><td>T5844</td><td>CTTCTTCTTGAGGTAGATCTCGG</td><td>1000</td><td></td><td></td><td></td><td></td><td></td><td></td><td></td><td></td><td></td></tr>
<tr><td>Oligo fwd</td><td>CACCCTTCTTCTTGAGGTAGATCT</td><td></td><td></td><td></td><td></td><td></td><td></td><td></td><td></td><td></td><td></td></tr>
<tr><td>Oligo rev</td><td>AAACAGATCTACCTCAAGAAGAAG</td><td></td><td></td><td></td><td></td><td></td><td></td><td></td><td></td><td></td><td></td></tr>
<tr><td>Chromosome</td><td>start</td><td>end</td><td>strand</td><td>MM</td><td>target_seq</td><td>PAM</td><td>alignment</td><td>distance</td><td>position</td><td>gene name</td><td>gene id</td></tr>
<tr><td>GU305914.1</td><td>81562</td><td>81584</td><td>+</td><td>0</td><td>CTTCTTCTTGAGGTAGATCT</td><td>CGG</td><td>||||||||[||||||||||||]PAM</td><td>NA</td><td>Intergenic</td><td>NA</td><td>NA</td></tr>
<tr><td>T5845</td><td>CTTGAGGTAGATCTCGGCCATGG</td><td>1000</td><td></td><td></td><td></td><td></td><td></td><td></td><td></td><td></td><td></td></tr>
<tr><td>Oligo fwd</td><td>CACCCTTGAGGTAGATCTCGGCCA</td><td></td><td></td><td></td><td></td><td></td><td></td><td></td><td></td><td></td><td></td></tr>
<tr><td>Oligo rev</td><td>AAACTGGCCGAGATCTACCTCAAG</td><td></td><td></td><td></td><td></td><td></td><td></td><td></td><td></td><td></td><td></td></tr>
<tr><td>Chromosome</td><td>start</td><td>end</td><td>strand</td><td>MM</td><td>target_seq</td><td>PAM</td><td>alignment</td><td>distance</td><td>position</td><td>gene name</td><td>gene id</td></tr>
<tr><td>GU305914.1</td><td>81568</td><td>81590</td><td>+</td><td>0</td><td>CTTGAGGTAGATCTCGGCCA</td><td>TGG</td><td>||||||||[||||||||||||]PAM</td><td>NA</td><td>Intergenic</td><td>NA</td><td>NA</td></tr>
<tr><td>T5846</td><td>CATGGTGTCGAGCTTGTAGTTGG</td><td>1000</td><td></td><td></td><td></td><td></td><td></td><td></td><td></td><td></td><td></td></tr>
<tr><td>Oligo fwd</td><td>CACCCATGGTGTCGAGCTTGTAGT</td><td></td><td></td><td></td><td></td><td></td><td></td><td></td><td></td><td></td><td></td></tr>
<tr><td>Oligo rev</td><td>AAACACTACAAGCTCGACACCATG</td><td></td><td></td><td></td><td></td><td></td><td></td><td></td><td></td><td></td><td></td></tr>
<tr><td>Chromosome</td><td>start</td><td>end</td><td>strand</td><td>MM</td><td>target_seq</td><td>PAM</td><td>alignment</td><td>distance</td><td>position</td><td>gene name</td><td>gene id</td></tr>
<tr><td>GU305914.1</td><td>81586</td><td>81608</td><td>+</td><td>0</td><td>CATGGTGTCGAGCTTGTAGT</td><td>TGG</td><td>||||||||[||||||||||||]PAM</td><td>NA</td><td>Intergenic</td><td>NA</td><td>NA</td></tr>
<tr><td>T5847</td><td>ATGGTGTCGAGCTTGTAGTTGGG</td><td>1000</td><td></td><td></td><td></td><td></td><td></td><td></td><td></td><td></td><td></td></tr>
<tr><td>Oligo fwd</td><td>CACCATGGTGTCGAGCTTGTAGTT</td><td></td><td></td><td></td><td></td><td></td><td></td><td></td><td></td><td></td><td></td></tr>
<tr><td>Oligo rev</td><td>AAACAACTACAAGCTCGACACCAT</td><td></td><td></td><td></td><td></td><td></td><td></td><td></td><td></td><td></td><td></td></tr>
<tr><td>Chromosome</td><td>start</td><td>end</td><td>strand</td><td>MM</td><td>target_seq</td><td>PAM</td><td>alignment</td><td>distance</td><td>position</td><td>gene name</td><td>gene id</td></tr>
<tr><td>GU305914.1</td><td>81587</td><td>81609</td><td>+</td><td>0</td><td>ATGGTGTCGAGCTTGTAGTT</td><td>GGG</td><td>||||||||[||||||||||||]PAM</td><td>NA</td><td>Intergenic</td><td>NA</td><td>NA</td></tr>
<tr><td>T5848</td><td>GAGCTTGTAGTTGGGCGAGCTGG</td><td>1000</td><td></td><td></td><td></td><td></td><td></td><td></td><td></td><td></td><td></td></tr>
<tr><td>Oligo fwd</td><td>CACCGAGCTTGTAGTTGGGCGAGC</td><td></td><td></td><td></td><td></td><td></td><td></td><td></td><td></td><td></td><td></td></tr>
<tr><td>Oligo rev</td><td>AAACGCTCGCCCAACTACAAGCTC</td><td></td><td></td><td></td><td></td><td></td><td></td><td></td><td></td><td></td><td></td></tr>
<tr><td>Chromosome</td><td>start</td><td>end</td><td>strand</td><td>MM</td><td>target_seq</td><td>PAM</td><td>alignment</td><td>distance</td><td>position</td><td>gene name</td><td>gene id</td></tr>
<tr><td>GU305914.1</td><td>81595</td><td>81617</td><td>+</td><td>0</td><td>GAGCTTGTAGTTGGGCGAGC</td><td>TGG</td><td>||||||||[||||||||||||]PAM</td><td>NA</td><td>Intergenic</td><td>NA</td><td>NA</td></tr>
<tr><td>T5849</td><td>GTAGTTGGGCGAGCTGGCCTTGG</td><td>1000</td><td></td><td></td><td></td><td></td><td></td><td></td><td></td><td></td><td></td></tr>
<tr><td>Oligo fwd</td><td>CACCGTAGTTGGGCGAGCTGGCCT</td><td></td><td></td><td></td><td></td><td></td><td></td><td></td><td></td><td></td><td></td></tr>
<tr><td>Oligo rev</td><td>AAACAGGCCAGCTCGCCCAACTAC</td><td></td><td></td><td></td><td></td><td></td><td></td><td></td><td></td><td></td><td></td></tr>
<tr><td>Chromosome</td><td>start</td><td>end</td><td>strand</td><td>MM</td><td>target_seq</td><td>PAM</td><td>alignment</td><td>distance</td><td>position</td><td>gene name</td><td>gene id</td></tr>
<tr><td>GU305914.1</td><td>81601</td><td>81623</td><td>+</td><td>0</td><td>GTAGTTGGGCGAGCTGGCCT</td><td>TGG</td><td>||||||||[||||||||||||]PAM</td><td>NA</td><td>Intergenic</td><td>NA</td><td>NA</td></tr>
<tr><td>T5850</td><td>GTTGGGCGAGCTGGCCTTGGCGG</td><td>1000</td><td></td><td></td><td></td><td></td><td></td><td></td><td></td><td></td><td></td></tr>
<tr><td>Oligo fwd</td><td>CACCGTTGGGCGAGCTGGCCTTGG</td><td></td><td></td><td></td><td></td><td></td><td></td><td></td><td></td><td></td><td></td></tr>
<tr><td>Oligo rev</td><td>AAACCCAAGGCCAGCTCGCCCAAC</td><td></td><td></td><td></td><td></td><td></td><td></td><td></td><td></td><td></td><td></td></tr>
<tr><td>Chromosome</td><td>start</td><td>end</td><td>strand</td><td>MM</td><td>target_seq</td><td>PAM</td><td>alignment</td><td>distance</td><td>position</td><td>gene name</td><td>gene id</td></tr>
<tr><td>GU305914.1</td><td>81604</td><td>81626</td><td>+</td><td>0</td><td>GTTGGGCGAGCTGGCCTTGG</td><td>CGG</td><td>||||||||[||||||||||||]PAM</td><td>NA</td><td>Intergenic</td><td>NA</td><td>NA</td></tr>
<tr><td>T5851</td><td>GCTGGCCTTGGCGGTACAGACGG</td><td>1000</td><td></td><td></td><td></td><td></td><td></td><td></td><td></td><td></td><td></td></tr>
<tr><td>Oligo fwd</td><td>CACCGCTGGCCTTGGCGGTACAGA</td><td></td><td></td><td></td><td></td><td></td><td></td><td></td><td></td><td></td><td></td></tr>
<tr><td>Oligo rev</td><td>AAACTCTGTACCGCCAAGGCCAGC</td><td></td><td></td><td></td><td></td><td></td><td></td><td></td><td></td><td></td><td></td></tr>
<tr><td>Chromosome</td><td>start</td><td>end</td><td>strand</td><td>MM</td><td>target_seq</td><td>PAM</td><td>alignment</td><td>distance</td><td>position</td><td>gene name</td><td>gene id</td></tr>
<tr><td>GU305914.1</td><td>81613</td><td>81635</td><td>+</td><td>0</td><td>GCTGGCCTTGGCGGTACAGA</td><td>CGG</td><td>||||||||[||||||||||||]PAM</td><td>NA</td><td>Intergenic</td><td>NA</td><td>NA</td></tr>
<tr><td>T5852</td><td>CTGGCCTTGGCGGTACAGACGGG</td><td>1000</td><td></td><td></td><td></td><td></td><td></td><td></td><td></td><td></td><td></td></tr>
<tr><td>Oligo fwd</td><td>CACCCTGGCCTTGGCGGTACAGAC</td><td></td><td></td><td></td><td></td><td></td><td></td><td></td><td></td><td></td><td></td></tr>
<tr><td>Oligo rev</td><td>AAACGTCTGTACCGCCAAGGCCAG</td><td></td><td></td><td></td><td></td><td></td><td></td><td></td><td></td><td></td><td></td></tr>
<tr><td>Chromosome</td><td>start</td><td>end</td><td>strand</td><td>MM</td><td>target_seq</td><td>PAM</td><td>alignment</td><td>distance</td><td>position</td><td>gene name</td><td>gene id</td></tr>
<tr><td>GU305914.1</td><td>81614</td><td>81636</td><td>+</td><td>0</td><td>CTGGCCTTGGCGGTACAGAC</td><td>GGG</td><td>||||||||[||||||||||||]PAM</td><td>NA</td><td>Intergenic</td><td>NA</td><td>NA</td></tr>
</table>

<table>
<tr><td>T5853</td><td>TGGCCTTGGCGGTACAGACGGGG</td><td>1000</td><td></td><td></td><td></td><td></td><td></td><td></td><td></td><td></td><td></td></tr>
<tr><td>Oligo fwd</td><td>CACCTGGCCTTGGCGGTACAGACG</td><td></td><td></td><td></td><td></td><td></td><td></td><td></td><td></td><td></td><td></td></tr>
<tr><td>Oligo rev</td><td>AAACCGTCTGTACCGCCAAGGCCA</td><td></td><td></td><td></td><td></td><td></td><td></td><td></td><td></td><td></td><td></td></tr>
<tr><td>Chromosome</td><td>start</td><td>end</td><td>strand</td><td>MM</td><td>target_seq</td><td>PAM</td><td>alignment</td><td>distance</td><td>position</td><td>gene name</td><td>gene id</td></tr>
<tr><td>GU305914.1</td><td>81615</td><td>81637</td><td>+</td><td>0</td><td>TGGCCTTGGCGGTACAGACG</td><td>GGG</td><td>||||||||[||||||||||||]PAM</td><td>NA</td><td>Intergenic</td><td>NA</td><td>NA</td></tr>
<tr><td>T5854</td><td>TACAGACGGGGTACATGTCGAGG</td><td>1000</td><td></td><td></td><td></td><td></td><td></td><td></td><td></td><td></td><td></td></tr>
<tr><td>Oligo fwd</td><td>CACCTACAGACGGGGTACATGTCG</td><td></td><td></td><td></td><td></td><td></td><td></td><td></td><td></td><td></td><td></td></tr>
<tr><td>Oligo rev</td><td>AAACCGACATGTACCCCGTCTGTA</td><td></td><td></td><td></td><td></td><td></td><td></td><td></td><td></td><td></td><td></td></tr>
<tr><td>Chromosome</td><td>start</td><td>end</td><td>strand</td><td>MM</td><td>target_seq</td><td>PAM</td><td>alignment</td><td>distance</td><td>position</td><td>gene name</td><td>gene id</td></tr>
<tr><td>GU305914.1</td><td>81627</td><td>81649</td><td>+</td><td>0</td><td>TACAGACGGGGTACATGTCG</td><td>AGG</td><td>||||||||[||||||||||||]PAM</td><td>NA</td><td>Intergenic</td><td>NA</td><td>NA</td></tr>
<tr><td>T5855</td><td>CCAGAAGACCTTGACCGTGTTGG</td><td>1000</td><td></td><td></td><td></td><td></td><td></td><td></td><td></td><td></td><td></td></tr>
<tr><td>Oligo fwd</td><td>CACCCCAGAAGACCTTGACCGTGT</td><td></td><td></td><td></td><td></td><td></td><td></td><td></td><td></td><td></td><td></td></tr>
<tr><td>Oligo rev</td><td>AAACACACGGTCAAGGTCTTCTGG</td><td></td><td></td><td></td><td></td><td></td><td></td><td></td><td></td><td></td><td></td></tr>
<tr><td>Chromosome</td><td>start</td><td>end</td><td>strand</td><td>MM</td><td>target_seq</td><td>PAM</td><td>alignment</td><td>distance</td><td>position</td><td>gene name</td><td>gene id</td></tr>
<tr><td>GU305914.1</td><td>81664</td><td>81686</td><td>+</td><td>0</td><td>CCAGAAGACCTTGACCGTGT</td><td>TGG</td><td>||||||||[||||||||||||]PAM</td><td>NA</td><td>Intergenic</td><td>NA</td><td>NA</td></tr>
<tr><td>T5856</td><td>GAAGACCTTGACCGTGTTGGAGG</td><td>1000</td><td></td><td></td><td></td><td></td><td></td><td></td><td></td><td></td><td></td></tr>
<tr><td>Oligo fwd</td><td>CACCGAAGACCTTGACCGTGTTGG</td><td></td><td></td><td></td><td></td><td></td><td></td><td></td><td></td><td></td><td></td></tr>
<tr><td>Oligo rev</td><td>AAACCCAACACGGTCAAGGTCTTC</td><td></td><td></td><td></td><td></td><td></td><td></td><td></td><td></td><td></td><td></td></tr>
<tr><td>Chromosome</td><td>start</td><td>end</td><td>strand</td><td>MM</td><td>target_seq</td><td>PAM</td><td>alignment</td><td>distance</td><td>position</td><td>gene name</td><td>gene id</td></tr>
<tr><td>GU305914.1</td><td>81667</td><td>81689</td><td>+</td><td>0</td><td>GAAGACCTTGACCGTGTTGG</td><td>AGG</td><td>||||||||[||||||||||||]PAM</td><td>NA</td><td>Intergenic</td><td>NA</td><td>NA</td></tr>
<tr><td>T5857</td><td>CGTGTTGGAGGTGCCGAAGCCGG</td><td>1000</td><td></td><td></td><td></td><td></td><td></td><td></td><td></td><td></td><td></td></tr>
<tr><td>Oligo fwd</td><td>CACCCGTGTTGGAGGTGCCGAAGC</td><td></td><td></td><td></td><td></td><td></td><td></td><td></td><td></td><td></td><td></td></tr>
<tr><td>Oligo rev</td><td>AAACGCTTCGGCACCTCCAACACG</td><td></td><td></td><td></td><td></td><td></td><td></td><td></td><td></td><td></td><td></td></tr>
<tr><td>Chromosome</td><td>start</td><td>end</td><td>strand</td><td>MM</td><td>target_seq</td><td>PAM</td><td>alignment</td><td>distance</td><td>position</td><td>gene name</td><td>gene id</td></tr>
<tr><td>GU305914.1</td><td>81679</td><td>81701</td><td>+</td><td>0</td><td>CGTGTTGGAGGTGCCGAAGC</td><td>CGG</td><td>||||||||[||||||||||||]PAM</td><td>NA</td><td>Intergenic</td><td>NA</td><td>NA</td></tr>
<tr><td>T5858</td><td>GAAGCCGGCCTTGCCCTTCTCGG</td><td>1000</td><td></td><td></td><td></td><td></td><td></td><td></td><td></td><td></td><td></td></tr>
<tr><td>Oligo fwd</td><td>CACCGAAGCCGGCCTTGCCCTTCT</td><td></td><td></td><td></td><td></td><td></td><td></td><td></td><td></td><td></td><td></td></tr>
<tr><td>Oligo rev</td><td>AAACAGAAGGGCAAGGCCGGCTTC</td><td></td><td></td><td></td><td></td><td></td><td></td><td></td><td></td><td></td><td></td></tr>
<tr><td>Chromosome</td><td>start</td><td>end</td><td>strand</td><td>MM</td><td>target_seq</td><td>PAM</td><td>alignment</td><td>distance</td><td>position</td><td>gene name</td><td>gene id</td></tr>
<tr><td>GU305914.1</td><td>81694</td><td>81716</td><td>+</td><td>0</td><td>GAAGCCGGCCTTGCCCTTCT</td><td>CGG</td><td>||||||||[||||||||||||]PAM</td><td>NA</td><td>Intergenic</td><td>NA</td><td>NA</td></tr>
<tr><td>T5859</td><td>AAGCCGGCCTTGCCCTTCTCGGG</td><td>1000</td><td></td><td></td><td></td><td></td><td></td><td></td><td></td><td></td><td></td></tr>
<tr><td>Oligo fwd</td><td>CACCAAGCCGGCCTTGCCCTTCTC</td><td></td><td></td><td></td><td></td><td></td><td></td><td></td><td></td><td></td><td></td></tr>
<tr><td>Oligo rev</td><td>AAACGAGAAGGGCAAGGCCGGCTT</td><td></td><td></td><td></td><td></td><td></td><td></td><td></td><td></td><td></td><td></td></tr>
<tr><td>Chromosome</td><td>start</td><td>end</td><td>strand</td><td>MM</td><td>target_seq</td><td>PAM</td><td>alignment</td><td>distance</td><td>position</td><td>gene name</td><td>gene id</td></tr>
<tr><td>GU305914.1</td><td>81695</td><td>81717</td><td>+</td><td>0</td><td>AAGCCGGCCTTGCCCTTCTC</td><td>GGG</td><td>||||||||[||||||||||||]PAM</td><td>NA</td><td>Intergenic</td><td>NA</td><td>NA</td></tr>
<tr><td>T5860</td><td>AGCCGGCCTTGCCCTTCTCGGGG</td><td>1000</td><td></td><td></td><td></td><td></td><td></td><td></td><td></td><td></td><td></td></tr>
<tr><td>Oligo fwd</td><td>CACCAGCCGGCCTTGCCCTTCTCG</td><td></td><td></td><td></td><td></td><td></td><td></td><td></td><td></td><td></td><td></td></tr>
<tr><td>Oligo rev</td><td>AAACCGAGAAGGGCAAGGCCGGCT</td><td></td><td></td><td></td><td></td><td></td><td></td><td></td><td></td><td></td><td></td></tr>
<tr><td>Chromosome</td><td>start</td><td>end</td><td>strand</td><td>MM</td><td>target_seq</td><td>PAM</td><td>alignment</td><td>distance</td><td>position</td><td>gene name</td><td>gene id</td></tr>
<tr><td>GU305914.1</td><td>81696</td><td>81718</td><td>+</td><td>0</td><td>AGCCGGCCTTGCCCTTCTCG</td><td>GGG</td><td>||||||||[||||||||||||]PAM</td><td>NA</td><td>Intergenic</td><td>NA</td><td>NA</td></tr>
<tr><td>T5861</td><td>CGGGGCTGAAGACGAACATTCGG</td><td>1000</td><td></td><td></td><td></td><td></td><td></td><td></td><td></td><td></td><td></td></tr>
<tr><td>Oligo fwd</td><td>CACCCGGGGCTGAAGACGAACATT</td><td></td><td></td><td></td><td></td><td></td><td></td><td></td><td></td><td></td><td></td></tr>
<tr><td>Oligo rev</td><td>AAACAATGTTCGTCTTCAGCCCCG</td><td></td><td></td><td></td><td></td><td></td><td></td><td></td><td></td><td></td><td></td></tr>
<tr><td>Chromosome</td><td>start</td><td>end</td><td>strand</td><td>MM</td><td>target_seq</td><td>PAM</td><td>alignment</td><td>distance</td><td>position</td><td>gene name</td><td>gene id</td></tr>
<tr><td>GU305914.1</td><td>81714</td><td>81736</td><td>+</td><td>0</td><td>CGGGGCTGAAGACGAACATT</td><td>CGG</td><td>||||||||[||||||||||||]PAM</td><td>NA</td><td>Intergenic</td><td>NA</td><td>NA</td></tr>
<tr><td>T5862</td><td>TCGGCCCCCCCTCCGCAGCTTGG</td><td>1000</td><td></td><td></td><td></td><td></td><td></td><td></td><td></td><td></td><td></td></tr>
<tr><td>Oligo fwd</td><td>CACCTCGGCCCCCCCTCCGCAGCT</td><td></td><td></td><td></td><td></td><td></td><td></td><td></td><td></td><td></td><td></td></tr>
<tr><td>Oligo rev</td><td>AAACAGCTGCGGAGGGGGGGCCGA</td><td></td><td></td><td></td><td></td><td></td><td></td><td></td><td></td><td></td><td></td></tr>
<tr><td>Chromosome</td><td>start</td><td>end</td><td>strand</td><td>MM</td><td>target_seq</td><td>PAM</td><td>alignment</td><td>distance</td><td>position</td><td>gene name</td><td>gene id</td></tr>
<tr><td>GU305914.1</td><td>81733</td><td>81755</td><td>+</td><td>0</td><td>TCGGCCCCCCCTCCGCAGCT</td><td>TGG</td><td>||||||||[||||||||||||]PAM</td><td>NA</td><td>Intergenic</td><td>NA</td><td>NA</td></tr>
<tr><td>T5863</td><td>CCCTCCGCAGCTTGGTGAATTGG</td><td>1000</td><td></td><td></td><td></td><td></td><td></td><td></td><td></td><td></td><td></td></tr>
<tr><td>Oligo fwd</td><td>CACCCCCTCCGCAGCTTGGTGAAT</td><td></td><td></td><td></td><td></td><td></td><td></td><td></td><td></td><td></td><td></td></tr>
<tr><td>Oligo rev</td><td>AAACATTCACCAAGCTGCGGAGGG</td><td></td><td></td><td></td><td></td><td></td><td></td><td></td><td></td><td></td><td></td></tr>
<tr><td>Chromosome</td><td>start</td><td>end</td><td>strand</td><td>MM</td><td>target_seq</td><td>PAM</td><td>alignment</td><td>distance</td><td>position</td><td>gene name</td><td>gene id</td></tr>
<tr><td>GU305914.1</td><td>81741</td><td>81763</td><td>+</td><td>0</td><td>CCCTCCGCAGCTTGGTGAAT</td><td>TGG</td><td>||||||||[||||||||||||]PAM</td><td>NA</td><td>Intergenic</td><td>NA</td><td>NA</td></tr>
<tr><td>T5864</td><td>ATCCATCTTGTAGAGTTTCTCGG</td><td>1000</td><td></td><td></td><td></td><td></td><td></td><td></td><td></td><td></td><td></td></tr>
<tr><td>Oligo fwd</td><td>CACCATCCATCTTGTAGAGTTTCT</td><td></td><td></td><td></td><td></td><td></td><td></td><td></td><td></td><td></td><td></td></tr>
<tr><td>Oligo rev</td><td>AAACAGAAACTCTACAAGATGGAT</td><td></td><td></td><td></td><td></td><td></td><td></td><td></td><td></td><td></td><td></td></tr>
<tr><td>Chromosome</td><td>start</td><td>end</td><td>strand</td><td>MM</td><td>target_seq</td><td>PAM</td><td>alignment</td><td>distance</td><td>position</td><td>gene name</td><td>gene id</td></tr>
<tr><td>GU305914.1</td><td>81769</td><td>81791</td><td>+</td><td>0</td><td>ATCCATCTTGTAGAGTTTCT</td><td>CGG</td><td>||||||||[||||||||||||]PAM</td><td>NA</td><td>Intergenic</td><td>NA</td><td>NA</td></tr>
<tr><td>T5865</td><td>TCTTGTAGAGTTTCTCGGCGCGG</td><td>1000</td><td></td><td></td><td></td><td></td><td></td><td></td><td></td><td></td><td></td></tr>
<tr><td>Oligo fwd</td><td>CACCTCTTGTAGAGTTTCTCGGCG</td><td></td><td></td><td></td><td></td><td></td><td></td><td></td><td></td><td></td><td></td></tr>
<tr><td>Oligo rev</td><td>AAACCGCCGAGAAACTCTACAAGA</td><td></td><td></td><td></td><td></td><td></td><td></td><td></td><td></td><td></td><td></td></tr>
<tr><td>Chromosome</td><td>start</td><td>end</td><td>strand</td><td>MM</td><td>target_seq</td><td>PAM</td><td>alignment</td><td>distance</td><td>position</td><td>gene name</td><td>gene id</td></tr>
<tr><td>GU305914.1</td><td>81774</td><td>81796</td><td>+</td><td>0</td><td>TCTTGTAGAGTTTCTCGGCG</td><td>CGG</td><td>||||||||[||||||||||||]PAM</td><td>NA</td><td>Intergenic</td><td>NA</td><td>NA</td></tr>
<tr><td>T5866</td><td>CTTGTAGAGTTTCTCGGCGCGGG</td><td>1000</td><td></td><td></td><td></td><td></td><td></td><td></td><td></td><td></td><td></td></tr>
<tr><td>Oligo fwd</td><td>CACCCTTGTAGAGTTTCTCGGCGC</td><td></td><td></td><td></td><td></td><td></td><td></td><td></td><td></td><td></td><td></td></tr>
<tr><td>Oligo rev</td><td>AAACGCGCCGAGAAACTCTACAAG</td><td></td><td></td><td></td><td></td><td></td><td></td><td></td><td></td><td></td><td></td></tr>
<tr><td>Chromosome</td><td>start</td><td>end</td><td>strand</td><td>MM</td><td>target_seq</td><td>PAM</td><td>alignment</td><td>distance</td><td>position</td><td>gene name</td><td>gene id</td></tr>
<tr><td>GU305914.1</td><td>81775</td><td>81797</td><td>+</td><td>0</td><td>CTTGTAGAGTTTCTCGGCGC</td><td>GGG</td><td>||||||||[||||||||||||]PAM</td><td>NA</td><td>Intergenic</td><td>NA</td><td>NA</td></tr>
<tr><td>T5867</td><td>AGAGTTTCTCGGCGCGGGTCAGG</td><td>1000</td><td></td><td></td><td></td><td></td><td></td><td></td><td></td><td></td><td></td></tr>
<tr><td>Oligo fwd</td><td>CACCAGAGTTTCTCGGCGCGGGTC</td><td></td><td></td><td></td><td></td><td></td><td></td><td></td><td></td><td></td><td></td></tr>
<tr><td>Oligo rev</td><td>AAACGACCCGCGCCGAGAAACTCT</td><td></td><td></td><td></td><td></td><td></td><td></td><td></td><td></td><td></td><td></td></tr>
<tr><td>Chromosome</td><td>start</td><td>end</td><td>strand</td><td>MM</td><td>target_seq</td><td>PAM</td><td>alignment</td><td>distance</td><td>position</td><td>gene name</td><td>gene id</td></tr>
</table>

<table>
<tr><td>GU305914.1</td><td>81780</td><td>81802</td><td>+</td><td>0</td><td>AGAGTTTCTCGGCGCGGGTC</td><td>AGG</td><td>||||||||[||||||||||||]PAM</td><td>NA</td><td>Intergenic</td><td>NA</td><td>NA</td></tr>
<tr><td>T5868</td><td>GAAGCCGTTGATGTTGTAGCCGG</td><td>1000</td><td></td><td></td><td></td><td></td><td></td><td></td><td></td><td></td><td></td></tr>
<tr><td>Oligo fwd</td><td>CACCGAAGCCGTTGATGTTGTAGC</td><td></td><td></td><td></td><td></td><td></td><td></td><td></td><td></td><td></td><td></td></tr>
<tr><td>Oligo rev</td><td>AAACGCTACAACATCAACGGCTTC</td><td></td><td></td><td></td><td></td><td></td><td></td><td></td><td></td><td></td><td></td></tr>
<tr><td>Chromosome</td><td>start</td><td>end</td><td>strand</td><td>MM</td><td>target_seq</td><td>PAM</td><td>alignment</td><td>distance</td><td>position</td><td>gene name</td><td>gene id</td></tr>
<tr><td>GU305914.1</td><td>81817</td><td>81839</td><td>+</td><td>0</td><td>GAAGCCGTTGATGTTGTAGC</td><td>CGG</td><td>||||||||[||||||||||||]PAM</td><td>NA</td><td>Intergenic</td><td>NA</td><td>NA</td></tr>
<tr><td>T5869</td><td>GTTGTAGCCGGTGATGAACTCGG</td><td>1000</td><td></td><td></td><td></td><td></td><td></td><td></td><td></td><td></td><td></td></tr>
<tr><td>Oligo fwd</td><td>CACCGTTGTAGCCGGTGATGAACT</td><td></td><td></td><td></td><td></td><td></td><td></td><td></td><td></td><td></td><td></td></tr>
<tr><td>Oligo rev</td><td>AAACAGTTCATCACCGGCTACAAC</td><td></td><td></td><td></td><td></td><td></td><td></td><td></td><td></td><td></td><td></td></tr>
<tr><td>Chromosome</td><td>start</td><td>end</td><td>strand</td><td>MM</td><td>target_seq</td><td>PAM</td><td>alignment</td><td>distance</td><td>position</td><td>gene name</td><td>gene id</td></tr>
<tr><td>GU305914.1</td><td>81829</td><td>81851</td><td>+</td><td>0</td><td>GTTGTAGCCGGTGATGAACT</td><td>CGG</td><td>||||||||[||||||||||||]PAM</td><td>NA</td><td>Intergenic</td><td>NA</td><td>NA</td></tr>
<tr><td>T5870</td><td>TTGTAGCCGGTGATGAACTCGGG</td><td>1000</td><td></td><td></td><td></td><td></td><td></td><td></td><td></td><td></td><td></td></tr>
<tr><td>Oligo fwd</td><td>CACCTTGTAGCCGGTGATGAACTC</td><td></td><td></td><td></td><td></td><td></td><td></td><td></td><td></td><td></td><td></td></tr>
<tr><td>Oligo rev</td><td>AAACGAGTTCATCACCGGCTACAA</td><td></td><td></td><td></td><td></td><td></td><td></td><td></td><td></td><td></td><td></td></tr>
<tr><td>Chromosome</td><td>start</td><td>end</td><td>strand</td><td>MM</td><td>target_seq</td><td>PAM</td><td>alignment</td><td>distance</td><td>position</td><td>gene name</td><td>gene id</td></tr>
<tr><td>GU305914.1</td><td>81830</td><td>81852</td><td>+</td><td>0</td><td>TTGTAGCCGGTGATGAACTC</td><td>GGG</td><td>||||||||[||||||||||||]PAM</td><td>NA</td><td>Intergenic</td><td>NA</td><td>NA</td></tr>
<tr><td>T5871</td><td>ACTCGGGCGAGTACCAGCTCAGG</td><td>1000</td><td></td><td></td><td></td><td></td><td></td><td></td><td></td><td></td><td></td></tr>
<tr><td>Oligo fwd</td><td>CACCACTCGGGCGAGTACCAGCTC</td><td></td><td></td><td></td><td></td><td></td><td></td><td></td><td></td><td></td><td></td></tr>
<tr><td>Oligo rev</td><td>AAACGAGCTGGTACTCGCCCGAGT</td><td></td><td></td><td></td><td></td><td></td><td></td><td></td><td></td><td></td><td></td></tr>
<tr><td>Chromosome</td><td>start</td><td>end</td><td>strand</td><td>MM</td><td>target_seq</td><td>PAM</td><td>alignment</td><td>distance</td><td>position</td><td>gene name</td><td>gene id</td></tr>
<tr><td>GU305914.1</td><td>81846</td><td>81868</td><td>+</td><td>0</td><td>ACTCGGGCGAGTACCAGCTC</td><td>AGG</td><td>||||||||[||||||||||||]PAM</td><td>NA</td><td>Intergenic</td><td>NA</td><td>NA</td></tr>
<tr><td>T5872</td><td>ACCCCGCAACATCTCGTACTCGG</td><td>1000</td><td></td><td></td><td></td><td></td><td></td><td></td><td></td><td></td><td></td></tr>
<tr><td>Oligo fwd</td><td>CACCACCCCGCAACATCTCGTACT</td><td></td><td></td><td></td><td></td><td></td><td></td><td></td><td></td><td></td><td></td></tr>
<tr><td>Oligo rev</td><td>AAACAGTACGAGATGTTGCGGGGT</td><td></td><td></td><td></td><td></td><td></td><td></td><td></td><td></td><td></td><td></td></tr>
<tr><td>Chromosome</td><td>start</td><td>end</td><td>strand</td><td>MM</td><td>target_seq</td><td>PAM</td><td>alignment</td><td>distance</td><td>position</td><td>gene name</td><td>gene id</td></tr>
<tr><td>GU305914.1</td><td>81880</td><td>81902</td><td>+</td><td>0</td><td>ACCCCGCAACATCTCGTACT</td><td>CGG</td><td>||||||||[||||||||||||]PAM</td><td>NA</td><td>Intergenic</td><td>NA</td><td>NA</td></tr>
<tr><td>T5873</td><td>CCGCAACATCTCGTACTCGGAGG</td><td>1000</td><td></td><td></td><td></td><td></td><td></td><td></td><td></td><td></td><td></td></tr>
<tr><td>Oligo fwd</td><td>CACCCCGCAACATCTCGTACTCGG</td><td></td><td></td><td></td><td></td><td></td><td></td><td></td><td></td><td></td><td></td></tr>
<tr><td>Oligo rev</td><td>AAACCCGAGTACGAGATGTTGCGG</td><td></td><td></td><td></td><td></td><td></td><td></td><td></td><td></td><td></td><td></td></tr>
<tr><td>Chromosome</td><td>start</td><td>end</td><td>strand</td><td>MM</td><td>target_seq</td><td>PAM</td><td>alignment</td><td>distance</td><td>position</td><td>gene name</td><td>gene id</td></tr>
<tr><td>GU305914.1</td><td>81883</td><td>81905</td><td>+</td><td>0</td><td>CCGCAACATCTCGTACTCGG</td><td>AGG</td><td>||||||||[||||||||||||]PAM</td><td>NA</td><td>Intergenic</td><td>NA</td><td>NA</td></tr>
<tr><td>T5874</td><td>CGCAACATCTCGTACTCGGAGGG</td><td>1000</td><td></td><td></td><td></td><td></td><td></td><td></td><td></td><td></td><td></td></tr>
<tr><td>Oligo fwd</td><td>CACCCGCAACATCTCGTACTCGGA</td><td></td><td></td><td></td><td></td><td></td><td></td><td></td><td></td><td></td><td></td></tr>
<tr><td>Oligo rev</td><td>AAACTCCGAGTACGAGATGTTGCG</td><td></td><td></td><td></td><td></td><td></td><td></td><td></td><td></td><td></td><td></td></tr>
<tr><td>Chromosome</td><td>start</td><td>end</td><td>strand</td><td>MM</td><td>target_seq</td><td>PAM</td><td>alignment</td><td>distance</td><td>position</td><td>gene name</td><td>gene id</td></tr>
<tr><td>GU305914.1</td><td>81884</td><td>81906</td><td>+</td><td>0</td><td>CGCAACATCTCGTACTCGGA</td><td>GGG</td><td>||||||||[||||||||||||]PAM</td><td>NA</td><td>Intergenic</td><td>NA</td><td>NA</td></tr>
<tr><td>T5875</td><td>GCAACATCTCGTACTCGGAGGGG</td><td>1000</td><td></td><td></td><td></td><td></td><td></td><td></td><td></td><td></td><td></td></tr>
<tr><td>Oligo fwd</td><td>CACCGCAACATCTCGTACTCGGAG</td><td></td><td></td><td></td><td></td><td></td><td></td><td></td><td></td><td></td><td></td></tr>
<tr><td>Oligo rev</td><td>AAACCTCCGAGTACGAGATGTTGC</td><td></td><td></td><td></td><td></td><td></td><td></td><td></td><td></td><td></td><td></td></tr>
<tr><td>Chromosome</td><td>start</td><td>end</td><td>strand</td><td>MM</td><td>target_seq</td><td>PAM</td><td>alignment</td><td>distance</td><td>position</td><td>gene name</td><td>gene id</td></tr>
<tr><td>GU305914.1</td><td>81885</td><td>81907</td><td>+</td><td>0</td><td>GCAACATCTCGTACTCGGAG</td><td>GGG</td><td>||||||||[||||||||||||]PAM</td><td>NA</td><td>Intergenic</td><td>NA</td><td>NA</td></tr>
<tr><td>T5876</td><td>CATCTCGTACTCGGAGGGGAAGG</td><td>1000</td><td></td><td></td><td></td><td></td><td></td><td></td><td></td><td></td><td></td></tr>
<tr><td>Oligo fwd</td><td>CACCCATCTCGTACTCGGAGGGGA</td><td></td><td></td><td></td><td></td><td></td><td></td><td></td><td></td><td></td><td></td></tr>
<tr><td>Oligo rev</td><td>AAACTCCCCTCCGAGTACGAGATG</td><td></td><td></td><td></td><td></td><td></td><td></td><td></td><td></td><td></td><td></td></tr>
<tr><td>Chromosome</td><td>start</td><td>end</td><td>strand</td><td>MM</td><td>target_seq</td><td>PAM</td><td>alignment</td><td>distance</td><td>position</td><td>gene name</td><td>gene id</td></tr>
<tr><td>GU305914.1</td><td>81889</td><td>81911</td><td>+</td><td>0</td><td>CATCTCGTACTCGGAGGGGA</td><td>AGG</td><td>||||||||[||||||||||||]PAM</td><td>NA</td><td>Intergenic</td><td>NA</td><td>NA</td></tr>
<tr><td>T5877</td><td>GAAGGTGTAGATCTTGACGTCGG</td><td>1000</td><td></td><td></td><td></td><td></td><td></td><td></td><td></td><td></td><td></td></tr>
<tr><td>Oligo fwd</td><td>CACCGAAGGTGTAGATCTTGACGT</td><td></td><td></td><td></td><td></td><td></td><td></td><td></td><td></td><td></td><td></td></tr>
<tr><td>Oligo rev</td><td>AAACACGTCAAGATCTACACCTTC</td><td></td><td></td><td></td><td></td><td></td><td></td><td></td><td></td><td></td><td></td></tr>
<tr><td>Chromosome</td><td>start</td><td>end</td><td>strand</td><td>MM</td><td>target_seq</td><td>PAM</td><td>alignment</td><td>distance</td><td>position</td><td>gene name</td><td>gene id</td></tr>
<tr><td>GU305914.1</td><td>81907</td><td>81929</td><td>+</td><td>0</td><td>GAAGGTGTAGATCTTGACGT</td><td>CGG</td><td>||||||||[||||||||||||]PAM</td><td>NA</td><td>Intergenic</td><td>NA</td><td>NA</td></tr>
<tr><td>T5878</td><td>AAGGTGTAGATCTTGACGTCGGG</td><td>1000</td><td></td><td></td><td></td><td></td><td></td><td></td><td></td><td></td><td></td></tr>
<tr><td>Oligo fwd</td><td>CACCAAGGTGTAGATCTTGACGTC</td><td></td><td></td><td></td><td></td><td></td><td></td><td></td><td></td><td></td><td></td></tr>
<tr><td>Oligo rev</td><td>AAACGACGTCAAGATCTACACCTT</td><td></td><td></td><td></td><td></td><td></td><td></td><td></td><td></td><td></td><td></td></tr>
<tr><td>Chromosome</td><td>start</td><td>end</td><td>strand</td><td>MM</td><td>target_seq</td><td>PAM</td><td>alignment</td><td>distance</td><td>position</td><td>gene name</td><td>gene id</td></tr>
<tr><td>GU305914.1</td><td>81908</td><td>81930</td><td>+</td><td>0</td><td>AAGGTGTAGATCTTGACGTC</td><td>GGG</td><td>||||||||[||||||||||||]PAM</td><td>NA</td><td>Intergenic</td><td>NA</td><td>NA</td></tr>
<tr><td>T5879</td><td>AGGTGTAGATCTTGACGTCGGGG</td><td>1000</td><td></td><td></td><td></td><td></td><td></td><td></td><td></td><td></td><td></td></tr>
<tr><td>Oligo fwd</td><td>CACCAGGTGTAGATCTTGACGTCG</td><td></td><td></td><td></td><td></td><td></td><td></td><td></td><td></td><td></td><td></td></tr>
<tr><td>Oligo rev</td><td>AAACCGACGTCAAGATCTACACCT</td><td></td><td></td><td></td><td></td><td></td><td></td><td></td><td></td><td></td><td></td></tr>
<tr><td>Chromosome</td><td>start</td><td>end</td><td>strand</td><td>MM</td><td>target_seq</td><td>PAM</td><td>alignment</td><td>distance</td><td>position</td><td>gene name</td><td>gene id</td></tr>
<tr><td>GU305914.1</td><td>81909</td><td>81931</td><td>+</td><td>0</td><td>AGGTGTAGATCTTGACGTCG</td><td>GGG</td><td>||||||||[||||||||||||]PAM</td><td>NA</td><td>Intergenic</td><td>NA</td><td>NA</td></tr>
<tr><td>T5880</td><td>TAGATCTTGACGTCGGGGATCGG</td><td>1000</td><td></td><td></td><td></td><td></td><td></td><td></td><td></td><td></td><td></td></tr>
<tr><td>Oligo fwd</td><td>CACCTAGATCTTGACGTCGGGGAT</td><td></td><td></td><td></td><td></td><td></td><td></td><td></td><td></td><td></td><td></td></tr>
<tr><td>Oligo rev</td><td>AAACATCCCCGACGTCAAGATCTA</td><td></td><td></td><td></td><td></td><td></td><td></td><td></td><td></td><td></td><td></td></tr>
<tr><td>Chromosome</td><td>start</td><td>end</td><td>strand</td><td>MM</td><td>target_seq</td><td>PAM</td><td>alignment</td><td>distance</td><td>position</td><td>gene name</td><td>gene id</td></tr>
<tr><td>GU305914.1</td><td>81914</td><td>81936</td><td>+</td><td>0</td><td>TAGATCTTGACGTCGGGGAT</td><td>CGG</td><td>||||||||[||||||||||||]PAM</td><td>NA</td><td>Intergenic</td><td>NA</td><td>NA</td></tr>
<tr><td>T5881</td><td>AGATCTTGACGTCGGGGATCGGG</td><td>1000</td><td></td><td></td><td></td><td></td><td></td><td></td><td></td><td></td><td></td></tr>
<tr><td>Oligo fwd</td><td>CACCAGATCTTGACGTCGGGGATC</td><td></td><td></td><td></td><td></td><td></td><td></td><td></td><td></td><td></td><td></td></tr>
<tr><td>Oligo rev</td><td>AAACGATCCCCGACGTCAAGATCT</td><td></td><td></td><td></td><td></td><td></td><td></td><td></td><td></td><td></td><td></td></tr>
<tr><td>Chromosome</td><td>start</td><td>end</td><td>strand</td><td>MM</td><td>target_seq</td><td>PAM</td><td>alignment</td><td>distance</td><td>position</td><td>gene name</td><td>gene id</td></tr>
<tr><td>GU305914.1</td><td>81915</td><td>81937</td><td>+</td><td>0</td><td>AGATCTTGACGTCGGGGATC</td><td>GGG</td><td>||||||||[||||||||||||]PAM</td><td>NA</td><td>Intergenic</td><td>NA</td><td>NA</td></tr>
<tr><td>T5882</td><td>GATCTTGACGTCGGGGATCGGGG</td><td>1000</td><td></td><td></td><td></td><td></td><td></td><td></td><td></td><td></td><td></td></tr>
<tr><td>Oligo fwd</td><td>CACCGATCTTGACGTCGGGGATCG</td><td></td><td></td><td></td><td></td><td></td><td></td><td></td><td></td><td></td><td></td></tr>
<tr><td>Oligo rev</td><td>AAACCGATCCCCGACGTCAAGATC</td><td></td><td></td><td></td><td></td><td></td><td></td><td></td><td></td><td></td><td></td></tr>
</table>

| | | | | | | | | | | | |
|---|---|---|---|---|---|---|---|---|---|---|---|
| Chromosome | start | end | strand | MM | target_seq | PAM | alignment | distance | position | gene name | gene id |
| GU305914.1 | 81916 | 81938 | + | 0 | GATCTTGACGTCGGGGATCG | GGG | \|\|\|\|\|\|\|\|[\|\|\|\|\|\|\|\|\|\|\|\|]PAM | NA | Intergenic | NA | NA |
| T5883 | ACGTCGGGGATCGGGGCGCACGG | 1000 | | | | | | | | | |
| Oligo fwd | CACCACGTCGGGGATCGGGGCGCA | | | | | | | | | | |
| Oligo rev | AAACTGCGCCCCGATCCCCGACGT | | | | | | | | | | |
| Chromosome | start | end | strand | MM | target_seq | PAM | alignment | distance | position | gene name | gene id |
| GU305914.1 | 81923 | 81945 | + | 0 | ACGTCGGGGATCGGGGCGCA | CGG | \|\|\|\|\|\|\|\|[\|\|\|\|\|\|\|\|\|\|\|\|]PAM | NA | Intergenic | NA | NA |
| T5884 | GATCGGGGCGCACGGCCCGATGG | 1000 | | | | | | | | | |
| Oligo fwd | CACCGATCGGGGCGCACGGCCCGA | | | | | | | | | | |
| Oligo rev | AAACTCGGGCCGTGCGCCCCGATC | | | | | | | | | | |
| Chromosome | start | end | strand | MM | target_seq | PAM | alignment | distance | position | gene name | gene id |
| GU305914.1 | 81931 | 81953 | + | 0 | GATCGGGGCGCACGGCCCGA | TGG | \|\|\|\|\|\|\|\|[\|\|\|\|\|\|\|\|\|\|\|\|]PAM | NA | Intergenic | NA | NA |
| T5885 | CGCACGGCCCGATGGTGAAGAGG | 1000 | | | | | | | | | |
| Oligo fwd | CACCCGCACGGCCCGATGGTGAAG | | | | | | | | | | |
| Oligo rev | AAACCTTCACCATCGGGCCGTGCG | | | | | | | | | | |
| Chromosome | start | end | strand | MM | target_seq | PAM | alignment | distance | position | gene name | gene id |
| GU305914.1 | 81939 | 81961 | + | 0 | CGCACGGCCCGATGGTGAAG | AGG | \|\|\|\|\|\|\|\|[\|\|\|\|\|\|\|\|\|\|\|\|]PAM | NA | Intergenic | NA | NA |
| T5886 | ACGGCCCGATGGTGAAGAGGTGG | 1000 | | | | | | | | | |
| Oligo fwd | CACCACGGCCCGATGGTGAAGAGG | | | | | | | | | | |
| Oligo rev | AAACCCTCTTCACCATCGGGCCGT | | | | | | | | | | |
| Chromosome | start | end | strand | MM | target_seq | PAM | alignment | distance | position | gene name | gene id |
| GU305914.1 | 81942 | 81964 | + | 0 | ACGGCCCGATGGTGAAGAGG | TGG | \|\|\|\|\|\|\|\|[\|\|\|\|\|\|\|\|\|\|\|\|]PAM | NA | Intergenic | NA | NA |
| T5887 | GATGGTGAAGAGGTGGTTCTCGG | 1000 | | | | | | | | | |
| Oligo fwd | CACCGATGGTGAAGAGGTGGTTCT | | | | | | | | | | |
| Oligo rev | AAACAGAACCACCTCTTCACCATC | | | | | | | | | | |
| Chromosome | start | end | strand | MM | target_seq | PAM | alignment | distance | position | gene name | gene id |
| GU305914.1 | 81949 | 81971 | + | 0 | GATGGTGAAGAGGTGGTTCT | CGG | \|\|\|\|\|\|\|\|[\|\|\|\|\|\|\|\|\|\|\|\|]PAM | NA | Intergenic | NA | NA |
| T5888 | ATGGTGAAGAGGTGGTTCTCGGG | 1000 | | | | | | | | | |
| Oligo fwd | CACCATGGTGAAGAGGTGGTTCTC | | | | | | | | | | |
| Oligo rev | AAACGAGAACCACCTCTTCACCAT | | | | | | | | | | |
| Chromosome | start | end | strand | MM | target_seq | PAM | alignment | distance | position | gene name | gene id |
| GU305914.1 | 81950 | 81972 | + | 0 | ATGGTGAAGAGGTGGTTCTC | GGG | \|\|\|\|\|\|\|\|[\|\|\|\|\|\|\|\|\|\|\|\|]PAM | NA | Intergenic | NA | NA |
| T5889 | GAAGAGGTGGTTCTCGGGCACGG | 1000 | | | | | | | | | |
| Oligo fwd | CACCGAAGAGGTGGTTCTCGGGCA | | | | | | | | | | |
| Oligo rev | AAACTGCCCGAGAACCACCTCTTC | | | | | | | | | | |
| Chromosome | start | end | strand | MM | target_seq | PAM | alignment | distance | position | gene name | gene id |
| GU305914.1 | 81955 | 81977 | + | 0 | GAAGAGGTGGTTCTCGGGCA | CGG | \|\|\|\|\|\|\|\|[\|\|\|\|\|\|\|\|\|\|\|\|]PAM | NA | Intergenic | NA | NA |
| T5890 | GAGGTGGTTCTCGGGCACGGCGG | 1000 | | | | | | | | | |
| Oligo fwd | CACCGAGGTGGTTCTCGGGCACGG | | | | | | | | | | |
| Oligo rev | AAACCCGTGCCCGAGAACCACCTC | | | | | | | | | | |
| Chromosome | start | end | strand | MM | target_seq | PAM | alignment | distance | position | gene name | gene id |
| GU305914.1 | 81958 | 81980 | + | 0 | GAGGTGGTTCTCGGGCACGG | CGG | \|\|\|\|\|\|\|\|[\|\|\|\|\|\|\|\|\|\|\|\|]PAM | NA | Intergenic | NA | NA |
| T5891 | AGGTGGTTCTCGGGCACGGCGGG | 1000 | | | | | | | | | |
| Oligo fwd | CACCAGGTGGTTCTCGGGCACGGC | | | | | | | | | | |
| Oligo rev | AAACGCCGTGCCCGAGAACCACCT | | | | | | | | | | |
| Chromosome | start | end | strand | MM | target_seq | PAM | alignment | distance | position | gene name | gene id |
| GU305914.1 | 81959 | 81981 | + | 0 | AGGTGGTTCTCGGGCACGGC | GGG | \|\|\|\|\|\|\|\|[\|\|\|\|\|\|\|\|\|\|\|\|]PAM | NA | Intergenic | NA | NA |
| T5892 | CGGGCACGGCGGGCGTGCTGAGG | 1000 | | | | | | | | | |
| Oligo fwd | CACCCGGGCACGGCGGGCGTGCTG | | | | | | | | | | |
| Oligo rev | AAACCAGCACGCCCGCCGTGCCCG | | | | | | | | | | |
| Chromosome | start | end | strand | MM | target_seq | PAM | alignment | distance | position | gene name | gene id |
| GU305914.1 | 81969 | 81991 | + | 0 | CGGGCACGGCGGGCGTGCTG | AGG | \|\|\|\|\|\|\|\|[\|\|\|\|\|\|\|\|\|\|\|\|]PAM | NA | Intergenic | NA | NA |
| T5893 | CACGGCGGGCGTGCTGAGGTCGG | 1000 | | | | | | | | | |
| Oligo fwd | CACCCACGGCGGGCGTGCTGAGGT | | | | | | | | | | |
| Oligo rev | AAACACCTCAGCACGCCCGCCGTG | | | | | | | | | | |
| Chromosome | start | end | strand | MM | target_seq | PAM | alignment | distance | position | gene name | gene id |
| GU305914.1 | 81973 | 81995 | + | 0 | CACGGCGGGCGTGCTGAGGT | CGG | \|\|\|\|\|\|\|\|[\|\|\|\|\|\|\|\|\|\|\|\|]PAM | NA | Intergenic | NA | NA |
| T5894 | CGGGCGTGCTGAGGTCGGCGTGG | 1000 | | | | | | | | | |
| Oligo fwd | CACCCGGGCGTGCTGAGGTCGGCG | | | | | | | | | | |
| Oligo rev | AAACCGCCGACCTCAGCACGCCCG | | | | | | | | | | |
| Chromosome | start | end | strand | MM | target_seq | PAM | alignment | distance | position | gene name | gene id |
| GU305914.1 | 81978 | 82000 | + | 0 | CGGGCGTGCTGAGGTCGGCG | TGG | \|\|\|\|\|\|\|\|[\|\|\|\|\|\|\|\|\|\|\|\|]PAM | NA | Intergenic | NA | NA |
| T5895 | GAGGTCGGCGTGGACGTCGTAGG | 1000 | | | | | | | | | |
| Oligo fwd | CACCGAGGTCGGCGTGGACGTCGT | | | | | | | | | | |
| Oligo rev | AAACACGACGTCCACGCCGACCTC | | | | | | | | | | |
| Chromosome | start | end | strand | MM | target_seq | PAM | alignment | distance | position | gene name | gene id |
| GU305914.1 | 81988 | 82010 | + | 0 | GAGGTCGGCGTGGACGTCGT | AGG | \|\|\|\|\|\|\|\|[\|\|\|\|\|\|\|\|\|\|\|\|]PAM | NA | Intergenic | NA | NA |
| T5896 | CGGCGTGGACGTCGTAGGCGTGG | 1000 | | | | | | | | | |
| Oligo fwd | CACCCGGCGTGGACGTCGTAGGCG | | | | | | | | | | |
| Oligo rev | AAACCGCCTACGACGTCCACGCCG | | | | | | | | | | |
| Chromosome | start | end | strand | MM | target_seq | PAM | alignment | distance | position | gene name | gene id |
| GU305914.1 | 81993 | 82015 | + | 0 | CGGCGTGGACGTCGTAGGCG | TGG | \|\|\|\|\|\|\|\|[\|\|\|\|\|\|\|\|\|\|\|\|]PAM | NA | Intergenic | NA | NA |
| T5897 | TCCGACGCCGAAACAGACGCAGG | 1000 | | | | | | | | | |
| Oligo fwd | CACCTCCGACGCCGAAACAGACGC | | | | | | | | | | |

Oligo rev AAACGCGTCTGTTTCGGCGTCGGA
Chromosome start end strand MM target_seq PAM alignment distance position gene name gene id
GU305914.1 82027 82049 + 0 TCCGACGCCGAAACAGACGC AGG ||||||||[||||||||||||]PAM NA Intergenic NA NA

T5898 CGAAACAGACGCAGGAGATCTGG 1000
Oligo fwd CACCCGAAACAGACGCAGGAGATC
Oligo rev AAACGATCTCCTGCGTCTGTTTCG
Chromosome start end strand MM target_seq PAM alignment distance position gene name gene id
GU305914.1 82035 82057 + 0 CGAAACAGACGCAGGAGATC TGG ||||||||[||||||||||||]PAM NA Intergenic NA NA

T5899 GATGATGTCGTCGACGTTTTCGG 1000
Oligo fwd CACCGATGATGTCGTCGACGTTTT
Oligo rev AAACAAAACGTCGACGACATCATC
Chromosome start end strand MM target_seq PAM alignment distance position gene name gene id
GU305914.1 82060 82082 + 0 GATGATGTCGTCGACGTTTT CGG ||||||||[||||||||||||]PAM NA Intergenic NA NA

T5900 GATGTCGTCGACGTTTTCGGCGG 1000
Oligo fwd CACCGATGTCGTCGACGTTTTCGG
Oligo rev AAACCCGAAAACGTCGACGACATC
Chromosome start end strand MM target_seq PAM alignment distance position gene name gene id
GU305914.1 82063 82085 + 0 GATGTCGTCGACGTTTTCGG CGG ||||||||[||||||||||||]PAM NA Intergenic NA NA

T5901 GTCGTCGACGTTTTCGGCGGCGG 1000
Oligo fwd CACCGTCGTCGACGTTTTCGGCGG
Oligo rev AAACCCGCCGAAAACGTCGACGAC
Chromosome start end strand MM target_seq PAM alignment distance position gene name gene id
GU305914.1 82066 82088 + 0 GTCGTCGACGTTTTCGGCGG CGG ||||||||[||||||||||||]PAM NA Intergenic NA NA

T5902 TCGTCGACGTTTTCGGCGGCGGG 1000
Oligo fwd CACCTCGTCGACGTTTTCGGCGGC
Oligo rev AAACGCCGCCGAAAACGTCGACGA
Chromosome start end strand MM target_seq PAM alignment distance position gene name gene id
GU305914.1 82067 82089 + 0 TCGTCGACGTTTTCGGCGGC GGG ||||||||[||||||||||||]PAM NA Intergenic NA NA

T5903 CGTCGACGTTTTCGGCGGCGGGG 1000
Oligo fwd CACCCGTCGACGTTTTCGGCGGCG
Oligo rev AAACCGCCGCCGAAAACGTCGACG
Chromosome start end strand MM target_seq PAM alignment distance position gene name gene id
GU305914.1 82068 82090 + 0 CGTCGACGTTTTCGGCGGCG GGG ||||||||[||||||||||||]PAM NA Intergenic NA NA

T5904 GACGTTTTCGGCGGCGGGGAAGG 1000
Oligo fwd CACCGACGTTTTCGGCGGCGGGGA
Oligo rev AAACTCCCCGCCGCCGAAAACGTC
Chromosome start end strand MM target_seq PAM alignment distance position gene name gene id
GU305914.1 82072 82094 + 0 GACGTTTTCGGCGGCGGGGA AGG ||||||||[||||||||||||]PAM NA Intergenic NA NA

T5905 GCTCATACACTCGATATCGAAGG 1000
Oligo fwd CACCGCTCATACACTCGATATCGA
Oligo rev AAACTCGATATCGAGTGTATGAGC
Chromosome start end strand MM target_seq PAM alignment distance position gene name gene id
GU305914.1 82105 82127 + 0 GCTCATACACTCGATATCGA AGG ||||||||[||||||||||||]PAM NA Intergenic NA NA

T5906 TACACTCGATATCGAAGGACAGG 1000
Oligo fwd CACCTACACTCGATATCGAAGGAC
Oligo rev AAACGTCCTTCGATATCGAGTGTA
Chromosome start end strand MM target_seq PAM alignment distance position gene name gene id
GU305914.1 82110 82132 + 0 TACACTCGATATCGAAGGAC AGG ||||||||[||||||||||||]PAM NA Intergenic NA NA

T5907 CGATATCGAAGGACAGGCAACGG 1000
Oligo fwd CACCCGATATCGAAGGACAGGCAA
Oligo rev AAACTTGCCTGTCCTTCGATATCG
Chromosome start end strand MM target_seq PAM alignment distance position gene name gene id
GU305914.1 82116 82138 + 0 CGATATCGAAGGACAGGCAA CGG ||||||||[||||||||||||]PAM NA Intergenic NA NA

T5908 AAGGACAGGCAACGGTAGAACGG 1000
Oligo fwd CACCAAGGACAGGCAACGGTAGAA
Oligo rev AAACTTCTACCGTTGCCTGTCCTT
Chromosome start end strand MM target_seq PAM alignment distance position gene name gene id
GU305914.1 82124 82146 + 0 AAGGACAGGCAACGGTAGAA CGG ||||||||[||||||||||||]PAM NA Intergenic NA NA

T5909 CAGTTGCTCTGCTCTTCGATCGG 1000
Oligo fwd CACCCAGTTGCTCTGCTCTTCGAT
Oligo rev AAACATCGAAGAGCAGAGCAACTG
Chromosome start end strand MM target_seq PAM alignment distance position gene name gene id
GU305914.1 82148 82170 + 0 CAGTTGCTCTGCTCTTCGAT CGG ||||||||[||||||||||||]PAM NA Intergenic NA NA

T5910 TGCTCTGCTCTTCGATCGGTAGG 1000
Oligo fwd CACCTGCTCTGCTCTTCGATCGGT
Oligo rev AAACACCGATCGAAGAGCAGAGCA
Chromosome start end strand MM target_seq PAM alignment distance position gene name gene id
GU305914.1 82152 82174 + 0 TGCTCTGCTCTTCGATCGGT AGG ||||||||[||||||||||||]PAM NA Intergenic NA NA

T5911 CTCTTCGATCGGTAGGATGTCGG 1000
Oligo fwd CACCCTCTTCGATCGGTAGGATGT
Oligo rev AAACACATCCTACCGATCGAAGAG
Chromosome start end strand MM target_seq PAM alignment distance position gene name gene id
GU305914.1 82159 82181 + 0 CTCTTCGATCGGTAGGATGT CGG ||||||||[||||||||||||]PAM NA Intergenic NA NA

T5912 AGTCGATCTCCAGCTGCGCCCGG 1000

<table>
<tr><td>Oligo fwd</td><td>CACCAGTCGATCTCCAGCTGCGCC</td><td></td><td></td><td></td><td></td><td></td><td></td><td></td><td></td><td></td><td></td></tr>
<tr><td>Oligo rev</td><td>AAACGGCGCAGCTGGAGATCGACT</td><td></td><td></td><td></td><td></td><td></td><td></td><td></td><td></td><td></td><td></td></tr>
<tr><td>Chromosome</td><td>start</td><td>end</td><td>strand</td><td>MM</td><td>target_seq</td><td>PAM</td><td>alignment</td><td>distance</td><td>position</td><td>gene name</td><td>gene id</td></tr>
<tr><td>GU305914.1</td><td>82191</td><td>82213</td><td>+</td><td>0</td><td>AGTCGATCTCCAGCTGCGCC</td><td>CGG</td><td>||||||||[||||||||||||]PAM</td><td>NA</td><td>Intergenic</td><td>NA</td><td>NA</td></tr>
<tr><td>T5913</td><td>CTGCGCCCGGCTGACGTATCCGG</td><td>1000</td><td></td><td></td><td></td><td></td><td></td><td></td><td></td><td></td><td></td></tr>
<tr><td>Oligo fwd</td><td>CACCCTGCGCCCGGCTGACGTATC</td><td></td><td></td><td></td><td></td><td></td><td></td><td></td><td></td><td></td><td></td></tr>
<tr><td>Oligo rev</td><td>AAACGATACGTCAGCCGGGCGCAG</td><td></td><td></td><td></td><td></td><td></td><td></td><td></td><td></td><td></td><td></td></tr>
<tr><td>Chromosome</td><td>start</td><td>end</td><td>strand</td><td>MM</td><td>target_seq</td><td>PAM</td><td>alignment</td><td>distance</td><td>position</td><td>gene name</td><td>gene id</td></tr>
<tr><td>GU305914.1</td><td>82204</td><td>82226</td><td>+</td><td>0</td><td>CTGCGCCCGGCTGACGTATC</td><td>CGG</td><td>||||||||[||||||||||||]PAM</td><td>NA</td><td>Intergenic</td><td>NA</td><td>NA</td></tr>
<tr><td>T5914</td><td>CGCCCGGCTGACGTATCCGGCGG</td><td>1000</td><td></td><td></td><td></td><td></td><td></td><td></td><td></td><td></td><td></td></tr>
<tr><td>Oligo fwd</td><td>CACCCGCCCGGCTGACGTATCCGG</td><td></td><td></td><td></td><td></td><td></td><td></td><td></td><td></td><td></td><td></td></tr>
<tr><td>Oligo rev</td><td>AAACCCGGATACGTCAGCCGGGCG</td><td></td><td></td><td></td><td></td><td></td><td></td><td></td><td></td><td></td><td></td></tr>
<tr><td>Chromosome</td><td>start</td><td>end</td><td>strand</td><td>MM</td><td>target_seq</td><td>PAM</td><td>alignment</td><td>distance</td><td>position</td><td>gene name</td><td>gene id</td></tr>
<tr><td>GU305914.1</td><td>82207</td><td>82229</td><td>+</td><td>0</td><td>CGCCCGGCTGACGTATCCGG</td><td>CGG</td><td>||||||||[||||||||||||]PAM</td><td>NA</td><td>Intergenic</td><td>NA</td><td>NA</td></tr>
<tr><td>T5915</td><td>GGCTGACGTATCCGGCGGCGCGG</td><td>1000</td><td></td><td></td><td></td><td></td><td></td><td></td><td></td><td></td><td></td></tr>
<tr><td>Oligo fwd</td><td>CACCGGCTGACGTATCCGGCGGCG</td><td></td><td></td><td></td><td></td><td></td><td></td><td></td><td></td><td></td><td></td></tr>
<tr><td>Oligo rev</td><td>AAACCGCCGCCGGATACGTCAGCC</td><td></td><td></td><td></td><td></td><td></td><td></td><td></td><td></td><td></td><td></td></tr>
<tr><td>Chromosome</td><td>start</td><td>end</td><td>strand</td><td>MM</td><td>target_seq</td><td>PAM</td><td>alignment</td><td>distance</td><td>position</td><td>gene name</td><td>gene id</td></tr>
<tr><td>GU305914.1</td><td>82212</td><td>82234</td><td>+</td><td>0</td><td>GGCTGACGTATCCGGCGGCG</td><td>CGG</td><td>||||||||[||||||||||||]PAM</td><td>NA</td><td>Intergenic</td><td>NA</td><td>NA</td></tr>
<tr><td>T5916</td><td>GGCGCGGACGCTGTACCTCCTGG</td><td>1000</td><td></td><td></td><td></td><td></td><td></td><td></td><td></td><td></td><td></td></tr>
<tr><td>Oligo fwd</td><td>CACCGGCGCGGACGCTGTACCTCC</td><td></td><td></td><td></td><td></td><td></td><td></td><td></td><td></td><td></td><td></td></tr>
<tr><td>Oligo rev</td><td>AAACGGAGGTACAGCGTCCGCGCC</td><td></td><td></td><td></td><td></td><td></td><td></td><td></td><td></td><td></td><td></td></tr>
<tr><td>Chromosome</td><td>start</td><td>end</td><td>strand</td><td>MM</td><td>target_seq</td><td>PAM</td><td>alignment</td><td>distance</td><td>position</td><td>gene name</td><td>gene id</td></tr>
<tr><td>GU305914.1</td><td>82228</td><td>82250</td><td>+</td><td>0</td><td>GGCGCGGACGCTGTACCTCC</td><td>TGG</td><td>||||||||[||||||||||||]PAM</td><td>NA</td><td>Intergenic</td><td>NA</td><td>NA</td></tr>
<tr><td>T5917</td><td>GGACGCTGTACCTCCTGGCGAGG</td><td>1000</td><td></td><td></td><td></td><td></td><td></td><td></td><td></td><td></td><td></td></tr>
<tr><td>Oligo fwd</td><td>CACCGGACGCTGTACCTCCTGGCG</td><td></td><td></td><td></td><td></td><td></td><td></td><td></td><td></td><td></td><td></td></tr>
<tr><td>Oligo rev</td><td>AAACCGCCAGGAGGTACAGCGTCC</td><td></td><td></td><td></td><td></td><td></td><td></td><td></td><td></td><td></td><td></td></tr>
<tr><td>Chromosome</td><td>start</td><td>end</td><td>strand</td><td>MM</td><td>target_seq</td><td>PAM</td><td>alignment</td><td>distance</td><td>position</td><td>gene name</td><td>gene id</td></tr>
<tr><td>GU305914.1</td><td>82233</td><td>82255</td><td>+</td><td>0</td><td>GGACGCTGTACCTCCTGGCG</td><td>AGG</td><td>||||||||[||||||||||||]PAM</td><td>NA</td><td>Intergenic</td><td>NA</td><td>NA</td></tr>
<tr><td>T5918</td><td>GGCGAGGCACCAGCCGAAGCTGG</td><td>1000</td><td></td><td></td><td></td><td></td><td></td><td></td><td></td><td></td><td></td></tr>
<tr><td>Oligo fwd</td><td>CACCGGCGAGGCACCAGCCGAAGC</td><td></td><td></td><td></td><td></td><td></td><td></td><td></td><td></td><td></td><td></td></tr>
<tr><td>Oligo rev</td><td>AAACGCTTCGGCTGGTGCCTCGCC</td><td></td><td></td><td></td><td></td><td></td><td></td><td></td><td></td><td></td><td></td></tr>
<tr><td>Chromosome</td><td>start</td><td>end</td><td>strand</td><td>MM</td><td>target_seq</td><td>PAM</td><td>alignment</td><td>distance</td><td>position</td><td>gene name</td><td>gene id</td></tr>
<tr><td>GU305914.1</td><td>82249</td><td>82271</td><td>+</td><td>0</td><td>GGCGAGGCACCAGCCGAAGC</td><td>TGG</td><td>||||||||[||||||||||||]PAM</td><td>NA</td><td>Intergenic</td><td>NA</td><td>NA</td></tr>
<tr><td>T5919</td><td>GCGAGGCACCAGCCGAAGCTGGG</td><td>1000</td><td></td><td></td><td></td><td></td><td></td><td></td><td></td><td></td><td></td></tr>
<tr><td>Oligo fwd</td><td>CACCGCGAGGCACCAGCCGAAGCT</td><td></td><td></td><td></td><td></td><td></td><td></td><td></td><td></td><td></td><td></td></tr>
<tr><td>Oligo rev</td><td>AAACAGCTTCGGCTGGTGCCTCGC</td><td></td><td></td><td></td><td></td><td></td><td></td><td></td><td></td><td></td><td></td></tr>
<tr><td>Chromosome</td><td>start</td><td>end</td><td>strand</td><td>MM</td><td>target_seq</td><td>PAM</td><td>alignment</td><td>distance</td><td>position</td><td>gene name</td><td>gene id</td></tr>
<tr><td>GU305914.1</td><td>82250</td><td>82272</td><td>+</td><td>0</td><td>GCGAGGCACCAGCCGAAGCT</td><td>GGG</td><td>||||||||[||||||||||||]PAM</td><td>NA</td><td>Intergenic</td><td>NA</td><td>NA</td></tr>
<tr><td>T5920</td><td>CGAGGCACCAGCCGAAGCTGGGG</td><td>1000</td><td></td><td></td><td></td><td></td><td></td><td></td><td></td><td></td><td></td></tr>
<tr><td>Oligo fwd</td><td>CACCCGAGGCACCAGCCGAAGCTG</td><td></td><td></td><td></td><td></td><td></td><td></td><td></td><td></td><td></td><td></td></tr>
<tr><td>Oligo rev</td><td>AAACCAGCTTCGGCTGGTGCCTCG</td><td></td><td></td><td></td><td></td><td></td><td></td><td></td><td></td><td></td><td></td></tr>
<tr><td>Chromosome</td><td>start</td><td>end</td><td>strand</td><td>MM</td><td>target_seq</td><td>PAM</td><td>alignment</td><td>distance</td><td>position</td><td>gene name</td><td>gene id</td></tr>
<tr><td>GU305914.1</td><td>82251</td><td>82273</td><td>+</td><td>0</td><td>CGAGGCACCAGCCGAAGCTG</td><td>GGG</td><td>||||||||[||||||||||||]PAM</td><td>NA</td><td>Intergenic</td><td>NA</td><td>NA</td></tr>
<tr><td>T5921</td><td>TGGGGATCTTCCTGTCGATCAGG</td><td>1000</td><td></td><td></td><td></td><td></td><td></td><td></td><td></td><td></td><td></td></tr>
<tr><td>Oligo fwd</td><td>CACCTGGGGATCTTCCTGTCGATC</td><td></td><td></td><td></td><td></td><td></td><td></td><td></td><td></td><td></td><td></td></tr>
<tr><td>Oligo rev</td><td>AAACGATCGACAGGAAGATCCCCA</td><td></td><td></td><td></td><td></td><td></td><td></td><td></td><td></td><td></td><td></td></tr>
<tr><td>Chromosome</td><td>start</td><td>end</td><td>strand</td><td>MM</td><td>target_seq</td><td>PAM</td><td>alignment</td><td>distance</td><td>position</td><td>gene name</td><td>gene id</td></tr>
<tr><td>GU305914.1</td><td>82269</td><td>82291</td><td>+</td><td>0</td><td>TGGGGATCTTCCTGTCGATC</td><td>AGG</td><td>||||||||[||||||||||||]PAM</td><td>NA</td><td>Intergenic</td><td>NA</td><td>NA</td></tr>
<tr><td>T5922</td><td>GTCGATCAGGAAGCGCGCCAAGG</td><td>1000</td><td></td><td></td><td></td><td></td><td></td><td></td><td></td><td></td><td></td></tr>
<tr><td>Oligo fwd</td><td>CACCGTCGATCAGGAAGCGCGCCA</td><td></td><td></td><td></td><td></td><td></td><td></td><td></td><td></td><td></td><td></td></tr>
<tr><td>Oligo rev</td><td>AAACTGGCGCGCTTCCTGATCGAC</td><td></td><td></td><td></td><td></td><td></td><td></td><td></td><td></td><td></td><td></td></tr>
<tr><td>Chromosome</td><td>start</td><td>end</td><td>strand</td><td>MM</td><td>target_seq</td><td>PAM</td><td>alignment</td><td>distance</td><td>position</td><td>gene name</td><td>gene id</td></tr>
<tr><td>GU305914.1</td><td>82282</td><td>82304</td><td>+</td><td>0</td><td>GTCGATCAGGAAGCGCGCCA</td><td>AGG</td><td>||||||||[||||||||||||]PAM</td><td>NA</td><td>Intergenic</td><td>NA</td><td>NA</td></tr>
<tr><td>T5923</td><td>TCGATCAGGAAGCGCGCCAAGGG</td><td>1000</td><td></td><td></td><td></td><td></td><td></td><td></td><td></td><td></td><td></td></tr>
<tr><td>Oligo fwd</td><td>CACCTCGATCAGGAAGCGCGCCAA</td><td></td><td></td><td></td><td></td><td></td><td></td><td></td><td></td><td></td><td></td></tr>
<tr><td>Oligo rev</td><td>AAACTTGGCGCGCTTCCTGATCGA</td><td></td><td></td><td></td><td></td><td></td><td></td><td></td><td></td><td></td><td></td></tr>
<tr><td>Chromosome</td><td>start</td><td>end</td><td>strand</td><td>MM</td><td>target_seq</td><td>PAM</td><td>alignment</td><td>distance</td><td>position</td><td>gene name</td><td>gene id</td></tr>
<tr><td>GU305914.1</td><td>82283</td><td>82305</td><td>+</td><td>0</td><td>TCGATCAGGAAGCGCGCCAA</td><td>GGG</td><td>||||||||[||||||||||||]PAM</td><td>NA</td><td>Intergenic</td><td>NA</td><td>NA</td></tr>
<tr><td>T5924</td><td>CGATCAGGAAGCGCGCCAAGGGG</td><td>1000</td><td></td><td></td><td></td><td></td><td></td><td></td><td></td><td></td><td></td></tr>
<tr><td>Oligo fwd</td><td>CACCCGATCAGGAAGCGCGCCAAG</td><td></td><td></td><td></td><td></td><td></td><td></td><td></td><td></td><td></td><td></td></tr>
<tr><td>Oligo rev</td><td>AAACCTTGGCGCGCTTCCTGATCG</td><td></td><td></td><td></td><td></td><td></td><td></td><td></td><td></td><td></td><td></td></tr>
<tr><td>Chromosome</td><td>start</td><td>end</td><td>strand</td><td>MM</td><td>target_seq</td><td>PAM</td><td>alignment</td><td>distance</td><td>position</td><td>gene name</td><td>gene id</td></tr>
<tr><td>GU305914.1</td><td>82284</td><td>82306</td><td>+</td><td>0</td><td>CGATCAGGAAGCGCGCCAAG</td><td>GGG</td><td>||||||||[||||||||||||]PAM</td><td>NA</td><td>Intergenic</td><td>NA</td><td>NA</td></tr>
<tr><td>T5925</td><td>GCATCCGCTTGCCGATCTTGCGG</td><td>1000</td><td></td><td></td><td></td><td></td><td></td><td></td><td></td><td></td><td></td></tr>
<tr><td>Oligo fwd</td><td>CACCGCATCCGCTTGCCGATCTTG</td><td></td><td></td><td></td><td></td><td></td><td></td><td></td><td></td><td></td><td></td></tr>
<tr><td>Oligo rev</td><td>AAACCAAGATCGGCAAGCGGATGC</td><td></td><td></td><td></td><td></td><td></td><td></td><td></td><td></td><td></td><td></td></tr>
<tr><td>Chromosome</td><td>start</td><td>end</td><td>strand</td><td>MM</td><td>target_seq</td><td>PAM</td><td>alignment</td><td>distance</td><td>position</td><td>gene name</td><td>gene id</td></tr>
<tr><td>GU305914.1</td><td>82344</td><td>82366</td><td>+</td><td>0</td><td>GCATCCGCTTGCCGATCTTG</td><td>CGG</td><td>||||||||[||||||||||||]PAM</td><td>NA</td><td>Intergenic</td><td>NA</td><td>NA</td></tr>
<tr><td>T5926</td><td>GCACATGTTCCAGTTGTTGAAGG</td><td>1000</td><td></td><td></td><td></td><td></td><td></td><td></td><td></td><td></td><td></td></tr>
<tr><td>Oligo fwd</td><td>CACCGCACATGTTCCAGTTGTTGA</td><td></td><td></td><td></td><td></td><td></td><td></td><td></td><td></td><td></td><td></td></tr>
<tr><td>Oligo rev</td><td>AAACTCAACAACTGGAACATGTGC</td><td></td><td></td><td></td><td></td><td></td><td></td><td></td><td></td><td></td><td></td></tr>
<tr><td>Chromosome</td><td>start</td><td>end</td><td>strand</td><td>MM</td><td>target_seq</td><td>PAM</td><td>alignment</td><td>distance</td><td>position</td><td>gene name</td><td>gene id</td></tr>
<tr><td>GU305914.1</td><td>82366</td><td>82388</td><td>+</td><td>0</td><td>GCACATGTTCCAGTTGTTGA</td><td>AGG</td><td>||||||||[||||||||||||]PAM</td><td>NA</td><td>Intergenic</td><td>NA</td><td>NA</td></tr>
</table>

<table>
<tr><td>T5927</td><td>TCCAGTTGTTGAAGGAGAGCCGG</td><td>1000</td><td></td><td></td><td></td><td></td><td></td><td></td><td></td><td></td><td></td></tr>
<tr><td>Oligo fwd</td><td>CACCTCCAGTTGTTGAAGGAGAGC</td><td></td><td></td><td></td><td></td><td></td><td></td><td></td><td></td><td></td><td></td></tr>
<tr><td>Oligo rev</td><td>AAACGCTCTCCTTCAACAACTGGA</td><td></td><td></td><td></td><td></td><td></td><td></td><td></td><td></td><td></td><td></td></tr>
<tr><td>Chromosome</td><td>start</td><td>end</td><td>strand</td><td>MM</td><td>target_seq</td><td>PAM</td><td>alignment</td><td>distance</td><td>position</td><td>gene name</td><td>gene id</td></tr>
<tr><td>GU305914.1</td><td>82374</td><td>82396</td><td>+</td><td>0</td><td>TCCAGTTGTTGAAGGAGAGC</td><td>CGG</td><td>||||||||[||||||||||||]PAM</td><td>NA</td><td>Intergenic</td><td>NA</td><td>NA</td></tr>
<tr><td>T5928</td><td>TGTTGAAGGAGAGCCGGTACAGG</td><td>1000</td><td></td><td></td><td></td><td></td><td></td><td></td><td></td><td></td><td></td></tr>
<tr><td>Oligo fwd</td><td>CACCTGTTGAAGGAGAGCCGGTAC</td><td></td><td></td><td></td><td></td><td></td><td></td><td></td><td></td><td></td><td></td></tr>
<tr><td>Oligo rev</td><td>AAACGTACCGGCTCTCCTTCAACA</td><td></td><td></td><td></td><td></td><td></td><td></td><td></td><td></td><td></td><td></td></tr>
<tr><td>Chromosome</td><td>start</td><td>end</td><td>strand</td><td>MM</td><td>target_seq</td><td>PAM</td><td>alignment</td><td>distance</td><td>position</td><td>gene name</td><td>gene id</td></tr>
<tr><td>GU305914.1</td><td>82380</td><td>82402</td><td>+</td><td>0</td><td>TGTTGAAGGAGAGCCGGTAC</td><td>AGG</td><td>||||||||[||||||||||||]PAM</td><td>NA</td><td>Intergenic</td><td>NA</td><td>NA</td></tr>
<tr><td>T5929</td><td>GAAGGAGAGCCGGTACAGGTTGG</td><td>1000</td><td></td><td></td><td></td><td></td><td></td><td></td><td></td><td></td><td></td></tr>
<tr><td>Oligo fwd</td><td>CACCGAAGGAGAGCCGGTACAGGT</td><td></td><td></td><td></td><td></td><td></td><td></td><td></td><td></td><td></td><td></td></tr>
<tr><td>Oligo rev</td><td>AAACACCTGTACCGGCTCTCCTTC</td><td></td><td></td><td></td><td></td><td></td><td></td><td></td><td></td><td></td><td></td></tr>
<tr><td>Chromosome</td><td>start</td><td>end</td><td>strand</td><td>MM</td><td>target_seq</td><td>PAM</td><td>alignment</td><td>distance</td><td>position</td><td>gene name</td><td>gene id</td></tr>
<tr><td>GU305914.1</td><td>82384</td><td>82406</td><td>+</td><td>0</td><td>GAAGGAGAGCCGGTACAGGT</td><td>TGG</td><td>||||||||[||||||||||||]PAM</td><td>NA</td><td>Intergenic</td><td>NA</td><td>NA</td></tr>
<tr><td>T5930</td><td>AAGGAGAGCCGGTACAGGTTGGG</td><td>1000</td><td></td><td></td><td></td><td></td><td></td><td></td><td></td><td></td><td></td></tr>
<tr><td>Oligo fwd</td><td>CACCAAGGAGAGCCGGTACAGGTT</td><td></td><td></td><td></td><td></td><td></td><td></td><td></td><td></td><td></td><td></td></tr>
<tr><td>Oligo rev</td><td>AAACAACCTGTACCGGCTCTCCTT</td><td></td><td></td><td></td><td></td><td></td><td></td><td></td><td></td><td></td><td></td></tr>
<tr><td>Chromosome</td><td>start</td><td>end</td><td>strand</td><td>MM</td><td>target_seq</td><td>PAM</td><td>alignment</td><td>distance</td><td>position</td><td>gene name</td><td>gene id</td></tr>
<tr><td>GU305914.1</td><td>82385</td><td>82407</td><td>+</td><td>0</td><td>AAGGAGAGCCGGTACAGGTT</td><td>GGG</td><td>||||||||[||||||||||||]PAM</td><td>NA</td><td>Intergenic</td><td>NA</td><td>NA</td></tr>
<tr><td>T5931</td><td>AGGAGAGCCGGTACAGGTTGGGG</td><td>1000</td><td></td><td></td><td></td><td></td><td></td><td></td><td></td><td></td><td></td></tr>
<tr><td>Oligo fwd</td><td>CACCAGGAGAGCCGGTACAGGTTG</td><td></td><td></td><td></td><td></td><td></td><td></td><td></td><td></td><td></td><td></td></tr>
<tr><td>Oligo rev</td><td>AAACCAACCTGTACCGGCTCTCCT</td><td></td><td></td><td></td><td></td><td></td><td></td><td></td><td></td><td></td><td></td></tr>
<tr><td>Chromosome</td><td>start</td><td>end</td><td>strand</td><td>MM</td><td>target_seq</td><td>PAM</td><td>alignment</td><td>distance</td><td>position</td><td>gene name</td><td>gene id</td></tr>
<tr><td>GU305914.1</td><td>82386</td><td>82408</td><td>+</td><td>0</td><td>AGGAGAGCCGGTACAGGTTG</td><td>GGG</td><td>||||||||[||||||||||||]PAM</td><td>NA</td><td>Intergenic</td><td>NA</td><td>NA</td></tr>
<tr><td>T5932</td><td>ACAGGTTGGGGATGTGCCCGAGG</td><td>1000</td><td></td><td></td><td></td><td></td><td></td><td></td><td></td><td></td><td></td></tr>
<tr><td>Oligo fwd</td><td>CACCACAGGTTGGGGATGTGCCCG</td><td></td><td></td><td></td><td></td><td></td><td></td><td></td><td></td><td></td><td></td></tr>
<tr><td>Oligo rev</td><td>AAACCGGGCACATCCCCAACCTGT</td><td></td><td></td><td></td><td></td><td></td><td></td><td></td><td></td><td></td><td></td></tr>
<tr><td>Chromosome</td><td>start</td><td>end</td><td>strand</td><td>MM</td><td>target_seq</td><td>PAM</td><td>alignment</td><td>distance</td><td>position</td><td>gene name</td><td>gene id</td></tr>
<tr><td>GU305914.1</td><td>82398</td><td>82420</td><td>+</td><td>0</td><td>ACAGGTTGGGGATGTGCCCG</td><td>AGG</td><td>||||||||[||||||||||||]PAM</td><td>NA</td><td>Intergenic</td><td>NA</td><td>NA</td></tr>
<tr><td>T5933</td><td>CCCGAGGCCGTAGCCGTAGAAGG</td><td>1000</td><td></td><td></td><td></td><td></td><td></td><td></td><td></td><td></td><td></td></tr>
<tr><td>Oligo fwd</td><td>CACCCCCGAGGCCGTAGCCGTAGA</td><td></td><td></td><td></td><td></td><td></td><td></td><td></td><td></td><td></td><td></td></tr>
<tr><td>Oligo rev</td><td>AAACTCTACGGCTACGGCCTCGGG</td><td></td><td></td><td></td><td></td><td></td><td></td><td></td><td></td><td></td><td></td></tr>
<tr><td>Chromosome</td><td>start</td><td>end</td><td>strand</td><td>MM</td><td>target_seq</td><td>PAM</td><td>alignment</td><td>distance</td><td>position</td><td>gene name</td><td>gene id</td></tr>
<tr><td>GU305914.1</td><td>82414</td><td>82436</td><td>+</td><td>0</td><td>CCCGAGGCCGTAGCCGTAGA</td><td>AGG</td><td>||||||||[||||||||||||]PAM</td><td>NA</td><td>Intergenic</td><td>NA</td><td>NA</td></tr>
<tr><td>T5934</td><td>GAGGCCGTAGCCGTAGAAGGAGG</td><td>1000</td><td></td><td></td><td></td><td></td><td></td><td></td><td></td><td></td><td></td></tr>
<tr><td>Oligo fwd</td><td>CACCGAGGCCGTAGCCGTAGAAGG</td><td></td><td></td><td></td><td></td><td></td><td></td><td></td><td></td><td></td><td></td></tr>
<tr><td>Oligo rev</td><td>AAACCCTTCTACGGCTACGGCCTC</td><td></td><td></td><td></td><td></td><td></td><td></td><td></td><td></td><td></td><td></td></tr>
<tr><td>Chromosome</td><td>start</td><td>end</td><td>strand</td><td>MM</td><td>target_seq</td><td>PAM</td><td>alignment</td><td>distance</td><td>position</td><td>gene name</td><td>gene id</td></tr>
<tr><td>GU305914.1</td><td>82417</td><td>82439</td><td>+</td><td>0</td><td>GAGGCCGTAGCCGTAGAAGG</td><td>AGG</td><td>||||||||[||||||||||||]PAM</td><td>NA</td><td>Intergenic</td><td>NA</td><td>NA</td></tr>
<tr><td>GU305914.1</td><td>32625</td><td>32647</td><td>+</td><td>4</td><td>GAGCCTGTCGCCGTAGAAGC</td><td>CGG</td><td>|||-|-||[-||||||||||-]PAM</td><td>NA</td><td>Intergenic</td><td>NA</td><td>NA</td></tr>
<tr><td>T5935</td><td>GTAGCCGTAGAAGGAGGATTTGG</td><td>1000</td><td></td><td></td><td></td><td></td><td></td><td></td><td></td><td></td><td></td></tr>
<tr><td>Oligo fwd</td><td>CACCGTAGCCGTAGAAGGAGGATT</td><td></td><td></td><td></td><td></td><td></td><td></td><td></td><td></td><td></td><td></td></tr>
<tr><td>Oligo rev</td><td>AAACAATCCTCCTTCTACGGCTAC</td><td></td><td></td><td></td><td></td><td></td><td></td><td></td><td></td><td></td><td></td></tr>
<tr><td>Chromosome</td><td>start</td><td>end</td><td>strand</td><td>MM</td><td>target_seq</td><td>PAM</td><td>alignment</td><td>distance</td><td>position</td><td>gene name</td><td>gene id</td></tr>
<tr><td>GU305914.1</td><td>82423</td><td>82445</td><td>+</td><td>0</td><td>GTAGCCGTAGAAGGAGGATT</td><td>TGG</td><td>||||||||[||||||||||||]PAM</td><td>NA</td><td>Intergenic</td><td>NA</td><td>NA</td></tr>
<tr><td>T5936</td><td>GTAGAAGGAGGATTTGGTGACGG</td><td>1000</td><td></td><td></td><td></td><td></td><td></td><td></td><td></td><td></td><td></td></tr>
<tr><td>Oligo fwd</td><td>CACCGTAGAAGGAGGATTTGGTGA</td><td></td><td></td><td></td><td></td><td></td><td></td><td></td><td></td><td></td><td></td></tr>
<tr><td>Oligo rev</td><td>AAACTCACCAAATCCTCCTTCTAC</td><td></td><td></td><td></td><td></td><td></td><td></td><td></td><td></td><td></td><td></td></tr>
<tr><td>Chromosome</td><td>start</td><td>end</td><td>strand</td><td>MM</td><td>target_seq</td><td>PAM</td><td>alignment</td><td>distance</td><td>position</td><td>gene name</td><td>gene id</td></tr>
<tr><td>GU305914.1</td><td>82429</td><td>82451</td><td>+</td><td>0</td><td>GTAGAAGGAGGATTTGGTGA</td><td>CGG</td><td>||||||||[||||||||||||]PAM</td><td>NA</td><td>Intergenic</td><td>NA</td><td>NA</td></tr>
<tr><td>T5937</td><td>TAGAAGGAGGATTTGGTGACGGG</td><td>1000</td><td></td><td></td><td></td><td></td><td></td><td></td><td></td><td></td><td></td></tr>
<tr><td>Oligo fwd</td><td>CACCTAGAAGGAGGATTTGGTGAC</td><td></td><td></td><td></td><td></td><td></td><td></td><td></td><td></td><td></td><td></td></tr>
<tr><td>Oligo rev</td><td>AAACGTCACCAAATCCTCCTTCTA</td><td></td><td></td><td></td><td></td><td></td><td></td><td></td><td></td><td></td><td></td></tr>
<tr><td>Chromosome</td><td>start</td><td>end</td><td>strand</td><td>MM</td><td>target_seq</td><td>PAM</td><td>alignment</td><td>distance</td><td>position</td><td>gene name</td><td>gene id</td></tr>
<tr><td>GU305914.1</td><td>82430</td><td>82452</td><td>+</td><td>0</td><td>TAGAAGGAGGATTTGGTGAC</td><td>GGG</td><td>||||||||[||||||||||||]PAM</td><td>NA</td><td>Intergenic</td><td>NA</td><td>NA</td></tr>
<tr><td>T5938</td><td>GGATTTGGTGACGGGAGAGATGG</td><td>1000</td><td></td><td></td><td></td><td></td><td></td><td></td><td></td><td></td><td></td></tr>
<tr><td>Oligo fwd</td><td>CACCGGATTTGGTGACGGGAGAGA</td><td></td><td></td><td></td><td></td><td></td><td></td><td></td><td></td><td></td><td></td></tr>
<tr><td>Oligo rev</td><td>AAACTCTCTCCCGTCACCAAATCC</td><td></td><td></td><td></td><td></td><td></td><td></td><td></td><td></td><td></td><td></td></tr>
<tr><td>Chromosome</td><td>start</td><td>end</td><td>strand</td><td>MM</td><td>target_seq</td><td>PAM</td><td>alignment</td><td>distance</td><td>position</td><td>gene name</td><td>gene id</td></tr>
<tr><td>GU305914.1</td><td>82438</td><td>82460</td><td>+</td><td>0</td><td>GGATTTGGTGACGGGAGAGA</td><td>TGG</td><td>||||||||[||||||||||||]PAM</td><td>NA</td><td>Intergenic</td><td>NA</td><td>NA</td></tr>
<tr><td>T5939</td><td>GGGAGAGATGGACACGCTGAAGG</td><td>1000</td><td></td><td></td><td></td><td></td><td></td><td></td><td></td><td></td><td></td></tr>
<tr><td>Oligo fwd</td><td>CACCGGGAGAGATGGACACGCTGA</td><td></td><td></td><td></td><td></td><td></td><td></td><td></td><td></td><td></td><td></td></tr>
<tr><td>Oligo rev</td><td>AAACTCAGCGTGTCCATCTCTCCC</td><td></td><td></td><td></td><td></td><td></td><td></td><td></td><td></td><td></td><td></td></tr>
<tr><td>Chromosome</td><td>start</td><td>end</td><td>strand</td><td>MM</td><td>target_seq</td><td>PAM</td><td>alignment</td><td>distance</td><td>position</td><td>gene name</td><td>gene id</td></tr>
<tr><td>GU305914.1</td><td>82450</td><td>82472</td><td>+</td><td>0</td><td>GGGAGAGATGGACACGCTGA</td><td>AGG</td><td>||||||||[||||||||||||]PAM</td><td>NA</td><td>Intergenic</td><td>NA</td><td>NA</td></tr>
<tr><td>T5940</td><td>GGAGAGATGGACACGCTGAAGGG</td><td>1000</td><td></td><td></td><td></td><td></td><td></td><td></td><td></td><td></td><td></td></tr>
<tr><td>Oligo fwd</td><td>CACCGGAGAGATGGACACGCTGAA</td><td></td><td></td><td></td><td></td><td></td><td></td><td></td><td></td><td></td><td></td></tr>
<tr><td>Oligo rev</td><td>AAACTTCAGCGTGTCCATCTCTCC</td><td></td><td></td><td></td><td></td><td></td><td></td><td></td><td></td><td></td><td></td></tr>
<tr><td>Chromosome</td><td>start</td><td>end</td><td>strand</td><td>MM</td><td>target_seq</td><td>PAM</td><td>alignment</td><td>distance</td><td>position</td><td>gene name</td><td>gene id</td></tr>
<tr><td>GU305914.1</td><td>82451</td><td>82473</td><td>+</td><td>0</td><td>GGAGAGATGGACACGCTGAA</td><td>GGG</td><td>||||||||[||||||||||||]PAM</td><td>NA</td><td>Intergenic</td><td>NA</td><td>NA</td></tr>
<tr><td>T5941</td><td>GAGAGATGGACACGCTGAAGGGG</td><td>1000</td><td></td><td></td><td></td><td></td><td></td><td></td><td></td><td></td><td></td></tr>
<tr><td>Oligo fwd</td><td>CACCGAGAGATGGACACGCTGAAG</td><td></td><td></td><td></td><td></td><td></td><td></td><td></td><td></td><td></td><td></td></tr>
<tr><td>Oligo rev</td><td>AAACCTTCAGCGTGTCCATCTCTC</td><td></td><td></td><td></td><td></td><td></td><td></td><td></td><td></td><td></td><td></td></tr>
<tr><td>Chromosome</td><td>start</td><td>end</td><td>strand</td><td>MM</td><td>target_seq</td><td>PAM</td><td>alignment</td><td>distance</td><td>position</td><td>gene name</td><td>gene id</td></tr>
</table>

| | | | | | | | | | | | |
|---|---|---|---|---|---|---|---|---|---|---|---|
| GU305914.1 | | 82452 | 82474 + | 0 | GAGAGATGGACACGCTGAAG | GGG | \|\|\|\|\|\|\|\|[\|\|\|\|\|\|\|\|\|\|\|\|]PAM | NA | Intergenic | NA | NA |

| | | | | | | | | | | | |
|---|---|---|---|---|---|---|---|---|---|---|---|
| T5942 | AGAGATGGACACGCTGAAGGGGG | | 1000 | | | | | | | | |
| Oligo fwd | CACCAGAGATGGACACGCTGAAGG | | | | | | | | | | |
| Oligo rev | AAACCCTTCAGCGTGTCCATCTCT | | | | | | | | | | |
| Chromosome | start | | end | strand | MM target_seq | PAM | alignment | distance | position | gene name | gene id |
| GU305914.1 | | 82453 | 82475 | + | 0 AGAGATGGACACGCTGAAGG | GGG | \|\|\|\|\|\|\|\|[\|\|\|\|\|\|\|\|\|\|\|\|]PAM | NA | Intergenic | NA | NA |
| GU305914.1 | | 126275 | 126297 | - | 4 GCAGATGGACGTGCTGAAGG | TGG | --\|\|\|\|\|\|[\|\|--\|\|\|\|\|\|\|\|]PAM | NA | Intergenic | NA | NA |

| | | | | | | | | | | | |
|---|---|---|---|---|---|---|---|---|---|---|---|
| T5943 | GGACACGCTGAAGGGGGAGCAGG | | 1000 | | | | | | | | |
| Oligo fwd | CACCGGACACGCTGAAGGGGGAGC | | | | | | | | | | |
| Oligo rev | AAACGCTCCCCCTTCAGCGTGTCC | | | | | | | | | | |
| Chromosome | start | | end | strand | MM target_seq | PAM | alignment | distance | position | gene name | gene id |
| GU305914.1 | | 82459 | 82481 | + | 0 GGACACGCTGAAGGGGGAGC | AGG | \|\|\|\|\|\|\|\|[\|\|\|\|\|\|\|\|\|\|\|\|]PAM | NA | Intergenic | NA | NA |

| | | | | | | | | | | | |
|---|---|---|---|---|---|---|---|---|---|---|---|
| T5944 | GACACGCTGAAGGGGGAGCAGGG | | 1000 | | | | | | | | |
| Oligo fwd | CACCGACACGCTGAAGGGGGAGCA | | | | | | | | | | |
| Oligo rev | AAACTGCTCCCCCTTCAGCGTGTC | | | | | | | | | | |
| Chromosome | start | | end | strand | MM target_seq | PAM | alignment | distance | position | gene name | gene id |
| GU305914.1 | | 82460 | 82482 | + | 0 GACACGCTGAAGGGGGAGCA | GGG | \|\|\|\|\|\|\|\|[\|\|\|\|\|\|\|\|\|\|\|\|]PAM | NA | Intergenic | NA | NA |

| | | | | | | | | | | | |
|---|---|---|---|---|---|---|---|---|---|---|---|
| T5945 | GCTGAAGGGGGAGCAGGGCTCGG | | 1000 | | | | | | | | |
| Oligo fwd | CACCGCTGAAGGGGGAGCAGGGCT | | | | | | | | | | |
| Oligo rev | AAACAGCCCTGCTCCCCCTTCAGC | | | | | | | | | | |
| Chromosome | start | | end | strand | MM target_seq | PAM | alignment | distance | position | gene name | gene id |
| GU305914.1 | | 82465 | 82487 | + | 0 GCTGAAGGGGGAGCAGGGCT | CGG | \|\|\|\|\|\|\|\|[\|\|\|\|\|\|\|\|\|\|\|\|]PAM | NA | Intergenic | NA | NA |

| | | | | | | | | | | | |
|---|---|---|---|---|---|---|---|---|---|---|---|
| T5946 | CTGAAGGGGGAGCAGGGCTCGGG | | 1000 | | | | | | | | |
| Oligo fwd | CACCCTGAAGGGGGAGCAGGGCTC | | | | | | | | | | |
| Oligo rev | AAACGAGCCCTGCTCCCCCTTCAG | | | | | | | | | | |
| Chromosome | start | | end | strand | MM target_seq | PAM | alignment | distance | position | gene name | gene id |
| GU305914.1 | | 82466 | 82488 | + | 0 CTGAAGGGGGAGCAGGGCTC | GGG | \|\|\|\|\|\|\|\|[\|\|\|\|\|\|\|\|\|\|\|\|]PAM | NA | Intergenic | NA | NA |

| | | | | | | | | | | | |
|---|---|---|---|---|---|---|---|---|---|---|---|
| T5947 | TGAAGGGGGAGCAGGGCTCGGGG | | 1000 | | | | | | | | |
| Oligo fwd | CACCTGAAGGGGGAGCAGGGCTCG | | | | | | | | | | |
| Oligo rev | AAACCGAGCCCTGCTCCCCCTTCA | | | | | | | | | | |
| Chromosome | start | | end | strand | MM target_seq | PAM | alignment | distance | position | gene name | gene id |
| GU305914.1 | | 82467 | 82489 | + | 0 TGAAGGGGGAGCAGGGCTCG | GGG | \|\|\|\|\|\|\|\|[\|\|\|\|\|\|\|\|\|\|\|\|]PAM | NA | Intergenic | NA | NA |

| | | | | | | | | | | | |
|---|---|---|---|---|---|---|---|---|---|---|---|
| T5948 | GCAGGGCTCGGGGACCTTATCGG | | 1000 | | | | | | | | |
| Oligo fwd | CACCGCAGGGCTCGGGGACCTTAT | | | | | | | | | | |
| Oligo rev | AAACATAAGGTCCCCGAGCCCTGC | | | | | | | | | | |
| Chromosome | start | | end | strand | MM target_seq | PAM | alignment | distance | position | gene name | gene id |
| GU305914.1 | | 82477 | 82499 | + | 0 GCAGGGCTCGGGGACCTTAT | CGG | \|\|\|\|\|\|\|\|[\|\|\|\|\|\|\|\|\|\|\|\|]PAM | NA | Intergenic | NA | NA |
| GU305914.1 | | 111364 | 111386 | - | 4 GTAGGGTTCGGTGACCTTAG | AGG | \|-\|\|\|\|-\|[\|\|\|-\|\|\|\|\|\|\|-]PAM | NA | Intergenic | NA | NA |

| | | | | | | | | | | | |
|---|---|---|---|---|---|---|---|---|---|---|---|
| T5949 | CTTATCGGCGATATCGTAGACGG | | 1000 | | | | | | | | |
| Oligo fwd | CACCCTTATCGGCGATATCGTAGA | | | | | | | | | | |
| Oligo rev | AAACTCTACGATATCGCCGATAAG | | | | | | | | | | |
| Chromosome | start | | end | strand | MM target_seq | PAM | alignment | distance | position | gene name | gene id |
| GU305914.1 | | 82492 | 82514 | + | 0 CTTATCGGCGATATCGTAGA | CGG | \|\|\|\|\|\|\|\|[\|\|\|\|\|\|\|\|\|\|\|\|]PAM | NA | Intergenic | NA | NA |

| | | | | | | | | | | | |
|---|---|---|---|---|---|---|---|---|---|---|---|
| T5950 | TATCGTAGACGGTGTTCTTGAGG | | 1000 | | | | | | | | |
| Oligo fwd | CACCTATCGTAGACGGTGTTCTTG | | | | | | | | | | |
| Oligo rev | AAACCAAGAACACCGTCTACGATA | | | | | | | | | | |
| Chromosome | start | | end | strand | MM target_seq | PAM | alignment | distance | position | gene name | gene id |
| GU305914.1 | | 82503 | 82525 | + | 0 TATCGTAGACGGTGTTCTTG | AGG | \|\|\|\|\|\|\|\|[\|\|\|\|\|\|\|\|\|\|\|\|]PAM | NA | Intergenic | NA | NA |

| | | | | | | | | | | | |
|---|---|---|---|---|---|---|---|---|---|---|---|
| T5951 | GACGGTGTTCTTGAGGTCCTCGG | | 1000 | | | | | | | | |
| Oligo fwd | CACCGACGGTGTTCTTGAGGTCCT | | | | | | | | | | |
| Oligo rev | AAACAGGACCTCAAGAACACCGTC | | | | | | | | | | |
| Chromosome | start | | end | strand | MM target_seq | PAM | alignment | distance | position | gene name | gene id |
| GU305914.1 | | 82510 | 82532 | + | 0 GACGGTGTTCTTGAGGTCCT | CGG | \|\|\|\|\|\|\|\|[\|\|\|\|\|\|\|\|\|\|\|\|]PAM | NA | Intergenic | NA | NA |

| | | | | | | | | | | | |
|---|---|---|---|---|---|---|---|---|---|---|---|
| T5952 | CATCCTCTCGCAGTAGAAGTAGG | | 1000 | | | | | | | | |
| Oligo fwd | CACCCATCCTCTCGCAGTAGAAGT | | | | | | | | | | |
| Oligo rev | AAACACTTCTACTGCGAGAGGATG | | | | | | | | | | |
| Chromosome | start | | end | strand | MM target_seq | PAM | alignment | distance | position | gene name | gene id |
| GU305914.1 | | 82537 | 82559 | + | 0 CATCCTCTCGCAGTAGAAGT | AGG | \|\|\|\|\|\|\|\|[\|\|\|\|\|\|\|\|\|\|\|\|]PAM | NA | Intergenic | NA | NA |

| | | | | | | | | | | | |
|---|---|---|---|---|---|---|---|---|---|---|---|
| T5953 | CGCAGTAGAAGTAGGCCTTCTGG | | 1000 | | | | | | | | |
| Oligo fwd | CACCCGCAGTAGAAGTAGGCCTTC | | | | | | | | | | |
| Oligo rev | AAACGAAGGCCTACTTCTACTGCG | | | | | | | | | | |
| Chromosome | start | | end | strand | MM target_seq | PAM | alignment | distance | position | gene name | gene id |
| GU305914.1 | | 82545 | 82567 | + | 0 CGCAGTAGAAGTAGGCCTTC | TGG | \|\|\|\|\|\|\|\|[\|\|\|\|\|\|\|\|\|\|\|\|]PAM | NA | Intergenic | NA | NA |

| | | | | | | | | | | | |
|---|---|---|---|---|---|---|---|---|---|---|---|
| T5954 | GCAGACCGAGACCCCCTCATCGG | | 1000 | | | | | | | | |
| Oligo fwd | CACCGCAGACCGAGACCCCCTCAT | | | | | | | | | | |
| Oligo rev | AAACATGAGGGGGTCTCGGTCTGC | | | | | | | | | | |
| Chromosome | start | | end | strand | MM target_seq | PAM | alignment | distance | position | gene name | gene id |
| GU305914.1 | | 82582 | 82604 | + | 0 GCAGACCGAGACCCCCTCAT | CGG | \|\|\|\|\|\|\|\|[\|\|\|\|\|\|\|\|\|\|\|\|]PAM | NA | Intergenic | NA | NA |

| | | | | | | | | | | | |
|---|---|---|---|---|---|---|---|---|---|---|---|
| T5955 | GACCGAGACCCCCTCATCGGTGG | | 1000 | | | | | | | | |
| Oligo fwd | CACCGACCGAGACCCCCTCATCGG | | | | | | | | | | |
| Oligo rev | AAACCCGATGAGGGGGTCTCGGTC | | | | | | | | | | |
| Chromosome | start | | end | strand | MM target_seq | PAM | alignment | distance | position | gene name | gene id |
| GU305914.1 | | 82585 | 82607 | + | 0 GACCGAGACCCCCTCATCGG | TGG | \|\|\|\|\|\|\|\|[\|\|\|\|\|\|\|\|\|\|\|\|]PAM | NA | Intergenic | NA | NA |

| | | | |
|---|---|---|---|
| T5956 | CATCGGTGGCCCCGAACATGCGG | | 1000 |

<table>
<tr><td>Oligo fwd</td><td>CACCCATCGGTGGCCCCGAACATG</td><td></td><td></td><td></td><td></td><td></td><td></td><td></td><td></td><td></td><td></td></tr>
<tr><td>Oligo rev</td><td>AAACCATGTTCGGGGCCACCGATG</td><td></td><td></td><td></td><td></td><td></td><td></td><td></td><td></td><td></td><td></td></tr>
<tr><td>Chromosome</td><td>start</td><td>end</td><td>strand</td><td>MM</td><td>target_seq</td><td>PAM</td><td>alignment</td><td>distance</td><td>position</td><td>gene name</td><td>gene id</td></tr>
<tr><td>GU305914.1</td><td>82599</td><td>82621</td><td>+</td><td>0</td><td>CATCGGTGGCCCCGAACATG</td><td>CGG</td><td>||||||||[||||||||||||]PAM</td><td>NA</td><td>Intergenic</td><td>NA</td><td>NA</td></tr>
<tr><td>T5957</td><td>ATGCGGATGACGTTCCCCGACGG</td><td>1000</td><td></td><td></td><td></td><td></td><td></td><td></td><td></td><td></td><td></td></tr>
<tr><td>Oligo fwd</td><td>CACCATGCGGATGACGTTCCCCGA</td><td></td><td></td><td></td><td></td><td></td><td></td><td></td><td></td><td></td><td></td></tr>
<tr><td>Oligo rev</td><td>AAACTCGGGGAACGTCATCCGCAT</td><td></td><td></td><td></td><td></td><td></td><td></td><td></td><td></td><td></td><td></td></tr>
<tr><td>Chromosome</td><td>start</td><td>end</td><td>strand</td><td>MM</td><td>target_seq</td><td>PAM</td><td>alignment</td><td>distance</td><td>position</td><td>gene name</td><td>gene id</td></tr>
<tr><td>GU305914.1</td><td>82616</td><td>82638</td><td>+</td><td>0</td><td>ATGCGGATGACGTTCCCCGA</td><td>CGG</td><td>||||||||[||||||||||||]PAM</td><td>NA</td><td>Intergenic</td><td>NA</td><td>NA</td></tr>
<tr><td>T5958</td><td>CGTTCCCCGACGGCACCACATGG</td><td>1000</td><td></td><td></td><td></td><td></td><td></td><td></td><td></td><td></td><td></td></tr>
<tr><td>Oligo fwd</td><td>CACCCGTTCCCCGACGGCACCACA</td><td></td><td></td><td></td><td></td><td></td><td></td><td></td><td></td><td></td><td></td></tr>
<tr><td>Oligo rev</td><td>AAACTGTGGTGCCGTCGGGGAACG</td><td></td><td></td><td></td><td></td><td></td><td></td><td></td><td></td><td></td><td></td></tr>
<tr><td>Chromosome</td><td>start</td><td>end</td><td>strand</td><td>MM</td><td>target_seq</td><td>PAM</td><td>alignment</td><td>distance</td><td>position</td><td>gene name</td><td>gene id</td></tr>
<tr><td>GU305914.1</td><td>82626</td><td>82648</td><td>+</td><td>0</td><td>CGTTCCCCGACGGCACCACA</td><td>TGG</td><td>||||||||[||||||||||||]PAM</td><td>NA</td><td>Intergenic</td><td>NA</td><td>NA</td></tr>
<tr><td>T5959</td><td>GCACCACATGGTGCTTCCAGCGG</td><td>1000</td><td></td><td></td><td></td><td></td><td></td><td></td><td></td><td></td><td></td></tr>
<tr><td>Oligo fwd</td><td>CACCGCACCACATGGTGCTTCCAG</td><td></td><td></td><td></td><td></td><td></td><td></td><td></td><td></td><td></td><td></td></tr>
<tr><td>Oligo rev</td><td>AAACCTGGAAGCACCATGTGGTGC</td><td></td><td></td><td></td><td></td><td></td><td></td><td></td><td></td><td></td><td></td></tr>
<tr><td>Chromosome</td><td>start</td><td>end</td><td>strand</td><td>MM</td><td>target_seq</td><td>PAM</td><td>alignment</td><td>distance</td><td>position</td><td>gene name</td><td>gene id</td></tr>
<tr><td>GU305914.1</td><td>82638</td><td>82660</td><td>+</td><td>0</td><td>GCACCACATGGTGCTTCCAG</td><td>CGG</td><td>||||||||[||||||||||||]PAM</td><td>NA</td><td>Intergenic</td><td>NA</td><td>NA</td></tr>
<tr><td>T5960</td><td>CACCACATGGTGCTTCCAGCGGG</td><td>1000</td><td></td><td></td><td></td><td></td><td></td><td></td><td></td><td></td><td></td></tr>
<tr><td>Oligo fwd</td><td>CACCCACCACATGGTGCTTCCAGC</td><td></td><td></td><td></td><td></td><td></td><td></td><td></td><td></td><td></td><td></td></tr>
<tr><td>Oligo rev</td><td>AAACGCTGGAAGCACCATGTGGTG</td><td></td><td></td><td></td><td></td><td></td><td></td><td></td><td></td><td></td><td></td></tr>
<tr><td>Chromosome</td><td>start</td><td>end</td><td>strand</td><td>MM</td><td>target_seq</td><td>PAM</td><td>alignment</td><td>distance</td><td>position</td><td>gene name</td><td>gene id</td></tr>
<tr><td>GU305914.1</td><td>82639</td><td>82661</td><td>+</td><td>0</td><td>CACCACATGGTGCTTCCAGC</td><td>GGG</td><td>||||||||[||||||||||||]PAM</td><td>NA</td><td>Intergenic</td><td>NA</td><td>NA</td></tr>
<tr><td>T5961</td><td>ACCACATGGTGCTTCCAGCGGGG</td><td>1000</td><td></td><td></td><td></td><td></td><td></td><td></td><td></td><td></td><td></td></tr>
<tr><td>Oligo fwd</td><td>CACCACCACATGGTGCTTCCAGCG</td><td></td><td></td><td></td><td></td><td></td><td></td><td></td><td></td><td></td><td></td></tr>
<tr><td>Oligo rev</td><td>AAACCGCTGGAAGCACCATGTGGT</td><td></td><td></td><td></td><td></td><td></td><td></td><td></td><td></td><td></td><td></td></tr>
<tr><td>Chromosome</td><td>start</td><td>end</td><td>strand</td><td>MM</td><td>target_seq</td><td>PAM</td><td>alignment</td><td>distance</td><td>position</td><td>gene name</td><td>gene id</td></tr>
<tr><td>GU305914.1</td><td>82640</td><td>82662</td><td>+</td><td>0</td><td>ACCACATGGTGCTTCCAGCG</td><td>GGG</td><td>||||||||[||||||||||||]PAM</td><td>NA</td><td>Intergenic</td><td>NA</td><td>NA</td></tr>
<tr><td>T5962</td><td>CGAGATCTGCTCGTCAGAGTCGG</td><td>1000</td><td></td><td></td><td></td><td></td><td></td><td></td><td></td><td></td><td></td></tr>
<tr><td>Oligo fwd</td><td>CACCCGAGATCTGCTCGTCAGAGT</td><td></td><td></td><td></td><td></td><td></td><td></td><td></td><td></td><td></td><td></td></tr>
<tr><td>Oligo rev</td><td>AAACACTCTGACGAGCAGATCTCG</td><td></td><td></td><td></td><td></td><td></td><td></td><td></td><td></td><td></td><td></td></tr>
<tr><td>Chromosome</td><td>start</td><td>end</td><td>strand</td><td>MM</td><td>target_seq</td><td>PAM</td><td>alignment</td><td>distance</td><td>position</td><td>gene name</td><td>gene id</td></tr>
<tr><td>GU305914.1</td><td>82663</td><td>82685</td><td>+</td><td>0</td><td>CGAGATCTGCTCGTCAGAGT</td><td>CGG</td><td>||||||||[||||||||||||]PAM</td><td>NA</td><td>Intergenic</td><td>NA</td><td>NA</td></tr>
<tr><td>T5963</td><td>CGTCAGAGTCGGCGAGCACCAGG</td><td>1000</td><td></td><td></td><td></td><td></td><td></td><td></td><td></td><td></td><td></td></tr>
<tr><td>Oligo fwd</td><td>CACCCGTCAGAGTCGGCGAGCACC</td><td></td><td></td><td></td><td></td><td></td><td></td><td></td><td></td><td></td><td></td></tr>
<tr><td>Oligo rev</td><td>AAACGGTGCTCGCCGACTCTGACG</td><td></td><td></td><td></td><td></td><td></td><td></td><td></td><td></td><td></td><td></td></tr>
<tr><td>Chromosome</td><td>start</td><td>end</td><td>strand</td><td>MM</td><td>target_seq</td><td>PAM</td><td>alignment</td><td>distance</td><td>position</td><td>gene name</td><td>gene id</td></tr>
<tr><td>GU305914.1</td><td>82674</td><td>82696</td><td>+</td><td>0</td><td>CGTCAGAGTCGGCGAGCACC</td><td>AGG</td><td>||||||||[||||||||||||]PAM</td><td>NA</td><td>Intergenic</td><td>NA</td><td>NA</td></tr>
<tr><td>T5964</td><td>CGAGCACCAGGTTGACGACCTGG</td><td>1000</td><td></td><td></td><td></td><td></td><td></td><td></td><td></td><td></td><td></td></tr>
<tr><td>Oligo fwd</td><td>CACCCGAGCACCAGGTTGACGACC</td><td></td><td></td><td></td><td></td><td></td><td></td><td></td><td></td><td></td><td></td></tr>
<tr><td>Oligo rev</td><td>AAACGGTCGTCAACCTGGTGCTCG</td><td></td><td></td><td></td><td></td><td></td><td></td><td></td><td></td><td></td><td></td></tr>
<tr><td>Chromosome</td><td>start</td><td>end</td><td>strand</td><td>MM</td><td>target_seq</td><td>PAM</td><td>alignment</td><td>distance</td><td>position</td><td>gene name</td><td>gene id</td></tr>
<tr><td>GU305914.1</td><td>82686</td><td>82708</td><td>+</td><td>0</td><td>CGAGCACCAGGTTGACGACC</td><td>TGG</td><td>||||||||[||||||||||||]PAM</td><td>NA</td><td>Intergenic</td><td>NA</td><td>NA</td></tr>
<tr><td>T5965</td><td>CAGGTTGACGACCTGGTCGTAGG</td><td>1000</td><td></td><td></td><td></td><td></td><td></td><td></td><td></td><td></td><td></td></tr>
<tr><td>Oligo fwd</td><td>CACCCAGGTTGACGACCTGGTCGT</td><td></td><td></td><td></td><td></td><td></td><td></td><td></td><td></td><td></td><td></td></tr>
<tr><td>Oligo rev</td><td>AAACACGACCAGGTCGTCAACCTG</td><td></td><td></td><td></td><td></td><td></td><td></td><td></td><td></td><td></td><td></td></tr>
<tr><td>Chromosome</td><td>start</td><td>end</td><td>strand</td><td>MM</td><td>target_seq</td><td>PAM</td><td>alignment</td><td>distance</td><td>position</td><td>gene name</td><td>gene id</td></tr>
<tr><td>GU305914.1</td><td>82693</td><td>82715</td><td>+</td><td>0</td><td>CAGGTTGACGACCTGGTCGT</td><td>AGG</td><td>||||||||[||||||||||||]PAM</td><td>NA</td><td>Intergenic</td><td>NA</td><td>NA</td></tr>
<tr><td>T5966</td><td>TGACGACCTGGTCGTAGGTGTGG</td><td>1000</td><td></td><td></td><td></td><td></td><td></td><td></td><td></td><td></td><td></td></tr>
<tr><td>Oligo fwd</td><td>CACCTGACGACCTGGTCGTAGGTG</td><td></td><td></td><td></td><td></td><td></td><td></td><td></td><td></td><td></td><td></td></tr>
<tr><td>Oligo rev</td><td>AAACCACCTACGACCAGGTCGTCA</td><td></td><td></td><td></td><td></td><td></td><td></td><td></td><td></td><td></td><td></td></tr>
<tr><td>Chromosome</td><td>start</td><td>end</td><td>strand</td><td>MM</td><td>target_seq</td><td>PAM</td><td>alignment</td><td>distance</td><td>position</td><td>gene name</td><td>gene id</td></tr>
<tr><td>GU305914.1</td><td>82698</td><td>82720</td><td>+</td><td>0</td><td>TGACGACCTGGTCGTAGGTG</td><td>TGG</td><td>||||||||[||||||||||||]PAM</td><td>NA</td><td>Intergenic</td><td>NA</td><td>NA</td></tr>
<tr><td>T5967</td><td>GGTGTGGAAGCGCAGCTCGTCGG</td><td>1000</td><td></td><td></td><td></td><td></td><td></td><td></td><td></td><td></td><td></td></tr>
<tr><td>Oligo fwd</td><td>CACCGGTGTGGAAGCGCAGCTCGT</td><td></td><td></td><td></td><td></td><td></td><td></td><td></td><td></td><td></td><td></td></tr>
<tr><td>Oligo rev</td><td>AAACACGAGCTGCGCTTCCACACC</td><td></td><td></td><td></td><td></td><td></td><td></td><td></td><td></td><td></td><td></td></tr>
<tr><td>Chromosome</td><td>start</td><td>end</td><td>strand</td><td>MM</td><td>target_seq</td><td>PAM</td><td>alignment</td><td>distance</td><td>position</td><td>gene name</td><td>gene id</td></tr>
<tr><td>GU305914.1</td><td>82714</td><td>82736</td><td>+</td><td>0</td><td>GGTGTGGAAGCGCAGCTCGT</td><td>CGG</td><td>||||||||[||||||||||||]PAM</td><td>NA</td><td>Intergenic</td><td>NA</td><td>NA</td></tr>
<tr><td>T5968</td><td>GAAGCGCAGCTCGTCGGAGTAGG</td><td>1000</td><td></td><td></td><td></td><td></td><td></td><td></td><td></td><td></td><td></td></tr>
<tr><td>Oligo fwd</td><td>CACCGAAGCGCAGCTCGTCGGAGT</td><td></td><td></td><td></td><td></td><td></td><td></td><td></td><td></td><td></td><td></td></tr>
<tr><td>Oligo rev</td><td>AAACACTCCGACGAGCTGCGCTTC</td><td></td><td></td><td></td><td></td><td></td><td></td><td></td><td></td><td></td><td></td></tr>
<tr><td>Chromosome</td><td>start</td><td>end</td><td>strand</td><td>MM</td><td>target_seq</td><td>PAM</td><td>alignment</td><td>distance</td><td>position</td><td>gene name</td><td>gene id</td></tr>
<tr><td>GU305914.1</td><td>82720</td><td>82742</td><td>+</td><td>0</td><td>GAAGCGCAGCTCGTCGGAGT</td><td>AGG</td><td>||||||||[||||||||||||]PAM</td><td>NA</td><td>Intergenic</td><td>NA</td><td>NA</td></tr>
<tr><td>T5969</td><td>AAGCGCAGCTCGTCGGAGTAGGG</td><td>1000</td><td></td><td></td><td></td><td></td><td></td><td></td><td></td><td></td><td></td></tr>
<tr><td>Oligo fwd</td><td>CACCAAGCGCAGCTCGTCGGAGTA</td><td></td><td></td><td></td><td></td><td></td><td></td><td></td><td></td><td></td><td></td></tr>
<tr><td>Oligo rev</td><td>AAACTACTCCGACGAGCTGCGCTT</td><td></td><td></td><td></td><td></td><td></td><td></td><td></td><td></td><td></td><td></td></tr>
<tr><td>Chromosome</td><td>start</td><td>end</td><td>strand</td><td>MM</td><td>target_seq</td><td>PAM</td><td>alignment</td><td>distance</td><td>position</td><td>gene name</td><td>gene id</td></tr>
<tr><td>GU305914.1</td><td>82721</td><td>82743</td><td>+</td><td>0</td><td>AAGCGCAGCTCGTCGGAGTA</td><td>GGG</td><td>||||||||[||||||||||||]PAM</td><td>NA</td><td>Intergenic</td><td>NA</td><td>NA</td></tr>
<tr><td>T5970</td><td>GCTCGTCGGAGTAGGGTCCCCGG</td><td>1000</td><td></td><td></td><td></td><td></td><td></td><td></td><td></td><td></td><td></td></tr>
<tr><td>Oligo fwd</td><td>CACCGCTCGTCGGAGTAGGGTCCC</td><td></td><td></td><td></td><td></td><td></td><td></td><td></td><td></td><td></td><td></td></tr>
<tr><td>Oligo rev</td><td>AAACGGGACCCTACTCCGACGAGC</td><td></td><td></td><td></td><td></td><td></td><td></td><td></td><td></td><td></td><td></td></tr>
<tr><td>Chromosome</td><td>start</td><td>end</td><td>strand</td><td>MM</td><td>target_seq</td><td>PAM</td><td>alignment</td><td>distance</td><td>position</td><td>gene name</td><td>gene id</td></tr>
<tr><td>GU305914.1</td><td>82728</td><td>82750</td><td>+</td><td>0</td><td>GCTCGTCGGAGTAGGGTCCC</td><td>CGG</td><td>||||||||[||||||||||||]PAM</td><td>NA</td><td>Intergenic</td><td>NA</td><td>NA</td></tr>
</table>

T5971 GGGTCCCCGGCACCAACCTCTGG 1000
Oligo fwd CACCGGGTCCCCGGCACCAACCTC
Oligo rev AAACGAGGTTGGTGCCGGGGACCC
Chromosome start end strand MM target_seq PAM alignment distance position gene name gene id
GU305914.1 82741 82763 + 0 GGGTCCCCGGCACCAACCTC TGG ||||||||[||||||||||||]PAM NA Intergenic NA NA

T5972 GGCACCAACCTCTGGTGCGCAGG 1000
Oligo fwd CACCGGCACCAACCTCTGGTGCGC
Oligo rev AAACGCGCACCAGAGGTTGGTGCC
Chromosome start end strand MM target_seq PAM alignment distance position gene name gene id
GU305914.1 82749 82771 + 0 GGCACCAACCTCTGGTGCGC AGG ||||||||[||||||||||||]PAM NA Intergenic NA NA

T5973 GCACCAACCTCTGGTGCGCAGGG 1000
Oligo fwd CACCGCACCAACCTCTGGTGCGCA
Oligo rev AAACTGCGCACCAGAGGTTGGTGC
Chromosome start end strand MM target_seq PAM alignment distance position gene name gene id
GU305914.1 82750 82772 + 0 GCACCAACCTCTGGTGCGCA GGG ||||||||[||||||||||||]PAM NA Intergenic NA NA

T5974 CAACCTCTGGTGCGCAGGGTCGG 1000
Oligo fwd CACCCAACCTCTGGTGCGCAGGGT
Oligo rev AAACACCCTGCGCACCAGAGGTTG
Chromosome start end strand MM target_seq PAM alignment distance position gene name gene id
GU305914.1 82754 82776 + 0 CAACCTCTGGTGCGCAGGGT CGG ||||||||[||||||||||||]PAM NA Intergenic NA NA

T5975 CTGGTGCGCAGGGTCGGCCACGG 1000
Oligo fwd CACCCTGGTGCGCAGGGTCGGCCA
Oligo rev AAACTGGCCGACCCTGCGCACCAG
Chromosome start end strand MM target_seq PAM alignment distance position gene name gene id
GU305914.1 82760 82782 + 0 CTGGTGCGCAGGGTCGGCCA CGG ||||||||[||||||||||||]PAM NA Intergenic NA NA

T5976 CAGGGTCGGCCACGGCTTGTCGG 1000
Oligo fwd CACCCAGGGTCGGCCACGGCTTGT
Oligo rev AAACACAAGCCGTGGCCGACCCTG
Chromosome start end strand MM target_seq PAM alignment distance position gene name gene id
GU305914.1 82768 82790 + 0 CAGGGTCGGCCACGGCTTGT CGG ||||||||[||||||||||||]PAM NA Intergenic NA NA

T5977 AGGGTCGGCCACGGCTTGTCGGG 1000
Oligo fwd CACCAGGGTCGGCCACGGCTTGTC
Oligo rev AAACGACAAGCCGTGGCCGACCCT
Chromosome start end strand MM target_seq PAM alignment distance position gene name gene id
GU305914.1 82769 82791 + 0 AGGGTCGGCCACGGCTTGTC GGG ||||||||[||||||||||||]PAM NA Intergenic NA NA

T5978 GGGTCGGCCACGGCTTGTCGGGG 1000
Oligo fwd CACCGGGTCGGCCACGGCTTGTCG
Oligo rev AAACCGACAAGCCGTGGCCGACCC
Chromosome start end strand MM target_seq PAM alignment distance position gene name gene id
GU305914.1 82770 82792 + 0 GGGTCGGCCACGGCTTGTCG GGG ||||||||[||||||||||||]PAM NA Intergenic NA NA

T5979 GTTGAGCACGTACTCTTTCTCGG 1000
Oligo fwd CACCGTTGAGCACGTACTCTTTCT
Oligo rev AAACAGAAAGAGTACGTGCTCAAC
Chromosome start end strand MM target_seq PAM alignment distance position gene name gene id
GU305914.1 82792 82814 + 0 GTTGAGCACGTACTCTTTCT CGG ||||||||[||||||||||||]PAM NA Intergenic NA NA

T5980 TTTCTCGGCGTAGAACATCCTGG 1000
Oligo fwd CACCTTTCTCGGCGTAGAACATCC
Oligo rev AAACGGATGTTCTACGCCGAGAAA
Chromosome start end strand MM target_seq PAM alignment distance position gene name gene id
GU305914.1 82807 82829 + 0 TTTCTCGGCGTAGAACATCC TGG ||||||||[||||||||||||]PAM NA Intergenic NA NA

T5981 TTCTCGGCGTAGAACATCCTGGG 1000
Oligo fwd CACCTTCTCGGCGTAGAACATCCT
Oligo rev AAACAGGATGTTCTACGCCGAGAA
Chromosome start end strand MM target_seq PAM alignment distance position gene name gene id
GU305914.1 82808 82830 + 0 TTCTCGGCGTAGAACATCCT GGG ||||||||[||||||||||||]PAM NA Intergenic NA NA

T5982 GAACATCCTGGGTTCCATGCCGG 1000
Oligo fwd CACCGAACATCCTGGGTTCCATGC
Oligo rev AAACGCATGGAACCCAGGATGTTC
Chromosome start end strand MM target_seq PAM alignment distance position gene name gene id
GU305914.1 82819 82841 + 0 GAACATCCTGGGTTCCATGC CGG ||||||||[||||||||||||]PAM NA Intergenic NA NA

T5983 CCATCATACAACACGCCACGCGG 1000
Oligo fwd CACCCCATCATACAACACGCCACG
Oligo rev AAACCGTGGCGTGTTGTATGATGG
Chromosome start end strand MM target_seq PAM alignment distance position gene name gene id
GU305914.1 82868 82890 + 0 CCATCATACAACACGCCACG CGG ||||||||[||||||||||||]PAM NA Intergenic NA NA

T5984 CGCAGCGTCTCCAATCGCGACGG 1000
Oligo fwd CACCCGCAGCGTCTCCAATCGCGA
Oligo rev AAACTCGCGATTGGAGACGCTGCG
Chromosome start end strand MM target_seq PAM alignment distance position gene name gene id
GU305914.1 82923 82945 + 0 CGCAGCGTCTCCAATCGCGA CGG ||||||||[||||||||||||]PAM NA Intergenic NA NA

T5985 GCAGCGTCTCCAATCGCGACGGG 1000
Oligo fwd CACCGCAGCGTCTCCAATCGCGAC
Oligo rev AAACGTCGCGATTGGAGACGCTGC
Chromosome start end strand MM target_seq PAM alignment distance position gene name gene id
GU305914.1 82924 82946 + 0 GCAGCGTCTCCAATCGCGAC GGG ||||||||[||||||||||||]PAM NA Intergenic NA NA

T5986 CAGCGTCTCCAATCGCGACGGGG 1000
Oligo fwd CACCCAGCGTCTCCAATCGCGACG
Oligo rev AAACCGTCGCGATTGGAGACGCTG
Chromosome start end strand MM target_seq PAM alignment distance position gene name gene id
GU305914.1 82925 82947 + 0 CAGCGTCTCCAATCGCGACG GGG ||||||||[||||||||||||]PAM NA Intergenic NA NA

T5987 CGCGACGGGGTTTCCGTCGCAGG 1000
Oligo fwd CACCCGCGACGGGGTTTCCGTCGC
Oligo rev AAACGCGACGGAAACCCCGTCGCG
Chromosome start end strand MM target_seq PAM alignment distance position gene name gene id
GU305914.1 82938 82960 + 0 CGCGACGGGGTTTCCGTCGC AGG ||||||||[||||||||||||]PAM NA Intergenic NA NA

T5988 ACGGGGTTTCCGTCGCAGGTAGG 1000
Oligo fwd CACCACGGGGTTTCCGTCGCAGGT
Oligo rev AAACACCTGCGACGGAAACCCCGT
Chromosome start end strand MM target_seq PAM alignment distance position gene name gene id
GU305914.1 82942 82964 + 0 ACGGGGTTTCCGTCGCAGGT AGG ||||||||[||||||||||||]PAM NA Intergenic NA NA

T5989 CGGGGTTTCCGTCGCAGGTAGGG 1000
Oligo fwd CACCCGGGGTTTCCGTCGCAGGTA
Oligo rev AAACTACCTGCGACGGAAACCCCG
Chromosome start end strand MM target_seq PAM alignment distance position gene name gene id
GU305914.1 82943 82965 + 0 CGGGGTTTCCGTCGCAGGTA GGG ||||||||[||||||||||||]PAM NA Intergenic NA NA

T5990 GGGGTTTCCGTCGCAGGTAGGGG 1000
Oligo fwd CACCGGGGTTTCCGTCGCAGGTAG
Oligo rev AAACCTACCTGCGACGGAAACCCC
Chromosome start end strand MM target_seq PAM alignment distance position gene name gene id
GU305914.1 82944 82966 + 0 GGGGTTTCCGTCGCAGGTAG GGG ||||||||[||||||||||||]PAM NA Intergenic NA NA

T5991 CTCCACGCAAGTGTCCATGTCGG 1000
Oligo fwd CACCCTCCACGCAAGTGTCCATGT
Oligo rev AAACACATGGACACTTGCGTGGAG
Chromosome start end strand MM target_seq PAM alignment distance position gene name gene id
GU305914.1 82978 83000 + 0 CTCCACGCAAGTGTCCATGT CGG ||||||||[||||||||||||]PAM NA Intergenic NA NA

T5992 CGGCCGTCAGTACTCGAAATCGG 1000
Oligo fwd CACCCGGCCGTCAGTACTCGAAAT
Oligo rev AAACATTTCGAGTACTGACGGCCG
Chromosome start end strand MM target_seq PAM alignment distance position gene name gene id
GU305914.1 82998 83020 + 0 CGGCCGTCAGTACTCGAAAT CGG ||||||||[||||||||||||]PAM NA Intergenic NA NA

T5993 TCGAAATCGGAGTCCTCCGCCGG 1000
Oligo fwd CACCTCGAAATCGGAGTCCTCCGC
Oligo rev AAACGCGGAGGACTCCGATTTCGA
Chromosome start end strand MM target_seq PAM alignment distance position gene name gene id
GU305914.1 83011 83033 + 0 TCGAAATCGGAGTCCTCCGC CGG ||||||||[||||||||||||]PAM NA Intergenic NA NA

T5994 AGTCCTCCGCCGGTAATCGCTGG 1000
Oligo fwd CACCAGTCCTCCGCCGGTAATCGC
Oligo rev AAACGCGATTACCGGCGGAGGACT
Chromosome start end strand MM target_seq PAM alignment distance position gene name gene id
GU305914.1 83021 83043 + 0 AGTCCTCCGCCGGTAATCGC TGG ||||||||[||||||||||||]PAM NA Intergenic NA NA

T5995 CGGTAATCGCTGGTAGCCACTGG 1000
Oligo fwd CACCCGGTAATCGCTGGTAGCCAC
Oligo rev AAACGTGGCTACCAGCGATTACCG
Chromosome start end strand MM target_seq PAM alignment distance position gene name gene id
GU305914.1 83031 83053 + 0 CGGTAATCGCTGGTAGCCAC TGG ||||||||[||||||||||||]PAM NA Intergenic NA NA

T5996 TGGTGTCGTTACCTTTGAGACGG 1000
Oligo fwd CACCTGGTGTCGTTACCTTTGAGA
Oligo rev AAACTCTCAAAGGTAACGACACCA
Chromosome start end strand MM target_seq PAM alignment distance position gene name gene id
GU305914.1 83051 83073 + 0 TGGTGTCGTTACCTTTGAGA CGG ||||||||[||||||||||||]PAM NA Intergenic NA NA

T5997 AGACGGTCCAAGATGCCCGTCGG 1000
Oligo fwd CACCAGACGGTCCAAGATGCCCGT
Oligo rev AAACACGGGCATCTTGGACCGTCT
Chromosome start end strand MM target_seq PAM alignment distance position gene name gene id
GU305914.1 83068 83090 + 0 AGACGGTCCAAGATGCCCGT CGG ||||||||[||||||||||||]PAM NA Intergenic NA NA

T5998 GCTGCGCTTTCTCGTTGCGTTGG 1000
Oligo fwd CACCGCTGCGCTTTCTCGTTGCGT
Oligo rev AAACACGCAACGAGAAAGCGCAGC
Chromosome start end strand MM target_seq PAM alignment distance position gene name gene id
GU305914.1 83108 83130 + 0 GCTGCGCTTTCTCGTTGCGT TGG ||||||||[||||||||||||]PAM NA Intergenic NA NA

T5999 CGTCTAGCTGTTTTAACGCGCGG 1000
Oligo fwd CACCCGTCTAGCTGTTTTAACGCG
Oligo rev AAACCGCGTTAAAACAGCTAGACG
Chromosome start end strand MM target_seq PAM alignment distance position gene name gene id
GU305914.1 83135 83157 + 0 CGTCTAGCTGTTTTAACGCG CGG ||||||||[||||||||||||]PAM NA Intergenic NA NA

T6000 TATCAATAGAGCATCTTGCTCGG 1000
Oligo fwd CACCTATCAATAGAGCATCTTGCT
Oligo rev AAACAGCAAGATGCTCTATTGATA
Chromosome start end strand MM target_seq PAM alignment distance position gene name gene id

<table>
<tr><td>GU305914.1</td><td></td><td>83160</td><td>83182</td><td>+</td><td>0</td><td>TATCAATAGAGCATCTTGCT</td><td>CGG</td><td>||||||||[||||||||||||]PAM</td><td>NA</td><td>Intergenic</td><td>NA</td><td>NA</td></tr>
<tr><td colspan="13"></td></tr>
<tr><td>T6001</td><td>TAGAGCATCTTGCTCGGTGTAGG</td><td></td><td>1000</td><td colspan="9"></td></tr>
<tr><td>Oligo fwd</td><td>CACCTAGAGCATCTTGCTCGGTGT</td><td colspan="11"></td></tr>
<tr><td>Oligo rev</td><td>AAACACACCGAGCAAGATGCTCTA</td><td colspan="11"></td></tr>
<tr><td>Chromosome</td><td>start</td><td></td><td>end</td><td>strand</td><td>MM</td><td>target_seq</td><td>PAM</td><td>alignment</td><td>distance</td><td>position</td><td>gene name</td><td>gene id</td></tr>
<tr><td>GU305914.1</td><td></td><td>83166</td><td>83188</td><td>+</td><td>0</td><td>TAGAGCATCTTGCTCGGTGT</td><td>AGG</td><td>||||||||[||||||||||||]PAM</td><td>NA</td><td>Intergenic</td><td>NA</td><td>NA</td></tr>
<tr><td colspan="13"></td></tr>
<tr><td>T6002</td><td>TTCACGCGCCGATGCAGACCAGG</td><td></td><td>1000</td><td colspan="9"></td></tr>
<tr><td>Oligo fwd</td><td>CACCTTCACGCGCCGATGCAGACC</td><td colspan="11"></td></tr>
<tr><td>Oligo rev</td><td>AAACGGTCTGCATCGGCGCGTGAA</td><td colspan="11"></td></tr>
<tr><td>Chromosome</td><td>start</td><td></td><td>end</td><td>strand</td><td>MM</td><td>target_seq</td><td>PAM</td><td>alignment</td><td>distance</td><td>position</td><td>gene name</td><td>gene id</td></tr>
<tr><td>GU305914.1</td><td></td><td>83251</td><td>83273</td><td>+</td><td>0</td><td>TTCACGCGCCGATGCAGACC</td><td>AGG</td><td>||||||||[||||||||||||]PAM</td><td>NA</td><td>Intergenic</td><td>NA</td><td>NA</td></tr>
<tr><td colspan="13"></td></tr>
<tr><td>T6003</td><td>CGATGCAGACCAGGCGCTCCCGG</td><td></td><td>1000</td><td colspan="9"></td></tr>
<tr><td>Oligo fwd</td><td>CACCCGATGCAGACCAGGCGCTCC</td><td colspan="11"></td></tr>
<tr><td>Oligo rev</td><td>AAACGGAGCGCCTGGTCTGCATCG</td><td colspan="11"></td></tr>
<tr><td>Chromosome</td><td>start</td><td></td><td>end</td><td>strand</td><td>MM</td><td>target_seq</td><td>PAM</td><td>alignment</td><td>distance</td><td>position</td><td>gene name</td><td>gene id</td></tr>
<tr><td>GU305914.1</td><td></td><td>83260</td><td>83282</td><td>+</td><td>0</td><td>CGATGCAGACCAGGCGCTCC</td><td>CGG</td><td>||||||||[||||||||||||]PAM</td><td>NA</td><td>Intergenic</td><td>NA</td><td>NA</td></tr>
<tr><td colspan="13"></td></tr>
<tr><td>T6004</td><td>TGCAGACCAGGCGCTCCCGGCGG</td><td></td><td>1000</td><td colspan="9"></td></tr>
<tr><td>Oligo fwd</td><td>CACCTGCAGACCAGGCGCTCCCGG</td><td colspan="11"></td></tr>
<tr><td>Oligo rev</td><td>AAACCCGGGAGCGCCTGGTCTGCA</td><td colspan="11"></td></tr>
<tr><td>Chromosome</td><td>start</td><td></td><td>end</td><td>strand</td><td>MM</td><td>target_seq</td><td>PAM</td><td>alignment</td><td>distance</td><td>position</td><td>gene name</td><td>gene id</td></tr>
<tr><td>GU305914.1</td><td></td><td>83263</td><td>83285</td><td>+</td><td>0</td><td>TGCAGACCAGGCGCTCCCGG</td><td>CGG</td><td>||||||||[||||||||||||]PAM</td><td>NA</td><td>Intergenic</td><td>NA</td><td>NA</td></tr>
<tr><td colspan="13"></td></tr>
<tr><td>T6005</td><td>GAGTCCCCCGTGCGCGCCGCCGG</td><td></td><td>1000</td><td colspan="9"></td></tr>
<tr><td>Oligo fwd</td><td>CACCGAGTCCCCCGTGCGCGCCGC</td><td colspan="11"></td></tr>
<tr><td>Oligo rev</td><td>AAACGCGGCGCGCACGGGGGACTC</td><td colspan="11"></td></tr>
<tr><td>Chromosome</td><td>start</td><td></td><td>end</td><td>strand</td><td>MM</td><td>target_seq</td><td>PAM</td><td>alignment</td><td>distance</td><td>position</td><td>gene name</td><td>gene id</td></tr>
<tr><td>GU305914.1</td><td></td><td>83295</td><td>83317</td><td>+</td><td>0</td><td>GAGTCCCCCGTGCGCGCCGC</td><td>CGG</td><td>||||||||[||||||||||||]PAM</td><td>NA</td><td>Intergenic</td><td>NA</td><td>NA</td></tr>
<tr><td colspan="13"></td></tr>
<tr><td>T6006</td><td>AGTCCCCCGTGCGCGCCGCCGGG</td><td></td><td>1000</td><td colspan="9"></td></tr>
<tr><td>Oligo fwd</td><td>CACCAGTCCCCCGTGCGCGCCGCC</td><td colspan="11"></td></tr>
<tr><td>Oligo rev</td><td>AAACGGCGGCGCGCACGGGGGACT</td><td colspan="11"></td></tr>
<tr><td>Chromosome</td><td>start</td><td></td><td>end</td><td>strand</td><td>MM</td><td>target_seq</td><td>PAM</td><td>alignment</td><td>distance</td><td>position</td><td>gene name</td><td>gene id</td></tr>
<tr><td>GU305914.1</td><td></td><td>83296</td><td>83318</td><td>+</td><td>0</td><td>AGTCCCCCGTGCGCGCCGCC</td><td>GGG</td><td>||||||||[||||||||||||]PAM</td><td>NA</td><td>Intergenic</td><td>NA</td><td>NA</td></tr>
<tr><td colspan="13"></td></tr>
<tr><td>T6007</td><td>CCCCCGTGCGCGCCGCCGGGAGG</td><td></td><td>1000</td><td colspan="9"></td></tr>
<tr><td>Oligo fwd</td><td>CACCCCCCCGTGCGCGCCGCCGGG</td><td colspan="11"></td></tr>
<tr><td>Oligo rev</td><td>AAACCCCGGCGGCGCGCACGGGGG</td><td colspan="11"></td></tr>
<tr><td>Chromosome</td><td>start</td><td></td><td>end</td><td>strand</td><td>MM</td><td>target_seq</td><td>PAM</td><td>alignment</td><td>distance</td><td>position</td><td>gene name</td><td>gene id</td></tr>
<tr><td>GU305914.1</td><td></td><td>83299</td><td>83321</td><td>+</td><td>0</td><td>CCCCCGTGCGCGCCGCCGGG</td><td>AGG</td><td>||||||||[||||||||||||]PAM</td><td>NA</td><td>Intergenic</td><td>NA</td><td>NA</td></tr>
<tr><td colspan="13"></td></tr>
<tr><td>T6008</td><td>GCCGGGAGGAGCATACTGCACGG</td><td></td><td>1000</td><td colspan="9"></td></tr>
<tr><td>Oligo fwd</td><td>CACCGCCGGGAGGAGCATACTGCA</td><td colspan="11"></td></tr>
<tr><td>Oligo rev</td><td>AAACTGCAGTATGCTCCTCCCGGC</td><td colspan="11"></td></tr>
<tr><td>Chromosome</td><td>start</td><td></td><td>end</td><td>strand</td><td>MM</td><td>target_seq</td><td>PAM</td><td>alignment</td><td>distance</td><td>position</td><td>gene name</td><td>gene id</td></tr>
<tr><td>GU305914.1</td><td></td><td>83313</td><td>83335</td><td>+</td><td>0</td><td>GCCGGGAGGAGCATACTGCA</td><td>CGG</td><td>||||||||[||||||||||||]PAM</td><td>NA</td><td>Intergenic</td><td>NA</td><td>NA</td></tr>
<tr><td colspan="13"></td></tr>
<tr><td>T6009</td><td>GGGAGGAGCATACTGCACGGCGG</td><td></td><td>1000</td><td colspan="9"></td></tr>
<tr><td>Oligo fwd</td><td>CACCGGGAGGAGCATACTGCACGG</td><td colspan="11"></td></tr>
<tr><td>Oligo rev</td><td>AAACCCGTGCAGTATGCTCCTCCC</td><td colspan="11"></td></tr>
<tr><td>Chromosome</td><td>start</td><td></td><td>end</td><td>strand</td><td>MM</td><td>target_seq</td><td>PAM</td><td>alignment</td><td>distance</td><td>position</td><td>gene name</td><td>gene id</td></tr>
<tr><td>GU305914.1</td><td></td><td>83316</td><td>83338</td><td>+</td><td>0</td><td>GGGAGGAGCATACTGCACGG</td><td>CGG</td><td>||||||||[||||||||||||]PAM</td><td>NA</td><td>Intergenic</td><td>NA</td><td>NA</td></tr>
<tr><td colspan="13"></td></tr>
<tr><td>T6010</td><td>GAGCATACTGCACGGCGGTCTGG</td><td></td><td>1000</td><td colspan="9"></td></tr>
<tr><td>Oligo fwd</td><td>CACCGAGCATACTGCACGGCGGTC</td><td colspan="11"></td></tr>
<tr><td>Oligo rev</td><td>AAACGACCGCCGTGCAGTATGCTC</td><td colspan="11"></td></tr>
<tr><td>Chromosome</td><td>start</td><td></td><td>end</td><td>strand</td><td>MM</td><td>target_seq</td><td>PAM</td><td>alignment</td><td>distance</td><td>position</td><td>gene name</td><td>gene id</td></tr>
<tr><td>GU305914.1</td><td></td><td>83321</td><td>83343</td><td>+</td><td>0</td><td>GAGCATACTGCACGGCGGTC</td><td>TGG</td><td>||||||||[||||||||||||]PAM</td><td>NA</td><td>Intergenic</td><td>NA</td><td>NA</td></tr>
<tr><td colspan="13"></td></tr>
<tr><td>T6011</td><td>AGCATACTGCACGGCGGTCTGGG</td><td></td><td>1000</td><td colspan="9"></td></tr>
<tr><td>Oligo fwd</td><td>CACCAGCATACTGCACGGCGGTCT</td><td colspan="11"></td></tr>
<tr><td>Oligo rev</td><td>AAACAGACCGCCGTGCAGTATGCT</td><td colspan="11"></td></tr>
<tr><td>Chromosome</td><td>start</td><td></td><td>end</td><td>strand</td><td>MM</td><td>target_seq</td><td>PAM</td><td>alignment</td><td>distance</td><td>position</td><td>gene name</td><td>gene id</td></tr>
<tr><td>GU305914.1</td><td></td><td>83322</td><td>83344</td><td>+</td><td>0</td><td>AGCATACTGCACGGCGGTCT</td><td>GGG</td><td>||||||||[||||||||||||]PAM</td><td>NA</td><td>Intergenic</td><td>NA</td><td>NA</td></tr>
<tr><td colspan="13"></td></tr>
<tr><td>T6012</td><td>ATACTGCACGGCGGTCTGGGTGG</td><td></td><td>1000</td><td colspan="9"></td></tr>
<tr><td>Oligo fwd</td><td>CACCATACTGCACGGCGGTCTGGG</td><td colspan="11"></td></tr>
<tr><td>Oligo rev</td><td>AAACCCCAGACCGCCGTGCAGTAT</td><td colspan="11"></td></tr>
<tr><td>Chromosome</td><td>start</td><td></td><td>end</td><td>strand</td><td>MM</td><td>target_seq</td><td>PAM</td><td>alignment</td><td>distance</td><td>position</td><td>gene name</td><td>gene id</td></tr>
<tr><td>GU305914.1</td><td></td><td>83325</td><td>83347</td><td>+</td><td>0</td><td>ATACTGCACGGCGGTCTGGG</td><td>TGG</td><td>||||||||[||||||||||||]PAM</td><td>NA</td><td>Intergenic</td><td>NA</td><td>NA</td></tr>
<tr><td colspan="13"></td></tr>
<tr><td>T6013</td><td>CACGGCGGTCTGGGTGGCGTAGG</td><td></td><td>1000</td><td colspan="9"></td></tr>
<tr><td>Oligo fwd</td><td>CACCCACGGCGGTCTGGGTGGCGT</td><td colspan="11"></td></tr>
<tr><td>Oligo rev</td><td>AAACACGCCACCCAGACCGCCGTG</td><td colspan="11"></td></tr>
<tr><td>Chromosome</td><td>start</td><td></td><td>end</td><td>strand</td><td>MM</td><td>target_seq</td><td>PAM</td><td>alignment</td><td>distance</td><td>position</td><td>gene name</td><td>gene id</td></tr>
<tr><td>GU305914.1</td><td></td><td>83331</td><td>83353</td><td>+</td><td>0</td><td>CACGGCGGTCTGGGTGGCGT</td><td>AGG</td><td>||||||||[||||||||||||]PAM</td><td>NA</td><td>Intergenic</td><td>NA</td><td>NA</td></tr>
<tr><td colspan="13"></td></tr>
<tr><td>T6014</td><td>ACGGCGGTCTGGGTGGCGTAGGG</td><td></td><td>1000</td><td colspan="9"></td></tr>
<tr><td>Oligo fwd</td><td>CACCACGGCGGTCTGGGTGGCGTA</td><td colspan="11"></td></tr>
<tr><td>Oligo rev</td><td>AAACTACGCCACCCAGACCGCCGT</td><td colspan="11"></td></tr>
<tr><td>Chromosome</td><td>start</td><td></td><td>end</td><td>strand</td><td>MM</td><td>target_seq</td><td>PAM</td><td>alignment</td><td>distance</td><td>position</td><td>gene name</td><td>gene id</td></tr>
<tr><td>GU305914.1</td><td></td><td>83332</td><td>83354</td><td>+</td><td>0</td><td>ACGGCGGTCTGGGTGGCGTA</td><td>GGG</td><td>||||||||[||||||||||||]PAM</td><td>NA</td><td>Intergenic</td><td>NA</td><td>NA</td></tr>
<tr><td colspan="13"></td></tr>
<tr><td>T6015</td><td>CGGCGGTCTGGGTGGCGTAGGGG</td><td></td><td>1000</td><td colspan="9"></td></tr>
<tr><td>Oligo fwd</td><td>CACCCGGCGGTCTGGGTGGCGTAG</td><td colspan="11"></td></tr>
<tr><td>Oligo rev</td><td>AAACCTACGCCACCCAGACCGCCG</td><td colspan="11"></td></tr>
</table>

| | | | | | | | | | | | |
|---|---|---|---|---|---|---|---|---|---|---|---|
| Chromosome | start | end | strand | MM | target_seq | PAM | alignment | distance | position | gene name | gene id |
| GU305914.1 | 83333 | 83355 | + | 0 | CGGCGGTCTGGGTGGCGTAG | GGG | \|\|\|\|\|\|\|\|[\|\|\|\|\|\|\|\|\|\|\|\|]PAM | NA | Intergenic | NA | NA |
| T6016 | TAGGGGAAGAAGTACTCGACCGG | 1000 | | | | | | | | | |
| Oligo fwd | CACCTAGGGGAAGAAGTACTCGAC | | | | | | | | | | |
| Oligo rev | AAACGTCGAGTACTTCTTCCCCTA | | | | | | | | | | |
| Chromosome | start | end | strand | MM | target_seq | PAM | alignment | distance | position | gene name | gene id |
| GU305914.1 | 83350 | 83372 | + | 0 | TAGGGGAAGAAGTACTCGAC | CGG | \|\|\|\|\|\|\|\|[\|\|\|\|\|\|\|\|\|\|\|\|]PAM | NA | Intergenic | NA | NA |
| T6017 | GAAGTACTCGACCGGCCTCGCGG | 1000 | | | | | | | | | |
| Oligo fwd | CACCGAAGTACTCGACCGGCCTCG | | | | | | | | | | |
| Oligo rev | AAACCGAGGCCGGTCGAGTACTTC | | | | | | | | | | |
| Chromosome | start | end | strand | MM | target_seq | PAM | alignment | distance | position | gene name | gene id |
| GU305914.1 | 83358 | 83380 | + | 0 | GAAGTACTCGACCGGCCTCG | CGG | \|\|\|\|\|\|\|\|[\|\|\|\|\|\|\|\|\|\|\|\|]PAM | NA | Intergenic | NA | NA |
| T6018 | GTACTCGACCGGCCTCGCGGCGG | 1000 | | | | | | | | | |
| Oligo fwd | CACCGTACTCGACCGGCCTCGCGG | | | | | | | | | | |
| Oligo rev | AAACCCGCGAGGCCGGTCGAGTAC | | | | | | | | | | |
| Chromosome | start | end | strand | MM | target_seq | PAM | alignment | distance | position | gene name | gene id |
| GU305914.1 | 83361 | 83383 | + | 0 | GTACTCGACCGGCCTCGCGG | CGG | \|\|\|\|\|\|\|\|[\|\|\|\|\|\|\|\|\|\|\|\|]PAM | NA | Intergenic | NA | NA |
| T6019 | CGTGAAGATGAGATACACGACGG | 1000 | | | | | | | | | |
| Oligo fwd | CACCCGTGAAGATGAGATACACGA | | | | | | | | | | |
| Oligo rev | AAACTCGTGTATCTCATCTTCACG | | | | | | | | | | |
| Chromosome | start | end | strand | MM | target_seq | PAM | alignment | distance | position | gene name | gene id |
| GU305914.1 | 83397 | 83419 | + | 0 | CGTGAAGATGAGATACACGA | CGG | \|\|\|\|\|\|\|\|[\|\|\|\|\|\|\|\|\|\|\|\|]PAM | NA | Intergenic | NA | NA |
| T6020 | CGGCTAGCACGCCGATCACCAGG | 1000 | | | | | | | | | |
| Oligo fwd | CACCCGGCTAGCACGCCGATCACC | | | | | | | | | | |
| Oligo rev | AAACGGTGATCGGCGTGCTAGCCG | | | | | | | | | | |
| Chromosome | start | end | strand | MM | target_seq | PAM | alignment | distance | position | gene name | gene id |
| GU305914.1 | 83417 | 83439 | + | 0 | CGGCTAGCACGCCGATCACC | AGG | \|\|\|\|\|\|\|\|[\|\|\|\|\|\|\|\|\|\|\|\|]PAM | NA | Intergenic | NA | NA |
| T6021 | CTAGCACGCCGATCACCAGGAGG | 1000 | | | | | | | | | |
| Oligo fwd | CACCCTAGCACGCCGATCACCAGG | | | | | | | | | | |
| Oligo rev | AAACCCTGGTGATCGGCGTGCTAG | | | | | | | | | | |
| Chromosome | start | end | strand | MM | target_seq | PAM | alignment | distance | position | gene name | gene id |
| GU305914.1 | 83420 | 83442 | + | 0 | CTAGCACGCCGATCACCAGG | AGG | \|\|\|\|\|\|\|\|[\|\|\|\|\|\|\|\|\|\|\|\|]PAM | NA | Intergenic | NA | NA |
| T6022 | CACGCCGATCACCAGGAGGATGG | 1000 | | | | | | | | | |
| Oligo fwd | CACCCACGCCGATCACCAGGAGGA | | | | | | | | | | |
| Oligo rev | AAACTCCTCCTGGTGATCGGCGTG | | | | | | | | | | |
| Chromosome | start | end | strand | MM | target_seq | PAM | alignment | distance | position | gene name | gene id |
| GU305914.1 | 83424 | 83446 | + | 0 | CACGCCGATCACCAGGAGGA | TGG | \|\|\|\|\|\|\|\|[\|\|\|\|\|\|\|\|\|\|\|\|]PAM | NA | Intergenic | NA | NA |
| T6023 | GCCGATCACCAGGAGGATGGTGG | 1000 | | | | | | | | | |
| Oligo fwd | CACCGCCGATCACCAGGAGGATGG | | | | | | | | | | |
| Oligo rev | AAACCCATCCTCCTGGTGATCGGC | | | | | | | | | | |
| Chromosome | start | end | strand | MM | target_seq | PAM | alignment | distance | position | gene name | gene id |
| GU305914.1 | 83427 | 83449 | + | 0 | GCCGATCACCAGGAGGATGG | TGG | \|\|\|\|\|\|\|\|[\|\|\|\|\|\|\|\|\|\|\|\|]PAM | NA | Intergenic | NA | NA |
| GU305914.1 | 70978 | 71000 | + | 4 | GGCGATCGCCAGGAAGATCG | AGG | \|-\|\|\|\|\|-[\|\|\|\|\|\|-\|\|\|-\|]PAM | NA | Intergenic | NA | NA |
| T6024 | ATGGTGGTGAAACCTCCAAACGG | 1000 | | | | | | | | | |
| Oligo fwd | CACCATGGTGGTGAAACCTCCAAA | | | | | | | | | | |
| Oligo rev | AAACTTTGGAGGTTTCACCACCAT | | | | | | | | | | |
| Chromosome | start | end | strand | MM | target_seq | PAM | alignment | distance | position | gene name | gene id |
| GU305914.1 | 83443 | 83465 | + | 0 | ATGGTGGTGAAACCTCCAAA | CGG | \|\|\|\|\|\|\|\|[\|\|\|\|\|\|\|\|\|\|\|\|]PAM | NA | Intergenic | NA | NA |
| T6025 | TGGTGGTGAAACCTCCAAACGGG | 1000 | | | | | | | | | |
| Oligo fwd | CACCTGGTGGTGAAACCTCCAAAC | | | | | | | | | | |
| Oligo rev | AAACGTTTGGAGGTTTCACCACCA | | | | | | | | | | |
| Chromosome | start | end | strand | MM | target_seq | PAM | alignment | distance | position | gene name | gene id |
| GU305914.1 | 83444 | 83466 | + | 0 | TGGTGGTGAAACCTCCAAAC | GGG | \|\|\|\|\|\|\|\|[\|\|\|\|\|\|\|\|\|\|\|\|]PAM | NA | Intergenic | NA | NA |
| T6026 | AACCTCCAAACGGGTTCGTCAGG | 1000 | | | | | | | | | |
| Oligo fwd | CACCAACCTCCAAACGGGTTCGTC | | | | | | | | | | |
| Oligo rev | AAACGACGAACCCGTTTGGAGGTT | | | | | | | | | | |
| Chromosome | start | end | strand | MM | target_seq | PAM | alignment | distance | position | gene name | gene id |
| GU305914.1 | 83453 | 83475 | + | 0 | AACCTCCAAACGGGTTCGTC | AGG | \|\|\|\|\|\|\|\|[\|\|\|\|\|\|\|\|\|\|\|\|]PAM | NA | Intergenic | NA | NA |
| T6027 | GTTCGTCAGGAAGCCCGTGACGG | 1000 | | | | | | | | | |
| Oligo fwd | CACCGTTCGTCAGGAAGCCCGTGA | | | | | | | | | | |
| Oligo rev | AAACTCACGGGCTTCCTGACGAAC | | | | | | | | | | |
| Chromosome | start | end | strand | MM | target_seq | PAM | alignment | distance | position | gene name | gene id |
| GU305914.1 | 83466 | 83488 | + | 0 | GTTCGTCAGGAAGCCCGTGA | CGG | \|\|\|\|\|\|\|\|[\|\|\|\|\|\|\|\|\|\|\|\|]PAM | NA | Intergenic | NA | NA |
| T6028 | GGCGTTCACCACCGAAGACACGG | 1000 | | | | | | | | | |
| Oligo fwd | CACCGGCGTTCACCACCGAAGACA | | | | | | | | | | |
| Oligo rev | AAACTGTCTTCGGTGGTGAACGCC | | | | | | | | | | |
| Chromosome | start | end | strand | MM | target_seq | PAM | alignment | distance | position | gene name | gene id |
| GU305914.1 | 83487 | 83509 | + | 0 | GGCGTTCACCACCGAAGACA | CGG | \|\|\|\|\|\|\|\|[\|\|\|\|\|\|\|\|\|\|\|\|]PAM | NA | Intergenic | NA | NA |
| T6029 | CGAAGACACGGCCCCGCCGACGG | 1000 | | | | | | | | | |
| Oligo fwd | CACCCGAAGACACGGCCCCGCCGA | | | | | | | | | | |
| Oligo rev | AAACTCGGCGGGGCCGTGTCTTCG | | | | | | | | | | |
| Chromosome | start | end | strand | MM | target_seq | PAM | alignment | distance | position | gene name | gene id |
| GU305914.1 | 83499 | 83521 | + | 0 | CGAAGACACGGCCCCGCCGA | CGG | \|\|\|\|\|\|\|\|[\|\|\|\|\|\|\|\|\|\|\|\|]PAM | NA | Intergenic | NA | NA |
| T6030 | GGCCCCGCCGACGGCCCCGATGG | 1000 | | | | | | | | | |

<table>
<tr><td>Oligo fwd</td><td>CACCGGCCCCGCCGACGGCCCCGA</td><td></td><td></td><td></td><td></td><td></td><td></td><td></td><td></td><td></td><td></td></tr>
<tr><td>Oligo rev</td><td>AAACTCGGGGCCGTCGGCGGGGCC</td><td></td><td></td><td></td><td></td><td></td><td></td><td></td><td></td><td></td><td></td></tr>
<tr><td>Chromosome</td><td>start</td><td>end</td><td>strand</td><td>MM</td><td>target_seq</td><td>PAM</td><td>alignment</td><td>distance</td><td>position</td><td>gene name</td><td>gene id</td></tr>
<tr><td>GU305914.1</td><td>83508</td><td>83530</td><td>+</td><td>0</td><td>GGCCCCGCCGACGGCCCCGA</td><td>TGG</td><td>||||||||[||||||||||||]PAM</td><td>NA</td><td>Intergenic</td><td>NA</td><td>NA</td></tr>
<tr><td>T6031</td><td>ATCCAACCCCAAGAGATAAGAGG</td><td>1000</td><td></td><td></td><td></td><td></td><td></td><td></td><td></td><td></td><td></td></tr>
<tr><td>Oligo fwd</td><td>CACCATCCAACCCCAAGAGATAAG</td><td></td><td></td><td></td><td></td><td></td><td></td><td></td><td></td><td></td><td></td></tr>
<tr><td>Oligo rev</td><td>AAACCTTATCTCTTGGGGTTGGAT</td><td></td><td></td><td></td><td></td><td></td><td></td><td></td><td></td><td></td><td></td></tr>
<tr><td>Chromosome</td><td>start</td><td>end</td><td>strand</td><td>MM</td><td>target_seq</td><td>PAM</td><td>alignment</td><td>distance</td><td>position</td><td>gene name</td><td>gene id</td></tr>
<tr><td>GU305914.1</td><td>83577</td><td>83599</td><td>+</td><td>0</td><td>ATCCAACCCCAAGAGATAAG</td><td>AGG</td><td>||||||||[||||||||||||]PAM</td><td>NA</td><td>Intergenic</td><td>NA</td><td>NA</td></tr>
<tr><td>T6032</td><td>TCCAACCCCAAGAGATAAGAGGG</td><td>1000</td><td></td><td></td><td></td><td></td><td></td><td></td><td></td><td></td><td></td></tr>
<tr><td>Oligo fwd</td><td>CACCTCCAACCCCAAGAGATAAGA</td><td></td><td></td><td></td><td></td><td></td><td></td><td></td><td></td><td></td><td></td></tr>
<tr><td>Oligo rev</td><td>AAACTCTTATCTCTTGGGGTTGGA</td><td></td><td></td><td></td><td></td><td></td><td></td><td></td><td></td><td></td><td></td></tr>
<tr><td>Chromosome</td><td>start</td><td>end</td><td>strand</td><td>MM</td><td>target_seq</td><td>PAM</td><td>alignment</td><td>distance</td><td>position</td><td>gene name</td><td>gene id</td></tr>
<tr><td>GU305914.1</td><td>83578</td><td>83600</td><td>+</td><td>0</td><td>TCCAACCCCAAGAGATAAGA</td><td>GGG</td><td>||||||||[||||||||||||]PAM</td><td>NA</td><td>Intergenic</td><td>NA</td><td>NA</td></tr>
<tr><td>T6033</td><td>GGGCGTGTTGTCGTTGACTTTGG</td><td>1000</td><td></td><td></td><td></td><td></td><td></td><td></td><td></td><td></td><td></td></tr>
<tr><td>Oligo fwd</td><td>CACCGGGCGTGTTGTCGTTGACTT</td><td></td><td></td><td></td><td></td><td></td><td></td><td></td><td></td><td></td><td></td></tr>
<tr><td>Oligo rev</td><td>AAACAAGTCAACGACAACACGCCC</td><td></td><td></td><td></td><td></td><td></td><td></td><td></td><td></td><td></td><td></td></tr>
<tr><td>Chromosome</td><td>start</td><td>end</td><td>strand</td><td>MM</td><td>target_seq</td><td>PAM</td><td>alignment</td><td>distance</td><td>position</td><td>gene name</td><td>gene id</td></tr>
<tr><td>GU305914.1</td><td>83598</td><td>83620</td><td>+</td><td>0</td><td>GGGCGTGTTGTCGTTGACTT</td><td>TGG</td><td>||||||||[||||||||||||]PAM</td><td>NA</td><td>Intergenic</td><td>NA</td><td>NA</td></tr>
<tr><td>T6034</td><td>TGTCGTTGACTTTGGCGCTCAGG</td><td>1000</td><td></td><td></td><td></td><td></td><td></td><td></td><td></td><td></td><td></td></tr>
<tr><td>Oligo fwd</td><td>CACCTGTCGTTGACTTTGGCGCTC</td><td></td><td></td><td></td><td></td><td></td><td></td><td></td><td></td><td></td><td></td></tr>
<tr><td>Oligo rev</td><td>AAACGAGCGCCAAAGTCAACGACA</td><td></td><td></td><td></td><td></td><td></td><td></td><td></td><td></td><td></td><td></td></tr>
<tr><td>Chromosome</td><td>start</td><td>end</td><td>strand</td><td>MM</td><td>target_seq</td><td>PAM</td><td>alignment</td><td>distance</td><td>position</td><td>gene name</td><td>gene id</td></tr>
<tr><td>GU305914.1</td><td>83606</td><td>83628</td><td>+</td><td>0</td><td>TGTCGTTGACTTTGGCGCTC</td><td>AGG</td><td>||||||||[||||||||||||]PAM</td><td>NA</td><td>Intergenic</td><td>NA</td><td>NA</td></tr>
<tr><td>T6035</td><td>GTCGTTGACTTTGGCGCTCAGGG</td><td>1000</td><td></td><td></td><td></td><td></td><td></td><td></td><td></td><td></td><td></td></tr>
<tr><td>Oligo fwd</td><td>CACCGTCGTTGACTTTGGCGCTCA</td><td></td><td></td><td></td><td></td><td></td><td></td><td></td><td></td><td></td><td></td></tr>
<tr><td>Oligo rev</td><td>AAACTGAGCGCCAAAGTCAACGAC</td><td></td><td></td><td></td><td></td><td></td><td></td><td></td><td></td><td></td><td></td></tr>
<tr><td>Chromosome</td><td>start</td><td>end</td><td>strand</td><td>MM</td><td>target_seq</td><td>PAM</td><td>alignment</td><td>distance</td><td>position</td><td>gene name</td><td>gene id</td></tr>
<tr><td>GU305914.1</td><td>83607</td><td>83629</td><td>+</td><td>0</td><td>GTCGTTGACTTTGGCGCTCA</td><td>GGG</td><td>||||||||[||||||||||||]PAM</td><td>NA</td><td>Intergenic</td><td>NA</td><td>NA</td></tr>
<tr><td>T6036</td><td>TGACTTTGGCGCTCAGGGTGCGG</td><td>1000</td><td></td><td></td><td></td><td></td><td></td><td></td><td></td><td></td><td></td></tr>
<tr><td>Oligo fwd</td><td>CACCTGACTTTGGCGCTCAGGGTG</td><td></td><td></td><td></td><td></td><td></td><td></td><td></td><td></td><td></td><td></td></tr>
<tr><td>Oligo rev</td><td>AAACCACCCTGAGCGCCAAAGTCA</td><td></td><td></td><td></td><td></td><td></td><td></td><td></td><td></td><td></td><td></td></tr>
<tr><td>Chromosome</td><td>start</td><td>end</td><td>strand</td><td>MM</td><td>target_seq</td><td>PAM</td><td>alignment</td><td>distance</td><td>position</td><td>gene name</td><td>gene id</td></tr>
<tr><td>GU305914.1</td><td>83612</td><td>83634</td><td>+</td><td>0</td><td>TGACTTTGGCGCTCAGGGTG</td><td>CGG</td><td>||||||||[||||||||||||]PAM</td><td>NA</td><td>Intergenic</td><td>NA</td><td>NA</td></tr>
<tr><td>T6037</td><td>ACTCCCGCATGATCTCGTCCAGG</td><td>1000</td><td></td><td></td><td></td><td></td><td></td><td></td><td></td><td></td><td></td></tr>
<tr><td>Oligo fwd</td><td>CACCACTCCCGCATGATCTCGTCC</td><td></td><td></td><td></td><td></td><td></td><td></td><td></td><td></td><td></td><td></td></tr>
<tr><td>Oligo rev</td><td>AAACGGACGAGATCATGCGGGAGT</td><td></td><td></td><td></td><td></td><td></td><td></td><td></td><td></td><td></td><td></td></tr>
<tr><td>Chromosome</td><td>start</td><td>end</td><td>strand</td><td>MM</td><td>target_seq</td><td>PAM</td><td>alignment</td><td>distance</td><td>position</td><td>gene name</td><td>gene id</td></tr>
<tr><td>GU305914.1</td><td>83657</td><td>83679</td><td>+</td><td>0</td><td>ACTCCCGCATGATCTCGTCC</td><td>AGG</td><td>||||||||[||||||||||||]PAM</td><td>NA</td><td>Intergenic</td><td>NA</td><td>NA</td></tr>
<tr><td>T6038</td><td>GTCCAGGCTGAACACGTTCGAGG</td><td>1000</td><td></td><td></td><td></td><td></td><td></td><td></td><td></td><td></td><td></td></tr>
<tr><td>Oligo fwd</td><td>CACCGTCCAGGCTGAACACGTTCG</td><td></td><td></td><td></td><td></td><td></td><td></td><td></td><td></td><td></td><td></td></tr>
<tr><td>Oligo rev</td><td>AAACCGAACGTGTTCAGCCTGGAC</td><td></td><td></td><td></td><td></td><td></td><td></td><td></td><td></td><td></td><td></td></tr>
<tr><td>Chromosome</td><td>start</td><td>end</td><td>strand</td><td>MM</td><td>target_seq</td><td>PAM</td><td>alignment</td><td>distance</td><td>position</td><td>gene name</td><td>gene id</td></tr>
<tr><td>GU305914.1</td><td>83673</td><td>83695</td><td>+</td><td>0</td><td>GTCCAGGCTGAACACGTTCG</td><td>AGG</td><td>||||||||[||||||||||||]PAM</td><td>NA</td><td>Intergenic</td><td>NA</td><td>NA</td></tr>
<tr><td>T6039</td><td>CAGCTCGAGCACCTTGAAGTCGG</td><td>1000</td><td></td><td></td><td></td><td></td><td></td><td></td><td></td><td></td><td></td></tr>
<tr><td>Oligo fwd</td><td>CACCCAGCTCGAGCACCTTGAAGT</td><td></td><td></td><td></td><td></td><td></td><td></td><td></td><td></td><td></td><td></td></tr>
<tr><td>Oligo rev</td><td>AAACACTTCAAGGTGCTCGAGCTG</td><td></td><td></td><td></td><td></td><td></td><td></td><td></td><td></td><td></td><td></td></tr>
<tr><td>Chromosome</td><td>start</td><td>end</td><td>strand</td><td>MM</td><td>target_seq</td><td>PAM</td><td>alignment</td><td>distance</td><td>position</td><td>gene name</td><td>gene id</td></tr>
<tr><td>GU305914.1</td><td>83718</td><td>83740</td><td>+</td><td>0</td><td>CAGCTCGAGCACCTTGAAGT</td><td>CGG</td><td>||||||||[||||||||||||]PAM</td><td>NA</td><td>Intergenic</td><td>NA</td><td>NA</td></tr>
<tr><td>T6040</td><td>CCTTGAAGTCGGTGTTCTCCAGG</td><td>1000</td><td></td><td></td><td></td><td></td><td></td><td></td><td></td><td></td><td></td></tr>
<tr><td>Oligo fwd</td><td>CACCCCTTGAAGTCGGTGTTCTCC</td><td></td><td></td><td></td><td></td><td></td><td></td><td></td><td></td><td></td><td></td></tr>
<tr><td>Oligo rev</td><td>AAACGGAGAACACCGACTTCAAGG</td><td></td><td></td><td></td><td></td><td></td><td></td><td></td><td></td><td></td><td></td></tr>
<tr><td>Chromosome</td><td>start</td><td>end</td><td>strand</td><td>MM</td><td>target_seq</td><td>PAM</td><td>alignment</td><td>distance</td><td>position</td><td>gene name</td><td>gene id</td></tr>
<tr><td>GU305914.1</td><td>83729</td><td>83751</td><td>+</td><td>0</td><td>CCTTGAAGTCGGTGTTCTCC</td><td>AGG</td><td>||||||||[||||||||||||]PAM</td><td>NA</td><td>Intergenic</td><td>NA</td><td>NA</td></tr>
<tr><td>T6041</td><td>CTTGAAGTCGGTGTTCTCCAGGG</td><td>1000</td><td></td><td></td><td></td><td></td><td></td><td></td><td></td><td></td><td></td></tr>
<tr><td>Oligo fwd</td><td>CACCCTTGAAGTCGGTGTTCTCCA</td><td></td><td></td><td></td><td></td><td></td><td></td><td></td><td></td><td></td><td></td></tr>
<tr><td>Oligo rev</td><td>AAACTGGAGAACACCGACTTCAAG</td><td></td><td></td><td></td><td></td><td></td><td></td><td></td><td></td><td></td><td></td></tr>
<tr><td>Chromosome</td><td>start</td><td>end</td><td>strand</td><td>MM</td><td>target_seq</td><td>PAM</td><td>alignment</td><td>distance</td><td>position</td><td>gene name</td><td>gene id</td></tr>
<tr><td>GU305914.1</td><td>83730</td><td>83752</td><td>+</td><td>0</td><td>CTTGAAGTCGGTGTTCTCCA</td><td>GGG</td><td>||||||||[||||||||||||]PAM</td><td>NA</td><td>Intergenic</td><td>NA</td><td>NA</td></tr>
<tr><td>T6042</td><td>TTGAAGTCGGTGTTCTCCAGGGG</td><td>1000</td><td></td><td></td><td></td><td></td><td></td><td></td><td></td><td></td><td></td></tr>
<tr><td>Oligo fwd</td><td>CACCTTGAAGTCGGTGTTCTCCAG</td><td></td><td></td><td></td><td></td><td></td><td></td><td></td><td></td><td></td><td></td></tr>
<tr><td>Oligo rev</td><td>AAACCTGGAGAACACCGACTTCAA</td><td></td><td></td><td></td><td></td><td></td><td></td><td></td><td></td><td></td><td></td></tr>
<tr><td>Chromosome</td><td>start</td><td>end</td><td>strand</td><td>MM</td><td>target_seq</td><td>PAM</td><td>alignment</td><td>distance</td><td>position</td><td>gene name</td><td>gene id</td></tr>
<tr><td>GU305914.1</td><td>83731</td><td>83753</td><td>+</td><td>0</td><td>TTGAAGTCGGTGTTCTCCAG</td><td>GGG</td><td>||||||||[||||||||||||]PAM</td><td>NA</td><td>Intergenic</td><td>NA</td><td>NA</td></tr>
<tr><td>T6043</td><td>GCTCGATGTCCAGCCCGATCAGG</td><td>1000</td><td></td><td></td><td></td><td></td><td></td><td></td><td></td><td></td><td></td></tr>
<tr><td>Oligo fwd</td><td>CACCGCTCGATGTCCAGCCCGATC</td><td></td><td></td><td></td><td></td><td></td><td></td><td></td><td></td><td></td><td></td></tr>
<tr><td>Oligo rev</td><td>AAACGATCGGGCTGGACATCGAGC</td><td></td><td></td><td></td><td></td><td></td><td></td><td></td><td></td><td></td><td></td></tr>
<tr><td>Chromosome</td><td>start</td><td>end</td><td>strand</td><td>MM</td><td>target_seq</td><td>PAM</td><td>alignment</td><td>distance</td><td>position</td><td>gene name</td><td>gene id</td></tr>
<tr><td>GU305914.1</td><td>83753</td><td>83775</td><td>+</td><td>0</td><td>GCTCGATGTCCAGCCCGATC</td><td>AGG</td><td>||||||||[||||||||||||]PAM</td><td>NA</td><td>Intergenic</td><td>NA</td><td>NA</td></tr>
<tr><td>T6044</td><td>CTCGATGTCCAGCCCGATCAGGG</td><td>1000</td><td></td><td></td><td></td><td></td><td></td><td></td><td></td><td></td><td></td></tr>
<tr><td>Oligo fwd</td><td>CACCCTCGATGTCCAGCCCGATCA</td><td></td><td></td><td></td><td></td><td></td><td></td><td></td><td></td><td></td><td></td></tr>
<tr><td>Oligo rev</td><td>AAACTGATCGGGCTGGACATCGAG</td><td></td><td></td><td></td><td></td><td></td><td></td><td></td><td></td><td></td><td></td></tr>
<tr><td>Chromosome</td><td>start</td><td>end</td><td>strand</td><td>MM</td><td>target_seq</td><td>PAM</td><td>alignment</td><td>distance</td><td>position</td><td>gene name</td><td>gene id</td></tr>
<tr><td>GU305914.1</td><td>83754</td><td>83776</td><td>+</td><td>0</td><td>CTCGATGTCCAGCCCGATCA</td><td>GGG</td><td>||||||||[||||||||||||]PAM</td><td>NA</td><td>Intergenic</td><td>NA</td><td>NA</td></tr>
</table>

T6045 GCCCGATCAGGGTGTCGACGAGG 1000
Oligo fwd CACCGCCCGATCAGGGTGTCGACG
Oligo rev AAACCGTCGACACCCTGATCGGGC

| Chromosome | start | end | strand | MM | target_seq | PAM | alignment | distance | position | gene name | gene id |
|---|---|---|---|---|---|---|---|---|---|---|---|
| GU305914.1 | 83765 | 83787 | + | 0 | GCCCGATCAGGGTGTCGACG | AGG | \|\|\|\|\|\|\|\|[\|\|\|\|\|\|\|\|\|\|\|\|]PAM | NA | Intergenic | NA | NA |

T6046 GAGGTCGATGTGCTCCAGACTGG 1000
Oligo fwd CACCGAGGTCGATGTGCTCCAGAC
Oligo rev AAACGTCTGGAGCACATCGACCTC

| Chromosome | start | end | strand | MM | target_seq | PAM | alignment | distance | position | gene name | gene id |
|---|---|---|---|---|---|---|---|---|---|---|---|
| GU305914.1 | 83784 | 83806 | + | 0 | GAGGTCGATGTGCTCCAGAC | TGG | \|\|\|\|\|\|\|\|[\|\|\|\|\|\|\|\|\|\|\|\|]PAM | NA | Intergenic | NA | NA |

T6047 GACTGGTCACGTTCCTGAAACGG 1000
Oligo fwd CACCGACTGGTCACGTTCCTGAAA
Oligo rev AAACTTTCAGGAACGTGACCAGTC

| Chromosome | start | end | strand | MM | target_seq | PAM | alignment | distance | position | gene name | gene id |
|---|---|---|---|---|---|---|---|---|---|---|---|
| GU305914.1 | 83801 | 83823 | + | 0 | GACTGGTCACGTTCCTGAAA | CGG | \|\|\|\|\|\|\|\|[\|\|\|\|\|\|\|\|\|\|\|\|]PAM | NA | Intergenic | NA | NA |

T6048 GTCGCAGTTCTCCGTCCGAAAGG 1000
Oligo fwd CACCGTCGCAGTTCTCCGTCCGAA
Oligo rev AAACTTCGGACGGAGAACTGCGAC

| Chromosome | start | end | strand | MM | target_seq | PAM | alignment | distance | position | gene name | gene id |
|---|---|---|---|---|---|---|---|---|---|---|---|
| GU305914.1 | 83880 | 83902 | + | 0 | GTCGCAGTTCTCCGTCCGAA | AGG | \|\|\|\|\|\|\|\|[\|\|\|\|\|\|\|\|\|\|\|\|]PAM | NA | Intergenic | NA | NA |

T6049 TCTCGTTGTTCTCGCCCAGCTGG 1000
Oligo fwd CACCTCTCGTTGTTCTCGCCCAGC
Oligo rev AAACGCTGGGCGAGAACAACGAGA

| Chromosome | start | end | strand | MM | target_seq | PAM | alignment | distance | position | gene name | gene id |
|---|---|---|---|---|---|---|---|---|---|---|---|
| GU305914.1 | 83915 | 83937 | + | 0 | TCTCGTTGTTCTCGCCCAGC | TGG | \|\|\|\|\|\|\|\|[\|\|\|\|\|\|\|\|\|\|\|\|]PAM | NA | Intergenic | NA | NA |
| GU305914.1 | 190006 | 190028 | + | 4 | TCTCGTAAATCTCGTCCAGC | GGG | \|\|\|\|\|\|--[-\|\|\|\|\|-\|\|\|\|\|]PAM | NA | Intergenic | NA | NA |

T6050 ATGAACTGAAAGACGACCAGCGG 1000
Oligo fwd CACCATGAACTGAAAGACGACCAG
Oligo rev AAACCTGGTCGTCTTTCAGTTCAT

| Chromosome | start | end | strand | MM | target_seq | PAM | alignment | distance | position | gene name | gene id |
|---|---|---|---|---|---|---|---|---|---|---|---|
| GU305914.1 | 83962 | 83984 | + | 0 | ATGAACTGAAAGACGACCAG | CGG | \|\|\|\|\|\|\|\|[\|\|\|\|\|\|\|\|\|\|\|\|]PAM | NA | Intergenic | NA | NA |

T6051 AGTTGACGACCTTCCCGTCCTGG 1000
Oligo fwd CACCAGTTGACGACCTTCCCGTCC
Oligo rev AAACGGACGGGAAGGTCGTCAACT

| Chromosome | start | end | strand | MM | target_seq | PAM | alignment | distance | position | gene name | gene id |
|---|---|---|---|---|---|---|---|---|---|---|---|
| GU305914.1 | 83996 | 84018 | + | 0 | AGTTGACGACCTTCCCGTCC | TGG | \|\|\|\|\|\|\|\|[\|\|\|\|\|\|\|\|\|\|\|\|]PAM | NA | Intergenic | NA | NA |

T6052 GGAAGATCCGCATGTCCTTCAGG 1000
Oligo fwd CACCGGAAGATCCGCATGTCCTTC
Oligo rev AAACGAAGGACATGCGGATCTTCC

| Chromosome | start | end | strand | MM | target_seq | PAM | alignment | distance | position | gene name | gene id |
|---|---|---|---|---|---|---|---|---|---|---|---|
| GU305914.1 | 84017 | 84039 | + | 0 | GGAAGATCCGCATGTCCTTC | AGG | \|\|\|\|\|\|\|\|[\|\|\|\|\|\|\|\|\|\|\|\|]PAM | NA | Intergenic | NA | NA |
| GU305914.1 | 46616 | 46638 | - | 4 | GGAGGAACAGCATGTCGTTC | AGG | \|\|\|-\|\|-\|[-\|\|\|\|\|\|\|-\|\|\|]PAM | NA | Intergenic | NA | NA |

T6053 CTGATCGACCTTGACACACTCGG 1000
Oligo fwd CACCCTGATCGACCTTGACACACT
Oligo rev AAACAGTGTGTCAAGGTCGATCAG

| Chromosome | start | end | strand | MM | target_seq | PAM | alignment | distance | position | gene name | gene id |
|---|---|---|---|---|---|---|---|---|---|---|---|
| GU305914.1 | 84054 | 84076 | + | 0 | CTGATCGACCTTGACACACT | CGG | \|\|\|\|\|\|\|\|[\|\|\|\|\|\|\|\|\|\|\|\|]PAM | NA | Intergenic | NA | NA |

T6054 GATGACGTCGCCCGCCAGCTTGG 1000
Oligo fwd CACCGATGACGTCGCCCGCCAGCT
Oligo rev AAACAGCTGGCGGGCGACGTCATC

| Chromosome | start | end | strand | MM | target_seq | PAM | alignment | distance | position | gene name | gene id |
|---|---|---|---|---|---|---|---|---|---|---|---|
| GU305914.1 | 84084 | 84106 | + | 0 | GATGACGTCGCCCGCCAGCT | TGG | \|\|\|\|\|\|\|\|[\|\|\|\|\|\|\|\|\|\|\|\|]PAM | NA | Intergenic | NA | NA |

T6055 GACGTCGCCCGCCAGCTTGGCGG 1000
Oligo fwd CACCGACGTCGCCCGCCAGCTTGG
Oligo rev AAACCCAAGCTGGCGGGCGACGTC

| Chromosome | start | end | strand | MM | target_seq | PAM | alignment | distance | position | gene name | gene id |
|---|---|---|---|---|---|---|---|---|---|---|---|
| GU305914.1 | 84087 | 84109 | + | 0 | GACGTCGCCCGCCAGCTTGG | CGG | \|\|\|\|\|\|\|\|[\|\|\|\|\|\|\|\|\|\|\|\|]PAM | NA | Intergenic | NA | NA |

T6056 CCCGCCAGCTTGGCGGTGACCGG 1000
Oligo fwd CACCCCCGCCAGCTTGGCGGTGAC
Oligo rev AAACGTCACCGCCAAGCTGGCGGG

| Chromosome | start | end | strand | MM | target_seq | PAM | alignment | distance | position | gene name | gene id |
|---|---|---|---|---|---|---|---|---|---|---|---|
| GU305914.1 | 84094 | 84116 | + | 0 | CCCGCCAGCTTGGCGGTGAC | CGG | \|\|\|\|\|\|\|\|[\|\|\|\|\|\|\|\|\|\|\|\|]PAM | NA | Intergenic | NA | NA |

T6057 GGTGACCGGTCTCTCGTAGATGG 1000
Oligo fwd CACCGGTGACCGGTCTCTCGTAGA
Oligo rev AAACTCTACGAGAGACCGGTCACC

| Chromosome | start | end | strand | MM | target_seq | PAM | alignment | distance | position | gene name | gene id |
|---|---|---|---|---|---|---|---|---|---|---|---|
| GU305914.1 | 84108 | 84130 | + | 0 | GGTGACCGGTCTCTCGTAGA | TGG | \|\|\|\|\|\|\|\|[\|\|\|\|\|\|\|\|\|\|\|\|]PAM | NA | Intergenic | NA | NA |

T6058 GTAGATGGCCGACAAGATGTTGG 1000
Oligo fwd CACCGTAGATGGCCGACAAGATGT
Oligo rev AAACACATCTTGTCGGCCATCTAC

| Chromosome | start | end | strand | MM | target_seq | PAM | alignment | distance | position | gene name | gene id |
|---|---|---|---|---|---|---|---|---|---|---|---|
| GU305914.1 | 84123 | 84145 | + | 0 | GTAGATGGCCGACAAGATGT | TGG | \|\|\|\|\|\|\|\|[\|\|\|\|\|\|\|\|\|\|\|\|]PAM | NA | Intergenic | NA | NA |

T6059 ATGGCCGACAAGATGTTGGACGG 1000
Oligo fwd CACCATGGCCGACAAGATGTTGGA
Oligo rev AAACTCCAACATCTTGTCGGCCAT

| | | | | | | | | | | | |
|---|---|---|---|---|---|---|---|---|---|---|---|
| Chromosome | start | end | strand | MM | target_seq | PAM | alignment | distance | position | gene name | gene id |
| GU305914.1 | 84127 | 84149 | + | 0 | ATGGCCGACAAGATGTTGGA | CGG | \|\|\|\|\|\|\|\|[\|\|\|\|\|\|\|\|\|\|\|\|]PAM | NA | Intergenic | NA | NA |
| T6060 | GCATCTCCGCCGTCCGTTTCTGG | 1000 | | | | | | | | | |
| Oligo fwd | CACCGCATCTCCGCCGTCCGTTTC | | | | | | | | | | |
| Oligo rev | AAACGAAACGGACGGCGGAGATGC | | | | | | | | | | |
| Chromosome | start | end | strand | MM | target_seq | PAM | alignment | distance | position | gene name | gene id |
| GU305914.1 | 84173 | 84195 | + | 0 | GCATCTCCGCCGTCCGTTTC | TGG | \|\|\|\|\|\|\|\|[\|\|\|\|\|\|\|\|\|\|\|\|]PAM | NA | Intergenic | NA | NA |
| T6061 | ACCAAGCGTCCATGATGTTTCGG | 1000 | | | | | | | | | |
| Oligo fwd | CACCACCAAGCGTCCATGATGTTT | | | | | | | | | | |
| Oligo rev | AAACAAACATCATGGACGCTTGGT | | | | | | | | | | |
| Chromosome | start | end | strand | MM | target_seq | PAM | alignment | distance | position | gene name | gene id |
| GU305914.1 | 84203 | 84225 | + | 0 | ACCAAGCGTCCATGATGTTT | CGG | \|\|\|\|\|\|\|\|[\|\|\|\|\|\|\|\|\|\|\|\|]PAM | NA | Intergenic | NA | NA |
| T6062 | GTAGTCCTTGAGCACGTCGTAGG | 1000 | | | | | | | | | |
| Oligo fwd | CACCGTAGTCCTTGAGCACGTCGT | | | | | | | | | | |
| Oligo rev | AAACACGACGTGCTCAAGGACTAC | | | | | | | | | | |
| Chromosome | start | end | strand | MM | target_seq | PAM | alignment | distance | position | gene name | gene id |
| GU305914.1 | 84240 | 84262 | + | 0 | GTAGTCCTTGAGCACGTCGT | AGG | \|\|\|\|\|\|\|\|[\|\|\|\|\|\|\|\|\|\|\|\|]PAM | NA | Intergenic | NA | NA |
| T6063 | TATCCGTCACCACATCATCTAGG | 1000 | | | | | | | | | |
| Oligo fwd | CACCTATCCGTCACCACATCATCT | | | | | | | | | | |
| Oligo rev | AAACAGATGATGTGGTGACGGATA | | | | | | | | | | |
| Chromosome | start | end | strand | MM | target_seq | PAM | alignment | distance | position | gene name | gene id |
| GU305914.1 | 84287 | 84309 | + | 0 | TATCCGTCACCACATCATCT | AGG | \|\|\|\|\|\|\|\|[\|\|\|\|\|\|\|\|\|\|\|\|]PAM | NA | Intergenic | NA | NA |
| T6064 | GGTTGTCAAAGCTGCGTTTTCGG | 1000 | | | | | | | | | |
| Oligo fwd | CACCGGTTGTCAAAGCTGCGTTTT | | | | | | | | | | |
| Oligo rev | AAACAAAACGCAGCTTTGACAACC | | | | | | | | | | |
| Chromosome | start | end | strand | MM | target_seq | PAM | alignment | distance | position | gene name | gene id |
| GU305914.1 | 84308 | 84330 | + | 0 | GGTTGTCAAAGCTGCGTTTT | CGG | \|\|\|\|\|\|\|\|[\|\|\|\|\|\|\|\|\|\|\|\|]PAM | NA | Intergenic | NA | NA |
| T6065 | AGTAGTACCATTCTCAGTCGCGG | 1000 | | | | | | | | | |
| Oligo fwd | CACCAGTAGTACCATTCTCAGTCG | | | | | | | | | | |
| Oligo rev | AAACCGACTGAGAATGGTACTACT | | | | | | | | | | |
| Chromosome | start | end | strand | MM | target_seq | PAM | alignment | distance | position | gene name | gene id |
| GU305914.1 | 84342 | 84364 | + | 0 | AGTAGTACCATTCTCAGTCG | CGG | \|\|\|\|\|\|\|\|[\|\|\|\|\|\|\|\|\|\|\|\|]PAM | NA | Intergenic | NA | NA |
| T6066 | ACCATTCTCAGTCGCGGCAGTGG | 1000 | | | | | | | | | |
| Oligo fwd | CACCACCATTCTCAGTCGCGGCAG | | | | | | | | | | |
| Oligo rev | AAACCTGCCGCGACTGAGAATGGT | | | | | | | | | | |
| Chromosome | start | end | strand | MM | target_seq | PAM | alignment | distance | position | gene name | gene id |
| GU305914.1 | 84348 | 84370 | + | 0 | ACCATTCTCAGTCGCGGCAG | TGG | \|\|\|\|\|\|\|\|[\|\|\|\|\|\|\|\|\|\|\|\|]PAM | NA | Intergenic | NA | NA |
| T6067 | TCAGTCGCGGCAGTGGTTACAGG | 1000 | | | | | | | | | |
| Oligo fwd | CACCTCAGTCGCGGCAGTGGTTAC | | | | | | | | | | |
| Oligo rev | AAACGTAACCACTGCCGCGACTGA | | | | | | | | | | |
| Chromosome | start | end | strand | MM | target_seq | PAM | alignment | distance | position | gene name | gene id |
| GU305914.1 | 84355 | 84377 | + | 0 | TCAGTCGCGGCAGTGGTTAC | AGG | \|\|\|\|\|\|\|\|[\|\|\|\|\|\|\|\|\|\|\|\|]PAM | NA | Intergenic | NA | NA |
| T6068 | GTCGCGGCAGTGGTTACAGGAGG | 1000 | | | | | | | | | |
| Oligo fwd | CACCGTCGCGGCAGTGGTTACAGG | | | | | | | | | | |
| Oligo rev | AAACCCTGTAACCACTGCCGCGAC | | | | | | | | | | |
| Chromosome | start | end | strand | MM | target_seq | PAM | alignment | distance | position | gene name | gene id |
| GU305914.1 | 84358 | 84380 | + | 0 | GTCGCGGCAGTGGTTACAGG | AGG | \|\|\|\|\|\|\|\|[\|\|\|\|\|\|\|\|\|\|\|\|]PAM | NA | Intergenic | NA | NA |
| T6069 | TCGCGGCAGTGGTTACAGGAGGG | 1000 | | | | | | | | | |
| Oligo fwd | CACCTCGCGGCAGTGGTTACAGGA | | | | | | | | | | |
| Oligo rev | AAACTCCTGTAACCACTGCCGCGA | | | | | | | | | | |
| Chromosome | start | end | strand | MM | target_seq | PAM | alignment | distance | position | gene name | gene id |
| GU305914.1 | 84359 | 84381 | + | 0 | TCGCGGCAGTGGTTACAGGA | GGG | \|\|\|\|\|\|\|\|[\|\|\|\|\|\|\|\|\|\|\|\|]PAM | NA | Intergenic | NA | NA |
| T6070 | CACTGAAACGTTATTCAACGCGG | 1000 | | | | | | | | | |
| Oligo fwd | CACCCACTGAAACGTTATTCAACG | | | | | | | | | | |
| Oligo rev | AAACCGTTGAATAACGTTTCAGTG | | | | | | | | | | |
| Chromosome | start | end | strand | MM | target_seq | PAM | alignment | distance | position | gene name | gene id |
| GU305914.1 | 84393 | 84415 | + | 0 | CACTGAAACGTTATTCAACG | CGG | \|\|\|\|\|\|\|\|[\|\|\|\|\|\|\|\|\|\|\|\|]PAM | NA | Intergenic | NA | NA |
| T6071 | CAAACTCATCATAAACACATCGG | 1000 | | | | | | | | | |
| Oligo fwd | CACCCAAACTCATCATAAACACAT | | | | | | | | | | |
| Oligo rev | AAACATGTGTTTATGATGAGTTTG | | | | | | | | | | |
| Chromosome | start | end | strand | MM | target_seq | PAM | alignment | distance | position | gene name | gene id |
| GU305914.1 | 84563 | 84585 | + | 0 | CAAACTCATCATAAACACAT | CGG | \|\|\|\|\|\|\|\|[\|\|\|\|\|\|\|\|\|\|\|\|]PAM | NA | Intergenic | NA | NA |
| T6072 | CTCATCATAAACACATCGGAAGG | 1000 | | | | | | | | | |
| Oligo fwd | CACCCTCATCATAAACACATCGGA | | | | | | | | | | |
| Oligo rev | AAACTCCGATGTGTTTATGATGAG | | | | | | | | | | |
| Chromosome | start | end | strand | MM | target_seq | PAM | alignment | distance | position | gene name | gene id |
| GU305914.1 | 84567 | 84589 | + | 0 | CTCATCATAAACACATCGGA | AGG | \|\|\|\|\|\|\|\|[\|\|\|\|\|\|\|\|\|\|\|\|]PAM | NA | Intergenic | NA | NA |
| T6073 | GGTGTTATAATTTTTACCTTCGG | 1000 | | | | | | | | | |
| Oligo fwd | CACCGGTGTTATAATTTTTACCTT | | | | | | | | | | |
| Oligo rev | AAACAAGGTAAAAATTATAACACC | | | | | | | | | | |
| Chromosome | start | end | strand | MM | target_seq | PAM | alignment | distance | position | gene name | gene id |
| GU305914.1 | 84588 | 84610 | + | 0 | GGTGTTATAATTTTTACCTT | CGG | \|\|\|\|\|\|\|\|[\|\|\|\|\|\|\|\|\|\|\|\|]PAM | NA | Intergenic | NA | NA |
| T6074 | ATAATCTATTTTGCGTTTACTGG | 1000 | | | | | | | | | |
| Oligo fwd | CACCATAATCTATTTTGCGTTTAC | | | | | | | | | | |

<table>
<tr><td>Oligo rev</td><td>AAACGTAAACGCAAAATAGATTAT</td><td></td><td></td><td></td><td></td><td></td><td></td><td></td><td></td><td></td><td></td></tr>
<tr><td>Chromosome</td><td>start</td><td>end</td><td>strand</td><td>MM</td><td>target_seq</td><td>PAM</td><td>alignment</td><td>distance</td><td>position</td><td>gene name</td><td>gene id</td></tr>
<tr><td>GU305914.1</td><td>84621</td><td>84643</td><td>+</td><td>0</td><td>ATAATCTATTTTGCGTTTAC</td><td>TGG</td><td>||||||||[||||||||||||]PAM</td><td>NA</td><td>Intergenic</td><td>NA</td><td>NA</td></tr>
<tr><td>T6075</td><td>AAAAGTAGCAGTTAAACTCTTGG</td><td>1000</td><td></td><td></td><td></td><td></td><td></td><td></td><td></td><td></td><td></td></tr>
<tr><td>Oligo fwd</td><td>CACCAAAAGTAGCAGTTAAACTCT</td><td></td><td></td><td></td><td></td><td></td><td></td><td></td><td></td><td></td><td></td></tr>
<tr><td>Oligo rev</td><td>AAACAGAGTTTAACTGCTACTTTT</td><td></td><td></td><td></td><td></td><td></td><td></td><td></td><td></td><td></td><td></td></tr>
<tr><td>Chromosome</td><td>start</td><td>end</td><td>strand</td><td>MM</td><td>target_seq</td><td>PAM</td><td>alignment</td><td>distance</td><td>position</td><td>gene name</td><td>gene id</td></tr>
<tr><td>GU305914.1</td><td>84648</td><td>84670</td><td>+</td><td>0</td><td>AAAAGTAGCAGTTAAACTCT</td><td>TGG</td><td>||||||||[||||||||||||]PAM</td><td>NA</td><td>Intergenic</td><td>NA</td><td>NA</td></tr>
<tr><td>T6076</td><td>AACAAAATGATAGCTAGTATTGG</td><td>1000</td><td></td><td></td><td></td><td></td><td></td><td></td><td></td><td></td><td></td></tr>
<tr><td>Oligo fwd</td><td>CACCAACAAAATGATAGCTAGTAT</td><td></td><td></td><td></td><td></td><td></td><td></td><td></td><td></td><td></td><td></td></tr>
<tr><td>Oligo rev</td><td>AAACATACTAGCTATCATTTTGTT</td><td></td><td></td><td></td><td></td><td></td><td></td><td></td><td></td><td></td><td></td></tr>
<tr><td>Chromosome</td><td>start</td><td>end</td><td>strand</td><td>MM</td><td>target_seq</td><td>PAM</td><td>alignment</td><td>distance</td><td>position</td><td>gene name</td><td>gene id</td></tr>
<tr><td>GU305914.1</td><td>84672</td><td>84694</td><td>+</td><td>0</td><td>AACAAAATGATAGCTAGTAT</td><td>TGG</td><td>||||||||[||||||||||||]PAM</td><td>NA</td><td>Intergenic</td><td>NA</td><td>NA</td></tr>
<tr><td>T6077</td><td>GTCGCACGTGACATTCTGTTGG</td><td>1000</td><td></td><td></td><td></td><td></td><td></td><td></td><td></td><td></td><td></td></tr>
<tr><td>Oligo fwd</td><td>CACCGTCGCACGTGACATTCTGTT</td><td></td><td></td><td></td><td></td><td></td><td></td><td></td><td></td><td></td><td></td></tr>
<tr><td>Oligo rev</td><td>AAACAACAGAATGTCACGTGCGAC</td><td></td><td></td><td></td><td></td><td></td><td></td><td></td><td></td><td></td><td></td></tr>
<tr><td>Chromosome</td><td>start</td><td>end</td><td>strand</td><td>MM</td><td>target_seq</td><td>PAM</td><td>alignment</td><td>distance</td><td>position</td><td>gene name</td><td>gene id</td></tr>
<tr><td>GU305914.1</td><td>84738</td><td>84760</td><td>+</td><td>0</td><td>GTCGCACGTGACATTCTGTT</td><td>TGG</td><td>||||||||[||||||||||||]PAM</td><td>NA</td><td>Intergenic</td><td>NA</td><td>NA</td></tr>
<tr><td>T6078</td><td>TCGCACGTGACATTCTGTTTGGG</td><td>1000</td><td></td><td></td><td></td><td></td><td></td><td></td><td></td><td></td><td></td></tr>
<tr><td>Oligo fwd</td><td>CACCTCGCACGTGACATTCTGTTT</td><td></td><td></td><td></td><td></td><td></td><td></td><td></td><td></td><td></td><td></td></tr>
<tr><td>Oligo rev</td><td>AAACAAACAGAATGTCACGTGCGA</td><td></td><td></td><td></td><td></td><td></td><td></td><td></td><td></td><td></td><td></td></tr>
<tr><td>Chromosome</td><td>start</td><td>end</td><td>strand</td><td>MM</td><td>target_seq</td><td>PAM</td><td>alignment</td><td>distance</td><td>position</td><td>gene name</td><td>gene id</td></tr>
<tr><td>GU305914.1</td><td>84739</td><td>84761</td><td>+</td><td>0</td><td>TCGCACGTGACATTCTGTTT</td><td>GGG</td><td>||||||||[||||||||||||]PAM</td><td>NA</td><td>Intergenic</td><td>NA</td><td>NA</td></tr>
<tr><td>T6079</td><td>GGGAAAGATCTCCCACCCGATGG</td><td>1000</td><td></td><td></td><td></td><td></td><td></td><td></td><td></td><td></td><td></td></tr>
<tr><td>Oligo fwd</td><td>CACCGGGAAAGATCTCCCACCCGA</td><td></td><td></td><td></td><td></td><td></td><td></td><td></td><td></td><td></td><td></td></tr>
<tr><td>Oligo rev</td><td>AAACTCGGGTGGGAGATCTTTCCC</td><td></td><td></td><td></td><td></td><td></td><td></td><td></td><td></td><td></td><td></td></tr>
<tr><td>Chromosome</td><td>start</td><td>end</td><td>strand</td><td>MM</td><td>target_seq</td><td>PAM</td><td>alignment</td><td>distance</td><td>position</td><td>gene name</td><td>gene id</td></tr>
<tr><td>GU305914.1</td><td>84759</td><td>84781</td><td>+</td><td>0</td><td>GGGAAAGATCTCCCACCCGA</td><td>TGG</td><td>||||||||[||||||||||||]PAM</td><td>NA</td><td>Intergenic</td><td>NA</td><td>NA</td></tr>
<tr><td>T6080</td><td>CTCTCTAGAAAAGCCATCAGAGG</td><td>1000</td><td></td><td></td><td></td><td></td><td></td><td></td><td></td><td></td><td></td></tr>
<tr><td>Oligo fwd</td><td>CACCCTCTCTAGAAAAGCCATCAG</td><td></td><td></td><td></td><td></td><td></td><td></td><td></td><td></td><td></td><td></td></tr>
<tr><td>Oligo rev</td><td>AAACCTGATGGCTTTTCTAGAGAG</td><td></td><td></td><td></td><td></td><td></td><td></td><td></td><td></td><td></td><td></td></tr>
<tr><td>Chromosome</td><td>start</td><td>end</td><td>strand</td><td>MM</td><td>target_seq</td><td>PAM</td><td>alignment</td><td>distance</td><td>position</td><td>gene name</td><td>gene id</td></tr>
<tr><td>GU305914.1</td><td>84793</td><td>84815</td><td>+</td><td>0</td><td>CTCTCTAGAAAAGCCATCAG</td><td>AGG</td><td>||||||||[||||||||||||]PAM</td><td>NA</td><td>Intergenic</td><td>NA</td><td>NA</td></tr>
<tr><td>T6081</td><td>CATCAGAGGTACTACTTTTTTGG</td><td>1000</td><td></td><td></td><td></td><td></td><td></td><td></td><td></td><td></td><td></td></tr>
<tr><td>Oligo fwd</td><td>CACCCATCAGAGGTACTACTTTTT</td><td></td><td></td><td></td><td></td><td></td><td></td><td></td><td></td><td></td><td></td></tr>
<tr><td>Oligo rev</td><td>AAACAAAAAGTAGTACCTCTGATG</td><td></td><td></td><td></td><td></td><td></td><td></td><td></td><td></td><td></td><td></td></tr>
<tr><td>Chromosome</td><td>start</td><td>end</td><td>strand</td><td>MM</td><td>target_seq</td><td>PAM</td><td>alignment</td><td>distance</td><td>position</td><td>gene name</td><td>gene id</td></tr>
<tr><td>GU305914.1</td><td>84807</td><td>84829</td><td>+</td><td>0</td><td>CATCAGAGGTACTACTTTTT</td><td>TGG</td><td>||||||||[||||||||||||]PAM</td><td>NA</td><td>Intergenic</td><td>NA</td><td>NA</td></tr>
<tr><td>T6082</td><td>ATCAGAGGTACTACTTTTTTGGG</td><td>1000</td><td></td><td></td><td></td><td></td><td></td><td></td><td></td><td></td><td></td></tr>
<tr><td>Oligo fwd</td><td>CACCATCAGAGGTACTACTTTTTT</td><td></td><td></td><td></td><td></td><td></td><td></td><td></td><td></td><td></td><td></td></tr>
<tr><td>Oligo rev</td><td>AAACAAAAAAGTAGTACCTCTGAT</td><td></td><td></td><td></td><td></td><td></td><td></td><td></td><td></td><td></td><td></td></tr>
<tr><td>Chromosome</td><td>start</td><td>end</td><td>strand</td><td>MM</td><td>target_seq</td><td>PAM</td><td>alignment</td><td>distance</td><td>position</td><td>gene name</td><td>gene id</td></tr>
<tr><td>GU305914.1</td><td>84808</td><td>84830</td><td>+</td><td>0</td><td>ATCAGAGGTACTACTTTTTT</td><td>GGG</td><td>||||||||[||||||||||||]PAM</td><td>NA</td><td>Intergenic</td><td>NA</td><td>NA</td></tr>
<tr><td>T6083</td><td>TTCACCGAAGTACGTCTTCATGG</td><td>1000</td><td></td><td></td><td></td><td></td><td></td><td></td><td></td><td></td><td></td></tr>
<tr><td>Oligo fwd</td><td>CACCTTCACCGAAGTACGTCTTCA</td><td></td><td></td><td></td><td></td><td></td><td></td><td></td><td></td><td></td><td></td></tr>
<tr><td>Oligo rev</td><td>AAACTGAAGACGTACTTCGGTGAA</td><td></td><td></td><td></td><td></td><td></td><td></td><td></td><td></td><td></td><td></td></tr>
<tr><td>Chromosome</td><td>start</td><td>end</td><td>strand</td><td>MM</td><td>target_seq</td><td>PAM</td><td>alignment</td><td>distance</td><td>position</td><td>gene name</td><td>gene id</td></tr>
<tr><td>GU305914.1</td><td>84840</td><td>84862</td><td>+</td><td>0</td><td>TTCACCGAAGTACGTCTTCA</td><td>TGG</td><td>||||||||[||||||||||||]PAM</td><td>NA</td><td>Intergenic</td><td>NA</td><td>NA</td></tr>
<tr><td>T6084</td><td>ATTTCTCGTGTCCTCTTCGAAGG</td><td>1000</td><td></td><td></td><td></td><td></td><td></td><td></td><td></td><td></td><td></td></tr>
<tr><td>Oligo fwd</td><td>CACCATTTCTCGTGTCCTCTTCGA</td><td></td><td></td><td></td><td></td><td></td><td></td><td></td><td></td><td></td><td></td></tr>
<tr><td>Oligo rev</td><td>AAACTCGAAGAGGACACGAGAAAT</td><td></td><td></td><td></td><td></td><td></td><td></td><td></td><td></td><td></td><td></td></tr>
<tr><td>Chromosome</td><td>start</td><td>end</td><td>strand</td><td>MM</td><td>target_seq</td><td>PAM</td><td>alignment</td><td>distance</td><td>position</td><td>gene name</td><td>gene id</td></tr>
<tr><td>GU305914.1</td><td>84885</td><td>84907</td><td>+</td><td>0</td><td>ATTTCTCGTGTCCTCTTCGA</td><td>AGG</td><td>||||||||[||||||||||||]PAM</td><td>NA</td><td>Intergenic</td><td>NA</td><td>NA</td></tr>
<tr><td>T6085</td><td>TTTCTCGTGTCCTCTTCGAAGGG</td><td>1000</td><td></td><td></td><td></td><td></td><td></td><td></td><td></td><td></td><td></td></tr>
<tr><td>Oligo fwd</td><td>CACCTTTCTCGTGTCCTCTTCGAA</td><td></td><td></td><td></td><td></td><td></td><td></td><td></td><td></td><td></td><td></td></tr>
<tr><td>Oligo rev</td><td>AAACTTCGAAGAGGACACGAGAAA</td><td></td><td></td><td></td><td></td><td></td><td></td><td></td><td></td><td></td><td></td></tr>
<tr><td>Chromosome</td><td>start</td><td>end</td><td>strand</td><td>MM</td><td>target_seq</td><td>PAM</td><td>alignment</td><td>distance</td><td>position</td><td>gene name</td><td>gene id</td></tr>
<tr><td>GU305914.1</td><td>84886</td><td>84908</td><td>+</td><td>0</td><td>TTTCTCGTGTCCTCTTCGAA</td><td>GGG</td><td>||||||||[||||||||||||]PAM</td><td>NA</td><td>Intergenic</td><td>NA</td><td>NA</td></tr>
<tr><td>T6086</td><td>CGTGTCCTCTTCGAAGGGCTCGG</td><td>1000</td><td></td><td></td><td></td><td></td><td></td><td></td><td></td><td></td><td></td></tr>
<tr><td>Oligo fwd</td><td>CACCCGTGTCCTCTTCGAAGGGCT</td><td></td><td></td><td></td><td></td><td></td><td></td><td></td><td></td><td></td><td></td></tr>
<tr><td>Oligo rev</td><td>AAACAGCCCTTCGAAGAGGACACG</td><td></td><td></td><td></td><td></td><td></td><td></td><td></td><td></td><td></td><td></td></tr>
<tr><td>Chromosome</td><td>start</td><td>end</td><td>strand</td><td>MM</td><td>target_seq</td><td>PAM</td><td>alignment</td><td>distance</td><td>position</td><td>gene name</td><td>gene id</td></tr>
<tr><td>GU305914.1</td><td>84891</td><td>84913</td><td>+</td><td>0</td><td>CGTGTCCTCTTCGAAGGGCT</td><td>CGG</td><td>||||||||[||||||||||||]PAM</td><td>NA</td><td>Intergenic</td><td>NA</td><td>NA</td></tr>
<tr><td>T6087</td><td>GAGTTTTCACCGTTGTAAAACGG</td><td>1000</td><td></td><td></td><td></td><td></td><td></td><td></td><td></td><td></td><td></td></tr>
<tr><td>Oligo fwd</td><td>CACCGAGTTTTCACCGTTGTAAAA</td><td></td><td></td><td></td><td></td><td></td><td></td><td></td><td></td><td></td><td></td></tr>
<tr><td>Oligo rev</td><td>AAACTTTTACAACGGTGAAAACTC</td><td></td><td></td><td></td><td></td><td></td><td></td><td></td><td></td><td></td><td></td></tr>
<tr><td>Chromosome</td><td>start</td><td>end</td><td>strand</td><td>MM</td><td>target_seq</td><td>PAM</td><td>alignment</td><td>distance</td><td>position</td><td>gene name</td><td>gene id</td></tr>
<tr><td>GU305914.1</td><td>84913</td><td>84935</td><td>+</td><td>0</td><td>GAGTTTTCACCGTTGTAAAA</td><td>CGG</td><td>||||||||[||||||||||||]PAM</td><td>NA</td><td>Intergenic</td><td>NA</td><td>NA</td></tr>
<tr><td>GU305914.1</td><td>84922</td><td>84944</td><td>-</td><td>4</td><td>GAGATATCACCGTTTTACAA</td><td>CGG</td><td>|||-|-||[||||||-||-||]PAM</td><td>NA</td><td>Intergenic</td><td>NA</td><td>NA</td></tr>
<tr><td>T6088</td><td>TGATATCTCGACGACTTCCCCGG</td><td>1000</td><td></td><td></td><td></td><td></td><td></td><td></td><td></td><td></td><td></td></tr>
<tr><td>Oligo fwd</td><td>CACCTGATATCTCGACGACTTCCC</td><td></td><td></td><td></td><td></td><td></td><td></td><td></td><td></td><td></td><td></td></tr>
<tr><td>Oligo rev</td><td>AAACGGGAAGTCGTCGAGATATCA</td><td></td><td></td><td></td><td></td><td></td><td></td><td></td><td></td><td></td><td></td></tr>
<tr><td>Chromosome</td><td>start</td><td>end</td><td>strand</td><td>MM</td><td>target_seq</td><td>PAM</td><td>alignment</td><td>distance</td><td>position</td><td>gene name</td><td>gene id</td></tr>
<tr><td>GU305914.1</td><td>84936</td><td>84958</td><td>+</td><td>0</td><td>TGATATCTCGACGACTTCCC</td><td>CGG</td><td>||||||||[||||||||||||]PAM</td><td>NA</td><td>Intergenic</td><td>NA</td><td>NA</td></tr>
</table>

T6089 GCTGAGCACGAAGAATTCGTAGG 1000
Oligo fwd CACCGCTGAGCACGAAGAATTCGT
Oligo rev AAACACGAATTCTTCGTGCTCAGC
Chromosome start end strand MM target_seq PAM alignment distance position gene name gene id
GU305914.1 84960 84982 + 0 GCTGAGCACGAAGAATTCGT AGG ||||||||[||||||||||||]PAM NA Intergenic NA NA

T6090 CTGAGCACGAAGAATTCGTAGGG 1000
Oligo fwd CACCCTGAGCACGAAGAATTCGTA
Oligo rev AAACTACGAATTCTTCGTGCTCAG
Chromosome start end strand MM target_seq PAM alignment distance position gene name gene id
GU305914.1 84961 84983 + 0 CTGAGCACGAAGAATTCGTA GGG ||||||||[||||||||||||]PAM NA Intergenic NA NA

T6091 TGAGCACGAAGAATTCGTAGGGG 1000
Oligo fwd CACCTGAGCACGAAGAATTCGTAG
Oligo rev AAACCTACGAATTCTTCGTGCTCA
Chromosome start end strand MM target_seq PAM alignment distance position gene name gene id
GU305914.1 84962 84984 + 0 TGAGCACGAAGAATTCGTAG GGG ||||||||[||||||||||||]PAM NA Intergenic NA NA

T6092 GAGCACGAAGAATTCGTAGGGGG 1000
Oligo fwd CACCGAGCACGAAGAATTCGTAGG
Oligo rev AAACCCTACGAATTCTTCGTGCTC
Chromosome start end strand MM target_seq PAM alignment distance position gene name gene id
GU305914.1 84963 84985 + 0 GAGCACGAAGAATTCGTAGG GGG ||||||||[||||||||||||]PAM NA Intergenic NA NA

T6093 GACGACGCAGTTCATGTTACAGG 1000
Oligo fwd CACCGACGACGCAGTTCATGTTAC
Oligo rev AAACGTAACATGAACTGCGTCGTC
Chromosome start end strand MM target_seq PAM alignment distance position gene name gene id
GU305914.1 85011 85033 + 0 GACGACGCAGTTCATGTTAC AGG ||||||||[||||||||||||]PAM NA Intergenic NA NA

T6094 GCAGTTCATGTTACAGGATTCGG 1000
Oligo fwd CACCGCAGTTCATGTTACAGGATT
Oligo rev AAACAATCCTGTAACATGAACTGC
Chromosome start end strand MM target_seq PAM alignment distance position gene name gene id
GU305914.1 85017 85039 + 0 GCAGTTCATGTTACAGGATT CGG ||||||||[||||||||||||]PAM NA Intergenic NA NA

T6095 GCCCTTATGATACCTACTCTTGG 1000
Oligo fwd CACCGCCCTTATGATACCTACTCT
Oligo rev AAACAGAGTAGGTATCATAAGGGC
Chromosome start end strand MM target_seq PAM alignment distance position gene name gene id
GU305914.1 85059 85081 + 0 GCCCTTATGATACCTACTCT TGG ||||||||[||||||||||||]PAM NA Intergenic NA NA

T6096 CTTATGATACCTACTCTTGGTGG 1000
Oligo fwd CACCCTTATGATACCTACTCTTGG
Oligo rev AAACCCAAGAGTAGGTATCATAAG
Chromosome start end strand MM target_seq PAM alignment distance position gene name gene id
GU305914.1 85062 85084 + 0 CTTATGATACCTACTCTTGG TGG ||||||||[||||||||||||]PAM NA Intergenic NA NA

T6097 AGTTCCTATAGCTATCTTCGTGG 1000
Oligo fwd CACCAGTTCCTATAGCTATCTTCG
Oligo rev AAACCGAAGATAGCTATAGGAACT
Chromosome start end strand MM target_seq PAM alignment distance position gene name gene id
GU305914.1 85142 85164 + 0 AGTTCCTATAGCTATCTTCG TGG ||||||||[||||||||||||]PAM NA Intergenic NA NA

T6098 CCCATTGAGTATCCTACTCGCGG 1000
Oligo fwd CACCCCCATTGAGTATCCTACTCG
Oligo rev AAACCGAGTAGGATACTCAATGGG
Chromosome start end strand MM target_seq PAM alignment distance position gene name gene id
GU305914.1 85182 85204 + 0 CCCATTGAGTATCCTACTCG CGG ||||||||[||||||||||||]PAM NA Intergenic NA NA

T6099 TACCTCATCCAGCTCCCAGATGG 1000
Oligo fwd CACCTACCTCATCCAGCTCCCAGA
Oligo rev AAACTCTGGGAGCTGGATGAGGTA
Chromosome start end strand MM target_seq PAM alignment distance position gene name gene id
GU305914.1 85230 85252 + 0 TACCTCATCCAGCTCCCAGA TGG ||||||||[||||||||||||]PAM NA Intergenic NA NA

T6100 ACCTCATCCAGCTCCCAGATGGG 1000
Oligo fwd CACCACCTCATCCAGCTCCCAGAT
Oligo rev AAACATCTGGGAGCTGGATGAGGT
Chromosome start end strand MM target_seq PAM alignment distance position gene name gene id
GU305914.1 85231 85253 + 0 ACCTCATCCAGCTCCCAGAT GGG ||||||||[||||||||||||]PAM NA Intergenic NA NA

T6101 CCTCATCCAGCTCCCAGATGGGG 1000
Oligo fwd CACCCCTCATCCAGCTCCCAGATG
Oligo rev AAACCATCTGGGAGCTGGATGAGG
Chromosome start end strand MM target_seq PAM alignment distance position gene name gene id
GU305914.1 85232 85254 + 0 CCTCATCCAGCTCCCAGATG GGG ||||||||[||||||||||||]PAM NA Intergenic NA NA

T6102 ATCCAGCTCCCAGATGGGGAAGG 1000
Oligo fwd CACCATCCAGCTCCCAGATGGGGA
Oligo rev AAACTCCCCATCTGGGAGCTGGAT
Chromosome start end strand MM target_seq PAM alignment distance position gene name gene id
GU305914.1 85236 85258 + 0 ATCCAGCTCCCAGATGGGGA AGG ||||||||[||||||||||||]PAM NA Intergenic NA NA

T6103 TACCATGATCCCCTCTTGTACGG 1000
Oligo fwd CACCTACCATGATCCCCTCTTGTA
Oligo rev AAACTACAAGAGGGGATCATGGTA
Chromosome start end strand MM target_seq PAM alignment distance position gene name gene id
GU305914.1 85386 85408 + 0 TACCATGATCCCCTCTTGTA CGG ||||||||[||||||||||||]PAM NA Intergenic NA NA

T6104 ACCATGATCCCCTCTTGTACGGG 1000
Oligo fwd CACCACCATGATCCCCTCTTGTAC
Oligo rev AAACGTACAAGAGGGGATCATGGT
Chromosome start end strand MM target_seq PAM alignment distance position gene name gene id
GU305914.1 85387 85409 + 0 ACCATGATCCCCTCTTGTAC GGG ||||||||[||||||||||||]PAM NA Intergenic NA NA

T6105 CCCTCTTGTACGGGTGTCTTCGG 1000
Oligo fwd CACCCCCTCTTGTACGGGTGTCTT
Oligo rev AAACAAGACACCCGTACAAGAGGG
Chromosome start end strand MM target_seq PAM alignment distance position gene name gene id
GU305914.1 85396 85418 + 0 CCCTCTTGTACGGGTGTCTT CGG ||||||||[||||||||||||]PAM NA Intergenic NA NA

T6106 CCTCTTGTACGGGTGTCTTCGGG 1000
Oligo fwd CACCCCTCTTGTACGGGTGTCTTC
Oligo rev AAACGAAGACACCCGTACAAGAGG
Chromosome start end strand MM target_seq PAM alignment distance position gene name gene id
GU305914.1 85397 85419 + 0 CCTCTTGTACGGGTGTCTTC GGG ||||||||[||||||||||||]PAM NA Intergenic NA NA

T6107 CTCTTGTACGGGTGTCTTCGGGG 1000
Oligo fwd CACCCTCTTGTACGGGTGTCTTCG
Oligo rev AAACCGAAGACACCCGTACAAGAG
Chromosome start end strand MM target_seq PAM alignment distance position gene name gene id
GU305914.1 85398 85420 + 0 CTCTTGTACGGGTGTCTTCG GGG ||||||||[||||||||||||]PAM NA Intergenic NA NA

T6108 ACAGCCATGCACACTCTGTAAGG 1000
Oligo fwd CACCACAGCCATGCACACTCTGTA
Oligo rev AAACTACAGAGTGTGCATGGCTGT
Chromosome start end strand MM target_seq PAM alignment distance position gene name gene id
GU305914.1 85471 85493 + 0 ACAGCCATGCACACTCTGTA AGG ||||||||[||||||||||||]PAM NA Intergenic NA NA

T6109 GTCCTCCGTCTCGTTTACGAAGG 1000
Oligo fwd CACCGTCCTCCGTCTCGTTTACGA
Oligo rev AAACTCGTAAACGAGACGGAGGAC
Chromosome start end strand MM target_seq PAM alignment distance position gene name gene id
GU305914.1 85521 85543 + 0 GTCCTCCGTCTCGTTTACGA AGG ||||||||[||||||||||||]PAM NA Intergenic NA NA

T6110 TCCTCCGTCTCGTTTACGAAGGG 1000
Oligo fwd CACCTCCTCCGTCTCGTTTACGAA
Oligo rev AAACTTCGTAAACGAGACGGAGGA
Chromosome start end strand MM target_seq PAM alignment distance position gene name gene id
GU305914.1 85522 85544 + 0 TCCTCCGTCTCGTTTACGAA GGG ||||||||[||||||||||||]PAM NA Intergenic NA NA

T6111 CTCGTTTACGAAGGGAACGATGG 1000
Oligo fwd CACCCTCGTTTACGAAGGGAACGA
Oligo rev AAACTCGTTCCCTTCGTAAACGAG
Chromosome start end strand MM target_seq PAM alignment distance position gene name gene id
GU305914.1 85530 85552 + 0 CTCGTTTACGAAGGGAACGA TGG ||||||||[||||||||||||]PAM NA Intergenic NA NA

T6112 GTTTACGAAGGGAACGATGGTGG 1000
Oligo fwd CACCGTTTACGAAGGGAACGATGG
Oligo rev AAACCCATCGTTCCCTTCGTAAAC
Chromosome start end strand MM target_seq PAM alignment distance position gene name gene id
GU305914.1 85533 85555 + 0 GTTTACGAAGGGAACGATGG TGG ||||||||[||||||||||||]PAM NA Intergenic NA NA

T6113 GGTGGTCCCCGTTATCTTCGAGG 1000
Oligo fwd CACCGGTGGTCCCCGTTATCTTCG
Oligo rev AAACCGAAGATAACGGGGACCACC
Chromosome start end strand MM target_seq PAM alignment distance position gene name gene id
GU305914.1 85551 85573 + 0 GGTGGTCCCCGTTATCTTCG AGG ||||||||[||||||||||||]PAM NA Intergenic NA NA

T6114 GTTATCTTCGAGGCCTGAGTCGG 1000
Oligo fwd CACCGTTATCTTCGAGGCCTGAGT
Oligo rev AAACACTCAGGCCTCGAAGATAAC
Chromosome start end strand MM target_seq PAM alignment distance position gene name gene id
GU305914.1 85561 85583 + 0 GTTATCTTCGAGGCCTGAGT CGG ||||||||[||||||||||||]PAM NA Intergenic NA NA

T6115 CTTCGAGGCCTGAGTCGGCGTGG 1000
Oligo fwd CACCCTTCGAGGCCTGAGTCGGCG
Oligo rev AAACCGCCGACTCAGGCCTCGAAG
Chromosome start end strand MM target_seq PAM alignment distance position gene name gene id
GU305914.1 85566 85588 + 0 CTTCGAGGCCTGAGTCGGCG TGG ||||||||[||||||||||||]PAM NA Intergenic NA NA

T6116 CGGCGTGGCTTCCTTCTTTCCGG 1000
Oligo fwd CACCCGGCGTGGCTTCCTTCTTTC
Oligo rev AAACGAAAGAAGGAAGCCACGCCG
Chromosome start end strand MM target_seq PAM alignment distance position gene name gene id
GU305914.1 85581 85603 + 0 CGGCGTGGCTTCCTTCTTTC CGG ||||||||[||||||||||||]PAM NA Intergenic NA NA

T6117 AGTCGCCGTCTCTGTTTCTGCGG 1000
Oligo fwd CACCAGTCGCCGTCTCTGTTTCTG
Oligo rev AAACCAGAAACAGAGACGGCGACT
Chromosome start end strand MM target_seq PAM alignment distance position gene name gene id
GU305914.1 85617 85639 + 0 AGTCGCCGTCTCTGTTTCTG CGG ||||||||[||||||||||||]PAM NA Intergenic NA NA

T6118 CGTCTCTGTTTCTGCGGACGCGG 1000
Oligo fwd CACCCGTCTCTGTTTCTGCGGACG
Oligo rev AAACCGTCCGCAGAAACAGAGACG
Chromosome start end strand MM target_seq PAM alignment distance position gene name gene id

<table>
<tr><td>GU305914.1</td><td>85623</td><td>85645</td><td>+</td><td>0</td><td>CGTCTCTGTTTCTGCGGACG</td><td>CGG</td><td>||||||||[||||||||||||]PAM</td><td>NA</td><td>Intergenic</td><td>NA</td><td>NA</td></tr>
<tr><td></td><td></td><td></td><td></td><td></td><td></td><td></td><td></td><td></td><td></td><td></td><td></td></tr>
<tr><td>T6119</td><td>GAAGTGTCAGTTTTTGCTTTCGG</td><td>1000</td><td></td><td></td><td></td><td></td><td></td><td></td><td></td><td></td><td></td></tr>
<tr><td>Oligo fwd</td><td>CACCGAAGTGTCAGTTTTTGCTTT</td><td></td><td></td><td></td><td></td><td></td><td></td><td></td><td></td><td></td><td></td></tr>
<tr><td>Oligo rev</td><td>AAACAAAGCAAAAACTGACACTTC</td><td></td><td></td><td></td><td></td><td></td><td></td><td></td><td></td><td></td><td></td></tr>
<tr><td>Chromosome</td><td>start</td><td>end</td><td>strand</td><td>MM</td><td>target_seq</td><td>PAM</td><td>alignment</td><td>distance</td><td>position</td><td>gene name</td><td>gene id</td></tr>
<tr><td>GU305914.1</td><td>85654</td><td>85676</td><td>+</td><td>0</td><td>GAAGTGTCAGTTTTTGCTTT</td><td>CGG</td><td>||||||||[||||||||||||]PAM</td><td>NA</td><td>Intergenic</td><td>NA</td><td>NA</td></tr>
<tr><td></td><td></td><td></td><td></td><td></td><td></td><td></td><td></td><td></td><td></td><td></td><td></td></tr>
<tr><td>T6120</td><td>TTTGCTTTCGGAGTAGTACCTGG</td><td>1000</td><td></td><td></td><td></td><td></td><td></td><td></td><td></td><td></td><td></td></tr>
<tr><td>Oligo fwd</td><td>CACCTTTGCTTTCGGAGTAGTACC</td><td></td><td></td><td></td><td></td><td></td><td></td><td></td><td></td><td></td><td></td></tr>
<tr><td>Oligo rev</td><td>AAACGGTACTACTCCGAAAGCAAA</td><td></td><td></td><td></td><td></td><td></td><td></td><td></td><td></td><td></td><td></td></tr>
<tr><td>Chromosome</td><td>start</td><td>end</td><td>strand</td><td>MM</td><td>target_seq</td><td>PAM</td><td>alignment</td><td>distance</td><td>position</td><td>gene name</td><td>gene id</td></tr>
<tr><td>GU305914.1</td><td>85666</td><td>85688</td><td>+</td><td>0</td><td>TTTGCTTTCGGAGTAGTACC</td><td>TGG</td><td>||||||||[||||||||||||]PAM</td><td>NA</td><td>Intergenic</td><td>NA</td><td>NA</td></tr>
<tr><td></td><td></td><td></td><td></td><td></td><td></td><td></td><td></td><td></td><td></td><td></td><td></td></tr>
<tr><td>T6121</td><td>CGCGATTACCTCACTTGTACAGG</td><td>1000</td><td></td><td></td><td></td><td></td><td></td><td></td><td></td><td></td><td></td></tr>
<tr><td>Oligo fwd</td><td>CACCCGCGATTACCTCACTTGTAC</td><td></td><td></td><td></td><td></td><td></td><td></td><td></td><td></td><td></td><td></td></tr>
<tr><td>Oligo rev</td><td>AAACGTACAAGTGAGGTAATCGCG</td><td></td><td></td><td></td><td></td><td></td><td></td><td></td><td></td><td></td><td></td></tr>
<tr><td>Chromosome</td><td>start</td><td>end</td><td>strand</td><td>MM</td><td>target_seq</td><td>PAM</td><td>alignment</td><td>distance</td><td>position</td><td>gene name</td><td>gene id</td></tr>
<tr><td>GU305914.1</td><td>85701</td><td>85723</td><td>+</td><td>0</td><td>CGCGATTACCTCACTTGTAC</td><td>AGG</td><td>||||||||[||||||||||||]PAM</td><td>NA</td><td>Intergenic</td><td>NA</td><td>NA</td></tr>
<tr><td></td><td></td><td></td><td></td><td></td><td></td><td></td><td></td><td></td><td></td><td></td><td></td></tr>
<tr><td>T6122</td><td>TTACCTCACTTGTACAGGCAAGG</td><td>1000</td><td></td><td></td><td></td><td></td><td></td><td></td><td></td><td></td><td></td></tr>
<tr><td>Oligo fwd</td><td>CACCTTACCTCACTTGTACAGGCA</td><td></td><td></td><td></td><td></td><td></td><td></td><td></td><td></td><td></td><td></td></tr>
<tr><td>Oligo rev</td><td>AAACTGCCTGTACAAGTGAGGTAA</td><td></td><td></td><td></td><td></td><td></td><td></td><td></td><td></td><td></td><td></td></tr>
<tr><td>Chromosome</td><td>start</td><td>end</td><td>strand</td><td>MM</td><td>target_seq</td><td>PAM</td><td>alignment</td><td>distance</td><td>position</td><td>gene name</td><td>gene id</td></tr>
<tr><td>GU305914.1</td><td>85706</td><td>85728</td><td>+</td><td>0</td><td>TTACCTCACTTGTACAGGCA</td><td>AGG</td><td>||||||||[||||||||||||]PAM</td><td>NA</td><td>Intergenic</td><td>NA</td><td>NA</td></tr>
<tr><td></td><td></td><td></td><td></td><td></td><td></td><td></td><td></td><td></td><td></td><td></td><td></td></tr>
<tr><td>T6123</td><td>CCTCACTTGTACAGGCAAGGAGG</td><td>1000</td><td></td><td></td><td></td><td></td><td></td><td></td><td></td><td></td><td></td></tr>
<tr><td>Oligo fwd</td><td>CACCCCTCACTTGTACAGGCAAGG</td><td></td><td></td><td></td><td></td><td></td><td></td><td></td><td></td><td></td><td></td></tr>
<tr><td>Oligo rev</td><td>AAACCCTTGCCTGTACAAGTGAGG</td><td></td><td></td><td></td><td></td><td></td><td></td><td></td><td></td><td></td><td></td></tr>
<tr><td>Chromosome</td><td>start</td><td>end</td><td>strand</td><td>MM</td><td>target_seq</td><td>PAM</td><td>alignment</td><td>distance</td><td>position</td><td>gene name</td><td>gene id</td></tr>
<tr><td>GU305914.1</td><td>85709</td><td>85731</td><td>+</td><td>0</td><td>CCTCACTTGTACAGGCAAGG</td><td>AGG</td><td>||||||||[||||||||||||]PAM</td><td>NA</td><td>Intergenic</td><td>NA</td><td>NA</td></tr>
<tr><td></td><td></td><td></td><td></td><td></td><td></td><td></td><td></td><td></td><td></td><td></td><td></td></tr>
<tr><td>T6124</td><td>CGACATCGCGTACCAGCTCGAGG</td><td>1000</td><td></td><td></td><td></td><td></td><td></td><td></td><td></td><td></td><td></td></tr>
<tr><td>Oligo fwd</td><td>CACCCGACATCGCGTACCAGCTCG</td><td></td><td></td><td></td><td></td><td></td><td></td><td></td><td></td><td></td><td></td></tr>
<tr><td>Oligo rev</td><td>AAACCGAGCTGGTACGCGATGTCG</td><td></td><td></td><td></td><td></td><td></td><td></td><td></td><td></td><td></td><td></td></tr>
<tr><td>Chromosome</td><td>start</td><td>end</td><td>strand</td><td>MM</td><td>target_seq</td><td>PAM</td><td>alignment</td><td>distance</td><td>position</td><td>gene name</td><td>gene id</td></tr>
<tr><td>GU305914.1</td><td>85758</td><td>85780</td><td>+</td><td>0</td><td>CGACATCGCGTACCAGCTCG</td><td>AGG</td><td>||||||||[||||||||||||]PAM</td><td>NA</td><td>Intergenic</td><td>NA</td><td>NA</td></tr>
<tr><td></td><td></td><td></td><td></td><td></td><td></td><td></td><td></td><td></td><td></td><td></td><td></td></tr>
<tr><td>T6125</td><td>GACGAGCGTTGTTCAGCGATCGG</td><td>1000</td><td></td><td></td><td></td><td></td><td></td><td></td><td></td><td></td><td></td></tr>
<tr><td>Oligo fwd</td><td>CACCGACGAGCGTTGTTCAGCGAT</td><td></td><td></td><td></td><td></td><td></td><td></td><td></td><td></td><td></td><td></td></tr>
<tr><td>Oligo rev</td><td>AAACATCGCTGAACAACGCTCGTC</td><td></td><td></td><td></td><td></td><td></td><td></td><td></td><td></td><td></td><td></td></tr>
<tr><td>Chromosome</td><td>start</td><td>end</td><td>strand</td><td>MM</td><td>target_seq</td><td>PAM</td><td>alignment</td><td>distance</td><td>position</td><td>gene name</td><td>gene id</td></tr>
<tr><td>GU305914.1</td><td>85820</td><td>85842</td><td>+</td><td>0</td><td>GACGAGCGTTGTTCAGCGAT</td><td>CGG</td><td>||||||||[||||||||||||]PAM</td><td>NA</td><td>Intergenic</td><td>NA</td><td>NA</td></tr>
<tr><td></td><td></td><td></td><td></td><td></td><td></td><td></td><td></td><td></td><td></td><td></td><td></td></tr>
<tr><td>T6126</td><td>GCGTTGTTCAGCGATCGGCCCGG</td><td>1000</td><td></td><td></td><td></td><td></td><td></td><td></td><td></td><td></td><td></td></tr>
<tr><td>Oligo fwd</td><td>CACCGCGTTGTTCAGCGATCGGCC</td><td></td><td></td><td></td><td></td><td></td><td></td><td></td><td></td><td></td><td></td></tr>
<tr><td>Oligo rev</td><td>AAACGGCCGATCGCTGAACAACGC</td><td></td><td></td><td></td><td></td><td></td><td></td><td></td><td></td><td></td><td></td></tr>
<tr><td>Chromosome</td><td>start</td><td>end</td><td>strand</td><td>MM</td><td>target_seq</td><td>PAM</td><td>alignment</td><td>distance</td><td>position</td><td>gene name</td><td>gene id</td></tr>
<tr><td>GU305914.1</td><td>85825</td><td>85847</td><td>+</td><td>0</td><td>GCGTTGTTCAGCGATCGGCC</td><td>CGG</td><td>||||||||[||||||||||||]PAM</td><td>NA</td><td>Intergenic</td><td>NA</td><td>NA</td></tr>
<tr><td></td><td></td><td></td><td></td><td></td><td></td><td></td><td></td><td></td><td></td><td></td><td></td></tr>
<tr><td>T6127</td><td>CGTTGTTCAGCGATCGGCCCGGG</td><td>1000</td><td></td><td></td><td></td><td></td><td></td><td></td><td></td><td></td><td></td></tr>
<tr><td>Oligo fwd</td><td>CACCCGTTGTTCAGCGATCGGCCC</td><td></td><td></td><td></td><td></td><td></td><td></td><td></td><td></td><td></td><td></td></tr>
<tr><td>Oligo rev</td><td>AAACGGGCCGATCGCTGAACAACG</td><td></td><td></td><td></td><td></td><td></td><td></td><td></td><td></td><td></td><td></td></tr>
<tr><td>Chromosome</td><td>start</td><td>end</td><td>strand</td><td>MM</td><td>target_seq</td><td>PAM</td><td>alignment</td><td>distance</td><td>position</td><td>gene name</td><td>gene id</td></tr>
<tr><td>GU305914.1</td><td>85826</td><td>85848</td><td>+</td><td>0</td><td>CGTTGTTCAGCGATCGGCCC</td><td>GGG</td><td>||||||||[||||||||||||]PAM</td><td>NA</td><td>Intergenic</td><td>NA</td><td>NA</td></tr>
<tr><td></td><td></td><td></td><td></td><td></td><td></td><td></td><td></td><td></td><td></td><td></td><td></td></tr>
<tr><td>T6128</td><td>TCTCTCGACCTGACATAAGTCGG</td><td>1000</td><td></td><td></td><td></td><td></td><td></td><td></td><td></td><td></td><td></td></tr>
<tr><td>Oligo fwd</td><td>CACCTCTCTCGACCTGACATAAGT</td><td></td><td></td><td></td><td></td><td></td><td></td><td></td><td></td><td></td><td></td></tr>
<tr><td>Oligo rev</td><td>AAACACTTATGTCAGGTCGAGAGA</td><td></td><td></td><td></td><td></td><td></td><td></td><td></td><td></td><td></td><td></td></tr>
<tr><td>Chromosome</td><td>start</td><td>end</td><td>strand</td><td>MM</td><td>target_seq</td><td>PAM</td><td>alignment</td><td>distance</td><td>position</td><td>gene name</td><td>gene id</td></tr>
<tr><td>GU305914.1</td><td>85849</td><td>85871</td><td>+</td><td>0</td><td>TCTCTCGACCTGACATAAGT</td><td>CGG</td><td>||||||||[||||||||||||]PAM</td><td>NA</td><td>Intergenic</td><td>NA</td><td>NA</td></tr>
<tr><td></td><td></td><td></td><td></td><td></td><td></td><td></td><td></td><td></td><td></td><td></td><td></td></tr>
<tr><td>T6129</td><td>CTGACATAAGTCGGCGACCGAGG</td><td>1000</td><td></td><td></td><td></td><td></td><td></td><td></td><td></td><td></td><td></td></tr>
<tr><td>Oligo fwd</td><td>CACCCTGACATAAGTCGGCGACCG</td><td></td><td></td><td></td><td></td><td></td><td></td><td></td><td></td><td></td><td></td></tr>
<tr><td>Oligo rev</td><td>AAACCGGTCGCCGACTTATGTCAG</td><td></td><td></td><td></td><td></td><td></td><td></td><td></td><td></td><td></td><td></td></tr>
<tr><td>Chromosome</td><td>start</td><td>end</td><td>strand</td><td>MM</td><td>target_seq</td><td>PAM</td><td>alignment</td><td>distance</td><td>position</td><td>gene name</td><td>gene id</td></tr>
<tr><td>GU305914.1</td><td>85858</td><td>85880</td><td>+</td><td>0</td><td>CTGACATAAGTCGGCGACCG</td><td>AGG</td><td>||||||||[||||||||||||]PAM</td><td>NA</td><td>Intergenic</td><td>NA</td><td>NA</td></tr>
<tr><td></td><td></td><td></td><td></td><td></td><td></td><td></td><td></td><td></td><td></td><td></td><td></td></tr>
<tr><td>T6130</td><td>GGCGACCGAGGCGCGAGAACCGG</td><td>1000</td><td></td><td></td><td></td><td></td><td></td><td></td><td></td><td></td><td></td></tr>
<tr><td>Oligo fwd</td><td>CACCGGCGACCGAGGCGCGAGAAC</td><td></td><td></td><td></td><td></td><td></td><td></td><td></td><td></td><td></td><td></td></tr>
<tr><td>Oligo rev</td><td>AAACGTTCTCGCGCCTCGGTCGCC</td><td></td><td></td><td></td><td></td><td></td><td></td><td></td><td></td><td></td><td></td></tr>
<tr><td>Chromosome</td><td>start</td><td>end</td><td>strand</td><td>MM</td><td>target_seq</td><td>PAM</td><td>alignment</td><td>distance</td><td>position</td><td>gene name</td><td>gene id</td></tr>
<tr><td>GU305914.1</td><td>85870</td><td>85892</td><td>+</td><td>0</td><td>GGCGACCGAGGCGCGAGAAC</td><td>CGG</td><td>||||||||[||||||||||||]PAM</td><td>NA</td><td>Intergenic</td><td>NA</td><td>NA</td></tr>
<tr><td></td><td></td><td></td><td></td><td></td><td></td><td></td><td></td><td></td><td></td><td></td><td></td></tr>
<tr><td>T6131</td><td>CGAGGCGCGAGAACCGGCGTCGG</td><td>1000</td><td></td><td></td><td></td><td></td><td></td><td></td><td></td><td></td><td></td></tr>
<tr><td>Oligo fwd</td><td>CACCCGAGGCGCGAGAACCGGCGT</td><td></td><td></td><td></td><td></td><td></td><td></td><td></td><td></td><td></td><td></td></tr>
<tr><td>Oligo rev</td><td>AAACACGCCGGTTCTCGCGCCTCG</td><td></td><td></td><td></td><td></td><td></td><td></td><td></td><td></td><td></td><td></td></tr>
<tr><td>Chromosome</td><td>start</td><td>end</td><td>strand</td><td>MM</td><td>target_seq</td><td>PAM</td><td>alignment</td><td>distance</td><td>position</td><td>gene name</td><td>gene id</td></tr>
<tr><td>GU305914.1</td><td>85876</td><td>85898</td><td>+</td><td>0</td><td>CGAGGCGCGAGAACCGGCGT</td><td>CGG</td><td>||||||||[||||||||||||]PAM</td><td>NA</td><td>Intergenic</td><td>NA</td><td>NA</td></tr>
<tr><td></td><td></td><td></td><td></td><td></td><td></td><td></td><td></td><td></td><td></td><td></td><td></td></tr>
<tr><td>T6132</td><td>ACCGGCGTCGGCGAGAGTCTCGG</td><td>1000</td><td></td><td></td><td></td><td></td><td></td><td></td><td></td><td></td><td></td></tr>
<tr><td>Oligo fwd</td><td>CACCACCGGCGTCGGCGAGAGTCT</td><td></td><td></td><td></td><td></td><td></td><td></td><td></td><td></td><td></td><td></td></tr>
<tr><td>Oligo rev</td><td>AAACAGACTCTCGCCGACGCCGGT</td><td></td><td></td><td></td><td></td><td></td><td></td><td></td><td></td><td></td><td></td></tr>
<tr><td>Chromosome</td><td>start</td><td>end</td><td>strand</td><td>MM</td><td>target_seq</td><td>PAM</td><td>alignment</td><td>distance</td><td>position</td><td>gene name</td><td>gene id</td></tr>
<tr><td>GU305914.1</td><td>85888</td><td>85910</td><td>+</td><td>0</td><td>ACCGGCGTCGGCGAGAGTCT</td><td>CGG</td><td>||||||||[||||||||||||]PAM</td><td>NA</td><td>Intergenic</td><td>NA</td><td>NA</td></tr>
<tr><td></td><td></td><td></td><td></td><td></td><td></td><td></td><td></td><td></td><td></td><td></td><td></td></tr>
<tr><td>T6133</td><td>GAGCAGCAAGTACAGATCCCTGG</td><td>1000</td><td></td><td></td><td></td><td></td><td></td><td></td><td></td><td></td><td></td></tr>
<tr><td>Oligo fwd</td><td>CACCGAGCAGCAAGTACAGATCCC</td><td></td><td></td><td></td><td></td><td></td><td></td><td></td><td></td><td></td><td></td></tr>
<tr><td>Oligo rev</td><td>AAACGGGATCTGTACTTGCTGCTC</td><td></td><td></td><td></td><td></td><td></td><td></td><td></td><td></td><td></td><td></td></tr>
</table>

<table>
<tr><td>Chromosome</td><td>start</td><td>end</td><td>strand</td><td>MM</td><td>target_seq</td><td>PAM</td><td>alignment</td><td>distance</td><td>position</td><td>gene name</td><td>gene id</td></tr>
<tr><td>GU305914.1</td><td>85933</td><td>85955</td><td>+</td><td>0</td><td>GAGCAGCAAGTACAGATCCC</td><td>TGG</td><td>||||||||[||||||||||||]PAM</td><td>NA</td><td>Intergenic</td><td>NA</td><td>NA</td></tr>
<tr><td></td><td></td><td></td><td></td><td></td><td></td><td></td><td></td><td></td><td></td><td></td><td></td></tr>
<tr><td>T6134</td><td>AACTTCGATCTCCGCGACGCAGG</td><td>1000</td><td></td><td></td><td></td><td></td><td></td><td></td><td></td><td></td><td></td></tr>
<tr><td>Oligo fwd</td><td>CACCAACTTCGATCTCCGCGACGC</td><td></td><td></td><td></td><td></td><td></td><td></td><td></td><td></td><td></td><td></td></tr>
<tr><td>Oligo rev</td><td>AAACGCGTCGCGGAGATCGAAGTT</td><td></td><td></td><td></td><td></td><td></td><td></td><td></td><td></td><td></td><td></td></tr>
<tr><td>Chromosome</td><td>start</td><td>end</td><td>strand</td><td>MM</td><td>target_seq</td><td>PAM</td><td>alignment</td><td>distance</td><td>position</td><td>gene name</td><td>gene id</td></tr>
<tr><td>GU305914.1</td><td>85962</td><td>85984</td><td>+</td><td>0</td><td>AACTTCGATCTCCGCGACGC</td><td>AGG</td><td>||||||||[||||||||||||]PAM</td><td>NA</td><td>Intergenic</td><td>NA</td><td>NA</td></tr>
<tr><td></td><td></td><td></td><td></td><td></td><td></td><td></td><td></td><td></td><td></td><td></td><td></td></tr>
<tr><td>T6135</td><td>TCTCCGCGACGCAGGATCAACGG</td><td>1000</td><td></td><td></td><td></td><td></td><td></td><td></td><td></td><td></td><td></td></tr>
<tr><td>Oligo fwd</td><td>CACCTCTCCGCGACGCAGGATCAA</td><td></td><td></td><td></td><td></td><td></td><td></td><td></td><td></td><td></td><td></td></tr>
<tr><td>Oligo rev</td><td>AAACTTGATCCTGCGTCGCGGAGA</td><td></td><td></td><td></td><td></td><td></td><td></td><td></td><td></td><td></td><td></td></tr>
<tr><td>Chromosome</td><td>start</td><td>end</td><td>strand</td><td>MM</td><td>target_seq</td><td>PAM</td><td>alignment</td><td>distance</td><td>position</td><td>gene name</td><td>gene id</td></tr>
<tr><td>GU305914.1</td><td>85970</td><td>85992</td><td>+</td><td>0</td><td>TCTCCGCGACGCAGGATCAA</td><td>CGG</td><td>||||||||[||||||||||||]PAM</td><td>NA</td><td>Intergenic</td><td>NA</td><td>NA</td></tr>
<tr><td></td><td></td><td></td><td></td><td></td><td></td><td></td><td></td><td></td><td></td><td></td><td></td></tr>
<tr><td>T6136</td><td>CGCAGGATCAACGGAGATTCTGG</td><td>1000</td><td></td><td></td><td></td><td></td><td></td><td></td><td></td><td></td><td></td></tr>
<tr><td>Oligo fwd</td><td>CACCCGCAGGATCAACGGAGATTC</td><td></td><td></td><td></td><td></td><td></td><td></td><td></td><td></td><td></td><td></td></tr>
<tr><td>Oligo rev</td><td>AAACGAATCTCCGTTGATCCTGCG</td><td></td><td></td><td></td><td></td><td></td><td></td><td></td><td></td><td></td><td></td></tr>
<tr><td>Chromosome</td><td>start</td><td>end</td><td>strand</td><td>MM</td><td>target_seq</td><td>PAM</td><td>alignment</td><td>distance</td><td>position</td><td>gene name</td><td>gene id</td></tr>
<tr><td>GU305914.1</td><td>85979</td><td>86001</td><td>+</td><td>0</td><td>CGCAGGATCAACGGAGATTC</td><td>TGG</td><td>||||||||[||||||||||||]PAM</td><td>NA</td><td>Intergenic</td><td>NA</td><td>NA</td></tr>
<tr><td></td><td></td><td></td><td></td><td></td><td></td><td></td><td></td><td></td><td></td><td></td><td></td></tr>
<tr><td>T6137</td><td>GCCGTCCTCGACGCGCCACAAGG</td><td>1000</td><td></td><td></td><td></td><td></td><td></td><td></td><td></td><td></td><td></td></tr>
<tr><td>Oligo fwd</td><td>CACCGCCGTCCTCGACGCGCCACA</td><td></td><td></td><td></td><td></td><td></td><td></td><td></td><td></td><td></td><td></td></tr>
<tr><td>Oligo rev</td><td>AAACTGTGGCGCGTCGAGGACGGC</td><td></td><td></td><td></td><td></td><td></td><td></td><td></td><td></td><td></td><td></td></tr>
<tr><td>Chromosome</td><td>start</td><td>end</td><td>strand</td><td>MM</td><td>target_seq</td><td>PAM</td><td>alignment</td><td>distance</td><td>position</td><td>gene name</td><td>gene id</td></tr>
<tr><td>GU305914.1</td><td>86038</td><td>86060</td><td>+</td><td>0</td><td>GCCGTCCTCGACGCGCCACA</td><td>AGG</td><td>||||||||[||||||||||||]PAM</td><td>NA</td><td>Intergenic</td><td>NA</td><td>NA</td></tr>
<tr><td></td><td></td><td></td><td></td><td></td><td></td><td></td><td></td><td></td><td></td><td></td><td></td></tr>
<tr><td>T6138</td><td>CGTCTCGTTATATAGTGCGACGG</td><td>1000</td><td></td><td></td><td></td><td></td><td></td><td></td><td></td><td></td><td></td></tr>
<tr><td>Oligo fwd</td><td>CACCCGTCTCGTTATATAGTGCGA</td><td></td><td></td><td></td><td></td><td></td><td></td><td></td><td></td><td></td><td></td></tr>
<tr><td>Oligo rev</td><td>AAACTCGCACTATATAACGAGACG</td><td></td><td></td><td></td><td></td><td></td><td></td><td></td><td></td><td></td><td></td></tr>
<tr><td>Chromosome</td><td>start</td><td>end</td><td>strand</td><td>MM</td><td>target_seq</td><td>PAM</td><td>alignment</td><td>distance</td><td>position</td><td>gene name</td><td>gene id</td></tr>
<tr><td>GU305914.1</td><td>86077</td><td>86099</td><td>+</td><td>0</td><td>CGTCTCGTTATATAGTGCGA</td><td>CGG</td><td>||||||||[||||||||||||]PAM</td><td>NA</td><td>Intergenic</td><td>NA</td><td>NA</td></tr>
<tr><td></td><td></td><td></td><td></td><td></td><td></td><td></td><td></td><td></td><td></td><td></td><td></td></tr>
<tr><td>T6139</td><td>CGTTATATAGTGCGACGGACAGG</td><td>1000</td><td></td><td></td><td></td><td></td><td></td><td></td><td></td><td></td><td></td></tr>
<tr><td>Oligo fwd</td><td>CACCCGTTATATAGTGCGACGGAC</td><td></td><td></td><td></td><td></td><td></td><td></td><td></td><td></td><td></td><td></td></tr>
<tr><td>Oligo rev</td><td>AAACGTCCGTCGCACTATATAACG</td><td></td><td></td><td></td><td></td><td></td><td></td><td></td><td></td><td></td><td></td></tr>
<tr><td>Chromosome</td><td>start</td><td>end</td><td>strand</td><td>MM</td><td>target_seq</td><td>PAM</td><td>alignment</td><td>distance</td><td>position</td><td>gene name</td><td>gene id</td></tr>
<tr><td>GU305914.1</td><td>86082</td><td>86104</td><td>+</td><td>0</td><td>CGTTATATAGTGCGACGGAC</td><td>AGG</td><td>||||||||[||||||||||||]PAM</td><td>NA</td><td>Intergenic</td><td>NA</td><td>NA</td></tr>
<tr><td></td><td></td><td></td><td></td><td></td><td></td><td></td><td></td><td></td><td></td><td></td><td></td></tr>
<tr><td>T6140</td><td>GCGCCCAGACCATCTTCTGCAGG</td><td>1000</td><td></td><td></td><td></td><td></td><td></td><td></td><td></td><td></td><td></td></tr>
<tr><td>Oligo fwd</td><td>CACCGCGCCCAGACCATCTTCTGC</td><td></td><td></td><td></td><td></td><td></td><td></td><td></td><td></td><td></td><td></td></tr>
<tr><td>Oligo rev</td><td>AAACGCAGAAGATGGTCTGGGCGC</td><td></td><td></td><td></td><td></td><td></td><td></td><td></td><td></td><td></td><td></td></tr>
<tr><td>Chromosome</td><td>start</td><td>end</td><td>strand</td><td>MM</td><td>target_seq</td><td>PAM</td><td>alignment</td><td>distance</td><td>position</td><td>gene name</td><td>gene id</td></tr>
<tr><td>GU305914.1</td><td>86121</td><td>86143</td><td>+</td><td>0</td><td>GCGCCCAGACCATCTTCTGC</td><td>AGG</td><td>||||||||[||||||||||||]PAM</td><td>NA</td><td>Intergenic</td><td>NA</td><td>NA</td></tr>
<tr><td></td><td></td><td></td><td></td><td></td><td></td><td></td><td></td><td></td><td></td><td></td><td></td></tr>
<tr><td>T6141</td><td>TCTGCAGGTCGTTGAGCTCGCGG</td><td>1000</td><td></td><td></td><td></td><td></td><td></td><td></td><td></td><td></td><td></td></tr>
<tr><td>Oligo fwd</td><td>CACCTCTGCAGGTCGTTGAGCTCG</td><td></td><td></td><td></td><td></td><td></td><td></td><td></td><td></td><td></td><td></td></tr>
<tr><td>Oligo rev</td><td>AAACCGAGCTCAACGACCTGCAGA</td><td></td><td></td><td></td><td></td><td></td><td></td><td></td><td></td><td></td><td></td></tr>
<tr><td>Chromosome</td><td>start</td><td>end</td><td>strand</td><td>MM</td><td>target_seq</td><td>PAM</td><td>alignment</td><td>distance</td><td>position</td><td>gene name</td><td>gene id</td></tr>
<tr><td>GU305914.1</td><td>86136</td><td>86158</td><td>+</td><td>0</td><td>TCTGCAGGTCGTTGAGCTCG</td><td>CGG</td><td>||||||||[||||||||||||]PAM</td><td>NA</td><td>Intergenic</td><td>NA</td><td>NA</td></tr>
<tr><td></td><td></td><td></td><td></td><td></td><td></td><td></td><td></td><td></td><td></td><td></td><td></td></tr>
<tr><td>T6142</td><td>GTCGTTGAGCTCGCGGATGTCGG</td><td>1000</td><td></td><td></td><td></td><td></td><td></td><td></td><td></td><td></td><td></td></tr>
<tr><td>Oligo fwd</td><td>CACCGTCGTTGAGCTCGCGGATGT</td><td></td><td></td><td></td><td></td><td></td><td></td><td></td><td></td><td></td><td></td></tr>
<tr><td>Oligo rev</td><td>AAACACATCCGCGAGCTCAACGAC</td><td></td><td></td><td></td><td></td><td></td><td></td><td></td><td></td><td></td><td></td></tr>
<tr><td>Chromosome</td><td>start</td><td>end</td><td>strand</td><td>MM</td><td>target_seq</td><td>PAM</td><td>alignment</td><td>distance</td><td>position</td><td>gene name</td><td>gene id</td></tr>
<tr><td>GU305914.1</td><td>86143</td><td>86165</td><td>+</td><td>0</td><td>GTCGTTGAGCTCGCGGATGT</td><td>CGG</td><td>||||||||[||||||||||||]PAM</td><td>NA</td><td>Intergenic</td><td>NA</td><td>NA</td></tr>
<tr><td></td><td></td><td></td><td></td><td></td><td></td><td></td><td></td><td></td><td></td><td></td><td></td></tr>
<tr><td>T6143</td><td>TCCGACTGTCATCCAGTCGCTGG</td><td>1000</td><td></td><td></td><td></td><td></td><td></td><td></td><td></td><td></td><td></td></tr>
<tr><td>Oligo fwd</td><td>CACCTCCGACTGTCATCCAGTCGC</td><td></td><td></td><td></td><td></td><td></td><td></td><td></td><td></td><td></td><td></td></tr>
<tr><td>Oligo rev</td><td>AAACGCGACTGGATGACAGTCGGA</td><td></td><td></td><td></td><td></td><td></td><td></td><td></td><td></td><td></td><td></td></tr>
<tr><td>Chromosome</td><td>start</td><td>end</td><td>strand</td><td>MM</td><td>target_seq</td><td>PAM</td><td>alignment</td><td>distance</td><td>position</td><td>gene name</td><td>gene id</td></tr>
<tr><td>GU305914.1</td><td>86188</td><td>86210</td><td>+</td><td>0</td><td>TCCGACTGTCATCCAGTCGC</td><td>TGG</td><td>||||||||[||||||||||||]PAM</td><td>NA</td><td>Intergenic</td><td>NA</td><td>NA</td></tr>
<tr><td></td><td></td><td></td><td></td><td></td><td></td><td></td><td></td><td></td><td></td><td></td><td></td></tr>
<tr><td>T6144</td><td>CCGACTGTCATCCAGTCGCTGGG</td><td>1000</td><td></td><td></td><td></td><td></td><td></td><td></td><td></td><td></td><td></td></tr>
<tr><td>Oligo fwd</td><td>CACCCCGACTGTCATCCAGTCGCT</td><td></td><td></td><td></td><td></td><td></td><td></td><td></td><td></td><td></td><td></td></tr>
<tr><td>Oligo rev</td><td>AAACAGCGACTGGATGACAGTCGG</td><td></td><td></td><td></td><td></td><td></td><td></td><td></td><td></td><td></td><td></td></tr>
<tr><td>Chromosome</td><td>start</td><td>end</td><td>strand</td><td>MM</td><td>target_seq</td><td>PAM</td><td>alignment</td><td>distance</td><td>position</td><td>gene name</td><td>gene id</td></tr>
<tr><td>GU305914.1</td><td>86189</td><td>86211</td><td>+</td><td>0</td><td>CCGACTGTCATCCAGTCGCT</td><td>GGG</td><td>||||||||[||||||||||||]PAM</td><td>NA</td><td>Intergenic</td><td>NA</td><td>NA</td></tr>
<tr><td></td><td></td><td></td><td></td><td></td><td></td><td></td><td></td><td></td><td></td><td></td><td></td></tr>
<tr><td>T6145</td><td>ACGAGATCCTCTTTGACGTGCGG</td><td>1000</td><td></td><td></td><td></td><td></td><td></td><td></td><td></td><td></td><td></td></tr>
<tr><td>Oligo fwd</td><td>CACCACGAGATCCTCTTTGACGTG</td><td></td><td></td><td></td><td></td><td></td><td></td><td></td><td></td><td></td><td></td></tr>
<tr><td>Oligo rev</td><td>AAACCACGTCAAAGAGGATCTCGT</td><td></td><td></td><td></td><td></td><td></td><td></td><td></td><td></td><td></td><td></td></tr>
<tr><td>Chromosome</td><td>start</td><td>end</td><td>strand</td><td>MM</td><td>target_seq</td><td>PAM</td><td>alignment</td><td>distance</td><td>position</td><td>gene name</td><td>gene id</td></tr>
<tr><td>GU305914.1</td><td>86237</td><td>86259</td><td>+</td><td>0</td><td>ACGAGATCCTCTTTGACGTG</td><td>CGG</td><td>||||||||[||||||||||||]PAM</td><td>NA</td><td>Intergenic</td><td>NA</td><td>NA</td></tr>
<tr><td></td><td></td><td></td><td></td><td></td><td></td><td></td><td></td><td></td><td></td><td></td><td></td></tr>
<tr><td>T6146</td><td>GATCCTCTTTGACGTGCGGCAGG</td><td>1000</td><td></td><td></td><td></td><td></td><td></td><td></td><td></td><td></td><td></td></tr>
<tr><td>Oligo fwd</td><td>CACCGATCCTCTTTGACGTGCGGC</td><td></td><td></td><td></td><td></td><td></td><td></td><td></td><td></td><td></td><td></td></tr>
<tr><td>Oligo rev</td><td>AAACGCCGCACGTCAAAGAGGATC</td><td></td><td></td><td></td><td></td><td></td><td></td><td></td><td></td><td></td><td></td></tr>
<tr><td>Chromosome</td><td>start</td><td>end</td><td>strand</td><td>MM</td><td>target_seq</td><td>PAM</td><td>alignment</td><td>distance</td><td>position</td><td>gene name</td><td>gene id</td></tr>
<tr><td>GU305914.1</td><td>86241</td><td>86263</td><td>+</td><td>0</td><td>GATCCTCTTTGACGTGCGGC</td><td>AGG</td><td>||||||||[||||||||||||]PAM</td><td>NA</td><td>Intergenic</td><td>NA</td><td>NA</td></tr>
<tr><td></td><td></td><td></td><td></td><td></td><td></td><td></td><td></td><td></td><td></td><td></td><td></td></tr>
<tr><td>T6147</td><td>TTTGACGTGCGGCAGGATGCCGG</td><td>1000</td><td></td><td></td><td></td><td></td><td></td><td></td><td></td><td></td><td></td></tr>
<tr><td>Oligo fwd</td><td>CACCTTTGACGTGCGGCAGGATGC</td><td></td><td></td><td></td><td></td><td></td><td></td><td></td><td></td><td></td><td></td></tr>
<tr><td>Oligo rev</td><td>AAACGCATCCTGCCGCACGTCAAA</td><td></td><td></td><td></td><td></td><td></td><td></td><td></td><td></td><td></td><td></td></tr>
<tr><td>Chromosome</td><td>start</td><td>end</td><td>strand</td><td>MM</td><td>target_seq</td><td>PAM</td><td>alignment</td><td>distance</td><td>position</td><td>gene name</td><td>gene id</td></tr>
<tr><td>GU305914.1</td><td>86248</td><td>86270</td><td>+</td><td>0</td><td>TTTGACGTGCGGCAGGATGC</td><td>CGG</td><td>||||||||[||||||||||||]PAM</td><td>NA</td><td>Intergenic</td><td>NA</td><td>NA</td></tr>
<tr><td></td><td></td><td></td><td></td><td></td><td></td><td></td><td></td><td></td><td></td><td></td><td></td></tr>
<tr><td>T6148</td><td>CAGGATGCCGGCGTTATCGCAGG</td><td>1000</td><td></td><td></td><td></td><td></td><td></td><td></td><td></td><td></td><td></td></tr>
<tr><td>Oligo fwd</td><td>CACCCAGGATGCCGGCGTTATCGC</td><td></td><td></td><td></td><td></td><td></td><td></td><td></td><td></td><td></td><td></td></tr>
</table>

Oligo rev AAACGCGATAACGCCGGCATCCTG
Chromosome start end strand MM target_seq PAM alignment distance position gene name gene id
GU305914.1 86260 86282 + 0 CAGGATGCCGGCGTTATCGC AGG ||||||||[||||||||||||]PAM NA Intergenic NA NA

T6149 GCCGGCGTTATCGCAGGCGTAGG 1000
Oligo fwd CACCGCCGGCGTTATCGCAGGCGT
Oligo rev AAACACGCCTGCGATAACGCCGGC
Chromosome start end strand MM target_seq PAM alignment distance position gene name gene id
GU305914.1 86266 86288 + 0 GCCGGCGTTATCGCAGGCGT AGG ||||||||[||||||||||||]PAM NA Intergenic NA NA

T6150 GTAGGCCATGTTAACGTTATAGG 1000
Oligo fwd CACCGTAGGCCATGTTAACGTTAT
Oligo rev AAACATAACGTTAACATGGCCTAC
Chromosome start end strand MM target_seq PAM alignment distance position gene name gene id
GU305914.1 86284 86306 + 0 GTAGGCCATGTTAACGTTAT AGG ||||||||[||||||||||||]PAM NA Intergenic NA NA

T6151 TAGGCCATGTTAACGTTATAGGG 1000
Oligo fwd CACCTAGGCCATGTTAACGTTATA
Oligo rev AAACTATAACGTTAACATGGCCTA
Chromosome start end strand MM target_seq PAM alignment distance position gene name gene id
GU305914.1 86285 86307 + 0 TAGGCCATGTTAACGTTATA GGG ||||||||[||||||||||||]PAM NA Intergenic NA NA

T6152 CCATGTTAACGTTATAGGGCAGG 1000
Oligo fwd CACCCCATGTTAACGTTATAGGGC
Oligo rev AAACGCCCTATAACGTTAACATGG
Chromosome start end strand MM target_seq PAM alignment distance position gene name gene id
GU305914.1 86289 86311 + 0 CCATGTTAACGTTATAGGGC AGG ||||||||[||||||||||||]PAM NA Intergenic NA NA

T6153 CATGTTAACGTTATAGGGCAGGG 1000
Oligo fwd CACCCATGTTAACGTTATAGGGCA
Oligo rev AAACTGCCCTATAACGTTAACATG
Chromosome start end strand MM target_seq PAM alignment distance position gene name gene id
GU305914.1 86290 86312 + 0 CATGTTAACGTTATAGGGCA GGG ||||||||[||||||||||||]PAM NA Intergenic NA NA

T6154 ATGTTAACGTTATAGGGCAGGGG 1000
Oligo fwd CACCATGTTAACGTTATAGGGCAG
Oligo rev AAACCTGCCCTATAACGTTAACAT
Chromosome start end strand MM target_seq PAM alignment distance position gene name gene id
GU305914.1 86291 86313 + 0 ATGTTAACGTTATAGGGCAG GGG ||||||||[||||||||||||]PAM NA Intergenic NA NA

T6155 TGTTAACGTTATAGGGCAGGGGG 1000
Oligo fwd CACCTGTTAACGTTATAGGGCAGG
Oligo rev AAACCCTGCCCTATAACGTTAACA
Chromosome start end strand MM target_seq PAM alignment distance position gene name gene id
GU305914.1 86292 86314 + 0 TGTTAACGTTATAGGGCAGG GGG ||||||||[||||||||||||]PAM NA Intergenic NA NA

T6156 GGGGGTGTCTGTCGAGATAGTGG 1000
Oligo fwd CACCGGGGGTGTCTGTCGAGATAG
Oligo rev AAACCTATCTCGACAGACACCCCC
Chromosome start end strand MM target_seq PAM alignment distance position gene name gene id
GU305914.1 86310 86332 + 0 GGGGGTGTCTGTCGAGATAG TGG ||||||||[||||||||||||]PAM NA Intergenic NA NA

T6157 GGGGTGTCTGTCGAGATAGTGGG 1000
Oligo fwd CACCGGGGTGTCTGTCGAGATAGT
Oligo rev AAACACTATCTCGACAGACACCCC
Chromosome start end strand MM target_seq PAM alignment distance position gene name gene id
GU305914.1 86311 86333 + 0 GGGGTGTCTGTCGAGATAGT GGG ||||||||[||||||||||||]PAM NA Intergenic NA NA

T6158 GTCGAGATAGTGGGTGAACATGG 1000
Oligo fwd CACCGTCGAGATAGTGGGTGAACA
Oligo rev AAACTGTTCACCCACTATCTCGAC
Chromosome start end strand MM target_seq PAM alignment distance position gene name gene id
GU305914.1 86320 86342 + 0 GTCGAGATAGTGGGTGAACA TGG ||||||||[||||||||||||]PAM NA Intergenic NA NA

T6159 TCGAGATAGTGGGTGAACATGGG 1000
Oligo fwd CACCTCGAGATAGTGGGTGAACAT
Oligo rev AAACATGTTCACCCACTATCTCGA
Chromosome start end strand MM target_seq PAM alignment distance position gene name gene id
GU305914.1 86321 86343 + 0 TCGAGATAGTGGGTGAACAT GGG ||||||||[||||||||||||]PAM NA Intergenic NA NA

T6160 ACATGGGACCGTTGATGAAACGG 1000
Oligo fwd CACCACATGGGACCGTTGATGAAA
Oligo rev AAACTTTCATCAACGGTCCCATGT
Chromosome start end strand MM target_seq PAM alignment distance position gene name gene id
GU305914.1 86337 86359 + 0 ACATGGGACCGTTGATGAAA CGG ||||||||[||||||||||||]PAM NA Intergenic NA NA

T6161 GGTAGATCTGCTGCCCGAGCTGG 1000
Oligo fwd CACCGGTAGATCTGCTGCCCGAGC
Oligo rev AAACGCTCGGGCAGCAGATCTACC
Chromosome start end strand MM target_seq PAM alignment distance position gene name gene id
GU305914.1 86358 86380 + 0 GGTAGATCTGCTGCCCGAGC TGG ||||||||[||||||||||||]PAM NA Intergenic NA NA

T6162 GTAGATCTGCTGCCCGAGCTGGG 1000
Oligo fwd CACCGTAGATCTGCTGCCCGAGCT
Oligo rev AAACAGCTCGGGCAGCAGATCTAC
Chromosome start end strand MM target_seq PAM alignment distance position gene name gene id
GU305914.1 86359 86381 + 0 GTAGATCTGCTGCCCGAGCT GGG ||||||||[||||||||||||]PAM NA Intergenic NA NA

T6163 TAGATCTGCTGCCCGAGCTGGGG 1000

Oligo fwd CACCTAGATCTGCTGCCCGAGCTG
Oligo rev AAACCAGCTCGGGCAGCAGATCTA
Chromosome start end strand MM target_seq PAM alignment distance position gene name gene id
GU305914.1 86360 86382 + 0 TAGATCTGCTGCCCGAGCTG GGG ||||||||[||||||||||||]PAM NA Intergenic NA NA

T6164 CCCGAGCTGGGGCAGCAGCTCGG 1000
Oligo fwd CACCCCCGAGCTGGGGCAGCAGCT
Oligo rev AAACAGCTGCTGCCCCAGCTCGGG
Chromosome start end strand MM target_seq PAM alignment distance position gene name gene id
GU305914.1 86371 86393 + 0 CCCGAGCTGGGGCAGCAGCT CGG ||||||||[||||||||||||]PAM NA Intergenic NA NA

T6165 GAGCTGGGGCAGCAGCTCGGAGG 1000
Oligo fwd CACCGAGCTGGGGCAGCAGCTCGG
Oligo rev AAACCCGAGCTGCTGCCCCAGCTC
Chromosome start end strand MM target_seq PAM alignment distance position gene name gene id
GU305914.1 86374 86396 + 0 GAGCTGGGGCAGCAGCTCGG AGG ||||||||[||||||||||||]PAM NA Intergenic NA NA

T6166 AGCTGGGGCAGCAGCTCGGAGGG 1000
Oligo fwd CACCAGCTGGGGCAGCAGCTCGGA
Oligo rev AAACTCCGAGCTGCTGCCCCAGCT
Chromosome start end strand MM target_seq PAM alignment distance position gene name gene id
GU305914.1 86375 86397 + 0 AGCTGGGGCAGCAGCTCGGA GGG ||||||||[||||||||||||]PAM NA Intergenic NA NA

T6167 GCTGGGGCAGCAGCTCGGAGGGG 1000
Oligo fwd CACCGCTGGGGCAGCAGCTCGGAG
Oligo rev AAACCTCCGAGCTGCTGCCCCAGC
Chromosome start end strand MM target_seq PAM alignment distance position gene name gene id
GU305914.1 86376 86398 + 0 GCTGGGGCAGCAGCTCGGAG GGG ||||||||[||||||||||||]PAM NA Intergenic NA NA

T6168 CGAGATTATTGATGACATAGAGG 1000
Oligo fwd CACCCGAGATTATTGATGACATAG
Oligo rev AAACCTATGTCATCAATAATCTCG
Chromosome start end strand MM target_seq PAM alignment distance position gene name gene id
GU305914.1 86412 86434 + 0 CGAGATTATTGATGACATAG AGG ||||||||[||||||||||||]PAM NA Intergenic NA NA

T6169 AGAGGTGTTCGTCGTAGCCGAGG 1000
Oligo fwd CACCAGAGGTGTTCGTCGTAGCCG
Oligo rev AAACCGGCTACGACGAACACCTCT
Chromosome start end strand MM target_seq PAM alignment distance position gene name gene id
GU305914.1 86430 86452 + 0 AGAGGTGTTCGTCGTAGCCG AGG ||||||||[||||||||||||]PAM NA Intergenic NA NA

T6170 TTCGTCGTAGCCGAGGCTCTCGG 1000
Oligo fwd CACCTTCGTCGTAGCCGAGGCTCT
Oligo rev AAACAGAGCCTCGGCTACGACGAA
Chromosome start end strand MM target_seq PAM alignment distance position gene name gene id
GU305914.1 86437 86459 + 0 TTCGTCGTAGCCGAGGCTCT CGG ||||||||[||||||||||||]PAM NA Intergenic NA NA

T6171 GCCGAGGCTCTCGGCCACGTCGG 1000
Oligo fwd CACCGCCGAGGCTCTCGGCCACGT
Oligo rev AAACACGTGGCCGAGAGCCTCGGC
Chromosome start end strand MM target_seq PAM alignment distance position gene name gene id
GU305914.1 86446 86468 + 0 GCCGAGGCTCTCGGCCACGT CGG ||||||||[||||||||||||]PAM NA Intergenic NA NA

T6172 CCGAGGCTCTCGGCCACGTCGGG 1000
Oligo fwd CACCCCGAGGCTCTCGGCCACGTC
Oligo rev AAACGACGTGGCCGAGAGCCTCGG
Chromosome start end strand MM target_seq PAM alignment distance position gene name gene id
GU305914.1 86447 86469 + 0 CCGAGGCTCTCGGCCACGTC GGG ||||||||[||||||||||||]PAM NA Intergenic NA NA

T6173 CGAGGCTCTCGGCCACGTCGGGG 1000
Oligo fwd CACCCGAGGCTCTCGGCCACGTCG
Oligo rev AAACCGACGTGGCCGAGAGCCTCG
Chromosome start end strand MM target_seq PAM alignment distance position gene name gene id
GU305914.1 86448 86470 + 0 CGAGGCTCTCGGCCACGTCG GGG ||||||||[||||||||||||]PAM NA Intergenic NA NA

T6174 GCTCTCGGCCACGTCGGGGTAGG 1000
Oligo fwd CACCGCTCTCGGCCACGTCGGGGT
Oligo rev AAACACCCCGACGTGGCCGAGAGC
Chromosome start end strand MM target_seq PAM alignment distance position gene name gene id
GU305914.1 86452 86474 + 0 GCTCTCGGCCACGTCGGGGT AGG ||||||||[||||||||||||]PAM NA Intergenic NA NA

T6175 GGCCACGTCGGGGTAGGTCTCGG 1000
Oligo fwd CACCGGCCACGTCGGGGTAGGTCT
Oligo rev AAACAGACCTACCCCGACGTGGCC
Chromosome start end strand MM target_seq PAM alignment distance position gene name gene id
GU305914.1 86458 86480 + 0 GGCCACGTCGGGGTAGGTCT CGG ||||||||[||||||||||||]PAM NA Intergenic NA NA

T6176 CCTCAGTTATAGACCGCTGTCGG 1000
Oligo fwd CACCCCTCAGTTATAGACCGCTGT
Oligo rev AAACACAGCGGTCTATAACTGAGG
Chromosome start end strand MM target_seq PAM alignment distance position gene name gene id
GU305914.1 86481 86503 + 0 CCTCAGTTATAGACCGCTGT CGG ||||||||[||||||||||||]PAM NA Intergenic NA NA

T6177 CTCAGTTATAGACCGCTGTCGGG 1000
Oligo fwd CACCCTCAGTTATAGACCGCTGTC
Oligo rev AAACGACAGCGGTCTATAACTGAG
Chromosome start end strand MM target_seq PAM alignment distance position gene name gene id
GU305914.1 86482 86504 + 0 CTCAGTTATAGACCGCTGTC GGG ||||||||[||||||||||||]PAM NA Intergenic NA NA

<table>
<tr><td>T6178</td><td>GAGAAAACCGTTCATAATTTTGG</td><td>1000</td><td></td><td></td><td></td><td></td><td></td><td></td><td></td><td></td><td></td><td></td></tr>
<tr><td>Oligo fwd</td><td>CACCGAGAAAACCGTTCATAATTT</td><td></td><td></td><td></td><td></td><td></td><td></td><td></td><td></td><td></td><td></td><td></td></tr>
<tr><td>Oligo rev</td><td>AAACAAATTATGAACGGTTTTCTC</td><td></td><td></td><td></td><td></td><td></td><td></td><td></td><td></td><td></td><td></td><td></td></tr>
<tr><td>Chromosome</td><td>start</td><td></td><td>end</td><td>strand</td><td>MM</td><td>target_seq</td><td>PAM</td><td>alignment</td><td>distance</td><td>position</td><td>gene name</td><td>gene id</td></tr>
<tr><td>GU305914.1</td><td></td><td>86506</td><td>86528</td><td>+</td><td>0</td><td>GAGAAAACCGTTCATAATTT</td><td>TGG</td><td>||||||||[||||||||||||]PAM</td><td>NA</td><td>Intergenic</td><td>NA</td><td>NA</td></tr>
<tr><td>T6179</td><td>CCAAACACACCGTTCCCACCAGG</td><td>1000</td><td></td><td></td><td></td><td></td><td></td><td></td><td></td><td></td><td></td><td></td></tr>
<tr><td>Oligo fwd</td><td>CACCCCAAACACACCGTTCCCACC</td><td></td><td></td><td></td><td></td><td></td><td></td><td></td><td></td><td></td><td></td><td></td></tr>
<tr><td>Oligo rev</td><td>AAACGGTGGGAACGGTGTGTTTGG</td><td></td><td></td><td></td><td></td><td></td><td></td><td></td><td></td><td></td><td></td><td></td></tr>
<tr><td>Chromosome</td><td>start</td><td></td><td>end</td><td>strand</td><td>MM</td><td>target_seq</td><td>PAM</td><td>alignment</td><td>distance</td><td>position</td><td>gene name</td><td>gene id</td></tr>
<tr><td>GU305914.1</td><td></td><td>86538</td><td>86560</td><td>+</td><td>0</td><td>CCAAACACACCGTTCCCACC</td><td>AGG</td><td>||||||||[||||||||||||]PAM</td><td>NA</td><td>Intergenic</td><td>NA</td><td>NA</td></tr>
<tr><td>T6180</td><td>GTTTCGTAACCAGATGCTCCTGG</td><td>1000</td><td></td><td></td><td></td><td></td><td></td><td></td><td></td><td></td><td></td><td></td></tr>
<tr><td>Oligo fwd</td><td>CACCGTTTCGTAACCAGATGCTCC</td><td></td><td></td><td></td><td></td><td></td><td></td><td></td><td></td><td></td><td></td><td></td></tr>
<tr><td>Oligo rev</td><td>AAACGGAGCATCTGGTTACGAAAC</td><td></td><td></td><td></td><td></td><td></td><td></td><td></td><td></td><td></td><td></td><td></td></tr>
<tr><td>Chromosome</td><td>start</td><td></td><td>end</td><td>strand</td><td>MM</td><td>target_seq</td><td>PAM</td><td>alignment</td><td>distance</td><td>position</td><td>gene name</td><td>gene id</td></tr>
<tr><td>GU305914.1</td><td></td><td>86574</td><td>86596</td><td>+</td><td>0</td><td>GTTTCGTAACCAGATGCTCC</td><td>TGG</td><td>||||||||[||||||||||||]PAM</td><td>NA</td><td>Intergenic</td><td>NA</td><td>NA</td></tr>
<tr><td>T6181</td><td>ATGCTCCTGGTTTCGAATACAGG</td><td>1000</td><td></td><td></td><td></td><td></td><td></td><td></td><td></td><td></td><td></td><td></td></tr>
<tr><td>Oligo fwd</td><td>CACCATGCTCCTGGTTTCGAATAC</td><td></td><td></td><td></td><td></td><td></td><td></td><td></td><td></td><td></td><td></td><td></td></tr>
<tr><td>Oligo rev</td><td>AAACGTATTCGAAACCAGGAGCAT</td><td></td><td></td><td></td><td></td><td></td><td></td><td></td><td></td><td></td><td></td><td></td></tr>
<tr><td>Chromosome</td><td>start</td><td></td><td>end</td><td>strand</td><td>MM</td><td>target_seq</td><td>PAM</td><td>alignment</td><td>distance</td><td>position</td><td>gene name</td><td>gene id</td></tr>
<tr><td>GU305914.1</td><td></td><td>86587</td><td>86609</td><td>+</td><td>0</td><td>ATGCTCCTGGTTTCGAATAC</td><td>AGG</td><td>||||||||[||||||||||||]PAM</td><td>NA</td><td>Intergenic</td><td>NA</td><td>NA</td></tr>
<tr><td>T6182</td><td>AACCTTGTTGTAGCCCATTTCGG</td><td>1000</td><td></td><td></td><td></td><td></td><td></td><td></td><td></td><td></td><td></td><td></td></tr>
<tr><td>Oligo fwd</td><td>CACCAACCTTGTTGTAGCCCATTT</td><td></td><td></td><td></td><td></td><td></td><td></td><td></td><td></td><td></td><td></td><td></td></tr>
<tr><td>Oligo rev</td><td>AAACAAATGGGCTACAACAAGGTT</td><td></td><td></td><td></td><td></td><td></td><td></td><td></td><td></td><td></td><td></td><td></td></tr>
<tr><td>Chromosome</td><td>start</td><td></td><td>end</td><td>strand</td><td>MM</td><td>target_seq</td><td>PAM</td><td>alignment</td><td>distance</td><td>position</td><td>gene name</td><td>gene id</td></tr>
<tr><td>GU305914.1</td><td></td><td>86614</td><td>86636</td><td>+</td><td>0</td><td>AACCTTGTTGTAGCCCATTT</td><td>CGG</td><td>||||||||[||||||||||||]PAM</td><td>NA</td><td>Intergenic</td><td>NA</td><td>NA</td></tr>
<tr><td>T6183</td><td>TTTTTAAGTATGCCCTTTTTCGG</td><td>1000</td><td></td><td></td><td></td><td></td><td></td><td></td><td></td><td></td><td></td><td></td></tr>
<tr><td>Oligo fwd</td><td>CACCTTTTTAAGTATGCCCTTTTT</td><td></td><td></td><td></td><td></td><td></td><td></td><td></td><td></td><td></td><td></td><td></td></tr>
<tr><td>Oligo rev</td><td>AAACAAAAAGGGCATACTTAAAAA</td><td></td><td></td><td></td><td></td><td></td><td></td><td></td><td></td><td></td><td></td><td></td></tr>
<tr><td>Chromosome</td><td>start</td><td></td><td>end</td><td>strand</td><td>MM</td><td>target_seq</td><td>PAM</td><td>alignment</td><td>distance</td><td>position</td><td>gene name</td><td>gene id</td></tr>
<tr><td>GU305914.1</td><td></td><td>86643</td><td>86665</td><td>+</td><td>0</td><td>TTTTTAAGTATGCCCTTTTT</td><td>CGG</td><td>||||||||[||||||||||||]PAM</td><td>NA</td><td>Intergenic</td><td>NA</td><td>NA</td></tr>
<tr><td>T6184</td><td>ACATCGTGCACCCTCAGCAGCGG</td><td>1000</td><td></td><td></td><td></td><td></td><td></td><td></td><td></td><td></td><td></td><td></td></tr>
<tr><td>Oligo fwd</td><td>CACCACATCGTGCACCCTCAGCAG</td><td></td><td></td><td></td><td></td><td></td><td></td><td></td><td></td><td></td><td></td><td></td></tr>
<tr><td>Oligo rev</td><td>AAACCTGCTGAGGGTGCACGATGT</td><td></td><td></td><td></td><td></td><td></td><td></td><td></td><td></td><td></td><td></td><td></td></tr>
<tr><td>Chromosome</td><td>start</td><td></td><td>end</td><td>strand</td><td>MM</td><td>target_seq</td><td>PAM</td><td>alignment</td><td>distance</td><td>position</td><td>gene name</td><td>gene id</td></tr>
<tr><td>GU305914.1</td><td></td><td>86684</td><td>86706</td><td>+</td><td>0</td><td>ACATCGTGCACCCTCAGCAG</td><td>CGG</td><td>||||||||[||||||||||||]PAM</td><td>NA</td><td>Intergenic</td><td>NA</td><td>NA</td></tr>
<tr><td>T6185</td><td>CGGATCTCCACACTCCAACGCGG</td><td>1000</td><td></td><td></td><td></td><td></td><td></td><td></td><td></td><td></td><td></td><td></td></tr>
<tr><td>Oligo fwd</td><td>CACCCGGATCTCCACACTCCAACG</td><td></td><td></td><td></td><td></td><td></td><td></td><td></td><td></td><td></td><td></td><td></td></tr>
<tr><td>Oligo rev</td><td>AAACCGTTGGAGTGTGGAGATCCG</td><td></td><td></td><td></td><td></td><td></td><td></td><td></td><td></td><td></td><td></td><td></td></tr>
<tr><td>Chromosome</td><td>start</td><td></td><td>end</td><td>strand</td><td>MM</td><td>target_seq</td><td>PAM</td><td>alignment</td><td>distance</td><td>position</td><td>gene name</td><td>gene id</td></tr>
<tr><td>GU305914.1</td><td></td><td>86704</td><td>86726</td><td>+</td><td>0</td><td>CGGATCTCCACACTCCAACG</td><td>CGG</td><td>||||||||[||||||||||||]PAM</td><td>NA</td><td>Intergenic</td><td>NA</td><td>NA</td></tr>
<tr><td>T6186</td><td>CCCCTCTTCCCCTCCTGCCCCGG</td><td>1000</td><td></td><td></td><td></td><td></td><td></td><td></td><td></td><td></td><td></td><td></td></tr>
<tr><td>Oligo fwd</td><td>CACCCCCCTCTTCCCCTCCTGCCC</td><td></td><td></td><td></td><td></td><td></td><td></td><td></td><td></td><td></td><td></td><td></td></tr>
<tr><td>Oligo rev</td><td>AAACGGGCAGGAGGGGAAGAGGGG</td><td></td><td></td><td></td><td></td><td></td><td></td><td></td><td></td><td></td><td></td><td></td></tr>
<tr><td>Chromosome</td><td>start</td><td></td><td>end</td><td>strand</td><td>MM</td><td>target_seq</td><td>PAM</td><td>alignment</td><td>distance</td><td>position</td><td>gene name</td><td>gene id</td></tr>
<tr><td>GU305914.1</td><td></td><td>86797</td><td>86819</td><td>+</td><td>0</td><td>CCCCTCTTCCCCTCCTGCCC</td><td>CGG</td><td>||||||||[||||||||||||]PAM</td><td>NA</td><td>Intergenic</td><td>NA</td><td>NA</td></tr>
<tr><td>T6187</td><td>CCCGGCCCCCGACGCCGCCTCGG</td><td>1000</td><td></td><td></td><td></td><td></td><td></td><td></td><td></td><td></td><td></td><td></td></tr>
<tr><td>Oligo fwd</td><td>CACCCCCGGCCCCCGACGCCGCCT</td><td></td><td></td><td></td><td></td><td></td><td></td><td></td><td></td><td></td><td></td><td></td></tr>
<tr><td>Oligo rev</td><td>AAACAGGCGGCGTCGGGGGCCGGG</td><td></td><td></td><td></td><td></td><td></td><td></td><td></td><td></td><td></td><td></td><td></td></tr>
<tr><td>Chromosome</td><td>start</td><td></td><td>end</td><td>strand</td><td>MM</td><td>target_seq</td><td>PAM</td><td>alignment</td><td>distance</td><td>position</td><td>gene name</td><td>gene id</td></tr>
<tr><td>GU305914.1</td><td></td><td>86815</td><td>86837</td><td>+</td><td>0</td><td>CCCGGCCCCCGACGCCGCCT</td><td>CGG</td><td>||||||||[||||||||||||]PAM</td><td>NA</td><td>Intergenic</td><td>NA</td><td>NA</td></tr>
<tr><td>GU305914.1</td><td></td><td>35655</td><td>35677</td><td>-</td><td>4</td><td>CCCCGCCGCCGCCGCCGCCG</td><td>TGG</td><td>|||-|||-[|||-|||||||-]PAM</td><td>NA</td><td>Intergenic</td><td>NA</td><td>NA</td></tr>
<tr><td>T6188</td><td>TGAAGCTGCCGCCCCCGCCGCGG</td><td>1000</td><td></td><td></td><td></td><td></td><td></td><td></td><td></td><td></td><td></td><td></td></tr>
<tr><td>Oligo fwd</td><td>CACCTGAAGCTGCCGCCCCCGCCG</td><td></td><td></td><td></td><td></td><td></td><td></td><td></td><td></td><td></td><td></td><td></td></tr>
<tr><td>Oligo rev</td><td>AAACCGGCGGGGGCGGCAGCTTCA</td><td></td><td></td><td></td><td></td><td></td><td></td><td></td><td></td><td></td><td></td><td></td></tr>
<tr><td>Chromosome</td><td>start</td><td></td><td>end</td><td>strand</td><td>MM</td><td>target_seq</td><td>PAM</td><td>alignment</td><td>distance</td><td>position</td><td>gene name</td><td>gene id</td></tr>
<tr><td>GU305914.1</td><td></td><td>86842</td><td>86864</td><td>+</td><td>0</td><td>TGAAGCTGCCGCCCCCGCCG</td><td>CGG</td><td>||||||||[||||||||||||]PAM</td><td>NA</td><td>Intergenic</td><td>NA</td><td>NA</td></tr>
<tr><td>GU305914.1</td><td></td><td>45592</td><td>45614</td><td>-</td><td>4</td><td>GGCGGCTGCCGCCGCCGCCG</td><td>CGG</td><td>-|--||||[|||||-||||||]PAM</td><td>NA</td><td>Intergenic</td><td>NA</td><td>NA</td></tr>
<tr><td>T6189</td><td>GAAGCTGCCGCCCCCGCCGCGGG</td><td>1000</td><td></td><td></td><td></td><td></td><td></td><td></td><td></td><td></td><td></td><td></td></tr>
<tr><td>Oligo fwd</td><td>CACCGAAGCTGCCGCCCCCGCCGC</td><td></td><td></td><td></td><td></td><td></td><td></td><td></td><td></td><td></td><td></td><td></td></tr>
<tr><td>Oligo rev</td><td>AAACGCGGCGGGGGCGGCAGCTTC</td><td></td><td></td><td></td><td></td><td></td><td></td><td></td><td></td><td></td><td></td><td></td></tr>
<tr><td>Chromosome</td><td>start</td><td></td><td>end</td><td>strand</td><td>MM</td><td>target_seq</td><td>PAM</td><td>alignment</td><td>distance</td><td>position</td><td>gene name</td><td>gene id</td></tr>
<tr><td>GU305914.1</td><td></td><td>86843</td><td>86865</td><td>+</td><td>0</td><td>GAAGCTGCCGCCCCCGCCGC</td><td>GGG</td><td>||||||||[||||||||||||]PAM</td><td>NA</td><td>Intergenic</td><td>NA</td><td>NA</td></tr>
<tr><td>GU305914.1</td><td></td><td>164964</td><td>164986</td><td>+</td><td>3</td><td>GCTGCTGCCGCCGCCGCCGC</td><td>CGG</td><td>|--|||||[||||-|||||||]PAM</td><td>NA</td><td>Intergenic</td><td>NA</td><td>NA</td></tr>
<tr><td>GU305914.1</td><td></td><td>164826</td><td>164848</td><td>-</td><td>4</td><td>GAAGCGGTCGGCGCCGCCGC</td><td>CGG</td><td>|||||-|-[||-|-|||||||]PAM</td><td>NA</td><td>Intergenic</td><td>NA</td><td>NA</td></tr>
<tr><td>GU305914.1</td><td></td><td>201386</td><td>201408</td><td>-</td><td>3</td><td>GAAGCCGCCGCCTCCGTCGC</td><td>CGG</td><td>|||||-||[||||-|||-|||]PAM</td><td>NA</td><td>Intergenic</td><td>NA</td><td>NA</td></tr>
<tr><td>T6190</td><td>CGGGTCCATCTCCCACCCCCCGG</td><td>1000</td><td></td><td></td><td></td><td></td><td></td><td></td><td></td><td></td><td></td><td></td></tr>
<tr><td>Oligo fwd</td><td>CACCCGGGTCCATCTCCCACCCCC</td><td></td><td></td><td></td><td></td><td></td><td></td><td></td><td></td><td></td><td></td><td></td></tr>
<tr><td>Oligo rev</td><td>AAACGGGGGTGGGAGATGGACCCG</td><td></td><td></td><td></td><td></td><td></td><td></td><td></td><td></td><td></td><td></td><td></td></tr>
<tr><td>Chromosome</td><td>start</td><td></td><td>end</td><td>strand</td><td>MM</td><td>target_seq</td><td>PAM</td><td>alignment</td><td>distance</td><td>position</td><td>gene name</td><td>gene id</td></tr>
<tr><td>GU305914.1</td><td></td><td>86862</td><td>86884</td><td>+</td><td>0</td><td>CGGGTCCATCTCCCACCCCC</td><td>CGG</td><td>||||||||[||||||||||||]PAM</td><td>NA</td><td>Intergenic</td><td>NA</td><td>NA</td></tr>
<tr><td>T6191</td><td>CTCGATCAGCGACTTGATCTTGG</td><td>1000</td><td></td><td></td><td></td><td></td><td></td><td></td><td></td><td></td><td></td><td></td></tr>
<tr><td>Oligo fwd</td><td>CACCCTCGATCAGCGACTTGATCT</td><td></td><td></td><td></td><td></td><td></td><td></td><td></td><td></td><td></td><td></td><td></td></tr>
<tr><td>Oligo rev</td><td>AAACAGATCAAGTCGCTGATCGAG</td><td></td><td></td><td></td><td></td><td></td><td></td><td></td><td></td><td></td><td></td><td></td></tr>
<tr><td>Chromosome</td><td>start</td><td></td><td>end</td><td>strand</td><td>MM</td><td>target_seq</td><td>PAM</td><td>alignment</td><td>distance</td><td>position</td><td>gene name</td><td>gene id</td></tr>
<tr><td>GU305914.1</td><td></td><td>86890</td><td>86912</td><td>+</td><td>0</td><td>CTCGATCAGCGACTTGATCT</td><td>TGG</td><td>||||||||[||||||||||||]PAM</td><td>NA</td><td>Intergenic</td><td>NA</td><td>NA</td></tr>
</table>

T6192 TCTCGTGCAAGCGATTGAAAAGG 1000
Oligo fwd CACCTCTCGTGCAAGCGATTGAAA
Oligo rev AAACTTTCAATCGCTTGCACGAGA
Chromosome start end strand MM target_seq PAM alignment distance position gene name gene id
GU305914.1 86928 86950 + 0 TCTCGTGCAAGCGATTGAAA AGG ||||||||[||||||||||||]PAM NA Intergenic NA NA

T6193 CTCGTGCAAGCGATTGAAAAGGG 1000
Oligo fwd CACCCTCGTGCAAGCGATTGAAAA
Oligo rev AAACTTTTCAATCGCTTGCACGAG
Chromosome start end strand MM target_seq PAM alignment distance position gene name gene id
GU305914.1 86929 86951 + 0 CTCGTGCAAGCGATTGAAAA GGG ||||||||[||||||||||||]PAM NA Intergenic NA NA

T6194 TCGTGCAAGCGATTGAAAAGGGG 1000
Oligo fwd CACCTCGTGCAAGCGATTGAAAAG
Oligo rev AAACCTTTTCAATCGCTTGCACGA
Chromosome start end strand MM target_seq PAM alignment distance position gene name gene id
GU305914.1 86930 86952 + 0 TCGTGCAAGCGATTGAAAAG GGG ||||||||[||||||||||||]PAM NA Intergenic NA NA

T6195 GAAAAGGGGATTGTTCTCGAAGG 1000
Oligo fwd CACCGAAAAGGGGATTGTTCTCGA
Oligo rev AAACTCGAGAACAATCCCCTTTTC
Chromosome start end strand MM target_seq PAM alignment distance position gene name gene id
GU305914.1 86944 86966 + 0 GAAAAGGGGATTGTTCTCGA AGG ||||||||[||||||||||||]PAM NA Intergenic NA NA

T6196 ATCATGTCGACGACCTTTCCCGG 1000
Oligo fwd CACCATCATGTCGACGACCTTTCC
Oligo rev AAACGGAAAGGTCGTCGACATGAT
Chromosome start end strand MM target_seq PAM alignment distance position gene name gene id
GU305914.1 86987 87009 + 0 ATCATGTCGACGACCTTTCC CGG ||||||||[||||||||||||]PAM NA Intergenic NA NA

T6197 CTTTCCCGGCGCCGCGAAGATGG 1000
Oligo fwd CACCCTTTCCCGGCGCCGCGAAGA
Oligo rev AAACTCTTCGCGGCGCCGGGAAAG
Chromosome start end strand MM target_seq PAM alignment distance position gene name gene id
GU305914.1 87001 87023 + 0 CTTTCCCGGCGCCGCGAAGA TGG ||||||||[||||||||||||]PAM NA Intergenic NA NA

T6198 CCAGAGTGAAAACGTCCTCGAGG 1000
Oligo fwd CACCCCAGAGTGAAAACGTCCTCG
Oligo rev AAACCGAGGACGTTTTCACTCTGG
Chromosome start end strand MM target_seq PAM alignment distance position gene name gene id
GU305914.1 87066 87088 + 0 CCAGAGTGAAAACGTCCTCG AGG ||||||||[||||||||||||]PAM NA Intergenic NA NA

T6199 GCGACTCAATTCCGTCCGCAAGG 1000
Oligo fwd CACCGCGACTCAATTCCGTCCGCA
Oligo rev AAACTGCGGACGGAATTGAGTCGC
Chromosome start end strand MM target_seq PAM alignment distance position gene name gene id
GU305914.1 87106 87128 + 0 GCGACTCAATTCCGTCCGCA AGG ||||||||[||||||||||||]PAM NA Intergenic NA NA

T6200 CCGTCCGCAAGGTTTTCCAGAGG 1000
Oligo fwd CACCCCGTCCGCAAGGTTTTCCAG
Oligo rev AAACCTGGAAAACCTTGCGGACGG
Chromosome start end strand MM target_seq PAM alignment distance position gene name gene id
GU305914.1 87117 87139 + 0 CCGTCCGCAAGGTTTTCCAG AGG ||||||||[||||||||||||]PAM NA Intergenic NA NA

T6201 GCAAGGTTTTCCAGAGGCTGTGG 1000
Oligo fwd CACCGCAAGGTTTTCCAGAGGCTG
Oligo rev AAACCAGCCTCTGGAAAACCTTGC
Chromosome start end strand MM target_seq PAM alignment distance position gene name gene id
GU305914.1 87123 87145 + 0 GCAAGGTTTTCCAGAGGCTG TGG ||||||||[||||||||||||]PAM NA Intergenic NA NA

T6202 CAAGGTTTTCCAGAGGCTGTGGG 1000
Oligo fwd CACCCAAGGTTTTCCAGAGGCTGT
Oligo rev AAACACAGCCTCTGGAAAACCTTG
Chromosome start end strand MM target_seq PAM alignment distance position gene name gene id
GU305914.1 87124 87146 + 0 CAAGGTTTTCCAGAGGCTGT GGG ||||||||[||||||||||||]PAM NA Intergenic NA NA

T6203 GCCTCATCGTAATCCCCACGTGG 1000
Oligo fwd CACCGCCTCATCGTAATCCCCACG
Oligo rev AAACCGTGGGGATTACGATGAGGC
Chromosome start end strand MM target_seq PAM alignment distance position gene name gene id
GU305914.1 87159 87181 + 0 GCCTCATCGTAATCCCCACG TGG ||||||||[||||||||||||]PAM NA Intergenic NA NA

T6204 TCATCGTAATCCCCACGTGGCGG 1000
Oligo fwd CACCTCATCGTAATCCCCACGTGG
Oligo rev AAACCCACGTGGGGATTACGATGA
Chromosome start end strand MM target_seq PAM alignment distance position gene name gene id
GU305914.1 87162 87184 + 0 TCATCGTAATCCCCACGTGG CGG ||||||||[||||||||||||]PAM NA Intergenic NA NA

T6205 GACAGTGAGTAGATCCGATCCGG 1000
Oligo fwd CACCGACAGTGAGTAGATCCGATC
Oligo rev AAACGATCGGATCTACTCACTGTC
Chromosome start end strand MM target_seq PAM alignment distance position gene name gene id
GU305914.1 87224 87246 + 0 GACAGTGAGTAGATCCGATC CGG ||||||||[||||||||||||]PAM NA Intergenic NA NA

T6206 ACAGTGAGTAGATCCGATCCGGG 1000
Oligo fwd CACCACAGTGAGTAGATCCGATCC
Oligo rev AAACGGATCGGATCTACTCACTGT
Chromosome start end strand MM target_seq PAM alignment distance position gene name gene id
GU305914.1 87225 87247 + 0 ACAGTGAGTAGATCCGATCC GGG ||||||||[||||||||||||]PAM NA Intergenic NA NA

T6207 TTGATGTACGCATACGACGCCGG 1000
Oligo fwd CACCTTGATGTACGCATACGACGC
Oligo rev AAACGCGTCGTATGCGTACATCAA
Chromosome start end strand MM target_seq PAM alignment distance position gene name gene id
GU305914.1 87293 87315 + 0 TTGATGTACGCATACGACGC CGG ||||||||[||||||||||||]PAM NA Intergenic NA NA

T6208 ACGCGAGACTCTTTATCGCCCGG 1000
Oligo fwd CACCACGCGAGACTCTTTATCGCC
Oligo rev AAACGGCGATAAAGAGTCTCGCGT
Chromosome start end strand MM target_seq PAM alignment distance position gene name gene id
GU305914.1 87324 87346 + 0 ACGCGAGACTCTTTATCGCC CGG ||||||||[||||||||||||]PAM NA Intergenic NA NA

T6209 CCGGATGATACCTTTCACAGAGG 1000
Oligo fwd CACCCCGGATGATACCTTTCACAG
Oligo rev AAACCTGTGAAAGGTATCATCCGG
Chromosome start end strand MM target_seq PAM alignment distance position gene name gene id
GU305914.1 87343 87365 + 0 CCGGATGATACCTTTCACAG AGG ||||||||[||||||||||||]PAM NA Intergenic NA NA

T6210 CGGATGATACCTTTCACAGAGGG 1000
Oligo fwd CACCCGGATGATACCTTTCACAGA
Oligo rev AAACTCTGTGAAAGGTATCATCCG
Chromosome start end strand MM target_seq PAM alignment distance position gene name gene id
GU305914.1 87344 87366 + 0 CGGATGATACCTTTCACAGA GGG ||||||||[||||||||||||]PAM NA Intergenic NA NA

T6211 CCATATCCTGCTCCACCGTCTGG 1000
Oligo fwd CACCCCATATCCTGCTCCACCGTC
Oligo rev AAACGACGGTGGAGCAGGATATGG
Chromosome start end strand MM target_seq PAM alignment distance position gene name gene id
GU305914.1 87387 87409 + 0 CCATATCCTGCTCCACCGTC TGG ||||||||[||||||||||||]PAM NA Intergenic NA NA

T6212 CATATCCTGCTCCACCGTCTGGG 1000
Oligo fwd CACCCATATCCTGCTCCACCGTCT
Oligo rev AAACAGACGGTGGAGCAGGATATG
Chromosome start end strand MM target_seq PAM alignment distance position gene name gene id
GU305914.1 87388 87410 + 0 CATATCCTGCTCCACCGTCT GGG ||||||||[||||||||||||]PAM NA Intergenic NA NA

T6213 AGACCTTTTTGATAGTGATATGG 1000
Oligo fwd CACCAGACCTTTTTGATAGTGATA
Oligo rev AAACTATCACTATCAAAAAGGTCT
Chromosome start end strand MM target_seq PAM alignment distance position gene name gene id
GU305914.1 87432 87454 + 0 AGACCTTTTTGATAGTGATA TGG ||||||||[||||||||||||]PAM NA Intergenic NA NA

T6214 TAGTGATATGGTCGCAGAGCAGG 1000
Oligo fwd CACCTAGTGATATGGTCGCAGAGC
Oligo rev AAACGCTCTGCGACCATATCACTA
Chromosome start end strand MM target_seq PAM alignment distance position gene name gene id
GU305914.1 87444 87466 + 0 TAGTGATATGGTCGCAGAGC AGG ||||||||[||||||||||||]PAM NA Intergenic NA NA

T6215 AGAGCAGGCCCTGCATCCGCCGG 1000
Oligo fwd CACCAGAGCAGGCCCTGCATCCGC
Oligo rev AAACGCGGATGCAGGGCCTGCTCT
Chromosome start end strand MM target_seq PAM alignment distance position gene name gene id
GU305914.1 87459 87481 + 0 AGAGCAGGCCCTGCATCCGC CGG ||||||||[||||||||||||]PAM NA Intergenic NA NA

T6216 GCATCCGCCGGTTGATCGAGCGG 1000
Oligo fwd CACCGCATCCGCCGGTTGATCGAG
Oligo rev AAACCTCGATCAACCGGCGGATGC
Chromosome start end strand MM target_seq PAM alignment distance position gene name gene id
GU305914.1 87471 87493 + 0 GCATCCGCCGGTTGATCGAG CGG ||||||||[||||||||||||]PAM NA Intergenic NA NA

T6217 GCCGGTTGATCGAGCGGCCTTGG 1000
Oligo fwd CACCGCCGGTTGATCGAGCGGCCT
Oligo rev AAACAGGCCGCTCGATCAACCGGC
Chromosome start end strand MM target_seq PAM alignment distance position gene name gene id
GU305914.1 87477 87499 + 0 GCCGGTTGATCGAGCGGCCT TGG ||||||||[||||||||||||]PAM NA Intergenic NA NA

T6218 TTGATCGAGCGGCCTTGGTTCGG 1000
Oligo fwd CACCTTGATCGAGCGGCCTTGGTT
Oligo rev AAACAACCAAGGCCGCTCGATCAA
Chromosome start end strand MM target_seq PAM alignment distance position gene name gene id
GU305914.1 87482 87504 + 0 TTGATCGAGCGGCCTTGGTT CGG ||||||||[||||||||||||]PAM NA Intergenic NA NA

T6219 CTCCTCGTAGCACTGAAAACAGG 1000
Oligo fwd CACCCTCCTCGTAGCACTGAAAAC
Oligo rev AAACGTTTTCAGTGCTACGAGGAG
Chromosome start end strand MM target_seq PAM alignment distance position gene name gene id
GU305914.1 87517 87539 + 0 CTCCTCGTAGCACTGAAAAC AGG ||||||||[||||||||||||]PAM NA Intergenic NA NA

T6220 ACCGTCTCCCGATAGATCTCCGG 1000
Oligo fwd CACCACCGTCTCCCGATAGATCTC
Oligo rev AAACGAGATCTATCGGGAGACGGT
Chromosome start end strand MM target_seq PAM alignment distance position gene name gene id
GU305914.1 87542 87564 + 0 ACCGTCTCCCGATAGATCTC CGG ||||||||[||||||||||||]PAM NA Intergenic NA NA

T6221 TAGATCTCCGGACCCGCCACCGG 1000
Oligo fwd CACCTAGATCTCCGGACCCGCCAC
Oligo rev AAACGTGGCGGGTCCGGAGATCTA
Chromosome start end strand MM target_seq PAM alignment distance position gene name gene id

<table>
<tr><td>GU305914.1</td><td>87554</td><td>87576</td><td>+</td><td>0</td><td>TAGATCTCCGGACCCGCCAC</td><td>CGG</td><td>||||||||[||||||||||||]PAM</td><td>NA</td><td>Intergenic</td><td>NA</td><td>NA</td></tr>
<tr><td>T6222</td><td>AGATCTCCGGACCCGCCACCGGG</td><td>1000</td><td></td><td></td><td></td><td></td><td></td><td></td><td></td><td></td><td></td></tr>
<tr><td>Oligo fwd</td><td>CACCAGATCTCCGGACCCGCCACC</td><td></td><td></td><td></td><td></td><td></td><td></td><td></td><td></td><td></td><td></td></tr>
<tr><td>Oligo rev</td><td>AAACGGTGGCGGGTCCGGAGATCT</td><td></td><td></td><td></td><td></td><td></td><td></td><td></td><td></td><td></td><td></td></tr>
<tr><td>Chromosome</td><td>start</td><td>end</td><td>strand</td><td>MM</td><td>target_seq</td><td>PAM</td><td>alignment</td><td>distance</td><td>position</td><td>gene name</td><td>gene id</td></tr>
<tr><td>GU305914.1</td><td>87555</td><td>87577</td><td>+</td><td>0</td><td>AGATCTCCGGACCCGCCACC</td><td>GGG</td><td>||||||||[||||||||||||]PAM</td><td>NA</td><td>Intergenic</td><td>NA</td><td>NA</td></tr>
<tr><td>T6223</td><td>TATCGCACAACCTCGCCAGCAGG</td><td>1000</td><td></td><td></td><td></td><td></td><td></td><td></td><td></td><td></td><td></td></tr>
<tr><td>Oligo fwd</td><td>CACCTATCGCACAACCTCGCCAGC</td><td></td><td></td><td></td><td></td><td></td><td></td><td></td><td></td><td></td><td></td></tr>
<tr><td>Oligo rev</td><td>AAACGCTGGCGAGGTTGTGCGATA</td><td></td><td></td><td></td><td></td><td></td><td></td><td></td><td></td><td></td><td></td></tr>
<tr><td>Chromosome</td><td>start</td><td>end</td><td>strand</td><td>MM</td><td>target_seq</td><td>PAM</td><td>alignment</td><td>distance</td><td>position</td><td>gene name</td><td>gene id</td></tr>
<tr><td>GU305914.1</td><td>87582</td><td>87604</td><td>+</td><td>0</td><td>TATCGCACAACCTCGCCAGC</td><td>AGG</td><td>||||||||[||||||||||||]PAM</td><td>NA</td><td>Intergenic</td><td>NA</td><td>NA</td></tr>
<tr><td>T6224</td><td>ATCGCACAACCTCGCCAGCAGGG</td><td>1000</td><td></td><td></td><td></td><td></td><td></td><td></td><td></td><td></td><td></td></tr>
<tr><td>Oligo fwd</td><td>CACCATCGCACAACCTCGCCAGCA</td><td></td><td></td><td></td><td></td><td></td><td></td><td></td><td></td><td></td><td></td></tr>
<tr><td>Oligo rev</td><td>AAACTGCTGGCGAGGTTGTGCGAT</td><td></td><td></td><td></td><td></td><td></td><td></td><td></td><td></td><td></td><td></td></tr>
<tr><td>Chromosome</td><td>start</td><td>end</td><td>strand</td><td>MM</td><td>target_seq</td><td>PAM</td><td>alignment</td><td>distance</td><td>position</td><td>gene name</td><td>gene id</td></tr>
<tr><td>GU305914.1</td><td>87583</td><td>87605</td><td>+</td><td>0</td><td>ATCGCACAACCTCGCCAGCA</td><td>GGG</td><td>||||||||[||||||||||||]PAM</td><td>NA</td><td>Intergenic</td><td>NA</td><td>NA</td></tr>
<tr><td>T6225</td><td>GTCGACAGACCGCCACCCCCTGG</td><td>1000</td><td></td><td></td><td></td><td></td><td></td><td></td><td></td><td></td><td></td></tr>
<tr><td>Oligo fwd</td><td>CACCGTCGACAGACCGCCACCCCC</td><td></td><td></td><td></td><td></td><td></td><td></td><td></td><td></td><td></td><td></td></tr>
<tr><td>Oligo rev</td><td>AAACGGGGGTGGCGGTCTGTCGAC</td><td></td><td></td><td></td><td></td><td></td><td></td><td></td><td></td><td></td><td></td></tr>
<tr><td>Chromosome</td><td>start</td><td>end</td><td>strand</td><td>MM</td><td>target_seq</td><td>PAM</td><td>alignment</td><td>distance</td><td>position</td><td>gene name</td><td>gene id</td></tr>
<tr><td>GU305914.1</td><td>87615</td><td>87637</td><td>+</td><td>0</td><td>GTCGACAGACCGCCACCCCC</td><td>TGG</td><td>||||||||[||||||||||||]PAM</td><td>NA</td><td>Intergenic</td><td>NA</td><td>NA</td></tr>
<tr><td>T6226</td><td>ACACACAGTTGATTTTGCACAGG</td><td>1000</td><td></td><td></td><td></td><td></td><td></td><td></td><td></td><td></td><td></td></tr>
<tr><td>Oligo fwd</td><td>CACCACACACAGTTGATTTTGCAC</td><td></td><td></td><td></td><td></td><td></td><td></td><td></td><td></td><td></td><td></td></tr>
<tr><td>Oligo rev</td><td>AAACGTGCAAAATCAACTGTGTGT</td><td></td><td></td><td></td><td></td><td></td><td></td><td></td><td></td><td></td><td></td></tr>
<tr><td>Chromosome</td><td>start</td><td>end</td><td>strand</td><td>MM</td><td>target_seq</td><td>PAM</td><td>alignment</td><td>distance</td><td>position</td><td>gene name</td><td>gene id</td></tr>
<tr><td>GU305914.1</td><td>87657</td><td>87679</td><td>+</td><td>0</td><td>ACACACAGTTGATTTTGCAC</td><td>AGG</td><td>||||||||[||||||||||||]PAM</td><td>NA</td><td>Intergenic</td><td>NA</td><td>NA</td></tr>
<tr><td>T6227</td><td>CCACGTCGTTCAGCGTCGCCAGG</td><td>1000</td><td></td><td></td><td></td><td></td><td></td><td></td><td></td><td></td><td></td></tr>
<tr><td>Oligo fwd</td><td>CACCCCACGTCGTTCAGCGTCGCC</td><td></td><td></td><td></td><td></td><td></td><td></td><td></td><td></td><td></td><td></td></tr>
<tr><td>Oligo rev</td><td>AAACGGCGACGCTGAACGACGTGG</td><td></td><td></td><td></td><td></td><td></td><td></td><td></td><td></td><td></td><td></td></tr>
<tr><td>Chromosome</td><td>start</td><td>end</td><td>strand</td><td>MM</td><td>target_seq</td><td>PAM</td><td>alignment</td><td>distance</td><td>position</td><td>gene name</td><td>gene id</td></tr>
<tr><td>GU305914.1</td><td>87687</td><td>87709</td><td>+</td><td>0</td><td>CCACGTCGTTCAGCGTCGCC</td><td>AGG</td><td>||||||||[||||||||||||]PAM</td><td>NA</td><td>Intergenic</td><td>NA</td><td>NA</td></tr>
<tr><td>T6228</td><td>GTCGATCGTCAGCGCCAGCTCGG</td><td>1000</td><td></td><td></td><td></td><td></td><td></td><td></td><td></td><td></td><td></td></tr>
<tr><td>Oligo fwd</td><td>CACCGTCGATCGTCAGCGCCAGCT</td><td></td><td></td><td></td><td></td><td></td><td></td><td></td><td></td><td></td><td></td></tr>
<tr><td>Oligo rev</td><td>AAACAGCTGGCGCTGACGATCGAC</td><td></td><td></td><td></td><td></td><td></td><td></td><td></td><td></td><td></td><td></td></tr>
<tr><td>Chromosome</td><td>start</td><td>end</td><td>strand</td><td>MM</td><td>target_seq</td><td>PAM</td><td>alignment</td><td>distance</td><td>position</td><td>gene name</td><td>gene id</td></tr>
<tr><td>GU305914.1</td><td>87709</td><td>87731</td><td>+</td><td>0</td><td>GTCGATCGTCAGCGCCAGCT</td><td>CGG</td><td>||||||||[||||||||||||]PAM</td><td>NA</td><td>Intergenic</td><td>NA</td><td>NA</td></tr>
<tr><td>T6229</td><td>TCGATCGTCAGCGCCAGCTCGGG</td><td>1000</td><td></td><td></td><td></td><td></td><td></td><td></td><td></td><td></td><td></td></tr>
<tr><td>Oligo fwd</td><td>CACCTCGATCGTCAGCGCCAGCTC</td><td></td><td></td><td></td><td></td><td></td><td></td><td></td><td></td><td></td><td></td></tr>
<tr><td>Oligo rev</td><td>AAACGAGCTGGCGCTGACGATCGA</td><td></td><td></td><td></td><td></td><td></td><td></td><td></td><td></td><td></td><td></td></tr>
<tr><td>Chromosome</td><td>start</td><td>end</td><td>strand</td><td>MM</td><td>target_seq</td><td>PAM</td><td>alignment</td><td>distance</td><td>position</td><td>gene name</td><td>gene id</td></tr>
<tr><td>GU305914.1</td><td>87710</td><td>87732</td><td>+</td><td>0</td><td>TCGATCGTCAGCGCCAGCTC</td><td>GGG</td><td>||||||||[||||||||||||]PAM</td><td>NA</td><td>Intergenic</td><td>NA</td><td>NA</td></tr>
<tr><td>GU305914.1</td><td>138217</td><td>138239</td><td>+</td><td>4</td><td>GCGTTCGTCAGGCCCAGCTC</td><td>CGG</td><td>-||-||||[|||--|||||||]PAM</td><td>NA</td><td>Intergenic</td><td>NA</td><td>NA</td></tr>
<tr><td>T6230</td><td>TTATCGTGATCGACTCGTGTCGG</td><td>1000</td><td></td><td></td><td></td><td></td><td></td><td></td><td></td><td></td><td></td></tr>
<tr><td>Oligo fwd</td><td>CACCTTATCGTGATCGACTCGTGT</td><td></td><td></td><td></td><td></td><td></td><td></td><td></td><td></td><td></td><td></td></tr>
<tr><td>Oligo rev</td><td>AAACACACGAGTCGATCACGATAA</td><td></td><td></td><td></td><td></td><td></td><td></td><td></td><td></td><td></td><td></td></tr>
<tr><td>Chromosome</td><td>start</td><td>end</td><td>strand</td><td>MM</td><td>target_seq</td><td>PAM</td><td>alignment</td><td>distance</td><td>position</td><td>gene name</td><td>gene id</td></tr>
<tr><td>GU305914.1</td><td>87741</td><td>87763</td><td>+</td><td>0</td><td>TTATCGTGATCGACTCGTGT</td><td>CGG</td><td>||||||||[||||||||||||]PAM</td><td>NA</td><td>Intergenic</td><td>NA</td><td>NA</td></tr>
<tr><td>T6231</td><td>AATCGAGTTTCAGCACCCGCCGG</td><td>1000</td><td></td><td></td><td></td><td></td><td></td><td></td><td></td><td></td><td></td></tr>
<tr><td>Oligo fwd</td><td>CACCAATCGAGTTTCAGCACCCGC</td><td></td><td></td><td></td><td></td><td></td><td></td><td></td><td></td><td></td><td></td></tr>
<tr><td>Oligo rev</td><td>AAACGCGGGTGCTGAAACTCGATT</td><td></td><td></td><td></td><td></td><td></td><td></td><td></td><td></td><td></td><td></td></tr>
<tr><td>Chromosome</td><td>start</td><td>end</td><td>strand</td><td>MM</td><td>target_seq</td><td>PAM</td><td>alignment</td><td>distance</td><td>position</td><td>gene name</td><td>gene id</td></tr>
<tr><td>GU305914.1</td><td>87771</td><td>87793</td><td>+</td><td>0</td><td>AATCGAGTTTCAGCACCCGC</td><td>CGG</td><td>||||||||[||||||||||||]PAM</td><td>NA</td><td>Intergenic</td><td>NA</td><td>NA</td></tr>
<tr><td>T6232</td><td>CCCGCCGGTAGTATCGCTGCCGG</td><td>1000</td><td></td><td></td><td></td><td></td><td></td><td></td><td></td><td></td><td></td></tr>
<tr><td>Oligo fwd</td><td>CACCCCCGCCGGTAGTATCGCTGC</td><td></td><td></td><td></td><td></td><td></td><td></td><td></td><td></td><td></td><td></td></tr>
<tr><td>Oligo rev</td><td>AAACGCAGCGATACTACCGGCGGG</td><td></td><td></td><td></td><td></td><td></td><td></td><td></td><td></td><td></td><td></td></tr>
<tr><td>Chromosome</td><td>start</td><td>end</td><td>strand</td><td>MM</td><td>target_seq</td><td>PAM</td><td>alignment</td><td>distance</td><td>position</td><td>gene name</td><td>gene id</td></tr>
<tr><td>GU305914.1</td><td>87786</td><td>87808</td><td>+</td><td>0</td><td>CCCGCCGGTAGTATCGCTGC</td><td>CGG</td><td>||||||||[||||||||||||]PAM</td><td>NA</td><td>Intergenic</td><td>NA</td><td>NA</td></tr>
<tr><td>T6233</td><td>GCAGCAGCATCTCCATGTAGAGG</td><td>1000</td><td></td><td></td><td></td><td></td><td></td><td></td><td></td><td></td><td></td></tr>
<tr><td>Oligo fwd</td><td>CACCGCAGCAGCATCTCCATGTAG</td><td></td><td></td><td></td><td></td><td></td><td></td><td></td><td></td><td></td><td></td></tr>
<tr><td>Oligo rev</td><td>AAACCTACATGGAGATGCTGCTGC</td><td></td><td></td><td></td><td></td><td></td><td></td><td></td><td></td><td></td><td></td></tr>
<tr><td>Chromosome</td><td>start</td><td>end</td><td>strand</td><td>MM</td><td>target_seq</td><td>PAM</td><td>alignment</td><td>distance</td><td>position</td><td>gene name</td><td>gene id</td></tr>
<tr><td>GU305914.1</td><td>87873</td><td>87895</td><td>+</td><td>0</td><td>GCAGCAGCATCTCCATGTAG</td><td>AGG</td><td>||||||||[||||||||||||]PAM</td><td>NA</td><td>Intergenic</td><td>NA</td><td>NA</td></tr>
<tr><td>T6234</td><td>CAGCAGCATCTCCATGTAGAGGG</td><td>1000</td><td></td><td></td><td></td><td></td><td></td><td></td><td></td><td></td><td></td></tr>
<tr><td>Oligo fwd</td><td>CACCCAGCAGCATCTCCATGTAGA</td><td></td><td></td><td></td><td></td><td></td><td></td><td></td><td></td><td></td><td></td></tr>
<tr><td>Oligo rev</td><td>AAACTCTACATGGAGATGCTGCTG</td><td></td><td></td><td></td><td></td><td></td><td></td><td></td><td></td><td></td><td></td></tr>
<tr><td>Chromosome</td><td>start</td><td>end</td><td>strand</td><td>MM</td><td>target_seq</td><td>PAM</td><td>alignment</td><td>distance</td><td>position</td><td>gene name</td><td>gene id</td></tr>
<tr><td>GU305914.1</td><td>87874</td><td>87896</td><td>+</td><td>0</td><td>CAGCAGCATCTCCATGTAGA</td><td>GGG</td><td>||||||||[||||||||||||]PAM</td><td>NA</td><td>Intergenic</td><td>NA</td><td>NA</td></tr>
<tr><td>T6235</td><td>CATGTAGAGGGTCAGCACCGAGG</td><td>1000</td><td></td><td></td><td></td><td></td><td></td><td></td><td></td><td></td><td></td></tr>
<tr><td>Oligo fwd</td><td>CACCCATGTAGAGGGTCAGCACCG</td><td></td><td></td><td></td><td></td><td></td><td></td><td></td><td></td><td></td><td></td></tr>
<tr><td>Oligo rev</td><td>AAACCGGTGCTGACCCTCTACATG</td><td></td><td></td><td></td><td></td><td></td><td></td><td></td><td></td><td></td><td></td></tr>
<tr><td>Chromosome</td><td>start</td><td>end</td><td>strand</td><td>MM</td><td>target_seq</td><td>PAM</td><td>alignment</td><td>distance</td><td>position</td><td>gene name</td><td>gene id</td></tr>
<tr><td>GU305914.1</td><td>87886</td><td>87908</td><td>+</td><td>0</td><td>CATGTAGAGGGTCAGCACCG</td><td>AGG</td><td>||||||||[||||||||||||]PAM</td><td>NA</td><td>Intergenic</td><td>NA</td><td>NA</td></tr>
<tr><td>T6236</td><td>GCGTTCTGCGAGACCAGCGCCGG</td><td>1000</td><td></td><td></td><td></td><td></td><td></td><td></td><td></td><td></td><td></td></tr>
<tr><td>Oligo fwd</td><td>CACCGCGTTCTGCGAGACCAGCGC</td><td></td><td></td><td></td><td></td><td></td><td></td><td></td><td></td><td></td><td></td></tr>
</table>

| | | | | | | | | | | | | |
|---|---|---|---|---|---|---|---|---|---|---|---|---|
| Oligo rev | AAACGCGCTGGTCTCGCAGAACGC | | | | | | | | | | | |
| Chromosome | start | | end | strand | MM | target_seq | PAM | alignment | distance | position | gene name | gene id |
| GU305914.1 | | 87917 | 87939 | + | 0 | GCGTTCTGCGAGACCAGCGC | CGG | \|\|\|\|\|\|\|\|[\|\|\|\|\|\|\|\|\|\|\|\|]PAM | NA | Intergenic | NA | NA |
| T6237 | GCGAGACCAGCGCCGGATAGAGG | | 1000 | | | | | | | | | |
| Oligo fwd | CACCGCGAGACCAGCGCCGGATAG | | | | | | | | | | | |
| Oligo rev | AAACCTATCCGGCGCTGGTCTCGC | | | | | | | | | | | |
| Chromosome | start | | end | strand | MM | target_seq | PAM | alignment | distance | position | gene name | gene id |
| GU305914.1 | | 87924 | 87946 | + | 0 | GCGAGACCAGCGCCGGATAG | AGG | \|\|\|\|\|\|\|\|[\|\|\|\|\|\|\|\|\|\|\|\|]PAM | NA | Intergenic | NA | NA |
| T6238 | CGAGACCAGCGCCGGATAGAGGG | | 1000 | | | | | | | | | |
| Oligo fwd | CACCCGAGACCAGCGCCGGATAGA | | | | | | | | | | | |
| Oligo rev | AAACTCTATCCGGCGCTGGTCTCG | | | | | | | | | | | |
| Chromosome | start | | end | strand | MM | target_seq | PAM | alignment | distance | position | gene name | gene id |
| GU305914.1 | | 87925 | 87947 | + | 0 | CGAGACCAGCGCCGGATAGA | GGG | \|\|\|\|\|\|\|\|[\|\|\|\|\|\|\|\|\|\|\|\|]PAM | NA | Intergenic | NA | NA |
| T6239 | CATGACCCCGTTCCTCCTGAAGG | | 1000 | | | | | | | | | |
| Oligo fwd | CACCCATGACCCCGTTCCTCCTGA | | | | | | | | | | | |
| Oligo rev | AAACTCAGGAGGAACGGGGTCATG | | | | | | | | | | | |
| Chromosome | start | | end | strand | MM | target_seq | PAM | alignment | distance | position | gene name | gene id |
| GU305914.1 | | 87967 | 87989 | + | 0 | CATGACCCCGTTCCTCCTGA | AGG | \|\|\|\|\|\|\|\|[\|\|\|\|\|\|\|\|\|\|\|\|]PAM | NA | Intergenic | NA | NA |
| T6240 | GAATCCATCGACACAATATTCGG | | 1000 | | | | | | | | | |
| Oligo fwd | CACCGAATCCATCGACACAATATT | | | | | | | | | | | |
| Oligo rev | AAACAATATTGTGTCGATGGATTC | | | | | | | | | | | |
| Chromosome | start | | end | strand | MM | target_seq | PAM | alignment | distance | position | gene name | gene id |
| GU305914.1 | | 87995 | 88017 | + | 0 | GAATCCATCGACACAATATT | CGG | \|\|\|\|\|\|\|\|[\|\|\|\|\|\|\|\|\|\|\|\|]PAM | NA | Intergenic | NA | NA |
| T6241 | CATCGCGCACTCGTTGCATTTGG | | 1000 | | | | | | | | | |
| Oligo fwd | CACCCATCGCGCACTCGTTGCATT | | | | | | | | | | | |
| Oligo rev | AAACAATGCAACGAGTGCGCGATG | | | | | | | | | | | |
| Chromosome | start | | end | strand | MM | target_seq | PAM | alignment | distance | position | gene name | gene id |
| GU305914.1 | | 88045 | 88067 | + | 0 | CATCGCGCACTCGTTGCATT | TGG | \|\|\|\|\|\|\|\|[\|\|\|\|\|\|\|\|\|\|\|\|]PAM | NA | Intergenic | NA | NA |
| T6242 | TGCCGACGTTGCGCGCGAGCAGG | | 1000 | | | | | | | | | |
| Oligo fwd | CACCTGCCGACGTTGCGCGCGAGC | | | | | | | | | | | |
| Oligo rev | AAACGCTCGCGCGCAACGTCGGCA | | | | | | | | | | | |
| Chromosome | start | | end | strand | MM | target_seq | PAM | alignment | distance | position | gene name | gene id |
| GU305914.1 | | 88111 | 88133 | + | 0 | TGCCGACGTTGCGCGCGAGC | AGG | \|\|\|\|\|\|\|\|[\|\|\|\|\|\|\|\|\|\|\|\|]PAM | NA | Intergenic | NA | NA |
| T6243 | CGCGCACAGACAACCCCGAACGG | | 1000 | | | | | | | | | |
| Oligo fwd | CACCCGCGCACAGACAACCCCGAA | | | | | | | | | | | |
| Oligo rev | AAACTTCGGGGTTGTCTGTGCGCG | | | | | | | | | | | |
| Chromosome | start | | end | strand | MM | target_seq | PAM | alignment | distance | position | gene name | gene id |
| GU305914.1 | | 88165 | 88187 | + | 0 | CGCGCACAGACAACCCCGAA | CGG | \|\|\|\|\|\|\|\|[\|\|\|\|\|\|\|\|\|\|\|\|]PAM | NA | Intergenic | NA | NA |
| T6244 | TTCAACGAGCTTTAATAGAAAGG | | 1000 | | | | | | | | | |
| Oligo fwd | CACCTTCAACGAGCTTTAATAGAA | | | | | | | | | | | |
| Oligo rev | AAACTTCTATTAAAGCTCGTTGAA | | | | | | | | | | | |
| Chromosome | start | | end | strand | MM | target_seq | PAM | alignment | distance | position | gene name | gene id |
| GU305914.1 | | 88219 | 88241 | + | 0 | TTCAACGAGCTTTAATAGAA | AGG | \|\|\|\|\|\|\|\|[\|\|\|\|\|\|\|\|\|\|\|\|]PAM | NA | Intergenic | NA | NA |
| T6245 | TCAACGAGCTTTAATAGAAAGGG | | 1000 | | | | | | | | | |
| Oligo fwd | CACCTCAACGAGCTTTAATAGAAA | | | | | | | | | | | |
| Oligo rev | AAACTTTCTATTAAAGCTCGTTGA | | | | | | | | | | | |
| Chromosome | start | | end | strand | MM | target_seq | PAM | alignment | distance | position | gene name | gene id |
| GU305914.1 | | 88220 | 88242 | + | 0 | TCAACGAGCTTTAATAGAAA | GGG | \|\|\|\|\|\|\|\|[\|\|\|\|\|\|\|\|\|\|\|\|]PAM | NA | Intergenic | NA | NA |
| T6246 | CAACGAGCTTTAATAGAAAGGGG | | 1000 | | | | | | | | | |
| Oligo fwd | CACCCAACGAGCTTTAATAGAAAG | | | | | | | | | | | |
| Oligo rev | AAACCTTTCTATTAAAGCTCGTTG | | | | | | | | | | | |
| Chromosome | start | | end | strand | MM | target_seq | PAM | alignment | distance | position | gene name | gene id |
| GU305914.1 | | 88221 | 88243 | + | 0 | CAACGAGCTTTAATAGAAAG | GGG | \|\|\|\|\|\|\|\|[\|\|\|\|\|\|\|\|\|\|\|\|]PAM | NA | Intergenic | NA | NA |
| T6247 | AGAAAGGGGCGCCCGCTCGCCGG | | 1000 | | | | | | | | | |
| Oligo fwd | CACCAGAAAGGGGCGCCCGCTCGC | | | | | | | | | | | |
| Oligo rev | AAACGCGAGCGGGCGCCCCTTTCT | | | | | | | | | | | |
| Chromosome | start | | end | strand | MM | target_seq | PAM | alignment | distance | position | gene name | gene id |
| GU305914.1 | | 88235 | 88257 | + | 0 | AGAAAGGGGCGCCCGCTCGC | CGG | \|\|\|\|\|\|\|\|[\|\|\|\|\|\|\|\|\|\|\|\|]PAM | NA | Intergenic | NA | NA |
| T6248 | GCGCCCGCTCGCCGGACGAGCGG | | 1000 | | | | | | | | | |
| Oligo fwd | CACCGCGCCCGCTCGCCGGACGAG | | | | | | | | | | | |
| Oligo rev | AAACCTCGTCCGGCGAGCGGGCGC | | | | | | | | | | | |
| Chromosome | start | | end | strand | MM | target_seq | PAM | alignment | distance | position | gene name | gene id |
| GU305914.1 | | 88243 | 88265 | + | 0 | GCGCCCGCTCGCCGGACGAG | CGG | \|\|\|\|\|\|\|\|[\|\|\|\|\|\|\|\|\|\|\|\|]PAM | NA | Intergenic | NA | NA |
| T6249 | CCCGCTCGCCGGACGAGCGGCGG | | 1000 | | | | | | | | | |
| Oligo fwd | CACCCCCGCTCGCCGGACGAGCGG | | | | | | | | | | | |
| Oligo rev | AAACCCGCTCGTCCGGCGAGCGGG | | | | | | | | | | | |
| Chromosome | start | | end | strand | MM | target_seq | PAM | alignment | distance | position | gene name | gene id |
| GU305914.1 | | 88246 | 88268 | + | 0 | CCCGCTCGCCGGACGAGCGG | CGG | \|\|\|\|\|\|\|\|[\|\|\|\|\|\|\|\|\|\|\|\|]PAM | NA | Intergenic | NA | NA |
| GU305914.1 | | 99540 | 99562 | + | 4 | CCTCCTCGCCCGACGAGCGT | TGG | \|\|--\|\|\|\|[\|\|-\|\|\|\|\|\|\|\|-]PAM | NA | Intergenic | NA | NA |
| T6250 | CTCGCCGGACGAGCGGCGGACGG | | 1000 | | | | | | | | | |
| Oligo fwd | CACCCTCGCCGGACGAGCGGCGGA | | | | | | | | | | | |
| Oligo rev | AAACTCCGCCGCTCGTCCGGCGAG | | | | | | | | | | | |
| Chromosome | start | | end | strand | MM | target_seq | PAM | alignment | distance | position | gene name | gene id |
| GU305914.1 | | 88250 | 88272 | + | 0 | CTCGCCGGACGAGCGGCGGA | CGG | \|\|\|\|\|\|\|\|[\|\|\|\|\|\|\|\|\|\|\|\|]PAM | NA | Intergenic | NA | NA |
| GU305914.1 | | 34403 | 34425 | - | 4 | CCCGGCAGCCGAGCGGCGGA | GGG | \|-\|\|-\|-\|[-\|\|\|\|\|\|\|\|\|\|\|]PAM | NA | Intergenic | NA | NA |

T6251 TCGCCGGACGAGCGGCGGACGGG 1000
Oligo fwd CACCTCGCCGGACGAGCGGCGGAC
Oligo rev AAACGTCCGCCGCTCGTCCGGCGA
Chromosome start end strand MM target_seq PAM alignment distance position gene name gene id
GU305914.1 88251 88273 + 0 TCGCCGGACGAGCGGCGGAC GGG ||||||||[||||||||||||]PAM NA Intergenic NA NA

T6252 GGACGAGCGGCGGACGGGAAAGG 1000
Oligo fwd CACCGGACGAGCGGCGGACGGGAA
Oligo rev AAACTTCCCGTCCGCCGCTCGTCC
Chromosome start end strand MM target_seq PAM alignment distance position gene name gene id
GU305914.1 88256 88278 + 0 GGACGAGCGGCGGACGGGAA AGG ||||||||[||||||||||||]PAM NA Intergenic NA NA

T6253 GCGGCGGACGGGAAAGGAGAAGG 1000
Oligo fwd CACCGCGGCGGACGGGAAAGGAGA
Oligo rev AAACTCTCCTTTCCCGTCCGCCGC
Chromosome start end strand MM target_seq PAM alignment distance position gene name gene id
GU305914.1 88262 88284 + 0 GCGGCGGACGGGAAAGGAGA AGG ||||||||[||||||||||||]PAM NA Intergenic NA NA

T6254 CGGGAAAGGAGAAGGAGTCGAGG 1000
Oligo fwd CACCCGGGAAAGGAGAAGGAGTCG
Oligo rev AAACCGACTCCTTCTCCTTTCCCG
Chromosome start end strand MM target_seq PAM alignment distance position gene name gene id
GU305914.1 88270 88292 + 0 CGGGAAAGGAGAAGGAGTCG AGG ||||||||[||||||||||||]PAM NA Intergenic NA NA

T6255 GGGAAAGGAGAAGGAGTCGAGGG 1000
Oligo fwd CACCGGGAAAGGAGAAGGAGTCGA
Oligo rev AAACTCGACTCCTTCTCCTTTCCC
Chromosome start end strand MM target_seq PAM alignment distance position gene name gene id
GU305914.1 88271 88293 + 0 GGGAAAGGAGAAGGAGTCGA GGG ||||||||[||||||||||||]PAM NA Intergenic NA NA

T6256 GGAAAGGAGAAGGAGTCGAGGGG 1000
Oligo fwd CACCGGAAAGGAGAAGGAGTCGAG
Oligo rev AAACCTCGACTCCTTCTCCTTTCC
Chromosome start end strand MM target_seq PAM alignment distance position gene name gene id
GU305914.1 88272 88294 + 0 GGAAAGGAGAAGGAGTCGAG GGG ||||||||[||||||||||||]PAM NA Intergenic NA NA

T6257 GAAAGGAGAAGGAGTCGAGGGGG 1000
Oligo fwd CACCGAAAGGAGAAGGAGTCGAGG
Oligo rev AAACCCTCGACTCCTTCTCCTTTC
Chromosome start end strand MM target_seq PAM alignment distance position gene name gene id
GU305914.1 88273 88295 + 0 GAAAGGAGAAGGAGTCGAGG GGG ||||||||[||||||||||||]PAM NA Intergenic NA NA

T6258 AGGAGAAGGAGTCGAGGGGGCGG 1000
Oligo fwd CACCAGGAGAAGGAGTCGAGGGGG
Oligo rev AAACCCCCCTCGACTCCTTCTCCT
Chromosome start end strand MM target_seq PAM alignment distance position gene name gene id
GU305914.1 88276 88298 + 0 AGGAGAAGGAGTCGAGGGGG CGG ||||||||[||||||||||||]PAM NA Intergenic NA NA

T6259 GGAGAAGGAGTCGAGGGGGCGGG 1000
Oligo fwd CACCGGAGAAGGAGTCGAGGGGGC
Oligo rev AAACGCCCCCTCGACTCCTTCTCC
Chromosome start end strand MM target_seq PAM alignment distance position gene name gene id
GU305914.1 88277 88299 + 0 GGAGAAGGAGTCGAGGGGGC GGG ||||||||[||||||||||||]PAM NA Intergenic NA NA

T6260 GAGAAGGAGTCGAGGGGGCGGGG 1000
Oligo fwd CACCGAGAAGGAGTCGAGGGGGCG
Oligo rev AAACCGCCCCCTCGACTCCTTCTC
Chromosome start end strand MM target_seq PAM alignment distance position gene name gene id
GU305914.1 88278 88300 + 0 GAGAAGGAGTCGAGGGGGCG GGG ||||||||[||||||||||||]PAM NA Intergenic NA NA

T6261 AAGGAGTCGAGGGGGCGGGGAGG 1000
Oligo fwd CACCAAGGAGTCGAGGGGGCGGGG
Oligo rev AAACCCCCGCCCCCTCGACTCCTT
Chromosome start end strand MM target_seq PAM alignment distance position gene name gene id
GU305914.1 88281 88303 + 0 AAGGAGTCGAGGGGGCGGGG AGG ||||||||[||||||||||||]PAM NA Intergenic NA NA

T6262 TCACAACCGAGACCGTTTGCTGG 1000
Oligo fwd CACCTCACAACCGAGACCGTTTGC
Oligo rev AAACGCAAACGGTCTCGGTTGTGA
Chromosome start end strand MM target_seq PAM alignment distance position gene name gene id
GU305914.1 88314 88336 + 0 TCACAACCGAGACCGTTTGC TGG ||||||||[||||||||||||]PAM NA Intergenic NA NA

T6263 CACAACCGAGACCGTTTGCTGGG 1000
Oligo fwd CACCCACAACCGAGACCGTTTGCT
Oligo rev AAACAGCAAACGGTCTCGGTTGTG
Chromosome start end strand MM target_seq PAM alignment distance position gene name gene id
GU305914.1 88315 88337 + 0 CACAACCGAGACCGTTTGCT GGG ||||||||[||||||||||||]PAM NA Intergenic NA NA

T6264 GCACGTCGTCGCCGTTGATGAGG 1000
Oligo fwd CACCGCACGTCGTCGCCGTTGATG
Oligo rev AAACCATCAACGGCGACGACGTGC
Chromosome start end strand MM target_seq PAM alignment distance position gene name gene id
GU305914.1 88361 88383 + 0 GCACGTCGTCGCCGTTGATG AGG ||||||||[||||||||||||]PAM NA Intergenic NA NA

T6265 GTCGTCGCCGTTGATGAGGCCGG 1000
Oligo fwd CACCGTCGTCGCCGTTGATGAGGC
Oligo rev AAACGCCTCATCAACGGCGACGAC
Chromosome start end strand MM target_seq PAM alignment distance position gene name gene id

<table>
<tr><td>GU305914.1</td><td>88365</td><td>88387</td><td>+</td><td>0</td><td>GTCGTCGCCGTTGATGAGGC</td><td>CGG</td><td>||||||||[||||||||||||]PAM</td><td>NA</td><td>Intergenic</td><td>NA</td><td>NA</td></tr>
<tr><td>T6266</td><td>GAGGCCGGCGTTCACGTCCTCGG</td><td>1000</td><td></td><td></td><td></td><td></td><td></td><td></td><td></td><td></td><td></td></tr>
<tr><td>Oligo fwd</td><td>CACCGAGGCCGGCGTTCACGTCCT</td><td></td><td></td><td></td><td></td><td></td><td></td><td></td><td></td><td></td><td></td></tr>
<tr><td>Oligo rev</td><td>AAACAGGACGTGAACGCCGGCCTC</td><td></td><td></td><td></td><td></td><td></td><td></td><td></td><td></td><td></td><td></td></tr>
<tr><td>Chromosome</td><td>start</td><td>end</td><td>strand</td><td>MM</td><td>target_seq</td><td>PAM</td><td>alignment</td><td>distance</td><td>position</td><td>gene name</td><td>gene id</td></tr>
<tr><td>GU305914.1</td><td>88380</td><td>88402</td><td>+</td><td>0</td><td>GAGGCCGGCGTTCACGTCCT</td><td>CGG</td><td>||||||||[||||||||||||]PAM</td><td>NA</td><td>Intergenic</td><td>NA</td><td>NA</td></tr>
<tr><td>T6267</td><td>GTCGCGCCTCGAAAACAGAGCGG</td><td>1000</td><td></td><td></td><td></td><td></td><td></td><td></td><td></td><td></td><td></td></tr>
<tr><td>Oligo fwd</td><td>CACCGTCGCGCCTCGAAAACAGAG</td><td></td><td></td><td></td><td></td><td></td><td></td><td></td><td></td><td></td><td></td></tr>
<tr><td>Oligo rev</td><td>AAACCTCTGTTTTCGAGGCGCGAC</td><td></td><td></td><td></td><td></td><td></td><td></td><td></td><td></td><td></td><td></td></tr>
<tr><td>Chromosome</td><td>start</td><td>end</td><td>strand</td><td>MM</td><td>target_seq</td><td>PAM</td><td>alignment</td><td>distance</td><td>position</td><td>gene name</td><td>gene id</td></tr>
<tr><td>GU305914.1</td><td>88407</td><td>88429</td><td>+</td><td>0</td><td>GTCGCGCCTCGAAAACAGAG</td><td>CGG</td><td>||||||||[||||||||||||]PAM</td><td>NA</td><td>Intergenic</td><td>NA</td><td>NA</td></tr>
<tr><td>T6268</td><td>CAGAGCGGAGAAATCGTACATGG</td><td>1000</td><td></td><td></td><td></td><td></td><td></td><td></td><td></td><td></td><td></td></tr>
<tr><td>Oligo fwd</td><td>CACCCAGAGCGGAGAAATCGTACA</td><td></td><td></td><td></td><td></td><td></td><td></td><td></td><td></td><td></td><td></td></tr>
<tr><td>Oligo rev</td><td>AAACTGTACGATTTCTCCGCTCTG</td><td></td><td></td><td></td><td></td><td></td><td></td><td></td><td></td><td></td><td></td></tr>
<tr><td>Chromosome</td><td>start</td><td>end</td><td>strand</td><td>MM</td><td>target_seq</td><td>PAM</td><td>alignment</td><td>distance</td><td>position</td><td>gene name</td><td>gene id</td></tr>
<tr><td>GU305914.1</td><td>88422</td><td>88444</td><td>+</td><td>0</td><td>CAGAGCGGAGAAATCGTACA</td><td>TGG</td><td>||||||||[||||||||||||]PAM</td><td>NA</td><td>Intergenic</td><td>NA</td><td>NA</td></tr>
<tr><td>T6269</td><td>AGAGCGGAGAAATCGTACATGGG</td><td>1000</td><td></td><td></td><td></td><td></td><td></td><td></td><td></td><td></td><td></td></tr>
<tr><td>Oligo fwd</td><td>CACCAGAGCGGAGAAATCGTACAT</td><td></td><td></td><td></td><td></td><td></td><td></td><td></td><td></td><td></td><td></td></tr>
<tr><td>Oligo rev</td><td>AAACATGTACGATTTCTCCGCTCT</td><td></td><td></td><td></td><td></td><td></td><td></td><td></td><td></td><td></td><td></td></tr>
<tr><td>Chromosome</td><td>start</td><td>end</td><td>strand</td><td>MM</td><td>target_seq</td><td>PAM</td><td>alignment</td><td>distance</td><td>position</td><td>gene name</td><td>gene id</td></tr>
<tr><td>GU305914.1</td><td>88423</td><td>88445</td><td>+</td><td>0</td><td>AGAGCGGAGAAATCGTACAT</td><td>GGG</td><td>||||||||[||||||||||||]PAM</td><td>NA</td><td>Intergenic</td><td>NA</td><td>NA</td></tr>
<tr><td>T6270</td><td>CATGGGACCGCTCTCGTCCGCGG</td><td>1000</td><td></td><td></td><td></td><td></td><td></td><td></td><td></td><td></td><td></td></tr>
<tr><td>Oligo fwd</td><td>CACCCATGGGACCGCTCTCGTCCG</td><td></td><td></td><td></td><td></td><td></td><td></td><td></td><td></td><td></td><td></td></tr>
<tr><td>Oligo rev</td><td>AAACCGGACGAGAGCGGTCCCATG</td><td></td><td></td><td></td><td></td><td></td><td></td><td></td><td></td><td></td><td></td></tr>
<tr><td>Chromosome</td><td>start</td><td>end</td><td>strand</td><td>MM</td><td>target_seq</td><td>PAM</td><td>alignment</td><td>distance</td><td>position</td><td>gene name</td><td>gene id</td></tr>
<tr><td>GU305914.1</td><td>88440</td><td>88462</td><td>+</td><td>0</td><td>CATGGGACCGCTCTCGTCCG</td><td>CGG</td><td>||||||||[||||||||||||]PAM</td><td>NA</td><td>Intergenic</td><td>NA</td><td>NA</td></tr>
<tr><td>T6271</td><td>CGAGCCGCCACCGCCAGCAGCGG</td><td>1000</td><td></td><td></td><td></td><td></td><td></td><td></td><td></td><td></td><td></td></tr>
<tr><td>Oligo fwd</td><td>CACCCGAGCCGCCACCGCCAGCAG</td><td></td><td></td><td></td><td></td><td></td><td></td><td></td><td></td><td></td><td></td></tr>
<tr><td>Oligo rev</td><td>AAACCTGCTGGCGGTGGCGGCTCG</td><td></td><td></td><td></td><td></td><td></td><td></td><td></td><td></td><td></td><td></td></tr>
<tr><td>Chromosome</td><td>start</td><td>end</td><td>strand</td><td>MM</td><td>target_seq</td><td>PAM</td><td>alignment</td><td>distance</td><td>position</td><td>gene name</td><td>gene id</td></tr>
<tr><td>GU305914.1</td><td>88491</td><td>88513</td><td>+</td><td>0</td><td>CGAGCCGCCACCGCCAGCAG</td><td>CGG</td><td>||||||||[||||||||||||]PAM</td><td>NA</td><td>Intergenic</td><td>NA</td><td>NA</td></tr>
<tr><td>T6272</td><td>GGAGCCCGCCGCCGTCGCGTTGG</td><td>1000</td><td></td><td></td><td></td><td></td><td></td><td></td><td></td><td></td><td></td></tr>
<tr><td>Oligo fwd</td><td>CACCGGAGCCCGCCGCCGTCGCGT</td><td></td><td></td><td></td><td></td><td></td><td></td><td></td><td></td><td></td><td></td></tr>
<tr><td>Oligo rev</td><td>AAACACGCGACGGCGGCGGGCTCC</td><td></td><td></td><td></td><td></td><td></td><td></td><td></td><td></td><td></td><td></td></tr>
<tr><td>Chromosome</td><td>start</td><td>end</td><td>strand</td><td>MM</td><td>target_seq</td><td>PAM</td><td>alignment</td><td>distance</td><td>position</td><td>gene name</td><td>gene id</td></tr>
<tr><td>GU305914.1</td><td>88512</td><td>88534</td><td>+</td><td>0</td><td>GGAGCCCGCCGCCGTCGCGT</td><td>TGG</td><td>||||||||[||||||||||||]PAM</td><td>NA</td><td>Intergenic</td><td>NA</td><td>NA</td></tr>
<tr><td>GU305914.1</td><td>138477</td><td>138499</td><td>+</td><td>4</td><td>GGCGCTCGCCGCCGTCGAGG</td><td>AGG</td><td>||-||-||[|||||||||-|-]PAM</td><td>NA</td><td>Intergenic</td><td>NA</td><td>NA</td></tr>
<tr><td>T6273</td><td>GCCCGCCGCCGTCGCGTTGGTGG</td><td>1000</td><td></td><td></td><td></td><td></td><td></td><td></td><td></td><td></td><td></td></tr>
<tr><td>Oligo fwd</td><td>CACCGCCCGCCGCCGTCGCGTTGG</td><td></td><td></td><td></td><td></td><td></td><td></td><td></td><td></td><td></td><td></td></tr>
<tr><td>Oligo rev</td><td>AAACCCAACGCGACGGCGGCGGGC</td><td></td><td></td><td></td><td></td><td></td><td></td><td></td><td></td><td></td><td></td></tr>
<tr><td>Chromosome</td><td>start</td><td>end</td><td>strand</td><td>MM</td><td>target_seq</td><td>PAM</td><td>alignment</td><td>distance</td><td>position</td><td>gene name</td><td>gene id</td></tr>
<tr><td>GU305914.1</td><td>88515</td><td>88537</td><td>+</td><td>0</td><td>GCCCGCCGCCGTCGCGTTGG</td><td>TGG</td><td>||||||||[||||||||||||]PAM</td><td>NA</td><td>Intergenic</td><td>NA</td><td>NA</td></tr>
<tr><td>T6274</td><td>CGCCGCCGTCGCGTTGGTGGCGG</td><td>1000</td><td></td><td></td><td></td><td></td><td></td><td></td><td></td><td></td><td></td></tr>
<tr><td>Oligo fwd</td><td>CACCCGCCGCCGTCGCGTTGGTGG</td><td></td><td></td><td></td><td></td><td></td><td></td><td></td><td></td><td></td><td></td></tr>
<tr><td>Oligo rev</td><td>AAACCCACCAACGCGACGGCGGCG</td><td></td><td></td><td></td><td></td><td></td><td></td><td></td><td></td><td></td><td></td></tr>
<tr><td>Chromosome</td><td>start</td><td>end</td><td>strand</td><td>MM</td><td>target_seq</td><td>PAM</td><td>alignment</td><td>distance</td><td>position</td><td>gene name</td><td>gene id</td></tr>
<tr><td>GU305914.1</td><td>88518</td><td>88540</td><td>+</td><td>0</td><td>CGCCGCCGTCGCGTTGGTGG</td><td>CGG</td><td>||||||||[||||||||||||]PAM</td><td>NA</td><td>Intergenic</td><td>NA</td><td>NA</td></tr>
<tr><td>T6275</td><td>GGCGGTCGCCTCGCGCAGAGAGG</td><td>1000</td><td></td><td></td><td></td><td></td><td></td><td></td><td></td><td></td><td></td></tr>
<tr><td>Oligo fwd</td><td>CACCGGCGGTCGCCTCGCGCAGAG</td><td></td><td></td><td></td><td></td><td></td><td></td><td></td><td></td><td></td><td></td></tr>
<tr><td>Oligo rev</td><td>AAACCTCTGCGCGAGGCGACCGCC</td><td></td><td></td><td></td><td></td><td></td><td></td><td></td><td></td><td></td><td></td></tr>
<tr><td>Chromosome</td><td>start</td><td>end</td><td>strand</td><td>MM</td><td>target_seq</td><td>PAM</td><td>alignment</td><td>distance</td><td>position</td><td>gene name</td><td>gene id</td></tr>
<tr><td>GU305914.1</td><td>88536</td><td>88558</td><td>+</td><td>0</td><td>GGCGGTCGCCTCGCGCAGAG</td><td>AGG</td><td>||||||||[||||||||||||]PAM</td><td>NA</td><td>Intergenic</td><td>NA</td><td>NA</td></tr>
<tr><td>T6276</td><td>AAAGTCCGTCACCCCCCTCTCGG</td><td>1000</td><td></td><td></td><td></td><td></td><td></td><td></td><td></td><td></td><td></td></tr>
<tr><td>Oligo fwd</td><td>CACCAAAGTCCGTCACCCCCCTCT</td><td></td><td></td><td></td><td></td><td></td><td></td><td></td><td></td><td></td><td></td></tr>
<tr><td>Oligo rev</td><td>AAACAGAGGGGGGTGACGGACTTT</td><td></td><td></td><td></td><td></td><td></td><td></td><td></td><td></td><td></td><td></td></tr>
<tr><td>Chromosome</td><td>start</td><td>end</td><td>strand</td><td>MM</td><td>target_seq</td><td>PAM</td><td>alignment</td><td>distance</td><td>position</td><td>gene name</td><td>gene id</td></tr>
<tr><td>GU305914.1</td><td>88566</td><td>88588</td><td>+</td><td>0</td><td>AAAGTCCGTCACCCCCCTCT</td><td>CGG</td><td>||||||||[||||||||||||]PAM</td><td>NA</td><td>Intergenic</td><td>NA</td><td>NA</td></tr>
<tr><td>T6277</td><td>ACTGCGCATCTTGCGATACATGG</td><td>1000</td><td></td><td></td><td></td><td></td><td></td><td></td><td></td><td></td><td></td></tr>
<tr><td>Oligo fwd</td><td>CACCACTGCGCATCTTGCGATACA</td><td></td><td></td><td></td><td></td><td></td><td></td><td></td><td></td><td></td><td></td></tr>
<tr><td>Oligo rev</td><td>AAACTGTATCGCAAGATGCGCAGT</td><td></td><td></td><td></td><td></td><td></td><td></td><td></td><td></td><td></td><td></td></tr>
<tr><td>Chromosome</td><td>start</td><td>end</td><td>strand</td><td>MM</td><td>target_seq</td><td>PAM</td><td>alignment</td><td>distance</td><td>position</td><td>gene name</td><td>gene id</td></tr>
<tr><td>GU305914.1</td><td>88593</td><td>88615</td><td>+</td><td>0</td><td>ACTGCGCATCTTGCGATACA</td><td>TGG</td><td>||||||||[||||||||||||]PAM</td><td>NA</td><td>Intergenic</td><td>NA</td><td>NA</td></tr>
<tr><td>T6278</td><td>CATCTTGCGATACATGGACCTGG</td><td>1000</td><td></td><td></td><td></td><td></td><td></td><td></td><td></td><td></td><td></td></tr>
<tr><td>Oligo fwd</td><td>CACCCATCTTGCGATACATGGACC</td><td></td><td></td><td></td><td></td><td></td><td></td><td></td><td></td><td></td><td></td></tr>
<tr><td>Oligo rev</td><td>AAACGGTCCATGTATCGCAAGATG</td><td></td><td></td><td></td><td></td><td></td><td></td><td></td><td></td><td></td><td></td></tr>
<tr><td>Chromosome</td><td>start</td><td>end</td><td>strand</td><td>MM</td><td>target_seq</td><td>PAM</td><td>alignment</td><td>distance</td><td>position</td><td>gene name</td><td>gene id</td></tr>
<tr><td>GU305914.1</td><td>88599</td><td>88621</td><td>+</td><td>0</td><td>CATCTTGCGATACATGGACC</td><td>TGG</td><td>||||||||[||||||||||||]PAM</td><td>NA</td><td>Intergenic</td><td>NA</td><td>NA</td></tr>
<tr><td>T6279</td><td>ACCTGGCGATGTACTCCTGCTGG</td><td>1000</td><td></td><td></td><td></td><td></td><td></td><td></td><td></td><td></td><td></td></tr>
<tr><td>Oligo fwd</td><td>CACCACCTGGCGATGTACTCCTGC</td><td></td><td></td><td></td><td></td><td></td><td></td><td></td><td></td><td></td><td></td></tr>
<tr><td>Oligo rev</td><td>AAACGCAGGAGTACATCGCCAGGT</td><td></td><td></td><td></td><td></td><td></td><td></td><td></td><td></td><td></td><td></td></tr>
<tr><td>Chromosome</td><td>start</td><td>end</td><td>strand</td><td>MM</td><td>target_seq</td><td>PAM</td><td>alignment</td><td>distance</td><td>position</td><td>gene name</td><td>gene id</td></tr>
<tr><td>GU305914.1</td><td>88616</td><td>88638</td><td>+</td><td>0</td><td>ACCTGGCGATGTACTCCTGC</td><td>TGG</td><td>||||||||[||||||||||||]PAM</td><td>NA</td><td>Intergenic</td><td>NA</td><td>NA</td></tr>
<tr><td>T6280</td><td>CCTGCTGGTCCACGTAGAAGAGG</td><td>1000</td><td></td><td></td><td></td><td></td><td></td><td></td><td></td><td></td><td></td></tr>
<tr><td>Oligo fwd</td><td>CACCCCTGCTGGTCCACGTAGAAG</td><td></td><td></td><td></td><td></td><td></td><td></td><td></td><td></td><td></td><td></td></tr>
</table>

| | | | | | | | | | | | | |
|---|---|---|---|---|---|---|---|---|---|---|---|---|
| Oligo rev | AAACCTTCTACGTGGACCAGCAGG | | | | | | | | | | | |
| Chromosome | start | | end | strand | MM | target_seq | PAM | alignment | distance | position | gene name | gene id |
| GU305914.1 | | 88631 | 88653 | + | 0 | CCTGCTGGTCCACGTAGAAG | AGG | \|\|\|\|\|\|\|\|[\|\|\|\|\|\|\|\|\|\|\|\|]PAM | NA | Intergenic | NA | NA |
| T6281 | GAAGAGGATGTCGTCGATCTCGG | | 1000 | | | | | | | | | |
| Oligo fwd | CACCGAAGAGGATGTCGTCGATCT | | | | | | | | | | | |
| Oligo rev | AAACAGATCGACGACATCCTCTTC | | | | | | | | | | | |
| Chromosome | start | | end | strand | MM | target_seq | PAM | alignment | distance | position | gene name | gene id |
| GU305914.1 | | 88647 | 88669 | + | 0 | GAAGAGGATGTCGTCGATCT | CGG | \|\|\|\|\|\|\|\|[\|\|\|\|\|\|\|\|\|\|\|\|]PAM | NA | Intergenic | NA | NA |
| T6282 | AAGAGGATGTCGTCGATCTCGGG | | 1000 | | | | | | | | | |
| Oligo fwd | CACCAAGAGGATGTCGTCGATCTC | | | | | | | | | | | |
| Oligo rev | AAACGAGATCGACGACATCCTCTT | | | | | | | | | | | |
| Chromosome | start | | end | strand | MM | target_seq | PAM | alignment | distance | position | gene name | gene id |
| GU305914.1 | | 88648 | 88670 | + | 0 | AAGAGGATGTCGTCGATCTC | GGG | \|\|\|\|\|\|\|\|[\|\|\|\|\|\|\|\|\|\|\|\|]PAM | NA | Intergenic | NA | NA |
| T6283 | AGAGGATGTCGTCGATCTCGGGG | | 1000 | | | | | | | | | |
| Oligo fwd | CACCAGAGGATGTCGTCGATCTCG | | | | | | | | | | | |
| Oligo rev | AAACCGAGATCGACGACATCCTCT | | | | | | | | | | | |
| Chromosome | start | | end | strand | MM | target_seq | PAM | alignment | distance | position | gene name | gene id |
| GU305914.1 | | 88649 | 88671 | + | 0 | AGAGGATGTCGTCGATCTCG | GGG | \|\|\|\|\|\|\|\|[\|\|\|\|\|\|\|\|\|\|\|\|]PAM | NA | Intergenic | NA | NA |
| T6284 | CGGGGATGTCGCCCTCGTCGCGG | | 1000 | | | | | | | | | |
| Oligo fwd | CACCCGGGGATGTCGCCCTCGTCG | | | | | | | | | | | |
| Oligo rev | AAACCGACGAGGGCGACATCCCCG | | | | | | | | | | | |
| Chromosome | start | | end | strand | MM | target_seq | PAM | alignment | distance | position | gene name | gene id |
| GU305914.1 | | 88667 | 88689 | + | 0 | CGGGGATGTCGCCCTCGTCG | CGG | \|\|\|\|\|\|\|\|[\|\|\|\|\|\|\|\|\|\|\|\|]PAM | NA | Intergenic | NA | NA |
| T6285 | GTCGCGGCCCTCCATGAGCTCGG | | 1000 | | | | | | | | | |
| Oligo fwd | CACCGTCGCGGCCCTCCATGAGCT | | | | | | | | | | | |
| Oligo rev | AAACAGCTCATGGAGGGCCGCGAC | | | | | | | | | | | |
| Chromosome | start | | end | strand | MM | target_seq | PAM | alignment | distance | position | gene name | gene id |
| GU305914.1 | | 88683 | 88705 | + | 0 | GTCGCGGCCCTCCATGAGCT | CGG | \|\|\|\|\|\|\|\|[\|\|\|\|\|\|\|\|\|\|\|\|]PAM | NA | Intergenic | NA | NA |
| T6286 | GCCCTCCATGAGCTCGGCGATGG | | 1000 | | | | | | | | | |
| Oligo fwd | CACCGCCCTCCATGAGCTCGGCGA | | | | | | | | | | | |
| Oligo rev | AAACTCGCCGAGCTCATGGAGGGC | | | | | | | | | | | |
| Chromosome | start | | end | strand | MM | target_seq | PAM | alignment | distance | position | gene name | gene id |
| GU305914.1 | | 88689 | 88711 | + | 0 | GCCCTCCATGAGCTCGGCGA | TGG | \|\|\|\|\|\|\|\|[\|\|\|\|\|\|\|\|\|\|\|\|]PAM | NA | Intergenic | NA | NA |
| T6287 | CTCGGCGATGGCCGCCAGCTCGG | | 1000 | | | | | | | | | |
| Oligo fwd | CACCCTCGGCGATGGCCGCCAGCT | | | | | | | | | | | |
| Oligo rev | AAACAGCTGGCGGCCATCGCCGAG | | | | | | | | | | | |
| Chromosome | start | | end | strand | MM | target_seq | PAM | alignment | distance | position | gene name | gene id |
| GU305914.1 | | 88701 | 88723 | + | 0 | CTCGGCGATGGCCGCCAGCT | CGG | \|\|\|\|\|\|\|\|[\|\|\|\|\|\|\|\|\|\|\|\|]PAM | NA | Intergenic | NA | NA |
| T6288 | TCGGCGATGGCCGCCAGCTCGGG | | 1000 | | | | | | | | | |
| Oligo fwd | CACCTCGGCGATGGCCGCCAGCTC | | | | | | | | | | | |
| Oligo rev | AAACGAGCTGGCGGCCATCGCCGA | | | | | | | | | | | |
| Chromosome | start | | end | strand | MM | target_seq | PAM | alignment | distance | position | gene name | gene id |
| GU305914.1 | | 88702 | 88724 | + | 0 | TCGGCGATGGCCGCCAGCTC | GGG | \|\|\|\|\|\|\|\|[\|\|\|\|\|\|\|\|\|\|\|\|]PAM | NA | Intergenic | NA | NA |
| T6289 | GCGCCGCCAGCACCCGCGAGCGG | | 1000 | | | | | | | | | |
| Oligo fwd | CACCGCGCCGCCAGCACCCGCGAG | | | | | | | | | | | |
| Oligo rev | AAACCTCGCGGGTGCTGGCGGCGC | | | | | | | | | | | |
| Chromosome | start | | end | strand | MM | target_seq | PAM | alignment | distance | position | gene name | gene id |
| GU305914.1 | | 88751 | 88773 | + | 0 | GCGCCGCCAGCACCCGCGAG | CGG | \|\|\|\|\|\|\|\|[\|\|\|\|\|\|\|\|\|\|\|\|]PAM | NA | Intergenic | NA | NA |
| GU305914.1 | | 177474 | 177496 | - | 4 | GCACCGTCAGCACGCTCGAG | TGG | \|\|-\|\|\|-\|[\|\|\|\|\|-\|-\|\|\|\|]PAM | NA | Intergenic | NA | NA |
| T6290 | CGTCGTACATGACGTTCTCGCGG | | 1000 | | | | | | | | | |
| Oligo fwd | CACCCGTCGTACATGACGTTCTCG | | | | | | | | | | | |
| Oligo rev | AAACCGAGAACGTCATGTACGACG | | | | | | | | | | | |
| Chromosome | start | | end | strand | MM | target_seq | PAM | alignment | distance | position | gene name | gene id |
| GU305914.1 | | 88784 | 88806 | + | 0 | CGTCGTACATGACGTTCTCG | CGG | \|\|\|\|\|\|\|\|[\|\|\|\|\|\|\|\|\|\|\|\|]PAM | NA | Intergenic | NA | NA |
| T6291 | TGACGTTCTCGCGGCGCGAGCGG | | 1000 | | | | | | | | | |
| Oligo fwd | CACCTGACGTTCTCGCGGCGCGAG | | | | | | | | | | | |
| Oligo rev | AAACCTCGCGCCGCGAGAACGTCA | | | | | | | | | | | |
| Chromosome | start | | end | strand | MM | target_seq | PAM | alignment | distance | position | gene name | gene id |
| GU305914.1 | | 88793 | 88815 | + | 0 | TGACGTTCTCGCGGCGCGAG | CGG | \|\|\|\|\|\|\|\|[\|\|\|\|\|\|\|\|\|\|\|\|]PAM | NA | Intergenic | NA | NA |
| T6292 | CGTTCTCGCGGCGCGAGCGGCGG | | 1000 | | | | | | | | | |
| Oligo fwd | CACCCGTTCTCGCGGCGCGAGCGG | | | | | | | | | | | |
| Oligo rev | AAACCCGCTCGCGCCGCGAGAACG | | | | | | | | | | | |
| Chromosome | start | | end | strand | MM | target_seq | PAM | alignment | distance | position | gene name | gene id |
| GU305914.1 | | 88796 | 88818 | + | 0 | CGTTCTCGCGGCGCGAGCGG | CGG | \|\|\|\|\|\|\|\|[\|\|\|\|\|\|\|\|\|\|\|\|]PAM | NA | Intergenic | NA | NA |
| T6293 | GATGAGCACGTCCACCAGCTTGG | | 1000 | | | | | | | | | |
| Oligo fwd | CACCGATGAGCACGTCCACCAGCT | | | | | | | | | | | |
| Oligo rev | AAACAGCTGGTGGACGTGCTCATC | | | | | | | | | | | |
| Chromosome | start | | end | strand | MM | target_seq | PAM | alignment | distance | position | gene name | gene id |
| GU305914.1 | | 88818 | 88840 | + | 0 | GATGAGCACGTCCACCAGCT | TGG | \|\|\|\|\|\|\|\|[\|\|\|\|\|\|\|\|\|\|\|\|]PAM | NA | Intergenic | NA | NA |
| T6294 | GAGCACGTCCACCAGCTTGGTGG | | 1000 | | | | | | | | | |
| Oligo fwd | CACCGAGCACGTCCACCAGCTTGG | | | | | | | | | | | |
| Oligo rev | AAACCCAAGCTGGTGGACGTGCTC | | | | | | | | | | | |
| Chromosome | start | | end | strand | MM | target_seq | PAM | alignment | distance | position | gene name | gene id |
| GU305914.1 | | 88821 | 88843 | + | 0 | GAGCACGTCCACCAGCTTGG | TGG | \|\|\|\|\|\|\|\|[\|\|\|\|\|\|\|\|\|\|\|\|]PAM | NA | Intergenic | NA | NA |

<table>
<tr><td>T6295</td><td>TGTACCTGTAGCGCATGAAGCGG</td><td></td><td>1000</td><td></td><td></td><td></td><td></td><td></td><td></td><td></td><td></td><td></td></tr>
<tr><td>Oligo fwd</td><td>CACCTGTACCTGTAGCGCATGAAG</td><td></td><td></td><td></td><td></td><td></td><td></td><td></td><td></td><td></td><td></td><td></td></tr>
<tr><td>Oligo rev</td><td>AAACCTTCATGCGCTACAGGTACA</td><td></td><td></td><td></td><td></td><td></td><td></td><td></td><td></td><td></td><td></td><td></td></tr>
<tr><td>Chromosome</td><td>start</td><td></td><td>end</td><td>strand</td><td>MM</td><td>target_seq</td><td>PAM</td><td>alignment</td><td>distance</td><td>position</td><td>gene name</td><td>gene id</td></tr>
<tr><td>GU305914.1</td><td></td><td>88850</td><td>88872</td><td>+</td><td>0</td><td>TGTACCTGTAGCGCATGAAG</td><td>CGG</td><td>||||||||[||||||||||||]PAM</td><td>NA</td><td>Intergenic</td><td>NA</td><td>NA</td></tr>
<tr><td>T6296</td><td>GCGGTAGTCCTGCGTGCCCATGG</td><td></td><td>1000</td><td></td><td></td><td></td><td></td><td></td><td></td><td></td><td></td><td></td></tr>
<tr><td>Oligo fwd</td><td>CACCGCGGTAGTCCTGCGTGCCCA</td><td></td><td></td><td></td><td></td><td></td><td></td><td></td><td></td><td></td><td></td><td></td></tr>
<tr><td>Oligo rev</td><td>AAACTGGGCACGCAGGACTACCGC</td><td></td><td></td><td></td><td></td><td></td><td></td><td></td><td></td><td></td><td></td><td></td></tr>
<tr><td>Chromosome</td><td>start</td><td></td><td>end</td><td>strand</td><td>MM</td><td>target_seq</td><td>PAM</td><td>alignment</td><td>distance</td><td>position</td><td>gene name</td><td>gene id</td></tr>
<tr><td>GU305914.1</td><td></td><td>88869</td><td>88891</td><td>+</td><td>0</td><td>GCGGTAGTCCTGCGTGCCCA</td><td>TGG</td><td>||||||||[||||||||||||]PAM</td><td>NA</td><td>Intergenic</td><td>NA</td><td>NA</td></tr>
<tr><td>T6297</td><td>TGCCCATGGAGCCCGTGTTGAGG</td><td></td><td>1000</td><td></td><td></td><td></td><td></td><td></td><td></td><td></td><td></td><td></td></tr>
<tr><td>Oligo fwd</td><td>CACCTGCCCATGGAGCCCGTGTTG</td><td></td><td></td><td></td><td></td><td></td><td></td><td></td><td></td><td></td><td></td><td></td></tr>
<tr><td>Oligo rev</td><td>AAACCAACACGGGCTCCATGGGCA</td><td></td><td></td><td></td><td></td><td></td><td></td><td></td><td></td><td></td><td></td><td></td></tr>
<tr><td>Chromosome</td><td>start</td><td></td><td>end</td><td>strand</td><td>MM</td><td>target_seq</td><td>PAM</td><td>alignment</td><td>distance</td><td>position</td><td>gene name</td><td>gene id</td></tr>
<tr><td>GU305914.1</td><td></td><td>88883</td><td>88905</td><td>+</td><td>0</td><td>TGCCCATGGAGCCCGTGTTG</td><td>AGG</td><td>||||||||[||||||||||||]PAM</td><td>NA</td><td>Intergenic</td><td>NA</td><td>NA</td></tr>
<tr><td>T6298</td><td>TGGAGCCCGTGTTGAGGCTGCGG</td><td></td><td>1000</td><td></td><td></td><td></td><td></td><td></td><td></td><td></td><td></td><td></td></tr>
<tr><td>Oligo fwd</td><td>CACCTGGAGCCCGTGTTGAGGCTG</td><td></td><td></td><td></td><td></td><td></td><td></td><td></td><td></td><td></td><td></td><td></td></tr>
<tr><td>Oligo rev</td><td>AAACCAGCCTCAACACGGGCTCCA</td><td></td><td></td><td></td><td></td><td></td><td></td><td></td><td></td><td></td><td></td><td></td></tr>
<tr><td>Chromosome</td><td>start</td><td></td><td>end</td><td>strand</td><td>MM</td><td>target_seq</td><td>PAM</td><td>alignment</td><td>distance</td><td>position</td><td>gene name</td><td>gene id</td></tr>
<tr><td>GU305914.1</td><td></td><td>88889</td><td>88911</td><td>+</td><td>0</td><td>TGGAGCCCGTGTTGAGGCTG</td><td>CGG</td><td>||||||||[||||||||||||]PAM</td><td>NA</td><td>Intergenic</td><td>NA</td><td>NA</td></tr>
<tr><td>T6299</td><td>GTCCGAACTGAAACACGTCCCGG</td><td></td><td>1000</td><td></td><td></td><td></td><td></td><td></td><td></td><td></td><td></td><td></td></tr>
<tr><td>Oligo fwd</td><td>CACCGTCCGAACTGAAACACGTCC</td><td></td><td></td><td></td><td></td><td></td><td></td><td></td><td></td><td></td><td></td><td></td></tr>
<tr><td>Oligo rev</td><td>AAACGGACGTGTTTCAGTTCGGAC</td><td></td><td></td><td></td><td></td><td></td><td></td><td></td><td></td><td></td><td></td><td></td></tr>
<tr><td>Chromosome</td><td>start</td><td></td><td>end</td><td>strand</td><td>MM</td><td>target_seq</td><td>PAM</td><td>alignment</td><td>distance</td><td>position</td><td>gene name</td><td>gene id</td></tr>
<tr><td>GU305914.1</td><td></td><td>88943</td><td>88965</td><td>+</td><td>0</td><td>GTCCGAACTGAAACACGTCC</td><td>CGG</td><td>||||||||[||||||||||||]PAM</td><td>NA</td><td>Intergenic</td><td>NA</td><td>NA</td></tr>
<tr><td>T6300</td><td>TCCGAACTGAAACACGTCCCGGG</td><td></td><td>1000</td><td></td><td></td><td></td><td></td><td></td><td></td><td></td><td></td><td></td></tr>
<tr><td>Oligo fwd</td><td>CACCTCCGAACTGAAACACGTCCC</td><td></td><td></td><td></td><td></td><td></td><td></td><td></td><td></td><td></td><td></td><td></td></tr>
<tr><td>Oligo rev</td><td>AAACGGGACGTGTTTCAGTTCGGA</td><td></td><td></td><td></td><td></td><td></td><td></td><td></td><td></td><td></td><td></td><td></td></tr>
<tr><td>Chromosome</td><td>start</td><td></td><td>end</td><td>strand</td><td>MM</td><td>target_seq</td><td>PAM</td><td>alignment</td><td>distance</td><td>position</td><td>gene name</td><td>gene id</td></tr>
<tr><td>GU305914.1</td><td></td><td>88944</td><td>88966</td><td>+</td><td>0</td><td>TCCGAACTGAAACACGTCCC</td><td>GGG</td><td>||||||||[||||||||||||]PAM</td><td>NA</td><td>Intergenic</td><td>NA</td><td>NA</td></tr>
<tr><td>T6301</td><td>AAACACGTCCCGGGTGCCCATGG</td><td></td><td>1000</td><td></td><td></td><td></td><td></td><td></td><td></td><td></td><td></td><td></td></tr>
<tr><td>Oligo fwd</td><td>CACCAAACACGTCCCGGGTGCCCA</td><td></td><td></td><td></td><td></td><td></td><td></td><td></td><td></td><td></td><td></td><td></td></tr>
<tr><td>Oligo rev</td><td>AAACTGGGCACCCGGGACGTGTTT</td><td></td><td></td><td></td><td></td><td></td><td></td><td></td><td></td><td></td><td></td><td></td></tr>
<tr><td>Chromosome</td><td>start</td><td></td><td>end</td><td>strand</td><td>MM</td><td>target_seq</td><td>PAM</td><td>alignment</td><td>distance</td><td>position</td><td>gene name</td><td>gene id</td></tr>
<tr><td>GU305914.1</td><td></td><td>88953</td><td>88975</td><td>+</td><td>0</td><td>AAACACGTCCCGGGTGCCCA</td><td>TGG</td><td>||||||||[||||||||||||]PAM</td><td>NA</td><td>Intergenic</td><td>NA</td><td>NA</td></tr>
<tr><td>T6302</td><td>GTCCCGGGTGCCCATGGAGCTGG</td><td></td><td>1000</td><td></td><td></td><td></td><td></td><td></td><td></td><td></td><td></td><td></td></tr>
<tr><td>Oligo fwd</td><td>CACCGTCCCGGGTGCCCATGGAGC</td><td></td><td></td><td></td><td></td><td></td><td></td><td></td><td></td><td></td><td></td><td></td></tr>
<tr><td>Oligo rev</td><td>AAACGCTCCATGGGCACCCGGGAC</td><td></td><td></td><td></td><td></td><td></td><td></td><td></td><td></td><td></td><td></td><td></td></tr>
<tr><td>Chromosome</td><td>start</td><td></td><td>end</td><td>strand</td><td>MM</td><td>target_seq</td><td>PAM</td><td>alignment</td><td>distance</td><td>position</td><td>gene name</td><td>gene id</td></tr>
<tr><td>GU305914.1</td><td></td><td>88959</td><td>88981</td><td>+</td><td>0</td><td>GTCCCGGGTGCCCATGGAGC</td><td>TGG</td><td>||||||||[||||||||||||]PAM</td><td>NA</td><td>Intergenic</td><td>NA</td><td>NA</td></tr>
<tr><td>T6303</td><td>GGAGCTGGAGTACTTCTCGACGG</td><td></td><td>1000</td><td></td><td></td><td></td><td></td><td></td><td></td><td></td><td></td><td></td></tr>
<tr><td>Oligo fwd</td><td>CACCGGAGCTGGAGTACTTCTCGA</td><td></td><td></td><td></td><td></td><td></td><td></td><td></td><td></td><td></td><td></td><td></td></tr>
<tr><td>Oligo rev</td><td>AAACTCGAGAAGTACTCCAGCTCC</td><td></td><td></td><td></td><td></td><td></td><td></td><td></td><td></td><td></td><td></td><td></td></tr>
<tr><td>Chromosome</td><td>start</td><td></td><td>end</td><td>strand</td><td>MM</td><td>target_seq</td><td>PAM</td><td>alignment</td><td>distance</td><td>position</td><td>gene name</td><td>gene id</td></tr>
<tr><td>GU305914.1</td><td></td><td>88974</td><td>88996</td><td>+</td><td>0</td><td>GGAGCTGGAGTACTTCTCGA</td><td>CGG</td><td>||||||||[||||||||||||]PAM</td><td>NA</td><td>Intergenic</td><td>NA</td><td>NA</td></tr>
<tr><td>GU305914.1</td><td></td><td>44435</td><td>44457</td><td>+</td><td>4</td><td>CGAGCTGCAGTACCACTCGA</td><td>CGG</td><td>-||||||-[|||||--|||||]PAM</td><td>NA</td><td>Intergenic</td><td>NA</td><td>NA</td></tr>
<tr><td>T6304</td><td>CTTCTCGACGGCGAAGCTGACGG</td><td></td><td>1000</td><td></td><td></td><td></td><td></td><td></td><td></td><td></td><td></td><td></td></tr>
<tr><td>Oligo fwd</td><td>CACCCTTCTCGACGGCGAAGCTGA</td><td></td><td></td><td></td><td></td><td></td><td></td><td></td><td></td><td></td><td></td><td></td></tr>
<tr><td>Oligo rev</td><td>AAACTCAGCTTCGCCGTCGAGAAG</td><td></td><td></td><td></td><td></td><td></td><td></td><td></td><td></td><td></td><td></td><td></td></tr>
<tr><td>Chromosome</td><td>start</td><td></td><td>end</td><td>strand</td><td>MM</td><td>target_seq</td><td>PAM</td><td>alignment</td><td>distance</td><td>position</td><td>gene name</td><td>gene id</td></tr>
<tr><td>GU305914.1</td><td></td><td>88986</td><td>89008</td><td>+</td><td>0</td><td>CTTCTCGACGGCGAAGCTGA</td><td>CGG</td><td>||||||||[||||||||||||]PAM</td><td>NA</td><td>Intergenic</td><td>NA</td><td>NA</td></tr>
<tr><td>T6305</td><td>AGCTGACGGAGACGAGCGAGCGG</td><td></td><td>1000</td><td></td><td></td><td></td><td></td><td></td><td></td><td></td><td></td><td></td></tr>
<tr><td>Oligo fwd</td><td>CACCAGCTGACGGAGACGAGCGAG</td><td></td><td></td><td></td><td></td><td></td><td></td><td></td><td></td><td></td><td></td><td></td></tr>
<tr><td>Oligo rev</td><td>AAACCTCGCTCGTCTCCGTCAGCT</td><td></td><td></td><td></td><td></td><td></td><td></td><td></td><td></td><td></td><td></td><td></td></tr>
<tr><td>Chromosome</td><td>start</td><td></td><td>end</td><td>strand</td><td>MM</td><td>target_seq</td><td>PAM</td><td>alignment</td><td>distance</td><td>position</td><td>gene name</td><td>gene id</td></tr>
<tr><td>GU305914.1</td><td></td><td>89000</td><td>89022</td><td>+</td><td>0</td><td>AGCTGACGGAGACGAGCGAG</td><td>CGG</td><td>||||||||[||||||||||||]PAM</td><td>NA</td><td>Intergenic</td><td>NA</td><td>NA</td></tr>
<tr><td>T6306</td><td>GTCGAAGACGTTCTTGAGCACGG</td><td></td><td>1000</td><td></td><td></td><td></td><td></td><td></td><td></td><td></td><td></td><td></td></tr>
<tr><td>Oligo fwd</td><td>CACCGTCGAAGACGTTCTTGAGCA</td><td></td><td></td><td></td><td></td><td></td><td></td><td></td><td></td><td></td><td></td><td></td></tr>
<tr><td>Oligo rev</td><td>AAACTGCTCAAGAACGTCTTCGAC</td><td></td><td></td><td></td><td></td><td></td><td></td><td></td><td></td><td></td><td></td><td></td></tr>
<tr><td>Chromosome</td><td>start</td><td></td><td>end</td><td>strand</td><td>MM</td><td>target_seq</td><td>PAM</td><td>alignment</td><td>distance</td><td>position</td><td>gene name</td><td>gene id</td></tr>
<tr><td>GU305914.1</td><td></td><td>89025</td><td>89047</td><td>+</td><td>0</td><td>GTCGAAGACGTTCTTGAGCA</td><td>CGG</td><td>||||||||[||||||||||||]PAM</td><td>NA</td><td>Intergenic</td><td>NA</td><td>NA</td></tr>
<tr><td>T6307</td><td>TCGAAGACGTTCTTGAGCACGGG</td><td></td><td>1000</td><td></td><td></td><td></td><td></td><td></td><td></td><td></td><td></td><td></td></tr>
<tr><td>Oligo fwd</td><td>CACCTCGAAGACGTTCTTGAGCAC</td><td></td><td></td><td></td><td></td><td></td><td></td><td></td><td></td><td></td><td></td><td></td></tr>
<tr><td>Oligo rev</td><td>AAACGTGCTCAAGAACGTCTTCGA</td><td></td><td></td><td></td><td></td><td></td><td></td><td></td><td></td><td></td><td></td><td></td></tr>
<tr><td>Chromosome</td><td>start</td><td></td><td>end</td><td>strand</td><td>MM</td><td>target_seq</td><td>PAM</td><td>alignment</td><td>distance</td><td>position</td><td>gene name</td><td>gene id</td></tr>
<tr><td>GU305914.1</td><td></td><td>89026</td><td>89048</td><td>+</td><td>0</td><td>TCGAAGACGTTCTTGAGCAC</td><td>GGG</td><td>||||||||[||||||||||||]PAM</td><td>NA</td><td>Intergenic</td><td>NA</td><td>NA</td></tr>
<tr><td>T6308</td><td>GAGCACGGGCTCGCGCGTGATGG</td><td></td><td>1000</td><td></td><td></td><td></td><td></td><td></td><td></td><td></td><td></td><td></td></tr>
<tr><td>Oligo fwd</td><td>CACCGAGCACGGGCTCGCGCGTGA</td><td></td><td></td><td></td><td></td><td></td><td></td><td></td><td></td><td></td><td></td><td></td></tr>
<tr><td>Oligo rev</td><td>AAACTCACGCGCGAGCCCGTGCTC</td><td></td><td></td><td></td><td></td><td></td><td></td><td></td><td></td><td></td><td></td><td></td></tr>
<tr><td>Chromosome</td><td>start</td><td></td><td>end</td><td>strand</td><td>MM</td><td>target_seq</td><td>PAM</td><td>alignment</td><td>distance</td><td>position</td><td>gene name</td><td>gene id</td></tr>
<tr><td>GU305914.1</td><td></td><td>89040</td><td>89062</td><td>+</td><td>0</td><td>GAGCACGGGCTCGCGCGTGA</td><td>TGG</td><td>||||||||[||||||||||||]PAM</td><td>NA</td><td>Intergenic</td><td>NA</td><td>NA</td></tr>
<tr><td>T6309</td><td>CACGGGCTCGCGCGTGATGGTGG</td><td></td><td>1000</td><td></td><td></td><td></td><td></td><td></td><td></td><td></td><td></td><td></td></tr>
<tr><td>Oligo fwd</td><td>CACCCACGGGCTCGCGCGTGATGG</td><td></td><td></td><td></td><td></td><td></td><td></td><td></td><td></td><td></td><td></td><td></td></tr>
<tr><td>Oligo rev</td><td>AAACCCATCACGCGCGAGCCCGTG</td><td></td><td></td><td></td><td></td><td></td><td></td><td></td><td></td><td></td><td></td><td></td></tr>
<tr><td>Chromosome</td><td>start</td><td></td><td>end</td><td>strand</td><td>MM</td><td>target_seq</td><td>PAM</td><td>alignment</td><td>distance</td><td>position</td><td>gene name</td><td>gene id</td></tr>
</table>

| | | | | | | | | | | | |
|---|---|---|---|---|---|---|---|---|---|---|---|
| GU305914.1 | | 89043 | 89065 + | | 0 CACGGGCTCGCGCGTGATGG | TGG | \|\|\|\|\|\|\|\|[\|\|\|\|\|\|\|\|\|\|\|\|]PAM | NA | Intergenic | NA | NA |
| GU305914.1 | | 187960 | 187982 + | | 3 CACGGGCACGAGCCTGATGG | TGG | \|\|\|\|\|\|\|-[\|\|-\|\|-\|\|\|\|\|\|]PAM | NA | Intergenic | NA | NA |
| T6310 | CGCGCGTGATGGTGGCCACCTGG | | 1000 | | | | | | | | |
| Oligo fwd | CACCCGCGCGTGATGGTGGCCACC | | | | | | | | | | |
| Oligo rev | AAACGGTGGCCACCATCACGCGCG | | | | | | | | | | |
| Chromosome | start | | end | strand | MM target_seq | PAM | alignment | distance | position | gene name | gene id |
| GU305914.1 | | 89051 | 89073 + | | 0 CGCGCGTGATGGTGGCCACC | TGG | \|\|\|\|\|\|\|\|[\|\|\|\|\|\|\|\|\|\|\|\|]PAM | NA | Intergenic | NA | NA |
| T6311 | GCGCGTGATGGTGGCCACCTGGG | | 1000 | | | | | | | | |
| Oligo fwd | CACCGCGCGTGATGGTGGCCACCT | | | | | | | | | | |
| Oligo rev | AAACAGGTGGCCACCATCACGCGC | | | | | | | | | | |
| Chromosome | start | | end | strand | MM target_seq | PAM | alignment | distance | position | gene name | gene id |
| GU305914.1 | | 89052 | 89074 + | | 0 GCGCGTGATGGTGGCCACCT | GGG | \|\|\|\|\|\|\|\|[\|\|\|\|\|\|\|\|\|\|\|\|]PAM | NA | Intergenic | NA | NA |
| T6312 | CGCGTGATGGTGGCCACCTGGGG | | 1000 | | | | | | | | |
| Oligo fwd | CACCCGCGTGATGGTGGCCACCTG | | | | | | | | | | |
| Oligo rev | AAACCAGGTGGCCACCATCACGCG | | | | | | | | | | |
| Chromosome | start | | end | strand | MM target_seq | PAM | alignment | distance | position | gene name | gene id |
| GU305914.1 | | 89053 | 89075 + | | 0 CGCGTGATGGTGGCCACCTG | GGG | \|\|\|\|\|\|\|\|[\|\|\|\|\|\|\|\|\|\|\|\|]PAM | NA | Intergenic | NA | NA |
| T6313 | CTTGAAGTGCAGCACGTACTCGG | | 1000 | | | | | | | | |
| Oligo fwd | CACCCTTGAAGTGCAGCACGTACT | | | | | | | | | | |
| Oligo rev | AAACAGTACGTGCTGCACTTCAAG | | | | | | | | | | |
| Chromosome | start | | end | strand | MM target_seq | PAM | alignment | distance | position | gene name | gene id |
| GU305914.1 | | 89103 | 89125 + | | 0 CTTGAAGTGCAGCACGTACT | CGG | \|\|\|\|\|\|\|\|[\|\|\|\|\|\|\|\|\|\|\|\|]PAM | NA | Intergenic | NA | NA |
| T6314 | GTGCAGCACGTACTCGGCGATGG | | 1000 | | | | | | | | |
| Oligo fwd | CACCGTGCAGCACGTACTCGGCGA | | | | | | | | | | |
| Oligo rev | AAACTCGCCGAGTACGTGCTGCAC | | | | | | | | | | |
| Chromosome | start | | end | strand | MM target_seq | PAM | alignment | distance | position | gene name | gene id |
| GU305914.1 | | 89109 | 89131 + | | 0 GTGCAGCACGTACTCGGCGA | TGG | \|\|\|\|\|\|\|\|[\|\|\|\|\|\|\|\|\|\|\|\|]PAM | NA | Intergenic | NA | NA |
| T6315 | CAGCACGTACTCGGCGATGGAGG | | 1000 | | | | | | | | |
| Oligo fwd | CACCCAGCACGTACTCGGCGATGG | | | | | | | | | | |
| Oligo rev | AAACCCATCGCCGAGTACGTGCTG | | | | | | | | | | |
| Chromosome | start | | end | strand | MM target_seq | PAM | alignment | distance | position | gene name | gene id |
| GU305914.1 | | 89112 | 89134 + | | 0 CAGCACGTACTCGGCGATGG | AGG | \|\|\|\|\|\|\|\|[\|\|\|\|\|\|\|\|\|\|\|\|]PAM | NA | Intergenic | NA | NA |
| T6316 | CTCGGCGATGGAGGCGAACTTGG | | 1000 | | | | | | | | |
| Oligo fwd | CACCCTCGGCGATGGAGGCGAACT | | | | | | | | | | |
| Oligo rev | AAACAGTTCGCCTCCATCGCCGAG | | | | | | | | | | |
| Chromosome | start | | end | strand | MM target_seq | PAM | alignment | distance | position | gene name | gene id |
| GU305914.1 | | 89121 | 89143 + | | 0 CTCGGCGATGGAGGCGAACT | TGG | \|\|\|\|\|\|\|\|[\|\|\|\|\|\|\|\|\|\|\|\|]PAM | NA | Intergenic | NA | NA |
| T6317 | TCGGCGATGGAGGCGAACTTGGG | | 1000 | | | | | | | | |
| Oligo fwd | CACCTCGGCGATGGAGGCGAACTT | | | | | | | | | | |
| Oligo rev | AAACAAGTTCGCCTCCATCGCCGA | | | | | | | | | | |
| Chromosome | start | | end | strand | MM target_seq | PAM | alignment | distance | position | gene name | gene id |
| GU305914.1 | | 89122 | 89144 + | | 0 TCGGCGATGGAGGCGAACTT | GGG | \|\|\|\|\|\|\|\|[\|\|\|\|\|\|\|\|\|\|\|\|]PAM | NA | Intergenic | NA | NA |
| T6318 | CGAACTTGGGCGCGTCCTTGAGG | | 1000 | | | | | | | | |
| Oligo fwd | CACCCGAACTTGGGCGCGTCCTTG | | | | | | | | | | |
| Oligo rev | AAACCAAGGACGCGCCCAAGTTCG | | | | | | | | | | |
| Chromosome | start | | end | strand | MM target_seq | PAM | alignment | distance | position | gene name | gene id |
| GU305914.1 | | 89135 | 89157 + | | 0 CGAACTTGGGCGCGTCCTTG | AGG | \|\|\|\|\|\|\|\|[\|\|\|\|\|\|\|\|\|\|\|\|]PAM | NA | Intergenic | NA | NA |
| T6319 | GGGCGCGTCCTTGAGGAGACAGG | | 1000 | | | | | | | | |
| Oligo fwd | CACCGGGCGCGTCCTTGAGGAGAC | | | | | | | | | | |
| Oligo rev | AAACGTCTCCTCAAGGACGCGCCC | | | | | | | | | | |
| Chromosome | start | | end | strand | MM target_seq | PAM | alignment | distance | position | gene name | gene id |
| GU305914.1 | | 89142 | 89164 + | | 0 GGGCGCGTCCTTGAGGAGAC | AGG | \|\|\|\|\|\|\|\|[\|\|\|\|\|\|\|\|\|\|\|\|]PAM | NA | Intergenic | NA | NA |
| T6320 | GGCGCGTCCTTGAGGAGACAGGG | | 1000 | | | | | | | | |
| Oligo fwd | CACCGGCGCGTCCTTGAGGAGACA | | | | | | | | | | |
| Oligo rev | AAACTGTCTCCTCAAGGACGCGCC | | | | | | | | | | |
| Chromosome | start | | end | strand | MM target_seq | PAM | alignment | distance | position | gene name | gene id |
| GU305914.1 | | 89143 | 89165 + | | 0 GGCGCGTCCTTGAGGAGACA | GGG | \|\|\|\|\|\|\|\|[\|\|\|\|\|\|\|\|\|\|\|\|]PAM | NA | Intergenic | NA | NA |
| T6321 | GCGCGTCCTTGAGGAGACAGGGG | | 1000 | | | | | | | | |
| Oligo fwd | CACCGCGCGTCCTTGAGGAGACAG | | | | | | | | | | |
| Oligo rev | AAACCTGTCTCCTCAAGGACGCGC | | | | | | | | | | |
| Chromosome | start | | end | strand | MM target_seq | PAM | alignment | distance | position | gene name | gene id |
| GU305914.1 | | 89144 | 89166 + | | 0 GCGCGTCCTTGAGGAGACAG | GGG | \|\|\|\|\|\|\|\|[\|\|\|\|\|\|\|\|\|\|\|\|]PAM | NA | Intergenic | NA | NA |
| T6322 | GGAGACAGGGGAACTGCGAGTGG | | 1000 | | | | | | | | |
| Oligo fwd | CACCGGAGACAGGGGAACTGCGAG | | | | | | | | | | |
| Oligo rev | AAACCTCGCAGTTCCCCTGTCTCC | | | | | | | | | | |
| Chromosome | start | | end | strand | MM target_seq | PAM | alignment | distance | position | gene name | gene id |
| GU305914.1 | | 89156 | 89178 + | | 0 GGAGACAGGGGAACTGCGAG | TGG | \|\|\|\|\|\|\|\|[\|\|\|\|\|\|\|\|\|\|\|\|]PAM | NA | Intergenic | NA | NA |
| T6323 | GGGAACTGCGAGTGGTTCTGCGG | | 1000 | | | | | | | | |
| Oligo fwd | CACCGGGAACTGCGAGTGGTTCTG | | | | | | | | | | |
| Oligo rev | AAACCAGAACCACTCGCAGTTCCC | | | | | | | | | | |
| Chromosome | start | | end | strand | MM target_seq | PAM | alignment | distance | position | gene name | gene id |
| GU305914.1 | | 89164 | 89186 + | | 0 GGGAACTGCGAGTGGTTCTG | CGG | \|\|\|\|\|\|\|\|[\|\|\|\|\|\|\|\|\|\|\|\|]PAM | NA | Intergenic | NA | NA |
| T6324 | GCGAGTGGTTCTGCGGCGTCAGG | | 1000 | | | | | | | | |
| Oligo fwd | CACCGCGAGTGGTTCTGCGGCGTC | | | | | | | | | | |

| | | | | | | | | | | | | |
|---|---|---|---|---|---|---|---|---|---|---|---|---|
| Oligo rev | AAACGACGCCGCAGAACCACTCGC | | | | | | | | | | | |
| Chromosome | start | | end | strand | MM | target_seq | PAM | alignment | distance | position | gene name | gene id |
| GU305914.1 | | 89171 | 89193 | + | 0 | GCGAGTGGTTCTGCGGCGTC | AGG | \|\|\|\|\|\|\|\|[\|\|\|\|\|\|\|\|\|\|\|\|]PAM | NA | Intergenic | NA | NA |
| T6325 | TCAGGCGCGTGCTCATGCCCTGG | | 1000 | | | | | | | | | |
| Oligo fwd | CACCTCAGGCGCGTGCTCATGCCC | | | | | | | | | | | |
| Oligo rev | AAACGGGCATGAGCACGCGCCTGA | | | | | | | | | | | |
| Chromosome | start | | end | strand | MM | target_seq | PAM | alignment | distance | position | gene name | gene id |
| GU305914.1 | | 89189 | 89211 | + | 0 | TCAGGCGCGTGCTCATGCCC | TGG | \|\|\|\|\|\|\|\|[\|\|\|\|\|\|\|\|\|\|\|\|]PAM | NA | Intergenic | NA | NA |
| T6326 | GCGCGTGCTCATGCCCTGGATGG | | 1000 | | | | | | | | | |
| Oligo fwd | CACCGCGCGTGCTCATGCCCTGGA | | | | | | | | | | | |
| Oligo rev | AAACTCCAGGGCATGAGCACGCGC | | | | | | | | | | | |
| Chromosome | start | | end | strand | MM | target_seq | PAM | alignment | distance | position | gene name | gene id |
| GU305914.1 | | 89193 | 89215 | + | 0 | GCGCGTGCTCATGCCCTGGA | TGG | \|\|\|\|\|\|\|\|[\|\|\|\|\|\|\|\|\|\|\|\|]PAM | NA | Intergenic | NA | NA |
| T6327 | CCCTGAAGCCGAACATGTAGAGG | | 1000 | | | | | | | | | |
| Oligo fwd | CACCCCCTGAAGCCGAACATGTAG | | | | | | | | | | | |
| Oligo rev | AAACCTACATGTTCGGCTTCAGGG | | | | | | | | | | | |
| Chromosome | start | | end | strand | MM | target_seq | PAM | alignment | distance | position | gene name | gene id |
| GU305914.1 | | 89228 | 89250 | + | 0 | CCCTGAAGCCGAACATGTAG | AGG | \|\|\|\|\|\|\|\|[\|\|\|\|\|\|\|\|\|\|\|\|]PAM | NA | Intergenic | NA | NA |
| T6328 | ACATGTAGAGGAACTTGTTGTGG | | 1000 | | | | | | | | | |
| Oligo fwd | CACCACATGTAGAGGAACTTGTTG | | | | | | | | | | | |
| Oligo rev | AAACCAACAAGTTCCTCTACATGT | | | | | | | | | | | |
| Chromosome | start | | end | strand | MM | target_seq | PAM | alignment | distance | position | gene name | gene id |
| GU305914.1 | | 89240 | 89262 | + | 0 | ACATGTAGAGGAACTTGTTG | TGG | \|\|\|\|\|\|\|\|[\|\|\|\|\|\|\|\|\|\|\|\|]PAM | NA | Intergenic | NA | NA |
| T6329 | AGAGGAACTTGTTGTGGAAGCGG | | 1000 | | | | | | | | | |
| Oligo fwd | CACCAGAGGAACTTGTTGTGGAAG | | | | | | | | | | | |
| Oligo rev | AAACCTTCCACAACAAGTTCCTCT | | | | | | | | | | | |
| Chromosome | start | | end | strand | MM | target_seq | PAM | alignment | distance | position | gene name | gene id |
| GU305914.1 | | 89246 | 89268 | + | 0 | AGAGGAACTTGTTGTGGAAG | CGG | \|\|\|\|\|\|\|\|[\|\|\|\|\|\|\|\|\|\|\|\|]PAM | NA | Intergenic | NA | NA |
| T6330 | GGTTCTCCACGTAGCTGAGCAGG | | 1000 | | | | | | | | | |
| Oligo fwd | CACCGGTTCTCCACGTAGCTGAGC | | | | | | | | | | | |
| Oligo rev | AAACGCTCAGCTACGTGGAGAACC | | | | | | | | | | | |
| Chromosome | start | | end | strand | MM | target_seq | PAM | alignment | distance | position | gene name | gene id |
| GU305914.1 | | 89267 | 89289 | + | 0 | GGTTCTCCACGTAGCTGAGC | AGG | \|\|\|\|\|\|\|\|[\|\|\|\|\|\|\|\|\|\|\|\|]PAM | NA | Intergenic | NA | NA |
| T6331 | ACGTAGCTGAGCAGGCAGTCCGG | | 1000 | | | | | | | | | |
| Oligo fwd | CACCACGTAGCTGAGCAGGCAGTC | | | | | | | | | | | |
| Oligo rev | AAACGACTGCCTGCTCAGCTACGT | | | | | | | | | | | |
| Chromosome | start | | end | strand | MM | target_seq | PAM | alignment | distance | position | gene name | gene id |
| GU305914.1 | | 89275 | 89297 | + | 0 | ACGTAGCTGAGCAGGCAGTC | CGG | \|\|\|\|\|\|\|\|[\|\|\|\|\|\|\|\|\|\|\|\|]PAM | NA | Intergenic | NA | NA |
| T6332 | AGCTGAGCAGGCAGTCCGGCTGG | | 1000 | | | | | | | | | |
| Oligo fwd | CACCAGCTGAGCAGGCAGTCCGGC | | | | | | | | | | | |
| Oligo rev | AAACGCCGGACTGCCTGCTCAGCT | | | | | | | | | | | |
| Chromosome | start | | end | strand | MM | target_seq | PAM | alignment | distance | position | gene name | gene id |
| GU305914.1 | | 89279 | 89301 | + | 0 | AGCTGAGCAGGCAGTCCGGC | TGG | \|\|\|\|\|\|\|\|[\|\|\|\|\|\|\|\|\|\|\|\|]PAM | NA | Intergenic | NA | NA |
| T6333 | GTCGCTCTCGTCGTAGTCGTCGG | | 1000 | | | | | | | | | |
| Oligo fwd | CACCGTCGCTCTCGTCGTAGTCGT | | | | | | | | | | | |
| Oligo rev | AAACACGACTACGACGAGAGCGAC | | | | | | | | | | | |
| Chromosome | start | | end | strand | MM | target_seq | PAM | alignment | distance | position | gene name | gene id |
| GU305914.1 | | 89313 | 89335 | + | 0 | GTCGCTCTCGTCGTAGTCGT | CGG | \|\|\|\|\|\|\|\|[\|\|\|\|\|\|\|\|\|\|\|\|]PAM | NA | Intergenic | NA | NA |
| T6334 | CTCGTCGTAGTCGTCGGTGACGG | | 1000 | | | | | | | | | |
| Oligo fwd | CACCCTCGTCGTAGTCGTCGGTGA | | | | | | | | | | | |
| Oligo rev | AAACTCACCGACGACTACGACGAG | | | | | | | | | | | |
| Chromosome | start | | end | strand | MM | target_seq | PAM | alignment | distance | position | gene name | gene id |
| GU305914.1 | | 89319 | 89341 | + | 0 | CTCGTCGTAGTCGTCGGTGA | CGG | \|\|\|\|\|\|\|\|[\|\|\|\|\|\|\|\|\|\|\|\|]PAM | NA | Intergenic | NA | NA |
| T6335 | CGTAGTCGTCGGTGACGGTGAGG | | 1000 | | | | | | | | | |
| Oligo fwd | CACCCGTAGTCGTCGGTGACGGTG | | | | | | | | | | | |
| Oligo rev | AAACCACCGTCACCGACGACTACG | | | | | | | | | | | |
| Chromosome | start | | end | strand | MM | target_seq | PAM | alignment | distance | position | gene name | gene id |
| GU305914.1 | | 89324 | 89346 | + | 0 | CGTAGTCGTCGGTGACGGTG | AGG | \|\|\|\|\|\|\|\|[\|\|\|\|\|\|\|\|\|\|\|\|]PAM | NA | Intergenic | NA | NA |
| GU305914.1 | | 142266 | 142288 | + | 3 | CGTCGTCGTCGACGACGGTG | GGG | \|\|\|-\|\|\|\|[\|\|\|--\|\|\|\|\|\|\|]PAM | NA | Intergenic | NA | NA |
| T6336 | AGTCGTCGGTGACGGTGAGGAGG | | 1000 | | | | | | | | | |
| Oligo fwd | CACCAGTCGTCGGTGACGGTGAGG | | | | | | | | | | | |
| Oligo rev | AAACCCTCACCGTCACCGACGACT | | | | | | | | | | | |
| Chromosome | start | | end | strand | MM | target_seq | PAM | alignment | distance | position | gene name | gene id |
| GU305914.1 | | 89327 | 89349 | + | 0 | AGTCGTCGGTGACGGTGAGG | AGG | \|\|\|\|\|\|\|\|[\|\|\|\|\|\|\|\|\|\|\|\|]PAM | NA | Intergenic | NA | NA |
| T6337 | CGGTGAGGAGGAAGCGCACCAGG | | 1000 | | | | | | | | | |
| Oligo fwd | CACCCGGTGAGGAGGAAGCGCACC | | | | | | | | | | | |
| Oligo rev | AAACGGTGCGCTTCCTCCTCACCG | | | | | | | | | | | |
| Chromosome | start | | end | strand | MM | target_seq | PAM | alignment | distance | position | gene name | gene id |
| GU305914.1 | | 89339 | 89361 | + | 0 | CGGTGAGGAGGAAGCGCACC | AGG | \|\|\|\|\|\|\|\|[\|\|\|\|\|\|\|\|\|\|\|\|]PAM | NA | Intergenic | NA | NA |
| T6338 | TTGCCCACGTTGACCGAGCGCGG | | 1000 | | | | | | | | | |
| Oligo fwd | CACCTTGCCCACGTTGACCGAGCG | | | | | | | | | | | |
| Oligo rev | AAACCGCTCGGTCAACGTGGGCAA | | | | | | | | | | | |
| Chromosome | start | | end | strand | MM | target_seq | PAM | alignment | distance | position | gene name | gene id |
| GU305914.1 | | 89380 | 89402 | + | 0 | TTGCCCACGTTGACCGAGCG | CGG | \|\|\|\|\|\|\|\|[\|\|\|\|\|\|\|\|\|\|\|\|]PAM | NA | Intergenic | NA | NA |

T6339 AGCGCGGCAGCGAGTTCTGCAGG 1000
Oligo fwd CACCAGCGCGGCAGCGAGTTCTGC
Oligo rev AAACGCAGAACTCGCTGCCGCGCT
Chromosome start end strand MM target_seq PAM alignment distance position gene name gene id
GU305914.1 89396 89418 + 0 AGCGCGGCAGCGAGTTCTGC AGG ||||||||[||||||||||||]PAM NA Intergenic NA NA

T6340 GGACGCGCTGCCAGAACTCGAGG 1000
Oligo fwd CACCGGACGCGCTGCCAGAACTCG
Oligo rev AAACCGAGTTCTGGCAGCGCGTCC
Chromosome start end strand MM target_seq PAM alignment distance position gene name gene id
GU305914.1 89417 89439 + 0 GGACGCGCTGCCAGAACTCG AGG ||||||||[||||||||||||]PAM NA Intergenic NA NA

T6341 GCCAGAACTCGAGGCACGACAGG 1000
Oligo fwd CACCGCCAGAACTCGAGGCACGAC
Oligo rev AAACGTCGTGCCTCGAGTTCTGGC
Chromosome start end strand MM target_seq PAM alignment distance position gene name gene id
GU305914.1 89426 89448 + 0 GCCAGAACTCGAGGCACGAC AGG ||||||||[||||||||||||]PAM NA Intergenic NA NA

T6342 CTCGAGGCACGACAGGCCGCAGG 1000
Oligo fwd CACCCTCGAGGCACGACAGGCCGC
Oligo rev AAACGCGGCCTGTCGTGCCTCGAG
Chromosome start end strand MM target_seq PAM alignment distance position gene name gene id
GU305914.1 89433 89455 + 0 CTCGAGGCACGACAGGCCGC AGG ||||||||[||||||||||||]PAM NA Intergenic NA NA

T6343 AGGCACGACAGGCCGCAGGCCGG 1000
Oligo fwd CACCAGGCACGACAGGCCGCAGGC
Oligo rev AAACGCCTGCGGCCTGTCGTGCCT
Chromosome start end strand MM target_seq PAM alignment distance position gene name gene id
GU305914.1 89437 89459 + 0 AGGCACGACAGGCCGCAGGC CGG ||||||||[||||||||||||]PAM NA Intergenic NA NA

T6344 GGCACGACAGGCCGCAGGCCGGG 1000
Oligo fwd CACCGGCACGACAGGCCGCAGGCC
Oligo rev AAACGGCCTGCGGCCTGTCGTGCC
Chromosome start end strand MM target_seq PAM alignment distance position gene name gene id
GU305914.1 89438 89460 + 0 GGCACGACAGGCCGCAGGCC GGG ||||||||[||||||||||||]PAM NA Intergenic NA NA

T6345 AGGCCGGGAAGAGCTTCTCGTGG 1000
Oligo fwd CACCAGGCCGGGAAGAGCTTCTCG
Oligo rev AAACCGAGAAGCTCTTCCCGGCCT
Chromosome start end strand MM target_seq PAM alignment distance position gene name gene id
GU305914.1 89453 89475 + 0 AGGCCGGGAAGAGCTTCTCG TGG ||||||||[||||||||||||]PAM NA Intergenic NA NA

T6346 GGAAGAGCTTCTCGTGGCAGCGG 1000
Oligo fwd CACCGGAAGAGCTTCTCGTGGCAG
Oligo rev AAACCTGCCACGAGAAGCTCTTCC
Chromosome start end strand MM target_seq PAM alignment distance position gene name gene id
GU305914.1 89459 89481 + 0 GGAAGAGCTTCTCGTGGCAG CGG ||||||||[||||||||||||]PAM NA Intergenic NA NA

T6347 TCTCGTGGCAGCGGAAGAGCAGG 1000
Oligo fwd CACCTCTCGTGGCAGCGGAAGAGC
Oligo rev AAACGCTCTTCCGCTGCCACGAGA
Chromosome start end strand MM target_seq PAM alignment distance position gene name gene id
GU305914.1 89468 89490 + 0 TCTCGTGGCAGCGGAAGAGC AGG ||||||||[||||||||||||]PAM NA Intergenic NA NA

T6348 GTGGCAGCGGAAGAGCAGGAAGG 1000
Oligo fwd CACCGTGGCAGCGGAAGAGCAGGA
Oligo rev AAACTCCTGCTCTTCCGCTGCCAC
Chromosome start end strand MM target_seq PAM alignment distance position gene name gene id
GU305914.1 89472 89494 + 0 GTGGCAGCGGAAGAGCAGGA AGG ||||||||[||||||||||||]PAM NA Intergenic NA NA

T6349 AGCGGAAGAGCAGGAAGGAGAGG 1000
Oligo fwd CACCAGCGGAAGAGCAGGAAGGAG
Oligo rev AAACCTCCTTCCTGCTCTTCCGCT
Chromosome start end strand MM target_seq PAM alignment distance position gene name gene id
GU305914.1 89477 89499 + 0 AGCGGAAGAGCAGGAAGGAG AGG ||||||||[||||||||||||]PAM NA Intergenic NA NA

T6350 GAAGGAGAGGCAGCCCTTGATGG 1000
Oligo fwd CACCGAAGGAGAGGCAGCCCTTGA
Oligo rev AAACTCAAGGGCTGCCTCTCCTTC
Chromosome start end strand MM target_seq PAM alignment distance position gene name gene id
GU305914.1 89490 89512 + 0 GAAGGAGAGGCAGCCCTTGA TGG ||||||||[||||||||||||]PAM NA Intergenic NA NA

T6351 AAGGAGAGGCAGCCCTTGATGGG 1000
Oligo fwd CACCAAGGAGAGGCAGCCCTTGAT
Oligo rev AAACATCAAGGGCTGCCTCTCCTT
Chromosome start end strand MM target_seq PAM alignment distance position gene name gene id
GU305914.1 89491 89513 + 0 AAGGAGAGGCAGCCCTTGAT GGG ||||||||[||||||||||||]PAM NA Intergenic NA NA

T6352 TGATGGGATTCTTCCTCTGCAGG 1000
Oligo fwd CACCTGATGGGATTCTTCCTCTGC
Oligo rev AAACGCAGAGGAAGAATCCCATCA
Chromosome start end strand MM target_seq PAM alignment distance position gene name gene id
GU305914.1 89507 89529 + 0 TGATGGGATTCTTCCTCTGC AGG ||||||||[||||||||||||]PAM NA Intergenic NA NA

T6353 GATGGGATTCTTCCTCTGCAGGG 1000
Oligo fwd CACCGATGGGATTCTTCCTCTGCA
Oligo rev AAACTGCAGAGGAAGAATCCCATC
Chromosome start end strand MM target_seq PAM alignment distance position gene name gene id
GU305914.1 89508 89530 + 0 GATGGGATTCTTCCTCTGCA GGG ||||||||[||||||||||||]PAM NA Intergenic NA NA

| | | | | | | | | | | | |
|---|---|---|---|---|---|---|---|---|---|---|---|
| T6354 | TTGAAGAAGACGCTGCTCTGCGG | 1000 | | | | | | | | | |
| Oligo fwd | CACCTTGAAGAAGACGCTGCTCTG | | | | | | | | | | |
| Oligo rev | AAACCAGAGCAGCGTCTTCTTCAA | | | | | | | | | | |
| Chromosome | start | end | strand | MM | target_seq | PAM | alignment | distance | position | gene name | gene id |
| GU305914.1 | 89539 | 89561 | + | 0 | TTGAAGAAGACGCTGCTCTG | CGG | \|\|\|\|\|\|\|\|[\|\|\|\|\|\|\|\|\|\|\|\|]PAM | NA | Intergenic | NA | NA |
| T6355 | GACGCTGCTCTGCGGCGTCTTGG | 1000 | | | | | | | | | |
| Oligo fwd | CACCGACGCTGCTCTGCGGCGTCT | | | | | | | | | | |
| Oligo rev | AAACAGACGCCGCAGAGCAGCGTC | | | | | | | | | | |
| Chromosome | start | end | strand | MM | target_seq | PAM | alignment | distance | position | gene name | gene id |
| GU305914.1 | 89547 | 89569 | + | 0 | GACGCTGCTCTGCGGCGTCT | TGG | \|\|\|\|\|\|\|\|[\|\|\|\|\|\|\|\|\|\|\|\|]PAM | NA | Intergenic | NA | NA |
| T6356 | GCTCTGCGGCGTCTTGGAGATGG | 1000 | | | | | | | | | |
| Oligo fwd | CACCGCTCTGCGGCGTCTTGGAGA | | | | | | | | | | |
| Oligo rev | AAACTCTCCAAGACGCCGCAGAGC | | | | | | | | | | |
| Chromosome | start | end | strand | MM | target_seq | PAM | alignment | distance | position | gene name | gene id |
| GU305914.1 | 89553 | 89575 | + | 0 | GCTCTGCGGCGTCTTGGAGA | TGG | \|\|\|\|\|\|\|\|[\|\|\|\|\|\|\|\|\|\|\|\|]PAM | NA | Intergenic | NA | NA |
| T6357 | CTCTGCGGCGTCTTGGAGATGGG | 1000 | | | | | | | | | |
| Oligo fwd | CACCCTCTGCGGCGTCTTGGAGAT | | | | | | | | | | |
| Oligo rev | AAACATCTCCAAGACGCCGCAGAG | | | | | | | | | | |
| Chromosome | start | end | strand | MM | target_seq | PAM | alignment | distance | position | gene name | gene id |
| GU305914.1 | 89554 | 89576 | + | 0 | CTCTGCGGCGTCTTGGAGAT | GGG | \|\|\|\|\|\|\|\|[\|\|\|\|\|\|\|\|\|\|\|\|]PAM | NA | Intergenic | NA | NA |
| T6358 | AGATGGGCCGATTCTTGATGCGG | 1000 | | | | | | | | | |
| Oligo fwd | CACCAGATGGGCCGATTCTTGATG | | | | | | | | | | |
| Oligo rev | AAACCATCAAGAATCGGCCCATCT | | | | | | | | | | |
| Chromosome | start | end | strand | MM | target_seq | PAM | alignment | distance | position | gene name | gene id |
| GU305914.1 | 89570 | 89592 | + | 0 | AGATGGGCCGATTCTTGATG | CGG | \|\|\|\|\|\|\|\|[\|\|\|\|\|\|\|\|\|\|\|\|]PAM | NA | Intergenic | NA | NA |
| T6359 | CATCTTACACGACTTGATCTCGG | 1000 | | | | | | | | | |
| Oligo fwd | CACCCATCTTACACGACTTGATCT | | | | | | | | | | |
| Oligo rev | AAACAGATCAAGTCGTGTAAGATG | | | | | | | | | | |
| Chromosome | start | end | strand | MM | target_seq | PAM | alignment | distance | position | gene name | gene id |
| GU305914.1 | 89616 | 89638 | + | 0 | CATCTTACACGACTTGATCT | CGG | \|\|\|\|\|\|\|\|[\|\|\|\|\|\|\|\|\|\|\|\|]PAM | NA | Intergenic | NA | NA |
| T6360 | CTTACACGACTTGATCTCGGTGG | 1000 | | | | | | | | | |
| Oligo fwd | CACCCTTACACGACTTGATCTCGG | | | | | | | | | | |
| Oligo rev | AAACCCGAGATCAAGTCGTGTAAG | | | | | | | | | | |
| Chromosome | start | end | strand | MM | target_seq | PAM | alignment | distance | position | gene name | gene id |
| GU305914.1 | 89619 | 89641 | + | 0 | CTTACACGACTTGATCTCGG | TGG | \|\|\|\|\|\|\|\|[\|\|\|\|\|\|\|\|\|\|\|\|]PAM | NA | Intergenic | NA | NA |
| T6361 | TGATCTCGGTGGAGATCTTGCGG | 1000 | | | | | | | | | |
| Oligo fwd | CACCTGATCTCGGTGGAGATCTTG | | | | | | | | | | |
| Oligo rev | AAACCAAGATCTCCACCGAGATCA | | | | | | | | | | |
| Chromosome | start | end | strand | MM | target_seq | PAM | alignment | distance | position | gene name | gene id |
| GU305914.1 | 89630 | 89652 | + | 0 | TGATCTCGGTGGAGATCTTG | CGG | \|\|\|\|\|\|\|\|[\|\|\|\|\|\|\|\|\|\|\|\|]PAM | NA | Intergenic | NA | NA |
| GU305914.1 | 113261 | 113283 | + | 4 | AGAACTCGGGCGAGATCTTG | CGG | -\|\|-\|\|\|\|[\|--\|\|\|\|\|\|\|\|\|]PAM | NA | Intergenic | NA | NA |
| T6362 | CGGTGGAGATCTTGCGGTAGCGG | 1000 | | | | | | | | | |
| Oligo fwd | CACCCGGTGGAGATCTTGCGGTAG | | | | | | | | | | |
| Oligo rev | AAACCTACCGCAAGATCTCCACCG | | | | | | | | | | |
| Chromosome | start | end | strand | MM | target_seq | PAM | alignment | distance | position | gene name | gene id |
| GU305914.1 | 89636 | 89658 | + | 0 | CGGTGGAGATCTTGCGGTAG | CGG | \|\|\|\|\|\|\|\|[\|\|\|\|\|\|\|\|\|\|\|\|]PAM | NA | Intergenic | NA | NA |
| GU305914.1 | 113267 | 113289 | + | 4 | CGGGCGAGATCTTGCGGTTC | AGG | \|\|\|--\|\|\|[\|\|\|\|\|\|\|\|\|\|--]PAM | NA | Intergenic | NA | NA |
| T6363 | TGCGGTAGCGGCCCGTCTCCTGG | 1000 | | | | | | | | | |
| Oligo fwd | CACCTGCGGTAGCGGCCCGTCTCC | | | | | | | | | | |
| Oligo rev | AAACGGAGACGGGCCGCTACCGCA | | | | | | | | | | |
| Chromosome | start | end | strand | MM | target_seq | PAM | alignment | distance | position | gene name | gene id |
| GU305914.1 | 89648 | 89670 | + | 0 | TGCGGTAGCGGCCCGTCTCC | TGG | \|\|\|\|\|\|\|\|[\|\|\|\|\|\|\|\|\|\|\|\|]PAM | NA | Intergenic | NA | NA |
| T6364 | GTAGCGGCCCGTCTCCTGGAAGG | 1000 | | | | | | | | | |
| Oligo fwd | CACCGTAGCGGCCCGTCTCCTGGA | | | | | | | | | | |
| Oligo rev | AAACTCCAGGAGACGGGCCGCTAC | | | | | | | | | | |
| Chromosome | start | end | strand | MM | target_seq | PAM | alignment | distance | position | gene name | gene id |
| GU305914.1 | 89652 | 89674 | + | 0 | GTAGCGGCCCGTCTCCTGGA | AGG | \|\|\|\|\|\|\|\|[\|\|\|\|\|\|\|\|\|\|\|\|]PAM | NA | Intergenic | NA | NA |
| T6365 | CATGATGCGCTCGAACTCGACGG | 1000 | | | | | | | | | |
| Oligo fwd | CACCCATGATGCGCTCGAACTCGA | | | | | | | | | | |
| Oligo rev | AAACTCGAGTTCGAGCGCATCATG | | | | | | | | | | |
| Chromosome | start | end | strand | MM | target_seq | PAM | alignment | distance | position | gene name | gene id |
| GU305914.1 | 89685 | 89707 | + | 0 | CATGATGCGCTCGAACTCGA | CGG | \|\|\|\|\|\|\|\|[\|\|\|\|\|\|\|\|\|\|\|\|]PAM | NA | Intergenic | NA | NA |
| T6366 | GCTCGAACTCGACGGATTTGAGG | 1000 | | | | | | | | | |
| Oligo fwd | CACCGCTCGAACTCGACGGATTTG | | | | | | | | | | |
| Oligo rev | AAACCAAATCCGTCGAGTTCGAGC | | | | | | | | | | |
| Chromosome | start | end | strand | MM | target_seq | PAM | alignment | distance | position | gene name | gene id |
| GU305914.1 | 89693 | 89715 | + | 0 | GCTCGAACTCGACGGATTTG | AGG | \|\|\|\|\|\|\|\|[\|\|\|\|\|\|\|\|\|\|\|\|]PAM | NA | Intergenic | NA | NA |
| T6367 | CTCGACGGATTTGAGGTTCTTGG | 1000 | | | | | | | | | |
| Oligo fwd | CACCCTCGACGGATTTGAGGTTCT | | | | | | | | | | |
| Oligo rev | AAACAGAACCTCAAATCCGTCGAG | | | | | | | | | | |
| Chromosome | start | end | strand | MM | target_seq | PAM | alignment | distance | position | gene name | gene id |
| GU305914.1 | 89700 | 89722 | + | 0 | CTCGACGGATTTGAGGTTCT | TGG | \|\|\|\|\|\|\|\|[\|\|\|\|\|\|\|\|\|\|\|\|]PAM | NA | Intergenic | NA | NA |
| T6368 | TCTTGGTGTCCTTGACGTTCAGG | 1000 | | | | | | | | | |
| Oligo fwd | CACCTCTTGGTGTCCTTGACGTTC | | | | | | | | | | |

<table>
<tr><td>Oligo rev</td><td>AAACGAACGTCAAGGACACCAAGA</td><td></td><td></td><td></td><td></td><td></td><td></td><td></td><td></td><td></td><td></td><td></td></tr>
<tr><td>Chromosome</td><td>start</td><td></td><td>end</td><td>strand</td><td>MM</td><td>target_seq</td><td>PAM</td><td>alignment</td><td>distance</td><td>position</td><td>gene name</td><td>gene id</td></tr>
<tr><td>GU305914.1</td><td></td><td>89717</td><td>89739</td><td>+</td><td>0</td><td>TCTTGGTGTCCTTGACGTTC</td><td>AGG</td><td>||||||||[||||||||||||]PAM</td><td>NA</td><td>Intergenic</td><td>NA</td><td>NA</td></tr>
<tr><td>T6369</td><td>CGTTCAGGAACCCCTTCTTGAGG</td><td></td><td>1000</td><td></td><td></td><td></td><td></td><td></td><td></td><td></td><td></td><td></td></tr>
<tr><td>Oligo fwd</td><td>CACCCGTTCAGGAACCCCTTCTTG</td><td></td><td></td><td></td><td></td><td></td><td></td><td></td><td></td><td></td><td></td><td></td></tr>
<tr><td>Oligo rev</td><td>AAACCAAGAAGGGGTTCCTGAACG</td><td></td><td></td><td></td><td></td><td></td><td></td><td></td><td></td><td></td><td></td><td></td></tr>
<tr><td>Chromosome</td><td>start</td><td></td><td>end</td><td>strand</td><td>MM</td><td>target_seq</td><td>PAM</td><td>alignment</td><td>distance</td><td>position</td><td>gene name</td><td>gene id</td></tr>
<tr><td>GU305914.1</td><td></td><td>89732</td><td>89754</td><td>+</td><td>0</td><td>CGTTCAGGAACCCCTTCTTG</td><td>AGG</td><td>||||||||[||||||||||||]PAM</td><td>NA</td><td>Intergenic</td><td>NA</td><td>NA</td></tr>
<tr><td>T6370</td><td>GTTCAGGAACCCCTTCTTGAGGG</td><td></td><td>1000</td><td></td><td></td><td></td><td></td><td></td><td></td><td></td><td></td><td></td></tr>
<tr><td>Oligo fwd</td><td>CACCGTTCAGGAACCCCTTCTTGA</td><td></td><td></td><td></td><td></td><td></td><td></td><td></td><td></td><td></td><td></td><td></td></tr>
<tr><td>Oligo rev</td><td>AAACTCAAGAAGGGGTTCCTGAAC</td><td></td><td></td><td></td><td></td><td></td><td></td><td></td><td></td><td></td><td></td><td></td></tr>
<tr><td>Chromosome</td><td>start</td><td></td><td>end</td><td>strand</td><td>MM</td><td>target_seq</td><td>PAM</td><td>alignment</td><td>distance</td><td>position</td><td>gene name</td><td>gene id</td></tr>
<tr><td>GU305914.1</td><td></td><td>89733</td><td>89755</td><td>+</td><td>0</td><td>GTTCAGGAACCCCTTCTTGA</td><td>GGG</td><td>||||||||[||||||||||||]PAM</td><td>NA</td><td>Intergenic</td><td>NA</td><td>NA</td></tr>
<tr><td>T6371</td><td>TTCAGGAACCCCTTCTTGAGGGG</td><td></td><td>1000</td><td></td><td></td><td></td><td></td><td></td><td></td><td></td><td></td><td></td></tr>
<tr><td>Oligo fwd</td><td>CACCTTCAGGAACCCCTTCTTGAG</td><td></td><td></td><td></td><td></td><td></td><td></td><td></td><td></td><td></td><td></td><td></td></tr>
<tr><td>Oligo rev</td><td>AAACCTCAAGAAGGGGTTCCTGAA</td><td></td><td></td><td></td><td></td><td></td><td></td><td></td><td></td><td></td><td></td><td></td></tr>
<tr><td>Chromosome</td><td>start</td><td></td><td>end</td><td>strand</td><td>MM</td><td>target_seq</td><td>PAM</td><td>alignment</td><td>distance</td><td>position</td><td>gene name</td><td>gene id</td></tr>
<tr><td>GU305914.1</td><td></td><td>89734</td><td>89756</td><td>+</td><td>0</td><td>TTCAGGAACCCCTTCTTGAG</td><td>GGG</td><td>||||||||[||||||||||||]PAM</td><td>NA</td><td>Intergenic</td><td>NA</td><td>NA</td></tr>
<tr><td>T6372</td><td>CGTCCCGAACGCGCCGCAGATGG</td><td></td><td>1000</td><td></td><td></td><td></td><td></td><td></td><td></td><td></td><td></td><td></td></tr>
<tr><td>Oligo fwd</td><td>CACCCGTCCCGAACGCGCCGCAGA</td><td></td><td></td><td></td><td></td><td></td><td></td><td></td><td></td><td></td><td></td><td></td></tr>
<tr><td>Oligo rev</td><td>AAACTCTGCGGCGCGTTCGGGACG</td><td></td><td></td><td></td><td></td><td></td><td></td><td></td><td></td><td></td><td></td><td></td></tr>
<tr><td>Chromosome</td><td>start</td><td></td><td>end</td><td>strand</td><td>MM</td><td>target_seq</td><td>PAM</td><td>alignment</td><td>distance</td><td>position</td><td>gene name</td><td>gene id</td></tr>
<tr><td>GU305914.1</td><td></td><td>89760</td><td>89782</td><td>+</td><td>0</td><td>CGTCCCGAACGCGCCGCAGA</td><td>TGG</td><td>||||||||[||||||||||||]PAM</td><td>NA</td><td>Intergenic</td><td>NA</td><td>NA</td></tr>
<tr><td>T6373</td><td>CGAACGCGCCGCAGATGGACTGG</td><td></td><td>1000</td><td></td><td></td><td></td><td></td><td></td><td></td><td></td><td></td><td></td></tr>
<tr><td>Oligo fwd</td><td>CACCCGAACGCGCCGCAGATGGAC</td><td></td><td></td><td></td><td></td><td></td><td></td><td></td><td></td><td></td><td></td><td></td></tr>
<tr><td>Oligo rev</td><td>AAACGTCCATCTGCGGCGCGTTCG</td><td></td><td></td><td></td><td></td><td></td><td></td><td></td><td></td><td></td><td></td><td></td></tr>
<tr><td>Chromosome</td><td>start</td><td></td><td>end</td><td>strand</td><td>MM</td><td>target_seq</td><td>PAM</td><td>alignment</td><td>distance</td><td>position</td><td>gene name</td><td>gene id</td></tr>
<tr><td>GU305914.1</td><td></td><td>89765</td><td>89787</td><td>+</td><td>0</td><td>CGAACGCGCCGCAGATGGAC</td><td>TGG</td><td>||||||||[||||||||||||]PAM</td><td>NA</td><td>Intergenic</td><td>NA</td><td>NA</td></tr>
<tr><td>T6374</td><td>GAAGTGCAGCGTCACCCACTTGG</td><td></td><td>1000</td><td></td><td></td><td></td><td></td><td></td><td></td><td></td><td></td><td></td></tr>
<tr><td>Oligo fwd</td><td>CACCGAAGTGCAGCGTCACCCACT</td><td></td><td></td><td></td><td></td><td></td><td></td><td></td><td></td><td></td><td></td><td></td></tr>
<tr><td>Oligo rev</td><td>AAACAGTGGGTGACGCTGCACTTC</td><td></td><td></td><td></td><td></td><td></td><td></td><td></td><td></td><td></td><td></td><td></td></tr>
<tr><td>Chromosome</td><td>start</td><td></td><td>end</td><td>strand</td><td>MM</td><td>target_seq</td><td>PAM</td><td>alignment</td><td>distance</td><td>position</td><td>gene name</td><td>gene id</td></tr>
<tr><td>GU305914.1</td><td></td><td>89787</td><td>89809</td><td>+</td><td>0</td><td>GAAGTGCAGCGTCACCCACT</td><td>TGG</td><td>||||||||[||||||||||||]PAM</td><td>NA</td><td>Intergenic</td><td>NA</td><td>NA</td></tr>
<tr><td>T6375</td><td>TGGAGATCTGCGCCGTCGTCAGG</td><td></td><td>1000</td><td></td><td></td><td></td><td></td><td></td><td></td><td></td><td></td><td></td></tr>
<tr><td>Oligo fwd</td><td>CACCTGGAGATCTGCGCCGTCGTC</td><td></td><td></td><td></td><td></td><td></td><td></td><td></td><td></td><td></td><td></td><td></td></tr>
<tr><td>Oligo rev</td><td>AAACGACGACGGCGCAGATCTCCA</td><td></td><td></td><td></td><td></td><td></td><td></td><td></td><td></td><td></td><td></td><td></td></tr>
<tr><td>Chromosome</td><td>start</td><td></td><td>end</td><td>strand</td><td>MM</td><td>target_seq</td><td>PAM</td><td>alignment</td><td>distance</td><td>position</td><td>gene name</td><td>gene id</td></tr>
<tr><td>GU305914.1</td><td></td><td>89807</td><td>89829</td><td>+</td><td>0</td><td>TGGAGATCTGCGCCGTCGTC</td><td>AGG</td><td>||||||||[||||||||||||]PAM</td><td>NA</td><td>Intergenic</td><td>NA</td><td>NA</td></tr>
<tr><td>T6376</td><td>GGAGATCTGCGCCGTCGTCAGGG</td><td></td><td>1000</td><td></td><td></td><td></td><td></td><td></td><td></td><td></td><td></td><td></td></tr>
<tr><td>Oligo fwd</td><td>CACCGGAGATCTGCGCCGTCGTCA</td><td></td><td></td><td></td><td></td><td></td><td></td><td></td><td></td><td></td><td></td><td></td></tr>
<tr><td>Oligo rev</td><td>AAACTGACGACGGCGCAGATCTCC</td><td></td><td></td><td></td><td></td><td></td><td></td><td></td><td></td><td></td><td></td><td></td></tr>
<tr><td>Chromosome</td><td>start</td><td></td><td>end</td><td>strand</td><td>MM</td><td>target_seq</td><td>PAM</td><td>alignment</td><td>distance</td><td>position</td><td>gene name</td><td>gene id</td></tr>
<tr><td>GU305914.1</td><td></td><td>89808</td><td>89830</td><td>+</td><td>0</td><td>GGAGATCTGCGCCGTCGTCA</td><td>GGG</td><td>||||||||[||||||||||||]PAM</td><td>NA</td><td>Intergenic</td><td>NA</td><td>NA</td></tr>
<tr><td>T6377</td><td>GAGATCTGCGCCGTCGTCAGGGG</td><td></td><td>1000</td><td></td><td></td><td></td><td></td><td></td><td></td><td></td><td></td><td></td></tr>
<tr><td>Oligo fwd</td><td>CACCGAGATCTGCGCCGTCGTCAG</td><td></td><td></td><td></td><td></td><td></td><td></td><td></td><td></td><td></td><td></td><td></td></tr>
<tr><td>Oligo rev</td><td>AAACCTGACGACGGCGCAGATCTC</td><td></td><td></td><td></td><td></td><td></td><td></td><td></td><td></td><td></td><td></td><td></td></tr>
<tr><td>Chromosome</td><td>start</td><td></td><td>end</td><td>strand</td><td>MM</td><td>target_seq</td><td>PAM</td><td>alignment</td><td>distance</td><td>position</td><td>gene name</td><td>gene id</td></tr>
<tr><td>GU305914.1</td><td></td><td>89809</td><td>89831</td><td>+</td><td>0</td><td>GAGATCTGCGCCGTCGTCAG</td><td>GGG</td><td>||||||||[||||||||||||]PAM</td><td>NA</td><td>Intergenic</td><td>NA</td><td>NA</td></tr>
<tr><td>T6378</td><td>AGATCTGCGCCGTCGTCAGGGGG</td><td></td><td>1000</td><td></td><td></td><td></td><td></td><td></td><td></td><td></td><td></td><td></td></tr>
<tr><td>Oligo fwd</td><td>CACCAGATCTGCGCCGTCGTCAGG</td><td></td><td></td><td></td><td></td><td></td><td></td><td></td><td></td><td></td><td></td><td></td></tr>
<tr><td>Oligo rev</td><td>AAACCCTGACGACGGCGCAGATCT</td><td></td><td></td><td></td><td></td><td></td><td></td><td></td><td></td><td></td><td></td><td></td></tr>
<tr><td>Chromosome</td><td>start</td><td></td><td>end</td><td>strand</td><td>MM</td><td>target_seq</td><td>PAM</td><td>alignment</td><td>distance</td><td>position</td><td>gene name</td><td>gene id</td></tr>
<tr><td>GU305914.1</td><td></td><td>89810</td><td>89832</td><td>+</td><td>0</td><td>AGATCTGCGCCGTCGTCAGG</td><td>GGG</td><td>||||||||[||||||||||||]PAM</td><td>NA</td><td>Intergenic</td><td>NA</td><td>NA</td></tr>
<tr><td>T6379</td><td>GTTGTCGACCACGTACGACGCGG</td><td></td><td>1000</td><td></td><td></td><td></td><td></td><td></td><td></td><td></td><td></td><td></td></tr>
<tr><td>Oligo fwd</td><td>CACCGTTGTCGACCACGTACGACG</td><td></td><td></td><td></td><td></td><td></td><td></td><td></td><td></td><td></td><td></td><td></td></tr>
<tr><td>Oligo rev</td><td>AAACCGTCGTACGTGGTCGACAAC</td><td></td><td></td><td></td><td></td><td></td><td></td><td></td><td></td><td></td><td></td><td></td></tr>
<tr><td>Chromosome</td><td>start</td><td></td><td>end</td><td>strand</td><td>MM</td><td>target_seq</td><td>PAM</td><td>alignment</td><td>distance</td><td>position</td><td>gene name</td><td>gene id</td></tr>
<tr><td>GU305914.1</td><td></td><td>89832</td><td>89854</td><td>+</td><td>0</td><td>GTTGTCGACCACGTACGACG</td><td>CGG</td><td>||||||||[||||||||||||]PAM</td><td>NA</td><td>Intergenic</td><td>NA</td><td>NA</td></tr>
<tr><td>T6380</td><td>CGTACGACGCGGCGTTGATGAGG</td><td></td><td>1000</td><td></td><td></td><td></td><td></td><td></td><td></td><td></td><td></td><td></td></tr>
<tr><td>Oligo fwd</td><td>CACCCGTACGACGCGGCGTTGATG</td><td></td><td></td><td></td><td></td><td></td><td></td><td></td><td></td><td></td><td></td><td></td></tr>
<tr><td>Oligo rev</td><td>AAACCATCAACGCCGCGTCGTACG</td><td></td><td></td><td></td><td></td><td></td><td></td><td></td><td></td><td></td><td></td><td></td></tr>
<tr><td>Chromosome</td><td>start</td><td></td><td>end</td><td>strand</td><td>MM</td><td>target_seq</td><td>PAM</td><td>alignment</td><td>distance</td><td>position</td><td>gene name</td><td>gene id</td></tr>
<tr><td>GU305914.1</td><td></td><td>89843</td><td>89865</td><td>+</td><td>0</td><td>CGTACGACGCGGCGTTGATG</td><td>AGG</td><td>||||||||[||||||||||||]PAM</td><td>NA</td><td>Intergenic</td><td>NA</td><td>NA</td></tr>
<tr><td>T6381</td><td>GTACGACGCGGCGTTGATGAGGG</td><td></td><td>1000</td><td></td><td></td><td></td><td></td><td></td><td></td><td></td><td></td><td></td></tr>
<tr><td>Oligo fwd</td><td>CACCGTACGACGCGGCGTTGATGA</td><td></td><td></td><td></td><td></td><td></td><td></td><td></td><td></td><td></td><td></td><td></td></tr>
<tr><td>Oligo rev</td><td>AAACTCATCAACGCCGCGTCGTAC</td><td></td><td></td><td></td><td></td><td></td><td></td><td></td><td></td><td></td><td></td><td></td></tr>
<tr><td>Chromosome</td><td>start</td><td></td><td>end</td><td>strand</td><td>MM</td><td>target_seq</td><td>PAM</td><td>alignment</td><td>distance</td><td>position</td><td>gene name</td><td>gene id</td></tr>
<tr><td>GU305914.1</td><td></td><td>89844</td><td>89866</td><td>+</td><td>0</td><td>GTACGACGCGGCGTTGATGA</td><td>GGG</td><td>||||||||[||||||||||||]PAM</td><td>NA</td><td>Intergenic</td><td>NA</td><td>NA</td></tr>
<tr><td>T6382</td><td>ACGCGGCGTTGATGAGGGCCAGG</td><td></td><td>1000</td><td></td><td></td><td></td><td></td><td></td><td></td><td></td><td></td><td></td></tr>
<tr><td>Oligo fwd</td><td>CACCACGCGGCGTTGATGAGGGCC</td><td></td><td></td><td></td><td></td><td></td><td></td><td></td><td></td><td></td><td></td><td></td></tr>
<tr><td>Oligo rev</td><td>AAACGGCCCTCATCAACGCCGCGT</td><td></td><td></td><td></td><td></td><td></td><td></td><td></td><td></td><td></td><td></td><td></td></tr>
<tr><td>Chromosome</td><td>start</td><td></td><td>end</td><td>strand</td><td>MM</td><td>target_seq</td><td>PAM</td><td>alignment</td><td>distance</td><td>position</td><td>gene name</td><td>gene id</td></tr>
<tr><td>GU305914.1</td><td></td><td>89849</td><td>89871</td><td>+</td><td>0</td><td>ACGCGGCGTTGATGAGGGCC</td><td>AGG</td><td>||||||||[||||||||||||]PAM</td><td>NA</td><td>Intergenic</td><td>NA</td><td>NA</td></tr>
<tr><td>T6383</td><td>CGTTGATGAGGGCCAGGTTCTGG</td><td></td><td>1000</td><td></td><td></td><td></td><td></td><td></td><td></td><td></td><td></td><td></td></tr>
</table>

<table>
<tr><td>Oligo fwd</td><td>CACCCGTTGATGAGGGCCAGGTTC</td><td></td><td></td><td></td><td></td><td></td><td></td><td></td><td></td><td></td><td></td></tr>
<tr><td>Oligo rev</td><td>AAACGAACCTGGCCCTCATCAACG</td><td></td><td></td><td></td><td></td><td></td><td></td><td></td><td></td><td></td><td></td></tr>
<tr><td>Chromosome</td><td>start</td><td>end</td><td>strand</td><td>MM</td><td>target_seq</td><td>PAM</td><td>alignment</td><td>distance</td><td>position</td><td>gene name</td><td>gene id</td></tr>
<tr><td>GU305914.1</td><td>89855</td><td>89877</td><td>+</td><td>0</td><td>CGTTGATGAGGGCCAGGTTC</td><td>TGG</td><td>||||||||[||||||||||||]PAM</td><td>NA</td><td>Intergenic</td><td>NA</td><td>NA</td></tr>
<tr><td>T6384</td><td>GAGGGCCAGGTTCTGGATGATGG</td><td>1000</td><td></td><td></td><td></td><td></td><td></td><td></td><td></td><td></td><td></td></tr>
<tr><td>Oligo fwd</td><td>CACCGAGGGCCAGGTTCTGGATGA</td><td></td><td></td><td></td><td></td><td></td><td></td><td></td><td></td><td></td><td></td></tr>
<tr><td>Oligo rev</td><td>AAACTCATCCAGAACCTGGCCCTC</td><td></td><td></td><td></td><td></td><td></td><td></td><td></td><td></td><td></td><td></td></tr>
<tr><td>Chromosome</td><td>start</td><td>end</td><td>strand</td><td>MM</td><td>target_seq</td><td>PAM</td><td>alignment</td><td>distance</td><td>position</td><td>gene name</td><td>gene id</td></tr>
<tr><td>GU305914.1</td><td>89862</td><td>89884</td><td>+</td><td>0</td><td>GAGGGCCAGGTTCTGGATGA</td><td>TGG</td><td>||||||||[||||||||||||]PAM</td><td>NA</td><td>Intergenic</td><td>NA</td><td>NA</td></tr>
<tr><td>T6385</td><td>GTTCTGGATGATGGAGAACATGG</td><td>1000</td><td></td><td></td><td></td><td></td><td></td><td></td><td></td><td></td><td></td></tr>
<tr><td>Oligo fwd</td><td>CACCGTTCTGGATGATGGAGAACA</td><td></td><td></td><td></td><td></td><td></td><td></td><td></td><td></td><td></td><td></td></tr>
<tr><td>Oligo rev</td><td>AAACTGTTCTCCATCATCCAGAAC</td><td></td><td></td><td></td><td></td><td></td><td></td><td></td><td></td><td></td><td></td></tr>
<tr><td>Chromosome</td><td>start</td><td>end</td><td>strand</td><td>MM</td><td>target_seq</td><td>PAM</td><td>alignment</td><td>distance</td><td>position</td><td>gene name</td><td>gene id</td></tr>
<tr><td>GU305914.1</td><td>89871</td><td>89893</td><td>+</td><td>0</td><td>GTTCTGGATGATGGAGAACA</td><td>TGG</td><td>||||||||[||||||||||||]PAM</td><td>NA</td><td>Intergenic</td><td>NA</td><td>NA</td></tr>
<tr><td>T6386</td><td>GGATGATGGAGAACATGGTCCGG</td><td>1000</td><td></td><td></td><td></td><td></td><td></td><td></td><td></td><td></td><td></td></tr>
<tr><td>Oligo fwd</td><td>CACCGGATGATGGAGAACATGGTC</td><td></td><td></td><td></td><td></td><td></td><td></td><td></td><td></td><td></td><td></td></tr>
<tr><td>Oligo rev</td><td>AAACGACCATGTTCTCCATCATCC</td><td></td><td></td><td></td><td></td><td></td><td></td><td></td><td></td><td></td><td></td></tr>
<tr><td>Chromosome</td><td>start</td><td>end</td><td>strand</td><td>MM</td><td>target_seq</td><td>PAM</td><td>alignment</td><td>distance</td><td>position</td><td>gene name</td><td>gene id</td></tr>
<tr><td>GU305914.1</td><td>89876</td><td>89898</td><td>+</td><td>0</td><td>GGATGATGGAGAACATGGTC</td><td>CGG</td><td>||||||||[||||||||||||]PAM</td><td>NA</td><td>Intergenic</td><td>NA</td><td>NA</td></tr>
<tr><td>T6387</td><td>AGAACATGGTCCGGTAATACTGG</td><td>1000</td><td></td><td></td><td></td><td></td><td></td><td></td><td></td><td></td><td></td></tr>
<tr><td>Oligo fwd</td><td>CACCAGAACATGGTCCGGTAATAC</td><td></td><td></td><td></td><td></td><td></td><td></td><td></td><td></td><td></td><td></td></tr>
<tr><td>Oligo rev</td><td>AAACGTATTACCGGACCATGTTCT</td><td></td><td></td><td></td><td></td><td></td><td></td><td></td><td></td><td></td><td></td></tr>
<tr><td>Chromosome</td><td>start</td><td>end</td><td>strand</td><td>MM</td><td>target_seq</td><td>PAM</td><td>alignment</td><td>distance</td><td>position</td><td>gene name</td><td>gene id</td></tr>
<tr><td>GU305914.1</td><td>89885</td><td>89907</td><td>+</td><td>0</td><td>AGAACATGGTCCGGTAATAC</td><td>TGG</td><td>||||||||[||||||||||||]PAM</td><td>NA</td><td>Intergenic</td><td>NA</td><td>NA</td></tr>
<tr><td>T6388</td><td>TCCGGTAATACTGGAACGACAGG</td><td>1000</td><td></td><td></td><td></td><td></td><td></td><td></td><td></td><td></td><td></td></tr>
<tr><td>Oligo fwd</td><td>CACCTCCGGTAATACTGGAACGAC</td><td></td><td></td><td></td><td></td><td></td><td></td><td></td><td></td><td></td><td></td></tr>
<tr><td>Oligo rev</td><td>AAACGTCGTTCCAGTATTACCGGA</td><td></td><td></td><td></td><td></td><td></td><td></td><td></td><td></td><td></td><td></td></tr>
<tr><td>Chromosome</td><td>start</td><td>end</td><td>strand</td><td>MM</td><td>target_seq</td><td>PAM</td><td>alignment</td><td>distance</td><td>position</td><td>gene name</td><td>gene id</td></tr>
<tr><td>GU305914.1</td><td>89894</td><td>89916</td><td>+</td><td>0</td><td>TCCGGTAATACTGGAACGAC</td><td>AGG</td><td>||||||||[||||||||||||]PAM</td><td>NA</td><td>Intergenic</td><td>NA</td><td>NA</td></tr>
<tr><td>T6389</td><td>GTAATACTGGAACGACAGGACGG</td><td>1000</td><td></td><td></td><td></td><td></td><td></td><td></td><td></td><td></td><td></td></tr>
<tr><td>Oligo fwd</td><td>CACCGTAATACTGGAACGACAGGA</td><td></td><td></td><td></td><td></td><td></td><td></td><td></td><td></td><td></td><td></td></tr>
<tr><td>Oligo rev</td><td>AAACTCCTGTCGTTCCAGTATTAC</td><td></td><td></td><td></td><td></td><td></td><td></td><td></td><td></td><td></td><td></td></tr>
<tr><td>Chromosome</td><td>start</td><td>end</td><td>strand</td><td>MM</td><td>target_seq</td><td>PAM</td><td>alignment</td><td>distance</td><td>position</td><td>gene name</td><td>gene id</td></tr>
<tr><td>GU305914.1</td><td>89898</td><td>89920</td><td>+</td><td>0</td><td>GTAATACTGGAACGACAGGA</td><td>CGG</td><td>||||||||[||||||||||||]PAM</td><td>NA</td><td>Intergenic</td><td>NA</td><td>NA</td></tr>
<tr><td>T6390</td><td>TAATACTGGAACGACAGGACGGG</td><td>1000</td><td></td><td></td><td></td><td></td><td></td><td></td><td></td><td></td><td></td></tr>
<tr><td>Oligo fwd</td><td>CACCTAATACTGGAACGACAGGAC</td><td></td><td></td><td></td><td></td><td></td><td></td><td></td><td></td><td></td><td></td></tr>
<tr><td>Oligo rev</td><td>AAACGTCCTGTCGTTCCAGTATTA</td><td></td><td></td><td></td><td></td><td></td><td></td><td></td><td></td><td></td><td></td></tr>
<tr><td>Chromosome</td><td>start</td><td>end</td><td>strand</td><td>MM</td><td>target_seq</td><td>PAM</td><td>alignment</td><td>distance</td><td>position</td><td>gene name</td><td>gene id</td></tr>
<tr><td>GU305914.1</td><td>89899</td><td>89921</td><td>+</td><td>0</td><td>TAATACTGGAACGACAGGAC</td><td>GGG</td><td>||||||||[||||||||||||]PAM</td><td>NA</td><td>Intergenic</td><td>NA</td><td>NA</td></tr>
<tr><td>T6391</td><td>AATACTGGAACGACAGGACGGGG</td><td>1000</td><td></td><td></td><td></td><td></td><td></td><td></td><td></td><td></td><td></td></tr>
<tr><td>Oligo fwd</td><td>CACCAATACTGGAACGACAGGACG</td><td></td><td></td><td></td><td></td><td></td><td></td><td></td><td></td><td></td><td></td></tr>
<tr><td>Oligo rev</td><td>AAACCGTCCTGTCGTTCCAGTATT</td><td></td><td></td><td></td><td></td><td></td><td></td><td></td><td></td><td></td><td></td></tr>
<tr><td>Chromosome</td><td>start</td><td>end</td><td>strand</td><td>MM</td><td>target_seq</td><td>PAM</td><td>alignment</td><td>distance</td><td>position</td><td>gene name</td><td>gene id</td></tr>
<tr><td>GU305914.1</td><td>89900</td><td>89922</td><td>+</td><td>0</td><td>AATACTGGAACGACAGGACG</td><td>GGG</td><td>||||||||[||||||||||||]PAM</td><td>NA</td><td>Intergenic</td><td>NA</td><td>NA</td></tr>
<tr><td>T6392</td><td>ATACTGGAACGACAGGACGGGGG</td><td>1000</td><td></td><td></td><td></td><td></td><td></td><td></td><td></td><td></td><td></td></tr>
<tr><td>Oligo fwd</td><td>CACCATACTGGAACGACAGGACGG</td><td></td><td></td><td></td><td></td><td></td><td></td><td></td><td></td><td></td><td></td></tr>
<tr><td>Oligo rev</td><td>AAACCCGTCCTGTCGTTCCAGTAT</td><td></td><td></td><td></td><td></td><td></td><td></td><td></td><td></td><td></td><td></td></tr>
<tr><td>Chromosome</td><td>start</td><td>end</td><td>strand</td><td>MM</td><td>target_seq</td><td>PAM</td><td>alignment</td><td>distance</td><td>position</td><td>gene name</td><td>gene id</td></tr>
<tr><td>GU305914.1</td><td>89901</td><td>89923</td><td>+</td><td>0</td><td>ATACTGGAACGACAGGACGG</td><td>GGG</td><td>||||||||[||||||||||||]PAM</td><td>NA</td><td>Intergenic</td><td>NA</td><td>NA</td></tr>
<tr><td>T6393</td><td>GAACGACAGGACGGGGGAGAAGG</td><td>1000</td><td></td><td></td><td></td><td></td><td></td><td></td><td></td><td></td><td></td></tr>
<tr><td>Oligo fwd</td><td>CACCGAACGACAGGACGGGGGAGA</td><td></td><td></td><td></td><td></td><td></td><td></td><td></td><td></td><td></td><td></td></tr>
<tr><td>Oligo rev</td><td>AAACTCTCCCCCGTCCTGTCGTTC</td><td></td><td></td><td></td><td></td><td></td><td></td><td></td><td></td><td></td><td></td></tr>
<tr><td>Chromosome</td><td>start</td><td>end</td><td>strand</td><td>MM</td><td>target_seq</td><td>PAM</td><td>alignment</td><td>distance</td><td>position</td><td>gene name</td><td>gene id</td></tr>
<tr><td>GU305914.1</td><td>89907</td><td>89929</td><td>+</td><td>0</td><td>GAACGACAGGACGGGGGAGA</td><td>AGG</td><td>||||||||[||||||||||||]PAM</td><td>NA</td><td>Intergenic</td><td>NA</td><td>NA</td></tr>
<tr><td>T6394</td><td>CAGGACGGGGGAGAAGGCCGTGG</td><td>1000</td><td></td><td></td><td></td><td></td><td></td><td></td><td></td><td></td><td></td></tr>
<tr><td>Oligo fwd</td><td>CACCCAGGACGGGGGAGAAGGCCG</td><td></td><td></td><td></td><td></td><td></td><td></td><td></td><td></td><td></td><td></td></tr>
<tr><td>Oligo rev</td><td>AAACCGGCCTTCTCCCCCGTCCTG</td><td></td><td></td><td></td><td></td><td></td><td></td><td></td><td></td><td></td><td></td></tr>
<tr><td>Chromosome</td><td>start</td><td>end</td><td>strand</td><td>MM</td><td>target_seq</td><td>PAM</td><td>alignment</td><td>distance</td><td>position</td><td>gene name</td><td>gene id</td></tr>
<tr><td>GU305914.1</td><td>89913</td><td>89935</td><td>+</td><td>0</td><td>CAGGACGGGGGAGAAGGCCG</td><td>TGG</td><td>||||||||[||||||||||||]PAM</td><td>NA</td><td>Intergenic</td><td>NA</td><td>NA</td></tr>
<tr><td>T6395</td><td>GGGGGAGAAGGCCGTGGCGTAGG</td><td>1000</td><td></td><td></td><td></td><td></td><td></td><td></td><td></td><td></td><td></td></tr>
<tr><td>Oligo fwd</td><td>CACCGGGGGAGAAGGCCGTGGCGT</td><td></td><td></td><td></td><td></td><td></td><td></td><td></td><td></td><td></td><td></td></tr>
<tr><td>Oligo rev</td><td>AAACACGCCACGGCCTTCTCCCCC</td><td></td><td></td><td></td><td></td><td></td><td></td><td></td><td></td><td></td><td></td></tr>
<tr><td>Chromosome</td><td>start</td><td>end</td><td>strand</td><td>MM</td><td>target_seq</td><td>PAM</td><td>alignment</td><td>distance</td><td>position</td><td>gene name</td><td>gene id</td></tr>
<tr><td>GU305914.1</td><td>89919</td><td>89941</td><td>+</td><td>0</td><td>GGGGGAGAAGGCCGTGGCGT</td><td>AGG</td><td>||||||||[||||||||||||]PAM</td><td>NA</td><td>Intergenic</td><td>NA</td><td>NA</td></tr>
<tr><td>T6396</td><td>GGGGAGAAGGCCGTGGCGTAGGG</td><td>1000</td><td></td><td></td><td></td><td></td><td></td><td></td><td></td><td></td><td></td></tr>
<tr><td>Oligo fwd</td><td>CACCGGGGAGAAGGCCGTGGCGTA</td><td></td><td></td><td></td><td></td><td></td><td></td><td></td><td></td><td></td><td></td></tr>
<tr><td>Oligo rev</td><td>AAACTACGCCACGGCCTTCTCCCC</td><td></td><td></td><td></td><td></td><td></td><td></td><td></td><td></td><td></td><td></td></tr>
<tr><td>Chromosome</td><td>start</td><td>end</td><td>strand</td><td>MM</td><td>target_seq</td><td>PAM</td><td>alignment</td><td>distance</td><td>position</td><td>gene name</td><td>gene id</td></tr>
<tr><td>GU305914.1</td><td>89920</td><td>89942</td><td>+</td><td>0</td><td>GGGGAGAAGGCCGTGGCGTA</td><td>GGG</td><td>||||||||[||||||||||||]PAM</td><td>NA</td><td>Intergenic</td><td>NA</td><td>NA</td></tr>
<tr><td>T6397</td><td>GGGAGAAGGCCGTGGCGTAGGGG</td><td>1000</td><td></td><td></td><td></td><td></td><td></td><td></td><td></td><td></td><td></td></tr>
<tr><td>Oligo fwd</td><td>CACCGGGAGAAGGCCGTGGCGTAG</td><td></td><td></td><td></td><td></td><td></td><td></td><td></td><td></td><td></td><td></td></tr>
<tr><td>Oligo rev</td><td>AAACCTACGCCACGGCCTTCTCCC</td><td></td><td></td><td></td><td></td><td></td><td></td><td></td><td></td><td></td><td></td></tr>
<tr><td>Chromosome</td><td>start</td><td>end</td><td>strand</td><td>MM</td><td>target_seq</td><td>PAM</td><td>alignment</td><td>distance</td><td>position</td><td>gene name</td><td>gene id</td></tr>
<tr><td>GU305914.1</td><td>89921</td><td>89943</td><td>+</td><td>0</td><td>GGGAGAAGGCCGTGGCGTAG</td><td>GGG</td><td>||||||||[||||||||||||]PAM</td><td>NA</td><td>Intergenic</td><td>NA</td><td>NA</td></tr>
</table>

<table>
<tr><td>T6398</td><td>AGGCCGTGGCGTAGGGGCTGAGG</td><td>1000</td><td></td><td></td><td></td><td></td><td></td><td></td><td></td><td></td><td></td></tr>
<tr><td>Oligo fwd</td><td>CACCAGGCCGTGGCGTAGGGGCTG</td><td></td><td></td><td></td><td></td><td></td><td></td><td></td><td></td><td></td><td></td></tr>
<tr><td>Oligo rev</td><td>AAACCAGCCCCTACGCCACGGCCT</td><td></td><td></td><td></td><td></td><td></td><td></td><td></td><td></td><td></td><td></td></tr>
<tr><td>Chromosome</td><td>start</td><td>end</td><td>strand</td><td>MM</td><td>target_seq</td><td>PAM</td><td>alignment</td><td>distance</td><td>position</td><td>gene name</td><td>gene id</td></tr>
<tr><td>GU305914.1</td><td>89927</td><td>89949</td><td>+</td><td>0</td><td>AGGCCGTGGCGTAGGGGCTG</td><td>AGG</td><td>||||||||[||||||||||||]PAM</td><td>NA</td><td>Intergenic</td><td>NA</td><td>NA</td></tr>
<tr><td>T6399</td><td>TGGCGTAGGGGCTGAGGTCCAGG</td><td>1000</td><td></td><td></td><td></td><td></td><td></td><td></td><td></td><td></td><td></td></tr>
<tr><td>Oligo fwd</td><td>CACCTGGCGTAGGGGCTGAGGTCC</td><td></td><td></td><td></td><td></td><td></td><td></td><td></td><td></td><td></td><td></td></tr>
<tr><td>Oligo rev</td><td>AAACGGACCTCAGCCCCTACGCCA</td><td></td><td></td><td></td><td></td><td></td><td></td><td></td><td></td><td></td><td></td></tr>
<tr><td>Chromosome</td><td>start</td><td>end</td><td>strand</td><td>MM</td><td>target_seq</td><td>PAM</td><td>alignment</td><td>distance</td><td>position</td><td>gene name</td><td>gene id</td></tr>
<tr><td>GU305914.1</td><td>89933</td><td>89955</td><td>+</td><td>0</td><td>TGGCGTAGGGGCTGAGGTCC</td><td>AGG</td><td>||||||||[||||||||||||]PAM</td><td>NA</td><td>Intergenic</td><td>NA</td><td>NA</td></tr>
<tr><td>T6400</td><td>GGGGCTGAGGTCCAGGTTGAAGG</td><td>1000</td><td></td><td></td><td></td><td></td><td></td><td></td><td></td><td></td><td></td></tr>
<tr><td>Oligo fwd</td><td>CACCGGGGCTGAGGTCCAGGTTGA</td><td></td><td></td><td></td><td></td><td></td><td></td><td></td><td></td><td></td><td></td></tr>
<tr><td>Oligo rev</td><td>AAACTCAACCTGGACCTCAGCCCC</td><td></td><td></td><td></td><td></td><td></td><td></td><td></td><td></td><td></td><td></td></tr>
<tr><td>Chromosome</td><td>start</td><td>end</td><td>strand</td><td>MM</td><td>target_seq</td><td>PAM</td><td>alignment</td><td>distance</td><td>position</td><td>gene name</td><td>gene id</td></tr>
<tr><td>GU305914.1</td><td>89940</td><td>89962</td><td>+</td><td>0</td><td>GGGGCTGAGGTCCAGGTTGA</td><td>AGG</td><td>||||||||[||||||||||||]PAM</td><td>NA</td><td>Intergenic</td><td>NA</td><td>NA</td></tr>
<tr><td>T6401</td><td>GATCTCGTCCCTCCCGTGCCCGG</td><td>1000</td><td></td><td></td><td></td><td></td><td></td><td></td><td></td><td></td><td></td></tr>
<tr><td>Oligo fwd</td><td>CACCGATCTCGTCCCTCCCGTGCC</td><td></td><td></td><td></td><td></td><td></td><td></td><td></td><td></td><td></td><td></td></tr>
<tr><td>Oligo rev</td><td>AAACGGCACGGGAGGGACGAGATC</td><td></td><td></td><td></td><td></td><td></td><td></td><td></td><td></td><td></td><td></td></tr>
<tr><td>Chromosome</td><td>start</td><td>end</td><td>strand</td><td>MM</td><td>target_seq</td><td>PAM</td><td>alignment</td><td>distance</td><td>position</td><td>gene name</td><td>gene id</td></tr>
<tr><td>GU305914.1</td><td>89979</td><td>90001</td><td>+</td><td>0</td><td>GATCTCGTCCCTCCCGTGCC</td><td>CGG</td><td>||||||||[||||||||||||]PAM</td><td>NA</td><td>Intergenic</td><td>NA</td><td>NA</td></tr>
<tr><td>T6402</td><td>CCCGGACCTCCGCACGTCCATGG</td><td>1000</td><td></td><td></td><td></td><td></td><td></td><td></td><td></td><td></td><td></td></tr>
<tr><td>Oligo fwd</td><td>CACCCCCGGACCTCCGCACGTCCA</td><td></td><td></td><td></td><td></td><td></td><td></td><td></td><td></td><td></td><td></td></tr>
<tr><td>Oligo rev</td><td>AAACTGGACGTGCGGAGGTCCGGG</td><td></td><td></td><td></td><td></td><td></td><td></td><td></td><td></td><td></td><td></td></tr>
<tr><td>Chromosome</td><td>start</td><td>end</td><td>strand</td><td>MM</td><td>target_seq</td><td>PAM</td><td>alignment</td><td>distance</td><td>position</td><td>gene name</td><td>gene id</td></tr>
<tr><td>GU305914.1</td><td>89997</td><td>90019</td><td>+</td><td>0</td><td>CCCGGACCTCCGCACGTCCA</td><td>TGG</td><td>||||||||[||||||||||||]PAM</td><td>NA</td><td>Intergenic</td><td>NA</td><td>NA</td></tr>
<tr><td>T6403</td><td>GTCCATGGCGAACCTGCACACGG</td><td>1000</td><td></td><td></td><td></td><td></td><td></td><td></td><td></td><td></td><td></td></tr>
<tr><td>Oligo fwd</td><td>CACCGTCCATGGCGAACCTGCACA</td><td></td><td></td><td></td><td></td><td></td><td></td><td></td><td></td><td></td><td></td></tr>
<tr><td>Oligo rev</td><td>AAACTGTGCAGGTTCGCCATGGAC</td><td></td><td></td><td></td><td></td><td></td><td></td><td></td><td></td><td></td><td></td></tr>
<tr><td>Chromosome</td><td>start</td><td>end</td><td>strand</td><td>MM</td><td>target_seq</td><td>PAM</td><td>alignment</td><td>distance</td><td>position</td><td>gene name</td><td>gene id</td></tr>
<tr><td>GU305914.1</td><td>90012</td><td>90034</td><td>+</td><td>0</td><td>GTCCATGGCGAACCTGCACA</td><td>CGG</td><td>||||||||[||||||||||||]PAM</td><td>NA</td><td>Intergenic</td><td>NA</td><td>NA</td></tr>
<tr><td>T6404</td><td>ACACGGCATCGTCGATCGTCTGG</td><td>1000</td><td></td><td></td><td></td><td></td><td></td><td></td><td></td><td></td><td></td></tr>
<tr><td>Oligo fwd</td><td>CACCACACGGCATCGTCGATCGTC</td><td></td><td></td><td></td><td></td><td></td><td></td><td></td><td></td><td></td><td></td></tr>
<tr><td>Oligo rev</td><td>AAACGACGATCGACGATGCCGTGT</td><td></td><td></td><td></td><td></td><td></td><td></td><td></td><td></td><td></td><td></td></tr>
<tr><td>Chromosome</td><td>start</td><td>end</td><td>strand</td><td>MM</td><td>target_seq</td><td>PAM</td><td>alignment</td><td>distance</td><td>position</td><td>gene name</td><td>gene id</td></tr>
<tr><td>GU305914.1</td><td>90029</td><td>90051</td><td>+</td><td>0</td><td>ACACGGCATCGTCGATCGTC</td><td>TGG</td><td>||||||||[||||||||||||]PAM</td><td>NA</td><td>Intergenic</td><td>NA</td><td>NA</td></tr>
<tr><td>T6405</td><td>CACGTTGATGATATCCTCGCCGG</td><td>1000</td><td></td><td></td><td></td><td></td><td></td><td></td><td></td><td></td><td></td></tr>
<tr><td>Oligo fwd</td><td>CACCCACGTTGATGATATCCTCGC</td><td></td><td></td><td></td><td></td><td></td><td></td><td></td><td></td><td></td><td></td></tr>
<tr><td>Oligo rev</td><td>AAACGCGAGGATATCATCAACGTG</td><td></td><td></td><td></td><td></td><td></td><td></td><td></td><td></td><td></td><td></td></tr>
<tr><td>Chromosome</td><td>start</td><td>end</td><td>strand</td><td>MM</td><td>target_seq</td><td>PAM</td><td>alignment</td><td>distance</td><td>position</td><td>gene name</td><td>gene id</td></tr>
<tr><td>GU305914.1</td><td>90093</td><td>90115</td><td>+</td><td>0</td><td>CACGTTGATGATATCCTCGC</td><td>CGG</td><td>||||||||[||||||||||||]PAM</td><td>NA</td><td>Intergenic</td><td>NA</td><td>NA</td></tr>
<tr><td>T6406</td><td>AATTCTTCTTACAGTAGTCCAGG</td><td>1000</td><td></td><td></td><td></td><td></td><td></td><td></td><td></td><td></td><td></td></tr>
<tr><td>Oligo fwd</td><td>CACCAATTCTTCTTACAGTAGTCC</td><td></td><td></td><td></td><td></td><td></td><td></td><td></td><td></td><td></td><td></td></tr>
<tr><td>Oligo rev</td><td>AAACGGACTACTGTAAGAAGAATT</td><td></td><td></td><td></td><td></td><td></td><td></td><td></td><td></td><td></td><td></td></tr>
<tr><td>Chromosome</td><td>start</td><td>end</td><td>strand</td><td>MM</td><td>target_seq</td><td>PAM</td><td>alignment</td><td>distance</td><td>position</td><td>gene name</td><td>gene id</td></tr>
<tr><td>GU305914.1</td><td>90134</td><td>90156</td><td>+</td><td>0</td><td>AATTCTTCTTACAGTAGTCC</td><td>AGG</td><td>||||||||[||||||||||||]PAM</td><td>NA</td><td>Intergenic</td><td>NA</td><td>NA</td></tr>
<tr><td>T6407</td><td>GAGATACTTCATCCTGTCGACGG</td><td>1000</td><td></td><td></td><td></td><td></td><td></td><td></td><td></td><td></td><td></td></tr>
<tr><td>Oligo fwd</td><td>CACCGAGATACTTCATCCTGTCGA</td><td></td><td></td><td></td><td></td><td></td><td></td><td></td><td></td><td></td><td></td></tr>
<tr><td>Oligo rev</td><td>AAACTCGACAGGATGAAGTATCTC</td><td></td><td></td><td></td><td></td><td></td><td></td><td></td><td></td><td></td><td></td></tr>
<tr><td>Chromosome</td><td>start</td><td>end</td><td>strand</td><td>MM</td><td>target_seq</td><td>PAM</td><td>alignment</td><td>distance</td><td>position</td><td>gene name</td><td>gene id</td></tr>
<tr><td>GU305914.1</td><td>90165</td><td>90187</td><td>+</td><td>0</td><td>GAGATACTTCATCCTGTCGA</td><td>CGG</td><td>||||||||[||||||||||||]PAM</td><td>NA</td><td>Intergenic</td><td>NA</td><td>NA</td></tr>
<tr><td>T6408</td><td>GGAGACGAAGTTCAACCCGCTGG</td><td>1000</td><td></td><td></td><td></td><td></td><td></td><td></td><td></td><td></td><td></td></tr>
<tr><td>Oligo fwd</td><td>CACCGGAGACGAAGTTCAACCCGC</td><td></td><td></td><td></td><td></td><td></td><td></td><td></td><td></td><td></td><td></td></tr>
<tr><td>Oligo rev</td><td>AAACGCGGGTTGAACTTCGTCTCC</td><td></td><td></td><td></td><td></td><td></td><td></td><td></td><td></td><td></td><td></td></tr>
<tr><td>Chromosome</td><td>start</td><td>end</td><td>strand</td><td>MM</td><td>target_seq</td><td>PAM</td><td>alignment</td><td>distance</td><td>position</td><td>gene name</td><td>gene id</td></tr>
<tr><td>GU305914.1</td><td>90186</td><td>90208</td><td>+</td><td>0</td><td>GGAGACGAAGTTCAACCCGC</td><td>TGG</td><td>||||||||[||||||||||||]PAM</td><td>NA</td><td>Intergenic</td><td>NA</td><td>NA</td></tr>
<tr><td>T6409</td><td>GTTCAACCCGCTGGCGCCGCCGG</td><td>1000</td><td></td><td></td><td></td><td></td><td></td><td></td><td></td><td></td><td></td></tr>
<tr><td>Oligo fwd</td><td>CACCGTTCAACCCGCTGGCGCCGC</td><td></td><td></td><td></td><td></td><td></td><td></td><td></td><td></td><td></td><td></td></tr>
<tr><td>Oligo rev</td><td>AAACGCGGCGCCAGCGGGTTGAAC</td><td></td><td></td><td></td><td></td><td></td><td></td><td></td><td></td><td></td><td></td></tr>
<tr><td>Chromosome</td><td>start</td><td>end</td><td>strand</td><td>MM</td><td>target_seq</td><td>PAM</td><td>alignment</td><td>distance</td><td>position</td><td>gene name</td><td>gene id</td></tr>
<tr><td>GU305914.1</td><td>90195</td><td>90217</td><td>+</td><td>0</td><td>GTTCAACCCGCTGGCGCCGC</td><td>CGG</td><td>||||||||[||||||||||||]PAM</td><td>NA</td><td>Intergenic</td><td>NA</td><td>NA</td></tr>
<tr><td>T6410</td><td>AACCCGCTGGCGCCGCCGGCCGG</td><td>1000</td><td></td><td></td><td></td><td></td><td></td><td></td><td></td><td></td><td></td></tr>
<tr><td>Oligo fwd</td><td>CACCAACCCGCTGGCGCCGCCGGC</td><td></td><td></td><td></td><td></td><td></td><td></td><td></td><td></td><td></td><td></td></tr>
<tr><td>Oligo rev</td><td>AAACGCCGGCGGCGCCAGCGGGTT</td><td></td><td></td><td></td><td></td><td></td><td></td><td></td><td></td><td></td><td></td></tr>
<tr><td>Chromosome</td><td>start</td><td>end</td><td>strand</td><td>MM</td><td>target_seq</td><td>PAM</td><td>alignment</td><td>distance</td><td>position</td><td>gene name</td><td>gene id</td></tr>
<tr><td>GU305914.1</td><td>90199</td><td>90221</td><td>+</td><td>0</td><td>AACCCGCTGGCGCCGCCGGC</td><td>CGG</td><td>||||||||[||||||||||||]PAM</td><td>NA</td><td>Intergenic</td><td>NA</td><td>NA</td></tr>
<tr><td>T6411</td><td>CCGCTGGCGCCGCCGGCCGGAGG</td><td>1000</td><td></td><td></td><td></td><td></td><td></td><td></td><td></td><td></td><td></td></tr>
<tr><td>Oligo fwd</td><td>CACCCCGCTGGCGCCGCCGGCCGG</td><td></td><td></td><td></td><td></td><td></td><td></td><td></td><td></td><td></td><td></td></tr>
<tr><td>Oligo rev</td><td>AAACCCGGCCGGCGGCGCCAGCGG</td><td></td><td></td><td></td><td></td><td></td><td></td><td></td><td></td><td></td><td></td></tr>
<tr><td>Chromosome</td><td>start</td><td>end</td><td>strand</td><td>MM</td><td>target_seq</td><td>PAM</td><td>alignment</td><td>distance</td><td>position</td><td>gene name</td><td>gene id</td></tr>
<tr><td>GU305914.1</td><td>90202</td><td>90224</td><td>+</td><td>0</td><td>CCGCTGGCGCCGCCGGCCGG</td><td>AGG</td><td>||||||||[||||||||||||]PAM</td><td>NA</td><td>Intergenic</td><td>NA</td><td>NA</td></tr>
<tr><td>GU305914.1</td><td>115372</td><td>115394</td><td>-</td><td>4</td><td>ACGACGGTGCCGCCGGCCGG</td><td>CGG</td><td>-||--||-[||||||||||||]PAM</td><td>NA</td><td>Intergenic</td><td>NA</td><td>NA</td></tr>
<tr><td>GU305914.1</td><td>121642</td><td>121664</td><td>-</td><td>3</td><td>CCGCTGAAGCCGCCGTCCGG</td><td>CGG</td><td>||||||--[|||||||-||||]PAM</td><td>NA</td><td>Intergenic</td><td>NA</td><td>NA</td></tr>
<tr><td>T6412</td><td>GCCGCCGGCCGGAGGCCCCGCGG</td><td>1000</td><td></td><td></td><td></td><td></td><td></td><td></td><td></td><td></td><td></td></tr>
<tr><td>Oligo fwd</td><td>CACCGCCGCCGGCCGGAGGCCCCG</td><td></td><td></td><td></td><td></td><td></td><td></td><td></td><td></td><td></td><td></td></tr>
<tr><td>Oligo rev</td><td>AAACCGGGGCCTCCGGCCGGCGGC</td><td></td><td></td><td></td><td></td><td></td><td></td><td></td><td></td><td></td><td></td></tr>
</table>

| | | | | | | | | | | | |
|---|---|---|---|---|---|---|---|---|---|---|---|
| Chromosome | start | end | strand | MM | target_seq | PAM | alignment | distance | position | gene name | gene id |
| GU305914.1 | 90210 | 90232 | + | 0 | GCCGCCGGCCGGAGGCCCCG | CGG | \|\|\|\|\|\|\|\|[\|\|\|\|\|\|\|\|\|\|\|\|]PAM | NA | Intergenic | NA | NA |
| T6413 | CACCTCGTCCGTCCTGCCGCCGG | 1000 | | | | | | | | | |
| Oligo fwd | CACCCACCTCGTCCGTCCTGCCGC | | | | | | | | | | |
| Oligo rev | AAACGCGGCAGGACGGACGAGGTG | | | | | | | | | | |
| Chromosome | start | end | strand | MM | target_seq | PAM | alignment | distance | position | gene name | gene id |
| GU305914.1 | 90234 | 90256 | + | 0 | CACCTCGTCCGTCCTGCCGC | CGG | \|\|\|\|\|\|\|\|[\|\|\|\|\|\|\|\|\|\|\|\|]PAM | NA | Intergenic | NA | NA |
| T6414 | CTCCGCCGTCGCCCGCCTCCCGG | 1000 | | | | | | | | | |
| Oligo fwd | CACCCTCCGCCGTCGCCCGCCTCC | | | | | | | | | | |
| Oligo rev | AAACGGAGGCGGGCGACGGCGGAG | | | | | | | | | | |
| Chromosome | start | end | strand | MM | target_seq | PAM | alignment | distance | position | gene name | gene id |
| GU305914.1 | 90266 | 90288 | + | 0 | CTCCGCCGTCGCCCGCCTCC | CGG | \|\|\|\|\|\|\|\|[\|\|\|\|\|\|\|\|\|\|\|\|]PAM | NA | Intergenic | NA | NA |
| T6415 | TCCGCCGTCGCCCGCCTCCCGGG | 1000 | | | | | | | | | |
| Oligo fwd | CACCTCCGCCGTCGCCCGCCTCCC | | | | | | | | | | |
| Oligo rev | AAACGGGAGGCGGGCGACGGCGGA | | | | | | | | | | |
| Chromosome | start | end | strand | MM | target_seq | PAM | alignment | distance | position | gene name | gene id |
| GU305914.1 | 90267 | 90289 | + | 0 | TCCGCCGTCGCCCGCCTCCC | GGG | \|\|\|\|\|\|\|\|[\|\|\|\|\|\|\|\|\|\|\|\|]PAM | NA | Intergenic | NA | NA |
| T6416 | CGCCCGCCTCCCGGGACTCCAGG | 1000 | | | | | | | | | |
| Oligo fwd | CACCCGCCCGCCTCCCGGGACTCC | | | | | | | | | | |
| Oligo rev | AAACGGAGTCCCGGGAGGCGGGCG | | | | | | | | | | |
| Chromosome | start | end | strand | MM | target_seq | PAM | alignment | distance | position | gene name | gene id |
| GU305914.1 | 90275 | 90297 | + | 0 | CGCCCGCCTCCCGGGACTCC | AGG | \|\|\|\|\|\|\|\|[\|\|\|\|\|\|\|\|\|\|\|\|]PAM | NA | Intergenic | NA | NA |
| T6417 | TCTTACCATAGTTTCCCAAGAGG | 1000 | | | | | | | | | |
| Oligo fwd | CACCTCTTACCATAGTTTCCCAAG | | | | | | | | | | |
| Oligo rev | AAACCTTGGGAAACTATGGTAAGA | | | | | | | | | | |
| Chromosome | start | end | strand | MM | target_seq | PAM | alignment | distance | position | gene name | gene id |
| GU305914.1 | 90305 | 90327 | + | 0 | TCTTACCATAGTTTCCCAAG | AGG | \|\|\|\|\|\|\|\|[\|\|\|\|\|\|\|\|\|\|\|\|]PAM | NA | Intergenic | NA | NA |
| T6418 | ACCATAGTTTCCCAAGAGGTCGG | 1000 | | | | | | | | | |
| Oligo fwd | CACCACCATAGTTTCCCAAGAGGT | | | | | | | | | | |
| Oligo rev | AAACACCTCTTGGGAAACTATGGT | | | | | | | | | | |
| Chromosome | start | end | strand | MM | target_seq | PAM | alignment | distance | position | gene name | gene id |
| GU305914.1 | 90309 | 90331 | + | 0 | ACCATAGTTTCCCAAGAGGT | CGG | \|\|\|\|\|\|\|\|[\|\|\|\|\|\|\|\|\|\|\|\|]PAM | NA | Intergenic | NA | NA |
| T6419 | GTTTCCCAAGAGGTCGGCGTCGG | 1000 | | | | | | | | | |
| Oligo fwd | CACCGTTTCCCAAGAGGTCGGCGT | | | | | | | | | | |
| Oligo rev | AAACACGCCGACCTCTTGGGAAAC | | | | | | | | | | |
| Chromosome | start | end | strand | MM | target_seq | PAM | alignment | distance | position | gene name | gene id |
| GU305914.1 | 90315 | 90337 | + | 0 | GTTTCCCAAGAGGTCGGCGT | CGG | \|\|\|\|\|\|\|\|[\|\|\|\|\|\|\|\|\|\|\|\|]PAM | NA | Intergenic | NA | NA |
| T6420 | CAAGAGGTCGGCGTCGGCGAAGG | 1000 | | | | | | | | | |
| Oligo fwd | CACCCAAGAGGTCGGCGTCGGCGA | | | | | | | | | | |
| Oligo rev | AAACTCGCCGACGCCGACCTCTTG | | | | | | | | | | |
| Chromosome | start | end | strand | MM | target_seq | PAM | alignment | distance | position | gene name | gene id |
| GU305914.1 | 90321 | 90343 | + | 0 | CAAGAGGTCGGCGTCGGCGA | AGG | \|\|\|\|\|\|\|\|[\|\|\|\|\|\|\|\|\|\|\|\|]PAM | NA | Intergenic | NA | NA |
| GU305914.1 | 113930 | 113952 | - | 4 | CGAGGTGTCGGCGTCGGCGG | CGG | \|-\|\|--\|\|[\|\|\|\|\|\|\|\|\|\|\|-]PAM | NA | Intergenic | NA | NA |
| GU305914.1 | 43632 | 43654 | + | 4 | CAACAGGGCGGCGGCGGCAA | CGG | \|\|\|-\|\|\|-[\|\|\|\|\|-\|\|\|\|-\|]PAM | NA | Intergenic | NA | NA |
| T6421 | CGAGAGCAGAGTGACCACGACGG | 1000 | | | | | | | | | |
| Oligo fwd | CACCCGAGAGCAGAGTGACCACGA | | | | | | | | | | |
| Oligo rev | AAACTCGTGGTCACTCTGCTCTCG | | | | | | | | | | |
| Chromosome | start | end | strand | MM | target_seq | PAM | alignment | distance | position | gene name | gene id |
| GU305914.1 | 90348 | 90370 | + | 0 | CGAGAGCAGAGTGACCACGA | CGG | \|\|\|\|\|\|\|\|[\|\|\|\|\|\|\|\|\|\|\|\|]PAM | NA | Intergenic | NA | NA |
| T6422 | GAGAGCAGAGTGACCACGACGGG | 1000 | | | | | | | | | |
| Oligo fwd | CACCGAGAGCAGAGTGACCACGAC | | | | | | | | | | |
| Oligo rev | AAACGTCGTGGTCACTCTGCTCTC | | | | | | | | | | |
| Chromosome | start | end | strand | MM | target_seq | PAM | alignment | distance | position | gene name | gene id |
| GU305914.1 | 90349 | 90371 | + | 0 | GAGAGCAGAGTGACCACGAC | GGG | \|\|\|\|\|\|\|\|[\|\|\|\|\|\|\|\|\|\|\|\|]PAM | NA | Intergenic | NA | NA |
| T6423 | GGGCTCCTTCTTGATCTGCTTGG | 1000 | | | | | | | | | |
| Oligo fwd | CACCGGGCTCCTTCTTGATCTGCT | | | | | | | | | | |
| Oligo rev | AAACAGCAGATCAAGAAGGAGCCC | | | | | | | | | | |
| Chromosome | start | end | strand | MM | target_seq | PAM | alignment | distance | position | gene name | gene id |
| GU305914.1 | 90369 | 90391 | + | 0 | GGGCTCCTTCTTGATCTGCT | TGG | \|\|\|\|\|\|\|\|[\|\|\|\|\|\|\|\|\|\|\|\|]PAM | NA | Intergenic | NA | NA |
| T6424 | GGCTCCTTCTTGATCTGCTTGGG | 1000 | | | | | | | | | |
| Oligo fwd | CACCGGCTCCTTCTTGATCTGCTT | | | | | | | | | | |
| Oligo rev | AAACAAGCAGATCAAGAAGGAGCC | | | | | | | | | | |
| Chromosome | start | end | strand | MM | target_seq | PAM | alignment | distance | position | gene name | gene id |
| GU305914.1 | 90370 | 90392 | + | 0 | GGCTCCTTCTTGATCTGCTT | GGG | \|\|\|\|\|\|\|\|[\|\|\|\|\|\|\|\|\|\|\|\|]PAM | NA | Intergenic | NA | NA |
| T6425 | GCTCCTTCTTGATCTGCTTGGGG | 1000 | | | | | | | | | |
| Oligo fwd | CACCGCTCCTTCTTGATCTGCTTG | | | | | | | | | | |
| Oligo rev | AAACCAAGCAGATCAAGAAGGAGC | | | | | | | | | | |
| Chromosome | start | end | strand | MM | target_seq | PAM | alignment | distance | position | gene name | gene id |
| GU305914.1 | 90371 | 90393 | + | 0 | GCTCCTTCTTGATCTGCTTG | GGG | \|\|\|\|\|\|\|\|[\|\|\|\|\|\|\|\|\|\|\|\|]PAM | NA | Intergenic | NA | NA |
| T6426 | TTGATCTGCTTGGGGATGCCCGG | 1000 | | | | | | | | | |
| Oligo fwd | CACCTTGATCTGCTTGGGGATGCC | | | | | | | | | | |
| Oligo rev | AAACGGCATCCCCAAGCAGATCAA | | | | | | | | | | |
| Chromosome | start | end | strand | MM | target_seq | PAM | alignment | distance | position | gene name | gene id |
| GU305914.1 | 90379 | 90401 | + | 0 | TTGATCTGCTTGGGGATGCC | CGG | \|\|\|\|\|\|\|\|[\|\|\|\|\|\|\|\|\|\|\|\|]PAM | NA | Intergenic | NA | NA |

<table>
<tr><td>T6427</td><td>CCGCGTGTTCACCCTGATGAAGG</td><td>1000</td><td></td><td></td><td></td><td></td><td></td><td></td><td></td><td></td><td></td></tr>
<tr><td>Oligo fwd</td><td>CACCCCGCGTGTTCACCCTGATGA</td><td></td><td></td><td></td><td></td><td></td><td></td><td></td><td></td><td></td><td></td></tr>
<tr><td>Oligo rev</td><td>AAACTCATCAGGGTGAACACGCGG</td><td></td><td></td><td></td><td></td><td></td><td></td><td></td><td></td><td></td><td></td></tr>
<tr><td>Chromosome</td><td>start</td><td>end</td><td>strand</td><td>MM</td><td>target_seq</td><td>PAM</td><td>alignment</td><td>distance</td><td>position</td><td>gene name</td><td>gene id</td></tr>
<tr><td>GU305914.1</td><td>90405</td><td>90427</td><td>+</td><td>0</td><td>CCGCGTGTTCACCCTGATGA</td><td>AGG</td><td>||||||||[||||||||||||]PAM</td><td>NA</td><td>Intergenic</td><td>NA</td><td>NA</td></tr>
<tr><td></td><td></td><td></td><td></td><td></td><td></td><td></td><td></td><td></td><td></td><td></td><td></td></tr>
<tr><td>T6428</td><td>CGTGTTCACCCTGATGAAGGAGG</td><td>1000</td><td></td><td></td><td></td><td></td><td></td><td></td><td></td><td></td><td></td></tr>
<tr><td>Oligo fwd</td><td>CACCCGTGTTCACCCTGATGAAGG</td><td></td><td></td><td></td><td></td><td></td><td></td><td></td><td></td><td></td><td></td></tr>
<tr><td>Oligo rev</td><td>AAACCCTTCATCAGGGTGAACACG</td><td></td><td></td><td></td><td></td><td></td><td></td><td></td><td></td><td></td><td></td></tr>
<tr><td>Chromosome</td><td>start</td><td>end</td><td>strand</td><td>MM</td><td>target_seq</td><td>PAM</td><td>alignment</td><td>distance</td><td>position</td><td>gene name</td><td>gene id</td></tr>
<tr><td>GU305914.1</td><td>90408</td><td>90430</td><td>+</td><td>0</td><td>CGTGTTCACCCTGATGAAGG</td><td>AGG</td><td>||||||||[||||||||||||]PAM</td><td>NA</td><td>Intergenic</td><td>NA</td><td>NA</td></tr>
<tr><td></td><td></td><td></td><td></td><td></td><td></td><td></td><td></td><td></td><td></td><td></td><td></td></tr>
<tr><td>T6429</td><td>GTTCACCCTGATGAAGGAGGTGG</td><td>1000</td><td></td><td></td><td></td><td></td><td></td><td></td><td></td><td></td><td></td></tr>
<tr><td>Oligo fwd</td><td>CACCGTTCACCCTGATGAAGGAGG</td><td></td><td></td><td></td><td></td><td></td><td></td><td></td><td></td><td></td><td></td></tr>
<tr><td>Oligo rev</td><td>AAACCCTCCTTCATCAGGGTGAAC</td><td></td><td></td><td></td><td></td><td></td><td></td><td></td><td></td><td></td><td></td></tr>
<tr><td>Chromosome</td><td>start</td><td>end</td><td>strand</td><td>MM</td><td>target_seq</td><td>PAM</td><td>alignment</td><td>distance</td><td>position</td><td>gene name</td><td>gene id</td></tr>
<tr><td>GU305914.1</td><td>90411</td><td>90433</td><td>+</td><td>0</td><td>GTTCACCCTGATGAAGGAGG</td><td>TGG</td><td>||||||||[||||||||||||]PAM</td><td>NA</td><td>Intergenic</td><td>NA</td><td>NA</td></tr>
<tr><td></td><td></td><td></td><td></td><td></td><td></td><td></td><td></td><td></td><td></td><td></td><td></td></tr>
<tr><td>T6430</td><td>CGTAGCACGTGTGACAGCAGCGG</td><td>1000</td><td></td><td></td><td></td><td></td><td></td><td></td><td></td><td></td><td></td></tr>
<tr><td>Oligo fwd</td><td>CACCCGTAGCACGTGTGACAGCAG</td><td></td><td></td><td></td><td></td><td></td><td></td><td></td><td></td><td></td><td></td></tr>
<tr><td>Oligo rev</td><td>AAACCTGCTGTCACACGTGCTACG</td><td></td><td></td><td></td><td></td><td></td><td></td><td></td><td></td><td></td><td></td></tr>
<tr><td>Chromosome</td><td>start</td><td>end</td><td>strand</td><td>MM</td><td>target_seq</td><td>PAM</td><td>alignment</td><td>distance</td><td>position</td><td>gene name</td><td>gene id</td></tr>
<tr><td>GU305914.1</td><td>90434</td><td>90456</td><td>+</td><td>0</td><td>CGTAGCACGTGTGACAGCAG</td><td>CGG</td><td>||||||||[||||||||||||]PAM</td><td>NA</td><td>Intergenic</td><td>NA</td><td>NA</td></tr>
<tr><td></td><td></td><td></td><td></td><td></td><td></td><td></td><td></td><td></td><td></td><td></td><td></td></tr>
<tr><td>T6431</td><td>GCCGCACGCGCCGCACAGCTTGG</td><td>1000</td><td></td><td></td><td></td><td></td><td></td><td></td><td></td><td></td><td></td></tr>
<tr><td>Oligo fwd</td><td>CACCGCCGCACGCGCCGCACAGCT</td><td></td><td></td><td></td><td></td><td></td><td></td><td></td><td></td><td></td><td></td></tr>
<tr><td>Oligo rev</td><td>AAACAGCTGTGCGGCGCGTGCGGC</td><td></td><td></td><td></td><td></td><td></td><td></td><td></td><td></td><td></td><td></td></tr>
<tr><td>Chromosome</td><td>start</td><td>end</td><td>strand</td><td>MM</td><td>target_seq</td><td>PAM</td><td>alignment</td><td>distance</td><td>position</td><td>gene name</td><td>gene id</td></tr>
<tr><td>GU305914.1</td><td>90459</td><td>90481</td><td>+</td><td>0</td><td>GCCGCACGCGCCGCACAGCT</td><td>TGG</td><td>||||||||[||||||||||||]PAM</td><td>NA</td><td>Intergenic</td><td>NA</td><td>NA</td></tr>
<tr><td></td><td></td><td></td><td></td><td></td><td></td><td></td><td></td><td></td><td></td><td></td><td></td></tr>
<tr><td>T6432</td><td>CGCGCCGCACAGCTTGGAGCCGG</td><td>1000</td><td></td><td></td><td></td><td></td><td></td><td></td><td></td><td></td><td></td></tr>
<tr><td>Oligo fwd</td><td>CACCCGCGCCGCACAGCTTGGAGC</td><td></td><td></td><td></td><td></td><td></td><td></td><td></td><td></td><td></td><td></td></tr>
<tr><td>Oligo rev</td><td>AAACGCTCCAAGCTGTGCGGCGCG</td><td></td><td></td><td></td><td></td><td></td><td></td><td></td><td></td><td></td><td></td></tr>
<tr><td>Chromosome</td><td>start</td><td>end</td><td>strand</td><td>MM</td><td>target_seq</td><td>PAM</td><td>alignment</td><td>distance</td><td>position</td><td>gene name</td><td>gene id</td></tr>
<tr><td>GU305914.1</td><td>90465</td><td>90487</td><td>+</td><td>0</td><td>CGCGCCGCACAGCTTGGAGC</td><td>CGG</td><td>||||||||[||||||||||||]PAM</td><td>NA</td><td>Intergenic</td><td>NA</td><td>NA</td></tr>
<tr><td></td><td></td><td></td><td></td><td></td><td></td><td></td><td></td><td></td><td></td><td></td><td></td></tr>
<tr><td>T6433</td><td>CGTGTCGCCGCTCGTGACCCCGG</td><td>1000</td><td></td><td></td><td></td><td></td><td></td><td></td><td></td><td></td><td></td></tr>
<tr><td>Oligo fwd</td><td>CACCCGTGTCGCCGCTCGTGACCC</td><td></td><td></td><td></td><td></td><td></td><td></td><td></td><td></td><td></td><td></td></tr>
<tr><td>Oligo rev</td><td>AAACGGGTCACGAGCGGCGACACG</td><td></td><td></td><td></td><td></td><td></td><td></td><td></td><td></td><td></td><td></td></tr>
<tr><td>Chromosome</td><td>start</td><td>end</td><td>strand</td><td>MM</td><td>target_seq</td><td>PAM</td><td>alignment</td><td>distance</td><td>position</td><td>gene name</td><td>gene id</td></tr>
<tr><td>GU305914.1</td><td>90510</td><td>90532</td><td>+</td><td>0</td><td>CGTGTCGCCGCTCGTGACCC</td><td>CGG</td><td>||||||||[||||||||||||]PAM</td><td>NA</td><td>Intergenic</td><td>NA</td><td>NA</td></tr>
<tr><td>GU305914.1</td><td>150625</td><td>150647</td><td>-</td><td>3</td><td>CGTGTCGGCGCTGATGACCC</td><td>GGG</td><td>|||||||-[||||--||||||]PAM</td><td>NA</td><td>Intergenic</td><td>NA</td><td>NA</td></tr>
<tr><td></td><td></td><td></td><td></td><td></td><td></td><td></td><td></td><td></td><td></td><td></td><td></td></tr>
<tr><td>T6434</td><td>CGTGACCCCGGTCGAGTAGATGG</td><td>1000</td><td></td><td></td><td></td><td></td><td></td><td></td><td></td><td></td><td></td></tr>
<tr><td>Oligo fwd</td><td>CACCCGTGACCCCGGTCGAGTAGA</td><td></td><td></td><td></td><td></td><td></td><td></td><td></td><td></td><td></td><td></td></tr>
<tr><td>Oligo rev</td><td>AAACTCTACTCGACCGGGGTCACG</td><td></td><td></td><td></td><td></td><td></td><td></td><td></td><td></td><td></td><td></td></tr>
<tr><td>Chromosome</td><td>start</td><td>end</td><td>strand</td><td>MM</td><td>target_seq</td><td>PAM</td><td>alignment</td><td>distance</td><td>position</td><td>gene name</td><td>gene id</td></tr>
<tr><td>GU305914.1</td><td>90522</td><td>90544</td><td>+</td><td>0</td><td>CGTGACCCCGGTCGAGTAGA</td><td>TGG</td><td>||||||||[||||||||||||]PAM</td><td>NA</td><td>Intergenic</td><td>NA</td><td>NA</td></tr>
<tr><td></td><td></td><td></td><td></td><td></td><td></td><td></td><td></td><td></td><td></td><td></td><td></td></tr>
<tr><td>T6435</td><td>CGGTCGAGTAGATGGACATGCGG</td><td>1000</td><td></td><td></td><td></td><td></td><td></td><td></td><td></td><td></td><td></td></tr>
<tr><td>Oligo fwd</td><td>CACCCGGTCGAGTAGATGGACATG</td><td></td><td></td><td></td><td></td><td></td><td></td><td></td><td></td><td></td><td></td></tr>
<tr><td>Oligo rev</td><td>AAACCATGTCCATCTACTCGACCG</td><td></td><td></td><td></td><td></td><td></td><td></td><td></td><td></td><td></td><td></td></tr>
<tr><td>Chromosome</td><td>start</td><td>end</td><td>strand</td><td>MM</td><td>target_seq</td><td>PAM</td><td>alignment</td><td>distance</td><td>position</td><td>gene name</td><td>gene id</td></tr>
<tr><td>GU305914.1</td><td>90530</td><td>90552</td><td>+</td><td>0</td><td>CGGTCGAGTAGATGGACATG</td><td>CGG</td><td>||||||||[||||||||||||]PAM</td><td>NA</td><td>Intergenic</td><td>NA</td><td>NA</td></tr>
<tr><td></td><td></td><td></td><td></td><td></td><td></td><td></td><td></td><td></td><td></td><td></td><td></td></tr>
<tr><td>T6436</td><td>TGCGGTTGAAGAACCACATGAGG</td><td>1000</td><td></td><td></td><td></td><td></td><td></td><td></td><td></td><td></td><td></td></tr>
<tr><td>Oligo fwd</td><td>CACCTGCGGTTGAAGAACCACATG</td><td></td><td></td><td></td><td></td><td></td><td></td><td></td><td></td><td></td><td></td></tr>
<tr><td>Oligo rev</td><td>AAACCATGTGGTTCTTCAACCGCA</td><td></td><td></td><td></td><td></td><td></td><td></td><td></td><td></td><td></td><td></td></tr>
<tr><td>Chromosome</td><td>start</td><td>end</td><td>strand</td><td>MM</td><td>target_seq</td><td>PAM</td><td>alignment</td><td>distance</td><td>position</td><td>gene name</td><td>gene id</td></tr>
<tr><td>GU305914.1</td><td>90548</td><td>90570</td><td>+</td><td>0</td><td>TGCGGTTGAAGAACCACATG</td><td>AGG</td><td>||||||||[||||||||||||]PAM</td><td>NA</td><td>Intergenic</td><td>NA</td><td>NA</td></tr>
<tr><td></td><td></td><td></td><td></td><td></td><td></td><td></td><td></td><td></td><td></td><td></td><td></td></tr>
<tr><td>T6437</td><td>GCGGTTGAAGAACCACATGAGGG</td><td>1000</td><td></td><td></td><td></td><td></td><td></td><td></td><td></td><td></td><td></td></tr>
<tr><td>Oligo fwd</td><td>CACCGCGGTTGAAGAACCACATGA</td><td></td><td></td><td></td><td></td><td></td><td></td><td></td><td></td><td></td><td></td></tr>
<tr><td>Oligo rev</td><td>AAACTCATGTGGTTCTTCAACCGC</td><td></td><td></td><td></td><td></td><td></td><td></td><td></td><td></td><td></td><td></td></tr>
<tr><td>Chromosome</td><td>start</td><td>end</td><td>strand</td><td>MM</td><td>target_seq</td><td>PAM</td><td>alignment</td><td>distance</td><td>position</td><td>gene name</td><td>gene id</td></tr>
<tr><td>GU305914.1</td><td>90549</td><td>90571</td><td>+</td><td>0</td><td>GCGGTTGAAGAACCACATGA</td><td>GGG</td><td>||||||||[||||||||||||]PAM</td><td>NA</td><td>Intergenic</td><td>NA</td><td>NA</td></tr>
<tr><td></td><td></td><td></td><td></td><td></td><td></td><td></td><td></td><td></td><td></td><td></td><td></td></tr>
<tr><td>T6438</td><td>GTTGAAGAACCACATGAGGGTGG</td><td>1000</td><td></td><td></td><td></td><td></td><td></td><td></td><td></td><td></td><td></td></tr>
<tr><td>Oligo fwd</td><td>CACCGTTGAAGAACCACATGAGGG</td><td></td><td></td><td></td><td></td><td></td><td></td><td></td><td></td><td></td><td></td></tr>
<tr><td>Oligo rev</td><td>AAACCCCTCATGTGGTTCTTCAAC</td><td></td><td></td><td></td><td></td><td></td><td></td><td></td><td></td><td></td><td></td></tr>
<tr><td>Chromosome</td><td>start</td><td>end</td><td>strand</td><td>MM</td><td>target_seq</td><td>PAM</td><td>alignment</td><td>distance</td><td>position</td><td>gene name</td><td>gene id</td></tr>
<tr><td>GU305914.1</td><td>90552</td><td>90574</td><td>+</td><td>0</td><td>GTTGAAGAACCACATGAGGG</td><td>TGG</td><td>||||||||[||||||||||||]PAM</td><td>NA</td><td>Intergenic</td><td>NA</td><td>NA</td></tr>
<tr><td></td><td></td><td></td><td></td><td></td><td></td><td></td><td></td><td></td><td></td><td></td><td></td></tr>
<tr><td>T6439</td><td>ATGAGGGTGGAGAAGAGCTGCGG</td><td>1000</td><td></td><td></td><td></td><td></td><td></td><td></td><td></td><td></td><td></td></tr>
<tr><td>Oligo fwd</td><td>CACCATGAGGGTGGAGAAGAGCTG</td><td></td><td></td><td></td><td></td><td></td><td></td><td></td><td></td><td></td><td></td></tr>
<tr><td>Oligo rev</td><td>AAACCAGCTCTTCTCCACCCTCAT</td><td></td><td></td><td></td><td></td><td></td><td></td><td></td><td></td><td></td><td></td></tr>
<tr><td>Chromosome</td><td>start</td><td>end</td><td>strand</td><td>MM</td><td>target_seq</td><td>PAM</td><td>alignment</td><td>distance</td><td>position</td><td>gene name</td><td>gene id</td></tr>
<tr><td>GU305914.1</td><td>90565</td><td>90587</td><td>+</td><td>0</td><td>ATGAGGGTGGAGAAGAGCTG</td><td>CGG</td><td>||||||||[||||||||||||]PAM</td><td>NA</td><td>Intergenic</td><td>NA</td><td>NA</td></tr>
<tr><td></td><td></td><td></td><td></td><td></td><td></td><td></td><td></td><td></td><td></td><td></td><td></td></tr>
<tr><td>T6440</td><td>GGTGGAGAAGAGCTGCGGCGAGG</td><td>1000</td><td></td><td></td><td></td><td></td><td></td><td></td><td></td><td></td><td></td></tr>
<tr><td>Oligo fwd</td><td>CACCGGTGGAGAAGAGCTGCGGCG</td><td></td><td></td><td></td><td></td><td></td><td></td><td></td><td></td><td></td><td></td></tr>
<tr><td>Oligo rev</td><td>AAACCGCCGCAGCTCTTCTCCACC</td><td></td><td></td><td></td><td></td><td></td><td></td><td></td><td></td><td></td><td></td></tr>
<tr><td>Chromosome</td><td>start</td><td>end</td><td>strand</td><td>MM</td><td>target_seq</td><td>PAM</td><td>alignment</td><td>distance</td><td>position</td><td>gene name</td><td>gene id</td></tr>
<tr><td>GU305914.1</td><td>90570</td><td>90592</td><td>+</td><td>0</td><td>GGTGGAGAAGAGCTGCGGCG</td><td>AGG</td><td>||||||||[||||||||||||]PAM</td><td>NA</td><td>Intergenic</td><td>NA</td><td>NA</td></tr>
<tr><td></td><td></td><td></td><td></td><td></td><td></td><td></td><td></td><td></td><td></td><td></td><td></td></tr>
<tr><td>T6441</td><td>CGGCGAGGTGCCGCAGAAGTAGG</td><td>1000</td><td></td><td></td><td></td><td></td><td></td><td></td><td></td><td></td><td></td></tr>
<tr><td>Oligo fwd</td><td>CACCCGGCGAGGTGCCGCAGAAGT</td><td></td><td></td><td></td><td></td><td></td><td></td><td></td><td></td><td></td><td></td></tr>
<tr><td>Oligo rev</td><td>AAACACTTCTGCGGCACCTCGCCG</td><td></td><td></td><td></td><td></td><td></td><td></td><td></td><td></td><td></td><td></td></tr>
<tr><td>Chromosome</td><td>start</td><td>end</td><td>strand</td><td>MM</td><td>target_seq</td><td>PAM</td><td>alignment</td><td>distance</td><td>position</td><td>gene name</td><td>gene id</td></tr>
</table>

<table>
<tr><td>GU305914.1</td><td>90585</td><td>90607</td><td>+</td><td>0</td><td>CGGCGAGGTGCCGCAGAAGT</td><td>AGG</td><td>||||||||[||||||||||||]PAM</td><td>NA</td><td>Intergenic</td><td>NA</td><td>NA</td></tr>
<tr><td></td><td></td><td></td><td></td><td></td><td></td><td></td><td></td><td></td><td></td><td></td><td></td></tr>
<tr><td>T6442</td><td>AGGTGCCGCAGAAGTAGGCCAGG</td><td>1000</td><td></td><td></td><td></td><td></td><td></td><td></td><td></td><td></td><td></td></tr>
<tr><td>Oligo fwd</td><td>CACCAGGTGCCGCAGAAGTAGGCC</td><td></td><td></td><td></td><td></td><td></td><td></td><td></td><td></td><td></td><td></td></tr>
<tr><td>Oligo rev</td><td>AAACGGCCTACTTCTGCGGCACCT</td><td></td><td></td><td></td><td></td><td></td><td></td><td></td><td></td><td></td><td></td></tr>
<tr><td>Chromosome</td><td>start</td><td>end</td><td>strand</td><td>MM</td><td>target_seq</td><td>PAM</td><td>alignment</td><td>distance</td><td>position</td><td>gene name</td><td>gene id</td></tr>
<tr><td>GU305914.1</td><td>90590</td><td>90612</td><td>+</td><td>0</td><td>AGGTGCCGCAGAAGTAGGCC</td><td>AGG</td><td>||||||||[||||||||||||]PAM</td><td>NA</td><td>Intergenic</td><td>NA</td><td>NA</td></tr>
<tr><td></td><td></td><td></td><td></td><td></td><td></td><td></td><td></td><td></td><td></td><td></td><td></td></tr>
<tr><td>T6443</td><td>TGCCGCAGAAGTAGGCCAGGTGG</td><td>1000</td><td></td><td></td><td></td><td></td><td></td><td></td><td></td><td></td><td></td></tr>
<tr><td>Oligo fwd</td><td>CACCTGCCGCAGAAGTAGGCCAGG</td><td></td><td></td><td></td><td></td><td></td><td></td><td></td><td></td><td></td><td></td></tr>
<tr><td>Oligo rev</td><td>AAACCCTGGCCTACTTCTGCGGCA</td><td></td><td></td><td></td><td></td><td></td><td></td><td></td><td></td><td></td><td></td></tr>
<tr><td>Chromosome</td><td>start</td><td>end</td><td>strand</td><td>MM</td><td>target_seq</td><td>PAM</td><td>alignment</td><td>distance</td><td>position</td><td>gene name</td><td>gene id</td></tr>
<tr><td>GU305914.1</td><td>90593</td><td>90615</td><td>+</td><td>0</td><td>TGCCGCAGAAGTAGGCCAGG</td><td>TGG</td><td>||||||||[||||||||||||]PAM</td><td>NA</td><td>Intergenic</td><td>NA</td><td>NA</td></tr>
<tr><td></td><td></td><td></td><td></td><td></td><td></td><td></td><td></td><td></td><td></td><td></td><td></td></tr>
<tr><td>T6444</td><td>CAGAAGTAGGCCAGGTGGTGCGG</td><td>1000</td><td></td><td></td><td></td><td></td><td></td><td></td><td></td><td></td><td></td></tr>
<tr><td>Oligo fwd</td><td>CACCCAGAAGTAGGCCAGGTGGTG</td><td></td><td></td><td></td><td></td><td></td><td></td><td></td><td></td><td></td><td></td></tr>
<tr><td>Oligo rev</td><td>AAACCACCACCTGGCCTACTTCTG</td><td></td><td></td><td></td><td></td><td></td><td></td><td></td><td></td><td></td><td></td></tr>
<tr><td>Chromosome</td><td>start</td><td>end</td><td>strand</td><td>MM</td><td>target_seq</td><td>PAM</td><td>alignment</td><td>distance</td><td>position</td><td>gene name</td><td>gene id</td></tr>
<tr><td>GU305914.1</td><td>90598</td><td>90620</td><td>+</td><td>0</td><td>CAGAAGTAGGCCAGGTGGTG</td><td>CGG</td><td>||||||||[||||||||||||]PAM</td><td>NA</td><td>Intergenic</td><td>NA</td><td>NA</td></tr>
<tr><td></td><td></td><td></td><td></td><td></td><td></td><td></td><td></td><td></td><td></td><td></td><td></td></tr>
<tr><td>T6445</td><td>AGAAGTAGGCCAGGTGGTGCGGG</td><td>1000</td><td></td><td></td><td></td><td></td><td></td><td></td><td></td><td></td><td></td></tr>
<tr><td>Oligo fwd</td><td>CACCAGAAGTAGGCCAGGTGGTGC</td><td></td><td></td><td></td><td></td><td></td><td></td><td></td><td></td><td></td><td></td></tr>
<tr><td>Oligo rev</td><td>AAACGCACCACCTGGCCTACTTCT</td><td></td><td></td><td></td><td></td><td></td><td></td><td></td><td></td><td></td><td></td></tr>
<tr><td>Chromosome</td><td>start</td><td>end</td><td>strand</td><td>MM</td><td>target_seq</td><td>PAM</td><td>alignment</td><td>distance</td><td>position</td><td>gene name</td><td>gene id</td></tr>
<tr><td>GU305914.1</td><td>90599</td><td>90621</td><td>+</td><td>0</td><td>AGAAGTAGGCCAGGTGGTGC</td><td>GGG</td><td>||||||||[||||||||||||]PAM</td><td>NA</td><td>Intergenic</td><td>NA</td><td>NA</td></tr>
<tr><td></td><td></td><td></td><td></td><td></td><td></td><td></td><td></td><td></td><td></td><td></td><td></td></tr>
<tr><td>T6446</td><td>GAAGTAGGCCAGGTGGTGCGGGG</td><td>1000</td><td></td><td></td><td></td><td></td><td></td><td></td><td></td><td></td><td></td></tr>
<tr><td>Oligo fwd</td><td>CACCGAAGTAGGCCAGGTGGTGCG</td><td></td><td></td><td></td><td></td><td></td><td></td><td></td><td></td><td></td><td></td></tr>
<tr><td>Oligo rev</td><td>AAACCGCACCACCTGGCCTACTTC</td><td></td><td></td><td></td><td></td><td></td><td></td><td></td><td></td><td></td><td></td></tr>
<tr><td>Chromosome</td><td>start</td><td>end</td><td>strand</td><td>MM</td><td>target_seq</td><td>PAM</td><td>alignment</td><td>distance</td><td>position</td><td>gene name</td><td>gene id</td></tr>
<tr><td>GU305914.1</td><td>90600</td><td>90622</td><td>+</td><td>0</td><td>GAAGTAGGCCAGGTGGTGCG</td><td>GGG</td><td>||||||||[||||||||||||]PAM</td><td>NA</td><td>Intergenic</td><td>NA</td><td>NA</td></tr>
<tr><td></td><td></td><td></td><td></td><td></td><td></td><td></td><td></td><td></td><td></td><td></td><td></td></tr>
<tr><td>T6447</td><td>GTACTTCTCGTCCCGCCCGACGG</td><td>1000</td><td></td><td></td><td></td><td></td><td></td><td></td><td></td><td></td><td></td></tr>
<tr><td>Oligo fwd</td><td>CACCGTACTTCTCGTCCCGCCCGA</td><td></td><td></td><td></td><td></td><td></td><td></td><td></td><td></td><td></td><td></td></tr>
<tr><td>Oligo rev</td><td>AAACTCGGGCGGGACGAGAAGTAC</td><td></td><td></td><td></td><td></td><td></td><td></td><td></td><td></td><td></td><td></td></tr>
<tr><td>Chromosome</td><td>start</td><td>end</td><td>strand</td><td>MM</td><td>target_seq</td><td>PAM</td><td>alignment</td><td>distance</td><td>position</td><td>gene name</td><td>gene id</td></tr>
<tr><td>GU305914.1</td><td>90624</td><td>90646</td><td>+</td><td>0</td><td>GTACTTCTCGTCCCGCCCGA</td><td>CGG</td><td>||||||||[||||||||||||]PAM</td><td>NA</td><td>Intergenic</td><td>NA</td><td>NA</td></tr>
<tr><td>GU305914.1</td><td>53490</td><td>53512</td><td>+</td><td>4</td><td>GTCGTTCTCGCCCCGACCGA</td><td>CGG</td><td>||--||||[||-||||-||||]PAM</td><td>NA</td><td>Intergenic</td><td>NA</td><td>NA</td></tr>
<tr><td></td><td></td><td></td><td></td><td></td><td></td><td></td><td></td><td></td><td></td><td></td><td></td></tr>
<tr><td>T6448</td><td>GACGGCCCCCTCGAACCTGATGG</td><td>1000</td><td></td><td></td><td></td><td></td><td></td><td></td><td></td><td></td><td></td></tr>
<tr><td>Oligo fwd</td><td>CACCGACGGCCCCCTCGAACCTGA</td><td></td><td></td><td></td><td></td><td></td><td></td><td></td><td></td><td></td><td></td></tr>
<tr><td>Oligo rev</td><td>AAACTCAGGTTCGAGGGGGCCGTC</td><td></td><td></td><td></td><td></td><td></td><td></td><td></td><td></td><td></td><td></td></tr>
<tr><td>Chromosome</td><td>start</td><td>end</td><td>strand</td><td>MM</td><td>target_seq</td><td>PAM</td><td>alignment</td><td>distance</td><td>position</td><td>gene name</td><td>gene id</td></tr>
<tr><td>GU305914.1</td><td>90642</td><td>90664</td><td>+</td><td>0</td><td>GACGGCCCCCTCGAACCTGA</td><td>TGG</td><td>||||||||[||||||||||||]PAM</td><td>NA</td><td>Intergenic</td><td>NA</td><td>NA</td></tr>
<tr><td></td><td></td><td></td><td></td><td></td><td></td><td></td><td></td><td></td><td></td><td></td><td></td></tr>
<tr><td>T6449</td><td>CTCGAACCTGATGGTCTTCTTGG</td><td>1000</td><td></td><td></td><td></td><td></td><td></td><td></td><td></td><td></td><td></td></tr>
<tr><td>Oligo fwd</td><td>CACCCTCGAACCTGATGGTCTTCT</td><td></td><td></td><td></td><td></td><td></td><td></td><td></td><td></td><td></td><td></td></tr>
<tr><td>Oligo rev</td><td>AAACAGAAGACCATCAGGTTCGAG</td><td></td><td></td><td></td><td></td><td></td><td></td><td></td><td></td><td></td><td></td></tr>
<tr><td>Chromosome</td><td>start</td><td>end</td><td>strand</td><td>MM</td><td>target_seq</td><td>PAM</td><td>alignment</td><td>distance</td><td>position</td><td>gene name</td><td>gene id</td></tr>
<tr><td>GU305914.1</td><td>90651</td><td>90673</td><td>+</td><td>0</td><td>CTCGAACCTGATGGTCTTCT</td><td>TGG</td><td>||||||||[||||||||||||]PAM</td><td>NA</td><td>Intergenic</td><td>NA</td><td>NA</td></tr>
<tr><td></td><td></td><td></td><td></td><td></td><td></td><td></td><td></td><td></td><td></td><td></td><td></td></tr>
<tr><td>T6450</td><td>TGGTCTTCTTGGCGTCGTTCAGG</td><td>1000</td><td></td><td></td><td></td><td></td><td></td><td></td><td></td><td></td><td></td></tr>
<tr><td>Oligo fwd</td><td>CACCTGGTCTTCTTGGCGTCGTTC</td><td></td><td></td><td></td><td></td><td></td><td></td><td></td><td></td><td></td><td></td></tr>
<tr><td>Oligo rev</td><td>AAACGAACGACGCCAAGAAGACCA</td><td></td><td></td><td></td><td></td><td></td><td></td><td></td><td></td><td></td><td></td></tr>
<tr><td>Chromosome</td><td>start</td><td>end</td><td>strand</td><td>MM</td><td>target_seq</td><td>PAM</td><td>alignment</td><td>distance</td><td>position</td><td>gene name</td><td>gene id</td></tr>
<tr><td>GU305914.1</td><td>90662</td><td>90684</td><td>+</td><td>0</td><td>TGGTCTTCTTGGCGTCGTTC</td><td>AGG</td><td>||||||||[||||||||||||]PAM</td><td>NA</td><td>Intergenic</td><td>NA</td><td>NA</td></tr>
<tr><td></td><td></td><td></td><td></td><td></td><td></td><td></td><td></td><td></td><td></td><td></td><td></td></tr>
<tr><td>T6451</td><td>TGTCATCGTACATGCCCGTGAGG</td><td>1000</td><td></td><td></td><td></td><td></td><td></td><td></td><td></td><td></td><td></td></tr>
<tr><td>Oligo fwd</td><td>CACCTGTCATCGTACATGCCCGTG</td><td></td><td></td><td></td><td></td><td></td><td></td><td></td><td></td><td></td><td></td></tr>
<tr><td>Oligo rev</td><td>AAACCACGGGCATGTACGATGACA</td><td></td><td></td><td></td><td></td><td></td><td></td><td></td><td></td><td></td><td></td></tr>
<tr><td>Chromosome</td><td>start</td><td>end</td><td>strand</td><td>MM</td><td>target_seq</td><td>PAM</td><td>alignment</td><td>distance</td><td>position</td><td>gene name</td><td>gene id</td></tr>
<tr><td>GU305914.1</td><td>90686</td><td>90708</td><td>+</td><td>0</td><td>TGTCATCGTACATGCCCGTG</td><td>AGG</td><td>||||||||[||||||||||||]PAM</td><td>NA</td><td>Intergenic</td><td>NA</td><td>NA</td></tr>
<tr><td></td><td></td><td></td><td></td><td></td><td></td><td></td><td></td><td></td><td></td><td></td><td></td></tr>
<tr><td>T6452</td><td>CCGTGAGGCCGTTGAAGAAGCGG</td><td>1000</td><td></td><td></td><td></td><td></td><td></td><td></td><td></td><td></td><td></td></tr>
<tr><td>Oligo fwd</td><td>CACCCCGTGAGGCCGTTGAAGAAG</td><td></td><td></td><td></td><td></td><td></td><td></td><td></td><td></td><td></td><td></td></tr>
<tr><td>Oligo rev</td><td>AAACCTTCTTCAACGGCCTCACGG</td><td></td><td></td><td></td><td></td><td></td><td></td><td></td><td></td><td></td><td></td></tr>
<tr><td>Chromosome</td><td>start</td><td>end</td><td>strand</td><td>MM</td><td>target_seq</td><td>PAM</td><td>alignment</td><td>distance</td><td>position</td><td>gene name</td><td>gene id</td></tr>
<tr><td>GU305914.1</td><td>90701</td><td>90723</td><td>+</td><td>0</td><td>CCGTGAGGCCGTTGAAGAAG</td><td>CGG</td><td>||||||||[||||||||||||]PAM</td><td>NA</td><td>Intergenic</td><td>NA</td><td>NA</td></tr>
<tr><td></td><td></td><td></td><td></td><td></td><td></td><td></td><td></td><td></td><td></td><td></td><td></td></tr>
<tr><td>T6453</td><td>GAGGCCGTTGAAGAAGCGGATGG</td><td>1000</td><td></td><td></td><td></td><td></td><td></td><td></td><td></td><td></td><td></td></tr>
<tr><td>Oligo fwd</td><td>CACCGAGGCCGTTGAAGAAGCGGA</td><td></td><td></td><td></td><td></td><td></td><td></td><td></td><td></td><td></td><td></td></tr>
<tr><td>Oligo rev</td><td>AAACTCCGCTTCTTCAACGGCCTC</td><td></td><td></td><td></td><td></td><td></td><td></td><td></td><td></td><td></td><td></td></tr>
<tr><td>Chromosome</td><td>start</td><td>end</td><td>strand</td><td>MM</td><td>target_seq</td><td>PAM</td><td>alignment</td><td>distance</td><td>position</td><td>gene name</td><td>gene id</td></tr>
<tr><td>GU305914.1</td><td>90705</td><td>90727</td><td>+</td><td>0</td><td>GAGGCCGTTGAAGAAGCGGA</td><td>TGG</td><td>||||||||[||||||||||||]PAM</td><td>NA</td><td>Intergenic</td><td>NA</td><td>NA</td></tr>
<tr><td></td><td></td><td></td><td></td><td></td><td></td><td></td><td></td><td></td><td></td><td></td><td></td></tr>
<tr><td>T6454</td><td>AGAAGCGGATGGACTTGAGCAGG</td><td>1000</td><td></td><td></td><td></td><td></td><td></td><td></td><td></td><td></td><td></td></tr>
<tr><td>Oligo fwd</td><td>CACCAGAAGCGGATGGACTTGAGC</td><td></td><td></td><td></td><td></td><td></td><td></td><td></td><td></td><td></td><td></td></tr>
<tr><td>Oligo rev</td><td>AAACGCTCAAGTCCATCCGCTTCT</td><td></td><td></td><td></td><td></td><td></td><td></td><td></td><td></td><td></td><td></td></tr>
<tr><td>Chromosome</td><td>start</td><td>end</td><td>strand</td><td>MM</td><td>target_seq</td><td>PAM</td><td>alignment</td><td>distance</td><td>position</td><td>gene name</td><td>gene id</td></tr>
<tr><td>GU305914.1</td><td>90716</td><td>90738</td><td>+</td><td>0</td><td>AGAAGCGGATGGACTTGAGC</td><td>AGG</td><td>||||||||[||||||||||||]PAM</td><td>NA</td><td>Intergenic</td><td>NA</td><td>NA</td></tr>
<tr><td></td><td></td><td></td><td></td><td></td><td></td><td></td><td></td><td></td><td></td><td></td><td></td></tr>
<tr><td>T6455</td><td>GAAGCGGATGGACTTGAGCAGGG</td><td>1000</td><td></td><td></td><td></td><td></td><td></td><td></td><td></td><td></td><td></td></tr>
<tr><td>Oligo fwd</td><td>CACCGAAGCGGATGGACTTGAGCA</td><td></td><td></td><td></td><td></td><td></td><td></td><td></td><td></td><td></td><td></td></tr>
<tr><td>Oligo rev</td><td>AAACTGCTCAAGTCCATCCGCTTC</td><td></td><td></td><td></td><td></td><td></td><td></td><td></td><td></td><td></td><td></td></tr>
<tr><td>Chromosome</td><td>start</td><td>end</td><td>strand</td><td>MM</td><td>target_seq</td><td>PAM</td><td>alignment</td><td>distance</td><td>position</td><td>gene name</td><td>gene id</td></tr>
<tr><td>GU305914.1</td><td>90717</td><td>90739</td><td>+</td><td>0</td><td>GAAGCGGATGGACTTGAGCA</td><td>GGG</td><td>||||||||[||||||||||||]PAM</td><td>NA</td><td>Intergenic</td><td>NA</td><td>NA</td></tr>
<tr><td></td><td></td><td></td><td></td><td></td><td></td><td></td><td></td><td></td><td></td><td></td><td></td></tr>
<tr><td>T6456</td><td>AGCAGGGAGTCCTGCGCCGACGG</td><td>1000</td><td></td><td></td><td></td><td></td><td></td><td></td><td></td><td></td><td></td></tr>
<tr><td>Oligo fwd</td><td>CACCAGCAGGGAGTCCTGCGCCGA</td><td></td><td></td><td></td><td></td><td></td><td></td><td></td><td></td><td></td><td></td></tr>
</table>

Oligo rev AAACTCGGCGCAGGACTCCCTGCT
Chromosome start end strand MM target_seq PAM alignment distance position gene name gene id
GU305914.1 90733 90755 + 0 AGCAGGGAGTCCTGCGCCGA CGG ||||||||[||||||||||||]PAM NA Intergenic NA NA

T6457 GCAGGGAGTCCTGCGCCGACGGG 1000
Oligo fwd CACCGCAGGGAGTCCTGCGCCGAC
Oligo rev AAACGTCGGCGCAGGACTCCCTGC
Chromosome start end strand MM target_seq PAM alignment distance position gene name gene id
GU305914.1 90734 90756 + 0 GCAGGGAGTCCTGCGCCGAC GGG ||||||||[||||||||||||]PAM NA Intergenic NA NA

T6458 CTGCGCCGACGGGTTGCTGTTGG 1000
Oligo fwd CACCCTGCGCCGACGGGTTGCTGT
Oligo rev AAACACAGCAACCCGTCGGCGCAG
Chromosome start end strand MM target_seq PAM alignment distance position gene name gene id
GU305914.1 90744 90766 + 0 CTGCGCCGACGGGTTGCTGT TGG ||||||||[||||||||||||]PAM NA Intergenic NA NA

T6459 GTTGCTGTTGGCCCCCTTGCCGG 1000
Oligo fwd CACCGTTGCTGTTGGCCCCCTTGC
Oligo rev AAACGCAAGGGGGCCAACAGCAAC
Chromosome start end strand MM target_seq PAM alignment distance position gene name gene id
GU305914.1 90756 90778 + 0 GTTGCTGTTGGCCCCCTTGC CGG ||||||||[||||||||||||]PAM NA Intergenic NA NA

T6460 GTTGGCCCCCTTGCCGGCGCTGG 1000
Oligo fwd CACCGTTGGCCCCCTTGCCGGCGC
Oligo rev AAACGCGCCGGCAAGGGGGCCAAC
Chromosome start end strand MM target_seq PAM alignment distance position gene name gene id
GU305914.1 90762 90784 + 0 GTTGGCCCCCTTGCCGGCGC TGG ||||||||[||||||||||||]PAM NA Intergenic NA NA

T6461 TTGGCCCCCTTGCCGGCGCTGGG 1000
Oligo fwd CACCTTGGCCCCCTTGCCGGCGCT
Oligo rev AAACAGCGCCGGCAAGGGGGCCAA
Chromosome start end strand MM target_seq PAM alignment distance position gene name gene id
GU305914.1 90763 90785 + 0 TTGGCCCCCTTGCCGGCGCT GGG ||||||||[||||||||||||]PAM NA Intergenic NA NA

T6462 GCACGAACACGATCCGCGTCTGG 1000
Oligo fwd CACCGCACGAACACGATCCGCGTC
Oligo rev AAACGACGCGGATCGTGTTCGTGC
Chromosome start end strand MM target_seq PAM alignment distance position gene name gene id
GU305914.1 90785 90807 + 0 GCACGAACACGATCCGCGTC TGG ||||||||[||||||||||||]PAM NA Intergenic NA NA

T6463 ACACGATCCGCGTCTGGTAGAGG 1000
Oligo fwd CACCACACGATCCGCGTCTGGTAG
Oligo rev AAACCTACCAGACGCGGATCGTGT
Chromosome start end strand MM target_seq PAM alignment distance position gene name gene id
GU305914.1 90791 90813 + 0 ACACGATCCGCGTCTGGTAG AGG ||||||||[||||||||||||]PAM NA Intergenic NA NA

T6464 GTTCGAGCTGAAGATGAGCGCGG 1000
Oligo fwd CACCGTTCGAGCTGAAGATGAGCG
Oligo rev AAACCGCTCATCTTCAGCTCGAAC
Chromosome start end strand MM target_seq PAM alignment distance position gene name gene id
GU305914.1 90819 90841 + 0 GTTCGAGCTGAAGATGAGCG CGG ||||||||[||||||||||||]PAM NA Intergenic NA NA

T6465 GCGCGGCGATGTGTTTAGCGAGG 1000
Oligo fwd CACCGCGCGGCGATGTGTTTAGCG
Oligo rev AAACCGCTAAACACATCGCCGCGC
Chromosome start end strand MM target_seq PAM alignment distance position gene name gene id
GU305914.1 90836 90858 + 0 GCGCGGCGATGTGTTTAGCG AGG ||||||||[||||||||||||]PAM NA Intergenic NA NA

T6466 TGTGTTTAGCGAGGTGCAACTGG 1000
Oligo fwd CACCTGTGTTTAGCGAGGTGCAAC
Oligo rev AAACGTTGCACCTCGCTAAACACA
Chromosome start end strand MM target_seq PAM alignment distance position gene name gene id
GU305914.1 90845 90867 + 0 TGTGTTTAGCGAGGTGCAAC TGG ||||||||[||||||||||||]PAM NA Intergenic NA NA

T6467 GGTGCAACTGGAAATCCCGCAGG 1000
Oligo fwd CACCGGTGCAACTGGAAATCCCGC
Oligo rev AAACGCGGGATTTCCAGTTGCACC
Chromosome start end strand MM target_seq PAM alignment distance position gene name gene id
GU305914.1 90857 90879 + 0 GGTGCAACTGGAAATCCCGC AGG ||||||||[||||||||||||]PAM NA Intergenic NA NA

T6468 CCGCAGGTTGTCCAGCAGATCGG 1000
Oligo fwd CACCCCGCAGGTTGTCCAGCAGAT
Oligo rev AAACATCTGCTGGACAACCTGCGG
Chromosome start end strand MM target_seq PAM alignment distance position gene name gene id
GU305914.1 90873 90895 + 0 CCGCAGGTTGTCCAGCAGAT CGG ||||||||[||||||||||||]PAM NA Intergenic NA NA

T6469 GGTTGTCCAGCAGATCGGCGTGG 1000
Oligo fwd CACCGGTTGTCCAGCAGATCGGCG
Oligo rev AAACCGCCGATCTGCTGGACAACC
Chromosome start end strand MM target_seq PAM alignment distance position gene name gene id
GU305914.1 90878 90900 + 0 GGTTGTCCAGCAGATCGGCG TGG ||||||||[||||||||||||]PAM NA Intergenic NA NA

T6470 GTTGTCCAGCAGATCGGCGTGGG 1000
Oligo fwd CACCGTTGTCCAGCAGATCGGCGT
Oligo rev AAACACGCCGATCTGCTGGACAAC
Chromosome start end strand MM target_seq PAM alignment distance position gene name gene id
GU305914.1 90879 90901 + 0 GTTGTCCAGCAGATCGGCGT GGG ||||||||[||||||||||||]PAM NA Intergenic NA NA

T6471 CAGCAGATCGGCGTGGGTCTTGG 1000

Oligo fwd CACCCAGCAGATCGGCGTGGGTCT
Oligo rev AAACAGACCCACGCCGATCTGCTG
Chromosome start end strand MM target_seq PAM alignment distance position gene name gene id
GU305914.1 90885 90907 + 0 CAGCAGATCGGCGTGGGTCT TGG ||||||||[||||||||||||]PAM NA Intergenic NA NA

T6472 GTCTTGGCGTTCTTGACGAGCGG 1000
Oligo fwd CACCGTCTTGGCGTTCTTGACGAG
Oligo rev AAACCTCGTCAAGAACGCCAAGAC
Chromosome start end strand MM target_seq PAM alignment distance position gene name gene id
GU305914.1 90901 90923 + 0 GTCTTGGCGTTCTTGACGAG CGG ||||||||[||||||||||||]PAM NA Intergenic NA NA

T6473 GAGCGGCCATTCGAGAAAGTTGG 1000
Oligo fwd CACCGAGCGGCCATTCGAGAAAGT
Oligo rev AAACACTTTCTCGAATGGCCGCTC
Chromosome start end strand MM target_seq PAM alignment distance position gene name gene id
GU305914.1 90918 90940 + 0 GAGCGGCCATTCGAGAAAGT TGG ||||||||[||||||||||||]PAM NA Intergenic NA NA

T6474 CGGCCATTCGAGAAAGTTGGAGG 1000
Oligo fwd CACCCGGCCATTCGAGAAAGTTGG
Oligo rev AAACCCAACTTTCTCGAATGGCCG
Chromosome start end strand MM target_seq PAM alignment distance position gene name gene id
GU305914.1 90921 90943 + 0 CGGCCATTCGAGAAAGTTGG AGG ||||||||[||||||||||||]PAM NA Intergenic NA NA

T6475 GGCCGTGCTGAATTCGTAGACGG 1000
Oligo fwd CACCGGCCGTGCTGAATTCGTAGA
Oligo rev AAACTCTACGAATTCAGCACGGCC
Chromosome start end strand MM target_seq PAM alignment distance position gene name gene id
GU305914.1 90942 90964 + 0 GGCCGTGCTGAATTCGTAGA CGG ||||||||[||||||||||||]PAM NA Intergenic NA NA

T6476 GTAGACGGAATCAAGATAGATGG 1000
Oligo fwd CACCGTAGACGGAATCAAGATAGA
Oligo rev AAACTCTATCTTGATTCCGTCTAC
Chromosome start end strand MM target_seq PAM alignment distance position gene name gene id
GU305914.1 90957 90979 + 0 GTAGACGGAATCAAGATAGA TGG ||||||||[||||||||||||]PAM NA Intergenic NA NA

T6477 CGCGTAGCTGAACGCGAGCTCGG 1000
Oligo fwd CACCCGCGTAGCTGAACGCGAGCT
Oligo rev AAACAGCTCGCGTTCAGCTACGCG
Chromosome start end strand MM target_seq PAM alignment distance position gene name gene id
GU305914.1 90981 91003 + 0 CGCGTAGCTGAACGCGAGCT CGG ||||||||[||||||||||||]PAM NA Intergenic NA NA

T6478 CCGCGTCGACCGTCATGAGCTGG 1000
Oligo fwd CACCCCGCGTCGACCGTCATGAGC
Oligo rev AAACGCTCATGACGGTCGACGCGG
Chromosome start end strand MM target_seq PAM alignment distance position gene name gene id
GU305914.1 91010 91032 + 0 CCGCGTCGACCGTCATGAGC TGG ||||||||[||||||||||||]PAM NA Intergenic NA NA

T6479 CATGAGCTGGTCCCTCTCGACGG 1000
Oligo fwd CACCCATGAGCTGGTCCCTCTCGA
Oligo rev AAACTCGAGAGGGACCAGCTCATG
Chromosome start end strand MM target_seq PAM alignment distance position gene name gene id
GU305914.1 91023 91045 + 0 CATGAGCTGGTCCCTCTCGA CGG ||||||||[||||||||||||]PAM NA Intergenic NA NA

T6480 GAGCTGGTCCCTCTCGACGGCGG 1000
Oligo fwd CACCGAGCTGGTCCCTCTCGACGG
Oligo rev AAACCCGTCGAGAGGGACCAGCTC
Chromosome start end strand MM target_seq PAM alignment distance position gene name gene id
GU305914.1 91026 91048 + 0 GAGCTGGTCCCTCTCGACGG CGG ||||||||[||||||||||||]PAM NA Intergenic NA NA

T6481 TCTCGACGGCGGTGAGCTTCTGG 1000
Oligo fwd CACCTCTCGACGGCGGTGAGCTTC
Oligo rev AAACGAAGCTCACCGCCGTCGAGA
Chromosome start end strand MM target_seq PAM alignment distance position gene name gene id
GU305914.1 91037 91059 + 0 TCTCGACGGCGGTGAGCTTC TGG ||||||||[||||||||||||]PAM NA Intergenic NA NA

T6482 GACGGCGGTGAGCTTCTGGCTGG 1000
Oligo fwd CACCGACGGCGGTGAGCTTCTGGC
Oligo rev AAACGCCAGAAGCTCACCGCCGTC
Chromosome start end strand MM target_seq PAM alignment distance position gene name gene id
GU305914.1 91041 91063 + 0 GACGGCGGTGAGCTTCTGGC TGG ||||||||[||||||||||||]PAM NA Intergenic NA NA

T6483 GCTTCTGGCTGGTGTAGCCGTGG 1000
Oligo fwd CACCGCTTCTGGCTGGTGTAGCCG
Oligo rev AAACCGGCTACACCAGCCAGAAGC
Chromosome start end strand MM target_seq PAM alignment distance position gene name gene id
GU305914.1 91052 91074 + 0 GCTTCTGGCTGGTGTAGCCG TGG ||||||||[||||||||||||]PAM NA Intergenic NA NA

T6484 TAGCCGTGGTACTTCTTGAACGG 1000
Oligo fwd CACCTAGCCGTGGTACTTCTTGAA
Oligo rev AAACTTCAAGAAGTACCACGGCTA
Chromosome start end strand MM target_seq PAM alignment distance position gene name gene id
GU305914.1 91066 91088 + 0 TAGCCGTGGTACTTCTTGAA CGG ||||||||[||||||||||||]PAM NA Intergenic NA NA

T6485 CTCGACGAACTGCTGCAGACCGG 1000
Oligo fwd CACCCTCGACGAACTGCTGCAGAC
Oligo rev AAACGTCTGCAGCAGTTCGTCGAG
Chromosome start end strand MM target_seq PAM alignment distance position gene name gene id
GU305914.1 91116 91138 + 0 CTCGACGAACTGCTGCAGAC CGG ||||||||[||||||||||||]PAM NA Intergenic NA NA

| | | | | | | | | | | | |
|---|---|---|---|---|---|---|---|---|---|---|---|
| T6486 | GCTGCAGACCGGCCTCGATGAGG | 1000 | | | | | | | | | |
| Oligo fwd | CACCGCTGCAGACCGGCCTCGATG | | | | | | | | | | |
| Oligo rev | AAACCATCGAGGCCGGTCTGCAGC | | | | | | | | | | |
| Chromosome | start | end | strand | MM | target_seq | PAM | alignment | distance | position | gene name | gene id |
| GU305914.1 | 91127 | 91149 | + | 0 | GCTGCAGACCGGCCTCGATG | AGG | \|\|\|\|\|\|\|\|[\|\|\|\|\|\|\|\|\|\|\|\|]PAM | NA | Intergenic | NA | NA |
| GU305914.1 | 25626 | 25648 | + | 4 | GCAGCAGGCCGATCTCGATG | GGG | \|\|-\|\|\|\|-[\|\|\|--\|\|\|\|\|\|\|]PAM | NA | Intergenic | NA | NA |
| T6487 | ACCGGCCTCGATGAGGCGACTGG | 1000 | | | | | | | | | |
| Oligo fwd | CACCACCGGCCTCGATGAGGCGAC | | | | | | | | | | |
| Oligo rev | AAACGTCGCCTCATCGAGGCCGGT | | | | | | | | | | |
| Chromosome | start | end | strand | MM | target_seq | PAM | alignment | distance | position | gene name | gene id |
| GU305914.1 | 91134 | 91156 | + | 0 | ACCGGCCTCGATGAGGCGAC | TGG | \|\|\|\|\|\|\|\|[\|\|\|\|\|\|\|\|\|\|\|\|]PAM | NA | Intergenic | NA | NA |
| T6488 | TCTCGCAGACGCGCAGCGACTGG | 1000 | | | | | | | | | |
| Oligo fwd | CACCTCTCGCAGACGCGCAGCGAC | | | | | | | | | | |
| Oligo rev | AAACGTCGCTGCGCGTCTGCGAGA | | | | | | | | | | |
| Chromosome | start | end | strand | MM | target_seq | PAM | alignment | distance | position | gene name | gene id |
| GU305914.1 | 91157 | 91179 | + | 0 | TCTCGCAGACGCGCAGCGAC | TGG | \|\|\|\|\|\|\|\|[\|\|\|\|\|\|\|\|\|\|\|\|]PAM | NA | Intergenic | NA | NA |
| T6489 | GACGCGCAGCGACTGGCCCCAGG | 1000 | | | | | | | | | |
| Oligo fwd | CACCGACGCGCAGCGACTGGCCCC | | | | | | | | | | |
| Oligo rev | AAACGGGGCCAGTCGCTGCGCGTC | | | | | | | | | | |
| Chromosome | start | end | strand | MM | target_seq | PAM | alignment | distance | position | gene name | gene id |
| GU305914.1 | 91164 | 91186 | + | 0 | GACGCGCAGCGACTGGCCCC | AGG | \|\|\|\|\|\|\|\|[\|\|\|\|\|\|\|\|\|\|\|\|]PAM | NA | Intergenic | NA | NA |
| T6490 | GCGCAGCGACTGGCCCCAGGAGG | 1000 | | | | | | | | | |
| Oligo fwd | CACCGCGCAGCGACTGGCCCCAGG | | | | | | | | | | |
| Oligo rev | AAACCCTGGGGCCAGTCGCTGCGC | | | | | | | | | | |
| Chromosome | start | end | strand | MM | target_seq | PAM | alignment | distance | position | gene name | gene id |
| GU305914.1 | 91167 | 91189 | + | 0 | GCGCAGCGACTGGCCCCAGG | AGG | \|\|\|\|\|\|\|\|[\|\|\|\|\|\|\|\|\|\|\|\|]PAM | NA | Intergenic | NA | NA |
| T6491 | GAAGAGCGCCTCGCTGAGCGCGG | 1000 | | | | | | | | | |
| Oligo fwd | CACCGAAGAGCGCCTCGCTGAGCG | | | | | | | | | | |
| Oligo rev | AAACCGCTCAGCGAGGCGCTCTTC | | | | | | | | | | |
| Chromosome | start | end | strand | MM | target_seq | PAM | alignment | distance | position | gene name | gene id |
| GU305914.1 | 91203 | 91225 | + | 0 | GAAGAGCGCCTCGCTGAGCG | CGG | \|\|\|\|\|\|\|\|[\|\|\|\|\|\|\|\|\|\|\|\|]PAM | NA | Intergenic | NA | NA |
| T6492 | AAGAGCGCCTCGCTGAGCGCGGG | 1000 | | | | | | | | | |
| Oligo fwd | CACCAAGAGCGCCTCGCTGAGCGC | | | | | | | | | | |
| Oligo rev | AAACGCGCTCAGCGAGGCGCTCTT | | | | | | | | | | |
| Chromosome | start | end | strand | MM | target_seq | PAM | alignment | distance | position | gene name | gene id |
| GU305914.1 | 91204 | 91226 | + | 0 | AAGAGCGCCTCGCTGAGCGC | GGG | \|\|\|\|\|\|\|\|[\|\|\|\|\|\|\|\|\|\|\|\|]PAM | NA | Intergenic | NA | NA |
| T6493 | AGAGCGCCTCGCTGAGCGCGGGG | 1000 | | | | | | | | | |
| Oligo fwd | CACCAGAGCGCCTCGCTGAGCGCG | | | | | | | | | | |
| Oligo rev | AAACCGCGCTCAGCGAGGCGCTCT | | | | | | | | | | |
| Chromosome | start | end | strand | MM | target_seq | PAM | alignment | distance | position | gene name | gene id |
| GU305914.1 | 91205 | 91227 | + | 0 | AGAGCGCCTCGCTGAGCGCG | GGG | \|\|\|\|\|\|\|\|[\|\|\|\|\|\|\|\|\|\|\|\|]PAM | NA | Intergenic | NA | NA |
| T6494 | GGTCGTAGAACCCGTGCTCGAGG | 1000 | | | | | | | | | |
| Oligo fwd | CACCGGTCGTAGAACCCGTGCTCG | | | | | | | | | | |
| Oligo rev | AAACCGAGCACGGGTTCTACGACC | | | | | | | | | | |
| Chromosome | start | end | strand | MM | target_seq | PAM | alignment | distance | position | gene name | gene id |
| GU305914.1 | 91226 | 91248 | + | 0 | GGTCGTAGAACCCGTGCTCG | AGG | \|\|\|\|\|\|\|\|[\|\|\|\|\|\|\|\|\|\|\|\|]PAM | NA | Intergenic | NA | NA |
| T6495 | AACCCGTGCTCGAGGACCCACGG | 1000 | | | | | | | | | |
| Oligo fwd | CACCAACCCGTGCTCGAGGACCCA | | | | | | | | | | |
| Oligo rev | AAACTGGGTCCTCGAGCACGGGTT | | | | | | | | | | |
| Chromosome | start | end | strand | MM | target_seq | PAM | alignment | distance | position | gene name | gene id |
| GU305914.1 | 91234 | 91256 | + | 0 | AACCCGTGCTCGAGGACCCA | CGG | \|\|\|\|\|\|\|\|[\|\|\|\|\|\|\|\|\|\|\|\|]PAM | NA | Intergenic | NA | NA |
| T6496 | GAGGACCCACGGATCGTCGCAGG | 1000 | | | | | | | | | |
| Oligo fwd | CACCGAGGACCCACGGATCGTCGC | | | | | | | | | | |
| Oligo rev | AAACGCGACGATCCGTGGGTCCTC | | | | | | | | | | |
| Chromosome | start | end | strand | MM | target_seq | PAM | alignment | distance | position | gene name | gene id |
| GU305914.1 | 91245 | 91267 | + | 0 | GAGGACCCACGGATCGTCGC | AGG | \|\|\|\|\|\|\|\|[\|\|\|\|\|\|\|\|\|\|\|\|]PAM | NA | Intergenic | NA | NA |
| T6497 | CCACGGATCGTCGCAGGCGACGG | 1000 | | | | | | | | | |
| Oligo fwd | CACCCCACGGATCGTCGCAGGCGA | | | | | | | | | | |
| Oligo rev | AAACTCGCCTGCGACGATCCGTGG | | | | | | | | | | |
| Chromosome | start | end | strand | MM | target_seq | PAM | alignment | distance | position | gene name | gene id |
| GU305914.1 | 91251 | 91273 | + | 0 | CCACGGATCGTCGCAGGCGA | CGG | \|\|\|\|\|\|\|\|[\|\|\|\|\|\|\|\|\|\|\|\|]PAM | NA | Intergenic | NA | NA |
| T6498 | ATCGTCGCAGGCGACGGCCGCGG | 1000 | | | | | | | | | |
| Oligo fwd | CACCATCGTCGCAGGCGACGGCCG | | | | | | | | | | |
| Oligo rev | AAACCGGCCGTCGCCTGCGACGAT | | | | | | | | | | |
| Chromosome | start | end | strand | MM | target_seq | PAM | alignment | distance | position | gene name | gene id |
| GU305914.1 | 91257 | 91279 | + | 0 | ATCGTCGCAGGCGACGGCCG | CGG | \|\|\|\|\|\|\|\|[\|\|\|\|\|\|\|\|\|\|\|\|]PAM | NA | Intergenic | NA | NA |
| GU305914.1 | 26382 | 26404 | + | 3 | ATCGGCGCTGGCGACGGCGG | CGG | \|\|\|\|-\|\|\|[-\|\|\|\|\|\|\|\|\|-\|]PAM | NA | Intergenic | NA | NA |
| T6499 | GACGGCCGCGGAAGCGCCGTCGG | 1000 | | | | | | | | | |
| Oligo fwd | CACCGACGGCCGCGGAAGCGCCGT | | | | | | | | | | |
| Oligo rev | AAACACGGCGCTTCCGCGGCCGTC | | | | | | | | | | |
| Chromosome | start | end | strand | MM | target_seq | PAM | alignment | distance | position | gene name | gene id |
| GU305914.1 | 91269 | 91291 | + | 0 | GACGGCCGCGGAAGCGCCGT | CGG | \|\|\|\|\|\|\|\|[\|\|\|\|\|\|\|\|\|\|\|\|]PAM | NA | Intergenic | NA | NA |
| GU305914.1 | 109443 | 109465 | + | 3 | GACGGCGGCGGTAGCGGCGT | CGG | \|\|\|\|\|\|-\|[\|\|\|-\|\|\|\|-\|\|\|]PAM | NA | Intergenic | NA | NA |
| T6500 | GGAAGCGCCGTCGGCGTGCGCGG | 1000 | | | | | | | | | |
| Oligo fwd | CACCGGAAGCGCCGTCGGCGTGCG | | | | | | | | | | |

<table>
<tr><td>Oligo rev</td><td>AAACCGCACGCCGACGGCGCTTCC</td><td></td><td></td><td></td><td></td><td></td><td></td><td></td><td></td><td></td><td></td><td></td></tr>
<tr><td>Chromosome</td><td>start</td><td></td><td>end</td><td>strand</td><td>MM</td><td>target_seq</td><td>PAM</td><td>alignment</td><td>distance</td><td>position</td><td>gene name</td><td>gene id</td></tr>
<tr><td>GU305914.1</td><td></td><td>91278</td><td>91300</td><td>+</td><td>0</td><td>GGAAGCGCCGTCGGCGTGCG</td><td>CGG</td><td>||||||||[||||||||||||]PAM</td><td>NA</td><td>Intergenic</td><td>NA</td><td>NA</td></tr>
<tr><td>T6501</td><td>GCCGTCGGCGTGCGCGGCATCGG</td><td></td><td>1000</td><td></td><td></td><td></td><td></td><td></td><td></td><td></td><td></td><td></td></tr>
<tr><td>Oligo fwd</td><td>CACCGCCGTCGGCGTGCGCGGCAT</td><td></td><td></td><td></td><td></td><td></td><td></td><td></td><td></td><td></td><td></td><td></td></tr>
<tr><td>Oligo rev</td><td>AAACATGCCGCGCACGCCGACGGC</td><td></td><td></td><td></td><td></td><td></td><td></td><td></td><td></td><td></td><td></td><td></td></tr>
<tr><td>Chromosome</td><td>start</td><td></td><td>end</td><td>strand</td><td>MM</td><td>target_seq</td><td>PAM</td><td>alignment</td><td>distance</td><td>position</td><td>gene name</td><td>gene id</td></tr>
<tr><td>GU305914.1</td><td></td><td>91284</td><td>91306</td><td>+</td><td>0</td><td>GCCGTCGGCGTGCGCGGCAT</td><td>CGG</td><td>||||||||[||||||||||||]PAM</td><td>NA</td><td>Intergenic</td><td>NA</td><td>NA</td></tr>
<tr><td>T6502</td><td>GTGCGCGGCATCGGCGCCGCCGG</td><td></td><td>1000</td><td></td><td></td><td></td><td></td><td></td><td></td><td></td><td></td><td></td></tr>
<tr><td>Oligo fwd</td><td>CACCGTGCGCGGCATCGGCGCCGC</td><td></td><td></td><td></td><td></td><td></td><td></td><td></td><td></td><td></td><td></td><td></td></tr>
<tr><td>Oligo rev</td><td>AAACGCGGCGCCGATGCCGCGCAC</td><td></td><td></td><td></td><td></td><td></td><td></td><td></td><td></td><td></td><td></td><td></td></tr>
<tr><td>Chromosome</td><td>start</td><td></td><td>end</td><td>strand</td><td>MM</td><td>target_seq</td><td>PAM</td><td>alignment</td><td>distance</td><td>position</td><td>gene name</td><td>gene id</td></tr>
<tr><td>GU305914.1</td><td></td><td>91293</td><td>91315</td><td>+</td><td>0</td><td>GTGCGCGGCATCGGCGCCGC</td><td>CGG</td><td>||||||||[||||||||||||]PAM</td><td>NA</td><td>Intergenic</td><td>NA</td><td>NA</td></tr>
<tr><td>T6503</td><td>GCCGCCGGAGAAGAGAGTGTAGG</td><td></td><td>1000</td><td></td><td></td><td></td><td></td><td></td><td></td><td></td><td></td><td></td></tr>
<tr><td>Oligo fwd</td><td>CACCGCCGCCGGAGAAGAGAGTGT</td><td></td><td></td><td></td><td></td><td></td><td></td><td></td><td></td><td></td><td></td><td></td></tr>
<tr><td>Oligo rev</td><td>AAACACACTCTCTTCTCCGGCGGC</td><td></td><td></td><td></td><td></td><td></td><td></td><td></td><td></td><td></td><td></td><td></td></tr>
<tr><td>Chromosome</td><td>start</td><td></td><td>end</td><td>strand</td><td>MM</td><td>target_seq</td><td>PAM</td><td>alignment</td><td>distance</td><td>position</td><td>gene name</td><td>gene id</td></tr>
<tr><td>GU305914.1</td><td></td><td>91308</td><td>91330</td><td>+</td><td>0</td><td>GCCGCCGGAGAAGAGAGTGT</td><td>AGG</td><td>||||||||[||||||||||||]PAM</td><td>NA</td><td>Intergenic</td><td>NA</td><td>NA</td></tr>
<tr><td>T6504</td><td>CCGCCGGAGAAGAGAGTGTAGGG</td><td></td><td>1000</td><td></td><td></td><td></td><td></td><td></td><td></td><td></td><td></td><td></td></tr>
<tr><td>Oligo fwd</td><td>CACCCCGCCGGAGAAGAGAGTGTA</td><td></td><td></td><td></td><td></td><td></td><td></td><td></td><td></td><td></td><td></td><td></td></tr>
<tr><td>Oligo rev</td><td>AAACTACACTCTCTTCTCCGGCGG</td><td></td><td></td><td></td><td></td><td></td><td></td><td></td><td></td><td></td><td></td><td></td></tr>
<tr><td>Chromosome</td><td>start</td><td></td><td>end</td><td>strand</td><td>MM</td><td>target_seq</td><td>PAM</td><td>alignment</td><td>distance</td><td>position</td><td>gene name</td><td>gene id</td></tr>
<tr><td>GU305914.1</td><td></td><td>91309</td><td>91331</td><td>+</td><td>0</td><td>CCGCCGGAGAAGAGAGTGTA</td><td>GGG</td><td>||||||||[||||||||||||]PAM</td><td>NA</td><td>Intergenic</td><td>NA</td><td>NA</td></tr>
<tr><td>T6505</td><td>AAGAGAGTGTAGGGATAGAGCGG</td><td></td><td>1000</td><td></td><td></td><td></td><td></td><td></td><td></td><td></td><td></td><td></td></tr>
<tr><td>Oligo fwd</td><td>CACCAAGAGAGTGTAGGGATAGAG</td><td></td><td></td><td></td><td></td><td></td><td></td><td></td><td></td><td></td><td></td><td></td></tr>
<tr><td>Oligo rev</td><td>AAACCTCTATCCCTACACTCTCTT</td><td></td><td></td><td></td><td></td><td></td><td></td><td></td><td></td><td></td><td></td><td></td></tr>
<tr><td>Chromosome</td><td>start</td><td></td><td>end</td><td>strand</td><td>MM</td><td>target_seq</td><td>PAM</td><td>alignment</td><td>distance</td><td>position</td><td>gene name</td><td>gene id</td></tr>
<tr><td>GU305914.1</td><td></td><td>91318</td><td>91340</td><td>+</td><td>0</td><td>AAGAGAGTGTAGGGATAGAG</td><td>CGG</td><td>||||||||[||||||||||||]PAM</td><td>NA</td><td>Intergenic</td><td>NA</td><td>NA</td></tr>
<tr><td>T6506</td><td>AGAGAGTGTAGGGATAGAGCGGG</td><td></td><td>1000</td><td></td><td></td><td></td><td></td><td></td><td></td><td></td><td></td><td></td></tr>
<tr><td>Oligo fwd</td><td>CACCAGAGAGTGTAGGGATAGAGC</td><td></td><td></td><td></td><td></td><td></td><td></td><td></td><td></td><td></td><td></td><td></td></tr>
<tr><td>Oligo rev</td><td>AAACGCTCTATCCCTACACTCTCT</td><td></td><td></td><td></td><td></td><td></td><td></td><td></td><td></td><td></td><td></td><td></td></tr>
<tr><td>Chromosome</td><td>start</td><td></td><td>end</td><td>strand</td><td>MM</td><td>target_seq</td><td>PAM</td><td>alignment</td><td>distance</td><td>position</td><td>gene name</td><td>gene id</td></tr>
<tr><td>GU305914.1</td><td></td><td>91319</td><td>91341</td><td>+</td><td>0</td><td>AGAGAGTGTAGGGATAGAGC</td><td>GGG</td><td>||||||||[||||||||||||]PAM</td><td>NA</td><td>Intergenic</td><td>NA</td><td>NA</td></tr>
<tr><td>T6507</td><td>CGCGCCGACCTGCACCTTCACGG</td><td></td><td>1000</td><td></td><td></td><td></td><td></td><td></td><td></td><td></td><td></td><td></td></tr>
<tr><td>Oligo fwd</td><td>CACCCGCGCCGACCTGCACCTTCA</td><td></td><td></td><td></td><td></td><td></td><td></td><td></td><td></td><td></td><td></td><td></td></tr>
<tr><td>Oligo rev</td><td>AAACTGAAGGTGCAGGTCGGCGCG</td><td></td><td></td><td></td><td></td><td></td><td></td><td></td><td></td><td></td><td></td><td></td></tr>
<tr><td>Chromosome</td><td>start</td><td></td><td>end</td><td>strand</td><td>MM</td><td>target_seq</td><td>PAM</td><td>alignment</td><td>distance</td><td>position</td><td>gene name</td><td>gene id</td></tr>
<tr><td>GU305914.1</td><td></td><td>91359</td><td>91381</td><td>+</td><td>0</td><td>CGCGCCGACCTGCACCTTCA</td><td>CGG</td><td>||||||||[||||||||||||]PAM</td><td>NA</td><td>Intergenic</td><td>NA</td><td>NA</td></tr>
<tr><td>T6508</td><td>CTGCACCTTCACGGCCTCTTCGG</td><td></td><td>1000</td><td></td><td></td><td></td><td></td><td></td><td></td><td></td><td></td><td></td></tr>
<tr><td>Oligo fwd</td><td>CACCCTGCACCTTCACGGCCTCTT</td><td></td><td></td><td></td><td></td><td></td><td></td><td></td><td></td><td></td><td></td><td></td></tr>
<tr><td>Oligo rev</td><td>AAACAAGAGGCCGTGAAGGTGCAG</td><td></td><td></td><td></td><td></td><td></td><td></td><td></td><td></td><td></td><td></td><td></td></tr>
<tr><td>Chromosome</td><td>start</td><td></td><td>end</td><td>strand</td><td>MM</td><td>target_seq</td><td>PAM</td><td>alignment</td><td>distance</td><td>position</td><td>gene name</td><td>gene id</td></tr>
<tr><td>GU305914.1</td><td></td><td>91368</td><td>91390</td><td>+</td><td>0</td><td>CTGCACCTTCACGGCCTCTT</td><td>CGG</td><td>||||||||[||||||||||||]PAM</td><td>NA</td><td>Intergenic</td><td>NA</td><td>NA</td></tr>
<tr><td>T6509</td><td>TGCACCTTCACGGCCTCTTCGGG</td><td></td><td>1000</td><td></td><td></td><td></td><td></td><td></td><td></td><td></td><td></td><td></td></tr>
<tr><td>Oligo fwd</td><td>CACCTGCACCTTCACGGCCTCTTC</td><td></td><td></td><td></td><td></td><td></td><td></td><td></td><td></td><td></td><td></td><td></td></tr>
<tr><td>Oligo rev</td><td>AAACGAAGAGGCCGTGAAGGTGCA</td><td></td><td></td><td></td><td></td><td></td><td></td><td></td><td></td><td></td><td></td><td></td></tr>
<tr><td>Chromosome</td><td>start</td><td></td><td>end</td><td>strand</td><td>MM</td><td>target_seq</td><td>PAM</td><td>alignment</td><td>distance</td><td>position</td><td>gene name</td><td>gene id</td></tr>
<tr><td>GU305914.1</td><td></td><td>91369</td><td>91391</td><td>+</td><td>0</td><td>TGCACCTTCACGGCCTCTTC</td><td>GGG</td><td>||||||||[||||||||||||]PAM</td><td>NA</td><td>Intergenic</td><td>NA</td><td>NA</td></tr>
<tr><td>T6510</td><td>CTTCACGGCCTCTTCGGGACAGG</td><td></td><td>1000</td><td></td><td></td><td></td><td></td><td></td><td></td><td></td><td></td><td></td></tr>
<tr><td>Oligo fwd</td><td>CACCCTTCACGGCCTCTTCGGGAC</td><td></td><td></td><td></td><td></td><td></td><td></td><td></td><td></td><td></td><td></td><td></td></tr>
<tr><td>Oligo rev</td><td>AAACGTCCCGAAGAGGCCGTGAAG</td><td></td><td></td><td></td><td></td><td></td><td></td><td></td><td></td><td></td><td></td><td></td></tr>
<tr><td>Chromosome</td><td>start</td><td></td><td>end</td><td>strand</td><td>MM</td><td>target_seq</td><td>PAM</td><td>alignment</td><td>distance</td><td>position</td><td>gene name</td><td>gene id</td></tr>
<tr><td>GU305914.1</td><td></td><td>91374</td><td>91396</td><td>+</td><td>0</td><td>CTTCACGGCCTCTTCGGGAC</td><td>AGG</td><td>||||||||[||||||||||||]PAM</td><td>NA</td><td>Intergenic</td><td>NA</td><td>NA</td></tr>
<tr><td>T6511</td><td>TTCACGGCCTCTTCGGGACAGGG</td><td></td><td>1000</td><td></td><td></td><td></td><td></td><td></td><td></td><td></td><td></td><td></td></tr>
<tr><td>Oligo fwd</td><td>CACCTTCACGGCCTCTTCGGGACA</td><td></td><td></td><td></td><td></td><td></td><td></td><td></td><td></td><td></td><td></td><td></td></tr>
<tr><td>Oligo rev</td><td>AAACTGTCCCGAAGAGGCCGTGAA</td><td></td><td></td><td></td><td></td><td></td><td></td><td></td><td></td><td></td><td></td><td></td></tr>
<tr><td>Chromosome</td><td>start</td><td></td><td>end</td><td>strand</td><td>MM</td><td>target_seq</td><td>PAM</td><td>alignment</td><td>distance</td><td>position</td><td>gene name</td><td>gene id</td></tr>
<tr><td>GU305914.1</td><td></td><td>91375</td><td>91397</td><td>+</td><td>0</td><td>TTCACGGCCTCTTCGGGACA</td><td>GGG</td><td>||||||||[||||||||||||]PAM</td><td>NA</td><td>Intergenic</td><td>NA</td><td>NA</td></tr>
<tr><td>T6512</td><td>TCACGGCCTCTTCGGGACAGGGG</td><td></td><td>1000</td><td></td><td></td><td></td><td></td><td></td><td></td><td></td><td></td><td></td></tr>
<tr><td>Oligo fwd</td><td>CACCTCACGGCCTCTTCGGGACAG</td><td></td><td></td><td></td><td></td><td></td><td></td><td></td><td></td><td></td><td></td><td></td></tr>
<tr><td>Oligo rev</td><td>AAACCTGTCCCGAAGAGGCCGTGA</td><td></td><td></td><td></td><td></td><td></td><td></td><td></td><td></td><td></td><td></td><td></td></tr>
<tr><td>Chromosome</td><td>start</td><td></td><td>end</td><td>strand</td><td>MM</td><td>target_seq</td><td>PAM</td><td>alignment</td><td>distance</td><td>position</td><td>gene name</td><td>gene id</td></tr>
<tr><td>GU305914.1</td><td></td><td>91376</td><td>91398</td><td>+</td><td>0</td><td>TCACGGCCTCTTCGGGACAG</td><td>GGG</td><td>||||||||[||||||||||||]PAM</td><td>NA</td><td>Intergenic</td><td>NA</td><td>NA</td></tr>
<tr><td>GU305914.1</td><td></td><td>50881</td><td>50903</td><td>-</td><td>4</td><td>TCACGTTCTCTTCGGGCCCG</td><td>TGG</td><td>|||||--|[||||||||-|-|]PAM</td><td>NA</td><td>Intergenic</td><td>NA</td><td>NA</td></tr>
<tr><td>T6513</td><td>GGATGAGCTGCCCCGCGTAGAGG</td><td></td><td>1000</td><td></td><td></td><td></td><td></td><td></td><td></td><td></td><td></td><td></td></tr>
<tr><td>Oligo fwd</td><td>CACCGGATGAGCTGCCCCGCGTAG</td><td></td><td></td><td></td><td></td><td></td><td></td><td></td><td></td><td></td><td></td><td></td></tr>
<tr><td>Oligo rev</td><td>AAACCTACGCGGGGCAGCTCATCC</td><td></td><td></td><td></td><td></td><td></td><td></td><td></td><td></td><td></td><td></td><td></td></tr>
<tr><td>Chromosome</td><td>start</td><td></td><td>end</td><td>strand</td><td>MM</td><td>target_seq</td><td>PAM</td><td>alignment</td><td>distance</td><td>position</td><td>gene name</td><td>gene id</td></tr>
<tr><td>GU305914.1</td><td></td><td>91397</td><td>91419</td><td>+</td><td>0</td><td>GGATGAGCTGCCCCGCGTAG</td><td>AGG</td><td>||||||||[||||||||||||]PAM</td><td>NA</td><td>Intergenic</td><td>NA</td><td>NA</td></tr>
<tr><td>T6514</td><td>CCGCGTAGAGGAGCTCCTTGAGG</td><td></td><td>1000</td><td></td><td></td><td></td><td></td><td></td><td></td><td></td><td></td><td></td></tr>
<tr><td>Oligo fwd</td><td>CACCCCGCGTAGAGGAGCTCCTTG</td><td></td><td></td><td></td><td></td><td></td><td></td><td></td><td></td><td></td><td></td><td></td></tr>
<tr><td>Oligo rev</td><td>AAACCAAGGAGCTCCTCTACGCGG</td><td></td><td></td><td></td><td></td><td></td><td></td><td></td><td></td><td></td><td></td><td></td></tr>
<tr><td>Chromosome</td><td>start</td><td></td><td>end</td><td>strand</td><td>MM</td><td>target_seq</td><td>PAM</td><td>alignment</td><td>distance</td><td>position</td><td>gene name</td><td>gene id</td></tr>
<tr><td>GU305914.1</td><td></td><td>91409</td><td>91431</td><td>+</td><td>0</td><td>CCGCGTAGAGGAGCTCCTTG</td><td>AGG</td><td>||||||||[||||||||||||]PAM</td><td>NA</td><td>Intergenic</td><td>NA</td><td>NA</td></tr>
</table>

| | | | | | | | | | | | |
|---|---|---|---|---|---|---|---|---|---|---|---|
| T6515 | CACGACGTGCCCCGTGTACTCGG | 1000 | | | | | | | | | |
| Oligo fwd | CACCCACGACGTGCCCCGTGTACT | | | | | | | | | | |
| Oligo rev | AAACAGTACACGGGGCACGTCGTG | | | | | | | | | | |
| Chromosome | start | end | strand | MM | target_seq | PAM | alignment | distance | position | gene name | gene id |
| GU305914.1 | 91443 | 91465 | + | 0 | CACGACGTGCCCCGTGTACT | CGG | \|\|\|\|\|\|\|\|[\|\|\|\|\|\|\|\|\|\|\|\|]PAM | NA | Intergenic | NA | NA |
| T6516 | GACGTGCCCCGTGTACTCGGCGG | 1000 | | | | | | | | | |
| Oligo fwd | CACCGACGTGCCCCGTGTACTCGG | | | | | | | | | | |
| Oligo rev | AAACCCGAGTACACGGGGCACGTC | | | | | | | | | | |
| Chromosome | start | end | strand | MM | target_seq | PAM | alignment | distance | position | gene name | gene id |
| GU305914.1 | 91446 | 91468 | + | 0 | GACGTGCCCCGTGTACTCGG | CGG | \|\|\|\|\|\|\|\|[\|\|\|\|\|\|\|\|\|\|\|\|]PAM | NA | Intergenic | NA | NA |
| T6517 | ACGTGCCCCGTGTACTCGGCGGG | 1000 | | | | | | | | | |
| Oligo fwd | CACCACGTGCCCCGTGTACTCGGC | | | | | | | | | | |
| Oligo rev | AAACGCCGAGTACACGGGGCACGT | | | | | | | | | | |
| Chromosome | start | end | strand | MM | target_seq | PAM | alignment | distance | position | gene name | gene id |
| GU305914.1 | 91447 | 91469 | + | 0 | ACGTGCCCCGTGTACTCGGC | GGG | \|\|\|\|\|\|\|\|[\|\|\|\|\|\|\|\|\|\|\|\|]PAM | NA | Intergenic | NA | NA |
| T6518 | CGTGCCCCGTGTACTCGGCGGGG | 1000 | | | | | | | | | |
| Oligo fwd | CACCCGTGCCCCGTGTACTCGGCG | | | | | | | | | | |
| Oligo rev | AAACCGCCGAGTACACGGGGCACG | | | | | | | | | | |
| Chromosome | start | end | strand | MM | target_seq | PAM | alignment | distance | position | gene name | gene id |
| GU305914.1 | 91448 | 91470 | + | 0 | CGTGCCCCGTGTACTCGGCG | GGG | \|\|\|\|\|\|\|\|[\|\|\|\|\|\|\|\|\|\|\|\|]PAM | NA | Intergenic | NA | NA |
| T6519 | GTGCCCCGTGTACTCGGCGGGGG | 1000 | | | | | | | | | |
| Oligo fwd | CACCGTGCCCCGTGTACTCGGCGG | | | | | | | | | | |
| Oligo rev | AAACCCGCCGAGTACACGGGGCAC | | | | | | | | | | |
| Chromosome | start | end | strand | MM | target_seq | PAM | alignment | distance | position | gene name | gene id |
| GU305914.1 | 91449 | 91471 | + | 0 | GTGCCCCGTGTACTCGGCGG | GGG | \|\|\|\|\|\|\|\|[\|\|\|\|\|\|\|\|\|\|\|\|]PAM | NA | Intergenic | NA | NA |
| T6520 | CGATACCTAGCTCGTCGCAGAGG | 1000 | | | | | | | | | |
| Oligo fwd | CACCCGATACCTAGCTCGTCGCAG | | | | | | | | | | |
| Oligo rev | AAACCTGCGACGAGCTAGGTATCG | | | | | | | | | | |
| Chromosome | start | end | strand | MM | target_seq | PAM | alignment | distance | position | gene name | gene id |
| GU305914.1 | 91472 | 91494 | + | 0 | CGATACCTAGCTCGTCGCAG | AGG | \|\|\|\|\|\|\|\|[\|\|\|\|\|\|\|\|\|\|\|\|]PAM | NA | Intergenic | NA | NA |
| T6521 | GTGCGATCTCGATTAGCGAGCGG | 1000 | | | | | | | | | |
| Oligo fwd | CACCGTGCGATCTCGATTAGCGAG | | | | | | | | | | |
| Oligo rev | AAACCTCGCTAATCGAGATCGCAC | | | | | | | | | | |
| Chromosome | start | end | strand | MM | target_seq | PAM | alignment | distance | position | gene name | gene id |
| GU305914.1 | 91507 | 91529 | + | 0 | GTGCGATCTCGATTAGCGAG | CGG | \|\|\|\|\|\|\|\|[\|\|\|\|\|\|\|\|\|\|\|\|]PAM | NA | Intergenic | NA | NA |
| T6522 | TCGATTAGCGAGCGGACTGAAGG | 1000 | | | | | | | | | |
| Oligo fwd | CACCTCGATTAGCGAGCGGACTGA | | | | | | | | | | |
| Oligo rev | AAACTCAGTCCGCTCGCTAATCGA | | | | | | | | | | |
| Chromosome | start | end | strand | MM | target_seq | PAM | alignment | distance | position | gene name | gene id |
| GU305914.1 | 91515 | 91537 | + | 0 | TCGATTAGCGAGCGGACTGA | AGG | \|\|\|\|\|\|\|\|[\|\|\|\|\|\|\|\|\|\|\|\|]PAM | NA | Intergenic | NA | NA |
| T6523 | TAGCGAGCGGACTGAAGGCCTGG | 1000 | | | | | | | | | |
| Oligo fwd | CACCTAGCGAGCGGACTGAAGGCC | | | | | | | | | | |
| Oligo rev | AAACGGCCTTCAGTCCGCTCGCTA | | | | | | | | | | |
| Chromosome | start | end | strand | MM | target_seq | PAM | alignment | distance | position | gene name | gene id |
| GU305914.1 | 91520 | 91542 | + | 0 | TAGCGAGCGGACTGAAGGCC | TGG | \|\|\|\|\|\|\|\|[\|\|\|\|\|\|\|\|\|\|\|\|]PAM | NA | Intergenic | NA | NA |
| T6524 | TGAAGGCCTGGACGCCGAAGCGG | 1000 | | | | | | | | | |
| Oligo fwd | CACCTGAAGGCCTGGACGCCGAAG | | | | | | | | | | |
| Oligo rev | AAACCTTCGGCGTCCAGGCCTTCA | | | | | | | | | | |
| Chromosome | start | end | strand | MM | target_seq | PAM | alignment | distance | position | gene name | gene id |
| GU305914.1 | 91532 | 91554 | + | 0 | TGAAGGCCTGGACGCCGAAG | CGG | \|\|\|\|\|\|\|\|[\|\|\|\|\|\|\|\|\|\|\|\|]PAM | NA | Intergenic | NA | NA |
| T6525 | CTGGACGCCGAAGCGGCGCCTGG | 1000 | | | | | | | | | |
| Oligo fwd | CACCCTGGACGCCGAAGCGGCGCC | | | | | | | | | | |
| Oligo rev | AAACGGCGCCGCTTCGGCGTCCAG | | | | | | | | | | |
| Chromosome | start | end | strand | MM | target_seq | PAM | alignment | distance | position | gene name | gene id |
| GU305914.1 | 91539 | 91561 | + | 0 | CTGGACGCCGAAGCGGCGCC | TGG | \|\|\|\|\|\|\|\|[\|\|\|\|\|\|\|\|\|\|\|\|]PAM | NA | Intergenic | NA | NA |
| T6526 | GGCGCCTGGCATCGTCACAGAGG | 1000 | | | | | | | | | |
| Oligo fwd | CACCGGCGCCTGGCATCGTCACAG | | | | | | | | | | |
| Oligo rev | AAACCTGTGACGATGCCAGGCGCC | | | | | | | | | | |
| Chromosome | start | end | strand | MM | target_seq | PAM | alignment | distance | position | gene name | gene id |
| GU305914.1 | 91553 | 91575 | + | 0 | GGCGCCTGGCATCGTCACAG | AGG | \|\|\|\|\|\|\|\|[\|\|\|\|\|\|\|\|\|\|\|\|]PAM | NA | Intergenic | NA | NA |
| T6527 | CGTGATCCTCGACGACGTCGAGG | 1000 | | | | | | | | | |
| Oligo fwd | CACCCGTGATCCTCGACGACGTCG | | | | | | | | | | |
| Oligo rev | AAACCGACGTCGTCGAGGATCACG | | | | | | | | | | |
| Chromosome | start | end | strand | MM | target_seq | PAM | alignment | distance | position | gene name | gene id |
| GU305914.1 | 91589 | 91611 | + | 0 | CGTGATCCTCGACGACGTCG | AGG | \|\|\|\|\|\|\|\|[\|\|\|\|\|\|\|\|\|\|\|\|]PAM | NA | Intergenic | NA | NA |
| GU305914.1 | 91595 | 91617 | - | 2 | CGAGATCCTCGACGTCGTCG | AGG | \|\|-\|\|\|\|\|[\|\|\|\|\|\|-\|\|\|\|\|]PAM | NA | Intergenic | NA | NA |
| T6528 | GACGACGTCGAGGATCTCGTCGG | 1000 | | | | | | | | | |
| Oligo fwd | CACCGACGACGTCGAGGATCTCGT | | | | | | | | | | |
| Oligo rev | AAACACGAGATCCTCGACGTCGTC | | | | | | | | | | |
| Chromosome | start | end | strand | MM | target_seq | PAM | alignment | distance | position | gene name | gene id |
| GU305914.1 | 91599 | 91621 | + | 0 | GACGACGTCGAGGATCTCGT | CGG | \|\|\|\|\|\|\|\|[\|\|\|\|\|\|\|\|\|\|\|\|]PAM | NA | Intergenic | NA | NA |
| T6529 | CGAGGATCTCGTCGGTGTTGTGG | 1000 | | | | | | | | | |
| Oligo fwd | CACCCGAGGATCTCGTCGGTGTTG | | | | | | | | | | |
| Oligo rev | AAACCAACACCGACGAGATCCTCG | | | | | | | | | | |
| Chromosome | start | end | strand | MM | target_seq | PAM | alignment | distance | position | gene name | gene id |

GU305914.1 91607 91629 + 0 CGAGGATCTCGTCGGTGTTG TGG ||||||||[||||||||||||]PAM NA Intergenic NA NA

T6530 GTCGGTGTTGTGGAAAAAGAAGG 1000
Oligo fwd CACCGTCGGTGTTGTGGAAAAAGA
Oligo rev AAACTCTTTTTCCACAACACCGAC
Chromosome start end strand MM target_seq PAM alignment distance position gene name gene id
GU305914.1 91617 91639 + 0 GTCGGTGTTGTGGAAAAAGA AGG ||||||||[||||||||||||]PAM NA Intergenic NA NA

T6531 GTTGTGGAAAAAGAAGGCGAAGG 1000
Oligo fwd CACCGTTGTGGAAAAAGAAGGCGA
Oligo rev AAACTCGCCTTCTTTTTCCACAAC
Chromosome start end strand MM target_seq PAM alignment distance position gene name gene id
GU305914.1 91623 91645 + 0 GTTGTGGAAAAAGAAGGCGA AGG ||||||||[||||||||||||]PAM NA Intergenic NA NA

T6532 TTGTGGAAAAAGAAGGCGAAGGG 1000
Oligo fwd CACCTTGTGGAAAAAGAAGGCGAA
Oligo rev AAACTTCGCCTTCTTTTTCCACAA
Chromosome start end strand MM target_seq PAM alignment distance position gene name gene id
GU305914.1 91624 91646 + 0 TTGTGGAAAAAGAAGGCGAA GGG ||||||||[||||||||||||]PAM NA Intergenic NA NA

T6533 GAAGGCGAAGGGACAGAAAGAGG 1000
Oligo fwd CACCGAAGGCGAAGGGACAGAAAG
Oligo rev AAACCTTTCTGTCCCTTCGCCTTC
Chromosome start end strand MM target_seq PAM alignment distance position gene name gene id
GU305914.1 91635 91657 + 0 GAAGGCGAAGGGACAGAAAG AGG ||||||||[||||||||||||]PAM NA Intergenic NA NA

T6534 GGGACAGAAAGAGGTGACTTTGG 1000
Oligo fwd CACCGGGACAGAAAGAGGTGACTT
Oligo rev AAACAAGTCACCTCTTTCTGTCCC
Chromosome start end strand MM target_seq PAM alignment distance position gene name gene id
GU305914.1 91644 91666 + 0 GGGACAGAAAGAGGTGACTT TGG ||||||||[||||||||||||]PAM NA Intergenic NA NA

T6535 AGAAAGAGGTGACTTTGGTGAGG 1000
Oligo fwd CACCAGAAAGAGGTGACTTTGGTG
Oligo rev AAACCACCAAAGTCACCTCTTTCT
Chromosome start end strand MM target_seq PAM alignment distance position gene name gene id
GU305914.1 91649 91671 + 0 AGAAAGAGGTGACTTTGGTG AGG ||||||||[||||||||||||]PAM NA Intergenic NA NA

T6536 CCCCCCTCGTATGTGCTCATCGG 1000
Oligo fwd CACCCCCCCCTCGTATGTGCTCAT
Oligo rev AAACATGAGCACATACGAGGGGGG
Chromosome start end strand MM target_seq PAM alignment distance position gene name gene id
GU305914.1 91675 91697 + 0 CCCCCCTCGTATGTGCTCAT CGG ||||||||[||||||||||||]PAM NA Intergenic NA NA

T6537 TGACCGCGCCGCAGAAGTCGCGG 1000
Oligo fwd CACCTGACCGCGCCGCAGAAGTCG
Oligo rev AAACCGACTTCTGCGGCGCGGTCA
Chromosome start end strand MM target_seq PAM alignment distance position gene name gene id
GU305914.1 91703 91725 + 0 TGACCGCGCCGCAGAAGTCG CGG ||||||||[||||||||||||]PAM NA Intergenic NA NA

T6538 GCCGCAGAAGTCGCGGTCGACGG 1000
Oligo fwd CACCGCCGCAGAAGTCGCGGTCGA
Oligo rev AAACTCGACCGCGACTTCTGCGGC
Chromosome start end strand MM target_seq PAM alignment distance position gene name gene id
GU305914.1 91710 91732 + 0 GCCGCAGAAGTCGCGGTCGA CGG ||||||||[||||||||||||]PAM NA Intergenic NA NA

T6539 AGAAGTCGCGGTCGACGGTCAGG 1000
Oligo fwd CACCAGAAGTCGCGGTCGACGGTC
Oligo rev AAACGACCGTCGACCGCGACTTCT
Chromosome start end strand MM target_seq PAM alignment distance position gene name gene id
GU305914.1 91715 91737 + 0 AGAAGTCGCGGTCGACGGTC AGG ||||||||[||||||||||||]PAM NA Intergenic NA NA

T6540 ACGGTCAGGTCTATCAACAGCGG 1000
Oligo fwd CACCACGGTCAGGTCTATCAACAG
Oligo rev AAACCTGTTGATAGACCTGACCGT
Chromosome start end strand MM target_seq PAM alignment distance position gene name gene id
GU305914.1 91729 91751 + 0 ACGGTCAGGTCTATCAACAG CGG ||||||||[||||||||||||]PAM NA Intergenic NA NA

T6541 CAACAGCGGCGCGATGACGACGG 1000
Oligo fwd CACCCAACAGCGGCGCGATGACGA
Oligo rev AAACTCGTCATCGCGCCGCTGTTG
Chromosome start end strand MM target_seq PAM alignment distance position gene name gene id
GU305914.1 91743 91765 + 0 CAACAGCGGCGCGATGACGA CGG ||||||||[||||||||||||]PAM NA Intergenic NA NA

T6542 AACAGCGGCGCGATGACGACGGG 1000
Oligo fwd CACCAACAGCGGCGCGATGACGAC
Oligo rev AAACGTCGTCATCGCGCCGCTGTT
Chromosome start end strand MM target_seq PAM alignment distance position gene name gene id
GU305914.1 91744 91766 + 0 AACAGCGGCGCGATGACGAC GGG ||||||||[||||||||||||]PAM NA Intergenic NA NA

T6543 ACAGCGGCGCGATGACGACGGGG 1000
Oligo fwd CACCACAGCGGCGCGATGACGACG
Oligo rev AAACCGTCGTCATCGCGCCGCTGT
Chromosome start end strand MM target_seq PAM alignment distance position gene name gene id
GU305914.1 91745 91767 + 0 ACAGCGGCGCGATGACGACG GGG ||||||||[||||||||||||]PAM NA Intergenic NA NA

T6544 CAGCGGCGCGATGACGACGGGGG 1000
Oligo fwd CACCCAGCGGCGCGATGACGACGG
Oligo rev AAACCCGTCGTCATCGCGCCGCTG

| | | | | | | | | | | | |
|---|---|---|---|---|---|---|---|---|---|---|---|
| Chromosome | start | end | strand | MM | target_seq | PAM | alignment | distance | position | gene name | gene id |
| GU305914.1 | 91746 | 91768 | + | 0 | CAGCGGCGCGATGACGACGG | GGG | \|\|\|\|\|\|\|\|[\|\|\|\|\|\|\|\|\|\|\|\|]PAM | NA | Intergenic | NA | NA |
| GU305914.1 | 19513 | 19535 | + | 4 | CATCGTCGCGATGGGGACGG | CGG | \|\|-\|\|-\|\|[\|\|\|\|\|--\|\|\|\|\|]PAM | NA | Intergenic | NA | NA |
| T6545 | CTTGTCGCAGAGCGACATCGCGG | 1000 | | | | | | | | | |
| Oligo fwd | CACCCTTGTCGCAGAGCGACATCG | | | | | | | | | | |
| Oligo rev | AAACCGATGTCGCTCTGCGACAAG | | | | | | | | | | |
| Chromosome | start | end | strand | MM | target_seq | PAM | alignment | distance | position | gene name | gene id |
| GU305914.1 | 91773 | 91795 | + | 0 | CTTGTCGCAGAGCGACATCG | CGG | \|\|\|\|\|\|\|\|[\|\|\|\|\|\|\|\|\|\|\|\|]PAM | NA | Intergenic | NA | NA |
| T6546 | CGAGCTCCCGCGTCTTTTTGAGG | 1000 | | | | | | | | | |
| Oligo fwd | CACCCGAGCTCCCGCGTCTTTTTG | | | | | | | | | | |
| Oligo rev | AAACCAAAAAGACGCGGGAGCTCG | | | | | | | | | | |
| Chromosome | start | end | strand | MM | target_seq | PAM | alignment | distance | position | gene name | gene id |
| GU305914.1 | 91808 | 91830 | + | 0 | CGAGCTCCCGCGTCTTTTTG | AGG | \|\|\|\|\|\|\|\|[\|\|\|\|\|\|\|\|\|\|\|\|]PAM | NA | Intergenic | NA | NA |
| T6547 | TTTGAGGAAGAACATCCAGACGG | 1000 | | | | | | | | | |
| Oligo fwd | CACCTTTGAGGAAGAACATCCAGA | | | | | | | | | | |
| Oligo rev | AAACTCTGGATGTTCTTCCTCAAA | | | | | | | | | | |
| Chromosome | start | end | strand | MM | target_seq | PAM | alignment | distance | position | gene name | gene id |
| GU305914.1 | 91824 | 91846 | + | 0 | TTTGAGGAAGAACATCCAGA | CGG | \|\|\|\|\|\|\|\|[\|\|\|\|\|\|\|\|\|\|\|\|]PAM | NA | Intergenic | NA | NA |
| T6548 | GAGGAAGAACATCCAGACGGCGG | 1000 | | | | | | | | | |
| Oligo fwd | CACCGAGGAAGAACATCCAGACGG | | | | | | | | | | |
| Oligo rev | AAACCCGTCTGGATGTTCTTCCTC | | | | | | | | | | |
| Chromosome | start | end | strand | MM | target_seq | PAM | alignment | distance | position | gene name | gene id |
| GU305914.1 | 91827 | 91849 | + | 0 | GAGGAAGAACATCCAGACGG | CGG | \|\|\|\|\|\|\|\|[\|\|\|\|\|\|\|\|\|\|\|\|]PAM | NA | Intergenic | NA | NA |
| GU305914.1 | 124160 | 124182 | - | 4 | GAGCCAGAACATCAACACGG | TGG | \|\|\|--\|\|\|[\|\|\|\|\|-\|-\|\|\|\|]PAM | NA | Intergenic | NA | NA |
| T6549 | AGGAAGAACATCCAGACGGCGGG | 1000 | | | | | | | | | |
| Oligo fwd | CACCAGGAAGAACATCCAGACGGC | | | | | | | | | | |
| Oligo rev | AAACGCCGTCTGGATGTTCTTCCT | | | | | | | | | | |
| Chromosome | start | end | strand | MM | target_seq | PAM | alignment | distance | position | gene name | gene id |
| GU305914.1 | 91828 | 91850 | + | 0 | AGGAAGAACATCCAGACGGC | GGG | \|\|\|\|\|\|\|\|[\|\|\|\|\|\|\|\|\|\|\|\|]PAM | NA | Intergenic | NA | NA |
| T6550 | CATCCAGACGGCGGGAGCGACGG | 1000 | | | | | | | | | |
| Oligo fwd | CACCCATCCAGACGGCGGGAGCGA | | | | | | | | | | |
| Oligo rev | AAACTCGCTCCCGCCGTCTGGATG | | | | | | | | | | |
| Chromosome | start | end | strand | MM | target_seq | PAM | alignment | distance | position | gene name | gene id |
| GU305914.1 | 91836 | 91858 | + | 0 | CATCCAGACGGCGGGAGCGA | CGG | \|\|\|\|\|\|\|\|[\|\|\|\|\|\|\|\|\|\|\|\|]PAM | NA | Intergenic | NA | NA |
| T6551 | ATCCAGACGGCGGGAGCGACGGG | 1000 | | | | | | | | | |
| Oligo fwd | CACCATCCAGACGGCGGGAGCGAC | | | | | | | | | | |
| Oligo rev | AAACGTCGCTCCCGCCGTCTGGAT | | | | | | | | | | |
| Chromosome | start | end | strand | MM | target_seq | PAM | alignment | distance | position | gene name | gene id |
| GU305914.1 | 91837 | 91859 | + | 0 | ATCCAGACGGCGGGAGCGAC | GGG | \|\|\|\|\|\|\|\|[\|\|\|\|\|\|\|\|\|\|\|\|]PAM | NA | Intergenic | NA | NA |
| T6552 | TCCAGACGGCGGGAGCGACGGGG | 1000 | | | | | | | | | |
| Oligo fwd | CACCTCCAGACGGCGGGAGCGACG | | | | | | | | | | |
| Oligo rev | AAACCGTCGCTCCCGCCGTCTGGA | | | | | | | | | | |
| Chromosome | start | end | strand | MM | target_seq | PAM | alignment | distance | position | gene name | gene id |
| GU305914.1 | 91838 | 91860 | + | 0 | TCCAGACGGCGGGAGCGACG | GGG | \|\|\|\|\|\|\|\|[\|\|\|\|\|\|\|\|\|\|\|\|]PAM | NA | Intergenic | NA | NA |
| T6553 | CCAGACGGCGGGAGCGACGGGGG | 1000 | | | | | | | | | |
| Oligo fwd | CACCCCAGACGGCGGGAGCGACGG | | | | | | | | | | |
| Oligo rev | AAACCCGTCGCTCCCGCCGTCTGG | | | | | | | | | | |
| Chromosome | start | end | strand | MM | target_seq | PAM | alignment | distance | position | gene name | gene id |
| GU305914.1 | 91839 | 91861 | + | 0 | CCAGACGGCGGGAGCGACGG | GGG | \|\|\|\|\|\|\|\|[\|\|\|\|\|\|\|\|\|\|\|\|]PAM | NA | Intergenic | NA | NA |
| T6554 | CGGCGGGAGCGACGGGGGCGAGG | 1000 | | | | | | | | | |
| Oligo fwd | CACCCGGCGGGAGCGACGGGGGCG | | | | | | | | | | |
| Oligo rev | AAACCGCCCCCGTCGCTCCCGCCG | | | | | | | | | | |
| Chromosome | start | end | strand | MM | target_seq | PAM | alignment | distance | position | gene name | gene id |
| GU305914.1 | 91844 | 91866 | + | 0 | CGGCGGGAGCGACGGGGGCG | AGG | \|\|\|\|\|\|\|\|[\|\|\|\|\|\|\|\|\|\|\|\|]PAM | NA | Intergenic | NA | NA |
| GU305914.1 | 26384 | 26406 | + | 4 | CGGCGCTGGCGACGGCGGCG | GGG | \|\|\|\|\|---[\|\|\|\|\|\|\|-\|\|\|\|]PAM | NA | Intergenic | NA | NA |
| T6555 | GGGAGCGACGGGGGCGAGGCTGG | 1000 | | | | | | | | | |
| Oligo fwd | CACCGGGAGCGACGGGGGCGAGGC | | | | | | | | | | |
| Oligo rev | AAACGCCTCGCCCCCGTCGCTCCC | | | | | | | | | | |
| Chromosome | start | end | strand | MM | target_seq | PAM | alignment | distance | position | gene name | gene id |
| GU305914.1 | 91848 | 91870 | + | 0 | GGGAGCGACGGGGGCGAGGC | TGG | \|\|\|\|\|\|\|\|[\|\|\|\|\|\|\|\|\|\|\|\|]PAM | NA | Intergenic | NA | NA |
| T6556 | GGAGAGATCGTCGTCCGCCATGG | 1000 | | | | | | | | | |
| Oligo fwd | CACCGGAGAGATCGTCGTCCGCCA | | | | | | | | | | |
| Oligo rev | AAACTGGCGGACGACGATCTCTCC | | | | | | | | | | |
| Chromosome | start | end | strand | MM | target_seq | PAM | alignment | distance | position | gene name | gene id |
| GU305914.1 | 91869 | 91891 | + | 0 | GGAGAGATCGTCGTCCGCCA | TGG | \|\|\|\|\|\|\|\|[\|\|\|\|\|\|\|\|\|\|\|\|]PAM | NA | Intergenic | NA | NA |
| T6557 | GTCCGCCATGGCGACCTCCCCGG | 1000 | | | | | | | | | |
| Oligo fwd | CACCGTCCGCCATGGCGACCTCCC | | | | | | | | | | |
| Oligo rev | AAACGGGAGGTCGCCATGGCGGAC | | | | | | | | | | |
| Chromosome | start | end | strand | MM | target_seq | PAM | alignment | distance | position | gene name | gene id |
| GU305914.1 | 91881 | 91903 | + | 0 | GTCCGCCATGGCGACCTCCC | CGG | \|\|\|\|\|\|\|\|[\|\|\|\|\|\|\|\|\|\|\|\|]PAM | NA | Intergenic | NA | NA |
| GU305914.1 | 125784 | 125806 | + | 4 | GTCCACCTTGGCGAACTCCT | TGG | \|\|\|\|-\|\|-[\|\|\|\|\|\|-\|\|\|\|-]PAM | NA | Intergenic | NA | NA |
| T6558 | CCCCGGCATCGAAATTATACAGG | 1000 | | | | | | | | | |
| Oligo fwd | CACCCCCCGGCATCGAAATTATAC | | | | | | | | | | |
| Oligo rev | AAACGTATAATTTCGATGCCGGGG | | | | | | | | | | |
| Chromosome | start | end | strand | MM | target_seq | PAM | alignment | distance | position | gene name | gene id |

| | | | | | | | | | | | |
|---|---|---|---|---|---|---|---|---|---|---|---|
| GU305914.1 | 91898 | 91920 | + | 0 | CCCCGGCATCGAAATTATAC | AGG | \|\|\|\|\|\|\|\|[\|\|\|\|\|\|\|\|\|\|\|\|]PAM | NA | Intergenic | NA | NA |
| T6559 | GCATCGAAATTATACAGGAGCGG | 1000 | | | | | | | | | |
| Oligo fwd | CACCGCATCGAAATTATACAGGAG | | | | | | | | | | |
| Oligo rev | AAACCTCCTGTATAATTTCGATGC | | | | | | | | | | |
| Chromosome | start | end | strand | MM | target_seq | PAM | alignment | distance | position | gene name | gene id |
| GU305914.1 | 91903 | 91925 | + | 0 | GCATCGAAATTATACAGGAG | CGG | \|\|\|\|\|\|\|\|[\|\|\|\|\|\|\|\|\|\|\|\|]PAM | NA | Intergenic | NA | NA |
| T6560 | CATCGAAATTATACAGGAGCGGG | 1000 | | | | | | | | | |
| Oligo fwd | CACCCATCGAAATTATACAGGAGC | | | | | | | | | | |
| Oligo rev | AAACGCTCCTGTATAATTTCGATG | | | | | | | | | | |
| Chromosome | start | end | strand | MM | target_seq | PAM | alignment | distance | position | gene name | gene id |
| GU305914.1 | 91904 | 91926 | + | 0 | CATCGAAATTATACAGGAGC | GGG | \|\|\|\|\|\|\|\|[\|\|\|\|\|\|\|\|\|\|\|\|]PAM | NA | Intergenic | NA | NA |
| T6561 | ATCGAAATTATACAGGAGCGGGG | 1000 | | | | | | | | | |
| Oligo fwd | CACCATCGAAATTATACAGGAGCG | | | | | | | | | | |
| Oligo rev | AAACCGCTCCTGTATAATTTCGAT | | | | | | | | | | |
| Chromosome | start | end | strand | MM | target_seq | PAM | alignment | distance | position | gene name | gene id |
| GU305914.1 | 91905 | 91927 | + | 0 | ATCGAAATTATACAGGAGCG | GGG | \|\|\|\|\|\|\|\|[\|\|\|\|\|\|\|\|\|\|\|\|]PAM | NA | Intergenic | NA | NA |
| T6562 | TCGAAATTATACAGGAGCGGGGG | 1000 | | | | | | | | | |
| Oligo fwd | CACCTCGAAATTATACAGGAGCGG | | | | | | | | | | |
| Oligo rev | AAACCCGCTCCTGTATAATTTCGA | | | | | | | | | | |
| Chromosome | start | end | strand | MM | target_seq | PAM | alignment | distance | position | gene name | gene id |
| GU305914.1 | 91906 | 91928 | + | 0 | TCGAAATTATACAGGAGCGG | GGG | \|\|\|\|\|\|\|\|[\|\|\|\|\|\|\|\|\|\|\|\|]PAM | NA | Intergenic | NA | NA |
| T6563 | TACAGGAGCGGGGGTCGTTCTGG | 1000 | | | | | | | | | |
| Oligo fwd | CACCTACAGGAGCGGGGGTCGTTC | | | | | | | | | | |
| Oligo rev | AAACGAACGACCCCCGCTCCTGTA | | | | | | | | | | |
| Chromosome | start | end | strand | MM | target_seq | PAM | alignment | distance | position | gene name | gene id |
| GU305914.1 | 91915 | 91937 | + | 0 | TACAGGAGCGGGGGTCGTTC | TGG | \|\|\|\|\|\|\|\|[\|\|\|\|\|\|\|\|\|\|\|\|]PAM | NA | Intergenic | NA | NA |
| T6564 | GGGGTCGTTCTGGTCGCCCGCGG | 1000 | | | | | | | | | |
| Oligo fwd | CACCGGGGTCGTTCTGGTCGCCCG | | | | | | | | | | |
| Oligo rev | AAACCGGGCGACCAGAACGACCCC | | | | | | | | | | |
| Chromosome | start | end | strand | MM | target_seq | PAM | alignment | distance | position | gene name | gene id |
| GU305914.1 | 91925 | 91947 | + | 0 | GGGGTCGTTCTGGTCGCCCG | CGG | \|\|\|\|\|\|\|\|[\|\|\|\|\|\|\|\|\|\|\|\|]PAM | NA | Intergenic | NA | NA |
| T6565 | CTTTATATACGTGAAGCCCCCGG | 1000 | | | | | | | | | |
| Oligo fwd | CACCCTTTATATACGTGAAGCCCC | | | | | | | | | | |
| Oligo rev | AAACGGGGCTTCACGTATATAAAG | | | | | | | | | | |
| Chromosome | start | end | strand | MM | target_seq | PAM | alignment | distance | position | gene name | gene id |
| GU305914.1 | 91957 | 91979 | + | 0 | CTTTATATACGTGAAGCCCC | CGG | \|\|\|\|\|\|\|\|[\|\|\|\|\|\|\|\|\|\|\|\|]PAM | NA | Intergenic | NA | NA |
| T6566 | TATATACGTGAAGCCCCCGGCGG | 1000 | | | | | | | | | |
| Oligo fwd | CACCTATATACGTGAAGCCCCCGG | | | | | | | | | | |
| Oligo rev | AAACCCGGGGGCTTCACGTATATA | | | | | | | | | | |
| Chromosome | start | end | strand | MM | target_seq | PAM | alignment | distance | position | gene name | gene id |
| GU305914.1 | 91960 | 91982 | + | 0 | TATATACGTGAAGCCCCCGG | CGG | \|\|\|\|\|\|\|\|[\|\|\|\|\|\|\|\|\|\|\|\|]PAM | NA | Intergenic | NA | NA |
| T6567 | ATATACGTGAAGCCCCCGGCGGG | 1000 | | | | | | | | | |
| Oligo fwd | CACCATATACGTGAAGCCCCCGGC | | | | | | | | | | |
| Oligo rev | AAACGCCGGGGGCTTCACGTATAT | | | | | | | | | | |
| Chromosome | start | end | strand | MM | target_seq | PAM | alignment | distance | position | gene name | gene id |
| GU305914.1 | 91961 | 91983 | + | 0 | ATATACGTGAAGCCCCCGGC | GGG | \|\|\|\|\|\|\|\|[\|\|\|\|\|\|\|\|\|\|\|\|]PAM | NA | Intergenic | NA | NA |
| T6568 | AAGCCTTTGTCACAACGTACTGG | 1000 | | | | | | | | | |
| Oligo fwd | CACCAAGCCTTTGTCACAACGTAC | | | | | | | | | | |
| Oligo rev | AAACGTACGTTGTGACAAAGGCTT | | | | | | | | | | |
| Chromosome | start | end | strand | MM | target_seq | PAM | alignment | distance | position | gene name | gene id |
| GU305914.1 | 92028 | 92050 | + | 0 | AAGCCTTTGTCACAACGTAC | TGG | \|\|\|\|\|\|\|\|[\|\|\|\|\|\|\|\|\|\|\|\|]PAM | NA | Intergenic | NA | NA |
| T6569 | TTGTCACAACGTACTGGACATGG | 1000 | | | | | | | | | |
| Oligo fwd | CACCTTGTCACAACGTACTGGACA | | | | | | | | | | |
| Oligo rev | AAACTGTCCAGTACGTTGTGACAA | | | | | | | | | | |
| Chromosome | start | end | strand | MM | target_seq | PAM | alignment | distance | position | gene name | gene id |
| GU305914.1 | 92034 | 92056 | + | 0 | TTGTCACAACGTACTGGACA | TGG | \|\|\|\|\|\|\|\|[\|\|\|\|\|\|\|\|\|\|\|\|]PAM | NA | Intergenic | NA | NA |
| T6570 | CGTACTGGACATGGTTTGCGCGG | 1000 | | | | | | | | | |
| Oligo fwd | CACCCGTACTGGACATGGTTTGCG | | | | | | | | | | |
| Oligo rev | AAACCGCAAACCATGTCCAGTACG | | | | | | | | | | |
| Chromosome | start | end | strand | MM | target_seq | PAM | alignment | distance | position | gene name | gene id |
| GU305914.1 | 92043 | 92065 | + | 0 | CGTACTGGACATGGTTTGCG | CGG | \|\|\|\|\|\|\|\|[\|\|\|\|\|\|\|\|\|\|\|\|]PAM | NA | Intergenic | NA | NA |
| T6571 | CACGAATCTGACCGCCCCGAAGG | 1000 | | | | | | | | | |
| Oligo fwd | CACCCACGAATCTGACCGCCCCGA | | | | | | | | | | |
| Oligo rev | AAACTCGGGGCGGTCAGATTCGTG | | | | | | | | | | |
| Chromosome | start | end | strand | MM | target_seq | PAM | alignment | distance | position | gene name | gene id |
| GU305914.1 | 92073 | 92095 | + | 0 | CACGAATCTGACCGCCCCGA | AGG | \|\|\|\|\|\|\|\|[\|\|\|\|\|\|\|\|\|\|\|\|]PAM | NA | Intergenic | NA | NA |
| T6572 | ACGAATCTGACCGCCCCGAAGGG | 1000 | | | | | | | | | |
| Oligo fwd | CACCACGAATCTGACCGCCCCGAA | | | | | | | | | | |
| Oligo rev | AAACTTCGGGGCGGTCAGATTCGT | | | | | | | | | | |
| Chromosome | start | end | strand | MM | target_seq | PAM | alignment | distance | position | gene name | gene id |
| GU305914.1 | 92074 | 92096 | + | 0 | ACGAATCTGACCGCCCCGAA | GGG | \|\|\|\|\|\|\|\|[\|\|\|\|\|\|\|\|\|\|\|\|]PAM | NA | Intergenic | NA | NA |
| T6573 | TCCGTGCGCCGCAATCGCAGAGG | 1000 | | | | | | | | | |
| Oligo fwd | CACCTCCGTGCGCCGCAATCGCAG | | | | | | | | | | |
| Oligo rev | AAACCTGCGATTGCGGCGCACGGA | | | | | | | | | | |

| | | | | | | | | | | | |
|---|---|---|---|---|---|---|---|---|---|---|---|
| Chromosome | start | end | strand | MM | target_seq | PAM | alignment | distance | position | gene name | gene id |
| GU305914.1 | 92098 | 92120 | + | 0 | TCCGTGCGCCGCAATCGCAG | AGG | \|\|\|\|\|\|\|\|[\|\|\|\|\|\|\|\|\|\|\|\|]PAM | NA | Intergenic | NA | NA |
| T6574 | GCCGCAATCGCAGAGGACCGCGG | 1000 | | | | | | | | | |
| Oligo fwd | CACCGCCGCAATCGCAGAGGACCG | | | | | | | | | | |
| Oligo rev | AAACCGGTCCTCTGCGATTGCGGC | | | | | | | | | | |
| Chromosome | start | end | strand | MM | target_seq | PAM | alignment | distance | position | gene name | gene id |
| GU305914.1 | 92105 | 92127 | + | 0 | GCCGCAATCGCAGAGGACCG | CGG | \|\|\|\|\|\|\|\|[\|\|\|\|\|\|\|\|\|\|\|\|]PAM | NA | Intergenic | NA | NA |
| T6575 | CCGCAATCGCAGAGGACCGCGGG | 1000 | | | | | | | | | |
| Oligo fwd | CACCCCGCAATCGCAGAGGACCGC | | | | | | | | | | |
| Oligo rev | AAACGCGGTCCTCTGCGATTGCGG | | | | | | | | | | |
| Chromosome | start | end | strand | MM | target_seq | PAM | alignment | distance | position | gene name | gene id |
| GU305914.1 | 92106 | 92128 | + | 0 | CCGCAATCGCAGAGGACCGC | GGG | \|\|\|\|\|\|\|\|[\|\|\|\|\|\|\|\|\|\|\|\|]PAM | NA | Intergenic | NA | NA |
| T6576 | CGCAGAGGACCGCGGGCTCTTGG | 1000 | | | | | | | | | |
| Oligo fwd | CACCCGCAGAGGACCGCGGGCTCT | | | | | | | | | | |
| Oligo rev | AAACAGAGCCCGCGGTCCTCTGCG | | | | | | | | | | |
| Chromosome | start | end | strand | MM | target_seq | PAM | alignment | distance | position | gene name | gene id |
| GU305914.1 | 92113 | 92135 | + | 0 | CGCAGAGGACCGCGGGCTCT | TGG | \|\|\|\|\|\|\|\|[\|\|\|\|\|\|\|\|\|\|\|\|]PAM | NA | Intergenic | NA | NA |
| T6577 | GCGGGCTCTTGGCGCGCTCGAGG | 1000 | | | | | | | | | |
| Oligo fwd | CACCGCGGGCTCTTGGCGCGCTCG | | | | | | | | | | |
| Oligo rev | AAACCGAGCGCGCCAAGAGCCCGC | | | | | | | | | | |
| Chromosome | start | end | strand | MM | target_seq | PAM | alignment | distance | position | gene name | gene id |
| GU305914.1 | 92124 | 92146 | + | 0 | GCGGGCTCTTGGCGCGCTCG | AGG | \|\|\|\|\|\|\|\|[\|\|\|\|\|\|\|\|\|\|\|\|]PAM | NA | Intergenic | NA | NA |
| T6578 | CGAGGCATCTCACGCGCGCTCGG | 1000 | | | | | | | | | |
| Oligo fwd | CACCCGAGGCATCTCACGCGCGCT | | | | | | | | | | |
| Oligo rev | AAACAGCGCGCGTGAGATGCCTCG | | | | | | | | | | |
| Chromosome | start | end | strand | MM | target_seq | PAM | alignment | distance | position | gene name | gene id |
| GU305914.1 | 92142 | 92164 | + | 0 | CGAGGCATCTCACGCGCGCT | CGG | \|\|\|\|\|\|\|\|[\|\|\|\|\|\|\|\|\|\|\|\|]PAM | NA | Intergenic | NA | NA |
| T6579 | GGCATCTCACGCGCGCTCGGAGG | 1000 | | | | | | | | | |
| Oligo fwd | CACCGGCATCTCACGCGCGCTCGG | | | | | | | | | | |
| Oligo rev | AAACCCGAGCGCGCGTGAGATGCC | | | | | | | | | | |
| Chromosome | start | end | strand | MM | target_seq | PAM | alignment | distance | position | gene name | gene id |
| GU305914.1 | 92145 | 92167 | + | 0 | GGCATCTCACGCGCGCTCGG | AGG | \|\|\|\|\|\|\|\|[\|\|\|\|\|\|\|\|\|\|\|\|]PAM | NA | Intergenic | NA | NA |
| T6580 | GCATCTCACGCGCGCTCGGAGGG | 1000 | | | | | | | | | |
| Oligo fwd | CACCGCATCTCACGCGCGCTCGGA | | | | | | | | | | |
| Oligo rev | AAACTCCGAGCGCGCGTGAGATGC | | | | | | | | | | |
| Chromosome | start | end | strand | MM | target_seq | PAM | alignment | distance | position | gene name | gene id |
| GU305914.1 | 92146 | 92168 | + | 0 | GCATCTCACGCGCGCTCGGA | GGG | \|\|\|\|\|\|\|\|[\|\|\|\|\|\|\|\|\|\|\|\|]PAM | NA | Intergenic | NA | NA |
| T6581 | CTCACGCGCGCTCGGAGGGCCGG | 1000 | | | | | | | | | |
| Oligo fwd | CACCCTCACGCGCGCTCGGAGGGC | | | | | | | | | | |
| Oligo rev | AAACGCCCTCCGAGCGCGCGTGAG | | | | | | | | | | |
| Chromosome | start | end | strand | MM | target_seq | PAM | alignment | distance | position | gene name | gene id |
| GU305914.1 | 92150 | 92172 | + | 0 | CTCACGCGCGCTCGGAGGGC | CGG | \|\|\|\|\|\|\|\|[\|\|\|\|\|\|\|\|\|\|\|\|]PAM | NA | Intergenic | NA | NA |
| T6582 | ACGCGCGCTCGGAGGGCCGGCGG | 1000 | | | | | | | | | |
| Oligo fwd | CACCACGCGCGCTCGGAGGGCCGG | | | | | | | | | | |
| Oligo rev | AAACCCGGCCCTCCGAGCGCGCGT | | | | | | | | | | |
| Chromosome | start | end | strand | MM | target_seq | PAM | alignment | distance | position | gene name | gene id |
| GU305914.1 | 92153 | 92175 | + | 0 | ACGCGCGCTCGGAGGGCCGG | CGG | \|\|\|\|\|\|\|\|[\|\|\|\|\|\|\|\|\|\|\|\|]PAM | NA | Intergenic | NA | NA |
| GU305914.1 | 230351 | 230373 | - | 4 | GCGCGCACTCAGACGGCCGG | GGG | -\|\|\|\|\|-\|[\|\|-\|\|-\|\|\|\|\|\|]PAM | NA | Intergenic | NA | NA |
| GU305914.1 | 36 | 58 | - | 4 | GCGCGCACTCAGACGGCCGG | GGG | -\|\|\|\|\|-\|[\|\|-\|\|-\|\|\|\|\|\|]PAM | NA | Intergenic | NA | NA |
| T6583 | TGCGCCAGCAGCTCCGAGCCCGG | 1000 | | | | | | | | | |
| Oligo fwd | CACCTGCGCCAGCAGCTCCGAGCC | | | | | | | | | | |
| Oligo rev | AAACGGCTCGGAGCTGCTGGCGCA | | | | | | | | | | |
| Chromosome | start | end | strand | MM | target_seq | PAM | alignment | distance | position | gene name | gene id |
| GU305914.1 | 92179 | 92201 | + | 0 | TGCGCCAGCAGCTCCGAGCC | CGG | \|\|\|\|\|\|\|\|[\|\|\|\|\|\|\|\|\|\|\|\|]PAM | NA | Intergenic | NA | NA |
| T6584 | GCGCCAGCAGCTCCGAGCCCGGG | 1000 | | | | | | | | | |
| Oligo fwd | CACCGCGCCAGCAGCTCCGAGCCC | | | | | | | | | | |
| Oligo rev | AAACGGGCTCGGAGCTGCTGGCGC | | | | | | | | | | |
| Chromosome | start | end | strand | MM | target_seq | PAM | alignment | distance | position | gene name | gene id |
| GU305914.1 | 92180 | 92202 | + | 0 | GCGCCAGCAGCTCCGAGCCC | GGG | \|\|\|\|\|\|\|\|[\|\|\|\|\|\|\|\|\|\|\|\|]PAM | NA | Intergenic | NA | NA |
| T6585 | TCCGAGCCCGGGCCGTCAGACGG | 1000 | | | | | | | | | |
| Oligo fwd | CACCTCCGAGCCCGGGCCGTCAGA | | | | | | | | | | |
| Oligo rev | AAACTCTGACGGCCCGGGCTCGGA | | | | | | | | | | |
| Chromosome | start | end | strand | MM | target_seq | PAM | alignment | distance | position | gene name | gene id |
| GU305914.1 | 92191 | 92213 | + | 0 | TCCGAGCCCGGGCCGTCAGA | CGG | \|\|\|\|\|\|\|\|[\|\|\|\|\|\|\|\|\|\|\|\|]PAM | NA | Intergenic | NA | NA |
| T6586 | GGCATTGACGAAAAAACGTACGG | 1000 | | | | | | | | | |
| Oligo fwd | CACCGGCATTGACGAAAAAACGTA | | | | | | | | | | |
| Oligo rev | AAACTACGTTTTTTCGTCAATGCC | | | | | | | | | | |
| Chromosome | start | end | strand | MM | target_seq | PAM | alignment | distance | position | gene name | gene id |
| GU305914.1 | 92212 | 92234 | + | 0 | GGCATTGACGAAAAAACGTA | CGG | \|\|\|\|\|\|\|\|[\|\|\|\|\|\|\|\|\|\|\|\|]PAM | NA | Intergenic | NA | NA |
| T6587 | CACACACGCGAAACGAGACGCGG | 1000 | | | | | | | | | |
| Oligo fwd | CACCCACACACGCGAAACGAGACG | | | | | | | | | | |
| Oligo rev | AAACCGTCTCGTTTCGCGTGTGTG | | | | | | | | | | |
| Chromosome | start | end | strand | MM | target_seq | PAM | alignment | distance | position | gene name | gene id |
| GU305914.1 | 92279 | 92301 | + | 0 | CACACACGCGAAACGAGACG | CGG | \|\|\|\|\|\|\|\|[\|\|\|\|\|\|\|\|\|\|\|\|]PAM | NA | Intergenic | NA | NA |

| | | | | | | | | | | | |
|---|---|---|---|---|---|---|---|---|---|---|---|
| T6588 | ACACACGCGAAACGAGACGCGGG | 1000 | | | | | | | | | |
| Oligo fwd | CACCACACACGCGAAACGAGACGC | | | | | | | | | | |
| Oligo rev | AAACGCGTCTCGTTTCGCGTGTGT | | | | | | | | | | |
| Chromosome | start | end | strand | MM | target_seq | PAM | alignment | distance | position | gene name | gene id |
| GU305914.1 | 92280 | 92302 | + | 0 | ACACACGCGAAACGAGACGC | GGG | \|\|\|\|\|\|\|\|[\|\|\|\|\|\|\|\|\|\|\|\|]PAM | NA | Intergenic | NA | NA |
| T6589 | ACGAGACGCGGGCCGCGAACAGG | 1000 | | | | | | | | | |
| Oligo fwd | CACCACGAGACGCGGGCCGCGAAC | | | | | | | | | | |
| Oligo rev | AAACGTTCGCGGCCCGCGTCTCGT | | | | | | | | | | |
| Chromosome | start | end | strand | MM | target_seq | PAM | alignment | distance | position | gene name | gene id |
| GU305914.1 | 92291 | 92313 | + | 0 | ACGAGACGCGGGCCGCGAAC | AGG | \|\|\|\|\|\|\|\|[\|\|\|\|\|\|\|\|\|\|\|\|]PAM | NA | Intergenic | NA | NA |
| T6590 | ACACTACGCAGACGAGCATGCGG | 1000 | | | | | | | | | |
| Oligo fwd | CACCACACTACGCAGACGAGCATG | | | | | | | | | | |
| Oligo rev | AAACCATGCTCGTCTGCGTAGTGT | | | | | | | | | | |
| Chromosome | start | end | strand | MM | target_seq | PAM | alignment | distance | position | gene name | gene id |
| GU305914.1 | 92330 | 92352 | + | 0 | ACACTACGCAGACGAGCATG | CGG | \|\|\|\|\|\|\|\|[\|\|\|\|\|\|\|\|\|\|\|\|]PAM | NA | Intergenic | NA | NA |
| T6591 | ACGAGCATGCGGTCGAAGCGTGG | 1000 | | | | | | | | | |
| Oligo fwd | CACCACGAGCATGCGGTCGAAGCG | | | | | | | | | | |
| Oligo rev | AAACCGCTTCGACCGCATGCTCGT | | | | | | | | | | |
| Chromosome | start | end | strand | MM | target_seq | PAM | alignment | distance | position | gene name | gene id |
| GU305914.1 | 92341 | 92363 | + | 0 | ACGAGCATGCGGTCGAAGCG | TGG | \|\|\|\|\|\|\|\|[\|\|\|\|\|\|\|\|\|\|\|\|]PAM | NA | Intergenic | NA | NA |
| T6592 | TGCGGTCGAAGCGTGGTCAGCGG | 1000 | | | | | | | | | |
| Oligo fwd | CACCTGCGGTCGAAGCGTGGTCAG | | | | | | | | | | |
| Oligo rev | AAACCTGACCACGCTTCGACCGCA | | | | | | | | | | |
| Chromosome | start | end | strand | MM | target_seq | PAM | alignment | distance | position | gene name | gene id |
| GU305914.1 | 92348 | 92370 | + | 0 | TGCGGTCGAAGCGTGGTCAG | CGG | \|\|\|\|\|\|\|\|[\|\|\|\|\|\|\|\|\|\|\|\|]PAM | NA | Intergenic | NA | NA |
| GU305914.1 | 30111 | 30133 | + | 4 | GGCGGTGGAAGCGAGGCCAG | CGG | -\|\|\|\|\|-\|[\|\|\|\|\|-\|\|-\|\|\|]PAM | NA | Intergenic | NA | NA |
| T6593 | GCGGTCGAAGCGTGGTCAGCGGG | 1000 | | | | | | | | | |
| Oligo fwd | CACCGCGGTCGAAGCGTGGTCAGC | | | | | | | | | | |
| Oligo rev | AAACGCTGACCACGCTTCGACCGC | | | | | | | | | | |
| Chromosome | start | end | strand | MM | target_seq | PAM | alignment | distance | position | gene name | gene id |
| GU305914.1 | 92349 | 92371 | + | 0 | GCGGTCGAAGCGTGGTCAGC | GGG | \|\|\|\|\|\|\|\|[\|\|\|\|\|\|\|\|\|\|\|\|]PAM | NA | Intergenic | NA | NA |
| T6594 | GGAGAGTTTGTTACGTGTGCCGG | 1000 | | | | | | | | | |
| Oligo fwd | CACCGGAGAGTTTGTTACGTGTGC | | | | | | | | | | |
| Oligo rev | AAACGCACACGTAACAAACTCTCC | | | | | | | | | | |
| Chromosome | start | end | strand | MM | target_seq | PAM | alignment | distance | position | gene name | gene id |
| GU305914.1 | 92370 | 92392 | + | 0 | GGAGAGTTTGTTACGTGTGC | CGG | \|\|\|\|\|\|\|\|[\|\|\|\|\|\|\|\|\|\|\|\|]PAM | NA | Intergenic | NA | NA |
| T6595 | GTGTTTGACACACCGTTAGACGG | 1000 | | | | | | | | | |
| Oligo fwd | CACCGTGTTTGACACACCGTTAGA | | | | | | | | | | |
| Oligo rev | AAACTCTAACGGTGTGTCAAACAC | | | | | | | | | | |
| Chromosome | start | end | strand | MM | target_seq | PAM | alignment | distance | position | gene name | gene id |
| GU305914.1 | 92392 | 92414 | + | 0 | GTGTTTGACACACCGTTAGA | CGG | \|\|\|\|\|\|\|\|[\|\|\|\|\|\|\|\|\|\|\|\|]PAM | NA | Intergenic | NA | NA |
| T6596 | CCCGCCCCGCAACCGATTTGTGG | 1000 | | | | | | | | | |
| Oligo fwd | CACCCCCGCCCCGCAACCGATTTG | | | | | | | | | | |
| Oligo rev | AAACCAAATCGGTTGCGGGGCGGG | | | | | | | | | | |
| Chromosome | start | end | strand | MM | target_seq | PAM | alignment | distance | position | gene name | gene id |
| GU305914.1 | 92430 | 92452 | + | 0 | CCCGCCCCGCAACCGATTTG | TGG | \|\|\|\|\|\|\|\|[\|\|\|\|\|\|\|\|\|\|\|\|]PAM | NA | Intergenic | NA | NA |
| T6597 | GGTCTCGTAGCACCCCCTAGCGG | 1000 | | | | | | | | | |
| Oligo fwd | CACCGGTCTCGTAGCACCCCCTAG | | | | | | | | | | |
| Oligo rev | AAACCTAGGGGGTGCTACGAGACC | | | | | | | | | | |
| Chromosome | start | end | strand | MM | target_seq | PAM | alignment | distance | position | gene name | gene id |
| GU305914.1 | 92451 | 92473 | + | 0 | GGTCTCGTAGCACCCCCTAG | CGG | \|\|\|\|\|\|\|\|[\|\|\|\|\|\|\|\|\|\|\|\|]PAM | NA | Intergenic | NA | NA |
| T6598 | CGCAGCACATGCTGTCGTCATGG | 1000 | | | | | | | | | |
| Oligo fwd | CACCCGCAGCACATGCTGTCGTCA | | | | | | | | | | |
| Oligo rev | AAACTGACGACAGCATGTGCTGCG | | | | | | | | | | |
| Chromosome | start | end | strand | MM | target_seq | PAM | alignment | distance | position | gene name | gene id |
| GU305914.1 | 92478 | 92500 | + | 0 | CGCAGCACATGCTGTCGTCA | TGG | \|\|\|\|\|\|\|\|[\|\|\|\|\|\|\|\|\|\|\|\|]PAM | NA | Intergenic | NA | NA |
| T6599 | TGGCAACATCCAACGCTTATCGG | 1000 | | | | | | | | | |
| Oligo fwd | CACCTGGCAACATCCAACGCTTAT | | | | | | | | | | |
| Oligo rev | AAACATAAGCGTTGGATGTTGCCA | | | | | | | | | | |
| Chromosome | start | end | strand | MM | target_seq | PAM | alignment | distance | position | gene name | gene id |
| GU305914.1 | 92498 | 92520 | + | 0 | TGGCAACATCCAACGCTTAT | CGG | \|\|\|\|\|\|\|\|[\|\|\|\|\|\|\|\|\|\|\|\|]PAM | NA | Intergenic | NA | NA |
| T6600 | GGCAACATCCAACGCTTATCGGG | 1000 | | | | | | | | | |
| Oligo fwd | CACCGGCAACATCCAACGCTTATC | | | | | | | | | | |
| Oligo rev | AAACGATAAGCGTTGGATGTTGCC | | | | | | | | | | |
| Chromosome | start | end | strand | MM | target_seq | PAM | alignment | distance | position | gene name | gene id |
| GU305914.1 | 92499 | 92521 | + | 0 | GGCAACATCCAACGCTTATC | GGG | \|\|\|\|\|\|\|\|[\|\|\|\|\|\|\|\|\|\|\|\|]PAM | NA | Intergenic | NA | NA |
| T6601 | AACATCCAACGCTTATCGGGCGG | 1000 | | | | | | | | | |
| Oligo fwd | CACCAACATCCAACGCTTATCGGG | | | | | | | | | | |
| Oligo rev | AAACCCCGATAAGCGTTGGATGTT | | | | | | | | | | |
| Chromosome | start | end | strand | MM | target_seq | PAM | alignment | distance | position | gene name | gene id |
| GU305914.1 | 92502 | 92524 | + | 0 | AACATCCAACGCTTATCGGG | CGG | \|\|\|\|\|\|\|\|[\|\|\|\|\|\|\|\|\|\|\|\|]PAM | NA | Intergenic | NA | NA |
| T6602 | ACATCCAACGCTTATCGGGCGGG | 1000 | | | | | | | | | |
| Oligo fwd | CACCACATCCAACGCTTATCGGGC | | | | | | | | | | |
| Oligo rev | AAACGCCCGATAAGCGTTGGATGT | | | | | | | | | | |
| Chromosome | start | end | strand | MM | target_seq | PAM | alignment | distance | position | gene name | gene id |

GU305914.1 92503 92525 + 0 ACATCCAACGCTTATCGGGC GGG ||||||||[||||||||||||]PAM NA Intergenic NA NA

T6603 CCAACGCTTATCGGGCGGGCAGG 1000
Oligo fwd CACCCCAACGCTTATCGGGCGGGC
Oligo rev AAACGCCCGCCCGATAAGCGTTGG
Chromosome start end strand MM target_seq PAM alignment distance position gene name gene id
GU305914.1 92507 92529 + 0 CCAACGCTTATCGGGCGGGC AGG ||||||||[||||||||||||]PAM NA Intergenic NA NA

T6604 CAACGCTTATCGGGCGGGCAGGG 1000
Oligo fwd CACCCAACGCTTATCGGGCGGGCA
Oligo rev AAACTGCCCGCCCGATAAGCGTTG
Chromosome start end strand MM target_seq PAM alignment distance position gene name gene id
GU305914.1 92508 92530 + 0 CAACGCTTATCGGGCGGGCA GGG ||||||||[||||||||||||]PAM NA Intergenic NA NA

T6605 GGGCGGGCAGGGCGTTGCCATGG 1000
Oligo fwd CACCGGGCGGGCAGGGCGTTGCCA
Oligo rev AAACTGGCAACGCCCTGCCCGCCC
Chromosome start end strand MM target_seq PAM alignment distance position gene name gene id
GU305914.1 92519 92541 + 0 GGGCGGGCAGGGCGTTGCCA TGG ||||||||[||||||||||||]PAM NA Intergenic NA NA

T6606 GTTACATGTCCCGCCTTCCGCGG 1000
Oligo fwd CACCGTTACATGTCCCGCCTTCCG
Oligo rev AAACCGGAAGGCGGGACATGTAAC
Chromosome start end strand MM target_seq PAM alignment distance position gene name gene id
GU305914.1 92541 92563 + 0 GTTACATGTCCCGCCTTCCG CGG ||||||||[||||||||||||]PAM NA Intergenic NA NA

T6607 CGCCTTCCGCGGCGATGTGCCGG 1000
Oligo fwd CACCCGCCTTCCGCGGCGATGTGC
Oligo rev AAACGCACATCGCCGCGGAAGGCG
Chromosome start end strand MM target_seq PAM alignment distance position gene name gene id
GU305914.1 92552 92574 + 0 CGCCTTCCGCGGCGATGTGC CGG ||||||||[||||||||||||]PAM NA Intergenic NA NA

T6608 GCCTTCCGCGGCGATGTGCCGGG 1000
Oligo fwd CACCGCCTTCCGCGGCGATGTGCC
Oligo rev AAACGGCACATCGCCGCGGAAGGC
Chromosome start end strand MM target_seq PAM alignment distance position gene name gene id
GU305914.1 92553 92575 + 0 GCCTTCCGCGGCGATGTGCC GGG ||||||||[||||||||||||]PAM NA Intergenic NA NA

T6609 GCGGCGATGTGCCGGGCCAGAGG 1000
Oligo fwd CACCGCGGCGATGTGCCGGGCCAG
Oligo rev AAACCTGGCCCGGCACATCGCCGC
Chromosome start end strand MM target_seq PAM alignment distance position gene name gene id
GU305914.1 92560 92582 + 0 GCGGCGATGTGCCGGGCCAG AGG ||||||||[||||||||||||]PAM NA Intergenic NA NA

T6610 GTGCCGGGCCAGAGGCTGACCGG 1000
Oligo fwd CACCGTGCCGGGCCAGAGGCTGAC
Oligo rev AAACGTCAGCCTCTGGCCCGGCAC
Chromosome start end strand MM target_seq PAM alignment distance position gene name gene id
GU305914.1 92568 92590 + 0 GTGCCGGGCCAGAGGCTGAC CGG ||||||||[||||||||||||]PAM NA Intergenic NA NA

T6611 TGACCGGAAGTCCCGCCTTTTGG 1000
Oligo fwd CACCTGACCGGAAGTCCCGCCTTT
Oligo rev AAACAAAGGCGGGACTTCCGGTCA
Chromosome start end strand MM target_seq PAM alignment distance position gene name gene id
GU305914.1 92584 92606 + 0 TGACCGGAAGTCCCGCCTTT TGG ||||||||[||||||||||||]PAM NA Intergenic NA NA

T6612 GGAAGTCCCGCCTTTTGGCCCGG 1000
Oligo fwd CACCGGAAGTCCCGCCTTTTGGCC
Oligo rev AAACGGCCAAAAGGCGGGACTTCC
Chromosome start end strand MM target_seq PAM alignment distance position gene name gene id
GU305914.1 92589 92611 + 0 GGAAGTCCCGCCTTTTGGCC CGG ||||||||[||||||||||||]PAM NA Intergenic NA NA

T6613 CGGTGCCTCGCGCGCGCGTACGG 1000
Oligo fwd CACCCGGTGCCTCGCGCGCGCGTA
Oligo rev AAACTACGCGCGCGCGAGGCACCG
Chromosome start end strand MM target_seq PAM alignment distance position gene name gene id
GU305914.1 92609 92631 + 0 CGGTGCCTCGCGCGCGCGTA CGG ||||||||[||||||||||||]PAM NA Intergenic NA NA

T6614 GGTGCCTCGCGCGCGCGTACGGG 1000
Oligo fwd CACCGGTGCCTCGCGCGCGCGTAC
Oligo rev AAACGTACGCGCGCGCGAGGCACC
Chromosome start end strand MM target_seq PAM alignment distance position gene name gene id
GU305914.1 92610 92632 + 0 GGTGCCTCGCGCGCGCGTAC GGG ||||||||[||||||||||||]PAM NA Intergenic NA NA

T6615 GCCTCGCGCGCGCGTACGGGCGG 1000
Oligo fwd CACCGCCTCGCGCGCGCGTACGGG
Oligo rev AAACCCCGTACGCGCGCGCGAGGC
Chromosome start end strand MM target_seq PAM alignment distance position gene name gene id
GU305914.1 92613 92635 + 0 GCCTCGCGCGCGCGTACGGG CGG ||||||||[||||||||||||]PAM NA Intergenic NA NA

T6616 CCTCGCGCGCGCGTACGGGCGGG 1000
Oligo fwd CACCCCTCGCGCGCGCGTACGGGC
Oligo rev AAACGCCCGTACGCGCGCGCGAGG
Chromosome start end strand MM target_seq PAM alignment distance position gene name gene id
GU305914.1 92614 92636 + 0 CCTCGCGCGCGCGTACGGGC GGG ||||||||[||||||||||||]PAM NA Intergenic NA NA

T6617 CGCGCGCGCGTACGGGCGGGCGG 1000
Oligo fwd CACCCGCGCGCGCGTACGGGCGGG
Oligo rev AAACCCCGCCCGTACGCGCGCGCG

Chromosome start end strand MM target_seq PAM alignment distance position gene name gene id
GU305914.1 92617 92639 + 0 CGCGCGCGCGTACGGGCGGG CGG ||||||||[||||||||||||]PAM NA Intergenic NA NA

T6618 GCGCGCGCGTACGGGCGGGCGGG 1000
Oligo fwd CACCGCGCGCGCGTACGGGCGGGC
Oligo rev AAACGCCCGCCCGTACGCGCGCGC
Chromosome start end strand MM target_seq PAM alignment distance position gene name gene id
GU305914.1 92618 92640 + 0 GCGCGCGCGTACGGGCGGGC GGG ||||||||[||||||||||||]PAM NA Intergenic NA NA

T6619 TATACCCGAAGCGCAACCCCCGG 1000
Oligo fwd CACCTATACCCGAAGCGCAACCCC
Oligo rev AAACGGGGTTGCGCTTCGGGTATA
Chromosome start end strand MM target_seq PAM alignment distance position gene name gene id
GU305914.1 92675 92697 + 0 TATACCCGAAGCGCAACCCC CGG ||||||||[||||||||||||]PAM NA Intergenic NA NA

T6620 GGAACCACGCCCCCTAAACTAGG 1000
Oligo fwd CACCGGAACCACGCCCCCTAAACT
Oligo rev AAACAGTTTAGGGGGCGTGGTTCC
Chromosome start end strand MM target_seq PAM alignment distance position gene name gene id
GU305914.1 92696 92718 + 0 GGAACCACGCCCCCTAAACT AGG ||||||||[||||||||||||]PAM NA Intergenic NA NA

T6621 AGGTACGCCCACCTAAACAATGG 1000
Oligo fwd CACCAGGTACGCCCACCTAAACAA
Oligo rev AAACTTGTTTAGGTGGGCGTACCT
Chromosome start end strand MM target_seq PAM alignment distance position gene name gene id
GU305914.1 92716 92738 + 0 AGGTACGCCCACCTAAACAA TGG ||||||||[||||||||||||]PAM NA Intergenic NA NA

T6622 GGTACGCCCACCTAAACAATGGG 1000
Oligo fwd CACCGGTACGCCCACCTAAACAAT
Oligo rev AAACATTGTTTAGGTGGGCGTACC
Chromosome start end strand MM target_seq PAM alignment distance position gene name gene id
GU305914.1 92717 92739 + 0 GGTACGCCCACCTAAACAAT GGG ||||||||[||||||||||||]PAM NA Intergenic NA NA

T6623 AACCCCTGTACAACTAATTATGG 1000
Oligo fwd CACCAACCCCTGTACAACTAATTA
Oligo rev AAACTAATTAGTTGTACAGGGGTT
Chromosome start end strand MM target_seq PAM alignment distance position gene name gene id
GU305914.1 92835 92857 + 0 AACCCCTGTACAACTAATTA TGG ||||||||[||||||||||||]PAM NA Intergenic NA NA

T6624 CTAATTATGGACCAGTCGACTGG 1000
Oligo fwd CACCCTAATTATGGACCAGTCGAC
Oligo rev AAACGTCGACTGGTCCATAATTAG
Chromosome start end strand MM target_seq PAM alignment distance position gene name gene id
GU305914.1 92848 92870 + 0 CTAATTATGGACCAGTCGAC TGG ||||||||[||||||||||||]PAM NA Intergenic NA NA

T6625 GGACCAGTCGACTGGTTCACCGG 1000
Oligo fwd CACCGGACCAGTCGACTGGTTCAC
Oligo rev AAACGTGAACCAGTCGACTGGTCC
Chromosome start end strand MM target_seq PAM alignment distance position gene name gene id
GU305914.1 92856 92878 + 0 GGACCAGTCGACTGGTTCAC CGG ||||||||[||||||||||||]PAM NA Intergenic NA NA

T6626 TCGACTGGTTCACCGGCCTACGG 1000
Oligo fwd CACCTCGACTGGTTCACCGGCCTA
Oligo rev AAACTAGGCCGGTGAACCAGTCGA
Chromosome start end strand MM target_seq PAM alignment distance position gene name gene id
GU305914.1 92863 92885 + 0 TCGACTGGTTCACCGGCCTA CGG ||||||||[||||||||||||]PAM NA Intergenic NA NA

T6627 CTGGTTCACCGGCCTACGGCCGG 1000
Oligo fwd CACCCTGGTTCACCGGCCTACGGC
Oligo rev AAACGCCGTAGGCCGGTGAACCAG
Chromosome start end strand MM target_seq PAM alignment distance position gene name gene id
GU305914.1 92867 92889 + 0 CTGGTTCACCGGCCTACGGC CGG ||||||||[||||||||||||]PAM NA Intergenic NA NA

T6628 CCGGTGAACCATCGAAACCTAGG 1000
Oligo fwd CACCCCGGTGAACCATCGAAACCT
Oligo rev AAACAGGTTTCGATGGTTCACCGG
Chromosome start end strand MM target_seq PAM alignment distance position gene name gene id
GU305914.1 92886 92908 + 0 CCGGTGAACCATCGAAACCT AGG ||||||||[||||||||||||]PAM NA Intergenic NA NA

T6629 CTCCCCCGTAACCAGCACCTAGG 1000
Oligo fwd CACCCTCCCCCGTAACCAGCACCT
Oligo rev AAACAGGTGCTGGTTACGGGGGAG
Chromosome start end strand MM target_seq PAM alignment distance position gene name gene id
GU305914.1 92931 92953 + 0 CTCCCCCGTAACCAGCACCT AGG ||||||||[||||||||||||]PAM NA Intergenic NA NA

T6630 TAGGTGACGTAATGCTCATTAGG 1000
Oligo fwd CACCTAGGTGACGTAATGCTCATT
Oligo rev AAACAATGAGCATTACGTCACCTA
Chromosome start end strand MM target_seq PAM alignment distance position gene name gene id
GU305914.1 92950 92972 + 0 TAGGTGACGTAATGCTCATT AGG ||||||||[||||||||||||]PAM NA Intergenic NA NA

T6631 AGGTGACGTAATGCTCATTAGGG 1000
Oligo fwd CACCAGGTGACGTAATGCTCATTA
Oligo rev AAACTAATGAGCATTACGTCACCT
Chromosome start end strand MM target_seq PAM alignment distance position gene name gene id
GU305914.1 92951 92973 + 0 AGGTGACGTAATGCTCATTA GGG ||||||||[||||||||||||]PAM NA Intergenic NA NA

T6632 ATCGATAAATTTTCGACATCCGG 1000
Oligo fwd CACCATCGATAAATTTTCGACATC

Oligo rev AAACGATGTCGAAAATTTATCGAT
Chromosome start end strand MM target_seq PAM alignment distance position gene name gene id
GU305914.1 93019 93041 + 0 ATCGATAAATTTTCGACATC CGG ||||||||[||||||||||||]PAM NA Intergenic NA NA

T6633 TTCGACATCCGGTTGACCGAAGG 1000
Oligo fwd CACCTTCGACATCCGGTTGACCGA
Oligo rev AAACTCGGTCAACCGGATGTCGAA
Chromosome start end strand MM target_seq PAM alignment distance position gene name gene id
GU305914.1 93030 93052 + 0 TTCGACATCCGGTTGACCGA AGG ||||||||[||||||||||||]PAM NA Intergenic NA NA
GU305914.1 93046 93068 - 3 TTCAACATCCGGTTGACCTT CGG |||-||||[||||||||||--]PAM NA Intergenic NA NA

T6634 CCGGTTGACCGAAGGTCAACCGG 1000
Oligo fwd CACCCCGGTTGACCGAAGGTCAAC
Oligo rev AAACGTTGACCTTCGGTCAACCGG
Chromosome start end strand MM target_seq PAM alignment distance position gene name gene id
GU305914.1 93038 93060 + 0 CCGGTTGACCGAAGGTCAAC CGG ||||||||[||||||||||||]PAM NA Intergenic NA NA
GU305914.1 164027 164049 - 4 ACGGTTCACCGAAGGTCGAG GGG -|||||-|[|||||||||-|-]PAM NA Intergenic NA NA

T6635 AAGGTCAACCGGATGTTGAAAGG 1000
Oligo fwd CACCAAGGTCAACCGGATGTTGAA
Oligo rev AAACTTCAACATCCGGTTGACCTT
Chromosome start end strand MM target_seq PAM alignment distance position gene name gene id
GU305914.1 93049 93071 + 0 AAGGTCAACCGGATGTTGAA AGG ||||||||[||||||||||||]PAM NA Intergenic NA NA

T6636 ACATTAAGCCATGTATACGTTGG 1000
Oligo fwd CACCACATTAAGCCATGTATACGT
Oligo rev AAACACGTATACATGGCTTAATGT
Chromosome start end strand MM target_seq PAM alignment distance position gene name gene id
GU305914.1 93086 93108 + 0 ACATTAAGCCATGTATACGT TGG ||||||||[||||||||||||]PAM NA Intergenic NA NA

T6637 ACGAATGACGTCACTCCCTGTGG 1000
Oligo fwd CACCACGAATGACGTCACTCCCTG
Oligo rev AAACCAGGGAGTGACGTCATTCGT
Chromosome start end strand MM target_seq PAM alignment distance position gene name gene id
GU305914.1 93112 93134 + 0 ACGAATGACGTCACTCCCTG TGG ||||||||[||||||||||||]PAM NA Intergenic NA NA

T6638 CCTGTGGAATAGTATACGTCTGG 1000
Oligo fwd CACCCCTGTGGAATAGTATACGTC
Oligo rev AAACGACGTATACTATTCCACAGG
Chromosome start end strand MM target_seq PAM alignment distance position gene name gene id
GU305914.1 93128 93150 + 0 CCTGTGGAATAGTATACGTC TGG ||||||||[||||||||||||]PAM NA Intergenic NA NA

T6639 TACGTCTGGATATTGCGCAATGG 1000
Oligo fwd CACCTACGTCTGGATATTGCGCAA
Oligo rev AAACTTGCGCAATATCCAGACGTA
Chromosome start end strand MM target_seq PAM alignment distance position gene name gene id
GU305914.1 93142 93164 + 0 TACGTCTGGATATTGCGCAA TGG ||||||||[||||||||||||]PAM NA Intergenic NA NA

T6640 ACGTCTGGATATTGCGCAATGGG 1000
Oligo fwd CACCACGTCTGGATATTGCGCAAT
Oligo rev AAACATTGCGCAATATCCAGACGT
Chromosome start end strand MM target_seq PAM alignment distance position gene name gene id
GU305914.1 93143 93165 + 0 ACGTCTGGATATTGCGCAAT GGG ||||||||[||||||||||||]PAM NA Intergenic NA NA

T6641 CAATGGGAACCTATATACATAGG 1000
Oligo fwd CACCCAATGGGAACCTATATACAT
Oligo rev AAACATGTATATAGGTTCCCATTG
Chromosome start end strand MM target_seq PAM alignment distance position gene name gene id
GU305914.1 93159 93181 + 0 CAATGGGAACCTATATACAT AGG ||||||||[||||||||||||]PAM NA Intergenic NA NA

T6642 AATGGGAACCTATATACATAGGG 1000
Oligo fwd CACCAATGGGAACCTATATACATA
Oligo rev AAACTATGTATATAGGTTCCCATT
Chromosome start end strand MM target_seq PAM alignment distance position gene name gene id
GU305914.1 93160 93182 + 0 AATGGGAACCTATATACATA GGG ||||||||[||||||||||||]PAM NA Intergenic NA NA

T6643 CTATATGTAACCCTATCCTATGG 1000
Oligo fwd CACCCTATATGTAACCCTATCCTA
Oligo rev AAACTAGGATAGGGTTACATATAG
Chromosome start end strand MM target_seq PAM alignment distance position gene name gene id
GU305914.1 93187 93209 + 0 CTATATGTAACCCTATCCTA TGG ||||||||[||||||||||||]PAM NA Intergenic NA NA

T6644 TATCCTATGGATCCTACTTAAGG 1000
Oligo fwd CACCTATCCTATGGATCCTACTTA
Oligo rev AAACTAAGTAGGATCCATAGGATA
Chromosome start end strand MM target_seq PAM alignment distance position gene name gene id
GU305914.1 93200 93222 + 0 TATCCTATGGATCCTACTTA AGG ||||||||[||||||||||||]PAM NA Intergenic NA NA

T6645 ATCCTATGGATCCTACTTAAGGG 1000
Oligo fwd CACCATCCTATGGATCCTACTTAA
Oligo rev AAACTTAAGTAGGATCCATAGGAT
Chromosome start end strand MM target_seq PAM alignment distance position gene name gene id
GU305914.1 93201 93223 + 0 ATCCTATGGATCCTACTTAA GGG ||||||||[||||||||||||]PAM NA Intergenic NA NA

T6646 TTAAGGGATAAAATAATATATGG 1000
Oligo fwd CACCTTAAGGGATAAAATAATATA
Oligo rev AAACTATATTATTTTATCCCTTAA
Chromosome start end strand MM target_seq PAM alignment distance position gene name gene id
GU305914.1 93217 93239 + 0 TTAAGGGATAAAATAATATA TGG ||||||||[||||||||||||]PAM NA Intergenic NA NA

<table>
<tr><td>T6647</td><td>TAAGGGATAAAATAATATATGGG</td><td>1000</td><td></td><td></td><td></td><td></td><td></td><td></td><td></td><td></td><td></td></tr>
<tr><td>Oligo fwd</td><td>CACCTAAGGGATAAAATAATATAT</td><td></td><td></td><td></td><td></td><td></td><td></td><td></td><td></td><td></td><td></td></tr>
<tr><td>Oligo rev</td><td>AAACATATATTATTTTATCCCTTA</td><td></td><td></td><td></td><td></td><td></td><td></td><td></td><td></td><td></td><td></td></tr>
<tr><td>Chromosome</td><td>start</td><td>end</td><td>strand</td><td>MM</td><td>target_seq</td><td>PAM</td><td>alignment</td><td>distance</td><td>position</td><td>gene name</td><td>gene id</td></tr>
<tr><td>GU305914.1</td><td>93218</td><td>93240</td><td>+</td><td>0</td><td>TAAGGGATAAAATAATATAT</td><td>GGG</td><td>||||||||[||||||||||||]PAM</td><td>NA</td><td>Intergenic</td><td>NA</td><td>NA</td></tr>
<tr><td></td><td></td><td></td><td></td><td></td><td></td><td></td><td></td><td></td><td></td><td></td><td></td></tr>
<tr><td>T6648</td><td>AAATAATATATGGGTTCACTAGG</td><td>1000</td><td></td><td></td><td></td><td></td><td></td><td></td><td></td><td></td><td></td></tr>
<tr><td>Oligo fwd</td><td>CACCAAATAATATATGGGTTCACT</td><td></td><td></td><td></td><td></td><td></td><td></td><td></td><td></td><td></td><td></td></tr>
<tr><td>Oligo rev</td><td>AAACAGTGAACCCATATATTATTT</td><td></td><td></td><td></td><td></td><td></td><td></td><td></td><td></td><td></td><td></td></tr>
<tr><td>Chromosome</td><td>start</td><td>end</td><td>strand</td><td>MM</td><td>target_seq</td><td>PAM</td><td>alignment</td><td>distance</td><td>position</td><td>gene name</td><td>gene id</td></tr>
<tr><td>GU305914.1</td><td>93227</td><td>93249</td><td>+</td><td>0</td><td>AAATAATATATGGGTTCACT</td><td>AGG</td><td>||||||||[||||||||||||]PAM</td><td>NA</td><td>Intergenic</td><td>NA</td><td>NA</td></tr>
<tr><td></td><td></td><td></td><td></td><td></td><td></td><td></td><td></td><td></td><td></td><td></td><td></td></tr>
<tr><td>T6649</td><td>AATAATATATGGGTTCACTAGGG</td><td>1000</td><td></td><td></td><td></td><td></td><td></td><td></td><td></td><td></td><td></td></tr>
<tr><td>Oligo fwd</td><td>CACCAATAATATATGGGTTCACTA</td><td></td><td></td><td></td><td></td><td></td><td></td><td></td><td></td><td></td><td></td></tr>
<tr><td>Oligo rev</td><td>AAACTAGTGAACCCATATATTATT</td><td></td><td></td><td></td><td></td><td></td><td></td><td></td><td></td><td></td><td></td></tr>
<tr><td>Chromosome</td><td>start</td><td>end</td><td>strand</td><td>MM</td><td>target_seq</td><td>PAM</td><td>alignment</td><td>distance</td><td>position</td><td>gene name</td><td>gene id</td></tr>
<tr><td>GU305914.1</td><td>93228</td><td>93250</td><td>+</td><td>0</td><td>AATAATATATGGGTTCACTA</td><td>GGG</td><td>||||||||[||||||||||||]PAM</td><td>NA</td><td>Intergenic</td><td>NA</td><td>NA</td></tr>
<tr><td></td><td></td><td></td><td></td><td></td><td></td><td></td><td></td><td></td><td></td><td></td><td></td></tr>
<tr><td>T6650</td><td>ATAATATATGGGTTCACTAGGGG</td><td>1000</td><td></td><td></td><td></td><td></td><td></td><td></td><td></td><td></td><td></td></tr>
<tr><td>Oligo fwd</td><td>CACCATAATATATGGGTTCACTAG</td><td></td><td></td><td></td><td></td><td></td><td></td><td></td><td></td><td></td><td></td></tr>
<tr><td>Oligo rev</td><td>AAACCTAGTGAACCCATATATTAT</td><td></td><td></td><td></td><td></td><td></td><td></td><td></td><td></td><td></td><td></td></tr>
<tr><td>Chromosome</td><td>start</td><td>end</td><td>strand</td><td>MM</td><td>target_seq</td><td>PAM</td><td>alignment</td><td>distance</td><td>position</td><td>gene name</td><td>gene id</td></tr>
<tr><td>GU305914.1</td><td>93229</td><td>93251</td><td>+</td><td>0</td><td>ATAATATATGGGTTCACTAG</td><td>GGG</td><td>||||||||[||||||||||||]PAM</td><td>NA</td><td>Intergenic</td><td>NA</td><td>NA</td></tr>
<tr><td></td><td></td><td></td><td></td><td></td><td></td><td></td><td></td><td></td><td></td><td></td><td></td></tr>
<tr><td>T6651</td><td>TCACTAGGGGTGCAATTCGAAGG</td><td>1000</td><td></td><td></td><td></td><td></td><td></td><td></td><td></td><td></td><td></td></tr>
<tr><td>Oligo fwd</td><td>CACCTCACTAGGGGTGCAATTCGA</td><td></td><td></td><td></td><td></td><td></td><td></td><td></td><td></td><td></td><td></td></tr>
<tr><td>Oligo rev</td><td>AAACTCGAATTGCACCCCTAGTGA</td><td></td><td></td><td></td><td></td><td></td><td></td><td></td><td></td><td></td><td></td></tr>
<tr><td>Chromosome</td><td>start</td><td>end</td><td>strand</td><td>MM</td><td>target_seq</td><td>PAM</td><td>alignment</td><td>distance</td><td>position</td><td>gene name</td><td>gene id</td></tr>
<tr><td>GU305914.1</td><td>93242</td><td>93264</td><td>+</td><td>0</td><td>TCACTAGGGGTGCAATTCGA</td><td>AGG</td><td>||||||||[||||||||||||]PAM</td><td>NA</td><td>Intergenic</td><td>NA</td><td>NA</td></tr>
<tr><td></td><td></td><td></td><td></td><td></td><td></td><td></td><td></td><td></td><td></td><td></td><td></td></tr>
<tr><td>T6652</td><td>CACTAGGGGTGCAATTCGAAGGG</td><td>1000</td><td></td><td></td><td></td><td></td><td></td><td></td><td></td><td></td><td></td></tr>
<tr><td>Oligo fwd</td><td>CACCCACTAGGGGTGCAATTCGAA</td><td></td><td></td><td></td><td></td><td></td><td></td><td></td><td></td><td></td><td></td></tr>
<tr><td>Oligo rev</td><td>AAACTTCGAATTGCACCCCTAGTG</td><td></td><td></td><td></td><td></td><td></td><td></td><td></td><td></td><td></td><td></td></tr>
<tr><td>Chromosome</td><td>start</td><td>end</td><td>strand</td><td>MM</td><td>target_seq</td><td>PAM</td><td>alignment</td><td>distance</td><td>position</td><td>gene name</td><td>gene id</td></tr>
<tr><td>GU305914.1</td><td>93243</td><td>93265</td><td>+</td><td>0</td><td>CACTAGGGGTGCAATTCGAA</td><td>GGG</td><td>||||||||[||||||||||||]PAM</td><td>NA</td><td>Intergenic</td><td>NA</td><td>NA</td></tr>
<tr><td></td><td></td><td></td><td></td><td></td><td></td><td></td><td></td><td></td><td></td><td></td><td></td></tr>
<tr><td>T6653</td><td>CCCGCCCTCCCAGCTCGAGCTGG</td><td>1000</td><td></td><td></td><td></td><td></td><td></td><td></td><td></td><td></td><td></td></tr>
<tr><td>Oligo fwd</td><td>CACCCCCGCCCTCCCAGCTCGAGC</td><td></td><td></td><td></td><td></td><td></td><td></td><td></td><td></td><td></td><td></td></tr>
<tr><td>Oligo rev</td><td>AAACGCTCGAGCTGGGAGGGCGGG</td><td></td><td></td><td></td><td></td><td></td><td></td><td></td><td></td><td></td><td></td></tr>
<tr><td>Chromosome</td><td>start</td><td>end</td><td>strand</td><td>MM</td><td>target_seq</td><td>PAM</td><td>alignment</td><td>distance</td><td>position</td><td>gene name</td><td>gene id</td></tr>
<tr><td>GU305914.1</td><td>93316</td><td>93338</td><td>+</td><td>0</td><td>CCCGCCCTCCCAGCTCGAGC</td><td>TGG</td><td>||||||||[||||||||||||]PAM</td><td>NA</td><td>Intergenic</td><td>NA</td><td>NA</td></tr>
<tr><td>GU305914.1</td><td>188848</td><td>188870</td><td>-</td><td>4</td><td>CCTGCTCTCCGGGCTCGAGC</td><td>TGG</td><td>||-||-||[||--||||||||]PAM</td><td>NA</td><td>Intergenic</td><td>NA</td><td>NA</td></tr>
<tr><td></td><td></td><td></td><td></td><td></td><td></td><td></td><td></td><td></td><td></td><td></td><td></td></tr>
<tr><td>T6654</td><td>CCGCGAAACCGCGCCCATTTTGG</td><td>1000</td><td></td><td></td><td></td><td></td><td></td><td></td><td></td><td></td><td></td></tr>
<tr><td>Oligo fwd</td><td>CACCCCGCGAAACCGCGCCCATTT</td><td></td><td></td><td></td><td></td><td></td><td></td><td></td><td></td><td></td><td></td></tr>
<tr><td>Oligo rev</td><td>AAACAAATGGGCGCGGTTTCGCGG</td><td></td><td></td><td></td><td></td><td></td><td></td><td></td><td></td><td></td><td></td></tr>
<tr><td>Chromosome</td><td>start</td><td>end</td><td>strand</td><td>MM</td><td>target_seq</td><td>PAM</td><td>alignment</td><td>distance</td><td>position</td><td>gene name</td><td>gene id</td></tr>
<tr><td>GU305914.1</td><td>93375</td><td>93397</td><td>+</td><td>0</td><td>CCGCGAAACCGCGCCCATTT</td><td>TGG</td><td>||||||||[||||||||||||]PAM</td><td>NA</td><td>Intergenic</td><td>NA</td><td>NA</td></tr>
<tr><td></td><td></td><td></td><td></td><td></td><td></td><td></td><td></td><td></td><td></td><td></td><td></td></tr>
<tr><td>T6655</td><td>AGCTGACCTATGCGTCATCGTGG</td><td>1000</td><td></td><td></td><td></td><td></td><td></td><td></td><td></td><td></td><td></td></tr>
<tr><td>Oligo fwd</td><td>CACCAGCTGACCTATGCGTCATCG</td><td></td><td></td><td></td><td></td><td></td><td></td><td></td><td></td><td></td><td></td></tr>
<tr><td>Oligo rev</td><td>AAACCGATGACGCATAGGTCAGCT</td><td></td><td></td><td></td><td></td><td></td><td></td><td></td><td></td><td></td><td></td></tr>
<tr><td>Chromosome</td><td>start</td><td>end</td><td>strand</td><td>MM</td><td>target_seq</td><td>PAM</td><td>alignment</td><td>distance</td><td>position</td><td>gene name</td><td>gene id</td></tr>
<tr><td>GU305914.1</td><td>93400</td><td>93422</td><td>+</td><td>0</td><td>AGCTGACCTATGCGTCATCG</td><td>TGG</td><td>||||||||[||||||||||||]PAM</td><td>NA</td><td>Intergenic</td><td>NA</td><td>NA</td></tr>
<tr><td></td><td></td><td></td><td></td><td></td><td></td><td></td><td></td><td></td><td></td><td></td><td></td></tr>
<tr><td>T6656</td><td>TGACCTATGCGTCATCGTGGCGG</td><td>1000</td><td></td><td></td><td></td><td></td><td></td><td></td><td></td><td></td><td></td></tr>
<tr><td>Oligo fwd</td><td>CACCTGACCTATGCGTCATCGTGG</td><td></td><td></td><td></td><td></td><td></td><td></td><td></td><td></td><td></td><td></td></tr>
<tr><td>Oligo rev</td><td>AAACCCACGATGACGCATAGGTCA</td><td></td><td></td><td></td><td></td><td></td><td></td><td></td><td></td><td></td><td></td></tr>
<tr><td>Chromosome</td><td>start</td><td>end</td><td>strand</td><td>MM</td><td>target_seq</td><td>PAM</td><td>alignment</td><td>distance</td><td>position</td><td>gene name</td><td>gene id</td></tr>
<tr><td>GU305914.1</td><td>93403</td><td>93425</td><td>+</td><td>0</td><td>TGACCTATGCGTCATCGTGG</td><td>CGG</td><td>||||||||[||||||||||||]PAM</td><td>NA</td><td>Intergenic</td><td>NA</td><td>NA</td></tr>
<tr><td></td><td></td><td></td><td></td><td></td><td></td><td></td><td></td><td></td><td></td><td></td><td></td></tr>
<tr><td>T6657</td><td>TGGCGGATAAAAGCTCGAGCCGG</td><td>1000</td><td></td><td></td><td></td><td></td><td></td><td></td><td></td><td></td><td></td></tr>
<tr><td>Oligo fwd</td><td>CACCTGGCGGATAAAAGCTCGAGC</td><td></td><td></td><td></td><td></td><td></td><td></td><td></td><td></td><td></td><td></td></tr>
<tr><td>Oligo rev</td><td>AAACGCTCGAGCTTTTATCCGCCA</td><td></td><td></td><td></td><td></td><td></td><td></td><td></td><td></td><td></td><td></td></tr>
<tr><td>Chromosome</td><td>start</td><td>end</td><td>strand</td><td>MM</td><td>target_seq</td><td>PAM</td><td>alignment</td><td>distance</td><td>position</td><td>gene name</td><td>gene id</td></tr>
<tr><td>GU305914.1</td><td>93420</td><td>93442</td><td>+</td><td>0</td><td>TGGCGGATAAAAGCTCGAGC</td><td>CGG</td><td>||||||||[||||||||||||]PAM</td><td>NA</td><td>Intergenic</td><td>NA</td><td>NA</td></tr>
<tr><td></td><td></td><td></td><td></td><td></td><td></td><td></td><td></td><td></td><td></td><td></td><td></td></tr>
<tr><td>T6658</td><td>AGCTCGAGCCGGCCCGCCGCCGG</td><td>1000</td><td></td><td></td><td></td><td></td><td></td><td></td><td></td><td></td><td></td></tr>
<tr><td>Oligo fwd</td><td>CACCAGCTCGAGCCGGCCCGCCGC</td><td></td><td></td><td></td><td></td><td></td><td></td><td></td><td></td><td></td><td></td></tr>
<tr><td>Oligo rev</td><td>AAACGCGGCGGGCCGGCTCGAGCT</td><td></td><td></td><td></td><td></td><td></td><td></td><td></td><td></td><td></td><td></td></tr>
<tr><td>Chromosome</td><td>start</td><td>end</td><td>strand</td><td>MM</td><td>target_seq</td><td>PAM</td><td>alignment</td><td>distance</td><td>position</td><td>gene name</td><td>gene id</td></tr>
<tr><td>GU305914.1</td><td>93431</td><td>93453</td><td>+</td><td>0</td><td>AGCTCGAGCCGGCCCGCCGC</td><td>CGG</td><td>||||||||[||||||||||||]PAM</td><td>NA</td><td>Intergenic</td><td>NA</td><td>NA</td></tr>
<tr><td></td><td></td><td></td><td></td><td></td><td></td><td></td><td></td><td></td><td></td><td></td><td></td></tr>
<tr><td>T6659</td><td>CCCGCCCGTCGCTCGCCACCAGG</td><td>1000</td><td></td><td></td><td></td><td></td><td></td><td></td><td></td><td></td><td></td></tr>
<tr><td>Oligo fwd</td><td>CACCCCCGCCCGTCGCTCGCCACC</td><td></td><td></td><td></td><td></td><td></td><td></td><td></td><td></td><td></td><td></td></tr>
<tr><td>Oligo rev</td><td>AAACGGTGGCGAGCGACGGGCGGG</td><td></td><td></td><td></td><td></td><td></td><td></td><td></td><td></td><td></td><td></td></tr>
<tr><td>Chromosome</td><td>start</td><td>end</td><td>strand</td><td>MM</td><td>target_seq</td><td>PAM</td><td>alignment</td><td>distance</td><td>position</td><td>gene name</td><td>gene id</td></tr>
<tr><td>GU305914.1</td><td>93459</td><td>93481</td><td>+</td><td>0</td><td>CCCGCCCGTCGCTCGCCACC</td><td>AGG</td><td>||||||||[||||||||||||]PAM</td><td>NA</td><td>Intergenic</td><td>NA</td><td>NA</td></tr>
<tr><td></td><td></td><td></td><td></td><td></td><td></td><td></td><td></td><td></td><td></td><td></td><td></td></tr>
<tr><td>T6660</td><td>CACCAGGATCCACGCCGCTCCGG</td><td>1000</td><td></td><td></td><td></td><td></td><td></td><td></td><td></td><td></td><td></td></tr>
<tr><td>Oligo fwd</td><td>CACCCACCAGGATCCACGCCGCTC</td><td></td><td></td><td></td><td></td><td></td><td></td><td></td><td></td><td></td><td></td></tr>
<tr><td>Oligo rev</td><td>AAACGAGCGGCGTGGATCCTGGTG</td><td></td><td></td><td></td><td></td><td></td><td></td><td></td><td></td><td></td><td></td></tr>
<tr><td>Chromosome</td><td>start</td><td>end</td><td>strand</td><td>MM</td><td>target_seq</td><td>PAM</td><td>alignment</td><td>distance</td><td>position</td><td>gene name</td><td>gene id</td></tr>
<tr><td>GU305914.1</td><td>93475</td><td>93497</td><td>+</td><td>0</td><td>CACCAGGATCCACGCCGCTC</td><td>CGG</td><td>||||||||[||||||||||||]PAM</td><td>NA</td><td>Intergenic</td><td>NA</td><td>NA</td></tr>
<tr><td></td><td></td><td></td><td></td><td></td><td></td><td></td><td></td><td></td><td></td><td></td><td></td></tr>
<tr><td>T6661</td><td>GCCGCTCCGGACCGCCGCCTCGG</td><td>1000</td><td></td><td></td><td></td><td></td><td></td><td></td><td></td><td></td><td></td></tr>
<tr><td>Oligo fwd</td><td>CACCGCCGCTCCGGACCGCCGCCT</td><td></td><td></td><td></td><td></td><td></td><td></td><td></td><td></td><td></td><td></td></tr>
<tr><td>Oligo rev</td><td>AAACAGGCGGCGGTCCGGAGCGGC</td><td></td><td></td><td></td><td></td><td></td><td></td><td></td><td></td><td></td><td></td></tr>
</table>

<table>
<tr><td>Chromosome</td><td>start</td><td>end</td><td>strand</td><td>MM</td><td>target_seq</td><td>PAM</td><td>alignment</td><td>distance</td><td>position</td><td>gene name</td><td>gene id</td></tr>
<tr><td>GU305914.1</td><td>93488</td><td>93510</td><td>+</td><td>0</td><td>GCCGCTCCGGACCGCCGCCT</td><td>CGG</td><td>||||||||[||||||||||||]PAM</td><td>NA</td><td>Intergenic</td><td>NA</td><td>NA</td></tr>
<tr><td></td><td></td><td></td><td></td><td></td><td></td><td></td><td></td><td></td><td></td><td></td><td></td></tr>
<tr><td>T6662</td><td>CCAGCGCCCCGCACCCCACCCGG</td><td>1000</td><td></td><td></td><td></td><td></td><td></td><td></td><td></td><td></td><td></td></tr>
<tr><td>Oligo fwd</td><td>CACCCCAGCGCCCCGCACCCCACC</td><td></td><td></td><td></td><td></td><td></td><td></td><td></td><td></td><td></td><td></td></tr>
<tr><td>Oligo rev</td><td>AAACGGTGGGGTGCGGGGCGCTGG</td><td></td><td></td><td></td><td></td><td></td><td></td><td></td><td></td><td></td><td></td></tr>
<tr><td>Chromosome</td><td>start</td><td>end</td><td>strand</td><td>MM</td><td>target_seq</td><td>PAM</td><td>alignment</td><td>distance</td><td>position</td><td>gene name</td><td>gene id</td></tr>
<tr><td>GU305914.1</td><td>93512</td><td>93534</td><td>+</td><td>0</td><td>CCAGCGCCCCGCACCCCACC</td><td>CGG</td><td>||||||||[||||||||||||]PAM</td><td>NA</td><td>Intergenic</td><td>NA</td><td>NA</td></tr>
<tr><td></td><td></td><td></td><td></td><td></td><td></td><td></td><td></td><td></td><td></td><td></td><td></td></tr>
<tr><td>T6663</td><td>CCGCACCCCACCCGGATCCCCGG</td><td>1000</td><td></td><td></td><td></td><td></td><td></td><td></td><td></td><td></td><td></td></tr>
<tr><td>Oligo fwd</td><td>CACCCCGCACCCCACCCGGATCCC</td><td></td><td></td><td></td><td></td><td></td><td></td><td></td><td></td><td></td><td></td></tr>
<tr><td>Oligo rev</td><td>AAACGGGATCCGGGTGGGGTGCGG</td><td></td><td></td><td></td><td></td><td></td><td></td><td></td><td></td><td></td><td></td></tr>
<tr><td>Chromosome</td><td>start</td><td>end</td><td>strand</td><td>MM</td><td>target_seq</td><td>PAM</td><td>alignment</td><td>distance</td><td>position</td><td>gene name</td><td>gene id</td></tr>
<tr><td>GU305914.1</td><td>93520</td><td>93542</td><td>+</td><td>0</td><td>CCGCACCCCACCCGGATCCC</td><td>CGG</td><td>||||||||[||||||||||||]PAM</td><td>NA</td><td>Intergenic</td><td>NA</td><td>NA</td></tr>
<tr><td></td><td></td><td></td><td></td><td></td><td></td><td></td><td></td><td></td><td></td><td></td><td></td></tr>
<tr><td>T6664</td><td>CACCCGGATCCCCGGACCGCAGG</td><td>1000</td><td></td><td></td><td></td><td></td><td></td><td></td><td></td><td></td><td></td></tr>
<tr><td>Oligo fwd</td><td>CACCCACCCGGATCCCCGGACCGC</td><td></td><td></td><td></td><td></td><td></td><td></td><td></td><td></td><td></td><td></td></tr>
<tr><td>Oligo rev</td><td>AAACGCGGTCCGGGGATCCGGGTG</td><td></td><td></td><td></td><td></td><td></td><td></td><td></td><td></td><td></td><td></td></tr>
<tr><td>Chromosome</td><td>start</td><td>end</td><td>strand</td><td>MM</td><td>target_seq</td><td>PAM</td><td>alignment</td><td>distance</td><td>position</td><td>gene name</td><td>gene id</td></tr>
<tr><td>GU305914.1</td><td>93528</td><td>93550</td><td>+</td><td>0</td><td>CACCCGGATCCCCGGACCGC</td><td>AGG</td><td>||||||||[||||||||||||]PAM</td><td>NA</td><td>Intergenic</td><td>NA</td><td>NA</td></tr>
<tr><td></td><td></td><td></td><td></td><td></td><td></td><td></td><td></td><td></td><td></td><td></td><td></td></tr>
<tr><td>T6665</td><td>GGACCGCAGGACCAGCTCCCCGG</td><td>1000</td><td></td><td></td><td></td><td></td><td></td><td></td><td></td><td></td><td></td></tr>
<tr><td>Oligo fwd</td><td>CACCGGACCGCAGGACCAGCTCCC</td><td></td><td></td><td></td><td></td><td></td><td></td><td></td><td></td><td></td><td></td></tr>
<tr><td>Oligo rev</td><td>AAACGGGAGCTGGTCCTGCGGTCC</td><td></td><td></td><td></td><td></td><td></td><td></td><td></td><td></td><td></td><td></td></tr>
<tr><td>Chromosome</td><td>start</td><td>end</td><td>strand</td><td>MM</td><td>target_seq</td><td>PAM</td><td>alignment</td><td>distance</td><td>position</td><td>gene name</td><td>gene id</td></tr>
<tr><td>GU305914.1</td><td>93541</td><td>93563</td><td>+</td><td>0</td><td>GGACCGCAGGACCAGCTCCC</td><td>CGG</td><td>||||||||[||||||||||||]PAM</td><td>NA</td><td>Intergenic</td><td>NA</td><td>NA</td></tr>
<tr><td>GU305914.1</td><td>93707</td><td>93729</td><td>+</td><td>2</td><td>GGCCCCCAGGACCAGCTCCC</td><td>CGG</td><td>||-||-||[||||||||||||]PAM</td><td>NA</td><td>Intergenic</td><td>NA</td><td>NA</td></tr>
<tr><td>GU305914.1</td><td>93811</td><td>93833</td><td>+</td><td>4</td><td>GACCTCCAGGACCAGCTCCC</td><td>CGG</td><td>|--|--||[||||||||||||]PAM</td><td>NA</td><td>Intergenic</td><td>NA</td><td>NA</td></tr>
<tr><td>GU305914.1</td><td>93583</td><td>93605</td><td>+</td><td>2</td><td>GGACCCCAGGACCAGCACCC</td><td>CGG</td><td>|||||-||[||||||||-|||]PAM</td><td>NA</td><td>Intergenic</td><td>NA</td><td>NA</td></tr>
<tr><td></td><td></td><td></td><td></td><td></td><td></td><td></td><td></td><td></td><td></td><td></td><td></td></tr>
<tr><td>T6666</td><td>AGGACCAGCTCCCCGGCTACCGG</td><td>1000</td><td></td><td></td><td></td><td></td><td></td><td></td><td></td><td></td><td></td></tr>
<tr><td>Oligo fwd</td><td>CACCAGGACCAGCTCCCCGGCTAC</td><td></td><td></td><td></td><td></td><td></td><td></td><td></td><td></td><td></td><td></td></tr>
<tr><td>Oligo rev</td><td>AAACGTAGCCGGGGAGCTGGTCCT</td><td></td><td></td><td></td><td></td><td></td><td></td><td></td><td></td><td></td><td></td></tr>
<tr><td>Chromosome</td><td>start</td><td>end</td><td>strand</td><td>MM</td><td>target_seq</td><td>PAM</td><td>alignment</td><td>distance</td><td>position</td><td>gene name</td><td>gene id</td></tr>
<tr><td>GU305914.1</td><td>93548</td><td>93570</td><td>+</td><td>0</td><td>AGGACCAGCTCCCCGGCTAC</td><td>CGG</td><td>||||||||[||||||||||||]PAM</td><td>NA</td><td>Intergenic</td><td>NA</td><td>NA</td></tr>
<tr><td>GU305914.1</td><td>93621</td><td>93643</td><td>+</td><td>1</td><td>CGGACCAGCTCCCCGGCTAC</td><td>CGG</td><td>-|||||||[||||||||||||]PAM</td><td>NA</td><td>Intergenic</td><td>NA</td><td>NA</td></tr>
<tr><td></td><td></td><td></td><td></td><td></td><td></td><td></td><td></td><td></td><td></td><td></td><td></td></tr>
<tr><td>T6667</td><td>AGCTCCCCGGCTACCGGCCTCGG</td><td>1000</td><td></td><td></td><td></td><td></td><td></td><td></td><td></td><td></td><td></td></tr>
<tr><td>Oligo fwd</td><td>CACCAGCTCCCCGGCTACCGGCCT</td><td></td><td></td><td></td><td></td><td></td><td></td><td></td><td></td><td></td><td></td></tr>
<tr><td>Oligo rev</td><td>AAACAGGCCGGTAGCCGGGGAGCT</td><td></td><td></td><td></td><td></td><td></td><td></td><td></td><td></td><td></td><td></td></tr>
<tr><td>Chromosome</td><td>start</td><td>end</td><td>strand</td><td>MM</td><td>target_seq</td><td>PAM</td><td>alignment</td><td>distance</td><td>position</td><td>gene name</td><td>gene id</td></tr>
<tr><td>GU305914.1</td><td>93627</td><td>93649</td><td>+</td><td>0</td><td>AGCTCCCCGGCTACCGGCCT</td><td>CGG</td><td>||||||||[||||||||||||]PAM</td><td>NA</td><td>Intergenic</td><td>NA</td><td>NA</td></tr>
<tr><td>GU305914.1</td><td>93554</td><td>93576</td><td>+</td><td>0</td><td>AGCTCCCCGGCTACCGGCCT</td><td>CGG</td><td>||||||||[||||||||||||]PAM</td><td>NA</td><td>Intergenic</td><td>NA</td><td>NA</td></tr>
<tr><td>GU305914.1</td><td>93720</td><td>93742</td><td>+</td><td>1</td><td>AGCTCCCCGGCTACCGACCT</td><td>CGG</td><td>||||||||[||||||||-|||]PAM</td><td>NA</td><td>Intergenic</td><td>NA</td><td>NA</td></tr>
<tr><td>GU305914.1</td><td>93677</td><td>93699</td><td>+</td><td>3</td><td>AGCTTCCCGCCTACCGACCT</td><td>CGG</td><td>||||-|||[|-||||||-|||]PAM</td><td>NA</td><td>Intergenic</td><td>NA</td><td>NA</td></tr>
<tr><td>GU305914.1</td><td>93781</td><td>93803</td><td>+</td><td>3</td><td>AGCTTCCCGGCTACCCTCCT</td><td>CGG</td><td>||||-|||[|||||||--|||]PAM</td><td>NA</td><td>Intergenic</td><td>NA</td><td>NA</td></tr>
<tr><td></td><td></td><td></td><td></td><td></td><td></td><td></td><td></td><td></td><td></td><td></td><td></td></tr>
<tr><td>T6668</td><td>GGCTACCGGCCTCGGCTCCCCGG</td><td>1000</td><td></td><td></td><td></td><td></td><td></td><td></td><td></td><td></td><td></td></tr>
<tr><td>Oligo fwd</td><td>CACCGGCTACCGGCCTCGGCTCCC</td><td></td><td></td><td></td><td></td><td></td><td></td><td></td><td></td><td></td><td></td></tr>
<tr><td>Oligo rev</td><td>AAACGGGAGCCGAGGCCGGTAGCC</td><td></td><td></td><td></td><td></td><td></td><td></td><td></td><td></td><td></td><td></td></tr>
<tr><td>Chromosome</td><td>start</td><td>end</td><td>strand</td><td>MM</td><td>target_seq</td><td>PAM</td><td>alignment</td><td>distance</td><td>position</td><td>gene name</td><td>gene id</td></tr>
<tr><td>GU305914.1</td><td>93562</td><td>93584</td><td>+</td><td>0</td><td>GGCTACCGGCCTCGGCTCCC</td><td>CGG</td><td>||||||||[||||||||||||]PAM</td><td>NA</td><td>Intergenic</td><td>NA</td><td>NA</td></tr>
<tr><td>GU305914.1</td><td>93635</td><td>93657</td><td>+</td><td>0</td><td>GGCTACCGGCCTCGGCTCCC</td><td>CGG</td><td>||||||||[||||||||||||]PAM</td><td>NA</td><td>Intergenic</td><td>NA</td><td>NA</td></tr>
<tr><td>GU305914.1</td><td>93685</td><td>93707</td><td>+</td><td>3</td><td>GCCTACCGACCTCGGCTCAC</td><td>CGG</td><td>|-||||||[-|||||||||-|]PAM</td><td>NA</td><td>Intergenic</td><td>NA</td><td>NA</td></tr>
<tr><td>GU305914.1</td><td>93789</td><td>93811</td><td>+</td><td>3</td><td>GGCTACCCTCCTCGGCTCAC</td><td>CGG</td><td>|||||||-[-|||||||||-|]PAM</td><td>NA</td><td>Intergenic</td><td>NA</td><td>NA</td></tr>
<tr><td></td><td></td><td></td><td></td><td></td><td></td><td></td><td></td><td></td><td></td><td></td><td></td></tr>
<tr><td>T6669</td><td>GCCTCGGCTCCCCGGACCCCAGG</td><td>1000</td><td></td><td></td><td></td><td></td><td></td><td></td><td></td><td></td><td></td></tr>
<tr><td>Oligo fwd</td><td>CACCGCCTCGGCTCCCCGGACCCC</td><td></td><td></td><td></td><td></td><td></td><td></td><td></td><td></td><td></td><td></td></tr>
<tr><td>Oligo rev</td><td>AAACGGGGTCCGGGGAGCCGAGGC</td><td></td><td></td><td></td><td></td><td></td><td></td><td></td><td></td><td></td><td></td></tr>
<tr><td>Chromosome</td><td>start</td><td>end</td><td>strand</td><td>MM</td><td>target_seq</td><td>PAM</td><td>alignment</td><td>distance</td><td>position</td><td>gene name</td><td>gene id</td></tr>
<tr><td>GU305914.1</td><td>93570</td><td>93592</td><td>+</td><td>0</td><td>GCCTCGGCTCCCCGGACCCC</td><td>AGG</td><td>||||||||[||||||||||||]PAM</td><td>NA</td><td>Intergenic</td><td>NA</td><td>NA</td></tr>
<tr><td></td><td></td><td></td><td></td><td></td><td></td><td></td><td></td><td></td><td></td><td></td><td></td></tr>
<tr><td>T6670</td><td>GGACCCCAGGACCAGCACCCCGG</td><td>1000</td><td></td><td></td><td></td><td></td><td></td><td></td><td></td><td></td><td></td></tr>
<tr><td>Oligo fwd</td><td>CACCGGACCCCAGGACCAGCACCC</td><td></td><td></td><td></td><td></td><td></td><td></td><td></td><td></td><td></td><td></td></tr>
<tr><td>Oligo rev</td><td>AAACGGGTGCTGGTCCTGGGGTCC</td><td></td><td></td><td></td><td></td><td></td><td></td><td></td><td></td><td></td><td></td></tr>
<tr><td>Chromosome</td><td>start</td><td>end</td><td>strand</td><td>MM</td><td>target_seq</td><td>PAM</td><td>alignment</td><td>distance</td><td>position</td><td>gene name</td><td>gene id</td></tr>
<tr><td>GU305914.1</td><td>93583</td><td>93605</td><td>+</td><td>0</td><td>GGACCCCAGGACCAGCACCC</td><td>CGG</td><td>||||||||[||||||||||||]PAM</td><td>NA</td><td>Intergenic</td><td>NA</td><td>NA</td></tr>
<tr><td>GU305914.1</td><td>93707</td><td>93729</td><td>+</td><td>2</td><td>GGCCCCCAGGACCAGCTCCC</td><td>CGG</td><td>||-|||||[||||||||-|||]PAM</td><td>NA</td><td>Intergenic</td><td>NA</td><td>NA</td></tr>
<tr><td>GU305914.1</td><td>93541</td><td>93563</td><td>+</td><td>2</td><td>GGACCGCAGGACCAGCTCCC</td><td>CGG</td><td>|||||-||[||||||||-|||]PAM</td><td>NA</td><td>Intergenic</td><td>NA</td><td>NA</td></tr>
<tr><td>GU305914.1</td><td>93811</td><td>93833</td><td>+</td><td>4</td><td>GACCTCCAGGACCAGCTCCC</td><td>CGG</td><td>|--|-|||[||||||||-|||]PAM</td><td>NA</td><td>Intergenic</td><td>NA</td><td>NA</td></tr>
<tr><td></td><td></td><td></td><td></td><td></td><td></td><td></td><td></td><td></td><td></td><td></td><td></td></tr>
<tr><td>T6671</td><td>ACCAGCACCCCGGACCCGCCTGG</td><td>1000</td><td></td><td></td><td></td><td></td><td></td><td></td><td></td><td></td><td></td></tr>
<tr><td>Oligo fwd</td><td>CACCACCAGCACCCCGGACCCGCC</td><td></td><td></td><td></td><td></td><td></td><td></td><td></td><td></td><td></td><td></td></tr>
<tr><td>Oligo rev</td><td>AAACGGCGGGTCCGGGGTGCTGGT</td><td></td><td></td><td></td><td></td><td></td><td></td><td></td><td></td><td></td><td></td></tr>
<tr><td>Chromosome</td><td>start</td><td>end</td><td>strand</td><td>MM</td><td>target_seq</td><td>PAM</td><td>alignment</td><td>distance</td><td>position</td><td>gene name</td><td>gene id</td></tr>
<tr><td>GU305914.1</td><td>93593</td><td>93615</td><td>+</td><td>0</td><td>ACCAGCACCCCGGACCCGCC</td><td>TGG</td><td>||||||||[||||||||||||]PAM</td><td>NA</td><td>Intergenic</td><td>NA</td><td>NA</td></tr>
<tr><td></td><td></td><td></td><td></td><td></td><td></td><td></td><td></td><td></td><td></td><td></td><td></td></tr>
<tr><td>T6672</td><td>CCCGGACCCGCCTGGCTTCCCGG</td><td>1000</td><td></td><td></td><td></td><td></td><td></td><td></td><td></td><td></td><td></td></tr>
<tr><td>Oligo fwd</td><td>CACCCCCGGACCCGCCTGGCTTCC</td><td></td><td></td><td></td><td></td><td></td><td></td><td></td><td></td><td></td><td></td></tr>
<tr><td>Oligo rev</td><td>AAACGGAAGCCAGGCGGGTCCGGG</td><td></td><td></td><td></td><td></td><td></td><td></td><td></td><td></td><td></td><td></td></tr>
<tr><td>Chromosome</td><td>start</td><td>end</td><td>strand</td><td>MM</td><td>target_seq</td><td>PAM</td><td>alignment</td><td>distance</td><td>position</td><td>gene name</td><td>gene id</td></tr>
<tr><td>GU305914.1</td><td>93601</td><td>93623</td><td>+</td><td>0</td><td>CCCGGACCCGCCTGGCTTCC</td><td>CGG</td><td>||||||||[||||||||||||]PAM</td><td>NA</td><td>Intergenic</td><td>NA</td><td>NA</td></tr>
<tr><td></td><td></td><td></td><td></td><td></td><td></td><td></td><td></td><td></td><td></td><td></td><td></td></tr>
<tr><td>T6673</td><td>GGCTTCCCGGACCAGCTCCCCGG</td><td>1000</td><td></td><td></td><td></td><td></td><td></td><td></td><td></td><td></td><td></td></tr>
<tr><td>Oligo fwd</td><td>CACCGGCTTCCCGGACCAGCTCCC</td><td></td><td></td><td></td><td></td><td></td><td></td><td></td><td></td><td></td><td></td></tr>
<tr><td>Oligo rev</td><td>AAACGGGAGCTGGTCCGGGAAGCC</td><td></td><td></td><td></td><td></td><td></td><td></td><td></td><td></td><td></td><td></td></tr>
<tr><td>Chromosome</td><td>start</td><td>end</td><td>strand</td><td>MM</td><td>target_seq</td><td>PAM</td><td>alignment</td><td>distance</td><td>position</td><td>gene name</td><td>gene id</td></tr>
<tr><td>GU305914.1</td><td>93614</td><td>93636</td><td>+</td><td>0</td><td>GGCTTCCCGGACCAGCTCCC</td><td>CGG</td><td>||||||||[||||||||||||]PAM</td><td>NA</td><td>Intergenic</td><td>NA</td><td>NA</td></tr>
<tr><td>GU305914.1</td><td>93811</td><td>93833</td><td>+</td><td>3</td><td>GACCTCCAGGACCAGCTCCC</td><td>CGG</td><td>|-|-|||-[||||||||||||]PAM</td><td>NA</td><td>Intergenic</td><td>NA</td><td>NA</td></tr>
</table>

```
GU305914.1	93707	93729	+	3	GGCCCCCAGGACCAGCTCCC	CGG	|||--||-[||||||||||||]PAM	NA	Intergenic	NA	NA

T6674	CGGACCAGCTCCCCGGCTACCGG	1000
Oligo fwd	CACCCGGACCAGCTCCCCGGCTAC
Oligo rev	AAACGTAGCCGGGGAGCTGGTCCG
Chromosome	start	end	strand	MM	target_seq	PAM	alignment	distance	position	gene name	gene id
GU305914.1	93621	93643	+	0	CGGACCAGCTCCCCGGCTAC	CGG	||||||||[||||||||||||]PAM	NA	Intergenic	NA	NA
GU305914.1	93548	93570	+	1	AGGACCAGCTCCCCGGCTAC	CGG	-|||||||[||||||||||||]PAM	NA	Intergenic	NA	NA

T6675	AGCTCCCCGGCTACCGGCCTCGG	1000
Oligo fwd	CACCAGCTCCCCGGCTACCGGCCT
Oligo rev	AAACAGGCCGGTAGCCGGGGAGCT
Chromosome	start	end	strand	MM	target_seq	PAM	alignment	distance	position	gene name	gene id
GU305914.1	93627	93649	+	0	AGCTCCCCGGCTACCGGCCT	CGG	||||||||[||||||||||||]PAM	NA	Intergenic	NA	NA
GU305914.1	93554	93576	+	0	AGCTCCCCGGCTACCGGCCT	CGG	||||||||[||||||||||||]PAM	NA	Intergenic	NA	NA
GU305914.1	93720	93742	+	1	AGCTCCCCGGCTACCGACCT	CGG	||||||||[||||||||-|||]PAM	NA	Intergenic	NA	NA
GU305914.1	93677	93699	+	3	AGCTTCCCGCCTACCGACCT	CGG	||||-|||[|-||||||-|||]PAM	NA	Intergenic	NA	NA
GU305914.1	93781	93803	+	3	AGCTTCCCGGCTACCCTCCT	CGG	||||-|||[|||||||--|||]PAM	NA	Intergenic	NA	NA

T6676	GGCTACCGGCCTCGGCTCCCCGG	1000
Oligo fwd	CACCGGCTACCGGCCTCGGCTCCC
Oligo rev	AAACGGGAGCCGAGGCCGGTAGCC
Chromosome	start	end	strand	MM	target_seq	PAM	alignment	distance	position	gene name	gene id
GU305914.1	93562	93584	+	0	GGCTACCGGCCTCGGCTCCC	CGG	||||||||[||||||||||||]PAM	NA	Intergenic	NA	NA
GU305914.1	93635	93657	+	0	GGCTACCGGCCTCGGCTCCC	CGG	||||||||[||||||||||||]PAM	NA	Intergenic	NA	NA
GU305914.1	93685	93707	+	3	GCCTACCGACCTCGGCTCAC	CGG	|-||||||[-|||||||||-|]PAM	NA	Intergenic	NA	NA
GU305914.1	93789	93811	+	3	GGCTACCCTCCTCGGCTCAC	CGG	|||||||-[-|||||||||-|]PAM	NA	Intergenic	NA	NA

T6677	GCCTCGGCTCCCCGGCTTTCCGG	1000
Oligo fwd	CACCGCCTCGGCTCCCCGGCTTTC
Oligo rev	AAACGAAAGCCGGGGAGCCGAGGC
Chromosome	start	end	strand	MM	target_seq	PAM	alignment	distance	position	gene name	gene id
GU305914.1	93643	93665	+	0	GCCTCGGCTCCCCGGCTTTC	CGG	||||||||[||||||||||||]PAM	NA	Intergenic	NA	NA
GU305914.1	140311	140333	-	4	GCGTCGCCTCAACGGCTTTC	TGG	||-|||-|[||--||||||||]PAM	NA	Intergenic	NA	NA
GU305914.1	93768	93790	+	3	GGCTCGGCTCCCCAGCTTCC	CGG	|-||||||[|||||-||||-|]PAM	NA	Intergenic	NA	NA

T6678	GGCTCCCCGGCTTTCCGGCTCGG	1000
Oligo fwd	CACCGGCTCCCCGGCTTTCCGGCT
Oligo rev	AAACAGCCGGAAAGCCGGGGAGCC
Chromosome	start	end	strand	MM	target_seq	PAM	alignment	distance	position	gene name	gene id
GU305914.1	93648	93670	+	0	GGCTCCCCGGCTTTCCGGCT	CGG	||||||||[||||||||||||]PAM	NA	Intergenic	NA	NA

T6679	AGCTTCCCGCCTACCGACCTCGG	1000
Oligo fwd	CACCAGCTTCCCGCCTACCGACCT
Oligo rev	AAACAGGTCGGTAGGCGGGAAGCT
Chromosome	start	end	strand	MM	target_seq	PAM	alignment	distance	position	gene name	gene id
GU305914.1	93677	93699	+	0	AGCTTCCCGCCTACCGACCT	CGG	||||||||[||||||||||||]PAM	NA	Intergenic	NA	NA
GU305914.1	93720	93742	+	2	AGCTCCCCGGCTACCGACCT	CGG	||||-|||[|-||||||||||]PAM	NA	Intergenic	NA	NA
GU305914.1	93554	93576	+	3	AGCTCCCCGGCTACCGGCCT	CGG	||||-|||[|-||||||-|||]PAM	NA	Intergenic	NA	NA
GU305914.1	93627	93649	+	3	AGCTCCCCGGCTACCGGCCT	CGG	||||-|||[|-||||||-|||]PAM	NA	Intergenic	NA	NA

T6680	GCCTACCGACCTCGGCTCACCGG	1000
Oligo fwd	CACCGCCTACCGACCTCGGCTCAC
Oligo rev	AAACGTGAGCCGAGGTCGGTAGGC
Chromosome	start	end	strand	MM	target_seq	PAM	alignment	distance	position	gene name	gene id
GU305914.1	93685	93707	+	0	GCCTACCGACCTCGGCTCAC	CGG	||||||||[||||||||||||]PAM	NA	Intergenic	NA	NA
GU305914.1	93789	93811	+	3	GGCTACCCTCCTCGGCTCAC	CGG	|-|||||-[-|||||||||||]PAM	NA	Intergenic	NA	NA
GU305914.1	93635	93657	+	3	GGCTACCGGCCTCGGCTCCC	CGG	|-||||||[-|||||||||-|]PAM	NA	Intergenic	NA	NA
GU305914.1	93562	93584	+	3	GGCTACCGGCCTCGGCTCCC	CGG	|-||||||[-|||||||||-|]PAM	NA	Intergenic	NA	NA

T6681	CCTACCGACCTCGGCTCACCGGG	1000
Oligo fwd	CACCCCTACCGACCTCGGCTCACC
Oligo rev	AAACGGTGAGCCGAGGTCGGTAGG
Chromosome	start	end	strand	MM	target_seq	PAM	alignment	distance	position	gene name	gene id
GU305914.1	93686	93708	+	0	CCTACCGACCTCGGCTCACC	GGG	||||||||[||||||||||||]PAM	NA	Intergenic	NA	NA

T6682	CCTCGGCTCACCGGGCCCCCAGG	1000
Oligo fwd	CACCCCTCGGCTCACCGGGCCCCC
Oligo rev	AAACGGGGGCCCGGTGAGCCGAGG
Chromosome	start	end	strand	MM	target_seq	PAM	alignment	distance	position	gene name	gene id
GU305914.1	93694	93716	+	0	CCTCGGCTCACCGGGCCCCC	AGG	||||||||[||||||||||||]PAM	NA	Intergenic	NA	NA
GU305914.1	93798	93820	+	2	CCTCGGCTCACCGGACCTCC	AGG	||||||||[||||||-||-||]PAM	NA	Intergenic	NA	NA

T6683	GGCCCCCAGGACCAGCTCCCCGG	1000
Oligo fwd	CACCGGCCCCCAGGACCAGCTCCC
Oligo rev	AAACGGGAGCTGGTCCTGGGGGCC
Chromosome	start	end	strand	MM	target_seq	PAM	alignment	distance	position	gene name	gene id
GU305914.1	93707	93729	+	0	GGCCCCCAGGACCAGCTCCC	CGG	||||||||[||||||||||||]PAM	NA	Intergenic	NA	NA
GU305914.1	93811	93833	+	2	GACCTCCAGGACCAGCTCCC	CGG	|-||-|||[||||||||||||]PAM	NA	Intergenic	NA	NA
GU305914.1	93541	93563	+	2	GGACCGCAGGACCAGCTCCC	CGG	||-||-||[||||||||||||]PAM	NA	Intergenic	NA	NA
GU305914.1	93614	93636	+	3	GGCTTCCCGGACCAGCTCCC	CGG	|||--||-[||||||||||||]PAM	NA	Intergenic	NA	NA
GU305914.1	93583	93605	+	2	GGACCCCAGGACCAGCACCC	CGG	||-|||||[||||||||-|||]PAM	NA	Intergenic	NA	NA

T6684	AGCTCCCCGGCTACCGACCTCGG	1000
Oligo fwd	CACCAGCTCCCCGGCTACCGACCT
Oligo rev	AAACAGGTCGGTAGCCGGGGAGCT
Chromosome	start	end	strand	MM	target_seq	PAM	alignment	distance	position	gene name	gene id
GU305914.1	93720	93742	+	0	AGCTCCCCGGCTACCGACCT	CGG	||||||||[||||||||||||]PAM	NA	Intergenic	NA	NA
GU305914.1	93677	93699	+	2	AGCTTCCCGCCTACCGACCT	CGG	||||-|||[|-||||||||||]PAM	NA	Intergenic	NA	NA
```

<table>
<tr><td>GU305914.1</td><td></td><td>93627</td><td>93649</td><td>+</td><td>1</td><td>AGCTCCCCGGCTACCGGCCT</td><td>CGG</td><td>||||||||[||||||||-|||]PAM</td><td>NA</td><td>Intergenic</td><td>NA</td><td>NA</td></tr>
<tr><td>GU305914.1</td><td></td><td>93554</td><td>93576</td><td>+</td><td>1</td><td>AGCTCCCCGGCTACCGGCCT</td><td>CGG</td><td>||||||||[||||||||-|||]PAM</td><td>NA</td><td>Intergenic</td><td>NA</td><td>NA</td></tr>
<tr><td>GU305914.1</td><td></td><td>93781</td><td>93803</td><td>+</td><td>3</td><td>AGCTTCCCGGCTACCCTCCT</td><td>CGG</td><td>||||-|||[|||||||--|||]PAM</td><td>NA</td><td>Intergenic</td><td>NA</td><td>NA</td></tr>
<tr><td></td><td></td><td></td><td></td><td></td><td></td><td></td><td></td><td></td><td></td><td></td><td></td><td></td></tr>
<tr><td>T6685</td><td>TCGGATCCCCGCACCCTTCCCGG</td><td></td><td>1000</td><td></td><td></td><td></td><td></td><td></td><td></td><td></td><td></td><td></td></tr>
<tr><td>Oligo fwd</td><td>CACCTCGGATCCCCGCACCCTTCC</td><td></td><td></td><td></td><td></td><td></td><td></td><td></td><td></td><td></td><td></td><td></td></tr>
<tr><td>Oligo rev</td><td>AAACGGAAGGGTGCGGGGATCCGA</td><td></td><td></td><td></td><td></td><td></td><td></td><td></td><td></td><td></td><td></td><td></td></tr>
<tr><td>Chromosome</td><td>start</td><td></td><td>end</td><td>strand</td><td>MM</td><td>target_seq</td><td>PAM</td><td>alignment</td><td>distance</td><td>position</td><td>gene name</td><td>gene id</td></tr>
<tr><td>GU305914.1</td><td></td><td>93739</td><td>93761</td><td>+</td><td>0</td><td>TCGGATCCCCGCACCCTTCC</td><td>CGG</td><td>||||||||[||||||||||||]PAM</td><td>NA</td><td>Intergenic</td><td>NA</td><td>NA</td></tr>
<tr><td></td><td></td><td></td><td></td><td></td><td></td><td></td><td></td><td></td><td></td><td></td><td></td><td></td></tr>
<tr><td>T6686</td><td>CCGCACCCTTCCCGGCTTCTCGG</td><td></td><td>1000</td><td></td><td></td><td></td><td></td><td></td><td></td><td></td><td></td><td></td></tr>
<tr><td>Oligo fwd</td><td>CACCCCGCACCCTTCCCGGCTTCT</td><td></td><td></td><td></td><td></td><td></td><td></td><td></td><td></td><td></td><td></td><td></td></tr>
<tr><td>Oligo rev</td><td>AAACAGAAGCCGGGAAGGGTGCGG</td><td></td><td></td><td></td><td></td><td></td><td></td><td></td><td></td><td></td><td></td><td></td></tr>
<tr><td>Chromosome</td><td>start</td><td></td><td>end</td><td>strand</td><td>MM</td><td>target_seq</td><td>PAM</td><td>alignment</td><td>distance</td><td>position</td><td>gene name</td><td>gene id</td></tr>
<tr><td>GU305914.1</td><td></td><td>93747</td><td>93769</td><td>+</td><td>0</td><td>CCGCACCCTTCCCGGCTTCT</td><td>CGG</td><td>||||||||[||||||||||||]PAM</td><td>NA</td><td>Intergenic</td><td>NA</td><td>NA</td></tr>
<tr><td></td><td></td><td></td><td></td><td></td><td></td><td></td><td></td><td></td><td></td><td></td><td></td><td></td></tr>
<tr><td>T6687</td><td>CCCTTCCCGGCTTCTCGGCTCGG</td><td></td><td>1000</td><td></td><td></td><td></td><td></td><td></td><td></td><td></td><td></td><td></td></tr>
<tr><td>Oligo fwd</td><td>CACCCCCTTCCCGGCTTCTCGGCT</td><td></td><td></td><td></td><td></td><td></td><td></td><td></td><td></td><td></td><td></td><td></td></tr>
<tr><td>Oligo rev</td><td>AAACAGCCGAGAAGCCGGGAAGGG</td><td></td><td></td><td></td><td></td><td></td><td></td><td></td><td></td><td></td><td></td><td></td></tr>
<tr><td>Chromosome</td><td>start</td><td></td><td>end</td><td>strand</td><td>MM</td><td>target_seq</td><td>PAM</td><td>alignment</td><td>distance</td><td>position</td><td>gene name</td><td>gene id</td></tr>
<tr><td>GU305914.1</td><td></td><td>93752</td><td>93774</td><td>+</td><td>0</td><td>CCCTTCCCGGCTTCTCGGCT</td><td>CGG</td><td>||||||||[||||||||||||]PAM</td><td>NA</td><td>Intergenic</td><td>NA</td><td>NA</td></tr>
<tr><td></td><td></td><td></td><td></td><td></td><td></td><td></td><td></td><td></td><td></td><td></td><td></td><td></td></tr>
<tr><td>T6688</td><td>GGCTCGGCTCCCCAGCTTCCCGG</td><td></td><td>1000</td><td></td><td></td><td></td><td></td><td></td><td></td><td></td><td></td><td></td></tr>
<tr><td>Oligo fwd</td><td>CACCGGCTCGGCTCCCCAGCTTCC</td><td></td><td></td><td></td><td></td><td></td><td></td><td></td><td></td><td></td><td></td><td></td></tr>
<tr><td>Oligo rev</td><td>AAACGGAAGCTGGGGAGCCGAGCC</td><td></td><td></td><td></td><td></td><td></td><td></td><td></td><td></td><td></td><td></td><td></td></tr>
<tr><td>Chromosome</td><td>start</td><td></td><td>end</td><td>strand</td><td>MM</td><td>target_seq</td><td>PAM</td><td>alignment</td><td>distance</td><td>position</td><td>gene name</td><td>gene id</td></tr>
<tr><td>GU305914.1</td><td></td><td>93768</td><td>93790</td><td>+</td><td>0</td><td>GGCTCGGCTCCCCAGCTTCC</td><td>CGG</td><td>||||||||[||||||||||||]PAM</td><td>NA</td><td>Intergenic</td><td>NA</td><td>NA</td></tr>
<tr><td>GU305914.1</td><td></td><td>93643</td><td>93665</td><td>+</td><td>3</td><td>GCCTCGGCTCCCCGGCTTTC</td><td>CGG</td><td>|-||||||[|||||-||||-|]PAM</td><td>NA</td><td>Intergenic</td><td>NA</td><td>NA</td></tr>
<tr><td></td><td></td><td></td><td></td><td></td><td></td><td></td><td></td><td></td><td></td><td></td><td></td><td></td></tr>
<tr><td>T6689</td><td>AGCTTCCCGGCTACCCTCCTCGG</td><td></td><td>1000</td><td></td><td></td><td></td><td></td><td></td><td></td><td></td><td></td><td></td></tr>
<tr><td>Oligo fwd</td><td>CACCAGCTTCCCGGCTACCCTCCT</td><td></td><td></td><td></td><td></td><td></td><td></td><td></td><td></td><td></td><td></td><td></td></tr>
<tr><td>Oligo rev</td><td>AAACAGGAGGGTAGCCGGGAAGCT</td><td></td><td></td><td></td><td></td><td></td><td></td><td></td><td></td><td></td><td></td><td></td></tr>
<tr><td>Chromosome</td><td>start</td><td></td><td>end</td><td>strand</td><td>MM</td><td>target_seq</td><td>PAM</td><td>alignment</td><td>distance</td><td>position</td><td>gene name</td><td>gene id</td></tr>
<tr><td>GU305914.1</td><td></td><td>93781</td><td>93803</td><td>+</td><td>0</td><td>AGCTTCCCGGCTACCCTCCT</td><td>CGG</td><td>||||||||[||||||||||||]PAM</td><td>NA</td><td>Intergenic</td><td>NA</td><td>NA</td></tr>
<tr><td>GU305914.1</td><td></td><td>93554</td><td>93576</td><td>+</td><td>3</td><td>AGCTCCCCGGCTACCGGCCT</td><td>CGG</td><td>||||-|||[|||||||--|||]PAM</td><td>NA</td><td>Intergenic</td><td>NA</td><td>NA</td></tr>
<tr><td>GU305914.1</td><td></td><td>93627</td><td>93649</td><td>+</td><td>3</td><td>AGCTCCCCGGCTACCGGCCT</td><td>CGG</td><td>||||-|||[|||||||--|||]PAM</td><td>NA</td><td>Intergenic</td><td>NA</td><td>NA</td></tr>
<tr><td>GU305914.1</td><td></td><td>93720</td><td>93742</td><td>+</td><td>3</td><td>AGCTCCCCGGCTACCGACCT</td><td>CGG</td><td>||||-|||[|||||||--|||]PAM</td><td>NA</td><td>Intergenic</td><td>NA</td><td>NA</td></tr>
<tr><td></td><td></td><td></td><td></td><td></td><td></td><td></td><td></td><td></td><td></td><td></td><td></td><td></td></tr>
<tr><td>T6690</td><td>GGCTACCCTCCTCGGCTCACCGG</td><td></td><td>1000</td><td></td><td></td><td></td><td></td><td></td><td></td><td></td><td></td><td></td></tr>
<tr><td>Oligo fwd</td><td>CACCGGCTACCCTCCTCGGCTCAC</td><td></td><td></td><td></td><td></td><td></td><td></td><td></td><td></td><td></td><td></td><td></td></tr>
<tr><td>Oligo rev</td><td>AAACGTGAGCCGAGGAGGGTAGCC</td><td></td><td></td><td></td><td></td><td></td><td></td><td></td><td></td><td></td><td></td><td></td></tr>
<tr><td>Chromosome</td><td>start</td><td></td><td>end</td><td>strand</td><td>MM</td><td>target_seq</td><td>PAM</td><td>alignment</td><td>distance</td><td>position</td><td>gene name</td><td>gene id</td></tr>
<tr><td>GU305914.1</td><td></td><td>93789</td><td>93811</td><td>+</td><td>0</td><td>GGCTACCCTCCTCGGCTCAC</td><td>CGG</td><td>||||||||[||||||||||||]PAM</td><td>NA</td><td>Intergenic</td><td>NA</td><td>NA</td></tr>
<tr><td>GU305914.1</td><td></td><td>93685</td><td>93707</td><td>+</td><td>3</td><td>GCCTACCGACCTCGGCTCAC</td><td>CGG</td><td>|-|||||-[-|||||||||||]PAM</td><td>NA</td><td>Intergenic</td><td>NA</td><td>NA</td></tr>
<tr><td>GU305914.1</td><td></td><td>93562</td><td>93584</td><td>+</td><td>3</td><td>GGCTACCGGCCTCGGCTCCC</td><td>CGG</td><td>|||||||-[-|||||||||-|]PAM</td><td>NA</td><td>Intergenic</td><td>NA</td><td>NA</td></tr>
<tr><td>GU305914.1</td><td></td><td>93635</td><td>93657</td><td>+</td><td>3</td><td>GGCTACCGGCCTCGGCTCCC</td><td>CGG</td><td>|||||||-[-|||||||||-|]PAM</td><td>NA</td><td>Intergenic</td><td>NA</td><td>NA</td></tr>
<tr><td></td><td></td><td></td><td></td><td></td><td></td><td></td><td></td><td></td><td></td><td></td><td></td><td></td></tr>
<tr><td>T6691</td><td>CCTCGGCTCACCGGACCTCCAGG</td><td></td><td>1000</td><td></td><td></td><td></td><td></td><td></td><td></td><td></td><td></td><td></td></tr>
<tr><td>Oligo fwd</td><td>CACCCCTCGGCTCACCGGACCTCC</td><td></td><td></td><td></td><td></td><td></td><td></td><td></td><td></td><td></td><td></td><td></td></tr>
<tr><td>Oligo rev</td><td>AAACGGAGGTCCGGTGAGCCGAGG</td><td></td><td></td><td></td><td></td><td></td><td></td><td></td><td></td><td></td><td></td><td></td></tr>
<tr><td>Chromosome</td><td>start</td><td></td><td>end</td><td>strand</td><td>MM</td><td>target_seq</td><td>PAM</td><td>alignment</td><td>distance</td><td>position</td><td>gene name</td><td>gene id</td></tr>
<tr><td>GU305914.1</td><td></td><td>93798</td><td>93820</td><td>+</td><td>0</td><td>CCTCGGCTCACCGGACCTCC</td><td>AGG</td><td>||||||||[||||||||||||]PAM</td><td>NA</td><td>Intergenic</td><td>NA</td><td>NA</td></tr>
<tr><td>GU305914.1</td><td></td><td>93694</td><td>93716</td><td>+</td><td>2</td><td>CCTCGGCTCACCGGGCCCCC</td><td>AGG</td><td>||||||||[||||||-||-||]PAM</td><td>NA</td><td>Intergenic</td><td>NA</td><td>NA</td></tr>
<tr><td></td><td></td><td></td><td></td><td></td><td></td><td></td><td></td><td></td><td></td><td></td><td></td><td></td></tr>
<tr><td>T6692</td><td>GACCTCCAGGACCAGCTCCCCGG</td><td></td><td>1000</td><td></td><td></td><td></td><td></td><td></td><td></td><td></td><td></td><td></td></tr>
<tr><td>Oligo fwd</td><td>CACCGACCTCCAGGACCAGCTCCC</td><td></td><td></td><td></td><td></td><td></td><td></td><td></td><td></td><td></td><td></td><td></td></tr>
<tr><td>Oligo rev</td><td>AAACGGGAGCTGGTCCTGGAGGTC</td><td></td><td></td><td></td><td></td><td></td><td></td><td></td><td></td><td></td><td></td><td></td></tr>
<tr><td>Chromosome</td><td>start</td><td></td><td>end</td><td>strand</td><td>MM</td><td>target_seq</td><td>PAM</td><td>alignment</td><td>distance</td><td>position</td><td>gene name</td><td>gene id</td></tr>
<tr><td>GU305914.1</td><td></td><td>93811</td><td>93833</td><td>+</td><td>0</td><td>GACCTCCAGGACCAGCTCCC</td><td>CGG</td><td>||||||||[||||||||||||]PAM</td><td>NA</td><td>Intergenic</td><td>NA</td><td>NA</td></tr>
<tr><td>GU305914.1</td><td></td><td>93707</td><td>93729</td><td>+</td><td>2</td><td>GGCCCCCAGGACCAGCTCCC</td><td>CGG</td><td>|-||-|||[||||||||||||]PAM</td><td>NA</td><td>Intergenic</td><td>NA</td><td>NA</td></tr>
<tr><td>GU305914.1</td><td></td><td>93614</td><td>93636</td><td>+</td><td>3</td><td>GGCTTCCCGGACCAGCTCCC</td><td>CGG</td><td>|-|-|||-[||||||||||||]PAM</td><td>NA</td><td>Intergenic</td><td>NA</td><td>NA</td></tr>
<tr><td>GU305914.1</td><td></td><td>93541</td><td>93563</td><td>+</td><td>4</td><td>GGACCGCAGGACCAGCTCCC</td><td>CGG</td><td>|--|--||[||||||||||||]PAM</td><td>NA</td><td>Intergenic</td><td>NA</td><td>NA</td></tr>
<tr><td>GU305914.1</td><td></td><td>93583</td><td>93605</td><td>+</td><td>4</td><td>GGACCCCAGGACCAGCACCC</td><td>CGG</td><td>|--|-|||[||||||||-|||]PAM</td><td>NA</td><td>Intergenic</td><td>NA</td><td>NA</td></tr>
<tr><td></td><td></td><td></td><td></td><td></td><td></td><td></td><td></td><td></td><td></td><td></td><td></td><td></td></tr>
<tr><td>T6693</td><td>TCGATTCCCCGCACCCCGCTTGG</td><td></td><td>1000</td><td></td><td></td><td></td><td></td><td></td><td></td><td></td><td></td><td></td></tr>
<tr><td>Oligo fwd</td><td>CACCTCGATTCCCCGCACCCCGCT</td><td></td><td></td><td></td><td></td><td></td><td></td><td></td><td></td><td></td><td></td><td></td></tr>
<tr><td>Oligo rev</td><td>AAACAGCGGGGTGCGGGGAATCGA</td><td></td><td></td><td></td><td></td><td></td><td></td><td></td><td></td><td></td><td></td><td></td></tr>
<tr><td>Chromosome</td><td>start</td><td></td><td>end</td><td>strand</td><td>MM</td><td>target_seq</td><td>PAM</td><td>alignment</td><td>distance</td><td>position</td><td>gene name</td><td>gene id</td></tr>
<tr><td>GU305914.1</td><td></td><td>93835</td><td>93857</td><td>+</td><td>0</td><td>TCGATTCCCCGCACCCCGCT</td><td>TGG</td><td>||||||||[||||||||||||]PAM</td><td>NA</td><td>Intergenic</td><td>NA</td><td>NA</td></tr>
<tr><td></td><td></td><td></td><td></td><td></td><td></td><td></td><td></td><td></td><td></td><td></td><td></td><td></td></tr>
<tr><td>T6694</td><td>CCAGACCAGCGCCCCGCTCCCGG</td><td></td><td>1000</td><td></td><td></td><td></td><td></td><td></td><td></td><td></td><td></td><td></td></tr>
<tr><td>Oligo fwd</td><td>CACCCCAGACCAGCGCCCCGCTCC</td><td></td><td></td><td></td><td></td><td></td><td></td><td></td><td></td><td></td><td></td><td></td></tr>
<tr><td>Oligo rev</td><td>AAACGGAGCGGGGCGCTGGTCTGG</td><td></td><td></td><td></td><td></td><td></td><td></td><td></td><td></td><td></td><td></td><td></td></tr>
<tr><td>Chromosome</td><td>start</td><td></td><td>end</td><td>strand</td><td>MM</td><td>target_seq</td><td>PAM</td><td>alignment</td><td>distance</td><td>position</td><td>gene name</td><td>gene id</td></tr>
<tr><td>GU305914.1</td><td></td><td>93862</td><td>93884</td><td>+</td><td>0</td><td>CCAGACCAGCGCCCCGCTCC</td><td>CGG</td><td>||||||||[||||||||||||]PAM</td><td>NA</td><td>Intergenic</td><td>NA</td><td>NA</td></tr>
<tr><td></td><td></td><td></td><td></td><td></td><td></td><td></td><td></td><td></td><td></td><td></td><td></td><td></td></tr>
<tr><td>T6695</td><td>CCCCGCTCCCGGTTCCTGTCCGG</td><td></td><td>1000</td><td></td><td></td><td></td><td></td><td></td><td></td><td></td><td></td><td></td></tr>
<tr><td>Oligo fwd</td><td>CACCCCCCGCTCCCGGTTCCTGTC</td><td></td><td></td><td></td><td></td><td></td><td></td><td></td><td></td><td></td><td></td><td></td></tr>
<tr><td>Oligo rev</td><td>AAACGACAGGAACCGGGAGCGGGG</td><td></td><td></td><td></td><td></td><td></td><td></td><td></td><td></td><td></td><td></td><td></td></tr>
<tr><td>Chromosome</td><td>start</td><td></td><td>end</td><td>strand</td><td>MM</td><td>target_seq</td><td>PAM</td><td>alignment</td><td>distance</td><td>position</td><td>gene name</td><td>gene id</td></tr>
<tr><td>GU305914.1</td><td></td><td>93873</td><td>93895</td><td>+</td><td>0</td><td>CCCCGCTCCCGGTTCCTGTC</td><td>CGG</td><td>||||||||[||||||||||||]PAM</td><td>NA</td><td>Intergenic</td><td>NA</td><td>NA</td></tr>
<tr><td>GU305914.1</td><td></td><td>93937</td><td>93959</td><td>+</td><td>3</td><td>CCGAGCTCCCGGTCCCTGTC</td><td>CGG</td><td>||--||||[|||||-||||||]PAM</td><td>NA</td><td>Intergenic</td><td>NA</td><td>NA</td></tr>
<tr><td>GU305914.1</td><td></td><td>93905</td><td>93927</td><td>+</td><td>3</td><td>CCGAGCTCCCGGTCCCTGTC</td><td>CGG</td><td>||--||||[|||||-||||||]PAM</td><td>NA</td><td>Intergenic</td><td>NA</td><td>NA</td></tr>
<tr><td>GU305914.1</td><td></td><td>94065</td><td>94087</td><td>+</td><td>3</td><td>CCGAGCTCCCGGTCCCTGTC</td><td>CGG</td><td>||--||||[|||||-||||||]PAM</td><td>NA</td><td>Intergenic</td><td>NA</td><td>NA</td></tr>
<tr><td>GU305914.1</td><td></td><td>94033</td><td>94055</td><td>+</td><td>3</td><td>CCGAGCTCCCGGTCCCTGTC</td><td>CGG</td><td>||--||||[|||||-||||||]PAM</td><td>NA</td><td>Intergenic</td><td>NA</td><td>NA</td></tr>
<tr><td>GU305914.1</td><td></td><td>94001</td><td>94023</td><td>+</td><td>3</td><td>CCGAGCTCCCGGTCCCTGTC</td><td>CGG</td><td>||--||||[|||||-||||||]PAM</td><td>NA</td><td>Intergenic</td><td>NA</td><td>NA</td></tr>
<tr><td>GU305914.1</td><td></td><td>93969</td><td>93991</td><td>+</td><td>3</td><td>CCGAGCTCCCGGTCCCTGTC</td><td>CGG</td><td>||--||||[|||||-||||||]PAM</td><td>NA</td><td>Intergenic</td><td>NA</td><td>NA</td></tr>
<tr><td>GU305914.1</td><td></td><td>74003</td><td>74025</td><td>-</td><td>4</td><td>CCACTCTCGCTGTTCCTGTC</td><td>TGG</td><td>||-|-|||[-|-|||||||||]PAM</td><td>NA</td><td>Intergenic</td><td>NA</td><td>NA</td></tr>
</table>

| | | | | | | | | | | | |
|---|---|---|---|---|---|---|---|---|---|---|---|
| T6696 | GGCTCCTCTCCCCGAGCTCCCGG | 1000 | | | | | | | | | |
| Oligo fwd | CACCGGCTCCTCTCCCCGAGCTCC | | | | | | | | | | |
| Oligo rev | AAACGGAGCTCGGGGAGAGGAGCC | | | | | | | | | | |
| Chromosome | start | end | strand | MM | target_seq | PAM | alignment | distance | position | gene name | gene id |
| GU305914.1 | 93894 | 93916 | + | 0 | GGCTCCTCTCCCCGAGCTCC | CGG | \|\|\|\|\|\|\|\|[\|\|\|\|\|\|\|\|\|\|\|\|]PAM | NA | Intergenic | NA | NA |
| GU305914.1 | 93958 | 93980 | + | 2 | GGCTTCCCTCCCCGAGCTCC | CGG | \|\|\|\|-\|-\|[\|\|\|\|\|\|\|\|\|\|\|\|]PAM | NA | Intergenic | NA | NA |
| GU305914.1 | 93926 | 93948 | + | 2 | GGCTTCCCTCCCCGAGCTCC | CGG | \|\|\|\|-\|-\|[\|\|\|\|\|\|\|\|\|\|\|\|]PAM | NA | Intergenic | NA | NA |
| GU305914.1 | 94086 | 94108 | + | 2 | GGCTTCCCTCCCCGAGCTCC | CGG | \|\|\|\|-\|-\|[\|\|\|\|\|\|\|\|\|\|\|\|]PAM | NA | Intergenic | NA | NA |
| GU305914.1 | 94054 | 94076 | + | 2 | GGCTTCCCTCCCCGAGCTCC | CGG | \|\|\|\|-\|-\|[\|\|\|\|\|\|\|\|\|\|\|\|]PAM | NA | Intergenic | NA | NA |
| GU305914.1 | 94022 | 94044 | + | 2 | GGCTTCCCTCCCCGAGCTCC | CGG | \|\|\|\|-\|-\|[\|\|\|\|\|\|\|\|\|\|\|\|]PAM | NA | Intergenic | NA | NA |
| GU305914.1 | 93990 | 94012 | + | 2 | GGCTTCCCTCCCCGAGCTCC | CGG | \|\|\|\|-\|-\|[\|\|\|\|\|\|\|\|\|\|\|\|]PAM | NA | Intergenic | NA | NA |
| T6697 | CCGAGCTCCCGGTCCCTGTCCGG | 1000 | | | | | | | | | |
| Oligo fwd | CACCCCGAGCTCCCGGTCCCTGTC | | | | | | | | | | |
| Oligo rev | AAACGACAGGGACCGGGAGCTCGG | | | | | | | | | | |
| Chromosome | start | end | strand | MM | target_seq | PAM | alignment | distance | position | gene name | gene id |
| GU305914.1 | 94001 | 94023 | + | 0 | CCGAGCTCCCGGTCCCTGTC | CGG | \|\|\|\|\|\|\|\|[\|\|\|\|\|\|\|\|\|\|\|\|]PAM | NA | Intergenic | NA | NA |
| GU305914.1 | 93969 | 93991 | + | 0 | CCGAGCTCCCGGTCCCTGTC | CGG | \|\|\|\|\|\|\|\|[\|\|\|\|\|\|\|\|\|\|\|\|]PAM | NA | Intergenic | NA | NA |
| GU305914.1 | 93937 | 93959 | + | 0 | CCGAGCTCCCGGTCCCTGTC | CGG | \|\|\|\|\|\|\|\|[\|\|\|\|\|\|\|\|\|\|\|\|]PAM | NA | Intergenic | NA | NA |
| GU305914.1 | 93905 | 93927 | + | 0 | CCGAGCTCCCGGTCCCTGTC | CGG | \|\|\|\|\|\|\|\|[\|\|\|\|\|\|\|\|\|\|\|\|]PAM | NA | Intergenic | NA | NA |
| GU305914.1 | 94065 | 94087 | + | 0 | CCGAGCTCCCGGTCCCTGTC | CGG | \|\|\|\|\|\|\|\|[\|\|\|\|\|\|\|\|\|\|\|\|]PAM | NA | Intergenic | NA | NA |
| GU305914.1 | 94033 | 94055 | + | 0 | CCGAGCTCCCGGTCCCTGTC | CGG | \|\|\|\|\|\|\|\|[\|\|\|\|\|\|\|\|\|\|\|\|]PAM | NA | Intergenic | NA | NA |
| GU305914.1 | 93873 | 93895 | + | 3 | CCCCGCTCCCGGTTCCTGTC | CGG | \|\|--\|\|\|\|[\|\|\|\|\|-\|\|\|\|\|\|]PAM | NA | Intergenic | NA | NA |
| T6698 | GGCTTCCCTCCCCGAGCTCCCGG | 1000 | | | | | | | | | |
| Oligo fwd | CACCGGCTTCCCTCCCCGAGCTCC | | | | | | | | | | |
| Oligo rev | AAACGGAGCTCGGGGAGGGAAGCC | | | | | | | | | | |
| Chromosome | start | end | strand | MM | target_seq | PAM | alignment | distance | position | gene name | gene id |
| GU305914.1 | 94086 | 94108 | + | 0 | GGCTTCCCTCCCCGAGCTCC | CGG | \|\|\|\|\|\|\|\|[\|\|\|\|\|\|\|\|\|\|\|\|]PAM | NA | Intergenic | NA | NA |
| GU305914.1 | 94054 | 94076 | + | 0 | GGCTTCCCTCCCCGAGCTCC | CGG | \|\|\|\|\|\|\|\|[\|\|\|\|\|\|\|\|\|\|\|\|]PAM | NA | Intergenic | NA | NA |
| GU305914.1 | 94022 | 94044 | + | 0 | GGCTTCCCTCCCCGAGCTCC | CGG | \|\|\|\|\|\|\|\|[\|\|\|\|\|\|\|\|\|\|\|\|]PAM | NA | Intergenic | NA | NA |
| GU305914.1 | 93990 | 94012 | + | 0 | GGCTTCCCTCCCCGAGCTCC | CGG | \|\|\|\|\|\|\|\|[\|\|\|\|\|\|\|\|\|\|\|\|]PAM | NA | Intergenic | NA | NA |
| GU305914.1 | 93958 | 93980 | + | 0 | GGCTTCCCTCCCCGAGCTCC | CGG | \|\|\|\|\|\|\|\|[\|\|\|\|\|\|\|\|\|\|\|\|]PAM | NA | Intergenic | NA | NA |
| GU305914.1 | 93926 | 93948 | + | 0 | GGCTTCCCTCCCCGAGCTCC | CGG | \|\|\|\|\|\|\|\|[\|\|\|\|\|\|\|\|\|\|\|\|]PAM | NA | Intergenic | NA | NA |
| GU305914.1 | 93894 | 93916 | + | 2 | GGCTCCTCTCCCCGAGCTCC | CGG | \|\|\|\|-\|-\|[\|\|\|\|\|\|\|\|\|\|\|\|]PAM | NA | Intergenic | NA | NA |
| T6699 | CCGAGCTCCCGGTCCCTGTCCGG | 1000 | | | | | | | | | |
| Oligo fwd | CACCCCGAGCTCCCGGTCCCTGTC | | | | | | | | | | |
| Oligo rev | AAACGACAGGGACCGGGAGCTCGG | | | | | | | | | | |
| Chromosome | start | end | strand | MM | target_seq | PAM | alignment | distance | position | gene name | gene id |
| GU305914.1 | 94001 | 94023 | + | 0 | CCGAGCTCCCGGTCCCTGTC | CGG | \|\|\|\|\|\|\|\|[\|\|\|\|\|\|\|\|\|\|\|\|]PAM | NA | Intergenic | NA | NA |
| GU305914.1 | 93969 | 93991 | + | 0 | CCGAGCTCCCGGTCCCTGTC | CGG | \|\|\|\|\|\|\|\|[\|\|\|\|\|\|\|\|\|\|\|\|]PAM | NA | Intergenic | NA | NA |
| GU305914.1 | 93937 | 93959 | + | 0 | CCGAGCTCCCGGTCCCTGTC | CGG | \|\|\|\|\|\|\|\|[\|\|\|\|\|\|\|\|\|\|\|\|]PAM | NA | Intergenic | NA | NA |
| GU305914.1 | 93905 | 93927 | + | 0 | CCGAGCTCCCGGTCCCTGTC | CGG | \|\|\|\|\|\|\|\|[\|\|\|\|\|\|\|\|\|\|\|\|]PAM | NA | Intergenic | NA | NA |
| GU305914.1 | 94065 | 94087 | + | 0 | CCGAGCTCCCGGTCCCTGTC | CGG | \|\|\|\|\|\|\|\|[\|\|\|\|\|\|\|\|\|\|\|\|]PAM | NA | Intergenic | NA | NA |
| GU305914.1 | 94033 | 94055 | + | 0 | CCGAGCTCCCGGTCCCTGTC | CGG | \|\|\|\|\|\|\|\|[\|\|\|\|\|\|\|\|\|\|\|\|]PAM | NA | Intergenic | NA | NA |
| GU305914.1 | 93873 | 93895 | + | 3 | CCCCGCTCCCGGTTCCTGTC | CGG | \|\|--\|\|\|\|[\|\|\|\|\|-\|\|\|\|\|\|]PAM | NA | Intergenic | NA | NA |
| T6700 | GGCTTCCCTCCCCGAGCTCCCGG | 1000 | | | | | | | | | |
| Oligo fwd | CACCGGCTTCCCTCCCCGAGCTCC | | | | | | | | | | |
| Oligo rev | AAACGGAGCTCGGGGAGGGAAGCC | | | | | | | | | | |
| Chromosome | start | end | strand | MM | target_seq | PAM | alignment | distance | position | gene name | gene id |
| GU305914.1 | 94086 | 94108 | + | 0 | GGCTTCCCTCCCCGAGCTCC | CGG | \|\|\|\|\|\|\|\|[\|\|\|\|\|\|\|\|\|\|\|\|]PAM | NA | Intergenic | NA | NA |
| GU305914.1 | 94054 | 94076 | + | 0 | GGCTTCCCTCCCCGAGCTCC | CGG | \|\|\|\|\|\|\|\|[\|\|\|\|\|\|\|\|\|\|\|\|]PAM | NA | Intergenic | NA | NA |
| GU305914.1 | 94022 | 94044 | + | 0 | GGCTTCCCTCCCCGAGCTCC | CGG | \|\|\|\|\|\|\|\|[\|\|\|\|\|\|\|\|\|\|\|\|]PAM | NA | Intergenic | NA | NA |
| GU305914.1 | 93990 | 94012 | + | 0 | GGCTTCCCTCCCCGAGCTCC | CGG | \|\|\|\|\|\|\|\|[\|\|\|\|\|\|\|\|\|\|\|\|]PAM | NA | Intergenic | NA | NA |
| GU305914.1 | 93958 | 93980 | + | 0 | GGCTTCCCTCCCCGAGCTCC | CGG | \|\|\|\|\|\|\|\|[\|\|\|\|\|\|\|\|\|\|\|\|]PAM | NA | Intergenic | NA | NA |
| GU305914.1 | 93926 | 93948 | + | 0 | GGCTTCCCTCCCCGAGCTCC | CGG | \|\|\|\|\|\|\|\|[\|\|\|\|\|\|\|\|\|\|\|\|]PAM | NA | Intergenic | NA | NA |
| GU305914.1 | 93894 | 93916 | + | 2 | GGCTCCTCTCCCCGAGCTCC | CGG | \|\|\|\|-\|-\|[\|\|\|\|\|\|\|\|\|\|\|\|]PAM | NA | Intergenic | NA | NA |
| T6701 | CCGAGCTCCCGGTCCCTGTCCGG | 1000 | | | | | | | | | |
| Oligo fwd | CACCCCGAGCTCCCGGTCCCTGTC | | | | | | | | | | |
| Oligo rev | AAACGACAGGGACCGGGAGCTCGG | | | | | | | | | | |
| Chromosome | start | end | strand | MM | target_seq | PAM | alignment | distance | position | gene name | gene id |
| GU305914.1 | 94001 | 94023 | + | 0 | CCGAGCTCCCGGTCCCTGTC | CGG | \|\|\|\|\|\|\|\|[\|\|\|\|\|\|\|\|\|\|\|\|]PAM | NA | Intergenic | NA | NA |
| GU305914.1 | 93969 | 93991 | + | 0 | CCGAGCTCCCGGTCCCTGTC | CGG | \|\|\|\|\|\|\|\|[\|\|\|\|\|\|\|\|\|\|\|\|]PAM | NA | Intergenic | NA | NA |
| GU305914.1 | 93937 | 93959 | + | 0 | CCGAGCTCCCGGTCCCTGTC | CGG | \|\|\|\|\|\|\|\|[\|\|\|\|\|\|\|\|\|\|\|\|]PAM | NA | Intergenic | NA | NA |
| GU305914.1 | 93905 | 93927 | + | 0 | CCGAGCTCCCGGTCCCTGTC | CGG | \|\|\|\|\|\|\|\|[\|\|\|\|\|\|\|\|\|\|\|\|]PAM | NA | Intergenic | NA | NA |
| GU305914.1 | 94065 | 94087 | + | 0 | CCGAGCTCCCGGTCCCTGTC | CGG | \|\|\|\|\|\|\|\|[\|\|\|\|\|\|\|\|\|\|\|\|]PAM | NA | Intergenic | NA | NA |
| GU305914.1 | 94033 | 94055 | + | 0 | CCGAGCTCCCGGTCCCTGTC | CGG | \|\|\|\|\|\|\|\|[\|\|\|\|\|\|\|\|\|\|\|\|]PAM | NA | Intergenic | NA | NA |
| GU305914.1 | 93873 | 93895 | + | 3 | CCCCGCTCCCGGTTCCTGTC | CGG | \|\|--\|\|\|\|[\|\|\|\|\|-\|\|\|\|\|\|]PAM | NA | Intergenic | NA | NA |
| T6702 | GGCTTCCCTCCCCGAGCTCCCGG | 1000 | | | | | | | | | |
| Oligo fwd | CACCGGCTTCCCTCCCCGAGCTCC | | | | | | | | | | |
| Oligo rev | AAACGGAGCTCGGGGAGGGAAGCC | | | | | | | | | | |
| Chromosome | start | end | strand | MM | target_seq | PAM | alignment | distance | position | gene name | gene id |
| GU305914.1 | 94086 | 94108 | + | 0 | GGCTTCCCTCCCCGAGCTCC | CGG | \|\|\|\|\|\|\|\|[\|\|\|\|\|\|\|\|\|\|\|\|]PAM | NA | Intergenic | NA | NA |
| GU305914.1 | 94054 | 94076 | + | 0 | GGCTTCCCTCCCCGAGCTCC | CGG | \|\|\|\|\|\|\|\|[\|\|\|\|\|\|\|\|\|\|\|\|]PAM | NA | Intergenic | NA | NA |
| GU305914.1 | 94022 | 94044 | + | 0 | GGCTTCCCTCCCCGAGCTCC | CGG | \|\|\|\|\|\|\|\|[\|\|\|\|\|\|\|\|\|\|\|\|]PAM | NA | Intergenic | NA | NA |
| GU305914.1 | 93990 | 94012 | + | 0 | GGCTTCCCTCCCCGAGCTCC | CGG | \|\|\|\|\|\|\|\|[\|\|\|\|\|\|\|\|\|\|\|\|]PAM | NA | Intergenic | NA | NA |
| GU305914.1 | 93958 | 93980 | + | 0 | GGCTTCCCTCCCCGAGCTCC | CGG | \|\|\|\|\|\|\|\|[\|\|\|\|\|\|\|\|\|\|\|\|]PAM | NA | Intergenic | NA | NA |
| GU305914.1 | 93926 | 93948 | + | 0 | GGCTTCCCTCCCCGAGCTCC | CGG | \|\|\|\|\|\|\|\|[\|\|\|\|\|\|\|\|\|\|\|\|]PAM | NA | Intergenic | NA | NA |
| GU305914.1 | 93894 | 93916 | + | 2 | GGCTCCTCTCCCCGAGCTCC | CGG | \|\|\|\|-\|-\|[\|\|\|\|\|\|\|\|\|\|\|\|]PAM | NA | Intergenic | NA | NA |
| T6703 | CCGAGCTCCCGGTCCCTGTCCGG | 1000 | | | | | | | | | |
| Oligo fwd | CACCCCGAGCTCCCGGTCCCTGTC | | | | | | | | | | |
| Oligo rev | AAACGACAGGGACCGGGAGCTCGG | | | | | | | | | | |
| Chromosome | start | end | strand | MM | target_seq | PAM | alignment | distance | position | gene name | gene id |
| GU305914.1 | 94001 | 94023 | + | 0 | CCGAGCTCCCGGTCCCTGTC | CGG | \|\|\|\|\|\|\|\|[\|\|\|\|\|\|\|\|\|\|\|\|]PAM | NA | Intergenic | NA | NA |

| | | | | | | | | | | | |
|---|---|---|---|---|---|---|---|---|---|---|---|
| GU305914.1 | 93969 | 93991 | + | 0 | CCGAGCTCCCGGTCCCTGTC | CGG | \|\|\|\|\|\|\|\|[\|\|\|\|\|\|\|\|\|\|\|\|]PAM | NA | Intergenic | NA | NA |
| GU305914.1 | 93937 | 93959 | + | 0 | CCGAGCTCCCGGTCCCTGTC | CGG | \|\|\|\|\|\|\|\|[\|\|\|\|\|\|\|\|\|\|\|\|]PAM | NA | Intergenic | NA | NA |
| GU305914.1 | 93905 | 93927 | + | 0 | CCGAGCTCCCGGTCCCTGTC | CGG | \|\|\|\|\|\|\|\|[\|\|\|\|\|\|\|\|\|\|\|\|]PAM | NA | Intergenic | NA | NA |
| GU305914.1 | 94065 | 94087 | + | 0 | CCGAGCTCCCGGTCCCTGTC | CGG | \|\|\|\|\|\|\|\|[\|\|\|\|\|\|\|\|\|\|\|\|]PAM | NA | Intergenic | NA | NA |
| GU305914.1 | 94033 | 94055 | + | 0 | CCGAGCTCCCGGTCCCTGTC | CGG | \|\|\|\|\|\|\|\|[\|\|\|\|\|\|\|\|\|\|\|\|]PAM | NA | Intergenic | NA | NA |
| GU305914.1 | 93873 | 93895 | + | 3 | CCCCGCTCCCGGTTCCTGTC | CGG | \|\|--\|\|\|\|[\|\|\|\|\|-\|\|\|\|\|\|]PAM | NA | Intergenic | NA | NA |

T6704 GGCTTCCCTCCCCGAGCTCCCGG 1000
Oligo fwd CACCGGCTTCCCTCCCCGAGCTCC
Oligo rev AAACGGAGCTCGGGGAGGGAAGCC

| Chromosome | start | end | strand | MM | target_seq | PAM | alignment | distance | position | gene name | gene id |
|---|---|---|---|---|---|---|---|---|---|---|---|
| GU305914.1 | 94086 | 94108 | + | 0 | GGCTTCCCTCCCCGAGCTCC | CGG | \|\|\|\|\|\|\|\|[\|\|\|\|\|\|\|\|\|\|\|\|]PAM | NA | Intergenic | NA | NA |
| GU305914.1 | 94054 | 94076 | + | 0 | GGCTTCCCTCCCCGAGCTCC | CGG | \|\|\|\|\|\|\|\|[\|\|\|\|\|\|\|\|\|\|\|\|]PAM | NA | Intergenic | NA | NA |
| GU305914.1 | 94022 | 94044 | + | 0 | GGCTTCCCTCCCCGAGCTCC | CGG | \|\|\|\|\|\|\|\|[\|\|\|\|\|\|\|\|\|\|\|\|]PAM | NA | Intergenic | NA | NA |
| GU305914.1 | 93990 | 94012 | + | 0 | GGCTTCCCTCCCCGAGCTCC | CGG | \|\|\|\|\|\|\|\|[\|\|\|\|\|\|\|\|\|\|\|\|]PAM | NA | Intergenic | NA | NA |
| GU305914.1 | 93958 | 93980 | + | 0 | GGCTTCCCTCCCCGAGCTCC | CGG | \|\|\|\|\|\|\|\|[\|\|\|\|\|\|\|\|\|\|\|\|]PAM | NA | Intergenic | NA | NA |
| GU305914.1 | 93926 | 93948 | + | 0 | GGCTTCCCTCCCCGAGCTCC | CGG | \|\|\|\|\|\|\|\|[\|\|\|\|\|\|\|\|\|\|\|\|]PAM | NA | Intergenic | NA | NA |
| GU305914.1 | 93894 | 93916 | + | 2 | GGCTCCTCTCCCCGAGCTCC | CGG | \|\|\|\|-\|-\|[\|\|\|\|\|\|\|\|\|\|\|\|]PAM | NA | Intergenic | NA | NA |

T6705 CCGAGCTCCCGGTCCCTGTCCGG 1000
Oligo fwd CACCCCGAGCTCCCGGTCCCTGTC
Oligo rev AAACGACAGGGACCGGGAGCTCGG

| Chromosome | start | end | strand | MM | target_seq | PAM | alignment | distance | position | gene name | gene id |
|---|---|---|---|---|---|---|---|---|---|---|---|
| GU305914.1 | 94001 | 94023 | + | 0 | CCGAGCTCCCGGTCCCTGTC | CGG | \|\|\|\|\|\|\|\|[\|\|\|\|\|\|\|\|\|\|\|\|]PAM | NA | Intergenic | NA | NA |
| GU305914.1 | 93969 | 93991 | + | 0 | CCGAGCTCCCGGTCCCTGTC | CGG | \|\|\|\|\|\|\|\|[\|\|\|\|\|\|\|\|\|\|\|\|]PAM | NA | Intergenic | NA | NA |
| GU305914.1 | 93937 | 93959 | + | 0 | CCGAGCTCCCGGTCCCTGTC | CGG | \|\|\|\|\|\|\|\|[\|\|\|\|\|\|\|\|\|\|\|\|]PAM | NA | Intergenic | NA | NA |
| GU305914.1 | 93905 | 93927 | + | 0 | CCGAGCTCCCGGTCCCTGTC | CGG | \|\|\|\|\|\|\|\|[\|\|\|\|\|\|\|\|\|\|\|\|]PAM | NA | Intergenic | NA | NA |
| GU305914.1 | 94065 | 94087 | + | 0 | CCGAGCTCCCGGTCCCTGTC | CGG | \|\|\|\|\|\|\|\|[\|\|\|\|\|\|\|\|\|\|\|\|]PAM | NA | Intergenic | NA | NA |
| GU305914.1 | 94033 | 94055 | + | 0 | CCGAGCTCCCGGTCCCTGTC | CGG | \|\|\|\|\|\|\|\|[\|\|\|\|\|\|\|\|\|\|\|\|]PAM | NA | Intergenic | NA | NA |
| GU305914.1 | 93873 | 93895 | + | 3 | CCCCGCTCCCGGTTCCTGTC | CGG | \|\|--\|\|\|\|[\|\|\|\|\|-\|\|\|\|\|\|]PAM | NA | Intergenic | NA | NA |

T6706 GGCTTCCCTCCCCGAGCTCCCGG 1000
Oligo fwd CACCGGCTTCCCTCCCCGAGCTCC
Oligo rev AAACGGAGCTCGGGGAGGGAAGCC

| Chromosome | start | end | strand | MM | target_seq | PAM | alignment | distance | position | gene name | gene id |
|---|---|---|---|---|---|---|---|---|---|---|---|
| GU305914.1 | 94086 | 94108 | + | 0 | GGCTTCCCTCCCCGAGCTCC | CGG | \|\|\|\|\|\|\|\|[\|\|\|\|\|\|\|\|\|\|\|\|]PAM | NA | Intergenic | NA | NA |
| GU305914.1 | 94054 | 94076 | + | 0 | GGCTTCCCTCCCCGAGCTCC | CGG | \|\|\|\|\|\|\|\|[\|\|\|\|\|\|\|\|\|\|\|\|]PAM | NA | Intergenic | NA | NA |
| GU305914.1 | 94022 | 94044 | + | 0 | GGCTTCCCTCCCCGAGCTCC | CGG | \|\|\|\|\|\|\|\|[\|\|\|\|\|\|\|\|\|\|\|\|]PAM | NA | Intergenic | NA | NA |
| GU305914.1 | 93990 | 94012 | + | 0 | GGCTTCCCTCCCCGAGCTCC | CGG | \|\|\|\|\|\|\|\|[\|\|\|\|\|\|\|\|\|\|\|\|]PAM | NA | Intergenic | NA | NA |
| GU305914.1 | 93958 | 93980 | + | 0 | GGCTTCCCTCCCCGAGCTCC | CGG | \|\|\|\|\|\|\|\|[\|\|\|\|\|\|\|\|\|\|\|\|]PAM | NA | Intergenic | NA | NA |
| GU305914.1 | 93926 | 93948 | + | 0 | GGCTTCCCTCCCCGAGCTCC | CGG | \|\|\|\|\|\|\|\|[\|\|\|\|\|\|\|\|\|\|\|\|]PAM | NA | Intergenic | NA | NA |
| GU305914.1 | 93894 | 93916 | + | 2 | GGCTCCTCTCCCCGAGCTCC | CGG | \|\|\|\|-\|-\|[\|\|\|\|\|\|\|\|\|\|\|\|]PAM | NA | Intergenic | NA | NA |

T6707 CCGAGCTCCCGGTCCCTGTCCGG 1000
Oligo fwd CACCCCGAGCTCCCGGTCCCTGTC
Oligo rev AAACGACAGGGACCGGGAGCTCGG

| Chromosome | start | end | strand | MM | target_seq | PAM | alignment | distance | position | gene name | gene id |
|---|---|---|---|---|---|---|---|---|---|---|---|
| GU305914.1 | 94001 | 94023 | + | 0 | CCGAGCTCCCGGTCCCTGTC | CGG | \|\|\|\|\|\|\|\|[\|\|\|\|\|\|\|\|\|\|\|\|]PAM | NA | Intergenic | NA | NA |
| GU305914.1 | 93969 | 93991 | + | 0 | CCGAGCTCCCGGTCCCTGTC | CGG | \|\|\|\|\|\|\|\|[\|\|\|\|\|\|\|\|\|\|\|\|]PAM | NA | Intergenic | NA | NA |
| GU305914.1 | 93937 | 93959 | + | 0 | CCGAGCTCCCGGTCCCTGTC | CGG | \|\|\|\|\|\|\|\|[\|\|\|\|\|\|\|\|\|\|\|\|]PAM | NA | Intergenic | NA | NA |
| GU305914.1 | 93905 | 93927 | + | 0 | CCGAGCTCCCGGTCCCTGTC | CGG | \|\|\|\|\|\|\|\|[\|\|\|\|\|\|\|\|\|\|\|\|]PAM | NA | Intergenic | NA | NA |
| GU305914.1 | 94065 | 94087 | + | 0 | CCGAGCTCCCGGTCCCTGTC | CGG | \|\|\|\|\|\|\|\|[\|\|\|\|\|\|\|\|\|\|\|\|]PAM | NA | Intergenic | NA | NA |
| GU305914.1 | 94033 | 94055 | + | 0 | CCGAGCTCCCGGTCCCTGTC | CGG | \|\|\|\|\|\|\|\|[\|\|\|\|\|\|\|\|\|\|\|\|]PAM | NA | Intergenic | NA | NA |
| GU305914.1 | 93873 | 93895 | + | 3 | CCCCGCTCCCGGTTCCTGTC | CGG | \|\|--\|\|\|\|[\|\|\|\|\|-\|\|\|\|\|\|]PAM | NA | Intergenic | NA | NA |

T6708 GGCTTCCCTCCCCGAGCTCCCGG 1000
Oligo fwd CACCGGCTTCCCTCCCCGAGCTCC
Oligo rev AAACGGAGCTCGGGGAGGGAAGCC

| Chromosome | start | end | strand | MM | target_seq | PAM | alignment | distance | position | gene name | gene id |
|---|---|---|---|---|---|---|---|---|---|---|---|
| GU305914.1 | 94086 | 94108 | + | 0 | GGCTTCCCTCCCCGAGCTCC | CGG | \|\|\|\|\|\|\|\|[\|\|\|\|\|\|\|\|\|\|\|\|]PAM | NA | Intergenic | NA | NA |
| GU305914.1 | 94054 | 94076 | + | 0 | GGCTTCCCTCCCCGAGCTCC | CGG | \|\|\|\|\|\|\|\|[\|\|\|\|\|\|\|\|\|\|\|\|]PAM | NA | Intergenic | NA | NA |
| GU305914.1 | 94022 | 94044 | + | 0 | GGCTTCCCTCCCCGAGCTCC | CGG | \|\|\|\|\|\|\|\|[\|\|\|\|\|\|\|\|\|\|\|\|]PAM | NA | Intergenic | NA | NA |
| GU305914.1 | 93990 | 94012 | + | 0 | GGCTTCCCTCCCCGAGCTCC | CGG | \|\|\|\|\|\|\|\|[\|\|\|\|\|\|\|\|\|\|\|\|]PAM | NA | Intergenic | NA | NA |
| GU305914.1 | 93958 | 93980 | + | 0 | GGCTTCCCTCCCCGAGCTCC | CGG | \|\|\|\|\|\|\|\|[\|\|\|\|\|\|\|\|\|\|\|\|]PAM | NA | Intergenic | NA | NA |
| GU305914.1 | 93926 | 93948 | + | 0 | GGCTTCCCTCCCCGAGCTCC | CGG | \|\|\|\|\|\|\|\|[\|\|\|\|\|\|\|\|\|\|\|\|]PAM | NA | Intergenic | NA | NA |
| GU305914.1 | 93894 | 93916 | + | 2 | GGCTCCTCTCCCCGAGCTCC | CGG | \|\|\|\|-\|-\|[\|\|\|\|\|\|\|\|\|\|\|\|]PAM | NA | Intergenic | NA | NA |

T6709 CGCTTCATTGCACCCTGCCCAGG 1000
Oligo fwd CACCCGCTTCATTGCACCCTGCCC
Oligo rev AAACGGGCAGGGTGCAATGAAGCG

| Chromosome | start | end | strand | MM | target_seq | PAM | alignment | distance | position | gene name | gene id |
|---|---|---|---|---|---|---|---|---|---|---|---|
| GU305914.1 | 94113 | 94135 | + | 0 | CGCTTCATTGCACCCTGCCC | AGG | \|\|\|\|\|\|\|\|[\|\|\|\|\|\|\|\|\|\|\|\|]PAM | NA | Intergenic | NA | NA |
| GU305914.1 | 94209 | 94231 | + | 3 | GGCTTCGTTGCTCCCTGCCC | AGG | -\|\|\|\|\|-\|[\|\|\|-\|\|\|\|\|\|\|\|]PAM | NA | Intergenic | NA | NA |

T6710 CCCGATTGCCGTTCTCCGAGCGG 1000
Oligo fwd CACCCCCGATTGCCGTTCTCCGAG
Oligo rev AAACCTCGGAGAACGGCAATCGGG

| Chromosome | start | end | strand | MM | target_seq | PAM | alignment | distance | position | gene name | gene id |
|---|---|---|---|---|---|---|---|---|---|---|---|
| GU305914.1 | 94137 | 94159 | + | 0 | CCCGATTGCCGTTCTCCGAG | CGG | \|\|\|\|\|\|\|\|[\|\|\|\|\|\|\|\|\|\|\|\|]PAM | NA | Intergenic | NA | NA |

T6711 GAGCGGCTCGATGCTCCGTTCGG 1000
Oligo fwd CACCGAGCGGCTCGATGCTCCGTT
Oligo rev AAACAACGGAGCATCGAGCCGCTC

| Chromosome | start | end | strand | MM | target_seq | PAM | alignment | distance | position | gene name | gene id |
|---|---|---|---|---|---|---|---|---|---|---|---|
| GU305914.1 | 94154 | 94176 | + | 0 | GAGCGGCTCGATGCTCCGTT | CGG | \|\|\|\|\|\|\|\|[\|\|\|\|\|\|\|\|\|\|\|\|]PAM | NA | Intergenic | NA | NA |

T6712 CGATGCTCCGTTCGGCACCCTGG 1000
Oligo fwd CACCCGATGCTCCGTTCGGCACCC
Oligo rev AAACGGGTGCCGAACGGAGCATCG

<table>
<tr><td>Chromosome</td><td>start</td><td>end</td><td>strand</td><td>MM</td><td>target_seq</td><td>PAM</td><td>alignment</td><td>distance</td><td>position</td><td>gene name</td><td>gene id</td></tr>
<tr><td>GU305914.1</td><td>94162</td><td>94184</td><td>+</td><td>0</td><td>CGATGCTCCGTTCGGCACCC</td><td>TGG</td><td>||||||||[||||||||||||]PAM</td><td>NA</td><td>Intergenic</td><td>NA</td><td>NA</td></tr>
<tr><td></td><td></td><td></td><td></td><td></td><td></td><td></td><td></td><td></td><td></td><td></td><td></td></tr>
<tr><td>T6713</td><td>GCTCCGTTCGGCACCCTGGCCGG</td><td>1000</td><td></td><td></td><td></td><td></td><td></td><td></td><td></td><td></td><td></td></tr>
<tr><td>Oligo fwd</td><td>CACCGCTCCGTTCGGCACCCTGGC</td><td></td><td></td><td></td><td></td><td></td><td></td><td></td><td></td><td></td><td></td></tr>
<tr><td>Oligo rev</td><td>AAACGCCAGGGTGCCGAACGGAGC</td><td></td><td></td><td></td><td></td><td></td><td></td><td></td><td></td><td></td><td></td></tr>
<tr><td>Chromosome</td><td>start</td><td>end</td><td>strand</td><td>MM</td><td>target_seq</td><td>PAM</td><td>alignment</td><td>distance</td><td>position</td><td>gene name</td><td>gene id</td></tr>
<tr><td>GU305914.1</td><td>94166</td><td>94188</td><td>+</td><td>0</td><td>GCTCCGTTCGGCACCCTGGC</td><td>CGG</td><td>||||||||[||||||||||||]PAM</td><td>NA</td><td>Intergenic</td><td>NA</td><td>NA</td></tr>
<tr><td></td><td></td><td></td><td></td><td></td><td></td><td></td><td></td><td></td><td></td><td></td><td></td></tr>
<tr><td>T6714</td><td>CGGCACCCTGGCCGGCTTTTCGG</td><td>1000</td><td></td><td></td><td></td><td></td><td></td><td></td><td></td><td></td><td></td></tr>
<tr><td>Oligo fwd</td><td>CACCCGGCACCCTGGCCGGCTTTT</td><td></td><td></td><td></td><td></td><td></td><td></td><td></td><td></td><td></td><td></td></tr>
<tr><td>Oligo rev</td><td>AAACAAAAGCCGGCCAGGGTGCCG</td><td></td><td></td><td></td><td></td><td></td><td></td><td></td><td></td><td></td><td></td></tr>
<tr><td>Chromosome</td><td>start</td><td>end</td><td>strand</td><td>MM</td><td>target_seq</td><td>PAM</td><td>alignment</td><td>distance</td><td>position</td><td>gene name</td><td>gene id</td></tr>
<tr><td>GU305914.1</td><td>94174</td><td>94196</td><td>+</td><td>0</td><td>CGGCACCCTGGCCGGCTTTT</td><td>CGG</td><td>||||||||[||||||||||||]PAM</td><td>NA</td><td>Intergenic</td><td>NA</td><td>NA</td></tr>
<tr><td></td><td></td><td></td><td></td><td></td><td></td><td></td><td></td><td></td><td></td><td></td><td></td></tr>
<tr><td>T6715</td><td>GGCACCCTGGCCGGCTTTTCGGG</td><td>1000</td><td></td><td></td><td></td><td></td><td></td><td></td><td></td><td></td><td></td></tr>
<tr><td>Oligo fwd</td><td>CACCGGCACCCTGGCCGGCTTTTC</td><td></td><td></td><td></td><td></td><td></td><td></td><td></td><td></td><td></td><td></td></tr>
<tr><td>Oligo rev</td><td>AAACGAAAAGCCGGCCAGGGTGCC</td><td></td><td></td><td></td><td></td><td></td><td></td><td></td><td></td><td></td><td></td></tr>
<tr><td>Chromosome</td><td>start</td><td>end</td><td>strand</td><td>MM</td><td>target_seq</td><td>PAM</td><td>alignment</td><td>distance</td><td>position</td><td>gene name</td><td>gene id</td></tr>
<tr><td>GU305914.1</td><td>94175</td><td>94197</td><td>+</td><td>0</td><td>GGCACCCTGGCCGGCTTTTC</td><td>GGG</td><td>||||||||[||||||||||||]PAM</td><td>NA</td><td>Intergenic</td><td>NA</td><td>NA</td></tr>
<tr><td></td><td></td><td></td><td></td><td></td><td></td><td></td><td></td><td></td><td></td><td></td><td></td></tr>
<tr><td>T6716</td><td>GCACCCTGGCCGGCTTTTCGGGG</td><td>1000</td><td></td><td></td><td></td><td></td><td></td><td></td><td></td><td></td><td></td></tr>
<tr><td>Oligo fwd</td><td>CACCGCACCCTGGCCGGCTTTTCG</td><td></td><td></td><td></td><td></td><td></td><td></td><td></td><td></td><td></td><td></td></tr>
<tr><td>Oligo rev</td><td>AAACCGAAAAGCCGGCCAGGGTGC</td><td></td><td></td><td></td><td></td><td></td><td></td><td></td><td></td><td></td><td></td></tr>
<tr><td>Chromosome</td><td>start</td><td>end</td><td>strand</td><td>MM</td><td>target_seq</td><td>PAM</td><td>alignment</td><td>distance</td><td>position</td><td>gene name</td><td>gene id</td></tr>
<tr><td>GU305914.1</td><td>94176</td><td>94198</td><td>+</td><td>0</td><td>GCACCCTGGCCGGCTTTTCG</td><td>GGG</td><td>||||||||[||||||||||||]PAM</td><td>NA</td><td>Intergenic</td><td>NA</td><td>NA</td></tr>
<tr><td></td><td></td><td></td><td></td><td></td><td></td><td></td><td></td><td></td><td></td><td></td><td></td></tr>
<tr><td>T6717</td><td>TGGCCGGCTTTTCGGGGTCCCGG</td><td>1000</td><td></td><td></td><td></td><td></td><td></td><td></td><td></td><td></td><td></td></tr>
<tr><td>Oligo fwd</td><td>CACCTGGCCGGCTTTTCGGGGTCC</td><td></td><td></td><td></td><td></td><td></td><td></td><td></td><td></td><td></td><td></td></tr>
<tr><td>Oligo rev</td><td>AAACGGACCCCGAAAAGCCGGCCA</td><td></td><td></td><td></td><td></td><td></td><td></td><td></td><td></td><td></td><td></td></tr>
<tr><td>Chromosome</td><td>start</td><td>end</td><td>strand</td><td>MM</td><td>target_seq</td><td>PAM</td><td>alignment</td><td>distance</td><td>position</td><td>gene name</td><td>gene id</td></tr>
<tr><td>GU305914.1</td><td>94182</td><td>94204</td><td>+</td><td>0</td><td>TGGCCGGCTTTTCGGGGTCC</td><td>CGG</td><td>||||||||[||||||||||||]PAM</td><td>NA</td><td>Intergenic</td><td>NA</td><td>NA</td></tr>
<tr><td></td><td></td><td></td><td></td><td></td><td></td><td></td><td></td><td></td><td></td><td></td><td></td></tr>
<tr><td>T6718</td><td>GCTTTTCGGGGTCCCGGTTCCGG</td><td>1000</td><td></td><td></td><td></td><td></td><td></td><td></td><td></td><td></td><td></td></tr>
<tr><td>Oligo fwd</td><td>CACCGCTTTTCGGGGTCCCGGTTC</td><td></td><td></td><td></td><td></td><td></td><td></td><td></td><td></td><td></td><td></td></tr>
<tr><td>Oligo rev</td><td>AAACGAACCGGGACCCCGAAAAGC</td><td></td><td></td><td></td><td></td><td></td><td></td><td></td><td></td><td></td><td></td></tr>
<tr><td>Chromosome</td><td>start</td><td>end</td><td>strand</td><td>MM</td><td>target_seq</td><td>PAM</td><td>alignment</td><td>distance</td><td>position</td><td>gene name</td><td>gene id</td></tr>
<tr><td>GU305914.1</td><td>94188</td><td>94210</td><td>+</td><td>0</td><td>GCTTTTCGGGGTCCCGGTTC</td><td>CGG</td><td>||||||||[||||||||||||]PAM</td><td>NA</td><td>Intergenic</td><td>NA</td><td>NA</td></tr>
<tr><td></td><td></td><td></td><td></td><td></td><td></td><td></td><td></td><td></td><td></td><td></td><td></td></tr>
<tr><td>T6719</td><td>GGCTTCGTTGCTCCCTGCCCAGG</td><td>1000</td><td></td><td></td><td></td><td></td><td></td><td></td><td></td><td></td><td></td></tr>
<tr><td>Oligo fwd</td><td>CACCGGCTTCGTTGCTCCCTGCCC</td><td></td><td></td><td></td><td></td><td></td><td></td><td></td><td></td><td></td><td></td></tr>
<tr><td>Oligo rev</td><td>AAACGGGCAGGGAGCAACGAAGCC</td><td></td><td></td><td></td><td></td><td></td><td></td><td></td><td></td><td></td><td></td></tr>
<tr><td>Chromosome</td><td>start</td><td>end</td><td>strand</td><td>MM</td><td>target_seq</td><td>PAM</td><td>alignment</td><td>distance</td><td>position</td><td>gene name</td><td>gene id</td></tr>
<tr><td>GU305914.1</td><td>94209</td><td>94231</td><td>+</td><td>0</td><td>GGCTTCGTTGCTCCCTGCCC</td><td>AGG</td><td>||||||||[||||||||||||]PAM</td><td>NA</td><td>Intergenic</td><td>NA</td><td>NA</td></tr>
<tr><td>GU305914.1</td><td>94113</td><td>94135</td><td>+</td><td>3</td><td>CGCTTCATTGCACCCTGCCC</td><td>AGG</td><td>-|||||-|[|||-||||||||]PAM</td><td>NA</td><td>Intergenic</td><td>NA</td><td>NA</td></tr>
<tr><td></td><td></td><td></td><td></td><td></td><td></td><td></td><td></td><td></td><td></td><td></td><td></td></tr>
<tr><td>T6720</td><td>GTTGCTCCCTGCCCAGGTCCCGG</td><td>1000</td><td></td><td></td><td></td><td></td><td></td><td></td><td></td><td></td><td></td></tr>
<tr><td>Oligo fwd</td><td>CACCGTTGCTCCCTGCCCAGGTCC</td><td></td><td></td><td></td><td></td><td></td><td></td><td></td><td></td><td></td><td></td></tr>
<tr><td>Oligo rev</td><td>AAACGGACCTGGGCAGGGAGCAAC</td><td></td><td></td><td></td><td></td><td></td><td></td><td></td><td></td><td></td><td></td></tr>
<tr><td>Chromosome</td><td>start</td><td>end</td><td>strand</td><td>MM</td><td>target_seq</td><td>PAM</td><td>alignment</td><td>distance</td><td>position</td><td>gene name</td><td>gene id</td></tr>
<tr><td>GU305914.1</td><td>94215</td><td>94237</td><td>+</td><td>0</td><td>GTTGCTCCCTGCCCAGGTCC</td><td>CGG</td><td>||||||||[||||||||||||]PAM</td><td>NA</td><td>Intergenic</td><td>NA</td><td>NA</td></tr>
<tr><td></td><td></td><td></td><td></td><td></td><td></td><td></td><td></td><td></td><td></td><td></td><td></td></tr>
<tr><td>T6721</td><td>TCTGCCTCTCGCCTTTCGCTCGG</td><td>1000</td><td></td><td></td><td></td><td></td><td></td><td></td><td></td><td></td><td></td></tr>
<tr><td>Oligo fwd</td><td>CACCTCTGCCTCTCGCCTTTCGCT</td><td></td><td></td><td></td><td></td><td></td><td></td><td></td><td></td><td></td><td></td></tr>
<tr><td>Oligo rev</td><td>AAACAGCGAAAGGCGAGAGGCAGA</td><td></td><td></td><td></td><td></td><td></td><td></td><td></td><td></td><td></td><td></td></tr>
<tr><td>Chromosome</td><td>start</td><td>end</td><td>strand</td><td>MM</td><td>target_seq</td><td>PAM</td><td>alignment</td><td>distance</td><td>position</td><td>gene name</td><td>gene id</td></tr>
<tr><td>GU305914.1</td><td>94270</td><td>94292</td><td>+</td><td>0</td><td>TCTGCCTCTCGCCTTTCGCT</td><td>CGG</td><td>||||||||[||||||||||||]PAM</td><td>NA</td><td>Intergenic</td><td>NA</td><td>NA</td></tr>
<tr><td></td><td></td><td></td><td></td><td></td><td></td><td></td><td></td><td></td><td></td><td></td><td></td></tr>
<tr><td>T6722</td><td>TTTGTGTTTTAGCTTGCCGTTGG</td><td>1000</td><td></td><td></td><td></td><td></td><td></td><td></td><td></td><td></td><td></td></tr>
<tr><td>Oligo fwd</td><td>CACCTTTGTGTTTTAGCTTGCCGT</td><td></td><td></td><td></td><td></td><td></td><td></td><td></td><td></td><td></td><td></td></tr>
<tr><td>Oligo rev</td><td>AAACACGGCAAGCTAAAACACAAA</td><td></td><td></td><td></td><td></td><td></td><td></td><td></td><td></td><td></td><td></td></tr>
<tr><td>Chromosome</td><td>start</td><td>end</td><td>strand</td><td>MM</td><td>target_seq</td><td>PAM</td><td>alignment</td><td>distance</td><td>position</td><td>gene name</td><td>gene id</td></tr>
<tr><td>GU305914.1</td><td>94344</td><td>94366</td><td>+</td><td>0</td><td>TTTGTGTTTTAGCTTGCCGT</td><td>TGG</td><td>||||||||[||||||||||||]PAM</td><td>NA</td><td>Intergenic</td><td>NA</td><td>NA</td></tr>
<tr><td></td><td></td><td></td><td></td><td></td><td></td><td></td><td></td><td></td><td></td><td></td><td></td></tr>
<tr><td>T6723</td><td>GTGTTTTAGCTTGCCGTTGGCGG</td><td>1000</td><td></td><td></td><td></td><td></td><td></td><td></td><td></td><td></td><td></td></tr>
<tr><td>Oligo fwd</td><td>CACCGTGTTTTAGCTTGCCGTTGG</td><td></td><td></td><td></td><td></td><td></td><td></td><td></td><td></td><td></td><td></td></tr>
<tr><td>Oligo rev</td><td>AAACCCAACGGCAAGCTAAAACAC</td><td></td><td></td><td></td><td></td><td></td><td></td><td></td><td></td><td></td><td></td></tr>
<tr><td>Chromosome</td><td>start</td><td>end</td><td>strand</td><td>MM</td><td>target_seq</td><td>PAM</td><td>alignment</td><td>distance</td><td>position</td><td>gene name</td><td>gene id</td></tr>
<tr><td>GU305914.1</td><td>94347</td><td>94369</td><td>+</td><td>0</td><td>GTGTTTTAGCTTGCCGTTGG</td><td>CGG</td><td>||||||||[||||||||||||]PAM</td><td>NA</td><td>Intergenic</td><td>NA</td><td>NA</td></tr>
<tr><td></td><td></td><td></td><td></td><td></td><td></td><td></td><td></td><td></td><td></td><td></td><td></td></tr>
<tr><td>T6724</td><td>ATTTTGTCGAGTTTTCGTTTAGG</td><td>1000</td><td></td><td></td><td></td><td></td><td></td><td></td><td></td><td></td><td></td></tr>
<tr><td>Oligo fwd</td><td>CACCATTTTGTCGAGTTTTCGTTT</td><td></td><td></td><td></td><td></td><td></td><td></td><td></td><td></td><td></td><td></td></tr>
<tr><td>Oligo rev</td><td>AAACAAACGAAAACTCGACAAAAT</td><td></td><td></td><td></td><td></td><td></td><td></td><td></td><td></td><td></td><td></td></tr>
<tr><td>Chromosome</td><td>start</td><td>end</td><td>strand</td><td>MM</td><td>target_seq</td><td>PAM</td><td>alignment</td><td>distance</td><td>position</td><td>gene name</td><td>gene id</td></tr>
<tr><td>GU305914.1</td><td>94372</td><td>94394</td><td>+</td><td>0</td><td>ATTTTGTCGAGTTTTCGTTT</td><td>AGG</td><td>||||||||[||||||||||||]PAM</td><td>NA</td><td>Intergenic</td><td>NA</td><td>NA</td></tr>
<tr><td></td><td></td><td></td><td></td><td></td><td></td><td></td><td></td><td></td><td></td><td></td><td></td></tr>
<tr><td>T6725</td><td>TTGTCGAGTTTTCGTTTAGGCGG</td><td>1000</td><td></td><td></td><td></td><td></td><td></td><td></td><td></td><td></td><td></td></tr>
<tr><td>Oligo fwd</td><td>CACCTTGTCGAGTTTTCGTTTAGG</td><td></td><td></td><td></td><td></td><td></td><td></td><td></td><td></td><td></td><td></td></tr>
<tr><td>Oligo rev</td><td>AAACCCTAAACGAAAACTCGACAA</td><td></td><td></td><td></td><td></td><td></td><td></td><td></td><td></td><td></td><td></td></tr>
<tr><td>Chromosome</td><td>start</td><td>end</td><td>strand</td><td>MM</td><td>target_seq</td><td>PAM</td><td>alignment</td><td>distance</td><td>position</td><td>gene name</td><td>gene id</td></tr>
<tr><td>GU305914.1</td><td>94375</td><td>94397</td><td>+</td><td>0</td><td>TTGTCGAGTTTTCGTTTAGG</td><td>CGG</td><td>||||||||[||||||||||||]PAM</td><td>NA</td><td>Intergenic</td><td>NA</td><td>NA</td></tr>
<tr><td></td><td></td><td></td><td></td><td></td><td></td><td></td><td></td><td></td><td></td><td></td><td></td></tr>
<tr><td>T6726</td><td>TGTCGAGTTTTCGTTTAGGCGGG</td><td>1000</td><td></td><td></td><td></td><td></td><td></td><td></td><td></td><td></td><td></td></tr>
<tr><td>Oligo fwd</td><td>CACCTGTCGAGTTTTCGTTTAGGC</td><td></td><td></td><td></td><td></td><td></td><td></td><td></td><td></td><td></td><td></td></tr>
<tr><td>Oligo rev</td><td>AAACGCCTAAACGAAAACTCGACA</td><td></td><td></td><td></td><td></td><td></td><td></td><td></td><td></td><td></td><td></td></tr>
<tr><td>Chromosome</td><td>start</td><td>end</td><td>strand</td><td>MM</td><td>target_seq</td><td>PAM</td><td>alignment</td><td>distance</td><td>position</td><td>gene name</td><td>gene id</td></tr>
<tr><td>GU305914.1</td><td>94376</td><td>94398</td><td>+</td><td>0</td><td>TGTCGAGTTTTCGTTTAGGC</td><td>GGG</td><td>||||||||[||||||||||||]PAM</td><td>NA</td><td>Intergenic</td><td>NA</td><td>NA</td></tr>
<tr><td></td><td></td><td></td><td></td><td></td><td></td><td></td><td></td><td></td><td></td><td></td><td></td></tr>
<tr><td>T6727</td><td>TTTCGTTTAGGCGGGAAAATTGG</td><td>1000</td><td></td><td></td><td></td><td></td><td></td><td></td><td></td><td></td><td></td></tr>
</table>

| | | | | | | | | | | | |
|---|---|---|---|---|---|---|---|---|---|---|---|
| Oligo fwd | CACCTTTCGTTTAGGCGGGAAAAT | | | | | | | | | | |
| Oligo rev | AAACATTTTCCCGCCTAAACGAAA | | | | | | | | | | |
| Chromosome | start | end | strand | MM | target_seq | PAM | alignment | distance | position | gene name | gene id |
| GU305914.1 | 94384 | 94406 | + | 0 | TTTCGTTTAGGCGGGAAAAT | TGG | \|\|\|\|\|\|\|\|[\|\|\|\|\|\|\|\|\|\|\|\|]PAM | NA | Intergenic | NA | NA |
| T6728 | GCGGGAAAATTGGCGCCATTTGG | 1000 | | | | | | | | | |
| Oligo fwd | CACCGCGGGAAAATTGGCGCCATT | | | | | | | | | | |
| Oligo rev | AAACAATGGCGCCAATTTTCCCGC | | | | | | | | | | |
| Chromosome | start | end | strand | MM | target_seq | PAM | alignment | distance | position | gene name | gene id |
| GU305914.1 | 94394 | 94416 | + | 0 | GCGGGAAAATTGGCGCCATT | TGG | \|\|\|\|\|\|\|\|[\|\|\|\|\|\|\|\|\|\|\|\|]PAM | NA | Intergenic | NA | NA |
| T6729 | ATTTGGTCCCGACGCCATTTTGG | 1000 | | | | | | | | | |
| Oligo fwd | CACCATTTGGTCCCGACGCCATTT | | | | | | | | | | |
| Oligo rev | AAACAAATGGCGTCGGGACCAAAT | | | | | | | | | | |
| Chromosome | start | end | strand | MM | target_seq | PAM | alignment | distance | position | gene name | gene id |
| GU305914.1 | 94411 | 94433 | + | 0 | ATTTGGTCCCGACGCCATTT | TGG | \|\|\|\|\|\|\|\|[\|\|\|\|\|\|\|\|\|\|\|\|]PAM | NA | Intergenic | NA | NA |
| T6730 | TGGTCCCGACGCCATTTTGGTGG | 1000 | | | | | | | | | |
| Oligo fwd | CACCTGGTCCCGACGCCATTTTGG | | | | | | | | | | |
| Oligo rev | AAACCCAAAATGGCGTCGGGACCA | | | | | | | | | | |
| Chromosome | start | end | strand | MM | target_seq | PAM | alignment | distance | position | gene name | gene id |
| GU305914.1 | 94414 | 94436 | + | 0 | TGGTCCCGACGCCATTTTGG | TGG | \|\|\|\|\|\|\|\|[\|\|\|\|\|\|\|\|\|\|\|\|]PAM | NA | Intergenic | NA | NA |
| T6731 | TTTAACGTTCGCACAAGCGCTGG | 1000 | | | | | | | | | |
| Oligo fwd | CACCTTTAACGTTCGCACAAGCGC | | | | | | | | | | |
| Oligo rev | AAACGCGCTTGTGCGAACGTTAAA | | | | | | | | | | |
| Chromosome | start | end | strand | MM | target_seq | PAM | alignment | distance | position | gene name | gene id |
| GU305914.1 | 94446 | 94468 | + | 0 | TTTAACGTTCGCACAAGCGC | TGG | \|\|\|\|\|\|\|\|[\|\|\|\|\|\|\|\|\|\|\|\|]PAM | NA | Intergenic | NA | NA |
| T6732 | ATCTCACTTTCCTCTTTCAAAGG | 1000 | | | | | | | | | |
| Oligo fwd | CACCATCTCACTTTCCTCTTTCAA | | | | | | | | | | |
| Oligo rev | AAACTTGAAAGAGGAAAGTGAGAT | | | | | | | | | | |
| Chromosome | start | end | strand | MM | target_seq | PAM | alignment | distance | position | gene name | gene id |
| GU305914.1 | 94533 | 94555 | + | 0 | ATCTCACTTTCCTCTTTCAA | AGG | \|\|\|\|\|\|\|\|[\|\|\|\|\|\|\|\|\|\|\|\|]PAM | NA | Intergenic | NA | NA |
| T6733 | TCTCACTTTCCTCTTTCAAAGGG | 1000 | | | | | | | | | |
| Oligo fwd | CACCTCTCACTTTCCTCTTTCAAA | | | | | | | | | | |
| Oligo rev | AAACTTTGAAAGAGGAAAGTGAGA | | | | | | | | | | |
| Chromosome | start | end | strand | MM | target_seq | PAM | alignment | distance | position | gene name | gene id |
| GU305914.1 | 94534 | 94556 | + | 0 | TCTCACTTTCCTCTTTCAAA | GGG | \|\|\|\|\|\|\|\|[\|\|\|\|\|\|\|\|\|\|\|\|]PAM | NA | Intergenic | NA | NA |
| T6734 | CTCACTTTCCTCTTTCAAAGGGG | 1000 | | | | | | | | | |
| Oligo fwd | CACCCTCACTTTCCTCTTTCAAAG | | | | | | | | | | |
| Oligo rev | AAACCTTTGAAAGAGGAAAGTGAG | | | | | | | | | | |
| Chromosome | start | end | strand | MM | target_seq | PAM | alignment | distance | position | gene name | gene id |
| GU305914.1 | 94535 | 94557 | + | 0 | CTCACTTTCCTCTTTCAAAG | GGG | \|\|\|\|\|\|\|\|[\|\|\|\|\|\|\|\|\|\|\|\|]PAM | NA | Intergenic | NA | NA |
| T6735 | TCACTTTCCTCTTTCAAAGGGGG | 1000 | | | | | | | | | |
| Oligo fwd | CACCTCACTTTCCTCTTTCAAAGG | | | | | | | | | | |
| Oligo rev | AAACCCTTTGAAAGAGGAAAGTGA | | | | | | | | | | |
| Chromosome | start | end | strand | MM | target_seq | PAM | alignment | distance | position | gene name | gene id |
| GU305914.1 | 94536 | 94558 | + | 0 | TCACTTTCCTCTTTCAAAGG | GGG | \|\|\|\|\|\|\|\|[\|\|\|\|\|\|\|\|\|\|\|\|]PAM | NA | Intergenic | NA | NA |
| T6736 | CACTTTCCTCTTTCAAAGGGGGG | 1000 | | | | | | | | | |
| Oligo fwd | CACCCACTTTCCTCTTTCAAAGGG | | | | | | | | | | |
| Oligo rev | AAACCCCTTTGAAAGAGGAAAGTG | | | | | | | | | | |
| Chromosome | start | end | strand | MM | target_seq | PAM | alignment | distance | position | gene name | gene id |
| GU305914.1 | 94537 | 94559 | + | 0 | CACTTTCCTCTTTCAAAGGG | GGG | \|\|\|\|\|\|\|\|[\|\|\|\|\|\|\|\|\|\|\|\|]PAM | NA | Intergenic | NA | NA |
| T6737 | TTTCAAAGGGGGGTCCGCAGAGG | 1000 | | | | | | | | | |
| Oligo fwd | CACCTTTCAAAGGGGGGTCCGCAG | | | | | | | | | | |
| Oligo rev | AAACCTGCGGACCCCCCTTTGAAA | | | | | | | | | | |
| Chromosome | start | end | strand | MM | target_seq | PAM | alignment | distance | position | gene name | gene id |
| GU305914.1 | 94547 | 94569 | + | 0 | TTTCAAAGGGGGGTCCGCAG | AGG | \|\|\|\|\|\|\|\|[\|\|\|\|\|\|\|\|\|\|\|\|]PAM | NA | Intergenic | NA | NA |
| T6738 | TTCAAAGGGGGGTCCGCAGAGGG | 1000 | | | | | | | | | |
| Oligo fwd | CACCTTCAAAGGGGGGTCCGCAGA | | | | | | | | | | |
| Oligo rev | AAACTCTGCGGACCCCCCTTTGAA | | | | | | | | | | |
| Chromosome | start | end | strand | MM | target_seq | PAM | alignment | distance | position | gene name | gene id |
| GU305914.1 | 94548 | 94570 | + | 0 | TTCAAAGGGGGGTCCGCAGA | GGG | \|\|\|\|\|\|\|\|[\|\|\|\|\|\|\|\|\|\|\|\|]PAM | NA | Intergenic | NA | NA |
| T6739 | GGGGGGTCCGCAGAGGGTCTAGG | 1000 | | | | | | | | | |
| Oligo fwd | CACCGGGGGGTCCGCAGAGGGTCT | | | | | | | | | | |
| Oligo rev | AAACAGACCCTCTGCGGACCCCCC | | | | | | | | | | |
| Chromosome | start | end | strand | MM | target_seq | PAM | alignment | distance | position | gene name | gene id |
| GU305914.1 | 94554 | 94576 | + | 0 | GGGGGGTCCGCAGAGGGTCT | AGG | \|\|\|\|\|\|\|\|[\|\|\|\|\|\|\|\|\|\|\|\|]PAM | NA | Intergenic | NA | NA |
| T6740 | CAGAGGGTCTAGGTCGCTGTCGG | 1000 | | | | | | | | | |
| Oligo fwd | CACCCAGAGGGTCTAGGTCGCTGT | | | | | | | | | | |
| Oligo rev | AAACACAGCGACCTAGACCCTCTG | | | | | | | | | | |
| Chromosome | start | end | strand | MM | target_seq | PAM | alignment | distance | position | gene name | gene id |
| GU305914.1 | 94564 | 94586 | + | 0 | CAGAGGGTCTAGGTCGCTGT | CGG | \|\|\|\|\|\|\|\|[\|\|\|\|\|\|\|\|\|\|\|\|]PAM | NA | Intergenic | NA | NA |
| T6741 | GTCTAGGTCGCTGTCGGCCGAGG | 1000 | | | | | | | | | |
| Oligo fwd | CACCGTCTAGGTCGCTGTCGGCCG | | | | | | | | | | |
| Oligo rev | AAACCGGCCGACAGCGACCTAGAC | | | | | | | | | | |
| Chromosome | start | end | strand | MM | target_seq | PAM | alignment | distance | position | gene name | gene id |
| GU305914.1 | 94570 | 94592 | + | 0 | GTCTAGGTCGCTGTCGGCCG | AGG | \|\|\|\|\|\|\|\|[\|\|\|\|\|\|\|\|\|\|\|\|]PAM | NA | Intergenic | NA | NA |

<table>
<tr><td>T6742</td><td>GAGGCACTGCTCCTTTTGAGCGG</td><td>1000</td><td></td><td></td><td></td><td></td><td></td><td></td><td></td><td></td><td></td></tr>
<tr><td>Oligo fwd</td><td>CACCGAGGCACTGCTCCTTTTGAG</td><td></td><td></td><td></td><td></td><td></td><td></td><td></td><td></td><td></td><td></td></tr>
<tr><td>Oligo rev</td><td>AAACCTCAAAAGGAGCAGTGCCTC</td><td></td><td></td><td></td><td></td><td></td><td></td><td></td><td></td><td></td><td></td></tr>
<tr><td>Chromosome</td><td>start</td><td>end</td><td>strand</td><td>MM</td><td>target_seq</td><td>PAM</td><td>alignment</td><td>distance</td><td>position</td><td>gene name</td><td>gene id</td></tr>
<tr><td>GU305914.1</td><td>94589</td><td>94611</td><td>+</td><td>0</td><td>GAGGCACTGCTCCTTTTGAG</td><td>CGG</td><td>||||||||[||||||||||||]PAM</td><td>NA</td><td>Intergenic</td><td>NA</td><td>NA</td></tr>
<tr><td>T6743</td><td>AAGATGACATACTCTCCCTGTGG</td><td>1000</td><td></td><td></td><td></td><td></td><td></td><td></td><td></td><td></td><td></td></tr>
<tr><td>Oligo fwd</td><td>CACCAAGATGACATACTCTCCCTG</td><td></td><td></td><td></td><td></td><td></td><td></td><td></td><td></td><td></td><td></td></tr>
<tr><td>Oligo rev</td><td>AAACCAGGGAGAGTATGTCATCTT</td><td></td><td></td><td></td><td></td><td></td><td></td><td></td><td></td><td></td><td></td></tr>
<tr><td>Chromosome</td><td>start</td><td>end</td><td>strand</td><td>MM</td><td>target_seq</td><td>PAM</td><td>alignment</td><td>distance</td><td>position</td><td>gene name</td><td>gene id</td></tr>
<tr><td>GU305914.1</td><td>94651</td><td>94673</td><td>+</td><td>0</td><td>AAGATGACATACTCTCCCTG</td><td>TGG</td><td>||||||||[||||||||||||]PAM</td><td>NA</td><td>Intergenic</td><td>NA</td><td>NA</td></tr>
<tr><td>T6744</td><td>CCTGTGGCTTTGAAAAGATCAGG</td><td>1000</td><td></td><td></td><td></td><td></td><td></td><td></td><td></td><td></td><td></td></tr>
<tr><td>Oligo fwd</td><td>CACCCCTGTGGCTTTGAAAAGATC</td><td></td><td></td><td></td><td></td><td></td><td></td><td></td><td></td><td></td><td></td></tr>
<tr><td>Oligo rev</td><td>AAACGATCTTTTCAAAGCCACAGG</td><td></td><td></td><td></td><td></td><td></td><td></td><td></td><td></td><td></td><td></td></tr>
<tr><td>Chromosome</td><td>start</td><td>end</td><td>strand</td><td>MM</td><td>target_seq</td><td>PAM</td><td>alignment</td><td>distance</td><td>position</td><td>gene name</td><td>gene id</td></tr>
<tr><td>GU305914.1</td><td>94667</td><td>94689</td><td>+</td><td>0</td><td>CCTGTGGCTTTGAAAAGATC</td><td>AGG</td><td>||||||||[||||||||||||]PAM</td><td>NA</td><td>Intergenic</td><td>NA</td><td>NA</td></tr>
<tr><td>GU305914.1</td><td>133427</td><td>133449</td><td>+</td><td>4</td><td>ACTGTCGTTTTGAAAACATC</td><td>TGG</td><td>-||||-|-[||||||||-|||]PAM</td><td>NA</td><td>Intergenic</td><td>NA</td><td>NA</td></tr>
<tr><td>T6745</td><td>GTGGCTTTGAAAAGATCAGGCGG</td><td>1000</td><td></td><td></td><td></td><td></td><td></td><td></td><td></td><td></td><td></td></tr>
<tr><td>Oligo fwd</td><td>CACCGTGGCTTTGAAAAGATCAGG</td><td></td><td></td><td></td><td></td><td></td><td></td><td></td><td></td><td></td><td></td></tr>
<tr><td>Oligo rev</td><td>AAACCCTGATCTTTTCAAAGCCAC</td><td></td><td></td><td></td><td></td><td></td><td></td><td></td><td></td><td></td><td></td></tr>
<tr><td>Chromosome</td><td>start</td><td>end</td><td>strand</td><td>MM</td><td>target_seq</td><td>PAM</td><td>alignment</td><td>distance</td><td>position</td><td>gene name</td><td>gene id</td></tr>
<tr><td>GU305914.1</td><td>94670</td><td>94692</td><td>+</td><td>0</td><td>GTGGCTTTGAAAAGATCAGG</td><td>CGG</td><td>||||||||[||||||||||||]PAM</td><td>NA</td><td>Intergenic</td><td>NA</td><td>NA</td></tr>
<tr><td>GU305914.1</td><td>133430</td><td>133452</td><td>+</td><td>4</td><td>GTCGTTTTGAAAACATCTGG</td><td>AGG</td><td>||-|-|||[|||||-|||-||]PAM</td><td>NA</td><td>Intergenic</td><td>NA</td><td>NA</td></tr>
<tr><td>T6746</td><td>ATCAGGCGGTTATCTCCGTTCGG</td><td>1000</td><td></td><td></td><td></td><td></td><td></td><td></td><td></td><td></td><td></td></tr>
<tr><td>Oligo fwd</td><td>CACCATCAGGCGGTTATCTCCGTT</td><td></td><td></td><td></td><td></td><td></td><td></td><td></td><td></td><td></td><td></td></tr>
<tr><td>Oligo rev</td><td>AAACAACGGAGATAACCGCCTGAT</td><td></td><td></td><td></td><td></td><td></td><td></td><td></td><td></td><td></td><td></td></tr>
<tr><td>Chromosome</td><td>start</td><td>end</td><td>strand</td><td>MM</td><td>target_seq</td><td>PAM</td><td>alignment</td><td>distance</td><td>position</td><td>gene name</td><td>gene id</td></tr>
<tr><td>GU305914.1</td><td>94684</td><td>94706</td><td>+</td><td>0</td><td>ATCAGGCGGTTATCTCCGTT</td><td>CGG</td><td>||||||||[||||||||||||]PAM</td><td>NA</td><td>Intergenic</td><td>NA</td><td>NA</td></tr>
<tr><td>T6747</td><td>AGCTAACCAGACCTCGAAAAAGG</td><td>1000</td><td></td><td></td><td></td><td></td><td></td><td></td><td></td><td></td><td></td></tr>
<tr><td>Oligo fwd</td><td>CACCAGCTAACCAGACCTCGAAAA</td><td></td><td></td><td></td><td></td><td></td><td></td><td></td><td></td><td></td><td></td></tr>
<tr><td>Oligo rev</td><td>AAACTTTTCGAGGTCTGGTTAGCT</td><td></td><td></td><td></td><td></td><td></td><td></td><td></td><td></td><td></td><td></td></tr>
<tr><td>Chromosome</td><td>start</td><td>end</td><td>strand</td><td>MM</td><td>target_seq</td><td>PAM</td><td>alignment</td><td>distance</td><td>position</td><td>gene name</td><td>gene id</td></tr>
<tr><td>GU305914.1</td><td>94733</td><td>94755</td><td>+</td><td>0</td><td>AGCTAACCAGACCTCGAAAA</td><td>AGG</td><td>||||||||[||||||||||||]PAM</td><td>NA</td><td>Intergenic</td><td>NA</td><td>NA</td></tr>
<tr><td>T6748</td><td>TGACTCTTTTGAGAAGCATATGG</td><td>1000</td><td></td><td></td><td></td><td></td><td></td><td></td><td></td><td></td><td></td></tr>
<tr><td>Oligo fwd</td><td>CACCTGACTCTTTTGAGAAGCATA</td><td></td><td></td><td></td><td></td><td></td><td></td><td></td><td></td><td></td><td></td></tr>
<tr><td>Oligo rev</td><td>AAACTATGCTTCTCAAAAGAGTCA</td><td></td><td></td><td></td><td></td><td></td><td></td><td></td><td></td><td></td><td></td></tr>
<tr><td>Chromosome</td><td>start</td><td>end</td><td>strand</td><td>MM</td><td>target_seq</td><td>PAM</td><td>alignment</td><td>distance</td><td>position</td><td>gene name</td><td>gene id</td></tr>
<tr><td>GU305914.1</td><td>94764</td><td>94786</td><td>+</td><td>0</td><td>TGACTCTTTTGAGAAGCATA</td><td>TGG</td><td>||||||||[||||||||||||]PAM</td><td>NA</td><td>Intergenic</td><td>NA</td><td>NA</td></tr>
<tr><td>T6749</td><td>GACTCTTTTGAGAAGCATATGGG</td><td>1000</td><td></td><td></td><td></td><td></td><td></td><td></td><td></td><td></td><td></td></tr>
<tr><td>Oligo fwd</td><td>CACCGACTCTTTTGAGAAGCATAT</td><td></td><td></td><td></td><td></td><td></td><td></td><td></td><td></td><td></td><td></td></tr>
<tr><td>Oligo rev</td><td>AAACATATGCTTCTCAAAAGAGTC</td><td></td><td></td><td></td><td></td><td></td><td></td><td></td><td></td><td></td><td></td></tr>
<tr><td>Chromosome</td><td>start</td><td>end</td><td>strand</td><td>MM</td><td>target_seq</td><td>PAM</td><td>alignment</td><td>distance</td><td>position</td><td>gene name</td><td>gene id</td></tr>
<tr><td>GU305914.1</td><td>94765</td><td>94787</td><td>+</td><td>0</td><td>GACTCTTTTGAGAAGCATAT</td><td>GGG</td><td>||||||||[||||||||||||]PAM</td><td>NA</td><td>Intergenic</td><td>NA</td><td>NA</td></tr>
<tr><td>T6750</td><td>TCTTTTGAGAAGCATATGGGTGG</td><td>1000</td><td></td><td></td><td></td><td></td><td></td><td></td><td></td><td></td><td></td></tr>
<tr><td>Oligo fwd</td><td>CACCTCTTTTGAGAAGCATATGGG</td><td></td><td></td><td></td><td></td><td></td><td></td><td></td><td></td><td></td><td></td></tr>
<tr><td>Oligo rev</td><td>AAACCCCATATGCTTCTCAAAAGA</td><td></td><td></td><td></td><td></td><td></td><td></td><td></td><td></td><td></td><td></td></tr>
<tr><td>Chromosome</td><td>start</td><td>end</td><td>strand</td><td>MM</td><td>target_seq</td><td>PAM</td><td>alignment</td><td>distance</td><td>position</td><td>gene name</td><td>gene id</td></tr>
<tr><td>GU305914.1</td><td>94768</td><td>94790</td><td>+</td><td>0</td><td>TCTTTTGAGAAGCATATGGG</td><td>TGG</td><td>||||||||[||||||||||||]PAM</td><td>NA</td><td>Intergenic</td><td>NA</td><td>NA</td></tr>
<tr><td>T6751</td><td>GGGTGGACGTGCCCGTTGATAGG</td><td>1000</td><td></td><td></td><td></td><td></td><td></td><td></td><td></td><td></td><td></td></tr>
<tr><td>Oligo fwd</td><td>CACCGGGTGGACGTGCCCGTTGAT</td><td></td><td></td><td></td><td></td><td></td><td></td><td></td><td></td><td></td><td></td></tr>
<tr><td>Oligo rev</td><td>AAACATCAACGGGCACGTCCACCC</td><td></td><td></td><td></td><td></td><td></td><td></td><td></td><td></td><td></td><td></td></tr>
<tr><td>Chromosome</td><td>start</td><td>end</td><td>strand</td><td>MM</td><td>target_seq</td><td>PAM</td><td>alignment</td><td>distance</td><td>position</td><td>gene name</td><td>gene id</td></tr>
<tr><td>GU305914.1</td><td>94785</td><td>94807</td><td>+</td><td>0</td><td>GGGTGGACGTGCCCGTTGAT</td><td>AGG</td><td>||||||||[||||||||||||]PAM</td><td>NA</td><td>Intergenic</td><td>NA</td><td>NA</td></tr>
<tr><td>T6752</td><td>ATAGAGCCTAAAGCAAACATTGG</td><td>1000</td><td></td><td></td><td></td><td></td><td></td><td></td><td></td><td></td><td></td></tr>
<tr><td>Oligo fwd</td><td>CACCATAGAGCCTAAAGCAAACAT</td><td></td><td></td><td></td><td></td><td></td><td></td><td></td><td></td><td></td><td></td></tr>
<tr><td>Oligo rev</td><td>AAACATGTTTGCTTTAGGCTCTAT</td><td></td><td></td><td></td><td></td><td></td><td></td><td></td><td></td><td></td><td></td></tr>
<tr><td>Chromosome</td><td>start</td><td>end</td><td>strand</td><td>MM</td><td>target_seq</td><td>PAM</td><td>alignment</td><td>distance</td><td>position</td><td>gene name</td><td>gene id</td></tr>
<tr><td>GU305914.1</td><td>94832</td><td>94854</td><td>+</td><td>0</td><td>ATAGAGCCTAAAGCAAACAT</td><td>TGG</td><td>||||||||[||||||||||||]PAM</td><td>NA</td><td>Intergenic</td><td>NA</td><td>NA</td></tr>
<tr><td>T6753</td><td>GCCATGTGCTTTATCTAAGATGG</td><td>1000</td><td></td><td></td><td></td><td></td><td></td><td></td><td></td><td></td><td></td></tr>
<tr><td>Oligo fwd</td><td>CACCGCCATGTGCTTTATCTAAGA</td><td></td><td></td><td></td><td></td><td></td><td></td><td></td><td></td><td></td><td></td></tr>
<tr><td>Oligo rev</td><td>AAACTCTTAGATAAAGCACATGGC</td><td></td><td></td><td></td><td></td><td></td><td></td><td></td><td></td><td></td><td></td></tr>
<tr><td>Chromosome</td><td>start</td><td>end</td><td>strand</td><td>MM</td><td>target_seq</td><td>PAM</td><td>alignment</td><td>distance</td><td>position</td><td>gene name</td><td>gene id</td></tr>
<tr><td>GU305914.1</td><td>94885</td><td>94907</td><td>+</td><td>0</td><td>GCCATGTGCTTTATCTAAGA</td><td>TGG</td><td>||||||||[||||||||||||]PAM</td><td>NA</td><td>Intergenic</td><td>NA</td><td>NA</td></tr>
<tr><td>T6754</td><td>ATGTGCTTTATCTAAGATGGCGG</td><td>1000</td><td></td><td></td><td></td><td></td><td></td><td></td><td></td><td></td><td></td></tr>
<tr><td>Oligo fwd</td><td>CACCATGTGCTTTATCTAAGATGG</td><td></td><td></td><td></td><td></td><td></td><td></td><td></td><td></td><td></td><td></td></tr>
<tr><td>Oligo rev</td><td>AAACCCATCTTAGATAAAGCACAT</td><td></td><td></td><td></td><td></td><td></td><td></td><td></td><td></td><td></td><td></td></tr>
<tr><td>Chromosome</td><td>start</td><td>end</td><td>strand</td><td>MM</td><td>target_seq</td><td>PAM</td><td>alignment</td><td>distance</td><td>position</td><td>gene name</td><td>gene id</td></tr>
<tr><td>GU305914.1</td><td>94888</td><td>94910</td><td>+</td><td>0</td><td>ATGTGCTTTATCTAAGATGG</td><td>CGG</td><td>||||||||[||||||||||||]PAM</td><td>NA</td><td>Intergenic</td><td>NA</td><td>NA</td></tr>
<tr><td>T6755</td><td>AAGATGGCGGACATAAAACATGG</td><td>1000</td><td></td><td></td><td></td><td></td><td></td><td></td><td></td><td></td><td></td></tr>
<tr><td>Oligo fwd</td><td>CACCAAGATGGCGGACATAAAACA</td><td></td><td></td><td></td><td></td><td></td><td></td><td></td><td></td><td></td><td></td></tr>
<tr><td>Oligo rev</td><td>AAACTGTTTTATGTCCGCCATCTT</td><td></td><td></td><td></td><td></td><td></td><td></td><td></td><td></td><td></td><td></td></tr>
<tr><td>Chromosome</td><td>start</td><td>end</td><td>strand</td><td>MM</td><td>target_seq</td><td>PAM</td><td>alignment</td><td>distance</td><td>position</td><td>gene name</td><td>gene id</td></tr>
<tr><td>GU305914.1</td><td>94901</td><td>94923</td><td>+</td><td>0</td><td>AAGATGGCGGACATAAAACA</td><td>TGG</td><td>||||||||[||||||||||||]PAM</td><td>NA</td><td>Intergenic</td><td>NA</td><td>NA</td></tr>
<tr><td>T6756</td><td>ACATAAAACATGGCCGTTTGCGG</td><td>1000</td><td></td><td></td><td></td><td></td><td></td><td></td><td></td><td></td><td></td></tr>
<tr><td>Oligo fwd</td><td>CACCACATAAAACATGGCCGTTTG</td><td></td><td></td><td></td><td></td><td></td><td></td><td></td><td></td><td></td><td></td></tr>
<tr><td>Oligo rev</td><td>AAACCAAACGGCCATGTTTTATGT</td><td></td><td></td><td></td><td></td><td></td><td></td><td></td><td></td><td></td><td></td></tr>
</table>

<table>
<tr><td>Chromosome</td><td>start</td><td>end</td><td>strand</td><td>MM</td><td>target_seq</td><td>PAM</td><td>alignment</td><td>distance</td><td>position</td><td>gene name</td><td>gene id</td></tr>
<tr><td>GU305914.1</td><td>94911</td><td>94933</td><td>+</td><td>0</td><td>ACATAAAACATGGCCGTTTG</td><td>CGG</td><td>||||||||[||||||||||||]PAM</td><td>NA</td><td>Intergenic</td><td>NA</td><td>NA</td></tr>
<tr><td></td><td></td><td></td><td></td><td></td><td></td><td></td><td></td><td></td><td></td><td></td><td></td></tr>
<tr><td>T6757</td><td>GGCCGTTTGCGGTTTCTTCTTGG</td><td>1000</td><td></td><td></td><td></td><td></td><td></td><td></td><td></td><td></td><td></td></tr>
<tr><td>Oligo fwd</td><td>CACCGGCCGTTTGCGGTTTCTTCT</td><td></td><td></td><td></td><td></td><td></td><td></td><td></td><td></td><td></td><td></td></tr>
<tr><td>Oligo rev</td><td>AAACAGAAGAAACCGCAAACGGCC</td><td></td><td></td><td></td><td></td><td></td><td></td><td></td><td></td><td></td><td></td></tr>
<tr><td>Chromosome</td><td>start</td><td>end</td><td>strand</td><td>MM</td><td>target_seq</td><td>PAM</td><td>alignment</td><td>distance</td><td>position</td><td>gene name</td><td>gene id</td></tr>
<tr><td>GU305914.1</td><td>94922</td><td>94944</td><td>+</td><td>0</td><td>GGCCGTTTGCGGTTTCTTCT</td><td>TGG</td><td>||||||||[||||||||||||]PAM</td><td>NA</td><td>Intergenic</td><td>NA</td><td>NA</td></tr>
<tr><td></td><td></td><td></td><td></td><td></td><td></td><td></td><td></td><td></td><td></td><td></td><td></td></tr>
<tr><td>T6758</td><td>TTTCTTCTTGGTTTGTTTTCTGG</td><td>1000</td><td></td><td></td><td></td><td></td><td></td><td></td><td></td><td></td><td></td></tr>
<tr><td>Oligo fwd</td><td>CACCTTTCTTCTTGGTTTGTTTTC</td><td></td><td></td><td></td><td></td><td></td><td></td><td></td><td></td><td></td><td></td></tr>
<tr><td>Oligo rev</td><td>AAACGAAAACAAACCAAGAAGAAA</td><td></td><td></td><td></td><td></td><td></td><td></td><td></td><td></td><td></td><td></td></tr>
<tr><td>Chromosome</td><td>start</td><td>end</td><td>strand</td><td>MM</td><td>target_seq</td><td>PAM</td><td>alignment</td><td>distance</td><td>position</td><td>gene name</td><td>gene id</td></tr>
<tr><td>GU305914.1</td><td>94934</td><td>94956</td><td>+</td><td>0</td><td>TTTCTTCTTGGTTTGTTTTC</td><td>TGG</td><td>||||||||[||||||||||||]PAM</td><td>NA</td><td>Intergenic</td><td>NA</td><td>NA</td></tr>
<tr><td></td><td></td><td></td><td></td><td></td><td></td><td></td><td></td><td></td><td></td><td></td><td></td></tr>
<tr><td>T6759</td><td>TTCTTCTTGGTTTGTTTTCTGGG</td><td>1000</td><td></td><td></td><td></td><td></td><td></td><td></td><td></td><td></td><td></td></tr>
<tr><td>Oligo fwd</td><td>CACCTTCTTCTTGGTTTGTTTTCT</td><td></td><td></td><td></td><td></td><td></td><td></td><td></td><td></td><td></td><td></td></tr>
<tr><td>Oligo rev</td><td>AAACAGAAAACAAACCAAGAAGAA</td><td></td><td></td><td></td><td></td><td></td><td></td><td></td><td></td><td></td><td></td></tr>
<tr><td>Chromosome</td><td>start</td><td>end</td><td>strand</td><td>MM</td><td>target_seq</td><td>PAM</td><td>alignment</td><td>distance</td><td>position</td><td>gene name</td><td>gene id</td></tr>
<tr><td>GU305914.1</td><td>94935</td><td>94957</td><td>+</td><td>0</td><td>TTCTTCTTGGTTTGTTTTCT</td><td>GGG</td><td>||||||||[||||||||||||]PAM</td><td>NA</td><td>Intergenic</td><td>NA</td><td>NA</td></tr>
<tr><td></td><td></td><td></td><td></td><td></td><td></td><td></td><td></td><td></td><td></td><td></td><td></td></tr>
<tr><td>T6760</td><td>CTTGGTTTGTTTTCTGGGTGCGG</td><td>1000</td><td></td><td></td><td></td><td></td><td></td><td></td><td></td><td></td><td></td></tr>
<tr><td>Oligo fwd</td><td>CACCCTTGGTTTGTTTTCTGGGTG</td><td></td><td></td><td></td><td></td><td></td><td></td><td></td><td></td><td></td><td></td></tr>
<tr><td>Oligo rev</td><td>AAACCACCCAGAAAACAAACCAAG</td><td></td><td></td><td></td><td></td><td></td><td></td><td></td><td></td><td></td><td></td></tr>
<tr><td>Chromosome</td><td>start</td><td>end</td><td>strand</td><td>MM</td><td>target_seq</td><td>PAM</td><td>alignment</td><td>distance</td><td>position</td><td>gene name</td><td>gene id</td></tr>
<tr><td>GU305914.1</td><td>94940</td><td>94962</td><td>+</td><td>0</td><td>CTTGGTTTGTTTTCTGGGTG</td><td>CGG</td><td>||||||||[||||||||||||]PAM</td><td>NA</td><td>Intergenic</td><td>NA</td><td>NA</td></tr>
<tr><td></td><td></td><td></td><td></td><td></td><td></td><td></td><td></td><td></td><td></td><td></td><td></td></tr>
<tr><td>T6761</td><td>TTTTCTGGGTGCGGTCTTCTTGG</td><td>1000</td><td></td><td></td><td></td><td></td><td></td><td></td><td></td><td></td><td></td></tr>
<tr><td>Oligo fwd</td><td>CACCTTTTCTGGGTGCGGTCTTCT</td><td></td><td></td><td></td><td></td><td></td><td></td><td></td><td></td><td></td><td></td></tr>
<tr><td>Oligo rev</td><td>AAACAGAAGACCGCACCCAGAAAA</td><td></td><td></td><td></td><td></td><td></td><td></td><td></td><td></td><td></td><td></td></tr>
<tr><td>Chromosome</td><td>start</td><td>end</td><td>strand</td><td>MM</td><td>target_seq</td><td>PAM</td><td>alignment</td><td>distance</td><td>position</td><td>gene name</td><td>gene id</td></tr>
<tr><td>GU305914.1</td><td>94949</td><td>94971</td><td>+</td><td>0</td><td>TTTTCTGGGTGCGGTCTTCT</td><td>TGG</td><td>||||||||[||||||||||||]PAM</td><td>NA</td><td>Intergenic</td><td>NA</td><td>NA</td></tr>
<tr><td></td><td></td><td></td><td></td><td></td><td></td><td></td><td></td><td></td><td></td><td></td><td></td></tr>
<tr><td>T6762</td><td>TTGGAAGTATGACTCATTAACGG</td><td>1000</td><td></td><td></td><td></td><td></td><td></td><td></td><td></td><td></td><td></td></tr>
<tr><td>Oligo fwd</td><td>CACCTTGGAAGTATGACTCATTAA</td><td></td><td></td><td></td><td></td><td></td><td></td><td></td><td></td><td></td><td></td></tr>
<tr><td>Oligo rev</td><td>AAACTTAATGAGTCATACTTCCAA</td><td></td><td></td><td></td><td></td><td></td><td></td><td></td><td></td><td></td><td></td></tr>
<tr><td>Chromosome</td><td>start</td><td>end</td><td>strand</td><td>MM</td><td>target_seq</td><td>PAM</td><td>alignment</td><td>distance</td><td>position</td><td>gene name</td><td>gene id</td></tr>
<tr><td>GU305914.1</td><td>94968</td><td>94990</td><td>+</td><td>0</td><td>TTGGAAGTATGACTCATTAA</td><td>CGG</td><td>||||||||[||||||||||||]PAM</td><td>NA</td><td>Intergenic</td><td>NA</td><td>NA</td></tr>
<tr><td></td><td></td><td></td><td></td><td></td><td></td><td></td><td></td><td></td><td></td><td></td><td></td></tr>
<tr><td>T6763</td><td>TCGCTGCTGTGTCATGCATCCGG</td><td>1000</td><td></td><td></td><td></td><td></td><td></td><td></td><td></td><td></td><td></td></tr>
<tr><td>Oligo fwd</td><td>CACCTCGCTGCTGTGTCATGCATC</td><td></td><td></td><td></td><td></td><td></td><td></td><td></td><td></td><td></td><td></td></tr>
<tr><td>Oligo rev</td><td>AAACGATGCATGACACAGCAGCGA</td><td></td><td></td><td></td><td></td><td></td><td></td><td></td><td></td><td></td><td></td></tr>
<tr><td>Chromosome</td><td>start</td><td>end</td><td>strand</td><td>MM</td><td>target_seq</td><td>PAM</td><td>alignment</td><td>distance</td><td>position</td><td>gene name</td><td>gene id</td></tr>
<tr><td>GU305914.1</td><td>94991</td><td>95013</td><td>+</td><td>0</td><td>TCGCTGCTGTGTCATGCATC</td><td>CGG</td><td>||||||||[||||||||||||]PAM</td><td>NA</td><td>Intergenic</td><td>NA</td><td>NA</td></tr>
<tr><td></td><td></td><td></td><td></td><td></td><td></td><td></td><td></td><td></td><td></td><td></td><td></td></tr>
<tr><td>T6764</td><td>TATGCCCGCCCCTTCCGCTTAGG</td><td>1000</td><td></td><td></td><td></td><td></td><td></td><td></td><td></td><td></td><td></td></tr>
<tr><td>Oligo fwd</td><td>CACCTATGCCCGCCCCTTCCGCTT</td><td></td><td></td><td></td><td></td><td></td><td></td><td></td><td></td><td></td><td></td></tr>
<tr><td>Oligo rev</td><td>AAACAAGCGGAAGGGGCGGGCATA</td><td></td><td></td><td></td><td></td><td></td><td></td><td></td><td></td><td></td><td></td></tr>
<tr><td>Chromosome</td><td>start</td><td>end</td><td>strand</td><td>MM</td><td>target_seq</td><td>PAM</td><td>alignment</td><td>distance</td><td>position</td><td>gene name</td><td>gene id</td></tr>
<tr><td>GU305914.1</td><td>95017</td><td>95039</td><td>+</td><td>0</td><td>TATGCCCGCCCCTTCCGCTT</td><td>AGG</td><td>||||||||[||||||||||||]PAM</td><td>NA</td><td>Intergenic</td><td>NA</td><td>NA</td></tr>
<tr><td></td><td></td><td></td><td></td><td></td><td></td><td></td><td></td><td></td><td></td><td></td><td></td></tr>
<tr><td>T6765</td><td>ATGCCCGCCCCTTCCGCTTAGGG</td><td>1000</td><td></td><td></td><td></td><td></td><td></td><td></td><td></td><td></td><td></td></tr>
<tr><td>Oligo fwd</td><td>CACCATGCCCGCCCCTTCCGCTTA</td><td></td><td></td><td></td><td></td><td></td><td></td><td></td><td></td><td></td><td></td></tr>
<tr><td>Oligo rev</td><td>AAACTAAGCGGAAGGGGCGGGCAT</td><td></td><td></td><td></td><td></td><td></td><td></td><td></td><td></td><td></td><td></td></tr>
<tr><td>Chromosome</td><td>start</td><td>end</td><td>strand</td><td>MM</td><td>target_seq</td><td>PAM</td><td>alignment</td><td>distance</td><td>position</td><td>gene name</td><td>gene id</td></tr>
<tr><td>GU305914.1</td><td>95018</td><td>95040</td><td>+</td><td>0</td><td>ATGCCCGCCCCTTCCGCTTA</td><td>GGG</td><td>||||||||[||||||||||||]PAM</td><td>NA</td><td>Intergenic</td><td>NA</td><td>NA</td></tr>
<tr><td></td><td></td><td></td><td></td><td></td><td></td><td></td><td></td><td></td><td></td><td></td><td></td></tr>
<tr><td>T6766</td><td>TGCCCGCCCCTTCCGCTTAGGGG</td><td>1000</td><td></td><td></td><td></td><td></td><td></td><td></td><td></td><td></td><td></td></tr>
<tr><td>Oligo fwd</td><td>CACCTGCCCGCCCCTTCCGCTTAG</td><td></td><td></td><td></td><td></td><td></td><td></td><td></td><td></td><td></td><td></td></tr>
<tr><td>Oligo rev</td><td>AAACCTAAGCGGAAGGGGCGGGCA</td><td></td><td></td><td></td><td></td><td></td><td></td><td></td><td></td><td></td><td></td></tr>
<tr><td>Chromosome</td><td>start</td><td>end</td><td>strand</td><td>MM</td><td>target_seq</td><td>PAM</td><td>alignment</td><td>distance</td><td>position</td><td>gene name</td><td>gene id</td></tr>
<tr><td>GU305914.1</td><td>95019</td><td>95041</td><td>+</td><td>0</td><td>TGCCCGCCCCTTCCGCTTAG</td><td>GGG</td><td>||||||||[||||||||||||]PAM</td><td>NA</td><td>Intergenic</td><td>NA</td><td>NA</td></tr>
<tr><td></td><td></td><td></td><td></td><td></td><td></td><td></td><td></td><td></td><td></td><td></td><td></td></tr>
<tr><td>T6767</td><td>TAGGGGCTGTCAATCATCTTAGG</td><td>1000</td><td></td><td></td><td></td><td></td><td></td><td></td><td></td><td></td><td></td></tr>
<tr><td>Oligo fwd</td><td>CACCTAGGGGCTGTCAATCATCTT</td><td></td><td></td><td></td><td></td><td></td><td></td><td></td><td></td><td></td><td></td></tr>
<tr><td>Oligo rev</td><td>AAACAAGATGATTGACAGCCCCTA</td><td></td><td></td><td></td><td></td><td></td><td></td><td></td><td></td><td></td><td></td></tr>
<tr><td>Chromosome</td><td>start</td><td>end</td><td>strand</td><td>MM</td><td>target_seq</td><td>PAM</td><td>alignment</td><td>distance</td><td>position</td><td>gene name</td><td>gene id</td></tr>
<tr><td>GU305914.1</td><td>95036</td><td>95058</td><td>+</td><td>0</td><td>TAGGGGCTGTCAATCATCTT</td><td>AGG</td><td>||||||||[||||||||||||]PAM</td><td>NA</td><td>Intergenic</td><td>NA</td><td>NA</td></tr>
<tr><td></td><td></td><td></td><td></td><td></td><td></td><td></td><td></td><td></td><td></td><td></td><td></td></tr>
<tr><td>T6768</td><td>CACAGAGCCTAAGTGTCCATCGG</td><td>1000</td><td></td><td></td><td></td><td></td><td></td><td></td><td></td><td></td><td></td></tr>
<tr><td>Oligo fwd</td><td>CACCCACAGAGCCTAAGTGTCCAT</td><td></td><td></td><td></td><td></td><td></td><td></td><td></td><td></td><td></td><td></td></tr>
<tr><td>Oligo rev</td><td>AAACATGGACACTTAGGCTCTGTG</td><td></td><td></td><td></td><td></td><td></td><td></td><td></td><td></td><td></td><td></td></tr>
<tr><td>Chromosome</td><td>start</td><td>end</td><td>strand</td><td>MM</td><td>target_seq</td><td>PAM</td><td>alignment</td><td>distance</td><td>position</td><td>gene name</td><td>gene id</td></tr>
<tr><td>GU305914.1</td><td>95073</td><td>95095</td><td>+</td><td>0</td><td>CACAGAGCCTAAGTGTCCAT</td><td>CGG</td><td>||||||||[||||||||||||]PAM</td><td>NA</td><td>Intergenic</td><td>NA</td><td>NA</td></tr>
<tr><td></td><td></td><td></td><td></td><td></td><td></td><td></td><td></td><td></td><td></td><td></td><td></td></tr>
<tr><td>T6769</td><td>CCATCGGACTTAGCCGCTGTCGG</td><td>1000</td><td></td><td></td><td></td><td></td><td></td><td></td><td></td><td></td><td></td></tr>
<tr><td>Oligo fwd</td><td>CACCCCATCGGACTTAGCCGCTGT</td><td></td><td></td><td></td><td></td><td></td><td></td><td></td><td></td><td></td><td></td></tr>
<tr><td>Oligo rev</td><td>AAACACAGCGGCTAAGTCCGATGG</td><td></td><td></td><td></td><td></td><td></td><td></td><td></td><td></td><td></td><td></td></tr>
<tr><td>Chromosome</td><td>start</td><td>end</td><td>strand</td><td>MM</td><td>target_seq</td><td>PAM</td><td>alignment</td><td>distance</td><td>position</td><td>gene name</td><td>gene id</td></tr>
<tr><td>GU305914.1</td><td>95089</td><td>95111</td><td>+</td><td>0</td><td>CCATCGGACTTAGCCGCTGT</td><td>CGG</td><td>||||||||[||||||||||||]PAM</td><td>NA</td><td>Intergenic</td><td>NA</td><td>NA</td></tr>
<tr><td></td><td></td><td></td><td></td><td></td><td></td><td></td><td></td><td></td><td></td><td></td><td></td></tr>
<tr><td>T6770</td><td>GACTTAGCCGCTGTCGGCCTCGG</td><td>1000</td><td></td><td></td><td></td><td></td><td></td><td></td><td></td><td></td><td></td></tr>
<tr><td>Oligo fwd</td><td>CACCGACTTAGCCGCTGTCGGCCT</td><td></td><td></td><td></td><td></td><td></td><td></td><td></td><td></td><td></td><td></td></tr>
<tr><td>Oligo rev</td><td>AAACAGGCCGACAGCGGCTAAGTC</td><td></td><td></td><td></td><td></td><td></td><td></td><td></td><td></td><td></td><td></td></tr>
<tr><td>Chromosome</td><td>start</td><td>end</td><td>strand</td><td>MM</td><td>target_seq</td><td>PAM</td><td>alignment</td><td>distance</td><td>position</td><td>gene name</td><td>gene id</td></tr>
<tr><td>GU305914.1</td><td>95095</td><td>95117</td><td>+</td><td>0</td><td>GACTTAGCCGCTGTCGGCCT</td><td>CGG</td><td>||||||||[||||||||||||]PAM</td><td>NA</td><td>Intergenic</td><td>NA</td><td>NA</td></tr>
<tr><td></td><td></td><td></td><td></td><td></td><td></td><td></td><td></td><td></td><td></td><td></td><td></td></tr>
<tr><td>T6771</td><td>AGCCGCTGTCGGCCTCGGAAAGG</td><td>1000</td><td></td><td></td><td></td><td></td><td></td><td></td><td></td><td></td><td></td></tr>
<tr><td>Oligo fwd</td><td>CACCAGCCGCTGTCGGCCTCGGAA</td><td></td><td></td><td></td><td></td><td></td><td></td><td></td><td></td><td></td><td></td></tr>
</table>

<table>
<tr><td>Oligo rev</td><td>AAACTTCCGAGGCCGACAGCGGCT</td><td></td><td></td><td></td><td></td><td></td><td></td><td></td><td></td><td></td><td></td><td></td></tr>
<tr><td>Chromosome</td><td>start</td><td></td><td>end</td><td>strand</td><td>MM</td><td>target_seq</td><td>PAM</td><td>alignment</td><td>distance</td><td>position</td><td>gene name</td><td>gene id</td></tr>
<tr><td>GU305914.1</td><td></td><td>95100</td><td>95122</td><td>+</td><td>0</td><td>AGCCGCTGTCGGCCTCGGAA</td><td>AGG</td><td>||||||||[||||||||||||]PAM</td><td>NA</td><td>Intergenic</td><td>NA</td><td>NA</td></tr>
<tr><td>T6772</td><td>GGCGTTCCTATGACCGTATATGG</td><td></td><td>1000</td><td></td><td></td><td></td><td></td><td></td><td></td><td></td><td></td><td></td></tr>
<tr><td>Oligo fwd</td><td>CACCGGCGTTCCTATGACCGTATA</td><td></td><td></td><td></td><td></td><td></td><td></td><td></td><td></td><td></td><td></td><td></td></tr>
<tr><td>Oligo rev</td><td>AAACTATACGGTCATAGGAACGCC</td><td></td><td></td><td></td><td></td><td></td><td></td><td></td><td></td><td></td><td></td><td></td></tr>
<tr><td>Chromosome</td><td>start</td><td></td><td>end</td><td>strand</td><td>MM</td><td>target_seq</td><td>PAM</td><td>alignment</td><td>distance</td><td>position</td><td>gene name</td><td>gene id</td></tr>
<tr><td>GU305914.1</td><td></td><td>95121</td><td>95143</td><td>+</td><td>0</td><td>GGCGTTCCTATGACCGTATA</td><td>TGG</td><td>||||||||[||||||||||||]PAM</td><td>NA</td><td>Intergenic</td><td>NA</td><td>NA</td></tr>
<tr><td>T6773</td><td>GCGTTCCTATGACCGTATATGGG</td><td></td><td>1000</td><td></td><td></td><td></td><td></td><td></td><td></td><td></td><td></td><td></td></tr>
<tr><td>Oligo fwd</td><td>CACCGCGTTCCTATGACCGTATAT</td><td></td><td></td><td></td><td></td><td></td><td></td><td></td><td></td><td></td><td></td><td></td></tr>
<tr><td>Oligo rev</td><td>AAACATATACGGTCATAGGAACGC</td><td></td><td></td><td></td><td></td><td></td><td></td><td></td><td></td><td></td><td></td><td></td></tr>
<tr><td>Chromosome</td><td>start</td><td></td><td>end</td><td>strand</td><td>MM</td><td>target_seq</td><td>PAM</td><td>alignment</td><td>distance</td><td>position</td><td>gene name</td><td>gene id</td></tr>
<tr><td>GU305914.1</td><td></td><td>95122</td><td>95144</td><td>+</td><td>0</td><td>GCGTTCCTATGACCGTATAT</td><td>GGG</td><td>||||||||[||||||||||||]PAM</td><td>NA</td><td>Intergenic</td><td>NA</td><td>NA</td></tr>
<tr><td>T6774</td><td>CGTTCCTATGACCGTATATGGGG</td><td></td><td>1000</td><td></td><td></td><td></td><td></td><td></td><td></td><td></td><td></td><td></td></tr>
<tr><td>Oligo fwd</td><td>CACCCGTTCCTATGACCGTATATG</td><td></td><td></td><td></td><td></td><td></td><td></td><td></td><td></td><td></td><td></td><td></td></tr>
<tr><td>Oligo rev</td><td>AAACCATATACGGTCATAGGAACG</td><td></td><td></td><td></td><td></td><td></td><td></td><td></td><td></td><td></td><td></td><td></td></tr>
<tr><td>Chromosome</td><td>start</td><td></td><td>end</td><td>strand</td><td>MM</td><td>target_seq</td><td>PAM</td><td>alignment</td><td>distance</td><td>position</td><td>gene name</td><td>gene id</td></tr>
<tr><td>GU305914.1</td><td></td><td>95123</td><td>95145</td><td>+</td><td>0</td><td>CGTTCCTATGACCGTATATG</td><td>GGG</td><td>||||||||[||||||||||||]PAM</td><td>NA</td><td>Intergenic</td><td>NA</td><td>NA</td></tr>
<tr><td>T6775</td><td>ACCGTATATGGGGCGTCACATGG</td><td></td><td>1000</td><td></td><td></td><td></td><td></td><td></td><td></td><td></td><td></td><td></td></tr>
<tr><td>Oligo fwd</td><td>CACCACCGTATATGGGGCGTCACA</td><td></td><td></td><td></td><td></td><td></td><td></td><td></td><td></td><td></td><td></td><td></td></tr>
<tr><td>Oligo rev</td><td>AAACTGTGACGCCCCATATACGGT</td><td></td><td></td><td></td><td></td><td></td><td></td><td></td><td></td><td></td><td></td><td></td></tr>
<tr><td>Chromosome</td><td>start</td><td></td><td>end</td><td>strand</td><td>MM</td><td>target_seq</td><td>PAM</td><td>alignment</td><td>distance</td><td>position</td><td>gene name</td><td>gene id</td></tr>
<tr><td>GU305914.1</td><td></td><td>95133</td><td>95155</td><td>+</td><td>0</td><td>ACCGTATATGGGGCGTCACA</td><td>TGG</td><td>||||||||[||||||||||||]PAM</td><td>NA</td><td>Intergenic</td><td>NA</td><td>NA</td></tr>
<tr><td>T6776</td><td>GGGGCGTCACATGGTAATAATGG</td><td></td><td>1000</td><td></td><td></td><td></td><td></td><td></td><td></td><td></td><td></td><td></td></tr>
<tr><td>Oligo fwd</td><td>CACCGGGGCGTCACATGGTAATAA</td><td></td><td></td><td></td><td></td><td></td><td></td><td></td><td></td><td></td><td></td><td></td></tr>
<tr><td>Oligo rev</td><td>AAACTTATTACCATGTGACGCCCC</td><td></td><td></td><td></td><td></td><td></td><td></td><td></td><td></td><td></td><td></td><td></td></tr>
<tr><td>Chromosome</td><td>start</td><td></td><td>end</td><td>strand</td><td>MM</td><td>target_seq</td><td>PAM</td><td>alignment</td><td>distance</td><td>position</td><td>gene name</td><td>gene id</td></tr>
<tr><td>GU305914.1</td><td></td><td>95142</td><td>95164</td><td>+</td><td>0</td><td>GGGGCGTCACATGGTAATAA</td><td>TGG</td><td>||||||||[||||||||||||]PAM</td><td>NA</td><td>Intergenic</td><td>NA</td><td>NA</td></tr>
<tr><td>T6777</td><td>GGGCGTCACATGGTAATAATGGG</td><td></td><td>1000</td><td></td><td></td><td></td><td></td><td></td><td></td><td></td><td></td><td></td></tr>
<tr><td>Oligo fwd</td><td>CACCGGGCGTCACATGGTAATAAT</td><td></td><td></td><td></td><td></td><td></td><td></td><td></td><td></td><td></td><td></td><td></td></tr>
<tr><td>Oligo rev</td><td>AAACATTATTACCATGTGACGCCC</td><td></td><td></td><td></td><td></td><td></td><td></td><td></td><td></td><td></td><td></td><td></td></tr>
<tr><td>Chromosome</td><td>start</td><td></td><td>end</td><td>strand</td><td>MM</td><td>target_seq</td><td>PAM</td><td>alignment</td><td>distance</td><td>position</td><td>gene name</td><td>gene id</td></tr>
<tr><td>GU305914.1</td><td></td><td>95143</td><td>95165</td><td>+</td><td>0</td><td>GGGCGTCACATGGTAATAAT</td><td>GGG</td><td>||||||||[||||||||||||]PAM</td><td>NA</td><td>Intergenic</td><td>NA</td><td>NA</td></tr>
<tr><td>T6778</td><td>TCACATGGTAATAATGGGCGTGG</td><td></td><td>1000</td><td></td><td></td><td></td><td></td><td></td><td></td><td></td><td></td><td></td></tr>
<tr><td>Oligo fwd</td><td>CACCTCACATGGTAATAATGGGCG</td><td></td><td></td><td></td><td></td><td></td><td></td><td></td><td></td><td></td><td></td><td></td></tr>
<tr><td>Oligo rev</td><td>AAACCGCCCATTATTACCATGTGA</td><td></td><td></td><td></td><td></td><td></td><td></td><td></td><td></td><td></td><td></td><td></td></tr>
<tr><td>Chromosome</td><td>start</td><td></td><td>end</td><td>strand</td><td>MM</td><td>target_seq</td><td>PAM</td><td>alignment</td><td>distance</td><td>position</td><td>gene name</td><td>gene id</td></tr>
<tr><td>GU305914.1</td><td></td><td>95148</td><td>95170</td><td>+</td><td>0</td><td>TCACATGGTAATAATGGGCG</td><td>TGG</td><td>||||||||[||||||||||||]PAM</td><td>NA</td><td>Intergenic</td><td>NA</td><td>NA</td></tr>
<tr><td>T6779</td><td>ATGGGCGTGGTTTCTGACTTAGG</td><td></td><td>1000</td><td></td><td></td><td></td><td></td><td></td><td></td><td></td><td></td><td></td></tr>
<tr><td>Oligo fwd</td><td>CACCATGGGCGTGGTTTCTGACTT</td><td></td><td></td><td></td><td></td><td></td><td></td><td></td><td></td><td></td><td></td><td></td></tr>
<tr><td>Oligo rev</td><td>AAACAAGTCAGAAACCACGCCCAT</td><td></td><td></td><td></td><td></td><td></td><td></td><td></td><td></td><td></td><td></td><td></td></tr>
<tr><td>Chromosome</td><td>start</td><td></td><td>end</td><td>strand</td><td>MM</td><td>target_seq</td><td>PAM</td><td>alignment</td><td>distance</td><td>position</td><td>gene name</td><td>gene id</td></tr>
<tr><td>GU305914.1</td><td></td><td>95161</td><td>95183</td><td>+</td><td>0</td><td>ATGGGCGTGGTTTCTGACTT</td><td>AGG</td><td>||||||||[||||||||||||]PAM</td><td>NA</td><td>Intergenic</td><td>NA</td><td>NA</td></tr>
<tr><td>T6780</td><td>TTCTGACTTAGGCTCTGCGAAGG</td><td></td><td>1000</td><td></td><td></td><td></td><td></td><td></td><td></td><td></td><td></td><td></td></tr>
<tr><td>Oligo fwd</td><td>CACCTTCTGACTTAGGCTCTGCGA</td><td></td><td></td><td></td><td></td><td></td><td></td><td></td><td></td><td></td><td></td><td></td></tr>
<tr><td>Oligo rev</td><td>AAACTCGCAGAGCCTAAGTCAGAA</td><td></td><td></td><td></td><td></td><td></td><td></td><td></td><td></td><td></td><td></td><td></td></tr>
<tr><td>Chromosome</td><td>start</td><td></td><td>end</td><td>strand</td><td>MM</td><td>target_seq</td><td>PAM</td><td>alignment</td><td>distance</td><td>position</td><td>gene name</td><td>gene id</td></tr>
<tr><td>GU305914.1</td><td></td><td>95172</td><td>95194</td><td>+</td><td>0</td><td>TTCTGACTTAGGCTCTGCGA</td><td>AGG</td><td>||||||||[||||||||||||]PAM</td><td>NA</td><td>Intergenic</td><td>NA</td><td>NA</td></tr>
<tr><td>T6781</td><td>AAGACTCGAGACTTAGATTCTGG</td><td></td><td>1000</td><td></td><td></td><td></td><td></td><td></td><td></td><td></td><td></td><td></td></tr>
<tr><td>Oligo fwd</td><td>CACCAAGACTCGAGACTTAGATTC</td><td></td><td></td><td></td><td></td><td></td><td></td><td></td><td></td><td></td><td></td><td></td></tr>
<tr><td>Oligo rev</td><td>AAACGAATCTAAGTCTCGAGTCTT</td><td></td><td></td><td></td><td></td><td></td><td></td><td></td><td></td><td></td><td></td><td></td></tr>
<tr><td>Chromosome</td><td>start</td><td></td><td>end</td><td>strand</td><td>MM</td><td>target_seq</td><td>PAM</td><td>alignment</td><td>distance</td><td>position</td><td>gene name</td><td>gene id</td></tr>
<tr><td>GU305914.1</td><td></td><td>95202</td><td>95224</td><td>+</td><td>0</td><td>AAGACTCGAGACTTAGATTC</td><td>TGG</td><td>||||||||[||||||||||||]PAM</td><td>NA</td><td>Intergenic</td><td>NA</td><td>NA</td></tr>
<tr><td>T6782</td><td>AGACTCGAGACTTAGATTCTGGG</td><td></td><td>1000</td><td></td><td></td><td></td><td></td><td></td><td></td><td></td><td></td><td></td></tr>
<tr><td>Oligo fwd</td><td>CACCAGACTCGAGACTTAGATTCT</td><td></td><td></td><td></td><td></td><td></td><td></td><td></td><td></td><td></td><td></td><td></td></tr>
<tr><td>Oligo rev</td><td>AAACAGAATCTAAGTCTCGAGTCT</td><td></td><td></td><td></td><td></td><td></td><td></td><td></td><td></td><td></td><td></td><td></td></tr>
<tr><td>Chromosome</td><td>start</td><td></td><td>end</td><td>strand</td><td>MM</td><td>target_seq</td><td>PAM</td><td>alignment</td><td>distance</td><td>position</td><td>gene name</td><td>gene id</td></tr>
<tr><td>GU305914.1</td><td></td><td>95203</td><td>95225</td><td>+</td><td>0</td><td>AGACTCGAGACTTAGATTCT</td><td>GGG</td><td>||||||||[||||||||||||]PAM</td><td>NA</td><td>Intergenic</td><td>NA</td><td>NA</td></tr>
<tr><td>T6783</td><td>GACTCGAGACTTAGATTCTGGGG</td><td></td><td>1000</td><td></td><td></td><td></td><td></td><td></td><td></td><td></td><td></td><td></td></tr>
<tr><td>Oligo fwd</td><td>CACCGACTCGAGACTTAGATTCTG</td><td></td><td></td><td></td><td></td><td></td><td></td><td></td><td></td><td></td><td></td><td></td></tr>
<tr><td>Oligo rev</td><td>AAACCAGAATCTAAGTCTCGAGTC</td><td></td><td></td><td></td><td></td><td></td><td></td><td></td><td></td><td></td><td></td><td></td></tr>
<tr><td>Chromosome</td><td>start</td><td></td><td>end</td><td>strand</td><td>MM</td><td>target_seq</td><td>PAM</td><td>alignment</td><td>distance</td><td>position</td><td>gene name</td><td>gene id</td></tr>
<tr><td>GU305914.1</td><td></td><td>95204</td><td>95226</td><td>+</td><td>0</td><td>GACTCGAGACTTAGATTCTG</td><td>GGG</td><td>||||||||[||||||||||||]PAM</td><td>NA</td><td>Intergenic</td><td>NA</td><td>NA</td></tr>
<tr><td>T6784</td><td>AGACTTAGATTCTGGGGACTCGG</td><td></td><td>1000</td><td></td><td></td><td></td><td></td><td></td><td></td><td></td><td></td><td></td></tr>
<tr><td>Oligo fwd</td><td>CACCAGACTTAGATTCTGGGGACT</td><td></td><td></td><td></td><td></td><td></td><td></td><td></td><td></td><td></td><td></td><td></td></tr>
<tr><td>Oligo rev</td><td>AAACAGTCCCCAGAATCTAAGTCT</td><td></td><td></td><td></td><td></td><td></td><td></td><td></td><td></td><td></td><td></td><td></td></tr>
<tr><td>Chromosome</td><td>start</td><td></td><td>end</td><td>strand</td><td>MM</td><td>target_seq</td><td>PAM</td><td>alignment</td><td>distance</td><td>position</td><td>gene name</td><td>gene id</td></tr>
<tr><td>GU305914.1</td><td></td><td>95210</td><td>95232</td><td>+</td><td>0</td><td>AGACTTAGATTCTGGGGACT</td><td>CGG</td><td>||||||||[||||||||||||]PAM</td><td>NA</td><td>Intergenic</td><td>NA</td><td>NA</td></tr>
<tr><td>T6785</td><td>GACTTAGATTCTGGGGACTCGGG</td><td></td><td>1000</td><td></td><td></td><td></td><td></td><td></td><td></td><td></td><td></td><td></td></tr>
<tr><td>Oligo fwd</td><td>CACCGACTTAGATTCTGGGGACTC</td><td></td><td></td><td></td><td></td><td></td><td></td><td></td><td></td><td></td><td></td><td></td></tr>
<tr><td>Oligo rev</td><td>AAACGAGTCCCCAGAATCTAAGTC</td><td></td><td></td><td></td><td></td><td></td><td></td><td></td><td></td><td></td><td></td><td></td></tr>
<tr><td>Chromosome</td><td>start</td><td></td><td>end</td><td>strand</td><td>MM</td><td>target_seq</td><td>PAM</td><td>alignment</td><td>distance</td><td>position</td><td>gene name</td><td>gene id</td></tr>
<tr><td>GU305914.1</td><td></td><td>95211</td><td>95233</td><td>+</td><td>0</td><td>GACTTAGATTCTGGGGACTC</td><td>GGG</td><td>||||||||[||||||||||||]PAM</td><td>NA</td><td>Intergenic</td><td>NA</td><td>NA</td></tr>
<tr><td>T6786</td><td>ACTTAGATTCTGGGGACTCGGGG</td><td></td><td>1000</td><td></td><td></td><td></td><td></td><td></td><td></td><td></td><td></td><td></td></tr>
</table>

<table>
<tr><td>Oligo fwd</td><td>CACCACTTAGATTCTGGGGACTCG</td><td></td><td></td><td></td><td></td><td></td><td></td><td></td><td></td><td></td><td></td></tr>
<tr><td>Oligo rev</td><td>AAACCGAGTCCCCAGAATCTAAGT</td><td></td><td></td><td></td><td></td><td></td><td></td><td></td><td></td><td></td><td></td></tr>
<tr><td>Chromosome</td><td>start</td><td>end</td><td>strand</td><td>MM</td><td>target_seq</td><td>PAM</td><td>alignment</td><td>distance</td><td>position</td><td>gene name</td><td>gene id</td></tr>
<tr><td>GU305914.1</td><td>95212</td><td>95234</td><td>+</td><td>0</td><td>ACTTAGATTCTGGGGACTCG</td><td>GGG</td><td>||||||||[||||||||||||]PAM</td><td>NA</td><td>Intergenic</td><td>NA</td><td>NA</td></tr>
<tr><td></td><td></td><td></td><td></td><td></td><td></td><td></td><td></td><td></td><td></td><td></td><td></td></tr>
<tr><td>T6787</td><td>TAGATTCTGGGGACTCGGGGTGG</td><td>1000</td><td></td><td></td><td></td><td></td><td></td><td></td><td></td><td></td><td></td></tr>
<tr><td>Oligo fwd</td><td>CACCTAGATTCTGGGGACTCGGGG</td><td></td><td></td><td></td><td></td><td></td><td></td><td></td><td></td><td></td><td></td></tr>
<tr><td>Oligo rev</td><td>AAACCCCCGAGTCCCCAGAATCTA</td><td></td><td></td><td></td><td></td><td></td><td></td><td></td><td></td><td></td><td></td></tr>
<tr><td>Chromosome</td><td>start</td><td>end</td><td>strand</td><td>MM</td><td>target_seq</td><td>PAM</td><td>alignment</td><td>distance</td><td>position</td><td>gene name</td><td>gene id</td></tr>
<tr><td>GU305914.1</td><td>95215</td><td>95237</td><td>+</td><td>0</td><td>TAGATTCTGGGGACTCGGGG</td><td>TGG</td><td>||||||||[||||||||||||]PAM</td><td>NA</td><td>Intergenic</td><td>NA</td><td>NA</td></tr>
<tr><td></td><td></td><td></td><td></td><td></td><td></td><td></td><td></td><td></td><td></td><td></td><td></td></tr>
<tr><td>T6788</td><td>AGATTCTGGGGACTCGGGGTGGG</td><td>1000</td><td></td><td></td><td></td><td></td><td></td><td></td><td></td><td></td><td></td></tr>
<tr><td>Oligo fwd</td><td>CACCAGATTCTGGGGACTCGGGGT</td><td></td><td></td><td></td><td></td><td></td><td></td><td></td><td></td><td></td><td></td></tr>
<tr><td>Oligo rev</td><td>AAACACCCCGAGTCCCCAGAATCT</td><td></td><td></td><td></td><td></td><td></td><td></td><td></td><td></td><td></td><td></td></tr>
<tr><td>Chromosome</td><td>start</td><td>end</td><td>strand</td><td>MM</td><td>target_seq</td><td>PAM</td><td>alignment</td><td>distance</td><td>position</td><td>gene name</td><td>gene id</td></tr>
<tr><td>GU305914.1</td><td>95216</td><td>95238</td><td>+</td><td>0</td><td>AGATTCTGGGGACTCGGGGT</td><td>GGG</td><td>||||||||[||||||||||||]PAM</td><td>NA</td><td>Intergenic</td><td>NA</td><td>NA</td></tr>
<tr><td></td><td></td><td></td><td></td><td></td><td></td><td></td><td></td><td></td><td></td><td></td><td></td></tr>
<tr><td>T6789</td><td>TCTGGGGACTCGGGGTGGGCTGG</td><td>1000</td><td></td><td></td><td></td><td></td><td></td><td></td><td></td><td></td><td></td></tr>
<tr><td>Oligo fwd</td><td>CACCTCTGGGGACTCGGGGTGGGC</td><td></td><td></td><td></td><td></td><td></td><td></td><td></td><td></td><td></td><td></td></tr>
<tr><td>Oligo rev</td><td>AAACGCCCACCCCGAGTCCCCAGA</td><td></td><td></td><td></td><td></td><td></td><td></td><td></td><td></td><td></td><td></td></tr>
<tr><td>Chromosome</td><td>start</td><td>end</td><td>strand</td><td>MM</td><td>target_seq</td><td>PAM</td><td>alignment</td><td>distance</td><td>position</td><td>gene name</td><td>gene id</td></tr>
<tr><td>GU305914.1</td><td>95220</td><td>95242</td><td>+</td><td>0</td><td>TCTGGGGACTCGGGGTGGGC</td><td>TGG</td><td>||||||||[||||||||||||]PAM</td><td>NA</td><td>Intergenic</td><td>NA</td><td>NA</td></tr>
<tr><td></td><td></td><td></td><td></td><td></td><td></td><td></td><td></td><td></td><td></td><td></td><td></td></tr>
<tr><td>T6790</td><td>CTGGGGACTCGGGGTGGGCTGGG</td><td>1000</td><td></td><td></td><td></td><td></td><td></td><td></td><td></td><td></td><td></td></tr>
<tr><td>Oligo fwd</td><td>CACCCTGGGGACTCGGGGTGGGCT</td><td></td><td></td><td></td><td></td><td></td><td></td><td></td><td></td><td></td><td></td></tr>
<tr><td>Oligo rev</td><td>AAACAGCCCACCCCGAGTCCCCAG</td><td></td><td></td><td></td><td></td><td></td><td></td><td></td><td></td><td></td><td></td></tr>
<tr><td>Chromosome</td><td>start</td><td>end</td><td>strand</td><td>MM</td><td>target_seq</td><td>PAM</td><td>alignment</td><td>distance</td><td>position</td><td>gene name</td><td>gene id</td></tr>
<tr><td>GU305914.1</td><td>95221</td><td>95243</td><td>+</td><td>0</td><td>CTGGGGACTCGGGGTGGGCT</td><td>GGG</td><td>||||||||[||||||||||||]PAM</td><td>NA</td><td>Intergenic</td><td>NA</td><td>NA</td></tr>
<tr><td></td><td></td><td></td><td></td><td></td><td></td><td></td><td></td><td></td><td></td><td></td><td></td></tr>
<tr><td>T6791</td><td>TGGGGACTCGGGGTGGGCTGGGG</td><td>1000</td><td></td><td></td><td></td><td></td><td></td><td></td><td></td><td></td><td></td></tr>
<tr><td>Oligo fwd</td><td>CACCTGGGGACTCGGGGTGGGCTG</td><td></td><td></td><td></td><td></td><td></td><td></td><td></td><td></td><td></td><td></td></tr>
<tr><td>Oligo rev</td><td>AAACCAGCCCACCCCGAGTCCCCA</td><td></td><td></td><td></td><td></td><td></td><td></td><td></td><td></td><td></td><td></td></tr>
<tr><td>Chromosome</td><td>start</td><td>end</td><td>strand</td><td>MM</td><td>target_seq</td><td>PAM</td><td>alignment</td><td>distance</td><td>position</td><td>gene name</td><td>gene id</td></tr>
<tr><td>GU305914.1</td><td>95222</td><td>95244</td><td>+</td><td>0</td><td>TGGGGACTCGGGGTGGGCTG</td><td>GGG</td><td>||||||||[||||||||||||]PAM</td><td>NA</td><td>Intergenic</td><td>NA</td><td>NA</td></tr>
<tr><td></td><td></td><td></td><td></td><td></td><td></td><td></td><td></td><td></td><td></td><td></td><td></td></tr>
<tr><td>T6792</td><td>GGACTCGGGGTGGGCTGGGGTGG</td><td>1000</td><td></td><td></td><td></td><td></td><td></td><td></td><td></td><td></td><td></td></tr>
<tr><td>Oligo fwd</td><td>CACCGGACTCGGGGTGGGCTGGGG</td><td></td><td></td><td></td><td></td><td></td><td></td><td></td><td></td><td></td><td></td></tr>
<tr><td>Oligo rev</td><td>AAACCCCCAGCCCACCCCGAGTCC</td><td></td><td></td><td></td><td></td><td></td><td></td><td></td><td></td><td></td><td></td></tr>
<tr><td>Chromosome</td><td>start</td><td>end</td><td>strand</td><td>MM</td><td>target_seq</td><td>PAM</td><td>alignment</td><td>distance</td><td>position</td><td>gene name</td><td>gene id</td></tr>
<tr><td>GU305914.1</td><td>95225</td><td>95247</td><td>+</td><td>0</td><td>GGACTCGGGGTGGGCTGGGG</td><td>TGG</td><td>||||||||[||||||||||||]PAM</td><td>NA</td><td>Intergenic</td><td>NA</td><td>NA</td></tr>
<tr><td></td><td></td><td></td><td></td><td></td><td></td><td></td><td></td><td></td><td></td><td></td><td></td></tr>
<tr><td>T6793</td><td>GACTCGGGGTGGGCTGGGGTGGG</td><td>1000</td><td></td><td></td><td></td><td></td><td></td><td></td><td></td><td></td><td></td></tr>
<tr><td>Oligo fwd</td><td>CACCGACTCGGGGTGGGCTGGGGT</td><td></td><td></td><td></td><td></td><td></td><td></td><td></td><td></td><td></td><td></td></tr>
<tr><td>Oligo rev</td><td>AAACACCCCAGCCCACCCCGAGTC</td><td></td><td></td><td></td><td></td><td></td><td></td><td></td><td></td><td></td><td></td></tr>
<tr><td>Chromosome</td><td>start</td><td>end</td><td>strand</td><td>MM</td><td>target_seq</td><td>PAM</td><td>alignment</td><td>distance</td><td>position</td><td>gene name</td><td>gene id</td></tr>
<tr><td>GU305914.1</td><td>95226</td><td>95248</td><td>+</td><td>0</td><td>GACTCGGGGTGGGCTGGGGT</td><td>GGG</td><td>||||||||[||||||||||||]PAM</td><td>NA</td><td>Intergenic</td><td>NA</td><td>NA</td></tr>
<tr><td></td><td></td><td></td><td></td><td></td><td></td><td></td><td></td><td></td><td></td><td></td><td></td></tr>
<tr><td>T6794</td><td>ACTCGGGGTGGGCTGGGGTGGGG</td><td>1000</td><td></td><td></td><td></td><td></td><td></td><td></td><td></td><td></td><td></td></tr>
<tr><td>Oligo fwd</td><td>CACCACTCGGGGTGGGCTGGGGTG</td><td></td><td></td><td></td><td></td><td></td><td></td><td></td><td></td><td></td><td></td></tr>
<tr><td>Oligo rev</td><td>AAACCACCCCAGCCCACCCCGAGT</td><td></td><td></td><td></td><td></td><td></td><td></td><td></td><td></td><td></td><td></td></tr>
<tr><td>Chromosome</td><td>start</td><td>end</td><td>strand</td><td>MM</td><td>target_seq</td><td>PAM</td><td>alignment</td><td>distance</td><td>position</td><td>gene name</td><td>gene id</td></tr>
<tr><td>GU305914.1</td><td>95227</td><td>95249</td><td>+</td><td>0</td><td>ACTCGGGGTGGGCTGGGGTG</td><td>GGG</td><td>||||||||[||||||||||||]PAM</td><td>NA</td><td>Intergenic</td><td>NA</td><td>NA</td></tr>
<tr><td></td><td></td><td></td><td></td><td></td><td></td><td></td><td></td><td></td><td></td><td></td><td></td></tr>
<tr><td>T6795</td><td>CTCGGGGTGGGCTGGGGTGGGGG</td><td>1000</td><td></td><td></td><td></td><td></td><td></td><td></td><td></td><td></td><td></td></tr>
<tr><td>Oligo fwd</td><td>CACCCTCGGGGTGGGCTGGGGTGG</td><td></td><td></td><td></td><td></td><td></td><td></td><td></td><td></td><td></td><td></td></tr>
<tr><td>Oligo rev</td><td>AAACCCACCCCAGCCCACCCCGAG</td><td></td><td></td><td></td><td></td><td></td><td></td><td></td><td></td><td></td><td></td></tr>
<tr><td>Chromosome</td><td>start</td><td>end</td><td>strand</td><td>MM</td><td>target_seq</td><td>PAM</td><td>alignment</td><td>distance</td><td>position</td><td>gene name</td><td>gene id</td></tr>
<tr><td>GU305914.1</td><td>95228</td><td>95250</td><td>+</td><td>0</td><td>CTCGGGGTGGGCTGGGGTGG</td><td>GGG</td><td>||||||||[||||||||||||]PAM</td><td>NA</td><td>Intergenic</td><td>NA</td><td>NA</td></tr>
<tr><td></td><td></td><td></td><td></td><td></td><td></td><td></td><td></td><td></td><td></td><td></td><td></td></tr>
<tr><td>T6796</td><td>GGGTGGGCTGGGGTGGGGGTTGG</td><td>1000</td><td></td><td></td><td></td><td></td><td></td><td></td><td></td><td></td><td></td></tr>
<tr><td>Oligo fwd</td><td>CACCGGGTGGGCTGGGGTGGGGGT</td><td></td><td></td><td></td><td></td><td></td><td></td><td></td><td></td><td></td><td></td></tr>
<tr><td>Oligo rev</td><td>AAACACCCCCACCCCAGCCCACCC</td><td></td><td></td><td></td><td></td><td></td><td></td><td></td><td></td><td></td><td></td></tr>
<tr><td>Chromosome</td><td>start</td><td>end</td><td>strand</td><td>MM</td><td>target_seq</td><td>PAM</td><td>alignment</td><td>distance</td><td>position</td><td>gene name</td><td>gene id</td></tr>
<tr><td>GU305914.1</td><td>95232</td><td>95254</td><td>+</td><td>0</td><td>GGGTGGGCTGGGGTGGGGGT</td><td>TGG</td><td>||||||||[||||||||||||]PAM</td><td>NA</td><td>Intergenic</td><td>NA</td><td>NA</td></tr>
<tr><td></td><td></td><td></td><td></td><td></td><td></td><td></td><td></td><td></td><td></td><td></td><td></td></tr>
<tr><td>T6797</td><td>GGTGGGCTGGGGTGGGGGTTGGG</td><td>1000</td><td></td><td></td><td></td><td></td><td></td><td></td><td></td><td></td><td></td></tr>
<tr><td>Oligo fwd</td><td>CACCGGTGGGCTGGGGTGGGGGTT</td><td></td><td></td><td></td><td></td><td></td><td></td><td></td><td></td><td></td><td></td></tr>
<tr><td>Oligo rev</td><td>AAACAACCCCCACCCCAGCCCACC</td><td></td><td></td><td></td><td></td><td></td><td></td><td></td><td></td><td></td><td></td></tr>
<tr><td>Chromosome</td><td>start</td><td>end</td><td>strand</td><td>MM</td><td>target_seq</td><td>PAM</td><td>alignment</td><td>distance</td><td>position</td><td>gene name</td><td>gene id</td></tr>
<tr><td>GU305914.1</td><td>95233</td><td>95255</td><td>+</td><td>0</td><td>GGTGGGCTGGGGTGGGGGTT</td><td>GGG</td><td>||||||||[||||||||||||]PAM</td><td>NA</td><td>Intergenic</td><td>NA</td><td>NA</td></tr>
<tr><td></td><td></td><td></td><td></td><td></td><td></td><td></td><td></td><td></td><td></td><td></td><td></td></tr>
<tr><td>T6798</td><td>GCTTGAATAATCTCCTTATAAGG</td><td>1000</td><td></td><td></td><td></td><td></td><td></td><td></td><td></td><td></td><td></td></tr>
<tr><td>Oligo fwd</td><td>CACCGCTTGAATAATCTCCTTATA</td><td></td><td></td><td></td><td></td><td></td><td></td><td></td><td></td><td></td><td></td></tr>
<tr><td>Oligo rev</td><td>AAACTATAAGGAGATTATTCAAGC</td><td></td><td></td><td></td><td></td><td></td><td></td><td></td><td></td><td></td><td></td></tr>
<tr><td>Chromosome</td><td>start</td><td>end</td><td>strand</td><td>MM</td><td>target_seq</td><td>PAM</td><td>alignment</td><td>distance</td><td>position</td><td>gene name</td><td>gene id</td></tr>
<tr><td>GU305914.1</td><td>95279</td><td>95301</td><td>+</td><td>0</td><td>GCTTGAATAATCTCCTTATA</td><td>AGG</td><td>||||||||[||||||||||||]PAM</td><td>NA</td><td>Intergenic</td><td>NA</td><td>NA</td></tr>
<tr><td></td><td></td><td></td><td></td><td></td><td></td><td></td><td></td><td></td><td></td><td></td><td></td></tr>
<tr><td>T6799</td><td>AATCTCCTTATAAGGACACAAGG</td><td>1000</td><td></td><td></td><td></td><td></td><td></td><td></td><td></td><td></td><td></td></tr>
<tr><td>Oligo fwd</td><td>CACCAATCTCCTTATAAGGACACA</td><td></td><td></td><td></td><td></td><td></td><td></td><td></td><td></td><td></td><td></td></tr>
<tr><td>Oligo rev</td><td>AAACTGTGTCCTTATAAGGAGATT</td><td></td><td></td><td></td><td></td><td></td><td></td><td></td><td></td><td></td><td></td></tr>
<tr><td>Chromosome</td><td>start</td><td>end</td><td>strand</td><td>MM</td><td>target_seq</td><td>PAM</td><td>alignment</td><td>distance</td><td>position</td><td>gene name</td><td>gene id</td></tr>
<tr><td>GU305914.1</td><td>95287</td><td>95309</td><td>+</td><td>0</td><td>AATCTCCTTATAAGGACACA</td><td>AGG</td><td>||||||||[||||||||||||]PAM</td><td>NA</td><td>Intergenic</td><td>NA</td><td>NA</td></tr>
<tr><td></td><td></td><td></td><td></td><td></td><td></td><td></td><td></td><td></td><td></td><td></td><td></td></tr>
<tr><td>T6800</td><td>CTCCTTATAAGGACACAAGGTGG</td><td>1000</td><td></td><td></td><td></td><td></td><td></td><td></td><td></td><td></td><td></td></tr>
<tr><td>Oligo fwd</td><td>CACCCTCCTTATAAGGACACAAGG</td><td></td><td></td><td></td><td></td><td></td><td></td><td></td><td></td><td></td><td></td></tr>
<tr><td>Oligo rev</td><td>AAACCCTTGTGTCCTTATAAGGAG</td><td></td><td></td><td></td><td></td><td></td><td></td><td></td><td></td><td></td><td></td></tr>
<tr><td>Chromosome</td><td>start</td><td>end</td><td>strand</td><td>MM</td><td>target_seq</td><td>PAM</td><td>alignment</td><td>distance</td><td>position</td><td>gene name</td><td>gene id</td></tr>
<tr><td>GU305914.1</td><td>95290</td><td>95312</td><td>+</td><td>0</td><td>CTCCTTATAAGGACACAAGG</td><td>TGG</td><td>||||||||[||||||||||||]PAM</td><td>NA</td><td>Intergenic</td><td>NA</td><td>NA</td></tr>
</table>

| | | | | | | | | | | | | |
|---|---|---|---|---|---|---|---|---|---|---|---|---|
| T6801 | CCCTCTCGACCACCACCTAGCGG | | 1000 | | | | | | | | | |
| Oligo fwd | CACCCCCTCTCGACCACCACCTAG | | | | | | | | | | | |
| Oligo rev | AAACCTAGGTGGTGGTCGAGAGGG | | | | | | | | | | | |
| Chromosome | start | | end | strand | MM | target_seq | PAM | alignment | distance | position | gene name | gene id |
| GU305914.1 | | 95323 | 95345 | + | 0 | CCCTCTCGACCACCACCTAG | CGG | \|\|\|\|\|\|\|\|[\|\|\|\|\|\|\|\|\|\|\|\|]PAM | NA | Intergenic | NA | NA |

| | | | | | | | | | | | | |
|---|---|---|---|---|---|---|---|---|---|---|---|---|
| T6802 | CACCTAGCGGACGACCTCGTTGG | | 1000 | | | | | | | | | |
| Oligo fwd | CACCCACCTAGCGGACGACCTCGT | | | | | | | | | | | |
| Oligo rev | AAACACGAGGTCGTCCGCTAGGTG | | | | | | | | | | | |
| Chromosome | start | | end | strand | MM | target_seq | PAM | alignment | distance | position | gene name | gene id |
| GU305914.1 | | 95336 | 95358 | + | 0 | CACCTAGCGGACGACCTCGT | TGG | \|\|\|\|\|\|\|\|[\|\|\|\|\|\|\|\|\|\|\|\|]PAM | NA | Intergenic | NA | NA |

| | | | | | | | | | | | | |
|---|---|---|---|---|---|---|---|---|---|---|---|---|
| T6803 | ACCTAGCGGACGACCTCGTTGGG | | 1000 | | | | | | | | | |
| Oligo fwd | CACCACCTAGCGGACGACCTCGTT | | | | | | | | | | | |
| Oligo rev | AAACAACGAGGTCGTCCGCTAGGT | | | | | | | | | | | |
| Chromosome | start | | end | strand | MM | target_seq | PAM | alignment | distance | position | gene name | gene id |
| GU305914.1 | | 95337 | 95359 | + | 0 | ACCTAGCGGACGACCTCGTT | GGG | \|\|\|\|\|\|\|\|[\|\|\|\|\|\|\|\|\|\|\|\|]PAM | NA | Intergenic | NA | NA |

| | | | | | | | | | | | | |
|---|---|---|---|---|---|---|---|---|---|---|---|---|
| T6804 | TAGCGGACGACCTCGTTGGGTGG | | 1000 | | | | | | | | | |
| Oligo fwd | CACCTAGCGGACGACCTCGTTGGG | | | | | | | | | | | |
| Oligo rev | AAACCCCAACGAGGTCGTCCGCTA | | | | | | | | | | | |
| Chromosome | start | | end | strand | MM | target_seq | PAM | alignment | distance | position | gene name | gene id |
| GU305914.1 | | 95340 | 95362 | + | 0 | TAGCGGACGACCTCGTTGGG | TGG | \|\|\|\|\|\|\|\|[\|\|\|\|\|\|\|\|\|\|\|\|]PAM | NA | Intergenic | NA | NA |

| | | | | | | | | | | | | |
|---|---|---|---|---|---|---|---|---|---|---|---|---|
| T6805 | AGCGGACGACCTCGTTGGGTGGG | | 1000 | | | | | | | | | |
| Oligo fwd | CACCAGCGGACGACCTCGTTGGGT | | | | | | | | | | | |
| Oligo rev | AAACACCCAACGAGGTCGTCCGCT | | | | | | | | | | | |
| Chromosome | start | | end | strand | MM | target_seq | PAM | alignment | distance | position | gene name | gene id |
| GU305914.1 | | 95341 | 95363 | + | 0 | AGCGGACGACCTCGTTGGGT | GGG | \|\|\|\|\|\|\|\|[\|\|\|\|\|\|\|\|\|\|\|\|]PAM | NA | Intergenic | NA | NA |

| | | | | | | | | | | | | |
|---|---|---|---|---|---|---|---|---|---|---|---|---|
| T6806 | GCGGACGACCTCGTTGGGTGGGG | | 1000 | | | | | | | | | |
| Oligo fwd | CACCGCGGACGACCTCGTTGGGTG | | | | | | | | | | | |
| Oligo rev | AAACCACCCAACGAGGTCGTCCGC | | | | | | | | | | | |
| Chromosome | start | | end | strand | MM | target_seq | PAM | alignment | distance | position | gene name | gene id |
| GU305914.1 | | 95342 | 95364 | + | 0 | GCGGACGACCTCGTTGGGTG | GGG | \|\|\|\|\|\|\|\|[\|\|\|\|\|\|\|\|\|\|\|\|]PAM | NA | Intergenic | NA | NA |

| | | | | | | | | | | | | |
|---|---|---|---|---|---|---|---|---|---|---|---|---|
| T6807 | CGGACGACCTCGTTGGGTGGGGG | | 1000 | | | | | | | | | |
| Oligo fwd | CACCCGGACGACCTCGTTGGGTGG | | | | | | | | | | | |
| Oligo rev | AAACCCACCCAACGAGGTCGTCCG | | | | | | | | | | | |
| Chromosome | start | | end | strand | MM | target_seq | PAM | alignment | distance | position | gene name | gene id |
| GU305914.1 | | 95343 | 95365 | + | 0 | CGGACGACCTCGTTGGGTGG | GGG | \|\|\|\|\|\|\|\|[\|\|\|\|\|\|\|\|\|\|\|\|]PAM | NA | Intergenic | NA | NA |

| | | | | | | | | | | | | |
|---|---|---|---|---|---|---|---|---|---|---|---|---|
| T6808 | GGACGACCTCGTTGGGTGGGGGG | | 1000 | | | | | | | | | |
| Oligo fwd | CACCGGACGACCTCGTTGGGTGGG | | | | | | | | | | | |
| Oligo rev | AAACCCCACCCAACGAGGTCGTCC | | | | | | | | | | | |
| Chromosome | start | | end | strand | MM | target_seq | PAM | alignment | distance | position | gene name | gene id |
| GU305914.1 | | 95344 | 95366 | + | 0 | GGACGACCTCGTTGGGTGGG | GGG | \|\|\|\|\|\|\|\|[\|\|\|\|\|\|\|\|\|\|\|\|]PAM | NA | Intergenic | NA | NA |

| | | | | | | | | | | | | |
|---|---|---|---|---|---|---|---|---|---|---|---|---|
| T6809 | GGGTAATAAGAAATAGTGTGTGG | | 1000 | | | | | | | | | |
| Oligo fwd | CACCGGGTAATAAGAAATAGTGTG | | | | | | | | | | | |
| Oligo rev | AAACCACACTATTTCTTATTACCC | | | | | | | | | | | |
| Chromosome | start | | end | strand | MM | target_seq | PAM | alignment | distance | position | gene name | gene id |
| GU305914.1 | | 95364 | 95386 | + | 0 | GGGTAATAAGAAATAGTGTG | TGG | \|\|\|\|\|\|\|\|[\|\|\|\|\|\|\|\|\|\|\|\|]PAM | NA | Intergenic | NA | NA |

| | | | | | | | | | | | | |
|---|---|---|---|---|---|---|---|---|---|---|---|---|
| T6810 | TAATAAGAAATAGTGTGTGGTGG | | 1000 | | | | | | | | | |
| Oligo fwd | CACCTAATAAGAAATAGTGTGTGG | | | | | | | | | | | |
| Oligo rev | AAACCCACACACTATTTCTTATTA | | | | | | | | | | | |
| Chromosome | start | | end | strand | MM | target_seq | PAM | alignment | distance | position | gene name | gene id |
| GU305914.1 | | 95367 | 95389 | + | 0 | TAATAAGAAATAGTGTGTGG | TGG | \|\|\|\|\|\|\|\|[\|\|\|\|\|\|\|\|\|\|\|\|]PAM | NA | Intergenic | NA | NA |

| | | | | | | | | | | | | |
|---|---|---|---|---|---|---|---|---|---|---|---|---|
| T6811 | AATAAGAAATAGTGTGTGGTGGG | | 1000 | | | | | | | | | |
| Oligo fwd | CACCAATAAGAAATAGTGTGTGGT | | | | | | | | | | | |
| Oligo rev | AAACACCACACACTATTTCTTATT | | | | | | | | | | | |
| Chromosome | start | | end | strand | MM | target_seq | PAM | alignment | distance | position | gene name | gene id |
| GU305914.1 | | 95368 | 95390 | + | 0 | AATAAGAAATAGTGTGTGGT | GGG | \|\|\|\|\|\|\|\|[\|\|\|\|\|\|\|\|\|\|\|\|]PAM | NA | Intergenic | NA | NA |

| | | | | | | | | | | | | |
|---|---|---|---|---|---|---|---|---|---|---|---|---|
| T6812 | ATAAGAAATAGTGTGTGGTGGGG | | 1000 | | | | | | | | | |
| Oligo fwd | CACCATAAGAAATAGTGTGTGGTG | | | | | | | | | | | |
| Oligo rev | AAACCACCACACACTATTTCTTAT | | | | | | | | | | | |
| Chromosome | start | | end | strand | MM | target_seq | PAM | alignment | distance | position | gene name | gene id |
| GU305914.1 | | 95369 | 95391 | + | 0 | ATAAGAAATAGTGTGTGGTG | GGG | \|\|\|\|\|\|\|\|[\|\|\|\|\|\|\|\|\|\|\|\|]PAM | NA | Intergenic | NA | NA |

| | | | | | | | | | | | | |
|---|---|---|---|---|---|---|---|---|---|---|---|---|
| T6813 | TAAGAAATAGTGTGTGGTGGGGG | | 1000 | | | | | | | | | |
| Oligo fwd | CACCTAAGAAATAGTGTGTGGTGG | | | | | | | | | | | |
| Oligo rev | AAACCCACCACACACTATTTCTTA | | | | | | | | | | | |
| Chromosome | start | | end | strand | MM | target_seq | PAM | alignment | distance | position | gene name | gene id |
| GU305914.1 | | 95370 | 95392 | + | 0 | TAAGAAATAGTGTGTGGTGG | GGG | \|\|\|\|\|\|\|\|[\|\|\|\|\|\|\|\|\|\|\|\|]PAM | NA | Intergenic | NA | NA |

| | | | | | | | | | | | | |
|---|---|---|---|---|---|---|---|---|---|---|---|---|
| T6814 | AAGAAATAGTGTGTGGTGGGGGG | | 1000 | | | | | | | | | |
| Oligo fwd | CACCAAGAAATAGTGTGTGGTGGG | | | | | | | | | | | |
| Oligo rev | AAACCCCACCACACACTATTTCTT | | | | | | | | | | | |
| Chromosome | start | | end | strand | MM | target_seq | PAM | alignment | distance | position | gene name | gene id |
| GU305914.1 | | 95371 | 95393 | + | 0 | AAGAAATAGTGTGTGGTGGG | GGG | \|\|\|\|\|\|\|\|[\|\|\|\|\|\|\|\|\|\|\|\|]PAM | NA | Intergenic | NA | NA |

| | | | | | | | | | | | | |
|---|---|---|---|---|---|---|---|---|---|---|---|---|
| T6815 | AATAGTGTGTGGTGGGGGGATGG | | 1000 | | | | | | | | | |
| Oligo fwd | CACCAATAGTGTGTGGTGGGGGGA | | | | | | | | | | | |
| Oligo rev | AAACTCCCCCCACCACACACTATT | | | | | | | | | | | |
| Chromosome | start | | end | strand | MM | target_seq | PAM | alignment | distance | position | gene name | gene id |
| GU305914.1 | | 95375 | 95397 | + | 0 | AATAGTGTGTGGTGGGGGGA | TGG | \|\|\|\|\|\|\|\|[\|\|\|\|\|\|\|\|\|\|\|\|]PAM | NA | Intergenic | NA | NA |

T6816 AGACGAAAAACTCATGTTGATGG 1000
Oligo fwd CACCAGACGAAAAACTCATGTTGA
Oligo rev AAACTCAACATGAGTTTTTCGTCT
Chromosome start end strand MM target_seq PAM alignment distance position gene name gene id
GU305914.1 95431 95453 + 0 AGACGAAAAACTCATGTTGA TGG ||||||||[||||||||||||]PAM NA Intergenic NA NA

T6817 GACGAAAAACTCATGTTGATGGG 1000
Oligo fwd CACCGACGAAAAACTCATGTTGAT
Oligo rev AAACATCAACATGAGTTTTTCGTC
Chromosome start end strand MM target_seq PAM alignment distance position gene name gene id
GU305914.1 95432 95454 + 0 GACGAAAAACTCATGTTGAT GGG ||||||||[||||||||||||]PAM NA Intergenic NA NA

T6818 ACGAAAAACTCATGTTGATGGGG 1000
Oligo fwd CACCACGAAAAACTCATGTTGATG
Oligo rev AAACCATCAACATGAGTTTTTCGT
Chromosome start end strand MM target_seq PAM alignment distance position gene name gene id
GU305914.1 95433 95455 + 0 ACGAAAAACTCATGTTGATG GGG ||||||||[||||||||||||]PAM NA Intergenic NA NA

T6819 CGAAAAACTCATGTTGATGGGGG 1000
Oligo fwd CACCCGAAAAACTCATGTTGATGG
Oligo rev AAACCCATCAACATGAGTTTTTCG
Chromosome start end strand MM target_seq PAM alignment distance position gene name gene id
GU305914.1 95434 95456 + 0 CGAAAAACTCATGTTGATGG GGG ||||||||[||||||||||||]PAM NA Intergenic NA NA

T6820 GAAAAACTCATGTTGATGGGGGG 1000
Oligo fwd CACCGAAAAACTCATGTTGATGGG
Oligo rev AAACCCCATCAACATGAGTTTTTC
Chromosome start end strand MM target_seq PAM alignment distance position gene name gene id
GU305914.1 95435 95457 + 0 GAAAAACTCATGTTGATGGG GGG ||||||||[||||||||||||]PAM NA Intergenic NA NA

T6821 GGTAATAAGAAAGCATGCAGTGG 1000
Oligo fwd CACCGGTAATAAGAAAGCATGCAG
Oligo rev AAACCTGCATGCTTTCTTATTACC
Chromosome start end strand MM target_seq PAM alignment distance position gene name gene id
GU305914.1 95456 95478 + 0 GGTAATAAGAAAGCATGCAG TGG ||||||||[||||||||||||]PAM NA Intergenic NA NA

T6822 GTAATAAGAAAGCATGCAGTGGG 1000
Oligo fwd CACCGTAATAAGAAAGCATGCAGT
Oligo rev AAACACTGCATGCTTTCTTATTAC
Chromosome start end strand MM target_seq PAM alignment distance position gene name gene id
GU305914.1 95457 95479 + 0 GTAATAAGAAAGCATGCAGT GGG ||||||||[||||||||||||]PAM NA Intergenic NA NA

T6823 TAATAAGAAAGCATGCAGTGGGG 1000
Oligo fwd CACCTAATAAGAAAGCATGCAGTG
Oligo rev AAACCACTGCATGCTTTCTTATTA
Chromosome start end strand MM target_seq PAM alignment distance position gene name gene id
GU305914.1 95458 95480 + 0 TAATAAGAAAGCATGCAGTG GGG ||||||||[||||||||||||]PAM NA Intergenic NA NA

T6824 AATAAGAAAGCATGCAGTGGGGG 1000
Oligo fwd CACCAATAAGAAAGCATGCAGTGG
Oligo rev AAACCCACTGCATGCTTTCTTATT
Chromosome start end strand MM target_seq PAM alignment distance position gene name gene id
GU305914.1 95459 95481 + 0 AATAAGAAAGCATGCAGTGG GGG ||||||||[||||||||||||]PAM NA Intergenic NA NA

T6825 AGAAAGCATGCAGTGGGGGTTGG 1000
Oligo fwd CACCAGAAAGCATGCAGTGGGGGT
Oligo rev AAACACCCCCACTGCATGCTTTCT
Chromosome start end strand MM target_seq PAM alignment distance position gene name gene id
GU305914.1 95463 95485 + 0 AGAAAGCATGCAGTGGGGGT TGG ||||||||[||||||||||||]PAM NA Intergenic NA NA

T6826 GAAAGCATGCAGTGGGGGTTGGG 1000
Oligo fwd CACCGAAAGCATGCAGTGGGGGTT
Oligo rev AAACAACCCCCACTGCATGCTTTC
Chromosome start end strand MM target_seq PAM alignment distance position gene name gene id
GU305914.1 95464 95486 + 0 GAAAGCATGCAGTGGGGGTT GGG ||||||||[||||||||||||]PAM NA Intergenic NA NA

T6827 CATGCAGTGGGGGTTGGGTGTGG 1000
Oligo fwd CACCCATGCAGTGGGGGTTGGGTG
Oligo rev AAACCACCCAACCCCCACTGCATG
Chromosome start end strand MM target_seq PAM alignment distance position gene name gene id
GU305914.1 95469 95491 + 0 CATGCAGTGGGGGTTGGGTG TGG ||||||||[||||||||||||]PAM NA Intergenic NA NA

T6828 AGTGGGGGTTGGGTGTGGAGTGG 1000
Oligo fwd CACCAGTGGGGGTTGGGTGTGGAG
Oligo rev AAACCTCCACACCCAACCCCCACT
Chromosome start end strand MM target_seq PAM alignment distance position gene name gene id
GU305914.1 95474 95496 + 0 AGTGGGGGTTGGGTGTGGAG TGG ||||||||[||||||||||||]PAM NA Intergenic NA NA

T6829 GTGGGGGTTGGGTGTGGAGTGGG 1000
Oligo fwd CACCGTGGGGGTTGGGTGTGGAGT
Oligo rev AAACACTCCACACCCAACCCCCAC
Chromosome start end strand MM target_seq PAM alignment distance position gene name gene id
GU305914.1 95475 95497 + 0 GTGGGGGTTGGGTGTGGAGT GGG ||||||||[||||||||||||]PAM NA Intergenic NA NA

T6830 TGGGGGTTGGGTGTGGAGTGGGG 1000
Oligo fwd CACCTGGGGGTTGGGTGTGGAGTG
Oligo rev AAACCACTCCACACCCAACCCCCA
Chromosome start end strand MM target_seq PAM alignment distance position gene name gene id

GU305914.1 95476 95498 + 0 TGGGGGTTGGGTGTGGAGTG GGG ||||||||[||||||||||||]PAM NA Intergenic NA NA

T6831 GGGGGTTGGGTGTGGAGTGGGGG 1000
Oligo fwd CACCGGGGGTTGGGTGTGGAGTGG
Oligo rev AAACCCACTCCACACCCAACCCCC
Chromosome start end strand MM target_seq PAM alignment distance position gene name gene id
GU305914.1 95477 95499 + 0 GGGGGTTGGGTGTGGAGTGG GGG ||||||||[||||||||||||]PAM NA Intergenic NA NA

T6832 GGTGTGGAGTGGGGGTAATAAGG 1000
Oligo fwd CACCGGTGTGGAGTGGGGGTAATA
Oligo rev AAACTATTACCCCCACTCCACACC
Chromosome start end strand MM target_seq PAM alignment distance position gene name gene id
GU305914.1 95485 95507 + 0 GGTGTGGAGTGGGGGTAATA AGG ||||||||[||||||||||||]PAM NA Intergenic NA NA

T6833 GTGGGGGTAATAAGGATTCATGG 1000
Oligo fwd CACCGTGGGGGTAATAAGGATTCA
Oligo rev AAACTGAATCCTTATTACCCCCAC
Chromosome start end strand MM target_seq PAM alignment distance position gene name gene id
GU305914.1 95493 95515 + 0 GTGGGGGTAATAAGGATTCA TGG ||||||||[||||||||||||]PAM NA Intergenic NA NA

T6834 TGGGGGTAATAAGGATTCATGGG 1000
Oligo fwd CACCTGGGGGTAATAAGGATTCAT
Oligo rev AAACATGAATCCTTATTACCCCCA
Chromosome start end strand MM target_seq PAM alignment distance position gene name gene id
GU305914.1 95494 95516 + 0 TGGGGGTAATAAGGATTCAT GGG ||||||||[||||||||||||]PAM NA Intergenic NA NA

T6835 GGGGGTAATAAGGATTCATGGGG 1000
Oligo fwd CACCGGGGGTAATAAGGATTCATG
Oligo rev AAACCATGAATCCTTATTACCCCC
Chromosome start end strand MM target_seq PAM alignment distance position gene name gene id
GU305914.1 95495 95517 + 0 GGGGGTAATAAGGATTCATG GGG ||||||||[||||||||||||]PAM NA Intergenic NA NA

T6836 TTAGCTAGATTTCAGACCCGAGG 1000
Oligo fwd CACCTTAGCTAGATTTCAGACCCG
Oligo rev AAACCGGGTCTGAAATCTAGCTAA
Chromosome start end strand MM target_seq PAM alignment distance position gene name gene id
GU305914.1 95524 95546 + 0 TTAGCTAGATTTCAGACCCG AGG ||||||||[||||||||||||]PAM NA Intergenic NA NA

T6837 TAGCTAGATTTCAGACCCGAGGG 1000
Oligo fwd CACCTAGCTAGATTTCAGACCCGA
Oligo rev AAACTCGGGTCTGAAATCTAGCTA
Chromosome start end strand MM target_seq PAM alignment distance position gene name gene id
GU305914.1 95525 95547 + 0 TAGCTAGATTTCAGACCCGA GGG ||||||||[||||||||||||]PAM NA Intergenic NA NA

T6838 AGCTAGATTTCAGACCCGAGGGG 1000
Oligo fwd CACCAGCTAGATTTCAGACCCGAG
Oligo rev AAACCTCGGGTCTGAAATCTAGCT
Chromosome start end strand MM target_seq PAM alignment distance position gene name gene id
GU305914.1 95526 95548 + 0 AGCTAGATTTCAGACCCGAG GGG ||||||||[||||||||||||]PAM NA Intergenic NA NA

T6839 GCTAGATTTCAGACCCGAGGGGG 1000
Oligo fwd CACCGCTAGATTTCAGACCCGAGG
Oligo rev AAACCCTCGGGTCTGAAATCTAGC
Chromosome start end strand MM target_seq PAM alignment distance position gene name gene id
GU305914.1 95527 95549 + 0 GCTAGATTTCAGACCCGAGG GGG ||||||||[||||||||||||]PAM NA Intergenic NA NA

T6840 TCAGACCCGAGGGGGTGTGTCGG 1000
Oligo fwd CACCTCAGACCCGAGGGGGTGTGT
Oligo rev AAACACACACCCCCTCGGGTCTGA
Chromosome start end strand MM target_seq PAM alignment distance position gene name gene id
GU305914.1 95535 95557 + 0 TCAGACCCGAGGGGGTGTGT CGG ||||||||[||||||||||||]PAM NA Intergenic NA NA

T6841 CCGAGGGGGTGTGTCGGTGTAGG 1000
Oligo fwd CACCCCGAGGGGGTGTGTCGGTGT
Oligo rev AAACACACCGACACACCCCCTCGG
Chromosome start end strand MM target_seq PAM alignment distance position gene name gene id
GU305914.1 95541 95563 + 0 CCGAGGGGGTGTGTCGGTGT AGG ||||||||[||||||||||||]PAM NA Intergenic NA NA
GU305914.1 32103 32125 - 4 CCGTGAGGGTCTGTCTGTGT CGG |||-|-||[||-||||-||||]PAM NA Intergenic NA NA

T6842 GTCCACTGCGCATGCGTGTGAGG 1000
Oligo fwd CACCGTCCACTGCGCATGCGTGTG
Oligo rev AAACCACACGCATGCGCAGTGGAC
Chromosome start end strand MM target_seq PAM alignment distance position gene name gene id
GU305914.1 95590 95612 + 0 GTCCACTGCGCATGCGTGTG AGG ||||||||[||||||||||||]PAM NA Intergenic NA NA

T6843 TCGTTCATCTATAGCATGATCGG 1000
Oligo fwd CACCTCGTTCATCTATAGCATGAT
Oligo rev AAACATCATGCTATAGATGAACGA
Chromosome start end strand MM target_seq PAM alignment distance position gene name gene id
GU305914.1 95637 95659 + 0 TCGTTCATCTATAGCATGAT CGG ||||||||[||||||||||||]PAM NA Intergenic NA NA

T6844 CATGATCGGCACATGCGCAGTGG 1000
Oligo fwd CACCCATGATCGGCACATGCGCAG
Oligo rev AAACCTGCGCATGTGCCGATCATG
Chromosome start end strand MM target_seq PAM alignment distance position gene name gene id
GU305914.1 95651 95673 + 0 CATGATCGGCACATGCGCAG TGG ||||||||[||||||||||||]PAM NA Intergenic NA NA

T6845 GATCAGATGCATGTGCAAAATGG 1000
Oligo fwd CACCGATCAGATGCATGTGCAAAA

Oligo rev AAACTTTTGCACATGCATCTGATC
Chromosome start end strand MM target_seq PAM alignment distance position gene name gene id
GU305914.1 95730 95752 + 0 GATCAGATGCATGTGCAAAA TGG ||||||||[||||||||||||]PAM NA Intergenic NA NA

T6846 CATATACTGACTCAACGTCCCGG 1000
Oligo fwd CACCCATATACTGACTCAACGTCC
Oligo rev AAACGGACGTTGAGTCAGTATATG
Chromosome start end strand MM target_seq PAM alignment distance position gene name gene id
GU305914.1 95786 95808 + 0 CATATACTGACTCAACGTCC CGG ||||||||[||||||||||||]PAM NA Intergenic NA NA

T6847 ATATACTGACTCAACGTCCCGGG 1000
Oligo fwd CACCATATACTGACTCAACGTCCC
Oligo rev AAACGGGACGTTGAGTCAGTATAT
Chromosome start end strand MM target_seq PAM alignment distance position gene name gene id
GU305914.1 95787 95809 + 0 ATATACTGACTCAACGTCCC GGG ||||||||[||||||||||||]PAM NA Intergenic NA NA

T6848 CGTCCCGGGAAGTGTGCCAGTGG 1000
Oligo fwd CACCCGTCCCGGGAAGTGTGCCAG
Oligo rev AAACCTGGCACACTTCCCGGGACG
Chromosome start end strand MM target_seq PAM alignment distance position gene name gene id
GU305914.1 95801 95823 + 0 CGTCCCGGGAAGTGTGCCAG TGG ||||||||[||||||||||||]PAM NA Intergenic NA NA

T6849 GTCCCGGGAAGTGTGCCAGTGGG 1000
Oligo fwd CACCGTCCCGGGAAGTGTGCCAGT
Oligo rev AAACACTGGCACACTTCCCGGGAC
Chromosome start end strand MM target_seq PAM alignment distance position gene name gene id
GU305914.1 95802 95824 + 0 GTCCCGGGAAGTGTGCCAGT GGG ||||||||[||||||||||||]PAM NA Intergenic NA NA

T6850 GCCAGTGGGCCTATGTGTATCGG 1000
Oligo fwd CACCGCCAGTGGGCCTATGTGTAT
Oligo rev AAACATACACATAGGCCCACTGGC
Chromosome start end strand MM target_seq PAM alignment distance position gene name gene id
GU305914.1 95816 95838 + 0 GCCAGTGGGCCTATGTGTAT CGG ||||||||[||||||||||||]PAM NA Intergenic NA NA

T6851 GCCTATGTGTATCGGTCGCTAGG 1000
Oligo fwd CACCGCCTATGTGTATCGGTCGCT
Oligo rev AAACAGCGACCGATACACATAGGC
Chromosome start end strand MM target_seq PAM alignment distance position gene name gene id
GU305914.1 95824 95846 + 0 GCCTATGTGTATCGGTCGCT AGG ||||||||[||||||||||||]PAM NA Intergenic NA NA

T6852 GTGTATCGGTCGCTAGGATACGG 1000
Oligo fwd CACCGTGTATCGGTCGCTAGGATA
Oligo rev AAACTATCCTAGCGACCGATACAC
Chromosome start end strand MM target_seq PAM alignment distance position gene name gene id
GU305914.1 95830 95852 + 0 GTGTATCGGTCGCTAGGATA CGG ||||||||[||||||||||||]PAM NA Intergenic NA NA

T6853 CAACCCAGAACTCCGATATTCGG 1000
Oligo fwd CACCCAACCCAGAACTCCGATATT
Oligo rev AAACAATATCGGAGTTCTGGGTTG
Chromosome start end strand MM target_seq PAM alignment distance position gene name gene id
GU305914.1 95899 95921 + 0 CAACCCAGAACTCCGATATT CGG ||||||||[||||||||||||]PAM NA Intergenic NA NA

T6854 CCAGAACTCCGATATTCGGATGG 1000
Oligo fwd CACCCCAGAACTCCGATATTCGGA
Oligo rev AAACTCCGAATATCGGAGTTCTGG
Chromosome start end strand MM target_seq PAM alignment distance position gene name gene id
GU305914.1 95903 95925 + 0 CCAGAACTCCGATATTCGGA TGG ||||||||[||||||||||||]PAM NA Intergenic NA NA

T6855 GGTAGCCAGCTTATCGTTGATGG 1000
Oligo fwd CACCGGTAGCCAGCTTATCGTTGA
Oligo rev AAACTCAACGATAAGCTGGCTACC
Chromosome start end strand MM target_seq PAM alignment distance position gene name gene id
GU305914.1 95924 95946 + 0 GGTAGCCAGCTTATCGTTGA TGG ||||||||[||||||||||||]PAM NA Intergenic NA NA

T6856 CCACCAATCGAAAGCTCGTACGG 1000
Oligo fwd CACCCCACCAATCGAAAGCTCGTA
Oligo rev AAACTACGAGCTTTCGATTGGTGG
Chromosome start end strand MM target_seq PAM alignment distance position gene name gene id
GU305914.1 95957 95979 + 0 CCACCAATCGAAAGCTCGTA CGG ||||||||[||||||||||||]PAM NA Intergenic NA NA

T6857 CCCCGTGCACGTTGAGTGACAGG 1000
Oligo fwd CACCCCCCGTGCACGTTGAGTGAC
Oligo rev AAACGTCACTCAACGTGCACGGGG
Chromosome start end strand MM target_seq PAM alignment distance position gene name gene id
GU305914.1 95981 96003 + 0 CCCCGTGCACGTTGAGTGAC AGG ||||||||[||||||||||||]PAM NA Intergenic NA NA

T6858 GTTGAGTGACAGGTCGAACCCGG 1000
Oligo fwd CACCGTTGAGTGACAGGTCGAACC
Oligo rev AAACGGTTCGACCTGTCACTCAAC
Chromosome start end strand MM target_seq PAM alignment distance position gene name gene id
GU305914.1 95991 96013 + 0 GTTGAGTGACAGGTCGAACC CGG ||||||||[||||||||||||]PAM NA Intergenic NA NA

T6859 GAACCCGGCAAGATTTCTAACGG 1000
Oligo fwd CACCGAACCCGGCAAGATTTCTAA
Oligo rev AAACTTAGAAATCTTGCCGGGTTC
Chromosome start end strand MM target_seq PAM alignment distance position gene name gene id
GU305914.1 96006 96028 + 0 GAACCCGGCAAGATTTCTAA CGG ||||||||[||||||||||||]PAM NA Intergenic NA NA

T6860 TAGCGCCATCTAGTCATCGACGG 1000

<table>
<tr><td>Oligo fwd</td><td>CACCTAGCGCCATCTAGTCATCGA</td></tr>
<tr><td>Oligo rev</td><td>AAACTCGATGACTAGATGGCGCTA</td></tr>
<tr><td>Chromosome</td><td>start</td><td>end</td><td>strand</td><td>MM</td><td>target_seq</td><td>PAM</td><td>alignment</td><td>distance</td><td>position</td><td>gene name</td><td>gene id</td></tr>
<tr><td>GU305914.1</td><td>96034</td><td>96056</td><td>+</td><td>0</td><td>TAGCGCCATCTAGTCATCGA</td><td>CGG</td><td>||||||||[||||||||||||]PAM</td><td>NA</td><td>Intergenic</td><td>NA</td><td>NA</td></tr>
<tr><td>T6861</td><td>TAGTCATCGACGGTGCACATCGG</td><td>1000</td></tr>
<tr><td>Oligo fwd</td><td>CACCTAGTCATCGACGGTGCACAT</td></tr>
<tr><td>Oligo rev</td><td>AAACATGTGCACCGTCGATGACTA</td></tr>
<tr><td>Chromosome</td><td>start</td><td>end</td><td>strand</td><td>MM</td><td>target_seq</td><td>PAM</td><td>alignment</td><td>distance</td><td>position</td><td>gene name</td><td>gene id</td></tr>
<tr><td>GU305914.1</td><td>96044</td><td>96066</td><td>+</td><td>0</td><td>TAGTCATCGACGGTGCACAT</td><td>CGG</td><td>||||||||[||||||||||||]PAM</td><td>NA</td><td>Intergenic</td><td>NA</td><td>NA</td></tr>
<tr><td>T6862</td><td>ACAGAATGACGCTCGTTTGACGG</td><td>1000</td></tr>
<tr><td>Oligo fwd</td><td>CACCACAGAATGACGCTCGTTTGA</td></tr>
<tr><td>Oligo rev</td><td>AAACTCAAACGAGCGTCATTCTGT</td></tr>
<tr><td>Chromosome</td><td>start</td><td>end</td><td>strand</td><td>MM</td><td>target_seq</td><td>PAM</td><td>alignment</td><td>distance</td><td>position</td><td>gene name</td><td>gene id</td></tr>
<tr><td>GU305914.1</td><td>96089</td><td>96111</td><td>+</td><td>0</td><td>ACAGAATGACGCTCGTTTGA</td><td>CGG</td><td>||||||||[||||||||||||]PAM</td><td>NA</td><td>Intergenic</td><td>NA</td><td>NA</td></tr>
<tr><td>T6863</td><td>CAGAATGACGCTCGTTTGACGGG</td><td>1000</td></tr>
<tr><td>Oligo fwd</td><td>CACCCAGAATGACGCTCGTTTGAC</td></tr>
<tr><td>Oligo rev</td><td>AAACGTCAAACGAGCGTCATTCTG</td></tr>
<tr><td>Chromosome</td><td>start</td><td>end</td><td>strand</td><td>MM</td><td>target_seq</td><td>PAM</td><td>alignment</td><td>distance</td><td>position</td><td>gene name</td><td>gene id</td></tr>
<tr><td>GU305914.1</td><td>96090</td><td>96112</td><td>+</td><td>0</td><td>CAGAATGACGCTCGTTTGAC</td><td>GGG</td><td>||||||||[||||||||||||]PAM</td><td>NA</td><td>Intergenic</td><td>NA</td><td>NA</td></tr>
<tr><td>T6864</td><td>TGACGCTCGTTTGACGGGCGCGG</td><td>1000</td></tr>
<tr><td>Oligo fwd</td><td>CACCTGACGCTCGTTTGACGGGCG</td></tr>
<tr><td>Oligo rev</td><td>AAACCGCCCGTCAAACGAGCGTCA</td></tr>
<tr><td>Chromosome</td><td>start</td><td>end</td><td>strand</td><td>MM</td><td>target_seq</td><td>PAM</td><td>alignment</td><td>distance</td><td>position</td><td>gene name</td><td>gene id</td></tr>
<tr><td>GU305914.1</td><td>96095</td><td>96117</td><td>+</td><td>0</td><td>TGACGCTCGTTTGACGGGCG</td><td>CGG</td><td>||||||||[||||||||||||]PAM</td><td>NA</td><td>Intergenic</td><td>NA</td><td>NA</td></tr>
<tr><td>T6865</td><td>ACGGGCGCGGACGCGATCTGAGG</td><td>1000</td></tr>
<tr><td>Oligo fwd</td><td>CACCACGGGCGCGGACGCGATCTG</td></tr>
<tr><td>Oligo rev</td><td>AAACCAGATCGCGTCCGCGCCCGT</td></tr>
<tr><td>Chromosome</td><td>start</td><td>end</td><td>strand</td><td>MM</td><td>target_seq</td><td>PAM</td><td>alignment</td><td>distance</td><td>position</td><td>gene name</td><td>gene id</td></tr>
<tr><td>GU305914.1</td><td>96108</td><td>96130</td><td>+</td><td>0</td><td>ACGGGCGCGGACGCGATCTG</td><td>AGG</td><td>||||||||[||||||||||||]PAM</td><td>NA</td><td>Intergenic</td><td>NA</td><td>NA</td></tr>
<tr><td>T6866</td><td>CGCGGACGCGATCTGAGGCTAGG</td><td>1000</td></tr>
<tr><td>Oligo fwd</td><td>CACCCGCGGACGCGATCTGAGGCT</td></tr>
<tr><td>Oligo rev</td><td>AAACAGCCTCAGATCGCGTCCGCG</td></tr>
<tr><td>Chromosome</td><td>start</td><td>end</td><td>strand</td><td>MM</td><td>target_seq</td><td>PAM</td><td>alignment</td><td>distance</td><td>position</td><td>gene name</td><td>gene id</td></tr>
<tr><td>GU305914.1</td><td>96113</td><td>96135</td><td>+</td><td>0</td><td>CGCGGACGCGATCTGAGGCT</td><td>AGG</td><td>||||||||[||||||||||||]PAM</td><td>NA</td><td>Intergenic</td><td>NA</td><td>NA</td></tr>
<tr><td>T6867</td><td>ACGCGATCTGAGGCTAGGAACGG</td><td>1000</td></tr>
<tr><td>Oligo fwd</td><td>CACCACGCGATCTGAGGCTAGGAA</td></tr>
<tr><td>Oligo rev</td><td>AAACTTCCTAGCCTCAGATCGCGT</td></tr>
<tr><td>Chromosome</td><td>start</td><td>end</td><td>strand</td><td>MM</td><td>target_seq</td><td>PAM</td><td>alignment</td><td>distance</td><td>position</td><td>gene name</td><td>gene id</td></tr>
<tr><td>GU305914.1</td><td>96118</td><td>96140</td><td>+</td><td>0</td><td>ACGCGATCTGAGGCTAGGAA</td><td>CGG</td><td>||||||||[||||||||||||]PAM</td><td>NA</td><td>Intergenic</td><td>NA</td><td>NA</td></tr>
<tr><td>T6868</td><td>ACGGACAACAGACGCTACAGCGG</td><td>1000</td></tr>
<tr><td>Oligo fwd</td><td>CACCACGGACAACAGACGCTACAG</td></tr>
<tr><td>Oligo rev</td><td>AAACCTGTAGCGTCTGTTGTCCGT</td></tr>
<tr><td>Chromosome</td><td>start</td><td>end</td><td>strand</td><td>MM</td><td>target_seq</td><td>PAM</td><td>alignment</td><td>distance</td><td>position</td><td>gene name</td><td>gene id</td></tr>
<tr><td>GU305914.1</td><td>96137</td><td>96159</td><td>+</td><td>0</td><td>ACGGACAACAGACGCTACAG</td><td>CGG</td><td>||||||||[||||||||||||]PAM</td><td>NA</td><td>Intergenic</td><td>NA</td><td>NA</td></tr>
<tr><td>T6869</td><td>GAAAATCGTACGAAAAACAACGG</td><td>1000</td></tr>
<tr><td>Oligo fwd</td><td>CACCGAAAATCGTACGAAAAACAA</td></tr>
<tr><td>Oligo rev</td><td>AAACTTGTTTTTCGTACGATTTTC</td></tr>
<tr><td>Chromosome</td><td>start</td><td>end</td><td>strand</td><td>MM</td><td>target_seq</td><td>PAM</td><td>alignment</td><td>distance</td><td>position</td><td>gene name</td><td>gene id</td></tr>
<tr><td>GU305914.1</td><td>96163</td><td>96185</td><td>+</td><td>0</td><td>GAAAATCGTACGAAAAACAA</td><td>CGG</td><td>||||||||[||||||||||||]PAM</td><td>NA</td><td>Intergenic</td><td>NA</td><td>NA</td></tr>
<tr><td>T6870</td><td>CAACGGTTTTTTATTAACGACGG</td><td>1000</td></tr>
<tr><td>Oligo fwd</td><td>CACCCAACGGTTTTTTATTAACGA</td></tr>
<tr><td>Oligo rev</td><td>AAACTCGTTAATAAAAAACCGTTG</td></tr>
<tr><td>Chromosome</td><td>start</td><td>end</td><td>strand</td><td>MM</td><td>target_seq</td><td>PAM</td><td>alignment</td><td>distance</td><td>position</td><td>gene name</td><td>gene id</td></tr>
<tr><td>GU305914.1</td><td>96180</td><td>96202</td><td>+</td><td>0</td><td>CAACGGTTTTTTATTAACGA</td><td>CGG</td><td>||||||||[||||||||||||]PAM</td><td>NA</td><td>Intergenic</td><td>NA</td><td>NA</td></tr>
<tr><td>T6871</td><td>CGGTTTTTTATTAACGACGGCGG</td><td>1000</td></tr>
<tr><td>Oligo fwd</td><td>CACCCGGTTTTTTATTAACGACGG</td></tr>
<tr><td>Oligo rev</td><td>AAACCCGTCGTTAATAAAAAACCG</td></tr>
<tr><td>Chromosome</td><td>start</td><td>end</td><td>strand</td><td>MM</td><td>target_seq</td><td>PAM</td><td>alignment</td><td>distance</td><td>position</td><td>gene name</td><td>gene id</td></tr>
<tr><td>GU305914.1</td><td>96183</td><td>96205</td><td>+</td><td>0</td><td>CGGTTTTTTATTAACGACGG</td><td>CGG</td><td>||||||||[||||||||||||]PAM</td><td>NA</td><td>Intergenic</td><td>NA</td><td>NA</td></tr>
<tr><td>T6872</td><td>GCAGCGTACCAACGTCTGAGAGG</td><td>1000</td></tr>
<tr><td>Oligo fwd</td><td>CACCGCAGCGTACCAACGTCTGAG</td></tr>
<tr><td>Oligo rev</td><td>AAACCTCAGACGTTGGTACGCTGC</td></tr>
<tr><td>Chromosome</td><td>start</td><td>end</td><td>strand</td><td>MM</td><td>target_seq</td><td>PAM</td><td>alignment</td><td>distance</td><td>position</td><td>gene name</td><td>gene id</td></tr>
<tr><td>GU305914.1</td><td>96210</td><td>96232</td><td>+</td><td>0</td><td>GCAGCGTACCAACGTCTGAG</td><td>AGG</td><td>||||||||[||||||||||||]PAM</td><td>NA</td><td>Intergenic</td><td>NA</td><td>NA</td></tr>
<tr><td>T6873</td><td>GTACCAACGTCTGAGAGGATAGG</td><td>1000</td></tr>
<tr><td>Oligo fwd</td><td>CACCGTACCAACGTCTGAGAGGAT</td></tr>
<tr><td>Oligo rev</td><td>AAACATCCTCTCAGACGTTGGTAC</td></tr>
<tr><td>Chromosome</td><td>start</td><td>end</td><td>strand</td><td>MM</td><td>target_seq</td><td>PAM</td><td>alignment</td><td>distance</td><td>position</td><td>gene name</td><td>gene id</td></tr>
<tr><td>GU305914.1</td><td>96215</td><td>96237</td><td>+</td><td>0</td><td>GTACCAACGTCTGAGAGGAT</td><td>AGG</td><td>||||||||[||||||||||||]PAM</td><td>NA</td><td>Intergenic</td><td>NA</td><td>NA</td></tr>
<tr><td>T6874</td><td>TACCAACGTCTGAGAGGATAGGG</td><td>1000</td></tr>
<tr><td>Oligo fwd</td><td>CACCTACCAACGTCTGAGAGGATA</td></tr>
<tr><td>Oligo rev</td><td>AAACTATCCTCTCAGACGTTGGTA</td></tr>
<tr><td>Chromosome</td><td>start</td><td>end</td><td>strand</td><td>MM</td><td>target_seq</td><td>PAM</td><td>alignment</td><td>distance</td><td>position</td><td>gene name</td><td>gene id</td></tr>
<tr><td>GU305914.1</td><td>96216</td><td>96238</td><td>+</td><td>0</td><td>TACCAACGTCTGAGAGGATA</td><td>GGG</td><td>||||||||[||||||||||||]PAM</td><td>NA</td><td>Intergenic</td><td>NA</td><td>NA</td></tr>
</table>

| | | | | | | | | | | | |
|---|---|---|---|---|---|---|---|---|---|---|---|
| T6875 | ACCAACGTCTGAGAGGATAGGGG | 1000 | | | | | | | | | |
| Oligo fwd | CACCACCAACGTCTGAGAGGATAG | | | | | | | | | | |
| Oligo rev | AAACCTATCCTCTCAGACGTTGGT | | | | | | | | | | |
| Chromosome | start | end | strand | MM | target_seq | PAM | alignment | distance | position | gene name | gene id |
| GU305914.1 | 96217 | 96239 | + | 0 | ACCAACGTCTGAGAGGATAG | GGG | \|\|\|\|\|\|\|\|[\|\|\|\|\|\|\|\|\|\|\|\|]PAM | NA | Intergenic | NA | NA |
| T6876 | CGTCTGAGAGGATAGGGGATCGG | 1000 | | | | | | | | | |
| Oligo fwd | CACCCGTCTGAGAGGATAGGGGAT | | | | | | | | | | |
| Oligo rev | AAACATCCCCTATCCTCTCAGACG | | | | | | | | | | |
| Chromosome | start | end | strand | MM | target_seq | PAM | alignment | distance | position | gene name | gene id |
| GU305914.1 | 96222 | 96244 | + | 0 | CGTCTGAGAGGATAGGGGAT | CGG | \|\|\|\|\|\|\|\|[\|\|\|\|\|\|\|\|\|\|\|\|]PAM | NA | Intergenic | NA | NA |
| T6877 | GTCTGAGAGGATAGGGGATCGGG | 1000 | | | | | | | | | |
| Oligo fwd | CACCGTCTGAGAGGATAGGGGATC | | | | | | | | | | |
| Oligo rev | AAACGATCCCCTATCCTCTCAGAC | | | | | | | | | | |
| Chromosome | start | end | strand | MM | target_seq | PAM | alignment | distance | position | gene name | gene id |
| GU305914.1 | 96223 | 96245 | + | 0 | GTCTGAGAGGATAGGGGATC | GGG | \|\|\|\|\|\|\|\|[\|\|\|\|\|\|\|\|\|\|\|\|]PAM | NA | Intergenic | NA | NA |
| T6878 | TCTGAGAGGATAGGGGATCGGGG | 1000 | | | | | | | | | |
| Oligo fwd | CACCTCTGAGAGGATAGGGGATCG | | | | | | | | | | |
| Oligo rev | AAACCGATCCCCTATCCTCTCAGA | | | | | | | | | | |
| Chromosome | start | end | strand | MM | target_seq | PAM | alignment | distance | position | gene name | gene id |
| GU305914.1 | 96224 | 96246 | + | 0 | TCTGAGAGGATAGGGGATCG | GGG | \|\|\|\|\|\|\|\|[\|\|\|\|\|\|\|\|\|\|\|\|]PAM | NA | Intergenic | NA | NA |
| T6879 | CGGGGAGTCACAGATATCGACGG | 1000 | | | | | | | | | |
| Oligo fwd | CACCCGGGGAGTCACAGATATCGA | | | | | | | | | | |
| Oligo rev | AAACTCGATATCTGTGACTCCCCG | | | | | | | | | | |
| Chromosome | start | end | strand | MM | target_seq | PAM | alignment | distance | position | gene name | gene id |
| GU305914.1 | 96242 | 96264 | + | 0 | CGGGGAGTCACAGATATCGA | CGG | \|\|\|\|\|\|\|\|[\|\|\|\|\|\|\|\|\|\|\|\|]PAM | NA | Intergenic | NA | NA |
| T6880 | ACAGATATCGACGGCGCGTTCGG | 1000 | | | | | | | | | |
| Oligo fwd | CACCACAGATATCGACGGCGCGTT | | | | | | | | | | |
| Oligo rev | AAACAACGCGCCGTCGATATCTGT | | | | | | | | | | |
| Chromosome | start | end | strand | MM | target_seq | PAM | alignment | distance | position | gene name | gene id |
| GU305914.1 | 96251 | 96273 | + | 0 | ACAGATATCGACGGCGCGTT | CGG | \|\|\|\|\|\|\|\|[\|\|\|\|\|\|\|\|\|\|\|\|]PAM | NA | Intergenic | NA | NA |
| T6881 | CAGATATCGACGGCGCGTTCGGG | 1000 | | | | | | | | | |
| Oligo fwd | CACCCAGATATCGACGGCGCGTTC | | | | | | | | | | |
| Oligo rev | AAACGAACGCGCCGTCGATATCTG | | | | | | | | | | |
| Chromosome | start | end | strand | MM | target_seq | PAM | alignment | distance | position | gene name | gene id |
| GU305914.1 | 96252 | 96274 | + | 0 | CAGATATCGACGGCGCGTTC | GGG | \|\|\|\|\|\|\|\|[\|\|\|\|\|\|\|\|\|\|\|\|]PAM | NA | Intergenic | NA | NA |
| T6882 | GTCAGAGTCCATCTCGCTGTAGG | 1000 | | | | | | | | | |
| Oligo fwd | CACCGTCAGAGTCCATCTCGCTGT | | | | | | | | | | |
| Oligo rev | AAACACAGCGAGATGGACTCTGAC | | | | | | | | | | |
| Chromosome | start | end | strand | MM | target_seq | PAM | alignment | distance | position | gene name | gene id |
| GU305914.1 | 96287 | 96309 | + | 0 | GTCAGAGTCCATCTCGCTGT | AGG | \|\|\|\|\|\|\|\|[\|\|\|\|\|\|\|\|\|\|\|\|]PAM | NA | Intergenic | NA | NA |
| T6883 | AGTCCATCTCGCTGTAGGAGAGG | 1000 | | | | | | | | | |
| Oligo fwd | CACCAGTCCATCTCGCTGTAGGAG | | | | | | | | | | |
| Oligo rev | AAACCTCCTACAGCGAGATGGACT | | | | | | | | | | |
| Chromosome | start | end | strand | MM | target_seq | PAM | alignment | distance | position | gene name | gene id |
| GU305914.1 | 96292 | 96314 | + | 0 | AGTCCATCTCGCTGTAGGAG | AGG | \|\|\|\|\|\|\|\|[\|\|\|\|\|\|\|\|\|\|\|\|]PAM | NA | Intergenic | NA | NA |
| T6884 | CTCCATGATCACGTCGTCCGTGG | 1000 | | | | | | | | | |
| Oligo fwd | CACCCTCCATGATCACGTCGTCCG | | | | | | | | | | |
| Oligo rev | AAACCGGACGACGTGATCATGGAG | | | | | | | | | | |
| Chromosome | start | end | strand | MM | target_seq | PAM | alignment | distance | position | gene name | gene id |
| GU305914.1 | 96320 | 96342 | + | 0 | CTCCATGATCACGTCGTCCG | TGG | \|\|\|\|\|\|\|\|[\|\|\|\|\|\|\|\|\|\|\|\|]PAM | NA | Intergenic | NA | NA |
| T6885 | TCCATGATCACGTCGTCCGTGGG | 1000 | | | | | | | | | |
| Oligo fwd | CACCTCCATGATCACGTCGTCCGT | | | | | | | | | | |
| Oligo rev | AAACACGGACGACGTGATCATGGA | | | | | | | | | | |
| Chromosome | start | end | strand | MM | target_seq | PAM | alignment | distance | position | gene name | gene id |
| GU305914.1 | 96321 | 96343 | + | 0 | TCCATGATCACGTCGTCCGT | GGG | \|\|\|\|\|\|\|\|[\|\|\|\|\|\|\|\|\|\|\|\|]PAM | NA | Intergenic | NA | NA |
| T6886 | CCATGATCACGTCGTCCGTGGGG | 1000 | | | | | | | | | |
| Oligo fwd | CACCCCATGATCACGTCGTCCGTG | | | | | | | | | | |
| Oligo rev | AAACCACGGACGACGTGATCATGG | | | | | | | | | | |
| Chromosome | start | end | strand | MM | target_seq | PAM | alignment | distance | position | gene name | gene id |
| GU305914.1 | 96322 | 96344 | + | 0 | CCATGATCACGTCGTCCGTG | GGG | \|\|\|\|\|\|\|\|[\|\|\|\|\|\|\|\|\|\|\|\|]PAM | NA | Intergenic | NA | NA |
| T6887 | CATGATCACGTCGTCCGTGGGGG | 1000 | | | | | | | | | |
| Oligo fwd | CACCCATGATCACGTCGTCCGTGG | | | | | | | | | | |
| Oligo rev | AAACCCACGGACGACGTGATCATG | | | | | | | | | | |
| Chromosome | start | end | strand | MM | target_seq | PAM | alignment | distance | position | gene name | gene id |
| GU305914.1 | 96323 | 96345 | + | 0 | CATGATCACGTCGTCCGTGG | GGG | \|\|\|\|\|\|\|\|[\|\|\|\|\|\|\|\|\|\|\|\|]PAM | NA | Intergenic | NA | NA |
| GU305914.1 | 224663 | 224685 | - | 4 | CGTCATCACGTGGTTCGTGG | CGG | \|-\|-\|\|\|\|[\|\|\|-\|\|-\|\|\|\|\|]PAM | NA | Intergenic | NA | NA |
| T6888 | TGGGGGTCGCGCTCCTCAAGTGG | 1000 | | | | | | | | | |
| Oligo fwd | CACCTGGGGGTCGCGCTCCTCAAG | | | | | | | | | | |
| Oligo rev | AAACCTTGAGGAGCGCGACCCCCA | | | | | | | | | | |
| Chromosome | start | end | strand | MM | target_seq | PAM | alignment | distance | position | gene name | gene id |
| GU305914.1 | 96340 | 96362 | + | 0 | TGGGGGTCGCGCTCCTCAAG | TGG | \|\|\|\|\|\|\|\|[\|\|\|\|\|\|\|\|\|\|\|\|]PAM | NA | Intergenic | NA | NA |
| T6889 | AAGTGGATGCGTTTGTTGAGCGG | 1000 | | | | | | | | | |
| Oligo fwd | CACCAAGTGGATGCGTTTGTTGAG | | | | | | | | | | |
| Oligo rev | AAACCTCAACAAACGCATCCACTT | | | | | | | | | | |
| Chromosome | start | end | strand | MM | target_seq | PAM | alignment | distance | position | gene name | gene id |

GU305914.1 96357 96379 + 0 AAGTGGATGCGTTTGTTGAG CGG ||||||||[||||||||||||]PAM NA Intergenic NA NA

T6890 GGATGCGTTTGTTGAGCGGCTGG 1000
Oligo fwd CACCGGATGCGTTTGTTGAGCGGC
Oligo rev AAACGCCGCTCAACAAACGCATCC
Chromosome start end strand MM target_seq PAM alignment distance position gene name gene id
GU305914.1 96361 96383 + 0 GGATGCGTTTGTTGAGCGGC TGG ||||||||[||||||||||||]PAM NA Intergenic NA NA

T6891 CTCGTCGTAGAAGTCCGTGTCGG 1000
Oligo fwd CACCCTCGTCGTAGAAGTCCGTGT
Oligo rev AAACACACGGACTTCTACGACGAG
Chromosome start end strand MM target_seq PAM alignment distance position gene name gene id
GU305914.1 96389 96411 + 0 CTCGTCGTAGAAGTCCGTGT CGG ||||||||[||||||||||||]PAM NA Intergenic NA NA

T6892 TCTTCTTCCTCCGCGTCTCCCGG 1000
Oligo fwd CACCTCTTCTTCCTCCGCGTCTCC
Oligo rev AAACGGAGACGCGGAGGAAGAAGA
Chromosome start end strand MM target_seq PAM alignment distance position gene name gene id
GU305914.1 96417 96439 + 0 TCTTCTTCCTCCGCGTCTCC CGG ||||||||[||||||||||||]PAM NA Intergenic NA NA

T6893 CCTCCGCGTCTCCCGGCTCGTGG 1000
Oligo fwd CACCCCTCCGCGTCTCCCGGCTCG
Oligo rev AAACCGAGCCGGGAGACGCGGAGG
Chromosome start end strand MM target_seq PAM alignment distance position gene name gene id
GU305914.1 96424 96446 + 0 CCTCCGCGTCTCCCGGCTCG TGG ||||||||[||||||||||||]PAM NA Intergenic NA NA

T6894 CGTCTCCCGGCTCGTGGTCGAGG 1000
Oligo fwd CACCCGTCTCCCGGCTCGTGGTCG
Oligo rev AAACCGACCACGAGCCGGGAGACG
Chromosome start end strand MM target_seq PAM alignment distance position gene name gene id
GU305914.1 96430 96452 + 0 CGTCTCCCGGCTCGTGGTCG AGG ||||||||[||||||||||||]PAM NA Intergenic NA NA

T6895 CCGGCTCGTGGTCGAGGATGCGG 1000
Oligo fwd CACCCCGGCTCGTGGTCGAGGATG
Oligo rev AAACCATCCTCGACCACGAGCCGG
Chromosome start end strand MM target_seq PAM alignment distance position gene name gene id
GU305914.1 96436 96458 + 0 CCGGCTCGTGGTCGAGGATG CGG ||||||||[||||||||||||]PAM NA Intergenic NA NA

T6896 CTCGTGGTCGAGGATGCGGATGG 1000
Oligo fwd CACCCTCGTGGTCGAGGATGCGGA
Oligo rev AAACTCCGCATCCTCGACCACGAG
Chromosome start end strand MM target_seq PAM alignment distance position gene name gene id
GU305914.1 96440 96462 + 0 CTCGTGGTCGAGGATGCGGA TGG ||||||||[||||||||||||]PAM NA Intergenic NA NA

T6897 TGGTCGAGGATGCGGATGGACGG 1000
Oligo fwd CACCTGGTCGAGGATGCGGATGGA
Oligo rev AAACTCCATCCGCATCCTCGACCA
Chromosome start end strand MM target_seq PAM alignment distance position gene name gene id
GU305914.1 96444 96466 + 0 TGGTCGAGGATGCGGATGGA CGG ||||||||[||||||||||||]PAM NA Intergenic NA NA

T6898 TCGAGGATGCGGATGGACGGCGG 1000
Oligo fwd CACCTCGAGGATGCGGATGGACGG
Oligo rev AAACCCGTCCATCCGCATCCTCGA
Chromosome start end strand MM target_seq PAM alignment distance position gene name gene id
GU305914.1 96447 96469 + 0 TCGAGGATGCGGATGGACGG CGG ||||||||[||||||||||||]PAM NA Intergenic NA NA

T6899 CGAGGATGCGGATGGACGGCGGG 1000
Oligo fwd CACCCGAGGATGCGGATGGACGGC
Oligo rev AAACGCCGTCCATCCGCATCCTCG
Chromosome start end strand MM target_seq PAM alignment distance position gene name gene id
GU305914.1 96448 96470 + 0 CGAGGATGCGGATGGACGGC GGG ||||||||[||||||||||||]PAM NA Intergenic NA NA

T6900 GAGGATGCGGATGGACGGCGGGG 1000
Oligo fwd CACCGAGGATGCGGATGGACGGCG
Oligo rev AAACCGCCGTCCATCCGCATCCTC
Chromosome start end strand MM target_seq PAM alignment distance position gene name gene id
GU305914.1 96449 96471 + 0 GAGGATGCGGATGGACGGCG GGG ||||||||[||||||||||||]PAM NA Intergenic NA NA

T6901 GATGCGGATGGACGGCGGGGCGG 1000
Oligo fwd CACCGATGCGGATGGACGGCGGGG
Oligo rev AAACCCCCGCCGTCCATCCGCATC
Chromosome start end strand MM target_seq PAM alignment distance position gene name gene id
GU305914.1 96452 96474 + 0 GATGCGGATGGACGGCGGGG CGG ||||||||[||||||||||||]PAM NA Intergenic NA NA

T6902 GGATGGACGGCGGGGCGGCCAGG 1000
Oligo fwd CACCGGATGGACGGCGGGGCGGCC
Oligo rev AAACGGCCGCCCCGCCGTCCATCC
Chromosome start end strand MM target_seq PAM alignment distance position gene name gene id
GU305914.1 96457 96479 + 0 GGATGGACGGCGGGGCGGCC AGG ||||||||[||||||||||||]PAM NA Intergenic NA NA
GU305914.1 52595 52617 - 4 GGGCGCACGGCGGGGCCGCC GGG ||--|-||[||||||||-|||]PAM NA Intergenic NA NA

T6903 GATGGACGGCGGGGCGGCCAGGG 1000
Oligo fwd CACCGATGGACGGCGGGGCGGCCA
Oligo rev AAACTGGCCGCCCCGCCGTCCATC
Chromosome start end strand MM target_seq PAM alignment distance position gene name gene id
GU305914.1 96458 96480 + 0 GATGGACGGCGGGGCGGCCA GGG ||||||||[||||||||||||]PAM NA Intergenic NA NA

T6904 ATGGACGGCGGGGCGGCCAGGGG 1000
Oligo fwd CACCATGGACGGCGGGGCGGCCAG

<table>
<tr><td>Oligo rev</td><td>AAACCTGGCCGCCCCGCCGTCCAT</td><td></td><td></td><td></td><td></td><td></td><td></td><td></td><td></td><td></td><td></td><td></td></tr>
<tr><td>Chromosome</td><td>start</td><td></td><td>end</td><td>strand</td><td>MM</td><td>target_seq</td><td>PAM</td><td>alignment</td><td>distance</td><td>position</td><td>gene name</td><td>gene id</td></tr>
<tr><td>GU305914.1</td><td></td><td>96459</td><td>96481</td><td>+</td><td>0</td><td>ATGGACGGCGGGGCGGCCAG</td><td>GGG</td><td>||||||||[||||||||||||]PAM</td><td>NA</td><td>Intergenic</td><td>NA</td><td>NA</td></tr>
<tr><td></td><td></td><td></td><td></td><td></td><td></td><td></td><td></td><td></td><td></td><td></td><td></td><td></td></tr>
<tr><td>T6905</td><td>CGGGGCGGCCAGGGGCGTCACGG</td><td></td><td>1000</td><td></td><td></td><td></td><td></td><td></td><td></td><td></td><td></td><td></td></tr>
<tr><td>Oligo fwd</td><td>CACCCGGGGCGGCCAGGGGCGTCA</td><td></td><td></td><td></td><td></td><td></td><td></td><td></td><td></td><td></td><td></td><td></td></tr>
<tr><td>Oligo rev</td><td>AAACTGACGCCCCTGGCCGCCCCG</td><td></td><td></td><td></td><td></td><td></td><td></td><td></td><td></td><td></td><td></td><td></td></tr>
<tr><td>Chromosome</td><td>start</td><td></td><td>end</td><td>strand</td><td>MM</td><td>target_seq</td><td>PAM</td><td>alignment</td><td>distance</td><td>position</td><td>gene name</td><td>gene id</td></tr>
<tr><td>GU305914.1</td><td></td><td>96467</td><td>96489</td><td>+</td><td>0</td><td>CGGGGCGGCCAGGGGCGTCA</td><td>CGG</td><td>||||||||[||||||||||||]PAM</td><td>NA</td><td>Intergenic</td><td>NA</td><td>NA</td></tr>
<tr><td></td><td></td><td></td><td></td><td></td><td></td><td></td><td></td><td></td><td></td><td></td><td></td><td></td></tr>
<tr><td>T6906</td><td>CGTCACGGACGCCGTCCACACGG</td><td></td><td>1000</td><td></td><td></td><td></td><td></td><td></td><td></td><td></td><td></td><td></td></tr>
<tr><td>Oligo fwd</td><td>CACCCGTCACGGACGCCGTCCACA</td><td></td><td></td><td></td><td></td><td></td><td></td><td></td><td></td><td></td><td></td><td></td></tr>
<tr><td>Oligo rev</td><td>AAACTGTGGACGGCGTCCGTGACG</td><td></td><td></td><td></td><td></td><td></td><td></td><td></td><td></td><td></td><td></td><td></td></tr>
<tr><td>Chromosome</td><td>start</td><td></td><td>end</td><td>strand</td><td>MM</td><td>target_seq</td><td>PAM</td><td>alignment</td><td>distance</td><td>position</td><td>gene name</td><td>gene id</td></tr>
<tr><td>GU305914.1</td><td></td><td>96482</td><td>96504</td><td>+</td><td>0</td><td>CGTCACGGACGCCGTCCACA</td><td>CGG</td><td>||||||||[||||||||||||]PAM</td><td>NA</td><td>Intergenic</td><td>NA</td><td>NA</td></tr>
<tr><td></td><td></td><td></td><td></td><td></td><td></td><td></td><td></td><td></td><td></td><td></td><td></td><td></td></tr>
<tr><td>T6907</td><td>CGTCCACACGGACTCCGTCTCGG</td><td></td><td>1000</td><td></td><td></td><td></td><td></td><td></td><td></td><td></td><td></td><td></td></tr>
<tr><td>Oligo fwd</td><td>CACCCGTCCACACGGACTCCGTCT</td><td></td><td></td><td></td><td></td><td></td><td></td><td></td><td></td><td></td><td></td><td></td></tr>
<tr><td>Oligo rev</td><td>AAACAGACGGAGTCCGTGTGGACG</td><td></td><td></td><td></td><td></td><td></td><td></td><td></td><td></td><td></td><td></td><td></td></tr>
<tr><td>Chromosome</td><td>start</td><td></td><td>end</td><td>strand</td><td>MM</td><td>target_seq</td><td>PAM</td><td>alignment</td><td>distance</td><td>position</td><td>gene name</td><td>gene id</td></tr>
<tr><td>GU305914.1</td><td></td><td>96494</td><td>96516</td><td>+</td><td>0</td><td>CGTCCACACGGACTCCGTCT</td><td>CGG</td><td>||||||||[||||||||||||]PAM</td><td>NA</td><td>Intergenic</td><td>NA</td><td>NA</td></tr>
<tr><td></td><td></td><td></td><td></td><td></td><td></td><td></td><td></td><td></td><td></td><td></td><td></td><td></td></tr>
<tr><td>T6908</td><td>GTCCACACGGACTCCGTCTCGGG</td><td></td><td>1000</td><td></td><td></td><td></td><td></td><td></td><td></td><td></td><td></td><td></td></tr>
<tr><td>Oligo fwd</td><td>CACCGTCCACACGGACTCCGTCTC</td><td></td><td></td><td></td><td></td><td></td><td></td><td></td><td></td><td></td><td></td><td></td></tr>
<tr><td>Oligo rev</td><td>AAACGAGACGGAGTCCGTGTGGAC</td><td></td><td></td><td></td><td></td><td></td><td></td><td></td><td></td><td></td><td></td><td></td></tr>
<tr><td>Chromosome</td><td>start</td><td></td><td>end</td><td>strand</td><td>MM</td><td>target_seq</td><td>PAM</td><td>alignment</td><td>distance</td><td>position</td><td>gene name</td><td>gene id</td></tr>
<tr><td>GU305914.1</td><td></td><td>96495</td><td>96517</td><td>+</td><td>0</td><td>GTCCACACGGACTCCGTCTC</td><td>GGG</td><td>||||||||[||||||||||||]PAM</td><td>NA</td><td>Intergenic</td><td>NA</td><td>NA</td></tr>
<tr><td></td><td></td><td></td><td></td><td></td><td></td><td></td><td></td><td></td><td></td><td></td><td></td><td></td></tr>
<tr><td>T6909</td><td>TCCACACGGACTCCGTCTCGGGG</td><td></td><td>1000</td><td></td><td></td><td></td><td></td><td></td><td></td><td></td><td></td><td></td></tr>
<tr><td>Oligo fwd</td><td>CACCTCCACACGGACTCCGTCTCG</td><td></td><td></td><td></td><td></td><td></td><td></td><td></td><td></td><td></td><td></td><td></td></tr>
<tr><td>Oligo rev</td><td>AAACCGAGACGGAGTCCGTGTGGA</td><td></td><td></td><td></td><td></td><td></td><td></td><td></td><td></td><td></td><td></td><td></td></tr>
<tr><td>Chromosome</td><td>start</td><td></td><td>end</td><td>strand</td><td>MM</td><td>target_seq</td><td>PAM</td><td>alignment</td><td>distance</td><td>position</td><td>gene name</td><td>gene id</td></tr>
<tr><td>GU305914.1</td><td></td><td>96496</td><td>96518</td><td>+</td><td>0</td><td>TCCACACGGACTCCGTCTCG</td><td>GGG</td><td>||||||||[||||||||||||]PAM</td><td>NA</td><td>Intergenic</td><td>NA</td><td>NA</td></tr>
<tr><td>GU305914.1</td><td></td><td>197897</td><td>197919</td><td>+</td><td>4</td><td>TCGTCATAGACTCCGTCTCG</td><td>TGG</td><td>||--||--[||||||||||||]PAM</td><td>NA</td><td>Intergenic</td><td>NA</td><td>NA</td></tr>
<tr><td></td><td></td><td></td><td></td><td></td><td></td><td></td><td></td><td></td><td></td><td></td><td></td><td></td></tr>
<tr><td>T6910</td><td>GGACTCCGTCTCGGGGTCCAAGG</td><td></td><td>1000</td><td></td><td></td><td></td><td></td><td></td><td></td><td></td><td></td><td></td></tr>
<tr><td>Oligo fwd</td><td>CACCGGACTCCGTCTCGGGGTCCA</td><td></td><td></td><td></td><td></td><td></td><td></td><td></td><td></td><td></td><td></td><td></td></tr>
<tr><td>Oligo rev</td><td>AAACTGGACCCCGAGACGGAGTCC</td><td></td><td></td><td></td><td></td><td></td><td></td><td></td><td></td><td></td><td></td><td></td></tr>
<tr><td>Chromosome</td><td>start</td><td></td><td>end</td><td>strand</td><td>MM</td><td>target_seq</td><td>PAM</td><td>alignment</td><td>distance</td><td>position</td><td>gene name</td><td>gene id</td></tr>
<tr><td>GU305914.1</td><td></td><td>96503</td><td>96525</td><td>+</td><td>0</td><td>GGACTCCGTCTCGGGGTCCA</td><td>AGG</td><td>||||||||[||||||||||||]PAM</td><td>NA</td><td>Intergenic</td><td>NA</td><td>NA</td></tr>
<tr><td></td><td></td><td></td><td></td><td></td><td></td><td></td><td></td><td></td><td></td><td></td><td></td><td></td></tr>
<tr><td>T6911</td><td>GACTCCGTCTCGGGGTCCAAGGG</td><td></td><td>1000</td><td></td><td></td><td></td><td></td><td></td><td></td><td></td><td></td><td></td></tr>
<tr><td>Oligo fwd</td><td>CACCGACTCCGTCTCGGGGTCCAA</td><td></td><td></td><td></td><td></td><td></td><td></td><td></td><td></td><td></td><td></td><td></td></tr>
<tr><td>Oligo rev</td><td>AAACTTGGACCCCGAGACGGAGTC</td><td></td><td></td><td></td><td></td><td></td><td></td><td></td><td></td><td></td><td></td><td></td></tr>
<tr><td>Chromosome</td><td>start</td><td></td><td>end</td><td>strand</td><td>MM</td><td>target_seq</td><td>PAM</td><td>alignment</td><td>distance</td><td>position</td><td>gene name</td><td>gene id</td></tr>
<tr><td>GU305914.1</td><td></td><td>96504</td><td>96526</td><td>+</td><td>0</td><td>GACTCCGTCTCGGGGTCCAA</td><td>GGG</td><td>||||||||[||||||||||||]PAM</td><td>NA</td><td>Intergenic</td><td>NA</td><td>NA</td></tr>
<tr><td></td><td></td><td></td><td></td><td></td><td></td><td></td><td></td><td></td><td></td><td></td><td></td><td></td></tr>
<tr><td>T6912</td><td>CCATGTCTAGCGAGCAGTGTAGG</td><td></td><td>1000</td><td></td><td></td><td></td><td></td><td></td><td></td><td></td><td></td><td></td></tr>
<tr><td>Oligo fwd</td><td>CACCCCATGTCTAGCGAGCAGTGT</td><td></td><td></td><td></td><td></td><td></td><td></td><td></td><td></td><td></td><td></td><td></td></tr>
<tr><td>Oligo rev</td><td>AAACACACTGCTCGCTAGACATGG</td><td></td><td></td><td></td><td></td><td></td><td></td><td></td><td></td><td></td><td></td><td></td></tr>
<tr><td>Chromosome</td><td>start</td><td></td><td>end</td><td>strand</td><td>MM</td><td>target_seq</td><td>PAM</td><td>alignment</td><td>distance</td><td>position</td><td>gene name</td><td>gene id</td></tr>
<tr><td>GU305914.1</td><td></td><td>96547</td><td>96569</td><td>+</td><td>0</td><td>CCATGTCTAGCGAGCAGTGT</td><td>AGG</td><td>||||||||[||||||||||||]PAM</td><td>NA</td><td>Intergenic</td><td>NA</td><td>NA</td></tr>
<tr><td></td><td></td><td></td><td></td><td></td><td></td><td></td><td></td><td></td><td></td><td></td><td></td><td></td></tr>
<tr><td>T6913</td><td>TGTAGGTCGTCTTCCTCGATCGG</td><td></td><td>1000</td><td></td><td></td><td></td><td></td><td></td><td></td><td></td><td></td><td></td></tr>
<tr><td>Oligo fwd</td><td>CACCTGTAGGTCGTCTTCCTCGAT</td><td></td><td></td><td></td><td></td><td></td><td></td><td></td><td></td><td></td><td></td><td></td></tr>
<tr><td>Oligo rev</td><td>AAACATCGAGGAAGACGACCTACA</td><td></td><td></td><td></td><td></td><td></td><td></td><td></td><td></td><td></td><td></td><td></td></tr>
<tr><td>Chromosome</td><td>start</td><td></td><td>end</td><td>strand</td><td>MM</td><td>target_seq</td><td>PAM</td><td>alignment</td><td>distance</td><td>position</td><td>gene name</td><td>gene id</td></tr>
<tr><td>GU305914.1</td><td></td><td>96564</td><td>96586</td><td>+</td><td>0</td><td>TGTAGGTCGTCTTCCTCGAT</td><td>CGG</td><td>||||||||[||||||||||||]PAM</td><td>NA</td><td>Intergenic</td><td>NA</td><td>NA</td></tr>
<tr><td></td><td></td><td></td><td></td><td></td><td></td><td></td><td></td><td></td><td></td><td></td><td></td><td></td></tr>
<tr><td>T6914</td><td>GTCGTCTTCCTCGATCGGCTCGG</td><td></td><td>1000</td><td></td><td></td><td></td><td></td><td></td><td></td><td></td><td></td><td></td></tr>
<tr><td>Oligo fwd</td><td>CACCGTCGTCTTCCTCGATCGGCT</td><td></td><td></td><td></td><td></td><td></td><td></td><td></td><td></td><td></td><td></td><td></td></tr>
<tr><td>Oligo rev</td><td>AAACAGCCGATCGAGGAAGACGAC</td><td></td><td></td><td></td><td></td><td></td><td></td><td></td><td></td><td></td><td></td><td></td></tr>
<tr><td>Chromosome</td><td>start</td><td></td><td>end</td><td>strand</td><td>MM</td><td>target_seq</td><td>PAM</td><td>alignment</td><td>distance</td><td>position</td><td>gene name</td><td>gene id</td></tr>
<tr><td>GU305914.1</td><td></td><td>96569</td><td>96591</td><td>+</td><td>0</td><td>GTCGTCTTCCTCGATCGGCT</td><td>CGG</td><td>||||||||[||||||||||||]PAM</td><td>NA</td><td>Intergenic</td><td>NA</td><td>NA</td></tr>
<tr><td></td><td></td><td></td><td></td><td></td><td></td><td></td><td></td><td></td><td></td><td></td><td></td><td></td></tr>
<tr><td>T6915</td><td>GACGTCAGATTCCTCCTCGTCGG</td><td></td><td>1000</td><td></td><td></td><td></td><td></td><td></td><td></td><td></td><td></td><td></td></tr>
<tr><td>Oligo fwd</td><td>CACCGACGTCAGATTCCTCCTCGT</td><td></td><td></td><td></td><td></td><td></td><td></td><td></td><td></td><td></td><td></td><td></td></tr>
<tr><td>Oligo rev</td><td>AAACACGAGGAGGAATCTGACGTC</td><td></td><td></td><td></td><td></td><td></td><td></td><td></td><td></td><td></td><td></td><td></td></tr>
<tr><td>Chromosome</td><td>start</td><td></td><td>end</td><td>strand</td><td>MM</td><td>target_seq</td><td>PAM</td><td>alignment</td><td>distance</td><td>position</td><td>gene name</td><td>gene id</td></tr>
<tr><td>GU305914.1</td><td></td><td>96689</td><td>96711</td><td>+</td><td>0</td><td>GACGTCAGATTCCTCCTCGT</td><td>CGG</td><td>||||||||[||||||||||||]PAM</td><td>NA</td><td>Intergenic</td><td>NA</td><td>NA</td></tr>
<tr><td></td><td></td><td></td><td></td><td></td><td></td><td></td><td></td><td></td><td></td><td></td><td></td><td></td></tr>
<tr><td>T6916</td><td>GTCAGATTCCTCCTCGTCGGAGG</td><td></td><td>1000</td><td></td><td></td><td></td><td></td><td></td><td></td><td></td><td></td><td></td></tr>
<tr><td>Oligo fwd</td><td>CACCGTCAGATTCCTCCTCGTCGG</td><td></td><td></td><td></td><td></td><td></td><td></td><td></td><td></td><td></td><td></td><td></td></tr>
<tr><td>Oligo rev</td><td>AAACCCGACGAGGAGGAATCTGAC</td><td></td><td></td><td></td><td></td><td></td><td></td><td></td><td></td><td></td><td></td><td></td></tr>
<tr><td>Chromosome</td><td>start</td><td></td><td>end</td><td>strand</td><td>MM</td><td>target_seq</td><td>PAM</td><td>alignment</td><td>distance</td><td>position</td><td>gene name</td><td>gene id</td></tr>
<tr><td>GU305914.1</td><td></td><td>96692</td><td>96714</td><td>+</td><td>0</td><td>GTCAGATTCCTCCTCGTCGG</td><td>AGG</td><td>||||||||[||||||||||||]PAM</td><td>NA</td><td>Intergenic</td><td>NA</td><td>NA</td></tr>
<tr><td></td><td></td><td></td><td></td><td></td><td></td><td></td><td></td><td></td><td></td><td></td><td></td><td></td></tr>
<tr><td>T6917</td><td>GGAGGACGACGACGAGTCCGAGG</td><td></td><td>1000</td><td></td><td></td><td></td><td></td><td></td><td></td><td></td><td></td><td></td></tr>
<tr><td>Oligo fwd</td><td>CACCGGAGGACGACGACGAGTCCG</td><td></td><td></td><td></td><td></td><td></td><td></td><td></td><td></td><td></td><td></td><td></td></tr>
<tr><td>Oligo rev</td><td>AAACCGGACTCGTCGTCGTCCTCC</td><td></td><td></td><td></td><td></td><td></td><td></td><td></td><td></td><td></td><td></td><td></td></tr>
<tr><td>Chromosome</td><td>start</td><td></td><td>end</td><td>strand</td><td>MM</td><td>target_seq</td><td>PAM</td><td>alignment</td><td>distance</td><td>position</td><td>gene name</td><td>gene id</td></tr>
<tr><td>GU305914.1</td><td></td><td>96710</td><td>96732</td><td>+</td><td>0</td><td>GGAGGACGACGACGAGTCCG</td><td>AGG</td><td>||||||||[||||||||||||]PAM</td><td>NA</td><td>Intergenic</td><td>NA</td><td>NA</td></tr>
<tr><td></td><td></td><td></td><td></td><td></td><td></td><td></td><td></td><td></td><td></td><td></td><td></td><td></td></tr>
<tr><td>T6918</td><td>CGACGACGAGTCCGAGGAGCTGG</td><td></td><td>1000</td><td></td><td></td><td></td><td></td><td></td><td></td><td></td><td></td><td></td></tr>
<tr><td>Oligo fwd</td><td>CACCCGACGACGAGTCCGAGGAGC</td><td></td><td></td><td></td><td></td><td></td><td></td><td></td><td></td><td></td><td></td><td></td></tr>
<tr><td>Oligo rev</td><td>AAACGCTCCTCGGACTCGTCGTCG</td><td></td><td></td><td></td><td></td><td></td><td></td><td></td><td></td><td></td><td></td><td></td></tr>
<tr><td>Chromosome</td><td>start</td><td></td><td>end</td><td>strand</td><td>MM</td><td>target_seq</td><td>PAM</td><td>alignment</td><td>distance</td><td>position</td><td>gene name</td><td>gene id</td></tr>
<tr><td>GU305914.1</td><td></td><td>96716</td><td>96738</td><td>+</td><td>0</td><td>CGACGACGAGTCCGAGGAGC</td><td>TGG</td><td>||||||||[||||||||||||]PAM</td><td>NA</td><td>Intergenic</td><td>NA</td><td>NA</td></tr>
</table>

<table>
<tr><td>T6919</td><td>CGAGTCCGAGGAGCTGGACGAGG</td><td></td><td>1000</td><td></td><td></td><td></td><td></td><td></td><td></td><td></td><td></td><td></td></tr>
<tr><td>Oligo fwd</td><td>CACCCGAGTCCGAGGAGCTGGACG</td><td></td><td></td><td></td><td></td><td></td><td></td><td></td><td></td><td></td><td></td><td></td></tr>
<tr><td>Oligo rev</td><td>AAACCGTCCAGCTCCTCGGACTCG</td><td></td><td></td><td></td><td></td><td></td><td></td><td></td><td></td><td></td><td></td><td></td></tr>
<tr><td>Chromosome</td><td>start</td><td></td><td>end</td><td>strand</td><td>MM</td><td>target_seq</td><td>PAM</td><td>alignment</td><td>distance</td><td>position</td><td>gene name</td><td>gene id</td></tr>
<tr><td>GU305914.1</td><td></td><td>96722</td><td>96744</td><td>+</td><td>0</td><td>CGAGTCCGAGGAGCTGGACG</td><td>AGG</td><td>||||||||[||||||||||||]PAM</td><td>NA</td><td>Intergenic</td><td>NA</td><td>NA</td></tr>
<tr><td>T6920</td><td>GGAGCTGGACGAGGAGTCGCTGG</td><td></td><td>1000</td><td></td><td></td><td></td><td></td><td></td><td></td><td></td><td></td><td></td></tr>
<tr><td>Oligo fwd</td><td>CACCGGAGCTGGACGAGGAGTCGC</td><td></td><td></td><td></td><td></td><td></td><td></td><td></td><td></td><td></td><td></td><td></td></tr>
<tr><td>Oligo rev</td><td>AAACGCGACTCCTCGTCCAGCTCC</td><td></td><td></td><td></td><td></td><td></td><td></td><td></td><td></td><td></td><td></td><td></td></tr>
<tr><td>Chromosome</td><td>start</td><td></td><td>end</td><td>strand</td><td>MM</td><td>target_seq</td><td>PAM</td><td>alignment</td><td>distance</td><td>position</td><td>gene name</td><td>gene id</td></tr>
<tr><td>GU305914.1</td><td></td><td>96731</td><td>96753</td><td>+</td><td>0</td><td>GGAGCTGGACGAGGAGTCGC</td><td>TGG</td><td>||||||||[||||||||||||]PAM</td><td>NA</td><td>Intergenic</td><td>NA</td><td>NA</td></tr>
<tr><td>T6921</td><td>CGCCGACGAGCCGCGACGCGTGG</td><td></td><td>1000</td><td></td><td></td><td></td><td></td><td></td><td></td><td></td><td></td><td></td></tr>
<tr><td>Oligo fwd</td><td>CACCCGCCGACGAGCCGCGACGCG</td><td></td><td></td><td></td><td></td><td></td><td></td><td></td><td></td><td></td><td></td><td></td></tr>
<tr><td>Oligo rev</td><td>AAACCGCGTCGCGGCTCGTCGGCG</td><td></td><td></td><td></td><td></td><td></td><td></td><td></td><td></td><td></td><td></td><td></td></tr>
<tr><td>Chromosome</td><td>start</td><td></td><td>end</td><td>strand</td><td>MM</td><td>target_seq</td><td>PAM</td><td>alignment</td><td>distance</td><td>position</td><td>gene name</td><td>gene id</td></tr>
<tr><td>GU305914.1</td><td></td><td>96761</td><td>96783</td><td>+</td><td>0</td><td>CGCCGACGAGCCGCGACGCG</td><td>TGG</td><td>||||||||[||||||||||||]PAM</td><td>NA</td><td>Intergenic</td><td>NA</td><td>NA</td></tr>
<tr><td>T6922</td><td>CGTGGCGACGCTGCACTCCCCGG</td><td></td><td>1000</td><td></td><td></td><td></td><td></td><td></td><td></td><td></td><td></td><td></td></tr>
<tr><td>Oligo fwd</td><td>CACCCGTGGCGACGCTGCACTCCC</td><td></td><td></td><td></td><td></td><td></td><td></td><td></td><td></td><td></td><td></td><td></td></tr>
<tr><td>Oligo rev</td><td>AAACGGGAGTGCAGCGTCGCCACG</td><td></td><td></td><td></td><td></td><td></td><td></td><td></td><td></td><td></td><td></td><td></td></tr>
<tr><td>Chromosome</td><td>start</td><td></td><td>end</td><td>strand</td><td>MM</td><td>target_seq</td><td>PAM</td><td>alignment</td><td>distance</td><td>position</td><td>gene name</td><td>gene id</td></tr>
<tr><td>GU305914.1</td><td></td><td>96779</td><td>96801</td><td>+</td><td>0</td><td>CGTGGCGACGCTGCACTCCC</td><td>CGG</td><td>||||||||[||||||||||||]PAM</td><td>NA</td><td>Intergenic</td><td>NA</td><td>NA</td></tr>
<tr><td>T6923</td><td>GTGGCGACGCTGCACTCCCCGGG</td><td></td><td>1000</td><td></td><td></td><td></td><td></td><td></td><td></td><td></td><td></td><td></td></tr>
<tr><td>Oligo fwd</td><td>CACCGTGGCGACGCTGCACTCCCC</td><td></td><td></td><td></td><td></td><td></td><td></td><td></td><td></td><td></td><td></td><td></td></tr>
<tr><td>Oligo rev</td><td>AAACGGGGAGTGCAGCGTCGCCAC</td><td></td><td></td><td></td><td></td><td></td><td></td><td></td><td></td><td></td><td></td><td></td></tr>
<tr><td>Chromosome</td><td>start</td><td></td><td>end</td><td>strand</td><td>MM</td><td>target_seq</td><td>PAM</td><td>alignment</td><td>distance</td><td>position</td><td>gene name</td><td>gene id</td></tr>
<tr><td>GU305914.1</td><td></td><td>96780</td><td>96802</td><td>+</td><td>0</td><td>GTGGCGACGCTGCACTCCCC</td><td>GGG</td><td>||||||||[||||||||||||]PAM</td><td>NA</td><td>Intergenic</td><td>NA</td><td>NA</td></tr>
<tr><td>T6924</td><td>GACCCCTTCGTCCTCGTCGCCGG</td><td></td><td>1000</td><td></td><td></td><td></td><td></td><td></td><td></td><td></td><td></td><td></td></tr>
<tr><td>Oligo fwd</td><td>CACCGACCCCTTCGTCCTCGTCGC</td><td></td><td></td><td></td><td></td><td></td><td></td><td></td><td></td><td></td><td></td><td></td></tr>
<tr><td>Oligo rev</td><td>AAACGCGACGAGGACGAAGGGGTC</td><td></td><td></td><td></td><td></td><td></td><td></td><td></td><td></td><td></td><td></td><td></td></tr>
<tr><td>Chromosome</td><td>start</td><td></td><td>end</td><td>strand</td><td>MM</td><td>target_seq</td><td>PAM</td><td>alignment</td><td>distance</td><td>position</td><td>gene name</td><td>gene id</td></tr>
<tr><td>GU305914.1</td><td></td><td>96809</td><td>96831</td><td>+</td><td>0</td><td>GACCCCTTCGTCCTCGTCGC</td><td>CGG</td><td>||||||||[||||||||||||]PAM</td><td>NA</td><td>Intergenic</td><td>NA</td><td>NA</td></tr>
<tr><td>T6925</td><td>CCCTTCGTCCTCGTCGCCGGCGG</td><td></td><td>1000</td><td></td><td></td><td></td><td></td><td></td><td></td><td></td><td></td><td></td></tr>
<tr><td>Oligo fwd</td><td>CACCCCCTTCGTCCTCGTCGCCGG</td><td></td><td></td><td></td><td></td><td></td><td></td><td></td><td></td><td></td><td></td><td></td></tr>
<tr><td>Oligo rev</td><td>AAACCCGGCGACGAGGACGAAGGG</td><td></td><td></td><td></td><td></td><td></td><td></td><td></td><td></td><td></td><td></td><td></td></tr>
<tr><td>Chromosome</td><td>start</td><td></td><td>end</td><td>strand</td><td>MM</td><td>target_seq</td><td>PAM</td><td>alignment</td><td>distance</td><td>position</td><td>gene name</td><td>gene id</td></tr>
<tr><td>GU305914.1</td><td></td><td>96812</td><td>96834</td><td>+</td><td>0</td><td>CCCTTCGTCCTCGTCGCCGG</td><td>CGG</td><td>||||||||[||||||||||||]PAM</td><td>NA</td><td>Intergenic</td><td>NA</td><td>NA</td></tr>
<tr><td>T6926</td><td>GGCGGTCGCGAGCTCTTTCTCGG</td><td></td><td>1000</td><td></td><td></td><td></td><td></td><td></td><td></td><td></td><td></td><td></td></tr>
<tr><td>Oligo fwd</td><td>CACCGGCGGTCGCGAGCTCTTTCT</td><td></td><td></td><td></td><td></td><td></td><td></td><td></td><td></td><td></td><td></td><td></td></tr>
<tr><td>Oligo rev</td><td>AAACAGAAAGAGCTCGCGACCGCC</td><td></td><td></td><td></td><td></td><td></td><td></td><td></td><td></td><td></td><td></td><td></td></tr>
<tr><td>Chromosome</td><td>start</td><td></td><td>end</td><td>strand</td><td>MM</td><td>target_seq</td><td>PAM</td><td>alignment</td><td>distance</td><td>position</td><td>gene name</td><td>gene id</td></tr>
<tr><td>GU305914.1</td><td></td><td>96830</td><td>96852</td><td>+</td><td>0</td><td>GGCGGTCGCGAGCTCTTTCT</td><td>CGG</td><td>||||||||[||||||||||||]PAM</td><td>NA</td><td>Intergenic</td><td>NA</td><td>NA</td></tr>
<tr><td>T6927</td><td>CGCGAGCTCTTTCTCGGCGTCGG</td><td></td><td>1000</td><td></td><td></td><td></td><td></td><td></td><td></td><td></td><td></td><td></td></tr>
<tr><td>Oligo fwd</td><td>CACCCGCGAGCTCTTTCTCGGCGT</td><td></td><td></td><td></td><td></td><td></td><td></td><td></td><td></td><td></td><td></td><td></td></tr>
<tr><td>Oligo rev</td><td>AAACACGCCGAGAAAGAGCTCGCG</td><td></td><td></td><td></td><td></td><td></td><td></td><td></td><td></td><td></td><td></td><td></td></tr>
<tr><td>Chromosome</td><td>start</td><td></td><td>end</td><td>strand</td><td>MM</td><td>target_seq</td><td>PAM</td><td>alignment</td><td>distance</td><td>position</td><td>gene name</td><td>gene id</td></tr>
<tr><td>GU305914.1</td><td></td><td>96836</td><td>96858</td><td>+</td><td>0</td><td>CGCGAGCTCTTTCTCGGCGT</td><td>CGG</td><td>||||||||[||||||||||||]PAM</td><td>NA</td><td>Intergenic</td><td>NA</td><td>NA</td></tr>
<tr><td>T6928</td><td>GAGCTCTTTCTCGGCGTCGGCGG</td><td></td><td>1000</td><td></td><td></td><td></td><td></td><td></td><td></td><td></td><td></td><td></td></tr>
<tr><td>Oligo fwd</td><td>CACCGAGCTCTTTCTCGGCGTCGG</td><td></td><td></td><td></td><td></td><td></td><td></td><td></td><td></td><td></td><td></td><td></td></tr>
<tr><td>Oligo rev</td><td>AAACCCGACGCCGAGAAAGAGCTC</td><td></td><td></td><td></td><td></td><td></td><td></td><td></td><td></td><td></td><td></td><td></td></tr>
<tr><td>Chromosome</td><td>start</td><td></td><td>end</td><td>strand</td><td>MM</td><td>target_seq</td><td>PAM</td><td>alignment</td><td>distance</td><td>position</td><td>gene name</td><td>gene id</td></tr>
<tr><td>GU305914.1</td><td></td><td>96839</td><td>96861</td><td>+</td><td>0</td><td>GAGCTCTTTCTCGGCGTCGG</td><td>CGG</td><td>||||||||[||||||||||||]PAM</td><td>NA</td><td>Intergenic</td><td>NA</td><td>NA</td></tr>
<tr><td>T6929</td><td>AGCTCTTTCTCGGCGTCGGCGGG</td><td></td><td>1000</td><td></td><td></td><td></td><td></td><td></td><td></td><td></td><td></td><td></td></tr>
<tr><td>Oligo fwd</td><td>CACCAGCTCTTTCTCGGCGTCGGC</td><td></td><td></td><td></td><td></td><td></td><td></td><td></td><td></td><td></td><td></td><td></td></tr>
<tr><td>Oligo rev</td><td>AAACGCCGACGCCGAGAAAGAGCT</td><td></td><td></td><td></td><td></td><td></td><td></td><td></td><td></td><td></td><td></td><td></td></tr>
<tr><td>Chromosome</td><td>start</td><td></td><td>end</td><td>strand</td><td>MM</td><td>target_seq</td><td>PAM</td><td>alignment</td><td>distance</td><td>position</td><td>gene name</td><td>gene id</td></tr>
<tr><td>GU305914.1</td><td></td><td>96840</td><td>96862</td><td>+</td><td>0</td><td>AGCTCTTTCTCGGCGTCGGC</td><td>GGG</td><td>||||||||[||||||||||||]PAM</td><td>NA</td><td>Intergenic</td><td>NA</td><td>NA</td></tr>
<tr><td>T6930</td><td>GCGTCGGCGGGCTCGCCCGACGG</td><td></td><td>1000</td><td></td><td></td><td></td><td></td><td></td><td></td><td></td><td></td><td></td></tr>
<tr><td>Oligo fwd</td><td>CACCGCGTCGGCGGGCTCGCCCGA</td><td></td><td></td><td></td><td></td><td></td><td></td><td></td><td></td><td></td><td></td><td></td></tr>
<tr><td>Oligo rev</td><td>AAACTCGGGCGAGCCCGCCGACGC</td><td></td><td></td><td></td><td></td><td></td><td></td><td></td><td></td><td></td><td></td><td></td></tr>
<tr><td>Chromosome</td><td>start</td><td></td><td>end</td><td>strand</td><td>MM</td><td>target_seq</td><td>PAM</td><td>alignment</td><td>distance</td><td>position</td><td>gene name</td><td>gene id</td></tr>
<tr><td>GU305914.1</td><td></td><td>96852</td><td>96874</td><td>+</td><td>0</td><td>GCGTCGGCGGGCTCGCCCGA</td><td>CGG</td><td>||||||||[||||||||||||]PAM</td><td>NA</td><td>Intergenic</td><td>NA</td><td>NA</td></tr>
<tr><td>GU305914.1</td><td></td><td>125947</td><td>125969</td><td>-</td><td>4</td><td>GCGCAGGCGGGCCCGCCCCA</td><td>GGG</td><td>|||--|||[||||-|||||-|]PAM</td><td>NA</td><td>Intergenic</td><td>NA</td><td>NA</td></tr>
<tr><td>T6931</td><td>TCGGCGGGCTCGCCCGACGGAGG</td><td></td><td>1000</td><td></td><td></td><td></td><td></td><td></td><td></td><td></td><td></td><td></td></tr>
<tr><td>Oligo fwd</td><td>CACCTCGGCGGGCTCGCCCGACGG</td><td></td><td></td><td></td><td></td><td></td><td></td><td></td><td></td><td></td><td></td><td></td></tr>
<tr><td>Oligo rev</td><td>AAACCCGTCGGGCGAGCCCGCCGA</td><td></td><td></td><td></td><td></td><td></td><td></td><td></td><td></td><td></td><td></td><td></td></tr>
<tr><td>Chromosome</td><td>start</td><td></td><td>end</td><td>strand</td><td>MM</td><td>target_seq</td><td>PAM</td><td>alignment</td><td>distance</td><td>position</td><td>gene name</td><td>gene id</td></tr>
<tr><td>GU305914.1</td><td></td><td>96855</td><td>96877</td><td>+</td><td>0</td><td>TCGGCGGGCTCGCCCGACGG</td><td>AGG</td><td>||||||||[||||||||||||]PAM</td><td>NA</td><td>Intergenic</td><td>NA</td><td>NA</td></tr>
<tr><td>T6932</td><td>TCGCCCGACGGAGGATCGTGCGG</td><td></td><td>1000</td><td></td><td></td><td></td><td></td><td></td><td></td><td></td><td></td><td></td></tr>
<tr><td>Oligo fwd</td><td>CACCTCGCCCGACGGAGGATCGTG</td><td></td><td></td><td></td><td></td><td></td><td></td><td></td><td></td><td></td><td></td><td></td></tr>
<tr><td>Oligo rev</td><td>AAACCACGATCCTCCGTCGGGCGA</td><td></td><td></td><td></td><td></td><td></td><td></td><td></td><td></td><td></td><td></td><td></td></tr>
<tr><td>Chromosome</td><td>start</td><td></td><td>end</td><td>strand</td><td>MM</td><td>target_seq</td><td>PAM</td><td>alignment</td><td>distance</td><td>position</td><td>gene name</td><td>gene id</td></tr>
<tr><td>GU305914.1</td><td></td><td>96864</td><td>96886</td><td>+</td><td>0</td><td>TCGCCCGACGGAGGATCGTG</td><td>CGG</td><td>||||||||[||||||||||||]PAM</td><td>NA</td><td>Intergenic</td><td>NA</td><td>NA</td></tr>
<tr><td>GU305914.1</td><td></td><td>118248</td><td>118270</td><td>+</td><td>4</td><td>CCTCCCGACGGAGGATCAGG</td><td>CGG</td><td>-|-|||||[|||||||||--|]PAM</td><td>NA</td><td>Intergenic</td><td>NA</td><td>NA</td></tr>
<tr><td>T6933</td><td>CGCCCGACGGAGGATCGTGCGGG</td><td></td><td>1000</td><td></td><td></td><td></td><td></td><td></td><td></td><td></td><td></td><td></td></tr>
<tr><td>Oligo fwd</td><td>CACCCGCCCGACGGAGGATCGTGC</td><td></td><td></td><td></td><td></td><td></td><td></td><td></td><td></td><td></td><td></td><td></td></tr>
<tr><td>Oligo rev</td><td>AAACGCACGATCCTCCGTCGGGCG</td><td></td><td></td><td></td><td></td><td></td><td></td><td></td><td></td><td></td><td></td><td></td></tr>
</table>

| | | | | | | | | | | | |
|---|---|---|---|---|---|---|---|---|---|---|---|
| Chromosome | start | end | strand | MM | target_seq | PAM | alignment | distance | position | gene name | gene id |
| GU305914.1 | 96865 | 96887 | + | 0 | CGCCCGACGGAGGATCGTGC | GGG | \|\|\|\|\|\|\|\|[\|\|\|\|\|\|\|\|\|\|\|\|]PAM | NA | Intergenic | NA | NA |
| T6934 | GACGGAGGATCGTGCGGGCGAGG | 1000 | | | | | | | | | |
| Oligo fwd | CACCGACGGAGGATCGTGCGGGCG | | | | | | | | | | |
| Oligo rev | AAACCGCCCGCACGATCCTCCGTC | | | | | | | | | | |
| Chromosome | start | end | strand | MM | target_seq | PAM | alignment | distance | position | gene name | gene id |
| GU305914.1 | 96870 | 96892 | + | 0 | GACGGAGGATCGTGCGGGCG | AGG | \|\|\|\|\|\|\|\|[\|\|\|\|\|\|\|\|\|\|\|\|]PAM | NA | Intergenic | NA | NA |
| T6935 | ACGGAGGATCGTGCGGGCGAGGG | 1000 | | | | | | | | | |
| Oligo fwd | CACCACGGAGGATCGTGCGGGCGA | | | | | | | | | | |
| Oligo rev | AAACTCGCCCGCACGATCCTCCGT | | | | | | | | | | |
| Chromosome | start | end | strand | MM | target_seq | PAM | alignment | distance | position | gene name | gene id |
| GU305914.1 | 96871 | 96893 | + | 0 | ACGGAGGATCGTGCGGGCGA | GGG | \|\|\|\|\|\|\|\|[\|\|\|\|\|\|\|\|\|\|\|\|]PAM | NA | Intergenic | NA | NA |
| T6936 | CGGAGGATCGTGCGGGCGAGGGG | 1000 | | | | | | | | | |
| Oligo fwd | CACCCGGAGGATCGTGCGGGCGAG | | | | | | | | | | |
| Oligo rev | AAACCTCGCCCGCACGATCCTCCG | | | | | | | | | | |
| Chromosome | start | end | strand | MM | target_seq | PAM | alignment | distance | position | gene name | gene id |
| GU305914.1 | 96872 | 96894 | + | 0 | CGGAGGATCGTGCGGGCGAG | GGG | \|\|\|\|\|\|\|\|[\|\|\|\|\|\|\|\|\|\|\|\|]PAM | NA | Intergenic | NA | NA |
| T6937 | TGCGGGCGAGGGGACGAAGACGG | 1000 | | | | | | | | | |
| Oligo fwd | CACCTGCGGGCGAGGGGACGAAGA | | | | | | | | | | |
| Oligo rev | AAACTCTTCGTCCCCTCGCCCGCA | | | | | | | | | | |
| Chromosome | start | end | strand | MM | target_seq | PAM | alignment | distance | position | gene name | gene id |
| GU305914.1 | 96882 | 96904 | + | 0 | TGCGGGCGAGGGGACGAAGA | CGG | \|\|\|\|\|\|\|\|[\|\|\|\|\|\|\|\|\|\|\|\|]PAM | NA | Intergenic | NA | NA |
| T6938 | GACGAAGACGGCGATTTCGACGG | 1000 | | | | | | | | | |
| Oligo fwd | CACCGACGAAGACGGCGATTTCGA | | | | | | | | | | |
| Oligo rev | AAACTCGAAATCGCCGTCTTCGTC | | | | | | | | | | |
| Chromosome | start | end | strand | MM | target_seq | PAM | alignment | distance | position | gene name | gene id |
| GU305914.1 | 96894 | 96916 | + | 0 | GACGAAGACGGCGATTTCGA | CGG | \|\|\|\|\|\|\|\|[\|\|\|\|\|\|\|\|\|\|\|\|]PAM | NA | Intergenic | NA | NA |
| T6939 | TTTCGACGGTCTCCTCTCTCTGG | 1000 | | | | | | | | | |
| Oligo fwd | CACCTTTCGACGGTCTCCTCTCTC | | | | | | | | | | |
| Oligo rev | AAACGAGAGAGGAGACCGTCGAAA | | | | | | | | | | |
| Chromosome | start | end | strand | MM | target_seq | PAM | alignment | distance | position | gene name | gene id |
| GU305914.1 | 96908 | 96930 | + | 0 | TTTCGACGGTCTCCTCTCTC | TGG | \|\|\|\|\|\|\|\|[\|\|\|\|\|\|\|\|\|\|\|\|]PAM | NA | Intergenic | NA | NA |
| T6940 | CGGTCTCCTCTCTCTGGACGCGG | 1000 | | | | | | | | | |
| Oligo fwd | CACCCGGTCTCCTCTCTCTGGACG | | | | | | | | | | |
| Oligo rev | AAACCGTCCAGAGAGAGGAGACCG | | | | | | | | | | |
| Chromosome | start | end | strand | MM | target_seq | PAM | alignment | distance | position | gene name | gene id |
| GU305914.1 | 96914 | 96936 | + | 0 | CGGTCTCCTCTCTCTGGACG | CGG | \|\|\|\|\|\|\|\|[\|\|\|\|\|\|\|\|\|\|\|\|]PAM | NA | Intergenic | NA | NA |
| T6941 | GGTCTCCTCTCTCTGGACGCGGG | 1000 | | | | | | | | | |
| Oligo fwd | CACCGGTCTCCTCTCTCTGGACGC | | | | | | | | | | |
| Oligo rev | AAACGCGTCCAGAGAGAGGAGACC | | | | | | | | | | |
| Chromosome | start | end | strand | MM | target_seq | PAM | alignment | distance | position | gene name | gene id |
| GU305914.1 | 96915 | 96937 | + | 0 | GGTCTCCTCTCTCTGGACGC | GGG | \|\|\|\|\|\|\|\|[\|\|\|\|\|\|\|\|\|\|\|\|]PAM | NA | Intergenic | NA | NA |
| T6942 | CCTCTCTCTGGACGCGGGCGAGG | 1000 | | | | | | | | | |
| Oligo fwd | CACCCCTCTCTCTGGACGCGGGCG | | | | | | | | | | |
| Oligo rev | AAACCGCCCGCGTCCAGAGAGAGG | | | | | | | | | | |
| Chromosome | start | end | strand | MM | target_seq | PAM | alignment | distance | position | gene name | gene id |
| GU305914.1 | 96920 | 96942 | + | 0 | CCTCTCTCTGGACGCGGGCG | AGG | \|\|\|\|\|\|\|\|[\|\|\|\|\|\|\|\|\|\|\|\|]PAM | NA | Intergenic | NA | NA |
| T6943 | CTCTCTCTGGACGCGGGCGAGGG | 1000 | | | | | | | | | |
| Oligo fwd | CACCCTCTCTCTGGACGCGGGCGA | | | | | | | | | | |
| Oligo rev | AAACTCGCCCGCGTCCAGAGAGAG | | | | | | | | | | |
| Chromosome | start | end | strand | MM | target_seq | PAM | alignment | distance | position | gene name | gene id |
| GU305914.1 | 96921 | 96943 | + | 0 | CTCTCTCTGGACGCGGGCGA | GGG | \|\|\|\|\|\|\|\|[\|\|\|\|\|\|\|\|\|\|\|\|]PAM | NA | Intergenic | NA | NA |
| T6944 | GGACGCGGGCGAGGGCGAGCAGG | 1000 | | | | | | | | | |
| Oligo fwd | CACCGGACGCGGGCGAGGGCGAGC | | | | | | | | | | |
| Oligo rev | AAACGCTCGCCCTCGCCCGCGTCC | | | | | | | | | | |
| Chromosome | start | end | strand | MM | target_seq | PAM | alignment | distance | position | gene name | gene id |
| GU305914.1 | 96929 | 96951 | + | 0 | GGACGCGGGCGAGGGCGAGC | AGG | \|\|\|\|\|\|\|\|[\|\|\|\|\|\|\|\|\|\|\|\|]PAM | NA | Intergenic | NA | NA |
| T6945 | GACGCGGGCGAGGGCGAGCAGGG | 1000 | | | | | | | | | |
| Oligo fwd | CACCGACGCGGGCGAGGGCGAGCA | | | | | | | | | | |
| Oligo rev | AAACTGCTCGCCCTCGCCCGCGTC | | | | | | | | | | |
| Chromosome | start | end | strand | MM | target_seq | PAM | alignment | distance | position | gene name | gene id |
| GU305914.1 | 96930 | 96952 | + | 0 | GACGCGGGCGAGGGCGAGCA | GGG | \|\|\|\|\|\|\|\|[\|\|\|\|\|\|\|\|\|\|\|\|]PAM | NA | Intergenic | NA | NA |
| T6946 | GGCGAGGGCGAGCAGGGCGACGG | 1000 | | | | | | | | | |
| Oligo fwd | CACCGGCGAGGGCGAGCAGGGCGA | | | | | | | | | | |
| Oligo rev | AAACTCGCCCTGCTCGCCCTCGCC | | | | | | | | | | |
| Chromosome | start | end | strand | MM | target_seq | PAM | alignment | distance | position | gene name | gene id |
| GU305914.1 | 96936 | 96958 | + | 0 | GGCGAGGGCGAGCAGGGCGA | CGG | \|\|\|\|\|\|\|\|[\|\|\|\|\|\|\|\|\|\|\|\|]PAM | NA | Intergenic | NA | NA |
| GU305914.1 | 20756 | 20778 | + | 4 | GACGACGGCGATCACGGCGA | GGG | \|-\|\|\|-\|\|[\|\|\|-\|\|-\|\|\|\|\|]PAM | NA | Intergenic | NA | NA |
| T6947 | GAGGGCGAGCAGGGCGACGGCGG | 1000 | | | | | | | | | |
| Oligo fwd | CACCGAGGGCGAGCAGGGCGACGG | | | | | | | | | | |
| Oligo rev | AAACCCGTCGCCCTGCTCGCCCTC | | | | | | | | | | |
| Chromosome | start | end | strand | MM | target_seq | PAM | alignment | distance | position | gene name | gene id |
| GU305914.1 | 96939 | 96961 | + | 0 | GAGGGCGAGCAGGGCGACGG | CGG | \|\|\|\|\|\|\|\|[\|\|\|\|\|\|\|\|\|\|\|\|]PAM | NA | Intergenic | NA | NA |
| GU305914.1 | 112392 | 112414 | + | 4 | GAGGGTGTGCAGGGCGATGA | AGG | \|\|\|\|\|-\|-[\|\|\|\|\|\|\|\|\|-\|-]PAM | NA | Intergenic | NA | NA |

| | | | | | | | | | | | |
|---|---|---|---|---|---|---|---|---|---|---|---|
| T6948 | GGCGAGCAGGGCGACGGCGGAGG | 1000 | | | | | | | | | |
| Oligo fwd | CACCGGCGAGCAGGGCGACGGCGG | | | | | | | | | | |
| Oligo rev | AAACCCGCCGTCGCCCTGCTCGCC | | | | | | | | | | |
| Chromosome | start | end | strand | MM | target_seq | PAM | alignment | distance | position | gene name | gene id |
| GU305914.1 | 96942 | 96964 | + | 0 | GGCGAGCAGGGCGACGGCGG | AGG | \|\|\|\|\|\|\|\|[\|\|\|\|\|\|\|\|\|\|\|\|]PAM | NA | Intergenic | NA | NA |
| GU305914.1 | 90265 | 90287 | - | 4 | CGGGAGGCGGGCGACGGCGG | AGG | -\|-\|\|\|--[\|\|\|\|\|\|\|\|\|\|\|\|]PAM | NA | Intergenic | NA | NA |
| GU305914.1 | 225987 | 226009 | - | 4 | CGCGTGGAGCGCGACGGCGG | CGG | -\|\|\|-\|-\|[\|-\|\|\|\|\|\|\|\|\|\|]PAM | NA | Intergenic | NA | NA |
| GU305914.1 | 110632 | 110654 | - | 4 | GACGTGCAGGACGCCGGCGG | CGG | \|-\|\|-\|\|\|[\|\|-\|\|-\|\|\|\|\|\|]PAM | NA | Intergenic | NA | NA |
| T6949 | GAGCAGGGCGACGGCGGAGGCGG | 1000 | | | | | | | | | |
| Oligo fwd | CACCGAGCAGGGCGACGGCGGAGG | | | | | | | | | | |
| Oligo rev | AAACCCTCCGCCGTCGCCCTGCTC | | | | | | | | | | |
| Chromosome | start | end | strand | MM | target_seq | PAM | alignment | distance | position | gene name | gene id |
| GU305914.1 | 96945 | 96967 | + | 0 | GAGCAGGGCGACGGCGGAGG | CGG | \|\|\|\|\|\|\|\|[\|\|\|\|\|\|\|\|\|\|\|\|]PAM | NA | Intergenic | NA | NA |
| GU305914.1 | 90262 | 90284 | - | 2 | GAGGCGGGCGACGGCGGAGG | CGG | \|\|\|--\|\|\|[\|\|\|\|\|\|\|\|\|\|\|\|]PAM | NA | Intergenic | NA | NA |
| GU305914.1 | 176454 | 176476 | + | 4 | GACGATGGCGACGGCGGCGG | AGG | \|\|--\|-\|\|[\|\|\|\|\|\|\|\|\|-\|\|]PAM | NA | Intergenic | NA | NA |
| GU305914.1 | 225984 | 226006 | - | 4 | GTGGAGCGCGACGGCGGCGG | CGG | \|-\|-\|\|-\|[\|\|\|\|\|\|\|\|\|-\|\|]PAM | NA | Intergenic | NA | NA |
| GU305914.1 | 110629 | 110651 | - | 4 | GTGCAGGACGCCGGCGGCGG | AGG | \|-\|\|\|\|\|-[\|\|-\|\|\|\|\|\|-\|\|]PAM | NA | Intergenic | NA | NA |
| T6950 | CGACGGCGGAGGCGGCATCGCGG | 1000 | | | | | | | | | |
| Oligo fwd | CACCCGACGGCGGAGGCGGCATCG | | | | | | | | | | |
| Oligo rev | AAACCGATGCCGCCTCCGCCGTCG | | | | | | | | | | |
| Chromosome | start | end | strand | MM | target_seq | PAM | alignment | distance | position | gene name | gene id |
| GU305914.1 | 96953 | 96975 | + | 0 | CGACGGCGGAGGCGGCATCG | CGG | \|\|\|\|\|\|\|\|[\|\|\|\|\|\|\|\|\|\|\|\|]PAM | NA | Intergenic | NA | NA |
| GU305914.1 | 61095 | 61117 | - | 4 | GGACCTCGGAGGCGACATCG | TGG | -\|\|\|--\|\|[\|\|\|\|\|\|-\|\|\|\|\|]PAM | NA | Intergenic | NA | NA |
| GU305914.1 | 86821 | 86843 | - | 4 | CAAGGGCCGAGGCGGCGTCG | GGG | \|-\|-\|\|\|-[\|\|\|\|\|\|\|\|-\|\|\|]PAM | NA | Intergenic | NA | NA |
| T6951 | GGCCGTCTTCGACGACTCTTTGG | 1000 | | | | | | | | | |
| Oligo fwd | CACCGGCCGTCTTCGACGACTCTT | | | | | | | | | | |
| Oligo rev | AAACAAGAGTCGTCGAAGACGGCC | | | | | | | | | | |
| Chromosome | start | end | strand | MM | target_seq | PAM | alignment | distance | position | gene name | gene id |
| GU305914.1 | 96974 | 96996 | + | 0 | GGCCGTCTTCGACGACTCTT | TGG | \|\|\|\|\|\|\|\|[\|\|\|\|\|\|\|\|\|\|\|\|]PAM | NA | Intergenic | NA | NA |
| T6952 | CTTCGACGACTCTTTGGACGTGG | 1000 | | | | | | | | | |
| Oligo fwd | CACCCTTCGACGACTCTTTGGACG | | | | | | | | | | |
| Oligo rev | AAACCGTCCAAAGAGTCGTCGAAG | | | | | | | | | | |
| Chromosome | start | end | strand | MM | target_seq | PAM | alignment | distance | position | gene name | gene id |
| GU305914.1 | 96980 | 97002 | + | 0 | CTTCGACGACTCTTTGGACG | TGG | \|\|\|\|\|\|\|\|[\|\|\|\|\|\|\|\|\|\|\|\|]PAM | NA | Intergenic | NA | NA |
| T6953 | CGACTCTTTGGACGTGGTCTCGG | 1000 | | | | | | | | | |
| Oligo fwd | CACCCGACTCTTTGGACGTGGTCT | | | | | | | | | | |
| Oligo rev | AAACAGACCACGTCCAAAGAGTCG | | | | | | | | | | |
| Chromosome | start | end | strand | MM | target_seq | PAM | alignment | distance | position | gene name | gene id |
| GU305914.1 | 96986 | 97008 | + | 0 | CGACTCTTTGGACGTGGTCT | CGG | \|\|\|\|\|\|\|\|[\|\|\|\|\|\|\|\|\|\|\|\|]PAM | NA | Intergenic | NA | NA |
| T6954 | GACTCTTTGGACGTGGTCTCGGG | 1000 | | | | | | | | | |
| Oligo fwd | CACCGACTCTTTGGACGTGGTCTC | | | | | | | | | | |
| Oligo rev | AAACGAGACCACGTCCAAAGAGTC | | | | | | | | | | |
| Chromosome | start | end | strand | MM | target_seq | PAM | alignment | distance | position | gene name | gene id |
| GU305914.1 | 96987 | 97009 | + | 0 | GACTCTTTGGACGTGGTCTC | GGG | \|\|\|\|\|\|\|\|[\|\|\|\|\|\|\|\|\|\|\|\|]PAM | NA | Intergenic | NA | NA |
| T6955 | GACTGTTGAGATTTTCGAGACGG | 1000 | | | | | | | | | |
| Oligo fwd | CACCGACTGTTGAGATTTTCGAGA | | | | | | | | | | |
| Oligo rev | AAACTCTCGAAAATCTCAACAGTC | | | | | | | | | | |
| Chromosome | start | end | strand | MM | target_seq | PAM | alignment | distance | position | gene name | gene id |
| GU305914.1 | 97020 | 97042 | + | 0 | GACTGTTGAGATTTTCGAGA | CGG | \|\|\|\|\|\|\|\|[\|\|\|\|\|\|\|\|\|\|\|\|]PAM | NA | Intergenic | NA | NA |
| T6956 | GAGACGGCGCGCTGTCGCCGCGG | 1000 | | | | | | | | | |
| Oligo fwd | CACCGAGACGGCGCGCTGTCGCCG | | | | | | | | | | |
| Oligo rev | AAACCGGCGACAGCGCGCCGTCTC | | | | | | | | | | |
| Chromosome | start | end | strand | MM | target_seq | PAM | alignment | distance | position | gene name | gene id |
| GU305914.1 | 97036 | 97058 | + | 0 | GAGACGGCGCGCTGTCGCCG | CGG | \|\|\|\|\|\|\|\|[\|\|\|\|\|\|\|\|\|\|\|\|]PAM | NA | Intergenic | NA | NA |
| T6957 | AGACGGCGCGCTGTCGCCGCGGG | 1000 | | | | | | | | | |
| Oligo fwd | CACCAGACGGCGCGCTGTCGCCGC | | | | | | | | | | |
| Oligo rev | AAACGCGGCGACAGCGCGCCGTCT | | | | | | | | | | |
| Chromosome | start | end | strand | MM | target_seq | PAM | alignment | distance | position | gene name | gene id |
| GU305914.1 | 97037 | 97059 | + | 0 | AGACGGCGCGCTGTCGCCGC | GGG | \|\|\|\|\|\|\|\|[\|\|\|\|\|\|\|\|\|\|\|\|]PAM | NA | Intergenic | NA | NA |
| GU305914.1 | 139479 | 139501 | - | 4 | AGGCGACGCGCTGTCCCGGC | AGG | \|\|-\|\|-\|\|[\|\|\|\|\|\|\|-\|-\|\|]PAM | NA | Intergenic | NA | NA |
| T6958 | TGTCGCCGCGGGCGCCGTTCTGG | 1000 | | | | | | | | | |
| Oligo fwd | CACCTGTCGCCGCGGGCGCCGTTC | | | | | | | | | | |
| Oligo rev | AAACGAACGGCGCCCGCGGCGACA | | | | | | | | | | |
| Chromosome | start | end | strand | MM | target_seq | PAM | alignment | distance | position | gene name | gene id |
| GU305914.1 | 97048 | 97070 | + | 0 | TGTCGCCGCGGGCGCCGTTC | TGG | \|\|\|\|\|\|\|\|[\|\|\|\|\|\|\|\|\|\|\|\|]PAM | NA | Intergenic | NA | NA |
| T6959 | CGCGGGCGCCGTTCTGGCCGTGG | 1000 | | | | | | | | | |
| Oligo fwd | CACCCGCGGGCGCCGTTCTGGCCG | | | | | | | | | | |
| Oligo rev | AAACCGGCCAGAACGGCGCCCGCG | | | | | | | | | | |
| Chromosome | start | end | strand | MM | target_seq | PAM | alignment | distance | position | gene name | gene id |
| GU305914.1 | 97054 | 97076 | + | 0 | CGCGGGCGCCGTTCTGGCCG | TGG | \|\|\|\|\|\|\|\|[\|\|\|\|\|\|\|\|\|\|\|\|]PAM | NA | Intergenic | NA | NA |
| T6960 | GCCGTTCTGGCCGTGGACCGTGG | 1000 | | | | | | | | | |
| Oligo fwd | CACCGCCGTTCTGGCCGTGGACCG | | | | | | | | | | |
| Oligo rev | AAACCGGTCCACGGCCAGAACGGC | | | | | | | | | | |
| Chromosome | start | end | strand | MM | target_seq | PAM | alignment | distance | position | gene name | gene id |
| GU305914.1 | 97061 | 97083 | + | 0 | GCCGTTCTGGCCGTGGACCG | TGG | \|\|\|\|\|\|\|\|[\|\|\|\|\|\|\|\|\|\|\|\|]PAM | NA | Intergenic | NA | NA |
| T6961 | GTTCTGGCCGTGGACCGTGGTGG | 1000 | | | | | | | | | |

<table>
<tr><td>Oligo fwd</td><td>CACCGTTCTGGCCGTGGACCGTGG</td><td></td><td></td><td></td><td></td><td></td><td></td><td></td><td></td><td></td><td></td></tr>
<tr><td>Oligo rev</td><td>AAACCCACGGTCCACGGCCAGAAC</td><td></td><td></td><td></td><td></td><td></td><td></td><td></td><td></td><td></td><td></td></tr>
<tr><td>Chromosome</td><td>start</td><td>end</td><td>strand</td><td>MM</td><td>target_seq</td><td>PAM</td><td>alignment</td><td>distance</td><td>position</td><td>gene name</td><td>gene id</td></tr>
<tr><td>GU305914.1</td><td>97064</td><td>97086</td><td>+</td><td>0</td><td>GTTCTGGCCGTGGACCGTGG</td><td>TGG</td><td>||||||||[||||||||||||]PAM</td><td>NA</td><td>Intergenic</td><td>NA</td><td>NA</td></tr>
<tr><td></td><td></td><td></td><td></td><td></td><td></td><td></td><td></td><td></td><td></td><td></td><td></td></tr>
<tr><td>T6962</td><td>GTGGACCGTGGTGGAGCTCACGG</td><td>1000</td><td></td><td></td><td></td><td></td><td></td><td></td><td></td><td></td><td></td></tr>
<tr><td>Oligo fwd</td><td>CACCGTGGACCGTGGTGGAGCTCA</td><td></td><td></td><td></td><td></td><td></td><td></td><td></td><td></td><td></td><td></td></tr>
<tr><td>Oligo rev</td><td>AAACTGAGCTCCACCACGGTCCAC</td><td></td><td></td><td></td><td></td><td></td><td></td><td></td><td></td><td></td><td></td></tr>
<tr><td>Chromosome</td><td>start</td><td>end</td><td>strand</td><td>MM</td><td>target_seq</td><td>PAM</td><td>alignment</td><td>distance</td><td>position</td><td>gene name</td><td>gene id</td></tr>
<tr><td>GU305914.1</td><td>97073</td><td>97095</td><td>+</td><td>0</td><td>GTGGACCGTGGTGGAGCTCA</td><td>CGG</td><td>||||||||[||||||||||||]PAM</td><td>NA</td><td>Intergenic</td><td>NA</td><td>NA</td></tr>
<tr><td></td><td></td><td></td><td></td><td></td><td></td><td></td><td></td><td></td><td></td><td></td><td></td></tr>
<tr><td>T6963</td><td>GTGGTGGAGCTCACGGTCGAAGG</td><td>1000</td><td></td><td></td><td></td><td></td><td></td><td></td><td></td><td></td><td></td></tr>
<tr><td>Oligo fwd</td><td>CACCGTGGTGGAGCTCACGGTCGA</td><td></td><td></td><td></td><td></td><td></td><td></td><td></td><td></td><td></td><td></td></tr>
<tr><td>Oligo rev</td><td>AAACTCGACCGTGAGCTCCACCAC</td><td></td><td></td><td></td><td></td><td></td><td></td><td></td><td></td><td></td><td></td></tr>
<tr><td>Chromosome</td><td>start</td><td>end</td><td>strand</td><td>MM</td><td>target_seq</td><td>PAM</td><td>alignment</td><td>distance</td><td>position</td><td>gene name</td><td>gene id</td></tr>
<tr><td>GU305914.1</td><td>97080</td><td>97102</td><td>+</td><td>0</td><td>GTGGTGGAGCTCACGGTCGA</td><td>AGG</td><td>||||||||[||||||||||||]PAM</td><td>NA</td><td>Intergenic</td><td>NA</td><td>NA</td></tr>
<tr><td></td><td></td><td></td><td></td><td></td><td></td><td></td><td></td><td></td><td></td><td></td><td></td></tr>
<tr><td>T6964</td><td>TCGAAGGCGATTTGCTGCTGTGG</td><td>1000</td><td></td><td></td><td></td><td></td><td></td><td></td><td></td><td></td><td></td></tr>
<tr><td>Oligo fwd</td><td>CACCTCGAAGGCGATTTGCTGCTG</td><td></td><td></td><td></td><td></td><td></td><td></td><td></td><td></td><td></td><td></td></tr>
<tr><td>Oligo rev</td><td>AAACCAGCAGCAAATCGCCTTCGA</td><td></td><td></td><td></td><td></td><td></td><td></td><td></td><td></td><td></td><td></td></tr>
<tr><td>Chromosome</td><td>start</td><td>end</td><td>strand</td><td>MM</td><td>target_seq</td><td>PAM</td><td>alignment</td><td>distance</td><td>position</td><td>gene name</td><td>gene id</td></tr>
<tr><td>GU305914.1</td><td>97096</td><td>97118</td><td>+</td><td>0</td><td>TCGAAGGCGATTTGCTGCTG</td><td>TGG</td><td>||||||||[||||||||||||]PAM</td><td>NA</td><td>Intergenic</td><td>NA</td><td>NA</td></tr>
<tr><td></td><td></td><td></td><td></td><td></td><td></td><td></td><td></td><td></td><td></td><td></td><td></td></tr>
<tr><td>T6965</td><td>TGCTGCCGCGCCTGCTGCGCCGG</td><td>1000</td><td></td><td></td><td></td><td></td><td></td><td></td><td></td><td></td><td></td></tr>
<tr><td>Oligo fwd</td><td>CACCTGCTGCCGCGCCTGCTGCGC</td><td></td><td></td><td></td><td></td><td></td><td></td><td></td><td></td><td></td><td></td></tr>
<tr><td>Oligo rev</td><td>AAACGCGCAGCAGGCGCGGCAGCA</td><td></td><td></td><td></td><td></td><td></td><td></td><td></td><td></td><td></td><td></td></tr>
<tr><td>Chromosome</td><td>start</td><td>end</td><td>strand</td><td>MM</td><td>target_seq</td><td>PAM</td><td>alignment</td><td>distance</td><td>position</td><td>gene name</td><td>gene id</td></tr>
<tr><td>GU305914.1</td><td>97153</td><td>97175</td><td>+</td><td>0</td><td>TGCTGCCGCGCCTGCTGCGC</td><td>CGG</td><td>||||||||[||||||||||||]PAM</td><td>NA</td><td>Intergenic</td><td>NA</td><td>NA</td></tr>
<tr><td></td><td></td><td></td><td></td><td></td><td></td><td></td><td></td><td></td><td></td><td></td><td></td></tr>
<tr><td>T6966</td><td>TGCCGCTGCTGCTACTGTCGTGG</td><td>1000</td><td></td><td></td><td></td><td></td><td></td><td></td><td></td><td></td><td></td></tr>
<tr><td>Oligo fwd</td><td>CACCTGCCGCTGCTGCTACTGTCG</td><td></td><td></td><td></td><td></td><td></td><td></td><td></td><td></td><td></td><td></td></tr>
<tr><td>Oligo rev</td><td>AAACCGACAGTAGCAGCAGCGGCA</td><td></td><td></td><td></td><td></td><td></td><td></td><td></td><td></td><td></td><td></td></tr>
<tr><td>Chromosome</td><td>start</td><td>end</td><td>strand</td><td>MM</td><td>target_seq</td><td>PAM</td><td>alignment</td><td>distance</td><td>position</td><td>gene name</td><td>gene id</td></tr>
<tr><td>GU305914.1</td><td>97204</td><td>97226</td><td>+</td><td>0</td><td>TGCCGCTGCTGCTACTGTCG</td><td>TGG</td><td>||||||||[||||||||||||]PAM</td><td>NA</td><td>Intergenic</td><td>NA</td><td>NA</td></tr>
<tr><td></td><td></td><td></td><td></td><td></td><td></td><td></td><td></td><td></td><td></td><td></td><td></td></tr>
<tr><td>T6967</td><td>GCCGCTGCTGCTACTGTCGTGGG</td><td>1000</td><td></td><td></td><td></td><td></td><td></td><td></td><td></td><td></td><td></td></tr>
<tr><td>Oligo fwd</td><td>CACCGCCGCTGCTGCTACTGTCGT</td><td></td><td></td><td></td><td></td><td></td><td></td><td></td><td></td><td></td><td></td></tr>
<tr><td>Oligo rev</td><td>AAACACGACAGTAGCAGCAGCGGC</td><td></td><td></td><td></td><td></td><td></td><td></td><td></td><td></td><td></td><td></td></tr>
<tr><td>Chromosome</td><td>start</td><td>end</td><td>strand</td><td>MM</td><td>target_seq</td><td>PAM</td><td>alignment</td><td>distance</td><td>position</td><td>gene name</td><td>gene id</td></tr>
<tr><td>GU305914.1</td><td>97205</td><td>97227</td><td>+</td><td>0</td><td>GCCGCTGCTGCTACTGTCGT</td><td>GGG</td><td>||||||||[||||||||||||]PAM</td><td>NA</td><td>Intergenic</td><td>NA</td><td>NA</td></tr>
<tr><td></td><td></td><td></td><td></td><td></td><td></td><td></td><td></td><td></td><td></td><td></td><td></td></tr>
<tr><td>T6968</td><td>GCTGCGCGTGCGAGCTCTCCTGG</td><td>1000</td><td></td><td></td><td></td><td></td><td></td><td></td><td></td><td></td><td></td></tr>
<tr><td>Oligo fwd</td><td>CACCGCTGCGCGTGCGAGCTCTCC</td><td></td><td></td><td></td><td></td><td></td><td></td><td></td><td></td><td></td><td></td></tr>
<tr><td>Oligo rev</td><td>AAACGGAGAGCTCGCACGCGCAGC</td><td></td><td></td><td></td><td></td><td></td><td></td><td></td><td></td><td></td><td></td></tr>
<tr><td>Chromosome</td><td>start</td><td>end</td><td>strand</td><td>MM</td><td>target_seq</td><td>PAM</td><td>alignment</td><td>distance</td><td>position</td><td>gene name</td><td>gene id</td></tr>
<tr><td>GU305914.1</td><td>97229</td><td>97251</td><td>+</td><td>0</td><td>GCTGCGCGTGCGAGCTCTCC</td><td>TGG</td><td>||||||||[||||||||||||]PAM</td><td>NA</td><td>Intergenic</td><td>NA</td><td>NA</td></tr>
<tr><td></td><td></td><td></td><td></td><td></td><td></td><td></td><td></td><td></td><td></td><td></td><td></td></tr>
<tr><td>T6969</td><td>GCGTGCGAGCTCTCCTGGTGTGG</td><td>1000</td><td></td><td></td><td></td><td></td><td></td><td></td><td></td><td></td><td></td></tr>
<tr><td>Oligo fwd</td><td>CACCGCGTGCGAGCTCTCCTGGTG</td><td></td><td></td><td></td><td></td><td></td><td></td><td></td><td></td><td></td><td></td></tr>
<tr><td>Oligo rev</td><td>AAACCACCAGGAGAGCTCGCACGC</td><td></td><td></td><td></td><td></td><td></td><td></td><td></td><td></td><td></td><td></td></tr>
<tr><td>Chromosome</td><td>start</td><td>end</td><td>strand</td><td>MM</td><td>target_seq</td><td>PAM</td><td>alignment</td><td>distance</td><td>position</td><td>gene name</td><td>gene id</td></tr>
<tr><td>GU305914.1</td><td>97234</td><td>97256</td><td>+</td><td>0</td><td>GCGTGCGAGCTCTCCTGGTG</td><td>TGG</td><td>||||||||[||||||||||||]PAM</td><td>NA</td><td>Intergenic</td><td>NA</td><td>NA</td></tr>
<tr><td></td><td></td><td></td><td></td><td></td><td></td><td></td><td></td><td></td><td></td><td></td><td></td></tr>
<tr><td>T6970</td><td>TGCGAGCTCTCCTGGTGTGGTGG</td><td>1000</td><td></td><td></td><td></td><td></td><td></td><td></td><td></td><td></td><td></td></tr>
<tr><td>Oligo fwd</td><td>CACCTGCGAGCTCTCCTGGTGTGG</td><td></td><td></td><td></td><td></td><td></td><td></td><td></td><td></td><td></td><td></td></tr>
<tr><td>Oligo rev</td><td>AAACCCACACCAGGAGAGCTCGCA</td><td></td><td></td><td></td><td></td><td></td><td></td><td></td><td></td><td></td><td></td></tr>
<tr><td>Chromosome</td><td>start</td><td>end</td><td>strand</td><td>MM</td><td>target_seq</td><td>PAM</td><td>alignment</td><td>distance</td><td>position</td><td>gene name</td><td>gene id</td></tr>
<tr><td>GU305914.1</td><td>97237</td><td>97259</td><td>+</td><td>0</td><td>TGCGAGCTCTCCTGGTGTGG</td><td>TGG</td><td>||||||||[||||||||||||]PAM</td><td>NA</td><td>Intergenic</td><td>NA</td><td>NA</td></tr>
<tr><td></td><td></td><td></td><td></td><td></td><td></td><td></td><td></td><td></td><td></td><td></td><td></td></tr>
<tr><td>T6971</td><td>CTCTCCTGGTGTGGTGGCGATGG</td><td>1000</td><td></td><td></td><td></td><td></td><td></td><td></td><td></td><td></td><td></td></tr>
<tr><td>Oligo fwd</td><td>CACCCTCTCCTGGTGTGGTGGCGA</td><td></td><td></td><td></td><td></td><td></td><td></td><td></td><td></td><td></td><td></td></tr>
<tr><td>Oligo rev</td><td>AAACTCGCCACCACACCAGGAGAG</td><td></td><td></td><td></td><td></td><td></td><td></td><td></td><td></td><td></td><td></td></tr>
<tr><td>Chromosome</td><td>start</td><td>end</td><td>strand</td><td>MM</td><td>target_seq</td><td>PAM</td><td>alignment</td><td>distance</td><td>position</td><td>gene name</td><td>gene id</td></tr>
<tr><td>GU305914.1</td><td>97243</td><td>97265</td><td>+</td><td>0</td><td>CTCTCCTGGTGTGGTGGCGA</td><td>TGG</td><td>||||||||[||||||||||||]PAM</td><td>NA</td><td>Intergenic</td><td>NA</td><td>NA</td></tr>
<tr><td></td><td></td><td></td><td></td><td></td><td></td><td></td><td></td><td></td><td></td><td></td><td></td></tr>
<tr><td>T6972</td><td>GTGGCGATGGTGATTGCTGCTGG</td><td>1000</td><td></td><td></td><td></td><td></td><td></td><td></td><td></td><td></td><td></td></tr>
<tr><td>Oligo fwd</td><td>CACCGTGGCGATGGTGATTGCTGC</td><td></td><td></td><td></td><td></td><td></td><td></td><td></td><td></td><td></td><td></td></tr>
<tr><td>Oligo rev</td><td>AAACGCAGCAATCACCATCGCCAC</td><td></td><td></td><td></td><td></td><td></td><td></td><td></td><td></td><td></td><td></td></tr>
<tr><td>Chromosome</td><td>start</td><td>end</td><td>strand</td><td>MM</td><td>target_seq</td><td>PAM</td><td>alignment</td><td>distance</td><td>position</td><td>gene name</td><td>gene id</td></tr>
<tr><td>GU305914.1</td><td>97256</td><td>97278</td><td>+</td><td>0</td><td>GTGGCGATGGTGATTGCTGC</td><td>TGG</td><td>||||||||[||||||||||||]PAM</td><td>NA</td><td>Intergenic</td><td>NA</td><td>NA</td></tr>
<tr><td></td><td></td><td></td><td></td><td></td><td></td><td></td><td></td><td></td><td></td><td></td><td></td></tr>
<tr><td>T6973</td><td>TGCTGCTGCTGTCTCCGCGACGG</td><td>1000</td><td></td><td></td><td></td><td></td><td></td><td></td><td></td><td></td><td></td></tr>
<tr><td>Oligo fwd</td><td>CACCTGCTGCTGCTGTCTCCGCGA</td><td></td><td></td><td></td><td></td><td></td><td></td><td></td><td></td><td></td><td></td></tr>
<tr><td>Oligo rev</td><td>AAACTCGCGGAGACAGCAGCAGCA</td><td></td><td></td><td></td><td></td><td></td><td></td><td></td><td></td><td></td><td></td></tr>
<tr><td>Chromosome</td><td>start</td><td>end</td><td>strand</td><td>MM</td><td>target_seq</td><td>PAM</td><td>alignment</td><td>distance</td><td>position</td><td>gene name</td><td>gene id</td></tr>
<tr><td>GU305914.1</td><td>97279</td><td>97301</td><td>+</td><td>0</td><td>TGCTGCTGCTGTCTCCGCGA</td><td>CGG</td><td>||||||||[||||||||||||]PAM</td><td>NA</td><td>Intergenic</td><td>NA</td><td>NA</td></tr>
<tr><td></td><td></td><td></td><td></td><td></td><td></td><td></td><td></td><td></td><td></td><td></td><td></td></tr>
<tr><td>T6974</td><td>CCTCCTCTGCTGTCCACACGAGG</td><td>1000</td><td></td><td></td><td></td><td></td><td></td><td></td><td></td><td></td><td></td></tr>
<tr><td>Oligo fwd</td><td>CACCCCTCCTCTGCTGTCCACACG</td><td></td><td></td><td></td><td></td><td></td><td></td><td></td><td></td><td></td><td></td></tr>
<tr><td>Oligo rev</td><td>AAACCGTGTGGACAGCAGAGGAGG</td><td></td><td></td><td></td><td></td><td></td><td></td><td></td><td></td><td></td><td></td></tr>
<tr><td>Chromosome</td><td>start</td><td>end</td><td>strand</td><td>MM</td><td>target_seq</td><td>PAM</td><td>alignment</td><td>distance</td><td>position</td><td>gene name</td><td>gene id</td></tr>
<tr><td>GU305914.1</td><td>97311</td><td>97333</td><td>+</td><td>0</td><td>CCTCCTCTGCTGTCCACACG</td><td>AGG</td><td>||||||||[||||||||||||]PAM</td><td>NA</td><td>Intergenic</td><td>NA</td><td>NA</td></tr>
<tr><td></td><td></td><td></td><td></td><td></td><td></td><td></td><td></td><td></td><td></td><td></td><td></td></tr>
<tr><td>T6975</td><td>GTCCACACGAGGACTGCTGCTGG</td><td>1000</td><td></td><td></td><td></td><td></td><td></td><td></td><td></td><td></td><td></td></tr>
<tr><td>Oligo fwd</td><td>CACCGTCCACACGAGGACTGCTGC</td><td></td><td></td><td></td><td></td><td></td><td></td><td></td><td></td><td></td><td></td></tr>
<tr><td>Oligo rev</td><td>AAACGCAGCAGTCCTCGTGTGGAC</td><td></td><td></td><td></td><td></td><td></td><td></td><td></td><td></td><td></td><td></td></tr>
<tr><td>Chromosome</td><td>start</td><td>end</td><td>strand</td><td>MM</td><td>target_seq</td><td>PAM</td><td>alignment</td><td>distance</td><td>position</td><td>gene name</td><td>gene id</td></tr>
<tr><td>GU305914.1</td><td>97322</td><td>97344</td><td>+</td><td>0</td><td>GTCCACACGAGGACTGCTGC</td><td>TGG</td><td>||||||||[||||||||||||]PAM</td><td>NA</td><td>Intergenic</td><td>NA</td><td>NA</td></tr>
</table>

| | | | | | | | | | | | |
|---|---|---|---|---|---|---|---|---|---|---|---|
| T6976 | CTGCTGCTGGTCATCCGACTCGG | 1000 | | | | | | | | | |
| Oligo fwd | CACCCTGCTGCTGGTCATCCGACT | | | | | | | | | | |
| Oligo rev | AAACAGTCGGATGACCAGCAGCAG | | | | | | | | | | |
| Chromosome | start | end | strand | MM | target_seq | PAM | alignment | distance | position | gene name | gene id |
| GU305914.1 | 97335 | 97357 | + | 0 | CTGCTGCTGGTCATCCGACT | CGG | \|\|\|\|\|\|\|\|[\|\|\|\|\|\|\|\|\|\|\|\|]PAM | NA | Intergenic | NA | NA |

| | | | | | | | | | | | |
|---|---|---|---|---|---|---|---|---|---|---|---|
| T6977 | CTGCTGGTCATCCGACTCGGAGG | 1000 | | | | | | | | | |
| Oligo fwd | CACCCTGCTGGTCATCCGACTCGG | | | | | | | | | | |
| Oligo rev | AAACCCGAGTCGGATGACCAGCAG | | | | | | | | | | |
| Chromosome | start | end | strand | MM | target_seq | PAM | alignment | distance | position | gene name | gene id |
| GU305914.1 | 97338 | 97360 | + | 0 | CTGCTGGTCATCCGACTCGG | AGG | \|\|\|\|\|\|\|\|[\|\|\|\|\|\|\|\|\|\|\|\|]PAM | NA | Intergenic | NA | NA |

| | | | | | | | | | | | |
|---|---|---|---|---|---|---|---|---|---|---|---|
| T6978 | TGCTGGTCATCCGACTCGGAGGG | 1000 | | | | | | | | | |
| Oligo fwd | CACCTGCTGGTCATCCGACTCGGA | | | | | | | | | | |
| Oligo rev | AAACTCCGAGTCGGATGACCAGCA | | | | | | | | | | |
| Chromosome | start | end | strand | MM | target_seq | PAM | alignment | distance | position | gene name | gene id |
| GU305914.1 | 97339 | 97361 | + | 0 | TGCTGGTCATCCGACTCGGA | GGG | \|\|\|\|\|\|\|\|[\|\|\|\|\|\|\|\|\|\|\|\|]PAM | NA | Intergenic | NA | NA |

| | | | | | | | | | | | |
|---|---|---|---|---|---|---|---|---|---|---|---|
| T6979 | GCTGGTCATCCGACTCGGAGGGG | 1000 | | | | | | | | | |
| Oligo fwd | CACCGCTGGTCATCCGACTCGGAG | | | | | | | | | | |
| Oligo rev | AAACCTCCGAGTCGGATGACCAGC | | | | | | | | | | |
| Chromosome | start | end | strand | MM | target_seq | PAM | alignment | distance | position | gene name | gene id |
| GU305914.1 | 97340 | 97362 | + | 0 | GCTGGTCATCCGACTCGGAG | GGG | \|\|\|\|\|\|\|\|[\|\|\|\|\|\|\|\|\|\|\|\|]PAM | NA | Intergenic | NA | NA |

| | | | | | | | | | | | |
|---|---|---|---|---|---|---|---|---|---|---|---|
| T6980 | GGTCATCCGACTCGGAGGGGCGG | 1000 | | | | | | | | | |
| Oligo fwd | CACCGGTCATCCGACTCGGAGGGG | | | | | | | | | | |
| Oligo rev | AAACCCCCTCCGAGTCGGATGACC | | | | | | | | | | |
| Chromosome | start | end | strand | MM | target_seq | PAM | alignment | distance | position | gene name | gene id |
| GU305914.1 | 97343 | 97365 | + | 0 | GGTCATCCGACTCGGAGGGG | CGG | \|\|\|\|\|\|\|\|[\|\|\|\|\|\|\|\|\|\|\|\|]PAM | NA | Intergenic | NA | NA |

| | | | | | | | | | | | |
|---|---|---|---|---|---|---|---|---|---|---|---|
| T6981 | GTCATCCGACTCGGAGGGGCGGG | 1000 | | | | | | | | | |
| Oligo fwd | CACCGTCATCCGACTCGGAGGGGC | | | | | | | | | | |
| Oligo rev | AAACGCCCCTCCGAGTCGGATGAC | | | | | | | | | | |
| Chromosome | start | end | strand | MM | target_seq | PAM | alignment | distance | position | gene name | gene id |
| GU305914.1 | 97344 | 97366 | + | 0 | GTCATCCGACTCGGAGGGGC | GGG | \|\|\|\|\|\|\|\|[\|\|\|\|\|\|\|\|\|\|\|\|]PAM | NA | Intergenic | NA | NA |

| | | | | | | | | | | | |
|---|---|---|---|---|---|---|---|---|---|---|---|
| T6982 | CGGAGGGGCGGGTTCCACTTCGG | 1000 | | | | | | | | | |
| Oligo fwd | CACCCGGAGGGGCGGGTTCCACTT | | | | | | | | | | |
| Oligo rev | AAACAAGTGGAACCCGCCCCTCCG | | | | | | | | | | |
| Chromosome | start | end | strand | MM | target_seq | PAM | alignment | distance | position | gene name | gene id |
| GU305914.1 | 97355 | 97377 | + | 0 | CGGAGGGGCGGGTTCCACTT | CGG | \|\|\|\|\|\|\|\|[\|\|\|\|\|\|\|\|\|\|\|\|]PAM | NA | Intergenic | NA | NA |

| | | | | | | | | | | | |
|---|---|---|---|---|---|---|---|---|---|---|---|
| T6983 | GGAGGGGCGGGTTCCACTTCGGG | 1000 | | | | | | | | | |
| Oligo fwd | CACCGGAGGGGCGGGTTCCACTTC | | | | | | | | | | |
| Oligo rev | AAACGAAGTGGAACCCGCCCCTCC | | | | | | | | | | |
| Chromosome | start | end | strand | MM | target_seq | PAM | alignment | distance | position | gene name | gene id |
| GU305914.1 | 97356 | 97378 | + | 0 | GGAGGGGCGGGTTCCACTTC | GGG | \|\|\|\|\|\|\|\|[\|\|\|\|\|\|\|\|\|\|\|\|]PAM | NA | Intergenic | NA | NA |
| GU305914.1 | 195015 | 195037 | + | 4 | AGAAGGGCGGCTTCCTCTTC | AGG | -\|\|-\|\|\|\|[\|\|-\|\|\|\|-\|\|\|\|]PAM | NA | Intergenic | NA | NA |

| | | | | | | | | | | | |
|---|---|---|---|---|---|---|---|---|---|---|---|
| T6984 | GAGGGGCGGGTTCCACTTCGGGG | 1000 | | | | | | | | | |
| Oligo fwd | CACCGAGGGGCGGGTTCCACTTCG | | | | | | | | | | |
| Oligo rev | AAACCGAAGTGGAACCCGCCCCTC | | | | | | | | | | |
| Chromosome | start | end | strand | MM | target_seq | PAM | alignment | distance | position | gene name | gene id |
| GU305914.1 | 97357 | 97379 | + | 0 | GAGGGGCGGGTTCCACTTCG | GGG | \|\|\|\|\|\|\|\|[\|\|\|\|\|\|\|\|\|\|\|\|]PAM | NA | Intergenic | NA | NA |

| | | | | | | | | | | | |
|---|---|---|---|---|---|---|---|---|---|---|---|
| T6985 | CGTCGTCGCGCTGCGCCTGTAGG | 1000 | | | | | | | | | |
| Oligo fwd | CACCCGTCGTCGCGCTGCGCCTGT | | | | | | | | | | |
| Oligo rev | AAACACAGGCGCAGCGCGACGACG | | | | | | | | | | |
| Chromosome | start | end | strand | MM | target_seq | PAM | alignment | distance | position | gene name | gene id |
| GU305914.1 | 97382 | 97404 | + | 0 | CGTCGTCGCGCTGCGCCTGT | AGG | \|\|\|\|\|\|\|\|[\|\|\|\|\|\|\|\|\|\|\|\|]PAM | NA | Intergenic | NA | NA |

| | | | | | | | | | | | |
|---|---|---|---|---|---|---|---|---|---|---|---|
| T6986 | GCTGCGCCTGTAGGCCAGCCTGG | 1000 | | | | | | | | | |
| Oligo fwd | CACCGCTGCGCCTGTAGGCCAGCC | | | | | | | | | | |
| Oligo rev | AAACGGCTGGCCTACAGGCGCAGC | | | | | | | | | | |
| Chromosome | start | end | strand | MM | target_seq | PAM | alignment | distance | position | gene name | gene id |
| GU305914.1 | 97391 | 97413 | + | 0 | GCTGCGCCTGTAGGCCAGCC | TGG | \|\|\|\|\|\|\|\|[\|\|\|\|\|\|\|\|\|\|\|\|]PAM | NA | Intergenic | NA | NA |

| | | | | | | | | | | | |
|---|---|---|---|---|---|---|---|---|---|---|---|
| T6987 | CTGCGCCTGTAGGCCAGCCTGGG | 1000 | | | | | | | | | |
| Oligo fwd | CACCCTGCGCCTGTAGGCCAGCCT | | | | | | | | | | |
| Oligo rev | AAACAGGCTGGCCTACAGGCGCAG | | | | | | | | | | |
| Chromosome | start | end | strand | MM | target_seq | PAM | alignment | distance | position | gene name | gene id |
| GU305914.1 | 97392 | 97414 | + | 0 | CTGCGCCTGTAGGCCAGCCT | GGG | \|\|\|\|\|\|\|\|[\|\|\|\|\|\|\|\|\|\|\|\|]PAM | NA | Intergenic | NA | NA |

| | | | | | | | | | | | |
|---|---|---|---|---|---|---|---|---|---|---|---|
| T6988 | TGCGCCTGTAGGCCAGCCTGGGG | 1000 | | | | | | | | | |
| Oligo fwd | CACCTGCGCCTGTAGGCCAGCCTG | | | | | | | | | | |
| Oligo rev | AAACCAGGCTGGCCTACAGGCGCA | | | | | | | | | | |
| Chromosome | start | end | strand | MM | target_seq | PAM | alignment | distance | position | gene name | gene id |
| GU305914.1 | 97393 | 97415 | + | 0 | TGCGCCTGTAGGCCAGCCTG | GGG | \|\|\|\|\|\|\|\|[\|\|\|\|\|\|\|\|\|\|\|\|]PAM | NA | Intergenic | NA | NA |

| | | | | | | | | | | | |
|---|---|---|---|---|---|---|---|---|---|---|---|
| T6989 | TAGGCCAGCCTGGGGATGTTCGG | 1000 | | | | | | | | | |
| Oligo fwd | CACCTAGGCCAGCCTGGGGATGTT | | | | | | | | | | |
| Oligo rev | AAACAACATCCCCAGGCTGGCCTA | | | | | | | | | | |
| Chromosome | start | end | strand | MM | target_seq | PAM | alignment | distance | position | gene name | gene id |
| GU305914.1 | 97401 | 97423 | + | 0 | TAGGCCAGCCTGGGGATGTT | CGG | \|\|\|\|\|\|\|\|[\|\|\|\|\|\|\|\|\|\|\|\|]PAM | NA | Intergenic | NA | NA |

| | | | | | | | | | | | |
|---|---|---|---|---|---|---|---|---|---|---|---|
| T6990 | AGGCCAGCCTGGGGATGTTCGGG | 1000 | | | | | | | | | |
| Oligo fwd | CACCAGGCCAGCCTGGGGATGTTC | | | | | | | | | | |
| Oligo rev | AAACGAACATCCCCAGGCTGGCCT | | | | | | | | | | |
| Chromosome | start | end | strand | MM | target_seq | PAM | alignment | distance | position | gene name | gene id |

| | | | | | | | | | | | |
|---|---|---|---|---|---|---|---|---|---|---|---|
| GU305914.1 | | 97402 | 97424 + | 0 | AGGCCAGCCTGGGGATGTTC | GGG | \|\|\|\|\|\|\|\|[\|\|\|\|\|\|\|\|\|\|\|\|]PAM | NA | Intergenic | NA | NA |
| T6991 | GGCCAGCCTGGGGATGTTCGGGG | 1000 | | | | | | | | | |
| Oligo fwd | CACCGGCCAGCCTGGGGATGTTCG | | | | | | | | | | |
| Oligo rev | AAACCGAACATCCCCAGGCTGGCC | | | | | | | | | | |
| Chromosome | start | end | strand | MM | target_seq | PAM | alignment | distance | position | gene name | gene id |
| GU305914.1 | 97403 | 97425 | + | 0 | GGCCAGCCTGGGGATGTTCG | GGG | \|\|\|\|\|\|\|\|[\|\|\|\|\|\|\|\|\|\|\|\|]PAM | NA | Intergenic | NA | NA |
| T6992 | GGGATGTTCGGGGTCGCGTCCGG | 1000 | | | | | | | | | |
| Oligo fwd | CACCGGGATGTTCGGGGTCGCGTC | | | | | | | | | | |
| Oligo rev | AAACGACGCGACCCCGAACATCCC | | | | | | | | | | |
| Chromosome | start | end | strand | MM | target_seq | PAM | alignment | distance | position | gene name | gene id |
| GU305914.1 | 97413 | 97435 | + | 0 | GGGATGTTCGGGGTCGCGTC | CGG | \|\|\|\|\|\|\|\|[\|\|\|\|\|\|\|\|\|\|\|\|]PAM | NA | Intergenic | NA | NA |
| T6993 | TCGGGGTCGCGTCCGGCGCACGG | 1000 | | | | | | | | | |
| Oligo fwd | CACCTCGGGGTCGCGTCCGGCGCA | | | | | | | | | | |
| Oligo rev | AAACTGCGCCGGACGCGACCCCGA | | | | | | | | | | |
| Chromosome | start | end | strand | MM | target_seq | PAM | alignment | distance | position | gene name | gene id |
| GU305914.1 | 97420 | 97442 | + | 0 | TCGGGGTCGCGTCCGGCGCA | CGG | \|\|\|\|\|\|\|\|[\|\|\|\|\|\|\|\|\|\|\|\|]PAM | NA | Intergenic | NA | NA |
| T6994 | CGTCCGGCGCACGGCGCTTCTGG | 1000 | | | | | | | | | |
| Oligo fwd | CACCCGTCCGGCGCACGGCGCTTC | | | | | | | | | | |
| Oligo rev | AAACGAAGCGCCGTGCGCCGGACG | | | | | | | | | | |
| Chromosome | start | end | strand | MM | target_seq | PAM | alignment | distance | position | gene name | gene id |
| GU305914.1 | 97429 | 97451 | + | 0 | CGTCCGGCGCACGGCGCTTC | TGG | \|\|\|\|\|\|\|\|[\|\|\|\|\|\|\|\|\|\|\|\|]PAM | NA | Intergenic | NA | NA |
| GU305914.1 | 307 | 329 | - | 4 | CGTCGCGCGCCCGGCGCTTA | CGG | \|\|\|\|--\|\|[\|\|-\|\|\|\|\|\|\|\|-]PAM | NA | Intergenic | NA | NA |
| T6995 | GCTTCTGGCGACCTTCGCGCCGG | 1000 | | | | | | | | | |
| Oligo fwd | CACCGCTTCTGGCGACCTTCGCGC | | | | | | | | | | |
| Oligo rev | AAACGCGCGAAGGTCGCCAGAAGC | | | | | | | | | | |
| Chromosome | start | end | strand | MM | target_seq | PAM | alignment | distance | position | gene name | gene id |
| GU305914.1 | 97444 | 97466 | + | 0 | GCTTCTGGCGACCTTCGCGC | CGG | \|\|\|\|\|\|\|\|[\|\|\|\|\|\|\|\|\|\|\|\|]PAM | NA | Intergenic | NA | NA |
| T6996 | CTGGCGACCTTCGCGCCGGTAGG | 1000 | | | | | | | | | |
| Oligo fwd | CACCCTGGCGACCTTCGCGCCGGT | | | | | | | | | | |
| Oligo rev | AAACACCGGCGCGAAGGTCGCCAG | | | | | | | | | | |
| Chromosome | start | end | strand | MM | target_seq | PAM | alignment | distance | position | gene name | gene id |
| GU305914.1 | 97448 | 97470 | + | 0 | CTGGCGACCTTCGCGCCGGT | AGG | \|\|\|\|\|\|\|\|[\|\|\|\|\|\|\|\|\|\|\|\|]PAM | NA | Intergenic | NA | NA |
| T6997 | TTCGCGCCGGTAGGTGCCGTTGG | 1000 | | | | | | | | | |
| Oligo fwd | CACCTTCGCGCCGGTAGGTGCCGT | | | | | | | | | | |
| Oligo rev | AAACACGGCACCTACCGGCGCGAA | | | | | | | | | | |
| Chromosome | start | end | strand | MM | target_seq | PAM | alignment | distance | position | gene name | gene id |
| GU305914.1 | 97457 | 97479 | + | 0 | TTCGCGCCGGTAGGTGCCGT | TGG | \|\|\|\|\|\|\|\|[\|\|\|\|\|\|\|\|\|\|\|\|]PAM | NA | Intergenic | NA | NA |
| T6998 | TCGCGCCGGTAGGTGCCGTTGGG | 1000 | | | | | | | | | |
| Oligo fwd | CACCTCGCGCCGGTAGGTGCCGTT | | | | | | | | | | |
| Oligo rev | AAACAACGGCACCTACCGGCGCGA | | | | | | | | | | |
| Chromosome | start | end | strand | MM | target_seq | PAM | alignment | distance | position | gene name | gene id |
| GU305914.1 | 97458 | 97480 | + | 0 | TCGCGCCGGTAGGTGCCGTT | GGG | \|\|\|\|\|\|\|\|[\|\|\|\|\|\|\|\|\|\|\|\|]PAM | NA | Intergenic | NA | NA |
| T6999 | CGCGCCGGTAGGTGCCGTTGGGG | 1000 | | | | | | | | | |
| Oligo fwd | CACCCGCGCCGGTAGGTGCCGTTG | | | | | | | | | | |
| Oligo rev | AAACCAACGGCACCTACCGGCGCG | | | | | | | | | | |
| Chromosome | start | end | strand | MM | target_seq | PAM | alignment | distance | position | gene name | gene id |
| GU305914.1 | 97459 | 97481 | + | 0 | CGCGCCGGTAGGTGCCGTTG | GGG | \|\|\|\|\|\|\|\|[\|\|\|\|\|\|\|\|\|\|\|\|]PAM | NA | Intergenic | NA | NA |
| T7000 | TGTTGTACGCGCAGAAGAACAGG | 1000 | | | | | | | | | |
| Oligo fwd | CACCTGTTGTACGCGCAGAAGAAC | | | | | | | | | | |
| Oligo rev | AAACGTTCTTCTGCGCGTACAACA | | | | | | | | | | |
| Chromosome | start | end | strand | MM | target_seq | PAM | alignment | distance | position | gene name | gene id |
| GU305914.1 | 97495 | 97517 | + | 0 | TGTTGTACGCGCAGAAGAAC | AGG | \|\|\|\|\|\|\|\|[\|\|\|\|\|\|\|\|\|\|\|\|]PAM | NA | Intergenic | NA | NA |
| T7001 | GTTGTACGCGCAGAAGAACAGGG | 1000 | | | | | | | | | |
| Oligo fwd | CACCGTTGTACGCGCAGAAGAACA | | | | | | | | | | |
| Oligo rev | AAACTGTTCTTCTGCGCGTACAAC | | | | | | | | | | |
| Chromosome | start | end | strand | MM | target_seq | PAM | alignment | distance | position | gene name | gene id |
| GU305914.1 | 97496 | 97518 | + | 0 | GTTGTACGCGCAGAAGAACA | GGG | \|\|\|\|\|\|\|\|[\|\|\|\|\|\|\|\|\|\|\|\|]PAM | NA | Intergenic | NA | NA |
| T7002 | GTACGCGCAGAAGAACAGGGTGG | 1000 | | | | | | | | | |
| Oligo fwd | CACCGTACGCGCAGAAGAACAGGG | | | | | | | | | | |
| Oligo rev | AAACCCCTGTTCTTCTGCGCGTAC | | | | | | | | | | |
| Chromosome | start | end | strand | MM | target_seq | PAM | alignment | distance | position | gene name | gene id |
| GU305914.1 | 97499 | 97521 | + | 0 | GTACGCGCAGAAGAACAGGG | TGG | \|\|\|\|\|\|\|\|[\|\|\|\|\|\|\|\|\|\|\|\|]PAM | NA | Intergenic | NA | NA |
| GU305914.1 | 116643 | 116665 | + | 4 | GTACTTGGGGAAGAACAGGG | CGG | \|\|\|\|--\|-[-\|\|\|\|\|\|\|\|\|\|\|]PAM | NA | Intergenic | NA | NA |
| T7003 | CGCGCAGAAGAACAGGGTGGAGG | 1000 | | | | | | | | | |
| Oligo fwd | CACCCGCGCAGAAGAACAGGGTGG | | | | | | | | | | |
| Oligo rev | AAACCCACCCTGTTCTTCTGCGCG | | | | | | | | | | |
| Chromosome | start | end | strand | MM | target_seq | PAM | alignment | distance | position | gene name | gene id |
| GU305914.1 | 97502 | 97524 | + | 0 | CGCGCAGAAGAACAGGGTGG | AGG | \|\|\|\|\|\|\|\|[\|\|\|\|\|\|\|\|\|\|\|\|]PAM | NA | Intergenic | NA | NA |
| T7004 | GAAGAACAGGGTGGAGGTGTAGG | 1000 | | | | | | | | | |
| Oligo fwd | CACCGAAGAACAGGGTGGAGGTGT | | | | | | | | | | |
| Oligo rev | AAACACACCTCCACCCTGTTCTTC | | | | | | | | | | |
| Chromosome | start | end | strand | MM | target_seq | PAM | alignment | distance | position | gene name | gene id |
| GU305914.1 | 97508 | 97530 | + | 0 | GAAGAACAGGGTGGAGGTGT | AGG | \|\|\|\|\|\|\|\|[\|\|\|\|\|\|\|\|\|\|\|\|]PAM | NA | Intergenic | NA | NA |
| T7005 | AAGAACAGGGTGGAGGTGTAGGG | 1000 | | | | | | | | | |

| | | | | | | | | | | | |
|---|---|---|---|---|---|---|---|---|---|---|---|
| Oligo fwd | CACCAAGAACAGGGTGGAGGTGTA | | | | | | | | | | |
| Oligo rev | AAACTACACCTCCACCCTGTTCTT | | | | | | | | | | |
| Chromosome | start | end | strand | MM | target_seq | PAM | alignment | distance | position | gene name | gene id |
| GU305914.1 | 97509 | 97531 | + | 0 | AAGAACAGGGTGGAGGTGTA | GGG | \|\|\|\|\|\|\|\|[\|\|\|\|\|\|\|\|\|\|\|\|]PAM | NA | Intergenic | NA | NA |
| T7006 | GACAGATCTCGTTCTGACAGCGG | 1000 | | | | | | | | | |
| Oligo fwd | CACCGACAGATCTCGTTCTGACAG | | | | | | | | | | |
| Oligo rev | AAACCTGTCAGAACGAGATCTGTC | | | | | | | | | | |
| Chromosome | start | end | strand | MM | target_seq | PAM | alignment | distance | position | gene name | gene id |
| GU305914.1 | 97558 | 97580 | + | 0 | GACAGATCTCGTTCTGACAG | CGG | \|\|\|\|\|\|\|\|[\|\|\|\|\|\|\|\|\|\|\|\|]PAM | NA | Intergenic | NA | NA |
| T7007 | AGATCTCGTTCTGACAGCGGTGG | 1000 | | | | | | | | | |
| Oligo fwd | CACCAGATCTCGTTCTGACAGCGG | | | | | | | | | | |
| Oligo rev | AAACCCGCTGTCAGAACGAGATCT | | | | | | | | | | |
| Chromosome | start | end | strand | MM | target_seq | PAM | alignment | distance | position | gene name | gene id |
| GU305914.1 | 97561 | 97583 | + | 0 | AGATCTCGTTCTGACAGCGG | TGG | \|\|\|\|\|\|\|\|[\|\|\|\|\|\|\|\|\|\|\|\|]PAM | NA | Intergenic | NA | NA |
| T7008 | TCTGACAGCGGTGGCCGCGCAGG | 1000 | | | | | | | | | |
| Oligo fwd | CACCTCTGACAGCGGTGGCCGCGC | | | | | | | | | | |
| Oligo rev | AAACGCGCGGCCACCGCTGTCAGA | | | | | | | | | | |
| Chromosome | start | end | strand | MM | target_seq | PAM | alignment | distance | position | gene name | gene id |
| GU305914.1 | 97570 | 97592 | + | 0 | TCTGACAGCGGTGGCCGCGC | AGG | \|\|\|\|\|\|\|\|[\|\|\|\|\|\|\|\|\|\|\|\|]PAM | NA | Intergenic | NA | NA |
| T7009 | CAGGTAGTCGAAGATCAGCTCGG | 1000 | | | | | | | | | |
| Oligo fwd | CACCCAGGTAGTCGAAGATCAGCT | | | | | | | | | | |
| Oligo rev | AAACAGCTGATCTTCGACTACCTG | | | | | | | | | | |
| Chromosome | start | end | strand | MM | target_seq | PAM | alignment | distance | position | gene name | gene id |
| GU305914.1 | 97589 | 97611 | + | 0 | CAGGTAGTCGAAGATCAGCT | CGG | \|\|\|\|\|\|\|\|[\|\|\|\|\|\|\|\|\|\|\|\|]PAM | NA | Intergenic | NA | NA |
| T7010 | GCTCGGAGATCATGCCGCAGCGG | 1000 | | | | | | | | | |
| Oligo fwd | CACCGCTCGGAGATCATGCCGCAG | | | | | | | | | | |
| Oligo rev | AAACCTGCGGCATGATCTCCGAGC | | | | | | | | | | |
| Chromosome | start | end | strand | MM | target_seq | PAM | alignment | distance | position | gene name | gene id |
| GU305914.1 | 97606 | 97628 | + | 0 | GCTCGGAGATCATGCCGCAG | CGG | \|\|\|\|\|\|\|\|[\|\|\|\|\|\|\|\|\|\|\|\|]PAM | NA | Intergenic | NA | NA |
| T7011 | GGAGATCATGCCGCAGCGGTAGG | 1000 | | | | | | | | | |
| Oligo fwd | CACCGGAGATCATGCCGCAGCGGT | | | | | | | | | | |
| Oligo rev | AAACACCGCTGCGGCATGATCTCC | | | | | | | | | | |
| Chromosome | start | end | strand | MM | target_seq | PAM | alignment | distance | position | gene name | gene id |
| GU305914.1 | 97610 | 97632 | + | 0 | GGAGATCATGCCGCAGCGGT | AGG | \|\|\|\|\|\|\|\|[\|\|\|\|\|\|\|\|\|\|\|\|]PAM | NA | Intergenic | NA | NA |
| T7012 | GCCGCAGCGGTAGGAATCCGTGG | 1000 | | | | | | | | | |
| Oligo fwd | CACCGCCGCAGCGGTAGGAATCCG | | | | | | | | | | |
| Oligo rev | AAACCGGATTCCTACCGCTGCGGC | | | | | | | | | | |
| Chromosome | start | end | strand | MM | target_seq | PAM | alignment | distance | position | gene name | gene id |
| GU305914.1 | 97619 | 97641 | + | 0 | GCCGCAGCGGTAGGAATCCG | TGG | \|\|\|\|\|\|\|\|[\|\|\|\|\|\|\|\|\|\|\|\|]PAM | NA | Intergenic | NA | NA |
| T7013 | CGCCCGTGTCGTCGAAGTCGCGG | 1000 | | | | | | | | | |
| Oligo fwd | CACCCGCCCGTGTCGTCGAAGTCG | | | | | | | | | | |
| Oligo rev | AAACCGACTTCGACGACACGGGCG | | | | | | | | | | |
| Chromosome | start | end | strand | MM | target_seq | PAM | alignment | distance | position | gene name | gene id |
| GU305914.1 | 97642 | 97664 | + | 0 | CGCCCGTGTCGTCGAAGTCG | CGG | \|\|\|\|\|\|\|\|[\|\|\|\|\|\|\|\|\|\|\|\|]PAM | NA | Intergenic | NA | NA |
| GU305914.1 | 201905 | 201927 | + | 4 | TGCCCGAGTCGTCGAAGACC | AGG | -\|\|\|\|\|-\|[\|\|\|\|\|\|\|\|\|-\|-]PAM | NA | Intergenic | NA | NA |
| T7014 | TCGAAGTCGCGGATGAAGAACGG | 1000 | | | | | | | | | |
| Oligo fwd | CACCTCGAAGTCGCGGATGAAGAA | | | | | | | | | | |
| Oligo rev | AAACTTCTTCATCCGCGACTTCGA | | | | | | | | | | |
| Chromosome | start | end | strand | MM | target_seq | PAM | alignment | distance | position | gene name | gene id |
| GU305914.1 | 97653 | 97675 | + | 0 | TCGAAGTCGCGGATGAAGAA | CGG | \|\|\|\|\|\|\|\|[\|\|\|\|\|\|\|\|\|\|\|\|]PAM | NA | Intergenic | NA | NA |
| T7015 | GTCGCGGATGAAGAACGGCCCGG | 1000 | | | | | | | | | |
| Oligo fwd | CACCGTCGCGGATGAAGAACGGCC | | | | | | | | | | |
| Oligo rev | AAACGGCCGTTCTTCATCCGCGAC | | | | | | | | | | |
| Chromosome | start | end | strand | MM | target_seq | PAM | alignment | distance | position | gene name | gene id |
| GU305914.1 | 97658 | 97680 | + | 0 | GTCGCGGATGAAGAACGGCC | CGG | \|\|\|\|\|\|\|\|[\|\|\|\|\|\|\|\|\|\|\|\|]PAM | NA | Intergenic | NA | NA |
| T7016 | CGGAGTGCCGCAGCGATCTGCGG | 1000 | | | | | | | | | |
| Oligo fwd | CACCCGGAGTGCCGCAGCGATCTG | | | | | | | | | | |
| Oligo rev | AAACCAGATCGCTGCGGCACTCCG | | | | | | | | | | |
| Chromosome | start | end | strand | MM | target_seq | PAM | alignment | distance | position | gene name | gene id |
| GU305914.1 | 97678 | 97700 | + | 0 | CGGAGTGCCGCAGCGATCTG | CGG | \|\|\|\|\|\|\|\|[\|\|\|\|\|\|\|\|\|\|\|\|]PAM | NA | Intergenic | NA | NA |
| T7017 | AGTGCCGCAGCGATCTGCGGCGG | 1000 | | | | | | | | | |
| Oligo fwd | CACCAGTGCCGCAGCGATCTGCGG | | | | | | | | | | |
| Oligo rev | AAACCCGCAGATCGCTGCGGCACT | | | | | | | | | | |
| Chromosome | start | end | strand | MM | target_seq | PAM | alignment | distance | position | gene name | gene id |
| GU305914.1 | 97681 | 97703 | + | 0 | AGTGCCGCAGCGATCTGCGG | CGG | \|\|\|\|\|\|\|\|[\|\|\|\|\|\|\|\|\|\|\|\|]PAM | NA | Intergenic | NA | NA |
| T7018 | AGCGCCAGCGCCTGCTGCAGCGG | 1000 | | | | | | | | | |
| Oligo fwd | CACCAGCGCCAGCGCCTGCTGCAG | | | | | | | | | | |
| Oligo rev | AAACCTGCAGCAGGCGCTGGCGCT | | | | | | | | | | |
| Chromosome | start | end | strand | MM | target_seq | PAM | alignment | distance | position | gene name | gene id |
| GU305914.1 | 97737 | 97759 | + | 0 | AGCGCCAGCGCCTGCTGCAG | CGG | \|\|\|\|\|\|\|\|[\|\|\|\|\|\|\|\|\|\|\|\|]PAM | NA | Intergenic | NA | NA |
| T7019 | GCGCCAGCGCCTGCTGCAGCGGG | 1000 | | | | | | | | | |
| Oligo fwd | CACCGCGCCAGCGCCTGCTGCAGC | | | | | | | | | | |
| Oligo rev | AAACGCTGCAGCAGGCGCTGGCGC | | | | | | | | | | |
| Chromosome | start | end | strand | MM | target_seq | PAM | alignment | distance | position | gene name | gene id |
| GU305914.1 | 97738 | 97760 | + | 0 | GCGCCAGCGCCTGCTGCAGC | GGG | \|\|\|\|\|\|\|\|[\|\|\|\|\|\|\|\|\|\|\|\|]PAM | NA | Intergenic | NA | NA |

| | | | | | | | | | | | |
|---|---|---|---|---|---|---|---|---|---|---|---|
| T7020 | GCAGCGGGTGCTTGAAAGACAGG | 1000 | | | | | | | | | |
| Oligo fwd | CACCGCAGCGGGTGCTTGAAAGAC | | | | | | | | | | |
| Oligo rev | AAACGTCTTTCAAGCACCCGCTGC | | | | | | | | | | |
| Chromosome | start | end | strand | MM | target_seq | PAM | alignment | distance | position | gene name | gene id |
| GU305914.1 | 97753 | 97775 | + | 0 | GCAGCGGGTGCTTGAAAGAC | AGG | \|\|\|\|\|\|\|\|[\|\|\|\|\|\|\|\|\|\|\|\|]PAM | NA | Intergenic | NA | NA |
| T7021 | CAGCGGGTGCTTGAAAGACAGGG | 1000 | | | | | | | | | |
| Oligo fwd | CACCCAGCGGGTGCTTGAAAGACA | | | | | | | | | | |
| Oligo rev | AAACTGTCTTTCAAGCACCCGCTG | | | | | | | | | | |
| Chromosome | start | end | strand | MM | target_seq | PAM | alignment | distance | position | gene name | gene id |
| GU305914.1 | 97754 | 97776 | + | 0 | CAGCGGGTGCTTGAAAGACA | GGG | \|\|\|\|\|\|\|\|[\|\|\|\|\|\|\|\|\|\|\|\|]PAM | NA | Intergenic | NA | NA |
| T7022 | AGCGGGTGCTTGAAAGACAGGGG | 1000 | | | | | | | | | |
| Oligo fwd | CACCAGCGGGTGCTTGAAAGACAG | | | | | | | | | | |
| Oligo rev | AAACCTGTCTTTCAAGCACCCGCT | | | | | | | | | | |
| Chromosome | start | end | strand | MM | target_seq | PAM | alignment | distance | position | gene name | gene id |
| GU305914.1 | 97755 | 97777 | + | 0 | AGCGGGTGCTTGAAAGACAG | GGG | \|\|\|\|\|\|\|\|[\|\|\|\|\|\|\|\|\|\|\|\|]PAM | NA | Intergenic | NA | NA |
| T7023 | GGGTGCTTGAAAGACAGGGGCGG | 1000 | | | | | | | | | |
| Oligo fwd | CACCGGGTGCTTGAAAGACAGGGG | | | | | | | | | | |
| Oligo rev | AAACCCCCTGTCTTTCAAGCACCC | | | | | | | | | | |
| Chromosome | start | end | strand | MM | target_seq | PAM | alignment | distance | position | gene name | gene id |
| GU305914.1 | 97758 | 97780 | + | 0 | GGGTGCTTGAAAGACAGGGG | CGG | \|\|\|\|\|\|\|\|[\|\|\|\|\|\|\|\|\|\|\|\|]PAM | NA | Intergenic | NA | NA |
| T7024 | GCTTGAAAGACAGGGGCGGACGG | 1000 | | | | | | | | | |
| Oligo fwd | CACCGCTTGAAAGACAGGGGCGGA | | | | | | | | | | |
| Oligo rev | AAACTCCGCCCCTGTCTTTCAAGC | | | | | | | | | | |
| Chromosome | start | end | strand | MM | target_seq | PAM | alignment | distance | position | gene name | gene id |
| GU305914.1 | 97762 | 97784 | + | 0 | GCTTGAAAGACAGGGGCGGA | CGG | \|\|\|\|\|\|\|\|[\|\|\|\|\|\|\|\|\|\|\|\|]PAM | NA | Intergenic | NA | NA |
| T7025 | CGGACGGCGCAACACGTCGAAGG | 1000 | | | | | | | | | |
| Oligo fwd | CACCCGGACGGCGCAACACGTCGA | | | | | | | | | | |
| Oligo rev | AAACTCGACGTGTTGCGCCGTCCG | | | | | | | | | | |
| Chromosome | start | end | strand | MM | target_seq | PAM | alignment | distance | position | gene name | gene id |
| GU305914.1 | 97778 | 97800 | + | 0 | CGGACGGCGCAACACGTCGA | AGG | \|\|\|\|\|\|\|\|[\|\|\|\|\|\|\|\|\|\|\|\|]PAM | NA | Intergenic | NA | NA |
| T7026 | ACGGCGCAACACGTCGAAGGAGG | 1000 | | | | | | | | | |
| Oligo fwd | CACCACGGCGCAACACGTCGAAGG | | | | | | | | | | |
| Oligo rev | AAACCCTTCGACGTGTTGCGCCGT | | | | | | | | | | |
| Chromosome | start | end | strand | MM | target_seq | PAM | alignment | distance | position | gene name | gene id |
| GU305914.1 | 97781 | 97803 | + | 0 | ACGGCGCAACACGTCGAAGG | AGG | \|\|\|\|\|\|\|\|[\|\|\|\|\|\|\|\|\|\|\|\|]PAM | NA | Intergenic | NA | NA |
| T7027 | CTGCCGAAGAGCCGCCCTTGCGG | 1000 | | | | | | | | | |
| Oligo fwd | CACCCTGCCGAAGAGCCGCCCTTG | | | | | | | | | | |
| Oligo rev | AAACCAAGGGCGGCTCTTCGGCAG | | | | | | | | | | |
| Chromosome | start | end | strand | MM | target_seq | PAM | alignment | distance | position | gene name | gene id |
| GU305914.1 | 97813 | 97835 | + | 0 | CTGCCGAAGAGCCGCCCTTG | CGG | \|\|\|\|\|\|\|\|[\|\|\|\|\|\|\|\|\|\|\|\|]PAM | NA | Intergenic | NA | NA |
| T7028 | GAAGAGCCGCCCTTGCGGCACGG | 1000 | | | | | | | | | |
| Oligo fwd | CACCGAAGAGCCGCCCTTGCGGCA | | | | | | | | | | |
| Oligo rev | AAACTGCCGCAAGGGCGGCTCTTC | | | | | | | | | | |
| Chromosome | start | end | strand | MM | target_seq | PAM | alignment | distance | position | gene name | gene id |
| GU305914.1 | 97818 | 97840 | + | 0 | GAAGAGCCGCCCTTGCGGCA | CGG | \|\|\|\|\|\|\|\|[\|\|\|\|\|\|\|\|\|\|\|\|]PAM | NA | Intergenic | NA | NA |
| T7029 | CCCTTGCGGCACGGAGACGCCGG | 1000 | | | | | | | | | |
| Oligo fwd | CACCCCCTTGCGGCACGGAGACGC | | | | | | | | | | |
| Oligo rev | AAACGCGTCTCCGTGCCGCAAGGG | | | | | | | | | | |
| Chromosome | start | end | strand | MM | target_seq | PAM | alignment | distance | position | gene name | gene id |
| GU305914.1 | 97827 | 97849 | + | 0 | CCCTTGCGGCACGGAGACGC | CGG | \|\|\|\|\|\|\|\|[\|\|\|\|\|\|\|\|\|\|\|\|]PAM | NA | Intergenic | NA | NA |
| T7030 | ACGCCGGTTTGTTCGCGATCTGG | 1000 | | | | | | | | | |
| Oligo fwd | CACCACGCCGGTTTGTTCGCGATC | | | | | | | | | | |
| Oligo rev | AAACGATCGCGAACAAACCGGCGT | | | | | | | | | | |
| Chromosome | start | end | strand | MM | target_seq | PAM | alignment | distance | position | gene name | gene id |
| GU305914.1 | 97843 | 97865 | + | 0 | ACGCCGGTTTGTTCGCGATC | TGG | \|\|\|\|\|\|\|\|[\|\|\|\|\|\|\|\|\|\|\|\|]PAM | NA | Intergenic | NA | NA |
| T7031 | GTTTGTTCGCGATCTGGCGCAGG | 1000 | | | | | | | | | |
| Oligo fwd | CACCGTTTGTTCGCGATCTGGCGC | | | | | | | | | | |
| Oligo rev | AAACGCGCCAGATCGCGAACAAAC | | | | | | | | | | |
| Chromosome | start | end | strand | MM | target_seq | PAM | alignment | distance | position | gene name | gene id |
| GU305914.1 | 97849 | 97871 | + | 0 | GTTTGTTCGCGATCTGGCGC | AGG | \|\|\|\|\|\|\|\|[\|\|\|\|\|\|\|\|\|\|\|\|]PAM | NA | Intergenic | NA | NA |
| T7032 | GGCGCAGGATGTGATACTCGCGG | 1000 | | | | | | | | | |
| Oligo fwd | CACCGGCGCAGGATGTGATACTCG | | | | | | | | | | |
| Oligo rev | AAACCGAGTATCACATCCTGCGCC | | | | | | | | | | |
| Chromosome | start | end | strand | MM | target_seq | PAM | alignment | distance | position | gene name | gene id |
| GU305914.1 | 97864 | 97886 | + | 0 | GGCGCAGGATGTGATACTCG | CGG | \|\|\|\|\|\|\|\|[\|\|\|\|\|\|\|\|\|\|\|\|]PAM | NA | Intergenic | NA | NA |
| T7033 | CGGTCGCTGCCGCCGCCGCCCGG | 1000 | | | | | | | | | |
| Oligo fwd | CACCCGGTCGCTGCCGCCGCCGCC | | | | | | | | | | |
| Oligo rev | AAACGGCGGCGGCGGCAGCGACCG | | | | | | | | | | |
| Chromosome | start | end | strand | MM | target_seq | PAM | alignment | distance | position | gene name | gene id |
| GU305914.1 | 97884 | 97906 | + | 0 | CGGTCGCTGCCGCCGCCGCC | CGG | \|\|\|\|\|\|\|\|[\|\|\|\|\|\|\|\|\|\|\|\|]PAM | NA | Intergenic | NA | NA |
| GU305914.1 | 154401 | 154423 | - | 4 | CCGCCGCCGCCGCCACCGCC | CGG | \|-\|-\|\|\|-[\|\|\|\|\|\|-\|\|\|\|\|]PAM | NA | Intergenic | NA | NA |
| T7034 | GGTCGCTGCCGCCGCCGCCCGGG | 1000 | | | | | | | | | |
| Oligo fwd | CACCGGTCGCTGCCGCCGCCGCCC | | | | | | | | | | |
| Oligo rev | AAACGGGCGGCGGCGGCAGCGACC | | | | | | | | | | |

<table>
<tr><td>Chromosome</td><td>start</td><td>end</td><td>strand</td><td>MM</td><td>target_seq</td><td>PAM</td><td>alignment</td><td>distance</td><td>position</td><td>gene name</td><td>gene id</td></tr>
<tr><td>GU305914.1</td><td>97885</td><td>97907</td><td>+</td><td>0</td><td>GGTCGCTGCCGCCGCCGCCC</td><td>GGG</td><td>||||||||[||||||||||||]PAM</td><td>NA</td><td>Intergenic</td><td>NA</td><td>NA</td></tr>
<tr><td>GU305914.1</td><td>154400</td><td>154422</td><td>-</td><td>4</td><td>CGCCGCCGCCGCCACCGCCC</td><td>GGG</td><td>-|-|||-|[|||||-||||||]PAM</td><td>NA</td><td>Intergenic</td><td>NA</td><td>NA</td></tr>
<tr><td>GU305914.1</td><td>45592</td><td>45614</td><td>-</td><td>3</td><td>GGCGGCTGCCGCCGCCGCCG</td><td>CGG</td><td>||--||||[|||||||||||-]PAM</td><td>NA</td><td>Intergenic</td><td>NA</td><td>NA</td></tr>
<tr><td>GU305914.1</td><td>135435</td><td>135457</td><td>+</td><td>4</td><td>GGGCGCCGCCGCCGACGGCC</td><td>CGG</td><td>||-|||-|[||||||-||-||]PAM</td><td>NA</td><td>Intergenic</td><td>NA</td><td>NA</td></tr>
<tr><td>T7035</td><td>CGCCGCCGCCCGGGCTGTCCAGG</td><td>1000</td><td></td><td></td><td></td><td></td><td></td><td></td><td></td><td></td><td></td></tr>
<tr><td>Oligo fwd</td><td>CACCCGCCGCCGCCCGGGCTGTCC</td><td></td><td></td><td></td><td></td><td></td><td></td><td></td><td></td><td></td><td></td></tr>
<tr><td>Oligo rev</td><td>AAACGGACAGCCCGGGCGGCGGCG</td><td></td><td></td><td></td><td></td><td></td><td></td><td></td><td></td><td></td><td></td></tr>
<tr><td>Chromosome</td><td>start</td><td>end</td><td>strand</td><td>MM</td><td>target_seq</td><td>PAM</td><td>alignment</td><td>distance</td><td>position</td><td>gene name</td><td>gene id</td></tr>
<tr><td>GU305914.1</td><td>97894</td><td>97916</td><td>+</td><td>0</td><td>CGCCGCCGCCCGGGCTGTCC</td><td>AGG</td><td>||||||||[||||||||||||]PAM</td><td>NA</td><td>Intergenic</td><td>NA</td><td>NA</td></tr>
<tr><td>T7036</td><td>CGCCGCCCGGGCTGTCCAGGAGG</td><td>1000</td><td></td><td></td><td></td><td></td><td></td><td></td><td></td><td></td><td></td></tr>
<tr><td>Oligo fwd</td><td>CACCCGCCGCCCGGGCTGTCCAGG</td><td></td><td></td><td></td><td></td><td></td><td></td><td></td><td></td><td></td><td></td></tr>
<tr><td>Oligo rev</td><td>AAACCCTGGACAGCCCGGGCGGCG</td><td></td><td></td><td></td><td></td><td></td><td></td><td></td><td></td><td></td><td></td></tr>
<tr><td>Chromosome</td><td>start</td><td>end</td><td>strand</td><td>MM</td><td>target_seq</td><td>PAM</td><td>alignment</td><td>distance</td><td>position</td><td>gene name</td><td>gene id</td></tr>
<tr><td>GU305914.1</td><td>97897</td><td>97919</td><td>+</td><td>0</td><td>CGCCGCCCGGGCTGTCCAGG</td><td>AGG</td><td>||||||||[||||||||||||]PAM</td><td>NA</td><td>Intergenic</td><td>NA</td><td>NA</td></tr>
<tr><td>T7037</td><td>GGAGGCACGACAGCTCGCCGAGG</td><td>1000</td><td></td><td></td><td></td><td></td><td></td><td></td><td></td><td></td><td></td></tr>
<tr><td>Oligo fwd</td><td>CACCGGAGGCACGACAGCTCGCCG</td><td></td><td></td><td></td><td></td><td></td><td></td><td></td><td></td><td></td><td></td></tr>
<tr><td>Oligo rev</td><td>AAACCGGCGAGCTGTCGTGCCTCC</td><td></td><td></td><td></td><td></td><td></td><td></td><td></td><td></td><td></td><td></td></tr>
<tr><td>Chromosome</td><td>start</td><td>end</td><td>strand</td><td>MM</td><td>target_seq</td><td>PAM</td><td>alignment</td><td>distance</td><td>position</td><td>gene name</td><td>gene id</td></tr>
<tr><td>GU305914.1</td><td>97915</td><td>97937</td><td>+</td><td>0</td><td>GGAGGCACGACAGCTCGCCG</td><td>AGG</td><td>||||||||[||||||||||||]PAM</td><td>NA</td><td>Intergenic</td><td>NA</td><td>NA</td></tr>
<tr><td>T7038</td><td>GAGGTTCAGTTTGTTGAACGTGG</td><td>1000</td><td></td><td></td><td></td><td></td><td></td><td></td><td></td><td></td><td></td></tr>
<tr><td>Oligo fwd</td><td>CACCGAGGTTCAGTTTGTTGAACG</td><td></td><td></td><td></td><td></td><td></td><td></td><td></td><td></td><td></td><td></td></tr>
<tr><td>Oligo rev</td><td>AAACCGTTCAACAAACTGAACCTC</td><td></td><td></td><td></td><td></td><td></td><td></td><td></td><td></td><td></td><td></td></tr>
<tr><td>Chromosome</td><td>start</td><td>end</td><td>strand</td><td>MM</td><td>target_seq</td><td>PAM</td><td>alignment</td><td>distance</td><td>position</td><td>gene name</td><td>gene id</td></tr>
<tr><td>GU305914.1</td><td>97934</td><td>97956</td><td>+</td><td>0</td><td>GAGGTTCAGTTTGTTGAACG</td><td>TGG</td><td>||||||||[||||||||||||]PAM</td><td>NA</td><td>Intergenic</td><td>NA</td><td>NA</td></tr>
<tr><td>T7039</td><td>TGAACGTGGCGTACTTTATCAGG</td><td>1000</td><td></td><td></td><td></td><td></td><td></td><td></td><td></td><td></td><td></td></tr>
<tr><td>Oligo fwd</td><td>CACCTGAACGTGGCGTACTTTATC</td><td></td><td></td><td></td><td></td><td></td><td></td><td></td><td></td><td></td><td></td></tr>
<tr><td>Oligo rev</td><td>AAACGATAAAGTACGCCACGTTCA</td><td></td><td></td><td></td><td></td><td></td><td></td><td></td><td></td><td></td><td></td></tr>
<tr><td>Chromosome</td><td>start</td><td>end</td><td>strand</td><td>MM</td><td>target_seq</td><td>PAM</td><td>alignment</td><td>distance</td><td>position</td><td>gene name</td><td>gene id</td></tr>
<tr><td>GU305914.1</td><td>97948</td><td>97970</td><td>+</td><td>0</td><td>TGAACGTGGCGTACTTTATC</td><td>AGG</td><td>||||||||[||||||||||||]PAM</td><td>NA</td><td>Intergenic</td><td>NA</td><td>NA</td></tr>
<tr><td>T7040</td><td>TGGCGTACTTTATCAGGCCCAGG</td><td>1000</td><td></td><td></td><td></td><td></td><td></td><td></td><td></td><td></td><td></td></tr>
<tr><td>Oligo fwd</td><td>CACCTGGCGTACTTTATCAGGCCC</td><td></td><td></td><td></td><td></td><td></td><td></td><td></td><td></td><td></td><td></td></tr>
<tr><td>Oligo rev</td><td>AAACGGGCCTGATAAAGTACGCCA</td><td></td><td></td><td></td><td></td><td></td><td></td><td></td><td></td><td></td><td></td></tr>
<tr><td>Chromosome</td><td>start</td><td>end</td><td>strand</td><td>MM</td><td>target_seq</td><td>PAM</td><td>alignment</td><td>distance</td><td>position</td><td>gene name</td><td>gene id</td></tr>
<tr><td>GU305914.1</td><td>97954</td><td>97976</td><td>+</td><td>0</td><td>TGGCGTACTTTATCAGGCCC</td><td>AGG</td><td>||||||||[||||||||||||]PAM</td><td>NA</td><td>Intergenic</td><td>NA</td><td>NA</td></tr>
<tr><td>T7041</td><td>GGCGTACTTTATCAGGCCCAGGG</td><td>1000</td><td></td><td></td><td></td><td></td><td></td><td></td><td></td><td></td><td></td></tr>
<tr><td>Oligo fwd</td><td>CACCGGCGTACTTTATCAGGCCCA</td><td></td><td></td><td></td><td></td><td></td><td></td><td></td><td></td><td></td><td></td></tr>
<tr><td>Oligo rev</td><td>AAACTGGGCCTGATAAAGTACGCC</td><td></td><td></td><td></td><td></td><td></td><td></td><td></td><td></td><td></td><td></td></tr>
<tr><td>Chromosome</td><td>start</td><td>end</td><td>strand</td><td>MM</td><td>target_seq</td><td>PAM</td><td>alignment</td><td>distance</td><td>position</td><td>gene name</td><td>gene id</td></tr>
<tr><td>GU305914.1</td><td>97955</td><td>97977</td><td>+</td><td>0</td><td>GGCGTACTTTATCAGGCCCA</td><td>GGG</td><td>||||||||[||||||||||||]PAM</td><td>NA</td><td>Intergenic</td><td>NA</td><td>NA</td></tr>
<tr><td>T7042</td><td>GGCCCAGGGTGCTCATGTACAGG</td><td>1000</td><td></td><td></td><td></td><td></td><td></td><td></td><td></td><td></td><td></td></tr>
<tr><td>Oligo fwd</td><td>CACCGGCCCAGGGTGCTCATGTAC</td><td></td><td></td><td></td><td></td><td></td><td></td><td></td><td></td><td></td><td></td></tr>
<tr><td>Oligo rev</td><td>AAACGTACATGAGCACCCTGGGCC</td><td></td><td></td><td></td><td></td><td></td><td></td><td></td><td></td><td></td><td></td></tr>
<tr><td>Chromosome</td><td>start</td><td>end</td><td>strand</td><td>MM</td><td>target_seq</td><td>PAM</td><td>alignment</td><td>distance</td><td>position</td><td>gene name</td><td>gene id</td></tr>
<tr><td>GU305914.1</td><td>97969</td><td>97991</td><td>+</td><td>0</td><td>GGCCCAGGGTGCTCATGTAC</td><td>AGG</td><td>||||||||[||||||||||||]PAM</td><td>NA</td><td>Intergenic</td><td>NA</td><td>NA</td></tr>
<tr><td>T7043</td><td>GGTGCTCATGTACAGGTCTTTGG</td><td>1000</td><td></td><td></td><td></td><td></td><td></td><td></td><td></td><td></td><td></td></tr>
<tr><td>Oligo fwd</td><td>CACCGGTGCTCATGTACAGGTCTT</td><td></td><td></td><td></td><td></td><td></td><td></td><td></td><td></td><td></td><td></td></tr>
<tr><td>Oligo rev</td><td>AAACAAGACCTGTACATGAGCACC</td><td></td><td></td><td></td><td></td><td></td><td></td><td></td><td></td><td></td><td></td></tr>
<tr><td>Chromosome</td><td>start</td><td>end</td><td>strand</td><td>MM</td><td>target_seq</td><td>PAM</td><td>alignment</td><td>distance</td><td>position</td><td>gene name</td><td>gene id</td></tr>
<tr><td>GU305914.1</td><td>97976</td><td>97998</td><td>+</td><td>0</td><td>GGTGCTCATGTACAGGTCTT</td><td>TGG</td><td>||||||||[||||||||||||]PAM</td><td>NA</td><td>Intergenic</td><td>NA</td><td>NA</td></tr>
<tr><td>T7044</td><td>GTGCTCATGTACAGGTCTTTGGG</td><td>1000</td><td></td><td></td><td></td><td></td><td></td><td></td><td></td><td></td><td></td></tr>
<tr><td>Oligo fwd</td><td>CACCGTGCTCATGTACAGGTCTTT</td><td></td><td></td><td></td><td></td><td></td><td></td><td></td><td></td><td></td><td></td></tr>
<tr><td>Oligo rev</td><td>AAACAAAGACCTGTACATGAGCAC</td><td></td><td></td><td></td><td></td><td></td><td></td><td></td><td></td><td></td><td></td></tr>
<tr><td>Chromosome</td><td>start</td><td>end</td><td>strand</td><td>MM</td><td>target_seq</td><td>PAM</td><td>alignment</td><td>distance</td><td>position</td><td>gene name</td><td>gene id</td></tr>
<tr><td>GU305914.1</td><td>97977</td><td>97999</td><td>+</td><td>0</td><td>GTGCTCATGTACAGGTCTTT</td><td>GGG</td><td>||||||||[||||||||||||]PAM</td><td>NA</td><td>Intergenic</td><td>NA</td><td>NA</td></tr>
<tr><td>T7045</td><td>TACAGGTCTTTGGGATTCACCGG</td><td>1000</td><td></td><td></td><td></td><td></td><td></td><td></td><td></td><td></td><td></td></tr>
<tr><td>Oligo fwd</td><td>CACCTACAGGTCTTTGGGATTCAC</td><td></td><td></td><td></td><td></td><td></td><td></td><td></td><td></td><td></td><td></td></tr>
<tr><td>Oligo rev</td><td>AAACGTGAATCCCAAAGACCTGTA</td><td></td><td></td><td></td><td></td><td></td><td></td><td></td><td></td><td></td><td></td></tr>
<tr><td>Chromosome</td><td>start</td><td>end</td><td>strand</td><td>MM</td><td>target_seq</td><td>PAM</td><td>alignment</td><td>distance</td><td>position</td><td>gene name</td><td>gene id</td></tr>
<tr><td>GU305914.1</td><td>97986</td><td>98008</td><td>+</td><td>0</td><td>TACAGGTCTTTGGGATTCAC</td><td>CGG</td><td>||||||||[||||||||||||]PAM</td><td>NA</td><td>Intergenic</td><td>NA</td><td>NA</td></tr>
<tr><td>T7046</td><td>GGTCTTTGGGATTCACCGGCAGG</td><td>1000</td><td></td><td></td><td></td><td></td><td></td><td></td><td></td><td></td><td></td></tr>
<tr><td>Oligo fwd</td><td>CACCGGTCTTTGGGATTCACCGGC</td><td></td><td></td><td></td><td></td><td></td><td></td><td></td><td></td><td></td><td></td></tr>
<tr><td>Oligo rev</td><td>AAACGCCGGTGAATCCCAAAGACC</td><td></td><td></td><td></td><td></td><td></td><td></td><td></td><td></td><td></td><td></td></tr>
<tr><td>Chromosome</td><td>start</td><td>end</td><td>strand</td><td>MM</td><td>target_seq</td><td>PAM</td><td>alignment</td><td>distance</td><td>position</td><td>gene name</td><td>gene id</td></tr>
<tr><td>GU305914.1</td><td>97990</td><td>98012</td><td>+</td><td>0</td><td>GGTCTTTGGGATTCACCGGC</td><td>AGG</td><td>||||||||[||||||||||||]PAM</td><td>NA</td><td>Intergenic</td><td>NA</td><td>NA</td></tr>
<tr><td>T7047</td><td>TCACCGGCAGGTTGTGTACGCGG</td><td>1000</td><td></td><td></td><td></td><td></td><td></td><td></td><td></td><td></td><td></td></tr>
<tr><td>Oligo fwd</td><td>CACCTCACCGGCAGGTTGTGTACG</td><td></td><td></td><td></td><td></td><td></td><td></td><td></td><td></td><td></td><td></td></tr>
<tr><td>Oligo rev</td><td>AAACCGTACACAACCTGCCGGTGA</td><td></td><td></td><td></td><td></td><td></td><td></td><td></td><td></td><td></td><td></td></tr>
<tr><td>Chromosome</td><td>start</td><td>end</td><td>strand</td><td>MM</td><td>target_seq</td><td>PAM</td><td>alignment</td><td>distance</td><td>position</td><td>gene name</td><td>gene id</td></tr>
<tr><td>GU305914.1</td><td>98002</td><td>98024</td><td>+</td><td>0</td><td>TCACCGGCAGGTTGTGTACG</td><td>CGG</td><td>||||||||[||||||||||||]PAM</td><td>NA</td><td>Intergenic</td><td>NA</td><td>NA</td></tr>
<tr><td>T7048</td><td>CCGGCAGGTTGTGTACGCGGTGG</td><td>1000</td><td></td><td></td><td></td><td></td><td></td><td></td><td></td><td></td><td></td></tr>
<tr><td>Oligo fwd</td><td>CACCCCGGCAGGTTGTGTACGCGG</td><td></td><td></td><td></td><td></td><td></td><td></td><td></td><td></td><td></td><td></td></tr>
<tr><td>Oligo rev</td><td>AAACCCGCGTACACAACCTGCCGG</td><td></td><td></td><td></td><td></td><td></td><td></td><td></td><td></td><td></td><td></td></tr>
<tr><td>Chromosome</td><td>start</td><td>end</td><td>strand</td><td>MM</td><td>target_seq</td><td>PAM</td><td>alignment</td><td>distance</td><td>position</td><td>gene name</td><td>gene id</td></tr>
<tr><td>GU305914.1</td><td>98005</td><td>98027</td><td>+</td><td>0</td><td>CCGGCAGGTTGTGTACGCGG</td><td>TGG</td><td>||||||||[||||||||||||]PAM</td><td>NA</td><td>Intergenic</td><td>NA</td><td>NA</td></tr>
</table>

| | | | | | | | | | | | | |
|---|---|---|---|---|---|---|---|---|---|---|---|---|
| T7049 | GGTGGAACTTGACGTACGTCAGG | | 1000 | | | | | | | | | |
| Oligo fwd | CACCGGTGGAACTTGACGTACGTC | | | | | | | | | | | |
| Oligo rev | AAACGACGTACGTCAAGTTCCACC | | | | | | | | | | | |
| Chromosome | start | | end | strand | MM | target_seq | PAM | alignment | distance | position | gene name | gene id |
| GU305914.1 | | 98023 | 98045 | + | 0 | GGTGGAACTTGACGTACGTC | AGG | ||||||||[||||||||||||]PAM | NA | Intergenic | NA | NA |
| T7050 | CCATCGCCGCCACCACGTCCAGG | | 1000 | | | | | | | | | |
| Oligo fwd | CACCCCATCGCCGCCACCACGTCC | | | | | | | | | | | |
| Oligo rev | AAACGGACGTGGTGGCGGCGATGG | | | | | | | | | | | |
| Chromosome | start | | end | strand | MM | target_seq | PAM | alignment | distance | position | gene name | gene id |
| GU305914.1 | | 98056 | 98078 | + | 0 | CCATCGCCGCCACCACGTCC | AGG | ||||||||[||||||||||||]PAM | NA | Intergenic | NA | NA |
| T7051 | GCCGCCACCACGTCCAGGCTCGG | | 1000 | | | | | | | | | |
| Oligo fwd | CACCGCCGCCACCACGTCCAGGCT | | | | | | | | | | | |
| Oligo rev | AAACAGCCTGGACGTGGTGGCGGC | | | | | | | | | | | |
| Chromosome | start | | end | strand | MM | target_seq | PAM | alignment | distance | position | gene name | gene id |
| GU305914.1 | | 98061 | 98083 | + | 0 | GCCGCCACCACGTCCAGGCT | CGG | ||||||||[||||||||||||]PAM | NA | Intergenic | NA | NA |
| T7052 | CAGGCTCGGCGCGTTGTTGATGG | | 1000 | | | | | | | | | |
| Oligo fwd | CACCCAGGCTCGGCGCGTTGTTGA | | | | | | | | | | | |
| Oligo rev | AAACTCAACAACGCGCCGAGCCTG | | | | | | | | | | | |
| Chromosome | start | | end | strand | MM | target_seq | PAM | alignment | distance | position | gene name | gene id |
| GU305914.1 | | 98075 | 98097 | + | 0 | CAGGCTCGGCGCGTTGTTGA | TGG | ||||||||[||||||||||||]PAM | NA | Intergenic | NA | NA |
| T7053 | CGAACGTGCGATCTTGTACCAGG | | 1000 | | | | | | | | | |
| Oligo fwd | CACCCGAACGTGCGATCTTGTACC | | | | | | | | | | | |
| Oligo rev | AAACGGTACAAGATCGCACGTTCG | | | | | | | | | | | |
| Chromosome | start | | end | strand | MM | target_seq | PAM | alignment | distance | position | gene name | gene id |
| GU305914.1 | | 98098 | 98120 | + | 0 | CGAACGTGCGATCTTGTACC | AGG | ||||||||[||||||||||||]PAM | NA | Intergenic | NA | NA |
| T7054 | ACCAGGACCCGCAGTTCCTCCGG | | 1000 | | | | | | | | | |
| Oligo fwd | CACCACCAGGACCCGCAGTTCCTC | | | | | | | | | | | |
| Oligo rev | AAACGAGGAACTGCGGGTCCTGGT | | | | | | | | | | | |
| Chromosome | start | | end | strand | MM | target_seq | PAM | alignment | distance | position | gene name | gene id |
| GU305914.1 | | 98115 | 98137 | + | 0 | ACCAGGACCCGCAGTTCCTC | CGG | ||||||||[||||||||||||]PAM | NA | Intergenic | NA | NA |
| GU305914.1 | | 77600 | 77622 | - | 4 | TCCAGCAGCCGCAGCTCCTC | GGG | -||||-|-[||||||-|||||]PAM | NA | Intergenic | NA | NA |
| T7055 | AGGACCCGCAGTTCCTCCGGCGG | | 1000 | | | | | | | | | |
| Oligo fwd | CACCAGGACCCGCAGTTCCTCCGG | | | | | | | | | | | |
| Oligo rev | AAACCCGGAGGAACTGCGGGTCCT | | | | | | | | | | | |
| Chromosome | start | | end | strand | MM | target_seq | PAM | alignment | distance | position | gene name | gene id |
| GU305914.1 | | 98118 | 98140 | + | 0 | AGGACCCGCAGTTCCTCCGG | CGG | ||||||||[||||||||||||]PAM | NA | Intergenic | NA | NA |
| T7056 | GCAGCGCGTCGAAGTGCTTGCGG | | 1000 | | | | | | | | | |
| Oligo fwd | CACCGCAGCGCGTCGAAGTGCTTG | | | | | | | | | | | |
| Oligo rev | AAACCAAGCACTTCGACGCGCTGC | | | | | | | | | | | |
| Chromosome | start | | end | strand | MM | target_seq | PAM | alignment | distance | position | gene name | gene id |
| GU305914.1 | | 98140 | 98162 | + | 0 | GCAGCGCGTCGAAGTGCTTG | CGG | ||||||||[||||||||||||]PAM | NA | Intergenic | NA | NA |
| T7057 | TAGTGTTGATCAGTTCGTCGTGG | | 1000 | | | | | | | | | |
| Oligo fwd | CACCTAGTGTTGATCAGTTCGTCG | | | | | | | | | | | |
| Oligo rev | AAACCGACGAACTGATCAACACTA | | | | | | | | | | | |
| Chromosome | start | | end | strand | MM | target_seq | PAM | alignment | distance | position | gene name | gene id |
| GU305914.1 | | 98179 | 98201 | + | 0 | TAGTGTTGATCAGTTCGTCG | TGG | ||||||||[||||||||||||]PAM | NA | Intergenic | NA | NA |
| T7058 | AGTGTTGATCAGTTCGTCGTGGG | | 1000 | | | | | | | | | |
| Oligo fwd | CACCAGTGTTGATCAGTTCGTCGT | | | | | | | | | | | |
| Oligo rev | AAACACGACGAACTGATCAACACT | | | | | | | | | | | |
| Chromosome | start | | end | strand | MM | target_seq | PAM | alignment | distance | position | gene name | gene id |
| GU305914.1 | | 98180 | 98202 | + | 0 | AGTGTTGATCAGTTCGTCGT | GGG | ||||||||[||||||||||||]PAM | NA | Intergenic | NA | NA |
| T7059 | TCGTCGTGGGTCACCACCGCCGG | | 1000 | | | | | | | | | |
| Oligo fwd | CACCTCGTCGTGGGTCACCACCGC | | | | | | | | | | | |
| Oligo rev | AAACGCGGTGGTGACCCACGACGA | | | | | | | | | | | |
| Chromosome | start | | end | strand | MM | target_seq | PAM | alignment | distance | position | gene name | gene id |
| GU305914.1 | | 98193 | 98215 | + | 0 | TCGTCGTGGGTCACCACCGC | CGG | ||||||||[||||||||||||]PAM | NA | Intergenic | NA | NA |
| T7060 | GGGTCACCACCGCCGGCGCGTGG | | 1000 | | | | | | | | | |
| Oligo fwd | CACCGGGTCACCACCGCCGGCGCG | | | | | | | | | | | |
| Oligo rev | AAACCGCGCCGGCGGTGGTGACCC | | | | | | | | | | | |
| Chromosome | start | | end | strand | MM | target_seq | PAM | alignment | distance | position | gene name | gene id |
| GU305914.1 | | 98200 | 98222 | + | 0 | GGGTCACCACCGCCGGCGCG | TGG | ||||||||[||||||||||||]PAM | NA | Intergenic | NA | NA |
| T7061 | TCACCACCGCCGGCGCGTGGCGG | | 1000 | | | | | | | | | |
| Oligo fwd | CACCTCACCACCGCCGGCGCGTGG | | | | | | | | | | | |
| Oligo rev | AAACCCACGCGCCGGCGGTGGTGA | | | | | | | | | | | |
| Chromosome | start | | end | strand | MM | target_seq | PAM | alignment | distance | position | gene name | gene id |
| GU305914.1 | | 98203 | 98225 | + | 0 | TCACCACCGCCGGCGCGTGG | CGG | ||||||||[||||||||||||]PAM | NA | Intergenic | NA | NA |
| T7062 | CGGCGCGTGGCGGTCGTTGTCGG | | 1000 | | | | | | | | | |
| Oligo fwd | CACCCGGCGCGTGGCGGTCGTTGT | | | | | | | | | | | |
| Oligo rev | AAACACAACGACCGCCACGCGCCG | | | | | | | | | | | |
| Chromosome | start | | end | strand | MM | target_seq | PAM | alignment | distance | position | gene name | gene id |
| GU305914.1 | | 98213 | 98235 | + | 0 | CGGCGCGTGGCGGTCGTTGT | CGG | ||||||||[||||||||||||]PAM | NA | Intergenic | NA | NA |
| T7063 | CCTGTTCTCTAGCAGCTTGATGG | | 1000 | | | | | | | | | |
| Oligo fwd | CACCCCTGTTCTCTAGCAGCTTGA | | | | | | | | | | | |
| Oligo rev | AAACTCAAGCTGCTAGAGAACAGG | | | | | | | | | | | |

<table>
<tr><td>Chromosome</td><td>start</td><td>end</td><td>strand</td><td>MM</td><td>target_seq</td><td>PAM</td><td>alignment</td><td>distance</td><td>position</td><td>gene name</td><td>gene id</td></tr>
<tr><td>GU305914.1</td><td>98240</td><td>98262</td><td>+</td><td>0</td><td>CCTGTTCTCTAGCAGCTTGA</td><td>TGG</td><td>||||||||[||||||||||||]PAM</td><td>NA</td><td>Intergenic</td><td>NA</td><td>NA</td></tr>
<tr><td>T7064</td><td>CTGTTCTCTAGCAGCTTGATGGG</td><td>1000</td><td></td><td></td><td></td><td></td><td></td><td></td><td></td><td></td><td></td></tr>
<tr><td>Oligo fwd</td><td>CACCCTGTTCTCTAGCAGCTTGAT</td><td></td><td></td><td></td><td></td><td></td><td></td><td></td><td></td><td></td><td></td></tr>
<tr><td>Oligo rev</td><td>AAACATCAAGCTGCTAGAGAACAG</td><td></td><td></td><td></td><td></td><td></td><td></td><td></td><td></td><td></td><td></td></tr>
<tr><td>Chromosome</td><td>start</td><td>end</td><td>strand</td><td>MM</td><td>target_seq</td><td>PAM</td><td>alignment</td><td>distance</td><td>position</td><td>gene name</td><td>gene id</td></tr>
<tr><td>GU305914.1</td><td>98241</td><td>98263</td><td>+</td><td>0</td><td>CTGTTCTCTAGCAGCTTGAT</td><td>GGG</td><td>||||||||[||||||||||||]PAM</td><td>NA</td><td>Intergenic</td><td>NA</td><td>NA</td></tr>
<tr><td>T7065</td><td>TGTTCTCTAGCAGCTTGATGGGG</td><td>1000</td><td></td><td></td><td></td><td></td><td></td><td></td><td></td><td></td><td></td></tr>
<tr><td>Oligo fwd</td><td>CACCTGTTCTCTAGCAGCTTGATG</td><td></td><td></td><td></td><td></td><td></td><td></td><td></td><td></td><td></td><td></td></tr>
<tr><td>Oligo rev</td><td>AAACCATCAAGCTGCTAGAGAACA</td><td></td><td></td><td></td><td></td><td></td><td></td><td></td><td></td><td></td><td></td></tr>
<tr><td>Chromosome</td><td>start</td><td>end</td><td>strand</td><td>MM</td><td>target_seq</td><td>PAM</td><td>alignment</td><td>distance</td><td>position</td><td>gene name</td><td>gene id</td></tr>
<tr><td>GU305914.1</td><td>98242</td><td>98264</td><td>+</td><td>0</td><td>TGTTCTCTAGCAGCTTGATG</td><td>GGG</td><td>||||||||[||||||||||||]PAM</td><td>NA</td><td>Intergenic</td><td>NA</td><td>NA</td></tr>
<tr><td>T7066</td><td>CTCTAGCAGCTTGATGGGGTCGG</td><td>1000</td><td></td><td></td><td></td><td></td><td></td><td></td><td></td><td></td><td></td></tr>
<tr><td>Oligo fwd</td><td>CACCCTCTAGCAGCTTGATGGGGT</td><td></td><td></td><td></td><td></td><td></td><td></td><td></td><td></td><td></td><td></td></tr>
<tr><td>Oligo rev</td><td>AAACACCCCATCAAGCTGCTAGAG</td><td></td><td></td><td></td><td></td><td></td><td></td><td></td><td></td><td></td><td></td></tr>
<tr><td>Chromosome</td><td>start</td><td>end</td><td>strand</td><td>MM</td><td>target_seq</td><td>PAM</td><td>alignment</td><td>distance</td><td>position</td><td>gene name</td><td>gene id</td></tr>
<tr><td>GU305914.1</td><td>98246</td><td>98268</td><td>+</td><td>0</td><td>CTCTAGCAGCTTGATGGGGT</td><td>CGG</td><td>||||||||[||||||||||||]PAM</td><td>NA</td><td>Intergenic</td><td>NA</td><td>NA</td></tr>
<tr><td>T7067</td><td>TCTAGCAGCTTGATGGGGTCGGG</td><td>1000</td><td></td><td></td><td></td><td></td><td></td><td></td><td></td><td></td><td></td></tr>
<tr><td>Oligo fwd</td><td>CACCTCTAGCAGCTTGATGGGGTC</td><td></td><td></td><td></td><td></td><td></td><td></td><td></td><td></td><td></td><td></td></tr>
<tr><td>Oligo rev</td><td>AAACGACCCCATCAAGCTGCTAGA</td><td></td><td></td><td></td><td></td><td></td><td></td><td></td><td></td><td></td><td></td></tr>
<tr><td>Chromosome</td><td>start</td><td>end</td><td>strand</td><td>MM</td><td>target_seq</td><td>PAM</td><td>alignment</td><td>distance</td><td>position</td><td>gene name</td><td>gene id</td></tr>
<tr><td>GU305914.1</td><td>98247</td><td>98269</td><td>+</td><td>0</td><td>TCTAGCAGCTTGATGGGGTC</td><td>GGG</td><td>||||||||[||||||||||||]PAM</td><td>NA</td><td>Intergenic</td><td>NA</td><td>NA</td></tr>
<tr><td>GU305914.1</td><td>170082</td><td>170104</td><td>+</td><td>4</td><td>TCTGGGAGCTTGACGGGCTC</td><td>TGG</td><td>|||-|-||[|||||-|||-||]PAM</td><td>NA</td><td>Intergenic</td><td>NA</td><td>NA</td></tr>
<tr><td>T7068</td><td>GCAGCTTGATGGGGTCGGGCAGG</td><td>1000</td><td></td><td></td><td></td><td></td><td></td><td></td><td></td><td></td><td></td></tr>
<tr><td>Oligo fwd</td><td>CACCGCAGCTTGATGGGGTCGGGC</td><td></td><td></td><td></td><td></td><td></td><td></td><td></td><td></td><td></td><td></td></tr>
<tr><td>Oligo rev</td><td>AAACGCCCGACCCCATCAAGCTGC</td><td></td><td></td><td></td><td></td><td></td><td></td><td></td><td></td><td></td><td></td></tr>
<tr><td>Chromosome</td><td>start</td><td>end</td><td>strand</td><td>MM</td><td>target_seq</td><td>PAM</td><td>alignment</td><td>distance</td><td>position</td><td>gene name</td><td>gene id</td></tr>
<tr><td>GU305914.1</td><td>98251</td><td>98273</td><td>+</td><td>0</td><td>GCAGCTTGATGGGGTCGGGC</td><td>AGG</td><td>||||||||[||||||||||||]PAM</td><td>NA</td><td>Intergenic</td><td>NA</td><td>NA</td></tr>
<tr><td>T7069</td><td>GATGGGGTCGGGCAGGTCCATGG</td><td>1000</td><td></td><td></td><td></td><td></td><td></td><td></td><td></td><td></td><td></td></tr>
<tr><td>Oligo fwd</td><td>CACCGATGGGGTCGGGCAGGTCCA</td><td></td><td></td><td></td><td></td><td></td><td></td><td></td><td></td><td></td><td></td></tr>
<tr><td>Oligo rev</td><td>AAACTGGACCTGCCCGACCCCATC</td><td></td><td></td><td></td><td></td><td></td><td></td><td></td><td></td><td></td><td></td></tr>
<tr><td>Chromosome</td><td>start</td><td>end</td><td>strand</td><td>MM</td><td>target_seq</td><td>PAM</td><td>alignment</td><td>distance</td><td>position</td><td>gene name</td><td>gene id</td></tr>
<tr><td>GU305914.1</td><td>98258</td><td>98280</td><td>+</td><td>0</td><td>GATGGGGTCGGGCAGGTCCA</td><td>TGG</td><td>||||||||[||||||||||||]PAM</td><td>NA</td><td>Intergenic</td><td>NA</td><td>NA</td></tr>
<tr><td>T7070</td><td>ATGGGGTCGGGCAGGTCCATGGG</td><td>1000</td><td></td><td></td><td></td><td></td><td></td><td></td><td></td><td></td><td></td></tr>
<tr><td>Oligo fwd</td><td>CACCATGGGGTCGGGCAGGTCCAT</td><td></td><td></td><td></td><td></td><td></td><td></td><td></td><td></td><td></td><td></td></tr>
<tr><td>Oligo rev</td><td>AAACATGGACCTGCCCGACCCCAT</td><td></td><td></td><td></td><td></td><td></td><td></td><td></td><td></td><td></td><td></td></tr>
<tr><td>Chromosome</td><td>start</td><td>end</td><td>strand</td><td>MM</td><td>target_seq</td><td>PAM</td><td>alignment</td><td>distance</td><td>position</td><td>gene name</td><td>gene id</td></tr>
<tr><td>GU305914.1</td><td>98259</td><td>98281</td><td>+</td><td>0</td><td>ATGGGGTCGGGCAGGTCCAT</td><td>GGG</td><td>||||||||[||||||||||||]PAM</td><td>NA</td><td>Intergenic</td><td>NA</td><td>NA</td></tr>
<tr><td>T7071</td><td>TGGGGTCGGGCAGGTCCATGGGG</td><td>1000</td><td></td><td></td><td></td><td></td><td></td><td></td><td></td><td></td><td></td></tr>
<tr><td>Oligo fwd</td><td>CACCTGGGGTCGGGCAGGTCCATG</td><td></td><td></td><td></td><td></td><td></td><td></td><td></td><td></td><td></td><td></td></tr>
<tr><td>Oligo rev</td><td>AAACCATGGACCTGCCCGACCCCA</td><td></td><td></td><td></td><td></td><td></td><td></td><td></td><td></td><td></td><td></td></tr>
<tr><td>Chromosome</td><td>start</td><td>end</td><td>strand</td><td>MM</td><td>target_seq</td><td>PAM</td><td>alignment</td><td>distance</td><td>position</td><td>gene name</td><td>gene id</td></tr>
<tr><td>GU305914.1</td><td>98260</td><td>98282</td><td>+</td><td>0</td><td>TGGGGTCGGGCAGGTCCATG</td><td>GGG</td><td>||||||||[||||||||||||]PAM</td><td>NA</td><td>Intergenic</td><td>NA</td><td>NA</td></tr>
<tr><td>T7072</td><td>GGTCCATGGGGAACATGCACAGG</td><td>1000</td><td></td><td></td><td></td><td></td><td></td><td></td><td></td><td></td><td></td></tr>
<tr><td>Oligo fwd</td><td>CACCGGTCCATGGGGAACATGCAC</td><td></td><td></td><td></td><td></td><td></td><td></td><td></td><td></td><td></td><td></td></tr>
<tr><td>Oligo rev</td><td>AAACGTGCATGTTCCCCATGGACC</td><td></td><td></td><td></td><td></td><td></td><td></td><td></td><td></td><td></td><td></td></tr>
<tr><td>Chromosome</td><td>start</td><td>end</td><td>strand</td><td>MM</td><td>target_seq</td><td>PAM</td><td>alignment</td><td>distance</td><td>position</td><td>gene name</td><td>gene id</td></tr>
<tr><td>GU305914.1</td><td>98272</td><td>98294</td><td>+</td><td>0</td><td>GGTCCATGGGGAACATGCAC</td><td>AGG</td><td>||||||||[||||||||||||]PAM</td><td>NA</td><td>Intergenic</td><td>NA</td><td>NA</td></tr>
<tr><td>T7073</td><td>GGGGAACATGCACAGGCCCAAGG</td><td>1000</td><td></td><td></td><td></td><td></td><td></td><td></td><td></td><td></td><td></td></tr>
<tr><td>Oligo fwd</td><td>CACCGGGGAACATGCACAGGCCCA</td><td></td><td></td><td></td><td></td><td></td><td></td><td></td><td></td><td></td><td></td></tr>
<tr><td>Oligo rev</td><td>AAACTGGGCCTGTGCATGTTCCCC</td><td></td><td></td><td></td><td></td><td></td><td></td><td></td><td></td><td></td><td></td></tr>
<tr><td>Chromosome</td><td>start</td><td>end</td><td>strand</td><td>MM</td><td>target_seq</td><td>PAM</td><td>alignment</td><td>distance</td><td>position</td><td>gene name</td><td>gene id</td></tr>
<tr><td>GU305914.1</td><td>98279</td><td>98301</td><td>+</td><td>0</td><td>GGGGAACATGCACAGGCCCA</td><td>AGG</td><td>||||||||[||||||||||||]PAM</td><td>NA</td><td>Intergenic</td><td>NA</td><td>NA</td></tr>
<tr><td>T7074</td><td>AGGAACCCGTCGTGTAGTCGCGG</td><td>1000</td><td></td><td></td><td></td><td></td><td></td><td></td><td></td><td></td><td></td></tr>
<tr><td>Oligo fwd</td><td>CACCAGGAACCCGTCGTGTAGTCG</td><td></td><td></td><td></td><td></td><td></td><td></td><td></td><td></td><td></td><td></td></tr>
<tr><td>Oligo rev</td><td>AAACCGACTACACGACGGGTTCCT</td><td></td><td></td><td></td><td></td><td></td><td></td><td></td><td></td><td></td><td></td></tr>
<tr><td>Chromosome</td><td>start</td><td>end</td><td>strand</td><td>MM</td><td>target_seq</td><td>PAM</td><td>alignment</td><td>distance</td><td>position</td><td>gene name</td><td>gene id</td></tr>
<tr><td>GU305914.1</td><td>98299</td><td>98321</td><td>+</td><td>0</td><td>AGGAACCCGTCGTGTAGTCG</td><td>CGG</td><td>||||||||[||||||||||||]PAM</td><td>NA</td><td>Intergenic</td><td>NA</td><td>NA</td></tr>
<tr><td>T7075</td><td>GTCGTGTAGTCGCGGATGTCCGG</td><td>1000</td><td></td><td></td><td></td><td></td><td></td><td></td><td></td><td></td><td></td></tr>
<tr><td>Oligo fwd</td><td>CACCGTCGTGTAGTCGCGGATGTC</td><td></td><td></td><td></td><td></td><td></td><td></td><td></td><td></td><td></td><td></td></tr>
<tr><td>Oligo rev</td><td>AAACGACATCCGCGACTACACGAC</td><td></td><td></td><td></td><td></td><td></td><td></td><td></td><td></td><td></td><td></td></tr>
<tr><td>Chromosome</td><td>start</td><td>end</td><td>strand</td><td>MM</td><td>target_seq</td><td>PAM</td><td>alignment</td><td>distance</td><td>position</td><td>gene name</td><td>gene id</td></tr>
<tr><td>GU305914.1</td><td>98307</td><td>98329</td><td>+</td><td>0</td><td>GTCGTGTAGTCGCGGATGTC</td><td>CGG</td><td>||||||||[||||||||||||]PAM</td><td>NA</td><td>Intergenic</td><td>NA</td><td>NA</td></tr>
<tr><td>T7076</td><td>GTAGTCGCGGATGTCCGGAGAGG</td><td>1000</td><td></td><td></td><td></td><td></td><td></td><td></td><td></td><td></td><td></td></tr>
<tr><td>Oligo fwd</td><td>CACCGTAGTCGCGGATGTCCGGAG</td><td></td><td></td><td></td><td></td><td></td><td></td><td></td><td></td><td></td><td></td></tr>
<tr><td>Oligo rev</td><td>AAACCTCCGGACATCCGCGACTAC</td><td></td><td></td><td></td><td></td><td></td><td></td><td></td><td></td><td></td><td></td></tr>
<tr><td>Chromosome</td><td>start</td><td>end</td><td>strand</td><td>MM</td><td>target_seq</td><td>PAM</td><td>alignment</td><td>distance</td><td>position</td><td>gene name</td><td>gene id</td></tr>
<tr><td>GU305914.1</td><td>98312</td><td>98334</td><td>+</td><td>0</td><td>GTAGTCGCGGATGTCCGGAG</td><td>AGG</td><td>||||||||[||||||||||||]PAM</td><td>NA</td><td>Intergenic</td><td>NA</td><td>NA</td></tr>
<tr><td>T7077</td><td>CGCGGATGTCCGGAGAGGTCTGG</td><td>1000</td><td></td><td></td><td></td><td></td><td></td><td></td><td></td><td></td><td></td></tr>
<tr><td>Oligo fwd</td><td>CACCCGCGGATGTCCGGAGAGGTC</td><td></td><td></td><td></td><td></td><td></td><td></td><td></td><td></td><td></td><td></td></tr>
<tr><td>Oligo rev</td><td>AAACGACCTCTCCGGACATCCGCG</td><td></td><td></td><td></td><td></td><td></td><td></td><td></td><td></td><td></td><td></td></tr>
<tr><td>Chromosome</td><td>start</td><td>end</td><td>strand</td><td>MM</td><td>target_seq</td><td>PAM</td><td>alignment</td><td>distance</td><td>position</td><td>gene name</td><td>gene id</td></tr>
<tr><td>GU305914.1</td><td>98317</td><td>98339</td><td>+</td><td>0</td><td>CGCGGATGTCCGGAGAGGTC</td><td>TGG</td><td>||||||||[||||||||||||]PAM</td><td>NA</td><td>Intergenic</td><td>NA</td><td>NA</td></tr>
<tr><td>T7078</td><td>GGATGTCCGGAGAGGTCTGGCGG</td><td>1000</td><td></td><td></td><td></td><td></td><td></td><td></td><td></td><td></td><td></td></tr>
</table>

| | | | | | | | | | | | |
|---|---|---|---|---|---|---|---|---|---|---|---|
| Oligo fwd | CACCGGATGTCCGGAGAGGTCTGG | | | | | | | | | | |
| Oligo rev | AAACCCAGACCTCTCCGGACATCC | | | | | | | | | | |
| Chromosome | start | end | strand | MM | target_seq | PAM | alignment | distance | position | gene name | gene id |
| GU305914.1 | 98320 | 98342 | + | 0 | GGATGTCCGGAGAGGTCTGG | CGG | ||||||||[||||||||||||]PAM | NA | Intergenic | NA | NA |
| T7079 | CGCCGCCGCCACCGCCGCCTAGG | 1000 | | | | | | | | | |
| Oligo fwd | CACCCGCCGCCGCCACCGCCGCCT | | | | | | | | | | |
| Oligo rev | AAACAGGCGGCGGTGGCGGCGGCG | | | | | | | | | | |
| Chromosome | start | end | strand | MM | target_seq | PAM | alignment | distance | position | gene name | gene id |
| GU305914.1 | 98350 | 98372 | + | 0 | CGCCGCCGCCACCGCCGCCT | AGG | ||||||||[||||||||||||]PAM | NA | Intergenic | NA | NA |
| GU305914.1 | 161752 | 161774 | + | 4 | CACCTCCGCCGCCGCCTCCT | CGG | |-||-|||[||-|||||-|||]PAM | NA | Intergenic | NA | NA |
| GU305914.1 | 35655 | 35677 | - | 3 | CCCCGCCGCCGCCGCCGCCG | TGG | |-||||||[||-||||||||-]PAM | NA | Intergenic | NA | NA |
| GU305914.1 | 74708 | 74730 | + | 4 | AACCGCCGCCACCGCCACCC | GGG | --||||||[||||||||-||-]PAM | NA | Intergenic | NA | NA |
| T7080 | ACCGCCGCCTAGGCTCAGCGTGG | 1000 | | | | | | | | | |
| Oligo fwd | CACCACCGCCGCCTAGGCTCAGCG | | | | | | | | | | |
| Oligo rev | AAACCGCTGAGCCTAGGCGGCGGT | | | | | | | | | | |
| Chromosome | start | end | strand | MM | target_seq | PAM | alignment | distance | position | gene name | gene id |
| GU305914.1 | 98360 | 98382 | + | 0 | ACCGCCGCCTAGGCTCAGCG | TGG | ||||||||[||||||||||||]PAM | NA | Intergenic | NA | NA |
| T7081 | GGTATTGTTGTTACTGTTGTTGG | 1000 | | | | | | | | | |
| Oligo fwd | CACCGGTATTGTTGTTACTGTTGT | | | | | | | | | | |
| Oligo rev | AAACACAACAGTAACAACAATACC | | | | | | | | | | |
| Chromosome | start | end | strand | MM | target_seq | PAM | alignment | distance | position | gene name | gene id |
| GU305914.1 | 98381 | 98403 | + | 0 | GGTATTGTTGTTACTGTTGT | TGG | ||||||||[||||||||||||]PAM | NA | Intergenic | NA | NA |
| T7082 | ATTGTTGTTACTGTTGTTGGAGG | 1000 | | | | | | | | | |
| Oligo fwd | CACCATTGTTGTTACTGTTGTTGG | | | | | | | | | | |
| Oligo rev | AAACCCAACAACAGTAACAACAAT | | | | | | | | | | |
| Chromosome | start | end | strand | MM | target_seq | PAM | alignment | distance | position | gene name | gene id |
| GU305914.1 | 98384 | 98406 | + | 0 | ATTGTTGTTACTGTTGTTGG | AGG | ||||||||[||||||||||||]PAM | NA | Intergenic | NA | NA |
| T7083 | GTTGTTGGAGGACACGAGCGAGG | 1000 | | | | | | | | | |
| Oligo fwd | CACCGTTGTTGGAGGACACGAGCG | | | | | | | | | | |
| Oligo rev | AAACCGCTCGTGTCCTCCAACAAC | | | | | | | | | | |
| Chromosome | start | end | strand | MM | target_seq | PAM | alignment | distance | position | gene name | gene id |
| GU305914.1 | 98396 | 98418 | + | 0 | GTTGTTGGAGGACACGAGCG | AGG | ||||||||[||||||||||||]PAM | NA | Intergenic | NA | NA |
| T7084 | AGGACACGAGCGAGGACTCGAGG | 1000 | | | | | | | | | |
| Oligo fwd | CACCAGGACACGAGCGAGGACTCG | | | | | | | | | | |
| Oligo rev | AAACCGAGTCCTCGCTCGTGTCCT | | | | | | | | | | |
| Chromosome | start | end | strand | MM | target_seq | PAM | alignment | distance | position | gene name | gene id |
| GU305914.1 | 98404 | 98426 | + | 0 | AGGACACGAGCGAGGACTCG | AGG | ||||||||[||||||||||||]PAM | NA | Intergenic | NA | NA |
| T7085 | GAGCGAGGACTCGAGGATCGAGG | 1000 | | | | | | | | | |
| Oligo fwd | CACCGAGCGAGGACTCGAGGATCG | | | | | | | | | | |
| Oligo rev | AAACCGATCCTCGAGTCCTCGCTC | | | | | | | | | | |
| Chromosome | start | end | strand | MM | target_seq | PAM | alignment | distance | position | gene name | gene id |
| GU305914.1 | 98411 | 98433 | + | 0 | GAGCGAGGACTCGAGGATCG | AGG | ||||||||[||||||||||||]PAM | NA | Intergenic | NA | NA |
| T7086 | AGCGAGGACTCGAGGATCGAGGG | 1000 | | | | | | | | | |
| Oligo fwd | CACCAGCGAGGACTCGAGGATCGA | | | | | | | | | | |
| Oligo rev | AAACTCGATCCTCGAGTCCTCGCT | | | | | | | | | | |
| Chromosome | start | end | strand | MM | target_seq | PAM | alignment | distance | position | gene name | gene id |
| GU305914.1 | 98412 | 98434 | + | 0 | AGCGAGGACTCGAGGATCGA | GGG | ||||||||[||||||||||||]PAM | NA | Intergenic | NA | NA |
| T7087 | AGGATCGAGGGTGCCGACGACGG | 1000 | | | | | | | | | |
| Oligo fwd | CACCAGGATCGAGGGTGCCGACGA | | | | | | | | | | |
| Oligo rev | AAACTCGTCGGCACCCTCGATCCT | | | | | | | | | | |
| Chromosome | start | end | strand | MM | target_seq | PAM | alignment | distance | position | gene name | gene id |
| GU305914.1 | 98424 | 98446 | + | 0 | AGGATCGAGGGTGCCGACGA | CGG | ||||||||[||||||||||||]PAM | NA | Intergenic | NA | NA |
| T7088 | CGAGGGTGCCGACGACGGCTCGG | 1000 | | | | | | | | | |
| Oligo fwd | CACCCGAGGGTGCCGACGACGGCT | | | | | | | | | | |
| Oligo rev | AAACAGCCGTCGTCGGCACCCTCG | | | | | | | | | | |
| Chromosome | start | end | strand | MM | target_seq | PAM | alignment | distance | position | gene name | gene id |
| GU305914.1 | 98429 | 98451 | + | 0 | CGAGGGTGCCGACGACGGCT | CGG | ||||||||[||||||||||||]PAM | NA | Intergenic | NA | NA |
| T7089 | GAGGGTGCCGACGACGGCTCGGG | 1000 | | | | | | | | | |
| Oligo fwd | CACCGAGGGTGCCGACGACGGCTC | | | | | | | | | | |
| Oligo rev | AAACGAGCCGTCGTCGGCACCCTC | | | | | | | | | | |
| Chromosome | start | end | strand | MM | target_seq | PAM | alignment | distance | position | gene name | gene id |
| GU305914.1 | 98430 | 98452 | + | 0 | GAGGGTGCCGACGACGGCTC | GGG | ||||||||[||||||||||||]PAM | NA | Intergenic | NA | NA |
| GU305914.1 | 151161 | 151183 | + | 4 | AAGGGGTCCGACGGCGGCTC | CGG | -||||--|[|||||-|||||||]PAM | NA | Intergenic | NA | NA |
| T7090 | GTGCCGACGACGGCTCGGGACGG | 1000 | | | | | | | | | |
| Oligo fwd | CACCGTGCCGACGACGGCTCGGGA | | | | | | | | | | |
| Oligo rev | AAACTCCCGAGCCGTCGTCGGCAC | | | | | | | | | | |
| Chromosome | start | end | strand | MM | target_seq | PAM | alignment | distance | position | gene name | gene id |
| GU305914.1 | 98434 | 98456 | + | 0 | GTGCCGACGACGGCTCGGGA | CGG | ||||||||[||||||||||||]PAM | NA | Intergenic | NA | NA |
| T7091 | CCGACGACGGCTCGGGACGGCGG | 1000 | | | | | | | | | |
| Oligo fwd | CACCCCGACGACGGCTCGGGACGG | | | | | | | | | | |
| Oligo rev | AAACCCGTCCCGAGCCGTCGTCGG | | | | | | | | | | |
| Chromosome | start | end | strand | MM | target_seq | PAM | alignment | distance | position | gene name | gene id |
| GU305914.1 | 98437 | 98459 | + | 0 | CCGACGACGGCTCGGGACGG | CGG | ||||||||[||||||||||||]PAM | NA | Intergenic | NA | NA |
| T7092 | ACGGCTCGGGACGGCGGCCCTGG | 1000 | | | | | | | | | |
| Oligo fwd | CACCACGGCTCGGGACGGCGGCCC | | | | | | | | | | |

| | | | | | | | | | | | | |
|---|---|---|---|---|---|---|---|---|---|---|---|---|
| Oligo rev | AAACGGGCCGCCGTCCCGAGCCGT | | | | | | | | | | | |
| Chromosome | start | | end | strand | MM | target_seq | PAM | alignment | distance | position | gene name | gene id |
| GU305914.1 | | 98443 | 98465 | + | 0 | ACGGCTCGGGACGGCGGCCC | TGG | \|\|\|\|\|\|\|\|[\|\|\|\|\|\|\|\|\|\|\|\|]PAM | NA | Intergenic | NA | NA |
| T7093 | CGGCTCGGGACGGCGGCCCTGGG | | 1000 | | | | | | | | | |
| Oligo fwd | CACCCGGCTCGGGACGGCGGCCCT | | | | | | | | | | | |
| Oligo rev | AAACAGGGCCGCCGTCCCGAGCCG | | | | | | | | | | | |
| Chromosome | start | | end | strand | MM | target_seq | PAM | alignment | distance | position | gene name | gene id |
| GU305914.1 | | 98444 | 98466 | + | 0 | CGGCTCGGGACGGCGGCCCT | GGG | \|\|\|\|\|\|\|\|[\|\|\|\|\|\|\|\|\|\|\|\|]PAM | NA | Intergenic | NA | NA |
| T7094 | GGCCCTGGGCGCGCGCCTCGCGG | | 1000 | | | | | | | | | |
| Oligo fwd | CACCGGCCCTGGGCGCGCGCCTCG | | | | | | | | | | | |
| Oligo rev | AAACCGAGGCGCGCGCCCAGGGCC | | | | | | | | | | | |
| Chromosome | start | | end | strand | MM | target_seq | PAM | alignment | distance | position | gene name | gene id |
| GU305914.1 | | 98458 | 98480 | + | 0 | GGCCCTGGGCGCGCGCCTCG | CGG | \|\|\|\|\|\|\|\|[\|\|\|\|\|\|\|\|\|\|\|\|]PAM | NA | Intergenic | NA | NA |
| T7095 | GGGCGCGCGCCTCGCGGATCCGG | | 1000 | | | | | | | | | |
| Oligo fwd | CACCGGGCGCGCGCCTCGCGGATC | | | | | | | | | | | |
| Oligo rev | AAACGATCCGCGAGGCGCGCGCCC | | | | | | | | | | | |
| Chromosome | start | | end | strand | MM | target_seq | PAM | alignment | distance | position | gene name | gene id |
| GU305914.1 | | 98464 | 98486 | + | 0 | GGGCGCGCGCCTCGCGGATC | CGG | \|\|\|\|\|\|\|\|[\|\|\|\|\|\|\|\|\|\|\|\|]PAM | NA | Intergenic | NA | NA |
| T7096 | CGCCTCGCGGATCCGGATCATGG | | 1000 | | | | | | | | | |
| Oligo fwd | CACCCGCCTCGCGGATCCGGATCA | | | | | | | | | | | |
| Oligo rev | AAACTGATCCGGATCCGCGAGGCG | | | | | | | | | | | |
| Chromosome | start | | end | strand | MM | target_seq | PAM | alignment | distance | position | gene name | gene id |
| GU305914.1 | | 98471 | 98493 | + | 0 | CGCCTCGCGGATCCGGATCA | TGG | \|\|\|\|\|\|\|\|[\|\|\|\|\|\|\|\|\|\|\|\|]PAM | NA | Intergenic | NA | NA |
| T7097 | TCATGGCTTTCAAGCATTCCAGG | | 1000 | | | | | | | | | |
| Oligo fwd | CACCTCATGGCTTTCAAGCATTCC | | | | | | | | | | | |
| Oligo rev | AAACGGAATGCTTGAAAGCCATGA | | | | | | | | | | | |
| Chromosome | start | | end | strand | MM | target_seq | PAM | alignment | distance | position | gene name | gene id |
| GU305914.1 | | 98488 | 98510 | + | 0 | TCATGGCTTTCAAGCATTCC | AGG | \|\|\|\|\|\|\|\|[\|\|\|\|\|\|\|\|\|\|\|\|]PAM | NA | Intergenic | NA | NA |
| T7098 | TTTCAAGCATTCCAGGTCGTCGG | | 1000 | | | | | | | | | |
| Oligo fwd | CACCTTTCAAGCATTCCAGGTCGT | | | | | | | | | | | |
| Oligo rev | AAACACGACCTGGAATGCTTGAAA | | | | | | | | | | | |
| Chromosome | start | | end | strand | MM | target_seq | PAM | alignment | distance | position | gene name | gene id |
| GU305914.1 | | 98495 | 98517 | + | 0 | TTTCAAGCATTCCAGGTCGT | CGG | \|\|\|\|\|\|\|\|[\|\|\|\|\|\|\|\|\|\|\|\|]PAM | NA | Intergenic | NA | NA |
| T7099 | TTCAAGCATTCCAGGTCGTCGGG | | 1000 | | | | | | | | | |
| Oligo fwd | CACCTTCAAGCATTCCAGGTCGTC | | | | | | | | | | | |
| Oligo rev | AAACGACGACCTGGAATGCTTGAA | | | | | | | | | | | |
| Chromosome | start | | end | strand | MM | target_seq | PAM | alignment | distance | position | gene name | gene id |
| GU305914.1 | | 98496 | 98518 | + | 0 | TTCAAGCATTCCAGGTCGTC | GGG | \|\|\|\|\|\|\|\|[\|\|\|\|\|\|\|\|\|\|\|\|]PAM | NA | Intergenic | NA | NA |
| T7100 | GTTTTTTATTCAAGTGAGTGAGG | | 1000 | | | | | | | | | |
| Oligo fwd | CACCGTTTTTTATTCAAGTGAGTG | | | | | | | | | | | |
| Oligo rev | AAACCACTCACTTGAATAAAAAAC | | | | | | | | | | | |
| Chromosome | start | | end | strand | MM | target_seq | PAM | alignment | distance | position | gene name | gene id |
| GU305914.1 | | 98533 | 98555 | + | 0 | GTTTTTTATTCAAGTGAGTG | AGG | \|\|\|\|\|\|\|\|[\|\|\|\|\|\|\|\|\|\|\|\|]PAM | NA | Intergenic | NA | NA |
| T7101 | TTTTTTATTCAAGTGAGTGAGGG | | 1000 | | | | | | | | | |
| Oligo fwd | CACCTTTTTTATTCAAGTGAGTGA | | | | | | | | | | | |
| Oligo rev | AAACTCACTCACTTGAATAAAAAA | | | | | | | | | | | |
| Chromosome | start | | end | strand | MM | target_seq | PAM | alignment | distance | position | gene name | gene id |
| GU305914.1 | | 98534 | 98556 | + | 0 | TTTTTTATTCAAGTGAGTGA | GGG | \|\|\|\|\|\|\|\|[\|\|\|\|\|\|\|\|\|\|\|\|]PAM | NA | Intergenic | NA | NA |
| T7102 | CCGCGATGTTTTTGTCATACTGG | | 1000 | | | | | | | | | |
| Oligo fwd | CACCCCGCGATGTTTTTGTCATAC | | | | | | | | | | | |
| Oligo rev | AAACGTATGACAAAAACATCGCGG | | | | | | | | | | | |
| Chromosome | start | | end | strand | MM | target_seq | PAM | alignment | distance | position | gene name | gene id |
| GU305914.1 | | 98557 | 98579 | + | 0 | CCGCGATGTTTTTGTCATAC | TGG | \|\|\|\|\|\|\|\|[\|\|\|\|\|\|\|\|\|\|\|\|]PAM | NA | Intergenic | NA | NA |
| T7103 | CGCGATGTTTTTGTCATACTGGG | | 1000 | | | | | | | | | |
| Oligo fwd | CACCCGCGATGTTTTTGTCATACT | | | | | | | | | | | |
| Oligo rev | AAACAGTATGACAAAAACATCGCG | | | | | | | | | | | |
| Chromosome | start | | end | strand | MM | target_seq | PAM | alignment | distance | position | gene name | gene id |
| GU305914.1 | | 98558 | 98580 | + | 0 | CGCGATGTTTTTGTCATACT | GGG | \|\|\|\|\|\|\|\|[\|\|\|\|\|\|\|\|\|\|\|\|]PAM | NA | Intergenic | NA | NA |
| T7104 | GCGATGTTTTTGTCATACTGGGG | | 1000 | | | | | | | | | |
| Oligo fwd | CACCGCGATGTTTTTGTCATACTG | | | | | | | | | | | |
| Oligo rev | AAACCAGTATGACAAAAACATCGC | | | | | | | | | | | |
| Chromosome | start | | end | strand | MM | target_seq | PAM | alignment | distance | position | gene name | gene id |
| GU305914.1 | | 98559 | 98581 | + | 0 | GCGATGTTTTTGTCATACTG | GGG | \|\|\|\|\|\|\|\|[\|\|\|\|\|\|\|\|\|\|\|\|]PAM | NA | Intergenic | NA | NA |
| T7105 | ATGTTTTTGTCATACTGGGGTGG | | 1000 | | | | | | | | | |
| Oligo fwd | CACCATGTTTTTGTCATACTGGGG | | | | | | | | | | | |
| Oligo rev | AAACCCCCAGTATGACAAAAACAT | | | | | | | | | | | |
| Chromosome | start | | end | strand | MM | target_seq | PAM | alignment | distance | position | gene name | gene id |
| GU305914.1 | | 98562 | 98584 | + | 0 | ATGTTTTTGTCATACTGGGG | TGG | \|\|\|\|\|\|\|\|[\|\|\|\|\|\|\|\|\|\|\|\|]PAM | NA | Intergenic | NA | NA |
| T7106 | TTTTTGTCATACTGGGGTGGTGG | | 1000 | | | | | | | | | |
| Oligo fwd | CACCTTTTTGTCATACTGGGGTGG | | | | | | | | | | | |
| Oligo rev | AAACCCACCCCAGTATGACAAAAA | | | | | | | | | | | |
| Chromosome | start | | end | strand | MM | target_seq | PAM | alignment | distance | position | gene name | gene id |
| GU305914.1 | | 98565 | 98587 | + | 0 | TTTTTGTCATACTGGGGTGG | TGG | \|\|\|\|\|\|\|\|[\|\|\|\|\|\|\|\|\|\|\|\|]PAM | NA | Intergenic | NA | NA |
| T7107 | GTCATACTGGGGTGGTGGCGAGG | | 1000 | | | | | | | | | |

Oligo fwd CACCGTCATACTGGGGTGGTGGCG
Oligo rev AAACCGCCACCACCCCAGTATGAC
Chromosome start end strand MM target_seq PAM alignment distance position gene name gene id
GU305914.1 98570 98592 + 0 GTCATACTGGGGTGGTGGCG AGG ||||||||[||||||||||||]PAM NA Intergenic NA NA

T7108 ATACTGGGGTGGTGGCGAGGAGG 1000
Oligo fwd CACCATACTGGGGTGGTGGCGAGG
Oligo rev AAACCCTCGCCACCACCCCAGTAT
Chromosome start end strand MM target_seq PAM alignment distance position gene name gene id
GU305914.1 98573 98595 + 0 ATACTGGGGTGGTGGCGAGG AGG ||||||||[||||||||||||]PAM NA Intergenic NA NA

T7109 TACTGGGGTGGTGGCGAGGAGGG 1000
Oligo fwd CACCTACTGGGGTGGTGGCGAGGA
Oligo rev AAACTCCTCGCCACCACCCCAGTA
Chromosome start end strand MM target_seq PAM alignment distance position gene name gene id
GU305914.1 98574 98596 + 0 TACTGGGGTGGTGGCGAGGA GGG ||||||||[||||||||||||]PAM NA Intergenic NA NA

T7110 GGCGAGGAGGGCGATGAGAGTGG 1000
Oligo fwd CACCGGCGAGGAGGGCGATGAGAG
Oligo rev AAACCTCTCATCGCCCTCCTCGCC
Chromosome start end strand MM target_seq PAM alignment distance position gene name gene id
GU305914.1 98586 98608 + 0 GGCGAGGAGGGCGATGAGAG TGG ||||||||[||||||||||||]PAM NA Intergenic NA NA
GU305914.1 130527 130549 + 3 CGCGAGGAGGACGACGAGAG GGG -|||||||[||-|||-|||||]PAM NA Intergenic NA NA
GU305914.1 86027 86049 - 3 GTCGAGGACGGCGATGAGAT CGG |-||||||[-||||||||||-]PAM NA Intergenic NA NA

T7111 GAGGAGGGCGATGAGAGTGGCGG 1000
Oligo fwd CACCGAGGAGGGCGATGAGAGTGG
Oligo rev AAACCCACTCTCATCGCCCTCCTC
Chromosome start end strand MM target_seq PAM alignment distance position gene name gene id
GU305914.1 98589 98611 + 0 GAGGAGGGCGATGAGAGTGG CGG ||||||||[||||||||||||]PAM NA Intergenic NA NA

T7112 GAGGGCGATGAGAGTGGCGGCGG 1000
Oligo fwd CACCGAGGGCGATGAGAGTGGCGG
Oligo rev AAACCCGCCACTCTCATCGCCCTC
Chromosome start end strand MM target_seq PAM alignment distance position gene name gene id
GU305914.1 98592 98614 + 0 GAGGGCGATGAGAGTGGCGG CGG ||||||||[||||||||||||]PAM NA Intergenic NA NA

T7113 ATGAGCATTTCGTTTCCAGTTGG 1000
Oligo fwd CACCATGAGCATTTCGTTTCCAGT
Oligo rev AAACACTGGAAACGAAATGCTCAT
Chromosome start end strand MM target_seq PAM alignment distance position gene name gene id
GU305914.1 98621 98643 + 0 ATGAGCATTTCGTTTCCAGT TGG ||||||||[||||||||||||]PAM NA Intergenic NA NA

T7114 TGAGCATTTCGTTTCCAGTTGGG 1000
Oligo fwd CACCTGAGCATTTCGTTTCCAGTT
Oligo rev AAACAACTGGAAACGAAATGCTCA
Chromosome start end strand MM target_seq PAM alignment distance position gene name gene id
GU305914.1 98622 98644 + 0 TGAGCATTTCGTTTCCAGTT GGG ||||||||[||||||||||||]PAM NA Intergenic NA NA

T7115 TCGTTTCCAGTTGGGATGAGAGG 1000
Oligo fwd CACCTCGTTTCCAGTTGGGATGAG
Oligo rev AAACCTCATCCCAACTGGAAACGA
Chromosome start end strand MM target_seq PAM alignment distance position gene name gene id
GU305914.1 98630 98652 + 0 TCGTTTCCAGTTGGGATGAG AGG ||||||||[||||||||||||]PAM NA Intergenic NA NA

T7116 GATGAGAGGCGCCCGAATCGCGG 1000
Oligo fwd CACCGATGAGAGGCGCCCGAATCG
Oligo rev AAACCGATTCGGGCGCCTCTCATC
Chromosome start end strand MM target_seq PAM alignment distance position gene name gene id
GU305914.1 98644 98666 + 0 GATGAGAGGCGCCCGAATCG CGG ||||||||[||||||||||||]PAM NA Intergenic NA NA

T7117 CCGAATCGCGGTAAGACGACTGG 1000
Oligo fwd CACCCCGAATCGCGGTAAGACGAC
Oligo rev AAACGTCGTCTTACCGCGATTCGG
Chromosome start end strand MM target_seq PAM alignment distance position gene name gene id
GU305914.1 98656 98678 + 0 CCGAATCGCGGTAAGACGAC TGG ||||||||[||||||||||||]PAM NA Intergenic NA NA

T7118 CGCGGTAAGACGACTGGTTGTGG 1000
Oligo fwd CACCCGCGGTAAGACGACTGGTTG
Oligo rev AAACCAACCAGTCGTCTTACCGCG
Chromosome start end strand MM target_seq PAM alignment distance position gene name gene id
GU305914.1 98662 98684 + 0 CGCGGTAAGACGACTGGTTG TGG ||||||||[||||||||||||]PAM NA Intergenic NA NA

T7119 GTTGCGAATATTTCGAACGTCGG 1000
Oligo fwd CACCGTTGCGAATATTTCGAACGT
Oligo rev AAACACGTTCGAAATATTCGCAAC
Chromosome start end strand MM target_seq PAM alignment distance position gene name gene id
GU305914.1 98710 98732 + 0 GTTGCGAATATTTCGAACGT CGG ||||||||[||||||||||||]PAM NA Intergenic NA NA

T7120 TTGCGAATATTTCGAACGTCGGG 1000
Oligo fwd CACCTTGCGAATATTTCGAACGTC
Oligo rev AAACGACGTTCGAAATATTCGCAA
Chromosome start end strand MM target_seq PAM alignment distance position gene name gene id
GU305914.1 98711 98733 + 0 TTGCGAATATTTCGAACGTC GGG ||||||||[||||||||||||]PAM NA Intergenic NA NA

T7121 TCGGGAGACGCCGCCCGTCGCGG 1000
Oligo fwd CACCTCGGGAGACGCCGCCCGTCG
Oligo rev AAACCGACGGGCGGCGTCTCCCGA
Chromosome start end strand MM target_seq PAM alignment distance position gene name gene id

| | | | | | | | | | | | | |
|---|---|---|---|---|---|---|---|---|---|---|---|---|
| GU305914.1 | | 98729 | 98751 | + | 0 | TCGGGAGACGCCGCCCGTCG | CGG | \|\|\|\|\|\|\|\|[\|\|\|\|\|\|\|\|\|\|\|\|]PAM | NA | Intergenic | NA | NA |
| T7122 | GGCATCGTCCTCGTCGTATCCGG | | 1000 | | | | | | | | | |
| Oligo fwd | CACCGGCATCGTCCTCGTCGTATC | | | | | | | | | | | |
| Oligo rev | AAACGATACGACGAGGACGATGCC | | | | | | | | | | | |
| Chromosome | start | | end | strand | MM | target_seq | PAM | alignment | distance | position | gene name | gene id |
| GU305914.1 | | 98750 | 98772 | + | 0 | GGCATCGTCCTCGTCGTATC | CGG | \|\|\|\|\|\|\|\|[\|\|\|\|\|\|\|\|\|\|\|\|]PAM | NA | Intergenic | NA | NA |
| T7123 | TCCTCGTCGTATCCGGCGAACGG | | 1000 | | | | | | | | | |
| Oligo fwd | CACCTCCTCGTCGTATCCGGCGAA | | | | | | | | | | | |
| Oligo rev | AAACTTCGCCGGATACGACGAGGA | | | | | | | | | | | |
| Chromosome | start | | end | strand | MM | target_seq | PAM | alignment | distance | position | gene name | gene id |
| GU305914.1 | | 98757 | 98779 | + | 0 | TCCTCGTCGTATCCGGCGAA | CGG | \|\|\|\|\|\|\|\|[\|\|\|\|\|\|\|\|\|\|\|\|]PAM | NA | Intergenic | NA | NA |
| T7124 | CCTCGTCGTATCCGGCGAACGGG | | 1000 | | | | | | | | | |
| Oligo fwd | CACCCCTCGTCGTATCCGGCGAAC | | | | | | | | | | | |
| Oligo rev | AAACGTTCGCCGGATACGACGAGG | | | | | | | | | | | |
| Chromosome | start | | end | strand | MM | target_seq | PAM | alignment | distance | position | gene name | gene id |
| GU305914.1 | | 98758 | 98780 | + | 0 | CCTCGTCGTATCCGGCGAAC | GGG | \|\|\|\|\|\|\|\|[\|\|\|\|\|\|\|\|\|\|\|\|]PAM | NA | Intergenic | NA | NA |
| T7125 | GCCCAGCCGTCTCTTGACACCGG | | 1000 | | | | | | | | | |
| Oligo fwd | CACCGCCCAGCCGTCTCTTGACAC | | | | | | | | | | | |
| Oligo rev | AAACGTGTCAAGAGACGGCTGGGC | | | | | | | | | | | |
| Chromosome | start | | end | strand | MM | target_seq | PAM | alignment | distance | position | gene name | gene id |
| GU305914.1 | | 98780 | 98802 | + | 0 | GCCCAGCCGTCTCTTGACAC | CGG | \|\|\|\|\|\|\|\|[\|\|\|\|\|\|\|\|\|\|\|\|]PAM | NA | Intergenic | NA | NA |
| T7126 | CTTGACACCGGTCCGCAGCATGG | | 1000 | | | | | | | | | |
| Oligo fwd | CACCCTTGACACCGGTCCGCAGCA | | | | | | | | | | | |
| Oligo rev | AAACTGCTGCGGACCGGTGTCAAG | | | | | | | | | | | |
| Chromosome | start | | end | strand | MM | target_seq | PAM | alignment | distance | position | gene name | gene id |
| GU305914.1 | | 98792 | 98814 | + | 0 | CTTGACACCGGTCCGCAGCA | TGG | \|\|\|\|\|\|\|\|[\|\|\|\|\|\|\|\|\|\|\|\|]PAM | NA | Intergenic | NA | NA |
| T7127 | CAGCATGGTTTCGCCGCCTTCGG | | 1000 | | | | | | | | | |
| Oligo fwd | CACCCAGCATGGTTTCGCCGCCTT | | | | | | | | | | | |
| Oligo rev | AAACAAGGCGGCGAAACCATGCTG | | | | | | | | | | | |
| Chromosome | start | | end | strand | MM | target_seq | PAM | alignment | distance | position | gene name | gene id |
| GU305914.1 | | 98807 | 98829 | + | 0 | CAGCATGGTTTCGCCGCCTT | CGG | \|\|\|\|\|\|\|\|[\|\|\|\|\|\|\|\|\|\|\|\|]PAM | NA | Intergenic | NA | NA |
| T7128 | TTCGCCGCCTTCGGTGCGCTCGG | | 1000 | | | | | | | | | |
| Oligo fwd | CACCTTCGCCGCCTTCGGTGCGCT | | | | | | | | | | | |
| Oligo rev | AAACAGCGCACCGAAGGCGGCGAA | | | | | | | | | | | |
| Chromosome | start | | end | strand | MM | target_seq | PAM | alignment | distance | position | gene name | gene id |
| GU305914.1 | | 98816 | 98838 | + | 0 | TTCGCCGCCTTCGGTGCGCT | CGG | \|\|\|\|\|\|\|\|[\|\|\|\|\|\|\|\|\|\|\|\|]PAM | NA | Intergenic | NA | NA |
| T7129 | CGGTCGCTCGTACGTACGCGCGG | | 1000 | | | | | | | | | |
| Oligo fwd | CACCCGGTCGCTCGTACGTACGCG | | | | | | | | | | | |
| Oligo rev | AAACCGCGTACGTACGAGCGACCG | | | | | | | | | | | |
| Chromosome | start | | end | strand | MM | target_seq | PAM | alignment | distance | position | gene name | gene id |
| GU305914.1 | | 98836 | 98858 | + | 0 | CGGTCGCTCGTACGTACGCG | CGG | \|\|\|\|\|\|\|\|[\|\|\|\|\|\|\|\|\|\|\|\|]PAM | NA | Intergenic | NA | NA |
| T7130 | TACGTACGCGCGGTTGCTGACGG | | 1000 | | | | | | | | | |
| Oligo fwd | CACCTACGTACGCGCGGTTGCTGA | | | | | | | | | | | |
| Oligo rev | AAACTCAGCAACCGCGCGTACGTA | | | | | | | | | | | |
| Chromosome | start | | end | strand | MM | target_seq | PAM | alignment | distance | position | gene name | gene id |
| GU305914.1 | | 98846 | 98868 | + | 0 | TACGTACGCGCGGTTGCTGA | CGG | \|\|\|\|\|\|\|\|[\|\|\|\|\|\|\|\|\|\|\|\|]PAM | NA | Intergenic | NA | NA |
| T7131 | ACGTACGCGCGGTTGCTGACGGG | | 1000 | | | | | | | | | |
| Oligo fwd | CACCACGTACGCGCGGTTGCTGAC | | | | | | | | | | | |
| Oligo rev | AAACGTCAGCAACCGCGCGTACGT | | | | | | | | | | | |
| Chromosome | start | | end | strand | MM | target_seq | PAM | alignment | distance | position | gene name | gene id |
| GU305914.1 | | 98847 | 98869 | + | 0 | ACGTACGCGCGGTTGCTGAC | GGG | \|\|\|\|\|\|\|\|[\|\|\|\|\|\|\|\|\|\|\|\|]PAM | NA | Intergenic | NA | NA |
| T7132 | CGTACGCGCGGTTGCTGACGGGG | | 1000 | | | | | | | | | |
| Oligo fwd | CACCCGTACGCGCGGTTGCTGACG | | | | | | | | | | | |
| Oligo rev | AAACCGTCAGCAACCGCGCGTACG | | | | | | | | | | | |
| Chromosome | start | | end | strand | MM | target_seq | PAM | alignment | distance | position | gene name | gene id |
| GU305914.1 | | 98848 | 98870 | + | 0 | CGTACGCGCGGTTGCTGACG | GGG | \|\|\|\|\|\|\|\|[\|\|\|\|\|\|\|\|\|\|\|\|]PAM | NA | Intergenic | NA | NA |
| T7133 | GTACGCGCGGTTGCTGACGGGGG | | 1000 | | | | | | | | | |
| Oligo fwd | CACCGTACGCGCGGTTGCTGACGG | | | | | | | | | | | |
| Oligo rev | AAACCCGTCAGCAACCGCGCGTAC | | | | | | | | | | | |
| Chromosome | start | | end | strand | MM | target_seq | PAM | alignment | distance | position | gene name | gene id |
| GU305914.1 | | 98849 | 98871 | + | 0 | GTACGCGCGGTTGCTGACGG | GGG | \|\|\|\|\|\|\|\|[\|\|\|\|\|\|\|\|\|\|\|\|]PAM | NA | Intergenic | NA | NA |
| T7134 | TACGCGCGGTTGCTGACGGGGGG | | 1000 | | | | | | | | | |
| Oligo fwd | CACCTACGCGCGGTTGCTGACGGG | | | | | | | | | | | |
| Oligo rev | AAACCCCGTCAGCAACCGCGCGTA | | | | | | | | | | | |
| Chromosome | start | | end | strand | MM | target_seq | PAM | alignment | distance | position | gene name | gene id |
| GU305914.1 | | 98850 | 98872 | + | 0 | TACGCGCGGTTGCTGACGGG | GGG | \|\|\|\|\|\|\|\|[\|\|\|\|\|\|\|\|\|\|\|\|]PAM | NA | Intergenic | NA | NA |
| T7135 | ACGCGCGGTTGCTGACGGGGGGG | | 1000 | | | | | | | | | |
| Oligo fwd | CACCACGCGCGGTTGCTGACGGGG | | | | | | | | | | | |
| Oligo rev | AAACCCCCGTCAGCAACCGCGCGT | | | | | | | | | | | |
| Chromosome | start | | end | strand | MM | target_seq | PAM | alignment | distance | position | gene name | gene id |
| GU305914.1 | | 98851 | 98873 | + | 0 | ACGCGCGGTTGCTGACGGGG | GGG | \|\|\|\|\|\|\|\|[\|\|\|\|\|\|\|\|\|\|\|\|]PAM | NA | Intergenic | NA | NA |
| T7136 | TGACGGGGGGGACGCCCGCGCGG | | 1000 | | | | | | | | | |
| Oligo fwd | CACCTGACGGGGGGGACGCCCGCG | | | | | | | | | | | |
| Oligo rev | AAACCGCGGGCGTCCCCCCCGTCA | | | | | | | | | | | |

<table>
<tr><td>Chromosome</td><td>start</td><td>end</td><td>strand</td><td>MM</td><td>target_seq</td><td>PAM</td><td>alignment</td><td>distance</td><td>position</td><td>gene name</td><td>gene id</td></tr>
<tr><td>GU305914.1</td><td>98863</td><td>98885</td><td>+</td><td>0</td><td>TGACGGGGGGGACGCCCGCG</td><td>CGG</td><td>||||||||[||||||||||||]PAM</td><td>NA</td><td>Intergenic</td><td>NA</td><td>NA</td></tr>
<tr><td>T7137</td><td>GGGACGCCCGCGCGGAAATGTGG</td><td>1000</td></tr>
<tr><td>Oligo fwd</td><td>CACCGGGACGCCCGCGCGGAAATG</td></tr>
<tr><td>Oligo rev</td><td>AAACCATTTCCGCGCGGGCGTCCC</td></tr>
<tr><td>Chromosome</td><td>start</td><td>end</td><td>strand</td><td>MM</td><td>target_seq</td><td>PAM</td><td>alignment</td><td>distance</td><td>position</td><td>gene name</td><td>gene id</td></tr>
<tr><td>GU305914.1</td><td>98871</td><td>98893</td><td>+</td><td>0</td><td>GGGACGCCCGCGCGGAAATG</td><td>TGG</td><td>||||||||[||||||||||||]PAM</td><td>NA</td><td>Intergenic</td><td>NA</td><td>NA</td></tr>
<tr><td>T7138</td><td>GCCCGCGCGGAAATGTGGCTAGG</td><td>1000</td></tr>
<tr><td>Oligo fwd</td><td>CACCGCCCGCGCGGAAATGTGGCT</td></tr>
<tr><td>Oligo rev</td><td>AAACAGCCACATTTCCGCGCGGGC</td></tr>
<tr><td>Chromosome</td><td>start</td><td>end</td><td>strand</td><td>MM</td><td>target_seq</td><td>PAM</td><td>alignment</td><td>distance</td><td>position</td><td>gene name</td><td>gene id</td></tr>
<tr><td>GU305914.1</td><td>98876</td><td>98898</td><td>+</td><td>0</td><td>GCCCGCGCGGAAATGTGGCT</td><td>AGG</td><td>||||||||[||||||||||||]PAM</td><td>NA</td><td>Intergenic</td><td>NA</td><td>NA</td></tr>
<tr><td>T7139</td><td>GACTGCGACTCCTTCCGCGCAGG</td><td>1000</td></tr>
<tr><td>Oligo fwd</td><td>CACCGACTGCGACTCCTTCCGCGC</td></tr>
<tr><td>Oligo rev</td><td>AAACGCGCGGAAGGAGTCGCAGTC</td></tr>
<tr><td>Chromosome</td><td>start</td><td>end</td><td>strand</td><td>MM</td><td>target_seq</td><td>PAM</td><td>alignment</td><td>distance</td><td>position</td><td>gene name</td><td>gene id</td></tr>
<tr><td>GU305914.1</td><td>98914</td><td>98936</td><td>+</td><td>0</td><td>GACTGCGACTCCTTCCGCGC</td><td>AGG</td><td>||||||||[||||||||||||]PAM</td><td>NA</td><td>Intergenic</td><td>NA</td><td>NA</td></tr>
<tr><td>T7140</td><td>CCCCCCGCTTTTTTATATCGCGG</td><td>1000</td></tr>
<tr><td>Oligo fwd</td><td>CACCCCCCCCGCTTTTTTATATCG</td></tr>
<tr><td>Oligo rev</td><td>AAACCGATATAAAAAAGCGGGGGG</td></tr>
<tr><td>Chromosome</td><td>start</td><td>end</td><td>strand</td><td>MM</td><td>target_seq</td><td>PAM</td><td>alignment</td><td>distance</td><td>position</td><td>gene name</td><td>gene id</td></tr>
<tr><td>GU305914.1</td><td>98940</td><td>98962</td><td>+</td><td>0</td><td>CCCCCCGCTTTTTTATATCG</td><td>CGG</td><td>||||||||[||||||||||||]PAM</td><td>NA</td><td>Intergenic</td><td>NA</td><td>NA</td></tr>
<tr><td>T7141</td><td>CCCCCGCTTTTTTATATCGCGGG</td><td>1000</td></tr>
<tr><td>Oligo fwd</td><td>CACCCCCCCGCTTTTTTATATCGC</td></tr>
<tr><td>Oligo rev</td><td>AAACGCGATATAAAAAAGCGGGGG</td></tr>
<tr><td>Chromosome</td><td>start</td><td>end</td><td>strand</td><td>MM</td><td>target_seq</td><td>PAM</td><td>alignment</td><td>distance</td><td>position</td><td>gene name</td><td>gene id</td></tr>
<tr><td>GU305914.1</td><td>98941</td><td>98963</td><td>+</td><td>0</td><td>CCCCCGCTTTTTTATATCGC</td><td>GGG</td><td>||||||||[||||||||||||]PAM</td><td>NA</td><td>Intergenic</td><td>NA</td><td>NA</td></tr>
<tr><td>T7142</td><td>CCGCTTTTTTATATCGCGGGAGG</td><td>1000</td></tr>
<tr><td>Oligo fwd</td><td>CACCCCGCTTTTTTATATCGCGGG</td></tr>
<tr><td>Oligo rev</td><td>AAACCCCGCGATATAAAAAAGCGG</td></tr>
<tr><td>Chromosome</td><td>start</td><td>end</td><td>strand</td><td>MM</td><td>target_seq</td><td>PAM</td><td>alignment</td><td>distance</td><td>position</td><td>gene name</td><td>gene id</td></tr>
<tr><td>GU305914.1</td><td>98944</td><td>98966</td><td>+</td><td>0</td><td>CCGCTTTTTTATATCGCGGG</td><td>AGG</td><td>||||||||[||||||||||||]PAM</td><td>NA</td><td>Intergenic</td><td>NA</td><td>NA</td></tr>
<tr><td>T7143</td><td>CGCTTTTTTATATCGCGGGAGGG</td><td>1000</td></tr>
<tr><td>Oligo fwd</td><td>CACCCGCTTTTTTATATCGCGGGA</td></tr>
<tr><td>Oligo rev</td><td>AAACTCCCGCGATATAAAAAAGCG</td></tr>
<tr><td>Chromosome</td><td>start</td><td>end</td><td>strand</td><td>MM</td><td>target_seq</td><td>PAM</td><td>alignment</td><td>distance</td><td>position</td><td>gene name</td><td>gene id</td></tr>
<tr><td>GU305914.1</td><td>98945</td><td>98967</td><td>+</td><td>0</td><td>CGCTTTTTTATATCGCGGGA</td><td>GGG</td><td>||||||||[||||||||||||]PAM</td><td>NA</td><td>Intergenic</td><td>NA</td><td>NA</td></tr>
<tr><td>T7144</td><td>GCTTTTTTATATCGCGGGAGGGG</td><td>1000</td></tr>
<tr><td>Oligo fwd</td><td>CACCGCTTTTTTATATCGCGGGAG</td></tr>
<tr><td>Oligo rev</td><td>AAACCTCCCGCGATATAAAAAAGC</td></tr>
<tr><td>Chromosome</td><td>start</td><td>end</td><td>strand</td><td>MM</td><td>target_seq</td><td>PAM</td><td>alignment</td><td>distance</td><td>position</td><td>gene name</td><td>gene id</td></tr>
<tr><td>GU305914.1</td><td>98946</td><td>98968</td><td>+</td><td>0</td><td>GCTTTTTTATATCGCGGGAG</td><td>GGG</td><td>||||||||[||||||||||||]PAM</td><td>NA</td><td>Intergenic</td><td>NA</td><td>NA</td></tr>
<tr><td>T7145</td><td>TTTTATATCGCGGGAGGGGAAGG</td><td>1000</td></tr>
<tr><td>Oligo fwd</td><td>CACCTTTTATATCGCGGGAGGGGA</td></tr>
<tr><td>Oligo rev</td><td>AAACTCCCCTCCCGCGATATAAAA</td></tr>
<tr><td>Chromosome</td><td>start</td><td>end</td><td>strand</td><td>MM</td><td>target_seq</td><td>PAM</td><td>alignment</td><td>distance</td><td>position</td><td>gene name</td><td>gene id</td></tr>
<tr><td>GU305914.1</td><td>98950</td><td>98972</td><td>+</td><td>0</td><td>TTTTATATCGCGGGAGGGGA</td><td>AGG</td><td>||||||||[||||||||||||]PAM</td><td>NA</td><td>Intergenic</td><td>NA</td><td>NA</td></tr>
<tr><td>T7146</td><td>TTTATATCGCGGGAGGGGAAGGG</td><td>1000</td></tr>
<tr><td>Oligo fwd</td><td>CACCTTTATATCGCGGGAGGGGAA</td></tr>
<tr><td>Oligo rev</td><td>AAACTTCCCCTCCCGCGATATAAA</td></tr>
<tr><td>Chromosome</td><td>start</td><td>end</td><td>strand</td><td>MM</td><td>target_seq</td><td>PAM</td><td>alignment</td><td>distance</td><td>position</td><td>gene name</td><td>gene id</td></tr>
<tr><td>GU305914.1</td><td>98951</td><td>98973</td><td>+</td><td>0</td><td>TTTATATCGCGGGAGGGGAA</td><td>GGG</td><td>||||||||[||||||||||||]PAM</td><td>NA</td><td>Intergenic</td><td>NA</td><td>NA</td></tr>
<tr><td>T7147</td><td>GGGTCGATGATACGTTAGTTAGG</td><td>1000</td></tr>
<tr><td>Oligo fwd</td><td>CACCGGGTCGATGATACGTTAGTT</td></tr>
<tr><td>Oligo rev</td><td>AAACAACTAACGTATCATCGACCC</td></tr>
<tr><td>Chromosome</td><td>start</td><td>end</td><td>strand</td><td>MM</td><td>target_seq</td><td>PAM</td><td>alignment</td><td>distance</td><td>position</td><td>gene name</td><td>gene id</td></tr>
<tr><td>GU305914.1</td><td>98971</td><td>98993</td><td>+</td><td>0</td><td>GGGTCGATGATACGTTAGTT</td><td>AGG</td><td>||||||||[||||||||||||]PAM</td><td>NA</td><td>Intergenic</td><td>NA</td><td>NA</td></tr>
<tr><td>T7148</td><td>GGTCGATGATACGTTAGTTAGGG</td><td>1000</td></tr>
<tr><td>Oligo fwd</td><td>CACCGGTCGATGATACGTTAGTTA</td></tr>
<tr><td>Oligo rev</td><td>AAACTAACTAACGTATCATCGACC</td></tr>
<tr><td>Chromosome</td><td>start</td><td>end</td><td>strand</td><td>MM</td><td>target_seq</td><td>PAM</td><td>alignment</td><td>distance</td><td>position</td><td>gene name</td><td>gene id</td></tr>
<tr><td>GU305914.1</td><td>98972</td><td>98994</td><td>+</td><td>0</td><td>GGTCGATGATACGTTAGTTA</td><td>GGG</td><td>||||||||[||||||||||||]PAM</td><td>NA</td><td>Intergenic</td><td>NA</td><td>NA</td></tr>
<tr><td>T7149</td><td>GATGATACGTTAGTTAGGGCCGG</td><td>1000</td></tr>
<tr><td>Oligo fwd</td><td>CACCGATGATACGTTAGTTAGGGC</td></tr>
<tr><td>Oligo rev</td><td>AAACGCCCTAACTAACGTATCATC</td></tr>
<tr><td>Chromosome</td><td>start</td><td>end</td><td>strand</td><td>MM</td><td>target_seq</td><td>PAM</td><td>alignment</td><td>distance</td><td>position</td><td>gene name</td><td>gene id</td></tr>
<tr><td>GU305914.1</td><td>98976</td><td>98998</td><td>+</td><td>0</td><td>GATGATACGTTAGTTAGGGC</td><td>CGG</td><td>||||||||[||||||||||||]PAM</td><td>NA</td><td>Intergenic</td><td>NA</td><td>NA</td></tr>
<tr><td>T7150</td><td>ATGATACGTTAGTTAGGGCCGGG</td><td>1000</td></tr>
<tr><td>Oligo fwd</td><td>CACCATGATACGTTAGTTAGGGCC</td></tr>
<tr><td>Oligo rev</td><td>AAACGGCCCTAACTAACGTATCAT</td></tr>
<tr><td>Chromosome</td><td>start</td><td>end</td><td>strand</td><td>MM</td><td>target_seq</td><td>PAM</td><td>alignment</td><td>distance</td><td>position</td><td>gene name</td><td>gene id</td></tr>
<tr><td>GU305914.1</td><td>98977</td><td>98999</td><td>+</td><td>0</td><td>ATGATACGTTAGTTAGGGCC</td><td>GGG</td><td>||||||||[||||||||||||]PAM</td><td>NA</td><td>Intergenic</td><td>NA</td><td>NA</td></tr>
<tr><td>T7151</td><td>GTTAGGGCCGGGTGCCAGCGTGG</td><td>1000</td></tr>
<tr><td>Oligo fwd</td><td>CACCGTTAGGGCCGGGTGCCAGCG</td></tr>
</table>

<table>
<tr><td>Oligo rev</td><td>AAACCGCTGGCACCCGGCCCTAAC</td><td></td><td></td><td></td><td></td><td></td><td></td><td></td><td></td><td></td><td></td></tr>
<tr><td>Chromosome</td><td>start</td><td>end</td><td>strand</td><td>MM</td><td>target_seq</td><td>PAM</td><td>alignment</td><td>distance</td><td>position</td><td>gene name</td><td>gene id</td></tr>
<tr><td>GU305914.1</td><td>98988</td><td>99010</td><td>+</td><td>0</td><td>GTTAGGGCCGGGTGCCAGCG</td><td>TGG</td><td>||||||||[||||||||||||]PAM</td><td>NA</td><td>Intergenic</td><td>NA</td><td>NA</td></tr>
<tr><td></td><td></td><td></td><td></td><td></td><td></td><td></td><td></td><td></td><td></td><td></td><td></td></tr>
<tr><td>T7152</td><td>TTAGGGCCGGGTGCCAGCGTGGG</td><td>1000</td><td></td><td></td><td></td><td></td><td></td><td></td><td></td><td></td><td></td></tr>
<tr><td>Oligo fwd</td><td>CACCTTAGGGCCGGGTGCCAGCGT</td><td></td><td></td><td></td><td></td><td></td><td></td><td></td><td></td><td></td><td></td></tr>
<tr><td>Oligo rev</td><td>AAACACGCTGGCACCCGGCCCTAA</td><td></td><td></td><td></td><td></td><td></td><td></td><td></td><td></td><td></td><td></td></tr>
<tr><td>Chromosome</td><td>start</td><td>end</td><td>strand</td><td>MM</td><td>target_seq</td><td>PAM</td><td>alignment</td><td>distance</td><td>position</td><td>gene name</td><td>gene id</td></tr>
<tr><td>GU305914.1</td><td>98989</td><td>99011</td><td>+</td><td>0</td><td>TTAGGGCCGGGTGCCAGCGT</td><td>GGG</td><td>||||||||[||||||||||||]PAM</td><td>NA</td><td>Intergenic</td><td>NA</td><td>NA</td></tr>
<tr><td></td><td></td><td></td><td></td><td></td><td></td><td></td><td></td><td></td><td></td><td></td><td></td></tr>
<tr><td>T7153</td><td>GGCCGGGTGCCAGCGTGGGCCGG</td><td>1000</td><td></td><td></td><td></td><td></td><td></td><td></td><td></td><td></td><td></td></tr>
<tr><td>Oligo fwd</td><td>CACCGGCCGGGTGCCAGCGTGGGC</td><td></td><td></td><td></td><td></td><td></td><td></td><td></td><td></td><td></td><td></td></tr>
<tr><td>Oligo rev</td><td>AAACGCCCACGCTGGCACCCGGCC</td><td></td><td></td><td></td><td></td><td></td><td></td><td></td><td></td><td></td><td></td></tr>
<tr><td>Chromosome</td><td>start</td><td>end</td><td>strand</td><td>MM</td><td>target_seq</td><td>PAM</td><td>alignment</td><td>distance</td><td>position</td><td>gene name</td><td>gene id</td></tr>
<tr><td>GU305914.1</td><td>98993</td><td>99015</td><td>+</td><td>0</td><td>GGCCGGGTGCCAGCGTGGGC</td><td>CGG</td><td>||||||||[||||||||||||]PAM</td><td>NA</td><td>Intergenic</td><td>NA</td><td>NA</td></tr>
<tr><td></td><td></td><td></td><td></td><td></td><td></td><td></td><td></td><td></td><td></td><td></td><td></td></tr>
<tr><td>T7154</td><td>GGGTGCCAGCGTGGGCCGGACGG</td><td>1000</td><td></td><td></td><td></td><td></td><td></td><td></td><td></td><td></td><td></td></tr>
<tr><td>Oligo fwd</td><td>CACCGGGTGCCAGCGTGGGCCGGA</td><td></td><td></td><td></td><td></td><td></td><td></td><td></td><td></td><td></td><td></td></tr>
<tr><td>Oligo rev</td><td>AAACTCCGGCCCACGCTGGCACCC</td><td></td><td></td><td></td><td></td><td></td><td></td><td></td><td></td><td></td><td></td></tr>
<tr><td>Chromosome</td><td>start</td><td>end</td><td>strand</td><td>MM</td><td>target_seq</td><td>PAM</td><td>alignment</td><td>distance</td><td>position</td><td>gene name</td><td>gene id</td></tr>
<tr><td>GU305914.1</td><td>98997</td><td>99019</td><td>+</td><td>0</td><td>GGGTGCCAGCGTGGGCCGGA</td><td>CGG</td><td>||||||||[||||||||||||]PAM</td><td>NA</td><td>Intergenic</td><td>NA</td><td>NA</td></tr>
<tr><td></td><td></td><td></td><td></td><td></td><td></td><td></td><td></td><td></td><td></td><td></td><td></td></tr>
<tr><td>T7155</td><td>ATTTATTCCTTGACGCAAGCAGG</td><td>1000</td><td></td><td></td><td></td><td></td><td></td><td></td><td></td><td></td><td></td></tr>
<tr><td>Oligo fwd</td><td>CACCATTTATTCCTTGACGCAAGC</td><td></td><td></td><td></td><td></td><td></td><td></td><td></td><td></td><td></td><td></td></tr>
<tr><td>Oligo rev</td><td>AAACGCTTGCGTCAAGGAATAAAT</td><td></td><td></td><td></td><td></td><td></td><td></td><td></td><td></td><td></td><td></td></tr>
<tr><td>Chromosome</td><td>start</td><td>end</td><td>strand</td><td>MM</td><td>target_seq</td><td>PAM</td><td>alignment</td><td>distance</td><td>position</td><td>gene name</td><td>gene id</td></tr>
<tr><td>GU305914.1</td><td>99020</td><td>99042</td><td>+</td><td>0</td><td>ATTTATTCCTTGACGCAAGC</td><td>AGG</td><td>||||||||[||||||||||||]PAM</td><td>NA</td><td>Intergenic</td><td>NA</td><td>NA</td></tr>
<tr><td></td><td></td><td></td><td></td><td></td><td></td><td></td><td></td><td></td><td></td><td></td><td></td></tr>
<tr><td>T7156</td><td>CTTGACGCAAGCAGGATGCGTGG</td><td>1000</td><td></td><td></td><td></td><td></td><td></td><td></td><td></td><td></td><td></td></tr>
<tr><td>Oligo fwd</td><td>CACCCTTGACGCAAGCAGGATGCG</td><td></td><td></td><td></td><td></td><td></td><td></td><td></td><td></td><td></td><td></td></tr>
<tr><td>Oligo rev</td><td>AAACCGCATCCTGCTTGCGTCAAG</td><td></td><td></td><td></td><td></td><td></td><td></td><td></td><td></td><td></td><td></td></tr>
<tr><td>Chromosome</td><td>start</td><td>end</td><td>strand</td><td>MM</td><td>target_seq</td><td>PAM</td><td>alignment</td><td>distance</td><td>position</td><td>gene name</td><td>gene id</td></tr>
<tr><td>GU305914.1</td><td>99028</td><td>99050</td><td>+</td><td>0</td><td>CTTGACGCAAGCAGGATGCG</td><td>TGG</td><td>||||||||[||||||||||||]PAM</td><td>NA</td><td>Intergenic</td><td>NA</td><td>NA</td></tr>
<tr><td></td><td></td><td></td><td></td><td></td><td></td><td></td><td></td><td></td><td></td><td></td><td></td></tr>
<tr><td>T7157</td><td>TTGACGCAAGCAGGATGCGTGGG</td><td>1000</td><td></td><td></td><td></td><td></td><td></td><td></td><td></td><td></td><td></td></tr>
<tr><td>Oligo fwd</td><td>CACCTTGACGCAAGCAGGATGCGT</td><td></td><td></td><td></td><td></td><td></td><td></td><td></td><td></td><td></td><td></td></tr>
<tr><td>Oligo rev</td><td>AAACACGCATCCTGCTTGCGTCAA</td><td></td><td></td><td></td><td></td><td></td><td></td><td></td><td></td><td></td><td></td></tr>
<tr><td>Chromosome</td><td>start</td><td>end</td><td>strand</td><td>MM</td><td>target_seq</td><td>PAM</td><td>alignment</td><td>distance</td><td>position</td><td>gene name</td><td>gene id</td></tr>
<tr><td>GU305914.1</td><td>99029</td><td>99051</td><td>+</td><td>0</td><td>TTGACGCAAGCAGGATGCGT</td><td>GGG</td><td>||||||||[||||||||||||]PAM</td><td>NA</td><td>Intergenic</td><td>NA</td><td>NA</td></tr>
<tr><td></td><td></td><td></td><td></td><td></td><td></td><td></td><td></td><td></td><td></td><td></td><td></td></tr>
<tr><td>T7158</td><td>AGGATGCGTGGGTGCGCTGCCGG</td><td>1000</td><td></td><td></td><td></td><td></td><td></td><td></td><td></td><td></td><td></td></tr>
<tr><td>Oligo fwd</td><td>CACCAGGATGCGTGGGTGCGCTGC</td><td></td><td></td><td></td><td></td><td></td><td></td><td></td><td></td><td></td><td></td></tr>
<tr><td>Oligo rev</td><td>AAACGCAGCGCACCCACGCATCCT</td><td></td><td></td><td></td><td></td><td></td><td></td><td></td><td></td><td></td><td></td></tr>
<tr><td>Chromosome</td><td>start</td><td>end</td><td>strand</td><td>MM</td><td>target_seq</td><td>PAM</td><td>alignment</td><td>distance</td><td>position</td><td>gene name</td><td>gene id</td></tr>
<tr><td>GU305914.1</td><td>99040</td><td>99062</td><td>+</td><td>0</td><td>AGGATGCGTGGGTGCGCTGC</td><td>CGG</td><td>||||||||[||||||||||||]PAM</td><td>NA</td><td>Intergenic</td><td>NA</td><td>NA</td></tr>
<tr><td></td><td></td><td></td><td></td><td></td><td></td><td></td><td></td><td></td><td></td><td></td><td></td></tr>
<tr><td>T7159</td><td>GCGTGGGTGCGCTGCCGGAACGG</td><td>1000</td><td></td><td></td><td></td><td></td><td></td><td></td><td></td><td></td><td></td></tr>
<tr><td>Oligo fwd</td><td>CACCGCGTGGGTGCGCTGCCGGAA</td><td></td><td></td><td></td><td></td><td></td><td></td><td></td><td></td><td></td><td></td></tr>
<tr><td>Oligo rev</td><td>AAACTTCCGGCAGCGCACCCACGC</td><td></td><td></td><td></td><td></td><td></td><td></td><td></td><td></td><td></td><td></td></tr>
<tr><td>Chromosome</td><td>start</td><td>end</td><td>strand</td><td>MM</td><td>target_seq</td><td>PAM</td><td>alignment</td><td>distance</td><td>position</td><td>gene name</td><td>gene id</td></tr>
<tr><td>GU305914.1</td><td>99045</td><td>99067</td><td>+</td><td>0</td><td>GCGTGGGTGCGCTGCCGGAA</td><td>CGG</td><td>||||||||[||||||||||||]PAM</td><td>NA</td><td>Intergenic</td><td>NA</td><td>NA</td></tr>
<tr><td></td><td></td><td></td><td></td><td></td><td></td><td></td><td></td><td></td><td></td><td></td><td></td></tr>
<tr><td>T7160</td><td>CGTGGGTGCGCTGCCGGAACGGG</td><td>1000</td><td></td><td></td><td></td><td></td><td></td><td></td><td></td><td></td><td></td></tr>
<tr><td>Oligo fwd</td><td>CACCCGTGGGTGCGCTGCCGGAAC</td><td></td><td></td><td></td><td></td><td></td><td></td><td></td><td></td><td></td><td></td></tr>
<tr><td>Oligo rev</td><td>AAACGTTCCGGCAGCGCACCCACG</td><td></td><td></td><td></td><td></td><td></td><td></td><td></td><td></td><td></td><td></td></tr>
<tr><td>Chromosome</td><td>start</td><td>end</td><td>strand</td><td>MM</td><td>target_seq</td><td>PAM</td><td>alignment</td><td>distance</td><td>position</td><td>gene name</td><td>gene id</td></tr>
<tr><td>GU305914.1</td><td>99046</td><td>99068</td><td>+</td><td>0</td><td>CGTGGGTGCGCTGCCGGAAC</td><td>GGG</td><td>||||||||[||||||||||||]PAM</td><td>NA</td><td>Intergenic</td><td>NA</td><td>NA</td></tr>
<tr><td></td><td></td><td></td><td></td><td></td><td></td><td></td><td></td><td></td><td></td><td></td><td></td></tr>
<tr><td>T7161</td><td>GTGGGTGCGCTGCCGGAACGGGG</td><td>1000</td><td></td><td></td><td></td><td></td><td></td><td></td><td></td><td></td><td></td></tr>
<tr><td>Oligo fwd</td><td>CACCGTGGGTGCGCTGCCGGAACG</td><td></td><td></td><td></td><td></td><td></td><td></td><td></td><td></td><td></td><td></td></tr>
<tr><td>Oligo rev</td><td>AAACCGTTCCGGCAGCGCACCCAC</td><td></td><td></td><td></td><td></td><td></td><td></td><td></td><td></td><td></td><td></td></tr>
<tr><td>Chromosome</td><td>start</td><td>end</td><td>strand</td><td>MM</td><td>target_seq</td><td>PAM</td><td>alignment</td><td>distance</td><td>position</td><td>gene name</td><td>gene id</td></tr>
<tr><td>GU305914.1</td><td>99047</td><td>99069</td><td>+</td><td>0</td><td>GTGGGTGCGCTGCCGGAACG</td><td>GGG</td><td>||||||||[||||||||||||]PAM</td><td>NA</td><td>Intergenic</td><td>NA</td><td>NA</td></tr>
<tr><td></td><td></td><td></td><td></td><td></td><td></td><td></td><td></td><td></td><td></td><td></td><td></td></tr>
<tr><td>T7162</td><td>TGGGTGCGCTGCCGGAACGGGGG</td><td>1000</td><td></td><td></td><td></td><td></td><td></td><td></td><td></td><td></td><td></td></tr>
<tr><td>Oligo fwd</td><td>CACCTGGGTGCGCTGCCGGAACGG</td><td></td><td></td><td></td><td></td><td></td><td></td><td></td><td></td><td></td><td></td></tr>
<tr><td>Oligo rev</td><td>AAACCCGTTCCGGCAGCGCACCCA</td><td></td><td></td><td></td><td></td><td></td><td></td><td></td><td></td><td></td><td></td></tr>
<tr><td>Chromosome</td><td>start</td><td>end</td><td>strand</td><td>MM</td><td>target_seq</td><td>PAM</td><td>alignment</td><td>distance</td><td>position</td><td>gene name</td><td>gene id</td></tr>
<tr><td>GU305914.1</td><td>99048</td><td>99070</td><td>+</td><td>0</td><td>TGGGTGCGCTGCCGGAACGG</td><td>GGG</td><td>||||||||[||||||||||||]PAM</td><td>NA</td><td>Intergenic</td><td>NA</td><td>NA</td></tr>
<tr><td></td><td></td><td></td><td></td><td></td><td></td><td></td><td></td><td></td><td></td><td></td><td></td></tr>
<tr><td>T7163</td><td>TCGACGCGTCTAATACTGCGAGG</td><td>1000</td><td></td><td></td><td></td><td></td><td></td><td></td><td></td><td></td><td></td></tr>
<tr><td>Oligo fwd</td><td>CACCTCGACGCGTCTAATACTGCG</td><td></td><td></td><td></td><td></td><td></td><td></td><td></td><td></td><td></td><td></td></tr>
<tr><td>Oligo rev</td><td>AAACCGCAGTATTAGACGCGTCGA</td><td></td><td></td><td></td><td></td><td></td><td></td><td></td><td></td><td></td><td></td></tr>
<tr><td>Chromosome</td><td>start</td><td>end</td><td>strand</td><td>MM</td><td>target_seq</td><td>PAM</td><td>alignment</td><td>distance</td><td>position</td><td>gene name</td><td>gene id</td></tr>
<tr><td>GU305914.1</td><td>99092</td><td>99114</td><td>+</td><td>0</td><td>TCGACGCGTCTAATACTGCG</td><td>AGG</td><td>||||||||[||||||||||||]PAM</td><td>NA</td><td>Intergenic</td><td>NA</td><td>NA</td></tr>
<tr><td>GU305914.1</td><td>114355</td><td>114377</td><td>-</td><td>4</td><td>GCGTCGCGTCTAATACTCTG</td><td>CGG</td><td>-||-||||[|||||||||--|]PAM</td><td>NA</td><td>Intergenic</td><td>NA</td><td>NA</td></tr>
<tr><td></td><td></td><td></td><td></td><td></td><td></td><td></td><td></td><td></td><td></td><td></td><td></td></tr>
<tr><td>T7164</td><td>CGACGCGTCTAATACTGCGAGGG</td><td>1000</td><td></td><td></td><td></td><td></td><td></td><td></td><td></td><td></td><td></td></tr>
<tr><td>Oligo fwd</td><td>CACCCGACGCGTCTAATACTGCGA</td><td></td><td></td><td></td><td></td><td></td><td></td><td></td><td></td><td></td><td></td></tr>
<tr><td>Oligo rev</td><td>AAACTCGCAGTATTAGACGCGTCG</td><td></td><td></td><td></td><td></td><td></td><td></td><td></td><td></td><td></td><td></td></tr>
<tr><td>Chromosome</td><td>start</td><td>end</td><td>strand</td><td>MM</td><td>target_seq</td><td>PAM</td><td>alignment</td><td>distance</td><td>position</td><td>gene name</td><td>gene id</td></tr>
<tr><td>GU305914.1</td><td>99093</td><td>99115</td><td>+</td><td>0</td><td>CGACGCGTCTAATACTGCGA</td><td>GGG</td><td>||||||||[||||||||||||]PAM</td><td>NA</td><td>Intergenic</td><td>NA</td><td>NA</td></tr>
<tr><td></td><td></td><td></td><td></td><td></td><td></td><td></td><td></td><td></td><td></td><td></td><td></td></tr>
<tr><td>T7165</td><td>GCGTCTAATACTGCGAGGGCAGG</td><td>1000</td><td></td><td></td><td></td><td></td><td></td><td></td><td></td><td></td><td></td></tr>
<tr><td>Oligo fwd</td><td>CACCGCGTCTAATACTGCGAGGGC</td><td></td><td></td><td></td><td></td><td></td><td></td><td></td><td></td><td></td><td></td></tr>
<tr><td>Oligo rev</td><td>AAACGCCCTCGCAGTATTAGACGC</td><td></td><td></td><td></td><td></td><td></td><td></td><td></td><td></td><td></td><td></td></tr>
<tr><td>Chromosome</td><td>start</td><td>end</td><td>strand</td><td>MM</td><td>target_seq</td><td>PAM</td><td>alignment</td><td>distance</td><td>position</td><td>gene name</td><td>gene id</td></tr>
<tr><td>GU305914.1</td><td>99097</td><td>99119</td><td>+</td><td>0</td><td>GCGTCTAATACTGCGAGGGC</td><td>AGG</td><td>||||||||[||||||||||||]PAM</td><td>NA</td><td>Intergenic</td><td>NA</td><td>NA</td></tr>
</table>

T7166 CGTCTAATACTGCGAGGGCAGGG 1000
Oligo fwd CACCCGTCTAATACTGCGAGGGCA
Oligo rev AAACTGCCCTCGCAGTATTAGACG
Chromosome start end strand MM target_seq PAM alignment distance position gene name gene id
GU305914.1 99098 99120 + 0 CGTCTAATACTGCGAGGGCA GGG ||||||||[||||||||||||]PAM NA Intergenic NA NA

T7167 GCAGGGAGATCTTCAGCGAGAGG 1000
Oligo fwd CACCGCAGGGAGATCTTCAGCGAG
Oligo rev AAACCTCGCTGAAGATCTCCCTGC
Chromosome start end strand MM target_seq PAM alignment distance position gene name gene id
GU305914.1 99115 99137 + 0 GCAGGGAGATCTTCAGCGAG AGG ||||||||[||||||||||||]PAM NA Intergenic NA NA

T7168 GGGAGATCTTCAGCGAGAGGTGG 1000
Oligo fwd CACCGGGAGATCTTCAGCGAGAGG
Oligo rev AAACCCTCTCGCTGAAGATCTCCC
Chromosome start end strand MM target_seq PAM alignment distance position gene name gene id
GU305914.1 99118 99140 + 0 GGGAGATCTTCAGCGAGAGG TGG ||||||||[||||||||||||]PAM NA Intergenic NA NA

T7169 GGAGATCTTCAGCGAGAGGTGGG 1000
Oligo fwd CACCGGAGATCTTCAGCGAGAGGT
Oligo rev AAACACCTCTCGCTGAAGATCTCC
Chromosome start end strand MM target_seq PAM alignment distance position gene name gene id
GU305914.1 99119 99141 + 0 GGAGATCTTCAGCGAGAGGT GGG ||||||||[||||||||||||]PAM NA Intergenic NA NA

T7170 TCGCGTTAGAATTGCATCGCCGG 1000
Oligo fwd CACCTCGCGTTAGAATTGCATCGC
Oligo rev AAACGCGATGCAATTCTAACGCGA
Chromosome start end strand MM target_seq PAM alignment distance position gene name gene id
GU305914.1 99148 99170 + 0 TCGCGTTAGAATTGCATCGC CGG ||||||||[||||||||||||]PAM NA Intergenic NA NA

T7171 CGCGTTAGAATTGCATCGCCGGG 1000
Oligo fwd CACCCGCGTTAGAATTGCATCGCC
Oligo rev AAACGGCGATGCAATTCTAACGCG
Chromosome start end strand MM target_seq PAM alignment distance position gene name gene id
GU305914.1 99149 99171 + 0 CGCGTTAGAATTGCATCGCC GGG ||||||||[||||||||||||]PAM NA Intergenic NA NA

T7172 GGGTGAAACACCTGCTCCACAGG 1000
Oligo fwd CACCGGGTGAAACACCTGCTCCAC
Oligo rev AAACGTGGAGCAGGTGTTTCACCC
Chromosome start end strand MM target_seq PAM alignment distance position gene name gene id
GU305914.1 99169 99191 + 0 GGGTGAAACACCTGCTCCAC AGG ||||||||[||||||||||||]PAM NA Intergenic NA NA

T7173 GGTGAAACACCTGCTCCACAGGG 1000
Oligo fwd CACCGGTGAAACACCTGCTCCACA
Oligo rev AAACTGTGGAGCAGGTGTTTCACC
Chromosome start end strand MM target_seq PAM alignment distance position gene name gene id
GU305914.1 99170 99192 + 0 GGTGAAACACCTGCTCCACA GGG ||||||||[||||||||||||]PAM NA Intergenic NA NA

T7174 TCGCCCACACGTTCCGCCGCTGG 1000
Oligo fwd CACCTCGCCCACACGTTCCGCCGC
Oligo rev AAACGCGGCGGAACGTGTGGGCGA
Chromosome start end strand MM target_seq PAM alignment distance position gene name gene id
GU305914.1 99193 99215 + 0 TCGCCCACACGTTCCGCCGC TGG ||||||||[||||||||||||]PAM NA Intergenic NA NA

T7175 CGTTCCGCCGCTGGTCTCCGCGG 1000
Oligo fwd CACCCGTTCCGCCGCTGGTCTCCG
Oligo rev AAACCGGAGACCAGCGGCGGAACG
Chromosome start end strand MM target_seq PAM alignment distance position gene name gene id
GU305914.1 99202 99224 + 0 CGTTCCGCCGCTGGTCTCCG CGG ||||||||[||||||||||||]PAM NA Intergenic NA NA

T7176 GCTGGTCTCCGCGGATGTCCAGG 1000
Oligo fwd CACCGCTGGTCTCCGCGGATGTCC
Oligo rev AAACGGACATCCGCGGAGACCAGC
Chromosome start end strand MM target_seq PAM alignment distance position gene name gene id
GU305914.1 99211 99233 + 0 GCTGGTCTCCGCGGATGTCC AGG ||||||||[||||||||||||]PAM NA Intergenic NA NA

T7177 GGAATACCTGCACCGTCTCCTGG 1000
Oligo fwd CACCGGAATACCTGCACCGTCTCC
Oligo rev AAACGGAGACGGTGCAGGTATTCC
Chromosome start end strand MM target_seq PAM alignment distance position gene name gene id
GU305914.1 99232 99254 + 0 GGAATACCTGCACCGTCTCC TGG ||||||||[||||||||||||]PAM NA Intergenic NA NA

T7178 GCGCTAGACACGAGAAGTCTCGG 1000
Oligo fwd CACCGCGCTAGACACGAGAAGTCT
Oligo rev AAACAGACTTCTCGTGTCTAGCGC
Chromosome start end strand MM target_seq PAM alignment distance position gene name gene id
GU305914.1 99271 99293 + 0 GCGCTAGACACGAGAAGTCT CGG ||||||||[||||||||||||]PAM NA Intergenic NA NA

T7179 CGCTAGACACGAGAAGTCTCGGG 1000
Oligo fwd CACCCGCTAGACACGAGAAGTCTC
Oligo rev AAACGAGACTTCTCGTGTCTAGCG
Chromosome start end strand MM target_seq PAM alignment distance position gene name gene id
GU305914.1 99272 99294 + 0 CGCTAGACACGAGAAGTCTC GGG ||||||||[||||||||||||]PAM NA Intergenic NA NA

T7180 GACACGAGAAGTCTCGGGCGCGG 1000
Oligo fwd CACCGACACGAGAAGTCTCGGGCG
Oligo rev AAACCGCCCGAGACTTCTCGTGTC
Chromosome start end strand MM target_seq PAM alignment distance position gene name gene id
GU305914.1 99277 99299 + 0 GACACGAGAAGTCTCGGGCG CGG ||||||||[||||||||||||]PAM NA Intergenic NA NA

| | | | | | | | | | | | |
|---|---|---|---|---|---|---|---|---|---|---|---|
| T7181 | GCCGTATAAACCCCGTTTGACGG | 1000 | | | | | | | | | |
| Oligo fwd | CACCGCCGTATAAACCCCGTTTGA | | | | | | | | | | |
| Oligo rev | AAACTCAAACGGGGTTTATACGGC | | | | | | | | | | |
| Chromosome | start | end | strand | MM | target_seq | PAM | alignment | distance | position | gene name | gene id |
| GU305914.1 | 99299 | 99321 | + | 0 | GCCGTATAAACCCCGTTTGA | CGG | \|\|\|\|\|\|\|\|[\|\|\|\|\|\|\|\|\|\|\|\|]PAM | NA | Intergenic | NA | NA |

| | | | | | | | | | | | |
|---|---|---|---|---|---|---|---|---|---|---|---|
| T7182 | ACCCCGTTTGACGGCAACGCAGG | 1000 | | | | | | | | | |
| Oligo fwd | CACCACCCCGTTTGACGGCAACGC | | | | | | | | | | |
| Oligo rev | AAACGCGTTGCCGTCAAACGGGGT | | | | | | | | | | |
| Chromosome | start | end | strand | MM | target_seq | PAM | alignment | distance | position | gene name | gene id |
| GU305914.1 | 99308 | 99330 | + | 0 | ACCCCGTTTGACGGCAACGC | AGG | \|\|\|\|\|\|\|\|[\|\|\|\|\|\|\|\|\|\|\|\|]PAM | NA | Intergenic | NA | NA |

| | | | | | | | | | | | |
|---|---|---|---|---|---|---|---|---|---|---|---|
| T7183 | GTTTGACGGCAACGCAGGTGCGG | 1000 | | | | | | | | | |
| Oligo fwd | CACCGTTTGACGGCAACGCAGGTG | | | | | | | | | | |
| Oligo rev | AAACCACCTGCGTTGCCGTCAAAC | | | | | | | | | | |
| Chromosome | start | end | strand | MM | target_seq | PAM | alignment | distance | position | gene name | gene id |
| GU305914.1 | 99313 | 99335 | + | 0 | GTTTGACGGCAACGCAGGTG | CGG | \|\|\|\|\|\|\|\|[\|\|\|\|\|\|\|\|\|\|\|\|]PAM | NA | Intergenic | NA | NA |

| | | | | | | | | | | | |
|---|---|---|---|---|---|---|---|---|---|---|---|
| T7184 | GGCAACGCAGGTGCGGCCGACGG | 1000 | | | | | | | | | |
| Oligo fwd | CACCGGCAACGCAGGTGCGGCCGA | | | | | | | | | | |
| Oligo rev | AAACTCGGCCGCACCTGCGTTGCC | | | | | | | | | | |
| Chromosome | start | end | strand | MM | target_seq | PAM | alignment | distance | position | gene name | gene id |
| GU305914.1 | 99320 | 99342 | + | 0 | GGCAACGCAGGTGCGGCCGA | CGG | \|\|\|\|\|\|\|\|[\|\|\|\|\|\|\|\|\|\|\|\|]PAM | NA | Intergenic | NA | NA |

| | | | | | | | | | | | |
|---|---|---|---|---|---|---|---|---|---|---|---|
| T7185 | AACGCAGGTGCGGCCGACGGCGG | 1000 | | | | | | | | | |
| Oligo fwd | CACCAACGCAGGTGCGGCCGACGG | | | | | | | | | | |
| Oligo rev | AAACCCGTCGGCCGCACCTGCGTT | | | | | | | | | | |
| Chromosome | start | end | strand | MM | target_seq | PAM | alignment | distance | position | gene name | gene id |
| GU305914.1 | 99323 | 99345 | + | 0 | AACGCAGGTGCGGCCGACGG | CGG | \|\|\|\|\|\|\|\|[\|\|\|\|\|\|\|\|\|\|\|\|]PAM | NA | Intergenic | NA | NA |

| | | | | | | | | | | | |
|---|---|---|---|---|---|---|---|---|---|---|---|
| T7186 | GCCGACGGCGGTAAACACCGTGG | 1000 | | | | | | | | | |
| Oligo fwd | CACCGCCGACGGCGGTAAACACCG | | | | | | | | | | |
| Oligo rev | AAACCGGTGTTTACCGCCGTCGGC | | | | | | | | | | |
| Chromosome | start | end | strand | MM | target_seq | PAM | alignment | distance | position | gene name | gene id |
| GU305914.1 | 99335 | 99357 | + | 0 | GCCGACGGCGGTAAACACCG | TGG | \|\|\|\|\|\|\|\|[\|\|\|\|\|\|\|\|\|\|\|\|]PAM | NA | Intergenic | NA | NA |

| | | | | | | | | | | | |
|---|---|---|---|---|---|---|---|---|---|---|---|
| T7187 | GCTGCTGCTGCACCTCCTCGCGG | 1000 | | | | | | | | | |
| Oligo fwd | CACCGCTGCTGCTGCACCTCCTCG | | | | | | | | | | |
| Oligo rev | AAACCGAGGAGGTGCAGCAGCAGC | | | | | | | | | | |
| Chromosome | start | end | strand | MM | target_seq | PAM | alignment | distance | position | gene name | gene id |
| GU305914.1 | 99367 | 99389 | + | 0 | GCTGCTGCTGCACCTCCTCG | CGG | \|\|\|\|\|\|\|\|[\|\|\|\|\|\|\|\|\|\|\|\|]PAM | NA | Intergenic | NA | NA |

| | | | | | | | | | | | |
|---|---|---|---|---|---|---|---|---|---|---|---|
| T7188 | GCTGCACCTCCTCGCGGTAGTGG | 1000 | | | | | | | | | |
| Oligo fwd | CACCGCTGCACCTCCTCGCGGTAG | | | | | | | | | | |
| Oligo rev | AAACCTACCGCGAGGAGGTGCAGC | | | | | | | | | | |
| Chromosome | start | end | strand | MM | target_seq | PAM | alignment | distance | position | gene name | gene id |
| GU305914.1 | 99373 | 99395 | + | 0 | GCTGCACCTCCTCGCGGTAG | TGG | \|\|\|\|\|\|\|\|[\|\|\|\|\|\|\|\|\|\|\|\|]PAM | NA | Intergenic | NA | NA |

| | | | | | | | | | | | |
|---|---|---|---|---|---|---|---|---|---|---|---|
| T7189 | CGCGTGTGTCTCAACAGCGCCGG | 1000 | | | | | | | | | |
| Oligo fwd | CACCCGCGTGTGTCTCAACAGCGC | | | | | | | | | | |
| Oligo rev | AAACGCGCTGTTGAGACACACGCG | | | | | | | | | | |
| Chromosome | start | end | strand | MM | target_seq | PAM | alignment | distance | position | gene name | gene id |
| GU305914.1 | 99399 | 99421 | + | 0 | CGCGTGTGTCTCAACAGCGC | CGG | \|\|\|\|\|\|\|\|[\|\|\|\|\|\|\|\|\|\|\|\|]PAM | NA | Intergenic | NA | NA |

| | | | | | | | | | | | |
|---|---|---|---|---|---|---|---|---|---|---|---|
| T7190 | GTGTCTCAACAGCGCCGGCCAGG | 1000 | | | | | | | | | |
| Oligo fwd | CACCGTGTCTCAACAGCGCCGGCC | | | | | | | | | | |
| Oligo rev | AAACGGCCGGCGCTGTTGAGACAC | | | | | | | | | | |
| Chromosome | start | end | strand | MM | target_seq | PAM | alignment | distance | position | gene name | gene id |
| GU305914.1 | 99404 | 99426 | + | 0 | GTGTCTCAACAGCGCCGGCC | AGG | \|\|\|\|\|\|\|\|[\|\|\|\|\|\|\|\|\|\|\|\|]PAM | NA | Intergenic | NA | NA |

| | | | | | | | | | | | |
|---|---|---|---|---|---|---|---|---|---|---|---|
| T7191 | TCCGCCTCGCGAACGCCTGCAGG | 1000 | | | | | | | | | |
| Oligo fwd | CACCTCCGCCTCGCGAACGCCTGC | | | | | | | | | | |
| Oligo rev | AAACGCAGGCGTTCGCGAGGCGGA | | | | | | | | | | |
| Chromosome | start | end | strand | MM | target_seq | PAM | alignment | distance | position | gene name | gene id |
| GU305914.1 | 99430 | 99452 | + | 0 | TCCGCCTCGCGAACGCCTGC | AGG | \|\|\|\|\|\|\|\|[\|\|\|\|\|\|\|\|\|\|\|\|]PAM | NA | Intergenic | NA | NA |

| | | | | | | | | | | | |
|---|---|---|---|---|---|---|---|---|---|---|---|
| T7192 | CCTCCTCGCCCGACGAGCGTTGG | 1000 | | | | | | | | | |
| Oligo fwd | CACCCCTCCTCGCCCGACGAGCGT | | | | | | | | | | |
| Oligo rev | AAACACGCTCGTCGGGCGAGGAGG | | | | | | | | | | |
| Chromosome | start | end | strand | MM | target_seq | PAM | alignment | distance | position | gene name | gene id |
| GU305914.1 | 99540 | 99562 | + | 0 | CCTCCTCGCCCGACGAGCGT | TGG | \|\|\|\|\|\|\|\|[\|\|\|\|\|\|\|\|\|\|\|\|]PAM | NA | Intergenic | NA | NA |
| GU305914.1 | 23540 | 23562 | + | 4 | GCGCCTCGATCGACGAGCGT | CGG | -\|-\|\|\|\|\|[--\|\|\|\|\|\|\|\|\|\|]PAM | NA | Intergenic | NA | NA |
| GU305914.1 | 92507 | 92529 | - | 4 | CCTGCCCGCCCGATAAGCGT | TGG | \|\|\|-\|-\|\|[\|\|\|\|\|--\|\|\|\|\|]PAM | NA | Intergenic | NA | NA |
| GU305914.1 | 88246 | 88268 | + | 4 | CCCGCTCGCCGGACGAGCGG | CGG | \|\|--\|\|\|\|[\|\|-\|\|\|\|\|\|\|\|-]PAM | NA | Intergenic | NA | NA |

| | | | | | | | | | | | |
|---|---|---|---|---|---|---|---|---|---|---|---|
| T7193 | CCGACGAGCGTTGGCAGCGACGG | 1000 | | | | | | | | | |
| Oligo fwd | CACCCCGACGAGCGTTGGCAGCGA | | | | | | | | | | |
| Oligo rev | AAACTCGCTGCCAACGCTCGTCGG | | | | | | | | | | |
| Chromosome | start | end | strand | MM | target_seq | PAM | alignment | distance | position | gene name | gene id |
| GU305914.1 | 99549 | 99571 | + | 0 | CCGACGAGCGTTGGCAGCGA | CGG | \|\|\|\|\|\|\|\|[\|\|\|\|\|\|\|\|\|\|\|\|]PAM | NA | Intergenic | NA | NA |

| | | | | | | | | | | | |
|---|---|---|---|---|---|---|---|---|---|---|---|
| T7194 | ACTGTCCCGACTCGAGCGACAGG | 1000 | | | | | | | | | |
| Oligo fwd | CACCACTGTCCCGACTCGAGCGAC | | | | | | | | | | |
| Oligo rev | AAACGTCGCTCGAGTCGGGACAGT | | | | | | | | | | |
| Chromosome | start | end | strand | MM | target_seq | PAM | alignment | distance | position | gene name | gene id |
| GU305914.1 | 99577 | 99599 | + | 0 | ACTGTCCCGACTCGAGCGAC | AGG | \|\|\|\|\|\|\|\|[\|\|\|\|\|\|\|\|\|\|\|\|]PAM | NA | Intergenic | NA | NA |

| | | |
|---|---|---|
| T7195 | GCGACAGGTGCGCTCGCAGCAGG | 1000 |

<table>
<tr><td>Oligo fwd</td><td>CACCGCGACAGGTGCGCTCGCAGC</td><td></td><td></td><td></td><td></td><td></td><td></td><td></td><td></td><td></td><td></td><td></td></tr>
<tr><td>Oligo rev</td><td>AAACGCTGCGAGCGCACCTGTCGC</td><td></td><td></td><td></td><td></td><td></td><td></td><td></td><td></td><td></td><td></td><td></td></tr>
<tr><td>Chromosome</td><td>start</td><td></td><td>end</td><td>strand</td><td>MM</td><td>target_seq</td><td>PAM</td><td>alignment</td><td>distance</td><td>position</td><td>gene name</td><td>gene id</td></tr>
<tr><td>GU305914.1</td><td></td><td>99592</td><td>99614</td><td>+</td><td>0</td><td>GCGACAGGTGCGCTCGCAGC</td><td>AGG</td><td>||||||||[||||||||||||]PAM</td><td>NA</td><td>Intergenic</td><td>NA</td><td>NA</td></tr>
<tr><td>T7196</td><td>CGACAGGTGCGCTCGCAGCAGGG</td><td></td><td>1000</td><td></td><td></td><td></td><td></td><td></td><td></td><td></td><td></td><td></td></tr>
<tr><td>Oligo fwd</td><td>CACCCGACAGGTGCGCTCGCAGCA</td><td></td><td></td><td></td><td></td><td></td><td></td><td></td><td></td><td></td><td></td><td></td></tr>
<tr><td>Oligo rev</td><td>AAACTGCTGCGAGCGCACCTGTCG</td><td></td><td></td><td></td><td></td><td></td><td></td><td></td><td></td><td></td><td></td><td></td></tr>
<tr><td>Chromosome</td><td>start</td><td></td><td>end</td><td>strand</td><td>MM</td><td>target_seq</td><td>PAM</td><td>alignment</td><td>distance</td><td>position</td><td>gene name</td><td>gene id</td></tr>
<tr><td>GU305914.1</td><td></td><td>99593</td><td>99615</td><td>+</td><td>0</td><td>CGACAGGTGCGCTCGCAGCA</td><td>GGG</td><td>||||||||[||||||||||||]PAM</td><td>NA</td><td>Intergenic</td><td>NA</td><td>NA</td></tr>
<tr><td>T7197</td><td>GACAGGTGCGCTCGCAGCAGGGG</td><td></td><td>1000</td><td></td><td></td><td></td><td></td><td></td><td></td><td></td><td></td><td></td></tr>
<tr><td>Oligo fwd</td><td>CACCGACAGGTGCGCTCGCAGCAG</td><td></td><td></td><td></td><td></td><td></td><td></td><td></td><td></td><td></td><td></td><td></td></tr>
<tr><td>Oligo rev</td><td>AAACCTGCTGCGAGCGCACCTGTC</td><td></td><td></td><td></td><td></td><td></td><td></td><td></td><td></td><td></td><td></td><td></td></tr>
<tr><td>Chromosome</td><td>start</td><td></td><td>end</td><td>strand</td><td>MM</td><td>target_seq</td><td>PAM</td><td>alignment</td><td>distance</td><td>position</td><td>gene name</td><td>gene id</td></tr>
<tr><td>GU305914.1</td><td></td><td>99594</td><td>99616</td><td>+</td><td>0</td><td>GACAGGTGCGCTCGCAGCAG</td><td>GGG</td><td>||||||||[||||||||||||]PAM</td><td>NA</td><td>Intergenic</td><td>NA</td><td>NA</td></tr>
<tr><td>T7198</td><td>GCTCGCAGCAGGGGCCCGAGCGG</td><td></td><td>1000</td><td></td><td></td><td></td><td></td><td></td><td></td><td></td><td></td><td></td></tr>
<tr><td>Oligo fwd</td><td>CACCGCTCGCAGCAGGGGCCCGAG</td><td></td><td></td><td></td><td></td><td></td><td></td><td></td><td></td><td></td><td></td><td></td></tr>
<tr><td>Oligo rev</td><td>AAACCTCGGGCCCCTGCTGCGAGC</td><td></td><td></td><td></td><td></td><td></td><td></td><td></td><td></td><td></td><td></td><td></td></tr>
<tr><td>Chromosome</td><td>start</td><td></td><td>end</td><td>strand</td><td>MM</td><td>target_seq</td><td>PAM</td><td>alignment</td><td>distance</td><td>position</td><td>gene name</td><td>gene id</td></tr>
<tr><td>GU305914.1</td><td></td><td>99603</td><td>99625</td><td>+</td><td>0</td><td>GCTCGCAGCAGGGGCCCGAG</td><td>CGG</td><td>||||||||[||||||||||||]PAM</td><td>NA</td><td>Intergenic</td><td>NA</td><td>NA</td></tr>
<tr><td>T7199</td><td>CAGAAAGTTGCCGTCGCTGTCGG</td><td></td><td>1000</td><td></td><td></td><td></td><td></td><td></td><td></td><td></td><td></td><td></td></tr>
<tr><td>Oligo fwd</td><td>CACCCAGAAAGTTGCCGTCGCTGT</td><td></td><td></td><td></td><td></td><td></td><td></td><td></td><td></td><td></td><td></td><td></td></tr>
<tr><td>Oligo rev</td><td>AAACACAGCGACGGCAACTTTCTG</td><td></td><td></td><td></td><td></td><td></td><td></td><td></td><td></td><td></td><td></td><td></td></tr>
<tr><td>Chromosome</td><td>start</td><td></td><td>end</td><td>strand</td><td>MM</td><td>target_seq</td><td>PAM</td><td>alignment</td><td>distance</td><td>position</td><td>gene name</td><td>gene id</td></tr>
<tr><td>GU305914.1</td><td></td><td>99665</td><td>99687</td><td>+</td><td>0</td><td>CAGAAAGTTGCCGTCGCTGT</td><td>CGG</td><td>||||||||[||||||||||||]PAM</td><td>NA</td><td>Intergenic</td><td>NA</td><td>NA</td></tr>
<tr><td>T7200</td><td>CGTCGCTGTCGGCGCATGCGCGG</td><td></td><td>1000</td><td></td><td></td><td></td><td></td><td></td><td></td><td></td><td></td><td></td></tr>
<tr><td>Oligo fwd</td><td>CACCCGTCGCTGTCGGCGCATGCG</td><td></td><td></td><td></td><td></td><td></td><td></td><td></td><td></td><td></td><td></td><td></td></tr>
<tr><td>Oligo rev</td><td>AAACCGCATGCGCCGACAGCGACG</td><td></td><td></td><td></td><td></td><td></td><td></td><td></td><td></td><td></td><td></td><td></td></tr>
<tr><td>Chromosome</td><td>start</td><td></td><td>end</td><td>strand</td><td>MM</td><td>target_seq</td><td>PAM</td><td>alignment</td><td>distance</td><td>position</td><td>gene name</td><td>gene id</td></tr>
<tr><td>GU305914.1</td><td></td><td>99676</td><td>99698</td><td>+</td><td>0</td><td>CGTCGCTGTCGGCGCATGCG</td><td>CGG</td><td>||||||||[||||||||||||]PAM</td><td>NA</td><td>Intergenic</td><td>NA</td><td>NA</td></tr>
<tr><td>T7201</td><td>GCTGTCGGCGCATGCGCGGTCGG</td><td></td><td>1000</td><td></td><td></td><td></td><td></td><td></td><td></td><td></td><td></td><td></td></tr>
<tr><td>Oligo fwd</td><td>CACCGCTGTCGGCGCATGCGCGGT</td><td></td><td></td><td></td><td></td><td></td><td></td><td></td><td></td><td></td><td></td><td></td></tr>
<tr><td>Oligo rev</td><td>AAACACCGCGCATGCGCCGACAGC</td><td></td><td></td><td></td><td></td><td></td><td></td><td></td><td></td><td></td><td></td><td></td></tr>
<tr><td>Chromosome</td><td>start</td><td></td><td>end</td><td>strand</td><td>MM</td><td>target_seq</td><td>PAM</td><td>alignment</td><td>distance</td><td>position</td><td>gene name</td><td>gene id</td></tr>
<tr><td>GU305914.1</td><td></td><td>99680</td><td>99702</td><td>+</td><td>0</td><td>GCTGTCGGCGCATGCGCGGT</td><td>CGG</td><td>||||||||[||||||||||||]PAM</td><td>NA</td><td>Intergenic</td><td>NA</td><td>NA</td></tr>
<tr><td>T7202</td><td>GCAACACCACCTCCGTCAGCTGG</td><td></td><td>1000</td><td></td><td></td><td></td><td></td><td></td><td></td><td></td><td></td><td></td></tr>
<tr><td>Oligo fwd</td><td>CACCGCAACACCACCTCCGTCAGC</td><td></td><td></td><td></td><td></td><td></td><td></td><td></td><td></td><td></td><td></td><td></td></tr>
<tr><td>Oligo rev</td><td>AAACGCTGACGGAGGTGGTGTTGC</td><td></td><td></td><td></td><td></td><td></td><td></td><td></td><td></td><td></td><td></td><td></td></tr>
<tr><td>Chromosome</td><td>start</td><td></td><td>end</td><td>strand</td><td>MM</td><td>target_seq</td><td>PAM</td><td>alignment</td><td>distance</td><td>position</td><td>gene name</td><td>gene id</td></tr>
<tr><td>GU305914.1</td><td></td><td>99709</td><td>99731</td><td>+</td><td>0</td><td>GCAACACCACCTCCGTCAGC</td><td>TGG</td><td>||||||||[||||||||||||]PAM</td><td>NA</td><td>Intergenic</td><td>NA</td><td>NA</td></tr>
<tr><td>GU305914.1</td><td></td><td>90417</td><td>90439</td><td>-</td><td>4</td><td>GCTACGCCACCTCCTTCATC</td><td>AGG</td><td>||-||-||[||||||-|||-|]PAM</td><td>NA</td><td>Intergenic</td><td>NA</td><td>NA</td></tr>
<tr><td>T7203</td><td>CACCACCTCCGTCAGCTGGTCGG</td><td></td><td>1000</td><td></td><td></td><td></td><td></td><td></td><td></td><td></td><td></td><td></td></tr>
<tr><td>Oligo fwd</td><td>CACCCACCACCTCCGTCAGCTGGT</td><td></td><td></td><td></td><td></td><td></td><td></td><td></td><td></td><td></td><td></td><td></td></tr>
<tr><td>Oligo rev</td><td>AAACACCAGCTGACGGAGGTGGTG</td><td></td><td></td><td></td><td></td><td></td><td></td><td></td><td></td><td></td><td></td><td></td></tr>
<tr><td>Chromosome</td><td>start</td><td></td><td>end</td><td>strand</td><td>MM</td><td>target_seq</td><td>PAM</td><td>alignment</td><td>distance</td><td>position</td><td>gene name</td><td>gene id</td></tr>
<tr><td>GU305914.1</td><td></td><td>99713</td><td>99735</td><td>+</td><td>0</td><td>CACCACCTCCGTCAGCTGGT</td><td>CGG</td><td>||||||||[||||||||||||]PAM</td><td>NA</td><td>Intergenic</td><td>NA</td><td>NA</td></tr>
<tr><td>T7204</td><td>TCGGCGCTCCCGTCTCGCCGCGG</td><td></td><td>1000</td><td></td><td></td><td></td><td></td><td></td><td></td><td></td><td></td><td></td></tr>
<tr><td>Oligo fwd</td><td>CACCTCGGCGCTCCCGTCTCGCCG</td><td></td><td></td><td></td><td></td><td></td><td></td><td></td><td></td><td></td><td></td><td></td></tr>
<tr><td>Oligo rev</td><td>AAACCGGCGAGACGGGAGCGCCGA</td><td></td><td></td><td></td><td></td><td></td><td></td><td></td><td></td><td></td><td></td><td></td></tr>
<tr><td>Chromosome</td><td>start</td><td></td><td>end</td><td>strand</td><td>MM</td><td>target_seq</td><td>PAM</td><td>alignment</td><td>distance</td><td>position</td><td>gene name</td><td>gene id</td></tr>
<tr><td>GU305914.1</td><td></td><td>99732</td><td>99754</td><td>+</td><td>0</td><td>TCGGCGCTCCCGTCTCGCCG</td><td>CGG</td><td>||||||||[||||||||||||]PAM</td><td>NA</td><td>Intergenic</td><td>NA</td><td>NA</td></tr>
<tr><td>T7205</td><td>GCGCTCCCGTCTCGCCGCGGCGG</td><td></td><td>1000</td><td></td><td></td><td></td><td></td><td></td><td></td><td></td><td></td><td></td></tr>
<tr><td>Oligo fwd</td><td>CACCGCGCTCCCGTCTCGCCGCGG</td><td></td><td></td><td></td><td></td><td></td><td></td><td></td><td></td><td></td><td></td><td></td></tr>
<tr><td>Oligo rev</td><td>AAACCCGCGGCGAGACGGGAGCGC</td><td></td><td></td><td></td><td></td><td></td><td></td><td></td><td></td><td></td><td></td><td></td></tr>
<tr><td>Chromosome</td><td>start</td><td></td><td>end</td><td>strand</td><td>MM</td><td>target_seq</td><td>PAM</td><td>alignment</td><td>distance</td><td>position</td><td>gene name</td><td>gene id</td></tr>
<tr><td>GU305914.1</td><td></td><td>99735</td><td>99757</td><td>+</td><td>0</td><td>GCGCTCCCGTCTCGCCGCGG</td><td>CGG</td><td>||||||||[||||||||||||]PAM</td><td>NA</td><td>Intergenic</td><td>NA</td><td>NA</td></tr>
<tr><td>GU305914.1</td><td></td><td>201076</td><td>201098</td><td>+</td><td>4</td><td>GCTCTCTCGTCGCGCGGCGG</td><td>TGG</td><td>||-|||-|[|||-|||-||||]PAM</td><td>NA</td><td>Intergenic</td><td>NA</td><td>NA</td></tr>
<tr><td>T7206</td><td>TCCCGTCTCGCCGCGGCGGACGG</td><td></td><td>1000</td><td></td><td></td><td></td><td></td><td></td><td></td><td></td><td></td><td></td></tr>
<tr><td>Oligo fwd</td><td>CACCTCCCGTCTCGCCGCGGCGGA</td><td></td><td></td><td></td><td></td><td></td><td></td><td></td><td></td><td></td><td></td><td></td></tr>
<tr><td>Oligo rev</td><td>AAACTCCGCCGCGGCGAGACGGGA</td><td></td><td></td><td></td><td></td><td></td><td></td><td></td><td></td><td></td><td></td><td></td></tr>
<tr><td>Chromosome</td><td>start</td><td></td><td>end</td><td>strand</td><td>MM</td><td>target_seq</td><td>PAM</td><td>alignment</td><td>distance</td><td>position</td><td>gene name</td><td>gene id</td></tr>
<tr><td>GU305914.1</td><td></td><td>99739</td><td>99761</td><td>+</td><td>0</td><td>TCCCGTCTCGCCGCGGCGGA</td><td>CGG</td><td>||||||||[||||||||||||]PAM</td><td>NA</td><td>Intergenic</td><td>NA</td><td>NA</td></tr>
<tr><td>T7207</td><td>CTCGCCGCGGCGGACGGCCGTGG</td><td></td><td>1000</td><td></td><td></td><td></td><td></td><td></td><td></td><td></td><td></td><td></td></tr>
<tr><td>Oligo fwd</td><td>CACCCTCGCCGCGGCGGACGGCCG</td><td></td><td></td><td></td><td></td><td></td><td></td><td></td><td></td><td></td><td></td><td></td></tr>
<tr><td>Oligo rev</td><td>AAACCGGCCGTCCGCCGCGGCGAG</td><td></td><td></td><td></td><td></td><td></td><td></td><td></td><td></td><td></td><td></td><td></td></tr>
<tr><td>Chromosome</td><td>start</td><td></td><td>end</td><td>strand</td><td>MM</td><td>target_seq</td><td>PAM</td><td>alignment</td><td>distance</td><td>position</td><td>gene name</td><td>gene id</td></tr>
<tr><td>GU305914.1</td><td></td><td>99745</td><td>99767</td><td>+</td><td>0</td><td>CTCGCCGCGGCGGACGGCCG</td><td>TGG</td><td>||||||||[||||||||||||]PAM</td><td>NA</td><td>Intergenic</td><td>NA</td><td>NA</td></tr>
<tr><td>T7208</td><td>GCGGACGGCCGTGGTGCGTCAGG</td><td></td><td>1000</td><td></td><td></td><td></td><td></td><td></td><td></td><td></td><td></td><td></td></tr>
<tr><td>Oligo fwd</td><td>CACCGCGGACGGCCGTGGTGCGTC</td><td></td><td></td><td></td><td></td><td></td><td></td><td></td><td></td><td></td><td></td><td></td></tr>
<tr><td>Oligo rev</td><td>AAACGACGCACCACGGCCGTCCGC</td><td></td><td></td><td></td><td></td><td></td><td></td><td></td><td></td><td></td><td></td><td></td></tr>
<tr><td>Chromosome</td><td>start</td><td></td><td>end</td><td>strand</td><td>MM</td><td>target_seq</td><td>PAM</td><td>alignment</td><td>distance</td><td>position</td><td>gene name</td><td>gene id</td></tr>
<tr><td>GU305914.1</td><td></td><td>99754</td><td>99776</td><td>+</td><td>0</td><td>GCGGACGGCCGTGGTGCGTC</td><td>AGG</td><td>||||||||[||||||||||||]PAM</td><td>NA</td><td>Intergenic</td><td>NA</td><td>NA</td></tr>
<tr><td>T7209</td><td>GGCCGTGGTGCGTCAGGTTCTGG</td><td></td><td>1000</td><td></td><td></td><td></td><td></td><td></td><td></td><td></td><td></td><td></td></tr>
<tr><td>Oligo fwd</td><td>CACCGGCCGTGGTGCGTCAGGTTC</td><td></td><td></td><td></td><td></td><td></td><td></td><td></td><td></td><td></td><td></td><td></td></tr>
<tr><td>Oligo rev</td><td>AAACGAACCTGACGCACCACGGCC</td><td></td><td></td><td></td><td></td><td></td><td></td><td></td><td></td><td></td><td></td><td></td></tr>
<tr><td>Chromosome</td><td>start</td><td></td><td>end</td><td>strand</td><td>MM</td><td>target_seq</td><td>PAM</td><td>alignment</td><td>distance</td><td>position</td><td>gene name</td><td>gene id</td></tr>
</table>

| | | | | | | | | | | | |
|---|---|---|---|---|---|---|---|---|---|---|---|
| GU305914.1 | 99760 | 99782 | + | 0 | GGCCGTGGTGCGTCAGGTTC | TGG | \|\|\|\|\|\|\|\|[\|\|\|\|\|\|\|\|\|\|\|\|]PAM | NA | Intergenic | NA | NA |
| T7210 | GCCGTGGTGCGTCAGGTTCTGGG | 1000 | | | | | | | | | |
| Oligo fwd | CACCGCCGTGGTGCGTCAGGTTCT | | | | | | | | | | |
| Oligo rev | AAACAGAACCTGACGCACCACGGC | | | | | | | | | | |
| Chromosome | start | end | strand | MM | target_seq | PAM | alignment | distance | position | gene name | gene id |
| GU305914.1 | 99761 | 99783 | + | 0 | GCCGTGGTGCGTCAGGTTCT | GGG | \|\|\|\|\|\|\|\|[\|\|\|\|\|\|\|\|\|\|\|\|]PAM | NA | Intergenic | NA | NA |
| T7211 | CCGTGGTGCGTCAGGTTCTGGGG | 1000 | | | | | | | | | |
| Oligo fwd | CACCCCGTGGTGCGTCAGGTTCTG | | | | | | | | | | |
| Oligo rev | AAACCAGAACCTGACGCACCACGG | | | | | | | | | | |
| Chromosome | start | end | strand | MM | target_seq | PAM | alignment | distance | position | gene name | gene id |
| GU305914.1 | 99762 | 99784 | + | 0 | CCGTGGTGCGTCAGGTTCTG | GGG | \|\|\|\|\|\|\|\|[\|\|\|\|\|\|\|\|\|\|\|\|]PAM | NA | Intergenic | NA | NA |
| T7212 | CGTGGTGCGTCAGGTTCTGGGGG | 1000 | | | | | | | | | |
| Oligo fwd | CACCCGTGGTGCGTCAGGTTCTGG | | | | | | | | | | |
| Oligo rev | AAACCCAGAACCTGACGCACCACG | | | | | | | | | | |
| Chromosome | start | end | strand | MM | target_seq | PAM | alignment | distance | position | gene name | gene id |
| GU305914.1 | 99763 | 99785 | + | 0 | CGTGGTGCGTCAGGTTCTGG | GGG | \|\|\|\|\|\|\|\|[\|\|\|\|\|\|\|\|\|\|\|\|]PAM | NA | Intergenic | NA | NA |
| T7213 | TGCTGCAGCACGAATTCCGCCGG | 1000 | | | | | | | | | |
| Oligo fwd | CACCTGCTGCAGCACGAATTCCGC | | | | | | | | | | |
| Oligo rev | AAACGCGGAATTCGTGCTGCAGCA | | | | | | | | | | |
| Chromosome | start | end | strand | MM | target_seq | PAM | alignment | distance | position | gene name | gene id |
| GU305914.1 | 99792 | 99814 | + | 0 | TGCTGCAGCACGAATTCCGC | CGG | \|\|\|\|\|\|\|\|[\|\|\|\|\|\|\|\|\|\|\|\|]PAM | NA | Intergenic | NA | NA |
| T7214 | GCAGCACGAATTCCGCCGGACGG | 1000 | | | | | | | | | |
| Oligo fwd | CACCGCAGCACGAATTCCGCCGGA | | | | | | | | | | |
| Oligo rev | AAACTCCGGCGGAATTCGTGCTGC | | | | | | | | | | |
| Chromosome | start | end | strand | MM | target_seq | PAM | alignment | distance | position | gene name | gene id |
| GU305914.1 | 99796 | 99818 | + | 0 | GCAGCACGAATTCCGCCGGA | CGG | \|\|\|\|\|\|\|\|[\|\|\|\|\|\|\|\|\|\|\|\|]PAM | NA | Intergenic | NA | NA |
| T7215 | CGCCGGACGGTCGCGATACCCGG | 1000 | | | | | | | | | |
| Oligo fwd | CACCCGCCGGACGGTCGCGATACC | | | | | | | | | | |
| Oligo rev | AAACGGTATCGCGACCGTCCGGCG | | | | | | | | | | |
| Chromosome | start | end | strand | MM | target_seq | PAM | alignment | distance | position | gene name | gene id |
| GU305914.1 | 99809 | 99831 | + | 0 | CGCCGGACGGTCGCGATACC | CGG | \|\|\|\|\|\|\|\|[\|\|\|\|\|\|\|\|\|\|\|\|]PAM | NA | Intergenic | NA | NA |
| T7216 | CGGACGGTCGCGATACCCGGCGG | 1000 | | | | | | | | | |
| Oligo fwd | CACCCGGACGGTCGCGATACCCGG | | | | | | | | | | |
| Oligo rev | AAACCCGGGTATCGCGACCGTCCG | | | | | | | | | | |
| Chromosome | start | end | strand | MM | target_seq | PAM | alignment | distance | position | gene name | gene id |
| GU305914.1 | 99812 | 99834 | + | 0 | CGGACGGTCGCGATACCCGG | CGG | \|\|\|\|\|\|\|\|[\|\|\|\|\|\|\|\|\|\|\|\|]PAM | NA | Intergenic | NA | NA |
| T7217 | GGACGGTCGCGATACCCGGCGGG | 1000 | | | | | | | | | |
| Oligo fwd | CACCGGACGGTCGCGATACCCGGC | | | | | | | | | | |
| Oligo rev | AAACGCCGGGTATCGCGACCGTCC | | | | | | | | | | |
| Chromosome | start | end | strand | MM | target_seq | PAM | alignment | distance | position | gene name | gene id |
| GU305914.1 | 99813 | 99835 | + | 0 | GGACGGTCGCGATACCCGGC | GGG | \|\|\|\|\|\|\|\|[\|\|\|\|\|\|\|\|\|\|\|\|]PAM | NA | Intergenic | NA | NA |
| T7218 | GACGGTCGCGATACCCGGCGGGG | 1000 | | | | | | | | | |
| Oligo fwd | CACCGACGGTCGCGATACCCGGCG | | | | | | | | | | |
| Oligo rev | AAACCGCCGGGTATCGCGACCGTC | | | | | | | | | | |
| Chromosome | start | end | strand | MM | target_seq | PAM | alignment | distance | position | gene name | gene id |
| GU305914.1 | 99814 | 99836 | + | 0 | GACGGTCGCGATACCCGGCG | GGG | \|\|\|\|\|\|\|\|[\|\|\|\|\|\|\|\|\|\|\|\|]PAM | NA | Intergenic | NA | NA |
| T7219 | CCCGGCGGGGACGATGAAGATGG | 1000 | | | | | | | | | |
| Oligo fwd | CACCCCCGGCGGGGACGATGAAGA | | | | | | | | | | |
| Oligo rev | AAACTCTTCATCGTCCCCGCCGGG | | | | | | | | | | |
| Chromosome | start | end | strand | MM | target_seq | PAM | alignment | distance | position | gene name | gene id |
| GU305914.1 | 99827 | 99849 | + | 0 | CCCGGCGGGGACGATGAAGA | TGG | \|\|\|\|\|\|\|\|[\|\|\|\|\|\|\|\|\|\|\|\|]PAM | NA | Intergenic | NA | NA |
| T7220 | CCGGCGGGGACGATGAAGATGGG | 1000 | | | | | | | | | |
| Oligo fwd | CACCCCGGCGGGGACGATGAAGAT | | | | | | | | | | |
| Oligo rev | AAACATCTTCATCGTCCCCGCCGG | | | | | | | | | | |
| Chromosome | start | end | strand | MM | target_seq | PAM | alignment | distance | position | gene name | gene id |
| GU305914.1 | 99828 | 99850 | + | 0 | CCGGCGGGGACGATGAAGAT | GGG | \|\|\|\|\|\|\|\|[\|\|\|\|\|\|\|\|\|\|\|\|]PAM | NA | Intergenic | NA | NA |
| T7221 | CGGCGGGGACGATGAAGATGGGG | 1000 | | | | | | | | | |
| Oligo fwd | CACCCGGCGGGGACGATGAAGATG | | | | | | | | | | |
| Oligo rev | AAACCATCTTCATCGTCCCCGCCG | | | | | | | | | | |
| Chromosome | start | end | strand | MM | target_seq | PAM | alignment | distance | position | gene name | gene id |
| GU305914.1 | 99829 | 99851 | + | 0 | CGGCGGGGACGATGAAGATG | GGG | \|\|\|\|\|\|\|\|[\|\|\|\|\|\|\|\|\|\|\|\|]PAM | NA | Intergenic | NA | NA |
| GU305914.1 | 130649 | 130671 | - | 4 | CGCCGGTGAGGACGAAGATG | AGG | \|\|-\|\|\|-\|[\|-\|\|-\|\|\|\|\|\|\|]PAM | NA | Intergenic | NA | NA |
| T7222 | CGATGAAGATGGGGTTTAATCGG | 1000 | | | | | | | | | |
| Oligo fwd | CACCCGATGAAGATGGGGTTTAAT | | | | | | | | | | |
| Oligo rev | AAACATTAAACCCCATCTTCATCG | | | | | | | | | | |
| Chromosome | start | end | strand | MM | target_seq | PAM | alignment | distance | position | gene name | gene id |
| GU305914.1 | 99838 | 99860 | + | 0 | CGATGAAGATGGGGTTTAAT | CGG | \|\|\|\|\|\|\|\|[\|\|\|\|\|\|\|\|\|\|\|\|]PAM | NA | Intergenic | NA | NA |
| T7223 | GCTGTGCAGAATCAGCCCGCTGG | 1000 | | | | | | | | | |
| Oligo fwd | CACCGCTGTGCAGAATCAGCCCGC | | | | | | | | | | |
| Oligo rev | AAACGCGGGCTGATTCTGCACAGC | | | | | | | | | | |
| Chromosome | start | end | strand | MM | target_seq | PAM | alignment | distance | position | gene name | gene id |
| GU305914.1 | 99860 | 99882 | + | 0 | GCTGTGCAGAATCAGCCCGC | TGG | \|\|\|\|\|\|\|\|[\|\|\|\|\|\|\|\|\|\|\|\|]PAM | NA | Intergenic | NA | NA |
| T7224 | TCGTCCAGCTTGTACATCAGCGG | 1000 | | | | | | | | | |
| Oligo fwd | CACCTCGTCCAGCTTGTACATCAG | | | | | | | | | | |

Oligo rev AAACCTGATGTACAAGCTGGACGA
Chromosome start end strand MM target_seq PAM alignment distance position gene name gene id
GU305914.1 99885 99907 + 0 TCGTCCAGCTTGTACATCAG CGG ||||||||[||||||||||||]PAM NA Intergenic NA NA

T7225 GCTTGTACATCAGCGGCATGCGG 1000
Oligo fwd CACCGCTTGTACATCAGCGGCATG
Oligo rev AAACCATGCCGCTGATGTACAAGC
Chromosome start end strand MM target_seq PAM alignment distance position gene name gene id
GU305914.1 99892 99914 + 0 GCTTGTACATCAGCGGCATG CGG ||||||||[||||||||||||]PAM NA Intergenic NA NA

T7226 GCATGCGGATCGAGCGCCCGTGG 1000
Oligo fwd CACCGCATGCGGATCGAGCGCCCG
Oligo rev AAACCGGGCGCTCGATCCGCATGC
Chromosome start end strand MM target_seq PAM alignment distance position gene name gene id
GU305914.1 99907 99929 + 0 GCATGCGGATCGAGCGCCCG TGG ||||||||[||||||||||||]PAM NA Intergenic NA NA

T7227 CCCCGTGTCGAAACACTCGCCGG 1000
Oligo fwd CACCCCCCGTGTCGAAACACTCGC
Oligo rev AAACGCGAGTGTTTCGACACGGGG
Chromosome start end strand MM target_seq PAM alignment distance position gene name gene id
GU305914.1 99938 99960 + 0 CCCCGTGTCGAAACACTCGC CGG ||||||||[||||||||||||]PAM NA Intergenic NA NA

T7228 CCCGTGTCGAAACACTCGCCGGG 1000
Oligo fwd CACCCCCGTGTCGAAACACTCGCC
Oligo rev AAACGGCGAGTGTTTCGACACGGG
Chromosome start end strand MM target_seq PAM alignment distance position gene name gene id
GU305914.1 99939 99961 + 0 CCCGTGTCGAAACACTCGCC GGG ||||||||[||||||||||||]PAM NA Intergenic NA NA

T7229 CCGTGTCGAAACACTCGCCGGGG 1000
Oligo fwd CACCCCGTGTCGAAACACTCGCCG
Oligo rev AAACCGGCGAGTGTTTCGACACGG
Chromosome start end strand MM target_seq PAM alignment distance position gene name gene id
GU305914.1 99940 99962 + 0 CCGTGTCGAAACACTCGCCG GGG ||||||||[||||||||||||]PAM NA Intergenic NA NA

T7230 ACACTCGCCGGGGTGCGAGATGG 1000
Oligo fwd CACCACACTCGCCGGGGTGCGAGA
Oligo rev AAACTCTCGCACCCCGGCGAGTGT
Chromosome start end strand MM target_seq PAM alignment distance position gene name gene id
GU305914.1 99950 99972 + 0 ACACTCGCCGGGGTGCGAGA TGG ||||||||[||||||||||||]PAM NA Intergenic NA NA

T7231 TGGCGTTGAGTTTGCACACCAGG 1000
Oligo fwd CACCTGGCGTTGAGTTTGCACACC
Oligo rev AAACGGTGTGCAAACTCAACGCCA
Chromosome start end strand MM target_seq PAM alignment distance position gene name gene id
GU305914.1 99970 99992 + 0 TGGCGTTGAGTTTGCACACC AGG ||||||||[||||||||||||]PAM NA Intergenic NA NA

T7232 TGCACACCAGGTCTCGATCGAGG 1000
Oligo fwd CACCTGCACACCAGGTCTCGATCG
Oligo rev AAACCGATCGAGACCTGGTGTGCA
Chromosome start end strand MM target_seq PAM alignment distance position gene name gene id
GU305914.1 99982 100004 + 0 TGCACACCAGGTCTCGATCG AGG ||||||||[||||||||||||]PAM NA Intergenic NA NA

T7233 CGAGGCACAGAGTGTGATCGAGG 1000
Oligo fwd CACCCGAGGCACAGAGTGTGATCG
Oligo rev AAACCGATCACACTCTGTGCCTCG
Chromosome start end strand MM target_seq PAM alignment distance position gene name gene id
GU305914.1 100000 100022 + 0 CGAGGCACAGAGTGTGATCG AGG ||||||||[||||||||||||]PAM NA Intergenic NA NA

T7234 CAGAGTGTGATCGAGGATGCCGG 1000
Oligo fwd CACCCAGAGTGTGATCGAGGATGC
Oligo rev AAACGCATCCTCGATCACACTCTG
Chromosome start end strand MM target_seq PAM alignment distance position gene name gene id
GU305914.1 100007 100029 + 0 CAGAGTGTGATCGAGGATGC CGG ||||||||[||||||||||||]PAM NA Intergenic NA NA

T7235 TGTGATCGAGGATGCCGGCCAGG 1000
Oligo fwd CACCTGTGATCGAGGATGCCGGCC
Oligo rev AAACGGCCGGCATCCTCGATCACA
Chromosome start end strand MM target_seq PAM alignment distance position gene name gene id
GU305914.1 100012 100034 + 0 TGTGATCGAGGATGCCGGCC AGG ||||||||[||||||||||||]PAM NA Intergenic NA NA

T7236 GCCCCGATCGCCGCCGTGCGAGG 1000
Oligo fwd CACCGCCCCGATCGCCGCCGTGCG
Oligo rev AAACCGCACGGCGGCGATCGGGGC
Chromosome start end strand MM target_seq PAM alignment distance position gene name gene id
GU305914.1 100050 100072 + 0 GCCCCGATCGCCGCCGTGCG AGG ||||||||[||||||||||||]PAM NA Intergenic NA NA

T7237 CCCCGATCGCCGCCGTGCGAGGG 1000
Oligo fwd CACCCCCCGATCGCCGCCGTGCGA
Oligo rev AAACTCGCACGGCGGCGATCGGGG
Chromosome start end strand MM target_seq PAM alignment distance position gene name gene id
GU305914.1 100051 100073 + 0 CCCCGATCGCCGCCGTGCGA GGG ||||||||[||||||||||||]PAM NA Intergenic NA NA
GU305914.1 167216 167238 - 4 CGCCGGTCGCCGGCGAGCGA CGG |-|||-||[||||-||-||||]PAM NA Intergenic NA NA

T7238 CCCGATCGCCGCCGTGCGAGGGG 1000
Oligo fwd CACCCCCGATCGCCGCCGTGCGAG
Oligo rev AAACCTCGCACGGCGGCGATCGGG
Chromosome start end strand MM target_seq PAM alignment distance position gene name gene id
GU305914.1 100052 100074 + 0 CCCGATCGCCGCCGTGCGAG GGG ||||||||[||||||||||||]PAM NA Intergenic NA NA

<table>
<tr><td>T7239</td><td>CCGATCGCCGCCGTGCGAGGGGG</td><td>1000</td><td></td><td></td><td></td><td></td><td></td><td></td><td></td><td></td><td></td></tr>
<tr><td>Oligo fwd</td><td>CACCCCGATCGCCGCCGTGCGAGG</td><td></td><td></td><td></td><td></td><td></td><td></td><td></td><td></td><td></td><td></td></tr>
<tr><td>Oligo rev</td><td>AAACCCTCGCACGGCGGCGATCGG</td><td></td><td></td><td></td><td></td><td></td><td></td><td></td><td></td><td></td><td></td></tr>
<tr><td>Chromosome</td><td>start</td><td>end</td><td>strand</td><td>MM</td><td>target_seq</td><td>PAM</td><td>alignment</td><td>distance</td><td>position</td><td>gene name</td><td>gene id</td></tr>
<tr><td>GU305914.1</td><td>100053</td><td>100075</td><td>+</td><td>0</td><td>CCGATCGCCGCCGTGCGAGG</td><td>GGG</td><td>||||||||[||||||||||||]PAM</td><td>NA</td><td>Intergenic</td><td>NA</td><td>NA</td></tr>
<tr><td>T7240</td><td>CGATCGCCGCCGTGCGAGGGGGG</td><td>1000</td><td></td><td></td><td></td><td></td><td></td><td></td><td></td><td></td><td></td></tr>
<tr><td>Oligo fwd</td><td>CACCCGATCGCCGCCGTGCGAGGG</td><td></td><td></td><td></td><td></td><td></td><td></td><td></td><td></td><td></td><td></td></tr>
<tr><td>Oligo rev</td><td>AAACCCCTCGCACGGCGGCGATCG</td><td></td><td></td><td></td><td></td><td></td><td></td><td></td><td></td><td></td><td></td></tr>
<tr><td>Chromosome</td><td>start</td><td>end</td><td>strand</td><td>MM</td><td>target_seq</td><td>PAM</td><td>alignment</td><td>distance</td><td>position</td><td>gene name</td><td>gene id</td></tr>
<tr><td>GU305914.1</td><td>100054</td><td>100076</td><td>+</td><td>0</td><td>CGATCGCCGCCGTGCGAGGG</td><td>GGG</td><td>||||||||[||||||||||||]PAM</td><td>NA</td><td>Intergenic</td><td>NA</td><td>NA</td></tr>
<tr><td>T7241</td><td>GCCGCCGTGCGAGGGGGGAAAGG</td><td>1000</td><td></td><td></td><td></td><td></td><td></td><td></td><td></td><td></td><td></td></tr>
<tr><td>Oligo fwd</td><td>CACCGCCGCCGTGCGAGGGGGGAA</td><td></td><td></td><td></td><td></td><td></td><td></td><td></td><td></td><td></td><td></td></tr>
<tr><td>Oligo rev</td><td>AAACTTCCCCCCTCGCACGGCGGC</td><td></td><td></td><td></td><td></td><td></td><td></td><td></td><td></td><td></td><td></td></tr>
<tr><td>Chromosome</td><td>start</td><td>end</td><td>strand</td><td>MM</td><td>target_seq</td><td>PAM</td><td>alignment</td><td>distance</td><td>position</td><td>gene name</td><td>gene id</td></tr>
<tr><td>GU305914.1</td><td>100059</td><td>100081</td><td>+</td><td>0</td><td>GCCGCCGTGCGAGGGGGGAA</td><td>AGG</td><td>||||||||[||||||||||||]PAM</td><td>NA</td><td>Intergenic</td><td>NA</td><td>NA</td></tr>
<tr><td>T7242</td><td>CCGCCGTGCGAGGGGGGAAAGGG</td><td>1000</td><td></td><td></td><td></td><td></td><td></td><td></td><td></td><td></td><td></td></tr>
<tr><td>Oligo fwd</td><td>CACCCCGCCGTGCGAGGGGGGAAA</td><td></td><td></td><td></td><td></td><td></td><td></td><td></td><td></td><td></td><td></td></tr>
<tr><td>Oligo rev</td><td>AAACTTTCCCCCCTCGCACGGCGG</td><td></td><td></td><td></td><td></td><td></td><td></td><td></td><td></td><td></td><td></td></tr>
<tr><td>Chromosome</td><td>start</td><td>end</td><td>strand</td><td>MM</td><td>target_seq</td><td>PAM</td><td>alignment</td><td>distance</td><td>position</td><td>gene name</td><td>gene id</td></tr>
<tr><td>GU305914.1</td><td>100060</td><td>100082</td><td>+</td><td>0</td><td>CCGCCGTGCGAGGGGGGAAA</td><td>GGG</td><td>||||||||[||||||||||||]PAM</td><td>NA</td><td>Intergenic</td><td>NA</td><td>NA</td></tr>
<tr><td>T7243</td><td>GGAAAGGGATGATCACCCTCAGG</td><td>1000</td><td></td><td></td><td></td><td></td><td></td><td></td><td></td><td></td><td></td></tr>
<tr><td>Oligo fwd</td><td>CACCGGAAAGGGATGATCACCCTC</td><td></td><td></td><td></td><td></td><td></td><td></td><td></td><td></td><td></td><td></td></tr>
<tr><td>Oligo rev</td><td>AAACGAGGGTGATCATCCCTTTCC</td><td></td><td></td><td></td><td></td><td></td><td></td><td></td><td></td><td></td><td></td></tr>
<tr><td>Chromosome</td><td>start</td><td>end</td><td>strand</td><td>MM</td><td>target_seq</td><td>PAM</td><td>alignment</td><td>distance</td><td>position</td><td>gene name</td><td>gene id</td></tr>
<tr><td>GU305914.1</td><td>100075</td><td>100097</td><td>+</td><td>0</td><td>GGAAAGGGATGATCACCCTC</td><td>AGG</td><td>||||||||[||||||||||||]PAM</td><td>NA</td><td>Intergenic</td><td>NA</td><td>NA</td></tr>
<tr><td>T7244</td><td>CTTGCGCCGACAGACACAGAAGG</td><td>1000</td><td></td><td></td><td></td><td></td><td></td><td></td><td></td><td></td><td></td></tr>
<tr><td>Oligo fwd</td><td>CACCCTTGCGCCGACAGACACAGA</td><td></td><td></td><td></td><td></td><td></td><td></td><td></td><td></td><td></td><td></td></tr>
<tr><td>Oligo rev</td><td>AAACTCTGTGTCTGTCGGCGCAAG</td><td></td><td></td><td></td><td></td><td></td><td></td><td></td><td></td><td></td><td></td></tr>
<tr><td>Chromosome</td><td>start</td><td>end</td><td>strand</td><td>MM</td><td>target_seq</td><td>PAM</td><td>alignment</td><td>distance</td><td>position</td><td>gene name</td><td>gene id</td></tr>
<tr><td>GU305914.1</td><td>100103</td><td>100125</td><td>+</td><td>0</td><td>CTTGCGCCGACAGACACAGA</td><td>AGG</td><td>||||||||[||||||||||||]PAM</td><td>NA</td><td>Intergenic</td><td>NA</td><td>NA</td></tr>
<tr><td>T7245</td><td>TTGCGCCGACAGACACAGAAGGG</td><td>1000</td><td></td><td></td><td></td><td></td><td></td><td></td><td></td><td></td><td></td></tr>
<tr><td>Oligo fwd</td><td>CACCTTGCGCCGACAGACACAGAA</td><td></td><td></td><td></td><td></td><td></td><td></td><td></td><td></td><td></td><td></td></tr>
<tr><td>Oligo rev</td><td>AAACTTCTGTGTCTGTCGGCGCAA</td><td></td><td></td><td></td><td></td><td></td><td></td><td></td><td></td><td></td><td></td></tr>
<tr><td>Chromosome</td><td>start</td><td>end</td><td>strand</td><td>MM</td><td>target_seq</td><td>PAM</td><td>alignment</td><td>distance</td><td>position</td><td>gene name</td><td>gene id</td></tr>
<tr><td>GU305914.1</td><td>100104</td><td>100126</td><td>+</td><td>0</td><td>TTGCGCCGACAGACACAGAA</td><td>GGG</td><td>||||||||[||||||||||||]PAM</td><td>NA</td><td>Intergenic</td><td>NA</td><td>NA</td></tr>
<tr><td>T7246</td><td>CGCCGACAGACACAGAAGGGCGG</td><td>1000</td><td></td><td></td><td></td><td></td><td></td><td></td><td></td><td></td><td></td></tr>
<tr><td>Oligo fwd</td><td>CACCCGCCGACAGACACAGAAGGG</td><td></td><td></td><td></td><td></td><td></td><td></td><td></td><td></td><td></td><td></td></tr>
<tr><td>Oligo rev</td><td>AAACCCCTTCTGTGTCTGTCGGCG</td><td></td><td></td><td></td><td></td><td></td><td></td><td></td><td></td><td></td><td></td></tr>
<tr><td>Chromosome</td><td>start</td><td>end</td><td>strand</td><td>MM</td><td>target_seq</td><td>PAM</td><td>alignment</td><td>distance</td><td>position</td><td>gene name</td><td>gene id</td></tr>
<tr><td>GU305914.1</td><td>100107</td><td>100129</td><td>+</td><td>0</td><td>CGCCGACAGACACAGAAGGG</td><td>CGG</td><td>||||||||[||||||||||||]PAM</td><td>NA</td><td>Intergenic</td><td>NA</td><td>NA</td></tr>
<tr><td>T7247</td><td>GCCGACAGACACAGAAGGGCGGG</td><td>1000</td><td></td><td></td><td></td><td></td><td></td><td></td><td></td><td></td><td></td></tr>
<tr><td>Oligo fwd</td><td>CACCGCCGACAGACACAGAAGGGC</td><td></td><td></td><td></td><td></td><td></td><td></td><td></td><td></td><td></td><td></td></tr>
<tr><td>Oligo rev</td><td>AAACGCCCTTCTGTGTCTGTCGGC</td><td></td><td></td><td></td><td></td><td></td><td></td><td></td><td></td><td></td><td></td></tr>
<tr><td>Chromosome</td><td>start</td><td>end</td><td>strand</td><td>MM</td><td>target_seq</td><td>PAM</td><td>alignment</td><td>distance</td><td>position</td><td>gene name</td><td>gene id</td></tr>
<tr><td>GU305914.1</td><td>100108</td><td>100130</td><td>+</td><td>0</td><td>GCCGACAGACACAGAAGGGC</td><td>GGG</td><td>||||||||[||||||||||||]PAM</td><td>NA</td><td>Intergenic</td><td>NA</td><td>NA</td></tr>
<tr><td>T7248</td><td>ACAGAAGGGCGGGCGCCGAACGG</td><td>1000</td><td></td><td></td><td></td><td></td><td></td><td></td><td></td><td></td><td></td></tr>
<tr><td>Oligo fwd</td><td>CACCACAGAAGGGCGGGCGCCGAA</td><td></td><td></td><td></td><td></td><td></td><td></td><td></td><td></td><td></td><td></td></tr>
<tr><td>Oligo rev</td><td>AAACTTCGGCGCCCGCCCTTCTGT</td><td></td><td></td><td></td><td></td><td></td><td></td><td></td><td></td><td></td><td></td></tr>
<tr><td>Chromosome</td><td>start</td><td>end</td><td>strand</td><td>MM</td><td>target_seq</td><td>PAM</td><td>alignment</td><td>distance</td><td>position</td><td>gene name</td><td>gene id</td></tr>
<tr><td>GU305914.1</td><td>100118</td><td>100140</td><td>+</td><td>0</td><td>ACAGAAGGGCGGGCGCCGAA</td><td>CGG</td><td>||||||||[||||||||||||]PAM</td><td>NA</td><td>Intergenic</td><td>NA</td><td>NA</td></tr>
<tr><td>T7249</td><td>GCCGTAGTCGACCATCGCCGCGG</td><td>1000</td><td></td><td></td><td></td><td></td><td></td><td></td><td></td><td></td><td></td></tr>
<tr><td>Oligo fwd</td><td>CACCGCCGTAGTCGACCATCGCCG</td><td></td><td></td><td></td><td></td><td></td><td></td><td></td><td></td><td></td><td></td></tr>
<tr><td>Oligo rev</td><td>AAACCGGCGATGGTCGACTACGGC</td><td></td><td></td><td></td><td></td><td></td><td></td><td></td><td></td><td></td><td></td></tr>
<tr><td>Chromosome</td><td>start</td><td>end</td><td>strand</td><td>MM</td><td>target_seq</td><td>PAM</td><td>alignment</td><td>distance</td><td>position</td><td>gene name</td><td>gene id</td></tr>
<tr><td>GU305914.1</td><td>100145</td><td>100167</td><td>+</td><td>0</td><td>GCCGTAGTCGACCATCGCCG</td><td>CGG</td><td>||||||||[||||||||||||]PAM</td><td>NA</td><td>Intergenic</td><td>NA</td><td>NA</td></tr>
<tr><td>T7250</td><td>GACCATCGCCGCGGCGTGCTCGG</td><td>1000</td><td></td><td></td><td></td><td></td><td></td><td></td><td></td><td></td><td></td></tr>
<tr><td>Oligo fwd</td><td>CACCGACCATCGCCGCGGCGTGCT</td><td></td><td></td><td></td><td></td><td></td><td></td><td></td><td></td><td></td><td></td></tr>
<tr><td>Oligo rev</td><td>AAACAGCACGCCGCGGCGATGGTC</td><td></td><td></td><td></td><td></td><td></td><td></td><td></td><td></td><td></td><td></td></tr>
<tr><td>Chromosome</td><td>start</td><td>end</td><td>strand</td><td>MM</td><td>target_seq</td><td>PAM</td><td>alignment</td><td>distance</td><td>position</td><td>gene name</td><td>gene id</td></tr>
<tr><td>GU305914.1</td><td>100154</td><td>100176</td><td>+</td><td>0</td><td>GACCATCGCCGCGGCGTGCT</td><td>CGG</td><td>||||||||[||||||||||||]PAM</td><td>NA</td><td>Intergenic</td><td>NA</td><td>NA</td></tr>
<tr><td>T7251</td><td>ACCATCGCCGCGGCGTGCTCGGG</td><td>1000</td><td></td><td></td><td></td><td></td><td></td><td></td><td></td><td></td><td></td></tr>
<tr><td>Oligo fwd</td><td>CACCACCATCGCCGCGGCGTGCTC</td><td></td><td></td><td></td><td></td><td></td><td></td><td></td><td></td><td></td><td></td></tr>
<tr><td>Oligo rev</td><td>AAACGAGCACGCCGCGGCGATGGT</td><td></td><td></td><td></td><td></td><td></td><td></td><td></td><td></td><td></td><td></td></tr>
<tr><td>Chromosome</td><td>start</td><td>end</td><td>strand</td><td>MM</td><td>target_seq</td><td>PAM</td><td>alignment</td><td>distance</td><td>position</td><td>gene name</td><td>gene id</td></tr>
<tr><td>GU305914.1</td><td>100155</td><td>100177</td><td>+</td><td>0</td><td>ACCATCGCCGCGGCGTGCTC</td><td>GGG</td><td>||||||||[||||||||||||]PAM</td><td>NA</td><td>Intergenic</td><td>NA</td><td>NA</td></tr>
<tr><td>GU305914.1</td><td>182443</td><td>182465</td><td>-</td><td>4</td><td>TCCATGGCCGCGGTGTGCTT</td><td>CGG</td><td>-||||-||[|||||-|||||-]PAM</td><td>NA</td><td>Intergenic</td><td>NA</td><td>NA</td></tr>
<tr><td>T7252</td><td>GCCGCGGCGTGCTCGGGCCGCGG</td><td>1000</td><td></td><td></td><td></td><td></td><td></td><td></td><td></td><td></td><td></td></tr>
<tr><td>Oligo fwd</td><td>CACCGCCGCGGCGTGCTCGGGCCG</td><td></td><td></td><td></td><td></td><td></td><td></td><td></td><td></td><td></td><td></td></tr>
<tr><td>Oligo rev</td><td>AAACCGGCCCGAGCACGCCGCGGC</td><td></td><td></td><td></td><td></td><td></td><td></td><td></td><td></td><td></td><td></td></tr>
<tr><td>Chromosome</td><td>start</td><td>end</td><td>strand</td><td>MM</td><td>target_seq</td><td>PAM</td><td>alignment</td><td>distance</td><td>position</td><td>gene name</td><td>gene id</td></tr>
<tr><td>GU305914.1</td><td>100161</td><td>100183</td><td>+</td><td>0</td><td>GCCGCGGCGTGCTCGGGCCG</td><td>CGG</td><td>||||||||[||||||||||||]PAM</td><td>NA</td><td>Intergenic</td><td>NA</td><td>NA</td></tr>
<tr><td>T7253</td><td>CCGCGGCGTGCTCGGGCCGCGGG</td><td>1000</td><td></td><td></td><td></td><td></td><td></td><td></td><td></td><td></td><td></td></tr>
<tr><td>Oligo fwd</td><td>CACCCCGCGGCGTGCTCGGGCCGC</td><td></td><td></td><td></td><td></td><td></td><td></td><td></td><td></td><td></td><td></td></tr>
<tr><td>Oligo rev</td><td>AAACGCGGCCCGAGCACGCCGCGG</td><td></td><td></td><td></td><td></td><td></td><td></td><td></td><td></td><td></td><td></td></tr>
<tr><td>Chromosome</td><td>start</td><td>end</td><td>strand</td><td>MM</td><td>target_seq</td><td>PAM</td><td>alignment</td><td>distance</td><td>position</td><td>gene name</td><td>gene id</td></tr>
</table>

<table>
<tr><td>GU305914.1</td><td>100162</td><td>100184</td><td>+</td><td>0</td><td>CCGCGGCGTGCTCGGGCCGC</td><td>GGG</td><td>||||||||[||||||||||||]PAM</td><td>NA</td><td>Intergenic</td><td>NA</td><td>NA</td></tr>
<tr><td>T7254</td><td>CGCCTCGTCCGCCGCCCCCGCGG</td><td>1000</td><td></td><td></td><td></td><td></td><td></td><td></td><td></td><td></td><td></td></tr>
<tr><td>Oligo fwd</td><td>CACCGCCTCGTCCGCCGCCCCCG</td><td></td><td></td><td></td><td></td><td></td><td></td><td></td><td></td><td></td><td></td></tr>
<tr><td>Oligo rev</td><td>AAACCGGGGGCGGCGGACGAGGCG</td><td></td><td></td><td></td><td></td><td></td><td></td><td></td><td></td><td></td><td></td></tr>
<tr><td>Chromosome</td><td>start</td><td>end</td><td>strand</td><td>MM</td><td>target_seq</td><td>PAM</td><td>alignment</td><td>distance</td><td>position</td><td>gene name</td><td>gene id</td></tr>
<tr><td>GU305914.1</td><td>100223</td><td>100245</td><td>+</td><td>0</td><td>CGCCTCGTCCGCCGCCCCCG</td><td>CGG</td><td>||||||||[||||||||||||]PAM</td><td>NA</td><td>Intergenic</td><td>NA</td><td>NA</td></tr>
<tr><td>T7255</td><td>GCCGCCCCCGCGGCGTTCTGCGG</td><td>1000</td><td></td><td></td><td></td><td></td><td></td><td></td><td></td><td></td><td></td></tr>
<tr><td>Oligo fwd</td><td>CACCGCCGCCCCCGCGGCGTTCTG</td><td></td><td></td><td></td><td></td><td></td><td></td><td></td><td></td><td></td><td></td></tr>
<tr><td>Oligo rev</td><td>AAACCAGAACGCCGCGGGGGCGGC</td><td></td><td></td><td></td><td></td><td></td><td></td><td></td><td></td><td></td><td></td></tr>
<tr><td>Chromosome</td><td>start</td><td>end</td><td>strand</td><td>MM</td><td>target_seq</td><td>PAM</td><td>alignment</td><td>distance</td><td>position</td><td>gene name</td><td>gene id</td></tr>
<tr><td>GU305914.1</td><td>100233</td><td>100255</td><td>+</td><td>0</td><td>GCCGCCCCCGCGGCGTTCTG</td><td>CGG</td><td>||||||||[||||||||||||]PAM</td><td>NA</td><td>Intergenic</td><td>NA</td><td>NA</td></tr>
<tr><td>GU305914.1</td><td>195107</td><td>195129</td><td>-</td><td>4</td><td>GCCACCGCCACGGCGTTCAG</td><td>GGG</td><td>|||-||-|[|-||||||||-|]PAM</td><td>NA</td><td>Intergenic</td><td>NA</td><td>NA</td></tr>
<tr><td>T7256</td><td>CGCGCTCTTGAAGAAGAAGACGG</td><td>1000</td><td></td><td></td><td></td><td></td><td></td><td></td><td></td><td></td><td></td></tr>
<tr><td>Oligo fwd</td><td>CACCCGCGCTCTTGAAGAAGAAGA</td><td></td><td></td><td></td><td></td><td></td><td></td><td></td><td></td><td></td><td></td></tr>
<tr><td>Oligo rev</td><td>AAACTCTTCTTCTTCAAGAGCGCG</td><td></td><td></td><td></td><td></td><td></td><td></td><td></td><td></td><td></td><td></td></tr>
<tr><td>Chromosome</td><td>start</td><td>end</td><td>strand</td><td>MM</td><td>target_seq</td><td>PAM</td><td>alignment</td><td>distance</td><td>position</td><td>gene name</td><td>gene id</td></tr>
<tr><td>GU305914.1</td><td>100259</td><td>100281</td><td>+</td><td>0</td><td>CGCGCTCTTGAAGAAGAAGA</td><td>CGG</td><td>||||||||[||||||||||||]PAM</td><td>NA</td><td>Intergenic</td><td>NA</td><td>NA</td></tr>
<tr><td>T7257</td><td>GCGCTCTTGAAGAAGAAGACGGG</td><td>1000</td><td></td><td></td><td></td><td></td><td></td><td></td><td></td><td></td><td></td></tr>
<tr><td>Oligo fwd</td><td>CACCGCGCTCTTGAAGAAGAAGAC</td><td></td><td></td><td></td><td></td><td></td><td></td><td></td><td></td><td></td><td></td></tr>
<tr><td>Oligo rev</td><td>AAACGTCTTCTTCTTCAAGAGCGC</td><td></td><td></td><td></td><td></td><td></td><td></td><td></td><td></td><td></td><td></td></tr>
<tr><td>Chromosome</td><td>start</td><td>end</td><td>strand</td><td>MM</td><td>target_seq</td><td>PAM</td><td>alignment</td><td>distance</td><td>position</td><td>gene name</td><td>gene id</td></tr>
<tr><td>GU305914.1</td><td>100260</td><td>100282</td><td>+</td><td>0</td><td>GCGCTCTTGAAGAAGAAGAC</td><td>GGG</td><td>||||||||[||||||||||||]PAM</td><td>NA</td><td>Intergenic</td><td>NA</td><td>NA</td></tr>
<tr><td>T7258</td><td>CGCTCTTGAAGAAGAAGACGGGG</td><td>1000</td><td></td><td></td><td></td><td></td><td></td><td></td><td></td><td></td><td></td></tr>
<tr><td>Oligo fwd</td><td>CACCCGCTCTTGAAGAAGAAGACG</td><td></td><td></td><td></td><td></td><td></td><td></td><td></td><td></td><td></td><td></td></tr>
<tr><td>Oligo rev</td><td>AAACCGTCTTCTTCTTCAAGAGCG</td><td></td><td></td><td></td><td></td><td></td><td></td><td></td><td></td><td></td><td></td></tr>
<tr><td>Chromosome</td><td>start</td><td>end</td><td>strand</td><td>MM</td><td>target_seq</td><td>PAM</td><td>alignment</td><td>distance</td><td>position</td><td>gene name</td><td>gene id</td></tr>
<tr><td>GU305914.1</td><td>100261</td><td>100283</td><td>+</td><td>0</td><td>CGCTCTTGAAGAAGAAGACG</td><td>GGG</td><td>||||||||[||||||||||||]PAM</td><td>NA</td><td>Intergenic</td><td>NA</td><td>NA</td></tr>
<tr><td>T7259</td><td>TCTTGAAGAAGAAGACGGGGTGG</td><td>1000</td><td></td><td></td><td></td><td></td><td></td><td></td><td></td><td></td><td></td></tr>
<tr><td>Oligo fwd</td><td>CACCTCTTGAAGAAGAAGACGGGG</td><td></td><td></td><td></td><td></td><td></td><td></td><td></td><td></td><td></td><td></td></tr>
<tr><td>Oligo rev</td><td>AAACCCCCGTCTTCTTCTTCAAGA</td><td></td><td></td><td></td><td></td><td></td><td></td><td></td><td></td><td></td><td></td></tr>
<tr><td>Chromosome</td><td>start</td><td>end</td><td>strand</td><td>MM</td><td>target_seq</td><td>PAM</td><td>alignment</td><td>distance</td><td>position</td><td>gene name</td><td>gene id</td></tr>
<tr><td>GU305914.1</td><td>100264</td><td>100286</td><td>+</td><td>0</td><td>TCTTGAAGAAGAAGACGGGG</td><td>TGG</td><td>||||||||[||||||||||||]PAM</td><td>NA</td><td>Intergenic</td><td>NA</td><td>NA</td></tr>
<tr><td>T7260</td><td>CTTGAAGAAGAAGACGGGGTGGG</td><td>1000</td><td></td><td></td><td></td><td></td><td></td><td></td><td></td><td></td><td></td></tr>
<tr><td>Oligo fwd</td><td>CACCCTTGAAGAAGAAGACGGGGT</td><td></td><td></td><td></td><td></td><td></td><td></td><td></td><td></td><td></td><td></td></tr>
<tr><td>Oligo rev</td><td>AAACACCCCGTCTTCTTCTTCAAG</td><td></td><td></td><td></td><td></td><td></td><td></td><td></td><td></td><td></td><td></td></tr>
<tr><td>Chromosome</td><td>start</td><td>end</td><td>strand</td><td>MM</td><td>target_seq</td><td>PAM</td><td>alignment</td><td>distance</td><td>position</td><td>gene name</td><td>gene id</td></tr>
<tr><td>GU305914.1</td><td>100265</td><td>100287</td><td>+</td><td>0</td><td>CTTGAAGAAGAAGACGGGGT</td><td>GGG</td><td>||||||||[||||||||||||]PAM</td><td>NA</td><td>Intergenic</td><td>NA</td><td>NA</td></tr>
<tr><td>T7261</td><td>AAGAAGACGGGGTGGGAGTCCGG</td><td>1000</td><td></td><td></td><td></td><td></td><td></td><td></td><td></td><td></td><td></td></tr>
<tr><td>Oligo fwd</td><td>CACCAAGAAGACGGGGTGGGAGTC</td><td></td><td></td><td></td><td></td><td></td><td></td><td></td><td></td><td></td><td></td></tr>
<tr><td>Oligo rev</td><td>AAACGACTCCCACCCCGTCTTCTT</td><td></td><td></td><td></td><td></td><td></td><td></td><td></td><td></td><td></td><td></td></tr>
<tr><td>Chromosome</td><td>start</td><td>end</td><td>strand</td><td>MM</td><td>target_seq</td><td>PAM</td><td>alignment</td><td>distance</td><td>position</td><td>gene name</td><td>gene id</td></tr>
<tr><td>GU305914.1</td><td>100272</td><td>100294</td><td>+</td><td>0</td><td>AAGAAGACGGGGTGGGAGTC</td><td>CGG</td><td>||||||||[||||||||||||]PAM</td><td>NA</td><td>Intergenic</td><td>NA</td><td>NA</td></tr>
<tr><td>T7262</td><td>AGAAGACGGGGTGGGAGTCCGGG</td><td>1000</td><td></td><td></td><td></td><td></td><td></td><td></td><td></td><td></td><td></td></tr>
<tr><td>Oligo fwd</td><td>CACCAGAAGACGGGGTGGGAGTCC</td><td></td><td></td><td></td><td></td><td></td><td></td><td></td><td></td><td></td><td></td></tr>
<tr><td>Oligo rev</td><td>AAACGGACTCCCACCCCGTCTTCT</td><td></td><td></td><td></td><td></td><td></td><td></td><td></td><td></td><td></td><td></td></tr>
<tr><td>Chromosome</td><td>start</td><td>end</td><td>strand</td><td>MM</td><td>target_seq</td><td>PAM</td><td>alignment</td><td>distance</td><td>position</td><td>gene name</td><td>gene id</td></tr>
<tr><td>GU305914.1</td><td>100273</td><td>100295</td><td>+</td><td>0</td><td>AGAAGACGGGGTGGGAGTCC</td><td>GGG</td><td>||||||||[||||||||||||]PAM</td><td>NA</td><td>Intergenic</td><td>NA</td><td>NA</td></tr>
<tr><td>T7263</td><td>GTGGGAGTCCGGGTCTACCTCGG</td><td>1000</td><td></td><td></td><td></td><td></td><td></td><td></td><td></td><td></td><td></td></tr>
<tr><td>Oligo fwd</td><td>CACCGTGGGAGTCCGGGTCTACCT</td><td></td><td></td><td></td><td></td><td></td><td></td><td></td><td></td><td></td><td></td></tr>
<tr><td>Oligo rev</td><td>AAACAGGTAGACCCGGACTCCCAC</td><td></td><td></td><td></td><td></td><td></td><td></td><td></td><td></td><td></td><td></td></tr>
<tr><td>Chromosome</td><td>start</td><td>end</td><td>strand</td><td>MM</td><td>target_seq</td><td>PAM</td><td>alignment</td><td>distance</td><td>position</td><td>gene name</td><td>gene id</td></tr>
<tr><td>GU305914.1</td><td>100283</td><td>100305</td><td>+</td><td>0</td><td>GTGGGAGTCCGGGTCTACCT</td><td>CGG</td><td>||||||||[||||||||||||]PAM</td><td>NA</td><td>Intergenic</td><td>NA</td><td>NA</td></tr>
<tr><td>T7264</td><td>TGGGAGTCCGGGTCTACCTCGGG</td><td>1000</td><td></td><td></td><td></td><td></td><td></td><td></td><td></td><td></td><td></td></tr>
<tr><td>Oligo fwd</td><td>CACCTGGGAGTCCGGGTCTACCTC</td><td></td><td></td><td></td><td></td><td></td><td></td><td></td><td></td><td></td><td></td></tr>
<tr><td>Oligo rev</td><td>AAACGAGGTAGACCCGGACTCCCA</td><td></td><td></td><td></td><td></td><td></td><td></td><td></td><td></td><td></td><td></td></tr>
<tr><td>Chromosome</td><td>start</td><td>end</td><td>strand</td><td>MM</td><td>target_seq</td><td>PAM</td><td>alignment</td><td>distance</td><td>position</td><td>gene name</td><td>gene id</td></tr>
<tr><td>GU305914.1</td><td>100284</td><td>100306</td><td>+</td><td>0</td><td>TGGGAGTCCGGGTCTACCTC</td><td>GGG</td><td>||||||||[||||||||||||]PAM</td><td>NA</td><td>Intergenic</td><td>NA</td><td>NA</td></tr>
<tr><td>T7265</td><td>CCGGGTCTACCTCGGGAAACAGG</td><td>1000</td><td></td><td></td><td></td><td></td><td></td><td></td><td></td><td></td><td></td></tr>
<tr><td>Oligo fwd</td><td>CACCCCGGGTCTACCTCGGGAAAC</td><td></td><td></td><td></td><td></td><td></td><td></td><td></td><td></td><td></td><td></td></tr>
<tr><td>Oligo rev</td><td>AAACGTTTCCCGAGGTAGACCCGG</td><td></td><td></td><td></td><td></td><td></td><td></td><td></td><td></td><td></td><td></td></tr>
<tr><td>Chromosome</td><td>start</td><td>end</td><td>strand</td><td>MM</td><td>target_seq</td><td>PAM</td><td>alignment</td><td>distance</td><td>position</td><td>gene name</td><td>gene id</td></tr>
<tr><td>GU305914.1</td><td>100291</td><td>100313</td><td>+</td><td>0</td><td>CCGGGTCTACCTCGGGAAAC</td><td>AGG</td><td>||||||||[||||||||||||]PAM</td><td>NA</td><td>Intergenic</td><td>NA</td><td>NA</td></tr>
<tr><td>T7266</td><td>CTCGGGAAACAGGTGCCGCCAGG</td><td>1000</td><td></td><td></td><td></td><td></td><td></td><td></td><td></td><td></td><td></td></tr>
<tr><td>Oligo fwd</td><td>CACCCTCGGGAAACAGGTGCCGCC</td><td></td><td></td><td></td><td></td><td></td><td></td><td></td><td></td><td></td><td></td></tr>
<tr><td>Oligo rev</td><td>AAACGGCGGCACCTGTTTCCCGAG</td><td></td><td></td><td></td><td></td><td></td><td></td><td></td><td></td><td></td><td></td></tr>
<tr><td>Chromosome</td><td>start</td><td>end</td><td>strand</td><td>MM</td><td>target_seq</td><td>PAM</td><td>alignment</td><td>distance</td><td>position</td><td>gene name</td><td>gene id</td></tr>
<tr><td>GU305914.1</td><td>100301</td><td>100323</td><td>+</td><td>0</td><td>CTCGGGAAACAGGTGCCGCC</td><td>AGG</td><td>||||||||[||||||||||||]PAM</td><td>NA</td><td>Intergenic</td><td>NA</td><td>NA</td></tr>
<tr><td>T7267</td><td>GGTGCCGCCAGGCCCCGATCAGG</td><td>1000</td><td></td><td></td><td></td><td></td><td></td><td></td><td></td><td></td><td></td></tr>
<tr><td>Oligo fwd</td><td>CACCGGTGCCGCCAGGCCCCGATC</td><td></td><td></td><td></td><td></td><td></td><td></td><td></td><td></td><td></td><td></td></tr>
<tr><td>Oligo rev</td><td>AAACGATCGGGGCCTGGCGGCACC</td><td></td><td></td><td></td><td></td><td></td><td></td><td></td><td></td><td></td><td></td></tr>
<tr><td>Chromosome</td><td>start</td><td>end</td><td>strand</td><td>MM</td><td>target_seq</td><td>PAM</td><td>alignment</td><td>distance</td><td>position</td><td>gene name</td><td>gene id</td></tr>
<tr><td>GU305914.1</td><td>100312</td><td>100334</td><td>+</td><td>0</td><td>GGTGCCGCCAGGCCCCGATC</td><td>AGG</td><td>||||||||[||||||||||||]PAM</td><td>NA</td><td>Intergenic</td><td>NA</td><td>NA</td></tr>
<tr><td>T7268</td><td>GTGCCGCCAGGCCCCGATCAGGG</td><td>1000</td><td></td><td></td><td></td><td></td><td></td><td></td><td></td><td></td><td></td></tr>
<tr><td>Oligo fwd</td><td>CACCGTGCCGCCAGGCCCCGATCA</td><td></td><td></td><td></td><td></td><td></td><td></td><td></td><td></td><td></td><td></td></tr>
</table>

<table>
<tr><td>Oligo rev</td><td>AAACTGATCGGGGCCTGGCGGCAC</td><td></td><td></td><td></td><td></td><td></td><td></td><td></td><td></td><td></td><td></td></tr>
<tr><td>Chromosome</td><td>start</td><td>end</td><td>strand</td><td>MM</td><td>target_seq</td><td>PAM</td><td>alignment</td><td>distance</td><td>position</td><td>gene name</td><td>gene id</td></tr>
<tr><td>GU305914.1</td><td>100313</td><td>100335</td><td>+</td><td>0</td><td>GTGCCGCCAGGCCCCGATCA</td><td>GGG</td><td>||||||||[||||||||||||]PAM</td><td>NA</td><td>Intergenic</td><td>NA</td><td>NA</td></tr>
<tr><td></td><td></td><td></td><td></td><td></td><td></td><td></td><td></td><td></td><td></td><td></td><td></td></tr>
<tr><td>T7269</td><td>TCAGGGTCCGCCGCAGCGCGCGG</td><td>1000</td><td></td><td></td><td></td><td></td><td></td><td></td><td></td><td></td><td></td></tr>
<tr><td>Oligo fwd</td><td>CACCTCAGGGTCCGCCGCAGCGCG</td><td></td><td></td><td></td><td></td><td></td><td></td><td></td><td></td><td></td><td></td></tr>
<tr><td>Oligo rev</td><td>AAACCGCGCTGCGGCGGACCCTGA</td><td></td><td></td><td></td><td></td><td></td><td></td><td></td><td></td><td></td><td></td></tr>
<tr><td>Chromosome</td><td>start</td><td>end</td><td>strand</td><td>MM</td><td>target_seq</td><td>PAM</td><td>alignment</td><td>distance</td><td>position</td><td>gene name</td><td>gene id</td></tr>
<tr><td>GU305914.1</td><td>100330</td><td>100352</td><td>+</td><td>0</td><td>TCAGGGTCCGCCGCAGCGCG</td><td>CGG</td><td>||||||||[||||||||||||]PAM</td><td>NA</td><td>Intergenic</td><td>NA</td><td>NA</td></tr>
<tr><td></td><td></td><td></td><td></td><td></td><td></td><td></td><td></td><td></td><td></td><td></td><td></td></tr>
<tr><td>T7270</td><td>TCCGCCGCAGCGCGCGGCACAGG</td><td>1000</td><td></td><td></td><td></td><td></td><td></td><td></td><td></td><td></td><td></td></tr>
<tr><td>Oligo fwd</td><td>CACCTCCGCCGCAGCGCGCGGCAC</td><td></td><td></td><td></td><td></td><td></td><td></td><td></td><td></td><td></td><td></td></tr>
<tr><td>Oligo rev</td><td>AAACGTGCCGCGCGCTGCGGCGGA</td><td></td><td></td><td></td><td></td><td></td><td></td><td></td><td></td><td></td><td></td></tr>
<tr><td>Chromosome</td><td>start</td><td>end</td><td>strand</td><td>MM</td><td>target_seq</td><td>PAM</td><td>alignment</td><td>distance</td><td>position</td><td>gene name</td><td>gene id</td></tr>
<tr><td>GU305914.1</td><td>100336</td><td>100358</td><td>+</td><td>0</td><td>TCCGCCGCAGCGCGCGGCAC</td><td>AGG</td><td>||||||||[||||||||||||]PAM</td><td>NA</td><td>Intergenic</td><td>NA</td><td>NA</td></tr>
<tr><td>GU305914.1</td><td>66689</td><td>66711</td><td>-</td><td>4</td><td>AGCGCCGCAGCCCGCGGCTC</td><td>CGG</td><td>--||||||[|||-||||||-|]PAM</td><td>NA</td><td>Intergenic</td><td>NA</td><td>NA</td></tr>
<tr><td></td><td></td><td></td><td></td><td></td><td></td><td></td><td></td><td></td><td></td><td></td><td></td></tr>
<tr><td>T7271</td><td>ACAGGTCGAACACCTCCTGACGG</td><td>1000</td><td></td><td></td><td></td><td></td><td></td><td></td><td></td><td></td><td></td></tr>
<tr><td>Oligo fwd</td><td>CACCACAGGTCGAACACCTCCTGA</td><td></td><td></td><td></td><td></td><td></td><td></td><td></td><td></td><td></td><td></td></tr>
<tr><td>Oligo rev</td><td>AAACTCAGGAGGTGTTCGACCTGT</td><td></td><td></td><td></td><td></td><td></td><td></td><td></td><td></td><td></td><td></td></tr>
<tr><td>Chromosome</td><td>start</td><td>end</td><td>strand</td><td>MM</td><td>target_seq</td><td>PAM</td><td>alignment</td><td>distance</td><td>position</td><td>gene name</td><td>gene id</td></tr>
<tr><td>GU305914.1</td><td>100354</td><td>100376</td><td>+</td><td>0</td><td>ACAGGTCGAACACCTCCTGA</td><td>CGG</td><td>||||||||[||||||||||||]PAM</td><td>NA</td><td>Intergenic</td><td>NA</td><td>NA</td></tr>
<tr><td></td><td></td><td></td><td></td><td></td><td></td><td></td><td></td><td></td><td></td><td></td><td></td></tr>
<tr><td>T7272</td><td>CCTGACGGCTCAGACCCTGCAGG</td><td>1000</td><td></td><td></td><td></td><td></td><td></td><td></td><td></td><td></td><td></td></tr>
<tr><td>Oligo fwd</td><td>CACCCCTGACGGCTCAGACCCTGC</td><td></td><td></td><td></td><td></td><td></td><td></td><td></td><td></td><td></td><td></td></tr>
<tr><td>Oligo rev</td><td>AAACGCAGGGTCTGAGCCGTCAGG</td><td></td><td></td><td></td><td></td><td></td><td></td><td></td><td></td><td></td><td></td></tr>
<tr><td>Chromosome</td><td>start</td><td>end</td><td>strand</td><td>MM</td><td>target_seq</td><td>PAM</td><td>alignment</td><td>distance</td><td>position</td><td>gene name</td><td>gene id</td></tr>
<tr><td>GU305914.1</td><td>100369</td><td>100391</td><td>+</td><td>0</td><td>CCTGACGGCTCAGACCCTGC</td><td>AGG</td><td>||||||||[||||||||||||]PAM</td><td>NA</td><td>Intergenic</td><td>NA</td><td>NA</td></tr>
<tr><td></td><td></td><td></td><td></td><td></td><td></td><td></td><td></td><td></td><td></td><td></td><td></td></tr>
<tr><td>T7273</td><td>GCAGGTCCTCGCGCAGCTTGAGG</td><td>1000</td><td></td><td></td><td></td><td></td><td></td><td></td><td></td><td></td><td></td></tr>
<tr><td>Oligo fwd</td><td>CACCGCAGGTCCTCGCGCAGCTTG</td><td></td><td></td><td></td><td></td><td></td><td></td><td></td><td></td><td></td><td></td></tr>
<tr><td>Oligo rev</td><td>AAACCAAGCTGCGCGAGGACCTGC</td><td></td><td></td><td></td><td></td><td></td><td></td><td></td><td></td><td></td><td></td></tr>
<tr><td>Chromosome</td><td>start</td><td>end</td><td>strand</td><td>MM</td><td>target_seq</td><td>PAM</td><td>alignment</td><td>distance</td><td>position</td><td>gene name</td><td>gene id</td></tr>
<tr><td>GU305914.1</td><td>100387</td><td>100409</td><td>+</td><td>0</td><td>GCAGGTCCTCGCGCAGCTTG</td><td>AGG</td><td>||||||||[||||||||||||]PAM</td><td>NA</td><td>Intergenic</td><td>NA</td><td>NA</td></tr>
<tr><td></td><td></td><td></td><td></td><td></td><td></td><td></td><td></td><td></td><td></td><td></td><td></td></tr>
<tr><td>T7274</td><td>GTCGCCCACGAAGTTGAACACGG</td><td>1000</td><td></td><td></td><td></td><td></td><td></td><td></td><td></td><td></td><td></td></tr>
<tr><td>Oligo fwd</td><td>CACCGTCGCCCACGAAGTTGAACA</td><td></td><td></td><td></td><td></td><td></td><td></td><td></td><td></td><td></td><td></td></tr>
<tr><td>Oligo rev</td><td>AAACTGTTCAACTTCGTGGGCGAC</td><td></td><td></td><td></td><td></td><td></td><td></td><td></td><td></td><td></td><td></td></tr>
<tr><td>Chromosome</td><td>start</td><td>end</td><td>strand</td><td>MM</td><td>target_seq</td><td>PAM</td><td>alignment</td><td>distance</td><td>position</td><td>gene name</td><td>gene id</td></tr>
<tr><td>GU305914.1</td><td>100415</td><td>100437</td><td>+</td><td>0</td><td>GTCGCCCACGAAGTTGAACA</td><td>CGG</td><td>||||||||[||||||||||||]PAM</td><td>NA</td><td>Intergenic</td><td>NA</td><td>NA</td></tr>
<tr><td></td><td></td><td></td><td></td><td></td><td></td><td></td><td></td><td></td><td></td><td></td><td></td></tr>
<tr><td>T7275</td><td>TCGCCCACGAAGTTGAACACGGG</td><td>1000</td><td></td><td></td><td></td><td></td><td></td><td></td><td></td><td></td><td></td></tr>
<tr><td>Oligo fwd</td><td>CACCTCGCCCACGAAGTTGAACAC</td><td></td><td></td><td></td><td></td><td></td><td></td><td></td><td></td><td></td><td></td></tr>
<tr><td>Oligo rev</td><td>AAACGTGTTCAACTTCGTGGGCGA</td><td></td><td></td><td></td><td></td><td></td><td></td><td></td><td></td><td></td><td></td></tr>
<tr><td>Chromosome</td><td>start</td><td>end</td><td>strand</td><td>MM</td><td>target_seq</td><td>PAM</td><td>alignment</td><td>distance</td><td>position</td><td>gene name</td><td>gene id</td></tr>
<tr><td>GU305914.1</td><td>100416</td><td>100438</td><td>+</td><td>0</td><td>TCGCCCACGAAGTTGAACAC</td><td>GGG</td><td>||||||||[||||||||||||]PAM</td><td>NA</td><td>Intergenic</td><td>NA</td><td>NA</td></tr>
<tr><td></td><td></td><td></td><td></td><td></td><td></td><td></td><td></td><td></td><td></td><td></td><td></td></tr>
<tr><td>T7276</td><td>CCACGAAGTTGAACACGGGCAGG</td><td>1000</td><td></td><td></td><td></td><td></td><td></td><td></td><td></td><td></td><td></td></tr>
<tr><td>Oligo fwd</td><td>CACCCCACGAAGTTGAACACGGGC</td><td></td><td></td><td></td><td></td><td></td><td></td><td></td><td></td><td></td><td></td></tr>
<tr><td>Oligo rev</td><td>AAACGCCCGTGTTCAACTTCGTGG</td><td></td><td></td><td></td><td></td><td></td><td></td><td></td><td></td><td></td><td></td></tr>
<tr><td>Chromosome</td><td>start</td><td>end</td><td>strand</td><td>MM</td><td>target_seq</td><td>PAM</td><td>alignment</td><td>distance</td><td>position</td><td>gene name</td><td>gene id</td></tr>
<tr><td>GU305914.1</td><td>100420</td><td>100442</td><td>+</td><td>0</td><td>CCACGAAGTTGAACACGGGC</td><td>AGG</td><td>||||||||[||||||||||||]PAM</td><td>NA</td><td>Intergenic</td><td>NA</td><td>NA</td></tr>
<tr><td></td><td></td><td></td><td></td><td></td><td></td><td></td><td></td><td></td><td></td><td></td><td></td></tr>
<tr><td>T7277</td><td>TGAACATCTCGTGACGCGTGCGG</td><td>1000</td><td></td><td></td><td></td><td></td><td></td><td></td><td></td><td></td><td></td></tr>
<tr><td>Oligo fwd</td><td>CACCTGAACATCTCGTGACGCGTG</td><td></td><td></td><td></td><td></td><td></td><td></td><td></td><td></td><td></td><td></td></tr>
<tr><td>Oligo rev</td><td>AAACCACGCGTCACGAGATGTTCA</td><td></td><td></td><td></td><td></td><td></td><td></td><td></td><td></td><td></td><td></td></tr>
<tr><td>Chromosome</td><td>start</td><td>end</td><td>strand</td><td>MM</td><td>target_seq</td><td>PAM</td><td>alignment</td><td>distance</td><td>position</td><td>gene name</td><td>gene id</td></tr>
<tr><td>GU305914.1</td><td>100450</td><td>100472</td><td>+</td><td>0</td><td>TGAACATCTCGTGACGCGTG</td><td>CGG</td><td>||||||||[||||||||||||]PAM</td><td>NA</td><td>Intergenic</td><td>NA</td><td>NA</td></tr>
<tr><td></td><td></td><td></td><td></td><td></td><td></td><td></td><td></td><td></td><td></td><td></td><td></td></tr>
<tr><td>T7278</td><td>CTGCTCGCCCACCTCGCACGAGG</td><td>1000</td><td></td><td></td><td></td><td></td><td></td><td></td><td></td><td></td><td></td></tr>
<tr><td>Oligo fwd</td><td>CACCCTGCTCGCCCACCTCGCACG</td><td></td><td></td><td></td><td></td><td></td><td></td><td></td><td></td><td></td><td></td></tr>
<tr><td>Oligo rev</td><td>AAACCGTGCGAGGTGGGCGAGCAG</td><td></td><td></td><td></td><td></td><td></td><td></td><td></td><td></td><td></td><td></td></tr>
<tr><td>Chromosome</td><td>start</td><td>end</td><td>strand</td><td>MM</td><td>target_seq</td><td>PAM</td><td>alignment</td><td>distance</td><td>position</td><td>gene name</td><td>gene id</td></tr>
<tr><td>GU305914.1</td><td>100478</td><td>100500</td><td>+</td><td>0</td><td>CTGCTCGCCCACCTCGCACG</td><td>AGG</td><td>||||||||[||||||||||||]PAM</td><td>NA</td><td>Intergenic</td><td>NA</td><td>NA</td></tr>
<tr><td></td><td></td><td></td><td></td><td></td><td></td><td></td><td></td><td></td><td></td><td></td><td></td></tr>
<tr><td>T7279</td><td>ACACTCGTCCGAGACGTACTCGG</td><td>1000</td><td></td><td></td><td></td><td></td><td></td><td></td><td></td><td></td><td></td></tr>
<tr><td>Oligo fwd</td><td>CACCACACTCGTCCGAGACGTACT</td><td></td><td></td><td></td><td></td><td></td><td></td><td></td><td></td><td></td><td></td></tr>
<tr><td>Oligo rev</td><td>AAACAGTACGTCTCGGACGAGTGT</td><td></td><td></td><td></td><td></td><td></td><td></td><td></td><td></td><td></td><td></td></tr>
<tr><td>Chromosome</td><td>start</td><td>end</td><td>strand</td><td>MM</td><td>target_seq</td><td>PAM</td><td>alignment</td><td>distance</td><td>position</td><td>gene name</td><td>gene id</td></tr>
<tr><td>GU305914.1</td><td>100538</td><td>100560</td><td>+</td><td>0</td><td>ACACTCGTCCGAGACGTACT</td><td>CGG</td><td>||||||||[||||||||||||]PAM</td><td>NA</td><td>Intergenic</td><td>NA</td><td>NA</td></tr>
<tr><td></td><td></td><td></td><td></td><td></td><td></td><td></td><td></td><td></td><td></td><td></td><td></td></tr>
<tr><td>T7280</td><td>TCGTCCGAGACGTACTCGGACGG</td><td>1000</td><td></td><td></td><td></td><td></td><td></td><td></td><td></td><td></td><td></td></tr>
<tr><td>Oligo fwd</td><td>CACCTCGTCCGAGACGTACTCGGA</td><td></td><td></td><td></td><td></td><td></td><td></td><td></td><td></td><td></td><td></td></tr>
<tr><td>Oligo rev</td><td>AAACTCCGAGTACGTCTCGGACGA</td><td></td><td></td><td></td><td></td><td></td><td></td><td></td><td></td><td></td><td></td></tr>
<tr><td>Chromosome</td><td>start</td><td>end</td><td>strand</td><td>MM</td><td>target_seq</td><td>PAM</td><td>alignment</td><td>distance</td><td>position</td><td>gene name</td><td>gene id</td></tr>
<tr><td>GU305914.1</td><td>100542</td><td>100564</td><td>+</td><td>0</td><td>TCGTCCGAGACGTACTCGGA</td><td>CGG</td><td>||||||||[||||||||||||]PAM</td><td>NA</td><td>Intergenic</td><td>NA</td><td>NA</td></tr>
<tr><td></td><td></td><td></td><td></td><td></td><td></td><td></td><td></td><td></td><td></td><td></td><td></td></tr>
<tr><td>T7281</td><td>CCGAGACGTACTCGGACGGCAGG</td><td>1000</td><td></td><td></td><td></td><td></td><td></td><td></td><td></td><td></td><td></td></tr>
<tr><td>Oligo fwd</td><td>CACCCCGAGACGTACTCGGACGGC</td><td></td><td></td><td></td><td></td><td></td><td></td><td></td><td></td><td></td><td></td></tr>
<tr><td>Oligo rev</td><td>AAACGCCGTCCGAGTACGTCTCGG</td><td></td><td></td><td></td><td></td><td></td><td></td><td></td><td></td><td></td><td></td></tr>
<tr><td>Chromosome</td><td>start</td><td>end</td><td>strand</td><td>MM</td><td>target_seq</td><td>PAM</td><td>alignment</td><td>distance</td><td>position</td><td>gene name</td><td>gene id</td></tr>
<tr><td>GU305914.1</td><td>100546</td><td>100568</td><td>+</td><td>0</td><td>CCGAGACGTACTCGGACGGC</td><td>AGG</td><td>||||||||[||||||||||||]PAM</td><td>NA</td><td>Intergenic</td><td>NA</td><td>NA</td></tr>
<tr><td></td><td></td><td></td><td></td><td></td><td></td><td></td><td></td><td></td><td></td><td></td><td></td></tr>
<tr><td>T7282</td><td>CGGACGGCAGGTTTTTCATCAGG</td><td>1000</td><td></td><td></td><td></td><td></td><td></td><td></td><td></td><td></td><td></td></tr>
<tr><td>Oligo fwd</td><td>CACCCGGACGGCAGGTTTTTCATC</td><td></td><td></td><td></td><td></td><td></td><td></td><td></td><td></td><td></td><td></td></tr>
<tr><td>Oligo rev</td><td>AAACGATGAAAAACCTGCCGTCCG</td><td></td><td></td><td></td><td></td><td></td><td></td><td></td><td></td><td></td><td></td></tr>
<tr><td>Chromosome</td><td>start</td><td>end</td><td>strand</td><td>MM</td><td>target_seq</td><td>PAM</td><td>alignment</td><td>distance</td><td>position</td><td>gene name</td><td>gene id</td></tr>
<tr><td>GU305914.1</td><td>100558</td><td>100580</td><td>+</td><td>0</td><td>CGGACGGCAGGTTTTTCATC</td><td>AGG</td><td>||||||||[||||||||||||]PAM</td><td>NA</td><td>Intergenic</td><td>NA</td><td>NA</td></tr>
</table>

<table>
<tr><td>T7283</td><td>CAGGTTTTTCATCAGGTCCTTGG</td><td>1000</td><td></td><td></td><td></td><td></td><td></td><td></td><td></td><td></td><td></td></tr>
<tr><td>Oligo fwd</td><td>CACCCAGGTTTTTCATCAGGTCCT</td><td></td><td></td><td></td><td></td><td></td><td></td><td></td><td></td><td></td><td></td></tr>
<tr><td>Oligo rev</td><td>AAACAGGACCTGATGAAAAACCTG</td><td></td><td></td><td></td><td></td><td></td><td></td><td></td><td></td><td></td><td></td></tr>
<tr><td>Chromosome</td><td>start</td><td>end</td><td>strand</td><td>MM</td><td>target_seq</td><td>PAM</td><td>alignment</td><td>distance</td><td>position</td><td>gene name</td><td>gene id</td></tr>
<tr><td>GU305914.1</td><td>100565</td><td>100587</td><td>+</td><td>0</td><td>CAGGTTTTTCATCAGGTCCT</td><td>TGG</td><td>||||||||[||||||||||||]PAM</td><td>NA</td><td>Intergenic</td><td>NA</td><td>NA</td></tr>
<tr><td>T7284</td><td>AGGTTTTTCATCAGGTCCTTGGG</td><td>1000</td><td></td><td></td><td></td><td></td><td></td><td></td><td></td><td></td><td></td></tr>
<tr><td>Oligo fwd</td><td>CACCAGGTTTTTCATCAGGTCCTT</td><td></td><td></td><td></td><td></td><td></td><td></td><td></td><td></td><td></td><td></td></tr>
<tr><td>Oligo rev</td><td>AAACAAGGACCTGATGAAAAACCT</td><td></td><td></td><td></td><td></td><td></td><td></td><td></td><td></td><td></td><td></td></tr>
<tr><td>Chromosome</td><td>start</td><td>end</td><td>strand</td><td>MM</td><td>target_seq</td><td>PAM</td><td>alignment</td><td>distance</td><td>position</td><td>gene name</td><td>gene id</td></tr>
<tr><td>GU305914.1</td><td>100566</td><td>100588</td><td>+</td><td>0</td><td>AGGTTTTTCATCAGGTCCTT</td><td>GGG</td><td>||||||||[||||||||||||]PAM</td><td>NA</td><td>Intergenic</td><td>NA</td><td>NA</td></tr>
<tr><td>T7285</td><td>TCAGGTCCTTGGGCAGCAGCCGG</td><td>1000</td><td></td><td></td><td></td><td></td><td></td><td></td><td></td><td></td><td></td></tr>
<tr><td>Oligo fwd</td><td>CACCTCAGGTCCTTGGGCAGCAGC</td><td></td><td></td><td></td><td></td><td></td><td></td><td></td><td></td><td></td><td></td></tr>
<tr><td>Oligo rev</td><td>AAACGCTGCTGCCCAAGGACCTGA</td><td></td><td></td><td></td><td></td><td></td><td></td><td></td><td></td><td></td><td></td></tr>
<tr><td>Chromosome</td><td>start</td><td>end</td><td>strand</td><td>MM</td><td>target_seq</td><td>PAM</td><td>alignment</td><td>distance</td><td>position</td><td>gene name</td><td>gene id</td></tr>
<tr><td>GU305914.1</td><td>100576</td><td>100598</td><td>+</td><td>0</td><td>TCAGGTCCTTGGGCAGCAGC</td><td>CGG</td><td>||||||||[||||||||||||]PAM</td><td>NA</td><td>Intergenic</td><td>NA</td><td>NA</td></tr>
<tr><td>T7286</td><td>GGTCCTTGGGCAGCAGCCGGCGG</td><td>1000</td><td></td><td></td><td></td><td></td><td></td><td></td><td></td><td></td><td></td></tr>
<tr><td>Oligo fwd</td><td>CACCGGTCCTTGGGCAGCAGCCGG</td><td></td><td></td><td></td><td></td><td></td><td></td><td></td><td></td><td></td><td></td></tr>
<tr><td>Oligo rev</td><td>AAACCCGGCTGCTGCCCAAGGACC</td><td></td><td></td><td></td><td></td><td></td><td></td><td></td><td></td><td></td><td></td></tr>
<tr><td>Chromosome</td><td>start</td><td>end</td><td>strand</td><td>MM</td><td>target_seq</td><td>PAM</td><td>alignment</td><td>distance</td><td>position</td><td>gene name</td><td>gene id</td></tr>
<tr><td>GU305914.1</td><td>100579</td><td>100601</td><td>+</td><td>0</td><td>GGTCCTTGGGCAGCAGCCGG</td><td>CGG</td><td>||||||||[||||||||||||]PAM</td><td>NA</td><td>Intergenic</td><td>NA</td><td>NA</td></tr>
<tr><td>T7287</td><td>GTCCTTGGGCAGCAGCCGGCGGG</td><td>1000</td><td></td><td></td><td></td><td></td><td></td><td></td><td></td><td></td><td></td></tr>
<tr><td>Oligo fwd</td><td>CACCGTCCTTGGGCAGCAGCCGGC</td><td></td><td></td><td></td><td></td><td></td><td></td><td></td><td></td><td></td><td></td></tr>
<tr><td>Oligo rev</td><td>AAACGCCGGCTGCTGCCCAAGGAC</td><td></td><td></td><td></td><td></td><td></td><td></td><td></td><td></td><td></td><td></td></tr>
<tr><td>Chromosome</td><td>start</td><td>end</td><td>strand</td><td>MM</td><td>target_seq</td><td>PAM</td><td>alignment</td><td>distance</td><td>position</td><td>gene name</td><td>gene id</td></tr>
<tr><td>GU305914.1</td><td>100580</td><td>100602</td><td>+</td><td>0</td><td>GTCCTTGGGCAGCAGCCGGC</td><td>GGG</td><td>||||||||[||||||||||||]PAM</td><td>NA</td><td>Intergenic</td><td>NA</td><td>NA</td></tr>
<tr><td>T7288</td><td>CAGAACACGTGCCGCTTCTCCGG</td><td>1000</td><td></td><td></td><td></td><td></td><td></td><td></td><td></td><td></td><td></td></tr>
<tr><td>Oligo fwd</td><td>CACCCAGAACACGTGCCGCTTCTC</td><td></td><td></td><td></td><td></td><td></td><td></td><td></td><td></td><td></td><td></td></tr>
<tr><td>Oligo rev</td><td>AAACGAGAAGCGGCACGTGTTCTG</td><td></td><td></td><td></td><td></td><td></td><td></td><td></td><td></td><td></td><td></td></tr>
<tr><td>Chromosome</td><td>start</td><td>end</td><td>strand</td><td>MM</td><td>target_seq</td><td>PAM</td><td>alignment</td><td>distance</td><td>position</td><td>gene name</td><td>gene id</td></tr>
<tr><td>GU305914.1</td><td>100626</td><td>100648</td><td>+</td><td>0</td><td>CAGAACACGTGCCGCTTCTC</td><td>CGG</td><td>||||||||[||||||||||||]PAM</td><td>NA</td><td>Intergenic</td><td>NA</td><td>NA</td></tr>
<tr><td>T7289</td><td>GCTTCTCCGGCATCTCCACCCGG</td><td>1000</td><td></td><td></td><td></td><td></td><td></td><td></td><td></td><td></td><td></td></tr>
<tr><td>Oligo fwd</td><td>CACCGCTTCTCCGGCATCTCCACC</td><td></td><td></td><td></td><td></td><td></td><td></td><td></td><td></td><td></td><td></td></tr>
<tr><td>Oligo rev</td><td>AAACGGTGGAGATGCCGGAGAAGC</td><td></td><td></td><td></td><td></td><td></td><td></td><td></td><td></td><td></td><td></td></tr>
<tr><td>Chromosome</td><td>start</td><td>end</td><td>strand</td><td>MM</td><td>target_seq</td><td>PAM</td><td>alignment</td><td>distance</td><td>position</td><td>gene name</td><td>gene id</td></tr>
<tr><td>GU305914.1</td><td>100639</td><td>100661</td><td>+</td><td>0</td><td>GCTTCTCCGGCATCTCCACC</td><td>CGG</td><td>||||||||[||||||||||||]PAM</td><td>NA</td><td>Intergenic</td><td>NA</td><td>NA</td></tr>
<tr><td>T7290</td><td>GGCATCTCCACCCGGAACACCGG</td><td>1000</td><td></td><td></td><td></td><td></td><td></td><td></td><td></td><td></td><td></td></tr>
<tr><td>Oligo fwd</td><td>CACCGGCATCTCCACCCGGAACAC</td><td></td><td></td><td></td><td></td><td></td><td></td><td></td><td></td><td></td><td></td></tr>
<tr><td>Oligo rev</td><td>AAACGTGTTCCGGGTGGAGATGCC</td><td></td><td></td><td></td><td></td><td></td><td></td><td></td><td></td><td></td><td></td></tr>
<tr><td>Chromosome</td><td>start</td><td>end</td><td>strand</td><td>MM</td><td>target_seq</td><td>PAM</td><td>alignment</td><td>distance</td><td>position</td><td>gene name</td><td>gene id</td></tr>
<tr><td>GU305914.1</td><td>100647</td><td>100669</td><td>+</td><td>0</td><td>GGCATCTCCACCCGGAACAC</td><td>CGG</td><td>||||||||[||||||||||||]PAM</td><td>NA</td><td>Intergenic</td><td>NA</td><td>NA</td></tr>
<tr><td>T7291</td><td>TCCACCCGGAACACCGGCATCGG</td><td>1000</td><td></td><td></td><td></td><td></td><td></td><td></td><td></td><td></td><td></td></tr>
<tr><td>Oligo fwd</td><td>CACCTCCACCCGGAACACCGGCAT</td><td></td><td></td><td></td><td></td><td></td><td></td><td></td><td></td><td></td><td></td></tr>
<tr><td>Oligo rev</td><td>AAACATGCCGGTGTTCCGGGTGGA</td><td></td><td></td><td></td><td></td><td></td><td></td><td></td><td></td><td></td><td></td></tr>
<tr><td>Chromosome</td><td>start</td><td>end</td><td>strand</td><td>MM</td><td>target_seq</td><td>PAM</td><td>alignment</td><td>distance</td><td>position</td><td>gene name</td><td>gene id</td></tr>
<tr><td>GU305914.1</td><td>100653</td><td>100675</td><td>+</td><td>0</td><td>TCCACCCGGAACACCGGCAT</td><td>CGG</td><td>||||||||[||||||||||||]PAM</td><td>NA</td><td>Intergenic</td><td>NA</td><td>NA</td></tr>
<tr><td>T7292</td><td>AACACCGGCATCGGCCCGATCGG</td><td>1000</td><td></td><td></td><td></td><td></td><td></td><td></td><td></td><td></td><td></td></tr>
<tr><td>Oligo fwd</td><td>CACCAACACCGGCATCGGCCCGAT</td><td></td><td></td><td></td><td></td><td></td><td></td><td></td><td></td><td></td><td></td></tr>
<tr><td>Oligo rev</td><td>AAACATCGGGCCGATGCCGGTGTT</td><td></td><td></td><td></td><td></td><td></td><td></td><td></td><td></td><td></td><td></td></tr>
<tr><td>Chromosome</td><td>start</td><td>end</td><td>strand</td><td>MM</td><td>target_seq</td><td>PAM</td><td>alignment</td><td>distance</td><td>position</td><td>gene name</td><td>gene id</td></tr>
<tr><td>GU305914.1</td><td>100662</td><td>100684</td><td>+</td><td>0</td><td>AACACCGGCATCGGCCCGAT</td><td>CGG</td><td>||||||||[||||||||||||]PAM</td><td>NA</td><td>Intergenic</td><td>NA</td><td>NA</td></tr>
<tr><td>T7293</td><td>CCGGCATCGGCCCGATCGGACGG</td><td>1000</td><td></td><td></td><td></td><td></td><td></td><td></td><td></td><td></td><td></td></tr>
<tr><td>Oligo fwd</td><td>CACCCCGGCATCGGCCCGATCGGA</td><td></td><td></td><td></td><td></td><td></td><td></td><td></td><td></td><td></td><td></td></tr>
<tr><td>Oligo rev</td><td>AAACTCCGATCGGGCCGATGCCGG</td><td></td><td></td><td></td><td></td><td></td><td></td><td></td><td></td><td></td><td></td></tr>
<tr><td>Chromosome</td><td>start</td><td>end</td><td>strand</td><td>MM</td><td>target_seq</td><td>PAM</td><td>alignment</td><td>distance</td><td>position</td><td>gene name</td><td>gene id</td></tr>
<tr><td>GU305914.1</td><td>100666</td><td>100688</td><td>+</td><td>0</td><td>CCGGCATCGGCCCGATCGGA</td><td>CGG</td><td>||||||||[||||||||||||]PAM</td><td>NA</td><td>Intergenic</td><td>NA</td><td>NA</td></tr>
<tr><td>T7294</td><td>GACGGCTCGTGTCGTTCAGCAGG</td><td>1000</td><td></td><td></td><td></td><td></td><td></td><td></td><td></td><td></td><td></td></tr>
<tr><td>Oligo fwd</td><td>CACCGACGGCTCGTGTCGTTCAGC</td><td></td><td></td><td></td><td></td><td></td><td></td><td></td><td></td><td></td><td></td></tr>
<tr><td>Oligo rev</td><td>AAACGCTGAACGACACGAGCCGTC</td><td></td><td></td><td></td><td></td><td></td><td></td><td></td><td></td><td></td><td></td></tr>
<tr><td>Chromosome</td><td>start</td><td>end</td><td>strand</td><td>MM</td><td>target_seq</td><td>PAM</td><td>alignment</td><td>distance</td><td>position</td><td>gene name</td><td>gene id</td></tr>
<tr><td>GU305914.1</td><td>100684</td><td>100706</td><td>+</td><td>0</td><td>GACGGCTCGTGTCGTTCAGC</td><td>AGG</td><td>||||||||[||||||||||||]PAM</td><td>NA</td><td>Intergenic</td><td>NA</td><td>NA</td></tr>
<tr><td>T7295</td><td>CGTTCAGCAGGTAGTTGCGCCGG</td><td>1000</td><td></td><td></td><td></td><td></td><td></td><td></td><td></td><td></td><td></td></tr>
<tr><td>Oligo fwd</td><td>CACCCGTTCAGCAGGTAGTTGCGC</td><td></td><td></td><td></td><td></td><td></td><td></td><td></td><td></td><td></td><td></td></tr>
<tr><td>Oligo rev</td><td>AAACGCGCAACTACCTGCTGAACG</td><td></td><td></td><td></td><td></td><td></td><td></td><td></td><td></td><td></td><td></td></tr>
<tr><td>Chromosome</td><td>start</td><td>end</td><td>strand</td><td>MM</td><td>target_seq</td><td>PAM</td><td>alignment</td><td>distance</td><td>position</td><td>gene name</td><td>gene id</td></tr>
<tr><td>GU305914.1</td><td>100696</td><td>100718</td><td>+</td><td>0</td><td>CGTTCAGCAGGTAGTTGCGC</td><td>CGG</td><td>||||||||[||||||||||||]PAM</td><td>NA</td><td>Intergenic</td><td>NA</td><td>NA</td></tr>
<tr><td>T7296</td><td>TTCTTTTTCTCTGCCGCCGCTGG</td><td>1000</td><td></td><td></td><td></td><td></td><td></td><td></td><td></td><td></td><td></td></tr>
<tr><td>Oligo fwd</td><td>CACCTTCTTTTTCTCTGCCGCCGC</td><td></td><td></td><td></td><td></td><td></td><td></td><td></td><td></td><td></td><td></td></tr>
<tr><td>Oligo rev</td><td>AAACGCGGCGGCAGAGAAAAAGAA</td><td></td><td></td><td></td><td></td><td></td><td></td><td></td><td></td><td></td><td></td></tr>
<tr><td>Chromosome</td><td>start</td><td>end</td><td>strand</td><td>MM</td><td>target_seq</td><td>PAM</td><td>alignment</td><td>distance</td><td>position</td><td>gene name</td><td>gene id</td></tr>
<tr><td>GU305914.1</td><td>100736</td><td>100758</td><td>+</td><td>0</td><td>TTCTTTTTCTCTGCCGCCGC</td><td>TGG</td><td>||||||||[||||||||||||]PAM</td><td>NA</td><td>Intergenic</td><td>NA</td><td>NA</td></tr>
<tr><td>T7297</td><td>TGCCGCCGCTGGTGCGCTTGCGG</td><td>1000</td><td></td><td></td><td></td><td></td><td></td><td></td><td></td><td></td><td></td></tr>
<tr><td>Oligo fwd</td><td>CACCTGCCGCCGCTGGTGCGCTTG</td><td></td><td></td><td></td><td></td><td></td><td></td><td></td><td></td><td></td><td></td></tr>
<tr><td>Oligo rev</td><td>AAACCAAGCGCACCAGCGGCGGCA</td><td></td><td></td><td></td><td></td><td></td><td></td><td></td><td></td><td></td><td></td></tr>
<tr><td>Chromosome</td><td>start</td><td>end</td><td>strand</td><td>MM</td><td>target_seq</td><td>PAM</td><td>alignment</td><td>distance</td><td>position</td><td>gene name</td><td>gene id</td></tr>
<tr><td>GU305914.1</td><td>100747</td><td>100769</td><td>+</td><td>0</td><td>TGCCGCCGCTGGTGCGCTTG</td><td>CGG</td><td>||||||||[||||||||||||]PAM</td><td>NA</td><td>Intergenic</td><td>NA</td><td>NA</td></tr>
</table>

| | | | | | | | | | | | |
|---|---|---|---|---|---|---|---|---|---|---|---|
| T7298 | TTGCGGCTTTTTTCCGCCCGCGG | 1000 | | | | | | | | | |
| Oligo fwd | CACCTTGCGGCTTTTTTCCGCCCG | | | | | | | | | | |
| Oligo rev | AAACCGGGCGGAAAAAAGCCGCAA | | | | | | | | | | |
| Chromosome | start | end | strand | MM | target_seq | PAM | alignment | distance | position | gene name | gene id |
| GU305914.1 | 100764 | 100786 | + | 0 | TTGCGGCTTTTTTCCGCCCG | CGG | ||||||||[||||||||||||]PAM | NA | Intergenic | NA | NA |
| T7299 | TTTCCGCCCGCGGCACGTACCGG | 1000 | | | | | | | | | |
| Oligo fwd | CACCTTTCCGCCCGCGGCACGTAC | | | | | | | | | | |
| Oligo rev | AAACGTACGTGCCGCGGGCGGAAA | | | | | | | | | | |
| Chromosome | start | end | strand | MM | target_seq | PAM | alignment | distance | position | gene name | gene id |
| GU305914.1 | 100774 | 100796 | + | 0 | TTTCCGCCCGCGGCACGTAC | CGG | ||||||||[||||||||||||]PAM | NA | Intergenic | NA | NA |
| T7300 | CCGCGGCACGTACCGGTCCGAGG | 1000 | | | | | | | | | |
| Oligo fwd | CACCCCGCGGCACGTACCGGTCCG | | | | | | | | | | |
| Oligo rev | AAACCGGACCGGTACGTGCCGCGG | | | | | | | | | | |
| Chromosome | start | end | strand | MM | target_seq | PAM | alignment | distance | position | gene name | gene id |
| GU305914.1 | 100781 | 100803 | + | 0 | CCGCGGCACGTACCGGTCCG | AGG | ||||||||[||||||||||||]PAM | NA | Intergenic | NA | NA |
| T7301 | TTCAGCAGCCCCATCAGCGTCGG | 1000 | | | | | | | | | |
| Oligo fwd | CACCTTCAGCAGCCCCATCAGCGT | | | | | | | | | | |
| Oligo rev | AAACACGCTGATGGGGCTGCTGAA | | | | | | | | | | |
| Chromosome | start | end | strand | MM | target_seq | PAM | alignment | distance | position | gene name | gene id |
| GU305914.1 | 100812 | 100834 | + | 0 | TTCAGCAGCCCCATCAGCGT | CGG | ||||||||[||||||||||||]PAM | NA | Intergenic | NA | NA |
| T7302 | AGCAGCCCCATCAGCGTCGGCGG | 1000 | | | | | | | | | |
| Oligo fwd | CACCAGCAGCCCCATCAGCGTCGG | | | | | | | | | | |
| Oligo rev | AAACCCGACGCTGATGGGGCTGCT | | | | | | | | | | |
| Chromosome | start | end | strand | MM | target_seq | PAM | alignment | distance | position | gene name | gene id |
| GU305914.1 | 100815 | 100837 | + | 0 | AGCAGCCCCATCAGCGTCGG | CGG | ||||||||[||||||||||||]PAM | NA | Intergenic | NA | NA |
| T7303 | CCCATCAGCGTCGGCGGCAGCGG | 1000 | | | | | | | | | |
| Oligo fwd | CACCCCCATCAGCGTCGGCGGCAG | | | | | | | | | | |
| Oligo rev | AAACCTGCCGCCGACGCTGATGGG | | | | | | | | | | |
| Chromosome | start | end | strand | MM | target_seq | PAM | alignment | distance | position | gene name | gene id |
| GU305914.1 | 100821 | 100843 | + | 0 | CCCATCAGCGTCGGCGGCAG | CGG | ||||||||[||||||||||||]PAM | NA | Intergenic | NA | NA |
| T7304 | TCGGCGGCAGCGGCGAGAACAGG | 1000 | | | | | | | | | |
| Oligo fwd | CACCTCGGCGGCAGCGGCGAGAAC | | | | | | | | | | |
| Oligo rev | AAACGTTCTCGCCGCTGCCGCCGA | | | | | | | | | | |
| Chromosome | start | end | strand | MM | target_seq | PAM | alignment | distance | position | gene name | gene id |
| GU305914.1 | 100831 | 100853 | + | 0 | TCGGCGGCAGCGGCGAGAAC | AGG | ||||||||[||||||||||||]PAM | NA | Intergenic | NA | NA |
| T7305 | CGGCGGCAGCGGCGAGAACAGGG | 1000 | | | | | | | | | |
| Oligo fwd | CACCCGGCGGCAGCGGCGAGAACA | | | | | | | | | | |
| Oligo rev | AAACTGTTCTCGCCGCTGCCGCCG | | | | | | | | | | |
| Chromosome | start | end | strand | MM | target_seq | PAM | alignment | distance | position | gene name | gene id |
| GU305914.1 | 100832 | 100854 | + | 0 | CGGCGGCAGCGGCGAGAACA | GGG | ||||||||[||||||||||||]PAM | NA | Intergenic | NA | NA |
| GU305914.1 | 42005 | 42027 | - | 4 | CAGCGCCAGCGTCGAGAACC | CGG | |-|||-||[|||-|||||||-]PAM | NA | Intergenic | NA | NA |
| T7306 | GTTCAGCTTGCGCAGCGAGATGG | 1000 | | | | | | | | | |
| Oligo fwd | CACCGTTCAGCTTGCGCAGCGAGA | | | | | | | | | | |
| Oligo rev | AAACTCTCGCTGCGCAAGCTGAAC | | | | | | | | | | |
| Chromosome | start | end | strand | MM | target_seq | PAM | alignment | distance | position | gene name | gene id |
| GU305914.1 | 100868 | 100890 | + | 0 | GTTCAGCTTGCGCAGCGAGA | TGG | ||||||||[||||||||||||]PAM | NA | Intergenic | NA | NA |
| GU305914.1 | 133107 | 133129 | + | 4 | GATCGGCTTGCGCGGCCAGA | GGG | |-||-|||[|||||-||-|||]PAM | NA | Intergenic | NA | NA |
| T7307 | GCTTGCGCAGCGAGATGGCCAGG | 1000 | | | | | | | | | |
| Oligo fwd | CACCGCTTGCGCAGCGAGATGGCC | | | | | | | | | | |
| Oligo rev | AAACGGCCATCTCGCTGCGCAAGC | | | | | | | | | | |
| Chromosome | start | end | strand | MM | target_seq | PAM | alignment | distance | position | gene name | gene id |
| GU305914.1 | 100873 | 100895 | + | 0 | GCTTGCGCAGCGAGATGGCC | AGG | ||||||||[||||||||||||]PAM | NA | Intergenic | NA | NA |
| T7308 | CGAGATGGCCAGGTTCGTCGCGG | 1000 | | | | | | | | | |
| Oligo fwd | CACCCGAGATGGCCAGGTTCGTCG | | | | | | | | | | |
| Oligo rev | AAACCGACGAACCTGGCCATCTCG | | | | | | | | | | |
| Chromosome | start | end | strand | MM | target_seq | PAM | alignment | distance | position | gene name | gene id |
| GU305914.1 | 100883 | 100905 | + | 0 | CGAGATGGCCAGGTTCGTCG | CGG | ||||||||[||||||||||||]PAM | NA | Intergenic | NA | NA |
| T7309 | GATGGCCAGGTTCGTCGCGGTGG | 1000 | | | | | | | | | |
| Oligo fwd | CACCGATGGCCAGGTTCGTCGCGG | | | | | | | | | | |
| Oligo rev | AAACCCGCGACGAACCTGGCCATC | | | | | | | | | | |
| Chromosome | start | end | strand | MM | target_seq | PAM | alignment | distance | position | gene name | gene id |
| GU305914.1 | 100886 | 100908 | + | 0 | GATGGCCAGGTTCGTCGCGG | TGG | ||||||||[||||||||||||]PAM | NA | Intergenic | NA | NA |
| T7310 | AGAGTAGCCCACCATCGTGTCGG | 1000 | | | | | | | | | |
| Oligo fwd | CACCAGAGTAGCCCACCATCGTGT | | | | | | | | | | |
| Oligo rev | AAACACACGATGGTGGGCTACTCT | | | | | | | | | | |
| Chromosome | start | end | strand | MM | target_seq | PAM | alignment | distance | position | gene name | gene id |
| GU305914.1 | 100910 | 100932 | + | 0 | AGAGTAGCCCACCATCGTGT | CGG | ||||||||[||||||||||||]PAM | NA | Intergenic | NA | NA |
| T7311 | CCGTGTTCCAGTCGTAGCCGCGG | 1000 | | | | | | | | | |
| Oligo fwd | CACCCCGTGTTCCAGTCGTAGCCG | | | | | | | | | | |
| Oligo rev | AAACCGGCTACGACTGGAACACGG | | | | | | | | | | |
| Chromosome | start | end | strand | MM | target_seq | PAM | alignment | distance | position | gene name | gene id |
| GU305914.1 | 100936 | 100958 | + | 0 | CCGTGTTCCAGTCGTAGCCG | CGG | ||||||||[||||||||||||]PAM | NA | Intergenic | NA | NA |
| T7312 | AGTCGTAGCCGCGGTAGCTGTGG | 1000 | | | | | | | | | |
| Oligo fwd | CACCAGTCGTAGCCGCGGTAGCTG | | | | | | | | | | |

Oligo rev AAACCAGCTACCGCGGCTACGACT
Chromosome start end strand MM target_seq PAM alignment distance position gene name gene id
GU305914.1 100945 100967 + 0 AGTCGTAGCCGCGGTAGCTG TGG ||||||||[||||||||||||]PAM NA Intergenic NA NA

T7313 GTCGTAGCCGCGGTAGCTGTGGG 1000
Oligo fwd CACCGTCGTAGCCGCGGTAGCTGT
Oligo rev AAACACAGCTACCGCGGCTACGAC
Chromosome start end strand MM target_seq PAM alignment distance position gene name gene id
GU305914.1 100946 100968 + 0 GTCGTAGCCGCGGTAGCTGT GGG ||||||||[||||||||||||]PAM NA Intergenic NA NA

T7314 TCGTAGCCGCGGTAGCTGTGGGG 1000
Oligo fwd CACCTCGTAGCCGCGGTAGCTGTG
Oligo rev AAACCACAGCTACCGCGGCTACGA
Chromosome start end strand MM target_seq PAM alignment distance position gene name gene id
GU305914.1 100947 100969 + 0 TCGTAGCCGCGGTAGCTGTG GGG ||||||||[||||||||||||]PAM NA Intergenic NA NA

T7315 TAGCCGCGGTAGCTGTGGGGCGG 1000
Oligo fwd CACCTAGCCGCGGTAGCTGTGGGG
Oligo rev AAACCCCCACAGCTACCGCGGCTA
Chromosome start end strand MM target_seq PAM alignment distance position gene name gene id
GU305914.1 100950 100972 + 0 TAGCCGCGGTAGCTGTGGGG CGG ||||||||[||||||||||||]PAM NA Intergenic NA NA

T7316 AGCCGCGGTAGCTGTGGGGCGGG 1000
Oligo fwd CACCAGCCGCGGTAGCTGTGGGGC
Oligo rev AAACGCCCCACAGCTACCGCGGCT
Chromosome start end strand MM target_seq PAM alignment distance position gene name gene id
GU305914.1 100951 100973 + 0 AGCCGCGGTAGCTGTGGGGC GGG ||||||||[||||||||||||]PAM NA Intergenic NA NA

T7317 GGGGCGGGCTCAGCTTCGCGCGG 1000
Oligo fwd CACCGGGGCGGGCTCAGCTTCGCG
Oligo rev AAACCGCGAAGCTGAGCCCGCCCC
Chromosome start end strand MM target_seq PAM alignment distance position gene name gene id
GU305914.1 100966 100988 + 0 GGGGCGGGCTCAGCTTCGCG CGG ||||||||[||||||||||||]PAM NA Intergenic NA NA

T7318 TCAGCTTCGCGCGGTCGACGTGG 1000
Oligo fwd CACCTCAGCTTCGCGCGGTCGACG
Oligo rev AAACCGTCGACCGCGCGAAGCTGA
Chromosome start end strand MM target_seq PAM alignment distance position gene name gene id
GU305914.1 100975 100997 + 0 TCAGCTTCGCGCGGTCGACG TGG ||||||||[||||||||||||]PAM NA Intergenic NA NA

T7319 ATAGCCCTCCACCGAAAAGTAGG 1000
Oligo fwd CACCATAGCCCTCCACCGAAAAGT
Oligo rev AAACACTTTTCGGTGGAGGGCTAT
Chromosome start end strand MM target_seq PAM alignment distance position gene name gene id
GU305914.1 101012 101034 + 0 ATAGCCCTCCACCGAAAAGT AGG ||||||||[||||||||||||]PAM NA Intergenic NA NA

T7320 CGAAAAGTAGGTATTCATGATGG 1000
Oligo fwd CACCCGAAAAGTAGGTATTCATGA
Oligo rev AAACTCATGAATACCTACTTTTCG
Chromosome start end strand MM target_seq PAM alignment distance position gene name gene id
GU305914.1 101024 101046 + 0 CGAAAAGTAGGTATTCATGA TGG ||||||||[||||||||||||]PAM NA Intergenic NA NA

T7321 ATTCATGATGGAGATCAGTTCGG 1000
Oligo fwd CACCATTCATGATGGAGATCAGTT
Oligo rev AAACAACTGATCTCCATCATGAAT
Chromosome start end strand MM target_seq PAM alignment distance position gene name gene id
GU305914.1 101036 101058 + 0 ATTCATGATGGAGATCAGTT CGG ||||||||[||||||||||||]PAM NA Intergenic NA NA

T7322 TGGAGATCAGTTCGGCGTCCAGG 1000
Oligo fwd CACCTGGAGATCAGTTCGGCGTCC
Oligo rev AAACGGACGCCGAACTGATCTCCA
Chromosome start end strand MM target_seq PAM alignment distance position gene name gene id
GU305914.1 101044 101066 + 0 TGGAGATCAGTTCGGCGTCC AGG ||||||||[||||||||||||]PAM NA Intergenic NA NA

T7323 CGGCGTCCAGGCTGCGCCCCAGG 1000
Oligo fwd CACCCGGCGTCCAGGCTGCGCCCC
Oligo rev AAACGGGGCGCAGCCTGGACGCCG
Chromosome start end strand MM target_seq PAM alignment distance position gene name gene id
GU305914.1 101056 101078 + 0 CGGCGTCCAGGCTGCGCCCC AGG ||||||||[||||||||||||]PAM NA Intergenic NA NA

T7324 GCCGCGCACTGCTGATCGTCCGG 1000
Oligo fwd CACCGCCGCGCACTGCTGATCGTC
Oligo rev AAACGACGATCAGCAGTGCGCGGC
Chromosome start end strand MM target_seq PAM alignment distance position gene name gene id
GU305914.1 101091 101113 + 0 GCCGCGCACTGCTGATCGTC CGG ||||||||[||||||||||||]PAM NA Intergenic NA NA

T7325 CGCACTGCTGATCGTCCGGCAGG 1000
Oligo fwd CACCCGCACTGCTGATCGTCCGGC
Oligo rev AAACGCCGGACGATCAGCAGTGCG
Chromosome start end strand MM target_seq PAM alignment distance position gene name gene id
GU305914.1 101095 101117 + 0 CGCACTGCTGATCGTCCGGC AGG ||||||||[||||||||||||]PAM NA Intergenic NA NA

T7326 CCGTGTGCTCGCAGTACTGCAGG 1000
Oligo fwd CACCCCGTGTGCTCGCAGTACTGC
Oligo rev AAACGCAGTACTGCGAGCACACGG
Chromosome start end strand MM target_seq PAM alignment distance position gene name gene id
GU305914.1 101131 101153 + 0 CCGTGTGCTCGCAGTACTGC AGG ||||||||[||||||||||||]PAM NA Intergenic NA NA

T7327 AGTACTGCAGGAACTTCTCCCGG 1000

| | | | | | | | | | | | |
|---|---|---|---|---|---|---|---|---|---|---|---|
| Oligo fwd | CACCAGTACTGCAGGAACTTCTCC | | | | | | | | | | |
| Oligo rev | AAACGGAGAAGTTCCTGCAGTACT | | | | | | | | | | |
| Chromosome | start | end | strand | MM | target_seq | PAM | alignment | distance | position | gene name | gene id |
| GU305914.1 | 101143 | 101165 | + | 0 | AGTACTGCAGGAACTTCTCC | CGG | \|\|\|\|\|\|\|\|[\|\|\|\|\|\|\|\|\|\|\|\|]PAM | NA | Intergenic | NA | NA |
| T7328 | TCTCCCGGCACAGCGCCGTCCGG | 1000 | | | | | | | | | |
| Oligo fwd | CACCTCTCCCGGCACAGCGCCGTC | | | | | | | | | | |
| Oligo rev | AAACGACGGCGCTGTGCCGGGAGA | | | | | | | | | | |
| Chromosome | start | end | strand | MM | target_seq | PAM | alignment | distance | position | gene name | gene id |
| GU305914.1 | 101158 | 101180 | + | 0 | TCTCCCGGCACAGCGCCGTC | CGG | \|\|\|\|\|\|\|\|[\|\|\|\|\|\|\|\|\|\|\|\|]PAM | NA | Intergenic | NA | NA |
| T7329 | CCGGCACAGCGCCGTCCGGTAGG | 1000 | | | | | | | | | |
| Oligo fwd | CACCCCGGCACAGCGCCGTCCGGT | | | | | | | | | | |
| Oligo rev | AAACACCGGACGGCGCTGTGCCGG | | | | | | | | | | |
| Chromosome | start | end | strand | MM | target_seq | PAM | alignment | distance | position | gene name | gene id |
| GU305914.1 | 101162 | 101184 | + | 0 | CCGGCACAGCGCCGTCCGGT | AGG | \|\|\|\|\|\|\|\|[\|\|\|\|\|\|\|\|\|\|\|\|]PAM | NA | Intergenic | NA | NA |
| T7330 | ACAGCGCCGTCCGGTAGGCCAGG | 1000 | | | | | | | | | |
| Oligo fwd | CACCACAGCGCCGTCCGGTAGGCC | | | | | | | | | | |
| Oligo rev | AAACGGCCTACCGGACGGCGCTGT | | | | | | | | | | |
| Chromosome | start | end | strand | MM | target_seq | PAM | alignment | distance | position | gene name | gene id |
| GU305914.1 | 101167 | 101189 | + | 0 | ACAGCGCCGTCCGGTAGGCC | AGG | \|\|\|\|\|\|\|\|[\|\|\|\|\|\|\|\|\|\|\|\|]PAM | NA | Intergenic | NA | NA |
| T7331 | CACGTAGTACACGCACGAGTCGG | 1000 | | | | | | | | | |
| Oligo fwd | CACCCACGTAGTACACGCACGAGT | | | | | | | | | | |
| Oligo rev | AAACACTCGTGCGTGTACTACGTG | | | | | | | | | | |
| Chromosome | start | end | strand | MM | target_seq | PAM | alignment | distance | position | gene name | gene id |
| GU305914.1 | 101192 | 101214 | + | 0 | CACGTAGTACACGCACGAGT | CGG | \|\|\|\|\|\|\|\|[\|\|\|\|\|\|\|\|\|\|\|\|]PAM | NA | Intergenic | NA | NA |
| T7332 | ACACGCACGAGTCGGCCGCCAGG | 1000 | | | | | | | | | |
| Oligo fwd | CACCACACGCACGAGTCGGCCGCC | | | | | | | | | | |
| Oligo rev | AAACGGCGGCCGACTCGTGCGTGT | | | | | | | | | | |
| Chromosome | start | end | strand | MM | target_seq | PAM | alignment | distance | position | gene name | gene id |
| GU305914.1 | 101200 | 101222 | + | 0 | ACACGCACGAGTCGGCCGCC | AGG | \|\|\|\|\|\|\|\|[\|\|\|\|\|\|\|\|\|\|\|\|]PAM | NA | Intergenic | NA | NA |
| T7333 | CCGCCAGGCGCGATTCCTCGCGG | 1000 | | | | | | | | | |
| Oligo fwd | CACCCCGCCAGGCGCGATTCCTCG | | | | | | | | | | |
| Oligo rev | AAACCGAGGAATCGCGCCTGGCGG | | | | | | | | | | |
| Chromosome | start | end | strand | MM | target_seq | PAM | alignment | distance | position | gene name | gene id |
| GU305914.1 | 101215 | 101237 | + | 0 | CCGCCAGGCGCGATTCCTCG | CGG | \|\|\|\|\|\|\|\|[\|\|\|\|\|\|\|\|\|\|\|\|]PAM | NA | Intergenic | NA | NA |
| T7334 | CTCGCGGAACGCGTTGATCTTGG | 1000 | | | | | | | | | |
| Oligo fwd | CACCCTCGCGGAACGCGTTGATCT | | | | | | | | | | |
| Oligo rev | AAACAGATCAACGCGTTCCGCGAG | | | | | | | | | | |
| Chromosome | start | end | strand | MM | target_seq | PAM | alignment | distance | position | gene name | gene id |
| GU305914.1 | 101231 | 101253 | + | 0 | CTCGCGGAACGCGTTGATCT | TGG | \|\|\|\|\|\|\|\|[\|\|\|\|\|\|\|\|\|\|\|\|]PAM | NA | Intergenic | NA | NA |
| T7335 | CCACGTATTCCAGCTCGCGCCGG | 1000 | | | | | | | | | |
| Oligo fwd | CACCCCACGTATTCCAGCTCGCGC | | | | | | | | | | |
| Oligo rev | AAACGCGCGAGCTGGAATACGTGG | | | | | | | | | | |
| Chromosome | start | end | strand | MM | target_seq | PAM | alignment | distance | position | gene name | gene id |
| GU305914.1 | 101257 | 101279 | + | 0 | CCACGTATTCCAGCTCGCGC | CGG | \|\|\|\|\|\|\|\|[\|\|\|\|\|\|\|\|\|\|\|\|]PAM | NA | Intergenic | NA | NA |
| T7336 | ATTCCAGCTCGCGCCGGTCTCGG | 1000 | | | | | | | | | |
| Oligo fwd | CACCATTCCAGCTCGCGCCGGTCT | | | | | | | | | | |
| Oligo rev | AAACAGACCGGCGCGAGCTGGAAT | | | | | | | | | | |
| Chromosome | start | end | strand | MM | target_seq | PAM | alignment | distance | position | gene name | gene id |
| GU305914.1 | 101263 | 101285 | + | 0 | ATTCCAGCTCGCGCCGGTCT | CGG | \|\|\|\|\|\|\|\|[\|\|\|\|\|\|\|\|\|\|\|\|]PAM | NA | Intergenic | NA | NA |
| T7337 | TTCCAGCTCGCGCCGGTCTCGGG | 1000 | | | | | | | | | |
| Oligo fwd | CACCTTCCAGCTCGCGCCGGTCTC | | | | | | | | | | |
| Oligo rev | AAACGAGACCGGCGCGAGCTGGAA | | | | | | | | | | |
| Chromosome | start | end | strand | MM | target_seq | PAM | alignment | distance | position | gene name | gene id |
| GU305914.1 | 101264 | 101286 | + | 0 | TTCCAGCTCGCGCCGGTCTC | GGG | \|\|\|\|\|\|\|\|[\|\|\|\|\|\|\|\|\|\|\|\|]PAM | NA | Intergenic | NA | NA |
| T7338 | GTCTCGGGCGTTTCTCGCCTTGG | 1000 | | | | | | | | | |
| Oligo fwd | CACCGTCTCGGGCGTTTCTCGCCT | | | | | | | | | | |
| Oligo rev | AAACAGGCGAGAAACGCCCGAGAC | | | | | | | | | | |
| Chromosome | start | end | strand | MM | target_seq | PAM | alignment | distance | position | gene name | gene id |
| GU305914.1 | 101279 | 101301 | + | 0 | GTCTCGGGCGTTTCTCGCCT | TGG | \|\|\|\|\|\|\|\|[\|\|\|\|\|\|\|\|\|\|\|\|]PAM | NA | Intergenic | NA | NA |
| T7339 | GGCGAACGCCGCGAACCGCGCGG | 1000 | | | | | | | | | |
| Oligo fwd | CACCGGCGAACGCCGCGAACCGCG | | | | | | | | | | |
| Oligo rev | AAACCGCGGTTCGCGGCGTTCGCC | | | | | | | | | | |
| Chromosome | start | end | strand | MM | target_seq | PAM | alignment | distance | position | gene name | gene id |
| GU305914.1 | 101300 | 101322 | + | 0 | GGCGAACGCCGCGAACCGCG | CGG | \|\|\|\|\|\|\|\|[\|\|\|\|\|\|\|\|\|\|\|\|]PAM | NA | Intergenic | NA | NA |
| T7340 | CGAACCGCGCGGTCTGCTTGCGG | 1000 | | | | | | | | | |
| Oligo fwd | CACCCGAACCGCGCGGTCTGCTTG | | | | | | | | | | |
| Oligo rev | AAACCAAGCAGACCGCGCGGTTCG | | | | | | | | | | |
| Chromosome | start | end | strand | MM | target_seq | PAM | alignment | distance | position | gene name | gene id |
| GU305914.1 | 101311 | 101333 | + | 0 | CGAACCGCGCGGTCTGCTTG | CGG | \|\|\|\|\|\|\|\|[\|\|\|\|\|\|\|\|\|\|\|\|]PAM | NA | Intergenic | NA | NA |
| T7341 | GCGGAACCGCGAGCCGTTCATGG | 1000 | | | | | | | | | |
| Oligo fwd | CACCGCGGAACCGCGAGCCGTTCA | | | | | | | | | | |
| Oligo rev | AAACTGAACGGCTCGCGGTTCCGC | | | | | | | | | | |
| Chromosome | start | end | strand | MM | target_seq | PAM | alignment | distance | position | gene name | gene id |
| GU305914.1 | 101330 | 101352 | + | 0 | GCGGAACCGCGAGCCGTTCA | TGG | \|\|\|\|\|\|\|\|[\|\|\|\|\|\|\|\|\|\|\|\|]PAM | NA | Intergenic | NA | NA |

<table>
<tr><td>T7342</td><td>GCATGTTGATGTACGTGAACAGG</td><td>1000</td><td></td><td></td><td></td><td></td><td></td><td></td><td></td><td></td><td></td></tr>
<tr><td>Oligo fwd</td><td>CACCGCATGTTGATGTACGTGAAC</td><td></td><td></td><td></td><td></td><td></td><td></td><td></td><td></td><td></td><td></td></tr>
<tr><td>Oligo rev</td><td>AAACGTTCACGTACATCAACATGC</td><td></td><td></td><td></td><td></td><td></td><td></td><td></td><td></td><td></td><td></td></tr>
<tr><td>Chromosome</td><td>start</td><td>end</td><td>strand</td><td>MM</td><td>target_seq</td><td>PAM</td><td>alignment</td><td>distance</td><td>position</td><td>gene name</td><td>gene id</td></tr>
<tr><td>GU305914.1</td><td>101368</td><td>101390</td><td>+</td><td>0</td><td>GCATGTTGATGTACGTGAAC</td><td>AGG</td><td>||||||||[||||||||||||]PAM</td><td>NA</td><td>Intergenic</td><td>NA</td><td>NA</td></tr>
<tr><td>T7343</td><td>TGATGTACGTGAACAGGTTGCGG</td><td>1000</td><td></td><td></td><td></td><td></td><td></td><td></td><td></td><td></td><td></td></tr>
<tr><td>Oligo fwd</td><td>CACCTGATGTACGTGAACAGGTTG</td><td></td><td></td><td></td><td></td><td></td><td></td><td></td><td></td><td></td><td></td></tr>
<tr><td>Oligo rev</td><td>AAACCAACCTGTTCACGTACATCA</td><td></td><td></td><td></td><td></td><td></td><td></td><td></td><td></td><td></td><td></td></tr>
<tr><td>Chromosome</td><td>start</td><td>end</td><td>strand</td><td>MM</td><td>target_seq</td><td>PAM</td><td>alignment</td><td>distance</td><td>position</td><td>gene name</td><td>gene id</td></tr>
<tr><td>GU305914.1</td><td>101374</td><td>101396</td><td>+</td><td>0</td><td>TGATGTACGTGAACAGGTTG</td><td>CGG</td><td>||||||||[||||||||||||]PAM</td><td>NA</td><td>Intergenic</td><td>NA</td><td>NA</td></tr>
<tr><td>T7344</td><td>GTACGTGAACAGGTTGCGGCCGG</td><td>1000</td><td></td><td></td><td></td><td></td><td></td><td></td><td></td><td></td><td></td></tr>
<tr><td>Oligo fwd</td><td>CACCGTACGTGAACAGGTTGCGGC</td><td></td><td></td><td></td><td></td><td></td><td></td><td></td><td></td><td></td><td></td></tr>
<tr><td>Oligo rev</td><td>AAACGCCGCAACCTGTTCACGTAC</td><td></td><td></td><td></td><td></td><td></td><td></td><td></td><td></td><td></td><td></td></tr>
<tr><td>Chromosome</td><td>start</td><td>end</td><td>strand</td><td>MM</td><td>target_seq</td><td>PAM</td><td>alignment</td><td>distance</td><td>position</td><td>gene name</td><td>gene id</td></tr>
<tr><td>GU305914.1</td><td>101378</td><td>101400</td><td>+</td><td>0</td><td>GTACGTGAACAGGTTGCGGC</td><td>CGG</td><td>||||||||[||||||||||||]PAM</td><td>NA</td><td>Intergenic</td><td>NA</td><td>NA</td></tr>
<tr><td>T7345</td><td>TACGTGAACAGGTTGCGGCCGGG</td><td>1000</td><td></td><td></td><td></td><td></td><td></td><td></td><td></td><td></td><td></td></tr>
<tr><td>Oligo fwd</td><td>CACCTACGTGAACAGGTTGCGGCC</td><td></td><td></td><td></td><td></td><td></td><td></td><td></td><td></td><td></td><td></td></tr>
<tr><td>Oligo rev</td><td>AAACGGCCGCAACCTGTTCACGTA</td><td></td><td></td><td></td><td></td><td></td><td></td><td></td><td></td><td></td><td></td></tr>
<tr><td>Chromosome</td><td>start</td><td>end</td><td>strand</td><td>MM</td><td>target_seq</td><td>PAM</td><td>alignment</td><td>distance</td><td>position</td><td>gene name</td><td>gene id</td></tr>
<tr><td>GU305914.1</td><td>101379</td><td>101401</td><td>+</td><td>0</td><td>TACGTGAACAGGTTGCGGCC</td><td>GGG</td><td>||||||||[||||||||||||]PAM</td><td>NA</td><td>Intergenic</td><td>NA</td><td>NA</td></tr>
<tr><td>T7346</td><td>ACGTGAACAGGTTGCGGCCGGGG</td><td>1000</td><td></td><td></td><td></td><td></td><td></td><td></td><td></td><td></td><td></td></tr>
<tr><td>Oligo fwd</td><td>CACCACGTGAACAGGTTGCGGCCG</td><td></td><td></td><td></td><td></td><td></td><td></td><td></td><td></td><td></td><td></td></tr>
<tr><td>Oligo rev</td><td>AAACCGGCCGCAACCTGTTCACGT</td><td></td><td></td><td></td><td></td><td></td><td></td><td></td><td></td><td></td><td></td></tr>
<tr><td>Chromosome</td><td>start</td><td>end</td><td>strand</td><td>MM</td><td>target_seq</td><td>PAM</td><td>alignment</td><td>distance</td><td>position</td><td>gene name</td><td>gene id</td></tr>
<tr><td>GU305914.1</td><td>101380</td><td>101402</td><td>+</td><td>0</td><td>ACGTGAACAGGTTGCGGCCG</td><td>GGG</td><td>||||||||[||||||||||||]PAM</td><td>NA</td><td>Intergenic</td><td>NA</td><td>NA</td></tr>
<tr><td>T7347</td><td>GAACAGGTTGCGGCCGGGGTCGG</td><td>1000</td><td></td><td></td><td></td><td></td><td></td><td></td><td></td><td></td><td></td></tr>
<tr><td>Oligo fwd</td><td>CACCGAACAGGTTGCGGCCGGGGT</td><td></td><td></td><td></td><td></td><td></td><td></td><td></td><td></td><td></td><td></td></tr>
<tr><td>Oligo rev</td><td>AAACACCCCGGCCGCAACCTGTTC</td><td></td><td></td><td></td><td></td><td></td><td></td><td></td><td></td><td></td><td></td></tr>
<tr><td>Chromosome</td><td>start</td><td>end</td><td>strand</td><td>MM</td><td>target_seq</td><td>PAM</td><td>alignment</td><td>distance</td><td>position</td><td>gene name</td><td>gene id</td></tr>
<tr><td>GU305914.1</td><td>101384</td><td>101406</td><td>+</td><td>0</td><td>GAACAGGTTGCGGCCGGGGT</td><td>CGG</td><td>||||||||[||||||||||||]PAM</td><td>NA</td><td>Intergenic</td><td>NA</td><td>NA</td></tr>
<tr><td>T7348</td><td>CAGGTTGCGGCCGGGGTCGGCGG</td><td>1000</td><td></td><td></td><td></td><td></td><td></td><td></td><td></td><td></td><td></td></tr>
<tr><td>Oligo fwd</td><td>CACCCAGGTTGCGGCCGGGGTCGG</td><td></td><td></td><td></td><td></td><td></td><td></td><td></td><td></td><td></td><td></td></tr>
<tr><td>Oligo rev</td><td>AAACCCGACCCCGGCCGCAACCTG</td><td></td><td></td><td></td><td></td><td></td><td></td><td></td><td></td><td></td><td></td></tr>
<tr><td>Chromosome</td><td>start</td><td>end</td><td>strand</td><td>MM</td><td>target_seq</td><td>PAM</td><td>alignment</td><td>distance</td><td>position</td><td>gene name</td><td>gene id</td></tr>
<tr><td>GU305914.1</td><td>101387</td><td>101409</td><td>+</td><td>0</td><td>CAGGTTGCGGCCGGGGTCGG</td><td>CGG</td><td>||||||||[||||||||||||]PAM</td><td>NA</td><td>Intergenic</td><td>NA</td><td>NA</td></tr>
<tr><td>T7349</td><td>TGAAACATTTCGTGTAATCGCGG</td><td>1000</td><td></td><td></td><td></td><td></td><td></td><td></td><td></td><td></td><td></td></tr>
<tr><td>Oligo fwd</td><td>CACCTGAAACATTTCGTGTAATCG</td><td></td><td></td><td></td><td></td><td></td><td></td><td></td><td></td><td></td><td></td></tr>
<tr><td>Oligo rev</td><td>AAACCGATTACACGAAATGTTTCA</td><td></td><td></td><td></td><td></td><td></td><td></td><td></td><td></td><td></td><td></td></tr>
<tr><td>Chromosome</td><td>start</td><td>end</td><td>strand</td><td>MM</td><td>target_seq</td><td>PAM</td><td>alignment</td><td>distance</td><td>position</td><td>gene name</td><td>gene id</td></tr>
<tr><td>GU305914.1</td><td>101410</td><td>101432</td><td>+</td><td>0</td><td>TGAAACATTTCGTGTAATCG</td><td>CGG</td><td>||||||||[||||||||||||]PAM</td><td>NA</td><td>Intergenic</td><td>NA</td><td>NA</td></tr>
<tr><td>T7350</td><td>GAAACATTTCGTGTAATCGCGGG</td><td>1000</td><td></td><td></td><td></td><td></td><td></td><td></td><td></td><td></td><td></td></tr>
<tr><td>Oligo fwd</td><td>CACCGAAACATTTCGTGTAATCGC</td><td></td><td></td><td></td><td></td><td></td><td></td><td></td><td></td><td></td><td></td></tr>
<tr><td>Oligo rev</td><td>AAACGCGATTACACGAAATGTTTC</td><td></td><td></td><td></td><td></td><td></td><td></td><td></td><td></td><td></td><td></td></tr>
<tr><td>Chromosome</td><td>start</td><td>end</td><td>strand</td><td>MM</td><td>target_seq</td><td>PAM</td><td>alignment</td><td>distance</td><td>position</td><td>gene name</td><td>gene id</td></tr>
<tr><td>GU305914.1</td><td>101411</td><td>101433</td><td>+</td><td>0</td><td>GAAACATTTCGTGTAATCGC</td><td>GGG</td><td>||||||||[||||||||||||]PAM</td><td>NA</td><td>Intergenic</td><td>NA</td><td>NA</td></tr>
<tr><td>T7351</td><td>ACATTTCGTGTAATCGCGGGTGG</td><td>1000</td><td></td><td></td><td></td><td></td><td></td><td></td><td></td><td></td><td></td></tr>
<tr><td>Oligo fwd</td><td>CACCACATTTCGTGTAATCGCGGG</td><td></td><td></td><td></td><td></td><td></td><td></td><td></td><td></td><td></td><td></td></tr>
<tr><td>Oligo rev</td><td>AAACCCCGCGATTACACGAAATGT</td><td></td><td></td><td></td><td></td><td></td><td></td><td></td><td></td><td></td><td></td></tr>
<tr><td>Chromosome</td><td>start</td><td>end</td><td>strand</td><td>MM</td><td>target_seq</td><td>PAM</td><td>alignment</td><td>distance</td><td>position</td><td>gene name</td><td>gene id</td></tr>
<tr><td>GU305914.1</td><td>101414</td><td>101436</td><td>+</td><td>0</td><td>ACATTTCGTGTAATCGCGGG</td><td>TGG</td><td>||||||||[||||||||||||]PAM</td><td>NA</td><td>Intergenic</td><td>NA</td><td>NA</td></tr>
<tr><td>T7352</td><td>CTGCCTCTCGAACAGATAGTAGG</td><td>1000</td><td></td><td></td><td></td><td></td><td></td><td></td><td></td><td></td><td></td></tr>
<tr><td>Oligo fwd</td><td>CACCCTGCCTCTCGAACAGATAGT</td><td></td><td></td><td></td><td></td><td></td><td></td><td></td><td></td><td></td><td></td></tr>
<tr><td>Oligo rev</td><td>AAACACTATCTGTTCGAGAGGCAG</td><td></td><td></td><td></td><td></td><td></td><td></td><td></td><td></td><td></td><td></td></tr>
<tr><td>Chromosome</td><td>start</td><td>end</td><td>strand</td><td>MM</td><td>target_seq</td><td>PAM</td><td>alignment</td><td>distance</td><td>position</td><td>gene name</td><td>gene id</td></tr>
<tr><td>GU305914.1</td><td>101453</td><td>101475</td><td>+</td><td>0</td><td>CTGCCTCTCGAACAGATAGT</td><td>AGG</td><td>||||||||[||||||||||||]PAM</td><td>NA</td><td>Intergenic</td><td>NA</td><td>NA</td></tr>
<tr><td>T7353</td><td>CTCGAACAGATAGTAGGACATGG</td><td>1000</td><td></td><td></td><td></td><td></td><td></td><td></td><td></td><td></td><td></td></tr>
<tr><td>Oligo fwd</td><td>CACCCTCGAACAGATAGTAGGACA</td><td></td><td></td><td></td><td></td><td></td><td></td><td></td><td></td><td></td><td></td></tr>
<tr><td>Oligo rev</td><td>AAACTGTCCTACTATCTGTTCGAG</td><td></td><td></td><td></td><td></td><td></td><td></td><td></td><td></td><td></td><td></td></tr>
<tr><td>Chromosome</td><td>start</td><td>end</td><td>strand</td><td>MM</td><td>target_seq</td><td>PAM</td><td>alignment</td><td>distance</td><td>position</td><td>gene name</td><td>gene id</td></tr>
<tr><td>GU305914.1</td><td>101459</td><td>101481</td><td>+</td><td>0</td><td>CTCGAACAGATAGTAGGACA</td><td>TGG</td><td>||||||||[||||||||||||]PAM</td><td>NA</td><td>Intergenic</td><td>NA</td><td>NA</td></tr>
<tr><td>T7354</td><td>ACAGATAGTAGGACATGGTCAGG</td><td>1000</td><td></td><td></td><td></td><td></td><td></td><td></td><td></td><td></td><td></td></tr>
<tr><td>Oligo fwd</td><td>CACCACAGATAGTAGGACATGGTC</td><td></td><td></td><td></td><td></td><td></td><td></td><td></td><td></td><td></td><td></td></tr>
<tr><td>Oligo rev</td><td>AAACGACCATGTCCTACTATCTGT</td><td></td><td></td><td></td><td></td><td></td><td></td><td></td><td></td><td></td><td></td></tr>
<tr><td>Chromosome</td><td>start</td><td>end</td><td>strand</td><td>MM</td><td>target_seq</td><td>PAM</td><td>alignment</td><td>distance</td><td>position</td><td>gene name</td><td>gene id</td></tr>
<tr><td>GU305914.1</td><td>101464</td><td>101486</td><td>+</td><td>0</td><td>ACAGATAGTAGGACATGGTC</td><td>AGG</td><td>||||||||[||||||||||||]PAM</td><td>NA</td><td>Intergenic</td><td>NA</td><td>NA</td></tr>
<tr><td>T7355</td><td>AGTAGGACATGGTCAGGATCAGG</td><td>1000</td><td></td><td></td><td></td><td></td><td></td><td></td><td></td><td></td><td></td></tr>
<tr><td>Oligo fwd</td><td>CACCAGTAGGACATGGTCAGGATC</td><td></td><td></td><td></td><td></td><td></td><td></td><td></td><td></td><td></td><td></td></tr>
<tr><td>Oligo rev</td><td>AAACGATCCTGACCATGTCCTACT</td><td></td><td></td><td></td><td></td><td></td><td></td><td></td><td></td><td></td><td></td></tr>
<tr><td>Chromosome</td><td>start</td><td>end</td><td>strand</td><td>MM</td><td>target_seq</td><td>PAM</td><td>alignment</td><td>distance</td><td>position</td><td>gene name</td><td>gene id</td></tr>
<tr><td>GU305914.1</td><td>101470</td><td>101492</td><td>+</td><td>0</td><td>AGTAGGACATGGTCAGGATC</td><td>AGG</td><td>||||||||[||||||||||||]PAM</td><td>NA</td><td>Intergenic</td><td>NA</td><td>NA</td></tr>
<tr><td>T7356</td><td>GTAGGACATGGTCAGGATCAGGG</td><td>1000</td><td></td><td></td><td></td><td></td><td></td><td></td><td></td><td></td><td></td></tr>
<tr><td>Oligo fwd</td><td>CACCGTAGGACATGGTCAGGATCA</td><td></td><td></td><td></td><td></td><td></td><td></td><td></td><td></td><td></td><td></td></tr>
<tr><td>Oligo rev</td><td>AAACTGATCCTGACCATGTCCTAC</td><td></td><td></td><td></td><td></td><td></td><td></td><td></td><td></td><td></td><td></td></tr>
<tr><td>Chromosome</td><td>start</td><td>end</td><td>strand</td><td>MM</td><td>target_seq</td><td>PAM</td><td>alignment</td><td>distance</td><td>position</td><td>gene name</td><td>gene id</td></tr>
<tr><td>GU305914.1</td><td>101471</td><td>101493</td><td>+</td><td>0</td><td>GTAGGACATGGTCAGGATCA</td><td>GGG</td><td>||||||||[||||||||||||]PAM</td><td>NA</td><td>Intergenic</td><td>NA</td><td>NA</td></tr>
</table>

| | | | | | | | | | | | |
|---|---|---|---|---|---|---|---|---|---|---|---|
| T7357 | TCAGGATCAGGGTACCCTCGTGG | 1000 | | | | | | | | | |
| Oligo fwd | CACCTCAGGATCAGGGTACCCTCG | | | | | | | | | | |
| Oligo rev | AAACCGAGGGTACCCTGATCCTGA | | | | | | | | | | |
| Chromosome | start | end | strand | MM | target_seq | PAM | alignment | distance | position | gene name | gene id |
| GU305914.1 | 101482 | 101504 | + | 0 | TCAGGATCAGGGTACCCTCG | TGG | ||||||||[||||||||||||]PAM | NA | Intergenic | NA | NA |
| T7358 | CAGGATCAGGGTACCCTCGTGGG | 1000 | | | | | | | | | |
| Oligo fwd | CACCCAGGATCAGGGTACCCTCGT | | | | | | | | | | |
| Oligo rev | AAACACGAGGGTACCCTGATCCTG | | | | | | | | | | |
| Chromosome | start | end | strand | MM | target_seq | PAM | alignment | distance | position | gene name | gene id |
| GU305914.1 | 101483 | 101505 | + | 0 | CAGGATCAGGGTACCCTCGT | GGG | ||||||||[||||||||||||]PAM | NA | Intergenic | NA | NA |
| T7359 | CGTGGGTCCCGTACGTGCTCAGG | 1000 | | | | | | | | | |
| Oligo fwd | CACCCGTGGGTCCCGTACGTGCTC | | | | | | | | | | |
| Oligo rev | AAACGAGCACGTACGGGACCCACG | | | | | | | | | | |
| Chromosome | start | end | strand | MM | target_seq | PAM | alignment | distance | position | gene name | gene id |
| GU305914.1 | 101500 | 101522 | + | 0 | CGTGGGTCCCGTACGTGCTC | AGG | ||||||||[||||||||||||]PAM | NA | Intergenic | NA | NA |
| T7360 | CCCGTACGTGCTCAGGAACCAGG | 1000 | | | | | | | | | |
| Oligo fwd | CACCCCCGTACGTGCTCAGGAACC | | | | | | | | | | |
| Oligo rev | AAACGGTTCCTGAGCACGTACGGG | | | | | | | | | | |
| Chromosome | start | end | strand | MM | target_seq | PAM | alignment | distance | position | gene name | gene id |
| GU305914.1 | 101507 | 101529 | + | 0 | CCCGTACGTGCTCAGGAACC | AGG | ||||||||[||||||||||||]PAM | NA | Intergenic | NA | NA |
| T7361 | GTACGTGCTCAGGAACCAGGAGG | 1000 | | | | | | | | | |
| Oligo fwd | CACCGTACGTGCTCAGGAACCAGG | | | | | | | | | | |
| Oligo rev | AAACCCTGGTTCCTGAGCACGTAC | | | | | | | | | | |
| Chromosome | start | end | strand | MM | target_seq | PAM | alignment | distance | position | gene name | gene id |
| GU305914.1 | 101510 | 101532 | + | 0 | GTACGTGCTCAGGAACCAGG | AGG | ||||||||[||||||||||||]PAM | NA | Intergenic | NA | NA |
| T7362 | TACGTGCTCAGGAACCAGGAGGG | 1000 | | | | | | | | | |
| Oligo fwd | CACCTACGTGCTCAGGAACCAGGA | | | | | | | | | | |
| Oligo rev | AAACTCCTGGTTCCTGAGCACGTA | | | | | | | | | | |
| Chromosome | start | end | strand | MM | target_seq | PAM | alignment | distance | position | gene name | gene id |
| GU305914.1 | 101511 | 101533 | + | 0 | TACGTGCTCAGGAACCAGGA | GGG | ||||||||[||||||||||||]PAM | NA | Intergenic | NA | NA |
| T7363 | ACGTGCTCAGGAACCAGGAGGGG | 1000 | | | | | | | | | |
| Oligo fwd | CACCACGTGCTCAGGAACCAGGAG | | | | | | | | | | |
| Oligo rev | AAACCTCCTGGTTCCTGAGCACGT | | | | | | | | | | |
| Chromosome | start | end | strand | MM | target_seq | PAM | alignment | distance | position | gene name | gene id |
| GU305914.1 | 101512 | 101534 | + | 0 | ACGTGCTCAGGAACCAGGAG | GGG | ||||||||[||||||||||||]PAM | NA | Intergenic | NA | NA |
| T7364 | CGTGCTCAGGAACCAGGAGGGGG | 1000 | | | | | | | | | |
| Oligo fwd | CACCCGTGCTCAGGAACCAGGAGG | | | | | | | | | | |
| Oligo rev | AAACCCTCCTGGTTCCTGAGCACG | | | | | | | | | | |
| Chromosome | start | end | strand | MM | target_seq | PAM | alignment | distance | position | gene name | gene id |
| GU305914.1 | 101513 | 101535 | + | 0 | CGTGCTCAGGAACCAGGAGG | GGG | ||||||||[||||||||||||]PAM | NA | Intergenic | NA | NA |
| T7365 | CAGGAACCAGGAGGGGGACGTGG | 1000 | | | | | | | | | |
| Oligo fwd | CACCCAGGAACCAGGAGGGGGACG | | | | | | | | | | |
| Oligo rev | AAACCGTCCCCCTCCTGGTTCCTG | | | | | | | | | | |
| Chromosome | start | end | strand | MM | target_seq | PAM | alignment | distance | position | gene name | gene id |
| GU305914.1 | 101519 | 101541 | + | 0 | CAGGAACCAGGAGGGGGACG | TGG | ||||||||[||||||||||||]PAM | NA | Intergenic | NA | NA |
| T7366 | AGGAACCAGGAGGGGGACGTGGG | 1000 | | | | | | | | | |
| Oligo fwd | CACCAGGAACCAGGAGGGGGACGT | | | | | | | | | | |
| Oligo rev | AAACACGTCCCCCTCCTGGTTCCT | | | | | | | | | | |
| Chromosome | start | end | strand | MM | target_seq | PAM | alignment | distance | position | gene name | gene id |
| GU305914.1 | 101520 | 101542 | + | 0 | AGGAACCAGGAGGGGGACGT | GGG | ||||||||[||||||||||||]PAM | NA | Intergenic | NA | NA |
| T7367 | GGAACCAGGAGGGGGACGTGGGG | 1000 | | | | | | | | | |
| Oligo fwd | CACCGGAACCAGGAGGGGGACGTG | | | | | | | | | | |
| Oligo rev | AAACCACGTCCCCCTCCTGGTTCC | | | | | | | | | | |
| Chromosome | start | end | strand | MM | target_seq | PAM | alignment | distance | position | gene name | gene id |
| GU305914.1 | 101521 | 101543 | + | 0 | GGAACCAGGAGGGGGACGTG | GGG | ||||||||[||||||||||||]PAM | NA | Intergenic | NA | NA |
| T7368 | GAACCAGGAGGGGGACGTGGGGG | 1000 | | | | | | | | | |
| Oligo fwd | CACCGAACCAGGAGGGGGACGTGG | | | | | | | | | | |
| Oligo rev | AAACCCACGTCCCCCTCCTGGTTC | | | | | | | | | | |
| Chromosome | start | end | strand | MM | target_seq | PAM | alignment | distance | position | gene name | gene id |
| GU305914.1 | 101522 | 101544 | + | 0 | GAACCAGGAGGGGGACGTGG | GGG | ||||||||[||||||||||||]PAM | NA | Intergenic | NA | NA |
| T7369 | AGGAGGGGGACGTGGGGGACAGG | 1000 | | | | | | | | | |
| Oligo fwd | CACCAGGAGGGGGACGTGGGGGAC | | | | | | | | | | |
| Oligo rev | AAACGTCCCCCACGTCCCCCTCCT | | | | | | | | | | |
| Chromosome | start | end | strand | MM | target_seq | PAM | alignment | distance | position | gene name | gene id |
| GU305914.1 | 101527 | 101549 | + | 0 | AGGAGGGGGACGTGGGGGAC | AGG | ||||||||[||||||||||||]PAM | NA | Intergenic | NA | NA |
| T7370 | GGGGGACGTGGGGGACAGGAAGG | 1000 | | | | | | | | | |
| Oligo fwd | CACCGGGGGACGTGGGGGACAGGA | | | | | | | | | | |
| Oligo rev | AAACTCCTGTCCCCCACGTCCCCC | | | | | | | | | | |
| Chromosome | start | end | strand | MM | target_seq | PAM | alignment | distance | position | gene name | gene id |
| GU305914.1 | 101531 | 101553 | + | 0 | GGGGGACGTGGGGGACAGGA | AGG | ||||||||[||||||||||||]PAM | NA | Intergenic | NA | NA |
| T7371 | AGATCGCGCTGCGATTGTAATGG | 1000 | | | | | | | | | |
| Oligo fwd | CACCAGATCGCGCTGCGATTGTAA | | | | | | | | | | |
| Oligo rev | AAACTTACAATCGCAGCGCGATCT | | | | | | | | | | |
| Chromosome | start | end | strand | MM | target_seq | PAM | alignment | distance | position | gene name | gene id |

<table>
<tr><td>GU305914.1</td><td>101572</td><td>101594</td><td>+</td><td>0</td><td>AGATCGCGCTGCGATTGTAA</td><td>TGG</td><td>||||||||[||||||||||||]PAM</td><td>NA</td><td>Intergenic</td><td>NA</td><td>NA</td></tr>
<tr><td>T7372</td><td>CGCTGCGATTGTAATGGACCAGG</td><td>1000</td><td></td><td></td><td></td><td></td><td></td><td></td><td></td><td></td><td></td></tr>
<tr><td>Oligo fwd</td><td>CACCCGCTGCGATTGTAATGGACC</td><td></td><td></td><td></td><td></td><td></td><td></td><td></td><td></td><td></td><td></td></tr>
<tr><td>Oligo rev</td><td>AAACGGTCCATTACAATCGCAGCG</td><td></td><td></td><td></td><td></td><td></td><td></td><td></td><td></td><td></td><td></td></tr>
<tr><td>Chromosome</td><td>start</td><td>end</td><td>strand</td><td>MM</td><td>target_seq</td><td>PAM</td><td>alignment</td><td>distance</td><td>position</td><td>gene name</td><td>gene id</td></tr>
<tr><td>GU305914.1</td><td>101578</td><td>101600</td><td>+</td><td>0</td><td>CGCTGCGATTGTAATGGACC</td><td>AGG</td><td>||||||||[||||||||||||]PAM</td><td>NA</td><td>Intergenic</td><td>NA</td><td>NA</td></tr>
<tr><td>T7373</td><td>TGTAATGGACCAGGCATTTGAGG</td><td>1000</td><td></td><td></td><td></td><td></td><td></td><td></td><td></td><td></td><td></td></tr>
<tr><td>Oligo fwd</td><td>CACCTGTAATGGACCAGGCATTTG</td><td></td><td></td><td></td><td></td><td></td><td></td><td></td><td></td><td></td><td></td></tr>
<tr><td>Oligo rev</td><td>AAACCAAATGCCTGGTCCATTACA</td><td></td><td></td><td></td><td></td><td></td><td></td><td></td><td></td><td></td><td></td></tr>
<tr><td>Chromosome</td><td>start</td><td>end</td><td>strand</td><td>MM</td><td>target_seq</td><td>PAM</td><td>alignment</td><td>distance</td><td>position</td><td>gene name</td><td>gene id</td></tr>
<tr><td>GU305914.1</td><td>101587</td><td>101609</td><td>+</td><td>0</td><td>TGTAATGGACCAGGCATTTG</td><td>AGG</td><td>||||||||[||||||||||||]PAM</td><td>NA</td><td>Intergenic</td><td>NA</td><td>NA</td></tr>
<tr><td>T7374</td><td>GTAATGGACCAGGCATTTGAGGG</td><td>1000</td><td></td><td></td><td></td><td></td><td></td><td></td><td></td><td></td><td></td></tr>
<tr><td>Oligo fwd</td><td>CACCGTAATGGACCAGGCATTTGA</td><td></td><td></td><td></td><td></td><td></td><td></td><td></td><td></td><td></td><td></td></tr>
<tr><td>Oligo rev</td><td>AAACTCAAATGCCTGGTCCATTAC</td><td></td><td></td><td></td><td></td><td></td><td></td><td></td><td></td><td></td><td></td></tr>
<tr><td>Chromosome</td><td>start</td><td>end</td><td>strand</td><td>MM</td><td>target_seq</td><td>PAM</td><td>alignment</td><td>distance</td><td>position</td><td>gene name</td><td>gene id</td></tr>
<tr><td>GU305914.1</td><td>101588</td><td>101610</td><td>+</td><td>0</td><td>GTAATGGACCAGGCATTTGA</td><td>GGG</td><td>||||||||[||||||||||||]PAM</td><td>NA</td><td>Intergenic</td><td>NA</td><td>NA</td></tr>
<tr><td>T7375</td><td>GACCAGGCATTTGAGGGTGCTGG</td><td>1000</td><td></td><td></td><td></td><td></td><td></td><td></td><td></td><td></td><td></td></tr>
<tr><td>Oligo fwd</td><td>CACCGACCAGGCATTTGAGGGTGC</td><td></td><td></td><td></td><td></td><td></td><td></td><td></td><td></td><td></td><td></td></tr>
<tr><td>Oligo rev</td><td>AAACGCACCCTCAAATGCCTGGTC</td><td></td><td></td><td></td><td></td><td></td><td></td><td></td><td></td><td></td><td></td></tr>
<tr><td>Chromosome</td><td>start</td><td>end</td><td>strand</td><td>MM</td><td>target_seq</td><td>PAM</td><td>alignment</td><td>distance</td><td>position</td><td>gene name</td><td>gene id</td></tr>
<tr><td>GU305914.1</td><td>101594</td><td>101616</td><td>+</td><td>0</td><td>GACCAGGCATTTGAGGGTGC</td><td>TGG</td><td>||||||||[||||||||||||]PAM</td><td>NA</td><td>Intergenic</td><td>NA</td><td>NA</td></tr>
<tr><td>T7376</td><td>GGCATTTGAGGGTGCTGGTGAGG</td><td>1000</td><td></td><td></td><td></td><td></td><td></td><td></td><td></td><td></td><td></td></tr>
<tr><td>Oligo fwd</td><td>CACCGGCATTTGAGGGTGCTGGTG</td><td></td><td></td><td></td><td></td><td></td><td></td><td></td><td></td><td></td><td></td></tr>
<tr><td>Oligo rev</td><td>AAACCACCAGCACCCTCAAATGCC</td><td></td><td></td><td></td><td></td><td></td><td></td><td></td><td></td><td></td><td></td></tr>
<tr><td>Chromosome</td><td>start</td><td>end</td><td>strand</td><td>MM</td><td>target_seq</td><td>PAM</td><td>alignment</td><td>distance</td><td>position</td><td>gene name</td><td>gene id</td></tr>
<tr><td>GU305914.1</td><td>101599</td><td>101621</td><td>+</td><td>0</td><td>GGCATTTGAGGGTGCTGGTG</td><td>AGG</td><td>||||||||[||||||||||||]PAM</td><td>NA</td><td>Intergenic</td><td>NA</td><td>NA</td></tr>
<tr><td>T7377</td><td>TTTGAGGGTGCTGGTGAGGTCGG</td><td>1000</td><td></td><td></td><td></td><td></td><td></td><td></td><td></td><td></td><td></td></tr>
<tr><td>Oligo fwd</td><td>CACCTTTGAGGGTGCTGGTGAGGT</td><td></td><td></td><td></td><td></td><td></td><td></td><td></td><td></td><td></td><td></td></tr>
<tr><td>Oligo rev</td><td>AAACACCTCACCAGCACCCTCAAA</td><td></td><td></td><td></td><td></td><td></td><td></td><td></td><td></td><td></td><td></td></tr>
<tr><td>Chromosome</td><td>start</td><td>end</td><td>strand</td><td>MM</td><td>target_seq</td><td>PAM</td><td>alignment</td><td>distance</td><td>position</td><td>gene name</td><td>gene id</td></tr>
<tr><td>GU305914.1</td><td>101603</td><td>101625</td><td>+</td><td>0</td><td>TTTGAGGGTGCTGGTGAGGT</td><td>CGG</td><td>||||||||[||||||||||||]PAM</td><td>NA</td><td>Intergenic</td><td>NA</td><td>NA</td></tr>
<tr><td>T7378</td><td>TGCTGGTGAGGTCGGAGAAGAGG</td><td>1000</td><td></td><td></td><td></td><td></td><td></td><td></td><td></td><td></td><td></td></tr>
<tr><td>Oligo fwd</td><td>CACCTGCTGGTGAGGTCGGAGAAG</td><td></td><td></td><td></td><td></td><td></td><td></td><td></td><td></td><td></td><td></td></tr>
<tr><td>Oligo rev</td><td>AAACCTTCTCCGACCTCACCAGCA</td><td></td><td></td><td></td><td></td><td></td><td></td><td></td><td></td><td></td><td></td></tr>
<tr><td>Chromosome</td><td>start</td><td>end</td><td>strand</td><td>MM</td><td>target_seq</td><td>PAM</td><td>alignment</td><td>distance</td><td>position</td><td>gene name</td><td>gene id</td></tr>
<tr><td>GU305914.1</td><td>101611</td><td>101633</td><td>+</td><td>0</td><td>TGCTGGTGAGGTCGGAGAAG</td><td>AGG</td><td>||||||||[||||||||||||]PAM</td><td>NA</td><td>Intergenic</td><td>NA</td><td>NA</td></tr>
<tr><td>T7379</td><td>GTCGGAGAAGAGGTCGCCGCCGG</td><td>1000</td><td></td><td></td><td></td><td></td><td></td><td></td><td></td><td></td><td></td></tr>
<tr><td>Oligo fwd</td><td>CACCGTCGGAGAAGAGGTCGCCGC</td><td></td><td></td><td></td><td></td><td></td><td></td><td></td><td></td><td></td><td></td></tr>
<tr><td>Oligo rev</td><td>AAACGCGGCGACCTCTTCTCCGAC</td><td></td><td></td><td></td><td></td><td></td><td></td><td></td><td></td><td></td><td></td></tr>
<tr><td>Chromosome</td><td>start</td><td>end</td><td>strand</td><td>MM</td><td>target_seq</td><td>PAM</td><td>alignment</td><td>distance</td><td>position</td><td>gene name</td><td>gene id</td></tr>
<tr><td>GU305914.1</td><td>101621</td><td>101643</td><td>+</td><td>0</td><td>GTCGGAGAAGAGGTCGCCGC</td><td>CGG</td><td>||||||||[||||||||||||]PAM</td><td>NA</td><td>Intergenic</td><td>NA</td><td>NA</td></tr>
<tr><td>T7380</td><td>GGAGAAGAGGTCGCCGCCGGCGG</td><td>1000</td><td></td><td></td><td></td><td></td><td></td><td></td><td></td><td></td><td></td></tr>
<tr><td>Oligo fwd</td><td>CACCGGAGAAGAGGTCGCCGCCGG</td><td></td><td></td><td></td><td></td><td></td><td></td><td></td><td></td><td></td><td></td></tr>
<tr><td>Oligo rev</td><td>AAACCCGGCGGCGACCTCTTCTCC</td><td></td><td></td><td></td><td></td><td></td><td></td><td></td><td></td><td></td><td></td></tr>
<tr><td>Chromosome</td><td>start</td><td>end</td><td>strand</td><td>MM</td><td>target_seq</td><td>PAM</td><td>alignment</td><td>distance</td><td>position</td><td>gene name</td><td>gene id</td></tr>
<tr><td>GU305914.1</td><td>101624</td><td>101646</td><td>+</td><td>0</td><td>GGAGAAGAGGTCGCCGCCGG</td><td>CGG</td><td>||||||||[||||||||||||]PAM</td><td>NA</td><td>Intergenic</td><td>NA</td><td>NA</td></tr>
<tr><td>T7381</td><td>CGTCTTTTTTACGATCGAACAGG</td><td>1000</td><td></td><td></td><td></td><td></td><td></td><td></td><td></td><td></td><td></td></tr>
<tr><td>Oligo fwd</td><td>CACCCGTCTTTTTTACGATCGAAC</td><td></td><td></td><td></td><td></td><td></td><td></td><td></td><td></td><td></td><td></td></tr>
<tr><td>Oligo rev</td><td>AAACGTTCGATCGTAAAAAAGACG</td><td></td><td></td><td></td><td></td><td></td><td></td><td></td><td></td><td></td><td></td></tr>
<tr><td>Chromosome</td><td>start</td><td>end</td><td>strand</td><td>MM</td><td>target_seq</td><td>PAM</td><td>alignment</td><td>distance</td><td>position</td><td>gene name</td><td>gene id</td></tr>
<tr><td>GU305914.1</td><td>101674</td><td>101696</td><td>+</td><td>0</td><td>CGTCTTTTTTACGATCGAAC</td><td>AGG</td><td>||||||||[||||||||||||]PAM</td><td>NA</td><td>Intergenic</td><td>NA</td><td>NA</td></tr>
<tr><td>T7382</td><td>GTACAACTCGCGAGTAGCGTCGG</td><td>1000</td><td></td><td></td><td></td><td></td><td></td><td></td><td></td><td></td><td></td></tr>
<tr><td>Oligo fwd</td><td>CACCGTACAACTCGCGAGTAGCGT</td><td></td><td></td><td></td><td></td><td></td><td></td><td></td><td></td><td></td><td></td></tr>
<tr><td>Oligo rev</td><td>AAACACGCTACTCGCGAGTTGTAC</td><td></td><td></td><td></td><td></td><td></td><td></td><td></td><td></td><td></td><td></td></tr>
<tr><td>Chromosome</td><td>start</td><td>end</td><td>strand</td><td>MM</td><td>target_seq</td><td>PAM</td><td>alignment</td><td>distance</td><td>position</td><td>gene name</td><td>gene id</td></tr>
<tr><td>GU305914.1</td><td>101702</td><td>101724</td><td>+</td><td>0</td><td>GTACAACTCGCGAGTAGCGT</td><td>CGG</td><td>||||||||[||||||||||||]PAM</td><td>NA</td><td>Intergenic</td><td>NA</td><td>NA</td></tr>
<tr><td>T7383</td><td>AGTAGCGTCGGCGTCTATGCCGG</td><td>1000</td><td></td><td></td><td></td><td></td><td></td><td></td><td></td><td></td><td></td></tr>
<tr><td>Oligo fwd</td><td>CACCAGTAGCGTCGGCGTCTATGC</td><td></td><td></td><td></td><td></td><td></td><td></td><td></td><td></td><td></td><td></td></tr>
<tr><td>Oligo rev</td><td>AAACGCATAGACGCCGACGCTACT</td><td></td><td></td><td></td><td></td><td></td><td></td><td></td><td></td><td></td><td></td></tr>
<tr><td>Chromosome</td><td>start</td><td>end</td><td>strand</td><td>MM</td><td>target_seq</td><td>PAM</td><td>alignment</td><td>distance</td><td>position</td><td>gene name</td><td>gene id</td></tr>
<tr><td>GU305914.1</td><td>101714</td><td>101736</td><td>+</td><td>0</td><td>AGTAGCGTCGGCGTCTATGC</td><td>CGG</td><td>||||||||[||||||||||||]PAM</td><td>NA</td><td>Intergenic</td><td>NA</td><td>NA</td></tr>
<tr><td>T7384</td><td>GCCGGCGCAGATAACGCGCGCGG</td><td>1000</td><td></td><td></td><td></td><td></td><td></td><td></td><td></td><td></td><td></td></tr>
<tr><td>Oligo fwd</td><td>CACCGCCGGCGCAGATAACGCGCG</td><td></td><td></td><td></td><td></td><td></td><td></td><td></td><td></td><td></td><td></td></tr>
<tr><td>Oligo rev</td><td>AAACCGCGCGTTATCTGCGCCGGC</td><td></td><td></td><td></td><td></td><td></td><td></td><td></td><td></td><td></td><td></td></tr>
<tr><td>Chromosome</td><td>start</td><td>end</td><td>strand</td><td>MM</td><td>target_seq</td><td>PAM</td><td>alignment</td><td>distance</td><td>position</td><td>gene name</td><td>gene id</td></tr>
<tr><td>GU305914.1</td><td>101732</td><td>101754</td><td>+</td><td>0</td><td>GCCGGCGCAGATAACGCGCG</td><td>CGG</td><td>||||||||[||||||||||||]PAM</td><td>NA</td><td>Intergenic</td><td>NA</td><td>NA</td></tr>
<tr><td>T7385</td><td>CCGGCGCAGATAACGCGCGCGGG</td><td>1000</td><td></td><td></td><td></td><td></td><td></td><td></td><td></td><td></td><td></td></tr>
<tr><td>Oligo fwd</td><td>CACCCCGGCGCAGATAACGCGCGC</td><td></td><td></td><td></td><td></td><td></td><td></td><td></td><td></td><td></td><td></td></tr>
<tr><td>Oligo rev</td><td>AAACGCGCGCGTTATCTGCGCCGG</td><td></td><td></td><td></td><td></td><td></td><td></td><td></td><td></td><td></td><td></td></tr>
<tr><td>Chromosome</td><td>start</td><td>end</td><td>strand</td><td>MM</td><td>target_seq</td><td>PAM</td><td>alignment</td><td>distance</td><td>position</td><td>gene name</td><td>gene id</td></tr>
<tr><td>GU305914.1</td><td>101733</td><td>101755</td><td>+</td><td>0</td><td>CCGGCGCAGATAACGCGCGC</td><td>GGG</td><td>||||||||[||||||||||||]PAM</td><td>NA</td><td>Intergenic</td><td>NA</td><td>NA</td></tr>
<tr><td>T7386</td><td>CAGATAACGCGCGCGGGTAGCGG</td><td>1000</td><td></td><td></td><td></td><td></td><td></td><td></td><td></td><td></td><td></td></tr>
<tr><td>Oligo fwd</td><td>CACCCAGATAACGCGCGCGGGTAG</td><td></td><td></td><td></td><td></td><td></td><td></td><td></td><td></td><td></td><td></td></tr>
<tr><td>Oligo rev</td><td>AAACCTACCCGCGCGCGTTATCTG</td><td></td><td></td><td></td><td></td><td></td><td></td><td></td><td></td><td></td><td></td></tr>
</table>

| | | | | | | | | | | | |
|---|---|---|---|---|---|---|---|---|---|---|---|
| Chromosome | start | end | strand | MM | target_seq | PAM | alignment | distance | position | gene name | gene id |
| GU305914.1 | 101739 | 101761 | + | 0 | CAGATAACGCGCGCGGGTAG | CGG | \|\|\|\|\|\|\|\|[\|\|\|\|\|\|\|\|\|\|\|\|]PAM | NA | Intergenic | NA | NA |
| T7387 | GCGCGCGGGTAGCGGTGTGTTGG | 1000 | | | | | | | | | |
| Oligo fwd | CACCGCGCGCGGGTAGCGGTGTGT | | | | | | | | | | |
| Oligo rev | AAACACACACCGCTACCCGCGCGC | | | | | | | | | | |
| Chromosome | start | end | strand | MM | target_seq | PAM | alignment | distance | position | gene name | gene id |
| GU305914.1 | 101747 | 101769 | + | 0 | GCGCGCGGGTAGCGGTGTGT | TGG | \|\|\|\|\|\|\|\|[\|\|\|\|\|\|\|\|\|\|\|\|]PAM | NA | Intergenic | NA | NA |
| T7388 | CGCGGGTAGCGGTGTGTTGGTGG | 1000 | | | | | | | | | |
| Oligo fwd | CACCCGCGGGTAGCGGTGTGTTGG | | | | | | | | | | |
| Oligo rev | AAACCCAACACACCGCTACCCGCG | | | | | | | | | | |
| Chromosome | start | end | strand | MM | target_seq | PAM | alignment | distance | position | gene name | gene id |
| GU305914.1 | 101750 | 101772 | + | 0 | CGCGGGTAGCGGTGTGTTGG | TGG | \|\|\|\|\|\|\|\|[\|\|\|\|\|\|\|\|\|\|\|\|]PAM | NA | Intergenic | NA | NA |
| T7389 | TGTGTTGGTGGACACGTATTCGG | 1000 | | | | | | | | | |
| Oligo fwd | CACCTGTGTTGGTGGACACGTATT | | | | | | | | | | |
| Oligo rev | AAACAATACGTGTCCACCAACACA | | | | | | | | | | |
| Chromosome | start | end | strand | MM | target_seq | PAM | alignment | distance | position | gene name | gene id |
| GU305914.1 | 101762 | 101784 | + | 0 | TGTGTTGGTGGACACGTATT | CGG | \|\|\|\|\|\|\|\|[\|\|\|\|\|\|\|\|\|\|\|\|]PAM | NA | Intergenic | NA | NA |
| T7390 | TGGTGGACACGTATTCGGACAGG | 1000 | | | | | | | | | |
| Oligo fwd | CACCTGGTGGACACGTATTCGGAC | | | | | | | | | | |
| Oligo rev | AAACGTCCGAATACGTGTCCACCA | | | | | | | | | | |
| Chromosome | start | end | strand | MM | target_seq | PAM | alignment | distance | position | gene name | gene id |
| GU305914.1 | 101767 | 101789 | + | 0 | TGGTGGACACGTATTCGGAC | AGG | \|\|\|\|\|\|\|\|[\|\|\|\|\|\|\|\|\|\|\|\|]PAM | NA | Intergenic | NA | NA |
| T7391 | GGTGGACACGTATTCGGACAGGG | 1000 | | | | | | | | | |
| Oligo fwd | CACCGGTGGACACGTATTCGGACA | | | | | | | | | | |
| Oligo rev | AAACTGTCCGAATACGTGTCCACC | | | | | | | | | | |
| Chromosome | start | end | strand | MM | target_seq | PAM | alignment | distance | position | gene name | gene id |
| GU305914.1 | 101768 | 101790 | + | 0 | GGTGGACACGTATTCGGACA | GGG | \|\|\|\|\|\|\|\|[\|\|\|\|\|\|\|\|\|\|\|\|]PAM | NA | Intergenic | NA | NA |
| T7392 | ACACGTATTCGGACAGGGAGAGG | 1000 | | | | | | | | | |
| Oligo fwd | CACCACACGTATTCGGACAGGGAG | | | | | | | | | | |
| Oligo rev | AAACCTCCCTGTCCGAATACGTGT | | | | | | | | | | |
| Chromosome | start | end | strand | MM | target_seq | PAM | alignment | distance | position | gene name | gene id |
| GU305914.1 | 101773 | 101795 | + | 0 | ACACGTATTCGGACAGGGAG | AGG | \|\|\|\|\|\|\|\|[\|\|\|\|\|\|\|\|\|\|\|\|]PAM | NA | Intergenic | NA | NA |
| GU305914.1 | 215731 | 215753 | + | 4 | ATACGTAGTCGGACAATGAG | GGG | \|-\|\|\|\|\|-[\|\|\|\|\|\|\|--\|\|\|]PAM | NA | Intergenic | NA | NA |
| T7393 | CGTATTCGGACAGGGAGAGGCGG | 1000 | | | | | | | | | |
| Oligo fwd | CACCCGTATTCGGACAGGGAGAGG | | | | | | | | | | |
| Oligo rev | AAACCCTCTCCCTGTCCGAATACG | | | | | | | | | | |
| Chromosome | start | end | strand | MM | target_seq | PAM | alignment | distance | position | gene name | gene id |
| GU305914.1 | 101776 | 101798 | + | 0 | CGTATTCGGACAGGGAGAGG | CGG | \|\|\|\|\|\|\|\|[\|\|\|\|\|\|\|\|\|\|\|\|]PAM | NA | Intergenic | NA | NA |
| T7394 | GCGCGCGTCGCATATGAAAACGG | 1000 | | | | | | | | | |
| Oligo fwd | CACCGCGCGCGTCGCATATGAAAA | | | | | | | | | | |
| Oligo rev | AAACTTTTCATATGCGACGCGCGC | | | | | | | | | | |
| Chromosome | start | end | strand | MM | target_seq | PAM | alignment | distance | position | gene name | gene id |
| GU305914.1 | 101798 | 101820 | + | 0 | GCGCGCGTCGCATATGAAAA | CGG | \|\|\|\|\|\|\|\|[\|\|\|\|\|\|\|\|\|\|\|\|]PAM | NA | Intergenic | NA | NA |
| T7395 | CGCGCGTCGCATATGAAAACGGG | 1000 | | | | | | | | | |
| Oligo fwd | CACCCGCGCGTCGCATATGAAAAC | | | | | | | | | | |
| Oligo rev | AAACGTTTTCATATGCGACGCGCG | | | | | | | | | | |
| Chromosome | start | end | strand | MM | target_seq | PAM | alignment | distance | position | gene name | gene id |
| GU305914.1 | 101799 | 101821 | + | 0 | CGCGCGTCGCATATGAAAAC | GGG | \|\|\|\|\|\|\|\|[\|\|\|\|\|\|\|\|\|\|\|\|]PAM | NA | Intergenic | NA | NA |
| T7396 | GCGCGTCGCATATGAAAACGGGG | 1000 | | | | | | | | | |
| Oligo fwd | CACCGCGCGTCGCATATGAAAACG | | | | | | | | | | |
| Oligo rev | AAACCGTTTTCATATGCGACGCGC | | | | | | | | | | |
| Chromosome | start | end | strand | MM | target_seq | PAM | alignment | distance | position | gene name | gene id |
| GU305914.1 | 101800 | 101822 | + | 0 | GCGCGTCGCATATGAAAACG | GGG | \|\|\|\|\|\|\|\|[\|\|\|\|\|\|\|\|\|\|\|\|]PAM | NA | Intergenic | NA | NA |
| T7397 | CGTCGCATATGAAAACGGGGCGG | 1000 | | | | | | | | | |
| Oligo fwd | CACCCGTCGCATATGAAAACGGGG | | | | | | | | | | |
| Oligo rev | AAACCCCCGTTTTCATATGCGACG | | | | | | | | | | |
| Chromosome | start | end | strand | MM | target_seq | PAM | alignment | distance | position | gene name | gene id |
| GU305914.1 | 101803 | 101825 | + | 0 | CGTCGCATATGAAAACGGGG | CGG | \|\|\|\|\|\|\|\|[\|\|\|\|\|\|\|\|\|\|\|\|]PAM | NA | Intergenic | NA | NA |
| T7398 | ATATGAAAACGGGGCGGACCCGG | 1000 | | | | | | | | | |
| Oligo fwd | CACCATATGAAAACGGGGCGGACC | | | | | | | | | | |
| Oligo rev | AAACGGTCCGCCCCGTTTTCATAT | | | | | | | | | | |
| Chromosome | start | end | strand | MM | target_seq | PAM | alignment | distance | position | gene name | gene id |
| GU305914.1 | 101809 | 101831 | + | 0 | ATATGAAAACGGGGCGGACC | CGG | \|\|\|\|\|\|\|\|[\|\|\|\|\|\|\|\|\|\|\|\|]PAM | NA | Intergenic | NA | NA |
| T7399 | TATGAAAACGGGGCGGACCCGGG | 1000 | | | | | | | | | |
| Oligo fwd | CACCTATGAAAACGGGGCGGACCC | | | | | | | | | | |
| Oligo rev | AAACGGGTCCGCCCCGTTTTCATA | | | | | | | | | | |
| Chromosome | start | end | strand | MM | target_seq | PAM | alignment | distance | position | gene name | gene id |
| GU305914.1 | 101810 | 101832 | + | 0 | TATGAAAACGGGGCGGACCC | GGG | \|\|\|\|\|\|\|\|[\|\|\|\|\|\|\|\|\|\|\|\|]PAM | NA | Intergenic | NA | NA |
| T7400 | GGGCGGACCCGGGTCGAATCCGG | 1000 | | | | | | | | | |
| Oligo fwd | CACCGGGCGGACCCGGGTCGAATC | | | | | | | | | | |
| Oligo rev | AAACGATTCGACCCGGGTCCGCCC | | | | | | | | | | |
| Chromosome | start | end | strand | MM | target_seq | PAM | alignment | distance | position | gene name | gene id |
| GU305914.1 | 101820 | 101842 | + | 0 | GGGCGGACCCGGGTCGAATC | CGG | \|\|\|\|\|\|\|\|[\|\|\|\|\|\|\|\|\|\|\|\|]PAM | NA | Intergenic | NA | NA |
| T7401 | GGCGGACCCGGGTCGAATCCGGG | 1000 | | | | | | | | | |

<table>
<tr><td>Oligo fwd</td><td>CACCGGCGGACCCGGGTCGAATCC</td><td></td><td></td><td></td><td></td><td></td><td></td><td></td><td></td><td></td><td></td></tr>
<tr><td>Oligo rev</td><td>AAACGGATTCGACCCGGGTCCGCC</td><td></td><td></td><td></td><td></td><td></td><td></td><td></td><td></td><td></td><td></td></tr>
<tr><td>Chromosome</td><td>start</td><td>end</td><td>strand</td><td>MM</td><td>target_seq</td><td>PAM</td><td>alignment</td><td>distance</td><td>position</td><td>gene name</td><td>gene id</td></tr>
<tr><td>GU305914.1</td><td>101821</td><td>101843</td><td>+</td><td>0</td><td>GGCGGACCCGGGTCGAATCC</td><td>GGG</td><td>||||||||[||||||||||||]PAM</td><td>NA</td><td>Intergenic</td><td>NA</td><td>NA</td></tr>
<tr><td>T7402</td><td>GGGTCGAATCCGGGCACCTCTGG</td><td>1000</td><td></td><td></td><td></td><td></td><td></td><td></td><td></td><td></td><td></td></tr>
<tr><td>Oligo fwd</td><td>CACCGGGTCGAATCCGGGCACCTC</td><td></td><td></td><td></td><td></td><td></td><td></td><td></td><td></td><td></td><td></td></tr>
<tr><td>Oligo rev</td><td>AAACGAGGTGCCCGGATTCGACCC</td><td></td><td></td><td></td><td></td><td></td><td></td><td></td><td></td><td></td><td></td></tr>
<tr><td>Chromosome</td><td>start</td><td>end</td><td>strand</td><td>MM</td><td>target_seq</td><td>PAM</td><td>alignment</td><td>distance</td><td>position</td><td>gene name</td><td>gene id</td></tr>
<tr><td>GU305914.1</td><td>101830</td><td>101852</td><td>+</td><td>0</td><td>GGGTCGAATCCGGGCACCTC</td><td>TGG</td><td>||||||||[||||||||||||]PAM</td><td>NA</td><td>Intergenic</td><td>NA</td><td>NA</td></tr>
<tr><td>T7403</td><td>GGTCGAATCCGGGCACCTCTGGG</td><td>1000</td><td></td><td></td><td></td><td></td><td></td><td></td><td></td><td></td><td></td></tr>
<tr><td>Oligo fwd</td><td>CACCGGTCGAATCCGGGCACCTCT</td><td></td><td></td><td></td><td></td><td></td><td></td><td></td><td></td><td></td><td></td></tr>
<tr><td>Oligo rev</td><td>AAACAGAGGTGCCCGGATTCGACC</td><td></td><td></td><td></td><td></td><td></td><td></td><td></td><td></td><td></td><td></td></tr>
<tr><td>Chromosome</td><td>start</td><td>end</td><td>strand</td><td>MM</td><td>target_seq</td><td>PAM</td><td>alignment</td><td>distance</td><td>position</td><td>gene name</td><td>gene id</td></tr>
<tr><td>GU305914.1</td><td>101831</td><td>101853</td><td>+</td><td>0</td><td>GGTCGAATCCGGGCACCTCT</td><td>GGG</td><td>||||||||[||||||||||||]PAM</td><td>NA</td><td>Intergenic</td><td>NA</td><td>NA</td></tr>
<tr><td>T7404</td><td>TCTGAAGTACGAATTCTATACGG</td><td>1000</td><td></td><td></td><td></td><td></td><td></td><td></td><td></td><td></td><td></td></tr>
<tr><td>Oligo fwd</td><td>CACCTCTGAAGTACGAATTCTATA</td><td></td><td></td><td></td><td></td><td></td><td></td><td></td><td></td><td></td><td></td></tr>
<tr><td>Oligo rev</td><td>AAACTATAGAATTCGTACTTCAGA</td><td></td><td></td><td></td><td></td><td></td><td></td><td></td><td></td><td></td><td></td></tr>
<tr><td>Chromosome</td><td>start</td><td>end</td><td>strand</td><td>MM</td><td>target_seq</td><td>PAM</td><td>alignment</td><td>distance</td><td>position</td><td>gene name</td><td>gene id</td></tr>
<tr><td>GU305914.1</td><td>101854</td><td>101876</td><td>+</td><td>0</td><td>TCTGAAGTACGAATTCTATA</td><td>CGG</td><td>||||||||[||||||||||||]PAM</td><td>NA</td><td>Intergenic</td><td>NA</td><td>NA</td></tr>
<tr><td>T7405</td><td>ACGAATTCTATACGGTTCGACGG</td><td>1000</td><td></td><td></td><td></td><td></td><td></td><td></td><td></td><td></td><td></td></tr>
<tr><td>Oligo fwd</td><td>CACCACGAATTCTATACGGTTCGA</td><td></td><td></td><td></td><td></td><td></td><td></td><td></td><td></td><td></td><td></td></tr>
<tr><td>Oligo rev</td><td>AAACTCGAACCGTATAGAATTCGT</td><td></td><td></td><td></td><td></td><td></td><td></td><td></td><td></td><td></td><td></td></tr>
<tr><td>Chromosome</td><td>start</td><td>end</td><td>strand</td><td>MM</td><td>target_seq</td><td>PAM</td><td>alignment</td><td>distance</td><td>position</td><td>gene name</td><td>gene id</td></tr>
<tr><td>GU305914.1</td><td>101862</td><td>101884</td><td>+</td><td>0</td><td>ACGAATTCTATACGGTTCGA</td><td>CGG</td><td>||||||||[||||||||||||]PAM</td><td>NA</td><td>Intergenic</td><td>NA</td><td>NA</td></tr>
<tr><td>T7406</td><td>ACGGTTTGTATTTAATCATCAGG</td><td>1000</td><td></td><td></td><td></td><td></td><td></td><td></td><td></td><td></td><td></td></tr>
<tr><td>Oligo fwd</td><td>CACCACGGTTTGTATTTAATCATC</td><td></td><td></td><td></td><td></td><td></td><td></td><td></td><td></td><td></td><td></td></tr>
<tr><td>Oligo rev</td><td>AAACGATGATTAAATACAAACCGT</td><td></td><td></td><td></td><td></td><td></td><td></td><td></td><td></td><td></td><td></td></tr>
<tr><td>Chromosome</td><td>start</td><td>end</td><td>strand</td><td>MM</td><td>target_seq</td><td>PAM</td><td>alignment</td><td>distance</td><td>position</td><td>gene name</td><td>gene id</td></tr>
<tr><td>GU305914.1</td><td>101881</td><td>101903</td><td>+</td><td>0</td><td>ACGGTTTGTATTTAATCATC</td><td>AGG</td><td>||||||||[||||||||||||]PAM</td><td>NA</td><td>Intergenic</td><td>NA</td><td>NA</td></tr>
<tr><td>T7407</td><td>CGGTTTGTATTTAATCATCAGGG</td><td>1000</td><td></td><td></td><td></td><td></td><td></td><td></td><td></td><td></td><td></td></tr>
<tr><td>Oligo fwd</td><td>CACCCGGTTTGTATTTAATCATCA</td><td></td><td></td><td></td><td></td><td></td><td></td><td></td><td></td><td></td><td></td></tr>
<tr><td>Oligo rev</td><td>AAACTGATGATTAAATACAAACCG</td><td></td><td></td><td></td><td></td><td></td><td></td><td></td><td></td><td></td><td></td></tr>
<tr><td>Chromosome</td><td>start</td><td>end</td><td>strand</td><td>MM</td><td>target_seq</td><td>PAM</td><td>alignment</td><td>distance</td><td>position</td><td>gene name</td><td>gene id</td></tr>
<tr><td>GU305914.1</td><td>101882</td><td>101904</td><td>+</td><td>0</td><td>CGGTTTGTATTTAATCATCA</td><td>GGG</td><td>||||||||[||||||||||||]PAM</td><td>NA</td><td>Intergenic</td><td>NA</td><td>NA</td></tr>
<tr><td>T7408</td><td>GGTTTGTATTTAATCATCAGGGG</td><td>1000</td><td></td><td></td><td></td><td></td><td></td><td></td><td></td><td></td><td></td></tr>
<tr><td>Oligo fwd</td><td>CACCGGTTTGTATTTAATCATCAG</td><td></td><td></td><td></td><td></td><td></td><td></td><td></td><td></td><td></td><td></td></tr>
<tr><td>Oligo rev</td><td>AAACCTGATGATTAAATACAAACC</td><td></td><td></td><td></td><td></td><td></td><td></td><td></td><td></td><td></td><td></td></tr>
<tr><td>Chromosome</td><td>start</td><td>end</td><td>strand</td><td>MM</td><td>target_seq</td><td>PAM</td><td>alignment</td><td>distance</td><td>position</td><td>gene name</td><td>gene id</td></tr>
<tr><td>GU305914.1</td><td>101883</td><td>101905</td><td>+</td><td>0</td><td>GGTTTGTATTTAATCATCAG</td><td>GGG</td><td>||||||||[||||||||||||]PAM</td><td>NA</td><td>Intergenic</td><td>NA</td><td>NA</td></tr>
<tr><td>T7409</td><td>GTTTGTATTTAATCATCAGGGGG</td><td>1000</td><td></td><td></td><td></td><td></td><td></td><td></td><td></td><td></td><td></td></tr>
<tr><td>Oligo fwd</td><td>CACCGTTTGTATTTAATCATCAGG</td><td></td><td></td><td></td><td></td><td></td><td></td><td></td><td></td><td></td><td></td></tr>
<tr><td>Oligo rev</td><td>AAACCCTGATGATTAAATACAAAC</td><td></td><td></td><td></td><td></td><td></td><td></td><td></td><td></td><td></td><td></td></tr>
<tr><td>Chromosome</td><td>start</td><td>end</td><td>strand</td><td>MM</td><td>target_seq</td><td>PAM</td><td>alignment</td><td>distance</td><td>position</td><td>gene name</td><td>gene id</td></tr>
<tr><td>GU305914.1</td><td>101884</td><td>101906</td><td>+</td><td>0</td><td>GTTTGTATTTAATCATCAGG</td><td>GGG</td><td>||||||||[||||||||||||]PAM</td><td>NA</td><td>Intergenic</td><td>NA</td><td>NA</td></tr>
<tr><td>T7410</td><td>ATTTAATCATCAGGGGGAACAGG</td><td>1000</td><td></td><td></td><td></td><td></td><td></td><td></td><td></td><td></td><td></td></tr>
<tr><td>Oligo fwd</td><td>CACCATTTAATCATCAGGGGGAAC</td><td></td><td></td><td></td><td></td><td></td><td></td><td></td><td></td><td></td><td></td></tr>
<tr><td>Oligo rev</td><td>AAACGTTCCCCCTGATGATTAAAT</td><td></td><td></td><td></td><td></td><td></td><td></td><td></td><td></td><td></td><td></td></tr>
<tr><td>Chromosome</td><td>start</td><td>end</td><td>strand</td><td>MM</td><td>target_seq</td><td>PAM</td><td>alignment</td><td>distance</td><td>position</td><td>gene name</td><td>gene id</td></tr>
<tr><td>GU305914.1</td><td>101890</td><td>101912</td><td>+</td><td>0</td><td>ATTTAATCATCAGGGGGAAC</td><td>AGG</td><td>||||||||[||||||||||||]PAM</td><td>NA</td><td>Intergenic</td><td>NA</td><td>NA</td></tr>
<tr><td>T7411</td><td>TAATCATCAGGGGGAACAGGTGG</td><td>1000</td><td></td><td></td><td></td><td></td><td></td><td></td><td></td><td></td><td></td></tr>
<tr><td>Oligo fwd</td><td>CACCTAATCATCAGGGGGAACAGG</td><td></td><td></td><td></td><td></td><td></td><td></td><td></td><td></td><td></td><td></td></tr>
<tr><td>Oligo rev</td><td>AAACCCTGTTCCCCCTGATGATTA</td><td></td><td></td><td></td><td></td><td></td><td></td><td></td><td></td><td></td><td></td></tr>
<tr><td>Chromosome</td><td>start</td><td>end</td><td>strand</td><td>MM</td><td>target_seq</td><td>PAM</td><td>alignment</td><td>distance</td><td>position</td><td>gene name</td><td>gene id</td></tr>
<tr><td>GU305914.1</td><td>101893</td><td>101915</td><td>+</td><td>0</td><td>TAATCATCAGGGGGAACAGG</td><td>TGG</td><td>||||||||[||||||||||||]PAM</td><td>NA</td><td>Intergenic</td><td>NA</td><td>NA</td></tr>
<tr><td>T7412</td><td>GGGAACAGGTGGTGCTCCGTCGG</td><td>1000</td><td></td><td></td><td></td><td></td><td></td><td></td><td></td><td></td><td></td></tr>
<tr><td>Oligo fwd</td><td>CACCGGGAACAGGTGGTGCTCCGT</td><td></td><td></td><td></td><td></td><td></td><td></td><td></td><td></td><td></td><td></td></tr>
<tr><td>Oligo rev</td><td>AAACACGGAGCACCACCTGTTCCC</td><td></td><td></td><td></td><td></td><td></td><td></td><td></td><td></td><td></td><td></td></tr>
<tr><td>Chromosome</td><td>start</td><td>end</td><td>strand</td><td>MM</td><td>target_seq</td><td>PAM</td><td>alignment</td><td>distance</td><td>position</td><td>gene name</td><td>gene id</td></tr>
<tr><td>GU305914.1</td><td>101904</td><td>101926</td><td>+</td><td>0</td><td>GGGAACAGGTGGTGCTCCGT</td><td>CGG</td><td>||||||||[||||||||||||]PAM</td><td>NA</td><td>Intergenic</td><td>NA</td><td>NA</td></tr>
<tr><td>T7413</td><td>AGCATATTGACTACGATATTTGG</td><td>1000</td><td></td><td></td><td></td><td></td><td></td><td></td><td></td><td></td><td></td></tr>
<tr><td>Oligo fwd</td><td>CACCAGCATATTGACTACGATATT</td><td></td><td></td><td></td><td></td><td></td><td></td><td></td><td></td><td></td><td></td></tr>
<tr><td>Oligo rev</td><td>AAACAATATCGTAGTCAATATGCT</td><td></td><td></td><td></td><td></td><td></td><td></td><td></td><td></td><td></td><td></td></tr>
<tr><td>Chromosome</td><td>start</td><td>end</td><td>strand</td><td>MM</td><td>target_seq</td><td>PAM</td><td>alignment</td><td>distance</td><td>position</td><td>gene name</td><td>gene id</td></tr>
<tr><td>GU305914.1</td><td>101937</td><td>101959</td><td>+</td><td>0</td><td>AGCATATTGACTACGATATT</td><td>TGG</td><td>||||||||[||||||||||||]PAM</td><td>NA</td><td>Intergenic</td><td>NA</td><td>NA</td></tr>
<tr><td>T7414</td><td>GCATATTGACTACGATATTTGGG</td><td>1000</td><td></td><td></td><td></td><td></td><td></td><td></td><td></td><td></td><td></td></tr>
<tr><td>Oligo fwd</td><td>CACCGCATATTGACTACGATATTT</td><td></td><td></td><td></td><td></td><td></td><td></td><td></td><td></td><td></td><td></td></tr>
<tr><td>Oligo rev</td><td>AAACAAATATCGTAGTCAATATGC</td><td></td><td></td><td></td><td></td><td></td><td></td><td></td><td></td><td></td><td></td></tr>
<tr><td>Chromosome</td><td>start</td><td>end</td><td>strand</td><td>MM</td><td>target_seq</td><td>PAM</td><td>alignment</td><td>distance</td><td>position</td><td>gene name</td><td>gene id</td></tr>
<tr><td>GU305914.1</td><td>101938</td><td>101960</td><td>+</td><td>0</td><td>GCATATTGACTACGATATTT</td><td>GGG</td><td>||||||||[||||||||||||]PAM</td><td>NA</td><td>Intergenic</td><td>NA</td><td>NA</td></tr>
<tr><td>T7415</td><td>CATATTGACTACGATATTTGGGG</td><td>1000</td><td></td><td></td><td></td><td></td><td></td><td></td><td></td><td></td><td></td></tr>
<tr><td>Oligo fwd</td><td>CACCCATATTGACTACGATATTTG</td><td></td><td></td><td></td><td></td><td></td><td></td><td></td><td></td><td></td><td></td></tr>
<tr><td>Oligo rev</td><td>AAACCAAATATCGTAGTCAATATG</td><td></td><td></td><td></td><td></td><td></td><td></td><td></td><td></td><td></td><td></td></tr>
<tr><td>Chromosome</td><td>start</td><td>end</td><td>strand</td><td>MM</td><td>target_seq</td><td>PAM</td><td>alignment</td><td>distance</td><td>position</td><td>gene name</td><td>gene id</td></tr>
<tr><td>GU305914.1</td><td>101939</td><td>101961</td><td>+</td><td>0</td><td>CATATTGACTACGATATTTG</td><td>GGG</td><td>||||||||[||||||||||||]PAM</td><td>NA</td><td>Intergenic</td><td>NA</td><td>NA</td></tr>
</table>

<table>
<tr><td>T7416</td><td>GATATTTGGGGTGTCGTATTCGG</td><td>1000</td><td></td><td></td><td></td><td></td><td></td><td></td><td></td><td></td><td></td></tr>
<tr><td>Oligo fwd</td><td>CACCGATATTTGGGGTGTCGTATT</td><td></td><td></td><td></td><td></td><td></td><td></td><td></td><td></td><td></td><td></td></tr>
<tr><td>Oligo rev</td><td>AAACAATACGACACCCCAAATATC</td><td></td><td></td><td></td><td></td><td></td><td></td><td></td><td></td><td></td><td></td></tr>
<tr><td>Chromosome</td><td>start</td><td>end</td><td>strand</td><td>MM</td><td>target_seq</td><td>PAM</td><td>alignment</td><td>distance</td><td>position</td><td>gene name</td><td>gene id</td></tr>
<tr><td>GU305914.1</td><td>101951</td><td>101973</td><td>+</td><td>0</td><td>GATATTTGGGGTGTCGTATT</td><td>CGG</td><td>||||||||[||||||||||||]PAM</td><td>NA</td><td>Intergenic</td><td>NA</td><td>NA</td></tr>
<tr><td>T7417</td><td>ATTTGGGGTGTCGTATTCGGTGG</td><td>1000</td><td></td><td></td><td></td><td></td><td></td><td></td><td></td><td></td><td></td></tr>
<tr><td>Oligo fwd</td><td>CACCATTTGGGGTGTCGTATTCGG</td><td></td><td></td><td></td><td></td><td></td><td></td><td></td><td></td><td></td><td></td></tr>
<tr><td>Oligo rev</td><td>AAACCCGAATACGACACCCCAAAT</td><td></td><td></td><td></td><td></td><td></td><td></td><td></td><td></td><td></td><td></td></tr>
<tr><td>Chromosome</td><td>start</td><td>end</td><td>strand</td><td>MM</td><td>target_seq</td><td>PAM</td><td>alignment</td><td>distance</td><td>position</td><td>gene name</td><td>gene id</td></tr>
<tr><td>GU305914.1</td><td>101954</td><td>101976</td><td>+</td><td>0</td><td>ATTTGGGGTGTCGTATTCGG</td><td>TGG</td><td>||||||||[||||||||||||]PAM</td><td>NA</td><td>Intergenic</td><td>NA</td><td>NA</td></tr>
<tr><td>T7418</td><td>TTCGGTGGCGAACAGCACGACGG</td><td>1000</td><td></td><td></td><td></td><td></td><td></td><td></td><td></td><td></td><td></td></tr>
<tr><td>Oligo fwd</td><td>CACCTTCGGTGGCGAACAGCACGA</td><td></td><td></td><td></td><td></td><td></td><td></td><td></td><td></td><td></td><td></td></tr>
<tr><td>Oligo rev</td><td>AAACTCGTGCTGTTCGCCACCGAA</td><td></td><td></td><td></td><td></td><td></td><td></td><td></td><td></td><td></td><td></td></tr>
<tr><td>Chromosome</td><td>start</td><td>end</td><td>strand</td><td>MM</td><td>target_seq</td><td>PAM</td><td>alignment</td><td>distance</td><td>position</td><td>gene name</td><td>gene id</td></tr>
<tr><td>GU305914.1</td><td>101969</td><td>101991</td><td>+</td><td>0</td><td>TTCGGTGGCGAACAGCACGA</td><td>CGG</td><td>||||||||[||||||||||||]PAM</td><td>NA</td><td>Intergenic</td><td>NA</td><td>NA</td></tr>
<tr><td>T7419</td><td>GTCATGATGATGACCGACTTCGG</td><td>1000</td><td></td><td></td><td></td><td></td><td></td><td></td><td></td><td></td><td></td></tr>
<tr><td>Oligo fwd</td><td>CACCGTCATGATGATGACCGACTT</td><td></td><td></td><td></td><td></td><td></td><td></td><td></td><td></td><td></td><td></td></tr>
<tr><td>Oligo rev</td><td>AAACAAGTCGGTCATCATCATGAC</td><td></td><td></td><td></td><td></td><td></td><td></td><td></td><td></td><td></td><td></td></tr>
<tr><td>Chromosome</td><td>start</td><td>end</td><td>strand</td><td>MM</td><td>target_seq</td><td>PAM</td><td>alignment</td><td>distance</td><td>position</td><td>gene name</td><td>gene id</td></tr>
<tr><td>GU305914.1</td><td>101991</td><td>102013</td><td>+</td><td>0</td><td>GTCATGATGATGACCGACTT</td><td>CGG</td><td>||||||||[||||||||||||]PAM</td><td>NA</td><td>Intergenic</td><td>NA</td><td>NA</td></tr>
<tr><td>T7420</td><td>ATGATGATGACCGACTTCGGCGG</td><td>1000</td><td></td><td></td><td></td><td></td><td></td><td></td><td></td><td></td><td></td></tr>
<tr><td>Oligo fwd</td><td>CACCATGATGATGACCGACTTCGG</td><td></td><td></td><td></td><td></td><td></td><td></td><td></td><td></td><td></td><td></td></tr>
<tr><td>Oligo rev</td><td>AAACCCGAAGTCGGTCATCATCAT</td><td></td><td></td><td></td><td></td><td></td><td></td><td></td><td></td><td></td><td></td></tr>
<tr><td>Chromosome</td><td>start</td><td>end</td><td>strand</td><td>MM</td><td>target_seq</td><td>PAM</td><td>alignment</td><td>distance</td><td>position</td><td>gene name</td><td>gene id</td></tr>
<tr><td>GU305914.1</td><td>101994</td><td>102016</td><td>+</td><td>0</td><td>ATGATGATGACCGACTTCGG</td><td>CGG</td><td>||||||||[||||||||||||]PAM</td><td>NA</td><td>Intergenic</td><td>NA</td><td>NA</td></tr>
<tr><td>T7421</td><td>ATGATGACCGACTTCGGCGGTGG</td><td>1000</td><td></td><td></td><td></td><td></td><td></td><td></td><td></td><td></td><td></td></tr>
<tr><td>Oligo fwd</td><td>CACCATGATGACCGACTTCGGCGG</td><td></td><td></td><td></td><td></td><td></td><td></td><td></td><td></td><td></td><td></td></tr>
<tr><td>Oligo rev</td><td>AAACCCGCCGAAGTCGGTCATCAT</td><td></td><td></td><td></td><td></td><td></td><td></td><td></td><td></td><td></td><td></td></tr>
<tr><td>Chromosome</td><td>start</td><td>end</td><td>strand</td><td>MM</td><td>target_seq</td><td>PAM</td><td>alignment</td><td>distance</td><td>position</td><td>gene name</td><td>gene id</td></tr>
<tr><td>GU305914.1</td><td>101997</td><td>102019</td><td>+</td><td>0</td><td>ATGATGACCGACTTCGGCGG</td><td>TGG</td><td>||||||||[||||||||||||]PAM</td><td>NA</td><td>Intergenic</td><td>NA</td><td>NA</td></tr>
<tr><td>T7422</td><td>ATGACCGACTTCGGCGGTGGCGG</td><td>1000</td><td></td><td></td><td></td><td></td><td></td><td></td><td></td><td></td><td></td></tr>
<tr><td>Oligo fwd</td><td>CACCATGACCGACTTCGGCGGTGG</td><td></td><td></td><td></td><td></td><td></td><td></td><td></td><td></td><td></td><td></td></tr>
<tr><td>Oligo rev</td><td>AAACCCACCGCCGAAGTCGGTCAT</td><td></td><td></td><td></td><td></td><td></td><td></td><td></td><td></td><td></td><td></td></tr>
<tr><td>Chromosome</td><td>start</td><td>end</td><td>strand</td><td>MM</td><td>target_seq</td><td>PAM</td><td>alignment</td><td>distance</td><td>position</td><td>gene name</td><td>gene id</td></tr>
<tr><td>GU305914.1</td><td>102000</td><td>102022</td><td>+</td><td>0</td><td>ATGACCGACTTCGGCGGTGG</td><td>CGG</td><td>||||||||[||||||||||||]PAM</td><td>NA</td><td>Intergenic</td><td>NA</td><td>NA</td></tr>
<tr><td>T7423</td><td>TGACCGACTTCGGCGGTGGCGGG</td><td>1000</td><td></td><td></td><td></td><td></td><td></td><td></td><td></td><td></td><td></td></tr>
<tr><td>Oligo fwd</td><td>CACCTGACCGACTTCGGCGGTGGC</td><td></td><td></td><td></td><td></td><td></td><td></td><td></td><td></td><td></td><td></td></tr>
<tr><td>Oligo rev</td><td>AAACGCCACCGCCGAAGTCGGTCA</td><td></td><td></td><td></td><td></td><td></td><td></td><td></td><td></td><td></td><td></td></tr>
<tr><td>Chromosome</td><td>start</td><td>end</td><td>strand</td><td>MM</td><td>target_seq</td><td>PAM</td><td>alignment</td><td>distance</td><td>position</td><td>gene name</td><td>gene id</td></tr>
<tr><td>GU305914.1</td><td>102001</td><td>102023</td><td>+</td><td>0</td><td>TGACCGACTTCGGCGGTGGC</td><td>GGG</td><td>||||||||[||||||||||||]PAM</td><td>NA</td><td>Intergenic</td><td>NA</td><td>NA</td></tr>
<tr><td>T7424</td><td>GACTTCGGCGGTGGCGGGACTGG</td><td>1000</td><td></td><td></td><td></td><td></td><td></td><td></td><td></td><td></td><td></td></tr>
<tr><td>Oligo fwd</td><td>CACCGACTTCGGCGGTGGCGGGAC</td><td></td><td></td><td></td><td></td><td></td><td></td><td></td><td></td><td></td><td></td></tr>
<tr><td>Oligo rev</td><td>AAACGTCCCGCCACCGCCGAAGTC</td><td></td><td></td><td></td><td></td><td></td><td></td><td></td><td></td><td></td><td></td></tr>
<tr><td>Chromosome</td><td>start</td><td>end</td><td>strand</td><td>MM</td><td>target_seq</td><td>PAM</td><td>alignment</td><td>distance</td><td>position</td><td>gene name</td><td>gene id</td></tr>
<tr><td>GU305914.1</td><td>102006</td><td>102028</td><td>+</td><td>0</td><td>GACTTCGGCGGTGGCGGGAC</td><td>TGG</td><td>||||||||[||||||||||||]PAM</td><td>NA</td><td>Intergenic</td><td>NA</td><td>NA</td></tr>
<tr><td>T7425</td><td>ACTGGCACTAGTAATGCTCGCGG</td><td>1000</td><td></td><td></td><td></td><td></td><td></td><td></td><td></td><td></td><td></td></tr>
<tr><td>Oligo fwd</td><td>CACCACTGGCACTAGTAATGCTCG</td><td></td><td></td><td></td><td></td><td></td><td></td><td></td><td></td><td></td><td></td></tr>
<tr><td>Oligo rev</td><td>AAACCGAGCATTACTAGTGCCAGT</td><td></td><td></td><td></td><td></td><td></td><td></td><td></td><td></td><td></td><td></td></tr>
<tr><td>Chromosome</td><td>start</td><td>end</td><td>strand</td><td>MM</td><td>target_seq</td><td>PAM</td><td>alignment</td><td>distance</td><td>position</td><td>gene name</td><td>gene id</td></tr>
<tr><td>GU305914.1</td><td>102024</td><td>102046</td><td>+</td><td>0</td><td>ACTGGCACTAGTAATGCTCG</td><td>CGG</td><td>||||||||[||||||||||||]PAM</td><td>NA</td><td>Intergenic</td><td>NA</td><td>NA</td></tr>
<tr><td>T7426</td><td>CTGGCACTAGTAATGCTCGCGGG</td><td>1000</td><td></td><td></td><td></td><td></td><td></td><td></td><td></td><td></td><td></td></tr>
<tr><td>Oligo fwd</td><td>CACCCTGGCACTAGTAATGCTCGC</td><td></td><td></td><td></td><td></td><td></td><td></td><td></td><td></td><td></td><td></td></tr>
<tr><td>Oligo rev</td><td>AAACGCGAGCATTACTAGTGCCAG</td><td></td><td></td><td></td><td></td><td></td><td></td><td></td><td></td><td></td><td></td></tr>
<tr><td>Chromosome</td><td>start</td><td>end</td><td>strand</td><td>MM</td><td>target_seq</td><td>PAM</td><td>alignment</td><td>distance</td><td>position</td><td>gene name</td><td>gene id</td></tr>
<tr><td>GU305914.1</td><td>102025</td><td>102047</td><td>+</td><td>0</td><td>CTGGCACTAGTAATGCTCGC</td><td>GGG</td><td>||||||||[||||||||||||]PAM</td><td>NA</td><td>Intergenic</td><td>NA</td><td>NA</td></tr>
<tr><td>T7427</td><td>TGGCACTAGTAATGCTCGCGGGG</td><td>1000</td><td></td><td></td><td></td><td></td><td></td><td></td><td></td><td></td><td></td></tr>
<tr><td>Oligo fwd</td><td>CACCTGGCACTAGTAATGCTCGCG</td><td></td><td></td><td></td><td></td><td></td><td></td><td></td><td></td><td></td><td></td></tr>
<tr><td>Oligo rev</td><td>AAACCGCGAGCATTACTAGTGCCA</td><td></td><td></td><td></td><td></td><td></td><td></td><td></td><td></td><td></td><td></td></tr>
<tr><td>Chromosome</td><td>start</td><td>end</td><td>strand</td><td>MM</td><td>target_seq</td><td>PAM</td><td>alignment</td><td>distance</td><td>position</td><td>gene name</td><td>gene id</td></tr>
<tr><td>GU305914.1</td><td>102026</td><td>102048</td><td>+</td><td>0</td><td>TGGCACTAGTAATGCTCGCG</td><td>GGG</td><td>||||||||[||||||||||||]PAM</td><td>NA</td><td>Intergenic</td><td>NA</td><td>NA</td></tr>
<tr><td>T7428</td><td>GGCACTAGTAATGCTCGCGGGGG</td><td>1000</td><td></td><td></td><td></td><td></td><td></td><td></td><td></td><td></td><td></td></tr>
<tr><td>Oligo fwd</td><td>CACCGGCACTAGTAATGCTCGCGG</td><td></td><td></td><td></td><td></td><td></td><td></td><td></td><td></td><td></td><td></td></tr>
<tr><td>Oligo rev</td><td>AAACCCGCGAGCATTACTAGTGCC</td><td></td><td></td><td></td><td></td><td></td><td></td><td></td><td></td><td></td><td></td></tr>
<tr><td>Chromosome</td><td>start</td><td>end</td><td>strand</td><td>MM</td><td>target_seq</td><td>PAM</td><td>alignment</td><td>distance</td><td>position</td><td>gene name</td><td>gene id</td></tr>
<tr><td>GU305914.1</td><td>102027</td><td>102049</td><td>+</td><td>0</td><td>GGCACTAGTAATGCTCGCGG</td><td>GGG</td><td>||||||||[||||||||||||]PAM</td><td>NA</td><td>Intergenic</td><td>NA</td><td>NA</td></tr>
<tr><td>T7429</td><td>AGTAATGCTCGCGGGGGCGTCGG</td><td>1000</td><td></td><td></td><td></td><td></td><td></td><td></td><td></td><td></td><td></td></tr>
<tr><td>Oligo fwd</td><td>CACCAGTAATGCTCGCGGGGGCGT</td><td></td><td></td><td></td><td></td><td></td><td></td><td></td><td></td><td></td><td></td></tr>
<tr><td>Oligo rev</td><td>AAACACGCCCCCGCGAGCATTACT</td><td></td><td></td><td></td><td></td><td></td><td></td><td></td><td></td><td></td><td></td></tr>
<tr><td>Chromosome</td><td>start</td><td>end</td><td>strand</td><td>MM</td><td>target_seq</td><td>PAM</td><td>alignment</td><td>distance</td><td>position</td><td>gene name</td><td>gene id</td></tr>
<tr><td>GU305914.1</td><td>102033</td><td>102055</td><td>+</td><td>0</td><td>AGTAATGCTCGCGGGGGCGT</td><td>CGG</td><td>||||||||[||||||||||||]PAM</td><td>NA</td><td>Intergenic</td><td>NA</td><td>NA</td></tr>
<tr><td>T7430</td><td>GTAATGCTCGCGGGGGCGTCGGG</td><td>1000</td><td></td><td></td><td></td><td></td><td></td><td></td><td></td><td></td><td></td></tr>
<tr><td>Oligo fwd</td><td>CACCGTAATGCTCGCGGGGGCGTC</td><td></td><td></td><td></td><td></td><td></td><td></td><td></td><td></td><td></td><td></td></tr>
<tr><td>Oligo rev</td><td>AAACGACGCCCCCGCGAGCATTAC</td><td></td><td></td><td></td><td></td><td></td><td></td><td></td><td></td><td></td><td></td></tr>
<tr><td>Chromosome</td><td>start</td><td>end</td><td>strand</td><td>MM</td><td>target_seq</td><td>PAM</td><td>alignment</td><td>distance</td><td>position</td><td>gene name</td><td>gene id</td></tr>
<tr><td>GU305914.1</td><td>102034</td><td>102056</td><td>+</td><td>0</td><td>GTAATGCTCGCGGGGGCGTC</td><td>GGG</td><td>||||||||[||||||||||||]PAM</td><td>NA</td><td>Intergenic</td><td>NA</td><td>NA</td></tr>
</table>

| | | | | | | | | | | | |
|---|---|---|---|---|---|---|---|---|---|---|---|
| T7431 | TAATGCTCGCGGGGGCGTCGGGG | 1000 | | | | | | | | | |
| Oligo fwd | CACCTAATGCTCGCGGGGGCGTCG | | | | | | | | | | |
| Oligo rev | AAACCGACGCCCCCGCGAGCATTA | | | | | | | | | | |
| Chromosome | start | end | strand | MM | target_seq | PAM | alignment | distance | position | gene name | gene id |
| GU305914.1 | 102035 | 102057 | + | 0 | TAATGCTCGCGGGGGCGTCG | GGG | \|\|\|\|\|\|\|\|[\|\|\|\|\|\|\|\|\|\|\|\|]PAM | NA | Intergenic | NA | NA |
| T7432 | AATGCTCGCGGGGGCGTCGGGGG | 1000 | | | | | | | | | |
| Oligo fwd | CACCAATGCTCGCGGGGGCGTCGG | | | | | | | | | | |
| Oligo rev | AAACCCGACGCCCCCGCGAGCATT | | | | | | | | | | |
| Chromosome | start | end | strand | MM | target_seq | PAM | alignment | distance | position | gene name | gene id |
| GU305914.1 | 102036 | 102058 | + | 0 | AATGCTCGCGGGGGCGTCGG | GGG | \|\|\|\|\|\|\|\|[\|\|\|\|\|\|\|\|\|\|\|\|]PAM | NA | Intergenic | NA | NA |
| T7433 | GCCGCCGCAGACGACGACGCCGG | 1000 | | | | | | | | | |
| Oligo fwd | CACCGCCGCCGCAGACGACGACGC | | | | | | | | | | |
| Oligo rev | AAACGCGTCGTCGTCTGCGGCGGC | | | | | | | | | | |
| Chromosome | start | end | strand | MM | target_seq | PAM | alignment | distance | position | gene name | gene id |
| GU305914.1 | 102066 | 102088 | + | 0 | GCCGCCGCAGACGACGACGC | CGG | \|\|\|\|\|\|\|\|[\|\|\|\|\|\|\|\|\|\|\|\|]PAM | NA | Intergenic | NA | NA |
| T7434 | CCGCCGCAGACGACGACGCCGGG | 1000 | | | | | | | | | |
| Oligo fwd | CACCCCGCCGCAGACGACGACGCC | | | | | | | | | | |
| Oligo rev | AAACGGCGTCGTCGTCTGCGGCGG | | | | | | | | | | |
| Chromosome | start | end | strand | MM | target_seq | PAM | alignment | distance | position | gene name | gene id |
| GU305914.1 | 102067 | 102089 | + | 0 | CCGCCGCAGACGACGACGCC | GGG | \|\|\|\|\|\|\|\|[\|\|\|\|\|\|\|\|\|\|\|\|]PAM | NA | Intergenic | NA | NA |
| T7435 | CGCCGCAGACGACGACGCCGGGG | 1000 | | | | | | | | | |
| Oligo fwd | CACCCGCCGCAGACGACGACGCCG | | | | | | | | | | |
| Oligo rev | AAACCGGCGTCGTCGTCTGCGGCG | | | | | | | | | | |
| Chromosome | start | end | strand | MM | target_seq | PAM | alignment | distance | position | gene name | gene id |
| GU305914.1 | 102068 | 102090 | + | 0 | CGCCGCAGACGACGACGCCG | GGG | \|\|\|\|\|\|\|\|[\|\|\|\|\|\|\|\|\|\|\|\|]PAM | NA | Intergenic | NA | NA |
| T7436 | CGCAGACGACGACGCCGGGGCGG | 1000 | | | | | | | | | |
| Oligo fwd | CACCCGCAGACGACGACGCCGGGG | | | | | | | | | | |
| Oligo rev | AAACCCCCGGCGTCGTCGTCTGCG | | | | | | | | | | |
| Chromosome | start | end | strand | MM | target_seq | PAM | alignment | distance | position | gene name | gene id |
| GU305914.1 | 102071 | 102093 | + | 0 | CGCAGACGACGACGCCGGGG | CGG | \|\|\|\|\|\|\|\|[\|\|\|\|\|\|\|\|\|\|\|\|]PAM | NA | Intergenic | NA | NA |
| T7437 | GGGCGGCTCCTCCGAGCTGCTGG | 1000 | | | | | | | | | |
| Oligo fwd | CACCGGGCGGCTCCTCCGAGCTGC | | | | | | | | | | |
| Oligo rev | AAACGCAGCTCGGAGGAGCCGCCC | | | | | | | | | | |
| Chromosome | start | end | strand | MM | target_seq | PAM | alignment | distance | position | gene name | gene id |
| GU305914.1 | 102088 | 102110 | + | 0 | GGGCGGCTCCTCCGAGCTGC | TGG | \|\|\|\|\|\|\|\|[\|\|\|\|\|\|\|\|\|\|\|\|]PAM | NA | Intergenic | NA | NA |
| T7438 | CGGCTCCTCCGAGCTGCTGGAGG | 1000 | | | | | | | | | |
| Oligo fwd | CACCCGGCTCCTCCGAGCTGCTGG | | | | | | | | | | |
| Oligo rev | AAACCCAGCAGCTCGGAGGAGCCG | | | | | | | | | | |
| Chromosome | start | end | strand | MM | target_seq | PAM | alignment | distance | position | gene name | gene id |
| GU305914.1 | 102091 | 102113 | + | 0 | CGGCTCCTCCGAGCTGCTGG | AGG | \|\|\|\|\|\|\|\|[\|\|\|\|\|\|\|\|\|\|\|\|]PAM | NA | Intergenic | NA | NA |
| T7439 | CTCCTCCGAGCTGCTGGAGGAGG | 1000 | | | | | | | | | |
| Oligo fwd | CACCCTCCTCCGAGCTGCTGGAGG | | | | | | | | | | |
| Oligo rev | AAACCCTCCAGCAGCTCGGAGGAG | | | | | | | | | | |
| Chromosome | start | end | strand | MM | target_seq | PAM | alignment | distance | position | gene name | gene id |
| GU305914.1 | 102094 | 102116 | + | 0 | CTCCTCCGAGCTGCTGGAGG | AGG | \|\|\|\|\|\|\|\|[\|\|\|\|\|\|\|\|\|\|\|\|]PAM | NA | Intergenic | NA | NA |
| GU305914.1 | 58575 | 58597 | + | 4 | CTGTACCGTGCTGCTGGAGG | TGG | \|\|---\|\|\|[-\|\|\|\|\|\|\|\|\|\|\|]PAM | NA | Intergenic | NA | NA |
| T7440 | CCCTGTGTCGACGCACGATTCGG | 1000 | | | | | | | | | |
| Oligo fwd | CACCCCCTGTGTCGACGCACGATT | | | | | | | | | | |
| Oligo rev | AAACAATCGTGCGTCGACACAGGG | | | | | | | | | | |
| Chromosome | start | end | strand | MM | target_seq | PAM | alignment | distance | position | gene name | gene id |
| GU305914.1 | 102130 | 102152 | + | 0 | CCCTGTGTCGACGCACGATT | CGG | \|\|\|\|\|\|\|\|[\|\|\|\|\|\|\|\|\|\|\|\|]PAM | NA | Intergenic | NA | NA |
| T7441 | CCTGTGTCGACGCACGATTCGGG | 1000 | | | | | | | | | |
| Oligo fwd | CACCCCTGTGTCGACGCACGATTC | | | | | | | | | | |
| Oligo rev | AAACGAATCGTGCGTCGACACAGG | | | | | | | | | | |
| Chromosome | start | end | strand | MM | target_seq | PAM | alignment | distance | position | gene name | gene id |
| GU305914.1 | 102131 | 102153 | + | 0 | CCTGTGTCGACGCACGATTC | GGG | \|\|\|\|\|\|\|\|[\|\|\|\|\|\|\|\|\|\|\|\|]PAM | NA | Intergenic | NA | NA |
| T7442 | TGTCGACGCACGATTCGGGACGG | 1000 | | | | | | | | | |
| Oligo fwd | CACCTGTCGACGCACGATTCGGGA | | | | | | | | | | |
| Oligo rev | AAACTCCCGAATCGTGCGTCGACA | | | | | | | | | | |
| Chromosome | start | end | strand | MM | target_seq | PAM | alignment | distance | position | gene name | gene id |
| GU305914.1 | 102135 | 102157 | + | 0 | TGTCGACGCACGATTCGGGA | CGG | \|\|\|\|\|\|\|\|[\|\|\|\|\|\|\|\|\|\|\|\|]PAM | NA | Intergenic | NA | NA |
| T7443 | GATCCGAATATATCGTCCTCAGG | 1000 | | | | | | | | | |
| Oligo fwd | CACCGATCCGAATATATCGTCCTC | | | | | | | | | | |
| Oligo rev | AAACGAGGACGATATATTCGGATC | | | | | | | | | | |
| Chromosome | start | end | strand | MM | target_seq | PAM | alignment | distance | position | gene name | gene id |
| GU305914.1 | 102157 | 102179 | + | 0 | GATCCGAATATATCGTCCTC | AGG | \|\|\|\|\|\|\|\|[\|\|\|\|\|\|\|\|\|\|\|\|]PAM | NA | Intergenic | NA | NA |
| T7444 | ATCCGAATATATCGTCCTCAGGG | 1000 | | | | | | | | | |
| Oligo fwd | CACCATCCGAATATATCGTCCTCA | | | | | | | | | | |
| Oligo rev | AAACTGAGGACGATATATTCGGAT | | | | | | | | | | |
| Chromosome | start | end | strand | MM | target_seq | PAM | alignment | distance | position | gene name | gene id |
| GU305914.1 | 102158 | 102180 | + | 0 | ATCCGAATATATCGTCCTCA | GGG | \|\|\|\|\|\|\|\|[\|\|\|\|\|\|\|\|\|\|\|\|]PAM | NA | Intergenic | NA | NA |
| T7445 | TATCGTCCTCAGGGCCGACGAGG | 1000 | | | | | | | | | |
| Oligo fwd | CACCTATCGTCCTCAGGGCCGACG | | | | | | | | | | |
| Oligo rev | AAACCGTCGGCCCTGAGGACGATA | | | | | | | | | | |

<table>
<tr><td>Chromosome</td><td>start</td><td>end</td><td>strand</td><td>MM</td><td>target_seq</td><td>PAM</td><td>alignment</td><td>distance</td><td>position</td><td>gene name</td><td>gene id</td></tr>
<tr><td>GU305914.1</td><td>102167</td><td>102189</td><td>+</td><td>0</td><td>TATCGTCCTCAGGGCCGACG</td><td>AGG</td><td>||||||||[||||||||||||]PAM</td><td>NA</td><td>Intergenic</td><td>NA</td><td>NA</td></tr>
<tr><td>T7446</td><td>GGCCGACGAGGACGTCGATATGG</td><td>1000</td><td></td><td></td><td></td><td></td><td></td><td></td><td></td><td></td><td></td></tr>
<tr><td>Oligo fwd</td><td>CACCGGCCGACGAGGACGTCGATA</td><td></td><td></td><td></td><td></td><td></td><td></td><td></td><td></td><td></td><td></td></tr>
<tr><td>Oligo rev</td><td>AAACTATCGACGTCCTCGTCGGCC</td><td></td><td></td><td></td><td></td><td></td><td></td><td></td><td></td><td></td><td></td></tr>
<tr><td>Chromosome</td><td>start</td><td>end</td><td>strand</td><td>MM</td><td>target_seq</td><td>PAM</td><td>alignment</td><td>distance</td><td>position</td><td>gene name</td><td>gene id</td></tr>
<tr><td>GU305914.1</td><td>102179</td><td>102201</td><td>+</td><td>0</td><td>GGCCGACGAGGACGTCGATA</td><td>TGG</td><td>||||||||[||||||||||||]PAM</td><td>NA</td><td>Intergenic</td><td>NA</td><td>NA</td></tr>
<tr><td>T7447</td><td>GTCGATATGGCCGAATTAGAAGG</td><td>1000</td><td></td><td></td><td></td><td></td><td></td><td></td><td></td><td></td><td></td></tr>
<tr><td>Oligo fwd</td><td>CACCGTCGATATGGCCGAATTAGA</td><td></td><td></td><td></td><td></td><td></td><td></td><td></td><td></td><td></td><td></td></tr>
<tr><td>Oligo rev</td><td>AAACTCTAATTCGGCCATATCGAC</td><td></td><td></td><td></td><td></td><td></td><td></td><td></td><td></td><td></td><td></td></tr>
<tr><td>Chromosome</td><td>start</td><td>end</td><td>strand</td><td>MM</td><td>target_seq</td><td>PAM</td><td>alignment</td><td>distance</td><td>position</td><td>gene name</td><td>gene id</td></tr>
<tr><td>GU305914.1</td><td>102192</td><td>102214</td><td>+</td><td>0</td><td>GTCGATATGGCCGAATTAGA</td><td>AGG</td><td>||||||||[||||||||||||]PAM</td><td>NA</td><td>Intergenic</td><td>NA</td><td>NA</td></tr>
<tr><td>T7448</td><td>TCGATATGGCCGAATTAGAAGGG</td><td>1000</td><td></td><td></td><td></td><td></td><td></td><td></td><td></td><td></td><td></td></tr>
<tr><td>Oligo fwd</td><td>CACCTCGATATGGCCGAATTAGAA</td><td></td><td></td><td></td><td></td><td></td><td></td><td></td><td></td><td></td><td></td></tr>
<tr><td>Oligo rev</td><td>AAACTTCTAATTCGGCCATATCGA</td><td></td><td></td><td></td><td></td><td></td><td></td><td></td><td></td><td></td><td></td></tr>
<tr><td>Chromosome</td><td>start</td><td>end</td><td>strand</td><td>MM</td><td>target_seq</td><td>PAM</td><td>alignment</td><td>distance</td><td>position</td><td>gene name</td><td>gene id</td></tr>
<tr><td>GU305914.1</td><td>102193</td><td>102215</td><td>+</td><td>0</td><td>TCGATATGGCCGAATTAGAA</td><td>GGG</td><td>||||||||[||||||||||||]PAM</td><td>NA</td><td>Intergenic</td><td>NA</td><td>NA</td></tr>
<tr><td>T7449</td><td>GGGTTCCTCAAAGATAATTTTGG</td><td>1000</td><td></td><td></td><td></td><td></td><td></td><td></td><td></td><td></td><td></td></tr>
<tr><td>Oligo fwd</td><td>CACCGGGTTCCTCAAAGATAATTT</td><td></td><td></td><td></td><td></td><td></td><td></td><td></td><td></td><td></td><td></td></tr>
<tr><td>Oligo rev</td><td>AAACAAATTATCTTTGAGGAACCC</td><td></td><td></td><td></td><td></td><td></td><td></td><td></td><td></td><td></td><td></td></tr>
<tr><td>Chromosome</td><td>start</td><td>end</td><td>strand</td><td>MM</td><td>target_seq</td><td>PAM</td><td>alignment</td><td>distance</td><td>position</td><td>gene name</td><td>gene id</td></tr>
<tr><td>GU305914.1</td><td>102213</td><td>102235</td><td>+</td><td>0</td><td>GGGTTCCTCAAAGATAATTT</td><td>TGG</td><td>||||||||[||||||||||||]PAM</td><td>NA</td><td>Intergenic</td><td>NA</td><td>NA</td></tr>
<tr><td>T7450</td><td>GGTTCCTCAAAGATAATTTTGGG</td><td>1000</td><td></td><td></td><td></td><td></td><td></td><td></td><td></td><td></td><td></td></tr>
<tr><td>Oligo fwd</td><td>CACCGGTTCCTCAAAGATAATTTT</td><td></td><td></td><td></td><td></td><td></td><td></td><td></td><td></td><td></td><td></td></tr>
<tr><td>Oligo rev</td><td>AAACAAAATTATCTTTGAGGAACC</td><td></td><td></td><td></td><td></td><td></td><td></td><td></td><td></td><td></td><td></td></tr>
<tr><td>Chromosome</td><td>start</td><td>end</td><td>strand</td><td>MM</td><td>target_seq</td><td>PAM</td><td>alignment</td><td>distance</td><td>position</td><td>gene name</td><td>gene id</td></tr>
<tr><td>GU305914.1</td><td>102214</td><td>102236</td><td>+</td><td>0</td><td>GGTTCCTCAAAGATAATTTT</td><td>GGG</td><td>||||||||[||||||||||||]PAM</td><td>NA</td><td>Intergenic</td><td>NA</td><td>NA</td></tr>
<tr><td>T7451</td><td>AAAGATAATTTTGGGAATCTCGG</td><td>1000</td><td></td><td></td><td></td><td></td><td></td><td></td><td></td><td></td><td></td></tr>
<tr><td>Oligo fwd</td><td>CACCAAAGATAATTTTGGGAATCT</td><td></td><td></td><td></td><td></td><td></td><td></td><td></td><td></td><td></td><td></td></tr>
<tr><td>Oligo rev</td><td>AAACAGATTCCCAAAATTATCTTT</td><td></td><td></td><td></td><td></td><td></td><td></td><td></td><td></td><td></td><td></td></tr>
<tr><td>Chromosome</td><td>start</td><td>end</td><td>strand</td><td>MM</td><td>target_seq</td><td>PAM</td><td>alignment</td><td>distance</td><td>position</td><td>gene name</td><td>gene id</td></tr>
<tr><td>GU305914.1</td><td>102222</td><td>102244</td><td>+</td><td>0</td><td>AAAGATAATTTTGGGAATCT</td><td>CGG</td><td>||||||||[||||||||||||]PAM</td><td>NA</td><td>Intergenic</td><td>NA</td><td>NA</td></tr>
<tr><td>T7452</td><td>AAGATAATTTTGGGAATCTCGGG</td><td>1000</td><td></td><td></td><td></td><td></td><td></td><td></td><td></td><td></td><td></td></tr>
<tr><td>Oligo fwd</td><td>CACCAAGATAATTTTGGGAATCTC</td><td></td><td></td><td></td><td></td><td></td><td></td><td></td><td></td><td></td><td></td></tr>
<tr><td>Oligo rev</td><td>AAACGAGATTCCCAAAATTATCTT</td><td></td><td></td><td></td><td></td><td></td><td></td><td></td><td></td><td></td><td></td></tr>
<tr><td>Chromosome</td><td>start</td><td>end</td><td>strand</td><td>MM</td><td>target_seq</td><td>PAM</td><td>alignment</td><td>distance</td><td>position</td><td>gene name</td><td>gene id</td></tr>
<tr><td>GU305914.1</td><td>102223</td><td>102245</td><td>+</td><td>0</td><td>AAGATAATTTTGGGAATCTC</td><td>GGG</td><td>||||||||[||||||||||||]PAM</td><td>NA</td><td>Intergenic</td><td>NA</td><td>NA</td></tr>
<tr><td>T7453</td><td>AGATAATTTTGGGAATCTCGGGG</td><td>1000</td><td></td><td></td><td></td><td></td><td></td><td></td><td></td><td></td><td></td></tr>
<tr><td>Oligo fwd</td><td>CACCAGATAATTTTGGGAATCTCG</td><td></td><td></td><td></td><td></td><td></td><td></td><td></td><td></td><td></td><td></td></tr>
<tr><td>Oligo rev</td><td>AAACCGAGATTCCCAAAATTATCT</td><td></td><td></td><td></td><td></td><td></td><td></td><td></td><td></td><td></td><td></td></tr>
<tr><td>Chromosome</td><td>start</td><td>end</td><td>strand</td><td>MM</td><td>target_seq</td><td>PAM</td><td>alignment</td><td>distance</td><td>position</td><td>gene name</td><td>gene id</td></tr>
<tr><td>GU305914.1</td><td>102224</td><td>102246</td><td>+</td><td>0</td><td>AGATAATTTTGGGAATCTCG</td><td>GGG</td><td>||||||||[||||||||||||]PAM</td><td>NA</td><td>Intergenic</td><td>NA</td><td>NA</td></tr>
<tr><td>T7454</td><td>TGGGAATCTCGGGGTCTCGTCGG</td><td>1000</td><td></td><td></td><td></td><td></td><td></td><td></td><td></td><td></td><td></td></tr>
<tr><td>Oligo fwd</td><td>CACCTGGGAATCTCGGGGTCTCGT</td><td></td><td></td><td></td><td></td><td></td><td></td><td></td><td></td><td></td><td></td></tr>
<tr><td>Oligo rev</td><td>AAACACGAGACCCCGAGATTCCCA</td><td></td><td></td><td></td><td></td><td></td><td></td><td></td><td></td><td></td><td></td></tr>
<tr><td>Chromosome</td><td>start</td><td>end</td><td>strand</td><td>MM</td><td>target_seq</td><td>PAM</td><td>alignment</td><td>distance</td><td>position</td><td>gene name</td><td>gene id</td></tr>
<tr><td>GU305914.1</td><td>102233</td><td>102255</td><td>+</td><td>0</td><td>TGGGAATCTCGGGGTCTCGT</td><td>CGG</td><td>||||||||[||||||||||||]PAM</td><td>NA</td><td>Intergenic</td><td>NA</td><td>NA</td></tr>
<tr><td>T7455</td><td>GAATCTCGGGGTCTCGTCGGCGG</td><td>1000</td><td></td><td></td><td></td><td></td><td></td><td></td><td></td><td></td><td></td></tr>
<tr><td>Oligo fwd</td><td>CACCGAATCTCGGGGTCTCGTCGG</td><td></td><td></td><td></td><td></td><td></td><td></td><td></td><td></td><td></td><td></td></tr>
<tr><td>Oligo rev</td><td>AAACCCGACGAGACCCCGAGATTC</td><td></td><td></td><td></td><td></td><td></td><td></td><td></td><td></td><td></td><td></td></tr>
<tr><td>Chromosome</td><td>start</td><td>end</td><td>strand</td><td>MM</td><td>target_seq</td><td>PAM</td><td>alignment</td><td>distance</td><td>position</td><td>gene name</td><td>gene id</td></tr>
<tr><td>GU305914.1</td><td>102236</td><td>102258</td><td>+</td><td>0</td><td>GAATCTCGGGGTCTCGTCGG</td><td>CGG</td><td>||||||||[||||||||||||]PAM</td><td>NA</td><td>Intergenic</td><td>NA</td><td>NA</td></tr>
<tr><td>T7456</td><td>CGGACCTGTCTGAAATAGATAGG</td><td>1000</td><td></td><td></td><td></td><td></td><td></td><td></td><td></td><td></td><td></td></tr>
<tr><td>Oligo fwd</td><td>CACCCGGACCTGTCTGAAATAGAT</td><td></td><td></td><td></td><td></td><td></td><td></td><td></td><td></td><td></td><td></td></tr>
<tr><td>Oligo rev</td><td>AAACATCTATTTCAGACAGGTCCG</td><td></td><td></td><td></td><td></td><td></td><td></td><td></td><td></td><td></td><td></td></tr>
<tr><td>Chromosome</td><td>start</td><td>end</td><td>strand</td><td>MM</td><td>target_seq</td><td>PAM</td><td>alignment</td><td>distance</td><td>position</td><td>gene name</td><td>gene id</td></tr>
<tr><td>GU305914.1</td><td>102256</td><td>102278</td><td>+</td><td>0</td><td>CGGACCTGTCTGAAATAGAT</td><td>AGG</td><td>||||||||[||||||||||||]PAM</td><td>NA</td><td>Intergenic</td><td>NA</td><td>NA</td></tr>
<tr><td>T7457</td><td>GGACCTGTCTGAAATAGATAGGG</td><td>1000</td><td></td><td></td><td></td><td></td><td></td><td></td><td></td><td></td><td></td></tr>
<tr><td>Oligo fwd</td><td>CACCGGACCTGTCTGAAATAGATA</td><td></td><td></td><td></td><td></td><td></td><td></td><td></td><td></td><td></td><td></td></tr>
<tr><td>Oligo rev</td><td>AAACTATCTATTTCAGACAGGTCC</td><td></td><td></td><td></td><td></td><td></td><td></td><td></td><td></td><td></td><td></td></tr>
<tr><td>Chromosome</td><td>start</td><td>end</td><td>strand</td><td>MM</td><td>target_seq</td><td>PAM</td><td>alignment</td><td>distance</td><td>position</td><td>gene name</td><td>gene id</td></tr>
<tr><td>GU305914.1</td><td>102257</td><td>102279</td><td>+</td><td>0</td><td>GGACCTGTCTGAAATAGATA</td><td>GGG</td><td>||||||||[||||||||||||]PAM</td><td>NA</td><td>Intergenic</td><td>NA</td><td>NA</td></tr>
<tr><td>T7458</td><td>GTCTGAAATAGATAGGGAGTCGG</td><td>1000</td><td></td><td></td><td></td><td></td><td></td><td></td><td></td><td></td><td></td></tr>
<tr><td>Oligo fwd</td><td>CACCGTCTGAAATAGATAGGGAGT</td><td></td><td></td><td></td><td></td><td></td><td></td><td></td><td></td><td></td><td></td></tr>
<tr><td>Oligo rev</td><td>AAACACTCCCTATCTATTTCAGAC</td><td></td><td></td><td></td><td></td><td></td><td></td><td></td><td></td><td></td><td></td></tr>
<tr><td>Chromosome</td><td>start</td><td>end</td><td>strand</td><td>MM</td><td>target_seq</td><td>PAM</td><td>alignment</td><td>distance</td><td>position</td><td>gene name</td><td>gene id</td></tr>
<tr><td>GU305914.1</td><td>102263</td><td>102285</td><td>+</td><td>0</td><td>GTCTGAAATAGATAGGGAGT</td><td>CGG</td><td>||||||||[||||||||||||]PAM</td><td>NA</td><td>Intergenic</td><td>NA</td><td>NA</td></tr>
<tr><td>T7459</td><td>AAGTCACCAAACACCTGCTCAGG</td><td>1000</td><td></td><td></td><td></td><td></td><td></td><td></td><td></td><td></td><td></td></tr>
<tr><td>Oligo fwd</td><td>CACCAAGTCACCAAACACCTGCTC</td><td></td><td></td><td></td><td></td><td></td><td></td><td></td><td></td><td></td><td></td></tr>
<tr><td>Oligo rev</td><td>AAACGAGCAGGTGTTTGGTGACTT</td><td></td><td></td><td></td><td></td><td></td><td></td><td></td><td></td><td></td><td></td></tr>
<tr><td>Chromosome</td><td>start</td><td>end</td><td>strand</td><td>MM</td><td>target_seq</td><td>PAM</td><td>alignment</td><td>distance</td><td>position</td><td>gene name</td><td>gene id</td></tr>
<tr><td>GU305914.1</td><td>102286</td><td>102308</td><td>+</td><td>0</td><td>AAGTCACCAAACACCTGCTC</td><td>AGG</td><td>||||||||[||||||||||||]PAM</td><td>NA</td><td>Intergenic</td><td>NA</td><td>NA</td></tr>
<tr><td>T7460</td><td>ACACCTGCTCAGGCTCCTTCCGG</td><td>1000</td><td></td><td></td><td></td><td></td><td></td><td></td><td></td><td></td><td></td></tr>
<tr><td>Oligo fwd</td><td>CACCACACCTGCTCAGGCTCCTTC</td><td></td><td></td><td></td><td></td><td></td><td></td><td></td><td></td><td></td><td></td></tr>
</table>

Oligo rev AAACGAAGGAGCCTGAGCAGGTGT
Chromosome start end strand MM target_seq PAM alignment distance position gene name gene id
GU305914.1 102296 102318 + 0 ACACCTGCTCAGGCTCCTTC CGG ||||||||[||||||||||||]PAM NA Intergenic NA NA

T7461 CGGTCTATAAGCGCTGCGTGCGG 1000
Oligo fwd CACCCGGTCTATAAGCGCTGCGTG
Oligo rev AAACCACGCAGCGCTTATAGACCG
Chromosome start end strand MM target_seq PAM alignment distance position gene name gene id
GU305914.1 102316 102338 + 0 CGGTCTATAAGCGCTGCGTG CGG ||||||||[||||||||||||]PAM NA Intergenic NA NA

T7462 TCTATAAGCGCTGCGTGCGGAGG 1000
Oligo fwd CACCTCTATAAGCGCTGCGTGCGG
Oligo rev AAACCCGCACGCAGCGCTTATAGA
Chromosome start end strand MM target_seq PAM alignment distance position gene name gene id
GU305914.1 102319 102341 + 0 TCTATAAGCGCTGCGTGCGG AGG ||||||||[||||||||||||]PAM NA Intergenic NA NA

T7463 GCTGCGTGCGGAGGCAGACCCGG 1000
Oligo fwd CACCGCTGCGTGCGGAGGCAGACC
Oligo rev AAACGGTCTGCCTCCGCACGCAGC
Chromosome start end strand MM target_seq PAM alignment distance position gene name gene id
GU305914.1 102328 102350 + 0 GCTGCGTGCGGAGGCAGACC CGG ||||||||[||||||||||||]PAM NA Intergenic NA NA

T7464 GACCCGGCTCGACCGTCTTCTGG 1000
Oligo fwd CACCGACCCGGCTCGACCGTCTTC
Oligo rev AAACGAAGACGGTCGAGCCGGGTC
Chromosome start end strand MM target_seq PAM alignment distance position gene name gene id
GU305914.1 102344 102366 + 0 GACCCGGCTCGACCGTCTTC TGG ||||||||[||||||||||||]PAM NA Intergenic NA NA

T7465 GAAATCCAAACGTGTCATGCAGG 1000
Oligo fwd CACCGAAATCCAAACGTGTCATGC
Oligo rev AAACGCATGACACGTTTGGATTTC
Chromosome start end strand MM target_seq PAM alignment distance position gene name gene id
GU305914.1 102416 102438 + 0 GAAATCCAAACGTGTCATGC AGG ||||||||[||||||||||||]PAM NA Intergenic NA NA

T7466 GTCATCCTCAAGCTCCTCGTCGG 1000
Oligo fwd CACCGTCATCCTCAAGCTCCTCGT
Oligo rev AAACACGAGGAGCTTGAGGATGAC
Chromosome start end strand MM target_seq PAM alignment distance position gene name gene id
GU305914.1 102450 102472 + 0 GTCATCCTCAAGCTCCTCGT CGG ||||||||[||||||||||||]PAM NA Intergenic NA NA

T7467 TCATCCTCAAGCTCCTCGTCGGG 1000
Oligo fwd CACCTCATCCTCAAGCTCCTCGTC
Oligo rev AAACGACGAGGAGCTTGAGGATGA
Chromosome start end strand MM target_seq PAM alignment distance position gene name gene id
GU305914.1 102451 102473 + 0 TCATCCTCAAGCTCCTCGTC GGG ||||||||[||||||||||||]PAM NA Intergenic NA NA

T7468 CATCCTCAAGCTCCTCGTCGGGG 1000
Oligo fwd CACCCATCCTCAAGCTCCTCGTCG
Oligo rev AAACCGACGAGGAGCTTGAGGATG
Chromosome start end strand MM target_seq PAM alignment distance position gene name gene id
GU305914.1 102452 102474 + 0 CATCCTCAAGCTCCTCGTCG GGG ||||||||[||||||||||||]PAM NA Intergenic NA NA

T7469 CGTCGGGGAGTTCACCCTCTCGG 1000
Oligo fwd CACCCGTCGGGGAGTTCACCCTCT
Oligo rev AAACAGAGGGTGAACTCCCCGACG
Chromosome start end strand MM target_seq PAM alignment distance position gene name gene id
GU305914.1 102467 102489 + 0 CGTCGGGGAGTTCACCCTCT CGG ||||||||[||||||||||||]PAM NA Intergenic NA NA

T7470 GGAGTTCACCCTCTCGGACGAGG 1000
Oligo fwd CACCGGAGTTCACCCTCTCGGACG
Oligo rev AAACCGTCCGAGAGGGTGAACTCC
Chromosome start end strand MM target_seq PAM alignment distance position gene name gene id
GU305914.1 102473 102495 + 0 GGAGTTCACCCTCTCGGACG AGG ||||||||[||||||||||||]PAM NA Intergenic NA NA

T7471 CGGACGAGGACAGCGTCGAGAGG 1000
Oligo fwd CACCCGGACGAGGACAGCGTCGAG
Oligo rev AAACCTCGACGCTGTCCTCGTCCG
Chromosome start end strand MM target_seq PAM alignment distance position gene name gene id
GU305914.1 102487 102509 + 0 CGGACGAGGACAGCGTCGAG AGG ||||||||[||||||||||||]PAM NA Intergenic NA NA

T7472 CGATCAGAGCACCCTCTGCGAGG 1000
Oligo fwd CACCCGATCAGAGCACCCTCTGCG
Oligo rev AAACCGCAGAGGGTGCTCTGATCG
Chromosome start end strand MM target_seq PAM alignment distance position gene name gene id
GU305914.1 102530 102552 + 0 CGATCAGAGCACCCTCTGCG AGG ||||||||[||||||||||||]PAM NA Intergenic NA NA

T7473 CAGAGCACCCTCTGCGAGGTCGG 1000
Oligo fwd CACCCAGAGCACCCTCTGCGAGGT
Oligo rev AAACACCTCGCAGAGGGTGCTCTG
Chromosome start end strand MM target_seq PAM alignment distance position gene name gene id
GU305914.1 102534 102556 + 0 CAGAGCACCCTCTGCGAGGT CGG ||||||||[||||||||||||]PAM NA Intergenic NA NA

T7474 GCACCCTCTGCGAGGTCGGCAGG 1000
Oligo fwd CACCGCACCCTCTGCGAGGTCGGC
Oligo rev AAACGCCGACCTCGCAGAGGGTGC
Chromosome start end strand MM target_seq PAM alignment distance position gene name gene id
GU305914.1 102538 102560 + 0 GCACCCTCTGCGAGGTCGGC AGG ||||||||[||||||||||||]PAM NA Intergenic NA NA

T7475 GATCGTGCGCCTCATAGACATGG 1000

<table>
<tr><td>Oligo fwd</td><td>CACCGATCGTGCGCCTCATAGACA</td><td></td><td></td><td></td><td></td><td></td><td></td><td></td><td></td><td></td><td></td></tr>
<tr><td>Oligo rev</td><td>AAACTGTCTATGAGGCGCACGATC</td><td></td><td></td><td></td><td></td><td></td><td></td><td></td><td></td><td></td><td></td></tr>
<tr><td>Chromosome</td><td>start</td><td>end</td><td>strand</td><td>MM</td><td>target_seq</td><td>PAM</td><td>alignment</td><td>distance</td><td>position</td><td>gene name</td><td>gene id</td></tr>
<tr><td>GU305914.1</td><td>102560</td><td>102582</td><td>+</td><td>0</td><td>GATCGTGCGCCTCATAGACA</td><td>TGG</td><td>||||||||[||||||||||||]PAM</td><td>NA</td><td>Intergenic</td><td>NA</td><td>NA</td></tr>
<tr><td></td><td></td><td></td><td></td><td></td><td></td><td></td><td></td><td></td><td></td><td></td><td></td></tr>
<tr><td>T7476</td><td>AACACTCAGCGCCTCGTCGACGG</td><td>1000</td><td></td><td></td><td></td><td></td><td></td><td></td><td></td><td></td><td></td></tr>
<tr><td>Oligo fwd</td><td>CACCAACACTCAGCGCCTCGTCGA</td><td></td><td></td><td></td><td></td><td></td><td></td><td></td><td></td><td></td><td></td></tr>
<tr><td>Oligo rev</td><td>AAACTCGACGAGGCGCTGAGTGTT</td><td></td><td></td><td></td><td></td><td></td><td></td><td></td><td></td><td></td><td></td></tr>
<tr><td>Chromosome</td><td>start</td><td>end</td><td>strand</td><td>MM</td><td>target_seq</td><td>PAM</td><td>alignment</td><td>distance</td><td>position</td><td>gene name</td><td>gene id</td></tr>
<tr><td>GU305914.1</td><td>102591</td><td>102613</td><td>+</td><td>0</td><td>AACACTCAGCGCCTCGTCGA</td><td>CGG</td><td>||||||||[||||||||||||]PAM</td><td>NA</td><td>Intergenic</td><td>NA</td><td>NA</td></tr>
<tr><td></td><td></td><td></td><td></td><td></td><td></td><td></td><td></td><td></td><td></td><td></td><td></td></tr>
<tr><td>T7477</td><td>CTCAGCGCCTCGTCGACGGACGG</td><td>1000</td><td></td><td></td><td></td><td></td><td></td><td></td><td></td><td></td><td></td></tr>
<tr><td>Oligo fwd</td><td>CACCCTCAGCGCCTCGTCGACGGA</td><td></td><td></td><td></td><td></td><td></td><td></td><td></td><td></td><td></td><td></td></tr>
<tr><td>Oligo rev</td><td>AAACTCCGTCGACGAGGCGCTGAG</td><td></td><td></td><td></td><td></td><td></td><td></td><td></td><td></td><td></td><td></td></tr>
<tr><td>Chromosome</td><td>start</td><td>end</td><td>strand</td><td>MM</td><td>target_seq</td><td>PAM</td><td>alignment</td><td>distance</td><td>position</td><td>gene name</td><td>gene id</td></tr>
<tr><td>GU305914.1</td><td>102595</td><td>102617</td><td>+</td><td>0</td><td>CTCAGCGCCTCGTCGACGGA</td><td>CGG</td><td>||||||||[||||||||||||]PAM</td><td>NA</td><td>Intergenic</td><td>NA</td><td>NA</td></tr>
<tr><td></td><td></td><td></td><td></td><td></td><td></td><td></td><td></td><td></td><td></td><td></td><td></td></tr>
<tr><td>T7478</td><td>TCAGCGCCTCGTCGACGGACGGG</td><td>1000</td><td></td><td></td><td></td><td></td><td></td><td></td><td></td><td></td><td></td></tr>
<tr><td>Oligo fwd</td><td>CACCTCAGCGCCTCGTCGACGGAC</td><td></td><td></td><td></td><td></td><td></td><td></td><td></td><td></td><td></td><td></td></tr>
<tr><td>Oligo rev</td><td>AAACGTCCGTCGACGAGGCGCTGA</td><td></td><td></td><td></td><td></td><td></td><td></td><td></td><td></td><td></td><td></td></tr>
<tr><td>Chromosome</td><td>start</td><td>end</td><td>strand</td><td>MM</td><td>target_seq</td><td>PAM</td><td>alignment</td><td>distance</td><td>position</td><td>gene name</td><td>gene id</td></tr>
<tr><td>GU305914.1</td><td>102596</td><td>102618</td><td>+</td><td>0</td><td>TCAGCGCCTCGTCGACGGAC</td><td>GGG</td><td>||||||||[||||||||||||]PAM</td><td>NA</td><td>Intergenic</td><td>NA</td><td>NA</td></tr>
<tr><td></td><td></td><td></td><td></td><td></td><td></td><td></td><td></td><td></td><td></td><td></td><td></td></tr>
<tr><td>T7479</td><td>GCGCCTCGTCGACGGACGGGAGG</td><td>1000</td><td></td><td></td><td></td><td></td><td></td><td></td><td></td><td></td><td></td></tr>
<tr><td>Oligo fwd</td><td>CACCGCGCCTCGTCGACGGACGGG</td><td></td><td></td><td></td><td></td><td></td><td></td><td></td><td></td><td></td><td></td></tr>
<tr><td>Oligo rev</td><td>AAACCCCGTCCGTCGACGAGGCGC</td><td></td><td></td><td></td><td></td><td></td><td></td><td></td><td></td><td></td><td></td></tr>
<tr><td>Chromosome</td><td>start</td><td>end</td><td>strand</td><td>MM</td><td>target_seq</td><td>PAM</td><td>alignment</td><td>distance</td><td>position</td><td>gene name</td><td>gene id</td></tr>
<tr><td>GU305914.1</td><td>102599</td><td>102621</td><td>+</td><td>0</td><td>GCGCCTCGTCGACGGACGGG</td><td>AGG</td><td>||||||||[||||||||||||]PAM</td><td>NA</td><td>Intergenic</td><td>NA</td><td>NA</td></tr>
<tr><td></td><td></td><td></td><td></td><td></td><td></td><td></td><td></td><td></td><td></td><td></td><td></td></tr>
<tr><td>T7480</td><td>CGTCGACGGACGGGAGGAGCCGG</td><td>1000</td><td></td><td></td><td></td><td></td><td></td><td></td><td></td><td></td><td></td></tr>
<tr><td>Oligo fwd</td><td>CACCCGTCGACGGACGGGAGGAGC</td><td></td><td></td><td></td><td></td><td></td><td></td><td></td><td></td><td></td><td></td></tr>
<tr><td>Oligo rev</td><td>AAACGCTCCTCCCGTCCGTCGACG</td><td></td><td></td><td></td><td></td><td></td><td></td><td></td><td></td><td></td><td></td></tr>
<tr><td>Chromosome</td><td>start</td><td>end</td><td>strand</td><td>MM</td><td>target_seq</td><td>PAM</td><td>alignment</td><td>distance</td><td>position</td><td>gene name</td><td>gene id</td></tr>
<tr><td>GU305914.1</td><td>102605</td><td>102627</td><td>+</td><td>0</td><td>CGTCGACGGACGGGAGGAGC</td><td>CGG</td><td>||||||||[||||||||||||]PAM</td><td>NA</td><td>Intergenic</td><td>NA</td><td>NA</td></tr>
<tr><td></td><td></td><td></td><td></td><td></td><td></td><td></td><td></td><td></td><td></td><td></td><td></td></tr>
<tr><td>T7481</td><td>CCTCCCCTCTGCGACCTGAACGG</td><td>1000</td><td></td><td></td><td></td><td></td><td></td><td></td><td></td><td></td><td></td></tr>
<tr><td>Oligo fwd</td><td>CACCCCTCCCCTCTGCGACCTGAA</td><td></td><td></td><td></td><td></td><td></td><td></td><td></td><td></td><td></td><td></td></tr>
<tr><td>Oligo rev</td><td>AAACTTCAGGTCGCAGAGGGGAGG</td><td></td><td></td><td></td><td></td><td></td><td></td><td></td><td></td><td></td><td></td></tr>
<tr><td>Chromosome</td><td>start</td><td>end</td><td>strand</td><td>MM</td><td>target_seq</td><td>PAM</td><td>alignment</td><td>distance</td><td>position</td><td>gene name</td><td>gene id</td></tr>
<tr><td>GU305914.1</td><td>102630</td><td>102652</td><td>+</td><td>0</td><td>CCTCCCCTCTGCGACCTGAA</td><td>CGG</td><td>||||||||[||||||||||||]PAM</td><td>NA</td><td>Intergenic</td><td>NA</td><td>NA</td></tr>
<tr><td></td><td></td><td></td><td></td><td></td><td></td><td></td><td></td><td></td><td></td><td></td><td></td></tr>
<tr><td>T7482</td><td>GTCTTCTTCCTCCTCCTACGCGG</td><td>1000</td><td></td><td></td><td></td><td></td><td></td><td></td><td></td><td></td><td></td></tr>
<tr><td>Oligo fwd</td><td>CACCGTCTTCTTCCTCCTCCTACG</td><td></td><td></td><td></td><td></td><td></td><td></td><td></td><td></td><td></td><td></td></tr>
<tr><td>Oligo rev</td><td>AAACCGTAGGAGGAGGAAGAAGAC</td><td></td><td></td><td></td><td></td><td></td><td></td><td></td><td></td><td></td><td></td></tr>
<tr><td>Chromosome</td><td>start</td><td>end</td><td>strand</td><td>MM</td><td>target_seq</td><td>PAM</td><td>alignment</td><td>distance</td><td>position</td><td>gene name</td><td>gene id</td></tr>
<tr><td>GU305914.1</td><td>102665</td><td>102687</td><td>+</td><td>0</td><td>GTCTTCTTCCTCCTCCTACG</td><td>CGG</td><td>||||||||[||||||||||||]PAM</td><td>NA</td><td>Intergenic</td><td>NA</td><td>NA</td></tr>
<tr><td></td><td></td><td></td><td></td><td></td><td></td><td></td><td></td><td></td><td></td><td></td><td></td></tr>
<tr><td>T7483</td><td>CTGCCTCCGACCTCTTACTCAGG</td><td>1000</td><td></td><td></td><td></td><td></td><td></td><td></td><td></td><td></td><td></td></tr>
<tr><td>Oligo fwd</td><td>CACCCTGCCTCCGACCTCTTACTC</td><td></td><td></td><td></td><td></td><td></td><td></td><td></td><td></td><td></td><td></td></tr>
<tr><td>Oligo rev</td><td>AAACGAGTAAGAGGTCGGAGGCAG</td><td></td><td></td><td></td><td></td><td></td><td></td><td></td><td></td><td></td><td></td></tr>
<tr><td>Chromosome</td><td>start</td><td>end</td><td>strand</td><td>MM</td><td>target_seq</td><td>PAM</td><td>alignment</td><td>distance</td><td>position</td><td>gene name</td><td>gene id</td></tr>
<tr><td>GU305914.1</td><td>102706</td><td>102728</td><td>+</td><td>0</td><td>CTGCCTCCGACCTCTTACTC</td><td>AGG</td><td>||||||||[||||||||||||]PAM</td><td>NA</td><td>Intergenic</td><td>NA</td><td>NA</td></tr>
<tr><td></td><td></td><td></td><td></td><td></td><td></td><td></td><td></td><td></td><td></td><td></td><td></td></tr>
<tr><td>T7484</td><td>TGCCTCCGACCTCTTACTCAGGG</td><td>1000</td><td></td><td></td><td></td><td></td><td></td><td></td><td></td><td></td><td></td></tr>
<tr><td>Oligo fwd</td><td>CACCTGCCTCCGACCTCTTACTCA</td><td></td><td></td><td></td><td></td><td></td><td></td><td></td><td></td><td></td><td></td></tr>
<tr><td>Oligo rev</td><td>AAACTGAGTAAGAGGTCGGAGGCA</td><td></td><td></td><td></td><td></td><td></td><td></td><td></td><td></td><td></td><td></td></tr>
<tr><td>Chromosome</td><td>start</td><td>end</td><td>strand</td><td>MM</td><td>target_seq</td><td>PAM</td><td>alignment</td><td>distance</td><td>position</td><td>gene name</td><td>gene id</td></tr>
<tr><td>GU305914.1</td><td>102707</td><td>102729</td><td>+</td><td>0</td><td>TGCCTCCGACCTCTTACTCA</td><td>GGG</td><td>||||||||[||||||||||||]PAM</td><td>NA</td><td>Intergenic</td><td>NA</td><td>NA</td></tr>
<tr><td></td><td></td><td></td><td></td><td></td><td></td><td></td><td></td><td></td><td></td><td></td><td></td></tr>
<tr><td>T7485</td><td>CGACCTCTTACTCAGGGAGCTGG</td><td>1000</td><td></td><td></td><td></td><td></td><td></td><td></td><td></td><td></td><td></td></tr>
<tr><td>Oligo fwd</td><td>CACCCGACCTCTTACTCAGGGAGC</td><td></td><td></td><td></td><td></td><td></td><td></td><td></td><td></td><td></td><td></td></tr>
<tr><td>Oligo rev</td><td>AAACGCTCCCTGAGTAAGAGGTCG</td><td></td><td></td><td></td><td></td><td></td><td></td><td></td><td></td><td></td><td></td></tr>
<tr><td>Chromosome</td><td>start</td><td>end</td><td>strand</td><td>MM</td><td>target_seq</td><td>PAM</td><td>alignment</td><td>distance</td><td>position</td><td>gene name</td><td>gene id</td></tr>
<tr><td>GU305914.1</td><td>102713</td><td>102735</td><td>+</td><td>0</td><td>CGACCTCTTACTCAGGGAGC</td><td>TGG</td><td>||||||||[||||||||||||]PAM</td><td>NA</td><td>Intergenic</td><td>NA</td><td>NA</td></tr>
<tr><td></td><td></td><td></td><td></td><td></td><td></td><td></td><td></td><td></td><td></td><td></td><td></td></tr>
<tr><td>T7486</td><td>GCCGCGCCCGACCACCTCCCCGG</td><td>1000</td><td></td><td></td><td></td><td></td><td></td><td></td><td></td><td></td><td></td></tr>
<tr><td>Oligo fwd</td><td>CACCGCCGCGCCCGACCACCTCCC</td><td></td><td></td><td></td><td></td><td></td><td></td><td></td><td></td><td></td><td></td></tr>
<tr><td>Oligo rev</td><td>AAACGGGAGGTGGTCGGGCGCGGC</td><td></td><td></td><td></td><td></td><td></td><td></td><td></td><td></td><td></td><td></td></tr>
<tr><td>Chromosome</td><td>start</td><td>end</td><td>strand</td><td>MM</td><td>target_seq</td><td>PAM</td><td>alignment</td><td>distance</td><td>position</td><td>gene name</td><td>gene id</td></tr>
<tr><td>GU305914.1</td><td>102747</td><td>102769</td><td>+</td><td>0</td><td>GCCGCGCCCGACCACCTCCC</td><td>CGG</td><td>||||||||[||||||||||||]PAM</td><td>NA</td><td>Intergenic</td><td>NA</td><td>NA</td></tr>
<tr><td></td><td></td><td></td><td></td><td></td><td></td><td></td><td></td><td></td><td></td><td></td><td></td></tr>
<tr><td>T7487</td><td>CCGCGCCCGACCACCTCCCCGGG</td><td>1000</td><td></td><td></td><td></td><td></td><td></td><td></td><td></td><td></td><td></td></tr>
<tr><td>Oligo fwd</td><td>CACCCCGCGCCCGACCACCTCCCC</td><td></td><td></td><td></td><td></td><td></td><td></td><td></td><td></td><td></td><td></td></tr>
<tr><td>Oligo rev</td><td>AAACGGGGAGGTGGTCGGGCGCGG</td><td></td><td></td><td></td><td></td><td></td><td></td><td></td><td></td><td></td><td></td></tr>
<tr><td>Chromosome</td><td>start</td><td>end</td><td>strand</td><td>MM</td><td>target_seq</td><td>PAM</td><td>alignment</td><td>distance</td><td>position</td><td>gene name</td><td>gene id</td></tr>
<tr><td>GU305914.1</td><td>102748</td><td>102770</td><td>+</td><td>0</td><td>CCGCGCCCGACCACCTCCCC</td><td>GGG</td><td>||||||||[||||||||||||]PAM</td><td>NA</td><td>Intergenic</td><td>NA</td><td>NA</td></tr>
<tr><td></td><td></td><td></td><td></td><td></td><td></td><td></td><td></td><td></td><td></td><td></td><td></td></tr>
<tr><td>T7488</td><td>CGCGCCCGACCACCTCCCCGGGG</td><td>1000</td><td></td><td></td><td></td><td></td><td></td><td></td><td></td><td></td><td></td></tr>
<tr><td>Oligo fwd</td><td>CACCCGCGCCCGACCACCTCCCCG</td><td></td><td></td><td></td><td></td><td></td><td></td><td></td><td></td><td></td><td></td></tr>
<tr><td>Oligo rev</td><td>AAACCGGGGAGGTGGTCGGGCGCG</td><td></td><td></td><td></td><td></td><td></td><td></td><td></td><td></td><td></td><td></td></tr>
<tr><td>Chromosome</td><td>start</td><td>end</td><td>strand</td><td>MM</td><td>target_seq</td><td>PAM</td><td>alignment</td><td>distance</td><td>position</td><td>gene name</td><td>gene id</td></tr>
<tr><td>GU305914.1</td><td>102749</td><td>102771</td><td>+</td><td>0</td><td>CGCGCCCGACCACCTCCCCG</td><td>GGG</td><td>||||||||[||||||||||||]PAM</td><td>NA</td><td>Intergenic</td><td>NA</td><td>NA</td></tr>
<tr><td></td><td></td><td></td><td></td><td></td><td></td><td></td><td></td><td></td><td></td><td></td><td></td></tr>
<tr><td>T7489</td><td>CTCCGCGACGAAGCGACGTCTGG</td><td>1000</td><td></td><td></td><td></td><td></td><td></td><td></td><td></td><td></td><td></td></tr>
<tr><td>Oligo fwd</td><td>CACCCTCCGCGACGAAGCGACGTC</td><td></td><td></td><td></td><td></td><td></td><td></td><td></td><td></td><td></td><td></td></tr>
<tr><td>Oligo rev</td><td>AAACGACGTCGCTTCGTCGCGGAG</td><td></td><td></td><td></td><td></td><td></td><td></td><td></td><td></td><td></td><td></td></tr>
<tr><td>Chromosome</td><td>start</td><td>end</td><td>strand</td><td>MM</td><td>target_seq</td><td>PAM</td><td>alignment</td><td>distance</td><td>position</td><td>gene name</td><td>gene id</td></tr>
<tr><td>GU305914.1</td><td>102789</td><td>102811</td><td>+</td><td>0</td><td>CTCCGCGACGAAGCGACGTC</td><td>TGG</td><td>||||||||[||||||||||||]PAM</td><td>NA</td><td>Intergenic</td><td>NA</td><td>NA</td></tr>
</table>

T7490 TCCGCGACGAAGCGACGTCTGGG 1000
Oligo fwd CACCTCCGCGACGAAGCGACGTCT
Oligo rev AAACAGACGTCGCTTCGTCGCGGA
Chromosome start end strand MM target_seq PAM alignment distance position gene name gene id
GU305914.1 102790 102812 + 0 TCCGCGACGAAGCGACGTCT GGG ||||||||[||||||||||||]PAM NA Intergenic NA NA

T7491 CGACGAAGCGACGTCTGGGACGG 1000
Oligo fwd CACCCGACGAAGCGACGTCTGGGA
Oligo rev AAACTCCCAGACGTCGCTTCGTCG
Chromosome start end strand MM target_seq PAM alignment distance position gene name gene id
GU305914.1 102794 102816 + 0 CGACGAAGCGACGTCTGGGA CGG ||||||||[||||||||||||]PAM NA Intergenic NA NA

T7492 GACGAAGCGACGTCTGGGACGGG 1000
Oligo fwd CACCGACGAAGCGACGTCTGGGAC
Oligo rev AAACGTCCCAGACGTCGCTTCGTC
Chromosome start end strand MM target_seq PAM alignment distance position gene name gene id
GU305914.1 102795 102817 + 0 GACGAAGCGACGTCTGGGAC GGG ||||||||[||||||||||||]PAM NA Intergenic NA NA

T7493 ACGGGACGCCTGCACAACGTCGG 1000
Oligo fwd CACCACGGGACGCCTGCACAACGT
Oligo rev AAACACGTTGTGCAGGCGTCCCGT
Chromosome start end strand MM target_seq PAM alignment distance position gene name gene id
GU305914.1 102813 102835 + 0 ACGGGACGCCTGCACAACGT CGG ||||||||[||||||||||||]PAM NA Intergenic NA NA

T7494 GACGCCTGCACAACGTCGGAAGG 1000
Oligo fwd CACCGACGCCTGCACAACGTCGGA
Oligo rev AAACTCCGACGTTGTGCAGGCGTC
Chromosome start end strand MM target_seq PAM alignment distance position gene name gene id
GU305914.1 102817 102839 + 0 GACGCCTGCACAACGTCGGA AGG ||||||||[||||||||||||]PAM NA Intergenic NA NA

T7495 CGGAAGGCGTCGCGACCTCGAGG 1000
Oligo fwd CACCCGGAAGGCGTCGCGACCTCG
Oligo rev AAACCGAGGTCGCGACGCCTTCCG
Chromosome start end strand MM target_seq PAM alignment distance position gene name gene id
GU305914.1 102833 102855 + 0 CGGAAGGCGTCGCGACCTCG AGG ||||||||[||||||||||||]PAM NA Intergenic NA NA

T7496 AAAGCAGCAGCACCAGCAGATGG 1000
Oligo fwd CACCAAAGCAGCAGCACCAGCAGA
Oligo rev AAACTCTGCTGGTGCTGCTGCTTT
Chromosome start end strand MM target_seq PAM alignment distance position gene name gene id
GU305914.1 102860 102882 + 0 AAAGCAGCAGCACCAGCAGA TGG ||||||||[||||||||||||]PAM NA Intergenic NA NA

T7497 AGATGGCCGCTCTGTGATCGAGG 1000
Oligo fwd CACCAGATGGCCGCTCTGTGATCG
Oligo rev AAACCGATCACAGAGCGGCCATCT
Chromosome start end strand MM target_seq PAM alignment distance position gene name gene id
GU305914.1 102877 102899 + 0 AGATGGCCGCTCTGTGATCG AGG ||||||||[||||||||||||]PAM NA Intergenic NA NA

T7498 TGATCGAGGCTTTTCTCGATCGG 1000
Oligo fwd CACCTGATCGAGGCTTTTCTCGAT
Oligo rev AAACATCGAGAAAAGCCTCGATCA
Chromosome start end strand MM target_seq PAM alignment distance position gene name gene id
GU305914.1 102891 102913 + 0 TGATCGAGGCTTTTCTCGAT CGG ||||||||[||||||||||||]PAM NA Intergenic NA NA

T7499 GATCGAGGCTTTTCTCGATCGGG 1000
Oligo fwd CACCGATCGAGGCTTTTCTCGATC
Oligo rev AAACGATCGAGAAAAGCCTCGATC
Chromosome start end strand MM target_seq PAM alignment distance position gene name gene id
GU305914.1 102892 102914 + 0 GATCGAGGCTTTTCTCGATC GGG ||||||||[||||||||||||]PAM NA Intergenic NA NA

T7500 ATCGAGGCTTTTCTCGATCGGGG 1000
Oligo fwd CACCATCGAGGCTTTTCTCGATCG
Oligo rev AAACCGATCGAGAAAAGCCTCGAT
Chromosome start end strand MM target_seq PAM alignment distance position gene name gene id
GU305914.1 102893 102915 + 0 ATCGAGGCTTTTCTCGATCG GGG ||||||||[||||||||||||]PAM NA Intergenic NA NA

T7501 TCGAGGCTTTTCTCGATCGGGGG 1000
Oligo fwd CACCTCGAGGCTTTTCTCGATCGG
Oligo rev AAACCCGATCGAGAAAAGCCTCGA
Chromosome start end strand MM target_seq PAM alignment distance position gene name gene id
GU305914.1 102894 102916 + 0 TCGAGGCTTTTCTCGATCGG GGG ||||||||[||||||||||||]PAM NA Intergenic NA NA

T7502 TCGGGGGACGTGTAAGTGTGTGG 1000
Oligo fwd CACCTCGGGGGACGTGTAAGTGTG
Oligo rev AAACCACACTTACACGTCCCCCGA
Chromosome start end strand MM target_seq PAM alignment distance position gene name gene id
GU305914.1 102910 102932 + 0 TCGGGGGACGTGTAAGTGTG TGG ||||||||[||||||||||||]PAM NA Intergenic NA NA

T7503 TGTGGATTGTTGTTATTTACCGG 1000
Oligo fwd CACCTGTGGATTGTTGTTATTTAC
Oligo rev AAACGTAAATAACAACAATCCACA
Chromosome start end strand MM target_seq PAM alignment distance position gene name gene id
GU305914.1 102928 102950 + 0 TGTGGATTGTTGTTATTTAC CGG ||||||||[||||||||||||]PAM NA Intergenic NA NA

T7504 GTGGATTGTTGTTATTTACCGGG 1000
Oligo fwd CACCGTGGATTGTTGTTATTTACC
Oligo rev AAACGGTAAATAACAACAATCCAC
Chromosome start end strand MM target_seq PAM alignment distance position gene name gene id
GU305914.1 102929 102951 + 0 GTGGATTGTTGTTATTTACC GGG ||||||||[||||||||||||]PAM NA Intergenic NA NA

<table>
<tr><td>T7505</td><td>TGGATTGTTGTTATTTACCGGGG</td><td>1000</td><td></td><td></td><td></td><td></td><td></td><td></td><td></td><td></td><td></td></tr>
<tr><td>Oligo fwd</td><td>CACCTGGATTGTTGTTATTTACCG</td><td></td><td></td><td></td><td></td><td></td><td></td><td></td><td></td><td></td><td></td></tr>
<tr><td>Oligo rev</td><td>AAACCGGTAAATAACAACAATCCA</td><td></td><td></td><td></td><td></td><td></td><td></td><td></td><td></td><td></td><td></td></tr>
<tr><td>Chromosome</td><td>start</td><td>end</td><td>strand</td><td>MM</td><td>target_seq</td><td>PAM</td><td>alignment</td><td>distance</td><td>position</td><td>gene name</td><td>gene id</td></tr>
<tr><td>GU305914.1</td><td>102930</td><td>102952</td><td>+</td><td>0</td><td>TGGATTGTTGTTATTTACCG</td><td>GGG</td><td>||||||||[||||||||||||]PAM</td><td>NA</td><td>Intergenic</td><td>NA</td><td>NA</td></tr>
<tr><td>T7506</td><td>ATTTACCGGGGCGTGTTTATTGG</td><td>1000</td><td></td><td></td><td></td><td></td><td></td><td></td><td></td><td></td><td></td></tr>
<tr><td>Oligo fwd</td><td>CACCATTTACCGGGGCGTGTTTAT</td><td></td><td></td><td></td><td></td><td></td><td></td><td></td><td></td><td></td><td></td></tr>
<tr><td>Oligo rev</td><td>AAACATAAACACGCCCCGGTAAAT</td><td></td><td></td><td></td><td></td><td></td><td></td><td></td><td></td><td></td><td></td></tr>
<tr><td>Chromosome</td><td>start</td><td>end</td><td>strand</td><td>MM</td><td>target_seq</td><td>PAM</td><td>alignment</td><td>distance</td><td>position</td><td>gene name</td><td>gene id</td></tr>
<tr><td>GU305914.1</td><td>102942</td><td>102964</td><td>+</td><td>0</td><td>ATTTACCGGGGCGTGTTTAT</td><td>TGG</td><td>||||||||[||||||||||||]PAM</td><td>NA</td><td>Intergenic</td><td>NA</td><td>NA</td></tr>
<tr><td>T7507</td><td>AAAATATACATACATCTCTGAGG</td><td>1000</td><td></td><td></td><td></td><td></td><td></td><td></td><td></td><td></td><td></td></tr>
<tr><td>Oligo fwd</td><td>CACCAAAATATACATACATCTCTG</td><td></td><td></td><td></td><td></td><td></td><td></td><td></td><td></td><td></td><td></td></tr>
<tr><td>Oligo rev</td><td>AAACCAGAGATGTATGTATATTTT</td><td></td><td></td><td></td><td></td><td></td><td></td><td></td><td></td><td></td><td></td></tr>
<tr><td>Chromosome</td><td>start</td><td>end</td><td>strand</td><td>MM</td><td>target_seq</td><td>PAM</td><td>alignment</td><td>distance</td><td>position</td><td>gene name</td><td>gene id</td></tr>
<tr><td>GU305914.1</td><td>103040</td><td>103062</td><td>+</td><td>0</td><td>AAAATATACATACATCTCTG</td><td>AGG</td><td>||||||||[||||||||||||]PAM</td><td>NA</td><td>Intergenic</td><td>NA</td><td>NA</td></tr>
<tr><td>T7508</td><td>AAATATACATACATCTCTGAGGG</td><td>1000</td><td></td><td></td><td></td><td></td><td></td><td></td><td></td><td></td><td></td></tr>
<tr><td>Oligo fwd</td><td>CACCAAATATACATACATCTCTGA</td><td></td><td></td><td></td><td></td><td></td><td></td><td></td><td></td><td></td><td></td></tr>
<tr><td>Oligo rev</td><td>AAACTCAGAGATGTATGTATATTT</td><td></td><td></td><td></td><td></td><td></td><td></td><td></td><td></td><td></td><td></td></tr>
<tr><td>Chromosome</td><td>start</td><td>end</td><td>strand</td><td>MM</td><td>target_seq</td><td>PAM</td><td>alignment</td><td>distance</td><td>position</td><td>gene name</td><td>gene id</td></tr>
<tr><td>GU305914.1</td><td>103041</td><td>103063</td><td>+</td><td>0</td><td>AAATATACATACATCTCTGA</td><td>GGG</td><td>||||||||[||||||||||||]PAM</td><td>NA</td><td>Intergenic</td><td>NA</td><td>NA</td></tr>
<tr><td>T7509</td><td>ATACATACATCTCTGAGGGTAGG</td><td>1000</td><td></td><td></td><td></td><td></td><td></td><td></td><td></td><td></td><td></td></tr>
<tr><td>Oligo fwd</td><td>CACCATACATACATCTCTGAGGGT</td><td></td><td></td><td></td><td></td><td></td><td></td><td></td><td></td><td></td><td></td></tr>
<tr><td>Oligo rev</td><td>AAACACCCTCAGAGATGTATGTAT</td><td></td><td></td><td></td><td></td><td></td><td></td><td></td><td></td><td></td><td></td></tr>
<tr><td>Chromosome</td><td>start</td><td>end</td><td>strand</td><td>MM</td><td>target_seq</td><td>PAM</td><td>alignment</td><td>distance</td><td>position</td><td>gene name</td><td>gene id</td></tr>
<tr><td>GU305914.1</td><td>103045</td><td>103067</td><td>+</td><td>0</td><td>ATACATACATCTCTGAGGGT</td><td>AGG</td><td>||||||||[||||||||||||]PAM</td><td>NA</td><td>Intergenic</td><td>NA</td><td>NA</td></tr>
<tr><td>T7510</td><td>ACATCTCTGAGGGTAGGTAGCGG</td><td>1000</td><td></td><td></td><td></td><td></td><td></td><td></td><td></td><td></td><td></td></tr>
<tr><td>Oligo fwd</td><td>CACCACATCTCTGAGGGTAGGTAG</td><td></td><td></td><td></td><td></td><td></td><td></td><td></td><td></td><td></td><td></td></tr>
<tr><td>Oligo rev</td><td>AAACCTACCTACCCTCAGAGATGT</td><td></td><td></td><td></td><td></td><td></td><td></td><td></td><td></td><td></td><td></td></tr>
<tr><td>Chromosome</td><td>start</td><td>end</td><td>strand</td><td>MM</td><td>target_seq</td><td>PAM</td><td>alignment</td><td>distance</td><td>position</td><td>gene name</td><td>gene id</td></tr>
<tr><td>GU305914.1</td><td>103051</td><td>103073</td><td>+</td><td>0</td><td>ACATCTCTGAGGGTAGGTAG</td><td>CGG</td><td>||||||||[||||||||||||]PAM</td><td>NA</td><td>Intergenic</td><td>NA</td><td>NA</td></tr>
<tr><td>T7511</td><td>TCCTGAGTACACTAGACAAGAGG</td><td>1000</td><td></td><td></td><td></td><td></td><td></td><td></td><td></td><td></td><td></td></tr>
<tr><td>Oligo fwd</td><td>CACCTCCTGAGTACACTAGACAAG</td><td></td><td></td><td></td><td></td><td></td><td></td><td></td><td></td><td></td><td></td></tr>
<tr><td>Oligo rev</td><td>AAACCTTGTCTAGTGTACTCAGGA</td><td></td><td></td><td></td><td></td><td></td><td></td><td></td><td></td><td></td><td></td></tr>
<tr><td>Chromosome</td><td>start</td><td>end</td><td>strand</td><td>MM</td><td>target_seq</td><td>PAM</td><td>alignment</td><td>distance</td><td>position</td><td>gene name</td><td>gene id</td></tr>
<tr><td>GU305914.1</td><td>103086</td><td>103108</td><td>+</td><td>0</td><td>TCCTGAGTACACTAGACAAG</td><td>AGG</td><td>||||||||[||||||||||||]PAM</td><td>NA</td><td>Intergenic</td><td>NA</td><td>NA</td></tr>
<tr><td>T7512</td><td>AGACAAGAGGTAATCTCTCTAGG</td><td>1000</td><td></td><td></td><td></td><td></td><td></td><td></td><td></td><td></td><td></td></tr>
<tr><td>Oligo fwd</td><td>CACCAGACAAGAGGTAATCTCTCT</td><td></td><td></td><td></td><td></td><td></td><td></td><td></td><td></td><td></td><td></td></tr>
<tr><td>Oligo rev</td><td>AAACAGAGAGATTACCTCTTGTCT</td><td></td><td></td><td></td><td></td><td></td><td></td><td></td><td></td><td></td><td></td></tr>
<tr><td>Chromosome</td><td>start</td><td>end</td><td>strand</td><td>MM</td><td>target_seq</td><td>PAM</td><td>alignment</td><td>distance</td><td>position</td><td>gene name</td><td>gene id</td></tr>
<tr><td>GU305914.1</td><td>103099</td><td>103121</td><td>+</td><td>0</td><td>AGACAAGAGGTAATCTCTCT</td><td>AGG</td><td>||||||||[||||||||||||]PAM</td><td>NA</td><td>Intergenic</td><td>NA</td><td>NA</td></tr>
<tr><td>T7513</td><td>TTCGCATTCTCTCTTCCGCGCGG</td><td>1000</td><td></td><td></td><td></td><td></td><td></td><td></td><td></td><td></td><td></td></tr>
<tr><td>Oligo fwd</td><td>CACCTTCGCATTCTCTCTTCCGCG</td><td></td><td></td><td></td><td></td><td></td><td></td><td></td><td></td><td></td><td></td></tr>
<tr><td>Oligo rev</td><td>AAACCGCGGAAGAGAGAATGCGAA</td><td></td><td></td><td></td><td></td><td></td><td></td><td></td><td></td><td></td><td></td></tr>
<tr><td>Chromosome</td><td>start</td><td>end</td><td>strand</td><td>MM</td><td>target_seq</td><td>PAM</td><td>alignment</td><td>distance</td><td>position</td><td>gene name</td><td>gene id</td></tr>
<tr><td>GU305914.1</td><td>103151</td><td>103173</td><td>+</td><td>0</td><td>TTCGCATTCTCTCTTCCGCG</td><td>CGG</td><td>||||||||[||||||||||||]PAM</td><td>NA</td><td>Intergenic</td><td>NA</td><td>NA</td></tr>
<tr><td>T7514</td><td>CTTCCGCGCGGTAGTAGTCGCGG</td><td>1000</td><td></td><td></td><td></td><td></td><td></td><td></td><td></td><td></td><td></td></tr>
<tr><td>Oligo fwd</td><td>CACCCTTCCGCGCGGTAGTAGTCG</td><td></td><td></td><td></td><td></td><td></td><td></td><td></td><td></td><td></td><td></td></tr>
<tr><td>Oligo rev</td><td>AAACCGACTACTACCGCGCGGAAG</td><td></td><td></td><td></td><td></td><td></td><td></td><td></td><td></td><td></td><td></td></tr>
<tr><td>Chromosome</td><td>start</td><td>end</td><td>strand</td><td>MM</td><td>target_seq</td><td>PAM</td><td>alignment</td><td>distance</td><td>position</td><td>gene name</td><td>gene id</td></tr>
<tr><td>GU305914.1</td><td>103163</td><td>103185</td><td>+</td><td>0</td><td>CTTCCGCGCGGTAGTAGTCG</td><td>CGG</td><td>||||||||[||||||||||||]PAM</td><td>NA</td><td>Intergenic</td><td>NA</td><td>NA</td></tr>
<tr><td>T7515</td><td>GCCCGCCCTTGACGAGCTTTAGG</td><td>1000</td><td></td><td></td><td></td><td></td><td></td><td></td><td></td><td></td><td></td></tr>
<tr><td>Oligo fwd</td><td>CACCGCCCGCCCTTGACGAGCTTT</td><td></td><td></td><td></td><td></td><td></td><td></td><td></td><td></td><td></td><td></td></tr>
<tr><td>Oligo rev</td><td>AAACAAAGCTCGTCAAGGGCGGGC</td><td></td><td></td><td></td><td></td><td></td><td></td><td></td><td></td><td></td><td></td></tr>
<tr><td>Chromosome</td><td>start</td><td>end</td><td>strand</td><td>MM</td><td>target_seq</td><td>PAM</td><td>alignment</td><td>distance</td><td>position</td><td>gene name</td><td>gene id</td></tr>
<tr><td>GU305914.1</td><td>103187</td><td>103209</td><td>+</td><td>0</td><td>GCCCGCCCTTGACGAGCTTT</td><td>AGG</td><td>||||||||[||||||||||||]PAM</td><td>NA</td><td>Intergenic</td><td>NA</td><td>NA</td></tr>
<tr><td>T7516</td><td>ACGAGCTTTAGGTCCGAGCGCGG</td><td>1000</td><td></td><td></td><td></td><td></td><td></td><td></td><td></td><td></td><td></td></tr>
<tr><td>Oligo fwd</td><td>CACCACGAGCTTTAGGTCCGAGCG</td><td></td><td></td><td></td><td></td><td></td><td></td><td></td><td></td><td></td><td></td></tr>
<tr><td>Oligo rev</td><td>AAACCGCTCGGACCTAAAGCTCGT</td><td></td><td></td><td></td><td></td><td></td><td></td><td></td><td></td><td></td><td></td></tr>
<tr><td>Chromosome</td><td>start</td><td>end</td><td>strand</td><td>MM</td><td>target_seq</td><td>PAM</td><td>alignment</td><td>distance</td><td>position</td><td>gene name</td><td>gene id</td></tr>
<tr><td>GU305914.1</td><td>103198</td><td>103220</td><td>+</td><td>0</td><td>ACGAGCTTTAGGTCCGAGCG</td><td>CGG</td><td>||||||||[||||||||||||]PAM</td><td>NA</td><td>Intergenic</td><td>NA</td><td>NA</td></tr>
<tr><td>T7517</td><td>ATCATGACGCTCGTCAGTGACGG</td><td>1000</td><td></td><td></td><td></td><td></td><td></td><td></td><td></td><td></td><td></td></tr>
<tr><td>Oligo fwd</td><td>CACCATCATGACGCTCGTCAGTGA</td><td></td><td></td><td></td><td></td><td></td><td></td><td></td><td></td><td></td><td></td></tr>
<tr><td>Oligo rev</td><td>AAACTCACTGACGAGCGTCATGAT</td><td></td><td></td><td></td><td></td><td></td><td></td><td></td><td></td><td></td><td></td></tr>
<tr><td>Chromosome</td><td>start</td><td>end</td><td>strand</td><td>MM</td><td>target_seq</td><td>PAM</td><td>alignment</td><td>distance</td><td>position</td><td>gene name</td><td>gene id</td></tr>
<tr><td>GU305914.1</td><td>103234</td><td>103256</td><td>+</td><td>0</td><td>ATCATGACGCTCGTCAGTGA</td><td>CGG</td><td>||||||||[||||||||||||]PAM</td><td>NA</td><td>Intergenic</td><td>NA</td><td>NA</td></tr>
<tr><td>T7518</td><td>CTCGTCAGTGACGGTTTGAACGG</td><td>1000</td><td></td><td></td><td></td><td></td><td></td><td></td><td></td><td></td><td></td></tr>
<tr><td>Oligo fwd</td><td>CACCCTCGTCAGTGACGGTTTGAA</td><td></td><td></td><td></td><td></td><td></td><td></td><td></td><td></td><td></td><td></td></tr>
<tr><td>Oligo rev</td><td>AAACTTCAAACCGTCACTGACGAG</td><td></td><td></td><td></td><td></td><td></td><td></td><td></td><td></td><td></td><td></td></tr>
<tr><td>Chromosome</td><td>start</td><td>end</td><td>strand</td><td>MM</td><td>target_seq</td><td>PAM</td><td>alignment</td><td>distance</td><td>position</td><td>gene name</td><td>gene id</td></tr>
<tr><td>GU305914.1</td><td>103243</td><td>103265</td><td>+</td><td>0</td><td>CTCGTCAGTGACGGTTTGAA</td><td>CGG</td><td>||||||||[||||||||||||]PAM</td><td>NA</td><td>Intergenic</td><td>NA</td><td>NA</td></tr>
<tr><td>T7519</td><td>GGTTTGAACGGCGTGATGCCCGG</td><td>1000</td><td></td><td></td><td></td><td></td><td></td><td></td><td></td><td></td><td></td></tr>
<tr><td>Oligo fwd</td><td>CACCGGTTTGAACGGCGTGATGCC</td><td></td><td></td><td></td><td></td><td></td><td></td><td></td><td></td><td></td><td></td></tr>
<tr><td>Oligo rev</td><td>AAACGGCATCACGCCGTTCAAACC</td><td></td><td></td><td></td><td></td><td></td><td></td><td></td><td></td><td></td><td></td></tr>
<tr><td>Chromosome</td><td>start</td><td>end</td><td>strand</td><td>MM</td><td>target_seq</td><td>PAM</td><td>alignment</td><td>distance</td><td>position</td><td>gene name</td><td>gene id</td></tr>
</table>

<table>
<tr><td>GU305914.1</td><td></td><td>103255</td><td>103277</td><td>+</td><td>0</td><td>GGTTTGAACGGCGTGATGCC</td><td>CGG</td><td>||||||||[||||||||||||]PAM</td><td>NA</td><td>Intergenic</td><td>NA</td><td>NA</td></tr>
<tr><td>T7520</td><td>CGGCGTCGTGAAGATGATCTTGG</td><td></td><td>1000</td><td></td><td></td><td></td><td></td><td></td><td></td><td></td><td></td><td></td></tr>
<tr><td>Oligo fwd</td><td>CACCGGCGTCGTGAAGATGATCT</td><td></td><td></td><td></td><td></td><td></td><td></td><td></td><td></td><td></td><td></td><td></td></tr>
<tr><td>Oligo rev</td><td>AAACAGATCATCTTCACGACGCCG</td><td></td><td></td><td></td><td></td><td></td><td></td><td></td><td></td><td></td><td></td><td></td></tr>
<tr><td>Chromosome</td><td>start</td><td></td><td>end</td><td>strand</td><td>MM</td><td>target_seq</td><td>PAM</td><td>alignment</td><td>distance</td><td>position</td><td>gene name</td><td>gene id</td></tr>
<tr><td>GU305914.1</td><td></td><td>103275</td><td>103297</td><td>+</td><td>0</td><td>CGGCGTCGTGAAGATGATCT</td><td>TGG</td><td>||||||||[||||||||||||]PAM</td><td>NA</td><td>Intergenic</td><td>NA</td><td>NA</td></tr>
<tr><td>T7521</td><td>GATCTTGGCCATCGCCATCCCGG</td><td></td><td>1000</td><td></td><td></td><td></td><td></td><td></td><td></td><td></td><td></td><td></td></tr>
<tr><td>Oligo fwd</td><td>CACCGATCTTGGCCATCGCCATCC</td><td></td><td></td><td></td><td></td><td></td><td></td><td></td><td></td><td></td><td></td><td></td></tr>
<tr><td>Oligo rev</td><td>AAACGGATGGCGATGGCCAAGATC</td><td></td><td></td><td></td><td></td><td></td><td></td><td></td><td></td><td></td><td></td><td></td></tr>
<tr><td>Chromosome</td><td>start</td><td></td><td>end</td><td>strand</td><td>MM</td><td>target_seq</td><td>PAM</td><td>alignment</td><td>distance</td><td>position</td><td>gene name</td><td>gene id</td></tr>
<tr><td>GU305914.1</td><td></td><td>103290</td><td>103312</td><td>+</td><td>0</td><td>GATCTTGGCCATCGCCATCC</td><td>CGG</td><td>||||||||[||||||||||||]PAM</td><td>NA</td><td>Intergenic</td><td>NA</td><td>NA</td></tr>
<tr><td>T7522</td><td>CAGATCGACGTGCTCGTCCGAGG</td><td></td><td>1000</td><td></td><td></td><td></td><td></td><td></td><td></td><td></td><td></td><td></td></tr>
<tr><td>Oligo fwd</td><td>CACCCAGATCGACGTGCTCGTCCG</td><td></td><td></td><td></td><td></td><td></td><td></td><td></td><td></td><td></td><td></td><td></td></tr>
<tr><td>Oligo rev</td><td>AAACCGGACGAGCACGTCGATCTG</td><td></td><td></td><td></td><td></td><td></td><td></td><td></td><td></td><td></td><td></td><td></td></tr>
<tr><td>Chromosome</td><td>start</td><td></td><td>end</td><td>strand</td><td>MM</td><td>target_seq</td><td>PAM</td><td>alignment</td><td>distance</td><td>position</td><td>gene name</td><td>gene id</td></tr>
<tr><td>GU305914.1</td><td></td><td>103317</td><td>103339</td><td>+</td><td>0</td><td>CAGATCGACGTGCTCGTCCG</td><td>AGG</td><td>||||||||[||||||||||||]PAM</td><td>NA</td><td>Intergenic</td><td>NA</td><td>NA</td></tr>
<tr><td>T7523</td><td>GCGTAGCACCGCCAGTGAGTTGG</td><td></td><td>1000</td><td></td><td></td><td></td><td></td><td></td><td></td><td></td><td></td><td></td></tr>
<tr><td>Oligo fwd</td><td>CACCGCGTAGCACCGCCAGTGAGT</td><td></td><td></td><td></td><td></td><td></td><td></td><td></td><td></td><td></td><td></td><td></td></tr>
<tr><td>Oligo rev</td><td>AAACACTCACTGGCGGTGCTACGC</td><td></td><td></td><td></td><td></td><td></td><td></td><td></td><td></td><td></td><td></td><td></td></tr>
<tr><td>Chromosome</td><td>start</td><td></td><td>end</td><td>strand</td><td>MM</td><td>target_seq</td><td>PAM</td><td>alignment</td><td>distance</td><td>position</td><td>gene name</td><td>gene id</td></tr>
<tr><td>GU305914.1</td><td></td><td>103350</td><td>103372</td><td>+</td><td>0</td><td>GCGTAGCACCGCCAGTGAGT</td><td>TGG</td><td>||||||||[||||||||||||]PAM</td><td>NA</td><td>Intergenic</td><td>NA</td><td>NA</td></tr>
<tr><td>T7524</td><td>CGTAGCACCGCCAGTGAGTTGGG</td><td></td><td>1000</td><td></td><td></td><td></td><td></td><td></td><td></td><td></td><td></td><td></td></tr>
<tr><td>Oligo fwd</td><td>CACCCGTAGCACCGCCAGTGAGTT</td><td></td><td></td><td></td><td></td><td></td><td></td><td></td><td></td><td></td><td></td><td></td></tr>
<tr><td>Oligo rev</td><td>AAACAACTCACTGGCGGTGCTACG</td><td></td><td></td><td></td><td></td><td></td><td></td><td></td><td></td><td></td><td></td><td></td></tr>
<tr><td>Chromosome</td><td>start</td><td></td><td>end</td><td>strand</td><td>MM</td><td>target_seq</td><td>PAM</td><td>alignment</td><td>distance</td><td>position</td><td>gene name</td><td>gene id</td></tr>
<tr><td>GU305914.1</td><td></td><td>103351</td><td>103373</td><td>+</td><td>0</td><td>CGTAGCACCGCCAGTGAGTT</td><td>GGG</td><td>||||||||[||||||||||||]PAM</td><td>NA</td><td>Intergenic</td><td>NA</td><td>NA</td></tr>
<tr><td>T7525</td><td>GTAGCACCGCCAGTGAGTTGGGG</td><td></td><td>1000</td><td></td><td></td><td></td><td></td><td></td><td></td><td></td><td></td><td></td></tr>
<tr><td>Oligo fwd</td><td>CACCGTAGCACCGCCAGTGAGTTG</td><td></td><td></td><td></td><td></td><td></td><td></td><td></td><td></td><td></td><td></td><td></td></tr>
<tr><td>Oligo rev</td><td>AAACCAACTCACTGGCGGTGCTAC</td><td></td><td></td><td></td><td></td><td></td><td></td><td></td><td></td><td></td><td></td><td></td></tr>
<tr><td>Chromosome</td><td>start</td><td></td><td>end</td><td>strand</td><td>MM</td><td>target_seq</td><td>PAM</td><td>alignment</td><td>distance</td><td>position</td><td>gene name</td><td>gene id</td></tr>
<tr><td>GU305914.1</td><td></td><td>103352</td><td>103374</td><td>+</td><td>0</td><td>GTAGCACCGCCAGTGAGTTG</td><td>GGG</td><td>||||||||[||||||||||||]PAM</td><td>NA</td><td>Intergenic</td><td>NA</td><td>NA</td></tr>
<tr><td>T7526</td><td>CAGTGAGTTGGGGCGCCACACGG</td><td></td><td>1000</td><td></td><td></td><td></td><td></td><td></td><td></td><td></td><td></td><td></td></tr>
<tr><td>Oligo fwd</td><td>CACCCAGTGAGTTGGGGCGCCACA</td><td></td><td></td><td></td><td></td><td></td><td></td><td></td><td></td><td></td><td></td><td></td></tr>
<tr><td>Oligo rev</td><td>AAACTGTGGCGCCCCAACTCACTG</td><td></td><td></td><td></td><td></td><td></td><td></td><td></td><td></td><td></td><td></td><td></td></tr>
<tr><td>Chromosome</td><td>start</td><td></td><td>end</td><td>strand</td><td>MM</td><td>target_seq</td><td>PAM</td><td>alignment</td><td>distance</td><td>position</td><td>gene name</td><td>gene id</td></tr>
<tr><td>GU305914.1</td><td></td><td>103362</td><td>103384</td><td>+</td><td>0</td><td>CAGTGAGTTGGGGCGCCACA</td><td>CGG</td><td>||||||||[||||||||||||]PAM</td><td>NA</td><td>Intergenic</td><td>NA</td><td>NA</td></tr>
<tr><td>T7527</td><td>GAGTTGGGGCGCCACACGGTCGG</td><td></td><td>1000</td><td></td><td></td><td></td><td></td><td></td><td></td><td></td><td></td><td></td></tr>
<tr><td>Oligo fwd</td><td>CACCGAGTTGGGGCGCCACACGGT</td><td></td><td></td><td></td><td></td><td></td><td></td><td></td><td></td><td></td><td></td><td></td></tr>
<tr><td>Oligo rev</td><td>AAACACCGTGTGGCGCCCCAACTC</td><td></td><td></td><td></td><td></td><td></td><td></td><td></td><td></td><td></td><td></td><td></td></tr>
<tr><td>Chromosome</td><td>start</td><td></td><td>end</td><td>strand</td><td>MM</td><td>target_seq</td><td>PAM</td><td>alignment</td><td>distance</td><td>position</td><td>gene name</td><td>gene id</td></tr>
<tr><td>GU305914.1</td><td></td><td>103366</td><td>103388</td><td>+</td><td>0</td><td>GAGTTGGGGCGCCACACGGT</td><td>CGG</td><td>||||||||[||||||||||||]PAM</td><td>NA</td><td>Intergenic</td><td>NA</td><td>NA</td></tr>
<tr><td>T7528</td><td>AGTTGGGGCGCCACACGGTCGGG</td><td></td><td>1000</td><td></td><td></td><td></td><td></td><td></td><td></td><td></td><td></td><td></td></tr>
<tr><td>Oligo fwd</td><td>CACCAGTTGGGGCGCCACACGGTC</td><td></td><td></td><td></td><td></td><td></td><td></td><td></td><td></td><td></td><td></td><td></td></tr>
<tr><td>Oligo rev</td><td>AAACGACCGTGTGGCGCCCCAACT</td><td></td><td></td><td></td><td></td><td></td><td></td><td></td><td></td><td></td><td></td><td></td></tr>
<tr><td>Chromosome</td><td>start</td><td></td><td>end</td><td>strand</td><td>MM</td><td>target_seq</td><td>PAM</td><td>alignment</td><td>distance</td><td>position</td><td>gene name</td><td>gene id</td></tr>
<tr><td>GU305914.1</td><td></td><td>103367</td><td>103389</td><td>+</td><td>0</td><td>AGTTGGGGCGCCACACGGTC</td><td>GGG</td><td>||||||||[||||||||||||]PAM</td><td>NA</td><td>Intergenic</td><td>NA</td><td>NA</td></tr>
<tr><td>T7529</td><td>GGCGCCACACGGTCGGGTCCAGG</td><td></td><td>1000</td><td></td><td></td><td></td><td></td><td></td><td></td><td></td><td></td><td></td></tr>
<tr><td>Oligo fwd</td><td>CACCGGCGCCACACGGTCGGGTCC</td><td></td><td></td><td></td><td></td><td></td><td></td><td></td><td></td><td></td><td></td><td></td></tr>
<tr><td>Oligo rev</td><td>AAACGGACCCGACCGTGTGGCGCC</td><td></td><td></td><td></td><td></td><td></td><td></td><td></td><td></td><td></td><td></td><td></td></tr>
<tr><td>Chromosome</td><td>start</td><td></td><td>end</td><td>strand</td><td>MM</td><td>target_seq</td><td>PAM</td><td>alignment</td><td>distance</td><td>position</td><td>gene name</td><td>gene id</td></tr>
<tr><td>GU305914.1</td><td></td><td>103373</td><td>103395</td><td>+</td><td>0</td><td>GGCGCCACACGGTCGGGTCC</td><td>AGG</td><td>||||||||[||||||||||||]PAM</td><td>NA</td><td>Intergenic</td><td>NA</td><td>NA</td></tr>
<tr><td>T7530</td><td>GGAGCACGCGCTTTGTCGCTAGG</td><td></td><td>1000</td><td></td><td></td><td></td><td></td><td></td><td></td><td></td><td></td><td></td></tr>
<tr><td>Oligo fwd</td><td>CACCGGAGCACGCGCTTTGTCGCT</td><td></td><td></td><td></td><td></td><td></td><td></td><td></td><td></td><td></td><td></td><td></td></tr>
<tr><td>Oligo rev</td><td>AAACAGCGACAAAGCGCGTGCTCC</td><td></td><td></td><td></td><td></td><td></td><td></td><td></td><td></td><td></td><td></td><td></td></tr>
<tr><td>Chromosome</td><td>start</td><td></td><td>end</td><td>strand</td><td>MM</td><td>target_seq</td><td>PAM</td><td>alignment</td><td>distance</td><td>position</td><td>gene name</td><td>gene id</td></tr>
<tr><td>GU305914.1</td><td></td><td>103394</td><td>103416</td><td>+</td><td>0</td><td>GGAGCACGCGCTTTGTCGCT</td><td>AGG</td><td>||||||||[||||||||||||]PAM</td><td>NA</td><td>Intergenic</td><td>NA</td><td>NA</td></tr>
<tr><td>T7531</td><td>GCACGCGCTTTGTCGCTAGGTGG</td><td></td><td>1000</td><td></td><td></td><td></td><td></td><td></td><td></td><td></td><td></td><td></td></tr>
<tr><td>Oligo fwd</td><td>CACCGCACGCGCTTTGTCGCTAGG</td><td></td><td></td><td></td><td></td><td></td><td></td><td></td><td></td><td></td><td></td><td></td></tr>
<tr><td>Oligo rev</td><td>AAACCCTAGCGACAAAGCGCGTGC</td><td></td><td></td><td></td><td></td><td></td><td></td><td></td><td></td><td></td><td></td><td></td></tr>
<tr><td>Chromosome</td><td>start</td><td></td><td>end</td><td>strand</td><td>MM</td><td>target_seq</td><td>PAM</td><td>alignment</td><td>distance</td><td>position</td><td>gene name</td><td>gene id</td></tr>
<tr><td>GU305914.1</td><td></td><td>103397</td><td>103419</td><td>+</td><td>0</td><td>GCACGCGCTTTGTCGCTAGG</td><td>TGG</td><td>||||||||[||||||||||||]PAM</td><td>NA</td><td>Intergenic</td><td>NA</td><td>NA</td></tr>
<tr><td>T7532</td><td>TTTGTCGCTAGGTGGCGACACGG</td><td></td><td>1000</td><td></td><td></td><td></td><td></td><td></td><td></td><td></td><td></td><td></td></tr>
<tr><td>Oligo fwd</td><td>CACCTTTGTCGCTAGGTGGCGACA</td><td></td><td></td><td></td><td></td><td></td><td></td><td></td><td></td><td></td><td></td><td></td></tr>
<tr><td>Oligo rev</td><td>AAACTGTCGCCACCTAGCGACAAA</td><td></td><td></td><td></td><td></td><td></td><td></td><td></td><td></td><td></td><td></td><td></td></tr>
<tr><td>Chromosome</td><td>start</td><td></td><td>end</td><td>strand</td><td>MM</td><td>target_seq</td><td>PAM</td><td>alignment</td><td>distance</td><td>position</td><td>gene name</td><td>gene id</td></tr>
<tr><td>GU305914.1</td><td></td><td>103405</td><td>103427</td><td>+</td><td>0</td><td>TTTGTCGCTAGGTGGCGACA</td><td>CGG</td><td>||||||||[||||||||||||]PAM</td><td>NA</td><td>Intergenic</td><td>NA</td><td>NA</td></tr>
<tr><td>T7533</td><td>TTGTCGCTAGGTGGCGACACGGG</td><td></td><td>1000</td><td></td><td></td><td></td><td></td><td></td><td></td><td></td><td></td><td></td></tr>
<tr><td>Oligo fwd</td><td>CACCTTGTCGCTAGGTGGCGACAC</td><td></td><td></td><td></td><td></td><td></td><td></td><td></td><td></td><td></td><td></td><td></td></tr>
<tr><td>Oligo rev</td><td>AAACGTGTCGCCACCTAGCGACAA</td><td></td><td></td><td></td><td></td><td></td><td></td><td></td><td></td><td></td><td></td><td></td></tr>
<tr><td>Chromosome</td><td>start</td><td></td><td>end</td><td>strand</td><td>MM</td><td>target_seq</td><td>PAM</td><td>alignment</td><td>distance</td><td>position</td><td>gene name</td><td>gene id</td></tr>
<tr><td>GU305914.1</td><td></td><td>103406</td><td>103428</td><td>+</td><td>0</td><td>TTGTCGCTAGGTGGCGACAC</td><td>GGG</td><td>||||||||[||||||||||||]PAM</td><td>NA</td><td>Intergenic</td><td>NA</td><td>NA</td></tr>
<tr><td>T7534</td><td>GGTGGCGACACGGGATGACCAGG</td><td></td><td>1000</td><td></td><td></td><td></td><td></td><td></td><td></td><td></td><td></td><td></td></tr>
<tr><td>Oligo fwd</td><td>CACCGGTGGCGACACGGGATGACC</td><td></td><td></td><td></td><td></td><td></td><td></td><td></td><td></td><td></td><td></td><td></td></tr>
<tr><td>Oligo rev</td><td>AAACGGTCATCCCGTGTCGCCACC</td><td></td><td></td><td></td><td></td><td></td><td></td><td></td><td></td><td></td><td></td><td></td></tr>
</table>

Chromosome start end strand MM target_seq PAM alignment distance position gene name gene id
GU305914.1 103415 103437 + 0 GGTGGCGACACGGGATGACC AGG ||||||||[||||||||||||]PAM NA Intergenic NA NA

T7535 TGCTCGCGTCCTCTTCACACAGG 1000
Oligo fwd CACCTGCTCGCGTCCTCTTCACAC
Oligo rev AAACGTGTGAAGAGGACGCGAGCA
Chromosome start end strand MM target_seq PAM alignment distance position gene name gene id
GU305914.1 103442 103464 + 0 TGCTCGCGTCCTCTTCACAC AGG ||||||||[||||||||||||]PAM NA Intergenic NA NA

T7536 GTCCTCTTCACACAGGTGTCCGG 1000
Oligo fwd CACCGTCCTCTTCACACAGGTGTC
Oligo rev AAACGACACCTGTGTGAAGAGGAC
Chromosome start end strand MM target_seq PAM alignment distance position gene name gene id
GU305914.1 103449 103471 + 0 GTCCTCTTCACACAGGTGTC CGG ||||||||[||||||||||||]PAM NA Intergenic NA NA

T7537 GGTGTCCGGCCTCGAACTTCAGG 1000
Oligo fwd CACCGGTGTCCGGCCTCGAACTTC
Oligo rev AAACGAAGTTCGAGGCCGGACACC
Chromosome start end strand MM target_seq PAM alignment distance position gene name gene id
GU305914.1 103463 103485 + 0 GGTGTCCGGCCTCGAACTTC AGG ||||||||[||||||||||||]PAM NA Intergenic NA NA

T7538 CTCGAACTTCAGGTTCTTCGAGG 1000
Oligo fwd CACCCTCGAACTTCAGGTTCTTCG
Oligo rev AAACCGAAGAACCTGAAGTTCGAG
Chromosome start end strand MM target_seq PAM alignment distance position gene name gene id
GU305914.1 103473 103495 + 0 CTCGAACTTCAGGTTCTTCG AGG ||||||||[||||||||||||]PAM NA Intergenic NA NA

T7539 TTCAGGTTCTTCGAGGCCCGCGG 1000
Oligo fwd CACCTTCAGGTTCTTCGAGGCCCG
Oligo rev AAACCGGGCCTCGAAGAACCTGAA
Chromosome start end strand MM target_seq PAM alignment distance position gene name gene id
GU305914.1 103480 103502 + 0 TTCAGGTTCTTCGAGGCCCG CGG ||||||||[||||||||||||]PAM NA Intergenic NA NA

T7540 AGGTTCTTCGAGGCCCGCGGCGG 1000
Oligo fwd CACCAGGTTCTTCGAGGCCCGCGG
Oligo rev AAACCCGCGGGCCTCGAAGAACCT
Chromosome start end strand MM target_seq PAM alignment distance position gene name gene id
GU305914.1 103483 103505 + 0 AGGTTCTTCGAGGCCCGCGG CGG ||||||||[||||||||||||]PAM NA Intergenic NA NA

T7541 TCTTCGAGGCCCGCGGCGGCAGG 1000
Oligo fwd CACCTCTTCGAGGCCCGCGGCGGC
Oligo rev AAACGCCGCCGCGGGCCTCGAAGA
Chromosome start end strand MM target_seq PAM alignment distance position gene name gene id
GU305914.1 103487 103509 + 0 TCTTCGAGGCCCGCGGCGGC AGG ||||||||[||||||||||||]PAM NA Intergenic NA NA

T7542 GCCCGCGGCGGCAGGCTCAGCGG 1000
Oligo fwd CACCGCCCGCGGCGGCAGGCTCAG
Oligo rev AAACCTGAGCCTGCCGCCGCGGGC
Chromosome start end strand MM target_seq PAM alignment distance position gene name gene id
GU305914.1 103495 103517 + 0 GCCCGCGGCGGCAGGCTCAG CGG ||||||||[||||||||||||]PAM NA Intergenic NA NA

T7543 GCAGGCTCAGCGGTCGCCGCAGG 1000
Oligo fwd CACCGCAGGCTCAGCGGTCGCCGC
Oligo rev AAACGCGGCGACCGCTGAGCCTGC
Chromosome start end strand MM target_seq PAM alignment distance position gene name gene id
GU305914.1 103505 103527 + 0 GCAGGCTCAGCGGTCGCCGC AGG ||||||||[||||||||||||]PAM NA Intergenic NA NA

T7544 TCAGCGGTCGCCGCAGGCGTAGG 1000
Oligo fwd CACCTCAGCGGTCGCCGCAGGCGT
Oligo rev AAACACGCCTGCGGCGACCGCTGA
Chromosome start end strand MM target_seq PAM alignment distance position gene name gene id
GU305914.1 103511 103533 + 0 TCAGCGGTCGCCGCAGGCGT AGG ||||||||[||||||||||||]PAM NA Intergenic NA NA

T7545 GGTCGAAGTACGTGTCGCCCTGG 1000
Oligo fwd CACCGGTCGAAGTACGTGTCGCCC
Oligo rev AAACGGGCGACACGTACTTCGACC
Chromosome start end strand MM target_seq PAM alignment distance position gene name gene id
GU305914.1 103532 103554 + 0 GGTCGAAGTACGTGTCGCCC TGG ||||||||[||||||||||||]PAM NA Intergenic NA NA

T7546 AAGTACGTGTCGCCCTGGTGCGG 1000
Oligo fwd CACCAAGTACGTGTCGCCCTGGTG
Oligo rev AAACCACCAGGGCGACACGTACTT
Chromosome start end strand MM target_seq PAM alignment distance position gene name gene id
GU305914.1 103537 103559 + 0 AAGTACGTGTCGCCCTGGTG CGG ||||||||[||||||||||||]PAM NA Intergenic NA NA

T7547 CGCCCTGGTGCGGCGCCATCAGG 1000
Oligo fwd CACCCGCCCTGGTGCGGCGCCATC
Oligo rev AAACGATGGCGCCGCACCAGGGCG
Chromosome start end strand MM target_seq PAM alignment distance position gene name gene id
GU305914.1 103547 103569 + 0 CGCCCTGGTGCGGCGCCATC AGG ||||||||[||||||||||||]PAM NA Intergenic NA NA

T7548 GTGCGGCGCCATCAGGTTGATGG 1000
Oligo fwd CACCGTGCGGCGCCATCAGGTTGA
Oligo rev AAACTCAACCTGATGGCGCCGCAC
Chromosome start end strand MM target_seq PAM alignment distance position gene name gene id
GU305914.1 103554 103576 + 0 GTGCGGCGCCATCAGGTTGA TGG ||||||||[||||||||||||]PAM NA Intergenic NA NA

T7549 ATCAGGTTGATGGTCTGCCACGG 1000
Oligo fwd CACCATCAGGTTGATGGTCTGCCA

Oligo rev AAACTGGCAGACCATCAACCTGAT
Chromosome start end strand MM target_seq PAM alignment distance position gene name gene id
GU305914.1 103564 103586 + 0 ATCAGGTTGATGGTCTGCCA CGG ||||||||[||||||||||||]PAM NA Intergenic NA NA

T7550 TTGATGGTCTGCCACGGCTCCGG 1000
Oligo fwd CACCTTGATGGTCTGCCACGGCTC
Oligo rev AAACGAGCCGTGGCAGACCATCAA
Chromosome start end strand MM target_seq PAM alignment distance position gene name gene id
GU305914.1 103570 103592 + 0 TTGATGGTCTGCCACGGCTC CGG ||||||||[||||||||||||]PAM NA Intergenic NA NA

T7551 CACGGCTCCGGCAGCAGCATCGG 1000
Oligo fwd CACCCACGGCTCCGGCAGCAGCAT
Oligo rev AAACATGCTGCTGCCGGAGCCGTG
Chromosome start end strand MM target_seq PAM alignment distance position gene name gene id
GU305914.1 103582 103604 + 0 CACGGCTCCGGCAGCAGCAT CGG ||||||||[||||||||||||]PAM NA Intergenic NA NA

T7552 GCAGCAGCATCGGCAACATGTGG 1000
Oligo fwd CACCGCAGCAGCATCGGCAACATG
Oligo rev AAACCATGTTGCCGATGCTGCTGC
Chromosome start end strand MM target_seq PAM alignment distance position gene name gene id
GU305914.1 103592 103614 + 0 GCAGCAGCATCGGCAACATG TGG ||||||||[||||||||||||]PAM NA Intergenic NA NA

T7553 ATGTGAATCTGCAGCATGTTCGG 1000
Oligo fwd CACCATGTGAATCTGCAGCATGTT
Oligo rev AAACAACATGCTGCAGATTCACAT
Chromosome start end strand MM target_seq PAM alignment distance position gene name gene id
GU305914.1 103615 103637 + 0 ATGTGAATCTGCAGCATGTT CGG ||||||||[||||||||||||]PAM NA Intergenic NA NA

T7554 TGAATCTGCAGCATGTTCGGCGG 1000
Oligo fwd CACCTGAATCTGCAGCATGTTCGG
Oligo rev AAACCCGAACATGCTGCAGATTCA
Chromosome start end strand MM target_seq PAM alignment distance position gene name gene id
GU305914.1 103618 103640 + 0 TGAATCTGCAGCATGTTCGG CGG ||||||||[||||||||||||]PAM NA Intergenic NA NA

T7555 ATGTTCGGCGGCAGCTCCAGCGG 1000
Oligo fwd CACCATGTTCGGCGGCAGCTCCAG
Oligo rev AAACCTGGAGCTGCCGCCGAACAT
Chromosome start end strand MM target_seq PAM alignment distance position gene name gene id
GU305914.1 103630 103652 + 0 ATGTTCGGCGGCAGCTCCAG CGG ||||||||[||||||||||||]PAM NA Intergenic NA NA

T7556 TGATGTTCACGATCTGCGCGCGG 1000
Oligo fwd CACCTGATGTTCACGATCTGCGCG
Oligo rev AAACCGCGCAGATCGTGAACATCA
Chromosome start end strand MM target_seq PAM alignment distance position gene name gene id
GU305914.1 103661 103683 + 0 TGATGTTCACGATCTGCGCG CGG ||||||||[||||||||||||]PAM NA Intergenic NA NA

T7557 GATCTGCGCGCGGATGTCGCTGG 1000
Oligo fwd CACCGATCTGCGCGCGGATGTCGC
Oligo rev AAACGCGACATCCGCGCGCAGATC
Chromosome start end strand MM target_seq PAM alignment distance position gene name gene id
GU305914.1 103671 103693 + 0 GATCTGCGCGCGGATGTCGC TGG ||||||||[||||||||||||]PAM NA Intergenic NA NA

T7558 ATCTGCGCGCGGATGTCGCTGGG 1000
Oligo fwd CACCATCTGCGCGCGGATGTCGCT
Oligo rev AAACAGCGACATCCGCGCGCAGAT
Chromosome start end strand MM target_seq PAM alignment distance position gene name gene id
GU305914.1 103672 103694 + 0 ATCTGCGCGCGGATGTCGCT GGG ||||||||[||||||||||||]PAM NA Intergenic NA NA

T7559 TCCATCACGCAGACGCACCCCGG 1000
Oligo fwd CACCTCCATCACGCAGACGCACCC
Oligo rev AAACGGGTGCGTCTGCGTGATGGA
Chromosome start end strand MM target_seq PAM alignment distance position gene name gene id
GU305914.1 103711 103733 + 0 TCCATCACGCAGACGCACCC CGG ||||||||[||||||||||||]PAM NA Intergenic NA NA

T7560 ATCACGCAGACGCACCCCGGTGG 1000
Oligo fwd CACCATCACGCAGACGCACCCCGG
Oligo rev AAACCCGGGGTGCGTCTGCGTGAT
Chromosome start end strand MM target_seq PAM alignment distance position gene name gene id
GU305914.1 103714 103736 + 0 ATCACGCAGACGCACCCCGG TGG ||||||||[||||||||||||]PAM NA Intergenic NA NA

T7561 TCACGCAGACGCACCCCGGTGGG 1000
Oligo fwd CACCTCACGCAGACGCACCCCGGT
Oligo rev AAACACCGGGGTGCGTCTGCGTGA
Chromosome start end strand MM target_seq PAM alignment distance position gene name gene id
GU305914.1 103715 103737 + 0 TCACGCAGACGCACCCCGGT GGG ||||||||[||||||||||||]PAM NA Intergenic NA NA

T7562 CGCAGACGCACCCCGGTGGGTGG 1000
Oligo fwd CACCCGCAGACGCACCCCGGTGGG
Oligo rev AAACCCCACCGGGGTGCGTCTGCG
Chromosome start end strand MM target_seq PAM alignment distance position gene name gene id
GU305914.1 103718 103740 + 0 CGCAGACGCACCCCGGTGGG TGG ||||||||[||||||||||||]PAM NA Intergenic NA NA

T7563 TGGGTGGAAACAGTTCGTGACGG 1000
Oligo fwd CACCTGGGTGGAAACAGTTCGTGA
Oligo rev AAACTCACGAACTGTTTCCACCCA
Chromosome start end strand MM target_seq PAM alignment distance position gene name gene id
GU305914.1 103734 103756 + 0 TGGGTGGAAACAGTTCGTGA CGG ||||||||[||||||||||||]PAM NA Intergenic NA NA

T7564 GTGACGGCGCAATAACCCATCGG 1000

Oligo fwd CACCGTGACGGCGCAATAACCCAT
Oligo rev AAACATGGGTTATTGCGCCGTCAC
Chromosome start end strand MM target_seq PAM alignment distance position gene name gene id
GU305914.1 103750 103772 + 0 GTGACGGCGCAATAACCCAT CGG ||||||||[||||||||||||]PAM NA Intergenic NA NA

T7565 TGACGGCGCAATAACCCATCGGG 1000
Oligo fwd CACCTGACGGCGCAATAACCCATC
Oligo rev AAACGATGGGTTATTGCGCCGTCA
Chromosome start end strand MM target_seq PAM alignment distance position gene name gene id
GU305914.1 103751 103773 + 0 TGACGGCGCAATAACCCATC GGG ||||||||[||||||||||||]PAM NA Intergenic NA NA

T7566 TCATCTTCACCCCCAGACCGAGG 1000
Oligo fwd CACCTCATCTTCACCCCCAGACCG
Oligo rev AAACCGGTCTGGGGGTGAAGATGA
Chromosome start end strand MM target_seq PAM alignment distance position gene name gene id
GU305914.1 103778 103800 + 0 TCATCTTCACCCCCAGACCG AGG ||||||||[||||||||||||]PAM NA Intergenic NA NA

T7567 TCACCCCCAGACCGAGGATGCGG 1000
Oligo fwd CACCTCACCCCCAGACCGAGGATG
Oligo rev AAACCATCCTCGGTCTGGGGGTGA
Chromosome start end strand MM target_seq PAM alignment distance position gene name gene id
GU305914.1 103784 103806 + 0 TCACCCCCAGACCGAGGATG CGG ||||||||[||||||||||||]PAM NA Intergenic NA NA

T7568 CGCGCTGATTCTGCGCGAGCAGG 1000
Oligo fwd CACCCGCGCTGATTCTGCGCGAGC
Oligo rev AAACGCTCGCGCAGAATCAGCGCG
Chromosome start end strand MM target_seq PAM alignment distance position gene name gene id
GU305914.1 103829 103851 + 0 CGCGCTGATTCTGCGCGAGC AGG ||||||||[||||||||||||]PAM NA Intergenic NA NA

T7569 GCTGATTCTGCGCGAGCAGGCGG 1000
Oligo fwd CACCGCTGATTCTGCGCGAGCAGG
Oligo rev AAACCCTGCTCGCGCAGAATCAGC
Chromosome start end strand MM target_seq PAM alignment distance position gene name gene id
GU305914.1 103832 103854 + 0 GCTGATTCTGCGCGAGCAGG CGG ||||||||[||||||||||||]PAM NA Intergenic NA NA

T7570 TGAAATCGTTGCCGTACGTCTGG 1000
Oligo fwd CACCTGAAATCGTTGCCGTACGTC
Oligo rev AAACGACGTACGGCAACGATTTCA
Chromosome start end strand MM target_seq PAM alignment distance position gene name gene id
GU305914.1 103874 103896 + 0 TGAAATCGTTGCCGTACGTC TGG ||||||||[||||||||||||]PAM NA Intergenic NA NA

T7571 GAAGATCTTCTGCATCTTCTCGG 1000
Oligo fwd CACCGAAGATCTTCTGCATCTTCT
Oligo rev AAACAGAAGATGCAGAAGATCTTC
Chromosome start end strand MM target_seq PAM alignment distance position gene name gene id
GU305914.1 103899 103921 + 0 GAAGATCTTCTGCATCTTCT CGG ||||||||[||||||||||||]PAM NA Intergenic NA NA

T7572 AAGATCTTCTGCATCTTCTCGGG 1000
Oligo fwd CACCAAGATCTTCTGCATCTTCTC
Oligo rev AAACGAGAAGATGCAGAAGATCTT
Chromosome start end strand MM target_seq PAM alignment distance position gene name gene id
GU305914.1 103900 103922 + 0 AAGATCTTCTGCATCTTCTC GGG ||||||||[||||||||||||]PAM NA Intergenic NA NA

T7573 TCGGGACGACGCGCTTCCTTCGG 1000
Oligo fwd CACCTCGGGACGACGCGCTTCCTT
Oligo rev AAACAAGGAAGCGCGTCGTCCCGA
Chromosome start end strand MM target_seq PAM alignment distance position gene name gene id
GU305914.1 103918 103940 + 0 TCGGGACGACGCGCTTCCTT CGG ||||||||[||||||||||||]PAM NA Intergenic NA NA

T7574 CGACGCGCTTCCTTCGGCGTCGG 1000
Oligo fwd CACCCGACGCGCTTCCTTCGGCGT
Oligo rev AAACACGCCGAAGGAAGCGCGTCG
Chromosome start end strand MM target_seq PAM alignment distance position gene name gene id
GU305914.1 103924 103946 + 0 CGACGCGCTTCCTTCGGCGT CGG ||||||||[||||||||||||]PAM NA Intergenic NA NA

T7575 TCCTTCGGCGTCGGTTCGCGCGG 1000
Oligo fwd CACCTCCTTCGGCGTCGGTTCGCG
Oligo rev AAACCGCGAACCGACGCCGAAGGA
Chromosome start end strand MM target_seq PAM alignment distance position gene name gene id
GU305914.1 103933 103955 + 0 TCCTTCGGCGTCGGTTCGCG CGG ||||||||[||||||||||||]PAM NA Intergenic NA NA

T7576 TTCGGCGTCGGTTCGCGCGGCGG 1000
Oligo fwd CACCTTCGGCGTCGGTTCGCGCGG
Oligo rev AAACCCGCGCGAACCGACGCCGAA
Chromosome start end strand MM target_seq PAM alignment distance position gene name gene id
GU305914.1 103936 103958 + 0 TTCGGCGTCGGTTCGCGCGG CGG ||||||||[||||||||||||]PAM NA Intergenic NA NA

T7577 TCGGCGTCGGTTCGCGCGGCGGG 1000
Oligo fwd CACCTCGGCGTCGGTTCGCGCGGC
Oligo rev AAACGCCGCGCGAACCGACGCCGA
Chromosome start end strand MM target_seq PAM alignment distance position gene name gene id
GU305914.1 103937 103959 + 0 TCGGCGTCGGTTCGCGCGGC GGG ||||||||[||||||||||||]PAM NA Intergenic NA NA

T7578 CGGCGTCGGTTCGCGCGGCGGGG 1000
Oligo fwd CACCCGGCGTCGGTTCGCGCGGCG
Oligo rev AAACCGCCGCGCGAACCGACGCCG
Chromosome start end strand MM target_seq PAM alignment distance position gene name gene id
GU305914.1 103938 103960 + 0 CGGCGTCGGTTCGCGCGGCG GGG ||||||||[||||||||||||]PAM NA Intergenic NA NA

| | | | | | | | | | | | |
|---|---|---|---|---|---|---|---|---|---|---|---|
| T7579 | GGTTCGCGCGGCGGGGCCTCCGG | 1000 | | | | | | | | | |
| Oligo fwd | CACCGGTTCGCGCGGCGGGGCCTC | | | | | | | | | | |
| Oligo rev | AAACGAGGCCCCGCCGCGCGAACC | | | | | | | | | | |
| Chromosome | start | end | strand | MM | target_seq | PAM | alignment | distance | position | gene name | gene id |
| GU305914.1 | 103945 | 103967 | + | 0 | GGTTCGCGCGGCGGGGCCTC | CGG | \|\|\|\|\|\|\|\|[\|\|\|\|\|\|\|\|\|\|\|\|]PAM | NA | Intergenic | NA | NA |
| GU305914.1 | 52596 | 52618 | - | 4 | GGGGCGCACGGCGGGGCCGC | CGG | \|\|--\|\|\|-[\|\|\|\|\|\|\|\|\|\|-\|]PAM | NA | Intergenic | NA | NA |

| | | | | | | | | | | | |
|---|---|---|---|---|---|---|---|---|---|---|---|
| T7580 | CCGGACCGTCGCGCAGCAGCTGG | 1000 | | | | | | | | | |
| Oligo fwd | CACCCCGGACCGTCGCGCAGCAGC | | | | | | | | | | |
| Oligo rev | AAACGCTGCTGCGCGACGGTCCGG | | | | | | | | | | |
| Chromosome | start | end | strand | MM | target_seq | PAM | alignment | distance | position | gene name | gene id |
| GU305914.1 | 103964 | 103986 | + | 0 | CCGGACCGTCGCGCAGCAGC | TGG | \|\|\|\|\|\|\|\|[\|\|\|\|\|\|\|\|\|\|\|\|]PAM | NA | Intergenic | NA | NA |

| | | | | | | | | | | | |
|---|---|---|---|---|---|---|---|---|---|---|---|
| T7581 | GAGCGCCGCGCCGTTCTCGCTGG | 1000 | | | | | | | | | |
| Oligo fwd | CACCGAGCGCCGCGCCGTTCTCGC | | | | | | | | | | |
| Oligo rev | AAACGCGAGAACGGCGCGGCGCTC | | | | | | | | | | |
| Chromosome | start | end | strand | MM | target_seq | PAM | alignment | distance | position | gene name | gene id |
| GU305914.1 | 103998 | 104020 | + | 0 | GAGCGCCGCGCCGTTCTCGC | TGG | \|\|\|\|\|\|\|\|[\|\|\|\|\|\|\|\|\|\|\|\|]PAM | NA | Intergenic | NA | NA |

| | | | | | | | | | | | |
|---|---|---|---|---|---|---|---|---|---|---|---|
| T7582 | TCGCTGGATTCGTCGTCCTCCGG | 1000 | | | | | | | | | |
| Oligo fwd | CACCTCGCTGGATTCGTCGTCCTC | | | | | | | | | | |
| Oligo rev | AAACGAGGACGACGAATCCAGCGA | | | | | | | | | | |
| Chromosome | start | end | strand | MM | target_seq | PAM | alignment | distance | position | gene name | gene id |
| GU305914.1 | 104014 | 104036 | + | 0 | TCGCTGGATTCGTCGTCCTC | CGG | \|\|\|\|\|\|\|\|[\|\|\|\|\|\|\|\|\|\|\|\|]PAM | NA | Intergenic | NA | NA |

| | | | | | | | | | | | |
|---|---|---|---|---|---|---|---|---|---|---|---|
| T7583 | TTCTCTCAGCGCGCAGCTGCTGG | 1000 | | | | | | | | | |
| Oligo fwd | CACCTTCTCTCAGCGCGCAGCTGC | | | | | | | | | | |
| Oligo rev | AAACGCAGCTGCGCGCTGAGAGAA | | | | | | | | | | |
| Chromosome | start | end | strand | MM | target_seq | PAM | alignment | distance | position | gene name | gene id |
| GU305914.1 | 104070 | 104092 | + | 0 | TTCTCTCAGCGCGCAGCTGC | TGG | \|\|\|\|\|\|\|\|[\|\|\|\|\|\|\|\|\|\|\|\|]PAM | NA | Intergenic | NA | NA |

| | | | | | | | | | | | |
|---|---|---|---|---|---|---|---|---|---|---|---|
| T7584 | CTGGAGATCATCATCTCACGAGG | 1000 | | | | | | | | | |
| Oligo fwd | CACCCTGGAGATCATCATCTCACG | | | | | | | | | | |
| Oligo rev | AAACCGTGAGATGATGATCTCCAG | | | | | | | | | | |
| Chromosome | start | end | strand | MM | target_seq | PAM | alignment | distance | position | gene name | gene id |
| GU305914.1 | 104089 | 104111 | + | 0 | CTGGAGATCATCATCTCACG | AGG | \|\|\|\|\|\|\|\|[\|\|\|\|\|\|\|\|\|\|\|\|]PAM | NA | Intergenic | NA | NA |

| | | | | | | | | | | | |
|---|---|---|---|---|---|---|---|---|---|---|---|
| T7585 | GAGATCATCATCTCACGAGGCGG | 1000 | | | | | | | | | |
| Oligo fwd | CACCGAGATCATCATCTCACGAGG | | | | | | | | | | |
| Oligo rev | AAACCCTCGTGAGATGATGATCTC | | | | | | | | | | |
| Chromosome | start | end | strand | MM | target_seq | PAM | alignment | distance | position | gene name | gene id |
| GU305914.1 | 104092 | 104114 | + | 0 | GAGATCATCATCTCACGAGG | CGG | \|\|\|\|\|\|\|\|[\|\|\|\|\|\|\|\|\|\|\|\|]PAM | NA | Intergenic | NA | NA |

| | | | | | | | | | | | |
|---|---|---|---|---|---|---|---|---|---|---|---|
| T7586 | ATCATCATCTCACGAGGCGGAGG | 1000 | | | | | | | | | |
| Oligo fwd | CACCATCATCATCTCACGAGGCGG | | | | | | | | | | |
| Oligo rev | AAACCCGCCTCGTGAGATGATGAT | | | | | | | | | | |
| Chromosome | start | end | strand | MM | target_seq | PAM | alignment | distance | position | gene name | gene id |
| GU305914.1 | 104095 | 104117 | + | 0 | ATCATCATCTCACGAGGCGG | AGG | \|\|\|\|\|\|\|\|[\|\|\|\|\|\|\|\|\|\|\|\|]PAM | NA | Intergenic | NA | NA |

| | | | | | | | | | | | |
|---|---|---|---|---|---|---|---|---|---|---|---|
| T7587 | TCGCGACGACTGCTCTGACGAGG | 1000 | | | | | | | | | |
| Oligo fwd | CACCTCGCGACGACTGCTCTGACG | | | | | | | | | | |
| Oligo rev | AAACCGTCAGAGCAGTCGTCGCGA | | | | | | | | | | |
| Chromosome | start | end | strand | MM | target_seq | PAM | alignment | distance | position | gene name | gene id |
| GU305914.1 | 104118 | 104140 | + | 0 | TCGCGACGACTGCTCTGACG | AGG | \|\|\|\|\|\|\|\|[\|\|\|\|\|\|\|\|\|\|\|\|]PAM | NA | Intergenic | NA | NA |

| | | | | | | | | | | | |
|---|---|---|---|---|---|---|---|---|---|---|---|
| T7588 | CGACGACTGCTCTGACGAGGAGG | 1000 | | | | | | | | | |
| Oligo fwd | CACCCGACGACTGCTCTGACGAGG | | | | | | | | | | |
| Oligo rev | AAACCCTCGTCAGAGCAGTCGTCG | | | | | | | | | | |
| Chromosome | start | end | strand | MM | target_seq | PAM | alignment | distance | position | gene name | gene id |
| GU305914.1 | 104121 | 104143 | + | 0 | CGACGACTGCTCTGACGAGG | AGG | \|\|\|\|\|\|\|\|[\|\|\|\|\|\|\|\|\|\|\|\|]PAM | NA | Intergenic | NA | NA |

| | | | | | | | | | | | |
|---|---|---|---|---|---|---|---|---|---|---|---|
| T7589 | CTCTGACGAGGAGGTATCTTCGG | 1000 | | | | | | | | | |
| Oligo fwd | CACCCTCTGACGAGGAGGTATCTT | | | | | | | | | | |
| Oligo rev | AAACAAGATACCTCCTCGTCAGAG | | | | | | | | | | |
| Chromosome | start | end | strand | MM | target_seq | PAM | alignment | distance | position | gene name | gene id |
| GU305914.1 | 104130 | 104152 | + | 0 | CTCTGACGAGGAGGTATCTT | CGG | \|\|\|\|\|\|\|\|[\|\|\|\|\|\|\|\|\|\|\|\|]PAM | NA | Intergenic | NA | NA |

| | | | | | | | | | | | |
|---|---|---|---|---|---|---|---|---|---|---|---|
| T7590 | GACGAGGAGGTATCTTCGGAAGG | 1000 | | | | | | | | | |
| Oligo fwd | CACCGACGAGGAGGTATCTTCGGA | | | | | | | | | | |
| Oligo rev | AAACTCCGAAGATACCTCCTCGTC | | | | | | | | | | |
| Chromosome | start | end | strand | MM | target_seq | PAM | alignment | distance | position | gene name | gene id |
| GU305914.1 | 104134 | 104156 | + | 0 | GACGAGGAGGTATCTTCGGA | AGG | \|\|\|\|\|\|\|\|[\|\|\|\|\|\|\|\|\|\|\|\|]PAM | NA | Intergenic | NA | NA |

| | | | | | | | | | | | |
|---|---|---|---|---|---|---|---|---|---|---|---|
| T7591 | GTCGTCGTCGAAGATATTAAAGG | 1000 | | | | | | | | | |
| Oligo fwd | CACCGTCGTCGTCGAAGATATTAA | | | | | | | | | | |
| Oligo rev | AAACTTAATATCTTCGACGACGAC | | | | | | | | | | |
| Chromosome | start | end | strand | MM | target_seq | PAM | alignment | distance | position | gene name | gene id |
| GU305914.1 | 104160 | 104182 | + | 0 | GTCGTCGTCGAAGATATTAA | AGG | \|\|\|\|\|\|\|\|[\|\|\|\|\|\|\|\|\|\|\|\|]PAM | NA | Intergenic | NA | NA |

| | | | | | | | | | | | |
|---|---|---|---|---|---|---|---|---|---|---|---|
| T7592 | TCGTCGTCGAAGATATTAAAGGG | 1000 | | | | | | | | | |
| Oligo fwd | CACCTCGTCGTCGAAGATATTAAA | | | | | | | | | | |
| Oligo rev | AAACTTTAATATCTTCGACGACGA | | | | | | | | | | |
| Chromosome | start | end | strand | MM | target_seq | PAM | alignment | distance | position | gene name | gene id |
| GU305914.1 | 104161 | 104183 | + | 0 | TCGTCGTCGAAGATATTAAA | GGG | \|\|\|\|\|\|\|\|[\|\|\|\|\|\|\|\|\|\|\|\|]PAM | NA | Intergenic | NA | NA |

| | | | | | | | | | | | |
|---|---|---|---|---|---|---|---|---|---|---|---|
| T7593 | GATATTAAAGGGATCCTTCATGG | 1000 | | | | | | | | | |
| Oligo fwd | CACCGATATTAAAGGGATCCTTCA | | | | | | | | | | |
| Oligo rev | AAACTGAAGGATCCCTTTAATATC | | | | | | | | | | |
| Chromosome | start | end | strand | MM | target_seq | PAM | alignment | distance | position | gene name | gene id |

<table>
<tr><td>GU305914.1</td><td>104172</td><td>104194</td><td>+</td><td>0</td><td>GATATTAAAGGGATCCTTCA</td><td>TGG</td><td>||||||||[||||||||||||]PAM</td><td>NA</td><td>Intergenic</td><td>NA</td><td>NA</td></tr>
<tr><td></td><td></td><td></td><td></td><td></td><td></td><td></td><td></td><td></td><td></td><td></td><td></td></tr>
<tr><td>T7594</td><td>AAGGGATCCTTCATGGCCTGCGG</td><td>1000</td><td></td><td></td><td></td><td></td><td></td><td></td><td></td><td></td><td></td></tr>
<tr><td>Oligo fwd</td><td>CACCAAGGGATCCTTCATGGCCTG</td><td></td><td></td><td></td><td></td><td></td><td></td><td></td><td></td><td></td><td></td></tr>
<tr><td>Oligo rev</td><td>AAACCAGGCCATGAAGGATCCCTT</td><td></td><td></td><td></td><td></td><td></td><td></td><td></td><td></td><td></td><td></td></tr>
<tr><td>Chromosome</td><td>start</td><td>end</td><td>strand</td><td>MM</td><td>target_seq</td><td>PAM</td><td>alignment</td><td>distance</td><td>position</td><td>gene name</td><td>gene id</td></tr>
<tr><td>GU305914.1</td><td>104179</td><td>104201</td><td>+</td><td>0</td><td>AAGGGATCCTTCATGGCCTG</td><td>CGG</td><td>||||||||[||||||||||||]PAM</td><td>NA</td><td>Intergenic</td><td>NA</td><td>NA</td></tr>
<tr><td></td><td></td><td></td><td></td><td></td><td></td><td></td><td></td><td></td><td></td><td></td><td></td></tr>
<tr><td>T7595</td><td>GCCTGCGGCAAGACTGAGTCTGG</td><td>1000</td><td></td><td></td><td></td><td></td><td></td><td></td><td></td><td></td><td></td></tr>
<tr><td>Oligo fwd</td><td>CACCGCCTGCGGCAAGACTGAGTC</td><td></td><td></td><td></td><td></td><td></td><td></td><td></td><td></td><td></td><td></td></tr>
<tr><td>Oligo rev</td><td>AAACGACTCAGTCTTGCCGCAGGC</td><td></td><td></td><td></td><td></td><td></td><td></td><td></td><td></td><td></td><td></td></tr>
<tr><td>Chromosome</td><td>start</td><td>end</td><td>strand</td><td>MM</td><td>target_seq</td><td>PAM</td><td>alignment</td><td>distance</td><td>position</td><td>gene name</td><td>gene id</td></tr>
<tr><td>GU305914.1</td><td>104194</td><td>104216</td><td>+</td><td>0</td><td>GCCTGCGGCAAGACTGAGTC</td><td>TGG</td><td>||||||||[||||||||||||]PAM</td><td>NA</td><td>Intergenic</td><td>NA</td><td>NA</td></tr>
<tr><td></td><td></td><td></td><td></td><td></td><td></td><td></td><td></td><td></td><td></td><td></td><td></td></tr>
<tr><td>T7596</td><td>ACTGAGTCTGGCGACGACAGTGG</td><td>1000</td><td></td><td></td><td></td><td></td><td></td><td></td><td></td><td></td><td></td></tr>
<tr><td>Oligo fwd</td><td>CACCACTGAGTCTGGCGACGACAG</td><td></td><td></td><td></td><td></td><td></td><td></td><td></td><td></td><td></td><td></td></tr>
<tr><td>Oligo rev</td><td>AAACCTGTCGTCGCCAGACTCAGT</td><td></td><td></td><td></td><td></td><td></td><td></td><td></td><td></td><td></td><td></td></tr>
<tr><td>Chromosome</td><td>start</td><td>end</td><td>strand</td><td>MM</td><td>target_seq</td><td>PAM</td><td>alignment</td><td>distance</td><td>position</td><td>gene name</td><td>gene id</td></tr>
<tr><td>GU305914.1</td><td>104206</td><td>104228</td><td>+</td><td>0</td><td>ACTGAGTCTGGCGACGACAG</td><td>TGG</td><td>||||||||[||||||||||||]PAM</td><td>NA</td><td>Intergenic</td><td>NA</td><td>NA</td></tr>
<tr><td></td><td></td><td></td><td></td><td></td><td></td><td></td><td></td><td></td><td></td><td></td><td></td></tr>
<tr><td>T7597</td><td>CTGAGTCTGGCGACGACAGTGGG</td><td>1000</td><td></td><td></td><td></td><td></td><td></td><td></td><td></td><td></td><td></td></tr>
<tr><td>Oligo fwd</td><td>CACCCTGAGTCTGGCGACGACAGT</td><td></td><td></td><td></td><td></td><td></td><td></td><td></td><td></td><td></td><td></td></tr>
<tr><td>Oligo rev</td><td>AAACACTGTCGTCGCCAGACTCAG</td><td></td><td></td><td></td><td></td><td></td><td></td><td></td><td></td><td></td><td></td></tr>
<tr><td>Chromosome</td><td>start</td><td>end</td><td>strand</td><td>MM</td><td>target_seq</td><td>PAM</td><td>alignment</td><td>distance</td><td>position</td><td>gene name</td><td>gene id</td></tr>
<tr><td>GU305914.1</td><td>104207</td><td>104229</td><td>+</td><td>0</td><td>CTGAGTCTGGCGACGACAGT</td><td>GGG</td><td>||||||||[||||||||||||]PAM</td><td>NA</td><td>Intergenic</td><td>NA</td><td>NA</td></tr>
<tr><td></td><td></td><td></td><td></td><td></td><td></td><td></td><td></td><td></td><td></td><td></td><td></td></tr>
<tr><td>T7598</td><td>GGCGACGACAGTGGGAGATTCGG</td><td>1000</td><td></td><td></td><td></td><td></td><td></td><td></td><td></td><td></td><td></td></tr>
<tr><td>Oligo fwd</td><td>CACCGGCGACGACAGTGGGAGATT</td><td></td><td></td><td></td><td></td><td></td><td></td><td></td><td></td><td></td><td></td></tr>
<tr><td>Oligo rev</td><td>AAACAATCTCCCACTGTCGTCGCC</td><td></td><td></td><td></td><td></td><td></td><td></td><td></td><td></td><td></td><td></td></tr>
<tr><td>Chromosome</td><td>start</td><td>end</td><td>strand</td><td>MM</td><td>target_seq</td><td>PAM</td><td>alignment</td><td>distance</td><td>position</td><td>gene name</td><td>gene id</td></tr>
<tr><td>GU305914.1</td><td>104215</td><td>104237</td><td>+</td><td>0</td><td>GGCGACGACAGTGGGAGATT</td><td>CGG</td><td>||||||||[||||||||||||]PAM</td><td>NA</td><td>Intergenic</td><td>NA</td><td>NA</td></tr>
<tr><td></td><td></td><td></td><td></td><td></td><td></td><td></td><td></td><td></td><td></td><td></td><td></td></tr>
<tr><td>T7599</td><td>GCGACGACAGTGGGAGATTCGGG</td><td>1000</td><td></td><td></td><td></td><td></td><td></td><td></td><td></td><td></td><td></td></tr>
<tr><td>Oligo fwd</td><td>CACCGCGACGACAGTGGGAGATTC</td><td></td><td></td><td></td><td></td><td></td><td></td><td></td><td></td><td></td><td></td></tr>
<tr><td>Oligo rev</td><td>AAACGAATCTCCCACTGTCGTCGC</td><td></td><td></td><td></td><td></td><td></td><td></td><td></td><td></td><td></td><td></td></tr>
<tr><td>Chromosome</td><td>start</td><td>end</td><td>strand</td><td>MM</td><td>target_seq</td><td>PAM</td><td>alignment</td><td>distance</td><td>position</td><td>gene name</td><td>gene id</td></tr>
<tr><td>GU305914.1</td><td>104216</td><td>104238</td><td>+</td><td>0</td><td>GCGACGACAGTGGGAGATTC</td><td>GGG</td><td>||||||||[||||||||||||]PAM</td><td>NA</td><td>Intergenic</td><td>NA</td><td>NA</td></tr>
<tr><td></td><td></td><td></td><td></td><td></td><td></td><td></td><td></td><td></td><td></td><td></td><td></td></tr>
<tr><td>T7600</td><td>ACGACAGTGGGAGATTCGGGCGG</td><td>1000</td><td></td><td></td><td></td><td></td><td></td><td></td><td></td><td></td><td></td></tr>
<tr><td>Oligo fwd</td><td>CACCACGACAGTGGGAGATTCGGG</td><td></td><td></td><td></td><td></td><td></td><td></td><td></td><td></td><td></td><td></td></tr>
<tr><td>Oligo rev</td><td>AAACCCCGAATCTCCCACTGTCGT</td><td></td><td></td><td></td><td></td><td></td><td></td><td></td><td></td><td></td><td></td></tr>
<tr><td>Chromosome</td><td>start</td><td>end</td><td>strand</td><td>MM</td><td>target_seq</td><td>PAM</td><td>alignment</td><td>distance</td><td>position</td><td>gene name</td><td>gene id</td></tr>
<tr><td>GU305914.1</td><td>104219</td><td>104241</td><td>+</td><td>0</td><td>ACGACAGTGGGAGATTCGGG</td><td>CGG</td><td>||||||||[||||||||||||]PAM</td><td>NA</td><td>Intergenic</td><td>NA</td><td>NA</td></tr>
<tr><td></td><td></td><td></td><td></td><td></td><td></td><td></td><td></td><td></td><td></td><td></td><td></td></tr>
<tr><td>T7601</td><td>AGTGGGAGATTCGGGCGGACCGG</td><td>1000</td><td></td><td></td><td></td><td></td><td></td><td></td><td></td><td></td><td></td></tr>
<tr><td>Oligo fwd</td><td>CACCAGTGGGAGATTCGGGCGGAC</td><td></td><td></td><td></td><td></td><td></td><td></td><td></td><td></td><td></td><td></td></tr>
<tr><td>Oligo rev</td><td>AAACGTCCGCCCGAATCTCCCACT</td><td></td><td></td><td></td><td></td><td></td><td></td><td></td><td></td><td></td><td></td></tr>
<tr><td>Chromosome</td><td>start</td><td>end</td><td>strand</td><td>MM</td><td>target_seq</td><td>PAM</td><td>alignment</td><td>distance</td><td>position</td><td>gene name</td><td>gene id</td></tr>
<tr><td>GU305914.1</td><td>104224</td><td>104246</td><td>+</td><td>0</td><td>AGTGGGAGATTCGGGCGGAC</td><td>CGG</td><td>||||||||[||||||||||||]PAM</td><td>NA</td><td>Intergenic</td><td>NA</td><td>NA</td></tr>
<tr><td></td><td></td><td></td><td></td><td></td><td></td><td></td><td></td><td></td><td></td><td></td><td></td></tr>
<tr><td>T7602</td><td>AGATTCGGGCGGACCGGAGCCGG</td><td>1000</td><td></td><td></td><td></td><td></td><td></td><td></td><td></td><td></td><td></td></tr>
<tr><td>Oligo fwd</td><td>CACCAGATTCGGGCGGACCGGAGC</td><td></td><td></td><td></td><td></td><td></td><td></td><td></td><td></td><td></td><td></td></tr>
<tr><td>Oligo rev</td><td>AAACGCTCCGGTCCGCCCGAATCT</td><td></td><td></td><td></td><td></td><td></td><td></td><td></td><td></td><td></td><td></td></tr>
<tr><td>Chromosome</td><td>start</td><td>end</td><td>strand</td><td>MM</td><td>target_seq</td><td>PAM</td><td>alignment</td><td>distance</td><td>position</td><td>gene name</td><td>gene id</td></tr>
<tr><td>GU305914.1</td><td>104230</td><td>104252</td><td>+</td><td>0</td><td>AGATTCGGGCGGACCGGAGC</td><td>CGG</td><td>||||||||[||||||||||||]PAM</td><td>NA</td><td>Intergenic</td><td>NA</td><td>NA</td></tr>
<tr><td></td><td></td><td></td><td></td><td></td><td></td><td></td><td></td><td></td><td></td><td></td><td></td></tr>
<tr><td>T7603</td><td>CGGACCGGAGCCGGTATGTTTGG</td><td>1000</td><td></td><td></td><td></td><td></td><td></td><td></td><td></td><td></td><td></td></tr>
<tr><td>Oligo fwd</td><td>CACCCGGACCGGAGCCGGTATGTT</td><td></td><td></td><td></td><td></td><td></td><td></td><td></td><td></td><td></td><td></td></tr>
<tr><td>Oligo rev</td><td>AAACAACATACCGGCTCCGGTCCG</td><td></td><td></td><td></td><td></td><td></td><td></td><td></td><td></td><td></td><td></td></tr>
<tr><td>Chromosome</td><td>start</td><td>end</td><td>strand</td><td>MM</td><td>target_seq</td><td>PAM</td><td>alignment</td><td>distance</td><td>position</td><td>gene name</td><td>gene id</td></tr>
<tr><td>GU305914.1</td><td>104239</td><td>104261</td><td>+</td><td>0</td><td>CGGACCGGAGCCGGTATGTT</td><td>TGG</td><td>||||||||[||||||||||||]PAM</td><td>NA</td><td>Intergenic</td><td>NA</td><td>NA</td></tr>
<tr><td></td><td></td><td></td><td></td><td></td><td></td><td></td><td></td><td></td><td></td><td></td><td></td></tr>
<tr><td>T7604</td><td>ATGTTTGGTTTTATCATGCCAGG</td><td>1000</td><td></td><td></td><td></td><td></td><td></td><td></td><td></td><td></td><td></td></tr>
<tr><td>Oligo fwd</td><td>CACCATGTTTGGTTTTATCATGCC</td><td></td><td></td><td></td><td></td><td></td><td></td><td></td><td></td><td></td><td></td></tr>
<tr><td>Oligo rev</td><td>AAACGGCATGATAAAACCAAACAT</td><td></td><td></td><td></td><td></td><td></td><td></td><td></td><td></td><td></td><td></td></tr>
<tr><td>Chromosome</td><td>start</td><td>end</td><td>strand</td><td>MM</td><td>target_seq</td><td>PAM</td><td>alignment</td><td>distance</td><td>position</td><td>gene name</td><td>gene id</td></tr>
<tr><td>GU305914.1</td><td>104254</td><td>104276</td><td>+</td><td>0</td><td>ATGTTTGGTTTTATCATGCC</td><td>AGG</td><td>||||||||[||||||||||||]PAM</td><td>NA</td><td>Intergenic</td><td>NA</td><td>NA</td></tr>
<tr><td></td><td></td><td></td><td></td><td></td><td></td><td></td><td></td><td></td><td></td><td></td><td></td></tr>
<tr><td>T7605</td><td>TTTATCATGCCAGGCTTTGTAGG</td><td>1000</td><td></td><td></td><td></td><td></td><td></td><td></td><td></td><td></td><td></td></tr>
<tr><td>Oligo fwd</td><td>CACCTTTATCATGCCAGGCTTTGT</td><td></td><td></td><td></td><td></td><td></td><td></td><td></td><td></td><td></td><td></td></tr>
<tr><td>Oligo rev</td><td>AAACACAAAGCCTGGCATGATAAA</td><td></td><td></td><td></td><td></td><td></td><td></td><td></td><td></td><td></td><td></td></tr>
<tr><td>Chromosome</td><td>start</td><td>end</td><td>strand</td><td>MM</td><td>target_seq</td><td>PAM</td><td>alignment</td><td>distance</td><td>position</td><td>gene name</td><td>gene id</td></tr>
<tr><td>GU305914.1</td><td>104263</td><td>104285</td><td>+</td><td>0</td><td>TTTATCATGCCAGGCTTTGT</td><td>AGG</td><td>||||||||[||||||||||||]PAM</td><td>NA</td><td>Intergenic</td><td>NA</td><td>NA</td></tr>
<tr><td></td><td></td><td></td><td></td><td></td><td></td><td></td><td></td><td></td><td></td><td></td><td></td></tr>
<tr><td>T7606</td><td>TTATCATGCCAGGCTTTGTAGGG</td><td>1000</td><td></td><td></td><td></td><td></td><td></td><td></td><td></td><td></td><td></td></tr>
<tr><td>Oligo fwd</td><td>CACCTTATCATGCCAGGCTTTGTA</td><td></td><td></td><td></td><td></td><td></td><td></td><td></td><td></td><td></td><td></td></tr>
<tr><td>Oligo rev</td><td>AAACTACAAAGCCTGGCATGATAA</td><td></td><td></td><td></td><td></td><td></td><td></td><td></td><td></td><td></td><td></td></tr>
<tr><td>Chromosome</td><td>start</td><td>end</td><td>strand</td><td>MM</td><td>target_seq</td><td>PAM</td><td>alignment</td><td>distance</td><td>position</td><td>gene name</td><td>gene id</td></tr>
<tr><td>GU305914.1</td><td>104264</td><td>104286</td><td>+</td><td>0</td><td>TTATCATGCCAGGCTTTGTA</td><td>GGG</td><td>||||||||[||||||||||||]PAM</td><td>NA</td><td>Intergenic</td><td>NA</td><td>NA</td></tr>
<tr><td></td><td></td><td></td><td></td><td></td><td></td><td></td><td></td><td></td><td></td><td></td><td></td></tr>
<tr><td>T7607</td><td>CCTTCTGTTGTGCTTCGCCATGG</td><td>1000</td><td></td><td></td><td></td><td></td><td></td><td></td><td></td><td></td><td></td></tr>
<tr><td>Oligo fwd</td><td>CACCCCTTCTGTTGTGCTTCGCCA</td><td></td><td></td><td></td><td></td><td></td><td></td><td></td><td></td><td></td><td></td></tr>
<tr><td>Oligo rev</td><td>AAACTGGCGAAGCACAACAGAAGG</td><td></td><td></td><td></td><td></td><td></td><td></td><td></td><td></td><td></td><td></td></tr>
<tr><td>Chromosome</td><td>start</td><td>end</td><td>strand</td><td>MM</td><td>target_seq</td><td>PAM</td><td>alignment</td><td>distance</td><td>position</td><td>gene name</td><td>gene id</td></tr>
<tr><td>GU305914.1</td><td>104307</td><td>104329</td><td>+</td><td>0</td><td>CCTTCTGTTGTGCTTCGCCA</td><td>TGG</td><td>||||||||[||||||||||||]PAM</td><td>NA</td><td>Intergenic</td><td>NA</td><td>NA</td></tr>
<tr><td></td><td></td><td></td><td></td><td></td><td></td><td></td><td></td><td></td><td></td><td></td><td></td></tr>
<tr><td>T7608</td><td>TTGTGCTTCGCCATGGCGAGCGG</td><td>1000</td><td></td><td></td><td></td><td></td><td></td><td></td><td></td><td></td><td></td></tr>
<tr><td>Oligo fwd</td><td>CACCTTGTGCTTCGCCATGGCGAG</td><td></td><td></td><td></td><td></td><td></td><td></td><td></td><td></td><td></td><td></td></tr>
<tr><td>Oligo rev</td><td>AAACCTCGCCATGGCGAAGCACAA</td><td></td><td></td><td></td><td></td><td></td><td></td><td></td><td></td><td></td><td></td></tr>
</table>

<table>
<tr><td>Chromosome</td><td>start</td><td>end</td><td>strand</td><td>MM</td><td>target_seq</td><td>PAM</td><td>alignment</td><td>distance</td><td>position</td><td>gene name</td><td>gene id</td></tr>
<tr><td>GU305914.1</td><td>104314</td><td>104336</td><td>+</td><td>0</td><td>TTGTGCTTCGCCATGGCGAG</td><td>CGG</td><td>||||||||[||||||||||||]PAM</td><td>NA</td><td>Intergenic</td><td>NA</td><td>NA</td></tr>
<tr><td></td><td></td><td></td><td></td><td></td><td></td><td></td><td></td><td></td><td></td><td></td><td></td></tr>
<tr><td>T7609</td><td>GGCGAGCGGATCCTCGTCGCCGG</td><td>1000</td><td></td><td></td><td></td><td></td><td></td><td></td><td></td><td></td><td></td></tr>
<tr><td>Oligo fwd</td><td>CACCGGCGAGCGGATCCTCGTCGC</td><td></td><td></td><td></td><td></td><td></td><td></td><td></td><td></td><td></td><td></td></tr>
<tr><td>Oligo rev</td><td>AAACGCGACGAGGATCCGCTCGCC</td><td></td><td></td><td></td><td></td><td></td><td></td><td></td><td></td><td></td><td></td></tr>
<tr><td>Chromosome</td><td>start</td><td>end</td><td>strand</td><td>MM</td><td>target_seq</td><td>PAM</td><td>alignment</td><td>distance</td><td>position</td><td>gene name</td><td>gene id</td></tr>
<tr><td>GU305914.1</td><td>104328</td><td>104350</td><td>+</td><td>0</td><td>GGCGAGCGGATCCTCGTCGC</td><td>CGG</td><td>||||||||[||||||||||||]PAM</td><td>NA</td><td>Intergenic</td><td>NA</td><td>NA</td></tr>
<tr><td></td><td></td><td></td><td></td><td></td><td></td><td></td><td></td><td></td><td></td><td></td><td></td></tr>
<tr><td>T7610</td><td>CTCGTCGCCGGCCTCCGTACCGG</td><td>1000</td><td></td><td></td><td></td><td></td><td></td><td></td><td></td><td></td><td></td></tr>
<tr><td>Oligo fwd</td><td>CACCCTCGTCGCCGGCCTCCGTAC</td><td></td><td></td><td></td><td></td><td></td><td></td><td></td><td></td><td></td><td></td></tr>
<tr><td>Oligo rev</td><td>AAACGTACGGAGGCCGGCGACGAG</td><td></td><td></td><td></td><td></td><td></td><td></td><td></td><td></td><td></td><td></td></tr>
<tr><td>Chromosome</td><td>start</td><td>end</td><td>strand</td><td>MM</td><td>target_seq</td><td>PAM</td><td>alignment</td><td>distance</td><td>position</td><td>gene name</td><td>gene id</td></tr>
<tr><td>GU305914.1</td><td>104340</td><td>104362</td><td>+</td><td>0</td><td>CTCGTCGCCGGCCTCCGTAC</td><td>CGG</td><td>||||||||[||||||||||||]PAM</td><td>NA</td><td>Intergenic</td><td>NA</td><td>NA</td></tr>
<tr><td></td><td></td><td></td><td></td><td></td><td></td><td></td><td></td><td></td><td></td><td></td><td></td></tr>
<tr><td>T7611</td><td>GCCGGCCTCCGTACCGGTGTCGG</td><td>1000</td><td></td><td></td><td></td><td></td><td></td><td></td><td></td><td></td><td></td></tr>
<tr><td>Oligo fwd</td><td>CACCGCCGGCCTCCGTACCGGTGT</td><td></td><td></td><td></td><td></td><td></td><td></td><td></td><td></td><td></td><td></td></tr>
<tr><td>Oligo rev</td><td>AAACACACCGGTACGGAGGCCGGC</td><td></td><td></td><td></td><td></td><td></td><td></td><td></td><td></td><td></td><td></td></tr>
<tr><td>Chromosome</td><td>start</td><td>end</td><td>strand</td><td>MM</td><td>target_seq</td><td>PAM</td><td>alignment</td><td>distance</td><td>position</td><td>gene name</td><td>gene id</td></tr>
<tr><td>GU305914.1</td><td>104346</td><td>104368</td><td>+</td><td>0</td><td>GCCGGCCTCCGTACCGGTGT</td><td>CGG</td><td>||||||||[||||||||||||]PAM</td><td>NA</td><td>Intergenic</td><td>NA</td><td>NA</td></tr>
<tr><td></td><td></td><td></td><td></td><td></td><td></td><td></td><td></td><td></td><td></td><td></td><td></td></tr>
<tr><td>T7612</td><td>ACCGGTGTCGGTCGCCGCTTCGG</td><td>1000</td><td></td><td></td><td></td><td></td><td></td><td></td><td></td><td></td><td></td></tr>
<tr><td>Oligo fwd</td><td>CACCACCGGTGTCGGTCGCCGCTT</td><td></td><td></td><td></td><td></td><td></td><td></td><td></td><td></td><td></td><td></td></tr>
<tr><td>Oligo rev</td><td>AAACAAGCGGCGACCGACACCGGT</td><td></td><td></td><td></td><td></td><td></td><td></td><td></td><td></td><td></td><td></td></tr>
<tr><td>Chromosome</td><td>start</td><td>end</td><td>strand</td><td>MM</td><td>target_seq</td><td>PAM</td><td>alignment</td><td>distance</td><td>position</td><td>gene name</td><td>gene id</td></tr>
<tr><td>GU305914.1</td><td>104358</td><td>104380</td><td>+</td><td>0</td><td>ACCGGTGTCGGTCGCCGCTT</td><td>CGG</td><td>||||||||[||||||||||||]PAM</td><td>NA</td><td>Intergenic</td><td>NA</td><td>NA</td></tr>
<tr><td></td><td></td><td></td><td></td><td></td><td></td><td></td><td></td><td></td><td></td><td></td><td></td></tr>
<tr><td>T7613</td><td>GTCGGTCGCCGCTTCGGTGCCGG</td><td>1000</td><td></td><td></td><td></td><td></td><td></td><td></td><td></td><td></td><td></td></tr>
<tr><td>Oligo fwd</td><td>CACCGTCGGTCGCCGCTTCGGTGC</td><td></td><td></td><td></td><td></td><td></td><td></td><td></td><td></td><td></td><td></td></tr>
<tr><td>Oligo rev</td><td>AAACGCACCGAAGCGGCGACCGAC</td><td></td><td></td><td></td><td></td><td></td><td></td><td></td><td></td><td></td><td></td></tr>
<tr><td>Chromosome</td><td>start</td><td>end</td><td>strand</td><td>MM</td><td>target_seq</td><td>PAM</td><td>alignment</td><td>distance</td><td>position</td><td>gene name</td><td>gene id</td></tr>
<tr><td>GU305914.1</td><td>104364</td><td>104386</td><td>+</td><td>0</td><td>GTCGGTCGCCGCTTCGGTGC</td><td>CGG</td><td>||||||||[||||||||||||]PAM</td><td>NA</td><td>Intergenic</td><td>NA</td><td>NA</td></tr>
<tr><td></td><td></td><td></td><td></td><td></td><td></td><td></td><td></td><td></td><td></td><td></td><td></td></tr>
<tr><td>T7614</td><td>AACAAGATCGTGATCAGCGACGG</td><td>1000</td><td></td><td></td><td></td><td></td><td></td><td></td><td></td><td></td><td></td></tr>
<tr><td>Oligo fwd</td><td>CACCAACAAGATCGTGATCAGCGA</td><td></td><td></td><td></td><td></td><td></td><td></td><td></td><td></td><td></td><td></td></tr>
<tr><td>Oligo rev</td><td>AAACTCGCTGATCACGATCTTGTT</td><td></td><td></td><td></td><td></td><td></td><td></td><td></td><td></td><td></td><td></td></tr>
<tr><td>Chromosome</td><td>start</td><td>end</td><td>strand</td><td>MM</td><td>target_seq</td><td>PAM</td><td>alignment</td><td>distance</td><td>position</td><td>gene name</td><td>gene id</td></tr>
<tr><td>GU305914.1</td><td>104398</td><td>104420</td><td>+</td><td>0</td><td>AACAAGATCGTGATCAGCGA</td><td>CGG</td><td>||||||||[||||||||||||]PAM</td><td>NA</td><td>Intergenic</td><td>NA</td><td>NA</td></tr>
<tr><td></td><td></td><td></td><td></td><td></td><td></td><td></td><td></td><td></td><td></td><td></td><td></td></tr>
<tr><td>T7615</td><td>CGTGATCAGCGACGGATCCGAGG</td><td>1000</td><td></td><td></td><td></td><td></td><td></td><td></td><td></td><td></td><td></td></tr>
<tr><td>Oligo fwd</td><td>CACCCGTGATCAGCGACGGATCCG</td><td></td><td></td><td></td><td></td><td></td><td></td><td></td><td></td><td></td><td></td></tr>
<tr><td>Oligo rev</td><td>AAACCGGATCCGTCGCTGATCACG</td><td></td><td></td><td></td><td></td><td></td><td></td><td></td><td></td><td></td><td></td></tr>
<tr><td>Chromosome</td><td>start</td><td>end</td><td>strand</td><td>MM</td><td>target_seq</td><td>PAM</td><td>alignment</td><td>distance</td><td>position</td><td>gene name</td><td>gene id</td></tr>
<tr><td>GU305914.1</td><td>104406</td><td>104428</td><td>+</td><td>0</td><td>CGTGATCAGCGACGGATCCG</td><td>AGG</td><td>||||||||[||||||||||||]PAM</td><td>NA</td><td>Intergenic</td><td>NA</td><td>NA</td></tr>
<tr><td>GU305914.1</td><td>76929</td><td>76951</td><td>-</td><td>4</td><td>CGTCGTCAGCGAGGGATCCC</td><td>AGG</td><td>|||--|||[||||-||||||-]PAM</td><td>NA</td><td>Intergenic</td><td>NA</td><td>NA</td></tr>
<tr><td></td><td></td><td></td><td></td><td></td><td></td><td></td><td></td><td></td><td></td><td></td><td></td></tr>
<tr><td>T7616</td><td>AAGTGTCACTCGCACTTCTACGG</td><td>1000</td><td></td><td></td><td></td><td></td><td></td><td></td><td></td><td></td><td></td></tr>
<tr><td>Oligo fwd</td><td>CACCAAGTGTCACTCGCACTTCTA</td><td></td><td></td><td></td><td></td><td></td><td></td><td></td><td></td><td></td><td></td></tr>
<tr><td>Oligo rev</td><td>AAACTAGAAGTGCGAGTGACACTT</td><td></td><td></td><td></td><td></td><td></td><td></td><td></td><td></td><td></td><td></td></tr>
<tr><td>Chromosome</td><td>start</td><td>end</td><td>strand</td><td>MM</td><td>target_seq</td><td>PAM</td><td>alignment</td><td>distance</td><td>position</td><td>gene name</td><td>gene id</td></tr>
<tr><td>GU305914.1</td><td>104458</td><td>104480</td><td>+</td><td>0</td><td>AAGTGTCACTCGCACTTCTA</td><td>CGG</td><td>||||||||[||||||||||||]PAM</td><td>NA</td><td>Intergenic</td><td>NA</td><td>NA</td></tr>
<tr><td></td><td></td><td></td><td></td><td></td><td></td><td></td><td></td><td></td><td></td><td></td><td></td></tr>
<tr><td>T7617</td><td>TCTCGTCGTTCGCCTCCATCTGG</td><td>1000</td><td></td><td></td><td></td><td></td><td></td><td></td><td></td><td></td><td></td></tr>
<tr><td>Oligo fwd</td><td>CACCTCTCGTCGTTCGCCTCCATC</td><td></td><td></td><td></td><td></td><td></td><td></td><td></td><td></td><td></td><td></td></tr>
<tr><td>Oligo rev</td><td>AAACGATGGAGGCGAACGACGAGA</td><td></td><td></td><td></td><td></td><td></td><td></td><td></td><td></td><td></td><td></td></tr>
<tr><td>Chromosome</td><td>start</td><td>end</td><td>strand</td><td>MM</td><td>target_seq</td><td>PAM</td><td>alignment</td><td>distance</td><td>position</td><td>gene name</td><td>gene id</td></tr>
<tr><td>GU305914.1</td><td>104489</td><td>104511</td><td>+</td><td>0</td><td>TCTCGTCGTTCGCCTCCATC</td><td>TGG</td><td>||||||||[||||||||||||]PAM</td><td>NA</td><td>Intergenic</td><td>NA</td><td>NA</td></tr>
<tr><td></td><td></td><td></td><td></td><td></td><td></td><td></td><td></td><td></td><td></td><td></td><td></td></tr>
<tr><td>T7618</td><td>GTTCGCCTCCATCTGGATGATGG</td><td>1000</td><td></td><td></td><td></td><td></td><td></td><td></td><td></td><td></td><td></td></tr>
<tr><td>Oligo fwd</td><td>CACCGTTCGCCTCCATCTGGATGA</td><td></td><td></td><td></td><td></td><td></td><td></td><td></td><td></td><td></td><td></td></tr>
<tr><td>Oligo rev</td><td>AAACTCATCCAGATGGAGGCGAAC</td><td></td><td></td><td></td><td></td><td></td><td></td><td></td><td></td><td></td><td></td></tr>
<tr><td>Chromosome</td><td>start</td><td>end</td><td>strand</td><td>MM</td><td>target_seq</td><td>PAM</td><td>alignment</td><td>distance</td><td>position</td><td>gene name</td><td>gene id</td></tr>
<tr><td>GU305914.1</td><td>104496</td><td>104518</td><td>+</td><td>0</td><td>GTTCGCCTCCATCTGGATGA</td><td>TGG</td><td>||||||||[||||||||||||]PAM</td><td>NA</td><td>Intergenic</td><td>NA</td><td>NA</td></tr>
<tr><td></td><td></td><td></td><td></td><td></td><td></td><td></td><td></td><td></td><td></td><td></td><td></td></tr>
<tr><td>T7619</td><td>ATCGTCTTTATTTGTGCGTTCGG</td><td>1000</td><td></td><td></td><td></td><td></td><td></td><td></td><td></td><td></td><td></td></tr>
<tr><td>Oligo fwd</td><td>CACCATCGTCTTTATTTGTGCGTT</td><td></td><td></td><td></td><td></td><td></td><td></td><td></td><td></td><td></td><td></td></tr>
<tr><td>Oligo rev</td><td>AAACAACGCACAAATAAAGACGAT</td><td></td><td></td><td></td><td></td><td></td><td></td><td></td><td></td><td></td><td></td></tr>
<tr><td>Chromosome</td><td>start</td><td>end</td><td>strand</td><td>MM</td><td>target_seq</td><td>PAM</td><td>alignment</td><td>distance</td><td>position</td><td>gene name</td><td>gene id</td></tr>
<tr><td>GU305914.1</td><td>104527</td><td>104549</td><td>+</td><td>0</td><td>ATCGTCTTTATTTGTGCGTT</td><td>CGG</td><td>||||||||[||||||||||||]PAM</td><td>NA</td><td>Intergenic</td><td>NA</td><td>NA</td></tr>
<tr><td></td><td></td><td></td><td></td><td></td><td></td><td></td><td></td><td></td><td></td><td></td><td></td></tr>
<tr><td>T7620</td><td>GTGCGTTCGGTGTTTTCATGAGG</td><td>1000</td><td></td><td></td><td></td><td></td><td></td><td></td><td></td><td></td><td></td></tr>
<tr><td>Oligo fwd</td><td>CACCGTGCGTTCGGTGTTTTCATG</td><td></td><td></td><td></td><td></td><td></td><td></td><td></td><td></td><td></td><td></td></tr>
<tr><td>Oligo rev</td><td>AAACCATGAAAACACCGAACGCAC</td><td></td><td></td><td></td><td></td><td></td><td></td><td></td><td></td><td></td><td></td></tr>
<tr><td>Chromosome</td><td>start</td><td>end</td><td>strand</td><td>MM</td><td>target_seq</td><td>PAM</td><td>alignment</td><td>distance</td><td>position</td><td>gene name</td><td>gene id</td></tr>
<tr><td>GU305914.1</td><td>104540</td><td>104562</td><td>+</td><td>0</td><td>GTGCGTTCGGTGTTTTCATG</td><td>AGG</td><td>||||||||[||||||||||||]PAM</td><td>NA</td><td>Intergenic</td><td>NA</td><td>NA</td></tr>
<tr><td></td><td></td><td></td><td></td><td></td><td></td><td></td><td></td><td></td><td></td><td></td><td></td></tr>
<tr><td>T7621</td><td>TCGGTGTTTTCATGAGGCACTGG</td><td>1000</td><td></td><td></td><td></td><td></td><td></td><td></td><td></td><td></td><td></td></tr>
<tr><td>Oligo fwd</td><td>CACCTCGGTGTTTTCATGAGGCAC</td><td></td><td></td><td></td><td></td><td></td><td></td><td></td><td></td><td></td><td></td></tr>
<tr><td>Oligo rev</td><td>AAACGTGCCTCATGAAAACACCGA</td><td></td><td></td><td></td><td></td><td></td><td></td><td></td><td></td><td></td><td></td></tr>
<tr><td>Chromosome</td><td>start</td><td>end</td><td>strand</td><td>MM</td><td>target_seq</td><td>PAM</td><td>alignment</td><td>distance</td><td>position</td><td>gene name</td><td>gene id</td></tr>
<tr><td>GU305914.1</td><td>104546</td><td>104568</td><td>+</td><td>0</td><td>TCGGTGTTTTCATGAGGCAC</td><td>TGG</td><td>||||||||[||||||||||||]PAM</td><td>NA</td><td>Intergenic</td><td>NA</td><td>NA</td></tr>
<tr><td></td><td></td><td></td><td></td><td></td><td></td><td></td><td></td><td></td><td></td><td></td><td></td></tr>
<tr><td>T7622</td><td>CATGAGGCACTGGTGTTACAAGG</td><td>1000</td><td></td><td></td><td></td><td></td><td></td><td></td><td></td><td></td><td></td></tr>
<tr><td>Oligo fwd</td><td>CACCCATGAGGCACTGGTGTTACA</td><td></td><td></td><td></td><td></td><td></td><td></td><td></td><td></td><td></td><td></td></tr>
<tr><td>Oligo rev</td><td>AAACTGTAACACCAGTGCCTCATG</td><td></td><td></td><td></td><td></td><td></td><td></td><td></td><td></td><td></td><td></td></tr>
<tr><td>Chromosome</td><td>start</td><td>end</td><td>strand</td><td>MM</td><td>target_seq</td><td>PAM</td><td>alignment</td><td>distance</td><td>position</td><td>gene name</td><td>gene id</td></tr>
<tr><td>GU305914.1</td><td>104556</td><td>104578</td><td>+</td><td>0</td><td>CATGAGGCACTGGTGTTACA</td><td>AGG</td><td>||||||||[||||||||||||]PAM</td><td>NA</td><td>Intergenic</td><td>NA</td><td>NA</td></tr>
<tr><td></td><td></td><td></td><td></td><td></td><td></td><td></td><td></td><td></td><td></td><td></td><td></td></tr>
<tr><td>T7623</td><td>CAAGGCCTTCACCTCAGACACGG</td><td>1000</td><td></td><td></td><td></td><td></td><td></td><td></td><td></td><td></td><td></td></tr>
</table>

<table>
<tr><td>Oligo fwd</td><td>CACCCAAGGCCTTCACCTCAGACA</td><td></td><td></td><td></td><td></td><td></td><td></td><td></td><td></td><td></td><td></td></tr>
<tr><td>Oligo rev</td><td>AAACTGTCTGAGGTGAAGGCCTTG</td><td></td><td></td><td></td><td></td><td></td><td></td><td></td><td></td><td></td><td></td></tr>
<tr><td>Chromosome</td><td>start</td><td>end</td><td>strand</td><td>MM</td><td>target_seq</td><td>PAM</td><td>alignment</td><td>distance</td><td>position</td><td>gene name</td><td>gene id</td></tr>
<tr><td>GU305914.1</td><td>104574</td><td>104596</td><td>+</td><td>0</td><td>CAAGGCCTTCACCTCAGACA</td><td>CGG</td><td>||||||||[||||||||||||]PAM</td><td>NA</td><td>Intergenic</td><td>NA</td><td>NA</td></tr>
<tr><td>T7624</td><td>TTCACCTCAGACACGGCTAAAGG</td><td>1000</td><td></td><td></td><td></td><td></td><td></td><td></td><td></td><td></td><td></td></tr>
<tr><td>Oligo fwd</td><td>CACCTTCACCTCAGACACGGCTAA</td><td></td><td></td><td></td><td></td><td></td><td></td><td></td><td></td><td></td><td></td></tr>
<tr><td>Oligo rev</td><td>AAACTTAGCCGTGTCTGAGGTGAA</td><td></td><td></td><td></td><td></td><td></td><td></td><td></td><td></td><td></td><td></td></tr>
<tr><td>Chromosome</td><td>start</td><td>end</td><td>strand</td><td>MM</td><td>target_seq</td><td>PAM</td><td>alignment</td><td>distance</td><td>position</td><td>gene name</td><td>gene id</td></tr>
<tr><td>GU305914.1</td><td>104581</td><td>104603</td><td>+</td><td>0</td><td>TTCACCTCAGACACGGCTAA</td><td>AGG</td><td>||||||||[||||||||||||]PAM</td><td>NA</td><td>Intergenic</td><td>NA</td><td>NA</td></tr>
<tr><td>T7625</td><td>ACGGCTAAAGGATATTGAGCCGG</td><td>1000</td><td></td><td></td><td></td><td></td><td></td><td></td><td></td><td></td><td></td></tr>
<tr><td>Oligo fwd</td><td>CACCACGGCTAAAGGATATTGAGC</td><td></td><td></td><td></td><td></td><td></td><td></td><td></td><td></td><td></td><td></td></tr>
<tr><td>Oligo rev</td><td>AAACGCTCAATATCCTTTAGCCGT</td><td></td><td></td><td></td><td></td><td></td><td></td><td></td><td></td><td></td><td></td></tr>
<tr><td>Chromosome</td><td>start</td><td>end</td><td>strand</td><td>MM</td><td>target_seq</td><td>PAM</td><td>alignment</td><td>distance</td><td>position</td><td>gene name</td><td>gene id</td></tr>
<tr><td>GU305914.1</td><td>104593</td><td>104615</td><td>+</td><td>0</td><td>ACGGCTAAAGGATATTGAGC</td><td>CGG</td><td>||||||||[||||||||||||]PAM</td><td>NA</td><td>Intergenic</td><td>NA</td><td>NA</td></tr>
<tr><td>T7626</td><td>TTGAGCCGGTGCGCCTTTTGTGG</td><td>1000</td><td></td><td></td><td></td><td></td><td></td><td></td><td></td><td></td><td></td></tr>
<tr><td>Oligo fwd</td><td>CACCTTGAGCCGGTGCGCCTTTTG</td><td></td><td></td><td></td><td></td><td></td><td></td><td></td><td></td><td></td><td></td></tr>
<tr><td>Oligo rev</td><td>AAACCAAAAGGCGCACCGGCTCAA</td><td></td><td></td><td></td><td></td><td></td><td></td><td></td><td></td><td></td><td></td></tr>
<tr><td>Chromosome</td><td>start</td><td>end</td><td>strand</td><td>MM</td><td>target_seq</td><td>PAM</td><td>alignment</td><td>distance</td><td>position</td><td>gene name</td><td>gene id</td></tr>
<tr><td>GU305914.1</td><td>104607</td><td>104629</td><td>+</td><td>0</td><td>TTGAGCCGGTGCGCCTTTTG</td><td>TGG</td><td>||||||||[||||||||||||]PAM</td><td>NA</td><td>Intergenic</td><td>NA</td><td>NA</td></tr>
<tr><td>T7627</td><td>TGAGCCGGTGCGCCTTTTGTGGG</td><td>1000</td><td></td><td></td><td></td><td></td><td></td><td></td><td></td><td></td><td></td></tr>
<tr><td>Oligo fwd</td><td>CACCTGAGCCGGTGCGCCTTTTGT</td><td></td><td></td><td></td><td></td><td></td><td></td><td></td><td></td><td></td><td></td></tr>
<tr><td>Oligo rev</td><td>AAACACAAAAGGCGCACCGGCTCA</td><td></td><td></td><td></td><td></td><td></td><td></td><td></td><td></td><td></td><td></td></tr>
<tr><td>Chromosome</td><td>start</td><td>end</td><td>strand</td><td>MM</td><td>target_seq</td><td>PAM</td><td>alignment</td><td>distance</td><td>position</td><td>gene name</td><td>gene id</td></tr>
<tr><td>GU305914.1</td><td>104608</td><td>104630</td><td>+</td><td>0</td><td>TGAGCCGGTGCGCCTTTTGT</td><td>GGG</td><td>||||||||[||||||||||||]PAM</td><td>NA</td><td>Intergenic</td><td>NA</td><td>NA</td></tr>
<tr><td>T7628</td><td>GAGCCGGTGCGCCTTTTGTGGGG</td><td>1000</td><td></td><td></td><td></td><td></td><td></td><td></td><td></td><td></td><td></td></tr>
<tr><td>Oligo fwd</td><td>CACCGAGCCGGTGCGCCTTTTGTG</td><td></td><td></td><td></td><td></td><td></td><td></td><td></td><td></td><td></td><td></td></tr>
<tr><td>Oligo rev</td><td>AAACCACAAAAGGCGCACCGGCTC</td><td></td><td></td><td></td><td></td><td></td><td></td><td></td><td></td><td></td><td></td></tr>
<tr><td>Chromosome</td><td>start</td><td>end</td><td>strand</td><td>MM</td><td>target_seq</td><td>PAM</td><td>alignment</td><td>distance</td><td>position</td><td>gene name</td><td>gene id</td></tr>
<tr><td>GU305914.1</td><td>104609</td><td>104631</td><td>+</td><td>0</td><td>GAGCCGGTGCGCCTTTTGTG</td><td>GGG</td><td>||||||||[||||||||||||]PAM</td><td>NA</td><td>Intergenic</td><td>NA</td><td>NA</td></tr>
<tr><td>T7629</td><td>CGCCTTTTGTGGGGCAGAGACGG</td><td>1000</td><td></td><td></td><td></td><td></td><td></td><td></td><td></td><td></td><td></td></tr>
<tr><td>Oligo fwd</td><td>CACCCGCCTTTTGTGGGGCAGAGA</td><td></td><td></td><td></td><td></td><td></td><td></td><td></td><td></td><td></td><td></td></tr>
<tr><td>Oligo rev</td><td>AAACTCTCTGCCCCACAAAAGGCG</td><td></td><td></td><td></td><td></td><td></td><td></td><td></td><td></td><td></td><td></td></tr>
<tr><td>Chromosome</td><td>start</td><td>end</td><td>strand</td><td>MM</td><td>target_seq</td><td>PAM</td><td>alignment</td><td>distance</td><td>position</td><td>gene name</td><td>gene id</td></tr>
<tr><td>GU305914.1</td><td>104618</td><td>104640</td><td>+</td><td>0</td><td>CGCCTTTTGTGGGGCAGAGA</td><td>CGG</td><td>||||||||[||||||||||||]PAM</td><td>NA</td><td>Intergenic</td><td>NA</td><td>NA</td></tr>
<tr><td>T7630</td><td>AGAGACGGTACGTGTTGTTGTGG</td><td>1000</td><td></td><td></td><td></td><td></td><td></td><td></td><td></td><td></td><td></td></tr>
<tr><td>Oligo fwd</td><td>CACCAGAGACGGTACGTGTTGTTG</td><td></td><td></td><td></td><td></td><td></td><td></td><td></td><td></td><td></td><td></td></tr>
<tr><td>Oligo rev</td><td>AAACCAACAACACGTACCGTCTCT</td><td></td><td></td><td></td><td></td><td></td><td></td><td></td><td></td><td></td><td></td></tr>
<tr><td>Chromosome</td><td>start</td><td>end</td><td>strand</td><td>MM</td><td>target_seq</td><td>PAM</td><td>alignment</td><td>distance</td><td>position</td><td>gene name</td><td>gene id</td></tr>
<tr><td>GU305914.1</td><td>104633</td><td>104655</td><td>+</td><td>0</td><td>AGAGACGGTACGTGTTGTTG</td><td>TGG</td><td>||||||||[||||||||||||]PAM</td><td>NA</td><td>Intergenic</td><td>NA</td><td>NA</td></tr>
<tr><td>GU305914.1</td><td>166945</td><td>166967</td><td>+</td><td>4</td><td>GGAGCTGGTCCGTGTTGTTG</td><td>TGG</td><td>-|||--||[|-||||||||||]PAM</td><td>NA</td><td>Intergenic</td><td>NA</td><td>NA</td></tr>
<tr><td>T7631</td><td>GTACGTGTTGTTGTGGTTTCTGG</td><td>1000</td><td></td><td></td><td></td><td></td><td></td><td></td><td></td><td></td><td></td></tr>
<tr><td>Oligo fwd</td><td>CACCGTACGTGTTGTTGTGGTTTC</td><td></td><td></td><td></td><td></td><td></td><td></td><td></td><td></td><td></td><td></td></tr>
<tr><td>Oligo rev</td><td>AAACGAAACCACAACAACACGTAC</td><td></td><td></td><td></td><td></td><td></td><td></td><td></td><td></td><td></td><td></td></tr>
<tr><td>Chromosome</td><td>start</td><td>end</td><td>strand</td><td>MM</td><td>target_seq</td><td>PAM</td><td>alignment</td><td>distance</td><td>position</td><td>gene name</td><td>gene id</td></tr>
<tr><td>GU305914.1</td><td>104640</td><td>104662</td><td>+</td><td>0</td><td>GTACGTGTTGTTGTGGTTTC</td><td>TGG</td><td>||||||||[||||||||||||]PAM</td><td>NA</td><td>Intergenic</td><td>NA</td><td>NA</td></tr>
<tr><td>T7632</td><td>TACGTGTTGTTGTGGTTTCTGGG</td><td>1000</td><td></td><td></td><td></td><td></td><td></td><td></td><td></td><td></td><td></td></tr>
<tr><td>Oligo fwd</td><td>CACCTACGTGTTGTTGTGGTTTCT</td><td></td><td></td><td></td><td></td><td></td><td></td><td></td><td></td><td></td><td></td></tr>
<tr><td>Oligo rev</td><td>AAACAGAAACCACAACAACACGTA</td><td></td><td></td><td></td><td></td><td></td><td></td><td></td><td></td><td></td><td></td></tr>
<tr><td>Chromosome</td><td>start</td><td>end</td><td>strand</td><td>MM</td><td>target_seq</td><td>PAM</td><td>alignment</td><td>distance</td><td>position</td><td>gene name</td><td>gene id</td></tr>
<tr><td>GU305914.1</td><td>104641</td><td>104663</td><td>+</td><td>0</td><td>TACGTGTTGTTGTGGTTTCT</td><td>GGG</td><td>||||||||[||||||||||||]PAM</td><td>NA</td><td>Intergenic</td><td>NA</td><td>NA</td></tr>
<tr><td>T7633</td><td>TTGTTGTGGTTTCTGGGAAATGG</td><td>1000</td><td></td><td></td><td></td><td></td><td></td><td></td><td></td><td></td><td></td></tr>
<tr><td>Oligo fwd</td><td>CACCTTGTTGTGGTTTCTGGGAAA</td><td></td><td></td><td></td><td></td><td></td><td></td><td></td><td></td><td></td><td></td></tr>
<tr><td>Oligo rev</td><td>AAACTTTCCCAGAAACCACAACAA</td><td></td><td></td><td></td><td></td><td></td><td></td><td></td><td></td><td></td><td></td></tr>
<tr><td>Chromosome</td><td>start</td><td>end</td><td>strand</td><td>MM</td><td>target_seq</td><td>PAM</td><td>alignment</td><td>distance</td><td>position</td><td>gene name</td><td>gene id</td></tr>
<tr><td>GU305914.1</td><td>104647</td><td>104669</td><td>+</td><td>0</td><td>TTGTTGTGGTTTCTGGGAAA</td><td>TGG</td><td>||||||||[||||||||||||]PAM</td><td>NA</td><td>Intergenic</td><td>NA</td><td>NA</td></tr>
<tr><td>T7634</td><td>TTCTGGGAAATGGTCCTCCAAGG</td><td>1000</td><td></td><td></td><td></td><td></td><td></td><td></td><td></td><td></td><td></td></tr>
<tr><td>Oligo fwd</td><td>CACCTTCTGGGAAATGGTCCTCCA</td><td></td><td></td><td></td><td></td><td></td><td></td><td></td><td></td><td></td><td></td></tr>
<tr><td>Oligo rev</td><td>AAACTGGAGGACCATTTCCCAGAA</td><td></td><td></td><td></td><td></td><td></td><td></td><td></td><td></td><td></td><td></td></tr>
<tr><td>Chromosome</td><td>start</td><td>end</td><td>strand</td><td>MM</td><td>target_seq</td><td>PAM</td><td>alignment</td><td>distance</td><td>position</td><td>gene name</td><td>gene id</td></tr>
<tr><td>GU305914.1</td><td>104657</td><td>104679</td><td>+</td><td>0</td><td>TTCTGGGAAATGGTCCTCCA</td><td>AGG</td><td>||||||||[||||||||||||]PAM</td><td>NA</td><td>Intergenic</td><td>NA</td><td>NA</td></tr>
<tr><td>T7635</td><td>TCTGGGAAATGGTCCTCCAAGGG</td><td>1000</td><td></td><td></td><td></td><td></td><td></td><td></td><td></td><td></td><td></td></tr>
<tr><td>Oligo fwd</td><td>CACCTCTGGGAAATGGTCCTCCAA</td><td></td><td></td><td></td><td></td><td></td><td></td><td></td><td></td><td></td><td></td></tr>
<tr><td>Oligo rev</td><td>AAACTTGGAGGACCATTTCCCAGA</td><td></td><td></td><td></td><td></td><td></td><td></td><td></td><td></td><td></td><td></td></tr>
<tr><td>Chromosome</td><td>start</td><td>end</td><td>strand</td><td>MM</td><td>target_seq</td><td>PAM</td><td>alignment</td><td>distance</td><td>position</td><td>gene name</td><td>gene id</td></tr>
<tr><td>GU305914.1</td><td>104658</td><td>104680</td><td>+</td><td>0</td><td>TCTGGGAAATGGTCCTCCAA</td><td>GGG</td><td>||||||||[||||||||||||]PAM</td><td>NA</td><td>Intergenic</td><td>NA</td><td>NA</td></tr>
<tr><td>T7636</td><td>AATGGTCCTCCAAGGGCTTGCGG</td><td>1000</td><td></td><td></td><td></td><td></td><td></td><td></td><td></td><td></td><td></td></tr>
<tr><td>Oligo fwd</td><td>CACCAATGGTCCTCCAAGGGCTTG</td><td></td><td></td><td></td><td></td><td></td><td></td><td></td><td></td><td></td><td></td></tr>
<tr><td>Oligo rev</td><td>AAACCAAGCCCTTGGAGGACCATT</td><td></td><td></td><td></td><td></td><td></td><td></td><td></td><td></td><td></td><td></td></tr>
<tr><td>Chromosome</td><td>start</td><td>end</td><td>strand</td><td>MM</td><td>target_seq</td><td>PAM</td><td>alignment</td><td>distance</td><td>position</td><td>gene name</td><td>gene id</td></tr>
<tr><td>GU305914.1</td><td>104665</td><td>104687</td><td>+</td><td>0</td><td>AATGGTCCTCCAAGGGCTTG</td><td>CGG</td><td>||||||||[||||||||||||]PAM</td><td>NA</td><td>Intergenic</td><td>NA</td><td>NA</td></tr>
<tr><td>T7637</td><td>AAGGGCTTGCGGTCAACGTCAGG</td><td>1000</td><td></td><td></td><td></td><td></td><td></td><td></td><td></td><td></td><td></td></tr>
<tr><td>Oligo fwd</td><td>CACCAAGGGCTTGCGGTCAACGTC</td><td></td><td></td><td></td><td></td><td></td><td></td><td></td><td></td><td></td><td></td></tr>
<tr><td>Oligo rev</td><td>AAACGACGTTGACCGCAAGCCCTT</td><td></td><td></td><td></td><td></td><td></td><td></td><td></td><td></td><td></td><td></td></tr>
<tr><td>Chromosome</td><td>start</td><td>end</td><td>strand</td><td>MM</td><td>target_seq</td><td>PAM</td><td>alignment</td><td>distance</td><td>position</td><td>gene name</td><td>gene id</td></tr>
<tr><td>GU305914.1</td><td>104676</td><td>104698</td><td>+</td><td>0</td><td>AAGGGCTTGCGGTCAACGTC</td><td>AGG</td><td>||||||||[||||||||||||]PAM</td><td>NA</td><td>Intergenic</td><td>NA</td><td>NA</td></tr>
</table>

<table>
<tr><td>T7638</td><td>AGGGCTTGCGGTCAACGTCAGGG</td><td>1000</td><td></td><td></td><td></td><td></td><td></td><td></td><td></td><td></td><td></td></tr>
<tr><td>Oligo fwd</td><td>CACCAGGGCTTGCGGTCAACGTCA</td><td></td><td></td><td></td><td></td><td></td><td></td><td></td><td></td><td></td><td></td></tr>
<tr><td>Oligo rev</td><td>AAACTGACGTTGACCGCAAGCCCT</td><td></td><td></td><td></td><td></td><td></td><td></td><td></td><td></td><td></td><td></td></tr>
<tr><td>Chromosome</td><td>start</td><td>end</td><td>strand</td><td>MM</td><td>target_seq</td><td>PAM</td><td>alignment</td><td>distance</td><td>position</td><td>gene name</td><td>gene id</td></tr>
<tr><td>GU305914.1</td><td>104677</td><td>104699</td><td>+</td><td>0</td><td>AGGGCTTGCGGTCAACGTCA</td><td>GGG</td><td>||||||||[||||||||||||]PAM</td><td>NA</td><td>Intergenic</td><td>NA</td><td>NA</td></tr>
<tr><td>T7639</td><td>CTAGAGTTTCGACGTCCTCCAGG</td><td>1000</td><td></td><td></td><td></td><td></td><td></td><td></td><td></td><td></td><td></td></tr>
<tr><td>Oligo fwd</td><td>CACCCTAGAGTTTCGACGTCCTCC</td><td></td><td></td><td></td><td></td><td></td><td></td><td></td><td></td><td></td><td></td></tr>
<tr><td>Oligo rev</td><td>AAACGGAGGACGTCGAAACTCTAG</td><td></td><td></td><td></td><td></td><td></td><td></td><td></td><td></td><td></td><td></td></tr>
<tr><td>Chromosome</td><td>start</td><td>end</td><td>strand</td><td>MM</td><td>target_seq</td><td>PAM</td><td>alignment</td><td>distance</td><td>position</td><td>gene name</td><td>gene id</td></tr>
<tr><td>GU305914.1</td><td>104727</td><td>104749</td><td>+</td><td>0</td><td>CTAGAGTTTCGACGTCCTCC</td><td>AGG</td><td>||||||||[||||||||||||]PAM</td><td>NA</td><td>Intergenic</td><td>NA</td><td>NA</td></tr>
<tr><td>T7640</td><td>GATCTTATTATTTTTGATTTCGG</td><td>1000</td><td></td><td></td><td></td><td></td><td></td><td></td><td></td><td></td><td></td></tr>
<tr><td>Oligo fwd</td><td>CACCGATCTTATTATTTTTGATTT</td><td></td><td></td><td></td><td></td><td></td><td></td><td></td><td></td><td></td><td></td></tr>
<tr><td>Oligo rev</td><td>AAACAAATCAAAAATAATAAGATC</td><td></td><td></td><td></td><td></td><td></td><td></td><td></td><td></td><td></td><td></td></tr>
<tr><td>Chromosome</td><td>start</td><td>end</td><td>strand</td><td>MM</td><td>target_seq</td><td>PAM</td><td>alignment</td><td>distance</td><td>position</td><td>gene name</td><td>gene id</td></tr>
<tr><td>GU305914.1</td><td>104752</td><td>104774</td><td>+</td><td>0</td><td>GATCTTATTATTTTTGATTT</td><td>CGG</td><td>||||||||[||||||||||||]PAM</td><td>NA</td><td>Intergenic</td><td>NA</td><td>NA</td></tr>
<tr><td>T7641</td><td>ATCTTATTATTTTTGATTTCGGG</td><td>1000</td><td></td><td></td><td></td><td></td><td></td><td></td><td></td><td></td><td></td></tr>
<tr><td>Oligo fwd</td><td>CACCATCTTATTATTTTTGATTTC</td><td></td><td></td><td></td><td></td><td></td><td></td><td></td><td></td><td></td><td></td></tr>
<tr><td>Oligo rev</td><td>AAACGAAATCAAAAATAATAAGAT</td><td></td><td></td><td></td><td></td><td></td><td></td><td></td><td></td><td></td><td></td></tr>
<tr><td>Chromosome</td><td>start</td><td>end</td><td>strand</td><td>MM</td><td>target_seq</td><td>PAM</td><td>alignment</td><td>distance</td><td>position</td><td>gene name</td><td>gene id</td></tr>
<tr><td>GU305914.1</td><td>104753</td><td>104775</td><td>+</td><td>0</td><td>ATCTTATTATTTTTGATTTC</td><td>GGG</td><td>||||||||[||||||||||||]PAM</td><td>NA</td><td>Intergenic</td><td>NA</td><td>NA</td></tr>
<tr><td>T7642</td><td>TATAACGTTGTAACGTTTACCGG</td><td>1000</td><td></td><td></td><td></td><td></td><td></td><td></td><td></td><td></td><td></td></tr>
<tr><td>Oligo fwd</td><td>CACCTATAACGTTGTAACGTTTAC</td><td></td><td></td><td></td><td></td><td></td><td></td><td></td><td></td><td></td><td></td></tr>
<tr><td>Oligo rev</td><td>AAACGTAAACGTTACAACGTTATA</td><td></td><td></td><td></td><td></td><td></td><td></td><td></td><td></td><td></td><td></td></tr>
<tr><td>Chromosome</td><td>start</td><td>end</td><td>strand</td><td>MM</td><td>target_seq</td><td>PAM</td><td>alignment</td><td>distance</td><td>position</td><td>gene name</td><td>gene id</td></tr>
<tr><td>GU305914.1</td><td>104798</td><td>104820</td><td>+</td><td>0</td><td>TATAACGTTGTAACGTTTAC</td><td>CGG</td><td>||||||||[||||||||||||]PAM</td><td>NA</td><td>Intergenic</td><td>NA</td><td>NA</td></tr>
<tr><td>T7643</td><td>CACCGTACCGTTCGTTAGTACGG</td><td>1000</td><td></td><td></td><td></td><td></td><td></td><td></td><td></td><td></td><td></td></tr>
<tr><td>Oligo fwd</td><td>CACCCACCGTACCGTTCGTTAGTA</td><td></td><td></td><td></td><td></td><td></td><td></td><td></td><td></td><td></td><td></td></tr>
<tr><td>Oligo rev</td><td>AAACTACTAACGAACGGTACGGTG</td><td></td><td></td><td></td><td></td><td></td><td></td><td></td><td></td><td></td><td></td></tr>
<tr><td>Chromosome</td><td>start</td><td>end</td><td>strand</td><td>MM</td><td>target_seq</td><td>PAM</td><td>alignment</td><td>distance</td><td>position</td><td>gene name</td><td>gene id</td></tr>
<tr><td>GU305914.1</td><td>104836</td><td>104858</td><td>+</td><td>0</td><td>CACCGTACCGTTCGTTAGTA</td><td>CGG</td><td>||||||||[||||||||||||]PAM</td><td>NA</td><td>Intergenic</td><td>NA</td><td>NA</td></tr>
<tr><td>T7644</td><td>GTACGGTTGCGATGTTAAACTGG</td><td>1000</td><td></td><td></td><td></td><td></td><td></td><td></td><td></td><td></td><td></td></tr>
<tr><td>Oligo fwd</td><td>CACCGTACGGTTGCGATGTTAAAC</td><td></td><td></td><td></td><td></td><td></td><td></td><td></td><td></td><td></td><td></td></tr>
<tr><td>Oligo rev</td><td>AAACGTTTAACATCGCAACCGTAC</td><td></td><td></td><td></td><td></td><td></td><td></td><td></td><td></td><td></td><td></td></tr>
<tr><td>Chromosome</td><td>start</td><td>end</td><td>strand</td><td>MM</td><td>target_seq</td><td>PAM</td><td>alignment</td><td>distance</td><td>position</td><td>gene name</td><td>gene id</td></tr>
<tr><td>GU305914.1</td><td>104853</td><td>104875</td><td>+</td><td>0</td><td>GTACGGTTGCGATGTTAAAC</td><td>TGG</td><td>||||||||[||||||||||||]PAM</td><td>NA</td><td>Intergenic</td><td>NA</td><td>NA</td></tr>
<tr><td>T7645</td><td>TACGGTTGCGATGTTAAACTGGG</td><td>1000</td><td></td><td></td><td></td><td></td><td></td><td></td><td></td><td></td><td></td></tr>
<tr><td>Oligo fwd</td><td>CACCTACGGTTGCGATGTTAAACT</td><td></td><td></td><td></td><td></td><td></td><td></td><td></td><td></td><td></td><td></td></tr>
<tr><td>Oligo rev</td><td>AAACAGTTTAACATCGCAACCGTA</td><td></td><td></td><td></td><td></td><td></td><td></td><td></td><td></td><td></td><td></td></tr>
<tr><td>Chromosome</td><td>start</td><td>end</td><td>strand</td><td>MM</td><td>target_seq</td><td>PAM</td><td>alignment</td><td>distance</td><td>position</td><td>gene name</td><td>gene id</td></tr>
<tr><td>GU305914.1</td><td>104854</td><td>104876</td><td>+</td><td>0</td><td>TACGGTTGCGATGTTAAACT</td><td>GGG</td><td>||||||||[||||||||||||]PAM</td><td>NA</td><td>Intergenic</td><td>NA</td><td>NA</td></tr>
<tr><td>T7646</td><td>ATATAGAGATGCGTGCATGAAGG</td><td>1000</td><td></td><td></td><td></td><td></td><td></td><td></td><td></td><td></td><td></td></tr>
<tr><td>Oligo fwd</td><td>CACCATATAGAGATGCGTGCATGA</td><td></td><td></td><td></td><td></td><td></td><td></td><td></td><td></td><td></td><td></td></tr>
<tr><td>Oligo rev</td><td>AAACTCATGCACGCATCTCTATAT</td><td></td><td></td><td></td><td></td><td></td><td></td><td></td><td></td><td></td><td></td></tr>
<tr><td>Chromosome</td><td>start</td><td>end</td><td>strand</td><td>MM</td><td>target_seq</td><td>PAM</td><td>alignment</td><td>distance</td><td>position</td><td>gene name</td><td>gene id</td></tr>
<tr><td>GU305914.1</td><td>104902</td><td>104924</td><td>+</td><td>0</td><td>ATATAGAGATGCGTGCATGA</td><td>AGG</td><td>||||||||[||||||||||||]PAM</td><td>NA</td><td>Intergenic</td><td>NA</td><td>NA</td></tr>
<tr><td>T7647</td><td>TAGAGATGCGTGCATGAAGGCGG</td><td>1000</td><td></td><td></td><td></td><td></td><td></td><td></td><td></td><td></td><td></td></tr>
<tr><td>Oligo fwd</td><td>CACCTAGAGATGCGTGCATGAAGG</td><td></td><td></td><td></td><td></td><td></td><td></td><td></td><td></td><td></td><td></td></tr>
<tr><td>Oligo rev</td><td>AAACCCTTCATGCACGCATCTCTA</td><td></td><td></td><td></td><td></td><td></td><td></td><td></td><td></td><td></td><td></td></tr>
<tr><td>Chromosome</td><td>start</td><td>end</td><td>strand</td><td>MM</td><td>target_seq</td><td>PAM</td><td>alignment</td><td>distance</td><td>position</td><td>gene name</td><td>gene id</td></tr>
<tr><td>GU305914.1</td><td>104905</td><td>104927</td><td>+</td><td>0</td><td>TAGAGATGCGTGCATGAAGG</td><td>CGG</td><td>||||||||[||||||||||||]PAM</td><td>NA</td><td>Intergenic</td><td>NA</td><td>NA</td></tr>
<tr><td>T7648</td><td>AAATCGTAGAAAAATTGTACCGG</td><td>1000</td><td></td><td></td><td></td><td></td><td></td><td></td><td></td><td></td><td></td></tr>
<tr><td>Oligo fwd</td><td>CACCAAATCGTAGAAAAATTGTAC</td><td></td><td></td><td></td><td></td><td></td><td></td><td></td><td></td><td></td><td></td></tr>
<tr><td>Oligo rev</td><td>AAACGTACAATTTTTCTACGATTT</td><td></td><td></td><td></td><td></td><td></td><td></td><td></td><td></td><td></td><td></td></tr>
<tr><td>Chromosome</td><td>start</td><td>end</td><td>strand</td><td>MM</td><td>target_seq</td><td>PAM</td><td>alignment</td><td>distance</td><td>position</td><td>gene name</td><td>gene id</td></tr>
<tr><td>GU305914.1</td><td>104930</td><td>104952</td><td>+</td><td>0</td><td>AAATCGTAGAAAAATTGTAC</td><td>CGG</td><td>||||||||[||||||||||||]PAM</td><td>NA</td><td>Intergenic</td><td>NA</td><td>NA</td></tr>
<tr><td>T7649</td><td>AATCGTAGAAAAATTGTACCGGG</td><td>1000</td><td></td><td></td><td></td><td></td><td></td><td></td><td></td><td></td><td></td></tr>
<tr><td>Oligo fwd</td><td>CACCAATCGTAGAAAAATTGTACC</td><td></td><td></td><td></td><td></td><td></td><td></td><td></td><td></td><td></td><td></td></tr>
<tr><td>Oligo rev</td><td>AAACGGTACAATTTTTCTACGATT</td><td></td><td></td><td></td><td></td><td></td><td></td><td></td><td></td><td></td><td></td></tr>
<tr><td>Chromosome</td><td>start</td><td>end</td><td>strand</td><td>MM</td><td>target_seq</td><td>PAM</td><td>alignment</td><td>distance</td><td>position</td><td>gene name</td><td>gene id</td></tr>
<tr><td>GU305914.1</td><td>104931</td><td>104953</td><td>+</td><td>0</td><td>AATCGTAGAAAAATTGTACC</td><td>GGG</td><td>||||||||[||||||||||||]PAM</td><td>NA</td><td>Intergenic</td><td>NA</td><td>NA</td></tr>
<tr><td>T7650</td><td>TGTACCGGGTGCATGTTTTTCGG</td><td>1000</td><td></td><td></td><td></td><td></td><td></td><td></td><td></td><td></td><td></td></tr>
<tr><td>Oligo fwd</td><td>CACCTGTACCGGGTGCATGTTTTT</td><td></td><td></td><td></td><td></td><td></td><td></td><td></td><td></td><td></td><td></td></tr>
<tr><td>Oligo rev</td><td>AAACAAAAACATGCACCCGGTACA</td><td></td><td></td><td></td><td></td><td></td><td></td><td></td><td></td><td></td><td></td></tr>
<tr><td>Chromosome</td><td>start</td><td>end</td><td>strand</td><td>MM</td><td>target_seq</td><td>PAM</td><td>alignment</td><td>distance</td><td>position</td><td>gene name</td><td>gene id</td></tr>
<tr><td>GU305914.1</td><td>104945</td><td>104967</td><td>+</td><td>0</td><td>TGTACCGGGTGCATGTTTTT</td><td>CGG</td><td>||||||||[||||||||||||]PAM</td><td>NA</td><td>Intergenic</td><td>NA</td><td>NA</td></tr>
<tr><td>T7651</td><td>CGGGTGCATGTTTTTCGGTATGG</td><td>1000</td><td></td><td></td><td></td><td></td><td></td><td></td><td></td><td></td><td></td></tr>
<tr><td>Oligo fwd</td><td>CACCCGGGTGCATGTTTTTCGGTA</td><td></td><td></td><td></td><td></td><td></td><td></td><td></td><td></td><td></td><td></td></tr>
<tr><td>Oligo rev</td><td>AAACTACCGAAAAACATGCACCCG</td><td></td><td></td><td></td><td></td><td></td><td></td><td></td><td></td><td></td><td></td></tr>
<tr><td>Chromosome</td><td>start</td><td>end</td><td>strand</td><td>MM</td><td>target_seq</td><td>PAM</td><td>alignment</td><td>distance</td><td>position</td><td>gene name</td><td>gene id</td></tr>
<tr><td>GU305914.1</td><td>104950</td><td>104972</td><td>+</td><td>0</td><td>CGGGTGCATGTTTTTCGGTA</td><td>TGG</td><td>||||||||[||||||||||||]PAM</td><td>NA</td><td>Intergenic</td><td>NA</td><td>NA</td></tr>
<tr><td>T7652</td><td>GGGTGCATGTTTTTCGGTATGGG</td><td>1000</td><td></td><td></td><td></td><td></td><td></td><td></td><td></td><td></td><td></td></tr>
<tr><td>Oligo fwd</td><td>CACCGGGTGCATGTTTTTCGGTAT</td><td></td><td></td><td></td><td></td><td></td><td></td><td></td><td></td><td></td><td></td></tr>
<tr><td>Oligo rev</td><td>AAACATACCGAAAAACATGCACCC</td><td></td><td></td><td></td><td></td><td></td><td></td><td></td><td></td><td></td><td></td></tr>
<tr><td>Chromosome</td><td>start</td><td>end</td><td>strand</td><td>MM</td><td>target_seq</td><td>PAM</td><td>alignment</td><td>distance</td><td>position</td><td>gene name</td><td>gene id</td></tr>
</table>

GU305914.1 104951 104973 + 0 GGGTGCATGTTTTTCGGTAT GGG ||||||||[||||||||||||]PAM NA Intergenic NA NA

T7653 GGTGCATGTTTTTCGGTATGGGG 1000
Oligo fwd CACCGGTGCATGTTTTTCGGTATG
Oligo rev AAACCATACCGAAAAACATGCACC
Chromosome start end strand MM target_seq PAM alignment distance position gene name gene id
GU305914.1 104952 104974 + 0 GGTGCATGTTTTTCGGTATG GGG ||||||||[||||||||||||]PAM NA Intergenic NA NA

T7654 GTGCATGTTTTTCGGTATGGGGG 1000
Oligo fwd CACCGTGCATGTTTTTCGGTATGG
Oligo rev AAACCCATACCGAAAAACATGCAC
Chromosome start end strand MM target_seq PAM alignment distance position gene name gene id
GU305914.1 104953 104975 + 0 GTGCATGTTTTTCGGTATGG GGG ||||||||[||||||||||||]PAM NA Intergenic NA NA

T7655 TGCATGTTTTTCGGTATGGGGGG 1000
Oligo fwd CACCTGCATGTTTTTCGGTATGGG
Oligo rev AAACCCCATACCGAAAAACATGCA
Chromosome start end strand MM target_seq PAM alignment distance position gene name gene id
GU305914.1 104954 104976 + 0 TGCATGTTTTTCGGTATGGG GGG ||||||||[||||||||||||]PAM NA Intergenic NA NA

T7656 ATGGGGGGCGTCACGACGATCGG 1000
Oligo fwd CACCATGGGGGGCGTCACGACGAT
Oligo rev AAACATCGTCGTGACGCCCCCCAT
Chromosome start end strand MM target_seq PAM alignment distance position gene name gene id
GU305914.1 104969 104991 + 0 ATGGGGGGCGTCACGACGAT CGG ||||||||[||||||||||||]PAM NA Intergenic NA NA

T7657 TGGGGGGCGTCACGACGATCGGG 1000
Oligo fwd CACCTGGGGGGCGTCACGACGATC
Oligo rev AAACGATCGTCGTGACGCCCCCCA
Chromosome start end strand MM target_seq PAM alignment distance position gene name gene id
GU305914.1 104970 104992 + 0 TGGGGGGCGTCACGACGATC GGG ||||||||[||||||||||||]PAM NA Intergenic NA NA

T7658 TGCGCACTTCGTTTTCTCCTCGG 1000
Oligo fwd CACCTGCGCACTTCGTTTTCTCCT
Oligo rev AAACAGGAGAAAACGAAGTGCGCA
Chromosome start end strand MM target_seq PAM alignment distance position gene name gene id
GU305914.1 104993 105015 + 0 TGCGCACTTCGTTTTCTCCT CGG ||||||||[||||||||||||]PAM NA Intergenic NA NA

T7659 GCGCACTTCGTTTTCTCCTCGGG 1000
Oligo fwd CACCGCGCACTTCGTTTTCTCCTC
Oligo rev AAACGAGGAGAAAACGAAGTGCGC
Chromosome start end strand MM target_seq PAM alignment distance position gene name gene id
GU305914.1 104994 105016 + 0 GCGCACTTCGTTTTCTCCTC GGG ||||||||[||||||||||||]PAM NA Intergenic NA NA

T7660 CGCACTTCGTTTTCTCCTCGGGG 1000
Oligo fwd CACCCGCACTTCGTTTTCTCCTCG
Oligo rev AAACCGAGGAGAAAACGAAGTGCG
Chromosome start end strand MM target_seq PAM alignment distance position gene name gene id
GU305914.1 104995 105017 + 0 CGCACTTCGTTTTCTCCTCG GGG ||||||||[||||||||||||]PAM NA Intergenic NA NA

T7661 GCACTTCGTTTTCTCCTCGGGGG 1000
Oligo fwd CACCGCACTTCGTTTTCTCCTCGG
Oligo rev AAACCCGAGGAGAAAACGAAGTGC
Chromosome start end strand MM target_seq PAM alignment distance position gene name gene id
GU305914.1 104996 105018 + 0 GCACTTCGTTTTCTCCTCGG GGG ||||||||[||||||||||||]PAM NA Intergenic NA NA

T7662 TCGTTTTCTCCTCGGGGGTAAGG 1000
Oligo fwd CACCTCGTTTTCTCCTCGGGGGTA
Oligo rev AAACTACCCCCGAGGAGAAAACGA
Chromosome start end strand MM target_seq PAM alignment distance position gene name gene id
GU305914.1 105001 105023 + 0 TCGTTTTCTCCTCGGGGGTA AGG ||||||||[||||||||||||]PAM NA Intergenic NA NA

T7663 CCTCGGGGGTAAGGTGACCGCGG 1000
Oligo fwd CACCCCTCGGGGGTAAGGTGACCG
Oligo rev AAACCGGTCACCTTACCCCCGAGG
Chromosome start end strand MM target_seq PAM alignment distance position gene name gene id
GU305914.1 105010 105032 + 0 CCTCGGGGGTAAGGTGACCG CGG ||||||||[||||||||||||]PAM NA Intergenic NA NA

T7664 CTCGGGGGTAAGGTGACCGCGGG 1000
Oligo fwd CACCCTCGGGGGTAAGGTGACCGC
Oligo rev AAACGCGGTCACCTTACCCCCGAG
Chromosome start end strand MM target_seq PAM alignment distance position gene name gene id
GU305914.1 105011 105033 + 0 CTCGGGGGTAAGGTGACCGC GGG ||||||||[||||||||||||]PAM NA Intergenic NA NA

T7665 TCGAAAAACGTTTTGATGTCCGG 1000
Oligo fwd CACCTCGAAAAACGTTTTGATGTC
Oligo rev AAACGACATCAAAACGTTTTTCGA
Chromosome start end strand MM target_seq PAM alignment distance position gene name gene id
GU305914.1 105068 105090 + 0 TCGAAAAACGTTTTGATGTC CGG ||||||||[||||||||||||]PAM NA Intergenic NA NA
GU305914.1 117187 117209 + 4 TCGAAATCCGTTTTGAAGTT GGG ||||||--[||||||||-||-]PAM NA Intergenic NA NA

T7666 AAACGTTTTGATGTCCGGCCAGG 1000
Oligo fwd CACCAAACGTTTTGATGTCCGGCC
Oligo rev AAACGGCCGGACATCAAAACGTTT
Chromosome start end strand MM target_seq PAM alignment distance position gene name gene id
GU305914.1 105073 105095 + 0 AAACGTTTTGATGTCCGGCC AGG ||||||||[||||||||||||]PAM NA Intergenic NA NA

T7667 CGTTTTGATGTCCGGCCAGGAGG 1000
Oligo fwd CACCCGTTTTGATGTCCGGCCAGG

<table>
<tr><td>Oligo rev</td><td>AAACCCTGGCCGGACATCAAAACG</td><td></td><td></td><td></td><td></td><td></td><td></td><td></td><td></td><td></td><td></td></tr>
<tr><td>Chromosome</td><td>start</td><td>end</td><td>strand</td><td>MM</td><td>target_seq</td><td>PAM</td><td>alignment</td><td>distance</td><td>position</td><td>gene name</td><td>gene id</td></tr>
<tr><td>GU305914.1</td><td>105076</td><td>105098</td><td>+</td><td>0</td><td>CGTTTTGATGTCCGGCCAGG</td><td>AGG</td><td>||||||||[||||||||||||]PAM</td><td>NA</td><td>Intergenic</td><td>NA</td><td>NA</td></tr>
<tr><td></td><td></td><td></td><td></td><td></td><td></td><td></td><td></td><td></td><td></td><td></td><td></td></tr>
<tr><td>T7668</td><td>CGTTTTTAGCTCTCGAGTAACGG</td><td>1000</td><td></td><td></td><td></td><td></td><td></td><td></td><td></td><td></td><td></td></tr>
<tr><td>Oligo fwd</td><td>CACCCGTTTTTAGCTCTCGAGTAA</td><td></td><td></td><td></td><td></td><td></td><td></td><td></td><td></td><td></td><td></td></tr>
<tr><td>Oligo rev</td><td>AAACTTACTCGAGAGCTAAAAACG</td><td></td><td></td><td></td><td></td><td></td><td></td><td></td><td></td><td></td><td></td></tr>
<tr><td>Chromosome</td><td>start</td><td>end</td><td>strand</td><td>MM</td><td>target_seq</td><td>PAM</td><td>alignment</td><td>distance</td><td>position</td><td>gene name</td><td>gene id</td></tr>
<tr><td>GU305914.1</td><td>105162</td><td>105184</td><td>+</td><td>0</td><td>CGTTTTTAGCTCTCGAGTAA</td><td>CGG</td><td>||||||||[||||||||||||]PAM</td><td>NA</td><td>Intergenic</td><td>NA</td><td>NA</td></tr>
<tr><td></td><td></td><td></td><td></td><td></td><td></td><td></td><td></td><td></td><td></td><td></td><td></td></tr>
<tr><td>T7669</td><td>ACGGCTGATGAGAGTCGCAACGG</td><td>1000</td><td></td><td></td><td></td><td></td><td></td><td></td><td></td><td></td><td></td></tr>
<tr><td>Oligo fwd</td><td>CACCACGGCTGATGAGAGTCGCAA</td><td></td><td></td><td></td><td></td><td></td><td></td><td></td><td></td><td></td><td></td></tr>
<tr><td>Oligo rev</td><td>AAACTTGCGACTCTCATCAGCCGT</td><td></td><td></td><td></td><td></td><td></td><td></td><td></td><td></td><td></td><td></td></tr>
<tr><td>Chromosome</td><td>start</td><td>end</td><td>strand</td><td>MM</td><td>target_seq</td><td>PAM</td><td>alignment</td><td>distance</td><td>position</td><td>gene name</td><td>gene id</td></tr>
<tr><td>GU305914.1</td><td>105181</td><td>105203</td><td>+</td><td>0</td><td>ACGGCTGATGAGAGTCGCAA</td><td>CGG</td><td>||||||||[||||||||||||]PAM</td><td>NA</td><td>Intergenic</td><td>NA</td><td>NA</td></tr>
<tr><td></td><td></td><td></td><td></td><td></td><td></td><td></td><td></td><td></td><td></td><td></td><td></td></tr>
<tr><td>T7670</td><td>GAGAGTCGCAACGGCTAACGTGG</td><td>1000</td><td></td><td></td><td></td><td></td><td></td><td></td><td></td><td></td><td></td></tr>
<tr><td>Oligo fwd</td><td>CACCGAGAGTCGCAACGGCTAACG</td><td></td><td></td><td></td><td></td><td></td><td></td><td></td><td></td><td></td><td></td></tr>
<tr><td>Oligo rev</td><td>AAACCGTTAGCCGTTGCGACTCTC</td><td></td><td></td><td></td><td></td><td></td><td></td><td></td><td></td><td></td><td></td></tr>
<tr><td>Chromosome</td><td>start</td><td>end</td><td>strand</td><td>MM</td><td>target_seq</td><td>PAM</td><td>alignment</td><td>distance</td><td>position</td><td>gene name</td><td>gene id</td></tr>
<tr><td>GU305914.1</td><td>105190</td><td>105212</td><td>+</td><td>0</td><td>GAGAGTCGCAACGGCTAACG</td><td>TGG</td><td>||||||||[||||||||||||]PAM</td><td>NA</td><td>Intergenic</td><td>NA</td><td>NA</td></tr>
<tr><td></td><td></td><td></td><td></td><td></td><td></td><td></td><td></td><td></td><td></td><td></td><td></td></tr>
<tr><td>T7671</td><td>CGTGGTGATAGTGAACGATTCGG</td><td>1000</td><td></td><td></td><td></td><td></td><td></td><td></td><td></td><td></td><td></td></tr>
<tr><td>Oligo fwd</td><td>CACCCGTGGTGATAGTGAACGATT</td><td></td><td></td><td></td><td></td><td></td><td></td><td></td><td></td><td></td><td></td></tr>
<tr><td>Oligo rev</td><td>AAACAATCGTTCACTATCACCACG</td><td></td><td></td><td></td><td></td><td></td><td></td><td></td><td></td><td></td><td></td></tr>
<tr><td>Chromosome</td><td>start</td><td>end</td><td>strand</td><td>MM</td><td>target_seq</td><td>PAM</td><td>alignment</td><td>distance</td><td>position</td><td>gene name</td><td>gene id</td></tr>
<tr><td>GU305914.1</td><td>105208</td><td>105230</td><td>+</td><td>0</td><td>CGTGGTGATAGTGAACGATT</td><td>CGG</td><td>||||||||[||||||||||||]PAM</td><td>NA</td><td>Intergenic</td><td>NA</td><td>NA</td></tr>
<tr><td></td><td></td><td></td><td></td><td></td><td></td><td></td><td></td><td></td><td></td><td></td><td></td></tr>
<tr><td>T7672</td><td>GTCACCAACACGCGTCTGTCCGG</td><td>1000</td><td></td><td></td><td></td><td></td><td></td><td></td><td></td><td></td><td></td></tr>
<tr><td>Oligo fwd</td><td>CACCGTCACCAACACGCGTCTGTC</td><td></td><td></td><td></td><td></td><td></td><td></td><td></td><td></td><td></td><td></td></tr>
<tr><td>Oligo rev</td><td>AAACGACAGACGCGTGTTGGTGAC</td><td></td><td></td><td></td><td></td><td></td><td></td><td></td><td></td><td></td><td></td></tr>
<tr><td>Chromosome</td><td>start</td><td>end</td><td>strand</td><td>MM</td><td>target_seq</td><td>PAM</td><td>alignment</td><td>distance</td><td>position</td><td>gene name</td><td>gene id</td></tr>
<tr><td>GU305914.1</td><td>105230</td><td>105252</td><td>+</td><td>0</td><td>GTCACCAACACGCGTCTGTC</td><td>CGG</td><td>||||||||[||||||||||||]PAM</td><td>NA</td><td>Intergenic</td><td>NA</td><td>NA</td></tr>
<tr><td></td><td></td><td></td><td></td><td></td><td></td><td></td><td></td><td></td><td></td><td></td><td></td></tr>
<tr><td>T7673</td><td>TCACCAACACGCGTCTGTCCGGG</td><td>1000</td><td></td><td></td><td></td><td></td><td></td><td></td><td></td><td></td><td></td></tr>
<tr><td>Oligo fwd</td><td>CACCTCACCAACACGCGTCTGTCC</td><td></td><td></td><td></td><td></td><td></td><td></td><td></td><td></td><td></td><td></td></tr>
<tr><td>Oligo rev</td><td>AAACGGACAGACGCGTGTTGGTGA</td><td></td><td></td><td></td><td></td><td></td><td></td><td></td><td></td><td></td><td></td></tr>
<tr><td>Chromosome</td><td>start</td><td>end</td><td>strand</td><td>MM</td><td>target_seq</td><td>PAM</td><td>alignment</td><td>distance</td><td>position</td><td>gene name</td><td>gene id</td></tr>
<tr><td>GU305914.1</td><td>105231</td><td>105253</td><td>+</td><td>0</td><td>TCACCAACACGCGTCTGTCC</td><td>GGG</td><td>||||||||[||||||||||||]PAM</td><td>NA</td><td>Intergenic</td><td>NA</td><td>NA</td></tr>
<tr><td></td><td></td><td></td><td></td><td></td><td></td><td></td><td></td><td></td><td></td><td></td><td></td></tr>
<tr><td>T7674</td><td>CACCAACACGCGTCTGTCCGGGG</td><td>1000</td><td></td><td></td><td></td><td></td><td></td><td></td><td></td><td></td><td></td></tr>
<tr><td>Oligo fwd</td><td>CACCCACCAACACGCGTCTGTCCG</td><td></td><td></td><td></td><td></td><td></td><td></td><td></td><td></td><td></td><td></td></tr>
<tr><td>Oligo rev</td><td>AAACCGGACAGACGCGTGTTGGTG</td><td></td><td></td><td></td><td></td><td></td><td></td><td></td><td></td><td></td><td></td></tr>
<tr><td>Chromosome</td><td>start</td><td>end</td><td>strand</td><td>MM</td><td>target_seq</td><td>PAM</td><td>alignment</td><td>distance</td><td>position</td><td>gene name</td><td>gene id</td></tr>
<tr><td>GU305914.1</td><td>105232</td><td>105254</td><td>+</td><td>0</td><td>CACCAACACGCGTCTGTCCG</td><td>GGG</td><td>||||||||[||||||||||||]PAM</td><td>NA</td><td>Intergenic</td><td>NA</td><td>NA</td></tr>
<tr><td></td><td></td><td></td><td></td><td></td><td></td><td></td><td></td><td></td><td></td><td></td><td></td></tr>
<tr><td>T7675</td><td>TCCTCAGTTCCACTTGCCAGCGG</td><td>1000</td><td></td><td></td><td></td><td></td><td></td><td></td><td></td><td></td><td></td></tr>
<tr><td>Oligo fwd</td><td>CACCTCCTCAGTTCCACTTGCCAG</td><td></td><td></td><td></td><td></td><td></td><td></td><td></td><td></td><td></td><td></td></tr>
<tr><td>Oligo rev</td><td>AAACCTGGCAAGTGGAACTGAGGA</td><td></td><td></td><td></td><td></td><td></td><td></td><td></td><td></td><td></td><td></td></tr>
<tr><td>Chromosome</td><td>start</td><td>end</td><td>strand</td><td>MM</td><td>target_seq</td><td>PAM</td><td>alignment</td><td>distance</td><td>position</td><td>gene name</td><td>gene id</td></tr>
<tr><td>GU305914.1</td><td>105255</td><td>105277</td><td>+</td><td>0</td><td>TCCTCAGTTCCACTTGCCAG</td><td>CGG</td><td>||||||||[||||||||||||]PAM</td><td>NA</td><td>Intergenic</td><td>NA</td><td>NA</td></tr>
<tr><td></td><td></td><td></td><td></td><td></td><td></td><td></td><td></td><td></td><td></td><td></td><td></td></tr>
<tr><td>T7676</td><td>CCTCAGTTCCACTTGCCAGCGGG</td><td>1000</td><td></td><td></td><td></td><td></td><td></td><td></td><td></td><td></td><td></td></tr>
<tr><td>Oligo fwd</td><td>CACCCCTCAGTTCCACTTGCCAGC</td><td></td><td></td><td></td><td></td><td></td><td></td><td></td><td></td><td></td><td></td></tr>
<tr><td>Oligo rev</td><td>AAACGCTGGCAAGTGGAACTGAGG</td><td></td><td></td><td></td><td></td><td></td><td></td><td></td><td></td><td></td><td></td></tr>
<tr><td>Chromosome</td><td>start</td><td>end</td><td>strand</td><td>MM</td><td>target_seq</td><td>PAM</td><td>alignment</td><td>distance</td><td>position</td><td>gene name</td><td>gene id</td></tr>
<tr><td>GU305914.1</td><td>105256</td><td>105278</td><td>+</td><td>0</td><td>CCTCAGTTCCACTTGCCAGC</td><td>GGG</td><td>||||||||[||||||||||||]PAM</td><td>NA</td><td>Intergenic</td><td>NA</td><td>NA</td></tr>
<tr><td></td><td></td><td></td><td></td><td></td><td></td><td></td><td></td><td></td><td></td><td></td><td></td></tr>
<tr><td>T7677</td><td>CTCAGTTCCACTTGCCAGCGGGG</td><td>1000</td><td></td><td></td><td></td><td></td><td></td><td></td><td></td><td></td><td></td></tr>
<tr><td>Oligo fwd</td><td>CACCCTCAGTTCCACTTGCCAGCG</td><td></td><td></td><td></td><td></td><td></td><td></td><td></td><td></td><td></td><td></td></tr>
<tr><td>Oligo rev</td><td>AAACCGCTGGCAAGTGGAACTGAG</td><td></td><td></td><td></td><td></td><td></td><td></td><td></td><td></td><td></td><td></td></tr>
<tr><td>Chromosome</td><td>start</td><td>end</td><td>strand</td><td>MM</td><td>target_seq</td><td>PAM</td><td>alignment</td><td>distance</td><td>position</td><td>gene name</td><td>gene id</td></tr>
<tr><td>GU305914.1</td><td>105257</td><td>105279</td><td>+</td><td>0</td><td>CTCAGTTCCACTTGCCAGCG</td><td>GGG</td><td>||||||||[||||||||||||]PAM</td><td>NA</td><td>Intergenic</td><td>NA</td><td>NA</td></tr>
<tr><td></td><td></td><td></td><td></td><td></td><td></td><td></td><td></td><td></td><td></td><td></td><td></td></tr>
<tr><td>T7678</td><td>GTTCCACTTGCCAGCGGGGCAGG</td><td>1000</td><td></td><td></td><td></td><td></td><td></td><td></td><td></td><td></td><td></td></tr>
<tr><td>Oligo fwd</td><td>CACCGTTCCACTTGCCAGCGGGGC</td><td></td><td></td><td></td><td></td><td></td><td></td><td></td><td></td><td></td><td></td></tr>
<tr><td>Oligo rev</td><td>AAACGCCCCGCTGGCAAGTGGAAC</td><td></td><td></td><td></td><td></td><td></td><td></td><td></td><td></td><td></td><td></td></tr>
<tr><td>Chromosome</td><td>start</td><td>end</td><td>strand</td><td>MM</td><td>target_seq</td><td>PAM</td><td>alignment</td><td>distance</td><td>position</td><td>gene name</td><td>gene id</td></tr>
<tr><td>GU305914.1</td><td>105261</td><td>105283</td><td>+</td><td>0</td><td>GTTCCACTTGCCAGCGGGGC</td><td>AGG</td><td>||||||||[||||||||||||]PAM</td><td>NA</td><td>Intergenic</td><td>NA</td><td>NA</td></tr>
<tr><td></td><td></td><td></td><td></td><td></td><td></td><td></td><td></td><td></td><td></td><td></td><td></td></tr>
<tr><td>T7679</td><td>GCCAGCGGGGCAGGTCGTACAGG</td><td>1000</td><td></td><td></td><td></td><td></td><td></td><td></td><td></td><td></td><td></td></tr>
<tr><td>Oligo fwd</td><td>CACCGCCAGCGGGGCAGGTCGTAC</td><td></td><td></td><td></td><td></td><td></td><td></td><td></td><td></td><td></td><td></td></tr>
<tr><td>Oligo rev</td><td>AAACGTACGACCTGCCCCGCTGGC</td><td></td><td></td><td></td><td></td><td></td><td></td><td></td><td></td><td></td><td></td></tr>
<tr><td>Chromosome</td><td>start</td><td>end</td><td>strand</td><td>MM</td><td>target_seq</td><td>PAM</td><td>alignment</td><td>distance</td><td>position</td><td>gene name</td><td>gene id</td></tr>
<tr><td>GU305914.1</td><td>105270</td><td>105292</td><td>+</td><td>0</td><td>GCCAGCGGGGCAGGTCGTAC</td><td>AGG</td><td>||||||||[||||||||||||]PAM</td><td>NA</td><td>Intergenic</td><td>NA</td><td>NA</td></tr>
<tr><td></td><td></td><td></td><td></td><td></td><td></td><td></td><td></td><td></td><td></td><td></td><td></td></tr>
<tr><td>T7680</td><td>CGGGGCAGGTCGTACAGGATAGG</td><td>1000</td><td></td><td></td><td></td><td></td><td></td><td></td><td></td><td></td><td></td></tr>
<tr><td>Oligo fwd</td><td>CACCCGGGGCAGGTCGTACAGGAT</td><td></td><td></td><td></td><td></td><td></td><td></td><td></td><td></td><td></td><td></td></tr>
<tr><td>Oligo rev</td><td>AAACATCCTGTACGACCTGCCCCG</td><td></td><td></td><td></td><td></td><td></td><td></td><td></td><td></td><td></td><td></td></tr>
<tr><td>Chromosome</td><td>start</td><td>end</td><td>strand</td><td>MM</td><td>target_seq</td><td>PAM</td><td>alignment</td><td>distance</td><td>position</td><td>gene name</td><td>gene id</td></tr>
<tr><td>GU305914.1</td><td>105275</td><td>105297</td><td>+</td><td>0</td><td>CGGGGCAGGTCGTACAGGAT</td><td>AGG</td><td>||||||||[||||||||||||]PAM</td><td>NA</td><td>Intergenic</td><td>NA</td><td>NA</td></tr>
<tr><td></td><td></td><td></td><td></td><td></td><td></td><td></td><td></td><td></td><td></td><td></td><td></td></tr>
<tr><td>T7681</td><td>CAGGTCGTACAGGATAGGCGCGG</td><td>1000</td><td></td><td></td><td></td><td></td><td></td><td></td><td></td><td></td><td></td></tr>
<tr><td>Oligo fwd</td><td>CACCCAGGTCGTACAGGATAGGCG</td><td></td><td></td><td></td><td></td><td></td><td></td><td></td><td></td><td></td><td></td></tr>
<tr><td>Oligo rev</td><td>AAACCGCCTATCCTGTACGACCTG</td><td></td><td></td><td></td><td></td><td></td><td></td><td></td><td></td><td></td><td></td></tr>
<tr><td>Chromosome</td><td>start</td><td>end</td><td>strand</td><td>MM</td><td>target_seq</td><td>PAM</td><td>alignment</td><td>distance</td><td>position</td><td>gene name</td><td>gene id</td></tr>
<tr><td>GU305914.1</td><td>105280</td><td>105302</td><td>+</td><td>0</td><td>CAGGTCGTACAGGATAGGCG</td><td>CGG</td><td>||||||||[||||||||||||]PAM</td><td>NA</td><td>Intergenic</td><td>NA</td><td>NA</td></tr>
<tr><td></td><td></td><td></td><td></td><td></td><td></td><td></td><td></td><td></td><td></td><td></td><td></td></tr>
<tr><td>T7682</td><td>ATCGCATTTTTTAAAATATTTGG</td><td>1000</td><td></td><td></td><td></td><td></td><td></td><td></td><td></td><td></td><td></td></tr>
</table>

<table>
<tr><td>Oligo fwd</td><td>CACCATCGCATTTTTTAAAATATT</td><td></td><td></td><td></td><td></td><td></td><td></td><td></td><td></td><td></td><td></td></tr>
<tr><td>Oligo rev</td><td>AAACAATATTTTAAAAAATGCGAT</td><td></td><td></td><td></td><td></td><td></td><td></td><td></td><td></td><td></td><td></td></tr>
<tr><td>Chromosome</td><td>start</td><td>end</td><td>strand</td><td>MM</td><td>target_seq</td><td>PAM</td><td>alignment</td><td>distance</td><td>position</td><td>gene name</td><td>gene id</td></tr>
<tr><td>GU305914.1</td><td>105313</td><td>105335</td><td>+</td><td>0</td><td>ATCGCATTTTTTAAAATATT</td><td>TGG</td><td>||||||||[||||||||||||]PAM</td><td>NA</td><td>Intergenic</td><td>NA</td><td>NA</td></tr>
<tr><td>T7683</td><td>GCATTTTTTAAAATATTTGGCGG</td><td>1000</td><td></td><td></td><td></td><td></td><td></td><td></td><td></td><td></td><td></td></tr>
<tr><td>Oligo fwd</td><td>CACCGCATTTTTTAAAATATTTGG</td><td></td><td></td><td></td><td></td><td></td><td></td><td></td><td></td><td></td><td></td></tr>
<tr><td>Oligo rev</td><td>AAACCCAAATATTTTAAAAAATGC</td><td></td><td></td><td></td><td></td><td></td><td></td><td></td><td></td><td></td><td></td></tr>
<tr><td>Chromosome</td><td>start</td><td>end</td><td>strand</td><td>MM</td><td>target_seq</td><td>PAM</td><td>alignment</td><td>distance</td><td>position</td><td>gene name</td><td>gene id</td></tr>
<tr><td>GU305914.1</td><td>105316</td><td>105338</td><td>+</td><td>0</td><td>GCATTTTTTAAAATATTTGG</td><td>CGG</td><td>||||||||[||||||||||||]PAM</td><td>NA</td><td>Intergenic</td><td>NA</td><td>NA</td></tr>
<tr><td>T7684</td><td>TTGGCGGTGATGTTACTTTTCGG</td><td>1000</td><td></td><td></td><td></td><td></td><td></td><td></td><td></td><td></td><td></td></tr>
<tr><td>Oligo fwd</td><td>CACCTTGGCGGTGATGTTACTTTT</td><td></td><td></td><td></td><td></td><td></td><td></td><td></td><td></td><td></td><td></td></tr>
<tr><td>Oligo rev</td><td>AAACAAAAGTAACATCACCGCCAA</td><td></td><td></td><td></td><td></td><td></td><td></td><td></td><td></td><td></td><td></td></tr>
<tr><td>Chromosome</td><td>start</td><td>end</td><td>strand</td><td>MM</td><td>target_seq</td><td>PAM</td><td>alignment</td><td>distance</td><td>position</td><td>gene name</td><td>gene id</td></tr>
<tr><td>GU305914.1</td><td>105332</td><td>105354</td><td>+</td><td>0</td><td>TTGGCGGTGATGTTACTTTT</td><td>CGG</td><td>||||||||[||||||||||||]PAM</td><td>NA</td><td>Intergenic</td><td>NA</td><td>NA</td></tr>
<tr><td>T7685</td><td>TGGCGGTGATGTTACTTTTCGGG</td><td>1000</td><td></td><td></td><td></td><td></td><td></td><td></td><td></td><td></td><td></td></tr>
<tr><td>Oligo fwd</td><td>CACCTGGCGGTGATGTTACTTTTC</td><td></td><td></td><td></td><td></td><td></td><td></td><td></td><td></td><td></td><td></td></tr>
<tr><td>Oligo rev</td><td>AAACGAAAAGTAACATCACCGCCA</td><td></td><td></td><td></td><td></td><td></td><td></td><td></td><td></td><td></td><td></td></tr>
<tr><td>Chromosome</td><td>start</td><td>end</td><td>strand</td><td>MM</td><td>target_seq</td><td>PAM</td><td>alignment</td><td>distance</td><td>position</td><td>gene name</td><td>gene id</td></tr>
<tr><td>GU305914.1</td><td>105333</td><td>105355</td><td>+</td><td>0</td><td>TGGCGGTGATGTTACTTTTC</td><td>GGG</td><td>||||||||[||||||||||||]PAM</td><td>NA</td><td>Intergenic</td><td>NA</td><td>NA</td></tr>
<tr><td>T7686</td><td>GGCGGTGATGTTACTTTTCGGGG</td><td>1000</td><td></td><td></td><td></td><td></td><td></td><td></td><td></td><td></td><td></td></tr>
<tr><td>Oligo fwd</td><td>CACCGGCGGTGATGTTACTTTTCG</td><td></td><td></td><td></td><td></td><td></td><td></td><td></td><td></td><td></td><td></td></tr>
<tr><td>Oligo rev</td><td>AAACCGAAAAGTAACATCACCGCC</td><td></td><td></td><td></td><td></td><td></td><td></td><td></td><td></td><td></td><td></td></tr>
<tr><td>Chromosome</td><td>start</td><td>end</td><td>strand</td><td>MM</td><td>target_seq</td><td>PAM</td><td>alignment</td><td>distance</td><td>position</td><td>gene name</td><td>gene id</td></tr>
<tr><td>GU305914.1</td><td>105334</td><td>105356</td><td>+</td><td>0</td><td>GGCGGTGATGTTACTTTTCG</td><td>GGG</td><td>||||||||[||||||||||||]PAM</td><td>NA</td><td>Intergenic</td><td>NA</td><td>NA</td></tr>
<tr><td>T7687</td><td>TGTTACTTTTCGGGGTGATGAGG</td><td>1000</td><td></td><td></td><td></td><td></td><td></td><td></td><td></td><td></td><td></td></tr>
<tr><td>Oligo fwd</td><td>CACCTGTTACTTTTCGGGGTGATG</td><td></td><td></td><td></td><td></td><td></td><td></td><td></td><td></td><td></td><td></td></tr>
<tr><td>Oligo rev</td><td>AAACCATCACCCCGAAAAGTAACA</td><td></td><td></td><td></td><td></td><td></td><td></td><td></td><td></td><td></td><td></td></tr>
<tr><td>Chromosome</td><td>start</td><td>end</td><td>strand</td><td>MM</td><td>target_seq</td><td>PAM</td><td>alignment</td><td>distance</td><td>position</td><td>gene name</td><td>gene id</td></tr>
<tr><td>GU305914.1</td><td>105342</td><td>105364</td><td>+</td><td>0</td><td>TGTTACTTTTCGGGGTGATG</td><td>AGG</td><td>||||||||[||||||||||||]PAM</td><td>NA</td><td>Intergenic</td><td>NA</td><td>NA</td></tr>
<tr><td>T7688</td><td>GGTGATGAGGTCTCTCCCCATGG</td><td>1000</td><td></td><td></td><td></td><td></td><td></td><td></td><td></td><td></td><td></td></tr>
<tr><td>Oligo fwd</td><td>CACCGGTGATGAGGTCTCTCCCCA</td><td></td><td></td><td></td><td></td><td></td><td></td><td></td><td></td><td></td><td></td></tr>
<tr><td>Oligo rev</td><td>AAACTGGGGAGAGACCTCATCACC</td><td></td><td></td><td></td><td></td><td></td><td></td><td></td><td></td><td></td><td></td></tr>
<tr><td>Chromosome</td><td>start</td><td>end</td><td>strand</td><td>MM</td><td>target_seq</td><td>PAM</td><td>alignment</td><td>distance</td><td>position</td><td>gene name</td><td>gene id</td></tr>
<tr><td>GU305914.1</td><td>105355</td><td>105377</td><td>+</td><td>0</td><td>GGTGATGAGGTCTCTCCCCA</td><td>TGG</td><td>||||||||[||||||||||||]PAM</td><td>NA</td><td>Intergenic</td><td>NA</td><td>NA</td></tr>
<tr><td>T7689</td><td>TGAGGTCTCTCCCCATGGATCGG</td><td>1000</td><td></td><td></td><td></td><td></td><td></td><td></td><td></td><td></td><td></td></tr>
<tr><td>Oligo fwd</td><td>CACCTGAGGTCTCTCCCCATGGAT</td><td></td><td></td><td></td><td></td><td></td><td></td><td></td><td></td><td></td><td></td></tr>
<tr><td>Oligo rev</td><td>AAACATCCATGGGGAGAGACCTCA</td><td></td><td></td><td></td><td></td><td></td><td></td><td></td><td></td><td></td><td></td></tr>
<tr><td>Chromosome</td><td>start</td><td>end</td><td>strand</td><td>MM</td><td>target_seq</td><td>PAM</td><td>alignment</td><td>distance</td><td>position</td><td>gene name</td><td>gene id</td></tr>
<tr><td>GU305914.1</td><td>105360</td><td>105382</td><td>+</td><td>0</td><td>TGAGGTCTCTCCCCATGGAT</td><td>CGG</td><td>||||||||[||||||||||||]PAM</td><td>NA</td><td>Intergenic</td><td>NA</td><td>NA</td></tr>
<tr><td>T7690</td><td>TGACACGAAAGCGACGAACTTGG</td><td>1000</td><td></td><td></td><td></td><td></td><td></td><td></td><td></td><td></td><td></td></tr>
<tr><td>Oligo fwd</td><td>CACCTGACACGAAAGCGACGAACT</td><td></td><td></td><td></td><td></td><td></td><td></td><td></td><td></td><td></td><td></td></tr>
<tr><td>Oligo rev</td><td>AAACAGTTCGTCGCTTTCGTGTCA</td><td></td><td></td><td></td><td></td><td></td><td></td><td></td><td></td><td></td><td></td></tr>
<tr><td>Chromosome</td><td>start</td><td>end</td><td>strand</td><td>MM</td><td>target_seq</td><td>PAM</td><td>alignment</td><td>distance</td><td>position</td><td>gene name</td><td>gene id</td></tr>
<tr><td>GU305914.1</td><td>105383</td><td>105405</td><td>+</td><td>0</td><td>TGACACGAAAGCGACGAACT</td><td>TGG</td><td>||||||||[||||||||||||]PAM</td><td>NA</td><td>Intergenic</td><td>NA</td><td>NA</td></tr>
<tr><td>T7691</td><td>AGCGACGAACTTGGTCCGCAAGG</td><td>1000</td><td></td><td></td><td></td><td></td><td></td><td></td><td></td><td></td><td></td></tr>
<tr><td>Oligo fwd</td><td>CACCAGCGACGAACTTGGTCCGCA</td><td></td><td></td><td></td><td></td><td></td><td></td><td></td><td></td><td></td><td></td></tr>
<tr><td>Oligo rev</td><td>AAACTGCGGACCAAGTTCGTCGCT</td><td></td><td></td><td></td><td></td><td></td><td></td><td></td><td></td><td></td><td></td></tr>
<tr><td>Chromosome</td><td>start</td><td>end</td><td>strand</td><td>MM</td><td>target_seq</td><td>PAM</td><td>alignment</td><td>distance</td><td>position</td><td>gene name</td><td>gene id</td></tr>
<tr><td>GU305914.1</td><td>105392</td><td>105414</td><td>+</td><td>0</td><td>AGCGACGAACTTGGTCCGCA</td><td>AGG</td><td>||||||||[||||||||||||]PAM</td><td>NA</td><td>Intergenic</td><td>NA</td><td>NA</td></tr>
<tr><td>T7692</td><td>AGGTACTCCGCTCACGATTACGG</td><td>1000</td><td></td><td></td><td></td><td></td><td></td><td></td><td></td><td></td><td></td></tr>
<tr><td>Oligo fwd</td><td>CACCAGGTACTCCGCTCACGATTA</td><td></td><td></td><td></td><td></td><td></td><td></td><td></td><td></td><td></td><td></td></tr>
<tr><td>Oligo rev</td><td>AAACTAATCGTGAGCGGAGTACCT</td><td></td><td></td><td></td><td></td><td></td><td></td><td></td><td></td><td></td><td></td></tr>
<tr><td>Chromosome</td><td>start</td><td>end</td><td>strand</td><td>MM</td><td>target_seq</td><td>PAM</td><td>alignment</td><td>distance</td><td>position</td><td>gene name</td><td>gene id</td></tr>
<tr><td>GU305914.1</td><td>105412</td><td>105434</td><td>+</td><td>0</td><td>AGGTACTCCGCTCACGATTA</td><td>CGG</td><td>||||||||[||||||||||||]PAM</td><td>NA</td><td>Intergenic</td><td>NA</td><td>NA</td></tr>
<tr><td>T7693</td><td>TTATTATTGTTATCTTTGATTGG</td><td>1000</td><td></td><td></td><td></td><td></td><td></td><td></td><td></td><td></td><td></td></tr>
<tr><td>Oligo fwd</td><td>CACCTTATTATTGTTATCTTTGAT</td><td></td><td></td><td></td><td></td><td></td><td></td><td></td><td></td><td></td><td></td></tr>
<tr><td>Oligo rev</td><td>AAACATCAAAGATAACAATAATAA</td><td></td><td></td><td></td><td></td><td></td><td></td><td></td><td></td><td></td><td></td></tr>
<tr><td>Chromosome</td><td>start</td><td>end</td><td>strand</td><td>MM</td><td>target_seq</td><td>PAM</td><td>alignment</td><td>distance</td><td>position</td><td>gene name</td><td>gene id</td></tr>
<tr><td>GU305914.1</td><td>105452</td><td>105474</td><td>+</td><td>0</td><td>TTATTATTGTTATCTTTGAT</td><td>TGG</td><td>||||||||[||||||||||||]PAM</td><td>NA</td><td>Intergenic</td><td>NA</td><td>NA</td></tr>
<tr><td>T7694</td><td>TATTATTGTTATCTTTGATTGGG</td><td>1000</td><td></td><td></td><td></td><td></td><td></td><td></td><td></td><td></td><td></td></tr>
<tr><td>Oligo fwd</td><td>CACCTATTATTGTTATCTTTGATT</td><td></td><td></td><td></td><td></td><td></td><td></td><td></td><td></td><td></td><td></td></tr>
<tr><td>Oligo rev</td><td>AAACAATCAAAGATAACAATAATA</td><td></td><td></td><td></td><td></td><td></td><td></td><td></td><td></td><td></td><td></td></tr>
<tr><td>Chromosome</td><td>start</td><td>end</td><td>strand</td><td>MM</td><td>target_seq</td><td>PAM</td><td>alignment</td><td>distance</td><td>position</td><td>gene name</td><td>gene id</td></tr>
<tr><td>GU305914.1</td><td>105453</td><td>105475</td><td>+</td><td>0</td><td>TATTATTGTTATCTTTGATT</td><td>GGG</td><td>||||||||[||||||||||||]PAM</td><td>NA</td><td>Intergenic</td><td>NA</td><td>NA</td></tr>
<tr><td>T7695</td><td>ATTATTGTTATCTTTGATTGGGG</td><td>1000</td><td></td><td></td><td></td><td></td><td></td><td></td><td></td><td></td><td></td></tr>
<tr><td>Oligo fwd</td><td>CACCATTATTGTTATCTTTGATTG</td><td></td><td></td><td></td><td></td><td></td><td></td><td></td><td></td><td></td><td></td></tr>
<tr><td>Oligo rev</td><td>AAACCAATCAAAGATAACAATAAT</td><td></td><td></td><td></td><td></td><td></td><td></td><td></td><td></td><td></td><td></td></tr>
<tr><td>Chromosome</td><td>start</td><td>end</td><td>strand</td><td>MM</td><td>target_seq</td><td>PAM</td><td>alignment</td><td>distance</td><td>position</td><td>gene name</td><td>gene id</td></tr>
<tr><td>GU305914.1</td><td>105454</td><td>105476</td><td>+</td><td>0</td><td>ATTATTGTTATCTTTGATTG</td><td>GGG</td><td>||||||||[||||||||||||]PAM</td><td>NA</td><td>Intergenic</td><td>NA</td><td>NA</td></tr>
<tr><td>T7696</td><td>GTTCACGTAATAGCCCGCGTAGG</td><td>1000</td><td></td><td></td><td></td><td></td><td></td><td></td><td></td><td></td><td></td></tr>
<tr><td>Oligo fwd</td><td>CACCGTTCACGTAATAGCCCGCGT</td><td></td><td></td><td></td><td></td><td></td><td></td><td></td><td></td><td></td><td></td></tr>
<tr><td>Oligo rev</td><td>AAACACGCGGGCTATTACGTGAAC</td><td></td><td></td><td></td><td></td><td></td><td></td><td></td><td></td><td></td><td></td></tr>
<tr><td>Chromosome</td><td>start</td><td>end</td><td>strand</td><td>MM</td><td>target_seq</td><td>PAM</td><td>alignment</td><td>distance</td><td>position</td><td>gene name</td><td>gene id</td></tr>
<tr><td>GU305914.1</td><td>105487</td><td>105509</td><td>+</td><td>0</td><td>GTTCACGTAATAGCCCGCGT</td><td>AGG</td><td>||||||||[||||||||||||]PAM</td><td>NA</td><td>Intergenic</td><td>NA</td><td>NA</td></tr>
</table>

<table>
<tr><td>T7697</td><td>TCCCTCGATGAGATACCAGATGG</td><td>1000</td><td></td><td></td><td></td><td></td><td></td><td></td><td></td><td></td><td></td></tr>
<tr><td>Oligo fwd</td><td>CACCTCCCTCGATGAGATACCAGA</td><td></td><td></td><td></td><td></td><td></td><td></td><td></td><td></td><td></td><td></td></tr>
<tr><td>Oligo rev</td><td>AAACTCTGGTATCTCATCGAGGGA</td><td></td><td></td><td></td><td></td><td></td><td></td><td></td><td></td><td></td><td></td></tr>
<tr><td>Chromosome</td><td>start</td><td>end</td><td>strand</td><td>MM</td><td>target_seq</td><td>PAM</td><td>alignment</td><td>distance</td><td>position</td><td>gene name</td><td>gene id</td></tr>
<tr><td>GU305914.1</td><td>105532</td><td>105554</td><td>+</td><td>0</td><td>TCCCTCGATGAGATACCAGA</td><td>TGG</td><td>||||||||[||||||||||||]PAM</td><td>NA</td><td>Intergenic</td><td>NA</td><td>NA</td></tr>
<tr><td>T7698</td><td>CCCTCGATGAGATACCAGATGGG</td><td>1000</td><td></td><td></td><td></td><td></td><td></td><td></td><td></td><td></td><td></td></tr>
<tr><td>Oligo fwd</td><td>CACCCCCTCGATGAGATACCAGAT</td><td></td><td></td><td></td><td></td><td></td><td></td><td></td><td></td><td></td><td></td></tr>
<tr><td>Oligo rev</td><td>AAACATCTGGTATCTCATCGAGGG</td><td></td><td></td><td></td><td></td><td></td><td></td><td></td><td></td><td></td><td></td></tr>
<tr><td>Chromosome</td><td>start</td><td>end</td><td>strand</td><td>MM</td><td>target_seq</td><td>PAM</td><td>alignment</td><td>distance</td><td>position</td><td>gene name</td><td>gene id</td></tr>
<tr><td>GU305914.1</td><td>105533</td><td>105555</td><td>+</td><td>0</td><td>CCCTCGATGAGATACCAGAT</td><td>GGG</td><td>||||||||[||||||||||||]PAM</td><td>NA</td><td>Intergenic</td><td>NA</td><td>NA</td></tr>
<tr><td>T7699</td><td>CAGATGGGAAAGACGTTGTCCGG</td><td>1000</td><td></td><td></td><td></td><td></td><td></td><td></td><td></td><td></td><td></td></tr>
<tr><td>Oligo fwd</td><td>CACCCAGATGGGAAAGACGTTGTC</td><td></td><td></td><td></td><td></td><td></td><td></td><td></td><td></td><td></td><td></td></tr>
<tr><td>Oligo rev</td><td>AAACGACAACGTCTTTCCCATCTG</td><td></td><td></td><td></td><td></td><td></td><td></td><td></td><td></td><td></td><td></td></tr>
<tr><td>Chromosome</td><td>start</td><td>end</td><td>strand</td><td>MM</td><td>target_seq</td><td>PAM</td><td>alignment</td><td>distance</td><td>position</td><td>gene name</td><td>gene id</td></tr>
<tr><td>GU305914.1</td><td>105548</td><td>105570</td><td>+</td><td>0</td><td>CAGATGGGAAAGACGTTGTC</td><td>CGG</td><td>||||||||[||||||||||||]PAM</td><td>NA</td><td>Intergenic</td><td>NA</td><td>NA</td></tr>
<tr><td>T7700</td><td>CCGGACGTCGATTTCTCCTTCGG</td><td>1000</td><td></td><td></td><td></td><td></td><td></td><td></td><td></td><td></td><td></td></tr>
<tr><td>Oligo fwd</td><td>CACCCCGGACGTCGATTTCTCCTT</td><td></td><td></td><td></td><td></td><td></td><td></td><td></td><td></td><td></td><td></td></tr>
<tr><td>Oligo rev</td><td>AAACAAGGAGAAATCGACGTCCGG</td><td></td><td></td><td></td><td></td><td></td><td></td><td></td><td></td><td></td><td></td></tr>
<tr><td>Chromosome</td><td>start</td><td>end</td><td>strand</td><td>MM</td><td>target_seq</td><td>PAM</td><td>alignment</td><td>distance</td><td>position</td><td>gene name</td><td>gene id</td></tr>
<tr><td>GU305914.1</td><td>105567</td><td>105589</td><td>+</td><td>0</td><td>CCGGACGTCGATTTCTCCTT</td><td>CGG</td><td>||||||||[||||||||||||]PAM</td><td>NA</td><td>Intergenic</td><td>NA</td><td>NA</td></tr>
<tr><td>T7701</td><td>CGTGAATGAGAGATAGTCTTTGG</td><td>1000</td><td></td><td></td><td></td><td></td><td></td><td></td><td></td><td></td><td></td></tr>
<tr><td>Oligo fwd</td><td>CACCCGTGAATGAGAGATAGTCTT</td><td></td><td></td><td></td><td></td><td></td><td></td><td></td><td></td><td></td><td></td></tr>
<tr><td>Oligo rev</td><td>AAACAAGACTATCTCTCATTCACG</td><td></td><td></td><td></td><td></td><td></td><td></td><td></td><td></td><td></td><td></td></tr>
<tr><td>Chromosome</td><td>start</td><td>end</td><td>strand</td><td>MM</td><td>target_seq</td><td>PAM</td><td>alignment</td><td>distance</td><td>position</td><td>gene name</td><td>gene id</td></tr>
<tr><td>GU305914.1</td><td>105622</td><td>105644</td><td>+</td><td>0</td><td>CGTGAATGAGAGATAGTCTT</td><td>TGG</td><td>||||||||[||||||||||||]PAM</td><td>NA</td><td>Intergenic</td><td>NA</td><td>NA</td></tr>
<tr><td>T7702</td><td>GTTCTCTTTTTCTTCGACGTCGG</td><td>1000</td><td></td><td></td><td></td><td></td><td></td><td></td><td></td><td></td><td></td></tr>
<tr><td>Oligo fwd</td><td>CACCGTTCTCTTTTTCTTCGACGT</td><td></td><td></td><td></td><td></td><td></td><td></td><td></td><td></td><td></td><td></td></tr>
<tr><td>Oligo rev</td><td>AAACACGTCGAAGAAAAAGAGAAC</td><td></td><td></td><td></td><td></td><td></td><td></td><td></td><td></td><td></td><td></td></tr>
<tr><td>Chromosome</td><td>start</td><td>end</td><td>strand</td><td>MM</td><td>target_seq</td><td>PAM</td><td>alignment</td><td>distance</td><td>position</td><td>gene name</td><td>gene id</td></tr>
<tr><td>GU305914.1</td><td>105652</td><td>105674</td><td>+</td><td>0</td><td>GTTCTCTTTTTCTTCGACGT</td><td>CGG</td><td>||||||||[||||||||||||]PAM</td><td>NA</td><td>Intergenic</td><td>NA</td><td>NA</td></tr>
<tr><td>T7703</td><td>TTCTCTTTTTCTTCGACGTCGGG</td><td>1000</td><td></td><td></td><td></td><td></td><td></td><td></td><td></td><td></td><td></td></tr>
<tr><td>Oligo fwd</td><td>CACCTTCTCTTTTTCTTCGACGTC</td><td></td><td></td><td></td><td></td><td></td><td></td><td></td><td></td><td></td><td></td></tr>
<tr><td>Oligo rev</td><td>AAACGACGTCGAAGAAAAAGAGAA</td><td></td><td></td><td></td><td></td><td></td><td></td><td></td><td></td><td></td><td></td></tr>
<tr><td>Chromosome</td><td>start</td><td>end</td><td>strand</td><td>MM</td><td>target_seq</td><td>PAM</td><td>alignment</td><td>distance</td><td>position</td><td>gene name</td><td>gene id</td></tr>
<tr><td>GU305914.1</td><td>105653</td><td>105675</td><td>+</td><td>0</td><td>TTCTCTTTTTCTTCGACGTC</td><td>GGG</td><td>||||||||[||||||||||||]PAM</td><td>NA</td><td>Intergenic</td><td>NA</td><td>NA</td></tr>
<tr><td>T7704</td><td>TCGACGTCGGGTTCTTTTTCTGG</td><td>1000</td><td></td><td></td><td></td><td></td><td></td><td></td><td></td><td></td><td></td></tr>
<tr><td>Oligo fwd</td><td>CACCTCGACGTCGGGTTCTTTTTC</td><td></td><td></td><td></td><td></td><td></td><td></td><td></td><td></td><td></td><td></td></tr>
<tr><td>Oligo rev</td><td>AAACGAAAAAGAACCCGACGTCGA</td><td></td><td></td><td></td><td></td><td></td><td></td><td></td><td></td><td></td><td></td></tr>
<tr><td>Chromosome</td><td>start</td><td>end</td><td>strand</td><td>MM</td><td>target_seq</td><td>PAM</td><td>alignment</td><td>distance</td><td>position</td><td>gene name</td><td>gene id</td></tr>
<tr><td>GU305914.1</td><td>105665</td><td>105687</td><td>+</td><td>0</td><td>TCGACGTCGGGTTCTTTTTC</td><td>TGG</td><td>||||||||[||||||||||||]PAM</td><td>NA</td><td>Intergenic</td><td>NA</td><td>NA</td></tr>
<tr><td>T7705</td><td>ACGTCGGGTTCTTTTTCTGGAGG</td><td>1000</td><td></td><td></td><td></td><td></td><td></td><td></td><td></td><td></td><td></td></tr>
<tr><td>Oligo fwd</td><td>CACCACGTCGGGTTCTTTTTCTGG</td><td></td><td></td><td></td><td></td><td></td><td></td><td></td><td></td><td></td><td></td></tr>
<tr><td>Oligo rev</td><td>AAACCCAGAAAAAGAACCCGACGT</td><td></td><td></td><td></td><td></td><td></td><td></td><td></td><td></td><td></td><td></td></tr>
<tr><td>Chromosome</td><td>start</td><td>end</td><td>strand</td><td>MM</td><td>target_seq</td><td>PAM</td><td>alignment</td><td>distance</td><td>position</td><td>gene name</td><td>gene id</td></tr>
<tr><td>GU305914.1</td><td>105668</td><td>105690</td><td>+</td><td>0</td><td>ACGTCGGGTTCTTTTTCTGG</td><td>AGG</td><td>||||||||[||||||||||||]PAM</td><td>NA</td><td>Intergenic</td><td>NA</td><td>NA</td></tr>
<tr><td>T7706</td><td>TCGGGTTCTTTTTCTGGAGGTGG</td><td>1000</td><td></td><td></td><td></td><td></td><td></td><td></td><td></td><td></td><td></td></tr>
<tr><td>Oligo fwd</td><td>CACCTCGGGTTCTTTTTCTGGAGG</td><td></td><td></td><td></td><td></td><td></td><td></td><td></td><td></td><td></td><td></td></tr>
<tr><td>Oligo rev</td><td>AAACCCTCCAGAAAAAGAACCCGA</td><td></td><td></td><td></td><td></td><td></td><td></td><td></td><td></td><td></td><td></td></tr>
<tr><td>Chromosome</td><td>start</td><td>end</td><td>strand</td><td>MM</td><td>target_seq</td><td>PAM</td><td>alignment</td><td>distance</td><td>position</td><td>gene name</td><td>gene id</td></tr>
<tr><td>GU305914.1</td><td>105671</td><td>105693</td><td>+</td><td>0</td><td>TCGGGTTCTTTTTCTGGAGG</td><td>TGG</td><td>||||||||[||||||||||||]PAM</td><td>NA</td><td>Intergenic</td><td>NA</td><td>NA</td></tr>
<tr><td>T7707</td><td>GGTTCTTTTTCTGGAGGTGGTGG</td><td>1000</td><td></td><td></td><td></td><td></td><td></td><td></td><td></td><td></td><td></td></tr>
<tr><td>Oligo fwd</td><td>CACCGGTTCTTTTTCTGGAGGTGG</td><td></td><td></td><td></td><td></td><td></td><td></td><td></td><td></td><td></td><td></td></tr>
<tr><td>Oligo rev</td><td>AAACCCACCTCCAGAAAAAGAACC</td><td></td><td></td><td></td><td></td><td></td><td></td><td></td><td></td><td></td><td></td></tr>
<tr><td>Chromosome</td><td>start</td><td>end</td><td>strand</td><td>MM</td><td>target_seq</td><td>PAM</td><td>alignment</td><td>distance</td><td>position</td><td>gene name</td><td>gene id</td></tr>
<tr><td>GU305914.1</td><td>105674</td><td>105696</td><td>+</td><td>0</td><td>GGTTCTTTTTCTGGAGGTGG</td><td>TGG</td><td>||||||||[||||||||||||]PAM</td><td>NA</td><td>Intergenic</td><td>NA</td><td>NA</td></tr>
<tr><td>T7708</td><td>CGAACCTATGTGTATTTCGCCGG</td><td>1000</td><td></td><td></td><td></td><td></td><td></td><td></td><td></td><td></td><td></td></tr>
<tr><td>Oligo fwd</td><td>CACCCGAACCTATGTGTATTTCGC</td><td></td><td></td><td></td><td></td><td></td><td></td><td></td><td></td><td></td><td></td></tr>
<tr><td>Oligo rev</td><td>AAACGCGAAATACACATAGGTTCG</td><td></td><td></td><td></td><td></td><td></td><td></td><td></td><td></td><td></td><td></td></tr>
<tr><td>Chromosome</td><td>start</td><td>end</td><td>strand</td><td>MM</td><td>target_seq</td><td>PAM</td><td>alignment</td><td>distance</td><td>position</td><td>gene name</td><td>gene id</td></tr>
<tr><td>GU305914.1</td><td>105712</td><td>105734</td><td>+</td><td>0</td><td>CGAACCTATGTGTATTTCGC</td><td>CGG</td><td>||||||||[||||||||||||]PAM</td><td>NA</td><td>Intergenic</td><td>NA</td><td>NA</td></tr>
<tr><td>T7709</td><td>GAACCTATGTGTATTTCGCCGGG</td><td>1000</td><td></td><td></td><td></td><td></td><td></td><td></td><td></td><td></td><td></td></tr>
<tr><td>Oligo fwd</td><td>CACCGAACCTATGTGTATTTCGCC</td><td></td><td></td><td></td><td></td><td></td><td></td><td></td><td></td><td></td><td></td></tr>
<tr><td>Oligo rev</td><td>AAACGGCGAAATACACATAGGTTC</td><td></td><td></td><td></td><td></td><td></td><td></td><td></td><td></td><td></td><td></td></tr>
<tr><td>Chromosome</td><td>start</td><td>end</td><td>strand</td><td>MM</td><td>target_seq</td><td>PAM</td><td>alignment</td><td>distance</td><td>position</td><td>gene name</td><td>gene id</td></tr>
<tr><td>GU305914.1</td><td>105713</td><td>105735</td><td>+</td><td>0</td><td>GAACCTATGTGTATTTCGCC</td><td>GGG</td><td>||||||||[||||||||||||]PAM</td><td>NA</td><td>Intergenic</td><td>NA</td><td>NA</td></tr>
<tr><td>T7710</td><td>CCTATGTGTATTTCGCCGGGAGG</td><td>1000</td><td></td><td></td><td></td><td></td><td></td><td></td><td></td><td></td><td></td></tr>
<tr><td>Oligo fwd</td><td>CACCCCTATGTGTATTTCGCCGGG</td><td></td><td></td><td></td><td></td><td></td><td></td><td></td><td></td><td></td><td></td></tr>
<tr><td>Oligo rev</td><td>AAACCCCGGCGAAATACACATAGG</td><td></td><td></td><td></td><td></td><td></td><td></td><td></td><td></td><td></td><td></td></tr>
<tr><td>Chromosome</td><td>start</td><td>end</td><td>strand</td><td>MM</td><td>target_seq</td><td>PAM</td><td>alignment</td><td>distance</td><td>position</td><td>gene name</td><td>gene id</td></tr>
<tr><td>GU305914.1</td><td>105716</td><td>105738</td><td>+</td><td>0</td><td>CCTATGTGTATTTCGCCGGG</td><td>AGG</td><td>||||||||[||||||||||||]PAM</td><td>NA</td><td>Intergenic</td><td>NA</td><td>NA</td></tr>
<tr><td>T7711</td><td>GTGTATTTCGCCGGGAGGTTCGG</td><td>1000</td><td></td><td></td><td></td><td></td><td></td><td></td><td></td><td></td><td></td></tr>
<tr><td>Oligo fwd</td><td>CACCGTGTATTTCGCCGGGAGGTT</td><td></td><td></td><td></td><td></td><td></td><td></td><td></td><td></td><td></td><td></td></tr>
<tr><td>Oligo rev</td><td>AAACAACCTCCCGGCGAAATACAC</td><td></td><td></td><td></td><td></td><td></td><td></td><td></td><td></td><td></td><td></td></tr>
<tr><td>Chromosome</td><td>start</td><td>end</td><td>strand</td><td>MM</td><td>target_seq</td><td>PAM</td><td>alignment</td><td>distance</td><td>position</td><td>gene name</td><td>gene id</td></tr>
<tr><td>GU305914.1</td><td>105721</td><td>105743</td><td>+</td><td>0</td><td>GTGTATTTCGCCGGGAGGTT</td><td>CGG</td><td>||||||||[||||||||||||]PAM</td><td>NA</td><td>Intergenic</td><td>NA</td><td>NA</td></tr>
</table>

| | | | | | | | | | | | |
|---|---|---|---|---|---|---|---|---|---|---|---|
| T7712 | TTCGCTCATCTCGATATAGCCGG | 1000 | | | | | | | | | |
| Oligo fwd | CACCTTCGCTCATCTCGATATAGC | | | | | | | | | | |
| Oligo rev | AAACGCTATATCGAGATGAGCGAA | | | | | | | | | | |
| Chromosome | start | end | strand | MM | target_seq | PAM | alignment | distance | position | gene name | gene id |
| GU305914.1 | 105784 | 105806 | + | 0 | TTCGCTCATCTCGATATAGC | CGG | \|\|\|\|\|\|\|\|[\|\|\|\|\|\|\|\|\|\|\|\|]PAM | NA | Intergenic | NA | NA |
| T7713 | CATCTCGATATAGCCGGTAACGG | 1000 | | | | | | | | | |
| Oligo fwd | CACCCATCTCGATATAGCCGGTAA | | | | | | | | | | |
| Oligo rev | AAACTTACCGGCTATATCGAGATG | | | | | | | | | | |
| Chromosome | start | end | strand | MM | target_seq | PAM | alignment | distance | position | gene name | gene id |
| GU305914.1 | 105790 | 105812 | + | 0 | CATCTCGATATAGCCGGTAA | CGG | \|\|\|\|\|\|\|\|[\|\|\|\|\|\|\|\|\|\|\|\|]PAM | NA | Intergenic | NA | NA |
| T7714 | TCGATATAGCCGGTAACGGCCGG | 1000 | | | | | | | | | |
| Oligo fwd | CACCTCGATATAGCCGGTAACGGC | | | | | | | | | | |
| Oligo rev | AAACGCCGTTACCGGCTATATCGA | | | | | | | | | | |
| Chromosome | start | end | strand | MM | target_seq | PAM | alignment | distance | position | gene name | gene id |
| GU305914.1 | 105794 | 105816 | + | 0 | TCGATATAGCCGGTAACGGC | CGG | \|\|\|\|\|\|\|\|[\|\|\|\|\|\|\|\|\|\|\|\|]PAM | NA | Intergenic | NA | NA |
| T7715 | CGATATAGCCGGTAACGGCCGGG | 1000 | | | | | | | | | |
| Oligo fwd | CACCCGATATAGCCGGTAACGGCC | | | | | | | | | | |
| Oligo rev | AAACGGCCGTTACCGGCTATATCG | | | | | | | | | | |
| Chromosome | start | end | strand | MM | target_seq | PAM | alignment | distance | position | gene name | gene id |
| GU305914.1 | 105795 | 105817 | + | 0 | CGATATAGCCGGTAACGGCC | GGG | \|\|\|\|\|\|\|\|[\|\|\|\|\|\|\|\|\|\|\|\|]PAM | NA | Intergenic | NA | NA |
| T7716 | TAGTGAGCTGTTCACTTTCGCGG | 1000 | | | | | | | | | |
| Oligo fwd | CACCTAGTGAGCTGTTCACTTTCG | | | | | | | | | | |
| Oligo rev | AAACCGAAAGTGAACAGCTCACTA | | | | | | | | | | |
| Chromosome | start | end | strand | MM | target_seq | PAM | alignment | distance | position | gene name | gene id |
| GU305914.1 | 105838 | 105860 | + | 0 | TAGTGAGCTGTTCACTTTCG | CGG | \|\|\|\|\|\|\|\|[\|\|\|\|\|\|\|\|\|\|\|\|]PAM | NA | Intergenic | NA | NA |
| T7717 | TGAGCTGTTCACTTTCGCGGCGG | 1000 | | | | | | | | | |
| Oligo fwd | CACCTGAGCTGTTCACTTTCGCGG | | | | | | | | | | |
| Oligo rev | AAACCCGCGAAAGTGAACAGCTCA | | | | | | | | | | |
| Chromosome | start | end | strand | MM | target_seq | PAM | alignment | distance | position | gene name | gene id |
| GU305914.1 | 105841 | 105863 | + | 0 | TGAGCTGTTCACTTTCGCGG | CGG | \|\|\|\|\|\|\|\|[\|\|\|\|\|\|\|\|\|\|\|\|]PAM | NA | Intergenic | NA | NA |
| T7718 | CACTTTCGCGGCGGATGCCGCGG | 1000 | | | | | | | | | |
| Oligo fwd | CACCCACTTTCGCGGCGGATGCCG | | | | | | | | | | |
| Oligo rev | AAACCGGCATCCGCCGCGAAAGTG | | | | | | | | | | |
| Chromosome | start | end | strand | MM | target_seq | PAM | alignment | distance | position | gene name | gene id |
| GU305914.1 | 105850 | 105872 | + | 0 | CACTTTCGCGGCGGATGCCG | CGG | \|\|\|\|\|\|\|\|[\|\|\|\|\|\|\|\|\|\|\|\|]PAM | NA | Intergenic | NA | NA |
| GU305914.1 | 93363 | 93385 | - | 4 | CGGTTTCGCGGGGGATGCGG | GGG | \|--\|\|\|\|\|[\|\|\|-\|\|\|\|\|\|-\|]PAM | NA | Intergenic | NA | NA |
| T7719 | GACGATCGACATGAGTAATAAGG | 1000 | | | | | | | | | |
| Oligo fwd | CACCGACGATCGACATGAGTAATA | | | | | | | | | | |
| Oligo rev | AAACTATTACTCATGTCGATCGTC | | | | | | | | | | |
| Chromosome | start | end | strand | MM | target_seq | PAM | alignment | distance | position | gene name | gene id |
| GU305914.1 | 105874 | 105896 | + | 0 | GACGATCGACATGAGTAATA | AGG | \|\|\|\|\|\|\|\|[\|\|\|\|\|\|\|\|\|\|\|\|]PAM | NA | Intergenic | NA | NA |
| T7720 | ACGATCGACATGAGTAATAAGGG | 1000 | | | | | | | | | |
| Oligo fwd | CACCACGATCGACATGAGTAATAA | | | | | | | | | | |
| Oligo rev | AAACTTATTACTCATGTCGATCGT | | | | | | | | | | |
| Chromosome | start | end | strand | MM | target_seq | PAM | alignment | distance | position | gene name | gene id |
| GU305914.1 | 105875 | 105897 | + | 0 | ACGATCGACATGAGTAATAA | GGG | \|\|\|\|\|\|\|\|[\|\|\|\|\|\|\|\|\|\|\|\|]PAM | NA | Intergenic | NA | NA |
| T7721 | CGATCGACATGAGTAATAAGGGG | 1000 | | | | | | | | | |
| Oligo fwd | CACCCGATCGACATGAGTAATAAG | | | | | | | | | | |
| Oligo rev | AAACCTTATTACTCATGTCGATCG | | | | | | | | | | |
| Chromosome | start | end | strand | MM | target_seq | PAM | alignment | distance | position | gene name | gene id |
| GU305914.1 | 105876 | 105898 | + | 0 | CGATCGACATGAGTAATAAG | GGG | \|\|\|\|\|\|\|\|[\|\|\|\|\|\|\|\|\|\|\|\|]PAM | NA | Intergenic | NA | NA |
| T7722 | ATGAGTAATAAGGGGTTCATCGG | 1000 | | | | | | | | | |
| Oligo fwd | CACCATGAGTAATAAGGGGTTCAT | | | | | | | | | | |
| Oligo rev | AAACATGAACCCCTTATTACTCAT | | | | | | | | | | |
| Chromosome | start | end | strand | MM | target_seq | PAM | alignment | distance | position | gene name | gene id |
| GU305914.1 | 105884 | 105906 | + | 0 | ATGAGTAATAAGGGGTTCAT | CGG | \|\|\|\|\|\|\|\|[\|\|\|\|\|\|\|\|\|\|\|\|]PAM | NA | Intergenic | NA | NA |
| T7723 | TGAGTAATAAGGGGTTCATCGGG | 1000 | | | | | | | | | |
| Oligo fwd | CACCTGAGTAATAAGGGGTTCATC | | | | | | | | | | |
| Oligo rev | AAACGATGAACCCCTTATTACTCA | | | | | | | | | | |
| Chromosome | start | end | strand | MM | target_seq | PAM | alignment | distance | position | gene name | gene id |
| GU305914.1 | 105885 | 105907 | + | 0 | TGAGTAATAAGGGGTTCATC | GGG | \|\|\|\|\|\|\|\|[\|\|\|\|\|\|\|\|\|\|\|\|]PAM | NA | Intergenic | NA | NA |
| T7724 | GTAATAAGGGGTTCATCGGGAGG | 1000 | | | | | | | | | |
| Oligo fwd | CACCGTAATAAGGGGTTCATCGGG | | | | | | | | | | |
| Oligo rev | AAACCCCGATGAACCCCTTATTAC | | | | | | | | | | |
| Chromosome | start | end | strand | MM | target_seq | PAM | alignment | distance | position | gene name | gene id |
| GU305914.1 | 105888 | 105910 | + | 0 | GTAATAAGGGGTTCATCGGG | AGG | \|\|\|\|\|\|\|\|[\|\|\|\|\|\|\|\|\|\|\|\|]PAM | NA | Intergenic | NA | NA |
| T7725 | TAATAAGGGGTTCATCGGGAGGG | 1000 | | | | | | | | | |
| Oligo fwd | CACCTAATAAGGGGTTCATCGGGA | | | | | | | | | | |
| Oligo rev | AAACTCCCGATGAACCCCTTATTA | | | | | | | | | | |
| Chromosome | start | end | strand | MM | target_seq | PAM | alignment | distance | position | gene name | gene id |
| GU305914.1 | 105889 | 105911 | + | 0 | TAATAAGGGGTTCATCGGGA | GGG | \|\|\|\|\|\|\|\|[\|\|\|\|\|\|\|\|\|\|\|\|]PAM | NA | Intergenic | NA | NA |
| T7726 | GATCTCCGCGTTGAATACAGCGG | 1000 | | | | | | | | | |
| Oligo fwd | CACCGATCTCCGCGTTGAATACAG | | | | | | | | | | |
| Oligo rev | AAACCTGTATTCAACGCGGAGATC | | | | | | | | | | |

| Chromosome | start | end | strand | MM | target_seq | PAM | alignment | distance | position | gene name | gene id |
|---|---|---|---|---|---|---|---|---|---|---|---|
| GU305914.1 | 105914 | 105936 | + | 0 | GATCTCCGCGTTGAATACAG | CGG | \|\|\|\|\|\|\|\|[\|\|\|\|\|\|\|\|\|\|\|\|]PAM | NA | Intergenic | NA | NA |
| T7727 | AATACAGCGGACATTAATCGCGG | 1000 | | | | | | | | | |
| Oligo fwd | CACCAATACAGCGGACATTAATCG | | | | | | | | | | |
| Oligo rev | AAACCGATTAATGTCCGCTGTATT | | | | | | | | | | |
| Chromosome | start | end | strand | MM | target_seq | PAM | alignment | distance | position | gene name | gene id |
| GU305914.1 | 105927 | 105949 | + | 0 | AATACAGCGGACATTAATCG | CGG | \|\|\|\|\|\|\|\|[\|\|\|\|\|\|\|\|\|\|\|\|]PAM | NA | Intergenic | NA | NA |
| T7728 | GGCTCTCTTTGTGTTTTAATAGG | 1000 | | | | | | | | | |
| Oligo fwd | CACCGGCTCTCTTTGTGTTTTAAT | | | | | | | | | | |
| Oligo rev | AAACATTAAAACACAAAGAGAGCC | | | | | | | | | | |
| Chromosome | start | end | strand | MM | target_seq | PAM | alignment | distance | position | gene name | gene id |
| GU305914.1 | 105948 | 105970 | + | 0 | GGCTCTCTTTGTGTTTTAAT | AGG | \|\|\|\|\|\|\|\|[\|\|\|\|\|\|\|\|\|\|\|\|]PAM | NA | Intergenic | NA | NA |
| T7729 | GTTTTAATAGGCAGTCTCTGCGG | 1000 | | | | | | | | | |
| Oligo fwd | CACCGTTTTAATAGGCAGTCTCTG | | | | | | | | | | |
| Oligo rev | AAACCAGAGACTGCCTATTAAAAC | | | | | | | | | | |
| Chromosome | start | end | strand | MM | target_seq | PAM | alignment | distance | position | gene name | gene id |
| GU305914.1 | 105960 | 105982 | + | 0 | GTTTTAATAGGCAGTCTCTG | CGG | \|\|\|\|\|\|\|\|[\|\|\|\|\|\|\|\|\|\|\|\|]PAM | NA | Intergenic | NA | NA |
| T7730 | GATAGTCAACACCGTGATCATGG | 1000 | | | | | | | | | |
| Oligo fwd | CACCGATAGTCAACACCGTGATCA | | | | | | | | | | |
| Oligo rev | AAACTGATCACGGTGTTGACTATC | | | | | | | | | | |
| Chromosome | start | end | strand | MM | target_seq | PAM | alignment | distance | position | gene name | gene id |
| GU305914.1 | 106004 | 106026 | + | 0 | GATAGTCAACACCGTGATCA | TGG | \|\|\|\|\|\|\|\|[\|\|\|\|\|\|\|\|\|\|\|\|]PAM | NA | Intergenic | NA | NA |
| T7731 | AGTCAACACCGTGATCATGGTGG | 1000 | | | | | | | | | |
| Oligo fwd | CACCAGTCAACACCGTGATCATGG | | | | | | | | | | |
| Oligo rev | AAACCCATGATCACGGTGTTGACT | | | | | | | | | | |
| Chromosome | start | end | strand | MM | target_seq | PAM | alignment | distance | position | gene name | gene id |
| GU305914.1 | 106007 | 106029 | + | 0 | AGTCAACACCGTGATCATGG | TGG | \|\|\|\|\|\|\|\|[\|\|\|\|\|\|\|\|\|\|\|\|]PAM | NA | Intergenic | NA | NA |
| T7732 | GATCATGGTGGCGATGATGTTGG | 1000 | | | | | | | | | |
| Oligo fwd | CACCGATCATGGTGGCGATGATGT | | | | | | | | | | |
| Oligo rev | AAACACATCATCGCCACCATGATC | | | | | | | | | | |
| Chromosome | start | end | strand | MM | target_seq | PAM | alignment | distance | position | gene name | gene id |
| GU305914.1 | 106019 | 106041 | + | 0 | GATCATGGTGGCGATGATGT | TGG | \|\|\|\|\|\|\|\|[\|\|\|\|\|\|\|\|\|\|\|\|]PAM | NA | Intergenic | NA | NA |
| GU305914.1 | 44666 | 44688 | - | 4 | GATGGTGGTGGGGATGATGG | TGG | \|\|\|--\|\|\|[\|\|\|-\|\|\|\|\|\|\|-]PAM | NA | Intergenic | NA | NA |
| T7733 | GTGTTGTACCGAGCCATCGTCGG | 1000 | | | | | | | | | |
| Oligo fwd | CACCGTGTTGTACCGAGCCATCGT | | | | | | | | | | |
| Oligo rev | AAACACGATGGCTCGGTACAACAC | | | | | | | | | | |
| Chromosome | start | end | strand | MM | target_seq | PAM | alignment | distance | position | gene name | gene id |
| GU305914.1 | 106041 | 106063 | + | 0 | GTGTTGTACCGAGCCATCGT | CGG | \|\|\|\|\|\|\|\|[\|\|\|\|\|\|\|\|\|\|\|\|]PAM | NA | Intergenic | NA | NA |
| T7734 | TGACATCGTCCTGAGAAGCACGG | 1000 | | | | | | | | | |
| Oligo fwd | CACCTGACATCGTCCTGAGAAGCA | | | | | | | | | | |
| Oligo rev | AAACTGCTTCTCAGGACGATGTCA | | | | | | | | | | |
| Chromosome | start | end | strand | MM | target_seq | PAM | alignment | distance | position | gene name | gene id |
| GU305914.1 | 106073 | 106095 | + | 0 | TGACATCGTCCTGAGAAGCA | CGG | \|\|\|\|\|\|\|\|[\|\|\|\|\|\|\|\|\|\|\|\|]PAM | NA | Intergenic | NA | NA |
| T7735 | CATCGTCCTGAGAAGCACGGAGG | 1000 | | | | | | | | | |
| Oligo fwd | CACCCATCGTCCTGAGAAGCACGG | | | | | | | | | | |
| Oligo rev | AAACCCGTGCTTCTCAGGACGATG | | | | | | | | | | |
| Chromosome | start | end | strand | MM | target_seq | PAM | alignment | distance | position | gene name | gene id |
| GU305914.1 | 106076 | 106098 | + | 0 | CATCGTCCTGAGAAGCACGG | AGG | \|\|\|\|\|\|\|\|[\|\|\|\|\|\|\|\|\|\|\|\|]PAM | NA | Intergenic | NA | NA |
| T7736 | GAAGCACGGAGGCTTTCAGACGG | 1000 | | | | | | | | | |
| Oligo fwd | CACCGAAGCACGGAGGCTTTCAGA | | | | | | | | | | |
| Oligo rev | AAACTCTGAAAGCCTCCGTGCTTC | | | | | | | | | | |
| Chromosome | start | end | strand | MM | target_seq | PAM | alignment | distance | position | gene name | gene id |
| GU305914.1 | 106087 | 106109 | + | 0 | GAAGCACGGAGGCTTTCAGA | CGG | \|\|\|\|\|\|\|\|[\|\|\|\|\|\|\|\|\|\|\|\|]PAM | NA | Intergenic | NA | NA |
| T7737 | CGGAGGCTTTCAGACGGATGTGG | 1000 | | | | | | | | | |
| Oligo fwd | CACCCGGAGGCTTTCAGACGGATG | | | | | | | | | | |
| Oligo rev | AAACCATCCGTCTGAAAGCCTCCG | | | | | | | | | | |
| Chromosome | start | end | strand | MM | target_seq | PAM | alignment | distance | position | gene name | gene id |
| GU305914.1 | 106093 | 106115 | + | 0 | CGGAGGCTTTCAGACGGATG | TGG | \|\|\|\|\|\|\|\|[\|\|\|\|\|\|\|\|\|\|\|\|]PAM | NA | Intergenic | NA | NA |
| T7738 | GGAGGCTTTCAGACGGATGTGGG | 1000 | | | | | | | | | |
| Oligo fwd | CACCGGAGGCTTTCAGACGGATGT | | | | | | | | | | |
| Oligo rev | AAACACATCCGTCTGAAAGCCTCC | | | | | | | | | | |
| Chromosome | start | end | strand | MM | target_seq | PAM | alignment | distance | position | gene name | gene id |
| GU305914.1 | 106094 | 106116 | + | 0 | GGAGGCTTTCAGACGGATGT | GGG | \|\|\|\|\|\|\|\|[\|\|\|\|\|\|\|\|\|\|\|\|]PAM | NA | Intergenic | NA | NA |
| T7739 | GCTTTCAGACGGATGTGGGCCGG | 1000 | | | | | | | | | |
| Oligo fwd | CACCGCTTTCAGACGGATGTGGGC | | | | | | | | | | |
| Oligo rev | AAACGCCCACATCCGTCTGAAAGC | | | | | | | | | | |
| Chromosome | start | end | strand | MM | target_seq | PAM | alignment | distance | position | gene name | gene id |
| GU305914.1 | 106098 | 106120 | + | 0 | GCTTTCAGACGGATGTGGGC | CGG | \|\|\|\|\|\|\|\|[\|\|\|\|\|\|\|\|\|\|\|\|]PAM | NA | Intergenic | NA | NA |
| T7740 | AAGAGCAACGCCACCGCGTGCGG | 1000 | | | | | | | | | |
| Oligo fwd | CACCAAGAGCAACGCCACCGCGTG | | | | | | | | | | |
| Oligo rev | AAACCACGCGGTGGCGTTGCTCTT | | | | | | | | | | |
| Chromosome | start | end | strand | MM | target_seq | PAM | alignment | distance | position | gene name | gene id |
| GU305914.1 | 106122 | 106144 | + | 0 | AAGAGCAACGCCACCGCGTG | CGG | \|\|\|\|\|\|\|\|[\|\|\|\|\|\|\|\|\|\|\|\|]PAM | NA | Intergenic | NA | NA |
| T7741 | AGAGCAACGCCACCGCGTGCGGG | 1000 | | | | | | | | | |

<table>
<tr><td>Oligo fwd</td><td>CACCAGAGCAACGCCACCGCGTGC</td><td></td><td></td><td></td><td></td><td></td><td></td><td></td><td></td><td></td><td></td></tr>
<tr><td>Oligo rev</td><td>AAACGCACGCGGTGGCGTTGCTCT</td><td></td><td></td><td></td><td></td><td></td><td></td><td></td><td></td><td></td><td></td></tr>
<tr><td>Chromosome</td><td>start</td><td>end</td><td>strand</td><td>MM</td><td>target_seq</td><td>PAM</td><td>alignment</td><td>distance</td><td>position</td><td>gene name</td><td>gene id</td></tr>
<tr><td>GU305914.1</td><td>106123</td><td>106145</td><td>+</td><td>0</td><td>AGAGCAACGCCACCGCGTGC</td><td>GGG</td><td>||||||||[||||||||||||]PAM</td><td>NA</td><td>Intergenic</td><td>NA</td><td>NA</td></tr>
<tr><td>T7742</td><td>GAGCAACGCCACCGCGTGCGGGG</td><td>1000</td><td></td><td></td><td></td><td></td><td></td><td></td><td></td><td></td><td></td></tr>
<tr><td>Oligo fwd</td><td>CACCGAGCAACGCCACCGCGTGCG</td><td></td><td></td><td></td><td></td><td></td><td></td><td></td><td></td><td></td><td></td></tr>
<tr><td>Oligo rev</td><td>AAACCGCACGCGGTGGCGTTGCTC</td><td></td><td></td><td></td><td></td><td></td><td></td><td></td><td></td><td></td><td></td></tr>
<tr><td>Chromosome</td><td>start</td><td>end</td><td>strand</td><td>MM</td><td>target_seq</td><td>PAM</td><td>alignment</td><td>distance</td><td>position</td><td>gene name</td><td>gene id</td></tr>
<tr><td>GU305914.1</td><td>106124</td><td>106146</td><td>+</td><td>0</td><td>GAGCAACGCCACCGCGTGCG</td><td>GGG</td><td>||||||||[||||||||||||]PAM</td><td>NA</td><td>Intergenic</td><td>NA</td><td>NA</td></tr>
<tr><td>T7743</td><td>AGCAACGCCACCGCGTGCGGGGG</td><td>1000</td><td></td><td></td><td></td><td></td><td></td><td></td><td></td><td></td><td></td></tr>
<tr><td>Oligo fwd</td><td>CACCAGCAACGCCACCGCGTGCGG</td><td></td><td></td><td></td><td></td><td></td><td></td><td></td><td></td><td></td><td></td></tr>
<tr><td>Oligo rev</td><td>AAACCCGCACGCGGTGGCGTTGCT</td><td></td><td></td><td></td><td></td><td></td><td></td><td></td><td></td><td></td><td></td></tr>
<tr><td>Chromosome</td><td>start</td><td>end</td><td>strand</td><td>MM</td><td>target_seq</td><td>PAM</td><td>alignment</td><td>distance</td><td>position</td><td>gene name</td><td>gene id</td></tr>
<tr><td>GU305914.1</td><td>106125</td><td>106147</td><td>+</td><td>0</td><td>AGCAACGCCACCGCGTGCGG</td><td>GGG</td><td>||||||||[||||||||||||]PAM</td><td>NA</td><td>Intergenic</td><td>NA</td><td>NA</td></tr>
<tr><td>T7744</td><td>GCAACGCCACCGCGTGCGGGGGG</td><td>1000</td><td></td><td></td><td></td><td></td><td></td><td></td><td></td><td></td><td></td></tr>
<tr><td>Oligo fwd</td><td>CACCGCAACGCCACCGCGTGCGGG</td><td></td><td></td><td></td><td></td><td></td><td></td><td></td><td></td><td></td><td></td></tr>
<tr><td>Oligo rev</td><td>AAACCCCGCACGCGGTGGCGTTGC</td><td></td><td></td><td></td><td></td><td></td><td></td><td></td><td></td><td></td><td></td></tr>
<tr><td>Chromosome</td><td>start</td><td>end</td><td>strand</td><td>MM</td><td>target_seq</td><td>PAM</td><td>alignment</td><td>distance</td><td>position</td><td>gene name</td><td>gene id</td></tr>
<tr><td>GU305914.1</td><td>106126</td><td>106148</td><td>+</td><td>0</td><td>GCAACGCCACCGCGTGCGGG</td><td>GGG</td><td>||||||||[||||||||||||]PAM</td><td>NA</td><td>Intergenic</td><td>NA</td><td>NA</td></tr>
<tr><td>T7745</td><td>TCCTTCCTTATCTTTTTTGCCGG</td><td>1000</td><td></td><td></td><td></td><td></td><td></td><td></td><td></td><td></td><td></td></tr>
<tr><td>Oligo fwd</td><td>CACCTCCTTCCTTATCTTTTTTGC</td><td></td><td></td><td></td><td></td><td></td><td></td><td></td><td></td><td></td><td></td></tr>
<tr><td>Oligo rev</td><td>AAACGCAAAAAAGATAAGGAAGGA</td><td></td><td></td><td></td><td></td><td></td><td></td><td></td><td></td><td></td><td></td></tr>
<tr><td>Chromosome</td><td>start</td><td>end</td><td>strand</td><td>MM</td><td>target_seq</td><td>PAM</td><td>alignment</td><td>distance</td><td>position</td><td>gene name</td><td>gene id</td></tr>
<tr><td>GU305914.1</td><td>106198</td><td>106220</td><td>+</td><td>0</td><td>TCCTTCCTTATCTTTTTTGC</td><td>CGG</td><td>||||||||[||||||||||||]PAM</td><td>NA</td><td>Intergenic</td><td>NA</td><td>NA</td></tr>
<tr><td>T7746</td><td>ATCTTTTTTGCCGGCACAACCGG</td><td>1000</td><td></td><td></td><td></td><td></td><td></td><td></td><td></td><td></td><td></td></tr>
<tr><td>Oligo fwd</td><td>CACCATCTTTTTTGCCGGCACAAC</td><td></td><td></td><td></td><td></td><td></td><td></td><td></td><td></td><td></td><td></td></tr>
<tr><td>Oligo rev</td><td>AAACGTTGTGCCGGCAAAAAAGAT</td><td></td><td></td><td></td><td></td><td></td><td></td><td></td><td></td><td></td><td></td></tr>
<tr><td>Chromosome</td><td>start</td><td>end</td><td>strand</td><td>MM</td><td>target_seq</td><td>PAM</td><td>alignment</td><td>distance</td><td>position</td><td>gene name</td><td>gene id</td></tr>
<tr><td>GU305914.1</td><td>106207</td><td>106229</td><td>+</td><td>0</td><td>ATCTTTTTTGCCGGCACAAC</td><td>CGG</td><td>||||||||[||||||||||||]PAM</td><td>NA</td><td>Intergenic</td><td>NA</td><td>NA</td></tr>
<tr><td>T7747</td><td>GGTACAGCCCTAGCGCCATGAGG</td><td>1000</td><td></td><td></td><td></td><td></td><td></td><td></td><td></td><td></td><td></td></tr>
<tr><td>Oligo fwd</td><td>CACCGGTACAGCCCTAGCGCCATG</td><td></td><td></td><td></td><td></td><td></td><td></td><td></td><td></td><td></td><td></td></tr>
<tr><td>Oligo rev</td><td>AAACCATGGCGCTAGGGCTGTACC</td><td></td><td></td><td></td><td></td><td></td><td></td><td></td><td></td><td></td><td></td></tr>
<tr><td>Chromosome</td><td>start</td><td>end</td><td>strand</td><td>MM</td><td>target_seq</td><td>PAM</td><td>alignment</td><td>distance</td><td>position</td><td>gene name</td><td>gene id</td></tr>
<tr><td>GU305914.1</td><td>106228</td><td>106250</td><td>+</td><td>0</td><td>GGTACAGCCCTAGCGCCATG</td><td>AGG</td><td>||||||||[||||||||||||]PAM</td><td>NA</td><td>Intergenic</td><td>NA</td><td>NA</td></tr>
<tr><td>T7748</td><td>GAGGATCGCTATGATCACCAAGG</td><td>1000</td><td></td><td></td><td></td><td></td><td></td><td></td><td></td><td></td><td></td></tr>
<tr><td>Oligo fwd</td><td>CACCGAGGATCGCTATGATCACCA</td><td></td><td></td><td></td><td></td><td></td><td></td><td></td><td></td><td></td><td></td></tr>
<tr><td>Oligo rev</td><td>AAACTGGTGATCATAGCGATCCTC</td><td></td><td></td><td></td><td></td><td></td><td></td><td></td><td></td><td></td><td></td></tr>
<tr><td>Chromosome</td><td>start</td><td>end</td><td>strand</td><td>MM</td><td>target_seq</td><td>PAM</td><td>alignment</td><td>distance</td><td>position</td><td>gene name</td><td>gene id</td></tr>
<tr><td>GU305914.1</td><td>106247</td><td>106269</td><td>+</td><td>0</td><td>GAGGATCGCTATGATCACCA</td><td>AGG</td><td>||||||||[||||||||||||]PAM</td><td>NA</td><td>Intergenic</td><td>NA</td><td>NA</td></tr>
<tr><td>T7749</td><td>GATCACCAAGGCGACGACCACGG</td><td>1000</td><td></td><td></td><td></td><td></td><td></td><td></td><td></td><td></td><td></td></tr>
<tr><td>Oligo fwd</td><td>CACCGATCACCAAGGCGACGACCA</td><td></td><td></td><td></td><td></td><td></td><td></td><td></td><td></td><td></td><td></td></tr>
<tr><td>Oligo rev</td><td>AAACTGGTCGTCGCCTTGGTGATC</td><td></td><td></td><td></td><td></td><td></td><td></td><td></td><td></td><td></td><td></td></tr>
<tr><td>Chromosome</td><td>start</td><td>end</td><td>strand</td><td>MM</td><td>target_seq</td><td>PAM</td><td>alignment</td><td>distance</td><td>position</td><td>gene name</td><td>gene id</td></tr>
<tr><td>GU305914.1</td><td>106259</td><td>106281</td><td>+</td><td>0</td><td>GATCACCAAGGCGACGACCA</td><td>CGG</td><td>||||||||[||||||||||||]PAM</td><td>NA</td><td>Intergenic</td><td>NA</td><td>NA</td></tr>
<tr><td>T7750</td><td>CAAGGCGACGACCACGGAGTAGG</td><td>1000</td><td></td><td></td><td></td><td></td><td></td><td></td><td></td><td></td><td></td></tr>
<tr><td>Oligo fwd</td><td>CACCCAAGGCGACGACCACGGAGT</td><td></td><td></td><td></td><td></td><td></td><td></td><td></td><td></td><td></td><td></td></tr>
<tr><td>Oligo rev</td><td>AAACACTCCGTGGTCGTCGCCTTG</td><td></td><td></td><td></td><td></td><td></td><td></td><td></td><td></td><td></td><td></td></tr>
<tr><td>Chromosome</td><td>start</td><td>end</td><td>strand</td><td>MM</td><td>target_seq</td><td>PAM</td><td>alignment</td><td>distance</td><td>position</td><td>gene name</td><td>gene id</td></tr>
<tr><td>GU305914.1</td><td>106265</td><td>106287</td><td>+</td><td>0</td><td>CAAGGCGACGACCACGGAGT</td><td>AGG</td><td>||||||||[||||||||||||]PAM</td><td>NA</td><td>Intergenic</td><td>NA</td><td>NA</td></tr>
<tr><td>T7751</td><td>AAGGCGACGACCACGGAGTAGGG</td><td>1000</td><td></td><td></td><td></td><td></td><td></td><td></td><td></td><td></td><td></td></tr>
<tr><td>Oligo fwd</td><td>CACCAAGGCGACGACCACGGAGTA</td><td></td><td></td><td></td><td></td><td></td><td></td><td></td><td></td><td></td><td></td></tr>
<tr><td>Oligo rev</td><td>AAACTACTCCGTGGTCGTCGCCTT</td><td></td><td></td><td></td><td></td><td></td><td></td><td></td><td></td><td></td><td></td></tr>
<tr><td>Chromosome</td><td>start</td><td>end</td><td>strand</td><td>MM</td><td>target_seq</td><td>PAM</td><td>alignment</td><td>distance</td><td>position</td><td>gene name</td><td>gene id</td></tr>
<tr><td>GU305914.1</td><td>106266</td><td>106288</td><td>+</td><td>0</td><td>AAGGCGACGACCACGGAGTA</td><td>GGG</td><td>||||||||[||||||||||||]PAM</td><td>NA</td><td>Intergenic</td><td>NA</td><td>NA</td></tr>
<tr><td>T7752</td><td>ACCACGGAGTAGGGCATCGATGG</td><td>1000</td><td></td><td></td><td></td><td></td><td></td><td></td><td></td><td></td><td></td></tr>
<tr><td>Oligo fwd</td><td>CACCACCACGGAGTAGGGCATCGA</td><td></td><td></td><td></td><td></td><td></td><td></td><td></td><td></td><td></td><td></td></tr>
<tr><td>Oligo rev</td><td>AAACTCGATGCCCTACTCCGTGGT</td><td></td><td></td><td></td><td></td><td></td><td></td><td></td><td></td><td></td><td></td></tr>
<tr><td>Chromosome</td><td>start</td><td>end</td><td>strand</td><td>MM</td><td>target_seq</td><td>PAM</td><td>alignment</td><td>distance</td><td>position</td><td>gene name</td><td>gene id</td></tr>
<tr><td>GU305914.1</td><td>106275</td><td>106297</td><td>+</td><td>0</td><td>ACCACGGAGTAGGGCATCGA</td><td>TGG</td><td>||||||||[||||||||||||]PAM</td><td>NA</td><td>Intergenic</td><td>NA</td><td>NA</td></tr>
<tr><td>T7753</td><td>CGGAGTAGGGCATCGATGGTCGG</td><td>1000</td><td></td><td></td><td></td><td></td><td></td><td></td><td></td><td></td><td></td></tr>
<tr><td>Oligo fwd</td><td>CACCCGGAGTAGGGCATCGATGGT</td><td></td><td></td><td></td><td></td><td></td><td></td><td></td><td></td><td></td><td></td></tr>
<tr><td>Oligo rev</td><td>AAACACCATCGATGCCCTACTCCG</td><td></td><td></td><td></td><td></td><td></td><td></td><td></td><td></td><td></td><td></td></tr>
<tr><td>Chromosome</td><td>start</td><td>end</td><td>strand</td><td>MM</td><td>target_seq</td><td>PAM</td><td>alignment</td><td>distance</td><td>position</td><td>gene name</td><td>gene id</td></tr>
<tr><td>GU305914.1</td><td>106279</td><td>106301</td><td>+</td><td>0</td><td>CGGAGTAGGGCATCGATGGT</td><td>CGG</td><td>||||||||[||||||||||||]PAM</td><td>NA</td><td>Intergenic</td><td>NA</td><td>NA</td></tr>
<tr><td>T7754</td><td>TCGATGGTCGGACGTTCACGAGG</td><td>1000</td><td></td><td></td><td></td><td></td><td></td><td></td><td></td><td></td><td></td></tr>
<tr><td>Oligo fwd</td><td>CACCTCGATGGTCGGACGTTCACG</td><td></td><td></td><td></td><td></td><td></td><td></td><td></td><td></td><td></td><td></td></tr>
<tr><td>Oligo rev</td><td>AAACCGTGAACGTCCGACCATCGA</td><td></td><td></td><td></td><td></td><td></td><td></td><td></td><td></td><td></td><td></td></tr>
<tr><td>Chromosome</td><td>start</td><td>end</td><td>strand</td><td>MM</td><td>target_seq</td><td>PAM</td><td>alignment</td><td>distance</td><td>position</td><td>gene name</td><td>gene id</td></tr>
<tr><td>GU305914.1</td><td>106291</td><td>106313</td><td>+</td><td>0</td><td>TCGATGGTCGGACGTTCACG</td><td>AGG</td><td>||||||||[||||||||||||]PAM</td><td>NA</td><td>Intergenic</td><td>NA</td><td>NA</td></tr>
<tr><td>T7755</td><td>TCGGACGTTCACGAGGATGTCGG</td><td>1000</td><td></td><td></td><td></td><td></td><td></td><td></td><td></td><td></td><td></td></tr>
<tr><td>Oligo fwd</td><td>CACCTCGGACGTTCACGAGGATGT</td><td></td><td></td><td></td><td></td><td></td><td></td><td></td><td></td><td></td><td></td></tr>
<tr><td>Oligo rev</td><td>AAACACATCCTCGTGAACGTCCGA</td><td></td><td></td><td></td><td></td><td></td><td></td><td></td><td></td><td></td><td></td></tr>
<tr><td>Chromosome</td><td>start</td><td>end</td><td>strand</td><td>MM</td><td>target_seq</td><td>PAM</td><td>alignment</td><td>distance</td><td>position</td><td>gene name</td><td>gene id</td></tr>
<tr><td>GU305914.1</td><td>106298</td><td>106320</td><td>+</td><td>0</td><td>TCGGACGTTCACGAGGATGT</td><td>CGG</td><td>||||||||[||||||||||||]PAM</td><td>NA</td><td>Intergenic</td><td>NA</td><td>NA</td></tr>
</table>

<table>
<tr><td>T7756</td><td>CTCGAACACCGTCCCGTTCGCGG</td><td>1000</td><td></td><td></td><td></td><td></td><td></td><td></td><td></td><td></td><td></td></tr>
<tr><td>Oligo fwd</td><td>CACCCTCGAACACCGTCCCGTTCG</td><td></td><td></td><td></td><td></td><td></td><td></td><td></td><td></td><td></td><td></td></tr>
<tr><td>Oligo rev</td><td>AAACCGAACGGGACGGTGTTCGAG</td><td></td><td></td><td></td><td></td><td></td><td></td><td></td><td></td><td></td><td></td></tr>
<tr><td>Chromosome</td><td>start</td><td>end</td><td>strand</td><td>MM</td><td>target_seq</td><td>PAM</td><td>alignment</td><td>distance</td><td>position</td><td>gene name</td><td>gene id</td></tr>
<tr><td>GU305914.1</td><td>106325</td><td>106347</td><td>+</td><td>0</td><td>CTCGAACACCGTCCCGTTCG</td><td>CGG</td><td>||||||||[||||||||||||]PAM</td><td>NA</td><td>Intergenic</td><td>NA</td><td>NA</td></tr>
<tr><td>T7757</td><td>TCCCGTTCGCGGTTAGTAATAGG</td><td>1000</td><td></td><td></td><td></td><td></td><td></td><td></td><td></td><td></td><td></td></tr>
<tr><td>Oligo fwd</td><td>CACCTCCCGTTCGCGGTTAGTAAT</td><td></td><td></td><td></td><td></td><td></td><td></td><td></td><td></td><td></td><td></td></tr>
<tr><td>Oligo rev</td><td>AAACATTACTAACCGCGAACGGGA</td><td></td><td></td><td></td><td></td><td></td><td></td><td></td><td></td><td></td><td></td></tr>
<tr><td>Chromosome</td><td>start</td><td>end</td><td>strand</td><td>MM</td><td>target_seq</td><td>PAM</td><td>alignment</td><td>distance</td><td>position</td><td>gene name</td><td>gene id</td></tr>
<tr><td>GU305914.1</td><td>106336</td><td>106358</td><td>+</td><td>0</td><td>TCCCGTTCGCGGTTAGTAAT</td><td>AGG</td><td>||||||||[||||||||||||]PAM</td><td>NA</td><td>Intergenic</td><td>NA</td><td>NA</td></tr>
<tr><td>T7758</td><td>TCGCGGTTAGTAATAGGTAGTGG</td><td>1000</td><td></td><td></td><td></td><td></td><td></td><td></td><td></td><td></td><td></td></tr>
<tr><td>Oligo fwd</td><td>CACCTCGCGGTTAGTAATAGGTAG</td><td></td><td></td><td></td><td></td><td></td><td></td><td></td><td></td><td></td><td></td></tr>
<tr><td>Oligo rev</td><td>AAACCTACCTATTACTAACCGCGA</td><td></td><td></td><td></td><td></td><td></td><td></td><td></td><td></td><td></td><td></td></tr>
<tr><td>Chromosome</td><td>start</td><td>end</td><td>strand</td><td>MM</td><td>target_seq</td><td>PAM</td><td>alignment</td><td>distance</td><td>position</td><td>gene name</td><td>gene id</td></tr>
<tr><td>GU305914.1</td><td>106342</td><td>106364</td><td>+</td><td>0</td><td>TCGCGGTTAGTAATAGGTAG</td><td>TGG</td><td>||||||||[||||||||||||]PAM</td><td>NA</td><td>Intergenic</td><td>NA</td><td>NA</td></tr>
<tr><td>T7759</td><td>AGTAATAGGTAGTGGACCCTAGG</td><td>1000</td><td></td><td></td><td></td><td></td><td></td><td></td><td></td><td></td><td></td></tr>
<tr><td>Oligo fwd</td><td>CACCAGTAATAGGTAGTGGACCCT</td><td></td><td></td><td></td><td></td><td></td><td></td><td></td><td></td><td></td><td></td></tr>
<tr><td>Oligo rev</td><td>AAACAGGGTCCACTACCTATTACT</td><td></td><td></td><td></td><td></td><td></td><td></td><td></td><td></td><td></td><td></td></tr>
<tr><td>Chromosome</td><td>start</td><td>end</td><td>strand</td><td>MM</td><td>target_seq</td><td>PAM</td><td>alignment</td><td>distance</td><td>position</td><td>gene name</td><td>gene id</td></tr>
<tr><td>GU305914.1</td><td>106350</td><td>106372</td><td>+</td><td>0</td><td>AGTAATAGGTAGTGGACCCT</td><td>AGG</td><td>||||||||[||||||||||||]PAM</td><td>NA</td><td>Intergenic</td><td>NA</td><td>NA</td></tr>
<tr><td>T7760</td><td>GGACCCTAGGCGTCGCCACCAGG</td><td>1000</td><td></td><td></td><td></td><td></td><td></td><td></td><td></td><td></td><td></td></tr>
<tr><td>Oligo fwd</td><td>CACCGGACCCTAGGCGTCGCCACC</td><td></td><td></td><td></td><td></td><td></td><td></td><td></td><td></td><td></td><td></td></tr>
<tr><td>Oligo rev</td><td>AAACGGTGGCGACGCCTAGGGTCC</td><td></td><td></td><td></td><td></td><td></td><td></td><td></td><td></td><td></td><td></td></tr>
<tr><td>Chromosome</td><td>start</td><td>end</td><td>strand</td><td>MM</td><td>target_seq</td><td>PAM</td><td>alignment</td><td>distance</td><td>position</td><td>gene name</td><td>gene id</td></tr>
<tr><td>GU305914.1</td><td>106363</td><td>106385</td><td>+</td><td>0</td><td>GGACCCTAGGCGTCGCCACC</td><td>AGG</td><td>||||||||[||||||||||||]PAM</td><td>NA</td><td>Intergenic</td><td>NA</td><td>NA</td></tr>
<tr><td>T7761</td><td>CGCCACCAGGATGTCGTTGTCGG</td><td>1000</td><td></td><td></td><td></td><td></td><td></td><td></td><td></td><td></td><td></td></tr>
<tr><td>Oligo fwd</td><td>CACCCGCCACCAGGATGTCGTTGT</td><td></td><td></td><td></td><td></td><td></td><td></td><td></td><td></td><td></td><td></td></tr>
<tr><td>Oligo rev</td><td>AAACACAACGACATCCTGGTGGCG</td><td></td><td></td><td></td><td></td><td></td><td></td><td></td><td></td><td></td><td></td></tr>
<tr><td>Chromosome</td><td>start</td><td>end</td><td>strand</td><td>MM</td><td>target_seq</td><td>PAM</td><td>alignment</td><td>distance</td><td>position</td><td>gene name</td><td>gene id</td></tr>
<tr><td>GU305914.1</td><td>106376</td><td>106398</td><td>+</td><td>0</td><td>CGCCACCAGGATGTCGTTGT</td><td>CGG</td><td>||||||||[||||||||||||]PAM</td><td>NA</td><td>Intergenic</td><td>NA</td><td>NA</td></tr>
<tr><td>GU305914.1</td><td>107642</td><td>107664</td><td>+</td><td>3</td><td>CACCACCAGGAGGTCGTGGT</td><td>CGG</td><td>|-||||||[|||-|||||-||]PAM</td><td>NA</td><td>Intergenic</td><td>NA</td><td>NA</td></tr>
<tr><td>T7762</td><td>GCCACCAGGATGTCGTTGTCGGG</td><td>1000</td><td></td><td></td><td></td><td></td><td></td><td></td><td></td><td></td><td></td></tr>
<tr><td>Oligo fwd</td><td>CACCGCCACCAGGATGTCGTTGTC</td><td></td><td></td><td></td><td></td><td></td><td></td><td></td><td></td><td></td><td></td></tr>
<tr><td>Oligo rev</td><td>AAACGACAACGACATCCTGGTGGC</td><td></td><td></td><td></td><td></td><td></td><td></td><td></td><td></td><td></td><td></td></tr>
<tr><td>Chromosome</td><td>start</td><td>end</td><td>strand</td><td>MM</td><td>target_seq</td><td>PAM</td><td>alignment</td><td>distance</td><td>position</td><td>gene name</td><td>gene id</td></tr>
<tr><td>GU305914.1</td><td>106377</td><td>106399</td><td>+</td><td>0</td><td>GCCACCAGGATGTCGTTGTC</td><td>GGG</td><td>||||||||[||||||||||||]PAM</td><td>NA</td><td>Intergenic</td><td>NA</td><td>NA</td></tr>
<tr><td>GU305914.1</td><td>107643</td><td>107665</td><td>+</td><td>3</td><td>ACCACCAGGAGGTCGTGGTC</td><td>GGG</td><td>-|||||||[||-|||||-|||]PAM</td><td>NA</td><td>Intergenic</td><td>NA</td><td>NA</td></tr>
<tr><td>T7763</td><td>CCACCAGGATGTCGTTGTCGGGG</td><td>1000</td><td></td><td></td><td></td><td></td><td></td><td></td><td></td><td></td><td></td></tr>
<tr><td>Oligo fwd</td><td>CACCCCACCAGGATGTCGTTGTCG</td><td></td><td></td><td></td><td></td><td></td><td></td><td></td><td></td><td></td><td></td></tr>
<tr><td>Oligo rev</td><td>AAACCGACAACGACATCCTGGTGG</td><td></td><td></td><td></td><td></td><td></td><td></td><td></td><td></td><td></td><td></td></tr>
<tr><td>Chromosome</td><td>start</td><td>end</td><td>strand</td><td>MM</td><td>target_seq</td><td>PAM</td><td>alignment</td><td>distance</td><td>position</td><td>gene name</td><td>gene id</td></tr>
<tr><td>GU305914.1</td><td>106378</td><td>106400</td><td>+</td><td>0</td><td>CCACCAGGATGTCGTTGTCG</td><td>GGG</td><td>||||||||[||||||||||||]PAM</td><td>NA</td><td>Intergenic</td><td>NA</td><td>NA</td></tr>
<tr><td>T7764</td><td>TGTCGTTGTCGGGGTCCGTCAGG</td><td>1000</td><td></td><td></td><td></td><td></td><td></td><td></td><td></td><td></td><td></td></tr>
<tr><td>Oligo fwd</td><td>CACCTGTCGTTGTCGGGGTCCGTC</td><td></td><td></td><td></td><td></td><td></td><td></td><td></td><td></td><td></td><td></td></tr>
<tr><td>Oligo rev</td><td>AAACGACGGACCCCGACAACGACA</td><td></td><td></td><td></td><td></td><td></td><td></td><td></td><td></td><td></td><td></td></tr>
<tr><td>Chromosome</td><td>start</td><td>end</td><td>strand</td><td>MM</td><td>target_seq</td><td>PAM</td><td>alignment</td><td>distance</td><td>position</td><td>gene name</td><td>gene id</td></tr>
<tr><td>GU305914.1</td><td>106387</td><td>106409</td><td>+</td><td>0</td><td>TGTCGTTGTCGGGGTCCGTC</td><td>AGG</td><td>||||||||[||||||||||||]PAM</td><td>NA</td><td>Intergenic</td><td>NA</td><td>NA</td></tr>
<tr><td>T7765</td><td>CGGGGTCCGTCAGGTATTTCAGG</td><td>1000</td><td></td><td></td><td></td><td></td><td></td><td></td><td></td><td></td><td></td></tr>
<tr><td>Oligo fwd</td><td>CACCCGGGGTCCGTCAGGTATTTC</td><td></td><td></td><td></td><td></td><td></td><td></td><td></td><td></td><td></td><td></td></tr>
<tr><td>Oligo rev</td><td>AAACGAAATACCTGACGGACCCCG</td><td></td><td></td><td></td><td></td><td></td><td></td><td></td><td></td><td></td><td></td></tr>
<tr><td>Chromosome</td><td>start</td><td>end</td><td>strand</td><td>MM</td><td>target_seq</td><td>PAM</td><td>alignment</td><td>distance</td><td>position</td><td>gene name</td><td>gene id</td></tr>
<tr><td>GU305914.1</td><td>106396</td><td>106418</td><td>+</td><td>0</td><td>CGGGGTCCGTCAGGTATTTC</td><td>AGG</td><td>||||||||[||||||||||||]PAM</td><td>NA</td><td>Intergenic</td><td>NA</td><td>NA</td></tr>
<tr><td>T7766</td><td>TATTTCAGGACCTCTATGCTCGG</td><td>1000</td><td></td><td></td><td></td><td></td><td></td><td></td><td></td><td></td><td></td></tr>
<tr><td>Oligo fwd</td><td>CACCTATTTCAGGACCTCTATGCT</td><td></td><td></td><td></td><td></td><td></td><td></td><td></td><td></td><td></td><td></td></tr>
<tr><td>Oligo rev</td><td>AAACAGCATAGAGGTCCTGAAATA</td><td></td><td></td><td></td><td></td><td></td><td></td><td></td><td></td><td></td><td></td></tr>
<tr><td>Chromosome</td><td>start</td><td>end</td><td>strand</td><td>MM</td><td>target_seq</td><td>PAM</td><td>alignment</td><td>distance</td><td>position</td><td>gene name</td><td>gene id</td></tr>
<tr><td>GU305914.1</td><td>106410</td><td>106432</td><td>+</td><td>0</td><td>TATTTCAGGACCTCTATGCT</td><td>CGG</td><td>||||||||[||||||||||||]PAM</td><td>NA</td><td>Intergenic</td><td>NA</td><td>NA</td></tr>
<tr><td>T7767</td><td>ATTTCAGGACCTCTATGCTCGGG</td><td>1000</td><td></td><td></td><td></td><td></td><td></td><td></td><td></td><td></td><td></td></tr>
<tr><td>Oligo fwd</td><td>CACCATTTCAGGACCTCTATGCTC</td><td></td><td></td><td></td><td></td><td></td><td></td><td></td><td></td><td></td><td></td></tr>
<tr><td>Oligo rev</td><td>AAACGAGCATAGAGGTCCTGAAAT</td><td></td><td></td><td></td><td></td><td></td><td></td><td></td><td></td><td></td><td></td></tr>
<tr><td>Chromosome</td><td>start</td><td>end</td><td>strand</td><td>MM</td><td>target_seq</td><td>PAM</td><td>alignment</td><td>distance</td><td>position</td><td>gene name</td><td>gene id</td></tr>
<tr><td>GU305914.1</td><td>106411</td><td>106433</td><td>+</td><td>0</td><td>ATTTCAGGACCTCTATGCTC</td><td>GGG</td><td>||||||||[||||||||||||]PAM</td><td>NA</td><td>Intergenic</td><td>NA</td><td>NA</td></tr>
<tr><td>T7768</td><td>TATGCTCGGGATGTGCATGATGG</td><td>1000</td><td></td><td></td><td></td><td></td><td></td><td></td><td></td><td></td><td></td></tr>
<tr><td>Oligo fwd</td><td>CACCTATGCTCGGGATGTGCATGA</td><td></td><td></td><td></td><td></td><td></td><td></td><td></td><td></td><td></td><td></td></tr>
<tr><td>Oligo rev</td><td>AAACTCATGCACATCCCGAGCATA</td><td></td><td></td><td></td><td></td><td></td><td></td><td></td><td></td><td></td><td></td></tr>
<tr><td>Chromosome</td><td>start</td><td>end</td><td>strand</td><td>MM</td><td>target_seq</td><td>PAM</td><td>alignment</td><td>distance</td><td>position</td><td>gene name</td><td>gene id</td></tr>
<tr><td>GU305914.1</td><td>106424</td><td>106446</td><td>+</td><td>0</td><td>TATGCTCGGGATGTGCATGA</td><td>TGG</td><td>||||||||[||||||||||||]PAM</td><td>NA</td><td>Intergenic</td><td>NA</td><td>NA</td></tr>
<tr><td>T7769</td><td>GCTCGGGATGTGCATGATGGAGG</td><td>1000</td><td></td><td></td><td></td><td></td><td></td><td></td><td></td><td></td><td></td></tr>
<tr><td>Oligo fwd</td><td>CACCGCTCGGGATGTGCATGATGG</td><td></td><td></td><td></td><td></td><td></td><td></td><td></td><td></td><td></td><td></td></tr>
<tr><td>Oligo rev</td><td>AAACCCATCATGCACATCCCGAGC</td><td></td><td></td><td></td><td></td><td></td><td></td><td></td><td></td><td></td><td></td></tr>
<tr><td>Chromosome</td><td>start</td><td>end</td><td>strand</td><td>MM</td><td>target_seq</td><td>PAM</td><td>alignment</td><td>distance</td><td>position</td><td>gene name</td><td>gene id</td></tr>
<tr><td>GU305914.1</td><td>106427</td><td>106449</td><td>+</td><td>0</td><td>GCTCGGGATGTGCATGATGG</td><td>AGG</td><td>||||||||[||||||||||||]PAM</td><td>NA</td><td>Intergenic</td><td>NA</td><td>NA</td></tr>
<tr><td>T7770</td><td>CCACCTCGTCGTACTCGACCAGG</td><td>1000</td><td></td><td></td><td></td><td></td><td></td><td></td><td></td><td></td><td></td></tr>
<tr><td>Oligo fwd</td><td>CACCCCACCTCGTCGTACTCGACC</td><td></td><td></td><td></td><td></td><td></td><td></td><td></td><td></td><td></td><td></td></tr>
<tr><td>Oligo rev</td><td>AAACGGTCGAGTACGACGAGGTGG</td><td></td><td></td><td></td><td></td><td></td><td></td><td></td><td></td><td></td><td></td></tr>
</table>

<table>
<tr><td>Chromosome</td><td>start</td><td>end</td><td>strand</td><td>MM</td><td>target_seq</td><td>PAM</td><td>alignment</td><td>distance</td><td>position</td><td>gene name</td><td>gene id</td></tr>
<tr><td>GU305914.1</td><td>106459</td><td>106481</td><td>+</td><td>0</td><td>CCACCTCGTCGTACTCGACC</td><td>AGG</td><td>||||||||[||||||||||||]PAM</td><td>NA</td><td>Intergenic</td><td>NA</td><td>NA</td></tr>
<tr><td></td><td></td><td></td><td></td><td></td><td></td><td></td><td></td><td></td><td></td><td></td><td></td></tr>
<tr><td>T7771</td><td>CACCTCGTCGTACTCGACCAGGG</td><td>1000</td><td></td><td></td><td></td><td></td><td></td><td></td><td></td><td></td><td></td></tr>
<tr><td>Oligo fwd</td><td>CACCCACCTCGTCGTACTCGACCA</td><td></td><td></td><td></td><td></td><td></td><td></td><td></td><td></td><td></td><td></td></tr>
<tr><td>Oligo rev</td><td>AAACTGGTCGAGTACGACGAGGTG</td><td></td><td></td><td></td><td></td><td></td><td></td><td></td><td></td><td></td><td></td></tr>
<tr><td>Chromosome</td><td>start</td><td>end</td><td>strand</td><td>MM</td><td>target_seq</td><td>PAM</td><td>alignment</td><td>distance</td><td>position</td><td>gene name</td><td>gene id</td></tr>
<tr><td>GU305914.1</td><td>106460</td><td>106482</td><td>+</td><td>0</td><td>CACCTCGTCGTACTCGACCA</td><td>GGG</td><td>||||||||[||||||||||||]PAM</td><td>NA</td><td>Intergenic</td><td>NA</td><td>NA</td></tr>
<tr><td>GU305914.1</td><td>45179</td><td>45201</td><td>+</td><td>4</td><td>CGACTCGTCGGACTCGTCCA</td><td>CGG</td><td>|--|||||[||-|||||-|||]PAM</td><td>NA</td><td>Intergenic</td><td>NA</td><td>NA</td></tr>
<tr><td>GU305914.1</td><td>108579</td><td>108601</td><td>-</td><td>4</td><td>CGCGTCGTCGTCCTCGAACA</td><td>CGG</td><td>|-|-||||[|||-|||||-||]PAM</td><td>NA</td><td>Intergenic</td><td>NA</td><td>NA</td></tr>
<tr><td></td><td></td><td></td><td></td><td></td><td></td><td></td><td></td><td></td><td></td><td></td><td></td></tr>
<tr><td>T7772</td><td>GTCGTACTCGACCAGGGTGCTGG</td><td>1000</td><td></td><td></td><td></td><td></td><td></td><td></td><td></td><td></td><td></td></tr>
<tr><td>Oligo fwd</td><td>CACCGTCGTACTCGACCAGGGTGC</td><td></td><td></td><td></td><td></td><td></td><td></td><td></td><td></td><td></td><td></td></tr>
<tr><td>Oligo rev</td><td>AAACGCACCCTGGTCGAGTACGAC</td><td></td><td></td><td></td><td></td><td></td><td></td><td></td><td></td><td></td><td></td></tr>
<tr><td>Chromosome</td><td>start</td><td>end</td><td>strand</td><td>MM</td><td>target_seq</td><td>PAM</td><td>alignment</td><td>distance</td><td>position</td><td>gene name</td><td>gene id</td></tr>
<tr><td>GU305914.1</td><td>106466</td><td>106488</td><td>+</td><td>0</td><td>GTCGTACTCGACCAGGGTGC</td><td>TGG</td><td>||||||||[||||||||||||]PAM</td><td>NA</td><td>Intergenic</td><td>NA</td><td>NA</td></tr>
<tr><td></td><td></td><td></td><td></td><td></td><td></td><td></td><td></td><td></td><td></td><td></td><td></td></tr>
<tr><td>T7773</td><td>GGCGCAGAACTCGCACCGTTCGG</td><td>1000</td><td></td><td></td><td></td><td></td><td></td><td></td><td></td><td></td><td></td></tr>
<tr><td>Oligo fwd</td><td>CACCGGCGCAGAACTCGCACCGTT</td><td></td><td></td><td></td><td></td><td></td><td></td><td></td><td></td><td></td><td></td></tr>
<tr><td>Oligo rev</td><td>AAACAACGGTGCGAGTTCTGCGCC</td><td></td><td></td><td></td><td></td><td></td><td></td><td></td><td></td><td></td><td></td></tr>
<tr><td>Chromosome</td><td>start</td><td>end</td><td>strand</td><td>MM</td><td>target_seq</td><td>PAM</td><td>alignment</td><td>distance</td><td>position</td><td>gene name</td><td>gene id</td></tr>
<tr><td>GU305914.1</td><td>106487</td><td>106509</td><td>+</td><td>0</td><td>GGCGCAGAACTCGCACCGTT</td><td>CGG</td><td>||||||||[||||||||||||]PAM</td><td>NA</td><td>Intergenic</td><td>NA</td><td>NA</td></tr>
<tr><td></td><td></td><td></td><td></td><td></td><td></td><td></td><td></td><td></td><td></td><td></td><td></td></tr>
<tr><td>T7774</td><td>CGATCACCGATCCGCGTTCGCGG</td><td>1000</td><td></td><td></td><td></td><td></td><td></td><td></td><td></td><td></td><td></td></tr>
<tr><td>Oligo fwd</td><td>CACCCGATCACCGATCCGCGTTCG</td><td></td><td></td><td></td><td></td><td></td><td></td><td></td><td></td><td></td><td></td></tr>
<tr><td>Oligo rev</td><td>AAACCGAACGCGGATCGGTGATCG</td><td></td><td></td><td></td><td></td><td></td><td></td><td></td><td></td><td></td><td></td></tr>
<tr><td>Chromosome</td><td>start</td><td>end</td><td>strand</td><td>MM</td><td>target_seq</td><td>PAM</td><td>alignment</td><td>distance</td><td>position</td><td>gene name</td><td>gene id</td></tr>
<tr><td>GU305914.1</td><td>106531</td><td>106553</td><td>+</td><td>0</td><td>CGATCACCGATCCGCGTTCG</td><td>CGG</td><td>||||||||[||||||||||||]PAM</td><td>NA</td><td>Intergenic</td><td>NA</td><td>NA</td></tr>
<tr><td></td><td></td><td></td><td></td><td></td><td></td><td></td><td></td><td></td><td></td><td></td><td></td></tr>
<tr><td>T7775</td><td>GTTCGCGGTAGCTCCTCGACAGG</td><td>1000</td><td></td><td></td><td></td><td></td><td></td><td></td><td></td><td></td><td></td></tr>
<tr><td>Oligo fwd</td><td>CACCGTTCGCGGTAGCTCCTCGAC</td><td></td><td></td><td></td><td></td><td></td><td></td><td></td><td></td><td></td><td></td></tr>
<tr><td>Oligo rev</td><td>AAACGTCGAGGAGCTACCGCGAAC</td><td></td><td></td><td></td><td></td><td></td><td></td><td></td><td></td><td></td><td></td></tr>
<tr><td>Chromosome</td><td>start</td><td>end</td><td>strand</td><td>MM</td><td>target_seq</td><td>PAM</td><td>alignment</td><td>distance</td><td>position</td><td>gene name</td><td>gene id</td></tr>
<tr><td>GU305914.1</td><td>106546</td><td>106568</td><td>+</td><td>0</td><td>GTTCGCGGTAGCTCCTCGAC</td><td>AGG</td><td>||||||||[||||||||||||]PAM</td><td>NA</td><td>Intergenic</td><td>NA</td><td>NA</td></tr>
<tr><td></td><td></td><td></td><td></td><td></td><td></td><td></td><td></td><td></td><td></td><td></td><td></td></tr>
<tr><td>T7776</td><td>GTAGCTCCTCGACAGGACGCAGG</td><td>1000</td><td></td><td></td><td></td><td></td><td></td><td></td><td></td><td></td><td></td></tr>
<tr><td>Oligo fwd</td><td>CACCGTAGCTCCTCGACAGGACGC</td><td></td><td></td><td></td><td></td><td></td><td></td><td></td><td></td><td></td><td></td></tr>
<tr><td>Oligo rev</td><td>AAACGCGTCCTGTCGAGGAGCTAC</td><td></td><td></td><td></td><td></td><td></td><td></td><td></td><td></td><td></td><td></td></tr>
<tr><td>Chromosome</td><td>start</td><td>end</td><td>strand</td><td>MM</td><td>target_seq</td><td>PAM</td><td>alignment</td><td>distance</td><td>position</td><td>gene name</td><td>gene id</td></tr>
<tr><td>GU305914.1</td><td>106553</td><td>106575</td><td>+</td><td>0</td><td>GTAGCTCCTCGACAGGACGC</td><td>AGG</td><td>||||||||[||||||||||||]PAM</td><td>NA</td><td>Intergenic</td><td>NA</td><td>NA</td></tr>
<tr><td></td><td></td><td></td><td></td><td></td><td></td><td></td><td></td><td></td><td></td><td></td><td></td></tr>
<tr><td>T7777</td><td>TAGCTCCTCGACAGGACGCAGGG</td><td>1000</td><td></td><td></td><td></td><td></td><td></td><td></td><td></td><td></td><td></td></tr>
<tr><td>Oligo fwd</td><td>CACCTAGCTCCTCGACAGGACGCA</td><td></td><td></td><td></td><td></td><td></td><td></td><td></td><td></td><td></td><td></td></tr>
<tr><td>Oligo rev</td><td>AAACTGCGTCCTGTCGAGGAGCTA</td><td></td><td></td><td></td><td></td><td></td><td></td><td></td><td></td><td></td><td></td></tr>
<tr><td>Chromosome</td><td>start</td><td>end</td><td>strand</td><td>MM</td><td>target_seq</td><td>PAM</td><td>alignment</td><td>distance</td><td>position</td><td>gene name</td><td>gene id</td></tr>
<tr><td>GU305914.1</td><td>106554</td><td>106576</td><td>+</td><td>0</td><td>TAGCTCCTCGACAGGACGCA</td><td>GGG</td><td>||||||||[||||||||||||]PAM</td><td>NA</td><td>Intergenic</td><td>NA</td><td>NA</td></tr>
<tr><td></td><td></td><td></td><td></td><td></td><td></td><td></td><td></td><td></td><td></td><td></td><td></td></tr>
<tr><td>T7778</td><td>AGCTCCTCGACAGGACGCAGGGG</td><td>1000</td><td></td><td></td><td></td><td></td><td></td><td></td><td></td><td></td><td></td></tr>
<tr><td>Oligo fwd</td><td>CACCAGCTCCTCGACAGGACGCAG</td><td></td><td></td><td></td><td></td><td></td><td></td><td></td><td></td><td></td><td></td></tr>
<tr><td>Oligo rev</td><td>AAACCTGCGTCCTGTCGAGGAGCT</td><td></td><td></td><td></td><td></td><td></td><td></td><td></td><td></td><td></td><td></td></tr>
<tr><td>Chromosome</td><td>start</td><td>end</td><td>strand</td><td>MM</td><td>target_seq</td><td>PAM</td><td>alignment</td><td>distance</td><td>position</td><td>gene name</td><td>gene id</td></tr>
<tr><td>GU305914.1</td><td>106555</td><td>106577</td><td>+</td><td>0</td><td>AGCTCCTCGACAGGACGCAG</td><td>GGG</td><td>||||||||[||||||||||||]PAM</td><td>NA</td><td>Intergenic</td><td>NA</td><td>NA</td></tr>
<tr><td></td><td></td><td></td><td></td><td></td><td></td><td></td><td></td><td></td><td></td><td></td><td></td></tr>
<tr><td>T7779</td><td>ACAGGACGCAGGGGCTCCTGCGG</td><td>1000</td><td></td><td></td><td></td><td></td><td></td><td></td><td></td><td></td><td></td></tr>
<tr><td>Oligo fwd</td><td>CACCACAGGACGCAGGGGCTCCTG</td><td></td><td></td><td></td><td></td><td></td><td></td><td></td><td></td><td></td><td></td></tr>
<tr><td>Oligo rev</td><td>AAACCAGGAGCCCCTGCGTCCTGT</td><td></td><td></td><td></td><td></td><td></td><td></td><td></td><td></td><td></td><td></td></tr>
<tr><td>Chromosome</td><td>start</td><td>end</td><td>strand</td><td>MM</td><td>target_seq</td><td>PAM</td><td>alignment</td><td>distance</td><td>position</td><td>gene name</td><td>gene id</td></tr>
<tr><td>GU305914.1</td><td>106564</td><td>106586</td><td>+</td><td>0</td><td>ACAGGACGCAGGGGCTCCTG</td><td>CGG</td><td>||||||||[||||||||||||]PAM</td><td>NA</td><td>Intergenic</td><td>NA</td><td>NA</td></tr>
<tr><td></td><td></td><td></td><td></td><td></td><td></td><td></td><td></td><td></td><td></td><td></td><td></td></tr>
<tr><td>T7780</td><td>CGCAGGGGCTCCTGCGGTCCAGG</td><td>1000</td><td></td><td></td><td></td><td></td><td></td><td></td><td></td><td></td><td></td></tr>
<tr><td>Oligo fwd</td><td>CACCCGCAGGGGCTCCTGCGGTCC</td><td></td><td></td><td></td><td></td><td></td><td></td><td></td><td></td><td></td><td></td></tr>
<tr><td>Oligo rev</td><td>AAACGGACCGCAGGAGCCCCTGCG</td><td></td><td></td><td></td><td></td><td></td><td></td><td></td><td></td><td></td><td></td></tr>
<tr><td>Chromosome</td><td>start</td><td>end</td><td>strand</td><td>MM</td><td>target_seq</td><td>PAM</td><td>alignment</td><td>distance</td><td>position</td><td>gene name</td><td>gene id</td></tr>
<tr><td>GU305914.1</td><td>106570</td><td>106592</td><td>+</td><td>0</td><td>CGCAGGGGCTCCTGCGGTCC</td><td>AGG</td><td>||||||||[||||||||||||]PAM</td><td>NA</td><td>Intergenic</td><td>NA</td><td>NA</td></tr>
<tr><td></td><td></td><td></td><td></td><td></td><td></td><td></td><td></td><td></td><td></td><td></td><td></td></tr>
<tr><td>T7781</td><td>GGTCCAGGACCGTGATCATCAGG</td><td>1000</td><td></td><td></td><td></td><td></td><td></td><td></td><td></td><td></td><td></td></tr>
<tr><td>Oligo fwd</td><td>CACCGGTCCAGGACCGTGATCATC</td><td></td><td></td><td></td><td></td><td></td><td></td><td></td><td></td><td></td><td></td></tr>
<tr><td>Oligo rev</td><td>AAACGATGATCACGGTCCTGGACC</td><td></td><td></td><td></td><td></td><td></td><td></td><td></td><td></td><td></td><td></td></tr>
<tr><td>Chromosome</td><td>start</td><td>end</td><td>strand</td><td>MM</td><td>target_seq</td><td>PAM</td><td>alignment</td><td>distance</td><td>position</td><td>gene name</td><td>gene id</td></tr>
<tr><td>GU305914.1</td><td>106585</td><td>106607</td><td>+</td><td>0</td><td>GGTCCAGGACCGTGATCATC</td><td>AGG</td><td>||||||||[||||||||||||]PAM</td><td>NA</td><td>Intergenic</td><td>NA</td><td>NA</td></tr>
<tr><td></td><td></td><td></td><td></td><td></td><td></td><td></td><td></td><td></td><td></td><td></td><td></td></tr>
<tr><td>T7782</td><td>CATCAGGTTCCTGTCCACCACGG</td><td>1000</td><td></td><td></td><td></td><td></td><td></td><td></td><td></td><td></td><td></td></tr>
<tr><td>Oligo fwd</td><td>CACCCATCAGGTTCCTGTCCACCA</td><td></td><td></td><td></td><td></td><td></td><td></td><td></td><td></td><td></td><td></td></tr>
<tr><td>Oligo rev</td><td>AAACTGGTGGACAGGAACCTGATG</td><td></td><td></td><td></td><td></td><td></td><td></td><td></td><td></td><td></td><td></td></tr>
<tr><td>Chromosome</td><td>start</td><td>end</td><td>strand</td><td>MM</td><td>target_seq</td><td>PAM</td><td>alignment</td><td>distance</td><td>position</td><td>gene name</td><td>gene id</td></tr>
<tr><td>GU305914.1</td><td>106601</td><td>106623</td><td>+</td><td>0</td><td>CATCAGGTTCCTGTCCACCA</td><td>CGG</td><td>||||||||[||||||||||||]PAM</td><td>NA</td><td>Intergenic</td><td>NA</td><td>NA</td></tr>
<tr><td></td><td></td><td></td><td></td><td></td><td></td><td></td><td></td><td></td><td></td><td></td><td></td></tr>
<tr><td>T7783</td><td>GTCCACCACGGTGTTGCTCACGG</td><td>1000</td><td></td><td></td><td></td><td></td><td></td><td></td><td></td><td></td><td></td></tr>
<tr><td>Oligo fwd</td><td>CACCGTCCACCACGGTGTTGCTCA</td><td></td><td></td><td></td><td></td><td></td><td></td><td></td><td></td><td></td><td></td></tr>
<tr><td>Oligo rev</td><td>AAACTGAGCAACACCGTGGTGGAC</td><td></td><td></td><td></td><td></td><td></td><td></td><td></td><td></td><td></td><td></td></tr>
<tr><td>Chromosome</td><td>start</td><td>end</td><td>strand</td><td>MM</td><td>target_seq</td><td>PAM</td><td>alignment</td><td>distance</td><td>position</td><td>gene name</td><td>gene id</td></tr>
<tr><td>GU305914.1</td><td>106613</td><td>106635</td><td>+</td><td>0</td><td>GTCCACCACGGTGTTGCTCA</td><td>CGG</td><td>||||||||[||||||||||||]PAM</td><td>NA</td><td>Intergenic</td><td>NA</td><td>NA</td></tr>
<tr><td></td><td></td><td></td><td></td><td></td><td></td><td></td><td></td><td></td><td></td><td></td><td></td></tr>
<tr><td>T7784</td><td>TCCACCACGGTGTTGCTCACGGG</td><td>1000</td><td></td><td></td><td></td><td></td><td></td><td></td><td></td><td></td><td></td></tr>
<tr><td>Oligo fwd</td><td>CACCTCCACCACGGTGTTGCTCAC</td><td></td><td></td><td></td><td></td><td></td><td></td><td></td><td></td><td></td><td></td></tr>
<tr><td>Oligo rev</td><td>AAACGTGAGCAACACCGTGGTGGA</td><td></td><td></td><td></td><td></td><td></td><td></td><td></td><td></td><td></td><td></td></tr>
<tr><td>Chromosome</td><td>start</td><td>end</td><td>strand</td><td>MM</td><td>target_seq</td><td>PAM</td><td>alignment</td><td>distance</td><td>position</td><td>gene name</td><td>gene id</td></tr>
<tr><td>GU305914.1</td><td>106614</td><td>106636</td><td>+</td><td>0</td><td>TCCACCACGGTGTTGCTCAC</td><td>GGG</td><td>||||||||[||||||||||||]PAM</td><td>NA</td><td>Intergenic</td><td>NA</td><td>NA</td></tr>
</table>

T7785 CCACCACGGTGTTGCTCACGGGG 1000
Oligo fwd CACCCCACCACGGTGTTGCTCACG
Oligo rev AAACCGTGAGCAACACCGTGGTGG
Chromosome start end strand MM target_seq PAM alignment distance position gene name gene id
GU305914.1 106615 106637 + 0 CCACCACGGTGTTGCTCACG GGG ||||||||[||||||||||||]PAM NA Intergenic NA NA

T7786 TGTTGCTCACGGGGATCGCCAGG 1000
Oligo fwd CACCTGTTGCTCACGGGGATCGCC
Oligo rev AAACGGCGATCCCCGTGAGCAACA
Chromosome start end strand MM target_seq PAM alignment distance position gene name gene id
GU305914.1 106624 106646 + 0 TGTTGCTCACGGGGATCGCC AGG ||||||||[||||||||||||]PAM NA Intergenic NA NA

T7787 CAGGCCCTTGACGATAAAATCGG 1000
Oligo fwd CACCCAGGCCCTTGACGATAAAAT
Oligo rev AAACATTTTATCGTCAAGGGCCTG
Chromosome start end strand MM target_seq PAM alignment distance position gene name gene id
GU305914.1 106643 106665 + 0 CAGGCCCTTGACGATAAAAT CGG ||||||||[||||||||||||]PAM NA Intergenic NA NA

T7788 CGATAAAATCGGACGAGATCAGG 1000
Oligo fwd CACCCGATAAAATCGGACGAGATC
Oligo rev AAACGATCTCGTCCGATTTTATCG
Chromosome start end strand MM target_seq PAM alignment distance position gene name gene id
GU305914.1 106654 106676 + 0 CGATAAAATCGGACGAGATC AGG ||||||||[||||||||||||]PAM NA Intergenic NA NA

T7789 ACCGCGCCCTGAGTCGAATCCGG 1000
Oligo fwd CACCACCGCGCCCTGAGTCGAATC
Oligo rev AAACGATTCGACTCAGGGCGCGGT
Chromosome start end strand MM target_seq PAM alignment distance position gene name gene id
GU305914.1 106701 106723 + 0 ACCGCGCCCTGAGTCGAATC CGG ||||||||[||||||||||||]PAM NA Intergenic NA NA

T7790 TCCGACATCTGCGAGCGCTTCGG 1000
Oligo fwd CACCTCCGACATCTGCGAGCGCTT
Oligo rev AAACAAGCGCTCGCAGATGTCGGA
Chromosome start end strand MM target_seq PAM alignment distance position gene name gene id
GU305914.1 106740 106762 + 0 TCCGACATCTGCGAGCGCTT CGG ||||||||[||||||||||||]PAM NA Intergenic NA NA

T7791 TCTGCGAGCGCTTCGGACGCAGG 1000
Oligo fwd CACCTCTGCGAGCGCTTCGGACGC
Oligo rev AAACGCGTCCGAAGCGCTCGCAGA
Chromosome start end strand MM target_seq PAM alignment distance position gene name gene id
GU305914.1 106747 106769 + 0 TCTGCGAGCGCTTCGGACGC AGG ||||||||[||||||||||||]PAM NA Intergenic NA NA

T7792 CTGCGAGCGCTTCGGACGCAGGG 1000
Oligo fwd CACCCTGCGAGCGCTTCGGACGCA
Oligo rev AAACTGCGTCCGAAGCGCTCGCAG
Chromosome start end strand MM target_seq PAM alignment distance position gene name gene id
GU305914.1 106748 106770 + 0 CTGCGAGCGCTTCGGACGCA GGG ||||||||[||||||||||||]PAM NA Intergenic NA NA

T7793 CGTCACCGACATCACCTGTTCGG 1000
Oligo fwd CACCCGTCACCGACATCACCTGTT
Oligo rev AAACAACAGGTGATGTCGGTGACG
Chromosome start end strand MM target_seq PAM alignment distance position gene name gene id
GU305914.1 106775 106797 + 0 CGTCACCGACATCACCTGTT CGG ||||||||[||||||||||||]PAM NA Intergenic NA NA

T7794 ACCTGTTCGGCCCGTCTCTCCGG 1000
Oligo fwd CACCACCTGTTCGGCCCGTCTCTC
Oligo rev AAACGAGAGACGGGCCGAACAGGT
Chromosome start end strand MM target_seq PAM alignment distance position gene name gene id
GU305914.1 106788 106810 + 0 ACCTGTTCGGCCCGTCTCTC CGG ||||||||[||||||||||||]PAM NA Intergenic NA NA

T7795 GTTCGGCCCGTCTCTCCGGACGG 1000
Oligo fwd CACCGTTCGGCCCGTCTCTCCGGA
Oligo rev AAACTCCGGAGAGACGGGCCGAAC
Chromosome start end strand MM target_seq PAM alignment distance position gene name gene id
GU305914.1 106792 106814 + 0 GTTCGGCCCGTCTCTCCGGA CGG ||||||||[||||||||||||]PAM NA Intergenic NA NA

T7796 GACGGTGCGCCGACTCCATCAGG 1000
Oligo fwd CACCGACGGTGCGCCGACTCCATC
Oligo rev AAACGATGGAGTCGGCGCACCGTC
Chromosome start end strand MM target_seq PAM alignment distance position gene name gene id
GU305914.1 106810 106832 + 0 GACGGTGCGCCGACTCCATC AGG ||||||||[||||||||||||]PAM NA Intergenic NA NA
GU305914.1 115046 115068 + 4 GAGGGAGCGCCGACTCCGGC AGG ||-||-||[|||||||||--|]PAM NA Intergenic NA NA

T7797 ACGGTGCGCCGACTCCATCAGGG 1000
Oligo fwd CACCACGGTGCGCCGACTCCATCA
Oligo rev AAACTGATGGAGTCGGCGCACCGT
Chromosome start end strand MM target_seq PAM alignment distance position gene name gene id
GU305914.1 106811 106833 + 0 ACGGTGCGCCGACTCCATCA GGG ||||||||[||||||||||||]PAM NA Intergenic NA NA

T7798 TCCATCAGGGCCCTGACCGATGG 1000
Oligo fwd CACCTCCATCAGGGCCCTGACCGA
Oligo rev AAACTCGGTCAGGGCCCTGATGGA
Chromosome start end strand MM target_seq PAM alignment distance position gene name gene id
GU305914.1 106824 106846 + 0 TCCATCAGGGCCCTGACCGA TGG ||||||||[||||||||||||]PAM NA Intergenic NA NA

T7799 CGCCTGCCTCCCGATACGCACGG 1000
Oligo fwd CACCCGCCTGCCTCCCGATACGCA
Oligo rev AAACTGCGTATCGGGAGGCAGGCG
Chromosome start end strand MM target_seq PAM alignment distance position gene name gene id

<table>
<tr><td>GU305914.1</td><td>106860</td><td>106882</td><td>+</td><td>0</td><td>CGCCTGCCTCCCGATACGCA</td><td>CGG</td><td>||||||||[||||||||||||]PAM</td><td>NA</td><td>Intergenic</td><td>NA</td><td>NA</td></tr>
<tr><td>T7800</td><td>GCCTGCCTCCCGATACGCACGGG</td><td>1000</td><td></td><td></td><td></td><td></td><td></td><td></td><td></td><td></td><td></td></tr>
<tr><td>Oligo fwd</td><td>CACCGCCTGCCTCCCGATACGCAC</td><td></td><td></td><td></td><td></td><td></td><td></td><td></td><td></td><td></td><td></td></tr>
<tr><td>Oligo rev</td><td>AAACGTGCGTATCGGGAGGCAGGC</td><td></td><td></td><td></td><td></td><td></td><td></td><td></td><td></td><td></td><td></td></tr>
<tr><td>Chromosome</td><td>start</td><td>end</td><td>strand</td><td>MM</td><td>target_seq</td><td>PAM</td><td>alignment</td><td>distance</td><td>position</td><td>gene name</td><td>gene id</td></tr>
<tr><td>GU305914.1</td><td>106861</td><td>106883</td><td>+</td><td>0</td><td>GCCTGCCTCCCGATACGCAC</td><td>GGG</td><td>||||||||[||||||||||||]PAM</td><td>NA</td><td>Intergenic</td><td>NA</td><td>NA</td></tr>
<tr><td>T7801</td><td>CACGGGCTGTAGATCTCCGACGG</td><td>1000</td><td></td><td></td><td></td><td></td><td></td><td></td><td></td><td></td><td></td></tr>
<tr><td>Oligo fwd</td><td>CACCCACGGGCTGTAGATCTCCGA</td><td></td><td></td><td></td><td></td><td></td><td></td><td></td><td></td><td></td><td></td></tr>
<tr><td>Oligo rev</td><td>AAACTCGGAGATCTACAGCCCGTG</td><td></td><td></td><td></td><td></td><td></td><td></td><td></td><td></td><td></td><td></td></tr>
<tr><td>Chromosome</td><td>start</td><td>end</td><td>strand</td><td>MM</td><td>target_seq</td><td>PAM</td><td>alignment</td><td>distance</td><td>position</td><td>gene name</td><td>gene id</td></tr>
<tr><td>GU305914.1</td><td>106878</td><td>106900</td><td>+</td><td>0</td><td>CACGGGCTGTAGATCTCCGA</td><td>CGG</td><td>||||||||[||||||||||||]PAM</td><td>NA</td><td>Intergenic</td><td>NA</td><td>NA</td></tr>
<tr><td>T7802</td><td>TCCGACGGTGCGCCCCGTTGTGG</td><td>1000</td><td></td><td></td><td></td><td></td><td></td><td></td><td></td><td></td><td></td></tr>
<tr><td>Oligo fwd</td><td>CACCTCCGACGGTGCGCCCCGTTG</td><td></td><td></td><td></td><td></td><td></td><td></td><td></td><td></td><td></td><td></td></tr>
<tr><td>Oligo rev</td><td>AAACCAACGGGGCGCACCGTCGGA</td><td></td><td></td><td></td><td></td><td></td><td></td><td></td><td></td><td></td><td></td></tr>
<tr><td>Chromosome</td><td>start</td><td>end</td><td>strand</td><td>MM</td><td>target_seq</td><td>PAM</td><td>alignment</td><td>distance</td><td>position</td><td>gene name</td><td>gene id</td></tr>
<tr><td>GU305914.1</td><td>106893</td><td>106915</td><td>+</td><td>0</td><td>TCCGACGGTGCGCCCCGTTG</td><td>TGG</td><td>||||||||[||||||||||||]PAM</td><td>NA</td><td>Intergenic</td><td>NA</td><td>NA</td></tr>
<tr><td>T7803</td><td>CCGACGGTGCGCCCCGTTGTGGG</td><td>1000</td><td></td><td></td><td></td><td></td><td></td><td></td><td></td><td></td><td></td></tr>
<tr><td>Oligo fwd</td><td>CACCCCGACGGTGCGCCCCGTTGT</td><td></td><td></td><td></td><td></td><td></td><td></td><td></td><td></td><td></td><td></td></tr>
<tr><td>Oligo rev</td><td>AAACACAACGGGGCGCACCGTCGG</td><td></td><td></td><td></td><td></td><td></td><td></td><td></td><td></td><td></td><td></td></tr>
<tr><td>Chromosome</td><td>start</td><td>end</td><td>strand</td><td>MM</td><td>target_seq</td><td>PAM</td><td>alignment</td><td>distance</td><td>position</td><td>gene name</td><td>gene id</td></tr>
<tr><td>GU305914.1</td><td>106894</td><td>106916</td><td>+</td><td>0</td><td>CCGACGGTGCGCCCCGTTGT</td><td>GGG</td><td>||||||||[||||||||||||]PAM</td><td>NA</td><td>Intergenic</td><td>NA</td><td>NA</td></tr>
<tr><td>T7804</td><td>CCCCATGAGATCTGTTCTCGCGG</td><td>1000</td><td></td><td></td><td></td><td></td><td></td><td></td><td></td><td></td><td></td></tr>
<tr><td>Oligo fwd</td><td>CACCCCCCATGAGATCTGTTCTCG</td><td></td><td></td><td></td><td></td><td></td><td></td><td></td><td></td><td></td><td></td></tr>
<tr><td>Oligo rev</td><td>AAACCGAGAACAGATCTCATGGGG</td><td></td><td></td><td></td><td></td><td></td><td></td><td></td><td></td><td></td><td></td></tr>
<tr><td>Chromosome</td><td>start</td><td>end</td><td>strand</td><td>MM</td><td>target_seq</td><td>PAM</td><td>alignment</td><td>distance</td><td>position</td><td>gene name</td><td>gene id</td></tr>
<tr><td>GU305914.1</td><td>106932</td><td>106954</td><td>+</td><td>0</td><td>CCCCATGAGATCTGTTCTCG</td><td>CGG</td><td>||||||||[||||||||||||]PAM</td><td>NA</td><td>Intergenic</td><td>NA</td><td>NA</td></tr>
<tr><td>T7805</td><td>CTGTTCTCGCGGCGAACACATGG</td><td>1000</td><td></td><td></td><td></td><td></td><td></td><td></td><td></td><td></td><td></td></tr>
<tr><td>Oligo fwd</td><td>CACCCTGTTCTCGCGGCGAACACA</td><td></td><td></td><td></td><td></td><td></td><td></td><td></td><td></td><td></td><td></td></tr>
<tr><td>Oligo rev</td><td>AAACTGTGTTCGCCGCGAGAACAG</td><td></td><td></td><td></td><td></td><td></td><td></td><td></td><td></td><td></td><td></td></tr>
<tr><td>Chromosome</td><td>start</td><td>end</td><td>strand</td><td>MM</td><td>target_seq</td><td>PAM</td><td>alignment</td><td>distance</td><td>position</td><td>gene name</td><td>gene id</td></tr>
<tr><td>GU305914.1</td><td>106943</td><td>106965</td><td>+</td><td>0</td><td>CTGTTCTCGCGGCGAACACA</td><td>TGG</td><td>||||||||[||||||||||||]PAM</td><td>NA</td><td>Intergenic</td><td>NA</td><td>NA</td></tr>
<tr><td>T7806</td><td>TTCTCGCGGCGAACACATGGAGG</td><td>1000</td><td></td><td></td><td></td><td></td><td></td><td></td><td></td><td></td><td></td></tr>
<tr><td>Oligo fwd</td><td>CACCTTCTCGCGGCGAACACATGG</td><td></td><td></td><td></td><td></td><td></td><td></td><td></td><td></td><td></td><td></td></tr>
<tr><td>Oligo rev</td><td>AAACCCATGTGTTCGCCGCGAGAA</td><td></td><td></td><td></td><td></td><td></td><td></td><td></td><td></td><td></td><td></td></tr>
<tr><td>Chromosome</td><td>start</td><td>end</td><td>strand</td><td>MM</td><td>target_seq</td><td>PAM</td><td>alignment</td><td>distance</td><td>position</td><td>gene name</td><td>gene id</td></tr>
<tr><td>GU305914.1</td><td>106946</td><td>106968</td><td>+</td><td>0</td><td>TTCTCGCGGCGAACACATGG</td><td>AGG</td><td>||||||||[||||||||||||]PAM</td><td>NA</td><td>Intergenic</td><td>NA</td><td>NA</td></tr>
<tr><td>T7807</td><td>TGCTCTCGTTCAGTTGTGTACGG</td><td>1000</td><td></td><td></td><td></td><td></td><td></td><td></td><td></td><td></td><td></td></tr>
<tr><td>Oligo fwd</td><td>CACCTGCTCTCGTTCAGTTGTGTA</td><td></td><td></td><td></td><td></td><td></td><td></td><td></td><td></td><td></td><td></td></tr>
<tr><td>Oligo rev</td><td>AAACTACACAACTGAACGAGAGCA</td><td></td><td></td><td></td><td></td><td></td><td></td><td></td><td></td><td></td><td></td></tr>
<tr><td>Chromosome</td><td>start</td><td>end</td><td>strand</td><td>MM</td><td>target_seq</td><td>PAM</td><td>alignment</td><td>distance</td><td>position</td><td>gene name</td><td>gene id</td></tr>
<tr><td>GU305914.1</td><td>106999</td><td>107021</td><td>+</td><td>0</td><td>TGCTCTCGTTCAGTTGTGTA</td><td>CGG</td><td>||||||||[||||||||||||]PAM</td><td>NA</td><td>Intergenic</td><td>NA</td><td>NA</td></tr>
<tr><td>T7808</td><td>TCAGTTGTGTACGGACCGTTAGG</td><td>1000</td><td></td><td></td><td></td><td></td><td></td><td></td><td></td><td></td><td></td></tr>
<tr><td>Oligo fwd</td><td>CACCTCAGTTGTGTACGGACCGTT</td><td></td><td></td><td></td><td></td><td></td><td></td><td></td><td></td><td></td><td></td></tr>
<tr><td>Oligo rev</td><td>AAACAACGGTCCGTACACAACTGA</td><td></td><td></td><td></td><td></td><td></td><td></td><td></td><td></td><td></td><td></td></tr>
<tr><td>Chromosome</td><td>start</td><td>end</td><td>strand</td><td>MM</td><td>target_seq</td><td>PAM</td><td>alignment</td><td>distance</td><td>position</td><td>gene name</td><td>gene id</td></tr>
<tr><td>GU305914.1</td><td>107008</td><td>107030</td><td>+</td><td>0</td><td>TCAGTTGTGTACGGACCGTT</td><td>AGG</td><td>||||||||[||||||||||||]PAM</td><td>NA</td><td>Intergenic</td><td>NA</td><td>NA</td></tr>
<tr><td>T7809</td><td>TACGGACCGTTAGGTCGACGCGG</td><td>1000</td><td></td><td></td><td></td><td></td><td></td><td></td><td></td><td></td><td></td></tr>
<tr><td>Oligo fwd</td><td>CACCTACGGACCGTTAGGTCGACG</td><td></td><td></td><td></td><td></td><td></td><td></td><td></td><td></td><td></td><td></td></tr>
<tr><td>Oligo rev</td><td>AAACCGTCGACCTAACGGTCCGTA</td><td></td><td></td><td></td><td></td><td></td><td></td><td></td><td></td><td></td><td></td></tr>
<tr><td>Chromosome</td><td>start</td><td>end</td><td>strand</td><td>MM</td><td>target_seq</td><td>PAM</td><td>alignment</td><td>distance</td><td>position</td><td>gene name</td><td>gene id</td></tr>
<tr><td>GU305914.1</td><td>107017</td><td>107039</td><td>+</td><td>0</td><td>TACGGACCGTTAGGTCGACG</td><td>CGG</td><td>||||||||[||||||||||||]PAM</td><td>NA</td><td>Intergenic</td><td>NA</td><td>NA</td></tr>
<tr><td>T7810</td><td>CAAGATGCGTCTCATCGATCCGG</td><td>1000</td><td></td><td></td><td></td><td></td><td></td><td></td><td></td><td></td><td></td></tr>
<tr><td>Oligo fwd</td><td>CACCCAAGATGCGTCTCATCGATC</td><td></td><td></td><td></td><td></td><td></td><td></td><td></td><td></td><td></td><td></td></tr>
<tr><td>Oligo rev</td><td>AAACGATCGATGAGACGCATCTTG</td><td></td><td></td><td></td><td></td><td></td><td></td><td></td><td></td><td></td><td></td></tr>
<tr><td>Chromosome</td><td>start</td><td>end</td><td>strand</td><td>MM</td><td>target_seq</td><td>PAM</td><td>alignment</td><td>distance</td><td>position</td><td>gene name</td><td>gene id</td></tr>
<tr><td>GU305914.1</td><td>107042</td><td>107064</td><td>+</td><td>0</td><td>CAAGATGCGTCTCATCGATC</td><td>CGG</td><td>||||||||[||||||||||||]PAM</td><td>NA</td><td>Intergenic</td><td>NA</td><td>NA</td></tr>
<tr><td>T7811</td><td>CGTCTCATCGATCCGGATAGCGG</td><td>1000</td><td></td><td></td><td></td><td></td><td></td><td></td><td></td><td></td><td></td></tr>
<tr><td>Oligo fwd</td><td>CACCCGTCTCATCGATCCGGATAG</td><td></td><td></td><td></td><td></td><td></td><td></td><td></td><td></td><td></td><td></td></tr>
<tr><td>Oligo rev</td><td>AAACCTATCCGGATCGATGAGACG</td><td></td><td></td><td></td><td></td><td></td><td></td><td></td><td></td><td></td><td></td></tr>
<tr><td>Chromosome</td><td>start</td><td>end</td><td>strand</td><td>MM</td><td>target_seq</td><td>PAM</td><td>alignment</td><td>distance</td><td>position</td><td>gene name</td><td>gene id</td></tr>
<tr><td>GU305914.1</td><td>107049</td><td>107071</td><td>+</td><td>0</td><td>CGTCTCATCGATCCGGATAG</td><td>CGG</td><td>||||||||[||||||||||||]PAM</td><td>NA</td><td>Intergenic</td><td>NA</td><td>NA</td></tr>
<tr><td>T7812</td><td>GTCTCATCGATCCGGATAGCGGG</td><td>1000</td><td></td><td></td><td></td><td></td><td></td><td></td><td></td><td></td><td></td></tr>
<tr><td>Oligo fwd</td><td>CACCGTCTCATCGATCCGGATAGC</td><td></td><td></td><td></td><td></td><td></td><td></td><td></td><td></td><td></td><td></td></tr>
<tr><td>Oligo rev</td><td>AAACGCTATCCGGATCGATGAGAC</td><td></td><td></td><td></td><td></td><td></td><td></td><td></td><td></td><td></td><td></td></tr>
<tr><td>Chromosome</td><td>start</td><td>end</td><td>strand</td><td>MM</td><td>target_seq</td><td>PAM</td><td>alignment</td><td>distance</td><td>position</td><td>gene name</td><td>gene id</td></tr>
<tr><td>GU305914.1</td><td>107050</td><td>107072</td><td>+</td><td>0</td><td>GTCTCATCGATCCGGATAGC</td><td>GGG</td><td>||||||||[||||||||||||]PAM</td><td>NA</td><td>Intergenic</td><td>NA</td><td>NA</td></tr>
<tr><td>T7813</td><td>GCGGGATCTTCAAGCTCATACGG</td><td>1000</td><td></td><td></td><td></td><td></td><td></td><td></td><td></td><td></td><td></td></tr>
<tr><td>Oligo fwd</td><td>CACCGCGGGATCTTCAAGCTCATA</td><td></td><td></td><td></td><td></td><td></td><td></td><td></td><td></td><td></td><td></td></tr>
<tr><td>Oligo rev</td><td>AAACTATGAGCTTGAAGATCCCGC</td><td></td><td></td><td></td><td></td><td></td><td></td><td></td><td></td><td></td><td></td></tr>
<tr><td>Chromosome</td><td>start</td><td>end</td><td>strand</td><td>MM</td><td>target_seq</td><td>PAM</td><td>alignment</td><td>distance</td><td>position</td><td>gene name</td><td>gene id</td></tr>
<tr><td>GU305914.1</td><td>107068</td><td>107090</td><td>+</td><td>0</td><td>GCGGGATCTTCAAGCTCATA</td><td>CGG</td><td>||||||||[||||||||||||]PAM</td><td>NA</td><td>Intergenic</td><td>NA</td><td>NA</td></tr>
<tr><td>T7814</td><td>CATACGGTAGTACGCGTCCGTGG</td><td>1000</td><td></td><td></td><td></td><td></td><td></td><td></td><td></td><td></td><td></td></tr>
<tr><td>Oligo fwd</td><td>CACCCATACGGTAGTACGCGTCCG</td><td></td><td></td><td></td><td></td><td></td><td></td><td></td><td></td><td></td><td></td></tr>
<tr><td>Oligo rev</td><td>AAACCGGACGCGTACTACCGTATG</td><td></td><td></td><td></td><td></td><td></td><td></td><td></td><td></td><td></td><td></td></tr>
</table>

<table>
<tr><td>Chromosome</td><td>start</td><td>end</td><td>strand</td><td>MM</td><td>target_seq</td><td>PAM</td><td>alignment</td><td>distance</td><td>position</td><td>gene name</td><td>gene id</td></tr>
<tr><td>GU305914.1</td><td>107084</td><td>107106</td><td>+</td><td>0</td><td>CATACGGTAGTACGCGTCCG</td><td>TGG</td><td>||||||||[||||||||||||]PAM</td><td>NA</td><td>Intergenic</td><td>NA</td><td>NA</td></tr>
<tr><td></td><td></td><td></td><td></td><td></td><td></td><td></td><td></td><td></td><td></td><td></td><td></td></tr>
<tr><td>T7815</td><td>TGAACAGTAACTTCTGTAACAGG</td><td>1000</td><td></td><td></td><td></td><td></td><td></td><td></td><td></td><td></td><td></td></tr>
<tr><td>Oligo fwd</td><td>CACCTGAACAGTAACTTCTGTAAC</td><td></td><td></td><td></td><td></td><td></td><td></td><td></td><td></td><td></td><td></td></tr>
<tr><td>Oligo rev</td><td>AAACGTTACAGAAGTTACTGTTCA</td><td></td><td></td><td></td><td></td><td></td><td></td><td></td><td></td><td></td><td></td></tr>
<tr><td>Chromosome</td><td>start</td><td>end</td><td>strand</td><td>MM</td><td>target_seq</td><td>PAM</td><td>alignment</td><td>distance</td><td>position</td><td>gene name</td><td>gene id</td></tr>
<tr><td>GU305914.1</td><td>107107</td><td>107129</td><td>+</td><td>0</td><td>TGAACAGTAACTTCTGTAAC</td><td>AGG</td><td>||||||||[||||||||||||]PAM</td><td>NA</td><td>Intergenic</td><td>NA</td><td>NA</td></tr>
<tr><td></td><td></td><td></td><td></td><td></td><td></td><td></td><td></td><td></td><td></td><td></td><td></td></tr>
<tr><td>T7816</td><td>GAACAGTAACTTCTGTAACAGGG</td><td>1000</td><td></td><td></td><td></td><td></td><td></td><td></td><td></td><td></td><td></td></tr>
<tr><td>Oligo fwd</td><td>CACCGAACAGTAACTTCTGTAACA</td><td></td><td></td><td></td><td></td><td></td><td></td><td></td><td></td><td></td><td></td></tr>
<tr><td>Oligo rev</td><td>AAACTGTTACAGAAGTTACTGTTC</td><td></td><td></td><td></td><td></td><td></td><td></td><td></td><td></td><td></td><td></td></tr>
<tr><td>Chromosome</td><td>start</td><td>end</td><td>strand</td><td>MM</td><td>target_seq</td><td>PAM</td><td>alignment</td><td>distance</td><td>position</td><td>gene name</td><td>gene id</td></tr>
<tr><td>GU305914.1</td><td>107108</td><td>107130</td><td>+</td><td>0</td><td>GAACAGTAACTTCTGTAACA</td><td>GGG</td><td>||||||||[||||||||||||]PAM</td><td>NA</td><td>Intergenic</td><td>NA</td><td>NA</td></tr>
<tr><td></td><td></td><td></td><td></td><td></td><td></td><td></td><td></td><td></td><td></td><td></td><td></td></tr>
<tr><td>T7817</td><td>AGGGCGTCCGTGTTGACATTCGG</td><td>1000</td><td></td><td></td><td></td><td></td><td></td><td></td><td></td><td></td><td></td></tr>
<tr><td>Oligo fwd</td><td>CACCAGGGCGTCCGTGTTGACATT</td><td></td><td></td><td></td><td></td><td></td><td></td><td></td><td></td><td></td><td></td></tr>
<tr><td>Oligo rev</td><td>AAACAATGTCAACACGGACGCCCT</td><td></td><td></td><td></td><td></td><td></td><td></td><td></td><td></td><td></td><td></td></tr>
<tr><td>Chromosome</td><td>start</td><td>end</td><td>strand</td><td>MM</td><td>target_seq</td><td>PAM</td><td>alignment</td><td>distance</td><td>position</td><td>gene name</td><td>gene id</td></tr>
<tr><td>GU305914.1</td><td>107127</td><td>107149</td><td>+</td><td>0</td><td>AGGGCGTCCGTGTTGACATT</td><td>CGG</td><td>||||||||[||||||||||||]PAM</td><td>NA</td><td>Intergenic</td><td>NA</td><td>NA</td></tr>
<tr><td></td><td></td><td></td><td></td><td></td><td></td><td></td><td></td><td></td><td></td><td></td><td></td></tr>
<tr><td>T7818</td><td>GCGTCCGTGTTGACATTCGGCGG</td><td>1000</td><td></td><td></td><td></td><td></td><td></td><td></td><td></td><td></td><td></td></tr>
<tr><td>Oligo fwd</td><td>CACCGCGTCCGTGTTGACATTCGG</td><td></td><td></td><td></td><td></td><td></td><td></td><td></td><td></td><td></td><td></td></tr>
<tr><td>Oligo rev</td><td>AAACCCGAATGTCAACACGGACGC</td><td></td><td></td><td></td><td></td><td></td><td></td><td></td><td></td><td></td><td></td></tr>
<tr><td>Chromosome</td><td>start</td><td>end</td><td>strand</td><td>MM</td><td>target_seq</td><td>PAM</td><td>alignment</td><td>distance</td><td>position</td><td>gene name</td><td>gene id</td></tr>
<tr><td>GU305914.1</td><td>107130</td><td>107152</td><td>+</td><td>0</td><td>GCGTCCGTGTTGACATTCGG</td><td>CGG</td><td>||||||||[||||||||||||]PAM</td><td>NA</td><td>Intergenic</td><td>NA</td><td>NA</td></tr>
<tr><td></td><td></td><td></td><td></td><td></td><td></td><td></td><td></td><td></td><td></td><td></td><td></td></tr>
<tr><td>T7819</td><td>CGTCCGTGTTGACATTCGGCGGG</td><td>1000</td><td></td><td></td><td></td><td></td><td></td><td></td><td></td><td></td><td></td></tr>
<tr><td>Oligo fwd</td><td>CACCCGTCCGTGTTGACATTCGGC</td><td></td><td></td><td></td><td></td><td></td><td></td><td></td><td></td><td></td><td></td></tr>
<tr><td>Oligo rev</td><td>AAACGCCGAATGTCAACACGGACG</td><td></td><td></td><td></td><td></td><td></td><td></td><td></td><td></td><td></td><td></td></tr>
<tr><td>Chromosome</td><td>start</td><td>end</td><td>strand</td><td>MM</td><td>target_seq</td><td>PAM</td><td>alignment</td><td>distance</td><td>position</td><td>gene name</td><td>gene id</td></tr>
<tr><td>GU305914.1</td><td>107131</td><td>107153</td><td>+</td><td>0</td><td>CGTCCGTGTTGACATTCGGC</td><td>GGG</td><td>||||||||[||||||||||||]PAM</td><td>NA</td><td>Intergenic</td><td>NA</td><td>NA</td></tr>
<tr><td></td><td></td><td></td><td></td><td></td><td></td><td></td><td></td><td></td><td></td><td></td><td></td></tr>
<tr><td>T7820</td><td>TCGGCGGGATGCGTGTCGAACGG</td><td>1000</td><td></td><td></td><td></td><td></td><td></td><td></td><td></td><td></td><td></td></tr>
<tr><td>Oligo fwd</td><td>CACCTCGGCGGGATGCGTGTCGAA</td><td></td><td></td><td></td><td></td><td></td><td></td><td></td><td></td><td></td><td></td></tr>
<tr><td>Oligo rev</td><td>AAACTTCGACACGCATCCCGCCGA</td><td></td><td></td><td></td><td></td><td></td><td></td><td></td><td></td><td></td><td></td></tr>
<tr><td>Chromosome</td><td>start</td><td>end</td><td>strand</td><td>MM</td><td>target_seq</td><td>PAM</td><td>alignment</td><td>distance</td><td>position</td><td>gene name</td><td>gene id</td></tr>
<tr><td>GU305914.1</td><td>107146</td><td>107168</td><td>+</td><td>0</td><td>TCGGCGGGATGCGTGTCGAA</td><td>CGG</td><td>||||||||[||||||||||||]PAM</td><td>NA</td><td>Intergenic</td><td>NA</td><td>NA</td></tr>
<tr><td></td><td></td><td></td><td></td><td></td><td></td><td></td><td></td><td></td><td></td><td></td><td></td></tr>
<tr><td>T7821</td><td>TCGAACGGAGCAAAAAAGCGAGG</td><td>1000</td><td></td><td></td><td></td><td></td><td></td><td></td><td></td><td></td><td></td></tr>
<tr><td>Oligo fwd</td><td>CACCTCGAACGGAGCAAAAAAGCG</td><td></td><td></td><td></td><td></td><td></td><td></td><td></td><td></td><td></td><td></td></tr>
<tr><td>Oligo rev</td><td>AAACCGCTTTTTTGCTCCGTTCGA</td><td></td><td></td><td></td><td></td><td></td><td></td><td></td><td></td><td></td><td></td></tr>
<tr><td>Chromosome</td><td>start</td><td>end</td><td>strand</td><td>MM</td><td>target_seq</td><td>PAM</td><td>alignment</td><td>distance</td><td>position</td><td>gene name</td><td>gene id</td></tr>
<tr><td>GU305914.1</td><td>107161</td><td>107183</td><td>+</td><td>0</td><td>TCGAACGGAGCAAAAAAGCG</td><td>AGG</td><td>||||||||[||||||||||||]PAM</td><td>NA</td><td>Intergenic</td><td>NA</td><td>NA</td></tr>
<tr><td></td><td></td><td></td><td></td><td></td><td></td><td></td><td></td><td></td><td></td><td></td><td></td></tr>
<tr><td>T7822</td><td>ACGGAGCAAAAAAGCGAGGTCGG</td><td>1000</td><td></td><td></td><td></td><td></td><td></td><td></td><td></td><td></td><td></td></tr>
<tr><td>Oligo fwd</td><td>CACCACGGAGCAAAAAAGCGAGGT</td><td></td><td></td><td></td><td></td><td></td><td></td><td></td><td></td><td></td><td></td></tr>
<tr><td>Oligo rev</td><td>AAACACCTCGCTTTTTTGCTCCGT</td><td></td><td></td><td></td><td></td><td></td><td></td><td></td><td></td><td></td><td></td></tr>
<tr><td>Chromosome</td><td>start</td><td>end</td><td>strand</td><td>MM</td><td>target_seq</td><td>PAM</td><td>alignment</td><td>distance</td><td>position</td><td>gene name</td><td>gene id</td></tr>
<tr><td>GU305914.1</td><td>107165</td><td>107187</td><td>+</td><td>0</td><td>ACGGAGCAAAAAAGCGAGGT</td><td>CGG</td><td>||||||||[||||||||||||]PAM</td><td>NA</td><td>Intergenic</td><td>NA</td><td>NA</td></tr>
<tr><td></td><td></td><td></td><td></td><td></td><td></td><td></td><td></td><td></td><td></td><td></td><td></td></tr>
<tr><td>T7823</td><td>GTGCATCCCTGACGAGTACGTGG</td><td>1000</td><td></td><td></td><td></td><td></td><td></td><td></td><td></td><td></td><td></td></tr>
<tr><td>Oligo fwd</td><td>CACCGTGCATCCCTGACGAGTACG</td><td></td><td></td><td></td><td></td><td></td><td></td><td></td><td></td><td></td><td></td></tr>
<tr><td>Oligo rev</td><td>AAACCGTACTCGTCAGGGATGCAC</td><td></td><td></td><td></td><td></td><td></td><td></td><td></td><td></td><td></td><td></td></tr>
<tr><td>Chromosome</td><td>start</td><td>end</td><td>strand</td><td>MM</td><td>target_seq</td><td>PAM</td><td>alignment</td><td>distance</td><td>position</td><td>gene name</td><td>gene id</td></tr>
<tr><td>GU305914.1</td><td>107201</td><td>107223</td><td>+</td><td>0</td><td>GTGCATCCCTGACGAGTACG</td><td>TGG</td><td>||||||||[||||||||||||]PAM</td><td>NA</td><td>Intergenic</td><td>NA</td><td>NA</td></tr>
<tr><td></td><td></td><td></td><td></td><td></td><td></td><td></td><td></td><td></td><td></td><td></td><td></td></tr>
<tr><td>T7824</td><td>AGTACGTGGCCGCCAATACCCGG</td><td>1000</td><td></td><td></td><td></td><td></td><td></td><td></td><td></td><td></td><td></td></tr>
<tr><td>Oligo fwd</td><td>CACCAGTACGTGGCCGCCAATACC</td><td></td><td></td><td></td><td></td><td></td><td></td><td></td><td></td><td></td><td></td></tr>
<tr><td>Oligo rev</td><td>AAACGGTATTGGCGGCCACGTACT</td><td></td><td></td><td></td><td></td><td></td><td></td><td></td><td></td><td></td><td></td></tr>
<tr><td>Chromosome</td><td>start</td><td>end</td><td>strand</td><td>MM</td><td>target_seq</td><td>PAM</td><td>alignment</td><td>distance</td><td>position</td><td>gene name</td><td>gene id</td></tr>
<tr><td>GU305914.1</td><td>107215</td><td>107237</td><td>+</td><td>0</td><td>AGTACGTGGCCGCCAATACC</td><td>CGG</td><td>||||||||[||||||||||||]PAM</td><td>NA</td><td>Intergenic</td><td>NA</td><td>NA</td></tr>
<tr><td></td><td></td><td></td><td></td><td></td><td></td><td></td><td></td><td></td><td></td><td></td><td></td></tr>
<tr><td>T7825</td><td>CGGACGTCGATCGAGTTGCTTGG</td><td>1000</td><td></td><td></td><td></td><td></td><td></td><td></td><td></td><td></td><td></td></tr>
<tr><td>Oligo fwd</td><td>CACCCGGACGTCGATCGAGTTGCT</td><td></td><td></td><td></td><td></td><td></td><td></td><td></td><td></td><td></td><td></td></tr>
<tr><td>Oligo rev</td><td>AAACAGCAACTCGATCGACGTCCG</td><td></td><td></td><td></td><td></td><td></td><td></td><td></td><td></td><td></td><td></td></tr>
<tr><td>Chromosome</td><td>start</td><td>end</td><td>strand</td><td>MM</td><td>target_seq</td><td>PAM</td><td>alignment</td><td>distance</td><td>position</td><td>gene name</td><td>gene id</td></tr>
<tr><td>GU305914.1</td><td>107235</td><td>107257</td><td>+</td><td>0</td><td>CGGACGTCGATCGAGTTGCT</td><td>TGG</td><td>||||||||[||||||||||||]PAM</td><td>NA</td><td>Intergenic</td><td>NA</td><td>NA</td></tr>
<tr><td></td><td></td><td></td><td></td><td></td><td></td><td></td><td></td><td></td><td></td><td></td><td></td></tr>
<tr><td>T7826</td><td>GATCGAGTTGCTTGGCTTTGTGG</td><td>1000</td><td></td><td></td><td></td><td></td><td></td><td></td><td></td><td></td><td></td></tr>
<tr><td>Oligo fwd</td><td>CACCGATCGAGTTGCTTGGCTTTG</td><td></td><td></td><td></td><td></td><td></td><td></td><td></td><td></td><td></td><td></td></tr>
<tr><td>Oligo rev</td><td>AAACCAAAGCCAAGCAACTCGATC</td><td></td><td></td><td></td><td></td><td></td><td></td><td></td><td></td><td></td><td></td></tr>
<tr><td>Chromosome</td><td>start</td><td>end</td><td>strand</td><td>MM</td><td>target_seq</td><td>PAM</td><td>alignment</td><td>distance</td><td>position</td><td>gene name</td><td>gene id</td></tr>
<tr><td>GU305914.1</td><td>107243</td><td>107265</td><td>+</td><td>0</td><td>GATCGAGTTGCTTGGCTTTG</td><td>TGG</td><td>||||||||[||||||||||||]PAM</td><td>NA</td><td>Intergenic</td><td>NA</td><td>NA</td></tr>
<tr><td></td><td></td><td></td><td></td><td></td><td></td><td></td><td></td><td></td><td></td><td></td><td></td></tr>
<tr><td>T7827</td><td>ATCGAGTTGCTTGGCTTTGTGGG</td><td>1000</td><td></td><td></td><td></td><td></td><td></td><td></td><td></td><td></td><td></td></tr>
<tr><td>Oligo fwd</td><td>CACCATCGAGTTGCTTGGCTTTGT</td><td></td><td></td><td></td><td></td><td></td><td></td><td></td><td></td><td></td><td></td></tr>
<tr><td>Oligo rev</td><td>AAACACAAAGCCAAGCAACTCGAT</td><td></td><td></td><td></td><td></td><td></td><td></td><td></td><td></td><td></td><td></td></tr>
<tr><td>Chromosome</td><td>start</td><td>end</td><td>strand</td><td>MM</td><td>target_seq</td><td>PAM</td><td>alignment</td><td>distance</td><td>position</td><td>gene name</td><td>gene id</td></tr>
<tr><td>GU305914.1</td><td>107244</td><td>107266</td><td>+</td><td>0</td><td>ATCGAGTTGCTTGGCTTTGT</td><td>GGG</td><td>||||||||[||||||||||||]PAM</td><td>NA</td><td>Intergenic</td><td>NA</td><td>NA</td></tr>
<tr><td></td><td></td><td></td><td></td><td></td><td></td><td></td><td></td><td></td><td></td><td></td><td></td></tr>
<tr><td>T7828</td><td>CTTTGTGGGTTTGTGATTAGCGG</td><td>1000</td><td></td><td></td><td></td><td></td><td></td><td></td><td></td><td></td><td></td></tr>
<tr><td>Oligo fwd</td><td>CACCCTTTGTGGGTTTGTGATTAG</td><td></td><td></td><td></td><td></td><td></td><td></td><td></td><td></td><td></td><td></td></tr>
<tr><td>Oligo rev</td><td>AAACCTAATCACAAACCCACAAAG</td><td></td><td></td><td></td><td></td><td></td><td></td><td></td><td></td><td></td><td></td></tr>
<tr><td>Chromosome</td><td>start</td><td>end</td><td>strand</td><td>MM</td><td>target_seq</td><td>PAM</td><td>alignment</td><td>distance</td><td>position</td><td>gene name</td><td>gene id</td></tr>
<tr><td>GU305914.1</td><td>107258</td><td>107280</td><td>+</td><td>0</td><td>CTTTGTGGGTTTGTGATTAG</td><td>CGG</td><td>||||||||[||||||||||||]PAM</td><td>NA</td><td>Intergenic</td><td>NA</td><td>NA</td></tr>
<tr><td></td><td></td><td></td><td></td><td></td><td></td><td></td><td></td><td></td><td></td><td></td><td></td></tr>
<tr><td>T7829</td><td>TGCATCCTCGTCTGAAACGCCGG</td><td>1000</td><td></td><td></td><td></td><td></td><td></td><td></td><td></td><td></td><td></td></tr>
<tr><td>Oligo fwd</td><td>CACCTGCATCCTCGTCTGAAACGC</td><td></td><td></td><td></td><td></td><td></td><td></td><td></td><td></td><td></td><td></td></tr>
</table>

Oligo rev AAACGCGTTTCAGACGAGGATGCA

| Chromosome | start | end | strand | MM | target_seq | PAM | alignment | distance | position | gene name | gene id |
|---|---|---|---|---|---|---|---|---|---|---|---|
| GU305914.1 | 107286 | 107308 | + | 0 | TGCATCCTCGTCTGAAACGC | CGG | ||||||||[||||||||||||]PAM | NA | Intergenic | NA | NA |

T7830 CTCGTCTGAAACGCCGGCGTCGG 1000
Oligo fwd CACCCTCGTCTGAAACGCCGGCGT
Oligo rev AAACACGCCGGCGTTTCAGACGAG

| Chromosome | start | end | strand | MM | target_seq | PAM | alignment | distance | position | gene name | gene id |
|---|---|---|---|---|---|---|---|---|---|---|---|
| GU305914.1 | 107292 | 107314 | + | 0 | CTCGTCTGAAACGCCGGCGT | CGG | ||||||||[||||||||||||]PAM | NA | Intergenic | NA | NA |

T7831 TCGTCTGAAACGCCGGCGTCGGG 1000
Oligo fwd CACCTCGTCTGAAACGCCGGCGTC
Oligo rev AAACGACGCCGGCGTTTCAGACGA

| Chromosome | start | end | strand | MM | target_seq | PAM | alignment | distance | position | gene name | gene id |
|---|---|---|---|---|---|---|---|---|---|---|---|
| GU305914.1 | 107293 | 107315 | + | 0 | TCGTCTGAAACGCCGGCGTC | GGG | ||||||||[||||||||||||]PAM | NA | Intergenic | NA | NA |

T7832 TGAAACGCCGGCGTCGGGTTCGG 1000
Oligo fwd CACCTGAAACGCCGGCGTCGGGTT
Oligo rev AAACAACCCGACGCCGGCGTTTCA

| Chromosome | start | end | strand | MM | target_seq | PAM | alignment | distance | position | gene name | gene id |
|---|---|---|---|---|---|---|---|---|---|---|---|
| GU305914.1 | 107298 | 107320 | + | 0 | TGAAACGCCGGCGTCGGGTT | CGG | ||||||||[||||||||||||]PAM | NA | Intergenic | NA | NA |

T7833 CGGCGTCGGGTTCGGTGTCGTGG 1000
Oligo fwd CACCCGGCGTCGGGTTCGGTGTCG
Oligo rev AAACCGACACCGAACCCGACGCCG

| Chromosome | start | end | strand | MM | target_seq | PAM | alignment | distance | position | gene name | gene id |
|---|---|---|---|---|---|---|---|---|---|---|---|
| GU305914.1 | 107306 | 107328 | + | 0 | CGGCGTCGGGTTCGGTGTCG | TGG | ||||||||[||||||||||||]PAM | NA | Intergenic | NA | NA |

T7834 GTGTCGTGGTTTTGAAACAGTGG 1000
Oligo fwd CACCGTGTCGTGGTTTTGAAACAG
Oligo rev AAACCTGTTTCAAAACCACGACAC

| Chromosome | start | end | strand | MM | target_seq | PAM | alignment | distance | position | gene name | gene id |
|---|---|---|---|---|---|---|---|---|---|---|---|
| GU305914.1 | 107320 | 107342 | + | 0 | GTGTCGTGGTTTTGAAACAG | TGG | ||||||||[||||||||||||]PAM | NA | Intergenic | NA | NA |

T7835 CCAATTCGCTCTGCCGCCAGAGG 1000
Oligo fwd CACCCCAATTCGCTCTGCCGCCAG
Oligo rev AAACCTGGCGGCAGAGCGAATTGG

| Chromosome | start | end | strand | MM | target_seq | PAM | alignment | distance | position | gene name | gene id |
|---|---|---|---|---|---|---|---|---|---|---|---|
| GU305914.1 | 107365 | 107387 | + | 0 | CCAATTCGCTCTGCCGCCAG | AGG | ||||||||[||||||||||||]PAM | NA | Intergenic | NA | NA |

T7836 CAATTCGCTCTGCCGCCAGAGGG 1000
Oligo fwd CACCCAATTCGCTCTGCCGCCAGA
Oligo rev AAACTCTGGCGGCAGAGCGAATTG

| Chromosome | start | end | strand | MM | target_seq | PAM | alignment | distance | position | gene name | gene id |
|---|---|---|---|---|---|---|---|---|---|---|---|
| GU305914.1 | 107366 | 107388 | + | 0 | CAATTCGCTCTGCCGCCAGA | GGG | ||||||||[||||||||||||]PAM | NA | Intergenic | NA | NA |

T7837 TTCGCTCTGCCGCCAGAGGGCGG 1000
Oligo fwd CACCTTCGCTCTGCCGCCAGAGGG
Oligo rev AAACCCCTCTGGCGGCAGAGCGAA

| Chromosome | start | end | strand | MM | target_seq | PAM | alignment | distance | position | gene name | gene id |
|---|---|---|---|---|---|---|---|---|---|---|---|
| GU305914.1 | 107369 | 107391 | + | 0 | TTCGCTCTGCCGCCAGAGGG | CGG | ||||||||[||||||||||||]PAM | NA | Intergenic | NA | NA |

T7838 CCGCCAGAGGGCGGTCGAGACGG 1000
Oligo fwd CACCCCGCCAGAGGGCGGTCGAGA
Oligo rev AAACTCTCGACCGCCCTCTGGCGG

| Chromosome | start | end | strand | MM | target_seq | PAM | alignment | distance | position | gene name | gene id |
|---|---|---|---|---|---|---|---|---|---|---|---|
| GU305914.1 | 107378 | 107400 | + | 0 | CCGCCAGAGGGCGGTCGAGA | CGG | ||||||||[||||||||||||]PAM | NA | Intergenic | NA | NA |
| GU305914.1 | 95323 | 95345 | - | 4 | CCGCTAGGTGGTGGTCGAGA | GGG | ||||-||-[-||-||||||||]PAM | NA | Intergenic | NA | NA |

T7839 CGCCAGAGGGCGGTCGAGACGGG 1000
Oligo fwd CACCCGCCAGAGGGCGGTCGAGAC
Oligo rev AAACGTCTCGACCGCCCTCTGGCG

| Chromosome | start | end | strand | MM | target_seq | PAM | alignment | distance | position | gene name | gene id |
|---|---|---|---|---|---|---|---|---|---|---|---|
| GU305914.1 | 107379 | 107401 | + | 0 | CGCCAGAGGGCGGTCGAGAC | GGG | ||||||||[||||||||||||]PAM | NA | Intergenic | NA | NA |
| GU305914.1 | 102346 | 102368 | - | 4 | AGCCAGAAGACGGTCGAGCC | GGG | -||||||-[|-||||||||-|]PAM | NA | Intergenic | NA | NA |

T7840 CATGAGCTGACACAGAGAATAGG 1000
Oligo fwd CACCCATGAGCTGACACAGAGAAT
Oligo rev AAACATTCTCTGTGTCAGCTCATG

| Chromosome | start | end | strand | MM | target_seq | PAM | alignment | distance | position | gene name | gene id |
|---|---|---|---|---|---|---|---|---|---|---|---|
| GU305914.1 | 107432 | 107454 | + | 0 | CATGAGCTGACACAGAGAAT | AGG | ||||||||[||||||||||||]PAM | NA | Intergenic | NA | NA |

T7841 CGACCGTGTCGCGCGTGATGAGG 1000
Oligo fwd CACCCGACCGTGTCGCGCGTGATG
Oligo rev AAACCATCACGCGCGACACGGTCG

| Chromosome | start | end | strand | MM | target_seq | PAM | alignment | distance | position | gene name | gene id |
|---|---|---|---|---|---|---|---|---|---|---|---|
| GU305914.1 | 107467 | 107489 | + | 0 | CGACCGTGTCGCGCGTGATG | AGG | ||||||||[||||||||||||]PAM | NA | Intergenic | NA | NA |

T7842 GAGGCCGCACTGCTTGCCTCTGG 1000
Oligo fwd CACCGAGGCCGCACTGCTTGCCTC
Oligo rev AAACGAGGCAAGCAGTGCGGCCTC

| Chromosome | start | end | strand | MM | target_seq | PAM | alignment | distance | position | gene name | gene id |
|---|---|---|---|---|---|---|---|---|---|---|---|
| GU305914.1 | 107486 | 107508 | + | 0 | GAGGCCGCACTGCTTGCCTC | TGG | ||||||||[||||||||||||]PAM | NA | Intergenic | NA | NA |

T7843 CACGCTCGTGTCCGCGTAGTCGG 1000
Oligo fwd CACCCACGCTCGTGTCCGCGTAGT
Oligo rev AAACACTACGCGGACACGAGCGTG

| Chromosome | start | end | strand | MM | target_seq | PAM | alignment | distance | position | gene name | gene id |
|---|---|---|---|---|---|---|---|---|---|---|---|
| GU305914.1 | 107546 | 107568 | + | 0 | CACGCTCGTGTCCGCGTAGT | CGG | ||||||||[||||||||||||]PAM | NA | Intergenic | NA | NA |

| | | | | | | | | | | | |
|---|---|---|---|---|---|---|---|---|---|---|---|
| T7844 | CGGCCGACAGCGTGCGTTGCAGG | 1000 | | | | | | | | | |
| Oligo fwd | CACCCGGCCGACAGCGTGCGTTGC | | | | | | | | | | |
| Oligo rev | AAACGCAACGCACGCTGTCGGCCG | | | | | | | | | | |
| Chromosome | start | end | strand | MM | target_seq | PAM | alignment | distance | position | gene name | gene id |
| GU305914.1 | 107566 | 107588 | + | 0 | CGGCCGACAGCGTGCGTTGC | AGG | \|\|\|\|\|\|\|\|[\|\|\|\|\|\|\|\|\|\|\|\|]PAM | NA | Intergenic | NA | NA |
| T7845 | GCGTTGCAGGAACACCACATCGG | 1000 | | | | | | | | | |
| Oligo fwd | CACCGCGTTGCAGGAACACCACAT | | | | | | | | | | |
| Oligo rev | AAACATGTGGTGTTCCTGCAACGC | | | | | | | | | | |
| Chromosome | start | end | strand | MM | target_seq | PAM | alignment | distance | position | gene name | gene id |
| GU305914.1 | 107579 | 107601 | + | 0 | GCGTTGCAGGAACACCACAT | CGG | \|\|\|\|\|\|\|\|[\|\|\|\|\|\|\|\|\|\|\|\|]PAM | NA | Intergenic | NA | NA |
| T7846 | GCAGGAACACCACATCGGTGAGG | 1000 | | | | | | | | | |
| Oligo fwd | CACCGCAGGAACACCACATCGGTG | | | | | | | | | | |
| Oligo rev | AAACCACCGATGTGGTGTTCCTGC | | | | | | | | | | |
| Chromosome | start | end | strand | MM | target_seq | PAM | alignment | distance | position | gene name | gene id |
| GU305914.1 | 107584 | 107606 | + | 0 | GCAGGAACACCACATCGGTG | AGG | \|\|\|\|\|\|\|\|[\|\|\|\|\|\|\|\|\|\|\|\|]PAM | NA | Intergenic | NA | NA |
| T7847 | CGGTGAGGAACATGTAGCGTTGG | 1000 | | | | | | | | | |
| Oligo fwd | CACCCGGTGAGGAACATGTAGCGT | | | | | | | | | | |
| Oligo rev | AAACACGCTACATGTTCCTCACCG | | | | | | | | | | |
| Chromosome | start | end | strand | MM | target_seq | PAM | alignment | distance | position | gene name | gene id |
| GU305914.1 | 107599 | 107621 | + | 0 | CGGTGAGGAACATGTAGCGT | TGG | \|\|\|\|\|\|\|\|[\|\|\|\|\|\|\|\|\|\|\|\|]PAM | NA | Intergenic | NA | NA |
| T7848 | CTGTCTGTTTGACCACCACCAGG | 1000 | | | | | | | | | |
| Oligo fwd | CACCCTGTCTGTTTGACCACCACC | | | | | | | | | | |
| Oligo rev | AAACGGTGGTGGTCAAACAGACAG | | | | | | | | | | |
| Chromosome | start | end | strand | MM | target_seq | PAM | alignment | distance | position | gene name | gene id |
| GU305914.1 | 107629 | 107651 | + | 0 | CTGTCTGTTTGACCACCACC | AGG | \|\|\|\|\|\|\|\|[\|\|\|\|\|\|\|\|\|\|\|\|]PAM | NA | Intergenic | NA | NA |
| T7849 | TCTGTTTGACCACCACCAGGAGG | 1000 | | | | | | | | | |
| Oligo fwd | CACCTCTGTTTGACCACCACCAGG | | | | | | | | | | |
| Oligo rev | AAACCCTGGTGGTGGTCAAACAGA | | | | | | | | | | |
| Chromosome | start | end | strand | MM | target_seq | PAM | alignment | distance | position | gene name | gene id |
| GU305914.1 | 107632 | 107654 | + | 0 | TCTGTTTGACCACCACCAGG | AGG | \|\|\|\|\|\|\|\|[\|\|\|\|\|\|\|\|\|\|\|\|]PAM | NA | Intergenic | NA | NA |
| T7850 | TGACCACCACCAGGAGGTCGTGG | 1000 | | | | | | | | | |
| Oligo fwd | CACCTGACCACCACCAGGAGGTCG | | | | | | | | | | |
| Oligo rev | AAACCGACCTCCTGGTGGTGGTCA | | | | | | | | | | |
| Chromosome | start | end | strand | MM | target_seq | PAM | alignment | distance | position | gene name | gene id |
| GU305914.1 | 107638 | 107660 | + | 0 | TGACCACCACCAGGAGGTCG | TGG | \|\|\|\|\|\|\|\|[\|\|\|\|\|\|\|\|\|\|\|\|]PAM | NA | Intergenic | NA | NA |
| GU305914.1 | 140440 | 140462 | + | 4 | GCAGCACCACCAGCAGGTCG | CGG | --\|-\|\|\|\|[\|\|\|\|\|-\|\|\|\|\|\|]PAM | NA | Intergenic | NA | NA |
| T7851 | CACCACCAGGAGGTCGTGGTCGG | 1000 | | | | | | | | | |
| Oligo fwd | CACCCACCACCAGGAGGTCGTGGT | | | | | | | | | | |
| Oligo rev | AAACACCACGACCTCCTGGTGGTG | | | | | | | | | | |
| Chromosome | start | end | strand | MM | target_seq | PAM | alignment | distance | position | gene name | gene id |
| GU305914.1 | 107642 | 107664 | + | 0 | CACCACCAGGAGGTCGTGGT | CGG | \|\|\|\|\|\|\|\|[\|\|\|\|\|\|\|\|\|\|\|\|]PAM | NA | Intergenic | NA | NA |
| GU305914.1 | 106376 | 106398 | + | 3 | CGCCACCAGGATGTCGTTGT | CGG | \|-\|\|\|\|\|\|[\|\|\|-\|\|\|\|\|-\|\|]PAM | NA | Intergenic | NA | NA |
| T7852 | ACCACCAGGAGGTCGTGGTCGGG | 1000 | | | | | | | | | |
| Oligo fwd | CACCACCACCAGGAGGTCGTGGTC | | | | | | | | | | |
| Oligo rev | AAACGACCACGACCTCCTGGTGGT | | | | | | | | | | |
| Chromosome | start | end | strand | MM | target_seq | PAM | alignment | distance | position | gene name | gene id |
| GU305914.1 | 107643 | 107665 | + | 0 | ACCACCAGGAGGTCGTGGTC | GGG | \|\|\|\|\|\|\|\|[\|\|\|\|\|\|\|\|\|\|\|\|]PAM | NA | Intergenic | NA | NA |
| GU305914.1 | 106377 | 106399 | + | 3 | GCCACCAGGATGTCGTTGTC | GGG | -\|\|\|\|\|\|\|[\|\|-\|\|\|\|\|-\|\|\|]PAM | NA | Intergenic | NA | NA |
| T7853 | CGTGGTCGGGCGTCTGCCGCAGG | 1000 | | | | | | | | | |
| Oligo fwd | CACCCGTGGTCGGGCGTCTGCCGC | | | | | | | | | | |
| Oligo rev | AAACGCGGCAGACGCCCGACCACG | | | | | | | | | | |
| Chromosome | start | end | strand | MM | target_seq | PAM | alignment | distance | position | gene name | gene id |
| GU305914.1 | 107656 | 107678 | + | 0 | CGTGGTCGGGCGTCTGCCGC | AGG | \|\|\|\|\|\|\|\|[\|\|\|\|\|\|\|\|\|\|\|\|]PAM | NA | Intergenic | NA | NA |
| GU305914.1 | 197436 | 197458 | - | 4 | CGCCGGCGAGCGTCTGCCGC | TGG | \|\|--\|-\|\|[-\|\|\|\|\|\|\|\|\|\|\|]PAM | NA | Intergenic | NA | NA |
| T7854 | GGTCGGGCGTCTGCCGCAGGAGG | 1000 | | | | | | | | | |
| Oligo fwd | CACCGGTCGGGCGTCTGCCGCAGG | | | | | | | | | | |
| Oligo rev | AAACCCTGCGGCAGACGCCCGACC | | | | | | | | | | |
| Chromosome | start | end | strand | MM | target_seq | PAM | alignment | distance | position | gene name | gene id |
| GU305914.1 | 107659 | 107681 | + | 0 | GGTCGGGCGTCTGCCGCAGG | AGG | \|\|\|\|\|\|\|\|[\|\|\|\|\|\|\|\|\|\|\|\|]PAM | NA | Intergenic | NA | NA |
| T7855 | GGGCGTCTGCCGCAGGAGGAAGG | 1000 | | | | | | | | | |
| Oligo fwd | CACCGGGCGTCTGCCGCAGGAGGA | | | | | | | | | | |
| Oligo rev | AAACTCCTCCTGCGGCAGACGCCC | | | | | | | | | | |
| Chromosome | start | end | strand | MM | target_seq | PAM | alignment | distance | position | gene name | gene id |
| GU305914.1 | 107663 | 107685 | + | 0 | GGGCGTCTGCCGCAGGAGGA | AGG | \|\|\|\|\|\|\|\|[\|\|\|\|\|\|\|\|\|\|\|\|]PAM | NA | Intergenic | NA | NA |
| T7856 | CCGCAGGAGGAAGGCTCCGACGG | 1000 | | | | | | | | | |
| Oligo fwd | CACCCCGCAGGAGGAAGGCTCCGA | | | | | | | | | | |
| Oligo rev | AAACTCGGAGCCTTCCTCCTGCGG | | | | | | | | | | |
| Chromosome | start | end | strand | MM | target_seq | PAM | alignment | distance | position | gene name | gene id |
| GU305914.1 | 107672 | 107694 | + | 0 | CCGCAGGAGGAAGGCTCCGA | CGG | \|\|\|\|\|\|\|\|[\|\|\|\|\|\|\|\|\|\|\|\|]PAM | NA | Intergenic | NA | NA |
| T7857 | AGGAAGGCTCCGACGGTGTACGG | 1000 | | | | | | | | | |
| Oligo fwd | CACCAGGAAGGCTCCGACGGTGTA | | | | | | | | | | |
| Oligo rev | AAACTACACCGTCGGAGCCTTCCT | | | | | | | | | | |
| Chromosome | start | end | strand | MM | target_seq | PAM | alignment | distance | position | gene name | gene id |
| GU305914.1 | 107679 | 107701 | + | 0 | AGGAAGGCTCCGACGGTGTA | CGG | \|\|\|\|\|\|\|\|[\|\|\|\|\|\|\|\|\|\|\|\|]PAM | NA | Intergenic | NA | NA |

T7858 TCTCACGTCTCCGAAGAACATGG 1000
Oligo fwd CACCTCTCACGTCTCCGAAGAACA
Oligo rev AAACTGTTCTTCGGAGACGTGAGA
Chromosome start end strand MM target_seq PAM alignment distance position gene name gene id
GU305914.1 107720 107742 + 0 TCTCACGTCTCCGAAGAACA TGG ||||||||[||||||||||||]PAM NA Intergenic NA NA

T7859 CATGGCCAACGTGTTCTCCGAGG 1000
Oligo fwd CACCCATGGCCAACGTGTTCTCCG
Oligo rev AAACCGGAGAACACGTTGGCCATG
Chromosome start end strand MM target_seq PAM alignment distance position gene name gene id
GU305914.1 107738 107760 + 0 CATGGCCAACGTGTTCTCCG AGG ||||||||[||||||||||||]PAM NA Intergenic NA NA

T7860 CCAACGTGTTCTCCGAGGTGTGG 1000
Oligo fwd CACCCCAACGTGTTCTCCGAGGTG
Oligo rev AAACCACCTCGGAGAACACGTTGG
Chromosome start end strand MM target_seq PAM alignment distance position gene name gene id
GU305914.1 107743 107765 + 0 CCAACGTGTTCTCCGAGGTG TGG ||||||||[||||||||||||]PAM NA Intergenic NA NA

T7861 TGTTCTCCGAGGTGTGGTCGCGG 1000
Oligo fwd CACCTGTTCTCCGAGGTGTGGTCG
Oligo rev AAACCGACCACACCTCGGAGAACA
Chromosome start end strand MM target_seq PAM alignment distance position gene name gene id
GU305914.1 107749 107771 + 0 TGTTCTCCGAGGTGTGGTCG CGG ||||||||[||||||||||||]PAM NA Intergenic NA NA

T7862 GTTCTCCGAGGTGTGGTCGCGGG 1000
Oligo fwd CACCGTTCTCCGAGGTGTGGTCGC
Oligo rev AAACGCGACCACACCTCGGAGAAC
Chromosome start end strand MM target_seq PAM alignment distance position gene name gene id
GU305914.1 107750 107772 + 0 GTTCTCCGAGGTGTGGTCGC GGG ||||||||[||||||||||||]PAM NA Intergenic NA NA

T7863 TCCGAGGTGTGGTCGCGGGTAGG 1000
Oligo fwd CACCTCCGAGGTGTGGTCGCGGGT
Oligo rev AAACACCCGCGACCACACCTCGGA
Chromosome start end strand MM target_seq PAM alignment distance position gene name gene id
GU305914.1 107754 107776 + 0 TCCGAGGTGTGGTCGCGGGT AGG ||||||||[||||||||||||]PAM NA Intergenic NA NA

T7864 GTGGTCGCGGGTAGGTTTGATGG 1000
Oligo fwd CACCGTGGTCGCGGGTAGGTTTGA
Oligo rev AAACTCAAACCTACCCGCGACCAC
Chromosome start end strand MM target_seq PAM alignment distance position gene name gene id
GU305914.1 107762 107784 + 0 GTGGTCGCGGGTAGGTTTGA TGG ||||||||[||||||||||||]PAM NA Intergenic NA NA

T7865 CGCGGGTAGGTTTGATGGTGAGG 1000
Oligo fwd CACCCGCGGGTAGGTTTGATGGTG
Oligo rev AAACCACCATCAAACCTACCCGCG
Chromosome start end strand MM target_seq PAM alignment distance position gene name gene id
GU305914.1 107767 107789 + 0 CGCGGGTAGGTTTGATGGTG AGG ||||||||[||||||||||||]PAM NA Intergenic NA NA

T7866 GGACGAAGAACTTCTGCGCGAGG 1000
Oligo fwd CACCGGACGAAGAACTTCTGCGCG
Oligo rev AAACCGCGCAGAAGTTCTTCGTCC
Chromosome start end strand MM target_seq PAM alignment distance position gene name gene id
GU305914.1 107788 107810 + 0 GGACGAAGAACTTCTGCGCG AGG ||||||||[||||||||||||]PAM NA Intergenic NA NA

T7867 CTGCGCGAGGCTCACCGTCACGG 1000
Oligo fwd CACCCTGCGCGAGGCTCACCGTCA
Oligo rev AAACTGACGGTGAGCCTCGCGCAG
Chromosome start end strand MM target_seq PAM alignment distance position gene name gene id
GU305914.1 107801 107823 + 0 CTGCGCGAGGCTCACCGTCA CGG ||||||||[||||||||||||]PAM NA Intergenic NA NA

T7868 CGTCACGGTATATCCCGTCCAGG 1000
Oligo fwd CACCCGTCACGGTATATCCCGTCC
Oligo rev AAACGGACGGGATATACCGTGACG
Chromosome start end strand MM target_seq PAM alignment distance position gene name gene id
GU305914.1 107816 107838 + 0 CGTCACGGTATATCCCGTCC AGG ||||||||[||||||||||||]PAM NA Intergenic NA NA

T7869 CCGTCCAGGTGATGACCTGATGG 1000
Oligo fwd CACCCCGTCCAGGTGATGACCTGA
Oligo rev AAACTCAGGTCATCACCTGGACGG
Chromosome start end strand MM target_seq PAM alignment distance position gene name gene id
GU305914.1 107830 107852 + 0 CCGTCCAGGTGATGACCTGA TGG ||||||||[||||||||||||]PAM NA Intergenic NA NA

T7870 GTGATGACCTGATGGCTGCACGG 1000
Oligo fwd CACCGTGATGACCTGATGGCTGCA
Oligo rev AAACTGCAGCCATCAGGTCATCAC
Chromosome start end strand MM target_seq PAM alignment distance position gene name gene id
GU305914.1 107838 107860 + 0 GTGATGACCTGATGGCTGCA CGG ||||||||[||||||||||||]PAM NA Intergenic NA NA

T7871 ATGGCTGCACGGTACCTTCATGG 1000
Oligo fwd CACCATGGCTGCACGGTACCTTCA
Oligo rev AAACTGAAGGTACCGTGCAGCCAT
Chromosome start end strand MM target_seq PAM alignment distance position gene name gene id
GU305914.1 107849 107871 + 0 ATGGCTGCACGGTACCTTCA TGG ||||||||[||||||||||||]PAM NA Intergenic NA NA

T7872 ACCTTCATGGAGAAGATCACCGG 1000
Oligo fwd CACCACCTTCATGGAGAAGATCAC
Oligo rev AAACGTGATCTTCTCCATGAAGGT
Chromosome start end strand MM target_seq PAM alignment distance position gene name gene id
GU305914.1 107862 107884 + 0 ACCTTCATGGAGAAGATCAC CGG ||||||||[||||||||||||]PAM NA Intergenic NA NA

<table>
<tr><td>T7873</td><td>CCTTCATGGAGAAGATCACCGGG</td><td>1000</td></tr>
<tr><td>Oligo fwd</td><td>CACCCCTTCATGGAGAAGATCACC</td></tr>
<tr><td>Oligo rev</td><td>AAACGGTGATCTTCTCCATGAAGG</td></tr>
<tr><td>Chromosome</td><td>start</td><td>end</td><td>strand</td><td>MM</td><td>target_seq</td><td>PAM</td><td>alignment</td><td>distance</td><td>position</td><td>gene name</td><td>gene id</td></tr>
<tr><td>GU305914.1</td><td>107863</td><td>107885</td><td>+</td><td>0</td><td>CCTTCATGGAGAAGATCACC</td><td>GGG</td><td>||||||||[||||||||||||]PAM</td><td>NA</td><td>Intergenic</td><td>NA</td><td>NA</td></tr>
<tr><td>T7874</td><td>GGAGAAGATCACCGGGTGCTCGG</td><td>1000</td></tr>
<tr><td>Oligo fwd</td><td>CACCGGAGAAGATCACCGGGTGCT</td></tr>
<tr><td>Oligo rev</td><td>AAACAGCACCCGGTGATCTTCTCC</td></tr>
<tr><td>Chromosome</td><td>start</td><td>end</td><td>strand</td><td>MM</td><td>target_seq</td><td>PAM</td><td>alignment</td><td>distance</td><td>position</td><td>gene name</td><td>gene id</td></tr>
<tr><td>GU305914.1</td><td>107870</td><td>107892</td><td>+</td><td>0</td><td>GGAGAAGATCACCGGGTGCT</td><td>CGG</td><td>||||||||[||||||||||||]PAM</td><td>NA</td><td>Intergenic</td><td>NA</td><td>NA</td></tr>
<tr><td>T7875</td><td>GTGATGACGCAGAGTTTATCCGG</td><td>1000</td></tr>
<tr><td>Oligo fwd</td><td>CACCGTGATGACGCAGAGTTTATC</td></tr>
<tr><td>Oligo rev</td><td>AAACGATAAACTCTGCGTCATCAC</td></tr>
<tr><td>Chromosome</td><td>start</td><td>end</td><td>strand</td><td>MM</td><td>target_seq</td><td>PAM</td><td>alignment</td><td>distance</td><td>position</td><td>gene name</td><td>gene id</td></tr>
<tr><td>GU305914.1</td><td>107892</td><td>107914</td><td>+</td><td>0</td><td>GTGATGACGCAGAGTTTATC</td><td>CGG</td><td>||||||||[||||||||||||]PAM</td><td>NA</td><td>Intergenic</td><td>NA</td><td>NA</td></tr>
<tr><td>T7876</td><td>GTTTATCCGGAACGACCGTGTGG</td><td>1000</td></tr>
<tr><td>Oligo fwd</td><td>CACCGTTTATCCGGAACGACCGTG</td></tr>
<tr><td>Oligo rev</td><td>AAACCACGGTCGTTCCGGATAAAC</td></tr>
<tr><td>Chromosome</td><td>start</td><td>end</td><td>strand</td><td>MM</td><td>target_seq</td><td>PAM</td><td>alignment</td><td>distance</td><td>position</td><td>gene name</td><td>gene id</td></tr>
<tr><td>GU305914.1</td><td>107905</td><td>107927</td><td>+</td><td>0</td><td>GTTTATCCGGAACGACCGTG</td><td>TGG</td><td>||||||||[||||||||||||]PAM</td><td>NA</td><td>Intergenic</td><td>NA</td><td>NA</td></tr>
<tr><td>T7877</td><td>TGTTCGTCGCGTCGACGATCAGG</td><td>1000</td></tr>
<tr><td>Oligo fwd</td><td>CACCTGTTCGTCGCGTCGACGATC</td></tr>
<tr><td>Oligo rev</td><td>AAACGATCGTCGACGCGACGAACA</td></tr>
<tr><td>Chromosome</td><td>start</td><td>end</td><td>strand</td><td>MM</td><td>target_seq</td><td>PAM</td><td>alignment</td><td>distance</td><td>position</td><td>gene name</td><td>gene id</td></tr>
<tr><td>GU305914.1</td><td>107929</td><td>107951</td><td>+</td><td>0</td><td>TGTTCGTCGCGTCGACGATC</td><td>AGG</td><td>||||||||[||||||||||||]PAM</td><td>NA</td><td>Intergenic</td><td>NA</td><td>NA</td></tr>
<tr><td>T7878</td><td>GTCGACGATCAGGTCCTGCATGG</td><td>1000</td></tr>
<tr><td>Oligo fwd</td><td>CACCGTCGACGATCAGGTCCTGCA</td></tr>
<tr><td>Oligo rev</td><td>AAACTGCAGGACCTGATCGTCGAC</td></tr>
<tr><td>Chromosome</td><td>start</td><td>end</td><td>strand</td><td>MM</td><td>target_seq</td><td>PAM</td><td>alignment</td><td>distance</td><td>position</td><td>gene name</td><td>gene id</td></tr>
<tr><td>GU305914.1</td><td>107939</td><td>107961</td><td>+</td><td>0</td><td>GTCGACGATCAGGTCCTGCA</td><td>TGG</td><td>||||||||[||||||||||||]PAM</td><td>NA</td><td>Intergenic</td><td>NA</td><td>NA</td></tr>
<tr><td>T7879</td><td>GTCCTGCATGGTCACTGAGCTGG</td><td>1000</td></tr>
<tr><td>Oligo fwd</td><td>CACCGTCCTGCATGGTCACTGAGC</td></tr>
<tr><td>Oligo rev</td><td>AAACGCTCAGTGACCATGCAGGAC</td></tr>
<tr><td>Chromosome</td><td>start</td><td>end</td><td>strand</td><td>MM</td><td>target_seq</td><td>PAM</td><td>alignment</td><td>distance</td><td>position</td><td>gene name</td><td>gene id</td></tr>
<tr><td>GU305914.1</td><td>107951</td><td>107973</td><td>+</td><td>0</td><td>GTCCTGCATGGTCACTGAGC</td><td>TGG</td><td>||||||||[||||||||||||]PAM</td><td>NA</td><td>Intergenic</td><td>NA</td><td>NA</td></tr>
<tr><td>T7880</td><td>TGCATGGTCACTGAGCTGGCCGG</td><td>1000</td></tr>
<tr><td>Oligo fwd</td><td>CACCTGCATGGTCACTGAGCTGGC</td></tr>
<tr><td>Oligo rev</td><td>AAACGCCAGCTCAGTGACCATGCA</td></tr>
<tr><td>Chromosome</td><td>start</td><td>end</td><td>strand</td><td>MM</td><td>target_seq</td><td>PAM</td><td>alignment</td><td>distance</td><td>position</td><td>gene name</td><td>gene id</td></tr>
<tr><td>GU305914.1</td><td>107955</td><td>107977</td><td>+</td><td>0</td><td>TGCATGGTCACTGAGCTGGC</td><td>CGG</td><td>||||||||[||||||||||||]PAM</td><td>NA</td><td>Intergenic</td><td>NA</td><td>NA</td></tr>
<tr><td>T7881</td><td>ATGGTCACTGAGCTGGCCGGCGG</td><td>1000</td></tr>
<tr><td>Oligo fwd</td><td>CACCATGGTCACTGAGCTGGCCGG</td></tr>
<tr><td>Oligo rev</td><td>AAACCCGGCCAGCTCAGTGACCAT</td></tr>
<tr><td>Chromosome</td><td>start</td><td>end</td><td>strand</td><td>MM</td><td>target_seq</td><td>PAM</td><td>alignment</td><td>distance</td><td>position</td><td>gene name</td><td>gene id</td></tr>
<tr><td>GU305914.1</td><td>107958</td><td>107980</td><td>+</td><td>0</td><td>ATGGTCACTGAGCTGGCCGG</td><td>CGG</td><td>||||||||[||||||||||||]PAM</td><td>NA</td><td>Intergenic</td><td>NA</td><td>NA</td></tr>
<tr><td>T7882</td><td>TGGTCACTGAGCTGGCCGGCGGG</td><td>1000</td></tr>
<tr><td>Oligo fwd</td><td>CACCTGGTCACTGAGCTGGCCGGC</td></tr>
<tr><td>Oligo rev</td><td>AAACGCCGGCCAGCTCAGTGACCA</td></tr>
<tr><td>Chromosome</td><td>start</td><td>end</td><td>strand</td><td>MM</td><td>target_seq</td><td>PAM</td><td>alignment</td><td>distance</td><td>position</td><td>gene name</td><td>gene id</td></tr>
<tr><td>GU305914.1</td><td>107959</td><td>107981</td><td>+</td><td>0</td><td>TGGTCACTGAGCTGGCCGGC</td><td>GGG</td><td>||||||||[||||||||||||]PAM</td><td>NA</td><td>Intergenic</td><td>NA</td><td>NA</td></tr>
<tr><td>GU305914.1</td><td>122448</td><td>122470</td><td>+</td><td>4</td><td>TGGTCCTCGAGCGGGCCGGC</td><td>GGG</td><td>|||||---[||||-|||||||]PAM</td><td>NA</td><td>Intergenic</td><td>NA</td><td>NA</td></tr>
<tr><td>T7883</td><td>GGTCACTGAGCTGGCCGGCGGGG</td><td>1000</td></tr>
<tr><td>Oligo fwd</td><td>CACCGGTCACTGAGCTGGCCGGCG</td></tr>
<tr><td>Oligo rev</td><td>AAACCGCCGGCCAGCTCAGTGACC</td></tr>
<tr><td>Chromosome</td><td>start</td><td>end</td><td>strand</td><td>MM</td><td>target_seq</td><td>PAM</td><td>alignment</td><td>distance</td><td>position</td><td>gene name</td><td>gene id</td></tr>
<tr><td>GU305914.1</td><td>107960</td><td>107982</td><td>+</td><td>0</td><td>GGTCACTGAGCTGGCCGGCG</td><td>GGG</td><td>||||||||[||||||||||||]PAM</td><td>NA</td><td>Intergenic</td><td>NA</td><td>NA</td></tr>
<tr><td>T7884</td><td>GTCACTGAGCTGGCCGGCGGGGG</td><td>1000</td></tr>
<tr><td>Oligo fwd</td><td>CACCGTCACTGAGCTGGCCGGCGG</td></tr>
<tr><td>Oligo rev</td><td>AAACCCGCCGGCCAGCTCAGTGAC</td></tr>
<tr><td>Chromosome</td><td>start</td><td>end</td><td>strand</td><td>MM</td><td>target_seq</td><td>PAM</td><td>alignment</td><td>distance</td><td>position</td><td>gene name</td><td>gene id</td></tr>
<tr><td>GU305914.1</td><td>107961</td><td>107983</td><td>+</td><td>0</td><td>GTCACTGAGCTGGCCGGCGG</td><td>GGG</td><td>||||||||[||||||||||||]PAM</td><td>NA</td><td>Intergenic</td><td>NA</td><td>NA</td></tr>
<tr><td>T7885</td><td>TCACTGAGCTGGCCGGCGGGGGG</td><td>1000</td></tr>
<tr><td>Oligo fwd</td><td>CACCTCACTGAGCTGGCCGGCGGG</td></tr>
<tr><td>Oligo rev</td><td>AAACCCCGCCGGCCAGCTCAGTGA</td></tr>
<tr><td>Chromosome</td><td>start</td><td>end</td><td>strand</td><td>MM</td><td>target_seq</td><td>PAM</td><td>alignment</td><td>distance</td><td>position</td><td>gene name</td><td>gene id</td></tr>
<tr><td>GU305914.1</td><td>107962</td><td>107984</td><td>+</td><td>0</td><td>TCACTGAGCTGGCCGGCGGG</td><td>GGG</td><td>||||||||[||||||||||||]PAM</td><td>NA</td><td>Intergenic</td><td>NA</td><td>NA</td></tr>
<tr><td>T7886</td><td>AGCTGGCCGGCGGGGGGTTCAGG</td><td>1000</td></tr>
<tr><td>Oligo fwd</td><td>CACCAGCTGGCCGGCGGGGGGTTC</td></tr>
<tr><td>Oligo rev</td><td>AAACGAACCCCCCGCCGGCCAGCT</td></tr>
<tr><td>Chromosome</td><td>start</td><td>end</td><td>strand</td><td>MM</td><td>target_seq</td><td>PAM</td><td>alignment</td><td>distance</td><td>position</td><td>gene name</td><td>gene id</td></tr>
<tr><td>GU305914.1</td><td>107968</td><td>107990</td><td>+</td><td>0</td><td>AGCTGGCCGGCGGGGGGTTC</td><td>AGG</td><td>||||||||[||||||||||||]PAM</td><td>NA</td><td>Intergenic</td><td>NA</td><td>NA</td></tr>
<tr><td>T7887</td><td>GGGGGGTTCAGGACGCCGCCCGG</td><td>1000</td></tr>
<tr><td>Oligo fwd</td><td>CACCGGGGGGTTCAGGACGCCGCC</td></tr>
<tr><td>Oligo rev</td><td>AAACGGCGGCGTCCTGAACCCCCC</td></tr>
</table>

<table>
<tr><td>Chromosome</td><td>start</td><td>end</td><td>strand</td><td>MM</td><td>target_seq</td><td>PAM</td><td>alignment</td><td>distance</td><td>position</td><td>gene name</td><td>gene id</td></tr>
<tr><td>GU305914.1</td><td>107979</td><td>108001</td><td>+</td><td>0</td><td>GGGGGGTTCAGGACGCCGCC</td><td>CGG</td><td>||||||||[||||||||||||]PAM</td><td>NA</td><td>Intergenic</td><td>NA</td><td>NA</td></tr>
<tr><td></td><td></td><td></td><td></td><td></td><td></td><td></td><td></td><td></td><td></td><td></td><td></td></tr>
<tr><td>T7888</td><td>CAGGACGCCGCCCGGATACTTGG</td><td>1000</td><td></td><td></td><td></td><td></td><td></td><td></td><td></td><td></td><td></td></tr>
<tr><td>Oligo fwd</td><td>CACCCAGGACGCCGCCCGGATACT</td><td></td><td></td><td></td><td></td><td></td><td></td><td></td><td></td><td></td><td></td></tr>
<tr><td>Oligo rev</td><td>AAACAGTATCCGGGCGGCGTCCTG</td><td></td><td></td><td></td><td></td><td></td><td></td><td></td><td></td><td></td><td></td></tr>
<tr><td>Chromosome</td><td>start</td><td>end</td><td>strand</td><td>MM</td><td>target_seq</td><td>PAM</td><td>alignment</td><td>distance</td><td>position</td><td>gene name</td><td>gene id</td></tr>
<tr><td>GU305914.1</td><td>107987</td><td>108009</td><td>+</td><td>0</td><td>CAGGACGCCGCCCGGATACT</td><td>TGG</td><td>||||||||[||||||||||||]PAM</td><td>NA</td><td>Intergenic</td><td>NA</td><td>NA</td></tr>
<tr><td></td><td></td><td></td><td></td><td></td><td></td><td></td><td></td><td></td><td></td><td></td><td></td></tr>
<tr><td>T7889</td><td>GCCGCCCGGATACTTGGCCGTGG</td><td>1000</td><td></td><td></td><td></td><td></td><td></td><td></td><td></td><td></td><td></td></tr>
<tr><td>Oligo fwd</td><td>CACCGCCGCCCGGATACTTGGCCG</td><td></td><td></td><td></td><td></td><td></td><td></td><td></td><td></td><td></td><td></td></tr>
<tr><td>Oligo rev</td><td>AAACCGGCCAAGTATCCGGGCGGC</td><td></td><td></td><td></td><td></td><td></td><td></td><td></td><td></td><td></td><td></td></tr>
<tr><td>Chromosome</td><td>start</td><td>end</td><td>strand</td><td>MM</td><td>target_seq</td><td>PAM</td><td>alignment</td><td>distance</td><td>position</td><td>gene name</td><td>gene id</td></tr>
<tr><td>GU305914.1</td><td>107993</td><td>108015</td><td>+</td><td>0</td><td>GCCGCCCGGATACTTGGCCG</td><td>TGG</td><td>||||||||[||||||||||||]PAM</td><td>NA</td><td>Intergenic</td><td>NA</td><td>NA</td></tr>
<tr><td></td><td></td><td></td><td></td><td></td><td></td><td></td><td></td><td></td><td></td><td></td><td></td></tr>
<tr><td>T7890</td><td>CCGCCCGGATACTTGGCCGTGGG</td><td>1000</td><td></td><td></td><td></td><td></td><td></td><td></td><td></td><td></td><td></td></tr>
<tr><td>Oligo fwd</td><td>CACCCCGCCCGGATACTTGGCCGT</td><td></td><td></td><td></td><td></td><td></td><td></td><td></td><td></td><td></td><td></td></tr>
<tr><td>Oligo rev</td><td>AAACACGGCCAAGTATCCGGGCGG</td><td></td><td></td><td></td><td></td><td></td><td></td><td></td><td></td><td></td><td></td></tr>
<tr><td>Chromosome</td><td>start</td><td>end</td><td>strand</td><td>MM</td><td>target_seq</td><td>PAM</td><td>alignment</td><td>distance</td><td>position</td><td>gene name</td><td>gene id</td></tr>
<tr><td>GU305914.1</td><td>107994</td><td>108016</td><td>+</td><td>0</td><td>CCGCCCGGATACTTGGCCGT</td><td>GGG</td><td>||||||||[||||||||||||]PAM</td><td>NA</td><td>Intergenic</td><td>NA</td><td>NA</td></tr>
<tr><td></td><td></td><td></td><td></td><td></td><td></td><td></td><td></td><td></td><td></td><td></td><td></td></tr>
<tr><td>T7891</td><td>GATACTTGGCCGTGGGCGAGCGG</td><td>1000</td><td></td><td></td><td></td><td></td><td></td><td></td><td></td><td></td><td></td></tr>
<tr><td>Oligo fwd</td><td>CACCGATACTTGGCCGTGGGCGAG</td><td></td><td></td><td></td><td></td><td></td><td></td><td></td><td></td><td></td><td></td></tr>
<tr><td>Oligo rev</td><td>AAACCTCGCCCACGGCCAAGTATC</td><td></td><td></td><td></td><td></td><td></td><td></td><td></td><td></td><td></td><td></td></tr>
<tr><td>Chromosome</td><td>start</td><td>end</td><td>strand</td><td>MM</td><td>target_seq</td><td>PAM</td><td>alignment</td><td>distance</td><td>position</td><td>gene name</td><td>gene id</td></tr>
<tr><td>GU305914.1</td><td>108001</td><td>108023</td><td>+</td><td>0</td><td>GATACTTGGCCGTGGGCGAG</td><td>CGG</td><td>||||||||[||||||||||||]PAM</td><td>NA</td><td>Intergenic</td><td>NA</td><td>NA</td></tr>
<tr><td></td><td></td><td></td><td></td><td></td><td></td><td></td><td></td><td></td><td></td><td></td><td></td></tr>
<tr><td>T7892</td><td>ATACTTGGCCGTGGGCGAGCGGG</td><td>1000</td><td></td><td></td><td></td><td></td><td></td><td></td><td></td><td></td><td></td></tr>
<tr><td>Oligo fwd</td><td>CACCATACTTGGCCGTGGGCGAGC</td><td></td><td></td><td></td><td></td><td></td><td></td><td></td><td></td><td></td><td></td></tr>
<tr><td>Oligo rev</td><td>AAACGCTCGCCCACGGCCAAGTAT</td><td></td><td></td><td></td><td></td><td></td><td></td><td></td><td></td><td></td><td></td></tr>
<tr><td>Chromosome</td><td>start</td><td>end</td><td>strand</td><td>MM</td><td>target_seq</td><td>PAM</td><td>alignment</td><td>distance</td><td>position</td><td>gene name</td><td>gene id</td></tr>
<tr><td>GU305914.1</td><td>108002</td><td>108024</td><td>+</td><td>0</td><td>ATACTTGGCCGTGGGCGAGC</td><td>GGG</td><td>||||||||[||||||||||||]PAM</td><td>NA</td><td>Intergenic</td><td>NA</td><td>NA</td></tr>
<tr><td></td><td></td><td></td><td></td><td></td><td></td><td></td><td></td><td></td><td></td><td></td><td></td></tr>
<tr><td>T7893</td><td>CGAGCGGGCCACCAGCCTGTAGG</td><td>1000</td><td></td><td></td><td></td><td></td><td></td><td></td><td></td><td></td><td></td></tr>
<tr><td>Oligo fwd</td><td>CACCCGAGCGGGCCACCAGCCTGT</td><td></td><td></td><td></td><td></td><td></td><td></td><td></td><td></td><td></td><td></td></tr>
<tr><td>Oligo rev</td><td>AAACACAGGCTGGTGGCCCGCTCG</td><td></td><td></td><td></td><td></td><td></td><td></td><td></td><td></td><td></td><td></td></tr>
<tr><td>Chromosome</td><td>start</td><td>end</td><td>strand</td><td>MM</td><td>target_seq</td><td>PAM</td><td>alignment</td><td>distance</td><td>position</td><td>gene name</td><td>gene id</td></tr>
<tr><td>GU305914.1</td><td>108017</td><td>108039</td><td>+</td><td>0</td><td>CGAGCGGGCCACCAGCCTGT</td><td>AGG</td><td>||||||||[||||||||||||]PAM</td><td>NA</td><td>Intergenic</td><td>NA</td><td>NA</td></tr>
<tr><td></td><td></td><td></td><td></td><td></td><td></td><td></td><td></td><td></td><td></td><td></td><td></td></tr>
<tr><td>T7894</td><td>CAGCCTGTAGGCGCCGTAGATGG</td><td>1000</td><td></td><td></td><td></td><td></td><td></td><td></td><td></td><td></td><td></td></tr>
<tr><td>Oligo fwd</td><td>CACCCAGCCTGTAGGCGCCGTAGA</td><td></td><td></td><td></td><td></td><td></td><td></td><td></td><td></td><td></td><td></td></tr>
<tr><td>Oligo rev</td><td>AAACTCTACGGCGCCTACAGGCTG</td><td></td><td></td><td></td><td></td><td></td><td></td><td></td><td></td><td></td><td></td></tr>
<tr><td>Chromosome</td><td>start</td><td>end</td><td>strand</td><td>MM</td><td>target_seq</td><td>PAM</td><td>alignment</td><td>distance</td><td>position</td><td>gene name</td><td>gene id</td></tr>
<tr><td>GU305914.1</td><td>108029</td><td>108051</td><td>+</td><td>0</td><td>CAGCCTGTAGGCGCCGTAGA</td><td>TGG</td><td>||||||||[||||||||||||]PAM</td><td>NA</td><td>Intergenic</td><td>NA</td><td>NA</td></tr>
<tr><td></td><td></td><td></td><td></td><td></td><td></td><td></td><td></td><td></td><td></td><td></td><td></td></tr>
<tr><td>T7895</td><td>AGCCTGTAGGCGCCGTAGATGGG</td><td>1000</td><td></td><td></td><td></td><td></td><td></td><td></td><td></td><td></td><td></td></tr>
<tr><td>Oligo fwd</td><td>CACCAGCCTGTAGGCGCCGTAGAT</td><td></td><td></td><td></td><td></td><td></td><td></td><td></td><td></td><td></td><td></td></tr>
<tr><td>Oligo rev</td><td>AAACATCTACGGCGCCTACAGGCT</td><td></td><td></td><td></td><td></td><td></td><td></td><td></td><td></td><td></td><td></td></tr>
<tr><td>Chromosome</td><td>start</td><td>end</td><td>strand</td><td>MM</td><td>target_seq</td><td>PAM</td><td>alignment</td><td>distance</td><td>position</td><td>gene name</td><td>gene id</td></tr>
<tr><td>GU305914.1</td><td>108030</td><td>108052</td><td>+</td><td>0</td><td>AGCCTGTAGGCGCCGTAGAT</td><td>GGG</td><td>||||||||[||||||||||||]PAM</td><td>NA</td><td>Intergenic</td><td>NA</td><td>NA</td></tr>
<tr><td></td><td></td><td></td><td></td><td></td><td></td><td></td><td></td><td></td><td></td><td></td><td></td></tr>
<tr><td>T7896</td><td>GCGATACGACTCTAGCGTCTCGG</td><td>1000</td><td></td><td></td><td></td><td></td><td></td><td></td><td></td><td></td><td></td></tr>
<tr><td>Oligo fwd</td><td>CACCGCGATACGACTCTAGCGTCT</td><td></td><td></td><td></td><td></td><td></td><td></td><td></td><td></td><td></td><td></td></tr>
<tr><td>Oligo rev</td><td>AAACAGACGCTAGAGTCGTATCGC</td><td></td><td></td><td></td><td></td><td></td><td></td><td></td><td></td><td></td><td></td></tr>
<tr><td>Chromosome</td><td>start</td><td>end</td><td>strand</td><td>MM</td><td>target_seq</td><td>PAM</td><td>alignment</td><td>distance</td><td>position</td><td>gene name</td><td>gene id</td></tr>
<tr><td>GU305914.1</td><td>108080</td><td>108102</td><td>+</td><td>0</td><td>GCGATACGACTCTAGCGTCT</td><td>CGG</td><td>||||||||[||||||||||||]PAM</td><td>NA</td><td>Intergenic</td><td>NA</td><td>NA</td></tr>
<tr><td></td><td></td><td></td><td></td><td></td><td></td><td></td><td></td><td></td><td></td><td></td><td></td></tr>
<tr><td>T7897</td><td>CGGTCATGTCGACGCTCGACAGG</td><td>1000</td><td></td><td></td><td></td><td></td><td></td><td></td><td></td><td></td><td></td></tr>
<tr><td>Oligo fwd</td><td>CACCCGGTCATGTCGACGCTCGAC</td><td></td><td></td><td></td><td></td><td></td><td></td><td></td><td></td><td></td><td></td></tr>
<tr><td>Oligo rev</td><td>AAACGTCGAGCGTCGACATGACCG</td><td></td><td></td><td></td><td></td><td></td><td></td><td></td><td></td><td></td><td></td></tr>
<tr><td>Chromosome</td><td>start</td><td>end</td><td>strand</td><td>MM</td><td>target_seq</td><td>PAM</td><td>alignment</td><td>distance</td><td>position</td><td>gene name</td><td>gene id</td></tr>
<tr><td>GU305914.1</td><td>108100</td><td>108122</td><td>+</td><td>0</td><td>CGGTCATGTCGACGCTCGAC</td><td>AGG</td><td>||||||||[||||||||||||]PAM</td><td>NA</td><td>Intergenic</td><td>NA</td><td>NA</td></tr>
<tr><td></td><td></td><td></td><td></td><td></td><td></td><td></td><td></td><td></td><td></td><td></td><td></td></tr>
<tr><td>T7898</td><td>CGTGCGTGTTGAAGAGACAGCGG</td><td>1000</td><td></td><td></td><td></td><td></td><td></td><td></td><td></td><td></td><td></td></tr>
<tr><td>Oligo fwd</td><td>CACCCGTGCGTGTTGAAGAGACAG</td><td></td><td></td><td></td><td></td><td></td><td></td><td></td><td></td><td></td><td></td></tr>
<tr><td>Oligo rev</td><td>AAACCTGTCTCTTCAACACGCACG</td><td></td><td></td><td></td><td></td><td></td><td></td><td></td><td></td><td></td><td></td></tr>
<tr><td>Chromosome</td><td>start</td><td>end</td><td>strand</td><td>MM</td><td>target_seq</td><td>PAM</td><td>alignment</td><td>distance</td><td>position</td><td>gene name</td><td>gene id</td></tr>
<tr><td>GU305914.1</td><td>108136</td><td>108158</td><td>+</td><td>0</td><td>CGTGCGTGTTGAAGAGACAG</td><td>CGG</td><td>||||||||[||||||||||||]PAM</td><td>NA</td><td>Intergenic</td><td>NA</td><td>NA</td></tr>
<tr><td></td><td></td><td></td><td></td><td></td><td></td><td></td><td></td><td></td><td></td><td></td><td></td></tr>
<tr><td>T7899</td><td>GTGCGTGTTGAAGAGACAGCGGG</td><td>1000</td><td></td><td></td><td></td><td></td><td></td><td></td><td></td><td></td><td></td></tr>
<tr><td>Oligo fwd</td><td>CACCGTGCGTGTTGAAGAGACAGC</td><td></td><td></td><td></td><td></td><td></td><td></td><td></td><td></td><td></td><td></td></tr>
<tr><td>Oligo rev</td><td>AAACGCTGTCTCTTCAACACGCAC</td><td></td><td></td><td></td><td></td><td></td><td></td><td></td><td></td><td></td><td></td></tr>
<tr><td>Chromosome</td><td>start</td><td>end</td><td>strand</td><td>MM</td><td>target_seq</td><td>PAM</td><td>alignment</td><td>distance</td><td>position</td><td>gene name</td><td>gene id</td></tr>
<tr><td>GU305914.1</td><td>108137</td><td>108159</td><td>+</td><td>0</td><td>GTGCGTGTTGAAGAGACAGC</td><td>GGG</td><td>||||||||[||||||||||||]PAM</td><td>NA</td><td>Intergenic</td><td>NA</td><td>NA</td></tr>
<tr><td></td><td></td><td></td><td></td><td></td><td></td><td></td><td></td><td></td><td></td><td></td><td></td></tr>
<tr><td>T7900</td><td>TGCGTGTTGAAGAGACAGCGGGG</td><td>1000</td><td></td><td></td><td></td><td></td><td></td><td></td><td></td><td></td><td></td></tr>
<tr><td>Oligo fwd</td><td>CACCTGCGTGTTGAAGAGACAGCG</td><td></td><td></td><td></td><td></td><td></td><td></td><td></td><td></td><td></td><td></td></tr>
<tr><td>Oligo rev</td><td>AAACCGCTGTCTCTTCAACACGCA</td><td></td><td></td><td></td><td></td><td></td><td></td><td></td><td></td><td></td><td></td></tr>
<tr><td>Chromosome</td><td>start</td><td>end</td><td>strand</td><td>MM</td><td>target_seq</td><td>PAM</td><td>alignment</td><td>distance</td><td>position</td><td>gene name</td><td>gene id</td></tr>
<tr><td>GU305914.1</td><td>108138</td><td>108160</td><td>+</td><td>0</td><td>TGCGTGTTGAAGAGACAGCG</td><td>GGG</td><td>||||||||[||||||||||||]PAM</td><td>NA</td><td>Intergenic</td><td>NA</td><td>NA</td></tr>
<tr><td></td><td></td><td></td><td></td><td></td><td></td><td></td><td></td><td></td><td></td><td></td><td></td></tr>
<tr><td>T7901</td><td>TGAAGAGACAGCGGGGCACCTGG</td><td>1000</td><td></td><td></td><td></td><td></td><td></td><td></td><td></td><td></td><td></td></tr>
<tr><td>Oligo fwd</td><td>CACCTGAAGAGACAGCGGGGCACC</td><td></td><td></td><td></td><td></td><td></td><td></td><td></td><td></td><td></td><td></td></tr>
<tr><td>Oligo rev</td><td>AAACGGTGCCCCGCTGTCTCTTCA</td><td></td><td></td><td></td><td></td><td></td><td></td><td></td><td></td><td></td><td></td></tr>
<tr><td>Chromosome</td><td>start</td><td>end</td><td>strand</td><td>MM</td><td>target_seq</td><td>PAM</td><td>alignment</td><td>distance</td><td>position</td><td>gene name</td><td>gene id</td></tr>
<tr><td>GU305914.1</td><td>108145</td><td>108167</td><td>+</td><td>0</td><td>TGAAGAGACAGCGGGGCACC</td><td>TGG</td><td>||||||||[||||||||||||]PAM</td><td>NA</td><td>Intergenic</td><td>NA</td><td>NA</td></tr>
<tr><td></td><td></td><td></td><td></td><td></td><td></td><td></td><td></td><td></td><td></td><td></td><td></td></tr>
<tr><td>T7902</td><td>AGCGGGGCACCTGGTAGACGCGG</td><td>1000</td><td></td><td></td><td></td><td></td><td></td><td></td><td></td><td></td><td></td></tr>
<tr><td>Oligo fwd</td><td>CACCAGCGGGGCACCTGGTAGACG</td><td></td><td></td><td></td><td></td><td></td><td></td><td></td><td></td><td></td><td></td></tr>
</table>

Oligo rev AAACCGTCTACCAGGTGCCCCGCT
Chromosome start end strand MM target_seq PAM alignment distance position gene name gene id
GU305914.1 108154 108176 + 0 AGCGGGGCACCTGGTAGACG CGG ||||||||[||||||||||||]PAM NA Intergenic NA NA

T7903 CGCGGTAGCCGTCCCCGTCCTGG 1000
Oligo fwd CACCCGCGGTAGCCGTCCCCGTCC
Oligo rev AAACGGACGGGGACGGCTACCGCG
Chromosome start end strand MM target_seq PAM alignment distance position gene name gene id
GU305914.1 108172 108194 + 0 CGCGGTAGCCGTCCCCGTCC TGG ||||||||[||||||||||||]PAM NA Intergenic NA NA

T7904 CTGGAAGTTGAACGTGAACATGG 1000
Oligo fwd CACCCTGGAAGTTGAACGTGAACA
Oligo rev AAACTGTTCACGTTCAACTTCCAG
Chromosome start end strand MM target_seq PAM alignment distance position gene name gene id
GU305914.1 108191 108213 + 0 CTGGAAGTTGAACGTGAACA TGG ||||||||[||||||||||||]PAM NA Intergenic NA NA

T7905 TGGCCGCGTGAGACACCGTCTGG 1000
Oligo fwd CACCTGGCCGCGTGAGACACCGTC
Oligo rev AAACGACGGTGTCTCACGCGGCCA
Chromosome start end strand MM target_seq PAM alignment distance position gene name gene id
GU305914.1 108211 108233 + 0 TGGCCGCGTGAGACACCGTC TGG ||||||||[||||||||||||]PAM NA Intergenic NA NA

T7906 GTGAGACACCGTCTGGTTAGAGG 1000
Oligo fwd CACCGTGAGACACCGTCTGGTTAG
Oligo rev AAACCTAACCAGACGGTGTCTCAC
Chromosome start end strand MM target_seq PAM alignment distance position gene name gene id
GU305914.1 108218 108240 + 0 GTGAGACACCGTCTGGTTAG AGG ||||||||[||||||||||||]PAM NA Intergenic NA NA

T7907 GGTTAGAGGATTCGTCGTGCCGG 1000
Oligo fwd CACCGGTTAGAGGATTCGTCGTGC
Oligo rev AAACGCACGACGAATCCTCTAACC
Chromosome start end strand MM target_seq PAM alignment distance position gene name gene id
GU305914.1 108232 108254 + 0 GGTTAGAGGATTCGTCGTGC CGG ||||||||[||||||||||||]PAM NA Intergenic NA NA

T7908 ATTCGTCGTGCCGGCACACCCGG 1000
Oligo fwd CACCATTCGTCGTGCCGGCACACC
Oligo rev AAACGGTGTGCCGGCACGACGAAT
Chromosome start end strand MM target_seq PAM alignment distance position gene name gene id
GU305914.1 108241 108263 + 0 ATTCGTCGTGCCGGCACACC CGG ||||||||[||||||||||||]PAM NA Intergenic NA NA

T7909 GTTGTACGAGTGCAACTTGACGG 1000
Oligo fwd CACCGTTGTACGAGTGCAACTTGA
Oligo rev AAACTCAAGTTGCACTCGTACAAC
Chromosome start end strand MM target_seq PAM alignment distance position gene name gene id
GU305914.1 108269 108291 + 0 GTTGTACGAGTGCAACTTGA CGG ||||||||[||||||||||||]PAM NA Intergenic NA NA

T7910 GTGCAACTTGACGGTGTTTCTGG 1000
Oligo fwd CACCGTGCAACTTGACGGTGTTTC
Oligo rev AAACGAAACACCGTCAAGTTGCAC
Chromosome start end strand MM target_seq PAM alignment distance position gene name gene id
GU305914.1 108278 108300 + 0 GTGCAACTTGACGGTGTTTC TGG ||||||||[||||||||||||]PAM NA Intergenic NA NA

T7911 ACGGTGTTTCTGGAGATCCGCGG 1000
Oligo fwd CACCACGGTGTTTCTGGAGATCCG
Oligo rev AAACCGGATCTCCAGAAACACCGT
Chromosome start end strand MM target_seq PAM alignment distance position gene name gene id
GU305914.1 108288 108310 + 0 ACGGTGTTTCTGGAGATCCG CGG ||||||||[||||||||||||]PAM NA Intergenic NA NA

T7912 TCTGGAGATCCGCGGCGCCTCGG 1000
Oligo fwd CACCTCTGGAGATCCGCGGCGCCT
Oligo rev AAACAGGCGCCGCGGATCTCCAGA
Chromosome start end strand MM target_seq PAM alignment distance position gene name gene id
GU305914.1 108296 108318 + 0 TCTGGAGATCCGCGGCGCCT CGG ||||||||[||||||||||||]PAM NA Intergenic NA NA

T7913 GCGCGTCGAACATAGCGCGATGG 1000
Oligo fwd CACCGCGCGTCGAACATAGCGCGA
Oligo rev AAACTCGCGCTATGTTCGACGCGC
Chromosome start end strand MM target_seq PAM alignment distance position gene name gene id
GU305914.1 108338 108360 + 0 GCGCGTCGAACATAGCGCGA TGG ||||||||[||||||||||||]PAM NA Intergenic NA NA

T7914 TTAATGACAACTTCATGTCGCGG 1000
Oligo fwd CACCTTAATGACAACTTCATGTCG
Oligo rev AAACCGACATGAAGTTGTCATTAA
Chromosome start end strand MM target_seq PAM alignment distance position gene name gene id
GU305914.1 108367 108389 + 0 TTAATGACAACTTCATGTCG CGG ||||||||[||||||||||||]PAM NA Intergenic NA NA

T7915 TAATGACAACTTCATGTCGCGGG 1000
Oligo fwd CACCTAATGACAACTTCATGTCGC
Oligo rev AAACGCGACATGAAGTTGTCATTA
Chromosome start end strand MM target_seq PAM alignment distance position gene name gene id
GU305914.1 108368 108390 + 0 TAATGACAACTTCATGTCGC GGG ||||||||[||||||||||||]PAM NA Intergenic NA NA

T7916 GACAACTTCATGTCGCGGGTCGG 1000
Oligo fwd CACCGACAACTTCATGTCGCGGGT
Oligo rev AAACACCCGCGACATGAAGTTGTC
Chromosome start end strand MM target_seq PAM alignment distance position gene name gene id
GU305914.1 108372 108394 + 0 GACAACTTCATGTCGCGGGT CGG ||||||||[||||||||||||]PAM NA Intergenic NA NA

T7917 ACAACTTCATGTCGCGGGTCGGG 1000

<table>
<tr><td>Oligo fwd</td><td>CACCACAACTTCATGTCGCGGGTC</td><td></td><td></td><td></td><td></td><td></td><td></td><td></td><td></td><td></td><td></td></tr>
<tr><td>Oligo rev</td><td>AAACGACCCGCGACATGAAGTTGT</td><td></td><td></td><td></td><td></td><td></td><td></td><td></td><td></td><td></td><td></td></tr>
<tr><td>Chromosome</td><td>start</td><td>end</td><td>strand</td><td>MM</td><td>target_seq</td><td>PAM</td><td>alignment</td><td>distance</td><td>position</td><td>gene name</td><td>gene id</td></tr>
<tr><td>GU305914.1</td><td>108373</td><td>108395</td><td>+</td><td>0</td><td>ACAACTTCATGTCGCGGGTC</td><td>GGG</td><td>||||||||[||||||||||||]PAM</td><td>NA</td><td>Intergenic</td><td>NA</td><td>NA</td></tr>
<tr><td>T7918</td><td>CAACTTCATGTCGCGGGTCGGGG</td><td>1000</td><td></td><td></td><td></td><td></td><td></td><td></td><td></td><td></td><td></td></tr>
<tr><td>Oligo fwd</td><td>CACCCAACTTCATGTCGCGGGTCG</td><td></td><td></td><td></td><td></td><td></td><td></td><td></td><td></td><td></td><td></td></tr>
<tr><td>Oligo rev</td><td>AAACCGACCCGCGACATGAAGTTG</td><td></td><td></td><td></td><td></td><td></td><td></td><td></td><td></td><td></td><td></td></tr>
<tr><td>Chromosome</td><td>start</td><td>end</td><td>strand</td><td>MM</td><td>target_seq</td><td>PAM</td><td>alignment</td><td>distance</td><td>position</td><td>gene name</td><td>gene id</td></tr>
<tr><td>GU305914.1</td><td>108374</td><td>108396</td><td>+</td><td>0</td><td>CAACTTCATGTCGCGGGTCG</td><td>GGG</td><td>||||||||[||||||||||||]PAM</td><td>NA</td><td>Intergenic</td><td>NA</td><td>NA</td></tr>
<tr><td>T7919</td><td>TCATGTCGCGGGTCGGGGTGCGG</td><td>1000</td><td></td><td></td><td></td><td></td><td></td><td></td><td></td><td></td><td></td></tr>
<tr><td>Oligo fwd</td><td>CACCTCATGTCGCGGGTCGGGGTG</td><td></td><td></td><td></td><td></td><td></td><td></td><td></td><td></td><td></td><td></td></tr>
<tr><td>Oligo rev</td><td>AAACCACCCCGACCCGCGACATGA</td><td></td><td></td><td></td><td></td><td></td><td></td><td></td><td></td><td></td><td></td></tr>
<tr><td>Chromosome</td><td>start</td><td>end</td><td>strand</td><td>MM</td><td>target_seq</td><td>PAM</td><td>alignment</td><td>distance</td><td>position</td><td>gene name</td><td>gene id</td></tr>
<tr><td>GU305914.1</td><td>108379</td><td>108401</td><td>+</td><td>0</td><td>TCATGTCGCGGGTCGGGGTG</td><td>CGG</td><td>||||||||[||||||||||||]PAM</td><td>NA</td><td>Intergenic</td><td>NA</td><td>NA</td></tr>
<tr><td>T7920</td><td>GGTCGGGGTGCGGACGTCGCAGG</td><td>1000</td><td></td><td></td><td></td><td></td><td></td><td></td><td></td><td></td><td></td></tr>
<tr><td>Oligo fwd</td><td>CACCGGTCGGGGTGCGGACGTCGC</td><td></td><td></td><td></td><td></td><td></td><td></td><td></td><td></td><td></td><td></td></tr>
<tr><td>Oligo rev</td><td>AAACGCGACGTCCGCACCCCGACC</td><td></td><td></td><td></td><td></td><td></td><td></td><td></td><td></td><td></td><td></td></tr>
<tr><td>Chromosome</td><td>start</td><td>end</td><td>strand</td><td>MM</td><td>target_seq</td><td>PAM</td><td>alignment</td><td>distance</td><td>position</td><td>gene name</td><td>gene id</td></tr>
<tr><td>GU305914.1</td><td>108389</td><td>108411</td><td>+</td><td>0</td><td>GGTCGGGGTGCGGACGTCGC</td><td>AGG</td><td>||||||||[||||||||||||]PAM</td><td>NA</td><td>Intergenic</td><td>NA</td><td>NA</td></tr>
<tr><td>GU305914.1</td><td>68896</td><td>68918</td><td>+</td><td>4</td><td>GGTCGAGCTGCTGAAGTCGC</td><td>AGG</td><td>|||||-|-[|||-||-|||||]PAM</td><td>NA</td><td>Intergenic</td><td>NA</td><td>NA</td></tr>
<tr><td>T7921</td><td>GTCGGGGTGCGGACGTCGCAGGG</td><td>1000</td><td></td><td></td><td></td><td></td><td></td><td></td><td></td><td></td><td></td></tr>
<tr><td>Oligo fwd</td><td>CACCGTCGGGGTGCGGACGTCGCA</td><td></td><td></td><td></td><td></td><td></td><td></td><td></td><td></td><td></td><td></td></tr>
<tr><td>Oligo rev</td><td>AAACTGCGACGTCCGCACCCCGAC</td><td></td><td></td><td></td><td></td><td></td><td></td><td></td><td></td><td></td><td></td></tr>
<tr><td>Chromosome</td><td>start</td><td>end</td><td>strand</td><td>MM</td><td>target_seq</td><td>PAM</td><td>alignment</td><td>distance</td><td>position</td><td>gene name</td><td>gene id</td></tr>
<tr><td>GU305914.1</td><td>108390</td><td>108412</td><td>+</td><td>0</td><td>GTCGGGGTGCGGACGTCGCA</td><td>GGG</td><td>||||||||[||||||||||||]PAM</td><td>NA</td><td>Intergenic</td><td>NA</td><td>NA</td></tr>
<tr><td>T7922</td><td>AATATCTTAATTACGCTCGTCGG</td><td>1000</td><td></td><td></td><td></td><td></td><td></td><td></td><td></td><td></td><td></td></tr>
<tr><td>Oligo fwd</td><td>CACCAATATCTTAATTACGCTCGT</td><td></td><td></td><td></td><td></td><td></td><td></td><td></td><td></td><td></td><td></td></tr>
<tr><td>Oligo rev</td><td>AAACACGAGCGTAATTAAGATATT</td><td></td><td></td><td></td><td></td><td></td><td></td><td></td><td></td><td></td><td></td></tr>
<tr><td>Chromosome</td><td>start</td><td>end</td><td>strand</td><td>MM</td><td>target_seq</td><td>PAM</td><td>alignment</td><td>distance</td><td>position</td><td>gene name</td><td>gene id</td></tr>
<tr><td>GU305914.1</td><td>108436</td><td>108458</td><td>+</td><td>0</td><td>AATATCTTAATTACGCTCGT</td><td>CGG</td><td>||||||||[||||||||||||]PAM</td><td>NA</td><td>Intergenic</td><td>NA</td><td>NA</td></tr>
<tr><td>T7923</td><td>GCTTTCCGTCCGACCCGTCATGG</td><td>1000</td><td></td><td></td><td></td><td></td><td></td><td></td><td></td><td></td><td></td></tr>
<tr><td>Oligo fwd</td><td>CACCGCTTTCCGTCCGACCCGTCA</td><td></td><td></td><td></td><td></td><td></td><td></td><td></td><td></td><td></td><td></td></tr>
<tr><td>Oligo rev</td><td>AAACTGACGGGTCGGACGGAAAGC</td><td></td><td></td><td></td><td></td><td></td><td></td><td></td><td></td><td></td><td></td></tr>
<tr><td>Chromosome</td><td>start</td><td>end</td><td>strand</td><td>MM</td><td>target_seq</td><td>PAM</td><td>alignment</td><td>distance</td><td>position</td><td>gene name</td><td>gene id</td></tr>
<tr><td>GU305914.1</td><td>108460</td><td>108482</td><td>+</td><td>0</td><td>GCTTTCCGTCCGACCCGTCA</td><td>TGG</td><td>||||||||[||||||||||||]PAM</td><td>NA</td><td>Intergenic</td><td>NA</td><td>NA</td></tr>
<tr><td>T7924</td><td>CTTTCCGTCCGACCCGTCATGGG</td><td>1000</td><td></td><td></td><td></td><td></td><td></td><td></td><td></td><td></td><td></td></tr>
<tr><td>Oligo fwd</td><td>CACCCTTTCCGTCCGACCCGTCAT</td><td></td><td></td><td></td><td></td><td></td><td></td><td></td><td></td><td></td><td></td></tr>
<tr><td>Oligo rev</td><td>AAACATGACGGGTCGGACGGAAAG</td><td></td><td></td><td></td><td></td><td></td><td></td><td></td><td></td><td></td><td></td></tr>
<tr><td>Chromosome</td><td>start</td><td>end</td><td>strand</td><td>MM</td><td>target_seq</td><td>PAM</td><td>alignment</td><td>distance</td><td>position</td><td>gene name</td><td>gene id</td></tr>
<tr><td>GU305914.1</td><td>108461</td><td>108483</td><td>+</td><td>0</td><td>CTTTCCGTCCGACCCGTCAT</td><td>GGG</td><td>||||||||[||||||||||||]PAM</td><td>NA</td><td>Intergenic</td><td>NA</td><td>NA</td></tr>
<tr><td>T7925</td><td>TTTCCGTCCGACCCGTCATGGGG</td><td>1000</td><td></td><td></td><td></td><td></td><td></td><td></td><td></td><td></td><td></td></tr>
<tr><td>Oligo fwd</td><td>CACCTTTCCGTCCGACCCGTCATG</td><td></td><td></td><td></td><td></td><td></td><td></td><td></td><td></td><td></td><td></td></tr>
<tr><td>Oligo rev</td><td>AAACCATGACGGGTCGGACGGAAA</td><td></td><td></td><td></td><td></td><td></td><td></td><td></td><td></td><td></td><td></td></tr>
<tr><td>Chromosome</td><td>start</td><td>end</td><td>strand</td><td>MM</td><td>target_seq</td><td>PAM</td><td>alignment</td><td>distance</td><td>position</td><td>gene name</td><td>gene id</td></tr>
<tr><td>GU305914.1</td><td>108462</td><td>108484</td><td>+</td><td>0</td><td>TTTCCGTCCGACCCGTCATG</td><td>GGG</td><td>||||||||[||||||||||||]PAM</td><td>NA</td><td>Intergenic</td><td>NA</td><td>NA</td></tr>
<tr><td>T7926</td><td>ACCCGTCATGGGGACGTCCGCGG</td><td>1000</td><td></td><td></td><td></td><td></td><td></td><td></td><td></td><td></td><td></td></tr>
<tr><td>Oligo fwd</td><td>CACCACCCGTCATGGGGACGTCCG</td><td></td><td></td><td></td><td></td><td></td><td></td><td></td><td></td><td></td><td></td></tr>
<tr><td>Oligo rev</td><td>AAACCGGACGTCCCCATGACGGGT</td><td></td><td></td><td></td><td></td><td></td><td></td><td></td><td></td><td></td><td></td></tr>
<tr><td>Chromosome</td><td>start</td><td>end</td><td>strand</td><td>MM</td><td>target_seq</td><td>PAM</td><td>alignment</td><td>distance</td><td>position</td><td>gene name</td><td>gene id</td></tr>
<tr><td>GU305914.1</td><td>108472</td><td>108494</td><td>+</td><td>0</td><td>ACCCGTCATGGGGACGTCCG</td><td>CGG</td><td>||||||||[||||||||||||]PAM</td><td>NA</td><td>Intergenic</td><td>NA</td><td>NA</td></tr>
<tr><td>T7927</td><td>CATGGGGACGTCCGCGGACACGG</td><td>1000</td><td></td><td></td><td></td><td></td><td></td><td></td><td></td><td></td><td></td></tr>
<tr><td>Oligo fwd</td><td>CACCCATGGGGACGTCCGCGGACA</td><td></td><td></td><td></td><td></td><td></td><td></td><td></td><td></td><td></td><td></td></tr>
<tr><td>Oligo rev</td><td>AAACTGTCCGCGGACGTCCCCATG</td><td></td><td></td><td></td><td></td><td></td><td></td><td></td><td></td><td></td><td></td></tr>
<tr><td>Chromosome</td><td>start</td><td>end</td><td>strand</td><td>MM</td><td>target_seq</td><td>PAM</td><td>alignment</td><td>distance</td><td>position</td><td>gene name</td><td>gene id</td></tr>
<tr><td>GU305914.1</td><td>108478</td><td>108500</td><td>+</td><td>0</td><td>CATGGGGACGTCCGCGGACA</td><td>CGG</td><td>||||||||[||||||||||||]PAM</td><td>NA</td><td>Intergenic</td><td>NA</td><td>NA</td></tr>
<tr><td>T7928</td><td>ACCTGACGCCCGCAAGCGTGCGG</td><td>1000</td><td></td><td></td><td></td><td></td><td></td><td></td><td></td><td></td><td></td></tr>
<tr><td>Oligo fwd</td><td>CACCACCTGACGCCCGCAAGCGTG</td><td></td><td></td><td></td><td></td><td></td><td></td><td></td><td></td><td></td><td></td></tr>
<tr><td>Oligo rev</td><td>AAACCACGCTTGCGGGCGTCAGGT</td><td></td><td></td><td></td><td></td><td></td><td></td><td></td><td></td><td></td><td></td></tr>
<tr><td>Chromosome</td><td>start</td><td>end</td><td>strand</td><td>MM</td><td>target_seq</td><td>PAM</td><td>alignment</td><td>distance</td><td>position</td><td>gene name</td><td>gene id</td></tr>
<tr><td>GU305914.1</td><td>108517</td><td>108539</td><td>+</td><td>0</td><td>ACCTGACGCCCGCAAGCGTG</td><td>CGG</td><td>||||||||[||||||||||||]PAM</td><td>NA</td><td>Intergenic</td><td>NA</td><td>NA</td></tr>
<tr><td>T7929</td><td>CCTGACGCCCGCAAGCGTGCGGG</td><td>1000</td><td></td><td></td><td></td><td></td><td></td><td></td><td></td><td></td><td></td></tr>
<tr><td>Oligo fwd</td><td>CACCCCTGACGCCCGCAAGCGTGC</td><td></td><td></td><td></td><td></td><td></td><td></td><td></td><td></td><td></td><td></td></tr>
<tr><td>Oligo rev</td><td>AAACGCACGCTTGCGGGCGTCAGG</td><td></td><td></td><td></td><td></td><td></td><td></td><td></td><td></td><td></td><td></td></tr>
<tr><td>Chromosome</td><td>start</td><td>end</td><td>strand</td><td>MM</td><td>target_seq</td><td>PAM</td><td>alignment</td><td>distance</td><td>position</td><td>gene name</td><td>gene id</td></tr>
<tr><td>GU305914.1</td><td>108518</td><td>108540</td><td>+</td><td>0</td><td>CCTGACGCCCGCAAGCGTGC</td><td>GGG</td><td>||||||||[||||||||||||]PAM</td><td>NA</td><td>Intergenic</td><td>NA</td><td>NA</td></tr>
<tr><td>T7930</td><td>CTGACGCCCGCAAGCGTGCGGGG</td><td>1000</td><td></td><td></td><td></td><td></td><td></td><td></td><td></td><td></td><td></td></tr>
<tr><td>Oligo fwd</td><td>CACCCTGACGCCCGCAAGCGTGCG</td><td></td><td></td><td></td><td></td><td></td><td></td><td></td><td></td><td></td><td></td></tr>
<tr><td>Oligo rev</td><td>AAACCGCACGCTTGCGGGCGTCAG</td><td></td><td></td><td></td><td></td><td></td><td></td><td></td><td></td><td></td><td></td></tr>
<tr><td>Chromosome</td><td>start</td><td>end</td><td>strand</td><td>MM</td><td>target_seq</td><td>PAM</td><td>alignment</td><td>distance</td><td>position</td><td>gene name</td><td>gene id</td></tr>
<tr><td>GU305914.1</td><td>108519</td><td>108541</td><td>+</td><td>0</td><td>CTGACGCCCGCAAGCGTGCG</td><td>GGG</td><td>||||||||[||||||||||||]PAM</td><td>NA</td><td>Intergenic</td><td>NA</td><td>NA</td></tr>
<tr><td>T7931</td><td>CCCGCAAGCGTGCGGGGCGACGG</td><td>1000</td><td></td><td></td><td></td><td></td><td></td><td></td><td></td><td></td><td></td></tr>
<tr><td>Oligo fwd</td><td>CACCCCCGCAAGCGTGCGGGGCGA</td><td></td><td></td><td></td><td></td><td></td><td></td><td></td><td></td><td></td><td></td></tr>
<tr><td>Oligo rev</td><td>AAACTCGCCCCGCACGCTTGCGGG</td><td></td><td></td><td></td><td></td><td></td><td></td><td></td><td></td><td></td><td></td></tr>
<tr><td>Chromosome</td><td>start</td><td>end</td><td>strand</td><td>MM</td><td>target_seq</td><td>PAM</td><td>alignment</td><td>distance</td><td>position</td><td>gene name</td><td>gene id</td></tr>
<tr><td>GU305914.1</td><td>108525</td><td>108547</td><td>+</td><td>0</td><td>CCCGCAAGCGTGCGGGGCGA</td><td>CGG</td><td>||||||||[||||||||||||]PAM</td><td>NA</td><td>Intergenic</td><td>NA</td><td>NA</td></tr>
</table>

T7932 CCGCAAGCGTGCGGGGCGACGGG 1000
Oligo fwd CACCCCGCAAGCGTGCGGGGCGAC
Oligo rev AAACGTCGCCCCGCACGCTTGCGG
Chromosome start end strand MM target_seq PAM alignment distance position gene name gene id
GU305914.1 108526 108548 + 0 CCGCAAGCGTGCGGGGCGAC GGG ||||||||[||||||||||||]PAM NA Intergenic NA NA

T7933 CGCAAGCGTGCGGGGCGACGGGG 1000
Oligo fwd CACCCGCAAGCGTGCGGGGCGACG
Oligo rev AAACCGTCGCCCCGCACGCTTGCG
Chromosome start end strand MM target_seq PAM alignment distance position gene name gene id
GU305914.1 108527 108549 + 0 CGCAAGCGTGCGGGGCGACG GGG ||||||||[||||||||||||]PAM NA Intergenic NA NA

T7934 GCAAGCGTGCGGGGCGACGGGGG 1000
Oligo fwd CACCGCAAGCGTGCGGGGCGACGG
Oligo rev AAACCCGTCGCCCCGCACGCTTGC
Chromosome start end strand MM target_seq PAM alignment distance position gene name gene id
GU305914.1 108528 108550 + 0 GCAAGCGTGCGGGGCGACGG GGG ||||||||[||||||||||||]PAM NA Intergenic NA NA

T7935 GAAAATCACTGCCGTGTTCGAGG 1000
Oligo fwd CACCGAAAATCACTGCCGTGTTCG
Oligo rev AAACCGAACACGGCAGTGATTTTC
Chromosome start end strand MM target_seq PAM alignment distance position gene name gene id
GU305914.1 108568 108590 + 0 GAAAATCACTGCCGTGTTCG AGG ||||||||[||||||||||||]PAM NA Intergenic NA NA

T7936 GACGCGCTCAACGCCATCCTCGG 1000
Oligo fwd CACCGACGCGCTCAACGCCATCCT
Oligo rev AAACAGGATGGCGTTGAGCGCGTC
Chromosome start end strand MM target_seq PAM alignment distance position gene name gene id
GU305914.1 108596 108618 + 0 GACGCGCTCAACGCCATCCT CGG ||||||||[||||||||||||]PAM NA Intergenic NA NA

T7937 ACGCGCTCAACGCCATCCTCGGG 1000
Oligo fwd CACCACGCGCTCAACGCCATCCTC
Oligo rev AAACGAGGATGGCGTTGAGCGCGT
Chromosome start end strand MM target_seq PAM alignment distance position gene name gene id
GU305914.1 108597 108619 + 0 ACGCGCTCAACGCCATCCTC GGG ||||||||[||||||||||||]PAM NA Intergenic NA NA

T7938 CGCGCTCAACGCCATCCTCGGGG 1000
Oligo fwd CACCCGCGCTCAACGCCATCCTCG
Oligo rev AAACCGAGGATGGCGTTGAGCGCG
Chromosome start end strand MM target_seq PAM alignment distance position gene name gene id
GU305914.1 108598 108620 + 0 CGCGCTCAACGCCATCCTCG GGG ||||||||[||||||||||||]PAM NA Intergenic NA NA

T7939 GCGCTCAACGCCATCCTCGGGGG 1000
Oligo fwd CACCGCGCTCAACGCCATCCTCGG
Oligo rev AAACCCGAGGATGGCGTTGAGCGC
Chromosome start end strand MM target_seq PAM alignment distance position gene name gene id
GU305914.1 108599 108621 + 0 GCGCTCAACGCCATCCTCGG GGG ||||||||[||||||||||||]PAM NA Intergenic NA NA

T7940 TCGGGGGCATCCTGCCCTACCGG 1000
Oligo fwd CACCTCGGGGGCATCCTGCCCTAC
Oligo rev AAACGTAGGGCAGGATGCCCCCGA
Chromosome start end strand MM target_seq PAM alignment distance position gene name gene id
GU305914.1 108615 108637 + 0 TCGGGGGCATCCTGCCCTAC CGG ||||||||[||||||||||||]PAM NA Intergenic NA NA

T7941 CGGGGGCATCCTGCCCTACCGGG 1000
Oligo fwd CACCCGGGGGCATCCTGCCCTACC
Oligo rev AAACGGTAGGGCAGGATGCCCCCG
Chromosome start end strand MM target_seq PAM alignment distance position gene name gene id
GU305914.1 108616 108638 + 0 CGGGGGCATCCTGCCCTACC GGG ||||||||[||||||||||||]PAM NA Intergenic NA NA

T7942 GGGGGCATCCTGCCCTACCGGGG 1000
Oligo fwd CACCGGGGGCATCCTGCCCTACCG
Oligo rev AAACCGGTAGGGCAGGATGCCCCC
Chromosome start end strand MM target_seq PAM alignment distance position gene name gene id
GU305914.1 108617 108639 + 0 GGGGGCATCCTGCCCTACCG GGG ||||||||[||||||||||||]PAM NA Intergenic NA NA

T7943 GGGGCATCCTGCCCTACCGGGGG 1000
Oligo fwd CACCGGGGCATCCTGCCCTACCGG
Oligo rev AAACCCGGTAGGGCAGGATGCCCC
Chromosome start end strand MM target_seq PAM alignment distance position gene name gene id
GU305914.1 108618 108640 + 0 GGGGCATCCTGCCCTACCGG GGG ||||||||[||||||||||||]PAM NA Intergenic NA NA

T7944 GGGCATCCTGCCCTACCGGGGGG 1000
Oligo fwd CACCGGGCATCCTGCCCTACCGGG
Oligo rev AAACCCCGGTAGGGCAGGATGCCC
Chromosome start end strand MM target_seq PAM alignment distance position gene name gene id
GU305914.1 108619 108641 + 0 GGGCATCCTGCCCTACCGGG GGG ||||||||[||||||||||||]PAM NA Intergenic NA NA

T7945 CCTGCCCTACCGGGGGGTCCCGG 1000
Oligo fwd CACCCCTGCCCTACCGGGGGGTCC
Oligo rev AAACGGACCCCCCGGTAGGGCAGG
Chromosome start end strand MM target_seq PAM alignment distance position gene name gene id
GU305914.1 108625 108647 + 0 CCTGCCCTACCGGGGGGTCC CGG ||||||||[||||||||||||]PAM NA Intergenic NA NA

T7946 CTGCCCTACCGGGGGGTCCCGGG 1000
Oligo fwd CACCCTGCCCTACCGGGGGGTCCC
Oligo rev AAACGGGACCCCCCGGTAGGGCAG
Chromosome start end strand MM target_seq PAM alignment distance position gene name gene id

<table>
<tr><td>GU305914.1</td><td>108626</td><td>108648</td><td>+</td><td>0</td><td>CTGCCCTACCGGGGGGTCCC</td><td>GGG</td><td>||||||||[||||||||||||]PAM</td><td>NA</td><td>Intergenic</td><td>NA</td><td>NA</td></tr>
<tr><td>T7947</td><td>GCGCATCCCCGACTGCATCGTGG</td><td>1000</td><td></td><td></td><td></td><td></td><td></td><td></td><td></td><td></td><td></td></tr>
<tr><td>Oligo fwd</td><td>CACCGCGCATCCCCGACTGCATCG</td><td></td><td></td><td></td><td></td><td></td><td></td><td></td><td></td><td></td><td></td></tr>
<tr><td>Oligo rev</td><td>AAACCGATGCAGTCGGGGATGCGC</td><td></td><td></td><td></td><td></td><td></td><td></td><td></td><td></td><td></td><td></td></tr>
<tr><td>Chromosome</td><td>start</td><td>end</td><td>strand</td><td>MM</td><td>target_seq</td><td>PAM</td><td>alignment</td><td>distance</td><td>position</td><td>gene name</td><td>gene id</td></tr>
<tr><td>GU305914.1</td><td>108685</td><td>108707</td><td>+</td><td>0</td><td>GCGCATCCCCGACTGCATCG</td><td>TGG</td><td>||||||||[||||||||||||]PAM</td><td>NA</td><td>Intergenic</td><td>NA</td><td>NA</td></tr>
<tr><td>T7948</td><td>GTCCTCGTCGACGACCGCGACGG</td><td>1000</td><td></td><td></td><td></td><td></td><td></td><td></td><td></td><td></td><td></td></tr>
<tr><td>Oligo fwd</td><td>CACCGTCCTCGTCGACGACCGCGA</td><td></td><td></td><td></td><td></td><td></td><td></td><td></td><td></td><td></td><td></td></tr>
<tr><td>Oligo rev</td><td>AAACTCGCGGTCGTCGACGAGGAC</td><td></td><td></td><td></td><td></td><td></td><td></td><td></td><td></td><td></td><td></td></tr>
<tr><td>Chromosome</td><td>start</td><td>end</td><td>strand</td><td>MM</td><td>target_seq</td><td>PAM</td><td>alignment</td><td>distance</td><td>position</td><td>gene name</td><td>gene id</td></tr>
<tr><td>GU305914.1</td><td>108707</td><td>108729</td><td>+</td><td>0</td><td>GTCCTCGTCGACGACCGCGA</td><td>CGG</td><td>||||||||[||||||||||||]PAM</td><td>NA</td><td>Intergenic</td><td>NA</td><td>NA</td></tr>
<tr><td>T7949</td><td>CGTCGACGACCGCGACGGCTCGG</td><td>1000</td><td></td><td></td><td></td><td></td><td></td><td></td><td></td><td></td><td></td></tr>
<tr><td>Oligo fwd</td><td>CACCCGTCGACGACCGCGACGGCT</td><td></td><td></td><td></td><td></td><td></td><td></td><td></td><td></td><td></td><td></td></tr>
<tr><td>Oligo rev</td><td>AAACAGCCGTCGCGGTCGTCGACG</td><td></td><td></td><td></td><td></td><td></td><td></td><td></td><td></td><td></td><td></td></tr>
<tr><td>Chromosome</td><td>start</td><td>end</td><td>strand</td><td>MM</td><td>target_seq</td><td>PAM</td><td>alignment</td><td>distance</td><td>position</td><td>gene name</td><td>gene id</td></tr>
<tr><td>GU305914.1</td><td>108712</td><td>108734</td><td>+</td><td>0</td><td>CGTCGACGACCGCGACGGCT</td><td>CGG</td><td>||||||||[||||||||||||]PAM</td><td>NA</td><td>Intergenic</td><td>NA</td><td>NA</td></tr>
<tr><td>T7950</td><td>ACTATGCGCTGCGCCGACACCGG</td><td>1000</td><td></td><td></td><td></td><td></td><td></td><td></td><td></td><td></td><td></td></tr>
<tr><td>Oligo fwd</td><td>CACCACTATGCGCTGCGCCGACAC</td><td></td><td></td><td></td><td></td><td></td><td></td><td></td><td></td><td></td><td></td></tr>
<tr><td>Oligo rev</td><td>AAACGTGTCGGCGCAGCGCATAGT</td><td></td><td></td><td></td><td></td><td></td><td></td><td></td><td></td><td></td><td></td></tr>
<tr><td>Chromosome</td><td>start</td><td>end</td><td>strand</td><td>MM</td><td>target_seq</td><td>PAM</td><td>alignment</td><td>distance</td><td>position</td><td>gene name</td><td>gene id</td></tr>
<tr><td>GU305914.1</td><td>108764</td><td>108786</td><td>+</td><td>0</td><td>ACTATGCGCTGCGCCGACAC</td><td>CGG</td><td>||||||||[||||||||||||]PAM</td><td>NA</td><td>Intergenic</td><td>NA</td><td>NA</td></tr>
<tr><td>T7951</td><td>GCATCGCCTCCAGTACCTGCAGG</td><td>1000</td><td></td><td></td><td></td><td></td><td></td><td></td><td></td><td></td><td></td></tr>
<tr><td>Oligo fwd</td><td>CACCGCATCGCCTCCAGTACCTGC</td><td></td><td></td><td></td><td></td><td></td><td></td><td></td><td></td><td></td><td></td></tr>
<tr><td>Oligo rev</td><td>AAACGCAGGTACTGGAGGCGATGC</td><td></td><td></td><td></td><td></td><td></td><td></td><td></td><td></td><td></td><td></td></tr>
<tr><td>Chromosome</td><td>start</td><td>end</td><td>strand</td><td>MM</td><td>target_seq</td><td>PAM</td><td>alignment</td><td>distance</td><td>position</td><td>gene name</td><td>gene id</td></tr>
<tr><td>GU305914.1</td><td>108808</td><td>108830</td><td>+</td><td>0</td><td>GCATCGCCTCCAGTACCTGC</td><td>AGG</td><td>||||||||[||||||||||||]PAM</td><td>NA</td><td>Intergenic</td><td>NA</td><td>NA</td></tr>
<tr><td>T7952</td><td>CATCGCCTCCAGTACCTGCAGGG</td><td>1000</td><td></td><td></td><td></td><td></td><td></td><td></td><td></td><td></td><td></td></tr>
<tr><td>Oligo fwd</td><td>CACCCATCGCCTCCAGTACCTGCA</td><td></td><td></td><td></td><td></td><td></td><td></td><td></td><td></td><td></td><td></td></tr>
<tr><td>Oligo rev</td><td>AAACTGCAGGTACTGGAGGCGATG</td><td></td><td></td><td></td><td></td><td></td><td></td><td></td><td></td><td></td><td></td></tr>
<tr><td>Chromosome</td><td>start</td><td>end</td><td>strand</td><td>MM</td><td>target_seq</td><td>PAM</td><td>alignment</td><td>distance</td><td>position</td><td>gene name</td><td>gene id</td></tr>
<tr><td>GU305914.1</td><td>108809</td><td>108831</td><td>+</td><td>0</td><td>CATCGCCTCCAGTACCTGCA</td><td>GGG</td><td>||||||||[||||||||||||]PAM</td><td>NA</td><td>Intergenic</td><td>NA</td><td>NA</td></tr>
<tr><td>T7953</td><td>GCGCATCTTCTCGCAGTTCACGG</td><td>1000</td><td></td><td></td><td></td><td></td><td></td><td></td><td></td><td></td><td></td></tr>
<tr><td>Oligo fwd</td><td>CACCGCGCATCTTCTCGCAGTTCA</td><td></td><td></td><td></td><td></td><td></td><td></td><td></td><td></td><td></td><td></td></tr>
<tr><td>Oligo rev</td><td>AAACTGAACTGCGAGAAGATGCGC</td><td></td><td></td><td></td><td></td><td></td><td></td><td></td><td></td><td></td><td></td></tr>
<tr><td>Chromosome</td><td>start</td><td>end</td><td>strand</td><td>MM</td><td>target_seq</td><td>PAM</td><td>alignment</td><td>distance</td><td>position</td><td>gene name</td><td>gene id</td></tr>
<tr><td>GU305914.1</td><td>108856</td><td>108878</td><td>+</td><td>0</td><td>GCGCATCTTCTCGCAGTTCA</td><td>CGG</td><td>||||||||[||||||||||||]PAM</td><td>NA</td><td>Intergenic</td><td>NA</td><td>NA</td></tr>
<tr><td>T7954</td><td>TCGCAGTTCACGGTCGCCGACGG</td><td>1000</td><td></td><td></td><td></td><td></td><td></td><td></td><td></td><td></td><td></td></tr>
<tr><td>Oligo fwd</td><td>CACCTCGCAGTTCACGGTCGCCGA</td><td></td><td></td><td></td><td></td><td></td><td></td><td></td><td></td><td></td><td></td></tr>
<tr><td>Oligo rev</td><td>AAACTCGGCGACCGTGAACTGCGA</td><td></td><td></td><td></td><td></td><td></td><td></td><td></td><td></td><td></td><td></td></tr>
<tr><td>Chromosome</td><td>start</td><td>end</td><td>strand</td><td>MM</td><td>target_seq</td><td>PAM</td><td>alignment</td><td>distance</td><td>position</td><td>gene name</td><td>gene id</td></tr>
<tr><td>GU305914.1</td><td>108866</td><td>108888</td><td>+</td><td>0</td><td>TCGCAGTTCACGGTCGCCGA</td><td>CGG</td><td>||||||||[||||||||||||]PAM</td><td>NA</td><td>Intergenic</td><td>NA</td><td>NA</td></tr>
<tr><td>T7955</td><td>CGGTCGCCGACGGTGTCTGTTGG</td><td>1000</td><td></td><td></td><td></td><td></td><td></td><td></td><td></td><td></td><td></td></tr>
<tr><td>Oligo fwd</td><td>CACCCGGTCGCCGACGGTGTCTGT</td><td></td><td></td><td></td><td></td><td></td><td></td><td></td><td></td><td></td><td></td></tr>
<tr><td>Oligo rev</td><td>AAACACAGACACCGTCGGCGACCG</td><td></td><td></td><td></td><td></td><td></td><td></td><td></td><td></td><td></td><td></td></tr>
<tr><td>Chromosome</td><td>start</td><td>end</td><td>strand</td><td>MM</td><td>target_seq</td><td>PAM</td><td>alignment</td><td>distance</td><td>position</td><td>gene name</td><td>gene id</td></tr>
<tr><td>GU305914.1</td><td>108876</td><td>108898</td><td>+</td><td>0</td><td>CGGTCGCCGACGGTGTCTGT</td><td>TGG</td><td>||||||||[||||||||||||]PAM</td><td>NA</td><td>Intergenic</td><td>NA</td><td>NA</td></tr>
<tr><td>T7956</td><td>CGCCGACGGTGTCTGTTGGAAGG</td><td>1000</td><td></td><td></td><td></td><td></td><td></td><td></td><td></td><td></td><td></td></tr>
<tr><td>Oligo fwd</td><td>CACCCGCCGACGGTGTCTGTTGGA</td><td></td><td></td><td></td><td></td><td></td><td></td><td></td><td></td><td></td><td></td></tr>
<tr><td>Oligo rev</td><td>AAACTCCAACAGACACCGTCGGCG</td><td></td><td></td><td></td><td></td><td></td><td></td><td></td><td></td><td></td><td></td></tr>
<tr><td>Chromosome</td><td>start</td><td>end</td><td>strand</td><td>MM</td><td>target_seq</td><td>PAM</td><td>alignment</td><td>distance</td><td>position</td><td>gene name</td><td>gene id</td></tr>
<tr><td>GU305914.1</td><td>108880</td><td>108902</td><td>+</td><td>0</td><td>CGCCGACGGTGTCTGTTGGA</td><td>AGG</td><td>||||||||[||||||||||||]PAM</td><td>NA</td><td>Intergenic</td><td>NA</td><td>NA</td></tr>
<tr><td>T7957</td><td>CATCTCTTTCTTTCGCCAGAAGG</td><td>1000</td><td></td><td></td><td></td><td></td><td></td><td></td><td></td><td></td><td></td></tr>
<tr><td>Oligo fwd</td><td>CACCCATCTCTTTCTTTCGCCAGA</td><td></td><td></td><td></td><td></td><td></td><td></td><td></td><td></td><td></td><td></td></tr>
<tr><td>Oligo rev</td><td>AAACTCTGGCGAAAGAAAGAGATG</td><td></td><td></td><td></td><td></td><td></td><td></td><td></td><td></td><td></td><td></td></tr>
<tr><td>Chromosome</td><td>start</td><td>end</td><td>strand</td><td>MM</td><td>target_seq</td><td>PAM</td><td>alignment</td><td>distance</td><td>position</td><td>gene name</td><td>gene id</td></tr>
<tr><td>GU305914.1</td><td>108913</td><td>108935</td><td>+</td><td>0</td><td>CATCTCTTTCTTTCGCCAGA</td><td>AGG</td><td>||||||||[||||||||||||]PAM</td><td>NA</td><td>Intergenic</td><td>NA</td><td>NA</td></tr>
<tr><td>T7958</td><td>ATCTCTTTCTTTCGCCAGAAGGG</td><td>1000</td><td></td><td></td><td></td><td></td><td></td><td></td><td></td><td></td><td></td></tr>
<tr><td>Oligo fwd</td><td>CACCATCTCTTTCTTTCGCCAGAA</td><td></td><td></td><td></td><td></td><td></td><td></td><td></td><td></td><td></td><td></td></tr>
<tr><td>Oligo rev</td><td>AAACTTCTGGCGAAAGAAAGAGAT</td><td></td><td></td><td></td><td></td><td></td><td></td><td></td><td></td><td></td><td></td></tr>
<tr><td>Chromosome</td><td>start</td><td>end</td><td>strand</td><td>MM</td><td>target_seq</td><td>PAM</td><td>alignment</td><td>distance</td><td>position</td><td>gene name</td><td>gene id</td></tr>
<tr><td>GU305914.1</td><td>108914</td><td>108936</td><td>+</td><td>0</td><td>ATCTCTTTCTTTCGCCAGAA</td><td>GGG</td><td>||||||||[||||||||||||]PAM</td><td>NA</td><td>Intergenic</td><td>NA</td><td>NA</td></tr>
<tr><td>T7959</td><td>TCTCTTTCTTTCGCCAGAAGGGG</td><td>1000</td><td></td><td></td><td></td><td></td><td></td><td></td><td></td><td></td><td></td></tr>
<tr><td>Oligo fwd</td><td>CACCTCTCTTTCTTTCGCCAGAAG</td><td></td><td></td><td></td><td></td><td></td><td></td><td></td><td></td><td></td><td></td></tr>
<tr><td>Oligo rev</td><td>AAACCTTCTGGCGAAAGAAAGAGA</td><td></td><td></td><td></td><td></td><td></td><td></td><td></td><td></td><td></td><td></td></tr>
<tr><td>Chromosome</td><td>start</td><td>end</td><td>strand</td><td>MM</td><td>target_seq</td><td>PAM</td><td>alignment</td><td>distance</td><td>position</td><td>gene name</td><td>gene id</td></tr>
<tr><td>GU305914.1</td><td>108915</td><td>108937</td><td>+</td><td>0</td><td>TCTCTTTCTTTCGCCAGAAG</td><td>GGG</td><td>||||||||[||||||||||||]PAM</td><td>NA</td><td>Intergenic</td><td>NA</td><td>NA</td></tr>
<tr><td>T7960</td><td>CTCTTTCTTTCGCCAGAAGGGGG</td><td>1000</td><td></td><td></td><td></td><td></td><td></td><td></td><td></td><td></td><td></td></tr>
<tr><td>Oligo fwd</td><td>CACCCTCTTTCTTTCGCCAGAAGG</td><td></td><td></td><td></td><td></td><td></td><td></td><td></td><td></td><td></td><td></td></tr>
<tr><td>Oligo rev</td><td>AAACCCTTCTGGCGAAAGAAAGAG</td><td></td><td></td><td></td><td></td><td></td><td></td><td></td><td></td><td></td><td></td></tr>
<tr><td>Chromosome</td><td>start</td><td>end</td><td>strand</td><td>MM</td><td>target_seq</td><td>PAM</td><td>alignment</td><td>distance</td><td>position</td><td>gene name</td><td>gene id</td></tr>
<tr><td>GU305914.1</td><td>108916</td><td>108938</td><td>+</td><td>0</td><td>CTCTTTCTTTCGCCAGAAGG</td><td>GGG</td><td>||||||||[||||||||||||]PAM</td><td>NA</td><td>Intergenic</td><td>NA</td><td>NA</td></tr>
<tr><td>T7961</td><td>GGGCGCGCGTCGCCCTCACGCGG</td><td>1000</td><td></td><td></td><td></td><td></td><td></td><td></td><td></td><td></td><td></td></tr>
<tr><td>Oligo fwd</td><td>CACCGGGCGCGCGTCGCCCTCACG</td><td></td><td></td><td></td><td></td><td></td><td></td><td></td><td></td><td></td><td></td></tr>
<tr><td>Oligo rev</td><td>AAACCGTGAGGGCGACGCGCGCCC</td><td></td><td></td><td></td><td></td><td></td><td></td><td></td><td></td><td></td><td></td></tr>
</table>

| | | | | | | | | | | | | |
|---|---|---|---|---|---|---|---|---|---|---|---|---|
| Chromosome | start | | end | strand | MM | target_seq | PAM | alignment | distance | position | gene name | gene id |
| GU305914.1 | | 108936 | 108958 | + | 0 | GGGCGCGCGTCGCCCTCACG | CGG | \|\|\|\|\|\|\|\|[\|\|\|\|\|\|\|\|\|\|\|\|]PAM | NA | Intergenic | NA | NA |
| T7962 | GGCGCGCGTCGCCCTCACGCGGG | | 1000 | | | | | | | | | |
| Oligo fwd | CACCGGCGCGCGTCGCCCTCACGC | | | | | | | | | | | |
| Oligo rev | AAACGCGTGAGGGCGACGCGCGCC | | | | | | | | | | | |
| Chromosome | start | | end | strand | MM | target_seq | PAM | alignment | distance | position | gene name | gene id |
| GU305914.1 | | 108937 | 108959 | + | 0 | GGCGCGCGTCGCCCTCACGC | GGG | \|\|\|\|\|\|\|\|[\|\|\|\|\|\|\|\|\|\|\|\|]PAM | NA | Intergenic | NA | NA |
| T7963 | TCACGCGGGTCTTCAAGCCCAGG | | 1000 | | | | | | | | | |
| Oligo fwd | CACCTCACGCGGGTCTTCAAGCCC | | | | | | | | | | | |
| Oligo rev | AAACGGGCTTGAAGACCCGCGTGA | | | | | | | | | | | |
| Chromosome | start | | end | strand | MM | target_seq | PAM | alignment | distance | position | gene name | gene id |
| GU305914.1 | | 108951 | 108973 | + | 0 | TCACGCGGGTCTTCAAGCCC | AGG | \|\|\|\|\|\|\|\|[\|\|\|\|\|\|\|\|\|\|\|\|]PAM | NA | Intergenic | NA | NA |
| T7964 | CACGCGGGTCTTCAAGCCCAGGG | | 1000 | | | | | | | | | |
| Oligo fwd | CACCCACGCGGGTCTTCAAGCCCA | | | | | | | | | | | |
| Oligo rev | AAACTGGGCTTGAAGACCCGCGTG | | | | | | | | | | | |
| Chromosome | start | | end | strand | MM | target_seq | PAM | alignment | distance | position | gene name | gene id |
| GU305914.1 | | 108952 | 108974 | + | 0 | CACGCGGGTCTTCAAGCCCA | GGG | \|\|\|\|\|\|\|\|[\|\|\|\|\|\|\|\|\|\|\|\|]PAM | NA | Intergenic | NA | NA |
| T7965 | CGTCCTCTCCTCCGTCGTCTTGG | | 1000 | | | | | | | | | |
| Oligo fwd | CACCCGTCCTCTCCTCCGTCGTCT | | | | | | | | | | | |
| Oligo rev | AAACAGACGACGGAGGAGAGGACG | | | | | | | | | | | |
| Chromosome | start | | end | strand | MM | target_seq | PAM | alignment | distance | position | gene name | gene id |
| GU305914.1 | | 108982 | 109004 | + | 0 | CGTCCTCTCCTCCGTCGTCT | TGG | \|\|\|\|\|\|\|\|[\|\|\|\|\|\|\|\|\|\|\|\|]PAM | NA | Intergenic | NA | NA |
| T7966 | GGATTTCCTGCGCCTGCACCAGG | | 1000 | | | | | | | | | |
| Oligo fwd | CACCGGATTTCCTGCGCCTGCACC | | | | | | | | | | | |
| Oligo rev | AAACGGTGCAGGCGCAGGAAATCC | | | | | | | | | | | |
| Chromosome | start | | end | strand | MM | target_seq | PAM | alignment | distance | position | gene name | gene id |
| GU305914.1 | | 109003 | 109025 | + | 0 | GGATTTCCTGCGCCTGCACC | AGG | \|\|\|\|\|\|\|\|[\|\|\|\|\|\|\|\|\|\|\|\|]PAM | NA | Intergenic | NA | NA |
| T7967 | GCGCCTGCACCAGGATGAGTCGG | | 1000 | | | | | | | | | |
| Oligo fwd | CACCGCGCCTGCACCAGGATGAGT | | | | | | | | | | | |
| Oligo rev | AAACACTCATCCTGGTGCAGGCGC | | | | | | | | | | | |
| Chromosome | start | | end | strand | MM | target_seq | PAM | alignment | distance | position | gene name | gene id |
| GU305914.1 | | 109012 | 109034 | + | 0 | GCGCCTGCACCAGGATGAGT | CGG | \|\|\|\|\|\|\|\|[\|\|\|\|\|\|\|\|\|\|\|\|]PAM | NA | Intergenic | NA | NA |
| T7968 | TCACGATGTGCCGTGTATAGTGG | | 1000 | | | | | | | | | |
| Oligo fwd | CACCTCACGATGTGCCGTGTATAG | | | | | | | | | | | |
| Oligo rev | AAACCTATACACGGCACATCGTGA | | | | | | | | | | | |
| Chromosome | start | | end | strand | MM | target_seq | PAM | alignment | distance | position | gene name | gene id |
| GU305914.1 | | 109046 | 109068 | + | 0 | TCACGATGTGCCGTGTATAG | TGG | \|\|\|\|\|\|\|\|[\|\|\|\|\|\|\|\|\|\|\|\|]PAM | NA | Intergenic | NA | NA |
| T7969 | GCCGTGTATAGTGGCCTTCGAGG | | 1000 | | | | | | | | | |
| Oligo fwd | CACCGCCGTGTATAGTGGCCTTCG | | | | | | | | | | | |
| Oligo rev | AAACCGAAGGCCACTATACACGGC | | | | | | | | | | | |
| Chromosome | start | | end | strand | MM | target_seq | PAM | alignment | distance | position | gene name | gene id |
| GU305914.1 | | 109055 | 109077 | + | 0 | GCCGTGTATAGTGGCCTTCG | AGG | \|\|\|\|\|\|\|\|[\|\|\|\|\|\|\|\|\|\|\|\|]PAM | NA | Intergenic | NA | NA |
| T7970 | TAGTGGCCTTCGAGGCGCATCGG | | 1000 | | | | | | | | | |
| Oligo fwd | CACCTAGTGGCCTTCGAGGCGCAT | | | | | | | | | | | |
| Oligo rev | AAACATGCGCCTCGAAGGCCACTA | | | | | | | | | | | |
| Chromosome | start | | end | strand | MM | target_seq | PAM | alignment | distance | position | gene name | gene id |
| GU305914.1 | | 109063 | 109085 | + | 0 | TAGTGGCCTTCGAGGCGCAT | CGG | \|\|\|\|\|\|\|\|[\|\|\|\|\|\|\|\|\|\|\|\|]PAM | NA | Intergenic | NA | NA |
| T7971 | AGTGGCCTTCGAGGCGCATCGGG | | 1000 | | | | | | | | | |
| Oligo fwd | CACCAGTGGCCTTCGAGGCGCATC | | | | | | | | | | | |
| Oligo rev | AAACGATGCGCCTCGAAGGCCACT | | | | | | | | | | | |
| Chromosome | start | | end | strand | MM | target_seq | PAM | alignment | distance | position | gene name | gene id |
| GU305914.1 | | 109064 | 109086 | + | 0 | AGTGGCCTTCGAGGCGCATC | GGG | \|\|\|\|\|\|\|\|[\|\|\|\|\|\|\|\|\|\|\|\|]PAM | NA | Intergenic | NA | NA |
| T7972 | GGCGCATCGGGAAAACGTCCTGG | | 1000 | | | | | | | | | |
| Oligo fwd | CACCGGCGCATCGGGAAAACGTCC | | | | | | | | | | | |
| Oligo rev | AAACGGACGTTTTCCCGATGCGCC | | | | | | | | | | | |
| Chromosome | start | | end | strand | MM | target_seq | PAM | alignment | distance | position | gene name | gene id |
| GU305914.1 | | 109076 | 109098 | + | 0 | GGCGCATCGGGAAAACGTCC | TGG | \|\|\|\|\|\|\|\|[\|\|\|\|\|\|\|\|\|\|\|\|]PAM | NA | Intergenic | NA | NA |
| T7973 | ACGAAAGCGCGCTGCGCCTGCGG | | 1000 | | | | | | | | | |
| Oligo fwd | CACCACGAAAGCGCGCTGCGCCTG | | | | | | | | | | | |
| Oligo rev | AAACCAGGCGCAGCGCGCTTTCGT | | | | | | | | | | | |
| Chromosome | start | | end | strand | MM | target_seq | PAM | alignment | distance | position | gene name | gene id |
| GU305914.1 | | 109132 | 109154 | + | 0 | ACGAAAGCGCGCTGCGCCTG | CGG | \|\|\|\|\|\|\|\|[\|\|\|\|\|\|\|\|\|\|\|\|]PAM | NA | Intergenic | NA | NA |
| T7974 | GCGGCAGTACGCCGACGACCTGG | | 1000 | | | | | | | | | |
| Oligo fwd | CACCGCGGCAGTACGCCGACGACC | | | | | | | | | | | |
| Oligo rev | AAACGGTCGTCGGCGTACTGCCGC | | | | | | | | | | | |
| Chromosome | start | | end | strand | MM | target_seq | PAM | alignment | distance | position | gene name | gene id |
| GU305914.1 | | 109151 | 109173 | + | 0 | GCGGCAGTACGCCGACGACC | TGG | \|\|\|\|\|\|\|\|[\|\|\|\|\|\|\|\|\|\|\|\|]PAM | NA | Intergenic | NA | NA |
| T7975 | CGGCAGTACGCCGACGACCTGGG | | 1000 | | | | | | | | | |
| Oligo fwd | CACCCGGCAGTACGCCGACGACCT | | | | | | | | | | | |
| Oligo rev | AAACAGGTCGTCGGCGTACTGCCG | | | | | | | | | | | |
| Chromosome | start | | end | strand | MM | target_seq | PAM | alignment | distance | position | gene name | gene id |
| GU305914.1 | | 109152 | 109174 | + | 0 | CGGCAGTACGCCGACGACCT | GGG | \|\|\|\|\|\|\|\|[\|\|\|\|\|\|\|\|\|\|\|\|]PAM | NA | Intergenic | NA | NA |
| T7976 | GGCAGTACGCCGACGACCTGGGG | | 1000 | | | | | | | | | |
| Oligo fwd | CACCGGCAGTACGCCGACGACCTG | | | | | | | | | | | |

<table>
<tr><td>Oligo rev</td><td>AAACCAGGTCGTCGGCGTACTGCC</td><td></td><td></td><td></td><td></td><td></td><td></td><td></td><td></td><td></td><td></td></tr>
<tr><td>Chromosome</td><td>start</td><td>end</td><td>strand</td><td>MM</td><td>target_seq</td><td>PAM</td><td>alignment</td><td>distance</td><td>position</td><td>gene name</td><td>gene id</td></tr>
<tr><td>GU305914.1</td><td>109153</td><td>109175</td><td>+</td><td>0</td><td>GGCAGTACGCCGACGACCTG</td><td>GGG</td><td>||||||||[||||||||||||]PAM</td><td>NA</td><td>Intergenic</td><td>NA</td><td>NA</td></tr>
<tr><td></td><td></td><td></td><td></td><td></td><td></td><td></td><td></td><td></td><td></td><td></td><td></td></tr>
<tr><td>T7977</td><td>ACGACCTGGGGCACGACGCCAGG</td><td>1000</td><td></td><td></td><td></td><td></td><td></td><td></td><td></td><td></td><td></td></tr>
<tr><td>Oligo fwd</td><td>CACCACGACCTGGGGCACGACGCC</td><td></td><td></td><td></td><td></td><td></td><td></td><td></td><td></td><td></td><td></td></tr>
<tr><td>Oligo rev</td><td>AAACGGCGTCGTGCCCCAGGTCGT</td><td></td><td></td><td></td><td></td><td></td><td></td><td></td><td></td><td></td><td></td></tr>
<tr><td>Chromosome</td><td>start</td><td>end</td><td>strand</td><td>MM</td><td>target_seq</td><td>PAM</td><td>alignment</td><td>distance</td><td>position</td><td>gene name</td><td>gene id</td></tr>
<tr><td>GU305914.1</td><td>109165</td><td>109187</td><td>+</td><td>0</td><td>ACGACCTGGGGCACGACGCC</td><td>AGG</td><td>||||||||[||||||||||||]PAM</td><td>NA</td><td>Intergenic</td><td>NA</td><td>NA</td></tr>
<tr><td></td><td></td><td></td><td></td><td></td><td></td><td></td><td></td><td></td><td></td><td></td><td></td></tr>
<tr><td>T7978</td><td>ACGACGCCAGGCTCCGACGAAGG</td><td>1000</td><td></td><td></td><td></td><td></td><td></td><td></td><td></td><td></td><td></td></tr>
<tr><td>Oligo fwd</td><td>CACCACGACGCCAGGCTCCGACGA</td><td></td><td></td><td></td><td></td><td></td><td></td><td></td><td></td><td></td><td></td></tr>
<tr><td>Oligo rev</td><td>AAACTCGTCGGAGCCTGGCGTCGT</td><td></td><td></td><td></td><td></td><td></td><td></td><td></td><td></td><td></td><td></td></tr>
<tr><td>Chromosome</td><td>start</td><td>end</td><td>strand</td><td>MM</td><td>target_seq</td><td>PAM</td><td>alignment</td><td>distance</td><td>position</td><td>gene name</td><td>gene id</td></tr>
<tr><td>GU305914.1</td><td>109177</td><td>109199</td><td>+</td><td>0</td><td>ACGACGCCAGGCTCCGACGA</td><td>AGG</td><td>||||||||[||||||||||||]PAM</td><td>NA</td><td>Intergenic</td><td>NA</td><td>NA</td></tr>
<tr><td></td><td></td><td></td><td></td><td></td><td></td><td></td><td></td><td></td><td></td><td></td><td></td></tr>
<tr><td>T7979</td><td>CGACGCCAGGCTCCGACGAAGGG</td><td>1000</td><td></td><td></td><td></td><td></td><td></td><td></td><td></td><td></td><td></td></tr>
<tr><td>Oligo fwd</td><td>CACCCGACGCCAGGCTCCGACGAA</td><td></td><td></td><td></td><td></td><td></td><td></td><td></td><td></td><td></td><td></td></tr>
<tr><td>Oligo rev</td><td>AAACTTCGTCGGAGCCTGGCGTCG</td><td></td><td></td><td></td><td></td><td></td><td></td><td></td><td></td><td></td><td></td></tr>
<tr><td>Chromosome</td><td>start</td><td>end</td><td>strand</td><td>MM</td><td>target_seq</td><td>PAM</td><td>alignment</td><td>distance</td><td>position</td><td>gene name</td><td>gene id</td></tr>
<tr><td>GU305914.1</td><td>109178</td><td>109200</td><td>+</td><td>0</td><td>CGACGCCAGGCTCCGACGAA</td><td>GGG</td><td>||||||||[||||||||||||]PAM</td><td>NA</td><td>Intergenic</td><td>NA</td><td>NA</td></tr>
<tr><td></td><td></td><td></td><td></td><td></td><td></td><td></td><td></td><td></td><td></td><td></td><td></td></tr>
<tr><td>T7980</td><td>CGCCAGGCTCCGACGAAGGGCGG</td><td>1000</td><td></td><td></td><td></td><td></td><td></td><td></td><td></td><td></td><td></td></tr>
<tr><td>Oligo fwd</td><td>CACCCGCCAGGCTCCGACGAAGGG</td><td></td><td></td><td></td><td></td><td></td><td></td><td></td><td></td><td></td><td></td></tr>
<tr><td>Oligo rev</td><td>AAACCCCTTCGTCGGAGCCTGGCG</td><td></td><td></td><td></td><td></td><td></td><td></td><td></td><td></td><td></td><td></td></tr>
<tr><td>Chromosome</td><td>start</td><td>end</td><td>strand</td><td>MM</td><td>target_seq</td><td>PAM</td><td>alignment</td><td>distance</td><td>position</td><td>gene name</td><td>gene id</td></tr>
<tr><td>GU305914.1</td><td>109181</td><td>109203</td><td>+</td><td>0</td><td>CGCCAGGCTCCGACGAAGGG</td><td>CGG</td><td>||||||||[||||||||||||]PAM</td><td>NA</td><td>Intergenic</td><td>NA</td><td>NA</td></tr>
<tr><td></td><td></td><td></td><td></td><td></td><td></td><td></td><td></td><td></td><td></td><td></td><td></td></tr>
<tr><td>T7981</td><td>GCCAGGCTCCGACGAAGGGCGGG</td><td>1000</td><td></td><td></td><td></td><td></td><td></td><td></td><td></td><td></td><td></td></tr>
<tr><td>Oligo fwd</td><td>CACCGCCAGGCTCCGACGAAGGGC</td><td></td><td></td><td></td><td></td><td></td><td></td><td></td><td></td><td></td><td></td></tr>
<tr><td>Oligo rev</td><td>AAACGCCCTTCGTCGGAGCCTGGC</td><td></td><td></td><td></td><td></td><td></td><td></td><td></td><td></td><td></td><td></td></tr>
<tr><td>Chromosome</td><td>start</td><td>end</td><td>strand</td><td>MM</td><td>target_seq</td><td>PAM</td><td>alignment</td><td>distance</td><td>position</td><td>gene name</td><td>gene id</td></tr>
<tr><td>GU305914.1</td><td>109182</td><td>109204</td><td>+</td><td>0</td><td>GCCAGGCTCCGACGAAGGGC</td><td>GGG</td><td>||||||||[||||||||||||]PAM</td><td>NA</td><td>Intergenic</td><td>NA</td><td>NA</td></tr>
<tr><td></td><td></td><td></td><td></td><td></td><td></td><td></td><td></td><td></td><td></td><td></td><td></td></tr>
<tr><td>T7982</td><td>GGCGAAGACCTAGAGTCTCTCGG</td><td>1000</td><td></td><td></td><td></td><td></td><td></td><td></td><td></td><td></td><td></td></tr>
<tr><td>Oligo fwd</td><td>CACCGGCGAAGACCTAGAGTCTCT</td><td></td><td></td><td></td><td></td><td></td><td></td><td></td><td></td><td></td><td></td></tr>
<tr><td>Oligo rev</td><td>AAACAGAGACTCTAGGTCTTCGCC</td><td></td><td></td><td></td><td></td><td></td><td></td><td></td><td></td><td></td><td></td></tr>
<tr><td>Chromosome</td><td>start</td><td>end</td><td>strand</td><td>MM</td><td>target_seq</td><td>PAM</td><td>alignment</td><td>distance</td><td>position</td><td>gene name</td><td>gene id</td></tr>
<tr><td>GU305914.1</td><td>109203</td><td>109225</td><td>+</td><td>0</td><td>GGCGAAGACCTAGAGTCTCT</td><td>CGG</td><td>||||||||[||||||||||||]PAM</td><td>NA</td><td>Intergenic</td><td>NA</td><td>NA</td></tr>
<tr><td></td><td></td><td></td><td></td><td></td><td></td><td></td><td></td><td></td><td></td><td></td><td></td></tr>
<tr><td>T7983</td><td>AATGAGTGCGAGCGCTTCCGCGG</td><td>1000</td><td></td><td></td><td></td><td></td><td></td><td></td><td></td><td></td><td></td></tr>
<tr><td>Oligo fwd</td><td>CACCAATGAGTGCGAGCGCTTCCG</td><td></td><td></td><td></td><td></td><td></td><td></td><td></td><td></td><td></td><td></td></tr>
<tr><td>Oligo rev</td><td>AAACCGGAAGCGCTCGCACTCATT</td><td></td><td></td><td></td><td></td><td></td><td></td><td></td><td></td><td></td><td></td></tr>
<tr><td>Chromosome</td><td>start</td><td>end</td><td>strand</td><td>MM</td><td>target_seq</td><td>PAM</td><td>alignment</td><td>distance</td><td>position</td><td>gene name</td><td>gene id</td></tr>
<tr><td>GU305914.1</td><td>109239</td><td>109261</td><td>+</td><td>0</td><td>AATGAGTGCGAGCGCTTCCG</td><td>CGG</td><td>||||||||[||||||||||||]PAM</td><td>NA</td><td>Intergenic</td><td>NA</td><td>NA</td></tr>
<tr><td></td><td></td><td></td><td></td><td></td><td></td><td></td><td></td><td></td><td></td><td></td><td></td></tr>
<tr><td>T7984</td><td>AGTGCGAGCGCTTCCGCGGCAGG</td><td>1000</td><td></td><td></td><td></td><td></td><td></td><td></td><td></td><td></td><td></td></tr>
<tr><td>Oligo fwd</td><td>CACCAGTGCGAGCGCTTCCGCGGC</td><td></td><td></td><td></td><td></td><td></td><td></td><td></td><td></td><td></td><td></td></tr>
<tr><td>Oligo rev</td><td>AAACGCCGCGGAAGCGCTCGCACT</td><td></td><td></td><td></td><td></td><td></td><td></td><td></td><td></td><td></td><td></td></tr>
<tr><td>Chromosome</td><td>start</td><td>end</td><td>strand</td><td>MM</td><td>target_seq</td><td>PAM</td><td>alignment</td><td>distance</td><td>position</td><td>gene name</td><td>gene id</td></tr>
<tr><td>GU305914.1</td><td>109243</td><td>109265</td><td>+</td><td>0</td><td>AGTGCGAGCGCTTCCGCGGC</td><td>AGG</td><td>||||||||[||||||||||||]PAM</td><td>NA</td><td>Intergenic</td><td>NA</td><td>NA</td></tr>
<tr><td></td><td></td><td></td><td></td><td></td><td></td><td></td><td></td><td></td><td></td><td></td><td></td></tr>
<tr><td>T7985</td><td>CTTCCGCGGCAGGATCGATCAGG</td><td>1000</td><td></td><td></td><td></td><td></td><td></td><td></td><td></td><td></td><td></td></tr>
<tr><td>Oligo fwd</td><td>CACCCTTCCGCGGCAGGATCGATC</td><td></td><td></td><td></td><td></td><td></td><td></td><td></td><td></td><td></td><td></td></tr>
<tr><td>Oligo rev</td><td>AAACGATCGATCCTGCCGCGGAAG</td><td></td><td></td><td></td><td></td><td></td><td></td><td></td><td></td><td></td><td></td></tr>
<tr><td>Chromosome</td><td>start</td><td>end</td><td>strand</td><td>MM</td><td>target_seq</td><td>PAM</td><td>alignment</td><td>distance</td><td>position</td><td>gene name</td><td>gene id</td></tr>
<tr><td>GU305914.1</td><td>109253</td><td>109275</td><td>+</td><td>0</td><td>CTTCCGCGGCAGGATCGATC</td><td>AGG</td><td>||||||||[||||||||||||]PAM</td><td>NA</td><td>Intergenic</td><td>NA</td><td>NA</td></tr>
<tr><td></td><td></td><td></td><td></td><td></td><td></td><td></td><td></td><td></td><td></td><td></td><td></td></tr>
<tr><td>T7986</td><td>CCGCGGCAGGATCGATCAGGCGG</td><td>1000</td><td></td><td></td><td></td><td></td><td></td><td></td><td></td><td></td><td></td></tr>
<tr><td>Oligo fwd</td><td>CACCCCGCGGCAGGATCGATCAGG</td><td></td><td></td><td></td><td></td><td></td><td></td><td></td><td></td><td></td><td></td></tr>
<tr><td>Oligo rev</td><td>AAACCCTGATCGATCCTGCCGCGG</td><td></td><td></td><td></td><td></td><td></td><td></td><td></td><td></td><td></td><td></td></tr>
<tr><td>Chromosome</td><td>start</td><td>end</td><td>strand</td><td>MM</td><td>target_seq</td><td>PAM</td><td>alignment</td><td>distance</td><td>position</td><td>gene name</td><td>gene id</td></tr>
<tr><td>GU305914.1</td><td>109256</td><td>109278</td><td>+</td><td>0</td><td>CCGCGGCAGGATCGATCAGG</td><td>CGG</td><td>||||||||[||||||||||||]PAM</td><td>NA</td><td>Intergenic</td><td>NA</td><td>NA</td></tr>
<tr><td></td><td></td><td></td><td></td><td></td><td></td><td></td><td></td><td></td><td></td><td></td><td></td></tr>
<tr><td>T7987</td><td>CAGGCGGAACAGCTGCTGTCCGG</td><td>1000</td><td></td><td></td><td></td><td></td><td></td><td></td><td></td><td></td><td></td></tr>
<tr><td>Oligo fwd</td><td>CACCCAGGCGGAACAGCTGCTGTC</td><td></td><td></td><td></td><td></td><td></td><td></td><td></td><td></td><td></td><td></td></tr>
<tr><td>Oligo rev</td><td>AAACGACAGCAGCTGTTCCGCCTG</td><td></td><td></td><td></td><td></td><td></td><td></td><td></td><td></td><td></td><td></td></tr>
<tr><td>Chromosome</td><td>start</td><td>end</td><td>strand</td><td>MM</td><td>target_seq</td><td>PAM</td><td>alignment</td><td>distance</td><td>position</td><td>gene name</td><td>gene id</td></tr>
<tr><td>GU305914.1</td><td>109272</td><td>109294</td><td>+</td><td>0</td><td>CAGGCGGAACAGCTGCTGTC</td><td>CGG</td><td>||||||||[||||||||||||]PAM</td><td>NA</td><td>Intergenic</td><td>NA</td><td>NA</td></tr>
<tr><td>GU305914.1</td><td>69470</td><td>69492</td><td>+</td><td>3</td><td>CTGGCGGAACAGCTGCCGAC</td><td>GGG</td><td>|-||||||[||||||||-|-|]PAM</td><td>NA</td><td>Intergenic</td><td>NA</td><td>NA</td></tr>
<tr><td></td><td></td><td></td><td></td><td></td><td></td><td></td><td></td><td></td><td></td><td></td><td></td></tr>
<tr><td>T7988</td><td>AGGCGGAACAGCTGCTGTCCGGG</td><td>1000</td><td></td><td></td><td></td><td></td><td></td><td></td><td></td><td></td><td></td></tr>
<tr><td>Oligo fwd</td><td>CACCAGGCGGAACAGCTGCTGTCC</td><td></td><td></td><td></td><td></td><td></td><td></td><td></td><td></td><td></td><td></td></tr>
<tr><td>Oligo rev</td><td>AAACGGACAGCAGCTGTTCCGCCT</td><td></td><td></td><td></td><td></td><td></td><td></td><td></td><td></td><td></td><td></td></tr>
<tr><td>Chromosome</td><td>start</td><td>end</td><td>strand</td><td>MM</td><td>target_seq</td><td>PAM</td><td>alignment</td><td>distance</td><td>position</td><td>gene name</td><td>gene id</td></tr>
<tr><td>GU305914.1</td><td>109273</td><td>109295</td><td>+</td><td>0</td><td>AGGCGGAACAGCTGCTGTCC</td><td>GGG</td><td>||||||||[||||||||||||]PAM</td><td>NA</td><td>Intergenic</td><td>NA</td><td>NA</td></tr>
<tr><td></td><td></td><td></td><td></td><td></td><td></td><td></td><td></td><td></td><td></td><td></td><td></td></tr>
<tr><td>T7989</td><td>GCTGCTGTCCGGGCCCATGTCGG</td><td>1000</td><td></td><td></td><td></td><td></td><td></td><td></td><td></td><td></td><td></td></tr>
<tr><td>Oligo fwd</td><td>CACCGCTGCTGTCCGGGCCCATGT</td><td></td><td></td><td></td><td></td><td></td><td></td><td></td><td></td><td></td><td></td></tr>
<tr><td>Oligo rev</td><td>AAACACATGGGCCCGGACAGCAGC</td><td></td><td></td><td></td><td></td><td></td><td></td><td></td><td></td><td></td><td></td></tr>
<tr><td>Chromosome</td><td>start</td><td>end</td><td>strand</td><td>MM</td><td>target_seq</td><td>PAM</td><td>alignment</td><td>distance</td><td>position</td><td>gene name</td><td>gene id</td></tr>
<tr><td>GU305914.1</td><td>109283</td><td>109305</td><td>+</td><td>0</td><td>GCTGCTGTCCGGGCCCATGT</td><td>CGG</td><td>||||||||[||||||||||||]PAM</td><td>NA</td><td>Intergenic</td><td>NA</td><td>NA</td></tr>
<tr><td></td><td></td><td></td><td></td><td></td><td></td><td></td><td></td><td></td><td></td><td></td><td></td></tr>
<tr><td>T7990</td><td>CGGGCCCATGTCGGACATTCTGG</td><td>1000</td><td></td><td></td><td></td><td></td><td></td><td></td><td></td><td></td><td></td></tr>
<tr><td>Oligo fwd</td><td>CACCCGGGCCCATGTCGGACATTC</td><td></td><td></td><td></td><td></td><td></td><td></td><td></td><td></td><td></td><td></td></tr>
<tr><td>Oligo rev</td><td>AAACGAATGTCCGACATGGGCCCG</td><td></td><td></td><td></td><td></td><td></td><td></td><td></td><td></td><td></td><td></td></tr>
<tr><td>Chromosome</td><td>start</td><td>end</td><td>strand</td><td>MM</td><td>target_seq</td><td>PAM</td><td>alignment</td><td>distance</td><td>position</td><td>gene name</td><td>gene id</td></tr>
<tr><td>GU305914.1</td><td>109292</td><td>109314</td><td>+</td><td>0</td><td>CGGGCCCATGTCGGACATTC</td><td>TGG</td><td>||||||||[||||||||||||]PAM</td><td>NA</td><td>Intergenic</td><td>NA</td><td>NA</td></tr>
</table>

<table>
<tr><td>T7991</td><td>GGGCCCATGTCGGACATTCTGGG</td><td>1000</td><td></td><td></td><td></td><td></td><td></td><td></td><td></td><td></td><td></td></tr>
<tr><td>Oligo fwd</td><td>CACCGGGCCCATGTCGGACATTCT</td><td></td><td></td><td></td><td></td><td></td><td></td><td></td><td></td><td></td><td></td></tr>
<tr><td>Oligo rev</td><td>AAACAGAATGTCCGACATGGGCCC</td><td></td><td></td><td></td><td></td><td></td><td></td><td></td><td></td><td></td><td></td></tr>
<tr><td>Chromosome</td><td>start</td><td>end</td><td>strand</td><td>MM</td><td>target_seq</td><td>PAM</td><td>alignment</td><td>distance</td><td>position</td><td>gene name</td><td>gene id</td></tr>
<tr><td>GU305914.1</td><td>109293</td><td>109315</td><td>+</td><td>0</td><td>GGGCCCATGTCGGACATTCT</td><td>GGG</td><td>||||||||[||||||||||||]PAM</td><td>NA</td><td>Intergenic</td><td>NA</td><td>NA</td></tr>
<tr><td>T7992</td><td>CCCATGTCGGACATTCTGGGCGG</td><td>1000</td><td></td><td></td><td></td><td></td><td></td><td></td><td></td><td></td><td></td></tr>
<tr><td>Oligo fwd</td><td>CACCCCCATGTCGGACATTCTGGG</td><td></td><td></td><td></td><td></td><td></td><td></td><td></td><td></td><td></td><td></td></tr>
<tr><td>Oligo rev</td><td>AAACCCCAGAATGTCCGACATGGG</td><td></td><td></td><td></td><td></td><td></td><td></td><td></td><td></td><td></td><td></td></tr>
<tr><td>Chromosome</td><td>start</td><td>end</td><td>strand</td><td>MM</td><td>target_seq</td><td>PAM</td><td>alignment</td><td>distance</td><td>position</td><td>gene name</td><td>gene id</td></tr>
<tr><td>GU305914.1</td><td>109296</td><td>109318</td><td>+</td><td>0</td><td>CCCATGTCGGACATTCTGGG</td><td>CGG</td><td>||||||||[||||||||||||]PAM</td><td>NA</td><td>Intergenic</td><td>NA</td><td>NA</td></tr>
<tr><td>T7993</td><td>ATGTCGGACATTCTGGGCGGAGG</td><td>1000</td><td></td><td></td><td></td><td></td><td></td><td></td><td></td><td></td><td></td></tr>
<tr><td>Oligo fwd</td><td>CACCATGTCGGACATTCTGGGCGG</td><td></td><td></td><td></td><td></td><td></td><td></td><td></td><td></td><td></td><td></td></tr>
<tr><td>Oligo rev</td><td>AAACCCGCCCAGAATGTCCGACAT</td><td></td><td></td><td></td><td></td><td></td><td></td><td></td><td></td><td></td><td></td></tr>
<tr><td>Chromosome</td><td>start</td><td>end</td><td>strand</td><td>MM</td><td>target_seq</td><td>PAM</td><td>alignment</td><td>distance</td><td>position</td><td>gene name</td><td>gene id</td></tr>
<tr><td>GU305914.1</td><td>109299</td><td>109321</td><td>+</td><td>0</td><td>ATGTCGGACATTCTGGGCGG</td><td>AGG</td><td>||||||||[||||||||||||]PAM</td><td>NA</td><td>Intergenic</td><td>NA</td><td>NA</td></tr>
<tr><td>T7994</td><td>GGACATTCTGGGCGGAGGAGAGG</td><td>1000</td><td></td><td></td><td></td><td></td><td></td><td></td><td></td><td></td><td></td></tr>
<tr><td>Oligo fwd</td><td>CACCGGACATTCTGGGCGGAGGAG</td><td></td><td></td><td></td><td></td><td></td><td></td><td></td><td></td><td></td><td></td></tr>
<tr><td>Oligo rev</td><td>AAACCTCCTCCGCCCAGAATGTCC</td><td></td><td></td><td></td><td></td><td></td><td></td><td></td><td></td><td></td><td></td></tr>
<tr><td>Chromosome</td><td>start</td><td>end</td><td>strand</td><td>MM</td><td>target_seq</td><td>PAM</td><td>alignment</td><td>distance</td><td>position</td><td>gene name</td><td>gene id</td></tr>
<tr><td>GU305914.1</td><td>109304</td><td>109326</td><td>+</td><td>0</td><td>GGACATTCTGGGCGGAGGAG</td><td>AGG</td><td>||||||||[||||||||||||]PAM</td><td>NA</td><td>Intergenic</td><td>NA</td><td>NA</td></tr>
<tr><td>T7995</td><td>GACATTCTGGGCGGAGGAGAGGG</td><td>1000</td><td></td><td></td><td></td><td></td><td></td><td></td><td></td><td></td><td></td></tr>
<tr><td>Oligo fwd</td><td>CACCGACATTCTGGGCGGAGGAGA</td><td></td><td></td><td></td><td></td><td></td><td></td><td></td><td></td><td></td><td></td></tr>
<tr><td>Oligo rev</td><td>AAACTCTCCTCCGCCCAGAATGTC</td><td></td><td></td><td></td><td></td><td></td><td></td><td></td><td></td><td></td><td></td></tr>
<tr><td>Chromosome</td><td>start</td><td>end</td><td>strand</td><td>MM</td><td>target_seq</td><td>PAM</td><td>alignment</td><td>distance</td><td>position</td><td>gene name</td><td>gene id</td></tr>
<tr><td>GU305914.1</td><td>109305</td><td>109327</td><td>+</td><td>0</td><td>GACATTCTGGGCGGAGGAGA</td><td>GGG</td><td>||||||||[||||||||||||]PAM</td><td>NA</td><td>Intergenic</td><td>NA</td><td>NA</td></tr>
<tr><td>T7996</td><td>ACATTCTGGGCGGAGGAGAGGGG</td><td>1000</td><td></td><td></td><td></td><td></td><td></td><td></td><td></td><td></td><td></td></tr>
<tr><td>Oligo fwd</td><td>CACCACATTCTGGGCGGAGGAGAG</td><td></td><td></td><td></td><td></td><td></td><td></td><td></td><td></td><td></td><td></td></tr>
<tr><td>Oligo rev</td><td>AAACCTCTCCTCCGCCCAGAATGT</td><td></td><td></td><td></td><td></td><td></td><td></td><td></td><td></td><td></td><td></td></tr>
<tr><td>Chromosome</td><td>start</td><td>end</td><td>strand</td><td>MM</td><td>target_seq</td><td>PAM</td><td>alignment</td><td>distance</td><td>position</td><td>gene name</td><td>gene id</td></tr>
<tr><td>GU305914.1</td><td>109306</td><td>109328</td><td>+</td><td>0</td><td>ACATTCTGGGCGGAGGAGAG</td><td>GGG</td><td>||||||||[||||||||||||]PAM</td><td>NA</td><td>Intergenic</td><td>NA</td><td>NA</td></tr>
<tr><td>T7997</td><td>TCTGGGCGGAGGAGAGGGGACGG</td><td>1000</td><td></td><td></td><td></td><td></td><td></td><td></td><td></td><td></td><td></td></tr>
<tr><td>Oligo fwd</td><td>CACCTCTGGGCGGAGGAGAGGGGA</td><td></td><td></td><td></td><td></td><td></td><td></td><td></td><td></td><td></td><td></td></tr>
<tr><td>Oligo rev</td><td>AAACTCCCCTCTCCTCCGCCCAGA</td><td></td><td></td><td></td><td></td><td></td><td></td><td></td><td></td><td></td><td></td></tr>
<tr><td>Chromosome</td><td>start</td><td>end</td><td>strand</td><td>MM</td><td>target_seq</td><td>PAM</td><td>alignment</td><td>distance</td><td>position</td><td>gene name</td><td>gene id</td></tr>
<tr><td>GU305914.1</td><td>109310</td><td>109332</td><td>+</td><td>0</td><td>TCTGGGCGGAGGAGAGGGGA</td><td>CGG</td><td>||||||||[||||||||||||]PAM</td><td>NA</td><td>Intergenic</td><td>NA</td><td>NA</td></tr>
<tr><td>T7998</td><td>GGGCGGAGGAGAGGGGACGGAGG</td><td>1000</td><td></td><td></td><td></td><td></td><td></td><td></td><td></td><td></td><td></td></tr>
<tr><td>Oligo fwd</td><td>CACCGGGCGGAGGAGAGGGGACGG</td><td></td><td></td><td></td><td></td><td></td><td></td><td></td><td></td><td></td><td></td></tr>
<tr><td>Oligo rev</td><td>AAACCCGTCCCCTCTCCTCCGCCC</td><td></td><td></td><td></td><td></td><td></td><td></td><td></td><td></td><td></td><td></td></tr>
<tr><td>Chromosome</td><td>start</td><td>end</td><td>strand</td><td>MM</td><td>target_seq</td><td>PAM</td><td>alignment</td><td>distance</td><td>position</td><td>gene name</td><td>gene id</td></tr>
<tr><td>GU305914.1</td><td>109313</td><td>109335</td><td>+</td><td>0</td><td>GGGCGGAGGAGAGGGGACGG</td><td>AGG</td><td>||||||||[||||||||||||]PAM</td><td>NA</td><td>Intergenic</td><td>NA</td><td>NA</td></tr>
<tr><td>T7999</td><td>CGGAGGAGAGGGGACGGAGGTGG</td><td>1000</td><td></td><td></td><td></td><td></td><td></td><td></td><td></td><td></td><td></td></tr>
<tr><td>Oligo fwd</td><td>CACCCGGAGGAGAGGGGACGGAGG</td><td></td><td></td><td></td><td></td><td></td><td></td><td></td><td></td><td></td><td></td></tr>
<tr><td>Oligo rev</td><td>AAACCCTCCGTCCCCTCTCCTCCG</td><td></td><td></td><td></td><td></td><td></td><td></td><td></td><td></td><td></td><td></td></tr>
<tr><td>Chromosome</td><td>start</td><td>end</td><td>strand</td><td>MM</td><td>target_seq</td><td>PAM</td><td>alignment</td><td>distance</td><td>position</td><td>gene name</td><td>gene id</td></tr>
<tr><td>GU305914.1</td><td>109316</td><td>109338</td><td>+</td><td>0</td><td>CGGAGGAGAGGGGACGGAGG</td><td>TGG</td><td>||||||||[||||||||||||]PAM</td><td>NA</td><td>Intergenic</td><td>NA</td><td>NA</td></tr>
<tr><td>T8000</td><td>AGGAGAGGGGACGGAGGTGGCGG</td><td>1000</td><td></td><td></td><td></td><td></td><td></td><td></td><td></td><td></td><td></td></tr>
<tr><td>Oligo fwd</td><td>CACCAGGAGAGGGGACGGAGGTGG</td><td></td><td></td><td></td><td></td><td></td><td></td><td></td><td></td><td></td><td></td></tr>
<tr><td>Oligo rev</td><td>AAACCCACCTCCGTCCCCTCTCCT</td><td></td><td></td><td></td><td></td><td></td><td></td><td></td><td></td><td></td><td></td></tr>
<tr><td>Chromosome</td><td>start</td><td>end</td><td>strand</td><td>MM</td><td>target_seq</td><td>PAM</td><td>alignment</td><td>distance</td><td>position</td><td>gene name</td><td>gene id</td></tr>
<tr><td>GU305914.1</td><td>109319</td><td>109341</td><td>+</td><td>0</td><td>AGGAGAGGGGACGGAGGTGG</td><td>CGG</td><td>||||||||[||||||||||||]PAM</td><td>NA</td><td>Intergenic</td><td>NA</td><td>NA</td></tr>
<tr><td>T8001</td><td>GGAGAGGGGACGGAGGTGGCGGG</td><td>1000</td><td></td><td></td><td></td><td></td><td></td><td></td><td></td><td></td><td></td></tr>
<tr><td>Oligo fwd</td><td>CACCGGAGAGGGGACGGAGGTGGC</td><td></td><td></td><td></td><td></td><td></td><td></td><td></td><td></td><td></td><td></td></tr>
<tr><td>Oligo rev</td><td>AAACGCCACCTCCGTCCCCTCTCC</td><td></td><td></td><td></td><td></td><td></td><td></td><td></td><td></td><td></td><td></td></tr>
<tr><td>Chromosome</td><td>start</td><td>end</td><td>strand</td><td>MM</td><td>target_seq</td><td>PAM</td><td>alignment</td><td>distance</td><td>position</td><td>gene name</td><td>gene id</td></tr>
<tr><td>GU305914.1</td><td>109320</td><td>109342</td><td>+</td><td>0</td><td>GGAGAGGGGACGGAGGTGGC</td><td>GGG</td><td>||||||||[||||||||||||]PAM</td><td>NA</td><td>Intergenic</td><td>NA</td><td>NA</td></tr>
<tr><td>T8002</td><td>GAGAGGGGACGGAGGTGGCGGGG</td><td>1000</td><td></td><td></td><td></td><td></td><td></td><td></td><td></td><td></td><td></td></tr>
<tr><td>Oligo fwd</td><td>CACCGAGAGGGGACGGAGGTGGCG</td><td></td><td></td><td></td><td></td><td></td><td></td><td></td><td></td><td></td><td></td></tr>
<tr><td>Oligo rev</td><td>AAACCGCCACCTCCGTCCCCTCTC</td><td></td><td></td><td></td><td></td><td></td><td></td><td></td><td></td><td></td><td></td></tr>
<tr><td>Chromosome</td><td>start</td><td>end</td><td>strand</td><td>MM</td><td>target_seq</td><td>PAM</td><td>alignment</td><td>distance</td><td>position</td><td>gene name</td><td>gene id</td></tr>
<tr><td>GU305914.1</td><td>109321</td><td>109343</td><td>+</td><td>0</td><td>GAGAGGGGACGGAGGTGGCG</td><td>GGG</td><td>||||||||[||||||||||||]PAM</td><td>NA</td><td>Intergenic</td><td>NA</td><td>NA</td></tr>
<tr><td>T8003</td><td>GGGGACGGAGGTGGCGGGGACGG</td><td>1000</td><td></td><td></td><td></td><td></td><td></td><td></td><td></td><td></td><td></td></tr>
<tr><td>Oligo fwd</td><td>CACCGGGGACGGAGGTGGCGGGGA</td><td></td><td></td><td></td><td></td><td></td><td></td><td></td><td></td><td></td><td></td></tr>
<tr><td>Oligo rev</td><td>AAACTCCCCGCCACCTCCGTCCCC</td><td></td><td></td><td></td><td></td><td></td><td></td><td></td><td></td><td></td><td></td></tr>
<tr><td>Chromosome</td><td>start</td><td>end</td><td>strand</td><td>MM</td><td>target_seq</td><td>PAM</td><td>alignment</td><td>distance</td><td>position</td><td>gene name</td><td>gene id</td></tr>
<tr><td>GU305914.1</td><td>109325</td><td>109347</td><td>+</td><td>0</td><td>GGGGACGGAGGTGGCGGGGA</td><td>CGG</td><td>||||||||[||||||||||||]PAM</td><td>NA</td><td>Intergenic</td><td>NA</td><td>NA</td></tr>
<tr><td>T8004</td><td>GACGGAGGTGGCGGGGACGGCGG</td><td>1000</td><td></td><td></td><td></td><td></td><td></td><td></td><td></td><td></td><td></td></tr>
<tr><td>Oligo fwd</td><td>CACCGACGGAGGTGGCGGGGACGG</td><td></td><td></td><td></td><td></td><td></td><td></td><td></td><td></td><td></td><td></td></tr>
<tr><td>Oligo rev</td><td>AAACCCGTCCCCGCCACCTCCGTC</td><td></td><td></td><td></td><td></td><td></td><td></td><td></td><td></td><td></td><td></td></tr>
<tr><td>Chromosome</td><td>start</td><td>end</td><td>strand</td><td>MM</td><td>target_seq</td><td>PAM</td><td>alignment</td><td>distance</td><td>position</td><td>gene name</td><td>gene id</td></tr>
<tr><td>GU305914.1</td><td>109328</td><td>109350</td><td>+</td><td>0</td><td>GACGGAGGTGGCGGGGACGG</td><td>CGG</td><td>||||||||[||||||||||||]PAM</td><td>NA</td><td>Intergenic</td><td>NA</td><td>NA</td></tr>
<tr><td>GU305914.1</td><td>18653</td><td>18675</td><td>-</td><td>4</td><td>GGCAGAGGTGGCGGAGGCGG</td><td>GGG</td><td>|-|-||||[||||||-|-|||]PAM</td><td>NA</td><td>Intergenic</td><td>NA</td><td>NA</td></tr>
<tr><td>GU305914.1</td><td>48159</td><td>48181</td><td>+</td><td>4</td><td>GGCGGCGGTGGCGGCGGCGG</td><td>TGG</td><td>|-|||-||[||||||-|-|||]PAM</td><td>NA</td><td>Intergenic</td><td>NA</td><td>NA</td></tr>
<tr><td>T8005</td><td>ACGGAGGTGGCGGGGACGGCGGG</td><td>1000</td><td></td><td></td><td></td><td></td><td></td><td></td><td></td><td></td><td></td></tr>
<tr><td>Oligo fwd</td><td>CACCACGGAGGTGGCGGGGACGGC</td><td></td><td></td><td></td><td></td><td></td><td></td><td></td><td></td><td></td><td></td></tr>
<tr><td>Oligo rev</td><td>AAACGCCGTCCCCGCCACCTCCGT</td><td></td><td></td><td></td><td></td><td></td><td></td><td></td><td></td><td></td><td></td></tr>
</table>

<table>
<tr><td>Chromosome</td><td>start</td><td>end</td><td>strand</td><td>MM</td><td>target_seq</td><td>PAM</td><td>alignment</td><td>distance</td><td>position</td><td>gene name</td><td>gene id</td></tr>
<tr><td>GU305914.1</td><td>109329</td><td>109351</td><td>+</td><td>0</td><td>ACGGAGGTGGCGGGGACGGC</td><td>GGG</td><td>||||||||[||||||||||||]PAM</td><td>NA</td><td>Intergenic</td><td>NA</td><td>NA</td></tr>
<tr><td>GU305914.1</td><td>146701</td><td>146723</td><td>-</td><td>4</td><td>ACGGAGTAGACGGCGACGGC</td><td>GGG</td><td>||||||--[|-|||-||||||]PAM</td><td>NA</td><td>Intergenic</td><td>NA</td><td>NA</td></tr>
<tr><td>GU305914.1</td><td>122056</td><td>122078</td><td>+</td><td>2</td><td>ACGGAGGTGGCGGGGATGTC</td><td>TGG</td><td>||||||||[||||||||-|-|]PAM</td><td>NA</td><td>Intergenic</td><td>NA</td><td>NA</td></tr>
<tr><td></td><td></td><td></td><td></td><td></td><td></td><td></td><td></td><td></td><td></td><td></td><td></td></tr>
<tr><td>T8006</td><td>GTGGCGGGGACGGCGGGTGACGG</td><td>1000</td><td></td><td></td><td></td><td></td><td></td><td></td><td></td><td></td><td></td></tr>
<tr><td>Oligo fwd</td><td>CACCGTGGCGGGGACGGCGGGTGA</td><td></td><td></td><td></td><td></td><td></td><td></td><td></td><td></td><td></td><td></td></tr>
<tr><td>Oligo rev</td><td>AAACTCACCCGCCGTCCCCGCCAC</td><td></td><td></td><td></td><td></td><td></td><td></td><td></td><td></td><td></td><td></td></tr>
<tr><td>Chromosome</td><td>start</td><td>end</td><td>strand</td><td>MM</td><td>target_seq</td><td>PAM</td><td>alignment</td><td>distance</td><td>position</td><td>gene name</td><td>gene id</td></tr>
<tr><td>GU305914.1</td><td>109335</td><td>109357</td><td>+</td><td>0</td><td>GTGGCGGGGACGGCGGGTGA</td><td>CGG</td><td>||||||||[||||||||||||]PAM</td><td>NA</td><td>Intergenic</td><td>NA</td><td>NA</td></tr>
<tr><td>GU305914.1</td><td>81239</td><td>81261</td><td>-</td><td>4</td><td>GAGGCGGCGGCGGGGGGTGA</td><td>GGG</td><td>|-|||||-[|-|||-||||||]PAM</td><td>NA</td><td>Intergenic</td><td>NA</td><td>NA</td></tr>
<tr><td></td><td></td><td></td><td></td><td></td><td></td><td></td><td></td><td></td><td></td><td></td><td></td></tr>
<tr><td>T8007</td><td>GCGGGGACGGCGGGTGACGGCGG</td><td>1000</td><td></td><td></td><td></td><td></td><td></td><td></td><td></td><td></td><td></td></tr>
<tr><td>Oligo fwd</td><td>CACCGCGGGGACGGCGGGTGACGG</td><td></td><td></td><td></td><td></td><td></td><td></td><td></td><td></td><td></td><td></td></tr>
<tr><td>Oligo rev</td><td>AAACCCGTCACCCGCCGTCCCCGC</td><td></td><td></td><td></td><td></td><td></td><td></td><td></td><td></td><td></td><td></td></tr>
<tr><td>Chromosome</td><td>start</td><td>end</td><td>strand</td><td>MM</td><td>target_seq</td><td>PAM</td><td>alignment</td><td>distance</td><td>position</td><td>gene name</td><td>gene id</td></tr>
<tr><td>GU305914.1</td><td>109338</td><td>109360</td><td>+</td><td>0</td><td>GCGGGGACGGCGGGTGACGG</td><td>CGG</td><td>||||||||[||||||||||||]PAM</td><td>NA</td><td>Intergenic</td><td>NA</td><td>NA</td></tr>
<tr><td></td><td></td><td></td><td></td><td></td><td></td><td></td><td></td><td></td><td></td><td></td><td></td></tr>
<tr><td>T8008</td><td>CCAGAAGAACAAGAGAAAGAAGG</td><td>1000</td><td></td><td></td><td></td><td></td><td></td><td></td><td></td><td></td><td></td></tr>
<tr><td>Oligo fwd</td><td>CACCCCAGAAGAACAAGAGAAAGA</td><td></td><td></td><td></td><td></td><td></td><td></td><td></td><td></td><td></td><td></td></tr>
<tr><td>Oligo rev</td><td>AAACTCTTTCTCTTGTTCTTCTGG</td><td></td><td></td><td></td><td></td><td></td><td></td><td></td><td></td><td></td><td></td></tr>
<tr><td>Chromosome</td><td>start</td><td>end</td><td>strand</td><td>MM</td><td>target_seq</td><td>PAM</td><td>alignment</td><td>distance</td><td>position</td><td>gene name</td><td>gene id</td></tr>
<tr><td>GU305914.1</td><td>109379</td><td>109401</td><td>+</td><td>0</td><td>CCAGAAGAACAAGAGAAAGA</td><td>AGG</td><td>||||||||[||||||||||||]PAM</td><td>NA</td><td>Intergenic</td><td>NA</td><td>NA</td></tr>
<tr><td></td><td></td><td></td><td></td><td></td><td></td><td></td><td></td><td></td><td></td><td></td><td></td></tr>
<tr><td>T8009</td><td>AAGAACAAGAGAAAGAAGGCCGG</td><td>1000</td><td></td><td></td><td></td><td></td><td></td><td></td><td></td><td></td><td></td></tr>
<tr><td>Oligo fwd</td><td>CACCAAGAACAAGAGAAAGAAGGC</td><td></td><td></td><td></td><td></td><td></td><td></td><td></td><td></td><td></td><td></td></tr>
<tr><td>Oligo rev</td><td>AAACGCCTTCTTTCTCTTGTTCTT</td><td></td><td></td><td></td><td></td><td></td><td></td><td></td><td></td><td></td><td></td></tr>
<tr><td>Chromosome</td><td>start</td><td>end</td><td>strand</td><td>MM</td><td>target_seq</td><td>PAM</td><td>alignment</td><td>distance</td><td>position</td><td>gene name</td><td>gene id</td></tr>
<tr><td>GU305914.1</td><td>109383</td><td>109405</td><td>+</td><td>0</td><td>AAGAACAAGAGAAAGAAGGC</td><td>CGG</td><td>||||||||[||||||||||||]PAM</td><td>NA</td><td>Intergenic</td><td>NA</td><td>NA</td></tr>
<tr><td></td><td></td><td></td><td></td><td></td><td></td><td></td><td></td><td></td><td></td><td></td><td></td></tr>
<tr><td>T8010</td><td>AACAAGAGAAAGAAGGCCGGCGG</td><td>1000</td><td></td><td></td><td></td><td></td><td></td><td></td><td></td><td></td><td></td></tr>
<tr><td>Oligo fwd</td><td>CACCAACAAGAGAAAGAAGGCCGG</td><td></td><td></td><td></td><td></td><td></td><td></td><td></td><td></td><td></td><td></td></tr>
<tr><td>Oligo rev</td><td>AAACCCGGCCTTCTTTCTCTTGTT</td><td></td><td></td><td></td><td></td><td></td><td></td><td></td><td></td><td></td><td></td></tr>
<tr><td>Chromosome</td><td>start</td><td>end</td><td>strand</td><td>MM</td><td>target_seq</td><td>PAM</td><td>alignment</td><td>distance</td><td>position</td><td>gene name</td><td>gene id</td></tr>
<tr><td>GU305914.1</td><td>109386</td><td>109408</td><td>+</td><td>0</td><td>AACAAGAGAAAGAAGGCCGG</td><td>CGG</td><td>||||||||[||||||||||||]PAM</td><td>NA</td><td>Intergenic</td><td>NA</td><td>NA</td></tr>
<tr><td></td><td></td><td></td><td></td><td></td><td></td><td></td><td></td><td></td><td></td><td></td><td></td></tr>
<tr><td>T8011</td><td>AAGAGAAAGAAGGCCGGCGGCGG</td><td>1000</td><td></td><td></td><td></td><td></td><td></td><td></td><td></td><td></td><td></td></tr>
<tr><td>Oligo fwd</td><td>CACCAAGAGAAAGAAGGCCGGCGG</td><td></td><td></td><td></td><td></td><td></td><td></td><td></td><td></td><td></td><td></td></tr>
<tr><td>Oligo rev</td><td>AAACCCGCCGGCCTTCTTTCTCTT</td><td></td><td></td><td></td><td></td><td></td><td></td><td></td><td></td><td></td><td></td></tr>
<tr><td>Chromosome</td><td>start</td><td>end</td><td>strand</td><td>MM</td><td>target_seq</td><td>PAM</td><td>alignment</td><td>distance</td><td>position</td><td>gene name</td><td>gene id</td></tr>
<tr><td>GU305914.1</td><td>109389</td><td>109411</td><td>+</td><td>0</td><td>AAGAGAAAGAAGGCCGGCGG</td><td>CGG</td><td>||||||||[||||||||||||]PAM</td><td>NA</td><td>Intergenic</td><td>NA</td><td>NA</td></tr>
<tr><td></td><td></td><td></td><td></td><td></td><td></td><td></td><td></td><td></td><td></td><td></td><td></td></tr>
<tr><td>T8012</td><td>GAAGGCCGGCGGCGGCGCCTCGG</td><td>1000</td><td></td><td></td><td></td><td></td><td></td><td></td><td></td><td></td><td></td></tr>
<tr><td>Oligo fwd</td><td>CACCGAAGGCCGGCGGCGGCGCCT</td><td></td><td></td><td></td><td></td><td></td><td></td><td></td><td></td><td></td><td></td></tr>
<tr><td>Oligo rev</td><td>AAACAGGCGCCGCCGCCGGCCTTC</td><td></td><td></td><td></td><td></td><td></td><td></td><td></td><td></td><td></td><td></td></tr>
<tr><td>Chromosome</td><td>start</td><td>end</td><td>strand</td><td>MM</td><td>target_seq</td><td>PAM</td><td>alignment</td><td>distance</td><td>position</td><td>gene name</td><td>gene id</td></tr>
<tr><td>GU305914.1</td><td>109397</td><td>109419</td><td>+</td><td>0</td><td>GAAGGCCGGCGGCGGCGCCT</td><td>CGG</td><td>||||||||[||||||||||||]PAM</td><td>NA</td><td>Intergenic</td><td>NA</td><td>NA</td></tr>
<tr><td>GU305914.1</td><td>125652</td><td>125674</td><td>+</td><td>4</td><td>GTCGGCCGGCGCCAGCGCCT</td><td>CGG</td><td>|--|||||[|||-|-||||||]PAM</td><td>NA</td><td>Intergenic</td><td>NA</td><td>NA</td></tr>
<tr><td></td><td></td><td></td><td></td><td></td><td></td><td></td><td></td><td></td><td></td><td></td><td></td></tr>
<tr><td>T8013</td><td>CGCCTCGGCCTCTTCCTGCTCGG</td><td>1000</td><td></td><td></td><td></td><td></td><td></td><td></td><td></td><td></td><td></td></tr>
<tr><td>Oligo fwd</td><td>CACCCGCCTCGGCCTCTTCCTGCT</td><td></td><td></td><td></td><td></td><td></td><td></td><td></td><td></td><td></td><td></td></tr>
<tr><td>Oligo rev</td><td>AAACAGCAGGAAGAGGCCGAGGCG</td><td></td><td></td><td></td><td></td><td></td><td></td><td></td><td></td><td></td><td></td></tr>
<tr><td>Chromosome</td><td>start</td><td>end</td><td>strand</td><td>MM</td><td>target_seq</td><td>PAM</td><td>alignment</td><td>distance</td><td>position</td><td>gene name</td><td>gene id</td></tr>
<tr><td>GU305914.1</td><td>109412</td><td>109434</td><td>+</td><td>0</td><td>CGCCTCGGCCTCTTCCTGCT</td><td>CGG</td><td>||||||||[||||||||||||]PAM</td><td>NA</td><td>Intergenic</td><td>NA</td><td>NA</td></tr>
<tr><td></td><td></td><td></td><td></td><td></td><td></td><td></td><td></td><td></td><td></td><td></td><td></td></tr>
<tr><td>T8014</td><td>GGCCTCTTCCTGCTCGGCGACGG</td><td>1000</td><td></td><td></td><td></td><td></td><td></td><td></td><td></td><td></td><td></td></tr>
<tr><td>Oligo fwd</td><td>CACCGGCCTCTTCCTGCTCGGCGA</td><td></td><td></td><td></td><td></td><td></td><td></td><td></td><td></td><td></td><td></td></tr>
<tr><td>Oligo rev</td><td>AAACTCGCCGAGCAGGAAGAGGCC</td><td></td><td></td><td></td><td></td><td></td><td></td><td></td><td></td><td></td><td></td></tr>
<tr><td>Chromosome</td><td>start</td><td>end</td><td>strand</td><td>MM</td><td>target_seq</td><td>PAM</td><td>alignment</td><td>distance</td><td>position</td><td>gene name</td><td>gene id</td></tr>
<tr><td>GU305914.1</td><td>109418</td><td>109440</td><td>+</td><td>0</td><td>GGCCTCTTCCTGCTCGGCGA</td><td>CGG</td><td>||||||||[||||||||||||]PAM</td><td>NA</td><td>Intergenic</td><td>NA</td><td>NA</td></tr>
<tr><td></td><td></td><td></td><td></td><td></td><td></td><td></td><td></td><td></td><td></td><td></td><td></td></tr>
<tr><td>T8015</td><td>GCCTCTTCCTGCTCGGCGACGGG</td><td>1000</td><td></td><td></td><td></td><td></td><td></td><td></td><td></td><td></td><td></td></tr>
<tr><td>Oligo fwd</td><td>CACCGCCTCTTCCTGCTCGGCGAC</td><td></td><td></td><td></td><td></td><td></td><td></td><td></td><td></td><td></td><td></td></tr>
<tr><td>Oligo rev</td><td>AAACGTCGCCGAGCAGGAAGAGGC</td><td></td><td></td><td></td><td></td><td></td><td></td><td></td><td></td><td></td><td></td></tr>
<tr><td>Chromosome</td><td>start</td><td>end</td><td>strand</td><td>MM</td><td>target_seq</td><td>PAM</td><td>alignment</td><td>distance</td><td>position</td><td>gene name</td><td>gene id</td></tr>
<tr><td>GU305914.1</td><td>109419</td><td>109441</td><td>+</td><td>0</td><td>GCCTCTTCCTGCTCGGCGAC</td><td>GGG</td><td>||||||||[||||||||||||]PAM</td><td>NA</td><td>Intergenic</td><td>NA</td><td>NA</td></tr>
<tr><td></td><td></td><td></td><td></td><td></td><td></td><td></td><td></td><td></td><td></td><td></td><td></td></tr>
<tr><td>T8016</td><td>TCCTGCTCGGCGACGGGAGACGG</td><td>1000</td><td></td><td></td><td></td><td></td><td></td><td></td><td></td><td></td><td></td></tr>
<tr><td>Oligo fwd</td><td>CACCTCCTGCTCGGCGACGGGAGA</td><td></td><td></td><td></td><td></td><td></td><td></td><td></td><td></td><td></td><td></td></tr>
<tr><td>Oligo rev</td><td>AAACTCTCCCGTCGCCGAGCAGGA</td><td></td><td></td><td></td><td></td><td></td><td></td><td></td><td></td><td></td><td></td></tr>
<tr><td>Chromosome</td><td>start</td><td>end</td><td>strand</td><td>MM</td><td>target_seq</td><td>PAM</td><td>alignment</td><td>distance</td><td>position</td><td>gene name</td><td>gene id</td></tr>
<tr><td>GU305914.1</td><td>109425</td><td>109447</td><td>+</td><td>0</td><td>TCCTGCTCGGCGACGGGAGA</td><td>CGG</td><td>||||||||[||||||||||||]PAM</td><td>NA</td><td>Intergenic</td><td>NA</td><td>NA</td></tr>
<tr><td></td><td></td><td></td><td></td><td></td><td></td><td></td><td></td><td></td><td></td><td></td><td></td></tr>
<tr><td>T8017</td><td>TGCTCGGCGACGGGAGACGGCGG</td><td>1000</td><td></td><td></td><td></td><td></td><td></td><td></td><td></td><td></td><td></td></tr>
<tr><td>Oligo fwd</td><td>CACCTGCTCGGCGACGGGAGACGG</td><td></td><td></td><td></td><td></td><td></td><td></td><td></td><td></td><td></td><td></td></tr>
<tr><td>Oligo rev</td><td>AAACCCGTCTCCCGTCGCCGAGCA</td><td></td><td></td><td></td><td></td><td></td><td></td><td></td><td></td><td></td><td></td></tr>
<tr><td>Chromosome</td><td>start</td><td>end</td><td>strand</td><td>MM</td><td>target_seq</td><td>PAM</td><td>alignment</td><td>distance</td><td>position</td><td>gene name</td><td>gene id</td></tr>
<tr><td>GU305914.1</td><td>109428</td><td>109450</td><td>+</td><td>0</td><td>TGCTCGGCGACGGGAGACGG</td><td>CGG</td><td>||||||||[||||||||||||]PAM</td><td>NA</td><td>Intergenic</td><td>NA</td><td>NA</td></tr>
<tr><td></td><td></td><td></td><td></td><td></td><td></td><td></td><td></td><td></td><td></td><td></td><td></td></tr>
<tr><td>T8018</td><td>TCGGCGACGGGAGACGGCGGCGG</td><td>1000</td><td></td><td></td><td></td><td></td><td></td><td></td><td></td><td></td><td></td></tr>
<tr><td>Oligo fwd</td><td>CACCTCGGCGACGGGAGACGGCGG</td><td></td><td></td><td></td><td></td><td></td><td></td><td></td><td></td><td></td><td></td></tr>
<tr><td>Oligo rev</td><td>AAACCCGCCGTCTCCCGTCGCCGA</td><td></td><td></td><td></td><td></td><td></td><td></td><td></td><td></td><td></td><td></td></tr>
<tr><td>Chromosome</td><td>start</td><td>end</td><td>strand</td><td>MM</td><td>target_seq</td><td>PAM</td><td>alignment</td><td>distance</td><td>position</td><td>gene name</td><td>gene id</td></tr>
<tr><td>GU305914.1</td><td>109431</td><td>109453</td><td>+</td><td>0</td><td>TCGGCGACGGGAGACGGCGG</td><td>CGG</td><td>||||||||[||||||||||||]PAM</td><td>NA</td><td>Intergenic</td><td>NA</td><td>NA</td></tr>
<tr><td></td><td></td><td></td><td></td><td></td><td></td><td></td><td></td><td></td><td></td><td></td><td></td></tr>
<tr><td>T8019</td><td>ACGGGAGACGGCGGCGGTAGCGG</td><td>1000</td><td></td><td></td><td></td><td></td><td></td><td></td><td></td><td></td><td></td></tr>
<tr><td>Oligo fwd</td><td>CACCACGGGAGACGGCGGCGGTAG</td><td></td><td></td><td></td><td></td><td></td><td></td><td></td><td></td><td></td><td></td></tr>
<tr><td>Oligo rev</td><td>AAACCTACCGCCGCCGTCTCCCGT</td><td></td><td></td><td></td><td></td><td></td><td></td><td></td><td></td><td></td><td></td></tr>
<tr><td>Chromosome</td><td>start</td><td>end</td><td>strand</td><td>MM</td><td>target_seq</td><td>PAM</td><td>alignment</td><td>distance</td><td>position</td><td>gene name</td><td>gene id</td></tr>
</table>

GU305914.1 109437 109459 + 0 ACGGGAGACGGCGGCGGTAG CGG ||||||||[||||||||||||]PAM NA Intergenic NA NA
GU305914.1 223537 223559 + 2 ACGGGAGTCGTCGGCGGTAG CGG |||||||-[||-|||||||||]PAM NA Intergenic NA NA
GU305914.1 62152 62174 + 4 AGGGGCGACGGCAGCGGCAG CGG |-|||-||[||||-||||-||]PAM NA Intergenic NA NA

T8020 GACGGCGGCGGTAGCGGCGTCGG 1000
Oligo fwd CACCGACGGCGGCGGTAGCGGCGT
Oligo rev AAACACGCCGCTACCGCCGCCGTC
Chromosome start end strand MM target_seq PAM alignment distance position gene name gene id
GU305914.1 109443 109465 + 0 GACGGCGGCGGTAGCGGCGT CGG ||||||||[||||||||||||]PAM NA Intergenic NA NA
GU305914.1 91269 91291 + 3 GACGGCCGCGGAAGCGCCGT CGG ||||||-|[|||-||||-|||]PAM NA Intergenic NA NA
GU305914.1 218521 218543 - 4 GACGGAGTCGGTGGCGGGGT CGG |||||-|-[||||-||||-||]PAM NA Intergenic NA NA
GU305914.1 113918 113940 - 3 GTCGGCGGCGGCAGCGGCGA CGG |-||||||[|||-|||||||-]PAM NA Intergenic NA NA
GU305914.1 48156 48178 + 3 GGCGGCGGCGGTGGCGGCGG CGG |-||||||[||||-||||||-]PAM NA Intergenic NA NA

T8021 TCGGTGACGATGACGACTCGCGG 1000
Oligo fwd CACCTCGGTGACGATGACGACTCG
Oligo rev AAACCGAGTCGTCATCGTCACCGA
Chromosome start end strand MM target_seq PAM alignment distance position gene name gene id
GU305914.1 109462 109484 + 0 TCGGTGACGATGACGACTCG CGG ||||||||[||||||||||||]PAM NA Intergenic NA NA

T8022 ACTCGCGGCAGCAGTGTCTCTGG 1000
Oligo fwd CACCACTCGCGGCAGCAGTGTCTC
Oligo rev AAACGAGACACTGCTGCCGCGAGT
Chromosome start end strand MM target_seq PAM alignment distance position gene name gene id
GU305914.1 109477 109499 + 0 ACTCGCGGCAGCAGTGTCTC TGG ||||||||[||||||||||||]PAM NA Intergenic NA NA

T8023 CTCGCGGCAGCAGTGTCTCTGGG 1000
Oligo fwd CACCCTCGCGGCAGCAGTGTCTCT
Oligo rev AAACAGAGACACTGCTGCCGCGAG
Chromosome start end strand MM target_seq PAM alignment distance position gene name gene id
GU305914.1 109478 109500 + 0 CTCGCGGCAGCAGTGTCTCT GGG ||||||||[||||||||||||]PAM NA Intergenic NA NA

T8024 TCCTACTCCACCGATTTCCGCGG 1000
Oligo fwd CACCTCCTACTCCACCGATTTCCG
Oligo rev AAACCGGAAATCGGTGGAGTAGGA
Chromosome start end strand MM target_seq PAM alignment distance position gene name gene id
GU305914.1 109524 109546 + 0 TCCTACTCCACCGATTTCCG CGG ||||||||[||||||||||||]PAM NA Intergenic NA NA

T8025 CCTACTCCACCGATTTCCGCGGG 1000
Oligo fwd CACCCCTACTCCACCGATTTCCGC
Oligo rev AAACGCGGAAATCGGTGGAGTAGG
Chromosome start end strand MM target_seq PAM alignment distance position gene name gene id
GU305914.1 109525 109547 + 0 CCTACTCCACCGATTTCCGC GGG ||||||||[||||||||||||]PAM NA Intergenic NA NA

T8026 CTACTCCACCGATTTCCGCGGGG 1000
Oligo fwd CACCCTACTCCACCGATTTCCGCG
Oligo rev AAACCGCGGAAATCGGTGGAGTAG
Chromosome start end strand MM target_seq PAM alignment distance position gene name gene id
GU305914.1 109526 109548 + 0 CTACTCCACCGATTTCCGCG GGG ||||||||[||||||||||||]PAM NA Intergenic NA NA

T8027 CACCGATTTCCGCGGGGAGCTGG 1000
Oligo fwd CACCCACCGATTTCCGCGGGGAGC
Oligo rev AAACGCTCCCCGCGGAAATCGGTG
Chromosome start end strand MM target_seq PAM alignment distance position gene name gene id
GU305914.1 109532 109554 + 0 CACCGATTTCCGCGGGGAGC TGG ||||||||[||||||||||||]PAM NA Intergenic NA NA

T8028 GACCATCTTCAACGTGAGTCAGG 1000
Oligo fwd CACCGACCATCTTCAACGTGAGTC
Oligo rev AAACGACTCACGTTGAAGATGGTC
Chromosome start end strand MM target_seq PAM alignment distance position gene name gene id
GU305914.1 109559 109581 + 0 GACCATCTTCAACGTGAGTC AGG ||||||||[||||||||||||]PAM NA Intergenic NA NA

T8029 TCTTCAACGTGAGTCAGGCCTGG 1000
Oligo fwd CACCTCTTCAACGTGAGTCAGGCC
Oligo rev AAACGGCCTGACTCACGTTGAAGA
Chromosome start end strand MM target_seq PAM alignment distance position gene name gene id
GU305914.1 109564 109586 + 0 TCTTCAACGTGAGTCAGGCC TGG ||||||||[||||||||||||]PAM NA Intergenic NA NA

T8030 CAGGCCTGGACTTTCTCCTTCGG 1000
Oligo fwd CACCCAGGCCTGGACTTTCTCCTT
Oligo rev AAACAAGGAGAAAGTCCAGGCCTG
Chromosome start end strand MM target_seq PAM alignment distance position gene name gene id
GU305914.1 109578 109600 + 0 CAGGCCTGGACTTTCTCCTT CGG ||||||||[||||||||||||]PAM NA Intergenic NA NA

T8031 GGACTTTCTCCTTCGGCTCCTGG 1000
Oligo fwd CACCGGACTTTCTCCTTCGGCTCC
Oligo rev AAACGGAGCCGAAGGAGAAAGTCC
Chromosome start end strand MM target_seq PAM alignment distance position gene name gene id
GU305914.1 109585 109607 + 0 GGACTTTCTCCTTCGGCTCC TGG ||||||||[||||||||||||]PAM NA Intergenic NA NA

T8032 CCTGGTACTACCGCCTCAAGAGG 1000
Oligo fwd CACCCCTGGTACTACCGCCTCAAG
Oligo rev AAACCTTGAGGCGGTAGTACCAGG
Chromosome start end strand MM target_seq PAM alignment distance position gene name gene id
GU305914.1 109603 109625 + 0 CCTGGTACTACCGCCTCAAG AGG ||||||||[||||||||||||]PAM NA Intergenic NA NA

T8033 GGTACTACCGCCTCAAGAGGTGG 1000
Oligo fwd CACCGGTACTACCGCCTCAAGAGG
Oligo rev AAACCCTCTTGAGGCGGTAGTACC

<table>
<tr><td>Chromosome</td><td>start</td><td>end</td><td>strand</td><td>MM</td><td>target_seq</td><td>PAM</td><td>alignment</td><td>distance</td><td>position</td><td>gene name</td><td>gene id</td></tr>
<tr><td>GU305914.1</td><td>109606</td><td>109628</td><td>+</td><td>0</td><td>GGTACTACCGCCTCAAGAGG</td><td>TGG</td><td>||||||||[||||||||||||]PAM</td><td>NA</td><td>Intergenic</td><td>NA</td><td>NA</td></tr>
<tr><td>T8034</td><td>GGCTCTACAACCAGCCGCGCTGG</td><td>1000</td><td></td><td></td><td></td><td></td><td></td><td></td><td></td><td></td><td></td></tr>
<tr><td>Oligo fwd</td><td>CACCGGCTCTACAACCAGCCGCGC</td><td></td><td></td><td></td><td></td><td></td><td></td><td></td><td></td><td></td><td></td></tr>
<tr><td>Oligo rev</td><td>AAACGCGCGGCTGGTTGTAGAGCC</td><td></td><td></td><td></td><td></td><td></td><td></td><td></td><td></td><td></td><td></td></tr>
<tr><td>Chromosome</td><td>start</td><td>end</td><td>strand</td><td>MM</td><td>target_seq</td><td>PAM</td><td>alignment</td><td>distance</td><td>position</td><td>gene name</td><td>gene id</td></tr>
<tr><td>GU305914.1</td><td>109627</td><td>109649</td><td>+</td><td>0</td><td>GGCTCTACAACCAGCCGCGC</td><td>TGG</td><td>||||||||[||||||||||||]PAM</td><td>NA</td><td>Intergenic</td><td>NA</td><td>NA</td></tr>
<tr><td>T8035</td><td>TCTACAACCAGCCGCGCTGGCGG</td><td>1000</td><td></td><td></td><td></td><td></td><td></td><td></td><td></td><td></td><td></td></tr>
<tr><td>Oligo fwd</td><td>CACCTCTACAACCAGCCGCGCTGG</td><td></td><td></td><td></td><td></td><td></td><td></td><td></td><td></td><td></td><td></td></tr>
<tr><td>Oligo rev</td><td>AAACCCAGCGCGGCTGGTTGTAGA</td><td></td><td></td><td></td><td></td><td></td><td></td><td></td><td></td><td></td><td></td></tr>
<tr><td>Chromosome</td><td>start</td><td>end</td><td>strand</td><td>MM</td><td>target_seq</td><td>PAM</td><td>alignment</td><td>distance</td><td>position</td><td>gene name</td><td>gene id</td></tr>
<tr><td>GU305914.1</td><td>109630</td><td>109652</td><td>+</td><td>0</td><td>TCTACAACCAGCCGCGCTGG</td><td>CGG</td><td>||||||||[||||||||||||]PAM</td><td>NA</td><td>Intergenic</td><td>NA</td><td>NA</td></tr>
<tr><td>T8036</td><td>CGAGAGCCTGTCCGTCTCGCAGG</td><td>1000</td><td></td><td></td><td></td><td></td><td></td><td></td><td></td><td></td><td></td></tr>
<tr><td>Oligo fwd</td><td>CACCCGAGAGCCTGTCCGTCTCGC</td><td></td><td></td><td></td><td></td><td></td><td></td><td></td><td></td><td></td><td></td></tr>
<tr><td>Oligo rev</td><td>AAACGCGAGACGGACAGGCTCTCG</td><td></td><td></td><td></td><td></td><td></td><td></td><td></td><td></td><td></td><td></td></tr>
<tr><td>Chromosome</td><td>start</td><td>end</td><td>strand</td><td>MM</td><td>target_seq</td><td>PAM</td><td>alignment</td><td>distance</td><td>position</td><td>gene name</td><td>gene id</td></tr>
<tr><td>GU305914.1</td><td>109676</td><td>109698</td><td>+</td><td>0</td><td>CGAGAGCCTGTCCGTCTCGC</td><td>AGG</td><td>||||||||[||||||||||||]PAM</td><td>NA</td><td>Intergenic</td><td>NA</td><td>NA</td></tr>
<tr><td>T8037</td><td>CGTCTCGCAGGAGCTCCTCATGG</td><td>1000</td><td></td><td></td><td></td><td></td><td></td><td></td><td></td><td></td><td></td></tr>
<tr><td>Oligo fwd</td><td>CACCCGTCTCGCAGGAGCTCCTCA</td><td></td><td></td><td></td><td></td><td></td><td></td><td></td><td></td><td></td><td></td></tr>
<tr><td>Oligo rev</td><td>AAACTGAGGAGCTCCTGCGAGACG</td><td></td><td></td><td></td><td></td><td></td><td></td><td></td><td></td><td></td><td></td></tr>
<tr><td>Chromosome</td><td>start</td><td>end</td><td>strand</td><td>MM</td><td>target_seq</td><td>PAM</td><td>alignment</td><td>distance</td><td>position</td><td>gene name</td><td>gene id</td></tr>
<tr><td>GU305914.1</td><td>109688</td><td>109710</td><td>+</td><td>0</td><td>CGTCTCGCAGGAGCTCCTCA</td><td>TGG</td><td>||||||||[||||||||||||]PAM</td><td>NA</td><td>Intergenic</td><td>NA</td><td>NA</td></tr>
<tr><td>T8038</td><td>GTCTCGCAGGAGCTCCTCATGGG</td><td>1000</td><td></td><td></td><td></td><td></td><td></td><td></td><td></td><td></td><td></td></tr>
<tr><td>Oligo fwd</td><td>CACCGTCTCGCAGGAGCTCCTCAT</td><td></td><td></td><td></td><td></td><td></td><td></td><td></td><td></td><td></td><td></td></tr>
<tr><td>Oligo rev</td><td>AAACATGAGGAGCTCCTGCGAGAC</td><td></td><td></td><td></td><td></td><td></td><td></td><td></td><td></td><td></td><td></td></tr>
<tr><td>Chromosome</td><td>start</td><td>end</td><td>strand</td><td>MM</td><td>target_seq</td><td>PAM</td><td>alignment</td><td>distance</td><td>position</td><td>gene name</td><td>gene id</td></tr>
<tr><td>GU305914.1</td><td>109689</td><td>109711</td><td>+</td><td>0</td><td>GTCTCGCAGGAGCTCCTCAT</td><td>GGG</td><td>||||||||[||||||||||||]PAM</td><td>NA</td><td>Intergenic</td><td>NA</td><td>NA</td></tr>
<tr><td>T8039</td><td>CATGGGCGTCCTCAACGCCGTGG</td><td>1000</td><td></td><td></td><td></td><td></td><td></td><td></td><td></td><td></td><td></td></tr>
<tr><td>Oligo fwd</td><td>CACCCATGGGCGTCCTCAACGCCG</td><td></td><td></td><td></td><td></td><td></td><td></td><td></td><td></td><td></td><td></td></tr>
<tr><td>Oligo rev</td><td>AAACCGGCGTTGAGGACGCCCATG</td><td></td><td></td><td></td><td></td><td></td><td></td><td></td><td></td><td></td><td></td></tr>
<tr><td>Chromosome</td><td>start</td><td>end</td><td>strand</td><td>MM</td><td>target_seq</td><td>PAM</td><td>alignment</td><td>distance</td><td>position</td><td>gene name</td><td>gene id</td></tr>
<tr><td>GU305914.1</td><td>109706</td><td>109728</td><td>+</td><td>0</td><td>CATGGGCGTCCTCAACGCCG</td><td>TGG</td><td>||||||||[||||||||||||]PAM</td><td>NA</td><td>Intergenic</td><td>NA</td><td>NA</td></tr>
<tr><td>T8040</td><td>CGTCCTCAACGCCGTGGAGCAGG</td><td>1000</td><td></td><td></td><td></td><td></td><td></td><td></td><td></td><td></td><td></td></tr>
<tr><td>Oligo fwd</td><td>CACCCGTCCTCAACGCCGTGGAGC</td><td></td><td></td><td></td><td></td><td></td><td></td><td></td><td></td><td></td><td></td></tr>
<tr><td>Oligo rev</td><td>AAACGCTCCACGGCGTTGAGGACG</td><td></td><td></td><td></td><td></td><td></td><td></td><td></td><td></td><td></td><td></td></tr>
<tr><td>Chromosome</td><td>start</td><td>end</td><td>strand</td><td>MM</td><td>target_seq</td><td>PAM</td><td>alignment</td><td>distance</td><td>position</td><td>gene name</td><td>gene id</td></tr>
<tr><td>GU305914.1</td><td>109712</td><td>109734</td><td>+</td><td>0</td><td>CGTCCTCAACGCCGTGGAGC</td><td>AGG</td><td>||||||||[||||||||||||]PAM</td><td>NA</td><td>Intergenic</td><td>NA</td><td>NA</td></tr>
<tr><td>GU305914.1</td><td>31678</td><td>31700</td><td>-</td><td>4</td><td>CGGCGTCAACGCCGTGGGCC</td><td>AGG</td><td>||-|-|||[|||||||||--|]PAM</td><td>NA</td><td>Intergenic</td><td>NA</td><td>NA</td></tr>
<tr><td>T8041</td><td>GAGCAGGTCACCGTCTACCCCGG</td><td>1000</td><td></td><td></td><td></td><td></td><td></td><td></td><td></td><td></td><td></td></tr>
<tr><td>Oligo fwd</td><td>CACCGAGCAGGTCACCGTCTACCC</td><td></td><td></td><td></td><td></td><td></td><td></td><td></td><td></td><td></td><td></td></tr>
<tr><td>Oligo rev</td><td>AAACGGGTAGACGGTGACCTGCTC</td><td></td><td></td><td></td><td></td><td></td><td></td><td></td><td></td><td></td><td></td></tr>
<tr><td>Chromosome</td><td>start</td><td>end</td><td>strand</td><td>MM</td><td>target_seq</td><td>PAM</td><td>alignment</td><td>distance</td><td>position</td><td>gene name</td><td>gene id</td></tr>
<tr><td>GU305914.1</td><td>109728</td><td>109750</td><td>+</td><td>0</td><td>GAGCAGGTCACCGTCTACCC</td><td>CGG</td><td>||||||||[||||||||||||]PAM</td><td>NA</td><td>Intergenic</td><td>NA</td><td>NA</td></tr>
<tr><td>T8042</td><td>AGCAGGTCACCGTCTACCCCGGG</td><td>1000</td><td></td><td></td><td></td><td></td><td></td><td></td><td></td><td></td><td></td></tr>
<tr><td>Oligo fwd</td><td>CACCAGCAGGTCACCGTCTACCCC</td><td></td><td></td><td></td><td></td><td></td><td></td><td></td><td></td><td></td><td></td></tr>
<tr><td>Oligo rev</td><td>AAACGGGGTAGACGGTGACCTGCT</td><td></td><td></td><td></td><td></td><td></td><td></td><td></td><td></td><td></td><td></td></tr>
<tr><td>Chromosome</td><td>start</td><td>end</td><td>strand</td><td>MM</td><td>target_seq</td><td>PAM</td><td>alignment</td><td>distance</td><td>position</td><td>gene name</td><td>gene id</td></tr>
<tr><td>GU305914.1</td><td>109729</td><td>109751</td><td>+</td><td>0</td><td>AGCAGGTCACCGTCTACCCC</td><td>GGG</td><td>||||||||[||||||||||||]PAM</td><td>NA</td><td>Intergenic</td><td>NA</td><td>NA</td></tr>
<tr><td>T8043</td><td>CACCGTCGTCAGCGACCTCGAGG</td><td>1000</td><td></td><td></td><td></td><td></td><td></td><td></td><td></td><td></td><td></td></tr>
<tr><td>Oligo fwd</td><td>CACCCACCGTCGTCAGCGACCTCG</td><td></td><td></td><td></td><td></td><td></td><td></td><td></td><td></td><td></td><td></td></tr>
<tr><td>Oligo rev</td><td>AAACCGAGGTCGCTGACGACGGTG</td><td></td><td></td><td></td><td></td><td></td><td></td><td></td><td></td><td></td><td></td></tr>
<tr><td>Chromosome</td><td>start</td><td>end</td><td>strand</td><td>MM</td><td>target_seq</td><td>PAM</td><td>alignment</td><td>distance</td><td>position</td><td>gene name</td><td>gene id</td></tr>
<tr><td>GU305914.1</td><td>109757</td><td>109779</td><td>+</td><td>0</td><td>CACCGTCGTCAGCGACCTCG</td><td>AGG</td><td>||||||||[||||||||||||]PAM</td><td>NA</td><td>Intergenic</td><td>NA</td><td>NA</td></tr>
<tr><td>T8044</td><td>CGTCGTCAGCGACCTCGAGGTGG</td><td>1000</td><td></td><td></td><td></td><td></td><td></td><td></td><td></td><td></td><td></td></tr>
<tr><td>Oligo fwd</td><td>CACCCGTCGTCAGCGACCTCGAGG</td><td></td><td></td><td></td><td></td><td></td><td></td><td></td><td></td><td></td><td></td></tr>
<tr><td>Oligo rev</td><td>AAACCCTCGAGGTCGCTGACGACG</td><td></td><td></td><td></td><td></td><td></td><td></td><td></td><td></td><td></td><td></td></tr>
<tr><td>Chromosome</td><td>start</td><td>end</td><td>strand</td><td>MM</td><td>target_seq</td><td>PAM</td><td>alignment</td><td>distance</td><td>position</td><td>gene name</td><td>gene id</td></tr>
<tr><td>GU305914.1</td><td>109760</td><td>109782</td><td>+</td><td>0</td><td>CGTCGTCAGCGACCTCGAGG</td><td>TGG</td><td>||||||||[||||||||||||]PAM</td><td>NA</td><td>Intergenic</td><td>NA</td><td>NA</td></tr>
<tr><td>T8045</td><td>CGAGGTGGCCGCCTGTCTCCTGG</td><td>1000</td><td></td><td></td><td></td><td></td><td></td><td></td><td></td><td></td><td></td></tr>
<tr><td>Oligo fwd</td><td>CACCCGAGGTGGCCGCCTGTCTCC</td><td></td><td></td><td></td><td></td><td></td><td></td><td></td><td></td><td></td><td></td></tr>
<tr><td>Oligo rev</td><td>AAACGGAGACAGGCGGCCACCTCG</td><td></td><td></td><td></td><td></td><td></td><td></td><td></td><td></td><td></td><td></td></tr>
<tr><td>Chromosome</td><td>start</td><td>end</td><td>strand</td><td>MM</td><td>target_seq</td><td>PAM</td><td>alignment</td><td>distance</td><td>position</td><td>gene name</td><td>gene id</td></tr>
<tr><td>GU305914.1</td><td>109775</td><td>109797</td><td>+</td><td>0</td><td>CGAGGTGGCCGCCTGTCTCC</td><td>TGG</td><td>||||||||[||||||||||||]PAM</td><td>NA</td><td>Intergenic</td><td>NA</td><td>NA</td></tr>
<tr><td>T8046</td><td>GGTGGCCGCCTGTCTCCTGGCGG</td><td>1000</td><td></td><td></td><td></td><td></td><td></td><td></td><td></td><td></td><td></td></tr>
<tr><td>Oligo fwd</td><td>CACCGGTGGCCGCCTGTCTCCTGG</td><td></td><td></td><td></td><td></td><td></td><td></td><td></td><td></td><td></td><td></td></tr>
<tr><td>Oligo rev</td><td>AAACCCAGGAGACAGGCGGCCACC</td><td></td><td></td><td></td><td></td><td></td><td></td><td></td><td></td><td></td><td></td></tr>
<tr><td>Chromosome</td><td>start</td><td>end</td><td>strand</td><td>MM</td><td>target_seq</td><td>PAM</td><td>alignment</td><td>distance</td><td>position</td><td>gene name</td><td>gene id</td></tr>
<tr><td>GU305914.1</td><td>109778</td><td>109800</td><td>+</td><td>0</td><td>GGTGGCCGCCTGTCTCCTGG</td><td>CGG</td><td>||||||||[||||||||||||]PAM</td><td>NA</td><td>Intergenic</td><td>NA</td><td>NA</td></tr>
<tr><td>T8047</td><td>CTGTCTCCTGGCGGCGTATCAGG</td><td>1000</td><td></td><td></td><td></td><td></td><td></td><td></td><td></td><td></td><td></td></tr>
<tr><td>Oligo fwd</td><td>CACCCTGTCTCCTGGCGGCGTATC</td><td></td><td></td><td></td><td></td><td></td><td></td><td></td><td></td><td></td><td></td></tr>
<tr><td>Oligo rev</td><td>AAACGATACGCCGCCAGGAGACAG</td><td></td><td></td><td></td><td></td><td></td><td></td><td></td><td></td><td></td><td></td></tr>
<tr><td>Chromosome</td><td>start</td><td>end</td><td>strand</td><td>MM</td><td>target_seq</td><td>PAM</td><td>alignment</td><td>distance</td><td>position</td><td>gene name</td><td>gene id</td></tr>
<tr><td>GU305914.1</td><td>109787</td><td>109809</td><td>+</td><td>0</td><td>CTGTCTCCTGGCGGCGTATC</td><td>AGG</td><td>||||||||[||||||||||||]PAM</td><td>NA</td><td>Intergenic</td><td>NA</td><td>NA</td></tr>
<tr><td>T8048</td><td>ATCAGGCCGCGCTCGACCCCCGG</td><td>1000</td><td></td><td></td><td></td><td></td><td></td><td></td><td></td><td></td><td></td></tr>
</table>

Oligo fwd CACCATCAGGCCGCGCTCGACCCC

Oligo rev AAACGGGGTCGAGCGCGGCCTGAT

Chromosome start end strand MM target_seq PAM alignment distance position gene name gene id

GU305914.1 109804 109826 + 0 ATCAGGCCGCGCTCGACCCC CGG ||||||||[||||||||||||]PAM NA Intergenic NA NA

T8049 TCAGGCCGCGCTCGACCCCCGGG 1000

Oligo fwd CACCTCAGGCCGCGCTCGACCCCC

Oligo rev AAACGGGGGTCGAGCGCGGCCTGA

Chromosome start end strand MM target_seq PAM alignment distance position gene name gene id

GU305914.1 109805 109827 + 0 TCAGGCCGCGCTCGACCCCC GGG ||||||||[||||||||||||]PAM NA Intergenic NA NA

T8050 CGCGCTCGACCCCCGGGCCGCGG 1000

Oligo fwd CACCCGCGCTCGACCCCCGGGCCG

Oligo rev AAACCGGCCCGGGGGTCGAGCGCG

Chromosome start end strand MM target_seq PAM alignment distance position gene name gene id

GU305914.1 109811 109833 + 0 CGCGCTCGACCCCCGGGCCG CGG ||||||||[||||||||||||]PAM NA Intergenic NA NA

T8051 CGCGGTGCCCACCACCGTCGAGG 1000

Oligo fwd CACCCGCGGTGCCCACCACCGTCG

Oligo rev AAACCGACGGTGGTGGGCACCGCG

Chromosome start end strand MM target_seq PAM alignment distance position gene name gene id

GU305914.1 109829 109851 + 0 CGCGGTGCCCACCACCGTCG AGG ||||||||[||||||||||||]PAM NA Intergenic NA NA

T8052 GCGGTGCCCACCACCGTCGAGGG 1000

Oligo fwd CACCGCGGTGCCCACCACCGTCGA

Oligo rev AAACTCGACGGTGGTGGGCACCGC

Chromosome start end strand MM target_seq PAM alignment distance position gene name gene id

GU305914.1 109830 109852 + 0 GCGGTGCCCACCACCGTCGA GGG ||||||||[||||||||||||]PAM NA Intergenic NA NA

T8053 GAGGGCGTCCTGCGCGACTCCGG 1000

Oligo fwd CACCGAGGGCGTCCTGCGCGACTC

Oligo rev AAACGAGTCGCGCAGGACGCCCTC

Chromosome start end strand MM target_seq PAM alignment distance position gene name gene id

GU305914.1 109848 109870 + 0 GAGGGCGTCCTGCGCGACTC CGG ||||||||[||||||||||||]PAM NA Intergenic NA NA

T8054 GCGTCCTGCGCGACTCCGGCAGG 1000

Oligo fwd CACCGCGTCCTGCGCGACTCCGGC

Oligo rev AAACGCCGGAGTCGCGCAGGACGC

Chromosome start end strand MM target_seq PAM alignment distance position gene name gene id

GU305914.1 109852 109874 + 0 GCGTCCTGCGCGACTCCGGC AGG ||||||||[||||||||||||]PAM NA Intergenic NA NA

T8055 CGTCCTGCGCGACTCCGGCAGGG 1000

Oligo fwd CACCCGTCCTGCGCGACTCCGGCA

Oligo rev AAACTGCCGGAGTCGCGCAGGACG

Chromosome start end strand MM target_seq PAM alignment distance position gene name gene id

GU305914.1 109853 109875 + 0 CGTCCTGCGCGACTCCGGCA GGG ||||||||[||||||||||||]PAM NA Intergenic NA NA

T8056 GCGACTCCGGCAGGGTCCTCCGG 1000

Oligo fwd CACCGCGACTCCGGCAGGGTCCTC

Oligo rev AAACGAGGACCCTGCCGGAGTCGC

Chromosome start end strand MM target_seq PAM alignment distance position gene name gene id

GU305914.1 109861 109883 + 0 GCGACTCCGGCAGGGTCCTC CGG ||||||||[||||||||||||]PAM NA Intergenic NA NA

T8057 CGACTCCGGCAGGGTCCTCCGGG 1000

Oligo fwd CACCCGACTCCGGCAGGGTCCTCC

Oligo rev AAACGGAGGACCCTGCCGGAGTCG

Chromosome start end strand MM target_seq PAM alignment distance position gene name gene id

GU305914.1 109862 109884 + 0 CGACTCCGGCAGGGTCCTCC GGG ||||||||[||||||||||||]PAM NA Intergenic NA NA

T8058 ACATCGCCGCCGAGATCGCGCGG 1000

Oligo fwd CACCACATCGCCGCCGAGATCGCG

Oligo rev AAACCGCGATCTCGGCGGCGATGT

Chromosome start end strand MM target_seq PAM alignment distance position gene name gene id

GU305914.1 109897 109919 + 0 ACATCGCCGCCGAGATCGCG CGG ||||||||[||||||||||||]PAM NA Intergenic NA NA

T8059 CGAGATCGCGCGGCGACCCTCGG 1000

Oligo fwd CACCCGAGATCGCGCGGCGACCCT

Oligo rev AAACAGGGTCGCCGCGCGATCTCG

Chromosome start end strand MM target_seq PAM alignment distance position gene name gene id

GU305914.1 109907 109929 + 0 CGAGATCGCGCGGCGACCCT CGG ||||||||[||||||||||||]PAM NA Intergenic NA NA

T8060 GAGATCGCGCGGCGACCCTCGGG 1000

Oligo fwd CACCGAGATCGCGCGGCGACCCTC

Oligo rev AAACGAGGGTCGCCGCGCGATCTC

Chromosome start end strand MM target_seq PAM alignment distance position gene name gene id

GU305914.1 109908 109930 + 0 GAGATCGCGCGGCGACCCTC GGG ||||||||[||||||||||||]PAM NA Intergenic NA NA

T8061 AGATCGCGCGGCGACCCTCGGGG 1000

Oligo fwd CACCAGATCGCGCGGCGACCCTCG

Oligo rev AAACCGAGGGTCGCCGCGCGATCT

Chromosome start end strand MM target_seq PAM alignment distance position gene name gene id

GU305914.1 109909 109931 + 0 AGATCGCGCGGCGACCCTCG GGG ||||||||[||||||||||||]PAM NA Intergenic NA NA

T8062 GATCGCGCGGCGACCCTCGGGGG 1000

Oligo fwd CACCGATCGCGCGGCGACCCTCGG

Oligo rev AAACCCGAGGGTCGCCGCGCGATC

Chromosome start end strand MM target_seq PAM alignment distance position gene name gene id

GU305914.1 109910 109932 + 0 GATCGCGCGGCGACCCTCGG GGG ||||||||[||||||||||||]PAM NA Intergenic NA NA

T8063 ATCGCGCGGCGACCCTCGGGGGG 1000
Oligo fwd CACCATCGCGCGGCGACCCTCGGG
Oligo rev AAACCCCGAGGGTCGCCGCGCGAT
Chromosome start end strand MM target_seq PAM alignment distance position gene name gene id
GU305914.1 109911 109933 + 0 ATCGCGCGGCGACCCTCGGG GGG ||||||||[||||||||||||]PAM NA Intergenic NA NA

T8064 GGGCAACGCTTTCGCCTACAAGG 1000
Oligo fwd CACCGGGCAACGCTTTCGCCTACA
Oligo rev AAACTGTAGGCGAAAGCGTTGCCC
Chromosome start end strand MM target_seq PAM alignment distance position gene name gene id
GU305914.1 109931 109953 + 0 GGGCAACGCTTTCGCCTACA AGG ||||||||[||||||||||||]PAM NA Intergenic NA NA

T8065 TTTCGCCTACAAGGACCCGCCGG 1000
Oligo fwd CACCTTTCGCCTACAAGGACCCGC
Oligo rev AAACGCGGGTCCTTGTAGGCGAAA
Chromosome start end strand MM target_seq PAM alignment distance position gene name gene id
GU305914.1 109940 109962 + 0 TTTCGCCTACAAGGACCCGC CGG ||||||||[||||||||||||]PAM NA Intergenic NA NA

T8066 TTCGCCTACAAGGACCCGCCGGG 1000
Oligo fwd CACCTTCGCCTACAAGGACCCGCC
Oligo rev AAACGGCGGGTCCTTGTAGGCGAA
Chromosome start end strand MM target_seq PAM alignment distance position gene name gene id
GU305914.1 109941 109963 + 0 TTCGCCTACAAGGACCCGCC GGG ||||||||[||||||||||||]PAM NA Intergenic NA NA

T8067 GGGCTTGCGTTTCTACGCGCCGG 1000
Oligo fwd CACCGGGCTTGCGTTTCTACGCGC
Oligo rev AAACGCGCGTAGAAACGCAAGCCC
Chromosome start end strand MM target_seq PAM alignment distance position gene name gene id
GU305914.1 109961 109983 + 0 GGGCTTGCGTTTCTACGCGC CGG ||||||||[||||||||||||]PAM NA Intergenic NA NA

T8068 TTTCTACGCGCCGGTCCAGCAGG 1000
Oligo fwd CACCTTTCTACGCGCCGGTCCAGC
Oligo rev AAACGCTGGACCGGCGCGTAGAAA
Chromosome start end strand MM target_seq PAM alignment distance position gene name gene id
GU305914.1 109970 109992 + 0 TTTCTACGCGCCGGTCCAGC AGG ||||||||[||||||||||||]PAM NA Intergenic NA NA

T8069 TTCTACGCGCCGGTCCAGCAGGG 1000
Oligo fwd CACCTTCTACGCGCCGGTCCAGCA
Oligo rev AAACTGCTGGACCGGCGCGTAGAA
Chromosome start end strand MM target_seq PAM alignment distance position gene name gene id
GU305914.1 109971 109993 + 0 TTCTACGCGCCGGTCCAGCA GGG ||||||||[||||||||||||]PAM NA Intergenic NA NA

T8070 ACGCGCCGGTCCAGCAGGGCAGG 1000
Oligo fwd CACCACGCGCCGGTCCAGCAGGGC
Oligo rev AAACGCCCTGCTGGACCGGCGCGT
Chromosome start end strand MM target_seq PAM alignment distance position gene name gene id
GU305914.1 109975 109997 + 0 ACGCGCCGGTCCAGCAGGGC AGG ||||||||[||||||||||||]PAM NA Intergenic NA NA

T8071 GCAGGGCAGGCGCTACGCCGCGG 1000
Oligo fwd CACCGCAGGGCAGGCGCTACGCCG
Oligo rev AAACCGGCGTAGCGCCTGCCCTGC
Chromosome start end strand MM target_seq PAM alignment distance position gene name gene id
GU305914.1 109988 110010 + 0 GCAGGGCAGGCGCTACGCCG CGG ||||||||[||||||||||||]PAM NA Intergenic NA NA

T8072 CAGGGCAGGCGCTACGCCGCGGG 1000
Oligo fwd CACCCAGGGCAGGCGCTACGCCGC
Oligo rev AAACGCGGCGTAGCGCCTGCCCTG
Chromosome start end strand MM target_seq PAM alignment distance position gene name gene id
GU305914.1 109989 110011 + 0 CAGGGCAGGCGCTACGCCGC GGG ||||||||[||||||||||||]PAM NA Intergenic NA NA

T8073 CCCTCGTCGCCGTCCTGCTGCGG 1000
Oligo fwd CACCCCCTCGTCGCCGTCCTGCTG
Oligo rev AAACCAGCAGGACGGCGACGAGGG
Chromosome start end strand MM target_seq PAM alignment distance position gene name gene id
GU305914.1 110029 110051 + 0 CCCTCGTCGCCGTCCTGCTG CGG ||||||||[||||||||||||]PAM NA Intergenic NA NA

T8074 TCGTCGCCGTCCTGCTGCGGCGG 1000
Oligo fwd CACCTCGTCGCCGTCCTGCTGCGG
Oligo rev AAACCCGCAGCAGGACGGCGACGA
Chromosome start end strand MM target_seq PAM alignment distance position gene name gene id
GU305914.1 110032 110054 + 0 TCGTCGCCGTCCTGCTGCGG CGG ||||||||[||||||||||||]PAM NA Intergenic NA NA

T8075 CGTCGCCGTCCTGCTGCGGCGGG 1000
Oligo fwd CACCCGTCGCCGTCCTGCTGCGGC
Oligo rev AAACGCCGCAGCAGGACGGCGACG
Chromosome start end strand MM target_seq PAM alignment distance position gene name gene id
GU305914.1 110033 110055 + 0 CGTCGCCGTCCTGCTGCGGC GGG ||||||||[||||||||||||]PAM NA Intergenic NA NA

T8076 GTCGCCGTCCTGCTGCGGCGGGG 1000
Oligo fwd CACCGTCGCCGTCCTGCTGCGGCG
Oligo rev AAACCGCCGCAGCAGGACGGCGAC
Chromosome start end strand MM target_seq PAM alignment distance position gene name gene id
GU305914.1 110034 110056 + 0 GTCGCCGTCCTGCTGCGGCG GGG ||||||||[||||||||||||]PAM NA Intergenic NA NA
GU305914.1 116358 116380 + 3 GTCGGCGTCCTGCGGCGGCT CGG ||||-|||[|||||-|||||-]PAM NA Intergenic NA NA

T8077 TCGCCGTCCTGCTGCGGCGGGGG 1000
Oligo fwd CACCTCGCCGTCCTGCTGCGGCGG
Oligo rev AAACCCGCCGCAGCAGGACGGCGA
Chromosome start end strand MM target_seq PAM alignment distance position gene name gene id

| | | | | | | | | | | | |
|---|---|---|---|---|---|---|---|---|---|---|---|
| GU305914.1 | 110035 | 110057 | + | 0 | TCGCCGTCCTGCTGCGGCGG | GGG | \|\|\|\|\|\|\|\|[\|\|\|\|\|\|\|\|\|\|\|\|]PAM | NA | Intergenic | NA | NA |
| GU305914.1 | 152544 | 152566 | + | 4 | GCCCAGTCCTGCGGCGGCGG | CGG | -\|-\|-\|\|\|[\|\|\|\|-\|\|\|\|\|\|\|]PAM | NA | Intergenic | NA | NA |
| GU305914.1 | 48144 | 48166 | + | 4 | TCGCGGACCTGTGGCGGCGG | CGG | \|\|\|\|-\|-\|[\|\|\|--\|\|\|\|\|\|\|]PAM | NA | Intergenic | NA | NA |
| T8078 | CGCCGTCCTGCTGCGGCGGGGGG | 1000 | | | | | | | | | |
| Oligo fwd | CACCCGCCGTCCTGCTGCGGCGGG | | | | | | | | | | |
| Oligo rev | AAACCCCGCCGCAGCAGGACGGCG | | | | | | | | | | |
| Chromosome | start | end | strand | MM | target_seq | PAM | alignment | distance | position | gene name | gene id |
| GU305914.1 | 110036 | 110058 | + | 0 | CGCCGTCCTGCTGCGGCGGG | GGG | \|\|\|\|\|\|\|\|[\|\|\|\|\|\|\|\|\|\|\|\|]PAM | NA | Intergenic | NA | NA |
| T8079 | GCGGCGGGGGGCCATCGCGCAGG | 1000 | | | | | | | | | |
| Oligo fwd | CACCGCGGCGGGGGGCCATCGCGC | | | | | | | | | | |
| Oligo rev | AAACGCGCGATGGCCCCCCGCCGC | | | | | | | | | | |
| Chromosome | start | end | strand | MM | target_seq | PAM | alignment | distance | position | gene name | gene id |
| GU305914.1 | 110048 | 110070 | + | 0 | GCGGCGGGGGGCCATCGCGC | AGG | \|\|\|\|\|\|\|\|[\|\|\|\|\|\|\|\|\|\|\|\|]PAM | NA | Intergenic | NA | NA |
| T8080 | GGGGGCCATCGCGCAGGTGCCGG | 1000 | | | | | | | | | |
| Oligo fwd | CACCGGGGGCCATCGCGCAGGTGC | | | | | | | | | | |
| Oligo rev | AAACGCACCTGCGCGATGGCCCCC | | | | | | | | | | |
| Chromosome | start | end | strand | MM | target_seq | PAM | alignment | distance | position | gene name | gene id |
| GU305914.1 | 110054 | 110076 | + | 0 | GGGGGCCATCGCGCAGGTGC | CGG | \|\|\|\|\|\|\|\|[\|\|\|\|\|\|\|\|\|\|\|\|]PAM | NA | Intergenic | NA | NA |
| GU305914.1 | 78394 | 78416 | + | 4 | GCGGGACTTCTCGCAGGTGC | TGG | \|-\|\|\|-\|-[\|\|-\|\|\|\|\|\|\|\|\|]PAM | NA | Intergenic | NA | NA |
| T8081 | GGGGCCATCGCGCAGGTGCCGGG | 1000 | | | | | | | | | |
| Oligo fwd | CACCGGGGCCATCGCGCAGGTGCC | | | | | | | | | | |
| Oligo rev | AAACGGCACCTGCGCGATGGCCCC | | | | | | | | | | |
| Chromosome | start | end | strand | MM | target_seq | PAM | alignment | distance | position | gene name | gene id |
| GU305914.1 | 110055 | 110077 | + | 0 | GGGGCCATCGCGCAGGTGCC | GGG | \|\|\|\|\|\|\|\|[\|\|\|\|\|\|\|\|\|\|\|\|]PAM | NA | Intergenic | NA | NA |
| T8082 | GCCATCGCGCAGGTGCCGGGCGG | 1000 | | | | | | | | | |
| Oligo fwd | CACCGCCATCGCGCAGGTGCCGGG | | | | | | | | | | |
| Oligo rev | AAACCCCGGCACCTGCGCGATGGC | | | | | | | | | | |
| Chromosome | start | end | strand | MM | target_seq | PAM | alignment | distance | position | gene name | gene id |
| GU305914.1 | 110058 | 110080 | + | 0 | GCCATCGCGCAGGTGCCGGG | CGG | \|\|\|\|\|\|\|\|[\|\|\|\|\|\|\|\|\|\|\|\|]PAM | NA | Intergenic | NA | NA |
| T8083 | CAGGTGCCGGGCGGTGCCACCGG | 1000 | | | | | | | | | |
| Oligo fwd | CACCCAGGTGCCGGGCGGTGCCAC | | | | | | | | | | |
| Oligo rev | AAACGTGGCACCGCCCGGCACCTG | | | | | | | | | | |
| Chromosome | start | end | strand | MM | target_seq | PAM | alignment | distance | position | gene name | gene id |
| GU305914.1 | 110067 | 110089 | + | 0 | CAGGTGCCGGGCGGTGCCAC | CGG | \|\|\|\|\|\|\|\|[\|\|\|\|\|\|\|\|\|\|\|\|]PAM | NA | Intergenic | NA | NA |
| T8084 | CGGTGCCACCGGCGTCGCCGCGG | 1000 | | | | | | | | | |
| Oligo fwd | CACCCGGTGCCACCGGCGTCGCCG | | | | | | | | | | |
| Oligo rev | AAACCGGCGACGCCGGTGGCACCG | | | | | | | | | | |
| Chromosome | start | end | strand | MM | target_seq | PAM | alignment | distance | position | gene name | gene id |
| GU305914.1 | 110078 | 110100 | + | 0 | CGGTGCCACCGGCGTCGCCG | CGG | \|\|\|\|\|\|\|\|[\|\|\|\|\|\|\|\|\|\|\|\|]PAM | NA | Intergenic | NA | NA |
| T8085 | CGGCGTCGCCGCGGCCGCGTCGG | 1000 | | | | | | | | | |
| Oligo fwd | CACCCGGCGTCGCCGCGGCCGCGT | | | | | | | | | | |
| Oligo rev | AAACACGCGGCCGCGGCGACGCCG | | | | | | | | | | |
| Chromosome | start | end | strand | MM | target_seq | PAM | alignment | distance | position | gene name | gene id |
| GU305914.1 | 110087 | 110109 | + | 0 | CGGCGTCGCCGCGGCCGCGT | CGG | \|\|\|\|\|\|\|\|[\|\|\|\|\|\|\|\|\|\|\|\|]PAM | NA | Intergenic | NA | NA |
| T8086 | GGCGTCGCCGCGGCCGCGTCGGG | 1000 | | | | | | | | | |
| Oligo fwd | CACCGGCGTCGCCGCGGCCGCGTC | | | | | | | | | | |
| Oligo rev | AAACGACGCGGCCGCGGCGACGCC | | | | | | | | | | |
| Chromosome | start | end | strand | MM | target_seq | PAM | alignment | distance | position | gene name | gene id |
| GU305914.1 | 110088 | 110110 | + | 0 | GGCGTCGCCGCGGCCGCGTC | GGG | \|\|\|\|\|\|\|\|[\|\|\|\|\|\|\|\|\|\|\|\|]PAM | NA | Intergenic | NA | NA |
| GU305914.1 | 198751 | 198773 | + | 3 | GACGTCGCCGCGTCCGCGTT | CGG | \|-\|\|\|\|\|\|[\|\|\|\|-\|\|\|\|\|\|-]PAM | NA | Intergenic | NA | NA |
| T8087 | GCGTCGCCGCGGCCGCGTCGGGG | 1000 | | | | | | | | | |
| Oligo fwd | CACCGCGTCGCCGCGGCCGCGTCG | | | | | | | | | | |
| Oligo rev | AAACCGACGCGGCCGCGGCGACGC | | | | | | | | | | |
| Chromosome | start | end | strand | MM | target_seq | PAM | alignment | distance | position | gene name | gene id |
| GU305914.1 | 110089 | 110111 | + | 0 | GCGTCGCCGCGGCCGCGTCG | GGG | \|\|\|\|\|\|\|\|[\|\|\|\|\|\|\|\|\|\|\|\|]PAM | NA | Intergenic | NA | NA |
| T8088 | CGTCGCCGCGGCCGCGTCGGGGG | 1000 | | | | | | | | | |
| Oligo fwd | CACCCGTCGCCGCGGCCGCGTCGG | | | | | | | | | | |
| Oligo rev | AAACCCGACGCGGCCGCGGCGACG | | | | | | | | | | |
| Chromosome | start | end | strand | MM | target_seq | PAM | alignment | distance | position | gene name | gene id |
| GU305914.1 | 110090 | 110112 | + | 0 | CGTCGCCGCGGCCGCGTCGG | GGG | \|\|\|\|\|\|\|\|[\|\|\|\|\|\|\|\|\|\|\|\|]PAM | NA | Intergenic | NA | NA |
| GU305914.1 | 56773 | 56795 | + | 4 | CGCCGCAGCGGCGGCGGCGG | CGG | \|\|-\|\|\|-\|[\|\|\|\|-\|\|\|-\|\|\|]PAM | NA | Intergenic | NA | NA |
| T8089 | GTCGCCGCGGCCGCGTCGGGGGG | 1000 | | | | | | | | | |
| Oligo fwd | CACCGTCGCCGCGGCCGCGTCGGG | | | | | | | | | | |
| Oligo rev | AAACCCCGACGCGGCCGCGGCGAC | | | | | | | | | | |
| Chromosome | start | end | strand | MM | target_seq | PAM | alignment | distance | position | gene name | gene id |
| GU305914.1 | 110091 | 110113 | + | 0 | GTCGCCGCGGCCGCGTCGGG | GGG | \|\|\|\|\|\|\|\|[\|\|\|\|\|\|\|\|\|\|\|\|]PAM | NA | Intergenic | NA | NA |
| T8090 | GCGGCCGCGTCGGGGGGTCCCGG | 1000 | | | | | | | | | |
| Oligo fwd | CACCGCGGCCGCGTCGGGGGGTCC | | | | | | | | | | |
| Oligo rev | AAACGGACCCCCCGACGCGGCCGC | | | | | | | | | | |
| Chromosome | start | end | strand | MM | target_seq | PAM | alignment | distance | position | gene name | gene id |
| GU305914.1 | 110097 | 110119 | + | 0 | GCGGCCGCGTCGGGGGGTCC | CGG | \|\|\|\|\|\|\|\|[\|\|\|\|\|\|\|\|\|\|\|\|]PAM | NA | Intergenic | NA | NA |
| T8091 | GGGTCCCGGCTCGTCCGCCGCGG | 1000 | | | | | | | | | |
| Oligo fwd | CACCGGGTCCCGGCTCGTCCGCCG | | | | | | | | | | |
| Oligo rev | AAACCGGCGGACGAGCCGGGACCC | | | | | | | | | | |
| Chromosome | start | end | strand | MM | target_seq | PAM | alignment | distance | position | gene name | gene id |

<table>
<tr><td>GU305914.1</td><td>110111</td><td>110133</td><td>+</td><td>0</td><td>GGGTCCCGGCTCGTCCGCCG</td><td>CGG</td><td>||||||||[||||||||||||]PAM</td><td>NA</td><td>Intergenic</td><td>NA</td><td>NA</td></tr>
<tr><td colspan="12"></td></tr>
<tr><td>T8092</td><td>CCGCGGCCGCCGTCGCGTCCCGG</td><td>1000</td></tr>
<tr><td>Oligo fwd</td><td>CACCCCGCGGCCGCCGTCGCGTCC</td></tr>
<tr><td>Oligo rev</td><td>AAACGGACGCGACGGCGGCCGCGG</td></tr>
<tr><td>Chromosome</td><td>start</td><td>end</td><td>strand</td><td>MM</td><td>target_seq</td><td>PAM</td><td>alignment</td><td>distance</td><td>position</td><td>gene name</td><td>gene id</td></tr>
<tr><td>GU305914.1</td><td>110128</td><td>110150</td><td>+</td><td>0</td><td>CCGCGGCCGCCGTCGCGTCC</td><td>CGG</td><td>||||||||[||||||||||||]PAM</td><td>NA</td><td>Intergenic</td><td>NA</td><td>NA</td></tr>
<tr><td colspan="12"></td></tr>
<tr><td>T8093</td><td>CGCGGCCGCCGTCGCGTCCCGGG</td><td>1000</td></tr>
<tr><td>Oligo fwd</td><td>CACCCGCGGCCGCCGTCGCGTCCC</td></tr>
<tr><td>Oligo rev</td><td>AAACGGGACGCGACGGCGGCCGCG</td></tr>
<tr><td>Chromosome</td><td>start</td><td>end</td><td>strand</td><td>MM</td><td>target_seq</td><td>PAM</td><td>alignment</td><td>distance</td><td>position</td><td>gene name</td><td>gene id</td></tr>
<tr><td>GU305914.1</td><td>110129</td><td>110151</td><td>+</td><td>0</td><td>CGCGGCCGCCGTCGCGTCCC</td><td>GGG</td><td>||||||||[||||||||||||]PAM</td><td>NA</td><td>Intergenic</td><td>NA</td><td>NA</td></tr>
<tr><td colspan="12"></td></tr>
<tr><td>T8094</td><td>GGCCGCCGTCGCGTCCCGGGAGG</td><td>1000</td></tr>
<tr><td>Oligo fwd</td><td>CACCGGCCGCCGTCGCGTCCCGGG</td></tr>
<tr><td>Oligo rev</td><td>AAACCCCGGGACGCGACGGCGGCC</td></tr>
<tr><td>Chromosome</td><td>start</td><td>end</td><td>strand</td><td>MM</td><td>target_seq</td><td>PAM</td><td>alignment</td><td>distance</td><td>position</td><td>gene name</td><td>gene id</td></tr>
<tr><td>GU305914.1</td><td>110132</td><td>110154</td><td>+</td><td>0</td><td>GGCCGCCGTCGCGTCCCGGG</td><td>AGG</td><td>||||||||[||||||||||||]PAM</td><td>NA</td><td>Intergenic</td><td>NA</td><td>NA</td></tr>
<tr><td>GU305914.1</td><td>45898</td><td>45920</td><td>-</td><td>4</td><td>AGCGGCCATCGCGTCCCGTG</td><td>CGG</td><td>-||-|||-[||||||||||-|]PAM</td><td>NA</td><td>Intergenic</td><td>NA</td><td>NA</td></tr>
<tr><td>GU305914.1</td><td>61920</td><td>61942</td><td>+</td><td>4</td><td>GCCCGCTGTCGCGCCCCGCG</td><td>CGG</td><td>|-||||-|[|||||-||||-|]PAM</td><td>NA</td><td>Intergenic</td><td>NA</td><td>NA</td></tr>
<tr><td colspan="12"></td></tr>
<tr><td>T8095</td><td>GAGGTCATGAGTCGCCTCTCCGG</td><td>1000</td></tr>
<tr><td>Oligo fwd</td><td>CACCGAGGTCATGAGTCGCCTCTC</td></tr>
<tr><td>Oligo rev</td><td>AAACGAGAGGCGACTCATGACCTC</td></tr>
<tr><td>Chromosome</td><td>start</td><td>end</td><td>strand</td><td>MM</td><td>target_seq</td><td>PAM</td><td>alignment</td><td>distance</td><td>position</td><td>gene name</td><td>gene id</td></tr>
<tr><td>GU305914.1</td><td>110151</td><td>110173</td><td>+</td><td>0</td><td>GAGGTCATGAGTCGCCTCTC</td><td>CGG</td><td>||||||||[||||||||||||]PAM</td><td>NA</td><td>Intergenic</td><td>NA</td><td>NA</td></tr>
<tr><td colspan="12"></td></tr>
<tr><td>T8096</td><td>CATGAGTCGCCTCTCCGGCGCGG</td><td>1000</td></tr>
<tr><td>Oligo fwd</td><td>CACCCATGAGTCGCCTCTCCGGCG</td></tr>
<tr><td>Oligo rev</td><td>AAACCGCCGGAGAGGCGACTCATG</td></tr>
<tr><td>Chromosome</td><td>start</td><td>end</td><td>strand</td><td>MM</td><td>target_seq</td><td>PAM</td><td>alignment</td><td>distance</td><td>position</td><td>gene name</td><td>gene id</td></tr>
<tr><td>GU305914.1</td><td>110156</td><td>110178</td><td>+</td><td>0</td><td>CATGAGTCGCCTCTCCGGCG</td><td>CGG</td><td>||||||||[||||||||||||]PAM</td><td>NA</td><td>Intergenic</td><td>NA</td><td>NA</td></tr>
<tr><td colspan="12"></td></tr>
<tr><td>T8097</td><td>CGCGGTCTCCGACGACGTCCTGG</td><td>1000</td></tr>
<tr><td>Oligo fwd</td><td>CACCCGCGGTCTCCGACGACGTCC</td></tr>
<tr><td>Oligo rev</td><td>AAACGGACGTCGTCGGAGACCGCG</td></tr>
<tr><td>Chromosome</td><td>start</td><td>end</td><td>strand</td><td>MM</td><td>target_seq</td><td>PAM</td><td>alignment</td><td>distance</td><td>position</td><td>gene name</td><td>gene id</td></tr>
<tr><td>GU305914.1</td><td>110174</td><td>110196</td><td>+</td><td>0</td><td>CGCGGTCTCCGACGACGTCC</td><td>TGG</td><td>||||||||[||||||||||||]PAM</td><td>NA</td><td>Intergenic</td><td>NA</td><td>NA</td></tr>
<tr><td>GU305914.1</td><td>120651</td><td>120673</td><td>+</td><td>4</td><td>CTCGCTCTCCGACCTCGTCC</td><td>GGG</td><td>|-||-|||[|||||--|||||]PAM</td><td>NA</td><td>Intergenic</td><td>NA</td><td>NA</td></tr>
<tr><td colspan="12"></td></tr>
<tr><td>T8098</td><td>CCGACGACGTCCTGGCGCTCTGG</td><td>1000</td></tr>
<tr><td>Oligo fwd</td><td>CACCCCGACGACGTCCTGGCGCTC</td></tr>
<tr><td>Oligo rev</td><td>AAACGAGCGCCAGGACGTCGTCGG</td></tr>
<tr><td>Chromosome</td><td>start</td><td>end</td><td>strand</td><td>MM</td><td>target_seq</td><td>PAM</td><td>alignment</td><td>distance</td><td>position</td><td>gene name</td><td>gene id</td></tr>
<tr><td>GU305914.1</td><td>110182</td><td>110204</td><td>+</td><td>0</td><td>CCGACGACGTCCTGGCGCTC</td><td>TGG</td><td>||||||||[||||||||||||]PAM</td><td>NA</td><td>Intergenic</td><td>NA</td><td>NA</td></tr>
<tr><td colspan="12"></td></tr>
<tr><td>T8099</td><td>CTCTGGACGCTGCGCCTCTTCGG</td><td>1000</td></tr>
<tr><td>Oligo fwd</td><td>CACCCTCTGGACGCTGCGCCTCTT</td></tr>
<tr><td>Oligo rev</td><td>AAACAAGAGGCGCAGCGTCCAGAG</td></tr>
<tr><td>Chromosome</td><td>start</td><td>end</td><td>strand</td><td>MM</td><td>target_seq</td><td>PAM</td><td>alignment</td><td>distance</td><td>position</td><td>gene name</td><td>gene id</td></tr>
<tr><td>GU305914.1</td><td>110199</td><td>110221</td><td>+</td><td>0</td><td>CTCTGGACGCTGCGCCTCTT</td><td>CGG</td><td>||||||||[||||||||||||]PAM</td><td>NA</td><td>Intergenic</td><td>NA</td><td>NA</td></tr>
<tr><td colspan="12"></td></tr>
<tr><td>T8100</td><td>CTCTTCGGCAAGCGCCTCAGCGG</td><td>1000</td></tr>
<tr><td>Oligo fwd</td><td>CACCCTCTTCGGCAAGCGCCTCAG</td></tr>
<tr><td>Oligo rev</td><td>AAACCTGAGGCGCTTGCCGAAGAG</td></tr>
<tr><td>Chromosome</td><td>start</td><td>end</td><td>strand</td><td>MM</td><td>target_seq</td><td>PAM</td><td>alignment</td><td>distance</td><td>position</td><td>gene name</td><td>gene id</td></tr>
<tr><td>GU305914.1</td><td>110214</td><td>110236</td><td>+</td><td>0</td><td>CTCTTCGGCAAGCGCCTCAG</td><td>CGG</td><td>||||||||[||||||||||||]PAM</td><td>NA</td><td>Intergenic</td><td>NA</td><td>NA</td></tr>
<tr><td colspan="12"></td></tr>
<tr><td>T8101</td><td>CGTCGTCCCCAACCTCCTCCAGG</td><td>1000</td></tr>
<tr><td>Oligo fwd</td><td>CACCCGTCGTCCCCAACCTCCTCC</td></tr>
<tr><td>Oligo rev</td><td>AAACGGAGGAGGTTGGGGACGACG</td></tr>
<tr><td>Chromosome</td><td>start</td><td>end</td><td>strand</td><td>MM</td><td>target_seq</td><td>PAM</td><td>alignment</td><td>distance</td><td>position</td><td>gene name</td><td>gene id</td></tr>
<tr><td>GU305914.1</td><td>110237</td><td>110259</td><td>+</td><td>0</td><td>CGTCGTCCCCAACCTCCTCC</td><td>AGG</td><td>||||||||[||||||||||||]PAM</td><td>NA</td><td>Intergenic</td><td>NA</td><td>NA</td></tr>
<tr><td colspan="12"></td></tr>
<tr><td>T8102</td><td>TCCAGGAGCAGCACTACCTCAGG</td><td>1000</td></tr>
<tr><td>Oligo fwd</td><td>CACCTCCAGGAGCAGCACTACCTC</td></tr>
<tr><td>Oligo rev</td><td>AAACGAGGTAGTGCTGCTCCTGGA</td></tr>
<tr><td>Chromosome</td><td>start</td><td>end</td><td>strand</td><td>MM</td><td>target_seq</td><td>PAM</td><td>alignment</td><td>distance</td><td>position</td><td>gene name</td><td>gene id</td></tr>
<tr><td>GU305914.1</td><td>110254</td><td>110276</td><td>+</td><td>0</td><td>TCCAGGAGCAGCACTACCTC</td><td>AGG</td><td>||||||||[||||||||||||]PAM</td><td>NA</td><td>Intergenic</td><td>NA</td><td>NA</td></tr>
<tr><td colspan="12"></td></tr>
<tr><td>T8103</td><td>GGAGCAGCACTACCTCAGGTCGG</td><td>1000</td></tr>
<tr><td>Oligo fwd</td><td>CACCGGAGCAGCACTACCTCAGGT</td></tr>
<tr><td>Oligo rev</td><td>AAACACCTGAGGTAGTGCTGCTCC</td></tr>
<tr><td>Chromosome</td><td>start</td><td>end</td><td>strand</td><td>MM</td><td>target_seq</td><td>PAM</td><td>alignment</td><td>distance</td><td>position</td><td>gene name</td><td>gene id</td></tr>
<tr><td>GU305914.1</td><td>110258</td><td>110280</td><td>+</td><td>0</td><td>GGAGCAGCACTACCTCAGGT</td><td>CGG</td><td>||||||||[||||||||||||]PAM</td><td>NA</td><td>Intergenic</td><td>NA</td><td>NA</td></tr>
<tr><td colspan="12"></td></tr>
<tr><td>T8104</td><td>GAGCAGCACTACCTCAGGTCGGG</td><td>1000</td></tr>
<tr><td>Oligo fwd</td><td>CACCGAGCAGCACTACCTCAGGTC</td></tr>
<tr><td>Oligo rev</td><td>AAACGACCTGAGGTAGTGCTGCTC</td></tr>
<tr><td>Chromosome</td><td>start</td><td>end</td><td>strand</td><td>MM</td><td>target_seq</td><td>PAM</td><td>alignment</td><td>distance</td><td>position</td><td>gene name</td><td>gene id</td></tr>
<tr><td>GU305914.1</td><td>110259</td><td>110281</td><td>+</td><td>0</td><td>GAGCAGCACTACCTCAGGTC</td><td>GGG</td><td>||||||||[||||||||||||]PAM</td><td>NA</td><td>Intergenic</td><td>NA</td><td>NA</td></tr>
<tr><td colspan="12"></td></tr>
<tr><td>T8105</td><td>AGCAGCACTACCTCAGGTCGGGG</td><td>1000</td></tr>
<tr><td>Oligo fwd</td><td>CACCAGCAGCACTACCTCAGGTCG</td></tr>
<tr><td>Oligo rev</td><td>AAACCGACCTGAGGTAGTGCTGCT</td></tr>
<tr><td>Chromosome</td><td>start</td><td>end</td><td>strand</td><td>MM</td><td>target_seq</td><td>PAM</td><td>alignment</td><td>distance</td><td>position</td><td>gene name</td><td>gene id</td></tr>
<tr><td>GU305914.1</td><td>110260</td><td>110282</td><td>+</td><td>0</td><td>AGCAGCACTACCTCAGGTCG</td><td>GGG</td><td>||||||||[||||||||||||]PAM</td><td>NA</td><td>Intergenic</td><td>NA</td><td>NA</td></tr>
</table>

<table>
<tr><td>T8106</td><td>CTACCTCAGGTCGGGGCTCACGG</td><td>1000</td><td></td><td></td><td></td><td></td><td></td><td></td><td></td><td></td><td></td></tr>
<tr><td>Oligo fwd</td><td>CACCCTACCTCAGGTCGGGGCTCA</td><td></td><td></td><td></td><td></td><td></td><td></td><td></td><td></td><td></td><td></td></tr>
<tr><td>Oligo rev</td><td>AAACTGAGCCCCGACCTGAGGTAG</td><td></td><td></td><td></td><td></td><td></td><td></td><td></td><td></td><td></td><td></td></tr>
<tr><td>Chromosome</td><td>start</td><td>end</td><td>strand</td><td>MM</td><td>target_seq</td><td>PAM</td><td>alignment</td><td>distance</td><td>position</td><td>gene name</td><td>gene id</td></tr>
<tr><td>GU305914.1</td><td>110267</td><td>110289</td><td>+</td><td>0</td><td>CTACCTCAGGTCGGGGCTCA</td><td>CGG</td><td>||||||||[||||||||||||]PAM</td><td>NA</td><td>Intergenic</td><td>NA</td><td>NA</td></tr>
<tr><td>T8107</td><td>TCCTCTGCCTGCTGTTCCTCTGG</td><td>1000</td><td></td><td></td><td></td><td></td><td></td><td></td><td></td><td></td><td></td></tr>
<tr><td>Oligo fwd</td><td>CACCTCCTCTGCCTGCTGTTCCTC</td><td></td><td></td><td></td><td></td><td></td><td></td><td></td><td></td><td></td><td></td></tr>
<tr><td>Oligo rev</td><td>AAACGAGGAACAGCAGGCAGAGGA</td><td></td><td></td><td></td><td></td><td></td><td></td><td></td><td></td><td></td><td></td></tr>
<tr><td>Chromosome</td><td>start</td><td>end</td><td>strand</td><td>MM</td><td>target_seq</td><td>PAM</td><td>alignment</td><td>distance</td><td>position</td><td>gene name</td><td>gene id</td></tr>
<tr><td>GU305914.1</td><td>110293</td><td>110315</td><td>+</td><td>0</td><td>TCCTCTGCCTGCTGTTCCTC</td><td>TGG</td><td>||||||||[||||||||||||]PAM</td><td>NA</td><td>Intergenic</td><td>NA</td><td>NA</td></tr>
<tr><td>T8108</td><td>AACTCCGAGAGCGTCTTCTCCGG</td><td>1000</td><td></td><td></td><td></td><td></td><td></td><td></td><td></td><td></td><td></td></tr>
<tr><td>Oligo fwd</td><td>CACCAACTCCGAGAGCGTCTTCTC</td><td></td><td></td><td></td><td></td><td></td><td></td><td></td><td></td><td></td><td></td></tr>
<tr><td>Oligo rev</td><td>AAACGAGAAGACGCTCTCGGAGTT</td><td></td><td></td><td></td><td></td><td></td><td></td><td></td><td></td><td></td><td></td></tr>
<tr><td>Chromosome</td><td>start</td><td>end</td><td>strand</td><td>MM</td><td>target_seq</td><td>PAM</td><td>alignment</td><td>distance</td><td>position</td><td>gene name</td><td>gene id</td></tr>
<tr><td>GU305914.1</td><td>110325</td><td>110347</td><td>+</td><td>0</td><td>AACTCCGAGAGCGTCTTCTC</td><td>CGG</td><td>||||||||[||||||||||||]PAM</td><td>NA</td><td>Intergenic</td><td>NA</td><td>NA</td></tr>
<tr><td>T8109</td><td>AGCGTCTTCTCCGGCCGCGCCGG</td><td>1000</td><td></td><td></td><td></td><td></td><td></td><td></td><td></td><td></td><td></td></tr>
<tr><td>Oligo fwd</td><td>CACCAGCGTCTTCTCCGGCCGCGC</td><td></td><td></td><td></td><td></td><td></td><td></td><td></td><td></td><td></td><td></td></tr>
<tr><td>Oligo rev</td><td>AAACGCGCGGCCGGAGAAGACGCT</td><td></td><td></td><td></td><td></td><td></td><td></td><td></td><td></td><td></td><td></td></tr>
<tr><td>Chromosome</td><td>start</td><td>end</td><td>strand</td><td>MM</td><td>target_seq</td><td>PAM</td><td>alignment</td><td>distance</td><td>position</td><td>gene name</td><td>gene id</td></tr>
<tr><td>GU305914.1</td><td>110334</td><td>110356</td><td>+</td><td>0</td><td>AGCGTCTTCTCCGGCCGCGC</td><td>CGG</td><td>||||||||[||||||||||||]PAM</td><td>NA</td><td>Intergenic</td><td>NA</td><td>NA</td></tr>
<tr><td>T8110</td><td>GCGCCGGCAAGTTCTCGCTCAGG</td><td>1000</td><td></td><td></td><td></td><td></td><td></td><td></td><td></td><td></td><td></td></tr>
<tr><td>Oligo fwd</td><td>CACCGCGCCGGCAAGTTCTCGCTC</td><td></td><td></td><td></td><td></td><td></td><td></td><td></td><td></td><td></td><td></td></tr>
<tr><td>Oligo rev</td><td>AAACGAGCGAGAACTTGCCGGCGC</td><td></td><td></td><td></td><td></td><td></td><td></td><td></td><td></td><td></td><td></td></tr>
<tr><td>Chromosome</td><td>start</td><td>end</td><td>strand</td><td>MM</td><td>target_seq</td><td>PAM</td><td>alignment</td><td>distance</td><td>position</td><td>gene name</td><td>gene id</td></tr>
<tr><td>GU305914.1</td><td>110350</td><td>110372</td><td>+</td><td>0</td><td>GCGCCGGCAAGTTCTCGCTC</td><td>AGG</td><td>||||||||[||||||||||||]PAM</td><td>NA</td><td>Intergenic</td><td>NA</td><td>NA</td></tr>
<tr><td>T8111</td><td>CGCCGGCAAGTTCTCGCTCAGGG</td><td>1000</td><td></td><td></td><td></td><td></td><td></td><td></td><td></td><td></td><td></td></tr>
<tr><td>Oligo fwd</td><td>CACCCGCCGGCAAGTTCTCGCTCA</td><td></td><td></td><td></td><td></td><td></td><td></td><td></td><td></td><td></td><td></td></tr>
<tr><td>Oligo rev</td><td>AAACTGAGCGAGAACTTGCCGGCG</td><td></td><td></td><td></td><td></td><td></td><td></td><td></td><td></td><td></td><td></td></tr>
<tr><td>Chromosome</td><td>start</td><td>end</td><td>strand</td><td>MM</td><td>target_seq</td><td>PAM</td><td>alignment</td><td>distance</td><td>position</td><td>gene name</td><td>gene id</td></tr>
<tr><td>GU305914.1</td><td>110351</td><td>110373</td><td>+</td><td>0</td><td>CGCCGGCAAGTTCTCGCTCA</td><td>GGG</td><td>||||||||[||||||||||||]PAM</td><td>NA</td><td>Intergenic</td><td>NA</td><td>NA</td></tr>
<tr><td>T8112</td><td>GACGTCTTCCCCGACCTCTGCGG</td><td>1000</td><td></td><td></td><td></td><td></td><td></td><td></td><td></td><td></td><td></td></tr>
<tr><td>Oligo fwd</td><td>CACCGACGTCTTCCCCGACCTCTG</td><td></td><td></td><td></td><td></td><td></td><td></td><td></td><td></td><td></td><td></td></tr>
<tr><td>Oligo rev</td><td>AAACCAGAGGTCGGGGAAGACGTC</td><td></td><td></td><td></td><td></td><td></td><td></td><td></td><td></td><td></td><td></td></tr>
<tr><td>Chromosome</td><td>start</td><td>end</td><td>strand</td><td>MM</td><td>target_seq</td><td>PAM</td><td>alignment</td><td>distance</td><td>position</td><td>gene name</td><td>gene id</td></tr>
<tr><td>GU305914.1</td><td>110373</td><td>110395</td><td>+</td><td>0</td><td>GACGTCTTCCCCGACCTCTG</td><td>CGG</td><td>||||||||[||||||||||||]PAM</td><td>NA</td><td>Intergenic</td><td>NA</td><td>NA</td></tr>
<tr><td>T8113</td><td>GTCTTCCCCGACCTCTGCGGCGG</td><td>1000</td><td></td><td></td><td></td><td></td><td></td><td></td><td></td><td></td><td></td></tr>
<tr><td>Oligo fwd</td><td>CACCGTCTTCCCCGACCTCTGCGG</td><td></td><td></td><td></td><td></td><td></td><td></td><td></td><td></td><td></td><td></td></tr>
<tr><td>Oligo rev</td><td>AAACCCGCAGAGGTCGGGGAAGAC</td><td></td><td></td><td></td><td></td><td></td><td></td><td></td><td></td><td></td><td></td></tr>
<tr><td>Chromosome</td><td>start</td><td>end</td><td>strand</td><td>MM</td><td>target_seq</td><td>PAM</td><td>alignment</td><td>distance</td><td>position</td><td>gene name</td><td>gene id</td></tr>
<tr><td>GU305914.1</td><td>110376</td><td>110398</td><td>+</td><td>0</td><td>GTCTTCCCCGACCTCTGCGG</td><td>CGG</td><td>||||||||[||||||||||||]PAM</td><td>NA</td><td>Intergenic</td><td>NA</td><td>NA</td></tr>
<tr><td>T8114</td><td>ACGCGCCGCCGCCCGTCGAGAGG</td><td>1000</td><td></td><td></td><td></td><td></td><td></td><td></td><td></td><td></td><td></td></tr>
<tr><td>Oligo fwd</td><td>CACCACGCGCCGCCGCCCGTCGAG</td><td></td><td></td><td></td><td></td><td></td><td></td><td></td><td></td><td></td><td></td></tr>
<tr><td>Oligo rev</td><td>AAACCTCGACGGGCGGCGGCGCGT</td><td></td><td></td><td></td><td></td><td></td><td></td><td></td><td></td><td></td><td></td></tr>
<tr><td>Chromosome</td><td>start</td><td>end</td><td>strand</td><td>MM</td><td>target_seq</td><td>PAM</td><td>alignment</td><td>distance</td><td>position</td><td>gene name</td><td>gene id</td></tr>
<tr><td>GU305914.1</td><td>110404</td><td>110426</td><td>+</td><td>0</td><td>ACGCGCCGCCGCCCGTCGAG</td><td>AGG</td><td>||||||||[||||||||||||]PAM</td><td>NA</td><td>Intergenic</td><td>NA</td><td>NA</td></tr>
<tr><td>T8115</td><td>CGCGCCGCCGCCCGTCGAGAGGG</td><td>1000</td><td></td><td></td><td></td><td></td><td></td><td></td><td></td><td></td><td></td></tr>
<tr><td>Oligo fwd</td><td>CACCCGCGCCGCCGCCCGTCGAGA</td><td></td><td></td><td></td><td></td><td></td><td></td><td></td><td></td><td></td><td></td></tr>
<tr><td>Oligo rev</td><td>AAACTCTCGACGGGCGGCGGCGCG</td><td></td><td></td><td></td><td></td><td></td><td></td><td></td><td></td><td></td><td></td></tr>
<tr><td>Chromosome</td><td>start</td><td>end</td><td>strand</td><td>MM</td><td>target_seq</td><td>PAM</td><td>alignment</td><td>distance</td><td>position</td><td>gene name</td><td>gene id</td></tr>
<tr><td>GU305914.1</td><td>110405</td><td>110427</td><td>+</td><td>0</td><td>CGCGCCGCCGCCCGTCGAGA</td><td>GGG</td><td>||||||||[||||||||||||]PAM</td><td>NA</td><td>Intergenic</td><td>NA</td><td>NA</td></tr>
<tr><td>T8116</td><td>GCCGCCCGTCGAGAGGGAAGAGG</td><td>1000</td><td></td><td></td><td></td><td></td><td></td><td></td><td></td><td></td><td></td></tr>
<tr><td>Oligo fwd</td><td>CACCGCCGCCCGTCGAGAGGGAAG</td><td></td><td></td><td></td><td></td><td></td><td></td><td></td><td></td><td></td><td></td></tr>
<tr><td>Oligo rev</td><td>AAACCTTCCCTCTCGACGGGCGGC</td><td></td><td></td><td></td><td></td><td></td><td></td><td></td><td></td><td></td><td></td></tr>
<tr><td>Chromosome</td><td>start</td><td>end</td><td>strand</td><td>MM</td><td>target_seq</td><td>PAM</td><td>alignment</td><td>distance</td><td>position</td><td>gene name</td><td>gene id</td></tr>
<tr><td>GU305914.1</td><td>110411</td><td>110433</td><td>+</td><td>0</td><td>GCCGCCCGTCGAGAGGGAAG</td><td>AGG</td><td>||||||||[||||||||||||]PAM</td><td>NA</td><td>Intergenic</td><td>NA</td><td>NA</td></tr>
<tr><td>T8117</td><td>CCGCCCGTCGAGAGGGAAGAGGG</td><td>1000</td><td></td><td></td><td></td><td></td><td></td><td></td><td></td><td></td><td></td></tr>
<tr><td>Oligo fwd</td><td>CACCCCGCCCGTCGAGAGGGAAGA</td><td></td><td></td><td></td><td></td><td></td><td></td><td></td><td></td><td></td><td></td></tr>
<tr><td>Oligo rev</td><td>AAACTCTTCCCTCTCGACGGGCGG</td><td></td><td></td><td></td><td></td><td></td><td></td><td></td><td></td><td></td><td></td></tr>
<tr><td>Chromosome</td><td>start</td><td>end</td><td>strand</td><td>MM</td><td>target_seq</td><td>PAM</td><td>alignment</td><td>distance</td><td>position</td><td>gene name</td><td>gene id</td></tr>
<tr><td>GU305914.1</td><td>110412</td><td>110434</td><td>+</td><td>0</td><td>CCGCCCGTCGAGAGGGAAGA</td><td>GGG</td><td>||||||||[||||||||||||]PAM</td><td>NA</td><td>Intergenic</td><td>NA</td><td>NA</td></tr>
<tr><td>T8118</td><td>GAGAGGGAAGAGGGCTTCGCCGG</td><td>1000</td><td></td><td></td><td></td><td></td><td></td><td></td><td></td><td></td><td></td></tr>
<tr><td>Oligo fwd</td><td>CACCGAGAGGGAAGAGGGCTTCGC</td><td></td><td></td><td></td><td></td><td></td><td></td><td></td><td></td><td></td><td></td></tr>
<tr><td>Oligo rev</td><td>AAACGCGAAGCCCTCTTCCCTCTC</td><td></td><td></td><td></td><td></td><td></td><td></td><td></td><td></td><td></td><td></td></tr>
<tr><td>Chromosome</td><td>start</td><td>end</td><td>strand</td><td>MM</td><td>target_seq</td><td>PAM</td><td>alignment</td><td>distance</td><td>position</td><td>gene name</td><td>gene id</td></tr>
<tr><td>GU305914.1</td><td>110421</td><td>110443</td><td>+</td><td>0</td><td>GAGAGGGAAGAGGGCTTCGC</td><td>CGG</td><td>||||||||[||||||||||||]PAM</td><td>NA</td><td>Intergenic</td><td>NA</td><td>NA</td></tr>
<tr><td>T8119</td><td>AGGGAAGAGGGCTTCGCCGGCGG</td><td>1000</td><td></td><td></td><td></td><td></td><td></td><td></td><td></td><td></td><td></td></tr>
<tr><td>Oligo fwd</td><td>CACCAGGGAAGAGGGCTTCGCCGG</td><td></td><td></td><td></td><td></td><td></td><td></td><td></td><td></td><td></td><td></td></tr>
<tr><td>Oligo rev</td><td>AAACCCGGCGAAGCCCTCTTCCCT</td><td></td><td></td><td></td><td></td><td></td><td></td><td></td><td></td><td></td><td></td></tr>
<tr><td>Chromosome</td><td>start</td><td>end</td><td>strand</td><td>MM</td><td>target_seq</td><td>PAM</td><td>alignment</td><td>distance</td><td>position</td><td>gene name</td><td>gene id</td></tr>
<tr><td>GU305914.1</td><td>110424</td><td>110446</td><td>+</td><td>0</td><td>AGGGAAGAGGGCTTCGCCGG</td><td>CGG</td><td>||||||||[||||||||||||]PAM</td><td>NA</td><td>Intergenic</td><td>NA</td><td>NA</td></tr>
<tr><td>T8120</td><td>CAAGAACTTCGAGTTCATGATGG</td><td>1000</td><td></td><td></td><td></td><td></td><td></td><td></td><td></td><td></td><td></td></tr>
<tr><td>Oligo fwd</td><td>CACCCAAGAACTTCGAGTTCATGA</td><td></td><td></td><td></td><td></td><td></td><td></td><td></td><td></td><td></td><td></td></tr>
<tr><td>Oligo rev</td><td>AAACTCATGAACTCGAAGTTCTTG</td><td></td><td></td><td></td><td></td><td></td><td></td><td></td><td></td><td></td><td></td></tr>
<tr><td>Chromosome</td><td>start</td><td>end</td><td>strand</td><td>MM</td><td>target_seq</td><td>PAM</td><td>alignment</td><td>distance</td><td>position</td><td>gene name</td><td>gene id</td></tr>
<tr><td>GU305914.1</td><td>110453</td><td>110475</td><td>+</td><td>0</td><td>CAAGAACTTCGAGTTCATGA</td><td>TGG</td><td>||||||||[||||||||||||]PAM</td><td>NA</td><td>Intergenic</td><td>NA</td><td>NA</td></tr>
</table>

| | | | | | | | | | | | |
|---|---|---|---|---|---|---|---|---|---|---|---|
| T8121 | TGGAGCGCTACGTCGTCCCGTGG | 1000 | | | | | | | | | |
| Oligo fwd | CACCTGGAGCGCTACGTCGTCCCG | | | | | | | | | | |
| Oligo rev | AAACCGGGACGACGTAGCGCTCCA | | | | | | | | | | |
| Chromosome | start | end | strand | MM | target_seq | PAM | alignment | distance | position | gene name | gene id |
| GU305914.1 | 110473 | 110495 | + | 0 | TGGAGCGCTACGTCGTCCCG | TGG | \|\|\|\|\|\|\|\|[\|\|\|\|\|\|\|\|\|\|\|\|]PAM | NA | Intergenic | NA | NA |

| | | | | | | | | | | | |
|---|---|---|---|---|---|---|---|---|---|---|---|
| T8122 | CCCGTGGTACTCGCGCGACCCGG | 1000 | | | | | | | | | |
| Oligo fwd | CACCCCCGTGGTACTCGCGCGACC | | | | | | | | | | |
| Oligo rev | AAACGGTCGCGCGAGTACCACGGG | | | | | | | | | | |
| Chromosome | start | end | strand | MM | target_seq | PAM | alignment | distance | position | gene name | gene id |
| GU305914.1 | 110489 | 110511 | + | 0 | CCCGTGGTACTCGCGCGACC | CGG | \|\|\|\|\|\|\|\|[\|\|\|\|\|\|\|\|\|\|\|\|]PAM | NA | Intergenic | NA | NA |

| | | | | | | | | | | | |
|---|---|---|---|---|---|---|---|---|---|---|---|
| T8123 | CTCGCGCGACCCGGCCGTCACGG | 1000 | | | | | | | | | |
| Oligo fwd | CACCCTCGCGCGACCCGGCCGTCA | | | | | | | | | | |
| Oligo rev | AAACTGACGGCCGGGTCGCGCGAG | | | | | | | | | | |
| Chromosome | start | end | strand | MM | target_seq | PAM | alignment | distance | position | gene name | gene id |
| GU305914.1 | 110498 | 110520 | + | 0 | CTCGCGCGACCCGGCCGTCA | CGG | \|\|\|\|\|\|\|\|[\|\|\|\|\|\|\|\|\|\|\|\|]PAM | NA | Intergenic | NA | NA |
| GU305914.1 | 224376 | 224398 | - | 4 | CTCGCGGCCACCGGCCGTCA | TGG | \|\|\|\|\|\|--[--\|\|\|\|\|\|\|\|\|\|]PAM | NA | Intergenic | NA | NA |

| | | | | | | | | | | | |
|---|---|---|---|---|---|---|---|---|---|---|---|
| T8124 | CCGTCACGGTCTCGCAGCTCTGG | 1000 | | | | | | | | | |
| Oligo fwd | CACCCCGTCACGGTCTCGCAGCTC | | | | | | | | | | |
| Oligo rev | AAACGAGCTGCGAGACCGTGACGG | | | | | | | | | | |
| Chromosome | start | end | strand | MM | target_seq | PAM | alignment | distance | position | gene name | gene id |
| GU305914.1 | 110512 | 110534 | + | 0 | CCGTCACGGTCTCGCAGCTC | TGG | \|\|\|\|\|\|\|\|[\|\|\|\|\|\|\|\|\|\|\|\|]PAM | NA | Intergenic | NA | NA |

| | | | | | | | | | | | |
|---|---|---|---|---|---|---|---|---|---|---|---|
| T8125 | CACGGTCTCGCAGCTCTGGCCGG | 1000 | | | | | | | | | |
| Oligo fwd | CACCCACGGTCTCGCAGCTCTGGC | | | | | | | | | | |
| Oligo rev | AAACGCCAGAGCTGCGAGACCGTG | | | | | | | | | | |
| Chromosome | start | end | strand | MM | target_seq | PAM | alignment | distance | position | gene name | gene id |
| GU305914.1 | 110516 | 110538 | + | 0 | CACGGTCTCGCAGCTCTGGC | CGG | \|\|\|\|\|\|\|\|[\|\|\|\|\|\|\|\|\|\|\|\|]PAM | NA | Intergenic | NA | NA |

| | | | | | | | | | | | |
|---|---|---|---|---|---|---|---|---|---|---|---|
| T8126 | ACGGTCTCGCAGCTCTGGCCGGG | 1000 | | | | | | | | | |
| Oligo fwd | CACCACGGTCTCGCAGCTCTGGCC | | | | | | | | | | |
| Oligo rev | AAACGGCCAGAGCTGCGAGACCGT | | | | | | | | | | |
| Chromosome | start | end | strand | MM | target_seq | PAM | alignment | distance | position | gene name | gene id |
| GU305914.1 | 110517 | 110539 | + | 0 | ACGGTCTCGCAGCTCTGGCC | GGG | \|\|\|\|\|\|\|\|[\|\|\|\|\|\|\|\|\|\|\|\|]PAM | NA | Intergenic | NA | NA |

| | | | | | | | | | | | |
|---|---|---|---|---|---|---|---|---|---|---|---|
| T8127 | CGGTCTCGCAGCTCTGGCCGGGG | 1000 | | | | | | | | | |
| Oligo fwd | CACCCGGTCTCGCAGCTCTGGCCG | | | | | | | | | | |
| Oligo rev | AAACCGGCCAGAGCTGCGAGACCG | | | | | | | | | | |
| Chromosome | start | end | strand | MM | target_seq | PAM | alignment | distance | position | gene name | gene id |
| GU305914.1 | 110518 | 110540 | + | 0 | CGGTCTCGCAGCTCTGGCCG | GGG | \|\|\|\|\|\|\|\|[\|\|\|\|\|\|\|\|\|\|\|\|]PAM | NA | Intergenic | NA | NA |

| | | | | | | | | | | | |
|---|---|---|---|---|---|---|---|---|---|---|---|
| T8128 | TACTGCGAGAGCCACCGCAGCGG | 1000 | | | | | | | | | |
| Oligo fwd | CACCTACTGCGAGAGCCACCGCAG | | | | | | | | | | |
| Oligo rev | AAACCTGCGGTGGCTCTCGCAGTA | | | | | | | | | | |
| Chromosome | start | end | strand | MM | target_seq | PAM | alignment | distance | position | gene name | gene id |
| GU305914.1 | 110556 | 110578 | + | 0 | TACTGCGAGAGCCACCGCAG | CGG | \|\|\|\|\|\|\|\|[\|\|\|\|\|\|\|\|\|\|\|\|]PAM | NA | Intergenic | NA | NA |

| | | | | | | | | | | | |
|---|---|---|---|---|---|---|---|---|---|---|---|
| T8129 | GCGAGAGCCACCGCAGCGGCTGG | 1000 | | | | | | | | | |
| Oligo fwd | CACCGCGAGAGCCACCGCAGCGGC | | | | | | | | | | |
| Oligo rev | AAACGCCGCTGCGGTGGCTCTCGC | | | | | | | | | | |
| Chromosome | start | end | strand | MM | target_seq | PAM | alignment | distance | position | gene name | gene id |
| GU305914.1 | 110560 | 110582 | + | 0 | GCGAGAGCCACCGCAGCGGC | TGG | \|\|\|\|\|\|\|\|[\|\|\|\|\|\|\|\|\|\|\|\|]PAM | NA | Intergenic | NA | NA |

| | | | | | | | | | | | |
|---|---|---|---|---|---|---|---|---|---|---|---|
| T8130 | CGAGAGCCACCGCAGCGGCTGGG | 1000 | | | | | | | | | |
| Oligo fwd | CACCCGAGAGCCACCGCAGCGGCT | | | | | | | | | | |
| Oligo rev | AAACAGCCGCTGCGGTGGCTCTCG | | | | | | | | | | |
| Chromosome | start | end | strand | MM | target_seq | PAM | alignment | distance | position | gene name | gene id |
| GU305914.1 | 110561 | 110583 | + | 0 | CGAGAGCCACCGCAGCGGCT | GGG | \|\|\|\|\|\|\|\|[\|\|\|\|\|\|\|\|\|\|\|\|]PAM | NA | Intergenic | NA | NA |

| | | | | | | | | | | | |
|---|---|---|---|---|---|---|---|---|---|---|---|
| T8131 | GCGGCTGGGACCTCTCGCGCAGG | 1000 | | | | | | | | | |
| Oligo fwd | CACCGCGGCTGGGACCTCTCGCGC | | | | | | | | | | |
| Oligo rev | AAACGCGCGAGAGGTCCCAGCCGC | | | | | | | | | | |
| Chromosome | start | end | strand | MM | target_seq | PAM | alignment | distance | position | gene name | gene id |
| GU305914.1 | 110575 | 110597 | + | 0 | GCGGCTGGGACCTCTCGCGC | AGG | \|\|\|\|\|\|\|\|[\|\|\|\|\|\|\|\|\|\|\|\|]PAM | NA | Intergenic | NA | NA |
| GU305914.1 | 120798 | 120820 | + | 4 | GCGGGCGGGACCTCCGGCGC | CGG | \|\|\|\|--\|\|[\|\|\|\|\|\|--\|\|\|\|]PAM | NA | Intergenic | NA | NA |

| | | | | | | | | | | | |
|---|---|---|---|---|---|---|---|---|---|---|---|
| T8132 | CAGGCCGTTCGAACACGCCACGG | 1000 | | | | | | | | | |
| Oligo fwd | CACCCAGGCCGTTCGAACACGCCA | | | | | | | | | | |
| Oligo rev | AAACTGGCGTGTTCGAACGGCCTG | | | | | | | | | | |
| Chromosome | start | end | strand | MM | target_seq | PAM | alignment | distance | position | gene name | gene id |
| GU305914.1 | 110594 | 110616 | + | 0 | CAGGCCGTTCGAACACGCCA | CGG | \|\|\|\|\|\|\|\|[\|\|\|\|\|\|\|\|\|\|\|\|]PAM | NA | Intergenic | NA | NA |

| | | | | | | | | | | | |
|---|---|---|---|---|---|---|---|---|---|---|---|
| T8133 | TTCGAACACGCCACGGCCGACGG | 1000 | | | | | | | | | |
| Oligo fwd | CACCTTCGAACACGCCACGGCCGA | | | | | | | | | | |
| Oligo rev | AAACTCGGCCGTGGCGTGTTCGAA | | | | | | | | | | |
| Chromosome | start | end | strand | MM | target_seq | PAM | alignment | distance | position | gene name | gene id |
| GU305914.1 | 110601 | 110623 | + | 0 | TTCGAACACGCCACGGCCGA | CGG | \|\|\|\|\|\|\|\|[\|\|\|\|\|\|\|\|\|\|\|\|]PAM | NA | Intergenic | NA | NA |

| | | | | | | | | | | | |
|---|---|---|---|---|---|---|---|---|---|---|---|
| T8134 | GACGGCGTCTCCTCCGCCGCCGG | 1000 | | | | | | | | | |
| Oligo fwd | CACCGACGGCGTCTCCTCCGCCGC | | | | | | | | | | |
| Oligo rev | AAACGCGGCGGAGGAGACGCCGTC | | | | | | | | | | |
| Chromosome | start | end | strand | MM | target_seq | PAM | alignment | distance | position | gene name | gene id |
| GU305914.1 | 110619 | 110641 | + | 0 | GACGGCGTCTCCTCCGCCGC | CGG | \|\|\|\|\|\|\|\|[\|\|\|\|\|\|\|\|\|\|\|\|]PAM | NA | Intergenic | NA | NA |

| | | |
|---|---|---|
| T8135 | CGCCGGCGTCCTGCACGTCCAGG | 1000 |
| Oligo fwd | CACCCGCCGGCGTCCTGCACGTCC | |

| | | | | | | | | | | | |
|---|---|---|---|---|---|---|---|---|---|---|---|
| Oligo rev | AAACGGACGTGCAGGACGCCGGCG | | | | | | | | | | |
| Chromosome | start | end | strand | MM | target_seq | PAM | alignment | distance | position | gene name | gene id |
| GU305914.1 | 110636 | 110658 | + | 0 | CGCCGGCGTCCTGCACGTCC | AGG | \|\|\|\|\|\|\|\|[\|\|\|\|\|\|\|\|\|\|\|\|]PAM | NA | Intergenic | NA | NA |
| T8136 | CTACATGCTGCTGCAGCAGACGG | 1000 | | | | | | | | | |
| Oligo fwd | CACCCTACATGCTGCTGCAGCAGA | | | | | | | | | | |
| Oligo rev | AAACTCTGCTGCAGCAGCATGTAG | | | | | | | | | | |
| Chromosome | start | end | strand | MM | target_seq | PAM | alignment | distance | position | gene name | gene id |
| GU305914.1 | 110684 | 110706 | + | 0 | CTACATGCTGCTGCAGCAGA | CGG | \|\|\|\|\|\|\|\|[\|\|\|\|\|\|\|\|\|\|\|\|]PAM | NA | Intergenic | NA | NA |
| T8137 | GCTGCAGCAGACGGCCGCCCCGG | 1000 | | | | | | | | | |
| Oligo fwd | CACCGCTGCAGCAGACGGCCGCCC | | | | | | | | | | |
| Oligo rev | AAACGGGCGGCCGTCTGCTGCAGC | | | | | | | | | | |
| Chromosome | start | end | strand | MM | target_seq | PAM | alignment | distance | position | gene name | gene id |
| GU305914.1 | 110693 | 110715 | + | 0 | GCTGCAGCAGACGGCCGCCC | CGG | \|\|\|\|\|\|\|\|[\|\|\|\|\|\|\|\|\|\|\|\|]PAM | NA | Intergenic | NA | NA |
| T8138 | GCAGACGGCCGCCCCGGACAAGG | 1000 | | | | | | | | | |
| Oligo fwd | CACCGCAGACGGCCGCCCCGGACA | | | | | | | | | | |
| Oligo rev | AAACTGTCCGGGGCGGCCGTCTGC | | | | | | | | | | |
| Chromosome | start | end | strand | MM | target_seq | PAM | alignment | distance | position | gene name | gene id |
| GU305914.1 | 110699 | 110721 | + | 0 | GCAGACGGCCGCCCCGGACA | AGG | \|\|\|\|\|\|\|\|[\|\|\|\|\|\|\|\|\|\|\|\|]PAM | NA | Intergenic | NA | NA |
| T8139 | CCCCGGACAAGGATGTCGACAGG | 1000 | | | | | | | | | |
| Oligo fwd | CACCCCCCGGACAAGGATGTCGAC | | | | | | | | | | |
| Oligo rev | AAACGTCGACATCCTTGTCCGGGG | | | | | | | | | | |
| Chromosome | start | end | strand | MM | target_seq | PAM | alignment | distance | position | gene name | gene id |
| GU305914.1 | 110710 | 110732 | + | 0 | CCCCGGACAAGGATGTCGAC | AGG | \|\|\|\|\|\|\|\|[\|\|\|\|\|\|\|\|\|\|\|\|]PAM | NA | Intergenic | NA | NA |
| T8140 | GCTCTCTTCCACTGCGAGAACGG | 1000 | | | | | | | | | |
| Oligo fwd | CACCGCTCTCTTCCACTGCGAGAA | | | | | | | | | | |
| Oligo rev | AAACTTCTCGCAGTGGAAGAGAGC | | | | | | | | | | |
| Chromosome | start | end | strand | MM | target_seq | PAM | alignment | distance | position | gene name | gene id |
| GU305914.1 | 110751 | 110773 | + | 0 | GCTCTCTTCCACTGCGAGAA | CGG | \|\|\|\|\|\|\|\|[\|\|\|\|\|\|\|\|\|\|\|\|]PAM | NA | Intergenic | NA | NA |
| T8141 | CTCTCTTCCACTGCGAGAACGGG | 1000 | | | | | | | | | |
| Oligo fwd | CACCCTCTCTTCCACTGCGAGAAC | | | | | | | | | | |
| Oligo rev | AAACGTTCTCGCAGTGGAAGAGAG | | | | | | | | | | |
| Chromosome | start | end | strand | MM | target_seq | PAM | alignment | distance | position | gene name | gene id |
| GU305914.1 | 110752 | 110774 | + | 0 | CTCTCTTCCACTGCGAGAAC | GGG | \|\|\|\|\|\|\|\|[\|\|\|\|\|\|\|\|\|\|\|\|]PAM | NA | Intergenic | NA | NA |
| T8142 | TTCCACTGCGAGAACGGGATCGG | 1000 | | | | | | | | | |
| Oligo fwd | CACCTTCCACTGCGAGAACGGGAT | | | | | | | | | | |
| Oligo rev | AAACATCCCGTTCTCGCAGTGGAA | | | | | | | | | | |
| Chromosome | start | end | strand | MM | target_seq | PAM | alignment | distance | position | gene name | gene id |
| GU305914.1 | 110757 | 110779 | + | 0 | TTCCACTGCGAGAACGGGAT | CGG | \|\|\|\|\|\|\|\|[\|\|\|\|\|\|\|\|\|\|\|\|]PAM | NA | Intergenic | NA | NA |
| T8143 | ACTGCGAGAACGGGATCGGCAGG | 1000 | | | | | | | | | |
| Oligo fwd | CACCACTGCGAGAACGGGATCGGC | | | | | | | | | | |
| Oligo rev | AAACGCCGATCCCGTTCTCGCAGT | | | | | | | | | | |
| Chromosome | start | end | strand | MM | target_seq | PAM | alignment | distance | position | gene name | gene id |
| GU305914.1 | 110761 | 110783 | + | 0 | ACTGCGAGAACGGGATCGGC | AGG | \|\|\|\|\|\|\|\|[\|\|\|\|\|\|\|\|\|\|\|\|]PAM | NA | Intergenic | NA | NA |
| T8144 | CCATCACGCTGCCCAAGCACAGG | 1000 | | | | | | | | | |
| Oligo fwd | CACCCCATCACGCTGCCCAAGCAC | | | | | | | | | | |
| Oligo rev | AAACGTGCTTGGGCAGCGTGATGG | | | | | | | | | | |
| Chromosome | start | end | strand | MM | target_seq | PAM | alignment | distance | position | gene name | gene id |
| GU305914.1 | 110791 | 110813 | + | 0 | CCATCACGCTGCCCAAGCAC | AGG | \|\|\|\|\|\|\|\|[\|\|\|\|\|\|\|\|\|\|\|\|]PAM | NA | Intergenic | NA | NA |
| GU305914.1 | 138632 | 138654 | + | 3 | CCAACGCGCTGCCCAAGGAC | CGG | \|\|\|-\|-\|\|[\|\|\|\|\|\|\|\|\|-\|\|]PAM | NA | Intergenic | NA | NA |
| T8145 | CATCACGCTGCCCAAGCACAGGG | 1000 | | | | | | | | | |
| Oligo fwd | CACCCATCACGCTGCCCAAGCACA | | | | | | | | | | |
| Oligo rev | AAACTGTGCTTGGGCAGCGTGATG | | | | | | | | | | |
| Chromosome | start | end | strand | MM | target_seq | PAM | alignment | distance | position | gene name | gene id |
| GU305914.1 | 110792 | 110814 | + | 0 | CATCACGCTGCCCAAGCACA | GGG | \|\|\|\|\|\|\|\|[\|\|\|\|\|\|\|\|\|\|\|\|]PAM | NA | Intergenic | NA | NA |
| T8146 | AAGCACAGGGTCCTGACCCTCGG | 1000 | | | | | | | | | |
| Oligo fwd | CACCAAGCACAGGGTCCTGACCCT | | | | | | | | | | |
| Oligo rev | AAACAGGGTCAGGACCCTGTGCTT | | | | | | | | | | |
| Chromosome | start | end | strand | MM | target_seq | PAM | alignment | distance | position | gene name | gene id |
| GU305914.1 | 110805 | 110827 | + | 0 | AAGCACAGGGTCCTGACCCT | CGG | \|\|\|\|\|\|\|\|[\|\|\|\|\|\|\|\|\|\|\|\|]PAM | NA | Intergenic | NA | NA |
| T8147 | TCCATGTATTTTTTTGTGCTCGG | 1000 | | | | | | | | | |
| Oligo fwd | CACCTCCATGTATTTTTTTGTGCT | | | | | | | | | | |
| Oligo rev | AAACAGCACAAAAAAATACATGGA | | | | | | | | | | |
| Chromosome | start | end | strand | MM | target_seq | PAM | alignment | distance | position | gene name | gene id |
| GU305914.1 | 110862 | 110884 | + | 0 | TCCATGTATTTTTTTGTGCT | CGG | \|\|\|\|\|\|\|\|[\|\|\|\|\|\|\|\|\|\|\|\|]PAM | NA | Intergenic | NA | NA |
| T8148 | TGTGCTCGGTTTCTTGCCTGTGG | 1000 | | | | | | | | | |
| Oligo fwd | CACCTGTGCTCGGTTTCTTGCCTG | | | | | | | | | | |
| Oligo rev | AAACCAGGCAAGAAACCGAGCACA | | | | | | | | | | |
| Chromosome | start | end | strand | MM | target_seq | PAM | alignment | distance | position | gene name | gene id |
| GU305914.1 | 110876 | 110898 | + | 0 | TGTGCTCGGTTTCTTGCCTG | TGG | \|\|\|\|\|\|\|\|[\|\|\|\|\|\|\|\|\|\|\|\|]PAM | NA | Intergenic | NA | NA |
| T8149 | CGTGAGCCGCCGCCGCGACTCGG | 1000 | | | | | | | | | |
| Oligo fwd | CACCCGTGAGCCGCCGCCGCGACT | | | | | | | | | | |
| Oligo rev | AAACAGTCGCGGCGGCGGCTCACG | | | | | | | | | | |
| Chromosome | start | end | strand | MM | target_seq | PAM | alignment | distance | position | gene name | gene id |
| GU305914.1 | 110908 | 110930 | + | 0 | CGTGAGCCGCCGCCGCGACT | CGG | \|\|\|\|\|\|\|\|[\|\|\|\|\|\|\|\|\|\|\|\|]PAM | NA | Intergenic | NA | NA |

<table>
<tr><td>T8150</td><td>CGACTCGGCCGAGTATTTTATGG</td><td>1000</td><td></td><td></td><td></td><td></td><td></td><td></td><td></td><td></td><td></td></tr>
<tr><td>Oligo fwd</td><td>CACCCGACTCGGCCGAGTATTTTA</td><td></td><td></td><td></td><td></td><td></td><td></td><td></td><td></td><td></td><td></td></tr>
<tr><td>Oligo rev</td><td>AAACTAAAATACTCGGCCGAGTCG</td><td></td><td></td><td></td><td></td><td></td><td></td><td></td><td></td><td></td><td></td></tr>
<tr><td>Chromosome</td><td>start</td><td>end</td><td>strand</td><td>MM</td><td>target_seq</td><td>PAM</td><td>alignment</td><td>distance</td><td>position</td><td>gene name</td><td>gene id</td></tr>
<tr><td>GU305914.1</td><td>110923</td><td>110945</td><td>+</td><td>0</td><td>CGACTCGGCCGAGTATTTTA</td><td>TGG</td><td>||||||||[||||||||||||]PAM</td><td>NA</td><td>Intergenic</td><td>NA</td><td>NA</td></tr>
<tr><td>T8151</td><td>CGGCCGAGTATTTTATGGATTGG</td><td>1000</td><td></td><td></td><td></td><td></td><td></td><td></td><td></td><td></td><td></td></tr>
<tr><td>Oligo fwd</td><td>CACCCGGCCGAGTATTTTATGGAT</td><td></td><td></td><td></td><td></td><td></td><td></td><td></td><td></td><td></td><td></td></tr>
<tr><td>Oligo rev</td><td>AAACATCCATAAAATACTCGGCCG</td><td></td><td></td><td></td><td></td><td></td><td></td><td></td><td></td><td></td><td></td></tr>
<tr><td>Chromosome</td><td>start</td><td>end</td><td>strand</td><td>MM</td><td>target_seq</td><td>PAM</td><td>alignment</td><td>distance</td><td>position</td><td>gene name</td><td>gene id</td></tr>
<tr><td>GU305914.1</td><td>110928</td><td>110950</td><td>+</td><td>0</td><td>CGGCCGAGTATTTTATGGAT</td><td>TGG</td><td>||||||||[||||||||||||]PAM</td><td>NA</td><td>Intergenic</td><td>NA</td><td>NA</td></tr>
<tr><td>T8152</td><td>CCGAGTATTTTATGGATTGGCGG</td><td>1000</td><td></td><td></td><td></td><td></td><td></td><td></td><td></td><td></td><td></td></tr>
<tr><td>Oligo fwd</td><td>CACCCCGAGTATTTTATGGATTGG</td><td></td><td></td><td></td><td></td><td></td><td></td><td></td><td></td><td></td><td></td></tr>
<tr><td>Oligo rev</td><td>AAACCCAATCCATAAAATACTCGG</td><td></td><td></td><td></td><td></td><td></td><td></td><td></td><td></td><td></td><td></td></tr>
<tr><td>Chromosome</td><td>start</td><td>end</td><td>strand</td><td>MM</td><td>target_seq</td><td>PAM</td><td>alignment</td><td>distance</td><td>position</td><td>gene name</td><td>gene id</td></tr>
<tr><td>GU305914.1</td><td>110931</td><td>110953</td><td>+</td><td>0</td><td>CCGAGTATTTTATGGATTGG</td><td>CGG</td><td>||||||||[||||||||||||]PAM</td><td>NA</td><td>Intergenic</td><td>NA</td><td>NA</td></tr>
<tr><td>T8153</td><td>GATTGGCGGACGTCCCCTTCTGG</td><td>1000</td><td></td><td></td><td></td><td></td><td></td><td></td><td></td><td></td><td></td></tr>
<tr><td>Oligo fwd</td><td>CACCGATTGGCGGACGTCCCCTTC</td><td></td><td></td><td></td><td></td><td></td><td></td><td></td><td></td><td></td><td></td></tr>
<tr><td>Oligo rev</td><td>AAACGAAGGGGACGTCCGCCAATC</td><td></td><td></td><td></td><td></td><td></td><td></td><td></td><td></td><td></td><td></td></tr>
<tr><td>Chromosome</td><td>start</td><td>end</td><td>strand</td><td>MM</td><td>target_seq</td><td>PAM</td><td>alignment</td><td>distance</td><td>position</td><td>gene name</td><td>gene id</td></tr>
<tr><td>GU305914.1</td><td>110945</td><td>110967</td><td>+</td><td>0</td><td>GATTGGCGGACGTCCCCTTC</td><td>TGG</td><td>||||||||[||||||||||||]PAM</td><td>NA</td><td>Intergenic</td><td>NA</td><td>NA</td></tr>
<tr><td>T8154</td><td>CTGGCGCCGCCTGACATATAAGG</td><td>1000</td><td></td><td></td><td></td><td></td><td></td><td></td><td></td><td></td><td></td></tr>
<tr><td>Oligo fwd</td><td>CACCCTGGCGCCGCCTGACATATA</td><td></td><td></td><td></td><td></td><td></td><td></td><td></td><td></td><td></td><td></td></tr>
<tr><td>Oligo rev</td><td>AAACTATATGTCAGGCGGCGCCAG</td><td></td><td></td><td></td><td></td><td></td><td></td><td></td><td></td><td></td><td></td></tr>
<tr><td>Chromosome</td><td>start</td><td>end</td><td>strand</td><td>MM</td><td>target_seq</td><td>PAM</td><td>alignment</td><td>distance</td><td>position</td><td>gene name</td><td>gene id</td></tr>
<tr><td>GU305914.1</td><td>110964</td><td>110986</td><td>+</td><td>0</td><td>CTGGCGCCGCCTGACATATA</td><td>AGG</td><td>||||||||[||||||||||||]PAM</td><td>NA</td><td>Intergenic</td><td>NA</td><td>NA</td></tr>
<tr><td>T8155</td><td>CGCCTGACATATAAGGCGCGCGG</td><td>1000</td><td></td><td></td><td></td><td></td><td></td><td></td><td></td><td></td><td></td></tr>
<tr><td>Oligo fwd</td><td>CACCCGCCTGACATATAAGGCGCG</td><td></td><td></td><td></td><td></td><td></td><td></td><td></td><td></td><td></td><td></td></tr>
<tr><td>Oligo rev</td><td>AAACCGCGCCTTATATGTCAGGCG</td><td></td><td></td><td></td><td></td><td></td><td></td><td></td><td></td><td></td><td></td></tr>
<tr><td>Chromosome</td><td>start</td><td>end</td><td>strand</td><td>MM</td><td>target_seq</td><td>PAM</td><td>alignment</td><td>distance</td><td>position</td><td>gene name</td><td>gene id</td></tr>
<tr><td>GU305914.1</td><td>110971</td><td>110993</td><td>+</td><td>0</td><td>CGCCTGACATATAAGGCGCG</td><td>CGG</td><td>||||||||[||||||||||||]PAM</td><td>NA</td><td>Intergenic</td><td>NA</td><td>NA</td></tr>
<tr><td>T8156</td><td>ATAAGGCGCGCGGCCCGTTTCGG</td><td>1000</td><td></td><td></td><td></td><td></td><td></td><td></td><td></td><td></td><td></td></tr>
<tr><td>Oligo fwd</td><td>CACCATAAGGCGCGCGGCCCGTTT</td><td></td><td></td><td></td><td></td><td></td><td></td><td></td><td></td><td></td><td></td></tr>
<tr><td>Oligo rev</td><td>AAACAAACGGGCCGCGCGCCTTAT</td><td></td><td></td><td></td><td></td><td></td><td></td><td></td><td></td><td></td><td></td></tr>
<tr><td>Chromosome</td><td>start</td><td>end</td><td>strand</td><td>MM</td><td>target_seq</td><td>PAM</td><td>alignment</td><td>distance</td><td>position</td><td>gene name</td><td>gene id</td></tr>
<tr><td>GU305914.1</td><td>110981</td><td>111003</td><td>+</td><td>0</td><td>ATAAGGCGCGCGGCCCGTTT</td><td>CGG</td><td>||||||||[||||||||||||]PAM</td><td>NA</td><td>Intergenic</td><td>NA</td><td>NA</td></tr>
<tr><td>T8157</td><td>CGGAGCGCACAGTCTCATACTGG</td><td>1000</td><td></td><td></td><td></td><td></td><td></td><td></td><td></td><td></td><td></td></tr>
<tr><td>Oligo fwd</td><td>CACCCGGAGCGCACAGTCTCATAC</td><td></td><td></td><td></td><td></td><td></td><td></td><td></td><td></td><td></td><td></td></tr>
<tr><td>Oligo rev</td><td>AAACGTATGAGACTGTGCGCTCCG</td><td></td><td></td><td></td><td></td><td></td><td></td><td></td><td></td><td></td><td></td></tr>
<tr><td>Chromosome</td><td>start</td><td>end</td><td>strand</td><td>MM</td><td>target_seq</td><td>PAM</td><td>alignment</td><td>distance</td><td>position</td><td>gene name</td><td>gene id</td></tr>
<tr><td>GU305914.1</td><td>111001</td><td>111023</td><td>+</td><td>0</td><td>CGGAGCGCACAGTCTCATAC</td><td>TGG</td><td>||||||||[||||||||||||]PAM</td><td>NA</td><td>Intergenic</td><td>NA</td><td>NA</td></tr>
<tr><td>T8158</td><td>CTGGTCCTTCACACAGTGCCAGG</td><td>1000</td><td></td><td></td><td></td><td></td><td></td><td></td><td></td><td></td><td></td></tr>
<tr><td>Oligo fwd</td><td>CACCCTGGTCCTTCACACAGTGCC</td><td></td><td></td><td></td><td></td><td></td><td></td><td></td><td></td><td></td><td></td></tr>
<tr><td>Oligo rev</td><td>AAACGGCACTGTGTGAAGGACCAG</td><td></td><td></td><td></td><td></td><td></td><td></td><td></td><td></td><td></td><td></td></tr>
<tr><td>Chromosome</td><td>start</td><td>end</td><td>strand</td><td>MM</td><td>target_seq</td><td>PAM</td><td>alignment</td><td>distance</td><td>position</td><td>gene name</td><td>gene id</td></tr>
<tr><td>GU305914.1</td><td>111020</td><td>111042</td><td>+</td><td>0</td><td>CTGGTCCTTCACACAGTGCC</td><td>AGG</td><td>||||||||[||||||||||||]PAM</td><td>NA</td><td>Intergenic</td><td>NA</td><td>NA</td></tr>
<tr><td>T8159</td><td>CACAGTGCCAGGCACCAGTGCGG</td><td>1000</td><td></td><td></td><td></td><td></td><td></td><td></td><td></td><td></td><td></td></tr>
<tr><td>Oligo fwd</td><td>CACCCACAGTGCCAGGCACCAGTG</td><td></td><td></td><td></td><td></td><td></td><td></td><td></td><td></td><td></td><td></td></tr>
<tr><td>Oligo rev</td><td>AAACCACTGGTGCCTGGCACTGTG</td><td></td><td></td><td></td><td></td><td></td><td></td><td></td><td></td><td></td><td></td></tr>
<tr><td>Chromosome</td><td>start</td><td>end</td><td>strand</td><td>MM</td><td>target_seq</td><td>PAM</td><td>alignment</td><td>distance</td><td>position</td><td>gene name</td><td>gene id</td></tr>
<tr><td>GU305914.1</td><td>111031</td><td>111053</td><td>+</td><td>0</td><td>CACAGTGCCAGGCACCAGTG</td><td>CGG</td><td>||||||||[||||||||||||]PAM</td><td>NA</td><td>Intergenic</td><td>NA</td><td>NA</td></tr>
<tr><td>T8160</td><td>ACAGTGCCAGGCACCAGTGCGGG</td><td>1000</td><td></td><td></td><td></td><td></td><td></td><td></td><td></td><td></td><td></td></tr>
<tr><td>Oligo fwd</td><td>CACCACAGTGCCAGGCACCAGTGC</td><td></td><td></td><td></td><td></td><td></td><td></td><td></td><td></td><td></td><td></td></tr>
<tr><td>Oligo rev</td><td>AAACGCACTGGTGCCTGGCACTGT</td><td></td><td></td><td></td><td></td><td></td><td></td><td></td><td></td><td></td><td></td></tr>
<tr><td>Chromosome</td><td>start</td><td>end</td><td>strand</td><td>MM</td><td>target_seq</td><td>PAM</td><td>alignment</td><td>distance</td><td>position</td><td>gene name</td><td>gene id</td></tr>
<tr><td>GU305914.1</td><td>111032</td><td>111054</td><td>+</td><td>0</td><td>ACAGTGCCAGGCACCAGTGC</td><td>GGG</td><td>||||||||[||||||||||||]PAM</td><td>NA</td><td>Intergenic</td><td>NA</td><td>NA</td></tr>
<tr><td>T8161</td><td>CAGTGCCAGGCACCAGTGCGGGG</td><td>1000</td><td></td><td></td><td></td><td></td><td></td><td></td><td></td><td></td><td></td></tr>
<tr><td>Oligo fwd</td><td>CACCCAGTGCCAGGCACCAGTGCG</td><td></td><td></td><td></td><td></td><td></td><td></td><td></td><td></td><td></td><td></td></tr>
<tr><td>Oligo rev</td><td>AAACCGCACTGGTGCCTGGCACTG</td><td></td><td></td><td></td><td></td><td></td><td></td><td></td><td></td><td></td><td></td></tr>
<tr><td>Chromosome</td><td>start</td><td>end</td><td>strand</td><td>MM</td><td>target_seq</td><td>PAM</td><td>alignment</td><td>distance</td><td>position</td><td>gene name</td><td>gene id</td></tr>
<tr><td>GU305914.1</td><td>111033</td><td>111055</td><td>+</td><td>0</td><td>CAGTGCCAGGCACCAGTGCG</td><td>GGG</td><td>||||||||[||||||||||||]PAM</td><td>NA</td><td>Intergenic</td><td>NA</td><td>NA</td></tr>
<tr><td>T8162</td><td>CCAGGCACCAGTGCGGGGCGCGG</td><td>1000</td><td></td><td></td><td></td><td></td><td></td><td></td><td></td><td></td><td></td></tr>
<tr><td>Oligo fwd</td><td>CACCCCAGGCACCAGTGCGGGGCG</td><td></td><td></td><td></td><td></td><td></td><td></td><td></td><td></td><td></td><td></td></tr>
<tr><td>Oligo rev</td><td>AAACCGCCCCGCACTGGTGCCTGG</td><td></td><td></td><td></td><td></td><td></td><td></td><td></td><td></td><td></td><td></td></tr>
<tr><td>Chromosome</td><td>start</td><td>end</td><td>strand</td><td>MM</td><td>target_seq</td><td>PAM</td><td>alignment</td><td>distance</td><td>position</td><td>gene name</td><td>gene id</td></tr>
<tr><td>GU305914.1</td><td>111038</td><td>111060</td><td>+</td><td>0</td><td>CCAGGCACCAGTGCGGGGCG</td><td>CGG</td><td>||||||||[||||||||||||]PAM</td><td>NA</td><td>Intergenic</td><td>NA</td><td>NA</td></tr>
<tr><td>T8163</td><td>CAGGCACCAGTGCGGGGCGCGGG</td><td>1000</td><td></td><td></td><td></td><td></td><td></td><td></td><td></td><td></td><td></td></tr>
<tr><td>Oligo fwd</td><td>CACCCAGGCACCAGTGCGGGGCGC</td><td></td><td></td><td></td><td></td><td></td><td></td><td></td><td></td><td></td><td></td></tr>
<tr><td>Oligo rev</td><td>AAACGCGCCCCGCACTGGTGCCTG</td><td></td><td></td><td></td><td></td><td></td><td></td><td></td><td></td><td></td><td></td></tr>
<tr><td>Chromosome</td><td>start</td><td>end</td><td>strand</td><td>MM</td><td>target_seq</td><td>PAM</td><td>alignment</td><td>distance</td><td>position</td><td>gene name</td><td>gene id</td></tr>
<tr><td>GU305914.1</td><td>111039</td><td>111061</td><td>+</td><td>0</td><td>CAGGCACCAGTGCGGGGCGC</td><td>GGG</td><td>||||||||[||||||||||||]PAM</td><td>NA</td><td>Intergenic</td><td>NA</td><td>NA</td></tr>
<tr><td>T8164</td><td>CGCCGTCGATTACTCCTACCCGG</td><td>1000</td><td></td><td></td><td></td><td></td><td></td><td></td><td></td><td></td><td></td></tr>
<tr><td>Oligo fwd</td><td>CACCCGCCGTCGATTACTCCTACC</td><td></td><td></td><td></td><td></td><td></td><td></td><td></td><td></td><td></td><td></td></tr>
<tr><td>Oligo rev</td><td>AAACGGTAGGAGTAATCGACGGCG</td><td></td><td></td><td></td><td></td><td></td><td></td><td></td><td></td><td></td><td></td></tr>
<tr><td>Chromosome</td><td>start</td><td>end</td><td>strand</td><td>MM</td><td>target_seq</td><td>PAM</td><td>alignment</td><td>distance</td><td>position</td><td>gene name</td><td>gene id</td></tr>
<tr><td>GU305914.1</td><td>111101</td><td>111123</td><td>+</td><td>0</td><td>CGCCGTCGATTACTCCTACC</td><td>CGG</td><td>||||||||[||||||||||||]PAM</td><td>NA</td><td>Intergenic</td><td>NA</td><td>NA</td></tr>
</table>

T8165 CGTCGATTACTCCTACCCGGAGG 1000
Oligo fwd CACCCGTCGATTACTCCTACCCGG
Oligo rev AAACCCGGGTAGGAGTAATCGACG
Chromosome start end strand MM target_seq PAM alignment distance position gene name gene id
GU305914.1 111104 111126 + 0 CGTCGATTACTCCTACCCGG AGG ||||||||[||||||||||||]PAM NA Intergenic NA NA

T8166 ACCTCCACCTTCGCCGTCGTCGG 1000
Oligo fwd CACCACCTCCACCTTCGCCGTCGT
Oligo rev AAACACGACGGCGAAGGTGGAGGT
Chromosome start end strand MM target_seq PAM alignment distance position gene name gene id
GU305914.1 111174 111196 + 0 ACCTCCACCTTCGCCGTCGT CGG ||||||||[||||||||||||]PAM NA Intergenic NA NA

T8167 ATCTCGATCGTTCTCATCCTAGG 1000
Oligo fwd CACCATCTCGATCGTTCTCATCCT
Oligo rev AAACAGGATGAGAACGATCGAGAT
Chromosome start end strand MM target_seq PAM alignment distance position gene name gene id
GU305914.1 111201 111223 + 0 ATCTCGATCGTTCTCATCCT AGG ||||||||[||||||||||||]PAM NA Intergenic NA NA

T8168 TCTCGATCGTTCTCATCCTAGGG 1000
Oligo fwd CACCTCTCGATCGTTCTCATCCTA
Oligo rev AAACTAGGATGAGAACGATCGAGA
Chromosome start end strand MM target_seq PAM alignment distance position gene name gene id
GU305914.1 111202 111224 + 0 TCTCGATCGTTCTCATCCTA GGG ||||||||[||||||||||||]PAM NA Intergenic NA NA

T8169 GGAGCCTCATCCTCTCGATGTGG 1000
Oligo fwd CACCGGAGCCTCATCCTCTCGATG
Oligo rev AAACCATCGAGAGGATGAGGCTCC
Chromosome start end strand MM target_seq PAM alignment distance position gene name gene id
GU305914.1 111223 111245 + 0 GGAGCCTCATCCTCTCGATG TGG ||||||||[||||||||||||]PAM NA Intergenic NA NA
GU305914.1 76587 76609 + 4 AGAGCTTCATCTTCTCGTTG AGG -||||-||[|||-|||||-||]PAM NA Intergenic NA NA

T8170 GCCTCATCCTCTCGATGTGGAGG 1000
Oligo fwd CACCGCCTCATCCTCTCGATGTGG
Oligo rev AAACCCACATCGAGAGGATGAGGC
Chromosome start end strand MM target_seq PAM alignment distance position gene name gene id
GU305914.1 111226 111248 + 0 GCCTCATCCTCTCGATGTGG AGG ||||||||[||||||||||||]PAM NA Intergenic NA NA

T8171 CATCCTCTCGATGTGGAGGAAGG 1000
Oligo fwd CACCCATCCTCTCGATGTGGAGGA
Oligo rev AAACTCCTCCACATCGAGAGGATG
Chromosome start end strand MM target_seq PAM alignment distance position gene name gene id
GU305914.1 111230 111252 + 0 CATCCTCTCGATGTGGAGGA AGG ||||||||[||||||||||||]PAM NA Intergenic NA NA

T8172 TCTCGATGTGGAGGAAGGTCTGG 1000
Oligo fwd CACCTCTCGATGTGGAGGAAGGTC
Oligo rev AAACGACCTTCCTCCACATCGAGA
Chromosome start end strand MM target_seq PAM alignment distance position gene name gene id
GU305914.1 111235 111257 + 0 TCTCGATGTGGAGGAAGGTC TGG ||||||||[||||||||||||]PAM NA Intergenic NA NA

T8173 AAGGTCTGGCACCGCCGTAACGG 1000
Oligo fwd CACCAAGGTCTGGCACCGCCGTAA
Oligo rev AAACTTACGGCGGTGCCAGACCTT
Chromosome start end strand MM target_seq PAM alignment distance position gene name gene id
GU305914.1 111249 111271 + 0 AAGGTCTGGCACCGCCGTAA CGG ||||||||[||||||||||||]PAM NA Intergenic NA NA

T8174 GTCTGGCACCGCCGTAACGGAGG 1000
Oligo fwd CACCGTCTGGCACCGCCGTAACGG
Oligo rev AAACCCGTTACGGCGGTGCCAGAC
Chromosome start end strand MM target_seq PAM alignment distance position gene name gene id
GU305914.1 111252 111274 + 0 GTCTGGCACCGCCGTAACGG AGG ||||||||[||||||||||||]PAM NA Intergenic NA NA

T8175 CTACGTTTTCAGCCTCTTCTTGG 1000
Oligo fwd CACCCTACGTTTTCAGCCTCTTCT
Oligo rev AAACAGAAGAGGCTGAAAACGTAG
Chromosome start end strand MM target_seq PAM alignment distance position gene name gene id
GU305914.1 111284 111306 + 0 CTACGTTTTCAGCCTCTTCT TGG ||||||||[||||||||||||]PAM NA Intergenic NA NA

T8176 TCGCCAGCTTCGTCGCCATCTGG 1000
Oligo fwd CACCTCGCCAGCTTCGTCGCCATC
Oligo rev AAACGATGGCGACGAAGCTGGCGA
Chromosome start end strand MM target_seq PAM alignment distance position gene name gene id
GU305914.1 111322 111344 + 0 TCGCCAGCTTCGTCGCCATC TGG ||||||||[||||||||||||]PAM NA Intergenic NA NA

T8177 GCGCGTCGTCTGCCCCTCTAAGG 1000
Oligo fwd CACCGCGCGTCGTCTGCCCCTCTA
Oligo rev AAACTAGAGGGGCAGACGACGCGC
Chromosome start end strand MM target_seq PAM alignment distance position gene name gene id
GU305914.1 111350 111372 + 0 GCGCGTCGTCTGCCCCTCTA AGG ||||||||[||||||||||||]PAM NA Intergenic NA NA

T8178 CACCGAACCCTACTGCCGCCTGG 1000
Oligo fwd CACCCACCGAACCCTACTGCCGCC
Oligo rev AAACGGCGGCAGTAGGGTTCGGTG
Chromosome start end strand MM target_seq PAM alignment distance position gene name gene id
GU305914.1 111374 111396 + 0 CACCGAACCCTACTGCCGCC TGG ||||||||[||||||||||||]PAM NA Intergenic NA NA

T8179 CTACTGCCGCCTGGCCATCATGG 1000
Oligo fwd CACCCTACTGCCGCCTGGCCATCA
Oligo rev AAACTGATGGCCAGGCGGCAGTAG

| | | | | | | | | | | | |
|---|---|---|---|---|---|---|---|---|---|---|---|
| Chromosome | start | end | strand | MM | target_seq | PAM | alignment | distance | position | gene name | gene id |
| GU305914.1 | 111383 | 111405 | + | 0 | CTACTGCCGCCTGGCCATCA | TGG | ||||||||[||||||||||||]PAM | NA | Intergenic | NA | NA |
| T8180 | CGCCTGGCCATCATGGTCAACGG | 1000 | | | | | | | | | |
| Oligo fwd | CACCCGCCTGGCCATCATGGTCAA | | | | | | | | | | |
| Oligo rev | AAACTTGACCATGATGGCCAGGCG | | | | | | | | | | |
| Chromosome | start | end | strand | MM | target_seq | PAM | alignment | distance | position | gene name | gene id |
| GU305914.1 | 111390 | 111412 | + | 0 | CGCCTGGCCATCATGGTCAA | CGG | ||||||||[||||||||||||]PAM | NA | Intergenic | NA | NA |
| T8181 | ATGGTCAACGGCATGTCCGAAGG | 1000 | | | | | | | | | |
| Oligo fwd | CACCATGGTCAACGGCATGTCCGA | | | | | | | | | | |
| Oligo rev | AAACTCGGACATGCCGTTGACCAT | | | | | | | | | | |
| Chromosome | start | end | strand | MM | target_seq | PAM | alignment | distance | position | gene name | gene id |
| GU305914.1 | 111402 | 111424 | + | 0 | ATGGTCAACGGCATGTCCGA | AGG | ||||||||[||||||||||||]PAM | NA | Intergenic | NA | NA |
| T8182 | CTCGATCGCCTCTGCCTCGTCGG | 1000 | | | | | | | | | |
| Oligo fwd | CACCCTCGATCGCCTCTGCCTCGT | | | | | | | | | | |
| Oligo rev | AAACACGAGGCAGAGGCGATCGAG | | | | | | | | | | |
| Chromosome | start | end | strand | MM | target_seq | PAM | alignment | distance | position | gene name | gene id |
| GU305914.1 | 111456 | 111478 | + | 0 | CTCGATCGCCTCTGCCTCGT | CGG | ||||||||[||||||||||||]PAM | NA | Intergenic | NA | NA |
| T8183 | ACCATCTTCAACACCTCCGCCGG | 1000 | | | | | | | | | |
| Oligo fwd | CACCACCATCTTCAACACCTCCGC | | | | | | | | | | |
| Oligo rev | AAACGCGGAGGTGTTGAAGATGGT | | | | | | | | | | |
| Chromosome | start | end | strand | MM | target_seq | PAM | alignment | distance | position | gene name | gene id |
| GU305914.1 | 111498 | 111520 | + | 0 | ACCATCTTCAACACCTCCGC | CGG | ||||||||[||||||||||||]PAM | NA | Intergenic | NA | NA |
| T8184 | CTTCAACACCTCCGCCGGCAAGG | 1000 | | | | | | | | | |
| Oligo fwd | CACCCTTCAACACCTCCGCCGGCA | | | | | | | | | | |
| Oligo rev | AAACTGCCGGCGGAGGTGTTGAAG | | | | | | | | | | |
| Chromosome | start | end | strand | MM | target_seq | PAM | alignment | distance | position | gene name | gene id |
| GU305914.1 | 111503 | 111525 | + | 0 | CTTCAACACCTCCGCCGGCA | AGG | ||||||||[||||||||||||]PAM | NA | Intergenic | NA | NA |
| T8185 | TCCGCCGGCAAGGATCTCGTCGG | 1000 | | | | | | | | | |
| Oligo fwd | CACCTCCGCCGGCAAGGATCTCGT | | | | | | | | | | |
| Oligo rev | AAACACGAGATCCTTGCCGGCGGA | | | | | | | | | | |
| Chromosome | start | end | strand | MM | target_seq | PAM | alignment | distance | position | gene name | gene id |
| GU305914.1 | 111513 | 111535 | + | 0 | TCCGCCGGCAAGGATCTCGT | CGG | ||||||||[||||||||||||]PAM | NA | Intergenic | NA | NA |
| T8186 | AGGATCTCGTCGGCGTCATCTGG | 1000 | | | | | | | | | |
| Oligo fwd | CACCAGGATCTCGTCGGCGTCATC | | | | | | | | | | |
| Oligo rev | AAACGATGACGCCGACGAGATCCT | | | | | | | | | | |
| Chromosome | start | end | strand | MM | target_seq | PAM | alignment | distance | position | gene name | gene id |
| GU305914.1 | 111523 | 111545 | + | 0 | AGGATCTCGTCGGCGTCATC | TGG | ||||||||[||||||||||||]PAM | NA | Intergenic | NA | NA |
| T8187 | GGATCTCGTCGGCGTCATCTGGG | 1000 | | | | | | | | | |
| Oligo fwd | CACCGGATCTCGTCGGCGTCATCT | | | | | | | | | | |
| Oligo rev | AAACAGATGACGCCGACGAGATCC | | | | | | | | | | |
| Chromosome | start | end | strand | MM | target_seq | PAM | alignment | distance | position | gene name | gene id |
| GU305914.1 | 111524 | 111546 | + | 0 | GGATCTCGTCGGCGTCATCT | GGG | ||||||||[||||||||||||]PAM | NA | Intergenic | NA | NA |
| T8188 | CGGCGTCATCTGGGTCACCGTGG | 1000 | | | | | | | | | |
| Oligo fwd | CACCCGGCGTCATCTGGGTCACCG | | | | | | | | | | |
| Oligo rev | AAACCGGTGACCCAGATGACGCCG | | | | | | | | | | |
| Chromosome | start | end | strand | MM | target_seq | PAM | alignment | distance | position | gene name | gene id |
| GU305914.1 | 111533 | 111555 | + | 0 | CGGCGTCATCTGGGTCACCG | TGG | ||||||||[||||||||||||]PAM | NA | Intergenic | NA | NA |
| T8189 | TCTGGGTCACCGTGGTCGCCTGG | 1000 | | | | | | | | | |
| Oligo fwd | CACCTCTGGGTCACCGTGGTCGCC | | | | | | | | | | |
| Oligo rev | AAACGGCGACCACGGTGACCCAGA | | | | | | | | | | |
| Chromosome | start | end | strand | MM | target_seq | PAM | alignment | distance | position | gene name | gene id |
| GU305914.1 | 111541 | 111563 | + | 0 | TCTGGGTCACCGTGGTCGCC | TGG | ||||||||[||||||||||||]PAM | NA | Intergenic | NA | NA |
| GU305914.1 | 148668 | 148690 | + | 4 | TCTCGGCCAGCATGGTCGCC | TGG | |||-||-|[|-|-||||||||]PAM | NA | Intergenic | NA | NA |
| T8190 | TGGATCTCCGCCTTCATCGTCGG | 1000 | | | | | | | | | |
| Oligo fwd | CACCTGGATCTCCGCCTTCATCGT | | | | | | | | | | |
| Oligo rev | AAACACGATGAAGGCGGAGATCCA | | | | | | | | | | |
| Chromosome | start | end | strand | MM | target_seq | PAM | alignment | distance | position | gene name | gene id |
| GU305914.1 | 111561 | 111583 | + | 0 | TGGATCTCCGCCTTCATCGT | CGG | ||||||||[||||||||||||]PAM | NA | Intergenic | NA | NA |
| T8191 | CGCTCCCGCTCTCACCAGCCAGG | 1000 | | | | | | | | | |
| Oligo fwd | CACCCGCTCCCGCTCTCACCAGCC | | | | | | | | | | |
| Oligo rev | AAACGGCTGGTGAGAGCGGGAGCG | | | | | | | | | | |
| Chromosome | start | end | strand | MM | target_seq | PAM | alignment | distance | position | gene name | gene id |
| GU305914.1 | 111584 | 111606 | + | 0 | CGCTCCCGCTCTCACCAGCC | AGG | ||||||||[||||||||||||]PAM | NA | Intergenic | NA | NA |
| T8192 | CCAGCCAGGTCATGCGCGACCGG | 1000 | | | | | | | | | |
| Oligo fwd | CACCCCAGCCAGGTCATGCGCGAC | | | | | | | | | | |
| Oligo rev | AAACGTCGCGCATGACCTGGCTGG | | | | | | | | | | |
| Chromosome | start | end | strand | MM | target_seq | PAM | alignment | distance | position | gene name | gene id |
| GU305914.1 | 111598 | 111620 | + | 0 | CCAGCCAGGTCATGCGCGAC | CGG | ||||||||[||||||||||||]PAM | NA | Intergenic | NA | NA |
| T8193 | CAGCCAGGTCATGCGCGACCGGG | 1000 | | | | | | | | | |
| Oligo fwd | CACCCAGCCAGGTCATGCGCGACC | | | | | | | | | | |
| Oligo rev | AAACGGTCGCGCATGACCTGGCTG | | | | | | | | | | |
| Chromosome | start | end | strand | MM | target_seq | PAM | alignment | distance | position | gene name | gene id |
| GU305914.1 | 111599 | 111621 | + | 0 | CAGCCAGGTCATGCGCGACC | GGG | ||||||||[||||||||||||]PAM | NA | Intergenic | NA | NA |
| T8194 | CAGGTCATGCGCGACCGGGCCGG | 1000 | | | | | | | | | |

| | | | | | | | | | | | |
|---|---|---|---|---|---|---|---|---|---|---|---|
| Oligo fwd | CACCCAGGTCATGCGCGACCGGGC | | | | | | | | | | |
| Oligo rev | AAACGCCCGGTCGCGCATGACCTG | | | | | | | | | | |
| Chromosome | start | end | strand | MM | target_seq | PAM | alignment | distance | position | gene name | gene id |
| GU305914.1 | 111603 | 111625 | + | 0 | CAGGTCATGCGCGACCGGGC | CGG | \|\|\|\|\|\|\|[\|\|\|\|\|\|\|\|\|\|\|\|\|]PAM | NA | Intergenic | NA | NA |
| T8195 | CCTCCCCGTCTGTGAGATGCAGG | 1000 | | | | | | | | | |
| Oligo fwd | CACCCCTCCCCGTCTGTGAGATGC | | | | | | | | | | |
| Oligo rev | AAACGCATCTCACAGACGGGGAGG | | | | | | | | | | |
| Chromosome | start | end | strand | MM | target_seq | PAM | alignment | distance | position | gene name | gene id |
| GU305914.1 | 111629 | 111651 | + | 0 | CCTCCCCGTCTGTGAGATGC | AGG | \|\|\|\|\|\|\|[\|\|\|\|\|\|\|\|\|\|\|\|\|]PAM | NA | Intergenic | NA | NA |
| T8196 | CTCCCCGTCTGTGAGATGCAGGG | 1000 | | | | | | | | | |
| Oligo fwd | CACCCTCCCCGTCTGTGAGATGCA | | | | | | | | | | |
| Oligo rev | AAACTGCATCTCACAGACGGGGAG | | | | | | | | | | |
| Chromosome | start | end | strand | MM | target_seq | PAM | alignment | distance | position | gene name | gene id |
| GU305914.1 | 111630 | 111652 | + | 0 | CTCCCCGTCTGTGAGATGCA | GGG | \|\|\|\|\|\|\|[\|\|\|\|\|\|\|\|\|\|\|\|\|]PAM | NA | Intergenic | NA | NA |
| T8197 | CTTTCCTCATCATCATCTCCAGG | 1000 | | | | | | | | | |
| Oligo fwd | CACCCTTTCCTCATCATCATCTCC | | | | | | | | | | |
| Oligo rev | AAACGGAGATGATGATGAGGAAAG | | | | | | | | | | |
| Chromosome | start | end | strand | MM | target_seq | PAM | alignment | distance | position | gene name | gene id |
| GU305914.1 | 111706 | 111728 | + | 0 | CTTTCCTCATCATCATCTCC | AGG | \|\|\|\|\|\|\|[\|\|\|\|\|\|\|\|\|\|\|\|\|]PAM | NA | Intergenic | NA | NA |
| T8198 | TCTCCAGGCTCTTCGAGATGCGG | 1000 | | | | | | | | | |
| Oligo fwd | CACCTCTCCAGGCTCTTCGAGATG | | | | | | | | | | |
| Oligo rev | AAACCATCTCGAAGAGCCTGGAGA | | | | | | | | | | |
| Chromosome | start | end | strand | MM | target_seq | PAM | alignment | distance | position | gene name | gene id |
| GU305914.1 | 111721 | 111743 | + | 0 | TCTCCAGGCTCTTCGAGATG | CGG | \|\|\|\|\|\|\|[\|\|\|\|\|\|\|\|\|\|\|\|\|]PAM | NA | Intergenic | NA | NA |
| T8199 | GCTCTTCGAGATGCGGTACTCGG | 1000 | | | | | | | | | |
| Oligo fwd | CACCGCTCTTCGAGATGCGGTACT | | | | | | | | | | |
| Oligo rev | AAACAGTACCGCATCTCGAAGAGC | | | | | | | | | | |
| Chromosome | start | end | strand | MM | target_seq | PAM | alignment | distance | position | gene name | gene id |
| GU305914.1 | 111728 | 111750 | + | 0 | GCTCTTCGAGATGCGGTACT | CGG | \|\|\|\|\|\|\|[\|\|\|\|\|\|\|\|\|\|\|\|\|]PAM | NA | Intergenic | NA | NA |
| T8200 | TCGAGATGCGGTACTCGGCCCGG | 1000 | | | | | | | | | |
| Oligo fwd | CACCTCGAGATGCGGTACTCGGCC | | | | | | | | | | |
| Oligo rev | AAACGGCCGAGTACCGCATCTCGA | | | | | | | | | | |
| Chromosome | start | end | strand | MM | target_seq | PAM | alignment | distance | position | gene name | gene id |
| GU305914.1 | 111733 | 111755 | + | 0 | TCGAGATGCGGTACTCGGCC | CGG | \|\|\|\|\|\|\|[\|\|\|\|\|\|\|\|\|\|\|\|\|]PAM | NA | Intergenic | NA | NA |
| T8201 | CGAGATGCGGTACTCGGCCCGGG | 1000 | | | | | | | | | |
| Oligo fwd | CACCCGAGATGCGGTACTCGGCCC | | | | | | | | | | |
| Oligo rev | AAACGGGCCGAGTACCGCATCTCG | | | | | | | | | | |
| Chromosome | start | end | strand | MM | target_seq | PAM | alignment | distance | position | gene name | gene id |
| GU305914.1 | 111734 | 111756 | + | 0 | CGAGATGCGGTACTCGGCCC | GGG | \|\|\|\|\|\|\|[\|\|\|\|\|\|\|\|\|\|\|\|\|]PAM | NA | Intergenic | NA | NA |
| T8202 | ATGCGGTACTCGGCCCGGGACGG | 1000 | | | | | | | | | |
| Oligo fwd | CACCATGCGGTACTCGGCCCGGGA | | | | | | | | | | |
| Oligo rev | AAACTCCCGGGCCGAGTACCGCAT | | | | | | | | | | |
| Chromosome | start | end | strand | MM | target_seq | PAM | alignment | distance | position | gene name | gene id |
| GU305914.1 | 111738 | 111760 | + | 0 | ATGCGGTACTCGGCCCGGGA | CGG | \|\|\|\|\|\|\|[\|\|\|\|\|\|\|\|\|\|\|\|\|]PAM | NA | Intergenic | NA | NA |
| T8203 | ACTCGGCCCGGGACGGCACCTGG | 1000 | | | | | | | | | |
| Oligo fwd | CACCACTCGGCCCGGGACGGCACC | | | | | | | | | | |
| Oligo rev | AAACGGTGCCGTCCCGGGCCGAGT | | | | | | | | | | |
| Chromosome | start | end | strand | MM | target_seq | PAM | alignment | distance | position | gene name | gene id |
| GU305914.1 | 111745 | 111767 | + | 0 | ACTCGGCCCGGGACGGCACC | TGG | \|\|\|\|\|\|\|[\|\|\|\|\|\|\|\|\|\|\|\|\|]PAM | NA | Intergenic | NA | NA |
| T8204 | GGCACCTGGCACTTCGTGAGCGG | 1000 | | | | | | | | | |
| Oligo fwd | CACCGGCACCTGGCACTTCGTGAG | | | | | | | | | | |
| Oligo rev | AAACCTCACGAAGTGCCAGGTGCC | | | | | | | | | | |
| Chromosome | start | end | strand | MM | target_seq | PAM | alignment | distance | position | gene name | gene id |
| GU305914.1 | 111759 | 111781 | + | 0 | GGCACCTGGCACTTCGTGAG | CGG | \|\|\|\|\|\|\|[\|\|\|\|\|\|\|\|\|\|\|\|\|]PAM | NA | Intergenic | NA | NA |
| T8205 | TCGTGCCCATCATCATCTGTCGG | 1000 | | | | | | | | | |
| Oligo fwd | CACCTCGTGCCCATCATCATCTGT | | | | | | | | | | |
| Oligo rev | AAACACAGATGATGATGGGCACGA | | | | | | | | | | |
| Chromosome | start | end | strand | MM | target_seq | PAM | alignment | distance | position | gene name | gene id |
| GU305914.1 | 111817 | 111839 | + | 0 | TCGTGCCCATCATCATCTGT | CGG | \|\|\|\|\|\|\|[\|\|\|\|\|\|\|\|\|\|\|\|\|]PAM | NA | Intergenic | NA | NA |
| T8206 | CGTGCCCATCATCATCTGTCGGG | 1000 | | | | | | | | | |
| Oligo fwd | CACCCGTGCCCATCATCATCTGTC | | | | | | | | | | |
| Oligo rev | AAACGACAGATGATGATGGGCACG | | | | | | | | | | |
| Chromosome | start | end | strand | MM | target_seq | PAM | alignment | distance | position | gene name | gene id |
| GU305914.1 | 111818 | 111840 | + | 0 | CGTGCCCATCATCATCTGTC | GGG | \|\|\|\|\|\|\|[\|\|\|\|\|\|\|\|\|\|\|\|\|]PAM | NA | Intergenic | NA | NA |
| T8207 | TGTCGGGTCGTCATCTCCTTCGG | 1000 | | | | | | | | | |
| Oligo fwd | CACCTGTCGGGTCGTCATCTCCTT | | | | | | | | | | |
| Oligo rev | AAACAAGGAGATGACGACCCGACA | | | | | | | | | | |
| Chromosome | start | end | strand | MM | target_seq | PAM | alignment | distance | position | gene name | gene id |
| GU305914.1 | 111834 | 111856 | + | 0 | TGTCGGGTCGTCATCTCCTT | CGG | \|\|\|\|\|\|\|[\|\|\|\|\|\|\|\|\|\|\|\|\|]PAM | NA | Intergenic | NA | NA |
| T8208 | GTCGGGTCGTCATCTCCTTCGGG | 1000 | | | | | | | | | |
| Oligo fwd | CACCGTCGGGTCGTCATCTCCTTC | | | | | | | | | | |
| Oligo rev | AAACGAAGGAGATGACGACCCGAC | | | | | | | | | | |
| Chromosome | start | end | strand | MM | target_seq | PAM | alignment | distance | position | gene name | gene id |
| GU305914.1 | 111835 | 111857 | + | 0 | GTCGGGTCGTCATCTCCTTC | GGG | \|\|\|\|\|\|\|[\|\|\|\|\|\|\|\|\|\|\|\|\|]PAM | NA | Intergenic | NA | NA |
| GU305914.1 | 112334 | 112356 | - | 4 | GTAGGGACGTCATCTTCTGC | TGG | \|\|-\|\|\|-\|[\|\|\|\|\|\|\|-\|\|-\|]PAM | NA | Intergenic | NA | NA |

<table>
<tr><td>T8209</td><td>TCGGGTCGTCATCTCCTTCGGGG</td><td>1000</td><td></td><td></td><td></td><td></td><td></td><td></td><td></td><td></td><td></td></tr>
<tr><td>Oligo fwd</td><td>CACCTCGGGTCGTCATCTCCTTCG</td><td></td><td></td><td></td><td></td><td></td><td></td><td></td><td></td><td></td><td></td></tr>
<tr><td>Oligo rev</td><td>AAACCGAAGGAGATGACGACCCGA</td><td></td><td></td><td></td><td></td><td></td><td></td><td></td><td></td><td></td><td></td></tr>
<tr><td>Chromosome</td><td>start</td><td>end</td><td>strand</td><td>MM</td><td>target_seq</td><td>PAM</td><td>alignment</td><td>distance</td><td>position</td><td>gene name</td><td>gene id</td></tr>
<tr><td>GU305914.1</td><td>111836</td><td>111858</td><td>+</td><td>0</td><td>TCGGGTCGTCATCTCCTTCG</td><td>GGG</td><td>||||||||[||||||||||||]PAM</td><td>NA</td><td>Intergenic</td><td>NA</td><td>NA</td></tr>
</table>

<table>
<tr><td>T8210</td><td>GTCATCTCCTTCGGGGAGATCGG</td><td>1000</td><td></td><td></td><td></td><td></td><td></td><td></td><td></td><td></td><td></td></tr>
<tr><td>Oligo fwd</td><td>CACCGTCATCTCCTTCGGGGAGAT</td><td></td><td></td><td></td><td></td><td></td><td></td><td></td><td></td><td></td><td></td></tr>
<tr><td>Oligo rev</td><td>AAACATCTCCCCGAAGGAGATGAC</td><td></td><td></td><td></td><td></td><td></td><td></td><td></td><td></td><td></td><td></td></tr>
<tr><td>Chromosome</td><td>start</td><td>end</td><td>strand</td><td>MM</td><td>target_seq</td><td>PAM</td><td>alignment</td><td>distance</td><td>position</td><td>gene name</td><td>gene id</td></tr>
<tr><td>GU305914.1</td><td>111843</td><td>111865</td><td>+</td><td>0</td><td>GTCATCTCCTTCGGGGAGAT</td><td>CGG</td><td>||||||||[||||||||||||]PAM</td><td>NA</td><td>Intergenic</td><td>NA</td><td>NA</td></tr>
</table>

<table>
<tr><td>T8211</td><td>GTCGATCTCATCTCTACCGCCGG</td><td>1000</td><td></td><td></td><td></td><td></td><td></td><td></td><td></td><td></td><td></td></tr>
<tr><td>Oligo fwd</td><td>CACCGTCGATCTCATCTCTACCGC</td><td></td><td></td><td></td><td></td><td></td><td></td><td></td><td></td><td></td><td></td></tr>
<tr><td>Oligo rev</td><td>AAACGCGGTAGAGATGAGATCGAC</td><td></td><td></td><td></td><td></td><td></td><td></td><td></td><td></td><td></td><td></td></tr>
<tr><td>Chromosome</td><td>start</td><td>end</td><td>strand</td><td>MM</td><td>target_seq</td><td>PAM</td><td>alignment</td><td>distance</td><td>position</td><td>gene name</td><td>gene id</td></tr>
<tr><td>GU305914.1</td><td>111897</td><td>111919</td><td>+</td><td>0</td><td>GTCGATCTCATCTCTACCGC</td><td>CGG</td><td>||||||||[||||||||||||]PAM</td><td>NA</td><td>Intergenic</td><td>NA</td><td>NA</td></tr>
</table>

<table>
<tr><td>T8212</td><td>ATCTCATCTCTACCGCCGGCTGG</td><td>1000</td><td></td><td></td><td></td><td></td><td></td><td></td><td></td><td></td><td></td></tr>
<tr><td>Oligo fwd</td><td>CACCATCTCATCTCTACCGCCGGC</td><td></td><td></td><td></td><td></td><td></td><td></td><td></td><td></td><td></td><td></td></tr>
<tr><td>Oligo rev</td><td>AAACGCCGGCGGTAGAGATGAGAT</td><td></td><td></td><td></td><td></td><td></td><td></td><td></td><td></td><td></td><td></td></tr>
<tr><td>Chromosome</td><td>start</td><td>end</td><td>strand</td><td>MM</td><td>target_seq</td><td>PAM</td><td>alignment</td><td>distance</td><td>position</td><td>gene name</td><td>gene id</td></tr>
<tr><td>GU305914.1</td><td>111901</td><td>111923</td><td>+</td><td>0</td><td>ATCTCATCTCTACCGCCGGC</td><td>TGG</td><td>||||||||[||||||||||||]PAM</td><td>NA</td><td>Intergenic</td><td>NA</td><td>NA</td></tr>
</table>

<table>
<tr><td>T8213</td><td>CATCTCTACCGCCGGCTGGCAGG</td><td>1000</td><td></td><td></td><td></td><td></td><td></td><td></td><td></td><td></td><td></td></tr>
<tr><td>Oligo fwd</td><td>CACCCATCTCTACCGCCGGCTGGC</td><td></td><td></td><td></td><td></td><td></td><td></td><td></td><td></td><td></td><td></td></tr>
<tr><td>Oligo rev</td><td>AAACGCCAGCCGGCGGTAGAGATG</td><td></td><td></td><td></td><td></td><td></td><td></td><td></td><td></td><td></td><td></td></tr>
<tr><td>Chromosome</td><td>start</td><td>end</td><td>strand</td><td>MM</td><td>target_seq</td><td>PAM</td><td>alignment</td><td>distance</td><td>position</td><td>gene name</td><td>gene id</td></tr>
<tr><td>GU305914.1</td><td>111905</td><td>111927</td><td>+</td><td>0</td><td>CATCTCTACCGCCGGCTGGC</td><td>AGG</td><td>||||||||[||||||||||||]PAM</td><td>NA</td><td>Intergenic</td><td>NA</td><td>NA</td></tr>
</table>

<table>
<tr><td>T8214</td><td>ATCTCTACCGCCGGCTGGCAGGG</td><td>1000</td><td></td><td></td><td></td><td></td><td></td><td></td><td></td><td></td><td></td></tr>
<tr><td>Oligo fwd</td><td>CACCATCTCTACCGCCGGCTGGCA</td><td></td><td></td><td></td><td></td><td></td><td></td><td></td><td></td><td></td><td></td></tr>
<tr><td>Oligo rev</td><td>AAACTGCCAGCCGGCGGTAGAGAT</td><td></td><td></td><td></td><td></td><td></td><td></td><td></td><td></td><td></td><td></td></tr>
<tr><td>Chromosome</td><td>start</td><td>end</td><td>strand</td><td>MM</td><td>target_seq</td><td>PAM</td><td>alignment</td><td>distance</td><td>position</td><td>gene name</td><td>gene id</td></tr>
<tr><td>GU305914.1</td><td>111906</td><td>111928</td><td>+</td><td>0</td><td>ATCTCTACCGCCGGCTGGCA</td><td>GGG</td><td>||||||||[||||||||||||]PAM</td><td>NA</td><td>Intergenic</td><td>NA</td><td>NA</td></tr>
</table>

<table>
<tr><td>T8215</td><td>CTACCGCCGGCTGGCAGGGAAGG</td><td>1000</td><td></td><td></td><td></td><td></td><td></td><td></td><td></td><td></td><td></td></tr>
<tr><td>Oligo fwd</td><td>CACCCTACCGCCGGCTGGCAGGGA</td><td></td><td></td><td></td><td></td><td></td><td></td><td></td><td></td><td></td><td></td></tr>
<tr><td>Oligo rev</td><td>AAACTCCCTGCCAGCCGGCGGTAG</td><td></td><td></td><td></td><td></td><td></td><td></td><td></td><td></td><td></td><td></td></tr>
<tr><td>Chromosome</td><td>start</td><td>end</td><td>strand</td><td>MM</td><td>target_seq</td><td>PAM</td><td>alignment</td><td>distance</td><td>position</td><td>gene name</td><td>gene id</td></tr>
<tr><td>GU305914.1</td><td>111910</td><td>111932</td><td>+</td><td>0</td><td>CTACCGCCGGCTGGCAGGGA</td><td>AGG</td><td>||||||||[||||||||||||]PAM</td><td>NA</td><td>Intergenic</td><td>NA</td><td>NA</td></tr>
</table>

<table>
<tr><td>T8216</td><td>TACCGCCGGCTGGCAGGGAAGGG</td><td>1000</td><td></td><td></td><td></td><td></td><td></td><td></td><td></td><td></td><td></td></tr>
<tr><td>Oligo fwd</td><td>CACCTACCGCCGGCTGGCAGGGAA</td><td></td><td></td><td></td><td></td><td></td><td></td><td></td><td></td><td></td><td></td></tr>
<tr><td>Oligo rev</td><td>AAACTTCCCTGCCAGCCGGCGGTA</td><td></td><td></td><td></td><td></td><td></td><td></td><td></td><td></td><td></td><td></td></tr>
<tr><td>Chromosome</td><td>start</td><td>end</td><td>strand</td><td>MM</td><td>target_seq</td><td>PAM</td><td>alignment</td><td>distance</td><td>position</td><td>gene name</td><td>gene id</td></tr>
<tr><td>GU305914.1</td><td>111911</td><td>111933</td><td>+</td><td>0</td><td>TACCGCCGGCTGGCAGGGAA</td><td>GGG</td><td>||||||||[||||||||||||]PAM</td><td>NA</td><td>Intergenic</td><td>NA</td><td>NA</td></tr>
</table>

<table>
<tr><td>T8217</td><td>ACCGCCGGCTGGCAGGGAAGGGG</td><td>1000</td><td></td><td></td><td></td><td></td><td></td><td></td><td></td><td></td><td></td></tr>
<tr><td>Oligo fwd</td><td>CACCACCGCCGGCTGGCAGGGAAG</td><td></td><td></td><td></td><td></td><td></td><td></td><td></td><td></td><td></td><td></td></tr>
<tr><td>Oligo rev</td><td>AAACCTTCCCTGCCAGCCGGCGGT</td><td></td><td></td><td></td><td></td><td></td><td></td><td></td><td></td><td></td><td></td></tr>
<tr><td>Chromosome</td><td>start</td><td>end</td><td>strand</td><td>MM</td><td>target_seq</td><td>PAM</td><td>alignment</td><td>distance</td><td>position</td><td>gene name</td><td>gene id</td></tr>
<tr><td>GU305914.1</td><td>111912</td><td>111934</td><td>+</td><td>0</td><td>ACCGCCGGCTGGCAGGGAAG</td><td>GGG</td><td>||||||||[||||||||||||]PAM</td><td>NA</td><td>Intergenic</td><td>NA</td><td>NA</td></tr>
</table>

<table>
<tr><td>T8218</td><td>CGTCGCCTTCGCCTGCGACGTGG</td><td>1000</td><td></td><td></td><td></td><td></td><td></td><td></td><td></td><td></td><td></td></tr>
<tr><td>Oligo fwd</td><td>CACCCGTCGCCTTCGCCTGCGACG</td><td></td><td></td><td></td><td></td><td></td><td></td><td></td><td></td><td></td><td></td></tr>
<tr><td>Oligo rev</td><td>AAACCGTCGCAGGCGAAGGCGACG</td><td></td><td></td><td></td><td></td><td></td><td></td><td></td><td></td><td></td><td></td></tr>
<tr><td>Chromosome</td><td>start</td><td>end</td><td>strand</td><td>MM</td><td>target_seq</td><td>PAM</td><td>alignment</td><td>distance</td><td>position</td><td>gene name</td><td>gene id</td></tr>
<tr><td>GU305914.1</td><td>111941</td><td>111963</td><td>+</td><td>0</td><td>CGTCGCCTTCGCCTGCGACG</td><td>TGG</td><td>||||||||[||||||||||||]PAM</td><td>NA</td><td>Intergenic</td><td>NA</td><td>NA</td></tr>
<tr><td>GU305914.1</td><td>29689</td><td>29711</td><td>-</td><td>4</td><td>CGTCCTCTTCGCCGGCGAGG</td><td>AGG</td><td>||||--||[|||||-||||-|]PAM</td><td>NA</td><td>Intergenic</td><td>NA</td><td>NA</td></tr>
</table>

<table>
<tr><td>T8219</td><td>CGTGGTCGTCAAGTACGTCGAGG</td><td>1000</td><td></td><td></td><td></td><td></td><td></td><td></td><td></td><td></td><td></td></tr>
<tr><td>Oligo fwd</td><td>CACCCGTGGTCGTCAAGTACGTCG</td><td></td><td></td><td></td><td></td><td></td><td></td><td></td><td></td><td></td><td></td></tr>
<tr><td>Oligo rev</td><td>AAACCGACGTACTTGACGACCACG</td><td></td><td></td><td></td><td></td><td></td><td></td><td></td><td></td><td></td><td></td></tr>
<tr><td>Chromosome</td><td>start</td><td>end</td><td>strand</td><td>MM</td><td>target_seq</td><td>PAM</td><td>alignment</td><td>distance</td><td>position</td><td>gene name</td><td>gene id</td></tr>
<tr><td>GU305914.1</td><td>111959</td><td>111981</td><td>+</td><td>0</td><td>CGTGGTCGTCAAGTACGTCG</td><td>AGG</td><td>||||||||[||||||||||||]PAM</td><td>NA</td><td>Intergenic</td><td>NA</td><td>NA</td></tr>
</table>

<table>
<tr><td>T8220</td><td>GTCAAGTACGTCGAGGCCGCCGG</td><td>1000</td><td></td><td></td><td></td><td></td><td></td><td></td><td></td><td></td><td></td></tr>
<tr><td>Oligo fwd</td><td>CACCGTCAAGTACGTCGAGGCCGC</td><td></td><td></td><td></td><td></td><td></td><td></td><td></td><td></td><td></td><td></td></tr>
<tr><td>Oligo rev</td><td>AAACGCGGCCTCGACGTACTTGAC</td><td></td><td></td><td></td><td></td><td></td><td></td><td></td><td></td><td></td><td></td></tr>
<tr><td>Chromosome</td><td>start</td><td>end</td><td>strand</td><td>MM</td><td>target_seq</td><td>PAM</td><td>alignment</td><td>distance</td><td>position</td><td>gene name</td><td>gene id</td></tr>
<tr><td>GU305914.1</td><td>111966</td><td>111988</td><td>+</td><td>0</td><td>GTCAAGTACGTCGAGGCCGC</td><td>CGG</td><td>||||||||[||||||||||||]PAM</td><td>NA</td><td>Intergenic</td><td>NA</td><td>NA</td></tr>
</table>

<table>
<tr><td>T8221</td><td>GAGCTCCATGCGTCCGTCGACGG</td><td>1000</td><td></td><td></td><td></td><td></td><td></td><td></td><td></td><td></td><td></td></tr>
<tr><td>Oligo fwd</td><td>CACCGAGCTCCATGCGTCCGTCGA</td><td></td><td></td><td></td><td></td><td></td><td></td><td></td><td></td><td></td><td></td></tr>
<tr><td>Oligo rev</td><td>AAACTCGACGGACGCATGGAGCTC</td><td></td><td></td><td></td><td></td><td></td><td></td><td></td><td></td><td></td><td></td></tr>
<tr><td>Chromosome</td><td>start</td><td>end</td><td>strand</td><td>MM</td><td>target_seq</td><td>PAM</td><td>alignment</td><td>distance</td><td>position</td><td>gene name</td><td>gene id</td></tr>
<tr><td>GU305914.1</td><td>111990</td><td>112012</td><td>+</td><td>0</td><td>GAGCTCCATGCGTCCGTCGA</td><td>CGG</td><td>||||||||[||||||||||||]PAM</td><td>NA</td><td>Intergenic</td><td>NA</td><td>NA</td></tr>
</table>

<table>
<tr><td>T8222</td><td>TCGACGGCTCCGATGACCGCCGG</td><td>1000</td><td></td><td></td><td></td><td></td><td></td><td></td><td></td><td></td><td></td></tr>
<tr><td>Oligo fwd</td><td>CACCTCGACGGCTCCGATGACCGC</td><td></td><td></td><td></td><td></td><td></td><td></td><td></td><td></td><td></td><td></td></tr>
<tr><td>Oligo rev</td><td>AAACGCGGTCATCGGAGCCGTCGA</td><td></td><td></td><td></td><td></td><td></td><td></td><td></td><td></td><td></td><td></td></tr>
<tr><td>Chromosome</td><td>start</td><td>end</td><td>strand</td><td>MM</td><td>target_seq</td><td>PAM</td><td>alignment</td><td>distance</td><td>position</td><td>gene name</td><td>gene id</td></tr>
<tr><td>GU305914.1</td><td>112006</td><td>112028</td><td>+</td><td>0</td><td>TCGACGGCTCCGATGACCGC</td><td>CGG</td><td>||||||||[||||||||||||]PAM</td><td>NA</td><td>Intergenic</td><td>NA</td><td>NA</td></tr>
</table>

<table>
<tr><td>T8223</td><td>TGACCGCCGGTCTTTGTGCGCGG</td><td>1000</td></tr>
<tr><td>Oligo fwd</td><td>CACCTGACCGCCGGTCTTTGTGCG</td><td></td></tr>
<tr><td>Oligo rev</td><td>AAACCGCACAAAGACCGGCGGTCA</td><td></td></tr>
</table>

<table>
<tr><td>Chromosome</td><td>start</td><td>end</td><td>strand</td><td>MM</td><td>target_seq</td><td>PAM</td><td>alignment</td><td>distance</td><td>position</td><td>gene name</td><td>gene id</td></tr>
<tr><td>GU305914.1</td><td>112019</td><td>112041</td><td>+</td><td>0</td><td>TGACCGCCGGTCTTTGTGCG</td><td>CGG</td><td>||||||||[||||||||||||]PAM</td><td>NA</td><td>Intergenic</td><td>NA</td><td>NA</td></tr>
<tr><td>T8224</td><td>TCTCCCCTCCTGCATCTACCCGG</td><td>1000</td><td></td><td></td><td></td><td></td><td></td><td></td><td></td><td></td><td></td></tr>
<tr><td>Oligo fwd</td><td>CACCTCTCCCCTCCTGCATCTACC</td><td></td><td></td><td></td><td></td><td></td><td></td><td></td><td></td><td></td><td></td></tr>
<tr><td>Oligo rev</td><td>AAACGGTAGATGCAGGAGGGGAGA</td><td></td><td></td><td></td><td></td><td></td><td></td><td></td><td></td><td></td><td></td></tr>
<tr><td>Chromosome</td><td>start</td><td>end</td><td>strand</td><td>MM</td><td>target_seq</td><td>PAM</td><td>alignment</td><td>distance</td><td>position</td><td>gene name</td><td>gene id</td></tr>
<tr><td>GU305914.1</td><td>112043</td><td>112065</td><td>+</td><td>0</td><td>TCTCCCCTCCTGCATCTACC</td><td>CGG</td><td>||||||||[||||||||||||]PAM</td><td>NA</td><td>Intergenic</td><td>NA</td><td>NA</td></tr>
<tr><td>T8225</td><td>CCCTCCTGCATCTACCCGGTCGG</td><td>1000</td><td></td><td></td><td></td><td></td><td></td><td></td><td></td><td></td><td></td></tr>
<tr><td>Oligo fwd</td><td>CACCCCCTCCTGCATCTACCCGGT</td><td></td><td></td><td></td><td></td><td></td><td></td><td></td><td></td><td></td><td></td></tr>
<tr><td>Oligo rev</td><td>AAACACCGGGTAGATGCAGGAGGG</td><td></td><td></td><td></td><td></td><td></td><td></td><td></td><td></td><td></td><td></td></tr>
<tr><td>Chromosome</td><td>start</td><td>end</td><td>strand</td><td>MM</td><td>target_seq</td><td>PAM</td><td>alignment</td><td>distance</td><td>position</td><td>gene name</td><td>gene id</td></tr>
<tr><td>GU305914.1</td><td>112047</td><td>112069</td><td>+</td><td>0</td><td>CCCTCCTGCATCTACCCGGT</td><td>CGG</td><td>||||||||[||||||||||||]PAM</td><td>NA</td><td>Intergenic</td><td>NA</td><td>NA</td></tr>
<tr><td>T8226</td><td>ACCCTCGTCAAGAGCACTTTCGG</td><td>1000</td><td></td><td></td><td></td><td></td><td></td><td></td><td></td><td></td><td></td></tr>
<tr><td>Oligo fwd</td><td>CACCACCCTCGTCAAGAGCACTTT</td><td></td><td></td><td></td><td></td><td></td><td></td><td></td><td></td><td></td><td></td></tr>
<tr><td>Oligo rev</td><td>AAACAAAGTGCTCTTGACGAGGGT</td><td></td><td></td><td></td><td></td><td></td><td></td><td></td><td></td><td></td><td></td></tr>
<tr><td>Chromosome</td><td>start</td><td>end</td><td>strand</td><td>MM</td><td>target_seq</td><td>PAM</td><td>alignment</td><td>distance</td><td>position</td><td>gene name</td><td>gene id</td></tr>
<tr><td>GU305914.1</td><td>112074</td><td>112096</td><td>+</td><td>0</td><td>ACCCTCGTCAAGAGCACTTT</td><td>CGG</td><td>||||||||[||||||||||||]PAM</td><td>NA</td><td>Intergenic</td><td>NA</td><td>NA</td></tr>
<tr><td>T8227</td><td>CACTTTCGGTTTCTTCCTCAAGG</td><td>1000</td><td></td><td></td><td></td><td></td><td></td><td></td><td></td><td></td><td></td></tr>
<tr><td>Oligo fwd</td><td>CACCCACTTTCGGTTTCTTCCTCA</td><td></td><td></td><td></td><td></td><td></td><td></td><td></td><td></td><td></td><td></td></tr>
<tr><td>Oligo rev</td><td>AAACTGAGGAAGAAACCGAAAGTG</td><td></td><td></td><td></td><td></td><td></td><td></td><td></td><td></td><td></td><td></td></tr>
<tr><td>Chromosome</td><td>start</td><td>end</td><td>strand</td><td>MM</td><td>target_seq</td><td>PAM</td><td>alignment</td><td>distance</td><td>position</td><td>gene name</td><td>gene id</td></tr>
<tr><td>GU305914.1</td><td>112088</td><td>112110</td><td>+</td><td>0</td><td>CACTTTCGGTTTCTTCCTCA</td><td>AGG</td><td>||||||||[||||||||||||]PAM</td><td>NA</td><td>Intergenic</td><td>NA</td><td>NA</td></tr>
<tr><td>T8228</td><td>ACTTTCGGTTTCTTCCTCAAGGG</td><td>1000</td><td></td><td></td><td></td><td></td><td></td><td></td><td></td><td></td><td></td></tr>
<tr><td>Oligo fwd</td><td>CACCACTTTCGGTTTCTTCCTCAA</td><td></td><td></td><td></td><td></td><td></td><td></td><td></td><td></td><td></td><td></td></tr>
<tr><td>Oligo rev</td><td>AAACTTGAGGAAGAAACCGAAAGT</td><td></td><td></td><td></td><td></td><td></td><td></td><td></td><td></td><td></td><td></td></tr>
<tr><td>Chromosome</td><td>start</td><td>end</td><td>strand</td><td>MM</td><td>target_seq</td><td>PAM</td><td>alignment</td><td>distance</td><td>position</td><td>gene name</td><td>gene id</td></tr>
<tr><td>GU305914.1</td><td>112089</td><td>112111</td><td>+</td><td>0</td><td>ACTTTCGGTTTCTTCCTCAA</td><td>GGG</td><td>||||||||[||||||||||||]PAM</td><td>NA</td><td>Intergenic</td><td>NA</td><td>NA</td></tr>
<tr><td>T8229</td><td>AGGGCGCCAAGACGATGCTGAGG</td><td>1000</td><td></td><td></td><td></td><td></td><td></td><td></td><td></td><td></td><td></td></tr>
<tr><td>Oligo fwd</td><td>CACCAGGGCGCCAAGACGATGCTG</td><td></td><td></td><td></td><td></td><td></td><td></td><td></td><td></td><td></td><td></td></tr>
<tr><td>Oligo rev</td><td>AAACCAGCATCGTCTTGGCGCCCT</td><td></td><td></td><td></td><td></td><td></td><td></td><td></td><td></td><td></td><td></td></tr>
<tr><td>Chromosome</td><td>start</td><td>end</td><td>strand</td><td>MM</td><td>target_seq</td><td>PAM</td><td>alignment</td><td>distance</td><td>position</td><td>gene name</td><td>gene id</td></tr>
<tr><td>GU305914.1</td><td>112108</td><td>112130</td><td>+</td><td>0</td><td>AGGGCGCCAAGACGATGCTG</td><td>AGG</td><td>||||||||[||||||||||||]PAM</td><td>NA</td><td>Intergenic</td><td>NA</td><td>NA</td></tr>
<tr><td>T8230</td><td>AGACGATGCTGAGGCTGTGTCGG</td><td>1000</td><td></td><td></td><td></td><td></td><td></td><td></td><td></td><td></td><td></td></tr>
<tr><td>Oligo fwd</td><td>CACCAGACGATGCTGAGGCTGTGT</td><td></td><td></td><td></td><td></td><td></td><td></td><td></td><td></td><td></td><td></td></tr>
<tr><td>Oligo rev</td><td>AAACACACAGCCTCAGCATCGTCT</td><td></td><td></td><td></td><td></td><td></td><td></td><td></td><td></td><td></td><td></td></tr>
<tr><td>Chromosome</td><td>start</td><td>end</td><td>strand</td><td>MM</td><td>target_seq</td><td>PAM</td><td>alignment</td><td>distance</td><td>position</td><td>gene name</td><td>gene id</td></tr>
<tr><td>GU305914.1</td><td>112117</td><td>112139</td><td>+</td><td>0</td><td>AGACGATGCTGAGGCTGTGT</td><td>CGG</td><td>||||||||[||||||||||||]PAM</td><td>NA</td><td>Intergenic</td><td>NA</td><td>NA</td></tr>
<tr><td>T8231</td><td>CGATGCTGAGGCTGTGTCGGAGG</td><td>1000</td><td></td><td></td><td></td><td></td><td></td><td></td><td></td><td></td><td></td></tr>
<tr><td>Oligo fwd</td><td>CACCCGATGCTGAGGCTGTGTCGG</td><td></td><td></td><td></td><td></td><td></td><td></td><td></td><td></td><td></td><td></td></tr>
<tr><td>Oligo rev</td><td>AAACCCGACACAGCCTCAGCATCG</td><td></td><td></td><td></td><td></td><td></td><td></td><td></td><td></td><td></td><td></td></tr>
<tr><td>Chromosome</td><td>start</td><td>end</td><td>strand</td><td>MM</td><td>target_seq</td><td>PAM</td><td>alignment</td><td>distance</td><td>position</td><td>gene name</td><td>gene id</td></tr>
<tr><td>GU305914.1</td><td>112120</td><td>112142</td><td>+</td><td>0</td><td>CGATGCTGAGGCTGTGTCGG</td><td>AGG</td><td>||||||||[||||||||||||]PAM</td><td>NA</td><td>Intergenic</td><td>NA</td><td>NA</td></tr>
<tr><td>T8232</td><td>GATGCTGAGGCTGTGTCGGAGGG</td><td>1000</td><td></td><td></td><td></td><td></td><td></td><td></td><td></td><td></td><td></td></tr>
<tr><td>Oligo fwd</td><td>CACCGATGCTGAGGCTGTGTCGGA</td><td></td><td></td><td></td><td></td><td></td><td></td><td></td><td></td><td></td><td></td></tr>
<tr><td>Oligo rev</td><td>AAACTCCGACACAGCCTCAGCATC</td><td></td><td></td><td></td><td></td><td></td><td></td><td></td><td></td><td></td><td></td></tr>
<tr><td>Chromosome</td><td>start</td><td>end</td><td>strand</td><td>MM</td><td>target_seq</td><td>PAM</td><td>alignment</td><td>distance</td><td>position</td><td>gene name</td><td>gene id</td></tr>
<tr><td>GU305914.1</td><td>112121</td><td>112143</td><td>+</td><td>0</td><td>GATGCTGAGGCTGTGTCGGA</td><td>GGG</td><td>||||||||[||||||||||||]PAM</td><td>NA</td><td>Intergenic</td><td>NA</td><td>NA</td></tr>
<tr><td>T8233</td><td>TGTCGGAGGGTCTCGCCTTTCGG</td><td>1000</td><td></td><td></td><td></td><td></td><td></td><td></td><td></td><td></td><td></td></tr>
<tr><td>Oligo fwd</td><td>CACCTGTCGGAGGGTCTCGCCTTT</td><td></td><td></td><td></td><td></td><td></td><td></td><td></td><td></td><td></td><td></td></tr>
<tr><td>Oligo rev</td><td>AAACAAAGGCGAGACCCTCCGACA</td><td></td><td></td><td></td><td></td><td></td><td></td><td></td><td></td><td></td><td></td></tr>
<tr><td>Chromosome</td><td>start</td><td>end</td><td>strand</td><td>MM</td><td>target_seq</td><td>PAM</td><td>alignment</td><td>distance</td><td>position</td><td>gene name</td><td>gene id</td></tr>
<tr><td>GU305914.1</td><td>112134</td><td>112156</td><td>+</td><td>0</td><td>TGTCGGAGGGTCTCGCCTTT</td><td>CGG</td><td>||||||||[||||||||||||]PAM</td><td>NA</td><td>Intergenic</td><td>NA</td><td>NA</td></tr>
<tr><td>T8234</td><td>GTCGGAGGGTCTCGCCTTTCGGG</td><td>1000</td><td></td><td></td><td></td><td></td><td></td><td></td><td></td><td></td><td></td></tr>
<tr><td>Oligo fwd</td><td>CACCGTCGGAGGGTCTCGCCTTTC</td><td></td><td></td><td></td><td></td><td></td><td></td><td></td><td></td><td></td><td></td></tr>
<tr><td>Oligo rev</td><td>AAACGAAAGGCGAGACCCTCCGAC</td><td></td><td></td><td></td><td></td><td></td><td></td><td></td><td></td><td></td><td></td></tr>
<tr><td>Chromosome</td><td>start</td><td>end</td><td>strand</td><td>MM</td><td>target_seq</td><td>PAM</td><td>alignment</td><td>distance</td><td>position</td><td>gene name</td><td>gene id</td></tr>
<tr><td>GU305914.1</td><td>112135</td><td>112157</td><td>+</td><td>0</td><td>GTCGGAGGGTCTCGCCTTTC</td><td>GGG</td><td>||||||||[||||||||||||]PAM</td><td>NA</td><td>Intergenic</td><td>NA</td><td>NA</td></tr>
<tr><td>T8235</td><td>CTCGCCTTTCGGGATCCAGAAGG</td><td>1000</td><td></td><td></td><td></td><td></td><td></td><td></td><td></td><td></td><td></td></tr>
<tr><td>Oligo fwd</td><td>CACCCTCGCCTTTCGGGATCCAGA</td><td></td><td></td><td></td><td></td><td></td><td></td><td></td><td></td><td></td><td></td></tr>
<tr><td>Oligo rev</td><td>AAACTCTGGATCCCGAAAGGCGAG</td><td></td><td></td><td></td><td></td><td></td><td></td><td></td><td></td><td></td><td></td></tr>
<tr><td>Chromosome</td><td>start</td><td>end</td><td>strand</td><td>MM</td><td>target_seq</td><td>PAM</td><td>alignment</td><td>distance</td><td>position</td><td>gene name</td><td>gene id</td></tr>
<tr><td>GU305914.1</td><td>112145</td><td>112167</td><td>+</td><td>0</td><td>CTCGCCTTTCGGGATCCAGA</td><td>AGG</td><td>||||||||[||||||||||||]PAM</td><td>NA</td><td>Intergenic</td><td>NA</td><td>NA</td></tr>
<tr><td>T8236</td><td>GAAGGACGCCCTCAAGTCACCGG</td><td>1000</td><td></td><td></td><td></td><td></td><td></td><td></td><td></td><td></td><td></td></tr>
<tr><td>Oligo fwd</td><td>CACCGAAGGACGCCCTCAAGTCAC</td><td></td><td></td><td></td><td></td><td></td><td></td><td></td><td></td><td></td><td></td></tr>
<tr><td>Oligo rev</td><td>AAACGTGACTTGAGGGCGTCCTTC</td><td></td><td></td><td></td><td></td><td></td><td></td><td></td><td></td><td></td><td></td></tr>
<tr><td>Chromosome</td><td>start</td><td>end</td><td>strand</td><td>MM</td><td>target_seq</td><td>PAM</td><td>alignment</td><td>distance</td><td>position</td><td>gene name</td><td>gene id</td></tr>
<tr><td>GU305914.1</td><td>112163</td><td>112185</td><td>+</td><td>0</td><td>GAAGGACGCCCTCAAGTCAC</td><td>CGG</td><td>||||||||[||||||||||||]PAM</td><td>NA</td><td>Intergenic</td><td>NA</td><td>NA</td></tr>
<tr><td>T8237</td><td>CAAGTCACCGGCCGCGATGAAGG</td><td>1000</td><td></td><td></td><td></td><td></td><td></td><td></td><td></td><td></td><td></td></tr>
<tr><td>Oligo fwd</td><td>CACCCAAGTCACCGGCCGCGATGA</td><td></td><td></td><td></td><td></td><td></td><td></td><td></td><td></td><td></td><td></td></tr>
<tr><td>Oligo rev</td><td>AAACTCATCGCGGCCGGTGACTTG</td><td></td><td></td><td></td><td></td><td></td><td></td><td></td><td></td><td></td><td></td></tr>
<tr><td>Chromosome</td><td>start</td><td>end</td><td>strand</td><td>MM</td><td>target_seq</td><td>PAM</td><td>alignment</td><td>distance</td><td>position</td><td>gene name</td><td>gene id</td></tr>
<tr><td>GU305914.1</td><td>112175</td><td>112197</td><td>+</td><td>0</td><td>CAAGTCACCGGCCGCGATGA</td><td>AGG</td><td>||||||||[||||||||||||]PAM</td><td>NA</td><td>Intergenic</td><td>NA</td><td>NA</td></tr>
<tr><td>GU305914.1</td><td>17547</td><td>17569</td><td>-</td><td>4</td><td>CAAATCGCCCGCTGCGATGA</td><td>CGG</td><td>|||-||-|[|-||-|||||||]PAM</td><td>NA</td><td>Intergenic</td><td>NA</td><td>NA</td></tr>
<tr><td>T8238</td><td>GGCCGCGATGAAGGCGCTCGAGG</td><td>1000</td><td></td><td></td><td></td><td></td><td></td><td></td><td></td><td></td><td></td></tr>
</table>

<table>
<tr><td>Oligo fwd</td><td>CACCGGCCGCGATGAAGGCGCTCG</td><td></td><td></td><td></td><td></td><td></td><td></td><td></td><td></td><td></td><td></td></tr>
<tr><td>Oligo rev</td><td>AAACCGAGCGCCTTCATCGCGGCC</td><td></td><td></td><td></td><td></td><td></td><td></td><td></td><td></td><td></td><td></td></tr>
<tr><td>Chromosome</td><td>start</td><td>end</td><td>strand</td><td>MM</td><td>target_seq</td><td>PAM</td><td>alignment</td><td>distance</td><td>position</td><td>gene name</td><td>gene id</td></tr>
<tr><td>GU305914.1</td><td>112184</td><td>112206</td><td>+</td><td>0</td><td>GGCCGCGATGAAGGCGCTCG</td><td>AGG</td><td>||||||||[||||||||||||]PAM</td><td>NA</td><td>Intergenic</td><td>NA</td><td>NA</td></tr>
<tr><td>T8239</td><td>GCCGCGATGAAGGCGCTCGAGGG</td><td>1000</td><td></td><td></td><td></td><td></td><td></td><td></td><td></td><td></td><td></td></tr>
<tr><td>Oligo fwd</td><td>CACCGCCGCGATGAAGGCGCTCGA</td><td></td><td></td><td></td><td></td><td></td><td></td><td></td><td></td><td></td><td></td></tr>
<tr><td>Oligo rev</td><td>AAACTCGAGCGCCTTCATCGCGGC</td><td></td><td></td><td></td><td></td><td></td><td></td><td></td><td></td><td></td><td></td></tr>
<tr><td>Chromosome</td><td>start</td><td>end</td><td>strand</td><td>MM</td><td>target_seq</td><td>PAM</td><td>alignment</td><td>distance</td><td>position</td><td>gene name</td><td>gene id</td></tr>
<tr><td>GU305914.1</td><td>112185</td><td>112207</td><td>+</td><td>0</td><td>GCCGCGATGAAGGCGCTCGA</td><td>GGG</td><td>||||||||[||||||||||||]PAM</td><td>NA</td><td>Intergenic</td><td>NA</td><td>NA</td></tr>
<tr><td>T8240</td><td>CCGCGATGAAGGCGCTCGAGGGG</td><td>1000</td><td></td><td></td><td></td><td></td><td></td><td></td><td></td><td></td><td></td></tr>
<tr><td>Oligo fwd</td><td>CACCCCGCGATGAAGGCGCTCGAG</td><td></td><td></td><td></td><td></td><td></td><td></td><td></td><td></td><td></td><td></td></tr>
<tr><td>Oligo rev</td><td>AAACCTCGAGCGCCTTCATCGCGG</td><td></td><td></td><td></td><td></td><td></td><td></td><td></td><td></td><td></td><td></td></tr>
<tr><td>Chromosome</td><td>start</td><td>end</td><td>strand</td><td>MM</td><td>target_seq</td><td>PAM</td><td>alignment</td><td>distance</td><td>position</td><td>gene name</td><td>gene id</td></tr>
<tr><td>GU305914.1</td><td>112186</td><td>112208</td><td>+</td><td>0</td><td>CCGCGATGAAGGCGCTCGAG</td><td>GGG</td><td>||||||||[||||||||||||]PAM</td><td>NA</td><td>Intergenic</td><td>NA</td><td>NA</td></tr>
<tr><td>T8241</td><td>CGCGATGAAGGCGCTCGAGGGGG</td><td>1000</td><td></td><td></td><td></td><td></td><td></td><td></td><td></td><td></td><td></td></tr>
<tr><td>Oligo fwd</td><td>CACCCGCGATGAAGGCGCTCGAGG</td><td></td><td></td><td></td><td></td><td></td><td></td><td></td><td></td><td></td><td></td></tr>
<tr><td>Oligo rev</td><td>AAACCCTCGAGCGCCTTCATCGCG</td><td></td><td></td><td></td><td></td><td></td><td></td><td></td><td></td><td></td><td></td></tr>
<tr><td>Chromosome</td><td>start</td><td>end</td><td>strand</td><td>MM</td><td>target_seq</td><td>PAM</td><td>alignment</td><td>distance</td><td>position</td><td>gene name</td><td>gene id</td></tr>
<tr><td>GU305914.1</td><td>112187</td><td>112209</td><td>+</td><td>0</td><td>CGCGATGAAGGCGCTCGAGG</td><td>GGG</td><td>||||||||[||||||||||||]PAM</td><td>NA</td><td>Intergenic</td><td>NA</td><td>NA</td></tr>
<tr><td>T8242</td><td>CAAGAGTTTCTCGCCCGACGAGG</td><td>1000</td><td></td><td></td><td></td><td></td><td></td><td></td><td></td><td></td><td></td></tr>
<tr><td>Oligo fwd</td><td>CACCCAAGAGTTTCTCGCCCGACG</td><td></td><td></td><td></td><td></td><td></td><td></td><td></td><td></td><td></td><td></td></tr>
<tr><td>Oligo rev</td><td>AAACCGTCGGGCGAGAAACTCTTG</td><td></td><td></td><td></td><td></td><td></td><td></td><td></td><td></td><td></td><td></td></tr>
<tr><td>Chromosome</td><td>start</td><td>end</td><td>strand</td><td>MM</td><td>target_seq</td><td>PAM</td><td>alignment</td><td>distance</td><td>position</td><td>gene name</td><td>gene id</td></tr>
<tr><td>GU305914.1</td><td>112223</td><td>112245</td><td>+</td><td>0</td><td>CAAGAGTTTCTCGCCCGACG</td><td>AGG</td><td>||||||||[||||||||||||]PAM</td><td>NA</td><td>Intergenic</td><td>NA</td><td>NA</td></tr>
<tr><td>T8243</td><td>CCCACCGTCTGTGAGAAGACCGG</td><td>1000</td><td></td><td></td><td></td><td></td><td></td><td></td><td></td><td></td><td></td></tr>
<tr><td>Oligo fwd</td><td>CACCCCCACCGTCTGTGAGAAGAC</td><td></td><td></td><td></td><td></td><td></td><td></td><td></td><td></td><td></td><td></td></tr>
<tr><td>Oligo rev</td><td>AAACGTCTTCTCACAGACGGTGGG</td><td></td><td></td><td></td><td></td><td></td><td></td><td></td><td></td><td></td><td></td></tr>
<tr><td>Chromosome</td><td>start</td><td>end</td><td>strand</td><td>MM</td><td>target_seq</td><td>PAM</td><td>alignment</td><td>distance</td><td>position</td><td>gene name</td><td>gene id</td></tr>
<tr><td>GU305914.1</td><td>112263</td><td>112285</td><td>+</td><td>0</td><td>CCCACCGTCTGTGAGAAGAC</td><td>CGG</td><td>||||||||[||||||||||||]PAM</td><td>NA</td><td>Intergenic</td><td>NA</td><td>NA</td></tr>
<tr><td>T8244</td><td>CGAGATCGCCATGACCCCTGTGG</td><td>1000</td><td></td><td></td><td></td><td></td><td></td><td></td><td></td><td></td><td></td></tr>
<tr><td>Oligo fwd</td><td>CACCCGAGATCGCCATGACCCCTG</td><td></td><td></td><td></td><td></td><td></td><td></td><td></td><td></td><td></td><td></td></tr>
<tr><td>Oligo rev</td><td>AAACCAGGGGTCATGGCGATCTCG</td><td></td><td></td><td></td><td></td><td></td><td></td><td></td><td></td><td></td><td></td></tr>
<tr><td>Chromosome</td><td>start</td><td>end</td><td>strand</td><td>MM</td><td>target_seq</td><td>PAM</td><td>alignment</td><td>distance</td><td>position</td><td>gene name</td><td>gene id</td></tr>
<tr><td>GU305914.1</td><td>112304</td><td>112326</td><td>+</td><td>0</td><td>CGAGATCGCCATGACCCCTG</td><td>TGG</td><td>||||||||[||||||||||||]PAM</td><td>NA</td><td>Intergenic</td><td>NA</td><td>NA</td></tr>
<tr><td>T8245</td><td>GATCGCCATGACCCCTGTGGTGG</td><td>1000</td><td></td><td></td><td></td><td></td><td></td><td></td><td></td><td></td><td></td></tr>
<tr><td>Oligo fwd</td><td>CACCGATCGCCATGACCCCTGTGG</td><td></td><td></td><td></td><td></td><td></td><td></td><td></td><td></td><td></td><td></td></tr>
<tr><td>Oligo rev</td><td>AAACCCACAGGGGTCATGGCGATC</td><td></td><td></td><td></td><td></td><td></td><td></td><td></td><td></td><td></td><td></td></tr>
<tr><td>Chromosome</td><td>start</td><td>end</td><td>strand</td><td>MM</td><td>target_seq</td><td>PAM</td><td>alignment</td><td>distance</td><td>position</td><td>gene name</td><td>gene id</td></tr>
<tr><td>GU305914.1</td><td>112307</td><td>112329</td><td>+</td><td>0</td><td>GATCGCCATGACCCCTGTGG</td><td>TGG</td><td>||||||||[||||||||||||]PAM</td><td>NA</td><td>Intergenic</td><td>NA</td><td>NA</td></tr>
<tr><td>T8246</td><td>ACCCATGTCTGTTTTTTACTCGG</td><td>1000</td><td></td><td></td><td></td><td></td><td></td><td></td><td></td><td></td><td></td></tr>
<tr><td>Oligo fwd</td><td>CACCACCCATGTCTGTTTTTTACT</td><td></td><td></td><td></td><td></td><td></td><td></td><td></td><td></td><td></td><td></td></tr>
<tr><td>Oligo rev</td><td>AAACAGTAAAAAACAGACATGGGT</td><td></td><td></td><td></td><td></td><td></td><td></td><td></td><td></td><td></td><td></td></tr>
<tr><td>Chromosome</td><td>start</td><td>end</td><td>strand</td><td>MM</td><td>target_seq</td><td>PAM</td><td>alignment</td><td>distance</td><td>position</td><td>gene name</td><td>gene id</td></tr>
<tr><td>GU305914.1</td><td>112355</td><td>112377</td><td>+</td><td>0</td><td>ACCCATGTCTGTTTTTTACT</td><td>CGG</td><td>||||||||[||||||||||||]PAM</td><td>NA</td><td>Intergenic</td><td>NA</td><td>NA</td></tr>
<tr><td>T8247</td><td>ATGTCTGTTTTTTACTCGGACGG</td><td>1000</td><td></td><td></td><td></td><td></td><td></td><td></td><td></td><td></td><td></td></tr>
<tr><td>Oligo fwd</td><td>CACCATGTCTGTTTTTTACTCGGA</td><td></td><td></td><td></td><td></td><td></td><td></td><td></td><td></td><td></td><td></td></tr>
<tr><td>Oligo rev</td><td>AAACTCCGAGTAAAAAACAGACAT</td><td></td><td></td><td></td><td></td><td></td><td></td><td></td><td></td><td></td><td></td></tr>
<tr><td>Chromosome</td><td>start</td><td>end</td><td>strand</td><td>MM</td><td>target_seq</td><td>PAM</td><td>alignment</td><td>distance</td><td>position</td><td>gene name</td><td>gene id</td></tr>
<tr><td>GU305914.1</td><td>112359</td><td>112381</td><td>+</td><td>0</td><td>ATGTCTGTTTTTTACTCGGA</td><td>CGG</td><td>||||||||[||||||||||||]PAM</td><td>NA</td><td>Intergenic</td><td>NA</td><td>NA</td></tr>
<tr><td>T8248</td><td>TCTGTTTTTTACTCGGACGGCGG</td><td>1000</td><td></td><td></td><td></td><td></td><td></td><td></td><td></td><td></td><td></td></tr>
<tr><td>Oligo fwd</td><td>CACCTCTGTTTTTTACTCGGACGG</td><td></td><td></td><td></td><td></td><td></td><td></td><td></td><td></td><td></td><td></td></tr>
<tr><td>Oligo rev</td><td>AAACCCGTCCGAGTAAAAAACAGA</td><td></td><td></td><td></td><td></td><td></td><td></td><td></td><td></td><td></td><td></td></tr>
<tr><td>Chromosome</td><td>start</td><td>end</td><td>strand</td><td>MM</td><td>target_seq</td><td>PAM</td><td>alignment</td><td>distance</td><td>position</td><td>gene name</td><td>gene id</td></tr>
<tr><td>GU305914.1</td><td>112362</td><td>112384</td><td>+</td><td>0</td><td>TCTGTTTTTTACTCGGACGG</td><td>CGG</td><td>||||||||[||||||||||||]PAM</td><td>NA</td><td>Intergenic</td><td>NA</td><td>NA</td></tr>
<tr><td>T8249</td><td>CTGTTTTTTACTCGGACGGCGGG</td><td>1000</td><td></td><td></td><td></td><td></td><td></td><td></td><td></td><td></td><td></td></tr>
<tr><td>Oligo fwd</td><td>CACCCTGTTTTTTACTCGGACGGC</td><td></td><td></td><td></td><td></td><td></td><td></td><td></td><td></td><td></td><td></td></tr>
<tr><td>Oligo rev</td><td>AAACGCCGTCCGAGTAAAAAACAG</td><td></td><td></td><td></td><td></td><td></td><td></td><td></td><td></td><td></td><td></td></tr>
<tr><td>Chromosome</td><td>start</td><td>end</td><td>strand</td><td>MM</td><td>target_seq</td><td>PAM</td><td>alignment</td><td>distance</td><td>position</td><td>gene name</td><td>gene id</td></tr>
<tr><td>GU305914.1</td><td>112363</td><td>112385</td><td>+</td><td>0</td><td>CTGTTTTTTACTCGGACGGC</td><td>GGG</td><td>||||||||[||||||||||||]PAM</td><td>NA</td><td>Intergenic</td><td>NA</td><td>NA</td></tr>
<tr><td>T8250</td><td>TGTTTTTTACTCGGACGGCGGGG</td><td>1000</td><td></td><td></td><td></td><td></td><td></td><td></td><td></td><td></td><td></td></tr>
<tr><td>Oligo fwd</td><td>CACCTGTTTTTTACTCGGACGGCG</td><td></td><td></td><td></td><td></td><td></td><td></td><td></td><td></td><td></td><td></td></tr>
<tr><td>Oligo rev</td><td>AAACCGCCGTCCGAGTAAAAAACA</td><td></td><td></td><td></td><td></td><td></td><td></td><td></td><td></td><td></td><td></td></tr>
<tr><td>Chromosome</td><td>start</td><td>end</td><td>strand</td><td>MM</td><td>target_seq</td><td>PAM</td><td>alignment</td><td>distance</td><td>position</td><td>gene name</td><td>gene id</td></tr>
<tr><td>GU305914.1</td><td>112364</td><td>112386</td><td>+</td><td>0</td><td>TGTTTTTTACTCGGACGGCG</td><td>GGG</td><td>||||||||[||||||||||||]PAM</td><td>NA</td><td>Intergenic</td><td>NA</td><td>NA</td></tr>
<tr><td>T8251</td><td>TTACTCGGACGGCGGGGAGAAGG</td><td>1000</td><td></td><td></td><td></td><td></td><td></td><td></td><td></td><td></td><td></td></tr>
<tr><td>Oligo fwd</td><td>CACCTTACTCGGACGGCGGGGAGA</td><td></td><td></td><td></td><td></td><td></td><td></td><td></td><td></td><td></td><td></td></tr>
<tr><td>Oligo rev</td><td>AAACTCTCCCCGCCGTCCGAGTAA</td><td></td><td></td><td></td><td></td><td></td><td></td><td></td><td></td><td></td><td></td></tr>
<tr><td>Chromosome</td><td>start</td><td>end</td><td>strand</td><td>MM</td><td>target_seq</td><td>PAM</td><td>alignment</td><td>distance</td><td>position</td><td>gene name</td><td>gene id</td></tr>
<tr><td>GU305914.1</td><td>112370</td><td>112392</td><td>+</td><td>0</td><td>TTACTCGGACGGCGGGGAGA</td><td>AGG</td><td>||||||||[||||||||||||]PAM</td><td>NA</td><td>Intergenic</td><td>NA</td><td>NA</td></tr>
<tr><td>T8252</td><td>CTCGGACGGCGGGGAGAAGGAGG</td><td>1000</td><td></td><td></td><td></td><td></td><td></td><td></td><td></td><td></td><td></td></tr>
<tr><td>Oligo fwd</td><td>CACCCTCGGACGGCGGGGAGAAGG</td><td></td><td></td><td></td><td></td><td></td><td></td><td></td><td></td><td></td><td></td></tr>
<tr><td>Oligo rev</td><td>AAACCCTTCTCCCCGCCGTCCGAG</td><td></td><td></td><td></td><td></td><td></td><td></td><td></td><td></td><td></td><td></td></tr>
<tr><td>Chromosome</td><td>start</td><td>end</td><td>strand</td><td>MM</td><td>target_seq</td><td>PAM</td><td>alignment</td><td>distance</td><td>position</td><td>gene name</td><td>gene id</td></tr>
<tr><td>GU305914.1</td><td>112373</td><td>112395</td><td>+</td><td>0</td><td>CTCGGACGGCGGGGAGAAGG</td><td>AGG</td><td>||||||||[||||||||||||]PAM</td><td>NA</td><td>Intergenic</td><td>NA</td><td>NA</td></tr>
</table>

| | | | | | | | | | | | |
|---|---|---|---|---|---|---|---|---|---|---|---|
| T8253 | TCGGACGGCGGGGAGAAGGAGGG | 1000 | | | | | | | | | |
| Oligo fwd | CACCTCGGACGGCGGGGAGAAGGA | | | | | | | | | | |
| Oligo rev | AAACTCCTTCTCCCCGCCGTCCGA | | | | | | | | | | |
| Chromosome | start | end | strand | MM | target_seq | PAM | alignment | distance | position | gene name | gene id |
| GU305914.1 | 112374 | 112396 | + | 0 | TCGGACGGCGGGGAGAAGGA | GGG | \|\|\|\|\|\|\|\|[\|\|\|\|\|\|\|\|\|\|\|\|]PAM | NA | Intergenic | NA | NA |
| T8254 | CGGGGAGAAGGAGGGTGTGCAGG | 1000 | | | | | | | | | |
| Oligo fwd | CACCCGGGGAGAAGGAGGGTGTGC | | | | | | | | | | |
| Oligo rev | AAACGCACACCCTCCTTCTCCCCG | | | | | | | | | | |
| Chromosome | start | end | strand | MM | target_seq | PAM | alignment | distance | position | gene name | gene id |
| GU305914.1 | 112382 | 112404 | + | 0 | CGGGGAGAAGGAGGGTGTGC | AGG | \|\|\|\|\|\|\|\|[\|\|\|\|\|\|\|\|\|\|\|\|]PAM | NA | Intergenic | NA | NA |
| T8255 | GGGGAGAAGGAGGGTGTGCAGGG | 1000 | | | | | | | | | |
| Oligo fwd | CACCGGGGAGAAGGAGGGTGTGCA | | | | | | | | | | |
| Oligo rev | AAACTGCACACCCTCCTTCTCCCC | | | | | | | | | | |
| Chromosome | start | end | strand | MM | target_seq | PAM | alignment | distance | position | gene name | gene id |
| GU305914.1 | 112383 | 112405 | + | 0 | GGGGAGAAGGAGGGTGTGCA | GGG | \|\|\|\|\|\|\|\|[\|\|\|\|\|\|\|\|\|\|\|\|]PAM | NA | Intergenic | NA | NA |
| T8256 | GAGGGTGTGCAGGGCGATGAAGG | 1000 | | | | | | | | | |
| Oligo fwd | CACCGAGGGTGTGCAGGGCGATGA | | | | | | | | | | |
| Oligo rev | AAACTCATCGCCCTGCACACCCTC | | | | | | | | | | |
| Chromosome | start | end | strand | MM | target_seq | PAM | alignment | distance | position | gene name | gene id |
| GU305914.1 | 112392 | 112414 | + | 0 | GAGGGTGTGCAGGGCGATGA | AGG | \|\|\|\|\|\|\|\|[\|\|\|\|\|\|\|\|\|\|\|\|]PAM | NA | Intergenic | NA | NA |
| GU305914.1 | 96939 | 96961 | + | 4 | GAGGGCGAGCAGGGCGACGG | CGG | \|\|\|\|\|-\|-[\|\|\|\|\|\|\|\|\|-\|-]PAM | NA | Intergenic | NA | NA |
| T8257 | GGTGTGCAGGGCGATGAAGGAGG | 1000 | | | | | | | | | |
| Oligo fwd | CACCGGTGTGCAGGGCGATGAAGG | | | | | | | | | | |
| Oligo rev | AAACCCTTCATCGCCCTGCACACC | | | | | | | | | | |
| Chromosome | start | end | strand | MM | target_seq | PAM | alignment | distance | position | gene name | gene id |
| GU305914.1 | 112395 | 112417 | + | 0 | GGTGTGCAGGGCGATGAAGG | AGG | \|\|\|\|\|\|\|\|[\|\|\|\|\|\|\|\|\|\|\|\|]PAM | NA | Intergenic | NA | NA |
| T8258 | ATACCGATACCGCCGTCCGACGG | 1000 | | | | | | | | | |
| Oligo fwd | CACCATACCGATACCGCCGTCCGA | | | | | | | | | | |
| Oligo rev | AAACTCGGACGGCGGTATCGGTAT | | | | | | | | | | |
| Chromosome | start | end | strand | MM | target_seq | PAM | alignment | distance | position | gene name | gene id |
| GU305914.1 | 112425 | 112447 | + | 0 | ATACCGATACCGCCGTCCGA | CGG | \|\|\|\|\|\|\|\|[\|\|\|\|\|\|\|\|\|\|\|\|]PAM | NA | Intergenic | NA | NA |
| T8259 | TACCGATACCGCCGTCCGACGGG | 1000 | | | | | | | | | |
| Oligo fwd | CACCTACCGATACCGCCGTCCGAC | | | | | | | | | | |
| Oligo rev | AAACGTCGGACGGCGGTATCGGTA | | | | | | | | | | |
| Chromosome | start | end | strand | MM | target_seq | PAM | alignment | distance | position | gene name | gene id |
| GU305914.1 | 112426 | 112448 | + | 0 | TACCGATACCGCCGTCCGAC | GGG | \|\|\|\|\|\|\|\|[\|\|\|\|\|\|\|\|\|\|\|\|]PAM | NA | Intergenic | NA | NA |
| T8260 | ACCGATACCGCCGTCCGACGGGG | 1000 | | | | | | | | | |
| Oligo fwd | CACCACCGATACCGCCGTCCGACG | | | | | | | | | | |
| Oligo rev | AAACCGTCGGACGGCGGTATCGGT | | | | | | | | | | |
| Chromosome | start | end | strand | MM | target_seq | PAM | alignment | distance | position | gene name | gene id |
| GU305914.1 | 112427 | 112449 | + | 0 | ACCGATACCGCCGTCCGACG | GGG | \|\|\|\|\|\|\|\|[\|\|\|\|\|\|\|\|\|\|\|\|]PAM | NA | Intergenic | NA | NA |
| T8261 | CGGGGTGACCGATGATGTCTCGG | 1000 | | | | | | | | | |
| Oligo fwd | CACCCGGGGTGACCGATGATGTCT | | | | | | | | | | |
| Oligo rev | AAACAGACATCATCGGTCACCCCG | | | | | | | | | | |
| Chromosome | start | end | strand | MM | target_seq | PAM | alignment | distance | position | gene name | gene id |
| GU305914.1 | 112445 | 112467 | + | 0 | CGGGGTGACCGATGATGTCT | CGG | \|\|\|\|\|\|\|\|[\|\|\|\|\|\|\|\|\|\|\|\|]PAM | NA | Intergenic | NA | NA |
| T8262 | TCTGTTGTCTGAAATAAAAACGG | 1000 | | | | | | | | | |
| Oligo fwd | CACCTCTGTTGTCTGAAATAAAAA | | | | | | | | | | |
| Oligo rev | AAACTTTTTATTTCAGACAACAGA | | | | | | | | | | |
| Chromosome | start | end | strand | MM | target_seq | PAM | alignment | distance | position | gene name | gene id |
| GU305914.1 | 112488 | 112510 | + | 0 | TCTGTTGTCTGAAATAAAAA | CGG | \|\|\|\|\|\|\|\|[\|\|\|\|\|\|\|\|\|\|\|\|]PAM | NA | Intergenic | NA | NA |
| T8263 | GTGCGCGCGCGAGAGACGCGAGG | 1000 | | | | | | | | | |
| Oligo fwd | CACCGTGCGCGCGCGAGAGACGCG | | | | | | | | | | |
| Oligo rev | AAACCGCGTCTCTCGCGCGCGCAC | | | | | | | | | | |
| Chromosome | start | end | strand | MM | target_seq | PAM | alignment | distance | position | gene name | gene id |
| GU305914.1 | 112510 | 112532 | + | 0 | GTGCGCGCGCGAGAGACGCG | AGG | \|\|\|\|\|\|\|\|[\|\|\|\|\|\|\|\|\|\|\|\|]PAM | NA | Intergenic | NA | NA |
| T8264 | TGCGCGCGCGAGAGACGCGAGGG | 1000 | | | | | | | | | |
| Oligo fwd | CACCTGCGCGCGCGAGAGACGCGA | | | | | | | | | | |
| Oligo rev | AAACTCGCGTCTCTCGCGCGCGCA | | | | | | | | | | |
| Chromosome | start | end | strand | MM | target_seq | PAM | alignment | distance | position | gene name | gene id |
| GU305914.1 | 112511 | 112533 | + | 0 | TGCGCGCGCGAGAGACGCGA | GGG | \|\|\|\|\|\|\|\|[\|\|\|\|\|\|\|\|\|\|\|\|]PAM | NA | Intergenic | NA | NA |
| GU305914.1 | 158800 | 158822 | - | 4 | TGAGCGTGCGAGAGATGCGC | CGG | \|\|-\|\|\|-\|[\|\|\|\|\|\|\|-\|\|\|-]PAM | NA | Intergenic | NA | NA |
| T8265 | CGCGAGAGACGCGAGGGTGACGG | 1000 | | | | | | | | | |
| Oligo fwd | CACCCGCGAGAGACGCGAGGGTGA | | | | | | | | | | |
| Oligo rev | AAACTCACCCTCGCGTCTCTCGCG | | | | | | | | | | |
| Chromosome | start | end | strand | MM | target_seq | PAM | alignment | distance | position | gene name | gene id |
| GU305914.1 | 112517 | 112539 | + | 0 | CGCGAGAGACGCGAGGGTGA | CGG | \|\|\|\|\|\|\|\|[\|\|\|\|\|\|\|\|\|\|\|\|]PAM | NA | Intergenic | NA | NA |
| GU305914.1 | 129134 | 129156 | + | 4 | GGCGTGCGACGCGAGGGCGA | GGG | -\|\|\|-\|-\|[\|\|\|\|\|\|\|\|\|-\|\|]PAM | NA | Intergenic | NA | NA |
| T8266 | GCGAGAGACGCGAGGGTGACGGG | 1000 | | | | | | | | | |
| Oligo fwd | CACCGCGAGAGACGCGAGGGTGAC | | | | | | | | | | |
| Oligo rev | AAACGTCACCCTCGCGTCTCTCGC | | | | | | | | | | |
| Chromosome | start | end | strand | MM | target_seq | PAM | alignment | distance | position | gene name | gene id |
| GU305914.1 | 112518 | 112540 | + | 0 | GCGAGAGACGCGAGGGTGAC | GGG | \|\|\|\|\|\|\|\|[\|\|\|\|\|\|\|\|\|\|\|\|]PAM | NA | Intergenic | NA | NA |
| T8267 | CTCAATCGATAAAGCATGCAAGG | 1000 | | | | | | | | | |
| Oligo fwd | CACCCTCAATCGATAAAGCATGCA | | | | | | | | | | |

Oligo rev AAACTGCATGCTTTATCGATTGAG
Chromosome start end strand MM target_seq PAM alignment distance position gene name gene id
GU305914.1 112564 112586 + 0 CTCAATCGATAAAGCATGCA AGG ||||||||[||||||||||||]PAM NA Intergenic NA NA

T8268 AGCATGCAAGGTGTTGTTGTCGG 1000
Oligo fwd CACCAGCATGCAAGGTGTTGTTGT
Oligo rev AAACACAACAACACCTTGCATGCT
Chromosome start end strand MM target_seq PAM alignment distance position gene name gene id
GU305914.1 112576 112598 + 0 AGCATGCAAGGTGTTGTTGT CGG ||||||||[||||||||||||]PAM NA Intergenic NA NA

T8269 TTGTCGGTCCGTCGTAACGACGG 1000
Oligo fwd CACCTTGTCGGTCCGTCGTAACGA
Oligo rev AAACTCGTTACGACGGACCGACAA
Chromosome start end strand MM target_seq PAM alignment distance position gene name gene id
GU305914.1 112592 112614 + 0 TTGTCGGTCCGTCGTAACGA CGG ||||||||[||||||||||||]PAM NA Intergenic NA NA

T8270 TCGGTCCGTCGTAACGACGGTGG 1000
Oligo fwd CACCTCGGTCCGTCGTAACGACGG
Oligo rev AAACCCGTCGTTACGACGGACCGA
Chromosome start end strand MM target_seq PAM alignment distance position gene name gene id
GU305914.1 112595 112617 + 0 TCGGTCCGTCGTAACGACGG TGG ||||||||[||||||||||||]PAM NA Intergenic NA NA

T8271 TTTCAGTAATAAAAAGAAAACGG 1000
Oligo fwd CACCTTTCAGTAATAAAAAGAAAA
Oligo rev AAACTTTTCTTTTTATTACTGAAA
Chromosome start end strand MM target_seq PAM alignment distance position gene name gene id
GU305914.1 112659 112681 + 0 TTTCAGTAATAAAAAGAAAA CGG ||||||||[||||||||||||]PAM NA Intergenic NA NA

T8272 AAAACGAGATATACCGTGTGTGG 1000
Oligo fwd CACCAAAACGAGATATACCGTGTG
Oligo rev AAACCACACGGTATATCTCGTTTT
Chromosome start end strand MM target_seq PAM alignment distance position gene name gene id
GU305914.1 112682 112704 + 0 AAAACGAGATATACCGTGTG TGG ||||||||[||||||||||||]PAM NA Intergenic NA NA

T8273 AAACGAGATATACCGTGTGTGGG 1000
Oligo fwd CACCAAACGAGATATACCGTGTGT
Oligo rev AAACACACACGGTATATCTCGTTT
Chromosome start end strand MM target_seq PAM alignment distance position gene name gene id
GU305914.1 112683 112705 + 0 AAACGAGATATACCGTGTGT GGG ||||||||[||||||||||||]PAM NA Intergenic NA NA

T8274 ATTCGAGCGCTCCGCTCAGCAGG 1000
Oligo fwd CACCATTCGAGCGCTCCGCTCAGC
Oligo rev AAACGCTGAGCGGAGCGCTCGAAT
Chromosome start end strand MM target_seq PAM alignment distance position gene name gene id
GU305914.1 112739 112761 + 0 ATTCGAGCGCTCCGCTCAGC AGG ||||||||[||||||||||||]PAM NA Intergenic NA NA

T8275 GAGCGCTCCGCTCAGCAGGTCGG 1000
Oligo fwd CACCGAGCGCTCCGCTCAGCAGGT
Oligo rev AAACACCTGCTGAGCGGAGCGCTC
Chromosome start end strand MM target_seq PAM alignment distance position gene name gene id
GU305914.1 112743 112765 + 0 GAGCGCTCCGCTCAGCAGGT CGG ||||||||[||||||||||||]PAM NA Intergenic NA NA

T8276 ACCTTCCTCCTTCTTGTCCATGG 1000
Oligo fwd CACCACCTTCCTCCTTCTTGTCCA
Oligo rev AAACTGGACAAGAAGGAGGAAGGT
Chromosome start end strand MM target_seq PAM alignment distance position gene name gene id
GU305914.1 112773 112795 + 0 ACCTTCCTCCTTCTTGTCCA TGG ||||||||[||||||||||||]PAM NA Intergenic NA NA

T8277 CCTTCCTCCTTCTTGTCCATGGG 1000
Oligo fwd CACCCCTTCCTCCTTCTTGTCCAT
Oligo rev AAACATGGACAAGAAGGAGGAAGG
Chromosome start end strand MM target_seq PAM alignment distance position gene name gene id
GU305914.1 112774 112796 + 0 CCTTCCTCCTTCTTGTCCAT GGG ||||||||[||||||||||||]PAM NA Intergenic NA NA

T8278 TCCTCCTTCTTGTCCATGGGCGG 1000
Oligo fwd CACCTCCTCCTTCTTGTCCATGGG
Oligo rev AAACCCCATGGACAAGAAGGAGGA
Chromosome start end strand MM target_seq PAM alignment distance position gene name gene id
GU305914.1 112777 112799 + 0 TCCTCCTTCTTGTCCATGGG CGG ||||||||[||||||||||||]PAM NA Intergenic NA NA

T8279 GCGTCGCAACGTGCCCGTGAAGG 1000
Oligo fwd CACCGCGTCGCAACGTGCCCGTGA
Oligo rev AAACTCACGGGCACGTTGCGACGC
Chromosome start end strand MM target_seq PAM alignment distance position gene name gene id
GU305914.1 112806 112828 + 0 GCGTCGCAACGTGCCCGTGA AGG ||||||||[||||||||||||]PAM NA Intergenic NA NA

T8280 GCAACGTGCCCGTGAAGGTCAGG 1000
Oligo fwd CACCGCAACGTGCCCGTGAAGGTC
Oligo rev AAACGACCTTCACGGGCACGTTGC
Chromosome start end strand MM target_seq PAM alignment distance position gene name gene id
GU305914.1 112811 112833 + 0 GCAACGTGCCCGTGAAGGTC AGG ||||||||[||||||||||||]PAM NA Intergenic NA NA

T8281 CCGTGAAGGTCAGGTAGTCCTGG 1000
Oligo fwd CACCCCGTGAAGGTCAGGTAGTCC
Oligo rev AAACGGACTACCTGACCTTCACGG
Chromosome start end strand MM target_seq PAM alignment distance position gene name gene id
GU305914.1 112820 112842 + 0 CCGTGAAGGTCAGGTAGTCC TGG ||||||||[||||||||||||]PAM NA Intergenic NA NA

T8282 AGGTCAGGTAGTCCTGGATCAGG 1000

<table>
<tr><td>Oligo fwd</td><td>CACCAGGTCAGGTAGTCCTGGATC</td><td></td><td></td><td></td><td></td><td></td><td></td><td></td><td></td><td></td><td></td></tr>
<tr><td>Oligo rev</td><td>AAACGATCCAGGACTACCTGACCT</td><td></td><td></td><td></td><td></td><td></td><td></td><td></td><td></td><td></td><td></td></tr>
<tr><td>Chromosome</td><td>start</td><td>end</td><td>strand</td><td>MM</td><td>target_seq</td><td>PAM</td><td>alignment</td><td>distance</td><td>position</td><td>gene name</td><td>gene id</td></tr>
<tr><td>GU305914.1</td><td>112826</td><td>112848</td><td>+</td><td>0</td><td>AGGTCAGGTAGTCCTGGATC</td><td>AGG</td><td>||||||||[||||||||||||]PAM</td><td>NA</td><td>Intergenic</td><td>NA</td><td>NA</td></tr>
<tr><td></td><td></td><td></td><td></td><td></td><td></td><td></td><td></td><td></td><td></td><td></td><td></td></tr>
<tr><td>T8283</td><td>CAGGTAGTCCTGGATCAGGAAGG</td><td>1000</td><td></td><td></td><td></td><td></td><td></td><td></td><td></td><td></td><td></td></tr>
<tr><td>Oligo fwd</td><td>CACCCAGGTAGTCCTGGATCAGGA</td><td></td><td></td><td></td><td></td><td></td><td></td><td></td><td></td><td></td><td></td></tr>
<tr><td>Oligo rev</td><td>AAACTCCTGATCCAGGACTACCTG</td><td></td><td></td><td></td><td></td><td></td><td></td><td></td><td></td><td></td><td></td></tr>
<tr><td>Chromosome</td><td>start</td><td>end</td><td>strand</td><td>MM</td><td>target_seq</td><td>PAM</td><td>alignment</td><td>distance</td><td>position</td><td>gene name</td><td>gene id</td></tr>
<tr><td>GU305914.1</td><td>112830</td><td>112852</td><td>+</td><td>0</td><td>CAGGTAGTCCTGGATCAGGA</td><td>AGG</td><td>||||||||[||||||||||||]PAM</td><td>NA</td><td>Intergenic</td><td>NA</td><td>NA</td></tr>
<tr><td></td><td></td><td></td><td></td><td></td><td></td><td></td><td></td><td></td><td></td><td></td><td></td></tr>
<tr><td>T8284</td><td>AGGAAAAGATCCCCAGCTCGTGG</td><td>1000</td><td></td><td></td><td></td><td></td><td></td><td></td><td></td><td></td><td></td></tr>
<tr><td>Oligo fwd</td><td>CACCAGGAAAAGATCCCCAGCTCG</td><td></td><td></td><td></td><td></td><td></td><td></td><td></td><td></td><td></td><td></td></tr>
<tr><td>Oligo rev</td><td>AAACCGAGCTGGGGATCTTTTCCT</td><td></td><td></td><td></td><td></td><td></td><td></td><td></td><td></td><td></td><td></td></tr>
<tr><td>Chromosome</td><td>start</td><td>end</td><td>strand</td><td>MM</td><td>target_seq</td><td>PAM</td><td>alignment</td><td>distance</td><td>position</td><td>gene name</td><td>gene id</td></tr>
<tr><td>GU305914.1</td><td>112850</td><td>112872</td><td>+</td><td>0</td><td>AGGAAAAGATCCCCAGCTCG</td><td>TGG</td><td>||||||||[||||||||||||]PAM</td><td>NA</td><td>Intergenic</td><td>NA</td><td>NA</td></tr>
<tr><td></td><td></td><td></td><td></td><td></td><td></td><td></td><td></td><td></td><td></td><td></td><td></td></tr>
<tr><td>T8285</td><td>GCGCTGCACGCTGAACGCCTCGG</td><td>1000</td><td></td><td></td><td></td><td></td><td></td><td></td><td></td><td></td><td></td></tr>
<tr><td>Oligo fwd</td><td>CACCGCGCTGCACGCTGAACGCCT</td><td></td><td></td><td></td><td></td><td></td><td></td><td></td><td></td><td></td><td></td></tr>
<tr><td>Oligo rev</td><td>AAACAGGCGTTCAGCGTGCAGCGC</td><td></td><td></td><td></td><td></td><td></td><td></td><td></td><td></td><td></td><td></td></tr>
<tr><td>Chromosome</td><td>start</td><td>end</td><td>strand</td><td>MM</td><td>target_seq</td><td>PAM</td><td>alignment</td><td>distance</td><td>position</td><td>gene name</td><td>gene id</td></tr>
<tr><td>GU305914.1</td><td>112878</td><td>112900</td><td>+</td><td>0</td><td>GCGCTGCACGCTGAACGCCT</td><td>CGG</td><td>||||||||[||||||||||||]PAM</td><td>NA</td><td>Intergenic</td><td>NA</td><td>NA</td></tr>
<tr><td></td><td></td><td></td><td></td><td></td><td></td><td></td><td></td><td></td><td></td><td></td><td></td></tr>
<tr><td>T8286</td><td>CGCTGCACGCTGAACGCCTCGGG</td><td>1000</td><td></td><td></td><td></td><td></td><td></td><td></td><td></td><td></td><td></td></tr>
<tr><td>Oligo fwd</td><td>CACCCGCTGCACGCTGAACGCCTC</td><td></td><td></td><td></td><td></td><td></td><td></td><td></td><td></td><td></td><td></td></tr>
<tr><td>Oligo rev</td><td>AAACGAGGCGTTCAGCGTGCAGCG</td><td></td><td></td><td></td><td></td><td></td><td></td><td></td><td></td><td></td><td></td></tr>
<tr><td>Chromosome</td><td>start</td><td>end</td><td>strand</td><td>MM</td><td>target_seq</td><td>PAM</td><td>alignment</td><td>distance</td><td>position</td><td>gene name</td><td>gene id</td></tr>
<tr><td>GU305914.1</td><td>112879</td><td>112901</td><td>+</td><td>0</td><td>CGCTGCACGCTGAACGCCTC</td><td>GGG</td><td>||||||||[||||||||||||]PAM</td><td>NA</td><td>Intergenic</td><td>NA</td><td>NA</td></tr>
<tr><td></td><td></td><td></td><td></td><td></td><td></td><td></td><td></td><td></td><td></td><td></td><td></td></tr>
<tr><td>T8287</td><td>CGACACCGTCAGCTTTTTCCAGG</td><td>1000</td><td></td><td></td><td></td><td></td><td></td><td></td><td></td><td></td><td></td></tr>
<tr><td>Oligo fwd</td><td>CACCCGACACCGTCAGCTTTTTCC</td><td></td><td></td><td></td><td></td><td></td><td></td><td></td><td></td><td></td><td></td></tr>
<tr><td>Oligo rev</td><td>AAACGGAAAAAGCTGACGGTGTCG</td><td></td><td></td><td></td><td></td><td></td><td></td><td></td><td></td><td></td><td></td></tr>
<tr><td>Chromosome</td><td>start</td><td>end</td><td>strand</td><td>MM</td><td>target_seq</td><td>PAM</td><td>alignment</td><td>distance</td><td>position</td><td>gene name</td><td>gene id</td></tr>
<tr><td>GU305914.1</td><td>112902</td><td>112924</td><td>+</td><td>0</td><td>CGACACCGTCAGCTTTTTCC</td><td>AGG</td><td>||||||||[||||||||||||]PAM</td><td>NA</td><td>Intergenic</td><td>NA</td><td>NA</td></tr>
<tr><td></td><td></td><td></td><td></td><td></td><td></td><td></td><td></td><td></td><td></td><td></td><td></td></tr>
<tr><td>T8288</td><td>TCAGCTTTTTCCAGGCGATCAGG</td><td>1000</td><td></td><td></td><td></td><td></td><td></td><td></td><td></td><td></td><td></td></tr>
<tr><td>Oligo fwd</td><td>CACCTCAGCTTTTTCCAGGCGATC</td><td></td><td></td><td></td><td></td><td></td><td></td><td></td><td></td><td></td><td></td></tr>
<tr><td>Oligo rev</td><td>AAACGATCGCCTGGAAAAAGCTGA</td><td></td><td></td><td></td><td></td><td></td><td></td><td></td><td></td><td></td><td></td></tr>
<tr><td>Chromosome</td><td>start</td><td>end</td><td>strand</td><td>MM</td><td>target_seq</td><td>PAM</td><td>alignment</td><td>distance</td><td>position</td><td>gene name</td><td>gene id</td></tr>
<tr><td>GU305914.1</td><td>112910</td><td>112932</td><td>+</td><td>0</td><td>TCAGCTTTTTCCAGGCGATC</td><td>AGG</td><td>||||||||[||||||||||||]PAM</td><td>NA</td><td>Intergenic</td><td>NA</td><td>NA</td></tr>
<tr><td></td><td></td><td></td><td></td><td></td><td></td><td></td><td></td><td></td><td></td><td></td><td></td></tr>
<tr><td>T8289</td><td>TGACAGATCGTCCTGATCGCCGG</td><td>1000</td><td></td><td></td><td></td><td></td><td></td><td></td><td></td><td></td><td></td></tr>
<tr><td>Oligo fwd</td><td>CACCTGACAGATCGTCCTGATCGC</td><td></td><td></td><td></td><td></td><td></td><td></td><td></td><td></td><td></td><td></td></tr>
<tr><td>Oligo rev</td><td>AAACGCGATCAGGACGATCTGTCA</td><td></td><td></td><td></td><td></td><td></td><td></td><td></td><td></td><td></td><td></td></tr>
<tr><td>Chromosome</td><td>start</td><td>end</td><td>strand</td><td>MM</td><td>target_seq</td><td>PAM</td><td>alignment</td><td>distance</td><td>position</td><td>gene name</td><td>gene id</td></tr>
<tr><td>GU305914.1</td><td>112951</td><td>112973</td><td>+</td><td>0</td><td>TGACAGATCGTCCTGATCGC</td><td>CGG</td><td>||||||||[||||||||||||]PAM</td><td>NA</td><td>Intergenic</td><td>NA</td><td>NA</td></tr>
<tr><td></td><td></td><td></td><td></td><td></td><td></td><td></td><td></td><td></td><td></td><td></td><td></td></tr>
<tr><td>T8290</td><td>GCGCCTCCTTGCCCCAGTGCCGG</td><td>1000</td><td></td><td></td><td></td><td></td><td></td><td></td><td></td><td></td><td></td></tr>
<tr><td>Oligo fwd</td><td>CACCGCGCCTCCTTGCCCCAGTGC</td><td></td><td></td><td></td><td></td><td></td><td></td><td></td><td></td><td></td><td></td></tr>
<tr><td>Oligo rev</td><td>AAACGCACTGGGGCAAGGAGGCGC</td><td></td><td></td><td></td><td></td><td></td><td></td><td></td><td></td><td></td><td></td></tr>
<tr><td>Chromosome</td><td>start</td><td>end</td><td>strand</td><td>MM</td><td>target_seq</td><td>PAM</td><td>alignment</td><td>distance</td><td>position</td><td>gene name</td><td>gene id</td></tr>
<tr><td>GU305914.1</td><td>112979</td><td>113001</td><td>+</td><td>0</td><td>GCGCCTCCTTGCCCCAGTGC</td><td>CGG</td><td>||||||||[||||||||||||]PAM</td><td>NA</td><td>Intergenic</td><td>NA</td><td>NA</td></tr>
<tr><td></td><td></td><td></td><td></td><td></td><td></td><td></td><td></td><td></td><td></td><td></td><td></td></tr>
<tr><td>T8291</td><td>GGTTGAAGAAGCGCAGCAGCCGG</td><td>1000</td><td></td><td></td><td></td><td></td><td></td><td></td><td></td><td></td><td></td></tr>
<tr><td>Oligo fwd</td><td>CACCGGTTGAAGAAGCGCAGCAGC</td><td></td><td></td><td></td><td></td><td></td><td></td><td></td><td></td><td></td><td></td></tr>
<tr><td>Oligo rev</td><td>AAACGCTGCTGCGCTTCTTCAACC</td><td></td><td></td><td></td><td></td><td></td><td></td><td></td><td></td><td></td><td></td></tr>
<tr><td>Chromosome</td><td>start</td><td>end</td><td>strand</td><td>MM</td><td>target_seq</td><td>PAM</td><td>alignment</td><td>distance</td><td>position</td><td>gene name</td><td>gene id</td></tr>
<tr><td>GU305914.1</td><td>113000</td><td>113022</td><td>+</td><td>0</td><td>GGTTGAAGAAGCGCAGCAGC</td><td>CGG</td><td>||||||||[||||||||||||]PAM</td><td>NA</td><td>Intergenic</td><td>NA</td><td>NA</td></tr>
<tr><td></td><td></td><td></td><td></td><td></td><td></td><td></td><td></td><td></td><td></td><td></td><td></td></tr>
<tr><td>T8292</td><td>AGCGCAGCAGCCGGAACAGACGG</td><td>1000</td><td></td><td></td><td></td><td></td><td></td><td></td><td></td><td></td><td></td></tr>
<tr><td>Oligo fwd</td><td>CACCAGCGCAGCAGCCGGAACAGA</td><td></td><td></td><td></td><td></td><td></td><td></td><td></td><td></td><td></td><td></td></tr>
<tr><td>Oligo rev</td><td>AAACTCTGTTCCGGCTGCTGCGCT</td><td></td><td></td><td></td><td></td><td></td><td></td><td></td><td></td><td></td><td></td></tr>
<tr><td>Chromosome</td><td>start</td><td>end</td><td>strand</td><td>MM</td><td>target_seq</td><td>PAM</td><td>alignment</td><td>distance</td><td>position</td><td>gene name</td><td>gene id</td></tr>
<tr><td>GU305914.1</td><td>113009</td><td>113031</td><td>+</td><td>0</td><td>AGCGCAGCAGCCGGAACAGA</td><td>CGG</td><td>||||||||[||||||||||||]PAM</td><td>NA</td><td>Intergenic</td><td>NA</td><td>NA</td></tr>
<tr><td></td><td></td><td></td><td></td><td></td><td></td><td></td><td></td><td></td><td></td><td></td><td></td></tr>
<tr><td>T8293</td><td>CGAGAAGAAGATGTTGCCGTCGG</td><td>1000</td><td></td><td></td><td></td><td></td><td></td><td></td><td></td><td></td><td></td></tr>
<tr><td>Oligo fwd</td><td>CACCCGAGAAGAAGATGTTGCCGT</td><td></td><td></td><td></td><td></td><td></td><td></td><td></td><td></td><td></td><td></td></tr>
<tr><td>Oligo rev</td><td>AAACACGGCAACATCTTCTTCTCG</td><td></td><td></td><td></td><td></td><td></td><td></td><td></td><td></td><td></td><td></td></tr>
<tr><td>Chromosome</td><td>start</td><td>end</td><td>strand</td><td>MM</td><td>target_seq</td><td>PAM</td><td>alignment</td><td>distance</td><td>position</td><td>gene name</td><td>gene id</td></tr>
<tr><td>GU305914.1</td><td>113052</td><td>113074</td><td>+</td><td>0</td><td>CGAGAAGAAGATGTTGCCGT</td><td>CGG</td><td>||||||||[||||||||||||]PAM</td><td>NA</td><td>Intergenic</td><td>NA</td><td>NA</td></tr>
<tr><td>GU305914.1</td><td>131194</td><td>131216</td><td>-</td><td>4</td><td>CGAGGCGAACATGTTGCCGA</td><td>TGG</td><td>||||--||[|-|||||||||-]PAM</td><td>NA</td><td>Intergenic</td><td>NA</td><td>NA</td></tr>
<tr><td></td><td></td><td></td><td></td><td></td><td></td><td></td><td></td><td></td><td></td><td></td><td></td></tr>
<tr><td>T8294</td><td>CCGTCGGACAGCAGCTCGTCCGG</td><td>1000</td><td></td><td></td><td></td><td></td><td></td><td></td><td></td><td></td><td></td></tr>
<tr><td>Oligo fwd</td><td>CACCCCGTCGGACAGCAGCTCGTC</td><td></td><td></td><td></td><td></td><td></td><td></td><td></td><td></td><td></td><td></td></tr>
<tr><td>Oligo rev</td><td>AAACGACGAGCTGCTGTCCGACGG</td><td></td><td></td><td></td><td></td><td></td><td></td><td></td><td></td><td></td><td></td></tr>
<tr><td>Chromosome</td><td>start</td><td>end</td><td>strand</td><td>MM</td><td>target_seq</td><td>PAM</td><td>alignment</td><td>distance</td><td>position</td><td>gene name</td><td>gene id</td></tr>
<tr><td>GU305914.1</td><td>113068</td><td>113090</td><td>+</td><td>0</td><td>CCGTCGGACAGCAGCTCGTC</td><td>CGG</td><td>||||||||[||||||||||||]PAM</td><td>NA</td><td>Intergenic</td><td>NA</td><td>NA</td></tr>
<tr><td></td><td></td><td></td><td></td><td></td><td></td><td></td><td></td><td></td><td></td><td></td><td></td></tr>
<tr><td>T8295</td><td>GCAGCTCGTCCGGCAGACTCAGG</td><td>1000</td><td></td><td></td><td></td><td></td><td></td><td></td><td></td><td></td><td></td></tr>
<tr><td>Oligo fwd</td><td>CACCGCAGCTCGTCCGGCAGACTC</td><td></td><td></td><td></td><td></td><td></td><td></td><td></td><td></td><td></td><td></td></tr>
<tr><td>Oligo rev</td><td>AAACGAGTCTGCCGGACGAGCTGC</td><td></td><td></td><td></td><td></td><td></td><td></td><td></td><td></td><td></td><td></td></tr>
<tr><td>Chromosome</td><td>start</td><td>end</td><td>strand</td><td>MM</td><td>target_seq</td><td>PAM</td><td>alignment</td><td>distance</td><td>position</td><td>gene name</td><td>gene id</td></tr>
<tr><td>GU305914.1</td><td>113078</td><td>113100</td><td>+</td><td>0</td><td>GCAGCTCGTCCGGCAGACTC</td><td>AGG</td><td>||||||||[||||||||||||]PAM</td><td>NA</td><td>Intergenic</td><td>NA</td><td>NA</td></tr>
<tr><td></td><td></td><td></td><td></td><td></td><td></td><td></td><td></td><td></td><td></td><td></td><td></td></tr>
<tr><td>T8296</td><td>CAGCTCGTCCGGCAGACTCAGGG</td><td>1000</td><td></td><td></td><td></td><td></td><td></td><td></td><td></td><td></td><td></td></tr>
<tr><td>Oligo fwd</td><td>CACCCAGCTCGTCCGGCAGACTCA</td><td></td><td></td><td></td><td></td><td></td><td></td><td></td><td></td><td></td><td></td></tr>
<tr><td>Oligo rev</td><td>AAACTGAGTCTGCCGGACGAGCTG</td><td></td><td></td><td></td><td></td><td></td><td></td><td></td><td></td><td></td><td></td></tr>
<tr><td>Chromosome</td><td>start</td><td>end</td><td>strand</td><td>MM</td><td>target_seq</td><td>PAM</td><td>alignment</td><td>distance</td><td>position</td><td>gene name</td><td>gene id</td></tr>
<tr><td>GU305914.1</td><td>113079</td><td>113101</td><td>+</td><td>0</td><td>CAGCTCGTCCGGCAGACTCA</td><td>GGG</td><td>||||||||[||||||||||||]PAM</td><td>NA</td><td>Intergenic</td><td>NA</td><td>NA</td></tr>
</table>

T8297 CGTCCGGCAGACTCAGGGCCAGG 1000
Oligo fwd CACCCGTCCGGCAGACTCAGGGCC
Oligo rev AAACGGCCCTGAGTCTGCCGGACG
Chromosome start end strand MM target_seq PAM alignment distance position gene name gene id
GU305914.1 113084 113106 + 0 CGTCCGGCAGACTCAGGGCC AGG ||||||||[||||||||||||]PAM NA Intergenic NA NA

T8298 GGGCCAGGCCCGTCTCCGTCCGG 1000
Oligo fwd CACCGGGCCAGGCCCGTCTCCGTC
Oligo rev AAACGACGGAGACGGGCCTGGCCC
Chromosome start end strand MM target_seq PAM alignment distance position gene name gene id
GU305914.1 113099 113121 + 0 GGGCCAGGCCCGTCTCCGTC CGG ||||||||[||||||||||||]PAM NA Intergenic NA NA

T8299 GGCCCGTCTCCGTCCGGATCAGG 1000
Oligo fwd CACCGGCCCGTCTCCGTCCGGATC
Oligo rev AAACGATCCGGACGGAGACGGGCC
Chromosome start end strand MM target_seq PAM alignment distance position gene name gene id
GU305914.1 113105 113127 + 0 GGCCCGTCTCCGTCCGGATC AGG ||||||||[||||||||||||]PAM NA Intergenic NA NA

T8300 TCCGGATCAGGTGCGACACCAGG 1000
Oligo fwd CACCTCCGGATCAGGTGCGACACC
Oligo rev AAACGGTGTCGCACCTGATCCGGA
Chromosome start end strand MM target_seq PAM alignment distance position gene name gene id
GU305914.1 113117 113139 + 0 TCCGGATCAGGTGCGACACC AGG ||||||||[||||||||||||]PAM NA Intergenic NA NA

T8301 CCGGATCAGGTGCGACACCAGGG 1000
Oligo fwd CACCCCGGATCAGGTGCGACACCA
Oligo rev AAACTGGTGTCGCACCTGATCCGG
Chromosome start end strand MM target_seq PAM alignment distance position gene name gene id
GU305914.1 113118 113140 + 0 CCGGATCAGGTGCGACACCA GGG ||||||||[||||||||||||]PAM NA Intergenic NA NA

T8302 TCAGGTGCGACACCAGGGCGCGG 1000
Oligo fwd CACCTCAGGTGCGACACCAGGGCG
Oligo rev AAACCGCCCTGGTGTCGCACCTGA
Chromosome start end strand MM target_seq PAM alignment distance position gene name gene id
GU305914.1 113123 113145 + 0 TCAGGTGCGACACCAGGGCG CGG ||||||||[||||||||||||]PAM NA Intergenic NA NA

T8303 CATCTCCACCGACAGCGCCGCGG 1000
Oligo fwd CACCCATCTCCACCGACAGCGCCG
Oligo rev AAACCGGCGCTGTCGGTGGAGATG
Chromosome start end strand MM target_seq PAM alignment distance position gene name gene id
GU305914.1 113163 113185 + 0 CATCTCCACCGACAGCGCCG CGG ||||||||[||||||||||||]PAM NA Intergenic NA NA
GU305914.1 138102 138124 + 4 GATCGCCGCCGACAGCGTCG GGG -|||-||-[|||||||||-||]PAM NA Intergenic NA NA

T8304 CGCGGCCGAACGCTCGCTGAAGG 1000
Oligo fwd CACCCGCGGCCGAACGCTCGCTGA
Oligo rev AAACTCAGCGAGCGTTCGGCCGCG
Chromosome start end strand MM target_seq PAM alignment distance position gene name gene id
GU305914.1 113181 113203 + 0 CGCGGCCGAACGCTCGCTGA AGG ||||||||[||||||||||||]PAM NA Intergenic NA NA

T8305 CCGAACGCTCGCTGAAGGCGCGG 1000
Oligo fwd CACCCCGAACGCTCGCTGAAGGCG
Oligo rev AAACCGCCTTCAGCGAGCGTTCGG
Chromosome start end strand MM target_seq PAM alignment distance position gene name gene id
GU305914.1 113186 113208 + 0 CCGAACGCTCGCTGAAGGCG CGG ||||||||[||||||||||||]PAM NA Intergenic NA NA

T8306 GAAGGCGCGGTGCAGCGTCTCGG 1000
Oligo fwd CACCGAAGGCGCGGTGCAGCGTCT
Oligo rev AAACAGACGCTGCACCGCGCCTTC
Chromosome start end strand MM target_seq PAM alignment distance position gene name gene id
GU305914.1 113199 113221 + 0 GAAGGCGCGGTGCAGCGTCT CGG ||||||||[||||||||||||]PAM NA Intergenic NA NA

T8307 GCGCGGTGCAGCGTCTCGGTCGG 1000
Oligo fwd CACCGCGCGGTGCAGCGTCTCGGT
Oligo rev AAACACCGAGACGCTGCACCGCGC
Chromosome start end strand MM target_seq PAM alignment distance position gene name gene id
GU305914.1 113203 113225 + 0 GCGCGGTGCAGCGTCTCGGT CGG ||||||||[||||||||||||]PAM NA Intergenic NA NA

T8308 GTACATCTTGCGCCAGAACTCGG 1000
Oligo fwd CACCGTACATCTTGCGCCAGAACT
Oligo rev AAACAGTTCTGGCGCAAGATGTAC
Chromosome start end strand MM target_seq PAM alignment distance position gene name gene id
GU305914.1 113247 113269 + 0 GTACATCTTGCGCCAGAACT CGG ||||||||[||||||||||||]PAM NA Intergenic NA NA

T8309 TACATCTTGCGCCAGAACTCGGG 1000
Oligo fwd CACCTACATCTTGCGCCAGAACTC
Oligo rev AAACGAGTTCTGGCGCAAGATGTA
Chromosome start end strand MM target_seq PAM alignment distance position gene name gene id
GU305914.1 113248 113270 + 0 TACATCTTGCGCCAGAACTC GGG ||||||||[||||||||||||]PAM NA Intergenic NA NA

T8310 AGAACTCGGGCGAGATCTTGCGG 1000
Oligo fwd CACCAGAACTCGGGCGAGATCTTG
Oligo rev AAACCAAGATCTCGCCCGAGTTCT
Chromosome start end strand MM target_seq PAM alignment distance position gene name gene id
GU305914.1 113261 113283 + 0 AGAACTCGGGCGAGATCTTG CGG ||||||||[||||||||||||]PAM NA Intergenic NA NA
GU305914.1 89630 89652 + 4 TGATCTCGGTGGAGATCTTG CGG -||-||||[|--|||||||||]PAM NA Intergenic NA NA

T8311 CGGGCGAGATCTTGCGGTTCAGG 1000
Oligo fwd CACCCGGGCGAGATCTTGCGGTTC

<table>
<tr><td>Oligo rev</td><td>AAACGAACCGCAAGATCTCGCCCG</td><td></td><td></td><td></td><td></td><td></td><td></td><td></td><td></td><td></td><td></td></tr>
<tr><td>Chromosome</td><td>start</td><td>end</td><td>strand</td><td>MM</td><td>target_seq</td><td>PAM</td><td>alignment</td><td>distance</td><td>position</td><td>gene name</td><td>gene id</td></tr>
<tr><td>GU305914.1</td><td>113267</td><td>113289</td><td>+</td><td>0</td><td>CGGGCGAGATCTTGCGGTTC</td><td>AGG</td><td>||||||||[||||||||||||]PAM</td><td>NA</td><td>Intergenic</td><td>NA</td><td>NA</td></tr>
<tr><td>GU305914.1</td><td>89636</td><td>89658</td><td>+</td><td>4</td><td>CGGTGGAGATCTTGCGGTAG</td><td>CGG</td><td>|||--|||[||||||||||--]PAM</td><td>NA</td><td>Intergenic</td><td>NA</td><td>NA</td></tr>
<tr><td></td><td></td><td></td><td></td><td></td><td></td><td></td><td></td><td></td><td></td><td></td><td></td></tr>
<tr><td>T8312</td><td>TGTTCGCCACCGTCTCCCTCAGG</td><td>1000</td><td></td><td></td><td></td><td></td><td></td><td></td><td></td><td></td><td></td></tr>
<tr><td>Oligo fwd</td><td>CACCTGTTCGCCACCGTCTCCCTC</td><td></td><td></td><td></td><td></td><td></td><td></td><td></td><td></td><td></td><td></td></tr>
<tr><td>Oligo rev</td><td>AAACGAGGGAGACGGTGGCGAACA</td><td></td><td></td><td></td><td></td><td></td><td></td><td></td><td></td><td></td><td></td></tr>
<tr><td>Chromosome</td><td>start</td><td>end</td><td>strand</td><td>MM</td><td>target_seq</td><td>PAM</td><td>alignment</td><td>distance</td><td>position</td><td>gene name</td><td>gene id</td></tr>
<tr><td>GU305914.1</td><td>113312</td><td>113334</td><td>+</td><td>0</td><td>TGTTCGCCACCGTCTCCCTC</td><td>AGG</td><td>||||||||[||||||||||||]PAM</td><td>NA</td><td>Intergenic</td><td>NA</td><td>NA</td></tr>
<tr><td></td><td></td><td></td><td></td><td></td><td></td><td></td><td></td><td></td><td></td><td></td><td></td></tr>
<tr><td>T8313</td><td>TCCCGCGCCCCAGCGTCAGCAGG</td><td>1000</td><td></td><td></td><td></td><td></td><td></td><td></td><td></td><td></td><td></td></tr>
<tr><td>Oligo fwd</td><td>CACCTCCCGCGCCCCAGCGTCAGC</td><td></td><td></td><td></td><td></td><td></td><td></td><td></td><td></td><td></td><td></td></tr>
<tr><td>Oligo rev</td><td>AAACGCTGACGCTGGGGCGCGGGA</td><td></td><td></td><td></td><td></td><td></td><td></td><td></td><td></td><td></td><td></td></tr>
<tr><td>Chromosome</td><td>start</td><td>end</td><td>strand</td><td>MM</td><td>target_seq</td><td>PAM</td><td>alignment</td><td>distance</td><td>position</td><td>gene name</td><td>gene id</td></tr>
<tr><td>GU305914.1</td><td>113348</td><td>113370</td><td>+</td><td>0</td><td>TCCCGCGCCCCAGCGTCAGC</td><td>AGG</td><td>||||||||[||||||||||||]PAM</td><td>NA</td><td>Intergenic</td><td>NA</td><td>NA</td></tr>
<tr><td></td><td></td><td></td><td></td><td></td><td></td><td></td><td></td><td></td><td></td><td></td><td></td></tr>
<tr><td>T8314</td><td>TGATGATCTTCAACTCCTCCAGG</td><td>1000</td><td></td><td></td><td></td><td></td><td></td><td></td><td></td><td></td><td></td></tr>
<tr><td>Oligo fwd</td><td>CACCTGATGATCTTCAACTCCTCC</td><td></td><td></td><td></td><td></td><td></td><td></td><td></td><td></td><td></td><td></td></tr>
<tr><td>Oligo rev</td><td>AAACGGAGGAGTTGAAGATCATCA</td><td></td><td></td><td></td><td></td><td></td><td></td><td></td><td></td><td></td><td></td></tr>
<tr><td>Chromosome</td><td>start</td><td>end</td><td>strand</td><td>MM</td><td>target_seq</td><td>PAM</td><td>alignment</td><td>distance</td><td>position</td><td>gene name</td><td>gene id</td></tr>
<tr><td>GU305914.1</td><td>113381</td><td>113403</td><td>+</td><td>0</td><td>TGATGATCTTCAACTCCTCC</td><td>AGG</td><td>||||||||[||||||||||||]PAM</td><td>NA</td><td>Intergenic</td><td>NA</td><td>NA</td></tr>
<tr><td></td><td></td><td></td><td></td><td></td><td></td><td></td><td></td><td></td><td></td><td></td><td></td></tr>
<tr><td>T8315</td><td>CTTCAACTCCTCCAGGCGCATGG</td><td>1000</td><td></td><td></td><td></td><td></td><td></td><td></td><td></td><td></td><td></td></tr>
<tr><td>Oligo fwd</td><td>CACCCTTCAACTCCTCCAGGCGCA</td><td></td><td></td><td></td><td></td><td></td><td></td><td></td><td></td><td></td><td></td></tr>
<tr><td>Oligo rev</td><td>AAACTGCGCCTGGAGGAGTTGAAG</td><td></td><td></td><td></td><td></td><td></td><td></td><td></td><td></td><td></td><td></td></tr>
<tr><td>Chromosome</td><td>start</td><td>end</td><td>strand</td><td>MM</td><td>target_seq</td><td>PAM</td><td>alignment</td><td>distance</td><td>position</td><td>gene name</td><td>gene id</td></tr>
<tr><td>GU305914.1</td><td>113388</td><td>113410</td><td>+</td><td>0</td><td>CTTCAACTCCTCCAGGCGCA</td><td>TGG</td><td>||||||||[||||||||||||]PAM</td><td>NA</td><td>Intergenic</td><td>NA</td><td>NA</td></tr>
<tr><td></td><td></td><td></td><td></td><td></td><td></td><td></td><td></td><td></td><td></td><td></td><td></td></tr>
<tr><td>T8316</td><td>TCCAGGCGCATGGACGACAGCGG</td><td>1000</td><td></td><td></td><td></td><td></td><td></td><td></td><td></td><td></td><td></td></tr>
<tr><td>Oligo fwd</td><td>CACCTCCAGGCGCATGGACGACAG</td><td></td><td></td><td></td><td></td><td></td><td></td><td></td><td></td><td></td><td></td></tr>
<tr><td>Oligo rev</td><td>AAACCTGTCGTCCATGCGCCTGGA</td><td></td><td></td><td></td><td></td><td></td><td></td><td></td><td></td><td></td><td></td></tr>
<tr><td>Chromosome</td><td>start</td><td>end</td><td>strand</td><td>MM</td><td>target_seq</td><td>PAM</td><td>alignment</td><td>distance</td><td>position</td><td>gene name</td><td>gene id</td></tr>
<tr><td>GU305914.1</td><td>113398</td><td>113420</td><td>+</td><td>0</td><td>TCCAGGCGCATGGACGACAG</td><td>CGG</td><td>||||||||[||||||||||||]PAM</td><td>NA</td><td>Intergenic</td><td>NA</td><td>NA</td></tr>
<tr><td></td><td></td><td></td><td></td><td></td><td></td><td></td><td></td><td></td><td></td><td></td><td></td></tr>
<tr><td>T8317</td><td>TTAGATGATTGTCGTCCGACAGG</td><td>1000</td><td></td><td></td><td></td><td></td><td></td><td></td><td></td><td></td><td></td></tr>
<tr><td>Oligo fwd</td><td>CACCTTAGATGATTGTCGTCCGAC</td><td></td><td></td><td></td><td></td><td></td><td></td><td></td><td></td><td></td><td></td></tr>
<tr><td>Oligo rev</td><td>AAACGTCGGACGACAATCATCTAA</td><td></td><td></td><td></td><td></td><td></td><td></td><td></td><td></td><td></td><td></td></tr>
<tr><td>Chromosome</td><td>start</td><td>end</td><td>strand</td><td>MM</td><td>target_seq</td><td>PAM</td><td>alignment</td><td>distance</td><td>position</td><td>gene name</td><td>gene id</td></tr>
<tr><td>GU305914.1</td><td>113456</td><td>113478</td><td>+</td><td>0</td><td>TTAGATGATTGTCGTCCGAC</td><td>AGG</td><td>||||||||[||||||||||||]PAM</td><td>NA</td><td>Intergenic</td><td>NA</td><td>NA</td></tr>
<tr><td></td><td></td><td></td><td></td><td></td><td></td><td></td><td></td><td></td><td></td><td></td><td></td></tr>
<tr><td>T8318</td><td>GGTGCAGAAACCGCCCGACGCGG</td><td>1000</td><td></td><td></td><td></td><td></td><td></td><td></td><td></td><td></td><td></td></tr>
<tr><td>Oligo fwd</td><td>CACCGGTGCAGAAACCGCCCGACG</td><td></td><td></td><td></td><td></td><td></td><td></td><td></td><td></td><td></td><td></td></tr>
<tr><td>Oligo rev</td><td>AAACCGTCGGGCGGTTTCTGCACC</td><td></td><td></td><td></td><td></td><td></td><td></td><td></td><td></td><td></td><td></td></tr>
<tr><td>Chromosome</td><td>start</td><td>end</td><td>strand</td><td>MM</td><td>target_seq</td><td>PAM</td><td>alignment</td><td>distance</td><td>position</td><td>gene name</td><td>gene id</td></tr>
<tr><td>GU305914.1</td><td>113477</td><td>113499</td><td>+</td><td>0</td><td>GGTGCAGAAACCGCCCGACG</td><td>CGG</td><td>||||||||[||||||||||||]PAM</td><td>NA</td><td>Intergenic</td><td>NA</td><td>NA</td></tr>
<tr><td></td><td></td><td></td><td></td><td></td><td></td><td></td><td></td><td></td><td></td><td></td><td></td></tr>
<tr><td>T8319</td><td>GCGGATCGTCTTCACCATGACGG</td><td>1000</td><td></td><td></td><td></td><td></td><td></td><td></td><td></td><td></td><td></td></tr>
<tr><td>Oligo fwd</td><td>CACCGCGGATCGTCTTCACCATGA</td><td></td><td></td><td></td><td></td><td></td><td></td><td></td><td></td><td></td><td></td></tr>
<tr><td>Oligo rev</td><td>AAACTCATGGTGAAGACGATCCGC</td><td></td><td></td><td></td><td></td><td></td><td></td><td></td><td></td><td></td><td></td></tr>
<tr><td>Chromosome</td><td>start</td><td>end</td><td>strand</td><td>MM</td><td>target_seq</td><td>PAM</td><td>alignment</td><td>distance</td><td>position</td><td>gene name</td><td>gene id</td></tr>
<tr><td>GU305914.1</td><td>113496</td><td>113518</td><td>+</td><td>0</td><td>GCGGATCGTCTTCACCATGA</td><td>CGG</td><td>||||||||[||||||||||||]PAM</td><td>NA</td><td>Intergenic</td><td>NA</td><td>NA</td></tr>
<tr><td></td><td></td><td></td><td></td><td></td><td></td><td></td><td></td><td></td><td></td><td></td><td></td></tr>
<tr><td>T8320</td><td>CGGATCGTCTTCACCATGACGGG</td><td>1000</td><td></td><td></td><td></td><td></td><td></td><td></td><td></td><td></td><td></td></tr>
<tr><td>Oligo fwd</td><td>CACCCGGATCGTCTTCACCATGAC</td><td></td><td></td><td></td><td></td><td></td><td></td><td></td><td></td><td></td><td></td></tr>
<tr><td>Oligo rev</td><td>AAACGTCATGGTGAAGACGATCCG</td><td></td><td></td><td></td><td></td><td></td><td></td><td></td><td></td><td></td><td></td></tr>
<tr><td>Chromosome</td><td>start</td><td>end</td><td>strand</td><td>MM</td><td>target_seq</td><td>PAM</td><td>alignment</td><td>distance</td><td>position</td><td>gene name</td><td>gene id</td></tr>
<tr><td>GU305914.1</td><td>113497</td><td>113519</td><td>+</td><td>0</td><td>CGGATCGTCTTCACCATGAC</td><td>GGG</td><td>||||||||[||||||||||||]PAM</td><td>NA</td><td>Intergenic</td><td>NA</td><td>NA</td></tr>
<tr><td></td><td></td><td></td><td></td><td></td><td></td><td></td><td></td><td></td><td></td><td></td><td></td></tr>
<tr><td>T8321</td><td>GGGCGACGCCGCTCGCGCGCCGG</td><td>1000</td><td></td><td></td><td></td><td></td><td></td><td></td><td></td><td></td><td></td></tr>
<tr><td>Oligo fwd</td><td>CACCGGGCGACGCCGCTCGCGCGC</td><td></td><td></td><td></td><td></td><td></td><td></td><td></td><td></td><td></td><td></td></tr>
<tr><td>Oligo rev</td><td>AAACGCGCGCGAGCGGCGTCGCCC</td><td></td><td></td><td></td><td></td><td></td><td></td><td></td><td></td><td></td><td></td></tr>
<tr><td>Chromosome</td><td>start</td><td>end</td><td>strand</td><td>MM</td><td>target_seq</td><td>PAM</td><td>alignment</td><td>distance</td><td>position</td><td>gene name</td><td>gene id</td></tr>
<tr><td>GU305914.1</td><td>113517</td><td>113539</td><td>+</td><td>0</td><td>GGGCGACGCCGCTCGCGCGC</td><td>CGG</td><td>||||||||[||||||||||||]PAM</td><td>NA</td><td>Intergenic</td><td>NA</td><td>NA</td></tr>
<tr><td></td><td></td><td></td><td></td><td></td><td></td><td></td><td></td><td></td><td></td><td></td><td></td></tr>
<tr><td>T8322</td><td>CGCCGCTCGCGCGCCGGACGCGG</td><td>1000</td><td></td><td></td><td></td><td></td><td></td><td></td><td></td><td></td><td></td></tr>
<tr><td>Oligo fwd</td><td>CACCCGCCGCTCGCGCGCCGGACG</td><td></td><td></td><td></td><td></td><td></td><td></td><td></td><td></td><td></td><td></td></tr>
<tr><td>Oligo rev</td><td>AAACCGTCCGGCGCGCGAGCGGCG</td><td></td><td></td><td></td><td></td><td></td><td></td><td></td><td></td><td></td><td></td></tr>
<tr><td>Chromosome</td><td>start</td><td>end</td><td>strand</td><td>MM</td><td>target_seq</td><td>PAM</td><td>alignment</td><td>distance</td><td>position</td><td>gene name</td><td>gene id</td></tr>
<tr><td>GU305914.1</td><td>113523</td><td>113545</td><td>+</td><td>0</td><td>CGCCGCTCGCGCGCCGGACG</td><td>CGG</td><td>||||||||[||||||||||||]PAM</td><td>NA</td><td>Intergenic</td><td>NA</td><td>NA</td></tr>
<tr><td>GU305914.1</td><td>100225</td><td>100247</td><td>-</td><td>4</td><td>CGCCGCGGGGGCGGCGGACG</td><td>AGG</td><td>||||||--[|-|||-||||||]PAM</td><td>NA</td><td>Intergenic</td><td>NA</td><td>NA</td></tr>
<tr><td></td><td></td><td></td><td></td><td></td><td></td><td></td><td></td><td></td><td></td><td></td><td></td></tr>
<tr><td>T8323</td><td>GCCGCTCGCGCGCCGGACGCGGG</td><td>1000</td><td></td><td></td><td></td><td></td><td></td><td></td><td></td><td></td><td></td></tr>
<tr><td>Oligo fwd</td><td>CACCGCCGCTCGCGCGCCGGACGC</td><td></td><td></td><td></td><td></td><td></td><td></td><td></td><td></td><td></td><td></td></tr>
<tr><td>Oligo rev</td><td>AAACGCGTCCGGCGCGCGAGCGGC</td><td></td><td></td><td></td><td></td><td></td><td></td><td></td><td></td><td></td><td></td></tr>
<tr><td>Chromosome</td><td>start</td><td>end</td><td>strand</td><td>MM</td><td>target_seq</td><td>PAM</td><td>alignment</td><td>distance</td><td>position</td><td>gene name</td><td>gene id</td></tr>
<tr><td>GU305914.1</td><td>113524</td><td>113546</td><td>+</td><td>0</td><td>GCCGCTCGCGCGCCGGACGC</td><td>GGG</td><td>||||||||[||||||||||||]PAM</td><td>NA</td><td>Intergenic</td><td>NA</td><td>NA</td></tr>
<tr><td></td><td></td><td></td><td></td><td></td><td></td><td></td><td></td><td></td><td></td><td></td><td></td></tr>
<tr><td>T8324</td><td>GCGGGATCGATGATCTACGTCGG</td><td>1000</td><td></td><td></td><td></td><td></td><td></td><td></td><td></td><td></td><td></td></tr>
<tr><td>Oligo fwd</td><td>CACCGCGGGATCGATGATCTACGT</td><td></td><td></td><td></td><td></td><td></td><td></td><td></td><td></td><td></td><td></td></tr>
<tr><td>Oligo rev</td><td>AAACACGTAGATCATCGATCCCGC</td><td></td><td></td><td></td><td></td><td></td><td></td><td></td><td></td><td></td><td></td></tr>
<tr><td>Chromosome</td><td>start</td><td>end</td><td>strand</td><td>MM</td><td>target_seq</td><td>PAM</td><td>alignment</td><td>distance</td><td>position</td><td>gene name</td><td>gene id</td></tr>
<tr><td>GU305914.1</td><td>113542</td><td>113564</td><td>+</td><td>0</td><td>GCGGGATCGATGATCTACGT</td><td>CGG</td><td>||||||||[||||||||||||]PAM</td><td>NA</td><td>Intergenic</td><td>NA</td><td>NA</td></tr>
<tr><td></td><td></td><td></td><td></td><td></td><td></td><td></td><td></td><td></td><td></td><td></td><td></td></tr>
<tr><td>T8325</td><td>CGGGATCGATGATCTACGTCGGG</td><td>1000</td><td></td><td></td><td></td><td></td><td></td><td></td><td></td><td></td><td></td></tr>
<tr><td>Oligo fwd</td><td>CACCCGGGATCGATGATCTACGTC</td><td></td><td></td><td></td><td></td><td></td><td></td><td></td><td></td><td></td><td></td></tr>
<tr><td>Oligo rev</td><td>AAACGACGTAGATCATCGATCCCG</td><td></td><td></td><td></td><td></td><td></td><td></td><td></td><td></td><td></td><td></td></tr>
<tr><td>Chromosome</td><td>start</td><td>end</td><td>strand</td><td>MM</td><td>target_seq</td><td>PAM</td><td>alignment</td><td>distance</td><td>position</td><td>gene name</td><td>gene id</td></tr>
<tr><td>GU305914.1</td><td>113543</td><td>113565</td><td>+</td><td>0</td><td>CGGGATCGATGATCTACGTC</td><td>GGG</td><td>||||||||[||||||||||||]PAM</td><td>NA</td><td>Intergenic</td><td>NA</td><td>NA</td></tr>
</table>

T8326 GGGATCGATGATCTACGTCGGGG 1000
Oligo fwd CACCGGGATCGATGATCTACGTCG
Oligo rev AAACCGACGTAGATCATCGATCCC
Chromosome start end strand MM target_seq PAM alignment distance position gene name gene id
GU305914.1 113544 113566 + 0 GGGATCGATGATCTACGTCG GGG ||||||||[||||||||||||]PAM NA Intergenic NA NA

T8327 GGATCGATGATCTACGTCGGGGG 1000
Oligo fwd CACCGGATCGATGATCTACGTCGG
Oligo rev AAACCCGACGTAGATCATCGATCC
Chromosome start end strand MM target_seq PAM alignment distance position gene name gene id
GU305914.1 113545 113567 + 0 GGATCGATGATCTACGTCGG GGG ||||||||[||||||||||||]PAM NA Intergenic NA NA

T8328 CTTCCTCACTCTGTATGACGAGG 1000
Oligo fwd CACCCTTCCTCACTCTGTATGACG
Oligo rev AAACCGTCATACAGAGTGAGGAAG
Chromosome start end strand MM target_seq PAM alignment distance position gene name gene id
GU305914.1 113568 113590 + 0 CTTCCTCACTCTGTATGACG AGG ||||||||[||||||||||||]PAM NA Intergenic NA NA

T8329 TCTGTATGACGAGGACCCTCAGG 1000
Oligo fwd CACCTCTGTATGACGAGGACCCTC
Oligo rev AAACGAGGGTCCTCGTCATACAGA
Chromosome start end strand MM target_seq PAM alignment distance position gene name gene id
GU305914.1 113577 113599 + 0 TCTGTATGACGAGGACCCTC AGG ||||||||[||||||||||||]PAM NA Intergenic NA NA

T8330 ACCCTCAGGACGAACGCCTGCGG 1000
Oligo fwd CACCACCCTCAGGACGAACGCCTG
Oligo rev AAACCAGGCGTTCGTCCTGAGGGT
Chromosome start end strand MM target_seq PAM alignment distance position gene name gene id
GU305914.1 113591 113613 + 0 ACCCTCAGGACGAACGCCTG CGG ||||||||[||||||||||||]PAM NA Intergenic NA NA

T8331 GCGGCTCCCTCGCGACGTCGTGG 1000
Oligo fwd CACCGCGGCTCCCTCGCGACGTCG
Oligo rev AAACCGACGTCGCGAGGGAGCCGC
Chromosome start end strand MM target_seq PAM alignment distance position gene name gene id
GU305914.1 113610 113632 + 0 GCGGCTCCCTCGCGACGTCG TGG ||||||||[||||||||||||]PAM NA Intergenic NA NA
GU305914.1 199633 199655 - 3 GCTGGTCCGTCGCGACGTCG AGG ||-|-|||[-|||||||||||]PAM NA Intergenic NA NA

T8332 TCCCTCGCGACGTCGTGGCCAGG 1000
Oligo fwd CACCTCCCTCGCGACGTCGTGGCC
Oligo rev AAACGGCCACGACGTCGCGAGGGA
Chromosome start end strand MM target_seq PAM alignment distance position gene name gene id
GU305914.1 113615 113637 + 0 TCCCTCGCGACGTCGTGGCC AGG ||||||||[||||||||||||]PAM NA Intergenic NA NA

T8333 CCCTCGCGACGTCGTGGCCAGGG 1000
Oligo fwd CACCCCCTCGCGACGTCGTGGCCA
Oligo rev AAACTGGCCACGACGTCGCGAGGG
Chromosome start end strand MM target_seq PAM alignment distance position gene name gene id
GU305914.1 113616 113638 + 0 CCCTCGCGACGTCGTGGCCA GGG ||||||||[||||||||||||]PAM NA Intergenic NA NA

T8334 GGCCAGGGAGCTGCGCAGAGCGG 1000
Oligo fwd CACCGGCCAGGGAGCTGCGCAGAG
Oligo rev AAACCTCTGCGCAGCTCCCTGGCC
Chromosome start end strand MM target_seq PAM alignment distance position gene name gene id
GU305914.1 113631 113653 + 0 GGCCAGGGAGCTGCGCAGAG CGG ||||||||[||||||||||||]PAM NA Intergenic NA NA

T8335 CAGGGAGCTGCGCAGAGCGGCGG 1000
Oligo fwd CACCCAGGGAGCTGCGCAGAGCGG
Oligo rev AAACCCGCTCTGCGCAGCTCCCTG
Chromosome start end strand MM target_seq PAM alignment distance position gene name gene id
GU305914.1 113634 113656 + 0 CAGGGAGCTGCGCAGAGCGG CGG ||||||||[||||||||||||]PAM NA Intergenic NA NA

T8336 GAGCTGCGCAGAGCGGCGGCCGG 1000
Oligo fwd CACCGAGCTGCGCAGAGCGGCGGC
Oligo rev AAACGCCGCCGCTCTGCGCAGCTC
Chromosome start end strand MM target_seq PAM alignment distance position gene name gene id
GU305914.1 113638 113660 + 0 GAGCTGCGCAGAGCGGCGGC CGG ||||||||[||||||||||||]PAM NA Intergenic NA NA

T8337 AGCTGCGCAGAGCGGCGGCCGGG 1000
Oligo fwd CACCAGCTGCGCAGAGCGGCGGCC
Oligo rev AAACGGCCGCCGCTCTGCGCAGCT
Chromosome start end strand MM target_seq PAM alignment distance position gene name gene id
GU305914.1 113639 113661 + 0 AGCTGCGCAGAGCGGCGGCC GGG ||||||||[||||||||||||]PAM NA Intergenic NA NA

T8338 GCTGCGCAGAGCGGCGGCCGGGG 1000
Oligo fwd CACCGCTGCGCAGAGCGGCGGCCG
Oligo rev AAACCGGCCGCCGCTCTGCGCAGC
Chromosome start end strand MM target_seq PAM alignment distance position gene name gene id
GU305914.1 113640 113662 + 0 GCTGCGCAGAGCGGCGGCCG GGG ||||||||[||||||||||||]PAM NA Intergenic NA NA
GU305914.1 31059 31081 - 4 GCTCGGCAGGGCCGCGGCCG AGG |||--|||[|-||-|||||||]PAM NA Intergenic NA NA

T8339 CTGCGCAGAGCGGCGGCCGGGGG 1000
Oligo fwd CACCCTGCGCAGAGCGGCGGCCGG
Oligo rev AAACCCGGCCGCCGCTCTGCGCAG
Chromosome start end strand MM target_seq PAM alignment distance position gene name gene id
GU305914.1 113641 113663 + 0 CTGCGCAGAGCGGCGGCCGG GGG ||||||||[||||||||||||]PAM NA Intergenic NA NA

T8340 TGCGCAGAGCGGCGGCCGGGGGG 1000
Oligo fwd CACCTGCGCAGAGCGGCGGCCGGG

| | | | | | | | | | | | |
|---|---|---|---|---|---|---|---|---|---|---|---|
| Oligo rev | AAACCCCGGCCGCCGCTCTGCGCA | | | | | | | | | | |
| Chromosome | start | end | strand | MM | target_seq | PAM | alignment | distance | position | gene name | gene id |
| GU305914.1 | 113642 | 113664 | + | 0 | TGCGCAGAGCGGCGGCCGGG | GGG | \|\|\|\|\|\|\|\|[\|\|\|\|\|\|\|\|\|\|\|\|]PAM | NA | Intergenic | NA | NA |
| T8341 | CAATCACGACGAATCCTCCACGG | 1000 | | | | | | | | | |
| Oligo fwd | CACCCAATCACGACGAATCCTCCA | | | | | | | | | | |
| Oligo rev | AAACTGGAGGATTCGTCGTGATTG | | | | | | | | | | |
| Chromosome | start | end | strand | MM | target_seq | PAM | alignment | distance | position | gene name | gene id |
| GU305914.1 | 113685 | 113707 | + | 0 | CAATCACGACGAATCCTCCA | CGG | \|\|\|\|\|\|\|\|[\|\|\|\|\|\|\|\|\|\|\|\|]PAM | NA | Intergenic | NA | NA |
| T8342 | CACGACGAATCCTCCACGGTCGG | 1000 | | | | | | | | | |
| Oligo fwd | CACCCACGACGAATCCTCCACGGT | | | | | | | | | | |
| Oligo rev | AAACACCGTGGAGGATTCGTCGTG | | | | | | | | | | |
| Chromosome | start | end | strand | MM | target_seq | PAM | alignment | distance | position | gene name | gene id |
| GU305914.1 | 113689 | 113711 | + | 0 | CACGACGAATCCTCCACGGT | CGG | \|\|\|\|\|\|\|\|[\|\|\|\|\|\|\|\|\|\|\|\|]PAM | NA | Intergenic | NA | NA |
| T8343 | CGTACGTCTCTTTGACGCCGAGG | 1000 | | | | | | | | | |
| Oligo fwd | CACCCGTACGTCTCTTTGACGCCG | | | | | | | | | | |
| Oligo rev | AAACCGGCGTCAAAGAGACGTACG | | | | | | | | | | |
| Chromosome | start | end | strand | MM | target_seq | PAM | alignment | distance | position | gene name | gene id |
| GU305914.1 | 113715 | 113737 | + | 0 | CGTACGTCTCTTTGACGCCG | AGG | \|\|\|\|\|\|\|\|[\|\|\|\|\|\|\|\|\|\|\|\|]PAM | NA | Intergenic | NA | NA |
| T8344 | CGTCTCTTTGACGCCGAGGCCGG | 1000 | | | | | | | | | |
| Oligo fwd | CACCCGTCTCTTTGACGCCGAGGC | | | | | | | | | | |
| Oligo rev | AAACGCCTCGGCGTCAAAGAGACG | | | | | | | | | | |
| Chromosome | start | end | strand | MM | target_seq | PAM | alignment | distance | position | gene name | gene id |
| GU305914.1 | 113719 | 113741 | + | 0 | CGTCTCTTTGACGCCGAGGC | CGG | \|\|\|\|\|\|\|\|[\|\|\|\|\|\|\|\|\|\|\|\|]PAM | NA | Intergenic | NA | NA |
| T8345 | CGAGGCCGGACTCTTCTGTCTGG | 1000 | | | | | | | | | |
| Oligo fwd | CACCCGAGGCCGGACTCTTCTGTC | | | | | | | | | | |
| Oligo rev | AAACGACAGAAGAGTCCGGCCTCG | | | | | | | | | | |
| Chromosome | start | end | strand | MM | target_seq | PAM | alignment | distance | position | gene name | gene id |
| GU305914.1 | 113733 | 113755 | + | 0 | CGAGGCCGGACTCTTCTGTC | TGG | \|\|\|\|\|\|\|\|[\|\|\|\|\|\|\|\|\|\|\|\|]PAM | NA | Intergenic | NA | NA |
| T8346 | GAGGCCGGACTCTTCTGTCTGGG | 1000 | | | | | | | | | |
| Oligo fwd | CACCGAGGCCGGACTCTTCTGTCT | | | | | | | | | | |
| Oligo rev | AAACAGACAGAAGAGTCCGGCCTC | | | | | | | | | | |
| Chromosome | start | end | strand | MM | target_seq | PAM | alignment | distance | position | gene name | gene id |
| GU305914.1 | 113734 | 113756 | + | 0 | GAGGCCGGACTCTTCTGTCT | GGG | \|\|\|\|\|\|\|\|[\|\|\|\|\|\|\|\|\|\|\|\|]PAM | NA | Intergenic | NA | NA |
| T8347 | CCGGACTCTTCTGTCTGGGCCGG | 1000 | | | | | | | | | |
| Oligo fwd | CACCCCGGACTCTTCTGTCTGGGC | | | | | | | | | | |
| Oligo rev | AAACGCCCAGACAGAAGAGTCCGG | | | | | | | | | | |
| Chromosome | start | end | strand | MM | target_seq | PAM | alignment | distance | position | gene name | gene id |
| GU305914.1 | 113738 | 113760 | + | 0 | CCGGACTCTTCTGTCTGGGC | CGG | \|\|\|\|\|\|\|\|[\|\|\|\|\|\|\|\|\|\|\|\|]PAM | NA | Intergenic | NA | NA |
| GU305914.1 | 188129 | 188151 | - | 4 | ACAGACTCTTGTGTCTGGTC | AGG | -\|-\|\|\|\|\|[\|\|-\|\|\|\|\|\|\|-\|]PAM | NA | Intergenic | NA | NA |
| T8348 | GCTCAGCTCGCCCGCCTTTCTGG | 1000 | | | | | | | | | |
| Oligo fwd | CACCGCTCAGCTCGCCCGCCTTTC | | | | | | | | | | |
| Oligo rev | AAACGAAAGGCGGGCGAGCTGAGC | | | | | | | | | | |
| Chromosome | start | end | strand | MM | target_seq | PAM | alignment | distance | position | gene name | gene id |
| GU305914.1 | 113760 | 113782 | + | 0 | GCTCAGCTCGCCCGCCTTTC | TGG | \|\|\|\|\|\|\|\|[\|\|\|\|\|\|\|\|\|\|\|\|]PAM | NA | Intergenic | NA | NA |
| T8349 | CTCAGCTCGCCCGCCTTTCTGGG | 1000 | | | | | | | | | |
| Oligo fwd | CACCCTCAGCTCGCCCGCCTTTCT | | | | | | | | | | |
| Oligo rev | AAACAGAAAGGCGGGCGAGCTGAG | | | | | | | | | | |
| Chromosome | start | end | strand | MM | target_seq | PAM | alignment | distance | position | gene name | gene id |
| GU305914.1 | 113761 | 113783 | + | 0 | CTCAGCTCGCCCGCCTTTCT | GGG | \|\|\|\|\|\|\|\|[\|\|\|\|\|\|\|\|\|\|\|\|]PAM | NA | Intergenic | NA | NA |
| T8350 | TCAGCTCGCCCGCCTTTCTGGGG | 1000 | | | | | | | | | |
| Oligo fwd | CACCTCAGCTCGCCCGCCTTTCTG | | | | | | | | | | |
| Oligo rev | AAACCAGAAAGGCGGGCGAGCTGA | | | | | | | | | | |
| Chromosome | start | end | strand | MM | target_seq | PAM | alignment | distance | position | gene name | gene id |
| GU305914.1 | 113762 | 113784 | + | 0 | TCAGCTCGCCCGCCTTTCTG | GGG | \|\|\|\|\|\|\|\|[\|\|\|\|\|\|\|\|\|\|\|\|]PAM | NA | Intergenic | NA | NA |
| T8351 | CTTTCTGGGGATCGTCGAGAAGG | 1000 | | | | | | | | | |
| Oligo fwd | CACCCTTTCTGGGGATCGTCGAGA | | | | | | | | | | |
| Oligo rev | AAACTCTCGACGATCCCCAGAAAG | | | | | | | | | | |
| Chromosome | start | end | strand | MM | target_seq | PAM | alignment | distance | position | gene name | gene id |
| GU305914.1 | 113775 | 113797 | + | 0 | CTTTCTGGGGATCGTCGAGA | AGG | \|\|\|\|\|\|\|\|[\|\|\|\|\|\|\|\|\|\|\|\|]PAM | NA | Intergenic | NA | NA |
| T8352 | GGGGATCGTCGAGAAGGCCGCGG | 1000 | | | | | | | | | |
| Oligo fwd | CACCGGGGATCGTCGAGAAGGCCG | | | | | | | | | | |
| Oligo rev | AAACCGGCCTTCTCGACGATCCCC | | | | | | | | | | |
| Chromosome | start | end | strand | MM | target_seq | PAM | alignment | distance | position | gene name | gene id |
| GU305914.1 | 113781 | 113803 | + | 0 | GGGGATCGTCGAGAAGGCCG | CGG | \|\|\|\|\|\|\|\|[\|\|\|\|\|\|\|\|\|\|\|\|]PAM | NA | Intergenic | NA | NA |
| T8353 | GGGATCGTCGAGAAGGCCGCGGG | 1000 | | | | | | | | | |
| Oligo fwd | CACCGGGATCGTCGAGAAGGCCGC | | | | | | | | | | |
| Oligo rev | AAACGCGGCCTTCTCGACGATCCC | | | | | | | | | | |
| Chromosome | start | end | strand | MM | target_seq | PAM | alignment | distance | position | gene name | gene id |
| GU305914.1 | 113782 | 113804 | + | 0 | GGGATCGTCGAGAAGGCCGC | GGG | \|\|\|\|\|\|\|\|[\|\|\|\|\|\|\|\|\|\|\|\|]PAM | NA | Intergenic | NA | NA |
| T8354 | AAATCCAAACTCGTCGCCCGCGG | 1000 | | | | | | | | | |
| Oligo fwd | CACCAAATCCAAACTCGTCGCCCG | | | | | | | | | | |
| Oligo rev | AAACCGGGCGACGAGTTTGGATTT | | | | | | | | | | |
| Chromosome | start | end | strand | MM | target_seq | PAM | alignment | distance | position | gene name | gene id |
| GU305914.1 | 113806 | 113828 | + | 0 | AAATCCAAACTCGTCGCCCG | CGG | \|\|\|\|\|\|\|\|[\|\|\|\|\|\|\|\|\|\|\|\|]PAM | NA | Intergenic | NA | NA |

<table>
<tr><td>T8355</td><td>CGTCGCCCGCGGTCCCGCCAAGG</td><td>1000</td><td></td><td></td><td></td><td></td><td></td><td></td><td></td><td></td><td></td></tr>
<tr><td>Oligo fwd</td><td>CACCCGTCGCCCGCGGTCCCGCCA</td><td></td><td></td><td></td><td></td><td></td><td></td><td></td><td></td><td></td><td></td></tr>
<tr><td>Oligo rev</td><td>AAACTGGCGGGACCGCGGGCGACG</td><td></td><td></td><td></td><td></td><td></td><td></td><td></td><td></td><td></td><td></td></tr>
<tr><td>Chromosome</td><td>start</td><td>end</td><td>strand</td><td>MM</td><td>target_seq</td><td>PAM</td><td>alignment</td><td>distance</td><td>position</td><td>gene name</td><td>gene id</td></tr>
<tr><td>GU305914.1</td><td>113817</td><td>113839</td><td>+</td><td>0</td><td>CGTCGCCCGCGGTCCCGCCA</td><td>AGG</td><td>||||||||[||||||||||||]PAM</td><td>NA</td><td>Intergenic</td><td>NA</td><td>NA</td></tr>
<tr><td>T8356</td><td>GTCGCCCGCGGTCCCGCCAAGGG</td><td>1000</td><td></td><td></td><td></td><td></td><td></td><td></td><td></td><td></td><td></td></tr>
<tr><td>Oligo fwd</td><td>CACCGTCGCCCGCGGTCCCGCCAA</td><td></td><td></td><td></td><td></td><td></td><td></td><td></td><td></td><td></td><td></td></tr>
<tr><td>Oligo rev</td><td>AAACTTGGCGGGACCGCGGGCGAC</td><td></td><td></td><td></td><td></td><td></td><td></td><td></td><td></td><td></td><td></td></tr>
<tr><td>Chromosome</td><td>start</td><td>end</td><td>strand</td><td>MM</td><td>target_seq</td><td>PAM</td><td>alignment</td><td>distance</td><td>position</td><td>gene name</td><td>gene id</td></tr>
<tr><td>GU305914.1</td><td>113818</td><td>113840</td><td>+</td><td>0</td><td>GTCGCCCGCGGTCCCGCCAA</td><td>GGG</td><td>||||||||[||||||||||||]PAM</td><td>NA</td><td>Intergenic</td><td>NA</td><td>NA</td></tr>
<tr><td>T8357</td><td>TCGCCCGCGGTCCCGCCAAGGGG</td><td>1000</td><td></td><td></td><td></td><td></td><td></td><td></td><td></td><td></td><td></td></tr>
<tr><td>Oligo fwd</td><td>CACCTCGCCCGCGGTCCCGCCAAG</td><td></td><td></td><td></td><td></td><td></td><td></td><td></td><td></td><td></td><td></td></tr>
<tr><td>Oligo rev</td><td>AAACCTTGGCGGGACCGCGGGCGA</td><td></td><td></td><td></td><td></td><td></td><td></td><td></td><td></td><td></td><td></td></tr>
<tr><td>Chromosome</td><td>start</td><td>end</td><td>strand</td><td>MM</td><td>target_seq</td><td>PAM</td><td>alignment</td><td>distance</td><td>position</td><td>gene name</td><td>gene id</td></tr>
<tr><td>GU305914.1</td><td>113819</td><td>113841</td><td>+</td><td>0</td><td>TCGCCCGCGGTCCCGCCAAG</td><td>GGG</td><td>||||||||[||||||||||||]PAM</td><td>NA</td><td>Intergenic</td><td>NA</td><td>NA</td></tr>
<tr><td>T8358</td><td>CCGCGGTCCCGCCAAGGGGCTGG</td><td>1000</td><td></td><td></td><td></td><td></td><td></td><td></td><td></td><td></td><td></td></tr>
<tr><td>Oligo fwd</td><td>CACCCCGCGGTCCCGCCAAGGGGC</td><td></td><td></td><td></td><td></td><td></td><td></td><td></td><td></td><td></td><td></td></tr>
<tr><td>Oligo rev</td><td>AAACGCCCCTTGGCGGGACCGCGG</td><td></td><td></td><td></td><td></td><td></td><td></td><td></td><td></td><td></td><td></td></tr>
<tr><td>Chromosome</td><td>start</td><td>end</td><td>strand</td><td>MM</td><td>target_seq</td><td>PAM</td><td>alignment</td><td>distance</td><td>position</td><td>gene name</td><td>gene id</td></tr>
<tr><td>GU305914.1</td><td>113823</td><td>113845</td><td>+</td><td>0</td><td>CCGCGGTCCCGCCAAGGGGC</td><td>TGG</td><td>||||||||[||||||||||||]PAM</td><td>NA</td><td>Intergenic</td><td>NA</td><td>NA</td></tr>
<tr><td>T8359</td><td>TCCCGCCAAGGGGCTGGAAGCGG</td><td>1000</td><td></td><td></td><td></td><td></td><td></td><td></td><td></td><td></td><td></td></tr>
<tr><td>Oligo fwd</td><td>CACCTCCCGCCAAGGGGCTGGAAG</td><td></td><td></td><td></td><td></td><td></td><td></td><td></td><td></td><td></td><td></td></tr>
<tr><td>Oligo rev</td><td>AAACCTTCCAGCCCCTTGGCGGGA</td><td></td><td></td><td></td><td></td><td></td><td></td><td></td><td></td><td></td><td></td></tr>
<tr><td>Chromosome</td><td>start</td><td>end</td><td>strand</td><td>MM</td><td>target_seq</td><td>PAM</td><td>alignment</td><td>distance</td><td>position</td><td>gene name</td><td>gene id</td></tr>
<tr><td>GU305914.1</td><td>113829</td><td>113851</td><td>+</td><td>0</td><td>TCCCGCCAAGGGGCTGGAAG</td><td>CGG</td><td>||||||||[||||||||||||]PAM</td><td>NA</td><td>Intergenic</td><td>NA</td><td>NA</td></tr>
<tr><td>T8360</td><td>GTCGTCGAGTATCTCAGCGCCGG</td><td>1000</td><td></td><td></td><td></td><td></td><td></td><td></td><td></td><td></td><td></td></tr>
<tr><td>Oligo fwd</td><td>CACCGTCGTCGAGTATCTCAGCGC</td><td></td><td></td><td></td><td></td><td></td><td></td><td></td><td></td><td></td><td></td></tr>
<tr><td>Oligo rev</td><td>AAACGCGCTGAGATACTCGACGAC</td><td></td><td></td><td></td><td></td><td></td><td></td><td></td><td></td><td></td><td></td></tr>
<tr><td>Chromosome</td><td>start</td><td>end</td><td>strand</td><td>MM</td><td>target_seq</td><td>PAM</td><td>alignment</td><td>distance</td><td>position</td><td>gene name</td><td>gene id</td></tr>
<tr><td>GU305914.1</td><td>113857</td><td>113879</td><td>+</td><td>0</td><td>GTCGTCGAGTATCTCAGCGC</td><td>CGG</td><td>||||||||[||||||||||||]PAM</td><td>NA</td><td>Intergenic</td><td>NA</td><td>NA</td></tr>
<tr><td>T8361</td><td>TCGTCGAGTATCTCAGCGCCGGG</td><td>1000</td><td></td><td></td><td></td><td></td><td></td><td></td><td></td><td></td><td></td></tr>
<tr><td>Oligo fwd</td><td>CACCTCGTCGAGTATCTCAGCGCC</td><td></td><td></td><td></td><td></td><td></td><td></td><td></td><td></td><td></td><td></td></tr>
<tr><td>Oligo rev</td><td>AAACGGCGCTGAGATACTCGACGA</td><td></td><td></td><td></td><td></td><td></td><td></td><td></td><td></td><td></td><td></td></tr>
<tr><td>Chromosome</td><td>start</td><td>end</td><td>strand</td><td>MM</td><td>target_seq</td><td>PAM</td><td>alignment</td><td>distance</td><td>position</td><td>gene name</td><td>gene id</td></tr>
<tr><td>GU305914.1</td><td>113858</td><td>113880</td><td>+</td><td>0</td><td>TCGTCGAGTATCTCAGCGCC</td><td>GGG</td><td>||||||||[||||||||||||]PAM</td><td>NA</td><td>Intergenic</td><td>NA</td><td>NA</td></tr>
<tr><td>T8362</td><td>CGCCGCCGACGCCGACACCTCGG</td><td>1000</td><td></td><td></td><td></td><td></td><td></td><td></td><td></td><td></td><td></td></tr>
<tr><td>Oligo fwd</td><td>CACCCGCCGCCGACGCCGACACCT</td><td></td><td></td><td></td><td></td><td></td><td></td><td></td><td></td><td></td><td></td></tr>
<tr><td>Oligo rev</td><td>AAACAGGTGTCGGCGTCGGCGGCG</td><td></td><td></td><td></td><td></td><td></td><td></td><td></td><td></td><td></td><td></td></tr>
<tr><td>Chromosome</td><td>start</td><td>end</td><td>strand</td><td>MM</td><td>target_seq</td><td>PAM</td><td>alignment</td><td>distance</td><td>position</td><td>gene name</td><td>gene id</td></tr>
<tr><td>GU305914.1</td><td>113931</td><td>113953</td><td>+</td><td>0</td><td>CGCCGCCGACGCCGACACCT</td><td>CGG</td><td>||||||||[||||||||||||]PAM</td><td>NA</td><td>Intergenic</td><td>NA</td><td>NA</td></tr>
<tr><td>GU305914.1</td><td>90320</td><td>90342</td><td>-</td><td>4</td><td>CTTCGCCGACGCCGACCTCT</td><td>TGG</td><td>|--|||||[||||||||--||]PAM</td><td>NA</td><td>Intergenic</td><td>NA</td><td>NA</td></tr>
<tr><td>GU305914.1</td><td>118569</td><td>118591</td><td>-</td><td>4</td><td>AGACGCCGACGCCGACGCCA</td><td>TGG</td><td>-|-|||||[||||||||-||-]PAM</td><td>NA</td><td>Intergenic</td><td>NA</td><td>NA</td></tr>
<tr><td>T8363</td><td>GCCGACACCTCGGAGAACTCCGG</td><td>1000</td><td></td><td></td><td></td><td></td><td></td><td></td><td></td><td></td><td></td></tr>
<tr><td>Oligo fwd</td><td>CACCGCCGACACCTCGGAGAACTC</td><td></td><td></td><td></td><td></td><td></td><td></td><td></td><td></td><td></td><td></td></tr>
<tr><td>Oligo rev</td><td>AAACGAGTTCTCCGAGGTGTCGGC</td><td></td><td></td><td></td><td></td><td></td><td></td><td></td><td></td><td></td><td></td></tr>
<tr><td>Chromosome</td><td>start</td><td>end</td><td>strand</td><td>MM</td><td>target_seq</td><td>PAM</td><td>alignment</td><td>distance</td><td>position</td><td>gene name</td><td>gene id</td></tr>
<tr><td>GU305914.1</td><td>113941</td><td>113963</td><td>+</td><td>0</td><td>GCCGACACCTCGGAGAACTC</td><td>CGG</td><td>||||||||[||||||||||||]PAM</td><td>NA</td><td>Intergenic</td><td>NA</td><td>NA</td></tr>
<tr><td>T8364</td><td>CACCTCGGAGAACTCCGGCGAGG</td><td>1000</td><td></td><td></td><td></td><td></td><td></td><td></td><td></td><td></td><td></td></tr>
<tr><td>Oligo fwd</td><td>CACCCACCTCGGAGAACTCCGGCG</td><td></td><td></td><td></td><td></td><td></td><td></td><td></td><td></td><td></td><td></td></tr>
<tr><td>Oligo rev</td><td>AAACCGCCGGAGTTCTCCGAGGTG</td><td></td><td></td><td></td><td></td><td></td><td></td><td></td><td></td><td></td><td></td></tr>
<tr><td>Chromosome</td><td>start</td><td>end</td><td>strand</td><td>MM</td><td>target_seq</td><td>PAM</td><td>alignment</td><td>distance</td><td>position</td><td>gene name</td><td>gene id</td></tr>
<tr><td>GU305914.1</td><td>113946</td><td>113968</td><td>+</td><td>0</td><td>CACCTCGGAGAACTCCGGCG</td><td>AGG</td><td>||||||||[||||||||||||]PAM</td><td>NA</td><td>Intergenic</td><td>NA</td><td>NA</td></tr>
<tr><td>T8365</td><td>CTCGGAGAACTCCGGCGAGGAGG</td><td>1000</td><td></td><td></td><td></td><td></td><td></td><td></td><td></td><td></td><td></td></tr>
<tr><td>Oligo fwd</td><td>CACCCTCGGAGAACTCCGGCGAGG</td><td></td><td></td><td></td><td></td><td></td><td></td><td></td><td></td><td></td><td></td></tr>
<tr><td>Oligo rev</td><td>AAACCCTCGCCGGAGTTCTCCGAG</td><td></td><td></td><td></td><td></td><td></td><td></td><td></td><td></td><td></td><td></td></tr>
<tr><td>Chromosome</td><td>start</td><td>end</td><td>strand</td><td>MM</td><td>target_seq</td><td>PAM</td><td>alignment</td><td>distance</td><td>position</td><td>gene name</td><td>gene id</td></tr>
<tr><td>GU305914.1</td><td>113949</td><td>113971</td><td>+</td><td>0</td><td>CTCGGAGAACTCCGGCGAGG</td><td>AGG</td><td>||||||||[||||||||||||]PAM</td><td>NA</td><td>Intergenic</td><td>NA</td><td>NA</td></tr>
<tr><td>T8366</td><td>GAACTCCGGCGAGGAGGCCGAGG</td><td>1000</td><td></td><td></td><td></td><td></td><td></td><td></td><td></td><td></td><td></td></tr>
<tr><td>Oligo fwd</td><td>CACCGAACTCCGGCGAGGAGGCCG</td><td></td><td></td><td></td><td></td><td></td><td></td><td></td><td></td><td></td><td></td></tr>
<tr><td>Oligo rev</td><td>AAACCGGCCTCCTCGCCGGAGTTC</td><td></td><td></td><td></td><td></td><td></td><td></td><td></td><td></td><td></td><td></td></tr>
<tr><td>Chromosome</td><td>start</td><td>end</td><td>strand</td><td>MM</td><td>target_seq</td><td>PAM</td><td>alignment</td><td>distance</td><td>position</td><td>gene name</td><td>gene id</td></tr>
<tr><td>GU305914.1</td><td>113955</td><td>113977</td><td>+</td><td>0</td><td>GAACTCCGGCGAGGAGGCCG</td><td>AGG</td><td>||||||||[||||||||||||]PAM</td><td>NA</td><td>Intergenic</td><td>NA</td><td>NA</td></tr>
<tr><td>GU305914.1</td><td>32327</td><td>32349</td><td>-</td><td>4</td><td>GGGCTCCGACGAGGAGACCG</td><td>AGG</td><td>|--|||||[-|||||||-|||]PAM</td><td>NA</td><td>Intergenic</td><td>NA</td><td>NA</td></tr>
<tr><td>T8367</td><td>AACTCCGGCGAGGAGGCCGAGGG</td><td>1000</td><td></td><td></td><td></td><td></td><td></td><td></td><td></td><td></td><td></td></tr>
<tr><td>Oligo fwd</td><td>CACCAACTCCGGCGAGGAGGCCGA</td><td></td><td></td><td></td><td></td><td></td><td></td><td></td><td></td><td></td><td></td></tr>
<tr><td>Oligo rev</td><td>AAACTCGGCCTCCTCGCCGGAGTT</td><td></td><td></td><td></td><td></td><td></td><td></td><td></td><td></td><td></td><td></td></tr>
<tr><td>Chromosome</td><td>start</td><td>end</td><td>strand</td><td>MM</td><td>target_seq</td><td>PAM</td><td>alignment</td><td>distance</td><td>position</td><td>gene name</td><td>gene id</td></tr>
<tr><td>GU305914.1</td><td>113956</td><td>113978</td><td>+</td><td>0</td><td>AACTCCGGCGAGGAGGCCGA</td><td>GGG</td><td>||||||||[||||||||||||]PAM</td><td>NA</td><td>Intergenic</td><td>NA</td><td>NA</td></tr>
<tr><td>GU305914.1</td><td>96813</td><td>96835</td><td>-</td><td>4</td><td>ACCGCCGGCGACGAGGACGA</td><td>AGG</td><td>|-|-||||[|||-||||-|||]PAM</td><td>NA</td><td>Intergenic</td><td>NA</td><td>NA</td></tr>
<tr><td>T8368</td><td>AGGAGGCCGAGGGACAGCCGCGG</td><td>1000</td><td></td><td></td><td></td><td></td><td></td><td></td><td></td><td></td><td></td></tr>
<tr><td>Oligo fwd</td><td>CACCAGGAGGCCGAGGGACAGCCG</td><td></td><td></td><td></td><td></td><td></td><td></td><td></td><td></td><td></td><td></td></tr>
<tr><td>Oligo rev</td><td>AAACCGGCTGTCCCTCGGCCTCCT</td><td></td><td></td><td></td><td></td><td></td><td></td><td></td><td></td><td></td><td></td></tr>
<tr><td>Chromosome</td><td>start</td><td>end</td><td>strand</td><td>MM</td><td>target_seq</td><td>PAM</td><td>alignment</td><td>distance</td><td>position</td><td>gene name</td><td>gene id</td></tr>
<tr><td>GU305914.1</td><td>113966</td><td>113988</td><td>+</td><td>0</td><td>AGGAGGCCGAGGGACAGCCG</td><td>CGG</td><td>||||||||[||||||||||||]PAM</td><td>NA</td><td>Intergenic</td><td>NA</td><td>NA</td></tr>
<tr><td>T8369</td><td>AGGCCGAGGGACAGCCGCGGCGG</td><td>1000</td><td></td><td></td><td></td><td></td><td></td><td></td><td></td><td></td><td></td></tr>
</table>

Oligo fwd CACCAGGCCGAGGGACAGCCGCGG
Oligo rev AAACCCGCGGCTGTCCCTCGGCCT
Chromosome start end strand MM target_seq PAM alignment distance position gene name gene id
GU305914.1 113969 113991 + 0 AGGCCGAGGGACAGCCGCGG CGG ||||||||[||||||||||||]PAM NA Intergenic NA NA

T8370 GCGGCGGCAGACGACCGACTCGG 1000
Oligo fwd CACCGCGGCGGCAGACGACCGACT
Oligo rev AAACAGTCGGTCGTCTGCCGCCGC
Chromosome start end strand MM target_seq PAM alignment distance position gene name gene id
GU305914.1 113985 114007 + 0 GCGGCGGCAGACGACCGACT CGG ||||||||[||||||||||||]PAM NA Intergenic NA NA

T8371 CGGCGGCAGACGACCGACTCGGG 1000
Oligo fwd CACCCGGCGGCAGACGACCGACTC
Oligo rev AAACGAGTCGGTCGTCTGCCGCCG
Chromosome start end strand MM target_seq PAM alignment distance position gene name gene id
GU305914.1 113986 114008 + 0 CGGCGGCAGACGACCGACTC GGG ||||||||[||||||||||||]PAM NA Intergenic NA NA

T8372 CGGCAGACGACCGACTCGGGCGG 1000
Oligo fwd CACCCGGCAGACGACCGACTCGGG
Oligo rev AAACCCCGAGTCGGTCGTCTGCCG
Chromosome start end strand MM target_seq PAM alignment distance position gene name gene id
GU305914.1 113989 114011 + 0 CGGCAGACGACCGACTCGGG CGG ||||||||[||||||||||||]PAM NA Intergenic NA NA

T8373 TTCAGACACGTTTCGCTCTGCGG 1000
Oligo fwd CACCTTCAGACACGTTTCGCTCTG
Oligo rev AAACCAGAGCGAAACGTGTCTGAA
Chromosome start end strand MM target_seq PAM alignment distance position gene name gene id
GU305914.1 114016 114038 + 0 TTCAGACACGTTTCGCTCTG CGG ||||||||[||||||||||||]PAM NA Intergenic NA NA

T8374 TCAGACACGTTTCGCTCTGCGGG 1000
Oligo fwd CACCTCAGACACGTTTCGCTCTGC
Oligo rev AAACGCAGAGCGAAACGTGTCTGA
Chromosome start end strand MM target_seq PAM alignment distance position gene name gene id
GU305914.1 114017 114039 + 0 TCAGACACGTTTCGCTCTGC GGG ||||||||[||||||||||||]PAM NA Intergenic NA NA

T8375 ACACGTTTCGCTCTGCGGGTTGG 1000
Oligo fwd CACCACACGTTTCGCTCTGCGGGT
Oligo rev AAACACCCGCAGAGCGAAACGTGT
Chromosome start end strand MM target_seq PAM alignment distance position gene name gene id
GU305914.1 114021 114043 + 0 ACACGTTTCGCTCTGCGGGT TGG ||||||||[||||||||||||]PAM NA Intergenic NA NA

T8376 CACGTTTCGCTCTGCGGGTTGGG 1000
Oligo fwd CACCCACGTTTCGCTCTGCGGGTT
Oligo rev AAACAACCCGCAGAGCGAAACGTG
Chromosome start end strand MM target_seq PAM alignment distance position gene name gene id
GU305914.1 114022 114044 + 0 CACGTTTCGCTCTGCGGGTT GGG ||||||||[||||||||||||]PAM NA Intergenic NA NA

T8377 ACGTTTCGCTCTGCGGGTTGGGG 1000
Oligo fwd CACCACGTTTCGCTCTGCGGGTTG
Oligo rev AAACCAACCCGCAGAGCGAAACGT
Chromosome start end strand MM target_seq PAM alignment distance position gene name gene id
GU305914.1 114023 114045 + 0 ACGTTTCGCTCTGCGGGTTG GGG ||||||||[||||||||||||]PAM NA Intergenic NA NA

T8378 TTTCGCTCTGCGGGTTGGGGAGG 1000
Oligo fwd CACCTTTCGCTCTGCGGGTTGGGG
Oligo rev AAACCCCCAACCCGCAGAGCGAAA
Chromosome start end strand MM target_seq PAM alignment distance position gene name gene id
GU305914.1 114026 114048 + 0 TTTCGCTCTGCGGGTTGGGG AGG ||||||||[||||||||||||]PAM NA Intergenic NA NA

T8379 CGCTCTGCGGGTTGGGGAGGCGG 1000
Oligo fwd CACCCGCTCTGCGGGTTGGGGAGG
Oligo rev AAACCCTCCCCAACCCGCAGAGCG
Chromosome start end strand MM target_seq PAM alignment distance position gene name gene id
GU305914.1 114029 114051 + 0 CGCTCTGCGGGTTGGGGAGG CGG ||||||||[||||||||||||]PAM NA Intergenic NA NA

T8380 TCTGCGGGTTGGGGAGGCGGAGG 1000
Oligo fwd CACCTCTGCGGGTTGGGGAGGCGG
Oligo rev AAACCCGCCTCCCCAACCCGCAGA
Chromosome start end strand MM target_seq PAM alignment distance position gene name gene id
GU305914.1 114032 114054 + 0 TCTGCGGGTTGGGGAGGCGG AGG ||||||||[||||||||||||]PAM NA Intergenic NA NA

T8381 CTGCGGGTTGGGGAGGCGGAGGG 1000
Oligo fwd CACCCTGCGGGTTGGGGAGGCGGA
Oligo rev AAACTCCGCCTCCCCAACCCGCAG
Chromosome start end strand MM target_seq PAM alignment distance position gene name gene id
GU305914.1 114033 114055 + 0 CTGCGGGTTGGGGAGGCGGA GGG ||||||||[||||||||||||]PAM NA Intergenic NA NA

T8382 TGCGGGTTGGGGAGGCGGAGGGG 1000
Oligo fwd CACCTGCGGGTTGGGGAGGCGGAG
Oligo rev AAACCTCCGCCTCCCCAACCCGCA
Chromosome start end strand MM target_seq PAM alignment distance position gene name gene id
GU305914.1 114034 114056 + 0 TGCGGGTTGGGGAGGCGGAG GGG ||||||||[||||||||||||]PAM NA Intergenic NA NA

T8383 GGGGAGGCGGAGGGGTACCTTGG 1000
Oligo fwd CACCGGGGAGGCGGAGGGGTACCT
Oligo rev AAACAGGTACCCCTCCGCCTCCCC
Chromosome start end strand MM target_seq PAM alignment distance position gene name gene id
GU305914.1 114042 114064 + 0 GGGGAGGCGGAGGGGTACCT TGG ||||||||[||||||||||||]PAM NA Intergenic NA NA

| | | | | | | | | | | | |
|---|---|---|---|---|---|---|---|---|---|---|---|
| T8384 | GAGGCGGAGGGGTACCTTGGCGG | 1000 | | | | | | | | | |
| Oligo fwd | CACCGAGGCGGAGGGGTACCTTGG | | | | | | | | | | |
| Oligo rev | AAACCCAAGGTACCCCTCCGCCTC | | | | | | | | | | |
| Chromosome | start | end | strand | MM | target_seq | PAM | alignment | distance | position | gene name | gene id |
| GU305914.1 | 114045 | 114067 | + | 0 | GAGGCGGAGGGGTACCTTGG | CGG | \|\|\|\|\|\|\|\|[\|\|\|\|\|\|\|\|\|\|\|\|]PAM | NA | Intergenic | NA | NA |
| T8385 | AGGGGTACCTTGGCGGTCTACGG | 1000 | | | | | | | | | |
| Oligo fwd | CACCAGGGGTACCTTGGCGGTCTA | | | | | | | | | | |
| Oligo rev | AAACTAGACCGCCAAGGTACCCCT | | | | | | | | | | |
| Chromosome | start | end | strand | MM | target_seq | PAM | alignment | distance | position | gene name | gene id |
| GU305914.1 | 114052 | 114074 | + | 0 | AGGGGTACCTTGGCGGTCTA | CGG | \|\|\|\|\|\|\|\|[\|\|\|\|\|\|\|\|\|\|\|\|]PAM | NA | Intergenic | NA | NA |
| T8386 | TCTACGGACGAGATCGCGACTGG | 1000 | | | | | | | | | |
| Oligo fwd | CACCTCTACGGACGAGATCGCGAC | | | | | | | | | | |
| Oligo rev | AAACGTCGCGATCTCGTCCGTAGA | | | | | | | | | | |
| Chromosome | start | end | strand | MM | target_seq | PAM | alignment | distance | position | gene name | gene id |
| GU305914.1 | 114068 | 114090 | + | 0 | TCTACGGACGAGATCGCGAC | TGG | \|\|\|\|\|\|\|\|[\|\|\|\|\|\|\|\|\|\|\|\|]PAM | NA | Intergenic | NA | NA |
| T8387 | ACGAGATCGCGACTGGATAGTGG | 1000 | | | | | | | | | |
| Oligo fwd | CACCACGAGATCGCGACTGGATAG | | | | | | | | | | |
| Oligo rev | AAACCTATCCAGTCGCGATCTCGT | | | | | | | | | | |
| Chromosome | start | end | strand | MM | target_seq | PAM | alignment | distance | position | gene name | gene id |
| GU305914.1 | 114075 | 114097 | + | 0 | ACGAGATCGCGACTGGATAG | TGG | \|\|\|\|\|\|\|\|[\|\|\|\|\|\|\|\|\|\|\|\|]PAM | NA | Intergenic | NA | NA |
| T8388 | CGAGATCGCGACTGGATAGTGGG | 1000 | | | | | | | | | |
| Oligo fwd | CACCCGAGATCGCGACTGGATAGT | | | | | | | | | | |
| Oligo rev | AAACACTATCCAGTCGCGATCTCG | | | | | | | | | | |
| Chromosome | start | end | strand | MM | target_seq | PAM | alignment | distance | position | gene name | gene id |
| GU305914.1 | 114076 | 114098 | + | 0 | CGAGATCGCGACTGGATAGT | GGG | \|\|\|\|\|\|\|\|[\|\|\|\|\|\|\|\|\|\|\|\|]PAM | NA | Intergenic | NA | NA |
| T8389 | GAGATCGCGACTGGATAGTGGGG | 1000 | | | | | | | | | |
| Oligo fwd | CACCGAGATCGCGACTGGATAGTG | | | | | | | | | | |
| Oligo rev | AAACCACTATCCAGTCGCGATCTC | | | | | | | | | | |
| Chromosome | start | end | strand | MM | target_seq | PAM | alignment | distance | position | gene name | gene id |
| GU305914.1 | 114077 | 114099 | + | 0 | GAGATCGCGACTGGATAGTG | GGG | \|\|\|\|\|\|\|\|[\|\|\|\|\|\|\|\|\|\|\|\|]PAM | NA | Intergenic | NA | NA |
| T8390 | ATCGCGACTGGATAGTGGGGAGG | 1000 | | | | | | | | | |
| Oligo fwd | CACCATCGCGACTGGATAGTGGGG | | | | | | | | | | |
| Oligo rev | AAACCCCCACTATCCAGTCGCGAT | | | | | | | | | | |
| Chromosome | start | end | strand | MM | target_seq | PAM | alignment | distance | position | gene name | gene id |
| GU305914.1 | 114080 | 114102 | + | 0 | ATCGCGACTGGATAGTGGGG | AGG | \|\|\|\|\|\|\|\|[\|\|\|\|\|\|\|\|\|\|\|\|]PAM | NA | Intergenic | NA | NA |
| T8391 | CTGGATAGTGGGGAGGTTCGCGG | 1000 | | | | | | | | | |
| Oligo fwd | CACCCTGGATAGTGGGGAGGTTCG | | | | | | | | | | |
| Oligo rev | AAACCGAACCTCCCCACTATCCAG | | | | | | | | | | |
| Chromosome | start | end | strand | MM | target_seq | PAM | alignment | distance | position | gene name | gene id |
| GU305914.1 | 114087 | 114109 | + | 0 | CTGGATAGTGGGGAGGTTCG | CGG | \|\|\|\|\|\|\|\|[\|\|\|\|\|\|\|\|\|\|\|\|]PAM | NA | Intergenic | NA | NA |
| T8392 | GAGGTTCGCGGCTCTTACGCCGG | 1000 | | | | | | | | | |
| Oligo fwd | CACCGAGGTTCGCGGCTCTTACGC | | | | | | | | | | |
| Oligo rev | AAACGCGTAAGAGCCGCGAACCTC | | | | | | | | | | |
| Chromosome | start | end | strand | MM | target_seq | PAM | alignment | distance | position | gene name | gene id |
| GU305914.1 | 114099 | 114121 | + | 0 | GAGGTTCGCGGCTCTTACGC | CGG | \|\|\|\|\|\|\|\|[\|\|\|\|\|\|\|\|\|\|\|\|]PAM | NA | Intergenic | NA | NA |
| T8393 | CGGCTCTTACGCCGGACGAGAGG | 1000 | | | | | | | | | |
| Oligo fwd | CACCCGGCTCTTACGCCGGACGAG | | | | | | | | | | |
| Oligo rev | AAACCTCGTCCGGCGTAAGAGCCG | | | | | | | | | | |
| Chromosome | start | end | strand | MM | target_seq | PAM | alignment | distance | position | gene name | gene id |
| GU305914.1 | 114107 | 114129 | + | 0 | CGGCTCTTACGCCGGACGAG | AGG | \|\|\|\|\|\|\|\|[\|\|\|\|\|\|\|\|\|\|\|\|]PAM | NA | Intergenic | NA | NA |
| T8394 | GGCTCTTACGCCGGACGAGAGGG | 1000 | | | | | | | | | |
| Oligo fwd | CACCGGCTCTTACGCCGGACGAGA | | | | | | | | | | |
| Oligo rev | AAACTCTCGTCCGGCGTAAGAGCC | | | | | | | | | | |
| Chromosome | start | end | strand | MM | target_seq | PAM | alignment | distance | position | gene name | gene id |
| GU305914.1 | 114108 | 114130 | + | 0 | GGCTCTTACGCCGGACGAGA | GGG | \|\|\|\|\|\|\|\|[\|\|\|\|\|\|\|\|\|\|\|\|]PAM | NA | Intergenic | NA | NA |
| T8395 | TCTTACGCCGGACGAGAGGGCGG | 1000 | | | | | | | | | |
| Oligo fwd | CACCTCTTACGCCGGACGAGAGGG | | | | | | | | | | |
| Oligo rev | AAACCCCTCTCGTCCGGCGTAAGA | | | | | | | | | | |
| Chromosome | start | end | strand | MM | target_seq | PAM | alignment | distance | position | gene name | gene id |
| GU305914.1 | 114111 | 114133 | + | 0 | TCTTACGCCGGACGAGAGGG | CGG | \|\|\|\|\|\|\|\|[\|\|\|\|\|\|\|\|\|\|\|\|]PAM | NA | Intergenic | NA | NA |
| T8396 | GATCGCGCGTGTCGACGACGCGG | 1000 | | | | | | | | | |
| Oligo fwd | CACCGATCGCGCGTGTCGACGACG | | | | | | | | | | |
| Oligo rev | AAACCGTCGTCGACACGCGCGATC | | | | | | | | | | |
| Chromosome | start | end | strand | MM | target_seq | PAM | alignment | distance | position | gene name | gene id |
| GU305914.1 | 114135 | 114157 | + | 0 | GATCGCGCGTGTCGACGACG | CGG | \|\|\|\|\|\|\|\|[\|\|\|\|\|\|\|\|\|\|\|\|]PAM | NA | Intergenic | NA | NA |
| T8397 | CGTGTCGACGACGCGGCGCTCGG | 1000 | | | | | | | | | |
| Oligo fwd | CACCCGTGTCGACGACGCGGCGCT | | | | | | | | | | |
| Oligo rev | AAACAGCGCCGCGTCGTCGACACG | | | | | | | | | | |
| Chromosome | start | end | strand | MM | target_seq | PAM | alignment | distance | position | gene name | gene id |
| GU305914.1 | 114142 | 114164 | + | 0 | CGTGTCGACGACGCGGCGCT | CGG | \|\|\|\|\|\|\|\|[\|\|\|\|\|\|\|\|\|\|\|\|]PAM | NA | Intergenic | NA | NA |
| GU305914.1 | 43394 | 43416 | - | 4 | AGTGTCGGGGACGCGGCGCG | TGG | -\|\|\|\|\|\|-[-\|\|\|\|\|\|\|\|\|\|-]PAM | NA | Intergenic | NA | NA |
| T8398 | GTGTCGACGACGCGGCGCTCGGG | 1000 | | | | | | | | | |
| Oligo fwd | CACCGTGTCGACGACGCGGCGCTC | | | | | | | | | | |
| Oligo rev | AAACGAGCGCCGCGTCGTCGACAC | | | | | | | | | | |
| Chromosome | start | end | strand | MM | target_seq | PAM | alignment | distance | position | gene name | gene id |

<table>
<tr><td>GU305914.1</td><td>114143</td><td>114165</td><td>+</td><td>0</td><td>GTGTCGACGACGCGGCGCTC</td><td>GGG</td><td>||||||||[||||||||||||]PAM</td><td>NA</td><td>Intergenic</td><td>NA</td><td>NA</td></tr>
<tr><td>GU305914.1</td><td>43393</td><td>43415</td><td>-</td><td>4</td><td>GTGTCGGGGACGCGGCGCGT</td><td>GGG</td><td>||||||--[||||||||||--]PAM</td><td>NA</td><td>Intergenic</td><td>NA</td><td>NA</td></tr>
<tr><td>T8399</td><td>TGTCGACGACGCGGCGCTCGGGG</td><td>1000</td><td></td><td></td><td></td><td></td><td></td><td></td><td></td><td></td><td></td></tr>
<tr><td>Oligo fwd</td><td>CACCTGTCGACGACGCGGCGCTCG</td><td></td><td></td><td></td><td></td><td></td><td></td><td></td><td></td><td></td><td></td></tr>
<tr><td>Oligo rev</td><td>AAACCGAGCGCCGCGTCGTCGACA</td><td></td><td></td><td></td><td></td><td></td><td></td><td></td><td></td><td></td><td></td></tr>
<tr><td>Chromosome</td><td>start</td><td>end</td><td>strand</td><td>MM</td><td>target_seq</td><td>PAM</td><td>alignment</td><td>distance</td><td>position</td><td>gene name</td><td>gene id</td></tr>
<tr><td>GU305914.1</td><td>114144</td><td>114166</td><td>+</td><td>0</td><td>TGTCGACGACGCGGCGCTCG</td><td>GGG</td><td>||||||||[||||||||||||]PAM</td><td>NA</td><td>Intergenic</td><td>NA</td><td>NA</td></tr>
<tr><td>GU305914.1</td><td>43392</td><td>43414</td><td>-</td><td>4</td><td>TGTCGGGGACGCGGCGCGTG</td><td>GGG</td><td>|||||--|[|||||||||--|]PAM</td><td>NA</td><td>Intergenic</td><td>NA</td><td>NA</td></tr>
<tr><td>T8400</td><td>GTCGACGACGCGGCGCTCGGGGG</td><td>1000</td><td></td><td></td><td></td><td></td><td></td><td></td><td></td><td></td><td></td></tr>
<tr><td>Oligo fwd</td><td>CACCGTCGACGACGCGGCGCTCGG</td><td></td><td></td><td></td><td></td><td></td><td></td><td></td><td></td><td></td><td></td></tr>
<tr><td>Oligo rev</td><td>AAACCCGAGCGCCGCGTCGTCGAC</td><td></td><td></td><td></td><td></td><td></td><td></td><td></td><td></td><td></td><td></td></tr>
<tr><td>Chromosome</td><td>start</td><td>end</td><td>strand</td><td>MM</td><td>target_seq</td><td>PAM</td><td>alignment</td><td>distance</td><td>position</td><td>gene name</td><td>gene id</td></tr>
<tr><td>GU305914.1</td><td>114145</td><td>114167</td><td>+</td><td>0</td><td>GTCGACGACGCGGCGCTCGG</td><td>GGG</td><td>||||||||[||||||||||||]PAM</td><td>NA</td><td>Intergenic</td><td>NA</td><td>NA</td></tr>
<tr><td>T8401</td><td>ACGACGCGGCGCTCGGGGGCTGG</td><td>1000</td><td></td><td></td><td></td><td></td><td></td><td></td><td></td><td></td><td></td></tr>
<tr><td>Oligo fwd</td><td>CACCACGACGCGGCGCTCGGGGGC</td><td></td><td></td><td></td><td></td><td></td><td></td><td></td><td></td><td></td><td></td></tr>
<tr><td>Oligo rev</td><td>AAACGCCCCCGAGCGCCGCGTCGT</td><td></td><td></td><td></td><td></td><td></td><td></td><td></td><td></td><td></td><td></td></tr>
<tr><td>Chromosome</td><td>start</td><td>end</td><td>strand</td><td>MM</td><td>target_seq</td><td>PAM</td><td>alignment</td><td>distance</td><td>position</td><td>gene name</td><td>gene id</td></tr>
<tr><td>GU305914.1</td><td>114149</td><td>114171</td><td>+</td><td>0</td><td>ACGACGCGGCGCTCGGGGGC</td><td>TGG</td><td>||||||||[||||||||||||]PAM</td><td>NA</td><td>Intergenic</td><td>NA</td><td>NA</td></tr>
<tr><td>GU305914.1</td><td>194661</td><td>194683</td><td>+</td><td>4</td><td>GCGACGTGGCGATCGGCGGC</td><td>CGG</td><td>-|||||-|[|||-||||-|||]PAM</td><td>NA</td><td>Intergenic</td><td>NA</td><td>NA</td></tr>
<tr><td>T8402</td><td>CGACGCGGCGCTCGGGGGCTGGG</td><td>1000</td><td></td><td></td><td></td><td></td><td></td><td></td><td></td><td></td><td></td></tr>
<tr><td>Oligo fwd</td><td>CACCCGACGCGGCGCTCGGGGGCT</td><td></td><td></td><td></td><td></td><td></td><td></td><td></td><td></td><td></td><td></td></tr>
<tr><td>Oligo rev</td><td>AAACAGCCCCCGAGCGCCGCGTCG</td><td></td><td></td><td></td><td></td><td></td><td></td><td></td><td></td><td></td><td></td></tr>
<tr><td>Chromosome</td><td>start</td><td>end</td><td>strand</td><td>MM</td><td>target_seq</td><td>PAM</td><td>alignment</td><td>distance</td><td>position</td><td>gene name</td><td>gene id</td></tr>
<tr><td>GU305914.1</td><td>114150</td><td>114172</td><td>+</td><td>0</td><td>CGACGCGGCGCTCGGGGGCT</td><td>GGG</td><td>||||||||[||||||||||||]PAM</td><td>NA</td><td>Intergenic</td><td>NA</td><td>NA</td></tr>
<tr><td>T8403</td><td>GCGGCGCTCGGGGGCTGGGACGG</td><td>1000</td><td></td><td></td><td></td><td></td><td></td><td></td><td></td><td></td><td></td></tr>
<tr><td>Oligo fwd</td><td>CACCGCGGCGCTCGGGGGCTGGGA</td><td></td><td></td><td></td><td></td><td></td><td></td><td></td><td></td><td></td><td></td></tr>
<tr><td>Oligo rev</td><td>AAACTCCCAGCCCCCGAGCGCCGC</td><td></td><td></td><td></td><td></td><td></td><td></td><td></td><td></td><td></td><td></td></tr>
<tr><td>Chromosome</td><td>start</td><td>end</td><td>strand</td><td>MM</td><td>target_seq</td><td>PAM</td><td>alignment</td><td>distance</td><td>position</td><td>gene name</td><td>gene id</td></tr>
<tr><td>GU305914.1</td><td>114154</td><td>114176</td><td>+</td><td>0</td><td>GCGGCGCTCGGGGGCTGGGA</td><td>CGG</td><td>||||||||[||||||||||||]PAM</td><td>NA</td><td>Intergenic</td><td>NA</td><td>NA</td></tr>
<tr><td>T8404</td><td>GACGGAGACGCCGATCCCTTCGG</td><td>1000</td><td></td><td></td><td></td><td></td><td></td><td></td><td></td><td></td><td></td></tr>
<tr><td>Oligo fwd</td><td>CACCGACGGAGACGCCGATCCCTT</td><td></td><td></td><td></td><td></td><td></td><td></td><td></td><td></td><td></td><td></td></tr>
<tr><td>Oligo rev</td><td>AAACAAGGGATCGGCGTCTCCGTC</td><td></td><td></td><td></td><td></td><td></td><td></td><td></td><td></td><td></td><td></td></tr>
<tr><td>Chromosome</td><td>start</td><td>end</td><td>strand</td><td>MM</td><td>target_seq</td><td>PAM</td><td>alignment</td><td>distance</td><td>position</td><td>gene name</td><td>gene id</td></tr>
<tr><td>GU305914.1</td><td>114172</td><td>114194</td><td>+</td><td>0</td><td>GACGGAGACGCCGATCCCTT</td><td>CGG</td><td>||||||||[||||||||||||]PAM</td><td>NA</td><td>Intergenic</td><td>NA</td><td>NA</td></tr>
<tr><td>GU305914.1</td><td>194506</td><td>194528</td><td>+</td><td>3</td><td>GATGGAGACGGGGATCCCTT</td><td>TGG</td><td>||-|||||[||--||||||||]PAM</td><td>NA</td><td>Intergenic</td><td>NA</td><td>NA</td></tr>
<tr><td>T8405</td><td>AGACGCCGATCCCTTCGGTTCGG</td><td>1000</td><td></td><td></td><td></td><td></td><td></td><td></td><td></td><td></td><td></td></tr>
<tr><td>Oligo fwd</td><td>CACCAGACGCCGATCCCTTCGGTT</td><td></td><td></td><td></td><td></td><td></td><td></td><td></td><td></td><td></td><td></td></tr>
<tr><td>Oligo rev</td><td>AAACAACCGAAGGGATCGGCGTCT</td><td></td><td></td><td></td><td></td><td></td><td></td><td></td><td></td><td></td><td></td></tr>
<tr><td>Chromosome</td><td>start</td><td>end</td><td>strand</td><td>MM</td><td>target_seq</td><td>PAM</td><td>alignment</td><td>distance</td><td>position</td><td>gene name</td><td>gene id</td></tr>
<tr><td>GU305914.1</td><td>114177</td><td>114199</td><td>+</td><td>0</td><td>AGACGCCGATCCCTTCGGTT</td><td>CGG</td><td>||||||||[||||||||||||]PAM</td><td>NA</td><td>Intergenic</td><td>NA</td><td>NA</td></tr>
<tr><td>T8406</td><td>CCCTTCGGTTCGGACTCATACGG</td><td>1000</td><td></td><td></td><td></td><td></td><td></td><td></td><td></td><td></td><td></td></tr>
<tr><td>Oligo fwd</td><td>CACCCCCTTCGGTTCGGACTCATA</td><td></td><td></td><td></td><td></td><td></td><td></td><td></td><td></td><td></td><td></td></tr>
<tr><td>Oligo rev</td><td>AAACTATGAGTCCGAACCGAAGGG</td><td></td><td></td><td></td><td></td><td></td><td></td><td></td><td></td><td></td><td></td></tr>
<tr><td>Chromosome</td><td>start</td><td>end</td><td>strand</td><td>MM</td><td>target_seq</td><td>PAM</td><td>alignment</td><td>distance</td><td>position</td><td>gene name</td><td>gene id</td></tr>
<tr><td>GU305914.1</td><td>114187</td><td>114209</td><td>+</td><td>0</td><td>CCCTTCGGTTCGGACTCATA</td><td>CGG</td><td>||||||||[||||||||||||]PAM</td><td>NA</td><td>Intergenic</td><td>NA</td><td>NA</td></tr>
<tr><td>T8407</td><td>ATACGGCCTCCTCGCCAGCACGG</td><td>1000</td><td></td><td></td><td></td><td></td><td></td><td></td><td></td><td></td><td></td></tr>
<tr><td>Oligo fwd</td><td>CACCATACGGCCTCCTCGCCAGCA</td><td></td><td></td><td></td><td></td><td></td><td></td><td></td><td></td><td></td><td></td></tr>
<tr><td>Oligo rev</td><td>AAACTGCTGGCGAGGAGGCCGTAT</td><td></td><td></td><td></td><td></td><td></td><td></td><td></td><td></td><td></td><td></td></tr>
<tr><td>Chromosome</td><td>start</td><td>end</td><td>strand</td><td>MM</td><td>target_seq</td><td>PAM</td><td>alignment</td><td>distance</td><td>position</td><td>gene name</td><td>gene id</td></tr>
<tr><td>GU305914.1</td><td>114204</td><td>114226</td><td>+</td><td>0</td><td>ATACGGCCTCCTCGCCAGCA</td><td>CGG</td><td>||||||||[||||||||||||]PAM</td><td>NA</td><td>Intergenic</td><td>NA</td><td>NA</td></tr>
<tr><td>T8408</td><td>CTCGCCAGCACGGTCGACGACGG</td><td>1000</td><td></td><td></td><td></td><td></td><td></td><td></td><td></td><td></td><td></td></tr>
<tr><td>Oligo fwd</td><td>CACCCTCGCCAGCACGGTCGACGA</td><td></td><td></td><td></td><td></td><td></td><td></td><td></td><td></td><td></td><td></td></tr>
<tr><td>Oligo rev</td><td>AAACTCGTCGACCGTGCTGGCGAG</td><td></td><td></td><td></td><td></td><td></td><td></td><td></td><td></td><td></td><td></td></tr>
<tr><td>Chromosome</td><td>start</td><td>end</td><td>strand</td><td>MM</td><td>target_seq</td><td>PAM</td><td>alignment</td><td>distance</td><td>position</td><td>gene name</td><td>gene id</td></tr>
<tr><td>GU305914.1</td><td>114214</td><td>114236</td><td>+</td><td>0</td><td>CTCGCCAGCACGGTCGACGA</td><td>CGG</td><td>||||||||[||||||||||||]PAM</td><td>NA</td><td>Intergenic</td><td>NA</td><td>NA</td></tr>
<tr><td>T8409</td><td>GGTCGACGACGGTTACATCGCGG</td><td>1000</td><td></td><td></td><td></td><td></td><td></td><td></td><td></td><td></td><td></td></tr>
<tr><td>Oligo fwd</td><td>CACCGGTCGACGACGGTTACATCG</td><td></td><td></td><td></td><td></td><td></td><td></td><td></td><td></td><td></td><td></td></tr>
<tr><td>Oligo rev</td><td>AAACCGATGTAACCGTCGTCGACC</td><td></td><td></td><td></td><td></td><td></td><td></td><td></td><td></td><td></td><td></td></tr>
<tr><td>Chromosome</td><td>start</td><td>end</td><td>strand</td><td>MM</td><td>target_seq</td><td>PAM</td><td>alignment</td><td>distance</td><td>position</td><td>gene name</td><td>gene id</td></tr>
<tr><td>GU305914.1</td><td>114225</td><td>114247</td><td>+</td><td>0</td><td>GGTCGACGACGGTTACATCG</td><td>CGG</td><td>||||||||[||||||||||||]PAM</td><td>NA</td><td>Intergenic</td><td>NA</td><td>NA</td></tr>
<tr><td>T8410</td><td>GTCGCCTCAGATACGACAAGCGG</td><td>1000</td><td></td><td></td><td></td><td></td><td></td><td></td><td></td><td></td><td></td></tr>
<tr><td>Oligo fwd</td><td>CACCGTCGCCTCAGATACGACAAG</td><td></td><td></td><td></td><td></td><td></td><td></td><td></td><td></td><td></td><td></td></tr>
<tr><td>Oligo rev</td><td>AAACCTTGTCGTATCTGAGGCGAC</td><td></td><td></td><td></td><td></td><td></td><td></td><td></td><td></td><td></td><td></td></tr>
<tr><td>Chromosome</td><td>start</td><td>end</td><td>strand</td><td>MM</td><td>target_seq</td><td>PAM</td><td>alignment</td><td>distance</td><td>position</td><td>gene name</td><td>gene id</td></tr>
<tr><td>GU305914.1</td><td>114257</td><td>114279</td><td>+</td><td>0</td><td>GTCGCCTCAGATACGACAAG</td><td>CGG</td><td>||||||||[||||||||||||]PAM</td><td>NA</td><td>Intergenic</td><td>NA</td><td>NA</td></tr>
<tr><td>T8411</td><td>CAGATACGACAAGCGGCTGCTGG</td><td>1000</td><td></td><td></td><td></td><td></td><td></td><td></td><td></td><td></td><td></td></tr>
<tr><td>Oligo fwd</td><td>CACCCAGATACGACAAGCGGCTGC</td><td></td><td></td><td></td><td></td><td></td><td></td><td></td><td></td><td></td><td></td></tr>
<tr><td>Oligo rev</td><td>AAACGCAGCCGCTTGTCGTATCTG</td><td></td><td></td><td></td><td></td><td></td><td></td><td></td><td></td><td></td><td></td></tr>
<tr><td>Chromosome</td><td>start</td><td>end</td><td>strand</td><td>MM</td><td>target_seq</td><td>PAM</td><td>alignment</td><td>distance</td><td>position</td><td>gene name</td><td>gene id</td></tr>
<tr><td>GU305914.1</td><td>114264</td><td>114286</td><td>+</td><td>0</td><td>CAGATACGACAAGCGGCTGC</td><td>TGG</td><td>||||||||[||||||||||||]PAM</td><td>NA</td><td>Intergenic</td><td>NA</td><td>NA</td></tr>
<tr><td>T8412</td><td>AGATACGACAAGCGGCTGCTGGG</td><td>1000</td><td></td><td></td><td></td><td></td><td></td><td></td><td></td><td></td><td></td></tr>
<tr><td>Oligo fwd</td><td>CACCAGATACGACAAGCGGCTGCT</td><td></td><td></td><td></td><td></td><td></td><td></td><td></td><td></td><td></td><td></td></tr>
<tr><td>Oligo rev</td><td>AAACAGCAGCCGCTTGTCGTATCT</td><td></td><td></td><td></td><td></td><td></td><td></td><td></td><td></td><td></td><td></td></tr>
<tr><td>Chromosome</td><td>start</td><td>end</td><td>strand</td><td>MM</td><td>target_seq</td><td>PAM</td><td>alignment</td><td>distance</td><td>position</td><td>gene name</td><td>gene id</td></tr>
<tr><td>GU305914.1</td><td>114265</td><td>114287</td><td>+</td><td>0</td><td>AGATACGACAAGCGGCTGCT</td><td>GGG</td><td>||||||||[||||||||||||]PAM</td><td>NA</td><td>Intergenic</td><td>NA</td><td>NA</td></tr>
</table>

| | | | | | | | | | | | |
|---|---|---|---|---|---|---|---|---|---|---|---|
| T8413 | GCTGCTGGGTTTGCAGTCCAAGG | 1000 | | | | | | | | | |
| Oligo fwd | CACCGCTGCTGGGTTTGCAGTCCA | | | | | | | | | | |
| Oligo rev | AAACTGGACTGCAAACCCAGCAGC | | | | | | | | | | |
| Chromosome | start | end | strand | MM | target_seq | PAM | alignment | distance | position | gene name | gene id |
| GU305914.1 | 114279 | 114301 | + | 0 | GCTGCTGGGTTTGCAGTCCA | AGG | \|\|\|\|\|\|\|\|[\|\|\|\|\|\|\|\|\|\|\|\|]PAM | NA | Intergenic | NA | NA |

| | | | | | | | | | | | |
|---|---|---|---|---|---|---|---|---|---|---|---|
| T8414 | CGTCAAAGCCAGCGAGCTGCCGG | 1000 | | | | | | | | | |
| Oligo fwd | CACCCGTCAAAGCCAGCGAGCTGC | | | | | | | | | | |
| Oligo rev | AAACGCAGCTCGCTGGCTTTGACG | | | | | | | | | | |
| Chromosome | start | end | strand | MM | target_seq | PAM | alignment | distance | position | gene name | gene id |
| GU305914.1 | 114309 | 114331 | + | 0 | CGTCAAAGCCAGCGAGCTGC | CGG | \|\|\|\|\|\|\|\|[\|\|\|\|\|\|\|\|\|\|\|\|]PAM | NA | Intergenic | NA | NA |

| | | | | | | | | | | | |
|---|---|---|---|---|---|---|---|---|---|---|---|
| T8415 | AGCCAGCGAGCTGCCGGCCGAGG | 1000 | | | | | | | | | |
| Oligo fwd | CACCAGCCAGCGAGCTGCCGGCCG | | | | | | | | | | |
| Oligo rev | AAACCGGCCGGCAGCTCGCTGGCT | | | | | | | | | | |
| Chromosome | start | end | strand | MM | target_seq | PAM | alignment | distance | position | gene name | gene id |
| GU305914.1 | 114315 | 114337 | + | 0 | AGCCAGCGAGCTGCCGGCCG | AGG | \|\|\|\|\|\|\|\|[\|\|\|\|\|\|\|\|\|\|\|\|]PAM | NA | Intergenic | NA | NA |

| | | | | | | | | | | | |
|---|---|---|---|---|---|---|---|---|---|---|---|
| T8416 | CTGCCGGCCGAGGCCGACGACGG | 1000 | | | | | | | | | |
| Oligo fwd | CACCCTGCCGGCCGAGGCCGACGA | | | | | | | | | | |
| Oligo rev | AAACTCGTCGGCCTCGGCCGGCAG | | | | | | | | | | |
| Chromosome | start | end | strand | MM | target_seq | PAM | alignment | distance | position | gene name | gene id |
| GU305914.1 | 114325 | 114347 | + | 0 | CTGCCGGCCGAGGCCGACGA | CGG | \|\|\|\|\|\|\|\|[\|\|\|\|\|\|\|\|\|\|\|\|]PAM | NA | Intergenic | NA | NA |

| | | | | | | | | | | | |
|---|---|---|---|---|---|---|---|---|---|---|---|
| T8417 | GTATTAGACGCGACGCTTCCCGG | 1000 | | | | | | | | | |
| Oligo fwd | CACCGTATTAGACGCGACGCTTCC | | | | | | | | | | |
| Oligo rev | AAACGGAAGCGTCGCGTCTAATAC | | | | | | | | | | |
| Chromosome | start | end | strand | MM | target_seq | PAM | alignment | distance | position | gene name | gene id |
| GU305914.1 | 114362 | 114384 | + | 0 | GTATTAGACGCGACGCTTCC | CGG | \|\|\|\|\|\|\|\|[\|\|\|\|\|\|\|\|\|\|\|\|]PAM | NA | Intergenic | NA | NA |

| | | | | | | | | | | | |
|---|---|---|---|---|---|---|---|---|---|---|---|
| T8418 | TATTAGACGCGACGCTTCCCGGG | 1000 | | | | | | | | | |
| Oligo fwd | CACCTATTAGACGCGACGCTTCCC | | | | | | | | | | |
| Oligo rev | AAACGGGAAGCGTCGCGTCTAATA | | | | | | | | | | |
| Chromosome | start | end | strand | MM | target_seq | PAM | alignment | distance | position | gene name | gene id |
| GU305914.1 | 114363 | 114385 | + | 0 | TATTAGACGCGACGCTTCCC | GGG | \|\|\|\|\|\|\|\|[\|\|\|\|\|\|\|\|\|\|\|\|]PAM | NA | Intergenic | NA | NA |

| | | | | | | | | | | | |
|---|---|---|---|---|---|---|---|---|---|---|---|
| T8419 | CGACGCTTCCCGGGACTCTGAGG | 1000 | | | | | | | | | |
| Oligo fwd | CACCCGACGCTTCCCGGGACTCTG | | | | | | | | | | |
| Oligo rev | AAACCAGAGTCCCGGGAAGCGTCG | | | | | | | | | | |
| Chromosome | start | end | strand | MM | target_seq | PAM | alignment | distance | position | gene name | gene id |
| GU305914.1 | 114372 | 114394 | + | 0 | CGACGCTTCCCGGGACTCTG | AGG | \|\|\|\|\|\|\|\|[\|\|\|\|\|\|\|\|\|\|\|\|]PAM | NA | Intergenic | NA | NA |

| | | | | | | | | | | | |
|---|---|---|---|---|---|---|---|---|---|---|---|
| T8420 | GACTCTGAGGAGCGCGCCGCCGG | 1000 | | | | | | | | | |
| Oligo fwd | CACCGACTCTGAGGAGCGCGCCGC | | | | | | | | | | |
| Oligo rev | AAACGCGGCGCGCTCCTCAGAGTC | | | | | | | | | | |
| Chromosome | start | end | strand | MM | target_seq | PAM | alignment | distance | position | gene name | gene id |
| GU305914.1 | 114385 | 114407 | + | 0 | GACTCTGAGGAGCGCGCCGC | CGG | \|\|\|\|\|\|\|\|[\|\|\|\|\|\|\|\|\|\|\|\|]PAM | NA | Intergenic | NA | NA |

| | | | | | | | | | | | |
|---|---|---|---|---|---|---|---|---|---|---|---|
| T8421 | GGAGCGCGCCGCCGGCCGCGAGG | 1000 | | | | | | | | | |
| Oligo fwd | CACCGGAGCGCGCCGCCGGCCGCG | | | | | | | | | | |
| Oligo rev | AAACCGCGGCCGGCGGCGCGCTCC | | | | | | | | | | |
| Chromosome | start | end | strand | MM | target_seq | PAM | alignment | distance | position | gene name | gene id |
| GU305914.1 | 114393 | 114415 | + | 0 | GGAGCGCGCCGCCGGCCGCG | AGG | \|\|\|\|\|\|\|\|[\|\|\|\|\|\|\|\|\|\|\|\|]PAM | NA | Intergenic | NA | NA |
| GU305914.1 | 76476 | 76498 | + | 4 | TGAGCTCGCCGCCGTCCGAG | TGG | -\|\|\|\|-\|\|[\|\|\|\|\|\|-\|\|\|-\|]PAM | NA | Intergenic | NA | NA |

| | | | | | | | | | | | |
|---|---|---|---|---|---|---|---|---|---|---|---|
| T8422 | CGCCGCCGGCCGCGAGGAGATGG | 1000 | | | | | | | | | |
| Oligo fwd | CACCCGCCGCCGGCCGCGAGGAGA | | | | | | | | | | |
| Oligo rev | AAACTCTCCTCGCGGCCGGCGGCG | | | | | | | | | | |
| Chromosome | start | end | strand | MM | target_seq | PAM | alignment | distance | position | gene name | gene id |
| GU305914.1 | 114399 | 114421 | + | 0 | CGCCGCCGGCCGCGAGGAGA | TGG | \|\|\|\|\|\|\|\|[\|\|\|\|\|\|\|\|\|\|\|\|]PAM | NA | Intergenic | NA | NA |

| | | | | | | | | | | | |
|---|---|---|---|---|---|---|---|---|---|---|---|
| T8423 | GAGATGGCTCAGTCGTCTCACGG | 1000 | | | | | | | | | |
| Oligo fwd | CACCGAGATGGCTCAGTCGTCTCA | | | | | | | | | | |
| Oligo rev | AAACTGAGACGACTGAGCCATCTC | | | | | | | | | | |
| Chromosome | start | end | strand | MM | target_seq | PAM | alignment | distance | position | gene name | gene id |
| GU305914.1 | 114415 | 114437 | + | 0 | GAGATGGCTCAGTCGTCTCA | CGG | \|\|\|\|\|\|\|\|[\|\|\|\|\|\|\|\|\|\|\|\|]PAM | NA | Intergenic | NA | NA |

| | | | | | | | | | | | |
|---|---|---|---|---|---|---|---|---|---|---|---|
| T8424 | GTCGTCTCACGGTCTGACACCGG | 1000 | | | | | | | | | |
| Oligo fwd | CACCGTCGTCTCACGGTCTGACAC | | | | | | | | | | |
| Oligo rev | AAACGTGTCAGACCGTGAGACGAC | | | | | | | | | | |
| Chromosome | start | end | strand | MM | target_seq | PAM | alignment | distance | position | gene name | gene id |
| GU305914.1 | 114426 | 114448 | + | 0 | GTCGTCTCACGGTCTGACAC | CGG | \|\|\|\|\|\|\|\|[\|\|\|\|\|\|\|\|\|\|\|\|]PAM | NA | Intergenic | NA | NA |

| | | | | | | | | | | | |
|---|---|---|---|---|---|---|---|---|---|---|---|
| T8425 | CCGGCCGCAGTTTCCGTGCCCGG | 1000 | | | | | | | | | |
| Oligo fwd | CACCCCGGCCGCAGTTTCCGTGCC | | | | | | | | | | |
| Oligo rev | AAACGGCACGGAAACTGCGGCCGG | | | | | | | | | | |
| Chromosome | start | end | strand | MM | target_seq | PAM | alignment | distance | position | gene name | gene id |
| GU305914.1 | 114445 | 114467 | + | 0 | CCGGCCGCAGTTTCCGTGCC | CGG | \|\|\|\|\|\|\|\|[\|\|\|\|\|\|\|\|\|\|\|\|]PAM | NA | Intergenic | NA | NA |

| | | | | | | | | | | | |
|---|---|---|---|---|---|---|---|---|---|---|---|
| T8426 | CGGCCGCAGTTTCCGTGCCCGGG | 1000 | | | | | | | | | |
| Oligo fwd | CACCCGGCCGCAGTTTCCGTGCCC | | | | | | | | | | |
| Oligo rev | AAACGGGCACGGAAACTGCGGCCG | | | | | | | | | | |
| Chromosome | start | end | strand | MM | target_seq | PAM | alignment | distance | position | gene name | gene id |
| GU305914.1 | 114446 | 114468 | + | 0 | CGGCCGCAGTTTCCGTGCCC | GGG | \|\|\|\|\|\|\|\|[\|\|\|\|\|\|\|\|\|\|\|\|]PAM | NA | Intergenic | NA | NA |

| | | |
|---|---|---|
| T8427 | CCCTGCCGCCTCCGCTTTCCCGG | 1000 |
| Oligo fwd | CACCCCCTGCCGCCTCCGCTTTCC | |
| Oligo rev | AAACGGAAAGCGGAGGCGGCAGGG | |

<table>
<tr><td>Chromosome</td><td>start</td><td>end</td><td>strand</td><td>MM</td><td>target_seq</td><td>PAM</td><td>alignment</td><td>distance</td><td>position</td><td>gene name</td><td>gene id</td></tr>
<tr><td>GU305914.1</td><td>114477</td><td>114499</td><td>+</td><td>0</td><td>CCCTGCCGCCTCCGCTTTCC</td><td>CGG</td><td>||||||||[||||||||||||]PAM</td><td>NA</td><td>Intergenic</td><td>NA</td><td>NA</td></tr>
<tr><td>T8428</td><td>CCGACTGCGTCTACCTCAGTCGG</td><td>1000</td><td></td><td></td><td></td><td></td><td></td><td></td><td></td><td></td><td></td></tr>
<tr><td>Oligo fwd</td><td>CACCCCGACTGCGTCTACCTCAGT</td><td></td><td></td><td></td><td></td><td></td><td></td><td></td><td></td><td></td><td></td></tr>
<tr><td>Oligo rev</td><td>AAACACTGAGGTAGACGCAGTCGG</td><td></td><td></td><td></td><td></td><td></td><td></td><td></td><td></td><td></td><td></td></tr>
<tr><td>Chromosome</td><td>start</td><td>end</td><td>strand</td><td>MM</td><td>target_seq</td><td>PAM</td><td>alignment</td><td>distance</td><td>position</td><td>gene name</td><td>gene id</td></tr>
<tr><td>GU305914.1</td><td>114500</td><td>114522</td><td>+</td><td>0</td><td>CCGACTGCGTCTACCTCAGT</td><td>CGG</td><td>||||||||[||||||||||||]PAM</td><td>NA</td><td>Intergenic</td><td>NA</td><td>NA</td></tr>
<tr><td>T8429</td><td>CGACTGCGTCTACCTCAGTCGGG</td><td>1000</td><td></td><td></td><td></td><td></td><td></td><td></td><td></td><td></td><td></td></tr>
<tr><td>Oligo fwd</td><td>CACCCGACTGCGTCTACCTCAGTC</td><td></td><td></td><td></td><td></td><td></td><td></td><td></td><td></td><td></td><td></td></tr>
<tr><td>Oligo rev</td><td>AAACGACTGAGGTAGACGCAGTCG</td><td></td><td></td><td></td><td></td><td></td><td></td><td></td><td></td><td></td><td></td></tr>
<tr><td>Chromosome</td><td>start</td><td>end</td><td>strand</td><td>MM</td><td>target_seq</td><td>PAM</td><td>alignment</td><td>distance</td><td>position</td><td>gene name</td><td>gene id</td></tr>
<tr><td>GU305914.1</td><td>114501</td><td>114523</td><td>+</td><td>0</td><td>CGACTGCGTCTACCTCAGTC</td><td>GGG</td><td>||||||||[||||||||||||]PAM</td><td>NA</td><td>Intergenic</td><td>NA</td><td>NA</td></tr>
<tr><td>T8430</td><td>GGATGCGCTGATGTCCATCCTGG</td><td>1000</td><td></td><td></td><td></td><td></td><td></td><td></td><td></td><td></td><td></td></tr>
<tr><td>Oligo fwd</td><td>CACCGGATGCGCTGATGTCCATCC</td><td></td><td></td><td></td><td></td><td></td><td></td><td></td><td></td><td></td><td></td></tr>
<tr><td>Oligo rev</td><td>AAACGGATGGACATCAGCGCATCC</td><td></td><td></td><td></td><td></td><td></td><td></td><td></td><td></td><td></td><td></td></tr>
<tr><td>Chromosome</td><td>start</td><td>end</td><td>strand</td><td>MM</td><td>target_seq</td><td>PAM</td><td>alignment</td><td>distance</td><td>position</td><td>gene name</td><td>gene id</td></tr>
<tr><td>GU305914.1</td><td>114522</td><td>114544</td><td>+</td><td>0</td><td>GGATGCGCTGATGTCCATCC</td><td>TGG</td><td>||||||||[||||||||||||]PAM</td><td>NA</td><td>Intergenic</td><td>NA</td><td>NA</td></tr>
<tr><td>T8431</td><td>TGCGCTGATGTCCATCCTGGCGG</td><td>1000</td><td></td><td></td><td></td><td></td><td></td><td></td><td></td><td></td><td></td></tr>
<tr><td>Oligo fwd</td><td>CACCTGCGCTGATGTCCATCCTGG</td><td></td><td></td><td></td><td></td><td></td><td></td><td></td><td></td><td></td><td></td></tr>
<tr><td>Oligo rev</td><td>AAACCCAGGATGGACATCAGCGCA</td><td></td><td></td><td></td><td></td><td></td><td></td><td></td><td></td><td></td><td></td></tr>
<tr><td>Chromosome</td><td>start</td><td>end</td><td>strand</td><td>MM</td><td>target_seq</td><td>PAM</td><td>alignment</td><td>distance</td><td>position</td><td>gene name</td><td>gene id</td></tr>
<tr><td>GU305914.1</td><td>114525</td><td>114547</td><td>+</td><td>0</td><td>TGCGCTGATGTCCATCCTGG</td><td>CGG</td><td>||||||||[||||||||||||]PAM</td><td>NA</td><td>Intergenic</td><td>NA</td><td>NA</td></tr>
<tr><td>T8432</td><td>GCGAAACAGAACGCCGTACCCGG</td><td>1000</td><td></td><td></td><td></td><td></td><td></td><td></td><td></td><td></td><td></td></tr>
<tr><td>Oligo fwd</td><td>CACCGCGAAACAGAACGCCGTACC</td><td></td><td></td><td></td><td></td><td></td><td></td><td></td><td></td><td></td><td></td></tr>
<tr><td>Oligo rev</td><td>AAACGGTACGGCGTTCTGTTTCGC</td><td></td><td></td><td></td><td></td><td></td><td></td><td></td><td></td><td></td><td></td></tr>
<tr><td>Chromosome</td><td>start</td><td>end</td><td>strand</td><td>MM</td><td>target_seq</td><td>PAM</td><td>alignment</td><td>distance</td><td>position</td><td>gene name</td><td>gene id</td></tr>
<tr><td>GU305914.1</td><td>114553</td><td>114575</td><td>+</td><td>0</td><td>GCGAAACAGAACGCCGTACC</td><td>CGG</td><td>||||||||[||||||||||||]PAM</td><td>NA</td><td>Intergenic</td><td>NA</td><td>NA</td></tr>
<tr><td>T8433</td><td>CGCTTTAACGTCGCCGCAGCAGG</td><td>1000</td><td></td><td></td><td></td><td></td><td></td><td></td><td></td><td></td><td></td></tr>
<tr><td>Oligo fwd</td><td>CACCCGCTTTAACGTCGCCGCAGC</td><td></td><td></td><td></td><td></td><td></td><td></td><td></td><td></td><td></td><td></td></tr>
<tr><td>Oligo rev</td><td>AAACGCTGCGGCGACGTTAAAGCG</td><td></td><td></td><td></td><td></td><td></td><td></td><td></td><td></td><td></td><td></td></tr>
<tr><td>Chromosome</td><td>start</td><td>end</td><td>strand</td><td>MM</td><td>target_seq</td><td>PAM</td><td>alignment</td><td>distance</td><td>position</td><td>gene name</td><td>gene id</td></tr>
<tr><td>GU305914.1</td><td>114576</td><td>114598</td><td>+</td><td>0</td><td>CGCTTTAACGTCGCCGCAGC</td><td>AGG</td><td>||||||||[||||||||||||]PAM</td><td>NA</td><td>Intergenic</td><td>NA</td><td>NA</td></tr>
<tr><td>T8434</td><td>GCCGCAGCAGGCCTTGCCTCAGG</td><td>1000</td><td></td><td></td><td></td><td></td><td></td><td></td><td></td><td></td><td></td></tr>
<tr><td>Oligo fwd</td><td>CACCGCCGCAGCAGGCCTTGCCTC</td><td></td><td></td><td></td><td></td><td></td><td></td><td></td><td></td><td></td><td></td></tr>
<tr><td>Oligo rev</td><td>AAACGAGGCAAGGCCTGCTGCGGC</td><td></td><td></td><td></td><td></td><td></td><td></td><td></td><td></td><td></td><td></td></tr>
<tr><td>Chromosome</td><td>start</td><td>end</td><td>strand</td><td>MM</td><td>target_seq</td><td>PAM</td><td>alignment</td><td>distance</td><td>position</td><td>gene name</td><td>gene id</td></tr>
<tr><td>GU305914.1</td><td>114588</td><td>114610</td><td>+</td><td>0</td><td>GCCGCAGCAGGCCTTGCCTC</td><td>AGG</td><td>||||||||[||||||||||||]PAM</td><td>NA</td><td>Intergenic</td><td>NA</td><td>NA</td></tr>
<tr><td>T8435</td><td>CCTCAGGTGCCCTCTTATTACGG</td><td>1000</td><td></td><td></td><td></td><td></td><td></td><td></td><td></td><td></td><td></td></tr>
<tr><td>Oligo fwd</td><td>CACCCCTCAGGTGCCCTCTTATTA</td><td></td><td></td><td></td><td></td><td></td><td></td><td></td><td></td><td></td><td></td></tr>
<tr><td>Oligo rev</td><td>AAACTAATAAGAGGGCACCTGAGG</td><td></td><td></td><td></td><td></td><td></td><td></td><td></td><td></td><td></td><td></td></tr>
<tr><td>Chromosome</td><td>start</td><td>end</td><td>strand</td><td>MM</td><td>target_seq</td><td>PAM</td><td>alignment</td><td>distance</td><td>position</td><td>gene name</td><td>gene id</td></tr>
<tr><td>GU305914.1</td><td>114604</td><td>114626</td><td>+</td><td>0</td><td>CCTCAGGTGCCCTCTTATTA</td><td>CGG</td><td>||||||||[||||||||||||]PAM</td><td>NA</td><td>Intergenic</td><td>NA</td><td>NA</td></tr>
<tr><td>T8436</td><td>CTCTTATTACGGAATGCCGCCGG</td><td>1000</td><td></td><td></td><td></td><td></td><td></td><td></td><td></td><td></td><td></td></tr>
<tr><td>Oligo fwd</td><td>CACCCTCTTATTACGGAATGCCGC</td><td></td><td></td><td></td><td></td><td></td><td></td><td></td><td></td><td></td><td></td></tr>
<tr><td>Oligo rev</td><td>AAACGCGGCATTCCGTAATAAGAG</td><td></td><td></td><td></td><td></td><td></td><td></td><td></td><td></td><td></td><td></td></tr>
<tr><td>Chromosome</td><td>start</td><td>end</td><td>strand</td><td>MM</td><td>target_seq</td><td>PAM</td><td>alignment</td><td>distance</td><td>position</td><td>gene name</td><td>gene id</td></tr>
<tr><td>GU305914.1</td><td>114615</td><td>114637</td><td>+</td><td>0</td><td>CTCTTATTACGGAATGCCGC</td><td>CGG</td><td>||||||||[||||||||||||]PAM</td><td>NA</td><td>Intergenic</td><td>NA</td><td>NA</td></tr>
<tr><td>T8437</td><td>TATTACGGAATGCCGCCGGACGG</td><td>1000</td><td></td><td></td><td></td><td></td><td></td><td></td><td></td><td></td><td></td></tr>
<tr><td>Oligo fwd</td><td>CACCTATTACGGAATGCCGCCGGA</td><td></td><td></td><td></td><td></td><td></td><td></td><td></td><td></td><td></td><td></td></tr>
<tr><td>Oligo rev</td><td>AAACTCCGGCGGCATTCCGTAATA</td><td></td><td></td><td></td><td></td><td></td><td></td><td></td><td></td><td></td><td></td></tr>
<tr><td>Chromosome</td><td>start</td><td>end</td><td>strand</td><td>MM</td><td>target_seq</td><td>PAM</td><td>alignment</td><td>distance</td><td>position</td><td>gene name</td><td>gene id</td></tr>
<tr><td>GU305914.1</td><td>114619</td><td>114641</td><td>+</td><td>0</td><td>TATTACGGAATGCCGCCGGA</td><td>CGG</td><td>||||||||[||||||||||||]PAM</td><td>NA</td><td>Intergenic</td><td>NA</td><td>NA</td></tr>
<tr><td>T8438</td><td>CCACCGCCGTCTCACCATCGCGG</td><td>1000</td><td></td><td></td><td></td><td></td><td></td><td></td><td></td><td></td><td></td></tr>
<tr><td>Oligo fwd</td><td>CACCCCACCGCCGTCTCACCATCG</td><td></td><td></td><td></td><td></td><td></td><td></td><td></td><td></td><td></td><td></td></tr>
<tr><td>Oligo rev</td><td>AAACCGATGGTGAGACGGCGGTGG</td><td></td><td></td><td></td><td></td><td></td><td></td><td></td><td></td><td></td><td></td></tr>
<tr><td>Chromosome</td><td>start</td><td>end</td><td>strand</td><td>MM</td><td>target_seq</td><td>PAM</td><td>alignment</td><td>distance</td><td>position</td><td>gene name</td><td>gene id</td></tr>
<tr><td>GU305914.1</td><td>114673</td><td>114695</td><td>+</td><td>0</td><td>CCACCGCCGTCTCACCATCG</td><td>CGG</td><td>||||||||[||||||||||||]PAM</td><td>NA</td><td>Intergenic</td><td>NA</td><td>NA</td></tr>
<tr><td>T8439</td><td>CCGCCGTCTCACCATCGCGGCGG</td><td>1000</td><td></td><td></td><td></td><td></td><td></td><td></td><td></td><td></td><td></td></tr>
<tr><td>Oligo fwd</td><td>CACCCCGCCGTCTCACCATCGCGG</td><td></td><td></td><td></td><td></td><td></td><td></td><td></td><td></td><td></td><td></td></tr>
<tr><td>Oligo rev</td><td>AAACCCGCGATGGTGAGACGGCGG</td><td></td><td></td><td></td><td></td><td></td><td></td><td></td><td></td><td></td><td></td></tr>
<tr><td>Chromosome</td><td>start</td><td>end</td><td>strand</td><td>MM</td><td>target_seq</td><td>PAM</td><td>alignment</td><td>distance</td><td>position</td><td>gene name</td><td>gene id</td></tr>
<tr><td>GU305914.1</td><td>114676</td><td>114698</td><td>+</td><td>0</td><td>CCGCCGTCTCACCATCGCGG</td><td>CGG</td><td>||||||||[||||||||||||]PAM</td><td>NA</td><td>Intergenic</td><td>NA</td><td>NA</td></tr>
<tr><td>T8440</td><td>CCGTCTCACCATCGCGGCGGTGG</td><td>1000</td><td></td><td></td><td></td><td></td><td></td><td></td><td></td><td></td><td></td></tr>
<tr><td>Oligo fwd</td><td>CACCCCGTCTCACCATCGCGGCGG</td><td></td><td></td><td></td><td></td><td></td><td></td><td></td><td></td><td></td><td></td></tr>
<tr><td>Oligo rev</td><td>AAACCCGCCGCGATGGTGAGACGG</td><td></td><td></td><td></td><td></td><td></td><td></td><td></td><td></td><td></td><td></td></tr>
<tr><td>Chromosome</td><td>start</td><td>end</td><td>strand</td><td>MM</td><td>target_seq</td><td>PAM</td><td>alignment</td><td>distance</td><td>position</td><td>gene name</td><td>gene id</td></tr>
<tr><td>GU305914.1</td><td>114679</td><td>114701</td><td>+</td><td>0</td><td>CCGTCTCACCATCGCGGCGG</td><td>TGG</td><td>||||||||[||||||||||||]PAM</td><td>NA</td><td>Intergenic</td><td>NA</td><td>NA</td></tr>
<tr><td>T8441</td><td>TCTCACCATCGCGGCGGTGGCGG</td><td>1000</td><td></td><td></td><td></td><td></td><td></td><td></td><td></td><td></td><td></td></tr>
<tr><td>Oligo fwd</td><td>CACCTCTCACCATCGCGGCGGTGG</td><td></td><td></td><td></td><td></td><td></td><td></td><td></td><td></td><td></td><td></td></tr>
<tr><td>Oligo rev</td><td>AAACCCACCGCCGCGATGGTGAGA</td><td></td><td></td><td></td><td></td><td></td><td></td><td></td><td></td><td></td><td></td></tr>
<tr><td>Chromosome</td><td>start</td><td>end</td><td>strand</td><td>MM</td><td>target_seq</td><td>PAM</td><td>alignment</td><td>distance</td><td>position</td><td>gene name</td><td>gene id</td></tr>
<tr><td>GU305914.1</td><td>114682</td><td>114704</td><td>+</td><td>0</td><td>TCTCACCATCGCGGCGGTGG</td><td>CGG</td><td>||||||||[||||||||||||]PAM</td><td>NA</td><td>Intergenic</td><td>NA</td><td>NA</td></tr>
<tr><td>T8442</td><td>TTCGATCCCCCTCTCCCACACGG</td><td>1000</td><td></td><td></td><td></td><td></td><td></td><td></td><td></td><td></td><td></td></tr>
<tr><td>Oligo fwd</td><td>CACCTTCGATCCCCCTCTCCCACA</td><td></td><td></td><td></td><td></td><td></td><td></td><td></td><td></td><td></td><td></td></tr>
</table>

<table>
<tr><td>Oligo rev</td><td>AAACTGTGGGAGAGGGGGATCGAA</td><td></td><td></td><td></td><td></td><td></td><td></td><td></td><td></td><td></td><td></td></tr>
<tr><td>Chromosome</td><td>start</td><td>end</td><td>strand</td><td>MM</td><td>target_seq</td><td>PAM</td><td>alignment</td><td>distance</td><td>position</td><td>gene name</td><td>gene id</td></tr>
<tr><td>GU305914.1</td><td>114709</td><td>114731</td><td>+</td><td>0</td><td>TTCGATCCCCCTCTCCCACA</td><td>CGG</td><td>||||||||[||||||||||||]PAM</td><td>NA</td><td>Intergenic</td><td>NA</td><td>NA</td></tr>
<tr><td>T8443</td><td>TCGATCCCCCTCTCCCACACGGG</td><td>1000</td><td></td><td></td><td></td><td></td><td></td><td></td><td></td><td></td><td></td></tr>
<tr><td>Oligo fwd</td><td>CACCTCGATCCCCCTCTCCCACAC</td><td></td><td></td><td></td><td></td><td></td><td></td><td></td><td></td><td></td><td></td></tr>
<tr><td>Oligo rev</td><td>AAACGTGTGGGAGAGGGGGATCGA</td><td></td><td></td><td></td><td></td><td></td><td></td><td></td><td></td><td></td><td></td></tr>
<tr><td>Chromosome</td><td>start</td><td>end</td><td>strand</td><td>MM</td><td>target_seq</td><td>PAM</td><td>alignment</td><td>distance</td><td>position</td><td>gene name</td><td>gene id</td></tr>
<tr><td>GU305914.1</td><td>114710</td><td>114732</td><td>+</td><td>0</td><td>TCGATCCCCCTCTCCCACAC</td><td>GGG</td><td>||||||||[||||||||||||]PAM</td><td>NA</td><td>Intergenic</td><td>NA</td><td>NA</td></tr>
<tr><td>T8444</td><td>CGATCCCCCTCTCCCACACGGGG</td><td>1000</td><td></td><td></td><td></td><td></td><td></td><td></td><td></td><td></td><td></td></tr>
<tr><td>Oligo fwd</td><td>CACCCGATCCCCCTCTCCCACACG</td><td></td><td></td><td></td><td></td><td></td><td></td><td></td><td></td><td></td><td></td></tr>
<tr><td>Oligo rev</td><td>AAACCGTGTGGGAGAGGGGGATCG</td><td></td><td></td><td></td><td></td><td></td><td></td><td></td><td></td><td></td><td></td></tr>
<tr><td>Chromosome</td><td>start</td><td>end</td><td>strand</td><td>MM</td><td>target_seq</td><td>PAM</td><td>alignment</td><td>distance</td><td>position</td><td>gene name</td><td>gene id</td></tr>
<tr><td>GU305914.1</td><td>114711</td><td>114733</td><td>+</td><td>0</td><td>CGATCCCCCTCTCCCACACG</td><td>GGG</td><td>||||||||[||||||||||||]PAM</td><td>NA</td><td>Intergenic</td><td>NA</td><td>NA</td></tr>
<tr><td>T8445</td><td>GATCCCCCTCTCCCACACGGGGG</td><td>1000</td><td></td><td></td><td></td><td></td><td></td><td></td><td></td><td></td><td></td></tr>
<tr><td>Oligo fwd</td><td>CACCGATCCCCCTCTCCCACACGG</td><td></td><td></td><td></td><td></td><td></td><td></td><td></td><td></td><td></td><td></td></tr>
<tr><td>Oligo rev</td><td>AAACCCGTGTGGGAGAGGGGGATC</td><td></td><td></td><td></td><td></td><td></td><td></td><td></td><td></td><td></td><td></td></tr>
<tr><td>Chromosome</td><td>start</td><td>end</td><td>strand</td><td>MM</td><td>target_seq</td><td>PAM</td><td>alignment</td><td>distance</td><td>position</td><td>gene name</td><td>gene id</td></tr>
<tr><td>GU305914.1</td><td>114712</td><td>114734</td><td>+</td><td>0</td><td>GATCCCCCTCTCCCACACGG</td><td>GGG</td><td>||||||||[||||||||||||]PAM</td><td>NA</td><td>Intergenic</td><td>NA</td><td>NA</td></tr>
<tr><td>T8446</td><td>CCTCTCCCACACGGGGGATACGG</td><td>1000</td><td></td><td></td><td></td><td></td><td></td><td></td><td></td><td></td><td></td></tr>
<tr><td>Oligo fwd</td><td>CACCCCTCTCCCACACGGGGGATA</td><td></td><td></td><td></td><td></td><td></td><td></td><td></td><td></td><td></td><td></td></tr>
<tr><td>Oligo rev</td><td>AAACTATCCCCCGTGTGGGAGAGG</td><td></td><td></td><td></td><td></td><td></td><td></td><td></td><td></td><td></td><td></td></tr>
<tr><td>Chromosome</td><td>start</td><td>end</td><td>strand</td><td>MM</td><td>target_seq</td><td>PAM</td><td>alignment</td><td>distance</td><td>position</td><td>gene name</td><td>gene id</td></tr>
<tr><td>GU305914.1</td><td>114718</td><td>114740</td><td>+</td><td>0</td><td>CCTCTCCCACACGGGGGATA</td><td>CGG</td><td>||||||||[||||||||||||]PAM</td><td>NA</td><td>Intergenic</td><td>NA</td><td>NA</td></tr>
<tr><td>T8447</td><td>CGGTCCTCCCTATCATCACCCGG</td><td>1000</td><td></td><td></td><td></td><td></td><td></td><td></td><td></td><td></td><td></td></tr>
<tr><td>Oligo fwd</td><td>CACCCGGTCCTCCCTATCATCACC</td><td></td><td></td><td></td><td></td><td></td><td></td><td></td><td></td><td></td><td></td></tr>
<tr><td>Oligo rev</td><td>AAACGGTGATGATAGGGAGGACCG</td><td></td><td></td><td></td><td></td><td></td><td></td><td></td><td></td><td></td><td></td></tr>
<tr><td>Chromosome</td><td>start</td><td>end</td><td>strand</td><td>MM</td><td>target_seq</td><td>PAM</td><td>alignment</td><td>distance</td><td>position</td><td>gene name</td><td>gene id</td></tr>
<tr><td>GU305914.1</td><td>114738</td><td>114760</td><td>+</td><td>0</td><td>CGGTCCTCCCTATCATCACC</td><td>CGG</td><td>||||||||[||||||||||||]PAM</td><td>NA</td><td>Intergenic</td><td>NA</td><td>NA</td></tr>
<tr><td>T8448</td><td>ATCATCACCCGGACGCCTATCGG</td><td>1000</td><td></td><td></td><td></td><td></td><td></td><td></td><td></td><td></td><td></td></tr>
<tr><td>Oligo fwd</td><td>CACCATCATCACCCGGACGCCTAT</td><td></td><td></td><td></td><td></td><td></td><td></td><td></td><td></td><td></td><td></td></tr>
<tr><td>Oligo rev</td><td>AAACATAGGCGTCCGGGTGATGAT</td><td></td><td></td><td></td><td></td><td></td><td></td><td></td><td></td><td></td><td></td></tr>
<tr><td>Chromosome</td><td>start</td><td>end</td><td>strand</td><td>MM</td><td>target_seq</td><td>PAM</td><td>alignment</td><td>distance</td><td>position</td><td>gene name</td><td>gene id</td></tr>
<tr><td>GU305914.1</td><td>114749</td><td>114771</td><td>+</td><td>0</td><td>ATCATCACCCGGACGCCTAT</td><td>CGG</td><td>||||||||[||||||||||||]PAM</td><td>NA</td><td>Intergenic</td><td>NA</td><td>NA</td></tr>
<tr><td>T8449</td><td>TCATCACCCGGACGCCTATCGGG</td><td>1000</td><td></td><td></td><td></td><td></td><td></td><td></td><td></td><td></td><td></td></tr>
<tr><td>Oligo fwd</td><td>CACCTCATCACCCGGACGCCTATC</td><td></td><td></td><td></td><td></td><td></td><td></td><td></td><td></td><td></td><td></td></tr>
<tr><td>Oligo rev</td><td>AAACGATAGGCGTCCGGGTGATGA</td><td></td><td></td><td></td><td></td><td></td><td></td><td></td><td></td><td></td><td></td></tr>
<tr><td>Chromosome</td><td>start</td><td>end</td><td>strand</td><td>MM</td><td>target_seq</td><td>PAM</td><td>alignment</td><td>distance</td><td>position</td><td>gene name</td><td>gene id</td></tr>
<tr><td>GU305914.1</td><td>114750</td><td>114772</td><td>+</td><td>0</td><td>TCATCACCCGGACGCCTATC</td><td>GGG</td><td>||||||||[||||||||||||]PAM</td><td>NA</td><td>Intergenic</td><td>NA</td><td>NA</td></tr>
<tr><td>T8450</td><td>CATCACCCGGACGCCTATCGGGG</td><td>1000</td><td></td><td></td><td></td><td></td><td></td><td></td><td></td><td></td><td></td></tr>
<tr><td>Oligo fwd</td><td>CACCCATCACCCGGACGCCTATCG</td><td></td><td></td><td></td><td></td><td></td><td></td><td></td><td></td><td></td><td></td></tr>
<tr><td>Oligo rev</td><td>AAACCGATAGGCGTCCGGGTGATG</td><td></td><td></td><td></td><td></td><td></td><td></td><td></td><td></td><td></td><td></td></tr>
<tr><td>Chromosome</td><td>start</td><td>end</td><td>strand</td><td>MM</td><td>target_seq</td><td>PAM</td><td>alignment</td><td>distance</td><td>position</td><td>gene name</td><td>gene id</td></tr>
<tr><td>GU305914.1</td><td>114751</td><td>114773</td><td>+</td><td>0</td><td>CATCACCCGGACGCCTATCG</td><td>GGG</td><td>||||||||[||||||||||||]PAM</td><td>NA</td><td>Intergenic</td><td>NA</td><td>NA</td></tr>
<tr><td>GU305914.1</td><td>51046</td><td>51068</td><td>-</td><td>4</td><td>CATCGCTCGGAGGGCTATCG</td><td>AGG</td><td>||||-|-|[|||-|-||||||]PAM</td><td>NA</td><td>Intergenic</td><td>NA</td><td>NA</td></tr>
<tr><td>T8451</td><td>ATCACCCGGACGCCTATCGGGGG</td><td>1000</td><td></td><td></td><td></td><td></td><td></td><td></td><td></td><td></td><td></td></tr>
<tr><td>Oligo fwd</td><td>CACCATCACCCGGACGCCTATCGG</td><td></td><td></td><td></td><td></td><td></td><td></td><td></td><td></td><td></td><td></td></tr>
<tr><td>Oligo rev</td><td>AAACCCGATAGGCGTCCGGGTGAT</td><td></td><td></td><td></td><td></td><td></td><td></td><td></td><td></td><td></td><td></td></tr>
<tr><td>Chromosome</td><td>start</td><td>end</td><td>strand</td><td>MM</td><td>target_seq</td><td>PAM</td><td>alignment</td><td>distance</td><td>position</td><td>gene name</td><td>gene id</td></tr>
<tr><td>GU305914.1</td><td>114752</td><td>114774</td><td>+</td><td>0</td><td>ATCACCCGGACGCCTATCGG</td><td>GGG</td><td>||||||||[||||||||||||]PAM</td><td>NA</td><td>Intergenic</td><td>NA</td><td>NA</td></tr>
<tr><td>T8452</td><td>TCACCCGGACGCCTATCGGGGGG</td><td>1000</td><td></td><td></td><td></td><td></td><td></td><td></td><td></td><td></td><td></td></tr>
<tr><td>Oligo fwd</td><td>CACCTCACCCGGACGCCTATCGGG</td><td></td><td></td><td></td><td></td><td></td><td></td><td></td><td></td><td></td><td></td></tr>
<tr><td>Oligo rev</td><td>AAACCCCGATAGGCGTCCGGGTGA</td><td></td><td></td><td></td><td></td><td></td><td></td><td></td><td></td><td></td><td></td></tr>
<tr><td>Chromosome</td><td>start</td><td>end</td><td>strand</td><td>MM</td><td>target_seq</td><td>PAM</td><td>alignment</td><td>distance</td><td>position</td><td>gene name</td><td>gene id</td></tr>
<tr><td>GU305914.1</td><td>114753</td><td>114775</td><td>+</td><td>0</td><td>TCACCCGGACGCCTATCGGG</td><td>GGG</td><td>||||||||[||||||||||||]PAM</td><td>NA</td><td>Intergenic</td><td>NA</td><td>NA</td></tr>
<tr><td>T8453</td><td>CACCCGGACGCCTATCGGGGGGG</td><td>1000</td><td></td><td></td><td></td><td></td><td></td><td></td><td></td><td></td><td></td></tr>
<tr><td>Oligo fwd</td><td>CACCCACCCGGACGCCTATCGGGG</td><td></td><td></td><td></td><td></td><td></td><td></td><td></td><td></td><td></td><td></td></tr>
<tr><td>Oligo rev</td><td>AAACCCCCGATAGGCGTCCGGGTG</td><td></td><td></td><td></td><td></td><td></td><td></td><td></td><td></td><td></td><td></td></tr>
<tr><td>Chromosome</td><td>start</td><td>end</td><td>strand</td><td>MM</td><td>target_seq</td><td>PAM</td><td>alignment</td><td>distance</td><td>position</td><td>gene name</td><td>gene id</td></tr>
<tr><td>GU305914.1</td><td>114754</td><td>114776</td><td>+</td><td>0</td><td>CACCCGGACGCCTATCGGGG</td><td>GGG</td><td>||||||||[||||||||||||]PAM</td><td>NA</td><td>Intergenic</td><td>NA</td><td>NA</td></tr>
<tr><td>T8454</td><td>CATCCTGATCGCGACCCGCGCGG</td><td>1000</td><td></td><td></td><td></td><td></td><td></td><td></td><td></td><td></td><td></td></tr>
<tr><td>Oligo fwd</td><td>CACCCATCCTGATCGCGACCCGCG</td><td></td><td></td><td></td><td></td><td></td><td></td><td></td><td></td><td></td><td></td></tr>
<tr><td>Oligo rev</td><td>AAACCGCGGGTCGCGATCAGGATG</td><td></td><td></td><td></td><td></td><td></td><td></td><td></td><td></td><td></td><td></td></tr>
<tr><td>Chromosome</td><td>start</td><td>end</td><td>strand</td><td>MM</td><td>target_seq</td><td>PAM</td><td>alignment</td><td>distance</td><td>position</td><td>gene name</td><td>gene id</td></tr>
<tr><td>GU305914.1</td><td>114784</td><td>114806</td><td>+</td><td>0</td><td>CATCCTGATCGCGACCCGCG</td><td>CGG</td><td>||||||||[||||||||||||]PAM</td><td>NA</td><td>Intergenic</td><td>NA</td><td>NA</td></tr>
<tr><td>T8455</td><td>CCTGATCGCGACCCGCGCGGCGG</td><td>1000</td><td></td><td></td><td></td><td></td><td></td><td></td><td></td><td></td><td></td></tr>
<tr><td>Oligo fwd</td><td>CACCCCTGATCGCGACCCGCGCGG</td><td></td><td></td><td></td><td></td><td></td><td></td><td></td><td></td><td></td><td></td></tr>
<tr><td>Oligo rev</td><td>AAACCCGCGCGGGTCGCGATCAGG</td><td></td><td></td><td></td><td></td><td></td><td></td><td></td><td></td><td></td><td></td></tr>
<tr><td>Chromosome</td><td>start</td><td>end</td><td>strand</td><td>MM</td><td>target_seq</td><td>PAM</td><td>alignment</td><td>distance</td><td>position</td><td>gene name</td><td>gene id</td></tr>
<tr><td>GU305914.1</td><td>114787</td><td>114809</td><td>+</td><td>0</td><td>CCTGATCGCGACCCGCGCGG</td><td>CGG</td><td>||||||||[||||||||||||]PAM</td><td>NA</td><td>Intergenic</td><td>NA</td><td>NA</td></tr>
<tr><td>T8456</td><td>GCGCGGCGGAGTGCCCTACGAGG</td><td>1000</td><td></td><td></td><td></td><td></td><td></td><td></td><td></td><td></td><td></td></tr>
<tr><td>Oligo fwd</td><td>CACCGCGCGGCGGAGTGCCCTACG</td><td></td><td></td><td></td><td></td><td></td><td></td><td></td><td></td><td></td><td></td></tr>
<tr><td>Oligo rev</td><td>AAACCGTAGGGCACTCCGCCGCGC</td><td></td><td></td><td></td><td></td><td></td><td></td><td></td><td></td><td></td><td></td></tr>
<tr><td>Chromosome</td><td>start</td><td>end</td><td>strand</td><td>MM</td><td>target_seq</td><td>PAM</td><td>alignment</td><td>distance</td><td>position</td><td>gene name</td><td>gene id</td></tr>
<tr><td>GU305914.1</td><td>114801</td><td>114823</td><td>+</td><td>0</td><td>GCGCGGCGGAGTGCCCTACG</td><td>AGG</td><td>||||||||[||||||||||||]PAM</td><td>NA</td><td>Intergenic</td><td>NA</td><td>NA</td></tr>
</table>

<table>
<tr><td>T8457</td><td>CGCGGCGGAGTGCCCTACGAGGG</td><td>1000</td><td></td><td></td><td></td><td></td><td></td><td></td><td></td><td></td><td></td></tr>
<tr><td>Oligo fwd</td><td>CACCCGCGGCGGAGTGCCCTACGA</td><td></td><td></td><td></td><td></td><td></td><td></td><td></td><td></td><td></td><td></td></tr>
<tr><td>Oligo rev</td><td>AAACTCGTAGGGCACTCCGCCGCG</td><td></td><td></td><td></td><td></td><td></td><td></td><td></td><td></td><td></td><td></td></tr>
<tr><td>Chromosome</td><td>start</td><td>end</td><td>strand</td><td>MM</td><td>target_seq</td><td>PAM</td><td>alignment</td><td>distance</td><td>position</td><td>gene name</td><td>gene id</td></tr>
<tr><td>GU305914.1</td><td>114802</td><td>114824</td><td>+</td><td>0</td><td>CGCGGCGGAGTGCCCTACGA</td><td>GGG</td><td>||||||||[||||||||||||]PAM</td><td>NA</td><td>Intergenic</td><td>NA</td><td>NA</td></tr>
<tr><td>T8458</td><td>GCGGAGTGCCCTACGAGGGCTGG</td><td>1000</td><td></td><td></td><td></td><td></td><td></td><td></td><td></td><td></td><td></td></tr>
<tr><td>Oligo fwd</td><td>CACCGCGGAGTGCCCTACGAGGGC</td><td></td><td></td><td></td><td></td><td></td><td></td><td></td><td></td><td></td><td></td></tr>
<tr><td>Oligo rev</td><td>AAACGCCCTCGTAGGGCACTCCGC</td><td></td><td></td><td></td><td></td><td></td><td></td><td></td><td></td><td></td><td></td></tr>
<tr><td>Chromosome</td><td>start</td><td>end</td><td>strand</td><td>MM</td><td>target_seq</td><td>PAM</td><td>alignment</td><td>distance</td><td>position</td><td>gene name</td><td>gene id</td></tr>
<tr><td>GU305914.1</td><td>114806</td><td>114828</td><td>+</td><td>0</td><td>GCGGAGTGCCCTACGAGGGC</td><td>TGG</td><td>||||||||[||||||||||||]PAM</td><td>NA</td><td>Intergenic</td><td>NA</td><td>NA</td></tr>
<tr><td>T8459</td><td>CGCCCTCGCTACGACCCCGCCGG</td><td>1000</td><td></td><td></td><td></td><td></td><td></td><td></td><td></td><td></td><td></td></tr>
<tr><td>Oligo fwd</td><td>CACCCGCCCTCGCTACGACCCCGC</td><td></td><td></td><td></td><td></td><td></td><td></td><td></td><td></td><td></td><td></td></tr>
<tr><td>Oligo rev</td><td>AAACGCGGGGTCGTAGCGAGGGCG</td><td></td><td></td><td></td><td></td><td></td><td></td><td></td><td></td><td></td><td></td></tr>
<tr><td>Chromosome</td><td>start</td><td>end</td><td>strand</td><td>MM</td><td>target_seq</td><td>PAM</td><td>alignment</td><td>distance</td><td>position</td><td>gene name</td><td>gene id</td></tr>
<tr><td>GU305914.1</td><td>114832</td><td>114854</td><td>+</td><td>0</td><td>CGCCCTCGCTACGACCCCGC</td><td>CGG</td><td>||||||||[||||||||||||]PAM</td><td>NA</td><td>Intergenic</td><td>NA</td><td>NA</td></tr>
<tr><td>T8460</td><td>GCCCTCGCTACGACCCCGCCGGG</td><td>1000</td><td></td><td></td><td></td><td></td><td></td><td></td><td></td><td></td><td></td></tr>
<tr><td>Oligo fwd</td><td>CACCGCCCTCGCTACGACCCCGCC</td><td></td><td></td><td></td><td></td><td></td><td></td><td></td><td></td><td></td><td></td></tr>
<tr><td>Oligo rev</td><td>AAACGGCGGGGTCGTAGCGAGGGC</td><td></td><td></td><td></td><td></td><td></td><td></td><td></td><td></td><td></td><td></td></tr>
<tr><td>Chromosome</td><td>start</td><td>end</td><td>strand</td><td>MM</td><td>target_seq</td><td>PAM</td><td>alignment</td><td>distance</td><td>position</td><td>gene name</td><td>gene id</td></tr>
<tr><td>GU305914.1</td><td>114833</td><td>114855</td><td>+</td><td>0</td><td>GCCCTCGCTACGACCCCGCC</td><td>GGG</td><td>||||||||[||||||||||||]PAM</td><td>NA</td><td>Intergenic</td><td>NA</td><td>NA</td></tr>
<tr><td>T8461</td><td>CCCTCGCTACGACCCCGCCGGGG</td><td>1000</td><td></td><td></td><td></td><td></td><td></td><td></td><td></td><td></td><td></td></tr>
<tr><td>Oligo fwd</td><td>CACCCCCTCGCTACGACCCCGCCG</td><td></td><td></td><td></td><td></td><td></td><td></td><td></td><td></td><td></td><td></td></tr>
<tr><td>Oligo rev</td><td>AAACCGGCGGGGTCGTAGCGAGGG</td><td></td><td></td><td></td><td></td><td></td><td></td><td></td><td></td><td></td><td></td></tr>
<tr><td>Chromosome</td><td>start</td><td>end</td><td>strand</td><td>MM</td><td>target_seq</td><td>PAM</td><td>alignment</td><td>distance</td><td>position</td><td>gene name</td><td>gene id</td></tr>
<tr><td>GU305914.1</td><td>114834</td><td>114856</td><td>+</td><td>0</td><td>CCCTCGCTACGACCCCGCCG</td><td>GGG</td><td>||||||||[||||||||||||]PAM</td><td>NA</td><td>Intergenic</td><td>NA</td><td>NA</td></tr>
<tr><td>T8462</td><td>ATCATCCCTCCTACAACAACAGG</td><td>1000</td><td></td><td></td><td></td><td></td><td></td><td></td><td></td><td></td><td></td></tr>
<tr><td>Oligo fwd</td><td>CACCATCATCCCTCCTACAACAAC</td><td></td><td></td><td></td><td></td><td></td><td></td><td></td><td></td><td></td><td></td></tr>
<tr><td>Oligo rev</td><td>AAACGTTGTTGTAGGAGGGATGAT</td><td></td><td></td><td></td><td></td><td></td><td></td><td></td><td></td><td></td><td></td></tr>
<tr><td>Chromosome</td><td>start</td><td>end</td><td>strand</td><td>MM</td><td>target_seq</td><td>PAM</td><td>alignment</td><td>distance</td><td>position</td><td>gene name</td><td>gene id</td></tr>
<tr><td>GU305914.1</td><td>114860</td><td>114882</td><td>+</td><td>0</td><td>ATCATCCCTCCTACAACAAC</td><td>AGG</td><td>||||||||[||||||||||||]PAM</td><td>NA</td><td>Intergenic</td><td>NA</td><td>NA</td></tr>
<tr><td>T8463</td><td>ATCCCTCCTACAACAACAGGCGG</td><td>1000</td><td></td><td></td><td></td><td></td><td></td><td></td><td></td><td></td><td></td></tr>
<tr><td>Oligo fwd</td><td>CACCATCCCTCCTACAACAACAGG</td><td></td><td></td><td></td><td></td><td></td><td></td><td></td><td></td><td></td><td></td></tr>
<tr><td>Oligo rev</td><td>AAACCCTGTTGTTGTAGGAGGGAT</td><td></td><td></td><td></td><td></td><td></td><td></td><td></td><td></td><td></td><td></td></tr>
<tr><td>Chromosome</td><td>start</td><td>end</td><td>strand</td><td>MM</td><td>target_seq</td><td>PAM</td><td>alignment</td><td>distance</td><td>position</td><td>gene name</td><td>gene id</td></tr>
<tr><td>GU305914.1</td><td>114863</td><td>114885</td><td>+</td><td>0</td><td>ATCCCTCCTACAACAACAGG</td><td>CGG</td><td>||||||||[||||||||||||]PAM</td><td>NA</td><td>Intergenic</td><td>NA</td><td>NA</td></tr>
<tr><td>T8464</td><td>TCCCTCCTACAACAACAGGCGGG</td><td>1000</td><td></td><td></td><td></td><td></td><td></td><td></td><td></td><td></td><td></td></tr>
<tr><td>Oligo fwd</td><td>CACCTCCCTCCTACAACAACAGGC</td><td></td><td></td><td></td><td></td><td></td><td></td><td></td><td></td><td></td><td></td></tr>
<tr><td>Oligo rev</td><td>AAACGCCTGTTGTTGTAGGAGGGA</td><td></td><td></td><td></td><td></td><td></td><td></td><td></td><td></td><td></td><td></td></tr>
<tr><td>Chromosome</td><td>start</td><td>end</td><td>strand</td><td>MM</td><td>target_seq</td><td>PAM</td><td>alignment</td><td>distance</td><td>position</td><td>gene name</td><td>gene id</td></tr>
<tr><td>GU305914.1</td><td>114864</td><td>114886</td><td>+</td><td>0</td><td>TCCCTCCTACAACAACAGGC</td><td>GGG</td><td>||||||||[||||||||||||]PAM</td><td>NA</td><td>Intergenic</td><td>NA</td><td>NA</td></tr>
<tr><td>T8465</td><td>CCCTCCTACAACAACAGGCGGGG</td><td>1000</td><td></td><td></td><td></td><td></td><td></td><td></td><td></td><td></td><td></td></tr>
<tr><td>Oligo fwd</td><td>CACCCCCTCCTACAACAACAGGCG</td><td></td><td></td><td></td><td></td><td></td><td></td><td></td><td></td><td></td><td></td></tr>
<tr><td>Oligo rev</td><td>AAACCGCCTGTTGTTGTAGGAGGG</td><td></td><td></td><td></td><td></td><td></td><td></td><td></td><td></td><td></td><td></td></tr>
<tr><td>Chromosome</td><td>start</td><td>end</td><td>strand</td><td>MM</td><td>target_seq</td><td>PAM</td><td>alignment</td><td>distance</td><td>position</td><td>gene name</td><td>gene id</td></tr>
<tr><td>GU305914.1</td><td>114865</td><td>114887</td><td>+</td><td>0</td><td>CCCTCCTACAACAACAGGCG</td><td>GGG</td><td>||||||||[||||||||||||]PAM</td><td>NA</td><td>Intergenic</td><td>NA</td><td>NA</td></tr>
<tr><td>T8466</td><td>ACAGGCGGGGAGACCGCTATCGG</td><td>1000</td><td></td><td></td><td></td><td></td><td></td><td></td><td></td><td></td><td></td></tr>
<tr><td>Oligo fwd</td><td>CACCACAGGCGGGGAGACCGCTAT</td><td></td><td></td><td></td><td></td><td></td><td></td><td></td><td></td><td></td><td></td></tr>
<tr><td>Oligo rev</td><td>AAACATAGCGGTCTCCCCGCCTGT</td><td></td><td></td><td></td><td></td><td></td><td></td><td></td><td></td><td></td><td></td></tr>
<tr><td>Chromosome</td><td>start</td><td>end</td><td>strand</td><td>MM</td><td>target_seq</td><td>PAM</td><td>alignment</td><td>distance</td><td>position</td><td>gene name</td><td>gene id</td></tr>
<tr><td>GU305914.1</td><td>114878</td><td>114900</td><td>+</td><td>0</td><td>ACAGGCGGGGAGACCGCTAT</td><td>CGG</td><td>||||||||[||||||||||||]PAM</td><td>NA</td><td>Intergenic</td><td>NA</td><td>NA</td></tr>
<tr><td>T8467</td><td>CAGGCGGGGAGACCGCTATCGGG</td><td>1000</td><td></td><td></td><td></td><td></td><td></td><td></td><td></td><td></td><td></td></tr>
<tr><td>Oligo fwd</td><td>CACCCAGGCGGGGAGACCGCTATC</td><td></td><td></td><td></td><td></td><td></td><td></td><td></td><td></td><td></td><td></td></tr>
<tr><td>Oligo rev</td><td>AAACGATAGCGGTCTCCCCGCCTG</td><td></td><td></td><td></td><td></td><td></td><td></td><td></td><td></td><td></td><td></td></tr>
<tr><td>Chromosome</td><td>start</td><td>end</td><td>strand</td><td>MM</td><td>target_seq</td><td>PAM</td><td>alignment</td><td>distance</td><td>position</td><td>gene name</td><td>gene id</td></tr>
<tr><td>GU305914.1</td><td>114879</td><td>114901</td><td>+</td><td>0</td><td>CAGGCGGGGAGACCGCTATC</td><td>GGG</td><td>||||||||[||||||||||||]PAM</td><td>NA</td><td>Intergenic</td><td>NA</td><td>NA</td></tr>
<tr><td>T8468</td><td>CTCAGCAGCAGCCCCTCTACAGG</td><td>1000</td><td></td><td></td><td></td><td></td><td></td><td></td><td></td><td></td><td></td></tr>
<tr><td>Oligo fwd</td><td>CACCCTCAGCAGCAGCCCCTCTAC</td><td></td><td></td><td></td><td></td><td></td><td></td><td></td><td></td><td></td><td></td></tr>
<tr><td>Oligo rev</td><td>AAACGTAGAGGGGCTGCTGCTGAG</td><td></td><td></td><td></td><td></td><td></td><td></td><td></td><td></td><td></td><td></td></tr>
<tr><td>Chromosome</td><td>start</td><td>end</td><td>strand</td><td>MM</td><td>target_seq</td><td>PAM</td><td>alignment</td><td>distance</td><td>position</td><td>gene name</td><td>gene id</td></tr>
<tr><td>GU305914.1</td><td>114914</td><td>114936</td><td>+</td><td>0</td><td>CTCAGCAGCAGCCCCTCTAC</td><td>AGG</td><td>||||||||[||||||||||||]PAM</td><td>NA</td><td>Intergenic</td><td>NA</td><td>NA</td></tr>
<tr><td>T8469</td><td>TCAGCAGCAGCCCCTCTACAGGG</td><td>1000</td><td></td><td></td><td></td><td></td><td></td><td></td><td></td><td></td><td></td></tr>
<tr><td>Oligo fwd</td><td>CACCTCAGCAGCAGCCCCTCTACA</td><td></td><td></td><td></td><td></td><td></td><td></td><td></td><td></td><td></td><td></td></tr>
<tr><td>Oligo rev</td><td>AAACTGTAGAGGGGCTGCTGCTGA</td><td></td><td></td><td></td><td></td><td></td><td></td><td></td><td></td><td></td><td></td></tr>
<tr><td>Chromosome</td><td>start</td><td>end</td><td>strand</td><td>MM</td><td>target_seq</td><td>PAM</td><td>alignment</td><td>distance</td><td>position</td><td>gene name</td><td>gene id</td></tr>
<tr><td>GU305914.1</td><td>114915</td><td>114937</td><td>+</td><td>0</td><td>TCAGCAGCAGCCCCTCTACA</td><td>GGG</td><td>||||||||[||||||||||||]PAM</td><td>NA</td><td>Intergenic</td><td>NA</td><td>NA</td></tr>
<tr><td>T8470</td><td>CAGCAGCAGCCCCTCTACAGGGG</td><td>1000</td><td></td><td></td><td></td><td></td><td></td><td></td><td></td><td></td><td></td></tr>
<tr><td>Oligo fwd</td><td>CACCCAGCAGCAGCCCCTCTACAG</td><td></td><td></td><td></td><td></td><td></td><td></td><td></td><td></td><td></td><td></td></tr>
<tr><td>Oligo rev</td><td>AAACCTGTAGAGGGGCTGCTGCTG</td><td></td><td></td><td></td><td></td><td></td><td></td><td></td><td></td><td></td><td></td></tr>
<tr><td>Chromosome</td><td>start</td><td>end</td><td>strand</td><td>MM</td><td>target_seq</td><td>PAM</td><td>alignment</td><td>distance</td><td>position</td><td>gene name</td><td>gene id</td></tr>
<tr><td>GU305914.1</td><td>114916</td><td>114938</td><td>+</td><td>0</td><td>CAGCAGCAGCCCCTCTACAG</td><td>GGG</td><td>||||||||[||||||||||||]PAM</td><td>NA</td><td>Intergenic</td><td>NA</td><td>NA</td></tr>
<tr><td>T8471</td><td>AGCCCCTCTACAGGGGAGAGAGG</td><td>1000</td><td></td><td></td><td></td><td></td><td></td><td></td><td></td><td></td><td></td></tr>
<tr><td>Oligo fwd</td><td>CACCAGCCCCTCTACAGGGGAGAG</td><td></td><td></td><td></td><td></td><td></td><td></td><td></td><td></td><td></td><td></td></tr>
<tr><td>Oligo rev</td><td>AAACCTCTCCCCTGTAGAGGGGCT</td><td></td><td></td><td></td><td></td><td></td><td></td><td></td><td></td><td></td><td></td></tr>
<tr><td>Chromosome</td><td>start</td><td>end</td><td>strand</td><td>MM</td><td>target_seq</td><td>PAM</td><td>alignment</td><td>distance</td><td>position</td><td>gene name</td><td>gene id</td></tr>
<tr><td>GU305914.1</td><td>114923</td><td>114945</td><td>+</td><td>0</td><td>AGCCCCTCTACAGGGGAGAG</td><td>AGG</td><td>||||||||[||||||||||||]PAM</td><td>NA</td><td>Intergenic</td><td>NA</td><td>NA</td></tr>
</table>

T8472 TCTACAGGGGAGAGAGGAATAGG 1000
Oligo fwd CACCTCTACAGGGGAGAGAGGAAT
Oligo rev AAACATTCCTCTCTCCCCTGTAGA
Chromosome start end strand MM target_seq PAM alignment distance position gene name gene id
GU305914.1 114929 114951 + 0 TCTACAGGGGAGAGAGGAAT AGG ||||||||[||||||||||||]PAM NA Intergenic NA NA

T8473 ACAGGGGAGAGAGGAATAGGCGG 1000
Oligo fwd CACCACAGGGGAGAGAGGAATAGG
Oligo rev AAACCCTATTCCTCTCTCCCCTGT
Chromosome start end strand MM target_seq PAM alignment distance position gene name gene id
GU305914.1 114932 114954 + 0 ACAGGGGAGAGAGGAATAGG CGG ||||||||[||||||||||||]PAM NA Intergenic NA NA

T8474 GGGGAGAGAGGAATAGGCGGCGG 1000
Oligo fwd CACCGGGGAGAGAGGAATAGGCGG
Oligo rev AAACCCGCCTATTCCTCTCTCCCC
Chromosome start end strand MM target_seq PAM alignment distance position gene name gene id
GU305914.1 114935 114957 + 0 GGGGAGAGAGGAATAGGCGG CGG ||||||||[||||||||||||]PAM NA Intergenic NA NA

T8475 GCGGCGGAGCCCTCCCGATTCGG 1000
Oligo fwd CACCGCGGCGGAGCCCTCCCGATT
Oligo rev AAACAATCGGGAGGGCTCCGCCGC
Chromosome start end strand MM target_seq PAM alignment distance position gene name gene id
GU305914.1 114951 114973 + 0 GCGGCGGAGCCCTCCCGATT CGG ||||||||[||||||||||||]PAM NA Intergenic NA NA

T8476 CGACGATGACGATGACGACGAGG 1000
Oligo fwd CACCCGACGATGACGATGACGACG
Oligo rev AAACCGTCGTCATCGTCATCGTCG
Chromosome start end strand MM target_seq PAM alignment distance position gene name gene id
GU305914.1 114978 115000 + 0 CGACGATGACGATGACGACG AGG ||||||||[||||||||||||]PAM NA Intergenic NA NA
GU305914.1 118812 118834 - 3 TGACGACGATGATGACGACG AGG -|||||-|[|-||||||||||]PAM NA Intergenic NA NA
GU305914.1 135393 135415 + 4 CGGGGACGTCGATGACGACG AGG ||--||-|[-|||||||||||]PAM NA Intergenic NA NA
GU305914.1 32834 32856 - 4 CTACGAGGACGACGACAACG AGG |-||||-|[||||-|||-|||]PAM NA Intergenic NA NA
GU305914.1 118515 118537 - 4 CGATGAGGACGATGATGAAG AGG |||-||-|[|||||||-||-|]PAM NA Intergenic NA NA
GU305914.1 174261 174283 + 4 CGACGGCGACGATGACGAAC AGG |||||--|[||||||||||--]PAM NA Intergenic NA NA

T8477 GACGACGAGGATCTCGAAGCCGG 1000
Oligo fwd CACCGACGACGAGGATCTCGAAGC
Oligo rev AAACGCTTCGAGATCCTCGTCGTC
Chromosome start end strand MM target_seq PAM alignment distance position gene name gene id
GU305914.1 114991 115013 + 0 GACGACGAGGATCTCGAAGC CGG ||||||||[||||||||||||]PAM NA Intergenic NA NA

T8478 TCTCGAAGCCGGCGAGCGTACGG 1000
Oligo fwd CACCTCTCGAAGCCGGCGAGCGTA
Oligo rev AAACTACGCTCGCCGGCTTCGAGA
Chromosome start end strand MM target_seq PAM alignment distance position gene name gene id
GU305914.1 115002 115024 + 0 TCTCGAAGCCGGCGAGCGTA CGG ||||||||[||||||||||||]PAM NA Intergenic NA NA

T8479 CTCGAAGCCGGCGAGCGTACGGG 1000
Oligo fwd CACCCTCGAAGCCGGCGAGCGTAC
Oligo rev AAACGTACGCTCGCCGGCTTCGAG
Chromosome start end strand MM target_seq PAM alignment distance position gene name gene id
GU305914.1 115003 115025 + 0 CTCGAAGCCGGCGAGCGTAC GGG ||||||||[||||||||||||]PAM NA Intergenic NA NA

T8480 TCGAAGCCGGCGAGCGTACGGGG 1000
Oligo fwd CACCTCGAAGCCGGCGAGCGTACG
Oligo rev AAACCGTACGCTCGCCGGCTTCGA
Chromosome start end strand MM target_seq PAM alignment distance position gene name gene id
GU305914.1 115004 115026 + 0 TCGAAGCCGGCGAGCGTACG GGG ||||||||[||||||||||||]PAM NA Intergenic NA NA

T8481 CGAAGCCGGCGAGCGTACGGGGG 1000
Oligo fwd CACCCGAAGCCGGCGAGCGTACGG
Oligo rev AAACCCGTACGCTCGCCGGCTTCG
Chromosome start end strand MM target_seq PAM alignment distance position gene name gene id
GU305914.1 115005 115027 + 0 CGAAGCCGGCGAGCGTACGG GGG ||||||||[||||||||||||]PAM NA Intergenic NA NA

T8482 GAAGCCGGCGAGCGTACGGGGGG 1000
Oligo fwd CACCGAAGCCGGCGAGCGTACGGG
Oligo rev AAACCCCGTACGCTCGCCGGCTTC
Chromosome start end strand MM target_seq PAM alignment distance position gene name gene id
GU305914.1 115006 115028 + 0 GAAGCCGGCGAGCGTACGGG GGG ||||||||[||||||||||||]PAM NA Intergenic NA NA

T8483 GTACGGGGGGCAAGAGAACGCGG 1000
Oligo fwd CACCGTACGGGGGGCAAGAGAACG
Oligo rev AAACCGTTCTCTTGCCCCCCGTAC
Chromosome start end strand MM target_seq PAM alignment distance position gene name gene id
GU305914.1 115019 115041 + 0 GTACGGGGGGCAAGAGAACG CGG ||||||||[||||||||||||]PAM NA Intergenic NA NA
GU305914.1 139107 139129 - 4 AGACGGGGGGCAGGAGCACG GGG --||||||[||||-|||-|||]PAM NA Intergenic NA NA

T8484 GGCAAGAGAACGCGGCAGAGAGG 1000
Oligo fwd CACCGGCAAGAGAACGCGGCAGAG
Oligo rev AAACCTCTGCCGCGTTCTCTTGCC
Chromosome start end strand MM target_seq PAM alignment distance position gene name gene id
GU305914.1 115027 115049 + 0 GGCAAGAGAACGCGGCAGAG AGG ||||||||[||||||||||||]PAM NA Intergenic NA NA

T8485 GCAAGAGAACGCGGCAGAGAGGG 1000
Oligo fwd CACCGCAAGAGAACGCGGCAGAGA
Oligo rev AAACTCTCTGCCGCGTTCTCTTGC
Chromosome start end strand MM target_seq PAM alignment distance position gene name gene id

GU305914.1 115028 115050 + 0 GCAAGAGAACGCGGCAGAGA GGG ||||||||[||||||||||||]PAM NA Intergenic NA NA

T8486 CAGAGAGGGAGCGCCGACTCCGG 1000
Oligo fwd CACCCAGAGAGGGAGCGCCGACTC
Oligo rev AAACGAGTCGGCGCTCCCTCTCTG
Chromosome start end strand MM target_seq PAM alignment distance position gene name gene id
GU305914.1 115042 115064 + 0 CAGAGAGGGAGCGCCGACTC CGG ||||||||[||||||||||||]PAM NA Intergenic NA NA

T8487 GAGGGAGCGCCGACTCCGGCAGG 1000
Oligo fwd CACCGAGGGAGCGCCGACTCCGGC
Oligo rev AAACGCCGGAGTCGGCGCTCCCTC
Chromosome start end strand MM target_seq PAM alignment distance position gene name gene id
GU305914.1 115046 115068 + 0 GAGGGAGCGCCGACTCCGGC AGG ||||||||[||||||||||||]PAM NA Intergenic NA NA
GU305914.1 106810 106832 + 4 GACGGTGCGCCGACTCCATC AGG ||-||-||[|||||||||--|]PAM NA Intergenic NA NA

T8488 GCGCCGACTCCGGCAGGAAGAGG 1000
Oligo fwd CACCGCGCCGACTCCGGCAGGAAG
Oligo rev AAACCTTCCTGCCGGAGTCGGCGC
Chromosome start end strand MM target_seq PAM alignment distance position gene name gene id
GU305914.1 115052 115074 + 0 GCGCCGACTCCGGCAGGAAG AGG ||||||||[||||||||||||]PAM NA Intergenic NA NA

T8489 CCGGCAGGAAGAGGAGACGTCGG 1000
Oligo fwd CACCCCGGCAGGAAGAGGAGACGT
Oligo rev AAACACGTCTCCTCTTCCTGCCGG
Chromosome start end strand MM target_seq PAM alignment distance position gene name gene id
GU305914.1 115061 115083 + 0 CCGGCAGGAAGAGGAGACGT CGG ||||||||[||||||||||||]PAM NA Intergenic NA NA

T8490 CGGCAGGAAGAGGAGACGTCGGG 1000
Oligo fwd CACCCGGCAGGAAGAGGAGACGTC
Oligo rev AAACGACGTCTCCTCTTCCTGCCG
Chromosome start end strand MM target_seq PAM alignment distance position gene name gene id
GU305914.1 115062 115084 + 0 CGGCAGGAAGAGGAGACGTC GGG ||||||||[||||||||||||]PAM NA Intergenic NA NA

T8491 GGCAGGAAGAGGAGACGTCGGGG 1000
Oligo fwd CACCGGCAGGAAGAGGAGACGTCG
Oligo rev AAACCGACGTCTCCTCTTCCTGCC
Chromosome start end strand MM target_seq PAM alignment distance position gene name gene id
GU305914.1 115063 115085 + 0 GGCAGGAAGAGGAGACGTCG GGG ||||||||[||||||||||||]PAM NA Intergenic NA NA

T8492 GAAGAGGAGACGTCGGGGCGCGG 1000
Oligo fwd CACCGAAGAGGAGACGTCGGGGCG
Oligo rev AAACCGCCCCGACGTCTCCTCTTC
Chromosome start end strand MM target_seq PAM alignment distance position gene name gene id
GU305914.1 115068 115090 + 0 GAAGAGGAGACGTCGGGGCG CGG ||||||||[||||||||||||]PAM NA Intergenic NA NA

T8493 GGCGCGGCGCCTGACGACGATGG 1000
Oligo fwd CACCGGCGCGGCGCCTGACGACGA
Oligo rev AAACTCGTCGTCAGGCGCCGCGCC
Chromosome start end strand MM target_seq PAM alignment distance position gene name gene id
GU305914.1 115084 115106 + 0 GGCGCGGCGCCTGACGACGA TGG ||||||||[||||||||||||]PAM NA Intergenic NA NA

T8494 GCGGCGCCTGACGACGATGGCGG 1000
Oligo fwd CACCGCGGCGCCTGACGACGATGG
Oligo rev AAACCCATCGTCGTCAGGCGCCGC
Chromosome start end strand MM target_seq PAM alignment distance position gene name gene id
GU305914.1 115087 115109 + 0 GCGGCGCCTGACGACGATGG CGG ||||||||[||||||||||||]PAM NA Intergenic NA NA

T8495 GGCGGTGACCTCTCCCTACCCGG 1000
Oligo fwd CACCGGCGGTGACCTCTCCCTACC
Oligo rev AAACGGTAGGGAGAGGTCACCGCC
Chromosome start end strand MM target_seq PAM alignment distance position gene name gene id
GU305914.1 115105 115127 + 0 GGCGGTGACCTCTCCCTACC CGG ||||||||[||||||||||||]PAM NA Intergenic NA NA

T8496 ACCTCTCCCTACCCGGAGAGCGG 1000
Oligo fwd CACCACCTCTCCCTACCCGGAGAG
Oligo rev AAACCTCTCCGGGTAGGGAGAGGT
Chromosome start end strand MM target_seq PAM alignment distance position gene name gene id
GU305914.1 115112 115134 + 0 ACCTCTCCCTACCCGGAGAG CGG ||||||||[||||||||||||]PAM NA Intergenic NA NA

T8497 CCTCTCCCTACCCGGAGAGCGGG 1000
Oligo fwd CACCCCTCTCCCTACCCGGAGAGC
Oligo rev AAACGCTCTCCGGGTAGGGAGAGG
Chromosome start end strand MM target_seq PAM alignment distance position gene name gene id
GU305914.1 115113 115135 + 0 CCTCTCCCTACCCGGAGAGC GGG ||||||||[||||||||||||]PAM NA Intergenic NA NA

T8498 CTCTCCCTACCCGGAGAGCGGGG 1000
Oligo fwd CACCCTCTCCCTACCCGGAGAGCG
Oligo rev AAACCGCTCTCCGGGTAGGGAGAG
Chromosome start end strand MM target_seq PAM alignment distance position gene name gene id
GU305914.1 115114 115136 + 0 CTCTCCCTACCCGGAGAGCG GGG ||||||||[||||||||||||]PAM NA Intergenic NA NA

T8499 GGATATCCCAAGCGCACCGCCGG 1000
Oligo fwd CACCGGATATCCCAAGCGCACCGC
Oligo rev AAACGCGGTGCGCTTGGGATATCC
Chromosome start end strand MM target_seq PAM alignment distance position gene name gene id
GU305914.1 115135 115157 + 0 GGATATCCCAAGCGCACCGC CGG ||||||||[||||||||||||]PAM NA Intergenic NA NA

T8500 CGCCGGTGATCATCACCAGTCGG 1000
Oligo fwd CACCCGCCGGTGATCATCACCAGT

Oligo rev AAACACTGGTGATGATCACCGGCG
Chromosome start end strand MM target_seq PAM alignment distance position gene name gene id
GU305914.1 115152 115174 + 0 CGCCGGTGATCATCACCAGT CGG ||||||||[||||||||||||]PAM NA Intergenic NA NA

T8501 TCATCACCAGTCGGCGCCGCCGG 1000
Oligo fwd CACCTCATCACCAGTCGGCGCCGC
Oligo rev AAACGCGGCGCCGACTGGTGATGA
Chromosome start end strand MM target_seq PAM alignment distance position gene name gene id
GU305914.1 115161 115183 + 0 TCATCACCAGTCGGCGCCGC CGG ||||||||[||||||||||||]PAM NA Intergenic NA NA

T8502 GCCTCCAGAACCGATGAGTTCGG 1000
Oligo fwd CACCGCCTCCAGAACCGATGAGTT
Oligo rev AAACAACTCATCGGTTCTGGAGGC
Chromosome start end strand MM target_seq PAM alignment distance position gene name gene id
GU305914.1 115183 115205 + 0 GCCTCCAGAACCGATGAGTT CGG ||||||||[||||||||||||]PAM NA Intergenic NA NA

T8503 CAGAACCGATGAGTTCGGAGAGG 1000
Oligo fwd CACCCAGAACCGATGAGTTCGGAG
Oligo rev AAACCTCCGAACTCATCGGTTCTG
Chromosome start end strand MM target_seq PAM alignment distance position gene name gene id
GU305914.1 115188 115210 + 0 CAGAACCGATGAGTTCGGAG AGG ||||||||[||||||||||||]PAM NA Intergenic NA NA

T8504 CCGATGAGTTCGGAGAGGTGAGG 1000
Oligo fwd CACCCCGATGAGTTCGGAGAGGTG
Oligo rev AAACCACCTCTCCGAACTCATCGG
Chromosome start end strand MM target_seq PAM alignment distance position gene name gene id
GU305914.1 115193 115215 + 0 CCGATGAGTTCGGAGAGGTG AGG ||||||||[||||||||||||]PAM NA Intergenic NA NA

T8505 CGATGAGTTCGGAGAGGTGAGGG 1000
Oligo fwd CACCCGATGAGTTCGGAGAGGTGA
Oligo rev AAACTCACCTCTCCGAACTCATCG
Chromosome start end strand MM target_seq PAM alignment distance position gene name gene id
GU305914.1 115194 115216 + 0 CGATGAGTTCGGAGAGGTGA GGG ||||||||[||||||||||||]PAM NA Intergenic NA NA

T8506 AGACATCTCGCAGATCCGCGCGG 1000
Oligo fwd CACCAGACATCTCGCAGATCCGCG
Oligo rev AAACCGCGGATCTGCGAGATGTCT
Chromosome start end strand MM target_seq PAM alignment distance position gene name gene id
GU305914.1 115239 115261 + 0 AGACATCTCGCAGATCCGCG CGG ||||||||[||||||||||||]PAM NA Intergenic NA NA

T8507 CGCAGATCCGCGCGGCCGCCAGG 1000
Oligo fwd CACCCGCAGATCCGCGCGGCCGCC
Oligo rev AAACGGCGGCCGCGCGGATCTGCG
Chromosome start end strand MM target_seq PAM alignment distance position gene name gene id
GU305914.1 115247 115269 + 0 CGCAGATCCGCGCGGCCGCC AGG ||||||||[||||||||||||]PAM NA Intergenic NA NA

T8508 GCAGATCCGCGCGGCCGCCAGGG 1000
Oligo fwd CACCGCAGATCCGCGCGGCCGCCA
Oligo rev AAACTGGCGGCCGCGCGGATCTGC
Chromosome start end strand MM target_seq PAM alignment distance position gene name gene id
GU305914.1 115248 115270 + 0 GCAGATCCGCGCGGCCGCCA GGG ||||||||[||||||||||||]PAM NA Intergenic NA NA

T8509 CCGCGCGGCCGCCAGGGCCGAGG 1000
Oligo fwd CACCCCGCGCGGCCGCCAGGGCCG
Oligo rev AAACCGGCCCTGGCGGCCGCGCGG
Chromosome start end strand MM target_seq PAM alignment distance position gene name gene id
GU305914.1 115254 115276 + 0 CCGCGCGGCCGCCAGGGCCG AGG ||||||||[||||||||||||]PAM NA Intergenic NA NA

T8510 CGCGCGGCCGCCAGGGCCGAGGG 1000
Oligo fwd CACCCGCGCGGCCGCCAGGGCCGA
Oligo rev AAACTCGGCCCTGGCGGCCGCGCG
Chromosome start end strand MM target_seq PAM alignment distance position gene name gene id
GU305914.1 115255 115277 + 0 CGCGCGGCCGCCAGGGCCGA GGG ||||||||[||||||||||||]PAM NA Intergenic NA NA

T8511 GCCGCCAGGGCCGAGGGCAACGG 1000
Oligo fwd CACCGCCGCCAGGGCCGAGGGCAA
Oligo rev AAACTTGCCCTCGGCCCTGGCGGC
Chromosome start end strand MM target_seq PAM alignment distance position gene name gene id
GU305914.1 115261 115283 + 0 GCCGCCAGGGCCGAGGGCAA CGG ||||||||[||||||||||||]PAM NA Intergenic NA NA

T8512 CGAGGGCAACGGCGCGCGCGAGG 1000
Oligo fwd CACCCGAGGGCAACGGCGCGCGCG
Oligo rev AAACCGCGCGCGCCGTTGCCCTCG
Chromosome start end strand MM target_seq PAM alignment distance position gene name gene id
GU305914.1 115272 115294 + 0 CGAGGGCAACGGCGCGCGCG AGG ||||||||[||||||||||||]PAM NA Intergenic NA NA

T8513 CAACGGCGCGCGCGAGGACGCGG 1000
Oligo fwd CACCCAACGGCGCGCGCGAGGACG
Oligo rev AAACCGTCCTCGCGCGCGCCGTTG
Chromosome start end strand MM target_seq PAM alignment distance position gene name gene id
GU305914.1 115278 115300 + 0 CAACGGCGCGCGCGAGGACG CGG ||||||||[||||||||||||]PAM NA Intergenic NA NA

T8514 CGCGCGCGAGGACGCGGCCTCGG 1000
Oligo fwd CACCCGCGCGCGAGGACGCGGCCT
Oligo rev AAACAGGCCGCGTCCTCGCGCGCG
Chromosome start end strand MM target_seq PAM alignment distance position gene name gene id
GU305914.1 115284 115306 + 0 CGCGCGCGAGGACGCGGCCT CGG ||||||||[||||||||||||]PAM NA Intergenic NA NA

T8515 CGCGAGGACGCGGCCTCGGTCGG 1000

| | | | | | | | | | | | |
|---|---|---|---|---|---|---|---|---|---|---|---|
| Oligo fwd | CACCCGCGAGGACGCGGCCTCGGT | | | | | | | | | | |
| Oligo rev | AAACACCGAGGCCGCGTCCTCGCG | | | | | | | | | | |
| Chromosome | start | end | strand | MM | target_seq | PAM | alignment | distance | position | gene name | gene id |
| GU305914.1 | 115288 | 115310 | + | 0 | CGCGAGGACGCGGCCTCGGT | CGG | \|\|\|\|\|\|\|\|[\|\|\|\|\|\|\|\|\|\|\|\|]PAM | NA | Intergenic | NA | NA |
| T8516 | AAATGTGCCGCCCCGCCCCCCGG | 1000 | | | | | | | | | |
| Oligo fwd | CACCAAATGTGCCGCCCCGCCCCC | | | | | | | | | | |
| Oligo rev | AAACGGGGGCGGGGCGGCACATTT | | | | | | | | | | |
| Chromosome | start | end | strand | MM | target_seq | PAM | alignment | distance | position | gene name | gene id |
| GU305914.1 | 115324 | 115346 | + | 0 | AAATGTGCCGCCCCGCCCCC | CGG | \|\|\|\|\|\|\|\|[\|\|\|\|\|\|\|\|\|\|\|\|]PAM | NA | Intergenic | NA | NA |
| T8517 | CCCCGGAGTCACCGAGATGATGG | 1000 | | | | | | | | | |
| Oligo fwd | CACCCCCCGGAGTCACCGAGATGA | | | | | | | | | | |
| Oligo rev | AAACTCATCTCGGTGACTCCGGGG | | | | | | | | | | |
| Chromosome | start | end | strand | MM | target_seq | PAM | alignment | distance | position | gene name | gene id |
| GU305914.1 | 115341 | 115363 | + | 0 | CCCCGGAGTCACCGAGATGA | TGG | \|\|\|\|\|\|\|\|[\|\|\|\|\|\|\|\|\|\|\|\|]PAM | NA | Intergenic | NA | NA |
| T8518 | AGTCACCGAGATGATGGCCTCGG | 1000 | | | | | | | | | |
| Oligo fwd | CACCAGTCACCGAGATGATGGCCT | | | | | | | | | | |
| Oligo rev | AAACAGGCCATCATCTCGGTGACT | | | | | | | | | | |
| Chromosome | start | end | strand | MM | target_seq | PAM | alignment | distance | position | gene name | gene id |
| GU305914.1 | 115347 | 115369 | + | 0 | AGTCACCGAGATGATGGCCT | CGG | \|\|\|\|\|\|\|\|[\|\|\|\|\|\|\|\|\|\|\|\|]PAM | NA | Intergenic | NA | NA |
| T8519 | GATGATGGCCTCGGAACCGCCGG | 1000 | | | | | | | | | |
| Oligo fwd | CACCGATGATGGCCTCGGAACCGC | | | | | | | | | | |
| Oligo rev | AAACGCGGTTCCGAGGCCATCATC | | | | | | | | | | |
| Chromosome | start | end | strand | MM | target_seq | PAM | alignment | distance | position | gene name | gene id |
| GU305914.1 | 115356 | 115378 | + | 0 | GATGATGGCCTCGGAACCGC | CGG | \|\|\|\|\|\|\|\|[\|\|\|\|\|\|\|\|\|\|\|\|]PAM | NA | Intergenic | NA | NA |
| T8520 | ATGGCCTCGGAACCGCCGGCCGG | 1000 | | | | | | | | | |
| Oligo fwd | CACCATGGCCTCGGAACCGCCGGC | | | | | | | | | | |
| Oligo rev | AAACGCCGGCGGTTCCGAGGCCAT | | | | | | | | | | |
| Chromosome | start | end | strand | MM | target_seq | PAM | alignment | distance | position | gene name | gene id |
| GU305914.1 | 115360 | 115382 | + | 0 | ATGGCCTCGGAACCGCCGGC | CGG | \|\|\|\|\|\|\|\|[\|\|\|\|\|\|\|\|\|\|\|\|]PAM | NA | Intergenic | NA | NA |
| T8521 | GCCTCGGAACCGCCGGCCGGCGG | 1000 | | | | | | | | | |
| Oligo fwd | CACCGCCTCGGAACCGCCGGCCGG | | | | | | | | | | |
| Oligo rev | AAACCCGGCCGGCGGTTCCGAGGC | | | | | | | | | | |
| Chromosome | start | end | strand | MM | target_seq | PAM | alignment | distance | position | gene name | gene id |
| GU305914.1 | 115363 | 115385 | + | 0 | GCCTCGGAACCGCCGGCCGG | CGG | \|\|\|\|\|\|\|\|[\|\|\|\|\|\|\|\|\|\|\|\|]PAM | NA | Intergenic | NA | NA |
| GU305914.1 | 90211 | 90233 | - | 4 | GCCGCGGGGCCTCCGGCCGG | CGG | \|\|\|-\|\|\|-[-\|\|-\|\|\|\|\|\|\|\|]PAM | NA | Intergenic | NA | NA |
| T8522 | CCGGCGGCACCGTCGTCGCCAGG | 1000 | | | | | | | | | |
| Oligo fwd | CACCCCGGCGGCACCGTCGTCGCC | | | | | | | | | | |
| Oligo rev | AAACGGCGACGACGGTGCCGCCGG | | | | | | | | | | |
| Chromosome | start | end | strand | MM | target_seq | PAM | alignment | distance | position | gene name | gene id |
| GU305914.1 | 115379 | 115401 | + | 0 | CCGGCGGCACCGTCGTCGCC | AGG | \|\|\|\|\|\|\|\|[\|\|\|\|\|\|\|\|\|\|\|\|]PAM | NA | Intergenic | NA | NA |
| GU305914.1 | 170560 | 170582 | + | 4 | CGGTCGGCACCATCGCCGCC | GGG | \|-\|-\|\|\|\|[\|\|\|-\|\|\|-\|\|\|\|]PAM | NA | Intergenic | NA | NA |
| T8523 | CGGCACCGTCGTCGCCAGGATGG | 1000 | | | | | | | | | |
| Oligo fwd | CACCCGGCACCGTCGTCGCCAGGA | | | | | | | | | | |
| Oligo rev | AAACTCCTGGCGACGACGGTGCCG | | | | | | | | | | |
| Chromosome | start | end | strand | MM | target_seq | PAM | alignment | distance | position | gene name | gene id |
| GU305914.1 | 115383 | 115405 | + | 0 | CGGCACCGTCGTCGCCAGGA | TGG | \|\|\|\|\|\|\|\|[\|\|\|\|\|\|\|\|\|\|\|\|]PAM | NA | Intergenic | NA | NA |
| GU305914.1 | 21983 | 22005 | + | 4 | CAGGGCCGTCGTCGGCAGGA | CGG | \|-\|--\|\|\|[\|\|\|\|\|\|-\|\|\|\|\|]PAM | NA | Intergenic | NA | NA |
| T8524 | CGTCGTCGCCAGGATGGCCCTGG | 1000 | | | | | | | | | |
| Oligo fwd | CACCCGTCGTCGCCAGGATGGCCC | | | | | | | | | | |
| Oligo rev | AAACGGGCCATCCTGGCGACGACG | | | | | | | | | | |
| Chromosome | start | end | strand | MM | target_seq | PAM | alignment | distance | position | gene name | gene id |
| GU305914.1 | 115389 | 115411 | + | 0 | CGTCGTCGCCAGGATGGCCC | TGG | \|\|\|\|\|\|\|\|[\|\|\|\|\|\|\|\|\|\|\|\|]PAM | NA | Intergenic | NA | NA |
| T8525 | CGCCAGGATGGCCCTGGATCCGG | 1000 | | | | | | | | | |
| Oligo fwd | CACCCGCCAGGATGGCCCTGGATC | | | | | | | | | | |
| Oligo rev | AAACGATCCAGGGCCATCCTGGCG | | | | | | | | | | |
| Chromosome | start | end | strand | MM | target_seq | PAM | alignment | distance | position | gene name | gene id |
| GU305914.1 | 115395 | 115417 | + | 0 | CGCCAGGATGGCCCTGGATC | CGG | \|\|\|\|\|\|\|\|[\|\|\|\|\|\|\|\|\|\|\|\|]PAM | NA | Intergenic | NA | NA |
| T8526 | GATGGCCCTGGATCCGGCCGTGG | 1000 | | | | | | | | | |
| Oligo fwd | CACCGATGGCCCTGGATCCGGCCG | | | | | | | | | | |
| Oligo rev | AAACCGGCCGGATCCAGGGCCATC | | | | | | | | | | |
| Chromosome | start | end | strand | MM | target_seq | PAM | alignment | distance | position | gene name | gene id |
| GU305914.1 | 115401 | 115423 | + | 0 | GATGGCCCTGGATCCGGCCG | TGG | \|\|\|\|\|\|\|\|[\|\|\|\|\|\|\|\|\|\|\|\|]PAM | NA | Intergenic | NA | NA |
| T8527 | TCCGGCCGTGGCCGCCGCGACGG | 1000 | | | | | | | | | |
| Oligo fwd | CACCTCCGGCCGTGGCCGCCGCGA | | | | | | | | | | |
| Oligo rev | AAACTCGCGGCGGCCACGGCCGGA | | | | | | | | | | |
| Chromosome | start | end | strand | MM | target_seq | PAM | alignment | distance | position | gene name | gene id |
| GU305914.1 | 115413 | 115435 | + | 0 | TCCGGCCGTGGCCGCCGCGA | CGG | \|\|\|\|\|\|\|\|[\|\|\|\|\|\|\|\|\|\|\|\|]PAM | NA | Intergenic | NA | NA |
| T8528 | CCGGCCGTGGCCGCCGCGACGGG | 1000 | | | | | | | | | |
| Oligo fwd | CACCCCGGCCGTGGCCGCCGCGAC | | | | | | | | | | |
| Oligo rev | AAACGTCGCGGCGGCCACGGCCGG | | | | | | | | | | |
| Chromosome | start | end | strand | MM | target_seq | PAM | alignment | distance | position | gene name | gene id |
| GU305914.1 | 115414 | 115436 | + | 0 | CCGGCCGTGGCCGCCGCGAC | GGG | \|\|\|\|\|\|\|\|[\|\|\|\|\|\|\|\|\|\|\|\|]PAM | NA | Intergenic | NA | NA |
| T8529 | GCCGCGACGGGACATACCGCCGG | 1000 | | | | | | | | | |
| Oligo fwd | CACCGCCGCGACGGGACATACCGC | | | | | | | | | | |
| Oligo rev | AAACGCGGTATGTCCCGTCGCGGC | | | | | | | | | | |

<table>
<tr><td>Chromosome</td><td>start</td><td>end</td><td>strand</td><td>MM</td><td>target_seq</td><td>PAM</td><td>alignment</td><td>distance</td><td>position</td><td>gene name</td><td>gene id</td></tr>
<tr><td>GU305914.1</td><td>115426</td><td>115448</td><td>+</td><td>0</td><td>GCCGCGACGGGACATACCGC</td><td>CGG</td><td>||||||||[||||||||||||]PAM</td><td>NA</td><td>Intergenic</td><td>NA</td><td>NA</td></tr>
<tr><td></td><td></td><td></td><td></td><td></td><td></td><td></td><td></td><td></td><td></td><td></td><td></td></tr>
<tr><td>T8530</td><td>CCGCGACGGGACATACCGCCGGG</td><td>1000</td><td></td><td></td><td></td><td></td><td></td><td></td><td></td><td></td><td></td></tr>
<tr><td>Oligo fwd</td><td>CACCCCGCGACGGGACATACCGCC</td><td></td><td></td><td></td><td></td><td></td><td></td><td></td><td></td><td></td><td></td></tr>
<tr><td>Oligo rev</td><td>AAACGGCGGTATGTCCCGTCGCGG</td><td></td><td></td><td></td><td></td><td></td><td></td><td></td><td></td><td></td><td></td></tr>
<tr><td>Chromosome</td><td>start</td><td>end</td><td>strand</td><td>MM</td><td>target_seq</td><td>PAM</td><td>alignment</td><td>distance</td><td>position</td><td>gene name</td><td>gene id</td></tr>
<tr><td>GU305914.1</td><td>115427</td><td>115449</td><td>+</td><td>0</td><td>CCGCGACGGGACATACCGCC</td><td>GGG</td><td>||||||||[||||||||||||]PAM</td><td>NA</td><td>Intergenic</td><td>NA</td><td>NA</td></tr>
<tr><td></td><td></td><td></td><td></td><td></td><td></td><td></td><td></td><td></td><td></td><td></td><td></td></tr>
<tr><td>T8531</td><td>ACCGCCGGGCTGCTTACCGCCGG</td><td>1000</td><td></td><td></td><td></td><td></td><td></td><td></td><td></td><td></td><td></td></tr>
<tr><td>Oligo fwd</td><td>CACCACCGCCGGGCTGCTTACCGC</td><td></td><td></td><td></td><td></td><td></td><td></td><td></td><td></td><td></td><td></td></tr>
<tr><td>Oligo rev</td><td>AAACGCGGTAAGCAGCCCGGCGGT</td><td></td><td></td><td></td><td></td><td></td><td></td><td></td><td></td><td></td><td></td></tr>
<tr><td>Chromosome</td><td>start</td><td>end</td><td>strand</td><td>MM</td><td>target_seq</td><td>PAM</td><td>alignment</td><td>distance</td><td>position</td><td>gene name</td><td>gene id</td></tr>
<tr><td>GU305914.1</td><td>115441</td><td>115463</td><td>+</td><td>0</td><td>ACCGCCGGGCTGCTTACCGC</td><td>CGG</td><td>||||||||[||||||||||||]PAM</td><td>NA</td><td>Intergenic</td><td>NA</td><td>NA</td></tr>
<tr><td></td><td></td><td></td><td></td><td></td><td></td><td></td><td></td><td></td><td></td><td></td><td></td></tr>
<tr><td>T8532</td><td>CCGCCGGGCTGCTTACCGCCGGG</td><td>1000</td><td></td><td></td><td></td><td></td><td></td><td></td><td></td><td></td><td></td></tr>
<tr><td>Oligo fwd</td><td>CACCCCGCCGGGCTGCTTACCGCC</td><td></td><td></td><td></td><td></td><td></td><td></td><td></td><td></td><td></td><td></td></tr>
<tr><td>Oligo rev</td><td>AAACGGCGGTAAGCAGCCCGGCGG</td><td></td><td></td><td></td><td></td><td></td><td></td><td></td><td></td><td></td><td></td></tr>
<tr><td>Chromosome</td><td>start</td><td>end</td><td>strand</td><td>MM</td><td>target_seq</td><td>PAM</td><td>alignment</td><td>distance</td><td>position</td><td>gene name</td><td>gene id</td></tr>
<tr><td>GU305914.1</td><td>115442</td><td>115464</td><td>+</td><td>0</td><td>CCGCCGGGCTGCTTACCGCC</td><td>GGG</td><td>||||||||[||||||||||||]PAM</td><td>NA</td><td>Intergenic</td><td>NA</td><td>NA</td></tr>
<tr><td></td><td></td><td></td><td></td><td></td><td></td><td></td><td></td><td></td><td></td><td></td><td></td></tr>
<tr><td>T8533</td><td>GCTGCTTACCGCCGGGAAATTGG</td><td>1000</td><td></td><td></td><td></td><td></td><td></td><td></td><td></td><td></td><td></td></tr>
<tr><td>Oligo fwd</td><td>CACCGCTGCTTACCGCCGGGAAAT</td><td></td><td></td><td></td><td></td><td></td><td></td><td></td><td></td><td></td><td></td></tr>
<tr><td>Oligo rev</td><td>AAACATTTCCCGGCGGTAAGCAGC</td><td></td><td></td><td></td><td></td><td></td><td></td><td></td><td></td><td></td><td></td></tr>
<tr><td>Chromosome</td><td>start</td><td>end</td><td>strand</td><td>MM</td><td>target_seq</td><td>PAM</td><td>alignment</td><td>distance</td><td>position</td><td>gene name</td><td>gene id</td></tr>
<tr><td>GU305914.1</td><td>115449</td><td>115471</td><td>+</td><td>0</td><td>GCTGCTTACCGCCGGGAAAT</td><td>TGG</td><td>||||||||[||||||||||||]PAM</td><td>NA</td><td>Intergenic</td><td>NA</td><td>NA</td></tr>
<tr><td></td><td></td><td></td><td></td><td></td><td></td><td></td><td></td><td></td><td></td><td></td><td></td></tr>
<tr><td>T8534</td><td>TTCGTGCGAACCGACTCCCATGG</td><td>1000</td><td></td><td></td><td></td><td></td><td></td><td></td><td></td><td></td><td></td></tr>
<tr><td>Oligo fwd</td><td>CACCTTCGTGCGAACCGACTCCCA</td><td></td><td></td><td></td><td></td><td></td><td></td><td></td><td></td><td></td><td></td></tr>
<tr><td>Oligo rev</td><td>AAACTGGGAGTCGGTTCGCACGAA</td><td></td><td></td><td></td><td></td><td></td><td></td><td></td><td></td><td></td><td></td></tr>
<tr><td>Chromosome</td><td>start</td><td>end</td><td>strand</td><td>MM</td><td>target_seq</td><td>PAM</td><td>alignment</td><td>distance</td><td>position</td><td>gene name</td><td>gene id</td></tr>
<tr><td>GU305914.1</td><td>115479</td><td>115501</td><td>+</td><td>0</td><td>TTCGTGCGAACCGACTCCCA</td><td>TGG</td><td>||||||||[||||||||||||]PAM</td><td>NA</td><td>Intergenic</td><td>NA</td><td>NA</td></tr>
<tr><td></td><td></td><td></td><td></td><td></td><td></td><td></td><td></td><td></td><td></td><td></td><td></td></tr>
<tr><td>T8535</td><td>GTGCGAACCGACTCCCATGGAGG</td><td>1000</td><td></td><td></td><td></td><td></td><td></td><td></td><td></td><td></td><td></td></tr>
<tr><td>Oligo fwd</td><td>CACCGTGCGAACCGACTCCCATGG</td><td></td><td></td><td></td><td></td><td></td><td></td><td></td><td></td><td></td><td></td></tr>
<tr><td>Oligo rev</td><td>AAACCCATGGGAGTCGGTTCGCAC</td><td></td><td></td><td></td><td></td><td></td><td></td><td></td><td></td><td></td><td></td></tr>
<tr><td>Chromosome</td><td>start</td><td>end</td><td>strand</td><td>MM</td><td>target_seq</td><td>PAM</td><td>alignment</td><td>distance</td><td>position</td><td>gene name</td><td>gene id</td></tr>
<tr><td>GU305914.1</td><td>115482</td><td>115504</td><td>+</td><td>0</td><td>GTGCGAACCGACTCCCATGG</td><td>AGG</td><td>||||||||[||||||||||||]PAM</td><td>NA</td><td>Intergenic</td><td>NA</td><td>NA</td></tr>
<tr><td></td><td></td><td></td><td></td><td></td><td></td><td></td><td></td><td></td><td></td><td></td><td></td></tr>
<tr><td>T8536</td><td>CGAACCGACTCCCATGGAGGTGG</td><td>1000</td><td></td><td></td><td></td><td></td><td></td><td></td><td></td><td></td><td></td></tr>
<tr><td>Oligo fwd</td><td>CACCCGAACCGACTCCCATGGAGG</td><td></td><td></td><td></td><td></td><td></td><td></td><td></td><td></td><td></td><td></td></tr>
<tr><td>Oligo rev</td><td>AAACCCTCCATGGGAGTCGGTTCG</td><td></td><td></td><td></td><td></td><td></td><td></td><td></td><td></td><td></td><td></td></tr>
<tr><td>Chromosome</td><td>start</td><td>end</td><td>strand</td><td>MM</td><td>target_seq</td><td>PAM</td><td>alignment</td><td>distance</td><td>position</td><td>gene name</td><td>gene id</td></tr>
<tr><td>GU305914.1</td><td>115485</td><td>115507</td><td>+</td><td>0</td><td>CGAACCGACTCCCATGGAGG</td><td>TGG</td><td>||||||||[||||||||||||]PAM</td><td>NA</td><td>Intergenic</td><td>NA</td><td>NA</td></tr>
<tr><td></td><td></td><td></td><td></td><td></td><td></td><td></td><td></td><td></td><td></td><td></td><td></td></tr>
<tr><td>T8537</td><td>GAACCGACTCCCATGGAGGTGGG</td><td>1000</td><td></td><td></td><td></td><td></td><td></td><td></td><td></td><td></td><td></td></tr>
<tr><td>Oligo fwd</td><td>CACCGAACCGACTCCCATGGAGGT</td><td></td><td></td><td></td><td></td><td></td><td></td><td></td><td></td><td></td><td></td></tr>
<tr><td>Oligo rev</td><td>AAACACCTCCATGGGAGTCGGTTC</td><td></td><td></td><td></td><td></td><td></td><td></td><td></td><td></td><td></td><td></td></tr>
<tr><td>Chromosome</td><td>start</td><td>end</td><td>strand</td><td>MM</td><td>target_seq</td><td>PAM</td><td>alignment</td><td>distance</td><td>position</td><td>gene name</td><td>gene id</td></tr>
<tr><td>GU305914.1</td><td>115486</td><td>115508</td><td>+</td><td>0</td><td>GAACCGACTCCCATGGAGGT</td><td>GGG</td><td>||||||||[||||||||||||]PAM</td><td>NA</td><td>Intergenic</td><td>NA</td><td>NA</td></tr>
<tr><td></td><td></td><td></td><td></td><td></td><td></td><td></td><td></td><td></td><td></td><td></td><td></td></tr>
<tr><td>T8538</td><td>CATGGAGGTGGGAGAACCCTCGG</td><td>1000</td><td></td><td></td><td></td><td></td><td></td><td></td><td></td><td></td><td></td></tr>
<tr><td>Oligo fwd</td><td>CACCCATGGAGGTGGGAGAACCCT</td><td></td><td></td><td></td><td></td><td></td><td></td><td></td><td></td><td></td><td></td></tr>
<tr><td>Oligo rev</td><td>AAACAGGGTTCTCCCACCTCCATG</td><td></td><td></td><td></td><td></td><td></td><td></td><td></td><td></td><td></td><td></td></tr>
<tr><td>Chromosome</td><td>start</td><td>end</td><td>strand</td><td>MM</td><td>target_seq</td><td>PAM</td><td>alignment</td><td>distance</td><td>position</td><td>gene name</td><td>gene id</td></tr>
<tr><td>GU305914.1</td><td>115497</td><td>115519</td><td>+</td><td>0</td><td>CATGGAGGTGGGAGAACCCT</td><td>CGG</td><td>||||||||[||||||||||||]PAM</td><td>NA</td><td>Intergenic</td><td>NA</td><td>NA</td></tr>
<tr><td></td><td></td><td></td><td></td><td></td><td></td><td></td><td></td><td></td><td></td><td></td><td></td></tr>
<tr><td>T8539</td><td>ATGGAGGTGGGAGAACCCTCGGG</td><td>1000</td><td></td><td></td><td></td><td></td><td></td><td></td><td></td><td></td><td></td></tr>
<tr><td>Oligo fwd</td><td>CACCATGGAGGTGGGAGAACCCTC</td><td></td><td></td><td></td><td></td><td></td><td></td><td></td><td></td><td></td><td></td></tr>
<tr><td>Oligo rev</td><td>AAACGAGGGTTCTCCCACCTCCAT</td><td></td><td></td><td></td><td></td><td></td><td></td><td></td><td></td><td></td><td></td></tr>
<tr><td>Chromosome</td><td>start</td><td>end</td><td>strand</td><td>MM</td><td>target_seq</td><td>PAM</td><td>alignment</td><td>distance</td><td>position</td><td>gene name</td><td>gene id</td></tr>
<tr><td>GU305914.1</td><td>115498</td><td>115520</td><td>+</td><td>0</td><td>ATGGAGGTGGGAGAACCCTC</td><td>GGG</td><td>||||||||[||||||||||||]PAM</td><td>NA</td><td>Intergenic</td><td>NA</td><td>NA</td></tr>
<tr><td></td><td></td><td></td><td></td><td></td><td></td><td></td><td></td><td></td><td></td><td></td><td></td></tr>
<tr><td>T8540</td><td>GAGGTGGGAGAACCCTCGGGAGG</td><td>1000</td><td></td><td></td><td></td><td></td><td></td><td></td><td></td><td></td><td></td></tr>
<tr><td>Oligo fwd</td><td>CACCGAGGTGGGAGAACCCTCGGG</td><td></td><td></td><td></td><td></td><td></td><td></td><td></td><td></td><td></td><td></td></tr>
<tr><td>Oligo rev</td><td>AAACCCCGAGGGTTCTCCCACCTC</td><td></td><td></td><td></td><td></td><td></td><td></td><td></td><td></td><td></td><td></td></tr>
<tr><td>Chromosome</td><td>start</td><td>end</td><td>strand</td><td>MM</td><td>target_seq</td><td>PAM</td><td>alignment</td><td>distance</td><td>position</td><td>gene name</td><td>gene id</td></tr>
<tr><td>GU305914.1</td><td>115501</td><td>115523</td><td>+</td><td>0</td><td>GAGGTGGGAGAACCCTCGGG</td><td>AGG</td><td>||||||||[||||||||||||]PAM</td><td>NA</td><td>Intergenic</td><td>NA</td><td>NA</td></tr>
<tr><td></td><td></td><td></td><td></td><td></td><td></td><td></td><td></td><td></td><td></td><td></td><td></td></tr>
<tr><td>T8541</td><td>GTGGGAGAACCCTCGGGAGGAGG</td><td>1000</td><td></td><td></td><td></td><td></td><td></td><td></td><td></td><td></td><td></td></tr>
<tr><td>Oligo fwd</td><td>CACCGTGGGAGAACCCTCGGGAGG</td><td></td><td></td><td></td><td></td><td></td><td></td><td></td><td></td><td></td><td></td></tr>
<tr><td>Oligo rev</td><td>AAACCCTCCCGAGGGTTCTCCCAC</td><td></td><td></td><td></td><td></td><td></td><td></td><td></td><td></td><td></td><td></td></tr>
<tr><td>Chromosome</td><td>start</td><td>end</td><td>strand</td><td>MM</td><td>target_seq</td><td>PAM</td><td>alignment</td><td>distance</td><td>position</td><td>gene name</td><td>gene id</td></tr>
<tr><td>GU305914.1</td><td>115504</td><td>115526</td><td>+</td><td>0</td><td>GTGGGAGAACCCTCGGGAGG</td><td>AGG</td><td>||||||||[||||||||||||]PAM</td><td>NA</td><td>Intergenic</td><td>NA</td><td>NA</td></tr>
<tr><td></td><td></td><td></td><td></td><td></td><td></td><td></td><td></td><td></td><td></td><td></td><td></td></tr>
<tr><td>T8542</td><td>CCTCGGGAGGAGGAACCTCGAGG</td><td>1000</td><td></td><td></td><td></td><td></td><td></td><td></td><td></td><td></td><td></td></tr>
<tr><td>Oligo fwd</td><td>CACCCCTCGGGAGGAGGAACCTCG</td><td></td><td></td><td></td><td></td><td></td><td></td><td></td><td></td><td></td><td></td></tr>
<tr><td>Oligo rev</td><td>AAACCGAGGTTCCTCCTCCCGAGG</td><td></td><td></td><td></td><td></td><td></td><td></td><td></td><td></td><td></td><td></td></tr>
<tr><td>Chromosome</td><td>start</td><td>end</td><td>strand</td><td>MM</td><td>target_seq</td><td>PAM</td><td>alignment</td><td>distance</td><td>position</td><td>gene name</td><td>gene id</td></tr>
<tr><td>GU305914.1</td><td>115514</td><td>115536</td><td>+</td><td>0</td><td>CCTCGGGAGGAGGAACCTCG</td><td>AGG</td><td>||||||||[||||||||||||]PAM</td><td>NA</td><td>Intergenic</td><td>NA</td><td>NA</td></tr>
<tr><td></td><td></td><td></td><td></td><td></td><td></td><td></td><td></td><td></td><td></td><td></td><td></td></tr>
<tr><td>T8543</td><td>GGGAGGAGGAACCTCGAGGAAGG</td><td>1000</td><td></td><td></td><td></td><td></td><td></td><td></td><td></td><td></td><td></td></tr>
<tr><td>Oligo fwd</td><td>CACCGGGAGGAGGAACCTCGAGGA</td><td></td><td></td><td></td><td></td><td></td><td></td><td></td><td></td><td></td><td></td></tr>
<tr><td>Oligo rev</td><td>AAACTCCTCGAGGTTCCTCCTCCC</td><td></td><td></td><td></td><td></td><td></td><td></td><td></td><td></td><td></td><td></td></tr>
<tr><td>Chromosome</td><td>start</td><td>end</td><td>strand</td><td>MM</td><td>target_seq</td><td>PAM</td><td>alignment</td><td>distance</td><td>position</td><td>gene name</td><td>gene id</td></tr>
<tr><td>GU305914.1</td><td>115518</td><td>115540</td><td>+</td><td>0</td><td>GGGAGGAGGAACCTCGAGGA</td><td>AGG</td><td>||||||||[||||||||||||]PAM</td><td>NA</td><td>Intergenic</td><td>NA</td><td>NA</td></tr>
<tr><td></td><td></td><td></td><td></td><td></td><td></td><td></td><td></td><td></td><td></td><td></td><td></td></tr>
<tr><td>T8544</td><td>GGAGGAGGAACCTCGAGGAAGGG</td><td>1000</td><td></td><td></td><td></td><td></td><td></td><td></td><td></td><td></td><td></td></tr>
<tr><td>Oligo fwd</td><td>CACCGGAGGAGGAACCTCGAGGAA</td><td></td><td></td><td></td><td></td><td></td><td></td><td></td><td></td><td></td><td></td></tr>
</table>

Oligo rev AAACTTCCTCGAGGTTCCTCCTCC
Chromosome start end strand MM target_seq PAM alignment distance position gene name gene id
GU305914.1 115519 115541 + 0 GGAGGAGGAACCTCGAGGAA GGG ||||||||[||||||||||||]PAM NA Intergenic NA NA

T8545 GGAGGAACCTCGAGGAAGGGAGG 1000
Oligo fwd CACCGGAGGAACCTCGAGGAAGGG
Oligo rev AAACCCCTTCCTCGAGGTTCCTCC
Chromosome start end strand MM target_seq PAM alignment distance position gene name gene id
GU305914.1 115522 115544 + 0 GGAGGAACCTCGAGGAAGGG AGG ||||||||[||||||||||||]PAM NA Intergenic NA NA

T8546 AACCTCGAGGAAGGGAGGTGAGG 1000
Oligo fwd CACCAACCTCGAGGAAGGGAGGTG
Oligo rev AAACCACCTCCCTTCCTCGAGGTT
Chromosome start end strand MM target_seq PAM alignment distance position gene name gene id
GU305914.1 115527 115549 + 0 AACCTCGAGGAAGGGAGGTG AGG ||||||||[||||||||||||]PAM NA Intergenic NA NA

T8547 GGGAGGTGAGGCCAGCATGTTGG 1000
Oligo fwd CACCGGGAGGTGAGGCCAGCATGT
Oligo rev AAACACATGCTGGCCTCACCTCCC
Chromosome start end strand MM target_seq PAM alignment distance position gene name gene id
GU305914.1 115539 115561 + 0 GGGAGGTGAGGCCAGCATGT TGG ||||||||[||||||||||||]PAM NA Intergenic NA NA

T8548 AGGTGAGGCCAGCATGTTGGAGG 1000
Oligo fwd CACCAGGTGAGGCCAGCATGTTGG
Oligo rev AAACCCAACATGCTGGCCTCACCT
Chromosome start end strand MM target_seq PAM alignment distance position gene name gene id
GU305914.1 115542 115564 + 0 AGGTGAGGCCAGCATGTTGG AGG ||||||||[||||||||||||]PAM NA Intergenic NA NA

T8549 GCATGTTGGAGGTGAACAAACGG 1000
Oligo fwd CACCGCATGTTGGAGGTGAACAAA
Oligo rev AAACTTTGTTCACCTCCAACATGC
Chromosome start end strand MM target_seq PAM alignment distance position gene name gene id
GU305914.1 115553 115575 + 0 GCATGTTGGAGGTGAACAAA CGG ||||||||[||||||||||||]PAM NA Intergenic NA NA

T8550 CGTGTCCCTCCTCAATAAAATGG 1000
Oligo fwd CACCCGTGTCCCTCCTCAATAAAA
Oligo rev AAACTTTTATTGAGGAGGGACACG
Chromosome start end strand MM target_seq PAM alignment distance position gene name gene id
GU305914.1 115581 115603 + 0 CGTGTCCCTCCTCAATAAAA TGG ||||||||[||||||||||||]PAM NA Intergenic NA NA
GU305914.1 26712 26734 - 4 CGTATCCGTCCTCATCAAAA AGG |||-|||-[||||||--||||]PAM NA Intergenic NA NA

T8551 GGAATGAGTGATTCAGACACCGG 1000
Oligo fwd CACCGGAATGAGTGATTCAGACAC
Oligo rev AAACGTGTCTGAATCACTCATTCC
Chromosome start end strand MM target_seq PAM alignment distance position gene name gene id
GU305914.1 115602 115624 + 0 GGAATGAGTGATTCAGACAC CGG ||||||||[||||||||||||]PAM NA Intergenic NA NA

T8552 TGATTCAGACACCGGACTAACGG 1000
Oligo fwd CACCTGATTCAGACACCGGACTAA
Oligo rev AAACTTAGTCCGGTGTCTGAATCA
Chromosome start end strand MM target_seq PAM alignment distance position gene name gene id
GU305914.1 115610 115632 + 0 TGATTCAGACACCGGACTAA CGG ||||||||[||||||||||||]PAM NA Intergenic NA NA

T8553 ACCTGTCTGTCACCGTGATGCGG 1000
Oligo fwd CACCACCTGTCTGTCACCGTGATG
Oligo rev AAACCATCACGGTGACAGACAGGT
Chromosome start end strand MM target_seq PAM alignment distance position gene name gene id
GU305914.1 115741 115763 + 0 ACCTGTCTGTCACCGTGATG CGG ||||||||[||||||||||||]PAM NA Intergenic NA NA

T8554 CCTGTCTGTCACCGTGATGCGGG 1000
Oligo fwd CACCCCTGTCTGTCACCGTGATGC
Oligo rev AAACGCATCACGGTGACAGACAGG
Chromosome start end strand MM target_seq PAM alignment distance position gene name gene id
GU305914.1 115742 115764 + 0 CCTGTCTGTCACCGTGATGC GGG ||||||||[||||||||||||]PAM NA Intergenic NA NA

T8555 GTCTGTCACCGTGATGCGGGCGG 1000
Oligo fwd CACCGTCTGTCACCGTGATGCGGG
Oligo rev AAACCCCGCATCACGGTGACAGAC
Chromosome start end strand MM target_seq PAM alignment distance position gene name gene id
GU305914.1 115745 115767 + 0 GTCTGTCACCGTGATGCGGG CGG ||||||||[||||||||||||]PAM NA Intergenic NA NA

T8556 TCTGTCACCGTGATGCGGGCGGG 1000
Oligo fwd CACCTCTGTCACCGTGATGCGGGC
Oligo rev AAACGCCCGCATCACGGTGACAGA
Chromosome start end strand MM target_seq PAM alignment distance position gene name gene id
GU305914.1 115746 115768 + 0 TCTGTCACCGTGATGCGGGC GGG ||||||||[||||||||||||]PAM NA Intergenic NA NA

T8557 GTCTGTCTGTTTGTCTCAGTCGG 1000
Oligo fwd CACCGTCTGTCTGTTTGTCTCAGT
Oligo rev AAACACTGAGACAAACAGACAGAC
Chromosome start end strand MM target_seq PAM alignment distance position gene name gene id
GU305914.1 115784 115806 + 0 GTCTGTCTGTTTGTCTCAGT CGG ||||||||[||||||||||||]PAM NA Intergenic NA NA

T8558 TCTGTCTGTTTGTCTCAGTCGGG 1000
Oligo fwd CACCTCTGTCTGTTTGTCTCAGTC
Oligo rev AAACGACTGAGACAAACAGACAGA
Chromosome start end strand MM target_seq PAM alignment distance position gene name gene id
GU305914.1 115785 115807 + 0 TCTGTCTGTTTGTCTCAGTC GGG ||||||||[||||||||||||]PAM NA Intergenic NA NA

T8559 CTGTCTGTTTGTCTCAGTCGGGG 1000
Oligo fwd CACCCTGTCTGTTTGTCTCAGTCG
Oligo rev AAACCGACTGAGACAAACAGACAG
Chromosome start end strand MM target_seq PAM alignment distance position gene name gene id
GU305914.1 115786 115808 + 0 CTGTCTGTTTGTCTCAGTCG GGG ||||||||[||||||||||||]PAM NA Intergenic NA NA

T8560 TTGTCTCAGTCGGGGAAATCAGG 1000
Oligo fwd CACCTTGTCTCAGTCGGGGAAATC
Oligo rev AAACGATTTCCCCGACTGAGACAA
Chromosome start end strand MM target_seq PAM alignment distance position gene name gene id
GU305914.1 115794 115816 + 0 TTGTCTCAGTCGGGGAAATC AGG ||||||||[||||||||||||]PAM NA Intergenic NA NA

T8561 TGTCTCAGTCGGGGAAATCAGGG 1000
Oligo fwd CACCTGTCTCAGTCGGGGAAATCA
Oligo rev AAACTGATTTCCCCGACTGAGACA
Chromosome start end strand MM target_seq PAM alignment distance position gene name gene id
GU305914.1 115795 115817 + 0 TGTCTCAGTCGGGGAAATCA GGG ||||||||[||||||||||||]PAM NA Intergenic NA NA

T8562 GGAAATCAGGGTTGTAGATGTGG 1000
Oligo fwd CACCGGAAATCAGGGTTGTAGATG
Oligo rev AAACCATCTACAACCCTGATTTCC
Chromosome start end strand MM target_seq PAM alignment distance position gene name gene id
GU305914.1 115807 115829 + 0 GGAAATCAGGGTTGTAGATG TGG ||||||||[||||||||||||]PAM NA Intergenic NA NA

T8563 GAAATCAGGGTTGTAGATGTGGG 1000
Oligo fwd CACCGAAATCAGGGTTGTAGATGT
Oligo rev AAACACATCTACAACCCTGATTTC
Chromosome start end strand MM target_seq PAM alignment distance position gene name gene id
GU305914.1 115808 115830 + 0 GAAATCAGGGTTGTAGATGT GGG ||||||||[||||||||||||]PAM NA Intergenic NA NA

T8564 AAATCAGGGTTGTAGATGTGGGG 1000
Oligo fwd CACCAAATCAGGGTTGTAGATGTG
Oligo rev AAACCACATCTACAACCCTGATTT
Chromosome start end strand MM target_seq PAM alignment distance position gene name gene id
GU305914.1 115809 115831 + 0 AAATCAGGGTTGTAGATGTG GGG ||||||||[||||||||||||]PAM NA Intergenic NA NA

T8565 AATCAGGGTTGTAGATGTGGGGG 1000
Oligo fwd CACCAATCAGGGTTGTAGATGTGG
Oligo rev AAACCCACATCTACAACCCTGATT
Chromosome start end strand MM target_seq PAM alignment distance position gene name gene id
GU305914.1 115810 115832 + 0 AATCAGGGTTGTAGATGTGG GGG ||||||||[||||||||||||]PAM NA Intergenic NA NA

T8566 ATCAGGGTTGTAGATGTGGGGGG 1000
Oligo fwd CACCATCAGGGTTGTAGATGTGGG
Oligo rev AAACCCCACATCTACAACCCTGAT
Chromosome start end strand MM target_seq PAM alignment distance position gene name gene id
GU305914.1 115811 115833 + 0 ATCAGGGTTGTAGATGTGGG GGG ||||||||[||||||||||||]PAM NA Intergenic NA NA

T8567 TTGTAGATGTGGGGGGATGATGG 1000
Oligo fwd CACCTTGTAGATGTGGGGGGATGA
Oligo rev AAACTCATCCCCCCACATCTACAA
Chromosome start end strand MM target_seq PAM alignment distance position gene name gene id
GU305914.1 115818 115840 + 0 TTGTAGATGTGGGGGGATGA TGG ||||||||[||||||||||||]PAM NA Intergenic NA NA

T8568 GGGGGATGATGGCTGTCTTCAGG 1000
Oligo fwd CACCGGGGGATGATGGCTGTCTTC
Oligo rev AAACGAAGACAGCCATCATCCCCC
Chromosome start end strand MM target_seq PAM alignment distance position gene name gene id
GU305914.1 115829 115851 + 0 GGGGGATGATGGCTGTCTTC AGG ||||||||[||||||||||||]PAM NA Intergenic NA NA

T8569 GGATGATGGCTGTCTTCAGGCGG 1000
Oligo fwd CACCGGATGATGGCTGTCTTCAGG
Oligo rev AAACCCTGAAGACAGCCATCATCC
Chromosome start end strand MM target_seq PAM alignment distance position gene name gene id
GU305914.1 115832 115854 + 0 GGATGATGGCTGTCTTCAGG CGG ||||||||[||||||||||||]PAM NA Intergenic NA NA

T8570 GATGATGGCTGTCTTCAGGCGGG 1000
Oligo fwd CACCGATGATGGCTGTCTTCAGGC
Oligo rev AAACGCCTGAAGACAGCCATCATC
Chromosome start end strand MM target_seq PAM alignment distance position gene name gene id
GU305914.1 115833 115855 + 0 GATGATGGCTGTCTTCAGGC GGG ||||||||[||||||||||||]PAM NA Intergenic NA NA

T8571 ATGATGGCTGTCTTCAGGCGGGG 1000
Oligo fwd CACCATGATGGCTGTCTTCAGGCG
Oligo rev AAACCGCCTGAAGACAGCCATCAT
Chromosome start end strand MM target_seq PAM alignment distance position gene name gene id
GU305914.1 115834 115856 + 0 ATGATGGCTGTCTTCAGGCG GGG ||||||||[||||||||||||]PAM NA Intergenic NA NA

T8572 TGGCTGTCTTCAGGCGGGGTAGG 1000
Oligo fwd CACCTGGCTGTCTTCAGGCGGGGT
Oligo rev AAACACCCCGCCTGAAGACAGCCA
Chromosome start end strand MM target_seq PAM alignment distance position gene name gene id
GU305914.1 115838 115860 + 0 TGGCTGTCTTCAGGCGGGGT AGG ||||||||[||||||||||||]PAM NA Intergenic NA NA

T8573 TGTCTTCAGGCGGGGTAGGCTGG 1000
Oligo fwd CACCTGTCTTCAGGCGGGGTAGGC
Oligo rev AAACGCCTACCCCGCCTGAAGACA
Chromosome start end strand MM target_seq PAM alignment distance position gene name gene id
GU305914.1 115842 115864 + 0 TGTCTTCAGGCGGGGTAGGC TGG ||||||||[||||||||||||]PAM NA Intergenic NA NA

<table>
<tr><td>T8574</td><td>TTCAGGCGGGGTAGGCTGGACGG</td><td>1000</td></tr>
<tr><td>Oligo fwd</td><td>CACCTTCAGGCGGGGTAGGCTGGA</td></tr>
<tr><td>Oligo rev</td><td>AAACTCCAGCCTACCCCGCCTGAA</td></tr>
<tr><td>Chromosome</td><td>start</td><td>end</td><td>strand</td><td>MM</td><td>target_seq</td><td>PAM</td><td>alignment</td><td>distance</td><td>position</td><td>gene name</td><td>gene id</td></tr>
<tr><td>GU305914.1</td><td>115846</td><td>115868</td><td>+</td><td>0</td><td>TTCAGGCGGGGTAGGCTGGA</td><td>CGG</td><td>||||||||[||||||||||||]PAM</td><td>NA</td><td>Intergenic</td><td>NA</td><td>NA</td></tr>
<tr><td>T8575</td><td>GCGGGGTAGGCTGGACGGCATGG</td><td>1000</td></tr>
<tr><td>Oligo fwd</td><td>CACCGCGGGGTAGGCTGGACGGCA</td></tr>
<tr><td>Oligo rev</td><td>AAACTGCCGTCCAGCCTACCCCGC</td></tr>
<tr><td>Chromosome</td><td>start</td><td>end</td><td>strand</td><td>MM</td><td>target_seq</td><td>PAM</td><td>alignment</td><td>distance</td><td>position</td><td>gene name</td><td>gene id</td></tr>
<tr><td>GU305914.1</td><td>115851</td><td>115873</td><td>+</td><td>0</td><td>GCGGGGTAGGCTGGACGGCA</td><td>TGG</td><td>||||||||[||||||||||||]PAM</td><td>NA</td><td>Intergenic</td><td>NA</td><td>NA</td></tr>
<tr><td>T8576</td><td>CGGGGTAGGCTGGACGGCATGGG</td><td>1000</td></tr>
<tr><td>Oligo fwd</td><td>CACCCGGGGTAGGCTGGACGGCAT</td></tr>
<tr><td>Oligo rev</td><td>AAACATGCCGTCCAGCCTACCCCG</td></tr>
<tr><td>Chromosome</td><td>start</td><td>end</td><td>strand</td><td>MM</td><td>target_seq</td><td>PAM</td><td>alignment</td><td>distance</td><td>position</td><td>gene name</td><td>gene id</td></tr>
<tr><td>GU305914.1</td><td>115852</td><td>115874</td><td>+</td><td>0</td><td>CGGGGTAGGCTGGACGGCAT</td><td>GGG</td><td>||||||||[||||||||||||]PAM</td><td>NA</td><td>Intergenic</td><td>NA</td><td>NA</td></tr>
<tr><td>T8577</td><td>CGCGCCGTATCTACCTTTCGTGG</td><td>1000</td></tr>
<tr><td>Oligo fwd</td><td>CACCCGCGCCGTATCTACCTTTCG</td></tr>
<tr><td>Oligo rev</td><td>AAACCGAAAGGTAGATACGGCGCG</td></tr>
<tr><td>Chromosome</td><td>start</td><td>end</td><td>strand</td><td>MM</td><td>target_seq</td><td>PAM</td><td>alignment</td><td>distance</td><td>position</td><td>gene name</td><td>gene id</td></tr>
<tr><td>GU305914.1</td><td>115878</td><td>115900</td><td>+</td><td>0</td><td>CGCGCCGTATCTACCTTTCG</td><td>TGG</td><td>||||||||[||||||||||||]PAM</td><td>NA</td><td>Intergenic</td><td>NA</td><td>NA</td></tr>
<tr><td>T8578</td><td>ACCTTTCGTGGTCAACTGCACGG</td><td>1000</td></tr>
<tr><td>Oligo fwd</td><td>CACCACCTTTCGTGGTCAACTGCA</td></tr>
<tr><td>Oligo rev</td><td>AAACTGCAGTTGACCACGAAAGGT</td></tr>
<tr><td>Chromosome</td><td>start</td><td>end</td><td>strand</td><td>MM</td><td>target_seq</td><td>PAM</td><td>alignment</td><td>distance</td><td>position</td><td>gene name</td><td>gene id</td></tr>
<tr><td>GU305914.1</td><td>115890</td><td>115912</td><td>+</td><td>0</td><td>ACCTTTCGTGGTCAACTGCA</td><td>CGG</td><td>||||||||[||||||||||||]PAM</td><td>NA</td><td>Intergenic</td><td>NA</td><td>NA</td></tr>
<tr><td>T8579</td><td>GGCGACCTTGTCGACTCCCATGG</td><td>1000</td></tr>
<tr><td>Oligo fwd</td><td>CACCGGCGACCTTGTCGACTCCCA</td></tr>
<tr><td>Oligo rev</td><td>AAACTGGGAGTCGACAAGGTCGCC</td></tr>
<tr><td>Chromosome</td><td>start</td><td>end</td><td>strand</td><td>MM</td><td>target_seq</td><td>PAM</td><td>alignment</td><td>distance</td><td>position</td><td>gene name</td><td>gene id</td></tr>
<tr><td>GU305914.1</td><td>115911</td><td>115933</td><td>+</td><td>0</td><td>GGCGACCTTGTCGACTCCCA</td><td>TGG</td><td>||||||||[||||||||||||]PAM</td><td>NA</td><td>Intergenic</td><td>NA</td><td>NA</td></tr>
<tr><td>T8580</td><td>TACACGCCTTCGTTGATGAACGG</td><td>1000</td></tr>
<tr><td>Oligo fwd</td><td>CACCTACACGCCTTCGTTGATGAA</td></tr>
<tr><td>Oligo rev</td><td>AAACTTCATCAACGAAGGCGTGTA</td></tr>
<tr><td>Chromosome</td><td>start</td><td>end</td><td>strand</td><td>MM</td><td>target_seq</td><td>PAM</td><td>alignment</td><td>distance</td><td>position</td><td>gene name</td><td>gene id</td></tr>
<tr><td>GU305914.1</td><td>115945</td><td>115967</td><td>+</td><td>0</td><td>TACACGCCTTCGTTGATGAA</td><td>CGG</td><td>||||||||[||||||||||||]PAM</td><td>NA</td><td>Intergenic</td><td>NA</td><td>NA</td></tr>
<tr><td>T8581</td><td>GTTGATGAACGGAGTGTACATGG</td><td>1000</td></tr>
<tr><td>Oligo fwd</td><td>CACCGTTGATGAACGGAGTGTACA</td></tr>
<tr><td>Oligo rev</td><td>AAACTGTACACTCCGTTCATCAAC</td></tr>
<tr><td>Chromosome</td><td>start</td><td>end</td><td>strand</td><td>MM</td><td>target_seq</td><td>PAM</td><td>alignment</td><td>distance</td><td>position</td><td>gene name</td><td>gene id</td></tr>
<tr><td>GU305914.1</td><td>115956</td><td>115978</td><td>+</td><td>0</td><td>GTTGATGAACGGAGTGTACA</td><td>TGG</td><td>||||||||[||||||||||||]PAM</td><td>NA</td><td>Intergenic</td><td>NA</td><td>NA</td></tr>
<tr><td>T8582</td><td>GAGTGTACATGGTGAACGTCAGG</td><td>1000</td></tr>
<tr><td>Oligo fwd</td><td>CACCGAGTGTACATGGTGAACGTC</td></tr>
<tr><td>Oligo rev</td><td>AAACGACGTTCACCATGTACACTC</td></tr>
<tr><td>Chromosome</td><td>start</td><td>end</td><td>strand</td><td>MM</td><td>target_seq</td><td>PAM</td><td>alignment</td><td>distance</td><td>position</td><td>gene name</td><td>gene id</td></tr>
<tr><td>GU305914.1</td><td>115967</td><td>115989</td><td>+</td><td>0</td><td>GAGTGTACATGGTGAACGTC</td><td>AGG</td><td>||||||||[||||||||||||]PAM</td><td>NA</td><td>Intergenic</td><td>NA</td><td>NA</td></tr>
<tr><td>T8583</td><td>AGTGTACATGGTGAACGTCAGGG</td><td>1000</td></tr>
<tr><td>Oligo fwd</td><td>CACCAGTGTACATGGTGAACGTCA</td></tr>
<tr><td>Oligo rev</td><td>AAACTGACGTTCACCATGTACACT</td></tr>
<tr><td>Chromosome</td><td>start</td><td>end</td><td>strand</td><td>MM</td><td>target_seq</td><td>PAM</td><td>alignment</td><td>distance</td><td>position</td><td>gene name</td><td>gene id</td></tr>
<tr><td>GU305914.1</td><td>115968</td><td>115990</td><td>+</td><td>0</td><td>AGTGTACATGGTGAACGTCA</td><td>GGG</td><td>||||||||[||||||||||||]PAM</td><td>NA</td><td>Intergenic</td><td>NA</td><td>NA</td></tr>
<tr><td>T8584</td><td>GTACATGGTGAACGTCAGGGTGG</td><td>1000</td></tr>
<tr><td>Oligo fwd</td><td>CACCGTACATGGTGAACGTCAGGG</td></tr>
<tr><td>Oligo rev</td><td>AAACCCCTGACGTTCACCATGTAC</td></tr>
<tr><td>Chromosome</td><td>start</td><td>end</td><td>strand</td><td>MM</td><td>target_seq</td><td>PAM</td><td>alignment</td><td>distance</td><td>position</td><td>gene name</td><td>gene id</td></tr>
<tr><td>GU305914.1</td><td>115971</td><td>115993</td><td>+</td><td>0</td><td>GTACATGGTGAACGTCAGGG</td><td>TGG</td><td>||||||||[||||||||||||]PAM</td><td>NA</td><td>Intergenic</td><td>NA</td><td>NA</td></tr>
<tr><td>T8585</td><td>CGTCAGGGTGGCCAGTCTCCCGG</td><td>1000</td></tr>
<tr><td>Oligo fwd</td><td>CACCCGTCAGGGTGGCCAGTCTCC</td></tr>
<tr><td>Oligo rev</td><td>AAACGGAGACTGGCCACCCTGACG</td></tr>
<tr><td>Chromosome</td><td>start</td><td>end</td><td>strand</td><td>MM</td><td>target_seq</td><td>PAM</td><td>alignment</td><td>distance</td><td>position</td><td>gene name</td><td>gene id</td></tr>
<tr><td>GU305914.1</td><td>115983</td><td>116005</td><td>+</td><td>0</td><td>CGTCAGGGTGGCCAGTCTCC</td><td>CGG</td><td>||||||||[||||||||||||]PAM</td><td>NA</td><td>Intergenic</td><td>NA</td><td>NA</td></tr>
<tr><td>T8586</td><td>GGGTGGCCAGTCTCCCGGCCCGG</td><td>1000</td></tr>
<tr><td>Oligo fwd</td><td>CACCGGGTGGCCAGTCTCCCGGCC</td></tr>
<tr><td>Oligo rev</td><td>AAACGGCCGGGAGACTGGCCACCC</td></tr>
<tr><td>Chromosome</td><td>start</td><td>end</td><td>strand</td><td>MM</td><td>target_seq</td><td>PAM</td><td>alignment</td><td>distance</td><td>position</td><td>gene name</td><td>gene id</td></tr>
<tr><td>GU305914.1</td><td>115988</td><td>116010</td><td>+</td><td>0</td><td>GGGTGGCCAGTCTCCCGGCC</td><td>CGG</td><td>||||||||[||||||||||||]PAM</td><td>NA</td><td>Intergenic</td><td>NA</td><td>NA</td></tr>
<tr><td>T8587</td><td>CCCGGATCCGTATCAGATCGTGG</td><td>1000</td></tr>
<tr><td>Oligo fwd</td><td>CACCCCCGGATCCGTATCAGATCG</td></tr>
<tr><td>Oligo rev</td><td>AAACCGATCTGATACGGATCCGGG</td></tr>
<tr><td>Chromosome</td><td>start</td><td>end</td><td>strand</td><td>MM</td><td>target_seq</td><td>PAM</td><td>alignment</td><td>distance</td><td>position</td><td>gene name</td><td>gene id</td></tr>
<tr><td>GU305914.1</td><td>116006</td><td>116028</td><td>+</td><td>0</td><td>CCCGGATCCGTATCAGATCG</td><td>TGG</td><td>||||||||[||||||||||||]PAM</td><td>NA</td><td>Intergenic</td><td>NA</td><td>NA</td></tr>
<tr><td>T8588</td><td>CCGGATCCGTATCAGATCGTGGG</td><td>1000</td></tr>
<tr><td>Oligo fwd</td><td>CACCCCGGATCCGTATCAGATCGT</td></tr>
<tr><td>Oligo rev</td><td>AAACACGATCTGATACGGATCCGG</td></tr>
<tr><td>Chromosome</td><td>start</td><td>end</td><td>strand</td><td>MM</td><td>target_seq</td><td>PAM</td><td>alignment</td><td>distance</td><td>position</td><td>gene name</td><td>gene id</td></tr>
</table>

| | | | | | | | | | | | |
|---|---|---|---|---|---|---|---|---|---|---|---|
| GU305914.1 | | 116007 | 116029 | + | 0 | CCGGATCCGTATCAGATCGT | GGG | \|\|\|\|\|\|\|\|[\|\|\|\|\|\|\|\|\|\|\|\|]PAM | NA | Intergenic | NA | NA |
| T8589 | CGGATCCGTATCAGATCGTGGGG | 1000 | | | | | | | | | |
| Oligo fwd | CACCCGGATCCGTATCAGATCGTG | | | | | | | | | | |
| Oligo rev | AAACCACGATCTGATACGGATCCG | | | | | | | | | | |
| Chromosome | start | end | strand | MM | target_seq | PAM | alignment | distance | position | gene name | gene id |
| GU305914.1 | 116008 | 116030 | + | 0 | CGGATCCGTATCAGATCGTG | GGG | \|\|\|\|\|\|\|\|[\|\|\|\|\|\|\|\|\|\|\|\|]PAM | NA | Intergenic | NA | NA |
| T8590 | GATCGTGGGGCTGCAGCGTCAGG | 1000 | | | | | | | | | |
| Oligo fwd | CACCGATCGTGGGGCTGCAGCGTC | | | | | | | | | | |
| Oligo rev | AAACGACGCTGCAGCCCCACGATC | | | | | | | | | | |
| Chromosome | start | end | strand | MM | target_seq | PAM | alignment | distance | position | gene name | gene id |
| GU305914.1 | 116021 | 116043 | + | 0 | GATCGTGGGGCTGCAGCGTC | AGG | \|\|\|\|\|\|\|\|[\|\|\|\|\|\|\|\|\|\|\|\|]PAM | NA | Intergenic | NA | NA |
| T8591 | CGTCAGGAGCTGTCTCTGAGCGG | 1000 | | | | | | | | | |
| Oligo fwd | CACCCGTCAGGAGCTGTCTCTGAG | | | | | | | | | | |
| Oligo rev | AAACCTCAGAGACAGCTCCTGACG | | | | | | | | | | |
| Chromosome | start | end | strand | MM | target_seq | PAM | alignment | distance | position | gene name | gene id |
| GU305914.1 | 116037 | 116059 | + | 0 | CGTCAGGAGCTGTCTCTGAG | CGG | \|\|\|\|\|\|\|\|[\|\|\|\|\|\|\|\|\|\|\|\|]PAM | NA | Intergenic | NA | NA |
| T8592 | CGCCTCCTCCTCGTACGAGTCGG | 1000 | | | | | | | | | |
| Oligo fwd | CACCCGCCTCCTCCTCGTACGAGT | | | | | | | | | | |
| Oligo rev | AAACACTCGTACGAGGAGGAGGCG | | | | | | | | | | |
| Chromosome | start | end | strand | MM | target_seq | PAM | alignment | distance | position | gene name | gene id |
| GU305914.1 | 116061 | 116083 | + | 0 | CGCCTCCTCCTCGTACGAGT | CGG | \|\|\|\|\|\|\|\|[\|\|\|\|\|\|\|\|\|\|\|\|]PAM | NA | Intergenic | NA | NA |
| T8593 | GCCTCCTCCTCGTACGAGTCGGG | 1000 | | | | | | | | | |
| Oligo fwd | CACCGCCTCCTCCTCGTACGAGTC | | | | | | | | | | |
| Oligo rev | AAACGACTCGTACGAGGAGGAGGC | | | | | | | | | | |
| Chromosome | start | end | strand | MM | target_seq | PAM | alignment | distance | position | gene name | gene id |
| GU305914.1 | 116062 | 116084 | + | 0 | GCCTCCTCCTCGTACGAGTC | GGG | \|\|\|\|\|\|\|\|[\|\|\|\|\|\|\|\|\|\|\|\|]PAM | NA | Intergenic | NA | NA |
| T8594 | CCTCCTCCTCGTACGAGTCGGGG | 1000 | | | | | | | | | |
| Oligo fwd | CACCCCTCCTCCTCGTACGAGTCG | | | | | | | | | | |
| Oligo rev | AAACCGACTCGTACGAGGAGGAGG | | | | | | | | | | |
| Chromosome | start | end | strand | MM | target_seq | PAM | alignment | distance | position | gene name | gene id |
| GU305914.1 | 116063 | 116085 | + | 0 | CCTCCTCCTCGTACGAGTCG | GGG | \|\|\|\|\|\|\|\|[\|\|\|\|\|\|\|\|\|\|\|\|]PAM | NA | Intergenic | NA | NA |
| T8595 | CTCCTCGTACGAGTCGGGGACGG | 1000 | | | | | | | | | |
| Oligo fwd | CACCCTCCTCGTACGAGTCGGGGA | | | | | | | | | | |
| Oligo rev | AAACTCCCCGACTCGTACGAGGAG | | | | | | | | | | |
| Chromosome | start | end | strand | MM | target_seq | PAM | alignment | distance | position | gene name | gene id |
| GU305914.1 | 116067 | 116089 | + | 0 | CTCCTCGTACGAGTCGGGGA | CGG | \|\|\|\|\|\|\|\|[\|\|\|\|\|\|\|\|\|\|\|\|]PAM | NA | Intergenic | NA | NA |
| T8596 | CGGCGTTGTAGACCCAGTTTCGG | 1000 | | | | | | | | | |
| Oligo fwd | CACCCGGCGTTGTAGACCCAGTTT | | | | | | | | | | |
| Oligo rev | AAACAAACTGGGTCTACAACGCCG | | | | | | | | | | |
| Chromosome | start | end | strand | MM | target_seq | PAM | alignment | distance | position | gene name | gene id |
| GU305914.1 | 116087 | 116109 | + | 0 | CGGCGTTGTAGACCCAGTTT | CGG | \|\|\|\|\|\|\|\|[\|\|\|\|\|\|\|\|\|\|\|\|]PAM | NA | Intergenic | NA | NA |
| T8597 | TTCGGTCTTCGTCGAAATCTTGG | 1000 | | | | | | | | | |
| Oligo fwd | CACCTTCGGTCTTCGTCGAAATCT | | | | | | | | | | |
| Oligo rev | AAACAGATTTCGACGAAGACCGAA | | | | | | | | | | |
| Chromosome | start | end | strand | MM | target_seq | PAM | alignment | distance | position | gene name | gene id |
| GU305914.1 | 116105 | 116127 | + | 0 | TTCGGTCTTCGTCGAAATCT | TGG | \|\|\|\|\|\|\|\|[\|\|\|\|\|\|\|\|\|\|\|\|]PAM | NA | Intergenic | NA | NA |
| T8598 | CGTCGCTATCATCCTCTTCCTGG | 1000 | | | | | | | | | |
| Oligo fwd | CACCCGTCGCTATCATCCTCTTCC | | | | | | | | | | |
| Oligo rev | AAACGGAAGAGGATGATAGCGACG | | | | | | | | | | |
| Chromosome | start | end | strand | MM | target_seq | PAM | alignment | distance | position | gene name | gene id |
| GU305914.1 | 116150 | 116172 | + | 0 | CGTCGCTATCATCCTCTTCC | TGG | \|\|\|\|\|\|\|\|[\|\|\|\|\|\|\|\|\|\|\|\|]PAM | NA | Intergenic | NA | NA |
| T8599 | TCATCCTCTTCCTGGTCCGACGG | 1000 | | | | | | | | | |
| Oligo fwd | CACCTCATCCTCTTCCTGGTCCGA | | | | | | | | | | |
| Oligo rev | AAACTCGGACCAGGAAGAGGATGA | | | | | | | | | | |
| Chromosome | start | end | strand | MM | target_seq | PAM | alignment | distance | position | gene name | gene id |
| GU305914.1 | 116158 | 116180 | + | 0 | TCATCCTCTTCCTGGTCCGA | CGG | \|\|\|\|\|\|\|\|[\|\|\|\|\|\|\|\|\|\|\|\|]PAM | NA | Intergenic | NA | NA |
| T8600 | CTGGTCCGACGGCACGTTGTCGG | 1000 | | | | | | | | | |
| Oligo fwd | CACCCTGGTCCGACGGCACGTTGT | | | | | | | | | | |
| Oligo rev | AAACACAACGTGCCGTCGGACCAG | | | | | | | | | | |
| Chromosome | start | end | strand | MM | target_seq | PAM | alignment | distance | position | gene name | gene id |
| GU305914.1 | 116169 | 116191 | + | 0 | CTGGTCCGACGGCACGTTGT | CGG | \|\|\|\|\|\|\|\|[\|\|\|\|\|\|\|\|\|\|\|\|]PAM | NA | Intergenic | NA | NA |
| T8601 | GCACGTTGTCGGCGTCCATCTGG | 1000 | | | | | | | | | |
| Oligo fwd | CACCGCACGTTGTCGGCGTCCATC | | | | | | | | | | |
| Oligo rev | AAACGATGGACGCCGACAACGTGC | | | | | | | | | | |
| Chromosome | start | end | strand | MM | target_seq | PAM | alignment | distance | position | gene name | gene id |
| GU305914.1 | 116180 | 116202 | + | 0 | GCACGTTGTCGGCGTCCATC | TGG | \|\|\|\|\|\|\|\|[\|\|\|\|\|\|\|\|\|\|\|\|]PAM | NA | Intergenic | NA | NA |
| T8602 | CACGTTGTCGGCGTCCATCTGGG | 1000 | | | | | | | | | |
| Oligo fwd | CACCCACGTTGTCGGCGTCCATCT | | | | | | | | | | |
| Oligo rev | AAACAGATGGACGCCGACAACGTG | | | | | | | | | | |
| Chromosome | start | end | strand | MM | target_seq | PAM | alignment | distance | position | gene name | gene id |
| GU305914.1 | 116181 | 116203 | + | 0 | CACGTTGTCGGCGTCCATCT | GGG | \|\|\|\|\|\|\|\|[\|\|\|\|\|\|\|\|\|\|\|\|]PAM | NA | Intergenic | NA | NA |
| T8603 | CGGCGTCCATCTGGGCGAAGAGG | 1000 | | | | | | | | | |
| Oligo fwd | CACCCGGCGTCCATCTGGGCGAAG | | | | | | | | | | |
| Oligo rev | AAACCTTCGCCCAGATGGACGCCG | | | | | | | | | | |

<table>
<tr><td>Chromosome</td><td>start</td><td>end</td><td>strand</td><td>MM</td><td>target_seq</td><td>PAM</td><td>alignment</td><td>distance</td><td>position</td><td>gene name</td><td>gene id</td></tr>
<tr><td>GU305914.1</td><td>116189</td><td>116211</td><td>+</td><td>0</td><td>CGGCGTCCATCTGGGCGAAG</td><td>AGG</td><td>||||||||[||||||||||||]PAM</td><td>NA</td><td>Intergenic</td><td>NA</td><td>NA</td></tr>
<tr><td></td><td></td><td></td><td></td><td></td><td></td><td></td><td></td><td></td><td></td><td></td><td></td></tr>
<tr><td>T8604</td><td>TCGTCCGCCGACGTGCGCCGCGG</td><td>1000</td><td></td><td></td><td></td><td></td><td></td><td></td><td></td><td></td><td></td></tr>
<tr><td>Oligo fwd</td><td>CACCTCGTCCGCCGACGTGCGCCG</td><td></td><td></td><td></td><td></td><td></td><td></td><td></td><td></td><td></td><td></td></tr>
<tr><td>Oligo rev</td><td>AAACCGGCGCACGTCGGCGGACGA</td><td></td><td></td><td></td><td></td><td></td><td></td><td></td><td></td><td></td><td></td></tr>
<tr><td>Chromosome</td><td>start</td><td>end</td><td>strand</td><td>MM</td><td>target_seq</td><td>PAM</td><td>alignment</td><td>distance</td><td>position</td><td>gene name</td><td>gene id</td></tr>
<tr><td>GU305914.1</td><td>116215</td><td>116237</td><td>+</td><td>0</td><td>TCGTCCGCCGACGTGCGCCG</td><td>CGG</td><td>||||||||[||||||||||||]PAM</td><td>NA</td><td>Intergenic</td><td>NA</td><td>NA</td></tr>
<tr><td></td><td></td><td></td><td></td><td></td><td></td><td></td><td></td><td></td><td></td><td></td><td></td></tr>
<tr><td>T8605</td><td>CGTCCGCCGACGTGCGCCGCGGG</td><td>1000</td><td></td><td></td><td></td><td></td><td></td><td></td><td></td><td></td><td></td></tr>
<tr><td>Oligo fwd</td><td>CACCCGTCCGCCGACGTGCGCCGC</td><td></td><td></td><td></td><td></td><td></td><td></td><td></td><td></td><td></td><td></td></tr>
<tr><td>Oligo rev</td><td>AAACGCGGCGCACGTCGGCGGACG</td><td></td><td></td><td></td><td></td><td></td><td></td><td></td><td></td><td></td><td></td></tr>
<tr><td>Chromosome</td><td>start</td><td>end</td><td>strand</td><td>MM</td><td>target_seq</td><td>PAM</td><td>alignment</td><td>distance</td><td>position</td><td>gene name</td><td>gene id</td></tr>
<tr><td>GU305914.1</td><td>116216</td><td>116238</td><td>+</td><td>0</td><td>CGTCCGCCGACGTGCGCCGC</td><td>GGG</td><td>||||||||[||||||||||||]PAM</td><td>NA</td><td>Intergenic</td><td>NA</td><td>NA</td></tr>
<tr><td></td><td></td><td></td><td></td><td></td><td></td><td></td><td></td><td></td><td></td><td></td><td></td></tr>
<tr><td>T8606</td><td>GTCCGCCGACGTGCGCCGCGGGG</td><td>1000</td><td></td><td></td><td></td><td></td><td></td><td></td><td></td><td></td><td></td></tr>
<tr><td>Oligo fwd</td><td>CACCGTCCGCCGACGTGCGCCGCG</td><td></td><td></td><td></td><td></td><td></td><td></td><td></td><td></td><td></td><td></td></tr>
<tr><td>Oligo rev</td><td>AAACCGCGGCGCACGTCGGCGGAC</td><td></td><td></td><td></td><td></td><td></td><td></td><td></td><td></td><td></td><td></td></tr>
<tr><td>Chromosome</td><td>start</td><td>end</td><td>strand</td><td>MM</td><td>target_seq</td><td>PAM</td><td>alignment</td><td>distance</td><td>position</td><td>gene name</td><td>gene id</td></tr>
<tr><td>GU305914.1</td><td>116217</td><td>116239</td><td>+</td><td>0</td><td>GTCCGCCGACGTGCGCCGCG</td><td>GGG</td><td>||||||||[||||||||||||]PAM</td><td>NA</td><td>Intergenic</td><td>NA</td><td>NA</td></tr>
<tr><td></td><td></td><td></td><td></td><td></td><td></td><td></td><td></td><td></td><td></td><td></td><td></td></tr>
<tr><td>T8607</td><td>TCCGCCGACGTGCGCCGCGGGGG</td><td>1000</td><td></td><td></td><td></td><td></td><td></td><td></td><td></td><td></td><td></td></tr>
<tr><td>Oligo fwd</td><td>CACCTCCGCCGACGTGCGCCGCGG</td><td></td><td></td><td></td><td></td><td></td><td></td><td></td><td></td><td></td><td></td></tr>
<tr><td>Oligo rev</td><td>AAACCCGCGGCGCACGTCGGCGGA</td><td></td><td></td><td></td><td></td><td></td><td></td><td></td><td></td><td></td><td></td></tr>
<tr><td>Chromosome</td><td>start</td><td>end</td><td>strand</td><td>MM</td><td>target_seq</td><td>PAM</td><td>alignment</td><td>distance</td><td>position</td><td>gene name</td><td>gene id</td></tr>
<tr><td>GU305914.1</td><td>116218</td><td>116240</td><td>+</td><td>0</td><td>TCCGCCGACGTGCGCCGCGG</td><td>GGG</td><td>||||||||[||||||||||||]PAM</td><td>NA</td><td>Intergenic</td><td>NA</td><td>NA</td></tr>
<tr><td></td><td></td><td></td><td></td><td></td><td></td><td></td><td></td><td></td><td></td><td></td><td></td></tr>
<tr><td>T8608</td><td>CGGGGGCAGCTCTCTGCCGACGG</td><td>1000</td><td></td><td></td><td></td><td></td><td></td><td></td><td></td><td></td><td></td></tr>
<tr><td>Oligo fwd</td><td>CACCCGGGGGCAGCTCTCTGCCGA</td><td></td><td></td><td></td><td></td><td></td><td></td><td></td><td></td><td></td><td></td></tr>
<tr><td>Oligo rev</td><td>AAACTCGGCAGAGAGCTGCCCCCG</td><td></td><td></td><td></td><td></td><td></td><td></td><td></td><td></td><td></td><td></td></tr>
<tr><td>Chromosome</td><td>start</td><td>end</td><td>strand</td><td>MM</td><td>target_seq</td><td>PAM</td><td>alignment</td><td>distance</td><td>position</td><td>gene name</td><td>gene id</td></tr>
<tr><td>GU305914.1</td><td>116235</td><td>116257</td><td>+</td><td>0</td><td>CGGGGGCAGCTCTCTGCCGA</td><td>CGG</td><td>||||||||[||||||||||||]PAM</td><td>NA</td><td>Intergenic</td><td>NA</td><td>NA</td></tr>
<tr><td></td><td></td><td></td><td></td><td></td><td></td><td></td><td></td><td></td><td></td><td></td><td></td></tr>
<tr><td>T8609</td><td>GGGGGCAGCTCTCTGCCGACGGG</td><td>1000</td><td></td><td></td><td></td><td></td><td></td><td></td><td></td><td></td><td></td></tr>
<tr><td>Oligo fwd</td><td>CACCGGGGGCAGCTCTCTGCCGAC</td><td></td><td></td><td></td><td></td><td></td><td></td><td></td><td></td><td></td><td></td></tr>
<tr><td>Oligo rev</td><td>AAACGTCGGCAGAGAGCTGCCCCC</td><td></td><td></td><td></td><td></td><td></td><td></td><td></td><td></td><td></td><td></td></tr>
<tr><td>Chromosome</td><td>start</td><td>end</td><td>strand</td><td>MM</td><td>target_seq</td><td>PAM</td><td>alignment</td><td>distance</td><td>position</td><td>gene name</td><td>gene id</td></tr>
<tr><td>GU305914.1</td><td>116236</td><td>116258</td><td>+</td><td>0</td><td>GGGGGCAGCTCTCTGCCGAC</td><td>GGG</td><td>||||||||[||||||||||||]PAM</td><td>NA</td><td>Intergenic</td><td>NA</td><td>NA</td></tr>
<tr><td></td><td></td><td></td><td></td><td></td><td></td><td></td><td></td><td></td><td></td><td></td><td></td></tr>
<tr><td>T8610</td><td>AGCTCTCTGCCGACGGGTCTCGG</td><td>1000</td><td></td><td></td><td></td><td></td><td></td><td></td><td></td><td></td><td></td></tr>
<tr><td>Oligo fwd</td><td>CACCAGCTCTCTGCCGACGGGTCT</td><td></td><td></td><td></td><td></td><td></td><td></td><td></td><td></td><td></td><td></td></tr>
<tr><td>Oligo rev</td><td>AAACAGACCCGTCGGCAGAGAGCT</td><td></td><td></td><td></td><td></td><td></td><td></td><td></td><td></td><td></td><td></td></tr>
<tr><td>Chromosome</td><td>start</td><td>end</td><td>strand</td><td>MM</td><td>target_seq</td><td>PAM</td><td>alignment</td><td>distance</td><td>position</td><td>gene name</td><td>gene id</td></tr>
<tr><td>GU305914.1</td><td>116242</td><td>116264</td><td>+</td><td>0</td><td>AGCTCTCTGCCGACGGGTCT</td><td>CGG</td><td>||||||||[||||||||||||]PAM</td><td>NA</td><td>Intergenic</td><td>NA</td><td>NA</td></tr>
<tr><td></td><td></td><td></td><td></td><td></td><td></td><td></td><td></td><td></td><td></td><td></td><td></td></tr>
<tr><td>T8611</td><td>GACGGGTCTCGGTGTGTTCAAGG</td><td>1000</td><td></td><td></td><td></td><td></td><td></td><td></td><td></td><td></td><td></td></tr>
<tr><td>Oligo fwd</td><td>CACCGACGGGTCTCGGTGTGTTCA</td><td></td><td></td><td></td><td></td><td></td><td></td><td></td><td></td><td></td><td></td></tr>
<tr><td>Oligo rev</td><td>AAACTGAACACACCGAGACCCGTC</td><td></td><td></td><td></td><td></td><td></td><td></td><td></td><td></td><td></td><td></td></tr>
<tr><td>Chromosome</td><td>start</td><td>end</td><td>strand</td><td>MM</td><td>target_seq</td><td>PAM</td><td>alignment</td><td>distance</td><td>position</td><td>gene name</td><td>gene id</td></tr>
<tr><td>GU305914.1</td><td>116253</td><td>116275</td><td>+</td><td>0</td><td>GACGGGTCTCGGTGTGTTCA</td><td>AGG</td><td>||||||||[||||||||||||]PAM</td><td>NA</td><td>Intergenic</td><td>NA</td><td>NA</td></tr>
<tr><td></td><td></td><td></td><td></td><td></td><td></td><td></td><td></td><td></td><td></td><td></td><td></td></tr>
<tr><td>T8612</td><td>TCAAGGAGAGTTGAGCGAATTGG</td><td>1000</td><td></td><td></td><td></td><td></td><td></td><td></td><td></td><td></td><td></td></tr>
<tr><td>Oligo fwd</td><td>CACCTCAAGGAGAGTTGAGCGAAT</td><td></td><td></td><td></td><td></td><td></td><td></td><td></td><td></td><td></td><td></td></tr>
<tr><td>Oligo rev</td><td>AAACATTCGCTCAACTCTCCTTGA</td><td></td><td></td><td></td><td></td><td></td><td></td><td></td><td></td><td></td><td></td></tr>
<tr><td>Chromosome</td><td>start</td><td>end</td><td>strand</td><td>MM</td><td>target_seq</td><td>PAM</td><td>alignment</td><td>distance</td><td>position</td><td>gene name</td><td>gene id</td></tr>
<tr><td>GU305914.1</td><td>116270</td><td>116292</td><td>+</td><td>0</td><td>TCAAGGAGAGTTGAGCGAAT</td><td>TGG</td><td>||||||||[||||||||||||]PAM</td><td>NA</td><td>Intergenic</td><td>NA</td><td>NA</td></tr>
<tr><td>GU305914.1</td><td>183945</td><td>183967</td><td>-</td><td>4</td><td>TCAATGGGAGGTGAGCCAAT</td><td>GGG</td><td>||||-|-|[||-|||||-|||]PAM</td><td>NA</td><td>Intergenic</td><td>NA</td><td>NA</td></tr>
<tr><td></td><td></td><td></td><td></td><td></td><td></td><td></td><td></td><td></td><td></td><td></td><td></td></tr>
<tr><td>T8613</td><td>GAGTTGAGCGAATTGGTCCTCGG</td><td>1000</td><td></td><td></td><td></td><td></td><td></td><td></td><td></td><td></td><td></td></tr>
<tr><td>Oligo fwd</td><td>CACCGAGTTGAGCGAATTGGTCCT</td><td></td><td></td><td></td><td></td><td></td><td></td><td></td><td></td><td></td><td></td></tr>
<tr><td>Oligo rev</td><td>AAACAGGACCAATTCGCTCAACTC</td><td></td><td></td><td></td><td></td><td></td><td></td><td></td><td></td><td></td><td></td></tr>
<tr><td>Chromosome</td><td>start</td><td>end</td><td>strand</td><td>MM</td><td>target_seq</td><td>PAM</td><td>alignment</td><td>distance</td><td>position</td><td>gene name</td><td>gene id</td></tr>
<tr><td>GU305914.1</td><td>116277</td><td>116299</td><td>+</td><td>0</td><td>GAGTTGAGCGAATTGGTCCT</td><td>CGG</td><td>||||||||[||||||||||||]PAM</td><td>NA</td><td>Intergenic</td><td>NA</td><td>NA</td></tr>
<tr><td></td><td></td><td></td><td></td><td></td><td></td><td></td><td></td><td></td><td></td><td></td><td></td></tr>
<tr><td>T8614</td><td>AGTTGAGCGAATTGGTCCTCGGG</td><td>1000</td><td></td><td></td><td></td><td></td><td></td><td></td><td></td><td></td><td></td></tr>
<tr><td>Oligo fwd</td><td>CACCAGTTGAGCGAATTGGTCCTC</td><td></td><td></td><td></td><td></td><td></td><td></td><td></td><td></td><td></td><td></td></tr>
<tr><td>Oligo rev</td><td>AAACGAGGACCAATTCGCTCAACT</td><td></td><td></td><td></td><td></td><td></td><td></td><td></td><td></td><td></td><td></td></tr>
<tr><td>Chromosome</td><td>start</td><td>end</td><td>strand</td><td>MM</td><td>target_seq</td><td>PAM</td><td>alignment</td><td>distance</td><td>position</td><td>gene name</td><td>gene id</td></tr>
<tr><td>GU305914.1</td><td>116278</td><td>116300</td><td>+</td><td>0</td><td>AGTTGAGCGAATTGGTCCTC</td><td>GGG</td><td>||||||||[||||||||||||]PAM</td><td>NA</td><td>Intergenic</td><td>NA</td><td>NA</td></tr>
<tr><td></td><td></td><td></td><td></td><td></td><td></td><td></td><td></td><td></td><td></td><td></td><td></td></tr>
<tr><td>T8615</td><td>CTCGGGTAGCTGCTGCTCGTCGG</td><td>1000</td><td></td><td></td><td></td><td></td><td></td><td></td><td></td><td></td><td></td></tr>
<tr><td>Oligo fwd</td><td>CACCCTCGGGTAGCTGCTGCTCGT</td><td></td><td></td><td></td><td></td><td></td><td></td><td></td><td></td><td></td><td></td></tr>
<tr><td>Oligo rev</td><td>AAACACGAGCAGCAGCTACCCGAG</td><td></td><td></td><td></td><td></td><td></td><td></td><td></td><td></td><td></td><td></td></tr>
<tr><td>Chromosome</td><td>start</td><td>end</td><td>strand</td><td>MM</td><td>target_seq</td><td>PAM</td><td>alignment</td><td>distance</td><td>position</td><td>gene name</td><td>gene id</td></tr>
<tr><td>GU305914.1</td><td>116295</td><td>116317</td><td>+</td><td>0</td><td>CTCGGGTAGCTGCTGCTCGT</td><td>CGG</td><td>||||||||[||||||||||||]PAM</td><td>NA</td><td>Intergenic</td><td>NA</td><td>NA</td></tr>
<tr><td></td><td></td><td></td><td></td><td></td><td></td><td></td><td></td><td></td><td></td><td></td><td></td></tr>
<tr><td>T8616</td><td>GCTCGTCGGCCGAGCTCATCCGG</td><td>1000</td><td></td><td></td><td></td><td></td><td></td><td></td><td></td><td></td><td></td></tr>
<tr><td>Oligo fwd</td><td>CACCGCTCGTCGGCCGAGCTCATC</td><td></td><td></td><td></td><td></td><td></td><td></td><td></td><td></td><td></td><td></td></tr>
<tr><td>Oligo rev</td><td>AAACGATGAGCTCGGCCGACGAGC</td><td></td><td></td><td></td><td></td><td></td><td></td><td></td><td></td><td></td><td></td></tr>
<tr><td>Chromosome</td><td>start</td><td>end</td><td>strand</td><td>MM</td><td>target_seq</td><td>PAM</td><td>alignment</td><td>distance</td><td>position</td><td>gene name</td><td>gene id</td></tr>
<tr><td>GU305914.1</td><td>116309</td><td>116331</td><td>+</td><td>0</td><td>GCTCGTCGGCCGAGCTCATC</td><td>CGG</td><td>||||||||[||||||||||||]PAM</td><td>NA</td><td>Intergenic</td><td>NA</td><td>NA</td></tr>
<tr><td></td><td></td><td></td><td></td><td></td><td></td><td></td><td></td><td></td><td></td><td></td><td></td></tr>
<tr><td>T8617</td><td>TCGGCCGAGCTCATCCGGCGAGG</td><td>1000</td><td></td><td></td><td></td><td></td><td></td><td></td><td></td><td></td><td></td></tr>
<tr><td>Oligo fwd</td><td>CACCTCGGCCGAGCTCATCCGGCG</td><td></td><td></td><td></td><td></td><td></td><td></td><td></td><td></td><td></td><td></td></tr>
<tr><td>Oligo rev</td><td>AAACCGCCGGATGAGCTCGGCCGA</td><td></td><td></td><td></td><td></td><td></td><td></td><td></td><td></td><td></td><td></td></tr>
<tr><td>Chromosome</td><td>start</td><td>end</td><td>strand</td><td>MM</td><td>target_seq</td><td>PAM</td><td>alignment</td><td>distance</td><td>position</td><td>gene name</td><td>gene id</td></tr>
<tr><td>GU305914.1</td><td>116314</td><td>116336</td><td>+</td><td>0</td><td>TCGGCCGAGCTCATCCGGCG</td><td>AGG</td><td>||||||||[||||||||||||]PAM</td><td>NA</td><td>Intergenic</td><td>NA</td><td>NA</td></tr>
<tr><td></td><td></td><td></td><td></td><td></td><td></td><td></td><td></td><td></td><td></td><td></td><td></td></tr>
<tr><td>T8618</td><td>CGGCCGAGCTCATCCGGCGAGGG</td><td>1000</td><td></td><td></td><td></td><td></td><td></td><td></td><td></td><td></td><td></td></tr>
</table>

| | | | | | | | | | | | |
|---|---|---|---|---|---|---|---|---|---|---|---|
| Oligo fwd | CACCCGGCCGAGCTCATCCGGCGA | | | | | | | | | | |
| Oligo rev | AAACTCGCCGGATGAGCTCGGCCG | | | | | | | | | | |
| Chromosome | start | end | strand | MM | target_seq | PAM | alignment | distance | position | gene name | gene id |
| GU305914.1 | 116315 | 116337 | + | 0 | CGGCCGAGCTCATCCGGCGA | GGG | \|\|\|\|\|\|\|\|[\|\|\|\|\|\|\|\|\|\|\|\|]PAM | NA | Intergenic | NA | NA |
| T8619 | GGCCGAGCTCATCCGGCGAGGGG | 1000 | | | | | | | | | |
| Oligo fwd | CACCGGCCGAGCTCATCCGGCGAG | | | | | | | | | | |
| Oligo rev | AAACCTCGCCGGATGAGCTCGGCC | | | | | | | | | | |
| Chromosome | start | end | strand | MM | target_seq | PAM | alignment | distance | position | gene name | gene id |
| GU305914.1 | 116316 | 116338 | + | 0 | GGCCGAGCTCATCCGGCGAG | GGG | \|\|\|\|\|\|\|\|[\|\|\|\|\|\|\|\|\|\|\|\|]PAM | NA | Intergenic | NA | NA |
| T8620 | CGAGCTCATCCGGCGAGGGGTGG | 1000 | | | | | | | | | |
| Oligo fwd | CACCCGAGCTCATCCGGCGAGGGG | | | | | | | | | | |
| Oligo rev | AAACCCCCTCGCCGGATGAGCTCG | | | | | | | | | | |
| Chromosome | start | end | strand | MM | target_seq | PAM | alignment | distance | position | gene name | gene id |
| GU305914.1 | 116319 | 116341 | + | 0 | CGAGCTCATCCGGCGAGGGG | TGG | \|\|\|\|\|\|\|\|[\|\|\|\|\|\|\|\|\|\|\|\|]PAM | NA | Intergenic | NA | NA |
| T8621 | ATCCGGCGAGGGGTGGTCGCTGG | 1000 | | | | | | | | | |
| Oligo fwd | CACCATCCGGCGAGGGGTGGTCGC | | | | | | | | | | |
| Oligo rev | AAACGCGACCACCCCTCGCCGGAT | | | | | | | | | | |
| Chromosome | start | end | strand | MM | target_seq | PAM | alignment | distance | position | gene name | gene id |
| GU305914.1 | 116326 | 116348 | + | 0 | ATCCGGCGAGGGGTGGTCGC | TGG | \|\|\|\|\|\|\|\|[\|\|\|\|\|\|\|\|\|\|\|\|]PAM | NA | Intergenic | NA | NA |
| T8622 | CGGCGAGGGGTGGTCGCTGGAGG | 1000 | | | | | | | | | |
| Oligo fwd | CACCCGGCGAGGGGTGGTCGCTGG | | | | | | | | | | |
| Oligo rev | AAACCCAGCGACCACCCCTCGCCG | | | | | | | | | | |
| Chromosome | start | end | strand | MM | target_seq | PAM | alignment | distance | position | gene name | gene id |
| GU305914.1 | 116329 | 116351 | + | 0 | CGGCGAGGGGTGGTCGCTGG | AGG | \|\|\|\|\|\|\|\|[\|\|\|\|\|\|\|\|\|\|\|\|]PAM | NA | Intergenic | NA | NA |
| T8623 | GGGTGGTCGCTGGAGGTAGCTGG | 1000 | | | | | | | | | |
| Oligo fwd | CACCGGGTGGTCGCTGGAGGTAGC | | | | | | | | | | |
| Oligo rev | AAACGCTACCTCCAGCGACCACCC | | | | | | | | | | |
| Chromosome | start | end | strand | MM | target_seq | PAM | alignment | distance | position | gene name | gene id |
| GU305914.1 | 116336 | 116358 | + | 0 | GGGTGGTCGCTGGAGGTAGC | TGG | \|\|\|\|\|\|\|\|[\|\|\|\|\|\|\|\|\|\|\|\|]PAM | NA | Intergenic | NA | NA |
| T8624 | GGTCGCTGGAGGTAGCTGGTCGG | 1000 | | | | | | | | | |
| Oligo fwd | CACCGGTCGCTGGAGGTAGCTGGT | | | | | | | | | | |
| Oligo rev | AAACACCAGCTACCTCCAGCGACC | | | | | | | | | | |
| Chromosome | start | end | strand | MM | target_seq | PAM | alignment | distance | position | gene name | gene id |
| GU305914.1 | 116340 | 116362 | + | 0 | GGTCGCTGGAGGTAGCTGGT | CGG | \|\|\|\|\|\|\|\|[\|\|\|\|\|\|\|\|\|\|\|\|]PAM | NA | Intergenic | NA | NA |
| T8625 | GGTAGCTGGTCGGCGTCCTGCGG | 1000 | | | | | | | | | |
| Oligo fwd | CACCGGTAGCTGGTCGGCGTCCTG | | | | | | | | | | |
| Oligo rev | AAACCAGGACGCCGACCAGCTACC | | | | | | | | | | |
| Chromosome | start | end | strand | MM | target_seq | PAM | alignment | distance | position | gene name | gene id |
| GU305914.1 | 116350 | 116372 | + | 0 | GGTAGCTGGTCGGCGTCCTG | CGG | \|\|\|\|\|\|\|\|[\|\|\|\|\|\|\|\|\|\|\|\|]PAM | NA | Intergenic | NA | NA |
| T8626 | AGCTGGTCGGCGTCCTGCGGCGG | 1000 | | | | | | | | | |
| Oligo fwd | CACCAGCTGGTCGGCGTCCTGCGG | | | | | | | | | | |
| Oligo rev | AAACCCGCAGGACGCCGACCAGCT | | | | | | | | | | |
| Chromosome | start | end | strand | MM | target_seq | PAM | alignment | distance | position | gene name | gene id |
| GU305914.1 | 116353 | 116375 | + | 0 | AGCTGGTCGGCGTCCTGCGG | CGG | \|\|\|\|\|\|\|\|[\|\|\|\|\|\|\|\|\|\|\|\|]PAM | NA | Intergenic | NA | NA |
| T8627 | GTCGGCGTCCTGCGGCGGCTCGG | 1000 | | | | | | | | | |
| Oligo fwd | CACCGTCGGCGTCCTGCGGCGGCT | | | | | | | | | | |
| Oligo rev | AAACAGCCGCCGCAGGACGCCGAC | | | | | | | | | | |
| Chromosome | start | end | strand | MM | target_seq | PAM | alignment | distance | position | gene name | gene id |
| GU305914.1 | 116358 | 116380 | + | 0 | GTCGGCGTCCTGCGGCGGCT | CGG | \|\|\|\|\|\|\|\|[\|\|\|\|\|\|\|\|\|\|\|\|]PAM | NA | Intergenic | NA | NA |
| GU305914.1 | 110034 | 110056 | + | 3 | GTCGCCGTCCTGCTGCGGCG | GGG | \|\|\|\|-\|\|\|[\|\|\|\|\|-\|\|\|\|\|-]PAM | NA | Intergenic | NA | NA |
| T8628 | TCGGCGTCCTGCGGCGGCTCGGG | 1000 | | | | | | | | | |
| Oligo fwd | CACCTCGGCGTCCTGCGGCGGCTC | | | | | | | | | | |
| Oligo rev | AAACGAGCCGCCGCAGGACGCCGA | | | | | | | | | | |
| Chromosome | start | end | strand | MM | target_seq | PAM | alignment | distance | position | gene name | gene id |
| GU305914.1 | 116359 | 116381 | + | 0 | TCGGCGTCCTGCGGCGGCTC | GGG | \|\|\|\|\|\|\|\|[\|\|\|\|\|\|\|\|\|\|\|\|]PAM | NA | Intergenic | NA | NA |
| T8629 | CGGCGTCCTGCGGCGGCTCGGGG | 1000 | | | | | | | | | |
| Oligo fwd | CACCCGGCGTCCTGCGGCGGCTCG | | | | | | | | | | |
| Oligo rev | AAACCGAGCCGCCGCAGGACGCCG | | | | | | | | | | |
| Chromosome | start | end | strand | MM | target_seq | PAM | alignment | distance | position | gene name | gene id |
| GU305914.1 | 116360 | 116382 | + | 0 | CGGCGTCCTGCGGCGGCTCG | GGG | \|\|\|\|\|\|\|\|[\|\|\|\|\|\|\|\|\|\|\|\|]PAM | NA | Intergenic | NA | NA |
| GU305914.1 | 40995 | 41017 | - | 3 | CGTCTTCCTGCGGGGGCTCG | AGG | \|\|-\|-\|\|\|[\|\|\|\|\|-\|\|\|\|\|\|]PAM | NA | Intergenic | NA | NA |
| GU305914.1 | 23667 | 23689 | - | 4 | CGGGCTCCTGCGTCGGCCCG | AGG | \|\|\|--\|\|\|[\|\|\|\|-\|\|\|\|-\|\|]PAM | NA | Intergenic | NA | NA |
| T8630 | GTCCTGCGGCGGCTCGGGGCCGG | 1000 | | | | | | | | | |
| Oligo fwd | CACCGTCCTGCGGCGGCTCGGGGC | | | | | | | | | | |
| Oligo rev | AAACGCCCCGAGCCGCCGCAGGAC | | | | | | | | | | |
| Chromosome | start | end | strand | MM | target_seq | PAM | alignment | distance | position | gene name | gene id |
| GU305914.1 | 116364 | 116386 | + | 0 | GTCCTGCGGCGGCTCGGGGC | CGG | \|\|\|\|\|\|\|\|[\|\|\|\|\|\|\|\|\|\|\|\|]PAM | NA | Intergenic | NA | NA |
| T8631 | TCCTGCGGCGGCTCGGGGCCGGG | 1000 | | | | | | | | | |
| Oligo fwd | CACCTCCTGCGGCGGCTCGGGGCC | | | | | | | | | | |
| Oligo rev | AAACGGCCCCGAGCCGCCGCAGGA | | | | | | | | | | |
| Chromosome | start | end | strand | MM | target_seq | PAM | alignment | distance | position | gene name | gene id |
| GU305914.1 | 116365 | 116387 | + | 0 | TCCTGCGGCGGCTCGGGGCC | GGG | \|\|\|\|\|\|\|\|[\|\|\|\|\|\|\|\|\|\|\|\|]PAM | NA | Intergenic | NA | NA |
| T8632 | GCGGCGGCTCGGGGCCGGGTCGG | 1000 | | | | | | | | | |
| Oligo fwd | CACCGCGGCGGCTCGGGGCCGGGT | | | | | | | | | | |
| Oligo rev | AAACACCCGGCCCCGAGCCGCCGC | | | | | | | | | | |

<table>
<tr><td>Chromosome</td><td>start</td><td>end</td><td>strand</td><td>MM</td><td>target_seq</td><td>PAM</td><td>alignment</td><td>distance</td><td>position</td><td>gene name</td><td>gene id</td></tr>
<tr><td>GU305914.1</td><td>116369</td><td>116391</td><td>+</td><td>0</td><td>GCGGCGGCTCGGGGCCGGGT</td><td>CGG</td><td>||||||||[||||||||||||]PAM</td><td>NA</td><td>Intergenic</td><td>NA</td><td>NA</td></tr>
<tr><td>GU305914.1</td><td>170176</td><td>170198</td><td>+</td><td>4</td><td>GCTCAGGCTCGGGGACGGGT</td><td>TGG</td><td>||---|||[||||||-|||||]PAM</td><td>NA</td><td>Intergenic</td><td>NA</td><td>NA</td></tr>
<tr><td>T8633</td><td>CGGCTCGGGGCCGGGTCGGTAGG</td><td>1000</td><td></td><td></td><td></td><td></td><td></td><td></td><td></td><td></td><td></td></tr>
<tr><td>Oligo fwd</td><td>CACCCGGCTCGGGGCCGGGTCGGT</td><td></td><td></td><td></td><td></td><td></td><td></td><td></td><td></td><td></td><td></td></tr>
<tr><td>Oligo rev</td><td>AAACACCGACCCGGCCCCGAGCCG</td><td></td><td></td><td></td><td></td><td></td><td></td><td></td><td></td><td></td><td></td></tr>
<tr><td>Chromosome</td><td>start</td><td>end</td><td>strand</td><td>MM</td><td>target_seq</td><td>PAM</td><td>alignment</td><td>distance</td><td>position</td><td>gene name</td><td>gene id</td></tr>
<tr><td>GU305914.1</td><td>116373</td><td>116395</td><td>+</td><td>0</td><td>CGGCTCGGGGCCGGGTCGGT</td><td>AGG</td><td>||||||||[||||||||||||]PAM</td><td>NA</td><td>Intergenic</td><td>NA</td><td>NA</td></tr>
<tr><td>T8634</td><td>GGGGCCGGGTCGGTAGGTCTCGG</td><td>1000</td><td></td><td></td><td></td><td></td><td></td><td></td><td></td><td></td><td></td></tr>
<tr><td>Oligo fwd</td><td>CACCGGGGCCGGGTCGGTAGGTCT</td><td></td><td></td><td></td><td></td><td></td><td></td><td></td><td></td><td></td><td></td></tr>
<tr><td>Oligo rev</td><td>AAACAGACCTACCGACCCGGCCCC</td><td></td><td></td><td></td><td></td><td></td><td></td><td></td><td></td><td></td><td></td></tr>
<tr><td>Chromosome</td><td>start</td><td>end</td><td>strand</td><td>MM</td><td>target_seq</td><td>PAM</td><td>alignment</td><td>distance</td><td>position</td><td>gene name</td><td>gene id</td></tr>
<tr><td>GU305914.1</td><td>116379</td><td>116401</td><td>+</td><td>0</td><td>GGGGCCGGGTCGGTAGGTCT</td><td>CGG</td><td>||||||||[||||||||||||]PAM</td><td>NA</td><td>Intergenic</td><td>NA</td><td>NA</td></tr>
<tr><td>T8635</td><td>TCGGTAGGTCTCGGATCCAACGG</td><td>1000</td><td></td><td></td><td></td><td></td><td></td><td></td><td></td><td></td><td></td></tr>
<tr><td>Oligo fwd</td><td>CACCTCGGTAGGTCTCGGATCCAA</td><td></td><td></td><td></td><td></td><td></td><td></td><td></td><td></td><td></td><td></td></tr>
<tr><td>Oligo rev</td><td>AAACTTGGATCCGAGACCTACCGA</td><td></td><td></td><td></td><td></td><td></td><td></td><td></td><td></td><td></td><td></td></tr>
<tr><td>Chromosome</td><td>start</td><td>end</td><td>strand</td><td>MM</td><td>target_seq</td><td>PAM</td><td>alignment</td><td>distance</td><td>position</td><td>gene name</td><td>gene id</td></tr>
<tr><td>GU305914.1</td><td>116388</td><td>116410</td><td>+</td><td>0</td><td>TCGGTAGGTCTCGGATCCAA</td><td>CGG</td><td>||||||||[||||||||||||]PAM</td><td>NA</td><td>Intergenic</td><td>NA</td><td>NA</td></tr>
<tr><td>T8636</td><td>CGGTAGGTCTCGGATCCAACGGG</td><td>1000</td><td></td><td></td><td></td><td></td><td></td><td></td><td></td><td></td><td></td></tr>
<tr><td>Oligo fwd</td><td>CACCCGGTAGGTCTCGGATCCAAC</td><td></td><td></td><td></td><td></td><td></td><td></td><td></td><td></td><td></td><td></td></tr>
<tr><td>Oligo rev</td><td>AAACGTTGGATCCGAGACCTACCG</td><td></td><td></td><td></td><td></td><td></td><td></td><td></td><td></td><td></td><td></td></tr>
<tr><td>Chromosome</td><td>start</td><td>end</td><td>strand</td><td>MM</td><td>target_seq</td><td>PAM</td><td>alignment</td><td>distance</td><td>position</td><td>gene name</td><td>gene id</td></tr>
<tr><td>GU305914.1</td><td>116389</td><td>116411</td><td>+</td><td>0</td><td>CGGTAGGTCTCGGATCCAAC</td><td>GGG</td><td>||||||||[||||||||||||]PAM</td><td>NA</td><td>Intergenic</td><td>NA</td><td>NA</td></tr>
<tr><td>T8637</td><td>GGTAGGTCTCGGATCCAACGGGG</td><td>1000</td><td></td><td></td><td></td><td></td><td></td><td></td><td></td><td></td><td></td></tr>
<tr><td>Oligo fwd</td><td>CACCGGTAGGTCTCGGATCCAACG</td><td></td><td></td><td></td><td></td><td></td><td></td><td></td><td></td><td></td><td></td></tr>
<tr><td>Oligo rev</td><td>AAACCGTTGGATCCGAGACCTACC</td><td></td><td></td><td></td><td></td><td></td><td></td><td></td><td></td><td></td><td></td></tr>
<tr><td>Chromosome</td><td>start</td><td>end</td><td>strand</td><td>MM</td><td>target_seq</td><td>PAM</td><td>alignment</td><td>distance</td><td>position</td><td>gene name</td><td>gene id</td></tr>
<tr><td>GU305914.1</td><td>116390</td><td>116412</td><td>+</td><td>0</td><td>GGTAGGTCTCGGATCCAACG</td><td>GGG</td><td>||||||||[||||||||||||]PAM</td><td>NA</td><td>Intergenic</td><td>NA</td><td>NA</td></tr>
<tr><td>T8638</td><td>GAGTTTGCATTCGTACTGTGAGG</td><td>1000</td><td></td><td></td><td></td><td></td><td></td><td></td><td></td><td></td><td></td></tr>
<tr><td>Oligo fwd</td><td>CACCGAGTTTGCATTCGTACTGTG</td><td></td><td></td><td></td><td></td><td></td><td></td><td></td><td></td><td></td><td></td></tr>
<tr><td>Oligo rev</td><td>AAACCACAGTACGAATGCAAACTC</td><td></td><td></td><td></td><td></td><td></td><td></td><td></td><td></td><td></td><td></td></tr>
<tr><td>Chromosome</td><td>start</td><td>end</td><td>strand</td><td>MM</td><td>target_seq</td><td>PAM</td><td>alignment</td><td>distance</td><td>position</td><td>gene name</td><td>gene id</td></tr>
<tr><td>GU305914.1</td><td>116418</td><td>116440</td><td>+</td><td>0</td><td>GAGTTTGCATTCGTACTGTG</td><td>AGG</td><td>||||||||[||||||||||||]PAM</td><td>NA</td><td>Intergenic</td><td>NA</td><td>NA</td></tr>
<tr><td>T8639</td><td>TTTGCATTCGTACTGTGAGGTGG</td><td>1000</td><td></td><td></td><td></td><td></td><td></td><td></td><td></td><td></td><td></td></tr>
<tr><td>Oligo fwd</td><td>CACCTTTGCATTCGTACTGTGAGG</td><td></td><td></td><td></td><td></td><td></td><td></td><td></td><td></td><td></td><td></td></tr>
<tr><td>Oligo rev</td><td>AAACCCTCACAGTACGAATGCAAA</td><td></td><td></td><td></td><td></td><td></td><td></td><td></td><td></td><td></td><td></td></tr>
<tr><td>Chromosome</td><td>start</td><td>end</td><td>strand</td><td>MM</td><td>target_seq</td><td>PAM</td><td>alignment</td><td>distance</td><td>position</td><td>gene name</td><td>gene id</td></tr>
<tr><td>GU305914.1</td><td>116421</td><td>116443</td><td>+</td><td>0</td><td>TTTGCATTCGTACTGTGAGG</td><td>TGG</td><td>||||||||[||||||||||||]PAM</td><td>NA</td><td>Intergenic</td><td>NA</td><td>NA</td></tr>
<tr><td>T8640</td><td>CTCTTTATTGATGATGCCGACGG</td><td>1000</td><td></td><td></td><td></td><td></td><td></td><td></td><td></td><td></td><td></td></tr>
<tr><td>Oligo fwd</td><td>CACCCTCTTTATTGATGATGCCGA</td><td></td><td></td><td></td><td></td><td></td><td></td><td></td><td></td><td></td><td></td></tr>
<tr><td>Oligo rev</td><td>AAACTCGGCATCATCAATAAAGAG</td><td></td><td></td><td></td><td></td><td></td><td></td><td></td><td></td><td></td><td></td></tr>
<tr><td>Chromosome</td><td>start</td><td>end</td><td>strand</td><td>MM</td><td>target_seq</td><td>PAM</td><td>alignment</td><td>distance</td><td>position</td><td>gene name</td><td>gene id</td></tr>
<tr><td>GU305914.1</td><td>116457</td><td>116479</td><td>+</td><td>0</td><td>CTCTTTATTGATGATGCCGA</td><td>CGG</td><td>||||||||[||||||||||||]PAM</td><td>NA</td><td>Intergenic</td><td>NA</td><td>NA</td></tr>
<tr><td>T8641</td><td>TCTTTATTGATGATGCCGACGGG</td><td>1000</td><td></td><td></td><td></td><td></td><td></td><td></td><td></td><td></td><td></td></tr>
<tr><td>Oligo fwd</td><td>CACCTCTTTATTGATGATGCCGAC</td><td></td><td></td><td></td><td></td><td></td><td></td><td></td><td></td><td></td><td></td></tr>
<tr><td>Oligo rev</td><td>AAACGTCGGCATCATCAATAAAGA</td><td></td><td></td><td></td><td></td><td></td><td></td><td></td><td></td><td></td><td></td></tr>
<tr><td>Chromosome</td><td>start</td><td>end</td><td>strand</td><td>MM</td><td>target_seq</td><td>PAM</td><td>alignment</td><td>distance</td><td>position</td><td>gene name</td><td>gene id</td></tr>
<tr><td>GU305914.1</td><td>116458</td><td>116480</td><td>+</td><td>0</td><td>TCTTTATTGATGATGCCGAC</td><td>GGG</td><td>||||||||[||||||||||||]PAM</td><td>NA</td><td>Intergenic</td><td>NA</td><td>NA</td></tr>
<tr><td>T8642</td><td>CTTTATTGATGATGCCGACGGGG</td><td>1000</td><td></td><td></td><td></td><td></td><td></td><td></td><td></td><td></td><td></td></tr>
<tr><td>Oligo fwd</td><td>CACCCTTTATTGATGATGCCGACG</td><td></td><td></td><td></td><td></td><td></td><td></td><td></td><td></td><td></td><td></td></tr>
<tr><td>Oligo rev</td><td>AAACCGTCGGCATCATCAATAAAG</td><td></td><td></td><td></td><td></td><td></td><td></td><td></td><td></td><td></td><td></td></tr>
<tr><td>Chromosome</td><td>start</td><td>end</td><td>strand</td><td>MM</td><td>target_seq</td><td>PAM</td><td>alignment</td><td>distance</td><td>position</td><td>gene name</td><td>gene id</td></tr>
<tr><td>GU305914.1</td><td>116459</td><td>116481</td><td>+</td><td>0</td><td>CTTTATTGATGATGCCGACG</td><td>GGG</td><td>||||||||[||||||||||||]PAM</td><td>NA</td><td>Intergenic</td><td>NA</td><td>NA</td></tr>
<tr><td>T8643</td><td>TTTATTGATGATGCCGACGGGGG</td><td>1000</td><td></td><td></td><td></td><td></td><td></td><td></td><td></td><td></td><td></td></tr>
<tr><td>Oligo fwd</td><td>CACCTTTATTGATGATGCCGACGG</td><td></td><td></td><td></td><td></td><td></td><td></td><td></td><td></td><td></td><td></td></tr>
<tr><td>Oligo rev</td><td>AAACCCGTCGGCATCATCAATAAA</td><td></td><td></td><td></td><td></td><td></td><td></td><td></td><td></td><td></td><td></td></tr>
<tr><td>Chromosome</td><td>start</td><td>end</td><td>strand</td><td>MM</td><td>target_seq</td><td>PAM</td><td>alignment</td><td>distance</td><td>position</td><td>gene name</td><td>gene id</td></tr>
<tr><td>GU305914.1</td><td>116460</td><td>116482</td><td>+</td><td>0</td><td>TTTATTGATGATGCCGACGG</td><td>GGG</td><td>||||||||[||||||||||||]PAM</td><td>NA</td><td>Intergenic</td><td>NA</td><td>NA</td></tr>
<tr><td>T8644</td><td>TGATGCCGACGGGGGTCAGCAGG</td><td>1000</td><td></td><td></td><td></td><td></td><td></td><td></td><td></td><td></td><td></td></tr>
<tr><td>Oligo fwd</td><td>CACCTGATGCCGACGGGGGTCAGC</td><td></td><td></td><td></td><td></td><td></td><td></td><td></td><td></td><td></td><td></td></tr>
<tr><td>Oligo rev</td><td>AAACGCTGACCCCCGTCGGCATCA</td><td></td><td></td><td></td><td></td><td></td><td></td><td></td><td></td><td></td><td></td></tr>
<tr><td>Chromosome</td><td>start</td><td>end</td><td>strand</td><td>MM</td><td>target_seq</td><td>PAM</td><td>alignment</td><td>distance</td><td>position</td><td>gene name</td><td>gene id</td></tr>
<tr><td>GU305914.1</td><td>116468</td><td>116490</td><td>+</td><td>0</td><td>TGATGCCGACGGGGGTCAGC</td><td>AGG</td><td>||||||||[||||||||||||]PAM</td><td>NA</td><td>Intergenic</td><td>NA</td><td>NA</td></tr>
<tr><td>T8645</td><td>CGACGAGCTCTTGTGTCAGTCGG</td><td>1000</td><td></td><td></td><td></td><td></td><td></td><td></td><td></td><td></td><td></td></tr>
<tr><td>Oligo fwd</td><td>CACCCGACGAGCTCTTGTGTCAGT</td><td></td><td></td><td></td><td></td><td></td><td></td><td></td><td></td><td></td><td></td></tr>
<tr><td>Oligo rev</td><td>AAACACTGACACAAGAGCTCGTCG</td><td></td><td></td><td></td><td></td><td></td><td></td><td></td><td></td><td></td><td></td></tr>
<tr><td>Chromosome</td><td>start</td><td>end</td><td>strand</td><td>MM</td><td>target_seq</td><td>PAM</td><td>alignment</td><td>distance</td><td>position</td><td>gene name</td><td>gene id</td></tr>
<tr><td>GU305914.1</td><td>116492</td><td>116514</td><td>+</td><td>0</td><td>CGACGAGCTCTTGTGTCAGT</td><td>CGG</td><td>||||||||[||||||||||||]PAM</td><td>NA</td><td>Intergenic</td><td>NA</td><td>NA</td></tr>
<tr><td>T8646</td><td>GAGCTCTTGTGTCAGTCGGTTGG</td><td>1000</td><td></td><td></td><td></td><td></td><td></td><td></td><td></td><td></td><td></td></tr>
<tr><td>Oligo fwd</td><td>CACCGAGCTCTTGTGTCAGTCGGT</td><td></td><td></td><td></td><td></td><td></td><td></td><td></td><td></td><td></td><td></td></tr>
<tr><td>Oligo rev</td><td>AAACACCGACTGACACAAGAGCTC</td><td></td><td></td><td></td><td></td><td></td><td></td><td></td><td></td><td></td><td></td></tr>
<tr><td>Chromosome</td><td>start</td><td>end</td><td>strand</td><td>MM</td><td>target_seq</td><td>PAM</td><td>alignment</td><td>distance</td><td>position</td><td>gene name</td><td>gene id</td></tr>
<tr><td>GU305914.1</td><td>116496</td><td>116518</td><td>+</td><td>0</td><td>GAGCTCTTGTGTCAGTCGGT</td><td>TGG</td><td>||||||||[||||||||||||]PAM</td><td>NA</td><td>Intergenic</td><td>NA</td><td>NA</td></tr>
<tr><td>T8647</td><td>AGCTCTTGTGTCAGTCGGTTGGG</td><td>1000</td><td></td><td></td><td></td><td></td><td></td><td></td><td></td><td></td><td></td></tr>
</table>

Oligo fwd CACCAGCTCTTGTGTCAGTCGGTT
Oligo rev AAACAACCGACTGACACAAGAGCT
Chromosome start end strand MM target_seq PAM alignment distance position gene name gene id
GU305914.1 116497 116519 + 0 AGCTCTTGTGTCAGTCGGTT GGG ||||||||[||||||||||||]PAM NA Intergenic NA NA

T8648 GCTCTTGTGTCAGTCGGTTGGGG 1000
Oligo fwd CACCGCTCTTGTGTCAGTCGGTTG
Oligo rev AAACCAACCGACTGACACAAGAGC
Chromosome start end strand MM target_seq PAM alignment distance position gene name gene id
GU305914.1 116498 116520 + 0 GCTCTTGTGTCAGTCGGTTG GGG ||||||||[||||||||||||]PAM NA Intergenic NA NA

T8649 GGTTGGGGAAGAAGTGCACTCGG 1000
Oligo fwd CACCGGTTGGGGAAGAAGTGCACT
Oligo rev AAACAGTGCACTTCTTCCCCAACC
Chromosome start end strand MM target_seq PAM alignment distance position gene name gene id
GU305914.1 116513 116535 + 0 GGTTGGGGAAGAAGTGCACT CGG ||||||||[||||||||||||]PAM NA Intergenic NA NA
GU305914.1 77416 77438 - 4 GGTTGATGAAGAAGTGCGCG TGG |||||--|[|||||||||-|-]PAM NA Intergenic NA NA

T8650 TCGGCCGAGCTCCATTCCTACGG 1000
Oligo fwd CACCTCGGCCGAGCTCCATTCCTA
Oligo rev AAACTAGGAATGGAGCTCGGCCGA
Chromosome start end strand MM target_seq PAM alignment distance position gene name gene id
GU305914.1 116532 116554 + 0 TCGGCCGAGCTCCATTCCTA CGG ||||||||[||||||||||||]PAM NA Intergenic NA NA

T8651 CGGCCGAGCTCCATTCCTACGGG 1000
Oligo fwd CACCCGGCCGAGCTCCATTCCTAC
Oligo rev AAACGTAGGAATGGAGCTCGGCCG
Chromosome start end strand MM target_seq PAM alignment distance position gene name gene id
GU305914.1 116533 116555 + 0 CGGCCGAGCTCCATTCCTAC GGG ||||||||[||||||||||||]PAM NA Intergenic NA NA

T8652 GAGCTCCATTCCTACGGGAATGG 1000
Oligo fwd CACCGAGCTCCATTCCTACGGGAA
Oligo rev AAACTTCCCGTAGGAATGGAGCTC
Chromosome start end strand MM target_seq PAM alignment distance position gene name gene id
GU305914.1 116538 116560 + 0 GAGCTCCATTCCTACGGGAA TGG ||||||||[||||||||||||]PAM NA Intergenic NA NA

T8653 TCCTACGGGAATGGTCACGTTGG 1000
Oligo fwd CACCTCCTACGGGAATGGTCACGT
Oligo rev AAACACGTGACCATTCCCGTAGGA
Chromosome start end strand MM target_seq PAM alignment distance position gene name gene id
GU305914.1 116547 116569 + 0 TCCTACGGGAATGGTCACGT TGG ||||||||[||||||||||||]PAM NA Intergenic NA NA

T8654 GGGAATGGTCACGTTGGCCATGG 1000
Oligo fwd CACCGGGAATGGTCACGTTGGCCA
Oligo rev AAACTGGCCAACGTGACCATTCCC
Chromosome start end strand MM target_seq PAM alignment distance position gene name gene id
GU305914.1 116553 116575 + 0 GGGAATGGTCACGTTGGCCA TGG ||||||||[||||||||||||]PAM NA Intergenic NA NA

T8655 GATGCAGATGTTGAGCGTCCTGG 1000
Oligo fwd CACCGATGCAGATGTTGAGCGTCC
Oligo rev AAACGGACGCTCAACATCTGCATC
Chromosome start end strand MM target_seq PAM alignment distance position gene name gene id
GU305914.1 116580 116602 + 0 GATGCAGATGTTGAGCGTCC TGG ||||||||[||||||||||||]PAM NA Intergenic NA NA

T8656 GCGTCCTGGTTTCGATCCATCGG 1000
Oligo fwd CACCGCGTCCTGGTTTCGATCCAT
Oligo rev AAACATGGATCGAAACCAGGACGC
Chromosome start end strand MM target_seq PAM alignment distance position gene name gene id
GU305914.1 116594 116616 + 0 GCGTCCTGGTTTCGATCCAT CGG ||||||||[||||||||||||]PAM NA Intergenic NA NA

T8657 TTTCGATCCATCGGCATACGTGG 1000
Oligo fwd CACCTTTCGATCCATCGGCATACG
Oligo rev AAACCGTATGCCGATGGATCGAAA
Chromosome start end strand MM target_seq PAM alignment distance position gene name gene id
GU305914.1 116603 116625 + 0 TTTCGATCCATCGGCATACG TGG ||||||||[||||||||||||]PAM NA Intergenic NA NA

T8658 CATACGTGGATGTCGAGCTGAGG 1000
Oligo fwd CACCCATACGTGGATGTCGAGCTG
Oligo rev AAACCAGCTCGACATCCACGTATG
Chromosome start end strand MM target_seq PAM alignment distance position gene name gene id
GU305914.1 116617 116639 + 0 CATACGTGGATGTCGAGCTG AGG ||||||||[||||||||||||]PAM NA Intergenic NA NA

T8659 ATACGTGGATGTCGAGCTGAGGG 1000
Oligo fwd CACCATACGTGGATGTCGAGCTGA
Oligo rev AAACTCAGCTCGACATCCACGTAT
Chromosome start end strand MM target_seq PAM alignment distance position gene name gene id
GU305914.1 116618 116640 + 0 ATACGTGGATGTCGAGCTGA GGG ||||||||[||||||||||||]PAM NA Intergenic NA NA

T8660 TACGTGGATGTCGAGCTGAGGGG 1000
Oligo fwd CACCTACGTGGATGTCGAGCTGAG
Oligo rev AAACCTCAGCTCGACATCCACGTA
Chromosome start end strand MM target_seq PAM alignment distance position gene name gene id
GU305914.1 116619 116641 + 0 TACGTGGATGTCGAGCTGAG GGG ||||||||[||||||||||||]PAM NA Intergenic NA NA

T8661 GTCGAGCTGAGGGGAGTACTTGG 1000
Oligo fwd CACCGTCGAGCTGAGGGGAGTACT
Oligo rev AAACAGTACTCCCCTCAGCTCGAC
Chromosome start end strand MM target_seq PAM alignment distance position gene name gene id
GU305914.1 116628 116650 + 0 GTCGAGCTGAGGGGAGTACT TGG ||||||||[||||||||||||]PAM NA Intergenic NA NA

T8662 TCGAGCTGAGGGGAGTACTTGGG 1000
Oligo fwd CACCTCGAGCTGAGGGGAGTACTT
Oligo rev AAACAAGTACTCCCCTCAGCTCGA

| Chromosome | start | end | strand | MM | target_seq | PAM | alignment | distance | position | gene name | gene id |
|---|---|---|---|---|---|---|---|---|---|---|---|
| GU305914.1 | 116629 | 116651 | + | 0 | TCGAGCTGAGGGGAGTACTT | GGG | ||||||||[||||||||||||]PAM | NA | Intergenic | NA | NA |

T8663 CGAGCTGAGGGGAGTACTTGGGG 1000
Oligo fwd CACCCGAGCTGAGGGGAGTACTTG
Oligo rev AAACCAAGTACTCCCCTCAGCTCG

| Chromosome | start | end | strand | MM | target_seq | PAM | alignment | distance | position | gene name | gene id |
|---|---|---|---|---|---|---|---|---|---|---|---|
| GU305914.1 | 116630 | 116652 | + | 0 | CGAGCTGAGGGGAGTACTTG | GGG | ||||||||[||||||||||||]PAM | NA | Intergenic | NA | NA |

T8664 GGGAGTACTTGGGGAAGAACAGG 1000
Oligo fwd CACCGGGAGTACTTGGGGAAGAAC
Oligo rev AAACGTTCTTCCCCAAGTACTCCC

| Chromosome | start | end | strand | MM | target_seq | PAM | alignment | distance | position | gene name | gene id |
|---|---|---|---|---|---|---|---|---|---|---|---|
| GU305914.1 | 116639 | 116661 | + | 0 | GGGAGTACTTGGGGAAGAAC | AGG | ||||||||[||||||||||||]PAM | NA | Intergenic | NA | NA |

T8665 GGAGTACTTGGGGAAGAACAGGG 1000
Oligo fwd CACCGGAGTACTTGGGGAAGAACA
Oligo rev AAACTGTTCTTCCCCAAGTACTCC

| Chromosome | start | end | strand | MM | target_seq | PAM | alignment | distance | position | gene name | gene id |
|---|---|---|---|---|---|---|---|---|---|---|---|
| GU305914.1 | 116640 | 116662 | + | 0 | GGAGTACTTGGGGAAGAACA | GGG | ||||||||[||||||||||||]PAM | NA | Intergenic | NA | NA |

T8666 GTACTTGGGGAAGAACAGGGCGG 1000
Oligo fwd CACCGTACTTGGGGAAGAACAGGG
Oligo rev AAACCCCTGTTCTTCCCCAAGTAC

| Chromosome | start | end | strand | MM | target_seq | PAM | alignment | distance | position | gene name | gene id |
|---|---|---|---|---|---|---|---|---|---|---|---|
| GU305914.1 | 116643 | 116665 | + | 0 | GTACTTGGGGAAGAACAGGG | CGG | ||||||||[||||||||||||]PAM | NA | Intergenic | NA | NA |
| GU305914.1 | 97499 | 97521 | + | 4 | GTACGCGCAGAAGAACAGGG | TGG | ||||--|-[-|||||||||||]PAM | NA | Intergenic | NA | NA |

T8667 GCACGAATAGCCGTTGTCGATGG 1000
Oligo fwd CACCGCACGAATAGCCGTTGTCGA
Oligo rev AAACTCGACAACGGCTATTCGTGC

| Chromosome | start | end | strand | MM | target_seq | PAM | alignment | distance | position | gene name | gene id |
|---|---|---|---|---|---|---|---|---|---|---|---|
| GU305914.1 | 116679 | 116701 | + | 0 | GCACGAATAGCCGTTGTCGA | TGG | ||||||||[||||||||||||]PAM | NA | Intergenic | NA | NA |

T8668 GTTGTCGATGGTGATCGTCCAGG 1000
Oligo fwd CACCGTTGTCGATGGTGATCGTCC
Oligo rev AAACGGACGATCACCATCGACAAC

| Chromosome | start | end | strand | MM | target_seq | PAM | alignment | distance | position | gene name | gene id |
|---|---|---|---|---|---|---|---|---|---|---|---|
| GU305914.1 | 116691 | 116713 | + | 0 | GTTGTCGATGGTGATCGTCC | AGG | ||||||||[||||||||||||]PAM | NA | Intergenic | NA | NA |

T8669 CGATGGTGATCGTCCAGGTCAGG 1000
Oligo fwd CACCCGATGGTGATCGTCCAGGTC
Oligo rev AAACGACCTGGACGATCACCATCG

| Chromosome | start | end | strand | MM | target_seq | PAM | alignment | distance | position | gene name | gene id |
|---|---|---|---|---|---|---|---|---|---|---|---|
| GU305914.1 | 116696 | 116718 | + | 0 | CGATGGTGATCGTCCAGGTC | AGG | ||||||||[||||||||||||]PAM | NA | Intergenic | NA | NA |

T8670 TTCCTGCTCTCGATCTTTCTCGG 1000
Oligo fwd CACCTTCCTGCTCTCGATCTTTCT
Oligo rev AAACAGAAAGATCGAGAGCAGGAA

| Chromosome | start | end | strand | MM | target_seq | PAM | alignment | distance | position | gene name | gene id |
|---|---|---|---|---|---|---|---|---|---|---|---|
| GU305914.1 | 116722 | 116744 | + | 0 | TTCCTGCTCTCGATCTTTCT | CGG | ||||||||[||||||||||||]PAM | NA | Intergenic | NA | NA |

T8671 TCCTGCTCTCGATCTTTCTCGGG 1000
Oligo fwd CACCTCCTGCTCTCGATCTTTCTC
Oligo rev AAACGAGAAAGATCGAGAGCAGGA

| Chromosome | start | end | strand | MM | target_seq | PAM | alignment | distance | position | gene name | gene id |
|---|---|---|---|---|---|---|---|---|---|---|---|
| GU305914.1 | 116723 | 116745 | + | 0 | TCCTGCTCTCGATCTTTCTC | GGG | ||||||||[||||||||||||]PAM | NA | Intergenic | NA | NA |

T8672 GAAGCGCTGCAACGACTTGTAGG 1000
Oligo fwd CACCGAAGCGCTGCAACGACTTGT
Oligo rev AAACACAAGTCGTTGCAGCGCTTC

| Chromosome | start | end | strand | MM | target_seq | PAM | alignment | distance | position | gene name | gene id |
|---|---|---|---|---|---|---|---|---|---|---|---|
| GU305914.1 | 116778 | 116800 | + | 0 | GAAGCGCTGCAACGACTTGT | AGG | ||||||||[||||||||||||]PAM | NA | Intergenic | NA | NA |

T8673 AAGCGCTGCAACGACTTGTAGGG 1000
Oligo fwd CACCAAGCGCTGCAACGACTTGTA
Oligo rev AAACTACAAGTCGTTGCAGCGCTT

| Chromosome | start | end | strand | MM | target_seq | PAM | alignment | distance | position | gene name | gene id |
|---|---|---|---|---|---|---|---|---|---|---|---|
| GU305914.1 | 116779 | 116801 | + | 0 | AAGCGCTGCAACGACTTGTA | GGG | ||||||||[||||||||||||]PAM | NA | Intergenic | NA | NA |

T8674 AGCGCTGCAACGACTTGTAGGGG 1000
Oligo fwd CACCAGCGCTGCAACGACTTGTAG
Oligo rev AAACCTACAAGTCGTTGCAGCGCT

| Chromosome | start | end | strand | MM | target_seq | PAM | alignment | distance | position | gene name | gene id |
|---|---|---|---|---|---|---|---|---|---|---|---|
| GU305914.1 | 116780 | 116802 | + | 0 | AGCGCTGCAACGACTTGTAG | GGG | ||||||||[||||||||||||]PAM | NA | Intergenic | NA | NA |

T8675 GCGCTGCAACGACTTGTAGGGGG 1000
Oligo fwd CACCGCGCTGCAACGACTTGTAGG
Oligo rev AAACCCTACAAGTCGTTGCAGCGC

| Chromosome | start | end | strand | MM | target_seq | PAM | alignment | distance | position | gene name | gene id |
|---|---|---|---|---|---|---|---|---|---|---|---|
| GU305914.1 | 116781 | 116803 | + | 0 | GCGCTGCAACGACTTGTAGG | GGG | ||||||||[||||||||||||]PAM | NA | Intergenic | NA | NA |

T8676 ACTTGTAGGGGGAATGTCGCAGG 1000
Oligo fwd CACCACTTGTAGGGGGAATGTCGC
Oligo rev AAACGCGACATTCCCCCTACAAGT

<table>
<tr><td>Chromosome</td><td>start</td><td>end</td><td>strand</td><td>MM</td><td>target_seq</td><td>PAM</td><td>alignment</td><td>distance</td><td>position</td><td>gene name</td><td>gene id</td></tr>
<tr><td>GU305914.1</td><td>116792</td><td>116814</td><td>+</td><td>0</td><td>ACTTGTAGGGGGAATGTCGC</td><td>AGG</td><td>||||||||[||||||||||||]PAM</td><td>NA</td><td>Intergenic</td><td>NA</td><td>NA</td></tr>
<tr><td></td><td></td><td></td><td></td><td></td><td></td><td></td><td></td><td></td><td></td><td></td><td></td></tr>
<tr><td>T8677</td><td>CTTGTAGGGGGAATGTCGCAGGG</td><td>1000</td><td></td><td></td><td></td><td></td><td></td><td></td><td></td><td></td><td></td></tr>
<tr><td>Oligo fwd</td><td>CACCCTTGTAGGGGGAATGTCGCA</td><td></td><td></td><td></td><td></td><td></td><td></td><td></td><td></td><td></td><td></td></tr>
<tr><td>Oligo rev</td><td>AAACTGCGACATTCCCCCTACAAG</td><td></td><td></td><td></td><td></td><td></td><td></td><td></td><td></td><td></td><td></td></tr>
<tr><td>Chromosome</td><td>start</td><td>end</td><td>strand</td><td>MM</td><td>target_seq</td><td>PAM</td><td>alignment</td><td>distance</td><td>position</td><td>gene name</td><td>gene id</td></tr>
<tr><td>GU305914.1</td><td>116793</td><td>116815</td><td>+</td><td>0</td><td>CTTGTAGGGGGAATGTCGCA</td><td>GGG</td><td>||||||||[||||||||||||]PAM</td><td>NA</td><td>Intergenic</td><td>NA</td><td>NA</td></tr>
<tr><td></td><td></td><td></td><td></td><td></td><td></td><td></td><td></td><td></td><td></td><td></td><td></td></tr>
<tr><td>T8678</td><td>CGAACGTGATCGCGTCTTTGTGG</td><td>1000</td><td></td><td></td><td></td><td></td><td></td><td></td><td></td><td></td><td></td></tr>
<tr><td>Oligo fwd</td><td>CACCCGAACGTGATCGCGTCTTTG</td><td></td><td></td><td></td><td></td><td></td><td></td><td></td><td></td><td></td><td></td></tr>
<tr><td>Oligo rev</td><td>AAACCAAAGACGCGATCACGTTCG</td><td></td><td></td><td></td><td></td><td></td><td></td><td></td><td></td><td></td><td></td></tr>
<tr><td>Chromosome</td><td>start</td><td>end</td><td>strand</td><td>MM</td><td>target_seq</td><td>PAM</td><td>alignment</td><td>distance</td><td>position</td><td>gene name</td><td>gene id</td></tr>
<tr><td>GU305914.1</td><td>116816</td><td>116838</td><td>+</td><td>0</td><td>CGAACGTGATCGCGTCTTTG</td><td>TGG</td><td>||||||||[||||||||||||]PAM</td><td>NA</td><td>Intergenic</td><td>NA</td><td>NA</td></tr>
<tr><td></td><td></td><td></td><td></td><td></td><td></td><td></td><td></td><td></td><td></td><td></td><td></td></tr>
<tr><td>T8679</td><td>ACGTGATCGCGTCTTTGTGGAGG</td><td>1000</td><td></td><td></td><td></td><td></td><td></td><td></td><td></td><td></td><td></td></tr>
<tr><td>Oligo fwd</td><td>CACCACGTGATCGCGTCTTTGTGG</td><td></td><td></td><td></td><td></td><td></td><td></td><td></td><td></td><td></td><td></td></tr>
<tr><td>Oligo rev</td><td>AAACCCACAAAGACGCGATCACGT</td><td></td><td></td><td></td><td></td><td></td><td></td><td></td><td></td><td></td><td></td></tr>
<tr><td>Chromosome</td><td>start</td><td>end</td><td>strand</td><td>MM</td><td>target_seq</td><td>PAM</td><td>alignment</td><td>distance</td><td>position</td><td>gene name</td><td>gene id</td></tr>
<tr><td>GU305914.1</td><td>116819</td><td>116841</td><td>+</td><td>0</td><td>ACGTGATCGCGTCTTTGTGG</td><td>AGG</td><td>||||||||[||||||||||||]PAM</td><td>NA</td><td>Intergenic</td><td>NA</td><td>NA</td></tr>
<tr><td></td><td></td><td></td><td></td><td></td><td></td><td></td><td></td><td></td><td></td><td></td><td></td></tr>
<tr><td>T8680</td><td>TGATCGCGTCTTTGTGGAGGAGG</td><td>1000</td><td></td><td></td><td></td><td></td><td></td><td></td><td></td><td></td><td></td></tr>
<tr><td>Oligo fwd</td><td>CACCTGATCGCGTCTTTGTGGAGG</td><td></td><td></td><td></td><td></td><td></td><td></td><td></td><td></td><td></td><td></td></tr>
<tr><td>Oligo rev</td><td>AAACCCTCCACAAAGACGCGATCA</td><td></td><td></td><td></td><td></td><td></td><td></td><td></td><td></td><td></td><td></td></tr>
<tr><td>Chromosome</td><td>start</td><td>end</td><td>strand</td><td>MM</td><td>target_seq</td><td>PAM</td><td>alignment</td><td>distance</td><td>position</td><td>gene name</td><td>gene id</td></tr>
<tr><td>GU305914.1</td><td>116822</td><td>116844</td><td>+</td><td>0</td><td>TGATCGCGTCTTTGTGGAGG</td><td>AGG</td><td>||||||||[||||||||||||]PAM</td><td>NA</td><td>Intergenic</td><td>NA</td><td>NA</td></tr>
<tr><td></td><td></td><td></td><td></td><td></td><td></td><td></td><td></td><td></td><td></td><td></td><td></td></tr>
<tr><td>T8681</td><td>GAGGATGTGCACCGTCAGCGAGG</td><td>1000</td><td></td><td></td><td></td><td></td><td></td><td></td><td></td><td></td><td></td></tr>
<tr><td>Oligo fwd</td><td>CACCGAGGATGTGCACCGTCAGCG</td><td></td><td></td><td></td><td></td><td></td><td></td><td></td><td></td><td></td><td></td></tr>
<tr><td>Oligo rev</td><td>AAACCGCTGACGGTGCACATCCTC</td><td></td><td></td><td></td><td></td><td></td><td></td><td></td><td></td><td></td><td></td></tr>
<tr><td>Chromosome</td><td>start</td><td>end</td><td>strand</td><td>MM</td><td>target_seq</td><td>PAM</td><td>alignment</td><td>distance</td><td>position</td><td>gene name</td><td>gene id</td></tr>
<tr><td>GU305914.1</td><td>116841</td><td>116863</td><td>+</td><td>0</td><td>GAGGATGTGCACCGTCAGCG</td><td>AGG</td><td>||||||||[||||||||||||]PAM</td><td>NA</td><td>Intergenic</td><td>NA</td><td>NA</td></tr>
<tr><td></td><td></td><td></td><td></td><td></td><td></td><td></td><td></td><td></td><td></td><td></td><td></td></tr>
<tr><td>T8682</td><td>ATGTGCACCGTCAGCGAGGTCGG</td><td>1000</td><td></td><td></td><td></td><td></td><td></td><td></td><td></td><td></td><td></td></tr>
<tr><td>Oligo fwd</td><td>CACCATGTGCACCGTCAGCGAGGT</td><td></td><td></td><td></td><td></td><td></td><td></td><td></td><td></td><td></td><td></td></tr>
<tr><td>Oligo rev</td><td>AAACACCTCGCTGACGGTGCACAT</td><td></td><td></td><td></td><td></td><td></td><td></td><td></td><td></td><td></td><td></td></tr>
<tr><td>Chromosome</td><td>start</td><td>end</td><td>strand</td><td>MM</td><td>target_seq</td><td>PAM</td><td>alignment</td><td>distance</td><td>position</td><td>gene name</td><td>gene id</td></tr>
<tr><td>GU305914.1</td><td>116845</td><td>116867</td><td>+</td><td>0</td><td>ATGTGCACCGTCAGCGAGGT</td><td>CGG</td><td>||||||||[||||||||||||]PAM</td><td>NA</td><td>Intergenic</td><td>NA</td><td>NA</td></tr>
<tr><td></td><td></td><td></td><td></td><td></td><td></td><td></td><td></td><td></td><td></td><td></td><td></td></tr>
<tr><td>T8683</td><td>TGCACCGTCAGCGAGGTCGGCGG</td><td>1000</td><td></td><td></td><td></td><td></td><td></td><td></td><td></td><td></td><td></td></tr>
<tr><td>Oligo fwd</td><td>CACCTGCACCGTCAGCGAGGTCGG</td><td></td><td></td><td></td><td></td><td></td><td></td><td></td><td></td><td></td><td></td></tr>
<tr><td>Oligo rev</td><td>AAACCCGACCTCGCTGACGGTGCA</td><td></td><td></td><td></td><td></td><td></td><td></td><td></td><td></td><td></td><td></td></tr>
<tr><td>Chromosome</td><td>start</td><td>end</td><td>strand</td><td>MM</td><td>target_seq</td><td>PAM</td><td>alignment</td><td>distance</td><td>position</td><td>gene name</td><td>gene id</td></tr>
<tr><td>GU305914.1</td><td>116848</td><td>116870</td><td>+</td><td>0</td><td>TGCACCGTCAGCGAGGTCGG</td><td>CGG</td><td>||||||||[||||||||||||]PAM</td><td>NA</td><td>Intergenic</td><td>NA</td><td>NA</td></tr>
<tr><td>GU305914.1</td><td>128803</td><td>128825</td><td>+</td><td>4</td><td>CGCAGCGGCAGCGACGTCGG</td><td>CGG</td><td>-|||-||-[||||||-|||||]PAM</td><td>NA</td><td>Intergenic</td><td>NA</td><td>NA</td></tr>
<tr><td></td><td></td><td></td><td></td><td></td><td></td><td></td><td></td><td></td><td></td><td></td><td></td></tr>
<tr><td>T8684</td><td>CGTGCCCGTCGAACACCAGCTGG</td><td>1000</td><td></td><td></td><td></td><td></td><td></td><td></td><td></td><td></td><td></td></tr>
<tr><td>Oligo fwd</td><td>CACCCGTGCCCGTCGAACACCAGC</td><td></td><td></td><td></td><td></td><td></td><td></td><td></td><td></td><td></td><td></td></tr>
<tr><td>Oligo rev</td><td>AAACGCTGGTGTTCGACGGGCACG</td><td></td><td></td><td></td><td></td><td></td><td></td><td></td><td></td><td></td><td></td></tr>
<tr><td>Chromosome</td><td>start</td><td>end</td><td>strand</td><td>MM</td><td>target_seq</td><td>PAM</td><td>alignment</td><td>distance</td><td>position</td><td>gene name</td><td>gene id</td></tr>
<tr><td>GU305914.1</td><td>116900</td><td>116922</td><td>+</td><td>0</td><td>CGTGCCCGTCGAACACCAGC</td><td>TGG</td><td>||||||||[||||||||||||]PAM</td><td>NA</td><td>Intergenic</td><td>NA</td><td>NA</td></tr>
<tr><td></td><td></td><td></td><td></td><td></td><td></td><td></td><td></td><td></td><td></td><td></td><td></td></tr>
<tr><td>T8685</td><td>CCAGCTGGATCCTCAGCTTGTGG</td><td>1000</td><td></td><td></td><td></td><td></td><td></td><td></td><td></td><td></td><td></td></tr>
<tr><td>Oligo fwd</td><td>CACCCCAGCTGGATCCTCAGCTTG</td><td></td><td></td><td></td><td></td><td></td><td></td><td></td><td></td><td></td><td></td></tr>
<tr><td>Oligo rev</td><td>AAACCAAGCTGAGGATCCAGCTGG</td><td></td><td></td><td></td><td></td><td></td><td></td><td></td><td></td><td></td><td></td></tr>
<tr><td>Chromosome</td><td>start</td><td>end</td><td>strand</td><td>MM</td><td>target_seq</td><td>PAM</td><td>alignment</td><td>distance</td><td>position</td><td>gene name</td><td>gene id</td></tr>
<tr><td>GU305914.1</td><td>116915</td><td>116937</td><td>+</td><td>0</td><td>CCAGCTGGATCCTCAGCTTG</td><td>TGG</td><td>||||||||[||||||||||||]PAM</td><td>NA</td><td>Intergenic</td><td>NA</td><td>NA</td></tr>
<tr><td></td><td></td><td></td><td></td><td></td><td></td><td></td><td></td><td></td><td></td><td></td><td></td></tr>
<tr><td>T8686</td><td>CTTGTGGTGCGCCGTCTCGCTGG</td><td>1000</td><td></td><td></td><td></td><td></td><td></td><td></td><td></td><td></td><td></td></tr>
<tr><td>Oligo fwd</td><td>CACCCTTGTGGTGCGCCGTCTCGC</td><td></td><td></td><td></td><td></td><td></td><td></td><td></td><td></td><td></td><td></td></tr>
<tr><td>Oligo rev</td><td>AAACGCGAGACGGCGCACCACAAG</td><td></td><td></td><td></td><td></td><td></td><td></td><td></td><td></td><td></td><td></td></tr>
<tr><td>Chromosome</td><td>start</td><td>end</td><td>strand</td><td>MM</td><td>target_seq</td><td>PAM</td><td>alignment</td><td>distance</td><td>position</td><td>gene name</td><td>gene id</td></tr>
<tr><td>GU305914.1</td><td>116931</td><td>116953</td><td>+</td><td>0</td><td>CTTGTGGTGCGCCGTCTCGC</td><td>TGG</td><td>||||||||[||||||||||||]PAM</td><td>NA</td><td>Intergenic</td><td>NA</td><td>NA</td></tr>
<tr><td></td><td></td><td></td><td></td><td></td><td></td><td></td><td></td><td></td><td></td><td></td><td></td></tr>
<tr><td>T8687</td><td>TTGTGGTGCGCCGTCTCGCTGGG</td><td>1000</td><td></td><td></td><td></td><td></td><td></td><td></td><td></td><td></td><td></td></tr>
<tr><td>Oligo fwd</td><td>CACCTTGTGGTGCGCCGTCTCGCT</td><td></td><td></td><td></td><td></td><td></td><td></td><td></td><td></td><td></td><td></td></tr>
<tr><td>Oligo rev</td><td>AAACAGCGAGACGGCGCACCACAA</td><td></td><td></td><td></td><td></td><td></td><td></td><td></td><td></td><td></td><td></td></tr>
<tr><td>Chromosome</td><td>start</td><td>end</td><td>strand</td><td>MM</td><td>target_seq</td><td>PAM</td><td>alignment</td><td>distance</td><td>position</td><td>gene name</td><td>gene id</td></tr>
<tr><td>GU305914.1</td><td>116932</td><td>116954</td><td>+</td><td>0</td><td>TTGTGGTGCGCCGTCTCGCT</td><td>GGG</td><td>||||||||[||||||||||||]PAM</td><td>NA</td><td>Intergenic</td><td>NA</td><td>NA</td></tr>
<tr><td></td><td></td><td></td><td></td><td></td><td></td><td></td><td></td><td></td><td></td><td></td><td></td></tr>
<tr><td>T8688</td><td>TGCGCCGTCTCGCTGGGTCGCGG</td><td>1000</td><td></td><td></td><td></td><td></td><td></td><td></td><td></td><td></td><td></td></tr>
<tr><td>Oligo fwd</td><td>CACCTGCGCCGTCTCGCTGGGTCG</td><td></td><td></td><td></td><td></td><td></td><td></td><td></td><td></td><td></td><td></td></tr>
<tr><td>Oligo rev</td><td>AAACCGACCCAGCGAGACGGCGCA</td><td></td><td></td><td></td><td></td><td></td><td></td><td></td><td></td><td></td><td></td></tr>
<tr><td>Chromosome</td><td>start</td><td>end</td><td>strand</td><td>MM</td><td>target_seq</td><td>PAM</td><td>alignment</td><td>distance</td><td>position</td><td>gene name</td><td>gene id</td></tr>
<tr><td>GU305914.1</td><td>116938</td><td>116960</td><td>+</td><td>0</td><td>TGCGCCGTCTCGCTGGGTCG</td><td>CGG</td><td>||||||||[||||||||||||]PAM</td><td>NA</td><td>Intergenic</td><td>NA</td><td>NA</td></tr>
<tr><td></td><td></td><td></td><td></td><td></td><td></td><td></td><td></td><td></td><td></td><td></td><td></td></tr>
<tr><td>T8689</td><td>GCGCCGTCTCGCTGGGTCGCGGG</td><td>1000</td><td></td><td></td><td></td><td></td><td></td><td></td><td></td><td></td><td></td></tr>
<tr><td>Oligo fwd</td><td>CACCGCGCCGTCTCGCTGGGTCGC</td><td></td><td></td><td></td><td></td><td></td><td></td><td></td><td></td><td></td><td></td></tr>
<tr><td>Oligo rev</td><td>AAACGCGACCCAGCGAGACGGCGC</td><td></td><td></td><td></td><td></td><td></td><td></td><td></td><td></td><td></td><td></td></tr>
<tr><td>Chromosome</td><td>start</td><td>end</td><td>strand</td><td>MM</td><td>target_seq</td><td>PAM</td><td>alignment</td><td>distance</td><td>position</td><td>gene name</td><td>gene id</td></tr>
<tr><td>GU305914.1</td><td>116939</td><td>116961</td><td>+</td><td>0</td><td>GCGCCGTCTCGCTGGGTCGC</td><td>GGG</td><td>||||||||[||||||||||||]PAM</td><td>NA</td><td>Intergenic</td><td>NA</td><td>NA</td></tr>
<tr><td></td><td></td><td></td><td></td><td></td><td></td><td></td><td></td><td></td><td></td><td></td><td></td></tr>
<tr><td>T8690</td><td>TCTCGCTGGGTCGCGGGATGAGG</td><td>1000</td><td></td><td></td><td></td><td></td><td></td><td></td><td></td><td></td><td></td></tr>
<tr><td>Oligo fwd</td><td>CACCTCTCGCTGGGTCGCGGGATG</td><td></td><td></td><td></td><td></td><td></td><td></td><td></td><td></td><td></td><td></td></tr>
<tr><td>Oligo rev</td><td>AAACCATCCCGCGACCCAGCGAGA</td><td></td><td></td><td></td><td></td><td></td><td></td><td></td><td></td><td></td><td></td></tr>
<tr><td>Chromosome</td><td>start</td><td>end</td><td>strand</td><td>MM</td><td>target_seq</td><td>PAM</td><td>alignment</td><td>distance</td><td>position</td><td>gene name</td><td>gene id</td></tr>
<tr><td>GU305914.1</td><td>116945</td><td>116967</td><td>+</td><td>0</td><td>TCTCGCTGGGTCGCGGGATG</td><td>AGG</td><td>||||||||[||||||||||||]PAM</td><td>NA</td><td>Intergenic</td><td>NA</td><td>NA</td></tr>
<tr><td></td><td></td><td></td><td></td><td></td><td></td><td></td><td></td><td></td><td></td><td></td><td></td></tr>
<tr><td>T8691</td><td>GCTGGGTCGCGGGATGAGGATGG</td><td>1000</td><td></td><td></td><td></td><td></td><td></td><td></td><td></td><td></td><td></td></tr>
</table>

Oligo fwd CACCGCTGGGTCGCGGGATGAGGA
Oligo rev AAACTCCTCATCCCGCGACCCAGC
Chromosome start end strand MM target_seq PAM alignment distance position gene name gene id
GU305914.1 116949 116971 + 0 GCTGGGTCGCGGGATGAGGA TGG ||||||||[||||||||||||]PAM NA Intergenic NA NA

T8692 GGGTCGCGGGATGAGGATGGAGG 1000
Oligo fwd CACCGGGTCGCGGGATGAGGATGG
Oligo rev AAACCCATCCTCATCCCGCGACCC
Chromosome start end strand MM target_seq PAM alignment distance position gene name gene id
GU305914.1 116952 116974 + 0 GGGTCGCGGGATGAGGATGG AGG ||||||||[||||||||||||]PAM NA Intergenic NA NA

T8693 GGATGAGGATGGAGGAATCGAGG 1000
Oligo fwd CACCGGATGAGGATGGAGGAATCG
Oligo rev AAACCGATTCCTCCATCCTCATCC
Chromosome start end strand MM target_seq PAM alignment distance position gene name gene id
GU305914.1 116960 116982 + 0 GGATGAGGATGGAGGAATCG AGG ||||||||[||||||||||||]PAM NA Intergenic NA NA

T8694 GAGGAATCGAGGCGTACCATCGG 1000
Oligo fwd CACCGAGGAATCGAGGCGTACCAT
Oligo rev AAACATGGTACGCCTCGATTCCTC
Chromosome start end strand MM target_seq PAM alignment distance position gene name gene id
GU305914.1 116971 116993 + 0 GAGGAATCGAGGCGTACCAT CGG ||||||||[||||||||||||]PAM NA Intergenic NA NA

T8695 AGGAATCGAGGCGTACCATCGGG 1000
Oligo fwd CACCAGGAATCGAGGCGTACCATC
Oligo rev AAACGATGGTACGCCTCGATTCCT
Chromosome start end strand MM target_seq PAM alignment distance position gene name gene id
GU305914.1 116972 116994 + 0 AGGAATCGAGGCGTACCATC GGG ||||||||[||||||||||||]PAM NA Intergenic NA NA

T8696 ATCGGGTCCGAGATCGCCTTCGG 1000
Oligo fwd CACCATCGGGTCCGAGATCGCCTT
Oligo rev AAACAAGGCGATCTCGGACCCGAT
Chromosome start end strand MM target_seq PAM alignment distance position gene name gene id
GU305914.1 116989 117011 + 0 ATCGGGTCCGAGATCGCCTT CGG ||||||||[||||||||||||]PAM NA Intergenic NA NA

T8697 CGAGATCGCCTTCGGTTGCGCGG 1000
Oligo fwd CACCCGAGATCGCCTTCGGTTGCG
Oligo rev AAACCGCAACCGAAGGCGATCTCG
Chromosome start end strand MM target_seq PAM alignment distance position gene name gene id
GU305914.1 116997 117019 + 0 CGAGATCGCCTTCGGTTGCG CGG ||||||||[||||||||||||]PAM NA Intergenic NA NA

T8698 TTGCGCGGCCACGTACGTGTTGG 1000
Oligo fwd CACCTTGCGCGGCCACGTACGTGT
Oligo rev AAACACACGTACGTGGCCGCGCAA
Chromosome start end strand MM target_seq PAM alignment distance position gene name gene id
GU305914.1 117012 117034 + 0 TTGCGCGGCCACGTACGTGT TGG ||||||||[||||||||||||]PAM NA Intergenic NA NA

T8699 GCGGCCACGTACGTGTTGGCCGG 1000
Oligo fwd CACCGCGGCCACGTACGTGTTGGC
Oligo rev AAACGCCAACACGTACGTGGCCGC
Chromosome start end strand MM target_seq PAM alignment distance position gene name gene id
GU305914.1 117016 117038 + 0 GCGGCCACGTACGTGTTGGC CGG ||||||||[||||||||||||]PAM NA Intergenic NA NA

T8700 ACGTGTTGGCCGGTACCGACCGG 1000
Oligo fwd CACCACGTGTTGGCCGGTACCGAC
Oligo rev AAACGTCGGTACCGGCCAACACGT
Chromosome start end strand MM target_seq PAM alignment distance position gene name gene id
GU305914.1 117026 117048 + 0 ACGTGTTGGCCGGTACCGAC CGG ||||||||[||||||||||||]PAM NA Intergenic NA NA

T8701 CGTGTTGGCCGGTACCGACCGGG 1000
Oligo fwd CACCCGTGTTGGCCGGTACCGACC
Oligo rev AAACGGTCGGTACCGGCCAACACG
Chromosome start end strand MM target_seq PAM alignment distance position gene name gene id
GU305914.1 117027 117049 + 0 CGTGTTGGCCGGTACCGACC GGG ||||||||[||||||||||||]PAM NA Intergenic NA NA

T8702 GTGTTGGCCGGTACCGACCGGGG 1000
Oligo fwd CACCGTGTTGGCCGGTACCGACCG
Oligo rev AAACCGGTCGGTACCGGCCAACAC
Chromosome start end strand MM target_seq PAM alignment distance position gene name gene id
GU305914.1 117028 117050 + 0 GTGTTGGCCGGTACCGACCG GGG ||||||||[||||||||||||]PAM NA Intergenic NA NA

T8703 CGGTACCGACCGGGGCAGCGTGG 1000
Oligo fwd CACCCGGTACCGACCGGGGCAGCG
Oligo rev AAACCGCTGCCCCGGTCGGTACCG
Chromosome start end strand MM target_seq PAM alignment distance position gene name gene id
GU305914.1 117036 117058 + 0 CGGTACCGACCGGGGCAGCG TGG ||||||||[||||||||||||]PAM NA Intergenic NA NA

T8704 CCGGGGCAGCGTGGCCGTGATGG 1000
Oligo fwd CACCCCGGGGCAGCGTGGCCGTGA
Oligo rev AAACTCACGGCCACGCTGCCCCGG
Chromosome start end strand MM target_seq PAM alignment distance position gene name gene id
GU305914.1 117045 117067 + 0 CCGGGGCAGCGTGGCCGTGA TGG ||||||||[||||||||||||]PAM NA Intergenic NA NA

T8705 GGGCAGCGTGGCCGTGATGGTGG 1000
Oligo fwd CACCGGGCAGCGTGGCCGTGATGG
Oligo rev AAACCCATCACGGCCACGCTGCCC
Chromosome start end strand MM target_seq PAM alignment distance position gene name gene id
GU305914.1 117048 117070 + 0 GGGCAGCGTGGCCGTGATGG TGG ||||||||[||||||||||||]PAM NA Intergenic NA NA

<table>
<tr><td>T8706</td><td>TGGCCGTGATGGTGGTGCAGTGG</td><td>1000</td><td></td><td></td><td></td><td></td><td></td><td></td><td></td><td></td><td></td></tr>
<tr><td>Oligo fwd</td><td>CACCTGGCCGTGATGGTGGTGCAG</td><td></td><td></td><td></td><td></td><td></td><td></td><td></td><td></td><td></td><td></td></tr>
<tr><td>Oligo rev</td><td>AAACCTGCACCACCATCACGGCCA</td><td></td><td></td><td></td><td></td><td></td><td></td><td></td><td></td><td></td><td></td></tr>
<tr><td>Chromosome</td><td>start</td><td>end</td><td>strand</td><td>MM</td><td>target_seq</td><td>PAM</td><td>alignment</td><td>distance</td><td>position</td><td>gene name</td><td>gene id</td></tr>
<tr><td>GU305914.1</td><td>117056</td><td>117078</td><td>+</td><td>0</td><td>TGGCCGTGATGGTGGTGCAG</td><td>TGG</td><td>||||||||[||||||||||||]PAM</td><td>NA</td><td>Intergenic</td><td>NA</td><td>NA</td></tr>
<tr><td>T8707</td><td>GGCCGTGATGGTGGTGCAGTGGG</td><td>1000</td><td></td><td></td><td></td><td></td><td></td><td></td><td></td><td></td><td></td></tr>
<tr><td>Oligo fwd</td><td>CACCGGCCGTGATGGTGGTGCAGT</td><td></td><td></td><td></td><td></td><td></td><td></td><td></td><td></td><td></td><td></td></tr>
<tr><td>Oligo rev</td><td>AAACACTGCACCACCATCACGGCC</td><td></td><td></td><td></td><td></td><td></td><td></td><td></td><td></td><td></td><td></td></tr>
<tr><td>Chromosome</td><td>start</td><td>end</td><td>strand</td><td>MM</td><td>target_seq</td><td>PAM</td><td>alignment</td><td>distance</td><td>position</td><td>gene name</td><td>gene id</td></tr>
<tr><td>GU305914.1</td><td>117057</td><td>117079</td><td>+</td><td>0</td><td>GGCCGTGATGGTGGTGCAGT</td><td>GGG</td><td>||||||||[||||||||||||]PAM</td><td>NA</td><td>Intergenic</td><td>NA</td><td>NA</td></tr>
<tr><td>T8708</td><td>TGGTGCAGTGGGTGCCGAACTGG</td><td>1000</td><td></td><td></td><td></td><td></td><td></td><td></td><td></td><td></td><td></td></tr>
<tr><td>Oligo fwd</td><td>CACCTGGTGCAGTGGGTGCCGAAC</td><td></td><td></td><td></td><td></td><td></td><td></td><td></td><td></td><td></td><td></td></tr>
<tr><td>Oligo rev</td><td>AAACGTTCGGCACCCACTGCACCA</td><td></td><td></td><td></td><td></td><td></td><td></td><td></td><td></td><td></td><td></td></tr>
<tr><td>Chromosome</td><td>start</td><td>end</td><td>strand</td><td>MM</td><td>target_seq</td><td>PAM</td><td>alignment</td><td>distance</td><td>position</td><td>gene name</td><td>gene id</td></tr>
<tr><td>GU305914.1</td><td>117068</td><td>117090</td><td>+</td><td>0</td><td>TGGTGCAGTGGGTGCCGAAC</td><td>TGG</td><td>||||||||[||||||||||||]PAM</td><td>NA</td><td>Intergenic</td><td>NA</td><td>NA</td></tr>
<tr><td>T8709</td><td>TGCTATGCACCCACCGCATGCGG</td><td>1000</td><td></td><td></td><td></td><td></td><td></td><td></td><td></td><td></td><td></td></tr>
<tr><td>Oligo fwd</td><td>CACCTGCTATGCACCCACCGCATG</td><td></td><td></td><td></td><td></td><td></td><td></td><td></td><td></td><td></td><td></td></tr>
<tr><td>Oligo rev</td><td>AAACCATGCGGTGGGTGCATAGCA</td><td></td><td></td><td></td><td></td><td></td><td></td><td></td><td></td><td></td><td></td></tr>
<tr><td>Chromosome</td><td>start</td><td>end</td><td>strand</td><td>MM</td><td>target_seq</td><td>PAM</td><td>alignment</td><td>distance</td><td>position</td><td>gene name</td><td>gene id</td></tr>
<tr><td>GU305914.1</td><td>117098</td><td>117120</td><td>+</td><td>0</td><td>TGCTATGCACCCACCGCATG</td><td>CGG</td><td>||||||||[||||||||||||]PAM</td><td>NA</td><td>Intergenic</td><td>NA</td><td>NA</td></tr>
<tr><td>T8710</td><td>TATGCACCCACCGCATGCGGCGG</td><td>1000</td><td></td><td></td><td></td><td></td><td></td><td></td><td></td><td></td><td></td></tr>
<tr><td>Oligo fwd</td><td>CACCTATGCACCCACCGCATGCGG</td><td></td><td></td><td></td><td></td><td></td><td></td><td></td><td></td><td></td><td></td></tr>
<tr><td>Oligo rev</td><td>AAACCCGCATGCGGTGGGTGCATA</td><td></td><td></td><td></td><td></td><td></td><td></td><td></td><td></td><td></td><td></td></tr>
<tr><td>Chromosome</td><td>start</td><td>end</td><td>strand</td><td>MM</td><td>target_seq</td><td>PAM</td><td>alignment</td><td>distance</td><td>position</td><td>gene name</td><td>gene id</td></tr>
<tr><td>GU305914.1</td><td>117101</td><td>117123</td><td>+</td><td>0</td><td>TATGCACCCACCGCATGCGG</td><td>CGG</td><td>||||||||[||||||||||||]PAM</td><td>NA</td><td>Intergenic</td><td>NA</td><td>NA</td></tr>
<tr><td>T8711</td><td>GCACCCACCGCATGCGGCGGTGG</td><td>1000</td><td></td><td></td><td></td><td></td><td></td><td></td><td></td><td></td><td></td></tr>
<tr><td>Oligo fwd</td><td>CACCGCACCCACCGCATGCGGCGG</td><td></td><td></td><td></td><td></td><td></td><td></td><td></td><td></td><td></td><td></td></tr>
<tr><td>Oligo rev</td><td>AAACCCGCCGCATGCGGTGGGTGC</td><td></td><td></td><td></td><td></td><td></td><td></td><td></td><td></td><td></td><td></td></tr>
<tr><td>Chromosome</td><td>start</td><td>end</td><td>strand</td><td>MM</td><td>target_seq</td><td>PAM</td><td>alignment</td><td>distance</td><td>position</td><td>gene name</td><td>gene id</td></tr>
<tr><td>GU305914.1</td><td>117104</td><td>117126</td><td>+</td><td>0</td><td>GCACCCACCGCATGCGGCGG</td><td>TGG</td><td>||||||||[||||||||||||]PAM</td><td>NA</td><td>Intergenic</td><td>NA</td><td>NA</td></tr>
<tr><td>T8712</td><td>CACCCACCGCATGCGGCGGTGGG</td><td>1000</td><td></td><td></td><td></td><td></td><td></td><td></td><td></td><td></td><td></td></tr>
<tr><td>Oligo fwd</td><td>CACCCACCCACCGCATGCGGCGGT</td><td></td><td></td><td></td><td></td><td></td><td></td><td></td><td></td><td></td><td></td></tr>
<tr><td>Oligo rev</td><td>AAACACCGCCGCATGCGGTGGGTG</td><td></td><td></td><td></td><td></td><td></td><td></td><td></td><td></td><td></td><td></td></tr>
<tr><td>Chromosome</td><td>start</td><td>end</td><td>strand</td><td>MM</td><td>target_seq</td><td>PAM</td><td>alignment</td><td>distance</td><td>position</td><td>gene name</td><td>gene id</td></tr>
<tr><td>GU305914.1</td><td>117105</td><td>117127</td><td>+</td><td>0</td><td>CACCCACCGCATGCGGCGGT</td><td>GGG</td><td>||||||||[||||||||||||]PAM</td><td>NA</td><td>Intergenic</td><td>NA</td><td>NA</td></tr>
<tr><td>T8713</td><td>GGCGGTGGGTGATCTTAATCAGG</td><td>1000</td><td></td><td></td><td></td><td></td><td></td><td></td><td></td><td></td><td></td></tr>
<tr><td>Oligo fwd</td><td>CACCGGCGGTGGGTGATCTTAATC</td><td></td><td></td><td></td><td></td><td></td><td></td><td></td><td></td><td></td><td></td></tr>
<tr><td>Oligo rev</td><td>AAACGATTAAGATCACCCACCGCC</td><td></td><td></td><td></td><td></td><td></td><td></td><td></td><td></td><td></td><td></td></tr>
<tr><td>Chromosome</td><td>start</td><td>end</td><td>strand</td><td>MM</td><td>target_seq</td><td>PAM</td><td>alignment</td><td>distance</td><td>position</td><td>gene name</td><td>gene id</td></tr>
<tr><td>GU305914.1</td><td>117119</td><td>117141</td><td>+</td><td>0</td><td>GGCGGTGGGTGATCTTAATC</td><td>AGG</td><td>||||||||[||||||||||||]PAM</td><td>NA</td><td>Intergenic</td><td>NA</td><td>NA</td></tr>
<tr><td>T8714</td><td>GGTGGGTGATCTTAATCAGGTGG</td><td>1000</td><td></td><td></td><td></td><td></td><td></td><td></td><td></td><td></td><td></td></tr>
<tr><td>Oligo fwd</td><td>CACCGGTGGGTGATCTTAATCAGG</td><td></td><td></td><td></td><td></td><td></td><td></td><td></td><td></td><td></td><td></td></tr>
<tr><td>Oligo rev</td><td>AAACCCTGATTAAGATCACCCACC</td><td></td><td></td><td></td><td></td><td></td><td></td><td></td><td></td><td></td><td></td></tr>
<tr><td>Chromosome</td><td>start</td><td>end</td><td>strand</td><td>MM</td><td>target_seq</td><td>PAM</td><td>alignment</td><td>distance</td><td>position</td><td>gene name</td><td>gene id</td></tr>
<tr><td>GU305914.1</td><td>117122</td><td>117144</td><td>+</td><td>0</td><td>GGTGGGTGATCTTAATCAGG</td><td>TGG</td><td>||||||||[||||||||||||]PAM</td><td>NA</td><td>Intergenic</td><td>NA</td><td>NA</td></tr>
<tr><td>T8715</td><td>GTGGCTGTCGTTCAGTTCTTCGG</td><td>1000</td><td></td><td></td><td></td><td></td><td></td><td></td><td></td><td></td><td></td></tr>
<tr><td>Oligo fwd</td><td>CACCGTGGCTGTCGTTCAGTTCTT</td><td></td><td></td><td></td><td></td><td></td><td></td><td></td><td></td><td></td><td></td></tr>
<tr><td>Oligo rev</td><td>AAACAAGAACTGAACGACAGCCAC</td><td></td><td></td><td></td><td></td><td></td><td></td><td></td><td></td><td></td><td></td></tr>
<tr><td>Chromosome</td><td>start</td><td>end</td><td>strand</td><td>MM</td><td>target_seq</td><td>PAM</td><td>alignment</td><td>distance</td><td>position</td><td>gene name</td><td>gene id</td></tr>
<tr><td>GU305914.1</td><td>117141</td><td>117163</td><td>+</td><td>0</td><td>GTGGCTGTCGTTCAGTTCTT</td><td>CGG</td><td>||||||||[||||||||||||]PAM</td><td>NA</td><td>Intergenic</td><td>NA</td><td>NA</td></tr>
<tr><td>T8716</td><td>GCTGTCGTTCAGTTCTTCGGCGG</td><td>1000</td><td></td><td></td><td></td><td></td><td></td><td></td><td></td><td></td><td></td></tr>
<tr><td>Oligo fwd</td><td>CACCGCTGTCGTTCAGTTCTTCGG</td><td></td><td></td><td></td><td></td><td></td><td></td><td></td><td></td><td></td><td></td></tr>
<tr><td>Oligo rev</td><td>AAACCCGAAGAACTGAACGACAGC</td><td></td><td></td><td></td><td></td><td></td><td></td><td></td><td></td><td></td><td></td></tr>
<tr><td>Chromosome</td><td>start</td><td>end</td><td>strand</td><td>MM</td><td>target_seq</td><td>PAM</td><td>alignment</td><td>distance</td><td>position</td><td>gene name</td><td>gene id</td></tr>
<tr><td>GU305914.1</td><td>117144</td><td>117166</td><td>+</td><td>0</td><td>GCTGTCGTTCAGTTCTTCGG</td><td>CGG</td><td>||||||||[||||||||||||]PAM</td><td>NA</td><td>Intergenic</td><td>NA</td><td>NA</td></tr>
<tr><td>T8717</td><td>GTCGTTCAGTTCTTCGGCGGTGG</td><td>1000</td><td></td><td></td><td></td><td></td><td></td><td></td><td></td><td></td><td></td></tr>
<tr><td>Oligo fwd</td><td>CACCGTCGTTCAGTTCTTCGGCGG</td><td></td><td></td><td></td><td></td><td></td><td></td><td></td><td></td><td></td><td></td></tr>
<tr><td>Oligo rev</td><td>AAACCCGCCGAAGAACTGAACGAC</td><td></td><td></td><td></td><td></td><td></td><td></td><td></td><td></td><td></td><td></td></tr>
<tr><td>Chromosome</td><td>start</td><td>end</td><td>strand</td><td>MM</td><td>target_seq</td><td>PAM</td><td>alignment</td><td>distance</td><td>position</td><td>gene name</td><td>gene id</td></tr>
<tr><td>GU305914.1</td><td>117147</td><td>117169</td><td>+</td><td>0</td><td>GTCGTTCAGTTCTTCGGCGG</td><td>TGG</td><td>||||||||[||||||||||||]PAM</td><td>NA</td><td>Intergenic</td><td>NA</td><td>NA</td></tr>
<tr><td>T8718</td><td>TCAGTTCTTCGGCGGTGGTCTGG</td><td>1000</td><td></td><td></td><td></td><td></td><td></td><td></td><td></td><td></td><td></td></tr>
<tr><td>Oligo fwd</td><td>CACCTCAGTTCTTCGGCGGTGGTC</td><td></td><td></td><td></td><td></td><td></td><td></td><td></td><td></td><td></td><td></td></tr>
<tr><td>Oligo rev</td><td>AAACGACCACCGCCGAAGAACTGA</td><td></td><td></td><td></td><td></td><td></td><td></td><td></td><td></td><td></td><td></td></tr>
<tr><td>Chromosome</td><td>start</td><td>end</td><td>strand</td><td>MM</td><td>target_seq</td><td>PAM</td><td>alignment</td><td>distance</td><td>position</td><td>gene name</td><td>gene id</td></tr>
<tr><td>GU305914.1</td><td>117152</td><td>117174</td><td>+</td><td>0</td><td>TCAGTTCTTCGGCGGTGGTC</td><td>TGG</td><td>||||||||[||||||||||||]PAM</td><td>NA</td><td>Intergenic</td><td>NA</td><td>NA</td></tr>
<tr><td>T8719</td><td>GTTCTTCGGCGGTGGTCTGGTGG</td><td>1000</td><td></td><td></td><td></td><td></td><td></td><td></td><td></td><td></td><td></td></tr>
<tr><td>Oligo fwd</td><td>CACCGTTCTTCGGCGGTGGTCTGG</td><td></td><td></td><td></td><td></td><td></td><td></td><td></td><td></td><td></td><td></td></tr>
<tr><td>Oligo rev</td><td>AAACCCAGACCACCGCCGAAGAAC</td><td></td><td></td><td></td><td></td><td></td><td></td><td></td><td></td><td></td><td></td></tr>
<tr><td>Chromosome</td><td>start</td><td>end</td><td>strand</td><td>MM</td><td>target_seq</td><td>PAM</td><td>alignment</td><td>distance</td><td>position</td><td>gene name</td><td>gene id</td></tr>
<tr><td>GU305914.1</td><td>117155</td><td>117177</td><td>+</td><td>0</td><td>GTTCTTCGGCGGTGGTCTGG</td><td>TGG</td><td>||||||||[||||||||||||]PAM</td><td>NA</td><td>Intergenic</td><td>NA</td><td>NA</td></tr>
<tr><td>T8720</td><td>TTCTTCGGCGGTGGTCTGGTGGG</td><td>1000</td><td></td><td></td><td></td><td></td><td></td><td></td><td></td><td></td><td></td></tr>
<tr><td>Oligo fwd</td><td>CACCTTCTTCGGCGGTGGTCTGGT</td><td></td><td></td><td></td><td></td><td></td><td></td><td></td><td></td><td></td><td></td></tr>
<tr><td>Oligo rev</td><td>AAACACCAGACCACCGCCGAAGAA</td><td></td><td></td><td></td><td></td><td></td><td></td><td></td><td></td><td></td><td></td></tr>
<tr><td>Chromosome</td><td>start</td><td>end</td><td>strand</td><td>MM</td><td>target_seq</td><td>PAM</td><td>alignment</td><td>distance</td><td>position</td><td>gene name</td><td>gene id</td></tr>
<tr><td>GU305914.1</td><td>117156</td><td>117178</td><td>+</td><td>0</td><td>TTCTTCGGCGGTGGTCTGGT</td><td>GGG</td><td>||||||||[||||||||||||]PAM</td><td>NA</td><td>Intergenic</td><td>NA</td><td>NA</td></tr>
</table>

<table>
<tr><td>T8721</td><td>CTCGAAATCCGTTTTGAAGTTGG</td><td></td><td>1000</td><td></td><td></td><td></td><td></td><td></td><td></td><td></td><td></td><td></td></tr>
<tr><td>Oligo fwd</td><td>CACCCTCGAAATCCGTTTTGAAGT</td><td></td><td></td><td></td><td></td><td></td><td></td><td></td><td></td><td></td><td></td><td></td></tr>
<tr><td>Oligo rev</td><td>AAACACTTCAAAACGGATTTCGAG</td><td></td><td></td><td></td><td></td><td></td><td></td><td></td><td></td><td></td><td></td><td></td></tr>
<tr><td>Chromosome</td><td>start</td><td></td><td>end</td><td>strand</td><td>MM</td><td>target_seq</td><td>PAM</td><td>alignment</td><td>distance</td><td>position</td><td>gene name</td><td>gene id</td></tr>
<tr><td>GU305914.1</td><td></td><td>117186</td><td>117208</td><td>+</td><td>0</td><td>CTCGAAATCCGTTTTGAAGT</td><td>TGG</td><td>||||||||[||||||||||||]PAM</td><td>NA</td><td>Intergenic</td><td>NA</td><td>NA</td></tr>
<tr><td>T8722</td><td>TCGAAATCCGTTTTGAAGTTGGG</td><td></td><td>1000</td><td></td><td></td><td></td><td></td><td></td><td></td><td></td><td></td><td></td></tr>
<tr><td>Oligo fwd</td><td>CACCTCGAAATCCGTTTTGAAGTT</td><td></td><td></td><td></td><td></td><td></td><td></td><td></td><td></td><td></td><td></td><td></td></tr>
<tr><td>Oligo rev</td><td>AAACAACTTCAAAACGGATTTCGA</td><td></td><td></td><td></td><td></td><td></td><td></td><td></td><td></td><td></td><td></td><td></td></tr>
<tr><td>Chromosome</td><td>start</td><td></td><td>end</td><td>strand</td><td>MM</td><td>target_seq</td><td>PAM</td><td>alignment</td><td>distance</td><td>position</td><td>gene name</td><td>gene id</td></tr>
<tr><td>GU305914.1</td><td></td><td>117187</td><td>117209</td><td>+</td><td>0</td><td>TCGAAATCCGTTTTGAAGTT</td><td>GGG</td><td>||||||||[||||||||||||]PAM</td><td>NA</td><td>Intergenic</td><td>NA</td><td>NA</td></tr>
<tr><td>GU305914.1</td><td></td><td>105068</td><td>105090</td><td>+</td><td>4</td><td>TCGAAAAACGTTTTGATGTC</td><td>CGG</td><td>||||||--[||||||||-||-]PAM</td><td>NA</td><td>Intergenic</td><td>NA</td><td>NA</td></tr>
<tr><td>T8723</td><td>CGAAATCCGTTTTGAAGTTGGGG</td><td></td><td>1000</td><td></td><td></td><td></td><td></td><td></td><td></td><td></td><td></td><td></td></tr>
<tr><td>Oligo fwd</td><td>CACCCGAAATCCGTTTTGAAGTTG</td><td></td><td></td><td></td><td></td><td></td><td></td><td></td><td></td><td></td><td></td><td></td></tr>
<tr><td>Oligo rev</td><td>AAACCAACTTCAAAACGGATTTCG</td><td></td><td></td><td></td><td></td><td></td><td></td><td></td><td></td><td></td><td></td><td></td></tr>
<tr><td>Chromosome</td><td>start</td><td></td><td>end</td><td>strand</td><td>MM</td><td>target_seq</td><td>PAM</td><td>alignment</td><td>distance</td><td>position</td><td>gene name</td><td>gene id</td></tr>
<tr><td>GU305914.1</td><td></td><td>117188</td><td>117210</td><td>+</td><td>0</td><td>CGAAATCCGTTTTGAAGTTG</td><td>GGG</td><td>||||||||[||||||||||||]PAM</td><td>NA</td><td>Intergenic</td><td>NA</td><td>NA</td></tr>
<tr><td>T8724</td><td>CCGTTTTGAAGTTGGGGTGCTGG</td><td></td><td>1000</td><td></td><td></td><td></td><td></td><td></td><td></td><td></td><td></td><td></td></tr>
<tr><td>Oligo fwd</td><td>CACCCCGTTTTGAAGTTGGGGTGC</td><td></td><td></td><td></td><td></td><td></td><td></td><td></td><td></td><td></td><td></td><td></td></tr>
<tr><td>Oligo rev</td><td>AAACGCACCCCAACTTCAAAACGG</td><td></td><td></td><td></td><td></td><td></td><td></td><td></td><td></td><td></td><td></td><td></td></tr>
<tr><td>Chromosome</td><td>start</td><td></td><td>end</td><td>strand</td><td>MM</td><td>target_seq</td><td>PAM</td><td>alignment</td><td>distance</td><td>position</td><td>gene name</td><td>gene id</td></tr>
<tr><td>GU305914.1</td><td></td><td>117194</td><td>117216</td><td>+</td><td>0</td><td>CCGTTTTGAAGTTGGGGTGC</td><td>TGG</td><td>||||||||[||||||||||||]PAM</td><td>NA</td><td>Intergenic</td><td>NA</td><td>NA</td></tr>
<tr><td>T8725</td><td>TTTTGAAGTTGGGGTGCTGGCGG</td><td></td><td>1000</td><td></td><td></td><td></td><td></td><td></td><td></td><td></td><td></td><td></td></tr>
<tr><td>Oligo fwd</td><td>CACCTTTTGAAGTTGGGGTGCTGG</td><td></td><td></td><td></td><td></td><td></td><td></td><td></td><td></td><td></td><td></td><td></td></tr>
<tr><td>Oligo rev</td><td>AAACCCAGCACCCCAACTTCAAAA</td><td></td><td></td><td></td><td></td><td></td><td></td><td></td><td></td><td></td><td></td><td></td></tr>
<tr><td>Chromosome</td><td>start</td><td></td><td>end</td><td>strand</td><td>MM</td><td>target_seq</td><td>PAM</td><td>alignment</td><td>distance</td><td>position</td><td>gene name</td><td>gene id</td></tr>
<tr><td>GU305914.1</td><td></td><td>117197</td><td>117219</td><td>+</td><td>0</td><td>TTTTGAAGTTGGGGTGCTGG</td><td>CGG</td><td>||||||||[||||||||||||]PAM</td><td>NA</td><td>Intergenic</td><td>NA</td><td>NA</td></tr>
<tr><td>T8726</td><td>TTGGGGTGCTGGCGGAACAGCGG</td><td></td><td>1000</td><td></td><td></td><td></td><td></td><td></td><td></td><td></td><td></td><td></td></tr>
<tr><td>Oligo fwd</td><td>CACCTTGGGGTGCTGGCGGAACAG</td><td></td><td></td><td></td><td></td><td></td><td></td><td></td><td></td><td></td><td></td><td></td></tr>
<tr><td>Oligo rev</td><td>AAACCTGTTCCGCCAGCACCCCAA</td><td></td><td></td><td></td><td></td><td></td><td></td><td></td><td></td><td></td><td></td><td></td></tr>
<tr><td>Chromosome</td><td>start</td><td></td><td>end</td><td>strand</td><td>MM</td><td>target_seq</td><td>PAM</td><td>alignment</td><td>distance</td><td>position</td><td>gene name</td><td>gene id</td></tr>
<tr><td>GU305914.1</td><td></td><td>117205</td><td>117227</td><td>+</td><td>0</td><td>TTGGGGTGCTGGCGGAACAG</td><td>CGG</td><td>||||||||[||||||||||||]PAM</td><td>NA</td><td>Intergenic</td><td>NA</td><td>NA</td></tr>
<tr><td>T8727</td><td>AACAGCGGATATTCGTTGATCGG</td><td></td><td>1000</td><td></td><td></td><td></td><td></td><td></td><td></td><td></td><td></td><td></td></tr>
<tr><td>Oligo fwd</td><td>CACCAACAGCGGATATTCGTTGAT</td><td></td><td></td><td></td><td></td><td></td><td></td><td></td><td></td><td></td><td></td><td></td></tr>
<tr><td>Oligo rev</td><td>AAACATCAACGAATATCCGCTGTT</td><td></td><td></td><td></td><td></td><td></td><td></td><td></td><td></td><td></td><td></td><td></td></tr>
<tr><td>Chromosome</td><td>start</td><td></td><td>end</td><td>strand</td><td>MM</td><td>target_seq</td><td>PAM</td><td>alignment</td><td>distance</td><td>position</td><td>gene name</td><td>gene id</td></tr>
<tr><td>GU305914.1</td><td></td><td>117220</td><td>117242</td><td>+</td><td>0</td><td>AACAGCGGATATTCGTTGAT</td><td>CGG</td><td>||||||||[||||||||||||]PAM</td><td>NA</td><td>Intergenic</td><td>NA</td><td>NA</td></tr>
<tr><td>T8728</td><td>TATTCGTTGATCGGCACGATCGG</td><td></td><td>1000</td><td></td><td></td><td></td><td></td><td></td><td></td><td></td><td></td><td></td></tr>
<tr><td>Oligo fwd</td><td>CACCTATTCGTTGATCGGCACGAT</td><td></td><td></td><td></td><td></td><td></td><td></td><td></td><td></td><td></td><td></td><td></td></tr>
<tr><td>Oligo rev</td><td>AAACATCGTGCCGATCAACGAATA</td><td></td><td></td><td></td><td></td><td></td><td></td><td></td><td></td><td></td><td></td><td></td></tr>
<tr><td>Chromosome</td><td>start</td><td></td><td>end</td><td>strand</td><td>MM</td><td>target_seq</td><td>PAM</td><td>alignment</td><td>distance</td><td>position</td><td>gene name</td><td>gene id</td></tr>
<tr><td>GU305914.1</td><td></td><td>117229</td><td>117251</td><td>+</td><td>0</td><td>TATTCGTTGATCGGCACGAT</td><td>CGG</td><td>||||||||[||||||||||||]PAM</td><td>NA</td><td>Intergenic</td><td>NA</td><td>NA</td></tr>
<tr><td>T8729</td><td>CGTTGATCGGCACGATCGGCAGG</td><td></td><td>1000</td><td></td><td></td><td></td><td></td><td></td><td></td><td></td><td></td><td></td></tr>
<tr><td>Oligo fwd</td><td>CACCCGTTGATCGGCACGATCGGC</td><td></td><td></td><td></td><td></td><td></td><td></td><td></td><td></td><td></td><td></td><td></td></tr>
<tr><td>Oligo rev</td><td>AAACGCCGATCGTGCCGATCAACG</td><td></td><td></td><td></td><td></td><td></td><td></td><td></td><td></td><td></td><td></td><td></td></tr>
<tr><td>Chromosome</td><td>start</td><td></td><td>end</td><td>strand</td><td>MM</td><td>target_seq</td><td>PAM</td><td>alignment</td><td>distance</td><td>position</td><td>gene name</td><td>gene id</td></tr>
<tr><td>GU305914.1</td><td></td><td>117233</td><td>117255</td><td>+</td><td>0</td><td>CGTTGATCGGCACGATCGGC</td><td>AGG</td><td>||||||||[||||||||||||]PAM</td><td>NA</td><td>Intergenic</td><td>NA</td><td>NA</td></tr>
<tr><td>T8730</td><td>GTTGATCGGCACGATCGGCAGGG</td><td></td><td>1000</td><td></td><td></td><td></td><td></td><td></td><td></td><td></td><td></td><td></td></tr>
<tr><td>Oligo fwd</td><td>CACCGTTGATCGGCACGATCGGCA</td><td></td><td></td><td></td><td></td><td></td><td></td><td></td><td></td><td></td><td></td><td></td></tr>
<tr><td>Oligo rev</td><td>AAACTGCCGATCGTGCCGATCAAC</td><td></td><td></td><td></td><td></td><td></td><td></td><td></td><td></td><td></td><td></td><td></td></tr>
<tr><td>Chromosome</td><td>start</td><td></td><td>end</td><td>strand</td><td>MM</td><td>target_seq</td><td>PAM</td><td>alignment</td><td>distance</td><td>position</td><td>gene name</td><td>gene id</td></tr>
<tr><td>GU305914.1</td><td></td><td>117234</td><td>117256</td><td>+</td><td>0</td><td>GTTGATCGGCACGATCGGCA</td><td>GGG</td><td>||||||||[||||||||||||]PAM</td><td>NA</td><td>Intergenic</td><td>NA</td><td>NA</td></tr>
<tr><td>T8731</td><td>TTGATCGGCACGATCGGCAGGGG</td><td></td><td>1000</td><td></td><td></td><td></td><td></td><td></td><td></td><td></td><td></td><td></td></tr>
<tr><td>Oligo fwd</td><td>CACCTTGATCGGCACGATCGGCAG</td><td></td><td></td><td></td><td></td><td></td><td></td><td></td><td></td><td></td><td></td><td></td></tr>
<tr><td>Oligo rev</td><td>AAACCTGCCGATCGTGCCGATCAA</td><td></td><td></td><td></td><td></td><td></td><td></td><td></td><td></td><td></td><td></td><td></td></tr>
<tr><td>Chromosome</td><td>start</td><td></td><td>end</td><td>strand</td><td>MM</td><td>target_seq</td><td>PAM</td><td>alignment</td><td>distance</td><td>position</td><td>gene name</td><td>gene id</td></tr>
<tr><td>GU305914.1</td><td></td><td>117235</td><td>117257</td><td>+</td><td>0</td><td>TTGATCGGCACGATCGGCAG</td><td>GGG</td><td>||||||||[||||||||||||]PAM</td><td>NA</td><td>Intergenic</td><td>NA</td><td>NA</td></tr>
<tr><td>T8732</td><td>TGATCGGCACGATCGGCAGGGGG</td><td></td><td>1000</td><td></td><td></td><td></td><td></td><td></td><td></td><td></td><td></td><td></td></tr>
<tr><td>Oligo fwd</td><td>CACCTGATCGGCACGATCGGCAGG</td><td></td><td></td><td></td><td></td><td></td><td></td><td></td><td></td><td></td><td></td><td></td></tr>
<tr><td>Oligo rev</td><td>AAACCCTGCCGATCGTGCCGATCA</td><td></td><td></td><td></td><td></td><td></td><td></td><td></td><td></td><td></td><td></td><td></td></tr>
<tr><td>Chromosome</td><td>start</td><td></td><td>end</td><td>strand</td><td>MM</td><td>target_seq</td><td>PAM</td><td>alignment</td><td>distance</td><td>position</td><td>gene name</td><td>gene id</td></tr>
<tr><td>GU305914.1</td><td></td><td>117236</td><td>117258</td><td>+</td><td>0</td><td>TGATCGGCACGATCGGCAGG</td><td>GGG</td><td>||||||||[||||||||||||]PAM</td><td>NA</td><td>Intergenic</td><td>NA</td><td>NA</td></tr>
<tr><td>T8733</td><td>GAAGATCTGATCTTGCGTCTCGG</td><td></td><td>1000</td><td></td><td></td><td></td><td></td><td></td><td></td><td></td><td></td><td></td></tr>
<tr><td>Oligo fwd</td><td>CACCGAAGATCTGATCTTGCGTCT</td><td></td><td></td><td></td><td></td><td></td><td></td><td></td><td></td><td></td><td></td><td></td></tr>
<tr><td>Oligo rev</td><td>AAACAGACGCAAGATCAGATCTTC</td><td></td><td></td><td></td><td></td><td></td><td></td><td></td><td></td><td></td><td></td><td></td></tr>
<tr><td>Chromosome</td><td>start</td><td></td><td>end</td><td>strand</td><td>MM</td><td>target_seq</td><td>PAM</td><td>alignment</td><td>distance</td><td>position</td><td>gene name</td><td>gene id</td></tr>
<tr><td>GU305914.1</td><td></td><td>117276</td><td>117298</td><td>+</td><td>0</td><td>GAAGATCTGATCTTGCGTCT</td><td>CGG</td><td>||||||||[||||||||||||]PAM</td><td>NA</td><td>Intergenic</td><td>NA</td><td>NA</td></tr>
<tr><td>T8734</td><td>CTCGGCCGAGTTTCTCGTGAAGG</td><td></td><td>1000</td><td></td><td></td><td></td><td></td><td></td><td></td><td></td><td></td><td></td></tr>
<tr><td>Oligo fwd</td><td>CACCCTCGGCCGAGTTTCTCGTGA</td><td></td><td></td><td></td><td></td><td></td><td></td><td></td><td></td><td></td><td></td><td></td></tr>
<tr><td>Oligo rev</td><td>AAACTCACGAGAAACTCGGCCGAG</td><td></td><td></td><td></td><td></td><td></td><td></td><td></td><td></td><td></td><td></td><td></td></tr>
<tr><td>Chromosome</td><td>start</td><td></td><td>end</td><td>strand</td><td>MM</td><td>target_seq</td><td>PAM</td><td>alignment</td><td>distance</td><td>position</td><td>gene name</td><td>gene id</td></tr>
<tr><td>GU305914.1</td><td></td><td>117294</td><td>117316</td><td>+</td><td>0</td><td>CTCGGCCGAGTTTCTCGTGA</td><td>AGG</td><td>||||||||[||||||||||||]PAM</td><td>NA</td><td>Intergenic</td><td>NA</td><td>NA</td></tr>
<tr><td>T8735</td><td>AAGGTTGACATGTTTCTCACAGG</td><td></td><td>1000</td><td></td><td></td><td></td><td></td><td></td><td></td><td></td><td></td><td></td></tr>
<tr><td>Oligo fwd</td><td>CACCAAGGTTGACATGTTTCTCAC</td><td></td><td></td><td></td><td></td><td></td><td></td><td></td><td></td><td></td><td></td><td></td></tr>
<tr><td>Oligo rev</td><td>AAACGTGAGAAACATGTCAACCTT</td><td></td><td></td><td></td><td></td><td></td><td></td><td></td><td></td><td></td><td></td><td></td></tr>
</table>

<table>
<tr><td>Chromosome</td><td>start</td><td>end</td><td>strand</td><td>MM</td><td>target_seq</td><td>PAM</td><td>alignment</td><td>distance</td><td>position</td><td>gene name</td><td>gene id</td></tr>
<tr><td>GU305914.1</td><td>117313</td><td>117335</td><td>+</td><td>0</td><td>AAGGTTGACATGTTTCTCAC</td><td>AGG</td><td>||||||||[||||||||||||]PAM</td><td>NA</td><td>Intergenic</td><td>NA</td><td>NA</td></tr>
<tr><td>T8736</td><td>AACCACGTTGATGAAGTCCTCGG</td><td>1000</td><td></td><td></td><td></td><td></td><td></td><td></td><td></td><td></td><td></td></tr>
<tr><td>Oligo fwd</td><td>CACCAACCACGTTGATGAAGTCCT</td><td></td><td></td><td></td><td></td><td></td><td></td><td></td><td></td><td></td><td></td></tr>
<tr><td>Oligo rev</td><td>AAACAGGACTTCATCAACGTGGTT</td><td></td><td></td><td></td><td></td><td></td><td></td><td></td><td></td><td></td><td></td></tr>
<tr><td>Chromosome</td><td>start</td><td>end</td><td>strand</td><td>MM</td><td>target_seq</td><td>PAM</td><td>alignment</td><td>distance</td><td>position</td><td>gene name</td><td>gene id</td></tr>
<tr><td>GU305914.1</td><td>117339</td><td>117361</td><td>+</td><td>0</td><td>AACCACGTTGATGAAGTCCT</td><td>CGG</td><td>||||||||[||||||||||||]PAM</td><td>NA</td><td>Intergenic</td><td>NA</td><td>NA</td></tr>
<tr><td>T8737</td><td>ACCACGTTGATGAAGTCCTCGGG</td><td>1000</td><td></td><td></td><td></td><td></td><td></td><td></td><td></td><td></td><td></td></tr>
<tr><td>Oligo fwd</td><td>CACCACCACGTTGATGAAGTCCTC</td><td></td><td></td><td></td><td></td><td></td><td></td><td></td><td></td><td></td><td></td></tr>
<tr><td>Oligo rev</td><td>AAACGAGGACTTCATCAACGTGGT</td><td></td><td></td><td></td><td></td><td></td><td></td><td></td><td></td><td></td><td></td></tr>
<tr><td>Chromosome</td><td>start</td><td>end</td><td>strand</td><td>MM</td><td>target_seq</td><td>PAM</td><td>alignment</td><td>distance</td><td>position</td><td>gene name</td><td>gene id</td></tr>
<tr><td>GU305914.1</td><td>117340</td><td>117362</td><td>+</td><td>0</td><td>ACCACGTTGATGAAGTCCTC</td><td>GGG</td><td>||||||||[||||||||||||]PAM</td><td>NA</td><td>Intergenic</td><td>NA</td><td>NA</td></tr>
<tr><td>T8738</td><td>TGAAGTCCTCGGGATCGCTCAGG</td><td>1000</td><td></td><td></td><td></td><td></td><td></td><td></td><td></td><td></td><td></td></tr>
<tr><td>Oligo fwd</td><td>CACCTGAAGTCCTCGGGATCGCTC</td><td></td><td></td><td></td><td></td><td></td><td></td><td></td><td></td><td></td><td></td></tr>
<tr><td>Oligo rev</td><td>AAACGAGCGATCCCGAGGACTTCA</td><td></td><td></td><td></td><td></td><td></td><td></td><td></td><td></td><td></td><td></td></tr>
<tr><td>Chromosome</td><td>start</td><td>end</td><td>strand</td><td>MM</td><td>target_seq</td><td>PAM</td><td>alignment</td><td>distance</td><td>position</td><td>gene name</td><td>gene id</td></tr>
<tr><td>GU305914.1</td><td>117350</td><td>117372</td><td>+</td><td>0</td><td>TGAAGTCCTCGGGATCGCTC</td><td>AGG</td><td>||||||||[||||||||||||]PAM</td><td>NA</td><td>Intergenic</td><td>NA</td><td>NA</td></tr>
<tr><td>T8739</td><td>GTCCTCGGGATCGCTCAGGACGG</td><td>1000</td><td></td><td></td><td></td><td></td><td></td><td></td><td></td><td></td><td></td></tr>
<tr><td>Oligo fwd</td><td>CACCGTCCTCGGGATCGCTCAGGA</td><td></td><td></td><td></td><td></td><td></td><td></td><td></td><td></td><td></td><td></td></tr>
<tr><td>Oligo rev</td><td>AAACTCCTGAGCGATCCCGAGGAC</td><td></td><td></td><td></td><td></td><td></td><td></td><td></td><td></td><td></td><td></td></tr>
<tr><td>Chromosome</td><td>start</td><td>end</td><td>strand</td><td>MM</td><td>target_seq</td><td>PAM</td><td>alignment</td><td>distance</td><td>position</td><td>gene name</td><td>gene id</td></tr>
<tr><td>GU305914.1</td><td>117354</td><td>117376</td><td>+</td><td>0</td><td>GTCCTCGGGATCGCTCAGGA</td><td>CGG</td><td>||||||||[||||||||||||]PAM</td><td>NA</td><td>Intergenic</td><td>NA</td><td>NA</td></tr>
<tr><td>T8740</td><td>GGATCGCTCAGGACGGTGAACGG</td><td>1000</td><td></td><td></td><td></td><td></td><td></td><td></td><td></td><td></td><td></td></tr>
<tr><td>Oligo fwd</td><td>CACCGGATCGCTCAGGACGGTGAA</td><td></td><td></td><td></td><td></td><td></td><td></td><td></td><td></td><td></td><td></td></tr>
<tr><td>Oligo rev</td><td>AAACTTCACCGTCCTGAGCGATCC</td><td></td><td></td><td></td><td></td><td></td><td></td><td></td><td></td><td></td><td></td></tr>
<tr><td>Chromosome</td><td>start</td><td>end</td><td>strand</td><td>MM</td><td>target_seq</td><td>PAM</td><td>alignment</td><td>distance</td><td>position</td><td>gene name</td><td>gene id</td></tr>
<tr><td>GU305914.1</td><td>117361</td><td>117383</td><td>+</td><td>0</td><td>GGATCGCTCAGGACGGTGAA</td><td>CGG</td><td>||||||||[||||||||||||]PAM</td><td>NA</td><td>Intergenic</td><td>NA</td><td>NA</td></tr>
<tr><td>T8741</td><td>AACGGCACCGACAGCGCCGACGG</td><td>1000</td><td></td><td></td><td></td><td></td><td></td><td></td><td></td><td></td><td></td></tr>
<tr><td>Oligo fwd</td><td>CACCAACGGCACCGACAGCGCCGA</td><td></td><td></td><td></td><td></td><td></td><td></td><td></td><td></td><td></td><td></td></tr>
<tr><td>Oligo rev</td><td>AAACTCGGCGCTGTCGGTGCCGTT</td><td></td><td></td><td></td><td></td><td></td><td></td><td></td><td></td><td></td><td></td></tr>
<tr><td>Chromosome</td><td>start</td><td>end</td><td>strand</td><td>MM</td><td>target_seq</td><td>PAM</td><td>alignment</td><td>distance</td><td>position</td><td>gene name</td><td>gene id</td></tr>
<tr><td>GU305914.1</td><td>117379</td><td>117401</td><td>+</td><td>0</td><td>AACGGCACCGACAGCGCCGA</td><td>CGG</td><td>||||||||[||||||||||||]PAM</td><td>NA</td><td>Intergenic</td><td>NA</td><td>NA</td></tr>
<tr><td>GU305914.1</td><td>20741</td><td>20763</td><td>+</td><td>3</td><td>GACGGCACCGACGGCGACGA</td><td>CGG</td><td>-|||||||[||||-|||-|||]PAM</td><td>NA</td><td>Intergenic</td><td>NA</td><td>NA</td></tr>
<tr><td>GU305914.1</td><td>70181</td><td>70203</td><td>+</td><td>4</td><td>AGCAGCACCGGCAGCGCCGG</td><td>CGG</td><td>|-|-||||[||-||||||||-]PAM</td><td>NA</td><td>Intergenic</td><td>NA</td><td>NA</td></tr>
<tr><td>T8742</td><td>ACGGCACCGACAGCGCCGACGGG</td><td>1000</td><td></td><td></td><td></td><td></td><td></td><td></td><td></td><td></td><td></td></tr>
<tr><td>Oligo fwd</td><td>CACCACGGCACCGACAGCGCCGAC</td><td></td><td></td><td></td><td></td><td></td><td></td><td></td><td></td><td></td><td></td></tr>
<tr><td>Oligo rev</td><td>AAACGTCGGCGCTGTCGGTGCCGT</td><td></td><td></td><td></td><td></td><td></td><td></td><td></td><td></td><td></td><td></td></tr>
<tr><td>Chromosome</td><td>start</td><td>end</td><td>strand</td><td>MM</td><td>target_seq</td><td>PAM</td><td>alignment</td><td>distance</td><td>position</td><td>gene name</td><td>gene id</td></tr>
<tr><td>GU305914.1</td><td>117380</td><td>117402</td><td>+</td><td>0</td><td>ACGGCACCGACAGCGCCGAC</td><td>GGG</td><td>||||||||[||||||||||||]PAM</td><td>NA</td><td>Intergenic</td><td>NA</td><td>NA</td></tr>
<tr><td>T8743</td><td>CGGCACCGACAGCGCCGACGGGG</td><td>1000</td><td></td><td></td><td></td><td></td><td></td><td></td><td></td><td></td><td></td></tr>
<tr><td>Oligo fwd</td><td>CACCCGGCACCGACAGCGCCGACG</td><td></td><td></td><td></td><td></td><td></td><td></td><td></td><td></td><td></td><td></td></tr>
<tr><td>Oligo rev</td><td>AAACCGTCGGCGCTGTCGGTGCCG</td><td></td><td></td><td></td><td></td><td></td><td></td><td></td><td></td><td></td><td></td></tr>
<tr><td>Chromosome</td><td>start</td><td>end</td><td>strand</td><td>MM</td><td>target_seq</td><td>PAM</td><td>alignment</td><td>distance</td><td>position</td><td>gene name</td><td>gene id</td></tr>
<tr><td>GU305914.1</td><td>117381</td><td>117403</td><td>+</td><td>0</td><td>CGGCACCGACAGCGCCGACG</td><td>GGG</td><td>||||||||[||||||||||||]PAM</td><td>NA</td><td>Intergenic</td><td>NA</td><td>NA</td></tr>
<tr><td>T8744</td><td>CCGACAGCGCCGACGGGGTCCGG</td><td>1000</td><td></td><td></td><td></td><td></td><td></td><td></td><td></td><td></td><td></td></tr>
<tr><td>Oligo fwd</td><td>CACCCCGACAGCGCCGACGGGGTC</td><td></td><td></td><td></td><td></td><td></td><td></td><td></td><td></td><td></td><td></td></tr>
<tr><td>Oligo rev</td><td>AAACGACCCCGTCGGCGCTGTCGG</td><td></td><td></td><td></td><td></td><td></td><td></td><td></td><td></td><td></td><td></td></tr>
<tr><td>Chromosome</td><td>start</td><td>end</td><td>strand</td><td>MM</td><td>target_seq</td><td>PAM</td><td>alignment</td><td>distance</td><td>position</td><td>gene name</td><td>gene id</td></tr>
<tr><td>GU305914.1</td><td>117386</td><td>117408</td><td>+</td><td>0</td><td>CCGACAGCGCCGACGGGGTC</td><td>CGG</td><td>||||||||[||||||||||||]PAM</td><td>NA</td><td>Intergenic</td><td>NA</td><td>NA</td></tr>
<tr><td>GU305914.1</td><td>113911</td><td>113933</td><td>-</td><td>4</td><td>GCGGCAGCGGCGACGGCGTC</td><td>GGG</td><td>-||-||||[|-||||||-|||]PAM</td><td>NA</td><td>Intergenic</td><td>NA</td><td>NA</td></tr>
<tr><td>T8745</td><td>GCGCCGACGGGGTCCGGCCCTGG</td><td>1000</td><td></td><td></td><td></td><td></td><td></td><td></td><td></td><td></td><td></td></tr>
<tr><td>Oligo fwd</td><td>CACCGCGCCGACGGGGTCCGGCCC</td><td></td><td></td><td></td><td></td><td></td><td></td><td></td><td></td><td></td><td></td></tr>
<tr><td>Oligo rev</td><td>AAACGGGCCGGACCCCGTCGGCGC</td><td></td><td></td><td></td><td></td><td></td><td></td><td></td><td></td><td></td><td></td></tr>
<tr><td>Chromosome</td><td>start</td><td>end</td><td>strand</td><td>MM</td><td>target_seq</td><td>PAM</td><td>alignment</td><td>distance</td><td>position</td><td>gene name</td><td>gene id</td></tr>
<tr><td>GU305914.1</td><td>117392</td><td>117414</td><td>+</td><td>0</td><td>GCGCCGACGGGGTCCGGCCC</td><td>TGG</td><td>||||||||[||||||||||||]PAM</td><td>NA</td><td>Intergenic</td><td>NA</td><td>NA</td></tr>
<tr><td>T8746</td><td>CGCCGACGGGGTCCGGCCCTGGG</td><td>1000</td><td></td><td></td><td></td><td></td><td></td><td></td><td></td><td></td><td></td></tr>
<tr><td>Oligo fwd</td><td>CACCCGCCGACGGGGTCCGGCCCT</td><td></td><td></td><td></td><td></td><td></td><td></td><td></td><td></td><td></td><td></td></tr>
<tr><td>Oligo rev</td><td>AAACAGGGCCGGACCCCGTCGGCG</td><td></td><td></td><td></td><td></td><td></td><td></td><td></td><td></td><td></td><td></td></tr>
<tr><td>Chromosome</td><td>start</td><td>end</td><td>strand</td><td>MM</td><td>target_seq</td><td>PAM</td><td>alignment</td><td>distance</td><td>position</td><td>gene name</td><td>gene id</td></tr>
<tr><td>GU305914.1</td><td>117393</td><td>117415</td><td>+</td><td>0</td><td>CGCCGACGGGGTCCGGCCCT</td><td>GGG</td><td>||||||||[||||||||||||]PAM</td><td>NA</td><td>Intergenic</td><td>NA</td><td>NA</td></tr>
<tr><td>T8747</td><td>GCCGACGGGGTCCGGCCCTGGGG</td><td>1000</td><td></td><td></td><td></td><td></td><td></td><td></td><td></td><td></td><td></td></tr>
<tr><td>Oligo fwd</td><td>CACCGCCGACGGGGTCCGGCCCTG</td><td></td><td></td><td></td><td></td><td></td><td></td><td></td><td></td><td></td><td></td></tr>
<tr><td>Oligo rev</td><td>AAACCAGGGCCGGACCCCGTCGGC</td><td></td><td></td><td></td><td></td><td></td><td></td><td></td><td></td><td></td><td></td></tr>
<tr><td>Chromosome</td><td>start</td><td>end</td><td>strand</td><td>MM</td><td>target_seq</td><td>PAM</td><td>alignment</td><td>distance</td><td>position</td><td>gene name</td><td>gene id</td></tr>
<tr><td>GU305914.1</td><td>117394</td><td>117416</td><td>+</td><td>0</td><td>GCCGACGGGGTCCGGCCCTG</td><td>GGG</td><td>||||||||[||||||||||||]PAM</td><td>NA</td><td>Intergenic</td><td>NA</td><td>NA</td></tr>
<tr><td>T8748</td><td>ACGGGGTCCGGCCCTGGGGCAGG</td><td>1000</td><td></td><td></td><td></td><td></td><td></td><td></td><td></td><td></td><td></td></tr>
<tr><td>Oligo fwd</td><td>CACCACGGGGTCCGGCCCTGGGGC</td><td></td><td></td><td></td><td></td><td></td><td></td><td></td><td></td><td></td><td></td></tr>
<tr><td>Oligo rev</td><td>AAACGCCCCAGGGCCGGACCCCGT</td><td></td><td></td><td></td><td></td><td></td><td></td><td></td><td></td><td></td><td></td></tr>
<tr><td>Chromosome</td><td>start</td><td>end</td><td>strand</td><td>MM</td><td>target_seq</td><td>PAM</td><td>alignment</td><td>distance</td><td>position</td><td>gene name</td><td>gene id</td></tr>
<tr><td>GU305914.1</td><td>117398</td><td>117420</td><td>+</td><td>0</td><td>ACGGGGTCCGGCCCTGGGGC</td><td>AGG</td><td>||||||||[||||||||||||]PAM</td><td>NA</td><td>Intergenic</td><td>NA</td><td>NA</td></tr>
<tr><td>T8749</td><td>CCTGGGGCAGGCGTTGCGCCAGG</td><td>1000</td><td></td><td></td><td></td><td></td><td></td><td></td><td></td><td></td><td></td></tr>
<tr><td>Oligo fwd</td><td>CACCCCTGGGGCAGGCGTTGCGCC</td><td></td><td></td><td></td><td></td><td></td><td></td><td></td><td></td><td></td><td></td></tr>
<tr><td>Oligo rev</td><td>AAACGGCGCAACGCCTGCCCCAGG</td><td></td><td></td><td></td><td></td><td></td><td></td><td></td><td></td><td></td><td></td></tr>
<tr><td>Chromosome</td><td>start</td><td>end</td><td>strand</td><td>MM</td><td>target_seq</td><td>PAM</td><td>alignment</td><td>distance</td><td>position</td><td>gene name</td><td>gene id</td></tr>
<tr><td>GU305914.1</td><td>117410</td><td>117432</td><td>+</td><td>0</td><td>CCTGGGGCAGGCGTTGCGCC</td><td>AGG</td><td>||||||||[||||||||||||]PAM</td><td>NA</td><td>Intergenic</td><td>NA</td><td>NA</td></tr>
</table>

<table>
<tr><td>T8750</td><td>GGGCAGGCGTTGCGCCAGGCAGG</td><td>1000</td><td></td><td></td><td></td><td></td><td></td><td></td><td></td><td></td><td></td></tr>
<tr><td>Oligo fwd</td><td>CACCGGGCAGGCGTTGCGCCAGGC</td><td></td><td></td><td></td><td></td><td></td><td></td><td></td><td></td><td></td><td></td></tr>
<tr><td>Oligo rev</td><td>AAACGCCTGGCGCAACGCCTGCCC</td><td></td><td></td><td></td><td></td><td></td><td></td><td></td><td></td><td></td><td></td></tr>
<tr><td>Chromosome</td><td>start</td><td>end</td><td>strand</td><td>MM</td><td>target_seq</td><td>PAM</td><td>alignment</td><td>distance</td><td>position</td><td>gene name</td><td>gene id</td></tr>
<tr><td>GU305914.1</td><td>117414</td><td>117436</td><td>+</td><td>0</td><td>GGGCAGGCGTTGCGCCAGGC</td><td>AGG</td><td>||||||||[||||||||||||]PAM</td><td>NA</td><td>Intergenic</td><td>NA</td><td>NA</td></tr>
<tr><td></td><td></td><td></td><td></td><td></td><td></td><td></td><td></td><td></td><td></td><td></td><td></td></tr>
<tr><td>T8751</td><td>GCGTTGCGCCAGGCAGGTGATGG</td><td>1000</td><td></td><td></td><td></td><td></td><td></td><td></td><td></td><td></td><td></td></tr>
<tr><td>Oligo fwd</td><td>CACCGCGTTGCGCCAGGCAGGTGA</td><td></td><td></td><td></td><td></td><td></td><td></td><td></td><td></td><td></td><td></td></tr>
<tr><td>Oligo rev</td><td>AAACTCACCTGCCTGGCGCAACGC</td><td></td><td></td><td></td><td></td><td></td><td></td><td></td><td></td><td></td><td></td></tr>
<tr><td>Chromosome</td><td>start</td><td>end</td><td>strand</td><td>MM</td><td>target_seq</td><td>PAM</td><td>alignment</td><td>distance</td><td>position</td><td>gene name</td><td>gene id</td></tr>
<tr><td>GU305914.1</td><td>117420</td><td>117442</td><td>+</td><td>0</td><td>GCGTTGCGCCAGGCAGGTGA</td><td>TGG</td><td>||||||||[||||||||||||]PAM</td><td>NA</td><td>Intergenic</td><td>NA</td><td>NA</td></tr>
<tr><td></td><td></td><td></td><td></td><td></td><td></td><td></td><td></td><td></td><td></td><td></td><td></td></tr>
<tr><td>T8752</td><td>TTGCGCCAGGCAGGTGATGGCGG</td><td>1000</td><td></td><td></td><td></td><td></td><td></td><td></td><td></td><td></td><td></td></tr>
<tr><td>Oligo fwd</td><td>CACCTTGCGCCAGGCAGGTGATGG</td><td></td><td></td><td></td><td></td><td></td><td></td><td></td><td></td><td></td><td></td></tr>
<tr><td>Oligo rev</td><td>AAACCCATCACCTGCCTGGCGCAA</td><td></td><td></td><td></td><td></td><td></td><td></td><td></td><td></td><td></td><td></td></tr>
<tr><td>Chromosome</td><td>start</td><td>end</td><td>strand</td><td>MM</td><td>target_seq</td><td>PAM</td><td>alignment</td><td>distance</td><td>position</td><td>gene name</td><td>gene id</td></tr>
<tr><td>GU305914.1</td><td>117423</td><td>117445</td><td>+</td><td>0</td><td>TTGCGCCAGGCAGGTGATGG</td><td>CGG</td><td>||||||||[||||||||||||]PAM</td><td>NA</td><td>Intergenic</td><td>NA</td><td>NA</td></tr>
<tr><td></td><td></td><td></td><td></td><td></td><td></td><td></td><td></td><td></td><td></td><td></td><td></td></tr>
<tr><td>T8753</td><td>TGCGCCAGGCAGGTGATGGCGGG</td><td>1000</td><td></td><td></td><td></td><td></td><td></td><td></td><td></td><td></td><td></td></tr>
<tr><td>Oligo fwd</td><td>CACCTGCGCCAGGCAGGTGATGGC</td><td></td><td></td><td></td><td></td><td></td><td></td><td></td><td></td><td></td><td></td></tr>
<tr><td>Oligo rev</td><td>AAACGCCATCACCTGCCTGGCGCA</td><td></td><td></td><td></td><td></td><td></td><td></td><td></td><td></td><td></td><td></td></tr>
<tr><td>Chromosome</td><td>start</td><td>end</td><td>strand</td><td>MM</td><td>target_seq</td><td>PAM</td><td>alignment</td><td>distance</td><td>position</td><td>gene name</td><td>gene id</td></tr>
<tr><td>GU305914.1</td><td>117424</td><td>117446</td><td>+</td><td>0</td><td>TGCGCCAGGCAGGTGATGGC</td><td>GGG</td><td>||||||||[||||||||||||]PAM</td><td>NA</td><td>Intergenic</td><td>NA</td><td>NA</td></tr>
<tr><td></td><td></td><td></td><td></td><td></td><td></td><td></td><td></td><td></td><td></td><td></td><td></td></tr>
<tr><td>T8754</td><td>GCGCCAGGCAGGTGATGGCGGGG</td><td>1000</td><td></td><td></td><td></td><td></td><td></td><td></td><td></td><td></td><td></td></tr>
<tr><td>Oligo fwd</td><td>CACCGCGCCAGGCAGGTGATGGCG</td><td></td><td></td><td></td><td></td><td></td><td></td><td></td><td></td><td></td><td></td></tr>
<tr><td>Oligo rev</td><td>AAACCGCCATCACCTGCCTGGCGC</td><td></td><td></td><td></td><td></td><td></td><td></td><td></td><td></td><td></td><td></td></tr>
<tr><td>Chromosome</td><td>start</td><td>end</td><td>strand</td><td>MM</td><td>target_seq</td><td>PAM</td><td>alignment</td><td>distance</td><td>position</td><td>gene name</td><td>gene id</td></tr>
<tr><td>GU305914.1</td><td>117425</td><td>117447</td><td>+</td><td>0</td><td>GCGCCAGGCAGGTGATGGCG</td><td>GGG</td><td>||||||||[||||||||||||]PAM</td><td>NA</td><td>Intergenic</td><td>NA</td><td>NA</td></tr>
<tr><td>GU305914.1</td><td>136441</td><td>136463</td><td>-</td><td>4</td><td>GCGACTGGCTGGTGACGGCG</td><td>CGG</td><td>|||-|-||[|-|||||-||||]PAM</td><td>NA</td><td>Intergenic</td><td>NA</td><td>NA</td></tr>
<tr><td>GU305914.1</td><td>135355</td><td>135377</td><td>-</td><td>4</td><td>GCGCCCGCCCGGTGATGGTG</td><td>AGG</td><td>|||||-|-[|-||||||||-|]PAM</td><td>NA</td><td>Intergenic</td><td>NA</td><td>NA</td></tr>
<tr><td></td><td></td><td></td><td></td><td></td><td></td><td></td><td></td><td></td><td></td><td></td><td></td></tr>
<tr><td>T8755</td><td>CAGGCAGGTGATGGCGGGGACGG</td><td>1000</td><td></td><td></td><td></td><td></td><td></td><td></td><td></td><td></td><td></td></tr>
<tr><td>Oligo fwd</td><td>CACCCAGGCAGGTGATGGCGGGGA</td><td></td><td></td><td></td><td></td><td></td><td></td><td></td><td></td><td></td><td></td></tr>
<tr><td>Oligo rev</td><td>AAACTCCCCGCCATCACCTGCCTG</td><td></td><td></td><td></td><td></td><td></td><td></td><td></td><td></td><td></td><td></td></tr>
<tr><td>Chromosome</td><td>start</td><td>end</td><td>strand</td><td>MM</td><td>target_seq</td><td>PAM</td><td>alignment</td><td>distance</td><td>position</td><td>gene name</td><td>gene id</td></tr>
<tr><td>GU305914.1</td><td>117429</td><td>117451</td><td>+</td><td>0</td><td>CAGGCAGGTGATGGCGGGGA</td><td>CGG</td><td>||||||||[||||||||||||]PAM</td><td>NA</td><td>Intergenic</td><td>NA</td><td>NA</td></tr>
<tr><td></td><td></td><td></td><td></td><td></td><td></td><td></td><td></td><td></td><td></td><td></td><td></td></tr>
<tr><td>T8756</td><td>GGACGTCGCGAGCGACAGACCGG</td><td>1000</td><td></td><td></td><td></td><td></td><td></td><td></td><td></td><td></td><td></td></tr>
<tr><td>Oligo fwd</td><td>CACCGGACGTCGCGAGCGACAGAC</td><td></td><td></td><td></td><td></td><td></td><td></td><td></td><td></td><td></td><td></td></tr>
<tr><td>Oligo rev</td><td>AAACGTCTGTCGCTCGCGACGTCC</td><td></td><td></td><td></td><td></td><td></td><td></td><td></td><td></td><td></td><td></td></tr>
<tr><td>Chromosome</td><td>start</td><td>end</td><td>strand</td><td>MM</td><td>target_seq</td><td>PAM</td><td>alignment</td><td>distance</td><td>position</td><td>gene name</td><td>gene id</td></tr>
<tr><td>GU305914.1</td><td>117450</td><td>117472</td><td>+</td><td>0</td><td>GGACGTCGCGAGCGACAGAC</td><td>CGG</td><td>||||||||[||||||||||||]PAM</td><td>NA</td><td>Intergenic</td><td>NA</td><td>NA</td></tr>
<tr><td></td><td></td><td></td><td></td><td></td><td></td><td></td><td></td><td></td><td></td><td></td><td></td></tr>
<tr><td>T8757</td><td>CGAGCGACAGACCGGTGTTCAGG</td><td>1000</td><td></td><td></td><td></td><td></td><td></td><td></td><td></td><td></td><td></td></tr>
<tr><td>Oligo fwd</td><td>CACCCGAGCGACAGACCGGTGTTC</td><td></td><td></td><td></td><td></td><td></td><td></td><td></td><td></td><td></td><td></td></tr>
<tr><td>Oligo rev</td><td>AAACGAACACCGGTCTGTCGCTCG</td><td></td><td></td><td></td><td></td><td></td><td></td><td></td><td></td><td></td><td></td></tr>
<tr><td>Chromosome</td><td>start</td><td>end</td><td>strand</td><td>MM</td><td>target_seq</td><td>PAM</td><td>alignment</td><td>distance</td><td>position</td><td>gene name</td><td>gene id</td></tr>
<tr><td>GU305914.1</td><td>117458</td><td>117480</td><td>+</td><td>0</td><td>CGAGCGACAGACCGGTGTTC</td><td>AGG</td><td>||||||||[||||||||||||]PAM</td><td>NA</td><td>Intergenic</td><td>NA</td><td>NA</td></tr>
<tr><td></td><td></td><td></td><td></td><td></td><td></td><td></td><td></td><td></td><td></td><td></td><td></td></tr>
<tr><td>T8758</td><td>GTCTCCTGCCCTCTGAAGAGCGG</td><td>1000</td><td></td><td></td><td></td><td></td><td></td><td></td><td></td><td></td><td></td></tr>
<tr><td>Oligo fwd</td><td>CACCGTCTCCTGCCCTCTGAAGAG</td><td></td><td></td><td></td><td></td><td></td><td></td><td></td><td></td><td></td><td></td></tr>
<tr><td>Oligo rev</td><td>AAACCTCTTCAGAGGGCAGGAGAC</td><td></td><td></td><td></td><td></td><td></td><td></td><td></td><td></td><td></td><td></td></tr>
<tr><td>Chromosome</td><td>start</td><td>end</td><td>strand</td><td>MM</td><td>target_seq</td><td>PAM</td><td>alignment</td><td>distance</td><td>position</td><td>gene name</td><td>gene id</td></tr>
<tr><td>GU305914.1</td><td>117487</td><td>117509</td><td>+</td><td>0</td><td>GTCTCCTGCCCTCTGAAGAG</td><td>CGG</td><td>||||||||[||||||||||||]PAM</td><td>NA</td><td>Intergenic</td><td>NA</td><td>NA</td></tr>
<tr><td></td><td></td><td></td><td></td><td></td><td></td><td></td><td></td><td></td><td></td><td></td><td></td></tr>
<tr><td>T8759</td><td>TCTCCTGCCCTCTGAAGAGCGGG</td><td>1000</td><td></td><td></td><td></td><td></td><td></td><td></td><td></td><td></td><td></td></tr>
<tr><td>Oligo fwd</td><td>CACCTCTCCTGCCCTCTGAAGAGC</td><td></td><td></td><td></td><td></td><td></td><td></td><td></td><td></td><td></td><td></td></tr>
<tr><td>Oligo rev</td><td>AAACGCTCTTCAGAGGGCAGGAGA</td><td></td><td></td><td></td><td></td><td></td><td></td><td></td><td></td><td></td><td></td></tr>
<tr><td>Chromosome</td><td>start</td><td>end</td><td>strand</td><td>MM</td><td>target_seq</td><td>PAM</td><td>alignment</td><td>distance</td><td>position</td><td>gene name</td><td>gene id</td></tr>
<tr><td>GU305914.1</td><td>117488</td><td>117510</td><td>+</td><td>0</td><td>TCTCCTGCCCTCTGAAGAGC</td><td>GGG</td><td>||||||||[||||||||||||]PAM</td><td>NA</td><td>Intergenic</td><td>NA</td><td>NA</td></tr>
<tr><td></td><td></td><td></td><td></td><td></td><td></td><td></td><td></td><td></td><td></td><td></td><td></td></tr>
<tr><td>T8760</td><td>CCCTCTGAAGAGCGGGCCCTTGG</td><td>1000</td><td></td><td></td><td></td><td></td><td></td><td></td><td></td><td></td><td></td></tr>
<tr><td>Oligo fwd</td><td>CACCCCCTCTGAAGAGCGGGCCCT</td><td></td><td></td><td></td><td></td><td></td><td></td><td></td><td></td><td></td><td></td></tr>
<tr><td>Oligo rev</td><td>AAACAGGGCCCGCTCTTCAGAGGG</td><td></td><td></td><td></td><td></td><td></td><td></td><td></td><td></td><td></td><td></td></tr>
<tr><td>Chromosome</td><td>start</td><td>end</td><td>strand</td><td>MM</td><td>target_seq</td><td>PAM</td><td>alignment</td><td>distance</td><td>position</td><td>gene name</td><td>gene id</td></tr>
<tr><td>GU305914.1</td><td>117495</td><td>117517</td><td>+</td><td>0</td><td>CCCTCTGAAGAGCGGGCCCT</td><td>TGG</td><td>||||||||[||||||||||||]PAM</td><td>NA</td><td>Intergenic</td><td>NA</td><td>NA</td></tr>
<tr><td></td><td></td><td></td><td></td><td></td><td></td><td></td><td></td><td></td><td></td><td></td><td></td></tr>
<tr><td>T8761</td><td>AGAGCGGGCCCTTGGTCACCCGG</td><td>1000</td><td></td><td></td><td></td><td></td><td></td><td></td><td></td><td></td><td></td></tr>
<tr><td>Oligo fwd</td><td>CACCAGAGCGGGCCCTTGGTCACC</td><td></td><td></td><td></td><td></td><td></td><td></td><td></td><td></td><td></td><td></td></tr>
<tr><td>Oligo rev</td><td>AAACGGTGACCAAGGGCCCGCTCT</td><td></td><td></td><td></td><td></td><td></td><td></td><td></td><td></td><td></td><td></td></tr>
<tr><td>Chromosome</td><td>start</td><td>end</td><td>strand</td><td>MM</td><td>target_seq</td><td>PAM</td><td>alignment</td><td>distance</td><td>position</td><td>gene name</td><td>gene id</td></tr>
<tr><td>GU305914.1</td><td>117503</td><td>117525</td><td>+</td><td>0</td><td>AGAGCGGGCCCTTGGTCACC</td><td>CGG</td><td>||||||||[||||||||||||]PAM</td><td>NA</td><td>Intergenic</td><td>NA</td><td>NA</td></tr>
<tr><td></td><td></td><td></td><td></td><td></td><td></td><td></td><td></td><td></td><td></td><td></td><td></td></tr>
<tr><td>T8762</td><td>GCGGGCCCTTGGTCACCCGGAGG</td><td>1000</td><td></td><td></td><td></td><td></td><td></td><td></td><td></td><td></td><td></td></tr>
<tr><td>Oligo fwd</td><td>CACCGCGGGCCCTTGGTCACCCGG</td><td></td><td></td><td></td><td></td><td></td><td></td><td></td><td></td><td></td><td></td></tr>
<tr><td>Oligo rev</td><td>AAACCCGGGTGACCAAGGGCCCGC</td><td></td><td></td><td></td><td></td><td></td><td></td><td></td><td></td><td></td><td></td></tr>
<tr><td>Chromosome</td><td>start</td><td>end</td><td>strand</td><td>MM</td><td>target_seq</td><td>PAM</td><td>alignment</td><td>distance</td><td>position</td><td>gene name</td><td>gene id</td></tr>
<tr><td>GU305914.1</td><td>117506</td><td>117528</td><td>+</td><td>0</td><td>GCGGGCCCTTGGTCACCCGG</td><td>AGG</td><td>||||||||[||||||||||||]PAM</td><td>NA</td><td>Intergenic</td><td>NA</td><td>NA</td></tr>
<tr><td></td><td></td><td></td><td></td><td></td><td></td><td></td><td></td><td></td><td></td><td></td><td></td></tr>
<tr><td>T8763</td><td>CCTTGGTCACCCGGAGGCTCAGG</td><td>1000</td><td></td><td></td><td></td><td></td><td></td><td></td><td></td><td></td><td></td></tr>
<tr><td>Oligo fwd</td><td>CACCCCTTGGTCACCCGGAGGCTC</td><td></td><td></td><td></td><td></td><td></td><td></td><td></td><td></td><td></td><td></td></tr>
<tr><td>Oligo rev</td><td>AAACGAGCCTCCGGGTGACCAAGG</td><td></td><td></td><td></td><td></td><td></td><td></td><td></td><td></td><td></td><td></td></tr>
<tr><td>Chromosome</td><td>start</td><td>end</td><td>strand</td><td>MM</td><td>target_seq</td><td>PAM</td><td>alignment</td><td>distance</td><td>position</td><td>gene name</td><td>gene id</td></tr>
<tr><td>GU305914.1</td><td>117512</td><td>117534</td><td>+</td><td>0</td><td>CCTTGGTCACCCGGAGGCTC</td><td>AGG</td><td>||||||||[||||||||||||]PAM</td><td>NA</td><td>Intergenic</td><td>NA</td><td>NA</td></tr>
<tr><td></td><td></td><td></td><td></td><td></td><td></td><td></td><td></td><td></td><td></td><td></td><td></td></tr>
<tr><td>T8764</td><td>GCACCATCCGTGAACGAACGTGG</td><td>1000</td><td></td><td></td><td></td><td></td><td></td><td></td><td></td><td></td><td></td></tr>
<tr><td>Oligo fwd</td><td>CACCGCACCATCCGTGAACGAACG</td><td></td><td></td><td></td><td></td><td></td><td></td><td></td><td></td><td></td><td></td></tr>
</table>

<table>
<tr><td>Oligo rev</td><td>AAACCGTTCGTTCACGGATGGTGC</td><td></td><td></td><td></td><td></td><td></td><td></td><td></td><td></td><td></td><td></td></tr>
<tr><td>Chromosome</td><td>start</td><td>end</td><td>strand</td><td>MM</td><td>target_seq</td><td>PAM</td><td>alignment</td><td>distance</td><td>position</td><td>gene name</td><td>gene id</td></tr>
<tr><td>GU305914.1</td><td>117540</td><td>117562</td><td>+</td><td>0</td><td>GCACCATCCGTGAACGAACG</td><td>TGG</td><td>||||||||[||||||||||||]PAM</td><td>NA</td><td>Intergenic</td><td>NA</td><td>NA</td></tr>
<tr><td>T8765</td><td>CCGTGAACGAACGTGGACCCCGG</td><td>1000</td><td></td><td></td><td></td><td></td><td></td><td></td><td></td><td></td><td></td></tr>
<tr><td>Oligo fwd</td><td>CACCCCGTGAACGAACGTGGACCC</td><td></td><td></td><td></td><td></td><td></td><td></td><td></td><td></td><td></td><td></td></tr>
<tr><td>Oligo rev</td><td>AAACGGGTCCACGTTCGTTCACGG</td><td></td><td></td><td></td><td></td><td></td><td></td><td></td><td></td><td></td><td></td></tr>
<tr><td>Chromosome</td><td>start</td><td>end</td><td>strand</td><td>MM</td><td>target_seq</td><td>PAM</td><td>alignment</td><td>distance</td><td>position</td><td>gene name</td><td>gene id</td></tr>
<tr><td>GU305914.1</td><td>117547</td><td>117569</td><td>+</td><td>0</td><td>CCGTGAACGAACGTGGACCC</td><td>CGG</td><td>||||||||[||||||||||||]PAM</td><td>NA</td><td>Intergenic</td><td>NA</td><td>NA</td></tr>
<tr><td>T8766</td><td>GAACGTGGACCCCGGTTTGTCGG</td><td>1000</td><td></td><td></td><td></td><td></td><td></td><td></td><td></td><td></td><td></td></tr>
<tr><td>Oligo fwd</td><td>CACCGAACGTGGACCCCGGTTTGT</td><td></td><td></td><td></td><td></td><td></td><td></td><td></td><td></td><td></td><td></td></tr>
<tr><td>Oligo rev</td><td>AAACACAAACCGGGGTCCACGTTC</td><td></td><td></td><td></td><td></td><td></td><td></td><td></td><td></td><td></td><td></td></tr>
<tr><td>Chromosome</td><td>start</td><td>end</td><td>strand</td><td>MM</td><td>target_seq</td><td>PAM</td><td>alignment</td><td>distance</td><td>position</td><td>gene name</td><td>gene id</td></tr>
<tr><td>GU305914.1</td><td>117555</td><td>117577</td><td>+</td><td>0</td><td>GAACGTGGACCCCGGTTTGT</td><td>CGG</td><td>||||||||[||||||||||||]PAM</td><td>NA</td><td>Intergenic</td><td>NA</td><td>NA</td></tr>
<tr><td>T8767</td><td>AACGTGGACCCCGGTTTGTCGGG</td><td>1000</td><td></td><td></td><td></td><td></td><td></td><td></td><td></td><td></td><td></td></tr>
<tr><td>Oligo fwd</td><td>CACCAACGTGGACCCCGGTTTGTC</td><td></td><td></td><td></td><td></td><td></td><td></td><td></td><td></td><td></td><td></td></tr>
<tr><td>Oligo rev</td><td>AAACGACAAACCGGGGTCCACGTT</td><td></td><td></td><td></td><td></td><td></td><td></td><td></td><td></td><td></td><td></td></tr>
<tr><td>Chromosome</td><td>start</td><td>end</td><td>strand</td><td>MM</td><td>target_seq</td><td>PAM</td><td>alignment</td><td>distance</td><td>position</td><td>gene name</td><td>gene id</td></tr>
<tr><td>GU305914.1</td><td>117556</td><td>117578</td><td>+</td><td>0</td><td>AACGTGGACCCCGGTTTGTC</td><td>GGG</td><td>||||||||[||||||||||||]PAM</td><td>NA</td><td>Intergenic</td><td>NA</td><td>NA</td></tr>
<tr><td>T8768</td><td>ACGTGGACCCCGGTTTGTCGGGG</td><td>1000</td><td></td><td></td><td></td><td></td><td></td><td></td><td></td><td></td><td></td></tr>
<tr><td>Oligo fwd</td><td>CACCACGTGGACCCCGGTTTGTCG</td><td></td><td></td><td></td><td></td><td></td><td></td><td></td><td></td><td></td><td></td></tr>
<tr><td>Oligo rev</td><td>AAACCGACAAACCGGGGTCCACGT</td><td></td><td></td><td></td><td></td><td></td><td></td><td></td><td></td><td></td><td></td></tr>
<tr><td>Chromosome</td><td>start</td><td>end</td><td>strand</td><td>MM</td><td>target_seq</td><td>PAM</td><td>alignment</td><td>distance</td><td>position</td><td>gene name</td><td>gene id</td></tr>
<tr><td>GU305914.1</td><td>117557</td><td>117579</td><td>+</td><td>0</td><td>ACGTGGACCCCGGTTTGTCG</td><td>GGG</td><td>||||||||[||||||||||||]PAM</td><td>NA</td><td>Intergenic</td><td>NA</td><td>NA</td></tr>
<tr><td>T8769</td><td>GTTTGTCGGGGTCCGCCGTCAGG</td><td>1000</td><td></td><td></td><td></td><td></td><td></td><td></td><td></td><td></td><td></td></tr>
<tr><td>Oligo fwd</td><td>CACCGTTTGTCGGGGTCCGCCGTC</td><td></td><td></td><td></td><td></td><td></td><td></td><td></td><td></td><td></td><td></td></tr>
<tr><td>Oligo rev</td><td>AAACGACGGCGGACCCCGACAAAC</td><td></td><td></td><td></td><td></td><td></td><td></td><td></td><td></td><td></td><td></td></tr>
<tr><td>Chromosome</td><td>start</td><td>end</td><td>strand</td><td>MM</td><td>target_seq</td><td>PAM</td><td>alignment</td><td>distance</td><td>position</td><td>gene name</td><td>gene id</td></tr>
<tr><td>GU305914.1</td><td>117569</td><td>117591</td><td>+</td><td>0</td><td>GTTTGTCGGGGTCCGCCGTC</td><td>AGG</td><td>||||||||[||||||||||||]PAM</td><td>NA</td><td>Intergenic</td><td>NA</td><td>NA</td></tr>
<tr><td>GU305914.1</td><td>124889</td><td>124911</td><td>+</td><td>4</td><td>GCGAGTCCGGGTCCGCCGTC</td><td>AGG</td><td>|---|||-[||||||||||||]PAM</td><td>NA</td><td>Intergenic</td><td>NA</td><td>NA</td></tr>
<tr><td>T8770</td><td>CGTCAGGAAGTTAAATTCCTCGG</td><td>1000</td><td></td><td></td><td></td><td></td><td></td><td></td><td></td><td></td><td></td></tr>
<tr><td>Oligo fwd</td><td>CACCCGTCAGGAAGTTAAATTCCT</td><td></td><td></td><td></td><td></td><td></td><td></td><td></td><td></td><td></td><td></td></tr>
<tr><td>Oligo rev</td><td>AAACAGGAATTTAACTTCCTGACG</td><td></td><td></td><td></td><td></td><td></td><td></td><td></td><td></td><td></td><td></td></tr>
<tr><td>Chromosome</td><td>start</td><td>end</td><td>strand</td><td>MM</td><td>target_seq</td><td>PAM</td><td>alignment</td><td>distance</td><td>position</td><td>gene name</td><td>gene id</td></tr>
<tr><td>GU305914.1</td><td>117585</td><td>117607</td><td>+</td><td>0</td><td>CGTCAGGAAGTTAAATTCCT</td><td>CGG</td><td>||||||||[||||||||||||]PAM</td><td>NA</td><td>Intergenic</td><td>NA</td><td>NA</td></tr>
<tr><td>T8771</td><td>TCCTCGGCCATATCGACGCTCGG</td><td>1000</td><td></td><td></td><td></td><td></td><td></td><td></td><td></td><td></td><td></td></tr>
<tr><td>Oligo fwd</td><td>CACCTCCTCGGCCATATCGACGCT</td><td></td><td></td><td></td><td></td><td></td><td></td><td></td><td></td><td></td><td></td></tr>
<tr><td>Oligo rev</td><td>AAACAGCGTCGATATGGCCGAGGA</td><td></td><td></td><td></td><td></td><td></td><td></td><td></td><td></td><td></td><td></td></tr>
<tr><td>Chromosome</td><td>start</td><td>end</td><td>strand</td><td>MM</td><td>target_seq</td><td>PAM</td><td>alignment</td><td>distance</td><td>position</td><td>gene name</td><td>gene id</td></tr>
<tr><td>GU305914.1</td><td>117601</td><td>117623</td><td>+</td><td>0</td><td>TCCTCGGCCATATCGACGCT</td><td>CGG</td><td>||||||||[||||||||||||]PAM</td><td>NA</td><td>Intergenic</td><td>NA</td><td>NA</td></tr>
<tr><td>T8772</td><td>CGACGCTCGGCCGCGCGCGCTGG</td><td>1000</td><td></td><td></td><td></td><td></td><td></td><td></td><td></td><td></td><td></td></tr>
<tr><td>Oligo fwd</td><td>CACCCGACGCTCGGCCGCGCGCGC</td><td></td><td></td><td></td><td></td><td></td><td></td><td></td><td></td><td></td><td></td></tr>
<tr><td>Oligo rev</td><td>AAACGCGCGCGCGGCCGAGCGTCG</td><td></td><td></td><td></td><td></td><td></td><td></td><td></td><td></td><td></td><td></td></tr>
<tr><td>Chromosome</td><td>start</td><td>end</td><td>strand</td><td>MM</td><td>target_seq</td><td>PAM</td><td>alignment</td><td>distance</td><td>position</td><td>gene name</td><td>gene id</td></tr>
<tr><td>GU305914.1</td><td>117614</td><td>117636</td><td>+</td><td>0</td><td>CGACGCTCGGCCGCGCGCGC</td><td>TGG</td><td>||||||||[||||||||||||]PAM</td><td>NA</td><td>Intergenic</td><td>NA</td><td>NA</td></tr>
<tr><td>T8773</td><td>GACGCTCGGCCGCGCGCGCTGGG</td><td>1000</td><td></td><td></td><td></td><td></td><td></td><td></td><td></td><td></td><td></td></tr>
<tr><td>Oligo fwd</td><td>CACCGACGCTCGGCCGCGCGCGCT</td><td></td><td></td><td></td><td></td><td></td><td></td><td></td><td></td><td></td><td></td></tr>
<tr><td>Oligo rev</td><td>AAACAGCGCGCGCGGCCGAGCGTC</td><td></td><td></td><td></td><td></td><td></td><td></td><td></td><td></td><td></td><td></td></tr>
<tr><td>Chromosome</td><td>start</td><td>end</td><td>strand</td><td>MM</td><td>target_seq</td><td>PAM</td><td>alignment</td><td>distance</td><td>position</td><td>gene name</td><td>gene id</td></tr>
<tr><td>GU305914.1</td><td>117615</td><td>117637</td><td>+</td><td>0</td><td>GACGCTCGGCCGCGCGCGCT</td><td>GGG</td><td>||||||||[||||||||||||]PAM</td><td>NA</td><td>Intergenic</td><td>NA</td><td>NA</td></tr>
<tr><td>T8774</td><td>GGCCGCGCGCGCTGGGCTCGCGG</td><td>1000</td><td></td><td></td><td></td><td></td><td></td><td></td><td></td><td></td><td></td></tr>
<tr><td>Oligo fwd</td><td>CACCGGCCGCGCGCGCTGGGCTCG</td><td></td><td></td><td></td><td></td><td></td><td></td><td></td><td></td><td></td><td></td></tr>
<tr><td>Oligo rev</td><td>AAACCGAGCCCAGCGCGCGCGGCC</td><td></td><td></td><td></td><td></td><td></td><td></td><td></td><td></td><td></td><td></td></tr>
<tr><td>Chromosome</td><td>start</td><td>end</td><td>strand</td><td>MM</td><td>target_seq</td><td>PAM</td><td>alignment</td><td>distance</td><td>position</td><td>gene name</td><td>gene id</td></tr>
<tr><td>GU305914.1</td><td>117622</td><td>117644</td><td>+</td><td>0</td><td>GGCCGCGCGCGCTGGGCTCG</td><td>CGG</td><td>||||||||[||||||||||||]PAM</td><td>NA</td><td>Intergenic</td><td>NA</td><td>NA</td></tr>
<tr><td>T8775</td><td>CGCGCGCTGGGCTCGCGGCGCGG</td><td>1000</td><td></td><td></td><td></td><td></td><td></td><td></td><td></td><td></td><td></td></tr>
<tr><td>Oligo fwd</td><td>CACCCGCGCGCTGGGCTCGCGGCG</td><td></td><td></td><td></td><td></td><td></td><td></td><td></td><td></td><td></td><td></td></tr>
<tr><td>Oligo rev</td><td>AAACCGCCGCGAGCCCAGCGCGCG</td><td></td><td></td><td></td><td></td><td></td><td></td><td></td><td></td><td></td><td></td></tr>
<tr><td>Chromosome</td><td>start</td><td>end</td><td>strand</td><td>MM</td><td>target_seq</td><td>PAM</td><td>alignment</td><td>distance</td><td>position</td><td>gene name</td><td>gene id</td></tr>
<tr><td>GU305914.1</td><td>117627</td><td>117649</td><td>+</td><td>0</td><td>CGCGCGCTGGGCTCGCGGCG</td><td>CGG</td><td>||||||||[||||||||||||]PAM</td><td>NA</td><td>Intergenic</td><td>NA</td><td>NA</td></tr>
<tr><td>T8776</td><td>GCTGGGCTCGCGGCGCGGAGTGG</td><td>1000</td><td></td><td></td><td></td><td></td><td></td><td></td><td></td><td></td><td></td></tr>
<tr><td>Oligo fwd</td><td>CACCGCTGGGCTCGCGGCGCGGAG</td><td></td><td></td><td></td><td></td><td></td><td></td><td></td><td></td><td></td><td></td></tr>
<tr><td>Oligo rev</td><td>AAACCTCCGCGCCGCGAGCCCAGC</td><td></td><td></td><td></td><td></td><td></td><td></td><td></td><td></td><td></td><td></td></tr>
<tr><td>Chromosome</td><td>start</td><td>end</td><td>strand</td><td>MM</td><td>target_seq</td><td>PAM</td><td>alignment</td><td>distance</td><td>position</td><td>gene name</td><td>gene id</td></tr>
<tr><td>GU305914.1</td><td>117632</td><td>117654</td><td>+</td><td>0</td><td>GCTGGGCTCGCGGCGCGGAG</td><td>TGG</td><td>||||||||[||||||||||||]PAM</td><td>NA</td><td>Intergenic</td><td>NA</td><td>NA</td></tr>
<tr><td>T8777</td><td>GAGTCAAATACTGCGTCGTCCGG</td><td>1000</td><td></td><td></td><td></td><td></td><td></td><td></td><td></td><td></td><td></td></tr>
<tr><td>Oligo fwd</td><td>CACCGAGTCAAATACTGCGTCGTC</td><td></td><td></td><td></td><td></td><td></td><td></td><td></td><td></td><td></td><td></td></tr>
<tr><td>Oligo rev</td><td>AAACGACGACGCAGTATTTGACTC</td><td></td><td></td><td></td><td></td><td></td><td></td><td></td><td></td><td></td><td></td></tr>
<tr><td>Chromosome</td><td>start</td><td>end</td><td>strand</td><td>MM</td><td>target_seq</td><td>PAM</td><td>alignment</td><td>distance</td><td>position</td><td>gene name</td><td>gene id</td></tr>
<tr><td>GU305914.1</td><td>117666</td><td>117688</td><td>+</td><td>0</td><td>GAGTCAAATACTGCGTCGTC</td><td>CGG</td><td>||||||||[||||||||||||]PAM</td><td>NA</td><td>Intergenic</td><td>NA</td><td>NA</td></tr>
<tr><td>T8778</td><td>TACTGCGTCGTCCGGCTCCGCGG</td><td>1000</td><td></td><td></td><td></td><td></td><td></td><td></td><td></td><td></td><td></td></tr>
<tr><td>Oligo fwd</td><td>CACCTACTGCGTCGTCCGGCTCCG</td><td></td><td></td><td></td><td></td><td></td><td></td><td></td><td></td><td></td><td></td></tr>
<tr><td>Oligo rev</td><td>AAACCGGAGCCGGACGACGCAGTA</td><td></td><td></td><td></td><td></td><td></td><td></td><td></td><td></td><td></td><td></td></tr>
<tr><td>Chromosome</td><td>start</td><td>end</td><td>strand</td><td>MM</td><td>target_seq</td><td>PAM</td><td>alignment</td><td>distance</td><td>position</td><td>gene name</td><td>gene id</td></tr>
<tr><td>GU305914.1</td><td>117674</td><td>117696</td><td>+</td><td>0</td><td>TACTGCGTCGTCCGGCTCCG</td><td>CGG</td><td>||||||||[||||||||||||]PAM</td><td>NA</td><td>Intergenic</td><td>NA</td><td>NA</td></tr>
</table>

<table>
<tr><td>T8779</td><td>CCGCGGCCGCTCGCCGCCACCGG</td><td></td><td>1000</td></tr>
<tr><td>Oligo fwd</td><td>CACCCCGCGGCCGCTCGCCGCCAC</td></tr>
<tr><td>Oligo rev</td><td>AAACGTGGCGGCGAGCGGCCGCGG</td></tr>
<tr><td>Chromosome</td><td>start</td><td></td><td>end</td><td>strand</td><td>MM</td><td>target_seq</td><td>PAM</td><td>alignment</td><td>distance</td><td>position</td><td>gene name</td><td>gene id</td></tr>
<tr><td>GU305914.1</td><td></td><td>117691</td><td>117713</td><td>+</td><td>0</td><td>CCGCGGCCGCTCGCCGCCAC</td><td>CGG</td><td>||||||||[||||||||||||]PAM</td><td>NA</td><td>Intergenic</td><td>NA</td><td>NA</td></tr>
<tr><td>T8780</td><td>TTCATGAACGCAGCAGTCTCCGG</td><td></td><td>1000</td></tr>
<tr><td>Oligo fwd</td><td>CACCTTCATGAACGCAGCAGTCTC</td></tr>
<tr><td>Oligo rev</td><td>AAACGAGACTGCTGCGTTCATGAA</td></tr>
<tr><td>Chromosome</td><td>start</td><td></td><td>end</td><td>strand</td><td>MM</td><td>target_seq</td><td>PAM</td><td>alignment</td><td>distance</td><td>position</td><td>gene name</td><td>gene id</td></tr>
<tr><td>GU305914.1</td><td></td><td>117717</td><td>117739</td><td>+</td><td>0</td><td>TTCATGAACGCAGCAGTCTC</td><td>CGG</td><td>||||||||[||||||||||||]PAM</td><td>NA</td><td>Intergenic</td><td>NA</td><td>NA</td></tr>
<tr><td>T8781</td><td>TCGTGTTTGAATCCCGCTCCTGG</td><td></td><td>1000</td></tr>
<tr><td>Oligo fwd</td><td>CACCTCGTGTTTGAATCCCGCTCC</td></tr>
<tr><td>Oligo rev</td><td>AAACGGAGCGGGATTCAAACACGA</td></tr>
<tr><td>Chromosome</td><td>start</td><td></td><td>end</td><td>strand</td><td>MM</td><td>target_seq</td><td>PAM</td><td>alignment</td><td>distance</td><td>position</td><td>gene name</td><td>gene id</td></tr>
<tr><td>GU305914.1</td><td></td><td>117758</td><td>117780</td><td>+</td><td>0</td><td>TCGTGTTTGAATCCCGCTCC</td><td>TGG</td><td>||||||||[||||||||||||]PAM</td><td>NA</td><td>Intergenic</td><td>NA</td><td>NA</td></tr>
<tr><td>T8782</td><td>CGTGTTTGAATCCCGCTCCTGGG</td><td></td><td>1000</td></tr>
<tr><td>Oligo fwd</td><td>CACCCGTGTTTGAATCCCGCTCCT</td></tr>
<tr><td>Oligo rev</td><td>AAACAGGAGCGGGATTCAAACACG</td></tr>
<tr><td>Chromosome</td><td>start</td><td></td><td>end</td><td>strand</td><td>MM</td><td>target_seq</td><td>PAM</td><td>alignment</td><td>distance</td><td>position</td><td>gene name</td><td>gene id</td></tr>
<tr><td>GU305914.1</td><td></td><td>117759</td><td>117781</td><td>+</td><td>0</td><td>CGTGTTTGAATCCCGCTCCT</td><td>GGG</td><td>||||||||[||||||||||||]PAM</td><td>NA</td><td>Intergenic</td><td>NA</td><td>NA</td></tr>
<tr><td>T8783</td><td>GGATCTCCTGTCCCGTCAGCTGG</td><td></td><td>1000</td></tr>
<tr><td>Oligo fwd</td><td>CACCGGATCTCCTGTCCCGTCAGC</td></tr>
<tr><td>Oligo rev</td><td>AAACGCTGACGGGACAGGAGATCC</td></tr>
<tr><td>Chromosome</td><td>start</td><td></td><td>end</td><td>strand</td><td>MM</td><td>target_seq</td><td>PAM</td><td>alignment</td><td>distance</td><td>position</td><td>gene name</td><td>gene id</td></tr>
<tr><td>GU305914.1</td><td></td><td>117780</td><td>117802</td><td>+</td><td>0</td><td>GGATCTCCTGTCCCGTCAGC</td><td>TGG</td><td>||||||||[||||||||||||]PAM</td><td>NA</td><td>Intergenic</td><td>NA</td><td>NA</td></tr>
<tr><td>T8784</td><td>CCTGTCCCGTCAGCTGGCTCAGG</td><td></td><td>1000</td></tr>
<tr><td>Oligo fwd</td><td>CACCCCTGTCCCGTCAGCTGGCTC</td></tr>
<tr><td>Oligo rev</td><td>AAACGAGCCAGCTGACGGGACAGG</td></tr>
<tr><td>Chromosome</td><td>start</td><td></td><td>end</td><td>strand</td><td>MM</td><td>target_seq</td><td>PAM</td><td>alignment</td><td>distance</td><td>position</td><td>gene name</td><td>gene id</td></tr>
<tr><td>GU305914.1</td><td></td><td>117786</td><td>117808</td><td>+</td><td>0</td><td>CCTGTCCCGTCAGCTGGCTC</td><td>AGG</td><td>||||||||[||||||||||||]PAM</td><td>NA</td><td>Intergenic</td><td>NA</td><td>NA</td></tr>
<tr><td>T8785</td><td>GTCCCGTCAGCTGGCTCAGGCGG</td><td></td><td>1000</td></tr>
<tr><td>Oligo fwd</td><td>CACCGTCCCGTCAGCTGGCTCAGG</td></tr>
<tr><td>Oligo rev</td><td>AAACCCTGAGCCAGCTGACGGGAC</td></tr>
<tr><td>Chromosome</td><td>start</td><td></td><td>end</td><td>strand</td><td>MM</td><td>target_seq</td><td>PAM</td><td>alignment</td><td>distance</td><td>position</td><td>gene name</td><td>gene id</td></tr>
<tr><td>GU305914.1</td><td></td><td>117789</td><td>117811</td><td>+</td><td>0</td><td>GTCCCGTCAGCTGGCTCAGG</td><td>CGG</td><td>||||||||[||||||||||||]PAM</td><td>NA</td><td>Intergenic</td><td>NA</td><td>NA</td></tr>
<tr><td>T8786</td><td>AGCTGGCTCAGGCGGTAAGTCGG</td><td></td><td>1000</td></tr>
<tr><td>Oligo fwd</td><td>CACCAGCTGGCTCAGGCGGTAAGT</td></tr>
<tr><td>Oligo rev</td><td>AAACACTTACCGCCTGAGCCAGCT</td></tr>
<tr><td>Chromosome</td><td>start</td><td></td><td>end</td><td>strand</td><td>MM</td><td>target_seq</td><td>PAM</td><td>alignment</td><td>distance</td><td>position</td><td>gene name</td><td>gene id</td></tr>
<tr><td>GU305914.1</td><td></td><td>117797</td><td>117819</td><td>+</td><td>0</td><td>AGCTGGCTCAGGCGGTAAGT</td><td>CGG</td><td>||||||||[||||||||||||]PAM</td><td>NA</td><td>Intergenic</td><td>NA</td><td>NA</td></tr>
<tr><td>T8787</td><td>GCTGGCTCAGGCGGTAAGTCGGG</td><td></td><td>1000</td></tr>
<tr><td>Oligo fwd</td><td>CACCGCTGGCTCAGGCGGTAAGTC</td></tr>
<tr><td>Oligo rev</td><td>AAACGACTTACCGCCTGAGCCAGC</td></tr>
<tr><td>Chromosome</td><td>start</td><td></td><td>end</td><td>strand</td><td>MM</td><td>target_seq</td><td>PAM</td><td>alignment</td><td>distance</td><td>position</td><td>gene name</td><td>gene id</td></tr>
<tr><td>GU305914.1</td><td></td><td>117798</td><td>117820</td><td>+</td><td>0</td><td>GCTGGCTCAGGCGGTAAGTC</td><td>GGG</td><td>||||||||[||||||||||||]PAM</td><td>NA</td><td>Intergenic</td><td>NA</td><td>NA</td></tr>
<tr><td>T8788</td><td>AAGTCGGGCGCGTTTTCGCCAGG</td><td></td><td>1000</td></tr>
<tr><td>Oligo fwd</td><td>CACCAAGTCGGGCGCGTTTTCGCC</td></tr>
<tr><td>Oligo rev</td><td>AAACGGCGAAAACGCGCCCGACTT</td></tr>
<tr><td>Chromosome</td><td>start</td><td></td><td>end</td><td>strand</td><td>MM</td><td>target_seq</td><td>PAM</td><td>alignment</td><td>distance</td><td>position</td><td>gene name</td><td>gene id</td></tr>
<tr><td>GU305914.1</td><td></td><td>117813</td><td>117835</td><td>+</td><td>0</td><td>AAGTCGGGCGCGTTTTCGCC</td><td>AGG</td><td>||||||||[||||||||||||]PAM</td><td>NA</td><td>Intergenic</td><td>NA</td><td>NA</td></tr>
<tr><td>T8789</td><td>GCCAGGAACGTGAAGTTGAACGG</td><td></td><td>1000</td></tr>
<tr><td>Oligo fwd</td><td>CACCGCCAGGAACGTGAAGTTGAA</td></tr>
<tr><td>Oligo rev</td><td>AAACTTCAACTTCACGTTCCTGGC</td></tr>
<tr><td>Chromosome</td><td>start</td><td></td><td>end</td><td>strand</td><td>MM</td><td>target_seq</td><td>PAM</td><td>alignment</td><td>distance</td><td>position</td><td>gene name</td><td>gene id</td></tr>
<tr><td>GU305914.1</td><td></td><td>117830</td><td>117852</td><td>+</td><td>0</td><td>GCCAGGAACGTGAAGTTGAA</td><td>CGG</td><td>||||||||[||||||||||||]PAM</td><td>NA</td><td>Intergenic</td><td>NA</td><td>NA</td></tr>
<tr><td>T8790</td><td>CGTGAAGTTGAACGGTTCTTTGG</td><td></td><td>1000</td></tr>
<tr><td>Oligo fwd</td><td>CACCCGTGAAGTTGAACGGTTCTT</td></tr>
<tr><td>Oligo rev</td><td>AAACAAGAACCGTTCAACTTCACG</td></tr>
<tr><td>Chromosome</td><td>start</td><td></td><td>end</td><td>strand</td><td>MM</td><td>target_seq</td><td>PAM</td><td>alignment</td><td>distance</td><td>position</td><td>gene name</td><td>gene id</td></tr>
<tr><td>GU305914.1</td><td></td><td>117838</td><td>117860</td><td>+</td><td>0</td><td>CGTGAAGTTGAACGGTTCTT</td><td>TGG</td><td>||||||||[||||||||||||]PAM</td><td>NA</td><td>Intergenic</td><td>NA</td><td>NA</td></tr>
<tr><td>T8791</td><td>AAGTTGAACGGTTCTTTGGACGG</td><td></td><td>1000</td></tr>
<tr><td>Oligo fwd</td><td>CACCAAGTTGAACGGTTCTTTGGA</td></tr>
<tr><td>Oligo rev</td><td>AAACTCCAAAGAACCGTTCAACTT</td></tr>
<tr><td>Chromosome</td><td>start</td><td></td><td>end</td><td>strand</td><td>MM</td><td>target_seq</td><td>PAM</td><td>alignment</td><td>distance</td><td>position</td><td>gene name</td><td>gene id</td></tr>
<tr><td>GU305914.1</td><td></td><td>117842</td><td>117864</td><td>+</td><td>0</td><td>AAGTTGAACGGTTCTTTGGA</td><td>CGG</td><td>||||||||[||||||||||||]PAM</td><td>NA</td><td>Intergenic</td><td>NA</td><td>NA</td></tr>
<tr><td>T8792</td><td>AGTTGAACGGTTCTTTGGACGGG</td><td></td><td>1000</td></tr>
<tr><td>Oligo fwd</td><td>CACCAGTTGAACGGTTCTTTGGAC</td></tr>
<tr><td>Oligo rev</td><td>AAACGTCCAAAGAACCGTTCAACT</td></tr>
<tr><td>Chromosome</td><td>start</td><td></td><td>end</td><td>strand</td><td>MM</td><td>target_seq</td><td>PAM</td><td>alignment</td><td>distance</td><td>position</td><td>gene name</td><td>gene id</td></tr>
<tr><td>GU305914.1</td><td></td><td>117843</td><td>117865</td><td>+</td><td>0</td><td>AGTTGAACGGTTCTTTGGAC</td><td>GGG</td><td>||||||||[||||||||||||]PAM</td><td>NA</td><td>Intergenic</td><td>NA</td><td>NA</td></tr>
<tr><td>T8793</td><td>GTTCTTTGGACGGGTAGAATTGG</td><td></td><td>1000</td></tr>
<tr><td>Oligo fwd</td><td>CACCGTTCTTTGGACGGGTAGAAT</td></tr>
<tr><td>Oligo rev</td><td>AAACATTCTACCCGTCCAAAGAAC</td></tr>
<tr><td>Chromosome</td><td>start</td><td></td><td>end</td><td>strand</td><td>MM</td><td>target_seq</td><td>PAM</td><td>alignment</td><td>distance</td><td>position</td><td>gene name</td><td>gene id</td></tr>
<tr><td>GU305914.1</td><td></td><td>117852</td><td>117874</td><td>+</td><td>0</td><td>GTTCTTTGGACGGGTAGAAT</td><td>TGG</td><td>||||||||[||||||||||||]PAM</td><td>NA</td><td>Intergenic</td><td>NA</td><td>NA</td></tr>
</table>

T8794 TTCTTTGGACGGGTAGAATTGGG 1000
Oligo fwd CACCTTCTTTGGACGGGTAGAATT
Oligo rev AAACAATTCTACCCGTCCAAAGAA
Chromosome start end strand MM target_seq PAM alignment distance position gene name gene id
GU305914.1 117853 117875 + 0 TTCTTTGGACGGGTAGAATT GGG ||||||||[||||||||||||]PAM NA Intergenic NA NA

T8795 TTGGACGGGTAGAATTGGGCCGG 1000
Oligo fwd CACCTTGGACGGGTAGAATTGGGC
Oligo rev AAACGCCCAATTCTACCCGTCCAA
Chromosome start end strand MM target_seq PAM alignment distance position gene name gene id
GU305914.1 117857 117879 + 0 TTGGACGGGTAGAATTGGGC CGG ||||||||[||||||||||||]PAM NA Intergenic NA NA

T8796 TGGACGGGTAGAATTGGGCCGGG 1000
Oligo fwd CACCTGGACGGGTAGAATTGGGCC
Oligo rev AAACGGCCCAATTCTACCCGTCCA
Chromosome start end strand MM target_seq PAM alignment distance position gene name gene id
GU305914.1 117858 117880 + 0 TGGACGGGTAGAATTGGGCC GGG ||||||||[||||||||||||]PAM NA Intergenic NA NA

T8797 GGACGGGTAGAATTGGGCCGGGG 1000
Oligo fwd CACCGGACGGGTAGAATTGGGCCG
Oligo rev AAACCGGCCCAATTCTACCCGTCC
Chromosome start end strand MM target_seq PAM alignment distance position gene name gene id
GU305914.1 117859 117881 + 0 GGACGGGTAGAATTGGGCCG GGG ||||||||[||||||||||||]PAM NA Intergenic NA NA

T8798 GACGGGTAGAATTGGGCCGGGGG 1000
Oligo fwd CACCGACGGGTAGAATTGGGCCGG
Oligo rev AAACCCGGCCCAATTCTACCCGTC
Chromosome start end strand MM target_seq PAM alignment distance position gene name gene id
GU305914.1 117860 117882 + 0 GACGGGTAGAATTGGGCCGG GGG ||||||||[||||||||||||]PAM NA Intergenic NA NA

T8799 ACGGGTAGAATTGGGCCGGGGGG 1000
Oligo fwd CACCACGGGTAGAATTGGGCCGGG
Oligo rev AAACCCCGGCCCAATTCTACCCGT
Chromosome start end strand MM target_seq PAM alignment distance position gene name gene id
GU305914.1 117861 117883 + 0 ACGGGTAGAATTGGGCCGGG GGG ||||||||[||||||||||||]PAM NA Intergenic NA NA

T8800 GGGCCGGGGGGATAATGAACAGG 1000
Oligo fwd CACCGGGCCGGGGGGATAATGAAC
Oligo rev AAACGTTCATTATCCCCCCGGCCC
Chromosome start end strand MM target_seq PAM alignment distance position gene name gene id
GU305914.1 117873 117895 + 0 GGGCCGGGGGGATAATGAAC AGG ||||||||[||||||||||||]PAM NA Intergenic NA NA

T8801 GGGGGGATAATGAACAGGAACGG 1000
Oligo fwd CACCGGGGGGATAATGAACAGGAA
Oligo rev AAACTTCCTGTTCATTATCCCCCC
Chromosome start end strand MM target_seq PAM alignment distance position gene name gene id
GU305914.1 117878 117900 + 0 GGGGGGATAATGAACAGGAA CGG ||||||||[||||||||||||]PAM NA Intergenic NA NA

T8802 GAGCCAGAACCGTTTCGTCCAGG 1000
Oligo fwd CACCGAGCCAGAACCGTTTCGTCC
Oligo rev AAACGGACGAAACGGTTCTGGCTC
Chromosome start end strand MM target_seq PAM alignment distance position gene name gene id
GU305914.1 117900 117922 + 0 GAGCCAGAACCGTTTCGTCC AGG ||||||||[||||||||||||]PAM NA Intergenic NA NA

T8803 TTTCGTCCAGGCACACCGACAGG 1000
Oligo fwd CACCTTTCGTCCAGGCACACCGAC
Oligo rev AAACGTCGGTGTGCCTGGACGAAA
Chromosome start end strand MM target_seq PAM alignment distance position gene name gene id
GU305914.1 117912 117934 + 0 TTTCGTCCAGGCACACCGAC AGG ||||||||[||||||||||||]PAM NA Intergenic NA NA

T8804 TTCGTCCAGGCACACCGACAGGG 1000
Oligo fwd CACCTTCGTCCAGGCACACCGACA
Oligo rev AAACTGTCGGTGTGCCTGGACGAA
Chromosome start end strand MM target_seq PAM alignment distance position gene name gene id
GU305914.1 117913 117935 + 0 TTCGTCCAGGCACACCGACA GGG ||||||||[||||||||||||]PAM NA Intergenic NA NA

T8805 TCGTCCAGGCACACCGACAGGGG 1000
Oligo fwd CACCTCGTCCAGGCACACCGACAG
Oligo rev AAACCTGTCGGTGTGCCTGGACGA
Chromosome start end strand MM target_seq PAM alignment distance position gene name gene id
GU305914.1 117914 117936 + 0 TCGTCCAGGCACACCGACAG GGG ||||||||[||||||||||||]PAM NA Intergenic NA NA

T8806 GGCACACCGACAGGGGAAAGAGG 1000
Oligo fwd CACCGGCACACCGACAGGGGAAAG
Oligo rev AAACCTTTCCCCTGTCGGTGTGCC
Chromosome start end strand MM target_seq PAM alignment distance position gene name gene id
GU305914.1 117921 117943 + 0 GGCACACCGACAGGGGAAAG AGG ||||||||[||||||||||||]PAM NA Intergenic NA NA

T8807 TGTCCAGATTGAACTGCTGCAGG 1000
Oligo fwd CACCTGTCCAGATTGAACTGCTGC
Oligo rev AAACGCAGCAGTTCAATCTGGACA
Chromosome start end strand MM target_seq PAM alignment distance position gene name gene id
GU305914.1 117960 117982 + 0 TGTCCAGATTGAACTGCTGC AGG ||||||||[||||||||||||]PAM NA Intergenic NA NA

T8808 GTTCATGTTTAGCGCCGTCGAGG 1000
Oligo fwd CACCGTTCATGTTTAGCGCCGTCG
Oligo rev AAACCGACGGCGCTAAACATGAAC
Chromosome start end strand MM target_seq PAM alignment distance position gene name gene id

<table>
<tr><td>GU305914.1</td><td>118036</td><td>118058</td><td>+</td><td>0</td><td>GTTCATGTTTAGCGCCGTCG</td><td>AGG</td><td>||||||||[||||||||||||]PAM</td><td>NA</td><td>Intergenic</td><td>NA</td><td>NA</td></tr>
<tr><td>T8809</td><td>TTCATGTTTAGCGCCGTCGAGGG</td><td>1000</td></tr>
<tr><td>Oligo fwd</td><td>CACCTTCATGTTTAGCGCCGTCGA</td></tr>
<tr><td>Oligo rev</td><td>AAACTCGACGGCGCTAAACATGAA</td></tr>
<tr><td>Chromosome</td><td>start</td><td>end</td><td>strand</td><td>MM</td><td>target_seq</td><td>PAM</td><td>alignment</td><td>distance</td><td>position</td><td>gene name</td><td>gene id</td></tr>
<tr><td>GU305914.1</td><td>118037</td><td>118059</td><td>+</td><td>0</td><td>TTCATGTTTAGCGCCGTCGA</td><td>GGG</td><td>||||||||[||||||||||||]PAM</td><td>NA</td><td>Intergenic</td><td>NA</td><td>NA</td></tr>
<tr><td>T8810</td><td>GCCGTCGAGGGCGACACCAGCGG</td><td>1000</td></tr>
<tr><td>Oligo fwd</td><td>CACCGCCGTCGAGGGCGACACCAG</td></tr>
<tr><td>Oligo rev</td><td>AAACCTGGTGTCGCCCTCGACGGC</td></tr>
<tr><td>Chromosome</td><td>start</td><td>end</td><td>strand</td><td>MM</td><td>target_seq</td><td>PAM</td><td>alignment</td><td>distance</td><td>position</td><td>gene name</td><td>gene id</td></tr>
<tr><td>GU305914.1</td><td>118049</td><td>118071</td><td>+</td><td>0</td><td>GCCGTCGAGGGCGACACCAG</td><td>CGG</td><td>||||||||[||||||||||||]PAM</td><td>NA</td><td>Intergenic</td><td>NA</td><td>NA</td></tr>
<tr><td>T8811</td><td>CCGTCGAGGGCGACACCAGCGGG</td><td>1000</td></tr>
<tr><td>Oligo fwd</td><td>CACCCCGTCGAGGGCGACACCAGC</td></tr>
<tr><td>Oligo rev</td><td>AAACGCTGGTGTCGCCCTCGACGG</td></tr>
<tr><td>Chromosome</td><td>start</td><td>end</td><td>strand</td><td>MM</td><td>target_seq</td><td>PAM</td><td>alignment</td><td>distance</td><td>position</td><td>gene name</td><td>gene id</td></tr>
<tr><td>GU305914.1</td><td>118050</td><td>118072</td><td>+</td><td>0</td><td>CCGTCGAGGGCGACACCAGC</td><td>GGG</td><td>||||||||[||||||||||||]PAM</td><td>NA</td><td>Intergenic</td><td>NA</td><td>NA</td></tr>
<tr><td>T8812</td><td>GCGGGTTCACTTGTCGTCCGTGG</td><td>1000</td></tr>
<tr><td>Oligo fwd</td><td>CACCGCGGGTTCACTTGTCGTCCG</td></tr>
<tr><td>Oligo rev</td><td>AAACCGGACGACAAGTGAACCCGC</td></tr>
<tr><td>Chromosome</td><td>start</td><td>end</td><td>strand</td><td>MM</td><td>target_seq</td><td>PAM</td><td>alignment</td><td>distance</td><td>position</td><td>gene name</td><td>gene id</td></tr>
<tr><td>GU305914.1</td><td>118068</td><td>118090</td><td>+</td><td>0</td><td>GCGGGTTCACTTGTCGTCCG</td><td>TGG</td><td>||||||||[||||||||||||]PAM</td><td>NA</td><td>Intergenic</td><td>NA</td><td>NA</td></tr>
<tr><td>T8813</td><td>TCCTGAAAAGAGATCCTGACCGG</td><td>1000</td></tr>
<tr><td>Oligo fwd</td><td>CACCTCCTGAAAAGAGATCCTGAC</td></tr>
<tr><td>Oligo rev</td><td>AAACGTCAGGATCTCTTTTCAGGA</td></tr>
<tr><td>Chromosome</td><td>start</td><td>end</td><td>strand</td><td>MM</td><td>target_seq</td><td>PAM</td><td>alignment</td><td>distance</td><td>position</td><td>gene name</td><td>gene id</td></tr>
<tr><td>GU305914.1</td><td>118103</td><td>118125</td><td>+</td><td>0</td><td>TCCTGAAAAGAGATCCTGAC</td><td>CGG</td><td>||||||||[||||||||||||]PAM</td><td>NA</td><td>Intergenic</td><td>NA</td><td>NA</td></tr>
<tr><td>T8814</td><td>CCGGTTCCCGCTGCAGAAAGAGG</td><td>1000</td></tr>
<tr><td>Oligo fwd</td><td>CACCCCGGTTCCCGCTGCAGAAAG</td></tr>
<tr><td>Oligo rev</td><td>AAACCTTTCTGCAGCGGGAACCGG</td></tr>
<tr><td>Chromosome</td><td>start</td><td>end</td><td>strand</td><td>MM</td><td>target_seq</td><td>PAM</td><td>alignment</td><td>distance</td><td>position</td><td>gene name</td><td>gene id</td></tr>
<tr><td>GU305914.1</td><td>118122</td><td>118144</td><td>+</td><td>0</td><td>CCGGTTCCCGCTGCAGAAAG</td><td>AGG</td><td>||||||||[||||||||||||]PAM</td><td>NA</td><td>Intergenic</td><td>NA</td><td>NA</td></tr>
<tr><td>T8815</td><td>CGGTTCCCGCTGCAGAAAGAGGG</td><td>1000</td></tr>
<tr><td>Oligo fwd</td><td>CACCCGGTTCCCGCTGCAGAAAGA</td></tr>
<tr><td>Oligo rev</td><td>AAACTCTTTCTGCAGCGGGAACCG</td></tr>
<tr><td>Chromosome</td><td>start</td><td>end</td><td>strand</td><td>MM</td><td>target_seq</td><td>PAM</td><td>alignment</td><td>distance</td><td>position</td><td>gene name</td><td>gene id</td></tr>
<tr><td>GU305914.1</td><td>118123</td><td>118145</td><td>+</td><td>0</td><td>CGGTTCCCGCTGCAGAAAGA</td><td>GGG</td><td>||||||||[||||||||||||]PAM</td><td>NA</td><td>Intergenic</td><td>NA</td><td>NA</td></tr>
<tr><td>T8816</td><td>CAGAAAGAGGGCTATCGCCTCGG</td><td>1000</td></tr>
<tr><td>Oligo fwd</td><td>CACCCAGAAAGAGGGCTATCGCCT</td></tr>
<tr><td>Oligo rev</td><td>AAACAGGCGATAGCCCTCTTTCTG</td></tr>
<tr><td>Chromosome</td><td>start</td><td>end</td><td>strand</td><td>MM</td><td>target_seq</td><td>PAM</td><td>alignment</td><td>distance</td><td>position</td><td>gene name</td><td>gene id</td></tr>
<tr><td>GU305914.1</td><td>118135</td><td>118157</td><td>+</td><td>0</td><td>CAGAAAGAGGGCTATCGCCT</td><td>CGG</td><td>||||||||[||||||||||||]PAM</td><td>NA</td><td>Intergenic</td><td>NA</td><td>NA</td></tr>
<tr><td>T8817</td><td>AGAAAGAGGGCTATCGCCTCGGG</td><td>1000</td></tr>
<tr><td>Oligo fwd</td><td>CACCAGAAAGAGGGCTATCGCCTC</td></tr>
<tr><td>Oligo rev</td><td>AAACGAGGCGATAGCCCTCTTTCT</td></tr>
<tr><td>Chromosome</td><td>start</td><td>end</td><td>strand</td><td>MM</td><td>target_seq</td><td>PAM</td><td>alignment</td><td>distance</td><td>position</td><td>gene name</td><td>gene id</td></tr>
<tr><td>GU305914.1</td><td>118136</td><td>118158</td><td>+</td><td>0</td><td>AGAAAGAGGGCTATCGCCTC</td><td>GGG</td><td>||||||||[||||||||||||]PAM</td><td>NA</td><td>Intergenic</td><td>NA</td><td>NA</td></tr>
<tr><td>T8818</td><td>AGGGCTATCGCCTCGGGATCCGG</td><td>1000</td></tr>
<tr><td>Oligo fwd</td><td>CACCAGGGCTATCGCCTCGGGATC</td></tr>
<tr><td>Oligo rev</td><td>AAACGATCCCGAGGCGATAGCCCT</td></tr>
<tr><td>Chromosome</td><td>start</td><td>end</td><td>strand</td><td>MM</td><td>target_seq</td><td>PAM</td><td>alignment</td><td>distance</td><td>position</td><td>gene name</td><td>gene id</td></tr>
<tr><td>GU305914.1</td><td>118142</td><td>118164</td><td>+</td><td>0</td><td>AGGGCTATCGCCTCGGGATC</td><td>CGG</td><td>||||||||[||||||||||||]PAM</td><td>NA</td><td>Intergenic</td><td>NA</td><td>NA</td></tr>
<tr><td>T8819</td><td>CGGTGTGCGCTCTTCTTCCGCGG</td><td>1000</td></tr>
<tr><td>Oligo fwd</td><td>CACCCGGTGTGCGCTCTTCTTCCG</td></tr>
<tr><td>Oligo rev</td><td>AAACCGGAAGAAGAGCGCACACCG</td></tr>
<tr><td>Chromosome</td><td>start</td><td>end</td><td>strand</td><td>MM</td><td>target_seq</td><td>PAM</td><td>alignment</td><td>distance</td><td>position</td><td>gene name</td><td>gene id</td></tr>
<tr><td>GU305914.1</td><td>118162</td><td>118184</td><td>+</td><td>0</td><td>CGGTGTGCGCTCTTCTTCCG</td><td>CGG</td><td>||||||||[||||||||||||]PAM</td><td>NA</td><td>Intergenic</td><td>NA</td><td>NA</td></tr>
<tr><td>T8820</td><td>CTTCCGCGGCTTCGTCTCTGCGG</td><td>1000</td></tr>
<tr><td>Oligo fwd</td><td>CACCCTTCCGCGGCTTCGTCTCTG</td></tr>
<tr><td>Oligo rev</td><td>AAACCAGAGACGAAGCCGCGGAAG</td></tr>
<tr><td>Chromosome</td><td>start</td><td>end</td><td>strand</td><td>MM</td><td>target_seq</td><td>PAM</td><td>alignment</td><td>distance</td><td>position</td><td>gene name</td><td>gene id</td></tr>
<tr><td>GU305914.1</td><td>118176</td><td>118198</td><td>+</td><td>0</td><td>CTTCCGCGGCTTCGTCTCTG</td><td>CGG</td><td>||||||||[||||||||||||]PAM</td><td>NA</td><td>Intergenic</td><td>NA</td><td>NA</td></tr>
<tr><td>T8821</td><td>CTGCGGTTACCGAACTCCGTCGG</td><td>1000</td></tr>
<tr><td>Oligo fwd</td><td>CACCCTGCGGTTACCGAACTCCGT</td></tr>
<tr><td>Oligo rev</td><td>AAACACGGAGTTCGGTAACCGCAG</td></tr>
<tr><td>Chromosome</td><td>start</td><td>end</td><td>strand</td><td>MM</td><td>target_seq</td><td>PAM</td><td>alignment</td><td>distance</td><td>position</td><td>gene name</td><td>gene id</td></tr>
<tr><td>GU305914.1</td><td>118193</td><td>118215</td><td>+</td><td>0</td><td>CTGCGGTTACCGAACTCCGT</td><td>CGG</td><td>||||||||[||||||||||||]PAM</td><td>NA</td><td>Intergenic</td><td>NA</td><td>NA</td></tr>
<tr><td>T8822</td><td>CCGAACTCCGTCGGCTGATTCGG</td><td>1000</td></tr>
<tr><td>Oligo fwd</td><td>CACCCCGAACTCCGTCGGCTGATT</td></tr>
<tr><td>Oligo rev</td><td>AAACAATCAGCCGACGGAGTTCGG</td></tr>
<tr><td>Chromosome</td><td>start</td><td>end</td><td>strand</td><td>MM</td><td>target_seq</td><td>PAM</td><td>alignment</td><td>distance</td><td>position</td><td>gene name</td><td>gene id</td></tr>
<tr><td>GU305914.1</td><td>118202</td><td>118224</td><td>+</td><td>0</td><td>CCGAACTCCGTCGGCTGATT</td><td>CGG</td><td>||||||||[||||||||||||]PAM</td><td>NA</td><td>Intergenic</td><td>NA</td><td>NA</td></tr>
<tr><td>T8823</td><td>TCCGTCGGCTGATTCGGCCGCGG</td><td>1000</td></tr>
<tr><td>Oligo fwd</td><td>CACCTCCGTCGGCTGATTCGGCCG</td></tr>
<tr><td>Oligo rev</td><td>AAACCGGCCGAATCAGCCGACGGA</td></tr>
</table>

| | | | | | | | | | | | |
|---|---|---|---|---|---|---|---|---|---|---|---|
| Chromosome | start | end | strand | MM | target_seq | PAM | alignment | distance | position | gene name | gene id |
| GU305914.1 | 118208 | 118230 | + | 0 | TCCGTCGGCTGATTCGGCCG | CGG | \|\|\|\|\|\|\|\|[\|\|\|\|\|\|\|\|\|\|\|\|]PAM | NA | Intergenic | NA | NA |
| T8824 | CGGCTGATTCGGCCGCGGTTCGG | 1000 | | | | | | | | | |
| Oligo fwd | CACCCGGCTGATTCGGCCGCGGTT | | | | | | | | | | |
| Oligo rev | AAACAACCGCGGCCGAATCAGCCG | | | | | | | | | | |
| Chromosome | start | end | strand | MM | target_seq | PAM | alignment | distance | position | gene name | gene id |
| GU305914.1 | 118213 | 118235 | + | 0 | CGGCTGATTCGGCCGCGGTT | CGG | \|\|\|\|\|\|\|\|[\|\|\|\|\|\|\|\|\|\|\|\|]PAM | NA | Intergenic | NA | NA |
| T8825 | GGCTGATTCGGCCGCGGTTCGGG | 1000 | | | | | | | | | |
| Oligo fwd | CACCGGCTGATTCGGCCGCGGTTC | | | | | | | | | | |
| Oligo rev | AAACGAACCGCGGCCGAATCAGCC | | | | | | | | | | |
| Chromosome | start | end | strand | MM | target_seq | PAM | alignment | distance | position | gene name | gene id |
| GU305914.1 | 118214 | 118236 | + | 0 | GGCTGATTCGGCCGCGGTTC | GGG | \|\|\|\|\|\|\|\|[\|\|\|\|\|\|\|\|\|\|\|\|]PAM | NA | Intergenic | NA | NA |
| T8826 | GCTCCTCGTCCTCCTCCCGACGG | 1000 | | | | | | | | | |
| Oligo fwd | CACCGCTCCTCGTCCTCCTCCCGA | | | | | | | | | | |
| Oligo rev | AAACTCGGGAGGAGGACGAGGAGC | | | | | | | | | | |
| Chromosome | start | end | strand | MM | target_seq | PAM | alignment | distance | position | gene name | gene id |
| GU305914.1 | 118236 | 118258 | + | 0 | GCTCCTCGTCCTCCTCCCGA | CGG | \|\|\|\|\|\|\|\|[\|\|\|\|\|\|\|\|\|\|\|\|]PAM | NA | Intergenic | NA | NA |
| T8827 | CCTCGTCCTCCTCCCGACGGAGG | 1000 | | | | | | | | | |
| Oligo fwd | CACCCCTCGTCCTCCTCCCGACGG | | | | | | | | | | |
| Oligo rev | AAACCCGTCGGGAGGAGGACGAGG | | | | | | | | | | |
| Chromosome | start | end | strand | MM | target_seq | PAM | alignment | distance | position | gene name | gene id |
| GU305914.1 | 118239 | 118261 | + | 0 | CCTCGTCCTCCTCCCGACGG | AGG | \|\|\|\|\|\|\|\|[\|\|\|\|\|\|\|\|\|\|\|\|]PAM | NA | Intergenic | NA | NA |
| GU305914.1 | 63649 | 63671 | - | 4 | CCTCTTCTTCCTCTCGATGG | TGG | \|\|\|\|-\|\|-[\|\|\|\|\|-\|\|\|-\|\|]PAM | NA | Intergenic | NA | NA |
| T8828 | CCTCCTCCCGACGGAGGATCAGG | 1000 | | | | | | | | | |
| Oligo fwd | CACCCCTCCTCCCGACGGAGGATC | | | | | | | | | | |
| Oligo rev | AAACGATCCTCCGTCGGGAGGAGG | | | | | | | | | | |
| Chromosome | start | end | strand | MM | target_seq | PAM | alignment | distance | position | gene name | gene id |
| GU305914.1 | 118245 | 118267 | + | 0 | CCTCCTCCCGACGGAGGATC | AGG | \|\|\|\|\|\|\|\|[\|\|\|\|\|\|\|\|\|\|\|\|]PAM | NA | Intergenic | NA | NA |
| T8829 | CCTCCCGACGGAGGATCAGGCGG | 1000 | | | | | | | | | |
| Oligo fwd | CACCCCTCCCGACGGAGGATCAGG | | | | | | | | | | |
| Oligo rev | AAACCCTGATCCTCCGTCGGGAGG | | | | | | | | | | |
| Chromosome | start | end | strand | MM | target_seq | PAM | alignment | distance | position | gene name | gene id |
| GU305914.1 | 118248 | 118270 | + | 0 | CCTCCCGACGGAGGATCAGG | CGG | \|\|\|\|\|\|\|\|[\|\|\|\|\|\|\|\|\|\|\|\|]PAM | NA | Intergenic | NA | NA |
| GU305914.1 | 96864 | 96886 | + | 4 | TCGCCCGACGGAGGATCGTG | CGG | -\|-\|\|\|\|\|[\|\|\|\|\|\|\|\|\|--\|]PAM | NA | Intergenic | NA | NA |
| T8830 | CCCGTAATCGTCCACCGCGACGG | 1000 | | | | | | | | | |
| Oligo fwd | CACCCCCGTAATCGTCCACCGCGA | | | | | | | | | | |
| Oligo rev | AAACTCGCGGTGGACGATTACGGG | | | | | | | | | | |
| Chromosome | start | end | strand | MM | target_seq | PAM | alignment | distance | position | gene name | gene id |
| GU305914.1 | 118303 | 118325 | + | 0 | CCCGTAATCGTCCACCGCGA | CGG | \|\|\|\|\|\|\|\|[\|\|\|\|\|\|\|\|\|\|\|\|]PAM | NA | Intergenic | NA | NA |
| T8831 | GTAATCGTCCACCGCGACGGCGG | 1000 | | | | | | | | | |
| Oligo fwd | CACCGTAATCGTCCACCGCGACGG | | | | | | | | | | |
| Oligo rev | AAACCCGTCGCGGTGGACGATTAC | | | | | | | | | | |
| Chromosome | start | end | strand | MM | target_seq | PAM | alignment | distance | position | gene name | gene id |
| GU305914.1 | 118306 | 118328 | + | 0 | GTAATCGTCCACCGCGACGG | CGG | \|\|\|\|\|\|\|\|[\|\|\|\|\|\|\|\|\|\|\|\|]PAM | NA | Intergenic | NA | NA |
| T8832 | CACCGCGACGGCGGCGCCCATGG | 1000 | | | | | | | | | |
| Oligo fwd | CACCCACCGCGACGGCGGCGCCCA | | | | | | | | | | |
| Oligo rev | AAACTGGGCGCCGCCGTCGCGGTG | | | | | | | | | | |
| Chromosome | start | end | strand | MM | target_seq | PAM | alignment | distance | position | gene name | gene id |
| GU305914.1 | 118315 | 118337 | + | 0 | CACCGCGACGGCGGCGCCCA | TGG | \|\|\|\|\|\|\|\|[\|\|\|\|\|\|\|\|\|\|\|\|]PAM | NA | Intergenic | NA | NA |
| T8833 | TGGCGTCTCTCGTCGCCGCGCGG | 1000 | | | | | | | | | |
| Oligo fwd | CACCTGGCGTCTCTCGTCGCCGCG | | | | | | | | | | |
| Oligo rev | AAACCGCGGCGACGAGAGACGCCA | | | | | | | | | | |
| Chromosome | start | end | strand | MM | target_seq | PAM | alignment | distance | position | gene name | gene id |
| GU305914.1 | 118335 | 118357 | + | 0 | TGGCGTCTCTCGTCGCCGCG | CGG | \|\|\|\|\|\|\|\|[\|\|\|\|\|\|\|\|\|\|\|\|]PAM | NA | Intergenic | NA | NA |
| T8834 | CTCTCGTCGCCGCGCGGACGAGG | 1000 | | | | | | | | | |
| Oligo fwd | CACCCTCTCGTCGCCGCGCGGACG | | | | | | | | | | |
| Oligo rev | AAACCGTCCGCGCGGCGACGAGAG | | | | | | | | | | |
| Chromosome | start | end | strand | MM | target_seq | PAM | alignment | distance | position | gene name | gene id |
| GU305914.1 | 118341 | 118363 | + | 0 | CTCTCGTCGCCGCGCGGACG | AGG | \|\|\|\|\|\|\|\|[\|\|\|\|\|\|\|\|\|\|\|\|]PAM | NA | Intergenic | NA | NA |
| T8835 | TCGCCGCGCGGACGAGGTTGCGG | 1000 | | | | | | | | | |
| Oligo fwd | CACCTCGCCGCGCGGACGAGGTTG | | | | | | | | | | |
| Oligo rev | AAACCAACCTCGTCCGCGCGGCGA | | | | | | | | | | |
| Chromosome | start | end | strand | MM | target_seq | PAM | alignment | distance | position | gene name | gene id |
| GU305914.1 | 118347 | 118369 | + | 0 | TCGCCGCGCGGACGAGGTTG | CGG | \|\|\|\|\|\|\|\|[\|\|\|\|\|\|\|\|\|\|\|\|]PAM | NA | Intergenic | NA | NA |
| GU305914.1 | 139922 | 139944 | + | 4 | TCGCGACGCGGACGAGTTCG | CGG | \|\|\|\|--\|\|[\|\|\|\|\|\|\|\|-\|-\|]PAM | NA | Intergenic | NA | NA |
| T8836 | CGGACGAGGTTGCGGTAGAATGG | 1000 | | | | | | | | | |
| Oligo fwd | CACCCGGACGAGGTTGCGGTAGAA | | | | | | | | | | |
| Oligo rev | AAACTTCTACCGCAACCTCGTCCG | | | | | | | | | | |
| Chromosome | start | end | strand | MM | target_seq | PAM | alignment | distance | position | gene name | gene id |
| GU305914.1 | 118355 | 118377 | + | 0 | CGGACGAGGTTGCGGTAGAA | TGG | \|\|\|\|\|\|\|\|[\|\|\|\|\|\|\|\|\|\|\|\|]PAM | NA | Intergenic | NA | NA |
| T8837 | ATGGATTGTCACCGAGACCGCGG | 1000 | | | | | | | | | |
| Oligo fwd | CACCATGGATTGTCACCGAGACCG | | | | | | | | | | |
| Oligo rev | AAACCGGTCTCGGTGACAATCCAT | | | | | | | | | | |
| Chromosome | start | end | strand | MM | target_seq | PAM | alignment | distance | position | gene name | gene id |
| GU305914.1 | 118374 | 118396 | + | 0 | ATGGATTGTCACCGAGACCG | CGG | \|\|\|\|\|\|\|\|[\|\|\|\|\|\|\|\|\|\|\|\|]PAM | NA | Intergenic | NA | NA |

T8838 GATTGTCACCGAGACCGCGGCGG 1000
Oligo fwd CACCGATTGTCACCGAGACCGCGG
Oligo rev AAACCCGCGGTCTCGGTGACAATC

<table><tr><th>Chromosome</th><th>start</th><th>end</th><th>strand</th><th>MM</th><th>target_seq</th><th>PAM</th><th>alignment</th><th>distance</th><th>position</th><th>gene name</th><th>gene id</th></tr>
<tr><td>GU305914.1</td><td>118377</td><td>118399</td><td>+</td><td>0</td><td>GATTGTCACCGAGACCGCGG</td><td>CGG</td><td>||||||||[||||||||||||]PAM</td><td>NA</td><td>Intergenic</td><td>NA</td><td>NA</td></tr></table>

T8839 TGTCACCGAGACCGCGGCGGCGG 1000
Oligo fwd CACCTGTCACCGAGACCGCGGCGG
Oligo rev AAACCCGCCGCGGTCTCGGTGACA

<table><tr><th>Chromosome</th><th>start</th><th>end</th><th>strand</th><th>MM</th><th>target_seq</th><th>PAM</th><th>alignment</th><th>distance</th><th>position</th><th>gene name</th><th>gene id</th></tr>
<tr><td>GU305914.1</td><td>118380</td><td>118402</td><td>+</td><td>0</td><td>TGTCACCGAGACCGCGGCGG</td><td>CGG</td><td>||||||||[||||||||||||]PAM</td><td>NA</td><td>Intergenic</td><td>NA</td><td>NA</td></tr></table>

T8840 GACCGCGGCGGCGGCCGTCACGG 1000
Oligo fwd CACCGACCGCGGCGGCGGCCGTCA
Oligo rev AAACTGACGGCCGCCGCCGCGGTC

<table><tr><th>Chromosome</th><th>start</th><th>end</th><th>strand</th><th>MM</th><th>target_seq</th><th>PAM</th><th>alignment</th><th>distance</th><th>position</th><th>gene name</th><th>gene id</th></tr>
<tr><td>GU305914.1</td><td>118389</td><td>118411</td><td>+</td><td>0</td><td>GACCGCGGCGGCGGCCGTCA</td><td>CGG</td><td>||||||||[||||||||||||]PAM</td><td>NA</td><td>Intergenic</td><td>NA</td><td>NA</td></tr></table>

T8841 CCGTCACGGCTGCCGCCGTTCGG 1000
Oligo fwd CACCCCGTCACGGCTGCCGCCGTT
Oligo rev AAACAACGGCGGCAGCCGTGACGG

<table><tr><th>Chromosome</th><th>start</th><th>end</th><th>strand</th><th>MM</th><th>target_seq</th><th>PAM</th><th>alignment</th><th>distance</th><th>position</th><th>gene name</th><th>gene id</th></tr>
<tr><td>GU305914.1</td><td>118403</td><td>118425</td><td>+</td><td>0</td><td>CCGTCACGGCTGCCGCCGTT</td><td>CGG</td><td>||||||||[||||||||||||]PAM</td><td>NA</td><td>Intergenic</td><td>NA</td><td>NA</td></tr></table>

T8842 CACGGCTGCCGCCGTTCGGTTGG 1000
Oligo fwd CACCCACGGCTGCCGCCGTTCGGT
Oligo rev AAACACCGAACGGCGGCAGCCGTG

<table><tr><th>Chromosome</th><th>start</th><th>end</th><th>strand</th><th>MM</th><th>target_seq</th><th>PAM</th><th>alignment</th><th>distance</th><th>position</th><th>gene name</th><th>gene id</th></tr>
<tr><td>GU305914.1</td><td>118407</td><td>118429</td><td>+</td><td>0</td><td>CACGGCTGCCGCCGTTCGGT</td><td>TGG</td><td>||||||||[||||||||||||]PAM</td><td>NA</td><td>Intergenic</td><td>NA</td><td>NA</td></tr></table>

T8843 CTGCCGCCGTTCGGTTGGCGCGG 1000
Oligo fwd CACCCTGCCGCCGTTCGGTTGGCG
Oligo rev AAACCGCCAACCGAACGGCGGCAG

<table><tr><th>Chromosome</th><th>start</th><th>end</th><th>strand</th><th>MM</th><th>target_seq</th><th>PAM</th><th>alignment</th><th>distance</th><th>position</th><th>gene name</th><th>gene id</th></tr>
<tr><td>GU305914.1</td><td>118412</td><td>118434</td><td>+</td><td>0</td><td>CTGCCGCCGTTCGGTTGGCG</td><td>CGG</td><td>||||||||[||||||||||||]PAM</td><td>NA</td><td>Intergenic</td><td>NA</td><td>NA</td></tr></table>

T8844 CTCTTCATCATCGTCCTCATCGG 1000
Oligo fwd CACCCTCTTCATCATCGTCCTCAT
Oligo rev AAACATGAGGACGATGATGAAGAG

<table><tr><th>Chromosome</th><th>start</th><th>end</th><th>strand</th><th>MM</th><th>target_seq</th><th>PAM</th><th>alignment</th><th>distance</th><th>position</th><th>gene name</th><th>gene id</th></tr>
<tr><td>GU305914.1</td><td>118516</td><td>118538</td><td>+</td><td>0</td><td>CTCTTCATCATCGTCCTCAT</td><td>CGG</td><td>||||||||[||||||||||||]PAM</td><td>NA</td><td>Intergenic</td><td>NA</td><td>NA</td></tr></table>

T8845 ATCGGCGTTCTCGTCGCCAGCGG 1000
Oligo fwd CACCATCGGCGTTCTCGTCGCCAG
Oligo rev AAACCTGGCGACGAGAACGCCGAT

<table><tr><th>Chromosome</th><th>start</th><th>end</th><th>strand</th><th>MM</th><th>target_seq</th><th>PAM</th><th>alignment</th><th>distance</th><th>position</th><th>gene name</th><th>gene id</th></tr>
<tr><td>GU305914.1</td><td>118534</td><td>118556</td><td>+</td><td>0</td><td>ATCGGCGTTCTCGTCGCCAG</td><td>CGG</td><td>||||||||[||||||||||||]PAM</td><td>NA</td><td>Intergenic</td><td>NA</td><td>NA</td></tr></table>

T8846 TCGGCGTTCTCGTCGCCAGCGGG 1000
Oligo fwd CACCTCGGCGTTCTCGTCGCCAGC
Oligo rev AAACGCTGGCGACGAGAACGCCGA

<table><tr><th>Chromosome</th><th>start</th><th>end</th><th>strand</th><th>MM</th><th>target_seq</th><th>PAM</th><th>alignment</th><th>distance</th><th>position</th><th>gene name</th><th>gene id</th></tr>
<tr><td>GU305914.1</td><td>118535</td><td>118557</td><td>+</td><td>0</td><td>TCGGCGTTCTCGTCGCCAGC</td><td>GGG</td><td>||||||||[||||||||||||]PAM</td><td>NA</td><td>Intergenic</td><td>NA</td><td>NA</td></tr></table>

T8847 AGCGGGACCGCCGACGTCCATGG 1000
Oligo fwd CACCAGCGGGACCGCCGACGTCCA
Oligo rev AAACTGGACGTCGGCGGTCCCGCT

<table><tr><th>Chromosome</th><th>start</th><th>end</th><th>strand</th><th>MM</th><th>target_seq</th><th>PAM</th><th>alignment</th><th>distance</th><th>position</th><th>gene name</th><th>gene id</th></tr>
<tr><td>GU305914.1</td><td>118552</td><td>118574</td><td>+</td><td>0</td><td>AGCGGGACCGCCGACGTCCA</td><td>TGG</td><td>||||||||[||||||||||||]PAM</td><td>NA</td><td>Intergenic</td><td>NA</td><td>NA</td></tr></table>

T8848 ACCGCCGACGTCCATGGCGTCGG 1000
Oligo fwd CACCACCGCCGACGTCCATGGCGT
Oligo rev AAACACGCCATGGACGTCGGCGGT

<table><tr><th>Chromosome</th><th>start</th><th>end</th><th>strand</th><th>MM</th><th>target_seq</th><th>PAM</th><th>alignment</th><th>distance</th><th>position</th><th>gene name</th><th>gene id</th></tr>
<tr><td>GU305914.1</td><td>118558</td><td>118580</td><td>+</td><td>0</td><td>ACCGCCGACGTCCATGGCGT</td><td>CGG</td><td>||||||||[||||||||||||]PAM</td><td>NA</td><td>Intergenic</td><td>NA</td><td>NA</td></tr></table>

T8849 GACGTCCATGGCGTCGGCGTCGG 1000
Oligo fwd CACCGACGTCCATGGCGTCGGCGT
Oligo rev AAACACGCCGACGCCATGGACGTC

<table><tr><th>Chromosome</th><th>start</th><th>end</th><th>strand</th><th>MM</th><th>target_seq</th><th>PAM</th><th>alignment</th><th>distance</th><th>position</th><th>gene name</th><th>gene id</th></tr>
<tr><td>GU305914.1</td><td>118564</td><td>118586</td><td>+</td><td>0</td><td>GACGTCCATGGCGTCGGCGT</td><td>CGG</td><td>||||||||[||||||||||||]PAM</td><td>NA</td><td>Intergenic</td><td>NA</td><td>NA</td></tr>
<tr><td>GU305914.1</td><td>24125</td><td>24147</td><td>+</td><td>2</td><td>GACGTCCATGGCGCCGACGT</td><td>CGG</td><td>||||||||[|||||-||-|||]PAM</td><td>NA</td><td>Intergenic</td><td>NA</td><td>NA</td></tr></table>

T8850 ATGGCGTCGGCGTCGGCGTCTGG 1000
Oligo fwd CACCATGGCGTCGGCGTCGGCGTC
Oligo rev AAACGACGCCGACGCCGACGCCAT

<table><tr><th>Chromosome</th><th>start</th><th>end</th><th>strand</th><th>MM</th><th>target_seq</th><th>PAM</th><th>alignment</th><th>distance</th><th>position</th><th>gene name</th><th>gene id</th></tr>
<tr><td>GU305914.1</td><td>118571</td><td>118593</td><td>+</td><td>0</td><td>ATGGCGTCGGCGTCGGCGTC</td><td>TGG</td><td>||||||||[||||||||||||]PAM</td><td>NA</td><td>Intergenic</td><td>NA</td><td>NA</td></tr></table>

T8851 GATTTCACCTGCGAATACAGCGG 1000
Oligo fwd CACCGATTTCACCTGCGAATACAG
Oligo rev AAACCTGTATTCGCAGGTGAAATC

<table><tr><th>Chromosome</th><th>start</th><th>end</th><th>strand</th><th>MM</th><th>target_seq</th><th>PAM</th><th>alignment</th><th>distance</th><th>position</th><th>gene name</th><th>gene id</th></tr>
<tr><td>GU305914.1</td><td>118597</td><td>118619</td><td>+</td><td>0</td><td>GATTTCACCTGCGAATACAG</td><td>CGG</td><td>||||||||[||||||||||||]PAM</td><td>NA</td><td>Intergenic</td><td>NA</td><td>NA</td></tr>
<tr><td>GU305914.1</td><td>118645</td><td>118667</td><td>+</td><td>4</td><td>GATCTCGCCGGCGAACACAG</td><td>CGG</td><td>|||-||-|[|-|||||-||||]PAM</td><td>NA</td><td>Intergenic</td><td>NA</td><td>NA</td></tr></table>

T8852 TACAGCGGCGTCATCATCTTCGG 1000
Oligo fwd CACCTACAGCGGCGTCATCATCTT

<table>
<tr><td>Oligo rev</td><td>AAACAAGATGATGACGCCGCTGTA</td><td></td><td></td><td></td><td></td><td></td><td></td><td></td><td></td><td></td><td></td></tr>
<tr><td>Chromosome</td><td>start</td><td>end</td><td>strand</td><td>MM</td><td>target_seq</td><td>PAM</td><td>alignment</td><td>distance</td><td>position</td><td>gene name</td><td>gene id</td></tr>
<tr><td>GU305914.1</td><td>118612</td><td>118634</td><td>+</td><td>0</td><td>TACAGCGGCGTCATCATCTT</td><td>CGG</td><td>||||||||[||||||||||||]PAM</td><td>NA</td><td>Intergenic</td><td>NA</td><td>NA</td></tr>
<tr><td></td><td></td><td></td><td></td><td></td><td></td><td></td><td></td><td></td><td></td><td></td><td></td></tr>
<tr><td>T8853</td><td>GGCGTCATCATCTTCGGCGTCGG</td><td>1000</td><td></td><td></td><td></td><td></td><td></td><td></td><td></td><td></td><td></td></tr>
<tr><td>Oligo fwd</td><td>CACCGGCGTCATCATCTTCGGCGT</td><td></td><td></td><td></td><td></td><td></td><td></td><td></td><td></td><td></td><td></td></tr>
<tr><td>Oligo rev</td><td>AAACACGCCGAAGATGATGACGCC</td><td></td><td></td><td></td><td></td><td></td><td></td><td></td><td></td><td></td><td></td></tr>
<tr><td>Chromosome</td><td>start</td><td>end</td><td>strand</td><td>MM</td><td>target_seq</td><td>PAM</td><td>alignment</td><td>distance</td><td>position</td><td>gene name</td><td>gene id</td></tr>
<tr><td>GU305914.1</td><td>118618</td><td>118640</td><td>+</td><td>0</td><td>GGCGTCATCATCTTCGGCGT</td><td>CGG</td><td>||||||||[||||||||||||]PAM</td><td>NA</td><td>Intergenic</td><td>NA</td><td>NA</td></tr>
<tr><td></td><td></td><td></td><td></td><td></td><td></td><td></td><td></td><td></td><td></td><td></td><td></td></tr>
<tr><td>T8854</td><td>GCGTCATCATCTTCGGCGTCGGG</td><td>1000</td><td></td><td></td><td></td><td></td><td></td><td></td><td></td><td></td><td></td></tr>
<tr><td>Oligo fwd</td><td>CACCGCGTCATCATCTTCGGCGTC</td><td></td><td></td><td></td><td></td><td></td><td></td><td></td><td></td><td></td><td></td></tr>
<tr><td>Oligo rev</td><td>AAACGACGCCGAAGATGATGACGC</td><td></td><td></td><td></td><td></td><td></td><td></td><td></td><td></td><td></td><td></td></tr>
<tr><td>Chromosome</td><td>start</td><td>end</td><td>strand</td><td>MM</td><td>target_seq</td><td>PAM</td><td>alignment</td><td>distance</td><td>position</td><td>gene name</td><td>gene id</td></tr>
<tr><td>GU305914.1</td><td>118619</td><td>118641</td><td>+</td><td>0</td><td>GCGTCATCATCTTCGGCGTC</td><td>GGG</td><td>||||||||[||||||||||||]PAM</td><td>NA</td><td>Intergenic</td><td>NA</td><td>NA</td></tr>
<tr><td></td><td></td><td></td><td></td><td></td><td></td><td></td><td></td><td></td><td></td><td></td><td></td></tr>
<tr><td>T8855</td><td>GGCGTCGGGATCGATCTCGCCGG</td><td>1000</td><td></td><td></td><td></td><td></td><td></td><td></td><td></td><td></td><td></td></tr>
<tr><td>Oligo fwd</td><td>CACCGGCGTCGGGATCGATCTCGC</td><td></td><td></td><td></td><td></td><td></td><td></td><td></td><td></td><td></td><td></td></tr>
<tr><td>Oligo rev</td><td>AAACGCGAGATCGATCCCGACGCC</td><td></td><td></td><td></td><td></td><td></td><td></td><td></td><td></td><td></td><td></td></tr>
<tr><td>Chromosome</td><td>start</td><td>end</td><td>strand</td><td>MM</td><td>target_seq</td><td>PAM</td><td>alignment</td><td>distance</td><td>position</td><td>gene name</td><td>gene id</td></tr>
<tr><td>GU305914.1</td><td>118633</td><td>118655</td><td>+</td><td>0</td><td>GGCGTCGGGATCGATCTCGC</td><td>CGG</td><td>||||||||[||||||||||||]PAM</td><td>NA</td><td>Intergenic</td><td>NA</td><td>NA</td></tr>
<tr><td></td><td></td><td></td><td></td><td></td><td></td><td></td><td></td><td></td><td></td><td></td><td></td></tr>
<tr><td>T8856</td><td>GATCTCGCCGGCGAACACAGCGG</td><td>1000</td><td></td><td></td><td></td><td></td><td></td><td></td><td></td><td></td><td></td></tr>
<tr><td>Oligo fwd</td><td>CACCGATCTCGCCGGCGAACACAG</td><td></td><td></td><td></td><td></td><td></td><td></td><td></td><td></td><td></td><td></td></tr>
<tr><td>Oligo rev</td><td>AAACCTGTGTTCGCCGGCGAGATC</td><td></td><td></td><td></td><td></td><td></td><td></td><td></td><td></td><td></td><td></td></tr>
<tr><td>Chromosome</td><td>start</td><td>end</td><td>strand</td><td>MM</td><td>target_seq</td><td>PAM</td><td>alignment</td><td>distance</td><td>position</td><td>gene name</td><td>gene id</td></tr>
<tr><td>GU305914.1</td><td>118645</td><td>118667</td><td>+</td><td>0</td><td>GATCTCGCCGGCGAACACAG</td><td>CGG</td><td>||||||||[||||||||||||]PAM</td><td>NA</td><td>Intergenic</td><td>NA</td><td>NA</td></tr>
<tr><td>GU305914.1</td><td>118597</td><td>118619</td><td>+</td><td>4</td><td>GATTTCACCTGCGAATACAG</td><td>CGG</td><td>|||-||-|[|-|||||-||||]PAM</td><td>NA</td><td>Intergenic</td><td>NA</td><td>NA</td></tr>
<tr><td></td><td></td><td></td><td></td><td></td><td></td><td></td><td></td><td></td><td></td><td></td><td></td></tr>
<tr><td>T8857</td><td>AGCGGCGTCCTCGTCTTCCTCGG</td><td>1000</td><td></td><td></td><td></td><td></td><td></td><td></td><td></td><td></td><td></td></tr>
<tr><td>Oligo fwd</td><td>CACCAGCGGCGTCCTCGTCTTCCT</td><td></td><td></td><td></td><td></td><td></td><td></td><td></td><td></td><td></td><td></td></tr>
<tr><td>Oligo rev</td><td>AAACAGGAAGACGAGGACGCCGCT</td><td></td><td></td><td></td><td></td><td></td><td></td><td></td><td></td><td></td><td></td></tr>
<tr><td>Chromosome</td><td>start</td><td>end</td><td>strand</td><td>MM</td><td>target_seq</td><td>PAM</td><td>alignment</td><td>distance</td><td>position</td><td>gene name</td><td>gene id</td></tr>
<tr><td>GU305914.1</td><td>118663</td><td>118685</td><td>+</td><td>0</td><td>AGCGGCGTCCTCGTCTTCCT</td><td>CGG</td><td>||||||||[||||||||||||]PAM</td><td>NA</td><td>Intergenic</td><td>NA</td><td>NA</td></tr>
<tr><td></td><td></td><td></td><td></td><td></td><td></td><td></td><td></td><td></td><td></td><td></td><td></td></tr>
<tr><td>T8858</td><td>GCGTCCTCGTCTTCCTCGGCCGG</td><td>1000</td><td></td><td></td><td></td><td></td><td></td><td></td><td></td><td></td><td></td></tr>
<tr><td>Oligo fwd</td><td>CACCGCGTCCTCGTCTTCCTCGGC</td><td></td><td></td><td></td><td></td><td></td><td></td><td></td><td></td><td></td><td></td></tr>
<tr><td>Oligo rev</td><td>AAACGCCGAGGAAGACGAGGACGC</td><td></td><td></td><td></td><td></td><td></td><td></td><td></td><td></td><td></td><td></td></tr>
<tr><td>Chromosome</td><td>start</td><td>end</td><td>strand</td><td>MM</td><td>target_seq</td><td>PAM</td><td>alignment</td><td>distance</td><td>position</td><td>gene name</td><td>gene id</td></tr>
<tr><td>GU305914.1</td><td>118667</td><td>118689</td><td>+</td><td>0</td><td>GCGTCCTCGTCTTCCTCGGC</td><td>CGG</td><td>||||||||[||||||||||||]PAM</td><td>NA</td><td>Intergenic</td><td>NA</td><td>NA</td></tr>
<tr><td></td><td></td><td></td><td></td><td></td><td></td><td></td><td></td><td></td><td></td><td></td><td></td></tr>
<tr><td>T8859</td><td>CGTCCTCGTCTTCCTCGGCCGGG</td><td>1000</td><td></td><td></td><td></td><td></td><td></td><td></td><td></td><td></td><td></td></tr>
<tr><td>Oligo fwd</td><td>CACCCGTCCTCGTCTTCCTCGGCC</td><td></td><td></td><td></td><td></td><td></td><td></td><td></td><td></td><td></td><td></td></tr>
<tr><td>Oligo rev</td><td>AAACGGCCGAGGAAGACGAGGACG</td><td></td><td></td><td></td><td></td><td></td><td></td><td></td><td></td><td></td><td></td></tr>
<tr><td>Chromosome</td><td>start</td><td>end</td><td>strand</td><td>MM</td><td>target_seq</td><td>PAM</td><td>alignment</td><td>distance</td><td>position</td><td>gene name</td><td>gene id</td></tr>
<tr><td>GU305914.1</td><td>118668</td><td>118690</td><td>+</td><td>0</td><td>CGTCCTCGTCTTCCTCGGCC</td><td>GGG</td><td>||||||||[||||||||||||]PAM</td><td>NA</td><td>Intergenic</td><td>NA</td><td>NA</td></tr>
<tr><td></td><td></td><td></td><td></td><td></td><td></td><td></td><td></td><td></td><td></td><td></td><td></td></tr>
<tr><td>T8860</td><td>CCTCGTCTTCCTCGGCCGGGCGG</td><td>1000</td><td></td><td></td><td></td><td></td><td></td><td></td><td></td><td></td><td></td></tr>
<tr><td>Oligo fwd</td><td>CACCCCTCGTCTTCCTCGGCCGGG</td><td></td><td></td><td></td><td></td><td></td><td></td><td></td><td></td><td></td><td></td></tr>
<tr><td>Oligo rev</td><td>AAACCCCGGCCGAGGAAGACGAGG</td><td></td><td></td><td></td><td></td><td></td><td></td><td></td><td></td><td></td><td></td></tr>
<tr><td>Chromosome</td><td>start</td><td>end</td><td>strand</td><td>MM</td><td>target_seq</td><td>PAM</td><td>alignment</td><td>distance</td><td>position</td><td>gene name</td><td>gene id</td></tr>
<tr><td>GU305914.1</td><td>118671</td><td>118693</td><td>+</td><td>0</td><td>CCTCGTCTTCCTCGGCCGGG</td><td>CGG</td><td>||||||||[||||||||||||]PAM</td><td>NA</td><td>Intergenic</td><td>NA</td><td>NA</td></tr>
<tr><td></td><td></td><td></td><td></td><td></td><td></td><td></td><td></td><td></td><td></td><td></td><td></td></tr>
<tr><td>T8861</td><td>GTCTTCCTCGGCCGGGCGGTTGG</td><td>1000</td><td></td><td></td><td></td><td></td><td></td><td></td><td></td><td></td><td></td></tr>
<tr><td>Oligo fwd</td><td>CACCGTCTTCCTCGGCCGGGCGGT</td><td></td><td></td><td></td><td></td><td></td><td></td><td></td><td></td><td></td><td></td></tr>
<tr><td>Oligo rev</td><td>AAACACCGCCCGGCCGAGGAAGAC</td><td></td><td></td><td></td><td></td><td></td><td></td><td></td><td></td><td></td><td></td></tr>
<tr><td>Chromosome</td><td>start</td><td>end</td><td>strand</td><td>MM</td><td>target_seq</td><td>PAM</td><td>alignment</td><td>distance</td><td>position</td><td>gene name</td><td>gene id</td></tr>
<tr><td>GU305914.1</td><td>118675</td><td>118697</td><td>+</td><td>0</td><td>GTCTTCCTCGGCCGGGCGGT</td><td>TGG</td><td>||||||||[||||||||||||]PAM</td><td>NA</td><td>Intergenic</td><td>NA</td><td>NA</td></tr>
<tr><td></td><td></td><td></td><td></td><td></td><td></td><td></td><td></td><td></td><td></td><td></td><td></td></tr>
<tr><td>T8862</td><td>TCGGCCGGGCGGTTGGCTCTCGG</td><td>1000</td><td></td><td></td><td></td><td></td><td></td><td></td><td></td><td></td><td></td></tr>
<tr><td>Oligo fwd</td><td>CACCTCGGCCGGGCGGTTGGCTCT</td><td></td><td></td><td></td><td></td><td></td><td></td><td></td><td></td><td></td><td></td></tr>
<tr><td>Oligo rev</td><td>AAACAGAGCCAACCGCCCGGCCGA</td><td></td><td></td><td></td><td></td><td></td><td></td><td></td><td></td><td></td><td></td></tr>
<tr><td>Chromosome</td><td>start</td><td>end</td><td>strand</td><td>MM</td><td>target_seq</td><td>PAM</td><td>alignment</td><td>distance</td><td>position</td><td>gene name</td><td>gene id</td></tr>
<tr><td>GU305914.1</td><td>118682</td><td>118704</td><td>+</td><td>0</td><td>TCGGCCGGGCGGTTGGCTCT</td><td>CGG</td><td>||||||||[||||||||||||]PAM</td><td>NA</td><td>Intergenic</td><td>NA</td><td>NA</td></tr>
<tr><td></td><td></td><td></td><td></td><td></td><td></td><td></td><td></td><td></td><td></td><td></td><td></td></tr>
<tr><td>T8863</td><td>GGGCGGTTGGCTCTCGGTGATGG</td><td>1000</td><td></td><td></td><td></td><td></td><td></td><td></td><td></td><td></td><td></td></tr>
<tr><td>Oligo fwd</td><td>CACCGGGCGGTTGGCTCTCGGTGA</td><td></td><td></td><td></td><td></td><td></td><td></td><td></td><td></td><td></td><td></td></tr>
<tr><td>Oligo rev</td><td>AAACTCACCGAGAGCCAACCGCCC</td><td></td><td></td><td></td><td></td><td></td><td></td><td></td><td></td><td></td><td></td></tr>
<tr><td>Chromosome</td><td>start</td><td>end</td><td>strand</td><td>MM</td><td>target_seq</td><td>PAM</td><td>alignment</td><td>distance</td><td>position</td><td>gene name</td><td>gene id</td></tr>
<tr><td>GU305914.1</td><td>118688</td><td>118710</td><td>+</td><td>0</td><td>GGGCGGTTGGCTCTCGGTGA</td><td>TGG</td><td>||||||||[||||||||||||]PAM</td><td>NA</td><td>Intergenic</td><td>NA</td><td>NA</td></tr>
<tr><td></td><td></td><td></td><td></td><td></td><td></td><td></td><td></td><td></td><td></td><td></td><td></td></tr>
<tr><td>T8864</td><td>GATGGTTGCCGCTGTTGCTGCGG</td><td>1000</td><td></td><td></td><td></td><td></td><td></td><td></td><td></td><td></td><td></td></tr>
<tr><td>Oligo fwd</td><td>CACCGATGGTTGCCGCTGTTGCTG</td><td></td><td></td><td></td><td></td><td></td><td></td><td></td><td></td><td></td><td></td></tr>
<tr><td>Oligo rev</td><td>AAACCAGCAACAGCGGCAACCATC</td><td></td><td></td><td></td><td></td><td></td><td></td><td></td><td></td><td></td><td></td></tr>
<tr><td>Chromosome</td><td>start</td><td>end</td><td>strand</td><td>MM</td><td>target_seq</td><td>PAM</td><td>alignment</td><td>distance</td><td>position</td><td>gene name</td><td>gene id</td></tr>
<tr><td>GU305914.1</td><td>118706</td><td>118728</td><td>+</td><td>0</td><td>GATGGTTGCCGCTGTTGCTG</td><td>CGG</td><td>||||||||[||||||||||||]PAM</td><td>NA</td><td>Intergenic</td><td>NA</td><td>NA</td></tr>
<tr><td></td><td></td><td></td><td></td><td></td><td></td><td></td><td></td><td></td><td></td><td></td><td></td></tr>
<tr><td>T8865</td><td>GGTTGCCGCTGTTGCTGCGGTGG</td><td>1000</td><td></td><td></td><td></td><td></td><td></td><td></td><td></td><td></td><td></td></tr>
<tr><td>Oligo fwd</td><td>CACCGGTTGCCGCTGTTGCTGCGG</td><td></td><td></td><td></td><td></td><td></td><td></td><td></td><td></td><td></td><td></td></tr>
<tr><td>Oligo rev</td><td>AAACCCGCAGCAACAGCGGCAACC</td><td></td><td></td><td></td><td></td><td></td><td></td><td></td><td></td><td></td><td></td></tr>
<tr><td>Chromosome</td><td>start</td><td>end</td><td>strand</td><td>MM</td><td>target_seq</td><td>PAM</td><td>alignment</td><td>distance</td><td>position</td><td>gene name</td><td>gene id</td></tr>
<tr><td>GU305914.1</td><td>118709</td><td>118731</td><td>+</td><td>0</td><td>GGTTGCCGCTGTTGCTGCGG</td><td>TGG</td><td>||||||||[||||||||||||]PAM</td><td>NA</td><td>Intergenic</td><td>NA</td><td>NA</td></tr>
<tr><td></td><td></td><td></td><td></td><td></td><td></td><td></td><td></td><td></td><td></td><td></td><td></td></tr>
<tr><td>T8866</td><td>GTTGCCGCTGTTGCTGCGGTGGG</td><td>1000</td><td></td><td></td><td></td><td></td><td></td><td></td><td></td><td></td><td></td></tr>
<tr><td>Oligo fwd</td><td>CACCGTTGCCGCTGTTGCTGCGGT</td><td></td><td></td><td></td><td></td><td></td><td></td><td></td><td></td><td></td><td></td></tr>
<tr><td>Oligo rev</td><td>AAACACCGCAGCAACAGCGGCAAC</td><td></td><td></td><td></td><td></td><td></td><td></td><td></td><td></td><td></td><td></td></tr>
<tr><td>Chromosome</td><td>start</td><td>end</td><td>strand</td><td>MM</td><td>target_seq</td><td>PAM</td><td>alignment</td><td>distance</td><td>position</td><td>gene name</td><td>gene id</td></tr>
<tr><td>GU305914.1</td><td>118710</td><td>118732</td><td>+</td><td>0</td><td>GTTGCCGCTGTTGCTGCGGT</td><td>GGG</td><td>||||||||[||||||||||||]PAM</td><td>NA</td><td>Intergenic</td><td>NA</td><td>NA</td></tr>
</table>

<table>
<tr><td>T8867</td><td>TTGCCGCTGTTGCTGCGGTGGGG</td><td>1000</td><td></td><td></td><td></td><td></td><td></td><td></td><td></td><td></td><td></td></tr>
<tr><td>Oligo fwd</td><td>CACCTTGCCGCTGTTGCTGCGGTG</td><td></td><td></td><td></td><td></td><td></td><td></td><td></td><td></td><td></td><td></td></tr>
<tr><td>Oligo rev</td><td>AAACCACCGCAGCAACAGCGGCAA</td><td></td><td></td><td></td><td></td><td></td><td></td><td></td><td></td><td></td><td></td></tr>
<tr><td>Chromosome</td><td>start</td><td>end</td><td>strand</td><td>MM</td><td>target_seq</td><td>PAM</td><td>alignment</td><td>distance</td><td>position</td><td>gene name</td><td>gene id</td></tr>
<tr><td>GU305914.1</td><td>118711</td><td>118733</td><td>+</td><td>0</td><td>TTGCCGCTGTTGCTGCGGTG</td><td>GGG</td><td>||||||||[||||||||||||]PAM</td><td>NA</td><td>Intergenic</td><td>NA</td><td>NA</td></tr>
<tr><td>T8868</td><td>CTGCGGTGGGGCCTGTCGCTCGG</td><td>1000</td><td></td><td></td><td></td><td></td><td></td><td></td><td></td><td></td><td></td></tr>
<tr><td>Oligo fwd</td><td>CACCCTGCGGTGGGGCCTGTCGCT</td><td></td><td></td><td></td><td></td><td></td><td></td><td></td><td></td><td></td><td></td></tr>
<tr><td>Oligo rev</td><td>AAACAGCGACAGGCCCCACCGCAG</td><td></td><td></td><td></td><td></td><td></td><td></td><td></td><td></td><td></td><td></td></tr>
<tr><td>Chromosome</td><td>start</td><td>end</td><td>strand</td><td>MM</td><td>target_seq</td><td>PAM</td><td>alignment</td><td>distance</td><td>position</td><td>gene name</td><td>gene id</td></tr>
<tr><td>GU305914.1</td><td>118723</td><td>118745</td><td>+</td><td>0</td><td>CTGCGGTGGGGCCTGTCGCT</td><td>CGG</td><td>||||||||[||||||||||||]PAM</td><td>NA</td><td>Intergenic</td><td>NA</td><td>NA</td></tr>
<tr><td>T8869</td><td>TGCGGTGGGGCCTGTCGCTCGGG</td><td>1000</td><td></td><td></td><td></td><td></td><td></td><td></td><td></td><td></td><td></td></tr>
<tr><td>Oligo fwd</td><td>CACCTGCGGTGGGGCCTGTCGCTC</td><td></td><td></td><td></td><td></td><td></td><td></td><td></td><td></td><td></td><td></td></tr>
<tr><td>Oligo rev</td><td>AAACGAGCGACAGGCCCCACCGCA</td><td></td><td></td><td></td><td></td><td></td><td></td><td></td><td></td><td></td><td></td></tr>
<tr><td>Chromosome</td><td>start</td><td>end</td><td>strand</td><td>MM</td><td>target_seq</td><td>PAM</td><td>alignment</td><td>distance</td><td>position</td><td>gene name</td><td>gene id</td></tr>
<tr><td>GU305914.1</td><td>118724</td><td>118746</td><td>+</td><td>0</td><td>TGCGGTGGGGCCTGTCGCTC</td><td>GGG</td><td>||||||||[||||||||||||]PAM</td><td>NA</td><td>Intergenic</td><td>NA</td><td>NA</td></tr>
<tr><td>T8870</td><td>TGGGGCCTGTCGCTCGGGTTCGG</td><td>1000</td><td></td><td></td><td></td><td></td><td></td><td></td><td></td><td></td><td></td></tr>
<tr><td>Oligo fwd</td><td>CACCTGGGGCCTGTCGCTCGGGTT</td><td></td><td></td><td></td><td></td><td></td><td></td><td></td><td></td><td></td><td></td></tr>
<tr><td>Oligo rev</td><td>AAACAACCCGAGCGACAGGCCCCA</td><td></td><td></td><td></td><td></td><td></td><td></td><td></td><td></td><td></td><td></td></tr>
<tr><td>Chromosome</td><td>start</td><td>end</td><td>strand</td><td>MM</td><td>target_seq</td><td>PAM</td><td>alignment</td><td>distance</td><td>position</td><td>gene name</td><td>gene id</td></tr>
<tr><td>GU305914.1</td><td>118729</td><td>118751</td><td>+</td><td>0</td><td>TGGGGCCTGTCGCTCGGGTT</td><td>CGG</td><td>||||||||[||||||||||||]PAM</td><td>NA</td><td>Intergenic</td><td>NA</td><td>NA</td></tr>
<tr><td>T8871</td><td>GGGTTCGGCATCTTCGTTCGAGG</td><td>1000</td><td></td><td></td><td></td><td></td><td></td><td></td><td></td><td></td><td></td></tr>
<tr><td>Oligo fwd</td><td>CACCGGGTTCGGCATCTTCGTTCG</td><td></td><td></td><td></td><td></td><td></td><td></td><td></td><td></td><td></td><td></td></tr>
<tr><td>Oligo rev</td><td>AAACCGAACGAAGATGCCGAACCC</td><td></td><td></td><td></td><td></td><td></td><td></td><td></td><td></td><td></td><td></td></tr>
<tr><td>Chromosome</td><td>start</td><td>end</td><td>strand</td><td>MM</td><td>target_seq</td><td>PAM</td><td>alignment</td><td>distance</td><td>position</td><td>gene name</td><td>gene id</td></tr>
<tr><td>GU305914.1</td><td>118744</td><td>118766</td><td>+</td><td>0</td><td>GGGTTCGGCATCTTCGTTCG</td><td>AGG</td><td>||||||||[||||||||||||]PAM</td><td>NA</td><td>Intergenic</td><td>NA</td><td>NA</td></tr>
<tr><td>T8872</td><td>TTCTCATCGTCCATCCCGTTTGG</td><td>1000</td><td></td><td></td><td></td><td></td><td></td><td></td><td></td><td></td><td></td></tr>
<tr><td>Oligo fwd</td><td>CACCTTCTCATCGTCCATCCCGTT</td><td></td><td></td><td></td><td></td><td></td><td></td><td></td><td></td><td></td><td></td></tr>
<tr><td>Oligo rev</td><td>AAACAACGGGATGGACGATGAGAA</td><td></td><td></td><td></td><td></td><td></td><td></td><td></td><td></td><td></td><td></td></tr>
<tr><td>Chromosome</td><td>start</td><td>end</td><td>strand</td><td>MM</td><td>target_seq</td><td>PAM</td><td>alignment</td><td>distance</td><td>position</td><td>gene name</td><td>gene id</td></tr>
<tr><td>GU305914.1</td><td>118772</td><td>118794</td><td>+</td><td>0</td><td>TTCTCATCGTCCATCCCGTT</td><td>TGG</td><td>||||||||[||||||||||||]PAM</td><td>NA</td><td>Intergenic</td><td>NA</td><td>NA</td></tr>
<tr><td>T8873</td><td>TCTCATCGTCCATCCCGTTTGGG</td><td>1000</td><td></td><td></td><td></td><td></td><td></td><td></td><td></td><td></td><td></td></tr>
<tr><td>Oligo fwd</td><td>CACCTCTCATCGTCCATCCCGTTT</td><td></td><td></td><td></td><td></td><td></td><td></td><td></td><td></td><td></td><td></td></tr>
<tr><td>Oligo rev</td><td>AAACAAACGGGATGGACGATGAGA</td><td></td><td></td><td></td><td></td><td></td><td></td><td></td><td></td><td></td><td></td></tr>
<tr><td>Chromosome</td><td>start</td><td>end</td><td>strand</td><td>MM</td><td>target_seq</td><td>PAM</td><td>alignment</td><td>distance</td><td>position</td><td>gene name</td><td>gene id</td></tr>
<tr><td>GU305914.1</td><td>118773</td><td>118795</td><td>+</td><td>0</td><td>TCTCATCGTCCATCCCGTTT</td><td>GGG</td><td>||||||||[||||||||||||]PAM</td><td>NA</td><td>Intergenic</td><td>NA</td><td>NA</td></tr>
<tr><td>T8874</td><td>TTCGTCTTCTTCCTCCTCCTCGG</td><td>1000</td><td></td><td></td><td></td><td></td><td></td><td></td><td></td><td></td><td></td></tr>
<tr><td>Oligo fwd</td><td>CACCTTCGTCTTCTTCCTCCTCCT</td><td></td><td></td><td></td><td></td><td></td><td></td><td></td><td></td><td></td><td></td></tr>
<tr><td>Oligo rev</td><td>AAACAGGAGGAGGAAGAAGACGAA</td><td></td><td></td><td></td><td></td><td></td><td></td><td></td><td></td><td></td><td></td></tr>
<tr><td>Chromosome</td><td>start</td><td>end</td><td>strand</td><td>MM</td><td>target_seq</td><td>PAM</td><td>alignment</td><td>distance</td><td>position</td><td>gene name</td><td>gene id</td></tr>
<tr><td>GU305914.1</td><td>118855</td><td>118877</td><td>+</td><td>0</td><td>TTCGTCTTCTTCCTCCTCCT</td><td>CGG</td><td>||||||||[||||||||||||]PAM</td><td>NA</td><td>Intergenic</td><td>NA</td><td>NA</td></tr>
<tr><td>T8875</td><td>TTCTTCCTCCTCCTCGGCGTCGG</td><td>1000</td><td></td><td></td><td></td><td></td><td></td><td></td><td></td><td></td><td></td></tr>
<tr><td>Oligo fwd</td><td>CACCTTCTTCCTCCTCCTCGGCGT</td><td></td><td></td><td></td><td></td><td></td><td></td><td></td><td></td><td></td><td></td></tr>
<tr><td>Oligo rev</td><td>AAACACGCCGAGGAGGAGGAAGAA</td><td></td><td></td><td></td><td></td><td></td><td></td><td></td><td></td><td></td><td></td></tr>
<tr><td>Chromosome</td><td>start</td><td>end</td><td>strand</td><td>MM</td><td>target_seq</td><td>PAM</td><td>alignment</td><td>distance</td><td>position</td><td>gene name</td><td>gene id</td></tr>
<tr><td>GU305914.1</td><td>118861</td><td>118883</td><td>+</td><td>0</td><td>TTCTTCCTCCTCCTCGGCGT</td><td>CGG</td><td>||||||||[||||||||||||]PAM</td><td>NA</td><td>Intergenic</td><td>NA</td><td>NA</td></tr>
<tr><td>T8876</td><td>CGGACGACGACGCCTCGTCCTGG</td><td>1000</td><td></td><td></td><td></td><td></td><td></td><td></td><td></td><td></td><td></td></tr>
<tr><td>Oligo fwd</td><td>CACCCGGACGACGACGCCTCGTCC</td><td></td><td></td><td></td><td></td><td></td><td></td><td></td><td></td><td></td><td></td></tr>
<tr><td>Oligo rev</td><td>AAACGGACGAGGCGTCGTCGTCCG</td><td></td><td></td><td></td><td></td><td></td><td></td><td></td><td></td><td></td><td></td></tr>
<tr><td>Chromosome</td><td>start</td><td>end</td><td>strand</td><td>MM</td><td>target_seq</td><td>PAM</td><td>alignment</td><td>distance</td><td>position</td><td>gene name</td><td>gene id</td></tr>
<tr><td>GU305914.1</td><td>118881</td><td>118903</td><td>+</td><td>0</td><td>CGGACGACGACGCCTCGTCC</td><td>TGG</td><td>||||||||[||||||||||||]PAM</td><td>NA</td><td>Intergenic</td><td>NA</td><td>NA</td></tr>
<tr><td>T8877</td><td>GGACGACGACGCCTCGTCCTGGG</td><td>1000</td><td></td><td></td><td></td><td></td><td></td><td></td><td></td><td></td><td></td></tr>
<tr><td>Oligo fwd</td><td>CACCGGACGACGACGCCTCGTCCT</td><td></td><td></td><td></td><td></td><td></td><td></td><td></td><td></td><td></td><td></td></tr>
<tr><td>Oligo rev</td><td>AAACAGGACGAGGCGTCGTCGTCC</td><td></td><td></td><td></td><td></td><td></td><td></td><td></td><td></td><td></td><td></td></tr>
<tr><td>Chromosome</td><td>start</td><td>end</td><td>strand</td><td>MM</td><td>target_seq</td><td>PAM</td><td>alignment</td><td>distance</td><td>position</td><td>gene name</td><td>gene id</td></tr>
<tr><td>GU305914.1</td><td>118882</td><td>118904</td><td>+</td><td>0</td><td>GGACGACGACGCCTCGTCCT</td><td>GGG</td><td>||||||||[||||||||||||]PAM</td><td>NA</td><td>Intergenic</td><td>NA</td><td>NA</td></tr>
<tr><td>GU305914.1</td><td>18510</td><td>18532</td><td>-</td><td>4</td><td>TGGCGACGACGCCCCGTCGT</td><td>TGG</td><td>-|-|||||[|||||-||||-|]PAM</td><td>NA</td><td>Intergenic</td><td>NA</td><td>NA</td></tr>
<tr><td>T8878</td><td>ACGACGCGTTGTCCTGCTCTAGG</td><td>1000</td><td></td><td></td><td></td><td></td><td></td><td></td><td></td><td></td><td></td></tr>
<tr><td>Oligo fwd</td><td>CACCACGACGCGTTGTCCTGCTCT</td><td></td><td></td><td></td><td></td><td></td><td></td><td></td><td></td><td></td><td></td></tr>
<tr><td>Oligo rev</td><td>AAACAGAGCAGGACAACGCGTCGT</td><td></td><td></td><td></td><td></td><td></td><td></td><td></td><td></td><td></td><td></td></tr>
<tr><td>Chromosome</td><td>start</td><td>end</td><td>strand</td><td>MM</td><td>target_seq</td><td>PAM</td><td>alignment</td><td>distance</td><td>position</td><td>gene name</td><td>gene id</td></tr>
<tr><td>GU305914.1</td><td>118905</td><td>118927</td><td>+</td><td>0</td><td>ACGACGCGTTGTCCTGCTCT</td><td>AGG</td><td>||||||||[||||||||||||]PAM</td><td>NA</td><td>Intergenic</td><td>NA</td><td>NA</td></tr>
<tr><td>T8879</td><td>GTCCTGCTCTAGGTCCTGTATGG</td><td>1000</td><td></td><td></td><td></td><td></td><td></td><td></td><td></td><td></td><td></td></tr>
<tr><td>Oligo fwd</td><td>CACCGTCCTGCTCTAGGTCCTGTA</td><td></td><td></td><td></td><td></td><td></td><td></td><td></td><td></td><td></td><td></td></tr>
<tr><td>Oligo rev</td><td>AAACTACAGGACCTAGAGCAGGAC</td><td></td><td></td><td></td><td></td><td></td><td></td><td></td><td></td><td></td><td></td></tr>
<tr><td>Chromosome</td><td>start</td><td>end</td><td>strand</td><td>MM</td><td>target_seq</td><td>PAM</td><td>alignment</td><td>distance</td><td>position</td><td>gene name</td><td>gene id</td></tr>
<tr><td>GU305914.1</td><td>118915</td><td>118937</td><td>+</td><td>0</td><td>GTCCTGCTCTAGGTCCTGTA</td><td>TGG</td><td>||||||||[||||||||||||]PAM</td><td>NA</td><td>Intergenic</td><td>NA</td><td>NA</td></tr>
<tr><td>T8880</td><td>TCCTGCTCTAGGTCCTGTATGGG</td><td>1000</td><td></td><td></td><td></td><td></td><td></td><td></td><td></td><td></td><td></td></tr>
<tr><td>Oligo fwd</td><td>CACCTCCTGCTCTAGGTCCTGTAT</td><td></td><td></td><td></td><td></td><td></td><td></td><td></td><td></td><td></td><td></td></tr>
<tr><td>Oligo rev</td><td>AAACATACAGGACCTAGAGCAGGA</td><td></td><td></td><td></td><td></td><td></td><td></td><td></td><td></td><td></td><td></td></tr>
<tr><td>Chromosome</td><td>start</td><td>end</td><td>strand</td><td>MM</td><td>target_seq</td><td>PAM</td><td>alignment</td><td>distance</td><td>position</td><td>gene name</td><td>gene id</td></tr>
<tr><td>GU305914.1</td><td>118916</td><td>118938</td><td>+</td><td>0</td><td>TCCTGCTCTAGGTCCTGTAT</td><td>GGG</td><td>||||||||[||||||||||||]PAM</td><td>NA</td><td>Intergenic</td><td>NA</td><td>NA</td></tr>
<tr><td>GU305914.1</td><td>74182</td><td>74204</td><td>-</td><td>4</td><td>TGCTCCTCTAGGACCTGTTT</td><td>CGG</td><td>|-||-|||[||||-|||||-|]PAM</td><td>NA</td><td>Intergenic</td><td>NA</td><td>NA</td></tr>
<tr><td>T8881</td><td>CCTGCTCTAGGTCCTGTATGGGG</td><td>1000</td><td></td><td></td><td></td><td></td><td></td><td></td><td></td><td></td><td></td></tr>
<tr><td>Oligo fwd</td><td>CACCCCTGCTCTAGGTCCTGTATG</td><td></td><td></td><td></td><td></td><td></td><td></td><td></td><td></td><td></td><td></td></tr>
<tr><td>Oligo rev</td><td>AAACCATACAGGACCTAGAGCAGG</td><td></td><td></td><td></td><td></td><td></td><td></td><td></td><td></td><td></td><td></td></tr>
</table>

<table>
<tr><td>Chromosome</td><td>start</td><td>end</td><td>strand</td><td>MM</td><td>target_seq</td><td>PAM</td><td>alignment</td><td>distance</td><td>position</td><td>gene name</td><td>gene id</td></tr>
<tr><td>GU305914.1</td><td>118917</td><td>118939</td><td>+</td><td>0</td><td>CCTGCTCTAGGTCCTGTATG</td><td>GGG</td><td>||||||||[||||||||||||]PAM</td><td>NA</td><td>Intergenic</td><td>NA</td><td>NA</td></tr>
<tr><td>T8882</td><td>GCTCTAGGTCCTGTATGGGGAGG</td><td>1000</td><td></td><td></td><td></td><td></td><td></td><td></td><td></td><td></td><td></td></tr>
<tr><td>Oligo fwd</td><td>CACCGCTCTAGGTCCTGTATGGGG</td><td></td><td></td><td></td><td></td><td></td><td></td><td></td><td></td><td></td><td></td></tr>
<tr><td>Oligo rev</td><td>AAACCCCCATACAGGACCTAGAGC</td><td></td><td></td><td></td><td></td><td></td><td></td><td></td><td></td><td></td><td></td></tr>
<tr><td>Chromosome</td><td>start</td><td>end</td><td>strand</td><td>MM</td><td>target_seq</td><td>PAM</td><td>alignment</td><td>distance</td><td>position</td><td>gene name</td><td>gene id</td></tr>
<tr><td>GU305914.1</td><td>118920</td><td>118942</td><td>+</td><td>0</td><td>GCTCTAGGTCCTGTATGGGG</td><td>AGG</td><td>||||||||[||||||||||||]PAM</td><td>NA</td><td>Intergenic</td><td>NA</td><td>NA</td></tr>
<tr><td>T8883</td><td>CTCTAGGTCCTGTATGGGGAGGG</td><td>1000</td><td></td><td></td><td></td><td></td><td></td><td></td><td></td><td></td><td></td></tr>
<tr><td>Oligo fwd</td><td>CACCCTCTAGGTCCTGTATGGGGA</td><td></td><td></td><td></td><td></td><td></td><td></td><td></td><td></td><td></td><td></td></tr>
<tr><td>Oligo rev</td><td>AAACTCCCCATACAGGACCTAGAG</td><td></td><td></td><td></td><td></td><td></td><td></td><td></td><td></td><td></td><td></td></tr>
<tr><td>Chromosome</td><td>start</td><td>end</td><td>strand</td><td>MM</td><td>target_seq</td><td>PAM</td><td>alignment</td><td>distance</td><td>position</td><td>gene name</td><td>gene id</td></tr>
<tr><td>GU305914.1</td><td>118921</td><td>118943</td><td>+</td><td>0</td><td>CTCTAGGTCCTGTATGGGGA</td><td>GGG</td><td>||||||||[||||||||||||]PAM</td><td>NA</td><td>Intergenic</td><td>NA</td><td>NA</td></tr>
<tr><td>T8884</td><td>TCTAGGTCCTGTATGGGGAGGGG</td><td>1000</td><td></td><td></td><td></td><td></td><td></td><td></td><td></td><td></td><td></td></tr>
<tr><td>Oligo fwd</td><td>CACCTCTAGGTCCTGTATGGGGAG</td><td></td><td></td><td></td><td></td><td></td><td></td><td></td><td></td><td></td><td></td></tr>
<tr><td>Oligo rev</td><td>AAACCTCCCCATACAGGACCTAGA</td><td></td><td></td><td></td><td></td><td></td><td></td><td></td><td></td><td></td><td></td></tr>
<tr><td>Chromosome</td><td>start</td><td>end</td><td>strand</td><td>MM</td><td>target_seq</td><td>PAM</td><td>alignment</td><td>distance</td><td>position</td><td>gene name</td><td>gene id</td></tr>
<tr><td>GU305914.1</td><td>118922</td><td>118944</td><td>+</td><td>0</td><td>TCTAGGTCCTGTATGGGGAG</td><td>GGG</td><td>||||||||[||||||||||||]PAM</td><td>NA</td><td>Intergenic</td><td>NA</td><td>NA</td></tr>
<tr><td>T8885</td><td>CTAGGTCCTGTATGGGGAGGGGG</td><td>1000</td><td></td><td></td><td></td><td></td><td></td><td></td><td></td><td></td><td></td></tr>
<tr><td>Oligo fwd</td><td>CACCCTAGGTCCTGTATGGGGAGG</td><td></td><td></td><td></td><td></td><td></td><td></td><td></td><td></td><td></td><td></td></tr>
<tr><td>Oligo rev</td><td>AAACCCTCCCCATACAGGACCTAG</td><td></td><td></td><td></td><td></td><td></td><td></td><td></td><td></td><td></td><td></td></tr>
<tr><td>Chromosome</td><td>start</td><td>end</td><td>strand</td><td>MM</td><td>target_seq</td><td>PAM</td><td>alignment</td><td>distance</td><td>position</td><td>gene name</td><td>gene id</td></tr>
<tr><td>GU305914.1</td><td>118923</td><td>118945</td><td>+</td><td>0</td><td>CTAGGTCCTGTATGGGGAGG</td><td>GGG</td><td>||||||||[||||||||||||]PAM</td><td>NA</td><td>Intergenic</td><td>NA</td><td>NA</td></tr>
<tr><td>T8886</td><td>GGGAGGGGGAGCGTCGTGTCCGG</td><td>1000</td><td></td><td></td><td></td><td></td><td></td><td></td><td></td><td></td><td></td></tr>
<tr><td>Oligo fwd</td><td>CACCGGGAGGGGGAGCGTCGTGTC</td><td></td><td></td><td></td><td></td><td></td><td></td><td></td><td></td><td></td><td></td></tr>
<tr><td>Oligo rev</td><td>AAACGACACGACGCTCCCCCTCCC</td><td></td><td></td><td></td><td></td><td></td><td></td><td></td><td></td><td></td><td></td></tr>
<tr><td>Chromosome</td><td>start</td><td>end</td><td>strand</td><td>MM</td><td>target_seq</td><td>PAM</td><td>alignment</td><td>distance</td><td>position</td><td>gene name</td><td>gene id</td></tr>
<tr><td>GU305914.1</td><td>118937</td><td>118959</td><td>+</td><td>0</td><td>GGGAGGGGGAGCGTCGTGTC</td><td>CGG</td><td>||||||||[||||||||||||]PAM</td><td>NA</td><td>Intergenic</td><td>NA</td><td>NA</td></tr>
<tr><td>T8887</td><td>GGGGGAGCGTCGTGTCCGGCAGG</td><td>1000</td><td></td><td></td><td></td><td></td><td></td><td></td><td></td><td></td><td></td></tr>
<tr><td>Oligo fwd</td><td>CACCGGGGGAGCGTCGTGTCCGGC</td><td></td><td></td><td></td><td></td><td></td><td></td><td></td><td></td><td></td><td></td></tr>
<tr><td>Oligo rev</td><td>AAACGCCGGACACGACGCTCCCCC</td><td></td><td></td><td></td><td></td><td></td><td></td><td></td><td></td><td></td><td></td></tr>
<tr><td>Chromosome</td><td>start</td><td>end</td><td>strand</td><td>MM</td><td>target_seq</td><td>PAM</td><td>alignment</td><td>distance</td><td>position</td><td>gene name</td><td>gene id</td></tr>
<tr><td>GU305914.1</td><td>118941</td><td>118963</td><td>+</td><td>0</td><td>GGGGGAGCGTCGTGTCCGGC</td><td>AGG</td><td>||||||||[||||||||||||]PAM</td><td>NA</td><td>Intergenic</td><td>NA</td><td>NA</td></tr>
<tr><td>T8888</td><td>TGTCCGGCAGGTCTCGCAGTCGG</td><td>1000</td><td></td><td></td><td></td><td></td><td></td><td></td><td></td><td></td><td></td></tr>
<tr><td>Oligo fwd</td><td>CACCTGTCCGGCAGGTCTCGCAGT</td><td></td><td></td><td></td><td></td><td></td><td></td><td></td><td></td><td></td><td></td></tr>
<tr><td>Oligo rev</td><td>AAACACTGCGAGACCTGCCGGACA</td><td></td><td></td><td></td><td></td><td></td><td></td><td></td><td></td><td></td><td></td></tr>
<tr><td>Chromosome</td><td>start</td><td>end</td><td>strand</td><td>MM</td><td>target_seq</td><td>PAM</td><td>alignment</td><td>distance</td><td>position</td><td>gene name</td><td>gene id</td></tr>
<tr><td>GU305914.1</td><td>118953</td><td>118975</td><td>+</td><td>0</td><td>TGTCCGGCAGGTCTCGCAGT</td><td>CGG</td><td>||||||||[||||||||||||]PAM</td><td>NA</td><td>Intergenic</td><td>NA</td><td>NA</td></tr>
<tr><td>T8889</td><td>GGCAGGTCTCGCAGTCGGCTCGG</td><td>1000</td><td></td><td></td><td></td><td></td><td></td><td></td><td></td><td></td><td></td></tr>
<tr><td>Oligo fwd</td><td>CACCGGCAGGTCTCGCAGTCGGCT</td><td></td><td></td><td></td><td></td><td></td><td></td><td></td><td></td><td></td><td></td></tr>
<tr><td>Oligo rev</td><td>AAACAGCCGACTGCGAGACCTGCC</td><td></td><td></td><td></td><td></td><td></td><td></td><td></td><td></td><td></td><td></td></tr>
<tr><td>Chromosome</td><td>start</td><td>end</td><td>strand</td><td>MM</td><td>target_seq</td><td>PAM</td><td>alignment</td><td>distance</td><td>position</td><td>gene name</td><td>gene id</td></tr>
<tr><td>GU305914.1</td><td>118958</td><td>118980</td><td>+</td><td>0</td><td>GGCAGGTCTCGCAGTCGGCT</td><td>CGG</td><td>||||||||[||||||||||||]PAM</td><td>NA</td><td>Intergenic</td><td>NA</td><td>NA</td></tr>
<tr><td>T8890</td><td>GCAGGTCTCGCAGTCGGCTCGGG</td><td>1000</td><td></td><td></td><td></td><td></td><td></td><td></td><td></td><td></td><td></td></tr>
<tr><td>Oligo fwd</td><td>CACCGCAGGTCTCGCAGTCGGCTC</td><td></td><td></td><td></td><td></td><td></td><td></td><td></td><td></td><td></td><td></td></tr>
<tr><td>Oligo rev</td><td>AAACGAGCCGACTGCGAGACCTGC</td><td></td><td></td><td></td><td></td><td></td><td></td><td></td><td></td><td></td><td></td></tr>
<tr><td>Chromosome</td><td>start</td><td>end</td><td>strand</td><td>MM</td><td>target_seq</td><td>PAM</td><td>alignment</td><td>distance</td><td>position</td><td>gene name</td><td>gene id</td></tr>
<tr><td>GU305914.1</td><td>118959</td><td>118981</td><td>+</td><td>0</td><td>GCAGGTCTCGCAGTCGGCTC</td><td>GGG</td><td>||||||||[||||||||||||]PAM</td><td>NA</td><td>Intergenic</td><td>NA</td><td>NA</td></tr>
<tr><td>T8891</td><td>CAGGTCTCGCAGTCGGCTCGGGG</td><td>1000</td><td></td><td></td><td></td><td></td><td></td><td></td><td></td><td></td><td></td></tr>
<tr><td>Oligo fwd</td><td>CACCCAGGTCTCGCAGTCGGCTCG</td><td></td><td></td><td></td><td></td><td></td><td></td><td></td><td></td><td></td><td></td></tr>
<tr><td>Oligo rev</td><td>AAACCGAGCCGACTGCGAGACCTG</td><td></td><td></td><td></td><td></td><td></td><td></td><td></td><td></td><td></td><td></td></tr>
<tr><td>Chromosome</td><td>start</td><td>end</td><td>strand</td><td>MM</td><td>target_seq</td><td>PAM</td><td>alignment</td><td>distance</td><td>position</td><td>gene name</td><td>gene id</td></tr>
<tr><td>GU305914.1</td><td>118960</td><td>118982</td><td>+</td><td>0</td><td>CAGGTCTCGCAGTCGGCTCG</td><td>GGG</td><td>||||||||[||||||||||||]PAM</td><td>NA</td><td>Intergenic</td><td>NA</td><td>NA</td></tr>
<tr><td>T8892</td><td>TTGTGTTTTCTGAAGACCTCCGG</td><td>1000</td><td></td><td></td><td></td><td></td><td></td><td></td><td></td><td></td><td></td></tr>
<tr><td>Oligo fwd</td><td>CACCTTGTGTTTTCTGAAGACCTC</td><td></td><td></td><td></td><td></td><td></td><td></td><td></td><td></td><td></td><td></td></tr>
<tr><td>Oligo rev</td><td>AAACGAGGTCTTCAGAAAACACAA</td><td></td><td></td><td></td><td></td><td></td><td></td><td></td><td></td><td></td><td></td></tr>
<tr><td>Chromosome</td><td>start</td><td>end</td><td>strand</td><td>MM</td><td>target_seq</td><td>PAM</td><td>alignment</td><td>distance</td><td>position</td><td>gene name</td><td>gene id</td></tr>
<tr><td>GU305914.1</td><td>118988</td><td>119010</td><td>+</td><td>0</td><td>TTGTGTTTTCTGAAGACCTC</td><td>CGG</td><td>||||||||[||||||||||||]PAM</td><td>NA</td><td>Intergenic</td><td>NA</td><td>NA</td></tr>
<tr><td>T8893</td><td>CGGCCCGTAGTTGATCACCTCGG</td><td>1000</td><td></td><td></td><td></td><td></td><td></td><td></td><td></td><td></td><td></td></tr>
<tr><td>Oligo fwd</td><td>CACCCGGCCCGTAGTTGATCACCT</td><td></td><td></td><td></td><td></td><td></td><td></td><td></td><td></td><td></td><td></td></tr>
<tr><td>Oligo rev</td><td>AAACAGGTGATCAACTACGGGCCG</td><td></td><td></td><td></td><td></td><td></td><td></td><td></td><td></td><td></td><td></td></tr>
<tr><td>Chromosome</td><td>start</td><td>end</td><td>strand</td><td>MM</td><td>target_seq</td><td>PAM</td><td>alignment</td><td>distance</td><td>position</td><td>gene name</td><td>gene id</td></tr>
<tr><td>GU305914.1</td><td>119008</td><td>119030</td><td>+</td><td>0</td><td>CGGCCCGTAGTTGATCACCT</td><td>CGG</td><td>||||||||[||||||||||||]PAM</td><td>NA</td><td>Intergenic</td><td>NA</td><td>NA</td></tr>
<tr><td>T8894</td><td>CCTCGGTCTGCGAACACGTGTGG</td><td>1000</td><td></td><td></td><td></td><td></td><td></td><td></td><td></td><td></td><td></td></tr>
<tr><td>Oligo fwd</td><td>CACCCCTCGGTCTGCGAACACGTG</td><td></td><td></td><td></td><td></td><td></td><td></td><td></td><td></td><td></td><td></td></tr>
<tr><td>Oligo rev</td><td>AAACCACGTGTTCGCAGACCGAGG</td><td></td><td></td><td></td><td></td><td></td><td></td><td></td><td></td><td></td><td></td></tr>
<tr><td>Chromosome</td><td>start</td><td>end</td><td>strand</td><td>MM</td><td>target_seq</td><td>PAM</td><td>alignment</td><td>distance</td><td>position</td><td>gene name</td><td>gene id</td></tr>
<tr><td>GU305914.1</td><td>119025</td><td>119047</td><td>+</td><td>0</td><td>CCTCGGTCTGCGAACACGTG</td><td>TGG</td><td>||||||||[||||||||||||]PAM</td><td>NA</td><td>Intergenic</td><td>NA</td><td>NA</td></tr>
<tr><td>T8895</td><td>GTGGCGAGCGTGTCGACTGTTGG</td><td>1000</td><td></td><td></td><td></td><td></td><td></td><td></td><td></td><td></td><td></td></tr>
<tr><td>Oligo fwd</td><td>CACCGTGGCGAGCGTGTCGACTGT</td><td></td><td></td><td></td><td></td><td></td><td></td><td></td><td></td><td></td><td></td></tr>
<tr><td>Oligo rev</td><td>AAACACAGTCGACACGCTCGCCAC</td><td></td><td></td><td></td><td></td><td></td><td></td><td></td><td></td><td></td><td></td></tr>
<tr><td>Chromosome</td><td>start</td><td>end</td><td>strand</td><td>MM</td><td>target_seq</td><td>PAM</td><td>alignment</td><td>distance</td><td>position</td><td>gene name</td><td>gene id</td></tr>
<tr><td>GU305914.1</td><td>119044</td><td>119066</td><td>+</td><td>0</td><td>GTGGCGAGCGTGTCGACTGT</td><td>TGG</td><td>||||||||[||||||||||||]PAM</td><td>NA</td><td>Intergenic</td><td>NA</td><td>NA</td></tr>
<tr><td>T8896</td><td>CACGAAGCGCAGCTCTCCGATGG</td><td>1000</td><td></td><td></td><td></td><td></td><td></td><td></td><td></td><td></td><td></td></tr>
<tr><td>Oligo fwd</td><td>CACCCACGAAGCGCAGCTCTCCGA</td><td></td><td></td><td></td><td></td><td></td><td></td><td></td><td></td><td></td><td></td></tr>
</table>

<table>
<tr><td>Oligo rev</td><td>AAACTCGGAGAGCTGCGCTTCGTG</td><td></td><td></td><td></td><td></td><td></td><td></td><td></td><td></td><td></td><td></td></tr>
<tr><td>Chromosome</td><td>start</td><td>end</td><td>strand</td><td>MM</td><td>target_seq</td><td>PAM</td><td>alignment</td><td>distance</td><td>position</td><td>gene name</td><td>gene id</td></tr>
<tr><td>GU305914.1</td><td>119092</td><td>119114</td><td>+</td><td>0</td><td>CACGAAGCGCAGCTCTCCGA</td><td>TGG</td><td>||||||||[||||||||||||]PAM</td><td>NA</td><td>Intergenic</td><td>NA</td><td>NA</td></tr>
<tr><td></td><td></td><td></td><td></td><td></td><td></td><td></td><td></td><td></td><td></td><td></td><td></td></tr>
<tr><td>T8897</td><td>GAAGCGCAGCTCTCCGATGGCGG</td><td>1000</td><td></td><td></td><td></td><td></td><td></td><td></td><td></td><td></td><td></td></tr>
<tr><td>Oligo fwd</td><td>CACCGAAGCGCAGCTCTCCGATGG</td><td></td><td></td><td></td><td></td><td></td><td></td><td></td><td></td><td></td><td></td></tr>
<tr><td>Oligo rev</td><td>AAACCCATCGGAGAGCTGCGCTTC</td><td></td><td></td><td></td><td></td><td></td><td></td><td></td><td></td><td></td><td></td></tr>
<tr><td>Chromosome</td><td>start</td><td>end</td><td>strand</td><td>MM</td><td>target_seq</td><td>PAM</td><td>alignment</td><td>distance</td><td>position</td><td>gene name</td><td>gene id</td></tr>
<tr><td>GU305914.1</td><td>119095</td><td>119117</td><td>+</td><td>0</td><td>GAAGCGCAGCTCTCCGATGG</td><td>CGG</td><td>||||||||[||||||||||||]PAM</td><td>NA</td><td>Intergenic</td><td>NA</td><td>NA</td></tr>
<tr><td></td><td></td><td></td><td></td><td></td><td></td><td></td><td></td><td></td><td></td><td></td><td></td></tr>
<tr><td>T8898</td><td>GACCTCCATGTCCTTCATGCCGG</td><td>1000</td><td></td><td></td><td></td><td></td><td></td><td></td><td></td><td></td><td></td></tr>
<tr><td>Oligo fwd</td><td>CACCGACCTCCATGTCCTTCATGC</td><td></td><td></td><td></td><td></td><td></td><td></td><td></td><td></td><td></td><td></td></tr>
<tr><td>Oligo rev</td><td>AAACGCATGAAGGACATGGAGGTC</td><td></td><td></td><td></td><td></td><td></td><td></td><td></td><td></td><td></td><td></td></tr>
<tr><td>Chromosome</td><td>start</td><td>end</td><td>strand</td><td>MM</td><td>target_seq</td><td>PAM</td><td>alignment</td><td>distance</td><td>position</td><td>gene name</td><td>gene id</td></tr>
<tr><td>GU305914.1</td><td>119128</td><td>119150</td><td>+</td><td>0</td><td>GACCTCCATGTCCTTCATGC</td><td>CGG</td><td>||||||||[||||||||||||]PAM</td><td>NA</td><td>Intergenic</td><td>NA</td><td>NA</td></tr>
<tr><td></td><td></td><td></td><td></td><td></td><td></td><td></td><td></td><td></td><td></td><td></td><td></td></tr>
<tr><td>T8899</td><td>TGCCGGTGATCGTGAACGACAGG</td><td>1000</td><td></td><td></td><td></td><td></td><td></td><td></td><td></td><td></td><td></td></tr>
<tr><td>Oligo fwd</td><td>CACCTGCCGGTGATCGTGAACGAC</td><td></td><td></td><td></td><td></td><td></td><td></td><td></td><td></td><td></td><td></td></tr>
<tr><td>Oligo rev</td><td>AAACGTCGTTCACGATCACCGGCA</td><td></td><td></td><td></td><td></td><td></td><td></td><td></td><td></td><td></td><td></td></tr>
<tr><td>Chromosome</td><td>start</td><td>end</td><td>strand</td><td>MM</td><td>target_seq</td><td>PAM</td><td>alignment</td><td>distance</td><td>position</td><td>gene name</td><td>gene id</td></tr>
<tr><td>GU305914.1</td><td>119145</td><td>119167</td><td>+</td><td>0</td><td>TGCCGGTGATCGTGAACGAC</td><td>AGG</td><td>||||||||[||||||||||||]PAM</td><td>NA</td><td>Intergenic</td><td>NA</td><td>NA</td></tr>
<tr><td></td><td></td><td></td><td></td><td></td><td></td><td></td><td></td><td></td><td></td><td></td><td></td></tr>
<tr><td>T8900</td><td>GTGAACGACAGGTTGCTGTTCGG</td><td>1000</td><td></td><td></td><td></td><td></td><td></td><td></td><td></td><td></td><td></td></tr>
<tr><td>Oligo fwd</td><td>CACCGTGAACGACAGGTTGCTGTT</td><td></td><td></td><td></td><td></td><td></td><td></td><td></td><td></td><td></td><td></td></tr>
<tr><td>Oligo rev</td><td>AAACAACAGCAACCTGTCGTTCAC</td><td></td><td></td><td></td><td></td><td></td><td></td><td></td><td></td><td></td><td></td></tr>
<tr><td>Chromosome</td><td>start</td><td>end</td><td>strand</td><td>MM</td><td>target_seq</td><td>PAM</td><td>alignment</td><td>distance</td><td>position</td><td>gene name</td><td>gene id</td></tr>
<tr><td>GU305914.1</td><td>119156</td><td>119178</td><td>+</td><td>0</td><td>GTGAACGACAGGTTGCTGTT</td><td>CGG</td><td>||||||||[||||||||||||]PAM</td><td>NA</td><td>Intergenic</td><td>NA</td><td>NA</td></tr>
<tr><td></td><td></td><td></td><td></td><td></td><td></td><td></td><td></td><td></td><td></td><td></td><td></td></tr>
<tr><td>T8901</td><td>CTGTTCGGCATCCACACGCTCGG</td><td>1000</td><td></td><td></td><td></td><td></td><td></td><td></td><td></td><td></td><td></td></tr>
<tr><td>Oligo fwd</td><td>CACCCTGTTCGGCATCCACACGCT</td><td></td><td></td><td></td><td></td><td></td><td></td><td></td><td></td><td></td><td></td></tr>
<tr><td>Oligo rev</td><td>AAACAGCGTGTGGATGCCGAACAG</td><td></td><td></td><td></td><td></td><td></td><td></td><td></td><td></td><td></td><td></td></tr>
<tr><td>Chromosome</td><td>start</td><td>end</td><td>strand</td><td>MM</td><td>target_seq</td><td>PAM</td><td>alignment</td><td>distance</td><td>position</td><td>gene name</td><td>gene id</td></tr>
<tr><td>GU305914.1</td><td>119171</td><td>119193</td><td>+</td><td>0</td><td>CTGTTCGGCATCCACACGCT</td><td>CGG</td><td>||||||||[||||||||||||]PAM</td><td>NA</td><td>Intergenic</td><td>NA</td><td>NA</td></tr>
<tr><td>GU305914.1</td><td>196799</td><td>196821</td><td>-</td><td>4</td><td>CTGCTCGTCATCGACACGTT</td><td>CGG</td><td>|||-|||-[||||-|||||-|]PAM</td><td>NA</td><td>Intergenic</td><td>NA</td><td>NA</td></tr>
<tr><td></td><td></td><td></td><td></td><td></td><td></td><td></td><td></td><td></td><td></td><td></td><td></td></tr>
<tr><td>T8902</td><td>TGTTCGGCATCCACACGCTCGGG</td><td>1000</td><td></td><td></td><td></td><td></td><td></td><td></td><td></td><td></td><td></td></tr>
<tr><td>Oligo fwd</td><td>CACCTGTTCGGCATCCACACGCTC</td><td></td><td></td><td></td><td></td><td></td><td></td><td></td><td></td><td></td><td></td></tr>
<tr><td>Oligo rev</td><td>AAACGAGCGTGTGGATGCCGAACA</td><td></td><td></td><td></td><td></td><td></td><td></td><td></td><td></td><td></td><td></td></tr>
<tr><td>Chromosome</td><td>start</td><td>end</td><td>strand</td><td>MM</td><td>target_seq</td><td>PAM</td><td>alignment</td><td>distance</td><td>position</td><td>gene name</td><td>gene id</td></tr>
<tr><td>GU305914.1</td><td>119172</td><td>119194</td><td>+</td><td>0</td><td>TGTTCGGCATCCACACGCTC</td><td>GGG</td><td>||||||||[||||||||||||]PAM</td><td>NA</td><td>Intergenic</td><td>NA</td><td>NA</td></tr>
<tr><td></td><td></td><td></td><td></td><td></td><td></td><td></td><td></td><td></td><td></td><td></td><td></td></tr>
<tr><td>T8903</td><td>TCGGCATCCACACGCTCGGGTGG</td><td>1000</td><td></td><td></td><td></td><td></td><td></td><td></td><td></td><td></td><td></td></tr>
<tr><td>Oligo fwd</td><td>CACCTCGGCATCCACACGCTCGGG</td><td></td><td></td><td></td><td></td><td></td><td></td><td></td><td></td><td></td><td></td></tr>
<tr><td>Oligo rev</td><td>AAACCCCGAGCGTGTGGATGCCGA</td><td></td><td></td><td></td><td></td><td></td><td></td><td></td><td></td><td></td><td></td></tr>
<tr><td>Chromosome</td><td>start</td><td>end</td><td>strand</td><td>MM</td><td>target_seq</td><td>PAM</td><td>alignment</td><td>distance</td><td>position</td><td>gene name</td><td>gene id</td></tr>
<tr><td>GU305914.1</td><td>119175</td><td>119197</td><td>+</td><td>0</td><td>TCGGCATCCACACGCTCGGG</td><td>TGG</td><td>||||||||[||||||||||||]PAM</td><td>NA</td><td>Intergenic</td><td>NA</td><td>NA</td></tr>
<tr><td></td><td></td><td></td><td></td><td></td><td></td><td></td><td></td><td></td><td></td><td></td><td></td></tr>
<tr><td>T8904</td><td>CTCGGGTGGATGTCCAGACCCGG</td><td>1000</td><td></td><td></td><td></td><td></td><td></td><td></td><td></td><td></td><td></td></tr>
<tr><td>Oligo fwd</td><td>CACCCTCGGGTGGATGTCCAGACC</td><td></td><td></td><td></td><td></td><td></td><td></td><td></td><td></td><td></td><td></td></tr>
<tr><td>Oligo rev</td><td>AAACGGTCTGGACATCCACCCGAG</td><td></td><td></td><td></td><td></td><td></td><td></td><td></td><td></td><td></td><td></td></tr>
<tr><td>Chromosome</td><td>start</td><td>end</td><td>strand</td><td>MM</td><td>target_seq</td><td>PAM</td><td>alignment</td><td>distance</td><td>position</td><td>gene name</td><td>gene id</td></tr>
<tr><td>GU305914.1</td><td>119189</td><td>119211</td><td>+</td><td>0</td><td>CTCGGGTGGATGTCCAGACC</td><td>CGG</td><td>||||||||[||||||||||||]PAM</td><td>NA</td><td>Intergenic</td><td>NA</td><td>NA</td></tr>
<tr><td></td><td></td><td></td><td></td><td></td><td></td><td></td><td></td><td></td><td></td><td></td><td></td></tr>
<tr><td>T8905</td><td>TCGGGTGGATGTCCAGACCCGGG</td><td>1000</td><td></td><td></td><td></td><td></td><td></td><td></td><td></td><td></td><td></td></tr>
<tr><td>Oligo fwd</td><td>CACCTCGGGTGGATGTCCAGACCC</td><td></td><td></td><td></td><td></td><td></td><td></td><td></td><td></td><td></td><td></td></tr>
<tr><td>Oligo rev</td><td>AAACGGGTCTGGACATCCACCCGA</td><td></td><td></td><td></td><td></td><td></td><td></td><td></td><td></td><td></td><td></td></tr>
<tr><td>Chromosome</td><td>start</td><td>end</td><td>strand</td><td>MM</td><td>target_seq</td><td>PAM</td><td>alignment</td><td>distance</td><td>position</td><td>gene name</td><td>gene id</td></tr>
<tr><td>GU305914.1</td><td>119190</td><td>119212</td><td>+</td><td>0</td><td>TCGGGTGGATGTCCAGACCC</td><td>GGG</td><td>||||||||[||||||||||||]PAM</td><td>NA</td><td>Intergenic</td><td>NA</td><td>NA</td></tr>
<tr><td></td><td></td><td></td><td></td><td></td><td></td><td></td><td></td><td></td><td></td><td></td><td></td></tr>
<tr><td>T8906</td><td>GGATGTCCAGACCCGGGATCAGG</td><td>1000</td><td></td><td></td><td></td><td></td><td></td><td></td><td></td><td></td><td></td></tr>
<tr><td>Oligo fwd</td><td>CACCGGATGTCCAGACCCGGGATC</td><td></td><td></td><td></td><td></td><td></td><td></td><td></td><td></td><td></td><td></td></tr>
<tr><td>Oligo rev</td><td>AAACGATCCCGGGTCTGGACATCC</td><td></td><td></td><td></td><td></td><td></td><td></td><td></td><td></td><td></td><td></td></tr>
<tr><td>Chromosome</td><td>start</td><td>end</td><td>strand</td><td>MM</td><td>target_seq</td><td>PAM</td><td>alignment</td><td>distance</td><td>position</td><td>gene name</td><td>gene id</td></tr>
<tr><td>GU305914.1</td><td>119196</td><td>119218</td><td>+</td><td>0</td><td>GGATGTCCAGACCCGGGATC</td><td>AGG</td><td>||||||||[||||||||||||]PAM</td><td>NA</td><td>Intergenic</td><td>NA</td><td>NA</td></tr>
<tr><td></td><td></td><td></td><td></td><td></td><td></td><td></td><td></td><td></td><td></td><td></td><td></td></tr>
<tr><td>T8907</td><td>TCCAGACCCGGGATCAGGTAAGG</td><td>1000</td><td></td><td></td><td></td><td></td><td></td><td></td><td></td><td></td><td></td></tr>
<tr><td>Oligo fwd</td><td>CACCTCCAGACCCGGGATCAGGTA</td><td></td><td></td><td></td><td></td><td></td><td></td><td></td><td></td><td></td><td></td></tr>
<tr><td>Oligo rev</td><td>AAACTACCTGATCCCGGGTCTGGA</td><td></td><td></td><td></td><td></td><td></td><td></td><td></td><td></td><td></td><td></td></tr>
<tr><td>Chromosome</td><td>start</td><td>end</td><td>strand</td><td>MM</td><td>target_seq</td><td>PAM</td><td>alignment</td><td>distance</td><td>position</td><td>gene name</td><td>gene id</td></tr>
<tr><td>GU305914.1</td><td>119201</td><td>119223</td><td>+</td><td>0</td><td>TCCAGACCCGGGATCAGGTA</td><td>AGG</td><td>||||||||[||||||||||||]PAM</td><td>NA</td><td>Intergenic</td><td>NA</td><td>NA</td></tr>
<tr><td></td><td></td><td></td><td></td><td></td><td></td><td></td><td></td><td></td><td></td><td></td><td></td></tr>
<tr><td>T8908</td><td>CCAGACCCGGGATCAGGTAAGGG</td><td>1000</td><td></td><td></td><td></td><td></td><td></td><td></td><td></td><td></td><td></td></tr>
<tr><td>Oligo fwd</td><td>CACCCCAGACCCGGGATCAGGTAA</td><td></td><td></td><td></td><td></td><td></td><td></td><td></td><td></td><td></td><td></td></tr>
<tr><td>Oligo rev</td><td>AAACTTACCTGATCCCGGGTCTGG</td><td></td><td></td><td></td><td></td><td></td><td></td><td></td><td></td><td></td><td></td></tr>
<tr><td>Chromosome</td><td>start</td><td>end</td><td>strand</td><td>MM</td><td>target_seq</td><td>PAM</td><td>alignment</td><td>distance</td><td>position</td><td>gene name</td><td>gene id</td></tr>
<tr><td>GU305914.1</td><td>119202</td><td>119224</td><td>+</td><td>0</td><td>CCAGACCCGGGATCAGGTAA</td><td>GGG</td><td>||||||||[||||||||||||]PAM</td><td>NA</td><td>Intergenic</td><td>NA</td><td>NA</td></tr>
<tr><td></td><td></td><td></td><td></td><td></td><td></td><td></td><td></td><td></td><td></td><td></td><td></td></tr>
<tr><td>T8909</td><td>GGATCAGGTAAGGGATGAAGAGG</td><td>1000</td><td></td><td></td><td></td><td></td><td></td><td></td><td></td><td></td><td></td></tr>
<tr><td>Oligo fwd</td><td>CACCGGATCAGGTAAGGGATGAAG</td><td></td><td></td><td></td><td></td><td></td><td></td><td></td><td></td><td></td><td></td></tr>
<tr><td>Oligo rev</td><td>AAACCTTCATCCCTTACCTGATCC</td><td></td><td></td><td></td><td></td><td></td><td></td><td></td><td></td><td></td><td></td></tr>
<tr><td>Chromosome</td><td>start</td><td>end</td><td>strand</td><td>MM</td><td>target_seq</td><td>PAM</td><td>alignment</td><td>distance</td><td>position</td><td>gene name</td><td>gene id</td></tr>
<tr><td>GU305914.1</td><td>119211</td><td>119233</td><td>+</td><td>0</td><td>GGATCAGGTAAGGGATGAAG</td><td>AGG</td><td>||||||||[||||||||||||]PAM</td><td>NA</td><td>Intergenic</td><td>NA</td><td>NA</td></tr>
<tr><td></td><td></td><td></td><td></td><td></td><td></td><td></td><td></td><td></td><td></td><td></td><td></td></tr>
<tr><td>T8910</td><td>GATCAGGTAAGGGATGAAGAGGG</td><td>1000</td><td></td><td></td><td></td><td></td><td></td><td></td><td></td><td></td><td></td></tr>
<tr><td>Oligo fwd</td><td>CACCGATCAGGTAAGGGATGAAGA</td><td></td><td></td><td></td><td></td><td></td><td></td><td></td><td></td><td></td><td></td></tr>
<tr><td>Oligo rev</td><td>AAACTCTTCATCCCTTACCTGATC</td><td></td><td></td><td></td><td></td><td></td><td></td><td></td><td></td><td></td><td></td></tr>
<tr><td>Chromosome</td><td>start</td><td>end</td><td>strand</td><td>MM</td><td>target_seq</td><td>PAM</td><td>alignment</td><td>distance</td><td>position</td><td>gene name</td><td>gene id</td></tr>
<tr><td>GU305914.1</td><td>119212</td><td>119234</td><td>+</td><td>0</td><td>GATCAGGTAAGGGATGAAGA</td><td>GGG</td><td>||||||||[||||||||||||]PAM</td><td>NA</td><td>Intergenic</td><td>NA</td><td>NA</td></tr>
</table>

T8911 AAGGGATGAAGAGGGCCGCGTGG 1000
Oligo fwd CACCAAGGGATGAAGAGGGCCGCG
Oligo rev AAACCGCGGCCCTCTTCATCCCTT
Chromosome start end strand MM target_seq PAM alignment distance position gene name gene id
GU305914.1 119220 119242 + 0 AAGGGATGAAGAGGGCCGCG TGG ||||||||[||||||||||||]PAM NA Intergenic NA NA

T8912 GATGAAGAGGGCCGCGTGGTTGG 1000
Oligo fwd CACCGATGAAGAGGGCCGCGTGGT
Oligo rev AAACACCACGCGGCCCTCTTCATC
Chromosome start end strand MM target_seq PAM alignment distance position gene name gene id
GU305914.1 119224 119246 + 0 GATGAAGAGGGCCGCGTGGT TGG ||||||||[||||||||||||]PAM NA Intergenic NA NA

T8913 GGGCCGCGTGGTTGGACGCGTGG 1000
Oligo fwd CACCGGGCCGCGTGGTTGGACGCG
Oligo rev AAACCGCGTCCAACCACGCGGCCC
Chromosome start end strand MM target_seq PAM alignment distance position gene name gene id
GU305914.1 119232 119254 + 0 GGGCCGCGTGGTTGGACGCG TGG ||||||||[||||||||||||]PAM NA Intergenic NA NA

T8914 CTGACCCGTCCTCAGCTGCACGG 1000
Oligo fwd CACCCTGACCCGTCCTCAGCTGCA
Oligo rev AAACTGCAGCTGAGGACGGGTCAG
Chromosome start end strand MM target_seq PAM alignment distance position gene name gene id
GU305914.1 119284 119306 + 0 CTGACCCGTCCTCAGCTGCA CGG ||||||||[||||||||||||]PAM NA Intergenic NA NA

T8915 GTCCTCAGCTGCACGGTGCGCGG 1000
Oligo fwd CACCGTCCTCAGCTGCACGGTGCG
Oligo rev AAACCGCACCGTGCAGCTGAGGAC
Chromosome start end strand MM target_seq PAM alignment distance position gene name gene id
GU305914.1 119291 119313 + 0 GTCCTCAGCTGCACGGTGCG CGG ||||||||[||||||||||||]PAM NA Intergenic NA NA

T8916 CGGTGCGCGGCATGACCACGCGG 1000
Oligo fwd CACCCGGTGCGCGGCATGACCACG
Oligo rev AAACCGTGGTCATGCCGCGCACCG
Chromosome start end strand MM target_seq PAM alignment distance position gene name gene id
GU305914.1 119304 119326 + 0 CGGTGCGCGGCATGACCACG CGG ||||||||[||||||||||||]PAM NA Intergenic NA NA

T8917 ACCACGCGGAACCCGTTCACCGG 1000
Oligo fwd CACCACCACGCGGAACCCGTTCAC
Oligo rev AAACGTGAACGGGTTCCGCGTGGT
Chromosome start end strand MM target_seq PAM alignment distance position gene name gene id
GU305914.1 119318 119340 + 0 ACCACGCGGAACCCGTTCAC CGG ||||||||[||||||||||||]PAM NA Intergenic NA NA

T8918 CCACGCGGAACCCGTTCACCGGG 1000
Oligo fwd CACCCCACGCGGAACCCGTTCACC
Oligo rev AAACGGTGAACGGGTTCCGCGTGG
Chromosome start end strand MM target_seq PAM alignment distance position gene name gene id
GU305914.1 119319 119341 + 0 CCACGCGGAACCCGTTCACC GGG ||||||||[||||||||||||]PAM NA Intergenic NA NA

T8919 CGGAACCCGTTCACCGGGAGCGG 1000
Oligo fwd CACCCGGAACCCGTTCACCGGGAG
Oligo rev AAACCTCCCGGTGAACGGGTTCCG
Chromosome start end strand MM target_seq PAM alignment distance position gene name gene id
GU305914.1 119324 119346 + 0 CGGAACCCGTTCACCGGGAG CGG ||||||||[||||||||||||]PAM NA Intergenic NA NA

T8920 ACCGGGAGCGGTTTCAGAAAAGG 1000
Oligo fwd CACCACCGGGAGCGGTTTCAGAAA
Oligo rev AAACTTTCTGAAACCGCTCCCGGT
Chromosome start end strand MM target_seq PAM alignment distance position gene name gene id
GU305914.1 119336 119358 + 0 ACCGGGAGCGGTTTCAGAAA AGG ||||||||[||||||||||||]PAM NA Intergenic NA NA

T8921 CCGGGAGCGGTTTCAGAAAAGGG 1000
Oligo fwd CACCCCGGGAGCGGTTTCAGAAAA
Oligo rev AAACTTTTCTGAAACCGCTCCCGG
Chromosome start end strand MM target_seq PAM alignment distance position gene name gene id
GU305914.1 119337 119359 + 0 CCGGGAGCGGTTTCAGAAAA GGG ||||||||[||||||||||||]PAM NA Intergenic NA NA

T8922 GCGGTTTCAGAAAAGGGTCCAGG 1000
Oligo fwd CACCGCGGTTTCAGAAAAGGGTCC
Oligo rev AAACGGACCCTTTTCTGAAACCGC
Chromosome start end strand MM target_seq PAM alignment distance position gene name gene id
GU305914.1 119343 119365 + 0 GCGGTTTCAGAAAAGGGTCC AGG ||||||||[||||||||||||]PAM NA Intergenic NA NA

T8923 AAGGGTCCAGGCTACGCTTCAGG 1000
Oligo fwd CACCAAGGGTCCAGGCTACGCTTC
Oligo rev AAACGAAGCGTAGCCTGGACCCTT
Chromosome start end strand MM target_seq PAM alignment distance position gene name gene id
GU305914.1 119355 119377 + 0 AAGGGTCCAGGCTACGCTTC AGG ||||||||[||||||||||||]PAM NA Intergenic NA NA

T8924 TCAGGTTCACCTTCTTCGCGCGG 1000
Oligo fwd CACCTCAGGTTCACCTTCTTCGCG
Oligo rev AAACCGCGAAGAAGGTGAACCTGA
Chromosome start end strand MM target_seq PAM alignment distance position gene name gene id
GU305914.1 119373 119395 + 0 TCAGGTTCACCTTCTTCGCG CGG ||||||||[||||||||||||]PAM NA Intergenic NA NA

T8925 TCGCGCGGTTACCCACCGCCAGG 1000
Oligo fwd CACCTCGCGCGGTTACCCACCGCC
Oligo rev AAACGGCGGTGGGTAACCGCGCGA
Chromosome start end strand MM target_seq PAM alignment distance position gene name gene id
GU305914.1 119388 119410 + 0 TCGCGCGGTTACCCACCGCC AGG ||||||||[||||||||||||]PAM NA Intergenic NA NA

<table>
<tr><td>T8926</td><td>GCGGTTACCCACCGCCAGGATGG</td><td>1000</td><td></td><td></td><td></td><td></td><td></td><td></td><td></td><td></td><td></td></tr>
<tr><td>Oligo fwd</td><td>CACCGCGGTTACCCACCGCCAGGA</td><td></td><td></td><td></td><td></td><td></td><td></td><td></td><td></td><td></td><td></td></tr>
<tr><td>Oligo rev</td><td>AAACTCCTGGCGGTGGGTAACCGC</td><td></td><td></td><td></td><td></td><td></td><td></td><td></td><td></td><td></td><td></td></tr>
<tr><td>Chromosome</td><td>start</td><td>end</td><td>strand</td><td>MM</td><td>target_seq</td><td>PAM</td><td>alignment</td><td>distance</td><td>position</td><td>gene name</td><td>gene id</td></tr>
<tr><td>GU305914.1</td><td>119392</td><td>119414</td><td>+</td><td>0</td><td>GCGGTTACCCACCGCCAGGA</td><td>TGG</td><td>||||||||[||||||||||||]PAM</td><td>NA</td><td>Intergenic</td><td>NA</td><td>NA</td></tr>
<tr><td>T8927</td><td>AGGATGGCCGTGAGCGAGTCTGG</td><td>1000</td><td></td><td></td><td></td><td></td><td></td><td></td><td></td><td></td><td></td></tr>
<tr><td>Oligo fwd</td><td>CACCAGGATGGCCGTGAGCGAGTC</td><td></td><td></td><td></td><td></td><td></td><td></td><td></td><td></td><td></td><td></td></tr>
<tr><td>Oligo rev</td><td>AAACGACTCGCTCACGGCCATCCT</td><td></td><td></td><td></td><td></td><td></td><td></td><td></td><td></td><td></td><td></td></tr>
<tr><td>Chromosome</td><td>start</td><td>end</td><td>strand</td><td>MM</td><td>target_seq</td><td>PAM</td><td>alignment</td><td>distance</td><td>position</td><td>gene name</td><td>gene id</td></tr>
<tr><td>GU305914.1</td><td>119408</td><td>119430</td><td>+</td><td>0</td><td>AGGATGGCCGTGAGCGAGTC</td><td>TGG</td><td>||||||||[||||||||||||]PAM</td><td>NA</td><td>Intergenic</td><td>NA</td><td>NA</td></tr>
<tr><td>T8928</td><td>ATGGCCGTGAGCGAGTCTGGAGG</td><td>1000</td><td></td><td></td><td></td><td></td><td></td><td></td><td></td><td></td><td></td></tr>
<tr><td>Oligo fwd</td><td>CACCATGGCCGTGAGCGAGTCTGG</td><td></td><td></td><td></td><td></td><td></td><td></td><td></td><td></td><td></td><td></td></tr>
<tr><td>Oligo rev</td><td>AAACCCAGACTCGCTCACGGCCAT</td><td></td><td></td><td></td><td></td><td></td><td></td><td></td><td></td><td></td><td></td></tr>
<tr><td>Chromosome</td><td>start</td><td>end</td><td>strand</td><td>MM</td><td>target_seq</td><td>PAM</td><td>alignment</td><td>distance</td><td>position</td><td>gene name</td><td>gene id</td></tr>
<tr><td>GU305914.1</td><td>119411</td><td>119433</td><td>+</td><td>0</td><td>ATGGCCGTGAGCGAGTCTGG</td><td>AGG</td><td>||||||||[||||||||||||]PAM</td><td>NA</td><td>Intergenic</td><td>NA</td><td>NA</td></tr>
<tr><td>T8929</td><td>CGAGTCTGGAGGCGTGCCGTTGG</td><td>1000</td><td></td><td></td><td></td><td></td><td></td><td></td><td></td><td></td><td></td></tr>
<tr><td>Oligo fwd</td><td>CACCCGAGTCTGGAGGCGTGCCGT</td><td></td><td></td><td></td><td></td><td></td><td></td><td></td><td></td><td></td><td></td></tr>
<tr><td>Oligo rev</td><td>AAACACGGCACGCCTCCAGACTCG</td><td></td><td></td><td></td><td></td><td></td><td></td><td></td><td></td><td></td><td></td></tr>
<tr><td>Chromosome</td><td>start</td><td>end</td><td>strand</td><td>MM</td><td>target_seq</td><td>PAM</td><td>alignment</td><td>distance</td><td>position</td><td>gene name</td><td>gene id</td></tr>
<tr><td>GU305914.1</td><td>119422</td><td>119444</td><td>+</td><td>0</td><td>CGAGTCTGGAGGCGTGCCGT</td><td>TGG</td><td>||||||||[||||||||||||]PAM</td><td>NA</td><td>Intergenic</td><td>NA</td><td>NA</td></tr>
<tr><td>T8930</td><td>CCTTCGCGTACCGCAGCGTCAGG</td><td>1000</td><td></td><td></td><td></td><td></td><td></td><td></td><td></td><td></td><td></td></tr>
<tr><td>Oligo fwd</td><td>CACCCCTTCGCGTACCGCAGCGTC</td><td></td><td></td><td></td><td></td><td></td><td></td><td></td><td></td><td></td><td></td></tr>
<tr><td>Oligo rev</td><td>AAACGACGCTGCGGTACGCGAAGG</td><td></td><td></td><td></td><td></td><td></td><td></td><td></td><td></td><td></td><td></td></tr>
<tr><td>Chromosome</td><td>start</td><td>end</td><td>strand</td><td>MM</td><td>target_seq</td><td>PAM</td><td>alignment</td><td>distance</td><td>position</td><td>gene name</td><td>gene id</td></tr>
<tr><td>GU305914.1</td><td>119445</td><td>119467</td><td>+</td><td>0</td><td>CCTTCGCGTACCGCAGCGTC</td><td>AGG</td><td>||||||||[||||||||||||]PAM</td><td>NA</td><td>Intergenic</td><td>NA</td><td>NA</td></tr>
<tr><td>T8931</td><td>CGCAGCGTCAGGTAGATCGTCGG</td><td>1000</td><td></td><td></td><td></td><td></td><td></td><td></td><td></td><td></td><td></td></tr>
<tr><td>Oligo fwd</td><td>CACCCGCAGCGTCAGGTAGATCGT</td><td></td><td></td><td></td><td></td><td></td><td></td><td></td><td></td><td></td><td></td></tr>
<tr><td>Oligo rev</td><td>AAACACGATCTACCTGACGCTGCG</td><td></td><td></td><td></td><td></td><td></td><td></td><td></td><td></td><td></td><td></td></tr>
<tr><td>Chromosome</td><td>start</td><td>end</td><td>strand</td><td>MM</td><td>target_seq</td><td>PAM</td><td>alignment</td><td>distance</td><td>position</td><td>gene name</td><td>gene id</td></tr>
<tr><td>GU305914.1</td><td>119456</td><td>119478</td><td>+</td><td>0</td><td>CGCAGCGTCAGGTAGATCGT</td><td>CGG</td><td>||||||||[||||||||||||]PAM</td><td>NA</td><td>Intergenic</td><td>NA</td><td>NA</td></tr>
<tr><td>T8932</td><td>AGTTTCTCTGCGCCGTCATCTGG</td><td>1000</td><td></td><td></td><td></td><td></td><td></td><td></td><td></td><td></td><td></td></tr>
<tr><td>Oligo fwd</td><td>CACCAGTTTCTCTGCGCCGTCATC</td><td></td><td></td><td></td><td></td><td></td><td></td><td></td><td></td><td></td><td></td></tr>
<tr><td>Oligo rev</td><td>AAACGATGACGGCGCAGAGAAACT</td><td></td><td></td><td></td><td></td><td></td><td></td><td></td><td></td><td></td><td></td></tr>
<tr><td>Chromosome</td><td>start</td><td>end</td><td>strand</td><td>MM</td><td>target_seq</td><td>PAM</td><td>alignment</td><td>distance</td><td>position</td><td>gene name</td><td>gene id</td></tr>
<tr><td>GU305914.1</td><td>119481</td><td>119503</td><td>+</td><td>0</td><td>AGTTTCTCTGCGCCGTCATC</td><td>TGG</td><td>||||||||[||||||||||||]PAM</td><td>NA</td><td>Intergenic</td><td>NA</td><td>NA</td></tr>
<tr><td>T8933</td><td>ATCTGGTAGAAATTGACGTGTGG</td><td>1000</td><td></td><td></td><td></td><td></td><td></td><td></td><td></td><td></td><td></td></tr>
<tr><td>Oligo fwd</td><td>CACCATCTGGTAGAAATTGACGTG</td><td></td><td></td><td></td><td></td><td></td><td></td><td></td><td></td><td></td><td></td></tr>
<tr><td>Oligo rev</td><td>AAACCACGTCAATTTCTACCAGAT</td><td></td><td></td><td></td><td></td><td></td><td></td><td></td><td></td><td></td><td></td></tr>
<tr><td>Chromosome</td><td>start</td><td>end</td><td>strand</td><td>MM</td><td>target_seq</td><td>PAM</td><td>alignment</td><td>distance</td><td>position</td><td>gene name</td><td>gene id</td></tr>
<tr><td>GU305914.1</td><td>119498</td><td>119520</td><td>+</td><td>0</td><td>ATCTGGTAGAAATTGACGTG</td><td>TGG</td><td>||||||||[||||||||||||]PAM</td><td>NA</td><td>Intergenic</td><td>NA</td><td>NA</td></tr>
<tr><td>T8934</td><td>ATTGACGTGTGGATCCGAGCAGG</td><td>1000</td><td></td><td></td><td></td><td></td><td></td><td></td><td></td><td></td><td></td></tr>
<tr><td>Oligo fwd</td><td>CACCATTGACGTGTGGATCCGAGC</td><td></td><td></td><td></td><td></td><td></td><td></td><td></td><td></td><td></td><td></td></tr>
<tr><td>Oligo rev</td><td>AAACGCTCGGATCCACACGTCAAT</td><td></td><td></td><td></td><td></td><td></td><td></td><td></td><td></td><td></td><td></td></tr>
<tr><td>Chromosome</td><td>start</td><td>end</td><td>strand</td><td>MM</td><td>target_seq</td><td>PAM</td><td>alignment</td><td>distance</td><td>position</td><td>gene name</td><td>gene id</td></tr>
<tr><td>GU305914.1</td><td>119509</td><td>119531</td><td>+</td><td>0</td><td>ATTGACGTGTGGATCCGAGC</td><td>AGG</td><td>||||||||[||||||||||||]PAM</td><td>NA</td><td>Intergenic</td><td>NA</td><td>NA</td></tr>
<tr><td>T8935</td><td>TGTGGATCCGAGCAGGTCTTCGG</td><td>1000</td><td></td><td></td><td></td><td></td><td></td><td></td><td></td><td></td><td></td></tr>
<tr><td>Oligo fwd</td><td>CACCTGTGGATCCGAGCAGGTCTT</td><td></td><td></td><td></td><td></td><td></td><td></td><td></td><td></td><td></td><td></td></tr>
<tr><td>Oligo rev</td><td>AAACAAGACCTGCTCGGATCCACA</td><td></td><td></td><td></td><td></td><td></td><td></td><td></td><td></td><td></td><td></td></tr>
<tr><td>Chromosome</td><td>start</td><td>end</td><td>strand</td><td>MM</td><td>target_seq</td><td>PAM</td><td>alignment</td><td>distance</td><td>position</td><td>gene name</td><td>gene id</td></tr>
<tr><td>GU305914.1</td><td>119516</td><td>119538</td><td>+</td><td>0</td><td>TGTGGATCCGAGCAGGTCTT</td><td>CGG</td><td>||||||||[||||||||||||]PAM</td><td>NA</td><td>Intergenic</td><td>NA</td><td>NA</td></tr>
<tr><td>T8936</td><td>GTGGATCCGAGCAGGTCTTCGGG</td><td>1000</td><td></td><td></td><td></td><td></td><td></td><td></td><td></td><td></td><td></td></tr>
<tr><td>Oligo fwd</td><td>CACCGTGGATCCGAGCAGGTCTTC</td><td></td><td></td><td></td><td></td><td></td><td></td><td></td><td></td><td></td><td></td></tr>
<tr><td>Oligo rev</td><td>AAACGAAGACCTGCTCGGATCCAC</td><td></td><td></td><td></td><td></td><td></td><td></td><td></td><td></td><td></td><td></td></tr>
<tr><td>Chromosome</td><td>start</td><td>end</td><td>strand</td><td>MM</td><td>target_seq</td><td>PAM</td><td>alignment</td><td>distance</td><td>position</td><td>gene name</td><td>gene id</td></tr>
<tr><td>GU305914.1</td><td>119517</td><td>119539</td><td>+</td><td>0</td><td>GTGGATCCGAGCAGGTCTTC</td><td>GGG</td><td>||||||||[||||||||||||]PAM</td><td>NA</td><td>Intergenic</td><td>NA</td><td>NA</td></tr>
<tr><td>T8937</td><td>CCGAGCAGGTCTTCGGGTCGAGG</td><td>1000</td><td></td><td></td><td></td><td></td><td></td><td></td><td></td><td></td><td></td></tr>
<tr><td>Oligo fwd</td><td>CACCCCGAGCAGGTCTTCGGGTCG</td><td></td><td></td><td></td><td></td><td></td><td></td><td></td><td></td><td></td><td></td></tr>
<tr><td>Oligo rev</td><td>AAACCGACCCGAAGACCTGCTCGG</td><td></td><td></td><td></td><td></td><td></td><td></td><td></td><td></td><td></td><td></td></tr>
<tr><td>Chromosome</td><td>start</td><td>end</td><td>strand</td><td>MM</td><td>target_seq</td><td>PAM</td><td>alignment</td><td>distance</td><td>position</td><td>gene name</td><td>gene id</td></tr>
<tr><td>GU305914.1</td><td>119523</td><td>119545</td><td>+</td><td>0</td><td>CCGAGCAGGTCTTCGGGTCG</td><td>AGG</td><td>||||||||[||||||||||||]PAM</td><td>NA</td><td>Intergenic</td><td>NA</td><td>NA</td></tr>
<tr><td>T8938</td><td>CGAGCAGGTCTTCGGGTCGAGGG</td><td>1000</td><td></td><td></td><td></td><td></td><td></td><td></td><td></td><td></td><td></td></tr>
<tr><td>Oligo fwd</td><td>CACCCGAGCAGGTCTTCGGGTCGA</td><td></td><td></td><td></td><td></td><td></td><td></td><td></td><td></td><td></td><td></td></tr>
<tr><td>Oligo rev</td><td>AAACTCGACCCGAAGACCTGCTCG</td><td></td><td></td><td></td><td></td><td></td><td></td><td></td><td></td><td></td><td></td></tr>
<tr><td>Chromosome</td><td>start</td><td>end</td><td>strand</td><td>MM</td><td>target_seq</td><td>PAM</td><td>alignment</td><td>distance</td><td>position</td><td>gene name</td><td>gene id</td></tr>
<tr><td>GU305914.1</td><td>119524</td><td>119546</td><td>+</td><td>0</td><td>CGAGCAGGTCTTCGGGTCGA</td><td>GGG</td><td>||||||||[||||||||||||]PAM</td><td>NA</td><td>Intergenic</td><td>NA</td><td>NA</td></tr>
<tr><td>T8939</td><td>GCAGGTCTTCGGGTCGAGGGTGG</td><td>1000</td><td></td><td></td><td></td><td></td><td></td><td></td><td></td><td></td><td></td></tr>
<tr><td>Oligo fwd</td><td>CACCGCAGGTCTTCGGGTCGAGGG</td><td></td><td></td><td></td><td></td><td></td><td></td><td></td><td></td><td></td><td></td></tr>
<tr><td>Oligo rev</td><td>AAACCCCTCGACCCGAAGACCTGC</td><td></td><td></td><td></td><td></td><td></td><td></td><td></td><td></td><td></td><td></td></tr>
<tr><td>Chromosome</td><td>start</td><td>end</td><td>strand</td><td>MM</td><td>target_seq</td><td>PAM</td><td>alignment</td><td>distance</td><td>position</td><td>gene name</td><td>gene id</td></tr>
<tr><td>GU305914.1</td><td>119527</td><td>119549</td><td>+</td><td>0</td><td>GCAGGTCTTCGGGTCGAGGG</td><td>TGG</td><td>||||||||[||||||||||||]PAM</td><td>NA</td><td>Intergenic</td><td>NA</td><td>NA</td></tr>
<tr><td>T8940</td><td>GGTCTTCGGGTCGAGGGTGGCGG</td><td>1000</td><td></td><td></td><td></td><td></td><td></td><td></td><td></td><td></td><td></td></tr>
<tr><td>Oligo fwd</td><td>CACCGGTCTTCGGGTCGAGGGTGG</td><td></td><td></td><td></td><td></td><td></td><td></td><td></td><td></td><td></td><td></td></tr>
<tr><td>Oligo rev</td><td>AAACCCACCCTCGACCCGAAGACC</td><td></td><td></td><td></td><td></td><td></td><td></td><td></td><td></td><td></td><td></td></tr>
<tr><td>Chromosome</td><td>start</td><td>end</td><td>strand</td><td>MM</td><td>target_seq</td><td>PAM</td><td>alignment</td><td>distance</td><td>position</td><td>gene name</td><td>gene id</td></tr>
</table>

GU305914.1 119530 119552 + 0 GGTCTTCGGGTCGAGGGTGG CGG ||||||||[||||||||||||]PAM NA Intergenic NA NA

T8941 GAGGGTGGCGGCTCTCGCCCTGG 1000
Oligo fwd CACCGAGGGTGGCGGCTCTCGCCC
Oligo rev AAACGGGCGAGAGCCGCCACCCTC
Chromosome start end strand MM target_seq PAM alignment distance position gene name gene id
GU305914.1 119542 119564 + 0 GAGGGTGGCGGCTCTCGCCC TGG ||||||||[||||||||||||]PAM NA Intergenic NA NA

T8942 AGGGTGGCGGCTCTCGCCCTGGG 1000
Oligo fwd CACCAGGGTGGCGGCTCTCGCCCT
Oligo rev AAACAGGGCGAGAGCCGCCACCCT
Chromosome start end strand MM target_seq PAM alignment distance position gene name gene id
GU305914.1 119543 119565 + 0 AGGGTGGCGGCTCTCGCCCT GGG ||||||||[||||||||||||]PAM NA Intergenic NA NA

T8943 GGGCAGCTCCAGAGACACGATGG 1000
Oligo fwd CACCGGGCAGCTCCAGAGACACGA
Oligo rev AAACTCGTGTCTCTGGAGCTGCCC
Chromosome start end strand MM target_seq PAM alignment distance position gene name gene id
GU305914.1 119563 119585 + 0 GGGCAGCTCCAGAGACACGA TGG ||||||||[||||||||||||]PAM NA Intergenic NA NA

T8944 GATGGCCTTCTTGAAGCGTTTGG 1000
Oligo fwd CACCGATGGCCTTCTTGAAGCGTT
Oligo rev AAACAACGCTTCAAGAAGGCCATC
Chromosome start end strand MM target_seq PAM alignment distance position gene name gene id
GU305914.1 119581 119603 + 0 GATGGCCTTCTTGAAGCGTT TGG ||||||||[||||||||||||]PAM NA Intergenic NA NA

T8945 TTCTTGAAGCGTTTGGAGTCCGG 1000
Oligo fwd CACCTTCTTGAAGCGTTTGGAGTC
Oligo rev AAACGACTCCAAACGCTTCAAGAA
Chromosome start end strand MM target_seq PAM alignment distance position gene name gene id
GU305914.1 119588 119610 + 0 TTCTTGAAGCGTTTGGAGTC CGG ||||||||[||||||||||||]PAM NA Intergenic NA NA

T8946 GCGTTTGGAGTCCGGATCGTTGG 1000
Oligo fwd CACCGCGTTTGGAGTCCGGATCGT
Oligo rev AAACACGATCCGGACTCCAAACGC
Chromosome start end strand MM target_seq PAM alignment distance position gene name gene id
GU305914.1 119596 119618 + 0 GCGTTTGGAGTCCGGATCGT TGG ||||||||[||||||||||||]PAM NA Intergenic NA NA

T8947 CGTTGGCCCAGCGCACCTTGTGG 1000
Oligo fwd CACCCGTTGGCCCAGCGCACCTTG
Oligo rev AAACCAAGGTGCGCTGGGCCAACG
Chromosome start end strand MM target_seq PAM alignment distance position gene name gene id
GU305914.1 119613 119635 + 0 CGTTGGCCCAGCGCACCTTG TGG ||||||||[||||||||||||]PAM NA Intergenic NA NA

T8948 CTTGTGGACCGTCATCCTGACGG 1000
Oligo fwd CACCCTTGTGGACCGTCATCCTGA
Oligo rev AAACTCAGGATGACGGTCCACAAG
Chromosome start end strand MM target_seq PAM alignment distance position gene name gene id
GU305914.1 119629 119651 + 0 CTTGTGGACCGTCATCCTGA CGG ||||||||[||||||||||||]PAM NA Intergenic NA NA

T8949 GGTCTTCATCCCCGTGCGCGTGG 1000
Oligo fwd CACCGGTCTTCATCCCCGTGCGCG
Oligo rev AAACCGCGCACGGGGATGAAGACC
Chromosome start end strand MM target_seq PAM alignment distance position gene name gene id
GU305914.1 119650 119672 + 0 GGTCTTCATCCCCGTGCGCG TGG ||||||||[||||||||||||]PAM NA Intergenic NA NA

T8950 CTTCATCCCCGTGCGCGTGGTGG 1000
Oligo fwd CACCCTTCATCCCCGTGCGCGTGG
Oligo rev AAACCCACGCGCACGGGGATGAAG
Chromosome start end strand MM target_seq PAM alignment distance position gene name gene id
GU305914.1 119653 119675 + 0 CTTCATCCCCGTGCGCGTGG TGG ||||||||[||||||||||||]PAM NA Intergenic NA NA

T8951 TGGTCGCCGAGACCGACGCCTGG 1000
Oligo fwd CACCTGGTCGCCGAGACCGACGCC
Oligo rev AAACGGCGTCGGTCTCGGCGACCA
Chromosome start end strand MM target_seq PAM alignment distance position gene name gene id
GU305914.1 119673 119695 + 0 TGGTCGCCGAGACCGACGCC TGG ||||||||[||||||||||||]PAM NA Intergenic NA NA

T8952 TCGCCGAGACCGACGCCTGGCGG 1000
Oligo fwd CACCTCGCCGAGACCGACGCCTGG
Oligo rev AAACCCAGGCGTCGGTCTCGGCGA
Chromosome start end strand MM target_seq PAM alignment distance position gene name gene id
GU305914.1 119676 119698 + 0 TCGCCGAGACCGACGCCTGG CGG ||||||||[||||||||||||]PAM NA Intergenic NA NA

T8953 CACCTTCACCGAGTTCACCCTGG 1000
Oligo fwd CACCCACCTTCACCGAGTTCACCC
Oligo rev AAACGGGTGAACTCGGTGAAGGTG
Chromosome start end strand MM target_seq PAM alignment distance position gene name gene id
GU305914.1 119713 119735 + 0 CACCTTCACCGAGTTCACCC TGG ||||||||[||||||||||||]PAM NA Intergenic NA NA

T8954 GTTCACCCTGGCGATCAGCATGG 1000
Oligo fwd CACCGTTCACCCTGGCGATCAGCA
Oligo rev AAACTGCTGATCGCCAGGGTGAAC
Chromosome start end strand MM target_seq PAM alignment distance position gene name gene id
GU305914.1 119725 119747 + 0 GTTCACCCTGGCGATCAGCA TGG ||||||||[||||||||||||]PAM NA Intergenic NA NA

T8955 TTCACCCTGGCGATCAGCATGGG 1000
Oligo fwd CACCTTCACCCTGGCGATCAGCAT
Oligo rev AAACATGCTGATCGCCAGGGTGAA

| | | | | | | | | | | | |
|---|---|---|---|---|---|---|---|---|---|---|---|
| Chromosome | start | end | strand | MM | target_seq | PAM | alignment | distance | position | gene name | gene id |
| GU305914.1 | 119726 | 119748 | + | 0 | TTCACCCTGGCGATCAGCAT | GGG | \|\|\|\|\|\|\|\|[\|\|\|\|\|\|\|\|\|\|\|\|]PAM | NA | Intergenic | NA | NA |
| T8956 | ACCCTGGCGATCAGCATGGGCGG | 1000 | | | | | | | | | |
| Oligo fwd | CACCACCCTGGCGATCAGCATGGG | | | | | | | | | | |
| Oligo rev | AAACCCCATGCTGATCGCCAGGGT | | | | | | | | | | |
| Chromosome | start | end | strand | MM | target_seq | PAM | alignment | distance | position | gene name | gene id |
| GU305914.1 | 119729 | 119751 | + | 0 | ACCCTGGCGATCAGCATGGG | CGG | \|\|\|\|\|\|\|\|[\|\|\|\|\|\|\|\|\|\|\|\|]PAM | NA | Intergenic | NA | NA |
| T8957 | GGCGATCAGCATGGGCGGCGCGG | 1000 | | | | | | | | | |
| Oligo fwd | CACCGGCGATCAGCATGGGCGGCG | | | | | | | | | | |
| Oligo rev | AAACCGCCGCCCATGCTGATCGCC | | | | | | | | | | |
| Chromosome | start | end | strand | MM | target_seq | PAM | alignment | distance | position | gene name | gene id |
| GU305914.1 | 119734 | 119756 | + | 0 | GGCGATCAGCATGGGCGGCG | CGG | \|\|\|\|\|\|\|\|[\|\|\|\|\|\|\|\|\|\|\|\|]PAM | NA | Intergenic | NA | NA |
| T8958 | GCGATCAGCATGGGCGGCGCGGG | 1000 | | | | | | | | | |
| Oligo fwd | CACCGCGATCAGCATGGGCGGCGC | | | | | | | | | | |
| Oligo rev | AAACGCGCCGCCCATGCTGATCGC | | | | | | | | | | |
| Chromosome | start | end | strand | MM | target_seq | PAM | alignment | distance | position | gene name | gene id |
| GU305914.1 | 119735 | 119757 | + | 0 | GCGATCAGCATGGGCGGCGC | GGG | \|\|\|\|\|\|\|\|[\|\|\|\|\|\|\|\|\|\|\|\|]PAM | NA | Intergenic | NA | NA |
| T8959 | CGATCAGCATGGGCGGCGCGGGG | 1000 | | | | | | | | | |
| Oligo fwd | CACCCGATCAGCATGGGCGGCGCG | | | | | | | | | | |
| Oligo rev | AAACCGCGCCGCCCATGCTGATCG | | | | | | | | | | |
| Chromosome | start | end | strand | MM | target_seq | PAM | alignment | distance | position | gene name | gene id |
| GU305914.1 | 119736 | 119758 | + | 0 | CGATCAGCATGGGCGGCGCG | GGG | \|\|\|\|\|\|\|\|[\|\|\|\|\|\|\|\|\|\|\|\|]PAM | NA | Intergenic | NA | NA |
| T8960 | GCATGGGCGGCGCGGGGACGTGG | 1000 | | | | | | | | | |
| Oligo fwd | CACCGCATGGGCGGCGCGGGGACG | | | | | | | | | | |
| Oligo rev | AAACCGTCCCCGCGCCGCCCATGC | | | | | | | | | | |
| Chromosome | start | end | strand | MM | target_seq | PAM | alignment | distance | position | gene name | gene id |
| GU305914.1 | 119742 | 119764 | + | 0 | GCATGGGCGGCGCGGGGACG | TGG | \|\|\|\|\|\|\|\|[\|\|\|\|\|\|\|\|\|\|\|\|]PAM | NA | Intergenic | NA | NA |
| T8961 | GCGGCGCGGGGACGTGGTCGAGG | 1000 | | | | | | | | | |
| Oligo fwd | CACCGCGGCGCGGGGACGTGGTCG | | | | | | | | | | |
| Oligo rev | AAACCGACCACGTCCCCGCGCCGC | | | | | | | | | | |
| Chromosome | start | end | strand | MM | target_seq | PAM | alignment | distance | position | gene name | gene id |
| GU305914.1 | 119748 | 119770 | + | 0 | GCGGCGCGGGGACGTGGTCG | AGG | \|\|\|\|\|\|\|\|[\|\|\|\|\|\|\|\|\|\|\|\|]PAM | NA | Intergenic | NA | NA |
| T8962 | CGGCGCGGGGACGTGGTCGAGGG | 1000 | | | | | | | | | |
| Oligo fwd | CACCCGGCGCGGGGACGTGGTCGA | | | | | | | | | | |
| Oligo rev | AAACTCGACCACGTCCCCGCGCCG | | | | | | | | | | |
| Chromosome | start | end | strand | MM | target_seq | PAM | alignment | distance | position | gene name | gene id |
| GU305914.1 | 119749 | 119771 | + | 0 | CGGCGCGGGGACGTGGTCGA | GGG | \|\|\|\|\|\|\|\|[\|\|\|\|\|\|\|\|\|\|\|\|]PAM | NA | Intergenic | NA | NA |
| T8963 | CGGGGACGTGGTCGAGGGCCAGG | 1000 | | | | | | | | | |
| Oligo fwd | CACCCGGGGACGTGGTCGAGGGCC | | | | | | | | | | |
| Oligo rev | AAACGGCCCTCGACCACGTCCCCG | | | | | | | | | | |
| Chromosome | start | end | strand | MM | target_seq | PAM | alignment | distance | position | gene name | gene id |
| GU305914.1 | 119754 | 119776 | + | 0 | CGGGGACGTGGTCGAGGGCC | AGG | \|\|\|\|\|\|\|\|[\|\|\|\|\|\|\|\|\|\|\|\|]PAM | NA | Intergenic | NA | NA |
| T8964 | GGGGACGTGGTCGAGGGCCAGGG | 1000 | | | | | | | | | |
| Oligo fwd | CACCGGGGACGTGGTCGAGGGCCA | | | | | | | | | | |
| Oligo rev | AAACTGGCCCTCGACCACGTCCCC | | | | | | | | | | |
| Chromosome | start | end | strand | MM | target_seq | PAM | alignment | distance | position | gene name | gene id |
| GU305914.1 | 119755 | 119777 | + | 0 | GGGGACGTGGTCGAGGGCCA | GGG | \|\|\|\|\|\|\|\|[\|\|\|\|\|\|\|\|\|\|\|\|]PAM | NA | Intergenic | NA | NA |
| T8965 | GGGCGAACACGCAGAAACCCAGG | 1000 | | | | | | | | | |
| Oligo fwd | CACCGGGCGAACACGCAGAAACCC | | | | | | | | | | |
| Oligo rev | AAACGGGTTTCTGCGTGTTCGCCC | | | | | | | | | | |
| Chromosome | start | end | strand | MM | target_seq | PAM | alignment | distance | position | gene name | gene id |
| GU305914.1 | 119775 | 119797 | + | 0 | GGGCGAACACGCAGAAACCC | AGG | \|\|\|\|\|\|\|\|[\|\|\|\|\|\|\|\|\|\|\|\|]PAM | NA | Intergenic | NA | NA |
| T8966 | GCAGAAACCCAGGTACTTGTTGG | 1000 | | | | | | | | | |
| Oligo fwd | CACCGCAGAAACCCAGGTACTTGT | | | | | | | | | | |
| Oligo rev | AAACACAAGTACCTGGGTTTCTGC | | | | | | | | | | |
| Chromosome | start | end | strand | MM | target_seq | PAM | alignment | distance | position | gene name | gene id |
| GU305914.1 | 119785 | 119807 | + | 0 | GCAGAAACCCAGGTACTTGT | TGG | \|\|\|\|\|\|\|\|[\|\|\|\|\|\|\|\|\|\|\|\|]PAM | NA | Intergenic | NA | NA |
| T8967 | AGCATCCGAGTCTGTTTCCGCGG | 1000 | | | | | | | | | |
| Oligo fwd | CACCAGCATCCGAGTCTGTTTCCG | | | | | | | | | | |
| Oligo rev | AAACCGGAAACAGACTCGGATGCT | | | | | | | | | | |
| Chromosome | start | end | strand | MM | target_seq | PAM | alignment | distance | position | gene name | gene id |
| GU305914.1 | 119813 | 119835 | + | 0 | AGCATCCGAGTCTGTTTCCG | CGG | \|\|\|\|\|\|\|\|[\|\|\|\|\|\|\|\|\|\|\|\|]PAM | NA | Intergenic | NA | NA |
| T8968 | TCCGAGTCTGTTTCCGCGGCTGG | 1000 | | | | | | | | | |
| Oligo fwd | CACCTCCGAGTCTGTTTCCGCGGC | | | | | | | | | | |
| Oligo rev | AAACGCCGCGGAAACAGACTCGGA | | | | | | | | | | |
| Chromosome | start | end | strand | MM | target_seq | PAM | alignment | distance | position | gene name | gene id |
| GU305914.1 | 119817 | 119839 | + | 0 | TCCGAGTCTGTTTCCGCGGC | TGG | \|\|\|\|\|\|\|\|[\|\|\|\|\|\|\|\|\|\|\|\|]PAM | NA | Intergenic | NA | NA |
| T8969 | CCGAGTCTGTTTCCGCGGCTGGG | 1000 | | | | | | | | | |
| Oligo fwd | CACCCCGAGTCTGTTTCCGCGGCT | | | | | | | | | | |
| Oligo rev | AAACAGCCGCGGAAACAGACTCGG | | | | | | | | | | |
| Chromosome | start | end | strand | MM | target_seq | PAM | alignment | distance | position | gene name | gene id |
| GU305914.1 | 119818 | 119840 | + | 0 | CCGAGTCTGTTTCCGCGGCT | GGG | \|\|\|\|\|\|\|\|[\|\|\|\|\|\|\|\|\|\|\|\|]PAM | NA | Intergenic | NA | NA |
| GU305914.1 | 174975 | 174997 | + | 3 | CCGTGTCCGTTTCCGCCGCT | CGG | \|\|\|-\|\|\|-[\|\|\|\|\|\|\|\|-\|\|\|]PAM | NA | Intergenic | NA | NA |
| T8970 | CGCGGCTGGGAGTTCTGTGCCGG | 1000 | | | | | | | | | |

<table>
<tr><td>Oligo fwd</td><td>CACCCGCGGCTGGGAGTTCTGTGC</td><td></td><td></td><td></td><td></td><td></td><td></td><td></td><td></td><td></td><td></td></tr>
<tr><td>Oligo rev</td><td>AAACGCACAGAACTCCCAGCCGCG</td><td></td><td></td><td></td><td></td><td></td><td></td><td></td><td></td><td></td><td></td></tr>
<tr><td>Chromosome</td><td>start</td><td>end</td><td>strand</td><td>MM</td><td>target_seq</td><td>PAM</td><td>alignment</td><td>distance</td><td>position</td><td>gene name</td><td>gene id</td></tr>
<tr><td>GU305914.1</td><td>119831</td><td>119853</td><td>+</td><td>0</td><td>CGCGGCTGGGAGTTCTGTGC</td><td>CGG</td><td>||||||||[||||||||||||]PAM</td><td>NA</td><td>Intergenic</td><td>NA</td><td>NA</td></tr>
<tr><td>T8971</td><td>GCGGCTGGGAGTTCTGTGCCGGG</td><td>1000</td><td></td><td></td><td></td><td></td><td></td><td></td><td></td><td></td><td></td></tr>
<tr><td>Oligo fwd</td><td>CACCGCGGCTGGGAGTTCTGTGCC</td><td></td><td></td><td></td><td></td><td></td><td></td><td></td><td></td><td></td><td></td></tr>
<tr><td>Oligo rev</td><td>AAACGGCACAGAACTCCCAGCCGC</td><td></td><td></td><td></td><td></td><td></td><td></td><td></td><td></td><td></td><td></td></tr>
<tr><td>Chromosome</td><td>start</td><td>end</td><td>strand</td><td>MM</td><td>target_seq</td><td>PAM</td><td>alignment</td><td>distance</td><td>position</td><td>gene name</td><td>gene id</td></tr>
<tr><td>GU305914.1</td><td>119832</td><td>119854</td><td>+</td><td>0</td><td>GCGGCTGGGAGTTCTGTGCC</td><td>GGG</td><td>||||||||[||||||||||||]PAM</td><td>NA</td><td>Intergenic</td><td>NA</td><td>NA</td></tr>
<tr><td>T8972</td><td>CTGGGAGTTCTGTGCCGGGACGG</td><td>1000</td><td></td><td></td><td></td><td></td><td></td><td></td><td></td><td></td><td></td></tr>
<tr><td>Oligo fwd</td><td>CACCCTGGGAGTTCTGTGCCGGGA</td><td></td><td></td><td></td><td></td><td></td><td></td><td></td><td></td><td></td><td></td></tr>
<tr><td>Oligo rev</td><td>AAACTCCCGGCACAGAACTCCCAG</td><td></td><td></td><td></td><td></td><td></td><td></td><td></td><td></td><td></td><td></td></tr>
<tr><td>Chromosome</td><td>start</td><td>end</td><td>strand</td><td>MM</td><td>target_seq</td><td>PAM</td><td>alignment</td><td>distance</td><td>position</td><td>gene name</td><td>gene id</td></tr>
<tr><td>GU305914.1</td><td>119836</td><td>119858</td><td>+</td><td>0</td><td>CTGGGAGTTCTGTGCCGGGA</td><td>CGG</td><td>||||||||[||||||||||||]PAM</td><td>NA</td><td>Intergenic</td><td>NA</td><td>NA</td></tr>
<tr><td>T8973</td><td>TGGGAGTTCTGTGCCGGGACGGG</td><td>1000</td><td></td><td></td><td></td><td></td><td></td><td></td><td></td><td></td><td></td></tr>
<tr><td>Oligo fwd</td><td>CACCTGGGAGTTCTGTGCCGGGAC</td><td></td><td></td><td></td><td></td><td></td><td></td><td></td><td></td><td></td><td></td></tr>
<tr><td>Oligo rev</td><td>AAACGTCCCGGCACAGAACTCCCA</td><td></td><td></td><td></td><td></td><td></td><td></td><td></td><td></td><td></td><td></td></tr>
<tr><td>Chromosome</td><td>start</td><td>end</td><td>strand</td><td>MM</td><td>target_seq</td><td>PAM</td><td>alignment</td><td>distance</td><td>position</td><td>gene name</td><td>gene id</td></tr>
<tr><td>GU305914.1</td><td>119837</td><td>119859</td><td>+</td><td>0</td><td>TGGGAGTTCTGTGCCGGGAC</td><td>GGG</td><td>||||||||[||||||||||||]PAM</td><td>NA</td><td>Intergenic</td><td>NA</td><td>NA</td></tr>
<tr><td>T8974</td><td>GGGAGTTCTGTGCCGGGACGGGG</td><td>1000</td><td></td><td></td><td></td><td></td><td></td><td></td><td></td><td></td><td></td></tr>
<tr><td>Oligo fwd</td><td>CACCGGGAGTTCTGTGCCGGGACG</td><td></td><td></td><td></td><td></td><td></td><td></td><td></td><td></td><td></td><td></td></tr>
<tr><td>Oligo rev</td><td>AAACCGTCCCGGCACAGAACTCCC</td><td></td><td></td><td></td><td></td><td></td><td></td><td></td><td></td><td></td><td></td></tr>
<tr><td>Chromosome</td><td>start</td><td>end</td><td>strand</td><td>MM</td><td>target_seq</td><td>PAM</td><td>alignment</td><td>distance</td><td>position</td><td>gene name</td><td>gene id</td></tr>
<tr><td>GU305914.1</td><td>119838</td><td>119860</td><td>+</td><td>0</td><td>GGGAGTTCTGTGCCGGGACG</td><td>GGG</td><td>||||||||[||||||||||||]PAM</td><td>NA</td><td>Intergenic</td><td>NA</td><td>NA</td></tr>
<tr><td>T8975</td><td>GGAGCGAGCCGATACCGAACTGG</td><td>1000</td><td></td><td></td><td></td><td></td><td></td><td></td><td></td><td></td><td></td></tr>
<tr><td>Oligo fwd</td><td>CACCGGAGCGAGCCGATACCGAAC</td><td></td><td></td><td></td><td></td><td></td><td></td><td></td><td></td><td></td><td></td></tr>
<tr><td>Oligo rev</td><td>AAACGTTCGGTATCGGCTCGCTCC</td><td></td><td></td><td></td><td></td><td></td><td></td><td></td><td></td><td></td><td></td></tr>
<tr><td>Chromosome</td><td>start</td><td>end</td><td>strand</td><td>MM</td><td>target_seq</td><td>PAM</td><td>alignment</td><td>distance</td><td>position</td><td>gene name</td><td>gene id</td></tr>
<tr><td>GU305914.1</td><td>119859</td><td>119881</td><td>+</td><td>0</td><td>GGAGCGAGCCGATACCGAAC</td><td>TGG</td><td>||||||||[||||||||||||]PAM</td><td>NA</td><td>Intergenic</td><td>NA</td><td>NA</td></tr>
<tr><td>T8976</td><td>GAGCGAGCCGATACCGAACTGGG</td><td>1000</td><td></td><td></td><td></td><td></td><td></td><td></td><td></td><td></td><td></td></tr>
<tr><td>Oligo fwd</td><td>CACCGAGCGAGCCGATACCGAACT</td><td></td><td></td><td></td><td></td><td></td><td></td><td></td><td></td><td></td><td></td></tr>
<tr><td>Oligo rev</td><td>AAACAGTTCGGTATCGGCTCGCTC</td><td></td><td></td><td></td><td></td><td></td><td></td><td></td><td></td><td></td><td></td></tr>
<tr><td>Chromosome</td><td>start</td><td>end</td><td>strand</td><td>MM</td><td>target_seq</td><td>PAM</td><td>alignment</td><td>distance</td><td>position</td><td>gene name</td><td>gene id</td></tr>
<tr><td>GU305914.1</td><td>119860</td><td>119882</td><td>+</td><td>0</td><td>GAGCGAGCCGATACCGAACT</td><td>GGG</td><td>||||||||[||||||||||||]PAM</td><td>NA</td><td>Intergenic</td><td>NA</td><td>NA</td></tr>
<tr><td>T8977</td><td>GATACCGAACTGGGTGTTCACGG</td><td>1000</td><td></td><td></td><td></td><td></td><td></td><td></td><td></td><td></td><td></td></tr>
<tr><td>Oligo fwd</td><td>CACCGATACCGAACTGGGTGTTCA</td><td></td><td></td><td></td><td></td><td></td><td></td><td></td><td></td><td></td><td></td></tr>
<tr><td>Oligo rev</td><td>AAACTGAACACCCAGTTCGGTATC</td><td></td><td></td><td></td><td></td><td></td><td></td><td></td><td></td><td></td><td></td></tr>
<tr><td>Chromosome</td><td>start</td><td>end</td><td>strand</td><td>MM</td><td>target_seq</td><td>PAM</td><td>alignment</td><td>distance</td><td>position</td><td>gene name</td><td>gene id</td></tr>
<tr><td>GU305914.1</td><td>119869</td><td>119891</td><td>+</td><td>0</td><td>GATACCGAACTGGGTGTTCA</td><td>CGG</td><td>||||||||[||||||||||||]PAM</td><td>NA</td><td>Intergenic</td><td>NA</td><td>NA</td></tr>
<tr><td>T8978</td><td>CGGCGTAAAACATCGTGTGCAGG</td><td>1000</td><td></td><td></td><td></td><td></td><td></td><td></td><td></td><td></td><td></td></tr>
<tr><td>Oligo fwd</td><td>CACCCGGCGTAAAACATCGTGTGC</td><td></td><td></td><td></td><td></td><td></td><td></td><td></td><td></td><td></td><td></td></tr>
<tr><td>Oligo rev</td><td>AAACGCACACGATGTTTTACGCCG</td><td></td><td></td><td></td><td></td><td></td><td></td><td></td><td></td><td></td><td></td></tr>
<tr><td>Chromosome</td><td>start</td><td>end</td><td>strand</td><td>MM</td><td>target_seq</td><td>PAM</td><td>alignment</td><td>distance</td><td>position</td><td>gene name</td><td>gene id</td></tr>
<tr><td>GU305914.1</td><td>119889</td><td>119911</td><td>+</td><td>0</td><td>CGGCGTAAAACATCGTGTGC</td><td>AGG</td><td>||||||||[||||||||||||]PAM</td><td>NA</td><td>Intergenic</td><td>NA</td><td>NA</td></tr>
<tr><td>T8979</td><td>GGAATCGCTTCTTGTTCTCCTGG</td><td>1000</td><td></td><td></td><td></td><td></td><td></td><td></td><td></td><td></td><td></td></tr>
<tr><td>Oligo fwd</td><td>CACCGGAATCGCTTCTTGTTCTCC</td><td></td><td></td><td></td><td></td><td></td><td></td><td></td><td></td><td></td><td></td></tr>
<tr><td>Oligo rev</td><td>AAACGGAGAACAAGAAGCGATTCC</td><td></td><td></td><td></td><td></td><td></td><td></td><td></td><td></td><td></td><td></td></tr>
<tr><td>Chromosome</td><td>start</td><td>end</td><td>strand</td><td>MM</td><td>target_seq</td><td>PAM</td><td>alignment</td><td>distance</td><td>position</td><td>gene name</td><td>gene id</td></tr>
<tr><td>GU305914.1</td><td>119910</td><td>119932</td><td>+</td><td>0</td><td>GGAATCGCTTCTTGTTCTCC</td><td>TGG</td><td>||||||||[||||||||||||]PAM</td><td>NA</td><td>Intergenic</td><td>NA</td><td>NA</td></tr>
<tr><td>T8980</td><td>TCGCTTCTTGTTCTCCTGGAAGG</td><td>1000</td><td></td><td></td><td></td><td></td><td></td><td></td><td></td><td></td><td></td></tr>
<tr><td>Oligo fwd</td><td>CACCTCGCTTCTTGTTCTCCTGGA</td><td></td><td></td><td></td><td></td><td></td><td></td><td></td><td></td><td></td><td></td></tr>
<tr><td>Oligo rev</td><td>AAACTCCAGGAGAACAAGAAGCGA</td><td></td><td></td><td></td><td></td><td></td><td></td><td></td><td></td><td></td><td></td></tr>
<tr><td>Chromosome</td><td>start</td><td>end</td><td>strand</td><td>MM</td><td>target_seq</td><td>PAM</td><td>alignment</td><td>distance</td><td>position</td><td>gene name</td><td>gene id</td></tr>
<tr><td>GU305914.1</td><td>119914</td><td>119936</td><td>+</td><td>0</td><td>TCGCTTCTTGTTCTCCTGGA</td><td>AGG</td><td>||||||||[||||||||||||]PAM</td><td>NA</td><td>Intergenic</td><td>NA</td><td>NA</td></tr>
<tr><td>T8981</td><td>CGCTTCTTGTTCTCCTGGAAGGG</td><td>1000</td><td></td><td></td><td></td><td></td><td></td><td></td><td></td><td></td><td></td></tr>
<tr><td>Oligo fwd</td><td>CACCCGCTTCTTGTTCTCCTGGAA</td><td></td><td></td><td></td><td></td><td></td><td></td><td></td><td></td><td></td><td></td></tr>
<tr><td>Oligo rev</td><td>AAACTTCCAGGAGAACAAGAAGCG</td><td></td><td></td><td></td><td></td><td></td><td></td><td></td><td></td><td></td><td></td></tr>
<tr><td>Chromosome</td><td>start</td><td>end</td><td>strand</td><td>MM</td><td>target_seq</td><td>PAM</td><td>alignment</td><td>distance</td><td>position</td><td>gene name</td><td>gene id</td></tr>
<tr><td>GU305914.1</td><td>119915</td><td>119937</td><td>+</td><td>0</td><td>CGCTTCTTGTTCTCCTGGAA</td><td>GGG</td><td>||||||||[||||||||||||]PAM</td><td>NA</td><td>Intergenic</td><td>NA</td><td>NA</td></tr>
<tr><td>T8982</td><td>CCTCGTCGCGACACACGCACAGG</td><td>1000</td><td></td><td></td><td></td><td></td><td></td><td></td><td></td><td></td><td></td></tr>
<tr><td>Oligo fwd</td><td>CACCCCTCGTCGCGACACACGCAC</td><td></td><td></td><td></td><td></td><td></td><td></td><td></td><td></td><td></td><td></td></tr>
<tr><td>Oligo rev</td><td>AAACGTGCGTGTGTCGCGACGAGG</td><td></td><td></td><td></td><td></td><td></td><td></td><td></td><td></td><td></td><td></td></tr>
<tr><td>Chromosome</td><td>start</td><td>end</td><td>strand</td><td>MM</td><td>target_seq</td><td>PAM</td><td>alignment</td><td>distance</td><td>position</td><td>gene name</td><td>gene id</td></tr>
<tr><td>GU305914.1</td><td>119949</td><td>119971</td><td>+</td><td>0</td><td>CCTCGTCGCGACACACGCAC</td><td>AGG</td><td>||||||||[||||||||||||]PAM</td><td>NA</td><td>Intergenic</td><td>NA</td><td>NA</td></tr>
<tr><td>T8983</td><td>GTCGCGACACACGCACAGGACGG</td><td>1000</td><td></td><td></td><td></td><td></td><td></td><td></td><td></td><td></td><td></td></tr>
<tr><td>Oligo fwd</td><td>CACCGTCGCGACACACGCACAGGA</td><td></td><td></td><td></td><td></td><td></td><td></td><td></td><td></td><td></td><td></td></tr>
<tr><td>Oligo rev</td><td>AAACTCCTGTGCGTGTGTCGCGAC</td><td></td><td></td><td></td><td></td><td></td><td></td><td></td><td></td><td></td><td></td></tr>
<tr><td>Chromosome</td><td>start</td><td>end</td><td>strand</td><td>MM</td><td>target_seq</td><td>PAM</td><td>alignment</td><td>distance</td><td>position</td><td>gene name</td><td>gene id</td></tr>
<tr><td>GU305914.1</td><td>119953</td><td>119975</td><td>+</td><td>0</td><td>GTCGCGACACACGCACAGGA</td><td>CGG</td><td>||||||||[||||||||||||]PAM</td><td>NA</td><td>Intergenic</td><td>NA</td><td>NA</td></tr>
<tr><td>T8984</td><td>GCGACACACGCACAGGACGGAGG</td><td>1000</td><td></td><td></td><td></td><td></td><td></td><td></td><td></td><td></td><td></td></tr>
<tr><td>Oligo fwd</td><td>CACCGCGACACACGCACAGGACGG</td><td></td><td></td><td></td><td></td><td></td><td></td><td></td><td></td><td></td><td></td></tr>
<tr><td>Oligo rev</td><td>AAACCCGTCCTGTGCGTGTGTCGC</td><td></td><td></td><td></td><td></td><td></td><td></td><td></td><td></td><td></td><td></td></tr>
<tr><td>Chromosome</td><td>start</td><td>end</td><td>strand</td><td>MM</td><td>target_seq</td><td>PAM</td><td>alignment</td><td>distance</td><td>position</td><td>gene name</td><td>gene id</td></tr>
<tr><td>GU305914.1</td><td>119956</td><td>119978</td><td>+</td><td>0</td><td>GCGACACACGCACAGGACGG</td><td>AGG</td><td>||||||||[||||||||||||]PAM</td><td>NA</td><td>Intergenic</td><td>NA</td><td>NA</td></tr>
</table>

<table>
<tr><td>T8985</td><td>CGACACACGCACAGGACGGAGGG</td><td>1000</td><td></td><td></td><td></td><td></td><td></td><td></td><td></td><td></td><td></td></tr>
<tr><td>Oligo fwd</td><td>CACCCGACACACGCACAGGACGGA</td><td></td><td></td><td></td><td></td><td></td><td></td><td></td><td></td><td></td><td></td></tr>
<tr><td>Oligo rev</td><td>AAACTCCGTCCTGTGCGTGTGTCG</td><td></td><td></td><td></td><td></td><td></td><td></td><td></td><td></td><td></td><td></td></tr>
<tr><td>Chromosome</td><td>start</td><td>end</td><td>strand</td><td>MM</td><td>target_seq</td><td>PAM</td><td>alignment</td><td>distance</td><td>position</td><td>gene name</td><td>gene id</td></tr>
<tr><td>GU305914.1</td><td>119957</td><td>119979</td><td>+</td><td>0</td><td>CGACACACGCACAGGACGGA</td><td>GGG</td><td>||||||||[||||||||||||]PAM</td><td>NA</td><td>Intergenic</td><td>NA</td><td>NA</td></tr>
<tr><td>T8986</td><td>ACAGGACGGAGGGTCGTTTCAGG</td><td>1000</td><td></td><td></td><td></td><td></td><td></td><td></td><td></td><td></td><td></td></tr>
<tr><td>Oligo fwd</td><td>CACCACAGGACGGAGGGTCGTTTC</td><td></td><td></td><td></td><td></td><td></td><td></td><td></td><td></td><td></td><td></td></tr>
<tr><td>Oligo rev</td><td>AAACGAAACGACCCTCCGTCCTGT</td><td></td><td></td><td></td><td></td><td></td><td></td><td></td><td></td><td></td><td></td></tr>
<tr><td>Chromosome</td><td>start</td><td>end</td><td>strand</td><td>MM</td><td>target_seq</td><td>PAM</td><td>alignment</td><td>distance</td><td>position</td><td>gene name</td><td>gene id</td></tr>
<tr><td>GU305914.1</td><td>119967</td><td>119989</td><td>+</td><td>0</td><td>ACAGGACGGAGGGTCGTTTC</td><td>AGG</td><td>||||||||[||||||||||||]PAM</td><td>NA</td><td>Intergenic</td><td>NA</td><td>NA</td></tr>
<tr><td>T8987</td><td>GGACGGAGGGTCGTTTCAGGAGG</td><td>1000</td><td></td><td></td><td></td><td></td><td></td><td></td><td></td><td></td><td></td></tr>
<tr><td>Oligo fwd</td><td>CACCGGACGGAGGGTCGTTTCAGG</td><td></td><td></td><td></td><td></td><td></td><td></td><td></td><td></td><td></td><td></td></tr>
<tr><td>Oligo rev</td><td>AAACCCTGAAACGACCCTCCGTCC</td><td></td><td></td><td></td><td></td><td></td><td></td><td></td><td></td><td></td><td></td></tr>
<tr><td>Chromosome</td><td>start</td><td>end</td><td>strand</td><td>MM</td><td>target_seq</td><td>PAM</td><td>alignment</td><td>distance</td><td>position</td><td>gene name</td><td>gene id</td></tr>
<tr><td>GU305914.1</td><td>119970</td><td>119992</td><td>+</td><td>0</td><td>GGACGGAGGGTCGTTTCAGG</td><td>AGG</td><td>||||||||[||||||||||||]PAM</td><td>NA</td><td>Intergenic</td><td>NA</td><td>NA</td></tr>
<tr><td>T8988</td><td>CGAGAACACTTTCAGCTCCTCGG</td><td>1000</td><td></td><td></td><td></td><td></td><td></td><td></td><td></td><td></td><td></td></tr>
<tr><td>Oligo fwd</td><td>CACCCGAGAACACTTTCAGCTCCT</td><td></td><td></td><td></td><td></td><td></td><td></td><td></td><td></td><td></td><td></td></tr>
<tr><td>Oligo rev</td><td>AAACAGGAGCTGAAAGTGTTCTCG</td><td></td><td></td><td></td><td></td><td></td><td></td><td></td><td></td><td></td><td></td></tr>
<tr><td>Chromosome</td><td>start</td><td>end</td><td>strand</td><td>MM</td><td>target_seq</td><td>PAM</td><td>alignment</td><td>distance</td><td>position</td><td>gene name</td><td>gene id</td></tr>
<tr><td>GU305914.1</td><td>120007</td><td>120029</td><td>+</td><td>0</td><td>CGAGAACACTTTCAGCTCCT</td><td>CGG</td><td>||||||||[||||||||||||]PAM</td><td>NA</td><td>Intergenic</td><td>NA</td><td>NA</td></tr>
<tr><td>T8989</td><td>GAGAACACTTTCAGCTCCTCGGG</td><td>1000</td><td></td><td></td><td></td><td></td><td></td><td></td><td></td><td></td><td></td></tr>
<tr><td>Oligo fwd</td><td>CACCGAGAACACTTTCAGCTCCTC</td><td></td><td></td><td></td><td></td><td></td><td></td><td></td><td></td><td></td><td></td></tr>
<tr><td>Oligo rev</td><td>AAACGAGGAGCTGAAAGTGTTCTC</td><td></td><td></td><td></td><td></td><td></td><td></td><td></td><td></td><td></td><td></td></tr>
<tr><td>Chromosome</td><td>start</td><td>end</td><td>strand</td><td>MM</td><td>target_seq</td><td>PAM</td><td>alignment</td><td>distance</td><td>position</td><td>gene name</td><td>gene id</td></tr>
<tr><td>GU305914.1</td><td>120008</td><td>120030</td><td>+</td><td>0</td><td>GAGAACACTTTCAGCTCCTC</td><td>GGG</td><td>||||||||[||||||||||||]PAM</td><td>NA</td><td>Intergenic</td><td>NA</td><td>NA</td></tr>
<tr><td>T8990</td><td>CGGGCTCGAAGAAGACCCGCTGG</td><td>1000</td><td></td><td></td><td></td><td></td><td></td><td></td><td></td><td></td><td></td></tr>
<tr><td>Oligo fwd</td><td>CACCCGGGCTCGAAGAAGACCCGC</td><td></td><td></td><td></td><td></td><td></td><td></td><td></td><td></td><td></td><td></td></tr>
<tr><td>Oligo rev</td><td>AAACGCGGGTCTTCTTCGAGCCCG</td><td></td><td></td><td></td><td></td><td></td><td></td><td></td><td></td><td></td><td></td></tr>
<tr><td>Chromosome</td><td>start</td><td>end</td><td>strand</td><td>MM</td><td>target_seq</td><td>PAM</td><td>alignment</td><td>distance</td><td>position</td><td>gene name</td><td>gene id</td></tr>
<tr><td>GU305914.1</td><td>120027</td><td>120049</td><td>+</td><td>0</td><td>CGGGCTCGAAGAAGACCCGC</td><td>TGG</td><td>||||||||[||||||||||||]PAM</td><td>NA</td><td>Intergenic</td><td>NA</td><td>NA</td></tr>
<tr><td>T8991</td><td>GCTCGAAGAAGACCCGCTGGAGG</td><td>1000</td><td></td><td></td><td></td><td></td><td></td><td></td><td></td><td></td><td></td></tr>
<tr><td>Oligo fwd</td><td>CACCGCTCGAAGAAGACCCGCTGG</td><td></td><td></td><td></td><td></td><td></td><td></td><td></td><td></td><td></td><td></td></tr>
<tr><td>Oligo rev</td><td>AAACCCAGCGGGTCTTCTTCGAGC</td><td></td><td></td><td></td><td></td><td></td><td></td><td></td><td></td><td></td><td></td></tr>
<tr><td>Chromosome</td><td>start</td><td>end</td><td>strand</td><td>MM</td><td>target_seq</td><td>PAM</td><td>alignment</td><td>distance</td><td>position</td><td>gene name</td><td>gene id</td></tr>
<tr><td>GU305914.1</td><td>120030</td><td>120052</td><td>+</td><td>0</td><td>GCTCGAAGAAGACCCGCTGG</td><td>AGG</td><td>||||||||[||||||||||||]PAM</td><td>NA</td><td>Intergenic</td><td>NA</td><td>NA</td></tr>
<tr><td>T8992</td><td>GCTGGAGGAAACTCGCCAGCAGG</td><td>1000</td><td></td><td></td><td></td><td></td><td></td><td></td><td></td><td></td><td></td></tr>
<tr><td>Oligo fwd</td><td>CACCGCTGGAGGAAACTCGCCAGC</td><td></td><td></td><td></td><td></td><td></td><td></td><td></td><td></td><td></td><td></td></tr>
<tr><td>Oligo rev</td><td>AAACGCTGGCGAGTTTCCTCCAGC</td><td></td><td></td><td></td><td></td><td></td><td></td><td></td><td></td><td></td><td></td></tr>
<tr><td>Chromosome</td><td>start</td><td>end</td><td>strand</td><td>MM</td><td>target_seq</td><td>PAM</td><td>alignment</td><td>distance</td><td>position</td><td>gene name</td><td>gene id</td></tr>
<tr><td>GU305914.1</td><td>120045</td><td>120067</td><td>+</td><td>0</td><td>GCTGGAGGAAACTCGCCAGC</td><td>AGG</td><td>||||||||[||||||||||||]PAM</td><td>NA</td><td>Intergenic</td><td>NA</td><td>NA</td></tr>
<tr><td>T8993</td><td>CGTCGAGCCCACCATCGACGAGG</td><td>1000</td><td></td><td></td><td></td><td></td><td></td><td></td><td></td><td></td><td></td></tr>
<tr><td>Oligo fwd</td><td>CACCCGTCGAGCCCACCATCGACG</td><td></td><td></td><td></td><td></td><td></td><td></td><td></td><td></td><td></td><td></td></tr>
<tr><td>Oligo rev</td><td>AAACCGTCGATGGTGGGCTCGACG</td><td></td><td></td><td></td><td></td><td></td><td></td><td></td><td></td><td></td><td></td></tr>
<tr><td>Chromosome</td><td>start</td><td>end</td><td>strand</td><td>MM</td><td>target_seq</td><td>PAM</td><td>alignment</td><td>distance</td><td>position</td><td>gene name</td><td>gene id</td></tr>
<tr><td>GU305914.1</td><td>120073</td><td>120095</td><td>+</td><td>0</td><td>CGTCGAGCCCACCATCGACG</td><td>AGG</td><td>||||||||[||||||||||||]PAM</td><td>NA</td><td>Intergenic</td><td>NA</td><td>NA</td></tr>
<tr><td>T8994</td><td>AGCCCACCATCGACGAGGTGCGG</td><td>1000</td><td></td><td></td><td></td><td></td><td></td><td></td><td></td><td></td><td></td></tr>
<tr><td>Oligo fwd</td><td>CACCAGCCCACCATCGACGAGGTG</td><td></td><td></td><td></td><td></td><td></td><td></td><td></td><td></td><td></td><td></td></tr>
<tr><td>Oligo rev</td><td>AAACCACCTCGTCGATGGTGGGCT</td><td></td><td></td><td></td><td></td><td></td><td></td><td></td><td></td><td></td><td></td></tr>
<tr><td>Chromosome</td><td>start</td><td>end</td><td>strand</td><td>MM</td><td>target_seq</td><td>PAM</td><td>alignment</td><td>distance</td><td>position</td><td>gene name</td><td>gene id</td></tr>
<tr><td>GU305914.1</td><td>120078</td><td>120100</td><td>+</td><td>0</td><td>AGCCCACCATCGACGAGGTG</td><td>CGG</td><td>||||||||[||||||||||||]PAM</td><td>NA</td><td>Intergenic</td><td>NA</td><td>NA</td></tr>
<tr><td>T8995</td><td>GCCCACCATCGACGAGGTGCGGG</td><td>1000</td><td></td><td></td><td></td><td></td><td></td><td></td><td></td><td></td><td></td></tr>
<tr><td>Oligo fwd</td><td>CACCGCCCACCATCGACGAGGTGC</td><td></td><td></td><td></td><td></td><td></td><td></td><td></td><td></td><td></td><td></td></tr>
<tr><td>Oligo rev</td><td>AAACGCACCTCGTCGATGGTGGGC</td><td></td><td></td><td></td><td></td><td></td><td></td><td></td><td></td><td></td><td></td></tr>
<tr><td>Chromosome</td><td>start</td><td>end</td><td>strand</td><td>MM</td><td>target_seq</td><td>PAM</td><td>alignment</td><td>distance</td><td>position</td><td>gene name</td><td>gene id</td></tr>
<tr><td>GU305914.1</td><td>120079</td><td>120101</td><td>+</td><td>0</td><td>GCCCACCATCGACGAGGTGC</td><td>GGG</td><td>||||||||[||||||||||||]PAM</td><td>NA</td><td>Intergenic</td><td>NA</td><td>NA</td></tr>
<tr><td>T8996</td><td>CGACGAGGTGCGGGAGTCCAAGG</td><td>1000</td><td></td><td></td><td></td><td></td><td></td><td></td><td></td><td></td><td></td></tr>
<tr><td>Oligo fwd</td><td>CACCCGACGAGGTGCGGGAGTCCA</td><td></td><td></td><td></td><td></td><td></td><td></td><td></td><td></td><td></td><td></td></tr>
<tr><td>Oligo rev</td><td>AAACTGGACTCCCGCACCTCGTCG</td><td></td><td></td><td></td><td></td><td></td><td></td><td></td><td></td><td></td><td></td></tr>
<tr><td>Chromosome</td><td>start</td><td>end</td><td>strand</td><td>MM</td><td>target_seq</td><td>PAM</td><td>alignment</td><td>distance</td><td>position</td><td>gene name</td><td>gene id</td></tr>
<tr><td>GU305914.1</td><td>120088</td><td>120110</td><td>+</td><td>0</td><td>CGACGAGGTGCGGGAGTCCA</td><td>AGG</td><td>||||||||[||||||||||||]PAM</td><td>NA</td><td>Intergenic</td><td>NA</td><td>NA</td></tr>
<tr><td>GU305914.1</td><td>141917</td><td>141939</td><td>+</td><td>4</td><td>CGACGACATGCGGGAGCCTA</td><td>TGG</td><td>||||||--[||||||||-|-|]PAM</td><td>NA</td><td>Intergenic</td><td>NA</td><td>NA</td></tr>
<tr><td>T8997</td><td>GACGAGGTGCGGGAGTCCAAGGG</td><td>1000</td><td></td><td></td><td></td><td></td><td></td><td></td><td></td><td></td><td></td></tr>
<tr><td>Oligo fwd</td><td>CACCGACGAGGTGCGGGAGTCCAA</td><td></td><td></td><td></td><td></td><td></td><td></td><td></td><td></td><td></td><td></td></tr>
<tr><td>Oligo rev</td><td>AAACTTGGACTCCCGCACCTCGTC</td><td></td><td></td><td></td><td></td><td></td><td></td><td></td><td></td><td></td><td></td></tr>
<tr><td>Chromosome</td><td>start</td><td>end</td><td>strand</td><td>MM</td><td>target_seq</td><td>PAM</td><td>alignment</td><td>distance</td><td>position</td><td>gene name</td><td>gene id</td></tr>
<tr><td>GU305914.1</td><td>120089</td><td>120111</td><td>+</td><td>0</td><td>GACGAGGTGCGGGAGTCCAA</td><td>GGG</td><td>||||||||[||||||||||||]PAM</td><td>NA</td><td>Intergenic</td><td>NA</td><td>NA</td></tr>
<tr><td>T8998</td><td>GTCCAAGGGATTGATCACTTCGG</td><td>1000</td><td></td><td></td><td></td><td></td><td></td><td></td><td></td><td></td><td></td></tr>
<tr><td>Oligo fwd</td><td>CACCGTCCAAGGGATTGATCACTT</td><td></td><td></td><td></td><td></td><td></td><td></td><td></td><td></td><td></td><td></td></tr>
<tr><td>Oligo rev</td><td>AAACAAGTGATCAATCCCTTGGAC</td><td></td><td></td><td></td><td></td><td></td><td></td><td></td><td></td><td></td><td></td></tr>
<tr><td>Chromosome</td><td>start</td><td>end</td><td>strand</td><td>MM</td><td>target_seq</td><td>PAM</td><td>alignment</td><td>distance</td><td>position</td><td>gene name</td><td>gene id</td></tr>
<tr><td>GU305914.1</td><td>120103</td><td>120125</td><td>+</td><td>0</td><td>GTCCAAGGGATTGATCACTT</td><td>CGG</td><td>||||||||[||||||||||||]PAM</td><td>NA</td><td>Intergenic</td><td>NA</td><td>NA</td></tr>
<tr><td>T8999</td><td>CACTTCGGTCCTGTTGTTCATGG</td><td>1000</td><td></td><td></td><td></td><td></td><td></td><td></td><td></td><td></td><td></td></tr>
<tr><td>Oligo fwd</td><td>CACCCACTTCGGTCCTGTTGTTCA</td><td></td><td></td><td></td><td></td><td></td><td></td><td></td><td></td><td></td><td></td></tr>
<tr><td>Oligo rev</td><td>AAACTGAACAACAGGACCGAAGTG</td><td></td><td></td><td></td><td></td><td></td><td></td><td></td><td></td><td></td><td></td></tr>
<tr><td>Chromosome</td><td>start</td><td>end</td><td>strand</td><td>MM</td><td>target_seq</td><td>PAM</td><td>alignment</td><td>distance</td><td>position</td><td>gene name</td><td>gene id</td></tr>
</table>

GU305914.1 120118 120140 + 0 CACTTCGGTCCTGTTGTTCA TGG ||||||||[||||||||||||]PAM NA Intergenic NA NA

T9000 GCGTTCATCTTGATTCCGATCGG 1000
Oligo fwd CACCGCGTTCATCTTGATTCCGAT
Oligo rev AAACATCGGAATCAAGATGAACGC
Chromosome start end strand MM target_seq PAM alignment distance position gene name gene id
GU305914.1 120140 120162 + 0 GCGTTCATCTTGATTCCGAT CGG ||||||||[||||||||||||]PAM NA Intergenic NA NA

T9001 CGTTCATCTTGATTCCGATCGGG 1000
Oligo fwd CACCCGTTCATCTTGATTCCGATC
Oligo rev AAACGATCGGAATCAAGATGAACG
Chromosome start end strand MM target_seq PAM alignment distance position gene name gene id
GU305914.1 120141 120163 + 0 CGTTCATCTTGATTCCGATC GGG ||||||||[||||||||||||]PAM NA Intergenic NA NA

T9002 AGATGTTCTGCTGCAGCCTCTGG 1000
Oligo fwd CACCAGATGTTCTGCTGCAGCCTC
Oligo rev AAACGAGGCTGCAGCAGAACATCT
Chromosome start end strand MM target_seq PAM alignment distance position gene name gene id
GU305914.1 120188 120210 + 0 AGATGTTCTGCTGCAGCCTC TGG ||||||||[||||||||||||]PAM NA Intergenic NA NA

T9003 AGCCTCTGGAGCGCGCCGAGCGG 1000
Oligo fwd CACCAGCCTCTGGAGCGCGCCGAG
Oligo rev AAACCTCGGCGCGCTCCAGAGGCT
Chromosome start end strand MM target_seq PAM alignment distance position gene name gene id
GU305914.1 120202 120224 + 0 AGCCTCTGGAGCGCGCCGAG CGG ||||||||[||||||||||||]PAM NA Intergenic NA NA

T9004 GCCTCTGGAGCGCGCCGAGCGGG 1000
Oligo fwd CACCGCCTCTGGAGCGCGCCGAGC
Oligo rev AAACGCTCGGCGCGCTCCAGAGGC
Chromosome start end strand MM target_seq PAM alignment distance position gene name gene id
GU305914.1 120203 120225 + 0 GCCTCTGGAGCGCGCCGAGC GGG ||||||||[||||||||||||]PAM NA Intergenic NA NA

T9005 TCTGGAGCGCGCCGAGCGGGTGG 1000
Oligo fwd CACCTCTGGAGCGCGCCGAGCGGG
Oligo rev AAACCCCGCTCGGCGCGCTCCAGA
Chromosome start end strand MM target_seq PAM alignment distance position gene name gene id
GU305914.1 120206 120228 + 0 TCTGGAGCGCGCCGAGCGGG TGG ||||||||[||||||||||||]PAM NA Intergenic NA NA

T9006 GCCGAGCGGGTGGAAGAACACGG 1000
Oligo fwd CACCGCCGAGCGGGTGGAAGAACA
Oligo rev AAACTGTTCTTCCACCCGCTCGGC
Chromosome start end strand MM target_seq PAM alignment distance position gene name gene id
GU305914.1 120216 120238 + 0 GCCGAGCGGGTGGAAGAACA CGG ||||||||[||||||||||||]PAM NA Intergenic NA NA

T9007 GAGCGGGTGGAAGAACACGGCGG 1000
Oligo fwd CACCGAGCGGGTGGAAGAACACGG
Oligo rev AAACCCGTGTTCTTCCACCCGCTC
Chromosome start end strand MM target_seq PAM alignment distance position gene name gene id
GU305914.1 120219 120241 + 0 GAGCGGGTGGAAGAACACGG CGG ||||||||[||||||||||||]PAM NA Intergenic NA NA

T9008 AGCGGGTGGAAGAACACGGCGGG 1000
Oligo fwd CACCAGCGGGTGGAAGAACACGGC
Oligo rev AAACGCCGTGTTCTTCCACCCGCT
Chromosome start end strand MM target_seq PAM alignment distance position gene name gene id
GU305914.1 120220 120242 + 0 AGCGGGTGGAAGAACACGGC GGG ||||||||[||||||||||||]PAM NA Intergenic NA NA

T9009 GGGTGGAAGAACACGGCGGGCGG 1000
Oligo fwd CACCGGGTGGAAGAACACGGCGGG
Oligo rev AAACCCCGCCGTGTTCTTCCACCC
Chromosome start end strand MM target_seq PAM alignment distance position gene name gene id
GU305914.1 120223 120245 + 0 GGGTGGAAGAACACGGCGGG CGG ||||||||[||||||||||||]PAM NA Intergenic NA NA

T9010 AAGAACACGGCGGGCGGTGCCGG 1000
Oligo fwd CACCAAGAACACGGCGGGCGGTGC
Oligo rev AAACGCACCGCCCGCCGTGTTCTT
Chromosome start end strand MM target_seq PAM alignment distance position gene name gene id
GU305914.1 120229 120251 + 0 AAGAACACGGCGGGCGGTGC CGG ||||||||[||||||||||||]PAM NA Intergenic NA NA

T9011 CGGGCGGTGCCGGCTGAAGCTGG 1000
Oligo fwd CACCCGGGCGGTGCCGGCTGAAGC
Oligo rev AAACGCTTCAGCCGGCACCGCCCG
Chromosome start end strand MM target_seq PAM alignment distance position gene name gene id
GU305914.1 120239 120261 + 0 CGGGCGGTGCCGGCTGAAGC TGG ||||||||[||||||||||||]PAM NA Intergenic NA NA

T9012 GCCGGCTGAAGCTGGAGCTGAGG 1000
Oligo fwd CACCGCCGGCTGAAGCTGGAGCTG
Oligo rev AAACCAGCTCCAGCTTCAGCCGGC
Chromosome start end strand MM target_seq PAM alignment distance position gene name gene id
GU305914.1 120247 120269 + 0 GCCGGCTGAAGCTGGAGCTG AGG ||||||||[||||||||||||]PAM NA Intergenic NA NA

T9013 GGCTGAAGCTGGAGCTGAGGAGG 1000
Oligo fwd CACCGGCTGAAGCTGGAGCTGAGG
Oligo rev AAACCCTCAGCTCCAGCTTCAGCC
Chromosome start end strand MM target_seq PAM alignment distance position gene name gene id
GU305914.1 120250 120272 + 0 GGCTGAAGCTGGAGCTGAGG AGG ||||||||[||||||||||||]PAM NA Intergenic NA NA

T9014 GCTGGAGCTGAGGAGGCGATTGG 1000
Oligo fwd CACCGCTGGAGCTGAGGAGGCGAT
Oligo rev AAACATCGCCTCCTCAGCTCCAGC

| | | | | | | | | | | | |
|---|---|---|---|---|---|---|---|---|---|---|---|
| Chromosome | start | end | strand | MM | target_seq | PAM | alignment | distance | position | gene name | gene id |
| GU305914.1 | 120257 | 120279 | + | 0 | GCTGGAGCTGAGGAGGCGAT | TGG | \|\|\|\|\|\|\|\|[\|\|\|\|\|\|\|\|\|\|\|\|]PAM | NA | Intergenic | NA | NA |
| T9015 | AGGAGGCGATTGGTAATGCTTGG | 1000 | | | | | | | | | |
| Oligo fwd | CACCAGGAGGCGATTGGTAATGCT | | | | | | | | | | |
| Oligo rev | AAACAGCATTACCAATCGCCTCCT | | | | | | | | | | |
| Chromosome | start | end | strand | MM | target_seq | PAM | alignment | distance | position | gene name | gene id |
| GU305914.1 | 120267 | 120289 | + | 0 | AGGAGGCGATTGGTAATGCT | TGG | \|\|\|\|\|\|\|\|[\|\|\|\|\|\|\|\|\|\|\|\|]PAM | NA | Intergenic | NA | NA |
| T9016 | TGCGCTGTGCCTGCAACGCCAGG | 1000 | | | | | | | | | |
| Oligo fwd | CACCTGCGCTGTGCCTGCAACGCC | | | | | | | | | | |
| Oligo rev | AAACGGCGTTGCAGGCACAGCGCA | | | | | | | | | | |
| Chromosome | start | end | strand | MM | target_seq | PAM | alignment | distance | position | gene name | gene id |
| GU305914.1 | 120290 | 120312 | + | 0 | TGCGCTGTGCCTGCAACGCC | AGG | \|\|\|\|\|\|\|\|[\|\|\|\|\|\|\|\|\|\|\|\|]PAM | NA | Intergenic | NA | NA |
| T9017 | TGCCTGCAACGCCAGGATGTTGG | 1000 | | | | | | | | | |
| Oligo fwd | CACCTGCCTGCAACGCCAGGATGT | | | | | | | | | | |
| Oligo rev | AAACACATCCTGGCGTTGCAGGCA | | | | | | | | | | |
| Chromosome | start | end | strand | MM | target_seq | PAM | alignment | distance | position | gene name | gene id |
| GU305914.1 | 120297 | 120319 | + | 0 | TGCCTGCAACGCCAGGATGT | TGG | \|\|\|\|\|\|\|\|[\|\|\|\|\|\|\|\|\|\|\|\|]PAM | NA | Intergenic | NA | NA |
| T9018 | AACGCCAGGATGTTGGAGAACGG | 1000 | | | | | | | | | |
| Oligo fwd | CACCAACGCCAGGATGTTGGAGAA | | | | | | | | | | |
| Oligo rev | AAACTTCTCCAACATCCTGGCGTT | | | | | | | | | | |
| Chromosome | start | end | strand | MM | target_seq | PAM | alignment | distance | position | gene name | gene id |
| GU305914.1 | 120304 | 120326 | + | 0 | AACGCCAGGATGTTGGAGAA | CGG | \|\|\|\|\|\|\|\|[\|\|\|\|\|\|\|\|\|\|\|\|]PAM | NA | Intergenic | NA | NA |
| T9019 | GCCAGGATGTTGGAGAACGGAGG | 1000 | | | | | | | | | |
| Oligo fwd | CACCGCCAGGATGTTGGAGAACGG | | | | | | | | | | |
| Oligo rev | AAACCCGTTCTCCAACATCCTGGC | | | | | | | | | | |
| Chromosome | start | end | strand | MM | target_seq | PAM | alignment | distance | position | gene name | gene id |
| GU305914.1 | 120307 | 120329 | + | 0 | GCCAGGATGTTGGAGAACGG | AGG | \|\|\|\|\|\|\|\|[\|\|\|\|\|\|\|\|\|\|\|\|]PAM | NA | Intergenic | NA | NA |
| T9020 | AACGGAGGCGAATGTCCGTACGG | 1000 | | | | | | | | | |
| Oligo fwd | CACCAACGGAGGCGAATGTCCGTA | | | | | | | | | | |
| Oligo rev | AAACTACGGACATTCGCCTCCGTT | | | | | | | | | | |
| Chromosome | start | end | strand | MM | target_seq | PAM | alignment | distance | position | gene name | gene id |
| GU305914.1 | 120322 | 120344 | + | 0 | AACGGAGGCGAATGTCCGTA | CGG | \|\|\|\|\|\|\|\|[\|\|\|\|\|\|\|\|\|\|\|\|]PAM | NA | Intergenic | NA | NA |
| T9021 | CCGTACGGCGCTTGACTCGCCGG | 1000 | | | | | | | | | |
| Oligo fwd | CACCCCGTACGGCGCTTGACTCGC | | | | | | | | | | |
| Oligo rev | AAACGCGAGTCAAGCGCCGTACGG | | | | | | | | | | |
| Chromosome | start | end | strand | MM | target_seq | PAM | alignment | distance | position | gene name | gene id |
| GU305914.1 | 120337 | 120359 | + | 0 | CCGTACGGCGCTTGACTCGC | CGG | \|\|\|\|\|\|\|\|[\|\|\|\|\|\|\|\|\|\|\|\|]PAM | NA | Intergenic | NA | NA |
| T9022 | GCACAATTCCGAGTGCCACCTGG | 1000 | | | | | | | | | |
| Oligo fwd | CACCGCACAATTCCGAGTGCCACC | | | | | | | | | | |
| Oligo rev | AAACGGTGGCACTCGGAATTGTGC | | | | | | | | | | |
| Chromosome | start | end | strand | MM | target_seq | PAM | alignment | distance | position | gene name | gene id |
| GU305914.1 | 120384 | 120406 | + | 0 | GCACAATTCCGAGTGCCACC | TGG | \|\|\|\|\|\|\|\|[\|\|\|\|\|\|\|\|\|\|\|\|]PAM | NA | Intergenic | NA | NA |
| T9023 | CACAATTCCGAGTGCCACCTGGG | 1000 | | | | | | | | | |
| Oligo fwd | CACCCACAATTCCGAGTGCCACCT | | | | | | | | | | |
| Oligo rev | AAACAGGTGGCACTCGGAATTGTG | | | | | | | | | | |
| Chromosome | start | end | strand | MM | target_seq | PAM | alignment | distance | position | gene name | gene id |
| GU305914.1 | 120385 | 120407 | + | 0 | CACAATTCCGAGTGCCACCT | GGG | \|\|\|\|\|\|\|\|[\|\|\|\|\|\|\|\|\|\|\|\|]PAM | NA | Intergenic | NA | NA |
| T9024 | TTCCGAGTGCCACCTGGGCGAGG | 1000 | | | | | | | | | |
| Oligo fwd | CACCTTCCGAGTGCCACCTGGGCG | | | | | | | | | | |
| Oligo rev | AAACCGCCCAGGTGGCACTCGGAA | | | | | | | | | | |
| Chromosome | start | end | strand | MM | target_seq | PAM | alignment | distance | position | gene name | gene id |
| GU305914.1 | 120390 | 120412 | + | 0 | TTCCGAGTGCCACCTGGGCG | AGG | \|\|\|\|\|\|\|\|[\|\|\|\|\|\|\|\|\|\|\|\|]PAM | NA | Intergenic | NA | NA |
| T9025 | TCCGAGTGCCACCTGGGCGAGGG | 1000 | | | | | | | | | |
| Oligo fwd | CACCTCCGAGTGCCACCTGGGCGA | | | | | | | | | | |
| Oligo rev | AAACTCGCCCAGGTGGCACTCGGA | | | | | | | | | | |
| Chromosome | start | end | strand | MM | target_seq | PAM | alignment | distance | position | gene name | gene id |
| GU305914.1 | 120391 | 120413 | + | 0 | TCCGAGTGCCACCTGGGCGA | GGG | \|\|\|\|\|\|\|\|[\|\|\|\|\|\|\|\|\|\|\|\|]PAM | NA | Intergenic | NA | NA |
| T9026 | CCGAGTGCCACCTGGGCGAGGGG | 1000 | | | | | | | | | |
| Oligo fwd | CACCCCGAGTGCCACCTGGGCGAG | | | | | | | | | | |
| Oligo rev | AAACCTCGCCCAGGTGGCACTCGG | | | | | | | | | | |
| Chromosome | start | end | strand | MM | target_seq | PAM | alignment | distance | position | gene name | gene id |
| GU305914.1 | 120392 | 120414 | + | 0 | CCGAGTGCCACCTGGGCGAG | GGG | \|\|\|\|\|\|\|\|[\|\|\|\|\|\|\|\|\|\|\|\|]PAM | NA | Intergenic | NA | NA |
| T9027 | GCTCGCCCTGCCGATCTTCTTGG | 1000 | | | | | | | | | |
| Oligo fwd | CACCGCTCGCCCTGCCGATCTTCT | | | | | | | | | | |
| Oligo rev | AAACAGAAGATCGGCAGGGCGAGC | | | | | | | | | | |
| Chromosome | start | end | strand | MM | target_seq | PAM | alignment | distance | position | gene name | gene id |
| GU305914.1 | 120426 | 120448 | + | 0 | GCTCGCCCTGCCGATCTTCT | TGG | \|\|\|\|\|\|\|\|[\|\|\|\|\|\|\|\|\|\|\|\|]PAM | NA | Intergenic | NA | NA |
| T9028 | CTCGCCCTGCCGATCTTCTTGGG | 1000 | | | | | | | | | |
| Oligo fwd | CACCCTCGCCCTGCCGATCTTCTT | | | | | | | | | | |
| Oligo rev | AAACAAGAAGATCGGCAGGGCGAG | | | | | | | | | | |
| Chromosome | start | end | strand | MM | target_seq | PAM | alignment | distance | position | gene name | gene id |
| GU305914.1 | 120427 | 120449 | + | 0 | CTCGCCCTGCCGATCTTCTT | GGG | \|\|\|\|\|\|\|\|[\|\|\|\|\|\|\|\|\|\|\|\|]PAM | NA | Intergenic | NA | NA |
| T9029 | GCCCTGCCGATCTTCTTGGGCGG | 1000 | | | | | | | | | |
| Oligo fwd | CACCGCCCTGCCGATCTTCTTGGG | | | | | | | | | | |

<table>
<tr><td>Oligo rev</td><td>AAACCCCAAGAAGATCGGCAGGGC</td><td></td><td></td><td></td><td></td><td></td><td></td><td></td><td></td><td></td><td></td><td></td></tr>
<tr><td>Chromosome</td><td>start</td><td></td><td>end</td><td>strand</td><td>MM</td><td>target_seq</td><td>PAM</td><td>alignment</td><td>distance</td><td>position</td><td>gene name</td><td>gene id</td></tr>
<tr><td>GU305914.1</td><td></td><td>120430</td><td>120452</td><td>+</td><td>0</td><td>GCCCTGCCGATCTTCTTGGG</td><td>CGG</td><td>||||||||[||||||||||||]PAM</td><td>NA</td><td>Intergenic</td><td>NA</td><td>NA</td></tr>
<tr><td>T9030</td><td>TTGGGCGGACCGCGCGTGATCGG</td><td>1000</td><td></td><td></td><td></td><td></td><td></td><td></td><td></td><td></td><td></td><td></td></tr>
<tr><td>Oligo fwd</td><td>CACCTTGGGCGGACCGCGCGTGAT</td><td></td><td></td><td></td><td></td><td></td><td></td><td></td><td></td><td></td><td></td><td></td></tr>
<tr><td>Oligo rev</td><td>AAACATCACGCGCGGTCCGCCCAA</td><td></td><td></td><td></td><td></td><td></td><td></td><td></td><td></td><td></td><td></td><td></td></tr>
<tr><td>Chromosome</td><td>start</td><td></td><td>end</td><td>strand</td><td>MM</td><td>target_seq</td><td>PAM</td><td>alignment</td><td>distance</td><td>position</td><td>gene name</td><td>gene id</td></tr>
<tr><td>GU305914.1</td><td></td><td>120445</td><td>120467</td><td>+</td><td>0</td><td>TTGGGCGGACCGCGCGTGAT</td><td>CGG</td><td>||||||||[||||||||||||]PAM</td><td>NA</td><td>Intergenic</td><td>NA</td><td>NA</td></tr>
<tr><td>T9031</td><td>CGCGCGTGATCGGCGAGTCTCGG</td><td>1000</td><td></td><td></td><td></td><td></td><td></td><td></td><td></td><td></td><td></td><td></td></tr>
<tr><td>Oligo fwd</td><td>CACCCGCGCGTGATCGGCGAGTCT</td><td></td><td></td><td></td><td></td><td></td><td></td><td></td><td></td><td></td><td></td><td></td></tr>
<tr><td>Oligo rev</td><td>AAACAGACTCGCCGATCACGCGCG</td><td></td><td></td><td></td><td></td><td></td><td></td><td></td><td></td><td></td><td></td><td></td></tr>
<tr><td>Chromosome</td><td>start</td><td></td><td>end</td><td>strand</td><td>MM</td><td>target_seq</td><td>PAM</td><td>alignment</td><td>distance</td><td>position</td><td>gene name</td><td>gene id</td></tr>
<tr><td>GU305914.1</td><td></td><td>120455</td><td>120477</td><td>+</td><td>0</td><td>CGCGCGTGATCGGCGAGTCT</td><td>CGG</td><td>||||||||[||||||||||||]PAM</td><td>NA</td><td>Intergenic</td><td>NA</td><td>NA</td></tr>
<tr><td>T9032</td><td>GCGAGTCTCGGTTCTTGCAGTGG</td><td>1000</td><td></td><td></td><td></td><td></td><td></td><td></td><td></td><td></td><td></td><td></td></tr>
<tr><td>Oligo fwd</td><td>CACCGCGAGTCTCGGTTCTTGCAG</td><td></td><td></td><td></td><td></td><td></td><td></td><td></td><td></td><td></td><td></td><td></td></tr>
<tr><td>Oligo rev</td><td>AAACCTGCAAGAACCGAGACTCGC</td><td></td><td></td><td></td><td></td><td></td><td></td><td></td><td></td><td></td><td></td><td></td></tr>
<tr><td>Chromosome</td><td>start</td><td></td><td>end</td><td>strand</td><td>MM</td><td>target_seq</td><td>PAM</td><td>alignment</td><td>distance</td><td>position</td><td>gene name</td><td>gene id</td></tr>
<tr><td>GU305914.1</td><td></td><td>120467</td><td>120489</td><td>+</td><td>0</td><td>GCGAGTCTCGGTTCTTGCAG</td><td>TGG</td><td>||||||||[||||||||||||]PAM</td><td>NA</td><td>Intergenic</td><td>NA</td><td>NA</td></tr>
<tr><td>T9033</td><td>CTCGGTTCTTGCAGTGGAGCAGG</td><td>1000</td><td></td><td></td><td></td><td></td><td></td><td></td><td></td><td></td><td></td><td></td></tr>
<tr><td>Oligo fwd</td><td>CACCCTCGGTTCTTGCAGTGGAGC</td><td></td><td></td><td></td><td></td><td></td><td></td><td></td><td></td><td></td><td></td><td></td></tr>
<tr><td>Oligo rev</td><td>AAACGCTCCACTGCAAGAACCGAG</td><td></td><td></td><td></td><td></td><td></td><td></td><td></td><td></td><td></td><td></td><td></td></tr>
<tr><td>Chromosome</td><td>start</td><td></td><td>end</td><td>strand</td><td>MM</td><td>target_seq</td><td>PAM</td><td>alignment</td><td>distance</td><td>position</td><td>gene name</td><td>gene id</td></tr>
<tr><td>GU305914.1</td><td></td><td>120473</td><td>120495</td><td>+</td><td>0</td><td>CTCGGTTCTTGCAGTGGAGC</td><td>AGG</td><td>||||||||[||||||||||||]PAM</td><td>NA</td><td>Intergenic</td><td>NA</td><td>NA</td></tr>
<tr><td>T9034</td><td>TTCTTGCAGTGGAGCAGGATCGG</td><td>1000</td><td></td><td></td><td></td><td></td><td></td><td></td><td></td><td></td><td></td><td></td></tr>
<tr><td>Oligo fwd</td><td>CACCTTCTTGCAGTGGAGCAGGAT</td><td></td><td></td><td></td><td></td><td></td><td></td><td></td><td></td><td></td><td></td><td></td></tr>
<tr><td>Oligo rev</td><td>AAACATCCTGCTCCACTGCAAGAA</td><td></td><td></td><td></td><td></td><td></td><td></td><td></td><td></td><td></td><td></td><td></td></tr>
<tr><td>Chromosome</td><td>start</td><td></td><td>end</td><td>strand</td><td>MM</td><td>target_seq</td><td>PAM</td><td>alignment</td><td>distance</td><td>position</td><td>gene name</td><td>gene id</td></tr>
<tr><td>GU305914.1</td><td></td><td>120478</td><td>120500</td><td>+</td><td>0</td><td>TTCTTGCAGTGGAGCAGGAT</td><td>CGG</td><td>||||||||[||||||||||||]PAM</td><td>NA</td><td>Intergenic</td><td>NA</td><td>NA</td></tr>
<tr><td>T9035</td><td>TCTTGCAGTGGAGCAGGATCGGG</td><td>1000</td><td></td><td></td><td></td><td></td><td></td><td></td><td></td><td></td><td></td><td></td></tr>
<tr><td>Oligo fwd</td><td>CACCTCTTGCAGTGGAGCAGGATC</td><td></td><td></td><td></td><td></td><td></td><td></td><td></td><td></td><td></td><td></td><td></td></tr>
<tr><td>Oligo rev</td><td>AAACGATCCTGCTCCACTGCAAGA</td><td></td><td></td><td></td><td></td><td></td><td></td><td></td><td></td><td></td><td></td><td></td></tr>
<tr><td>Chromosome</td><td>start</td><td></td><td>end</td><td>strand</td><td>MM</td><td>target_seq</td><td>PAM</td><td>alignment</td><td>distance</td><td>position</td><td>gene name</td><td>gene id</td></tr>
<tr><td>GU305914.1</td><td></td><td>120479</td><td>120501</td><td>+</td><td>0</td><td>TCTTGCAGTGGAGCAGGATC</td><td>GGG</td><td>||||||||[||||||||||||]PAM</td><td>NA</td><td>Intergenic</td><td>NA</td><td>NA</td></tr>
<tr><td>T9036</td><td>AGCTTAAATACAGCTCCCGTCGG</td><td>1000</td><td></td><td></td><td></td><td></td><td></td><td></td><td></td><td></td><td></td><td></td></tr>
<tr><td>Oligo fwd</td><td>CACCAGCTTAAATACAGCTCCCGT</td><td></td><td></td><td></td><td></td><td></td><td></td><td></td><td></td><td></td><td></td><td></td></tr>
<tr><td>Oligo rev</td><td>AAACACGGGAGCTGTATTTAAGCT</td><td></td><td></td><td></td><td></td><td></td><td></td><td></td><td></td><td></td><td></td><td></td></tr>
<tr><td>Chromosome</td><td>start</td><td></td><td>end</td><td>strand</td><td>MM</td><td>target_seq</td><td>PAM</td><td>alignment</td><td>distance</td><td>position</td><td>gene name</td><td>gene id</td></tr>
<tr><td>GU305914.1</td><td></td><td>120514</td><td>120536</td><td>+</td><td>0</td><td>AGCTTAAATACAGCTCCCGT</td><td>CGG</td><td>||||||||[||||||||||||]PAM</td><td>NA</td><td>Intergenic</td><td>NA</td><td>NA</td></tr>
<tr><td>T9037</td><td>CTCCCGTCGGCACGATCTCTAGG</td><td>1000</td><td></td><td></td><td></td><td></td><td></td><td></td><td></td><td></td><td></td><td></td></tr>
<tr><td>Oligo fwd</td><td>CACCCTCCCGTCGGCACGATCTCT</td><td></td><td></td><td></td><td></td><td></td><td></td><td></td><td></td><td></td><td></td><td></td></tr>
<tr><td>Oligo rev</td><td>AAACAGAGATCGTGCCGACGGGAG</td><td></td><td></td><td></td><td></td><td></td><td></td><td></td><td></td><td></td><td></td><td></td></tr>
<tr><td>Chromosome</td><td>start</td><td></td><td>end</td><td>strand</td><td>MM</td><td>target_seq</td><td>PAM</td><td>alignment</td><td>distance</td><td>position</td><td>gene name</td><td>gene id</td></tr>
<tr><td>GU305914.1</td><td></td><td>120527</td><td>120549</td><td>+</td><td>0</td><td>CTCCCGTCGGCACGATCTCT</td><td>AGG</td><td>||||||||[||||||||||||]PAM</td><td>NA</td><td>Intergenic</td><td>NA</td><td>NA</td></tr>
<tr><td>T9038</td><td>TCCCGTCGGCACGATCTCTAGGG</td><td>1000</td><td></td><td></td><td></td><td></td><td></td><td></td><td></td><td></td><td></td><td></td></tr>
<tr><td>Oligo fwd</td><td>CACCTCCCGTCGGCACGATCTCTA</td><td></td><td></td><td></td><td></td><td></td><td></td><td></td><td></td><td></td><td></td><td></td></tr>
<tr><td>Oligo rev</td><td>AAACTAGAGATCGTGCCGACGGGA</td><td></td><td></td><td></td><td></td><td></td><td></td><td></td><td></td><td></td><td></td><td></td></tr>
<tr><td>Chromosome</td><td>start</td><td></td><td>end</td><td>strand</td><td>MM</td><td>target_seq</td><td>PAM</td><td>alignment</td><td>distance</td><td>position</td><td>gene name</td><td>gene id</td></tr>
<tr><td>GU305914.1</td><td></td><td>120528</td><td>120550</td><td>+</td><td>0</td><td>TCCCGTCGGCACGATCTCTA</td><td>GGG</td><td>||||||||[||||||||||||]PAM</td><td>NA</td><td>Intergenic</td><td>NA</td><td>NA</td></tr>
<tr><td>T9039</td><td>CTCCGAGAATCTCTCCACCTCGG</td><td>1000</td><td></td><td></td><td></td><td></td><td></td><td></td><td></td><td></td><td></td><td></td></tr>
<tr><td>Oligo fwd</td><td>CACCCTCCGAGAATCTCTCCACCT</td><td></td><td></td><td></td><td></td><td></td><td></td><td></td><td></td><td></td><td></td><td></td></tr>
<tr><td>Oligo rev</td><td>AAACAGGTGGAGAGATTCTCGGAG</td><td></td><td></td><td></td><td></td><td></td><td></td><td></td><td></td><td></td><td></td><td></td></tr>
<tr><td>Chromosome</td><td>start</td><td></td><td>end</td><td>strand</td><td>MM</td><td>target_seq</td><td>PAM</td><td>alignment</td><td>distance</td><td>position</td><td>gene name</td><td>gene id</td></tr>
<tr><td>GU305914.1</td><td></td><td>120561</td><td>120583</td><td>+</td><td>0</td><td>CTCCGAGAATCTCTCCACCT</td><td>CGG</td><td>||||||||[||||||||||||]PAM</td><td>NA</td><td>Intergenic</td><td>NA</td><td>NA</td></tr>
<tr><td>T9040</td><td>TCCGAGAATCTCTCCACCTCGGG</td><td>1000</td><td></td><td></td><td></td><td></td><td></td><td></td><td></td><td></td><td></td><td></td></tr>
<tr><td>Oligo fwd</td><td>CACCTCCGAGAATCTCTCCACCTC</td><td></td><td></td><td></td><td></td><td></td><td></td><td></td><td></td><td></td><td></td><td></td></tr>
<tr><td>Oligo rev</td><td>AAACGAGGTGGAGAGATTCTCGGA</td><td></td><td></td><td></td><td></td><td></td><td></td><td></td><td></td><td></td><td></td><td></td></tr>
<tr><td>Chromosome</td><td>start</td><td></td><td>end</td><td>strand</td><td>MM</td><td>target_seq</td><td>PAM</td><td>alignment</td><td>distance</td><td>position</td><td>gene name</td><td>gene id</td></tr>
<tr><td>GU305914.1</td><td></td><td>120562</td><td>120584</td><td>+</td><td>0</td><td>TCCGAGAATCTCTCCACCTC</td><td>GGG</td><td>||||||||[||||||||||||]PAM</td><td>NA</td><td>Intergenic</td><td>NA</td><td>NA</td></tr>
<tr><td>T9041</td><td>CCGAGAATCTCTCCACCTCGGGG</td><td>1000</td><td></td><td></td><td></td><td></td><td></td><td></td><td></td><td></td><td></td><td></td></tr>
<tr><td>Oligo fwd</td><td>CACCCCGAGAATCTCTCCACCTCG</td><td></td><td></td><td></td><td></td><td></td><td></td><td></td><td></td><td></td><td></td><td></td></tr>
<tr><td>Oligo rev</td><td>AAACCGAGGTGGAGAGATTCTCGG</td><td></td><td></td><td></td><td></td><td></td><td></td><td></td><td></td><td></td><td></td><td></td></tr>
<tr><td>Chromosome</td><td>start</td><td></td><td>end</td><td>strand</td><td>MM</td><td>target_seq</td><td>PAM</td><td>alignment</td><td>distance</td><td>position</td><td>gene name</td><td>gene id</td></tr>
<tr><td>GU305914.1</td><td></td><td>120563</td><td>120585</td><td>+</td><td>0</td><td>CCGAGAATCTCTCCACCTCG</td><td>GGG</td><td>||||||||[||||||||||||]PAM</td><td>NA</td><td>Intergenic</td><td>NA</td><td>NA</td></tr>
<tr><td>T9042</td><td>CTCTCCACCTCGGGGTACGTCGG</td><td>1000</td><td></td><td></td><td></td><td></td><td></td><td></td><td></td><td></td><td></td><td></td></tr>
<tr><td>Oligo fwd</td><td>CACCCTCTCCACCTCGGGGTACGT</td><td></td><td></td><td></td><td></td><td></td><td></td><td></td><td></td><td></td><td></td><td></td></tr>
<tr><td>Oligo rev</td><td>AAACACGTACCCCGAGGTGGAGAG</td><td></td><td></td><td></td><td></td><td></td><td></td><td></td><td></td><td></td><td></td><td></td></tr>
<tr><td>Chromosome</td><td>start</td><td></td><td>end</td><td>strand</td><td>MM</td><td>target_seq</td><td>PAM</td><td>alignment</td><td>distance</td><td>position</td><td>gene name</td><td>gene id</td></tr>
<tr><td>GU305914.1</td><td></td><td>120571</td><td>120593</td><td>+</td><td>0</td><td>CTCTCCACCTCGGGGTACGT</td><td>CGG</td><td>||||||||[||||||||||||]PAM</td><td>NA</td><td>Intergenic</td><td>NA</td><td>NA</td></tr>
<tr><td>T9043</td><td>GGCGCCTGACGTTCGCTCTCCGG</td><td>1000</td><td></td><td></td><td></td><td></td><td></td><td></td><td></td><td></td><td></td><td></td></tr>
<tr><td>Oligo fwd</td><td>CACCGGCGCCTGACGTTCGCTCTC</td><td></td><td></td><td></td><td></td><td></td><td></td><td></td><td></td><td></td><td></td><td></td></tr>
<tr><td>Oligo rev</td><td>AAACGAGAGCGAACGTCAGGCGCC</td><td></td><td></td><td></td><td></td><td></td><td></td><td></td><td></td><td></td><td></td><td></td></tr>
<tr><td>Chromosome</td><td>start</td><td></td><td>end</td><td>strand</td><td>MM</td><td>target_seq</td><td>PAM</td><td>alignment</td><td>distance</td><td>position</td><td>gene name</td><td>gene id</td></tr>
<tr><td>GU305914.1</td><td></td><td>120592</td><td>120614</td><td>+</td><td>0</td><td>GGCGCCTGACGTTCGCTCTC</td><td>CGG</td><td>||||||||[||||||||||||]PAM</td><td>NA</td><td>Intergenic</td><td>NA</td><td>NA</td></tr>
<tr><td>T9044</td><td>GCCTGACGTTCGCTCTCCGGCGG</td><td>1000</td><td></td><td></td><td></td><td></td><td></td><td></td><td></td><td></td><td></td><td></td></tr>
</table>

<table>
<tr><td>Oligo fwd</td><td>CACCGCCTGACGTTCGCTCTCCGG</td><td></td><td></td><td></td><td></td><td></td><td></td><td></td><td></td><td></td><td></td></tr>
<tr><td>Oligo rev</td><td>AAACCCGGAGAGCGAACGTCAGGC</td><td></td><td></td><td></td><td></td><td></td><td></td><td></td><td></td><td></td><td></td></tr>
<tr><td>Chromosome</td><td>start</td><td>end</td><td>strand</td><td>MM</td><td>target_seq</td><td>PAM</td><td>alignment</td><td>distance</td><td>position</td><td>gene name</td><td>gene id</td></tr>
<tr><td>GU305914.1</td><td>120595</td><td>120617</td><td>+</td><td>0</td><td>GCCTGACGTTCGCTCTCCGG</td><td>CGG</td><td>||||||||[||||||||||||]PAM</td><td>NA</td><td>Intergenic</td><td>NA</td><td>NA</td></tr>
<tr><td>T9045</td><td>ACGTTCGCTCTCCGGCGGATCGG</td><td>1000</td><td></td><td></td><td></td><td></td><td></td><td></td><td></td><td></td><td></td></tr>
<tr><td>Oligo fwd</td><td>CACCACGTTCGCTCTCCGGCGGAT</td><td></td><td></td><td></td><td></td><td></td><td></td><td></td><td></td><td></td><td></td></tr>
<tr><td>Oligo rev</td><td>AAACATCCGCCGGAGAGCGAACGT</td><td></td><td></td><td></td><td></td><td></td><td></td><td></td><td></td><td></td><td></td></tr>
<tr><td>Chromosome</td><td>start</td><td>end</td><td>strand</td><td>MM</td><td>target_seq</td><td>PAM</td><td>alignment</td><td>distance</td><td>position</td><td>gene name</td><td>gene id</td></tr>
<tr><td>GU305914.1</td><td>120600</td><td>120622</td><td>+</td><td>0</td><td>ACGTTCGCTCTCCGGCGGAT</td><td>CGG</td><td>||||||||[||||||||||||]PAM</td><td>NA</td><td>Intergenic</td><td>NA</td><td>NA</td></tr>
<tr><td>T9046</td><td>CGTTCGCTCTCCGGCGGATCGGG</td><td>1000</td><td></td><td></td><td></td><td></td><td></td><td></td><td></td><td></td><td></td></tr>
<tr><td>Oligo fwd</td><td>CACCCGTTCGCTCTCCGGCGGATC</td><td></td><td></td><td></td><td></td><td></td><td></td><td></td><td></td><td></td><td></td></tr>
<tr><td>Oligo rev</td><td>AAACGATCCGCCGGAGAGCGAACG</td><td></td><td></td><td></td><td></td><td></td><td></td><td></td><td></td><td></td><td></td></tr>
<tr><td>Chromosome</td><td>start</td><td>end</td><td>strand</td><td>MM</td><td>target_seq</td><td>PAM</td><td>alignment</td><td>distance</td><td>position</td><td>gene name</td><td>gene id</td></tr>
<tr><td>GU305914.1</td><td>120601</td><td>120623</td><td>+</td><td>0</td><td>CGTTCGCTCTCCGGCGGATC</td><td>GGG</td><td>||||||||[||||||||||||]PAM</td><td>NA</td><td>Intergenic</td><td>NA</td><td>NA</td></tr>
<tr><td>T9047</td><td>TCGCTCTCCGGCGGATCGGGTGG</td><td>1000</td><td></td><td></td><td></td><td></td><td></td><td></td><td></td><td></td><td></td></tr>
<tr><td>Oligo fwd</td><td>CACCTCGCTCTCCGGCGGATCGGG</td><td></td><td></td><td></td><td></td><td></td><td></td><td></td><td></td><td></td><td></td></tr>
<tr><td>Oligo rev</td><td>AAACCCCGATCCGCCGGAGAGCGA</td><td></td><td></td><td></td><td></td><td></td><td></td><td></td><td></td><td></td><td></td></tr>
<tr><td>Chromosome</td><td>start</td><td>end</td><td>strand</td><td>MM</td><td>target_seq</td><td>PAM</td><td>alignment</td><td>distance</td><td>position</td><td>gene name</td><td>gene id</td></tr>
<tr><td>GU305914.1</td><td>120604</td><td>120626</td><td>+</td><td>0</td><td>TCGCTCTCCGGCGGATCGGG</td><td>TGG</td><td>||||||||[||||||||||||]PAM</td><td>NA</td><td>Intergenic</td><td>NA</td><td>NA</td></tr>
<tr><td>T9048</td><td>TCTCGCTCTCCGACCTCGTCCGG</td><td>1000</td><td></td><td></td><td></td><td></td><td></td><td></td><td></td><td></td><td></td></tr>
<tr><td>Oligo fwd</td><td>CACCTCTCGCTCTCCGACCTCGTC</td><td></td><td></td><td></td><td></td><td></td><td></td><td></td><td></td><td></td><td></td></tr>
<tr><td>Oligo rev</td><td>AAACGACGAGGTCGGAGAGCGAGA</td><td></td><td></td><td></td><td></td><td></td><td></td><td></td><td></td><td></td><td></td></tr>
<tr><td>Chromosome</td><td>start</td><td>end</td><td>strand</td><td>MM</td><td>target_seq</td><td>PAM</td><td>alignment</td><td>distance</td><td>position</td><td>gene name</td><td>gene id</td></tr>
<tr><td>GU305914.1</td><td>120650</td><td>120672</td><td>+</td><td>0</td><td>TCTCGCTCTCCGACCTCGTC</td><td>CGG</td><td>||||||||[||||||||||||]PAM</td><td>NA</td><td>Intergenic</td><td>NA</td><td>NA</td></tr>
<tr><td>T9049</td><td>CTCGCTCTCCGACCTCGTCCGGG</td><td>1000</td><td></td><td></td><td></td><td></td><td></td><td></td><td></td><td></td><td></td></tr>
<tr><td>Oligo fwd</td><td>CACCCTCGCTCTCCGACCTCGTCC</td><td></td><td></td><td></td><td></td><td></td><td></td><td></td><td></td><td></td><td></td></tr>
<tr><td>Oligo rev</td><td>AAACGGACGAGGTCGGAGAGCGAG</td><td></td><td></td><td></td><td></td><td></td><td></td><td></td><td></td><td></td><td></td></tr>
<tr><td>Chromosome</td><td>start</td><td>end</td><td>strand</td><td>MM</td><td>target_seq</td><td>PAM</td><td>alignment</td><td>distance</td><td>position</td><td>gene name</td><td>gene id</td></tr>
<tr><td>GU305914.1</td><td>120651</td><td>120673</td><td>+</td><td>0</td><td>CTCGCTCTCCGACCTCGTCC</td><td>GGG</td><td>||||||||[||||||||||||]PAM</td><td>NA</td><td>Intergenic</td><td>NA</td><td>NA</td></tr>
<tr><td>GU305914.1</td><td>110174</td><td>110196</td><td>+</td><td>4</td><td>CGCGGTCTCCGACGACGTCC</td><td>TGG</td><td>|-||-|||[|||||--|||||]PAM</td><td>NA</td><td>Intergenic</td><td>NA</td><td>NA</td></tr>
<tr><td>T9050</td><td>CGTCCGGGCCCTCTTGCTGTCGG</td><td>1000</td><td></td><td></td><td></td><td></td><td></td><td></td><td></td><td></td><td></td></tr>
<tr><td>Oligo fwd</td><td>CACCCGTCCGGGCCCTCTTGCTGT</td><td></td><td></td><td></td><td></td><td></td><td></td><td></td><td></td><td></td><td></td></tr>
<tr><td>Oligo rev</td><td>AAACACAGCAAGAGGGCCCGGACG</td><td></td><td></td><td></td><td></td><td></td><td></td><td></td><td></td><td></td><td></td></tr>
<tr><td>Chromosome</td><td>start</td><td>end</td><td>strand</td><td>MM</td><td>target_seq</td><td>PAM</td><td>alignment</td><td>distance</td><td>position</td><td>gene name</td><td>gene id</td></tr>
<tr><td>GU305914.1</td><td>120666</td><td>120688</td><td>+</td><td>0</td><td>CGTCCGGGCCCTCTTGCTGT</td><td>CGG</td><td>||||||||[||||||||||||]PAM</td><td>NA</td><td>Intergenic</td><td>NA</td><td>NA</td></tr>
<tr><td>T9051</td><td>GTCCGGGCCCTCTTGCTGTCGGG</td><td>1000</td><td></td><td></td><td></td><td></td><td></td><td></td><td></td><td></td><td></td></tr>
<tr><td>Oligo fwd</td><td>CACCGTCCGGGCCCTCTTGCTGTC</td><td></td><td></td><td></td><td></td><td></td><td></td><td></td><td></td><td></td><td></td></tr>
<tr><td>Oligo rev</td><td>AAACGACAGCAAGAGGGCCCGGAC</td><td></td><td></td><td></td><td></td><td></td><td></td><td></td><td></td><td></td><td></td></tr>
<tr><td>Chromosome</td><td>start</td><td>end</td><td>strand</td><td>MM</td><td>target_seq</td><td>PAM</td><td>alignment</td><td>distance</td><td>position</td><td>gene name</td><td>gene id</td></tr>
<tr><td>GU305914.1</td><td>120667</td><td>120689</td><td>+</td><td>0</td><td>GTCCGGGCCCTCTTGCTGTC</td><td>GGG</td><td>||||||||[||||||||||||]PAM</td><td>NA</td><td>Intergenic</td><td>NA</td><td>NA</td></tr>
<tr><td>T9052</td><td>CGGGCCCTCTTGCTGTCGGGCGG</td><td>1000</td><td></td><td></td><td></td><td></td><td></td><td></td><td></td><td></td><td></td></tr>
<tr><td>Oligo fwd</td><td>CACCCGGGCCCTCTTGCTGTCGGG</td><td></td><td></td><td></td><td></td><td></td><td></td><td></td><td></td><td></td><td></td></tr>
<tr><td>Oligo rev</td><td>AAACCCCGACAGCAAGAGGGCCCG</td><td></td><td></td><td></td><td></td><td></td><td></td><td></td><td></td><td></td><td></td></tr>
<tr><td>Chromosome</td><td>start</td><td>end</td><td>strand</td><td>MM</td><td>target_seq</td><td>PAM</td><td>alignment</td><td>distance</td><td>position</td><td>gene name</td><td>gene id</td></tr>
<tr><td>GU305914.1</td><td>120670</td><td>120692</td><td>+</td><td>0</td><td>CGGGCCCTCTTGCTGTCGGG</td><td>CGG</td><td>||||||||[||||||||||||]PAM</td><td>NA</td><td>Intergenic</td><td>NA</td><td>NA</td></tr>
<tr><td>T9053</td><td>GCCCTCTTGCTGTCGGGCGGCGG</td><td>1000</td><td></td><td></td><td></td><td></td><td></td><td></td><td></td><td></td><td></td></tr>
<tr><td>Oligo fwd</td><td>CACCGCCCTCTTGCTGTCGGGCGG</td><td></td><td></td><td></td><td></td><td></td><td></td><td></td><td></td><td></td><td></td></tr>
<tr><td>Oligo rev</td><td>AAACCCGCCCGACAGCAAGAGGGC</td><td></td><td></td><td></td><td></td><td></td><td></td><td></td><td></td><td></td><td></td></tr>
<tr><td>Chromosome</td><td>start</td><td>end</td><td>strand</td><td>MM</td><td>target_seq</td><td>PAM</td><td>alignment</td><td>distance</td><td>position</td><td>gene name</td><td>gene id</td></tr>
<tr><td>GU305914.1</td><td>120673</td><td>120695</td><td>+</td><td>0</td><td>GCCCTCTTGCTGTCGGGCGG</td><td>CGG</td><td>||||||||[||||||||||||]PAM</td><td>NA</td><td>Intergenic</td><td>NA</td><td>NA</td></tr>
<tr><td>T9054</td><td>CCCTCTTGCTGTCGGGCGGCGGG</td><td>1000</td><td></td><td></td><td></td><td></td><td></td><td></td><td></td><td></td><td></td></tr>
<tr><td>Oligo fwd</td><td>CACCCCCTCTTGCTGTCGGGCGGC</td><td></td><td></td><td></td><td></td><td></td><td></td><td></td><td></td><td></td><td></td></tr>
<tr><td>Oligo rev</td><td>AAACGCCGCCCGACAGCAAGAGGG</td><td></td><td></td><td></td><td></td><td></td><td></td><td></td><td></td><td></td><td></td></tr>
<tr><td>Chromosome</td><td>start</td><td>end</td><td>strand</td><td>MM</td><td>target_seq</td><td>PAM</td><td>alignment</td><td>distance</td><td>position</td><td>gene name</td><td>gene id</td></tr>
<tr><td>GU305914.1</td><td>120674</td><td>120696</td><td>+</td><td>0</td><td>CCCTCTTGCTGTCGGGCGGC</td><td>GGG</td><td>||||||||[||||||||||||]PAM</td><td>NA</td><td>Intergenic</td><td>NA</td><td>NA</td></tr>
<tr><td>T9055</td><td>AGACCAGAACTCGCTAAATCCGG</td><td>1000</td><td></td><td></td><td></td><td></td><td></td><td></td><td></td><td></td><td></td></tr>
<tr><td>Oligo fwd</td><td>CACCAGACCAGAACTCGCTAAATC</td><td></td><td></td><td></td><td></td><td></td><td></td><td></td><td></td><td></td><td></td></tr>
<tr><td>Oligo rev</td><td>AAACGATTTAGCGAGTTCTGGTCT</td><td></td><td></td><td></td><td></td><td></td><td></td><td></td><td></td><td></td><td></td></tr>
<tr><td>Chromosome</td><td>start</td><td>end</td><td>strand</td><td>MM</td><td>target_seq</td><td>PAM</td><td>alignment</td><td>distance</td><td>position</td><td>gene name</td><td>gene id</td></tr>
<tr><td>GU305914.1</td><td>120702</td><td>120724</td><td>+</td><td>0</td><td>AGACCAGAACTCGCTAAATC</td><td>CGG</td><td>||||||||[||||||||||||]PAM</td><td>NA</td><td>Intergenic</td><td>NA</td><td>NA</td></tr>
<tr><td>T9056</td><td>GACCAGAACTCGCTAAATCCGGG</td><td>1000</td><td></td><td></td><td></td><td></td><td></td><td></td><td></td><td></td><td></td></tr>
<tr><td>Oligo fwd</td><td>CACCGACCAGAACTCGCTAAATCC</td><td></td><td></td><td></td><td></td><td></td><td></td><td></td><td></td><td></td><td></td></tr>
<tr><td>Oligo rev</td><td>AAACGGATTTAGCGAGTTCTGGTC</td><td></td><td></td><td></td><td></td><td></td><td></td><td></td><td></td><td></td><td></td></tr>
<tr><td>Chromosome</td><td>start</td><td>end</td><td>strand</td><td>MM</td><td>target_seq</td><td>PAM</td><td>alignment</td><td>distance</td><td>position</td><td>gene name</td><td>gene id</td></tr>
<tr><td>GU305914.1</td><td>120703</td><td>120725</td><td>+</td><td>0</td><td>GACCAGAACTCGCTAAATCC</td><td>GGG</td><td>||||||||[||||||||||||]PAM</td><td>NA</td><td>Intergenic</td><td>NA</td><td>NA</td></tr>
<tr><td>T9057</td><td>ACCAGAACTCGCTAAATCCGGGG</td><td>1000</td><td></td><td></td><td></td><td></td><td></td><td></td><td></td><td></td><td></td></tr>
<tr><td>Oligo fwd</td><td>CACCACCAGAACTCGCTAAATCCG</td><td></td><td></td><td></td><td></td><td></td><td></td><td></td><td></td><td></td><td></td></tr>
<tr><td>Oligo rev</td><td>AAACCGGATTTAGCGAGTTCTGGT</td><td></td><td></td><td></td><td></td><td></td><td></td><td></td><td></td><td></td><td></td></tr>
<tr><td>Chromosome</td><td>start</td><td>end</td><td>strand</td><td>MM</td><td>target_seq</td><td>PAM</td><td>alignment</td><td>distance</td><td>position</td><td>gene name</td><td>gene id</td></tr>
<tr><td>GU305914.1</td><td>120704</td><td>120726</td><td>+</td><td>0</td><td>ACCAGAACTCGCTAAATCCG</td><td>GGG</td><td>||||||||[||||||||||||]PAM</td><td>NA</td><td>Intergenic</td><td>NA</td><td>NA</td></tr>
<tr><td>T9058</td><td>GGGGTTCATCGACCCCCTCTTGG</td><td>1000</td><td></td><td></td><td></td><td></td><td></td><td></td><td></td><td></td><td></td></tr>
<tr><td>Oligo fwd</td><td>CACCGGGGTTCATCGACCCCCTCT</td><td></td><td></td><td></td><td></td><td></td><td></td><td></td><td></td><td></td><td></td></tr>
<tr><td>Oligo rev</td><td>AAACAGAGGGGGTCGATGAACCCC</td><td></td><td></td><td></td><td></td><td></td><td></td><td></td><td></td><td></td><td></td></tr>
<tr><td>Chromosome</td><td>start</td><td>end</td><td>strand</td><td>MM</td><td>target_seq</td><td>PAM</td><td>alignment</td><td>distance</td><td>position</td><td>gene name</td><td>gene id</td></tr>
<tr><td>GU305914.1</td><td>120723</td><td>120745</td><td>+</td><td>0</td><td>GGGGTTCATCGACCCCCTCT</td><td>TGG</td><td>||||||||[||||||||||||]PAM</td><td>NA</td><td>Intergenic</td><td>NA</td><td>NA</td></tr>
</table>

T9059 GGCGCCGCTCGACGATGTCGAGG 1000
Oligo fwd CACCGGCGCCGCTCGACGATGTCG
Oligo rev AAACCGACATCGTCGAGCGGCGCC
Chromosome start end strand MM target_seq PAM alignment distance position gene name gene id
GU305914.1 120744 120766 + 0 GGCGCCGCTCGACGATGTCG AGG ||||||||[||||||||||||]PAM NA Intergenic NA NA

T9060 GCGCCGCTCGACGATGTCGAGGG 1000
Oligo fwd CACCGCGCCGCTCGACGATGTCGA
Oligo rev AAACTCGACATCGTCGAGCGGCGC
Chromosome start end strand MM target_seq PAM alignment distance position gene name gene id
GU305914.1 120745 120767 + 0 GCGCCGCTCGACGATGTCGA GGG ||||||||[||||||||||||]PAM NA Intergenic NA NA

T9061 TCGACGATGTCGAGGGTCTCCGG 1000
Oligo fwd CACCTCGACGATGTCGAGGGTCTC
Oligo rev AAACGAGACCCTCGACATCGTCGA
Chromosome start end strand MM target_seq PAM alignment distance position gene name gene id
GU305914.1 120752 120774 + 0 TCGACGATGTCGAGGGTCTC CGG ||||||||[||||||||||||]PAM NA Intergenic NA NA
GU305914.1 62394 62416 + 4 TCGAGGCTGTAGAGGGGCTC GGG ||||-|-|[||-|||||-|||]PAM NA Intergenic NA NA

T9062 GTCGAGGGTCTCCGGTGTGTCGG 1000
Oligo fwd CACCGTCGAGGGTCTCCGGTGTGT
Oligo rev AAACACACACCGGAGACCCTCGAC
Chromosome start end strand MM target_seq PAM alignment distance position gene name gene id
GU305914.1 120760 120782 + 0 GTCGAGGGTCTCCGGTGTGT CGG ||||||||[||||||||||||]PAM NA Intergenic NA NA

T9063 GAGGGTCTCCGGTGTGTCGGCGG 1000
Oligo fwd CACCGAGGGTCTCCGGTGTGTCGG
Oligo rev AAACCCGACACACCGGAGACCCTC
Chromosome start end strand MM target_seq PAM alignment distance position gene name gene id
GU305914.1 120763 120785 + 0 GAGGGTCTCCGGTGTGTCGG CGG ||||||||[||||||||||||]PAM NA Intergenic NA NA

T9064 GGTCTCCGGTGTGTCGGCGGTGG 1000
Oligo fwd CACCGGTCTCCGGTGTGTCGGCGG
Oligo rev AAACCCGCCGACACACCGGAGACC
Chromosome start end strand MM target_seq PAM alignment distance position gene name gene id
GU305914.1 120766 120788 + 0 GGTCTCCGGTGTGTCGGCGG TGG ||||||||[||||||||||||]PAM NA Intergenic NA NA

T9065 TCCGGTGTGTCGGCGGTGGTCGG 1000
Oligo fwd CACCTCCGGTGTGTCGGCGGTGGT
Oligo rev AAACACCACCGCCGACACACCGGA
Chromosome start end strand MM target_seq PAM alignment distance position gene name gene id
GU305914.1 120770 120792 + 0 TCCGGTGTGTCGGCGGTGGT CGG ||||||||[||||||||||||]PAM NA Intergenic NA NA

T9066 TCGGCGGTGGTCGGCGCGAGCGG 1000
Oligo fwd CACCTCGGCGGTGGTCGGCGCGAG
Oligo rev AAACCTCGCGCCGACCACCGCCGA
Chromosome start end strand MM target_seq PAM alignment distance position gene name gene id
GU305914.1 120779 120801 + 0 TCGGCGGTGGTCGGCGCGAG CGG ||||||||[||||||||||||]PAM NA Intergenic NA NA

T9067 CGGCGGTGGTCGGCGCGAGCGGG 1000
Oligo fwd CACCCGGCGGTGGTCGGCGCGAGC
Oligo rev AAACGCTCGCGCCGACCACCGCCG
Chromosome start end strand MM target_seq PAM alignment distance position gene name gene id
GU305914.1 120780 120802 + 0 CGGCGGTGGTCGGCGCGAGC GGG ||||||||[||||||||||||]PAM NA Intergenic NA NA

T9068 CGGTGGTCGGCGCGAGCGGGCGG 1000
Oligo fwd CACCCGGTGGTCGGCGCGAGCGGG
Oligo rev AAACCCCGCTCGCGCCGACCACCG
Chromosome start end strand MM target_seq PAM alignment distance position gene name gene id
GU305914.1 120783 120805 + 0 CGGTGGTCGGCGCGAGCGGG CGG ||||||||[||||||||||||]PAM NA Intergenic NA NA

T9069 GGTGGTCGGCGCGAGCGGGCGGG 1000
Oligo fwd CACCGGTGGTCGGCGCGAGCGGGC
Oligo rev AAACGCCCGCTCGCGCCGACCACC
Chromosome start end strand MM target_seq PAM alignment distance position gene name gene id
GU305914.1 120784 120806 + 0 GGTGGTCGGCGCGAGCGGGC GGG ||||||||[||||||||||||]PAM NA Intergenic NA NA

T9070 GCGCGAGCGGGCGGGACCTCCGG 1000
Oligo fwd CACCGCGCGAGCGGGCGGGACCTC
Oligo rev AAACGAGGTCCCGCCCGCTCGCGC
Chromosome start end strand MM target_seq PAM alignment distance position gene name gene id
GU305914.1 120792 120814 + 0 GCGCGAGCGGGCGGGACCTC CGG ||||||||[||||||||||||]PAM NA Intergenic NA NA

T9071 GCGGGCGGGACCTCCGGCGCCGG 1000
Oligo fwd CACCGCGGGCGGGACCTCCGGCGC
Oligo rev AAACGCGCCGGAGGTCCCGCCCGC
Chromosome start end strand MM target_seq PAM alignment distance position gene name gene id
GU305914.1 120798 120820 + 0 GCGGGCGGGACCTCCGGCGC CGG ||||||||[||||||||||||]PAM NA Intergenic NA NA
GU305914.1 110575 110597 + 4 GCGGCTGGGACCTCTCGCGC AGG ||||--||[||||||--||||]PAM NA Intergenic NA NA

T9072 GGGACCTCCGGCGCCGGTCTCGG 1000
Oligo fwd CACCGGGACCTCCGGCGCCGGTCT
Oligo rev AAACAGACCGGCGCCGGAGGTCCC
Chromosome start end strand MM target_seq PAM alignment distance position gene name gene id
GU305914.1 120804 120826 + 0 GGGACCTCCGGCGCCGGTCT CGG ||||||||[||||||||||||]PAM NA Intergenic NA NA

T9073 CCTCCGGCGCCGGTCTCGGCCGG 1000
Oligo fwd CACCCCTCCGGCGCCGGTCTCGGC

<table>
<tr><td>Oligo rev</td><td>AAACGCCGAGACCGGCGCCGGAGG</td><td></td><td></td><td></td><td></td><td></td><td></td><td></td><td></td><td></td><td></td></tr>
<tr><td>Chromosome</td><td>start</td><td>end</td><td>strand</td><td>MM</td><td>target_seq</td><td>PAM</td><td>alignment</td><td>distance</td><td>position</td><td>gene name</td><td>gene id</td></tr>
<tr><td>GU305914.1</td><td>120808</td><td>120830</td><td>+</td><td>0</td><td>CCTCCGGCGCCGGTCTCGGC</td><td>CGG</td><td>||||||||[||||||||||||]PAM</td><td>NA</td><td>Intergenic</td><td>NA</td><td>NA</td></tr>
<tr><td>T9074</td><td>CTCCGGCGCCGGTCTCGGCCGGG</td><td>1000</td><td></td><td></td><td></td><td></td><td></td><td></td><td></td><td></td><td></td></tr>
<tr><td>Oligo fwd</td><td>CACCCTCCGGCGCCGGTCTCGGCC</td><td></td><td></td><td></td><td></td><td></td><td></td><td></td><td></td><td></td><td></td></tr>
<tr><td>Oligo rev</td><td>AAACGGCCGAGACCGGCGCCGGAG</td><td></td><td></td><td></td><td></td><td></td><td></td><td></td><td></td><td></td><td></td></tr>
<tr><td>Chromosome</td><td>start</td><td>end</td><td>strand</td><td>MM</td><td>target_seq</td><td>PAM</td><td>alignment</td><td>distance</td><td>position</td><td>gene name</td><td>gene id</td></tr>
<tr><td>GU305914.1</td><td>120809</td><td>120831</td><td>+</td><td>0</td><td>CTCCGGCGCCGGTCTCGGCC</td><td>GGG</td><td>||||||||[||||||||||||]PAM</td><td>NA</td><td>Intergenic</td><td>NA</td><td>NA</td></tr>
<tr><td>T9075</td><td>CGGTCTCGGCCGGGTGCACGAGG</td><td>1000</td><td></td><td></td><td></td><td></td><td></td><td></td><td></td><td></td><td></td></tr>
<tr><td>Oligo fwd</td><td>CACCCGGTCTCGGCCGGGTGCACG</td><td></td><td></td><td></td><td></td><td></td><td></td><td></td><td></td><td></td><td></td></tr>
<tr><td>Oligo rev</td><td>AAACCGTGCACCCGGCCGAGACCG</td><td></td><td></td><td></td><td></td><td></td><td></td><td></td><td></td><td></td><td></td></tr>
<tr><td>Chromosome</td><td>start</td><td>end</td><td>strand</td><td>MM</td><td>target_seq</td><td>PAM</td><td>alignment</td><td>distance</td><td>position</td><td>gene name</td><td>gene id</td></tr>
<tr><td>GU305914.1</td><td>120818</td><td>120840</td><td>+</td><td>0</td><td>CGGTCTCGGCCGGGTGCACG</td><td>AGG</td><td>||||||||[||||||||||||]PAM</td><td>NA</td><td>Intergenic</td><td>NA</td><td>NA</td></tr>
<tr><td>T9076</td><td>CACGAGGATCGAGTAGCTCGTGG</td><td>1000</td><td></td><td></td><td></td><td></td><td></td><td></td><td></td><td></td><td></td></tr>
<tr><td>Oligo fwd</td><td>CACCCACGAGGATCGAGTAGCTCG</td><td></td><td></td><td></td><td></td><td></td><td></td><td></td><td></td><td></td><td></td></tr>
<tr><td>Oligo rev</td><td>AAACCGAGCTACTCGATCCTCGTG</td><td></td><td></td><td></td><td></td><td></td><td></td><td></td><td></td><td></td><td></td></tr>
<tr><td>Chromosome</td><td>start</td><td>end</td><td>strand</td><td>MM</td><td>target_seq</td><td>PAM</td><td>alignment</td><td>distance</td><td>position</td><td>gene name</td><td>gene id</td></tr>
<tr><td>GU305914.1</td><td>120834</td><td>120856</td><td>+</td><td>0</td><td>CACGAGGATCGAGTAGCTCG</td><td>TGG</td><td>||||||||[||||||||||||]PAM</td><td>NA</td><td>Intergenic</td><td>NA</td><td>NA</td></tr>
<tr><td>T9077</td><td>GTAGCTCGTGGCGTCCGCCGCGG</td><td>1000</td><td></td><td></td><td></td><td></td><td></td><td></td><td></td><td></td><td></td></tr>
<tr><td>Oligo fwd</td><td>CACCGTAGCTCGTGGCGTCCGCCG</td><td></td><td></td><td></td><td></td><td></td><td></td><td></td><td></td><td></td><td></td></tr>
<tr><td>Oligo rev</td><td>AAACCGGCGGACGCCACGAGCTAC</td><td></td><td></td><td></td><td></td><td></td><td></td><td></td><td></td><td></td><td></td></tr>
<tr><td>Chromosome</td><td>start</td><td>end</td><td>strand</td><td>MM</td><td>target_seq</td><td>PAM</td><td>alignment</td><td>distance</td><td>position</td><td>gene name</td><td>gene id</td></tr>
<tr><td>GU305914.1</td><td>120846</td><td>120868</td><td>+</td><td>0</td><td>GTAGCTCGTGGCGTCCGCCG</td><td>CGG</td><td>||||||||[||||||||||||]PAM</td><td>NA</td><td>Intergenic</td><td>NA</td><td>NA</td></tr>
<tr><td>T9078</td><td>GTGGCGTCCGCCGCGGCCGACGG</td><td>1000</td><td></td><td></td><td></td><td></td><td></td><td></td><td></td><td></td><td></td></tr>
<tr><td>Oligo fwd</td><td>CACCGTGGCGTCCGCCGCGGCCGA</td><td></td><td></td><td></td><td></td><td></td><td></td><td></td><td></td><td></td><td></td></tr>
<tr><td>Oligo rev</td><td>AAACTCGGCCGCGGCGGACGCCAC</td><td></td><td></td><td></td><td></td><td></td><td></td><td></td><td></td><td></td><td></td></tr>
<tr><td>Chromosome</td><td>start</td><td>end</td><td>strand</td><td>MM</td><td>target_seq</td><td>PAM</td><td>alignment</td><td>distance</td><td>position</td><td>gene name</td><td>gene id</td></tr>
<tr><td>GU305914.1</td><td>120853</td><td>120875</td><td>+</td><td>0</td><td>GTGGCGTCCGCCGCGGCCGA</td><td>CGG</td><td>||||||||[||||||||||||]PAM</td><td>NA</td><td>Intergenic</td><td>NA</td><td>NA</td></tr>
<tr><td>T9079</td><td>TGGCGTCCGCCGCGGCCGACGGG</td><td>1000</td><td></td><td></td><td></td><td></td><td></td><td></td><td></td><td></td><td></td></tr>
<tr><td>Oligo fwd</td><td>CACCTGGCGTCCGCCGCGGCCGAC</td><td></td><td></td><td></td><td></td><td></td><td></td><td></td><td></td><td></td><td></td></tr>
<tr><td>Oligo rev</td><td>AAACGTCGGCCGCGGCGGACGCCA</td><td></td><td></td><td></td><td></td><td></td><td></td><td></td><td></td><td></td><td></td></tr>
<tr><td>Chromosome</td><td>start</td><td>end</td><td>strand</td><td>MM</td><td>target_seq</td><td>PAM</td><td>alignment</td><td>distance</td><td>position</td><td>gene name</td><td>gene id</td></tr>
<tr><td>GU305914.1</td><td>120854</td><td>120876</td><td>+</td><td>0</td><td>TGGCGTCCGCCGCGGCCGAC</td><td>GGG</td><td>||||||||[||||||||||||]PAM</td><td>NA</td><td>Intergenic</td><td>NA</td><td>NA</td></tr>
<tr><td>T9080</td><td>ACGAAGTACATGTCTCCCAGCGG</td><td>1000</td><td></td><td></td><td></td><td></td><td></td><td></td><td></td><td></td><td></td></tr>
<tr><td>Oligo fwd</td><td>CACCACGAAGTACATGTCTCCCAG</td><td></td><td></td><td></td><td></td><td></td><td></td><td></td><td></td><td></td><td></td></tr>
<tr><td>Oligo rev</td><td>AAACCTGGGAGACATGTACTTCGT</td><td></td><td></td><td></td><td></td><td></td><td></td><td></td><td></td><td></td><td></td></tr>
<tr><td>Chromosome</td><td>start</td><td>end</td><td>strand</td><td>MM</td><td>target_seq</td><td>PAM</td><td>alignment</td><td>distance</td><td>position</td><td>gene name</td><td>gene id</td></tr>
<tr><td>GU305914.1</td><td>120910</td><td>120932</td><td>+</td><td>0</td><td>ACGAAGTACATGTCTCCCAG</td><td>CGG</td><td>||||||||[||||||||||||]PAM</td><td>NA</td><td>Intergenic</td><td>NA</td><td>NA</td></tr>
<tr><td>T9081</td><td>AGCACCTGACGCTCCTTCCACGG</td><td>1000</td><td></td><td></td><td></td><td></td><td></td><td></td><td></td><td></td><td></td></tr>
<tr><td>Oligo fwd</td><td>CACCAGCACCTGACGCTCCTTCCA</td><td></td><td></td><td></td><td></td><td></td><td></td><td></td><td></td><td></td><td></td></tr>
<tr><td>Oligo rev</td><td>AAACTGGAAGGAGCGTCAGGTGCT</td><td></td><td></td><td></td><td></td><td></td><td></td><td></td><td></td><td></td><td></td></tr>
<tr><td>Chromosome</td><td>start</td><td>end</td><td>strand</td><td>MM</td><td>target_seq</td><td>PAM</td><td>alignment</td><td>distance</td><td>position</td><td>gene name</td><td>gene id</td></tr>
<tr><td>GU305914.1</td><td>120982</td><td>121004</td><td>+</td><td>0</td><td>AGCACCTGACGCTCCTTCCA</td><td>CGG</td><td>||||||||[||||||||||||]PAM</td><td>NA</td><td>Intergenic</td><td>NA</td><td>NA</td></tr>
<tr><td>T9082</td><td>GCTCCTTCCACGGTACGATCTGG</td><td>1000</td><td></td><td></td><td></td><td></td><td></td><td></td><td></td><td></td><td></td></tr>
<tr><td>Oligo fwd</td><td>CACCGCTCCTTCCACGGTACGATC</td><td></td><td></td><td></td><td></td><td></td><td></td><td></td><td></td><td></td><td></td></tr>
<tr><td>Oligo rev</td><td>AAACGATCGTACCGTGGAAGGAGC</td><td></td><td></td><td></td><td></td><td></td><td></td><td></td><td></td><td></td><td></td></tr>
<tr><td>Chromosome</td><td>start</td><td>end</td><td>strand</td><td>MM</td><td>target_seq</td><td>PAM</td><td>alignment</td><td>distance</td><td>position</td><td>gene name</td><td>gene id</td></tr>
<tr><td>GU305914.1</td><td>120992</td><td>121014</td><td>+</td><td>0</td><td>GCTCCTTCCACGGTACGATC</td><td>TGG</td><td>||||||||[||||||||||||]PAM</td><td>NA</td><td>Intergenic</td><td>NA</td><td>NA</td></tr>
<tr><td>T9083</td><td>ACGATCTGGCAGCTCAACCCCGG</td><td>1000</td><td></td><td></td><td></td><td></td><td></td><td></td><td></td><td></td><td></td></tr>
<tr><td>Oligo fwd</td><td>CACCACGATCTGGCAGCTCAACCC</td><td></td><td></td><td></td><td></td><td></td><td></td><td></td><td></td><td></td><td></td></tr>
<tr><td>Oligo rev</td><td>AAACGGGTTGAGCTGCCAGATCGT</td><td></td><td></td><td></td><td></td><td></td><td></td><td></td><td></td><td></td><td></td></tr>
<tr><td>Chromosome</td><td>start</td><td>end</td><td>strand</td><td>MM</td><td>target_seq</td><td>PAM</td><td>alignment</td><td>distance</td><td>position</td><td>gene name</td><td>gene id</td></tr>
<tr><td>GU305914.1</td><td>121006</td><td>121028</td><td>+</td><td>0</td><td>ACGATCTGGCAGCTCAACCC</td><td>CGG</td><td>||||||||[||||||||||||]PAM</td><td>NA</td><td>Intergenic</td><td>NA</td><td>NA</td></tr>
<tr><td>T9084</td><td>TCTGGCAGCTCAACCCCGGACGG</td><td>1000</td><td></td><td></td><td></td><td></td><td></td><td></td><td></td><td></td><td></td></tr>
<tr><td>Oligo fwd</td><td>CACCTCTGGCAGCTCAACCCCGGA</td><td></td><td></td><td></td><td></td><td></td><td></td><td></td><td></td><td></td><td></td></tr>
<tr><td>Oligo rev</td><td>AAACTCCGGGGTTGAGCTGCCAGA</td><td></td><td></td><td></td><td></td><td></td><td></td><td></td><td></td><td></td><td></td></tr>
<tr><td>Chromosome</td><td>start</td><td>end</td><td>strand</td><td>MM</td><td>target_seq</td><td>PAM</td><td>alignment</td><td>distance</td><td>position</td><td>gene name</td><td>gene id</td></tr>
<tr><td>GU305914.1</td><td>121010</td><td>121032</td><td>+</td><td>0</td><td>TCTGGCAGCTCAACCCCGGA</td><td>CGG</td><td>||||||||[||||||||||||]PAM</td><td>NA</td><td>Intergenic</td><td>NA</td><td>NA</td></tr>
<tr><td>T9085</td><td>CTCAACCCCGGACGGTACAGCGG</td><td>1000</td><td></td><td></td><td></td><td></td><td></td><td></td><td></td><td></td><td></td></tr>
<tr><td>Oligo fwd</td><td>CACCCTCAACCCCGGACGGTACAG</td><td></td><td></td><td></td><td></td><td></td><td></td><td></td><td></td><td></td><td></td></tr>
<tr><td>Oligo rev</td><td>AAACCTGTACCGTCCGGGGTTGAG</td><td></td><td></td><td></td><td></td><td></td><td></td><td></td><td></td><td></td><td></td></tr>
<tr><td>Chromosome</td><td>start</td><td>end</td><td>strand</td><td>MM</td><td>target_seq</td><td>PAM</td><td>alignment</td><td>distance</td><td>position</td><td>gene name</td><td>gene id</td></tr>
<tr><td>GU305914.1</td><td>121018</td><td>121040</td><td>+</td><td>0</td><td>CTCAACCCCGGACGGTACAG</td><td>CGG</td><td>||||||||[||||||||||||]PAM</td><td>NA</td><td>Intergenic</td><td>NA</td><td>NA</td></tr>
<tr><td>T9086</td><td>GACGGTACAGCGGCGTGAACAGG</td><td>1000</td><td></td><td></td><td></td><td></td><td></td><td></td><td></td><td></td><td></td></tr>
<tr><td>Oligo fwd</td><td>CACCGACGGTACAGCGGCGTGAAC</td><td></td><td></td><td></td><td></td><td></td><td></td><td></td><td></td><td></td><td></td></tr>
<tr><td>Oligo rev</td><td>AAACGTTCACGCCGCTGTACCGTC</td><td></td><td></td><td></td><td></td><td></td><td></td><td></td><td></td><td></td><td></td></tr>
<tr><td>Chromosome</td><td>start</td><td>end</td><td>strand</td><td>MM</td><td>target_seq</td><td>PAM</td><td>alignment</td><td>distance</td><td>position</td><td>gene name</td><td>gene id</td></tr>
<tr><td>GU305914.1</td><td>121028</td><td>121050</td><td>+</td><td>0</td><td>GACGGTACAGCGGCGTGAAC</td><td>AGG</td><td>||||||||[||||||||||||]PAM</td><td>NA</td><td>Intergenic</td><td>NA</td><td>NA</td></tr>
<tr><td>T9087</td><td>CGTGAACAGGCCCGCGTACGCGG</td><td>1000</td><td></td><td></td><td></td><td></td><td></td><td></td><td></td><td></td><td></td></tr>
<tr><td>Oligo fwd</td><td>CACCCGTGAACAGGCCCGCGTACG</td><td></td><td></td><td></td><td></td><td></td><td></td><td></td><td></td><td></td><td></td></tr>
<tr><td>Oligo rev</td><td>AAACCGTACGCGGGCCTGTTCACG</td><td></td><td></td><td></td><td></td><td></td><td></td><td></td><td></td><td></td><td></td></tr>
<tr><td>Chromosome</td><td>start</td><td>end</td><td>strand</td><td>MM</td><td>target_seq</td><td>PAM</td><td>alignment</td><td>distance</td><td>position</td><td>gene name</td><td>gene id</td></tr>
<tr><td>GU305914.1</td><td>121041</td><td>121063</td><td>+</td><td>0</td><td>CGTGAACAGGCCCGCGTACG</td><td>CGG</td><td>||||||||[||||||||||||]PAM</td><td>NA</td><td>Intergenic</td><td>NA</td><td>NA</td></tr>
<tr><td>T9088</td><td>ATCGACACGTACATGCGCACCGG</td><td>1000</td><td></td><td></td><td></td><td></td><td></td><td></td><td></td><td></td><td></td></tr>
</table>

<table>
<tr><td>Oligo fwd</td><td>CACCATCGACACGTACATGCGCAC</td><td></td><td></td><td></td><td></td><td></td><td></td><td></td><td></td><td></td><td></td></tr>
<tr><td>Oligo rev</td><td>AAACGTGCGCATGTACGTGTCGAT</td><td></td><td></td><td></td><td></td><td></td><td></td><td></td><td></td><td></td><td></td></tr>
<tr><td>Chromosome</td><td>start</td><td>end</td><td>strand</td><td>MM</td><td>target_seq</td><td>PAM</td><td>alignment</td><td>distance</td><td>position</td><td>gene name</td><td>gene id</td></tr>
<tr><td>GU305914.1</td><td>121081</td><td>121103</td><td>+</td><td>0</td><td>ATCGACACGTACATGCGCAC</td><td>CGG</td><td>||||||||[||||||||||||]PAM</td><td>NA</td><td>Intergenic</td><td>NA</td><td>NA</td></tr>
<tr><td></td><td></td><td></td><td></td><td></td><td></td><td></td><td></td><td></td><td></td><td></td><td></td></tr>
<tr><td>T9089</td><td>TACATGCGCACCGGCTTGCCCGG</td><td>1000</td><td></td><td></td><td></td><td></td><td></td><td></td><td></td><td></td><td></td></tr>
<tr><td>Oligo fwd</td><td>CACCTACATGCGCACCGGCTTGCC</td><td></td><td></td><td></td><td></td><td></td><td></td><td></td><td></td><td></td><td></td></tr>
<tr><td>Oligo rev</td><td>AAACGGCAAGCCGGTGCGCATGTA</td><td></td><td></td><td></td><td></td><td></td><td></td><td></td><td></td><td></td><td></td></tr>
<tr><td>Chromosome</td><td>start</td><td>end</td><td>strand</td><td>MM</td><td>target_seq</td><td>PAM</td><td>alignment</td><td>distance</td><td>position</td><td>gene name</td><td>gene id</td></tr>
<tr><td>GU305914.1</td><td>121090</td><td>121112</td><td>+</td><td>0</td><td>TACATGCGCACCGGCTTGCC</td><td>CGG</td><td>||||||||[||||||||||||]PAM</td><td>NA</td><td>Intergenic</td><td>NA</td><td>NA</td></tr>
<tr><td></td><td></td><td></td><td></td><td></td><td></td><td></td><td></td><td></td><td></td><td></td><td></td></tr>
<tr><td>T9090</td><td>ACATGCGCACCGGCTTGCCCGGG</td><td>1000</td><td></td><td></td><td></td><td></td><td></td><td></td><td></td><td></td><td></td></tr>
<tr><td>Oligo fwd</td><td>CACCACATGCGCACCGGCTTGCCC</td><td></td><td></td><td></td><td></td><td></td><td></td><td></td><td></td><td></td><td></td></tr>
<tr><td>Oligo rev</td><td>AAACGGGCAAGCCGGTGCGCATGT</td><td></td><td></td><td></td><td></td><td></td><td></td><td></td><td></td><td></td><td></td></tr>
<tr><td>Chromosome</td><td>start</td><td>end</td><td>strand</td><td>MM</td><td>target_seq</td><td>PAM</td><td>alignment</td><td>distance</td><td>position</td><td>gene name</td><td>gene id</td></tr>
<tr><td>GU305914.1</td><td>121091</td><td>121113</td><td>+</td><td>0</td><td>ACATGCGCACCGGCTTGCCC</td><td>GGG</td><td>||||||||[||||||||||||]PAM</td><td>NA</td><td>Intergenic</td><td>NA</td><td>NA</td></tr>
<tr><td></td><td></td><td></td><td></td><td></td><td></td><td></td><td></td><td></td><td></td><td></td><td></td></tr>
<tr><td>T9091</td><td>CATGCGCACCGGCTTGCCCGGGG</td><td>1000</td><td></td><td></td><td></td><td></td><td></td><td></td><td></td><td></td><td></td></tr>
<tr><td>Oligo fwd</td><td>CACCCATGCGCACCGGCTTGCCCG</td><td></td><td></td><td></td><td></td><td></td><td></td><td></td><td></td><td></td><td></td></tr>
<tr><td>Oligo rev</td><td>AAACCGGGCAAGCCGGTGCGCATG</td><td></td><td></td><td></td><td></td><td></td><td></td><td></td><td></td><td></td><td></td></tr>
<tr><td>Chromosome</td><td>start</td><td>end</td><td>strand</td><td>MM</td><td>target_seq</td><td>PAM</td><td>alignment</td><td>distance</td><td>position</td><td>gene name</td><td>gene id</td></tr>
<tr><td>GU305914.1</td><td>121092</td><td>121114</td><td>+</td><td>0</td><td>CATGCGCACCGGCTTGCCCG</td><td>GGG</td><td>||||||||[||||||||||||]PAM</td><td>NA</td><td>Intergenic</td><td>NA</td><td>NA</td></tr>
<tr><td></td><td></td><td></td><td></td><td></td><td></td><td></td><td></td><td></td><td></td><td></td><td></td></tr>
<tr><td>T9092</td><td>GCACCGGCTTGCCCGGGGTCAGG</td><td>1000</td><td></td><td></td><td></td><td></td><td></td><td></td><td></td><td></td><td></td></tr>
<tr><td>Oligo fwd</td><td>CACCGCACCGGCTTGCCCGGGGTC</td><td></td><td></td><td></td><td></td><td></td><td></td><td></td><td></td><td></td><td></td></tr>
<tr><td>Oligo rev</td><td>AAACGACCCCGGGCAAGCCGGTGC</td><td></td><td></td><td></td><td></td><td></td><td></td><td></td><td></td><td></td><td></td></tr>
<tr><td>Chromosome</td><td>start</td><td>end</td><td>strand</td><td>MM</td><td>target_seq</td><td>PAM</td><td>alignment</td><td>distance</td><td>position</td><td>gene name</td><td>gene id</td></tr>
<tr><td>GU305914.1</td><td>121097</td><td>121119</td><td>+</td><td>0</td><td>GCACCGGCTTGCCCGGGGTC</td><td>AGG</td><td>||||||||[||||||||||||]PAM</td><td>NA</td><td>Intergenic</td><td>NA</td><td>NA</td></tr>
<tr><td></td><td></td><td></td><td></td><td></td><td></td><td></td><td></td><td></td><td></td><td></td><td></td></tr>
<tr><td>T9093</td><td>CACCGGCTTGCCCGGGGTCAGGG</td><td>1000</td><td></td><td></td><td></td><td></td><td></td><td></td><td></td><td></td><td></td></tr>
<tr><td>Oligo fwd</td><td>CACCCACCGGCTTGCCCGGGGTCA</td><td></td><td></td><td></td><td></td><td></td><td></td><td></td><td></td><td></td><td></td></tr>
<tr><td>Oligo rev</td><td>AAACTGACCCCGGGCAAGCCGGTG</td><td></td><td></td><td></td><td></td><td></td><td></td><td></td><td></td><td></td><td></td></tr>
<tr><td>Chromosome</td><td>start</td><td>end</td><td>strand</td><td>MM</td><td>target_seq</td><td>PAM</td><td>alignment</td><td>distance</td><td>position</td><td>gene name</td><td>gene id</td></tr>
<tr><td>GU305914.1</td><td>121098</td><td>121120</td><td>+</td><td>0</td><td>CACCGGCTTGCCCGGGGTCA</td><td>GGG</td><td>||||||||[||||||||||||]PAM</td><td>NA</td><td>Intergenic</td><td>NA</td><td>NA</td></tr>
<tr><td></td><td></td><td></td><td></td><td></td><td></td><td></td><td></td><td></td><td></td><td></td><td></td></tr>
<tr><td>T9094</td><td>CGGCTTGCCCGGGGTCAGGGCGG</td><td>1000</td><td></td><td></td><td></td><td></td><td></td><td></td><td></td><td></td><td></td></tr>
<tr><td>Oligo fwd</td><td>CACCCGGCTTGCCCGGGGTCAGGG</td><td></td><td></td><td></td><td></td><td></td><td></td><td></td><td></td><td></td><td></td></tr>
<tr><td>Oligo rev</td><td>AAACCCCTGACCCCGGGCAAGCCG</td><td></td><td></td><td></td><td></td><td></td><td></td><td></td><td></td><td></td><td></td></tr>
<tr><td>Chromosome</td><td>start</td><td>end</td><td>strand</td><td>MM</td><td>target_seq</td><td>PAM</td><td>alignment</td><td>distance</td><td>position</td><td>gene name</td><td>gene id</td></tr>
<tr><td>GU305914.1</td><td>121101</td><td>121123</td><td>+</td><td>0</td><td>CGGCTTGCCCGGGGTCAGGG</td><td>CGG</td><td>||||||||[||||||||||||]PAM</td><td>NA</td><td>Intergenic</td><td>NA</td><td>NA</td></tr>
<tr><td></td><td></td><td></td><td></td><td></td><td></td><td></td><td></td><td></td><td></td><td></td><td></td></tr>
<tr><td>T9095</td><td>GGGGTCAGGGCGGCGTCTCGCGG</td><td>1000</td><td></td><td></td><td></td><td></td><td></td><td></td><td></td><td></td><td></td></tr>
<tr><td>Oligo fwd</td><td>CACCGGGGTCAGGGCGGCGTCTCG</td><td></td><td></td><td></td><td></td><td></td><td></td><td></td><td></td><td></td><td></td></tr>
<tr><td>Oligo rev</td><td>AAACCGAGACGCCGCCCTGACCCC</td><td></td><td></td><td></td><td></td><td></td><td></td><td></td><td></td><td></td><td></td></tr>
<tr><td>Chromosome</td><td>start</td><td>end</td><td>strand</td><td>MM</td><td>target_seq</td><td>PAM</td><td>alignment</td><td>distance</td><td>position</td><td>gene name</td><td>gene id</td></tr>
<tr><td>GU305914.1</td><td>121111</td><td>121133</td><td>+</td><td>0</td><td>GGGGTCAGGGCGGCGTCTCG</td><td>CGG</td><td>||||||||[||||||||||||]PAM</td><td>NA</td><td>Intergenic</td><td>NA</td><td>NA</td></tr>
<tr><td></td><td></td><td></td><td></td><td></td><td></td><td></td><td></td><td></td><td></td><td></td><td></td></tr>
<tr><td>T9096</td><td>ACCCGCAGTCCGTACTTGTGCGG</td><td>1000</td><td></td><td></td><td></td><td></td><td></td><td></td><td></td><td></td><td></td></tr>
<tr><td>Oligo fwd</td><td>CACCACCCGCAGTCCGTACTTGTG</td><td></td><td></td><td></td><td></td><td></td><td></td><td></td><td></td><td></td><td></td></tr>
<tr><td>Oligo rev</td><td>AAACCACAAGTACGGACTGCGGGT</td><td></td><td></td><td></td><td></td><td></td><td></td><td></td><td></td><td></td><td></td></tr>
<tr><td>Chromosome</td><td>start</td><td>end</td><td>strand</td><td>MM</td><td>target_seq</td><td>PAM</td><td>alignment</td><td>distance</td><td>position</td><td>gene name</td><td>gene id</td></tr>
<tr><td>GU305914.1</td><td>121141</td><td>121163</td><td>+</td><td>0</td><td>ACCCGCAGTCCGTACTTGTG</td><td>CGG</td><td>||||||||[||||||||||||]PAM</td><td>NA</td><td>Intergenic</td><td>NA</td><td>NA</td></tr>
<tr><td></td><td></td><td></td><td></td><td></td><td></td><td></td><td></td><td></td><td></td><td></td><td></td></tr>
<tr><td>T9097</td><td>GTACTTGTGCGGCGTGCACACGG</td><td>1000</td><td></td><td></td><td></td><td></td><td></td><td></td><td></td><td></td><td></td></tr>
<tr><td>Oligo fwd</td><td>CACCGTACTTGTGCGGCGTGCACA</td><td></td><td></td><td></td><td></td><td></td><td></td><td></td><td></td><td></td><td></td></tr>
<tr><td>Oligo rev</td><td>AAACTGTGCACGCCGCACAAGTAC</td><td></td><td></td><td></td><td></td><td></td><td></td><td></td><td></td><td></td><td></td></tr>
<tr><td>Chromosome</td><td>start</td><td>end</td><td>strand</td><td>MM</td><td>target_seq</td><td>PAM</td><td>alignment</td><td>distance</td><td>position</td><td>gene name</td><td>gene id</td></tr>
<tr><td>GU305914.1</td><td>121152</td><td>121174</td><td>+</td><td>0</td><td>GTACTTGTGCGGCGTGCACA</td><td>CGG</td><td>||||||||[||||||||||||]PAM</td><td>NA</td><td>Intergenic</td><td>NA</td><td>NA</td></tr>
<tr><td></td><td></td><td></td><td></td><td></td><td></td><td></td><td></td><td></td><td></td><td></td><td></td></tr>
<tr><td>T9098</td><td>TTGTGCGGCGTGCACACGGTCGG</td><td>1000</td><td></td><td></td><td></td><td></td><td></td><td></td><td></td><td></td><td></td></tr>
<tr><td>Oligo fwd</td><td>CACCTTGTGCGGCGTGCACACGGT</td><td></td><td></td><td></td><td></td><td></td><td></td><td></td><td></td><td></td><td></td></tr>
<tr><td>Oligo rev</td><td>AAACACCGTGTGCACGCCGCACAA</td><td></td><td></td><td></td><td></td><td></td><td></td><td></td><td></td><td></td><td></td></tr>
<tr><td>Chromosome</td><td>start</td><td>end</td><td>strand</td><td>MM</td><td>target_seq</td><td>PAM</td><td>alignment</td><td>distance</td><td>position</td><td>gene name</td><td>gene id</td></tr>
<tr><td>GU305914.1</td><td>121156</td><td>121178</td><td>+</td><td>0</td><td>TTGTGCGGCGTGCACACGGT</td><td>CGG</td><td>||||||||[||||||||||||]PAM</td><td>NA</td><td>Intergenic</td><td>NA</td><td>NA</td></tr>
<tr><td></td><td></td><td></td><td></td><td></td><td></td><td></td><td></td><td></td><td></td><td></td><td></td></tr>
<tr><td>T9099</td><td>TGTGCGGCGTGCACACGGTCGGG</td><td>1000</td><td></td><td></td><td></td><td></td><td></td><td></td><td></td><td></td><td></td></tr>
<tr><td>Oligo fwd</td><td>CACCTGTGCGGCGTGCACACGGTC</td><td></td><td></td><td></td><td></td><td></td><td></td><td></td><td></td><td></td><td></td></tr>
<tr><td>Oligo rev</td><td>AAACGACCGTGTGCACGCCGCACA</td><td></td><td></td><td></td><td></td><td></td><td></td><td></td><td></td><td></td><td></td></tr>
<tr><td>Chromosome</td><td>start</td><td>end</td><td>strand</td><td>MM</td><td>target_seq</td><td>PAM</td><td>alignment</td><td>distance</td><td>position</td><td>gene name</td><td>gene id</td></tr>
<tr><td>GU305914.1</td><td>121157</td><td>121179</td><td>+</td><td>0</td><td>TGTGCGGCGTGCACACGGTC</td><td>GGG</td><td>||||||||[||||||||||||]PAM</td><td>NA</td><td>Intergenic</td><td>NA</td><td>NA</td></tr>
<tr><td></td><td></td><td></td><td></td><td></td><td></td><td></td><td></td><td></td><td></td><td></td><td></td></tr>
<tr><td>T9100</td><td>GGCGTGCACACGGTCGGGTCCGG</td><td>1000</td><td></td><td></td><td></td><td></td><td></td><td></td><td></td><td></td><td></td></tr>
<tr><td>Oligo fwd</td><td>CACCGGCGTGCACACGGTCGGGTC</td><td></td><td></td><td></td><td></td><td></td><td></td><td></td><td></td><td></td><td></td></tr>
<tr><td>Oligo rev</td><td>AAACGACCCGACCGTGTGCACGCC</td><td></td><td></td><td></td><td></td><td></td><td></td><td></td><td></td><td></td><td></td></tr>
<tr><td>Chromosome</td><td>start</td><td>end</td><td>strand</td><td>MM</td><td>target_seq</td><td>PAM</td><td>alignment</td><td>distance</td><td>position</td><td>gene name</td><td>gene id</td></tr>
<tr><td>GU305914.1</td><td>121162</td><td>121184</td><td>+</td><td>0</td><td>GGCGTGCACACGGTCGGGTC</td><td>CGG</td><td>||||||||[||||||||||||]PAM</td><td>NA</td><td>Intergenic</td><td>NA</td><td>NA</td></tr>
<tr><td></td><td></td><td></td><td></td><td></td><td></td><td></td><td></td><td></td><td></td><td></td><td></td></tr>
<tr><td>T9101</td><td>ACACGGTCGGGTCCGGATAGCGG</td><td>1000</td><td></td><td></td><td></td><td></td><td></td><td></td><td></td><td></td><td></td></tr>
<tr><td>Oligo fwd</td><td>CACCACACGGTCGGGTCCGGATAG</td><td></td><td></td><td></td><td></td><td></td><td></td><td></td><td></td><td></td><td></td></tr>
<tr><td>Oligo rev</td><td>AAACCTATCCGGACCCGACCGTGT</td><td></td><td></td><td></td><td></td><td></td><td></td><td></td><td></td><td></td><td></td></tr>
<tr><td>Chromosome</td><td>start</td><td>end</td><td>strand</td><td>MM</td><td>target_seq</td><td>PAM</td><td>alignment</td><td>distance</td><td>position</td><td>gene name</td><td>gene id</td></tr>
<tr><td>GU305914.1</td><td>121169</td><td>121191</td><td>+</td><td>0</td><td>ACACGGTCGGGTCCGGATAG</td><td>CGG</td><td>||||||||[||||||||||||]PAM</td><td>NA</td><td>Intergenic</td><td>NA</td><td>NA</td></tr>
<tr><td></td><td></td><td></td><td></td><td></td><td></td><td></td><td></td><td></td><td></td><td></td><td></td></tr>
<tr><td>T9102</td><td>CACGGTCGGGTCCGGATAGCGGG</td><td>1000</td><td></td><td></td><td></td><td></td><td></td><td></td><td></td><td></td><td></td></tr>
<tr><td>Oligo fwd</td><td>CACCCACGGTCGGGTCCGGATAGC</td><td></td><td></td><td></td><td></td><td></td><td></td><td></td><td></td><td></td><td></td></tr>
<tr><td>Oligo rev</td><td>AAACGCTATCCGGACCCGACCGTG</td><td></td><td></td><td></td><td></td><td></td><td></td><td></td><td></td><td></td><td></td></tr>
<tr><td>Chromosome</td><td>start</td><td>end</td><td>strand</td><td>MM</td><td>target_seq</td><td>PAM</td><td>alignment</td><td>distance</td><td>position</td><td>gene name</td><td>gene id</td></tr>
<tr><td>GU305914.1</td><td>121170</td><td>121192</td><td>+</td><td>0</td><td>CACGGTCGGGTCCGGATAGC</td><td>GGG</td><td>||||||||[||||||||||||]PAM</td><td>NA</td><td>Intergenic</td><td>NA</td><td>NA</td></tr>
</table>

<table>
<tr><td>T9103</td><td>ACGGTCGGGTCCGGATAGCGGGG</td><td>1000</td></tr>
<tr><td>Oligo fwd</td><td>CACCACGGTCGGGTCCGGATAGCG</td></tr>
<tr><td>Oligo rev</td><td>AAACCGCTATCCGGACCCGACCGT</td></tr>
<tr><td>Chromosome</td><td>start</td><td>end</td><td>strand</td><td>MM</td><td>target_seq</td><td>PAM</td><td>alignment</td><td>distance</td><td>position</td><td>gene name</td><td>gene id</td></tr>
<tr><td>GU305914.1</td><td>121171</td><td>121193</td><td>+</td><td>0</td><td>ACGGTCGGGTCCGGATAGCG</td><td>GGG</td><td>||||||||[||||||||||||]PAM</td><td>NA</td><td>Intergenic</td><td>NA</td><td>NA</td></tr>
<tr><td></td></tr>
<tr><td>T9104</td><td>CGGTCGGGTCCGGATAGCGGGGG</td><td>1000</td></tr>
<tr><td>Oligo fwd</td><td>CACCCGGTCGGGTCCGGATAGCGG</td></tr>
<tr><td>Oligo rev</td><td>AAACCCGCTATCCGGACCCGACCG</td></tr>
<tr><td>Chromosome</td><td>start</td><td>end</td><td>strand</td><td>MM</td><td>target_seq</td><td>PAM</td><td>alignment</td><td>distance</td><td>position</td><td>gene name</td><td>gene id</td></tr>
<tr><td>GU305914.1</td><td>121172</td><td>121194</td><td>+</td><td>0</td><td>CGGTCGGGTCCGGATAGCGG</td><td>GGG</td><td>||||||||[||||||||||||]PAM</td><td>NA</td><td>Intergenic</td><td>NA</td><td>NA</td></tr>
<tr><td></td></tr>
<tr><td>T9105</td><td>GCTGCGCGTCGTCCTCCTCGCGG</td><td>1000</td></tr>
<tr><td>Oligo fwd</td><td>CACCGCTGCGCGTCGTCCTCCTCG</td></tr>
<tr><td>Oligo rev</td><td>AAACCGAGGAGGACGACGCGCAGC</td></tr>
<tr><td>Chromosome</td><td>start</td><td>end</td><td>strand</td><td>MM</td><td>target_seq</td><td>PAM</td><td>alignment</td><td>distance</td><td>position</td><td>gene name</td><td>gene id</td></tr>
<tr><td>GU305914.1</td><td>121217</td><td>121239</td><td>+</td><td>0</td><td>GCTGCGCGTCGTCCTCCTCG</td><td>CGG</td><td>||||||||[||||||||||||]PAM</td><td>NA</td><td>Intergenic</td><td>NA</td><td>NA</td></tr>
<tr><td></td></tr>
<tr><td>T9106</td><td>CGTCGTCCTCCTCGCGGAACAGG</td><td>1000</td></tr>
<tr><td>Oligo fwd</td><td>CACCCGTCGTCCTCCTCGCGGAAC</td></tr>
<tr><td>Oligo rev</td><td>AAACGTTCCGCGAGGAGGACGACG</td></tr>
<tr><td>Chromosome</td><td>start</td><td>end</td><td>strand</td><td>MM</td><td>target_seq</td><td>PAM</td><td>alignment</td><td>distance</td><td>position</td><td>gene name</td><td>gene id</td></tr>
<tr><td>GU305914.1</td><td>121223</td><td>121245</td><td>+</td><td>0</td><td>CGTCGTCCTCCTCGCGGAAC</td><td>AGG</td><td>||||||||[||||||||||||]PAM</td><td>NA</td><td>Intergenic</td><td>NA</td><td>NA</td></tr>
<tr><td></td></tr>
<tr><td>T9107</td><td>CTCGCGGAACAGGATCTTGAAGG</td><td>1000</td></tr>
<tr><td>Oligo fwd</td><td>CACCCTCGCGGAACAGGATCTTGA</td></tr>
<tr><td>Oligo rev</td><td>AAACTCAAGATCCTGTTCCGCGAG</td></tr>
<tr><td>Chromosome</td><td>start</td><td>end</td><td>strand</td><td>MM</td><td>target_seq</td><td>PAM</td><td>alignment</td><td>distance</td><td>position</td><td>gene name</td><td>gene id</td></tr>
<tr><td>GU305914.1</td><td>121233</td><td>121255</td><td>+</td><td>0</td><td>CTCGCGGAACAGGATCTTGA</td><td>AGG</td><td>||||||||[||||||||||||]PAM</td><td>NA</td><td>Intergenic</td><td>NA</td><td>NA</td></tr>
<tr><td></td></tr>
<tr><td>T9108</td><td>ATCTTGAAGGCGATGTTGCTCGG</td><td>1000</td></tr>
<tr><td>Oligo fwd</td><td>CACCATCTTGAAGGCGATGTTGCT</td></tr>
<tr><td>Oligo rev</td><td>AAACAGCAACATCGCCTTCAAGAT</td></tr>
<tr><td>Chromosome</td><td>start</td><td>end</td><td>strand</td><td>MM</td><td>target_seq</td><td>PAM</td><td>alignment</td><td>distance</td><td>position</td><td>gene name</td><td>gene id</td></tr>
<tr><td>GU305914.1</td><td>121246</td><td>121268</td><td>+</td><td>0</td><td>ATCTTGAAGGCGATGTTGCT</td><td>CGG</td><td>||||||||[||||||||||||]PAM</td><td>NA</td><td>Intergenic</td><td>NA</td><td>NA</td></tr>
<tr><td></td></tr>
<tr><td>T9109</td><td>TCTTGAAGGCGATGTTGCTCGGG</td><td>1000</td></tr>
<tr><td>Oligo fwd</td><td>CACCTCTTGAAGGCGATGTTGCTC</td></tr>
<tr><td>Oligo rev</td><td>AAACGAGCAACATCGCCTTCAAGA</td></tr>
<tr><td>Chromosome</td><td>start</td><td>end</td><td>strand</td><td>MM</td><td>target_seq</td><td>PAM</td><td>alignment</td><td>distance</td><td>position</td><td>gene name</td><td>gene id</td></tr>
<tr><td>GU305914.1</td><td>121247</td><td>121269</td><td>+</td><td>0</td><td>TCTTGAAGGCGATGTTGCTC</td><td>GGG</td><td>||||||||[||||||||||||]PAM</td><td>NA</td><td>Intergenic</td><td>NA</td><td>NA</td></tr>
<tr><td></td></tr>
<tr><td>T9110</td><td>CTTGAAGGCGATGTTGCTCGGGG</td><td>1000</td></tr>
<tr><td>Oligo fwd</td><td>CACCCTTGAAGGCGATGTTGCTCG</td></tr>
<tr><td>Oligo rev</td><td>AAACCGAGCAACATCGCCTTCAAG</td></tr>
<tr><td>Chromosome</td><td>start</td><td>end</td><td>strand</td><td>MM</td><td>target_seq</td><td>PAM</td><td>alignment</td><td>distance</td><td>position</td><td>gene name</td><td>gene id</td></tr>
<tr><td>GU305914.1</td><td>121248</td><td>121270</td><td>+</td><td>0</td><td>CTTGAAGGCGATGTTGCTCG</td><td>GGG</td><td>||||||||[||||||||||||]PAM</td><td>NA</td><td>Intergenic</td><td>NA</td><td>NA</td></tr>
<tr><td></td></tr>
<tr><td>T9111</td><td>GGCCGCGATCGTCGCGTACTTGG</td><td>1000</td></tr>
<tr><td>Oligo fwd</td><td>CACCGGCCGCGATCGTCGCGTACT</td></tr>
<tr><td>Oligo rev</td><td>AAACAGTACGCGACGATCGCGGCC</td></tr>
<tr><td>Chromosome</td><td>start</td><td>end</td><td>strand</td><td>MM</td><td>target_seq</td><td>PAM</td><td>alignment</td><td>distance</td><td>position</td><td>gene name</td><td>gene id</td></tr>
<tr><td>GU305914.1</td><td>121269</td><td>121291</td><td>+</td><td>0</td><td>GGCCGCGATCGTCGCGTACT</td><td>TGG</td><td>||||||||[||||||||||||]PAM</td><td>NA</td><td>Intergenic</td><td>NA</td><td>NA</td></tr>
<tr><td></td></tr>
<tr><td>T9112</td><td>CGTACACCACCCCCGAGCTCTGG</td><td>1000</td></tr>
<tr><td>Oligo fwd</td><td>CACCCGTACACCACCCCCGAGCTC</td></tr>
<tr><td>Oligo rev</td><td>AAACGAGCTCGGGGGTGGTGTACG</td></tr>
<tr><td>Chromosome</td><td>start</td><td>end</td><td>strand</td><td>MM</td><td>target_seq</td><td>PAM</td><td>alignment</td><td>distance</td><td>position</td><td>gene name</td><td>gene id</td></tr>
<tr><td>GU305914.1</td><td>121304</td><td>121326</td><td>+</td><td>0</td><td>CGTACACCACCCCCGAGCTC</td><td>TGG</td><td>||||||||[||||||||||||]PAM</td><td>NA</td><td>Intergenic</td><td>NA</td><td>NA</td></tr>
<tr><td></td></tr>
<tr><td>T9113</td><td>AGCTCTGGACCACCGTCACCTGG</td><td>1000</td></tr>
<tr><td>Oligo fwd</td><td>CACCAGCTCTGGACCACCGTCACC</td></tr>
<tr><td>Oligo rev</td><td>AAACGGTGACGGTGGTCCAGAGCT</td></tr>
<tr><td>Chromosome</td><td>start</td><td>end</td><td>strand</td><td>MM</td><td>target_seq</td><td>PAM</td><td>alignment</td><td>distance</td><td>position</td><td>gene name</td><td>gene id</td></tr>
<tr><td>GU305914.1</td><td>121319</td><td>121341</td><td>+</td><td>0</td><td>AGCTCTGGACCACCGTCACC</td><td>TGG</td><td>||||||||[||||||||||||]PAM</td><td>NA</td><td>Intergenic</td><td>NA</td><td>NA</td></tr>
<tr><td></td></tr>
<tr><td>T9114</td><td>GTAGATGTCCGAGTGCGAGCAGG</td><td>1000</td></tr>
<tr><td>Oligo fwd</td><td>CACCGTAGATGTCCGAGTGCGAGC</td></tr>
<tr><td>Oligo rev</td><td>AAACGCTCGCACTCGGACATCTAC</td></tr>
<tr><td>Chromosome</td><td>start</td><td>end</td><td>strand</td><td>MM</td><td>target_seq</td><td>PAM</td><td>alignment</td><td>distance</td><td>position</td><td>gene name</td><td>gene id</td></tr>
<tr><td>GU305914.1</td><td>121347</td><td>121369</td><td>+</td><td>0</td><td>GTAGATGTCCGAGTGCGAGC</td><td>AGG</td><td>||||||||[||||||||||||]PAM</td><td>NA</td><td>Intergenic</td><td>NA</td><td>NA</td></tr>
<tr><td></td></tr>
<tr><td>T9115</td><td>TGTCCGAGTGCGAGCAGGAGAGG</td><td>1000</td></tr>
<tr><td>Oligo fwd</td><td>CACCTGTCCGAGTGCGAGCAGGAG</td></tr>
<tr><td>Oligo rev</td><td>AAACCTCCTGCTCGCACTCGGACA</td></tr>
<tr><td>Chromosome</td><td>start</td><td>end</td><td>strand</td><td>MM</td><td>target_seq</td><td>PAM</td><td>alignment</td><td>distance</td><td>position</td><td>gene name</td><td>gene id</td></tr>
<tr><td>GU305914.1</td><td>121352</td><td>121374</td><td>+</td><td>0</td><td>TGTCCGAGTGCGAGCAGGAG</td><td>AGG</td><td>||||||||[||||||||||||]PAM</td><td>NA</td><td>Intergenic</td><td>NA</td><td>NA</td></tr>
<tr><td></td></tr>
<tr><td>T9116</td><td>AGGAGAGGACGTGCGCTGTGTGG</td><td>1000</td></tr>
<tr><td>Oligo fwd</td><td>CACCAGGAGAGGACGTGCGCTGTG</td></tr>
<tr><td>Oligo rev</td><td>AAACCACAGCGCACGTCCTCTCCT</td></tr>
<tr><td>Chromosome</td><td>start</td><td>end</td><td>strand</td><td>MM</td><td>target_seq</td><td>PAM</td><td>alignment</td><td>distance</td><td>position</td><td>gene name</td><td>gene id</td></tr>
<tr><td>GU305914.1</td><td>121367</td><td>121389</td><td>+</td><td>0</td><td>AGGAGAGGACGTGCGCTGTG</td><td>TGG</td><td>||||||||[||||||||||||]PAM</td><td>NA</td><td>Intergenic</td><td>NA</td><td>NA</td></tr>
<tr><td></td></tr>
<tr><td>T9117</td><td>TGTTCGTCGACGACACGTCCAGG</td><td>1000</td></tr>
<tr><td>Oligo fwd</td><td>CACCTGTTCGTCGACGACACGTCC</td></tr>
<tr><td>Oligo rev</td><td>AAACGGACGTGTCGTCGACGAACA</td></tr>
<tr><td>Chromosome</td><td>start</td><td>end</td><td>strand</td><td>MM</td><td>target_seq</td><td>PAM</td><td>alignment</td><td>distance</td><td>position</td><td>gene name</td><td>gene id</td></tr>
<tr><td>GU305914.1</td><td>121400</td><td>121422</td><td>+</td><td>0</td><td>TGTTCGTCGACGACACGTCC</td><td>AGG</td><td>||||||||[||||||||||||]PAM</td><td>NA</td><td>Intergenic</td><td>NA</td><td>NA</td></tr>
</table>

T9118 CGACACGTCCAGGCTCACCGAGG 1000
Oligo fwd CACCCGACACGTCCAGGCTCACCG
Oligo rev AAACCGGTGAGCCTGGACGTGTCG
Chromosome start end strand MM target_seq PAM alignment distance position gene name gene id
GU305914.1 121410 121432 + 0 CGACACGTCCAGGCTCACCG AGG ||||||||[||||||||||||]PAM NA Intergenic NA NA

T9119 AGGCCGTCAGCCTGTTGCCGCGG 1000
Oligo fwd CACCAGGCCGTCAGCCTGTTGCCG
Oligo rev AAACCGGCAACAGGCTGACGGCCT
Chromosome start end strand MM target_seq PAM alignment distance position gene name gene id
GU305914.1 121430 121452 + 0 AGGCCGTCAGCCTGTTGCCG CGG ||||||||[||||||||||||]PAM NA Intergenic NA NA

T9120 AGCCTGTTGCCGCGGCTGAACGG 1000
Oligo fwd CACCAGCCTGTTGCCGCGGCTGAA
Oligo rev AAACTTCAGCCGCGGCAACAGGCT
Chromosome start end strand MM target_seq PAM alignment distance position gene name gene id
GU305914.1 121438 121460 + 0 AGCCTGTTGCCGCGGCTGAA CGG ||||||||[||||||||||||]PAM NA Intergenic NA NA

T9121 GGCTGAACGGCCGCCACGCCAGG 1000
Oligo fwd CACCGGCTGAACGGCCGCCACGCC
Oligo rev AAACGGCGTGGCGGCCGTTCAGCC
Chromosome start end strand MM target_seq PAM alignment distance position gene name gene id
GU305914.1 121451 121473 + 0 GGCTGAACGGCCGCCACGCC AGG ||||||||[||||||||||||]PAM NA Intergenic NA NA

T9122 GGTTCGACGCCGCCACCTCTAGG 1000
Oligo fwd CACCGGTTCGACGCCGCCACCTCT
Oligo rev AAACAGAGGTGGCGGCGTCGAACC
Chromosome start end strand MM target_seq PAM alignment distance position gene name gene id
GU305914.1 121472 121494 + 0 GGTTCGACGCCGCCACCTCT AGG ||||||||[||||||||||||]PAM NA Intergenic NA NA

T9123 TCTTGCTCGTCCGCATCGCCAGG 1000
Oligo fwd CACCTCTTGCTCGTCCGCATCGCC
Oligo rev AAACGGCGATGCGGACGAGCAAGA
Chromosome start end strand MM target_seq PAM alignment distance position gene name gene id
GU305914.1 121505 121527 + 0 TCTTGCTCGTCCGCATCGCC AGG ||||||||[||||||||||||]PAM NA Intergenic NA NA

T9124 CGCATCGCCAGGATGCTCACCGG 1000
Oligo fwd CACCCGCATCGCCAGGATGCTCAC
Oligo rev AAACGTGAGCATCCTGGCGATGCG
Chromosome start end strand MM target_seq PAM alignment distance position gene name gene id
GU305914.1 121516 121538 + 0 CGCATCGCCAGGATGCTCAC CGG ||||||||[||||||||||||]PAM NA Intergenic NA NA

T9125 GCATCGCCAGGATGCTCACCGGG 1000
Oligo fwd CACCGCATCGCCAGGATGCTCACC
Oligo rev AAACGGTGAGCATCCTGGCGATGC
Chromosome start end strand MM target_seq PAM alignment distance position gene name gene id
GU305914.1 121517 121539 + 0 GCATCGCCAGGATGCTCACC GGG ||||||||[||||||||||||]PAM NA Intergenic NA NA

T9126 CCAGGATGCTCACCGGGCAGCGG 1000
Oligo fwd CACCCCAGGATGCTCACCGGGCAG
Oligo rev AAACCTGCCCGGTGAGCATCCTGG
Chromosome start end strand MM target_seq PAM alignment distance position gene name gene id
GU305914.1 121523 121545 + 0 CCAGGATGCTCACCGGGCAG CGG ||||||||[||||||||||||]PAM NA Intergenic NA NA

T9127 CCGGGCAGCGGCCCCGCACTCGG 1000
Oligo fwd CACCCCGGGCAGCGGCCCCGCACT
Oligo rev AAACAGTGCGGGGCCGCTGCCCGG
Chromosome start end strand MM target_seq PAM alignment distance position gene name gene id
GU305914.1 121535 121557 + 0 CCGGGCAGCGGCCCCGCACT CGG ||||||||[||||||||||||]PAM NA Intergenic NA NA

T9128 CGGGCAGCGGCCCCGCACTCGGG 1000
Oligo fwd CACCCGGGCAGCGGCCCCGCACTC
Oligo rev AAACGAGTGCGGGGCCGCTGCCCG
Chromosome start end strand MM target_seq PAM alignment distance position gene name gene id
GU305914.1 121536 121558 + 0 CGGGCAGCGGCCCCGCACTC GGG ||||||||[||||||||||||]PAM NA Intergenic NA NA

T9129 GGGCAGCGGCCCCGCACTCGGGG 1000
Oligo fwd CACCGGGCAGCGGCCCCGCACTCG
Oligo rev AAACCGAGTGCGGGGCCGCTGCCC
Chromosome start end strand MM target_seq PAM alignment distance position gene name gene id
GU305914.1 121537 121559 + 0 GGGCAGCGGCCCCGCACTCG GGG ||||||||[||||||||||||]PAM NA Intergenic NA NA

T9130 GTCTCCTCTGCACGCACAGCAGG 1000
Oligo fwd CACCGTCTCCTCTGCACGCACAGC
Oligo rev AAACGCTGTGCGTGCAGAGGAGAC
Chromosome start end strand MM target_seq PAM alignment distance position gene name gene id
GU305914.1 121568 121590 + 0 GTCTCCTCTGCACGCACAGC AGG ||||||||[||||||||||||]PAM NA Intergenic NA NA

T9131 GCACAGCAGGTCCATCCGCACGG 1000
Oligo fwd CACCGCACAGCAGGTCCATCCGCA
Oligo rev AAACTGCGGATGGACCTGCTGTGC
Chromosome start end strand MM target_seq PAM alignment distance position gene name gene id
GU305914.1 121581 121603 + 0 GCACAGCAGGTCCATCCGCA CGG ||||||||[||||||||||||]PAM NA Intergenic NA NA

T9132 CACAGCAGGTCCATCCGCACGGG 1000
Oligo fwd CACCCACAGCAGGTCCATCCGCAC
Oligo rev AAACGTGCGGATGGACCTGCTGTG
Chromosome start end strand MM target_seq PAM alignment distance position gene name gene id

GU305914.1 121582 121604 + 0 CACAGCAGGTCCATCCGCAC GGG ||||||||[||||||||||||]PAM NA Intergenic NA NA

T9133 TCCATCCGCACGGGCGCCAGCGG 1000
Oligo fwd CACCTCCATCCGCACGGGCGCCAG
Oligo rev AAACCTGGCGCCCGTGCGGATGGA
Chromosome start end strand MM target_seq PAM alignment distance position gene name gene id
GU305914.1 121591 121613 + 0 TCCATCCGCACGGGCGCCAG CGG ||||||||[||||||||||||]PAM NA Intergenic NA NA

T9134 GATCAGCACCAGCTGCCCGCCGG 1000
Oligo fwd CACCGATCAGCACCAGCTGCCCGC
Oligo rev AAACGCGGGCAGCTGGTGCTGATC
Chromosome start end strand MM target_seq PAM alignment distance position gene name gene id
GU305914.1 121626 121648 + 0 GATCAGCACCAGCTGCCCGC CGG ||||||||[||||||||||||]PAM NA Intergenic NA NA

T9135 AGCACCAGCTGCCCGCCGGACGG 1000
Oligo fwd CACCAGCACCAGCTGCCCGCCGGA
Oligo rev AAACTCCGGCGGGCAGCTGGTGCT
Chromosome start end strand MM target_seq PAM alignment distance position gene name gene id
GU305914.1 121630 121652 + 0 AGCACCAGCTGCCCGCCGGA CGG ||||||||[||||||||||||]PAM NA Intergenic NA NA

T9136 ACCAGCTGCCCGCCGGACGGCGG 1000
Oligo fwd CACCACCAGCTGCCCGCCGGACGG
Oligo rev AAACCCGTCCGGCGGGCAGCTGGT
Chromosome start end strand MM target_seq PAM alignment distance position gene name gene id
GU305914.1 121633 121655 + 0 ACCAGCTGCCCGCCGGACGG CGG ||||||||[||||||||||||]PAM NA Intergenic NA NA

T9137 CCGCCGGACGGCGGCTTCAGCGG 1000
Oligo fwd CACCCCGCCGGACGGCGGCTTCAG
Oligo rev AAACCTGAAGCCGCCGTCCGGCGG
Chromosome start end strand MM target_seq PAM alignment distance position gene name gene id
GU305914.1 121642 121664 + 0 CCGCCGGACGGCGGCTTCAG CGG ||||||||[||||||||||||]PAM NA Intergenic NA NA
GU305914.1 90202 90224 - 4 CCTCCGGCCGGCGGCGCCAG CGG ||-||||-[|||||||--|||]PAM NA Intergenic NA NA

T9138 CGGACGGCGGCTTCAGCGGATGG 1000
Oligo fwd CACCCGGACGGCGGCTTCAGCGGA
Oligo rev AAACTCCGCTGAAGCCGCCGTCCG
Chromosome start end strand MM target_seq PAM alignment distance position gene name gene id
GU305914.1 121646 121668 + 0 CGGACGGCGGCTTCAGCGGA TGG ||||||||[||||||||||||]PAM NA Intergenic NA NA

T9139 GGACGGCGGCTTCAGCGGATGGG 1000
Oligo fwd CACCGGACGGCGGCTTCAGCGGAT
Oligo rev AAACATCCGCTGAAGCCGCCGTCC
Chromosome start end strand MM target_seq PAM alignment distance position gene name gene id
GU305914.1 121647 121669 + 0 GGACGGCGGCTTCAGCGGAT GGG ||||||||[||||||||||||]PAM NA Intergenic NA NA

T9140 GGCTTCAGCGGATGGGCGCTCGG 1000
Oligo fwd CACCGGCTTCAGCGGATGGGCGCT
Oligo rev AAACAGCGCCCATCCGCTGAAGCC
Chromosome start end strand MM target_seq PAM alignment distance position gene name gene id
GU305914.1 121654 121676 + 0 GGCTTCAGCGGATGGGCGCT CGG ||||||||[||||||||||||]PAM NA Intergenic NA NA

T9141 GCTTCAGCGGATGGGCGCTCGGG 1000
Oligo fwd CACCGCTTCAGCGGATGGGCGCTC
Oligo rev AAACGAGCGCCCATCCGCTGAAGC
Chromosome start end strand MM target_seq PAM alignment distance position gene name gene id
GU305914.1 121655 121677 + 0 GCTTCAGCGGATGGGCGCTC GGG ||||||||[||||||||||||]PAM NA Intergenic NA NA

T9142 GGTTCGACACTTCGACGACCAGG 1000
Oligo fwd CACCGGTTCGACACTTCGACGACC
Oligo rev AAACGGTCGTCGAAGTGTCGAACC
Chromosome start end strand MM target_seq PAM alignment distance position gene name gene id
GU305914.1 121676 121698 + 0 GGTTCGACACTTCGACGACC AGG ||||||||[||||||||||||]PAM NA Intergenic NA NA

T9143 CGACACTTCGACGACCAGGTCGG 1000
Oligo fwd CACCCGACACTTCGACGACCAGGT
Oligo rev AAACACCTGGTCGTCGAAGTGTCG
Chromosome start end strand MM target_seq PAM alignment distance position gene name gene id
GU305914.1 121680 121702 + 0 CGACACTTCGACGACCAGGT CGG ||||||||[||||||||||||]PAM NA Intergenic NA NA

T9144 ACTTCGACGACCAGGTCGGTCGG 1000
Oligo fwd CACCACTTCGACGACCAGGTCGGT
Oligo rev AAACACCGACCTGGTCGTCGAAGT
Chromosome start end strand MM target_seq PAM alignment distance position gene name gene id
GU305914.1 121684 121706 + 0 ACTTCGACGACCAGGTCGGT CGG ||||||||[||||||||||||]PAM NA Intergenic NA NA

T9145 CAGGTCGGTCGGCGCGTCCGCGG 1000
Oligo fwd CACCCAGGTCGGTCGGCGCGTCCG
Oligo rev AAACCGGACGCGCCGACCGACCTG
Chromosome start end strand MM target_seq PAM alignment distance position gene name gene id
GU305914.1 121695 121717 + 0 CAGGTCGGTCGGCGCGTCCG CGG ||||||||[||||||||||||]PAM NA Intergenic NA NA

T9146 AGGTCGGTCGGCGCGTCCGCGGG 1000
Oligo fwd CACCAGGTCGGTCGGCGCGTCCGC
Oligo rev AAACGCGGACGCGCCGACCGACCT
Chromosome start end strand MM target_seq PAM alignment distance position gene name gene id
GU305914.1 121696 121718 + 0 AGGTCGGTCGGCGCGTCCGC GGG ||||||||[||||||||||||]PAM NA Intergenic NA NA

T9147 CGGCGCGTCCGCGGGCGTGAAGG 1000
Oligo fwd CACCCGGCGCGTCCGCGGGCGTGA

| | | | | | | | | | | | |
|---|---|---|---|---|---|---|---|---|---|---|---|
| Oligo rev | AAACTCACGCCCGCGGACGCGCCG | | | | | | | | | | |
| Chromosome | start | end | strand | MM | target_seq | PAM | alignment | distance | position | gene name | gene id |
| GU305914.1 | 121704 | 121726 | + | 0 | CGGCGCGTCCGCGGGCGTGA | AGG | \|\|\|\|\|\|\|\|[\|\|\|\|\|\|\|\|\|\|\|\|]PAM | NA | Intergenic | NA | NA |
| T9148 | CGCGTCCGCGGGCGTGAAGGTGG | 1000 | | | | | | | | | |
| Oligo fwd | CACCCGCGTCCGCGGGCGTGAAGG | | | | | | | | | | |
| Oligo rev | AAACCCTTCACGCCCGCGGACGCG | | | | | | | | | | |
| Chromosome | start | end | strand | MM | target_seq | PAM | alignment | distance | position | gene name | gene id |
| GU305914.1 | 121707 | 121729 | + | 0 | CGCGTCCGCGGGCGTGAAGG | TGG | \|\|\|\|\|\|\|\|[\|\|\|\|\|\|\|\|\|\|\|\|]PAM | NA | Intergenic | NA | NA |
| GU305914.1 | 20502 | 20524 | - | 4 | CCCGACCGCGGGACTGAAGG | AGG | \|-\|\|-\|\|\|[\|\|\|\|--\|\|\|\|\|\|]PAM | NA | Intergenic | NA | NA |
| T9149 | TGAAGGTGGCGAAGTTCACCAGG | 1000 | | | | | | | | | |
| Oligo fwd | CACCTGAAGGTGGCGAAGTTCACC | | | | | | | | | | |
| Oligo rev | AAACGGTGAACTTCGCCACCTTCA | | | | | | | | | | |
| Chromosome | start | end | strand | MM | target_seq | PAM | alignment | distance | position | gene name | gene id |
| GU305914.1 | 121721 | 121743 | + | 0 | TGAAGGTGGCGAAGTTCACC | AGG | \|\|\|\|\|\|\|\|[\|\|\|\|\|\|\|\|\|\|\|\|]PAM | NA | Intergenic | NA | NA |
| GU305914.1 | 147306 | 147328 | - | 4 | CGATGGTGTCGAAGTTGACC | TGG | -\|\|-\|\|\|\|[-\|\|\|\|\|\|\|-\|\|\|]PAM | NA | Intergenic | NA | NA |
| T9150 | GTGGCGAAGTTCACCAGGTGTGG | 1000 | | | | | | | | | |
| Oligo fwd | CACCGTGGCGAAGTTCACCAGGTG | | | | | | | | | | |
| Oligo rev | AAACCACCTGGTGAACTTCGCCAC | | | | | | | | | | |
| Chromosome | start | end | strand | MM | target_seq | PAM | alignment | distance | position | gene name | gene id |
| GU305914.1 | 121726 | 121748 | + | 0 | GTGGCGAAGTTCACCAGGTG | TGG | \|\|\|\|\|\|\|\|[\|\|\|\|\|\|\|\|\|\|\|\|]PAM | NA | Intergenic | NA | NA |
| T9151 | TGGCGAAGTTCACCAGGTGTGGG | 1000 | | | | | | | | | |
| Oligo fwd | CACCTGGCGAAGTTCACCAGGTGT | | | | | | | | | | |
| Oligo rev | AAACACACCTGGTGAACTTCGCCA | | | | | | | | | | |
| Chromosome | start | end | strand | MM | target_seq | PAM | alignment | distance | position | gene name | gene id |
| GU305914.1 | 121727 | 121749 | + | 0 | TGGCGAAGTTCACCAGGTGT | GGG | \|\|\|\|\|\|\|\|[\|\|\|\|\|\|\|\|\|\|\|\|]PAM | NA | Intergenic | NA | NA |
| T9152 | CGAAGTTCACCAGGTGTGGGTGG | 1000 | | | | | | | | | |
| Oligo fwd | CACCCGAAGTTCACCAGGTGTGGG | | | | | | | | | | |
| Oligo rev | AAACCCCACACCTGGTGAACTTCG | | | | | | | | | | |
| Chromosome | start | end | strand | MM | target_seq | PAM | alignment | distance | position | gene name | gene id |
| GU305914.1 | 121730 | 121752 | + | 0 | CGAAGTTCACCAGGTGTGGG | TGG | \|\|\|\|\|\|\|\|[\|\|\|\|\|\|\|\|\|\|\|\|]PAM | NA | Intergenic | NA | NA |
| T9153 | ACCAGGTGTGGGTGGTTGTCCGG | 1000 | | | | | | | | | |
| Oligo fwd | CACCACCAGGTGTGGGTGGTTGTC | | | | | | | | | | |
| Oligo rev | AAACGACAACCACCCACACCTGGT | | | | | | | | | | |
| Chromosome | start | end | strand | MM | target_seq | PAM | alignment | distance | position | gene name | gene id |
| GU305914.1 | 121738 | 121760 | + | 0 | ACCAGGTGTGGGTGGTTGTC | CGG | \|\|\|\|\|\|\|\|[\|\|\|\|\|\|\|\|\|\|\|\|]PAM | NA | Intergenic | NA | NA |
| T9154 | CCAGGTGTGGGTGGTTGTCCGGG | 1000 | | | | | | | | | |
| Oligo fwd | CACCCCAGGTGTGGGTGGTTGTCC | | | | | | | | | | |
| Oligo rev | AAACGGACAACCACCCACACCTGG | | | | | | | | | | |
| Chromosome | start | end | strand | MM | target_seq | PAM | alignment | distance | position | gene name | gene id |
| GU305914.1 | 121739 | 121761 | + | 0 | CCAGGTGTGGGTGGTTGTCC | GGG | \|\|\|\|\|\|\|\|[\|\|\|\|\|\|\|\|\|\|\|\|]PAM | NA | Intergenic | NA | NA |
| T9155 | GTGGGTGGTTGTCCGGGTGCCGG | 1000 | | | | | | | | | |
| Oligo fwd | CACCGTGGGTGGTTGTCCGGGTGC | | | | | | | | | | |
| Oligo rev | AAACGCACCCGGACAACCACCCAC | | | | | | | | | | |
| Chromosome | start | end | strand | MM | target_seq | PAM | alignment | distance | position | gene name | gene id |
| GU305914.1 | 121745 | 121767 | + | 0 | GTGGGTGGTTGTCCGGGTGC | CGG | \|\|\|\|\|\|\|\|[\|\|\|\|\|\|\|\|\|\|\|\|]PAM | NA | Intergenic | NA | NA |
| T9156 | GGTACATCTTCCCCACACACAGG | 1000 | | | | | | | | | |
| Oligo fwd | CACCGGTACATCTTCCCCACACAC | | | | | | | | | | |
| Oligo rev | AAACGTGTGTGGGGAAGATGTACC | | | | | | | | | | |
| Chromosome | start | end | strand | MM | target_seq | PAM | alignment | distance | position | gene name | gene id |
| GU305914.1 | 121766 | 121788 | + | 0 | GGTACATCTTCCCCACACAC | AGG | \|\|\|\|\|\|\|\|[\|\|\|\|\|\|\|\|\|\|\|\|]PAM | NA | Intergenic | NA | NA |
| T9157 | CTTCCCCACACACAGGACGCAGG | 1000 | | | | | | | | | |
| Oligo fwd | CACCCTTCCCCACACACAGGACGC | | | | | | | | | | |
| Oligo rev | AAACGCGTCCTGTGTGTGGGGAAG | | | | | | | | | | |
| Chromosome | start | end | strand | MM | target_seq | PAM | alignment | distance | position | gene name | gene id |
| GU305914.1 | 121773 | 121795 | + | 0 | CTTCCCCACACACAGGACGC | AGG | \|\|\|\|\|\|\|\|[\|\|\|\|\|\|\|\|\|\|\|\|]PAM | NA | Intergenic | NA | NA |
| T9158 | TTCCCCACACACAGGACGCAGGG | 1000 | | | | | | | | | |
| Oligo fwd | CACCTTCCCCACACACAGGACGCA | | | | | | | | | | |
| Oligo rev | AAACTGCGTCCTGTGTGTGGGGAA | | | | | | | | | | |
| Chromosome | start | end | strand | MM | target_seq | PAM | alignment | distance | position | gene name | gene id |
| GU305914.1 | 121774 | 121796 | + | 0 | TTCCCCACACACAGGACGCA | GGG | \|\|\|\|\|\|\|\|[\|\|\|\|\|\|\|\|\|\|\|\|]PAM | NA | Intergenic | NA | NA |
| T9159 | CACAGGACGCAGGGCGCCTGCGG | 1000 | | | | | | | | | |
| Oligo fwd | CACCCACAGGACGCAGGGCGCCTG | | | | | | | | | | |
| Oligo rev | AAACCAGGCGCCCTGCGTCCTGTG | | | | | | | | | | |
| Chromosome | start | end | strand | MM | target_seq | PAM | alignment | distance | position | gene name | gene id |
| GU305914.1 | 121783 | 121805 | + | 0 | CACAGGACGCAGGGCGCCTG | CGG | \|\|\|\|\|\|\|\|[\|\|\|\|\|\|\|\|\|\|\|\|]PAM | NA | Intergenic | NA | NA |
| T9160 | AGGGCGCCTGCGGACTCAGCCGG | 1000 | | | | | | | | | |
| Oligo fwd | CACCAGGGCGCCTGCGGACTCAGC | | | | | | | | | | |
| Oligo rev | AAACGCTGAGTCCGCAGGCGCCCT | | | | | | | | | | |
| Chromosome | start | end | strand | MM | target_seq | PAM | alignment | distance | position | gene name | gene id |
| GU305914.1 | 121793 | 121815 | + | 0 | AGGGCGCCTGCGGACTCAGC | CGG | \|\|\|\|\|\|\|\|[\|\|\|\|\|\|\|\|\|\|\|\|]PAM | NA | Intergenic | NA | NA |
| T9161 | CTGCGGACTCAGCCGGATCCCGG | 1000 | | | | | | | | | |
| Oligo fwd | CACCCTGCGGACTCAGCCGGATCC | | | | | | | | | | |
| Oligo rev | AAACGGATCCGGCTGAGTCCGCAG | | | | | | | | | | |
| Chromosome | start | end | strand | MM | target_seq | PAM | alignment | distance | position | gene name | gene id |
| GU305914.1 | 121800 | 121822 | + | 0 | CTGCGGACTCAGCCGGATCC | CGG | \|\|\|\|\|\|\|\|[\|\|\|\|\|\|\|\|\|\|\|\|]PAM | NA | Intergenic | NA | NA |

| | | | | | | | | | | | |
|---|---|---|---|---|---|---|---|---|---|---|---|
| GU305914.1 | 115407 | 115429 | - | 4 | CGGCGGCCACGGCCGGATCC | AGG | \|-\|\|\|\|-\|[-\|-\|\|\|\|\|\|\|\|\|]PAM | NA | Intergenic | NA | NA |
| T9162 | GACTCAGCCGGATCCCGGTCTGG | 1000 | | | | | | | | | |
| Oligo fwd | CACCGACTCAGCCGGATCCCGGTC | | | | | | | | | | |
| Oligo rev | AAACGACCGGGATCCGGCTGAGTC | | | | | | | | | | |
| Chromosome | start | end | strand | MM | target_seq | PAM | alignment | distance | position | gene name | gene id |
| GU305914.1 | 121805 | 121827 | + | 0 | GACTCAGCCGGATCCCGGTC | TGG | \|\|\|\|\|\|\|\|[\|\|\|\|\|\|\|\|\|\|\|\|]PAM | NA | Intergenic | NA | NA |
| T9163 | TCAGCCGGATCCCGGTCTGGAGG | 1000 | | | | | | | | | |
| Oligo fwd | CACCTCAGCCGGATCCCGGTCTGG | | | | | | | | | | |
| Oligo rev | AAACCCAGACCGGGATCCGGCTGA | | | | | | | | | | |
| Chromosome | start | end | strand | MM | target_seq | PAM | alignment | distance | position | gene name | gene id |
| GU305914.1 | 121808 | 121830 | + | 0 | TCAGCCGGATCCCGGTCTGG | AGG | \|\|\|\|\|\|\|\|[\|\|\|\|\|\|\|\|\|\|\|\|]PAM | NA | Intergenic | NA | NA |
| T9164 | GATCCCGGTCTGGAGGAATTTGG | 1000 | | | | | | | | | |
| Oligo fwd | CACCGATCCCGGTCTGGAGGAATT | | | | | | | | | | |
| Oligo rev | AAACAATTCCTCCAGACCGGGATC | | | | | | | | | | |
| Chromosome | start | end | strand | MM | target_seq | PAM | alignment | distance | position | gene name | gene id |
| GU305914.1 | 121815 | 121837 | + | 0 | GATCCCGGTCTGGAGGAATT | TGG | \|\|\|\|\|\|\|\|[\|\|\|\|\|\|\|\|\|\|\|\|]PAM | NA | Intergenic | NA | NA |
| T9165 | CTGGAGGAATTTGGTCTCGTAGG | 1000 | | | | | | | | | |
| Oligo fwd | CACCCTGGAGGAATTTGGTCTCGT | | | | | | | | | | |
| Oligo rev | AAACACGAGACCAAATTCCTCCAG | | | | | | | | | | |
| Chromosome | start | end | strand | MM | target_seq | PAM | alignment | distance | position | gene name | gene id |
| GU305914.1 | 121824 | 121846 | + | 0 | CTGGAGGAATTTGGTCTCGT | AGG | \|\|\|\|\|\|\|\|[\|\|\|\|\|\|\|\|\|\|\|\|]PAM | NA | Intergenic | NA | NA |
| T9166 | TGGAGGAATTTGGTCTCGTAGGG | 1000 | | | | | | | | | |
| Oligo fwd | CACCTGGAGGAATTTGGTCTCGTA | | | | | | | | | | |
| Oligo rev | AAACTACGAGACCAAATTCCTCCA | | | | | | | | | | |
| Chromosome | start | end | strand | MM | target_seq | PAM | alignment | distance | position | gene name | gene id |
| GU305914.1 | 121825 | 121847 | + | 0 | TGGAGGAATTTGGTCTCGTA | GGG | \|\|\|\|\|\|\|\|[\|\|\|\|\|\|\|\|\|\|\|\|]PAM | NA | Intergenic | NA | NA |
| T9167 | GGAGGAATTTGGTCTCGTAGGGG | 1000 | | | | | | | | | |
| Oligo fwd | CACCGGAGGAATTTGGTCTCGTAG | | | | | | | | | | |
| Oligo rev | AAACCTACGAGACCAAATTCCTCC | | | | | | | | | | |
| Chromosome | start | end | strand | MM | target_seq | PAM | alignment | distance | position | gene name | gene id |
| GU305914.1 | 121826 | 121848 | + | 0 | GGAGGAATTTGGTCTCGTAG | GGG | \|\|\|\|\|\|\|\|[\|\|\|\|\|\|\|\|\|\|\|\|]PAM | NA | Intergenic | NA | NA |
| T9168 | GGGGCCGAAATCTACAGTCACGG | 1000 | | | | | | | | | |
| Oligo fwd | CACCGGGGCCGAAATCTACAGTCA | | | | | | | | | | |
| Oligo rev | AAACTGACTGTAGATTTCGGCCCC | | | | | | | | | | |
| Chromosome | start | end | strand | MM | target_seq | PAM | alignment | distance | position | gene name | gene id |
| GU305914.1 | 121845 | 121867 | + | 0 | GGGGCCGAAATCTACAGTCA | CGG | \|\|\|\|\|\|\|\|[\|\|\|\|\|\|\|\|\|\|\|\|]PAM | NA | Intergenic | NA | NA |
| T9169 | GGGCCGAAATCTACAGTCACGGG | 1000 | | | | | | | | | |
| Oligo fwd | CACCGGGCCGAAATCTACAGTCAC | | | | | | | | | | |
| Oligo rev | AAACGTGACTGTAGATTTCGGCCC | | | | | | | | | | |
| Chromosome | start | end | strand | MM | target_seq | PAM | alignment | distance | position | gene name | gene id |
| GU305914.1 | 121846 | 121868 | + | 0 | GGGCCGAAATCTACAGTCAC | GGG | \|\|\|\|\|\|\|\|[\|\|\|\|\|\|\|\|\|\|\|\|]PAM | NA | Intergenic | NA | NA |
| T9170 | GGCCGAAATCTACAGTCACGGGG | 1000 | | | | | | | | | |
| Oligo fwd | CACCGGCCGAAATCTACAGTCACG | | | | | | | | | | |
| Oligo rev | AAACCGTGACTGTAGATTTCGGCC | | | | | | | | | | |
| Chromosome | start | end | strand | MM | target_seq | PAM | alignment | distance | position | gene name | gene id |
| GU305914.1 | 121847 | 121869 | + | 0 | GGCCGAAATCTACAGTCACG | GGG | \|\|\|\|\|\|\|\|[\|\|\|\|\|\|\|\|\|\|\|\|]PAM | NA | Intergenic | NA | NA |
| T9171 | CGGGGACGTGTAGCTCCAGCAGG | 1000 | | | | | | | | | |
| Oligo fwd | CACCCGGGGACGTGTAGCTCCAGC | | | | | | | | | | |
| Oligo rev | AAACGCTGGAGCTACACGTCCCCG | | | | | | | | | | |
| Chromosome | start | end | strand | MM | target_seq | PAM | alignment | distance | position | gene name | gene id |
| GU305914.1 | 121865 | 121887 | + | 0 | CGGGGACGTGTAGCTCCAGC | AGG | \|\|\|\|\|\|\|\|[\|\|\|\|\|\|\|\|\|\|\|\|]PAM | NA | Intergenic | NA | NA |
| T9172 | AGACAGACTCCCCGCGAACAAGG | 1000 | | | | | | | | | |
| Oligo fwd | CACCAGACAGACTCCCCGCGAACA | | | | | | | | | | |
| Oligo rev | AAACTGTTCGCGGGGAGTCTGTCT | | | | | | | | | | |
| Chromosome | start | end | strand | MM | target_seq | PAM | alignment | distance | position | gene name | gene id |
| GU305914.1 | 121890 | 121912 | + | 0 | AGACAGACTCCCCGCGAACA | AGG | \|\|\|\|\|\|\|\|[\|\|\|\|\|\|\|\|\|\|\|\|]PAM | NA | Intergenic | NA | NA |
| T9173 | AACAAGGAGCTCGTGCCGCTCGG | 1000 | | | | | | | | | |
| Oligo fwd | CACCAACAAGGAGCTCGTGCCGCT | | | | | | | | | | |
| Oligo rev | AAACAGCGGCACGAGCTCCTTGTT | | | | | | | | | | |
| Chromosome | start | end | strand | MM | target_seq | PAM | alignment | distance | position | gene name | gene id |
| GU305914.1 | 121906 | 121928 | + | 0 | AACAAGGAGCTCGTGCCGCT | CGG | \|\|\|\|\|\|\|\|[\|\|\|\|\|\|\|\|\|\|\|\|]PAM | NA | Intergenic | NA | NA |
| T9174 | TTGACCGACATGTCGAACGAAGG | 1000 | | | | | | | | | |
| Oligo fwd | CACCTTGACCGACATGTCGAACGA | | | | | | | | | | |
| Oligo rev | AAACTCGTTCGACATGTCGGTCAA | | | | | | | | | | |
| Chromosome | start | end | strand | MM | target_seq | PAM | alignment | distance | position | gene name | gene id |
| GU305914.1 | 121939 | 121961 | + | 0 | TTGACCGACATGTCGAACGA | AGG | \|\|\|\|\|\|\|\|[\|\|\|\|\|\|\|\|\|\|\|\|]PAM | NA | Intergenic | NA | NA |
| T9175 | ACATGTCGAACGAAGGAAGAAGG | 1000 | | | | | | | | | |
| Oligo fwd | CACCACATGTCGAACGAAGGAAGA | | | | | | | | | | |
| Oligo rev | AAACTCTTCCTTCGTTCGACATGT | | | | | | | | | | |
| Chromosome | start | end | strand | MM | target_seq | PAM | alignment | distance | position | gene name | gene id |
| GU305914.1 | 121946 | 121968 | + | 0 | ACATGTCGAACGAAGGAAGA | AGG | \|\|\|\|\|\|\|\|[\|\|\|\|\|\|\|\|\|\|\|\|]PAM | NA | Intergenic | NA | NA |
| T9176 | GTCGAACGAAGGAAGAAGGAAGG | 1000 | | | | | | | | | |
| Oligo fwd | CACCGTCGAACGAAGGAAGAAGGA | | | | | | | | | | |
| Oligo rev | AAACTCCTTCTTCCTTCGTTCGAC | | | | | | | | | | |

| Chromosome | start | end | strand | MM | target_seq | PAM | alignment | distance | position | gene name | gene id |
|---|---|---|---|---|---|---|---|---|---|---|---|
| GU305914.1 | 121950 | 121972 | + | 0 | GTCGAACGAAGGAAGAAGGA | AGG | \|\|\|\|\|\|\|\|[\|\|\|\|\|\|\|\|\|\|\|\|]PAM | NA | Intergenic | NA | NA |
| T9177 | AACGAAGGAAGAAGGAAGGACGG | 1000 | | | | | | | | | |
| Oligo fwd | CACCAACGAAGGAAGAAGGAAGGA | | | | | | | | | | |
| Oligo rev | AAACTCCTTCCTTCTTCCTTCGTT | | | | | | | | | | |
| Chromosome | start | end | strand | MM | target_seq | PAM | alignment | distance | position | gene name | gene id |
| GU305914.1 | 121954 | 121976 | + | 0 | AACGAAGGAAGAAGGAAGGA | CGG | \|\|\|\|\|\|\|\|[\|\|\|\|\|\|\|\|\|\|\|\|]PAM | NA | Intergenic | NA | NA |
| GU305914.1 | 121958 | 121980 | + | 4 | AAGGAAGAAGGAAGGACGGA | CGG | \|\|-\|\|\|\|-[\|-\|\|\|\|\|\|-\|\|\|]PAM | NA | Intergenic | NA | NA |
| T9178 | AAGGAAGAAGGAAGGACGGACGG | 1000 | | | | | | | | | |
| Oligo fwd | CACCAAGGAAGAAGGAAGGACGGA | | | | | | | | | | |
| Oligo rev | AAACTCCGTCCTTCCTTCTTCCTT | | | | | | | | | | |
| Chromosome | start | end | strand | MM | target_seq | PAM | alignment | distance | position | gene name | gene id |
| GU305914.1 | 121958 | 121980 | + | 0 | AAGGAAGAAGGAAGGACGGA | CGG | \|\|\|\|\|\|\|\|[\|\|\|\|\|\|\|\|\|\|\|\|]PAM | NA | Intergenic | NA | NA |
| GU305914.1 | 121954 | 121976 | + | 4 | AACGAAGGAAGAAGGAAGGA | CGG | \|\|-\|\|\|\|-[\|-\|\|\|\|\|\|-\|\|\|]PAM | NA | Intergenic | NA | NA |
| T9179 | TCTATCCCTCTCTCCTTCTGCGG | 1000 | | | | | | | | | |
| Oligo fwd | CACCTCTATCCCTCTCTCCTTCTG | | | | | | | | | | |
| Oligo rev | AAACCAGAAGGAGAGAGGGATAGA | | | | | | | | | | |
| Chromosome | start | end | strand | MM | target_seq | PAM | alignment | distance | position | gene name | gene id |
| GU305914.1 | 122033 | 122055 | + | 0 | TCTATCCCTCTCTCCTTCTG | CGG | \|\|\|\|\|\|\|\|[\|\|\|\|\|\|\|\|\|\|\|\|]PAM | NA | Intergenic | NA | NA |
| T9180 | TCCCTCTCTCCTTCTGCGGACGG | 1000 | | | | | | | | | |
| Oligo fwd | CACCTCCCTCTCTCCTTCTGCGGA | | | | | | | | | | |
| Oligo rev | AAACTCCGCAGAAGGAGAGAGGGA | | | | | | | | | | |
| Chromosome | start | end | strand | MM | target_seq | PAM | alignment | distance | position | gene name | gene id |
| GU305914.1 | 122037 | 122059 | + | 0 | TCCCTCTCTCCTTCTGCGGA | CGG | \|\|\|\|\|\|\|\|[\|\|\|\|\|\|\|\|\|\|\|\|]PAM | NA | Intergenic | NA | NA |
| GU305914.1 | 212878 | 212900 | - | 4 | TCCCCTTCTCCTTCTGCCGT | TGG | \|\|\|\|--\|\|[\|\|\|\|\|\|\|\|\|-\|-]PAM | NA | Intergenic | NA | NA |
| T9181 | CTCTCTCCTTCTGCGGACGGAGG | 1000 | | | | | | | | | |
| Oligo fwd | CACCCTCTCTCCTTCTGCGGACGG | | | | | | | | | | |
| Oligo rev | AAACCCGTCCGCAGAAGGAGAGAG | | | | | | | | | | |
| Chromosome | start | end | strand | MM | target_seq | PAM | alignment | distance | position | gene name | gene id |
| GU305914.1 | 122040 | 122062 | + | 0 | CTCTCTCCTTCTGCGGACGG | AGG | \|\|\|\|\|\|\|\|[\|\|\|\|\|\|\|\|\|\|\|\|]PAM | NA | Intergenic | NA | NA |
| T9182 | TCTCCTTCTGCGGACGGAGGTGG | 1000 | | | | | | | | | |
| Oligo fwd | CACCTCTCCTTCTGCGGACGGAGG | | | | | | | | | | |
| Oligo rev | AAACCCTCCGTCCGCAGAAGGAGA | | | | | | | | | | |
| Chromosome | start | end | strand | MM | target_seq | PAM | alignment | distance | position | gene name | gene id |
| GU305914.1 | 122043 | 122065 | + | 0 | TCTCCTTCTGCGGACGGAGG | TGG | \|\|\|\|\|\|\|\|[\|\|\|\|\|\|\|\|\|\|\|\|]PAM | NA | Intergenic | NA | NA |
| T9183 | CCTTCTGCGGACGGAGGTGGCGG | 1000 | | | | | | | | | |
| Oligo fwd | CACCCCTTCTGCGGACGGAGGTGG | | | | | | | | | | |
| Oligo rev | AAACCCACCTCCGTCCGCAGAAGG | | | | | | | | | | |
| Chromosome | start | end | strand | MM | target_seq | PAM | alignment | distance | position | gene name | gene id |
| GU305914.1 | 122046 | 122068 | + | 0 | CCTTCTGCGGACGGAGGTGG | CGG | \|\|\|\|\|\|\|\|[\|\|\|\|\|\|\|\|\|\|\|\|]PAM | NA | Intergenic | NA | NA |
| T9184 | CTTCTGCGGACGGAGGTGGCGGG | 1000 | | | | | | | | | |
| Oligo fwd | CACCCTTCTGCGGACGGAGGTGGC | | | | | | | | | | |
| Oligo rev | AAACGCCACCTCCGTCCGCAGAAG | | | | | | | | | | |
| Chromosome | start | end | strand | MM | target_seq | PAM | alignment | distance | position | gene name | gene id |
| GU305914.1 | 122047 | 122069 | + | 0 | CTTCTGCGGACGGAGGTGGC | GGG | \|\|\|\|\|\|\|\|[\|\|\|\|\|\|\|\|\|\|\|\|]PAM | NA | Intergenic | NA | NA |
| T9185 | TTCTGCGGACGGAGGTGGCGGGG | 1000 | | | | | | | | | |
| Oligo fwd | CACCTTCTGCGGACGGAGGTGGCG | | | | | | | | | | |
| Oligo rev | AAACCGCCACCTCCGTCCGCAGAA | | | | | | | | | | |
| Chromosome | start | end | strand | MM | target_seq | PAM | alignment | distance | position | gene name | gene id |
| GU305914.1 | 122048 | 122070 | + | 0 | TTCTGCGGACGGAGGTGGCG | GGG | \|\|\|\|\|\|\|\|[\|\|\|\|\|\|\|\|\|\|\|\|]PAM | NA | Intergenic | NA | NA |
| T9186 | ACGGAGGTGGCGGGGATGTCTGG | 1000 | | | | | | | | | |
| Oligo fwd | CACCACGGAGGTGGCGGGGATGTC | | | | | | | | | | |
| Oligo rev | AAACGACATCCCCGCCACCTCCGT | | | | | | | | | | |
| Chromosome | start | end | strand | MM | target_seq | PAM | alignment | distance | position | gene name | gene id |
| GU305914.1 | 122056 | 122078 | + | 0 | ACGGAGGTGGCGGGGATGTC | TGG | \|\|\|\|\|\|\|\|[\|\|\|\|\|\|\|\|\|\|\|\|]PAM | NA | Intergenic | NA | NA |
| GU305914.1 | 109329 | 109351 | + | 2 | ACGGAGGTGGCGGGGACGGC | GGG | \|\|\|\|\|\|\|\|[\|\|\|\|\|\|\|\|-\|-\|]PAM | NA | Intergenic | NA | NA |
| T9187 | CGGAGGTGGCGGGGATGTCTGGG | 1000 | | | | | | | | | |
| Oligo fwd | CACCCGGAGGTGGCGGGGATGTCT | | | | | | | | | | |
| Oligo rev | AAACAGACATCCCCGCCACCTCCG | | | | | | | | | | |
| Chromosome | start | end | strand | MM | target_seq | PAM | alignment | distance | position | gene name | gene id |
| GU305914.1 | 122057 | 122079 | + | 0 | CGGAGGTGGCGGGGATGTCT | GGG | \|\|\|\|\|\|\|\|[\|\|\|\|\|\|\|\|\|\|\|\|]PAM | NA | Intergenic | NA | NA |
| T9188 | GGAGGTGGCGGGGATGTCTGGGG | 1000 | | | | | | | | | |
| Oligo fwd | CACCGGAGGTGGCGGGGATGTCTG | | | | | | | | | | |
| Oligo rev | AAACCAGACATCCCCGCCACCTCC | | | | | | | | | | |
| Chromosome | start | end | strand | MM | target_seq | PAM | alignment | distance | position | gene name | gene id |
| GU305914.1 | 122058 | 122080 | + | 0 | GGAGGTGGCGGGGATGTCTG | GGG | \|\|\|\|\|\|\|\|[\|\|\|\|\|\|\|\|\|\|\|\|]PAM | NA | Intergenic | NA | NA |
| GU305914.1 | 35665 | 35687 | + | 4 | GGCGGCGGCGGGGATGTCAT | CGG | \|\|-\|\|-\|\|[\|\|\|\|\|\|\|\|\|\|--]PAM | NA | Intergenic | NA | NA |
| T9189 | GGCGGGGATGTCTGGGGTTTCGG | 1000 | | | | | | | | | |
| Oligo fwd | CACCGGCGGGGATGTCTGGGGTTT | | | | | | | | | | |
| Oligo rev | AAACAAACCCCAGACATCCCCGCC | | | | | | | | | | |
| Chromosome | start | end | strand | MM | target_seq | PAM | alignment | distance | position | gene name | gene id |
| GU305914.1 | 122064 | 122086 | + | 0 | GGCGGGGATGTCTGGGGTTT | CGG | \|\|\|\|\|\|\|\|[\|\|\|\|\|\|\|\|\|\|\|\|]PAM | NA | Intergenic | NA | NA |
| T9190 | CGCGCACTTTCTCTTATAAGTGG | 1000 | | | | | | | | | |
| Oligo fwd | CACCCGCGCACTTTCTCTTATAAG | | | | | | | | | | |
| Oligo rev | AAACCTTATAAGAGAAAGTGCGCG | | | | | | | | | | |

Chromosome start end strand MM target_seq PAM alignment distance position gene name gene id
GU305914.1 122093 122115 + 0 CGCGCACTTTCTCTTATAAG TGG ||||||||[||||||||||||]PAM NA Intergenic NA NA

T9191 ATAAGTGGTTAGCGCTACGCTGG 1000
Oligo fwd CACCATAAGTGGTTAGCGCTACGC
Oligo rev AAACGCGTAGCGCTAACCACTTAT
Chromosome start end strand MM target_seq PAM alignment distance position gene name gene id
GU305914.1 122108 122130 + 0 ATAAGTGGTTAGCGCTACGC TGG ||||||||[||||||||||||]PAM NA Intergenic NA NA

T9192 TAAGTGGTTAGCGCTACGCTGGG 1000
Oligo fwd CACCTAAGTGGTTAGCGCTACGCT
Oligo rev AAACAGCGTAGCGCTAACCACTTA
Chromosome start end strand MM target_seq PAM alignment distance position gene name gene id
GU305914.1 122109 122131 + 0 TAAGTGGTTAGCGCTACGCT GGG ||||||||[||||||||||||]PAM NA Intergenic NA NA

T9193 GTGGTTAGCGCTACGCTGGGCGG 1000
Oligo fwd CACCGTGGTTAGCGCTACGCTGGG
Oligo rev AAACCCCAGCGTAGCGCTAACCAC
Chromosome start end strand MM target_seq PAM alignment distance position gene name gene id
GU305914.1 122112 122134 + 0 GTGGTTAGCGCTACGCTGGG CGG ||||||||[||||||||||||]PAM NA Intergenic NA NA

T9194 TGGTTAGCGCTACGCTGGGCGGG 1000
Oligo fwd CACCTGGTTAGCGCTACGCTGGGC
Oligo rev AAACGCCCAGCGTAGCGCTAACCA
Chromosome start end strand MM target_seq PAM alignment distance position gene name gene id
GU305914.1 122113 122135 + 0 TGGTTAGCGCTACGCTGGGC GGG ||||||||[||||||||||||]PAM NA Intergenic NA NA

T9195 TCTCAGACATACATAGATATAGG 1000
Oligo fwd CACCTCTCAGACATACATAGATAT
Oligo rev AAACATATCTATGTATGTCTGAGA
Chromosome start end strand MM target_seq PAM alignment distance position gene name gene id
GU305914.1 122143 122165 + 0 TCTCAGACATACATAGATAT AGG ||||||||[||||||||||||]PAM NA Intergenic NA NA

T9196 TACATAGATATAGGTGCATGAGG 1000
Oligo fwd CACCTACATAGATATAGGTGCATG
Oligo rev AAACCATGCACCTATATCTATGTA
Chromosome start end strand MM target_seq PAM alignment distance position gene name gene id
GU305914.1 122152 122174 + 0 TACATAGATATAGGTGCATG AGG ||||||||[||||||||||||]PAM NA Intergenic NA NA

T9197 GTGCATGAGGAAAACGTAGACGG 1000
Oligo fwd CACCGTGCATGAGGAAAACGTAGA
Oligo rev AAACTCTACGTTTTCCTCATGCAC
Chromosome start end strand MM target_seq PAM alignment distance position gene name gene id
GU305914.1 122165 122187 + 0 GTGCATGAGGAAAACGTAGA CGG ||||||||[||||||||||||]PAM NA Intergenic NA NA

T9198 GAAAACGTAGACGGTCCGCTCGG 1000
Oligo fwd CACCGAAAACGTAGACGGTCCGCT
Oligo rev AAACAGCGGACCGTCTACGTTTTC
Chromosome start end strand MM target_seq PAM alignment distance position gene name gene id
GU305914.1 122174 122196 + 0 GAAAACGTAGACGGTCCGCT CGG ||||||||[||||||||||||]PAM NA Intergenic NA NA

T9199 AAAACGTAGACGGTCCGCTCGGG 1000
Oligo fwd CACCAAAACGTAGACGGTCCGCTC
Oligo rev AAACGAGCGGACCGTCTACGTTTT
Chromosome start end strand MM target_seq PAM alignment distance position gene name gene id
GU305914.1 122175 122197 + 0 AAAACGTAGACGGTCCGCTC GGG ||||||||[||||||||||||]PAM NA Intergenic NA NA

T9200 AAACGTAGACGGTCCGCTCGGGG 1000
Oligo fwd CACCAAACGTAGACGGTCCGCTCG
Oligo rev AAACCGAGCGGACCGTCTACGTTT
Chromosome start end strand MM target_seq PAM alignment distance position gene name gene id
GU305914.1 122176 122198 + 0 AAACGTAGACGGTCCGCTCG GGG ||||||||[||||||||||||]PAM NA Intergenic NA NA

T9201 AACGTAGACGGTCCGCTCGGGGG 1000
Oligo fwd CACCAACGTAGACGGTCCGCTCGG
Oligo rev AAACCCGAGCGGACCGTCTACGTT
Chromosome start end strand MM target_seq PAM alignment distance position gene name gene id
GU305914.1 122177 122199 + 0 AACGTAGACGGTCCGCTCGG GGG ||||||||[||||||||||||]PAM NA Intergenic NA NA

T9202 TAGACGGTCCGCTCGGGGGTTGG 1000
Oligo fwd CACCTAGACGGTCCGCTCGGGGGT
Oligo rev AAACACCCCCGAGCGGACCGTCTA
Chromosome start end strand MM target_seq PAM alignment distance position gene name gene id
GU305914.1 122181 122203 + 0 TAGACGGTCCGCTCGGGGGT TGG ||||||||[||||||||||||]PAM NA Intergenic NA NA

T9203 AGACGGTCCGCTCGGGGGTTGGG 1000
Oligo fwd CACCAGACGGTCCGCTCGGGGGTT
Oligo rev AAACAACCCCCGAGCGGACCGTCT
Chromosome start end strand MM target_seq PAM alignment distance position gene name gene id
GU305914.1 122182 122204 + 0 AGACGGTCCGCTCGGGGGTT GGG ||||||||[||||||||||||]PAM NA Intergenic NA NA

T9204 TACGACGTCGTTTTTCCTCGCGG 1000
Oligo fwd CACCTACGACGTCGTTTTTCCTCG
Oligo rev AAACCGAGGAAAAACGACGTCGTA
Chromosome start end strand MM target_seq PAM alignment distance position gene name gene id
GU305914.1 122206 122228 + 0 TACGACGTCGTTTTTCCTCG CGG ||||||||[||||||||||||]PAM NA Intergenic NA NA

T9205 TTTCCTCGCGGTGTTGTCGAAGG 1000
Oligo fwd CACCTTTCCTCGCGGTGTTGTCGA

Oligo rev AAACTCGACAACACCGCGAGGAAA
Chromosome start end strand MM target_seq PAM alignment distance position gene name gene id
GU305914.1 122218 122240 + 0 TTTCCTCGCGGTGTTGTCGA AGG ||||||||[||||||||||||]PAM NA Intergenic NA NA

T9206 CTCGCGGTGTTGTCGAAGGTCGG 1000
Oligo fwd CACCCTCGCGGTGTTGTCGAAGGT
Oligo rev AAACACCTTCGACAACACCGCGAG
Chromosome start end strand MM target_seq PAM alignment distance position gene name gene id
GU305914.1 122222 122244 + 0 CTCGCGGTGTTGTCGAAGGT CGG ||||||||[||||||||||||]PAM NA Intergenic NA NA

T9207 TCGCGGTGTTGTCGAAGGTCGGG 1000
Oligo fwd CACCTCGCGGTGTTGTCGAAGGTC
Oligo rev AAACGACCTTCGACAACACCGCGA
Chromosome start end strand MM target_seq PAM alignment distance position gene name gene id
GU305914.1 122223 122245 + 0 TCGCGGTGTTGTCGAAGGTC GGG ||||||||[||||||||||||]PAM NA Intergenic NA NA

T9208 CGGTGTTGTCGAAGGTCGGGAGG 1000
Oligo fwd CACCCGGTGTTGTCGAAGGTCGGG
Oligo rev AAACCCCGACCTTCGACAACACCG
Chromosome start end strand MM target_seq PAM alignment distance position gene name gene id
GU305914.1 122226 122248 + 0 CGGTGTTGTCGAAGGTCGGG AGG ||||||||[||||||||||||]PAM NA Intergenic NA NA

T9209 GTTGTCGAAGGTCGGGAGGTCGG 1000
Oligo fwd CACCGTTGTCGAAGGTCGGGAGGT
Oligo rev AAACACCTCCCGACCTTCGACAAC
Chromosome start end strand MM target_seq PAM alignment distance position gene name gene id
GU305914.1 122230 122252 + 0 GTTGTCGAAGGTCGGGAGGT CGG ||||||||[||||||||||||]PAM NA Intergenic NA NA

T9210 TTGTCGAAGGTCGGGAGGTCGGG 1000
Oligo fwd CACCTTGTCGAAGGTCGGGAGGTC
Oligo rev AAACGACCTCCCGACCTTCGACAA
Chromosome start end strand MM target_seq PAM alignment distance position gene name gene id
GU305914.1 122231 122253 + 0 TTGTCGAAGGTCGGGAGGTC GGG ||||||||[||||||||||||]PAM NA Intergenic NA NA

T9211 TCGAAGGTCGGGAGGTCGGGAGG 1000
Oligo fwd CACCTCGAAGGTCGGGAGGTCGGG
Oligo rev AAACCCCGACCTCCCGACCTTCGA
Chromosome start end strand MM target_seq PAM alignment distance position gene name gene id
GU305914.1 122234 122256 + 0 TCGAAGGTCGGGAGGTCGGG AGG ||||||||[||||||||||||]PAM NA Intergenic NA NA

T9212 CGAAGGTCGGGAGGTCGGGAGGG 1000
Oligo fwd CACCCGAAGGTCGGGAGGTCGGGA
Oligo rev AAACTCCCGACCTCCCGACCTTCG
Chromosome start end strand MM target_seq PAM alignment distance position gene name gene id
GU305914.1 122235 122257 + 0 CGAAGGTCGGGAGGTCGGGA GGG ||||||||[||||||||||||]PAM NA Intergenic NA NA

T9213 GAAGGTCGGGAGGTCGGGAGGGG 1000
Oligo fwd CACCGAAGGTCGGGAGGTCGGGAG
Oligo rev AAACCTCCCGACCTCCCGACCTTC
Chromosome start end strand MM target_seq PAM alignment distance position gene name gene id
GU305914.1 122236 122258 + 0 GAAGGTCGGGAGGTCGGGAG GGG ||||||||[||||||||||||]PAM NA Intergenic NA NA

T9214 AAGGTCGGGAGGTCGGGAGGGGG 1000
Oligo fwd CACCAAGGTCGGGAGGTCGGGAGG
Oligo rev AAACCCTCCCGACCTCCCGACCTT
Chromosome start end strand MM target_seq PAM alignment distance position gene name gene id
GU305914.1 122237 122259 + 0 AAGGTCGGGAGGTCGGGAGG GGG ||||||||[||||||||||||]PAM NA Intergenic NA NA

T9215 TCTGTCTGTCCGTCCGCTGCCGG 1000
Oligo fwd CACCTCTGTCTGTCCGTCCGCTGC
Oligo rev AAACGCAGCGGACGGACAGACAGA
Chromosome start end strand MM target_seq PAM alignment distance position gene name gene id
GU305914.1 122268 122290 + 0 TCTGTCTGTCCGTCCGCTGC CGG ||||||||[||||||||||||]PAM NA Intergenic NA NA

T9216 CTCAACCCTTGTAGATGCACAGG 1000
Oligo fwd CACCCTCAACCCTTGTAGATGCAC
Oligo rev AAACGTGCATCTACAAGGGTTGAG
Chromosome start end strand MM target_seq PAM alignment distance position gene name gene id
GU305914.1 122292 122314 + 0 CTCAACCCTTGTAGATGCAC AGG ||||||||[||||||||||||]PAM NA Intergenic NA NA

T9217 TCAACCCTTGTAGATGCACAGGG 1000
Oligo fwd CACCTCAACCCTTGTAGATGCACA
Oligo rev AAACTGTGCATCTACAAGGGTTGA
Chromosome start end strand MM target_seq PAM alignment distance position gene name gene id
GU305914.1 122293 122315 + 0 TCAACCCTTGTAGATGCACA GGG ||||||||[||||||||||||]PAM NA Intergenic NA NA

T9218 GCACAGGGCCGATTTGTCGTCGG 1000
Oligo fwd CACCGCACAGGGCCGATTTGTCGT
Oligo rev AAACACGACAAATCGGCCCTGTGC
Chromosome start end strand MM target_seq PAM alignment distance position gene name gene id
GU305914.1 122308 122330 + 0 GCACAGGGCCGATTTGTCGT CGG ||||||||[||||||||||||]PAM NA Intergenic NA NA

T9219 CACAGGGCCGATTTGTCGTCGGG 1000
Oligo fwd CACCCACAGGGCCGATTTGTCGTC
Oligo rev AAACGACGACAAATCGGCCCTGTG
Chromosome start end strand MM target_seq PAM alignment distance position gene name gene id
GU305914.1 122309 122331 + 0 CACAGGGCCGATTTGTCGTC GGG ||||||||[||||||||||||]PAM NA Intergenic NA NA

T9220 ACAGGGCCGATTTGTCGTCGGGG 1000

<table>
<tr><td>Oligo fwd</td><td>CACCACAGGGCCGATTTGTCGTCG</td><td></td><td></td><td></td><td></td><td></td><td></td><td></td><td></td><td></td><td></td></tr>
<tr><td>Oligo rev</td><td>AAACCGACGACAAATCGGCCCTGT</td><td></td><td></td><td></td><td></td><td></td><td></td><td></td><td></td><td></td><td></td></tr>
<tr><td>Chromosome</td><td>start</td><td>end</td><td>strand</td><td>MM</td><td>target_seq</td><td>PAM</td><td>alignment</td><td>distance</td><td>position</td><td>gene name</td><td>gene id</td></tr>
<tr><td>GU305914.1</td><td>122310</td><td>122332</td><td>+</td><td>0</td><td>ACAGGGCCGATTTGTCGTCG</td><td>GGG</td><td>||||||||[||||||||||||]PAM</td><td>NA</td><td>Intergenic</td><td>NA</td><td>NA</td></tr>
<tr><td>T9221</td><td>CAGGGCCGATTTGTCGTCGGGGG</td><td>1000</td><td></td><td></td><td></td><td></td><td></td><td></td><td></td><td></td><td></td></tr>
<tr><td>Oligo fwd</td><td>CACCCAGGGCCGATTTGTCGTCGG</td><td></td><td></td><td></td><td></td><td></td><td></td><td></td><td></td><td></td><td></td></tr>
<tr><td>Oligo rev</td><td>AAACCCGACGACAAATCGGCCCTG</td><td></td><td></td><td></td><td></td><td></td><td></td><td></td><td></td><td></td><td></td></tr>
<tr><td>Chromosome</td><td>start</td><td>end</td><td>strand</td><td>MM</td><td>target_seq</td><td>PAM</td><td>alignment</td><td>distance</td><td>position</td><td>gene name</td><td>gene id</td></tr>
<tr><td>GU305914.1</td><td>122311</td><td>122333</td><td>+</td><td>0</td><td>CAGGGCCGATTTGTCGTCGG</td><td>GGG</td><td>||||||||[||||||||||||]PAM</td><td>NA</td><td>Intergenic</td><td>NA</td><td>NA</td></tr>
<tr><td>T9222</td><td>CGGGGGACACGTCGCAGACCAGG</td><td>1000</td><td></td><td></td><td></td><td></td><td></td><td></td><td></td><td></td><td></td></tr>
<tr><td>Oligo fwd</td><td>CACCCGGGGGACACGTCGCAGACC</td><td></td><td></td><td></td><td></td><td></td><td></td><td></td><td></td><td></td><td></td></tr>
<tr><td>Oligo rev</td><td>AAACGGTCTGCGACGTGTCCCCCG</td><td></td><td></td><td></td><td></td><td></td><td></td><td></td><td></td><td></td><td></td></tr>
<tr><td>Chromosome</td><td>start</td><td>end</td><td>strand</td><td>MM</td><td>target_seq</td><td>PAM</td><td>alignment</td><td>distance</td><td>position</td><td>gene name</td><td>gene id</td></tr>
<tr><td>GU305914.1</td><td>122328</td><td>122350</td><td>+</td><td>0</td><td>CGGGGGACACGTCGCAGACC</td><td>AGG</td><td>||||||||[||||||||||||]PAM</td><td>NA</td><td>Intergenic</td><td>NA</td><td>NA</td></tr>
<tr><td>T9223</td><td>CACGTCGCAGACCAGGAACGCGG</td><td>1000</td><td></td><td></td><td></td><td></td><td></td><td></td><td></td><td></td><td></td></tr>
<tr><td>Oligo fwd</td><td>CACCCACGTCGCAGACCAGGAACG</td><td></td><td></td><td></td><td></td><td></td><td></td><td></td><td></td><td></td><td></td></tr>
<tr><td>Oligo rev</td><td>AAACCGTTCCTGGTCTGCGACGTG</td><td></td><td></td><td></td><td></td><td></td><td></td><td></td><td></td><td></td><td></td></tr>
<tr><td>Chromosome</td><td>start</td><td>end</td><td>strand</td><td>MM</td><td>target_seq</td><td>PAM</td><td>alignment</td><td>distance</td><td>position</td><td>gene name</td><td>gene id</td></tr>
<tr><td>GU305914.1</td><td>122335</td><td>122357</td><td>+</td><td>0</td><td>CACGTCGCAGACCAGGAACG</td><td>CGG</td><td>||||||||[||||||||||||]PAM</td><td>NA</td><td>Intergenic</td><td>NA</td><td>NA</td></tr>
<tr><td>T9224</td><td>ACGTCGCAGACCAGGAACGCGGG</td><td>1000</td><td></td><td></td><td></td><td></td><td></td><td></td><td></td><td></td><td></td></tr>
<tr><td>Oligo fwd</td><td>CACCACGTCGCAGACCAGGAACGC</td><td></td><td></td><td></td><td></td><td></td><td></td><td></td><td></td><td></td><td></td></tr>
<tr><td>Oligo rev</td><td>AAACGCGTTCCTGGTCTGCGACGT</td><td></td><td></td><td></td><td></td><td></td><td></td><td></td><td></td><td></td><td></td></tr>
<tr><td>Chromosome</td><td>start</td><td>end</td><td>strand</td><td>MM</td><td>target_seq</td><td>PAM</td><td>alignment</td><td>distance</td><td>position</td><td>gene name</td><td>gene id</td></tr>
<tr><td>GU305914.1</td><td>122336</td><td>122358</td><td>+</td><td>0</td><td>ACGTCGCAGACCAGGAACGC</td><td>GGG</td><td>||||||||[||||||||||||]PAM</td><td>NA</td><td>Intergenic</td><td>NA</td><td>NA</td></tr>
<tr><td>T9225</td><td>GACCAGGAACGCGGGCTGCTCGG</td><td>1000</td><td></td><td></td><td></td><td></td><td></td><td></td><td></td><td></td><td></td></tr>
<tr><td>Oligo fwd</td><td>CACCGACCAGGAACGCGGGCTGCT</td><td></td><td></td><td></td><td></td><td></td><td></td><td></td><td></td><td></td><td></td></tr>
<tr><td>Oligo rev</td><td>AAACAGCAGCCCGCGTTCCTGGTC</td><td></td><td></td><td></td><td></td><td></td><td></td><td></td><td></td><td></td><td></td></tr>
<tr><td>Chromosome</td><td>start</td><td>end</td><td>strand</td><td>MM</td><td>target_seq</td><td>PAM</td><td>alignment</td><td>distance</td><td>position</td><td>gene name</td><td>gene id</td></tr>
<tr><td>GU305914.1</td><td>122344</td><td>122366</td><td>+</td><td>0</td><td>GACCAGGAACGCGGGCTGCT</td><td>CGG</td><td>||||||||[||||||||||||]PAM</td><td>NA</td><td>Intergenic</td><td>NA</td><td>NA</td></tr>
<tr><td>T9226</td><td>AGATCACGTCCGTGATCTGCCGG</td><td>1000</td><td></td><td></td><td></td><td></td><td></td><td></td><td></td><td></td><td></td></tr>
<tr><td>Oligo fwd</td><td>CACCAGATCACGTCCGTGATCTGC</td><td></td><td></td><td></td><td></td><td></td><td></td><td></td><td></td><td></td><td></td></tr>
<tr><td>Oligo rev</td><td>AAACGCAGATCACGGACGTGATCT</td><td></td><td></td><td></td><td></td><td></td><td></td><td></td><td></td><td></td><td></td></tr>
<tr><td>Chromosome</td><td>start</td><td>end</td><td>strand</td><td>MM</td><td>target_seq</td><td>PAM</td><td>alignment</td><td>distance</td><td>position</td><td>gene name</td><td>gene id</td></tr>
<tr><td>GU305914.1</td><td>122367</td><td>122389</td><td>+</td><td>0</td><td>AGATCACGTCCGTGATCTGC</td><td>CGG</td><td>||||||||[||||||||||||]PAM</td><td>NA</td><td>Intergenic</td><td>NA</td><td>NA</td></tr>
<tr><td>T9227</td><td>TCTGCCGGATCATGACGAACAGG</td><td>1000</td><td></td><td></td><td></td><td></td><td></td><td></td><td></td><td></td><td></td></tr>
<tr><td>Oligo fwd</td><td>CACCTCTGCCGGATCATGACGAAC</td><td></td><td></td><td></td><td></td><td></td><td></td><td></td><td></td><td></td><td></td></tr>
<tr><td>Oligo rev</td><td>AAACGTTCGTCATGATCCGGCAGA</td><td></td><td></td><td></td><td></td><td></td><td></td><td></td><td></td><td></td><td></td></tr>
<tr><td>Chromosome</td><td>start</td><td>end</td><td>strand</td><td>MM</td><td>target_seq</td><td>PAM</td><td>alignment</td><td>distance</td><td>position</td><td>gene name</td><td>gene id</td></tr>
<tr><td>GU305914.1</td><td>122382</td><td>122404</td><td>+</td><td>0</td><td>TCTGCCGGATCATGACGAAC</td><td>AGG</td><td>||||||||[||||||||||||]PAM</td><td>NA</td><td>Intergenic</td><td>NA</td><td>NA</td></tr>
<tr><td>T9228</td><td>CTGCCGGATCATGACGAACAGGG</td><td>1000</td><td></td><td></td><td></td><td></td><td></td><td></td><td></td><td></td><td></td></tr>
<tr><td>Oligo fwd</td><td>CACCCTGCCGGATCATGACGAACA</td><td></td><td></td><td></td><td></td><td></td><td></td><td></td><td></td><td></td><td></td></tr>
<tr><td>Oligo rev</td><td>AAACTGTTCGTCATGATCCGGCAG</td><td></td><td></td><td></td><td></td><td></td><td></td><td></td><td></td><td></td><td></td></tr>
<tr><td>Chromosome</td><td>start</td><td>end</td><td>strand</td><td>MM</td><td>target_seq</td><td>PAM</td><td>alignment</td><td>distance</td><td>position</td><td>gene name</td><td>gene id</td></tr>
<tr><td>GU305914.1</td><td>122383</td><td>122405</td><td>+</td><td>0</td><td>CTGCCGGATCATGACGAACA</td><td>GGG</td><td>||||||||[||||||||||||]PAM</td><td>NA</td><td>Intergenic</td><td>NA</td><td>NA</td></tr>
<tr><td>T9229</td><td>CCGGATCATGACGAACAGGGCGG</td><td>1000</td><td></td><td></td><td></td><td></td><td></td><td></td><td></td><td></td><td></td></tr>
<tr><td>Oligo fwd</td><td>CACCCCGGATCATGACGAACAGGG</td><td></td><td></td><td></td><td></td><td></td><td></td><td></td><td></td><td></td><td></td></tr>
<tr><td>Oligo rev</td><td>AAACCCCTGTTCGTCATGATCCGG</td><td></td><td></td><td></td><td></td><td></td><td></td><td></td><td></td><td></td><td></td></tr>
<tr><td>Chromosome</td><td>start</td><td>end</td><td>strand</td><td>MM</td><td>target_seq</td><td>PAM</td><td>alignment</td><td>distance</td><td>position</td><td>gene name</td><td>gene id</td></tr>
<tr><td>GU305914.1</td><td>122386</td><td>122408</td><td>+</td><td>0</td><td>CCGGATCATGACGAACAGGG</td><td>CGG</td><td>||||||||[||||||||||||]PAM</td><td>NA</td><td>Intergenic</td><td>NA</td><td>NA</td></tr>
<tr><td>T9230</td><td>GGTGAGTTTGTTGAGCTCCTCGG</td><td>1000</td><td></td><td></td><td></td><td></td><td></td><td></td><td></td><td></td><td></td></tr>
<tr><td>Oligo fwd</td><td>CACCGGTGAGTTTGTTGAGCTCCT</td><td></td><td></td><td></td><td></td><td></td><td></td><td></td><td></td><td></td><td></td></tr>
<tr><td>Oligo rev</td><td>AAACAGGAGCTCAACAAACTCACC</td><td></td><td></td><td></td><td></td><td></td><td></td><td></td><td></td><td></td><td></td></tr>
<tr><td>Chromosome</td><td>start</td><td>end</td><td>strand</td><td>MM</td><td>target_seq</td><td>PAM</td><td>alignment</td><td>distance</td><td>position</td><td>gene name</td><td>gene id</td></tr>
<tr><td>GU305914.1</td><td>122407</td><td>122429</td><td>+</td><td>0</td><td>GGTGAGTTTGTTGAGCTCCT</td><td>CGG</td><td>||||||||[||||||||||||]PAM</td><td>NA</td><td>Intergenic</td><td>NA</td><td>NA</td></tr>
<tr><td>T9231</td><td>GAGTTTGTTGAGCTCCTCGGTGG</td><td>1000</td><td></td><td></td><td></td><td></td><td></td><td></td><td></td><td></td><td></td></tr>
<tr><td>Oligo fwd</td><td>CACCGAGTTTGTTGAGCTCCTCGG</td><td></td><td></td><td></td><td></td><td></td><td></td><td></td><td></td><td></td><td></td></tr>
<tr><td>Oligo rev</td><td>AAACCCGAGGAGCTCAACAAACTC</td><td></td><td></td><td></td><td></td><td></td><td></td><td></td><td></td><td></td><td></td></tr>
<tr><td>Chromosome</td><td>start</td><td>end</td><td>strand</td><td>MM</td><td>target_seq</td><td>PAM</td><td>alignment</td><td>distance</td><td>position</td><td>gene name</td><td>gene id</td></tr>
<tr><td>GU305914.1</td><td>122410</td><td>122432</td><td>+</td><td>0</td><td>GAGTTTGTTGAGCTCCTCGG</td><td>TGG</td><td>||||||||[||||||||||||]PAM</td><td>NA</td><td>Intergenic</td><td>NA</td><td>NA</td></tr>
<tr><td>T9232</td><td>AGTTTGTTGAGCTCCTCGGTGGG</td><td>1000</td><td></td><td></td><td></td><td></td><td></td><td></td><td></td><td></td><td></td></tr>
<tr><td>Oligo fwd</td><td>CACCAGTTTGTTGAGCTCCTCGGT</td><td></td><td></td><td></td><td></td><td></td><td></td><td></td><td></td><td></td><td></td></tr>
<tr><td>Oligo rev</td><td>AAACACCGAGGAGCTCAACAAACT</td><td></td><td></td><td></td><td></td><td></td><td></td><td></td><td></td><td></td><td></td></tr>
<tr><td>Chromosome</td><td>start</td><td>end</td><td>strand</td><td>MM</td><td>target_seq</td><td>PAM</td><td>alignment</td><td>distance</td><td>position</td><td>gene name</td><td>gene id</td></tr>
<tr><td>GU305914.1</td><td>122411</td><td>122433</td><td>+</td><td>0</td><td>AGTTTGTTGAGCTCCTCGGT</td><td>GGG</td><td>||||||||[||||||||||||]PAM</td><td>NA</td><td>Intergenic</td><td>NA</td><td>NA</td></tr>
<tr><td>T9233</td><td>GTTTGTTGAGCTCCTCGGTGGGG</td><td>1000</td><td></td><td></td><td></td><td></td><td></td><td></td><td></td><td></td><td></td></tr>
<tr><td>Oligo fwd</td><td>CACCGTTTGTTGAGCTCCTCGGTG</td><td></td><td></td><td></td><td></td><td></td><td></td><td></td><td></td><td></td><td></td></tr>
<tr><td>Oligo rev</td><td>AAACCACCGAGGAGCTCAACAAAC</td><td></td><td></td><td></td><td></td><td></td><td></td><td></td><td></td><td></td><td></td></tr>
<tr><td>Chromosome</td><td>start</td><td>end</td><td>strand</td><td>MM</td><td>target_seq</td><td>PAM</td><td>alignment</td><td>distance</td><td>position</td><td>gene name</td><td>gene id</td></tr>
<tr><td>GU305914.1</td><td>122412</td><td>122434</td><td>+</td><td>0</td><td>GTTTGTTGAGCTCCTCGGTG</td><td>GGG</td><td>||||||||[||||||||||||]PAM</td><td>NA</td><td>Intergenic</td><td>NA</td><td>NA</td></tr>
<tr><td>T9234</td><td>TTTGTTGAGCTCCTCGGTGGGGG</td><td>1000</td><td></td><td></td><td></td><td></td><td></td><td></td><td></td><td></td><td></td></tr>
<tr><td>Oligo fwd</td><td>CACCTTTGTTGAGCTCCTCGGTGG</td><td></td><td></td><td></td><td></td><td></td><td></td><td></td><td></td><td></td><td></td></tr>
<tr><td>Oligo rev</td><td>AAACCCACCGAGGAGCTCAACAAA</td><td></td><td></td><td></td><td></td><td></td><td></td><td></td><td></td><td></td><td></td></tr>
<tr><td>Chromosome</td><td>start</td><td>end</td><td>strand</td><td>MM</td><td>target_seq</td><td>PAM</td><td>alignment</td><td>distance</td><td>position</td><td>gene name</td><td>gene id</td></tr>
<tr><td>GU305914.1</td><td>122413</td><td>122435</td><td>+</td><td>0</td><td>TTTGTTGAGCTCCTCGGTGG</td><td>GGG</td><td>||||||||[||||||||||||]PAM</td><td>NA</td><td>Intergenic</td><td>NA</td><td>NA</td></tr>
</table>

<table>
<tr><td>T9235</td><td>GGTGGGGGTGATGTTGTAGTTGG</td><td>1000</td><td></td><td></td><td></td><td></td><td></td><td></td><td></td><td></td><td></td><td></td></tr>
<tr><td>Oligo fwd</td><td>CACCGGTGGGGGTGATGTTGTAGT</td><td></td><td></td><td></td><td></td><td></td><td></td><td></td><td></td><td></td><td></td><td></td></tr>
<tr><td>Oligo rev</td><td>AAACACTACAACATCACCCCCACC</td><td></td><td></td><td></td><td></td><td></td><td></td><td></td><td></td><td></td><td></td><td></td></tr>
<tr><td>Chromosome</td><td>start</td><td>end</td><td>strand</td><td>MM</td><td>target_seq</td><td>PAM</td><td>alignment</td><td>distance</td><td>position</td><td>gene name</td><td>gene id</td></tr>
<tr><td>GU305914.1</td><td>122428</td><td>122450</td><td>+</td><td>0</td><td>GGTGGGGGTGATGTTGTAGT</td><td>TGG</td><td>||||||||[||||||||||||]PAM</td><td>NA</td><td>Intergenic</td><td>NA</td><td>NA</td></tr>
<tr><td>T9236</td><td>TGTTGTAGTTGGTCCTCGAGCGG</td><td>1000</td></tr>
<tr><td>Oligo fwd</td><td>CACCTGTTGTAGTTGGTCCTCGAG</td></tr>
<tr><td>Oligo rev</td><td>AAACCTCGAGGACCAACTACAACA</td></tr>
<tr><td>Chromosome</td><td>start</td><td>end</td><td>strand</td><td>MM</td><td>target_seq</td><td>PAM</td><td>alignment</td><td>distance</td><td>position</td><td>gene name</td><td>gene id</td></tr>
<tr><td>GU305914.1</td><td>122439</td><td>122461</td><td>+</td><td>0</td><td>TGTTGTAGTTGGTCCTCGAG</td><td>CGG</td><td>||||||||[||||||||||||]PAM</td><td>NA</td><td>Intergenic</td><td>NA</td><td>NA</td></tr>
<tr><td>T9237</td><td>GTTGTAGTTGGTCCTCGAGCGGG</td><td>1000</td></tr>
<tr><td>Oligo fwd</td><td>CACCGTTGTAGTTGGTCCTCGAGC</td></tr>
<tr><td>Oligo rev</td><td>AAACGCTCGAGGACCAACTACAAC</td></tr>
<tr><td>Chromosome</td><td>start</td><td>end</td><td>strand</td><td>MM</td><td>target_seq</td><td>PAM</td><td>alignment</td><td>distance</td><td>position</td><td>gene name</td><td>gene id</td></tr>
<tr><td>GU305914.1</td><td>122440</td><td>122462</td><td>+</td><td>0</td><td>GTTGTAGTTGGTCCTCGAGC</td><td>GGG</td><td>||||||||[||||||||||||]PAM</td><td>NA</td><td>Intergenic</td><td>NA</td><td>NA</td></tr>
<tr><td>T9238</td><td>TAGTTGGTCCTCGAGCGGGCCGG</td><td>1000</td></tr>
<tr><td>Oligo fwd</td><td>CACCTAGTTGGTCCTCGAGCGGGC</td></tr>
<tr><td>Oligo rev</td><td>AAACGCCCGCTCGAGGACCAACTA</td></tr>
<tr><td>Chromosome</td><td>start</td><td>end</td><td>strand</td><td>MM</td><td>target_seq</td><td>PAM</td><td>alignment</td><td>distance</td><td>position</td><td>gene name</td><td>gene id</td></tr>
<tr><td>GU305914.1</td><td>122444</td><td>122466</td><td>+</td><td>0</td><td>TAGTTGGTCCTCGAGCGGGC</td><td>CGG</td><td>||||||||[||||||||||||]PAM</td><td>NA</td><td>Intergenic</td><td>NA</td><td>NA</td></tr>
<tr><td>T9239</td><td>TTGGTCCTCGAGCGGGCCGGCGG</td><td>1000</td></tr>
<tr><td>Oligo fwd</td><td>CACCTTGGTCCTCGAGCGGGCCGG</td></tr>
<tr><td>Oligo rev</td><td>AAACCCGGCCCGCTCGAGGACCAA</td></tr>
<tr><td>Chromosome</td><td>start</td><td>end</td><td>strand</td><td>MM</td><td>target_seq</td><td>PAM</td><td>alignment</td><td>distance</td><td>position</td><td>gene name</td><td>gene id</td></tr>
<tr><td>GU305914.1</td><td>122447</td><td>122469</td><td>+</td><td>0</td><td>TTGGTCCTCGAGCGGGCCGG</td><td>CGG</td><td>||||||||[||||||||||||]PAM</td><td>NA</td><td>Intergenic</td><td>NA</td><td>NA</td></tr>
<tr><td>T9240</td><td>TGGTCCTCGAGCGGGCCGGCGGG</td><td>1000</td></tr>
<tr><td>Oligo fwd</td><td>CACCTGGTCCTCGAGCGGGCCGGC</td></tr>
<tr><td>Oligo rev</td><td>AAACGCCGGCCCGCTCGAGGACCA</td></tr>
<tr><td>Chromosome</td><td>start</td><td>end</td><td>strand</td><td>MM</td><td>target_seq</td><td>PAM</td><td>alignment</td><td>distance</td><td>position</td><td>gene name</td><td>gene id</td></tr>
<tr><td>GU305914.1</td><td>122448</td><td>122470</td><td>+</td><td>0</td><td>TGGTCCTCGAGCGGGCCGGC</td><td>GGG</td><td>||||||||[||||||||||||]PAM</td><td>NA</td><td>Intergenic</td><td>NA</td><td>NA</td></tr>
<tr><td>GU305914.1</td><td>107959</td><td>107981</td><td>+</td><td>4</td><td>TGGTCACTGAGCTGGCCGGC</td><td>GGG</td><td>|||||---[||||-|||||||]PAM</td><td>NA</td><td>Intergenic</td><td>NA</td><td>NA</td></tr>
<tr><td>T9241</td><td>TGCCCGCGCCGACACCGACCTGG</td><td>1000</td></tr>
<tr><td>Oligo fwd</td><td>CACCTGCCCGCGCCGACACCGACC</td></tr>
<tr><td>Oligo rev</td><td>AAACGGTCGGTGTCGGCGCGGGCA</td></tr>
<tr><td>Chromosome</td><td>start</td><td>end</td><td>strand</td><td>MM</td><td>target_seq</td><td>PAM</td><td>alignment</td><td>distance</td><td>position</td><td>gene name</td><td>gene id</td></tr>
<tr><td>GU305914.1</td><td>122490</td><td>122512</td><td>+</td><td>0</td><td>TGCCCGCGCCGACACCGACC</td><td>TGG</td><td>||||||||[||||||||||||]PAM</td><td>NA</td><td>Intergenic</td><td>NA</td><td>NA</td></tr>
<tr><td>T9242</td><td>GCCCGCGCCGACACCGACCTGGG</td><td>1000</td></tr>
<tr><td>Oligo fwd</td><td>CACCGCCCGCGCCGACACCGACCT</td></tr>
<tr><td>Oligo rev</td><td>AAACAGGTCGGTGTCGGCGCGGGC</td></tr>
<tr><td>Chromosome</td><td>start</td><td>end</td><td>strand</td><td>MM</td><td>target_seq</td><td>PAM</td><td>alignment</td><td>distance</td><td>position</td><td>gene name</td><td>gene id</td></tr>
<tr><td>GU305914.1</td><td>122491</td><td>122513</td><td>+</td><td>0</td><td>GCCCGCGCCGACACCGACCT</td><td>GGG</td><td>||||||||[||||||||||||]PAM</td><td>NA</td><td>Intergenic</td><td>NA</td><td>NA</td></tr>
<tr><td>T9243</td><td>CCCGCGCCGACACCGACCTGGGG</td><td>1000</td></tr>
<tr><td>Oligo fwd</td><td>CACCCCCGCGCCGACACCGACCTG</td></tr>
<tr><td>Oligo rev</td><td>AAACCAGGTCGGTGTCGGCGCGGG</td></tr>
<tr><td>Chromosome</td><td>start</td><td>end</td><td>strand</td><td>MM</td><td>target_seq</td><td>PAM</td><td>alignment</td><td>distance</td><td>position</td><td>gene name</td><td>gene id</td></tr>
<tr><td>GU305914.1</td><td>122492</td><td>122514</td><td>+</td><td>0</td><td>CCCGCGCCGACACCGACCTG</td><td>GGG</td><td>||||||||[||||||||||||]PAM</td><td>NA</td><td>Intergenic</td><td>NA</td><td>NA</td></tr>
<tr><td>T9244</td><td>CACCGACCTGGGGCATCCGCCGG</td><td>1000</td></tr>
<tr><td>Oligo fwd</td><td>CACCCACCGACCTGGGGCATCCGC</td></tr>
<tr><td>Oligo rev</td><td>AAACGCGGATGCCCCAGGTCGGTG</td></tr>
<tr><td>Chromosome</td><td>start</td><td>end</td><td>strand</td><td>MM</td><td>target_seq</td><td>PAM</td><td>alignment</td><td>distance</td><td>position</td><td>gene name</td><td>gene id</td></tr>
<tr><td>GU305914.1</td><td>122502</td><td>122524</td><td>+</td><td>0</td><td>CACCGACCTGGGGCATCCGC</td><td>CGG</td><td>||||||||[||||||||||||]PAM</td><td>NA</td><td>Intergenic</td><td>NA</td><td>NA</td></tr>
<tr><td>T9245</td><td>CCTGGGGCATCCGCCGGCCGAGG</td><td>1000</td></tr>
<tr><td>Oligo fwd</td><td>CACCCCTGGGGCATCCGCCGGCCG</td></tr>
<tr><td>Oligo rev</td><td>AAACCGGCCGGCGGATGCCCCAGG</td></tr>
<tr><td>Chromosome</td><td>start</td><td>end</td><td>strand</td><td>MM</td><td>target_seq</td><td>PAM</td><td>alignment</td><td>distance</td><td>position</td><td>gene name</td><td>gene id</td></tr>
<tr><td>GU305914.1</td><td>122508</td><td>122530</td><td>+</td><td>0</td><td>CCTGGGGCATCCGCCGGCCG</td><td>AGG</td><td>||||||||[||||||||||||]PAM</td><td>NA</td><td>Intergenic</td><td>NA</td><td>NA</td></tr>
<tr><td>T9246</td><td>GGATCTCCTGACATTTCATGAGG</td><td>1000</td></tr>
<tr><td>Oligo fwd</td><td>CACCGGATCTCCTGACATTTCATG</td></tr>
<tr><td>Oligo rev</td><td>AAACCATGAAATGTCAGGAGATCC</td></tr>
<tr><td>Chromosome</td><td>start</td><td>end</td><td>strand</td><td>MM</td><td>target_seq</td><td>PAM</td><td>alignment</td><td>distance</td><td>position</td><td>gene name</td><td>gene id</td></tr>
<tr><td>GU305914.1</td><td>122529</td><td>122551</td><td>+</td><td>0</td><td>GGATCTCCTGACATTTCATG</td><td>AGG</td><td>||||||||[||||||||||||]PAM</td><td>NA</td><td>Intergenic</td><td>NA</td><td>NA</td></tr>
<tr><td>T9247</td><td>GGAGCATGTTGCTGTCCTCCAGG</td><td>1000</td></tr>
<tr><td>Oligo fwd</td><td>CACCGGAGCATGTTGCTGTCCTCC</td></tr>
<tr><td>Oligo rev</td><td>AAACGGAGGACAGCAACATGCTCC</td></tr>
<tr><td>Chromosome</td><td>start</td><td>end</td><td>strand</td><td>MM</td><td>target_seq</td><td>PAM</td><td>alignment</td><td>distance</td><td>position</td><td>gene name</td><td>gene id</td></tr>
<tr><td>GU305914.1</td><td>122550</td><td>122572</td><td>+</td><td>0</td><td>GGAGCATGTTGCTGTCCTCC</td><td>AGG</td><td>||||||||[||||||||||||]PAM</td><td>NA</td><td>Intergenic</td><td>NA</td><td>NA</td></tr>
<tr><td>T9248</td><td>GAGCATGTTGCTGTCCTCCAGGG</td><td>1000</td></tr>
<tr><td>Oligo fwd</td><td>CACCGAGCATGTTGCTGTCCTCCA</td></tr>
<tr><td>Oligo rev</td><td>AAACTGGAGGACAGCAACATGCTC</td></tr>
<tr><td>Chromosome</td><td>start</td><td>end</td><td>strand</td><td>MM</td><td>target_seq</td><td>PAM</td><td>alignment</td><td>distance</td><td>position</td><td>gene name</td><td>gene id</td></tr>
<tr><td>GU305914.1</td><td>122551</td><td>122573</td><td>+</td><td>0</td><td>GAGCATGTTGCTGTCCTCCA</td><td>GGG</td><td>||||||||[||||||||||||]PAM</td><td>NA</td><td>Intergenic</td><td>NA</td><td>NA</td></tr>
<tr><td>T9249</td><td>CAGGGCCAGCTTCCGCACGTAGG</td><td>1000</td></tr>
<tr><td>Oligo fwd</td><td>CACCCAGGGCCAGCTTCCGCACGT</td></tr>
<tr><td>Oligo rev</td><td>AAACACGTGCGGAAGCTGGCCCTG</td></tr>
<tr><td>Chromosome</td><td>start</td><td>end</td><td>strand</td><td>MM</td><td>target_seq</td><td>PAM</td><td>alignment</td><td>distance</td><td>position</td><td>gene name</td><td>gene id</td></tr>
</table>

<table>
<tr><td>GU305914.1</td><td>122569</td><td>122591</td><td>+</td><td>0</td><td>CAGGGCCAGCTTCCGCACGT</td><td>AGG</td><td>||||||||[||||||||||||]PAM</td><td>NA</td><td>Intergenic</td><td>NA</td><td>NA</td></tr>
<tr><td></td><td></td><td></td><td></td><td></td><td></td><td></td><td></td><td></td><td></td><td></td><td></td></tr>
<tr><td>T9250</td><td>CACGTAGGCCAGCGTCACCTCGG</td><td>1000</td><td></td><td></td><td></td><td></td><td></td><td></td><td></td><td></td><td></td></tr>
<tr><td>Oligo fwd</td><td>CACCCACGTAGGCCAGCGTCACCT</td><td></td><td></td><td></td><td></td><td></td><td></td><td></td><td></td><td></td><td></td></tr>
<tr><td>Oligo rev</td><td>AAACAGGTGACGCTGGCCTACGTG</td><td></td><td></td><td></td><td></td><td></td><td></td><td></td><td></td><td></td><td></td></tr>
<tr><td>Chromosome</td><td>start</td><td>end</td><td>strand</td><td>MM</td><td>target_seq</td><td>PAM</td><td>alignment</td><td>distance</td><td>position</td><td>gene name</td><td>gene id</td></tr>
<tr><td>GU305914.1</td><td>122584</td><td>122606</td><td>+</td><td>0</td><td>CACGTAGGCCAGCGTCACCT</td><td>CGG</td><td>||||||||[||||||||||||]PAM</td><td>NA</td><td>Intergenic</td><td>NA</td><td>NA</td></tr>
<tr><td></td><td></td><td></td><td></td><td></td><td></td><td></td><td></td><td></td><td></td><td></td><td></td></tr>
<tr><td>T9251</td><td>GTAGGCCAGCGTCACCTCGGAGG</td><td>1000</td><td></td><td></td><td></td><td></td><td></td><td></td><td></td><td></td><td></td></tr>
<tr><td>Oligo fwd</td><td>CACCGTAGGCCAGCGTCACCTCGG</td><td></td><td></td><td></td><td></td><td></td><td></td><td></td><td></td><td></td><td></td></tr>
<tr><td>Oligo rev</td><td>AAACCCGAGGTGACGCTGGCCTAC</td><td></td><td></td><td></td><td></td><td></td><td></td><td></td><td></td><td></td><td></td></tr>
<tr><td>Chromosome</td><td>start</td><td>end</td><td>strand</td><td>MM</td><td>target_seq</td><td>PAM</td><td>alignment</td><td>distance</td><td>position</td><td>gene name</td><td>gene id</td></tr>
<tr><td>GU305914.1</td><td>122587</td><td>122609</td><td>+</td><td>0</td><td>GTAGGCCAGCGTCACCTCGG</td><td>AGG</td><td>||||||||[||||||||||||]PAM</td><td>NA</td><td>Intergenic</td><td>NA</td><td>NA</td></tr>
<tr><td></td><td></td><td></td><td></td><td></td><td></td><td></td><td></td><td></td><td></td><td></td><td></td></tr>
<tr><td>T9252</td><td>CCAGCGTCACCTCGGAGGATAGG</td><td>1000</td><td></td><td></td><td></td><td></td><td></td><td></td><td></td><td></td><td></td></tr>
<tr><td>Oligo fwd</td><td>CACCCCAGCGTCACCTCGGAGGAT</td><td></td><td></td><td></td><td></td><td></td><td></td><td></td><td></td><td></td><td></td></tr>
<tr><td>Oligo rev</td><td>AAACATCCTCCGAGGTGACGCTGG</td><td></td><td></td><td></td><td></td><td></td><td></td><td></td><td></td><td></td><td></td></tr>
<tr><td>Chromosome</td><td>start</td><td>end</td><td>strand</td><td>MM</td><td>target_seq</td><td>PAM</td><td>alignment</td><td>distance</td><td>position</td><td>gene name</td><td>gene id</td></tr>
<tr><td>GU305914.1</td><td>122592</td><td>122614</td><td>+</td><td>0</td><td>CCAGCGTCACCTCGGAGGAT</td><td>AGG</td><td>||||||||[||||||||||||]PAM</td><td>NA</td><td>Intergenic</td><td>NA</td><td>NA</td></tr>
<tr><td></td><td></td><td></td><td></td><td></td><td></td><td></td><td></td><td></td><td></td><td></td><td></td></tr>
<tr><td>T9253</td><td>CGTCACCTCGGAGGATAGGTTGG</td><td>1000</td><td></td><td></td><td></td><td></td><td></td><td></td><td></td><td></td><td></td></tr>
<tr><td>Oligo fwd</td><td>CACCCGTCACCTCGGAGGATAGGT</td><td></td><td></td><td></td><td></td><td></td><td></td><td></td><td></td><td></td><td></td></tr>
<tr><td>Oligo rev</td><td>AAACACCTATCCTCCGAGGTGACG</td><td></td><td></td><td></td><td></td><td></td><td></td><td></td><td></td><td></td><td></td></tr>
<tr><td>Chromosome</td><td>start</td><td>end</td><td>strand</td><td>MM</td><td>target_seq</td><td>PAM</td><td>alignment</td><td>distance</td><td>position</td><td>gene name</td><td>gene id</td></tr>
<tr><td>GU305914.1</td><td>122596</td><td>122618</td><td>+</td><td>0</td><td>CGTCACCTCGGAGGATAGGT</td><td>TGG</td><td>||||||||[||||||||||||]PAM</td><td>NA</td><td>Intergenic</td><td>NA</td><td>NA</td></tr>
<tr><td></td><td></td><td></td><td></td><td></td><td></td><td></td><td></td><td></td><td></td><td></td><td></td></tr>
<tr><td>T9254</td><td>GGAGGATAGGTTGGCGATCATGG</td><td>1000</td><td></td><td></td><td></td><td></td><td></td><td></td><td></td><td></td><td></td></tr>
<tr><td>Oligo fwd</td><td>CACCGGAGGATAGGTTGGCGATCA</td><td></td><td></td><td></td><td></td><td></td><td></td><td></td><td></td><td></td><td></td></tr>
<tr><td>Oligo rev</td><td>AAACTGATCGCCAACCTATCCTCC</td><td></td><td></td><td></td><td></td><td></td><td></td><td></td><td></td><td></td><td></td></tr>
<tr><td>Chromosome</td><td>start</td><td>end</td><td>strand</td><td>MM</td><td>target_seq</td><td>PAM</td><td>alignment</td><td>distance</td><td>position</td><td>gene name</td><td>gene id</td></tr>
<tr><td>GU305914.1</td><td>122605</td><td>122627</td><td>+</td><td>0</td><td>GGAGGATAGGTTGGCGATCA</td><td>TGG</td><td>||||||||[||||||||||||]PAM</td><td>NA</td><td>Intergenic</td><td>NA</td><td>NA</td></tr>
<tr><td></td><td></td><td></td><td></td><td></td><td></td><td></td><td></td><td></td><td></td><td></td><td></td></tr>
<tr><td>T9255</td><td>CGTGTCGTCGATCGTGATCTCGG</td><td>1000</td><td></td><td></td><td></td><td></td><td></td><td></td><td></td><td></td><td></td></tr>
<tr><td>Oligo fwd</td><td>CACCCGTGTCGTCGATCGTGATCT</td><td></td><td></td><td></td><td></td><td></td><td></td><td></td><td></td><td></td><td></td></tr>
<tr><td>Oligo rev</td><td>AAACAGATCACGATCGACGACACG</td><td></td><td></td><td></td><td></td><td></td><td></td><td></td><td></td><td></td><td></td></tr>
<tr><td>Chromosome</td><td>start</td><td>end</td><td>strand</td><td>MM</td><td>target_seq</td><td>PAM</td><td>alignment</td><td>distance</td><td>position</td><td>gene name</td><td>gene id</td></tr>
<tr><td>GU305914.1</td><td>122665</td><td>122687</td><td>+</td><td>0</td><td>CGTGTCGTCGATCGTGATCT</td><td>CGG</td><td>||||||||[||||||||||||]PAM</td><td>NA</td><td>Intergenic</td><td>NA</td><td>NA</td></tr>
<tr><td>GU305914.1</td><td>61174</td><td>61196</td><td>+</td><td>4</td><td>GGTGCCGTTGATCGTGCTCT</td><td>CGG</td><td>-|||-|||[-|||||||-|||]PAM</td><td>NA</td><td>Intergenic</td><td>NA</td><td>NA</td></tr>
<tr><td></td><td></td><td></td><td></td><td></td><td></td><td></td><td></td><td></td><td></td><td></td><td></td></tr>
<tr><td>T9256</td><td>GTGTCGTCGATCGTGATCTCGGG</td><td>1000</td><td></td><td></td><td></td><td></td><td></td><td></td><td></td><td></td><td></td></tr>
<tr><td>Oligo fwd</td><td>CACCGTGTCGTCGATCGTGATCTC</td><td></td><td></td><td></td><td></td><td></td><td></td><td></td><td></td><td></td><td></td></tr>
<tr><td>Oligo rev</td><td>AAACGAGATCACGATCGACGACAC</td><td></td><td></td><td></td><td></td><td></td><td></td><td></td><td></td><td></td><td></td></tr>
<tr><td>Chromosome</td><td>start</td><td>end</td><td>strand</td><td>MM</td><td>target_seq</td><td>PAM</td><td>alignment</td><td>distance</td><td>position</td><td>gene name</td><td>gene id</td></tr>
<tr><td>GU305914.1</td><td>122666</td><td>122688</td><td>+</td><td>0</td><td>GTGTCGTCGATCGTGATCTC</td><td>GGG</td><td>||||||||[||||||||||||]PAM</td><td>NA</td><td>Intergenic</td><td>NA</td><td>NA</td></tr>
<tr><td></td><td></td><td></td><td></td><td></td><td></td><td></td><td></td><td></td><td></td><td></td><td></td></tr>
<tr><td>T9257</td><td>CGTCGATCGTGATCTCGGGCAGG</td><td>1000</td><td></td><td></td><td></td><td></td><td></td><td></td><td></td><td></td><td></td></tr>
<tr><td>Oligo fwd</td><td>CACCCGTCGATCGTGATCTCGGGC</td><td></td><td></td><td></td><td></td><td></td><td></td><td></td><td></td><td></td><td></td></tr>
<tr><td>Oligo rev</td><td>AAACGCCCGAGATCACGATCGACG</td><td></td><td></td><td></td><td></td><td></td><td></td><td></td><td></td><td></td><td></td></tr>
<tr><td>Chromosome</td><td>start</td><td>end</td><td>strand</td><td>MM</td><td>target_seq</td><td>PAM</td><td>alignment</td><td>distance</td><td>position</td><td>gene name</td><td>gene id</td></tr>
<tr><td>GU305914.1</td><td>122670</td><td>122692</td><td>+</td><td>0</td><td>CGTCGATCGTGATCTCGGGC</td><td>AGG</td><td>||||||||[||||||||||||]PAM</td><td>NA</td><td>Intergenic</td><td>NA</td><td>NA</td></tr>
<tr><td></td><td></td><td></td><td></td><td></td><td></td><td></td><td></td><td></td><td></td><td></td><td></td></tr>
<tr><td>T9258</td><td>GTCGATCGTGATCTCGGGCAGGG</td><td>1000</td><td></td><td></td><td></td><td></td><td></td><td></td><td></td><td></td><td></td></tr>
<tr><td>Oligo fwd</td><td>CACCGTCGATCGTGATCTCGGGCA</td><td></td><td></td><td></td><td></td><td></td><td></td><td></td><td></td><td></td><td></td></tr>
<tr><td>Oligo rev</td><td>AAACTGCCCGAGATCACGATCGAC</td><td></td><td></td><td></td><td></td><td></td><td></td><td></td><td></td><td></td><td></td></tr>
<tr><td>Chromosome</td><td>start</td><td>end</td><td>strand</td><td>MM</td><td>target_seq</td><td>PAM</td><td>alignment</td><td>distance</td><td>position</td><td>gene name</td><td>gene id</td></tr>
<tr><td>GU305914.1</td><td>122671</td><td>122693</td><td>+</td><td>0</td><td>GTCGATCGTGATCTCGGGCA</td><td>GGG</td><td>||||||||[||||||||||||]PAM</td><td>NA</td><td>Intergenic</td><td>NA</td><td>NA</td></tr>
<tr><td></td><td></td><td></td><td></td><td></td><td></td><td></td><td></td><td></td><td></td><td></td><td></td></tr>
<tr><td>T9259</td><td>TCGTGATCTCGGGCAGGGTCAGG</td><td>1000</td><td></td><td></td><td></td><td></td><td></td><td></td><td></td><td></td><td></td></tr>
<tr><td>Oligo fwd</td><td>CACCTCGTGATCTCGGGCAGGGTC</td><td></td><td></td><td></td><td></td><td></td><td></td><td></td><td></td><td></td><td></td></tr>
<tr><td>Oligo rev</td><td>AAACGACCCTGCCCGAGATCACGA</td><td></td><td></td><td></td><td></td><td></td><td></td><td></td><td></td><td></td><td></td></tr>
<tr><td>Chromosome</td><td>start</td><td>end</td><td>strand</td><td>MM</td><td>target_seq</td><td>PAM</td><td>alignment</td><td>distance</td><td>position</td><td>gene name</td><td>gene id</td></tr>
<tr><td>GU305914.1</td><td>122676</td><td>122698</td><td>+</td><td>0</td><td>TCGTGATCTCGGGCAGGGTC</td><td>AGG</td><td>||||||||[||||||||||||]PAM</td><td>NA</td><td>Intergenic</td><td>NA</td><td>NA</td></tr>
<tr><td></td><td></td><td></td><td></td><td></td><td></td><td></td><td></td><td></td><td></td><td></td><td></td></tr>
<tr><td>T9260</td><td>CGTGATCTCGGGCAGGGTCAGGG</td><td>1000</td><td></td><td></td><td></td><td></td><td></td><td></td><td></td><td></td><td></td></tr>
<tr><td>Oligo fwd</td><td>CACCCGTGATCTCGGGCAGGGTCA</td><td></td><td></td><td></td><td></td><td></td><td></td><td></td><td></td><td></td><td></td></tr>
<tr><td>Oligo rev</td><td>AAACTGACCCTGCCCGAGATCACG</td><td></td><td></td><td></td><td></td><td></td><td></td><td></td><td></td><td></td><td></td></tr>
<tr><td>Chromosome</td><td>start</td><td>end</td><td>strand</td><td>MM</td><td>target_seq</td><td>PAM</td><td>alignment</td><td>distance</td><td>position</td><td>gene name</td><td>gene id</td></tr>
<tr><td>GU305914.1</td><td>122677</td><td>122699</td><td>+</td><td>0</td><td>CGTGATCTCGGGCAGGGTCA</td><td>GGG</td><td>||||||||[||||||||||||]PAM</td><td>NA</td><td>Intergenic</td><td>NA</td><td>NA</td></tr>
<tr><td></td><td></td><td></td><td></td><td></td><td></td><td></td><td></td><td></td><td></td><td></td><td></td></tr>
<tr><td>T9261</td><td>GATCTCGGGCAGGGTCAGGGTGG</td><td>1000</td><td></td><td></td><td></td><td></td><td></td><td></td><td></td><td></td><td></td></tr>
<tr><td>Oligo fwd</td><td>CACCGATCTCGGGCAGGGTCAGGG</td><td></td><td></td><td></td><td></td><td></td><td></td><td></td><td></td><td></td><td></td></tr>
<tr><td>Oligo rev</td><td>AAACCCCTGACCCTGCCCGAGATC</td><td></td><td></td><td></td><td></td><td></td><td></td><td></td><td></td><td></td><td></td></tr>
<tr><td>Chromosome</td><td>start</td><td>end</td><td>strand</td><td>MM</td><td>target_seq</td><td>PAM</td><td>alignment</td><td>distance</td><td>position</td><td>gene name</td><td>gene id</td></tr>
<tr><td>GU305914.1</td><td>122680</td><td>122702</td><td>+</td><td>0</td><td>GATCTCGGGCAGGGTCAGGG</td><td>TGG</td><td>||||||||[||||||||||||]PAM</td><td>NA</td><td>Intergenic</td><td>NA</td><td>NA</td></tr>
<tr><td></td><td></td><td></td><td></td><td></td><td></td><td></td><td></td><td></td><td></td><td></td><td></td></tr>
<tr><td>T9262</td><td>GCAGGGTCAGGGTGGCGTCCCGG</td><td>1000</td><td></td><td></td><td></td><td></td><td></td><td></td><td></td><td></td><td></td></tr>
<tr><td>Oligo fwd</td><td>CACCGCAGGGTCAGGGTGGCGTCC</td><td></td><td></td><td></td><td></td><td></td><td></td><td></td><td></td><td></td><td></td></tr>
<tr><td>Oligo rev</td><td>AAACGGACGCCACCCTGACCCTGC</td><td></td><td></td><td></td><td></td><td></td><td></td><td></td><td></td><td></td><td></td></tr>
<tr><td>Chromosome</td><td>start</td><td>end</td><td>strand</td><td>MM</td><td>target_seq</td><td>PAM</td><td>alignment</td><td>distance</td><td>position</td><td>gene name</td><td>gene id</td></tr>
<tr><td>GU305914.1</td><td>122688</td><td>122710</td><td>+</td><td>0</td><td>GCAGGGTCAGGGTGGCGTCC</td><td>CGG</td><td>||||||||[||||||||||||]PAM</td><td>NA</td><td>Intergenic</td><td>NA</td><td>NA</td></tr>
<tr><td></td><td></td><td></td><td></td><td></td><td></td><td></td><td></td><td></td><td></td><td></td><td></td></tr>
<tr><td>T9263</td><td>GGTAGCGCACGACCCGCAGCTGG</td><td>1000</td><td></td><td></td><td></td><td></td><td></td><td></td><td></td><td></td><td></td></tr>
<tr><td>Oligo fwd</td><td>CACCGGTAGCGCACGACCCGCAGC</td><td></td><td></td><td></td><td></td><td></td><td></td><td></td><td></td><td></td><td></td></tr>
<tr><td>Oligo rev</td><td>AAACGCTGCGGGTCGTGCGCTACC</td><td></td><td></td><td></td><td></td><td></td><td></td><td></td><td></td><td></td><td></td></tr>
<tr><td>Chromosome</td><td>start</td><td>end</td><td>strand</td><td>MM</td><td>target_seq</td><td>PAM</td><td>alignment</td><td>distance</td><td>position</td><td>gene name</td><td>gene id</td></tr>
<tr><td>GU305914.1</td><td>122709</td><td>122731</td><td>+</td><td>0</td><td>GGTAGCGCACGACCCGCAGC</td><td>TGG</td><td>||||||||[||||||||||||]PAM</td><td>NA</td><td>Intergenic</td><td>NA</td><td>NA</td></tr>
<tr><td></td><td></td><td></td><td></td><td></td><td></td><td></td><td></td><td></td><td></td><td></td><td></td></tr>
<tr><td>T9264</td><td>GTAGCGCACGACCCGCAGCTGGG</td><td>1000</td><td></td><td></td><td></td><td></td><td></td><td></td><td></td><td></td><td></td></tr>
<tr><td>Oligo fwd</td><td>CACCGTAGCGCACGACCCGCAGCT</td><td></td><td></td><td></td><td></td><td></td><td></td><td></td><td></td><td></td><td></td></tr>
</table>

Oligo rev AAACAGCTGCGGGTCGTGCGCTAC
Chromosome start end strand MM target_seq PAM alignment distance position gene name gene id
GU305914.1 122710 122732 + 0 GTAGCGCACGACCCGCAGCT GGG ||||||||[||||||||||||]PAM NA Intergenic NA NA

T9265 GCGCACGACCCGCAGCTGGGCGG 1000
Oligo fwd CACCGCGCACGACCCGCAGCTGGG
Oligo rev AAACCCCAGCTGCGGGTCGTGCGC
Chromosome start end strand MM target_seq PAM alignment distance position gene name gene id
GU305914.1 122713 122735 + 0 GCGCACGACCCGCAGCTGGG CGG ||||||||[||||||||||||]PAM NA Intergenic NA NA

T9266 CGCGCACCTCCTCGTGCGAGCGG 1000
Oligo fwd CACCCGCGCACCTCCTCGTGCGAG
Oligo rev AAACCTCGCACGAGGAGGTGCGCG
Chromosome start end strand MM target_seq PAM alignment distance position gene name gene id
GU305914.1 122742 122764 + 0 CGCGCACCTCCTCGTGCGAG CGG ||||||||[||||||||||||]PAM NA Intergenic NA NA

T9267 GCGAGCGGTCGAGCGACAGCAGG 1000
Oligo fwd CACCGCGAGCGGTCGAGCGACAGC
Oligo rev AAACGCTGTCGCTCGACCGCTCGC
Chromosome start end strand MM target_seq PAM alignment distance position gene name gene id
GU305914.1 122757 122779 + 0 GCGAGCGGTCGAGCGACAGC AGG ||||||||[||||||||||||]PAM NA Intergenic NA NA

T9268 CGAGCGGTCGAGCGACAGCAGGG 1000
Oligo fwd CACCCGAGCGGTCGAGCGACAGCA
Oligo rev AAACTGCTGTCGCTCGACCGCTCG
Chromosome start end strand MM target_seq PAM alignment distance position gene name gene id
GU305914.1 122758 122780 + 0 CGAGCGGTCGAGCGACAGCA GGG ||||||||[||||||||||||]PAM NA Intergenic NA NA

T9269 GGGCCAGCATCATCATGCGCTGG 1000
Oligo fwd CACCGGGCCAGCATCATCATGCGC
Oligo rev AAACGCGCATGATGATGCTGGCCC
Chromosome start end strand MM target_seq PAM alignment distance position gene name gene id
GU305914.1 122778 122800 + 0 GGGCCAGCATCATCATGCGC TGG ||||||||[||||||||||||]PAM NA Intergenic NA NA

T9270 CATCATGCGCTGGTTGATCTCGG 1000
Oligo fwd CACCCATCATGCGCTGGTTGATCT
Oligo rev AAACAGATCAACCAGCGCATGATG
Chromosome start end strand MM target_seq PAM alignment distance position gene name gene id
GU305914.1 122788 122810 + 0 CATCATGCGCTGGTTGATCT CGG ||||||||[||||||||||||]PAM NA Intergenic NA NA

T9271 GATCTCGGTCGCCAGCGCCATGG 1000
Oligo fwd CACCGATCTCGGTCGCCAGCGCCA
Oligo rev AAACTGGCGCTGGCGACCGAGATC
Chromosome start end strand MM target_seq PAM alignment distance position gene name gene id
GU305914.1 122803 122825 + 0 GATCTCGGTCGCCAGCGCCA TGG ||||||||[||||||||||||]PAM NA Intergenic NA NA

T9272 ATCTCGGTCGCCAGCGCCATGGG 1000
Oligo fwd CACCATCTCGGTCGCCAGCGCCAT
Oligo rev AAACATGGCGCTGGCGACCGAGAT
Chromosome start end strand MM target_seq PAM alignment distance position gene name gene id
GU305914.1 122804 122826 + 0 ATCTCGGTCGCCAGCGCCAT GGG ||||||||[||||||||||||]PAM NA Intergenic NA NA

T9273 CAGCGCCATGGGCACGATCCAGG 1000
Oligo fwd CACCCAGCGCCATGGGCACGATCC
Oligo rev AAACGGATCGTGCCCATGGCGCTG
Chromosome start end strand MM target_seq PAM alignment distance position gene name gene id
GU305914.1 122815 122837 + 0 CAGCGCCATGGGCACGATCC AGG ||||||||[||||||||||||]PAM NA Intergenic NA NA

T9274 AGCGCCATGGGCACGATCCAGGG 1000
Oligo fwd CACCAGCGCCATGGGCACGATCCA
Oligo rev AAACTGGATCGTGCCCATGGCGCT
Chromosome start end strand MM target_seq PAM alignment distance position gene name gene id
GU305914.1 122816 122838 + 0 AGCGCCATGGGCACGATCCA GGG ||||||||[||||||||||||]PAM NA Intergenic NA NA

T9275 GCACGATCCAGGGCAGCACGAGG 1000
Oligo fwd CACCGCACGATCCAGGGCAGCACG
Oligo rev AAACCGTGCTGCCCTGGATCGTGC
Chromosome start end strand MM target_seq PAM alignment distance position gene name gene id
GU305914.1 122826 122848 + 0 GCACGATCCAGGGCAGCACG AGG ||||||||[||||||||||||]PAM NA Intergenic NA NA

T9276 CACGAGGTCCCAGTCCGAGACGG 1000
Oligo fwd CACCCACGAGGTCCCAGTCCGAGA
Oligo rev AAACTCTCGGACTGGGACCTCGTG
Chromosome start end strand MM target_seq PAM alignment distance position gene name gene id
GU305914.1 122842 122864 + 0 CACGAGGTCCCAGTCCGAGA CGG ||||||||[||||||||||||]PAM NA Intergenic NA NA

T9277 CCCAGTCCGAGACGGAGACGAGG 1000
Oligo fwd CACCCCCAGTCCGAGACGGAGACG
Oligo rev AAACCGTCTCCGTCTCGGACTGGG
Chromosome start end strand MM target_seq PAM alignment distance position gene name gene id
GU305914.1 122850 122872 + 0 CCCAGTCCGAGACGGAGACG AGG ||||||||[||||||||||||]PAM NA Intergenic NA NA

T9278 CGTCACCTCTCCCGAGAAGACGG 1000
Oligo fwd CACCCGTCACCTCTCCCGAGAAGA
Oligo rev AAACTCTTCTCGGGAGAGGTGACG
Chromosome start end strand MM target_seq PAM alignment distance position gene name gene id
GU305914.1 122875 122897 + 0 CGTCACCTCTCCCGAGAAGA CGG ||||||||[||||||||||||]PAM NA Intergenic NA NA

T9279 GTCACCTCTCCCGAGAAGACGGG 1000

<table>
<tr><td>Oligo fwd</td><td>CACCGTCACCTCTCCCGAGAAGAC</td><td></td><td></td><td></td><td></td><td></td><td></td><td></td><td></td><td></td><td></td></tr>
<tr><td>Oligo rev</td><td>AAACGTCTTCTCGGGAGAGGTGAC</td><td></td><td></td><td></td><td></td><td></td><td></td><td></td><td></td><td></td><td></td></tr>
<tr><td>Chromosome</td><td>start</td><td>end</td><td>strand</td><td>MM</td><td>target_seq</td><td>PAM</td><td>alignment</td><td>distance</td><td>position</td><td>gene name</td><td>gene id</td></tr>
<tr><td>GU305914.1</td><td>122876</td><td>122898</td><td>+</td><td>0</td><td>GTCACCTCTCCCGAGAAGAC</td><td>GGG</td><td>||||||||[||||||||||||]PAM</td><td>NA</td><td>Intergenic</td><td>NA</td><td>NA</td></tr>
<tr><td>GU305914.1</td><td>178270</td><td>178292</td><td>+</td><td>4</td><td>TTCACCGCTCCCAAGAAGTC</td><td>TGG</td><td>-|||||-|[||||-|||||-|]PAM</td><td>NA</td><td>Intergenic</td><td>NA</td><td>NA</td></tr>
<tr><td></td><td></td><td></td><td></td><td></td><td></td><td></td><td></td><td></td><td></td><td></td><td></td></tr>
<tr><td>T9280</td><td>ACCTCTCCCGAGAAGACGGGCGG</td><td>1000</td><td></td><td></td><td></td><td></td><td></td><td></td><td></td><td></td><td></td></tr>
<tr><td>Oligo fwd</td><td>CACCACCTCTCCCGAGAAGACGGG</td><td></td><td></td><td></td><td></td><td></td><td></td><td></td><td></td><td></td><td></td></tr>
<tr><td>Oligo rev</td><td>AAACCCCGTCTTCTCGGGAGAGGT</td><td></td><td></td><td></td><td></td><td></td><td></td><td></td><td></td><td></td><td></td></tr>
<tr><td>Chromosome</td><td>start</td><td>end</td><td>strand</td><td>MM</td><td>target_seq</td><td>PAM</td><td>alignment</td><td>distance</td><td>position</td><td>gene name</td><td>gene id</td></tr>
<tr><td>GU305914.1</td><td>122879</td><td>122901</td><td>+</td><td>0</td><td>ACCTCTCCCGAGAAGACGGG</td><td>CGG</td><td>||||||||[||||||||||||]PAM</td><td>NA</td><td>Intergenic</td><td>NA</td><td>NA</td></tr>
<tr><td></td><td></td><td></td><td></td><td></td><td></td><td></td><td></td><td></td><td></td><td></td><td></td></tr>
<tr><td>T9281</td><td>CTCCCGAGAAGACGGGCGGCAGG</td><td>1000</td><td></td><td></td><td></td><td></td><td></td><td></td><td></td><td></td><td></td></tr>
<tr><td>Oligo fwd</td><td>CACCCTCCCGAGAAGACGGGCGGC</td><td></td><td></td><td></td><td></td><td></td><td></td><td></td><td></td><td></td><td></td></tr>
<tr><td>Oligo rev</td><td>AAACGCCGCCCGTCTTCTCGGGAG</td><td></td><td></td><td></td><td></td><td></td><td></td><td></td><td></td><td></td><td></td></tr>
<tr><td>Chromosome</td><td>start</td><td>end</td><td>strand</td><td>MM</td><td>target_seq</td><td>PAM</td><td>alignment</td><td>distance</td><td>position</td><td>gene name</td><td>gene id</td></tr>
<tr><td>GU305914.1</td><td>122883</td><td>122905</td><td>+</td><td>0</td><td>CTCCCGAGAAGACGGGCGGC</td><td>AGG</td><td>||||||||[||||||||||||]PAM</td><td>NA</td><td>Intergenic</td><td>NA</td><td>NA</td></tr>
<tr><td></td><td></td><td></td><td></td><td></td><td></td><td></td><td></td><td></td><td></td><td></td><td></td></tr>
<tr><td>T9282</td><td>TCCCGAGAAGACGGGCGGCAGGG</td><td>1000</td><td></td><td></td><td></td><td></td><td></td><td></td><td></td><td></td><td></td></tr>
<tr><td>Oligo fwd</td><td>CACCTCCCGAGAAGACGGGCGGCA</td><td></td><td></td><td></td><td></td><td></td><td></td><td></td><td></td><td></td><td></td></tr>
<tr><td>Oligo rev</td><td>AAACTGCCGCCCGTCTTCTCGGGA</td><td></td><td></td><td></td><td></td><td></td><td></td><td></td><td></td><td></td><td></td></tr>
<tr><td>Chromosome</td><td>start</td><td>end</td><td>strand</td><td>MM</td><td>target_seq</td><td>PAM</td><td>alignment</td><td>distance</td><td>position</td><td>gene name</td><td>gene id</td></tr>
<tr><td>GU305914.1</td><td>122884</td><td>122906</td><td>+</td><td>0</td><td>TCCCGAGAAGACGGGCGGCA</td><td>GGG</td><td>||||||||[||||||||||||]PAM</td><td>NA</td><td>Intergenic</td><td>NA</td><td>NA</td></tr>
<tr><td></td><td></td><td></td><td></td><td></td><td></td><td></td><td></td><td></td><td></td><td></td><td></td></tr>
<tr><td>T9283</td><td>CGAGAAGACGGGCGGCAGGGTGG</td><td>1000</td><td></td><td></td><td></td><td></td><td></td><td></td><td></td><td></td><td></td></tr>
<tr><td>Oligo fwd</td><td>CACCCGAGAAGACGGGCGGCAGGG</td><td></td><td></td><td></td><td></td><td></td><td></td><td></td><td></td><td></td><td></td></tr>
<tr><td>Oligo rev</td><td>AAACCCCTGCCGCCCGTCTTCTCG</td><td></td><td></td><td></td><td></td><td></td><td></td><td></td><td></td><td></td><td></td></tr>
<tr><td>Chromosome</td><td>start</td><td>end</td><td>strand</td><td>MM</td><td>target_seq</td><td>PAM</td><td>alignment</td><td>distance</td><td>position</td><td>gene name</td><td>gene id</td></tr>
<tr><td>GU305914.1</td><td>122887</td><td>122909</td><td>+</td><td>0</td><td>CGAGAAGACGGGCGGCAGGG</td><td>TGG</td><td>||||||||[||||||||||||]PAM</td><td>NA</td><td>Intergenic</td><td>NA</td><td>NA</td></tr>
<tr><td></td><td></td><td></td><td></td><td></td><td></td><td></td><td></td><td></td><td></td><td></td><td></td></tr>
<tr><td>T9284</td><td>GTCTCCCTTCTCCCAGCTCACGG</td><td>1000</td><td></td><td></td><td></td><td></td><td></td><td></td><td></td><td></td><td></td></tr>
<tr><td>Oligo fwd</td><td>CACCGTCTCCCTTCTCCCAGCTCA</td><td></td><td></td><td></td><td></td><td></td><td></td><td></td><td></td><td></td><td></td></tr>
<tr><td>Oligo rev</td><td>AAACTGAGCTGGGAGAAGGGAGAC</td><td></td><td></td><td></td><td></td><td></td><td></td><td></td><td></td><td></td><td></td></tr>
<tr><td>Chromosome</td><td>start</td><td>end</td><td>strand</td><td>MM</td><td>target_seq</td><td>PAM</td><td>alignment</td><td>distance</td><td>position</td><td>gene name</td><td>gene id</td></tr>
<tr><td>GU305914.1</td><td>122917</td><td>122939</td><td>+</td><td>0</td><td>GTCTCCCTTCTCCCAGCTCA</td><td>CGG</td><td>||||||||[||||||||||||]PAM</td><td>NA</td><td>Intergenic</td><td>NA</td><td>NA</td></tr>
<tr><td></td><td></td><td></td><td></td><td></td><td></td><td></td><td></td><td></td><td></td><td></td><td></td></tr>
<tr><td>T9285</td><td>TCTCCCTTCTCCCAGCTCACGGG</td><td>1000</td><td></td><td></td><td></td><td></td><td></td><td></td><td></td><td></td><td></td></tr>
<tr><td>Oligo fwd</td><td>CACCTCTCCCTTCTCCCAGCTCAC</td><td></td><td></td><td></td><td></td><td></td><td></td><td></td><td></td><td></td><td></td></tr>
<tr><td>Oligo rev</td><td>AAACGTGAGCTGGGAGAAGGGAGA</td><td></td><td></td><td></td><td></td><td></td><td></td><td></td><td></td><td></td><td></td></tr>
<tr><td>Chromosome</td><td>start</td><td>end</td><td>strand</td><td>MM</td><td>target_seq</td><td>PAM</td><td>alignment</td><td>distance</td><td>position</td><td>gene name</td><td>gene id</td></tr>
<tr><td>GU305914.1</td><td>122918</td><td>122940</td><td>+</td><td>0</td><td>TCTCCCTTCTCCCAGCTCAC</td><td>GGG</td><td>||||||||[||||||||||||]PAM</td><td>NA</td><td>Intergenic</td><td>NA</td><td>NA</td></tr>
<tr><td></td><td></td><td></td><td></td><td></td><td></td><td></td><td></td><td></td><td></td><td></td><td></td></tr>
<tr><td>T9286</td><td>AGCTCACGGGACCCGTGTTGCGG</td><td>1000</td><td></td><td></td><td></td><td></td><td></td><td></td><td></td><td></td><td></td></tr>
<tr><td>Oligo fwd</td><td>CACCAGCTCACGGGACCCGTGTTG</td><td></td><td></td><td></td><td></td><td></td><td></td><td></td><td></td><td></td><td></td></tr>
<tr><td>Oligo rev</td><td>AAACCAACACGGGTCCCGTGAGCT</td><td></td><td></td><td></td><td></td><td></td><td></td><td></td><td></td><td></td><td></td></tr>
<tr><td>Chromosome</td><td>start</td><td>end</td><td>strand</td><td>MM</td><td>target_seq</td><td>PAM</td><td>alignment</td><td>distance</td><td>position</td><td>gene name</td><td>gene id</td></tr>
<tr><td>GU305914.1</td><td>122931</td><td>122953</td><td>+</td><td>0</td><td>AGCTCACGGGACCCGTGTTG</td><td>CGG</td><td>||||||||[||||||||||||]PAM</td><td>NA</td><td>Intergenic</td><td>NA</td><td>NA</td></tr>
<tr><td></td><td></td><td></td><td></td><td></td><td></td><td></td><td></td><td></td><td></td><td></td><td></td></tr>
<tr><td>T9287</td><td>CACGGGACCCGTGTTGCGGATGG</td><td>1000</td><td></td><td></td><td></td><td></td><td></td><td></td><td></td><td></td><td></td></tr>
<tr><td>Oligo fwd</td><td>CACCCACGGGACCCGTGTTGCGGA</td><td></td><td></td><td></td><td></td><td></td><td></td><td></td><td></td><td></td><td></td></tr>
<tr><td>Oligo rev</td><td>AAACTCCGCAACACGGGTCCCGTG</td><td></td><td></td><td></td><td></td><td></td><td></td><td></td><td></td><td></td><td></td></tr>
<tr><td>Chromosome</td><td>start</td><td>end</td><td>strand</td><td>MM</td><td>target_seq</td><td>PAM</td><td>alignment</td><td>distance</td><td>position</td><td>gene name</td><td>gene id</td></tr>
<tr><td>GU305914.1</td><td>122935</td><td>122957</td><td>+</td><td>0</td><td>CACGGGACCCGTGTTGCGGA</td><td>TGG</td><td>||||||||[||||||||||||]PAM</td><td>NA</td><td>Intergenic</td><td>NA</td><td>NA</td></tr>
<tr><td></td><td></td><td></td><td></td><td></td><td></td><td></td><td></td><td></td><td></td><td></td><td></td></tr>
<tr><td>T9288</td><td>TGTTGCGGATGGTGTAGCACTGG</td><td>1000</td><td></td><td></td><td></td><td></td><td></td><td></td><td></td><td></td><td></td></tr>
<tr><td>Oligo fwd</td><td>CACCTGTTGCGGATGGTGTAGCAC</td><td></td><td></td><td></td><td></td><td></td><td></td><td></td><td></td><td></td><td></td></tr>
<tr><td>Oligo rev</td><td>AAACGTGCTACACCATCCGCAACA</td><td></td><td></td><td></td><td></td><td></td><td></td><td></td><td></td><td></td><td></td></tr>
<tr><td>Chromosome</td><td>start</td><td>end</td><td>strand</td><td>MM</td><td>target_seq</td><td>PAM</td><td>alignment</td><td>distance</td><td>position</td><td>gene name</td><td>gene id</td></tr>
<tr><td>GU305914.1</td><td>122946</td><td>122968</td><td>+</td><td>0</td><td>TGTTGCGGATGGTGTAGCAC</td><td>TGG</td><td>||||||||[||||||||||||]PAM</td><td>NA</td><td>Intergenic</td><td>NA</td><td>NA</td></tr>
<tr><td></td><td></td><td></td><td></td><td></td><td></td><td></td><td></td><td></td><td></td><td></td><td></td></tr>
<tr><td>T9289</td><td>GTGTAGCACTGGCCGTGCGTCGG</td><td>1000</td><td></td><td></td><td></td><td></td><td></td><td></td><td></td><td></td><td></td></tr>
<tr><td>Oligo fwd</td><td>CACCGTGTAGCACTGGCCGTGCGT</td><td></td><td></td><td></td><td></td><td></td><td></td><td></td><td></td><td></td><td></td></tr>
<tr><td>Oligo rev</td><td>AAACACGCACGGCCAGTGCTACAC</td><td></td><td></td><td></td><td></td><td></td><td></td><td></td><td></td><td></td><td></td></tr>
<tr><td>Chromosome</td><td>start</td><td>end</td><td>strand</td><td>MM</td><td>target_seq</td><td>PAM</td><td>alignment</td><td>distance</td><td>position</td><td>gene name</td><td>gene id</td></tr>
<tr><td>GU305914.1</td><td>122957</td><td>122979</td><td>+</td><td>0</td><td>GTGTAGCACTGGCCGTGCGT</td><td>CGG</td><td>||||||||[||||||||||||]PAM</td><td>NA</td><td>Intergenic</td><td>NA</td><td>NA</td></tr>
<tr><td></td><td></td><td></td><td></td><td></td><td></td><td></td><td></td><td></td><td></td><td></td><td></td></tr>
<tr><td>T9290</td><td>TGTAGCACTGGCCGTGCGTCGGG</td><td>1000</td><td></td><td></td><td></td><td></td><td></td><td></td><td></td><td></td><td></td></tr>
<tr><td>Oligo fwd</td><td>CACCTGTAGCACTGGCCGTGCGTC</td><td></td><td></td><td></td><td></td><td></td><td></td><td></td><td></td><td></td><td></td></tr>
<tr><td>Oligo rev</td><td>AAACGACGCACGGCCAGTGCTACA</td><td></td><td></td><td></td><td></td><td></td><td></td><td></td><td></td><td></td><td></td></tr>
<tr><td>Chromosome</td><td>start</td><td>end</td><td>strand</td><td>MM</td><td>target_seq</td><td>PAM</td><td>alignment</td><td>distance</td><td>position</td><td>gene name</td><td>gene id</td></tr>
<tr><td>GU305914.1</td><td>122958</td><td>122980</td><td>+</td><td>0</td><td>TGTAGCACTGGCCGTGCGTC</td><td>GGG</td><td>||||||||[||||||||||||]PAM</td><td>NA</td><td>Intergenic</td><td>NA</td><td>NA</td></tr>
<tr><td></td><td></td><td></td><td></td><td></td><td></td><td></td><td></td><td></td><td></td><td></td><td></td></tr>
<tr><td>T9291</td><td>GGCCGTGCGTCGGGACGCCCAGG</td><td>1000</td><td></td><td></td><td></td><td></td><td></td><td></td><td></td><td></td><td></td></tr>
<tr><td>Oligo fwd</td><td>CACCGGCCGTGCGTCGGGACGCCC</td><td></td><td></td><td></td><td></td><td></td><td></td><td></td><td></td><td></td><td></td></tr>
<tr><td>Oligo rev</td><td>AAACGGGCGTCCCGACGCACGGCC</td><td></td><td></td><td></td><td></td><td></td><td></td><td></td><td></td><td></td><td></td></tr>
<tr><td>Chromosome</td><td>start</td><td>end</td><td>strand</td><td>MM</td><td>target_seq</td><td>PAM</td><td>alignment</td><td>distance</td><td>position</td><td>gene name</td><td>gene id</td></tr>
<tr><td>GU305914.1</td><td>122967</td><td>122989</td><td>+</td><td>0</td><td>GGCCGTGCGTCGGGACGCCC</td><td>AGG</td><td>||||||||[||||||||||||]PAM</td><td>NA</td><td>Intergenic</td><td>NA</td><td>NA</td></tr>
<tr><td></td><td></td><td></td><td></td><td></td><td></td><td></td><td></td><td></td><td></td><td></td><td></td></tr>
<tr><td>T9292</td><td>TCGGGACGCCCAGGATCATCTGG</td><td>1000</td><td></td><td></td><td></td><td></td><td></td><td></td><td></td><td></td><td></td></tr>
<tr><td>Oligo fwd</td><td>CACCTCGGGACGCCCAGGATCATC</td><td></td><td></td><td></td><td></td><td></td><td></td><td></td><td></td><td></td><td></td></tr>
<tr><td>Oligo rev</td><td>AAACGATGATCCTGGGCGTCCCGA</td><td></td><td></td><td></td><td></td><td></td><td></td><td></td><td></td><td></td><td></td></tr>
<tr><td>Chromosome</td><td>start</td><td>end</td><td>strand</td><td>MM</td><td>target_seq</td><td>PAM</td><td>alignment</td><td>distance</td><td>position</td><td>gene name</td><td>gene id</td></tr>
<tr><td>GU305914.1</td><td>122976</td><td>122998</td><td>+</td><td>0</td><td>TCGGGACGCCCAGGATCATC</td><td>TGG</td><td>||||||||[||||||||||||]PAM</td><td>NA</td><td>Intergenic</td><td>NA</td><td>NA</td></tr>
<tr><td>GU305914.1</td><td>144797</td><td>144819</td><td>+</td><td>4</td><td>TCGCGAAGCCCAAGATCAGC</td><td>TGG</td><td>|||-||-|[||||-|||||-|]PAM</td><td>NA</td><td>Intergenic</td><td>NA</td><td>NA</td></tr>
<tr><td></td><td></td><td></td><td></td><td></td><td></td><td></td><td></td><td></td><td></td><td></td><td></td></tr>
<tr><td>T9293</td><td>ACCCTCGACGCGCCGCATGATGG</td><td>1000</td><td></td><td></td><td></td><td></td><td></td><td></td><td></td><td></td><td></td></tr>
<tr><td>Oligo fwd</td><td>CACCACCCTCGACGCGCCGCATGA</td><td></td><td></td><td></td><td></td><td></td><td></td><td></td><td></td><td></td><td></td></tr>
<tr><td>Oligo rev</td><td>AAACTCATGCGGCGCGTCGAGGGT</td><td></td><td></td><td></td><td></td><td></td><td></td><td></td><td></td><td></td><td></td></tr>
<tr><td>Chromosome</td><td>start</td><td>end</td><td>strand</td><td>MM</td><td>target_seq</td><td>PAM</td><td>alignment</td><td>distance</td><td>position</td><td>gene name</td><td>gene id</td></tr>
</table>

GU305914.1 123001 123023 + 0 ACCCTCGACGCGCCGCATGA TGG ||||||||[||||||||||||]PAM NA Intergenic NA NA

T9294 CCTGCGAGCTGCCCAGCAGATGG 1000
Oligo fwd CACCCCTGCGAGCTGCCCAGCAGA
Oligo rev AAACTCTGCTGGGCAGCTCGCAGG
Chromosome start end strand MM target_seq PAM alignment distance position gene name gene id
GU305914.1 123102 123124 + 0 CCTGCGAGCTGCCCAGCAGA TGG ||||||||[||||||||||||]PAM NA Intergenic NA NA

T9295 GCGAGCTGCCCAGCAGATGGTGG 1000
Oligo fwd CACCGCGAGCTGCCCAGCAGATGG
Oligo rev AAACCCATCTGCTGGGCAGCTCGC
Chromosome start end strand MM target_seq PAM alignment distance position gene name gene id
GU305914.1 123105 123127 + 0 GCGAGCTGCCCAGCAGATGG TGG ||||||||[||||||||||||]PAM NA Intergenic NA NA

T9296 AGCTGCCCAGCAGATGGTGGCGG 1000
Oligo fwd CACCAGCTGCCCAGCAGATGGTGG
Oligo rev AAACCCACCATCTGCTGGGCAGCT
Chromosome start end strand MM target_seq PAM alignment distance position gene name gene id
GU305914.1 123108 123130 + 0 AGCTGCCCAGCAGATGGTGG CGG ||||||||[||||||||||||]PAM NA Intergenic NA NA

T9297 GCCCAGCAGATGGTGGCGGTAGG 1000
Oligo fwd CACCGCCCAGCAGATGGTGGCGGT
Oligo rev AAACACCGCCACCATCTGCTGGGC
Chromosome start end strand MM target_seq PAM alignment distance position gene name gene id
GU305914.1 123112 123134 + 0 GCCCAGCAGATGGTGGCGGT AGG ||||||||[||||||||||||]PAM NA Intergenic NA NA

T9298 CCCAGCAGATGGTGGCGGTAGGG 1000
Oligo fwd CACCCCCAGCAGATGGTGGCGGTA
Oligo rev AAACTACCGCCACCATCTGCTGGG
Chromosome start end strand MM target_seq PAM alignment distance position gene name gene id
GU305914.1 123113 123135 + 0 CCCAGCAGATGGTGGCGGTA GGG ||||||||[||||||||||||]PAM NA Intergenic NA NA

T9299 CCAGCAGATGGTGGCGGTAGGGG 1000
Oligo fwd CACCCCAGCAGATGGTGGCGGTAG
Oligo rev AAACCTACCGCCACCATCTGCTGG
Chromosome start end strand MM target_seq PAM alignment distance position gene name gene id
GU305914.1 123114 123136 + 0 CCAGCAGATGGTGGCGGTAG GGG ||||||||[||||||||||||]PAM NA Intergenic NA NA

T9300 CAGATGGTGGCGGTAGGGGATGG 1000
Oligo fwd CACCCAGATGGTGGCGGTAGGGGA
Oligo rev AAACTCCCCTACCGCCACCATCTG
Chromosome start end strand MM target_seq PAM alignment distance position gene name gene id
GU305914.1 123118 123140 + 0 CAGATGGTGGCGGTAGGGGA TGG ||||||||[||||||||||||]PAM NA Intergenic NA NA

T9301 AGATGGTGGCGGTAGGGGATGGG 1000
Oligo fwd CACCAGATGGTGGCGGTAGGGGAT
Oligo rev AAACATCCCCTACCGCCACCATCT
Chromosome start end strand MM target_seq PAM alignment distance position gene name gene id
GU305914.1 123119 123141 + 0 AGATGGTGGCGGTAGGGGAT GGG ||||||||[||||||||||||]PAM NA Intergenic NA NA

T9302 GATGGTGGCGGTAGGGGATGGGG 1000
Oligo fwd CACCGATGGTGGCGGTAGGGGATG
Oligo rev AAACCATCCCCTACCGCCACCATC
Chromosome start end strand MM target_seq PAM alignment distance position gene name gene id
GU305914.1 123120 123142 + 0 GATGGTGGCGGTAGGGGATG GGG ||||||||[||||||||||||]PAM NA Intergenic NA NA

T9303 CGACAGCTTCCCCACGTCCCCGG 1000
Oligo fwd CACCCGACAGCTTCCCCACGTCCC
Oligo rev AAACGGGACGTGGGGAAGCTGTCG
Chromosome start end strand MM target_seq PAM alignment distance position gene name gene id
GU305914.1 123163 123185 + 0 CGACAGCTTCCCCACGTCCC CGG ||||||||[||||||||||||]PAM NA Intergenic NA NA

T9304 TCCCCACGTCCCCGGTCGTGAGG 1000
Oligo fwd CACCTCCCCACGTCCCCGGTCGTG
Oligo rev AAACCACGACCGGGGACGTGGGGA
Chromosome start end strand MM target_seq PAM alignment distance position gene name gene id
GU305914.1 123171 123193 + 0 TCCCCACGTCCCCGGTCGTG AGG ||||||||[||||||||||||]PAM NA Intergenic NA NA

T9305 GACCAAAACCGTCGTTTCCATGG 1000
Oligo fwd CACCGACCAAAACCGTCGTTTCCA
Oligo rev AAACTGGAAACGACGGTTTTGGTC
Chromosome start end strand MM target_seq PAM alignment distance position gene name gene id
GU305914.1 123208 123230 + 0 GACCAAAACCGTCGTTTCCA TGG ||||||||[||||||||||||]PAM NA Intergenic NA NA

T9306 AACCGTCGTTTCCATGGTCGTGG 1000
Oligo fwd CACCAACCGTCGTTTCCATGGTCG
Oligo rev AAACCGACCATGGAAACGACGGTT
Chromosome start end strand MM target_seq PAM alignment distance position gene name gene id
GU305914.1 123214 123236 + 0 AACCGTCGTTTCCATGGTCG TGG ||||||||[||||||||||||]PAM NA Intergenic NA NA

T9307 ACCGTCGTTTCCATGGTCGTGGG 1000
Oligo fwd CACCACCGTCGTTTCCATGGTCGT
Oligo rev AAACACGACCATGGAAACGACGGT
Chromosome start end strand MM target_seq PAM alignment distance position gene name gene id
GU305914.1 123215 123237 + 0 ACCGTCGTTTCCATGGTCGT GGG ||||||||[||||||||||||]PAM NA Intergenic NA NA

T9308 GTTTCCATGGTCGTGGGTGTCGG 1000
Oligo fwd CACCGTTTCCATGGTCGTGGGTGT
Oligo rev AAACACACCCACGACCATGGAAAC

<table>
<tr><td>Chromosome</td><td>start</td><td>end</td><td>strand</td><td>MM</td><td>target_seq</td><td>PAM</td><td>alignment</td><td>distance</td><td>position</td><td>gene name</td><td>gene id</td></tr>
<tr><td>GU305914.1</td><td>123221</td><td>123243</td><td>+</td><td>0</td><td>GTTTCCATGGTCGTGGGTGT</td><td>CGG</td><td>||||||||[||||||||||||]PAM</td><td>NA</td><td>Intergenic</td><td>NA</td><td>NA</td></tr>
<tr><td></td><td></td><td></td><td></td><td></td><td></td><td></td><td></td><td></td><td></td><td></td><td></td></tr>
<tr><td>T9309</td><td>TGGTCGTGGGTGTCGGTCTCCGG</td><td>1000</td><td></td><td></td><td></td><td></td><td></td><td></td><td></td><td></td><td></td></tr>
<tr><td>Oligo fwd</td><td>CACCTGGTCGTGGGTGTCGGTCTC</td><td></td><td></td><td></td><td></td><td></td><td></td><td></td><td></td><td></td><td></td></tr>
<tr><td>Oligo rev</td><td>AAACGAGACCGACACCCACGACCA</td><td></td><td></td><td></td><td></td><td></td><td></td><td></td><td></td><td></td><td></td></tr>
<tr><td>Chromosome</td><td>start</td><td>end</td><td>strand</td><td>MM</td><td>target_seq</td><td>PAM</td><td>alignment</td><td>distance</td><td>position</td><td>gene name</td><td>gene id</td></tr>
<tr><td>GU305914.1</td><td>123228</td><td>123250</td><td>+</td><td>0</td><td>TGGTCGTGGGTGTCGGTCTC</td><td>CGG</td><td>||||||||[||||||||||||]PAM</td><td>NA</td><td>Intergenic</td><td>NA</td><td>NA</td></tr>
<tr><td></td><td></td><td></td><td></td><td></td><td></td><td></td><td></td><td></td><td></td><td></td><td></td></tr>
<tr><td>T9310</td><td>CCGGCGAGCGTGACGTCCGACGG</td><td>1000</td><td></td><td></td><td></td><td></td><td></td><td></td><td></td><td></td><td></td></tr>
<tr><td>Oligo fwd</td><td>CACCCCGGCGAGCGTGACGTCCGA</td><td></td><td></td><td></td><td></td><td></td><td></td><td></td><td></td><td></td><td></td></tr>
<tr><td>Oligo rev</td><td>AAACTCGGACGTCACGCTCGCCGG</td><td></td><td></td><td></td><td></td><td></td><td></td><td></td><td></td><td></td><td></td></tr>
<tr><td>Chromosome</td><td>start</td><td>end</td><td>strand</td><td>MM</td><td>target_seq</td><td>PAM</td><td>alignment</td><td>distance</td><td>position</td><td>gene name</td><td>gene id</td></tr>
<tr><td>GU305914.1</td><td>123247</td><td>123269</td><td>+</td><td>0</td><td>CCGGCGAGCGTGACGTCCGA</td><td>CGG</td><td>||||||||[||||||||||||]PAM</td><td>NA</td><td>Intergenic</td><td>NA</td><td>NA</td></tr>
<tr><td></td><td></td><td></td><td></td><td></td><td></td><td></td><td></td><td></td><td></td><td></td><td></td></tr>
<tr><td>T9311</td><td>ACGGCGCTGCGCCTTCCGCTCGG</td><td>1000</td><td></td><td></td><td></td><td></td><td></td><td></td><td></td><td></td><td></td></tr>
<tr><td>Oligo fwd</td><td>CACCACGGCGCTGCGCCTTCCGCT</td><td></td><td></td><td></td><td></td><td></td><td></td><td></td><td></td><td></td><td></td></tr>
<tr><td>Oligo rev</td><td>AAACAGCGGAAGGCGCAGCGCCGT</td><td></td><td></td><td></td><td></td><td></td><td></td><td></td><td></td><td></td><td></td></tr>
<tr><td>Chromosome</td><td>start</td><td>end</td><td>strand</td><td>MM</td><td>target_seq</td><td>PAM</td><td>alignment</td><td>distance</td><td>position</td><td>gene name</td><td>gene id</td></tr>
<tr><td>GU305914.1</td><td>123266</td><td>123288</td><td>+</td><td>0</td><td>ACGGCGCTGCGCCTTCCGCT</td><td>CGG</td><td>||||||||[||||||||||||]PAM</td><td>NA</td><td>Intergenic</td><td>NA</td><td>NA</td></tr>
<tr><td></td><td></td><td></td><td></td><td></td><td></td><td></td><td></td><td></td><td></td><td></td><td></td></tr>
<tr><td>T9312</td><td>CGGCGCTGCGCCTTCCGCTCGGG</td><td>1000</td><td></td><td></td><td></td><td></td><td></td><td></td><td></td><td></td><td></td></tr>
<tr><td>Oligo fwd</td><td>CACCCGGCGCTGCGCCTTCCGCTC</td><td></td><td></td><td></td><td></td><td></td><td></td><td></td><td></td><td></td><td></td></tr>
<tr><td>Oligo rev</td><td>AAACGAGCGGAAGGCGCAGCGCCG</td><td></td><td></td><td></td><td></td><td></td><td></td><td></td><td></td><td></td><td></td></tr>
<tr><td>Chromosome</td><td>start</td><td>end</td><td>strand</td><td>MM</td><td>target_seq</td><td>PAM</td><td>alignment</td><td>distance</td><td>position</td><td>gene name</td><td>gene id</td></tr>
<tr><td>GU305914.1</td><td>123267</td><td>123289</td><td>+</td><td>0</td><td>CGGCGCTGCGCCTTCCGCTC</td><td>GGG</td><td>||||||||[||||||||||||]PAM</td><td>NA</td><td>Intergenic</td><td>NA</td><td>NA</td></tr>
<tr><td></td><td></td><td></td><td></td><td></td><td></td><td></td><td></td><td></td><td></td><td></td><td></td></tr>
<tr><td>T9313</td><td>CGCTGCGCCTTCCGCTCGGGAGG</td><td>1000</td><td></td><td></td><td></td><td></td><td></td><td></td><td></td><td></td><td></td></tr>
<tr><td>Oligo fwd</td><td>CACCCGCTGCGCCTTCCGCTCGGG</td><td></td><td></td><td></td><td></td><td></td><td></td><td></td><td></td><td></td><td></td></tr>
<tr><td>Oligo rev</td><td>AAACCCCGAGCGGAAGGCGCAGCG</td><td></td><td></td><td></td><td></td><td></td><td></td><td></td><td></td><td></td><td></td></tr>
<tr><td>Chromosome</td><td>start</td><td>end</td><td>strand</td><td>MM</td><td>target_seq</td><td>PAM</td><td>alignment</td><td>distance</td><td>position</td><td>gene name</td><td>gene id</td></tr>
<tr><td>GU305914.1</td><td>123270</td><td>123292</td><td>+</td><td>0</td><td>CGCTGCGCCTTCCGCTCGGG</td><td>AGG</td><td>||||||||[||||||||||||]PAM</td><td>NA</td><td>Intergenic</td><td>NA</td><td>NA</td></tr>
<tr><td></td><td></td><td></td><td></td><td></td><td></td><td></td><td></td><td></td><td></td><td></td><td></td></tr>
<tr><td>T9314</td><td>AGTTAAATAAGATGCTTTGCTGG</td><td>1000</td><td></td><td></td><td></td><td></td><td></td><td></td><td></td><td></td><td></td></tr>
<tr><td>Oligo fwd</td><td>CACCAGTTAAATAAGATGCTTTGC</td><td></td><td></td><td></td><td></td><td></td><td></td><td></td><td></td><td></td><td></td></tr>
<tr><td>Oligo rev</td><td>AAACGCAAAGCATCTTATTTAACT</td><td></td><td></td><td></td><td></td><td></td><td></td><td></td><td></td><td></td><td></td></tr>
<tr><td>Chromosome</td><td>start</td><td>end</td><td>strand</td><td>MM</td><td>target_seq</td><td>PAM</td><td>alignment</td><td>distance</td><td>position</td><td>gene name</td><td>gene id</td></tr>
<tr><td>GU305914.1</td><td>123308</td><td>123330</td><td>+</td><td>0</td><td>AGTTAAATAAGATGCTTTGC</td><td>TGG</td><td>||||||||[||||||||||||]PAM</td><td>NA</td><td>Intergenic</td><td>NA</td><td>NA</td></tr>
<tr><td></td><td></td><td></td><td></td><td></td><td></td><td></td><td></td><td></td><td></td><td></td><td></td></tr>
<tr><td>T9315</td><td>TAAATAAGATGCTTTGCTGGAGG</td><td>1000</td><td></td><td></td><td></td><td></td><td></td><td></td><td></td><td></td><td></td></tr>
<tr><td>Oligo fwd</td><td>CACCTAAATAAGATGCTTTGCTGG</td><td></td><td></td><td></td><td></td><td></td><td></td><td></td><td></td><td></td><td></td></tr>
<tr><td>Oligo rev</td><td>AAACCCAGCAAAGCATCTTATTTA</td><td></td><td></td><td></td><td></td><td></td><td></td><td></td><td></td><td></td><td></td></tr>
<tr><td>Chromosome</td><td>start</td><td>end</td><td>strand</td><td>MM</td><td>target_seq</td><td>PAM</td><td>alignment</td><td>distance</td><td>position</td><td>gene name</td><td>gene id</td></tr>
<tr><td>GU305914.1</td><td>123311</td><td>123333</td><td>+</td><td>0</td><td>TAAATAAGATGCTTTGCTGG</td><td>AGG</td><td>||||||||[||||||||||||]PAM</td><td>NA</td><td>Intergenic</td><td>NA</td><td>NA</td></tr>
<tr><td></td><td></td><td></td><td></td><td></td><td></td><td></td><td></td><td></td><td></td><td></td><td></td></tr>
<tr><td>T9316</td><td>AAATAAGATGCTTTGCTGGAGGG</td><td>1000</td><td></td><td></td><td></td><td></td><td></td><td></td><td></td><td></td><td></td></tr>
<tr><td>Oligo fwd</td><td>CACCAAATAAGATGCTTTGCTGGA</td><td></td><td></td><td></td><td></td><td></td><td></td><td></td><td></td><td></td><td></td></tr>
<tr><td>Oligo rev</td><td>AAACTCCAGCAAAGCATCTTATTT</td><td></td><td></td><td></td><td></td><td></td><td></td><td></td><td></td><td></td><td></td></tr>
<tr><td>Chromosome</td><td>start</td><td>end</td><td>strand</td><td>MM</td><td>target_seq</td><td>PAM</td><td>alignment</td><td>distance</td><td>position</td><td>gene name</td><td>gene id</td></tr>
<tr><td>GU305914.1</td><td>123312</td><td>123334</td><td>+</td><td>0</td><td>AAATAAGATGCTTTGCTGGA</td><td>GGG</td><td>||||||||[||||||||||||]PAM</td><td>NA</td><td>Intergenic</td><td>NA</td><td>NA</td></tr>
<tr><td></td><td></td><td></td><td></td><td></td><td></td><td></td><td></td><td></td><td></td><td></td><td></td></tr>
<tr><td>T9317</td><td>AATAAGATGCTTTGCTGGAGGGG</td><td>1000</td><td></td><td></td><td></td><td></td><td></td><td></td><td></td><td></td><td></td></tr>
<tr><td>Oligo fwd</td><td>CACCAATAAGATGCTTTGCTGGAG</td><td></td><td></td><td></td><td></td><td></td><td></td><td></td><td></td><td></td><td></td></tr>
<tr><td>Oligo rev</td><td>AAACCTCCAGCAAAGCATCTTATT</td><td></td><td></td><td></td><td></td><td></td><td></td><td></td><td></td><td></td><td></td></tr>
<tr><td>Chromosome</td><td>start</td><td>end</td><td>strand</td><td>MM</td><td>target_seq</td><td>PAM</td><td>alignment</td><td>distance</td><td>position</td><td>gene name</td><td>gene id</td></tr>
<tr><td>GU305914.1</td><td>123313</td><td>123335</td><td>+</td><td>0</td><td>AATAAGATGCTTTGCTGGAG</td><td>GGG</td><td>||||||||[||||||||||||]PAM</td><td>NA</td><td>Intergenic</td><td>NA</td><td>NA</td></tr>
<tr><td></td><td></td><td></td><td></td><td></td><td></td><td></td><td></td><td></td><td></td><td></td><td></td></tr>
<tr><td>T9318</td><td>GATGCTTTGCTGGAGGGGAATGG</td><td>1000</td><td></td><td></td><td></td><td></td><td></td><td></td><td></td><td></td><td></td></tr>
<tr><td>Oligo fwd</td><td>CACCGATGCTTTGCTGGAGGGGAA</td><td></td><td></td><td></td><td></td><td></td><td></td><td></td><td></td><td></td><td></td></tr>
<tr><td>Oligo rev</td><td>AAACTTCCCCTCCAGCAAAGCATC</td><td></td><td></td><td></td><td></td><td></td><td></td><td></td><td></td><td></td><td></td></tr>
<tr><td>Chromosome</td><td>start</td><td>end</td><td>strand</td><td>MM</td><td>target_seq</td><td>PAM</td><td>alignment</td><td>distance</td><td>position</td><td>gene name</td><td>gene id</td></tr>
<tr><td>GU305914.1</td><td>123318</td><td>123340</td><td>+</td><td>0</td><td>GATGCTTTGCTGGAGGGGAA</td><td>TGG</td><td>||||||||[||||||||||||]PAM</td><td>NA</td><td>Intergenic</td><td>NA</td><td>NA</td></tr>
<tr><td></td><td></td><td></td><td></td><td></td><td></td><td></td><td></td><td></td><td></td><td></td><td></td></tr>
<tr><td>T9319</td><td>CGATGCTGTAGTTGCCGTAGTGG</td><td>1000</td><td></td><td></td><td></td><td></td><td></td><td></td><td></td><td></td><td></td></tr>
<tr><td>Oligo fwd</td><td>CACCCGATGCTGTAGTTGCCGTAG</td><td></td><td></td><td></td><td></td><td></td><td></td><td></td><td></td><td></td><td></td></tr>
<tr><td>Oligo rev</td><td>AAACCTACGGCAACTACAGCATCG</td><td></td><td></td><td></td><td></td><td></td><td></td><td></td><td></td><td></td><td></td></tr>
<tr><td>Chromosome</td><td>start</td><td>end</td><td>strand</td><td>MM</td><td>target_seq</td><td>PAM</td><td>alignment</td><td>distance</td><td>position</td><td>gene name</td><td>gene id</td></tr>
<tr><td>GU305914.1</td><td>123347</td><td>123369</td><td>+</td><td>0</td><td>CGATGCTGTAGTTGCCGTAG</td><td>TGG</td><td>||||||||[||||||||||||]PAM</td><td>NA</td><td>Intergenic</td><td>NA</td><td>NA</td></tr>
<tr><td></td><td></td><td></td><td></td><td></td><td></td><td></td><td></td><td></td><td></td><td></td><td></td></tr>
<tr><td>T9320</td><td>GATGCTGTAGTTGCCGTAGTGGG</td><td>1000</td><td></td><td></td><td></td><td></td><td></td><td></td><td></td><td></td><td></td></tr>
<tr><td>Oligo fwd</td><td>CACCGATGCTGTAGTTGCCGTAGT</td><td></td><td></td><td></td><td></td><td></td><td></td><td></td><td></td><td></td><td></td></tr>
<tr><td>Oligo rev</td><td>AAACACTACGGCAACTACAGCATC</td><td></td><td></td><td></td><td></td><td></td><td></td><td></td><td></td><td></td><td></td></tr>
<tr><td>Chromosome</td><td>start</td><td>end</td><td>strand</td><td>MM</td><td>target_seq</td><td>PAM</td><td>alignment</td><td>distance</td><td>position</td><td>gene name</td><td>gene id</td></tr>
<tr><td>GU305914.1</td><td>123348</td><td>123370</td><td>+</td><td>0</td><td>GATGCTGTAGTTGCCGTAGT</td><td>GGG</td><td>||||||||[||||||||||||]PAM</td><td>NA</td><td>Intergenic</td><td>NA</td><td>NA</td></tr>
<tr><td></td><td></td><td></td><td></td><td></td><td></td><td></td><td></td><td></td><td></td><td></td><td></td></tr>
<tr><td>T9321</td><td>GTTTGCTCGTGCCCTTCAGCTGG</td><td>1000</td><td></td><td></td><td></td><td></td><td></td><td></td><td></td><td></td><td></td></tr>
<tr><td>Oligo fwd</td><td>CACCGTTTGCTCGTGCCCTTCAGC</td><td></td><td></td><td></td><td></td><td></td><td></td><td></td><td></td><td></td><td></td></tr>
<tr><td>Oligo rev</td><td>AAACGCTGAAGGGCACGAGCAAAC</td><td></td><td></td><td></td><td></td><td></td><td></td><td></td><td></td><td></td><td></td></tr>
<tr><td>Chromosome</td><td>start</td><td>end</td><td>strand</td><td>MM</td><td>target_seq</td><td>PAM</td><td>alignment</td><td>distance</td><td>position</td><td>gene name</td><td>gene id</td></tr>
<tr><td>GU305914.1</td><td>123386</td><td>123408</td><td>+</td><td>0</td><td>GTTTGCTCGTGCCCTTCAGC</td><td>TGG</td><td>||||||||[||||||||||||]PAM</td><td>NA</td><td>Intergenic</td><td>NA</td><td>NA</td></tr>
<tr><td></td><td></td><td></td><td></td><td></td><td></td><td></td><td></td><td></td><td></td><td></td><td></td></tr>
<tr><td>T9322</td><td>CCTTCAGCTGGCTCTCCATCAGG</td><td>1000</td><td></td><td></td><td></td><td></td><td></td><td></td><td></td><td></td><td></td></tr>
<tr><td>Oligo fwd</td><td>CACCCCTTCAGCTGGCTCTCCATC</td><td></td><td></td><td></td><td></td><td></td><td></td><td></td><td></td><td></td><td></td></tr>
<tr><td>Oligo rev</td><td>AAACGATGGAGAGCCAGCTGAAGG</td><td></td><td></td><td></td><td></td><td></td><td></td><td></td><td></td><td></td><td></td></tr>
<tr><td>Chromosome</td><td>start</td><td>end</td><td>strand</td><td>MM</td><td>target_seq</td><td>PAM</td><td>alignment</td><td>distance</td><td>position</td><td>gene name</td><td>gene id</td></tr>
<tr><td>GU305914.1</td><td>123398</td><td>123420</td><td>+</td><td>0</td><td>CCTTCAGCTGGCTCTCCATC</td><td>AGG</td><td>||||||||[||||||||||||]PAM</td><td>NA</td><td>Intergenic</td><td>NA</td><td>NA</td></tr>
<tr><td></td><td></td><td></td><td></td><td></td><td></td><td></td><td></td><td></td><td></td><td></td><td></td></tr>
<tr><td>T9323</td><td>CTTCAGCTGGCTCTCCATCAGGG</td><td>1000</td><td></td><td></td><td></td><td></td><td></td><td></td><td></td><td></td><td></td></tr>
<tr><td>Oligo fwd</td><td>CACCCTTCAGCTGGCTCTCCATCA</td><td></td><td></td><td></td><td></td><td></td><td></td><td></td><td></td><td></td><td></td></tr>
</table>

| | | | | | | | | | | | |
|---|---|---|---|---|---|---|---|---|---|---|---|
| Oligo rev | AAACTGATGGAGAGCCAGCTGAAG | | | | | | | | | | |
| Chromosome | start | end | strand | MM | target_seq | PAM | alignment | distance | position | gene name | gene id |
| GU305914.1 | 123399 | 123421 | + | 0 | CTTCAGCTGGCTCTCCATCA | GGG | \|\|\|\|\|\|\|\|[\|\|\|\|\|\|\|\|\|\|\|\|]PAM | NA | Intergenic | NA | NA |
| T9324 | TCGTGCCCGTGTGTTGCGTGAGG | 1000 | | | | | | | | | |
| Oligo fwd | CACCTCGTGCCCGTGTGTTGCGTG | | | | | | | | | | |
| Oligo rev | AAACCACGCAACACACGGGCACGA | | | | | | | | | | |
| Chromosome | start | end | strand | MM | target_seq | PAM | alignment | distance | position | gene name | gene id |
| GU305914.1 | 123428 | 123450 | + | 0 | TCGTGCCCGTGTGTTGCGTG | AGG | \|\|\|\|\|\|\|\|[\|\|\|\|\|\|\|\|\|\|\|\|]PAM | NA | Intergenic | NA | NA |
| T9325 | CGTGCCCGTGTGTTGCGTGAGGG | 1000 | | | | | | | | | |
| Oligo fwd | CACCCGTGCCCGTGTGTTGCGTGA | | | | | | | | | | |
| Oligo rev | AAACTCACGCAACACACGGGCACG | | | | | | | | | | |
| Chromosome | start | end | strand | MM | target_seq | PAM | alignment | distance | position | gene name | gene id |
| GU305914.1 | 123429 | 123451 | + | 0 | CGTGCCCGTGTGTTGCGTGA | GGG | \|\|\|\|\|\|\|\|[\|\|\|\|\|\|\|\|\|\|\|\|]PAM | NA | Intergenic | NA | NA |
| T9326 | TGTGTTGCGTGAGGGCTTCCTGG | 1000 | | | | | | | | | |
| Oligo fwd | CACCTGTGTTGCGTGAGGGCTTCC | | | | | | | | | | |
| Oligo rev | AAACGGAAGCCCTCACGCAACACA | | | | | | | | | | |
| Chromosome | start | end | strand | MM | target_seq | PAM | alignment | distance | position | gene name | gene id |
| GU305914.1 | 123437 | 123459 | + | 0 | TGTGTTGCGTGAGGGCTTCC | TGG | \|\|\|\|\|\|\|\|[\|\|\|\|\|\|\|\|\|\|\|\|]PAM | NA | Intergenic | NA | NA |
| T9327 | GTTGCGTGAGGGCTTCCTGGAGG | 1000 | | | | | | | | | |
| Oligo fwd | CACCGTTGCGTGAGGGCTTCCTGG | | | | | | | | | | |
| Oligo rev | AAACCCAGGAAGCCCTCACGCAAC | | | | | | | | | | |
| Chromosome | start | end | strand | MM | target_seq | PAM | alignment | distance | position | gene name | gene id |
| GU305914.1 | 123440 | 123462 | + | 0 | GTTGCGTGAGGGCTTCCTGG | AGG | \|\|\|\|\|\|\|\|[\|\|\|\|\|\|\|\|\|\|\|\|]PAM | NA | Intergenic | NA | NA |
| T9328 | GGCTTCCTGGAGGAAGCGACAGG | 1000 | | | | | | | | | |
| Oligo fwd | CACCGGCTTCCTGGAGGAAGCGAC | | | | | | | | | | |
| Oligo rev | AAACGTCGCTTCCTCCAGGAAGCC | | | | | | | | | | |
| Chromosome | start | end | strand | MM | target_seq | PAM | alignment | distance | position | gene name | gene id |
| GU305914.1 | 123450 | 123472 | + | 0 | GGCTTCCTGGAGGAAGCGAC | AGG | \|\|\|\|\|\|\|\|[\|\|\|\|\|\|\|\|\|\|\|\|]PAM | NA | Intergenic | NA | NA |
| T9329 | GCTTCCTGGAGGAAGCGACAGGG | 1000 | | | | | | | | | |
| Oligo fwd | CACCGCTTCCTGGAGGAAGCGACA | | | | | | | | | | |
| Oligo rev | AAACTGTCGCTTCCTCCAGGAAGC | | | | | | | | | | |
| Chromosome | start | end | strand | MM | target_seq | PAM | alignment | distance | position | gene name | gene id |
| GU305914.1 | 123451 | 123473 | + | 0 | GCTTCCTGGAGGAAGCGACA | GGG | \|\|\|\|\|\|\|\|[\|\|\|\|\|\|\|\|\|\|\|\|]PAM | NA | Intergenic | NA | NA |
| T9330 | GGGCTTCTCCACGATCGAGTTGG | 1000 | | | | | | | | | |
| Oligo fwd | CACCGGGCTTCTCCACGATCGAGT | | | | | | | | | | |
| Oligo rev | AAACACTCGATCGTGGAGAAGCCC | | | | | | | | | | |
| Chromosome | start | end | strand | MM | target_seq | PAM | alignment | distance | position | gene name | gene id |
| GU305914.1 | 123471 | 123493 | + | 0 | GGGCTTCTCCACGATCGAGT | TGG | \|\|\|\|\|\|\|\|[\|\|\|\|\|\|\|\|\|\|\|\|]PAM | NA | Intergenic | NA | NA |
| GU305914.1 | 168235 | 168257 | + | 3 | CGGCTTGTCCACGATCGAGG | CGG | -\|\|\|\|\|-\|[\|\|\|\|\|\|\|\|\|\|\|-]PAM | NA | Intergenic | NA | NA |
| T9331 | CCTCCACGCAGATGTACTGCAGG | 1000 | | | | | | | | | |
| Oligo fwd | CACCCCTCCACGCAGATGTACTGC | | | | | | | | | | |
| Oligo rev | AAACGCAGTACATCTGCGTGGAGG | | | | | | | | | | |
| Chromosome | start | end | strand | MM | target_seq | PAM | alignment | distance | position | gene name | gene id |
| GU305914.1 | 123497 | 123519 | + | 0 | CCTCCACGCAGATGTACTGC | AGG | \|\|\|\|\|\|\|\|[\|\|\|\|\|\|\|\|\|\|\|\|]PAM | NA | Intergenic | NA | NA |
| T9332 | ACTGCAGGTCCGTCTCGCCGTGG | 1000 | | | | | | | | | |
| Oligo fwd | CACCACTGCAGGTCCGTCTCGCCG | | | | | | | | | | |
| Oligo rev | AAACCGGCGAGACGGACCTGCAGT | | | | | | | | | | |
| Chromosome | start | end | strand | MM | target_seq | PAM | alignment | distance | position | gene name | gene id |
| GU305914.1 | 123512 | 123534 | + | 0 | ACTGCAGGTCCGTCTCGCCG | TGG | \|\|\|\|\|\|\|\|[\|\|\|\|\|\|\|\|\|\|\|\|]PAM | NA | Intergenic | NA | NA |
| T9333 | CGTGGATGCAGTCCTGAGAGCGG | 1000 | | | | | | | | | |
| Oligo fwd | CACCCGTGGATGCAGTCCTGAGAG | | | | | | | | | | |
| Oligo rev | AAACCTCTCAGGACTGCATCCACG | | | | | | | | | | |
| Chromosome | start | end | strand | MM | target_seq | PAM | alignment | distance | position | gene name | gene id |
| GU305914.1 | 123530 | 123552 | + | 0 | CGTGGATGCAGTCCTGAGAG | CGG | \|\|\|\|\|\|\|\|[\|\|\|\|\|\|\|\|\|\|\|\|]PAM | NA | Intergenic | NA | NA |
| T9334 | TGCAGTCCTGAGAGCGGTTCAGG | 1000 | | | | | | | | | |
| Oligo fwd | CACCTGCAGTCCTGAGAGCGGTTC | | | | | | | | | | |
| Oligo rev | AAACGAACCGCTCTCAGGACTGCA | | | | | | | | | | |
| Chromosome | start | end | strand | MM | target_seq | PAM | alignment | distance | position | gene name | gene id |
| GU305914.1 | 123536 | 123558 | + | 0 | TGCAGTCCTGAGAGCGGTTC | AGG | \|\|\|\|\|\|\|\|[\|\|\|\|\|\|\|\|\|\|\|\|]PAM | NA | Intergenic | NA | NA |
| T9335 | ACTCCTCCACCGTCTTGAACAGG | 1000 | | | | | | | | | |
| Oligo fwd | CACCACTCCTCCACCGTCTTGAAC | | | | | | | | | | |
| Oligo rev | AAACGTTCAAGACGGTGGAGGAGT | | | | | | | | | | |
| Chromosome | start | end | strand | MM | target_seq | PAM | alignment | distance | position | gene name | gene id |
| GU305914.1 | 123560 | 123582 | + | 0 | ACTCCTCCACCGTCTTGAAC | AGG | \|\|\|\|\|\|\|\|[\|\|\|\|\|\|\|\|\|\|\|\|]PAM | NA | Intergenic | NA | NA |
| T9336 | CTCCTCCACCGTCTTGAACAGGG | 1000 | | | | | | | | | |
| Oligo fwd | CACCCTCCTCCACCGTCTTGAACA | | | | | | | | | | |
| Oligo rev | AAACTGTTCAAGACGGTGGAGGAG | | | | | | | | | | |
| Chromosome | start | end | strand | MM | target_seq | PAM | alignment | distance | position | gene name | gene id |
| GU305914.1 | 123561 | 123583 | + | 0 | CTCCTCCACCGTCTTGAACA | GGG | \|\|\|\|\|\|\|\|[\|\|\|\|\|\|\|\|\|\|\|\|]PAM | NA | Intergenic | NA | NA |
| T9337 | GTCGTGAAGAACTGCGCGCACGG | 1000 | | | | | | | | | |
| Oligo fwd | CACCGTCGTGAAGAACTGCGCGCA | | | | | | | | | | |
| Oligo rev | AAACTGCGCGCAGTTCTTCACGAC | | | | | | | | | | |
| Chromosome | start | end | strand | MM | target_seq | PAM | alignment | distance | position | gene name | gene id |
| GU305914.1 | 123610 | 123632 | + | 0 | GTCGTGAAGAACTGCGCGCA | CGG | \|\|\|\|\|\|\|\|[\|\|\|\|\|\|\|\|\|\|\|\|]PAM | NA | Intergenic | NA | NA |

T9338 TCGTGAAGAACTGCGCGCACGGG 1000
Oligo fwd CACCTCGTGAAGAACTGCGCGCAC
Oligo rev AAACGTGCGCGCAGTTCTTCACGA
Chromosome start end strand MM target_seq PAM alignment distance position gene name gene id
GU305914.1 123611 123633 + 0 TCGTGAAGAACTGCGCGCAC GGG ||||||||[||||||||||||]PAM NA Intergenic NA NA

T9339 GCGCACGGGCTGTAGAAGCTCGG 1000
Oligo fwd CACCGCGCACGGGCTGTAGAAGCT
Oligo rev AAACAGCTTCTACAGCCCGTGCGC
Chromosome start end strand MM target_seq PAM alignment distance position gene name gene id
GU305914.1 123625 123647 + 0 GCGCACGGGCTGTAGAAGCT CGG ||||||||[||||||||||||]PAM NA Intergenic NA NA

T9340 CGCACGGGCTGTAGAAGCTCGGG 1000
Oligo fwd CACCCGCACGGGCTGTAGAAGCTC
Oligo rev AAACGAGCTTCTACAGCCCGTGCG
Chromosome start end strand MM target_seq PAM alignment distance position gene name gene id
GU305914.1 123626 123648 + 0 CGCACGGGCTGTAGAAGCTC GGG ||||||||[||||||||||||]PAM NA Intergenic NA NA
GU305914.1 128274 128296 + 4 TGCTCGAGCTGCAGAAGCTC TGG -||-||-|[|||-||||||||]PAM NA Intergenic NA NA

T9341 GGCTGTAGAAGCTCGGGTTGTGG 1000
Oligo fwd CACCGGCTGTAGAAGCTCGGGTTG
Oligo rev AAACCAACCCGAGCTTCTACAGCC
Chromosome start end strand MM target_seq PAM alignment distance position gene name gene id
GU305914.1 123632 123654 + 0 GGCTGTAGAAGCTCGGGTTG TGG ||||||||[||||||||||||]PAM NA Intergenic NA NA

T9342 GGTTGTGGCCCAGCTTCTCGCGG 1000
Oligo fwd CACCGGTTGTGGCCCAGCTTCTCG
Oligo rev AAACCGAGAAGCTGGGCCACAACC
Chromosome start end strand MM target_seq PAM alignment distance position gene name gene id
GU305914.1 123647 123669 + 0 GGTTGTGGCCCAGCTTCTCG CGG ||||||||[||||||||||||]PAM NA Intergenic NA NA

T9343 GGCCCAGCTTCTCGCGGTTGCGG 1000
Oligo fwd CACCGGCCCAGCTTCTCGCGGTTG
Oligo rev AAACCAACCGCGAGAAGCTGGGCC
Chromosome start end strand MM target_seq PAM alignment distance position gene name gene id
GU305914.1 123653 123675 + 0 GGCCCAGCTTCTCGCGGTTG CGG ||||||||[||||||||||||]PAM NA Intergenic NA NA

T9344 GCCCAGCTTCTCGCGGTTGCGGG 1000
Oligo fwd CACCGCCCAGCTTCTCGCGGTTGC
Oligo rev AAACGCAACCGCGAGAAGCTGGGC
Chromosome start end strand MM target_seq PAM alignment distance position gene name gene id
GU305914.1 123654 123676 + 0 GCCCAGCTTCTCGCGGTTGC GGG ||||||||[||||||||||||]PAM NA Intergenic NA NA

T9345 GACCCGCGCTGCGAGCCCCACGG 1000
Oligo fwd CACCGACCCGCGCTGCGAGCCCCA
Oligo rev AAACTGGGGCTCGCAGCGCGGGTC
Chromosome start end strand MM target_seq PAM alignment distance position gene name gene id
GU305914.1 123700 123722 + 0 GACCCGCGCTGCGAGCCCCA CGG ||||||||[||||||||||||]PAM NA Intergenic NA NA

T9346 ACCCGCGCTGCGAGCCCCACGGG 1000
Oligo fwd CACCACCCGCGCTGCGAGCCCCAC
Oligo rev AAACGTGGGGCTCGCAGCGCGGGT
Chromosome start end strand MM target_seq PAM alignment distance position gene name gene id
GU305914.1 123701 123723 + 0 ACCCGCGCTGCGAGCCCCAC GGG ||||||||[||||||||||||]PAM NA Intergenic NA NA

T9347 CTGCGAGCCCCACGGGTTGTTGG 1000
Oligo fwd CACCCTGCGAGCCCCACGGGTTGT
Oligo rev AAACACAACCCGTGGGGCTCGCAG
Chromosome start end strand MM target_seq PAM alignment distance position gene name gene id
GU305914.1 123708 123730 + 0 CTGCGAGCCCCACGGGTTGT TGG ||||||||[||||||||||||]PAM NA Intergenic NA NA

T9348 TTGGTCGACACGAACGTCTGCGG 1000
Oligo fwd CACCTTGGTCGACACGAACGTCTG
Oligo rev AAACCAGACGTTCGTGTCGACCAA
Chromosome start end strand MM target_seq PAM alignment distance position gene name gene id
GU305914.1 123727 123749 + 0 TTGGTCGACACGAACGTCTG CGG ||||||||[||||||||||||]PAM NA Intergenic NA NA

T9349 TGGTCGACACGAACGTCTGCGGG 1000
Oligo fwd CACCTGGTCGACACGAACGTCTGC
Oligo rev AAACGCAGACGTTCGTGTCGACCA
Chromosome start end strand MM target_seq PAM alignment distance position gene name gene id
GU305914.1 123728 123750 + 0 TGGTCGACACGAACGTCTGC GGG ||||||||[||||||||||||]PAM NA Intergenic NA NA

T9350 GACACGAACGTCTGCGGGTCCGG 1000
Oligo fwd CACCGACACGAACGTCTGCGGGTC
Oligo rev AAACGACCCGCAGACGTTCGTGTC
Chromosome start end strand MM target_seq PAM alignment distance position gene name gene id
GU305914.1 123733 123755 + 0 GACACGAACGTCTGCGGGTC CGG ||||||||[||||||||||||]PAM NA Intergenic NA NA

T9351 ACACGAACGTCTGCGGGTCCGGG 1000
Oligo fwd CACCACACGAACGTCTGCGGGTCC
Oligo rev AAACGGACCCGCAGACGTTCGTGT
Chromosome start end strand MM target_seq PAM alignment distance position gene name gene id
GU305914.1 123734 123756 + 0 ACACGAACGTCTGCGGGTCC GGG ||||||||[||||||||||||]PAM NA Intergenic NA NA

T9352 GAACGTCTGCGGGTCCGGGCTGG 1000
Oligo fwd CACCGAACGTCTGCGGGTCCGGGC
Oligo rev AAACGCCCGGACCCGCAGACGTTC
Chromosome start end strand MM target_seq PAM alignment distance position gene name gene id

<table>
<tr><td>GU305914.1</td><td></td><td>123738</td><td>123760</td><td>+</td><td>0</td><td>GAACGTCTGCGGGTCCGGGC</td><td>TGG</td><td>||||||||[||||||||||||]PAM</td><td>NA</td><td>Intergenic</td><td>NA</td><td>NA</td></tr>
<tr><td>T9353</td><td>TCTGCGGGTCCGGGCTGGTGTGG</td><td></td><td>1000</td><td></td><td></td><td></td><td></td><td></td><td></td><td></td><td></td><td></td></tr>
<tr><td>Oligo fwd</td><td>CACCTCTGCGGGTCCGGGCTGGTG</td><td></td><td></td><td></td><td></td><td></td><td></td><td></td><td></td><td></td><td></td><td></td></tr>
<tr><td>Oligo rev</td><td>AAACCACCAGCCCGGACCCGCAGA</td><td></td><td></td><td></td><td></td><td></td><td></td><td></td><td></td><td></td><td></td><td></td></tr>
<tr><td>Chromosome</td><td>start</td><td></td><td>end</td><td>strand</td><td>MM</td><td>target_seq</td><td>PAM</td><td>alignment</td><td>distance</td><td>position</td><td>gene name</td><td>gene id</td></tr>
<tr><td>GU305914.1</td><td></td><td>123743</td><td>123765</td><td>+</td><td>0</td><td>TCTGCGGGTCCGGGCTGGTG</td><td>TGG</td><td>||||||||[||||||||||||]PAM</td><td>NA</td><td>Intergenic</td><td>NA</td><td>NA</td></tr>
<tr><td>T9354</td><td>CCGGGCTGGTGTGGTCGTACAGG</td><td></td><td>1000</td><td></td><td></td><td></td><td></td><td></td><td></td><td></td><td></td><td></td></tr>
<tr><td>Oligo fwd</td><td>CACCCCGGGCTGGTGTGGTCGTAC</td><td></td><td></td><td></td><td></td><td></td><td></td><td></td><td></td><td></td><td></td><td></td></tr>
<tr><td>Oligo rev</td><td>AAACGTACGACCACACCAGCCCGG</td><td></td><td></td><td></td><td></td><td></td><td></td><td></td><td></td><td></td><td></td><td></td></tr>
<tr><td>Chromosome</td><td>start</td><td></td><td>end</td><td>strand</td><td>MM</td><td>target_seq</td><td>PAM</td><td>alignment</td><td>distance</td><td>position</td><td>gene name</td><td>gene id</td></tr>
<tr><td>GU305914.1</td><td></td><td>123752</td><td>123774</td><td>+</td><td>0</td><td>CCGGGCTGGTGTGGTCGTAC</td><td>AGG</td><td>||||||||[||||||||||||]PAM</td><td>NA</td><td>Intergenic</td><td>NA</td><td>NA</td></tr>
<tr><td>T9355</td><td>CGGGCTGGTGTGGTCGTACAGGG</td><td></td><td>1000</td><td></td><td></td><td></td><td></td><td></td><td></td><td></td><td></td><td></td></tr>
<tr><td>Oligo fwd</td><td>CACCCGGGCTGGTGTGGTCGTACA</td><td></td><td></td><td></td><td></td><td></td><td></td><td></td><td></td><td></td><td></td><td></td></tr>
<tr><td>Oligo rev</td><td>AAACTGTACGACCACACCAGCCCG</td><td></td><td></td><td></td><td></td><td></td><td></td><td></td><td></td><td></td><td></td><td></td></tr>
<tr><td>Chromosome</td><td>start</td><td></td><td>end</td><td>strand</td><td>MM</td><td>target_seq</td><td>PAM</td><td>alignment</td><td>distance</td><td>position</td><td>gene name</td><td>gene id</td></tr>
<tr><td>GU305914.1</td><td></td><td>123753</td><td>123775</td><td>+</td><td>0</td><td>CGGGCTGGTGTGGTCGTACA</td><td>GGG</td><td>||||||||[||||||||||||]PAM</td><td>NA</td><td>Intergenic</td><td>NA</td><td>NA</td></tr>
<tr><td>T9356</td><td>GCTGGTGTGGTCGTACAGGGTGG</td><td></td><td>1000</td><td></td><td></td><td></td><td></td><td></td><td></td><td></td><td></td><td></td></tr>
<tr><td>Oligo fwd</td><td>CACCGCTGGTGTGGTCGTACAGGG</td><td></td><td></td><td></td><td></td><td></td><td></td><td></td><td></td><td></td><td></td><td></td></tr>
<tr><td>Oligo rev</td><td>AAACCCCTGTACGACCACACCAGC</td><td></td><td></td><td></td><td></td><td></td><td></td><td></td><td></td><td></td><td></td><td></td></tr>
<tr><td>Chromosome</td><td>start</td><td></td><td>end</td><td>strand</td><td>MM</td><td>target_seq</td><td>PAM</td><td>alignment</td><td>distance</td><td>position</td><td>gene name</td><td>gene id</td></tr>
<tr><td>GU305914.1</td><td></td><td>123756</td><td>123778</td><td>+</td><td>0</td><td>GCTGGTGTGGTCGTACAGGG</td><td>TGG</td><td>||||||||[||||||||||||]PAM</td><td>NA</td><td>Intergenic</td><td>NA</td><td>NA</td></tr>
<tr><td>T9357</td><td>AGCGCCGCGTCCTCGTCGTACGG</td><td></td><td>1000</td><td></td><td></td><td></td><td></td><td></td><td></td><td></td><td></td><td></td></tr>
<tr><td>Oligo fwd</td><td>CACCAGCGCCGCGTCCTCGTCGTA</td><td></td><td></td><td></td><td></td><td></td><td></td><td></td><td></td><td></td><td></td><td></td></tr>
<tr><td>Oligo rev</td><td>AAACTACGACGAGGACGCGGCGCT</td><td></td><td></td><td></td><td></td><td></td><td></td><td></td><td></td><td></td><td></td><td></td></tr>
<tr><td>Chromosome</td><td>start</td><td></td><td>end</td><td>strand</td><td>MM</td><td>target_seq</td><td>PAM</td><td>alignment</td><td>distance</td><td>position</td><td>gene name</td><td>gene id</td></tr>
<tr><td>GU305914.1</td><td></td><td>123781</td><td>123803</td><td>+</td><td>0</td><td>AGCGCCGCGTCCTCGTCGTA</td><td>CGG</td><td>||||||||[||||||||||||]PAM</td><td>NA</td><td>Intergenic</td><td>NA</td><td>NA</td></tr>
<tr><td>T9358</td><td>GCGCCGCGTCCTCGTCGTACGGG</td><td></td><td>1000</td><td></td><td></td><td></td><td></td><td></td><td></td><td></td><td></td><td></td></tr>
<tr><td>Oligo fwd</td><td>CACCGCGCCGCGTCCTCGTCGTAC</td><td></td><td></td><td></td><td></td><td></td><td></td><td></td><td></td><td></td><td></td><td></td></tr>
<tr><td>Oligo rev</td><td>AAACGTACGACGAGGACGCGGCGC</td><td></td><td></td><td></td><td></td><td></td><td></td><td></td><td></td><td></td><td></td><td></td></tr>
<tr><td>Chromosome</td><td>start</td><td></td><td>end</td><td>strand</td><td>MM</td><td>target_seq</td><td>PAM</td><td>alignment</td><td>distance</td><td>position</td><td>gene name</td><td>gene id</td></tr>
<tr><td>GU305914.1</td><td></td><td>123782</td><td>123804</td><td>+</td><td>0</td><td>GCGCCGCGTCCTCGTCGTAC</td><td>GGG</td><td>||||||||[||||||||||||]PAM</td><td>NA</td><td>Intergenic</td><td>NA</td><td>NA</td></tr>
<tr><td>T9359</td><td>ATCGACGACGAGCGCCCGCGCGG</td><td></td><td>1000</td><td></td><td></td><td></td><td></td><td></td><td></td><td></td><td></td><td></td></tr>
<tr><td>Oligo fwd</td><td>CACCATCGACGACGAGCGCCCGCG</td><td></td><td></td><td></td><td></td><td></td><td></td><td></td><td></td><td></td><td></td><td></td></tr>
<tr><td>Oligo rev</td><td>AAACCGCGGGCGCTCGTCGTCGAT</td><td></td><td></td><td></td><td></td><td></td><td></td><td></td><td></td><td></td><td></td><td></td></tr>
<tr><td>Chromosome</td><td>start</td><td></td><td>end</td><td>strand</td><td>MM</td><td>target_seq</td><td>PAM</td><td>alignment</td><td>distance</td><td>position</td><td>gene name</td><td>gene id</td></tr>
<tr><td>GU305914.1</td><td></td><td>123817</td><td>123839</td><td>+</td><td>0</td><td>ATCGACGACGAGCGCCCGCG</td><td>CGG</td><td>||||||||[||||||||||||]PAM</td><td>NA</td><td>Intergenic</td><td>NA</td><td>NA</td></tr>
<tr><td>T9360</td><td>TCGACGACGAGCGCCCGCGCGGG</td><td></td><td>1000</td><td></td><td></td><td></td><td></td><td></td><td></td><td></td><td></td><td></td></tr>
<tr><td>Oligo fwd</td><td>CACCTCGACGACGAGCGCCCGCGC</td><td></td><td></td><td></td><td></td><td></td><td></td><td></td><td></td><td></td><td></td><td></td></tr>
<tr><td>Oligo rev</td><td>AAACGCGCGGGCGCTCGTCGTCGA</td><td></td><td></td><td></td><td></td><td></td><td></td><td></td><td></td><td></td><td></td><td></td></tr>
<tr><td>Chromosome</td><td>start</td><td></td><td>end</td><td>strand</td><td>MM</td><td>target_seq</td><td>PAM</td><td>alignment</td><td>distance</td><td>position</td><td>gene name</td><td>gene id</td></tr>
<tr><td>GU305914.1</td><td></td><td>123818</td><td>123840</td><td>+</td><td>0</td><td>TCGACGACGAGCGCCCGCGC</td><td>GGG</td><td>||||||||[||||||||||||]PAM</td><td>NA</td><td>Intergenic</td><td>NA</td><td>NA</td></tr>
<tr><td>T9361</td><td>GAGCGCCCGCGCGGGTTGTTCGG</td><td></td><td>1000</td><td></td><td></td><td></td><td></td><td></td><td></td><td></td><td></td><td></td></tr>
<tr><td>Oligo fwd</td><td>CACCGAGCGCCCGCGCGGGTTGTT</td><td></td><td></td><td></td><td></td><td></td><td></td><td></td><td></td><td></td><td></td><td></td></tr>
<tr><td>Oligo rev</td><td>AAACAACAACCCGCGCGGGCGCTC</td><td></td><td></td><td></td><td></td><td></td><td></td><td></td><td></td><td></td><td></td><td></td></tr>
<tr><td>Chromosome</td><td>start</td><td></td><td>end</td><td>strand</td><td>MM</td><td>target_seq</td><td>PAM</td><td>alignment</td><td>distance</td><td>position</td><td>gene name</td><td>gene id</td></tr>
<tr><td>GU305914.1</td><td></td><td>123826</td><td>123848</td><td>+</td><td>0</td><td>GAGCGCCCGCGCGGGTTGTT</td><td>CGG</td><td>||||||||[||||||||||||]PAM</td><td>NA</td><td>Intergenic</td><td>NA</td><td>NA</td></tr>
<tr><td>T9362</td><td>AGCGCCCGCGCGGGTTGTTCGGG</td><td></td><td>1000</td><td></td><td></td><td></td><td></td><td></td><td></td><td></td><td></td><td></td></tr>
<tr><td>Oligo fwd</td><td>CACCAGCGCCCGCGCGGGTTGTTC</td><td></td><td></td><td></td><td></td><td></td><td></td><td></td><td></td><td></td><td></td><td></td></tr>
<tr><td>Oligo rev</td><td>AAACGAACAACCCGCGCGGGCGCT</td><td></td><td></td><td></td><td></td><td></td><td></td><td></td><td></td><td></td><td></td><td></td></tr>
<tr><td>Chromosome</td><td>start</td><td></td><td>end</td><td>strand</td><td>MM</td><td>target_seq</td><td>PAM</td><td>alignment</td><td>distance</td><td>position</td><td>gene name</td><td>gene id</td></tr>
<tr><td>GU305914.1</td><td></td><td>123827</td><td>123849</td><td>+</td><td>0</td><td>AGCGCCCGCGCGGGTTGTTC</td><td>GGG</td><td>||||||||[||||||||||||]PAM</td><td>NA</td><td>Intergenic</td><td>NA</td><td>NA</td></tr>
<tr><td>T9363</td><td>CGCGCGGGTTGTTCGGGTAGCGG</td><td></td><td>1000</td><td></td><td></td><td></td><td></td><td></td><td></td><td></td><td></td><td></td></tr>
<tr><td>Oligo fwd</td><td>CACCCGCGCGGGTTGTTCGGGTAG</td><td></td><td></td><td></td><td></td><td></td><td></td><td></td><td></td><td></td><td></td><td></td></tr>
<tr><td>Oligo rev</td><td>AAACCTACCCGAACAACCCGCGCG</td><td></td><td></td><td></td><td></td><td></td><td></td><td></td><td></td><td></td><td></td><td></td></tr>
<tr><td>Chromosome</td><td>start</td><td></td><td>end</td><td>strand</td><td>MM</td><td>target_seq</td><td>PAM</td><td>alignment</td><td>distance</td><td>position</td><td>gene name</td><td>gene id</td></tr>
<tr><td>GU305914.1</td><td></td><td>123833</td><td>123855</td><td>+</td><td>0</td><td>CGCGCGGGTTGTTCGGGTAG</td><td>CGG</td><td>||||||||[||||||||||||]PAM</td><td>NA</td><td>Intergenic</td><td>NA</td><td>NA</td></tr>
<tr><td>T9364</td><td>TCGGGTAGCGGAAGAACTCCAGG</td><td></td><td>1000</td><td></td><td></td><td></td><td></td><td></td><td></td><td></td><td></td><td></td></tr>
<tr><td>Oligo fwd</td><td>CACCTCGGGTAGCGGAAGAACTCC</td><td></td><td></td><td></td><td></td><td></td><td></td><td></td><td></td><td></td><td></td><td></td></tr>
<tr><td>Oligo rev</td><td>AAACGGAGTTCTTCCGCTACCCGA</td><td></td><td></td><td></td><td></td><td></td><td></td><td></td><td></td><td></td><td></td><td></td></tr>
<tr><td>Chromosome</td><td>start</td><td></td><td>end</td><td>strand</td><td>MM</td><td>target_seq</td><td>PAM</td><td>alignment</td><td>distance</td><td>position</td><td>gene name</td><td>gene id</td></tr>
<tr><td>GU305914.1</td><td></td><td>123845</td><td>123867</td><td>+</td><td>0</td><td>TCGGGTAGCGGAAGAACTCC</td><td>AGG</td><td>||||||||[||||||||||||]PAM</td><td>NA</td><td>Intergenic</td><td>NA</td><td>NA</td></tr>
<tr><td>T9365</td><td>GTAGCGGAAGAACTCCAGGTCGG</td><td></td><td>1000</td><td></td><td></td><td></td><td></td><td></td><td></td><td></td><td></td><td></td></tr>
<tr><td>Oligo fwd</td><td>CACCGTAGCGGAAGAACTCCAGGT</td><td></td><td></td><td></td><td></td><td></td><td></td><td></td><td></td><td></td><td></td><td></td></tr>
<tr><td>Oligo rev</td><td>AAACACCTGGAGTTCTTCCGCTAC</td><td></td><td></td><td></td><td></td><td></td><td></td><td></td><td></td><td></td><td></td><td></td></tr>
<tr><td>Chromosome</td><td>start</td><td></td><td>end</td><td>strand</td><td>MM</td><td>target_seq</td><td>PAM</td><td>alignment</td><td>distance</td><td>position</td><td>gene name</td><td>gene id</td></tr>
<tr><td>GU305914.1</td><td></td><td>123849</td><td>123871</td><td>+</td><td>0</td><td>GTAGCGGAAGAACTCCAGGT</td><td>CGG</td><td>||||||||[||||||||||||]PAM</td><td>NA</td><td>Intergenic</td><td>NA</td><td>NA</td></tr>
<tr><td>T9366</td><td>AACTCCAGGTCGGCCGACACCGG</td><td></td><td>1000</td><td></td><td></td><td></td><td></td><td></td><td></td><td></td><td></td><td></td></tr>
<tr><td>Oligo fwd</td><td>CACCAACTCCAGGTCGGCCGACAC</td><td></td><td></td><td></td><td></td><td></td><td></td><td></td><td></td><td></td><td></td><td></td></tr>
<tr><td>Oligo rev</td><td>AAACGTGTCGGCCGACCTGGAGTT</td><td></td><td></td><td></td><td></td><td></td><td></td><td></td><td></td><td></td><td></td><td></td></tr>
<tr><td>Chromosome</td><td>start</td><td></td><td>end</td><td>strand</td><td>MM</td><td>target_seq</td><td>PAM</td><td>alignment</td><td>distance</td><td>position</td><td>gene name</td><td>gene id</td></tr>
<tr><td>GU305914.1</td><td></td><td>123859</td><td>123881</td><td>+</td><td>0</td><td>AACTCCAGGTCGGCCGACAC</td><td>CGG</td><td>||||||||[||||||||||||]PAM</td><td>NA</td><td>Intergenic</td><td>NA</td><td>NA</td></tr>
<tr><td>T9367</td><td>GGTCGGCCGACACCGGCGTCAGG</td><td></td><td>1000</td><td></td><td></td><td></td><td></td><td></td><td></td><td></td><td></td><td></td></tr>
<tr><td>Oligo fwd</td><td>CACCGGTCGGCCGACACCGGCGTC</td><td></td><td></td><td></td><td></td><td></td><td></td><td></td><td></td><td></td><td></td><td></td></tr>
<tr><td>Oligo rev</td><td>AAACGACGCCGGTGTCGGCCGACC</td><td></td><td></td><td></td><td></td><td></td><td></td><td></td><td></td><td></td><td></td><td></td></tr>
</table>

<table>
<tr><td>Chromosome</td><td>start</td><td>end</td><td>strand</td><td>MM</td><td>target_seq</td><td>PAM</td><td>alignment</td><td>distance</td><td>position</td><td>gene name</td><td>gene id</td></tr>
<tr><td>GU305914.1</td><td>123866</td><td>123888</td><td>+</td><td>0</td><td>GGTCGGCCGACACCGGCGTC</td><td>AGG</td><td>||||||||[||||||||||||]PAM</td><td>NA</td><td>Intergenic</td><td>NA</td><td>NA</td></tr>
<tr><td></td><td></td><td></td><td></td><td></td><td></td><td></td><td></td><td></td><td></td><td></td><td></td></tr>
<tr><td>T9368</td><td>CGTCAGGATCACCTCGCACGTGG</td><td>1000</td><td></td><td></td><td></td><td></td><td></td><td></td><td></td><td></td><td></td></tr>
<tr><td>Oligo fwd</td><td>CACCCGTCAGGATCACCTCGCACG</td><td></td><td></td><td></td><td></td><td></td><td></td><td></td><td></td><td></td><td></td></tr>
<tr><td>Oligo rev</td><td>AAACCGTGCGAGGTGATCCTGACG</td><td></td><td></td><td></td><td></td><td></td><td></td><td></td><td></td><td></td><td></td></tr>
<tr><td>Chromosome</td><td>start</td><td>end</td><td>strand</td><td>MM</td><td>target_seq</td><td>PAM</td><td>alignment</td><td>distance</td><td>position</td><td>gene name</td><td>gene id</td></tr>
<tr><td>GU305914.1</td><td>123882</td><td>123904</td><td>+</td><td>0</td><td>CGTCAGGATCACCTCGCACG</td><td>TGG</td><td>||||||||[||||||||||||]PAM</td><td>NA</td><td>Intergenic</td><td>NA</td><td>NA</td></tr>
<tr><td></td><td></td><td></td><td></td><td></td><td></td><td></td><td></td><td></td><td></td><td></td><td></td></tr>
<tr><td>T9369</td><td>CGCTGCCCGTGCAGCAGCGCCGG</td><td>1000</td><td></td><td></td><td></td><td></td><td></td><td></td><td></td><td></td><td></td></tr>
<tr><td>Oligo fwd</td><td>CACCCGCTGCCCGTGCAGCAGCGC</td><td></td><td></td><td></td><td></td><td></td><td></td><td></td><td></td><td></td><td></td></tr>
<tr><td>Oligo rev</td><td>AAACGCGCTGCTGCACGGGCAGCG</td><td></td><td></td><td></td><td></td><td></td><td></td><td></td><td></td><td></td><td></td></tr>
<tr><td>Chromosome</td><td>start</td><td>end</td><td>strand</td><td>MM</td><td>target_seq</td><td>PAM</td><td>alignment</td><td>distance</td><td>position</td><td>gene name</td><td>gene id</td></tr>
<tr><td>GU305914.1</td><td>123907</td><td>123929</td><td>+</td><td>0</td><td>CGCTGCCCGTGCAGCAGCGC</td><td>CGG</td><td>||||||||[||||||||||||]PAM</td><td>NA</td><td>Intergenic</td><td>NA</td><td>NA</td></tr>
<tr><td></td><td></td><td></td><td></td><td></td><td></td><td></td><td></td><td></td><td></td><td></td><td></td></tr>
<tr><td>T9370</td><td>GCTGCCCGTGCAGCAGCGCCGGG</td><td>1000</td><td></td><td></td><td></td><td></td><td></td><td></td><td></td><td></td><td></td></tr>
<tr><td>Oligo fwd</td><td>CACCGCTGCCCGTGCAGCAGCGCC</td><td></td><td></td><td></td><td></td><td></td><td></td><td></td><td></td><td></td><td></td></tr>
<tr><td>Oligo rev</td><td>AAACGGCGCTGCTGCACGGGCAGC</td><td></td><td></td><td></td><td></td><td></td><td></td><td></td><td></td><td></td><td></td></tr>
<tr><td>Chromosome</td><td>start</td><td>end</td><td>strand</td><td>MM</td><td>target_seq</td><td>PAM</td><td>alignment</td><td>distance</td><td>position</td><td>gene name</td><td>gene id</td></tr>
<tr><td>GU305914.1</td><td>123908</td><td>123930</td><td>+</td><td>0</td><td>GCTGCCCGTGCAGCAGCGCC</td><td>GGG</td><td>||||||||[||||||||||||]PAM</td><td>NA</td><td>Intergenic</td><td>NA</td><td>NA</td></tr>
<tr><td></td><td></td><td></td><td></td><td></td><td></td><td></td><td></td><td></td><td></td><td></td><td></td></tr>
<tr><td>T9371</td><td>CTTGTCCGTCTTGCGCCCGAAGG</td><td>1000</td><td></td><td></td><td></td><td></td><td></td><td></td><td></td><td></td><td></td></tr>
<tr><td>Oligo fwd</td><td>CACCCTTGTCCGTCTTGCGCCCGA</td><td></td><td></td><td></td><td></td><td></td><td></td><td></td><td></td><td></td><td></td></tr>
<tr><td>Oligo rev</td><td>AAACTCGGGCGCAAGACGGACAAG</td><td></td><td></td><td></td><td></td><td></td><td></td><td></td><td></td><td></td><td></td></tr>
<tr><td>Chromosome</td><td>start</td><td>end</td><td>strand</td><td>MM</td><td>target_seq</td><td>PAM</td><td>alignment</td><td>distance</td><td>position</td><td>gene name</td><td>gene id</td></tr>
<tr><td>GU305914.1</td><td>123936</td><td>123958</td><td>+</td><td>0</td><td>CTTGTCCGTCTTGCGCCCGA</td><td>AGG</td><td>||||||||[||||||||||||]PAM</td><td>NA</td><td>Intergenic</td><td>NA</td><td>NA</td></tr>
<tr><td></td><td></td><td></td><td></td><td></td><td></td><td></td><td></td><td></td><td></td><td></td><td></td></tr>
<tr><td>T9372</td><td>CCGTCTTGCGCCCGAAGGTCAGG</td><td>1000</td><td></td><td></td><td></td><td></td><td></td><td></td><td></td><td></td><td></td></tr>
<tr><td>Oligo fwd</td><td>CACCCCGTCTTGCGCCCGAAGGTC</td><td></td><td></td><td></td><td></td><td></td><td></td><td></td><td></td><td></td><td></td></tr>
<tr><td>Oligo rev</td><td>AAACGACCTTCGGGCGCAAGACGG</td><td></td><td></td><td></td><td></td><td></td><td></td><td></td><td></td><td></td><td></td></tr>
<tr><td>Chromosome</td><td>start</td><td>end</td><td>strand</td><td>MM</td><td>target_seq</td><td>PAM</td><td>alignment</td><td>distance</td><td>position</td><td>gene name</td><td>gene id</td></tr>
<tr><td>GU305914.1</td><td>123941</td><td>123963</td><td>+</td><td>0</td><td>CCGTCTTGCGCCCGAAGGTC</td><td>AGG</td><td>||||||||[||||||||||||]PAM</td><td>NA</td><td>Intergenic</td><td>NA</td><td>NA</td></tr>
<tr><td></td><td></td><td></td><td></td><td></td><td></td><td></td><td></td><td></td><td></td><td></td><td></td></tr>
<tr><td>T9373</td><td>CCCGAAGGTCAGGATCGAGATGG</td><td>1000</td><td></td><td></td><td></td><td></td><td></td><td></td><td></td><td></td><td></td></tr>
<tr><td>Oligo fwd</td><td>CACCCCCGAAGGTCAGGATCGAGA</td><td></td><td></td><td></td><td></td><td></td><td></td><td></td><td></td><td></td><td></td></tr>
<tr><td>Oligo rev</td><td>AAACTCTCGATCCTGACCTTCGGG</td><td></td><td></td><td></td><td></td><td></td><td></td><td></td><td></td><td></td><td></td></tr>
<tr><td>Chromosome</td><td>start</td><td>end</td><td>strand</td><td>MM</td><td>target_seq</td><td>PAM</td><td>alignment</td><td>distance</td><td>position</td><td>gene name</td><td>gene id</td></tr>
<tr><td>GU305914.1</td><td>123951</td><td>123973</td><td>+</td><td>0</td><td>CCCGAAGGTCAGGATCGAGA</td><td>TGG</td><td>||||||||[||||||||||||]PAM</td><td>NA</td><td>Intergenic</td><td>NA</td><td>NA</td></tr>
<tr><td></td><td></td><td></td><td></td><td></td><td></td><td></td><td></td><td></td><td></td><td></td><td></td></tr>
<tr><td>T9374</td><td>GGCCTCCGAGTCCAGCACCTCGG</td><td>1000</td><td></td><td></td><td></td><td></td><td></td><td></td><td></td><td></td><td></td></tr>
<tr><td>Oligo fwd</td><td>CACCGGCCTCCGAGTCCAGCACCT</td><td></td><td></td><td></td><td></td><td></td><td></td><td></td><td></td><td></td><td></td></tr>
<tr><td>Oligo rev</td><td>AAACAGGTGCTGGACTCGGAGGCC</td><td></td><td></td><td></td><td></td><td></td><td></td><td></td><td></td><td></td><td></td></tr>
<tr><td>Chromosome</td><td>start</td><td>end</td><td>strand</td><td>MM</td><td>target_seq</td><td>PAM</td><td>alignment</td><td>distance</td><td>position</td><td>gene name</td><td>gene id</td></tr>
<tr><td>GU305914.1</td><td>123972</td><td>123994</td><td>+</td><td>0</td><td>GGCCTCCGAGTCCAGCACCT</td><td>CGG</td><td>||||||||[||||||||||||]PAM</td><td>NA</td><td>Intergenic</td><td>NA</td><td>NA</td></tr>
<tr><td></td><td></td><td></td><td></td><td></td><td></td><td></td><td></td><td></td><td></td><td></td><td></td></tr>
<tr><td>T9375</td><td>CACCTCGGTGCGCCGCGCGCCGG</td><td>1000</td><td></td><td></td><td></td><td></td><td></td><td></td><td></td><td></td><td></td></tr>
<tr><td>Oligo fwd</td><td>CACCCACCTCGGTGCGCCGCGCGC</td><td></td><td></td><td></td><td></td><td></td><td></td><td></td><td></td><td></td><td></td></tr>
<tr><td>Oligo rev</td><td>AAACGCGCGCGGCGCACCGAGGTG</td><td></td><td></td><td></td><td></td><td></td><td></td><td></td><td></td><td></td><td></td></tr>
<tr><td>Chromosome</td><td>start</td><td>end</td><td>strand</td><td>MM</td><td>target_seq</td><td>PAM</td><td>alignment</td><td>distance</td><td>position</td><td>gene name</td><td>gene id</td></tr>
<tr><td>GU305914.1</td><td>123987</td><td>124009</td><td>+</td><td>0</td><td>CACCTCGGTGCGCCGCGCGC</td><td>CGG</td><td>||||||||[||||||||||||]PAM</td><td>NA</td><td>Intergenic</td><td>NA</td><td>NA</td></tr>
<tr><td></td><td></td><td></td><td></td><td></td><td></td><td></td><td></td><td></td><td></td><td></td><td></td></tr>
<tr><td>T9376</td><td>GCCGCGCGCCGGTCGCCTGTCGG</td><td>1000</td><td></td><td></td><td></td><td></td><td></td><td></td><td></td><td></td><td></td></tr>
<tr><td>Oligo fwd</td><td>CACCGCCGCGCGCCGGTCGCCTGT</td><td></td><td></td><td></td><td></td><td></td><td></td><td></td><td></td><td></td><td></td></tr>
<tr><td>Oligo rev</td><td>AAACACAGGCGACCGGCGCGCGGC</td><td></td><td></td><td></td><td></td><td></td><td></td><td></td><td></td><td></td><td></td></tr>
<tr><td>Chromosome</td><td>start</td><td>end</td><td>strand</td><td>MM</td><td>target_seq</td><td>PAM</td><td>alignment</td><td>distance</td><td>position</td><td>gene name</td><td>gene id</td></tr>
<tr><td>GU305914.1</td><td>123998</td><td>124020</td><td>+</td><td>0</td><td>GCCGCGCGCCGGTCGCCTGT</td><td>CGG</td><td>||||||||[||||||||||||]PAM</td><td>NA</td><td>Intergenic</td><td>NA</td><td>NA</td></tr>
<tr><td></td><td></td><td></td><td></td><td></td><td></td><td></td><td></td><td></td><td></td><td></td><td></td></tr>
<tr><td>T9377</td><td>GCCGGTCGCCTGTCGGACCCAGG</td><td>1000</td><td></td><td></td><td></td><td></td><td></td><td></td><td></td><td></td><td></td></tr>
<tr><td>Oligo fwd</td><td>CACCGCCGGTCGCCTGTCGGACCC</td><td></td><td></td><td></td><td></td><td></td><td></td><td></td><td></td><td></td><td></td></tr>
<tr><td>Oligo rev</td><td>AAACGGGTCCGACAGGCGACCGGC</td><td></td><td></td><td></td><td></td><td></td><td></td><td></td><td></td><td></td><td></td></tr>
<tr><td>Chromosome</td><td>start</td><td>end</td><td>strand</td><td>MM</td><td>target_seq</td><td>PAM</td><td>alignment</td><td>distance</td><td>position</td><td>gene name</td><td>gene id</td></tr>
<tr><td>GU305914.1</td><td>124005</td><td>124027</td><td>+</td><td>0</td><td>GCCGGTCGCCTGTCGGACCC</td><td>AGG</td><td>||||||||[||||||||||||]PAM</td><td>NA</td><td>Intergenic</td><td>NA</td><td>NA</td></tr>
<tr><td></td><td></td><td></td><td></td><td></td><td></td><td></td><td></td><td></td><td></td><td></td><td></td></tr>
<tr><td>T9378</td><td>TCGTTGCGATACACGTGCATCGG</td><td>1000</td><td></td><td></td><td></td><td></td><td></td><td></td><td></td><td></td><td></td></tr>
<tr><td>Oligo fwd</td><td>CACCTCGTTGCGATACACGTGCAT</td><td></td><td></td><td></td><td></td><td></td><td></td><td></td><td></td><td></td><td></td></tr>
<tr><td>Oligo rev</td><td>AAACATGCACGTGTATCGCAACGA</td><td></td><td></td><td></td><td></td><td></td><td></td><td></td><td></td><td></td><td></td></tr>
<tr><td>Chromosome</td><td>start</td><td>end</td><td>strand</td><td>MM</td><td>target_seq</td><td>PAM</td><td>alignment</td><td>distance</td><td>position</td><td>gene name</td><td>gene id</td></tr>
<tr><td>GU305914.1</td><td>124039</td><td>124061</td><td>+</td><td>0</td><td>TCGTTGCGATACACGTGCAT</td><td>CGG</td><td>||||||||[||||||||||||]PAM</td><td>NA</td><td>Intergenic</td><td>NA</td><td>NA</td></tr>
<tr><td></td><td></td><td></td><td></td><td></td><td></td><td></td><td></td><td></td><td></td><td></td><td></td></tr>
<tr><td>T9379</td><td>CGTTGCGATACACGTGCATCGGG</td><td>1000</td><td></td><td></td><td></td><td></td><td></td><td></td><td></td><td></td><td></td></tr>
<tr><td>Oligo fwd</td><td>CACCCGTTGCGATACACGTGCATC</td><td></td><td></td><td></td><td></td><td></td><td></td><td></td><td></td><td></td><td></td></tr>
<tr><td>Oligo rev</td><td>AAACGATGCACGTGTATCGCAACG</td><td></td><td></td><td></td><td></td><td></td><td></td><td></td><td></td><td></td><td></td></tr>
<tr><td>Chromosome</td><td>start</td><td>end</td><td>strand</td><td>MM</td><td>target_seq</td><td>PAM</td><td>alignment</td><td>distance</td><td>position</td><td>gene name</td><td>gene id</td></tr>
<tr><td>GU305914.1</td><td>124040</td><td>124062</td><td>+</td><td>0</td><td>CGTTGCGATACACGTGCATC</td><td>GGG</td><td>||||||||[||||||||||||]PAM</td><td>NA</td><td>Intergenic</td><td>NA</td><td>NA</td></tr>
<tr><td></td><td></td><td></td><td></td><td></td><td></td><td></td><td></td><td></td><td></td><td></td><td></td></tr>
<tr><td>T9380</td><td>ACACGTGCATCGGGAAGACCTGG</td><td>1000</td><td></td><td></td><td></td><td></td><td></td><td></td><td></td><td></td><td></td></tr>
<tr><td>Oligo fwd</td><td>CACCACACGTGCATCGGGAAGACC</td><td></td><td></td><td></td><td></td><td></td><td></td><td></td><td></td><td></td><td></td></tr>
<tr><td>Oligo rev</td><td>AAACGGTCTTCCCGATGCACGTGT</td><td></td><td></td><td></td><td></td><td></td><td></td><td></td><td></td><td></td><td></td></tr>
<tr><td>Chromosome</td><td>start</td><td>end</td><td>strand</td><td>MM</td><td>target_seq</td><td>PAM</td><td>alignment</td><td>distance</td><td>position</td><td>gene name</td><td>gene id</td></tr>
<tr><td>GU305914.1</td><td>124049</td><td>124071</td><td>+</td><td>0</td><td>ACACGTGCATCGGGAAGACC</td><td>TGG</td><td>||||||||[||||||||||||]PAM</td><td>NA</td><td>Intergenic</td><td>NA</td><td>NA</td></tr>
<tr><td></td><td></td><td></td><td></td><td></td><td></td><td></td><td></td><td></td><td></td><td></td><td></td></tr>
<tr><td>T9381</td><td>GCATCGGGAAGACCTGGAACAGG</td><td>1000</td><td></td><td></td><td></td><td></td><td></td><td></td><td></td><td></td><td></td></tr>
<tr><td>Oligo fwd</td><td>CACCGCATCGGGAAGACCTGGAAC</td><td></td><td></td><td></td><td></td><td></td><td></td><td></td><td></td><td></td><td></td></tr>
<tr><td>Oligo rev</td><td>AAACGTTCCAGGTCTTCCCGATGC</td><td></td><td></td><td></td><td></td><td></td><td></td><td></td><td></td><td></td><td></td></tr>
<tr><td>Chromosome</td><td>start</td><td>end</td><td>strand</td><td>MM</td><td>target_seq</td><td>PAM</td><td>alignment</td><td>distance</td><td>position</td><td>gene name</td><td>gene id</td></tr>
<tr><td>GU305914.1</td><td>124055</td><td>124077</td><td>+</td><td>0</td><td>GCATCGGGAAGACCTGGAAC</td><td>AGG</td><td>||||||||[||||||||||||]PAM</td><td>NA</td><td>Intergenic</td><td>NA</td><td>NA</td></tr>
<tr><td></td><td></td><td></td><td></td><td></td><td></td><td></td><td></td><td></td><td></td><td></td><td></td></tr>
<tr><td>T9382</td><td>TGTCGCTCCGCACGCGCCGCAGG</td><td>1000</td><td></td><td></td><td></td><td></td><td></td><td></td><td></td><td></td><td></td></tr>
<tr><td>Oligo fwd</td><td>CACCTGTCGCTCCGCACGCGCCGC</td><td></td><td></td><td></td><td></td><td></td><td></td><td></td><td></td><td></td><td></td></tr>
</table>

<table>
<tr><td>Oligo rev</td><td>AAACGCGGCGCGTGCGGAGCGACA</td><td></td><td></td><td></td><td></td><td></td><td></td><td></td><td></td><td></td><td></td></tr>
<tr><td>Chromosome</td><td>start</td><td>end</td><td>strand</td><td>MM</td><td>target_seq</td><td>PAM</td><td>alignment</td><td>distance</td><td>position</td><td>gene name</td><td>gene id</td></tr>
<tr><td>GU305914.1</td><td>124097</td><td>124119</td><td>+</td><td>0</td><td>TGTCGCTCCGCACGCGCCGC</td><td>AGG</td><td>||||||||[||||||||||||]PAM</td><td>NA</td><td>Intergenic</td><td>NA</td><td>NA</td></tr>
<tr><td></td><td></td><td></td><td></td><td></td><td></td><td></td><td></td><td></td><td></td><td></td><td></td></tr>
<tr><td>T9383</td><td>ACGTCGTCGCCCCGTACCCCAGG</td><td>1000</td><td></td><td></td><td></td><td></td><td></td><td></td><td></td><td></td><td></td></tr>
<tr><td>Oligo fwd</td><td>CACCACGTCGTCGCCCCGTACCCC</td><td></td><td></td><td></td><td></td><td></td><td></td><td></td><td></td><td></td><td></td></tr>
<tr><td>Oligo rev</td><td>AAACGGGGTACGGGGCGACGACGT</td><td></td><td></td><td></td><td></td><td></td><td></td><td></td><td></td><td></td><td></td></tr>
<tr><td>Chromosome</td><td>start</td><td>end</td><td>strand</td><td>MM</td><td>target_seq</td><td>PAM</td><td>alignment</td><td>distance</td><td>position</td><td>gene name</td><td>gene id</td></tr>
<tr><td>GU305914.1</td><td>124130</td><td>124152</td><td>+</td><td>0</td><td>ACGTCGTCGCCCCGTACCCC</td><td>AGG</td><td>||||||||[||||||||||||]PAM</td><td>NA</td><td>Intergenic</td><td>NA</td><td>NA</td></tr>
<tr><td></td><td></td><td></td><td></td><td></td><td></td><td></td><td></td><td></td><td></td><td></td><td></td></tr>
<tr><td>T9384</td><td>TCTCCACCGTGTTGATGTTCTGG</td><td>1000</td><td></td><td></td><td></td><td></td><td></td><td></td><td></td><td></td><td></td></tr>
<tr><td>Oligo fwd</td><td>CACCTCTCCACCGTGTTGATGTTC</td><td></td><td></td><td></td><td></td><td></td><td></td><td></td><td></td><td></td><td></td></tr>
<tr><td>Oligo rev</td><td>AAACGAACATCAACACGGTGGAGA</td><td></td><td></td><td></td><td></td><td></td><td></td><td></td><td></td><td></td><td></td></tr>
<tr><td>Chromosome</td><td>start</td><td>end</td><td>strand</td><td>MM</td><td>target_seq</td><td>PAM</td><td>alignment</td><td>distance</td><td>position</td><td>gene name</td><td>gene id</td></tr>
<tr><td>GU305914.1</td><td>124157</td><td>124179</td><td>+</td><td>0</td><td>TCTCCACCGTGTTGATGTTC</td><td>TGG</td><td>||||||||[||||||||||||]PAM</td><td>NA</td><td>Intergenic</td><td>NA</td><td>NA</td></tr>
<tr><td></td><td></td><td></td><td></td><td></td><td></td><td></td><td></td><td></td><td></td><td></td><td></td></tr>
<tr><td>T9385</td><td>TGTTGATGTTCTGGCTCACGTGG</td><td>1000</td><td></td><td></td><td></td><td></td><td></td><td></td><td></td><td></td><td></td></tr>
<tr><td>Oligo fwd</td><td>CACCTGTTGATGTTCTGGCTCACG</td><td></td><td></td><td></td><td></td><td></td><td></td><td></td><td></td><td></td><td></td></tr>
<tr><td>Oligo rev</td><td>AAACCGTGAGCCAGAACATCAACA</td><td></td><td></td><td></td><td></td><td></td><td></td><td></td><td></td><td></td><td></td></tr>
<tr><td>Chromosome</td><td>start</td><td>end</td><td>strand</td><td>MM</td><td>target_seq</td><td>PAM</td><td>alignment</td><td>distance</td><td>position</td><td>gene name</td><td>gene id</td></tr>
<tr><td>GU305914.1</td><td>124166</td><td>124188</td><td>+</td><td>0</td><td>TGTTGATGTTCTGGCTCACG</td><td>TGG</td><td>||||||||[||||||||||||]PAM</td><td>NA</td><td>Intergenic</td><td>NA</td><td>NA</td></tr>
<tr><td></td><td></td><td></td><td></td><td></td><td></td><td></td><td></td><td></td><td></td><td></td><td></td></tr>
<tr><td>T9386</td><td>GGAACACCGTGTTGATGTCCCGG</td><td>1000</td><td></td><td></td><td></td><td></td><td></td><td></td><td></td><td></td><td></td></tr>
<tr><td>Oligo fwd</td><td>CACCGGAACACCGTGTTGATGTCC</td><td></td><td></td><td></td><td></td><td></td><td></td><td></td><td></td><td></td><td></td></tr>
<tr><td>Oligo rev</td><td>AAACGGACATCAACACGGTGTTCC</td><td></td><td></td><td></td><td></td><td></td><td></td><td></td><td></td><td></td><td></td></tr>
<tr><td>Chromosome</td><td>start</td><td>end</td><td>strand</td><td>MM</td><td>target_seq</td><td>PAM</td><td>alignment</td><td>distance</td><td>position</td><td>gene name</td><td>gene id</td></tr>
<tr><td>GU305914.1</td><td>124187</td><td>124209</td><td>+</td><td>0</td><td>GGAACACCGTGTTGATGTCC</td><td>CGG</td><td>||||||||[||||||||||||]PAM</td><td>NA</td><td>Intergenic</td><td>NA</td><td>NA</td></tr>
<tr><td></td><td></td><td></td><td></td><td></td><td></td><td></td><td></td><td></td><td></td><td></td><td></td></tr>
<tr><td>T9387</td><td>GTTGATGTCCCGGTCCTCCTTGG</td><td>1000</td><td></td><td></td><td></td><td></td><td></td><td></td><td></td><td></td><td></td></tr>
<tr><td>Oligo fwd</td><td>CACCGTTGATGTCCCGGTCCTCCT</td><td></td><td></td><td></td><td></td><td></td><td></td><td></td><td></td><td></td><td></td></tr>
<tr><td>Oligo rev</td><td>AAACAGGAGGACCGGGACATCAAC</td><td></td><td></td><td></td><td></td><td></td><td></td><td></td><td></td><td></td><td></td></tr>
<tr><td>Chromosome</td><td>start</td><td>end</td><td>strand</td><td>MM</td><td>target_seq</td><td>PAM</td><td>alignment</td><td>distance</td><td>position</td><td>gene name</td><td>gene id</td></tr>
<tr><td>GU305914.1</td><td>124197</td><td>124219</td><td>+</td><td>0</td><td>GTTGATGTCCCGGTCCTCCT</td><td>TGG</td><td>||||||||[||||||||||||]PAM</td><td>NA</td><td>Intergenic</td><td>NA</td><td>NA</td></tr>
<tr><td></td><td></td><td></td><td></td><td></td><td></td><td></td><td></td><td></td><td></td><td></td><td></td></tr>
<tr><td>T9388</td><td>CGGTCCTCCTTGGTCACGATCGG</td><td>1000</td><td></td><td></td><td></td><td></td><td></td><td></td><td></td><td></td><td></td></tr>
<tr><td>Oligo fwd</td><td>CACCCGGTCCTCCTTGGTCACGAT</td><td></td><td></td><td></td><td></td><td></td><td></td><td></td><td></td><td></td><td></td></tr>
<tr><td>Oligo rev</td><td>AAACATCGTGACCAAGGAGGACCG</td><td></td><td></td><td></td><td></td><td></td><td></td><td></td><td></td><td></td><td></td></tr>
<tr><td>Chromosome</td><td>start</td><td>end</td><td>strand</td><td>MM</td><td>target_seq</td><td>PAM</td><td>alignment</td><td>distance</td><td>position</td><td>gene name</td><td>gene id</td></tr>
<tr><td>GU305914.1</td><td>124207</td><td>124229</td><td>+</td><td>0</td><td>CGGTCCTCCTTGGTCACGAT</td><td>CGG</td><td>||||||||[||||||||||||]PAM</td><td>NA</td><td>Intergenic</td><td>NA</td><td>NA</td></tr>
<tr><td></td><td></td><td></td><td></td><td></td><td></td><td></td><td></td><td></td><td></td><td></td><td></td></tr>
<tr><td>T9389</td><td>GATCACCGCCGTCGACGCCTTGG</td><td>1000</td><td></td><td></td><td></td><td></td><td></td><td></td><td></td><td></td><td></td></tr>
<tr><td>Oligo fwd</td><td>CACCGATCACCGCCGTCGACGCCT</td><td></td><td></td><td></td><td></td><td></td><td></td><td></td><td></td><td></td><td></td></tr>
<tr><td>Oligo rev</td><td>AAACAGGCGTCGACGGCGGTGATC</td><td></td><td></td><td></td><td></td><td></td><td></td><td></td><td></td><td></td><td></td></tr>
<tr><td>Chromosome</td><td>start</td><td>end</td><td>strand</td><td>MM</td><td>target_seq</td><td>PAM</td><td>alignment</td><td>distance</td><td>position</td><td>gene name</td><td>gene id</td></tr>
<tr><td>GU305914.1</td><td>124239</td><td>124261</td><td>+</td><td>0</td><td>GATCACCGCCGTCGACGCCT</td><td>TGG</td><td>||||||||[||||||||||||]PAM</td><td>NA</td><td>Intergenic</td><td>NA</td><td>NA</td></tr>
<tr><td></td><td></td><td></td><td></td><td></td><td></td><td></td><td></td><td></td><td></td><td></td><td></td></tr>
<tr><td>T9390</td><td>GAACGTGTCCGTCCGCACCACGG</td><td>1000</td><td></td><td></td><td></td><td></td><td></td><td></td><td></td><td></td><td></td></tr>
<tr><td>Oligo fwd</td><td>CACCGAACGTGTCCGTCCGCACCA</td><td></td><td></td><td></td><td></td><td></td><td></td><td></td><td></td><td></td><td></td></tr>
<tr><td>Oligo rev</td><td>AAACTGGTGCGGACGGACACGTTC</td><td></td><td></td><td></td><td></td><td></td><td></td><td></td><td></td><td></td><td></td></tr>
<tr><td>Chromosome</td><td>start</td><td>end</td><td>strand</td><td>MM</td><td>target_seq</td><td>PAM</td><td>alignment</td><td>distance</td><td>position</td><td>gene name</td><td>gene id</td></tr>
<tr><td>GU305914.1</td><td>124287</td><td>124309</td><td>+</td><td>0</td><td>GAACGTGTCCGTCCGCACCA</td><td>CGG</td><td>||||||||[||||||||||||]PAM</td><td>NA</td><td>Intergenic</td><td>NA</td><td>NA</td></tr>
<tr><td></td><td></td><td></td><td></td><td></td><td></td><td></td><td></td><td></td><td></td><td></td><td></td></tr>
<tr><td>T9391</td><td>ACCACGGTCAGCGCGAAGCCCGG</td><td>1000</td><td></td><td></td><td></td><td></td><td></td><td></td><td></td><td></td><td></td></tr>
<tr><td>Oligo fwd</td><td>CACCACCACGGTCAGCGCGAAGCC</td><td></td><td></td><td></td><td></td><td></td><td></td><td></td><td></td><td></td><td></td></tr>
<tr><td>Oligo rev</td><td>AAACGGCTTCGCGCTGACCGTGGT</td><td></td><td></td><td></td><td></td><td></td><td></td><td></td><td></td><td></td><td></td></tr>
<tr><td>Chromosome</td><td>start</td><td>end</td><td>strand</td><td>MM</td><td>target_seq</td><td>PAM</td><td>alignment</td><td>distance</td><td>position</td><td>gene name</td><td>gene id</td></tr>
<tr><td>GU305914.1</td><td>124303</td><td>124325</td><td>+</td><td>0</td><td>ACCACGGTCAGCGCGAAGCC</td><td>CGG</td><td>||||||||[||||||||||||]PAM</td><td>NA</td><td>Intergenic</td><td>NA</td><td>NA</td></tr>
<tr><td></td><td></td><td></td><td></td><td></td><td></td><td></td><td></td><td></td><td></td><td></td><td></td></tr>
<tr><td>T9392</td><td>CCACGGTCAGCGCGAAGCCCGGG</td><td>1000</td><td></td><td></td><td></td><td></td><td></td><td></td><td></td><td></td><td></td></tr>
<tr><td>Oligo fwd</td><td>CACCCCACGGTCAGCGCGAAGCCC</td><td></td><td></td><td></td><td></td><td></td><td></td><td></td><td></td><td></td><td></td></tr>
<tr><td>Oligo rev</td><td>AAACGGGCTTCGCGCTGACCGTGG</td><td></td><td></td><td></td><td></td><td></td><td></td><td></td><td></td><td></td><td></td></tr>
<tr><td>Chromosome</td><td>start</td><td>end</td><td>strand</td><td>MM</td><td>target_seq</td><td>PAM</td><td>alignment</td><td>distance</td><td>position</td><td>gene name</td><td>gene id</td></tr>
<tr><td>GU305914.1</td><td>124304</td><td>124326</td><td>+</td><td>0</td><td>CCACGGTCAGCGCGAAGCCC</td><td>GGG</td><td>||||||||[||||||||||||]PAM</td><td>NA</td><td>Intergenic</td><td>NA</td><td>NA</td></tr>
<tr><td></td><td></td><td></td><td></td><td></td><td></td><td></td><td></td><td></td><td></td><td></td><td></td></tr>
<tr><td>T9393</td><td>TCAGCGCGAAGCCCGGGTGCAGG</td><td>1000</td><td></td><td></td><td></td><td></td><td></td><td></td><td></td><td></td><td></td></tr>
<tr><td>Oligo fwd</td><td>CACCTCAGCGCGAAGCCCGGGTGC</td><td></td><td></td><td></td><td></td><td></td><td></td><td></td><td></td><td></td><td></td></tr>
<tr><td>Oligo rev</td><td>AAACGCACCCGGGCTTCGCGCTGA</td><td></td><td></td><td></td><td></td><td></td><td></td><td></td><td></td><td></td><td></td></tr>
<tr><td>Chromosome</td><td>start</td><td>end</td><td>strand</td><td>MM</td><td>target_seq</td><td>PAM</td><td>alignment</td><td>distance</td><td>position</td><td>gene name</td><td>gene id</td></tr>
<tr><td>GU305914.1</td><td>124310</td><td>124332</td><td>+</td><td>0</td><td>TCAGCGCGAAGCCCGGGTGC</td><td>AGG</td><td>||||||||[||||||||||||]PAM</td><td>NA</td><td>Intergenic</td><td>NA</td><td>NA</td></tr>
<tr><td></td><td></td><td></td><td></td><td></td><td></td><td></td><td></td><td></td><td></td><td></td><td></td></tr>
<tr><td>T9394</td><td>AGCCCGGGTGCAGGCCCATCCGG</td><td>1000</td><td></td><td></td><td></td><td></td><td></td><td></td><td></td><td></td><td></td></tr>
<tr><td>Oligo fwd</td><td>CACCAGCCCGGGTGCAGGCCCATC</td><td></td><td></td><td></td><td></td><td></td><td></td><td></td><td></td><td></td><td></td></tr>
<tr><td>Oligo rev</td><td>AAACGATGGGCCTGCACCCGGGCT</td><td></td><td></td><td></td><td></td><td></td><td></td><td></td><td></td><td></td><td></td></tr>
<tr><td>Chromosome</td><td>start</td><td>end</td><td>strand</td><td>MM</td><td>target_seq</td><td>PAM</td><td>alignment</td><td>distance</td><td>position</td><td>gene name</td><td>gene id</td></tr>
<tr><td>GU305914.1</td><td>124319</td><td>124341</td><td>+</td><td>0</td><td>AGCCCGGGTGCAGGCCCATC</td><td>CGG</td><td>||||||||[||||||||||||]PAM</td><td>NA</td><td>Intergenic</td><td>NA</td><td>NA</td></tr>
<tr><td></td><td></td><td></td><td></td><td></td><td></td><td></td><td></td><td></td><td></td><td></td><td></td></tr>
<tr><td>T9395</td><td>CTCAGCGCCGCGATCGACACCGG</td><td>1000</td><td></td><td></td><td></td><td></td><td></td><td></td><td></td><td></td><td></td></tr>
<tr><td>Oligo fwd</td><td>CACCCTCAGCGCCGCGATCGACAC</td><td></td><td></td><td></td><td></td><td></td><td></td><td></td><td></td><td></td><td></td></tr>
<tr><td>Oligo rev</td><td>AAACGTGTCGATCGCGGCGCTGAG</td><td></td><td></td><td></td><td></td><td></td><td></td><td></td><td></td><td></td><td></td></tr>
<tr><td>Chromosome</td><td>start</td><td>end</td><td>strand</td><td>MM</td><td>target_seq</td><td>PAM</td><td>alignment</td><td>distance</td><td>position</td><td>gene name</td><td>gene id</td></tr>
<tr><td>GU305914.1</td><td>124342</td><td>124364</td><td>+</td><td>0</td><td>CTCAGCGCCGCGATCGACAC</td><td>CGG</td><td>||||||||[||||||||||||]PAM</td><td>NA</td><td>Intergenic</td><td>NA</td><td>NA</td></tr>
<tr><td></td><td></td><td></td><td></td><td></td><td></td><td></td><td></td><td></td><td></td><td></td><td></td></tr>
<tr><td>T9396</td><td>TCAGCGCCGCGATCGACACCGGG</td><td>1000</td><td></td><td></td><td></td><td></td><td></td><td></td><td></td><td></td><td></td></tr>
<tr><td>Oligo fwd</td><td>CACCTCAGCGCCGCGATCGACACC</td><td></td><td></td><td></td><td></td><td></td><td></td><td></td><td></td><td></td><td></td></tr>
<tr><td>Oligo rev</td><td>AAACGGTGTCGATCGCGGCGCTGA</td><td></td><td></td><td></td><td></td><td></td><td></td><td></td><td></td><td></td><td></td></tr>
<tr><td>Chromosome</td><td>start</td><td>end</td><td>strand</td><td>MM</td><td>target_seq</td><td>PAM</td><td>alignment</td><td>distance</td><td>position</td><td>gene name</td><td>gene id</td></tr>
<tr><td>GU305914.1</td><td>124343</td><td>124365</td><td>+</td><td>0</td><td>TCAGCGCCGCGATCGACACC</td><td>GGG</td><td>||||||||[||||||||||||]PAM</td><td>NA</td><td>Intergenic</td><td>NA</td><td>NA</td></tr>
<tr><td>GU305914.1</td><td>99988</td><td>100010</td><td>-</td><td>4</td><td>TCTGTGCCTCGATCGAGACC</td><td>TGG</td><td>||-|-|||[-|||||||-|||]PAM</td><td>NA</td><td>Intergenic</td><td>NA</td><td>NA</td></tr>
</table>

<table>
<tr><td>T9397</td><td>CAGCGCCGCGATCGACACCGGGG</td><td>1000</td><td></td><td></td><td></td><td></td><td></td><td></td><td></td><td></td><td></td></tr>
<tr><td>Oligo fwd</td><td>CACCCAGCGCCGCGATCGACACCG</td><td></td><td></td><td></td><td></td><td></td><td></td><td></td><td></td><td></td><td></td></tr>
<tr><td>Oligo rev</td><td>AAACCGGTGTCGATCGCGGCGCTG</td><td></td><td></td><td></td><td></td><td></td><td></td><td></td><td></td><td></td><td></td></tr>
<tr><td>Chromosome</td><td>start</td><td>end</td><td>strand</td><td>MM</td><td>target_seq</td><td>PAM</td><td>alignment</td><td>distance</td><td>position</td><td>gene name</td><td>gene id</td></tr>
<tr><td>GU305914.1</td><td>124344</td><td>124366</td><td>+</td><td>0</td><td>CAGCGCCGCGATCGACACCG</td><td>GGG</td><td>||||||||[||||||||||||]PAM</td><td>NA</td><td>Intergenic</td><td>NA</td><td>NA</td></tr>
<tr><td>T9398</td><td>CCAGCGTCAACACCGACTCCAGG</td><td>1000</td><td></td><td></td><td></td><td></td><td></td><td></td><td></td><td></td><td></td></tr>
<tr><td>Oligo fwd</td><td>CACCCCAGCGTCAACACCGACTCC</td><td></td><td></td><td></td><td></td><td></td><td></td><td></td><td></td><td></td><td></td></tr>
<tr><td>Oligo rev</td><td>AAACGGAGTCGGTGTTGACGCTGG</td><td></td><td></td><td></td><td></td><td></td><td></td><td></td><td></td><td></td><td></td></tr>
<tr><td>Chromosome</td><td>start</td><td>end</td><td>strand</td><td>MM</td><td>target_seq</td><td>PAM</td><td>alignment</td><td>distance</td><td>position</td><td>gene name</td><td>gene id</td></tr>
<tr><td>GU305914.1</td><td>124388</td><td>124410</td><td>+</td><td>0</td><td>CCAGCGTCAACACCGACTCC</td><td>AGG</td><td>||||||||[||||||||||||]PAM</td><td>NA</td><td>Intergenic</td><td>NA</td><td>NA</td></tr>
<tr><td>T9399</td><td>CAGCGTCAACACCGACTCCAGGG</td><td>1000</td><td></td><td></td><td></td><td></td><td></td><td></td><td></td><td></td><td></td></tr>
<tr><td>Oligo fwd</td><td>CACCCAGCGTCAACACCGACTCCA</td><td></td><td></td><td></td><td></td><td></td><td></td><td></td><td></td><td></td><td></td></tr>
<tr><td>Oligo rev</td><td>AAACTGGAGTCGGTGTTGACGCTG</td><td></td><td></td><td></td><td></td><td></td><td></td><td></td><td></td><td></td><td></td></tr>
<tr><td>Chromosome</td><td>start</td><td>end</td><td>strand</td><td>MM</td><td>target_seq</td><td>PAM</td><td>alignment</td><td>distance</td><td>position</td><td>gene name</td><td>gene id</td></tr>
<tr><td>GU305914.1</td><td>124389</td><td>124411</td><td>+</td><td>0</td><td>CAGCGTCAACACCGACTCCA</td><td>GGG</td><td>||||||||[||||||||||||]PAM</td><td>NA</td><td>Intergenic</td><td>NA</td><td>NA</td></tr>
<tr><td>GU305914.1</td><td>88886</td><td>88908</td><td>-</td><td>3</td><td>CAGCCTCAACACGGGCTCCA</td><td>TGG</td><td>||||-|||[||||-|-|||||]PAM</td><td>NA</td><td>Intergenic</td><td>NA</td><td>NA</td></tr>
<tr><td>T9400</td><td>CGACTCCAGGGTGTGCGCGCAGG</td><td>1000</td><td></td><td></td><td></td><td></td><td></td><td></td><td></td><td></td><td></td></tr>
<tr><td>Oligo fwd</td><td>CACCCGACTCCAGGGTGTGCGCGC</td><td></td><td></td><td></td><td></td><td></td><td></td><td></td><td></td><td></td><td></td></tr>
<tr><td>Oligo rev</td><td>AAACGCGCGCACACCCTGGAGTCG</td><td></td><td></td><td></td><td></td><td></td><td></td><td></td><td></td><td></td><td></td></tr>
<tr><td>Chromosome</td><td>start</td><td>end</td><td>strand</td><td>MM</td><td>target_seq</td><td>PAM</td><td>alignment</td><td>distance</td><td>position</td><td>gene name</td><td>gene id</td></tr>
<tr><td>GU305914.1</td><td>124401</td><td>124423</td><td>+</td><td>0</td><td>CGACTCCAGGGTGTGCGCGC</td><td>AGG</td><td>||||||||[||||||||||||]PAM</td><td>NA</td><td>Intergenic</td><td>NA</td><td>NA</td></tr>
<tr><td>T9401</td><td>GCCGCCGAGTAGCACGAGAACGG</td><td>1000</td><td></td><td></td><td></td><td></td><td></td><td></td><td></td><td></td><td></td></tr>
<tr><td>Oligo fwd</td><td>CACCGCCGCCGAGTAGCACGAGAA</td><td></td><td></td><td></td><td></td><td></td><td></td><td></td><td></td><td></td><td></td></tr>
<tr><td>Oligo rev</td><td>AAACTTCTCGTGCTACTCGGCGGC</td><td></td><td></td><td></td><td></td><td></td><td></td><td></td><td></td><td></td><td></td></tr>
<tr><td>Chromosome</td><td>start</td><td>end</td><td>strand</td><td>MM</td><td>target_seq</td><td>PAM</td><td>alignment</td><td>distance</td><td>position</td><td>gene name</td><td>gene id</td></tr>
<tr><td>GU305914.1</td><td>124423</td><td>124445</td><td>+</td><td>0</td><td>GCCGCCGAGTAGCACGAGAA</td><td>CGG</td><td>||||||||[||||||||||||]PAM</td><td>NA</td><td>Intergenic</td><td>NA</td><td>NA</td></tr>
<tr><td>T9402</td><td>AGTAGCACGAGAACGGCGTGCGG</td><td>1000</td><td></td><td></td><td></td><td></td><td></td><td></td><td></td><td></td><td></td></tr>
<tr><td>Oligo fwd</td><td>CACCAGTAGCACGAGAACGGCGTG</td><td></td><td></td><td></td><td></td><td></td><td></td><td></td><td></td><td></td><td></td></tr>
<tr><td>Oligo rev</td><td>AAACCACGCCGTTCTCGTGCTACT</td><td></td><td></td><td></td><td></td><td></td><td></td><td></td><td></td><td></td><td></td></tr>
<tr><td>Chromosome</td><td>start</td><td>end</td><td>strand</td><td>MM</td><td>target_seq</td><td>PAM</td><td>alignment</td><td>distance</td><td>position</td><td>gene name</td><td>gene id</td></tr>
<tr><td>GU305914.1</td><td>124430</td><td>124452</td><td>+</td><td>0</td><td>AGTAGCACGAGAACGGCGTG</td><td>CGG</td><td>||||||||[||||||||||||]PAM</td><td>NA</td><td>Intergenic</td><td>NA</td><td>NA</td></tr>
<tr><td>T9403</td><td>AGAACGGCGTGCGGTGCCAGTGG</td><td>1000</td><td></td><td></td><td></td><td></td><td></td><td></td><td></td><td></td><td></td></tr>
<tr><td>Oligo fwd</td><td>CACCAGAACGGCGTGCGGTGCCAG</td><td></td><td></td><td></td><td></td><td></td><td></td><td></td><td></td><td></td><td></td></tr>
<tr><td>Oligo rev</td><td>AAACCTGGCACCGCACGCCGTTCT</td><td></td><td></td><td></td><td></td><td></td><td></td><td></td><td></td><td></td><td></td></tr>
<tr><td>Chromosome</td><td>start</td><td>end</td><td>strand</td><td>MM</td><td>target_seq</td><td>PAM</td><td>alignment</td><td>distance</td><td>position</td><td>gene name</td><td>gene id</td></tr>
<tr><td>GU305914.1</td><td>124439</td><td>124461</td><td>+</td><td>0</td><td>AGAACGGCGTGCGGTGCCAG</td><td>TGG</td><td>||||||||[||||||||||||]PAM</td><td>NA</td><td>Intergenic</td><td>NA</td><td>NA</td></tr>
<tr><td>T9404</td><td>ACGGCGTGCGGTGCCAGTGGTGG</td><td>1000</td><td></td><td></td><td></td><td></td><td></td><td></td><td></td><td></td><td></td></tr>
<tr><td>Oligo fwd</td><td>CACCACGGCGTGCGGTGCCAGTGG</td><td></td><td></td><td></td><td></td><td></td><td></td><td></td><td></td><td></td><td></td></tr>
<tr><td>Oligo rev</td><td>AAACCCACTGGCACCGCACGCCGT</td><td></td><td></td><td></td><td></td><td></td><td></td><td></td><td></td><td></td><td></td></tr>
<tr><td>Chromosome</td><td>start</td><td>end</td><td>strand</td><td>MM</td><td>target_seq</td><td>PAM</td><td>alignment</td><td>distance</td><td>position</td><td>gene name</td><td>gene id</td></tr>
<tr><td>GU305914.1</td><td>124442</td><td>124464</td><td>+</td><td>0</td><td>ACGGCGTGCGGTGCCAGTGG</td><td>TGG</td><td>||||||||[||||||||||||]PAM</td><td>NA</td><td>Intergenic</td><td>NA</td><td>NA</td></tr>
<tr><td>T9405</td><td>GTGGTACTCGTGCGCGAAGATGG</td><td>1000</td><td></td><td></td><td></td><td></td><td></td><td></td><td></td><td></td><td></td></tr>
<tr><td>Oligo fwd</td><td>CACCGTGGTACTCGTGCGCGAAGA</td><td></td><td></td><td></td><td></td><td></td><td></td><td></td><td></td><td></td><td></td></tr>
<tr><td>Oligo rev</td><td>AAACTCTTCGCGCACGAGTACCAC</td><td></td><td></td><td></td><td></td><td></td><td></td><td></td><td></td><td></td><td></td></tr>
<tr><td>Chromosome</td><td>start</td><td>end</td><td>strand</td><td>MM</td><td>target_seq</td><td>PAM</td><td>alignment</td><td>distance</td><td>position</td><td>gene name</td><td>gene id</td></tr>
<tr><td>GU305914.1</td><td>124461</td><td>124483</td><td>+</td><td>0</td><td>GTGGTACTCGTGCGCGAAGA</td><td>TGG</td><td>||||||||[||||||||||||]PAM</td><td>NA</td><td>Intergenic</td><td>NA</td><td>NA</td></tr>
<tr><td>T9406</td><td>TGGTACTCGTGCGCGAAGATGGG</td><td>1000</td><td></td><td></td><td></td><td></td><td></td><td></td><td></td><td></td><td></td></tr>
<tr><td>Oligo fwd</td><td>CACCTGGTACTCGTGCGCGAAGAT</td><td></td><td></td><td></td><td></td><td></td><td></td><td></td><td></td><td></td><td></td></tr>
<tr><td>Oligo rev</td><td>AAACATCTTCGCGCACGAGTACCA</td><td></td><td></td><td></td><td></td><td></td><td></td><td></td><td></td><td></td><td></td></tr>
<tr><td>Chromosome</td><td>start</td><td>end</td><td>strand</td><td>MM</td><td>target_seq</td><td>PAM</td><td>alignment</td><td>distance</td><td>position</td><td>gene name</td><td>gene id</td></tr>
<tr><td>GU305914.1</td><td>124462</td><td>124484</td><td>+</td><td>0</td><td>TGGTACTCGTGCGCGAAGAT</td><td>GGG</td><td>||||||||[||||||||||||]PAM</td><td>NA</td><td>Intergenic</td><td>NA</td><td>NA</td></tr>
<tr><td>T9407</td><td>TGCGCGAAGATGGGCGACAGCGG</td><td>1000</td><td></td><td></td><td></td><td></td><td></td><td></td><td></td><td></td><td></td></tr>
<tr><td>Oligo fwd</td><td>CACCTGCGCGAAGATGGGCGACAG</td><td></td><td></td><td></td><td></td><td></td><td></td><td></td><td></td><td></td><td></td></tr>
<tr><td>Oligo rev</td><td>AAACCTGTCGCCCATCTTCGCGCA</td><td></td><td></td><td></td><td></td><td></td><td></td><td></td><td></td><td></td><td></td></tr>
<tr><td>Chromosome</td><td>start</td><td>end</td><td>strand</td><td>MM</td><td>target_seq</td><td>PAM</td><td>alignment</td><td>distance</td><td>position</td><td>gene name</td><td>gene id</td></tr>
<tr><td>GU305914.1</td><td>124471</td><td>124493</td><td>+</td><td>0</td><td>TGCGCGAAGATGGGCGACAG</td><td>CGG</td><td>||||||||[||||||||||||]PAM</td><td>NA</td><td>Intergenic</td><td>NA</td><td>NA</td></tr>
<tr><td>T9408</td><td>GCGCGAAGATGGGCGACAGCGGG</td><td>1000</td><td></td><td></td><td></td><td></td><td></td><td></td><td></td><td></td><td></td></tr>
<tr><td>Oligo fwd</td><td>CACCGCGCGAAGATGGGCGACAGC</td><td></td><td></td><td></td><td></td><td></td><td></td><td></td><td></td><td></td><td></td></tr>
<tr><td>Oligo rev</td><td>AAACGCTGTCGCCCATCTTCGCGC</td><td></td><td></td><td></td><td></td><td></td><td></td><td></td><td></td><td></td><td></td></tr>
<tr><td>Chromosome</td><td>start</td><td>end</td><td>strand</td><td>MM</td><td>target_seq</td><td>PAM</td><td>alignment</td><td>distance</td><td>position</td><td>gene name</td><td>gene id</td></tr>
<tr><td>GU305914.1</td><td>124472</td><td>124494</td><td>+</td><td>0</td><td>GCGCGAAGATGGGCGACAGC</td><td>GGG</td><td>||||||||[||||||||||||]PAM</td><td>NA</td><td>Intergenic</td><td>NA</td><td>NA</td></tr>
<tr><td>T9409</td><td>GGAAGCCGCCGTCGCACCGCTGG</td><td>1000</td><td></td><td></td><td></td><td></td><td></td><td></td><td></td><td></td><td></td></tr>
<tr><td>Oligo fwd</td><td>CACCGGAAGCCGCCGTCGCACCGC</td><td></td><td></td><td></td><td></td><td></td><td></td><td></td><td></td><td></td><td></td></tr>
<tr><td>Oligo rev</td><td>AAACGCGGTGCGACGGCGGCTTCC</td><td></td><td></td><td></td><td></td><td></td><td></td><td></td><td></td><td></td><td></td></tr>
<tr><td>Chromosome</td><td>start</td><td>end</td><td>strand</td><td>MM</td><td>target_seq</td><td>PAM</td><td>alignment</td><td>distance</td><td>position</td><td>gene name</td><td>gene id</td></tr>
<tr><td>GU305914.1</td><td>124493</td><td>124515</td><td>+</td><td>0</td><td>GGAAGCCGCCGTCGCACCGC</td><td>TGG</td><td>||||||||[||||||||||||]PAM</td><td>NA</td><td>Intergenic</td><td>NA</td><td>NA</td></tr>
<tr><td>T9410</td><td>CCGTCGCACCGCTGGTACTGCGG</td><td>1000</td><td></td><td></td><td></td><td></td><td></td><td></td><td></td><td></td><td></td></tr>
<tr><td>Oligo fwd</td><td>CACCCCGTCGCACCGCTGGTACTG</td><td></td><td></td><td></td><td></td><td></td><td></td><td></td><td></td><td></td><td></td></tr>
<tr><td>Oligo rev</td><td>AAACCAGTACCAGCGGTGCGACGG</td><td></td><td></td><td></td><td></td><td></td><td></td><td></td><td></td><td></td><td></td></tr>
<tr><td>Chromosome</td><td>start</td><td>end</td><td>strand</td><td>MM</td><td>target_seq</td><td>PAM</td><td>alignment</td><td>distance</td><td>position</td><td>gene name</td><td>gene id</td></tr>
<tr><td>GU305914.1</td><td>124501</td><td>124523</td><td>+</td><td>0</td><td>CCGTCGCACCGCTGGTACTG</td><td>CGG</td><td>||||||||[||||||||||||]PAM</td><td>NA</td><td>Intergenic</td><td>NA</td><td>NA</td></tr>
<tr><td>T9411</td><td>CGTCGCACCGCTGGTACTGCGGG</td><td>1000</td><td></td><td></td><td></td><td></td><td></td><td></td><td></td><td></td><td></td></tr>
<tr><td>Oligo fwd</td><td>CACCCGTCGCACCGCTGGTACTGC</td><td></td><td></td><td></td><td></td><td></td><td></td><td></td><td></td><td></td><td></td></tr>
<tr><td>Oligo rev</td><td>AAACGCAGTACCAGCGGTGCGACG</td><td></td><td></td><td></td><td></td><td></td><td></td><td></td><td></td><td></td><td></td></tr>
<tr><td>Chromosome</td><td>start</td><td>end</td><td>strand</td><td>MM</td><td>target_seq</td><td>PAM</td><td>alignment</td><td>distance</td><td>position</td><td>gene name</td><td>gene id</td></tr>
</table>

| | | | | | | | | | | | |
|---|---|---|---|---|---|---|---|---|---|---|---|
| GU305914.1 | 124502 | 124524 | + | 0 | CGTCGCACCGCTGGTACTGC | GGG | \|\|\|\|\|\|\|\|[\|\|\|\|\|\|\|\|\|\|\|\|]PAM | NA | Intergenic | NA | NA |
| T9412 | GGTACTGCGGGAAGTCGTTCAGG | 1000 | | | | | | | | | |
| Oligo fwd | CACCGGTACTGCGGGAAGTCGTTC | | | | | | | | | | |
| Oligo rev | AAACGAACGACTTCCCGCAGTACC | | | | | | | | | | |
| Chromosome | start | end | strand | MM | target_seq | PAM | alignment | distance | position | gene name | gene id |
| GU305914.1 | 124514 | 124536 | + | 0 | GGTACTGCGGGAAGTCGTTC | AGG | \|\|\|\|\|\|\|\|[\|\|\|\|\|\|\|\|\|\|\|\|]PAM | NA | Intergenic | NA | NA |
| T9413 | GGAAGTCGTTCAGGAACACGCGG | 1000 | | | | | | | | | |
| Oligo fwd | CACCGGAAGTCGTTCAGGAACACG | | | | | | | | | | |
| Oligo rev | AAACCGTGTTCCTGAACGACTTCC | | | | | | | | | | |
| Chromosome | start | end | strand | MM | target_seq | PAM | alignment | distance | position | gene name | gene id |
| GU305914.1 | 124523 | 124545 | + | 0 | GGAAGTCGTTCAGGAACACG | CGG | \|\|\|\|\|\|\|\|[\|\|\|\|\|\|\|\|\|\|\|\|]PAM | NA | Intergenic | NA | NA |
| T9414 | TCAGGAACACGCGGATGTCGTGG | 1000 | | | | | | | | | |
| Oligo fwd | CACCTCAGGAACACGCGGATGTCG | | | | | | | | | | |
| Oligo rev | AAACCGACATCCGCGTGTTCCTGA | | | | | | | | | | |
| Chromosome | start | end | strand | MM | target_seq | PAM | alignment | distance | position | gene name | gene id |
| GU305914.1 | 124532 | 124554 | + | 0 | TCAGGAACACGCGGATGTCG | TGG | \|\|\|\|\|\|\|\|[\|\|\|\|\|\|\|\|\|\|\|\|]PAM | NA | Intergenic | NA | NA |
| T9415 | TTCGCCGTCGCCGCCAGTCCCGG | 1000 | | | | | | | | | |
| Oligo fwd | CACCTTCGCCGTCGCCGCCAGTCC | | | | | | | | | | |
| Oligo rev | AAACGGACTGGCGGCGACGGCGAA | | | | | | | | | | |
| Chromosome | start | end | strand | MM | target_seq | PAM | alignment | distance | position | gene name | gene id |
| GU305914.1 | 124555 | 124577 | + | 0 | TTCGCCGTCGCCGCCAGTCC | CGG | \|\|\|\|\|\|\|\|[\|\|\|\|\|\|\|\|\|\|\|\|]PAM | NA | Intergenic | NA | NA |
| T9416 | TCGCCGTCGCCGCCAGTCCCGGG | 1000 | | | | | | | | | |
| Oligo fwd | CACCTCGCCGTCGCCGCCAGTCCC | | | | | | | | | | |
| Oligo rev | AAACGGGACTGGCGGCGACGGCGA | | | | | | | | | | |
| Chromosome | start | end | strand | MM | target_seq | PAM | alignment | distance | position | gene name | gene id |
| GU305914.1 | 124556 | 124578 | + | 0 | TCGCCGTCGCCGCCAGTCCC | GGG | \|\|\|\|\|\|\|\|[\|\|\|\|\|\|\|\|\|\|\|\|]PAM | NA | Intergenic | NA | NA |
| T9417 | GTCCCGGGTCCGAGTAGAACCGG | 1000 | | | | | | | | | |
| Oligo fwd | CACCGTCCCGGGTCCGAGTAGAAC | | | | | | | | | | |
| Oligo rev | AAACGTTCTACTCGGACCCGGGAC | | | | | | | | | | |
| Chromosome | start | end | strand | MM | target_seq | PAM | alignment | distance | position | gene name | gene id |
| GU305914.1 | 124571 | 124593 | + | 0 | GTCCCGGGTCCGAGTAGAAC | CGG | \|\|\|\|\|\|\|\|[\|\|\|\|\|\|\|\|\|\|\|\|]PAM | NA | Intergenic | NA | NA |
| T9418 | CCGGGTCCGAGTAGAACCGGTGG | 1000 | | | | | | | | | |
| Oligo fwd | CACCCCGGGTCCGAGTAGAACCGG | | | | | | | | | | |
| Oligo rev | AAACCCGGTTCTACTCGGACCCGG | | | | | | | | | | |
| Chromosome | start | end | strand | MM | target_seq | PAM | alignment | distance | position | gene name | gene id |
| GU305914.1 | 124574 | 124596 | + | 0 | CCGGGTCCGAGTAGAACCGG | TGG | \|\|\|\|\|\|\|\|[\|\|\|\|\|\|\|\|\|\|\|\|]PAM | NA | Intergenic | NA | NA |
| GU305914.1 | 208125 | 208147 | + | 4 | CCGTATCCGAAAAGAACCGG | CGG | \|\|\|--\|\|\|[\|\|--\|\|\|\|\|\|\|\|]PAM | NA | Intergenic | NA | NA |
| T9419 | GTCCGAGTAGAACCGGTGGAAGG | 1000 | | | | | | | | | |
| Oligo fwd | CACCGTCCGAGTAGAACCGGTGGA | | | | | | | | | | |
| Oligo rev | AAACTCCACCGGTTCTACTCGGAC | | | | | | | | | | |
| Chromosome | start | end | strand | MM | target_seq | PAM | alignment | distance | position | gene name | gene id |
| GU305914.1 | 124578 | 124600 | + | 0 | GTCCGAGTAGAACCGGTGGA | AGG | \|\|\|\|\|\|\|\|[\|\|\|\|\|\|\|\|\|\|\|\|]PAM | NA | Intergenic | NA | NA |
| T9420 | TCCGAGTAGAACCGGTGGAAGGG | 1000 | | | | | | | | | |
| Oligo fwd | CACCTCCGAGTAGAACCGGTGGAA | | | | | | | | | | |
| Oligo rev | AAACTTCCACCGGTTCTACTCGGA | | | | | | | | | | |
| Chromosome | start | end | strand | MM | target_seq | PAM | alignment | distance | position | gene name | gene id |
| GU305914.1 | 124579 | 124601 | + | 0 | TCCGAGTAGAACCGGTGGAA | GGG | \|\|\|\|\|\|\|\|[\|\|\|\|\|\|\|\|\|\|\|\|]PAM | NA | Intergenic | NA | NA |
| GU305914.1 | 209035 | 209057 | + | 4 | GACGAGTAGAACCGCGGGAA | CGG | --\|\|\|\|\|\|[\|\|\|\|\|\|--\|\|\|\|]PAM | NA | Intergenic | NA | NA |
| T9421 | CCGAGTAGAACCGGTGGAAGGGG | 1000 | | | | | | | | | |
| Oligo fwd | CACCCCGAGTAGAACCGGTGGAAG | | | | | | | | | | |
| Oligo rev | AAACCTTCCACCGGTTCTACTCGG | | | | | | | | | | |
| Chromosome | start | end | strand | MM | target_seq | PAM | alignment | distance | position | gene name | gene id |
| GU305914.1 | 124580 | 124602 | + | 0 | CCGAGTAGAACCGGTGGAAG | GGG | \|\|\|\|\|\|\|\|[\|\|\|\|\|\|\|\|\|\|\|\|]PAM | NA | Intergenic | NA | NA |
| T9422 | GGTGGAAGGGGATCACCGTCAGG | 1000 | | | | | | | | | |
| Oligo fwd | CACCGGTGGAAGGGGATCACCGTC | | | | | | | | | | |
| Oligo rev | AAACGACGGTGATCCCCTTCCACC | | | | | | | | | | |
| Chromosome | start | end | strand | MM | target_seq | PAM | alignment | distance | position | gene name | gene id |
| GU305914.1 | 124592 | 124614 | + | 0 | GGTGGAAGGGGATCACCGTC | AGG | \|\|\|\|\|\|\|\|[\|\|\|\|\|\|\|\|\|\|\|\|]PAM | NA | Intergenic | NA | NA |
| T9423 | AGGTACTCCTGCAGCAGCTTCGG | 1000 | | | | | | | | | |
| Oligo fwd | CACCAGGTACTCCTGCAGCAGCTT | | | | | | | | | | |
| Oligo rev | AAACAAGCTGCTGCAGGAGTACCT | | | | | | | | | | |
| Chromosome | start | end | strand | MM | target_seq | PAM | alignment | distance | position | gene name | gene id |
| GU305914.1 | 124612 | 124634 | + | 0 | AGGTACTCCTGCAGCAGCTT | CGG | \|\|\|\|\|\|\|\|[\|\|\|\|\|\|\|\|\|\|\|\|]PAM | NA | Intergenic | NA | NA |
| GU305914.1 | 77886 | 77908 | + | 4 | ACGTGTTCCTGCAGCAGCTG | CGG | \|-\|\|--\|\|[\|\|\|\|\|\|\|\|\|\|\|-]PAM | NA | Intergenic | NA | NA |
| T9424 | CTCCTGCAGCAGCTTCGGCACGG | 1000 | | | | | | | | | |
| Oligo fwd | CACCCTCCTGCAGCAGCTTCGGCA | | | | | | | | | | |
| Oligo rev | AAACTGCCGAAGCTGCTGCAGGAG | | | | | | | | | | |
| Chromosome | start | end | strand | MM | target_seq | PAM | alignment | distance | position | gene name | gene id |
| GU305914.1 | 124617 | 124639 | + | 0 | CTCCTGCAGCAGCTTCGGCA | CGG | \|\|\|\|\|\|\|\|[\|\|\|\|\|\|\|\|\|\|\|\|]PAM | NA | Intergenic | NA | NA |
| T9425 | TCATGCAGAACCCGTTGTACAGG | 1000 | | | | | | | | | |
| Oligo fwd | CACCTCATGCAGAACCCGTTGTAC | | | | | | | | | | |
| Oligo rev | AAACGTACAACGGGTTCTGCATGA | | | | | | | | | | |
| Chromosome | start | end | strand | MM | target_seq | PAM | alignment | distance | position | gene name | gene id |
| GU305914.1 | 124643 | 124665 | + | 0 | TCATGCAGAACCCGTTGTAC | AGG | \|\|\|\|\|\|\|\|[\|\|\|\|\|\|\|\|\|\|\|\|]PAM | NA | Intergenic | NA | NA |

<table>
<tr><td>T9426</td><td>AGCGACTCGAAGCCCGCCGACGG</td><td>1000</td><td></td><td></td><td></td><td></td><td></td><td></td><td></td><td></td><td></td></tr>
<tr><td>Oligo fwd</td><td>CACCAGCGACTCGAAGCCCGCCGA</td><td></td><td></td><td></td><td></td><td></td><td></td><td></td><td></td><td></td><td></td></tr>
<tr><td>Oligo rev</td><td>AAACTCGGCGGGCTTCGAGTCGCT</td><td></td><td></td><td></td><td></td><td></td><td></td><td></td><td></td><td></td><td></td></tr>
<tr><td>Chromosome</td><td>start</td><td>end</td><td>strand</td><td>MM</td><td>target_seq</td><td>PAM</td><td>alignment</td><td>distance</td><td>position</td><td>gene name</td><td>gene id</td></tr>
<tr><td>GU305914.1</td><td>124675</td><td>124697</td><td>+</td><td>0</td><td>AGCGACTCGAAGCCCGCCGA</td><td>CGG</td><td>||||||||[||||||||||||]PAM</td><td>NA</td><td>Intergenic</td><td>NA</td><td>NA</td></tr>
<tr><td>T9427</td><td>GGCGCGCCGTGCCGCTGCGCCGG</td><td>1000</td><td></td><td></td><td></td><td></td><td></td><td></td><td></td><td></td><td></td></tr>
<tr><td>Oligo fwd</td><td>CACCGGCGCGCCGTGCCGCTGCGC</td><td></td><td></td><td></td><td></td><td></td><td></td><td></td><td></td><td></td><td></td></tr>
<tr><td>Oligo rev</td><td>AAACGCGCAGCGGCACGGCGCGCC</td><td></td><td></td><td></td><td></td><td></td><td></td><td></td><td></td><td></td><td></td></tr>
<tr><td>Chromosome</td><td>start</td><td>end</td><td>strand</td><td>MM</td><td>target_seq</td><td>PAM</td><td>alignment</td><td>distance</td><td>position</td><td>gene name</td><td>gene id</td></tr>
<tr><td>GU305914.1</td><td>124696</td><td>124718</td><td>+</td><td>0</td><td>GGCGCGCCGTGCCGCTGCGC</td><td>CGG</td><td>||||||||[||||||||||||]PAM</td><td>NA</td><td>Intergenic</td><td>NA</td><td>NA</td></tr>
<tr><td>T9428</td><td>GCGCGCCGTGCCGCTGCGCCGGG</td><td>1000</td><td></td><td></td><td></td><td></td><td></td><td></td><td></td><td></td><td></td></tr>
<tr><td>Oligo fwd</td><td>CACCGCGCGCCGTGCCGCTGCGCC</td><td></td><td></td><td></td><td></td><td></td><td></td><td></td><td></td><td></td><td></td></tr>
<tr><td>Oligo rev</td><td>AAACGGCGCAGCGGCACGGCGCGC</td><td></td><td></td><td></td><td></td><td></td><td></td><td></td><td></td><td></td><td></td></tr>
<tr><td>Chromosome</td><td>start</td><td>end</td><td>strand</td><td>MM</td><td>target_seq</td><td>PAM</td><td>alignment</td><td>distance</td><td>position</td><td>gene name</td><td>gene id</td></tr>
<tr><td>GU305914.1</td><td>124697</td><td>124719</td><td>+</td><td>0</td><td>GCGCGCCGTGCCGCTGCGCC</td><td>GGG</td><td>||||||||[||||||||||||]PAM</td><td>NA</td><td>Intergenic</td><td>NA</td><td>NA</td></tr>
<tr><td>T9429</td><td>CGTGCCGCTGCGCCGGGTCCAGG</td><td>1000</td><td></td><td></td><td></td><td></td><td></td><td></td><td></td><td></td><td></td></tr>
<tr><td>Oligo fwd</td><td>CACCCGTGCCGCTGCGCCGGGTCC</td><td></td><td></td><td></td><td></td><td></td><td></td><td></td><td></td><td></td><td></td></tr>
<tr><td>Oligo rev</td><td>AAACGGACCCGGCGCAGCGGCACG</td><td></td><td></td><td></td><td></td><td></td><td></td><td></td><td></td><td></td><td></td></tr>
<tr><td>Chromosome</td><td>start</td><td>end</td><td>strand</td><td>MM</td><td>target_seq</td><td>PAM</td><td>alignment</td><td>distance</td><td>position</td><td>gene name</td><td>gene id</td></tr>
<tr><td>GU305914.1</td><td>124703</td><td>124725</td><td>+</td><td>0</td><td>CGTGCCGCTGCGCCGGGTCC</td><td>AGG</td><td>||||||||[||||||||||||]PAM</td><td>NA</td><td>Intergenic</td><td>NA</td><td>NA</td></tr>
<tr><td>T9430</td><td>GTGCCGCTGCGCCGGGTCCAGGG</td><td>1000</td><td></td><td></td><td></td><td></td><td></td><td></td><td></td><td></td><td></td></tr>
<tr><td>Oligo fwd</td><td>CACCGTGCCGCTGCGCCGGGTCCA</td><td></td><td></td><td></td><td></td><td></td><td></td><td></td><td></td><td></td><td></td></tr>
<tr><td>Oligo rev</td><td>AAACTGGACCCGGCGCAGCGGCAC</td><td></td><td></td><td></td><td></td><td></td><td></td><td></td><td></td><td></td><td></td></tr>
<tr><td>Chromosome</td><td>start</td><td>end</td><td>strand</td><td>MM</td><td>target_seq</td><td>PAM</td><td>alignment</td><td>distance</td><td>position</td><td>gene name</td><td>gene id</td></tr>
<tr><td>GU305914.1</td><td>124704</td><td>124726</td><td>+</td><td>0</td><td>GTGCCGCTGCGCCGGGTCCA</td><td>GGG</td><td>||||||||[||||||||||||]PAM</td><td>NA</td><td>Intergenic</td><td>NA</td><td>NA</td></tr>
<tr><td>T9431</td><td>GCGCCGGGTCCAGGGCCGCGCGG</td><td>1000</td><td></td><td></td><td></td><td></td><td></td><td></td><td></td><td></td><td></td></tr>
<tr><td>Oligo fwd</td><td>CACCGCGCCGGGTCCAGGGCCGCG</td><td></td><td></td><td></td><td></td><td></td><td></td><td></td><td></td><td></td><td></td></tr>
<tr><td>Oligo rev</td><td>AAACCGCGGCCCTGGACCCGGCGC</td><td></td><td></td><td></td><td></td><td></td><td></td><td></td><td></td><td></td><td></td></tr>
<tr><td>Chromosome</td><td>start</td><td>end</td><td>strand</td><td>MM</td><td>target_seq</td><td>PAM</td><td>alignment</td><td>distance</td><td>position</td><td>gene name</td><td>gene id</td></tr>
<tr><td>GU305914.1</td><td>124712</td><td>124734</td><td>+</td><td>0</td><td>GCGCCGGGTCCAGGGCCGCG</td><td>CGG</td><td>||||||||[||||||||||||]PAM</td><td>NA</td><td>Intergenic</td><td>NA</td><td>NA</td></tr>
<tr><td>T9432</td><td>CTCCAGCATCTGCGCGTTCTCGG</td><td>1000</td><td></td><td></td><td></td><td></td><td></td><td></td><td></td><td></td><td></td></tr>
<tr><td>Oligo fwd</td><td>CACCCTCCAGCATCTGCGCGTTCT</td><td></td><td></td><td></td><td></td><td></td><td></td><td></td><td></td><td></td><td></td></tr>
<tr><td>Oligo rev</td><td>AAACAGAACGCGCAGATGCTGGAG</td><td></td><td></td><td></td><td></td><td></td><td></td><td></td><td></td><td></td><td></td></tr>
<tr><td>Chromosome</td><td>start</td><td>end</td><td>strand</td><td>MM</td><td>target_seq</td><td>PAM</td><td>alignment</td><td>distance</td><td>position</td><td>gene name</td><td>gene id</td></tr>
<tr><td>GU305914.1</td><td>124737</td><td>124759</td><td>+</td><td>0</td><td>CTCCAGCATCTGCGCGTTCT</td><td>CGG</td><td>||||||||[||||||||||||]PAM</td><td>NA</td><td>Intergenic</td><td>NA</td><td>NA</td></tr>
<tr><td>T9433</td><td>TCCAGCATCTGCGCGTTCTCGGG</td><td>1000</td><td></td><td></td><td></td><td></td><td></td><td></td><td></td><td></td><td></td></tr>
<tr><td>Oligo fwd</td><td>CACCTCCAGCATCTGCGCGTTCTC</td><td></td><td></td><td></td><td></td><td></td><td></td><td></td><td></td><td></td><td></td></tr>
<tr><td>Oligo rev</td><td>AAACGAGAACGCGCAGATGCTGGA</td><td></td><td></td><td></td><td></td><td></td><td></td><td></td><td></td><td></td><td></td></tr>
<tr><td>Chromosome</td><td>start</td><td>end</td><td>strand</td><td>MM</td><td>target_seq</td><td>PAM</td><td>alignment</td><td>distance</td><td>position</td><td>gene name</td><td>gene id</td></tr>
<tr><td>GU305914.1</td><td>124738</td><td>124760</td><td>+</td><td>0</td><td>TCCAGCATCTGCGCGTTCTC</td><td>GGG</td><td>||||||||[||||||||||||]PAM</td><td>NA</td><td>Intergenic</td><td>NA</td><td>NA</td></tr>
<tr><td>T9434</td><td>TCTCGGGCACGTGCGCCACCAGG</td><td>1000</td><td></td><td></td><td></td><td></td><td></td><td></td><td></td><td></td><td></td></tr>
<tr><td>Oligo fwd</td><td>CACCTCTCGGGCACGTGCGCCACC</td><td></td><td></td><td></td><td></td><td></td><td></td><td></td><td></td><td></td><td></td></tr>
<tr><td>Oligo rev</td><td>AAACGGTGGCGCACGTGCCCGAGA</td><td></td><td></td><td></td><td></td><td></td><td></td><td></td><td></td><td></td><td></td></tr>
<tr><td>Chromosome</td><td>start</td><td>end</td><td>strand</td><td>MM</td><td>target_seq</td><td>PAM</td><td>alignment</td><td>distance</td><td>position</td><td>gene name</td><td>gene id</td></tr>
<tr><td>GU305914.1</td><td>124754</td><td>124776</td><td>+</td><td>0</td><td>TCTCGGGCACGTGCGCCACC</td><td>AGG</td><td>||||||||[||||||||||||]PAM</td><td>NA</td><td>Intergenic</td><td>NA</td><td>NA</td></tr>
<tr><td>T9435</td><td>CGGGCACGTGCGCCACCAGGTGG</td><td>1000</td><td></td><td></td><td></td><td></td><td></td><td></td><td></td><td></td><td></td></tr>
<tr><td>Oligo fwd</td><td>CACCCGGGCACGTGCGCCACCAGG</td><td></td><td></td><td></td><td></td><td></td><td></td><td></td><td></td><td></td><td></td></tr>
<tr><td>Oligo rev</td><td>AAACCCTGGTGGCGCACGTGCCCG</td><td></td><td></td><td></td><td></td><td></td><td></td><td></td><td></td><td></td><td></td></tr>
<tr><td>Chromosome</td><td>start</td><td>end</td><td>strand</td><td>MM</td><td>target_seq</td><td>PAM</td><td>alignment</td><td>distance</td><td>position</td><td>gene name</td><td>gene id</td></tr>
<tr><td>GU305914.1</td><td>124757</td><td>124779</td><td>+</td><td>0</td><td>CGGGCACGTGCGCCACCAGG</td><td>TGG</td><td>||||||||[||||||||||||]PAM</td><td>NA</td><td>Intergenic</td><td>NA</td><td>NA</td></tr>
<tr><td>T9436</td><td>CACCAGGTGGAACAGCTCCTCGG</td><td>1000</td><td></td><td></td><td></td><td></td><td></td><td></td><td></td><td></td><td></td></tr>
<tr><td>Oligo fwd</td><td>CACCCACCAGGTGGAACAGCTCCT</td><td></td><td></td><td></td><td></td><td></td><td></td><td></td><td></td><td></td><td></td></tr>
<tr><td>Oligo rev</td><td>AAACAGGAGCTGTTCCACCTGGTG</td><td></td><td></td><td></td><td></td><td></td><td></td><td></td><td></td><td></td><td></td></tr>
<tr><td>Chromosome</td><td>start</td><td>end</td><td>strand</td><td>MM</td><td>target_seq</td><td>PAM</td><td>alignment</td><td>distance</td><td>position</td><td>gene name</td><td>gene id</td></tr>
<tr><td>GU305914.1</td><td>124770</td><td>124792</td><td>+</td><td>0</td><td>CACCAGGTGGAACAGCTCCT</td><td>CGG</td><td>||||||||[||||||||||||]PAM</td><td>NA</td><td>Intergenic</td><td>NA</td><td>NA</td></tr>
<tr><td>T9437</td><td>GGAACAGCTCCTCGGCCGCCTGG</td><td>1000</td><td></td><td></td><td></td><td></td><td></td><td></td><td></td><td></td><td></td></tr>
<tr><td>Oligo fwd</td><td>CACCGGAACAGCTCCTCGGCCGCC</td><td></td><td></td><td></td><td></td><td></td><td></td><td></td><td></td><td></td><td></td></tr>
<tr><td>Oligo rev</td><td>AAACGGCGGCCGAGGAGCTGTTCC</td><td></td><td></td><td></td><td></td><td></td><td></td><td></td><td></td><td></td><td></td></tr>
<tr><td>Chromosome</td><td>start</td><td>end</td><td>strand</td><td>MM</td><td>target_seq</td><td>PAM</td><td>alignment</td><td>distance</td><td>position</td><td>gene name</td><td>gene id</td></tr>
<tr><td>GU305914.1</td><td>124778</td><td>124800</td><td>+</td><td>0</td><td>GGAACAGCTCCTCGGCCGCC</td><td>TGG</td><td>||||||||[||||||||||||]PAM</td><td>NA</td><td>Intergenic</td><td>NA</td><td>NA</td></tr>
<tr><td>GU305914.1</td><td>148554</td><td>148576</td><td>+</td><td>4</td><td>TGTTCAGCACCTCGGCCGCC</td><td>AGG</td><td>-|--||||[-|||||||||||]PAM</td><td>NA</td><td>Intergenic</td><td>NA</td><td>NA</td></tr>
<tr><td>T9438</td><td>CGCCACGTCCCGCACCAGCGCGG</td><td>1000</td><td></td><td></td><td></td><td></td><td></td><td></td><td></td><td></td><td></td></tr>
<tr><td>Oligo fwd</td><td>CACCCGCCACGTCCCGCACCAGCG</td><td></td><td></td><td></td><td></td><td></td><td></td><td></td><td></td><td></td><td></td></tr>
<tr><td>Oligo rev</td><td>AAACCGCTGGTGCGGGACGTGGCG</td><td></td><td></td><td></td><td></td><td></td><td></td><td></td><td></td><td></td><td></td></tr>
<tr><td>Chromosome</td><td>start</td><td>end</td><td>strand</td><td>MM</td><td>target_seq</td><td>PAM</td><td>alignment</td><td>distance</td><td>position</td><td>gene name</td><td>gene id</td></tr>
<tr><td>GU305914.1</td><td>124824</td><td>124846</td><td>+</td><td>0</td><td>CGCCACGTCCCGCACCAGCG</td><td>CGG</td><td>||||||||[||||||||||||]PAM</td><td>NA</td><td>Intergenic</td><td>NA</td><td>NA</td></tr>
<tr><td>T9439</td><td>CGCACCAGCGCGGCCAGCAGCGG</td><td>1000</td><td></td><td></td><td></td><td></td><td></td><td></td><td></td><td></td><td></td></tr>
<tr><td>Oligo fwd</td><td>CACCCGCACCAGCGCGGCCAGCAG</td><td></td><td></td><td></td><td></td><td></td><td></td><td></td><td></td><td></td><td></td></tr>
<tr><td>Oligo rev</td><td>AAACCTGCTGGCCGCGCTGGTGCG</td><td></td><td></td><td></td><td></td><td></td><td></td><td></td><td></td><td></td><td></td></tr>
<tr><td>Chromosome</td><td>start</td><td>end</td><td>strand</td><td>MM</td><td>target_seq</td><td>PAM</td><td>alignment</td><td>distance</td><td>position</td><td>gene name</td><td>gene id</td></tr>
<tr><td>GU305914.1</td><td>124834</td><td>124856</td><td>+</td><td>0</td><td>CGCACCAGCGCGGCCAGCAG</td><td>CGG</td><td>||||||||[||||||||||||]PAM</td><td>NA</td><td>Intergenic</td><td>NA</td><td>NA</td></tr>
<tr><td>T9440</td><td>GCACCAGCGCGGCCAGCAGCGGG</td><td>1000</td><td></td><td></td><td></td><td></td><td></td><td></td><td></td><td></td><td></td></tr>
<tr><td>Oligo fwd</td><td>CACCGCACCAGCGCGGCCAGCAGC</td><td></td><td></td><td></td><td></td><td></td><td></td><td></td><td></td><td></td><td></td></tr>
<tr><td>Oligo rev</td><td>AAACGCTGCTGGCCGCGCTGGTGC</td><td></td><td></td><td></td><td></td><td></td><td></td><td></td><td></td><td></td><td></td></tr>
<tr><td>Chromosome</td><td>start</td><td>end</td><td>strand</td><td>MM</td><td>target_seq</td><td>PAM</td><td>alignment</td><td>distance</td><td>position</td><td>gene name</td><td>gene id</td></tr>
</table>

GU305914.1 124835 124857 + 0 GCACCAGCGCGGCCAGCAGC GGG ||||||||[||||||||||||]PAM NA Intergenic NA NA

T9441 CGCGGCCAGCAGCGGGTTGTTGG 1000
Oligo fwd CACCCGCGGCCAGCAGCGGGTTGT
Oligo rev AAACACAACCCGCTGCTGGCCGCG
Chromosome start end strand MM target_seq PAM alignment distance position gene name gene id
GU305914.1 124842 124864 + 0 CGCGGCCAGCAGCGGGTTGT TGG ||||||||[||||||||||||]PAM NA Intergenic NA NA

T9442 GTTGGTCAGCGCCTCCTCGCCGG 1000
Oligo fwd CACCGTTGGTCAGCGCCTCCTCGC
Oligo rev AAACGCGAGGAGGCGCTGACCAAC
Chromosome start end strand MM target_seq PAM alignment distance position gene name gene id
GU305914.1 124860 124882 + 0 GTTGGTCAGCGCCTCCTCGC CGG ||||||||[||||||||||||]PAM NA Intergenic NA NA

T9443 TCGCCGGCCGTCAGCGAGTCCGG 1000
Oligo fwd CACCTCGCCGGCCGTCAGCGAGTC
Oligo rev AAACGACTCGCTGACGGCCGGCGA
Chromosome start end strand MM target_seq PAM alignment distance position gene name gene id
GU305914.1 124876 124898 + 0 TCGCCGGCCGTCAGCGAGTC CGG ||||||||[||||||||||||]PAM NA Intergenic NA NA

T9444 CGCCGGCCGTCAGCGAGTCCGGG 1000
Oligo fwd CACCCGCCGGCCGTCAGCGAGTCC
Oligo rev AAACGGACTCGCTGACGGCCGGCG
Chromosome start end strand MM target_seq PAM alignment distance position gene name gene id
GU305914.1 124877 124899 + 0 CGCCGGCCGTCAGCGAGTCC GGG ||||||||[||||||||||||]PAM NA Intergenic NA NA
GU305914.1 126614 126636 + 4 CGCAGTCCGTCAGCATGTCC AGG |||-|-||[||||||--||||]PAM NA Intergenic NA NA

T9445 GCGAGTCCGGGTCCGCCGTCAGG 1000
Oligo fwd CACCGCGAGTCCGGGTCCGCCGTC
Oligo rev AAACGACGGCGGACCCGGACTCGC
Chromosome start end strand MM target_seq PAM alignment distance position gene name gene id
GU305914.1 124889 124911 + 0 GCGAGTCCGGGTCCGCCGTC AGG ||||||||[||||||||||||]PAM NA Intergenic NA NA
GU305914.1 117569 117591 + 4 GTTTGTCGGGGTCCGCCGTC AGG |---|||-[||||||||||||]PAM NA Intergenic NA NA

T9446 GTCCGCCGTCAGGAAGAACTTGG 1000
Oligo fwd CACCGTCCGCCGTCAGGAAGAACT
Oligo rev AAACAGTTCTTCCTGACGGCGGAC
Chromosome start end strand MM target_seq PAM alignment distance position gene name gene id
GU305914.1 124899 124921 + 0 GTCCGCCGTCAGGAAGAACT TGG ||||||||[||||||||||||]PAM NA Intergenic NA NA

T9447 TGTAGAACAGCTTCACCAGCAGG 1000
Oligo fwd CACCTGTAGAACAGCTTCACCAGC
Oligo rev AAACGCTGGTGAAGCTGTTCTACA
Chromosome start end strand MM target_seq PAM alignment distance position gene name gene id
GU305914.1 124922 124944 + 0 TGTAGAACAGCTTCACCAGC AGG ||||||||[||||||||||||]PAM NA Intergenic NA NA

T9448 GCTTCACCAGCAGGTGCTTCAGG 1000
Oligo fwd CACCGCTTCACCAGCAGGTGCTTC
Oligo rev AAACGAAGCACCTGCTGGTGAAGC
Chromosome start end strand MM target_seq PAM alignment distance position gene name gene id
GU305914.1 124931 124953 + 0 GCTTCACCAGCAGGTGCTTC AGG ||||||||[||||||||||||]PAM NA Intergenic NA NA

T9449 CCAGCAGGTGCTTCAGGTTCAGG 1000
Oligo fwd CACCCCAGCAGGTGCTTCAGGTTC
Oligo rev AAACGAACCTGAAGCACCTGCTGG
Chromosome start end strand MM target_seq PAM alignment distance position gene name gene id
GU305914.1 124937 124959 + 0 CCAGCAGGTGCTTCAGGTTC AGG ||||||||[||||||||||||]PAM NA Intergenic NA NA

T9450 TCAGGCCCGCGCCGCACGCGTGG 1000
Oligo fwd CACCTCAGGCCCGCGCCGCACGCG
Oligo rev AAACCGCGTGCGGCGCGGGCCTGA
Chromosome start end strand MM target_seq PAM alignment distance position gene name gene id
GU305914.1 124955 124977 + 0 TCAGGCCCGCGCCGCACGCG TGG ||||||||[||||||||||||]PAM NA Intergenic NA NA

T9451 GCACGCGTGGTTGTTCAGCACGG 1000
Oligo fwd CACCGCACGCGTGGTTGTTCAGCA
Oligo rev AAACTGCTGAACAACCACGCGTGC
Chromosome start end strand MM target_seq PAM alignment distance position gene name gene id
GU305914.1 124968 124990 + 0 GCACGCGTGGTTGTTCAGCA CGG ||||||||[||||||||||||]PAM NA Intergenic NA NA

T9452 CGCGTGGTTGTTCAGCACGGCGG 1000
Oligo fwd CACCCGCGTGGTTGTTCAGCACGG
Oligo rev AAACCCGTGCTGAACAACCACGCG
Chromosome start end strand MM target_seq PAM alignment distance position gene name gene id
GU305914.1 124971 124993 + 0 CGCGTGGTTGTTCAGCACGG CGG ||||||||[||||||||||||]PAM NA Intergenic NA NA

T9453 GCGTGGTTGTTCAGCACGGCGGG 1000
Oligo fwd CACCGCGTGGTTGTTCAGCACGGC
Oligo rev AAACGCCGTGCTGAACAACCACGC
Chromosome start end strand MM target_seq PAM alignment distance position gene name gene id
GU305914.1 124972 124994 + 0 GCGTGGTTGTTCAGCACGGC GGG ||||||||[||||||||||||]PAM NA Intergenic NA NA

T9454 TGTTCAGCACGGCGGGCACCAGG 1000
Oligo fwd CACCTGTTCAGCACGGCGGGCACC
Oligo rev AAACGGTGCCCGCCGTGCTGAACA
Chromosome start end strand MM target_seq PAM alignment distance position gene name gene id
GU305914.1 124979 125001 + 0 TGTTCAGCACGGCGGGCACC AGG ||||||||[||||||||||||]PAM NA Intergenic NA NA

T9455 GCACGGCGGGCACCAGGCACAGG 1000

<table>
<tr><td>Oligo fwd</td><td>CACCGCACGGCGGGCACCAGGCAC</td><td></td><td></td><td></td><td></td><td></td><td></td><td></td><td></td><td></td><td></td></tr>
<tr><td>Oligo rev</td><td>AAACGTGCCTGGTGCCCGCCGTGC</td><td></td><td></td><td></td><td></td><td></td><td></td><td></td><td></td><td></td><td></td></tr>
<tr><td>Chromosome</td><td>start</td><td>end</td><td>strand</td><td>MM</td><td>target_seq</td><td>PAM</td><td>alignment</td><td>distance</td><td>position</td><td>gene name</td><td>gene id</td></tr>
<tr><td>GU305914.1</td><td>124985</td><td>125007</td><td>+</td><td>0</td><td>GCACGGCGGGCACCAGGCAC</td><td>AGG</td><td>||||||||[||||||||||||]PAM</td><td>NA</td><td>Intergenic</td><td>NA</td><td>NA</td></tr>
<tr><td></td><td></td><td></td><td></td><td></td><td></td><td></td><td></td><td></td><td></td><td></td><td></td></tr>
<tr><td>T9456</td><td>TCTCCTCGTCCGACGTCCGCCGG</td><td>1000</td><td></td><td></td><td></td><td></td><td></td><td></td><td></td><td></td><td></td></tr>
<tr><td>Oligo fwd</td><td>CACCTCTCCTCGTCCGACGTCCGC</td><td></td><td></td><td></td><td></td><td></td><td></td><td></td><td></td><td></td><td></td></tr>
<tr><td>Oligo rev</td><td>AAACGCGGACGTCGGACGAGGAGA</td><td></td><td></td><td></td><td></td><td></td><td></td><td></td><td></td><td></td><td></td></tr>
<tr><td>Chromosome</td><td>start</td><td>end</td><td>strand</td><td>MM</td><td>target_seq</td><td>PAM</td><td>alignment</td><td>distance</td><td>position</td><td>gene name</td><td>gene id</td></tr>
<tr><td>GU305914.1</td><td>125030</td><td>125052</td><td>+</td><td>0</td><td>TCTCCTCGTCCGACGTCCGC</td><td>CGG</td><td>||||||||[||||||||||||]PAM</td><td>NA</td><td>Intergenic</td><td>NA</td><td>NA</td></tr>
<tr><td></td><td></td><td></td><td></td><td></td><td></td><td></td><td></td><td></td><td></td><td></td><td></td></tr>
<tr><td>T9457</td><td>CTCCTCGTCCGACGTCCGCCGGG</td><td>1000</td><td></td><td></td><td></td><td></td><td></td><td></td><td></td><td></td><td></td></tr>
<tr><td>Oligo fwd</td><td>CACCCTCCTCGTCCGACGTCCGCC</td><td></td><td></td><td></td><td></td><td></td><td></td><td></td><td></td><td></td><td></td></tr>
<tr><td>Oligo rev</td><td>AAACGGCGGACGTCGGACGAGGAG</td><td></td><td></td><td></td><td></td><td></td><td></td><td></td><td></td><td></td><td></td></tr>
<tr><td>Chromosome</td><td>start</td><td>end</td><td>strand</td><td>MM</td><td>target_seq</td><td>PAM</td><td>alignment</td><td>distance</td><td>position</td><td>gene name</td><td>gene id</td></tr>
<tr><td>GU305914.1</td><td>125031</td><td>125053</td><td>+</td><td>0</td><td>CTCCTCGTCCGACGTCCGCC</td><td>GGG</td><td>||||||||[||||||||||||]PAM</td><td>NA</td><td>Intergenic</td><td>NA</td><td>NA</td></tr>
<tr><td></td><td></td><td></td><td></td><td></td><td></td><td></td><td></td><td></td><td></td><td></td><td></td></tr>
<tr><td>T9458</td><td>CGCCGGGCGTCCACGAACAGCGG</td><td>1000</td><td></td><td></td><td></td><td></td><td></td><td></td><td></td><td></td><td></td></tr>
<tr><td>Oligo fwd</td><td>CACCCGCCGGGCGTCCACGAACAG</td><td></td><td></td><td></td><td></td><td></td><td></td><td></td><td></td><td></td><td></td></tr>
<tr><td>Oligo rev</td><td>AAACCTGTTCGTGGACGCCCGGCG</td><td></td><td></td><td></td><td></td><td></td><td></td><td></td><td></td><td></td><td></td></tr>
<tr><td>Chromosome</td><td>start</td><td>end</td><td>strand</td><td>MM</td><td>target_seq</td><td>PAM</td><td>alignment</td><td>distance</td><td>position</td><td>gene name</td><td>gene id</td></tr>
<tr><td>GU305914.1</td><td>125047</td><td>125069</td><td>+</td><td>0</td><td>CGCCGGGCGTCCACGAACAG</td><td>CGG</td><td>||||||||[||||||||||||]PAM</td><td>NA</td><td>Intergenic</td><td>NA</td><td>NA</td></tr>
<tr><td></td><td></td><td></td><td></td><td></td><td></td><td></td><td></td><td></td><td></td><td></td><td></td></tr>
<tr><td>T9459</td><td>CCACGAACAGCGGCTCCGCCTGG</td><td>1000</td><td></td><td></td><td></td><td></td><td></td><td></td><td></td><td></td><td></td></tr>
<tr><td>Oligo fwd</td><td>CACCCCACGAACAGCGGCTCCGCC</td><td></td><td></td><td></td><td></td><td></td><td></td><td></td><td></td><td></td><td></td></tr>
<tr><td>Oligo rev</td><td>AAACGGCGGAGCCGCTGTTCGTGG</td><td></td><td></td><td></td><td></td><td></td><td></td><td></td><td></td><td></td><td></td></tr>
<tr><td>Chromosome</td><td>start</td><td>end</td><td>strand</td><td>MM</td><td>target_seq</td><td>PAM</td><td>alignment</td><td>distance</td><td>position</td><td>gene name</td><td>gene id</td></tr>
<tr><td>GU305914.1</td><td>125057</td><td>125079</td><td>+</td><td>0</td><td>CCACGAACAGCGGCTCCGCC</td><td>TGG</td><td>||||||||[||||||||||||]PAM</td><td>NA</td><td>Intergenic</td><td>NA</td><td>NA</td></tr>
<tr><td></td><td></td><td></td><td></td><td></td><td></td><td></td><td></td><td></td><td></td><td></td><td></td></tr>
<tr><td>T9460</td><td>GATCATCCGCGCCACGTCCGAGG</td><td>1000</td><td></td><td></td><td></td><td></td><td></td><td></td><td></td><td></td><td></td></tr>
<tr><td>Oligo fwd</td><td>CACCGATCATCCGCGCCACGTCCG</td><td></td><td></td><td></td><td></td><td></td><td></td><td></td><td></td><td></td><td></td></tr>
<tr><td>Oligo rev</td><td>AAACCGGACGTGGCGCGGATGATC</td><td></td><td></td><td></td><td></td><td></td><td></td><td></td><td></td><td></td><td></td></tr>
<tr><td>Chromosome</td><td>start</td><td>end</td><td>strand</td><td>MM</td><td>target_seq</td><td>PAM</td><td>alignment</td><td>distance</td><td>position</td><td>gene name</td><td>gene id</td></tr>
<tr><td>GU305914.1</td><td>125088</td><td>125110</td><td>+</td><td>0</td><td>GATCATCCGCGCCACGTCCG</td><td>AGG</td><td>||||||||[||||||||||||]PAM</td><td>NA</td><td>Intergenic</td><td>NA</td><td>NA</td></tr>
<tr><td></td><td></td><td></td><td></td><td></td><td></td><td></td><td></td><td></td><td></td><td></td><td></td></tr>
<tr><td>T9461</td><td>CGCGCCACGTCCGAGGTGCCCGG</td><td>1000</td><td></td><td></td><td></td><td></td><td></td><td></td><td></td><td></td><td></td></tr>
<tr><td>Oligo fwd</td><td>CACCCGCGCCACGTCCGAGGTGCC</td><td></td><td></td><td></td><td></td><td></td><td></td><td></td><td></td><td></td><td></td></tr>
<tr><td>Oligo rev</td><td>AAACGGCACCTCGGACGTGGCGCG</td><td></td><td></td><td></td><td></td><td></td><td></td><td></td><td></td><td></td><td></td></tr>
<tr><td>Chromosome</td><td>start</td><td>end</td><td>strand</td><td>MM</td><td>target_seq</td><td>PAM</td><td>alignment</td><td>distance</td><td>position</td><td>gene name</td><td>gene id</td></tr>
<tr><td>GU305914.1</td><td>125095</td><td>125117</td><td>+</td><td>0</td><td>CGCGCCACGTCCGAGGTGCC</td><td>CGG</td><td>||||||||[||||||||||||]PAM</td><td>NA</td><td>Intergenic</td><td>NA</td><td>NA</td></tr>
<tr><td></td><td></td><td></td><td></td><td></td><td></td><td></td><td></td><td></td><td></td><td></td><td></td></tr>
<tr><td>T9462</td><td>GCGCCACGTCCGAGGTGCCCGGG</td><td>1000</td><td></td><td></td><td></td><td></td><td></td><td></td><td></td><td></td><td></td></tr>
<tr><td>Oligo fwd</td><td>CACCGCGCCACGTCCGAGGTGCCC</td><td></td><td></td><td></td><td></td><td></td><td></td><td></td><td></td><td></td><td></td></tr>
<tr><td>Oligo rev</td><td>AAACGGGCACCTCGGACGTGGCGC</td><td></td><td></td><td></td><td></td><td></td><td></td><td></td><td></td><td></td><td></td></tr>
<tr><td>Chromosome</td><td>start</td><td>end</td><td>strand</td><td>MM</td><td>target_seq</td><td>PAM</td><td>alignment</td><td>distance</td><td>position</td><td>gene name</td><td>gene id</td></tr>
<tr><td>GU305914.1</td><td>125096</td><td>125118</td><td>+</td><td>0</td><td>GCGCCACGTCCGAGGTGCCC</td><td>GGG</td><td>||||||||[||||||||||||]PAM</td><td>NA</td><td>Intergenic</td><td>NA</td><td>NA</td></tr>
<tr><td></td><td></td><td></td><td></td><td></td><td></td><td></td><td></td><td></td><td></td><td></td><td></td></tr>
<tr><td>T9463</td><td>TCGATGTGCGCCGCGTCCAGCGG</td><td>1000</td><td></td><td></td><td></td><td></td><td></td><td></td><td></td><td></td><td></td></tr>
<tr><td>Oligo fwd</td><td>CACCTCGATGTGCGCCGCGTCCAG</td><td></td><td></td><td></td><td></td><td></td><td></td><td></td><td></td><td></td><td></td></tr>
<tr><td>Oligo rev</td><td>AAACCTGGACGCGGCGCACATCGA</td><td></td><td></td><td></td><td></td><td></td><td></td><td></td><td></td><td></td><td></td></tr>
<tr><td>Chromosome</td><td>start</td><td>end</td><td>strand</td><td>MM</td><td>target_seq</td><td>PAM</td><td>alignment</td><td>distance</td><td>position</td><td>gene name</td><td>gene id</td></tr>
<tr><td>GU305914.1</td><td>125128</td><td>125150</td><td>+</td><td>0</td><td>TCGATGTGCGCCGCGTCCAG</td><td>CGG</td><td>||||||||[||||||||||||]PAM</td><td>NA</td><td>Intergenic</td><td>NA</td><td>NA</td></tr>
<tr><td></td><td></td><td></td><td></td><td></td><td></td><td></td><td></td><td></td><td></td><td></td><td></td></tr>
<tr><td>T9464</td><td>CGCGTCCAGCGGCACGTCGTCGG</td><td>1000</td><td></td><td></td><td></td><td></td><td></td><td></td><td></td><td></td><td></td></tr>
<tr><td>Oligo fwd</td><td>CACCCGCGTCCAGCGGCACGTCGT</td><td></td><td></td><td></td><td></td><td></td><td></td><td></td><td></td><td></td><td></td></tr>
<tr><td>Oligo rev</td><td>AAACACGACGTGCCGCTGGACGCG</td><td></td><td></td><td></td><td></td><td></td><td></td><td></td><td></td><td></td><td></td></tr>
<tr><td>Chromosome</td><td>start</td><td>end</td><td>strand</td><td>MM</td><td>target_seq</td><td>PAM</td><td>alignment</td><td>distance</td><td>position</td><td>gene name</td><td>gene id</td></tr>
<tr><td>GU305914.1</td><td>125139</td><td>125161</td><td>+</td><td>0</td><td>CGCGTCCAGCGGCACGTCGT</td><td>CGG</td><td>||||||||[||||||||||||]PAM</td><td>NA</td><td>Intergenic</td><td>NA</td><td>NA</td></tr>
<tr><td></td><td></td><td></td><td></td><td></td><td></td><td></td><td></td><td></td><td></td><td></td><td></td></tr>
<tr><td>T9465</td><td>CGTCGGCCACCACGCGCACCAGG</td><td>1000</td><td></td><td></td><td></td><td></td><td></td><td></td><td></td><td></td><td></td></tr>
<tr><td>Oligo fwd</td><td>CACCCGTCGGCCACCACGCGCACC</td><td></td><td></td><td></td><td></td><td></td><td></td><td></td><td></td><td></td><td></td></tr>
<tr><td>Oligo rev</td><td>AAACGGTGCGCGTGGTGGCCGACG</td><td></td><td></td><td></td><td></td><td></td><td></td><td></td><td></td><td></td><td></td></tr>
<tr><td>Chromosome</td><td>start</td><td>end</td><td>strand</td><td>MM</td><td>target_seq</td><td>PAM</td><td>alignment</td><td>distance</td><td>position</td><td>gene name</td><td>gene id</td></tr>
<tr><td>GU305914.1</td><td>125156</td><td>125178</td><td>+</td><td>0</td><td>CGTCGGCCACCACGCGCACC</td><td>AGG</td><td>||||||||[||||||||||||]PAM</td><td>NA</td><td>Intergenic</td><td>NA</td><td>NA</td></tr>
<tr><td>GU305914.1</td><td>119659</td><td>119681</td><td>-</td><td>3</td><td>CGGCGACCACCACGCGCACG</td><td>GGG</td><td>||-||-||[|||||||||||-]PAM</td><td>NA</td><td>Intergenic</td><td>NA</td><td>NA</td></tr>
<tr><td></td><td></td><td></td><td></td><td></td><td></td><td></td><td></td><td></td><td></td><td></td><td></td></tr>
<tr><td>T9466</td><td>CGCACCAGGTTCGCGTTGCGCGG</td><td>1000</td><td></td><td></td><td></td><td></td><td></td><td></td><td></td><td></td><td></td></tr>
<tr><td>Oligo fwd</td><td>CACCCGCACCAGGTTCGCGTTGCG</td><td></td><td></td><td></td><td></td><td></td><td></td><td></td><td></td><td></td><td></td></tr>
<tr><td>Oligo rev</td><td>AAACCGCAACGCGAACCTGGTGCG</td><td></td><td></td><td></td><td></td><td></td><td></td><td></td><td></td><td></td><td></td></tr>
<tr><td>Chromosome</td><td>start</td><td>end</td><td>strand</td><td>MM</td><td>target_seq</td><td>PAM</td><td>alignment</td><td>distance</td><td>position</td><td>gene name</td><td>gene id</td></tr>
<tr><td>GU305914.1</td><td>125170</td><td>125192</td><td>+</td><td>0</td><td>CGCACCAGGTTCGCGTTGCG</td><td>CGG</td><td>||||||||[||||||||||||]PAM</td><td>NA</td><td>Intergenic</td><td>NA</td><td>NA</td></tr>
<tr><td></td><td></td><td></td><td></td><td></td><td></td><td></td><td></td><td></td><td></td><td></td><td></td></tr>
<tr><td>T9467</td><td>GCGCGGCAGCGCCTGCACGAAGG</td><td>1000</td><td></td><td></td><td></td><td></td><td></td><td></td><td></td><td></td><td></td></tr>
<tr><td>Oligo fwd</td><td>CACCGCGCGGCAGCGCCTGCACGA</td><td></td><td></td><td></td><td></td><td></td><td></td><td></td><td></td><td></td><td></td></tr>
<tr><td>Oligo rev</td><td>AAACTCGTGCAGGCGCTGCCGCGC</td><td></td><td></td><td></td><td></td><td></td><td></td><td></td><td></td><td></td><td></td></tr>
<tr><td>Chromosome</td><td>start</td><td>end</td><td>strand</td><td>MM</td><td>target_seq</td><td>PAM</td><td>alignment</td><td>distance</td><td>position</td><td>gene name</td><td>gene id</td></tr>
<tr><td>GU305914.1</td><td>125187</td><td>125209</td><td>+</td><td>0</td><td>GCGCGGCAGCGCCTGCACGA</td><td>AGG</td><td>||||||||[||||||||||||]PAM</td><td>NA</td><td>Intergenic</td><td>NA</td><td>NA</td></tr>
<tr><td>GU305914.1</td><td>126507</td><td>126529</td><td>+</td><td>4</td><td>GTTCAGCAGCGCCTGCACCA</td><td>CGG</td><td>|--|-|||[||||||||||-|]PAM</td><td>NA</td><td>Intergenic</td><td>NA</td><td>NA</td></tr>
<tr><td>GU305914.1</td><td>76452</td><td>76474</td><td>-</td><td>4</td><td>CCGCGGCGGCGCCGGCGCGA</td><td>TGG</td><td>-||||||-[|||||-||-|||]PAM</td><td>NA</td><td>Intergenic</td><td>NA</td><td>NA</td></tr>
<tr><td></td><td></td><td></td><td></td><td></td><td></td><td></td><td></td><td></td><td></td><td></td><td></td></tr>
<tr><td>T9468</td><td>CGCGGCAGCGCCTGCACGAAGGG</td><td>1000</td><td></td><td></td><td></td><td></td><td></td><td></td><td></td><td></td><td></td></tr>
<tr><td>Oligo fwd</td><td>CACCCGCGGCAGCGCCTGCACGAA</td><td></td><td></td><td></td><td></td><td></td><td></td><td></td><td></td><td></td><td></td></tr>
<tr><td>Oligo rev</td><td>AAACTTCGTGCAGGCGCTGCCGCG</td><td></td><td></td><td></td><td></td><td></td><td></td><td></td><td></td><td></td><td></td></tr>
<tr><td>Chromosome</td><td>start</td><td>end</td><td>strand</td><td>MM</td><td>target_seq</td><td>PAM</td><td>alignment</td><td>distance</td><td>position</td><td>gene name</td><td>gene id</td></tr>
<tr><td>GU305914.1</td><td>125188</td><td>125210</td><td>+</td><td>0</td><td>CGCGGCAGCGCCTGCACGAA</td><td>GGG</td><td>||||||||[||||||||||||]PAM</td><td>NA</td><td>Intergenic</td><td>NA</td><td>NA</td></tr>
<tr><td></td><td></td><td></td><td></td><td></td><td></td><td></td><td></td><td></td><td></td><td></td><td></td></tr>
<tr><td>T9469</td><td>GGCAGCGCCTGCACGAAGGGCGG</td><td>1000</td><td></td><td></td><td></td><td></td><td></td><td></td><td></td><td></td><td></td></tr>
<tr><td>Oligo fwd</td><td>CACCGGCAGCGCCTGCACGAAGGG</td><td></td><td></td><td></td><td></td><td></td><td></td><td></td><td></td><td></td><td></td></tr>
<tr><td>Oligo rev</td><td>AAACCCCTTCGTGCAGGCGCTGCC</td><td></td><td></td><td></td><td></td><td></td><td></td><td></td><td></td><td></td><td></td></tr>
</table>

<table>
<tr><td>Chromosome</td><td>start</td><td>end</td><td>strand</td><td>MM</td><td>target_seq</td><td>PAM</td><td>alignment</td><td>distance</td><td>position</td><td>gene name</td><td>gene id</td></tr>
<tr><td>GU305914.1</td><td>125191</td><td>125213</td><td>+</td><td>0</td><td>GGCAGCGCCTGCACGAAGGG</td><td>CGG</td><td>||||||||[||||||||||||]PAM</td><td>NA</td><td>Intergenic</td><td>NA</td><td>NA</td></tr>
<tr><td></td><td></td><td></td><td></td><td></td><td></td><td></td><td></td><td></td><td></td><td></td><td></td></tr>
<tr><td>T9470</td><td>CCTGCACGAAGGGCGGCCAGAGG</td><td>1000</td><td></td><td></td><td></td><td></td><td></td><td></td><td></td><td></td><td></td></tr>
<tr><td>Oligo fwd</td><td>CACCCCTGCACGAAGGGCGGCCAG</td><td></td><td></td><td></td><td></td><td></td><td></td><td></td><td></td><td></td><td></td></tr>
<tr><td>Oligo rev</td><td>AAACCTGGCCGCCCTTCGTGCAGG</td><td></td><td></td><td></td><td></td><td></td><td></td><td></td><td></td><td></td><td></td></tr>
<tr><td>Chromosome</td><td>start</td><td>end</td><td>strand</td><td>MM</td><td>target_seq</td><td>PAM</td><td>alignment</td><td>distance</td><td>position</td><td>gene name</td><td>gene id</td></tr>
<tr><td>GU305914.1</td><td>125198</td><td>125220</td><td>+</td><td>0</td><td>CCTGCACGAAGGGCGGCCAG</td><td>AGG</td><td>||||||||[||||||||||||]PAM</td><td>NA</td><td>Intergenic</td><td>NA</td><td>NA</td></tr>
<tr><td></td><td></td><td></td><td></td><td></td><td></td><td></td><td></td><td></td><td></td><td></td><td></td></tr>
<tr><td>T9471</td><td>ACGAAGGGCGGCCAGAGGCGCGG</td><td>1000</td><td></td><td></td><td></td><td></td><td></td><td></td><td></td><td></td><td></td></tr>
<tr><td>Oligo fwd</td><td>CACCACGAAGGGCGGCCAGAGGCG</td><td></td><td></td><td></td><td></td><td></td><td></td><td></td><td></td><td></td><td></td></tr>
<tr><td>Oligo rev</td><td>AAACCGCCTCTGGCCGCCCTTCGT</td><td></td><td></td><td></td><td></td><td></td><td></td><td></td><td></td><td></td><td></td></tr>
<tr><td>Chromosome</td><td>start</td><td>end</td><td>strand</td><td>MM</td><td>target_seq</td><td>PAM</td><td>alignment</td><td>distance</td><td>position</td><td>gene name</td><td>gene id</td></tr>
<tr><td>GU305914.1</td><td>125203</td><td>125225</td><td>+</td><td>0</td><td>ACGAAGGGCGGCCAGAGGCG</td><td>CGG</td><td>||||||||[||||||||||||]PAM</td><td>NA</td><td>Intergenic</td><td>NA</td><td>NA</td></tr>
<tr><td></td><td></td><td></td><td></td><td></td><td></td><td></td><td></td><td></td><td></td><td></td><td></td></tr>
<tr><td>T9472</td><td>CGAAGGGCGGCCAGAGGCGCGGG</td><td>1000</td><td></td><td></td><td></td><td></td><td></td><td></td><td></td><td></td><td></td></tr>
<tr><td>Oligo fwd</td><td>CACCCGAAGGGCGGCCAGAGGCGC</td><td></td><td></td><td></td><td></td><td></td><td></td><td></td><td></td><td></td><td></td></tr>
<tr><td>Oligo rev</td><td>AAACGCGCCTCTGGCCGCCCTTCG</td><td></td><td></td><td></td><td></td><td></td><td></td><td></td><td></td><td></td><td></td></tr>
<tr><td>Chromosome</td><td>start</td><td>end</td><td>strand</td><td>MM</td><td>target_seq</td><td>PAM</td><td>alignment</td><td>distance</td><td>position</td><td>gene name</td><td>gene id</td></tr>
<tr><td>GU305914.1</td><td>125204</td><td>125226</td><td>+</td><td>0</td><td>CGAAGGGCGGCCAGAGGCGC</td><td>GGG</td><td>||||||||[||||||||||||]PAM</td><td>NA</td><td>Intergenic</td><td>NA</td><td>NA</td></tr>
<tr><td></td><td></td><td></td><td></td><td></td><td></td><td></td><td></td><td></td><td></td><td></td><td></td></tr>
<tr><td>T9473</td><td>GCGCTCACGTACGCCAGCAGCGG</td><td>1000</td><td></td><td></td><td></td><td></td><td></td><td></td><td></td><td></td><td></td></tr>
<tr><td>Oligo fwd</td><td>CACCGCGCTCACGTACGCCAGCAG</td><td></td><td></td><td></td><td></td><td></td><td></td><td></td><td></td><td></td><td></td></tr>
<tr><td>Oligo rev</td><td>AAACCTGCTGGCGTACGTGAGCGC</td><td></td><td></td><td></td><td></td><td></td><td></td><td></td><td></td><td></td><td></td></tr>
<tr><td>Chromosome</td><td>start</td><td>end</td><td>strand</td><td>MM</td><td>target_seq</td><td>PAM</td><td>alignment</td><td>distance</td><td>position</td><td>gene name</td><td>gene id</td></tr>
<tr><td>GU305914.1</td><td>125236</td><td>125258</td><td>+</td><td>0</td><td>GCGCTCACGTACGCCAGCAG</td><td>CGG</td><td>||||||||[||||||||||||]PAM</td><td>NA</td><td>Intergenic</td><td>NA</td><td>NA</td></tr>
<tr><td></td><td></td><td></td><td></td><td></td><td></td><td></td><td></td><td></td><td></td><td></td><td></td></tr>
<tr><td>T9474</td><td>CGTCACACAGCCGCCCGTTCAGG</td><td>1000</td><td></td><td></td><td></td><td></td><td></td><td></td><td></td><td></td><td></td></tr>
<tr><td>Oligo fwd</td><td>CACCCGTCACACAGCCGCCCGTTC</td><td></td><td></td><td></td><td></td><td></td><td></td><td></td><td></td><td></td><td></td></tr>
<tr><td>Oligo rev</td><td>AAACGAACGGGCGGCTGTGTGACG</td><td></td><td></td><td></td><td></td><td></td><td></td><td></td><td></td><td></td><td></td></tr>
<tr><td>Chromosome</td><td>start</td><td>end</td><td>strand</td><td>MM</td><td>target_seq</td><td>PAM</td><td>alignment</td><td>distance</td><td>position</td><td>gene name</td><td>gene id</td></tr>
<tr><td>GU305914.1</td><td>125261</td><td>125283</td><td>+</td><td>0</td><td>CGTCACACAGCCGCCCGTTC</td><td>AGG</td><td>||||||||[||||||||||||]PAM</td><td>NA</td><td>Intergenic</td><td>NA</td><td>NA</td></tr>
<tr><td></td><td></td><td></td><td></td><td></td><td></td><td></td><td></td><td></td><td></td><td></td><td></td></tr>
<tr><td>T9475</td><td>GTTCAGGCCCGTCAGTTCGCAGG</td><td>1000</td><td></td><td></td><td></td><td></td><td></td><td></td><td></td><td></td><td></td></tr>
<tr><td>Oligo fwd</td><td>CACCGTTCAGGCCCGTCAGTTCGC</td><td></td><td></td><td></td><td></td><td></td><td></td><td></td><td></td><td></td><td></td></tr>
<tr><td>Oligo rev</td><td>AAACGCGAACTGACGGGCCTGAAC</td><td></td><td></td><td></td><td></td><td></td><td></td><td></td><td></td><td></td><td></td></tr>
<tr><td>Chromosome</td><td>start</td><td>end</td><td>strand</td><td>MM</td><td>target_seq</td><td>PAM</td><td>alignment</td><td>distance</td><td>position</td><td>gene name</td><td>gene id</td></tr>
<tr><td>GU305914.1</td><td>125277</td><td>125299</td><td>+</td><td>0</td><td>GTTCAGGCCCGTCAGTTCGC</td><td>AGG</td><td>||||||||[||||||||||||]PAM</td><td>NA</td><td>Intergenic</td><td>NA</td><td>NA</td></tr>
<tr><td></td><td></td><td></td><td></td><td></td><td></td><td></td><td></td><td></td><td></td><td></td><td></td></tr>
<tr><td>T9476</td><td>CCGCGCCACGAACCGTGTGATGG</td><td>1000</td><td></td><td></td><td></td><td></td><td></td><td></td><td></td><td></td><td></td></tr>
<tr><td>Oligo fwd</td><td>CACCCCGCGCCACGAACCGTGTGA</td><td></td><td></td><td></td><td></td><td></td><td></td><td></td><td></td><td></td><td></td></tr>
<tr><td>Oligo rev</td><td>AAACTCACACGGTTCGTGGCGCGG</td><td></td><td></td><td></td><td></td><td></td><td></td><td></td><td></td><td></td><td></td></tr>
<tr><td>Chromosome</td><td>start</td><td>end</td><td>strand</td><td>MM</td><td>target_seq</td><td>PAM</td><td>alignment</td><td>distance</td><td>position</td><td>gene name</td><td>gene id</td></tr>
<tr><td>GU305914.1</td><td>125301</td><td>125323</td><td>+</td><td>0</td><td>CCGCGCCACGAACCGTGTGA</td><td>TGG</td><td>||||||||[||||||||||||]PAM</td><td>NA</td><td>Intergenic</td><td>NA</td><td>NA</td></tr>
<tr><td></td><td></td><td></td><td></td><td></td><td></td><td></td><td></td><td></td><td></td><td></td><td></td></tr>
<tr><td>T9477</td><td>CGAACCGTGTGATGGACAGCAGG</td><td>1000</td><td></td><td></td><td></td><td></td><td></td><td></td><td></td><td></td><td></td></tr>
<tr><td>Oligo fwd</td><td>CACCCGAACCGTGTGATGGACAGC</td><td></td><td></td><td></td><td></td><td></td><td></td><td></td><td></td><td></td><td></td></tr>
<tr><td>Oligo rev</td><td>AAACGCTGTCCATCACACGGTTCG</td><td></td><td></td><td></td><td></td><td></td><td></td><td></td><td></td><td></td><td></td></tr>
<tr><td>Chromosome</td><td>start</td><td>end</td><td>strand</td><td>MM</td><td>target_seq</td><td>PAM</td><td>alignment</td><td>distance</td><td>position</td><td>gene name</td><td>gene id</td></tr>
<tr><td>GU305914.1</td><td>125309</td><td>125331</td><td>+</td><td>0</td><td>CGAACCGTGTGATGGACAGC</td><td>AGG</td><td>||||||||[||||||||||||]PAM</td><td>NA</td><td>Intergenic</td><td>NA</td><td>NA</td></tr>
<tr><td></td><td></td><td></td><td></td><td></td><td></td><td></td><td></td><td></td><td></td><td></td><td></td></tr>
<tr><td>T9478</td><td>GTGTGATGGACAGCAGGTTGCGG</td><td>1000</td><td></td><td></td><td></td><td></td><td></td><td></td><td></td><td></td><td></td></tr>
<tr><td>Oligo fwd</td><td>CACCGTGTGATGGACAGCAGGTTG</td><td></td><td></td><td></td><td></td><td></td><td></td><td></td><td></td><td></td><td></td></tr>
<tr><td>Oligo rev</td><td>AAACCAACCTGCTGTCCATCACAC</td><td></td><td></td><td></td><td></td><td></td><td></td><td></td><td></td><td></td><td></td></tr>
<tr><td>Chromosome</td><td>start</td><td>end</td><td>strand</td><td>MM</td><td>target_seq</td><td>PAM</td><td>alignment</td><td>distance</td><td>position</td><td>gene name</td><td>gene id</td></tr>
<tr><td>GU305914.1</td><td>125315</td><td>125337</td><td>+</td><td>0</td><td>GTGTGATGGACAGCAGGTTG</td><td>CGG</td><td>||||||||[||||||||||||]PAM</td><td>NA</td><td>Intergenic</td><td>NA</td><td>NA</td></tr>
<tr><td></td><td></td><td></td><td></td><td></td><td></td><td></td><td></td><td></td><td></td><td></td><td></td></tr>
<tr><td>T9479</td><td>CGGTAGTGCGTGTACGCCGCCGG</td><td>1000</td><td></td><td></td><td></td><td></td><td></td><td></td><td></td><td></td><td></td></tr>
<tr><td>Oligo fwd</td><td>CACCCGGTAGTGCGTGTACGCCGC</td><td></td><td></td><td></td><td></td><td></td><td></td><td></td><td></td><td></td><td></td></tr>
<tr><td>Oligo rev</td><td>AAACGCGGCGTACACGCACTACCG</td><td></td><td></td><td></td><td></td><td></td><td></td><td></td><td></td><td></td><td></td></tr>
<tr><td>Chromosome</td><td>start</td><td>end</td><td>strand</td><td>MM</td><td>target_seq</td><td>PAM</td><td>alignment</td><td>distance</td><td>position</td><td>gene name</td><td>gene id</td></tr>
<tr><td>GU305914.1</td><td>125335</td><td>125357</td><td>+</td><td>0</td><td>CGGTAGTGCGTGTACGCCGC</td><td>CGG</td><td>||||||||[||||||||||||]PAM</td><td>NA</td><td>Intergenic</td><td>NA</td><td>NA</td></tr>
<tr><td></td><td></td><td></td><td></td><td></td><td></td><td></td><td></td><td></td><td></td><td></td><td></td></tr>
<tr><td>T9480</td><td>CGCGCCGTCGCCCAGCCGCGTGG</td><td>1000</td><td></td><td></td><td></td><td></td><td></td><td></td><td></td><td></td><td></td></tr>
<tr><td>Oligo fwd</td><td>CACCCGCGCCGTCGCCCAGCCGCG</td><td></td><td></td><td></td><td></td><td></td><td></td><td></td><td></td><td></td><td></td></tr>
<tr><td>Oligo rev</td><td>AAACCGCGGCTGGGCGACGGCGCG</td><td></td><td></td><td></td><td></td><td></td><td></td><td></td><td></td><td></td><td></td></tr>
<tr><td>Chromosome</td><td>start</td><td>end</td><td>strand</td><td>MM</td><td>target_seq</td><td>PAM</td><td>alignment</td><td>distance</td><td>position</td><td>gene name</td><td>gene id</td></tr>
<tr><td>GU305914.1</td><td>125364</td><td>125386</td><td>+</td><td>0</td><td>CGCGCCGTCGCCCAGCCGCG</td><td>TGG</td><td>||||||||[||||||||||||]PAM</td><td>NA</td><td>Intergenic</td><td>NA</td><td>NA</td></tr>
<tr><td></td><td></td><td></td><td></td><td></td><td></td><td></td><td></td><td></td><td></td><td></td><td></td></tr>
<tr><td>T9481</td><td>CCGCACCATGTCGTAGCTGTTGG</td><td>1000</td><td></td><td></td><td></td><td></td><td></td><td></td><td></td><td></td><td></td></tr>
<tr><td>Oligo fwd</td><td>CACCCCGCACCATGTCGTAGCTGT</td><td></td><td></td><td></td><td></td><td></td><td></td><td></td><td></td><td></td><td></td></tr>
<tr><td>Oligo rev</td><td>AAACACAGCTACGACATGGTGCGG</td><td></td><td></td><td></td><td></td><td></td><td></td><td></td><td></td><td></td><td></td></tr>
<tr><td>Chromosome</td><td>start</td><td>end</td><td>strand</td><td>MM</td><td>target_seq</td><td>PAM</td><td>alignment</td><td>distance</td><td>position</td><td>gene name</td><td>gene id</td></tr>
<tr><td>GU305914.1</td><td>125394</td><td>125416</td><td>+</td><td>0</td><td>CCGCACCATGTCGTAGCTGT</td><td>TGG</td><td>||||||||[||||||||||||]PAM</td><td>NA</td><td>Intergenic</td><td>NA</td><td>NA</td></tr>
<tr><td></td><td></td><td></td><td></td><td></td><td></td><td></td><td></td><td></td><td></td><td></td><td></td></tr>
<tr><td>T9482</td><td>AGCTGTTGGCGAACGCCAGCAGG</td><td>1000</td><td></td><td></td><td></td><td></td><td></td><td></td><td></td><td></td><td></td></tr>
<tr><td>Oligo fwd</td><td>CACCAGCTGTTGGCGAACGCCAGC</td><td></td><td></td><td></td><td></td><td></td><td></td><td></td><td></td><td></td><td></td></tr>
<tr><td>Oligo rev</td><td>AAACGCTGGCGTTCGCCAACAGCT</td><td></td><td></td><td></td><td></td><td></td><td></td><td></td><td></td><td></td><td></td></tr>
<tr><td>Chromosome</td><td>start</td><td>end</td><td>strand</td><td>MM</td><td>target_seq</td><td>PAM</td><td>alignment</td><td>distance</td><td>position</td><td>gene name</td><td>gene id</td></tr>
<tr><td>GU305914.1</td><td>125408</td><td>125430</td><td>+</td><td>0</td><td>AGCTGTTGGCGAACGCCAGC</td><td>AGG</td><td>||||||||[||||||||||||]PAM</td><td>NA</td><td>Intergenic</td><td>NA</td><td>NA</td></tr>
<tr><td></td><td></td><td></td><td></td><td></td><td></td><td></td><td></td><td></td><td></td><td></td><td></td></tr>
<tr><td>T9483</td><td>CAGCTCGATGCCCGTCTCGAAGG</td><td>1000</td><td></td><td></td><td></td><td></td><td></td><td></td><td></td><td></td><td></td></tr>
<tr><td>Oligo fwd</td><td>CACCCAGCTCGATGCCCGTCTCGA</td><td></td><td></td><td></td><td></td><td></td><td></td><td></td><td></td><td></td><td></td></tr>
<tr><td>Oligo rev</td><td>AAACTCGAGACGGGCATCGAGCTG</td><td></td><td></td><td></td><td></td><td></td><td></td><td></td><td></td><td></td><td></td></tr>
<tr><td>Chromosome</td><td>start</td><td>end</td><td>strand</td><td>MM</td><td>target_seq</td><td>PAM</td><td>alignment</td><td>distance</td><td>position</td><td>gene name</td><td>gene id</td></tr>
<tr><td>GU305914.1</td><td>125469</td><td>125491</td><td>+</td><td>0</td><td>CAGCTCGATGCCCGTCTCGA</td><td>AGG</td><td>||||||||[||||||||||||]PAM</td><td>NA</td><td>Intergenic</td><td>NA</td><td>NA</td></tr>
<tr><td></td><td></td><td></td><td></td><td></td><td></td><td></td><td></td><td></td><td></td><td></td><td></td></tr>
<tr><td>T9484</td><td>ATGCCCGTCTCGAAGGCCGCCGG</td><td>1000</td><td></td><td></td><td></td><td></td><td></td><td></td><td></td><td></td><td></td></tr>
<tr><td>Oligo fwd</td><td>CACCATGCCCGTCTCGAAGGCCGC</td><td></td><td></td><td></td><td></td><td></td><td></td><td></td><td></td><td></td><td></td></tr>
</table>

Oligo rev AAACGCGGCCTTCGAGACGGGCAT
Chromosome start end strand MM target_seq PAM alignment distance position gene name gene id
GU305914.1 125476 125498 + 0 ATGCCCGTCTCGAAGGCCGC CGG ||||||||[||||||||||||]PAM NA Intergenic NA NA

T9485 TGCCCGTCTCGAAGGCCGCCGGG 1000
Oligo fwd CACCTGCCCGTCTCGAAGGCCGCC
Oligo rev AAACGGCGGCCTTCGAGACGGGCA
Chromosome start end strand MM target_seq PAM alignment distance position gene name gene id
GU305914.1 125477 125499 + 0 TGCCCGTCTCGAAGGCCGCC GGG ||||||||[||||||||||||]PAM NA Intergenic NA NA

T9486 CGAAGGCCGCCGGGTCGCCGTGG 1000
Oligo fwd CACCCGAAGGCCGCCGGGTCGCCG
Oligo rev AAACCGGCGACCCGGCGGCCTTCG
Chromosome start end strand MM target_seq PAM alignment distance position gene name gene id
GU305914.1 125486 125508 + 0 CGAAGGCCGCCGGGTCGCCG TGG ||||||||[||||||||||||]PAM NA Intergenic NA NA

T9487 CGTGGATCAGAGACTCTATCAGG 1000
Oligo fwd CACCCGTGGATCAGAGACTCTATC
Oligo rev AAACGATAGAGTCTCTGATCCACG
Chromosome start end strand MM target_seq PAM alignment distance position gene name gene id
GU305914.1 125504 125526 + 0 CGTGGATCAGAGACTCTATC AGG ||||||||[||||||||||||]PAM NA Intergenic NA NA

T9488 TCTATCAGGTAGAACAGCTCCGG 1000
Oligo fwd CACCTCTATCAGGTAGAACAGCTC
Oligo rev AAACGAGCTGTTCTACCTGATAGA
Chromosome start end strand MM target_seq PAM alignment distance position gene name gene id
GU305914.1 125518 125540 + 0 TCTATCAGGTAGAACAGCTC CGG ||||||||[||||||||||||]PAM NA Intergenic NA NA

T9489 CTATCAGGTAGAACAGCTCCGGG 1000
Oligo fwd CACCCTATCAGGTAGAACAGCTCC
Oligo rev AAACGGAGCTGTTCTACCTGATAG
Chromosome start end strand MM target_seq PAM alignment distance position gene name gene id
GU305914.1 125519 125541 + 0 CTATCAGGTAGAACAGCTCC GGG ||||||||[||||||||||||]PAM NA Intergenic NA NA

T9490 GGTAGAACAGCTCCGGGTACTGG 1000
Oligo fwd CACCGGTAGAACAGCTCCGGGTAC
Oligo rev AAACGTACCCGGAGCTGTTCTACC
Chromosome start end strand MM target_seq PAM alignment distance position gene name gene id
GU305914.1 125525 125547 + 0 GGTAGAACAGCTCCGGGTAC TGG ||||||||[||||||||||||]PAM NA Intergenic NA NA

T9491 AGAACAGCTCCGGGTACTGGTGG 1000
Oligo fwd CACCAGAACAGCTCCGGGTACTGG
Oligo rev AAACCCAGTACCCGGAGCTGTTCT
Chromosome start end strand MM target_seq PAM alignment distance position gene name gene id
GU305914.1 125528 125550 + 0 AGAACAGCTCCGGGTACTGG TGG ||||||||[||||||||||||]PAM NA Intergenic NA NA

T9492 GTACTGGTGGTCCGTCAGCGAGG 1000
Oligo fwd CACCGTACTGGTGGTCCGTCAGCG
Oligo rev AAACCGCTGACGGACCACCAGTAC
Chromosome start end strand MM target_seq PAM alignment distance position gene name gene id
GU305914.1 125541 125563 + 0 GTACTGGTGGTCCGTCAGCG AGG ||||||||[||||||||||||]PAM NA Intergenic NA NA

T9493 AGCGTCGCCTCGTGACCCTCCGG 1000
Oligo fwd CACCAGCGTCGCCTCGTGACCCTC
Oligo rev AAACGAGGGTCACGAGGCGACGCT
Chromosome start end strand MM target_seq PAM alignment distance position gene name gene id
GU305914.1 125578 125600 + 0 AGCGTCGCCTCGTGACCCTC CGG ||||||||[||||||||||||]PAM NA Intergenic NA NA

T9494 ACCCTCCGGCGCGCGCACCTTGG 1000
Oligo fwd CACCACCCTCCGGCGCGCGCACCT
Oligo rev AAACAGGTGCGCGCGCCGGAGGGT
Chromosome start end strand MM target_seq PAM alignment distance position gene name gene id
GU305914.1 125592 125614 + 0 ACCCTCCGGCGCGCGCACCT TGG ||||||||[||||||||||||]PAM NA Intergenic NA NA

T9495 CCAGCGCCTGCTTCGCCCGCAGG 1000
Oligo fwd CACCCCAGCGCCTGCTTCGCCCGC
Oligo rev AAACGCGGGCGAAGCAGGCGCTGG
Chromosome start end strand MM target_seq PAM alignment distance position gene name gene id
GU305914.1 125621 125643 + 0 CCAGCGCCTGCTTCGCCCGC AGG ||||||||[||||||||||||]PAM NA Intergenic NA NA

T9496 CCTGCTTCGCCCGCAGGTCGTGG 1000
Oligo fwd CACCCCTGCTTCGCCCGCAGGTCG
Oligo rev AAACCGACCTGCGGGCGAAGCAGG
Chromosome start end strand MM target_seq PAM alignment distance position gene name gene id
GU305914.1 125627 125649 + 0 CCTGCTTCGCCCGCAGGTCG TGG ||||||||[||||||||||||]PAM NA Intergenic NA NA

T9497 CGCCCGCAGGTCGTGGAAGTCGG 1000
Oligo fwd CACCCGCCCGCAGGTCGTGGAAGT
Oligo rev AAACACTTCCACGACCTGCGGGCG
Chromosome start end strand MM target_seq PAM alignment distance position gene name gene id
GU305914.1 125634 125656 + 0 CGCCCGCAGGTCGTGGAAGT CGG ||||||||[||||||||||||]PAM NA Intergenic NA NA

T9498 CGCAGGTCGTGGAAGTCGGCCGG 1000
Oligo fwd CACCCGCAGGTCGTGGAAGTCGGC
Oligo rev AAACGCCGACTTCCACGACCTGCG
Chromosome start end strand MM target_seq PAM alignment distance position gene name gene id
GU305914.1 125638 125660 + 0 CGCAGGTCGTGGAAGTCGGC CGG ||||||||[||||||||||||]PAM NA Intergenic NA NA

T9499 GTCGGCCGGCGCCAGCGCCTCGG 1000

| | | | | | | | | | | | |
|---|---|---|---|---|---|---|---|---|---|---|---|
| Oligo fwd | CACCGTCGGCCGGCGCCAGCGCCT | | | | | | | | | | |
| Oligo rev | AAACAGGCGCTGGCGCCGGCCGAC | | | | | | | | | | |
| Chromosome | start | end | strand | MM | target_seq | PAM | alignment | distance | position | gene name | gene id |
| GU305914.1 | 125652 | 125674 | + | 0 | GTCGGCCGGCGCCAGCGCCT | CGG | \|\|\|\|\|\|\|\|[\|\|\|\|\|\|\|\|\|\|\|\|]PAM | NA | Intergenic | NA | NA |
| GU305914.1 | 109397 | 109419 | + | 4 | GAAGGCCGGCGGCGGCGCCT | CGG | \|--\|\|\|\|\|[\|\|\|-\|-\|\|\|\|\|\|]PAM | NA | Intergenic | NA | NA |

| | | | | | | | | | | | |
|---|---|---|---|---|---|---|---|---|---|---|---|
| T9500 | TCGGCCGGCGCCAGCGCCTCGGG | 1000 | | | | | | | | | |
| Oligo fwd | CACCTCGGCCGGCGCCAGCGCCTC | | | | | | | | | | |
| Oligo rev | AAACGAGGCGCTGGCGCCGGCCGA | | | | | | | | | | |
| Chromosome | start | end | strand | MM | target_seq | PAM | alignment | distance | position | gene name | gene id |
| GU305914.1 | 125653 | 125675 | + | 0 | TCGGCCGGCGCCAGCGCCTC | GGG | \|\|\|\|\|\|\|\|[\|\|\|\|\|\|\|\|\|\|\|\|]PAM | NA | Intergenic | NA | NA |

| | | | | | | | | | | | |
|---|---|---|---|---|---|---|---|---|---|---|---|
| T9501 | CCGGCGCCAGCGCCTCGGGCAGG | 1000 | | | | | | | | | |
| Oligo fwd | CACCCCGGCGCCAGCGCCTCGGGC | | | | | | | | | | |
| Oligo rev | AAACGCCCGAGGCGCTGGCGCCGG | | | | | | | | | | |
| Chromosome | start | end | strand | MM | target_seq | PAM | alignment | distance | position | gene name | gene id |
| GU305914.1 | 125657 | 125679 | + | 0 | CCGGCGCCAGCGCCTCGGGC | AGG | \|\|\|\|\|\|\|\|[\|\|\|\|\|\|\|\|\|\|\|\|]PAM | NA | Intergenic | NA | NA |

| | | | | | | | | | | | |
|---|---|---|---|---|---|---|---|---|---|---|---|
| T9502 | AGGTTACCCACCATGATGCGCGG | 1000 | | | | | | | | | |
| Oligo fwd | CACCAGGTTACCCACCATGATGCG | | | | | | | | | | |
| Oligo rev | AAACCGCATCATGGTGGGTAACCT | | | | | | | | | | |
| Chromosome | start | end | strand | MM | target_seq | PAM | alignment | distance | position | gene name | gene id |
| GU305914.1 | 125677 | 125699 | + | 0 | AGGTTACCCACCATGATGCG | CGG | \|\|\|\|\|\|\|\|[\|\|\|\|\|\|\|\|\|\|\|\|]PAM | NA | Intergenic | NA | NA |

| | | | | | | | | | | | |
|---|---|---|---|---|---|---|---|---|---|---|---|
| T9503 | ATGATGCGCGGCGTGCACAGCGG | 1000 | | | | | | | | | |
| Oligo fwd | CACCATGATGCGCGGCGTGCACAG | | | | | | | | | | |
| Oligo rev | AAACCTGTGCACGCCGCGCATCAT | | | | | | | | | | |
| Chromosome | start | end | strand | MM | target_seq | PAM | alignment | distance | position | gene name | gene id |
| GU305914.1 | 125689 | 125711 | + | 0 | ATGATGCGCGGCGTGCACAG | CGG | \|\|\|\|\|\|\|\|[\|\|\|\|\|\|\|\|\|\|\|\|]PAM | NA | Intergenic | NA | NA |

| | | | | | | | | | | | |
|---|---|---|---|---|---|---|---|---|---|---|---|
| T9504 | GCACCTGCCCGTTCTCCGTGTGG | 1000 | | | | | | | | | |
| Oligo fwd | CACCGCACCTGCCCGTTCTCCGTG | | | | | | | | | | |
| Oligo rev | AAACCACGGAGAACGGGCAGGTGC | | | | | | | | | | |
| Chromosome | start | end | strand | MM | target_seq | PAM | alignment | distance | position | gene name | gene id |
| GU305914.1 | 125714 | 125736 | + | 0 | GCACCTGCCCGTTCTCCGTG | TGG | \|\|\|\|\|\|\|\|[\|\|\|\|\|\|\|\|\|\|\|\|]PAM | NA | Intergenic | NA | NA |

| | | | | | | | | | | | |
|---|---|---|---|---|---|---|---|---|---|---|---|
| T9505 | GTGGAAGCAGAAGTCGAAGAAGG | 1000 | | | | | | | | | |
| Oligo fwd | CACCGTGGAAGCAGAAGTCGAAGA | | | | | | | | | | |
| Oligo rev | AAACTCTTCGACTTCTGCTTCCAC | | | | | | | | | | |
| Chromosome | start | end | strand | MM | target_seq | PAM | alignment | distance | position | gene name | gene id |
| GU305914.1 | 125733 | 125755 | + | 0 | GTGGAAGCAGAAGTCGAAGA | AGG | \|\|\|\|\|\|\|\|[\|\|\|\|\|\|\|\|\|\|\|\|]PAM | NA | Intergenic | NA | NA |

| | | | | | | | | | | | |
|---|---|---|---|---|---|---|---|---|---|---|---|
| T9506 | TGGAAGCAGAAGTCGAAGAAGGG | 1000 | | | | | | | | | |
| Oligo fwd | CACCTGGAAGCAGAAGTCGAAGAA | | | | | | | | | | |
| Oligo rev | AAACTTCTTCGACTTCTGCTTCCA | | | | | | | | | | |
| Chromosome | start | end | strand | MM | target_seq | PAM | alignment | distance | position | gene name | gene id |
| GU305914.1 | 125734 | 125756 | + | 0 | TGGAAGCAGAAGTCGAAGAA | GGG | \|\|\|\|\|\|\|\|[\|\|\|\|\|\|\|\|\|\|\|\|]PAM | NA | Intergenic | NA | NA |

| | | | | | | | | | | | |
|---|---|---|---|---|---|---|---|---|---|---|---|
| T9507 | GGAAGCAGAAGTCGAAGAAGGGG | 1000 | | | | | | | | | |
| Oligo fwd | CACCGGAAGCAGAAGTCGAAGAAG | | | | | | | | | | |
| Oligo rev | AAACCTTCTTCGACTTCTGCTTCC | | | | | | | | | | |
| Chromosome | start | end | strand | MM | target_seq | PAM | alignment | distance | position | gene name | gene id |
| GU305914.1 | 125735 | 125757 | + | 0 | GGAAGCAGAAGTCGAAGAAG | GGG | \|\|\|\|\|\|\|\|[\|\|\|\|\|\|\|\|\|\|\|\|]PAM | NA | Intergenic | NA | NA |

| | | | | | | | | | | | |
|---|---|---|---|---|---|---|---|---|---|---|---|
| T9508 | GTCGAAGAAGGGGTGCAGCTCGG | 1000 | | | | | | | | | |
| Oligo fwd | CACCGTCGAAGAAGGGGTGCAGCT | | | | | | | | | | |
| Oligo rev | AAACAGCTGCACCCCTTCTTCGAC | | | | | | | | | | |
| Chromosome | start | end | strand | MM | target_seq | PAM | alignment | distance | position | gene name | gene id |
| GU305914.1 | 125745 | 125767 | + | 0 | GTCGAAGAAGGGGTGCAGCT | CGG | \|\|\|\|\|\|\|\|[\|\|\|\|\|\|\|\|\|\|\|\|]PAM | NA | Intergenic | NA | NA |

| | | | | | | | | | | | |
|---|---|---|---|---|---|---|---|---|---|---|---|
| T9509 | GGTGCAGCTCGGTGAAGAGCAGG | 1000 | | | | | | | | | |
| Oligo fwd | CACCGGTGCAGCTCGGTGAAGAGC | | | | | | | | | | |
| Oligo rev | AAACGCTCTTCACCGAGCTGCACC | | | | | | | | | | |
| Chromosome | start | end | strand | MM | target_seq | PAM | alignment | distance | position | gene name | gene id |
| GU305914.1 | 125756 | 125778 | + | 0 | GGTGCAGCTCGGTGAAGAGC | AGG | \|\|\|\|\|\|\|\|[\|\|\|\|\|\|\|\|\|\|\|\|]PAM | NA | Intergenic | NA | NA |

| | | | | | | | | | | | |
|---|---|---|---|---|---|---|---|---|---|---|---|
| T9510 | GAGCAGGTAGTTGTCCACCTTGG | 1000 | | | | | | | | | |
| Oligo fwd | CACCGAGCAGGTAGTTGTCCACCT | | | | | | | | | | |
| Oligo rev | AAACAGGTGGACAACTACCTGCTC | | | | | | | | | | |
| Chromosome | start | end | strand | MM | target_seq | PAM | alignment | distance | position | gene name | gene id |
| GU305914.1 | 125772 | 125794 | + | 0 | GAGCAGGTAGTTGTCCACCT | TGG | \|\|\|\|\|\|\|\|[\|\|\|\|\|\|\|\|\|\|\|\|]PAM | NA | Intergenic | NA | NA |

| | | | | | | | | | | | |
|---|---|---|---|---|---|---|---|---|---|---|---|
| T9511 | GTCCACCTTGGCGAACTCCTTGG | 1000 | | | | | | | | | |
| Oligo fwd | CACCGTCCACCTTGGCGAACTCCT | | | | | | | | | | |
| Oligo rev | AAACAGGAGTTCGCCAAGGTGGAC | | | | | | | | | | |
| Chromosome | start | end | strand | MM | target_seq | PAM | alignment | distance | position | gene name | gene id |
| GU305914.1 | 125784 | 125806 | + | 0 | GTCCACCTTGGCGAACTCCT | TGG | \|\|\|\|\|\|\|\|[\|\|\|\|\|\|\|\|\|\|\|\|]PAM | NA | Intergenic | NA | NA |
| GU305914.1 | 91881 | 91903 | + | 4 | GTCCGCCATGGCGACCTCCC | CGG | \|\|\|\|-\|\|-[\|\|\|\|\|\|-\|\|\|\|-]PAM | NA | Intergenic | NA | NA |

| | | | | | | | | | | | |
|---|---|---|---|---|---|---|---|---|---|---|---|
| T9512 | GAAATCGTGCTTCGCCTCGTTGG | 1000 | | | | | | | | | |
| Oligo fwd | CACCGAAATCGTGCTTCGCCTCGT | | | | | | | | | | |
| Oligo rev | AAACACGAGGCGAAGCACGATTTC | | | | | | | | | | |
| Chromosome | start | end | strand | MM | target_seq | PAM | alignment | distance | position | gene name | gene id |
| GU305914.1 | 125811 | 125833 | + | 0 | GAAATCGTGCTTCGCCTCGT | TGG | \|\|\|\|\|\|\|\|[\|\|\|\|\|\|\|\|\|\|\|\|]PAM | NA | Intergenic | NA | NA |

| | | | | | | | | | | | |
|---|---|---|---|---|---|---|---|---|---|---|---|
| T9513 | TTCGCCTCGTTGGCCGTGCGCGG | 1000 | | | | | | | | | |
| Oligo fwd | CACCTTCGCCTCGTTGGCCGTGCG | | | | | | | | | | |
| Oligo rev | AAACCGCACGGCCAACGAGGCGAA | | | | | | | | | | |
| Chromosome | start | end | strand | MM | target_seq | PAM | alignment | distance | position | gene name | gene id |

<table>
<tr><td>GU305914.1</td><td>125821</td><td>125843</td><td>+</td><td>0</td><td>TTCGCCTCGTTGGCCGTGCG</td><td>CGG</td><td>||||||||[||||||||||||]PAM</td><td>NA</td><td>Intergenic</td><td>NA</td><td>NA</td></tr>
<tr><td>T9514</td><td>GCCTCGTTGGCCGTGCGCGGCGG</td><td>1000</td><td></td><td></td><td></td><td></td><td></td><td></td><td></td><td></td><td></td></tr>
<tr><td>Oligo fwd</td><td>CACCGCCTCGTTGGCCGTGCGCGG</td><td></td><td></td><td></td><td></td><td></td><td></td><td></td><td></td><td></td><td></td></tr>
<tr><td>Oligo rev</td><td>AAACCCGCGCACGGCCAACGAGGC</td><td></td><td></td><td></td><td></td><td></td><td></td><td></td><td></td><td></td><td></td></tr>
<tr><td>Chromosome</td><td>start</td><td>end</td><td>strand</td><td>MM</td><td>target_seq</td><td>PAM</td><td>alignment</td><td>distance</td><td>position</td><td>gene name</td><td>gene id</td></tr>
<tr><td>GU305914.1</td><td>125824</td><td>125846</td><td>+</td><td>0</td><td>GCCTCGTTGGCCGTGCGCGG</td><td>CGG</td><td>||||||||[||||||||||||]PAM</td><td>NA</td><td>Intergenic</td><td>NA</td><td>NA</td></tr>
<tr><td>T9515</td><td>CCTCGTTGGCCGTGCGCGGCGGG</td><td>1000</td><td></td><td></td><td></td><td></td><td></td><td></td><td></td><td></td><td></td></tr>
<tr><td>Oligo fwd</td><td>CACCCCTCGTTGGCCGTGCGCGGC</td><td></td><td></td><td></td><td></td><td></td><td></td><td></td><td></td><td></td><td></td></tr>
<tr><td>Oligo rev</td><td>AAACGCCGCGCACGGCCAACGAGG</td><td></td><td></td><td></td><td></td><td></td><td></td><td></td><td></td><td></td><td></td></tr>
<tr><td>Chromosome</td><td>start</td><td>end</td><td>strand</td><td>MM</td><td>target_seq</td><td>PAM</td><td>alignment</td><td>distance</td><td>position</td><td>gene name</td><td>gene id</td></tr>
<tr><td>GU305914.1</td><td>125825</td><td>125847</td><td>+</td><td>0</td><td>CCTCGTTGGCCGTGCGCGGC</td><td>GGG</td><td>||||||||[||||||||||||]PAM</td><td>NA</td><td>Intergenic</td><td>NA</td><td>NA</td></tr>
<tr><td>T9516</td><td>GCGGCGGGTCGATGCGCGCCTGG</td><td>1000</td><td></td><td></td><td></td><td></td><td></td><td></td><td></td><td></td><td></td></tr>
<tr><td>Oligo fwd</td><td>CACCGCGGCGGGTCGATGCGCGCC</td><td></td><td></td><td></td><td></td><td></td><td></td><td></td><td></td><td></td><td></td></tr>
<tr><td>Oligo rev</td><td>AAACGGCGCGCATCGACCCGCCGC</td><td></td><td></td><td></td><td></td><td></td><td></td><td></td><td></td><td></td><td></td></tr>
<tr><td>Chromosome</td><td>start</td><td>end</td><td>strand</td><td>MM</td><td>target_seq</td><td>PAM</td><td>alignment</td><td>distance</td><td>position</td><td>gene name</td><td>gene id</td></tr>
<tr><td>GU305914.1</td><td>125840</td><td>125862</td><td>+</td><td>0</td><td>GCGGCGGGTCGATGCGCGCC</td><td>TGG</td><td>||||||||[||||||||||||]PAM</td><td>NA</td><td>Intergenic</td><td>NA</td><td>NA</td></tr>
<tr><td>GU305914.1</td><td>27783</td><td>27805</td><td>-</td><td>4</td><td>GCAGCGTCTCGATGCGCGGC</td><td>GGG</td><td>||-|||--[||||||||||-|]PAM</td><td>NA</td><td>Intergenic</td><td>NA</td><td>NA</td></tr>
<tr><td>T9517</td><td>GAAGTGCACGAGCCGCCGCGTGG</td><td>1000</td><td></td><td></td><td></td><td></td><td></td><td></td><td></td><td></td><td></td></tr>
<tr><td>Oligo fwd</td><td>CACCGAAGTGCACGAGCCGCCGCG</td><td></td><td></td><td></td><td></td><td></td><td></td><td></td><td></td><td></td><td></td></tr>
<tr><td>Oligo rev</td><td>AAACCGCGGCGGCTCGTGCACTTC</td><td></td><td></td><td></td><td></td><td></td><td></td><td></td><td></td><td></td><td></td></tr>
<tr><td>Chromosome</td><td>start</td><td>end</td><td>strand</td><td>MM</td><td>target_seq</td><td>PAM</td><td>alignment</td><td>distance</td><td>position</td><td>gene name</td><td>gene id</td></tr>
<tr><td>GU305914.1</td><td>125865</td><td>125887</td><td>+</td><td>0</td><td>GAAGTGCACGAGCCGCCGCG</td><td>TGG</td><td>||||||||[||||||||||||]PAM</td><td>NA</td><td>Intergenic</td><td>NA</td><td>NA</td></tr>
<tr><td>T9518</td><td>GCTCCCTGACGAACTCTCGCCGG</td><td>1000</td><td></td><td></td><td></td><td></td><td></td><td></td><td></td><td></td><td></td></tr>
<tr><td>Oligo fwd</td><td>CACCGCTCCCTGACGAACTCTCGC</td><td></td><td></td><td></td><td></td><td></td><td></td><td></td><td></td><td></td><td></td></tr>
<tr><td>Oligo rev</td><td>AAACGCGAGAGTTCGTCAGGGAGC</td><td></td><td></td><td></td><td></td><td></td><td></td><td></td><td></td><td></td><td></td></tr>
<tr><td>Chromosome</td><td>start</td><td>end</td><td>strand</td><td>MM</td><td>target_seq</td><td>PAM</td><td>alignment</td><td>distance</td><td>position</td><td>gene name</td><td>gene id</td></tr>
<tr><td>GU305914.1</td><td>125900</td><td>125922</td><td>+</td><td>0</td><td>GCTCCCTGACGAACTCTCGC</td><td>CGG</td><td>||||||||[||||||||||||]PAM</td><td>NA</td><td>Intergenic</td><td>NA</td><td>NA</td></tr>
<tr><td>T9519</td><td>CCCTGACCCGATCATCCCCCTGG</td><td>1000</td><td></td><td></td><td></td><td></td><td></td><td></td><td></td><td></td><td></td></tr>
<tr><td>Oligo fwd</td><td>CACCCCCTGACCCGATCATCCCCC</td><td></td><td></td><td></td><td></td><td></td><td></td><td></td><td></td><td></td><td></td></tr>
<tr><td>Oligo rev</td><td>AAACGGGGGATGATCGGGTCAGGG</td><td></td><td></td><td></td><td></td><td></td><td></td><td></td><td></td><td></td><td></td></tr>
<tr><td>Chromosome</td><td>start</td><td>end</td><td>strand</td><td>MM</td><td>target_seq</td><td>PAM</td><td>alignment</td><td>distance</td><td>position</td><td>gene name</td><td>gene id</td></tr>
<tr><td>GU305914.1</td><td>125930</td><td>125952</td><td>+</td><td>0</td><td>CCCTGACCCGATCATCCCCC</td><td>TGG</td><td>||||||||[||||||||||||]PAM</td><td>NA</td><td>Intergenic</td><td>NA</td><td>NA</td></tr>
<tr><td>T9520</td><td>CCTGACCCGATCATCCCCCTGGG</td><td>1000</td><td></td><td></td><td></td><td></td><td></td><td></td><td></td><td></td><td></td></tr>
<tr><td>Oligo fwd</td><td>CACCCCTGACCCGATCATCCCCCT</td><td></td><td></td><td></td><td></td><td></td><td></td><td></td><td></td><td></td><td></td></tr>
<tr><td>Oligo rev</td><td>AAACAGGGGGATGATCGGGTCAGG</td><td></td><td></td><td></td><td></td><td></td><td></td><td></td><td></td><td></td><td></td></tr>
<tr><td>Chromosome</td><td>start</td><td>end</td><td>strand</td><td>MM</td><td>target_seq</td><td>PAM</td><td>alignment</td><td>distance</td><td>position</td><td>gene name</td><td>gene id</td></tr>
<tr><td>GU305914.1</td><td>125931</td><td>125953</td><td>+</td><td>0</td><td>CCTGACCCGATCATCCCCCT</td><td>GGG</td><td>||||||||[||||||||||||]PAM</td><td>NA</td><td>Intergenic</td><td>NA</td><td>NA</td></tr>
<tr><td>T9521</td><td>CTGACCCGATCATCCCCCTGGGG</td><td>1000</td><td></td><td></td><td></td><td></td><td></td><td></td><td></td><td></td><td></td></tr>
<tr><td>Oligo fwd</td><td>CACCCTGACCCGATCATCCCCCTG</td><td></td><td></td><td></td><td></td><td></td><td></td><td></td><td></td><td></td><td></td></tr>
<tr><td>Oligo rev</td><td>AAACCAGGGGGATGATCGGGTCAG</td><td></td><td></td><td></td><td></td><td></td><td></td><td></td><td></td><td></td><td></td></tr>
<tr><td>Chromosome</td><td>start</td><td>end</td><td>strand</td><td>MM</td><td>target_seq</td><td>PAM</td><td>alignment</td><td>distance</td><td>position</td><td>gene name</td><td>gene id</td></tr>
<tr><td>GU305914.1</td><td>125932</td><td>125954</td><td>+</td><td>0</td><td>CTGACCCGATCATCCCCCTG</td><td>GGG</td><td>||||||||[||||||||||||]PAM</td><td>NA</td><td>Intergenic</td><td>NA</td><td>NA</td></tr>
<tr><td>T9522</td><td>ACCCGATCATCCCCCTGGGGCGG</td><td>1000</td><td></td><td></td><td></td><td></td><td></td><td></td><td></td><td></td><td></td></tr>
<tr><td>Oligo fwd</td><td>CACCACCCGATCATCCCCCTGGGG</td><td></td><td></td><td></td><td></td><td></td><td></td><td></td><td></td><td></td><td></td></tr>
<tr><td>Oligo rev</td><td>AAACCCCCAGGGGGATGATCGGGT</td><td></td><td></td><td></td><td></td><td></td><td></td><td></td><td></td><td></td><td></td></tr>
<tr><td>Chromosome</td><td>start</td><td>end</td><td>strand</td><td>MM</td><td>target_seq</td><td>PAM</td><td>alignment</td><td>distance</td><td>position</td><td>gene name</td><td>gene id</td></tr>
<tr><td>GU305914.1</td><td>125935</td><td>125957</td><td>+</td><td>0</td><td>ACCCGATCATCCCCCTGGGG</td><td>CGG</td><td>||||||||[||||||||||||]PAM</td><td>NA</td><td>Intergenic</td><td>NA</td><td>NA</td></tr>
<tr><td>T9523</td><td>CCCGATCATCCCCCTGGGGCGGG</td><td>1000</td><td></td><td></td><td></td><td></td><td></td><td></td><td></td><td></td><td></td></tr>
<tr><td>Oligo fwd</td><td>CACCCCCGATCATCCCCCTGGGGC</td><td></td><td></td><td></td><td></td><td></td><td></td><td></td><td></td><td></td><td></td></tr>
<tr><td>Oligo rev</td><td>AAACGCCCCAGGGGGATGATCGGG</td><td></td><td></td><td></td><td></td><td></td><td></td><td></td><td></td><td></td><td></td></tr>
<tr><td>Chromosome</td><td>start</td><td>end</td><td>strand</td><td>MM</td><td>target_seq</td><td>PAM</td><td>alignment</td><td>distance</td><td>position</td><td>gene name</td><td>gene id</td></tr>
<tr><td>GU305914.1</td><td>125936</td><td>125958</td><td>+</td><td>0</td><td>CCCGATCATCCCCCTGGGGC</td><td>GGG</td><td>||||||||[||||||||||||]PAM</td><td>NA</td><td>Intergenic</td><td>NA</td><td>NA</td></tr>
<tr><td>T9524</td><td>CTGCGCGAACGCCTCCAGACAGG</td><td>1000</td><td></td><td></td><td></td><td></td><td></td><td></td><td></td><td></td><td></td></tr>
<tr><td>Oligo fwd</td><td>CACCCTGCGCGAACGCCTCCAGAC</td><td></td><td></td><td></td><td></td><td></td><td></td><td></td><td></td><td></td><td></td></tr>
<tr><td>Oligo rev</td><td>AAACGTCTGGAGGCGTTCGCGCAG</td><td></td><td></td><td></td><td></td><td></td><td></td><td></td><td></td><td></td><td></td></tr>
<tr><td>Chromosome</td><td>start</td><td>end</td><td>strand</td><td>MM</td><td>target_seq</td><td>PAM</td><td>alignment</td><td>distance</td><td>position</td><td>gene name</td><td>gene id</td></tr>
<tr><td>GU305914.1</td><td>125964</td><td>125986</td><td>+</td><td>0</td><td>CTGCGCGAACGCCTCCAGAC</td><td>AGG</td><td>||||||||[||||||||||||]PAM</td><td>NA</td><td>Intergenic</td><td>NA</td><td>NA</td></tr>
<tr><td>T9525</td><td>TGCGCGAACGCCTCCAGACAGGG</td><td>1000</td><td></td><td></td><td></td><td></td><td></td><td></td><td></td><td></td><td></td></tr>
<tr><td>Oligo fwd</td><td>CACCTGCGCGAACGCCTCCAGACA</td><td></td><td></td><td></td><td></td><td></td><td></td><td></td><td></td><td></td><td></td></tr>
<tr><td>Oligo rev</td><td>AAACTGTCTGGAGGCGTTCGCGCA</td><td></td><td></td><td></td><td></td><td></td><td></td><td></td><td></td><td></td><td></td></tr>
<tr><td>Chromosome</td><td>start</td><td>end</td><td>strand</td><td>MM</td><td>target_seq</td><td>PAM</td><td>alignment</td><td>distance</td><td>position</td><td>gene name</td><td>gene id</td></tr>
<tr><td>GU305914.1</td><td>125965</td><td>125987</td><td>+</td><td>0</td><td>TGCGCGAACGCCTCCAGACA</td><td>GGG</td><td>||||||||[||||||||||||]PAM</td><td>NA</td><td>Intergenic</td><td>NA</td><td>NA</td></tr>
<tr><td>T9526</td><td>GAACGCCTCCAGACAGGGCCCGG</td><td>1000</td><td></td><td></td><td></td><td></td><td></td><td></td><td></td><td></td><td></td></tr>
<tr><td>Oligo fwd</td><td>CACCGAACGCCTCCAGACAGGGCC</td><td></td><td></td><td></td><td></td><td></td><td></td><td></td><td></td><td></td><td></td></tr>
<tr><td>Oligo rev</td><td>AAACGGCCCTGTCTGGAGGCGTTC</td><td></td><td></td><td></td><td></td><td></td><td></td><td></td><td></td><td></td><td></td></tr>
<tr><td>Chromosome</td><td>start</td><td>end</td><td>strand</td><td>MM</td><td>target_seq</td><td>PAM</td><td>alignment</td><td>distance</td><td>position</td><td>gene name</td><td>gene id</td></tr>
<tr><td>GU305914.1</td><td>125970</td><td>125992</td><td>+</td><td>0</td><td>GAACGCCTCCAGACAGGGCC</td><td>CGG</td><td>||||||||[||||||||||||]PAM</td><td>NA</td><td>Intergenic</td><td>NA</td><td>NA</td></tr>
<tr><td>T9527</td><td>GGGCCCGGCGTCGTGTATCGCGG</td><td>1000</td><td></td><td></td><td></td><td></td><td></td><td></td><td></td><td></td><td></td></tr>
<tr><td>Oligo fwd</td><td>CACCGGGCCCGGCGTCGTGTATCG</td><td></td><td></td><td></td><td></td><td></td><td></td><td></td><td></td><td></td><td></td></tr>
<tr><td>Oligo rev</td><td>AAACCGATACACGACGCCGGGCCC</td><td></td><td></td><td></td><td></td><td></td><td></td><td></td><td></td><td></td><td></td></tr>
<tr><td>Chromosome</td><td>start</td><td>end</td><td>strand</td><td>MM</td><td>target_seq</td><td>PAM</td><td>alignment</td><td>distance</td><td>position</td><td>gene name</td><td>gene id</td></tr>
<tr><td>GU305914.1</td><td>125985</td><td>126007</td><td>+</td><td>0</td><td>GGGCCCGGCGTCGTGTATCG</td><td>CGG</td><td>||||||||[||||||||||||]PAM</td><td>NA</td><td>Intergenic</td><td>NA</td><td>NA</td></tr>
<tr><td>T9528</td><td>GGCCCGGCGTCGTGTATCGCGGG</td><td>1000</td><td></td><td></td><td></td><td></td><td></td><td></td><td></td><td></td><td></td></tr>
<tr><td>Oligo fwd</td><td>CACCGGCCCGGCGTCGTGTATCGC</td><td></td><td></td><td></td><td></td><td></td><td></td><td></td><td></td><td></td><td></td></tr>
</table>

Oligo rev AAACGCGATACACGACGCCGGGCC
Chromosome start end strand MM target_seq PAM alignment distance position gene name gene id
GU305914.1 125986 126008 + 0 GGCCCGGCGTCGTGTATCGC GGG ||||||||[||||||||||||]PAM NA Intergenic NA NA

T9529 GCCCGGCGTCGTGTATCGCGGGG 1000
Oligo fwd CACCGCCCGGCGTCGTGTATCGCG
Oligo rev AAACCGCGATACACGACGCCGGGC
Chromosome start end strand MM target_seq PAM alignment distance position gene name gene id
GU305914.1 125987 126009 + 0 GCCCGGCGTCGTGTATCGCG GGG ||||||||[||||||||||||]PAM NA Intergenic NA NA

T9530 CGGCGTCGTGTATCGCGGGGTGG 1000
Oligo fwd CACCCGGCGTCGTGTATCGCGGGG
Oligo rev AAACCCCCGCGATACACGACGCCG
Chromosome start end strand MM target_seq PAM alignment distance position gene name gene id
GU305914.1 125990 126012 + 0 CGGCGTCGTGTATCGCGGGG TGG ||||||||[||||||||||||]PAM NA Intergenic NA NA

T9531 GCAGAGAGTCTTGAGCGCCTCGG 1000
Oligo fwd CACCGCAGAGAGTCTTGAGCGCCT
Oligo rev AAACAGGCGCTCAAGACTCTCTGC
Chromosome start end strand MM target_seq PAM alignment distance position gene name gene id
GU305914.1 126012 126034 + 0 GCAGAGAGTCTTGAGCGCCT CGG ||||||||[||||||||||||]PAM NA Intergenic NA NA

T9532 CCTCGGCGTGCGAGATGCGCTGG 1000
Oligo fwd CACCCCTCGGCGTGCGAGATGCGC
Oligo rev AAACGCGCATCTCGCACGCCGAGG
Chromosome start end strand MM target_seq PAM alignment distance position gene name gene id
GU305914.1 126029 126051 + 0 CCTCGGCGTGCGAGATGCGC TGG ||||||||[||||||||||||]PAM NA Intergenic NA NA

T9533 CTCGGCGTGCGAGATGCGCTGGG 1000
Oligo fwd CACCCTCGGCGTGCGAGATGCGCT
Oligo rev AAACAGCGCATCTCGCACGCCGAG
Chromosome start end strand MM target_seq PAM alignment distance position gene name gene id
GU305914.1 126030 126052 + 0 CTCGGCGTGCGAGATGCGCT GGG ||||||||[||||||||||||]PAM NA Intergenic NA NA

T9534 TCGGCGTGCGAGATGCGCTGGGG 1000
Oligo fwd CACCTCGGCGTGCGAGATGCGCTG
Oligo rev AAACCAGCGCATCTCGCACGCCGA
Chromosome start end strand MM target_seq PAM alignment distance position gene name gene id
GU305914.1 126031 126053 + 0 TCGGCGTGCGAGATGCGCTG GGG ||||||||[||||||||||||]PAM NA Intergenic NA NA

T9535 CGGCGTGCGAGATGCGCTGGGGG 1000
Oligo fwd CACCCGGCGTGCGAGATGCGCTGG
Oligo rev AAACCCAGCGCATCTCGCACGCCG
Chromosome start end strand MM target_seq PAM alignment distance position gene name gene id
GU305914.1 126032 126054 + 0 CGGCGTGCGAGATGCGCTGG GGG ||||||||[||||||||||||]PAM NA Intergenic NA NA

T9536 GTCCTTGTTGTAGAAGTACACGG 1000
Oligo fwd CACCGTCCTTGTTGTAGAAGTACA
Oligo rev AAACTGTACTTCTACAACAAGGAC
Chromosome start end strand MM target_seq PAM alignment distance position gene name gene id
GU305914.1 126060 126082 + 0 GTCCTTGTTGTAGAAGTACA CGG ||||||||[||||||||||||]PAM NA Intergenic NA NA

T9537 GTTGTAGAAGTACACGGTCGTGG 1000
Oligo fwd CACCGTTGTAGAAGTACACGGTCG
Oligo rev AAACCGACCGTGTACTTCTACAAC
Chromosome start end strand MM target_seq PAM alignment distance position gene name gene id
GU305914.1 126066 126088 + 0 GTTGTAGAAGTACACGGTCG TGG ||||||||[||||||||||||]PAM NA Intergenic NA NA

T9538 TTGTAGAAGTACACGGTCGTGGG 1000
Oligo fwd CACCTTGTAGAAGTACACGGTCGT
Oligo rev AAACACGACCGTGTACTTCTACAA
Chromosome start end strand MM target_seq PAM alignment distance position gene name gene id
GU305914.1 126067 126089 + 0 TTGTAGAAGTACACGGTCGT GGG ||||||||[||||||||||||]PAM NA Intergenic NA NA

T9539 AGTACACGGTCGTGGGCAGCTGG 1000
Oligo fwd CACCAGTACACGGTCGTGGGCAGC
Oligo rev AAACGCTGCCCACGACCGTGTACT
Chromosome start end strand MM target_seq PAM alignment distance position gene name gene id
GU305914.1 126074 126096 + 0 AGTACACGGTCGTGGGCAGC TGG ||||||||[||||||||||||]PAM NA Intergenic NA NA

T9540 GGTCGTGGGCAGCTGGTTCTCGG 1000
Oligo fwd CACCGGTCGTGGGCAGCTGGTTCT
Oligo rev AAACAGAACCAGCTGCCCACGACC
Chromosome start end strand MM target_seq PAM alignment distance position gene name gene id
GU305914.1 126081 126103 + 0 GGTCGTGGGCAGCTGGTTCT CGG ||||||||[||||||||||||]PAM NA Intergenic NA NA

T9541 CTTGTTCTCCGCCGTGCTGTAGG 1000
Oligo fwd CACCCTTGTTCTCCGCCGTGCTGT
Oligo rev AAACACAGCACGGCGGAGAACAAG
Chromosome start end strand MM target_seq PAM alignment distance position gene name gene id
GU305914.1 126123 126145 + 0 CTTGTTCTCCGCCGTGCTGT AGG ||||||||[||||||||||||]PAM NA Intergenic NA NA

T9542 TCTCCGCCGTGCTGTAGGCCCGG 1000
Oligo fwd CACCTCTCCGCCGTGCTGTAGGCC
Oligo rev AAACGGCCTACAGCACGGCGGAGA
Chromosome start end strand MM target_seq PAM alignment distance position gene name gene id
GU305914.1 126128 126150 + 0 TCTCCGCCGTGCTGTAGGCC CGG ||||||||[||||||||||||]PAM NA Intergenic NA NA
GU305914.1 70849 70871 - 4 TCTTGGCCGTGATGTAGGGC AGG |||--|||[|||-||||||-|]PAM NA Intergenic NA NA

| | | | | | | | | | | | |
|---|---|---|---|---|---|---|---|---|---|---|---|
| T9543 | GTGCTGTAGGCCCGGCCGCTCGG | 1000 | | | | | | | | | |
| Oligo fwd | CACCGTGCTGTAGGCCCGGCCGCT | | | | | | | | | | |
| Oligo rev | AAACAGCGGCCGGGCCTACAGCAC | | | | | | | | | | |
| Chromosome | start | end | strand | MM | target_seq | PAM | alignment | distance | position | gene name | gene id |
| GU305914.1 | 126136 | 126158 | + | 0 | GTGCTGTAGGCCCGGCCGCT | CGG | \|\|\|\|\|\|\|\|[\|\|\|\|\|\|\|\|\|\|\|\|]PAM | NA | Intergenic | NA | NA |
| T9544 | CCCGGCCGCTCGGCACGTGCAGG | 1000 | | | | | | | | | |
| Oligo fwd | CACCCCCGGCCGCTCGGCACGTGC | | | | | | | | | | |
| Oligo rev | AAACGCACGTGCCGAGCGGCCGGG | | | | | | | | | | |
| Chromosome | start | end | strand | MM | target_seq | PAM | alignment | distance | position | gene name | gene id |
| GU305914.1 | 126146 | 126168 | + | 0 | CCCGGCCGCTCGGCACGTGC | AGG | \|\|\|\|\|\|\|\|[\|\|\|\|\|\|\|\|\|\|\|\|]PAM | NA | Intergenic | NA | NA |
| T9545 | CTCGGCACGTGCAGGCCGACCGG | 1000 | | | | | | | | | |
| Oligo fwd | CACCCTCGGCACGTGCAGGCCGAC | | | | | | | | | | |
| Oligo rev | AAACGTCGGCCTGCACGTGCCGAG | | | | | | | | | | |
| Chromosome | start | end | strand | MM | target_seq | PAM | alignment | distance | position | gene name | gene id |
| GU305914.1 | 126154 | 126176 | + | 0 | CTCGGCACGTGCAGGCCGAC | CGG | \|\|\|\|\|\|\|\|[\|\|\|\|\|\|\|\|\|\|\|\|]PAM | NA | Intergenic | NA | NA |
| T9546 | CAGGCCGACCGGAAAGTAGAAGG | 1000 | | | | | | | | | |
| Oligo fwd | CACCCAGGCCGACCGGAAAGTAGA | | | | | | | | | | |
| Oligo rev | AAACTCTACTTTCCGGTCGGCCTG | | | | | | | | | | |
| Chromosome | start | end | strand | MM | target_seq | PAM | alignment | distance | position | gene name | gene id |
| GU305914.1 | 126165 | 126187 | + | 0 | CAGGCCGACCGGAAAGTAGA | AGG | \|\|\|\|\|\|\|\|[\|\|\|\|\|\|\|\|\|\|\|\|]PAM | NA | Intergenic | NA | NA |
| T9547 | AGCTCGACGTCCCGCTCCAGCGG | 1000 | | | | | | | | | |
| Oligo fwd | CACCAGCTCGACGTCCCGCTCCAG | | | | | | | | | | |
| Oligo rev | AAACCTGGAGCGGGACGTCGAGCT | | | | | | | | | | |
| Chromosome | start | end | strand | MM | target_seq | PAM | alignment | distance | position | gene name | gene id |
| GU305914.1 | 126190 | 126212 | + | 0 | AGCTCGACGTCCCGCTCCAG | CGG | \|\|\|\|\|\|\|\|[\|\|\|\|\|\|\|\|\|\|\|\|]PAM | NA | Intergenic | NA | NA |
| T9548 | GCTCGACGTCCCGCTCCAGCGGG | 1000 | | | | | | | | | |
| Oligo fwd | CACCGCTCGACGTCCCGCTCCAGC | | | | | | | | | | |
| Oligo rev | AAACGCTGGAGCGGGACGTCGAGC | | | | | | | | | | |
| Chromosome | start | end | strand | MM | target_seq | PAM | alignment | distance | position | gene name | gene id |
| GU305914.1 | 126191 | 126213 | + | 0 | GCTCGACGTCCCGCTCCAGC | GGG | \|\|\|\|\|\|\|\|[\|\|\|\|\|\|\|\|\|\|\|\|]PAM | NA | Intergenic | NA | NA |
| T9549 | TGTTCTTGTACACCTTTCGCAGG | 1000 | | | | | | | | | |
| Oligo fwd | CACCTGTTCTTGTACACCTTTCGC | | | | | | | | | | |
| Oligo rev | AAACGCGAAAGGTGTACAAGAACA | | | | | | | | | | |
| Chromosome | start | end | strand | MM | target_seq | PAM | alignment | distance | position | gene name | gene id |
| GU305914.1 | 126227 | 126249 | + | 0 | TGTTCTTGTACACCTTTCGC | AGG | \|\|\|\|\|\|\|\|[\|\|\|\|\|\|\|\|\|\|\|\|]PAM | NA | Intergenic | NA | NA |
| T9550 | CAGCACGTCCATCTGCGTCTCGG | 1000 | | | | | | | | | |
| Oligo fwd | CACCCAGCACGTCCATCTGCGTCT | | | | | | | | | | |
| Oligo rev | AAACAGACGCAGATGGACGTGCTG | | | | | | | | | | |
| Chromosome | start | end | strand | MM | target_seq | PAM | alignment | distance | position | gene name | gene id |
| GU305914.1 | 126282 | 126304 | + | 0 | CAGCACGTCCATCTGCGTCT | CGG | \|\|\|\|\|\|\|\|[\|\|\|\|\|\|\|\|\|\|\|\|]PAM | NA | Intergenic | NA | NA |
| T9551 | GTCCATCTGCGTCTCGGTGAAGG | 1000 | | | | | | | | | |
| Oligo fwd | CACCGTCCATCTGCGTCTCGGTGA | | | | | | | | | | |
| Oligo rev | AAACTCACCGAGACGCAGATGGAC | | | | | | | | | | |
| Chromosome | start | end | strand | MM | target_seq | PAM | alignment | distance | position | gene name | gene id |
| GU305914.1 | 126288 | 126310 | + | 0 | GTCCATCTGCGTCTCGGTGA | AGG | \|\|\|\|\|\|\|\|[\|\|\|\|\|\|\|\|\|\|\|\|]PAM | NA | Intergenic | NA | NA |
| GU305914.1 | 134488 | 134510 | - | 4 | GTCGACGTGCGGCTCGGTGA | CGG | \|\|\|-\|--\|[\|\|\|-\|\|\|\|\|\|\|\|]PAM | NA | Intergenic | NA | NA |
| T9552 | TCTCGGTGAAGGTCAGCCGCTGG | 1000 | | | | | | | | | |
| Oligo fwd | CACCTCTCGGTGAAGGTCAGCCGC | | | | | | | | | | |
| Oligo rev | AAACGCGGCTGACCTTCACCGAGA | | | | | | | | | | |
| Chromosome | start | end | strand | MM | target_seq | PAM | alignment | distance | position | gene name | gene id |
| GU305914.1 | 126299 | 126321 | + | 0 | TCTCGGTGAAGGTCAGCCGC | TGG | \|\|\|\|\|\|\|\|[\|\|\|\|\|\|\|\|\|\|\|\|]PAM | NA | Intergenic | NA | NA |
| T9553 | TGAAGGTCAGCCGCTGGTCCAGG | 1000 | | | | | | | | | |
| Oligo fwd | CACCTGAAGGTCAGCCGCTGGTCC | | | | | | | | | | |
| Oligo rev | AAACGGACCAGCGGCTGACCTTCA | | | | | | | | | | |
| Chromosome | start | end | strand | MM | target_seq | PAM | alignment | distance | position | gene name | gene id |
| GU305914.1 | 126305 | 126327 | + | 0 | TGAAGGTCAGCCGCTGGTCC | AGG | \|\|\|\|\|\|\|\|[\|\|\|\|\|\|\|\|\|\|\|\|]PAM | NA | Intergenic | NA | NA |
| T9554 | CTGGTCCAGGAACTGCGACTCGG | 1000 | | | | | | | | | |
| Oligo fwd | CACCCTGGTCCAGGAACTGCGACT | | | | | | | | | | |
| Oligo rev | AAACAGTCGCAGTTCCTGGACCAG | | | | | | | | | | |
| Chromosome | start | end | strand | MM | target_seq | PAM | alignment | distance | position | gene name | gene id |
| GU305914.1 | 126318 | 126340 | + | 0 | CTGGTCCAGGAACTGCGACT | CGG | \|\|\|\|\|\|\|\|[\|\|\|\|\|\|\|\|\|\|\|\|]PAM | NA | Intergenic | NA | NA |
| T9555 | TGGTCCAGGAACTGCGACTCGGG | 1000 | | | | | | | | | |
| Oligo fwd | CACCTGGTCCAGGAACTGCGACTC | | | | | | | | | | |
| Oligo rev | AAACGAGTCGCAGTTCCTGGACCA | | | | | | | | | | |
| Chromosome | start | end | strand | MM | target_seq | PAM | alignment | distance | position | gene name | gene id |
| GU305914.1 | 126319 | 126341 | + | 0 | TGGTCCAGGAACTGCGACTC | GGG | \|\|\|\|\|\|\|\|[\|\|\|\|\|\|\|\|\|\|\|\|]PAM | NA | Intergenic | NA | NA |
| T9556 | AGTCCTTCTGCGAGCTCTGCTGG | 1000 | | | | | | | | | |
| Oligo fwd | CACCAGTCCTTCTGCGAGCTCTGC | | | | | | | | | | |
| Oligo rev | AAACGCAGAGCTCGCAGAAGGACT | | | | | | | | | | |
| Chromosome | start | end | strand | MM | target_seq | PAM | alignment | distance | position | gene name | gene id |
| GU305914.1 | 126344 | 126366 | + | 0 | AGTCCTTCTGCGAGCTCTGC | TGG | \|\|\|\|\|\|\|\|[\|\|\|\|\|\|\|\|\|\|\|\|]PAM | NA | Intergenic | NA | NA |
| T9557 | TCTGCGAGCTCTGCTGGATGCGG | 1000 | | | | | | | | | |
| Oligo fwd | CACCTCTGCGAGCTCTGCTGGATG | | | | | | | | | | |
| Oligo rev | AAACCATCCAGCAGAGCTCGCAGA | | | | | | | | | | |
| Chromosome | start | end | strand | MM | target_seq | PAM | alignment | distance | position | gene name | gene id |

| | | | | | | | | | | | |
|---|---|---|---|---|---|---|---|---|---|---|---|
| GU305914.1 | 126350 | 126372 | + | 0 | TCTGCGAGCTCTGCTGGATG | CGG | \|\|\|\|\|\|\|\|[\|\|\|\|\|\|\|\|\|\|\|\|]PAM | NA | Intergenic | NA | NA |
| T9558 | CGAGCTCTGCTGGATGCGGTCGG | 1000 | | | | | | | | | |
| Oligo fwd | CACCCGAGCTCTGCTGGATGCGGT | | | | | | | | | | |
| Oligo rev | AAACACCGCATCCAGCAGAGCTCG | | | | | | | | | | |
| Chromosome | start | end | strand | MM | target_seq | PAM | alignment | distance | position | gene name | gene id |
| GU305914.1 | 126354 | 126376 | + | 0 | CGAGCTCTGCTGGATGCGGT | CGG | \|\|\|\|\|\|\|\|[\|\|\|\|\|\|\|\|\|\|\|\|]PAM | NA | Intergenic | NA | NA |
| T9559 | AGAAGTCCGCCATGATCGCGTGG | 1000 | | | | | | | | | |
| Oligo fwd | CACCAGAAGTCCGCCATGATCGCG | | | | | | | | | | |
| Oligo rev | AAACCGCGATCATGGCGGACTTCT | | | | | | | | | | |
| Chromosome | start | end | strand | MM | target_seq | PAM | alignment | distance | position | gene name | gene id |
| GU305914.1 | 126386 | 126408 | + | 0 | AGAAGTCCGCCATGATCGCG | TGG | \|\|\|\|\|\|\|\|[\|\|\|\|\|\|\|\|\|\|\|\|]PAM | NA | Intergenic | NA | NA |
| T9560 | GTGGTGCGCGATCGCCGTCACGG | 1000 | | | | | | | | | |
| Oligo fwd | CACCGTGGTGCGCGATCGCCGTCA | | | | | | | | | | |
| Oligo rev | AAACTGACGGCGATCGCGCACCAC | | | | | | | | | | |
| Chromosome | start | end | strand | MM | target_seq | PAM | alignment | distance | position | gene name | gene id |
| GU305914.1 | 126405 | 126427 | + | 0 | GTGGTGCGCGATCGCCGTCA | CGG | \|\|\|\|\|\|\|\|[\|\|\|\|\|\|\|\|\|\|\|\|]PAM | NA | Intergenic | NA | NA |
| T9561 | CAGCACGAACTTGCCGTACGAGG | 1000 | | | | | | | | | |
| Oligo fwd | CACCCAGCACGAACTTGCCGTACG | | | | | | | | | | |
| Oligo rev | AAACCGTACGGCAAGTTCGTGCTG | | | | | | | | | | |
| Chromosome | start | end | strand | MM | target_seq | PAM | alignment | distance | position | gene name | gene id |
| GU305914.1 | 126441 | 126463 | + | 0 | CAGCACGAACTTGCCGTACG | AGG | \|\|\|\|\|\|\|\|[\|\|\|\|\|\|\|\|\|\|\|\|]PAM | NA | Intergenic | NA | NA |
| T9562 | CACGAACTTGCCGTACGAGGCGG | 1000 | | | | | | | | | |
| Oligo fwd | CACCCACGAACTTGCCGTACGAGG | | | | | | | | | | |
| Oligo rev | AAACCCTCGTACGGCAAGTTCGTG | | | | | | | | | | |
| Chromosome | start | end | strand | MM | target_seq | PAM | alignment | distance | position | gene name | gene id |
| GU305914.1 | 126444 | 126466 | + | 0 | CACGAACTTGCCGTACGAGG | CGG | \|\|\|\|\|\|\|\|[\|\|\|\|\|\|\|\|\|\|\|\|]PAM | NA | Intergenic | NA | NA |
| T9563 | ACGAACTTGCCGTACGAGGCGGG | 1000 | | | | | | | | | |
| Oligo fwd | CACCACGAACTTGCCGTACGAGGC | | | | | | | | | | |
| Oligo rev | AAACGCCTCGTACGGCAAGTTCGT | | | | | | | | | | |
| Chromosome | start | end | strand | MM | target_seq | PAM | alignment | distance | position | gene name | gene id |
| GU305914.1 | 126445 | 126467 | + | 0 | ACGAACTTGCCGTACGAGGC | GGG | \|\|\|\|\|\|\|\|[\|\|\|\|\|\|\|\|\|\|\|\|]PAM | NA | Intergenic | NA | NA |
| T9564 | CGGGCGTCTTCACGCCCGTGCGG | 1000 | | | | | | | | | |
| Oligo fwd | CACCCGGGCGTCTTCACGCCCGTG | | | | | | | | | | |
| Oligo rev | AAACCACGGGCGTGAAGACGCCCG | | | | | | | | | | |
| Chromosome | start | end | strand | MM | target_seq | PAM | alignment | distance | position | gene name | gene id |
| GU305914.1 | 126464 | 126486 | + | 0 | CGGGCGTCTTCACGCCCGTG | CGG | \|\|\|\|\|\|\|\|[\|\|\|\|\|\|\|\|\|\|\|\|]PAM | NA | Intergenic | NA | NA |
| T9565 | GCGTCTTCACGCCCGTGCGGCGG | 1000 | | | | | | | | | |
| Oligo fwd | CACCGCGTCTTCACGCCCGTGCGG | | | | | | | | | | |
| Oligo rev | AAACCCGCACGGGCGTGAAGACGC | | | | | | | | | | |
| Chromosome | start | end | strand | MM | target_seq | PAM | alignment | distance | position | gene name | gene id |
| GU305914.1 | 126467 | 126489 | + | 0 | GCGTCTTCACGCCCGTGCGG | CGG | \|\|\|\|\|\|\|\|[\|\|\|\|\|\|\|\|\|\|\|\|]PAM | NA | Intergenic | NA | NA |
| GU305914.1 | 38242 | 38264 | + | 4 | GCATCATGACCCCCGTGCGG | AGG | \|\|-\|\|-\|-[\|\|-\|\|\|\|\|\|\|\|\|]PAM | NA | Intergenic | NA | NA |
| T9566 | GTTCAGCAGCGCCTGCACCACGG | 1000 | | | | | | | | | |
| Oligo fwd | CACCGTTCAGCAGCGCCTGCACCA | | | | | | | | | | |
| Oligo rev | AAACTGGTGCAGGCGCTGCTGAAC | | | | | | | | | | |
| Chromosome | start | end | strand | MM | target_seq | PAM | alignment | distance | position | gene name | gene id |
| GU305914.1 | 126507 | 126529 | + | 0 | GTTCAGCAGCGCCTGCACCA | CGG | \|\|\|\|\|\|\|\|[\|\|\|\|\|\|\|\|\|\|\|\|]PAM | NA | Intergenic | NA | NA |
| GU305914.1 | 125187 | 125209 | + | 4 | GCGCGGCAGCGCCTGCACGA | AGG | \|--\|-\|\|\|[\|\|\|\|\|\|\|\|\|\|-\|]PAM | NA | Intergenic | NA | NA |
| T9567 | TGCTCACGATCACGCCCGCCAGG | 1000 | | | | | | | | | |
| Oligo fwd | CACCTGCTCACGATCACGCCCGCC | | | | | | | | | | |
| Oligo rev | AAACGGCGGGCGTGATCGTGAGCA | | | | | | | | | | |
| Chromosome | start | end | strand | MM | target_seq | PAM | alignment | distance | position | gene name | gene id |
| GU305914.1 | 126536 | 126558 | + | 0 | TGCTCACGATCACGCCCGCC | AGG | \|\|\|\|\|\|\|\|[\|\|\|\|\|\|\|\|\|\|\|\|]PAM | NA | Intergenic | NA | NA |
| T9568 | CTCCCCGCCGCCCACCGTGTAGG | 1000 | | | | | | | | | |
| Oligo fwd | CACCCTCCCCGCCGCCCACCGTGT | | | | | | | | | | |
| Oligo rev | AAACACACGGTGGGCGGCGGGGAG | | | | | | | | | | |
| Chromosome | start | end | strand | MM | target_seq | PAM | alignment | distance | position | gene name | gene id |
| GU305914.1 | 126561 | 126583 | + | 0 | CTCCCCGCCGCCCACCGTGT | AGG | \|\|\|\|\|\|\|\|[\|\|\|\|\|\|\|\|\|\|\|\|]PAM | NA | Intergenic | NA | NA |
| T9569 | CCGCCCACCGTGTAGGTCTCCGG | 1000 | | | | | | | | | |
| Oligo fwd | CACCCCGCCCACCGTGTAGGTCTC | | | | | | | | | | |
| Oligo rev | AAACGAGACCTACACGGTGGGCGG | | | | | | | | | | |
| Chromosome | start | end | strand | MM | target_seq | PAM | alignment | distance | position | gene name | gene id |
| GU305914.1 | 126568 | 126590 | + | 0 | CCGCCCACCGTGTAGGTCTC | CGG | \|\|\|\|\|\|\|\|[\|\|\|\|\|\|\|\|\|\|\|\|]PAM | NA | Intergenic | NA | NA |
| T9570 | CGCCCACCGTGTAGGTCTCCGGG | 1000 | | | | | | | | | |
| Oligo fwd | CACCCGCCCACCGTGTAGGTCTCC | | | | | | | | | | |
| Oligo rev | AAACGGAGACCTACACGGTGGGCG | | | | | | | | | | |
| Chromosome | start | end | strand | MM | target_seq | PAM | alignment | distance | position | gene name | gene id |
| GU305914.1 | 126569 | 126591 | + | 0 | CGCCCACCGTGTAGGTCTCC | GGG | \|\|\|\|\|\|\|\|[\|\|\|\|\|\|\|\|\|\|\|\|]PAM | NA | Intergenic | NA | NA |
| T9571 | TGTAGGTCTCCGGGTTGTCCAGG | 1000 | | | | | | | | | |
| Oligo fwd | CACCTGTAGGTCTCCGGGTTGTCC | | | | | | | | | | |
| Oligo rev | AAACGGACAACCCGGAGACCTACA | | | | | | | | | | |
| Chromosome | start | end | strand | MM | target_seq | PAM | alignment | distance | position | gene name | gene id |
| GU305914.1 | 126578 | 126600 | + | 0 | TGTAGGTCTCCGGGTTGTCC | AGG | \|\|\|\|\|\|\|\|[\|\|\|\|\|\|\|\|\|\|\|\|]PAM | NA | Intergenic | NA | NA |
| T9572 | CGCAGTCCGTCAGCATGTCCAGG | 1000 | | | | | | | | | |

<table>
<tr><td>Oligo fwd</td><td>CACCCGCAGTCCGTCAGCATGTCC</td><td></td><td></td><td></td><td></td><td></td><td></td><td></td><td></td><td></td><td></td></tr>
<tr><td>Oligo rev</td><td>AAACGGACATGCTGACGGACTGCG</td><td></td><td></td><td></td><td></td><td></td><td></td><td></td><td></td><td></td><td></td></tr>
<tr><td>Chromosome</td><td>start</td><td>end</td><td>strand</td><td>MM</td><td>target_seq</td><td>PAM</td><td>alignment</td><td>distance</td><td>position</td><td>gene name</td><td>gene id</td></tr>
<tr><td>GU305914.1</td><td>126614</td><td>126636</td><td>+</td><td>0</td><td>CGCAGTCCGTCAGCATGTCC</td><td>AGG</td><td>||||||||[||||||||||||]PAM</td><td>NA</td><td>Intergenic</td><td>NA</td><td>NA</td></tr>
<tr><td>GU305914.1</td><td>124877</td><td>124899</td><td>+</td><td>4</td><td>CGCCGGCCGTCAGCGAGTCC</td><td>GGG</td><td>|||-|-||[||||||--||||]PAM</td><td>NA</td><td>Intergenic</td><td>NA</td><td>NA</td></tr>
<tr><td>T9573</td><td>GGTAACGCGTCAGCTGCCCGCGG</td><td>1000</td><td></td><td></td><td></td><td></td><td></td><td></td><td></td><td></td><td></td></tr>
<tr><td>Oligo fwd</td><td>CACCGGTAACGCGTCAGCTGCCCG</td><td></td><td></td><td></td><td></td><td></td><td></td><td></td><td></td><td></td><td></td></tr>
<tr><td>Oligo rev</td><td>AAACCGGGCAGCTGACGCGTTACC</td><td></td><td></td><td></td><td></td><td></td><td></td><td></td><td></td><td></td><td></td></tr>
<tr><td>Chromosome</td><td>start</td><td>end</td><td>strand</td><td>MM</td><td>target_seq</td><td>PAM</td><td>alignment</td><td>distance</td><td>position</td><td>gene name</td><td>gene id</td></tr>
<tr><td>GU305914.1</td><td>126635</td><td>126657</td><td>+</td><td>0</td><td>GGTAACGCGTCAGCTGCCCG</td><td>CGG</td><td>||||||||[||||||||||||]PAM</td><td>NA</td><td>Intergenic</td><td>NA</td><td>NA</td></tr>
<tr><td>T9574</td><td>GCGTCAGCTGCCCGCGGTCCCGG</td><td>1000</td><td></td><td></td><td></td><td></td><td></td><td></td><td></td><td></td><td></td></tr>
<tr><td>Oligo fwd</td><td>CACCGCGTCAGCTGCCCGCGGTCC</td><td></td><td></td><td></td><td></td><td></td><td></td><td></td><td></td><td></td><td></td></tr>
<tr><td>Oligo rev</td><td>AAACGGACCGCGGGCAGCTGACGC</td><td></td><td></td><td></td><td></td><td></td><td></td><td></td><td></td><td></td><td></td></tr>
<tr><td>Chromosome</td><td>start</td><td>end</td><td>strand</td><td>MM</td><td>target_seq</td><td>PAM</td><td>alignment</td><td>distance</td><td>position</td><td>gene name</td><td>gene id</td></tr>
<tr><td>GU305914.1</td><td>126641</td><td>126663</td><td>+</td><td>0</td><td>GCGTCAGCTGCCCGCGGTCC</td><td>CGG</td><td>||||||||[||||||||||||]PAM</td><td>NA</td><td>Intergenic</td><td>NA</td><td>NA</td></tr>
<tr><td>GU305914.1</td><td>68692</td><td>68714</td><td>-</td><td>4</td><td>ACGCCAGCTGCTCGCTGTCC</td><td>AGG</td><td>-||-||||[|||-|||-||||]PAM</td><td>NA</td><td>Intergenic</td><td>NA</td><td>NA</td></tr>
<tr><td>T9575</td><td>GCTGCCCGCGGTCCCGGCTGCGG</td><td>1000</td><td></td><td></td><td></td><td></td><td></td><td></td><td></td><td></td><td></td></tr>
<tr><td>Oligo fwd</td><td>CACCGCTGCCCGCGGTCCCGGCTG</td><td></td><td></td><td></td><td></td><td></td><td></td><td></td><td></td><td></td><td></td></tr>
<tr><td>Oligo rev</td><td>AAACCAGCCGGGACCGCGGGCAGC</td><td></td><td></td><td></td><td></td><td></td><td></td><td></td><td></td><td></td><td></td></tr>
<tr><td>Chromosome</td><td>start</td><td>end</td><td>strand</td><td>MM</td><td>target_seq</td><td>PAM</td><td>alignment</td><td>distance</td><td>position</td><td>gene name</td><td>gene id</td></tr>
<tr><td>GU305914.1</td><td>126647</td><td>126669</td><td>+</td><td>0</td><td>GCTGCCCGCGGTCCCGGCTG</td><td>CGG</td><td>||||||||[||||||||||||]PAM</td><td>NA</td><td>Intergenic</td><td>NA</td><td>NA</td></tr>
<tr><td>T9576</td><td>GGTCCCGGCTGCGGTCCAGCAGG</td><td>1000</td><td></td><td></td><td></td><td></td><td></td><td></td><td></td><td></td><td></td></tr>
<tr><td>Oligo fwd</td><td>CACCGGTCCCGGCTGCGGTCCAGC</td><td></td><td></td><td></td><td></td><td></td><td></td><td></td><td></td><td></td><td></td></tr>
<tr><td>Oligo rev</td><td>AAACGCTGGACCGCAGCCGGGACC</td><td></td><td></td><td></td><td></td><td></td><td></td><td></td><td></td><td></td><td></td></tr>
<tr><td>Chromosome</td><td>start</td><td>end</td><td>strand</td><td>MM</td><td>target_seq</td><td>PAM</td><td>alignment</td><td>distance</td><td>position</td><td>gene name</td><td>gene id</td></tr>
<tr><td>GU305914.1</td><td>126656</td><td>126678</td><td>+</td><td>0</td><td>GGTCCCGGCTGCGGTCCAGC</td><td>AGG</td><td>||||||||[||||||||||||]PAM</td><td>NA</td><td>Intergenic</td><td>NA</td><td>NA</td></tr>
<tr><td>T9577</td><td>GCTGCGGTCCAGCAGGAAGATGG</td><td>1000</td><td></td><td></td><td></td><td></td><td></td><td></td><td></td><td></td><td></td></tr>
<tr><td>Oligo fwd</td><td>CACCGCTGCGGTCCAGCAGGAAGA</td><td></td><td></td><td></td><td></td><td></td><td></td><td></td><td></td><td></td><td></td></tr>
<tr><td>Oligo rev</td><td>AAACTCTTCCTGCTGGACCGCAGC</td><td></td><td></td><td></td><td></td><td></td><td></td><td></td><td></td><td></td><td></td></tr>
<tr><td>Chromosome</td><td>start</td><td>end</td><td>strand</td><td>MM</td><td>target_seq</td><td>PAM</td><td>alignment</td><td>distance</td><td>position</td><td>gene name</td><td>gene id</td></tr>
<tr><td>GU305914.1</td><td>126663</td><td>126685</td><td>+</td><td>0</td><td>GCTGCGGTCCAGCAGGAAGA</td><td>TGG</td><td>||||||||[||||||||||||]PAM</td><td>NA</td><td>Intergenic</td><td>NA</td><td>NA</td></tr>
<tr><td>T9578</td><td>GCGGTCCAGCAGGAAGATGGTGG</td><td>1000</td><td></td><td></td><td></td><td></td><td></td><td></td><td></td><td></td><td></td></tr>
<tr><td>Oligo fwd</td><td>CACCGCGGTCCAGCAGGAAGATGG</td><td></td><td></td><td></td><td></td><td></td><td></td><td></td><td></td><td></td><td></td></tr>
<tr><td>Oligo rev</td><td>AAACCCATCTTCCTGCTGGACCGC</td><td></td><td></td><td></td><td></td><td></td><td></td><td></td><td></td><td></td><td></td></tr>
<tr><td>Chromosome</td><td>start</td><td>end</td><td>strand</td><td>MM</td><td>target_seq</td><td>PAM</td><td>alignment</td><td>distance</td><td>position</td><td>gene name</td><td>gene id</td></tr>
<tr><td>GU305914.1</td><td>126666</td><td>126688</td><td>+</td><td>0</td><td>GCGGTCCAGCAGGAAGATGG</td><td>TGG</td><td>||||||||[||||||||||||]PAM</td><td>NA</td><td>Intergenic</td><td>NA</td><td>NA</td></tr>
<tr><td>T9579</td><td>CTTGAAGCCCTGCGCGATGTTGG</td><td>1000</td><td></td><td></td><td></td><td></td><td></td><td></td><td></td><td></td><td></td></tr>
<tr><td>Oligo fwd</td><td>CACCCTTGAAGCCCTGCGCGATGT</td><td></td><td></td><td></td><td></td><td></td><td></td><td></td><td></td><td></td><td></td></tr>
<tr><td>Oligo rev</td><td>AAACACATCGCGCAGGGCTTCAAG</td><td></td><td></td><td></td><td></td><td></td><td></td><td></td><td></td><td></td><td></td></tr>
<tr><td>Chromosome</td><td>start</td><td>end</td><td>strand</td><td>MM</td><td>target_seq</td><td>PAM</td><td>alignment</td><td>distance</td><td>position</td><td>gene name</td><td>gene id</td></tr>
<tr><td>GU305914.1</td><td>126702</td><td>126724</td><td>+</td><td>0</td><td>CTTGAAGCCCTGCGCGATGT</td><td>TGG</td><td>||||||||[||||||||||||]PAM</td><td>NA</td><td>Intergenic</td><td>NA</td><td>NA</td></tr>
<tr><td>T9580</td><td>AGCGTACGCAACACGAAGCACGG</td><td>1000</td><td></td><td></td><td></td><td></td><td></td><td></td><td></td><td></td><td></td></tr>
<tr><td>Oligo fwd</td><td>CACCAGCGTACGCAACACGAAGCA</td><td></td><td></td><td></td><td></td><td></td><td></td><td></td><td></td><td></td><td></td></tr>
<tr><td>Oligo rev</td><td>AAACTGCTTCGTGTTGCGTACGCT</td><td></td><td></td><td></td><td></td><td></td><td></td><td></td><td></td><td></td><td></td></tr>
<tr><td>Chromosome</td><td>start</td><td>end</td><td>strand</td><td>MM</td><td>target_seq</td><td>PAM</td><td>alignment</td><td>distance</td><td>position</td><td>gene name</td><td>gene id</td></tr>
<tr><td>GU305914.1</td><td>126775</td><td>126797</td><td>+</td><td>0</td><td>AGCGTACGCAACACGAAGCA</td><td>CGG</td><td>||||||||[||||||||||||]PAM</td><td>NA</td><td>Intergenic</td><td>NA</td><td>NA</td></tr>
<tr><td>T9581</td><td>GCGTACGCAACACGAAGCACGGG</td><td>1000</td><td></td><td></td><td></td><td></td><td></td><td></td><td></td><td></td><td></td></tr>
<tr><td>Oligo fwd</td><td>CACCGCGTACGCAACACGAAGCAC</td><td></td><td></td><td></td><td></td><td></td><td></td><td></td><td></td><td></td><td></td></tr>
<tr><td>Oligo rev</td><td>AAACGTGCTTCGTGTTGCGTACGC</td><td></td><td></td><td></td><td></td><td></td><td></td><td></td><td></td><td></td><td></td></tr>
<tr><td>Chromosome</td><td>start</td><td>end</td><td>strand</td><td>MM</td><td>target_seq</td><td>PAM</td><td>alignment</td><td>distance</td><td>position</td><td>gene name</td><td>gene id</td></tr>
<tr><td>GU305914.1</td><td>126776</td><td>126798</td><td>+</td><td>0</td><td>GCGTACGCAACACGAAGCAC</td><td>GGG</td><td>||||||||[||||||||||||]PAM</td><td>NA</td><td>Intergenic</td><td>NA</td><td>NA</td></tr>
<tr><td>T9582</td><td>CGTACGCAACACGAAGCACGGGG</td><td>1000</td><td></td><td></td><td></td><td></td><td></td><td></td><td></td><td></td><td></td></tr>
<tr><td>Oligo fwd</td><td>CACCCGTACGCAACACGAAGCACG</td><td></td><td></td><td></td><td></td><td></td><td></td><td></td><td></td><td></td><td></td></tr>
<tr><td>Oligo rev</td><td>AAACCGTGCTTCGTGTTGCGTACG</td><td></td><td></td><td></td><td></td><td></td><td></td><td></td><td></td><td></td><td></td></tr>
<tr><td>Chromosome</td><td>start</td><td>end</td><td>strand</td><td>MM</td><td>target_seq</td><td>PAM</td><td>alignment</td><td>distance</td><td>position</td><td>gene name</td><td>gene id</td></tr>
<tr><td>GU305914.1</td><td>126777</td><td>126799</td><td>+</td><td>0</td><td>CGTACGCAACACGAAGCACG</td><td>GGG</td><td>||||||||[||||||||||||]PAM</td><td>NA</td><td>Intergenic</td><td>NA</td><td>NA</td></tr>
<tr><td>T9583</td><td>GTACGCAACACGAAGCACGGGGG</td><td>1000</td><td></td><td></td><td></td><td></td><td></td><td></td><td></td><td></td><td></td></tr>
<tr><td>Oligo fwd</td><td>CACCGTACGCAACACGAAGCACGG</td><td></td><td></td><td></td><td></td><td></td><td></td><td></td><td></td><td></td><td></td></tr>
<tr><td>Oligo rev</td><td>AAACCCGTGCTTCGTGTTGCGTAC</td><td></td><td></td><td></td><td></td><td></td><td></td><td></td><td></td><td></td><td></td></tr>
<tr><td>Chromosome</td><td>start</td><td>end</td><td>strand</td><td>MM</td><td>target_seq</td><td>PAM</td><td>alignment</td><td>distance</td><td>position</td><td>gene name</td><td>gene id</td></tr>
<tr><td>GU305914.1</td><td>126778</td><td>126800</td><td>+</td><td>0</td><td>GTACGCAACACGAAGCACGG</td><td>GGG</td><td>||||||||[||||||||||||]PAM</td><td>NA</td><td>Intergenic</td><td>NA</td><td>NA</td></tr>
<tr><td>T9584</td><td>ACACGAAGCACGGGGGCGCGTGG</td><td>1000</td><td></td><td></td><td></td><td></td><td></td><td></td><td></td><td></td><td></td></tr>
<tr><td>Oligo fwd</td><td>CACCACACGAAGCACGGGGGCGCG</td><td></td><td></td><td></td><td></td><td></td><td></td><td></td><td></td><td></td><td></td></tr>
<tr><td>Oligo rev</td><td>AAACCGCGCCCCCGTGCTTCGTGT</td><td></td><td></td><td></td><td></td><td></td><td></td><td></td><td></td><td></td><td></td></tr>
<tr><td>Chromosome</td><td>start</td><td>end</td><td>strand</td><td>MM</td><td>target_seq</td><td>PAM</td><td>alignment</td><td>distance</td><td>position</td><td>gene name</td><td>gene id</td></tr>
<tr><td>GU305914.1</td><td>126785</td><td>126807</td><td>+</td><td>0</td><td>ACACGAAGCACGGGGGCGCG</td><td>TGG</td><td>||||||||[||||||||||||]PAM</td><td>NA</td><td>Intergenic</td><td>NA</td><td>NA</td></tr>
<tr><td>T9585</td><td>CGTGGCGCAGCAGCACGCGCAGG</td><td>1000</td><td></td><td></td><td></td><td></td><td></td><td></td><td></td><td></td><td></td></tr>
<tr><td>Oligo fwd</td><td>CACCCGTGGCGCAGCAGCACGCGC</td><td></td><td></td><td></td><td></td><td></td><td></td><td></td><td></td><td></td><td></td></tr>
<tr><td>Oligo rev</td><td>AAACGCGCGTGCTGCTGCGCCACG</td><td></td><td></td><td></td><td></td><td></td><td></td><td></td><td></td><td></td><td></td></tr>
<tr><td>Chromosome</td><td>start</td><td>end</td><td>strand</td><td>MM</td><td>target_seq</td><td>PAM</td><td>alignment</td><td>distance</td><td>position</td><td>gene name</td><td>gene id</td></tr>
<tr><td>GU305914.1</td><td>126803</td><td>126825</td><td>+</td><td>0</td><td>CGTGGCGCAGCAGCACGCGC</td><td>AGG</td><td>||||||||[||||||||||||]PAM</td><td>NA</td><td>Intergenic</td><td>NA</td><td>NA</td></tr>
<tr><td>T9586</td><td>GCGCAGCAGCACGCGCAGGAAGG</td><td>1000</td><td></td><td></td><td></td><td></td><td></td><td></td><td></td><td></td><td></td></tr>
<tr><td>Oligo fwd</td><td>CACCGCGCAGCAGCACGCGCAGGA</td><td></td><td></td><td></td><td></td><td></td><td></td><td></td><td></td><td></td><td></td></tr>
<tr><td>Oligo rev</td><td>AAACTCCTGCGCGTGCTGCTGCGC</td><td></td><td></td><td></td><td></td><td></td><td></td><td></td><td></td><td></td><td></td></tr>
<tr><td>Chromosome</td><td>start</td><td>end</td><td>strand</td><td>MM</td><td>target_seq</td><td>PAM</td><td>alignment</td><td>distance</td><td>position</td><td>gene name</td><td>gene id</td></tr>
</table>

<table>
<tr><td>GU305914.1</td><td>126807</td><td>126829</td><td>+</td><td>0</td><td>GCGCAGCAGCACGCGCAGGA</td><td>AGG</td><td>||||||||[||||||||||||]PAM</td><td>NA</td><td>Intergenic</td><td>NA</td><td>NA</td></tr>
<tr><td></td><td></td><td></td><td></td><td></td><td></td><td></td><td></td><td></td><td></td><td></td><td></td></tr>
<tr><td>T9587</td><td>CGCGCAGGAAGGTGTCGATCAGG</td><td>1000</td><td></td><td></td><td></td><td></td><td></td><td></td><td></td><td></td><td></td></tr>
<tr><td>Oligo fwd</td><td>CACCCGCGCAGGAAGGTGTCGATC</td><td></td><td></td><td></td><td></td><td></td><td></td><td></td><td></td><td></td><td></td></tr>
<tr><td>Oligo rev</td><td>AAACGATCGACACCTTCCTGCGCG</td><td></td><td></td><td></td><td></td><td></td><td></td><td></td><td></td><td></td><td></td></tr>
<tr><td>Chromosome</td><td>start</td><td>end</td><td>strand</td><td>MM</td><td>target_seq</td><td>PAM</td><td>alignment</td><td>distance</td><td>position</td><td>gene name</td><td>gene id</td></tr>
<tr><td>GU305914.1</td><td>126818</td><td>126840</td><td>+</td><td>0</td><td>CGCGCAGGAAGGTGTCGATC</td><td>AGG</td><td>||||||||[||||||||||||]PAM</td><td>NA</td><td>Intergenic</td><td>NA</td><td>NA</td></tr>
<tr><td>GU305914.1</td><td>81336</td><td>81358</td><td>+</td><td>4</td><td>CGAGCAGACAGGTGTAGATC</td><td>CGG</td><td>||-||||-[-||||||-||||]PAM</td><td>NA</td><td>Intergenic</td><td>NA</td><td>NA</td></tr>
<tr><td></td><td></td><td></td><td></td><td></td><td></td><td></td><td></td><td></td><td></td><td></td><td></td></tr>
<tr><td>T9588</td><td>CAGGCCGCGCTCCATCGCGTCGG</td><td>1000</td><td></td><td></td><td></td><td></td><td></td><td></td><td></td><td></td><td></td></tr>
<tr><td>Oligo fwd</td><td>CACCCAGGCCGCGCTCCATCGCGT</td><td></td><td></td><td></td><td></td><td></td><td></td><td></td><td></td><td></td><td></td></tr>
<tr><td>Oligo rev</td><td>AAACACGCGATGGAGCGCGGCCTG</td><td></td><td></td><td></td><td></td><td></td><td></td><td></td><td></td><td></td><td></td></tr>
<tr><td>Chromosome</td><td>start</td><td>end</td><td>strand</td><td>MM</td><td>target_seq</td><td>PAM</td><td>alignment</td><td>distance</td><td>position</td><td>gene name</td><td>gene id</td></tr>
<tr><td>GU305914.1</td><td>126837</td><td>126859</td><td>+</td><td>0</td><td>CAGGCCGCGCTCCATCGCGT</td><td>CGG</td><td>||||||||[||||||||||||]PAM</td><td>NA</td><td>Intergenic</td><td>NA</td><td>NA</td></tr>
<tr><td></td><td></td><td></td><td></td><td></td><td></td><td></td><td></td><td></td><td></td><td></td><td></td></tr>
<tr><td>T9589</td><td>CGCACGCAACACCGTGTTGACGG</td><td>1000</td><td></td><td></td><td></td><td></td><td></td><td></td><td></td><td></td><td></td></tr>
<tr><td>Oligo fwd</td><td>CACCCGCACGCAACACCGTGTTGA</td><td></td><td></td><td></td><td></td><td></td><td></td><td></td><td></td><td></td><td></td></tr>
<tr><td>Oligo rev</td><td>AAACTCAACACGGTGTTGCGTGCG</td><td></td><td></td><td></td><td></td><td></td><td></td><td></td><td></td><td></td><td></td></tr>
<tr><td>Chromosome</td><td>start</td><td>end</td><td>strand</td><td>MM</td><td>target_seq</td><td>PAM</td><td>alignment</td><td>distance</td><td>position</td><td>gene name</td><td>gene id</td></tr>
<tr><td>GU305914.1</td><td>126873</td><td>126895</td><td>+</td><td>0</td><td>CGCACGCAACACCGTGTTGA</td><td>CGG</td><td>||||||||[||||||||||||]PAM</td><td>NA</td><td>Intergenic</td><td>NA</td><td>NA</td></tr>
<tr><td></td><td></td><td></td><td></td><td></td><td></td><td></td><td></td><td></td><td></td><td></td><td></td></tr>
<tr><td>T9590</td><td>CAACACCGTGTTGACGGCGTCGG</td><td>1000</td><td></td><td></td><td></td><td></td><td></td><td></td><td></td><td></td><td></td></tr>
<tr><td>Oligo fwd</td><td>CACCCAACACCGTGTTGACGGCGT</td><td></td><td></td><td></td><td></td><td></td><td></td><td></td><td></td><td></td><td></td></tr>
<tr><td>Oligo rev</td><td>AAACACGCCGTCAACACGGTGTTG</td><td></td><td></td><td></td><td></td><td></td><td></td><td></td><td></td><td></td><td></td></tr>
<tr><td>Chromosome</td><td>start</td><td>end</td><td>strand</td><td>MM</td><td>target_seq</td><td>PAM</td><td>alignment</td><td>distance</td><td>position</td><td>gene name</td><td>gene id</td></tr>
<tr><td>GU305914.1</td><td>126879</td><td>126901</td><td>+</td><td>0</td><td>CAACACCGTGTTGACGGCGT</td><td>CGG</td><td>||||||||[||||||||||||]PAM</td><td>NA</td><td>Intergenic</td><td>NA</td><td>NA</td></tr>
<tr><td></td><td></td><td></td><td></td><td></td><td></td><td></td><td></td><td></td><td></td><td></td><td></td></tr>
<tr><td>T9591</td><td>GTCCAGCAGCGTGTCCTCGAAGG</td><td>1000</td><td></td><td></td><td></td><td></td><td></td><td></td><td></td><td></td><td></td></tr>
<tr><td>Oligo fwd</td><td>CACCGTCCAGCAGCGTGTCCTCGA</td><td></td><td></td><td></td><td></td><td></td><td></td><td></td><td></td><td></td><td></td></tr>
<tr><td>Oligo rev</td><td>AAACTCGAGGACACGCTGCTGGAC</td><td></td><td></td><td></td><td></td><td></td><td></td><td></td><td></td><td></td><td></td></tr>
<tr><td>Chromosome</td><td>start</td><td>end</td><td>strand</td><td>MM</td><td>target_seq</td><td>PAM</td><td>alignment</td><td>distance</td><td>position</td><td>gene name</td><td>gene id</td></tr>
<tr><td>GU305914.1</td><td>126915</td><td>126937</td><td>+</td><td>0</td><td>GTCCAGCAGCGTGTCCTCGA</td><td>AGG</td><td>||||||||[||||||||||||]PAM</td><td>NA</td><td>Intergenic</td><td>NA</td><td>NA</td></tr>
<tr><td></td><td></td><td></td><td></td><td></td><td></td><td></td><td></td><td></td><td></td><td></td><td></td></tr>
<tr><td>T9592</td><td>TGTCCTCGAAGGTCTCCTTCAGG</td><td>1000</td><td></td><td></td><td></td><td></td><td></td><td></td><td></td><td></td><td></td></tr>
<tr><td>Oligo fwd</td><td>CACCTGTCCTCGAAGGTCTCCTTC</td><td></td><td></td><td></td><td></td><td></td><td></td><td></td><td></td><td></td><td></td></tr>
<tr><td>Oligo rev</td><td>AAACGAAGGAGACCTTCGAGGACA</td><td></td><td></td><td></td><td></td><td></td><td></td><td></td><td></td><td></td><td></td></tr>
<tr><td>Chromosome</td><td>start</td><td>end</td><td>strand</td><td>MM</td><td>target_seq</td><td>PAM</td><td>alignment</td><td>distance</td><td>position</td><td>gene name</td><td>gene id</td></tr>
<tr><td>GU305914.1</td><td>126926</td><td>126948</td><td>+</td><td>0</td><td>TGTCCTCGAAGGTCTCCTTC</td><td>AGG</td><td>||||||||[||||||||||||]PAM</td><td>NA</td><td>Intergenic</td><td>NA</td><td>NA</td></tr>
<tr><td></td><td></td><td></td><td></td><td></td><td></td><td></td><td></td><td></td><td></td><td></td><td></td></tr>
<tr><td>T9593</td><td>CCTCGAAGGTCTCCTTCAGGTGG</td><td>1000</td><td></td><td></td><td></td><td></td><td></td><td></td><td></td><td></td><td></td></tr>
<tr><td>Oligo fwd</td><td>CACCCCTCGAAGGTCTCCTTCAGG</td><td></td><td></td><td></td><td></td><td></td><td></td><td></td><td></td><td></td><td></td></tr>
<tr><td>Oligo rev</td><td>AAACCCTGAAGGAGACCTTCGAGG</td><td></td><td></td><td></td><td></td><td></td><td></td><td></td><td></td><td></td><td></td></tr>
<tr><td>Chromosome</td><td>start</td><td>end</td><td>strand</td><td>MM</td><td>target_seq</td><td>PAM</td><td>alignment</td><td>distance</td><td>position</td><td>gene name</td><td>gene id</td></tr>
<tr><td>GU305914.1</td><td>126929</td><td>126951</td><td>+</td><td>0</td><td>CCTCGAAGGTCTCCTTCAGG</td><td>TGG</td><td>||||||||[||||||||||||]PAM</td><td>NA</td><td>Intergenic</td><td>NA</td><td>NA</td></tr>
<tr><td></td><td></td><td></td><td></td><td></td><td></td><td></td><td></td><td></td><td></td><td></td><td></td></tr>
<tr><td>T9594</td><td>CTCGAAGGTCTCCTTCAGGTGGG</td><td>1000</td><td></td><td></td><td></td><td></td><td></td><td></td><td></td><td></td><td></td></tr>
<tr><td>Oligo fwd</td><td>CACCCTCGAAGGTCTCCTTCAGGT</td><td></td><td></td><td></td><td></td><td></td><td></td><td></td><td></td><td></td><td></td></tr>
<tr><td>Oligo rev</td><td>AAACACCTGAAGGAGACCTTCGAG</td><td></td><td></td><td></td><td></td><td></td><td></td><td></td><td></td><td></td><td></td></tr>
<tr><td>Chromosome</td><td>start</td><td>end</td><td>strand</td><td>MM</td><td>target_seq</td><td>PAM</td><td>alignment</td><td>distance</td><td>position</td><td>gene name</td><td>gene id</td></tr>
<tr><td>GU305914.1</td><td>126930</td><td>126952</td><td>+</td><td>0</td><td>CTCGAAGGTCTCCTTCAGGT</td><td>GGG</td><td>||||||||[||||||||||||]PAM</td><td>NA</td><td>Intergenic</td><td>NA</td><td>NA</td></tr>
<tr><td></td><td></td><td></td><td></td><td></td><td></td><td></td><td></td><td></td><td></td><td></td><td></td></tr>
<tr><td>T9595</td><td>CTCCTTCAGGTGGGTCAGACAGG</td><td>1000</td><td></td><td></td><td></td><td></td><td></td><td></td><td></td><td></td><td></td></tr>
<tr><td>Oligo fwd</td><td>CACCCTCCTTCAGGTGGGTCAGAC</td><td></td><td></td><td></td><td></td><td></td><td></td><td></td><td></td><td></td><td></td></tr>
<tr><td>Oligo rev</td><td>AAACGTCTGACCCACCTGAAGGAG</td><td></td><td></td><td></td><td></td><td></td><td></td><td></td><td></td><td></td><td></td></tr>
<tr><td>Chromosome</td><td>start</td><td>end</td><td>strand</td><td>MM</td><td>target_seq</td><td>PAM</td><td>alignment</td><td>distance</td><td>position</td><td>gene name</td><td>gene id</td></tr>
<tr><td>GU305914.1</td><td>126939</td><td>126961</td><td>+</td><td>0</td><td>CTCCTTCAGGTGGGTCAGAC</td><td>AGG</td><td>||||||||[||||||||||||]PAM</td><td>NA</td><td>Intergenic</td><td>NA</td><td>NA</td></tr>
<tr><td></td><td></td><td></td><td></td><td></td><td></td><td></td><td></td><td></td><td></td><td></td><td></td></tr>
<tr><td>T9596</td><td>CTTCAGGTGGGTCAGACAGGCGG</td><td>1000</td><td></td><td></td><td></td><td></td><td></td><td></td><td></td><td></td><td></td></tr>
<tr><td>Oligo fwd</td><td>CACCCTTCAGGTGGGTCAGACAGG</td><td></td><td></td><td></td><td></td><td></td><td></td><td></td><td></td><td></td><td></td></tr>
<tr><td>Oligo rev</td><td>AAACCCTGTCTGACCCACCTGAAG</td><td></td><td></td><td></td><td></td><td></td><td></td><td></td><td></td><td></td><td></td></tr>
<tr><td>Chromosome</td><td>start</td><td>end</td><td>strand</td><td>MM</td><td>target_seq</td><td>PAM</td><td>alignment</td><td>distance</td><td>position</td><td>gene name</td><td>gene id</td></tr>
<tr><td>GU305914.1</td><td>126942</td><td>126964</td><td>+</td><td>0</td><td>CTTCAGGTGGGTCAGACAGG</td><td>CGG</td><td>||||||||[||||||||||||]PAM</td><td>NA</td><td>Intergenic</td><td>NA</td><td>NA</td></tr>
<tr><td></td><td></td><td></td><td></td><td></td><td></td><td></td><td></td><td></td><td></td><td></td><td></td></tr>
<tr><td>T9597</td><td>CAGGCGGCGCTGATCTCGAAAGG</td><td>1000</td><td></td><td></td><td></td><td></td><td></td><td></td><td></td><td></td><td></td></tr>
<tr><td>Oligo fwd</td><td>CACCCAGGCGGCGCTGATCTCGAA</td><td></td><td></td><td></td><td></td><td></td><td></td><td></td><td></td><td></td><td></td></tr>
<tr><td>Oligo rev</td><td>AAACTTCGAGATCAGCGCCGCCTG</td><td></td><td></td><td></td><td></td><td></td><td></td><td></td><td></td><td></td><td></td></tr>
<tr><td>Chromosome</td><td>start</td><td>end</td><td>strand</td><td>MM</td><td>target_seq</td><td>PAM</td><td>alignment</td><td>distance</td><td>position</td><td>gene name</td><td>gene id</td></tr>
<tr><td>GU305914.1</td><td>126958</td><td>126980</td><td>+</td><td>0</td><td>CAGGCGGCGCTGATCTCGAA</td><td>AGG</td><td>||||||||[||||||||||||]PAM</td><td>NA</td><td>Intergenic</td><td>NA</td><td>NA</td></tr>
<tr><td></td><td></td><td></td><td></td><td></td><td></td><td></td><td></td><td></td><td></td><td></td><td></td></tr>
<tr><td>T9598</td><td>AGGCGGCGCTGATCTCGAAAGGG</td><td>1000</td><td></td><td></td><td></td><td></td><td></td><td></td><td></td><td></td><td></td></tr>
<tr><td>Oligo fwd</td><td>CACCAGGCGGCGCTGATCTCGAAA</td><td></td><td></td><td></td><td></td><td></td><td></td><td></td><td></td><td></td><td></td></tr>
<tr><td>Oligo rev</td><td>AAACTTTCGAGATCAGCGCCGCCT</td><td></td><td></td><td></td><td></td><td></td><td></td><td></td><td></td><td></td><td></td></tr>
<tr><td>Chromosome</td><td>start</td><td>end</td><td>strand</td><td>MM</td><td>target_seq</td><td>PAM</td><td>alignment</td><td>distance</td><td>position</td><td>gene name</td><td>gene id</td></tr>
<tr><td>GU305914.1</td><td>126959</td><td>126981</td><td>+</td><td>0</td><td>AGGCGGCGCTGATCTCGAAA</td><td>GGG</td><td>||||||||[||||||||||||]PAM</td><td>NA</td><td>Intergenic</td><td>NA</td><td>NA</td></tr>
<tr><td></td><td></td><td></td><td></td><td></td><td></td><td></td><td></td><td></td><td></td><td></td><td></td></tr>
<tr><td>T9599</td><td>GGCGCTGATCTCGAAAGGGATGG</td><td>1000</td><td></td><td></td><td></td><td></td><td></td><td></td><td></td><td></td><td></td></tr>
<tr><td>Oligo fwd</td><td>CACCGGCGCTGATCTCGAAAGGGA</td><td></td><td></td><td></td><td></td><td></td><td></td><td></td><td></td><td></td><td></td></tr>
<tr><td>Oligo rev</td><td>AAACTCCCTTTCGAGATCAGCGCC</td><td></td><td></td><td></td><td></td><td></td><td></td><td></td><td></td><td></td><td></td></tr>
<tr><td>Chromosome</td><td>start</td><td>end</td><td>strand</td><td>MM</td><td>target_seq</td><td>PAM</td><td>alignment</td><td>distance</td><td>position</td><td>gene name</td><td>gene id</td></tr>
<tr><td>GU305914.1</td><td>126963</td><td>126985</td><td>+</td><td>0</td><td>GGCGCTGATCTCGAAAGGGA</td><td>TGG</td><td>||||||||[||||||||||||]PAM</td><td>NA</td><td>Intergenic</td><td>NA</td><td>NA</td></tr>
<tr><td></td><td></td><td></td><td></td><td></td><td></td><td></td><td></td><td></td><td></td><td></td><td></td></tr>
<tr><td>T9600</td><td>GATCTCGAAAGGGATGGTGATGG</td><td>1000</td><td></td><td></td><td></td><td></td><td></td><td></td><td></td><td></td><td></td></tr>
<tr><td>Oligo fwd</td><td>CACCGATCTCGAAAGGGATGGTGA</td><td></td><td></td><td></td><td></td><td></td><td></td><td></td><td></td><td></td><td></td></tr>
<tr><td>Oligo rev</td><td>AAACTCACCATCCCTTTCGAGATC</td><td></td><td></td><td></td><td></td><td></td><td></td><td></td><td></td><td></td><td></td></tr>
<tr><td>Chromosome</td><td>start</td><td>end</td><td>strand</td><td>MM</td><td>target_seq</td><td>PAM</td><td>alignment</td><td>distance</td><td>position</td><td>gene name</td><td>gene id</td></tr>
<tr><td>GU305914.1</td><td>126969</td><td>126991</td><td>+</td><td>0</td><td>GATCTCGAAAGGGATGGTGA</td><td>TGG</td><td>||||||||[||||||||||||]PAM</td><td>NA</td><td>Intergenic</td><td>NA</td><td>NA</td></tr>
<tr><td></td><td></td><td></td><td></td><td></td><td></td><td></td><td></td><td></td><td></td><td></td><td></td></tr>
<tr><td>T9601</td><td>ATCTCGAAAGGGATGGTGATGGG</td><td>1000</td><td></td><td></td><td></td><td></td><td></td><td></td><td></td><td></td><td></td></tr>
<tr><td>Oligo fwd</td><td>CACCATCTCGAAAGGGATGGTGAT</td><td></td><td></td><td></td><td></td><td></td><td></td><td></td><td></td><td></td><td></td></tr>
</table>

<table>
<tr><td>Oligo rev</td><td>AAACATCACCATCCCTTTCGAGAT</td><td></td><td></td><td></td><td></td><td></td><td></td><td></td><td></td><td></td><td></td><td></td></tr>
<tr><td>Chromosome</td><td>start</td><td></td><td>end</td><td>strand</td><td>MM</td><td>target_seq</td><td>PAM</td><td>alignment</td><td>distance</td><td>position</td><td>gene name</td><td>gene id</td></tr>
<tr><td>GU305914.1</td><td></td><td>126970</td><td>126992</td><td>+</td><td>0</td><td>ATCTCGAAAGGGATGGTGAT</td><td>GGG</td><td>||||||||[||||||||||||]PAM</td><td>NA</td><td>Intergenic</td><td>NA</td><td>NA</td></tr>
<tr><td>T9602</td><td>TCTCGAAAGGGATGGTGATGGGG</td><td></td><td>1000</td><td></td><td></td><td></td><td></td><td></td><td></td><td></td><td></td><td></td></tr>
<tr><td>Oligo fwd</td><td>CACCTCTCGAAAGGGATGGTGATG</td><td></td><td></td><td></td><td></td><td></td><td></td><td></td><td></td><td></td><td></td><td></td></tr>
<tr><td>Oligo rev</td><td>AAACCATCACCATCCCTTTCGAGA</td><td></td><td></td><td></td><td></td><td></td><td></td><td></td><td></td><td></td><td></td><td></td></tr>
<tr><td>Chromosome</td><td>start</td><td></td><td>end</td><td>strand</td><td>MM</td><td>target_seq</td><td>PAM</td><td>alignment</td><td>distance</td><td>position</td><td>gene name</td><td>gene id</td></tr>
<tr><td>GU305914.1</td><td></td><td>126971</td><td>126993</td><td>+</td><td>0</td><td>TCTCGAAAGGGATGGTGATG</td><td>GGG</td><td>||||||||[||||||||||||]PAM</td><td>NA</td><td>Intergenic</td><td>NA</td><td>NA</td></tr>
<tr><td>T9603</td><td>GGGGCTCTTCTCGCTGTACTTGG</td><td></td><td>1000</td><td></td><td></td><td></td><td></td><td></td><td></td><td></td><td></td><td></td></tr>
<tr><td>Oligo fwd</td><td>CACCGGGGCTCTTCTCGCTGTACT</td><td></td><td></td><td></td><td></td><td></td><td></td><td></td><td></td><td></td><td></td><td></td></tr>
<tr><td>Oligo rev</td><td>AAACAGTACAGCGAGAAGAGCCCC</td><td></td><td></td><td></td><td></td><td></td><td></td><td></td><td></td><td></td><td></td><td></td></tr>
<tr><td>Chromosome</td><td>start</td><td></td><td>end</td><td>strand</td><td>MM</td><td>target_seq</td><td>PAM</td><td>alignment</td><td>distance</td><td>position</td><td>gene name</td><td>gene id</td></tr>
<tr><td>GU305914.1</td><td></td><td>126990</td><td>127012</td><td>+</td><td>0</td><td>GGGGCTCTTCTCGCTGTACT</td><td>TGG</td><td>||||||||[||||||||||||]PAM</td><td>NA</td><td>Intergenic</td><td>NA</td><td>NA</td></tr>
<tr><td>T9604</td><td>CATCACCGTCGTCTGTCGCGAGG</td><td></td><td>1000</td><td></td><td></td><td></td><td></td><td></td><td></td><td></td><td></td><td></td></tr>
<tr><td>Oligo fwd</td><td>CACCCATCACCGTCGTCTGTCGCG</td><td></td><td></td><td></td><td></td><td></td><td></td><td></td><td></td><td></td><td></td><td></td></tr>
<tr><td>Oligo rev</td><td>AAACCGCGACAGACGACGGTGATG</td><td></td><td></td><td></td><td></td><td></td><td></td><td></td><td></td><td></td><td></td><td></td></tr>
<tr><td>Chromosome</td><td>start</td><td></td><td>end</td><td>strand</td><td>MM</td><td>target_seq</td><td>PAM</td><td>alignment</td><td>distance</td><td>position</td><td>gene name</td><td>gene id</td></tr>
<tr><td>GU305914.1</td><td></td><td>127017</td><td>127039</td><td>+</td><td>0</td><td>CATCACCGTCGTCTGTCGCG</td><td>AGG</td><td>||||||||[||||||||||||]PAM</td><td>NA</td><td>Intergenic</td><td>NA</td><td>NA</td></tr>
<tr><td>T9605</td><td>CACCGTCGTCTGTCGCGAGGTGG</td><td></td><td>1000</td><td></td><td></td><td></td><td></td><td></td><td></td><td></td><td></td><td></td></tr>
<tr><td>Oligo fwd</td><td>CACCCACCGTCGTCTGTCGCGAGG</td><td></td><td></td><td></td><td></td><td></td><td></td><td></td><td></td><td></td><td></td><td></td></tr>
<tr><td>Oligo rev</td><td>AAACCCTCGCGACAGACGACGGTG</td><td></td><td></td><td></td><td></td><td></td><td></td><td></td><td></td><td></td><td></td><td></td></tr>
<tr><td>Chromosome</td><td>start</td><td></td><td>end</td><td>strand</td><td>MM</td><td>target_seq</td><td>PAM</td><td>alignment</td><td>distance</td><td>position</td><td>gene name</td><td>gene id</td></tr>
<tr><td>GU305914.1</td><td></td><td>127020</td><td>127042</td><td>+</td><td>0</td><td>CACCGTCGTCTGTCGCGAGG</td><td>TGG</td><td>||||||||[||||||||||||]PAM</td><td>NA</td><td>Intergenic</td><td>NA</td><td>NA</td></tr>
<tr><td>T9606</td><td>ACCGTCGTCTGTCGCGAGGTGGG</td><td></td><td>1000</td><td></td><td></td><td></td><td></td><td></td><td></td><td></td><td></td><td></td></tr>
<tr><td>Oligo fwd</td><td>CACCACCGTCGTCTGTCGCGAGGT</td><td></td><td></td><td></td><td></td><td></td><td></td><td></td><td></td><td></td><td></td><td></td></tr>
<tr><td>Oligo rev</td><td>AAACACCTCGCGACAGACGACGGT</td><td></td><td></td><td></td><td></td><td></td><td></td><td></td><td></td><td></td><td></td><td></td></tr>
<tr><td>Chromosome</td><td>start</td><td></td><td>end</td><td>strand</td><td>MM</td><td>target_seq</td><td>PAM</td><td>alignment</td><td>distance</td><td>position</td><td>gene name</td><td>gene id</td></tr>
<tr><td>GU305914.1</td><td></td><td>127021</td><td>127043</td><td>+</td><td>0</td><td>ACCGTCGTCTGTCGCGAGGT</td><td>GGG</td><td>||||||||[||||||||||||]PAM</td><td>NA</td><td>Intergenic</td><td>NA</td><td>NA</td></tr>
<tr><td>T9607</td><td>CCGTCGTCTGTCGCGAGGTGGGG</td><td></td><td>1000</td><td></td><td></td><td></td><td></td><td></td><td></td><td></td><td></td><td></td></tr>
<tr><td>Oligo fwd</td><td>CACCCCGTCGTCTGTCGCGAGGTG</td><td></td><td></td><td></td><td></td><td></td><td></td><td></td><td></td><td></td><td></td><td></td></tr>
<tr><td>Oligo rev</td><td>AAACCACCTCGCGACAGACGACGG</td><td></td><td></td><td></td><td></td><td></td><td></td><td></td><td></td><td></td><td></td><td></td></tr>
<tr><td>Chromosome</td><td>start</td><td></td><td>end</td><td>strand</td><td>MM</td><td>target_seq</td><td>PAM</td><td>alignment</td><td>distance</td><td>position</td><td>gene name</td><td>gene id</td></tr>
<tr><td>GU305914.1</td><td></td><td>127022</td><td>127044</td><td>+</td><td>0</td><td>CCGTCGTCTGTCGCGAGGTG</td><td>GGG</td><td>||||||||[||||||||||||]PAM</td><td>NA</td><td>Intergenic</td><td>NA</td><td>NA</td></tr>
<tr><td>T9608</td><td>CTGTCGCGAGGTGGGGATCCCGG</td><td></td><td>1000</td><td></td><td></td><td></td><td></td><td></td><td></td><td></td><td></td><td></td></tr>
<tr><td>Oligo fwd</td><td>CACCCTGTCGCGAGGTGGGGATCC</td><td></td><td></td><td></td><td></td><td></td><td></td><td></td><td></td><td></td><td></td><td></td></tr>
<tr><td>Oligo rev</td><td>AAACGGATCCCCACCTCGCGACAG</td><td></td><td></td><td></td><td></td><td></td><td></td><td></td><td></td><td></td><td></td><td></td></tr>
<tr><td>Chromosome</td><td>start</td><td></td><td>end</td><td>strand</td><td>MM</td><td>target_seq</td><td>PAM</td><td>alignment</td><td>distance</td><td>position</td><td>gene name</td><td>gene id</td></tr>
<tr><td>GU305914.1</td><td></td><td>127029</td><td>127051</td><td>+</td><td>0</td><td>CTGTCGCGAGGTGGGGATCC</td><td>CGG</td><td>||||||||[||||||||||||]PAM</td><td>NA</td><td>Intergenic</td><td>NA</td><td>NA</td></tr>
<tr><td>T9609</td><td>CGAGGTGGGGATCCCGGCCCCGG</td><td></td><td>1000</td><td></td><td></td><td></td><td></td><td></td><td></td><td></td><td></td><td></td></tr>
<tr><td>Oligo fwd</td><td>CACCCGAGGTGGGGATCCCGGCCC</td><td></td><td></td><td></td><td></td><td></td><td></td><td></td><td></td><td></td><td></td><td></td></tr>
<tr><td>Oligo rev</td><td>AAACGGGCCGGGATCCCCACCTCG</td><td></td><td></td><td></td><td></td><td></td><td></td><td></td><td></td><td></td><td></td><td></td></tr>
<tr><td>Chromosome</td><td>start</td><td></td><td>end</td><td>strand</td><td>MM</td><td>target_seq</td><td>PAM</td><td>alignment</td><td>distance</td><td>position</td><td>gene name</td><td>gene id</td></tr>
<tr><td>GU305914.1</td><td></td><td>127035</td><td>127057</td><td>+</td><td>0</td><td>CGAGGTGGGGATCCCGGCCC</td><td>CGG</td><td>||||||||[||||||||||||]PAM</td><td>NA</td><td>Intergenic</td><td>NA</td><td>NA</td></tr>
<tr><td>T9610</td><td>GGTGGGGATCCCGGCCCCGGTGG</td><td></td><td>1000</td><td></td><td></td><td></td><td></td><td></td><td></td><td></td><td></td><td></td></tr>
<tr><td>Oligo fwd</td><td>CACCGGTGGGGATCCCGGCCCCGG</td><td></td><td></td><td></td><td></td><td></td><td></td><td></td><td></td><td></td><td></td><td></td></tr>
<tr><td>Oligo rev</td><td>AAACCCGGGGCCGGGATCCCCACC</td><td></td><td></td><td></td><td></td><td></td><td></td><td></td><td></td><td></td><td></td><td></td></tr>
<tr><td>Chromosome</td><td>start</td><td></td><td>end</td><td>strand</td><td>MM</td><td>target_seq</td><td>PAM</td><td>alignment</td><td>distance</td><td>position</td><td>gene name</td><td>gene id</td></tr>
<tr><td>GU305914.1</td><td></td><td>127038</td><td>127060</td><td>+</td><td>0</td><td>GGTGGGGATCCCGGCCCCGG</td><td>TGG</td><td>||||||||[||||||||||||]PAM</td><td>NA</td><td>Intergenic</td><td>NA</td><td>NA</td></tr>
<tr><td>T9611</td><td>CCGGCCCCGGTGGCCACTCGAGG</td><td></td><td>1000</td><td></td><td></td><td></td><td></td><td></td><td></td><td></td><td></td><td></td></tr>
<tr><td>Oligo fwd</td><td>CACCCCGGCCCCGGTGGCCACTCG</td><td></td><td></td><td></td><td></td><td></td><td></td><td></td><td></td><td></td><td></td><td></td></tr>
<tr><td>Oligo rev</td><td>AAACCGAGTGGCCACCGGGGCCGG</td><td></td><td></td><td></td><td></td><td></td><td></td><td></td><td></td><td></td><td></td><td></td></tr>
<tr><td>Chromosome</td><td>start</td><td></td><td>end</td><td>strand</td><td>MM</td><td>target_seq</td><td>PAM</td><td>alignment</td><td>distance</td><td>position</td><td>gene name</td><td>gene id</td></tr>
<tr><td>GU305914.1</td><td></td><td>127048</td><td>127070</td><td>+</td><td>0</td><td>CCGGCCCCGGTGGCCACTCG</td><td>AGG</td><td>||||||||[||||||||||||]PAM</td><td>NA</td><td>Intergenic</td><td>NA</td><td>NA</td></tr>
<tr><td>T9612</td><td>CGGCCCCGGTGGCCACTCGAGGG</td><td></td><td>1000</td><td></td><td></td><td></td><td></td><td></td><td></td><td></td><td></td><td></td></tr>
<tr><td>Oligo fwd</td><td>CACCCGGCCCCGGTGGCCACTCGA</td><td></td><td></td><td></td><td></td><td></td><td></td><td></td><td></td><td></td><td></td><td></td></tr>
<tr><td>Oligo rev</td><td>AAACTCGAGTGGCCACCGGGGCCG</td><td></td><td></td><td></td><td></td><td></td><td></td><td></td><td></td><td></td><td></td><td></td></tr>
<tr><td>Chromosome</td><td>start</td><td></td><td>end</td><td>strand</td><td>MM</td><td>target_seq</td><td>PAM</td><td>alignment</td><td>distance</td><td>position</td><td>gene name</td><td>gene id</td></tr>
<tr><td>GU305914.1</td><td></td><td>127049</td><td>127071</td><td>+</td><td>0</td><td>CGGCCCCGGTGGCCACTCGA</td><td>GGG</td><td>||||||||[||||||||||||]PAM</td><td>NA</td><td>Intergenic</td><td>NA</td><td>NA</td></tr>
<tr><td>T9613</td><td>TGCCCGTAGTCATTTTGTTCAGG</td><td></td><td>1000</td><td></td><td></td><td></td><td></td><td></td><td></td><td></td><td></td><td></td></tr>
<tr><td>Oligo fwd</td><td>CACCTGCCCGTAGTCATTTTGTTC</td><td></td><td></td><td></td><td></td><td></td><td></td><td></td><td></td><td></td><td></td><td></td></tr>
<tr><td>Oligo rev</td><td>AAACGAACAAAATGACTACGGGCA</td><td></td><td></td><td></td><td></td><td></td><td></td><td></td><td></td><td></td><td></td><td></td></tr>
<tr><td>Chromosome</td><td>start</td><td></td><td>end</td><td>strand</td><td>MM</td><td>target_seq</td><td>PAM</td><td>alignment</td><td>distance</td><td>position</td><td>gene name</td><td>gene id</td></tr>
<tr><td>GU305914.1</td><td></td><td>127094</td><td>127116</td><td>+</td><td>0</td><td>TGCCCGTAGTCATTTTGTTC</td><td>AGG</td><td>||||||||[||||||||||||]PAM</td><td>NA</td><td>Intergenic</td><td>NA</td><td>NA</td></tr>
<tr><td>T9614</td><td>TAGTCATTTTGTTCAGGTCGTGG</td><td></td><td>1000</td><td></td><td></td><td></td><td></td><td></td><td></td><td></td><td></td><td></td></tr>
<tr><td>Oligo fwd</td><td>CACCTAGTCATTTTGTTCAGGTCG</td><td></td><td></td><td></td><td></td><td></td><td></td><td></td><td></td><td></td><td></td><td></td></tr>
<tr><td>Oligo rev</td><td>AAACCGACCTGAACAAAATGACTA</td><td></td><td></td><td></td><td></td><td></td><td></td><td></td><td></td><td></td><td></td><td></td></tr>
<tr><td>Chromosome</td><td>start</td><td></td><td>end</td><td>strand</td><td>MM</td><td>target_seq</td><td>PAM</td><td>alignment</td><td>distance</td><td>position</td><td>gene name</td><td>gene id</td></tr>
<tr><td>GU305914.1</td><td></td><td>127100</td><td>127122</td><td>+</td><td>0</td><td>TAGTCATTTTGTTCAGGTCG</td><td>TGG</td><td>||||||||[||||||||||||]PAM</td><td>NA</td><td>Intergenic</td><td>NA</td><td>NA</td></tr>
<tr><td>T9615</td><td>CCGTCGCCAACGCCGTTCCCAGG</td><td></td><td>1000</td><td></td><td></td><td></td><td></td><td></td><td></td><td></td><td></td><td></td></tr>
<tr><td>Oligo fwd</td><td>CACCCCGTCGCCAACGCCGTTCCC</td><td></td><td></td><td></td><td></td><td></td><td></td><td></td><td></td><td></td><td></td><td></td></tr>
<tr><td>Oligo rev</td><td>AAACGGGAACGGCGTTGGCGACGG</td><td></td><td></td><td></td><td></td><td></td><td></td><td></td><td></td><td></td><td></td><td></td></tr>
<tr><td>Chromosome</td><td>start</td><td></td><td>end</td><td>strand</td><td>MM</td><td>target_seq</td><td>PAM</td><td>alignment</td><td>distance</td><td>position</td><td>gene name</td><td>gene id</td></tr>
<tr><td>GU305914.1</td><td></td><td>127142</td><td>127164</td><td>+</td><td>0</td><td>CCGTCGCCAACGCCGTTCCC</td><td>AGG</td><td>||||||||[||||||||||||]PAM</td><td>NA</td><td>Intergenic</td><td>NA</td><td>NA</td></tr>
<tr><td>T9616</td><td>CGCCAACGCCGTTCCCAGGAAGG</td><td></td><td>1000</td><td></td><td></td><td></td><td></td><td></td><td></td><td></td><td></td><td></td></tr>
</table>

<table>
<tr><td>Oligo fwd</td><td>CACCCGCCAACGCCGTTCCCAGGA</td><td></td><td></td><td></td><td></td><td></td><td></td><td></td><td></td><td></td><td></td></tr>
<tr><td>Oligo rev</td><td>AAACTCCTGGGAACGGCGTTGGCG</td><td></td><td></td><td></td><td></td><td></td><td></td><td></td><td></td><td></td><td></td></tr>
<tr><td>Chromosome</td><td>start</td><td>end</td><td>strand</td><td>MM</td><td>target_seq</td><td>PAM</td><td>alignment</td><td>distance</td><td>position</td><td>gene name</td><td>gene id</td></tr>
<tr><td>GU305914.1</td><td>127146</td><td>127168</td><td>+</td><td>0</td><td>CGCCAACGCCGTTCCCAGGA</td><td>AGG</td><td>||||||||[||||||||||||]PAM</td><td>NA</td><td>Intergenic</td><td>NA</td><td>NA</td></tr>
<tr><td></td><td></td><td></td><td></td><td></td><td></td><td></td><td></td><td></td><td></td><td></td><td></td></tr>
<tr><td>T9617</td><td>CGAAGATCGCCTCGAAGCTCAGG</td><td>1000</td><td></td><td></td><td></td><td></td><td></td><td></td><td></td><td></td><td></td></tr>
<tr><td>Oligo fwd</td><td>CACCCGAAGATCGCCTCGAAGCTC</td><td></td><td></td><td></td><td></td><td></td><td></td><td></td><td></td><td></td><td></td></tr>
<tr><td>Oligo rev</td><td>AAACGAGCTTCGAGGCGATCTTCG</td><td></td><td></td><td></td><td></td><td></td><td></td><td></td><td></td><td></td><td></td></tr>
<tr><td>Chromosome</td><td>start</td><td>end</td><td>strand</td><td>MM</td><td>target_seq</td><td>PAM</td><td>alignment</td><td>distance</td><td>position</td><td>gene name</td><td>gene id</td></tr>
<tr><td>GU305914.1</td><td>127199</td><td>127221</td><td>+</td><td>0</td><td>CGAAGATCGCCTCGAAGCTC</td><td>AGG</td><td>||||||||[||||||||||||]PAM</td><td>NA</td><td>Intergenic</td><td>NA</td><td>NA</td></tr>
<tr><td></td><td></td><td></td><td></td><td></td><td></td><td></td><td></td><td></td><td></td><td></td><td></td></tr>
<tr><td>T9618</td><td>GAAGCTCAGGTTGTACCGTTCGG</td><td>1000</td><td></td><td></td><td></td><td></td><td></td><td></td><td></td><td></td><td></td></tr>
<tr><td>Oligo fwd</td><td>CACCGAAGCTCAGGTTGTACCGTT</td><td></td><td></td><td></td><td></td><td></td><td></td><td></td><td></td><td></td><td></td></tr>
<tr><td>Oligo rev</td><td>AAACAACGGTACAACCTGAGCTTC</td><td></td><td></td><td></td><td></td><td></td><td></td><td></td><td></td><td></td><td></td></tr>
<tr><td>Chromosome</td><td>start</td><td>end</td><td>strand</td><td>MM</td><td>target_seq</td><td>PAM</td><td>alignment</td><td>distance</td><td>position</td><td>gene name</td><td>gene id</td></tr>
<tr><td>GU305914.1</td><td>127212</td><td>127234</td><td>+</td><td>0</td><td>GAAGCTCAGGTTGTACCGTT</td><td>CGG</td><td>||||||||[||||||||||||]PAM</td><td>NA</td><td>Intergenic</td><td>NA</td><td>NA</td></tr>
<tr><td></td><td></td><td></td><td></td><td></td><td></td><td></td><td></td><td></td><td></td><td></td><td></td></tr>
<tr><td>T9619</td><td>AAGCTCAGGTTGTACCGTTCGGG</td><td>1000</td><td></td><td></td><td></td><td></td><td></td><td></td><td></td><td></td><td></td></tr>
<tr><td>Oligo fwd</td><td>CACCAAGCTCAGGTTGTACCGTTC</td><td></td><td></td><td></td><td></td><td></td><td></td><td></td><td></td><td></td><td></td></tr>
<tr><td>Oligo rev</td><td>AAACGAACGGTACAACCTGAGCTT</td><td></td><td></td><td></td><td></td><td></td><td></td><td></td><td></td><td></td><td></td></tr>
<tr><td>Chromosome</td><td>start</td><td>end</td><td>strand</td><td>MM</td><td>target_seq</td><td>PAM</td><td>alignment</td><td>distance</td><td>position</td><td>gene name</td><td>gene id</td></tr>
<tr><td>GU305914.1</td><td>127213</td><td>127235</td><td>+</td><td>0</td><td>AAGCTCAGGTTGTACCGTTC</td><td>GGG</td><td>||||||||[||||||||||||]PAM</td><td>NA</td><td>Intergenic</td><td>NA</td><td>NA</td></tr>
<tr><td></td><td></td><td></td><td></td><td></td><td></td><td></td><td></td><td></td><td></td><td></td><td></td></tr>
<tr><td>T9620</td><td>AGCTCAGGTTGTACCGTTCGGGG</td><td>1000</td><td></td><td></td><td></td><td></td><td></td><td></td><td></td><td></td><td></td></tr>
<tr><td>Oligo fwd</td><td>CACCAGCTCAGGTTGTACCGTTCG</td><td></td><td></td><td></td><td></td><td></td><td></td><td></td><td></td><td></td><td></td></tr>
<tr><td>Oligo rev</td><td>AAACCGAACGGTACAACCTGAGCT</td><td></td><td></td><td></td><td></td><td></td><td></td><td></td><td></td><td></td><td></td></tr>
<tr><td>Chromosome</td><td>start</td><td>end</td><td>strand</td><td>MM</td><td>target_seq</td><td>PAM</td><td>alignment</td><td>distance</td><td>position</td><td>gene name</td><td>gene id</td></tr>
<tr><td>GU305914.1</td><td>127214</td><td>127236</td><td>+</td><td>0</td><td>AGCTCAGGTTGTACCGTTCG</td><td>GGG</td><td>||||||||[||||||||||||]PAM</td><td>NA</td><td>Intergenic</td><td>NA</td><td>NA</td></tr>
<tr><td></td><td></td><td></td><td></td><td></td><td></td><td></td><td></td><td></td><td></td><td></td><td></td></tr>
<tr><td>T9621</td><td>CGGGGTCGTCCCCGTAGTACAGG</td><td>1000</td><td></td><td></td><td></td><td></td><td></td><td></td><td></td><td></td><td></td></tr>
<tr><td>Oligo fwd</td><td>CACCCGGGGTCGTCCCCGTAGTAC</td><td></td><td></td><td></td><td></td><td></td><td></td><td></td><td></td><td></td><td></td></tr>
<tr><td>Oligo rev</td><td>AAACGTACTACGGGGACGACCCCG</td><td></td><td></td><td></td><td></td><td></td><td></td><td></td><td></td><td></td><td></td></tr>
<tr><td>Chromosome</td><td>start</td><td>end</td><td>strand</td><td>MM</td><td>target_seq</td><td>PAM</td><td>alignment</td><td>distance</td><td>position</td><td>gene name</td><td>gene id</td></tr>
<tr><td>GU305914.1</td><td>127232</td><td>127254</td><td>+</td><td>0</td><td>CGGGGTCGTCCCCGTAGTAC</td><td>AGG</td><td>||||||||[||||||||||||]PAM</td><td>NA</td><td>Intergenic</td><td>NA</td><td>NA</td></tr>
<tr><td></td><td></td><td></td><td></td><td></td><td></td><td></td><td></td><td></td><td></td><td></td><td></td></tr>
<tr><td>T9622</td><td>ATGTGCGTCAGCAGATCGATCGG</td><td>1000</td><td></td><td></td><td></td><td></td><td></td><td></td><td></td><td></td><td></td></tr>
<tr><td>Oligo fwd</td><td>CACCATGTGCGTCAGCAGATCGAT</td><td></td><td></td><td></td><td></td><td></td><td></td><td></td><td></td><td></td><td></td></tr>
<tr><td>Oligo rev</td><td>AAACATCGATCTGCTGACGCACAT</td><td></td><td></td><td></td><td></td><td></td><td></td><td></td><td></td><td></td><td></td></tr>
<tr><td>Chromosome</td><td>start</td><td>end</td><td>strand</td><td>MM</td><td>target_seq</td><td>PAM</td><td>alignment</td><td>distance</td><td>position</td><td>gene name</td><td>gene id</td></tr>
<tr><td>GU305914.1</td><td>127294</td><td>127316</td><td>+</td><td>0</td><td>ATGTGCGTCAGCAGATCGAT</td><td>CGG</td><td>||||||||[||||||||||||]PAM</td><td>NA</td><td>Intergenic</td><td>NA</td><td>NA</td></tr>
<tr><td></td><td></td><td></td><td></td><td></td><td></td><td></td><td></td><td></td><td></td><td></td><td></td></tr>
<tr><td>T9623</td><td>TCGATCGGCACATCGAGCTTAGG</td><td>1000</td><td></td><td></td><td></td><td></td><td></td><td></td><td></td><td></td><td></td></tr>
<tr><td>Oligo fwd</td><td>CACCTCGATCGGCACATCGAGCTT</td><td></td><td></td><td></td><td></td><td></td><td></td><td></td><td></td><td></td><td></td></tr>
<tr><td>Oligo rev</td><td>AAACAAGCTCGATGTGCCGATCGA</td><td></td><td></td><td></td><td></td><td></td><td></td><td></td><td></td><td></td><td></td></tr>
<tr><td>Chromosome</td><td>start</td><td>end</td><td>strand</td><td>MM</td><td>target_seq</td><td>PAM</td><td>alignment</td><td>distance</td><td>position</td><td>gene name</td><td>gene id</td></tr>
<tr><td>GU305914.1</td><td>127309</td><td>127331</td><td>+</td><td>0</td><td>TCGATCGGCACATCGAGCTT</td><td>AGG</td><td>||||||||[||||||||||||]PAM</td><td>NA</td><td>Intergenic</td><td>NA</td><td>NA</td></tr>
<tr><td></td><td></td><td></td><td></td><td></td><td></td><td></td><td></td><td></td><td></td><td></td><td></td></tr>
<tr><td>T9624</td><td>CGTCCAGTTCTCCCCCATCGCGG</td><td>1000</td><td></td><td></td><td></td><td></td><td></td><td></td><td></td><td></td><td></td></tr>
<tr><td>Oligo fwd</td><td>CACCCGTCCAGTTCTCCCCCATCG</td><td></td><td></td><td></td><td></td><td></td><td></td><td></td><td></td><td></td><td></td></tr>
<tr><td>Oligo rev</td><td>AAACCGATGGGGGAGAACTGGACG</td><td></td><td></td><td></td><td></td><td></td><td></td><td></td><td></td><td></td><td></td></tr>
<tr><td>Chromosome</td><td>start</td><td>end</td><td>strand</td><td>MM</td><td>target_seq</td><td>PAM</td><td>alignment</td><td>distance</td><td>position</td><td>gene name</td><td>gene id</td></tr>
<tr><td>GU305914.1</td><td>127347</td><td>127369</td><td>+</td><td>0</td><td>CGTCCAGTTCTCCCCCATCG</td><td>CGG</td><td>||||||||[||||||||||||]PAM</td><td>NA</td><td>Intergenic</td><td>NA</td><td>NA</td></tr>
<tr><td></td><td></td><td></td><td></td><td></td><td></td><td></td><td></td><td></td><td></td><td></td><td></td></tr>
<tr><td>T9625</td><td>CCAGTTCTCCCCCATCGCGGCGG</td><td>1000</td><td></td><td></td><td></td><td></td><td></td><td></td><td></td><td></td><td></td></tr>
<tr><td>Oligo fwd</td><td>CACCCCAGTTCTCCCCCATCGCGG</td><td></td><td></td><td></td><td></td><td></td><td></td><td></td><td></td><td></td><td></td></tr>
<tr><td>Oligo rev</td><td>AAACCCGCGATGGGGGAGAACTGG</td><td></td><td></td><td></td><td></td><td></td><td></td><td></td><td></td><td></td><td></td></tr>
<tr><td>Chromosome</td><td>start</td><td>end</td><td>strand</td><td>MM</td><td>target_seq</td><td>PAM</td><td>alignment</td><td>distance</td><td>position</td><td>gene name</td><td>gene id</td></tr>
<tr><td>GU305914.1</td><td>127350</td><td>127372</td><td>+</td><td>0</td><td>CCAGTTCTCCCCCATCGCGG</td><td>CGG</td><td>||||||||[||||||||||||]PAM</td><td>NA</td><td>Intergenic</td><td>NA</td><td>NA</td></tr>
<tr><td></td><td></td><td></td><td></td><td></td><td></td><td></td><td></td><td></td><td></td><td></td><td></td></tr>
<tr><td>T9626</td><td>GTTCTCCCCCATCGCGGCGGCGG</td><td>1000</td><td></td><td></td><td></td><td></td><td></td><td></td><td></td><td></td><td></td></tr>
<tr><td>Oligo fwd</td><td>CACCGTTCTCCCCCATCGCGGCGG</td><td></td><td></td><td></td><td></td><td></td><td></td><td></td><td></td><td></td><td></td></tr>
<tr><td>Oligo rev</td><td>AAACCCGCCGCGATGGGGGAGAAC</td><td></td><td></td><td></td><td></td><td></td><td></td><td></td><td></td><td></td><td></td></tr>
<tr><td>Chromosome</td><td>start</td><td>end</td><td>strand</td><td>MM</td><td>target_seq</td><td>PAM</td><td>alignment</td><td>distance</td><td>position</td><td>gene name</td><td>gene id</td></tr>
<tr><td>GU305914.1</td><td>127353</td><td>127375</td><td>+</td><td>0</td><td>GTTCTCCCCCATCGCGGCGG</td><td>CGG</td><td>||||||||[||||||||||||]PAM</td><td>NA</td><td>Intergenic</td><td>NA</td><td>NA</td></tr>
<tr><td></td><td></td><td></td><td></td><td></td><td></td><td></td><td></td><td></td><td></td><td></td><td></td></tr>
<tr><td>T9627</td><td>CATCGCGGCGGCGGCGCTACCGG</td><td>1000</td><td></td><td></td><td></td><td></td><td></td><td></td><td></td><td></td><td></td></tr>
<tr><td>Oligo fwd</td><td>CACCCATCGCGGCGGCGGCGCTAC</td><td></td><td></td><td></td><td></td><td></td><td></td><td></td><td></td><td></td><td></td></tr>
<tr><td>Oligo rev</td><td>AAACGTAGCGCCGCCGCCGCGATG</td><td></td><td></td><td></td><td></td><td></td><td></td><td></td><td></td><td></td><td></td></tr>
<tr><td>Chromosome</td><td>start</td><td>end</td><td>strand</td><td>MM</td><td>target_seq</td><td>PAM</td><td>alignment</td><td>distance</td><td>position</td><td>gene name</td><td>gene id</td></tr>
<tr><td>GU305914.1</td><td>127362</td><td>127384</td><td>+</td><td>0</td><td>CATCGCGGCGGCGGCGCTAC</td><td>CGG</td><td>||||||||[||||||||||||]PAM</td><td>NA</td><td>Intergenic</td><td>NA</td><td>NA</td></tr>
<tr><td></td><td></td><td></td><td></td><td></td><td></td><td></td><td></td><td></td><td></td><td></td><td></td></tr>
<tr><td>T9628</td><td>CGCGGCGGCGGCGCTACCGGCGG</td><td>1000</td><td></td><td></td><td></td><td></td><td></td><td></td><td></td><td></td><td></td></tr>
<tr><td>Oligo fwd</td><td>CACCCGCGGCGGCGGCGCTACCGG</td><td></td><td></td><td></td><td></td><td></td><td></td><td></td><td></td><td></td><td></td></tr>
<tr><td>Oligo rev</td><td>AAACCCGGTAGCGCCGCCGCCGCG</td><td></td><td></td><td></td><td></td><td></td><td></td><td></td><td></td><td></td><td></td></tr>
<tr><td>Chromosome</td><td>start</td><td>end</td><td>strand</td><td>MM</td><td>target_seq</td><td>PAM</td><td>alignment</td><td>distance</td><td>position</td><td>gene name</td><td>gene id</td></tr>
<tr><td>GU305914.1</td><td>127365</td><td>127387</td><td>+</td><td>0</td><td>CGCGGCGGCGGCGCTACCGG</td><td>CGG</td><td>||||||||[||||||||||||]PAM</td><td>NA</td><td>Intergenic</td><td>NA</td><td>NA</td></tr>
<tr><td>GU305914.1</td><td>152553</td><td>152575</td><td>+</td><td>3</td><td>TGCGGCGGCGGCGGTGCCGG</td><td>CGG</td><td>-|||||||[|||||-|-||||]PAM</td><td>NA</td><td>Intergenic</td><td>NA</td><td>NA</td></tr>
<tr><td></td><td></td><td></td><td></td><td></td><td></td><td></td><td></td><td></td><td></td><td></td><td></td></tr>
<tr><td>T9629</td><td>GGCGGCGGCGCTACCGGCGGCGG</td><td>1000</td><td></td><td></td><td></td><td></td><td></td><td></td><td></td><td></td><td></td></tr>
<tr><td>Oligo fwd</td><td>CACCGGCGGCGGCGCTACCGGCGG</td><td></td><td></td><td></td><td></td><td></td><td></td><td></td><td></td><td></td><td></td></tr>
<tr><td>Oligo rev</td><td>AAACCCGCCGGTAGCGCCGCCGCC</td><td></td><td></td><td></td><td></td><td></td><td></td><td></td><td></td><td></td><td></td></tr>
<tr><td>Chromosome</td><td>start</td><td>end</td><td>strand</td><td>MM</td><td>target_seq</td><td>PAM</td><td>alignment</td><td>distance</td><td>position</td><td>gene name</td><td>gene id</td></tr>
<tr><td>GU305914.1</td><td>127368</td><td>127390</td><td>+</td><td>0</td><td>GGCGGCGGCGCTACCGGCGG</td><td>CGG</td><td>||||||||[||||||||||||]PAM</td><td>NA</td><td>Intergenic</td><td>NA</td><td>NA</td></tr>
<tr><td></td><td></td><td></td><td></td><td></td><td></td><td></td><td></td><td></td><td></td><td></td><td></td></tr>
<tr><td>T9630</td><td>GGCGGCGGCGATCGCCCTCTCGG</td><td>1000</td><td></td><td></td><td></td><td></td><td></td><td></td><td></td><td></td><td></td></tr>
<tr><td>Oligo fwd</td><td>CACCGGCGGCGGCGATCGCCCTCT</td><td></td><td></td><td></td><td></td><td></td><td></td><td></td><td></td><td></td><td></td></tr>
<tr><td>Oligo rev</td><td>AAACAGAGGGCGATCGCCGCCGCC</td><td></td><td></td><td></td><td></td><td></td><td></td><td></td><td></td><td></td><td></td></tr>
<tr><td>Chromosome</td><td>start</td><td>end</td><td>strand</td><td>MM</td><td>target_seq</td><td>PAM</td><td>alignment</td><td>distance</td><td>position</td><td>gene name</td><td>gene id</td></tr>
<tr><td>GU305914.1</td><td>127383</td><td>127405</td><td>+</td><td>0</td><td>GGCGGCGGCGATCGCCCTCT</td><td>CGG</td><td>||||||||[||||||||||||]PAM</td><td>NA</td><td>Intergenic</td><td>NA</td><td>NA</td></tr>
</table>

| | | | | | | | | | | | | |
|---|---|---|---|---|---|---|---|---|---|---|---|---|
| GU305914.1 | | 189049 | 189071 | - | 4 | GGCGGTCGCGATCGCCCTGG | TGG | \|\|\|\|\|--\|[\|\|\|\|\|\|\|\|\|\|--]PAM | NA | Intergenic | NA | NA |
| T9631 | GGTCTCTCTCTCGCTCTTTCAGG | | 1000 | | | | | | | | | |
| Oligo fwd | CACCGGTCTCTCTCTCGCTCTTTC | | | | | | | | | | | |
| Oligo rev | AAACGAAAGAGCGAGAGAGAGACC | | | | | | | | | | | |
| Chromosome | start | | end | strand | MM | target_seq | PAM | alignment | distance | position | gene name | gene id |
| GU305914.1 | | 127404 | 127426 | + | 0 | GGTCTCTCTCTCGCTCTTTC | AGG | \|\|\|\|\|\|\|\|[\|\|\|\|\|\|\|\|\|\|\|\|]PAM | NA | Intergenic | NA | NA |
| T9632 | CTCTCTCTCGCTCTTTCAGGCGG | | 1000 | | | | | | | | | |
| Oligo fwd | CACCCTCTCTCTCGCTCTTTCAGG | | | | | | | | | | | |
| Oligo rev | AAACCCTGAAAGAGCGAGAGAGAG | | | | | | | | | | | |
| Chromosome | start | | end | strand | MM | target_seq | PAM | alignment | distance | position | gene name | gene id |
| GU305914.1 | | 127407 | 127429 | + | 0 | CTCTCTCTCGCTCTTTCAGG | CGG | \|\|\|\|\|\|\|\|[\|\|\|\|\|\|\|\|\|\|\|\|]PAM | NA | Intergenic | NA | NA |
| T9633 | TCTCTCTCGCTCTTTCAGGCGGG | | 1000 | | | | | | | | | |
| Oligo fwd | CACCTCTCTCTCGCTCTTTCAGGC | | | | | | | | | | | |
| Oligo rev | AAACGCCTGAAAGAGCGAGAGAGA | | | | | | | | | | | |
| Chromosome | start | | end | strand | MM | target_seq | PAM | alignment | distance | position | gene name | gene id |
| GU305914.1 | | 127408 | 127430 | + | 0 | TCTCTCTCGCTCTTTCAGGC | GGG | \|\|\|\|\|\|\|\|[\|\|\|\|\|\|\|\|\|\|\|\|]PAM | NA | Intergenic | NA | NA |
| T9634 | CGCTCTTTCAGGCGGGCCCTCGG | | 1000 | | | | | | | | | |
| Oligo fwd | CACCCGCTCTTTCAGGCGGGCCCT | | | | | | | | | | | |
| Oligo rev | AAACAGGGCCCGCCTGAAAGAGCG | | | | | | | | | | | |
| Chromosome | start | | end | strand | MM | target_seq | PAM | alignment | distance | position | gene name | gene id |
| GU305914.1 | | 127415 | 127437 | + | 0 | CGCTCTTTCAGGCGGGCCCT | CGG | \|\|\|\|\|\|\|\|[\|\|\|\|\|\|\|\|\|\|\|\|]PAM | NA | Intergenic | NA | NA |
| T9635 | GCTCTTTCAGGCGGGCCCTCGGG | | 1000 | | | | | | | | | |
| Oligo fwd | CACCGCTCTTTCAGGCGGGCCCTC | | | | | | | | | | | |
| Oligo rev | AAACGAGGGCCCGCCTGAAAGAGC | | | | | | | | | | | |
| Chromosome | start | | end | strand | MM | target_seq | PAM | alignment | distance | position | gene name | gene id |
| GU305914.1 | | 127416 | 127438 | + | 0 | GCTCTTTCAGGCGGGCCCTC | GGG | \|\|\|\|\|\|\|\|[\|\|\|\|\|\|\|\|\|\|\|\|]PAM | NA | Intergenic | NA | NA |
| T9636 | CTCTTTCAGGCGGGCCCTCGGGG | | 1000 | | | | | | | | | |
| Oligo fwd | CACCCTCTTTCAGGCGGGCCCTCG | | | | | | | | | | | |
| Oligo rev | AAACCGAGGGCCCGCCTGAAAGAG | | | | | | | | | | | |
| Chromosome | start | | end | strand | MM | target_seq | PAM | alignment | distance | position | gene name | gene id |
| GU305914.1 | | 127417 | 127439 | + | 0 | CTCTTTCAGGCGGGCCCTCG | GGG | \|\|\|\|\|\|\|\|[\|\|\|\|\|\|\|\|\|\|\|\|]PAM | NA | Intergenic | NA | NA |
| T9637 | CAGGCGGGCCCTCGGGGACGTGG | | 1000 | | | | | | | | | |
| Oligo fwd | CACCCAGGCGGGCCCTCGGGGACG | | | | | | | | | | | |
| Oligo rev | AAACCGTCCCCGAGGGCCCGCCTG | | | | | | | | | | | |
| Chromosome | start | | end | strand | MM | target_seq | PAM | alignment | distance | position | gene name | gene id |
| GU305914.1 | | 127423 | 127445 | + | 0 | CAGGCGGGCCCTCGGGGACG | TGG | \|\|\|\|\|\|\|\|[\|\|\|\|\|\|\|\|\|\|\|\|]PAM | NA | Intergenic | NA | NA |
| T9638 | GGGCCCTCGGGGACGTGGTTCGG | | 1000 | | | | | | | | | |
| Oligo fwd | CACCGGGCCCTCGGGGACGTGGTT | | | | | | | | | | | |
| Oligo rev | AAACAACCACGTCCCCGAGGGCCC | | | | | | | | | | | |
| Chromosome | start | | end | strand | MM | target_seq | PAM | alignment | distance | position | gene name | gene id |
| GU305914.1 | | 127428 | 127450 | + | 0 | GGGCCCTCGGGGACGTGGTT | CGG | \|\|\|\|\|\|\|\|[\|\|\|\|\|\|\|\|\|\|\|\|]PAM | NA | Intergenic | NA | NA |
| T9639 | GGCCCTCGGGGACGTGGTTCGGG | | 1000 | | | | | | | | | |
| Oligo fwd | CACCGGCCCTCGGGGACGTGGTTC | | | | | | | | | | | |
| Oligo rev | AAACGAACCACGTCCCCGAGGGCC | | | | | | | | | | | |
| Chromosome | start | | end | strand | MM | target_seq | PAM | alignment | distance | position | gene name | gene id |
| GU305914.1 | | 127429 | 127451 | + | 0 | GGCCCTCGGGGACGTGGTTC | GGG | \|\|\|\|\|\|\|\|[\|\|\|\|\|\|\|\|\|\|\|\|]PAM | NA | Intergenic | NA | NA |
| T9640 | TCGGGGACGTGGTTCGGGAGCGG | | 1000 | | | | | | | | | |
| Oligo fwd | CACCTCGGGGACGTGGTTCGGGAG | | | | | | | | | | | |
| Oligo rev | AAACCTCCCGAACCACGTCCCCGA | | | | | | | | | | | |
| Chromosome | start | | end | strand | MM | target_seq | PAM | alignment | distance | position | gene name | gene id |
| GU305914.1 | | 127434 | 127456 | + | 0 | TCGGGGACGTGGTTCGGGAG | CGG | \|\|\|\|\|\|\|\|[\|\|\|\|\|\|\|\|\|\|\|\|]PAM | NA | Intergenic | NA | NA |
| T9641 | GGGACGTGGTTCGGGAGCGGCGG | | 1000 | | | | | | | | | |
| Oligo fwd | CACCGGGACGTGGTTCGGGAGCGG | | | | | | | | | | | |
| Oligo rev | AAACCCGCTCCCGAACCACGTCCC | | | | | | | | | | | |
| Chromosome | start | | end | strand | MM | target_seq | PAM | alignment | distance | position | gene name | gene id |
| GU305914.1 | | 127437 | 127459 | + | 0 | GGGACGTGGTTCGGGAGCGG | CGG | \|\|\|\|\|\|\|\|[\|\|\|\|\|\|\|\|\|\|\|\|]PAM | NA | Intergenic | NA | NA |
| T9642 | GTTCGGGAGCGGCGGCCGCGAGG | | 1000 | | | | | | | | | |
| Oligo fwd | CACCGTTCGGGAGCGGCGGCCGCG | | | | | | | | | | | |
| Oligo rev | AAACCGCGGCCGCCGCTCCCGAAC | | | | | | | | | | | |
| Chromosome | start | | end | strand | MM | target_seq | PAM | alignment | distance | position | gene name | gene id |
| GU305914.1 | | 127445 | 127467 | + | 0 | GTTCGGGAGCGGCGGCCGCG | AGG | \|\|\|\|\|\|\|\|[\|\|\|\|\|\|\|\|\|\|\|\|]PAM | NA | Intergenic | NA | NA |
| T9643 | GGAGCGGCGGCCGCGAGGATCGG | | 1000 | | | | | | | | | |
| Oligo fwd | CACCGGAGCGGCGGCCGCGAGGAT | | | | | | | | | | | |
| Oligo rev | AAACATCCTCGCGGCCGCCGCTCC | | | | | | | | | | | |
| Chromosome | start | | end | strand | MM | target_seq | PAM | alignment | distance | position | gene name | gene id |
| GU305914.1 | | 127450 | 127472 | + | 0 | GGAGCGGCGGCCGCGAGGAT | CGG | \|\|\|\|\|\|\|\|[\|\|\|\|\|\|\|\|\|\|\|\|]PAM | NA | Intergenic | NA | NA |
| T9644 | GAGCGGCGGCCGCGAGGATCGGG | | 1000 | | | | | | | | | |
| Oligo fwd | CACCGAGCGGCGGCCGCGAGGATC | | | | | | | | | | | |
| Oligo rev | AAACGATCCTCGCGGCCGCCGCTC | | | | | | | | | | | |
| Chromosome | start | | end | strand | MM | target_seq | PAM | alignment | distance | position | gene name | gene id |
| GU305914.1 | | 127451 | 127473 | + | 0 | GAGCGGCGGCCGCGAGGATC | GGG | \|\|\|\|\|\|\|\|[\|\|\|\|\|\|\|\|\|\|\|\|]PAM | NA | Intergenic | NA | NA |
| T9645 | GCGGCCGCGAGGATCGGGCGCGG | | 1000 | | | | | | | | | |
| Oligo fwd | CACCGCGGCCGCGAGGATCGGGCG | | | | | | | | | | | |
| Oligo rev | AAACCGCCCGATCCTCGCGGCCGC | | | | | | | | | | | |

Chromosome start end strand MM target_seq PAM alignment distance position gene name gene id
GU305914.1 127456 127478 + 0 GCGGCCGCGAGGATCGGGCG CGG ||||||||[||||||||||||]PAM NA Intergenic NA NA

T9646 GCCGCGAGGATCGGGCGCGGTGG 1000
Oligo fwd CACCGCCGCGAGGATCGGGCGCGG
Oligo rev AAACCCGCGCCCGATCCTCGCGGC
Chromosome start end strand MM target_seq PAM alignment distance position gene name gene id
GU305914.1 127459 127481 + 0 GCCGCGAGGATCGGGCGCGG TGG ||||||||[||||||||||||]PAM NA Intergenic NA NA

T9647 GCGAGGATCGGGCGCGGTGGCGG 1000
Oligo fwd CACCGCGAGGATCGGGCGCGGTGG
Oligo rev AAACCCACCGCGCCCGATCCTCGC
Chromosome start end strand MM target_seq PAM alignment distance position gene name gene id
GU305914.1 127462 127484 + 0 GCGAGGATCGGGCGCGGTGG CGG ||||||||[||||||||||||]PAM NA Intergenic NA NA

T9648 CGAGGATCGGGCGCGGTGGCGGG 1000
Oligo fwd CACCCGAGGATCGGGCGCGGTGGC
Oligo rev AAACGCCACCGCGCCCGATCCTCG
Chromosome start end strand MM target_seq PAM alignment distance position gene name gene id
GU305914.1 127463 127485 + 0 CGAGGATCGGGCGCGGTGGC GGG ||||||||[||||||||||||]PAM NA Intergenic NA NA

T9649 GAGGATCGGGCGCGGTGGCGGGG 1000
Oligo fwd CACCGAGGATCGGGCGCGGTGGCG
Oligo rev AAACCGCCACCGCGCCCGATCCTC
Chromosome start end strand MM target_seq PAM alignment distance position gene name gene id
GU305914.1 127464 127486 + 0 GAGGATCGGGCGCGGTGGCG GGG ||||||||[||||||||||||]PAM NA Intergenic NA NA

T9650 GGGATGATGAGCGCGAGCGATGG 1000
Oligo fwd CACCGGGATGATGAGCGCGAGCGA
Oligo rev AAACTCGCTCGCGCTCATCATCCC
Chromosome start end strand MM target_seq PAM alignment distance position gene name gene id
GU305914.1 127484 127506 + 0 GGGATGATGAGCGCGAGCGA TGG ||||||||[||||||||||||]PAM NA Intergenic NA NA

T9651 GGATGATGAGCGCGAGCGATGGG 1000
Oligo fwd CACCGGATGATGAGCGCGAGCGAT
Oligo rev AAACATCGCTCGCGCTCATCATCC
Chromosome start end strand MM target_seq PAM alignment distance position gene name gene id
GU305914.1 127485 127507 + 0 GGATGATGAGCGCGAGCGAT GGG ||||||||[||||||||||||]PAM NA Intergenic NA NA

T9652 GAGCGCGAGCGATGGGTCGCCGG 1000
Oligo fwd CACCGAGCGCGAGCGATGGGTCGC
Oligo rev AAACGCGACCCATCGCTCGCGCTC
Chromosome start end strand MM target_seq PAM alignment distance position gene name gene id
GU305914.1 127492 127514 + 0 GAGCGCGAGCGATGGGTCGC CGG ||||||||[||||||||||||]PAM NA Intergenic NA NA

T9653 GTCTCTGTCCTGTCTCGATCCGG 1000
Oligo fwd CACCGTCTCTGTCCTGTCTCGATC
Oligo rev AAACGATCGAGACAGGACAGAGAC
Chromosome start end strand MM target_seq PAM alignment distance position gene name gene id
GU305914.1 127516 127538 + 0 GTCTCTGTCCTGTCTCGATC CGG ||||||||[||||||||||||]PAM NA Intergenic NA NA

T9654 TGCGCGCGTCAAGAGCGCTCCGG 1000
Oligo fwd CACCTGCGCGCGTCAAGAGCGCTC
Oligo rev AAACGAGCGCTCTTGACGCGCGCA
Chromosome start end strand MM target_seq PAM alignment distance position gene name gene id
GU305914.1 127561 127583 + 0 TGCGCGCGTCAAGAGCGCTC CGG ||||||||[||||||||||||]PAM NA Intergenic NA NA

T9655 CCGAGAGATCTCGCCGTTCGAGG 1000
Oligo fwd CACCCCGAGAGATCTCGCCGTTCG
Oligo rev AAACCGAACGGCGAGATCTCTCGG
Chromosome start end strand MM target_seq PAM alignment distance position gene name gene id
GU305914.1 127609 127631 + 0 CCGAGAGATCTCGCCGTTCG AGG ||||||||[||||||||||||]PAM NA Intergenic NA NA

T9656 CGAGGACTCCCGCCTCTCCCAGG 1000
Oligo fwd CACCCGAGGACTCCCGCCTCTCCC
Oligo rev AAACGGGAGAGGCGGGAGTCCTCG
Chromosome start end strand MM target_seq PAM alignment distance position gene name gene id
GU305914.1 127627 127649 + 0 CGAGGACTCCCGCCTCTCCC AGG ||||||||[||||||||||||]PAM NA Intergenic NA NA

T9657 CCAGGCCGTCACCGTCGACGAGG 1000
Oligo fwd CACCCCAGGCCGTCACCGTCGACG
Oligo rev AAACCGTCGACGGTGACGGCCTGG
Chromosome start end strand MM target_seq PAM alignment distance position gene name gene id
GU305914.1 127645 127667 + 0 CCAGGCCGTCACCGTCGACG AGG ||||||||[||||||||||||]PAM NA Intergenic NA NA

T9658 AGCACATCTCCTCGATCTTCAGG 1000
Oligo fwd CACCAGCACATCTCCTCGATCTTC
Oligo rev AAACGAAGATCGAGGAGATGTGCT
Chromosome start end strand MM target_seq PAM alignment distance position gene name gene id
GU305914.1 127668 127690 + 0 AGCACATCTCCTCGATCTTC AGG ||||||||[||||||||||||]PAM NA Intergenic NA NA

T9659 CTCGATCTTCAGGACGCTGATGG 1000
Oligo fwd CACCCTCGATCTTCAGGACGCTGA
Oligo rev AAACTCAGCGTCCTGAAGATCGAG
Chromosome start end strand MM target_seq PAM alignment distance position gene name gene id
GU305914.1 127678 127700 + 0 CTCGATCTTCAGGACGCTGA TGG ||||||||[||||||||||||]PAM NA Intergenic NA NA

T9660 CTTCAGGACGCTGATGGCCGCGG 1000
Oligo fwd CACCCTTCAGGACGCTGATGGCCG

<table>
<tr><td>Oligo rev</td><td>AAACCGGCCATCAGCGTCCTGAAG</td><td></td><td></td><td></td><td></td><td></td><td></td><td></td><td></td><td></td><td></td></tr>
<tr><td>Chromosome</td><td>start</td><td>end</td><td>strand</td><td>MM</td><td>target_seq</td><td>PAM</td><td>alignment</td><td>distance</td><td>position</td><td>gene name</td><td>gene id</td></tr>
<tr><td>GU305914.1</td><td>127684</td><td>127706</td><td>+</td><td>0</td><td>CTTCAGGACGCTGATGGCCG</td><td>CGG</td><td>||||||||[||||||||||||]PAM</td><td>NA</td><td>Intergenic</td><td>NA</td><td>NA</td></tr>
<tr><td>GU305914.1</td><td>173322</td><td>173344</td><td>-</td><td>4</td><td>CTCCGGGATCCTGATGGCCG</td><td>TGG</td><td>||-|-|||[--||||||||||]PAM</td><td>NA</td><td>Intergenic</td><td>NA</td><td>NA</td></tr>
<tr><td>T9661</td><td>TTCAGGACGCTGATGGCCGCGGG</td><td>1000</td><td></td><td></td><td></td><td></td><td></td><td></td><td></td><td></td><td></td></tr>
<tr><td>Oligo fwd</td><td>CACCTTCAGGACGCTGATGGCCGC</td><td></td><td></td><td></td><td></td><td></td><td></td><td></td><td></td><td></td><td></td></tr>
<tr><td>Oligo rev</td><td>AAACGCGGCCATCAGCGTCCTGAA</td><td></td><td></td><td></td><td></td><td></td><td></td><td></td><td></td><td></td><td></td></tr>
<tr><td>Chromosome</td><td>start</td><td>end</td><td>strand</td><td>MM</td><td>target_seq</td><td>PAM</td><td>alignment</td><td>distance</td><td>position</td><td>gene name</td><td>gene id</td></tr>
<tr><td>GU305914.1</td><td>127685</td><td>127707</td><td>+</td><td>0</td><td>TTCAGGACGCTGATGGCCGC</td><td>GGG</td><td>||||||||[||||||||||||]PAM</td><td>NA</td><td>Intergenic</td><td>NA</td><td>NA</td></tr>
<tr><td>T9662</td><td>ATGGCCGCGGGACCCGACCCCGG</td><td>1000</td><td></td><td></td><td></td><td></td><td></td><td></td><td></td><td></td><td></td></tr>
<tr><td>Oligo fwd</td><td>CACCATGGCCGCGGGACCCGACCC</td><td></td><td></td><td></td><td></td><td></td><td></td><td></td><td></td><td></td><td></td></tr>
<tr><td>Oligo rev</td><td>AAACGGGTCGGGTCCCGCGGCCAT</td><td></td><td></td><td></td><td></td><td></td><td></td><td></td><td></td><td></td><td></td></tr>
<tr><td>Chromosome</td><td>start</td><td>end</td><td>strand</td><td>MM</td><td>target_seq</td><td>PAM</td><td>alignment</td><td>distance</td><td>position</td><td>gene name</td><td>gene id</td></tr>
<tr><td>GU305914.1</td><td>127697</td><td>127719</td><td>+</td><td>0</td><td>ATGGCCGCGGGACCCGACCC</td><td>CGG</td><td>||||||||[||||||||||||]PAM</td><td>NA</td><td>Intergenic</td><td>NA</td><td>NA</td></tr>
<tr><td>T9663</td><td>CGACCCCGGCGCCACCGACGAGG</td><td>1000</td><td></td><td></td><td></td><td></td><td></td><td></td><td></td><td></td><td></td></tr>
<tr><td>Oligo fwd</td><td>CACCCGACCCCGGCGCCACCGACG</td><td></td><td></td><td></td><td></td><td></td><td></td><td></td><td></td><td></td><td></td></tr>
<tr><td>Oligo rev</td><td>AAACCGTCGGTGGCGCCGGGGTCG</td><td></td><td></td><td></td><td></td><td></td><td></td><td></td><td></td><td></td><td></td></tr>
<tr><td>Chromosome</td><td>start</td><td>end</td><td>strand</td><td>MM</td><td>target_seq</td><td>PAM</td><td>alignment</td><td>distance</td><td>position</td><td>gene name</td><td>gene id</td></tr>
<tr><td>GU305914.1</td><td>127711</td><td>127733</td><td>+</td><td>0</td><td>CGACCCCGGCGCCACCGACG</td><td>AGG</td><td>||||||||[||||||||||||]PAM</td><td>NA</td><td>Intergenic</td><td>NA</td><td>NA</td></tr>
<tr><td>T9664</td><td>CGGCGCCACCGACGAGGACAAGG</td><td>1000</td><td></td><td></td><td></td><td></td><td></td><td></td><td></td><td></td><td></td></tr>
<tr><td>Oligo fwd</td><td>CACCCGGCGCCACCGACGAGGACA</td><td></td><td></td><td></td><td></td><td></td><td></td><td></td><td></td><td></td><td></td></tr>
<tr><td>Oligo rev</td><td>AAACTGTCCTCGTCGGTGGCGCCG</td><td></td><td></td><td></td><td></td><td></td><td></td><td></td><td></td><td></td><td></td></tr>
<tr><td>Chromosome</td><td>start</td><td>end</td><td>strand</td><td>MM</td><td>target_seq</td><td>PAM</td><td>alignment</td><td>distance</td><td>position</td><td>gene name</td><td>gene id</td></tr>
<tr><td>GU305914.1</td><td>127717</td><td>127739</td><td>+</td><td>0</td><td>CGGCGCCACCGACGAGGACA</td><td>AGG</td><td>||||||||[||||||||||||]PAM</td><td>NA</td><td>Intergenic</td><td>NA</td><td>NA</td></tr>
<tr><td>T9665</td><td>CGTCCTCTGCCGCCTGCTCATGG</td><td>1000</td><td></td><td></td><td></td><td></td><td></td><td></td><td></td><td></td><td></td></tr>
<tr><td>Oligo fwd</td><td>CACCCGTCCTCTGCCGCCTGCTCA</td><td></td><td></td><td></td><td></td><td></td><td></td><td></td><td></td><td></td><td></td></tr>
<tr><td>Oligo rev</td><td>AAACTGAGCAGGCGGCAGAGGACG</td><td></td><td></td><td></td><td></td><td></td><td></td><td></td><td></td><td></td><td></td></tr>
<tr><td>Chromosome</td><td>start</td><td>end</td><td>strand</td><td>MM</td><td>target_seq</td><td>PAM</td><td>alignment</td><td>distance</td><td>position</td><td>gene name</td><td>gene id</td></tr>
<tr><td>GU305914.1</td><td>127747</td><td>127769</td><td>+</td><td>0</td><td>CGTCCTCTGCCGCCTGCTCA</td><td>TGG</td><td>||||||||[||||||||||||]PAM</td><td>NA</td><td>Intergenic</td><td>NA</td><td>NA</td></tr>
<tr><td>T9666</td><td>GTCCTCTGCCGCCTGCTCATGGG</td><td>1000</td><td></td><td></td><td></td><td></td><td></td><td></td><td></td><td></td><td></td></tr>
<tr><td>Oligo fwd</td><td>CACCGTCCTCTGCCGCCTGCTCAT</td><td></td><td></td><td></td><td></td><td></td><td></td><td></td><td></td><td></td><td></td></tr>
<tr><td>Oligo rev</td><td>AAACATGAGCAGGCGGCAGAGGAC</td><td></td><td></td><td></td><td></td><td></td><td></td><td></td><td></td><td></td><td></td></tr>
<tr><td>Chromosome</td><td>start</td><td>end</td><td>strand</td><td>MM</td><td>target_seq</td><td>PAM</td><td>alignment</td><td>distance</td><td>position</td><td>gene name</td><td>gene id</td></tr>
<tr><td>GU305914.1</td><td>127748</td><td>127770</td><td>+</td><td>0</td><td>GTCCTCTGCCGCCTGCTCAT</td><td>GGG</td><td>||||||||[||||||||||||]PAM</td><td>NA</td><td>Intergenic</td><td>NA</td><td>NA</td></tr>
<tr><td>T9667</td><td>TGCCCTGCTACTGCGACGAGTGG</td><td>1000</td><td></td><td></td><td></td><td></td><td></td><td></td><td></td><td></td><td></td></tr>
<tr><td>Oligo fwd</td><td>CACCTGCCCTGCTACTGCGACGAG</td><td></td><td></td><td></td><td></td><td></td><td></td><td></td><td></td><td></td><td></td></tr>
<tr><td>Oligo rev</td><td>AAACCTCGTCGCAGTAGCAGGGCA</td><td></td><td></td><td></td><td></td><td></td><td></td><td></td><td></td><td></td><td></td></tr>
<tr><td>Chromosome</td><td>start</td><td>end</td><td>strand</td><td>MM</td><td>target_seq</td><td>PAM</td><td>alignment</td><td>distance</td><td>position</td><td>gene name</td><td>gene id</td></tr>
<tr><td>GU305914.1</td><td>127782</td><td>127804</td><td>+</td><td>0</td><td>TGCCCTGCTACTGCGACGAG</td><td>TGG</td><td>||||||||[||||||||||||]PAM</td><td>NA</td><td>Intergenic</td><td>NA</td><td>NA</td></tr>
<tr><td>T9668</td><td>GCCCTGCTACTGCGACGAGTGGG</td><td>1000</td><td></td><td></td><td></td><td></td><td></td><td></td><td></td><td></td><td></td></tr>
<tr><td>Oligo fwd</td><td>CACCGCCCTGCTACTGCGACGAGT</td><td></td><td></td><td></td><td></td><td></td><td></td><td></td><td></td><td></td><td></td></tr>
<tr><td>Oligo rev</td><td>AAACACTCGTCGCAGTAGCAGGGC</td><td></td><td></td><td></td><td></td><td></td><td></td><td></td><td></td><td></td><td></td></tr>
<tr><td>Chromosome</td><td>start</td><td>end</td><td>strand</td><td>MM</td><td>target_seq</td><td>PAM</td><td>alignment</td><td>distance</td><td>position</td><td>gene name</td><td>gene id</td></tr>
<tr><td>GU305914.1</td><td>127783</td><td>127805</td><td>+</td><td>0</td><td>GCCCTGCTACTGCGACGAGT</td><td>GGG</td><td>||||||||[||||||||||||]PAM</td><td>NA</td><td>Intergenic</td><td>NA</td><td>NA</td></tr>
<tr><td>T9669</td><td>CAAGTGCGCCTACCGCTGCTCGG</td><td>1000</td><td></td><td></td><td></td><td></td><td></td><td></td><td></td><td></td><td></td></tr>
<tr><td>Oligo fwd</td><td>CACCCAAGTGCGCCTACCGCTGCT</td><td></td><td></td><td></td><td></td><td></td><td></td><td></td><td></td><td></td><td></td></tr>
<tr><td>Oligo rev</td><td>AAACAGCAGCGGTAGGCGCACTTG</td><td></td><td></td><td></td><td></td><td></td><td></td><td></td><td></td><td></td><td></td></tr>
<tr><td>Chromosome</td><td>start</td><td>end</td><td>strand</td><td>MM</td><td>target_seq</td><td>PAM</td><td>alignment</td><td>distance</td><td>position</td><td>gene name</td><td>gene id</td></tr>
<tr><td>GU305914.1</td><td>127825</td><td>127847</td><td>+</td><td>0</td><td>CAAGTGCGCCTACCGCTGCT</td><td>CGG</td><td>||||||||[||||||||||||]PAM</td><td>NA</td><td>Intergenic</td><td>NA</td><td>NA</td></tr>
<tr><td>T9670</td><td>AAGTGCGCCTACCGCTGCTCGGG</td><td>1000</td><td></td><td></td><td></td><td></td><td></td><td></td><td></td><td></td><td></td></tr>
<tr><td>Oligo fwd</td><td>CACCAAGTGCGCCTACCGCTGCTC</td><td></td><td></td><td></td><td></td><td></td><td></td><td></td><td></td><td></td><td></td></tr>
<tr><td>Oligo rev</td><td>AAACGAGCAGCGGTAGGCGCACTT</td><td></td><td></td><td></td><td></td><td></td><td></td><td></td><td></td><td></td><td></td></tr>
<tr><td>Chromosome</td><td>start</td><td>end</td><td>strand</td><td>MM</td><td>target_seq</td><td>PAM</td><td>alignment</td><td>distance</td><td>position</td><td>gene name</td><td>gene id</td></tr>
<tr><td>GU305914.1</td><td>127826</td><td>127848</td><td>+</td><td>0</td><td>AAGTGCGCCTACCGCTGCTC</td><td>GGG</td><td>||||||||[||||||||||||]PAM</td><td>NA</td><td>Intergenic</td><td>NA</td><td>NA</td></tr>
<tr><td>T9671</td><td>CGCCTACCGCTGCTCGGGACCGG</td><td>1000</td><td></td><td></td><td></td><td></td><td></td><td></td><td></td><td></td><td></td></tr>
<tr><td>Oligo fwd</td><td>CACCCGCCTACCGCTGCTCGGGAC</td><td></td><td></td><td></td><td></td><td></td><td></td><td></td><td></td><td></td><td></td></tr>
<tr><td>Oligo rev</td><td>AAACGTCCCGAGCAGCGGTAGGCG</td><td></td><td></td><td></td><td></td><td></td><td></td><td></td><td></td><td></td><td></td></tr>
<tr><td>Chromosome</td><td>start</td><td>end</td><td>strand</td><td>MM</td><td>target_seq</td><td>PAM</td><td>alignment</td><td>distance</td><td>position</td><td>gene name</td><td>gene id</td></tr>
<tr><td>GU305914.1</td><td>127831</td><td>127853</td><td>+</td><td>0</td><td>CGCCTACCGCTGCTCGGGAC</td><td>CGG</td><td>||||||||[||||||||||||]PAM</td><td>NA</td><td>Intergenic</td><td>NA</td><td>NA</td></tr>
<tr><td>T9672</td><td>CACCGCTCGCGCTGCCGCTGCGG</td><td>1000</td><td></td><td></td><td></td><td></td><td></td><td></td><td></td><td></td><td></td></tr>
<tr><td>Oligo fwd</td><td>CACCCACCGCTCGCGCTGCCGCTG</td><td></td><td></td><td></td><td></td><td></td><td></td><td></td><td></td><td></td><td></td></tr>
<tr><td>Oligo rev</td><td>AAACCAGCGGCAGCGCGAGCGGTG</td><td></td><td></td><td></td><td></td><td></td><td></td><td></td><td></td><td></td><td></td></tr>
<tr><td>Chromosome</td><td>start</td><td>end</td><td>strand</td><td>MM</td><td>target_seq</td><td>PAM</td><td>alignment</td><td>distance</td><td>position</td><td>gene name</td><td>gene id</td></tr>
<tr><td>GU305914.1</td><td>127865</td><td>127887</td><td>+</td><td>0</td><td>CACCGCTCGCGCTGCCGCTG</td><td>CGG</td><td>||||||||[||||||||||||]PAM</td><td>NA</td><td>Intergenic</td><td>NA</td><td>NA</td></tr>
<tr><td>T9673</td><td>GCGCTGCCGCTGCGGCGCCGAGG</td><td>1000</td><td></td><td></td><td></td><td></td><td></td><td></td><td></td><td></td><td></td></tr>
<tr><td>Oligo fwd</td><td>CACCGCGCTGCCGCTGCGGCGCCG</td><td></td><td></td><td></td><td></td><td></td><td></td><td></td><td></td><td></td><td></td></tr>
<tr><td>Oligo rev</td><td>AAACCGGCGCCGCAGCGGCAGCGC</td><td></td><td></td><td></td><td></td><td></td><td></td><td></td><td></td><td></td><td></td></tr>
<tr><td>Chromosome</td><td>start</td><td>end</td><td>strand</td><td>MM</td><td>target_seq</td><td>PAM</td><td>alignment</td><td>distance</td><td>position</td><td>gene name</td><td>gene id</td></tr>
<tr><td>GU305914.1</td><td>127873</td><td>127895</td><td>+</td><td>0</td><td>GCGCTGCCGCTGCGGCGCCG</td><td>AGG</td><td>||||||||[||||||||||||]PAM</td><td>NA</td><td>Intergenic</td><td>NA</td><td>NA</td></tr>
<tr><td>T9674</td><td>CGCTGCCGCTGCGGCGCCGAGGG</td><td>1000</td><td></td><td></td><td></td><td></td><td></td><td></td><td></td><td></td><td></td></tr>
<tr><td>Oligo fwd</td><td>CACCCGCTGCCGCTGCGGCGCCGA</td><td></td><td></td><td></td><td></td><td></td><td></td><td></td><td></td><td></td><td></td></tr>
<tr><td>Oligo rev</td><td>AAACTCGGCGCCGCAGCGGCAGCG</td><td></td><td></td><td></td><td></td><td></td><td></td><td></td><td></td><td></td><td></td></tr>
<tr><td>Chromosome</td><td>start</td><td>end</td><td>strand</td><td>MM</td><td>target_seq</td><td>PAM</td><td>alignment</td><td>distance</td><td>position</td><td>gene name</td><td>gene id</td></tr>
<tr><td>GU305914.1</td><td>127874</td><td>127896</td><td>+</td><td>0</td><td>CGCTGCCGCTGCGGCGCCGA</td><td>GGG</td><td>||||||||[||||||||||||]PAM</td><td>NA</td><td>Intergenic</td><td>NA</td><td>NA</td></tr>
<tr><td>GU305914.1</td><td>128755</td><td>128777</td><td>+</td><td>4</td><td>CGATGCCTCCGCGGCGTCGA</td><td>CGG</td><td>||-||||-[|-||||||-|||]PAM</td><td>NA</td><td>Intergenic</td><td>NA</td><td>NA</td></tr>
</table>

| | | | | | | | | | | | |
|---|---|---|---|---|---|---|---|---|---|---|---|
| T9675 | TGCCGCTGCGGCGCCGAGGGCGG | 1000 | | | | | | | | | |
| Oligo fwd | CACCTGCCGCTGCGGCGCCGAGGG | | | | | | | | | | |
| Oligo rev | AAACCCCTCGGCGCCGCAGCGGCA | | | | | | | | | | |
| Chromosome | start | end | strand | MM | target_seq | PAM | alignment | distance | position | gene name | gene id |
| GU305914.1 | 127877 | 127899 | + | 0 | TGCCGCTGCGGCGCCGAGGG | CGG | \|\|\|\|\|\|\|\|[\|\|\|\|\|\|\|\|\|\|\|\|]PAM | NA | Intergenic | NA | NA |
| T9676 | CGCTGCGGCGCCGAGGGCGGCGG | 1000 | | | | | | | | | |
| Oligo fwd | CACCCGCTGCGGCGCCGAGGGCGG | | | | | | | | | | |
| Oligo rev | AAACCCGCCCTCGGCGCCGCAGCG | | | | | | | | | | |
| Chromosome | start | end | strand | MM | target_seq | PAM | alignment | distance | position | gene name | gene id |
| GU305914.1 | 127880 | 127902 | + | 0 | CGCTGCGGCGCCGAGGGCGG | CGG | \|\|\|\|\|\|\|\|[\|\|\|\|\|\|\|\|\|\|\|\|]PAM | NA | Intergenic | NA | NA |
| T9677 | GCTGCGGCGCCGAGGGCGGCGGG | 1000 | | | | | | | | | |
| Oligo fwd | CACCGCTGCGGCGCCGAGGGCGGC | | | | | | | | | | |
| Oligo rev | AAACGCCGCCCTCGGCGCCGCAGC | | | | | | | | | | |
| Chromosome | start | end | strand | MM | target_seq | PAM | alignment | distance | position | gene name | gene id |
| GU305914.1 | 127881 | 127903 | + | 0 | GCTGCGGCGCCGAGGGCGGC | GGG | \|\|\|\|\|\|\|\|[\|\|\|\|\|\|\|\|\|\|\|\|]PAM | NA | Intergenic | NA | NA |
| T9678 | ACTACACCACGCACGTGTTTCGG | 1000 | | | | | | | | | |
| Oligo fwd | CACCACTACACCACGCACGTGTTT | | | | | | | | | | |
| Oligo rev | AAACAAACACGTGCGTGGTGTAGT | | | | | | | | | | |
| Chromosome | start | end | strand | MM | target_seq | PAM | alignment | distance | position | gene name | gene id |
| GU305914.1 | 127929 | 127951 | + | 0 | ACTACACCACGCACGTGTTT | CGG | \|\|\|\|\|\|\|\|[\|\|\|\|\|\|\|\|\|\|\|\|]PAM | NA | Intergenic | NA | NA |
| T9679 | CTACACCACGCACGTGTTTCGGG | 1000 | | | | | | | | | |
| Oligo fwd | CACCCTACACCACGCACGTGTTTC | | | | | | | | | | |
| Oligo rev | AAACGAAACACGTGCGTGGTGTAG | | | | | | | | | | |
| Chromosome | start | end | strand | MM | target_seq | PAM | alignment | distance | position | gene name | gene id |
| GU305914.1 | 127930 | 127952 | + | 0 | CTACACCACGCACGTGTTTC | GGG | \|\|\|\|\|\|\|\|[\|\|\|\|\|\|\|\|\|\|\|\|]PAM | NA | Intergenic | NA | NA |
| T9680 | TACACCACGCACGTGTTTCGGGG | 1000 | | | | | | | | | |
| Oligo fwd | CACCTACACCACGCACGTGTTTCG | | | | | | | | | | |
| Oligo rev | AAACCGAAACACGTGCGTGGTGTA | | | | | | | | | | |
| Chromosome | start | end | strand | MM | target_seq | PAM | alignment | distance | position | gene name | gene id |
| GU305914.1 | 127931 | 127953 | + | 0 | TACACCACGCACGTGTTTCG | GGG | \|\|\|\|\|\|\|\|[\|\|\|\|\|\|\|\|\|\|\|\|]PAM | NA | Intergenic | NA | NA |
| T9681 | ACACCACGCACGTGTTTCGGGGG | 1000 | | | | | | | | | |
| Oligo fwd | CACCACACCACGCACGTGTTTCGG | | | | | | | | | | |
| Oligo rev | AAACCCGAAACACGTGCGTGGTGT | | | | | | | | | | |
| Chromosome | start | end | strand | MM | target_seq | PAM | alignment | distance | position | gene name | gene id |
| GU305914.1 | 127932 | 127954 | + | 0 | ACACCACGCACGTGTTTCGG | GGG | \|\|\|\|\|\|\|\|[\|\|\|\|\|\|\|\|\|\|\|\|]PAM | NA | Intergenic | NA | NA |
| T9682 | GGCTGCTCTCGCTCTCCGAGTGG | 1000 | | | | | | | | | |
| Oligo fwd | CACCGGCTGCTCTCGCTCTCCGAG | | | | | | | | | | |
| Oligo rev | AAACCTCGGAGAGCGAGAGCAGCC | | | | | | | | | | |
| Chromosome | start | end | strand | MM | target_seq | PAM | alignment | distance | position | gene name | gene id |
| GU305914.1 | 127953 | 127975 | + | 0 | GGCTGCTCTCGCTCTCCGAG | TGG | \|\|\|\|\|\|\|\|[\|\|\|\|\|\|\|\|\|\|\|\|]PAM | NA | Intergenic | NA | NA |
| T9683 | GTAACGCCTTTCGCTCCGACCGG | 1000 | | | | | | | | | |
| Oligo fwd | CACCGTAACGCCTTTCGCTCCGAC | | | | | | | | | | |
| Oligo rev | AAACGTCGGAGCGAAAGGCGTTAC | | | | | | | | | | |
| Chromosome | start | end | strand | MM | target_seq | PAM | alignment | distance | position | gene name | gene id |
| GU305914.1 | 128007 | 128029 | + | 0 | GTAACGCCTTTCGCTCCGAC | CGG | \|\|\|\|\|\|\|\|[\|\|\|\|\|\|\|\|\|\|\|\|]PAM | NA | Intergenic | NA | NA |
| T9684 | TCGCTCCGACCGGTATGTCATGG | 1000 | | | | | | | | | |
| Oligo fwd | CACCTCGCTCCGACCGGTATGTCA | | | | | | | | | | |
| Oligo rev | AAACTGACATACCGGTCGGAGCGA | | | | | | | | | | |
| Chromosome | start | end | strand | MM | target_seq | PAM | alignment | distance | position | gene name | gene id |
| GU305914.1 | 128017 | 128039 | + | 0 | TCGCTCCGACCGGTATGTCA | TGG | \|\|\|\|\|\|\|\|[\|\|\|\|\|\|\|\|\|\|\|\|]PAM | NA | Intergenic | NA | NA |
| T9685 | ATTTCCTCGTCTGCCTCGCCAGG | 1000 | | | | | | | | | |
| Oligo fwd | CACCATTTCCTCGTCTGCCTCGCC | | | | | | | | | | |
| Oligo rev | AAACGGCGAGGCAGACGAGGAAAT | | | | | | | | | | |
| Chromosome | start | end | strand | MM | target_seq | PAM | alignment | distance | position | gene name | gene id |
| GU305914.1 | 128085 | 128107 | + | 0 | ATTTCCTCGTCTGCCTCGCC | AGG | \|\|\|\|\|\|\|\|[\|\|\|\|\|\|\|\|\|\|\|\|]PAM | NA | Intergenic | NA | NA |
| T9686 | AACTCCATGACCGCGCACCTCGG | 1000 | | | | | | | | | |
| Oligo fwd | CACCAACTCCATGACCGCGCACCT | | | | | | | | | | |
| Oligo rev | AAACAGGTGCGCGGTCATGGAGTT | | | | | | | | | | |
| Chromosome | start | end | strand | MM | target_seq | PAM | alignment | distance | position | gene name | gene id |
| GU305914.1 | 128141 | 128163 | + | 0 | AACTCCATGACCGCGCACCT | CGG | \|\|\|\|\|\|\|\|[\|\|\|\|\|\|\|\|\|\|\|\|]PAM | NA | Intergenic | NA | NA |
| T9687 | CATGACCGCGCACCTCGGTCCGG | 1000 | | | | | | | | | |
| Oligo fwd | CACCCATGACCGCGCACCTCGGTC | | | | | | | | | | |
| Oligo rev | AAACGACCGAGGTGCGCGGTCATG | | | | | | | | | | |
| Chromosome | start | end | strand | MM | target_seq | PAM | alignment | distance | position | gene name | gene id |
| GU305914.1 | 128146 | 128168 | + | 0 | CATGACCGCGCACCTCGGTC | CGG | \|\|\|\|\|\|\|\|[\|\|\|\|\|\|\|\|\|\|\|\|]PAM | NA | Intergenic | NA | NA |
| T9688 | CCGGCCATCGCCGCCCGCGTCGG | 1000 | | | | | | | | | |
| Oligo fwd | CACCCCGGCCATCGCCGCCCGCGT | | | | | | | | | | |
| Oligo rev | AAACACGCGGGCGGCGATGGCCGG | | | | | | | | | | |
| Chromosome | start | end | strand | MM | target_seq | PAM | alignment | distance | position | gene name | gene id |
| GU305914.1 | 128165 | 128187 | + | 0 | CCGGCCATCGCCGCCCGCGT | CGG | \|\|\|\|\|\|\|\|[\|\|\|\|\|\|\|\|\|\|\|\|]PAM | NA | Intergenic | NA | NA |
| T9689 | GTGCACTACAAGATGCTCTTCGG | 1000 | | | | | | | | | |
| Oligo fwd | CACCGTGCACTACAAGATGCTCTT | | | | | | | | | | |
| Oligo rev | AAACAAGAGCATCTTGTAGTGCAC | | | | | | | | | | |
| Chromosome | start | end | strand | MM | target_seq | PAM | alignment | distance | position | gene name | gene id |

GU305914.1 128189 128211 + 0 GTGCACTACAAGATGCTCTT CGG ||||||||[||||||||||||]PAM NA Intergenic NA NA

T9690 TGCTCTTCGGCGCTAACGCCCGG 1000
Oligo fwd CACCTGCTCTTCGGCGCTAACGCC
Oligo rev AAACGGCGTTAGCGCCGAAGAGCA
Chromosome start end strand MM target_seq PAM alignment distance position gene name gene id
GU305914.1 128202 128224 + 0 TGCTCTTCGGCGCTAACGCC CGG ||||||||[||||||||||||]PAM NA Intergenic NA NA

T9691 GCGCTAACGCCCGGCCGCCGCGG 1000
Oligo fwd CACCGCGCTAACGCCCGGCCGCCG
Oligo rev AAACCGGCGGCCGGGCGTTAGCGC
Chromosome start end strand MM target_seq PAM alignment distance position gene name gene id
GU305914.1 128211 128233 + 0 GCGCTAACGCCCGGCCGCCG CGG ||||||||[||||||||||||]PAM NA Intergenic NA NA

T9692 CGCTAACGCCCGGCCGCCGCGGG 1000
Oligo fwd CACCCGCTAACGCCCGGCCGCCGC
Oligo rev AAACGCGGCGGCCGGGCGTTAGCG
Chromosome start end strand MM target_seq PAM alignment distance position gene name gene id
GU305914.1 128212 128234 + 0 CGCTAACGCCCGGCCGCCGC GGG ||||||||[||||||||||||]PAM NA Intergenic NA NA

T9693 CCGGCCGCCGCGGGTCACCGAGG 1000
Oligo fwd CACCCCGGCCGCCGCGGGTCACCG
Oligo rev AAACCGGTGACCCGCGGCGGCCGG
Chromosome start end strand MM target_seq PAM alignment distance position gene name gene id
GU305914.1 128221 128243 + 0 CCGGCCGCCGCGGGTCACCG AGG ||||||||[||||||||||||]PAM NA Intergenic NA NA

T9694 GCCGCCGCGGGTCACCGAGGTGG 1000
Oligo fwd CACCGCCGCCGCGGGTCACCGAGG
Oligo rev AAACCCTCGGTGACCCGCGGCGGC
Chromosome start end strand MM target_seq PAM alignment distance position gene name gene id
GU305914.1 128224 128246 + 0 GCCGCCGCGGGTCACCGAGG TGG ||||||||[||||||||||||]PAM NA Intergenic NA NA

T9695 TGCTCGAGCTGCAGAAGCTCTGG 1000
Oligo fwd CACCTGCTCGAGCTGCAGAAGCTC
Oligo rev AAACGAGCTTCTGCAGCTCGAGCA
Chromosome start end strand MM target_seq PAM alignment distance position gene name gene id
GU305914.1 128274 128296 + 0 TGCTCGAGCTGCAGAAGCTC TGG ||||||||[||||||||||||]PAM NA Intergenic NA NA
GU305914.1 123626 123648 + 4 CGCACGGGCTGTAGAAGCTC GGG -||-||-|[|||-||||||||]PAM NA Intergenic NA NA

T9696 CAACGTCTCCTACCGTAATGCGG 1000
Oligo fwd CACCCAACGTCTCCTACCGTAATG
Oligo rev AAACCATTACGGTAGGAGACGTTG
Chromosome start end strand MM target_seq PAM alignment distance position gene name gene id
GU305914.1 128299 128321 + 0 CAACGTCTCCTACCGTAATG CGG ||||||||[||||||||||||]PAM NA Intergenic NA NA

T9697 TCCGCCTTCCACCACGAGACAGG 1000
Oligo fwd CACCTCCGCCTTCCACCACGAGAC
Oligo rev AAACGTCTCGTGGTGGAAGGCGGA
Chromosome start end strand MM target_seq PAM alignment distance position gene name gene id
GU305914.1 128351 128373 + 0 TCCGCCTTCCACCACGAGAC AGG ||||||||[||||||||||||]PAM NA Intergenic NA NA

T9698 CTTCCACCACGAGACAGGTAAGG 1000
Oligo fwd CACCCTTCCACCACGAGACAGGTA
Oligo rev AAACTACCTGTCTCGTGGTGGAAG
Chromosome start end strand MM target_seq PAM alignment distance position gene name gene id
GU305914.1 128356 128378 + 0 CTTCCACCACGAGACAGGTA AGG ||||||||[||||||||||||]PAM NA Intergenic NA NA

T9699 CGAGACAGGTAAGGTCATGCTGG 1000
Oligo fwd CACCCGAGACAGGTAAGGTCATGC
Oligo rev AAACGCATGACCTTACCTGTCTCG
Chromosome start end strand MM target_seq PAM alignment distance position gene name gene id
GU305914.1 128365 128387 + 0 CGAGACAGGTAAGGTCATGC TGG ||||||||[||||||||||||]PAM NA Intergenic NA NA

T9700 CATGCTGGCGTTACGCTCGCCGG 1000
Oligo fwd CACCCATGCTGGCGTTACGCTCGC
Oligo rev AAACGCGAGCGTAACGCCAGCATG
Chromosome start end strand MM target_seq PAM alignment distance position gene name gene id
GU305914.1 128380 128402 + 0 CATGCTGGCGTTACGCTCGC CGG ||||||||[||||||||||||]PAM NA Intergenic NA NA

T9701 ATGCTGGCGTTACGCTCGCCGGG 1000
Oligo fwd CACCATGCTGGCGTTACGCTCGCC
Oligo rev AAACGGCGAGCGTAACGCCAGCAT
Chromosome start end strand MM target_seq PAM alignment distance position gene name gene id
GU305914.1 128381 128403 + 0 ATGCTGGCGTTACGCTCGCC GGG ||||||||[||||||||||||]PAM NA Intergenic NA NA

T9702 GCCAGCCGCTGTTTCCGCGATGG 1000
Oligo fwd CACCGCCAGCCGCTGTTTCCGCGA
Oligo rev AAACTCGCGGAAACAGCGGCTGGC
Chromosome start end strand MM target_seq PAM alignment distance position gene name gene id
GU305914.1 128406 128428 + 0 GCCAGCCGCTGTTTCCGCGA TGG ||||||||[||||||||||||]PAM NA Intergenic NA NA

T9703 CGCGATGGATATCGATGTCGCGG 1000
Oligo fwd CACCCGCGATGGATATCGATGTCG
Oligo rev AAACCGACATCGATATCCATCGCG
Chromosome start end strand MM target_seq PAM alignment distance position gene name gene id
GU305914.1 128421 128443 + 0 CGCGATGGATATCGATGTCG CGG ||||||||[||||||||||||]PAM NA Intergenic NA NA

T9704 GATGTCGCGGTTCAAGAAACAGG 1000
Oligo fwd CACCGATGTCGCGGTTCAAGAAAC

| | | | | | | | | | | | |
|---|---|---|---|---|---|---|---|---|---|---|---|
| Oligo rev | AAACGTTTCTTGAACCGCGACATC | | | | | | | | | | |
| Chromosome | start | end | strand | MM | target_seq | PAM | alignment | distance | position | gene name | gene id |
| GU305914.1 | 128434 | 128456 | + | 0 | GATGTCGCGGTTCAAGAAAC | AGG | ||||||||[||||||||||||]PAM | NA | Intergenic | NA | NA |
| T9705 | AAGTCCGATGTACTAAGAGCCGG | 1000 | | | | | | | | | |
| Oligo fwd | CACCAAGTCCGATGTACTAAGAGC | | | | | | | | | | |
| Oligo rev | AAACGCTCTTAGTACATCGGACTT | | | | | | | | | | |
| Chromosome | start | end | strand | MM | target_seq | PAM | alignment | distance | position | gene name | gene id |
| GU305914.1 | 128475 | 128497 | + | 0 | AAGTCCGATGTACTAAGAGC | CGG | ||||||||[||||||||||||]PAM | NA | Intergenic | NA | NA |
| T9706 | CCGATGTACTAAGAGCCGGAAGG | 1000 | | | | | | | | | |
| Oligo fwd | CACCCCGATGTACTAAGAGCCGGA | | | | | | | | | | |
| Oligo rev | AAACTCCGGCTCTTAGTACATCGG | | | | | | | | | | |
| Chromosome | start | end | strand | MM | target_seq | PAM | alignment | distance | position | gene name | gene id |
| GU305914.1 | 128479 | 128501 | + | 0 | CCGATGTACTAAGAGCCGGA | AGG | ||||||||[||||||||||||]PAM | NA | Intergenic | NA | NA |
| T9707 | TCAAAAATGCGTTGATTTTCAGG | 1000 | | | | | | | | | |
| Oligo fwd | CACCTCAAAAATGCGTTGATTTTC | | | | | | | | | | |
| Oligo rev | AAACGAAAATCAACGCATTTTTGA | | | | | | | | | | |
| Chromosome | start | end | strand | MM | target_seq | PAM | alignment | distance | position | gene name | gene id |
| GU305914.1 | 128508 | 128530 | + | 0 | TCAAAAATGCGTTGATTTTC | AGG | ||||||||[||||||||||||]PAM | NA | Intergenic | NA | NA |
| T9708 | GAAGACCAGCGTTCTGTTCGCGG | 1000 | | | | | | | | | |
| Oligo fwd | CACCGAAGACCAGCGTTCTGTTCG | | | | | | | | | | |
| Oligo rev | AAACCGAACAGAACGCTGGTCTTC | | | | | | | | | | |
| Chromosome | start | end | strand | MM | target_seq | PAM | alignment | distance | position | gene name | gene id |
| GU305914.1 | 128530 | 128552 | + | 0 | GAAGACCAGCGTTCTGTTCG | CGG | ||||||||[||||||||||||]PAM | NA | Intergenic | NA | NA |
| T9709 | TCGCGGACCACGACGTGATCTGG | 1000 | | | | | | | | | |
| Oligo fwd | CACCTCGCGGACCACGACGTGATC | | | | | | | | | | |
| Oligo rev | AAACGATCACGTCGTGGTCCGCGA | | | | | | | | | | |
| Chromosome | start | end | strand | MM | target_seq | PAM | alignment | distance | position | gene name | gene id |
| GU305914.1 | 128547 | 128569 | + | 0 | TCGCGGACCACGACGTGATC | TGG | ||||||||[||||||||||||]PAM | NA | Intergenic | NA | NA |
| T9710 | CGGACCACGACGTGATCTGGAGG | 1000 | | | | | | | | | |
| Oligo fwd | CACCCGGACCACGACGTGATCTGG | | | | | | | | | | |
| Oligo rev | AAACCCAGATCACGTCGTGGTCCG | | | | | | | | | | |
| Chromosome | start | end | strand | MM | target_seq | PAM | alignment | distance | position | gene name | gene id |
| GU305914.1 | 128550 | 128572 | + | 0 | CGGACCACGACGTGATCTGG | AGG | ||||||||[||||||||||||]PAM | NA | Intergenic | NA | NA |
| GU305914.1 | 199949 | 199971 | - | 4 | CGGACATGGACGTGACCTGG | CGG | |||||---[|||||||-||||]PAM | NA | Intergenic | NA | NA |
| T9711 | TGTTCTACACCTACTATGCCTGG | 1000 | | | | | | | | | |
| Oligo fwd | CACCTGTTCTACACCTACTATGCC | | | | | | | | | | |
| Oligo rev | AAACGGCATAGTAGGTGTAGAACA | | | | | | | | | | |
| Chromosome | start | end | strand | MM | target_seq | PAM | alignment | distance | position | gene name | gene id |
| GU305914.1 | 128577 | 128599 | + | 0 | TGTTCTACACCTACTATGCC | TGG | ||||||||[||||||||||||]PAM | NA | Intergenic | NA | NA |
| T9712 | ACTATGCCTGGTGCGCCCATCGG | 1000 | | | | | | | | | |
| Oligo fwd | CACCACTATGCCTGGTGCGCCCAT | | | | | | | | | | |
| Oligo rev | AAACATGGGCGCACCAGGCATAGT | | | | | | | | | | |
| Chromosome | start | end | strand | MM | target_seq | PAM | alignment | distance | position | gene name | gene id |
| GU305914.1 | 128589 | 128611 | + | 0 | ACTATGCCTGGTGCGCCCAT | CGG | ||||||||[||||||||||||]PAM | NA | Intergenic | NA | NA |
| T9713 | CTATGCCTGGTGCGCCCATCGGG | 1000 | | | | | | | | | |
| Oligo fwd | CACCCTATGCCTGGTGCGCCCATC | | | | | | | | | | |
| Oligo rev | AAACGATGGGCGCACCAGGCATAG | | | | | | | | | | |
| Chromosome | start | end | strand | MM | target_seq | PAM | alignment | distance | position | gene name | gene id |
| GU305914.1 | 128590 | 128612 | + | 0 | CTATGCCTGGTGCGCCCATC | GGG | ||||||||[||||||||||||]PAM | NA | Intergenic | NA | NA |
| T9714 | TATGCCTGGTGCGCCCATCGGGG | 1000 | | | | | | | | | |
| Oligo fwd | CACCTATGCCTGGTGCGCCCATCG | | | | | | | | | | |
| Oligo rev | AAACCGATGGGCGCACCAGGCATA | | | | | | | | | | |
| Chromosome | start | end | strand | MM | target_seq | PAM | alignment | distance | position | gene name | gene id |
| GU305914.1 | 128591 | 128613 | + | 0 | TATGCCTGGTGCGCCCATCG | GGG | ||||||||[||||||||||||]PAM | NA | Intergenic | NA | NA |
| T9715 | TGGTGCGCCCATCGGGGCTTCGG | 1000 | | | | | | | | | |
| Oligo fwd | CACCTGGTGCGCCCATCGGGGCTT | | | | | | | | | | |
| Oligo rev | AAACAAGCCCCGATGGGCGCACCA | | | | | | | | | | |
| Chromosome | start | end | strand | MM | target_seq | PAM | alignment | distance | position | gene name | gene id |
| GU305914.1 | 128597 | 128619 | + | 0 | TGGTGCGCCCATCGGGGCTT | CGG | ||||||||[||||||||||||]PAM | NA | Intergenic | NA | NA |
| T9716 | TGCGCCCATCGGGGCTTCGGCGG | 1000 | | | | | | | | | |
| Oligo fwd | CACCTGCGCCCATCGGGGCTTCGG | | | | | | | | | | |
| Oligo rev | AAACCCGAAGCCCCGATGGGCGCA | | | | | | | | | | |
| Chromosome | start | end | strand | MM | target_seq | PAM | alignment | distance | position | gene name | gene id |
| GU305914.1 | 128600 | 128622 | + | 0 | TGCGCCCATCGGGGCTTCGG | CGG | ||||||||[||||||||||||]PAM | NA | Intergenic | NA | NA |
| T9717 | GGGGCTTCGGCGGCGAGAGCCGG | 1000 | | | | | | | | | |
| Oligo fwd | CACCGGGGCTTCGGCGGCGAGAGC | | | | | | | | | | |
| Oligo rev | AAACGCTCTCGCCGCCGAAGCCCC | | | | | | | | | | |
| Chromosome | start | end | strand | MM | target_seq | PAM | alignment | distance | position | gene name | gene id |
| GU305914.1 | 128610 | 128632 | + | 0 | GGGGCTTCGGCGGCGAGAGC | CGG | ||||||||[||||||||||||]PAM | NA | Intergenic | NA | NA |
| GU305914.1 | 168830 | 168852 | + | 4 | GCGGCGGCGGCGGCGACAGC | GGG | |-|||--|[||||||||-|||]PAM | NA | Intergenic | NA | NA |
| T9718 | TCGGCGGCGAGAGCCGGCTGTGG | 1000 | | | | | | | | | |
| Oligo fwd | CACCTCGGCGGCGAGAGCCGGCTG | | | | | | | | | | |
| Oligo rev | AAACCAGCCGGCTCTCGCCGCCGA | | | | | | | | | | |
| Chromosome | start | end | strand | MM | target_seq | PAM | alignment | distance | position | gene name | gene id |
| GU305914.1 | 128616 | 128638 | + | 0 | TCGGCGGCGAGAGCCGGCTG | TGG | ||||||||[||||||||||||]PAM | NA | Intergenic | NA | NA |

<table>
<tr><td>T9719</td><td>CGGCGGCGAGAGCCGGCTGTGGG</td><td></td><td>1000</td><td></td><td></td><td></td><td></td><td></td><td></td><td></td><td></td></tr>
<tr><td>Oligo fwd</td><td>CACCCGGCGGCGAGAGCCGGCTGT</td><td></td><td></td><td></td><td></td><td></td><td></td><td></td><td></td><td></td><td></td></tr>
<tr><td>Oligo rev</td><td>AAACACAGCCGGCTCTCGCCGCCG</td><td></td><td></td><td></td><td></td><td></td><td></td><td></td><td></td><td></td><td></td></tr>
<tr><td>Chromosome</td><td>start</td><td>end</td><td></td><td>strand</td><td>MM</td><td>target_seq</td><td>PAM</td><td>alignment</td><td>distance</td><td>position</td><td>gene name</td><td>gene id</td></tr>
<tr><td>GU305914.1</td><td>128617</td><td>128639</td><td></td><td>+</td><td>0</td><td>CGGCGGCGAGAGCCGGCTGT</td><td>GGG</td><td>||||||||[||||||||||||]PAM</td><td>NA</td><td>Intergenic</td><td>NA</td><td>NA</td></tr>
<tr><td>T9720</td><td>GGCGGCGAGAGCCGGCTGTGGGG</td><td></td><td>1000</td><td></td><td></td><td></td><td></td><td></td><td></td><td></td><td></td></tr>
<tr><td>Oligo fwd</td><td>CACCGGCGGCGAGAGCCGGCTGTG</td><td></td><td></td><td></td><td></td><td></td><td></td><td></td><td></td><td></td><td></td></tr>
<tr><td>Oligo rev</td><td>AAACCACAGCCGGCTCTCGCCGCC</td><td></td><td></td><td></td><td></td><td></td><td></td><td></td><td></td><td></td><td></td></tr>
<tr><td>Chromosome</td><td>start</td><td>end</td><td></td><td>strand</td><td>MM</td><td>target_seq</td><td>PAM</td><td>alignment</td><td>distance</td><td>position</td><td>gene name</td><td>gene id</td></tr>
<tr><td>GU305914.1</td><td>128618</td><td>128640</td><td></td><td>+</td><td>0</td><td>GGCGGCGAGAGCCGGCTGTG</td><td>GGG</td><td>||||||||[||||||||||||]PAM</td><td>NA</td><td>Intergenic</td><td>NA</td><td>NA</td></tr>
<tr><td>T9721</td><td>GCGGCGAGAGCCGGCTGTGGGGG</td><td></td><td>1000</td><td></td><td></td><td></td><td></td><td></td><td></td><td></td><td></td></tr>
<tr><td>Oligo fwd</td><td>CACCGCGGCGAGAGCCGGCTGTGG</td><td></td><td></td><td></td><td></td><td></td><td></td><td></td><td></td><td></td><td></td></tr>
<tr><td>Oligo rev</td><td>AAACCCACAGCCGGCTCTCGCCGC</td><td></td><td></td><td></td><td></td><td></td><td></td><td></td><td></td><td></td><td></td></tr>
<tr><td>Chromosome</td><td>start</td><td>end</td><td></td><td>strand</td><td>MM</td><td>target_seq</td><td>PAM</td><td>alignment</td><td>distance</td><td>position</td><td>gene name</td><td>gene id</td></tr>
<tr><td>GU305914.1</td><td>128619</td><td>128641</td><td></td><td>+</td><td>0</td><td>GCGGCGAGAGCCGGCTGTGG</td><td>GGG</td><td>||||||||[||||||||||||]PAM</td><td>NA</td><td>Intergenic</td><td>NA</td><td>NA</td></tr>
<tr><td>T9722</td><td>AGCCGGCTGTGGGGGCCGAGCGG</td><td></td><td>1000</td><td></td><td></td><td></td><td></td><td></td><td></td><td></td><td></td></tr>
<tr><td>Oligo fwd</td><td>CACCAGCCGGCTGTGGGGGCCGAG</td><td></td><td></td><td></td><td></td><td></td><td></td><td></td><td></td><td></td><td></td></tr>
<tr><td>Oligo rev</td><td>AAACCTCGGCCCCCACAGCCGGCT</td><td></td><td></td><td></td><td></td><td></td><td></td><td></td><td></td><td></td><td></td></tr>
<tr><td>Chromosome</td><td>start</td><td>end</td><td></td><td>strand</td><td>MM</td><td>target_seq</td><td>PAM</td><td>alignment</td><td>distance</td><td>position</td><td>gene name</td><td>gene id</td></tr>
<tr><td>GU305914.1</td><td>128627</td><td>128649</td><td></td><td>+</td><td>0</td><td>AGCCGGCTGTGGGGGCCGAG</td><td>CGG</td><td>||||||||[||||||||||||]PAM</td><td>NA</td><td>Intergenic</td><td>NA</td><td>NA</td></tr>
<tr><td>T9723</td><td>CTGTGGGGGCCGAGCGGCAGCGG</td><td></td><td>1000</td><td></td><td></td><td></td><td></td><td></td><td></td><td></td><td></td></tr>
<tr><td>Oligo fwd</td><td>CACCCTGTGGGGGCCGAGCGGCAG</td><td></td><td></td><td></td><td></td><td></td><td></td><td></td><td></td><td></td><td></td></tr>
<tr><td>Oligo rev</td><td>AAACCTGCCGCTCGGCCCCCACAG</td><td></td><td></td><td></td><td></td><td></td><td></td><td></td><td></td><td></td><td></td></tr>
<tr><td>Chromosome</td><td>start</td><td>end</td><td></td><td>strand</td><td>MM</td><td>target_seq</td><td>PAM</td><td>alignment</td><td>distance</td><td>position</td><td>gene name</td><td>gene id</td></tr>
<tr><td>GU305914.1</td><td>128633</td><td>128655</td><td></td><td>+</td><td>0</td><td>CTGTGGGGGCCGAGCGGCAG</td><td>CGG</td><td>||||||||[||||||||||||]PAM</td><td>NA</td><td>Intergenic</td><td>NA</td><td>NA</td></tr>
<tr><td>T9724</td><td>TGGGGGCCGAGCGGCAGCGGCGG</td><td></td><td>1000</td><td></td><td></td><td></td><td></td><td></td><td></td><td></td><td></td></tr>
<tr><td>Oligo fwd</td><td>CACCTGGGGGCCGAGCGGCAGCGG</td><td></td><td></td><td></td><td></td><td></td><td></td><td></td><td></td><td></td><td></td></tr>
<tr><td>Oligo rev</td><td>AAACCCGCTGCCGCTCGGCCCCCA</td><td></td><td></td><td></td><td></td><td></td><td></td><td></td><td></td><td></td><td></td></tr>
<tr><td>Chromosome</td><td>start</td><td>end</td><td></td><td>strand</td><td>MM</td><td>target_seq</td><td>PAM</td><td>alignment</td><td>distance</td><td>position</td><td>gene name</td><td>gene id</td></tr>
<tr><td>GU305914.1</td><td>128636</td><td>128658</td><td></td><td>+</td><td>0</td><td>TGGGGGCCGAGCGGCAGCGG</td><td>CGG</td><td>||||||||[||||||||||||]PAM</td><td>NA</td><td>Intergenic</td><td>NA</td><td>NA</td></tr>
<tr><td>T9725</td><td>GGGCCGAGCGGCAGCGGCGGCGG</td><td></td><td>1000</td><td></td><td></td><td></td><td></td><td></td><td></td><td></td><td></td></tr>
<tr><td>Oligo fwd</td><td>CACCGGGCCGAGCGGCAGCGGCGG</td><td></td><td></td><td></td><td></td><td></td><td></td><td></td><td></td><td></td><td></td></tr>
<tr><td>Oligo rev</td><td>AAACCCGCCGCTGCCGCTCGGCCC</td><td></td><td></td><td></td><td></td><td></td><td></td><td></td><td></td><td></td><td></td></tr>
<tr><td>Chromosome</td><td>start</td><td>end</td><td></td><td>strand</td><td>MM</td><td>target_seq</td><td>PAM</td><td>alignment</td><td>distance</td><td>position</td><td>gene name</td><td>gene id</td></tr>
<tr><td>GU305914.1</td><td>128639</td><td>128661</td><td></td><td>+</td><td>0</td><td>GGGCCGAGCGGCAGCGGCGG</td><td>CGG</td><td>||||||||[||||||||||||]PAM</td><td>NA</td><td>Intergenic</td><td>NA</td><td>NA</td></tr>
<tr><td>GU305914.1</td><td>163896</td><td>163918</td><td></td><td>+</td><td>3</td><td>GGATCGAGCGGCGGCGGCGG</td><td>CGG</td><td>||--||||[||||-|||||||]PAM</td><td>NA</td><td>Intergenic</td><td>NA</td><td>NA</td></tr>
<tr><td>T9726</td><td>GCCGCCACCGAGTCTGAGCGCGG</td><td></td><td>1000</td><td></td><td></td><td></td><td></td><td></td><td></td><td></td><td></td></tr>
<tr><td>Oligo fwd</td><td>CACCGCCGCCACCGAGTCTGAGCG</td><td></td><td></td><td></td><td></td><td></td><td></td><td></td><td></td><td></td><td></td></tr>
<tr><td>Oligo rev</td><td>AAACCGCTCAGACTCGGTGGCGGC</td><td></td><td></td><td></td><td></td><td></td><td></td><td></td><td></td><td></td><td></td></tr>
<tr><td>Chromosome</td><td>start</td><td>end</td><td></td><td>strand</td><td>MM</td><td>target_seq</td><td>PAM</td><td>alignment</td><td>distance</td><td>position</td><td>gene name</td><td>gene id</td></tr>
<tr><td>GU305914.1</td><td>128663</td><td>128685</td><td></td><td>+</td><td>0</td><td>GCCGCCACCGAGTCTGAGCG</td><td>CGG</td><td>||||||||[||||||||||||]PAM</td><td>NA</td><td>Intergenic</td><td>NA</td><td>NA</td></tr>
<tr><td>T9727</td><td>GCCACCGAGTCTGAGCGCGGCGG</td><td></td><td>1000</td><td></td><td></td><td></td><td></td><td></td><td></td><td></td><td></td></tr>
<tr><td>Oligo fwd</td><td>CACCGCCACCGAGTCTGAGCGCGG</td><td></td><td></td><td></td><td></td><td></td><td></td><td></td><td></td><td></td><td></td></tr>
<tr><td>Oligo rev</td><td>AAACCCGCGCTCAGACTCGGTGGC</td><td></td><td></td><td></td><td></td><td></td><td></td><td></td><td></td><td></td><td></td></tr>
<tr><td>Chromosome</td><td>start</td><td>end</td><td></td><td>strand</td><td>MM</td><td>target_seq</td><td>PAM</td><td>alignment</td><td>distance</td><td>position</td><td>gene name</td><td>gene id</td></tr>
<tr><td>GU305914.1</td><td>128666</td><td>128688</td><td></td><td>+</td><td>0</td><td>GCCACCGAGTCTGAGCGCGG</td><td>CGG</td><td>||||||||[||||||||||||]PAM</td><td>NA</td><td>Intergenic</td><td>NA</td><td>NA</td></tr>
<tr><td>T9728</td><td>ACCGAGTCTGAGCGCGGCGGAGG</td><td></td><td>1000</td><td></td><td></td><td></td><td></td><td></td><td></td><td></td><td></td></tr>
<tr><td>Oligo fwd</td><td>CACCACCGAGTCTGAGCGCGGCGG</td><td></td><td></td><td></td><td></td><td></td><td></td><td></td><td></td><td></td><td></td></tr>
<tr><td>Oligo rev</td><td>AAACCCGCCGCGCTCAGACTCGGT</td><td></td><td></td><td></td><td></td><td></td><td></td><td></td><td></td><td></td><td></td></tr>
<tr><td>Chromosome</td><td>start</td><td>end</td><td></td><td>strand</td><td>MM</td><td>target_seq</td><td>PAM</td><td>alignment</td><td>distance</td><td>position</td><td>gene name</td><td>gene id</td></tr>
<tr><td>GU305914.1</td><td>128669</td><td>128691</td><td></td><td>+</td><td>0</td><td>ACCGAGTCTGAGCGCGGCGG</td><td>AGG</td><td>||||||||[||||||||||||]PAM</td><td>NA</td><td>Intergenic</td><td>NA</td><td>NA</td></tr>
<tr><td>T9729</td><td>GAGCGCGGCGGAGGCGTCCGCGG</td><td></td><td>1000</td><td></td><td></td><td></td><td></td><td></td><td></td><td></td><td></td></tr>
<tr><td>Oligo fwd</td><td>CACCGAGCGCGGCGGAGGCGTCCG</td><td></td><td></td><td></td><td></td><td></td><td></td><td></td><td></td><td></td><td></td></tr>
<tr><td>Oligo rev</td><td>AAACCGGACGCCTCCGCCGCGCTC</td><td></td><td></td><td></td><td></td><td></td><td></td><td></td><td></td><td></td><td></td></tr>
<tr><td>Chromosome</td><td>start</td><td>end</td><td></td><td>strand</td><td>MM</td><td>target_seq</td><td>PAM</td><td>alignment</td><td>distance</td><td>position</td><td>gene name</td><td>gene id</td></tr>
<tr><td>GU305914.1</td><td>128678</td><td>128700</td><td></td><td>+</td><td>0</td><td>GAGCGCGGCGGAGGCGTCCG</td><td>CGG</td><td>||||||||[||||||||||||]PAM</td><td>NA</td><td>Intergenic</td><td>NA</td><td>NA</td></tr>
<tr><td>T9730</td><td>AGCGCGGCGGAGGCGTCCGCGGG</td><td></td><td>1000</td><td></td><td></td><td></td><td></td><td></td><td></td><td></td><td></td></tr>
<tr><td>Oligo fwd</td><td>CACCAGCGCGGCGGAGGCGTCCGC</td><td></td><td></td><td></td><td></td><td></td><td></td><td></td><td></td><td></td><td></td></tr>
<tr><td>Oligo rev</td><td>AAACGCGGACGCCTCCGCCGCGCT</td><td></td><td></td><td></td><td></td><td></td><td></td><td></td><td></td><td></td><td></td></tr>
<tr><td>Chromosome</td><td>start</td><td>end</td><td></td><td>strand</td><td>MM</td><td>target_seq</td><td>PAM</td><td>alignment</td><td>distance</td><td>position</td><td>gene name</td><td>gene id</td></tr>
<tr><td>GU305914.1</td><td>128679</td><td>128701</td><td></td><td>+</td><td>0</td><td>AGCGCGGCGGAGGCGTCCGC</td><td>GGG</td><td>||||||||[||||||||||||]PAM</td><td>NA</td><td>Intergenic</td><td>NA</td><td>NA</td></tr>
<tr><td>T9731</td><td>AGGCGTCCGCGGGCGCGATCAGG</td><td></td><td>1000</td><td></td><td></td><td></td><td></td><td></td><td></td><td></td><td></td></tr>
<tr><td>Oligo fwd</td><td>CACCAGGCGTCCGCGGGCGCGATC</td><td></td><td></td><td></td><td></td><td></td><td></td><td></td><td></td><td></td><td></td></tr>
<tr><td>Oligo rev</td><td>AAACGATCGCGCCCGCGGACGCCT</td><td></td><td></td><td></td><td></td><td></td><td></td><td></td><td></td><td></td><td></td></tr>
<tr><td>Chromosome</td><td>start</td><td>end</td><td></td><td>strand</td><td>MM</td><td>target_seq</td><td>PAM</td><td>alignment</td><td>distance</td><td>position</td><td>gene name</td><td>gene id</td></tr>
<tr><td>GU305914.1</td><td>128689</td><td>128711</td><td></td><td>+</td><td>0</td><td>AGGCGTCCGCGGGCGCGATC</td><td>AGG</td><td>||||||||[||||||||||||]PAM</td><td>NA</td><td>Intergenic</td><td>NA</td><td>NA</td></tr>
<tr><td>T9732</td><td>CACCGACGACGATGCCTCCGCGG</td><td></td><td>1000</td><td></td><td></td><td></td><td></td><td></td><td></td><td></td><td></td></tr>
<tr><td>Oligo fwd</td><td>CACCCACCGACGACGATGCCTCCG</td><td></td><td></td><td></td><td></td><td></td><td></td><td></td><td></td><td></td><td></td></tr>
<tr><td>Oligo rev</td><td>AAACCGGAGGCATCGTCGTCGGTG</td><td></td><td></td><td></td><td></td><td></td><td></td><td></td><td></td><td></td><td></td></tr>
<tr><td>Chromosome</td><td>start</td><td>end</td><td></td><td>strand</td><td>MM</td><td>target_seq</td><td>PAM</td><td>alignment</td><td>distance</td><td>position</td><td>gene name</td><td>gene id</td></tr>
<tr><td>GU305914.1</td><td>128746</td><td>128768</td><td></td><td>+</td><td>0</td><td>CACCGACGACGATGCCTCCG</td><td>CGG</td><td>||||||||[||||||||||||]PAM</td><td>NA</td><td>Intergenic</td><td>NA</td><td>NA</td></tr>
<tr><td>T9733</td><td>CGATGCCTCCGCGGCGTCGACGG</td><td></td><td>1000</td><td></td><td></td><td></td><td></td><td></td><td></td><td></td><td></td></tr>
<tr><td>Oligo fwd</td><td>CACCCGATGCCTCCGCGGCGTCGA</td><td></td><td></td><td></td><td></td><td></td><td></td><td></td><td></td><td></td><td></td></tr>
<tr><td>Oligo rev</td><td>AAACTCGACGCCGCGGAGGCATCG</td><td></td><td></td><td></td><td></td><td></td><td></td><td></td><td></td><td></td><td></td></tr>
</table>

<table>
<tr><td>Chromosome</td><td>start</td><td>end</td><td>strand</td><td>MM</td><td>target_seq</td><td>PAM</td><td>alignment</td><td>distance</td><td>position</td><td>gene name</td><td>gene id</td></tr>
<tr><td>GU305914.1</td><td>128755</td><td>128777</td><td>+</td><td>0</td><td>CGATGCCTCCGCGGCGTCGA</td><td>CGG</td><td>||||||||[||||||||||||]PAM</td><td>NA</td><td>Intergenic</td><td>NA</td><td>NA</td></tr>
<tr><td>GU305914.1</td><td>61324</td><td>61346</td><td>+</td><td>4</td><td>GAAGGCCTGCGCGGCGTCGA</td><td>CGG</td><td>--|-||||[-|||||||||||]PAM</td><td>NA</td><td>Intergenic</td><td>NA</td><td>NA</td></tr>
<tr><td>GU305914.1</td><td>127874</td><td>127896</td><td>+</td><td>4</td><td>CGCTGCCGCTGCGGCGCCGA</td><td>GGG</td><td>||-||||-[|-||||||-|||]PAM</td><td>NA</td><td>Intergenic</td><td>NA</td><td>NA</td></tr>
<tr><td></td><td></td><td></td><td></td><td></td><td></td><td></td><td></td><td></td><td></td><td></td><td></td></tr>
<tr><td>T9734</td><td>TCCGCGGCGTCGACGGAGAGTGG</td><td>1000</td><td></td><td></td><td></td><td></td><td></td><td></td><td></td><td></td><td></td></tr>
<tr><td>Oligo fwd</td><td>CACCTCCGCGGCGTCGACGGAGAG</td><td></td><td></td><td></td><td></td><td></td><td></td><td></td><td></td><td></td><td></td></tr>
<tr><td>Oligo rev</td><td>AAACCTCTCCGTCGACGCCGCGGA</td><td></td><td></td><td></td><td></td><td></td><td></td><td></td><td></td><td></td><td></td></tr>
<tr><td>Chromosome</td><td>start</td><td>end</td><td>strand</td><td>MM</td><td>target_seq</td><td>PAM</td><td>alignment</td><td>distance</td><td>position</td><td>gene name</td><td>gene id</td></tr>
<tr><td>GU305914.1</td><td>128762</td><td>128784</td><td>+</td><td>0</td><td>TCCGCGGCGTCGACGGAGAG</td><td>TGG</td><td>||||||||[||||||||||||]PAM</td><td>NA</td><td>Intergenic</td><td>NA</td><td>NA</td></tr>
<tr><td>GU305914.1</td><td>225226</td><td>225248</td><td>+</td><td>4</td><td>TCCACGGTGTCGTCGGTGAG</td><td>CGG</td><td>|||-|||-[||||-|||-|||]PAM</td><td>NA</td><td>Intergenic</td><td>NA</td><td>NA</td></tr>
<tr><td></td><td></td><td></td><td></td><td></td><td></td><td></td><td></td><td></td><td></td><td></td><td></td></tr>
<tr><td>T9735</td><td>GGCGTCGACGGAGAGTGGTGTGG</td><td>1000</td><td></td><td></td><td></td><td></td><td></td><td></td><td></td><td></td><td></td></tr>
<tr><td>Oligo fwd</td><td>CACCGGCGTCGACGGAGAGTGGTG</td><td></td><td></td><td></td><td></td><td></td><td></td><td></td><td></td><td></td><td></td></tr>
<tr><td>Oligo rev</td><td>AAACCACCACTCTCCGTCGACGCC</td><td></td><td></td><td></td><td></td><td></td><td></td><td></td><td></td><td></td><td></td></tr>
<tr><td>Chromosome</td><td>start</td><td>end</td><td>strand</td><td>MM</td><td>target_seq</td><td>PAM</td><td>alignment</td><td>distance</td><td>position</td><td>gene name</td><td>gene id</td></tr>
<tr><td>GU305914.1</td><td>128767</td><td>128789</td><td>+</td><td>0</td><td>GGCGTCGACGGAGAGTGGTG</td><td>TGG</td><td>||||||||[||||||||||||]PAM</td><td>NA</td><td>Intergenic</td><td>NA</td><td>NA</td></tr>
<tr><td></td><td></td><td></td><td></td><td></td><td></td><td></td><td></td><td></td><td></td><td></td><td></td></tr>
<tr><td>T9736</td><td>GGCTGTGTCTGCGCCCGCAGCGG</td><td>1000</td><td></td><td></td><td></td><td></td><td></td><td></td><td></td><td></td><td></td></tr>
<tr><td>Oligo fwd</td><td>CACCGGCTGTGTCTGCGCCCGCAG</td><td></td><td></td><td></td><td></td><td></td><td></td><td></td><td></td><td></td><td></td></tr>
<tr><td>Oligo rev</td><td>AAACCTGCGGGCGCAGACACAGCC</td><td></td><td></td><td></td><td></td><td></td><td></td><td></td><td></td><td></td><td></td></tr>
<tr><td>Chromosome</td><td>start</td><td>end</td><td>strand</td><td>MM</td><td>target_seq</td><td>PAM</td><td>alignment</td><td>distance</td><td>position</td><td>gene name</td><td>gene id</td></tr>
<tr><td>GU305914.1</td><td>128788</td><td>128810</td><td>+</td><td>0</td><td>GGCTGTGTCTGCGCCCGCAG</td><td>CGG</td><td>||||||||[||||||||||||]PAM</td><td>NA</td><td>Intergenic</td><td>NA</td><td>NA</td></tr>
<tr><td></td><td></td><td></td><td></td><td></td><td></td><td></td><td></td><td></td><td></td><td></td><td></td></tr>
<tr><td>T9737</td><td>GCCCGCAGCGGCAGCGACGTCGG</td><td>1000</td><td></td><td></td><td></td><td></td><td></td><td></td><td></td><td></td><td></td></tr>
<tr><td>Oligo fwd</td><td>CACCGCCCGCAGCGGCAGCGACGT</td><td></td><td></td><td></td><td></td><td></td><td></td><td></td><td></td><td></td><td></td></tr>
<tr><td>Oligo rev</td><td>AAACACGTCGCTGCCGCTGCGGGC</td><td></td><td></td><td></td><td></td><td></td><td></td><td></td><td></td><td></td><td></td></tr>
<tr><td>Chromosome</td><td>start</td><td>end</td><td>strand</td><td>MM</td><td>target_seq</td><td>PAM</td><td>alignment</td><td>distance</td><td>position</td><td>gene name</td><td>gene id</td></tr>
<tr><td>GU305914.1</td><td>128800</td><td>128822</td><td>+</td><td>0</td><td>GCCCGCAGCGGCAGCGACGT</td><td>CGG</td><td>||||||||[||||||||||||]PAM</td><td>NA</td><td>Intergenic</td><td>NA</td><td>NA</td></tr>
<tr><td></td><td></td><td></td><td></td><td></td><td></td><td></td><td></td><td></td><td></td><td></td><td></td></tr>
<tr><td>T9738</td><td>CGCAGCGGCAGCGACGTCGGCGG</td><td>1000</td><td></td><td></td><td></td><td></td><td></td><td></td><td></td><td></td><td></td></tr>
<tr><td>Oligo fwd</td><td>CACCCGCAGCGGCAGCGACGTCGG</td><td></td><td></td><td></td><td></td><td></td><td></td><td></td><td></td><td></td><td></td></tr>
<tr><td>Oligo rev</td><td>AAACCCGACGTCGCTGCCGCTGCG</td><td></td><td></td><td></td><td></td><td></td><td></td><td></td><td></td><td></td><td></td></tr>
<tr><td>Chromosome</td><td>start</td><td>end</td><td>strand</td><td>MM</td><td>target_seq</td><td>PAM</td><td>alignment</td><td>distance</td><td>position</td><td>gene name</td><td>gene id</td></tr>
<tr><td>GU305914.1</td><td>128803</td><td>128825</td><td>+</td><td>0</td><td>CGCAGCGGCAGCGACGTCGG</td><td>CGG</td><td>||||||||[||||||||||||]PAM</td><td>NA</td><td>Intergenic</td><td>NA</td><td>NA</td></tr>
<tr><td>GU305914.1</td><td>198746</td><td>198768</td><td>-</td><td>4</td><td>CGCGGACGCGGCGACGTCGG</td><td>TGG</td><td>|||-|--|[|-||||||||||]PAM</td><td>NA</td><td>Intergenic</td><td>NA</td><td>NA</td></tr>
<tr><td>GU305914.1</td><td>116848</td><td>116870</td><td>+</td><td>4</td><td>TGCACCGTCAGCGAGGTCGG</td><td>CGG</td><td>-|||-||-[||||||-|||||]PAM</td><td>NA</td><td>Intergenic</td><td>NA</td><td>NA</td></tr>
<tr><td>GU305914.1</td><td>128745</td><td>128767</td><td>-</td><td>4</td><td>CGCGGAGGCATCGTCGTCGG</td><td>TGG</td><td>|||-|-||[||-||-||||||]PAM</td><td>NA</td><td>Intergenic</td><td>NA</td><td>NA</td></tr>
<tr><td>GU305914.1</td><td>110083</td><td>110105</td><td>-</td><td>4</td><td>CGCGGCCGCGGCGACGCCGG</td><td>TGG</td><td>|||-||-|[|-||||||-|||]PAM</td><td>NA</td><td>Intergenic</td><td>NA</td><td>NA</td></tr>
<tr><td>GU305914.1</td><td>70184</td><td>70206</td><td>+</td><td>4</td><td>AGCACCGGCAGCGCCGGCGG</td><td>CGG</td><td>-|||-|||[|||||-||-|||]PAM</td><td>NA</td><td>Intergenic</td><td>NA</td><td>NA</td></tr>
<tr><td></td><td></td><td></td><td></td><td></td><td></td><td></td><td></td><td></td><td></td><td></td><td></td></tr>
<tr><td>T9739</td><td>GGCAGCGACGTCGGCGGCGAAGG</td><td>1000</td><td></td><td></td><td></td><td></td><td></td><td></td><td></td><td></td><td></td></tr>
<tr><td>Oligo fwd</td><td>CACCGGCAGCGACGTCGGCGGCGA</td><td></td><td></td><td></td><td></td><td></td><td></td><td></td><td></td><td></td><td></td></tr>
<tr><td>Oligo rev</td><td>AAACTCGCCGCCGACGTCGCTGCC</td><td></td><td></td><td></td><td></td><td></td><td></td><td></td><td></td><td></td><td></td></tr>
<tr><td>Chromosome</td><td>start</td><td>end</td><td>strand</td><td>MM</td><td>target_seq</td><td>PAM</td><td>alignment</td><td>distance</td><td>position</td><td>gene name</td><td>gene id</td></tr>
<tr><td>GU305914.1</td><td>128809</td><td>128831</td><td>+</td><td>0</td><td>GGCAGCGACGTCGGCGGCGA</td><td>AGG</td><td>||||||||[||||||||||||]PAM</td><td>NA</td><td>Intergenic</td><td>NA</td><td>NA</td></tr>
<tr><td>GU305914.1</td><td>67742</td><td>67764</td><td>+</td><td>4</td><td>GGGAGCGGCGGCGGCGGCGG</td><td>CGG</td><td>||-||||-[||-||||||||-]PAM</td><td>NA</td><td>Intergenic</td><td>NA</td><td>NA</td></tr>
<tr><td>GU305914.1</td><td>166404</td><td>166426</td><td>+</td><td>3</td><td>GGCAGAGACGTCGGCGCCGG</td><td>CGG</td><td>|||||-||[||||||||-||-]PAM</td><td>NA</td><td>Intergenic</td><td>NA</td><td>NA</td></tr>
<tr><td></td><td></td><td></td><td></td><td></td><td></td><td></td><td></td><td></td><td></td><td></td><td></td></tr>
<tr><td>T9740</td><td>GCACGCGCCAGCCGCCCCCACGG</td><td>1000</td><td></td><td></td><td></td><td></td><td></td><td></td><td></td><td></td><td></td></tr>
<tr><td>Oligo fwd</td><td>CACCGCACGCGCCAGCCGCCCCCA</td><td></td><td></td><td></td><td></td><td></td><td></td><td></td><td></td><td></td><td></td></tr>
<tr><td>Oligo rev</td><td>AAACTGGGGGCGGCTGGCGCGTGC</td><td></td><td></td><td></td><td></td><td></td><td></td><td></td><td></td><td></td><td></td></tr>
<tr><td>Chromosome</td><td>start</td><td>end</td><td>strand</td><td>MM</td><td>target_seq</td><td>PAM</td><td>alignment</td><td>distance</td><td>position</td><td>gene name</td><td>gene id</td></tr>
<tr><td>GU305914.1</td><td>128839</td><td>128861</td><td>+</td><td>0</td><td>GCACGCGCCAGCCGCCCCCA</td><td>CGG</td><td>||||||||[||||||||||||]PAM</td><td>NA</td><td>Intergenic</td><td>NA</td><td>NA</td></tr>
<tr><td></td><td></td><td></td><td></td><td></td><td></td><td></td><td></td><td></td><td></td><td></td><td></td></tr>
<tr><td>T9741</td><td>CCAGCCGCCCCCACGGTCGTTGG</td><td>1000</td><td></td><td></td><td></td><td></td><td></td><td></td><td></td><td></td><td></td></tr>
<tr><td>Oligo fwd</td><td>CACCCCAGCCGCCCCCACGGTCGT</td><td></td><td></td><td></td><td></td><td></td><td></td><td></td><td></td><td></td><td></td></tr>
<tr><td>Oligo rev</td><td>AAACACGACCGTGGGGGCGGCTGG</td><td></td><td></td><td></td><td></td><td></td><td></td><td></td><td></td><td></td><td></td></tr>
<tr><td>Chromosome</td><td>start</td><td>end</td><td>strand</td><td>MM</td><td>target_seq</td><td>PAM</td><td>alignment</td><td>distance</td><td>position</td><td>gene name</td><td>gene id</td></tr>
<tr><td>GU305914.1</td><td>128846</td><td>128868</td><td>+</td><td>0</td><td>CCAGCCGCCCCCACGGTCGT</td><td>TGG</td><td>||||||||[||||||||||||]PAM</td><td>NA</td><td>Intergenic</td><td>NA</td><td>NA</td></tr>
<tr><td></td><td></td><td></td><td></td><td></td><td></td><td></td><td></td><td></td><td></td><td></td><td></td></tr>
<tr><td>T9742</td><td>GCCGCCCCCACGGTCGTTGGCGG</td><td>1000</td><td></td><td></td><td></td><td></td><td></td><td></td><td></td><td></td><td></td></tr>
<tr><td>Oligo fwd</td><td>CACCGCCGCCCCCACGGTCGTTGG</td><td></td><td></td><td></td><td></td><td></td><td></td><td></td><td></td><td></td><td></td></tr>
<tr><td>Oligo rev</td><td>AAACCCAACGACCGTGGGGGCGGC</td><td></td><td></td><td></td><td></td><td></td><td></td><td></td><td></td><td></td><td></td></tr>
<tr><td>Chromosome</td><td>start</td><td>end</td><td>strand</td><td>MM</td><td>target_seq</td><td>PAM</td><td>alignment</td><td>distance</td><td>position</td><td>gene name</td><td>gene id</td></tr>
<tr><td>GU305914.1</td><td>128849</td><td>128871</td><td>+</td><td>0</td><td>GCCGCCCCCACGGTCGTTGG</td><td>CGG</td><td>||||||||[||||||||||||]PAM</td><td>NA</td><td>Intergenic</td><td>NA</td><td>NA</td></tr>
<tr><td></td><td></td><td></td><td></td><td></td><td></td><td></td><td></td><td></td><td></td><td></td><td></td></tr>
<tr><td>T9743</td><td>GCCCCCACGGTCGTTGGCGGAGG</td><td>1000</td><td></td><td></td><td></td><td></td><td></td><td></td><td></td><td></td><td></td></tr>
<tr><td>Oligo fwd</td><td>CACCGCCCCCACGGTCGTTGGCGG</td><td></td><td></td><td></td><td></td><td></td><td></td><td></td><td></td><td></td><td></td></tr>
<tr><td>Oligo rev</td><td>AAACCCGCCAACGACCGTGGGGGC</td><td></td><td></td><td></td><td></td><td></td><td></td><td></td><td></td><td></td><td></td></tr>
<tr><td>Chromosome</td><td>start</td><td>end</td><td>strand</td><td>MM</td><td>target_seq</td><td>PAM</td><td>alignment</td><td>distance</td><td>position</td><td>gene name</td><td>gene id</td></tr>
<tr><td>GU305914.1</td><td>128852</td><td>128874</td><td>+</td><td>0</td><td>GCCCCCACGGTCGTTGGCGG</td><td>AGG</td><td>||||||||[||||||||||||]PAM</td><td>NA</td><td>Intergenic</td><td>NA</td><td>NA</td></tr>
<tr><td></td><td></td><td></td><td></td><td></td><td></td><td></td><td></td><td></td><td></td><td></td><td></td></tr>
<tr><td>T9744</td><td>CACGGTCGTTGGCGGAGGTCAGG</td><td>1000</td><td></td><td></td><td></td><td></td><td></td><td></td><td></td><td></td><td></td></tr>
<tr><td>Oligo fwd</td><td>CACCCACGGTCGTTGGCGGAGGTC</td><td></td><td></td><td></td><td></td><td></td><td></td><td></td><td></td><td></td><td></td></tr>
<tr><td>Oligo rev</td><td>AAACGACCTCCGCCAACGACCGTG</td><td></td><td></td><td></td><td></td><td></td><td></td><td></td><td></td><td></td><td></td></tr>
<tr><td>Chromosome</td><td>start</td><td>end</td><td>strand</td><td>MM</td><td>target_seq</td><td>PAM</td><td>alignment</td><td>distance</td><td>position</td><td>gene name</td><td>gene id</td></tr>
<tr><td>GU305914.1</td><td>128857</td><td>128879</td><td>+</td><td>0</td><td>CACGGTCGTTGGCGGAGGTC</td><td>AGG</td><td>||||||||[||||||||||||]PAM</td><td>NA</td><td>Intergenic</td><td>NA</td><td>NA</td></tr>
<tr><td></td><td></td><td></td><td></td><td></td><td></td><td></td><td></td><td></td><td></td><td></td><td></td></tr>
<tr><td>T9745</td><td>ACGGTCGTTGGCGGAGGTCAGGG</td><td>1000</td><td></td><td></td><td></td><td></td><td></td><td></td><td></td><td></td><td></td></tr>
<tr><td>Oligo fwd</td><td>CACCACGGTCGTTGGCGGAGGTCA</td><td></td><td></td><td></td><td></td><td></td><td></td><td></td><td></td><td></td><td></td></tr>
<tr><td>Oligo rev</td><td>AAACTGACCTCCGCCAACGACCGT</td><td></td><td></td><td></td><td></td><td></td><td></td><td></td><td></td><td></td><td></td></tr>
<tr><td>Chromosome</td><td>start</td><td>end</td><td>strand</td><td>MM</td><td>target_seq</td><td>PAM</td><td>alignment</td><td>distance</td><td>position</td><td>gene name</td><td>gene id</td></tr>
<tr><td>GU305914.1</td><td>128858</td><td>128880</td><td>+</td><td>0</td><td>ACGGTCGTTGGCGGAGGTCA</td><td>GGG</td><td>||||||||[||||||||||||]PAM</td><td>NA</td><td>Intergenic</td><td>NA</td><td>NA</td></tr>
<tr><td></td><td></td><td></td><td></td><td></td><td></td><td></td><td></td><td></td><td></td><td></td><td></td></tr>
<tr><td>T9746</td><td>CGGTCGTTGGCGGAGGTCAGGGG</td><td>1000</td><td></td><td></td><td></td><td></td><td></td><td></td><td></td><td></td><td></td></tr>
<tr><td>Oligo fwd</td><td>CACCCGGTCGTTGGCGGAGGTCAG</td><td></td><td></td><td></td><td></td><td></td><td></td><td></td><td></td><td></td><td></td></tr>
<tr><td>Oligo rev</td><td>AAACCTGACCTCCGCCAACGACCG</td><td></td><td></td><td></td><td></td><td></td><td></td><td></td><td></td><td></td><td></td></tr>
<tr><td>Chromosome</td><td>start</td><td>end</td><td>strand</td><td>MM</td><td>target_seq</td><td>PAM</td><td>alignment</td><td>distance</td><td>position</td><td>gene name</td><td>gene id</td></tr>
</table>

GU305914.1 128859 128881 + 0 CGGTCGTTGGCGGAGGTCAG GGG ||||||||[||||||||||||]PAM NA Intergenic NA NA
GU305914.1 22933 22955 + 4 CGGTCAGTGGCGGCGGTCGG CGG |||||--|[|||||-||||-|]PAM NA Intergenic NA NA

T9747 CGTAAAGATCGTAGATCGCCTGG 1000
Oligo fwd CACCCGTAAAGATCGTAGATCGCC
Oligo rev AAACGGCGATCTACGATCTTTACG
Chromosome start end strand MM target_seq PAM alignment distance position gene name gene id
GU305914.1 128914 128936 + 0 CGTAAAGATCGTAGATCGCC TGG ||||||||[||||||||||||]PAM NA Intergenic NA NA

T9748 GATCGTAGATCGCCTGGCTCTGG 1000
Oligo fwd CACCGATCGTAGATCGCCTGGCTC
Oligo rev AAACGAGCCAGGCGATCTACGATC
Chromosome start end strand MM target_seq PAM alignment distance position gene name gene id
GU305914.1 128920 128942 + 0 GATCGTAGATCGCCTGGCTC TGG ||||||||[||||||||||||]PAM NA Intergenic NA NA

T9749 TAGATCGCCTGGCTCTGGTTCGG 1000
Oligo fwd CACCTAGATCGCCTGGCTCTGGTT
Oligo rev AAACAACCAGAGCCAGGCGATCTA
Chromosome start end strand MM target_seq PAM alignment distance position gene name gene id
GU305914.1 128925 128947 + 0 TAGATCGCCTGGCTCTGGTT CGG ||||||||[||||||||||||]PAM NA Intergenic NA NA

T9750 CTCTGGTTCGGCTCCGCTTCAGG 1000
Oligo fwd CACCCTCTGGTTCGGCTCCGCTTC
Oligo rev AAACGAAGCGGAGCCGAACCAGAG
Chromosome start end strand MM target_seq PAM alignment distance position gene name gene id
GU305914.1 128937 128959 + 0 CTCTGGTTCGGCTCCGCTTC AGG ||||||||[||||||||||||]PAM NA Intergenic NA NA

T9751 TCTGGTTCGGCTCCGCTTCAGGG 1000
Oligo fwd CACCTCTGGTTCGGCTCCGCTTCA
Oligo rev AAACTGAAGCGGAGCCGAACCAGA
Chromosome start end strand MM target_seq PAM alignment distance position gene name gene id
GU305914.1 128938 128960 + 0 TCTGGTTCGGCTCCGCTTCA GGG ||||||||[||||||||||||]PAM NA Intergenic NA NA

T9752 TTCGGCTCCGCTTCAGGGAGAGG 1000
Oligo fwd CACCTTCGGCTCCGCTTCAGGGAG
Oligo rev AAACCTCCCTGAAGCGGAGCCGAA
Chromosome start end strand MM target_seq PAM alignment distance position gene name gene id
GU305914.1 128943 128965 + 0 TTCGGCTCCGCTTCAGGGAG AGG ||||||||[||||||||||||]PAM NA Intergenic NA NA

T9753 GGCTCCGCTTCAGGGAGAGGCGG 1000
Oligo fwd CACCGGCTCCGCTTCAGGGAGAGG
Oligo rev AAACCCTCTCCCTGAAGCGGAGCC
Chromosome start end strand MM target_seq PAM alignment distance position gene name gene id
GU305914.1 128946 128968 + 0 GGCTCCGCTTCAGGGAGAGG CGG ||||||||[||||||||||||]PAM NA Intergenic NA NA

T9754 CCGCTTCAGGGAGAGGCGGCTGG 1000
Oligo fwd CACCCCGCTTCAGGGAGAGGCGGC
Oligo rev AAACGCCGCCTCTCCCTGAAGCGG
Chromosome start end strand MM target_seq PAM alignment distance position gene name gene id
GU305914.1 128950 128972 + 0 CCGCTTCAGGGAGAGGCGGC TGG ||||||||[||||||||||||]PAM NA Intergenic NA NA

T9755 GGAGAGGCGGCTGGCCGCCGAGG 1000
Oligo fwd CACCGGAGAGGCGGCTGGCCGCCG
Oligo rev AAACCGGCGGCCAGCCGCCTCTCC
Chromosome start end strand MM target_seq PAM alignment distance position gene name gene id
GU305914.1 128959 128981 + 0 GGAGAGGCGGCTGGCCGCCG AGG ||||||||[||||||||||||]PAM NA Intergenic NA NA

T9756 CTGGCCGCCGAGGCAGCCGTCGG 1000
Oligo fwd CACCCTGGCCGCCGAGGCAGCCGT
Oligo rev AAACACGGCTGCCTCGGCGGCCAG
Chromosome start end strand MM target_seq PAM alignment distance position gene name gene id
GU305914.1 128969 128991 + 0 CTGGCCGCCGAGGCAGCCGT CGG ||||||||[||||||||||||]PAM NA Intergenic NA NA
GU305914.1 74028 74050 + 3 CTGGCAGCCGAGACAGGCGT CGG |||||-||[||||-|||-|||]PAM NA Intergenic NA NA
GU305914.1 24138 24160 - 4 TGGGCCGCCGAGGCCGACGT CGG --||||||[||||||-|-|||]PAM NA Intergenic NA NA

T9757 GCCGAGGCAGCCGTCGGCAGCGG 1000
Oligo fwd CACCGCCGAGGCAGCCGTCGGCAG
Oligo rev AAACCTGCCGACGGCTGCCTCGGC
Chromosome start end strand MM target_seq PAM alignment distance position gene name gene id
GU305914.1 128975 128997 + 0 GCCGAGGCAGCCGTCGGCAG CGG ||||||||[||||||||||||]PAM NA Intergenic NA NA

T9758 GCAGCCGTCGGCAGCGGCAGCGG 1000
Oligo fwd CACCGCAGCCGTCGGCAGCGGCAG
Oligo rev AAACCTGCCGCTGCCGACGGCTGC
Chromosome start end strand MM target_seq PAM alignment distance position gene name gene id
GU305914.1 128981 129003 + 0 GCAGCCGTCGGCAGCGGCAG CGG ||||||||[||||||||||||]PAM NA Intergenic NA NA
GU305914.1 163908 163930 + 4 GGCGGCGGCGGCAGCGGCAG CGG |--|-||-[||||||||||||]PAM NA Intergenic NA NA
GU305914.1 113924 113946 - 4 GTCGGCGTCGGCGGCGGCAG CGG |--|-|||[||||-|||||||]PAM NA Intergenic NA NA
GU305914.1 149485 149507 - 4 GCAGGCGACGGCGGCGGCGG CGG ||||-||-[||||-|||||-|]PAM NA Intergenic NA NA

T9759 CAGCGGCAGCGGCAGCGTTGAGG 1000
Oligo fwd CACCCAGCGGCAGCGGCAGCGTTG
Oligo rev AAACCAACGCTGCCGCTGCCGCTG
Chromosome start end strand MM target_seq PAM alignment distance position gene name gene id
GU305914.1 128992 129014 + 0 CAGCGGCAGCGGCAGCGTTG AGG ||||||||[||||||||||||]PAM NA Intergenic NA NA

T9760 AGCGGCAGCGGCAGCGTTGAGGG 1000
Oligo fwd CACCAGCGGCAGCGGCAGCGTTGA
Oligo rev AAACTCAACGCTGCCGCTGCCGCT

<table>
<tr><td>Chromosome</td><td>start</td><td>end</td><td>strand</td><td>MM</td><td>target_seq</td><td>PAM</td><td>alignment</td><td>distance</td><td>position</td><td>gene name</td><td>gene id</td></tr>
<tr><td>GU305914.1</td><td>128993</td><td>129015</td><td>+</td><td>0</td><td>AGCGGCAGCGGCAGCGTTGA</td><td>GGG</td><td>||||||||[||||||||||||]PAM</td><td>NA</td><td>Intergenic</td><td>NA</td><td>NA</td></tr>
<tr><td>GU305914.1</td><td>114470</td><td>114492</td><td>-</td><td>4</td><td>AGCGGAGGCGGCAGGGTTGG</td><td>CGG</td><td>|||||--|[||||||-||||-]PAM</td><td>NA</td><td>Intergenic</td><td>NA</td><td>NA</td></tr>
<tr><td>T9761</td><td>AGCGTTGAGGGCGCCGTCGCCGG</td><td>1000</td><td></td><td></td><td></td><td></td><td></td><td></td><td></td><td></td><td></td></tr>
<tr><td>Oligo fwd</td><td>CACCAGCGTTGAGGGCGCCGTCGC</td><td></td><td></td><td></td><td></td><td></td><td></td><td></td><td></td><td></td><td></td></tr>
<tr><td>Oligo rev</td><td>AAACGCGACGGCGCCCTCAACGCT</td><td></td><td></td><td></td><td></td><td></td><td></td><td></td><td></td><td></td><td></td></tr>
<tr><td>Chromosome</td><td>start</td><td>end</td><td>strand</td><td>MM</td><td>target_seq</td><td>PAM</td><td>alignment</td><td>distance</td><td>position</td><td>gene name</td><td>gene id</td></tr>
<tr><td>GU305914.1</td><td>129005</td><td>129027</td><td>+</td><td>0</td><td>AGCGTTGAGGGCGCCGTCGC</td><td>CGG</td><td>||||||||[||||||||||||]PAM</td><td>NA</td><td>Intergenic</td><td>NA</td><td>NA</td></tr>
<tr><td>T9762</td><td>GTTGAGGGCGCCGTCGCCGGCGG</td><td>1000</td><td></td><td></td><td></td><td></td><td></td><td></td><td></td><td></td><td></td></tr>
<tr><td>Oligo fwd</td><td>CACCGTTGAGGGCGCCGTCGCCGG</td><td></td><td></td><td></td><td></td><td></td><td></td><td></td><td></td><td></td><td></td></tr>
<tr><td>Oligo rev</td><td>AAACCCGGCGACGGCGCCCTCAAC</td><td></td><td></td><td></td><td></td><td></td><td></td><td></td><td></td><td></td><td></td></tr>
<tr><td>Chromosome</td><td>start</td><td>end</td><td>strand</td><td>MM</td><td>target_seq</td><td>PAM</td><td>alignment</td><td>distance</td><td>position</td><td>gene name</td><td>gene id</td></tr>
<tr><td>GU305914.1</td><td>129008</td><td>129030</td><td>+</td><td>0</td><td>GTTGAGGGCGCCGTCGCCGG</td><td>CGG</td><td>||||||||[||||||||||||]PAM</td><td>NA</td><td>Intergenic</td><td>NA</td><td>NA</td></tr>
<tr><td>T9763</td><td>GCCGTCGCCGGCGGCACCTCTGG</td><td>1000</td><td></td><td></td><td></td><td></td><td></td><td></td><td></td><td></td><td></td></tr>
<tr><td>Oligo fwd</td><td>CACCGCCGTCGCCGGCGGCACCTC</td><td></td><td></td><td></td><td></td><td></td><td></td><td></td><td></td><td></td><td></td></tr>
<tr><td>Oligo rev</td><td>AAACGAGGTGCCGCCGGCGACGGC</td><td></td><td></td><td></td><td></td><td></td><td></td><td></td><td></td><td></td><td></td></tr>
<tr><td>Chromosome</td><td>start</td><td>end</td><td>strand</td><td>MM</td><td>target_seq</td><td>PAM</td><td>alignment</td><td>distance</td><td>position</td><td>gene name</td><td>gene id</td></tr>
<tr><td>GU305914.1</td><td>129017</td><td>129039</td><td>+</td><td>0</td><td>GCCGTCGCCGGCGGCACCTC</td><td>TGG</td><td>||||||||[||||||||||||]PAM</td><td>NA</td><td>Intergenic</td><td>NA</td><td>NA</td></tr>
<tr><td>T9764</td><td>TGGCGTTCAGACCGACGTGAAGG</td><td>1000</td><td></td><td></td><td></td><td></td><td></td><td></td><td></td><td></td><td></td></tr>
<tr><td>Oligo fwd</td><td>CACCTGGCGTTCAGACCGACGTGA</td><td></td><td></td><td></td><td></td><td></td><td></td><td></td><td></td><td></td><td></td></tr>
<tr><td>Oligo rev</td><td>AAACTCACGTCGGTCTGAACGCCA</td><td></td><td></td><td></td><td></td><td></td><td></td><td></td><td></td><td></td><td></td></tr>
<tr><td>Chromosome</td><td>start</td><td>end</td><td>strand</td><td>MM</td><td>target_seq</td><td>PAM</td><td>alignment</td><td>distance</td><td>position</td><td>gene name</td><td>gene id</td></tr>
<tr><td>GU305914.1</td><td>129037</td><td>129059</td><td>+</td><td>0</td><td>TGGCGTTCAGACCGACGTGA</td><td>AGG</td><td>||||||||[||||||||||||]PAM</td><td>NA</td><td>Intergenic</td><td>NA</td><td>NA</td></tr>
<tr><td>T9765</td><td>CGTTCAGACCGACGTGAAGGCGG</td><td>1000</td><td></td><td></td><td></td><td></td><td></td><td></td><td></td><td></td><td></td></tr>
<tr><td>Oligo fwd</td><td>CACCCGTTCAGACCGACGTGAAGG</td><td></td><td></td><td></td><td></td><td></td><td></td><td></td><td></td><td></td><td></td></tr>
<tr><td>Oligo rev</td><td>AAACCCTTCACGTCGGTCTGAACG</td><td></td><td></td><td></td><td></td><td></td><td></td><td></td><td></td><td></td><td></td></tr>
<tr><td>Chromosome</td><td>start</td><td>end</td><td>strand</td><td>MM</td><td>target_seq</td><td>PAM</td><td>alignment</td><td>distance</td><td>position</td><td>gene name</td><td>gene id</td></tr>
<tr><td>GU305914.1</td><td>129040</td><td>129062</td><td>+</td><td>0</td><td>CGTTCAGACCGACGTGAAGG</td><td>CGG</td><td>||||||||[||||||||||||]PAM</td><td>NA</td><td>Intergenic</td><td>NA</td><td>NA</td></tr>
<tr><td>T9766</td><td>ATGCCTCGTCAGACATCGTGCGG</td><td>1000</td><td></td><td></td><td></td><td></td><td></td><td></td><td></td><td></td><td></td></tr>
<tr><td>Oligo fwd</td><td>CACCATGCCTCGTCAGACATCGTG</td><td></td><td></td><td></td><td></td><td></td><td></td><td></td><td></td><td></td><td></td></tr>
<tr><td>Oligo rev</td><td>AAACCACGATGTCTGACGAGGCAT</td><td></td><td></td><td></td><td></td><td></td><td></td><td></td><td></td><td></td><td></td></tr>
<tr><td>Chromosome</td><td>start</td><td>end</td><td>strand</td><td>MM</td><td>target_seq</td><td>PAM</td><td>alignment</td><td>distance</td><td>position</td><td>gene name</td><td>gene id</td></tr>
<tr><td>GU305914.1</td><td>129106</td><td>129128</td><td>+</td><td>0</td><td>ATGCCTCGTCAGACATCGTG</td><td>CGG</td><td>||||||||[||||||||||||]PAM</td><td>NA</td><td>Intergenic</td><td>NA</td><td>NA</td></tr>
<tr><td>T9767</td><td>CGTCAGACATCGTGCGGCCGAGG</td><td>1000</td><td></td><td></td><td></td><td></td><td></td><td></td><td></td><td></td><td></td></tr>
<tr><td>Oligo fwd</td><td>CACCCGTCAGACATCGTGCGGCCG</td><td></td><td></td><td></td><td></td><td></td><td></td><td></td><td></td><td></td><td></td></tr>
<tr><td>Oligo rev</td><td>AAACCGGCCGCACGATGTCTGACG</td><td></td><td></td><td></td><td></td><td></td><td></td><td></td><td></td><td></td><td></td></tr>
<tr><td>Chromosome</td><td>start</td><td>end</td><td>strand</td><td>MM</td><td>target_seq</td><td>PAM</td><td>alignment</td><td>distance</td><td>position</td><td>gene name</td><td>gene id</td></tr>
<tr><td>GU305914.1</td><td>129112</td><td>129134</td><td>+</td><td>0</td><td>CGTCAGACATCGTGCGGCCG</td><td>AGG</td><td>||||||||[||||||||||||]PAM</td><td>NA</td><td>Intergenic</td><td>NA</td><td>NA</td></tr>
<tr><td>T9768</td><td>GTCAGACATCGTGCGGCCGAGGG</td><td>1000</td><td></td><td></td><td></td><td></td><td></td><td></td><td></td><td></td><td></td></tr>
<tr><td>Oligo fwd</td><td>CACCGTCAGACATCGTGCGGCCGA</td><td></td><td></td><td></td><td></td><td></td><td></td><td></td><td></td><td></td><td></td></tr>
<tr><td>Oligo rev</td><td>AAACTCGGCCGCACGATGTCTGAC</td><td></td><td></td><td></td><td></td><td></td><td></td><td></td><td></td><td></td><td></td></tr>
<tr><td>Chromosome</td><td>start</td><td>end</td><td>strand</td><td>MM</td><td>target_seq</td><td>PAM</td><td>alignment</td><td>distance</td><td>position</td><td>gene name</td><td>gene id</td></tr>
<tr><td>GU305914.1</td><td>129113</td><td>129135</td><td>+</td><td>0</td><td>GTCAGACATCGTGCGGCCGA</td><td>GGG</td><td>||||||||[||||||||||||]PAM</td><td>NA</td><td>Intergenic</td><td>NA</td><td>NA</td></tr>
<tr><td>T9769</td><td>GGCCGAGGGCGTGCGACGCGAGG</td><td>1000</td><td></td><td></td><td></td><td></td><td></td><td></td><td></td><td></td><td></td></tr>
<tr><td>Oligo fwd</td><td>CACCGGCCGAGGGCGTGCGACGCG</td><td></td><td></td><td></td><td></td><td></td><td></td><td></td><td></td><td></td><td></td></tr>
<tr><td>Oligo rev</td><td>AAACCGCGTCGCACGCCCTCGGCC</td><td></td><td></td><td></td><td></td><td></td><td></td><td></td><td></td><td></td><td></td></tr>
<tr><td>Chromosome</td><td>start</td><td>end</td><td>strand</td><td>MM</td><td>target_seq</td><td>PAM</td><td>alignment</td><td>distance</td><td>position</td><td>gene name</td><td>gene id</td></tr>
<tr><td>GU305914.1</td><td>129127</td><td>129149</td><td>+</td><td>0</td><td>GGCCGAGGGCGTGCGACGCG</td><td>AGG</td><td>||||||||[||||||||||||]PAM</td><td>NA</td><td>Intergenic</td><td>NA</td><td>NA</td></tr>
<tr><td>T9770</td><td>GCCGAGGGCGTGCGACGCGAGGG</td><td>1000</td><td></td><td></td><td></td><td></td><td></td><td></td><td></td><td></td><td></td></tr>
<tr><td>Oligo fwd</td><td>CACCGCCGAGGGCGTGCGACGCGA</td><td></td><td></td><td></td><td></td><td></td><td></td><td></td><td></td><td></td><td></td></tr>
<tr><td>Oligo rev</td><td>AAACTCGCGTCGCACGCCCTCGGC</td><td></td><td></td><td></td><td></td><td></td><td></td><td></td><td></td><td></td><td></td></tr>
<tr><td>Chromosome</td><td>start</td><td>end</td><td>strand</td><td>MM</td><td>target_seq</td><td>PAM</td><td>alignment</td><td>distance</td><td>position</td><td>gene name</td><td>gene id</td></tr>
<tr><td>GU305914.1</td><td>129128</td><td>129150</td><td>+</td><td>0</td><td>GCCGAGGGCGTGCGACGCGA</td><td>GGG</td><td>||||||||[||||||||||||]PAM</td><td>NA</td><td>Intergenic</td><td>NA</td><td>NA</td></tr>
<tr><td>T9771</td><td>GGGCGTGCGACGCGAGGGCGAGG</td><td>1000</td><td></td><td></td><td></td><td></td><td></td><td></td><td></td><td></td><td></td></tr>
<tr><td>Oligo fwd</td><td>CACCGGGCGTGCGACGCGAGGGCG</td><td></td><td></td><td></td><td></td><td></td><td></td><td></td><td></td><td></td><td></td></tr>
<tr><td>Oligo rev</td><td>AAACCGCCCTCGCGTCGCACGCCC</td><td></td><td></td><td></td><td></td><td></td><td></td><td></td><td></td><td></td><td></td></tr>
<tr><td>Chromosome</td><td>start</td><td>end</td><td>strand</td><td>MM</td><td>target_seq</td><td>PAM</td><td>alignment</td><td>distance</td><td>position</td><td>gene name</td><td>gene id</td></tr>
<tr><td>GU305914.1</td><td>129133</td><td>129155</td><td>+</td><td>0</td><td>GGGCGTGCGACGCGAGGGCG</td><td>AGG</td><td>||||||||[||||||||||||]PAM</td><td>NA</td><td>Intergenic</td><td>NA</td><td>NA</td></tr>
<tr><td>T9772</td><td>GGCGTGCGACGCGAGGGCGAGGG</td><td>1000</td><td></td><td></td><td></td><td></td><td></td><td></td><td></td><td></td><td></td></tr>
<tr><td>Oligo fwd</td><td>CACCGGCGTGCGACGCGAGGGCGA</td><td></td><td></td><td></td><td></td><td></td><td></td><td></td><td></td><td></td><td></td></tr>
<tr><td>Oligo rev</td><td>AAACTCGCCCTCGCGTCGCACGCC</td><td></td><td></td><td></td><td></td><td></td><td></td><td></td><td></td><td></td><td></td></tr>
<tr><td>Chromosome</td><td>start</td><td>end</td><td>strand</td><td>MM</td><td>target_seq</td><td>PAM</td><td>alignment</td><td>distance</td><td>position</td><td>gene name</td><td>gene id</td></tr>
<tr><td>GU305914.1</td><td>129134</td><td>129156</td><td>+</td><td>0</td><td>GGCGTGCGACGCGAGGGCGA</td><td>GGG</td><td>||||||||[||||||||||||]PAM</td><td>NA</td><td>Intergenic</td><td>NA</td><td>NA</td></tr>
<tr><td>GU305914.1</td><td>200840</td><td>200862</td><td>+</td><td>4</td><td>GGCGTCCATGGCGAGGGCGA</td><td>CGG</td><td>|||||-|-[--||||||||||]PAM</td><td>NA</td><td>Intergenic</td><td>NA</td><td>NA</td></tr>
<tr><td>GU305914.1</td><td>112517</td><td>112539</td><td>+</td><td>4</td><td>CGCGAGAGACGCGAGGGTGA</td><td>CGG</td><td>-|||-|-|[|||||||||-||]PAM</td><td>NA</td><td>Intergenic</td><td>NA</td><td>NA</td></tr>
<tr><td>T9773</td><td>GAGGGCGAGGGCTCCGTTTACGG</td><td>1000</td><td></td><td></td><td></td><td></td><td></td><td></td><td></td><td></td><td></td></tr>
<tr><td>Oligo fwd</td><td>CACCGAGGGCGAGGGCTCCGTTTA</td><td></td><td></td><td></td><td></td><td></td><td></td><td></td><td></td><td></td><td></td></tr>
<tr><td>Oligo rev</td><td>AAACTAAACGGAGCCCTCGCCCTC</td><td></td><td></td><td></td><td></td><td></td><td></td><td></td><td></td><td></td><td></td></tr>
<tr><td>Chromosome</td><td>start</td><td>end</td><td>strand</td><td>MM</td><td>target_seq</td><td>PAM</td><td>alignment</td><td>distance</td><td>position</td><td>gene name</td><td>gene id</td></tr>
<tr><td>GU305914.1</td><td>129146</td><td>129168</td><td>+</td><td>0</td><td>GAGGGCGAGGGCTCCGTTTA</td><td>CGG</td><td>||||||||[||||||||||||]PAM</td><td>NA</td><td>Intergenic</td><td>NA</td><td>NA</td></tr>
<tr><td>T9774</td><td>TACGGCGCACGTAAGATCATCGG</td><td>1000</td><td></td><td></td><td></td><td></td><td></td><td></td><td></td><td></td><td></td></tr>
<tr><td>Oligo fwd</td><td>CACCTACGGCGCACGTAAGATCAT</td><td></td><td></td><td></td><td></td><td></td><td></td><td></td><td></td><td></td><td></td></tr>
<tr><td>Oligo rev</td><td>AAACATGATCTTACGTGCGCCGTA</td><td></td><td></td><td></td><td></td><td></td><td></td><td></td><td></td><td></td><td></td></tr>
<tr><td>Chromosome</td><td>start</td><td>end</td><td>strand</td><td>MM</td><td>target_seq</td><td>PAM</td><td>alignment</td><td>distance</td><td>position</td><td>gene name</td><td>gene id</td></tr>
<tr><td>GU305914.1</td><td>129164</td><td>129186</td><td>+</td><td>0</td><td>TACGGCGCACGTAAGATCAT</td><td>CGG</td><td>||||||||[||||||||||||]PAM</td><td>NA</td><td>Intergenic</td><td>NA</td><td>NA</td></tr>
</table>

<table>
<tr><td>T9775</td><td>GGCGCACGTAAGATCATCGGCGG</td><td>1000</td><td></td><td></td><td></td><td></td><td></td><td></td><td></td><td></td><td></td></tr>
<tr><td>Oligo fwd</td><td>CACCGGCGCACGTAAGATCATCGG</td><td></td><td></td><td></td><td></td><td></td><td></td><td></td><td></td><td></td><td></td></tr>
<tr><td>Oligo rev</td><td>AAACCCGATGATCTTACGTGCGCC</td><td></td><td></td><td></td><td></td><td></td><td></td><td></td><td></td><td></td><td></td></tr>
<tr><td>Chromosome</td><td>start</td><td>end</td><td>strand</td><td>MM</td><td>target_seq</td><td>PAM</td><td>alignment</td><td>distance</td><td>position</td><td>gene name</td><td>gene id</td></tr>
<tr><td>GU305914.1</td><td>129167</td><td>129189</td><td>+</td><td>0</td><td>GGCGCACGTAAGATCATCGG</td><td>CGG</td><td>||||||||[||||||||||||]PAM</td><td>NA</td><td>Intergenic</td><td>NA</td><td>NA</td></tr>
<tr><td>T9776</td><td>GCGCACGTAAGATCATCGGCGGG</td><td>1000</td><td></td><td></td><td></td><td></td><td></td><td></td><td></td><td></td><td></td></tr>
<tr><td>Oligo fwd</td><td>CACCGCGCACGTAAGATCATCGGC</td><td></td><td></td><td></td><td></td><td></td><td></td><td></td><td></td><td></td><td></td></tr>
<tr><td>Oligo rev</td><td>AAACGCCGATGATCTTACGTGCGC</td><td></td><td></td><td></td><td></td><td></td><td></td><td></td><td></td><td></td><td></td></tr>
<tr><td>Chromosome</td><td>start</td><td>end</td><td>strand</td><td>MM</td><td>target_seq</td><td>PAM</td><td>alignment</td><td>distance</td><td>position</td><td>gene name</td><td>gene id</td></tr>
<tr><td>GU305914.1</td><td>129168</td><td>129190</td><td>+</td><td>0</td><td>GCGCACGTAAGATCATCGGC</td><td>GGG</td><td>||||||||[||||||||||||]PAM</td><td>NA</td><td>Intergenic</td><td>NA</td><td>NA</td></tr>
<tr><td>T9777</td><td>CATCGGCGGGCGCGAGTTCTCGG</td><td>1000</td><td></td><td></td><td></td><td></td><td></td><td></td><td></td><td></td><td></td></tr>
<tr><td>Oligo fwd</td><td>CACCCATCGGCGGGCGCGAGTTCT</td><td></td><td></td><td></td><td></td><td></td><td></td><td></td><td></td><td></td><td></td></tr>
<tr><td>Oligo rev</td><td>AAACAGAACTCGCGCCCGCCGATG</td><td></td><td></td><td></td><td></td><td></td><td></td><td></td><td></td><td></td><td></td></tr>
<tr><td>Chromosome</td><td>start</td><td>end</td><td>strand</td><td>MM</td><td>target_seq</td><td>PAM</td><td>alignment</td><td>distance</td><td>position</td><td>gene name</td><td>gene id</td></tr>
<tr><td>GU305914.1</td><td>129181</td><td>129203</td><td>+</td><td>0</td><td>CATCGGCGGGCGCGAGTTCT</td><td>CGG</td><td>||||||||[||||||||||||]PAM</td><td>NA</td><td>Intergenic</td><td>NA</td><td>NA</td></tr>
<tr><td>T9778</td><td>TGACTGCCGTCTCGCTCAATCGG</td><td>1000</td><td></td><td></td><td></td><td></td><td></td><td></td><td></td><td></td><td></td></tr>
<tr><td>Oligo fwd</td><td>CACCTGACTGCCGTCTCGCTCAAT</td><td></td><td></td><td></td><td></td><td></td><td></td><td></td><td></td><td></td><td></td></tr>
<tr><td>Oligo rev</td><td>AAACATTGAGCGAGACGGCAGTCA</td><td></td><td></td><td></td><td></td><td></td><td></td><td></td><td></td><td></td><td></td></tr>
<tr><td>Chromosome</td><td>start</td><td>end</td><td>strand</td><td>MM</td><td>target_seq</td><td>PAM</td><td>alignment</td><td>distance</td><td>position</td><td>gene name</td><td>gene id</td></tr>
<tr><td>GU305914.1</td><td>129207</td><td>129229</td><td>+</td><td>0</td><td>TGACTGCCGTCTCGCTCAAT</td><td>CGG</td><td>||||||||[||||||||||||]PAM</td><td>NA</td><td>Intergenic</td><td>NA</td><td>NA</td></tr>
<tr><td>T9779</td><td>GACTGCCGTCTCGCTCAATCGGG</td><td>1000</td><td></td><td></td><td></td><td></td><td></td><td></td><td></td><td></td><td></td></tr>
<tr><td>Oligo fwd</td><td>CACCGACTGCCGTCTCGCTCAATC</td><td></td><td></td><td></td><td></td><td></td><td></td><td></td><td></td><td></td><td></td></tr>
<tr><td>Oligo rev</td><td>AAACGATTGAGCGAGACGGCAGTC</td><td></td><td></td><td></td><td></td><td></td><td></td><td></td><td></td><td></td><td></td></tr>
<tr><td>Chromosome</td><td>start</td><td>end</td><td>strand</td><td>MM</td><td>target_seq</td><td>PAM</td><td>alignment</td><td>distance</td><td>position</td><td>gene name</td><td>gene id</td></tr>
<tr><td>GU305914.1</td><td>129208</td><td>129230</td><td>+</td><td>0</td><td>GACTGCCGTCTCGCTCAATC</td><td>GGG</td><td>||||||||[||||||||||||]PAM</td><td>NA</td><td>Intergenic</td><td>NA</td><td>NA</td></tr>
<tr><td>T9780</td><td>TGCCGTCTCGCTCAATCGGGTGG</td><td>1000</td><td></td><td></td><td></td><td></td><td></td><td></td><td></td><td></td><td></td></tr>
<tr><td>Oligo fwd</td><td>CACCTGCCGTCTCGCTCAATCGGG</td><td></td><td></td><td></td><td></td><td></td><td></td><td></td><td></td><td></td><td></td></tr>
<tr><td>Oligo rev</td><td>AAACCCCGATTGAGCGAGACGGCA</td><td></td><td></td><td></td><td></td><td></td><td></td><td></td><td></td><td></td><td></td></tr>
<tr><td>Chromosome</td><td>start</td><td>end</td><td>strand</td><td>MM</td><td>target_seq</td><td>PAM</td><td>alignment</td><td>distance</td><td>position</td><td>gene name</td><td>gene id</td></tr>
<tr><td>GU305914.1</td><td>129211</td><td>129233</td><td>+</td><td>0</td><td>TGCCGTCTCGCTCAATCGGG</td><td>TGG</td><td>||||||||[||||||||||||]PAM</td><td>NA</td><td>Intergenic</td><td>NA</td><td>NA</td></tr>
<tr><td>T9781</td><td>CGACCATCGTGCAGACGCCCAGG</td><td>1000</td><td></td><td></td><td></td><td></td><td></td><td></td><td></td><td></td><td></td></tr>
<tr><td>Oligo fwd</td><td>CACCCGACCATCGTGCAGACGCCC</td><td></td><td></td><td></td><td></td><td></td><td></td><td></td><td></td><td></td><td></td></tr>
<tr><td>Oligo rev</td><td>AAACGGGCGTCTGCACGATGGTCG</td><td></td><td></td><td></td><td></td><td></td><td></td><td></td><td></td><td></td><td></td></tr>
<tr><td>Chromosome</td><td>start</td><td>end</td><td>strand</td><td>MM</td><td>target_seq</td><td>PAM</td><td>alignment</td><td>distance</td><td>position</td><td>gene name</td><td>gene id</td></tr>
<tr><td>GU305914.1</td><td>129276</td><td>129298</td><td>+</td><td>0</td><td>CGACCATCGTGCAGACGCCC</td><td>AGG</td><td>||||||||[||||||||||||]PAM</td><td>NA</td><td>Intergenic</td><td>NA</td><td>NA</td></tr>
<tr><td>GU305914.1</td><td>74894</td><td>74916</td><td>+</td><td>4</td><td>TGACGAGCGTGGAGACGCCC</td><td>GGG</td><td>-|||-|-|[|||-||||||||]PAM</td><td>NA</td><td>Intergenic</td><td>NA</td><td>NA</td></tr>
<tr><td>T9782</td><td>CTTCAAAGAACCCCCATATACGG</td><td>1000</td><td></td><td></td><td></td><td></td><td></td><td></td><td></td><td></td><td></td></tr>
<tr><td>Oligo fwd</td><td>CACCCTTCAAAGAACCCCCATATA</td><td></td><td></td><td></td><td></td><td></td><td></td><td></td><td></td><td></td><td></td></tr>
<tr><td>Oligo rev</td><td>AAACTATATGGGGGTTCTTTGAAG</td><td></td><td></td><td></td><td></td><td></td><td></td><td></td><td></td><td></td><td></td></tr>
<tr><td>Chromosome</td><td>start</td><td>end</td><td>strand</td><td>MM</td><td>target_seq</td><td>PAM</td><td>alignment</td><td>distance</td><td>position</td><td>gene name</td><td>gene id</td></tr>
<tr><td>GU305914.1</td><td>129358</td><td>129380</td><td>+</td><td>0</td><td>CTTCAAAGAACCCCCATATA</td><td>CGG</td><td>||||||||[||||||||||||]PAM</td><td>NA</td><td>Intergenic</td><td>NA</td><td>NA</td></tr>
<tr><td>T9783</td><td>CACATTCGTCTCTAACGACGCGG</td><td>1000</td><td></td><td></td><td></td><td></td><td></td><td></td><td></td><td></td><td></td></tr>
<tr><td>Oligo fwd</td><td>CACCCACATTCGTCTCTAACGACG</td><td></td><td></td><td></td><td></td><td></td><td></td><td></td><td></td><td></td><td></td></tr>
<tr><td>Oligo rev</td><td>AAACCGTCGTTAGAGACGAATGTG</td><td></td><td></td><td></td><td></td><td></td><td></td><td></td><td></td><td></td><td></td></tr>
<tr><td>Chromosome</td><td>start</td><td>end</td><td>strand</td><td>MM</td><td>target_seq</td><td>PAM</td><td>alignment</td><td>distance</td><td>position</td><td>gene name</td><td>gene id</td></tr>
<tr><td>GU305914.1</td><td>129385</td><td>129407</td><td>+</td><td>0</td><td>CACATTCGTCTCTAACGACG</td><td>CGG</td><td>||||||||[||||||||||||]PAM</td><td>NA</td><td>Intergenic</td><td>NA</td><td>NA</td></tr>
<tr><td>GU305914.1</td><td>44498</td><td>44520</td><td>+</td><td>4</td><td>CGCGTTCGTCTTGAACGACG</td><td>CGG</td><td>|-|-||||[|||--|||||||]PAM</td><td>NA</td><td>Intergenic</td><td>NA</td><td>NA</td></tr>
<tr><td>T9784</td><td>ACTCTCTAAACGTGAATATCAGG</td><td>1000</td><td></td><td></td><td></td><td></td><td></td><td></td><td></td><td></td><td></td></tr>
<tr><td>Oligo fwd</td><td>CACCACTCTCTAAACGTGAATATC</td><td></td><td></td><td></td><td></td><td></td><td></td><td></td><td></td><td></td><td></td></tr>
<tr><td>Oligo rev</td><td>AAACGATATTCACGTTTAGAGAGT</td><td></td><td></td><td></td><td></td><td></td><td></td><td></td><td></td><td></td><td></td></tr>
<tr><td>Chromosome</td><td>start</td><td>end</td><td>strand</td><td>MM</td><td>target_seq</td><td>PAM</td><td>alignment</td><td>distance</td><td>position</td><td>gene name</td><td>gene id</td></tr>
<tr><td>GU305914.1</td><td>129417</td><td>129439</td><td>+</td><td>0</td><td>ACTCTCTAAACGTGAATATC</td><td>AGG</td><td>||||||||[||||||||||||]PAM</td><td>NA</td><td>Intergenic</td><td>NA</td><td>NA</td></tr>
<tr><td>T9785</td><td>CTCTCTAAACGTGAATATCAGGG</td><td>1000</td><td></td><td></td><td></td><td></td><td></td><td></td><td></td><td></td><td></td></tr>
<tr><td>Oligo fwd</td><td>CACCCTCTCTAAACGTGAATATCA</td><td></td><td></td><td></td><td></td><td></td><td></td><td></td><td></td><td></td><td></td></tr>
<tr><td>Oligo rev</td><td>AAACTGATATTCACGTTTAGAGAG</td><td></td><td></td><td></td><td></td><td></td><td></td><td></td><td></td><td></td><td></td></tr>
<tr><td>Chromosome</td><td>start</td><td>end</td><td>strand</td><td>MM</td><td>target_seq</td><td>PAM</td><td>alignment</td><td>distance</td><td>position</td><td>gene name</td><td>gene id</td></tr>
<tr><td>GU305914.1</td><td>129418</td><td>129440</td><td>+</td><td>0</td><td>CTCTCTAAACGTGAATATCA</td><td>GGG</td><td>||||||||[||||||||||||]PAM</td><td>NA</td><td>Intergenic</td><td>NA</td><td>NA</td></tr>
<tr><td>T9786</td><td>TCTCTAAACGTGAATATCAGGGG</td><td>1000</td><td></td><td></td><td></td><td></td><td></td><td></td><td></td><td></td><td></td></tr>
<tr><td>Oligo fwd</td><td>CACCTCTCTAAACGTGAATATCAG</td><td></td><td></td><td></td><td></td><td></td><td></td><td></td><td></td><td></td><td></td></tr>
<tr><td>Oligo rev</td><td>AAACCTGATATTCACGTTTAGAGA</td><td></td><td></td><td></td><td></td><td></td><td></td><td></td><td></td><td></td><td></td></tr>
<tr><td>Chromosome</td><td>start</td><td>end</td><td>strand</td><td>MM</td><td>target_seq</td><td>PAM</td><td>alignment</td><td>distance</td><td>position</td><td>gene name</td><td>gene id</td></tr>
<tr><td>GU305914.1</td><td>129419</td><td>129441</td><td>+</td><td>0</td><td>TCTCTAAACGTGAATATCAG</td><td>GGG</td><td>||||||||[||||||||||||]PAM</td><td>NA</td><td>Intergenic</td><td>NA</td><td>NA</td></tr>
<tr><td>T9787</td><td>GAATATCAGGGGCTCCTACCAGG</td><td>1000</td><td></td><td></td><td></td><td></td><td></td><td></td><td></td><td></td><td></td></tr>
<tr><td>Oligo fwd</td><td>CACCGAATATCAGGGGCTCCTACC</td><td></td><td></td><td></td><td></td><td></td><td></td><td></td><td></td><td></td><td></td></tr>
<tr><td>Oligo rev</td><td>AAACGGTAGGAGCCCCTGATATTC</td><td></td><td></td><td></td><td></td><td></td><td></td><td></td><td></td><td></td><td></td></tr>
<tr><td>Chromosome</td><td>start</td><td>end</td><td>strand</td><td>MM</td><td>target_seq</td><td>PAM</td><td>alignment</td><td>distance</td><td>position</td><td>gene name</td><td>gene id</td></tr>
<tr><td>GU305914.1</td><td>129430</td><td>129452</td><td>+</td><td>0</td><td>GAATATCAGGGGCTCCTACC</td><td>AGG</td><td>||||||||[||||||||||||]PAM</td><td>NA</td><td>Intergenic</td><td>NA</td><td>NA</td></tr>
<tr><td>T9788</td><td>AACTCCTCGCTAGACGTACACGG</td><td>1000</td><td></td><td></td><td></td><td></td><td></td><td></td><td></td><td></td><td></td></tr>
<tr><td>Oligo fwd</td><td>CACCAACTCCTCGCTAGACGTACA</td><td></td><td></td><td></td><td></td><td></td><td></td><td></td><td></td><td></td><td></td></tr>
<tr><td>Oligo rev</td><td>AAACTGTACGTCTAGCGAGGAGTT</td><td></td><td></td><td></td><td></td><td></td><td></td><td></td><td></td><td></td><td></td></tr>
<tr><td>Chromosome</td><td>start</td><td>end</td><td>strand</td><td>MM</td><td>target_seq</td><td>PAM</td><td>alignment</td><td>distance</td><td>position</td><td>gene name</td><td>gene id</td></tr>
<tr><td>GU305914.1</td><td>129533</td><td>129555</td><td>+</td><td>0</td><td>AACTCCTCGCTAGACGTACA</td><td>CGG</td><td>||||||||[||||||||||||]PAM</td><td>NA</td><td>Intergenic</td><td>NA</td><td>NA</td></tr>
<tr><td>T9789</td><td>ACTCCTCGCTAGACGTACACGGG</td><td>1000</td><td></td><td></td><td></td><td></td><td></td><td></td><td></td><td></td><td></td></tr>
<tr><td>Oligo fwd</td><td>CACCACTCCTCGCTAGACGTACAC</td><td></td><td></td><td></td><td></td><td></td><td></td><td></td><td></td><td></td><td></td></tr>
</table>

<table>
<tr><td>Oligo rev</td><td>AAACGTGTACGTCTAGCGAGGAGT</td><td></td><td></td><td></td><td></td><td></td><td></td><td></td><td></td><td></td><td></td><td></td></tr>
<tr><td>Chromosome</td><td>start</td><td></td><td>end</td><td>strand</td><td>MM</td><td>target_seq</td><td>PAM</td><td>alignment</td><td>distance</td><td>position</td><td>gene name</td><td>gene id</td></tr>
<tr><td>GU305914.1</td><td></td><td>129534</td><td>129556</td><td>+</td><td>0</td><td>ACTCCTCGCTAGACGTACAC</td><td>GGG</td><td>||||||||[||||||||||||]PAM</td><td>NA</td><td>Intergenic</td><td>NA</td><td>NA</td></tr>
<tr><td>T9790</td><td>GCTAGACGTACACGGGATCGAGG</td><td></td><td>1000</td><td></td><td></td><td></td><td></td><td></td><td></td><td></td><td></td><td></td></tr>
<tr><td>Oligo fwd</td><td>CACCGCTAGACGTACACGGGATCG</td><td></td><td></td><td></td><td></td><td></td><td></td><td></td><td></td><td></td><td></td><td></td></tr>
<tr><td>Oligo rev</td><td>AAACCGATCCCGTGTACGTCTAGC</td><td></td><td></td><td></td><td></td><td></td><td></td><td></td><td></td><td></td><td></td><td></td></tr>
<tr><td>Chromosome</td><td>start</td><td></td><td>end</td><td>strand</td><td>MM</td><td>target_seq</td><td>PAM</td><td>alignment</td><td>distance</td><td>position</td><td>gene name</td><td>gene id</td></tr>
<tr><td>GU305914.1</td><td></td><td>129541</td><td>129563</td><td>+</td><td>0</td><td>GCTAGACGTACACGGGATCG</td><td>AGG</td><td>||||||||[||||||||||||]PAM</td><td>NA</td><td>Intergenic</td><td>NA</td><td>NA</td></tr>
<tr><td>T9791</td><td>CGTACACGGGATCGAGGACCAGG</td><td></td><td>1000</td><td></td><td></td><td></td><td></td><td></td><td></td><td></td><td></td><td></td></tr>
<tr><td>Oligo fwd</td><td>CACCCGTACACGGGATCGAGGACC</td><td></td><td></td><td></td><td></td><td></td><td></td><td></td><td></td><td></td><td></td><td></td></tr>
<tr><td>Oligo rev</td><td>AAACGGTCCTCGATCCCGTGTACG</td><td></td><td></td><td></td><td></td><td></td><td></td><td></td><td></td><td></td><td></td><td></td></tr>
<tr><td>Chromosome</td><td>start</td><td></td><td>end</td><td>strand</td><td>MM</td><td>target_seq</td><td>PAM</td><td>alignment</td><td>distance</td><td>position</td><td>gene name</td><td>gene id</td></tr>
<tr><td>GU305914.1</td><td></td><td>129547</td><td>129569</td><td>+</td><td>0</td><td>CGTACACGGGATCGAGGACC</td><td>AGG</td><td>||||||||[||||||||||||]PAM</td><td>NA</td><td>Intergenic</td><td>NA</td><td>NA</td></tr>
<tr><td>T9792</td><td>GTACACGGGATCGAGGACCAGGG</td><td></td><td>1000</td><td></td><td></td><td></td><td></td><td></td><td></td><td></td><td></td><td></td></tr>
<tr><td>Oligo fwd</td><td>CACCGTACACGGGATCGAGGACCA</td><td></td><td></td><td></td><td></td><td></td><td></td><td></td><td></td><td></td><td></td><td></td></tr>
<tr><td>Oligo rev</td><td>AAACTGGTCCTCGATCCCGTGTAC</td><td></td><td></td><td></td><td></td><td></td><td></td><td></td><td></td><td></td><td></td><td></td></tr>
<tr><td>Chromosome</td><td>start</td><td></td><td>end</td><td>strand</td><td>MM</td><td>target_seq</td><td>PAM</td><td>alignment</td><td>distance</td><td>position</td><td>gene name</td><td>gene id</td></tr>
<tr><td>GU305914.1</td><td></td><td>129548</td><td>129570</td><td>+</td><td>0</td><td>GTACACGGGATCGAGGACCA</td><td>GGG</td><td>||||||||[||||||||||||]PAM</td><td>NA</td><td>Intergenic</td><td>NA</td><td>NA</td></tr>
<tr><td>T9793</td><td>TACACGGGATCGAGGACCAGGGG</td><td></td><td>1000</td><td></td><td></td><td></td><td></td><td></td><td></td><td></td><td></td><td></td></tr>
<tr><td>Oligo fwd</td><td>CACCTACACGGGATCGAGGACCAG</td><td></td><td></td><td></td><td></td><td></td><td></td><td></td><td></td><td></td><td></td><td></td></tr>
<tr><td>Oligo rev</td><td>AAACCTGGTCCTCGATCCCGTGTA</td><td></td><td></td><td></td><td></td><td></td><td></td><td></td><td></td><td></td><td></td><td></td></tr>
<tr><td>Chromosome</td><td>start</td><td></td><td>end</td><td>strand</td><td>MM</td><td>target_seq</td><td>PAM</td><td>alignment</td><td>distance</td><td>position</td><td>gene name</td><td>gene id</td></tr>
<tr><td>GU305914.1</td><td></td><td>129549</td><td>129571</td><td>+</td><td>0</td><td>TACACGGGATCGAGGACCAG</td><td>GGG</td><td>||||||||[||||||||||||]PAM</td><td>NA</td><td>Intergenic</td><td>NA</td><td>NA</td></tr>
<tr><td>T9794</td><td>ACACGGGATCGAGGACCAGGGGG</td><td></td><td>1000</td><td></td><td></td><td></td><td></td><td></td><td></td><td></td><td></td><td></td></tr>
<tr><td>Oligo fwd</td><td>CACCACACGGGATCGAGGACCAGG</td><td></td><td></td><td></td><td></td><td></td><td></td><td></td><td></td><td></td><td></td><td></td></tr>
<tr><td>Oligo rev</td><td>AAACCCTGGTCCTCGATCCCGTGT</td><td></td><td></td><td></td><td></td><td></td><td></td><td></td><td></td><td></td><td></td><td></td></tr>
<tr><td>Chromosome</td><td>start</td><td></td><td>end</td><td>strand</td><td>MM</td><td>target_seq</td><td>PAM</td><td>alignment</td><td>distance</td><td>position</td><td>gene name</td><td>gene id</td></tr>
<tr><td>GU305914.1</td><td></td><td>129550</td><td>129572</td><td>+</td><td>0</td><td>ACACGGGATCGAGGACCAGG</td><td>GGG</td><td>||||||||[||||||||||||]PAM</td><td>NA</td><td>Intergenic</td><td>NA</td><td>NA</td></tr>
<tr><td>T9795</td><td>AGGGGGTCATCCGATCCGAGCGG</td><td></td><td>1000</td><td></td><td></td><td></td><td></td><td></td><td></td><td></td><td></td><td></td></tr>
<tr><td>Oligo fwd</td><td>CACCAGGGGGTCATCCGATCCGAG</td><td></td><td></td><td></td><td></td><td></td><td></td><td></td><td></td><td></td><td></td><td></td></tr>
<tr><td>Oligo rev</td><td>AAACCTCGGATCGGATGACCCCCT</td><td></td><td></td><td></td><td></td><td></td><td></td><td></td><td></td><td></td><td></td><td></td></tr>
<tr><td>Chromosome</td><td>start</td><td></td><td>end</td><td>strand</td><td>MM</td><td>target_seq</td><td>PAM</td><td>alignment</td><td>distance</td><td>position</td><td>gene name</td><td>gene id</td></tr>
<tr><td>GU305914.1</td><td></td><td>129567</td><td>129589</td><td>+</td><td>0</td><td>AGGGGGTCATCCGATCCGAG</td><td>CGG</td><td>||||||||[||||||||||||]PAM</td><td>NA</td><td>Intergenic</td><td>NA</td><td>NA</td></tr>
<tr><td>T9796</td><td>GGGGGTCATCCGATCCGAGCGGG</td><td></td><td>1000</td><td></td><td></td><td></td><td></td><td></td><td></td><td></td><td></td><td></td></tr>
<tr><td>Oligo fwd</td><td>CACCGGGGGTCATCCGATCCGAGC</td><td></td><td></td><td></td><td></td><td></td><td></td><td></td><td></td><td></td><td></td><td></td></tr>
<tr><td>Oligo rev</td><td>AAACGCTCGGATCGGATGACCCCC</td><td></td><td></td><td></td><td></td><td></td><td></td><td></td><td></td><td></td><td></td><td></td></tr>
<tr><td>Chromosome</td><td>start</td><td></td><td>end</td><td>strand</td><td>MM</td><td>target_seq</td><td>PAM</td><td>alignment</td><td>distance</td><td>position</td><td>gene name</td><td>gene id</td></tr>
<tr><td>GU305914.1</td><td></td><td>129568</td><td>129590</td><td>+</td><td>0</td><td>GGGGGTCATCCGATCCGAGC</td><td>GGG</td><td>||||||||[||||||||||||]PAM</td><td>NA</td><td>Intergenic</td><td>NA</td><td>NA</td></tr>
<tr><td>T9797</td><td>CATCCGATCCGAGCGGGATAAGG</td><td></td><td>1000</td><td></td><td></td><td></td><td></td><td></td><td></td><td></td><td></td><td></td></tr>
<tr><td>Oligo fwd</td><td>CACCCATCCGATCCGAGCGGGATA</td><td></td><td></td><td></td><td></td><td></td><td></td><td></td><td></td><td></td><td></td><td></td></tr>
<tr><td>Oligo rev</td><td>AAACTATCCCGCTCGGATCGGATG</td><td></td><td></td><td></td><td></td><td></td><td></td><td></td><td></td><td></td><td></td><td></td></tr>
<tr><td>Chromosome</td><td>start</td><td></td><td>end</td><td>strand</td><td>MM</td><td>target_seq</td><td>PAM</td><td>alignment</td><td>distance</td><td>position</td><td>gene name</td><td>gene id</td></tr>
<tr><td>GU305914.1</td><td></td><td>129574</td><td>129596</td><td>+</td><td>0</td><td>CATCCGATCCGAGCGGGATA</td><td>AGG</td><td>||||||||[||||||||||||]PAM</td><td>NA</td><td>Intergenic</td><td>NA</td><td>NA</td></tr>
<tr><td>T9798</td><td>CCGAGCGGGATAAGGTCTACTGG</td><td></td><td>1000</td><td></td><td></td><td></td><td></td><td></td><td></td><td></td><td></td><td></td></tr>
<tr><td>Oligo fwd</td><td>CACCCCGAGCGGGATAAGGTCTAC</td><td></td><td></td><td></td><td></td><td></td><td></td><td></td><td></td><td></td><td></td><td></td></tr>
<tr><td>Oligo rev</td><td>AAACGTAGACCTTATCCCGCTCGG</td><td></td><td></td><td></td><td></td><td></td><td></td><td></td><td></td><td></td><td></td><td></td></tr>
<tr><td>Chromosome</td><td>start</td><td></td><td>end</td><td>strand</td><td>MM</td><td>target_seq</td><td>PAM</td><td>alignment</td><td>distance</td><td>position</td><td>gene name</td><td>gene id</td></tr>
<tr><td>GU305914.1</td><td></td><td>129582</td><td>129604</td><td>+</td><td>0</td><td>CCGAGCGGGATAAGGTCTAC</td><td>TGG</td><td>||||||||[||||||||||||]PAM</td><td>NA</td><td>Intergenic</td><td>NA</td><td>NA</td></tr>
<tr><td>T9799</td><td>AACACCGACAACATTAACGTCGG</td><td></td><td>1000</td><td></td><td></td><td></td><td></td><td></td><td></td><td></td><td></td><td></td></tr>
<tr><td>Oligo fwd</td><td>CACCAACACCGACAACATTAACGT</td><td></td><td></td><td></td><td></td><td></td><td></td><td></td><td></td><td></td><td></td><td></td></tr>
<tr><td>Oligo rev</td><td>AAACACGTTAATGTTGTCGGTGTT</td><td></td><td></td><td></td><td></td><td></td><td></td><td></td><td></td><td></td><td></td><td></td></tr>
<tr><td>Chromosome</td><td>start</td><td></td><td>end</td><td>strand</td><td>MM</td><td>target_seq</td><td>PAM</td><td>alignment</td><td>distance</td><td>position</td><td>gene name</td><td>gene id</td></tr>
<tr><td>GU305914.1</td><td></td><td>129632</td><td>129654</td><td>+</td><td>0</td><td>AACACCGACAACATTAACGT</td><td>CGG</td><td>||||||||[||||||||||||]PAM</td><td>NA</td><td>Intergenic</td><td>NA</td><td>NA</td></tr>
<tr><td>T9800</td><td>CCGACAACATTAACGTCGGCTGG</td><td></td><td>1000</td><td></td><td></td><td></td><td></td><td></td><td></td><td></td><td></td><td></td></tr>
<tr><td>Oligo fwd</td><td>CACCCCGACAACATTAACGTCGGC</td><td></td><td></td><td></td><td></td><td></td><td></td><td></td><td></td><td></td><td></td><td></td></tr>
<tr><td>Oligo rev</td><td>AAACGCCGACGTTAATGTTGTCGG</td><td></td><td></td><td></td><td></td><td></td><td></td><td></td><td></td><td></td><td></td><td></td></tr>
<tr><td>Chromosome</td><td>start</td><td></td><td>end</td><td>strand</td><td>MM</td><td>target_seq</td><td>PAM</td><td>alignment</td><td>distance</td><td>position</td><td>gene name</td><td>gene id</td></tr>
<tr><td>GU305914.1</td><td></td><td>129636</td><td>129658</td><td>+</td><td>0</td><td>CCGACAACATTAACGTCGGC</td><td>TGG</td><td>||||||||[||||||||||||]PAM</td><td>NA</td><td>Intergenic</td><td>NA</td><td>NA</td></tr>
<tr><td>T9801</td><td>CATTAACGTCGGCTGGTTCAAGG</td><td></td><td>1000</td><td></td><td></td><td></td><td></td><td></td><td></td><td></td><td></td><td></td></tr>
<tr><td>Oligo fwd</td><td>CACCCATTAACGTCGGCTGGTTCA</td><td></td><td></td><td></td><td></td><td></td><td></td><td></td><td></td><td></td><td></td><td></td></tr>
<tr><td>Oligo rev</td><td>AAACTGAACCAGCCGACGTTAATG</td><td></td><td></td><td></td><td></td><td></td><td></td><td></td><td></td><td></td><td></td><td></td></tr>
<tr><td>Chromosome</td><td>start</td><td></td><td>end</td><td>strand</td><td>MM</td><td>target_seq</td><td>PAM</td><td>alignment</td><td>distance</td><td>position</td><td>gene name</td><td>gene id</td></tr>
<tr><td>GU305914.1</td><td></td><td>129643</td><td>129665</td><td>+</td><td>0</td><td>CATTAACGTCGGCTGGTTCA</td><td>AGG</td><td>||||||||[||||||||||||]PAM</td><td>NA</td><td>Intergenic</td><td>NA</td><td>NA</td></tr>
<tr><td>T9802</td><td>CGCGACCGCCATCATACCTAAGG</td><td></td><td>1000</td><td></td><td></td><td></td><td></td><td></td><td></td><td></td><td></td><td></td></tr>
<tr><td>Oligo fwd</td><td>CACCCGCGACCGCCATCATACCTA</td><td></td><td></td><td></td><td></td><td></td><td></td><td></td><td></td><td></td><td></td><td></td></tr>
<tr><td>Oligo rev</td><td>AAACTAGGTATGATGGCGGTCGCG</td><td></td><td></td><td></td><td></td><td></td><td></td><td></td><td></td><td></td><td></td><td></td></tr>
<tr><td>Chromosome</td><td>start</td><td></td><td>end</td><td>strand</td><td>MM</td><td>target_seq</td><td>PAM</td><td>alignment</td><td>distance</td><td>position</td><td>gene name</td><td>gene id</td></tr>
<tr><td>GU305914.1</td><td></td><td>129667</td><td>129689</td><td>+</td><td>0</td><td>CGCGACCGCCATCATACCTA</td><td>AGG</td><td>||||||||[||||||||||||]PAM</td><td>NA</td><td>Intergenic</td><td>NA</td><td>NA</td></tr>
<tr><td>T9803</td><td>GCCATCATACCTAAGGTCTCCGG</td><td></td><td>1000</td><td></td><td></td><td></td><td></td><td></td><td></td><td></td><td></td><td></td></tr>
<tr><td>Oligo fwd</td><td>CACCGCCATCATACCTAAGGTCTC</td><td></td><td></td><td></td><td></td><td></td><td></td><td></td><td></td><td></td><td></td><td></td></tr>
<tr><td>Oligo rev</td><td>AAACGAGACCTTAGGTATGATGGC</td><td></td><td></td><td></td><td></td><td></td><td></td><td></td><td></td><td></td><td></td><td></td></tr>
<tr><td>Chromosome</td><td>start</td><td></td><td>end</td><td>strand</td><td>MM</td><td>target_seq</td><td>PAM</td><td>alignment</td><td>distance</td><td>position</td><td>gene name</td><td>gene id</td></tr>
<tr><td>GU305914.1</td><td></td><td>129674</td><td>129696</td><td>+</td><td>0</td><td>GCCATCATACCTAAGGTCTC</td><td>CGG</td><td>||||||||[||||||||||||]PAM</td><td>NA</td><td>Intergenic</td><td>NA</td><td>NA</td></tr>
<tr><td>T9804</td><td>GAACGTGCTGCTCAAAGAGCTGG</td><td></td><td>1000</td><td></td><td></td><td></td><td></td><td></td><td></td><td></td><td></td><td></td></tr>
</table>

<table>
<tr><td>Oligo fwd</td><td>CACCGAACGTGCTGCTCAAAGAGC</td><td></td><td></td><td></td><td></td><td></td><td></td><td></td><td></td><td></td><td></td></tr>
<tr><td>Oligo rev</td><td>AAACGCTCTTTGAGCAGCACGTTC</td><td></td><td></td><td></td><td></td><td></td><td></td><td></td><td></td><td></td><td></td></tr>
<tr><td>Chromosome</td><td>start</td><td>end</td><td>strand</td><td>MM</td><td>target_seq</td><td>PAM</td><td>alignment</td><td>distance</td><td>position</td><td>gene name</td><td>gene id</td></tr>
<tr><td>GU305914.1</td><td>129709</td><td>129731</td><td>+</td><td>0</td><td>GAACGTGCTGCTCAAAGAGC</td><td>TGG</td><td>||||||||[||||||||||||]PAM</td><td>NA</td><td>Intergenic</td><td>NA</td><td>NA</td></tr>
<tr><td></td><td></td><td></td><td></td><td></td><td></td><td></td><td></td><td></td><td></td><td></td><td></td></tr>
<tr><td>T9805</td><td>AGCAGTTTATTTTATTCGTGCGG</td><td>1000</td><td></td><td></td><td></td><td></td><td></td><td></td><td></td><td></td><td></td></tr>
<tr><td>Oligo fwd</td><td>CACCAGCAGTTTATTTTATTCGTG</td><td></td><td></td><td></td><td></td><td></td><td></td><td></td><td></td><td></td><td></td></tr>
<tr><td>Oligo rev</td><td>AAACCACGAATAAAATAAACTGCT</td><td></td><td></td><td></td><td></td><td></td><td></td><td></td><td></td><td></td><td></td></tr>
<tr><td>Chromosome</td><td>start</td><td>end</td><td>strand</td><td>MM</td><td>target_seq</td><td>PAM</td><td>alignment</td><td>distance</td><td>position</td><td>gene name</td><td>gene id</td></tr>
<tr><td>GU305914.1</td><td>129834</td><td>129856</td><td>+</td><td>0</td><td>AGCAGTTTATTTTATTCGTG</td><td>CGG</td><td>||||||||[||||||||||||]PAM</td><td>NA</td><td>Intergenic</td><td>NA</td><td>NA</td></tr>
<tr><td></td><td></td><td></td><td></td><td></td><td></td><td></td><td></td><td></td><td></td><td></td><td></td></tr>
<tr><td>T9806</td><td>ATCATGATGCTCCGCATCTGCGG</td><td>1000</td><td></td><td></td><td></td><td></td><td></td><td></td><td></td><td></td><td></td></tr>
<tr><td>Oligo fwd</td><td>CACCATCATGATGCTCCGCATCTG</td><td></td><td></td><td></td><td></td><td></td><td></td><td></td><td></td><td></td><td></td></tr>
<tr><td>Oligo rev</td><td>AAACCAGATGCGGAGCATCATGAT</td><td></td><td></td><td></td><td></td><td></td><td></td><td></td><td></td><td></td><td></td></tr>
<tr><td>Chromosome</td><td>start</td><td>end</td><td>strand</td><td>MM</td><td>target_seq</td><td>PAM</td><td>alignment</td><td>distance</td><td>position</td><td>gene name</td><td>gene id</td></tr>
<tr><td>GU305914.1</td><td>129857</td><td>129879</td><td>+</td><td>0</td><td>ATCATGATGCTCCGCATCTG</td><td>CGG</td><td>||||||||[||||||||||||]PAM</td><td>NA</td><td>Intergenic</td><td>NA</td><td>NA</td></tr>
<tr><td></td><td></td><td></td><td></td><td></td><td></td><td></td><td></td><td></td><td></td><td></td><td></td></tr>
<tr><td>T9807</td><td>GATGCTCCGCATCTGCGGTCTGG</td><td>1000</td><td></td><td></td><td></td><td></td><td></td><td></td><td></td><td></td><td></td></tr>
<tr><td>Oligo fwd</td><td>CACCGATGCTCCGCATCTGCGGTC</td><td></td><td></td><td></td><td></td><td></td><td></td><td></td><td></td><td></td><td></td></tr>
<tr><td>Oligo rev</td><td>AAACGACCGCAGATGCGGAGCATC</td><td></td><td></td><td></td><td></td><td></td><td></td><td></td><td></td><td></td><td></td></tr>
<tr><td>Chromosome</td><td>start</td><td>end</td><td>strand</td><td>MM</td><td>target_seq</td><td>PAM</td><td>alignment</td><td>distance</td><td>position</td><td>gene name</td><td>gene id</td></tr>
<tr><td>GU305914.1</td><td>129862</td><td>129884</td><td>+</td><td>0</td><td>GATGCTCCGCATCTGCGGTC</td><td>TGG</td><td>||||||||[||||||||||||]PAM</td><td>NA</td><td>Intergenic</td><td>NA</td><td>NA</td></tr>
<tr><td></td><td></td><td></td><td></td><td></td><td></td><td></td><td></td><td></td><td></td><td></td><td></td></tr>
<tr><td>T9808</td><td>CTGCGGTCTGGAACACAGACTGG</td><td>1000</td><td></td><td></td><td></td><td></td><td></td><td></td><td></td><td></td><td></td></tr>
<tr><td>Oligo fwd</td><td>CACCCTGCGGTCTGGAACACAGAC</td><td></td><td></td><td></td><td></td><td></td><td></td><td></td><td></td><td></td><td></td></tr>
<tr><td>Oligo rev</td><td>AAACGTCTGTGTTCCAGACCGCAG</td><td></td><td></td><td></td><td></td><td></td><td></td><td></td><td></td><td></td><td></td></tr>
<tr><td>Chromosome</td><td>start</td><td>end</td><td>strand</td><td>MM</td><td>target_seq</td><td>PAM</td><td>alignment</td><td>distance</td><td>position</td><td>gene name</td><td>gene id</td></tr>
<tr><td>GU305914.1</td><td>129874</td><td>129896</td><td>+</td><td>0</td><td>CTGCGGTCTGGAACACAGAC</td><td>TGG</td><td>||||||||[||||||||||||]PAM</td><td>NA</td><td>Intergenic</td><td>NA</td><td>NA</td></tr>
<tr><td></td><td></td><td></td><td></td><td></td><td></td><td></td><td></td><td></td><td></td><td></td><td></td></tr>
<tr><td>T9809</td><td>AACACAGACTGGCCGTCGACCGG</td><td>1000</td><td></td><td></td><td></td><td></td><td></td><td></td><td></td><td></td><td></td></tr>
<tr><td>Oligo fwd</td><td>CACCAACACAGACTGGCCGTCGAC</td><td></td><td></td><td></td><td></td><td></td><td></td><td></td><td></td><td></td><td></td></tr>
<tr><td>Oligo rev</td><td>AAACGTCGACGGCCAGTCTGTGTT</td><td></td><td></td><td></td><td></td><td></td><td></td><td></td><td></td><td></td><td></td></tr>
<tr><td>Chromosome</td><td>start</td><td>end</td><td>strand</td><td>MM</td><td>target_seq</td><td>PAM</td><td>alignment</td><td>distance</td><td>position</td><td>gene name</td><td>gene id</td></tr>
<tr><td>GU305914.1</td><td>129885</td><td>129907</td><td>+</td><td>0</td><td>AACACAGACTGGCCGTCGAC</td><td>CGG</td><td>||||||||[||||||||||||]PAM</td><td>NA</td><td>Intergenic</td><td>NA</td><td>NA</td></tr>
<tr><td></td><td></td><td></td><td></td><td></td><td></td><td></td><td></td><td></td><td></td><td></td><td></td></tr>
<tr><td>T9810</td><td>CATCTTCCGCGCTATACGCCAGG</td><td>1000</td><td></td><td></td><td></td><td></td><td></td><td></td><td></td><td></td><td></td></tr>
<tr><td>Oligo fwd</td><td>CACCCATCTTCCGCGCTATACGCC</td><td></td><td></td><td></td><td></td><td></td><td></td><td></td><td></td><td></td><td></td></tr>
<tr><td>Oligo rev</td><td>AAACGGCGTATAGCGCGGAAGATG</td><td></td><td></td><td></td><td></td><td></td><td></td><td></td><td></td><td></td><td></td></tr>
<tr><td>Chromosome</td><td>start</td><td>end</td><td>strand</td><td>MM</td><td>target_seq</td><td>PAM</td><td>alignment</td><td>distance</td><td>position</td><td>gene name</td><td>gene id</td></tr>
<tr><td>GU305914.1</td><td>129910</td><td>129932</td><td>+</td><td>0</td><td>CATCTTCCGCGCTATACGCC</td><td>AGG</td><td>||||||||[||||||||||||]PAM</td><td>NA</td><td>Intergenic</td><td>NA</td><td>NA</td></tr>
<tr><td></td><td></td><td></td><td></td><td></td><td></td><td></td><td></td><td></td><td></td><td></td><td></td></tr>
<tr><td>T9811</td><td>ATCTTCCGCGCTATACGCCAGGG</td><td>1000</td><td></td><td></td><td></td><td></td><td></td><td></td><td></td><td></td><td></td></tr>
<tr><td>Oligo fwd</td><td>CACCATCTTCCGCGCTATACGCCA</td><td></td><td></td><td></td><td></td><td></td><td></td><td></td><td></td><td></td><td></td></tr>
<tr><td>Oligo rev</td><td>AAACTGGCGTATAGCGCGGAAGAT</td><td></td><td></td><td></td><td></td><td></td><td></td><td></td><td></td><td></td><td></td></tr>
<tr><td>Chromosome</td><td>start</td><td>end</td><td>strand</td><td>MM</td><td>target_seq</td><td>PAM</td><td>alignment</td><td>distance</td><td>position</td><td>gene name</td><td>gene id</td></tr>
<tr><td>GU305914.1</td><td>129911</td><td>129933</td><td>+</td><td>0</td><td>ATCTTCCGCGCTATACGCCA</td><td>GGG</td><td>||||||||[||||||||||||]PAM</td><td>NA</td><td>Intergenic</td><td>NA</td><td>NA</td></tr>
<tr><td></td><td></td><td></td><td></td><td></td><td></td><td></td><td></td><td></td><td></td><td></td><td></td></tr>
<tr><td>T9812</td><td>CTTCGATTATCACAAGAATACGG</td><td>1000</td><td></td><td></td><td></td><td></td><td></td><td></td><td></td><td></td><td></td></tr>
<tr><td>Oligo fwd</td><td>CACCCTTCGATTATCACAAGAATA</td><td></td><td></td><td></td><td></td><td></td><td></td><td></td><td></td><td></td><td></td></tr>
<tr><td>Oligo rev</td><td>AAACTATTCTTGTGATAATCGAAG</td><td></td><td></td><td></td><td></td><td></td><td></td><td></td><td></td><td></td><td></td></tr>
<tr><td>Chromosome</td><td>start</td><td>end</td><td>strand</td><td>MM</td><td>target_seq</td><td>PAM</td><td>alignment</td><td>distance</td><td>position</td><td>gene name</td><td>gene id</td></tr>
<tr><td>GU305914.1</td><td>129937</td><td>129959</td><td>+</td><td>0</td><td>CTTCGATTATCACAAGAATA</td><td>CGG</td><td>||||||||[||||||||||||]PAM</td><td>NA</td><td>Intergenic</td><td>NA</td><td>NA</td></tr>
<tr><td></td><td></td><td></td><td></td><td></td><td></td><td></td><td></td><td></td><td></td><td></td><td></td></tr>
<tr><td>T9813</td><td>GATCAAACACACATGCGCTCTGG</td><td>1000</td><td></td><td></td><td></td><td></td><td></td><td></td><td></td><td></td><td></td></tr>
<tr><td>Oligo fwd</td><td>CACCGATCAAACACACATGCGCTC</td><td></td><td></td><td></td><td></td><td></td><td></td><td></td><td></td><td></td><td></td></tr>
<tr><td>Oligo rev</td><td>AAACGAGCGCATGTGTGTTTGATC</td><td></td><td></td><td></td><td></td><td></td><td></td><td></td><td></td><td></td><td></td></tr>
<tr><td>Chromosome</td><td>start</td><td>end</td><td>strand</td><td>MM</td><td>target_seq</td><td>PAM</td><td>alignment</td><td>distance</td><td>position</td><td>gene name</td><td>gene id</td></tr>
<tr><td>GU305914.1</td><td>129973</td><td>129995</td><td>+</td><td>0</td><td>GATCAAACACACATGCGCTC</td><td>TGG</td><td>||||||||[||||||||||||]PAM</td><td>NA</td><td>Intergenic</td><td>NA</td><td>NA</td></tr>
<tr><td></td><td></td><td></td><td></td><td></td><td></td><td></td><td></td><td></td><td></td><td></td><td></td></tr>
<tr><td>T9814</td><td>AAACACACATGCGCTCTGGTCGG</td><td>1000</td><td></td><td></td><td></td><td></td><td></td><td></td><td></td><td></td><td></td></tr>
<tr><td>Oligo fwd</td><td>CACCAAACACACATGCGCTCTGGT</td><td></td><td></td><td></td><td></td><td></td><td></td><td></td><td></td><td></td><td></td></tr>
<tr><td>Oligo rev</td><td>AAACACCAGAGCGCATGTGTGTTT</td><td></td><td></td><td></td><td></td><td></td><td></td><td></td><td></td><td></td><td></td></tr>
<tr><td>Chromosome</td><td>start</td><td>end</td><td>strand</td><td>MM</td><td>target_seq</td><td>PAM</td><td>alignment</td><td>distance</td><td>position</td><td>gene name</td><td>gene id</td></tr>
<tr><td>GU305914.1</td><td>129977</td><td>129999</td><td>+</td><td>0</td><td>AAACACACATGCGCTCTGGT</td><td>CGG</td><td>||||||||[||||||||||||]PAM</td><td>NA</td><td>Intergenic</td><td>NA</td><td>NA</td></tr>
<tr><td></td><td></td><td></td><td></td><td></td><td></td><td></td><td></td><td></td><td></td><td></td><td></td></tr>
<tr><td>T9815</td><td>CGCTCTGGTCGGCACAAGATTGG</td><td>1000</td><td></td><td></td><td></td><td></td><td></td><td></td><td></td><td></td><td></td></tr>
<tr><td>Oligo fwd</td><td>CACCCGCTCTGGTCGGCACAAGAT</td><td></td><td></td><td></td><td></td><td></td><td></td><td></td><td></td><td></td><td></td></tr>
<tr><td>Oligo rev</td><td>AAACATCTTGTGCCGACCAGAGCG</td><td></td><td></td><td></td><td></td><td></td><td></td><td></td><td></td><td></td><td></td></tr>
<tr><td>Chromosome</td><td>start</td><td>end</td><td>strand</td><td>MM</td><td>target_seq</td><td>PAM</td><td>alignment</td><td>distance</td><td>position</td><td>gene name</td><td>gene id</td></tr>
<tr><td>GU305914.1</td><td>129988</td><td>130010</td><td>+</td><td>0</td><td>CGCTCTGGTCGGCACAAGAT</td><td>TGG</td><td>||||||||[||||||||||||]PAM</td><td>NA</td><td>Intergenic</td><td>NA</td><td>NA</td></tr>
<tr><td></td><td></td><td></td><td></td><td></td><td></td><td></td><td></td><td></td><td></td><td></td><td></td></tr>
<tr><td>T9816</td><td>CAAGAACAAAAAAATCAAACTGG</td><td>1000</td><td></td><td></td><td></td><td></td><td></td><td></td><td></td><td></td><td></td></tr>
<tr><td>Oligo fwd</td><td>CACCCAAGAACAAAAAAATCAAAC</td><td></td><td></td><td></td><td></td><td></td><td></td><td></td><td></td><td></td><td></td></tr>
<tr><td>Oligo rev</td><td>AAACGTTTGATTTTTTTGTTCTTG</td><td></td><td></td><td></td><td></td><td></td><td></td><td></td><td></td><td></td><td></td></tr>
<tr><td>Chromosome</td><td>start</td><td>end</td><td>strand</td><td>MM</td><td>target_seq</td><td>PAM</td><td>alignment</td><td>distance</td><td>position</td><td>gene name</td><td>gene id</td></tr>
<tr><td>GU305914.1</td><td>130036</td><td>130058</td><td>+</td><td>0</td><td>CAAGAACAAAAAAATCAAAC</td><td>TGG</td><td>||||||||[||||||||||||]PAM</td><td>NA</td><td>Intergenic</td><td>NA</td><td>NA</td></tr>
<tr><td></td><td></td><td></td><td></td><td></td><td></td><td></td><td></td><td></td><td></td><td></td><td></td></tr>
<tr><td>T9817</td><td>AAAATCAAACTGGACTACTTAGG</td><td>1000</td><td></td><td></td><td></td><td></td><td></td><td></td><td></td><td></td><td></td></tr>
<tr><td>Oligo fwd</td><td>CACCAAAATCAAACTGGACTACTT</td><td></td><td></td><td></td><td></td><td></td><td></td><td></td><td></td><td></td><td></td></tr>
<tr><td>Oligo rev</td><td>AAACAAGTAGTCCAGTTTGATTTT</td><td></td><td></td><td></td><td></td><td></td><td></td><td></td><td></td><td></td><td></td></tr>
<tr><td>Chromosome</td><td>start</td><td>end</td><td>strand</td><td>MM</td><td>target_seq</td><td>PAM</td><td>alignment</td><td>distance</td><td>position</td><td>gene name</td><td>gene id</td></tr>
<tr><td>GU305914.1</td><td>130046</td><td>130068</td><td>+</td><td>0</td><td>AAAATCAAACTGGACTACTT</td><td>AGG</td><td>||||||||[||||||||||||]PAM</td><td>NA</td><td>Intergenic</td><td>NA</td><td>NA</td></tr>
<tr><td></td><td></td><td></td><td></td><td></td><td></td><td></td><td></td><td></td><td></td><td></td><td></td></tr>
<tr><td>T9818</td><td>AAATCAAACTGGACTACTTAGGG</td><td>1000</td><td></td><td></td><td></td><td></td><td></td><td></td><td></td><td></td><td></td></tr>
<tr><td>Oligo fwd</td><td>CACCAAATCAAACTGGACTACTTA</td><td></td><td></td><td></td><td></td><td></td><td></td><td></td><td></td><td></td><td></td></tr>
<tr><td>Oligo rev</td><td>AAACTAAGTAGTCCAGTTTGATTT</td><td></td><td></td><td></td><td></td><td></td><td></td><td></td><td></td><td></td><td></td></tr>
<tr><td>Chromosome</td><td>start</td><td>end</td><td>strand</td><td>MM</td><td>target_seq</td><td>PAM</td><td>alignment</td><td>distance</td><td>position</td><td>gene name</td><td>gene id</td></tr>
<tr><td>GU305914.1</td><td>130047</td><td>130069</td><td>+</td><td>0</td><td>AAATCAAACTGGACTACTTA</td><td>GGG</td><td>||||||||[||||||||||||]PAM</td><td>NA</td><td>Intergenic</td><td>NA</td><td>NA</td></tr>
</table>

<table>
<tr><td>T9819</td><td>TCAAACTGGACTACTTAGGGAGG</td><td>1000</td><td></td><td></td><td></td><td></td><td></td><td></td><td></td><td></td><td></td></tr>
<tr><td>Oligo fwd</td><td>CACCTCAAACTGGACTACTTAGGG</td><td></td><td></td><td></td><td></td><td></td><td></td><td></td><td></td><td></td><td></td></tr>
<tr><td>Oligo rev</td><td>AAACCCCTAAGTAGTCCAGTTTGA</td><td></td><td></td><td></td><td></td><td></td><td></td><td></td><td></td><td></td><td></td></tr>
<tr><td>Chromosome</td><td>start</td><td>end</td><td>strand</td><td>MM</td><td>target_seq</td><td>PAM</td><td>alignment</td><td>distance</td><td>position</td><td>gene name</td><td>gene id</td></tr>
<tr><td>GU305914.1</td><td>130050</td><td>130072</td><td>+</td><td>0</td><td>TCAAACTGGACTACTTAGGG</td><td>AGG</td><td>||||||||[||||||||||||]PAM</td><td>NA</td><td>Intergenic</td><td>NA</td><td>NA</td></tr>
<tr><td>T9820</td><td>ACGCCTGCGTCGATGCTCATCGG</td><td>1000</td><td></td><td></td><td></td><td></td><td></td><td></td><td></td><td></td><td></td></tr>
<tr><td>Oligo fwd</td><td>CACCACGCCTGCGTCGATGCTCAT</td><td></td><td></td><td></td><td></td><td></td><td></td><td></td><td></td><td></td><td></td></tr>
<tr><td>Oligo rev</td><td>AAACATGAGCATCGACGCAGGCGT</td><td></td><td></td><td></td><td></td><td></td><td></td><td></td><td></td><td></td><td></td></tr>
<tr><td>Chromosome</td><td>start</td><td>end</td><td>strand</td><td>MM</td><td>target_seq</td><td>PAM</td><td>alignment</td><td>distance</td><td>position</td><td>gene name</td><td>gene id</td></tr>
<tr><td>GU305914.1</td><td>130110</td><td>130132</td><td>+</td><td>0</td><td>ACGCCTGCGTCGATGCTCAT</td><td>CGG</td><td>||||||||[||||||||||||]PAM</td><td>NA</td><td>Intergenic</td><td>NA</td><td>NA</td></tr>
<tr><td>T9821</td><td>CTGCGTCGATGCTCATCGGCTGG</td><td>1000</td><td></td><td></td><td></td><td></td><td></td><td></td><td></td><td></td><td></td></tr>
<tr><td>Oligo fwd</td><td>CACCCTGCGTCGATGCTCATCGGC</td><td></td><td></td><td></td><td></td><td></td><td></td><td></td><td></td><td></td><td></td></tr>
<tr><td>Oligo rev</td><td>AAACGCCGATGAGCATCGACGCAG</td><td></td><td></td><td></td><td></td><td></td><td></td><td></td><td></td><td></td><td></td></tr>
<tr><td>Chromosome</td><td>start</td><td>end</td><td>strand</td><td>MM</td><td>target_seq</td><td>PAM</td><td>alignment</td><td>distance</td><td>position</td><td>gene name</td><td>gene id</td></tr>
<tr><td>GU305914.1</td><td>130114</td><td>130136</td><td>+</td><td>0</td><td>CTGCGTCGATGCTCATCGGC</td><td>TGG</td><td>||||||||[||||||||||||]PAM</td><td>NA</td><td>Intergenic</td><td>NA</td><td>NA</td></tr>
<tr><td>T9822</td><td>CGTCGATGCTCATCGGCTGGAGG</td><td>1000</td><td></td><td></td><td></td><td></td><td></td><td></td><td></td><td></td><td></td></tr>
<tr><td>Oligo fwd</td><td>CACCCGTCGATGCTCATCGGCTGG</td><td></td><td></td><td></td><td></td><td></td><td></td><td></td><td></td><td></td><td></td></tr>
<tr><td>Oligo rev</td><td>AAACCCAGCCGATGAGCATCGACG</td><td></td><td></td><td></td><td></td><td></td><td></td><td></td><td></td><td></td><td></td></tr>
<tr><td>Chromosome</td><td>start</td><td>end</td><td>strand</td><td>MM</td><td>target_seq</td><td>PAM</td><td>alignment</td><td>distance</td><td>position</td><td>gene name</td><td>gene id</td></tr>
<tr><td>GU305914.1</td><td>130117</td><td>130139</td><td>+</td><td>0</td><td>CGTCGATGCTCATCGGCTGG</td><td>AGG</td><td>||||||||[||||||||||||]PAM</td><td>NA</td><td>Intergenic</td><td>NA</td><td>NA</td></tr>
<tr><td>T9823</td><td>CATCGGCTGGAGGCGCTGATCGG</td><td>1000</td><td></td><td></td><td></td><td></td><td></td><td></td><td></td><td></td><td></td></tr>
<tr><td>Oligo fwd</td><td>CACCCATCGGCTGGAGGCGCTGAT</td><td></td><td></td><td></td><td></td><td></td><td></td><td></td><td></td><td></td><td></td></tr>
<tr><td>Oligo rev</td><td>AAACATCAGCGCCTCCAGCCGATG</td><td></td><td></td><td></td><td></td><td></td><td></td><td></td><td></td><td></td><td></td></tr>
<tr><td>Chromosome</td><td>start</td><td>end</td><td>strand</td><td>MM</td><td>target_seq</td><td>PAM</td><td>alignment</td><td>distance</td><td>position</td><td>gene name</td><td>gene id</td></tr>
<tr><td>GU305914.1</td><td>130127</td><td>130149</td><td>+</td><td>0</td><td>CATCGGCTGGAGGCGCTGAT</td><td>CGG</td><td>||||||||[||||||||||||]PAM</td><td>NA</td><td>Intergenic</td><td>NA</td><td>NA</td></tr>
<tr><td>T9824</td><td>ATCGGCTGGAGGCGCTGATCGGG</td><td>1000</td><td></td><td></td><td></td><td></td><td></td><td></td><td></td><td></td><td></td></tr>
<tr><td>Oligo fwd</td><td>CACCATCGGCTGGAGGCGCTGATC</td><td></td><td></td><td></td><td></td><td></td><td></td><td></td><td></td><td></td><td></td></tr>
<tr><td>Oligo rev</td><td>AAACGATCAGCGCCTCCAGCCGAT</td><td></td><td></td><td></td><td></td><td></td><td></td><td></td><td></td><td></td><td></td></tr>
<tr><td>Chromosome</td><td>start</td><td>end</td><td>strand</td><td>MM</td><td>target_seq</td><td>PAM</td><td>alignment</td><td>distance</td><td>position</td><td>gene name</td><td>gene id</td></tr>
<tr><td>GU305914.1</td><td>130128</td><td>130150</td><td>+</td><td>0</td><td>ATCGGCTGGAGGCGCTGATC</td><td>GGG</td><td>||||||||[||||||||||||]PAM</td><td>NA</td><td>Intergenic</td><td>NA</td><td>NA</td></tr>
<tr><td>T9825</td><td>TCGGCTGGAGGCGCTGATCGGGG</td><td>1000</td><td></td><td></td><td></td><td></td><td></td><td></td><td></td><td></td><td></td></tr>
<tr><td>Oligo fwd</td><td>CACCTCGGCTGGAGGCGCTGATCG</td><td></td><td></td><td></td><td></td><td></td><td></td><td></td><td></td><td></td><td></td></tr>
<tr><td>Oligo rev</td><td>AAACCGATCAGCGCCTCCAGCCGA</td><td></td><td></td><td></td><td></td><td></td><td></td><td></td><td></td><td></td><td></td></tr>
<tr><td>Chromosome</td><td>start</td><td>end</td><td>strand</td><td>MM</td><td>target_seq</td><td>PAM</td><td>alignment</td><td>distance</td><td>position</td><td>gene name</td><td>gene id</td></tr>
<tr><td>GU305914.1</td><td>130129</td><td>130151</td><td>+</td><td>0</td><td>TCGGCTGGAGGCGCTGATCG</td><td>GGG</td><td>||||||||[||||||||||||]PAM</td><td>NA</td><td>Intergenic</td><td>NA</td><td>NA</td></tr>
<tr><td>T9826</td><td>AGAAGCTCACGTCCATCGACAGG</td><td>1000</td><td></td><td></td><td></td><td></td><td></td><td></td><td></td><td></td><td></td></tr>
<tr><td>Oligo fwd</td><td>CACCAGAAGCTCACGTCCATCGAC</td><td></td><td></td><td></td><td></td><td></td><td></td><td></td><td></td><td></td><td></td></tr>
<tr><td>Oligo rev</td><td>AAACGTCGATGGACGTGAGCTTCT</td><td></td><td></td><td></td><td></td><td></td><td></td><td></td><td></td><td></td><td></td></tr>
<tr><td>Chromosome</td><td>start</td><td>end</td><td>strand</td><td>MM</td><td>target_seq</td><td>PAM</td><td>alignment</td><td>distance</td><td>position</td><td>gene name</td><td>gene id</td></tr>
<tr><td>GU305914.1</td><td>130167</td><td>130189</td><td>+</td><td>0</td><td>AGAAGCTCACGTCCATCGAC</td><td>AGG</td><td>||||||||[||||||||||||]PAM</td><td>NA</td><td>Intergenic</td><td>NA</td><td>NA</td></tr>
<tr><td>T9827</td><td>GTCCATCGACAGGACCAAAGAGG</td><td>1000</td><td></td><td></td><td></td><td></td><td></td><td></td><td></td><td></td><td></td></tr>
<tr><td>Oligo fwd</td><td>CACCGTCCATCGACAGGACCAAAG</td><td></td><td></td><td></td><td></td><td></td><td></td><td></td><td></td><td></td><td></td></tr>
<tr><td>Oligo rev</td><td>AAACCTTTGGTCCTGTCGATGGAC</td><td></td><td></td><td></td><td></td><td></td><td></td><td></td><td></td><td></td><td></td></tr>
<tr><td>Chromosome</td><td>start</td><td>end</td><td>strand</td><td>MM</td><td>target_seq</td><td>PAM</td><td>alignment</td><td>distance</td><td>position</td><td>gene name</td><td>gene id</td></tr>
<tr><td>GU305914.1</td><td>130177</td><td>130199</td><td>+</td><td>0</td><td>GTCCATCGACAGGACCAAAG</td><td>AGG</td><td>||||||||[||||||||||||]PAM</td><td>NA</td><td>Intergenic</td><td>NA</td><td>NA</td></tr>
<tr><td>T9828</td><td>AAGAGGCGCTCACGCGAGCGAGG</td><td>1000</td><td></td><td></td><td></td><td></td><td></td><td></td><td></td><td></td><td></td></tr>
<tr><td>Oligo fwd</td><td>CACCAAGAGGCGCTCACGCGAGCG</td><td></td><td></td><td></td><td></td><td></td><td></td><td></td><td></td><td></td><td></td></tr>
<tr><td>Oligo rev</td><td>AAACCGCTCGCGTGAGCGCCTCTT</td><td></td><td></td><td></td><td></td><td></td><td></td><td></td><td></td><td></td><td></td></tr>
<tr><td>Chromosome</td><td>start</td><td>end</td><td>strand</td><td>MM</td><td>target_seq</td><td>PAM</td><td>alignment</td><td>distance</td><td>position</td><td>gene name</td><td>gene id</td></tr>
<tr><td>GU305914.1</td><td>130194</td><td>130216</td><td>+</td><td>0</td><td>AAGAGGCGCTCACGCGAGCG</td><td>AGG</td><td>||||||||[||||||||||||]PAM</td><td>NA</td><td>Intergenic</td><td>NA</td><td>NA</td></tr>
<tr><td>T9829</td><td>AGAGGCGCTCACGCGAGCGAGGG</td><td>1000</td><td></td><td></td><td></td><td></td><td></td><td></td><td></td><td></td><td></td></tr>
<tr><td>Oligo fwd</td><td>CACCAGAGGCGCTCACGCGAGCGA</td><td></td><td></td><td></td><td></td><td></td><td></td><td></td><td></td><td></td><td></td></tr>
<tr><td>Oligo rev</td><td>AAACTCGCTCGCGTGAGCGCCTCT</td><td></td><td></td><td></td><td></td><td></td><td></td><td></td><td></td><td></td><td></td></tr>
<tr><td>Chromosome</td><td>start</td><td>end</td><td>strand</td><td>MM</td><td>target_seq</td><td>PAM</td><td>alignment</td><td>distance</td><td>position</td><td>gene name</td><td>gene id</td></tr>
<tr><td>GU305914.1</td><td>130195</td><td>130217</td><td>+</td><td>0</td><td>AGAGGCGCTCACGCGAGCGA</td><td>GGG</td><td>||||||||[||||||||||||]PAM</td><td>NA</td><td>Intergenic</td><td>NA</td><td>NA</td></tr>
<tr><td>T9830</td><td>CGCTCACGCGAGCGAGGGTCAGG</td><td>1000</td><td></td><td></td><td></td><td></td><td></td><td></td><td></td><td></td><td></td></tr>
<tr><td>Oligo fwd</td><td>CACCCGCTCACGCGAGCGAGGGTC</td><td></td><td></td><td></td><td></td><td></td><td></td><td></td><td></td><td></td><td></td></tr>
<tr><td>Oligo rev</td><td>AAACGACCCTCGCTCGCGTGAGCG</td><td></td><td></td><td></td><td></td><td></td><td></td><td></td><td></td><td></td><td></td></tr>
<tr><td>Chromosome</td><td>start</td><td>end</td><td>strand</td><td>MM</td><td>target_seq</td><td>PAM</td><td>alignment</td><td>distance</td><td>position</td><td>gene name</td><td>gene id</td></tr>
<tr><td>GU305914.1</td><td>130200</td><td>130222</td><td>+</td><td>0</td><td>CGCTCACGCGAGCGAGGGTC</td><td>AGG</td><td>||||||||[||||||||||||]PAM</td><td>NA</td><td>Intergenic</td><td>NA</td><td>NA</td></tr>
<tr><td>T9831</td><td>CGAGCGAGGGTCAGGCTCTGCGG</td><td>1000</td><td></td><td></td><td></td><td></td><td></td><td></td><td></td><td></td><td></td></tr>
<tr><td>Oligo fwd</td><td>CACCCGAGCGAGGGTCAGGCTCTG</td><td></td><td></td><td></td><td></td><td></td><td></td><td></td><td></td><td></td><td></td></tr>
<tr><td>Oligo rev</td><td>AAACCAGAGCCTGACCCTCGCTCG</td><td></td><td></td><td></td><td></td><td></td><td></td><td></td><td></td><td></td><td></td></tr>
<tr><td>Chromosome</td><td>start</td><td>end</td><td>strand</td><td>MM</td><td>target_seq</td><td>PAM</td><td>alignment</td><td>distance</td><td>position</td><td>gene name</td><td>gene id</td></tr>
<tr><td>GU305914.1</td><td>130208</td><td>130230</td><td>+</td><td>0</td><td>CGAGCGAGGGTCAGGCTCTG</td><td>CGG</td><td>||||||||[||||||||||||]PAM</td><td>NA</td><td>Intergenic</td><td>NA</td><td>NA</td></tr>
<tr><td>T9832</td><td>GCGAGGGTCAGGCTCTGCGGAGG</td><td>1000</td><td></td><td></td><td></td><td></td><td></td><td></td><td></td><td></td><td></td></tr>
<tr><td>Oligo fwd</td><td>CACCGCGAGGGTCAGGCTCTGCGG</td><td></td><td></td><td></td><td></td><td></td><td></td><td></td><td></td><td></td><td></td></tr>
<tr><td>Oligo rev</td><td>AAACCCGCAGAGCCTGACCCTCGC</td><td></td><td></td><td></td><td></td><td></td><td></td><td></td><td></td><td></td><td></td></tr>
<tr><td>Chromosome</td><td>start</td><td>end</td><td>strand</td><td>MM</td><td>target_seq</td><td>PAM</td><td>alignment</td><td>distance</td><td>position</td><td>gene name</td><td>gene id</td></tr>
<tr><td>GU305914.1</td><td>130211</td><td>130233</td><td>+</td><td>0</td><td>GCGAGGGTCAGGCTCTGCGG</td><td>AGG</td><td>||||||||[||||||||||||]PAM</td><td>NA</td><td>Intergenic</td><td>NA</td><td>NA</td></tr>
<tr><td>T9833</td><td>CTGCGGAGGATACCGCCCCGAGG</td><td>1000</td><td></td><td></td><td></td><td></td><td></td><td></td><td></td><td></td><td></td></tr>
<tr><td>Oligo fwd</td><td>CACCCTGCGGAGGATACCGCCCCG</td><td></td><td></td><td></td><td></td><td></td><td></td><td></td><td></td><td></td><td></td></tr>
<tr><td>Oligo rev</td><td>AAACCGGGGCGGTATCCTCCGCAG</td><td></td><td></td><td></td><td></td><td></td><td></td><td></td><td></td><td></td><td></td></tr>
<tr><td>Chromosome</td><td>start</td><td>end</td><td>strand</td><td>MM</td><td>target_seq</td><td>PAM</td><td>alignment</td><td>distance</td><td>position</td><td>gene name</td><td>gene id</td></tr>
<tr><td>GU305914.1</td><td>130225</td><td>130247</td><td>+</td><td>0</td><td>CTGCGGAGGATACCGCCCCG</td><td>AGG</td><td>||||||||[||||||||||||]PAM</td><td>NA</td><td>Intergenic</td><td>NA</td><td>NA</td></tr>
</table>

<table>
<tr><td>T9834</td><td>GCCCCGAGGCAGCGACGAGCCGG</td><td>1000</td><td></td><td></td><td></td><td></td><td></td><td></td><td></td><td></td><td></td></tr>
<tr><td>Oligo fwd</td><td>CACCGCCCCGAGGCAGCGACGAGC</td><td></td><td></td><td></td><td></td><td></td><td></td><td></td><td></td><td></td><td></td></tr>
<tr><td>Oligo rev</td><td>AAACGCTCGTCGCTGCCTCGGGGC</td><td></td><td></td><td></td><td></td><td></td><td></td><td></td><td></td><td></td><td></td></tr>
<tr><td>Chromosome</td><td>start</td><td>end</td><td>strand</td><td>MM</td><td>target_seq</td><td>PAM</td><td>alignment</td><td>distance</td><td>position</td><td>gene name</td><td>gene id</td></tr>
<tr><td>GU305914.1</td><td>130239</td><td>130261</td><td>+</td><td>0</td><td>GCCCCGAGGCAGCGACGAGC</td><td>CGG</td><td>||||||||[||||||||||||]PAM</td><td>NA</td><td>Intergenic</td><td>NA</td><td>NA</td></tr>
<tr><td></td><td></td><td></td><td></td><td></td><td></td><td></td><td></td><td></td><td></td><td></td><td></td></tr>
<tr><td>T9835</td><td>GTCACTCACTCACGCTCACTTGG</td><td>1000</td><td></td><td></td><td></td><td></td><td></td><td></td><td></td><td></td><td></td></tr>
<tr><td>Oligo fwd</td><td>CACCGTCACTCACTCACGCTCACT</td><td></td><td></td><td></td><td></td><td></td><td></td><td></td><td></td><td></td><td></td></tr>
<tr><td>Oligo rev</td><td>AAACAGTGAGCGTGAGTGAGTGAC</td><td></td><td></td><td></td><td></td><td></td><td></td><td></td><td></td><td></td><td></td></tr>
<tr><td>Chromosome</td><td>start</td><td>end</td><td>strand</td><td>MM</td><td>target_seq</td><td>PAM</td><td>alignment</td><td>distance</td><td>position</td><td>gene name</td><td>gene id</td></tr>
<tr><td>GU305914.1</td><td>130277</td><td>130299</td><td>+</td><td>0</td><td>GTCACTCACTCACGCTCACT</td><td>TGG</td><td>||||||||[||||||||||||]PAM</td><td>NA</td><td>Intergenic</td><td>NA</td><td>NA</td></tr>
<tr><td></td><td></td><td></td><td></td><td></td><td></td><td></td><td></td><td></td><td></td><td></td><td></td></tr>
<tr><td>T9836</td><td>ACTCACTCACGCTCACTTGGCGG</td><td>1000</td><td></td><td></td><td></td><td></td><td></td><td></td><td></td><td></td><td></td></tr>
<tr><td>Oligo fwd</td><td>CACCACTCACTCACGCTCACTTGG</td><td></td><td></td><td></td><td></td><td></td><td></td><td></td><td></td><td></td><td></td></tr>
<tr><td>Oligo rev</td><td>AAACCCAAGTGAGCGTGAGTGAGT</td><td></td><td></td><td></td><td></td><td></td><td></td><td></td><td></td><td></td><td></td></tr>
<tr><td>Chromosome</td><td>start</td><td>end</td><td>strand</td><td>MM</td><td>target_seq</td><td>PAM</td><td>alignment</td><td>distance</td><td>position</td><td>gene name</td><td>gene id</td></tr>
<tr><td>GU305914.1</td><td>130280</td><td>130302</td><td>+</td><td>0</td><td>ACTCACTCACGCTCACTTGG</td><td>CGG</td><td>||||||||[||||||||||||]PAM</td><td>NA</td><td>Intergenic</td><td>NA</td><td>NA</td></tr>
<tr><td></td><td></td><td></td><td></td><td></td><td></td><td></td><td></td><td></td><td></td><td></td><td></td></tr>
<tr><td>T9837</td><td>ACTCACGCTCACTTGGCGGACGG</td><td>1000</td><td></td><td></td><td></td><td></td><td></td><td></td><td></td><td></td><td></td></tr>
<tr><td>Oligo fwd</td><td>CACCACTCACGCTCACTTGGCGGA</td><td></td><td></td><td></td><td></td><td></td><td></td><td></td><td></td><td></td><td></td></tr>
<tr><td>Oligo rev</td><td>AAACTCCGCCAAGTGAGCGTGAGT</td><td></td><td></td><td></td><td></td><td></td><td></td><td></td><td></td><td></td><td></td></tr>
<tr><td>Chromosome</td><td>start</td><td>end</td><td>strand</td><td>MM</td><td>target_seq</td><td>PAM</td><td>alignment</td><td>distance</td><td>position</td><td>gene name</td><td>gene id</td></tr>
<tr><td>GU305914.1</td><td>130284</td><td>130306</td><td>+</td><td>0</td><td>ACTCACGCTCACTTGGCGGA</td><td>CGG</td><td>||||||||[||||||||||||]PAM</td><td>NA</td><td>Intergenic</td><td>NA</td><td>NA</td></tr>
<tr><td></td><td></td><td></td><td></td><td></td><td></td><td></td><td></td><td></td><td></td><td></td><td></td></tr>
<tr><td>T9838</td><td>CGCAGCAGCAGCCGCCGCCGCGG</td><td>1000</td><td></td><td></td><td></td><td></td><td></td><td></td><td></td><td></td><td></td></tr>
<tr><td>Oligo fwd</td><td>CACCCGCAGCAGCAGCCGCCGCCG</td><td></td><td></td><td></td><td></td><td></td><td></td><td></td><td></td><td></td><td></td></tr>
<tr><td>Oligo rev</td><td>AAACCGGCGGCGGCTGCTGCTGCG</td><td></td><td></td><td></td><td></td><td></td><td></td><td></td><td></td><td></td><td></td></tr>
<tr><td>Chromosome</td><td>start</td><td>end</td><td>strand</td><td>MM</td><td>target_seq</td><td>PAM</td><td>alignment</td><td>distance</td><td>position</td><td>gene name</td><td>gene id</td></tr>
<tr><td>GU305914.1</td><td>130312</td><td>130334</td><td>+</td><td>0</td><td>CGCAGCAGCAGCCGCCGCCG</td><td>CGG</td><td>||||||||[||||||||||||]PAM</td><td>NA</td><td>Intergenic</td><td>NA</td><td>NA</td></tr>
<tr><td>GU305914.1</td><td>45592</td><td>45614</td><td>-</td><td>4</td><td>GGCGGCTGCCGCCGCCGCCG</td><td>CGG</td><td>-||-||-|[|-||||||||||]PAM</td><td>NA</td><td>Intergenic</td><td>NA</td><td>NA</td></tr>
<tr><td>GU305914.1</td><td>35655</td><td>35677</td><td>-</td><td>4</td><td>CCCCGCCGCCGCCGCCGCCG</td><td>TGG</td><td>|-|-||-|[|-||||||||||]PAM</td><td>NA</td><td>Intergenic</td><td>NA</td><td>NA</td></tr>
<tr><td>GU305914.1</td><td>62679</td><td>62701</td><td>-</td><td>4</td><td>CGCGGCTGCTGCCGCCGGCG</td><td>GGG</td><td>|||-||-|[|-|||||||-||]PAM</td><td>NA</td><td>Intergenic</td><td>NA</td><td>NA</td></tr>
<tr><td>GU305914.1</td><td>130315</td><td>130337</td><td>+</td><td>3</td><td>AGCAGCAGCCGCCGCCGCGG</td><td>CGG</td><td>-|||||||[|-||||||||-|]PAM</td><td>NA</td><td>Intergenic</td><td>NA</td><td>NA</td></tr>
<tr><td></td><td></td><td></td><td></td><td></td><td></td><td></td><td></td><td></td><td></td><td></td><td></td></tr>
<tr><td>T9839</td><td>AGCAGCAGCCGCCGCCGCGGCGG</td><td>1000</td><td></td><td></td><td></td><td></td><td></td><td></td><td></td><td></td><td></td></tr>
<tr><td>Oligo fwd</td><td>CACCAGCAGCAGCCGCCGCCGCGG</td><td></td><td></td><td></td><td></td><td></td><td></td><td></td><td></td><td></td><td></td></tr>
<tr><td>Oligo rev</td><td>AAACCCGCGGCGGCGGCTGCTGCT</td><td></td><td></td><td></td><td></td><td></td><td></td><td></td><td></td><td></td><td></td></tr>
<tr><td>Chromosome</td><td>start</td><td>end</td><td>strand</td><td>MM</td><td>target_seq</td><td>PAM</td><td>alignment</td><td>distance</td><td>position</td><td>gene name</td><td>gene id</td></tr>
<tr><td>GU305914.1</td><td>130315</td><td>130337</td><td>+</td><td>0</td><td>AGCAGCAGCCGCCGCCGCGG</td><td>CGG</td><td>||||||||[||||||||||||]PAM</td><td>NA</td><td>Intergenic</td><td>NA</td><td>NA</td></tr>
<tr><td>GU305914.1</td><td>56772</td><td>56794</td><td>-</td><td>4</td><td>CGCCGCCGCCGCCGCTGCGG</td><td>CGG</td><td>-||-||-|[|||||||-||||]PAM</td><td>NA</td><td>Intergenic</td><td>NA</td><td>NA</td></tr>
<tr><td>GU305914.1</td><td>45592</td><td>45614</td><td>-</td><td>4</td><td>GGCGGCTGCCGCCGCCGCCG</td><td>CGG</td><td>-||-||-|[||||||||||-|]PAM</td><td>NA</td><td>Intergenic</td><td>NA</td><td>NA</td></tr>
<tr><td>GU305914.1</td><td>56775</td><td>56797</td><td>-</td><td>4</td><td>TGCCGCCGCCGCCGCCGCTG</td><td>CGG</td><td>-||-||-|[||||||||||-|]PAM</td><td>NA</td><td>Intergenic</td><td>NA</td><td>NA</td></tr>
<tr><td>GU305914.1</td><td>130312</td><td>130334</td><td>+</td><td>3</td><td>CGCAGCAGCAGCCGCCGCCG</td><td>CGG</td><td>-|||||||[|-||||||||-|]PAM</td><td>NA</td><td>Intergenic</td><td>NA</td><td>NA</td></tr>
<tr><td></td><td></td><td></td><td></td><td></td><td></td><td></td><td></td><td></td><td></td><td></td><td></td></tr>
<tr><td>T9840</td><td>CAGCCGCCGCCGCGGCGGTCCGG</td><td>1000</td><td></td><td></td><td></td><td></td><td></td><td></td><td></td><td></td><td></td></tr>
<tr><td>Oligo fwd</td><td>CACCCAGCCGCCGCCGCGGCGGTC</td><td></td><td></td><td></td><td></td><td></td><td></td><td></td><td></td><td></td><td></td></tr>
<tr><td>Oligo rev</td><td>AAACGACCGCCGCGGCGGCGGCTG</td><td></td><td></td><td></td><td></td><td></td><td></td><td></td><td></td><td></td><td></td></tr>
<tr><td>Chromosome</td><td>start</td><td>end</td><td>strand</td><td>MM</td><td>target_seq</td><td>PAM</td><td>alignment</td><td>distance</td><td>position</td><td>gene name</td><td>gene id</td></tr>
<tr><td>GU305914.1</td><td>130320</td><td>130342</td><td>+</td><td>0</td><td>CAGCCGCCGCCGCGGCGGTC</td><td>CGG</td><td>||||||||[||||||||||||]PAM</td><td>NA</td><td>Intergenic</td><td>NA</td><td>NA</td></tr>
<tr><td>GU305914.1</td><td>195109</td><td>195131</td><td>-</td><td>4</td><td>CGGCCACCGCCACGGCGTTC</td><td>AGG</td><td>|-|||-||[|||-|||||-||]PAM</td><td>NA</td><td>Intergenic</td><td>NA</td><td>NA</td></tr>
<tr><td></td><td></td><td></td><td></td><td></td><td></td><td></td><td></td><td></td><td></td><td></td><td></td></tr>
<tr><td>T9841</td><td>GCCGCGGCGGTCCGGCGTCATGG</td><td>1000</td><td></td><td></td><td></td><td></td><td></td><td></td><td></td><td></td><td></td></tr>
<tr><td>Oligo fwd</td><td>CACCGCCGCGGCGGTCCGGCGTCA</td><td></td><td></td><td></td><td></td><td></td><td></td><td></td><td></td><td></td><td></td></tr>
<tr><td>Oligo rev</td><td>AAACTGACGCCGGACCGCCGCGGC</td><td></td><td></td><td></td><td></td><td></td><td></td><td></td><td></td><td></td><td></td></tr>
<tr><td>Chromosome</td><td>start</td><td>end</td><td>strand</td><td>MM</td><td>target_seq</td><td>PAM</td><td>alignment</td><td>distance</td><td>position</td><td>gene name</td><td>gene id</td></tr>
<tr><td>GU305914.1</td><td>130328</td><td>130350</td><td>+</td><td>0</td><td>GCCGCGGCGGTCCGGCGTCA</td><td>TGG</td><td>||||||||[||||||||||||]PAM</td><td>NA</td><td>Intergenic</td><td>NA</td><td>NA</td></tr>
<tr><td>GU305914.1</td><td>163556</td><td>163578</td><td>+</td><td>4</td><td>GCCGCCGGGGTCCGACGGCA</td><td>CGG</td><td>|||||-|-[||||||-||-||]PAM</td><td>NA</td><td>Intergenic</td><td>NA</td><td>NA</td></tr>
<tr><td></td><td></td><td></td><td></td><td></td><td></td><td></td><td></td><td></td><td></td><td></td><td></td></tr>
<tr><td>T9842</td><td>GTCATGGACGAAGAAGACGCCGG</td><td>1000</td><td></td><td></td><td></td><td></td><td></td><td></td><td></td><td></td><td></td></tr>
<tr><td>Oligo fwd</td><td>CACCGTCATGGACGAAGAAGACGC</td><td></td><td></td><td></td><td></td><td></td><td></td><td></td><td></td><td></td><td></td></tr>
<tr><td>Oligo rev</td><td>AAACGCGTCTTCTTCGTCCATGAC</td><td></td><td></td><td></td><td></td><td></td><td></td><td></td><td></td><td></td><td></td></tr>
<tr><td>Chromosome</td><td>start</td><td>end</td><td>strand</td><td>MM</td><td>target_seq</td><td>PAM</td><td>alignment</td><td>distance</td><td>position</td><td>gene name</td><td>gene id</td></tr>
<tr><td>GU305914.1</td><td>130344</td><td>130366</td><td>+</td><td>0</td><td>GTCATGGACGAAGAAGACGC</td><td>CGG</td><td>||||||||[||||||||||||]PAM</td><td>NA</td><td>Intergenic</td><td>NA</td><td>NA</td></tr>
<tr><td></td><td></td><td></td><td></td><td></td><td></td><td></td><td></td><td></td><td></td><td></td><td></td></tr>
<tr><td>T9843</td><td>GAAGACGCCGGCAGCATCAACGG</td><td>1000</td><td></td><td></td><td></td><td></td><td></td><td></td><td></td><td></td><td></td></tr>
<tr><td>Oligo fwd</td><td>CACCGAAGACGCCGGCAGCATCAA</td><td></td><td></td><td></td><td></td><td></td><td></td><td></td><td></td><td></td><td></td></tr>
<tr><td>Oligo rev</td><td>AAACTTGATGCTGCCGGCGTCTTC</td><td></td><td></td><td></td><td></td><td></td><td></td><td></td><td></td><td></td><td></td></tr>
<tr><td>Chromosome</td><td>start</td><td>end</td><td>strand</td><td>MM</td><td>target_seq</td><td>PAM</td><td>alignment</td><td>distance</td><td>position</td><td>gene name</td><td>gene id</td></tr>
<tr><td>GU305914.1</td><td>130356</td><td>130378</td><td>+</td><td>0</td><td>GAAGACGCCGGCAGCATCAA</td><td>CGG</td><td>||||||||[||||||||||||]PAM</td><td>NA</td><td>Intergenic</td><td>NA</td><td>NA</td></tr>
<tr><td></td><td></td><td></td><td></td><td></td><td></td><td></td><td></td><td></td><td></td><td></td><td></td></tr>
<tr><td>T9844</td><td>CCGGCAGCATCAACGGCGAGAGG</td><td>1000</td><td></td><td></td><td></td><td></td><td></td><td></td><td></td><td></td><td></td></tr>
<tr><td>Oligo fwd</td><td>CACCCCGGCAGCATCAACGGCGAG</td><td></td><td></td><td></td><td></td><td></td><td></td><td></td><td></td><td></td><td></td></tr>
<tr><td>Oligo rev</td><td>AAACCTCGCCGTTGATGCTGCCGG</td><td></td><td></td><td></td><td></td><td></td><td></td><td></td><td></td><td></td><td></td></tr>
<tr><td>Chromosome</td><td>start</td><td>end</td><td>strand</td><td>MM</td><td>target_seq</td><td>PAM</td><td>alignment</td><td>distance</td><td>position</td><td>gene name</td><td>gene id</td></tr>
<tr><td>GU305914.1</td><td>130363</td><td>130385</td><td>+</td><td>0</td><td>CCGGCAGCATCAACGGCGAG</td><td>AGG</td><td>||||||||[||||||||||||]PAM</td><td>NA</td><td>Intergenic</td><td>NA</td><td>NA</td></tr>
<tr><td></td><td></td><td></td><td></td><td></td><td></td><td></td><td></td><td></td><td></td><td></td><td></td></tr>
<tr><td>T9845</td><td>CAGCATCAACGGCGAGAGGAAGG</td><td>1000</td><td></td><td></td><td></td><td></td><td></td><td></td><td></td><td></td><td></td></tr>
<tr><td>Oligo fwd</td><td>CACCCAGCATCAACGGCGAGAGGA</td><td></td><td></td><td></td><td></td><td></td><td></td><td></td><td></td><td></td><td></td></tr>
<tr><td>Oligo rev</td><td>AAACTCCTCTCGCCGTTGATGCTG</td><td></td><td></td><td></td><td></td><td></td><td></td><td></td><td></td><td></td><td></td></tr>
<tr><td>Chromosome</td><td>start</td><td>end</td><td>strand</td><td>MM</td><td>target_seq</td><td>PAM</td><td>alignment</td><td>distance</td><td>position</td><td>gene name</td><td>gene id</td></tr>
<tr><td>GU305914.1</td><td>130367</td><td>130389</td><td>+</td><td>0</td><td>CAGCATCAACGGCGAGAGGA</td><td>AGG</td><td>||||||||[||||||||||||]PAM</td><td>NA</td><td>Intergenic</td><td>NA</td><td>NA</td></tr>
<tr><td></td><td></td><td></td><td></td><td></td><td></td><td></td><td></td><td></td><td></td><td></td><td></td></tr>
<tr><td>T9846</td><td>CAACGGCGAGAGGAAGGACGAGG</td><td>1000</td><td></td><td></td><td></td><td></td><td></td><td></td><td></td><td></td><td></td></tr>
<tr><td>Oligo fwd</td><td>CACCCAACGGCGAGAGGAAGGACG</td><td></td><td></td><td></td><td></td><td></td><td></td><td></td><td></td><td></td><td></td></tr>
<tr><td>Oligo rev</td><td>AAACCGTCCTTCCTCTCGCCGTTG</td><td></td><td></td><td></td><td></td><td></td><td></td><td></td><td></td><td></td><td></td></tr>
<tr><td>Chromosome</td><td>start</td><td>end</td><td>strand</td><td>MM</td><td>target_seq</td><td>PAM</td><td>alignment</td><td>distance</td><td>position</td><td>gene name</td><td>gene id</td></tr>
<tr><td>GU305914.1</td><td>130373</td><td>130395</td><td>+</td><td>0</td><td>CAACGGCGAGAGGAAGGACG</td><td>AGG</td><td>||||||||[||||||||||||]PAM</td><td>NA</td><td>Intergenic</td><td>NA</td><td>NA</td></tr>
</table>

<table>
<tr><td>T9847</td><td>GCCGTCGCTATCCGCGACCCCGG</td><td>1000</td><td></td><td></td><td></td><td></td><td></td><td></td><td></td><td></td><td></td></tr>
<tr><td>Oligo fwd</td><td>CACCGCCGTCGCTATCCGCGACCC</td><td></td><td></td><td></td><td></td><td></td><td></td><td></td><td></td><td></td><td></td></tr>
<tr><td>Oligo rev</td><td>AAACGGGTCGCGGATAGCGACGGC</td><td></td><td></td><td></td><td></td><td></td><td></td><td></td><td></td><td></td><td></td></tr>
<tr><td>Chromosome</td><td>start</td><td>end</td><td>strand</td><td>MM</td><td>target_seq</td><td>PAM</td><td>alignment</td><td>distance</td><td>position</td><td>gene name</td><td>gene id</td></tr>
<tr><td>GU305914.1</td><td>130395</td><td>130417</td><td>+</td><td>0</td><td>GCCGTCGCTATCCGCGACCC</td><td>CGG</td><td>||||||||[||||||||||||]PAM</td><td>NA</td><td>Intergenic</td><td>NA</td><td>NA</td></tr>
<tr><td>T9848</td><td>TCCGCGACCCCGGCGCCATCTGG</td><td>1000</td><td></td><td></td><td></td><td></td><td></td><td></td><td></td><td></td><td></td></tr>
<tr><td>Oligo fwd</td><td>CACCTCCGCGACCCCGGCGCCATC</td><td></td><td></td><td></td><td></td><td></td><td></td><td></td><td></td><td></td><td></td></tr>
<tr><td>Oligo rev</td><td>AAACGATGGCGCCGGGGTCGCGGA</td><td></td><td></td><td></td><td></td><td></td><td></td><td></td><td></td><td></td><td></td></tr>
<tr><td>Chromosome</td><td>start</td><td>end</td><td>strand</td><td>MM</td><td>target_seq</td><td>PAM</td><td>alignment</td><td>distance</td><td>position</td><td>gene name</td><td>gene id</td></tr>
<tr><td>GU305914.1</td><td>130405</td><td>130427</td><td>+</td><td>0</td><td>TCCGCGACCCCGGCGCCATC</td><td>TGG</td><td>||||||||[||||||||||||]PAM</td><td>NA</td><td>Intergenic</td><td>NA</td><td>NA</td></tr>
<tr><td>T9849</td><td>CCCGGCGCCATCTGGTGCGACGG</td><td>1000</td><td></td><td></td><td></td><td></td><td></td><td></td><td></td><td></td><td></td></tr>
<tr><td>Oligo fwd</td><td>CACCCCCGGCGCCATCTGGTGCGA</td><td></td><td></td><td></td><td></td><td></td><td></td><td></td><td></td><td></td><td></td></tr>
<tr><td>Oligo rev</td><td>AAACTCGCACCAGATGGCGCCGGG</td><td></td><td></td><td></td><td></td><td></td><td></td><td></td><td></td><td></td><td></td></tr>
<tr><td>Chromosome</td><td>start</td><td>end</td><td>strand</td><td>MM</td><td>target_seq</td><td>PAM</td><td>alignment</td><td>distance</td><td>position</td><td>gene name</td><td>gene id</td></tr>
<tr><td>GU305914.1</td><td>130413</td><td>130435</td><td>+</td><td>0</td><td>CCCGGCGCCATCTGGTGCGA</td><td>CGG</td><td>||||||||[||||||||||||]PAM</td><td>NA</td><td>Intergenic</td><td>NA</td><td>NA</td></tr>
<tr><td>T9850</td><td>CCGGCGCCATCTGGTGCGACGGG</td><td>1000</td><td></td><td></td><td></td><td></td><td></td><td></td><td></td><td></td><td></td></tr>
<tr><td>Oligo fwd</td><td>CACCCCGGCGCCATCTGGTGCGAC</td><td></td><td></td><td></td><td></td><td></td><td></td><td></td><td></td><td></td><td></td></tr>
<tr><td>Oligo rev</td><td>AAACGTCGCACCAGATGGCGCCGG</td><td></td><td></td><td></td><td></td><td></td><td></td><td></td><td></td><td></td><td></td></tr>
<tr><td>Chromosome</td><td>start</td><td>end</td><td>strand</td><td>MM</td><td>target_seq</td><td>PAM</td><td>alignment</td><td>distance</td><td>position</td><td>gene name</td><td>gene id</td></tr>
<tr><td>GU305914.1</td><td>130414</td><td>130436</td><td>+</td><td>0</td><td>CCGGCGCCATCTGGTGCGAC</td><td>GGG</td><td>||||||||[||||||||||||]PAM</td><td>NA</td><td>Intergenic</td><td>NA</td><td>NA</td></tr>
<tr><td>T9851</td><td>CGGCGCCATCTGGTGCGACGGGG</td><td>1000</td><td></td><td></td><td></td><td></td><td></td><td></td><td></td><td></td><td></td></tr>
<tr><td>Oligo fwd</td><td>CACCCGGCGCCATCTGGTGCGACG</td><td></td><td></td><td></td><td></td><td></td><td></td><td></td><td></td><td></td><td></td></tr>
<tr><td>Oligo rev</td><td>AAACCGTCGCACCAGATGGCGCCG</td><td></td><td></td><td></td><td></td><td></td><td></td><td></td><td></td><td></td><td></td></tr>
<tr><td>Chromosome</td><td>start</td><td>end</td><td>strand</td><td>MM</td><td>target_seq</td><td>PAM</td><td>alignment</td><td>distance</td><td>position</td><td>gene name</td><td>gene id</td></tr>
<tr><td>GU305914.1</td><td>130415</td><td>130437</td><td>+</td><td>0</td><td>CGGCGCCATCTGGTGCGACG</td><td>GGG</td><td>||||||||[||||||||||||]PAM</td><td>NA</td><td>Intergenic</td><td>NA</td><td>NA</td></tr>
<tr><td>T9852</td><td>GGGGCCGTGCTCATGAGCGACGG</td><td>1000</td><td></td><td></td><td></td><td></td><td></td><td></td><td></td><td></td><td></td></tr>
<tr><td>Oligo fwd</td><td>CACCGGGGCCGTGCTCATGAGCGA</td><td></td><td></td><td></td><td></td><td></td><td></td><td></td><td></td><td></td><td></td></tr>
<tr><td>Oligo rev</td><td>AAACTCGCTCATGAGCACGGCCCC</td><td></td><td></td><td></td><td></td><td></td><td></td><td></td><td></td><td></td><td></td></tr>
<tr><td>Chromosome</td><td>start</td><td>end</td><td>strand</td><td>MM</td><td>target_seq</td><td>PAM</td><td>alignment</td><td>distance</td><td>position</td><td>gene name</td><td>gene id</td></tr>
<tr><td>GU305914.1</td><td>130434</td><td>130456</td><td>+</td><td>0</td><td>GGGGCCGTGCTCATGAGCGA</td><td>CGG</td><td>||||||||[||||||||||||]PAM</td><td>NA</td><td>Intergenic</td><td>NA</td><td>NA</td></tr>
<tr><td>T9853</td><td>GGGCCGTGCTCATGAGCGACGGG</td><td>1000</td><td></td><td></td><td></td><td></td><td></td><td></td><td></td><td></td><td></td></tr>
<tr><td>Oligo fwd</td><td>CACCGGGCCGTGCTCATGAGCGAC</td><td></td><td></td><td></td><td></td><td></td><td></td><td></td><td></td><td></td><td></td></tr>
<tr><td>Oligo rev</td><td>AAACGTCGCTCATGAGCACGGCCC</td><td></td><td></td><td></td><td></td><td></td><td></td><td></td><td></td><td></td><td></td></tr>
<tr><td>Chromosome</td><td>start</td><td>end</td><td>strand</td><td>MM</td><td>target_seq</td><td>PAM</td><td>alignment</td><td>distance</td><td>position</td><td>gene name</td><td>gene id</td></tr>
<tr><td>GU305914.1</td><td>130435</td><td>130457</td><td>+</td><td>0</td><td>GGGCCGTGCTCATGAGCGAC</td><td>GGG</td><td>||||||||[||||||||||||]PAM</td><td>NA</td><td>Intergenic</td><td>NA</td><td>NA</td></tr>
<tr><td>T9854</td><td>CATGAGCGACGGGACCATCATGG</td><td>1000</td><td></td><td></td><td></td><td></td><td></td><td></td><td></td><td></td><td></td></tr>
<tr><td>Oligo fwd</td><td>CACCCATGAGCGACGGGACCATCA</td><td></td><td></td><td></td><td></td><td></td><td></td><td></td><td></td><td></td><td></td></tr>
<tr><td>Oligo rev</td><td>AAACTGATGGTCCCGTCGCTCATG</td><td></td><td></td><td></td><td></td><td></td><td></td><td></td><td></td><td></td><td></td></tr>
<tr><td>Chromosome</td><td>start</td><td>end</td><td>strand</td><td>MM</td><td>target_seq</td><td>PAM</td><td>alignment</td><td>distance</td><td>position</td><td>gene name</td><td>gene id</td></tr>
<tr><td>GU305914.1</td><td>130445</td><td>130467</td><td>+</td><td>0</td><td>CATGAGCGACGGGACCATCA</td><td>TGG</td><td>||||||||[||||||||||||]PAM</td><td>NA</td><td>Intergenic</td><td>NA</td><td>NA</td></tr>
<tr><td>T9855</td><td>CGAGTTCTACTCCAAACCCCTGG</td><td>1000</td><td></td><td></td><td></td><td></td><td></td><td></td><td></td><td></td><td></td></tr>
<tr><td>Oligo fwd</td><td>CACCCGAGTTCTACTCCAAACCCC</td><td></td><td></td><td></td><td></td><td></td><td></td><td></td><td></td><td></td><td></td></tr>
<tr><td>Oligo rev</td><td>AAACGGGGTTTGGAGTAGAACTCG</td><td></td><td></td><td></td><td></td><td></td><td></td><td></td><td></td><td></td><td></td></tr>
<tr><td>Chromosome</td><td>start</td><td>end</td><td>strand</td><td>MM</td><td>target_seq</td><td>PAM</td><td>alignment</td><td>distance</td><td>position</td><td>gene name</td><td>gene id</td></tr>
<tr><td>GU305914.1</td><td>130472</td><td>130494</td><td>+</td><td>0</td><td>CGAGTTCTACTCCAAACCCC</td><td>TGG</td><td>||||||||[||||||||||||]PAM</td><td>NA</td><td>Intergenic</td><td>NA</td><td>NA</td></tr>
<tr><td>T9856</td><td>CCCTGGTCTCTCTGATCAGAAGG</td><td>1000</td><td></td><td></td><td></td><td></td><td></td><td></td><td></td><td></td><td></td></tr>
<tr><td>Oligo fwd</td><td>CACCCCCTGGTCTCTCTGATCAGA</td><td></td><td></td><td></td><td></td><td></td><td></td><td></td><td></td><td></td><td></td></tr>
<tr><td>Oligo rev</td><td>AAACTCTGATCAGAGAGACCAGGG</td><td></td><td></td><td></td><td></td><td></td><td></td><td></td><td></td><td></td><td></td></tr>
<tr><td>Chromosome</td><td>start</td><td>end</td><td>strand</td><td>MM</td><td>target_seq</td><td>PAM</td><td>alignment</td><td>distance</td><td>position</td><td>gene name</td><td>gene id</td></tr>
<tr><td>GU305914.1</td><td>130489</td><td>130511</td><td>+</td><td>0</td><td>CCCTGGTCTCTCTGATCAGA</td><td>AGG</td><td>||||||||[||||||||||||]PAM</td><td>NA</td><td>Intergenic</td><td>NA</td><td>NA</td></tr>
<tr><td>T9857</td><td>CTCTGATCAGAAGGAAGATCAGG</td><td>1000</td><td></td><td></td><td></td><td></td><td></td><td></td><td></td><td></td><td></td></tr>
<tr><td>Oligo fwd</td><td>CACCCTCTGATCAGAAGGAAGATC</td><td></td><td></td><td></td><td></td><td></td><td></td><td></td><td></td><td></td><td></td></tr>
<tr><td>Oligo rev</td><td>AAACGATCTTCCTTCTGATCAGAG</td><td></td><td></td><td></td><td></td><td></td><td></td><td></td><td></td><td></td><td></td></tr>
<tr><td>Chromosome</td><td>start</td><td>end</td><td>strand</td><td>MM</td><td>target_seq</td><td>PAM</td><td>alignment</td><td>distance</td><td>position</td><td>gene name</td><td>gene id</td></tr>
<tr><td>GU305914.1</td><td>130498</td><td>130520</td><td>+</td><td>0</td><td>CTCTGATCAGAAGGAAGATC</td><td>AGG</td><td>||||||||[||||||||||||]PAM</td><td>NA</td><td>Intergenic</td><td>NA</td><td>NA</td></tr>
<tr><td>T9858</td><td>TCTGATCAGAAGGAAGATCAGGG</td><td>1000</td><td></td><td></td><td></td><td></td><td></td><td></td><td></td><td></td><td></td></tr>
<tr><td>Oligo fwd</td><td>CACCTCTGATCAGAAGGAAGATCA</td><td></td><td></td><td></td><td></td><td></td><td></td><td></td><td></td><td></td><td></td></tr>
<tr><td>Oligo rev</td><td>AAACTGATCTTCCTTCTGATCAGA</td><td></td><td></td><td></td><td></td><td></td><td></td><td></td><td></td><td></td><td></td></tr>
<tr><td>Chromosome</td><td>start</td><td>end</td><td>strand</td><td>MM</td><td>target_seq</td><td>PAM</td><td>alignment</td><td>distance</td><td>position</td><td>gene name</td><td>gene id</td></tr>
<tr><td>GU305914.1</td><td>130499</td><td>130521</td><td>+</td><td>0</td><td>TCTGATCAGAAGGAAGATCA</td><td>GGG</td><td>||||||||[||||||||||||]PAM</td><td>NA</td><td>Intergenic</td><td>NA</td><td>NA</td></tr>
<tr><td>T9859</td><td>GATCAGAAGGAAGATCAGGGTGG</td><td>1000</td><td></td><td></td><td></td><td></td><td></td><td></td><td></td><td></td><td></td></tr>
<tr><td>Oligo fwd</td><td>CACCGATCAGAAGGAAGATCAGGG</td><td></td><td></td><td></td><td></td><td></td><td></td><td></td><td></td><td></td><td></td></tr>
<tr><td>Oligo rev</td><td>AAACCCCTGATCTTCCTTCTGATC</td><td></td><td></td><td></td><td></td><td></td><td></td><td></td><td></td><td></td><td></td></tr>
<tr><td>Chromosome</td><td>start</td><td>end</td><td>strand</td><td>MM</td><td>target_seq</td><td>PAM</td><td>alignment</td><td>distance</td><td>position</td><td>gene name</td><td>gene id</td></tr>
<tr><td>GU305914.1</td><td>130502</td><td>130524</td><td>+</td><td>0</td><td>GATCAGAAGGAAGATCAGGG</td><td>TGG</td><td>||||||||[||||||||||||]PAM</td><td>NA</td><td>Intergenic</td><td>NA</td><td>NA</td></tr>
<tr><td>GU305914.1</td><td>171176</td><td>171198</td><td>+</td><td>4</td><td>GACCAGCAGGACGGTCAGGG</td><td>CGG</td><td>||-|||-|[|||-|-||||||]PAM</td><td>NA</td><td>Intergenic</td><td>NA</td><td>NA</td></tr>
<tr><td>T9860</td><td>GAAGATCAGGGTGGAGCGCGAGG</td><td>1000</td><td></td><td></td><td></td><td></td><td></td><td></td><td></td><td></td><td></td></tr>
<tr><td>Oligo fwd</td><td>CACCGAAGATCAGGGTGGAGCGCG</td><td></td><td></td><td></td><td></td><td></td><td></td><td></td><td></td><td></td><td></td></tr>
<tr><td>Oligo rev</td><td>AAACCGCGCTCCACCCTGATCTTC</td><td></td><td></td><td></td><td></td><td></td><td></td><td></td><td></td><td></td><td></td></tr>
<tr><td>Chromosome</td><td>start</td><td>end</td><td>strand</td><td>MM</td><td>target_seq</td><td>PAM</td><td>alignment</td><td>distance</td><td>position</td><td>gene name</td><td>gene id</td></tr>
<tr><td>GU305914.1</td><td>130511</td><td>130533</td><td>+</td><td>0</td><td>GAAGATCAGGGTGGAGCGCG</td><td>AGG</td><td>||||||||[||||||||||||]PAM</td><td>NA</td><td>Intergenic</td><td>NA</td><td>NA</td></tr>
<tr><td>T9861</td><td>GATCAGGGTGGAGCGCGAGGAGG</td><td>1000</td><td></td><td></td><td></td><td></td><td></td><td></td><td></td><td></td><td></td></tr>
<tr><td>Oligo fwd</td><td>CACCGATCAGGGTGGAGCGCGAGG</td><td></td><td></td><td></td><td></td><td></td><td></td><td></td><td></td><td></td><td></td></tr>
<tr><td>Oligo rev</td><td>AAACCCTCGCGCTCCACCCTGATC</td><td></td><td></td><td></td><td></td><td></td><td></td><td></td><td></td><td></td><td></td></tr>
<tr><td>Chromosome</td><td>start</td><td>end</td><td>strand</td><td>MM</td><td>target_seq</td><td>PAM</td><td>alignment</td><td>distance</td><td>position</td><td>gene name</td><td>gene id</td></tr>
</table>

<table>
<tr><td>GU305914.1</td><td>130514</td><td>130536</td><td>+</td><td>0</td><td>GATCAGGGTGGAGCGCGAGG</td><td>AGG</td><td>||||||||[||||||||||||]PAM</td><td>NA</td><td>Intergenic</td><td>NA</td><td>NA</td></tr>
<tr><td>T9862</td><td>AGCGCGAGGAGGACGACGAGAGG</td><td>1000</td><td></td><td></td><td></td><td></td><td></td><td></td><td></td><td></td><td></td></tr>
<tr><td>Oligo fwd</td><td>CACCAGCGCGAGGAGGACGACGAG</td><td></td><td></td><td></td><td></td><td></td><td></td><td></td><td></td><td></td><td></td></tr>
<tr><td>Oligo rev</td><td>AAACCTCGTCGTCCTCCTCGCGCT</td><td></td><td></td><td></td><td></td><td></td><td></td><td></td><td></td><td></td><td></td></tr>
<tr><td>Chromosome</td><td>start</td><td>end</td><td>strand</td><td>MM</td><td>target_seq</td><td>PAM</td><td>alignment</td><td>distance</td><td>position</td><td>gene name</td><td>gene id</td></tr>
<tr><td>GU305914.1</td><td>130525</td><td>130547</td><td>+</td><td>0</td><td>AGCGCGAGGAGGACGACGAG</td><td>AGG</td><td>||||||||[||||||||||||]PAM</td><td>NA</td><td>Intergenic</td><td>NA</td><td>NA</td></tr>
<tr><td>T9863</td><td>GCGCGAGGAGGACGACGAGAGGG</td><td>1000</td><td></td><td></td><td></td><td></td><td></td><td></td><td></td><td></td><td></td></tr>
<tr><td>Oligo fwd</td><td>CACCGCGCGAGGAGGACGACGAGA</td><td></td><td></td><td></td><td></td><td></td><td></td><td></td><td></td><td></td><td></td></tr>
<tr><td>Oligo rev</td><td>AAACTCTCGTCGTCCTCCTCGCGC</td><td></td><td></td><td></td><td></td><td></td><td></td><td></td><td></td><td></td><td></td></tr>
<tr><td>Chromosome</td><td>start</td><td>end</td><td>strand</td><td>MM</td><td>target_seq</td><td>PAM</td><td>alignment</td><td>distance</td><td>position</td><td>gene name</td><td>gene id</td></tr>
<tr><td>GU305914.1</td><td>130526</td><td>130548</td><td>+</td><td>0</td><td>GCGCGAGGAGGACGACGAGA</td><td>GGG</td><td>||||||||[||||||||||||]PAM</td><td>NA</td><td>Intergenic</td><td>NA</td><td>NA</td></tr>
<tr><td>T9864</td><td>CGCGAGGAGGACGACGAGAGGGG</td><td>1000</td><td></td><td></td><td></td><td></td><td></td><td></td><td></td><td></td><td></td></tr>
<tr><td>Oligo fwd</td><td>CACCCGCGAGGAGGACGACGAGAG</td><td></td><td></td><td></td><td></td><td></td><td></td><td></td><td></td><td></td><td></td></tr>
<tr><td>Oligo rev</td><td>AAACCTCTCGTCGTCCTCCTCGCG</td><td></td><td></td><td></td><td></td><td></td><td></td><td></td><td></td><td></td><td></td></tr>
<tr><td>Chromosome</td><td>start</td><td>end</td><td>strand</td><td>MM</td><td>target_seq</td><td>PAM</td><td>alignment</td><td>distance</td><td>position</td><td>gene name</td><td>gene id</td></tr>
<tr><td>GU305914.1</td><td>130527</td><td>130549</td><td>+</td><td>0</td><td>CGCGAGGAGGACGACGAGAG</td><td>GGG</td><td>||||||||[||||||||||||]PAM</td><td>NA</td><td>Intergenic</td><td>NA</td><td>NA</td></tr>
<tr><td>GU305914.1</td><td>98586</td><td>98608</td><td>+</td><td>3</td><td>GGCGAGGAGGGCGATGAGAG</td><td>TGG</td><td>-|||||||[||-|||-|||||]PAM</td><td>NA</td><td>Intergenic</td><td>NA</td><td>NA</td></tr>
<tr><td>T9865</td><td>GAACCGCCTTCGTCAGCTCCCGG</td><td>1000</td><td></td><td></td><td></td><td></td><td></td><td></td><td></td><td></td><td></td></tr>
<tr><td>Oligo fwd</td><td>CACCGAACCGCCTTCGTCAGCTCC</td><td></td><td></td><td></td><td></td><td></td><td></td><td></td><td></td><td></td><td></td></tr>
<tr><td>Oligo rev</td><td>AAACGGAGCTGACGAAGGCGGTTC</td><td></td><td></td><td></td><td></td><td></td><td></td><td></td><td></td><td></td><td></td></tr>
<tr><td>Chromosome</td><td>start</td><td>end</td><td>strand</td><td>MM</td><td>target_seq</td><td>PAM</td><td>alignment</td><td>distance</td><td>position</td><td>gene name</td><td>gene id</td></tr>
<tr><td>GU305914.1</td><td>130549</td><td>130571</td><td>+</td><td>0</td><td>GAACCGCCTTCGTCAGCTCC</td><td>CGG</td><td>||||||||[||||||||||||]PAM</td><td>NA</td><td>Intergenic</td><td>NA</td><td>NA</td></tr>
<tr><td>T9866</td><td>AACCGCCTTCGTCAGCTCCCGGG</td><td>1000</td><td></td><td></td><td></td><td></td><td></td><td></td><td></td><td></td><td></td></tr>
<tr><td>Oligo fwd</td><td>CACCAACCGCCTTCGTCAGCTCCC</td><td></td><td></td><td></td><td></td><td></td><td></td><td></td><td></td><td></td><td></td></tr>
<tr><td>Oligo rev</td><td>AAACGGGAGCTGACGAAGGCGGTT</td><td></td><td></td><td></td><td></td><td></td><td></td><td></td><td></td><td></td><td></td></tr>
<tr><td>Chromosome</td><td>start</td><td>end</td><td>strand</td><td>MM</td><td>target_seq</td><td>PAM</td><td>alignment</td><td>distance</td><td>position</td><td>gene name</td><td>gene id</td></tr>
<tr><td>GU305914.1</td><td>130550</td><td>130572</td><td>+</td><td>0</td><td>AACCGCCTTCGTCAGCTCCC</td><td>GGG</td><td>||||||||[||||||||||||]PAM</td><td>NA</td><td>Intergenic</td><td>NA</td><td>NA</td></tr>
<tr><td>T9867</td><td>GAGCTCTCGCTCTTTGTCACCGG</td><td>1000</td><td></td><td></td><td></td><td></td><td></td><td></td><td></td><td></td><td></td></tr>
<tr><td>Oligo fwd</td><td>CACCGAGCTCTCGCTCTTTGTCAC</td><td></td><td></td><td></td><td></td><td></td><td></td><td></td><td></td><td></td><td></td></tr>
<tr><td>Oligo rev</td><td>AAACGTGACAAAGAGCGAGAGCTC</td><td></td><td></td><td></td><td></td><td></td><td></td><td></td><td></td><td></td><td></td></tr>
<tr><td>Chromosome</td><td>start</td><td>end</td><td>strand</td><td>MM</td><td>target_seq</td><td>PAM</td><td>alignment</td><td>distance</td><td>position</td><td>gene name</td><td>gene id</td></tr>
<tr><td>GU305914.1</td><td>130572</td><td>130594</td><td>+</td><td>0</td><td>GAGCTCTCGCTCTTTGTCAC</td><td>CGG</td><td>||||||||[||||||||||||]PAM</td><td>NA</td><td>Intergenic</td><td>NA</td><td>NA</td></tr>
<tr><td>T9868</td><td>TCTTTGTCACCGGACGCTGTCGG</td><td>1000</td><td></td><td></td><td></td><td></td><td></td><td></td><td></td><td></td><td></td></tr>
<tr><td>Oligo fwd</td><td>CACCTCTTTGTCACCGGACGCTGT</td><td></td><td></td><td></td><td></td><td></td><td></td><td></td><td></td><td></td><td></td></tr>
<tr><td>Oligo rev</td><td>AAACACAGCGTCCGGTGACAAAGA</td><td></td><td></td><td></td><td></td><td></td><td></td><td></td><td></td><td></td><td></td></tr>
<tr><td>Chromosome</td><td>start</td><td>end</td><td>strand</td><td>MM</td><td>target_seq</td><td>PAM</td><td>alignment</td><td>distance</td><td>position</td><td>gene name</td><td>gene id</td></tr>
<tr><td>GU305914.1</td><td>130582</td><td>130604</td><td>+</td><td>0</td><td>TCTTTGTCACCGGACGCTGT</td><td>CGG</td><td>||||||||[||||||||||||]PAM</td><td>NA</td><td>Intergenic</td><td>NA</td><td>NA</td></tr>
<tr><td>T9869</td><td>CGCTGTCGGCGCCGCGCCGCCGG</td><td>1000</td><td></td><td></td><td></td><td></td><td></td><td></td><td></td><td></td><td></td></tr>
<tr><td>Oligo fwd</td><td>CACCCGCTGTCGGCGCCGCGCCGC</td><td></td><td></td><td></td><td></td><td></td><td></td><td></td><td></td><td></td><td></td></tr>
<tr><td>Oligo rev</td><td>AAACGCGGCGCGGCGCCGACAGCG</td><td></td><td></td><td></td><td></td><td></td><td></td><td></td><td></td><td></td><td></td></tr>
<tr><td>Chromosome</td><td>start</td><td>end</td><td>strand</td><td>MM</td><td>target_seq</td><td>PAM</td><td>alignment</td><td>distance</td><td>position</td><td>gene name</td><td>gene id</td></tr>
<tr><td>GU305914.1</td><td>130596</td><td>130618</td><td>+</td><td>0</td><td>CGCTGTCGGCGCCGCGCCGC</td><td>CGG</td><td>||||||||[||||||||||||]PAM</td><td>NA</td><td>Intergenic</td><td>NA</td><td>NA</td></tr>
<tr><td>T9870</td><td>CCGCCGGCTTCTCCATCTACTGG</td><td>1000</td><td></td><td></td><td></td><td></td><td></td><td></td><td></td><td></td><td></td></tr>
<tr><td>Oligo fwd</td><td>CACCCCGCCGGCTTCTCCATCTAC</td><td></td><td></td><td></td><td></td><td></td><td></td><td></td><td></td><td></td><td></td></tr>
<tr><td>Oligo rev</td><td>AAACGTAGATGGAGAAGCCGGCGG</td><td></td><td></td><td></td><td></td><td></td><td></td><td></td><td></td><td></td><td></td></tr>
<tr><td>Chromosome</td><td>start</td><td>end</td><td>strand</td><td>MM</td><td>target_seq</td><td>PAM</td><td>alignment</td><td>distance</td><td>position</td><td>gene name</td><td>gene id</td></tr>
<tr><td>GU305914.1</td><td>130612</td><td>130634</td><td>+</td><td>0</td><td>CCGCCGGCTTCTCCATCTAC</td><td>TGG</td><td>||||||||[||||||||||||]PAM</td><td>NA</td><td>Intergenic</td><td>NA</td><td>NA</td></tr>
<tr><td>T9871</td><td>GACCTCATCTTCGTCCTCACCGG</td><td>1000</td><td></td><td></td><td></td><td></td><td></td><td></td><td></td><td></td><td></td></tr>
<tr><td>Oligo fwd</td><td>CACCGACCTCATCTTCGTCCTCAC</td><td></td><td></td><td></td><td></td><td></td><td></td><td></td><td></td><td></td><td></td></tr>
<tr><td>Oligo rev</td><td>AAACGTGAGGACGAAGATGAGGTC</td><td></td><td></td><td></td><td></td><td></td><td></td><td></td><td></td><td></td><td></td></tr>
<tr><td>Chromosome</td><td>start</td><td>end</td><td>strand</td><td>MM</td><td>target_seq</td><td>PAM</td><td>alignment</td><td>distance</td><td>position</td><td>gene name</td><td>gene id</td></tr>
<tr><td>GU305914.1</td><td>130647</td><td>130669</td><td>+</td><td>0</td><td>GACCTCATCTTCGTCCTCAC</td><td>CGG</td><td>||||||||[||||||||||||]PAM</td><td>NA</td><td>Intergenic</td><td>NA</td><td>NA</td></tr>
<tr><td>T9872</td><td>GCGCAGATCTACGTCGAGTGCGG</td><td>1000</td><td></td><td></td><td></td><td></td><td></td><td></td><td></td><td></td><td></td></tr>
<tr><td>Oligo fwd</td><td>CACCGCGCAGATCTACGTCGAGTG</td><td></td><td></td><td></td><td></td><td></td><td></td><td></td><td></td><td></td><td></td></tr>
<tr><td>Oligo rev</td><td>AAACCACTCGACGTAGATCTGCGC</td><td></td><td></td><td></td><td></td><td></td><td></td><td></td><td></td><td></td><td></td></tr>
<tr><td>Chromosome</td><td>start</td><td>end</td><td>strand</td><td>MM</td><td>target_seq</td><td>PAM</td><td>alignment</td><td>distance</td><td>position</td><td>gene name</td><td>gene id</td></tr>
<tr><td>GU305914.1</td><td>130683</td><td>130705</td><td>+</td><td>0</td><td>GCGCAGATCTACGTCGAGTG</td><td>CGG</td><td>||||||||[||||||||||||]PAM</td><td>NA</td><td>Intergenic</td><td>NA</td><td>NA</td></tr>
<tr><td>T9873</td><td>CGCAGATCTACGTCGAGTGCGGG</td><td>1000</td><td></td><td></td><td></td><td></td><td></td><td></td><td></td><td></td><td></td></tr>
<tr><td>Oligo fwd</td><td>CACCCGCAGATCTACGTCGAGTGC</td><td></td><td></td><td></td><td></td><td></td><td></td><td></td><td></td><td></td><td></td></tr>
<tr><td>Oligo rev</td><td>AAACGCACTCGACGTAGATCTGCG</td><td></td><td></td><td></td><td></td><td></td><td></td><td></td><td></td><td></td><td></td></tr>
<tr><td>Chromosome</td><td>start</td><td>end</td><td>strand</td><td>MM</td><td>target_seq</td><td>PAM</td><td>alignment</td><td>distance</td><td>position</td><td>gene name</td><td>gene id</td></tr>
<tr><td>GU305914.1</td><td>130684</td><td>130706</td><td>+</td><td>0</td><td>CGCAGATCTACGTCGAGTGC</td><td>GGG</td><td>||||||||[||||||||||||]PAM</td><td>NA</td><td>Intergenic</td><td>NA</td><td>NA</td></tr>
<tr><td>T9874</td><td>AGATCTACGTCGAGTGCGGGAGG</td><td>1000</td><td></td><td></td><td></td><td></td><td></td><td></td><td></td><td></td><td></td></tr>
<tr><td>Oligo fwd</td><td>CACCAGATCTACGTCGAGTGCGGG</td><td></td><td></td><td></td><td></td><td></td><td></td><td></td><td></td><td></td><td></td></tr>
<tr><td>Oligo rev</td><td>AAACCCCGCACTCGACGTAGATCT</td><td></td><td></td><td></td><td></td><td></td><td></td><td></td><td></td><td></td><td></td></tr>
<tr><td>Chromosome</td><td>start</td><td>end</td><td>strand</td><td>MM</td><td>target_seq</td><td>PAM</td><td>alignment</td><td>distance</td><td>position</td><td>gene name</td><td>gene id</td></tr>
<tr><td>GU305914.1</td><td>130687</td><td>130709</td><td>+</td><td>0</td><td>AGATCTACGTCGAGTGCGGG</td><td>AGG</td><td>||||||||[||||||||||||]PAM</td><td>NA</td><td>Intergenic</td><td>NA</td><td>NA</td></tr>
<tr><td>T9875</td><td>CTACGTCGAGTGCGGGAGGATGG</td><td>1000</td><td></td><td></td><td></td><td></td><td></td><td></td><td></td><td></td><td></td></tr>
<tr><td>Oligo fwd</td><td>CACCCTACGTCGAGTGCGGGAGGA</td><td></td><td></td><td></td><td></td><td></td><td></td><td></td><td></td><td></td><td></td></tr>
<tr><td>Oligo rev</td><td>AAACTCCTCCCGCACTCGACGTAG</td><td></td><td></td><td></td><td></td><td></td><td></td><td></td><td></td><td></td><td></td></tr>
<tr><td>Chromosome</td><td>start</td><td>end</td><td>strand</td><td>MM</td><td>target_seq</td><td>PAM</td><td>alignment</td><td>distance</td><td>position</td><td>gene name</td><td>gene id</td></tr>
<tr><td>GU305914.1</td><td>130691</td><td>130713</td><td>+</td><td>0</td><td>CTACGTCGAGTGCGGGAGGA</td><td>TGG</td><td>||||||||[||||||||||||]PAM</td><td>NA</td><td>Intergenic</td><td>NA</td><td>NA</td></tr>
<tr><td>T9876</td><td>CGAGTGCGGGAGGATGGCGCCGG</td><td>1000</td><td></td><td></td><td></td><td></td><td></td><td></td><td></td><td></td><td></td></tr>
<tr><td>Oligo fwd</td><td>CACCCGAGTGCGGGAGGATGGCGC</td><td></td><td></td><td></td><td></td><td></td><td></td><td></td><td></td><td></td><td></td></tr>
</table>

<table>
<tr><td>Oligo rev</td><td>AAACGCGCCATCCTCCCGCACTCG</td></tr>
<tr><td>Chromosome</td><td>start</td><td>end</td><td>strand</td><td>MM</td><td>target_seq</td><td>PAM</td><td>alignment</td><td>distance</td><td>position</td><td>gene name</td><td>gene id</td></tr>
<tr><td>GU305914.1</td><td>130697</td><td>130719</td><td>+</td><td>0</td><td>CGAGTGCGGGAGGATGGCGC</td><td>CGG</td><td>||||||||[||||||||||||]PAM</td><td>NA</td><td>Intergenic</td><td>NA</td><td>NA</td></tr>
<tr><td>T9877</td><td>TGCGGGAGGATGGCGCCGGACGG</td><td>1000</td></tr>
<tr><td>Oligo fwd</td><td>CACCTGCGGGAGGATGGCGCCGGA</td></tr>
<tr><td>Oligo rev</td><td>AAACTCCGGCGCCATCCTCCCGCA</td></tr>
<tr><td>Chromosome</td><td>start</td><td>end</td><td>strand</td><td>MM</td><td>target_seq</td><td>PAM</td><td>alignment</td><td>distance</td><td>position</td><td>gene name</td><td>gene id</td></tr>
<tr><td>GU305914.1</td><td>130701</td><td>130723</td><td>+</td><td>0</td><td>TGCGGGAGGATGGCGCCGGA</td><td>CGG</td><td>||||||||[||||||||||||]PAM</td><td>NA</td><td>Intergenic</td><td>NA</td><td>NA</td></tr>
<tr><td>T9878</td><td>GATGGCGCCGGACGGCACCATGG</td><td>1000</td></tr>
<tr><td>Oligo fwd</td><td>CACCGATGGCGCCGGACGGCACCA</td></tr>
<tr><td>Oligo rev</td><td>AAACTGGTGCCGTCCGGCGCCATC</td></tr>
<tr><td>Chromosome</td><td>start</td><td>end</td><td>strand</td><td>MM</td><td>target_seq</td><td>PAM</td><td>alignment</td><td>distance</td><td>position</td><td>gene name</td><td>gene id</td></tr>
<tr><td>GU305914.1</td><td>130709</td><td>130731</td><td>+</td><td>0</td><td>GATGGCGCCGGACGGCACCA</td><td>TGG</td><td>||||||||[||||||||||||]PAM</td><td>NA</td><td>Intergenic</td><td>NA</td><td>NA</td></tr>
<tr><td>T9879</td><td>GCGTCCTCCTCGTGCGCGCCAGG</td><td>1000</td></tr>
<tr><td>Oligo fwd</td><td>CACCGCGTCCTCCTCGTGCGCGCC</td></tr>
<tr><td>Oligo rev</td><td>AAACGGCGCGCACGAGGAGGACGC</td></tr>
<tr><td>Chromosome</td><td>start</td><td>end</td><td>strand</td><td>MM</td><td>target_seq</td><td>PAM</td><td>alignment</td><td>distance</td><td>position</td><td>gene name</td><td>gene id</td></tr>
<tr><td>GU305914.1</td><td>130750</td><td>130772</td><td>+</td><td>0</td><td>GCGTCCTCCTCGTGCGCGCC</td><td>AGG</td><td>||||||||[||||||||||||]PAM</td><td>NA</td><td>Intergenic</td><td>NA</td><td>NA</td></tr>
<tr><td>T9880</td><td>CGTCCTCCTCGTGCGCGCCAGGG</td><td>1000</td></tr>
<tr><td>Oligo fwd</td><td>CACCCGTCCTCCTCGTGCGCGCCA</td></tr>
<tr><td>Oligo rev</td><td>AAACTGGCGCGCACGAGGAGGACG</td></tr>
<tr><td>Chromosome</td><td>start</td><td>end</td><td>strand</td><td>MM</td><td>target_seq</td><td>PAM</td><td>alignment</td><td>distance</td><td>position</td><td>gene name</td><td>gene id</td></tr>
<tr><td>GU305914.1</td><td>130751</td><td>130773</td><td>+</td><td>0</td><td>CGTCCTCCTCGTGCGCGCCA</td><td>GGG</td><td>||||||||[||||||||||||]PAM</td><td>NA</td><td>Intergenic</td><td>NA</td><td>NA</td></tr>
<tr><td>T9881</td><td>GTGCGCGCCAGGGACCCCAACGG</td><td>1000</td></tr>
<tr><td>Oligo fwd</td><td>CACCGTGCGCGCCAGGGACCCCAA</td></tr>
<tr><td>Oligo rev</td><td>AAACTTGGGGTCCCTGGCGCGCAC</td></tr>
<tr><td>Chromosome</td><td>start</td><td>end</td><td>strand</td><td>MM</td><td>target_seq</td><td>PAM</td><td>alignment</td><td>distance</td><td>position</td><td>gene name</td><td>gene id</td></tr>
<tr><td>GU305914.1</td><td>130761</td><td>130783</td><td>+</td><td>0</td><td>GTGCGCGCCAGGGACCCCAA</td><td>CGG</td><td>||||||||[||||||||||||]PAM</td><td>NA</td><td>Intergenic</td><td>NA</td><td>NA</td></tr>
<tr><td>T9882</td><td>CGCGCCAGGGACCCCAACGGCGG</td><td>1000</td></tr>
<tr><td>Oligo fwd</td><td>CACCCGCGCCAGGGACCCCAACGG</td></tr>
<tr><td>Oligo rev</td><td>AAACCCGTTGGGGTCCCTGGCGCG</td></tr>
<tr><td>Chromosome</td><td>start</td><td>end</td><td>strand</td><td>MM</td><td>target_seq</td><td>PAM</td><td>alignment</td><td>distance</td><td>position</td><td>gene name</td><td>gene id</td></tr>
<tr><td>GU305914.1</td><td>130764</td><td>130786</td><td>+</td><td>0</td><td>CGCGCCAGGGACCCCAACGG</td><td>CGG</td><td>||||||||[||||||||||||]PAM</td><td>NA</td><td>Intergenic</td><td>NA</td><td>NA</td></tr>
<tr><td>T9883</td><td>GCGGCATCTCGCACGTCTCCTGG</td><td>1000</td></tr>
<tr><td>Oligo fwd</td><td>CACCGCGGCATCTCGCACGTCTCC</td></tr>
<tr><td>Oligo rev</td><td>AAACGGAGACGTGCGAGATGCCGC</td></tr>
<tr><td>Chromosome</td><td>start</td><td>end</td><td>strand</td><td>MM</td><td>target_seq</td><td>PAM</td><td>alignment</td><td>distance</td><td>position</td><td>gene name</td><td>gene id</td></tr>
<tr><td>GU305914.1</td><td>130783</td><td>130805</td><td>+</td><td>0</td><td>GCGGCATCTCGCACGTCTCC</td><td>TGG</td><td>||||||||[||||||||||||]PAM</td><td>NA</td><td>Intergenic</td><td>NA</td><td>NA</td></tr>
<tr><td>T9884</td><td>CGTCTCCTGGTCCAAGACTCCGG</td><td>1000</td></tr>
<tr><td>Oligo fwd</td><td>CACCCGTCTCCTGGTCCAAGACTC</td></tr>
<tr><td>Oligo rev</td><td>AAACGAGTCTTGGACCAGGAGACG</td></tr>
<tr><td>Chromosome</td><td>start</td><td>end</td><td>strand</td><td>MM</td><td>target_seq</td><td>PAM</td><td>alignment</td><td>distance</td><td>position</td><td>gene name</td><td>gene id</td></tr>
<tr><td>GU305914.1</td><td>130796</td><td>130818</td><td>+</td><td>0</td><td>CGTCTCCTGGTCCAAGACTC</td><td>CGG</td><td>||||||||[||||||||||||]PAM</td><td>NA</td><td>Intergenic</td><td>NA</td><td>NA</td></tr>
<tr><td>T9885</td><td>GTCTCCTGGTCCAAGACTCCGGG</td><td>1000</td></tr>
<tr><td>Oligo fwd</td><td>CACCGTCTCCTGGTCCAAGACTCC</td></tr>
<tr><td>Oligo rev</td><td>AAACGGAGTCTTGGACCAGGAGAC</td></tr>
<tr><td>Chromosome</td><td>start</td><td>end</td><td>strand</td><td>MM</td><td>target_seq</td><td>PAM</td><td>alignment</td><td>distance</td><td>position</td><td>gene name</td><td>gene id</td></tr>
<tr><td>GU305914.1</td><td>130797</td><td>130819</td><td>+</td><td>0</td><td>GTCTCCTGGTCCAAGACTCC</td><td>GGG</td><td>||||||||[||||||||||||]PAM</td><td>NA</td><td>Intergenic</td><td>NA</td><td>NA</td></tr>
<tr><td>T9886</td><td>GGTCCAAGACTCCGGGCATCTGG</td><td>1000</td></tr>
<tr><td>Oligo fwd</td><td>CACCGGTCCAAGACTCCGGGCATC</td></tr>
<tr><td>Oligo rev</td><td>AAACGATGCCCGGAGTCTTGGACC</td></tr>
<tr><td>Chromosome</td><td>start</td><td>end</td><td>strand</td><td>MM</td><td>target_seq</td><td>PAM</td><td>alignment</td><td>distance</td><td>position</td><td>gene name</td><td>gene id</td></tr>
<tr><td>GU305914.1</td><td>130804</td><td>130826</td><td>+</td><td>0</td><td>GGTCCAAGACTCCGGGCATC</td><td>TGG</td><td>||||||||[||||||||||||]PAM</td><td>NA</td><td>Intergenic</td><td>NA</td><td>NA</td></tr>
<tr><td>T9887</td><td>GACTCCGGGCATCTGGCCGAAGG</td><td>1000</td></tr>
<tr><td>Oligo fwd</td><td>CACCGACTCCGGGCATCTGGCCGA</td></tr>
<tr><td>Oligo rev</td><td>AAACTCGGCCAGATGCCCGGAGTC</td></tr>
<tr><td>Chromosome</td><td>start</td><td>end</td><td>strand</td><td>MM</td><td>target_seq</td><td>PAM</td><td>alignment</td><td>distance</td><td>position</td><td>gene name</td><td>gene id</td></tr>
<tr><td>GU305914.1</td><td>130811</td><td>130833</td><td>+</td><td>0</td><td>GACTCCGGGCATCTGGCCGA</td><td>AGG</td><td>||||||||[||||||||||||]PAM</td><td>NA</td><td>Intergenic</td><td>NA</td><td>NA</td></tr>
<tr><td>T9888</td><td>GGCATCTGGCCGAAGGACGTCGG</td><td>1000</td></tr>
<tr><td>Oligo fwd</td><td>CACCGGCATCTGGCCGAAGGACGT</td></tr>
<tr><td>Oligo rev</td><td>AAACACGTCCTTCGGCCAGATGCC</td></tr>
<tr><td>Chromosome</td><td>start</td><td>end</td><td>strand</td><td>MM</td><td>target_seq</td><td>PAM</td><td>alignment</td><td>distance</td><td>position</td><td>gene name</td><td>gene id</td></tr>
<tr><td>GU305914.1</td><td>130818</td><td>130840</td><td>+</td><td>0</td><td>GGCATCTGGCCGAAGGACGT</td><td>CGG</td><td>||||||||[||||||||||||]PAM</td><td>NA</td><td>Intergenic</td><td>NA</td><td>NA</td></tr>
<tr><td>T9889</td><td>CCAGCAACGCGCCCGCCGCCAGG</td><td>1000</td></tr>
<tr><td>Oligo fwd</td><td>CACCCCAGCAACGCGCCCGCCGCC</td></tr>
<tr><td>Oligo rev</td><td>AAACGGCGGCGGGCGCGTTGCTGG</td></tr>
<tr><td>Chromosome</td><td>start</td><td>end</td><td>strand</td><td>MM</td><td>target_seq</td><td>PAM</td><td>alignment</td><td>distance</td><td>position</td><td>gene name</td><td>gene id</td></tr>
<tr><td>GU305914.1</td><td>130861</td><td>130883</td><td>+</td><td>0</td><td>CCAGCAACGCGCCCGCCGCC</td><td>AGG</td><td>||||||||[||||||||||||]PAM</td><td>NA</td><td>Intergenic</td><td>NA</td><td>NA</td></tr>
<tr><td>T9890</td><td>ACGCCCGTCATGCCCAACGTCGG</td><td>1000</td></tr>
<tr><td>Oligo fwd</td><td>CACCACGCCCGTCATGCCCAACGT</td></tr>
<tr><td>Oligo rev</td><td>AAACACGTTGGGCATGACGGGCGT</td></tr>
<tr><td>Chromosome</td><td>start</td><td>end</td><td>strand</td><td>MM</td><td>target_seq</td><td>PAM</td><td>alignment</td><td>distance</td><td>position</td><td>gene name</td><td>gene id</td></tr>
<tr><td>GU305914.1</td><td>130890</td><td>130912</td><td>+</td><td>0</td><td>ACGCCCGTCATGCCCAACGT</td><td>CGG</td><td>||||||||[||||||||||||]PAM</td><td>NA</td><td>Intergenic</td><td>NA</td><td>NA</td></tr>
<tr><td>T9891</td><td>ATGCCCAACGTCGGCATGATCGG</td><td>1000</td></tr>
</table>

Oligo fwd CACCATGCCCAACGTCGGCATGAT
Oligo rev AAACATCATGCCGACGTTGGGCAT
Chromosome start end strand MM target_seq PAM alignment distance position gene name gene id
GU305914.1 130899 130921 + 0 ATGCCCAACGTCGGCATGAT CGG ||||||||[||||||||||||]PAM NA Intergenic NA NA

T9892 TTCCCTTGCGTCAGACTCTTCGG 1000
Oligo fwd CACCTTCCCTTGCGTCAGACTCTT
Oligo rev AAACAAGAGTCTGACGCAAGGGAA
Chromosome start end strand MM target_seq PAM alignment distance position gene name gene id
GU305914.1 130923 130945 + 0 TTCCCTTGCGTCAGACTCTT CGG ||||||||[||||||||||||]PAM NA Intergenic NA NA

T9893 GAGCCCACCAACATCGACATCGG 1000
Oligo fwd CACCGAGCCCACCAACATCGACAT
Oligo rev AAACATGTCGATGTTGGTGGGCTC
Chromosome start end strand MM target_seq PAM alignment distance position gene name gene id
GU305914.1 130947 130969 + 0 GAGCCCACCAACATCGACAT CGG ||||||||[||||||||||||]PAM NA Intergenic NA NA

T9894 AGCCCACCAACATCGACATCGGG 1000
Oligo fwd CACCAGCCCACCAACATCGACATC
Oligo rev AAACGATGTCGATGTTGGTGGGCT
Chromosome start end strand MM target_seq PAM alignment distance position gene name gene id
GU305914.1 130948 130970 + 0 AGCCCACCAACATCGACATC GGG ||||||||[||||||||||||]PAM NA Intergenic NA NA

T9895 GCCCACCAACATCGACATCGGGG 1000
Oligo fwd CACCGCCCACCAACATCGACATCG
Oligo rev AAACCGATGTCGATGTTGGTGGGC
Chromosome start end strand MM target_seq PAM alignment distance position gene name gene id
GU305914.1 130949 130971 + 0 GCCCACCAACATCGACATCG GGG ||||||||[||||||||||||]PAM NA Intergenic NA NA

T9896 GGCCTATGTGCCCCTCGTCGTGG 1000
Oligo fwd CACCGGCCTATGTGCCCCTCGTCG
Oligo rev AAACCGACGAGGGGCACATAGGCC
Chromosome start end strand MM target_seq PAM alignment distance position gene name gene id
GU305914.1 130970 130992 + 0 GGCCTATGTGCCCCTCGTCG TGG ||||||||[||||||||||||]PAM NA Intergenic NA NA

T9897 GCCTATGTGCCCCTCGTCGTGGG 1000
Oligo fwd CACCGCCTATGTGCCCCTCGTCGT
Oligo rev AAACACGACGAGGGGCACATAGGC
Chromosome start end strand MM target_seq PAM alignment distance position gene name gene id
GU305914.1 130971 130993 + 0 GCCTATGTGCCCCTCGTCGT GGG ||||||||[||||||||||||]PAM NA Intergenic NA NA

T9898 GCGAGCAGTACACCGACGAGAGG 1000
Oligo fwd CACCGCGAGCAGTACACCGACGAG
Oligo rev AAACCTCGTCGGTGTACTGCTCGC
Chromosome start end strand MM target_seq PAM alignment distance position gene name gene id
GU305914.1 130999 131021 + 0 GCGAGCAGTACACCGACGAG AGG ||||||||[||||||||||||]PAM NA Intergenic NA NA

T9899 AGTACACCGACGAGAGGCTACGG 1000
Oligo fwd CACCAGTACACCGACGAGAGGCTA
Oligo rev AAACTAGCCTCTCGTCGGTGTACT
Chromosome start end strand MM target_seq PAM alignment distance position gene name gene id
GU305914.1 131005 131027 + 0 AGTACACCGACGAGAGGCTA CGG ||||||||[||||||||||||]PAM NA Intergenic NA NA

T9900 CTGCGAGACGTCCATCTTCCTGG 1000
Oligo fwd CACCCTGCGAGACGTCCATCTTCC
Oligo rev AAACGGAAGATGGACGTCTCGCAG
Chromosome start end strand MM target_seq PAM alignment distance position gene name gene id
GU305914.1 131033 131055 + 0 CTGCGAGACGTCCATCTTCC TGG ||||||||[||||||||||||]PAM NA Intergenic NA NA

T9901 CATCTTCCTGGAGCGCGCCTCGG 1000
Oligo fwd CACCCATCTTCCTGGAGCGCGCCT
Oligo rev AAACAGGCGCGCTCCAGGAAGATG
Chromosome start end strand MM target_seq PAM alignment distance position gene name gene id
GU305914.1 131045 131067 + 0 CATCTTCCTGGAGCGCGCCT CGG ||||||||[||||||||||||]PAM NA Intergenic NA NA

T9902 GCTCGCCATCGAGCGCGAACTGG 1000
Oligo fwd CACCGCTCGCCATCGAGCGCGAAC
Oligo rev AAACGTTCGCGCTCGATGGCGAGC
Chromosome start end strand MM target_seq PAM alignment distance position gene name gene id
GU305914.1 131084 131106 + 0 GCTCGCCATCGAGCGCGAAC TGG ||||||||[||||||||||||]PAM NA Intergenic NA NA

T9903 CGCGAACTGGAGACCAGCCTCGG 1000
Oligo fwd CACCCGCGAACTGGAGACCAGCCT
Oligo rev AAACAGGCTGGTCTCCAGTTCGCG
Chromosome start end strand MM target_seq PAM alignment distance position gene name gene id
GU305914.1 131097 131119 + 0 CGCGAACTGGAGACCAGCCT CGG ||||||||[||||||||||||]PAM NA Intergenic NA NA

T9904 GCGAACTGGAGACCAGCCTCGGG 1000
Oligo fwd CACCGCGAACTGGAGACCAGCCTC
Oligo rev AAACGAGGCTGGTCTCCAGTTCGC
Chromosome start end strand MM target_seq PAM alignment distance position gene name gene id
GU305914.1 131098 131120 + 0 GCGAACTGGAGACCAGCCTC GGG ||||||||[||||||||||||]PAM NA Intergenic NA NA

T9905 TGGAGACCAGCCTCGGGCGCCGG 1000
Oligo fwd CACCTGGAGACCAGCCTCGGGCGC
Oligo rev AAACGCGCCCGAGGCTGGTCTCCA
Chromosome start end strand MM target_seq PAM alignment distance position gene name gene id
GU305914.1 131104 131126 + 0 TGGAGACCAGCCTCGGGCGC CGG ||||||||[||||||||||||]PAM NA Intergenic NA NA

| | | | | | | | | | | | |
|---|---|---|---|---|---|---|---|---|---|---|---|
| T9906 | CAGCCTCGGGCGCCGGCGTGAGG | 1000 | | | | | | | | | |
| Oligo fwd | CACCCAGCCTCGGGCGCCGGCGTG | | | | | | | | | | |
| Oligo rev | AAACCACGCCGGCGCCCGAGGCTG | | | | | | | | | | |
| Chromosome | start | end | strand | MM | target_seq | PAM | alignment | distance | position | gene name | gene id |
| GU305914.1 | 131111 | 131133 | + | 0 | CAGCCTCGGGCGCCGGCGTG | AGG | ||||||||[||||||||||||]PAM | NA | Intergenic | NA | NA |
| T9907 | GCCGGCGTGAGGTCCTCATGAGG | 1000 | | | | | | | | | |
| Oligo fwd | CACCGCCGGCGTGAGGTCCTCATG | | | | | | | | | | |
| Oligo rev | AAACCATGAGGACCTCACGCCGGC | | | | | | | | | | |
| Chromosome | start | end | strand | MM | target_seq | PAM | alignment | distance | position | gene name | gene id |
| GU305914.1 | 131122 | 131144 | + | 0 | GCCGGCGTGAGGTCCTCATG | AGG | ||||||||[||||||||||||]PAM | NA | Intergenic | NA | NA |
| T9908 | TGAGGTCCTCATGAGGTGTGTGG | 1000 | | | | | | | | | |
| Oligo fwd | CACCTGAGGTCCTCATGAGGTGTG | | | | | | | | | | |
| Oligo rev | AAACCACACCTCATGAGGACCTCA | | | | | | | | | | |
| Chromosome | start | end | strand | MM | target_seq | PAM | alignment | distance | position | gene name | gene id |
| GU305914.1 | 131129 | 131151 | + | 0 | TGAGGTCCTCATGAGGTGTG | TGG | ||||||||[||||||||||||]PAM | NA | Intergenic | NA | NA |
| T9909 | GGTGTGTGGAGCTCGCCGCCAGG | 1000 | | | | | | | | | |
| Oligo fwd | CACCGGTGTGTGGAGCTCGCCGCC | | | | | | | | | | |
| Oligo rev | AAACGGCGGCGAGCTCCACACACC | | | | | | | | | | |
| Chromosome | start | end | strand | MM | target_seq | PAM | alignment | distance | position | gene name | gene id |
| GU305914.1 | 131143 | 131165 | + | 0 | GGTGTGTGGAGCTCGCCGCC | AGG | ||||||||[||||||||||||]PAM | NA | Intergenic | NA | NA |
| T9910 | CGCCAGGAAGAGACTCTTGTTGG | 1000 | | | | | | | | | |
| Oligo fwd | CACCCGCCAGGAAGAGACTCTTGT | | | | | | | | | | |
| Oligo rev | AAACACAAGAGTCTCTTCCTGGCG | | | | | | | | | | |
| Chromosome | start | end | strand | MM | target_seq | PAM | alignment | distance | position | gene name | gene id |
| GU305914.1 | 131159 | 131181 | + | 0 | CGCCAGGAAGAGACTCTTGT | TGG | ||||||||[||||||||||||]PAM | NA | Intergenic | NA | NA |
| T9911 | AAGAGACTCTTGTTGGTCGACGG | 1000 | | | | | | | | | |
| Oligo fwd | CACCAAGAGACTCTTGTTGGTCGA | | | | | | | | | | |
| Oligo rev | AAACTCGACCAACAAGAGTCTCTT | | | | | | | | | | |
| Chromosome | start | end | strand | MM | target_seq | PAM | alignment | distance | position | gene name | gene id |
| GU305914.1 | 131166 | 131188 | + | 0 | AAGAGACTCTTGTTGGTCGA | CGG | ||||||||[||||||||||||]PAM | NA | Intergenic | NA | NA |
| T9912 | TTGGTCGACGGCACCCCCATCGG | 1000 | | | | | | | | | |
| Oligo fwd | CACCTTGGTCGACGGCACCCCCAT | | | | | | | | | | |
| Oligo rev | AAACATGGGGGTGCCGTCGACCAA | | | | | | | | | | |
| Chromosome | start | end | strand | MM | target_seq | PAM | alignment | distance | position | gene name | gene id |
| GU305914.1 | 131178 | 131200 | + | 0 | TTGGTCGACGGCACCCCCAT | CGG | ||||||||[||||||||||||]PAM | NA | Intergenic | NA | NA |
| T9913 | CATCGGCAACATGTTCGCCTCGG | 1000 | | | | | | | | | |
| Oligo fwd | CACCCATCGGCAACATGTTCGCCT | | | | | | | | | | |
| Oligo rev | AAACAGGCGAACATGTTGCCGATG | | | | | | | | | | |
| Chromosome | start | end | strand | MM | target_seq | PAM | alignment | distance | position | gene name | gene id |
| GU305914.1 | 131195 | 131217 | + | 0 | CATCGGCAACATGTTCGCCT | CGG | ||||||||[||||||||||||]PAM | NA | Intergenic | NA | NA |
| T9914 | CGGCAACATGTTCGCCTCGGAGG | 1000 | | | | | | | | | |
| Oligo fwd | CACCCGGCAACATGTTCGCCTCGG | | | | | | | | | | |
| Oligo rev | AAACCCGAGGCGAACATGTTGCCG | | | | | | | | | | |
| Chromosome | start | end | strand | MM | target_seq | PAM | alignment | distance | position | gene name | gene id |
| GU305914.1 | 131198 | 131220 | + | 0 | CGGCAACATGTTCGCCTCGG | AGG | ||||||||[||||||||||||]PAM | NA | Intergenic | NA | NA |
| T9915 | GCCTCGGAGGTCTGCCTCTGTGG | 1000 | | | | | | | | | |
| Oligo fwd | CACCGCCTCGGAGGTCTGCCTCTG | | | | | | | | | | |
| Oligo rev | AAACCAGAGGCAGACCTCCGAGGC | | | | | | | | | | |
| Chromosome | start | end | strand | MM | target_seq | PAM | alignment | distance | position | gene name | gene id |
| GU305914.1 | 131211 | 131233 | + | 0 | GCCTCGGAGGTCTGCCTCTG | TGG | ||||||||[||||||||||||]PAM | NA | Intergenic | NA | NA |
| GU305914.1 | 37395 | 37417 | - | 4 | GCGTCGGGGTTCTTCCTCTG | CGG | ||-||||-[|-|||-||||||]PAM | NA | Intergenic | NA | NA |
| T9916 | GAGGTCTGCCTCTGTGGACTCGG | 1000 | | | | | | | | | |
| Oligo fwd | CACCGAGGTCTGCCTCTGTGGACT | | | | | | | | | | |
| Oligo rev | AAACAGTCCACAGAGGCAGACCTC | | | | | | | | | | |
| Chromosome | start | end | strand | MM | target_seq | PAM | alignment | distance | position | gene name | gene id |
| GU305914.1 | 131217 | 131239 | + | 0 | GAGGTCTGCCTCTGTGGACT | CGG | ||||||||[||||||||||||]PAM | NA | Intergenic | NA | NA |
| T9917 | CTGCCTCTGTGGACTCGGAGAGG | 1000 | | | | | | | | | |
| Oligo fwd | CACCCTGCCTCTGTGGACTCGGAG | | | | | | | | | | |
| Oligo rev | AAACCTCCGAGTCCACAGAGGCAG | | | | | | | | | | |
| Chromosome | start | end | strand | MM | target_seq | PAM | alignment | distance | position | gene name | gene id |
| GU305914.1 | 131222 | 131244 | + | 0 | CTGCCTCTGTGGACTCGGAG | AGG | ||||||||[||||||||||||]PAM | NA | Intergenic | NA | NA |
| T9918 | CAGTTACGCCAATGAGATAGTGG | 1000 | | | | | | | | | |
| Oligo fwd | CACCCAGTTACGCCAATGAGATAG | | | | | | | | | | |
| Oligo rev | AAACCTATCTCATTGGCGTAACTG | | | | | | | | | | |
| Chromosome | start | end | strand | MM | target_seq | PAM | alignment | distance | position | gene name | gene id |
| GU305914.1 | 131246 | 131268 | + | 0 | CAGTTACGCCAATGAGATAG | TGG | ||||||||[||||||||||||]PAM | NA | Intergenic | NA | NA |
| T9919 | AGTTACGCCAATGAGATAGTGGG | 1000 | | | | | | | | | |
| Oligo fwd | CACCAGTTACGCCAATGAGATAGT | | | | | | | | | | |
| Oligo rev | AAACACTATCTCATTGGCGTAACT | | | | | | | | | | |
| Chromosome | start | end | strand | MM | target_seq | PAM | alignment | distance | position | gene name | gene id |
| GU305914.1 | 131247 | 131269 | + | 0 | AGTTACGCCAATGAGATAGT | GGG | ||||||||[||||||||||||]PAM | NA | Intergenic | NA | NA |
| T9920 | GTTACGCCAATGAGATAGTGGGG | 1000 | | | | | | | | | |
| Oligo fwd | CACCGTTACGCCAATGAGATAGTG | | | | | | | | | | |
| Oligo rev | AAACCACTATCTCATTGGCGTAAC | | | | | | | | | | |
| Chromosome | start | end | strand | MM | target_seq | PAM | alignment | distance | position | gene name | gene id |

GU305914.1 131248 131270 + 0 GTTACGCCAATGAGATAGTG GGG ||||||||[||||||||||||]PAM NA Intergenic NA NA

T9921 TTACGCCAATGAGATAGTGGGGG 1000
Oligo fwd CACCTTACGCCAATGAGATAGTGG
Oligo rev AAACCCACTATCTCATTGGCGTAA
Chromosome start end strand MM target_seq PAM alignment distance position gene name gene id
GU305914.1 131249 131271 + 0 TTACGCCAATGAGATAGTGG GGG ||||||||[||||||||||||]PAM NA Intergenic NA NA

T9922 ATGAGATAGTGGGGGTGCTCTGG 1000
Oligo fwd CACCATGAGATAGTGGGGGTGCTC
Oligo rev AAACGAGCACCCCCACTATCTCAT
Chromosome start end strand MM target_seq PAM alignment distance position gene name gene id
GU305914.1 131257 131279 + 0 ATGAGATAGTGGGGGTGCTC TGG ||||||||[||||||||||||]PAM NA Intergenic NA NA

T9923 AGATAGTGGGGGTGCTCTGGCGG 1000
Oligo fwd CACCAGATAGTGGGGGTGCTCTGG
Oligo rev AAACCCAGAGCACCCCCACTATCT
Chromosome start end strand MM target_seq PAM alignment distance position gene name gene id
GU305914.1 131260 131282 + 0 AGATAGTGGGGGTGCTCTGG CGG ||||||||[||||||||||||]PAM NA Intergenic NA NA

T9924 TAGTGGGGGTGCTCTGGCGGCGG 1000
Oligo fwd CACCTAGTGGGGGTGCTCTGGCGG
Oligo rev AAACCCGCCAGAGCACCCCCACTA
Chromosome start end strand MM target_seq PAM alignment distance position gene name gene id
GU305914.1 131263 131285 + 0 TAGTGGGGGTGCTCTGGCGG CGG ||||||||[||||||||||||]PAM NA Intergenic NA NA

T9925 TGGGGGTGCTCTGGCGGCGGCGG 1000
Oligo fwd CACCTGGGGGTGCTCTGGCGGCGG
Oligo rev AAACCCGCCGCCAGAGCACCCCCA
Chromosome start end strand MM target_seq PAM alignment distance position gene name gene id
GU305914.1 131266 131288 + 0 TGGGGGTGCTCTGGCGGCGG CGG ||||||||[||||||||||||]PAM NA Intergenic NA NA

T9926 GGGGGTGCTCTGGCGGCGGCGGG 1000
Oligo fwd CACCGGGGGTGCTCTGGCGGCGGC
Oligo rev AAACGCCGCCGCCAGAGCACCCCC
Chromosome start end strand MM target_seq PAM alignment distance position gene name gene id
GU305914.1 131267 131289 + 0 GGGGGTGCTCTGGCGGCGGC GGG ||||||||[||||||||||||]PAM NA Intergenic NA NA

T9927 CGGCGGCGGGATACTAAGAGCGG 1000
Oligo fwd CACCCGGCGGCGGGATACTAAGAG
Oligo rev AAACCTCTTAGTATCCCGCCGCCG
Chromosome start end strand MM target_seq PAM alignment distance position gene name gene id
GU305914.1 131280 131302 + 0 CGGCGGCGGGATACTAAGAG CGG ||||||||[||||||||||||]PAM NA Intergenic NA NA

T9928 GCGGGATACTAAGAGCGGCGCGG 1000
Oligo fwd CACCGCGGGATACTAAGAGCGGCG
Oligo rev AAACCGCCGCTCTTAGTATCCCGC
Chromosome start end strand MM target_seq PAM alignment distance position gene name gene id
GU305914.1 131285 131307 + 0 GCGGGATACTAAGAGCGGCG CGG ||||||||[||||||||||||]PAM NA Intergenic NA NA

T9929 TAACTTGCACAACGTCGCTCTGG 1000
Oligo fwd CACCTAACTTGCACAACGTCGCTC
Oligo rev AAACGAGCGACGTTGTGCAAGTTA
Chromosome start end strand MM target_seq PAM alignment distance position gene name gene id
GU305914.1 131312 131334 + 0 TAACTTGCACAACGTCGCTC TGG ||||||||[||||||||||||]PAM NA Intergenic NA NA

T9930 CACAACGTCGCTCTGGCCAACGG 1000
Oligo fwd CACCCACAACGTCGCTCTGGCCAA
Oligo rev AAACTTGGCCAGAGCGACGTTGTG
Chromosome start end strand MM target_seq PAM alignment distance position gene name gene id
GU305914.1 131319 131341 + 0 CACAACGTCGCTCTGGCCAA CGG ||||||||[||||||||||||]PAM NA Intergenic NA NA

T9931 CGCAGCTCGCCCTGATCCTCAGG 1000
Oligo fwd CACCCGCAGCTCGCCCTGATCCTC
Oligo rev AAACGAGGATCAGGGCGAGCTGCG
Chromosome start end strand MM target_seq PAM alignment distance position gene name gene id
GU305914.1 131344 131366 + 0 CGCAGCTCGCCCTGATCCTC AGG ||||||||[||||||||||||]PAM NA Intergenic NA NA

T9932 AGCTCGCCCTGATCCTCAGGAGG 1000
Oligo fwd CACCAGCTCGCCCTGATCCTCAGG
Oligo rev AAACCCTGAGGATCAGGGCGAGCT
Chromosome start end strand MM target_seq PAM alignment distance position gene name gene id
GU305914.1 131347 131369 + 0 AGCTCGCCCTGATCCTCAGG AGG ||||||||[||||||||||||]PAM NA Intergenic NA NA

T9933 CCTGATCCTCAGGAGGATTCAGG 1000
Oligo fwd CACCCCTGATCCTCAGGAGGATTC
Oligo rev AAACGAATCCTCCTGAGGATCAGG
Chromosome start end strand MM target_seq PAM alignment distance position gene name gene id
GU305914.1 131354 131376 + 0 CCTGATCCTCAGGAGGATTC AGG ||||||||[||||||||||||]PAM NA Intergenic NA NA

T9934 CAGGAGGATTCAGGAGTACTCGG 1000
Oligo fwd CACCCAGGAGGATTCAGGAGTACT
Oligo rev AAACAGTACTCCTGAATCCTCCTG
Chromosome start end strand MM target_seq PAM alignment distance position gene name gene id
GU305914.1 131363 131385 + 0 CAGGAGGATTCAGGAGTACT CGG ||||||||[||||||||||||]PAM NA Intergenic NA NA

T9935 CCTACCTGCCGCCCGTAGAGCGG 1000
Oligo fwd CACCCCTACCTGCCGCCCGTAGAG
Oligo rev AAACCTCTACGGGCGGCAGGTAGG

Chromosome start end strand MM target_seq PAM alignment distance position gene name gene id
GU305914.1 131386 131408 + 0 CCTACCTGCCGCCCGTAGAG CGG ||||||||[||||||||||||]PAM NA Intergenic NA NA

T9936 GCCGCCCGTAGAGCGGCGTCTGG 1000
Oligo fwd CACCGCCGCCCGTAGAGCGGCGTC
Oligo rev AAACGACGCCGCTCTACGGGCGGC
Chromosome start end strand MM target_seq PAM alignment distance position gene name gene id
GU305914.1 131393 131415 + 0 GCCGCCCGTAGAGCGGCGTC TGG ||||||||[||||||||||||]PAM NA Intergenic NA NA

T9937 TATGACCGACCCCCTGACATCGG 1000
Oligo fwd CACCTATGACCGACCCCCTGACAT
Oligo rev AAACATGTCAGGGGGTCGGTCATA
Chromosome start end strand MM target_seq PAM alignment distance position gene name gene id
GU305914.1 131417 131439 + 0 TATGACCGACCCCCTGACAT CGG ||||||||[||||||||||||]PAM NA Intergenic NA NA

T9938 GACCGACCCCCTGACATCGGCGG 1000
Oligo fwd CACCGACCGACCCCCTGACATCGG
Oligo rev AAACCCGATGTCAGGGGGTCGGTC
Chromosome start end strand MM target_seq PAM alignment distance position gene name gene id
GU305914.1 131420 131442 + 0 GACCGACCCCCTGACATCGG CGG ||||||||[||||||||||||]PAM NA Intergenic NA NA

T9939 TATGCCGCAGAAGTCGATGTCGG 1000
Oligo fwd CACCTATGCCGCAGAAGTCGATGT
Oligo rev AAACACATCGACTTCTGCGGCATA
Chromosome start end strand MM target_seq PAM alignment distance position gene name gene id
GU305914.1 131454 131476 + 0 TATGCCGCAGAAGTCGATGT CGG ||||||||[||||||||||||]PAM NA Intergenic NA NA

T9940 ATGCCGCAGAAGTCGATGTCGGG 1000
Oligo fwd CACCATGCCGCAGAAGTCGATGTC
Oligo rev AAACGACATCGACTTCTGCGGCAT
Chromosome start end strand MM target_seq PAM alignment distance position gene name gene id
GU305914.1 131455 131477 + 0 ATGCCGCAGAAGTCGATGTC GGG ||||||||[||||||||||||]PAM NA Intergenic NA NA

T9941 TGCCGCAGAAGTCGATGTCGGGG 1000
Oligo fwd CACCTGCCGCAGAAGTCGATGTCG
Oligo rev AAACCGACATCGACTTCTGCGGCA
Chromosome start end strand MM target_seq PAM alignment distance position gene name gene id
GU305914.1 131456 131478 + 0 TGCCGCAGAAGTCGATGTCG GGG ||||||||[||||||||||||]PAM NA Intergenic NA NA

T9942 AGAAGTCGATGTCGGGGTCTCGG 1000
Oligo fwd CACCAGAAGTCGATGTCGGGGTCT
Oligo rev AAACAGACCCCGACATCGACTTCT
Chromosome start end strand MM target_seq PAM alignment distance position gene name gene id
GU305914.1 131462 131484 + 0 AGAAGTCGATGTCGGGGTCT CGG ||||||||[||||||||||||]PAM NA Intergenic NA NA

T9943 CGGGGTCTCGGTCGTCTCCGTGG 1000
Oligo fwd CACCCGGGGTCTCGGTCGTCTCCG
Oligo rev AAACCGGAGACGACCGAGACCCCG
Chromosome start end strand MM target_seq PAM alignment distance position gene name gene id
GU305914.1 131474 131496 + 0 CGGGGTCTCGGTCGTCTCCG TGG ||||||||[||||||||||||]PAM NA Intergenic NA NA

T9944 GGTCTCGGTCGTCTCCGTGGCGG 1000
Oligo fwd CACCGGTCTCGGTCGTCTCCGTGG
Oligo rev AAACCCACGGAGACGACCGAGACC
Chromosome start end strand MM target_seq PAM alignment distance position gene name gene id
GU305914.1 131477 131499 + 0 GGTCTCGGTCGTCTCCGTGG CGG ||||||||[||||||||||||]PAM NA Intergenic NA NA

T9945 CTCGGTCGTCTCCGTGGCGGCGG 1000
Oligo fwd CACCCTCGGTCGTCTCCGTGGCGG
Oligo rev AAACCCGCCACGGAGACGACCGAG
Chromosome start end strand MM target_seq PAM alignment distance position gene name gene id
GU305914.1 131480 131502 + 0 CTCGGTCGTCTCCGTGGCGG CGG ||||||||[||||||||||||]PAM NA Intergenic NA NA

T9946 TTCTGCGCGCCTTCTCCCTCCGG 1000
Oligo fwd CACCTTCTGCGCGCCTTCTCCCTC
Oligo rev AAACGAGGGAGAAGGCGCGCAGAA
Chromosome start end strand MM target_seq PAM alignment distance position gene name gene id
GU305914.1 131506 131528 + 0 TTCTGCGCGCCTTCTCCCTC CGG ||||||||[||||||||||||]PAM NA Intergenic NA NA

T9947 TCTGCGCGCCTTCTCCCTCCGGG 1000
Oligo fwd CACCTCTGCGCGCCTTCTCCCTCC
Oligo rev AAACGGAGGGAGAAGGCGCGCAGA
Chromosome start end strand MM target_seq PAM alignment distance position gene name gene id
GU305914.1 131507 131529 + 0 TCTGCGCGCCTTCTCCCTCC GGG ||||||||[||||||||||||]PAM NA Intergenic NA NA

T9948 CTGCGCGCCTTCTCCCTCCGGGG 1000
Oligo fwd CACCCTGCGCGCCTTCTCCCTCCG
Oligo rev AAACCGGAGGGAGAAGGCGCGCAG
Chromosome start end strand MM target_seq PAM alignment distance position gene name gene id
GU305914.1 131508 131530 + 0 CTGCGCGCCTTCTCCCTCCG GGG ||||||||[||||||||||||]PAM NA Intergenic NA NA

T9949 TGCGCGCCTTCTCCCTCCGGGGG 1000
Oligo fwd CACCTGCGCGCCTTCTCCCTCCGG
Oligo rev AAACCCGGAGGGAGAAGGCGCGCA
Chromosome start end strand MM target_seq PAM alignment distance position gene name gene id
GU305914.1 131509 131531 + 0 TGCGCGCCTTCTCCCTCCGG GGG ||||||||[||||||||||||]PAM NA Intergenic NA NA

T9950 CGCCTTCTCCCTCCGGGGGTTGG 1000
Oligo fwd CACCCGCCTTCTCCCTCCGGGGGT

| | | | | | | | | | | | |
|---|---|---|---|---|---|---|---|---|---|---|---|
| Oligo rev | AAACACCCCCGGAGGGAGAAGGCG | | | | | | | | | | |
| Chromosome | start | end | strand | MM | target_seq | PAM | alignment | distance | position | gene name | gene id |
| GU305914.1 | 131513 | 131535 | + | 0 | CGCCTTCTCCCTCCGGGGGT | TGG | \|\|\|\|\|\|\|\|[\|\|\|\|\|\|\|\|\|\|\|\|]PAM | NA | Intergenic | NA | NA |
| T9951 | CTCCGGGGGTTGGTGAGCGACGG | 1000 | | | | | | | | | |
| Oligo fwd | CACCCTCCGGGGGTTGGTGAGCGA | | | | | | | | | | |
| Oligo rev | AAACTCGCTCACCAACCCCCGGAG | | | | | | | | | | |
| Chromosome | start | end | strand | MM | target_seq | PAM | alignment | distance | position | gene name | gene id |
| GU305914.1 | 131523 | 131545 | + | 0 | CTCCGGGGGTTGGTGAGCGA | CGG | \|\|\|\|\|\|\|\|[\|\|\|\|\|\|\|\|\|\|\|\|]PAM | NA | Intergenic | NA | NA |
| T9952 | GTTGGTGAGCGACGGCGTCAAGG | 1000 | | | | | | | | | |
| Oligo fwd | CACCGTTGGTGAGCGACGGCGTCA | | | | | | | | | | |
| Oligo rev | AAACTGACGCCGTCGCTCACCAAC | | | | | | | | | | |
| Chromosome | start | end | strand | MM | target_seq | PAM | alignment | distance | position | gene name | gene id |
| GU305914.1 | 131531 | 131553 | + | 0 | GTTGGTGAGCGACGGCGTCA | AGG | \|\|\|\|\|\|\|\|[\|\|\|\|\|\|\|\|\|\|\|\|]PAM | NA | Intergenic | NA | NA |
| GU305914.1 | 201094 | 201116 | + | 4 | GGTGGCGGGCGACGGCGTCT | CGG | \|-\|\|\|-\|-[\|\|\|\|\|\|\|\|\|\|\|-]PAM | NA | Intergenic | NA | NA |
| T9953 | GGTGAGCGACGGCGTCAAGGTGG | 1000 | | | | | | | | | |
| Oligo fwd | CACCGGTGAGCGACGGCGTCAAGG | | | | | | | | | | |
| Oligo rev | AAACCCTTGACGCCGTCGCTCACC | | | | | | | | | | |
| Chromosome | start | end | strand | MM | target_seq | PAM | alignment | distance | position | gene name | gene id |
| GU305914.1 | 131534 | 131556 | + | 0 | GGTGAGCGACGGCGTCAAGG | TGG | \|\|\|\|\|\|\|\|[\|\|\|\|\|\|\|\|\|\|\|\|]PAM | NA | Intergenic | NA | NA |
| T9954 | CGTCAAGGTGGTCGAGAAGCTGG | 1000 | | | | | | | | | |
| Oligo fwd | CACCCGTCAAGGTGGTCGAGAAGC | | | | | | | | | | |
| Oligo rev | AAACGCTTCTCGACCACCTTGACG | | | | | | | | | | |
| Chromosome | start | end | strand | MM | target_seq | PAM | alignment | distance | position | gene name | gene id |
| GU305914.1 | 131546 | 131568 | + | 0 | CGTCAAGGTGGTCGAGAAGC | TGG | \|\|\|\|\|\|\|\|[\|\|\|\|\|\|\|\|\|\|\|\|]PAM | NA | Intergenic | NA | NA |
| GU305914.1 | 64478 | 64500 | + | 4 | CCTAAAGACGGTCGAGAAGC | TGG | \|-\|-\|\|\|-[-\|\|\|\|\|\|\|\|\|\|\|]PAM | NA | Intergenic | NA | NA |
| T9955 | GTCAAGGTGGTCGAGAAGCTGGG | 1000 | | | | | | | | | |
| Oligo fwd | CACCGTCAAGGTGGTCGAGAAGCT | | | | | | | | | | |
| Oligo rev | AAACAGCTTCTCGACCACCTTGAC | | | | | | | | | | |
| Chromosome | start | end | strand | MM | target_seq | PAM | alignment | distance | position | gene name | gene id |
| GU305914.1 | 131547 | 131569 | + | 0 | GTCAAGGTGGTCGAGAAGCT | GGG | \|\|\|\|\|\|\|\|[\|\|\|\|\|\|\|\|\|\|\|\|]PAM | NA | Intergenic | NA | NA |
| T9956 | GAAGCTGGGCGTCCCCATGACGG | 1000 | | | | | | | | | |
| Oligo fwd | CACCGAAGCTGGGCGTCCCCATGA | | | | | | | | | | |
| Oligo rev | AAACTCATGGGGACGCCCAGCTTC | | | | | | | | | | |
| Chromosome | start | end | strand | MM | target_seq | PAM | alignment | distance | position | gene name | gene id |
| GU305914.1 | 131561 | 131583 | + | 0 | GAAGCTGGGCGTCCCCATGA | CGG | \|\|\|\|\|\|\|\|[\|\|\|\|\|\|\|\|\|\|\|\|]PAM | NA | Intergenic | NA | NA |
| T9957 | AAGCTGGGCGTCCCCATGACGGG | 1000 | | | | | | | | | |
| Oligo fwd | CACCAAGCTGGGCGTCCCCATGAC | | | | | | | | | | |
| Oligo rev | AAACGTCATGGGGACGCCCAGCTT | | | | | | | | | | |
| Chromosome | start | end | strand | MM | target_seq | PAM | alignment | distance | position | gene name | gene id |
| GU305914.1 | 131562 | 131584 | + | 0 | AAGCTGGGCGTCCCCATGAC | GGG | \|\|\|\|\|\|\|\|[\|\|\|\|\|\|\|\|\|\|\|\|]PAM | NA | Intergenic | NA | NA |
| T9958 | CCATGACGGGTCCCGTCGATCGG | 1000 | | | | | | | | | |
| Oligo fwd | CACCCCATGACGGGTCCCGTCGAT | | | | | | | | | | |
| Oligo rev | AAACATCGACGGGACCCGTCATGG | | | | | | | | | | |
| Chromosome | start | end | strand | MM | target_seq | PAM | alignment | distance | position | gene name | gene id |
| GU305914.1 | 131575 | 131597 | + | 0 | CCATGACGGGTCCCGTCGAT | CGG | \|\|\|\|\|\|\|\|[\|\|\|\|\|\|\|\|\|\|\|\|]PAM | NA | Intergenic | NA | NA |
| T9959 | CATGACGGGTCCCGTCGATCGGG | 1000 | | | | | | | | | |
| Oligo fwd | CACCCATGACGGGTCCCGTCGATC | | | | | | | | | | |
| Oligo rev | AAACGATCGACGGGACCCGTCATG | | | | | | | | | | |
| Chromosome | start | end | strand | MM | target_seq | PAM | alignment | distance | position | gene name | gene id |
| GU305914.1 | 131576 | 131598 | + | 0 | CATGACGGGTCCCGTCGATC | GGG | \|\|\|\|\|\|\|\|[\|\|\|\|\|\|\|\|\|\|\|\|]PAM | NA | Intergenic | NA | NA |
| T9960 | TCGATCGGGACGACGTGATCAGG | 1000 | | | | | | | | | |
| Oligo fwd | CACCTCGATCGGGACGACGTGATC | | | | | | | | | | |
| Oligo rev | AAACGATCACGTCGTCCCGATCGA | | | | | | | | | | |
| Chromosome | start | end | strand | MM | target_seq | PAM | alignment | distance | position | gene name | gene id |
| GU305914.1 | 131590 | 131612 | + | 0 | TCGATCGGGACGACGTGATC | AGG | \|\|\|\|\|\|\|\|[\|\|\|\|\|\|\|\|\|\|\|\|]PAM | NA | Intergenic | NA | NA |
| GU305914.1 | 136103 | 136125 | + | 4 | TCGGTTGGGCCGATGTGATC | TGG | \|\|\|-\|-\|\|[\|-\|\|\|-\|\|\|\|\|\|]PAM | NA | Intergenic | NA | NA |
| T9961 | CGATCGGGACGACGTGATCAGGG | 1000 | | | | | | | | | |
| Oligo fwd | CACCCGATCGGGACGACGTGATCA | | | | | | | | | | |
| Oligo rev | AAACTGATCACGTCGTCCCGATCG | | | | | | | | | | |
| Chromosome | start | end | strand | MM | target_seq | PAM | alignment | distance | position | gene name | gene id |
| GU305914.1 | 131591 | 131613 | + | 0 | CGATCGGGACGACGTGATCA | GGG | \|\|\|\|\|\|\|\|[\|\|\|\|\|\|\|\|\|\|\|\|]PAM | NA | Intergenic | NA | NA |
| T9962 | TCAGGGTGTTACAGCTTTAAAGG | 1000 | | | | | | | | | |
| Oligo fwd | CACCTCAGGGTGTTACAGCTTTAA | | | | | | | | | | |
| Oligo rev | AAACTTAAAGCTGTAACACCCTGA | | | | | | | | | | |
| Chromosome | start | end | strand | MM | target_seq | PAM | alignment | distance | position | gene name | gene id |
| GU305914.1 | 131608 | 131630 | + | 0 | TCAGGGTGTTACAGCTTTAA | AGG | \|\|\|\|\|\|\|\|[\|\|\|\|\|\|\|\|\|\|\|\|]PAM | NA | Intergenic | NA | NA |
| T9963 | CAGGGTGTTACAGCTTTAAAGGG | 1000 | | | | | | | | | |
| Oligo fwd | CACCCAGGGTGTTACAGCTTTAAA | | | | | | | | | | |
| Oligo rev | AAACTTTAAAGCTGTAACACCCTG | | | | | | | | | | |
| Chromosome | start | end | strand | MM | target_seq | PAM | alignment | distance | position | gene name | gene id |
| GU305914.1 | 131609 | 131631 | + | 0 | CAGGGTGTTACAGCTTTAAA | GGG | \|\|\|\|\|\|\|\|[\|\|\|\|\|\|\|\|\|\|\|\|]PAM | NA | Intergenic | NA | NA |
| T9964 | GTCAGTTAGACAGACAAGAGAGG | 1000 | | | | | | | | | |
| Oligo fwd | CACCGTCAGTTAGACAGACAAGAG | | | | | | | | | | |
| Oligo rev | AAACCTCTTGTCTGTCTAACTGAC | | | | | | | | | | |
| Chromosome | start | end | strand | MM | target_seq | PAM | alignment | distance | position | gene name | gene id |

GU305914.1 131649 131671 + 0 GTCAGTTAGACAGACAAGAG AGG ||||||||[||||||||||||]PAM NA Intergenic NA NA

T9965 TCAGTTAGACAGACAAGAGAGGG 1000
Oligo fwd CACCTCAGTTAGACAGACAAGAGA
Oligo rev AAACTCTCTTGTCTGTCTAACTGA
Chromosome start end strand MM target_seq PAM alignment distance position gene name gene id
GU305914.1 131650 131672 + 0 TCAGTTAGACAGACAAGAGA GGG ||||||||[||||||||||||]PAM NA Intergenic NA NA

T9966 GTTAGACAGACAAGAGAGGGAGG 1000
Oligo fwd CACCGTTAGACAGACAAGAGAGGG
Oligo rev AAACCCCTCTCTTGTCTGTCTAAC
Chromosome start end strand MM target_seq PAM alignment distance position gene name gene id
GU305914.1 131653 131675 + 0 GTTAGACAGACAAGAGAGGG AGG ||||||||[||||||||||||]PAM NA Intergenic NA NA

T9967 GACAGACAAGAGAGGGAGGAAGG 1000
Oligo fwd CACCGACAGACAAGAGAGGGAGGA
Oligo rev AAACTCCTCCCTCTCTTGTCTGTC
Chromosome start end strand MM target_seq PAM alignment distance position gene name gene id
GU305914.1 131657 131679 + 0 GACAGACAAGAGAGGGAGGA AGG ||||||||[||||||||||||]PAM NA Intergenic NA NA

T9968 AGAGGGAGGAAGGAAGAAGTAGG 1000
Oligo fwd CACCAGAGGGAGGAAGGAAGAAGT
Oligo rev AAACACTTCTTCCTTCCTCCCTCT
Chromosome start end strand MM target_seq PAM alignment distance position gene name gene id
GU305914.1 131667 131689 + 0 AGAGGGAGGAAGGAAGAAGT AGG ||||||||[||||||||||||]PAM NA Intergenic NA NA

T9969 AGGAAGAAGTAGGTACGAGTTGG 1000
Oligo fwd CACCAGGAAGAAGTAGGTACGAGT
Oligo rev AAACACTCGTACCTACTTCTTCCT
Chromosome start end strand MM target_seq PAM alignment distance position gene name gene id
GU305914.1 131677 131699 + 0 AGGAAGAAGTAGGTACGAGT TGG ||||||||[||||||||||||]PAM NA Intergenic NA NA

T9970 AAGAAGTAGGTACGAGTTGGCGG 1000
Oligo fwd CACCAAGAAGTAGGTACGAGTTGG
Oligo rev AAACCCAACTCGTACCTACTTCTT
Chromosome start end strand MM target_seq PAM alignment distance position gene name gene id
GU305914.1 131680 131702 + 0 AAGAAGTAGGTACGAGTTGG CGG ||||||||[||||||||||||]PAM NA Intergenic NA NA

T9971 AGAAGTAGGTACGAGTTGGCGGG 1000
Oligo fwd CACCAGAAGTAGGTACGAGTTGGC
Oligo rev AAACGCCAACTCGTACCTACTTCT
Chromosome start end strand MM target_seq PAM alignment distance position gene name gene id
GU305914.1 131681 131703 + 0 AGAAGTAGGTACGAGTTGGC GGG ||||||||[||||||||||||]PAM NA Intergenic NA NA

T9972 GATCGATCGATTGATTAGATCGG 1000
Oligo fwd CACCGATCGATCGATTGATTAGAT
Oligo rev AAACATCTAATCAATCGATCGATC
Chromosome start end strand MM target_seq PAM alignment distance position gene name gene id
GU305914.1 131732 131754 + 0 GATCGATCGATTGATTAGAT CGG ||||||||[||||||||||||]PAM NA Intergenic NA NA

T9973 TTGAAGTTGAAACTCCTCTCCGG 1000
Oligo fwd CACCTTGAAGTTGAAACTCCTCTC
Oligo rev AAACGAGAGGAGTTTCAACTTCAA
Chromosome start end strand MM target_seq PAM alignment distance position gene name gene id
GU305914.1 131756 131778 + 0 TTGAAGTTGAAACTCCTCTC CGG ||||||||[||||||||||||]PAM NA Intergenic NA NA

T9974 CTCCGGCGAGCACATGTACACGG 1000
Oligo fwd CACCCTCCGGCGAGCACATGTACA
Oligo rev AAACTGTACATGTGCTCGCCGGAG
Chromosome start end strand MM target_seq PAM alignment distance position gene name gene id
GU305914.1 131773 131795 + 0 CTCCGGCGAGCACATGTACA CGG ||||||||[||||||||||||]PAM NA Intergenic NA NA

T9975 ACATGTACACGGCCATGATCAGG 1000
Oligo fwd CACCACATGTACACGGCCATGATC
Oligo rev AAACGATCATGGCCGTGTACATGT
Chromosome start end strand MM target_seq PAM alignment distance position gene name gene id
GU305914.1 131784 131806 + 0 ACATGTACACGGCCATGATC AGG ||||||||[||||||||||||]PAM NA Intergenic NA NA

T9976 CATGTACACGGCCATGATCAGGG 1000
Oligo fwd CACCCATGTACACGGCCATGATCA
Oligo rev AAACTGATCATGGCCGTGTACATG
Chromosome start end strand MM target_seq PAM alignment distance position gene name gene id
GU305914.1 131785 131807 + 0 CATGTACACGGCCATGATCA GGG ||||||||[||||||||||||]PAM NA Intergenic NA NA

T9977 CCATGATCAGGGCGACCACCAGG 1000
Oligo fwd CACCCCATGATCAGGGCGACCACC
Oligo rev AAACGGTGGTCGCCCTGATCATGG
Chromosome start end strand MM target_seq PAM alignment distance position gene name gene id
GU305914.1 131796 131818 + 0 CCATGATCAGGGCGACCACC AGG ||||||||[||||||||||||]PAM NA Intergenic NA NA

T9978 CGACCACCAGGTCGTCCGACTGG 1000
Oligo fwd CACCCGACCACCAGGTCGTCCGAC
Oligo rev AAACGTCGGACGACCTGGTGGTCG
Chromosome start end strand MM target_seq PAM alignment distance position gene name gene id
GU305914.1 131808 131830 + 0 CGACCACCAGGTCGTCCGAC TGG ||||||||[||||||||||||]PAM NA Intergenic NA NA

T9979 TGTATGTCACGTACTCGTTCAGG 1000
Oligo fwd CACCTGTATGTCACGTACTCGTTC
Oligo rev AAACGAACGAGTACGTGACATACA

<table>
<tr><td>Chromosome</td><td>start</td><td>end</td><td>strand</td><td>MM</td><td>target_seq</td><td>PAM</td><td>alignment</td><td>distance</td><td>position</td><td>gene name</td><td>gene id</td></tr>
<tr><td>GU305914.1</td><td>131847</td><td>131869</td><td>+</td><td>0</td><td>TGTATGTCACGTACTCGTTC</td><td>AGG</td><td>||||||||[||||||||||||]PAM</td><td>NA</td><td>Intergenic</td><td>NA</td><td>NA</td></tr>
<tr><td>T9980</td><td>GTATGTCACGTACTCGTTCAGGG</td><td>1000</td><td></td><td></td><td></td><td></td><td></td><td></td><td></td><td></td><td></td></tr>
<tr><td>Oligo fwd</td><td>CACCGTATGTCACGTACTCGTTCA</td><td></td><td></td><td></td><td></td><td></td><td></td><td></td><td></td><td></td><td></td></tr>
<tr><td>Oligo rev</td><td>AAACTGAACGAGTACGTGACATAC</td><td></td><td></td><td></td><td></td><td></td><td></td><td></td><td></td><td></td><td></td></tr>
<tr><td>Chromosome</td><td>start</td><td>end</td><td>strand</td><td>MM</td><td>target_seq</td><td>PAM</td><td>alignment</td><td>distance</td><td>position</td><td>gene name</td><td>gene id</td></tr>
<tr><td>GU305914.1</td><td>131848</td><td>131870</td><td>+</td><td>0</td><td>GTATGTCACGTACTCGTTCA</td><td>GGG</td><td>||||||||[||||||||||||]PAM</td><td>NA</td><td>Intergenic</td><td>NA</td><td>NA</td></tr>
<tr><td>T9981</td><td>ACTCGTTCAGGGTGATGCGCTGG</td><td>1000</td><td></td><td></td><td></td><td></td><td></td><td></td><td></td><td></td><td></td></tr>
<tr><td>Oligo fwd</td><td>CACCACTCGTTCAGGGTGATGCGC</td><td></td><td></td><td></td><td></td><td></td><td></td><td></td><td></td><td></td><td></td></tr>
<tr><td>Oligo rev</td><td>AAACGCGCATCACCCTGAACGAGT</td><td></td><td></td><td></td><td></td><td></td><td></td><td></td><td></td><td></td><td></td></tr>
<tr><td>Chromosome</td><td>start</td><td>end</td><td>strand</td><td>MM</td><td>target_seq</td><td>PAM</td><td>alignment</td><td>distance</td><td>position</td><td>gene name</td><td>gene id</td></tr>
<tr><td>GU305914.1</td><td>131859</td><td>131881</td><td>+</td><td>0</td><td>ACTCGTTCAGGGTGATGCGC</td><td>TGG</td><td>||||||||[||||||||||||]PAM</td><td>NA</td><td>Intergenic</td><td>NA</td><td>NA</td></tr>
<tr><td>T9982</td><td>TCAGGGTGATGCGCTGGATGTGG</td><td>1000</td><td></td><td></td><td></td><td></td><td></td><td></td><td></td><td></td><td></td></tr>
<tr><td>Oligo fwd</td><td>CACCTCAGGGTGATGCGCTGGATG</td><td></td><td></td><td></td><td></td><td></td><td></td><td></td><td></td><td></td><td></td></tr>
<tr><td>Oligo rev</td><td>AAACCATCCAGCGCATCACCCTGA</td><td></td><td></td><td></td><td></td><td></td><td></td><td></td><td></td><td></td><td></td></tr>
<tr><td>Chromosome</td><td>start</td><td>end</td><td>strand</td><td>MM</td><td>target_seq</td><td>PAM</td><td>alignment</td><td>distance</td><td>position</td><td>gene name</td><td>gene id</td></tr>
<tr><td>GU305914.1</td><td>131865</td><td>131887</td><td>+</td><td>0</td><td>TCAGGGTGATGCGCTGGATG</td><td>TGG</td><td>||||||||[||||||||||||]PAM</td><td>NA</td><td>Intergenic</td><td>NA</td><td>NA</td></tr>
<tr><td>T9983</td><td>GGGTGATGCGCTGGATGTGGCGG</td><td>1000</td><td></td><td></td><td></td><td></td><td></td><td></td><td></td><td></td><td></td></tr>
<tr><td>Oligo fwd</td><td>CACCGGGTGATGCGCTGGATGTGG</td><td></td><td></td><td></td><td></td><td></td><td></td><td></td><td></td><td></td><td></td></tr>
<tr><td>Oligo rev</td><td>AAACCCACATCCAGCGCATCACCC</td><td></td><td></td><td></td><td></td><td></td><td></td><td></td><td></td><td></td><td></td></tr>
<tr><td>Chromosome</td><td>start</td><td>end</td><td>strand</td><td>MM</td><td>target_seq</td><td>PAM</td><td>alignment</td><td>distance</td><td>position</td><td>gene name</td><td>gene id</td></tr>
<tr><td>GU305914.1</td><td>131868</td><td>131890</td><td>+</td><td>0</td><td>GGGTGATGCGCTGGATGTGG</td><td>CGG</td><td>||||||||[||||||||||||]PAM</td><td>NA</td><td>Intergenic</td><td>NA</td><td>NA</td></tr>
<tr><td>T9984</td><td>TGCGCTGGATGTGGCGGAGCTGG</td><td>1000</td><td></td><td></td><td></td><td></td><td></td><td></td><td></td><td></td><td></td></tr>
<tr><td>Oligo fwd</td><td>CACCTGCGCTGGATGTGGCGGAGC</td><td></td><td></td><td></td><td></td><td></td><td></td><td></td><td></td><td></td><td></td></tr>
<tr><td>Oligo rev</td><td>AAACGCTCCGCCACATCCAGCGCA</td><td></td><td></td><td></td><td></td><td></td><td></td><td></td><td></td><td></td><td></td></tr>
<tr><td>Chromosome</td><td>start</td><td>end</td><td>strand</td><td>MM</td><td>target_seq</td><td>PAM</td><td>alignment</td><td>distance</td><td>position</td><td>gene name</td><td>gene id</td></tr>
<tr><td>GU305914.1</td><td>131874</td><td>131896</td><td>+</td><td>0</td><td>TGCGCTGGATGTGGCGGAGC</td><td>TGG</td><td>||||||||[||||||||||||]PAM</td><td>NA</td><td>Intergenic</td><td>NA</td><td>NA</td></tr>
<tr><td>T9985</td><td>TGTGGCGGAGCTGGTCGAGCAGG</td><td>1000</td><td></td><td></td><td></td><td></td><td></td><td></td><td></td><td></td><td></td></tr>
<tr><td>Oligo fwd</td><td>CACCTGTGGCGGAGCTGGTCGAGC</td><td></td><td></td><td></td><td></td><td></td><td></td><td></td><td></td><td></td><td></td></tr>
<tr><td>Oligo rev</td><td>AAACGCTCGACCAGCTCCGCCACA</td><td></td><td></td><td></td><td></td><td></td><td></td><td></td><td></td><td></td><td></td></tr>
<tr><td>Chromosome</td><td>start</td><td>end</td><td>strand</td><td>MM</td><td>target_seq</td><td>PAM</td><td>alignment</td><td>distance</td><td>position</td><td>gene name</td><td>gene id</td></tr>
<tr><td>GU305914.1</td><td>131883</td><td>131905</td><td>+</td><td>0</td><td>TGTGGCGGAGCTGGTCGAGC</td><td>AGG</td><td>||||||||[||||||||||||]PAM</td><td>NA</td><td>Intergenic</td><td>NA</td><td>NA</td></tr>
<tr><td>T9986</td><td>TGGTCGAGCAGGTACTCGACCGG</td><td>1000</td><td></td><td></td><td></td><td></td><td></td><td></td><td></td><td></td><td></td></tr>
<tr><td>Oligo fwd</td><td>CACCTGGTCGAGCAGGTACTCGAC</td><td></td><td></td><td></td><td></td><td></td><td></td><td></td><td></td><td></td><td></td></tr>
<tr><td>Oligo rev</td><td>AAACGTCGAGTACCTGCTCGACCA</td><td></td><td></td><td></td><td></td><td></td><td></td><td></td><td></td><td></td><td></td></tr>
<tr><td>Chromosome</td><td>start</td><td>end</td><td>strand</td><td>MM</td><td>target_seq</td><td>PAM</td><td>alignment</td><td>distance</td><td>position</td><td>gene name</td><td>gene id</td></tr>
<tr><td>GU305914.1</td><td>131894</td><td>131916</td><td>+</td><td>0</td><td>TGGTCGAGCAGGTACTCGAC</td><td>CGG</td><td>||||||||[||||||||||||]PAM</td><td>NA</td><td>Intergenic</td><td>NA</td><td>NA</td></tr>
<tr><td>T9987</td><td>GGTCGAGCAGGTACTCGACCGGG</td><td>1000</td><td></td><td></td><td></td><td></td><td></td><td></td><td></td><td></td><td></td></tr>
<tr><td>Oligo fwd</td><td>CACCGGTCGAGCAGGTACTCGACC</td><td></td><td></td><td></td><td></td><td></td><td></td><td></td><td></td><td></td><td></td></tr>
<tr><td>Oligo rev</td><td>AAACGGTCGAGTACCTGCTCGACC</td><td></td><td></td><td></td><td></td><td></td><td></td><td></td><td></td><td></td><td></td></tr>
<tr><td>Chromosome</td><td>start</td><td>end</td><td>strand</td><td>MM</td><td>target_seq</td><td>PAM</td><td>alignment</td><td>distance</td><td>position</td><td>gene name</td><td>gene id</td></tr>
<tr><td>GU305914.1</td><td>131895</td><td>131917</td><td>+</td><td>0</td><td>GGTCGAGCAGGTACTCGACC</td><td>GGG</td><td>||||||||[||||||||||||]PAM</td><td>NA</td><td>Intergenic</td><td>NA</td><td>NA</td></tr>
<tr><td>T9988</td><td>CGGGTCGTAACTCAGCTTGATGG</td><td>1000</td><td></td><td></td><td></td><td></td><td></td><td></td><td></td><td></td><td></td></tr>
<tr><td>Oligo fwd</td><td>CACCCGGGTCGTAACTCAGCTTGA</td><td></td><td></td><td></td><td></td><td></td><td></td><td></td><td></td><td></td><td></td></tr>
<tr><td>Oligo rev</td><td>AAACTCAAGCTGAGTTACGACCCG</td><td></td><td></td><td></td><td></td><td></td><td></td><td></td><td></td><td></td><td></td></tr>
<tr><td>Chromosome</td><td>start</td><td>end</td><td>strand</td><td>MM</td><td>target_seq</td><td>PAM</td><td>alignment</td><td>distance</td><td>position</td><td>gene name</td><td>gene id</td></tr>
<tr><td>GU305914.1</td><td>131914</td><td>131936</td><td>+</td><td>0</td><td>CGGGTCGTAACTCAGCTTGA</td><td>TGG</td><td>||||||||[||||||||||||]PAM</td><td>NA</td><td>Intergenic</td><td>NA</td><td>NA</td></tr>
<tr><td>T9989</td><td>TGGTGTGCGAGATCAGCTCCTGG</td><td>1000</td><td></td><td></td><td></td><td></td><td></td><td></td><td></td><td></td><td></td></tr>
<tr><td>Oligo fwd</td><td>CACCTGGTGTGCGAGATCAGCTCC</td><td></td><td></td><td></td><td></td><td></td><td></td><td></td><td></td><td></td><td></td></tr>
<tr><td>Oligo rev</td><td>AAACGGAGCTGATCTCGCACACCA</td><td></td><td></td><td></td><td></td><td></td><td></td><td></td><td></td><td></td><td></td></tr>
<tr><td>Chromosome</td><td>start</td><td>end</td><td>strand</td><td>MM</td><td>target_seq</td><td>PAM</td><td>alignment</td><td>distance</td><td>position</td><td>gene name</td><td>gene id</td></tr>
<tr><td>GU305914.1</td><td>131934</td><td>131956</td><td>+</td><td>0</td><td>TGGTGTGCGAGATCAGCTCC</td><td>TGG</td><td>||||||||[||||||||||||]PAM</td><td>NA</td><td>Intergenic</td><td>NA</td><td>NA</td></tr>
<tr><td>T9990</td><td>GGTGTGCGAGATCAGCTCCTGGG</td><td>1000</td><td></td><td></td><td></td><td></td><td></td><td></td><td></td><td></td><td></td></tr>
<tr><td>Oligo fwd</td><td>CACCGGTGTGCGAGATCAGCTCCT</td><td></td><td></td><td></td><td></td><td></td><td></td><td></td><td></td><td></td><td></td></tr>
<tr><td>Oligo rev</td><td>AAACAGGAGCTGATCTCGCACACC</td><td></td><td></td><td></td><td></td><td></td><td></td><td></td><td></td><td></td><td></td></tr>
<tr><td>Chromosome</td><td>start</td><td>end</td><td>strand</td><td>MM</td><td>target_seq</td><td>PAM</td><td>alignment</td><td>distance</td><td>position</td><td>gene name</td><td>gene id</td></tr>
<tr><td>GU305914.1</td><td>131935</td><td>131957</td><td>+</td><td>0</td><td>GGTGTGCGAGATCAGCTCCT</td><td>GGG</td><td>||||||||[||||||||||||]PAM</td><td>NA</td><td>Intergenic</td><td>NA</td><td>NA</td></tr>
<tr><td>T9991</td><td>GTGCGAGATCAGCTCCTGGGAGG</td><td>1000</td><td></td><td></td><td></td><td></td><td></td><td></td><td></td><td></td><td></td></tr>
<tr><td>Oligo fwd</td><td>CACCGTGCGAGATCAGCTCCTGGG</td><td></td><td></td><td></td><td></td><td></td><td></td><td></td><td></td><td></td><td></td></tr>
<tr><td>Oligo rev</td><td>AAACCCCAGGAGCTGATCTCGCAC</td><td></td><td></td><td></td><td></td><td></td><td></td><td></td><td></td><td></td><td></td></tr>
<tr><td>Chromosome</td><td>start</td><td>end</td><td>strand</td><td>MM</td><td>target_seq</td><td>PAM</td><td>alignment</td><td>distance</td><td>position</td><td>gene name</td><td>gene id</td></tr>
<tr><td>GU305914.1</td><td>131938</td><td>131960</td><td>+</td><td>0</td><td>GTGCGAGATCAGCTCCTGGG</td><td>AGG</td><td>||||||||[||||||||||||]PAM</td><td>NA</td><td>Intergenic</td><td>NA</td><td>NA</td></tr>
<tr><td>T9992</td><td>GATGTACGACGAATTAAACTTGG</td><td>1000</td><td></td><td></td><td></td><td></td><td></td><td></td><td></td><td></td><td></td></tr>
<tr><td>Oligo fwd</td><td>CACCGATGTACGACGAATTAAACT</td><td></td><td></td><td></td><td></td><td></td><td></td><td></td><td></td><td></td><td></td></tr>
<tr><td>Oligo rev</td><td>AAACAGTTTAATTCGTCGTACATC</td><td></td><td></td><td></td><td></td><td></td><td></td><td></td><td></td><td></td><td></td></tr>
<tr><td>Chromosome</td><td>start</td><td>end</td><td>strand</td><td>MM</td><td>target_seq</td><td>PAM</td><td>alignment</td><td>distance</td><td>position</td><td>gene name</td><td>gene id</td></tr>
<tr><td>GU305914.1</td><td>131965</td><td>131987</td><td>+</td><td>0</td><td>GATGTACGACGAATTAAACT</td><td>TGG</td><td>||||||||[||||||||||||]PAM</td><td>NA</td><td>Intergenic</td><td>NA</td><td>NA</td></tr>
<tr><td>T9993</td><td>GTCTCTTGTCTCTGTTCATGAGG</td><td>1000</td><td></td><td></td><td></td><td></td><td></td><td></td><td></td><td></td><td></td></tr>
<tr><td>Oligo fwd</td><td>CACCGTCTCTTGTCTCTGTTCATG</td><td></td><td></td><td></td><td></td><td></td><td></td><td></td><td></td><td></td><td></td></tr>
<tr><td>Oligo rev</td><td>AAACCATGAACAGAGACAAGAGAC</td><td></td><td></td><td></td><td></td><td></td><td></td><td></td><td></td><td></td><td></td></tr>
<tr><td>Chromosome</td><td>start</td><td>end</td><td>strand</td><td>MM</td><td>target_seq</td><td>PAM</td><td>alignment</td><td>distance</td><td>position</td><td>gene name</td><td>gene id</td></tr>
<tr><td>GU305914.1</td><td>132009</td><td>132031</td><td>+</td><td>0</td><td>GTCTCTTGTCTCTGTTCATG</td><td>AGG</td><td>||||||||[||||||||||||]PAM</td><td>NA</td><td>Intergenic</td><td>NA</td><td>NA</td></tr>
<tr><td>T9994</td><td>TCTCTGTTCATGAGGTAGAACGG</td><td>1000</td><td></td><td></td><td></td><td></td><td></td><td></td><td></td><td></td><td></td></tr>
<tr><td>Oligo fwd</td><td>CACCTCTCTGTTCATGAGGTAGAA</td><td></td><td></td><td></td><td></td><td></td><td></td><td></td><td></td><td></td><td></td></tr>
</table>

<table>
<tr><td>Oligo rev</td><td>AAACTTCTACCTCATGAACAGAGA</td><td></td><td></td><td></td><td></td><td></td><td></td><td></td><td></td><td></td><td></td></tr>
<tr><td>Chromosome</td><td>start</td><td>end</td><td>strand</td><td>MM</td><td>target_seq</td><td>PAM</td><td>alignment</td><td>distance</td><td>position</td><td>gene name</td><td>gene id</td></tr>
<tr><td>GU305914.1</td><td>132017</td><td>132039</td><td>+</td><td>0</td><td>TCTCTGTTCATGAGGTAGAA</td><td>CGG</td><td>||||||||[||||||||||||]PAM</td><td>NA</td><td>Intergenic</td><td>NA</td><td>NA</td></tr>
<tr><td></td><td></td><td></td><td></td><td></td><td></td><td></td><td></td><td></td><td></td><td></td><td></td></tr>
<tr><td>T9995</td><td>TCATGAGGTAGAACGGTTGTTGG</td><td>1000</td><td></td><td></td><td></td><td></td><td></td><td></td><td></td><td></td><td></td></tr>
<tr><td>Oligo fwd</td><td>CACCTCATGAGGTAGAACGGTTGT</td><td></td><td></td><td></td><td></td><td></td><td></td><td></td><td></td><td></td><td></td></tr>
<tr><td>Oligo rev</td><td>AAACACAACCGTTCTACCTCATGA</td><td></td><td></td><td></td><td></td><td></td><td></td><td></td><td></td><td></td><td></td></tr>
<tr><td>Chromosome</td><td>start</td><td>end</td><td>strand</td><td>MM</td><td>target_seq</td><td>PAM</td><td>alignment</td><td>distance</td><td>position</td><td>gene name</td><td>gene id</td></tr>
<tr><td>GU305914.1</td><td>132024</td><td>132046</td><td>+</td><td>0</td><td>TCATGAGGTAGAACGGTTGT</td><td>TGG</td><td>||||||||[||||||||||||]PAM</td><td>NA</td><td>Intergenic</td><td>NA</td><td>NA</td></tr>
<tr><td></td><td></td><td></td><td></td><td></td><td></td><td></td><td></td><td></td><td></td><td></td><td></td></tr>
<tr><td>T9996</td><td>ACGGTTGTTGGATCTCGTTCTGG</td><td>1000</td><td></td><td></td><td></td><td></td><td></td><td></td><td></td><td></td><td></td></tr>
<tr><td>Oligo fwd</td><td>CACCACGGTTGTTGGATCTCGTTC</td><td></td><td></td><td></td><td></td><td></td><td></td><td></td><td></td><td></td><td></td></tr>
<tr><td>Oligo rev</td><td>AAACGAACGAGATCCAACAACCGT</td><td></td><td></td><td></td><td></td><td></td><td></td><td></td><td></td><td></td><td></td></tr>
<tr><td>Chromosome</td><td>start</td><td>end</td><td>strand</td><td>MM</td><td>target_seq</td><td>PAM</td><td>alignment</td><td>distance</td><td>position</td><td>gene name</td><td>gene id</td></tr>
<tr><td>GU305914.1</td><td>132036</td><td>132058</td><td>+</td><td>0</td><td>ACGGTTGTTGGATCTCGTTC</td><td>TGG</td><td>||||||||[||||||||||||]PAM</td><td>NA</td><td>Intergenic</td><td>NA</td><td>NA</td></tr>
<tr><td></td><td></td><td></td><td></td><td></td><td></td><td></td><td></td><td></td><td></td><td></td><td></td></tr>
<tr><td>T9997</td><td>TGTTGGATCTCGTTCTGGTCCGG</td><td>1000</td><td></td><td></td><td></td><td></td><td></td><td></td><td></td><td></td><td></td></tr>
<tr><td>Oligo fwd</td><td>CACCTGTTGGATCTCGTTCTGGTC</td><td></td><td></td><td></td><td></td><td></td><td></td><td></td><td></td><td></td><td></td></tr>
<tr><td>Oligo rev</td><td>AAACGACCAGAACGAGATCCAACA</td><td></td><td></td><td></td><td></td><td></td><td></td><td></td><td></td><td></td><td></td></tr>
<tr><td>Chromosome</td><td>start</td><td>end</td><td>strand</td><td>MM</td><td>target_seq</td><td>PAM</td><td>alignment</td><td>distance</td><td>position</td><td>gene name</td><td>gene id</td></tr>
<tr><td>GU305914.1</td><td>132041</td><td>132063</td><td>+</td><td>0</td><td>TGTTGGATCTCGTTCTGGTC</td><td>CGG</td><td>||||||||[||||||||||||]PAM</td><td>NA</td><td>Intergenic</td><td>NA</td><td>NA</td></tr>
<tr><td></td><td></td><td></td><td></td><td></td><td></td><td></td><td></td><td></td><td></td><td></td><td></td></tr>
<tr><td>T9998</td><td>GTTGGATCTCGTTCTGGTCCGGG</td><td>1000</td><td></td><td></td><td></td><td></td><td></td><td></td><td></td><td></td><td></td></tr>
<tr><td>Oligo fwd</td><td>CACCGTTGGATCTCGTTCTGGTCC</td><td></td><td></td><td></td><td></td><td></td><td></td><td></td><td></td><td></td><td></td></tr>
<tr><td>Oligo rev</td><td>AAACGGACCAGAACGAGATCCAAC</td><td></td><td></td><td></td><td></td><td></td><td></td><td></td><td></td><td></td><td></td></tr>
<tr><td>Chromosome</td><td>start</td><td>end</td><td>strand</td><td>MM</td><td>target_seq</td><td>PAM</td><td>alignment</td><td>distance</td><td>position</td><td>gene name</td><td>gene id</td></tr>
<tr><td>GU305914.1</td><td>132042</td><td>132064</td><td>+</td><td>0</td><td>GTTGGATCTCGTTCTGGTCC</td><td>GGG</td><td>||||||||[||||||||||||]PAM</td><td>NA</td><td>Intergenic</td><td>NA</td><td>NA</td></tr>
<tr><td>GU305914.1</td><td>225416</td><td>225438</td><td>+</td><td>4</td><td>ACTGGATCACGTTCTGGTCT</td><td>TGG</td><td>--||||||[-||||||||||-]PAM</td><td>NA</td><td>Intergenic</td><td>NA</td><td>NA</td></tr>
<tr><td></td><td></td><td></td><td></td><td></td><td></td><td></td><td></td><td></td><td></td><td></td><td></td></tr>
<tr><td>T9999</td><td>TCTCGTTCTGGTCCGGGCTGTGG</td><td>1000</td><td></td><td></td><td></td><td></td><td></td><td></td><td></td><td></td><td></td></tr>
<tr><td>Oligo fwd</td><td>CACCTCTCGTTCTGGTCCGGGCTG</td><td></td><td></td><td></td><td></td><td></td><td></td><td></td><td></td><td></td><td></td></tr>
<tr><td>Oligo rev</td><td>AAACCAGCCCGGACCAGAACGAGA</td><td></td><td></td><td></td><td></td><td></td><td></td><td></td><td></td><td></td><td></td></tr>
<tr><td>Chromosome</td><td>start</td><td>end</td><td>strand</td><td>MM</td><td>target_seq</td><td>PAM</td><td>alignment</td><td>distance</td><td>position</td><td>gene name</td><td>gene id</td></tr>
<tr><td>GU305914.1</td><td>132048</td><td>132070</td><td>+</td><td>0</td><td>TCTCGTTCTGGTCCGGGCTG</td><td>TGG</td><td>||||||||[||||||||||||]PAM</td><td>NA</td><td>Intergenic</td><td>NA</td><td>NA</td></tr>
<tr><td></td><td></td><td></td><td></td><td></td><td></td><td></td><td></td><td></td><td></td><td></td><td></td></tr>
<tr><td>T10000</td><td>GCTGTGGTAGAACAGCGTCTCGG</td><td>1000</td><td></td><td></td><td></td><td></td><td></td><td></td><td></td><td></td><td></td></tr>
<tr><td>Oligo fwd</td><td>CACCGCTGTGGTAGAACAGCGTCT</td><td></td><td></td><td></td><td></td><td></td><td></td><td></td><td></td><td></td><td></td></tr>
<tr><td>Oligo rev</td><td>AAACAGACGCTGTTCTACCACAGC</td><td></td><td></td><td></td><td></td><td></td><td></td><td></td><td></td><td></td><td></td></tr>
<tr><td>Chromosome</td><td>start</td><td>end</td><td>strand</td><td>MM</td><td>target_seq</td><td>PAM</td><td>alignment</td><td>distance</td><td>position</td><td>gene name</td><td>gene id</td></tr>
<tr><td>GU305914.1</td><td>132064</td><td>132086</td><td>+</td><td>0</td><td>GCTGTGGTAGAACAGCGTCT</td><td>CGG</td><td>||||||||[||||||||||||]PAM</td><td>NA</td><td>Intergenic</td><td>NA</td><td>NA</td></tr>
<tr><td></td><td></td><td></td><td></td><td></td><td></td><td></td><td></td><td></td><td></td><td></td><td></td></tr>
<tr><td>T10001</td><td>TCTCGGCGTGCGCCGACAGCAGG</td><td>1000</td><td></td><td></td><td></td><td></td><td></td><td></td><td></td><td></td><td></td></tr>
<tr><td>Oligo fwd</td><td>CACCTCTCGGCGTGCGCCGACAGC</td><td></td><td></td><td></td><td></td><td></td><td></td><td></td><td></td><td></td><td></td></tr>
<tr><td>Oligo rev</td><td>AAACGCTGTCGGCGCACGCCGAGA</td><td></td><td></td><td></td><td></td><td></td><td></td><td></td><td></td><td></td><td></td></tr>
<tr><td>Chromosome</td><td>start</td><td>end</td><td>strand</td><td>MM</td><td>target_seq</td><td>PAM</td><td>alignment</td><td>distance</td><td>position</td><td>gene name</td><td>gene id</td></tr>
<tr><td>GU305914.1</td><td>132081</td><td>132103</td><td>+</td><td>0</td><td>TCTCGGCGTGCGCCGACAGC</td><td>AGG</td><td>||||||||[||||||||||||]PAM</td><td>NA</td><td>Intergenic</td><td>NA</td><td>NA</td></tr>
<tr><td>GU305914.1</td><td>26647</td><td>26669</td><td>+</td><td>4</td><td>AGTCGGAGTGCGGCGACAGC</td><td>GGG</td><td>--||||-|[||||-|||||||]PAM</td><td>NA</td><td>Intergenic</td><td>NA</td><td>NA</td></tr>
<tr><td></td><td></td><td></td><td></td><td></td><td></td><td></td><td></td><td></td><td></td><td></td><td></td></tr>
<tr><td>T10002</td><td>CGATGCGCACCGCCGCCGCCTGG</td><td>1000</td><td></td><td></td><td></td><td></td><td></td><td></td><td></td><td></td><td></td></tr>
<tr><td>Oligo fwd</td><td>CACCCGATGCGCACCGCCGCCGCC</td><td></td><td></td><td></td><td></td><td></td><td></td><td></td><td></td><td></td><td></td></tr>
<tr><td>Oligo rev</td><td>AAACGGCGGCGGCGGTGCGCATCG</td><td></td><td></td><td></td><td></td><td></td><td></td><td></td><td></td><td></td><td></td></tr>
<tr><td>Chromosome</td><td>start</td><td>end</td><td>strand</td><td>MM</td><td>target_seq</td><td>PAM</td><td>alignment</td><td>distance</td><td>position</td><td>gene name</td><td>gene id</td></tr>
<tr><td>GU305914.1</td><td>132123</td><td>132145</td><td>+</td><td>0</td><td>CGATGCGCACCGCCGCCGCC</td><td>TGG</td><td>||||||||[||||||||||||]PAM</td><td>NA</td><td>Intergenic</td><td>NA</td><td>NA</td></tr>
<tr><td></td><td></td><td></td><td></td><td></td><td></td><td></td><td></td><td></td><td></td><td></td><td></td></tr>
<tr><td>T10003</td><td>TCGTCACCCGCACCTGTGACAGG</td><td>1000</td><td></td><td></td><td></td><td></td><td></td><td></td><td></td><td></td><td></td></tr>
<tr><td>Oligo fwd</td><td>CACCTCGTCACCCGCACCTGTGAC</td><td></td><td></td><td></td><td></td><td></td><td></td><td></td><td></td><td></td><td></td></tr>
<tr><td>Oligo rev</td><td>AAACGTCACAGGTGCGGGTGACGA</td><td></td><td></td><td></td><td></td><td></td><td></td><td></td><td></td><td></td><td></td></tr>
<tr><td>Chromosome</td><td>start</td><td>end</td><td>strand</td><td>MM</td><td>target_seq</td><td>PAM</td><td>alignment</td><td>distance</td><td>position</td><td>gene name</td><td>gene id</td></tr>
<tr><td>GU305914.1</td><td>132162</td><td>132184</td><td>+</td><td>0</td><td>TCGTCACCCGCACCTGTGAC</td><td>AGG</td><td>||||||||[||||||||||||]PAM</td><td>NA</td><td>Intergenic</td><td>NA</td><td>NA</td></tr>
<tr><td></td><td></td><td></td><td></td><td></td><td></td><td></td><td></td><td></td><td></td><td></td><td></td></tr>
<tr><td>T10004</td><td>ACCCGCACCTGTGACAGGAACGG</td><td>1000</td><td></td><td></td><td></td><td></td><td></td><td></td><td></td><td></td><td></td></tr>
<tr><td>Oligo fwd</td><td>CACCACCCGCACCTGTGACAGGAA</td><td></td><td></td><td></td><td></td><td></td><td></td><td></td><td></td><td></td><td></td></tr>
<tr><td>Oligo rev</td><td>AAACTTCCTGTCACAGGTGCGGGT</td><td></td><td></td><td></td><td></td><td></td><td></td><td></td><td></td><td></td><td></td></tr>
<tr><td>Chromosome</td><td>start</td><td>end</td><td>strand</td><td>MM</td><td>target_seq</td><td>PAM</td><td>alignment</td><td>distance</td><td>position</td><td>gene name</td><td>gene id</td></tr>
<tr><td>GU305914.1</td><td>132167</td><td>132189</td><td>+</td><td>0</td><td>ACCCGCACCTGTGACAGGAA</td><td>CGG</td><td>||||||||[||||||||||||]PAM</td><td>NA</td><td>Intergenic</td><td>NA</td><td>NA</td></tr>
<tr><td></td><td></td><td></td><td></td><td></td><td></td><td></td><td></td><td></td><td></td><td></td><td></td></tr>
<tr><td>T10005</td><td>CCCGCACCTGTGACAGGAACGGG</td><td>1000</td><td></td><td></td><td></td><td></td><td></td><td></td><td></td><td></td><td></td></tr>
<tr><td>Oligo fwd</td><td>CACCCCCGCACCTGTGACAGGAAC</td><td></td><td></td><td></td><td></td><td></td><td></td><td></td><td></td><td></td><td></td></tr>
<tr><td>Oligo rev</td><td>AAACGTTCCTGTCACAGGTGCGGG</td><td></td><td></td><td></td><td></td><td></td><td></td><td></td><td></td><td></td><td></td></tr>
<tr><td>Chromosome</td><td>start</td><td>end</td><td>strand</td><td>MM</td><td>target_seq</td><td>PAM</td><td>alignment</td><td>distance</td><td>position</td><td>gene name</td><td>gene id</td></tr>
<tr><td>GU305914.1</td><td>132168</td><td>132190</td><td>+</td><td>0</td><td>CCCGCACCTGTGACAGGAAC</td><td>GGG</td><td>||||||||[||||||||||||]PAM</td><td>NA</td><td>Intergenic</td><td>NA</td><td>NA</td></tr>
<tr><td></td><td></td><td></td><td></td><td></td><td></td><td></td><td></td><td></td><td></td><td></td><td></td></tr>
<tr><td>T10006</td><td>CCTGTGACAGGAACGGGTGCAGG</td><td>1000</td><td></td><td></td><td></td><td></td><td></td><td></td><td></td><td></td><td></td></tr>
<tr><td>Oligo fwd</td><td>CACCCCTGTGACAGGAACGGGTGC</td><td></td><td></td><td></td><td></td><td></td><td></td><td></td><td></td><td></td><td></td></tr>
<tr><td>Oligo rev</td><td>AAACGCACCCGTTCCTGTCACAGG</td><td></td><td></td><td></td><td></td><td></td><td></td><td></td><td></td><td></td><td></td></tr>
<tr><td>Chromosome</td><td>start</td><td>end</td><td>strand</td><td>MM</td><td>target_seq</td><td>PAM</td><td>alignment</td><td>distance</td><td>position</td><td>gene name</td><td>gene id</td></tr>
<tr><td>GU305914.1</td><td>132174</td><td>132196</td><td>+</td><td>0</td><td>CCTGTGACAGGAACGGGTGC</td><td>AGG</td><td>||||||||[||||||||||||]PAM</td><td>NA</td><td>Intergenic</td><td>NA</td><td>NA</td></tr>
<tr><td></td><td></td><td></td><td></td><td></td><td></td><td></td><td></td><td></td><td></td><td></td><td></td></tr>
<tr><td>T10007</td><td>GCAAGATCGACAGCAGCATGTGG</td><td>1000</td><td></td><td></td><td></td><td></td><td></td><td></td><td></td><td></td><td></td></tr>
<tr><td>Oligo fwd</td><td>CACCGCAAGATCGACAGCAGCATG</td><td></td><td></td><td></td><td></td><td></td><td></td><td></td><td></td><td></td><td></td></tr>
<tr><td>Oligo rev</td><td>AAACCATGCTGCTGTCGATCTTGC</td><td></td><td></td><td></td><td></td><td></td><td></td><td></td><td></td><td></td><td></td></tr>
<tr><td>Chromosome</td><td>start</td><td>end</td><td>strand</td><td>MM</td><td>target_seq</td><td>PAM</td><td>alignment</td><td>distance</td><td>position</td><td>gene name</td><td>gene id</td></tr>
<tr><td>GU305914.1</td><td>132198</td><td>132220</td><td>+</td><td>0</td><td>GCAAGATCGACAGCAGCATG</td><td>TGG</td><td>||||||||[||||||||||||]PAM</td><td>NA</td><td>Intergenic</td><td>NA</td><td>NA</td></tr>
<tr><td></td><td></td><td></td><td></td><td></td><td></td><td></td><td></td><td></td><td></td><td></td><td></td></tr>
<tr><td>T10008</td><td>CAAGATCGACAGCAGCATGTGGG</td><td>1000</td><td></td><td></td><td></td><td></td><td></td><td></td><td></td><td></td><td></td></tr>
<tr><td>Oligo fwd</td><td>CACCCAAGATCGACAGCAGCATGT</td><td></td><td></td><td></td><td></td><td></td><td></td><td></td><td></td><td></td><td></td></tr>
<tr><td>Oligo rev</td><td>AAACACATGCTGCTGTCGATCTTG</td><td></td><td></td><td></td><td></td><td></td><td></td><td></td><td></td><td></td><td></td></tr>
<tr><td>Chromosome</td><td>start</td><td>end</td><td>strand</td><td>MM</td><td>target_seq</td><td>PAM</td><td>alignment</td><td>distance</td><td>position</td><td>gene name</td><td>gene id</td></tr>
<tr><td>GU305914.1</td><td>132199</td><td>132221</td><td>+</td><td>0</td><td>CAAGATCGACAGCAGCATGT</td><td>GGG</td><td>||||||||[||||||||||||]PAM</td><td>NA</td><td>Intergenic</td><td>NA</td><td>NA</td></tr>
</table>

| | | | | | | | | | | | |
|---|---|---|---|---|---|---|---|---|---|---|---|
| T10009 | ACGTCTCCGAGCTGTCCAGCAGG | 1000 | | | | | | | | | |
| Oligo fwd | CACCACGTCTCCGAGCTGTCCAGC | | | | | | | | | | |
| Oligo rev | AAACGCTGGACAGCTCGGAGACGT | | | | | | | | | | |
| Chromosome | start | end | strand | MM | target_seq | PAM | alignment | distance | position | gene name | gene id |
| GU305914.1 | 132240 | 132262 | + | 0 | ACGTCTCCGAGCTGTCCAGC | AGG | \|\|\|\|\|\|\|\|[\|\|\|\|\|\|\|\|\|\|\|\|]PAM | NA | Intergenic | NA | NA |
| GU305914.1 | 70960 | 70982 | - | 4 | TCGCCTGCGAGTTGTCCAGC | AGG | -\|\|-\|\|-\|[\|\|\|-\|\|\|\|\|\|\|\|]PAM | NA | Intergenic | NA | NA |
| T10010 | AGCTGTCCAGCAGGTCCTTCAGG | 1000 | | | | | | | | | |
| Oligo fwd | CACCAGCTGTCCAGCAGGTCCTTC | | | | | | | | | | |
| Oligo rev | AAACGAAGGACCTGCTGGACAGCT | | | | | | | | | | |
| Chromosome | start | end | strand | MM | target_seq | PAM | alignment | distance | position | gene name | gene id |
| GU305914.1 | 132249 | 132271 | + | 0 | AGCTGTCCAGCAGGTCCTTC | AGG | \|\|\|\|\|\|\|\|[\|\|\|\|\|\|\|\|\|\|\|\|]PAM | NA | Intergenic | NA | NA |
| T10011 | CCAACCCGTACACGATAAACTGG | 1000 | | | | | | | | | |
| Oligo fwd | CACCCCAACCCGTACACGATAAAC | | | | | | | | | | |
| Oligo rev | AAACGTTTATCGTGTACGGGTTGG | | | | | | | | | | |
| Chromosome | start | end | strand | MM | target_seq | PAM | alignment | distance | position | gene name | gene id |
| GU305914.1 | 132282 | 132304 | + | 0 | CCAACCCGTACACGATAAAC | TGG | \|\|\|\|\|\|\|\|[\|\|\|\|\|\|\|\|\|\|\|\|]PAM | NA | Intergenic | NA | NA |
| T10012 | CGTACACGATAAACTGGTCGCGG | 1000 | | | | | | | | | |
| Oligo fwd | CACCCGTACACGATAAACTGGTCG | | | | | | | | | | |
| Oligo rev | AAACCGACCAGTTTATCGTGTACG | | | | | | | | | | |
| Chromosome | start | end | strand | MM | target_seq | PAM | alignment | distance | position | gene name | gene id |
| GU305914.1 | 132288 | 132310 | + | 0 | CGTACACGATAAACTGGTCG | CGG | \|\|\|\|\|\|\|\|[\|\|\|\|\|\|\|\|\|\|\|\|]PAM | NA | Intergenic | NA | NA |
| T10013 | CACGATAAACTGGTCGCGGTAGG | 1000 | | | | | | | | | |
| Oligo fwd | CACCCACGATAAACTGGTCGCGGT | | | | | | | | | | |
| Oligo rev | AAACACCGCGACCAGTTTATCGTG | | | | | | | | | | |
| Chromosome | start | end | strand | MM | target_seq | PAM | alignment | distance | position | gene name | gene id |
| GU305914.1 | 132292 | 132314 | + | 0 | CACGATAAACTGGTCGCGGT | AGG | \|\|\|\|\|\|\|\|[\|\|\|\|\|\|\|\|\|\|\|\|]PAM | NA | Intergenic | NA | NA |
| T10014 | GGTCCCCACCGCTGCCACGCCGG | 1000 | | | | | | | | | |
| Oligo fwd | CACCGGTCCCCACCGCTGCCACGC | | | | | | | | | | |
| Oligo rev | AAACGCGTGGCAGCGGTGGGGACC | | | | | | | | | | |
| Chromosome | start | end | strand | MM | target_seq | PAM | alignment | distance | position | gene name | gene id |
| GU305914.1 | 132313 | 132335 | + | 0 | GGTCCCCACCGCTGCCACGC | CGG | \|\|\|\|\|\|\|\|[\|\|\|\|\|\|\|\|\|\|\|\|]PAM | NA | Intergenic | NA | NA |
| T10015 | CACCGCTGCCACGCCGGTGCCGG | 1000 | | | | | | | | | |
| Oligo fwd | CACCCACCGCTGCCACGCCGGTGC | | | | | | | | | | |
| Oligo rev | AAACGCACCGGCGTGGCAGCGGTG | | | | | | | | | | |
| Chromosome | start | end | strand | MM | target_seq | PAM | alignment | distance | position | gene name | gene id |
| GU305914.1 | 132319 | 132341 | + | 0 | CACCGCTGCCACGCCGGTGC | CGG | \|\|\|\|\|\|\|\|[\|\|\|\|\|\|\|\|\|\|\|\|]PAM | NA | Intergenic | NA | NA |
| T10016 | CCACGCCGGTGCCGGACGCGCGG | 1000 | | | | | | | | | |
| Oligo fwd | CACCCCACGCCGGTGCCGGACGCG | | | | | | | | | | |
| Oligo rev | AAACCGCGTCCGGCACCGGCGTGG | | | | | | | | | | |
| Chromosome | start | end | strand | MM | target_seq | PAM | alignment | distance | position | gene name | gene id |
| GU305914.1 | 132327 | 132349 | + | 0 | CCACGCCGGTGCCGGACGCG | CGG | \|\|\|\|\|\|\|\|[\|\|\|\|\|\|\|\|\|\|\|\|]PAM | NA | Intergenic | NA | NA |
| T10017 | CGCCGGTGCCGGACGCGCGGCGG | 1000 | | | | | | | | | |
| Oligo fwd | CACCCGCCGGTGCCGGACGCGCGG | | | | | | | | | | |
| Oligo rev | AAACCCGCGCGTCCGGCACCGGCG | | | | | | | | | | |
| Chromosome | start | end | strand | MM | target_seq | PAM | alignment | distance | position | gene name | gene id |
| GU305914.1 | 132330 | 132352 | + | 0 | CGCCGGTGCCGGACGCGCGG | CGG | \|\|\|\|\|\|\|\|[\|\|\|\|\|\|\|\|\|\|\|\|]PAM | NA | Intergenic | NA | NA |
| T10018 | GCCGGACGCGCGGCGGTTCGAGG | 1000 | | | | | | | | | |
| Oligo fwd | CACCGCCGGACGCGCGGCGGTTCG | | | | | | | | | | |
| Oligo rev | AAACCGAACCGCCGCGCGTCCGGC | | | | | | | | | | |
| Chromosome | start | end | strand | MM | target_seq | PAM | alignment | distance | position | gene name | gene id |
| GU305914.1 | 132337 | 132359 | + | 0 | GCCGGACGCGCGGCGGTTCG | AGG | \|\|\|\|\|\|\|\|[\|\|\|\|\|\|\|\|\|\|\|\|]PAM | NA | Intergenic | NA | NA |
| T10019 | CGGCGGTTCGAGGTGAACGCCGG | 1000 | | | | | | | | | |
| Oligo fwd | CACCCGGCGGTTCGAGGTGAACGC | | | | | | | | | | |
| Oligo rev | AAACGCGTTCACCTCGAACCGCCG | | | | | | | | | | |
| Chromosome | start | end | strand | MM | target_seq | PAM | alignment | distance | position | gene name | gene id |
| GU305914.1 | 132347 | 132369 | + | 0 | CGGCGGTTCGAGGTGAACGC | CGG | \|\|\|\|\|\|\|\|[\|\|\|\|\|\|\|\|\|\|\|\|]PAM | NA | Intergenic | NA | NA |
| T10020 | GGCGGTTCGAGGTGAACGCCGGG | 1000 | | | | | | | | | |
| Oligo fwd | CACCGGCGGTTCGAGGTGAACGCC | | | | | | | | | | |
| Oligo rev | AAACGGCGTTCACCTCGAACCGCC | | | | | | | | | | |
| Chromosome | start | end | strand | MM | target_seq | PAM | alignment | distance | position | gene name | gene id |
| GU305914.1 | 132348 | 132370 | + | 0 | GGCGGTTCGAGGTGAACGCC | GGG | \|\|\|\|\|\|\|\|[\|\|\|\|\|\|\|\|\|\|\|\|]PAM | NA | Intergenic | NA | NA |
| T10021 | CCGGGTCCACGTAGACGCTCAGG | 1000 | | | | | | | | | |
| Oligo fwd | CACCCCGGGTCCACGTAGACGCTC | | | | | | | | | | |
| Oligo rev | AAACGAGCGTCTACGTGGACCCGG | | | | | | | | | | |
| Chromosome | start | end | strand | MM | target_seq | PAM | alignment | distance | position | gene name | gene id |
| GU305914.1 | 132366 | 132388 | + | 0 | CCGGGTCCACGTAGACGCTC | AGG | \|\|\|\|\|\|\|\|[\|\|\|\|\|\|\|\|\|\|\|\|]PAM | NA | Intergenic | NA | NA |
| T10022 | AGACGCTCAGGATCGCGCCCAGG | 1000 | | | | | | | | | |
| Oligo fwd | CACCAGACGCTCAGGATCGCGCCC | | | | | | | | | | |
| Oligo rev | AAACGGGCGCGATCCTGAGCGTCT | | | | | | | | | | |
| Chromosome | start | end | strand | MM | target_seq | PAM | alignment | distance | position | gene name | gene id |
| GU305914.1 | 132378 | 132400 | + | 0 | AGACGCTCAGGATCGCGCCC | AGG | \|\|\|\|\|\|\|\|[\|\|\|\|\|\|\|\|\|\|\|\|]PAM | NA | Intergenic | NA | NA |
| GU305914.1 | 77200 | 77222 | - | 4 | AGATGCTGACGATCGCGCAC | AGG | \|\|\|-\|\|\|-[\|-\|\|\|\|\|\|\|\|-\|]PAM | NA | Intergenic | NA | NA |
| T10023 | CTCAGGATCGCGCCCAGGTGAGG | 1000 | | | | | | | | | |
| Oligo fwd | CACCCTCAGGATCGCGCCCAGGTG | | | | | | | | | | |

<table>
<tr><td>Oligo rev</td><td>AAACCACCTGGGCGCGATCCTGAG</td><td></td><td></td><td></td><td></td><td></td><td></td><td></td><td></td><td></td><td></td></tr>
<tr><td>Chromosome</td><td>start</td><td>end</td><td>strand</td><td>MM</td><td>target_seq</td><td>PAM</td><td>alignment</td><td>distance</td><td>position</td><td>gene name</td><td>gene id</td></tr>
<tr><td>GU305914.1</td><td>132383</td><td>132405</td><td>+</td><td>0</td><td>CTCAGGATCGCGCCCAGGTG</td><td>AGG</td><td>||||||||[||||||||||||]PAM</td><td>NA</td><td>Intergenic</td><td>NA</td><td>NA</td></tr>
<tr><td>T10024</td><td>TCAGGATCGCGCCCAGGTGAGGG</td><td>1000</td><td></td><td></td><td></td><td></td><td></td><td></td><td></td><td></td><td></td></tr>
<tr><td>Oligo fwd</td><td>CACCTCAGGATCGCGCCCAGGTGA</td><td></td><td></td><td></td><td></td><td></td><td></td><td></td><td></td><td></td><td></td></tr>
<tr><td>Oligo rev</td><td>AAACTCACCTGGGCGCGATCCTGA</td><td></td><td></td><td></td><td></td><td></td><td></td><td></td><td></td><td></td><td></td></tr>
<tr><td>Chromosome</td><td>start</td><td>end</td><td>strand</td><td>MM</td><td>target_seq</td><td>PAM</td><td>alignment</td><td>distance</td><td>position</td><td>gene name</td><td>gene id</td></tr>
<tr><td>GU305914.1</td><td>132384</td><td>132406</td><td>+</td><td>0</td><td>TCAGGATCGCGCCCAGGTGA</td><td>GGG</td><td>||||||||[||||||||||||]PAM</td><td>NA</td><td>Intergenic</td><td>NA</td><td>NA</td></tr>
<tr><td>T10025</td><td>GGATCGCGCCCAGGTGAGGGTGG</td><td>1000</td><td></td><td></td><td></td><td></td><td></td><td></td><td></td><td></td><td></td></tr>
<tr><td>Oligo fwd</td><td>CACCGGATCGCGCCCAGGTGAGGG</td><td></td><td></td><td></td><td></td><td></td><td></td><td></td><td></td><td></td><td></td></tr>
<tr><td>Oligo rev</td><td>AAACCCCTCACCTGGGCGCGATCC</td><td></td><td></td><td></td><td></td><td></td><td></td><td></td><td></td><td></td><td></td></tr>
<tr><td>Chromosome</td><td>start</td><td>end</td><td>strand</td><td>MM</td><td>target_seq</td><td>PAM</td><td>alignment</td><td>distance</td><td>position</td><td>gene name</td><td>gene id</td></tr>
<tr><td>GU305914.1</td><td>132387</td><td>132409</td><td>+</td><td>0</td><td>GGATCGCGCCCAGGTGAGGG</td><td>TGG</td><td>||||||||[||||||||||||]PAM</td><td>NA</td><td>Intergenic</td><td>NA</td><td>NA</td></tr>
<tr><td>T10026</td><td>CCCAGGTGAGGGTGGAACTGCGG</td><td>1000</td><td></td><td></td><td></td><td></td><td></td><td></td><td></td><td></td><td></td></tr>
<tr><td>Oligo fwd</td><td>CACCCCCAGGTGAGGGTGGAACTG</td><td></td><td></td><td></td><td></td><td></td><td></td><td></td><td></td><td></td><td></td></tr>
<tr><td>Oligo rev</td><td>AAACCAGTTCCACCCTCACCTGGG</td><td></td><td></td><td></td><td></td><td></td><td></td><td></td><td></td><td></td><td></td></tr>
<tr><td>Chromosome</td><td>start</td><td>end</td><td>strand</td><td>MM</td><td>target_seq</td><td>PAM</td><td>alignment</td><td>distance</td><td>position</td><td>gene name</td><td>gene id</td></tr>
<tr><td>GU305914.1</td><td>132395</td><td>132417</td><td>+</td><td>0</td><td>CCCAGGTGAGGGTGGAACTG</td><td>CGG</td><td>||||||||[||||||||||||]PAM</td><td>NA</td><td>Intergenic</td><td>NA</td><td>NA</td></tr>
<tr><td>T10027</td><td>CCAGGTGAGGGTGGAACTGCGGG</td><td>1000</td><td></td><td></td><td></td><td></td><td></td><td></td><td></td><td></td><td></td></tr>
<tr><td>Oligo fwd</td><td>CACCCCAGGTGAGGGTGGAACTGC</td><td></td><td></td><td></td><td></td><td></td><td></td><td></td><td></td><td></td><td></td></tr>
<tr><td>Oligo rev</td><td>AAACGCAGTTCCACCCTCACCTGG</td><td></td><td></td><td></td><td></td><td></td><td></td><td></td><td></td><td></td><td></td></tr>
<tr><td>Chromosome</td><td>start</td><td>end</td><td>strand</td><td>MM</td><td>target_seq</td><td>PAM</td><td>alignment</td><td>distance</td><td>position</td><td>gene name</td><td>gene id</td></tr>
<tr><td>GU305914.1</td><td>132396</td><td>132418</td><td>+</td><td>0</td><td>CCAGGTGAGGGTGGAACTGC</td><td>GGG</td><td>||||||||[||||||||||||]PAM</td><td>NA</td><td>Intergenic</td><td>NA</td><td>NA</td></tr>
<tr><td>T10028</td><td>GCGGGTTCGTCGTACTGTAGCGG</td><td>1000</td><td></td><td></td><td></td><td></td><td></td><td></td><td></td><td></td><td></td></tr>
<tr><td>Oligo fwd</td><td>CACCGCGGGTTCGTCGTACTGTAG</td><td></td><td></td><td></td><td></td><td></td><td></td><td></td><td></td><td></td><td></td></tr>
<tr><td>Oligo rev</td><td>AAACCTACAGTACGACGAACCCGC</td><td></td><td></td><td></td><td></td><td></td><td></td><td></td><td></td><td></td><td></td></tr>
<tr><td>Chromosome</td><td>start</td><td>end</td><td>strand</td><td>MM</td><td>target_seq</td><td>PAM</td><td>alignment</td><td>distance</td><td>position</td><td>gene name</td><td>gene id</td></tr>
<tr><td>GU305914.1</td><td>132414</td><td>132436</td><td>+</td><td>0</td><td>GCGGGTTCGTCGTACTGTAG</td><td>CGG</td><td>||||||||[||||||||||||]PAM</td><td>NA</td><td>Intergenic</td><td>NA</td><td>NA</td></tr>
<tr><td>T10029</td><td>TCGTCGTACTGTAGCGGAACAGG</td><td>1000</td><td></td><td></td><td></td><td></td><td></td><td></td><td></td><td></td><td></td></tr>
<tr><td>Oligo fwd</td><td>CACCTCGTCGTACTGTAGCGGAAC</td><td></td><td></td><td></td><td></td><td></td><td></td><td></td><td></td><td></td><td></td></tr>
<tr><td>Oligo rev</td><td>AAACGTTCCGCTACAGTACGACGA</td><td></td><td></td><td></td><td></td><td></td><td></td><td></td><td></td><td></td><td></td></tr>
<tr><td>Chromosome</td><td>start</td><td>end</td><td>strand</td><td>MM</td><td>target_seq</td><td>PAM</td><td>alignment</td><td>distance</td><td>position</td><td>gene name</td><td>gene id</td></tr>
<tr><td>GU305914.1</td><td>132420</td><td>132442</td><td>+</td><td>0</td><td>TCGTCGTACTGTAGCGGAAC</td><td>AGG</td><td>||||||||[||||||||||||]PAM</td><td>NA</td><td>Intergenic</td><td>NA</td><td>NA</td></tr>
<tr><td>T10030</td><td>GGAACAGGTCGAACTCCTCGCGG</td><td>1000</td><td></td><td></td><td></td><td></td><td></td><td></td><td></td><td></td><td></td></tr>
<tr><td>Oligo fwd</td><td>CACCGGAACAGGTCGAACTCCTCG</td><td></td><td></td><td></td><td></td><td></td><td></td><td></td><td></td><td></td><td></td></tr>
<tr><td>Oligo rev</td><td>AAACCGAGGAGTTCGACCTGTTCC</td><td></td><td></td><td></td><td></td><td></td><td></td><td></td><td></td><td></td><td></td></tr>
<tr><td>Chromosome</td><td>start</td><td>end</td><td>strand</td><td>MM</td><td>target_seq</td><td>PAM</td><td>alignment</td><td>distance</td><td>position</td><td>gene name</td><td>gene id</td></tr>
<tr><td>GU305914.1</td><td>132435</td><td>132457</td><td>+</td><td>0</td><td>GGAACAGGTCGAACTCCTCG</td><td>CGG</td><td>||||||||[||||||||||||]PAM</td><td>NA</td><td>Intergenic</td><td>NA</td><td>NA</td></tr>
<tr><td>T10031</td><td>GAACTCCTCGCGGCTCTCGTCGG</td><td>1000</td><td></td><td></td><td></td><td></td><td></td><td></td><td></td><td></td><td></td></tr>
<tr><td>Oligo fwd</td><td>CACCGAACTCCTCGCGGCTCTCGT</td><td></td><td></td><td></td><td></td><td></td><td></td><td></td><td></td><td></td><td></td></tr>
<tr><td>Oligo rev</td><td>AAACACGAGAGCCGCGAGGAGTTC</td><td></td><td></td><td></td><td></td><td></td><td></td><td></td><td></td><td></td><td></td></tr>
<tr><td>Chromosome</td><td>start</td><td>end</td><td>strand</td><td>MM</td><td>target_seq</td><td>PAM</td><td>alignment</td><td>distance</td><td>position</td><td>gene name</td><td>gene id</td></tr>
<tr><td>GU305914.1</td><td>132445</td><td>132467</td><td>+</td><td>0</td><td>GAACTCCTCGCGGCTCTCGT</td><td>CGG</td><td>||||||||[||||||||||||]PAM</td><td>NA</td><td>Intergenic</td><td>NA</td><td>NA</td></tr>
<tr><td>T10032</td><td>GATCTCGTCCATGAACGAGCCGG</td><td>1000</td><td></td><td></td><td></td><td></td><td></td><td></td><td></td><td></td><td></td></tr>
<tr><td>Oligo fwd</td><td>CACCGATCTCGTCCATGAACGAGC</td><td></td><td></td><td></td><td></td><td></td><td></td><td></td><td></td><td></td><td></td></tr>
<tr><td>Oligo rev</td><td>AAACGCTCGTTCATGGACGAGATC</td><td></td><td></td><td></td><td></td><td></td><td></td><td></td><td></td><td></td><td></td></tr>
<tr><td>Chromosome</td><td>start</td><td>end</td><td>strand</td><td>MM</td><td>target_seq</td><td>PAM</td><td>alignment</td><td>distance</td><td>position</td><td>gene name</td><td>gene id</td></tr>
<tr><td>GU305914.1</td><td>132511</td><td>132533</td><td>+</td><td>0</td><td>GATCTCGTCCATGAACGAGC</td><td>CGG</td><td>||||||||[||||||||||||]PAM</td><td>NA</td><td>Intergenic</td><td>NA</td><td>NA</td></tr>
<tr><td>T10033</td><td>ATCTCGTCCATGAACGAGCCGGG</td><td>1000</td><td></td><td></td><td></td><td></td><td></td><td></td><td></td><td></td><td></td></tr>
<tr><td>Oligo fwd</td><td>CACCATCTCGTCCATGAACGAGCC</td><td></td><td></td><td></td><td></td><td></td><td></td><td></td><td></td><td></td><td></td></tr>
<tr><td>Oligo rev</td><td>AAACGGCTCGTTCATGGACGAGAT</td><td></td><td></td><td></td><td></td><td></td><td></td><td></td><td></td><td></td><td></td></tr>
<tr><td>Chromosome</td><td>start</td><td>end</td><td>strand</td><td>MM</td><td>target_seq</td><td>PAM</td><td>alignment</td><td>distance</td><td>position</td><td>gene name</td><td>gene id</td></tr>
<tr><td>GU305914.1</td><td>132512</td><td>132534</td><td>+</td><td>0</td><td>ATCTCGTCCATGAACGAGCC</td><td>GGG</td><td>||||||||[||||||||||||]PAM</td><td>NA</td><td>Intergenic</td><td>NA</td><td>NA</td></tr>
<tr><td>T10034</td><td>GTCCGAGTTCAGCGTGATGAAGG</td><td>1000</td><td></td><td></td><td></td><td></td><td></td><td></td><td></td><td></td><td></td></tr>
<tr><td>Oligo fwd</td><td>CACCGTCCGAGTTCAGCGTGATGA</td><td></td><td></td><td></td><td></td><td></td><td></td><td></td><td></td><td></td><td></td></tr>
<tr><td>Oligo rev</td><td>AAACTCATCACGCTGAACTCGGAC</td><td></td><td></td><td></td><td></td><td></td><td></td><td></td><td></td><td></td><td></td></tr>
<tr><td>Chromosome</td><td>start</td><td>end</td><td>strand</td><td>MM</td><td>target_seq</td><td>PAM</td><td>alignment</td><td>distance</td><td>position</td><td>gene name</td><td>gene id</td></tr>
<tr><td>GU305914.1</td><td>132562</td><td>132584</td><td>+</td><td>0</td><td>GTCCGAGTTCAGCGTGATGA</td><td>AGG</td><td>||||||||[||||||||||||]PAM</td><td>NA</td><td>Intergenic</td><td>NA</td><td>NA</td></tr>
<tr><td>T10035</td><td>GAGTTCAGCGTGATGAAGGTCGG</td><td>1000</td><td></td><td></td><td></td><td></td><td></td><td></td><td></td><td></td><td></td></tr>
<tr><td>Oligo fwd</td><td>CACCGAGTTCAGCGTGATGAAGGT</td><td></td><td></td><td></td><td></td><td></td><td></td><td></td><td></td><td></td><td></td></tr>
<tr><td>Oligo rev</td><td>AAACACCTTCATCACGCTGAACTC</td><td></td><td></td><td></td><td></td><td></td><td></td><td></td><td></td><td></td><td></td></tr>
<tr><td>Chromosome</td><td>start</td><td>end</td><td>strand</td><td>MM</td><td>target_seq</td><td>PAM</td><td>alignment</td><td>distance</td><td>position</td><td>gene name</td><td>gene id</td></tr>
<tr><td>GU305914.1</td><td>132566</td><td>132588</td><td>+</td><td>0</td><td>GAGTTCAGCGTGATGAAGGT</td><td>CGG</td><td>||||||||[||||||||||||]PAM</td><td>NA</td><td>Intergenic</td><td>NA</td><td>NA</td></tr>
<tr><td>T10036</td><td>GCGTGATGAAGGTCGGCTTGTGG</td><td>1000</td><td></td><td></td><td></td><td></td><td></td><td></td><td></td><td></td><td></td></tr>
<tr><td>Oligo fwd</td><td>CACCGCGTGATGAAGGTCGGCTTG</td><td></td><td></td><td></td><td></td><td></td><td></td><td></td><td></td><td></td><td></td></tr>
<tr><td>Oligo rev</td><td>AAACCAAGCCGACCTTCATCACGC</td><td></td><td></td><td></td><td></td><td></td><td></td><td></td><td></td><td></td><td></td></tr>
<tr><td>Chromosome</td><td>start</td><td>end</td><td>strand</td><td>MM</td><td>target_seq</td><td>PAM</td><td>alignment</td><td>distance</td><td>position</td><td>gene name</td><td>gene id</td></tr>
<tr><td>GU305914.1</td><td>132573</td><td>132595</td><td>+</td><td>0</td><td>GCGTGATGAAGGTCGGCTTG</td><td>TGG</td><td>||||||||[||||||||||||]PAM</td><td>NA</td><td>Intergenic</td><td>NA</td><td>NA</td></tr>
<tr><td>T10037</td><td>TGATGAAGGTCGGCTTGTGGAGG</td><td>1000</td><td></td><td></td><td></td><td></td><td></td><td></td><td></td><td></td><td></td></tr>
<tr><td>Oligo fwd</td><td>CACCTGATGAAGGTCGGCTTGTGG</td><td></td><td></td><td></td><td></td><td></td><td></td><td></td><td></td><td></td><td></td></tr>
<tr><td>Oligo rev</td><td>AAACCCACAAGCCGACCTTCATCA</td><td></td><td></td><td></td><td></td><td></td><td></td><td></td><td></td><td></td><td></td></tr>
<tr><td>Chromosome</td><td>start</td><td>end</td><td>strand</td><td>MM</td><td>target_seq</td><td>PAM</td><td>alignment</td><td>distance</td><td>position</td><td>gene name</td><td>gene id</td></tr>
<tr><td>GU305914.1</td><td>132576</td><td>132598</td><td>+</td><td>0</td><td>TGATGAAGGTCGGCTTGTGG</td><td>AGG</td><td>||||||||[||||||||||||]PAM</td><td>NA</td><td>Intergenic</td><td>NA</td><td>NA</td></tr>
<tr><td>T10038</td><td>TGAAGGTCGGCTTGTGGAGGCGG</td><td>1000</td><td></td><td></td><td></td><td></td><td></td><td></td><td></td><td></td><td></td></tr>
</table>

<table>
<tr><td>Oligo fwd</td><td>CACCTGAAGGTCGGCTTGTGGAGG</td><td></td><td></td><td></td><td></td><td></td><td></td><td></td><td></td><td></td><td></td></tr>
<tr><td>Oligo rev</td><td>AAACCCTCCACAAGCCGACCTTCA</td><td></td><td></td><td></td><td></td><td></td><td></td><td></td><td></td><td></td><td></td></tr>
<tr><td>Chromosome</td><td>start</td><td>end</td><td>strand</td><td>MM</td><td>target_seq</td><td>PAM</td><td>alignment</td><td>distance</td><td>position</td><td>gene name</td><td>gene id</td></tr>
<tr><td>GU305914.1</td><td>132579</td><td>132601</td><td>+</td><td>0</td><td>TGAAGGTCGGCTTGTGGAGG</td><td>CGG</td><td>||||||||[||||||||||||]PAM</td><td>NA</td><td>Intergenic</td><td>NA</td><td>NA</td></tr>
<tr><td>T10039</td><td>GGCTTGTGGAGGCGGTAGCACGG</td><td>1000</td><td></td><td></td><td></td><td></td><td></td><td></td><td></td><td></td><td></td></tr>
<tr><td>Oligo fwd</td><td>CACCGGCTTGTGGAGGCGGTAGCA</td><td></td><td></td><td></td><td></td><td></td><td></td><td></td><td></td><td></td><td></td></tr>
<tr><td>Oligo rev</td><td>AAACTGCTACCGCCTCCACAAGCC</td><td></td><td></td><td></td><td></td><td></td><td></td><td></td><td></td><td></td><td></td></tr>
<tr><td>Chromosome</td><td>start</td><td>end</td><td>strand</td><td>MM</td><td>target_seq</td><td>PAM</td><td>alignment</td><td>distance</td><td>position</td><td>gene name</td><td>gene id</td></tr>
<tr><td>GU305914.1</td><td>132587</td><td>132609</td><td>+</td><td>0</td><td>GGCTTGTGGAGGCGGTAGCA</td><td>CGG</td><td>||||||||[||||||||||||]PAM</td><td>NA</td><td>Intergenic</td><td>NA</td><td>NA</td></tr>
<tr><td>T10040</td><td>GCTTGTGGAGGCGGTAGCACGGG</td><td>1000</td><td></td><td></td><td></td><td></td><td></td><td></td><td></td><td></td><td></td></tr>
<tr><td>Oligo fwd</td><td>CACCGCTTGTGGAGGCGGTAGCAC</td><td></td><td></td><td></td><td></td><td></td><td></td><td></td><td></td><td></td><td></td></tr>
<tr><td>Oligo rev</td><td>AAACGTGCTACCGCCTCCACAAGC</td><td></td><td></td><td></td><td></td><td></td><td></td><td></td><td></td><td></td><td></td></tr>
<tr><td>Chromosome</td><td>start</td><td>end</td><td>strand</td><td>MM</td><td>target_seq</td><td>PAM</td><td>alignment</td><td>distance</td><td>position</td><td>gene name</td><td>gene id</td></tr>
<tr><td>GU305914.1</td><td>132588</td><td>132610</td><td>+</td><td>0</td><td>GCTTGTGGAGGCGGTAGCAC</td><td>GGG</td><td>||||||||[||||||||||||]PAM</td><td>NA</td><td>Intergenic</td><td>NA</td><td>NA</td></tr>
<tr><td>T10041</td><td>GTGGAGGCGGTAGCACGGGCAGG</td><td>1000</td><td></td><td></td><td></td><td></td><td></td><td></td><td></td><td></td><td></td></tr>
<tr><td>Oligo fwd</td><td>CACCGTGGAGGCGGTAGCACGGGC</td><td></td><td></td><td></td><td></td><td></td><td></td><td></td><td></td><td></td><td></td></tr>
<tr><td>Oligo rev</td><td>AAACGCCCGTGCTACCGCCTCCAC</td><td></td><td></td><td></td><td></td><td></td><td></td><td></td><td></td><td></td><td></td></tr>
<tr><td>Chromosome</td><td>start</td><td>end</td><td>strand</td><td>MM</td><td>target_seq</td><td>PAM</td><td>alignment</td><td>distance</td><td>position</td><td>gene name</td><td>gene id</td></tr>
<tr><td>GU305914.1</td><td>132592</td><td>132614</td><td>+</td><td>0</td><td>GTGGAGGCGGTAGCACGGGC</td><td>AGG</td><td>||||||||[||||||||||||]PAM</td><td>NA</td><td>Intergenic</td><td>NA</td><td>NA</td></tr>
<tr><td>T10042</td><td>CGCCCTTCTCCGAGAACGCCTGG</td><td>1000</td><td></td><td></td><td></td><td></td><td></td><td></td><td></td><td></td><td></td></tr>
<tr><td>Oligo fwd</td><td>CACCCGCCCTTCTCCGAGAACGCC</td><td></td><td></td><td></td><td></td><td></td><td></td><td></td><td></td><td></td><td></td></tr>
<tr><td>Oligo rev</td><td>AAACGGCGTTCTCGGAGAAGGGCG</td><td></td><td></td><td></td><td></td><td></td><td></td><td></td><td></td><td></td><td></td></tr>
<tr><td>Chromosome</td><td>start</td><td>end</td><td>strand</td><td>MM</td><td>target_seq</td><td>PAM</td><td>alignment</td><td>distance</td><td>position</td><td>gene name</td><td>gene id</td></tr>
<tr><td>GU305914.1</td><td>132624</td><td>132646</td><td>+</td><td>0</td><td>CGCCCTTCTCCGAGAACGCC</td><td>TGG</td><td>||||||||[||||||||||||]PAM</td><td>NA</td><td>Intergenic</td><td>NA</td><td>NA</td></tr>
<tr><td>T10043</td><td>GATGTGCTCCTCGCACACGTAGG</td><td>1000</td><td></td><td></td><td></td><td></td><td></td><td></td><td></td><td></td><td></td></tr>
<tr><td>Oligo fwd</td><td>CACCGATGTGCTCCTCGCACACGT</td><td></td><td></td><td></td><td></td><td></td><td></td><td></td><td></td><td></td><td></td></tr>
<tr><td>Oligo rev</td><td>AAACACGTGTGCGAGGAGCACATC</td><td></td><td></td><td></td><td></td><td></td><td></td><td></td><td></td><td></td><td></td></tr>
<tr><td>Chromosome</td><td>start</td><td>end</td><td>strand</td><td>MM</td><td>target_seq</td><td>PAM</td><td>alignment</td><td>distance</td><td>position</td><td>gene name</td><td>gene id</td></tr>
<tr><td>GU305914.1</td><td>132646</td><td>132668</td><td>+</td><td>0</td><td>GATGTGCTCCTCGCACACGT</td><td>AGG</td><td>||||||||[||||||||||||]PAM</td><td>NA</td><td>Intergenic</td><td>NA</td><td>NA</td></tr>
<tr><td>GU305914.1</td><td>210984</td><td>211006</td><td>+</td><td>4</td><td>GATGTCATCCTCGGATACGT</td><td>CGG</td><td>|||||--|[|||||-|-||||]PAM</td><td>NA</td><td>Intergenic</td><td>NA</td><td>NA</td></tr>
<tr><td>T10044</td><td>TACCACGTTCAGCATGTCGAAGG</td><td>1000</td><td></td><td></td><td></td><td></td><td></td><td></td><td></td><td></td><td></td></tr>
<tr><td>Oligo fwd</td><td>CACCTACCACGTTCAGCATGTCGA</td><td></td><td></td><td></td><td></td><td></td><td></td><td></td><td></td><td></td><td></td></tr>
<tr><td>Oligo rev</td><td>AAACTCGACATGCTGAACGTGGTA</td><td></td><td></td><td></td><td></td><td></td><td></td><td></td><td></td><td></td><td></td></tr>
<tr><td>Chromosome</td><td>start</td><td>end</td><td>strand</td><td>MM</td><td>target_seq</td><td>PAM</td><td>alignment</td><td>distance</td><td>position</td><td>gene name</td><td>gene id</td></tr>
<tr><td>GU305914.1</td><td>132670</td><td>132692</td><td>+</td><td>0</td><td>TACCACGTTCAGCATGTCGA</td><td>AGG</td><td>||||||||[||||||||||||]PAM</td><td>NA</td><td>Intergenic</td><td>NA</td><td>NA</td></tr>
<tr><td>T10045</td><td>ACCACGTTCAGCATGTCGAAGGG</td><td>1000</td><td></td><td></td><td></td><td></td><td></td><td></td><td></td><td></td><td></td></tr>
<tr><td>Oligo fwd</td><td>CACCACCACGTTCAGCATGTCGAA</td><td></td><td></td><td></td><td></td><td></td><td></td><td></td><td></td><td></td><td></td></tr>
<tr><td>Oligo rev</td><td>AAACTTCGACATGCTGAACGTGGT</td><td></td><td></td><td></td><td></td><td></td><td></td><td></td><td></td><td></td><td></td></tr>
<tr><td>Chromosome</td><td>start</td><td>end</td><td>strand</td><td>MM</td><td>target_seq</td><td>PAM</td><td>alignment</td><td>distance</td><td>position</td><td>gene name</td><td>gene id</td></tr>
<tr><td>GU305914.1</td><td>132671</td><td>132693</td><td>+</td><td>0</td><td>ACCACGTTCAGCATGTCGAA</td><td>GGG</td><td>||||||||[||||||||||||]PAM</td><td>NA</td><td>Intergenic</td><td>NA</td><td>NA</td></tr>
<tr><td>T10046</td><td>CAGCATGTCGAAGGGCGCCGAGG</td><td>1000</td><td></td><td></td><td></td><td></td><td></td><td></td><td></td><td></td><td></td></tr>
<tr><td>Oligo fwd</td><td>CACCCAGCATGTCGAAGGGCGCCG</td><td></td><td></td><td></td><td></td><td></td><td></td><td></td><td></td><td></td><td></td></tr>
<tr><td>Oligo rev</td><td>AAACCGGCGCCCTTCGACATGCTG</td><td></td><td></td><td></td><td></td><td></td><td></td><td></td><td></td><td></td><td></td></tr>
<tr><td>Chromosome</td><td>start</td><td>end</td><td>strand</td><td>MM</td><td>target_seq</td><td>PAM</td><td>alignment</td><td>distance</td><td>position</td><td>gene name</td><td>gene id</td></tr>
<tr><td>GU305914.1</td><td>132679</td><td>132701</td><td>+</td><td>0</td><td>CAGCATGTCGAAGGGCGCCG</td><td>AGG</td><td>||||||||[||||||||||||]PAM</td><td>NA</td><td>Intergenic</td><td>NA</td><td>NA</td></tr>
<tr><td>T10047</td><td>GCGCCGAGGTCAGCTTCGTCAGG</td><td>1000</td><td></td><td></td><td></td><td></td><td></td><td></td><td></td><td></td><td></td></tr>
<tr><td>Oligo fwd</td><td>CACCGCGCCGAGGTCAGCTTCGTC</td><td></td><td></td><td></td><td></td><td></td><td></td><td></td><td></td><td></td><td></td></tr>
<tr><td>Oligo rev</td><td>AAACGACGAAGCTGACCTCGGCGC</td><td></td><td></td><td></td><td></td><td></td><td></td><td></td><td></td><td></td><td></td></tr>
<tr><td>Chromosome</td><td>start</td><td>end</td><td>strand</td><td>MM</td><td>target_seq</td><td>PAM</td><td>alignment</td><td>distance</td><td>position</td><td>gene name</td><td>gene id</td></tr>
<tr><td>GU305914.1</td><td>132693</td><td>132715</td><td>+</td><td>0</td><td>GCGCCGAGGTCAGCTTCGTC</td><td>AGG</td><td>||||||||[||||||||||||]PAM</td><td>NA</td><td>Intergenic</td><td>NA</td><td>NA</td></tr>
<tr><td>GU305914.1</td><td>77953</td><td>77975</td><td>-</td><td>4</td><td>CCGCCATGGTCAGCTGCGTC</td><td>CGG</td><td>-||||--|[|||||||-||||]PAM</td><td>NA</td><td>Intergenic</td><td>NA</td><td>NA</td></tr>
<tr><td>T10048</td><td>GGTCAGCTTCGTCAGGAAGCAGG</td><td>1000</td><td></td><td></td><td></td><td></td><td></td><td></td><td></td><td></td><td></td></tr>
<tr><td>Oligo fwd</td><td>CACCGGTCAGCTTCGTCAGGAAGC</td><td></td><td></td><td></td><td></td><td></td><td></td><td></td><td></td><td></td><td></td></tr>
<tr><td>Oligo rev</td><td>AAACGCTTCCTGACGAAGCTGACC</td><td></td><td></td><td></td><td></td><td></td><td></td><td></td><td></td><td></td><td></td></tr>
<tr><td>Chromosome</td><td>start</td><td>end</td><td>strand</td><td>MM</td><td>target_seq</td><td>PAM</td><td>alignment</td><td>distance</td><td>position</td><td>gene name</td><td>gene id</td></tr>
<tr><td>GU305914.1</td><td>132700</td><td>132722</td><td>+</td><td>0</td><td>GGTCAGCTTCGTCAGGAAGC</td><td>AGG</td><td>||||||||[||||||||||||]PAM</td><td>NA</td><td>Intergenic</td><td>NA</td><td>NA</td></tr>
<tr><td>T10049</td><td>AGATGATCTTCGTCGTGTTCTGG</td><td>1000</td><td></td><td></td><td></td><td></td><td></td><td></td><td></td><td></td><td></td></tr>
<tr><td>Oligo fwd</td><td>CACCAGATGATCTTCGTCGTGTTC</td><td></td><td></td><td></td><td></td><td></td><td></td><td></td><td></td><td></td><td></td></tr>
<tr><td>Oligo rev</td><td>AAACGAACACGACGAAGATCATCT</td><td></td><td></td><td></td><td></td><td></td><td></td><td></td><td></td><td></td><td></td></tr>
<tr><td>Chromosome</td><td>start</td><td>end</td><td>strand</td><td>MM</td><td>target_seq</td><td>PAM</td><td>alignment</td><td>distance</td><td>position</td><td>gene name</td><td>gene id</td></tr>
<tr><td>GU305914.1</td><td>132756</td><td>132778</td><td>+</td><td>0</td><td>AGATGATCTTCGTCGTGTTC</td><td>TGG</td><td>||||||||[||||||||||||]PAM</td><td>NA</td><td>Intergenic</td><td>NA</td><td>NA</td></tr>
<tr><td>T10050</td><td>GATGATCTTCGTCGTGTTCTGGG</td><td>1000</td><td></td><td></td><td></td><td></td><td></td><td></td><td></td><td></td><td></td></tr>
<tr><td>Oligo fwd</td><td>CACCGATGATCTTCGTCGTGTTCT</td><td></td><td></td><td></td><td></td><td></td><td></td><td></td><td></td><td></td><td></td></tr>
<tr><td>Oligo rev</td><td>AAACAGAACACGACGAAGATCATC</td><td></td><td></td><td></td><td></td><td></td><td></td><td></td><td></td><td></td><td></td></tr>
<tr><td>Chromosome</td><td>start</td><td>end</td><td>strand</td><td>MM</td><td>target_seq</td><td>PAM</td><td>alignment</td><td>distance</td><td>position</td><td>gene name</td><td>gene id</td></tr>
<tr><td>GU305914.1</td><td>132757</td><td>132779</td><td>+</td><td>0</td><td>GATGATCTTCGTCGTGTTCT</td><td>GGG</td><td>||||||||[||||||||||||]PAM</td><td>NA</td><td>Intergenic</td><td>NA</td><td>NA</td></tr>
<tr><td>T10051</td><td>TCTTCGTCGTGTTCTGGGCCAGG</td><td>1000</td><td></td><td></td><td></td><td></td><td></td><td></td><td></td><td></td><td></td></tr>
<tr><td>Oligo fwd</td><td>CACCTCTTCGTCGTGTTCTGGGCC</td><td></td><td></td><td></td><td></td><td></td><td></td><td></td><td></td><td></td><td></td></tr>
<tr><td>Oligo rev</td><td>AAACGGCCCAGAACACGACGAAGA</td><td></td><td></td><td></td><td></td><td></td><td></td><td></td><td></td><td></td><td></td></tr>
<tr><td>Chromosome</td><td>start</td><td>end</td><td>strand</td><td>MM</td><td>target_seq</td><td>PAM</td><td>alignment</td><td>distance</td><td>position</td><td>gene name</td><td>gene id</td></tr>
<tr><td>GU305914.1</td><td>132762</td><td>132784</td><td>+</td><td>0</td><td>TCTTCGTCGTGTTCTGGGCC</td><td>AGG</td><td>||||||||[||||||||||||]PAM</td><td>NA</td><td>Intergenic</td><td>NA</td><td>NA</td></tr>
<tr><td>T10052</td><td>TGTTCTGGGCCAGGAAGCCCAGG</td><td>1000</td><td></td><td></td><td></td><td></td><td></td><td></td><td></td><td></td><td></td></tr>
<tr><td>Oligo fwd</td><td>CACCTGTTCTGGGCCAGGAAGCCC</td><td></td><td></td><td></td><td></td><td></td><td></td><td></td><td></td><td></td><td></td></tr>
<tr><td>Oligo rev</td><td>AAACGGGCTTCCTGGCCCAGAACA</td><td></td><td></td><td></td><td></td><td></td><td></td><td></td><td></td><td></td><td></td></tr>
<tr><td>Chromosome</td><td>start</td><td>end</td><td>strand</td><td>MM</td><td>target_seq</td><td>PAM</td><td>alignment</td><td>distance</td><td>position</td><td>gene name</td><td>gene id</td></tr>
</table>

<table>
<tr><td>GU305914.1</td><td>132771</td><td>132793</td><td>+</td><td>0</td><td>TGTTCTGGGCCAGGAAGCCC</td><td>AGG</td><td>||||||||[||||||||||||]PAM</td><td>NA</td><td>Intergenic</td><td>NA</td><td>NA</td></tr>
<tr><td>T10053</td><td>GAAGCCCAGGATCGTGTTAAAGG</td><td>1000</td><td></td><td></td><td></td><td></td><td></td><td></td><td></td><td></td><td></td></tr>
<tr><td>Oligo fwd</td><td>CACCGAAGCCCAGGATCGTGTTAA</td><td></td><td></td><td></td><td></td><td></td><td></td><td></td><td></td><td></td><td></td></tr>
<tr><td>Oligo rev</td><td>AAACTTAACACGATCCTGGGCTTC</td><td></td><td></td><td></td><td></td><td></td><td></td><td></td><td></td><td></td><td></td></tr>
<tr><td>Chromosome</td><td>start</td><td>end</td><td>strand</td><td>MM</td><td>target_seq</td><td>PAM</td><td>alignment</td><td>distance</td><td>position</td><td>gene name</td><td>gene id</td></tr>
<tr><td>GU305914.1</td><td>132784</td><td>132806</td><td>+</td><td>0</td><td>GAAGCCCAGGATCGTGTTAA</td><td>AGG</td><td>||||||||[||||||||||||]PAM</td><td>NA</td><td>Intergenic</td><td>NA</td><td>NA</td></tr>
<tr><td>T10054</td><td>GTGCCTCGTCCACGATCAAGAGG</td><td>1000</td><td></td><td></td><td></td><td></td><td></td><td></td><td></td><td></td><td></td></tr>
<tr><td>Oligo fwd</td><td>CACCGTGCCTCGTCCACGATCAAG</td><td></td><td></td><td></td><td></td><td></td><td></td><td></td><td></td><td></td><td></td></tr>
<tr><td>Oligo rev</td><td>AAACCTTGATCGTGGACGAGGCAC</td><td></td><td></td><td></td><td></td><td></td><td></td><td></td><td></td><td></td><td></td></tr>
<tr><td>Chromosome</td><td>start</td><td>end</td><td>strand</td><td>MM</td><td>target_seq</td><td>PAM</td><td>alignment</td><td>distance</td><td>position</td><td>gene name</td><td>gene id</td></tr>
<tr><td>GU305914.1</td><td>132825</td><td>132847</td><td>+</td><td>0</td><td>GTGCCTCGTCCACGATCAAG</td><td>AGG</td><td>||||||||[||||||||||||]PAM</td><td>NA</td><td>Intergenic</td><td>NA</td><td>NA</td></tr>
<tr><td>T10055</td><td>AGAGGTTGAAGTTCTGCCCGCGG</td><td>1000</td><td></td><td></td><td></td><td></td><td></td><td></td><td></td><td></td><td></td></tr>
<tr><td>Oligo fwd</td><td>CACCAGAGGTTGAAGTTCTGCCCG</td><td></td><td></td><td></td><td></td><td></td><td></td><td></td><td></td><td></td><td></td></tr>
<tr><td>Oligo rev</td><td>AAACCGGGCAGAACTTCAACCTCT</td><td></td><td></td><td></td><td></td><td></td><td></td><td></td><td></td><td></td><td></td></tr>
<tr><td>Chromosome</td><td>start</td><td>end</td><td>strand</td><td>MM</td><td>target_seq</td><td>PAM</td><td>alignment</td><td>distance</td><td>position</td><td>gene name</td><td>gene id</td></tr>
<tr><td>GU305914.1</td><td>132843</td><td>132865</td><td>+</td><td>0</td><td>AGAGGTTGAAGTTCTGCCCG</td><td>CGG</td><td>||||||||[||||||||||||]PAM</td><td>NA</td><td>Intergenic</td><td>NA</td><td>NA</td></tr>
<tr><td>T10056</td><td>GCCCGCGGATGCTCTGCGTGAGG</td><td>1000</td><td></td><td></td><td></td><td></td><td></td><td></td><td></td><td></td><td></td></tr>
<tr><td>Oligo fwd</td><td>CACCGCCCGCGGATGCTCTGCGTG</td><td></td><td></td><td></td><td></td><td></td><td></td><td></td><td></td><td></td><td></td></tr>
<tr><td>Oligo rev</td><td>AAACCACGCAGAGCATCCGCGGGC</td><td></td><td></td><td></td><td></td><td></td><td></td><td></td><td></td><td></td><td></td></tr>
<tr><td>Chromosome</td><td>start</td><td>end</td><td>strand</td><td>MM</td><td>target_seq</td><td>PAM</td><td>alignment</td><td>distance</td><td>position</td><td>gene name</td><td>gene id</td></tr>
<tr><td>GU305914.1</td><td>132858</td><td>132880</td><td>+</td><td>0</td><td>GCCCGCGGATGCTCTGCGTG</td><td>AGG</td><td>||||||||[||||||||||||]PAM</td><td>NA</td><td>Intergenic</td><td>NA</td><td>NA</td></tr>
<tr><td>T10057</td><td>AACACATATAAAACAGCGTACGG</td><td>1000</td><td></td><td></td><td></td><td></td><td></td><td></td><td></td><td></td><td></td></tr>
<tr><td>Oligo fwd</td><td>CACCAACACATATAAAACAGCGTA</td><td></td><td></td><td></td><td></td><td></td><td></td><td></td><td></td><td></td><td></td></tr>
<tr><td>Oligo rev</td><td>AAACTACGCTGTTTTATATGTGTT</td><td></td><td></td><td></td><td></td><td></td><td></td><td></td><td></td><td></td><td></td></tr>
<tr><td>Chromosome</td><td>start</td><td>end</td><td>strand</td><td>MM</td><td>target_seq</td><td>PAM</td><td>alignment</td><td>distance</td><td>position</td><td>gene name</td><td>gene id</td></tr>
<tr><td>GU305914.1</td><td>132893</td><td>132915</td><td>+</td><td>0</td><td>AACACATATAAAACAGCGTA</td><td>CGG</td><td>||||||||[||||||||||||]PAM</td><td>NA</td><td>Intergenic</td><td>NA</td><td>NA</td></tr>
<tr><td>T10058</td><td>AACAGCGTACGGTATACGAGAGG</td><td>1000</td><td></td><td></td><td></td><td></td><td></td><td></td><td></td><td></td><td></td></tr>
<tr><td>Oligo fwd</td><td>CACCAACAGCGTACGGTATACGAG</td><td></td><td></td><td></td><td></td><td></td><td></td><td></td><td></td><td></td><td></td></tr>
<tr><td>Oligo rev</td><td>AAACCTCGTATACCGTACGCTGTT</td><td></td><td></td><td></td><td></td><td></td><td></td><td></td><td></td><td></td><td></td></tr>
<tr><td>Chromosome</td><td>start</td><td>end</td><td>strand</td><td>MM</td><td>target_seq</td><td>PAM</td><td>alignment</td><td>distance</td><td>position</td><td>gene name</td><td>gene id</td></tr>
<tr><td>GU305914.1</td><td>132904</td><td>132926</td><td>+</td><td>0</td><td>AACAGCGTACGGTATACGAG</td><td>AGG</td><td>||||||||[||||||||||||]PAM</td><td>NA</td><td>Intergenic</td><td>NA</td><td>NA</td></tr>
<tr><td>T10059</td><td>CGAGAGGTAGAAGAACCCGACGG</td><td>1000</td><td></td><td></td><td></td><td></td><td></td><td></td><td></td><td></td><td></td></tr>
<tr><td>Oligo fwd</td><td>CACCCGAGAGGTAGAAGAACCCGA</td><td></td><td></td><td></td><td></td><td></td><td></td><td></td><td></td><td></td><td></td></tr>
<tr><td>Oligo rev</td><td>AAACTCGGGTTCTTCTACCTCTCG</td><td></td><td></td><td></td><td></td><td></td><td></td><td></td><td></td><td></td><td></td></tr>
<tr><td>Chromosome</td><td>start</td><td>end</td><td>strand</td><td>MM</td><td>target_seq</td><td>PAM</td><td>alignment</td><td>distance</td><td>position</td><td>gene name</td><td>gene id</td></tr>
<tr><td>GU305914.1</td><td>132920</td><td>132942</td><td>+</td><td>0</td><td>CGAGAGGTAGAAGAACCCGA</td><td>CGG</td><td>||||||||[||||||||||||]PAM</td><td>NA</td><td>Intergenic</td><td>NA</td><td>NA</td></tr>
<tr><td>T10060</td><td>GAGAGGTAGAAGAACCCGACGGG</td><td>1000</td><td></td><td></td><td></td><td></td><td></td><td></td><td></td><td></td><td></td></tr>
<tr><td>Oligo fwd</td><td>CACCGAGAGGTAGAAGAACCCGAC</td><td></td><td></td><td></td><td></td><td></td><td></td><td></td><td></td><td></td><td></td></tr>
<tr><td>Oligo rev</td><td>AAACGTCGGGTTCTTCTACCTCTC</td><td></td><td></td><td></td><td></td><td></td><td></td><td></td><td></td><td></td><td></td></tr>
<tr><td>Chromosome</td><td>start</td><td>end</td><td>strand</td><td>MM</td><td>target_seq</td><td>PAM</td><td>alignment</td><td>distance</td><td>position</td><td>gene name</td><td>gene id</td></tr>
<tr><td>GU305914.1</td><td>132921</td><td>132943</td><td>+</td><td>0</td><td>GAGAGGTAGAAGAACCCGAC</td><td>GGG</td><td>||||||||[||||||||||||]PAM</td><td>NA</td><td>Intergenic</td><td>NA</td><td>NA</td></tr>
<tr><td>T10061</td><td>AGAGGTAGAAGAACCCGACGGGG</td><td>1000</td><td></td><td></td><td></td><td></td><td></td><td></td><td></td><td></td><td></td></tr>
<tr><td>Oligo fwd</td><td>CACCAGAGGTAGAAGAACCCGACG</td><td></td><td></td><td></td><td></td><td></td><td></td><td></td><td></td><td></td><td></td></tr>
<tr><td>Oligo rev</td><td>AAACCGTCGGGTTCTTCTACCTCT</td><td></td><td></td><td></td><td></td><td></td><td></td><td></td><td></td><td></td><td></td></tr>
<tr><td>Chromosome</td><td>start</td><td>end</td><td>strand</td><td>MM</td><td>target_seq</td><td>PAM</td><td>alignment</td><td>distance</td><td>position</td><td>gene name</td><td>gene id</td></tr>
<tr><td>GU305914.1</td><td>132922</td><td>132944</td><td>+</td><td>0</td><td>AGAGGTAGAAGAACCCGACG</td><td>GGG</td><td>||||||||[||||||||||||]PAM</td><td>NA</td><td>Intergenic</td><td>NA</td><td>NA</td></tr>
<tr><td>T10062</td><td>GTAGAAGAACCCGACGGGGTCGG</td><td>1000</td><td></td><td></td><td></td><td></td><td></td><td></td><td></td><td></td><td></td></tr>
<tr><td>Oligo fwd</td><td>CACCGTAGAAGAACCCGACGGGGT</td><td></td><td></td><td></td><td></td><td></td><td></td><td></td><td></td><td></td><td></td></tr>
<tr><td>Oligo rev</td><td>AAACACCCCGTCGGGTTCTTCTAC</td><td></td><td></td><td></td><td></td><td></td><td></td><td></td><td></td><td></td><td></td></tr>
<tr><td>Chromosome</td><td>start</td><td>end</td><td>strand</td><td>MM</td><td>target_seq</td><td>PAM</td><td>alignment</td><td>distance</td><td>position</td><td>gene name</td><td>gene id</td></tr>
<tr><td>GU305914.1</td><td>132926</td><td>132948</td><td>+</td><td>0</td><td>GTAGAAGAACCCGACGGGGT</td><td>CGG</td><td>||||||||[||||||||||||]PAM</td><td>NA</td><td>Intergenic</td><td>NA</td><td>NA</td></tr>
<tr><td>T10063</td><td>TAGAAGAACCCGACGGGGTCGGG</td><td>1000</td><td></td><td></td><td></td><td></td><td></td><td></td><td></td><td></td><td></td></tr>
<tr><td>Oligo fwd</td><td>CACCTAGAAGAACCCGACGGGGTC</td><td></td><td></td><td></td><td></td><td></td><td></td><td></td><td></td><td></td><td></td></tr>
<tr><td>Oligo rev</td><td>AAACGACCCCGTCGGGTTCTTCTA</td><td></td><td></td><td></td><td></td><td></td><td></td><td></td><td></td><td></td><td></td></tr>
<tr><td>Chromosome</td><td>start</td><td>end</td><td>strand</td><td>MM</td><td>target_seq</td><td>PAM</td><td>alignment</td><td>distance</td><td>position</td><td>gene name</td><td>gene id</td></tr>
<tr><td>GU305914.1</td><td>132927</td><td>132949</td><td>+</td><td>0</td><td>TAGAAGAACCCGACGGGGTC</td><td>GGG</td><td>||||||||[||||||||||||]PAM</td><td>NA</td><td>Intergenic</td><td>NA</td><td>NA</td></tr>
<tr><td>T10064</td><td>AGAAGAACCCGACGGGGTCGGGG</td><td>1000</td><td></td><td></td><td></td><td></td><td></td><td></td><td></td><td></td><td></td></tr>
<tr><td>Oligo fwd</td><td>CACCAGAAGAACCCGACGGGGTCG</td><td></td><td></td><td></td><td></td><td></td><td></td><td></td><td></td><td></td><td></td></tr>
<tr><td>Oligo rev</td><td>AAACCGACCCCGTCGGGTTCTTCT</td><td></td><td></td><td></td><td></td><td></td><td></td><td></td><td></td><td></td><td></td></tr>
<tr><td>Chromosome</td><td>start</td><td>end</td><td>strand</td><td>MM</td><td>target_seq</td><td>PAM</td><td>alignment</td><td>distance</td><td>position</td><td>gene name</td><td>gene id</td></tr>
<tr><td>GU305914.1</td><td>132928</td><td>132950</td><td>+</td><td>0</td><td>AGAAGAACCCGACGGGGTCG</td><td>GGG</td><td>||||||||[||||||||||||]PAM</td><td>NA</td><td>Intergenic</td><td>NA</td><td>NA</td></tr>
<tr><td>T10065</td><td>AGAACCCGACGGGGTCGGGGCGG</td><td>1000</td><td></td><td></td><td></td><td></td><td></td><td></td><td></td><td></td><td></td></tr>
<tr><td>Oligo fwd</td><td>CACCAGAACCCGACGGGGTCGGGG</td><td></td><td></td><td></td><td></td><td></td><td></td><td></td><td></td><td></td><td></td></tr>
<tr><td>Oligo rev</td><td>AAACCCCCGACCCCGTCGGGTTCT</td><td></td><td></td><td></td><td></td><td></td><td></td><td></td><td></td><td></td><td></td></tr>
<tr><td>Chromosome</td><td>start</td><td>end</td><td>strand</td><td>MM</td><td>target_seq</td><td>PAM</td><td>alignment</td><td>distance</td><td>position</td><td>gene name</td><td>gene id</td></tr>
<tr><td>GU305914.1</td><td>132931</td><td>132953</td><td>+</td><td>0</td><td>AGAACCCGACGGGGTCGGGG</td><td>CGG</td><td>||||||||[||||||||||||]PAM</td><td>NA</td><td>Intergenic</td><td>NA</td><td>NA</td></tr>
<tr><td>T10066</td><td>CCCGACGGGGTCGGGGCGGCTGG</td><td>1000</td><td></td><td></td><td></td><td></td><td></td><td></td><td></td><td></td><td></td></tr>
<tr><td>Oligo fwd</td><td>CACCCCCGACGGGGTCGGGGCGGC</td><td></td><td></td><td></td><td></td><td></td><td></td><td></td><td></td><td></td><td></td></tr>
<tr><td>Oligo rev</td><td>AAACGCCGCCCCGACCCCGTCGGG</td><td></td><td></td><td></td><td></td><td></td><td></td><td></td><td></td><td></td><td></td></tr>
<tr><td>Chromosome</td><td>start</td><td>end</td><td>strand</td><td>MM</td><td>target_seq</td><td>PAM</td><td>alignment</td><td>distance</td><td>position</td><td>gene name</td><td>gene id</td></tr>
<tr><td>GU305914.1</td><td>132935</td><td>132957</td><td>+</td><td>0</td><td>CCCGACGGGGTCGGGGCGGC</td><td>TGG</td><td>||||||||[||||||||||||]PAM</td><td>NA</td><td>Intergenic</td><td>NA</td><td>NA</td></tr>
<tr><td>GU305914.1</td><td>48960</td><td>48982</td><td>-</td><td>4</td><td>CATGACGGGGTGGCGGCGGC</td><td>GGG</td><td>|--|||||[|||-|-||||||]PAM</td><td>NA</td><td>Intergenic</td><td>NA</td><td>NA</td></tr>
<tr><td>T10067</td><td>TTACGTATATACATACACACAGG</td><td>1000</td><td></td><td></td><td></td><td></td><td></td><td></td><td></td><td></td><td></td></tr>
<tr><td>Oligo fwd</td><td>CACCTTACGTATATACATACACAC</td><td></td><td></td><td></td><td></td><td></td><td></td><td></td><td></td><td></td><td></td></tr>
</table>

<table>
<tr><td>Oligo rev</td><td>AAACGTGTGTATGTATATACGTAA</td><td></td><td></td><td></td><td></td><td></td><td></td><td></td><td></td><td></td><td></td></tr>
<tr><td>Chromosome</td><td>start</td><td>end</td><td>strand</td><td>MM</td><td>target_seq</td><td>PAM</td><td>alignment</td><td>distance</td><td>position</td><td>gene name</td><td>gene id</td></tr>
<tr><td>GU305914.1</td><td>132958</td><td>132980</td><td>+</td><td>0</td><td>TTACGTATATACATACACAC</td><td>AGG</td><td>||||||||[||||||||||||]PAM</td><td>NA</td><td>Intergenic</td><td>NA</td><td>NA</td></tr>
<tr><td>T10068</td><td>TACGTATATACATACACACAGGG</td><td>1000</td><td></td><td></td><td></td><td></td><td></td><td></td><td></td><td></td><td></td></tr>
<tr><td>Oligo fwd</td><td>CACCTACGTATATACATACACACA</td><td></td><td></td><td></td><td></td><td></td><td></td><td></td><td></td><td></td><td></td></tr>
<tr><td>Oligo rev</td><td>AAACTGTGTGTATGTATATACGTA</td><td></td><td></td><td></td><td></td><td></td><td></td><td></td><td></td><td></td><td></td></tr>
<tr><td>Chromosome</td><td>start</td><td>end</td><td>strand</td><td>MM</td><td>target_seq</td><td>PAM</td><td>alignment</td><td>distance</td><td>position</td><td>gene name</td><td>gene id</td></tr>
<tr><td>GU305914.1</td><td>132959</td><td>132981</td><td>+</td><td>0</td><td>TACGTATATACATACACACA</td><td>GGG</td><td>||||||||[||||||||||||]PAM</td><td>NA</td><td>Intergenic</td><td>NA</td><td>NA</td></tr>
<tr><td>T10069</td><td>ACGACATAAAACATCGAAATCGG</td><td>1000</td><td></td><td></td><td></td><td></td><td></td><td></td><td></td><td></td><td></td></tr>
<tr><td>Oligo fwd</td><td>CACCACGACATAAAACATCGAAAT</td><td></td><td></td><td></td><td></td><td></td><td></td><td></td><td></td><td></td><td></td></tr>
<tr><td>Oligo rev</td><td>AAACATTTCGATGTTTTATGTCGT</td><td></td><td></td><td></td><td></td><td></td><td></td><td></td><td></td><td></td><td></td></tr>
<tr><td>Chromosome</td><td>start</td><td>end</td><td>strand</td><td>MM</td><td>target_seq</td><td>PAM</td><td>alignment</td><td>distance</td><td>position</td><td>gene name</td><td>gene id</td></tr>
<tr><td>GU305914.1</td><td>132984</td><td>133006</td><td>+</td><td>0</td><td>ACGACATAAAACATCGAAAT</td><td>CGG</td><td>||||||||[||||||||||||]PAM</td><td>NA</td><td>Intergenic</td><td>NA</td><td>NA</td></tr>
<tr><td>T10070</td><td>ACATAAAACATCGAAATCGGCGG</td><td>1000</td><td></td><td></td><td></td><td></td><td></td><td></td><td></td><td></td><td></td></tr>
<tr><td>Oligo fwd</td><td>CACCACATAAAACATCGAAATCGG</td><td></td><td></td><td></td><td></td><td></td><td></td><td></td><td></td><td></td><td></td></tr>
<tr><td>Oligo rev</td><td>AAACCCGATTTCGATGTTTTATGT</td><td></td><td></td><td></td><td></td><td></td><td></td><td></td><td></td><td></td><td></td></tr>
<tr><td>Chromosome</td><td>start</td><td>end</td><td>strand</td><td>MM</td><td>target_seq</td><td>PAM</td><td>alignment</td><td>distance</td><td>position</td><td>gene name</td><td>gene id</td></tr>
<tr><td>GU305914.1</td><td>132987</td><td>133009</td><td>+</td><td>0</td><td>ACATAAAACATCGAAATCGG</td><td>CGG</td><td>||||||||[||||||||||||]PAM</td><td>NA</td><td>Intergenic</td><td>NA</td><td>NA</td></tr>
<tr><td>T10071</td><td>CAGTCTTTGTACATGAACCCCGG</td><td>1000</td><td></td><td></td><td></td><td></td><td></td><td></td><td></td><td></td><td></td></tr>
<tr><td>Oligo fwd</td><td>CACCCAGTCTTTGTACATGAACCC</td><td></td><td></td><td></td><td></td><td></td><td></td><td></td><td></td><td></td><td></td></tr>
<tr><td>Oligo rev</td><td>AAACGGGTTCATGTACAAAGACTG</td><td></td><td></td><td></td><td></td><td></td><td></td><td></td><td></td><td></td><td></td></tr>
<tr><td>Chromosome</td><td>start</td><td>end</td><td>strand</td><td>MM</td><td>target_seq</td><td>PAM</td><td>alignment</td><td>distance</td><td>position</td><td>gene name</td><td>gene id</td></tr>
<tr><td>GU305914.1</td><td>133021</td><td>133043</td><td>+</td><td>0</td><td>CAGTCTTTGTACATGAACCC</td><td>CGG</td><td>||||||||[||||||||||||]PAM</td><td>NA</td><td>Intergenic</td><td>NA</td><td>NA</td></tr>
<tr><td>T10072</td><td>AGTCTTTGTACATGAACCCCGGG</td><td>1000</td><td></td><td></td><td></td><td></td><td></td><td></td><td></td><td></td><td></td></tr>
<tr><td>Oligo fwd</td><td>CACCAGTCTTTGTACATGAACCCC</td><td></td><td></td><td></td><td></td><td></td><td></td><td></td><td></td><td></td><td></td></tr>
<tr><td>Oligo rev</td><td>AAACGGGGTTCATGTACAAAGACT</td><td></td><td></td><td></td><td></td><td></td><td></td><td></td><td></td><td></td><td></td></tr>
<tr><td>Chromosome</td><td>start</td><td>end</td><td>strand</td><td>MM</td><td>target_seq</td><td>PAM</td><td>alignment</td><td>distance</td><td>position</td><td>gene name</td><td>gene id</td></tr>
<tr><td>GU305914.1</td><td>133022</td><td>133044</td><td>+</td><td>0</td><td>AGTCTTTGTACATGAACCCC</td><td>GGG</td><td>||||||||[||||||||||||]PAM</td><td>NA</td><td>Intergenic</td><td>NA</td><td>NA</td></tr>
<tr><td>T10073</td><td>CATGAACCCCGGGTTGACCACGG</td><td>1000</td><td></td><td></td><td></td><td></td><td></td><td></td><td></td><td></td><td></td></tr>
<tr><td>Oligo fwd</td><td>CACCCATGAACCCCGGGTTGACCA</td><td></td><td></td><td></td><td></td><td></td><td></td><td></td><td></td><td></td><td></td></tr>
<tr><td>Oligo rev</td><td>AAACTGGTCAACCCGGGGTTCATG</td><td></td><td></td><td></td><td></td><td></td><td></td><td></td><td></td><td></td><td></td></tr>
<tr><td>Chromosome</td><td>start</td><td>end</td><td>strand</td><td>MM</td><td>target_seq</td><td>PAM</td><td>alignment</td><td>distance</td><td>position</td><td>gene name</td><td>gene id</td></tr>
<tr><td>GU305914.1</td><td>133032</td><td>133054</td><td>+</td><td>0</td><td>CATGAACCCCGGGTTGACCA</td><td>CGG</td><td>||||||||[||||||||||||]PAM</td><td>NA</td><td>Intergenic</td><td>NA</td><td>NA</td></tr>
<tr><td>T10074</td><td>CCCCGGGTTGACCACGGCGTAGG</td><td>1000</td><td></td><td></td><td></td><td></td><td></td><td></td><td></td><td></td><td></td></tr>
<tr><td>Oligo fwd</td><td>CACCCCCCGGGTTGACCACGGCGT</td><td></td><td></td><td></td><td></td><td></td><td></td><td></td><td></td><td></td><td></td></tr>
<tr><td>Oligo rev</td><td>AAACACGCCGTGGTCAACCCGGGG</td><td></td><td></td><td></td><td></td><td></td><td></td><td></td><td></td><td></td><td></td></tr>
<tr><td>Chromosome</td><td>start</td><td>end</td><td>strand</td><td>MM</td><td>target_seq</td><td>PAM</td><td>alignment</td><td>distance</td><td>position</td><td>gene name</td><td>gene id</td></tr>
<tr><td>GU305914.1</td><td>133038</td><td>133060</td><td>+</td><td>0</td><td>CCCCGGGTTGACCACGGCGT</td><td>AGG</td><td>||||||||[||||||||||||]PAM</td><td>NA</td><td>Intergenic</td><td>NA</td><td>NA</td></tr>
<tr><td>T10075</td><td>CCCGGGTTGACCACGGCGTAGGG</td><td>1000</td><td></td><td></td><td></td><td></td><td></td><td></td><td></td><td></td><td></td></tr>
<tr><td>Oligo fwd</td><td>CACCCCCGGGTTGACCACGGCGTA</td><td></td><td></td><td></td><td></td><td></td><td></td><td></td><td></td><td></td><td></td></tr>
<tr><td>Oligo rev</td><td>AAACTACGCCGTGGTCAACCCGGG</td><td></td><td></td><td></td><td></td><td></td><td></td><td></td><td></td><td></td><td></td></tr>
<tr><td>Chromosome</td><td>start</td><td>end</td><td>strand</td><td>MM</td><td>target_seq</td><td>PAM</td><td>alignment</td><td>distance</td><td>position</td><td>gene name</td><td>gene id</td></tr>
<tr><td>GU305914.1</td><td>133039</td><td>133061</td><td>+</td><td>0</td><td>CCCGGGTTGACCACGGCGTA</td><td>GGG</td><td>||||||||[||||||||||||]PAM</td><td>NA</td><td>Intergenic</td><td>NA</td><td>NA</td></tr>
<tr><td>T10076</td><td>GTAGGGCTTCTTGATGCTCCCGG</td><td>1000</td><td></td><td></td><td></td><td></td><td></td><td></td><td></td><td></td><td></td></tr>
<tr><td>Oligo fwd</td><td>CACCGTAGGGCTTCTTGATGCTCC</td><td></td><td></td><td></td><td></td><td></td><td></td><td></td><td></td><td></td><td></td></tr>
<tr><td>Oligo rev</td><td>AAACGGAGCATCAAGAAGCCCTAC</td><td></td><td></td><td></td><td></td><td></td><td></td><td></td><td></td><td></td><td></td></tr>
<tr><td>Chromosome</td><td>start</td><td>end</td><td>strand</td><td>MM</td><td>target_seq</td><td>PAM</td><td>alignment</td><td>distance</td><td>position</td><td>gene name</td><td>gene id</td></tr>
<tr><td>GU305914.1</td><td>133056</td><td>133078</td><td>+</td><td>0</td><td>GTAGGGCTTCTTGATGCTCC</td><td>CGG</td><td>||||||||[||||||||||||]PAM</td><td>NA</td><td>Intergenic</td><td>NA</td><td>NA</td></tr>
<tr><td>T10077</td><td>TAGGGCTTCTTGATGCTCCCGGG</td><td>1000</td><td></td><td></td><td></td><td></td><td></td><td></td><td></td><td></td><td></td></tr>
<tr><td>Oligo fwd</td><td>CACCTAGGGCTTCTTGATGCTCCC</td><td></td><td></td><td></td><td></td><td></td><td></td><td></td><td></td><td></td><td></td></tr>
<tr><td>Oligo rev</td><td>AAACGGGAGCATCAAGAAGCCCTA</td><td></td><td></td><td></td><td></td><td></td><td></td><td></td><td></td><td></td><td></td></tr>
<tr><td>Chromosome</td><td>start</td><td>end</td><td>strand</td><td>MM</td><td>target_seq</td><td>PAM</td><td>alignment</td><td>distance</td><td>position</td><td>gene name</td><td>gene id</td></tr>
<tr><td>GU305914.1</td><td>133057</td><td>133079</td><td>+</td><td>0</td><td>TAGGGCTTCTTGATGCTCCC</td><td>GGG</td><td>||||||||[||||||||||||]PAM</td><td>NA</td><td>Intergenic</td><td>NA</td><td>NA</td></tr>
<tr><td>T10078</td><td>GGCTTCTTGATGCTCCCGGGCGG</td><td>1000</td><td></td><td></td><td></td><td></td><td></td><td></td><td></td><td></td><td></td></tr>
<tr><td>Oligo fwd</td><td>CACCGGCTTCTTGATGCTCCCGGG</td><td></td><td></td><td></td><td></td><td></td><td></td><td></td><td></td><td></td><td></td></tr>
<tr><td>Oligo rev</td><td>AAACCCCGGGAGCATCAAGAAGCC</td><td></td><td></td><td></td><td></td><td></td><td></td><td></td><td></td><td></td><td></td></tr>
<tr><td>Chromosome</td><td>start</td><td>end</td><td>strand</td><td>MM</td><td>target_seq</td><td>PAM</td><td>alignment</td><td>distance</td><td>position</td><td>gene name</td><td>gene id</td></tr>
<tr><td>GU305914.1</td><td>133060</td><td>133082</td><td>+</td><td>0</td><td>GGCTTCTTGATGCTCCCGGG</td><td>CGG</td><td>||||||||[||||||||||||]PAM</td><td>NA</td><td>Intergenic</td><td>NA</td><td>NA</td></tr>
<tr><td>T10079</td><td>TAATATTCTTCCGTAAAGATCGG</td><td>1000</td><td></td><td></td><td></td><td></td><td></td><td></td><td></td><td></td><td></td></tr>
<tr><td>Oligo fwd</td><td>CACCTAATATTCTTCCGTAAAGAT</td><td></td><td></td><td></td><td></td><td></td><td></td><td></td><td></td><td></td><td></td></tr>
<tr><td>Oligo rev</td><td>AAACATCTTTACGGAAGAATATTA</td><td></td><td></td><td></td><td></td><td></td><td></td><td></td><td></td><td></td><td></td></tr>
<tr><td>Chromosome</td><td>start</td><td>end</td><td>strand</td><td>MM</td><td>target_seq</td><td>PAM</td><td>alignment</td><td>distance</td><td>position</td><td>gene name</td><td>gene id</td></tr>
<tr><td>GU305914.1</td><td>133090</td><td>133112</td><td>+</td><td>0</td><td>TAATATTCTTCCGTAAAGAT</td><td>CGG</td><td>||||||||[||||||||||||]PAM</td><td>NA</td><td>Intergenic</td><td>NA</td><td>NA</td></tr>
<tr><td>T10080</td><td>TCCGTAAAGATCGGCTTGCGCGG</td><td>1000</td><td></td><td></td><td></td><td></td><td></td><td></td><td></td><td></td><td></td></tr>
<tr><td>Oligo fwd</td><td>CACCTCCGTAAAGATCGGCTTGCG</td><td></td><td></td><td></td><td></td><td></td><td></td><td></td><td></td><td></td><td></td></tr>
<tr><td>Oligo rev</td><td>AAACCGCAAGCCGATCTTTACGGA</td><td></td><td></td><td></td><td></td><td></td><td></td><td></td><td></td><td></td><td></td></tr>
<tr><td>Chromosome</td><td>start</td><td>end</td><td>strand</td><td>MM</td><td>target_seq</td><td>PAM</td><td>alignment</td><td>distance</td><td>position</td><td>gene name</td><td>gene id</td></tr>
<tr><td>GU305914.1</td><td>133099</td><td>133121</td><td>+</td><td>0</td><td>TCCGTAAAGATCGGCTTGCG</td><td>CGG</td><td>||||||||[||||||||||||]PAM</td><td>NA</td><td>Intergenic</td><td>NA</td><td>NA</td></tr>
<tr><td>T10081</td><td>AGATCGGCTTGCGCGGCCAGAGG</td><td>1000</td><td></td><td></td><td></td><td></td><td></td><td></td><td></td><td></td><td></td></tr>
<tr><td>Oligo fwd</td><td>CACCAGATCGGCTTGCGCGGCCAG</td><td></td><td></td><td></td><td></td><td></td><td></td><td></td><td></td><td></td><td></td></tr>
<tr><td>Oligo rev</td><td>AAACCTGGCCGCGCAAGCCGATCT</td><td></td><td></td><td></td><td></td><td></td><td></td><td></td><td></td><td></td><td></td></tr>
<tr><td>Chromosome</td><td>start</td><td>end</td><td>strand</td><td>MM</td><td>target_seq</td><td>PAM</td><td>alignment</td><td>distance</td><td>position</td><td>gene name</td><td>gene id</td></tr>
<tr><td>GU305914.1</td><td>133106</td><td>133128</td><td>+</td><td>0</td><td>AGATCGGCTTGCGCGGCCAG</td><td>AGG</td><td>||||||||[||||||||||||]PAM</td><td>NA</td><td>Intergenic</td><td>NA</td><td>NA</td></tr>
<tr><td>T10082</td><td>GATCGGCTTGCGCGGCCAGAGGG</td><td>1000</td><td></td><td></td><td></td><td></td><td></td><td></td><td></td><td></td><td></td></tr>
</table>

| | | | | | | | | | | | |
|---|---|---|---|---|---|---|---|---|---|---|---|
| Oligo fwd | CACCGATCGGCTTGCGCGGCCAGA | | | | | | | | | | |
| Oligo rev | AAACTCTGGCCGCGCAAGCCGATC | | | | | | | | | | |
| Chromosome | start | end | strand | MM | target_seq | PAM | alignment | distance | position | gene name | gene id |
| GU305914.1 | 133107 | 133129 | + | 0 | GATCGGCTTGCGCGGCCAGA | GGG | \|\|\|\|\|\|\|\|[\|\|\|\|\|\|\|\|\|\|\|\|]PAM | NA | Intergenic | NA | NA |
| GU305914.1 | 100868 | 100890 | + | 4 | GTTCAGCTTGCGCAGCGAGA | TGG | \|-\|\|-\|\|\|[\|\|\|\|\|-\|\|-\|\|\|]PAM | NA | Intergenic | NA | NA |
| GU305914.1 | 141643 | 141665 | - | 4 | GAACTGCTTGCGCGGCTCGA | TGG | \|\|-\|-\|\|\|[\|\|\|\|\|\|\|\|--\|\|]PAM | NA | Intergenic | NA | NA |
| T10083 | GTCTCTGTTCTTTTTCTTCTTGG | 1000 | | | | | | | | | |
| Oligo fwd | CACCGTCTCTGTTCTTTTTCTTCT | | | | | | | | | | |
| Oligo rev | AAACAGAAGAAAAAGAACAGAGAC | | | | | | | | | | |
| Chromosome | start | end | strand | MM | target_seq | PAM | alignment | distance | position | gene name | gene id |
| GU305914.1 | 133140 | 133162 | + | 0 | GTCTCTGTTCTTTTTCTTCT | TGG | \|\|\|\|\|\|\|\|[\|\|\|\|\|\|\|\|\|\|\|\|]PAM | NA | Intergenic | NA | NA |
| T10084 | TCTCTGTTCTTTTTCTTCTTGGG | 1000 | | | | | | | | | |
| Oligo fwd | CACCTCTCTGTTCTTTTTCTTCTT | | | | | | | | | | |
| Oligo rev | AAACAAGAAGAAAAAGAACAGAGA | | | | | | | | | | |
| Chromosome | start | end | strand | MM | target_seq | PAM | alignment | distance | position | gene name | gene id |
| GU305914.1 | 133141 | 133163 | + | 0 | TCTCTGTTCTTTTTCTTCTT | GGG | \|\|\|\|\|\|\|\|[\|\|\|\|\|\|\|\|\|\|\|\|]PAM | NA | Intergenic | NA | NA |
| T10085 | ATGCGTTCGCCGACTTCTTCTGG | 1000 | | | | | | | | | |
| Oligo fwd | CACCATGCGTTCGCCGACTTCTTC | | | | | | | | | | |
| Oligo rev | AAACGAAGAAGTCGGCGAACGCAT | | | | | | | | | | |
| Chromosome | start | end | strand | MM | target_seq | PAM | alignment | distance | position | gene name | gene id |
| GU305914.1 | 133166 | 133188 | + | 0 | ATGCGTTCGCCGACTTCTTC | TGG | \|\|\|\|\|\|\|\|[\|\|\|\|\|\|\|\|\|\|\|\|]PAM | NA | Intergenic | NA | NA |
| T10086 | GCCGACTTCTTCTGGTAGTACGG | 1000 | | | | | | | | | |
| Oligo fwd | CACCGCCGACTTCTTCTGGTAGTA | | | | | | | | | | |
| Oligo rev | AAACTACTACCAGAAGAAGTCGGC | | | | | | | | | | |
| Chromosome | start | end | strand | MM | target_seq | PAM | alignment | distance | position | gene name | gene id |
| GU305914.1 | 133174 | 133196 | + | 0 | GCCGACTTCTTCTGGTAGTA | CGG | \|\|\|\|\|\|\|\|[\|\|\|\|\|\|\|\|\|\|\|\|]PAM | NA | Intergenic | NA | NA |
| T10087 | CCGACTTCTTCTGGTAGTACGGG | 1000 | | | | | | | | | |
| Oligo fwd | CACCCCGACTTCTTCTGGTAGTAC | | | | | | | | | | |
| Oligo rev | AAACGTACTACCAGAAGAAGTCGG | | | | | | | | | | |
| Chromosome | start | end | strand | MM | target_seq | PAM | alignment | distance | position | gene name | gene id |
| GU305914.1 | 133175 | 133197 | + | 0 | CCGACTTCTTCTGGTAGTAC | GGG | \|\|\|\|\|\|\|\|[\|\|\|\|\|\|\|\|\|\|\|\|]PAM | NA | Intergenic | NA | NA |
| T10088 | CTGGTAGTACGGGCGCAGTATGG | 1000 | | | | | | | | | |
| Oligo fwd | CACCCTGGTAGTACGGGCGCAGTA | | | | | | | | | | |
| Oligo rev | AAACTACTGCGCCCGTACTACCAG | | | | | | | | | | |
| Chromosome | start | end | strand | MM | target_seq | PAM | alignment | distance | position | gene name | gene id |
| GU305914.1 | 133185 | 133207 | + | 0 | CTGGTAGTACGGGCGCAGTA | TGG | \|\|\|\|\|\|\|\|[\|\|\|\|\|\|\|\|\|\|\|\|]PAM | NA | Intergenic | NA | NA |
| T10089 | ACGGGCGCAGTATGGCGTACAGG | 1000 | | | | | | | | | |
| Oligo fwd | CACCACGGGCGCAGTATGGCGTAC | | | | | | | | | | |
| Oligo rev | AAACGTACGCCATACTGCGCCCGT | | | | | | | | | | |
| Chromosome | start | end | strand | MM | target_seq | PAM | alignment | distance | position | gene name | gene id |
| GU305914.1 | 133193 | 133215 | + | 0 | ACGGGCGCAGTATGGCGTAC | AGG | \|\|\|\|\|\|\|\|[\|\|\|\|\|\|\|\|\|\|\|\|]PAM | NA | Intergenic | NA | NA |
| T10090 | TGGCGTACAGGCACACGATCAGG | 1000 | | | | | | | | | |
| Oligo fwd | CACCTGGCGTACAGGCACACGATC | | | | | | | | | | |
| Oligo rev | AAACGATCGTGTGCCTGTACGCCA | | | | | | | | | | |
| Chromosome | start | end | strand | MM | target_seq | PAM | alignment | distance | position | gene name | gene id |
| GU305914.1 | 133205 | 133227 | + | 0 | TGGCGTACAGGCACACGATC | AGG | \|\|\|\|\|\|\|\|[\|\|\|\|\|\|\|\|\|\|\|\|]PAM | NA | Intergenic | NA | NA |
| T10091 | GGCCCGTGCCGATGCAGAAGAGG | 1000 | | | | | | | | | |
| Oligo fwd | CACCGGCCCGTGCCGATGCAGAAG | | | | | | | | | | |
| Oligo rev | AAACCTTCTGCATCGGCACGGGCC | | | | | | | | | | |
| Chromosome | start | end | strand | MM | target_seq | PAM | alignment | distance | position | gene name | gene id |
| GU305914.1 | 133226 | 133248 | + | 0 | GGCCCGTGCCGATGCAGAAG | AGG | \|\|\|\|\|\|\|\|[\|\|\|\|\|\|\|\|\|\|\|\|]PAM | NA | Intergenic | NA | NA |
| T10092 | CGTGCCGATGCAGAAGAGGAAGG | 1000 | | | | | | | | | |
| Oligo fwd | CACCCGTGCCGATGCAGAAGAGGA | | | | | | | | | | |
| Oligo rev | AAACTCCTCTTCTGCATCGGCACG | | | | | | | | | | |
| Chromosome | start | end | strand | MM | target_seq | PAM | alignment | distance | position | gene name | gene id |
| GU305914.1 | 133230 | 133252 | + | 0 | CGTGCCGATGCAGAAGAGGA | AGG | \|\|\|\|\|\|\|\|[\|\|\|\|\|\|\|\|\|\|\|\|]PAM | NA | Intergenic | NA | NA |
| GU305914.1 | 79382 | 79404 | + | 3 | CGTGCTGATGCTGCAGAGGA | AGG | \|\|\|\|\|-\|\|[\|\|\|-\|-\|\|\|\|\|\|]PAM | NA | Intergenic | NA | NA |
| T10093 | GGAAGGCGATCAGCACGTGCAGG | 1000 | | | | | | | | | |
| Oligo fwd | CACCGGAAGGCGATCAGCACGTGC | | | | | | | | | | |
| Oligo rev | AAACGCACGTGCTGATCGCCTTCC | | | | | | | | | | |
| Chromosome | start | end | strand | MM | target_seq | PAM | alignment | distance | position | gene name | gene id |
| GU305914.1 | 133247 | 133269 | + | 0 | GGAAGGCGATCAGCACGTGC | AGG | \|\|\|\|\|\|\|\|[\|\|\|\|\|\|\|\|\|\|\|\|]PAM | NA | Intergenic | NA | NA |
| T10094 | GGCGATCAGCACGTGCAGGAAGG | 1000 | | | | | | | | | |
| Oligo fwd | CACCGGCGATCAGCACGTGCAGGA | | | | | | | | | | |
| Oligo rev | AAACTCCTGCACGTGCTGATCGCC | | | | | | | | | | |
| Chromosome | start | end | strand | MM | target_seq | PAM | alignment | distance | position | gene name | gene id |
| GU305914.1 | 133251 | 133273 | + | 0 | GGCGATCAGCACGTGCAGGA | AGG | \|\|\|\|\|\|\|\|[\|\|\|\|\|\|\|\|\|\|\|\|]PAM | NA | Intergenic | NA | NA |
| T10095 | GCAGGAAGGTGTAAGCCTCCTGG | 1000 | | | | | | | | | |
| Oligo fwd | CACCGCAGGAAGGTGTAAGCCTCC | | | | | | | | | | |
| Oligo rev | AAACGGAGGCTTACACCTTCCTGC | | | | | | | | | | |
| Chromosome | start | end | strand | MM | target_seq | PAM | alignment | distance | position | gene name | gene id |
| GU305914.1 | 133265 | 133287 | + | 0 | GCAGGAAGGTGTAAGCCTCC | TGG | \|\|\|\|\|\|\|\|[\|\|\|\|\|\|\|\|\|\|\|\|]PAM | NA | Intergenic | NA | NA |
| T10096 | CAGGAAGGTGTAAGCCTCCTGGG | 1000 | | | | | | | | | |
| Oligo fwd | CACCCAGGAAGGTGTAAGCCTCCT | | | | | | | | | | |
| Oligo rev | AAACAGGAGGCTTACACCTTCCTG | | | | | | | | | | |

<table>
<tr><td>Chromosome</td><td>start</td><td>end</td><td>strand</td><td>MM</td><td>target_seq</td><td>PAM</td><td>alignment</td><td>distance</td><td>position</td><td>gene name</td><td>gene id</td></tr>
<tr><td>GU305914.1</td><td>133266</td><td>133288</td><td>+</td><td>0</td><td>CAGGAAGGTGTAAGCCTCCT</td><td>GGG</td><td>||||||||[||||||||||||]PAM</td><td>NA</td><td>Intergenic</td><td>NA</td><td>NA</td></tr>
<tr><td>T10097</td><td>AGGAAGGTGTAAGCCTCCTGGGG</td><td>1000</td><td></td><td></td><td></td><td></td><td></td><td></td><td></td><td></td><td></td></tr>
<tr><td>Oligo fwd</td><td>CACCAGGAAGGTGTAAGCCTCCTG</td><td></td><td></td><td></td><td></td><td></td><td></td><td></td><td></td><td></td><td></td></tr>
<tr><td>Oligo rev</td><td>AAACCAGGAGGCTTACACCTTCCT</td><td></td><td></td><td></td><td></td><td></td><td></td><td></td><td></td><td></td><td></td></tr>
<tr><td>Chromosome</td><td>start</td><td>end</td><td>strand</td><td>MM</td><td>target_seq</td><td>PAM</td><td>alignment</td><td>distance</td><td>position</td><td>gene name</td><td>gene id</td></tr>
<tr><td>GU305914.1</td><td>133267</td><td>133289</td><td>+</td><td>0</td><td>AGGAAGGTGTAAGCCTCCTG</td><td>GGG</td><td>||||||||[||||||||||||]PAM</td><td>NA</td><td>Intergenic</td><td>NA</td><td>NA</td></tr>
<tr><td>T10098</td><td>CTCCTGGGGTTCTCTGCCCACGG</td><td>1000</td><td></td><td></td><td></td><td></td><td></td><td></td><td></td><td></td><td></td></tr>
<tr><td>Oligo fwd</td><td>CACCCTCCTGGGGTTCTCTGCCCA</td><td></td><td></td><td></td><td></td><td></td><td></td><td></td><td></td><td></td><td></td></tr>
<tr><td>Oligo rev</td><td>AAACTGGGCAGAGAACCCCAGGAG</td><td></td><td></td><td></td><td></td><td></td><td></td><td></td><td></td><td></td><td></td></tr>
<tr><td>Chromosome</td><td>start</td><td>end</td><td>strand</td><td>MM</td><td>target_seq</td><td>PAM</td><td>alignment</td><td>distance</td><td>position</td><td>gene name</td><td>gene id</td></tr>
<tr><td>GU305914.1</td><td>133281</td><td>133303</td><td>+</td><td>0</td><td>CTCCTGGGGTTCTCTGCCCA</td><td>CGG</td><td>||||||||[||||||||||||]PAM</td><td>NA</td><td>Intergenic</td><td>NA</td><td>NA</td></tr>
<tr><td>T10099</td><td>TTCTCTGCCCACGGTTCCCTCGG</td><td>1000</td><td></td><td></td><td></td><td></td><td></td><td></td><td></td><td></td><td></td></tr>
<tr><td>Oligo fwd</td><td>CACCTTCTCTGCCCACGGTTCCCT</td><td></td><td></td><td></td><td></td><td></td><td></td><td></td><td></td><td></td><td></td></tr>
<tr><td>Oligo rev</td><td>AAACAGGGAACCGTGGGCAGAGAA</td><td></td><td></td><td></td><td></td><td></td><td></td><td></td><td></td><td></td><td></td></tr>
<tr><td>Chromosome</td><td>start</td><td>end</td><td>strand</td><td>MM</td><td>target_seq</td><td>PAM</td><td>alignment</td><td>distance</td><td>position</td><td>gene name</td><td>gene id</td></tr>
<tr><td>GU305914.1</td><td>133290</td><td>133312</td><td>+</td><td>0</td><td>TTCTCTGCCCACGGTTCCCT</td><td>CGG</td><td>||||||||[||||||||||||]PAM</td><td>NA</td><td>Intergenic</td><td>NA</td><td>NA</td></tr>
<tr><td>T10100</td><td>CCACGGTTCCCTCGGCGAGCTGG</td><td>1000</td><td></td><td></td><td></td><td></td><td></td><td></td><td></td><td></td><td></td></tr>
<tr><td>Oligo fwd</td><td>CACCCCACGGTTCCCTCGGCGAGC</td><td></td><td></td><td></td><td></td><td></td><td></td><td></td><td></td><td></td><td></td></tr>
<tr><td>Oligo rev</td><td>AAACGCTCGCCGAGGGAACCGTGG</td><td></td><td></td><td></td><td></td><td></td><td></td><td></td><td></td><td></td><td></td></tr>
<tr><td>Chromosome</td><td>start</td><td>end</td><td>strand</td><td>MM</td><td>target_seq</td><td>PAM</td><td>alignment</td><td>distance</td><td>position</td><td>gene name</td><td>gene id</td></tr>
<tr><td>GU305914.1</td><td>133298</td><td>133320</td><td>+</td><td>0</td><td>CCACGGTTCCCTCGGCGAGC</td><td>TGG</td><td>||||||||[||||||||||||]PAM</td><td>NA</td><td>Intergenic</td><td>NA</td><td>NA</td></tr>
<tr><td>T10101</td><td>TCCCTCGGCGAGCTGGACCTCGG</td><td>1000</td><td></td><td></td><td></td><td></td><td></td><td></td><td></td><td></td><td></td></tr>
<tr><td>Oligo fwd</td><td>CACCTCCCTCGGCGAGCTGGACCT</td><td></td><td></td><td></td><td></td><td></td><td></td><td></td><td></td><td></td><td></td></tr>
<tr><td>Oligo rev</td><td>AAACAGGTCCAGCTCGCCGAGGGA</td><td></td><td></td><td></td><td></td><td></td><td></td><td></td><td></td><td></td><td></td></tr>
<tr><td>Chromosome</td><td>start</td><td>end</td><td>strand</td><td>MM</td><td>target_seq</td><td>PAM</td><td>alignment</td><td>distance</td><td>position</td><td>gene name</td><td>gene id</td></tr>
<tr><td>GU305914.1</td><td>133305</td><td>133327</td><td>+</td><td>0</td><td>TCCCTCGGCGAGCTGGACCT</td><td>CGG</td><td>||||||||[||||||||||||]PAM</td><td>NA</td><td>Intergenic</td><td>NA</td><td>NA</td></tr>
<tr><td>T10102</td><td>CCCTCGGCGAGCTGGACCTCGGG</td><td>1000</td><td></td><td></td><td></td><td></td><td></td><td></td><td></td><td></td><td></td></tr>
<tr><td>Oligo fwd</td><td>CACCCCCTCGGCGAGCTGGACCTC</td><td></td><td></td><td></td><td></td><td></td><td></td><td></td><td></td><td></td><td></td></tr>
<tr><td>Oligo rev</td><td>AAACGAGGTCCAGCTCGCCGAGGG</td><td></td><td></td><td></td><td></td><td></td><td></td><td></td><td></td><td></td><td></td></tr>
<tr><td>Chromosome</td><td>start</td><td>end</td><td>strand</td><td>MM</td><td>target_seq</td><td>PAM</td><td>alignment</td><td>distance</td><td>position</td><td>gene name</td><td>gene id</td></tr>
<tr><td>GU305914.1</td><td>133306</td><td>133328</td><td>+</td><td>0</td><td>CCCTCGGCGAGCTGGACCTC</td><td>GGG</td><td>||||||||[||||||||||||]PAM</td><td>NA</td><td>Intergenic</td><td>NA</td><td>NA</td></tr>
<tr><td>T10103</td><td>GGCGAGCTGGACCTCGGGATCGG</td><td>1000</td><td></td><td></td><td></td><td></td><td></td><td></td><td></td><td></td><td></td></tr>
<tr><td>Oligo fwd</td><td>CACCGGCGAGCTGGACCTCGGGAT</td><td></td><td></td><td></td><td></td><td></td><td></td><td></td><td></td><td></td><td></td></tr>
<tr><td>Oligo rev</td><td>AAACATCCCGAGGTCCAGCTCGCC</td><td></td><td></td><td></td><td></td><td></td><td></td><td></td><td></td><td></td><td></td></tr>
<tr><td>Chromosome</td><td>start</td><td>end</td><td>strand</td><td>MM</td><td>target_seq</td><td>PAM</td><td>alignment</td><td>distance</td><td>position</td><td>gene name</td><td>gene id</td></tr>
<tr><td>GU305914.1</td><td>133311</td><td>133333</td><td>+</td><td>0</td><td>GGCGAGCTGGACCTCGGGAT</td><td>CGG</td><td>||||||||[||||||||||||]PAM</td><td>NA</td><td>Intergenic</td><td>NA</td><td>NA</td></tr>
<tr><td>T10104</td><td>CCTCGGGATCGGCCGCGCGACGG</td><td>1000</td><td></td><td></td><td></td><td></td><td></td><td></td><td></td><td></td><td></td></tr>
<tr><td>Oligo fwd</td><td>CACCCCTCGGGATCGGCCGCGCGA</td><td></td><td></td><td></td><td></td><td></td><td></td><td></td><td></td><td></td><td></td></tr>
<tr><td>Oligo rev</td><td>AAACTCGCGCGGCCGATCCCGAGG</td><td></td><td></td><td></td><td></td><td></td><td></td><td></td><td></td><td></td><td></td></tr>
<tr><td>Chromosome</td><td>start</td><td>end</td><td>strand</td><td>MM</td><td>target_seq</td><td>PAM</td><td>alignment</td><td>distance</td><td>position</td><td>gene name</td><td>gene id</td></tr>
<tr><td>GU305914.1</td><td>133322</td><td>133344</td><td>+</td><td>0</td><td>CCTCGGGATCGGCCGCGCGA</td><td>CGG</td><td>||||||||[||||||||||||]PAM</td><td>NA</td><td>Intergenic</td><td>NA</td><td>NA</td></tr>
<tr><td>T10105</td><td>CGGCCGCGCGACGGTAAGTGAGG</td><td>1000</td><td></td><td></td><td></td><td></td><td></td><td></td><td></td><td></td><td></td></tr>
<tr><td>Oligo fwd</td><td>CACCCGGCCGCGCGACGGTAAGTG</td><td></td><td></td><td></td><td></td><td></td><td></td><td></td><td></td><td></td><td></td></tr>
<tr><td>Oligo rev</td><td>AAACCACTTACCGTCGCGCGGCCG</td><td></td><td></td><td></td><td></td><td></td><td></td><td></td><td></td><td></td><td></td></tr>
<tr><td>Chromosome</td><td>start</td><td>end</td><td>strand</td><td>MM</td><td>target_seq</td><td>PAM</td><td>alignment</td><td>distance</td><td>position</td><td>gene name</td><td>gene id</td></tr>
<tr><td>GU305914.1</td><td>133331</td><td>133353</td><td>+</td><td>0</td><td>CGGCCGCGCGACGGTAAGTG</td><td>AGG</td><td>||||||||[||||||||||||]PAM</td><td>NA</td><td>Intergenic</td><td>NA</td><td>NA</td></tr>
<tr><td>T10106</td><td>CGCGCGACGGTAAGTGAGGCAGG</td><td>1000</td><td></td><td></td><td></td><td></td><td></td><td></td><td></td><td></td><td></td></tr>
<tr><td>Oligo fwd</td><td>CACCCGCGCGACGGTAAGTGAGGC</td><td></td><td></td><td></td><td></td><td></td><td></td><td></td><td></td><td></td><td></td></tr>
<tr><td>Oligo rev</td><td>AAACGCCTCACTTACCGTCGCGCG</td><td></td><td></td><td></td><td></td><td></td><td></td><td></td><td></td><td></td><td></td></tr>
<tr><td>Chromosome</td><td>start</td><td>end</td><td>strand</td><td>MM</td><td>target_seq</td><td>PAM</td><td>alignment</td><td>distance</td><td>position</td><td>gene name</td><td>gene id</td></tr>
<tr><td>GU305914.1</td><td>133335</td><td>133357</td><td>+</td><td>0</td><td>CGCGCGACGGTAAGTGAGGC</td><td>AGG</td><td>||||||||[||||||||||||]PAM</td><td>NA</td><td>Intergenic</td><td>NA</td><td>NA</td></tr>
<tr><td>T10107</td><td>AAGTGAGGCAGGTTTTGCTCAGG</td><td>1000</td><td></td><td></td><td></td><td></td><td></td><td></td><td></td><td></td><td></td></tr>
<tr><td>Oligo fwd</td><td>CACCAAGTGAGGCAGGTTTTGCTC</td><td></td><td></td><td></td><td></td><td></td><td></td><td></td><td></td><td></td><td></td></tr>
<tr><td>Oligo rev</td><td>AAACGAGCAAAACCTGCCTCACTT</td><td></td><td></td><td></td><td></td><td></td><td></td><td></td><td></td><td></td><td></td></tr>
<tr><td>Chromosome</td><td>start</td><td>end</td><td>strand</td><td>MM</td><td>target_seq</td><td>PAM</td><td>alignment</td><td>distance</td><td>position</td><td>gene name</td><td>gene id</td></tr>
<tr><td>GU305914.1</td><td>133346</td><td>133368</td><td>+</td><td>0</td><td>AAGTGAGGCAGGTTTTGCTC</td><td>AGG</td><td>||||||||[||||||||||||]PAM</td><td>NA</td><td>Intergenic</td><td>NA</td><td>NA</td></tr>
<tr><td>T10108</td><td>GGCAGGTTTTGCTCAGGATACGG</td><td>1000</td><td></td><td></td><td></td><td></td><td></td><td></td><td></td><td></td><td></td></tr>
<tr><td>Oligo fwd</td><td>CACCGGCAGGTTTTGCTCAGGATA</td><td></td><td></td><td></td><td></td><td></td><td></td><td></td><td></td><td></td><td></td></tr>
<tr><td>Oligo rev</td><td>AAACTATCCTGAGCAAAACCTGCC</td><td></td><td></td><td></td><td></td><td></td><td></td><td></td><td></td><td></td><td></td></tr>
<tr><td>Chromosome</td><td>start</td><td>end</td><td>strand</td><td>MM</td><td>target_seq</td><td>PAM</td><td>alignment</td><td>distance</td><td>position</td><td>gene name</td><td>gene id</td></tr>
<tr><td>GU305914.1</td><td>133352</td><td>133374</td><td>+</td><td>0</td><td>GGCAGGTTTTGCTCAGGATA</td><td>CGG</td><td>||||||||[||||||||||||]PAM</td><td>NA</td><td>Intergenic</td><td>NA</td><td>NA</td></tr>
<tr><td>T10109</td><td>GTGAGAACGCTTTGTGATGATGG</td><td>1000</td><td></td><td></td><td></td><td></td><td></td><td></td><td></td><td></td><td></td></tr>
<tr><td>Oligo fwd</td><td>CACCGTGAGAACGCTTTGTGATGA</td><td></td><td></td><td></td><td></td><td></td><td></td><td></td><td></td><td></td><td></td></tr>
<tr><td>Oligo rev</td><td>AAACTCATCACAAAGCGTTCTCAC</td><td></td><td></td><td></td><td></td><td></td><td></td><td></td><td></td><td></td><td></td></tr>
<tr><td>Chromosome</td><td>start</td><td>end</td><td>strand</td><td>MM</td><td>target_seq</td><td>PAM</td><td>alignment</td><td>distance</td><td>position</td><td>gene name</td><td>gene id</td></tr>
<tr><td>GU305914.1</td><td>133386</td><td>133408</td><td>+</td><td>0</td><td>GTGAGAACGCTTTGTGATGA</td><td>TGG</td><td>||||||||[||||||||||||]PAM</td><td>NA</td><td>Intergenic</td><td>NA</td><td>NA</td></tr>
<tr><td>T10110</td><td>TGAGAACGCTTTGTGATGATGGG</td><td>1000</td><td></td><td></td><td></td><td></td><td></td><td></td><td></td><td></td><td></td></tr>
<tr><td>Oligo fwd</td><td>CACCTGAGAACGCTTTGTGATGAT</td><td></td><td></td><td></td><td></td><td></td><td></td><td></td><td></td><td></td><td></td></tr>
<tr><td>Oligo rev</td><td>AAACATCATCACAAAGCGTTCTCA</td><td></td><td></td><td></td><td></td><td></td><td></td><td></td><td></td><td></td><td></td></tr>
<tr><td>Chromosome</td><td>start</td><td>end</td><td>strand</td><td>MM</td><td>target_seq</td><td>PAM</td><td>alignment</td><td>distance</td><td>position</td><td>gene name</td><td>gene id</td></tr>
<tr><td>GU305914.1</td><td>133387</td><td>133409</td><td>+</td><td>0</td><td>TGAGAACGCTTTGTGATGAT</td><td>GGG</td><td>||||||||[||||||||||||]PAM</td><td>NA</td><td>Intergenic</td><td>NA</td><td>NA</td></tr>
<tr><td>T10111</td><td>ACTGTCGTTTTGAAAACATCTGG</td><td>1000</td><td></td><td></td><td></td><td></td><td></td><td></td><td></td><td></td><td></td></tr>
<tr><td>Oligo fwd</td><td>CACCACTGTCGTTTTGAAAACATC</td><td></td><td></td><td></td><td></td><td></td><td></td><td></td><td></td><td></td><td></td></tr>
</table>

| | | | | | | | | | | | |
|---|---|---|---|---|---|---|---|---|---|---|---|
| Oligo rev | AAACGATGTTTTCAAAACGACAGT | | | | | | | | | | |
| Chromosome | start | end | strand | MM | target_seq | PAM | alignment | distance | position | gene name | gene id |
| GU305914.1 | 133427 | 133449 | + | 0 | ACTGTCGTTTTGAAAACATC | TGG | \|\|\|\|\|\|\|\|[\|\|\|\|\|\|\|\|\|\|\|\|]PAM | NA | Intergenic | NA | NA |
| GU305914.1 | 94667 | 94689 | + | 4 | CCTGTGGCTTTGAAAAGATC | AGG | -\|\|\|\|-\|-[\|\|\|\|\|\|\|\|-\|\|\|]PAM | NA | Intergenic | NA | NA |
| T10112 | GTCGTTTTGAAAACATCTGGAGG | 1000 | | | | | | | | | |
| Oligo fwd | CACCGTCGTTTTGAAAACATCTGG | | | | | | | | | | |
| Oligo rev | AAACCCAGATGTTTTCAAAACGAC | | | | | | | | | | |
| Chromosome | start | end | strand | MM | target_seq | PAM | alignment | distance | position | gene name | gene id |
| GU305914.1 | 133430 | 133452 | + | 0 | GTCGTTTTGAAAACATCTGG | AGG | \|\|\|\|\|\|\|\|[\|\|\|\|\|\|\|\|\|\|\|\|]PAM | NA | Intergenic | NA | NA |
| GU305914.1 | 94670 | 94692 | + | 4 | GTGGCTTTGAAAAGATCAGG | CGG | \|\|-\|-\|\|\|[\|\|\|\|\|-\|\|\|-\|\|]PAM | NA | Intergenic | NA | NA |
| T10113 | GAAAACATCTGGAGGTAGAGCGG | 1000 | | | | | | | | | |
| Oligo fwd | CACCGAAAACATCTGGAGGTAGAG | | | | | | | | | | |
| Oligo rev | AAACCTCTACCTCCAGATGTTTTC | | | | | | | | | | |
| Chromosome | start | end | strand | MM | target_seq | PAM | alignment | distance | position | gene name | gene id |
| GU305914.1 | 133438 | 133460 | + | 0 | GAAAACATCTGGAGGTAGAG | CGG | \|\|\|\|\|\|\|\|[\|\|\|\|\|\|\|\|\|\|\|\|]PAM | NA | Intergenic | NA | NA |
| T10114 | CTGGAGGTAGAGCGGTTGTGTGG | 1000 | | | | | | | | | |
| Oligo fwd | CACCCTGGAGGTAGAGCGGTTGTG | | | | | | | | | | |
| Oligo rev | AAACCACAACCGCTCTACCTCCAG | | | | | | | | | | |
| Chromosome | start | end | strand | MM | target_seq | PAM | alignment | distance | position | gene name | gene id |
| GU305914.1 | 133446 | 133468 | + | 0 | CTGGAGGTAGAGCGGTTGTG | TGG | \|\|\|\|\|\|\|\|[\|\|\|\|\|\|\|\|\|\|\|\|]PAM | NA | Intergenic | NA | NA |
| T10115 | GTGTGGCGTTAGTGTTGTCGAGG | 1000 | | | | | | | | | |
| Oligo fwd | CACCGTGTGGCGTTAGTGTTGTCG | | | | | | | | | | |
| Oligo rev | AAACCGACAACACTAACGCCACAC | | | | | | | | | | |
| Chromosome | start | end | strand | MM | target_seq | PAM | alignment | distance | position | gene name | gene id |
| GU305914.1 | 133463 | 133485 | + | 0 | GTGTGGCGTTAGTGTTGTCG | AGG | \|\|\|\|\|\|\|\|[\|\|\|\|\|\|\|\|\|\|\|\|]PAM | NA | Intergenic | NA | NA |
| T10116 | GTTGTCGAGGTCGATTCTCATGG | 1000 | | | | | | | | | |
| Oligo fwd | CACCGTTGTCGAGGTCGATTCTCA | | | | | | | | | | |
| Oligo rev | AAACTGAGAATCGACCTCGACAAC | | | | | | | | | | |
| Chromosome | start | end | strand | MM | target_seq | PAM | alignment | distance | position | gene name | gene id |
| GU305914.1 | 133476 | 133498 | + | 0 | GTTGTCGAGGTCGATTCTCA | TGG | \|\|\|\|\|\|\|\|[\|\|\|\|\|\|\|\|\|\|\|\|]PAM | NA | Intergenic | NA | NA |
| T10117 | GTCGAGGTCGATTCTCATGGCGG | 1000 | | | | | | | | | |
| Oligo fwd | CACCGTCGAGGTCGATTCTCATGG | | | | | | | | | | |
| Oligo rev | AAACCCATGAGAATCGACCTCGAC | | | | | | | | | | |
| Chromosome | start | end | strand | MM | target_seq | PAM | alignment | distance | position | gene name | gene id |
| GU305914.1 | 133479 | 133501 | + | 0 | GTCGAGGTCGATTCTCATGG | CGG | \|\|\|\|\|\|\|\|[\|\|\|\|\|\|\|\|\|\|\|\|]PAM | NA | Intergenic | NA | NA |
| T10118 | ATTCTCATGGCGGTGAATCTCGG | 1000 | | | | | | | | | |
| Oligo fwd | CACCATTCTCATGGCGGTGAATCT | | | | | | | | | | |
| Oligo rev | AAACAGATTCACCGCCATGAGAAT | | | | | | | | | | |
| Chromosome | start | end | strand | MM | target_seq | PAM | alignment | distance | position | gene name | gene id |
| GU305914.1 | 133489 | 133511 | + | 0 | ATTCTCATGGCGGTGAATCT | CGG | \|\|\|\|\|\|\|\|[\|\|\|\|\|\|\|\|\|\|\|\|]PAM | NA | Intergenic | NA | NA |
| T10119 | CTCATGGCGGTGAATCTCGGTGG | 1000 | | | | | | | | | |
| Oligo fwd | CACCCTCATGGCGGTGAATCTCGG | | | | | | | | | | |
| Oligo rev | AAACCCGAGATTCACCGCCATGAG | | | | | | | | | | |
| Chromosome | start | end | strand | MM | target_seq | PAM | alignment | distance | position | gene name | gene id |
| GU305914.1 | 133492 | 133514 | + | 0 | CTCATGGCGGTGAATCTCGG | TGG | \|\|\|\|\|\|\|\|[\|\|\|\|\|\|\|\|\|\|\|\|]PAM | NA | Intergenic | NA | NA |
| T10120 | ATCTCGGTGGTGAGTTCATCAGG | 1000 | | | | | | | | | |
| Oligo fwd | CACCATCTCGGTGGTGAGTTCATC | | | | | | | | | | |
| Oligo rev | AAACGATGAACTCACCACCGAGAT | | | | | | | | | | |
| Chromosome | start | end | strand | MM | target_seq | PAM | alignment | distance | position | gene name | gene id |
| GU305914.1 | 133505 | 133527 | + | 0 | ATCTCGGTGGTGAGTTCATC | AGG | \|\|\|\|\|\|\|\|[\|\|\|\|\|\|\|\|\|\|\|\|]PAM | NA | Intergenic | NA | NA |
| T10121 | AGTTCATCAGGAGCGAGCTGTGG | 1000 | | | | | | | | | |
| Oligo fwd | CACCAGTTCATCAGGAGCGAGCTG | | | | | | | | | | |
| Oligo rev | AAACCAGCTCGCTCCTGATGAACT | | | | | | | | | | |
| Chromosome | start | end | strand | MM | target_seq | PAM | alignment | distance | position | gene name | gene id |
| GU305914.1 | 133517 | 133539 | + | 0 | AGTTCATCAGGAGCGAGCTG | TGG | \|\|\|\|\|\|\|\|[\|\|\|\|\|\|\|\|\|\|\|\|]PAM | NA | Intergenic | NA | NA |
| T10122 | GTTCATCAGGAGCGAGCTGTGGG | 1000 | | | | | | | | | |
| Oligo fwd | CACCGTTCATCAGGAGCGAGCTGT | | | | | | | | | | |
| Oligo rev | AAACACAGCTCGCTCCTGATGAAC | | | | | | | | | | |
| Chromosome | start | end | strand | MM | target_seq | PAM | alignment | distance | position | gene name | gene id |
| GU305914.1 | 133518 | 133540 | + | 0 | GTTCATCAGGAGCGAGCTGT | GGG | \|\|\|\|\|\|\|\|[\|\|\|\|\|\|\|\|\|\|\|\|]PAM | NA | Intergenic | NA | NA |
| T10123 | TGTGGGTCTCGTTGCACGTCAGG | 1000 | | | | | | | | | |
| Oligo fwd | CACCTGTGGGTCTCGTTGCACGTC | | | | | | | | | | |
| Oligo rev | AAACGACGTGCAACGAGACCCACA | | | | | | | | | | |
| Chromosome | start | end | strand | MM | target_seq | PAM | alignment | distance | position | gene name | gene id |
| GU305914.1 | 133535 | 133557 | + | 0 | TGTGGGTCTCGTTGCACGTC | AGG | \|\|\|\|\|\|\|\|[\|\|\|\|\|\|\|\|\|\|\|\|]PAM | NA | Intergenic | NA | NA |
| T10124 | AAAGAGCACCGTCCTGAACCTGG | 1000 | | | | | | | | | |
| Oligo fwd | CACCAAAGAGCACCGTCCTGAACC | | | | | | | | | | |
| Oligo rev | AAACGGTTCAGGACGGTGCTCTTT | | | | | | | | | | |
| Chromosome | start | end | strand | MM | target_seq | PAM | alignment | distance | position | gene name | gene id |
| GU305914.1 | 133560 | 133582 | + | 0 | AAAGAGCACCGTCCTGAACC | TGG | \|\|\|\|\|\|\|\|[\|\|\|\|\|\|\|\|\|\|\|\|]PAM | NA | Intergenic | NA | NA |
| T10125 | GAGCACCGTCCTGAACCTGGCGG | 1000 | | | | | | | | | |
| Oligo fwd | CACCGAGCACCGTCCTGAACCTGG | | | | | | | | | | |
| Oligo rev | AAACCCAGGTTCAGGACGGTGCTC | | | | | | | | | | |
| Chromosome | start | end | strand | MM | target_seq | PAM | alignment | distance | position | gene name | gene id |
| GU305914.1 | 133563 | 133585 | + | 0 | GAGCACCGTCCTGAACCTGG | CGG | \|\|\|\|\|\|\|\|[\|\|\|\|\|\|\|\|\|\|\|\|]PAM | NA | Intergenic | NA | NA |

<table>
<tr><td>T10126</td><td>CCTGGCGGTGTTGTTCTTGCCGG</td><td>1000</td><td></td><td></td><td></td><td></td><td></td><td></td><td></td><td></td><td></td></tr>
<tr><td>Oligo fwd</td><td>CACCCCTGGCGGTGTTGTTCTTGC</td><td></td><td></td><td></td><td></td><td></td><td></td><td></td><td></td><td></td><td></td></tr>
<tr><td>Oligo rev</td><td>AAACGCAAGAACAACACCGCCAGG</td><td></td><td></td><td></td><td></td><td></td><td></td><td></td><td></td><td></td><td></td></tr>
<tr><td>Chromosome</td><td>start</td><td>end</td><td>strand</td><td>MM</td><td>target_seq</td><td>PAM</td><td>alignment</td><td>distance</td><td>position</td><td>gene name</td><td>gene id</td></tr>
<tr><td>GU305914.1</td><td>133578</td><td>133600</td><td>+</td><td>0</td><td>CCTGGCGGTGTTGTTCTTGC</td><td>CGG</td><td>||||||||[||||||||||||]PAM</td><td>NA</td><td>Intergenic</td><td>NA</td><td>NA</td></tr>
<tr><td>T10127</td><td>GCGAAACCAGTTGAAGAGTACGG</td><td>1000</td><td></td><td></td><td></td><td></td><td></td><td></td><td></td><td></td><td></td></tr>
<tr><td>Oligo fwd</td><td>CACCGCGAAACCAGTTGAAGAGTA</td><td></td><td></td><td></td><td></td><td></td><td></td><td></td><td></td><td></td><td></td></tr>
<tr><td>Oligo rev</td><td>AAACTACTCTTCAACTGGTTTCGC</td><td></td><td></td><td></td><td></td><td></td><td></td><td></td><td></td><td></td><td></td></tr>
<tr><td>Chromosome</td><td>start</td><td>end</td><td>strand</td><td>MM</td><td>target_seq</td><td>PAM</td><td>alignment</td><td>distance</td><td>position</td><td>gene name</td><td>gene id</td></tr>
<tr><td>GU305914.1</td><td>133605</td><td>133627</td><td>+</td><td>0</td><td>GCGAAACCAGTTGAAGAGTA</td><td>CGG</td><td>||||||||[||||||||||||]PAM</td><td>NA</td><td>Intergenic</td><td>NA</td><td>NA</td></tr>
<tr><td>T10128</td><td>CGAAACCAGTTGAAGAGTACGGG</td><td>1000</td><td></td><td></td><td></td><td></td><td></td><td></td><td></td><td></td><td></td></tr>
<tr><td>Oligo fwd</td><td>CACCCGAAACCAGTTGAAGAGTAC</td><td></td><td></td><td></td><td></td><td></td><td></td><td></td><td></td><td></td><td></td></tr>
<tr><td>Oligo rev</td><td>AAACGTACTCTTCAACTGGTTTCG</td><td></td><td></td><td></td><td></td><td></td><td></td><td></td><td></td><td></td><td></td></tr>
<tr><td>Chromosome</td><td>start</td><td>end</td><td>strand</td><td>MM</td><td>target_seq</td><td>PAM</td><td>alignment</td><td>distance</td><td>position</td><td>gene name</td><td>gene id</td></tr>
<tr><td>GU305914.1</td><td>133606</td><td>133628</td><td>+</td><td>0</td><td>CGAAACCAGTTGAAGAGTAC</td><td>GGG</td><td>||||||||[||||||||||||]PAM</td><td>NA</td><td>Intergenic</td><td>NA</td><td>NA</td></tr>
<tr><td>T10129</td><td>GAAGAGTACGGGCGCGAACGTGG</td><td>1000</td><td></td><td></td><td></td><td></td><td></td><td></td><td></td><td></td><td></td></tr>
<tr><td>Oligo fwd</td><td>CACCGAAGAGTACGGGCGCGAACG</td><td></td><td></td><td></td><td></td><td></td><td></td><td></td><td></td><td></td><td></td></tr>
<tr><td>Oligo rev</td><td>AAACCGTTCGCGCCCGTACTCTTC</td><td></td><td></td><td></td><td></td><td></td><td></td><td></td><td></td><td></td><td></td></tr>
<tr><td>Chromosome</td><td>start</td><td>end</td><td>strand</td><td>MM</td><td>target_seq</td><td>PAM</td><td>alignment</td><td>distance</td><td>position</td><td>gene name</td><td>gene id</td></tr>
<tr><td>GU305914.1</td><td>133617</td><td>133639</td><td>+</td><td>0</td><td>GAAGAGTACGGGCGCGAACG</td><td>TGG</td><td>||||||||[||||||||||||]PAM</td><td>NA</td><td>Intergenic</td><td>NA</td><td>NA</td></tr>
<tr><td>T10130</td><td>GTACGGGCGCGAACGTGGTGTGG</td><td>1000</td><td></td><td></td><td></td><td></td><td></td><td></td><td></td><td></td><td></td></tr>
<tr><td>Oligo fwd</td><td>CACCGTACGGGCGCGAACGTGGTG</td><td></td><td></td><td></td><td></td><td></td><td></td><td></td><td></td><td></td><td></td></tr>
<tr><td>Oligo rev</td><td>AAACCACCACGTTCGCGCCCGTAC</td><td></td><td></td><td></td><td></td><td></td><td></td><td></td><td></td><td></td><td></td></tr>
<tr><td>Chromosome</td><td>start</td><td>end</td><td>strand</td><td>MM</td><td>target_seq</td><td>PAM</td><td>alignment</td><td>distance</td><td>position</td><td>gene name</td><td>gene id</td></tr>
<tr><td>GU305914.1</td><td>133622</td><td>133644</td><td>+</td><td>0</td><td>GTACGGGCGCGAACGTGGTG</td><td>TGG</td><td>||||||||[||||||||||||]PAM</td><td>NA</td><td>Intergenic</td><td>NA</td><td>NA</td></tr>
<tr><td>T10131</td><td>GAACGTGGTGTGGATTTGTTTGG</td><td>1000</td><td></td><td></td><td></td><td></td><td></td><td></td><td></td><td></td><td></td></tr>
<tr><td>Oligo fwd</td><td>CACCGAACGTGGTGTGGATTTGTT</td><td></td><td></td><td></td><td></td><td></td><td></td><td></td><td></td><td></td><td></td></tr>
<tr><td>Oligo rev</td><td>AAACAACAAATCCACACCACGTTC</td><td></td><td></td><td></td><td></td><td></td><td></td><td></td><td></td><td></td><td></td></tr>
<tr><td>Chromosome</td><td>start</td><td>end</td><td>strand</td><td>MM</td><td>target_seq</td><td>PAM</td><td>alignment</td><td>distance</td><td>position</td><td>gene name</td><td>gene id</td></tr>
<tr><td>GU305914.1</td><td>133632</td><td>133654</td><td>+</td><td>0</td><td>GAACGTGGTGTGGATTTGTT</td><td>TGG</td><td>||||||||[||||||||||||]PAM</td><td>NA</td><td>Intergenic</td><td>NA</td><td>NA</td></tr>
<tr><td>T10132</td><td>TTTGGACGCCGTGTCGTATTCGG</td><td>1000</td><td></td><td></td><td></td><td></td><td></td><td></td><td></td><td></td><td></td></tr>
<tr><td>Oligo fwd</td><td>CACCTTTGGACGCCGTGTCGTATT</td><td></td><td></td><td></td><td></td><td></td><td></td><td></td><td></td><td></td><td></td></tr>
<tr><td>Oligo rev</td><td>AAACAATACGACACGGCGTCCAAA</td><td></td><td></td><td></td><td></td><td></td><td></td><td></td><td></td><td></td><td></td></tr>
<tr><td>Chromosome</td><td>start</td><td>end</td><td>strand</td><td>MM</td><td>target_seq</td><td>PAM</td><td>alignment</td><td>distance</td><td>position</td><td>gene name</td><td>gene id</td></tr>
<tr><td>GU305914.1</td><td>133650</td><td>133672</td><td>+</td><td>0</td><td>TTTGGACGCCGTGTCGTATT</td><td>CGG</td><td>||||||||[||||||||||||]PAM</td><td>NA</td><td>Intergenic</td><td>NA</td><td>NA</td></tr>
<tr><td>T10133</td><td>TTGGACGCCGTGTCGTATTCGGG</td><td>1000</td><td></td><td></td><td></td><td></td><td></td><td></td><td></td><td></td><td></td></tr>
<tr><td>Oligo fwd</td><td>CACCTTGGACGCCGTGTCGTATTC</td><td></td><td></td><td></td><td></td><td></td><td></td><td></td><td></td><td></td><td></td></tr>
<tr><td>Oligo rev</td><td>AAACGAATACGACACGGCGTCCAA</td><td></td><td></td><td></td><td></td><td></td><td></td><td></td><td></td><td></td><td></td></tr>
<tr><td>Chromosome</td><td>start</td><td>end</td><td>strand</td><td>MM</td><td>target_seq</td><td>PAM</td><td>alignment</td><td>distance</td><td>position</td><td>gene name</td><td>gene id</td></tr>
<tr><td>GU305914.1</td><td>133651</td><td>133673</td><td>+</td><td>0</td><td>TTGGACGCCGTGTCGTATTC</td><td>GGG</td><td>||||||||[||||||||||||]PAM</td><td>NA</td><td>Intergenic</td><td>NA</td><td>NA</td></tr>
<tr><td>T10134</td><td>CGTGTCGTATTCGGGCGCGTCGG</td><td>1000</td><td></td><td></td><td></td><td></td><td></td><td></td><td></td><td></td><td></td></tr>
<tr><td>Oligo fwd</td><td>CACCCGTGTCGTATTCGGGCGCGT</td><td></td><td></td><td></td><td></td><td></td><td></td><td></td><td></td><td></td><td></td></tr>
<tr><td>Oligo rev</td><td>AAACACGCGCCCGAATACGACACG</td><td></td><td></td><td></td><td></td><td></td><td></td><td></td><td></td><td></td><td></td></tr>
<tr><td>Chromosome</td><td>start</td><td>end</td><td>strand</td><td>MM</td><td>target_seq</td><td>PAM</td><td>alignment</td><td>distance</td><td>position</td><td>gene name</td><td>gene id</td></tr>
<tr><td>GU305914.1</td><td>133659</td><td>133681</td><td>+</td><td>0</td><td>CGTGTCGTATTCGGGCGCGT</td><td>CGG</td><td>||||||||[||||||||||||]PAM</td><td>NA</td><td>Intergenic</td><td>NA</td><td>NA</td></tr>
<tr><td>T10135</td><td>GTTCTGCGATTGTTCCGCCTCGG</td><td>1000</td><td></td><td></td><td></td><td></td><td></td><td></td><td></td><td></td><td></td></tr>
<tr><td>Oligo fwd</td><td>CACCGTTCTGCGATTGTTCCGCCT</td><td></td><td></td><td></td><td></td><td></td><td></td><td></td><td></td><td></td><td></td></tr>
<tr><td>Oligo rev</td><td>AAACAGGCGGAACAATCGCAGAAC</td><td></td><td></td><td></td><td></td><td></td><td></td><td></td><td></td><td></td><td></td></tr>
<tr><td>Chromosome</td><td>start</td><td>end</td><td>strand</td><td>MM</td><td>target_seq</td><td>PAM</td><td>alignment</td><td>distance</td><td>position</td><td>gene name</td><td>gene id</td></tr>
<tr><td>GU305914.1</td><td>133686</td><td>133708</td><td>+</td><td>0</td><td>GTTCTGCGATTGTTCCGCCT</td><td>CGG</td><td>||||||||[||||||||||||]PAM</td><td>NA</td><td>Intergenic</td><td>NA</td><td>NA</td></tr>
<tr><td>T10136</td><td>TTGTTCCGCCTCGGTCACGTAGG</td><td>1000</td><td></td><td></td><td></td><td></td><td></td><td></td><td></td><td></td><td></td></tr>
<tr><td>Oligo fwd</td><td>CACCTTGTTCCGCCTCGGTCACGT</td><td></td><td></td><td></td><td></td><td></td><td></td><td></td><td></td><td></td><td></td></tr>
<tr><td>Oligo rev</td><td>AAACACGTGACCGAGGCGGAACAA</td><td></td><td></td><td></td><td></td><td></td><td></td><td></td><td></td><td></td><td></td></tr>
<tr><td>Chromosome</td><td>start</td><td>end</td><td>strand</td><td>MM</td><td>target_seq</td><td>PAM</td><td>alignment</td><td>distance</td><td>position</td><td>gene name</td><td>gene id</td></tr>
<tr><td>GU305914.1</td><td>133695</td><td>133717</td><td>+</td><td>0</td><td>TTGTTCCGCCTCGGTCACGT</td><td>AGG</td><td>||||||||[||||||||||||]PAM</td><td>NA</td><td>Intergenic</td><td>NA</td><td>NA</td></tr>
<tr><td>T10137</td><td>GGTCACGTAGGCGAAAGTGCTGG</td><td>1000</td><td></td><td></td><td></td><td></td><td></td><td></td><td></td><td></td><td></td></tr>
<tr><td>Oligo fwd</td><td>CACCGGTCACGTAGGCGAAAGTGC</td><td></td><td></td><td></td><td></td><td></td><td></td><td></td><td></td><td></td><td></td></tr>
<tr><td>Oligo rev</td><td>AAACGCACTTTCGCCTACGTGACC</td><td></td><td></td><td></td><td></td><td></td><td></td><td></td><td></td><td></td><td></td></tr>
<tr><td>Chromosome</td><td>start</td><td>end</td><td>strand</td><td>MM</td><td>target_seq</td><td>PAM</td><td>alignment</td><td>distance</td><td>position</td><td>gene name</td><td>gene id</td></tr>
<tr><td>GU305914.1</td><td>133707</td><td>133729</td><td>+</td><td>0</td><td>GGTCACGTAGGCGAAAGTGC</td><td>TGG</td><td>||||||||[||||||||||||]PAM</td><td>NA</td><td>Intergenic</td><td>NA</td><td>NA</td></tr>
<tr><td>T10138</td><td>TGGCGTTGATACGTCCGCTACGG</td><td>1000</td><td></td><td></td><td></td><td></td><td></td><td></td><td></td><td></td><td></td></tr>
<tr><td>Oligo fwd</td><td>CACCTGGCGTTGATACGTCCGCTA</td><td></td><td></td><td></td><td></td><td></td><td></td><td></td><td></td><td></td><td></td></tr>
<tr><td>Oligo rev</td><td>AAACTAGCGGACGTATCAACGCCA</td><td></td><td></td><td></td><td></td><td></td><td></td><td></td><td></td><td></td><td></td></tr>
<tr><td>Chromosome</td><td>start</td><td>end</td><td>strand</td><td>MM</td><td>target_seq</td><td>PAM</td><td>alignment</td><td>distance</td><td>position</td><td>gene name</td><td>gene id</td></tr>
<tr><td>GU305914.1</td><td>133727</td><td>133749</td><td>+</td><td>0</td><td>TGGCGTTGATACGTCCGCTA</td><td>CGG</td><td>||||||||[||||||||||||]PAM</td><td>NA</td><td>Intergenic</td><td>NA</td><td>NA</td></tr>
<tr><td>T10139</td><td>ACGTCCGCTACGGTCCGAGTCGG</td><td>1000</td><td></td><td></td><td></td><td></td><td></td><td></td><td></td><td></td><td></td></tr>
<tr><td>Oligo fwd</td><td>CACCACGTCCGCTACGGTCCGAGT</td><td></td><td></td><td></td><td></td><td></td><td></td><td></td><td></td><td></td><td></td></tr>
<tr><td>Oligo rev</td><td>AAACACTCGGACCGTAGCGGACGT</td><td></td><td></td><td></td><td></td><td></td><td></td><td></td><td></td><td></td><td></td></tr>
<tr><td>Chromosome</td><td>start</td><td>end</td><td>strand</td><td>MM</td><td>target_seq</td><td>PAM</td><td>alignment</td><td>distance</td><td>position</td><td>gene name</td><td>gene id</td></tr>
<tr><td>GU305914.1</td><td>133737</td><td>133759</td><td>+</td><td>0</td><td>ACGTCCGCTACGGTCCGAGT</td><td>CGG</td><td>||||||||[||||||||||||]PAM</td><td>NA</td><td>Intergenic</td><td>NA</td><td>NA</td></tr>
<tr><td>T10140</td><td>GTCCGAGTCGGTCGCCGAGATGG</td><td>1000</td><td></td><td></td><td></td><td></td><td></td><td></td><td></td><td></td><td></td></tr>
<tr><td>Oligo fwd</td><td>CACCGTCCGAGTCGGTCGCCGAGA</td><td></td><td></td><td></td><td></td><td></td><td></td><td></td><td></td><td></td><td></td></tr>
<tr><td>Oligo rev</td><td>AAACTCTCGGCGACCGACTCGGAC</td><td></td><td></td><td></td><td></td><td></td><td></td><td></td><td></td><td></td><td></td></tr>
<tr><td>Chromosome</td><td>start</td><td>end</td><td>strand</td><td>MM</td><td>target_seq</td><td>PAM</td><td>alignment</td><td>distance</td><td>position</td><td>gene name</td><td>gene id</td></tr>
</table>

GU305914.1 133749 133771 + 0 GTCCGAGTCGGTCGCCGAGA TGG ||||||||[||||||||||||]PAM NA Intergenic NA NA

T10141 CGCCGAGATGGTGACGTTCATGG 1000
Oligo fwd CACCGCCGAGATGGTGACGTTCA
Oligo rev AAACTGAACGTCACCATCTCGGCG
Chromosome start end strand MM target_seq PAM alignment distance position gene name gene id
GU305914.1 133761 133783 + 0 CGCCGAGATGGTGACGTTCA TGG ||||||||[||||||||||||]PAM NA Intergenic NA NA

T10142 GTTCATGGTAAACACGTCCGAGG 1000
Oligo fwd CACCGTTCATGGTAAACACGTCCG
Oligo rev AAACCGGACGTGTTTACCATGAAC
Chromosome start end strand MM target_seq PAM alignment distance position gene name gene id
GU305914.1 133776 133798 + 0 GTTCATGGTAAACACGTCCG AGG ||||||||[||||||||||||]PAM NA Intergenic NA NA

T10143 TTCATGGTAAACACGTCCGAGGG 1000
Oligo fwd CACCTTCATGGTAAACACGTCCGA
Oligo rev AAACTCGGACGTGTTTACCATGAA
Chromosome start end strand MM target_seq PAM alignment distance position gene name gene id
GU305914.1 133777 133799 + 0 TTCATGGTAAACACGTCCGA GGG ||||||||[||||||||||||]PAM NA Intergenic NA NA

T10144 GAGGGCGACGTGATGTTGCTCGG 1000
Oligo fwd CACCGAGGGCGACGTGATGTTGCT
Oligo rev AAACAGCAACATCACGTCGCCCTC
Chromosome start end strand MM target_seq PAM alignment distance position gene name gene id
GU305914.1 133795 133817 + 0 GAGGGCGACGTGATGTTGCT CGG ||||||||[||||||||||||]PAM NA Intergenic NA NA

T10145 AGGGCGACGTGATGTTGCTCGGG 1000
Oligo fwd CACCAGGGCGACGTGATGTTGCTC
Oligo rev AAACGAGCAACATCACGTCGCCCT
Chromosome start end strand MM target_seq PAM alignment distance position gene name gene id
GU305914.1 133796 133818 + 0 AGGGCGACGTGATGTTGCTC GGG ||||||||[||||||||||||]PAM NA Intergenic NA NA

T10146 GGGCGACGTGATGTTGCTCGGGG 1000
Oligo fwd CACCGGGCGACGTGATGTTGCTCG
Oligo rev AAACCGAGCAACATCACGTCGCCC
Chromosome start end strand MM target_seq PAM alignment distance position gene name gene id
GU305914.1 133797 133819 + 0 GGGCGACGTGATGTTGCTCG GGG ||||||||[||||||||||||]PAM NA Intergenic NA NA

T10147 GCTCGGGGTCGCCTGTATCCAGG 1000
Oligo fwd CACCGCTCGGGGTCGCCTGTATCC
Oligo rev AAACGGATACAGGCGACCCCGAGC
Chromosome start end strand MM target_seq PAM alignment distance position gene name gene id
GU305914.1 133812 133834 + 0 GCTCGGGGTCGCCTGTATCC AGG ||||||||[||||||||||||]PAM NA Intergenic NA NA

T10148 AGCAGCATGAATGTGATCCTCGG 1000
Oligo fwd CACCAGCAGCATGAATGTGATCCT
Oligo rev AAACAGGATCACATTCATGCTGCT
Chromosome start end strand MM target_seq PAM alignment distance position gene name gene id
GU305914.1 133888 133910 + 0 AGCAGCATGAATGTGATCCT CGG ||||||||[||||||||||||]PAM NA Intergenic NA NA

T10149 ATGTGATCCTCGGCGAGCTCAGG 1000
Oligo fwd CACCATGTGATCCTCGGCGAGCTC
Oligo rev AAACGAGCTCGCCGAGGATCACAT
Chromosome start end strand MM target_seq PAM alignment distance position gene name gene id
GU305914.1 133898 133920 + 0 ATGTGATCCTCGGCGAGCTC AGG ||||||||[||||||||||||]PAM NA Intergenic NA NA

T10150 GATCCTCGGCGAGCTCAGGAAGG 1000
Oligo fwd CACCGATCCTCGGCGAGCTCAGGA
Oligo rev AAACTCCTGAGCTCGCCGAGGATC
Chromosome start end strand MM target_seq PAM alignment distance position gene name gene id
GU305914.1 133902 133924 + 0 GATCCTCGGCGAGCTCAGGA AGG ||||||||[||||||||||||]PAM NA Intergenic NA NA

T10151 CTCGGCGAGCTCAGGAAGGCCGG 1000
Oligo fwd CACCCTCGGCGAGCTCAGGAAGGC
Oligo rev AAACGCCTTCCTGAGCTCGCCGAG
Chromosome start end strand MM target_seq PAM alignment distance position gene name gene id
GU305914.1 133906 133928 + 0 CTCGGCGAGCTCAGGAAGGC CGG ||||||||[||||||||||||]PAM NA Intergenic NA NA

T10152 CGGCATCGAGCACGAGACCCTGG 1000
Oligo fwd CACCCGGCATCGAGCACGAGACCC
Oligo rev AAACGGGTCTCGTGCTCGATGCCG
Chromosome start end strand MM target_seq PAM alignment distance position gene name gene id
GU305914.1 133926 133948 + 0 CGGCATCGAGCACGAGACCC TGG ||||||||[||||||||||||]PAM NA Intergenic NA NA

T10153 TTTCGTCTCGCCGAGTCCATCGG 1000
Oligo fwd CACCTTTCGTCTCGCCGAGTCCAT
Oligo rev AAACATGGACTCGGCGAGACGAAA
Chromosome start end strand MM target_seq PAM alignment distance position gene name gene id
GU305914.1 133957 133979 + 0 TTTCGTCTCGCCGAGTCCAT CGG ||||||||[||||||||||||]PAM NA Intergenic NA NA

T10154 TTCGTCTCGCCGAGTCCATCGGG 1000
Oligo fwd CACCTTCGTCTCGCCGAGTCCATC
Oligo rev AAACGATGGACTCGGCGAGACGAA
Chromosome start end strand MM target_seq PAM alignment distance position gene name gene id
GU305914.1 133958 133980 + 0 TTCGTCTCGCCGAGTCCATC GGG ||||||||[||||||||||||]PAM NA Intergenic NA NA

T10155 TCGTCTCGCCGAGTCCATCGGGG 1000
Oligo fwd CACCTCGTCTCGCCGAGTCCATCG
Oligo rev AAACCGATGGACTCGGCGAGACGA

<table>
<tr><td>Chromosome</td><td>start</td><td>end</td><td>strand</td><td>MM</td><td>target_seq</td><td>PAM</td><td>alignment</td><td>distance</td><td>position</td><td>gene name</td><td>gene id</td></tr>
<tr><td>GU305914.1</td><td>133959</td><td>133981</td><td>+</td><td>0</td><td>TCGTCTCGCCGAGTCCATCG</td><td>GGG</td><td>||||||||[||||||||||||]PAM</td><td>NA</td><td>Intergenic</td><td>NA</td><td>NA</td></tr>
<tr><td></td><td></td><td></td><td></td><td></td><td></td><td></td><td></td><td></td><td></td><td></td><td></td></tr>
<tr><td>T10156</td><td>TCTCGCCGAGTCCATCGGGGAGG</td><td>1000</td><td></td><td></td><td></td><td></td><td></td><td></td><td></td><td></td><td></td></tr>
<tr><td>Oligo fwd</td><td>CACCTCTCGCCGAGTCCATCGGGG</td><td></td><td></td><td></td><td></td><td></td><td></td><td></td><td></td><td></td><td></td></tr>
<tr><td>Oligo rev</td><td>AAACCCCCGATGGACTCGGCGAGA</td><td></td><td></td><td></td><td></td><td></td><td></td><td></td><td></td><td></td><td></td></tr>
<tr><td>Chromosome</td><td>start</td><td>end</td><td>strand</td><td>MM</td><td>target_seq</td><td>PAM</td><td>alignment</td><td>distance</td><td>position</td><td>gene name</td><td>gene id</td></tr>
<tr><td>GU305914.1</td><td>133962</td><td>133984</td><td>+</td><td>0</td><td>TCTCGCCGAGTCCATCGGGG</td><td>AGG</td><td>||||||||[||||||||||||]PAM</td><td>NA</td><td>Intergenic</td><td>NA</td><td>NA</td></tr>
<tr><td></td><td></td><td></td><td></td><td></td><td></td><td></td><td></td><td></td><td></td><td></td><td></td></tr>
<tr><td>T10157</td><td>GGGAGGCGTGCGACTTCTTCAGG</td><td>1000</td><td></td><td></td><td></td><td></td><td></td><td></td><td></td><td></td><td></td></tr>
<tr><td>Oligo fwd</td><td>CACCGGGAGGCGTGCGACTTCTTC</td><td></td><td></td><td></td><td></td><td></td><td></td><td></td><td></td><td></td><td></td></tr>
<tr><td>Oligo rev</td><td>AAACGAAGAAGTCGCACGCCTCCC</td><td></td><td></td><td></td><td></td><td></td><td></td><td></td><td></td><td></td><td></td></tr>
<tr><td>Chromosome</td><td>start</td><td>end</td><td>strand</td><td>MM</td><td>target_seq</td><td>PAM</td><td>alignment</td><td>distance</td><td>position</td><td>gene name</td><td>gene id</td></tr>
<tr><td>GU305914.1</td><td>133979</td><td>134001</td><td>+</td><td>0</td><td>GGGAGGCGTGCGACTTCTTC</td><td>AGG</td><td>||||||||[||||||||||||]PAM</td><td>NA</td><td>Intergenic</td><td>NA</td><td>NA</td></tr>
<tr><td></td><td></td><td></td><td></td><td></td><td></td><td></td><td></td><td></td><td></td><td></td><td></td></tr>
<tr><td>T10158</td><td>TGCGACTTCTTCAGGCAACCCGG</td><td>1000</td><td></td><td></td><td></td><td></td><td></td><td></td><td></td><td></td><td></td></tr>
<tr><td>Oligo fwd</td><td>CACCTGCGACTTCTTCAGGCAACC</td><td></td><td></td><td></td><td></td><td></td><td></td><td></td><td></td><td></td><td></td></tr>
<tr><td>Oligo rev</td><td>AAACGGTTGCCTGAAGAAGTCGCA</td><td></td><td></td><td></td><td></td><td></td><td></td><td></td><td></td><td></td><td></td></tr>
<tr><td>Chromosome</td><td>start</td><td>end</td><td>strand</td><td>MM</td><td>target_seq</td><td>PAM</td><td>alignment</td><td>distance</td><td>position</td><td>gene name</td><td>gene id</td></tr>
<tr><td>GU305914.1</td><td>133987</td><td>134009</td><td>+</td><td>0</td><td>TGCGACTTCTTCAGGCAACC</td><td>CGG</td><td>||||||||[||||||||||||]PAM</td><td>NA</td><td>Intergenic</td><td>NA</td><td>NA</td></tr>
<tr><td></td><td></td><td></td><td></td><td></td><td></td><td></td><td></td><td></td><td></td><td></td><td></td></tr>
<tr><td>T10159</td><td>GCGACTTCTTCAGGCAACCCGGG</td><td>1000</td><td></td><td></td><td></td><td></td><td></td><td></td><td></td><td></td><td></td></tr>
<tr><td>Oligo fwd</td><td>CACCGCGACTTCTTCAGGCAACCC</td><td></td><td></td><td></td><td></td><td></td><td></td><td></td><td></td><td></td><td></td></tr>
<tr><td>Oligo rev</td><td>AAACGGGTTGCCTGAAGAAGTCGC</td><td></td><td></td><td></td><td></td><td></td><td></td><td></td><td></td><td></td><td></td></tr>
<tr><td>Chromosome</td><td>start</td><td>end</td><td>strand</td><td>MM</td><td>target_seq</td><td>PAM</td><td>alignment</td><td>distance</td><td>position</td><td>gene name</td><td>gene id</td></tr>
<tr><td>GU305914.1</td><td>133988</td><td>134010</td><td>+</td><td>0</td><td>GCGACTTCTTCAGGCAACCC</td><td>GGG</td><td>||||||||[||||||||||||]PAM</td><td>NA</td><td>Intergenic</td><td>NA</td><td>NA</td></tr>
<tr><td></td><td></td><td></td><td></td><td></td><td></td><td></td><td></td><td></td><td></td><td></td><td></td></tr>
<tr><td>T10160</td><td>CGACTTCTTCAGGCAACCCGGGG</td><td>1000</td><td></td><td></td><td></td><td></td><td></td><td></td><td></td><td></td><td></td></tr>
<tr><td>Oligo fwd</td><td>CACCCGACTTCTTCAGGCAACCCG</td><td></td><td></td><td></td><td></td><td></td><td></td><td></td><td></td><td></td><td></td></tr>
<tr><td>Oligo rev</td><td>AAACCGGGTTGCCTGAAGAAGTCG</td><td></td><td></td><td></td><td></td><td></td><td></td><td></td><td></td><td></td><td></td></tr>
<tr><td>Chromosome</td><td>start</td><td>end</td><td>strand</td><td>MM</td><td>target_seq</td><td>PAM</td><td>alignment</td><td>distance</td><td>position</td><td>gene name</td><td>gene id</td></tr>
<tr><td>GU305914.1</td><td>133989</td><td>134011</td><td>+</td><td>0</td><td>CGACTTCTTCAGGCAACCCG</td><td>GGG</td><td>||||||||[||||||||||||]PAM</td><td>NA</td><td>Intergenic</td><td>NA</td><td>NA</td></tr>
<tr><td></td><td></td><td></td><td></td><td></td><td></td><td></td><td></td><td></td><td></td><td></td><td></td></tr>
<tr><td>T10161</td><td>AACCCGGGGAATCCCGACTGCGG</td><td>1000</td><td></td><td></td><td></td><td></td><td></td><td></td><td></td><td></td><td></td></tr>
<tr><td>Oligo fwd</td><td>CACCAACCCGGGGAATCCCGACTG</td><td></td><td></td><td></td><td></td><td></td><td></td><td></td><td></td><td></td><td></td></tr>
<tr><td>Oligo rev</td><td>AAACCAGTCGGGATTCCCCGGGTT</td><td></td><td></td><td></td><td></td><td></td><td></td><td></td><td></td><td></td><td></td></tr>
<tr><td>Chromosome</td><td>start</td><td>end</td><td>strand</td><td>MM</td><td>target_seq</td><td>PAM</td><td>alignment</td><td>distance</td><td>position</td><td>gene name</td><td>gene id</td></tr>
<tr><td>GU305914.1</td><td>134003</td><td>134025</td><td>+</td><td>0</td><td>AACCCGGGGAATCCCGACTG</td><td>CGG</td><td>||||||||[||||||||||||]PAM</td><td>NA</td><td>Intergenic</td><td>NA</td><td>NA</td></tr>
<tr><td></td><td></td><td></td><td></td><td></td><td></td><td></td><td></td><td></td><td></td><td></td><td></td></tr>
<tr><td>T10162</td><td>ACCCGGGGAATCCCGACTGCGGG</td><td>1000</td><td></td><td></td><td></td><td></td><td></td><td></td><td></td><td></td><td></td></tr>
<tr><td>Oligo fwd</td><td>CACCACCCGGGGAATCCCGACTGC</td><td></td><td></td><td></td><td></td><td></td><td></td><td></td><td></td><td></td><td></td></tr>
<tr><td>Oligo rev</td><td>AAACGCAGTCGGGATTCCCCGGGT</td><td></td><td></td><td></td><td></td><td></td><td></td><td></td><td></td><td></td><td></td></tr>
<tr><td>Chromosome</td><td>start</td><td>end</td><td>strand</td><td>MM</td><td>target_seq</td><td>PAM</td><td>alignment</td><td>distance</td><td>position</td><td>gene name</td><td>gene id</td></tr>
<tr><td>GU305914.1</td><td>134004</td><td>134026</td><td>+</td><td>0</td><td>ACCCGGGGAATCCCGACTGC</td><td>GGG</td><td>||||||||[||||||||||||]PAM</td><td>NA</td><td>Intergenic</td><td>NA</td><td>NA</td></tr>
<tr><td></td><td></td><td></td><td></td><td></td><td></td><td></td><td></td><td></td><td></td><td></td><td></td></tr>
<tr><td>T10163</td><td>GGAATCCCGACTGCGGGTCTTGG</td><td>1000</td><td></td><td></td><td></td><td></td><td></td><td></td><td></td><td></td><td></td></tr>
<tr><td>Oligo fwd</td><td>CACCGGAATCCCGACTGCGGGTCT</td><td></td><td></td><td></td><td></td><td></td><td></td><td></td><td></td><td></td><td></td></tr>
<tr><td>Oligo rev</td><td>AAACAGACCCGCAGTCGGGATTCC</td><td></td><td></td><td></td><td></td><td></td><td></td><td></td><td></td><td></td><td></td></tr>
<tr><td>Chromosome</td><td>start</td><td>end</td><td>strand</td><td>MM</td><td>target_seq</td><td>PAM</td><td>alignment</td><td>distance</td><td>position</td><td>gene name</td><td>gene id</td></tr>
<tr><td>GU305914.1</td><td>134010</td><td>134032</td><td>+</td><td>0</td><td>GGAATCCCGACTGCGGGTCT</td><td>TGG</td><td>||||||||[||||||||||||]PAM</td><td>NA</td><td>Intergenic</td><td>NA</td><td>NA</td></tr>
<tr><td></td><td></td><td></td><td></td><td></td><td></td><td></td><td></td><td></td><td></td><td></td><td></td></tr>
<tr><td>T10164</td><td>ACTGCGGGTCTTGGATCTCGCGG</td><td>1000</td><td></td><td></td><td></td><td></td><td></td><td></td><td></td><td></td><td></td></tr>
<tr><td>Oligo fwd</td><td>CACCACTGCGGGTCTTGGATCTCG</td><td></td><td></td><td></td><td></td><td></td><td></td><td></td><td></td><td></td><td></td></tr>
<tr><td>Oligo rev</td><td>AAACCGAGATCCAAGACCCGCAGT</td><td></td><td></td><td></td><td></td><td></td><td></td><td></td><td></td><td></td><td></td></tr>
<tr><td>Chromosome</td><td>start</td><td>end</td><td>strand</td><td>MM</td><td>target_seq</td><td>PAM</td><td>alignment</td><td>distance</td><td>position</td><td>gene name</td><td>gene id</td></tr>
<tr><td>GU305914.1</td><td>134019</td><td>134041</td><td>+</td><td>0</td><td>ACTGCGGGTCTTGGATCTCG</td><td>CGG</td><td>||||||||[||||||||||||]PAM</td><td>NA</td><td>Intergenic</td><td>NA</td><td>NA</td></tr>
<tr><td></td><td></td><td></td><td></td><td></td><td></td><td></td><td></td><td></td><td></td><td></td><td></td></tr>
<tr><td>T10165</td><td>GTCCTTATTTGATCACGTTGCGG</td><td>1000</td><td></td><td></td><td></td><td></td><td></td><td></td><td></td><td></td><td></td></tr>
<tr><td>Oligo fwd</td><td>CACCGTCCTTATTTGATCACGTTG</td><td></td><td></td><td></td><td></td><td></td><td></td><td></td><td></td><td></td><td></td></tr>
<tr><td>Oligo rev</td><td>AAACCAACGTGATCAAATAAGGAC</td><td></td><td></td><td></td><td></td><td></td><td></td><td></td><td></td><td></td><td></td></tr>
<tr><td>Chromosome</td><td>start</td><td>end</td><td>strand</td><td>MM</td><td>target_seq</td><td>PAM</td><td>alignment</td><td>distance</td><td>position</td><td>gene name</td><td>gene id</td></tr>
<tr><td>GU305914.1</td><td>134043</td><td>134065</td><td>+</td><td>0</td><td>GTCCTTATTTGATCACGTTG</td><td>CGG</td><td>||||||||[||||||||||||]PAM</td><td>NA</td><td>Intergenic</td><td>NA</td><td>NA</td></tr>
<tr><td></td><td></td><td></td><td></td><td></td><td></td><td></td><td></td><td></td><td></td><td></td><td></td></tr>
<tr><td>T10166</td><td>CTTATTTGATCACGTTGCGGCGG</td><td>1000</td><td></td><td></td><td></td><td></td><td></td><td></td><td></td><td></td><td></td></tr>
<tr><td>Oligo fwd</td><td>CACCCTTATTTGATCACGTTGCGG</td><td></td><td></td><td></td><td></td><td></td><td></td><td></td><td></td><td></td><td></td></tr>
<tr><td>Oligo rev</td><td>AAACCCGCAACGTGATCAAATAAG</td><td></td><td></td><td></td><td></td><td></td><td></td><td></td><td></td><td></td><td></td></tr>
<tr><td>Chromosome</td><td>start</td><td>end</td><td>strand</td><td>MM</td><td>target_seq</td><td>PAM</td><td>alignment</td><td>distance</td><td>position</td><td>gene name</td><td>gene id</td></tr>
<tr><td>GU305914.1</td><td>134046</td><td>134068</td><td>+</td><td>0</td><td>CTTATTTGATCACGTTGCGG</td><td>CGG</td><td>||||||||[||||||||||||]PAM</td><td>NA</td><td>Intergenic</td><td>NA</td><td>NA</td></tr>
<tr><td></td><td></td><td></td><td></td><td></td><td></td><td></td><td></td><td></td><td></td><td></td><td></td></tr>
<tr><td>T10167</td><td>CACGTTGCGGCGGAGTGTATAGG</td><td>1000</td><td></td><td></td><td></td><td></td><td></td><td></td><td></td><td></td><td></td></tr>
<tr><td>Oligo fwd</td><td>CACCCACGTTGCGGCGGAGTGTAT</td><td></td><td></td><td></td><td></td><td></td><td></td><td></td><td></td><td></td><td></td></tr>
<tr><td>Oligo rev</td><td>AAACATACACTCCGCCGCAACGTG</td><td></td><td></td><td></td><td></td><td></td><td></td><td></td><td></td><td></td><td></td></tr>
<tr><td>Chromosome</td><td>start</td><td>end</td><td>strand</td><td>MM</td><td>target_seq</td><td>PAM</td><td>alignment</td><td>distance</td><td>position</td><td>gene name</td><td>gene id</td></tr>
<tr><td>GU305914.1</td><td>134056</td><td>134078</td><td>+</td><td>0</td><td>CACGTTGCGGCGGAGTGTAT</td><td>AGG</td><td>||||||||[||||||||||||]PAM</td><td>NA</td><td>Intergenic</td><td>NA</td><td>NA</td></tr>
<tr><td></td><td></td><td></td><td></td><td></td><td></td><td></td><td></td><td></td><td></td><td></td><td></td></tr>
<tr><td>T10168</td><td>GGCGGAGTGTATAGGAGACGTGG</td><td>1000</td><td></td><td></td><td></td><td></td><td></td><td></td><td></td><td></td><td></td></tr>
<tr><td>Oligo fwd</td><td>CACCGGCGGAGTGTATAGGAGACG</td><td></td><td></td><td></td><td></td><td></td><td></td><td></td><td></td><td></td><td></td></tr>
<tr><td>Oligo rev</td><td>AAACCGTCTCCTATACACTCCGCC</td><td></td><td></td><td></td><td></td><td></td><td></td><td></td><td></td><td></td><td></td></tr>
<tr><td>Chromosome</td><td>start</td><td>end</td><td>strand</td><td>MM</td><td>target_seq</td><td>PAM</td><td>alignment</td><td>distance</td><td>position</td><td>gene name</td><td>gene id</td></tr>
<tr><td>GU305914.1</td><td>134064</td><td>134086</td><td>+</td><td>0</td><td>GGCGGAGTGTATAGGAGACG</td><td>TGG</td><td>||||||||[||||||||||||]PAM</td><td>NA</td><td>Intergenic</td><td>NA</td><td>NA</td></tr>
<tr><td></td><td></td><td></td><td></td><td></td><td></td><td></td><td></td><td></td><td></td><td></td><td></td></tr>
<tr><td>T10169</td><td>TATAGGAGACGTGGTTTCGCTGG</td><td>1000</td><td></td><td></td><td></td><td></td><td></td><td></td><td></td><td></td><td></td></tr>
<tr><td>Oligo fwd</td><td>CACCTATAGGAGACGTGGTTTCGC</td><td></td><td></td><td></td><td></td><td></td><td></td><td></td><td></td><td></td><td></td></tr>
<tr><td>Oligo rev</td><td>AAACGCGAAACCACGTCTCCTATA</td><td></td><td></td><td></td><td></td><td></td><td></td><td></td><td></td><td></td><td></td></tr>
<tr><td>Chromosome</td><td>start</td><td>end</td><td>strand</td><td>MM</td><td>target_seq</td><td>PAM</td><td>alignment</td><td>distance</td><td>position</td><td>gene name</td><td>gene id</td></tr>
<tr><td>GU305914.1</td><td>134073</td><td>134095</td><td>+</td><td>0</td><td>TATAGGAGACGTGGTTTCGC</td><td>TGG</td><td>||||||||[||||||||||||]PAM</td><td>NA</td><td>Intergenic</td><td>NA</td><td>NA</td></tr>
<tr><td></td><td></td><td></td><td></td><td></td><td></td><td></td><td></td><td></td><td></td><td></td><td></td></tr>
<tr><td>T10170</td><td>ATAGGAGACGTGGTTTCGCTGGG</td><td>1000</td><td></td><td></td><td></td><td></td><td></td><td></td><td></td><td></td><td></td></tr>
<tr><td>Oligo fwd</td><td>CACCATAGGAGACGTGGTTTCGCT</td><td></td><td></td><td></td><td></td><td></td><td></td><td></td><td></td><td></td><td></td></tr>
</table>

Oligo rev AAACAGCGAAACCACGTCTCCTAT
Chromosome start end strand MM target_seq PAM alignment distance position gene name gene id
GU305914.1 134074 134096 + 0 ATAGGAGACGTGGTTTCGCT GGG ||||||||[||||||||||||]PAM NA Intergenic NA NA

T10171 GGCTTCGTCGAGTCGCAAGACGG 1000
Oligo fwd CACCGGCTTCGTCGAGTCGCAAGA
Oligo rev AAACTCTTGCGACTCGACGAAGCC
Chromosome start end strand MM target_seq PAM alignment distance position gene name gene id
GU305914.1 134095 134117 + 0 GGCTTCGTCGAGTCGCAAGA CGG ||||||||[||||||||||||]PAM NA Intergenic NA NA

T10172 CGAGTCGCAAGACGGCGCGCCGG 1000
Oligo fwd CACCCGAGTCGCAAGACGGCGCGC
Oligo rev AAACGCGCGCCGTCTTGCGACTCG
Chromosome start end strand MM target_seq PAM alignment distance position gene name gene id
GU305914.1 134103 134125 + 0 CGAGTCGCAAGACGGCGCGC CGG ||||||||[||||||||||||]PAM NA Intergenic NA NA

T10173 GAGTCGCAAGACGGCGCGCCGGG 1000
Oligo fwd CACCGAGTCGCAAGACGGCGCGCC
Oligo rev AAACGGCGCGCCGTCTTGCGACTC
Chromosome start end strand MM target_seq PAM alignment distance position gene name gene id
GU305914.1 134104 134126 + 0 GAGTCGCAAGACGGCGCGCC GGG ||||||||[||||||||||||]PAM NA Intergenic NA NA

T10174 TCGCAAGACGGCGCGCCGGGCGG 1000
Oligo fwd CACCTCGCAAGACGGCGCGCCGGG
Oligo rev AAACCCCGGCGCGCCGTCTTGCGA
Chromosome start end strand MM target_seq PAM alignment distance position gene name gene id
GU305914.1 134107 134129 + 0 TCGCAAGACGGCGCGCCGGG CGG ||||||||[||||||||||||]PAM NA Intergenic NA NA

T10175 CGGCGCGCCGGGCGGAACGACGG 1000
Oligo fwd CACCCGGCGCGCCGGGCGGAACGA
Oligo rev AAACTCGTTCCGCCCGGCGCGCCG
Chromosome start end strand MM target_seq PAM alignment distance position gene name gene id
GU305914.1 134115 134137 + 0 CGGCGCGCCGGGCGGAACGA CGG ||||||||[||||||||||||]PAM NA Intergenic NA NA

T10176 GGCGCGCCGGGCGGAACGACGGG 1000
Oligo fwd CACCGGCGCGCCGGGCGGAACGAC
Oligo rev AAACGTCGTTCCGCCCGGCGCGCC
Chromosome start end strand MM target_seq PAM alignment distance position gene name gene id
GU305914.1 134116 134138 + 0 GGCGCGCCGGGCGGAACGAC GGG ||||||||[||||||||||||]PAM NA Intergenic NA NA

T10177 GGAACGACGGGCCACCCTCGCGG 1000
Oligo fwd CACCGGAACGACGGGCCACCCTCG
Oligo rev AAACCGAGGGTGGCCCGTCGTTCC
Chromosome start end strand MM target_seq PAM alignment distance position gene name gene id
GU305914.1 134128 134150 + 0 GGAACGACGGGCCACCCTCG CGG ||||||||[||||||||||||]PAM NA Intergenic NA NA

T10178 TCGCGGAGCGACCGCCTCGAAGG 1000
Oligo fwd CACCTCGCGGAGCGACCGCCTCGA
Oligo rev AAACTCGAGGCGGTCGCTCCGCGA
Chromosome start end strand MM target_seq PAM alignment distance position gene name gene id
GU305914.1 134145 134167 + 0 TCGCGGAGCGACCGCCTCGA AGG ||||||||[||||||||||||]PAM NA Intergenic NA NA

T10179 GGACTGACTATGTTCTCACACGG 1000
Oligo fwd CACCGGACTGACTATGTTCTCACA
Oligo rev AAACTGTGAGAACATAGTCAGTCC
Chromosome start end strand MM target_seq PAM alignment distance position gene name gene id
GU305914.1 134166 134188 + 0 GGACTGACTATGTTCTCACA CGG ||||||||[||||||||||||]PAM NA Intergenic NA NA

T10180 TCTCACACGGCCGAGACCCGCGG 1000
Oligo fwd CACCTCTCACACGGCCGAGACCCG
Oligo rev AAACCGGGTCTCGGCCGTGTGAGA
Chromosome start end strand MM target_seq PAM alignment distance position gene name gene id
GU305914.1 134179 134201 + 0 TCTCACACGGCCGAGACCCG CGG ||||||||[||||||||||||]PAM NA Intergenic NA NA

T10181 ACACGGCCGAGACCCGCGGACGG 1000
Oligo fwd CACCACACGGCCGAGACCCGCGGA
Oligo rev AAACTCCGCGGGTCTCGGCCGTGT
Chromosome start end strand MM target_seq PAM alignment distance position gene name gene id
GU305914.1 134183 134205 + 0 ACACGGCCGAGACCCGCGGA CGG ||||||||[||||||||||||]PAM NA Intergenic NA NA

T10182 CACGGCCGAGACCCGCGGACGGG 1000
Oligo fwd CACCCACGGCCGAGACCCGCGGAC
Oligo rev AAACGTCCGCGGGTCTCGGCCGTG
Chromosome start end strand MM target_seq PAM alignment distance position gene name gene id
GU305914.1 134184 134206 + 0 CACGGCCGAGACCCGCGGAC GGG ||||||||[||||||||||||]PAM NA Intergenic NA NA

T10183 GCGGACGGGCAAGCCTATATCGG 1000
Oligo fwd CACCGCGGACGGGCAAGCCTATAT
Oligo rev AAACATATAGGCTTGCCCGTCCGC
Chromosome start end strand MM target_seq PAM alignment distance position gene name gene id
GU305914.1 134198 134220 + 0 GCGGACGGGCAAGCCTATAT CGG ||||||||[||||||||||||]PAM NA Intergenic NA NA

T10184 GACGGGCAAGCCTATATCGGCGG 1000
Oligo fwd CACCGACGGGCAAGCCTATATCGG
Oligo rev AAACCCGATATAGGCTTGCCCGTC
Chromosome start end strand MM target_seq PAM alignment distance position gene name gene id
GU305914.1 134201 134223 + 0 GACGGGCAAGCCTATATCGG CGG ||||||||[||||||||||||]PAM NA Intergenic NA NA

T10185 GGCAAGCCTATATCGGCGGCCGG 1000

<table>
<tr><td>Oligo fwd</td><td>CACCGGCAAGCCTATATCGGCGGC</td><td></td><td></td><td></td><td></td><td></td><td></td><td></td><td></td><td></td><td></td></tr>
<tr><td>Oligo rev</td><td>AAACGCCGCCGATATAGGCTTGCC</td><td></td><td></td><td></td><td></td><td></td><td></td><td></td><td></td><td></td><td></td></tr>
<tr><td>Chromosome</td><td>start</td><td>end</td><td>strand</td><td>MM</td><td>target_seq</td><td>PAM</td><td>alignment</td><td>distance</td><td>position</td><td>gene name</td><td>gene id</td></tr>
<tr><td>GU305914.1</td><td>134205</td><td>134227</td><td>+</td><td>0</td><td>GGCAAGCCTATATCGGCGGC</td><td>CGG</td><td>||||||||[||||||||||||]PAM</td><td>NA</td><td>Intergenic</td><td>NA</td><td>NA</td></tr>
<tr><td>T10186</td><td>GCAAGCCTATATCGGCGGCCGGG</td><td>1000</td><td></td><td></td><td></td><td></td><td></td><td></td><td></td><td></td><td></td></tr>
<tr><td>Oligo fwd</td><td>CACCGCAAGCCTATATCGGCGGCC</td><td></td><td></td><td></td><td></td><td></td><td></td><td></td><td></td><td></td><td></td></tr>
<tr><td>Oligo rev</td><td>AAACGGCCGCCGATATAGGCTTGC</td><td></td><td></td><td></td><td></td><td></td><td></td><td></td><td></td><td></td><td></td></tr>
<tr><td>Chromosome</td><td>start</td><td>end</td><td>strand</td><td>MM</td><td>target_seq</td><td>PAM</td><td>alignment</td><td>distance</td><td>position</td><td>gene name</td><td>gene id</td></tr>
<tr><td>GU305914.1</td><td>134206</td><td>134228</td><td>+</td><td>0</td><td>GCAAGCCTATATCGGCGGCC</td><td>GGG</td><td>||||||||[||||||||||||]PAM</td><td>NA</td><td>Intergenic</td><td>NA</td><td>NA</td></tr>
<tr><td>T10187</td><td>CAAGCCTATATCGGCGGCCGGGG</td><td>1000</td><td></td><td></td><td></td><td></td><td></td><td></td><td></td><td></td><td></td></tr>
<tr><td>Oligo fwd</td><td>CACCCAAGCCTATATCGGCGGCCG</td><td></td><td></td><td></td><td></td><td></td><td></td><td></td><td></td><td></td><td></td></tr>
<tr><td>Oligo rev</td><td>AAACCGGCCGCCGATATAGGCTTG</td><td></td><td></td><td></td><td></td><td></td><td></td><td></td><td></td><td></td><td></td></tr>
<tr><td>Chromosome</td><td>start</td><td>end</td><td>strand</td><td>MM</td><td>target_seq</td><td>PAM</td><td>alignment</td><td>distance</td><td>position</td><td>gene name</td><td>gene id</td></tr>
<tr><td>GU305914.1</td><td>134207</td><td>134229</td><td>+</td><td>0</td><td>CAAGCCTATATCGGCGGCCG</td><td>GGG</td><td>||||||||[||||||||||||]PAM</td><td>NA</td><td>Intergenic</td><td>NA</td><td>NA</td></tr>
<tr><td>T10188</td><td>GGGGCCCGTGCCCACGCACGCGG</td><td>1000</td><td></td><td></td><td></td><td></td><td></td><td></td><td></td><td></td><td></td></tr>
<tr><td>Oligo fwd</td><td>CACCGGGGCCCGTGCCCACGCACG</td><td></td><td></td><td></td><td></td><td></td><td></td><td></td><td></td><td></td><td></td></tr>
<tr><td>Oligo rev</td><td>AAACCGTGCGTGGGCACGGGCCCC</td><td></td><td></td><td></td><td></td><td></td><td></td><td></td><td></td><td></td><td></td></tr>
<tr><td>Chromosome</td><td>start</td><td>end</td><td>strand</td><td>MM</td><td>target_seq</td><td>PAM</td><td>alignment</td><td>distance</td><td>position</td><td>gene name</td><td>gene id</td></tr>
<tr><td>GU305914.1</td><td>134226</td><td>134248</td><td>+</td><td>0</td><td>GGGGCCCGTGCCCACGCACG</td><td>CGG</td><td>||||||||[||||||||||||]PAM</td><td>NA</td><td>Intergenic</td><td>NA</td><td>NA</td></tr>
<tr><td>T10189</td><td>GGGCCCGTGCCCACGCACGCGGG</td><td>1000</td><td></td><td></td><td></td><td></td><td></td><td></td><td></td><td></td><td></td></tr>
<tr><td>Oligo fwd</td><td>CACCGGGCCCGTGCCCACGCACGC</td><td></td><td></td><td></td><td></td><td></td><td></td><td></td><td></td><td></td><td></td></tr>
<tr><td>Oligo rev</td><td>AAACGCGTGCGTGGGCACGGGCCC</td><td></td><td></td><td></td><td></td><td></td><td></td><td></td><td></td><td></td><td></td></tr>
<tr><td>Chromosome</td><td>start</td><td>end</td><td>strand</td><td>MM</td><td>target_seq</td><td>PAM</td><td>alignment</td><td>distance</td><td>position</td><td>gene name</td><td>gene id</td></tr>
<tr><td>GU305914.1</td><td>134227</td><td>134249</td><td>+</td><td>0</td><td>GGGCCCGTGCCCACGCACGC</td><td>GGG</td><td>||||||||[||||||||||||]PAM</td><td>NA</td><td>Intergenic</td><td>NA</td><td>NA</td></tr>
<tr><td>T10190</td><td>GGCCCGTGCCCACGCACGCGGGG</td><td>1000</td><td></td><td></td><td></td><td></td><td></td><td></td><td></td><td></td><td></td></tr>
<tr><td>Oligo fwd</td><td>CACCGGCCCGTGCCCACGCACGCG</td><td></td><td></td><td></td><td></td><td></td><td></td><td></td><td></td><td></td><td></td></tr>
<tr><td>Oligo rev</td><td>AAACCGCGTGCGTGGGCACGGGCC</td><td></td><td></td><td></td><td></td><td></td><td></td><td></td><td></td><td></td><td></td></tr>
<tr><td>Chromosome</td><td>start</td><td>end</td><td>strand</td><td>MM</td><td>target_seq</td><td>PAM</td><td>alignment</td><td>distance</td><td>position</td><td>gene name</td><td>gene id</td></tr>
<tr><td>GU305914.1</td><td>134228</td><td>134250</td><td>+</td><td>0</td><td>GGCCCGTGCCCACGCACGCG</td><td>GGG</td><td>||||||||[||||||||||||]PAM</td><td>NA</td><td>Intergenic</td><td>NA</td><td>NA</td></tr>
<tr><td>T10191</td><td>GCCCGTGCCCACGCACGCGGGGG</td><td>1000</td><td></td><td></td><td></td><td></td><td></td><td></td><td></td><td></td><td></td></tr>
<tr><td>Oligo fwd</td><td>CACCGCCCGTGCCCACGCACGCGG</td><td></td><td></td><td></td><td></td><td></td><td></td><td></td><td></td><td></td><td></td></tr>
<tr><td>Oligo rev</td><td>AAACCCGCGTGCGTGGGCACGGGC</td><td></td><td></td><td></td><td></td><td></td><td></td><td></td><td></td><td></td><td></td></tr>
<tr><td>Chromosome</td><td>start</td><td>end</td><td>strand</td><td>MM</td><td>target_seq</td><td>PAM</td><td>alignment</td><td>distance</td><td>position</td><td>gene name</td><td>gene id</td></tr>
<tr><td>GU305914.1</td><td>134229</td><td>134251</td><td>+</td><td>0</td><td>GCCCGTGCCCACGCACGCGG</td><td>GGG</td><td>||||||||[||||||||||||]PAM</td><td>NA</td><td>Intergenic</td><td>NA</td><td>NA</td></tr>
<tr><td>T10192</td><td>CGTGCCCACGCACGCGGGGGCGG</td><td>1000</td><td></td><td></td><td></td><td></td><td></td><td></td><td></td><td></td><td></td></tr>
<tr><td>Oligo fwd</td><td>CACCCGTGCCCACGCACGCGGGGG</td><td></td><td></td><td></td><td></td><td></td><td></td><td></td><td></td><td></td><td></td></tr>
<tr><td>Oligo rev</td><td>AAACCCCCCGCGTGCGTGGGCACG</td><td></td><td></td><td></td><td></td><td></td><td></td><td></td><td></td><td></td><td></td></tr>
<tr><td>Chromosome</td><td>start</td><td>end</td><td>strand</td><td>MM</td><td>target_seq</td><td>PAM</td><td>alignment</td><td>distance</td><td>position</td><td>gene name</td><td>gene id</td></tr>
<tr><td>GU305914.1</td><td>134232</td><td>134254</td><td>+</td><td>0</td><td>CGTGCCCACGCACGCGGGGG</td><td>CGG</td><td>||||||||[||||||||||||]PAM</td><td>NA</td><td>Intergenic</td><td>NA</td><td>NA</td></tr>
<tr><td>T10193</td><td>CACGCACGCGGGGGCGGTAGTGG</td><td>1000</td><td></td><td></td><td></td><td></td><td></td><td></td><td></td><td></td><td></td></tr>
<tr><td>Oligo fwd</td><td>CACCCACGCACGCGGGGGCGGTAG</td><td></td><td></td><td></td><td></td><td></td><td></td><td></td><td></td><td></td><td></td></tr>
<tr><td>Oligo rev</td><td>AAACCTACCGCCCCCGCGTGCGTG</td><td></td><td></td><td></td><td></td><td></td><td></td><td></td><td></td><td></td><td></td></tr>
<tr><td>Chromosome</td><td>start</td><td>end</td><td>strand</td><td>MM</td><td>target_seq</td><td>PAM</td><td>alignment</td><td>distance</td><td>position</td><td>gene name</td><td>gene id</td></tr>
<tr><td>GU305914.1</td><td>134238</td><td>134260</td><td>+</td><td>0</td><td>CACGCACGCGGGGGCGGTAG</td><td>TGG</td><td>||||||||[||||||||||||]PAM</td><td>NA</td><td>Intergenic</td><td>NA</td><td>NA</td></tr>
<tr><td>T10194</td><td>ACGCACGCGGGGGCGGTAGTGGG</td><td>1000</td><td></td><td></td><td></td><td></td><td></td><td></td><td></td><td></td><td></td></tr>
<tr><td>Oligo fwd</td><td>CACCACGCACGCGGGGGCGGTAGT</td><td></td><td></td><td></td><td></td><td></td><td></td><td></td><td></td><td></td><td></td></tr>
<tr><td>Oligo rev</td><td>AAACACTACCGCCCCCGCGTGCGT</td><td></td><td></td><td></td><td></td><td></td><td></td><td></td><td></td><td></td><td></td></tr>
<tr><td>Chromosome</td><td>start</td><td>end</td><td>strand</td><td>MM</td><td>target_seq</td><td>PAM</td><td>alignment</td><td>distance</td><td>position</td><td>gene name</td><td>gene id</td></tr>
<tr><td>GU305914.1</td><td>134239</td><td>134261</td><td>+</td><td>0</td><td>ACGCACGCGGGGGCGGTAGT</td><td>GGG</td><td>||||||||[||||||||||||]PAM</td><td>NA</td><td>Intergenic</td><td>NA</td><td>NA</td></tr>
<tr><td>T10195</td><td>GGCGGTAGTGGGAAGTGCGATGG</td><td>1000</td><td></td><td></td><td></td><td></td><td></td><td></td><td></td><td></td><td></td></tr>
<tr><td>Oligo fwd</td><td>CACCGGCGGTAGTGGGAAGTGCGA</td><td></td><td></td><td></td><td></td><td></td><td></td><td></td><td></td><td></td><td></td></tr>
<tr><td>Oligo rev</td><td>AAACTCGCACTTCCCACTACCGCC</td><td></td><td></td><td></td><td></td><td></td><td></td><td></td><td></td><td></td><td></td></tr>
<tr><td>Chromosome</td><td>start</td><td>end</td><td>strand</td><td>MM</td><td>target_seq</td><td>PAM</td><td>alignment</td><td>distance</td><td>position</td><td>gene name</td><td>gene id</td></tr>
<tr><td>GU305914.1</td><td>134250</td><td>134272</td><td>+</td><td>0</td><td>GGCGGTAGTGGGAAGTGCGA</td><td>TGG</td><td>||||||||[||||||||||||]PAM</td><td>NA</td><td>Intergenic</td><td>NA</td><td>NA</td></tr>
<tr><td>T10196</td><td>GGTAGTGGGAAGTGCGATGGCGG</td><td>1000</td><td></td><td></td><td></td><td></td><td></td><td></td><td></td><td></td><td></td></tr>
<tr><td>Oligo fwd</td><td>CACCGGTAGTGGGAAGTGCGATGG</td><td></td><td></td><td></td><td></td><td></td><td></td><td></td><td></td><td></td><td></td></tr>
<tr><td>Oligo rev</td><td>AAACCCATCGCACTTCCCACTACC</td><td></td><td></td><td></td><td></td><td></td><td></td><td></td><td></td><td></td><td></td></tr>
<tr><td>Chromosome</td><td>start</td><td>end</td><td>strand</td><td>MM</td><td>target_seq</td><td>PAM</td><td>alignment</td><td>distance</td><td>position</td><td>gene name</td><td>gene id</td></tr>
<tr><td>GU305914.1</td><td>134253</td><td>134275</td><td>+</td><td>0</td><td>GGTAGTGGGAAGTGCGATGG</td><td>CGG</td><td>||||||||[||||||||||||]PAM</td><td>NA</td><td>Intergenic</td><td>NA</td><td>NA</td></tr>
<tr><td>T10197</td><td>AGTGGGAAGTGCGATGGCGGCGG</td><td>1000</td><td></td><td></td><td></td><td></td><td></td><td></td><td></td><td></td><td></td></tr>
<tr><td>Oligo fwd</td><td>CACCAGTGGGAAGTGCGATGGCGG</td><td></td><td></td><td></td><td></td><td></td><td></td><td></td><td></td><td></td><td></td></tr>
<tr><td>Oligo rev</td><td>AAACCCGCCATCGCACTTCCCACT</td><td></td><td></td><td></td><td></td><td></td><td></td><td></td><td></td><td></td><td></td></tr>
<tr><td>Chromosome</td><td>start</td><td>end</td><td>strand</td><td>MM</td><td>target_seq</td><td>PAM</td><td>alignment</td><td>distance</td><td>position</td><td>gene name</td><td>gene id</td></tr>
<tr><td>GU305914.1</td><td>134256</td><td>134278</td><td>+</td><td>0</td><td>AGTGGGAAGTGCGATGGCGG</td><td>CGG</td><td>||||||||[||||||||||||]PAM</td><td>NA</td><td>Intergenic</td><td>NA</td><td>NA</td></tr>
<tr><td>T10198</td><td>GGGAAGTGCGATGGCGGCGGAGG</td><td>1000</td><td></td><td></td><td></td><td></td><td></td><td></td><td></td><td></td><td></td></tr>
<tr><td>Oligo fwd</td><td>CACCGGGAAGTGCGATGGCGGCGG</td><td></td><td></td><td></td><td></td><td></td><td></td><td></td><td></td><td></td><td></td></tr>
<tr><td>Oligo rev</td><td>AAACCCGCCGCCATCGCACTTCCC</td><td></td><td></td><td></td><td></td><td></td><td></td><td></td><td></td><td></td><td></td></tr>
<tr><td>Chromosome</td><td>start</td><td>end</td><td>strand</td><td>MM</td><td>target_seq</td><td>PAM</td><td>alignment</td><td>distance</td><td>position</td><td>gene name</td><td>gene id</td></tr>
<tr><td>GU305914.1</td><td>134259</td><td>134281</td><td>+</td><td>0</td><td>GGGAAGTGCGATGGCGGCGG</td><td>AGG</td><td>||||||||[||||||||||||]PAM</td><td>NA</td><td>Intergenic</td><td>NA</td><td>NA</td></tr>
<tr><td>GU305914.1</td><td>225984</td><td>226006</td><td>-</td><td>4</td><td>GTGGAGCGCGACGGCGGCGG</td><td>CGG</td><td>|-|-||-|[|||-||||||||]PAM</td><td>NA</td><td>Intergenic</td><td>NA</td><td>NA</td></tr>
<tr><td>GU305914.1</td><td>127139</td><td>127161</td><td>-</td><td>4</td><td>GGGAACGGCGTTGGCGACGG</td><td>CGG</td><td>|||||--|[||-|||||-|||]PAM</td><td>NA</td><td>Intergenic</td><td>NA</td><td>NA</td></tr>
<tr><td>T10199</td><td>GCGGCGGAGGCTGCGATATGAGG</td><td>1000</td><td></td><td></td><td></td><td></td><td></td><td></td><td></td><td></td><td></td></tr>
<tr><td>Oligo fwd</td><td>CACCGCGGCGGAGGCTGCGATATG</td><td></td><td></td><td></td><td></td><td></td><td></td><td></td><td></td><td></td><td></td></tr>
<tr><td>Oligo rev</td><td>AAACCATATCGCAGCCTCCGCCGC</td><td></td><td></td><td></td><td></td><td></td><td></td><td></td><td></td><td></td><td></td></tr>
<tr><td>Chromosome</td><td>start</td><td>end</td><td>strand</td><td>MM</td><td>target_seq</td><td>PAM</td><td>alignment</td><td>distance</td><td>position</td><td>gene name</td><td>gene id</td></tr>
</table>

GU305914.1 134272 134294 + 0 GCGGCGGAGGCTGCGATATG AGG ||||||||[||||||||||||]PAM NA Intergenic NA NA

T10200 CTTGTGCAACCCGCTGACGCAGG 1000
Oligo fwd CACCCTTGTGCAACCCGCTGACGC
Oligo rev AAACGCGTCAGCGGGTTGCACAAG
Chromosome start end strand MM target_seq PAM alignment distance position gene name gene id
GU305914.1 134297 134319 + 0 CTTGTGCAACCCGCTGACGC AGG ||||||||[||||||||||||]PAM NA Intergenic NA NA

T10201 TGACGCAGGAGCTCAACCTCAGG 1000
Oligo fwd CACCTGACGCAGGAGCTCAACCTC
Oligo rev AAACGAGGTTGAGCTCCTGCGTCA
Chromosome start end strand MM target_seq PAM alignment distance position gene name gene id
GU305914.1 134311 134333 + 0 TGACGCAGGAGCTCAACCTC AGG ||||||||[||||||||||||]PAM NA Intergenic NA NA

T10202 GCACTCACCTCTGCGATCTACGG 1000
Oligo fwd CACCGCACTCACCTCTGCGATCTA
Oligo rev AAACTAGATCGCAGAGGTGAGTGC
Chromosome start end strand MM target_seq PAM alignment distance position gene name gene id
GU305914.1 134362 134384 + 0 GCACTCACCTCTGCGATCTA CGG ||||||||[||||||||||||]PAM NA Intergenic NA NA

T10203 CGATCTACGGCACGACTGTGTGG 1000
Oligo fwd CACCCGATCTACGGCACGACTGTG
Oligo rev AAACCACAGTCGTGCCGTAGATCG
Chromosome start end strand MM target_seq PAM alignment distance position gene name gene id
GU305914.1 134375 134397 + 0 CGATCTACGGCACGACTGTG TGG ||||||||[||||||||||||]PAM NA Intergenic NA NA

T10204 TCTACGGCACGACTGTGTGGTGG 1000
Oligo fwd CACCTCTACGGCACGACTGTGTGG
Oligo rev AAACCCACACAGTCGTGCCGTAGA
Chromosome start end strand MM target_seq PAM alignment distance position gene name gene id
GU305914.1 134378 134400 + 0 TCTACGGCACGACTGTGTGG TGG ||||||||[||||||||||||]PAM NA Intergenic NA NA

T10205 CTGTGTGGTGGTCCACACGCAGG 1000
Oligo fwd CACCCTGTGTGGTGGTCCACACGC
Oligo rev AAACGCGTGTGGACCACCACACAG
Chromosome start end strand MM target_seq PAM alignment distance position gene name gene id
GU305914.1 134390 134412 + 0 CTGTGTGGTGGTCCACACGC AGG ||||||||[||||||||||||]PAM NA Intergenic NA NA

T10206 GTGGTGGTCCACACGCAGGACGG 1000
Oligo fwd CACCGTGGTGGTCCACACGCAGGA
Oligo rev AAACTCCTGCGTGTGGACCACCAC
Chromosome start end strand MM target_seq PAM alignment distance position gene name gene id
GU305914.1 134394 134416 + 0 GTGGTGGTCCACACGCAGGA CGG ||||||||[||||||||||||]PAM NA Intergenic NA NA

T10207 CGGCTCCGTCTGCATCAAGACGG 1000
Oligo fwd CACCCGGCTCCGTCTGCATCAAGA
Oligo rev AAACTCTTGATGCAGACGGAGCCG
Chromosome start end strand MM target_seq PAM alignment distance position gene name gene id
GU305914.1 134414 134436 + 0 CGGCTCCGTCTGCATCAAGA CGG ||||||||[||||||||||||]PAM NA Intergenic NA NA

T10208 GGCTCCGTCTGCATCAAGACGGG 1000
Oligo fwd CACCGGCTCCGTCTGCATCAAGAC
Oligo rev AAACGTCTTGATGCAGACGGAGCC
Chromosome start end strand MM target_seq PAM alignment distance position gene name gene id
GU305914.1 134415 134437 + 0 GGCTCCGTCTGCATCAAGAC GGG ||||||||[||||||||||||]PAM NA Intergenic NA NA

T10209 ATCAAGACGGGCCTCACCTACGG 1000
Oligo fwd CACCATCAAGACGGGCCTCACCTA
Oligo rev AAACTAGGTGAGGCCCGTCTTGAT
Chromosome start end strand MM target_seq PAM alignment distance position gene name gene id
GU305914.1 134427 134449 + 0 ATCAAGACGGGCCTCACCTA CGG ||||||||[||||||||||||]PAM NA Intergenic NA NA

T10210 ACCTACGGCTCCGTTTTCCCCGG 1000
Oligo fwd CACCACCTACGGCTCCGTTTTCCC
Oligo rev AAACGGGAAAACGGAGCCGTAGGT
Chromosome start end strand MM target_seq PAM alignment distance position gene name gene id
GU305914.1 134442 134464 + 0 ACCTACGGCTCCGTTTTCCC CGG ||||||||[||||||||||||]PAM NA Intergenic NA NA

T10211 CCTACGGCTCCGTTTTCCCCGGG 1000
Oligo fwd CACCCCTACGGCTCCGTTTTCCCC
Oligo rev AAACGGGGAAAACGGAGCCGTAGG
Chromosome start end strand MM target_seq PAM alignment distance position gene name gene id
GU305914.1 134443 134465 + 0 CCTACGGCTCCGTTTTCCCC GGG ||||||||[||||||||||||]PAM NA Intergenic NA NA

T10212 CTACGGCTCCGTTTTCCCCGGGG 1000
Oligo fwd CACCCTACGGCTCCGTTTTCCCCG
Oligo rev AAACCGGGGAAAACGGAGCCGTAG
Chromosome start end strand MM target_seq PAM alignment distance position gene name gene id
GU305914.1 134444 134466 + 0 CTACGGCTCCGTTTTCCCCG GGG ||||||||[||||||||||||]PAM NA Intergenic NA NA

T10213 TACGGCTCCGTTTTCCCCGGGGG 1000
Oligo fwd CACCTACGGCTCCGTTTTCCCCGG
Oligo rev AAACCCGGGGAAAACGGAGCCGTA
Chromosome start end strand MM target_seq PAM alignment distance position gene name gene id
GU305914.1 134445 134467 + 0 TACGGCTCCGTTTTCCCCGG GGG ||||||||[||||||||||||]PAM NA Intergenic NA NA

T10214 CGGGGGCTGCGTGAGCGCCCTGG 1000
Oligo fwd CACCCGGGGGCTGCGTGAGCGCCC
Oligo rev AAACGGGCGCTCACGCAGCCCCCG

Chromosome start end strand MM target_seq PAM alignment distance position gene name gene id
GU305914.1 134462 134484 + 0 CGGGGGCTGCGTGAGCGCCC TGG ||||||||[||||||||||||]PAM NA Intergenic NA NA

T10215 GCACGTCGACGAGATCAATGTGG 1000
Oligo fwd CACCGCACGTCGACGAGATCAATG
Oligo rev AAACCATTGATCTCGTCGACGTGC
Chromosome start end strand MM target_seq PAM alignment distance position gene name gene id
GU305914.1 134501 134523 + 0 GCACGTCGACGAGATCAATG TGG ||||||||[||||||||||||]PAM NA Intergenic NA NA

T10216 GTCGACGAGATCAATGTGGTCGG 1000
Oligo fwd CACCGTCGACGAGATCAATGTGGT
Oligo rev AAACACCACATTGATCTCGTCGAC
Chromosome start end strand MM target_seq PAM alignment distance position gene name gene id
GU305914.1 134505 134527 + 0 GTCGACGAGATCAATGTGGT CGG ||||||||[||||||||||||]PAM NA Intergenic NA NA

T10217 ACGTCTACACCTATCTCACCAGG 1000
Oligo fwd CACCACGTCTACACCTATCTCACC
Oligo rev AAACGGTGAGATAGGTGTAGACGT
Chromosome start end strand MM target_seq PAM alignment distance position gene name gene id
GU305914.1 134542 134564 + 0 ACGTCTACACCTATCTCACC AGG ||||||||[||||||||||||]PAM NA Intergenic NA NA

T10218 CAGGAACGCTGACCACTACGCGG 1000
Oligo fwd CACCCAGGAACGCTGACCACTACG
Oligo rev AAACCGTAGTGGTCAGCGTTCCTG
Chromosome start end strand MM target_seq PAM alignment distance position gene name gene id
GU305914.1 134561 134583 + 0 CAGGAACGCTGACCACTACG CGG ||||||||[||||||||||||]PAM NA Intergenic NA NA

T10219 GACCACTACGCGGATGTCATAGG 1000
Oligo fwd CACCGACCACTACGCGGATGTCAT
Oligo rev AAACATGACATCCGCGTAGTGGTC
Chromosome start end strand MM target_seq PAM alignment distance position gene name gene id
GU305914.1 134571 134593 + 0 GACCACTACGCGGATGTCAT AGG ||||||||[||||||||||||]PAM NA Intergenic NA NA

T10220 ACCACTACGCGGATGTCATAGGG 1000
Oligo fwd CACCACCACTACGCGGATGTCATA
Oligo rev AAACTATGACATCCGCGTAGTGGT
Chromosome start end strand MM target_seq PAM alignment distance position gene name gene id
GU305914.1 134572 134594 + 0 ACCACTACGCGGATGTCATA GGG ||||||||[||||||||||||]PAM NA Intergenic NA NA

T10221 TGTCATAGGGAGCGTGATCGAGG 1000
Oligo fwd CACCTGTCATAGGGAGCGTGATCG
Oligo rev AAACCGATCACGCTCCCTATGACA
Chromosome start end strand MM target_seq PAM alignment distance position gene name gene id
GU305914.1 134585 134607 + 0 TGTCATAGGGAGCGTGATCG AGG ||||||||[||||||||||||]PAM NA Intergenic NA NA

T10222 GTCATAGGGAGCGTGATCGAGGG 1000
Oligo fwd CACCGTCATAGGGAGCGTGATCGA
Oligo rev AAACTCGATCACGCTCCCTATGAC
Chromosome start end strand MM target_seq PAM alignment distance position gene name gene id
GU305914.1 134586 134608 + 0 GTCATAGGGAGCGTGATCGA GGG ||||||||[||||||||||||]PAM NA Intergenic NA NA

T10223 TCATAGGGAGCGTGATCGAGGGG 1000
Oligo fwd CACCTCATAGGGAGCGTGATCGAG
Oligo rev AAACCTCGATCACGCTCCCTATGA
Chromosome start end strand MM target_seq PAM alignment distance position gene name gene id
GU305914.1 134587 134609 + 0 TCATAGGGAGCGTGATCGAG GGG ||||||||[||||||||||||]PAM NA Intergenic NA NA

T10224 CATAGGGAGCGTGATCGAGGGGG 1000
Oligo fwd CACCCATAGGGAGCGTGATCGAGG
Oligo rev AAACCCTCGATCACGCTCCCTATG
Chromosome start end strand MM target_seq PAM alignment distance position gene name gene id
GU305914.1 134588 134610 + 0 CATAGGGAGCGTGATCGAGG GGG ||||||||[||||||||||||]PAM NA Intergenic NA NA

T10225 ATAGGGAGCGTGATCGAGGGGGG 1000
Oligo fwd CACCATAGGGAGCGTGATCGAGGG
Oligo rev AAACCCCTCGATCACGCTCCCTAT
Chromosome start end strand MM target_seq PAM alignment distance position gene name gene id
GU305914.1 134589 134611 + 0 ATAGGGAGCGTGATCGAGGG GGG ||||||||[||||||||||||]PAM NA Intergenic NA NA

T10226 GGAGCGTGATCGAGGGGGGTTGG 1000
Oligo fwd CACCGGAGCGTGATCGAGGGGGGT
Oligo rev AAACACCCCCCTCGATCACGCTCC
Chromosome start end strand MM target_seq PAM alignment distance position gene name gene id
GU305914.1 134593 134615 + 0 GGAGCGTGATCGAGGGGGGT TGG ||||||||[||||||||||||]PAM NA Intergenic NA NA

T10227 GGGTTGGTTTAACAAACCCACGG 1000
Oligo fwd CACCGGGTTGGTTTAACAAACCCA
Oligo rev AAACTGGGTTTGTTAAACCAACCC
Chromosome start end strand MM target_seq PAM alignment distance position gene name gene id
GU305914.1 134609 134631 + 0 GGGTTGGTTTAACAAACCCA CGG ||||||||[||||||||||||]PAM NA Intergenic NA NA

T10228 GCAGCATAACGCGCTGCAGAAGG 1000
Oligo fwd CACCGCAGCATAACGCGCTGCAGA
Oligo rev AAACTCTGCAGCGCGTTATGCTGC
Chromosome start end strand MM target_seq PAM alignment distance position gene name gene id
GU305914.1 134669 134691 + 0 GCAGCATAACGCGCTGCAGA AGG ||||||||[||||||||||||]PAM NA Intergenic NA NA

T10229 AAGGTGCCCATCAGCGTCATAGG 1000
Oligo fwd CACCAAGGTGCCCATCAGCGTCAT

Oligo rev AAACATGACGCTGATGGGCACCTT
Chromosome start end strand MM target_seq PAM alignment distance position gene name gene id
GU305914.1 134688 134710 + 0 AAGGTGCCCATCAGCGTCAT AGG ||||||||[||||||||||||]PAM NA Intergenic NA NA

T10230 AGGTCAGCTGTTCGTGCAGCTGG 1000
Oligo fwd CACCAGGTCAGCTGTTCGTGCAGC
Oligo rev AAACGCTGCACGAACAGCTGACCT
Chromosome start end strand MM target_seq PAM alignment distance position gene name gene id
GU305914.1 134708 134730 + 0 AGGTCAGCTGTTCGTGCAGC TGG ||||||||[||||||||||||]PAM NA Intergenic NA NA

T10231 CTGTTCGTGCAGCTGGTCATCGG 1000
Oligo fwd CACCCTGTTCGTGCAGCTGGTCAT
Oligo rev AAACATGACCAGCTGCACGAACAG
Chromosome start end strand MM target_seq PAM alignment distance position gene name gene id
GU305914.1 134715 134737 + 0 CTGTTCGTGCAGCTGGTCAT CGG ||||||||[||||||||||||]PAM NA Intergenic NA NA

T10232 TGGTCATCGGAGTGCATGCCAGG 1000
Oligo fwd CACCTGGTCATCGGAGTGCATGCC
Oligo rev AAACGGCATGCACTCCGATGACCA
Chromosome start end strand MM target_seq PAM alignment distance position gene name gene id
GU305914.1 134728 134750 + 0 TGGTCATCGGAGTGCATGCC AGG ||||||||[||||||||||||]PAM NA Intergenic NA NA

T10233 GGTCATCGGAGTGCATGCCAGGG 1000
Oligo fwd CACCGGTCATCGGAGTGCATGCCA
Oligo rev AAACTGGCATGCACTCCGATGACC
Chromosome start end strand MM target_seq PAM alignment distance position gene name gene id
GU305914.1 134729 134751 + 0 GGTCATCGGAGTGCATGCCA GGG ||||||||[||||||||||||]PAM NA Intergenic NA NA

T10234 CGACCGTGATCAAAGTCAGTCGG 1000
Oligo fwd CACCCGACCGTGATCAAAGTCAGT
Oligo rev AAACACTGACTTTGATCACGGTCG
Chromosome start end strand MM target_seq PAM alignment distance position gene name gene id
GU305914.1 134767 134789 + 0 CGACCGTGATCAAAGTCAGT CGG ||||||||[||||||||||||]PAM NA Intergenic NA NA

T10235 CCGTGATCAAAGTCAGTCGGAGG 1000
Oligo fwd CACCCCGTGATCAAAGTCAGTCGG
Oligo rev AAACCCGACTGACTTTGATCACGG
Chromosome start end strand MM target_seq PAM alignment distance position gene name gene id
GU305914.1 134770 134792 + 0 CCGTGATCAAAGTCAGTCGG AGG ||||||||[||||||||||||]PAM NA Intergenic NA NA

T10236 TCAAAGTCAGTCGGAGGAAGAGG 1000
Oligo fwd CACCTCAAAGTCAGTCGGAGGAAG
Oligo rev AAACCTTCCTCCGACTGACTTTGA
Chromosome start end strand MM target_seq PAM alignment distance position gene name gene id
GU305914.1 134776 134798 + 0 TCAAAGTCAGTCGGAGGAAG AGG ||||||||[||||||||||||]PAM NA Intergenic NA NA

T10237 CAAAGTCAGTCGGAGGAAGAGGG 1000
Oligo fwd CACCCAAAGTCAGTCGGAGGAAGA
Oligo rev AAACTCTTCCTCCGACTGACTTTG
Chromosome start end strand MM target_seq PAM alignment distance position gene name gene id
GU305914.1 134777 134799 + 0 CAAAGTCAGTCGGAGGAAGA GGG ||||||||[||||||||||||]PAM NA Intergenic NA NA

T10238 AGTCAGTCGGAGGAAGAGGGAGG 1000
Oligo fwd CACCAGTCAGTCGGAGGAAGAGGG
Oligo rev AAACCCCTCTTCCTCCGACTGACT
Chromosome start end strand MM target_seq PAM alignment distance position gene name gene id
GU305914.1 134780 134802 + 0 AGTCAGTCGGAGGAAGAGGG AGG ||||||||[||||||||||||]PAM NA Intergenic NA NA

T10239 AGTCGGAGGAAGAGGGAGGACGG 1000
Oligo fwd CACCAGTCGGAGGAAGAGGGAGGA
Oligo rev AAACTCCTCCCTCTTCCTCCGACT
Chromosome start end strand MM target_seq PAM alignment distance position gene name gene id
GU305914.1 134784 134806 + 0 AGTCGGAGGAAGAGGGAGGA CGG ||||||||[||||||||||||]PAM NA Intergenic NA NA

T10240 GTCGGAGGAAGAGGGAGGACGGG 1000
Oligo fwd CACCGTCGGAGGAAGAGGGAGGAC
Oligo rev AAACGTCCTCCCTCTTCCTCCGAC
Chromosome start end strand MM target_seq PAM alignment distance position gene name gene id
GU305914.1 134785 134807 + 0 GTCGGAGGAAGAGGGAGGAC GGG ||||||||[||||||||||||]PAM NA Intergenic NA NA

T10241 GGGAGGACGGGCTCCTGAAACGG 1000
Oligo fwd CACCGGGAGGACGGGCTCCTGAAA
Oligo rev AAACTTTCAGGAGCCCGTCCTCCC
Chromosome start end strand MM target_seq PAM alignment distance position gene name gene id
GU305914.1 134797 134819 + 0 GGGAGGACGGGCTCCTGAAA CGG ||||||||[||||||||||||]PAM NA Intergenic NA NA

T10242 AAACGGATGCGTTTCGAATATGG 1000
Oligo fwd CACCAAACGGATGCGTTTCGAATA
Oligo rev AAACTATTCGAAACGCATCCGTTT
Chromosome start end strand MM target_seq PAM alignment distance position gene name gene id
GU305914.1 134814 134836 + 0 AAACGGATGCGTTTCGAATA TGG ||||||||[||||||||||||]PAM NA Intergenic NA NA

T10243 AACGCACCTTCGTTTCGAACAGG 1000
Oligo fwd CACCAACGCACCTTCGTTTCGAAC
Oligo rev AAACGTTCGAAACGAAGGTGCGTT
Chromosome start end strand MM target_seq PAM alignment distance position gene name gene id
GU305914.1 134838 134860 + 0 AACGCACCTTCGTTTCGAAC AGG ||||||||[||||||||||||]PAM NA Intergenic NA NA

T10244 TTTCGAACAGGCCGCTAGCGAGG 1000

Oligo fwd CACCTTTCGAACAGGCCGCTAGCG
Oligo rev AAACCGCTAGCGGCCTGTTCGAAA
Chromosome start end strand MM target_seq PAM alignment distance position gene name gene id
GU305914.1 134850 134872 + 0 TTTCGAACAGGCCGCTAGCG AGG ||||||||[||||||||||||]PAM NA Intergenic NA NA

T10245 CTAGCGAGGACGACCGCGACAGG 1000
Oligo fwd CACCCTAGCGAGGACGACCGCGAC
Oligo rev AAACGTCGCGGTCGTCCTCGCTAG
Chromosome start end strand MM target_seq PAM alignment distance position gene name gene id
GU305914.1 134864 134886 + 0 CTAGCGAGGACGACCGCGAC AGG ||||||||[||||||||||||]PAM NA Intergenic NA NA

T10246 CGACCGCGACAGGCTCGTGCCGG 1000
Oligo fwd CACCCGACCGCGACAGGCTCGTGC
Oligo rev AAACGCACGAGCCTGTCGCGGTCG
Chromosome start end strand MM target_seq PAM alignment distance position gene name gene id
GU305914.1 134874 134896 + 0 CGACCGCGACAGGCTCGTGC CGG ||||||||[||||||||||||]PAM NA Intergenic NA NA

T10247 GGTACACGTCGTTTTCGCTGAGG 1000
Oligo fwd CACCGGTACACGTCGTTTTCGCTG
Oligo rev AAACCAGCGAAAACGACGTGTACC
Chromosome start end strand MM target_seq PAM alignment distance position gene name gene id
GU305914.1 134895 134917 + 0 GGTACACGTCGTTTTCGCTG AGG ||||||||[||||||||||||]PAM NA Intergenic NA NA

T10248 CTGAGGATGCGCTCTCCTACCGG 1000
Oligo fwd CACCCTGAGGATGCGCTCTCCTAC
Oligo rev AAACGTAGGAGAGCGCATCCTCAG
Chromosome start end strand MM target_seq PAM alignment distance position gene name gene id
GU305914.1 134912 134934 + 0 CTGAGGATGCGCTCTCCTAC CGG ||||||||[||||||||||||]PAM NA Intergenic NA NA

T10249 TGAGGATGCGCTCTCCTACCGGG 1000
Oligo fwd CACCTGAGGATGCGCTCTCCTACC
Oligo rev AAACGGTAGGAGAGCGCATCCTCA
Chromosome start end strand MM target_seq PAM alignment distance position gene name gene id
GU305914.1 134913 134935 + 0 TGAGGATGCGCTCTCCTACC GGG ||||||||[||||||||||||]PAM NA Intergenic NA NA

T10250 GCTCTCCTACCGGGAGTTCGAGG 1000
Oligo fwd CACCGCTCTCCTACCGGGAGTTCG
Oligo rev AAACCGAACTCCCGGTAGGAGAGC
Chromosome start end strand MM target_seq PAM alignment distance position gene name gene id
GU305914.1 134922 134944 + 0 GCTCTCCTACCGGGAGTTCG AGG ||||||||[||||||||||||]PAM NA Intergenic NA NA

T10251 CTCTCCTACCGGGAGTTCGAGGG 1000
Oligo fwd CACCCTCTCCTACCGGGAGTTCGA
Oligo rev AAACTCGAACTCCCGGTAGGAGAG
Chromosome start end strand MM target_seq PAM alignment distance position gene name gene id
GU305914.1 134923 134945 + 0 CTCTCCTACCGGGAGTTCGA GGG ||||||||[||||||||||||]PAM NA Intergenic NA NA

T10252 TCTCCAGACCCGACGCGCGCTGG 1000
Oligo fwd CACCTCTCCAGACCCGACGCGCGC
Oligo rev AAACGCGCGCGTCGGGTCTGGAGA
Chromosome start end strand MM target_seq PAM alignment distance position gene name gene id
GU305914.1 134972 134994 + 0 TCTCCAGACCCGACGCGCGC TGG ||||||||[||||||||||||]PAM NA Intergenic NA NA

T10253 CTCCAGACCCGACGCGCGCTGGG 1000
Oligo fwd CACCCTCCAGACCCGACGCGCGCT
Oligo rev AAACAGCGCGCGTCGGGTCTGGAG
Chromosome start end strand MM target_seq PAM alignment distance position gene name gene id
GU305914.1 134973 134995 + 0 CTCCAGACCCGACGCGCGCT GGG ||||||||[||||||||||||]PAM NA Intergenic NA NA

T10254 CATCCCCTACTTCGAGCGTCTGG 1000
Oligo fwd CACCCATCCCCTACTTCGAGCGTC
Oligo rev AAACGACGCTCGAAGTAGGGGATG
Chromosome start end strand MM target_seq PAM alignment distance position gene name gene id
GU305914.1 135000 135022 + 0 CATCCCCTACTTCGAGCGTC TGG ||||||||[||||||||||||]PAM NA Intergenic NA NA

T10255 GTCTCTTCCACGACAAGTGCAGG 1000
Oligo fwd CACCGTCTCTTCCACGACAAGTGC
Oligo rev AAACGCACTTGTCGTGGAAGAGAC
Chromosome start end strand MM target_seq PAM alignment distance position gene name gene id
GU305914.1 135044 135066 + 0 GTCTCTTCCACGACAAGTGC AGG ||||||||[||||||||||||]PAM NA Intergenic NA NA

T10256 CGACAAGTGCAGGCACCTCTCGG 1000
Oligo fwd CACCCGACAAGTGCAGGCACCTCT
Oligo rev AAACAGAGGTGCCTGCACTTGTCG
Chromosome start end strand MM target_seq PAM alignment distance position gene name gene id
GU305914.1 135054 135076 + 0 CGACAAGTGCAGGCACCTCT CGG ||||||||[||||||||||||]PAM NA Intergenic NA NA

T10257 AGGCACCTCTCGGCCTCCCCCGG 1000
Oligo fwd CACCAGGCACCTCTCGGCCTCCCC
Oligo rev AAACGGGGAGGCCGAGAGGTGCCT
Chromosome start end strand MM target_seq PAM alignment distance position gene name gene id
GU305914.1 135064 135086 + 0 AGGCACCTCTCGGCCTCCCC CGG ||||||||[||||||||||||]PAM NA Intergenic NA NA

T10258 CTCTCGGCCTCCCCCGGAGACGG 1000
Oligo fwd CACCCTCTCGGCCTCCCCCGGAGA
Oligo rev AAACTCTCCGGGGGAGGCCGAGAG
Chromosome start end strand MM target_seq PAM alignment distance position gene name gene id
GU305914.1 135070 135092 + 0 CTCTCGGCCTCCCCCGGAGA CGG ||||||||[||||||||||||]PAM NA Intergenic NA NA

<table>
<tr><td>T10259</td><td>TCTCGGCCTCCCCCGGAGACGGG</td><td></td><td>1000</td><td></td><td></td><td></td><td></td><td></td><td></td><td></td><td></td><td></td></tr>
<tr><td>Oligo fwd</td><td>CACCTCTCGGCCTCCCCCGGAGAC</td><td></td><td></td><td></td><td></td><td></td><td></td><td></td><td></td><td></td><td></td><td></td></tr>
<tr><td>Oligo rev</td><td>AAACGTCTCCGGGGGAGGCCGAGA</td><td></td><td></td><td></td><td></td><td></td><td></td><td></td><td></td><td></td><td></td><td></td></tr>
<tr><td>Chromosome</td><td>start</td><td></td><td>end</td><td>strand</td><td>MM</td><td>target_seq</td><td>PAM</td><td>alignment</td><td>distance</td><td>position</td><td>gene name</td><td>gene id</td></tr>
<tr><td>GU305914.1</td><td></td><td>135071</td><td>135093</td><td>+</td><td>0</td><td>TCTCGGCCTCCCCCGGAGAC</td><td>GGG</td><td>||||||||[||||||||||||]PAM</td><td>NA</td><td>Intergenic</td><td>NA</td><td>NA</td></tr>
<tr><td>T10260</td><td>CCTCCCCCGGAGACGGGCCCTGG</td><td></td><td>1000</td><td></td><td></td><td></td><td></td><td></td><td></td><td></td><td></td><td></td></tr>
<tr><td>Oligo fwd</td><td>CACCCCTCCCCCGGAGACGGGCCC</td><td></td><td></td><td></td><td></td><td></td><td></td><td></td><td></td><td></td><td></td><td></td></tr>
<tr><td>Oligo rev</td><td>AAACGGGCCCGTCTCCGGGGGAGG</td><td></td><td></td><td></td><td></td><td></td><td></td><td></td><td></td><td></td><td></td><td></td></tr>
<tr><td>Chromosome</td><td>start</td><td></td><td>end</td><td>strand</td><td>MM</td><td>target_seq</td><td>PAM</td><td>alignment</td><td>distance</td><td>position</td><td>gene name</td><td>gene id</td></tr>
<tr><td>GU305914.1</td><td></td><td>135077</td><td>135099</td><td>+</td><td>0</td><td>CCTCCCCCGGAGACGGGCCC</td><td>TGG</td><td>||||||||[||||||||||||]PAM</td><td>NA</td><td>Intergenic</td><td>NA</td><td>NA</td></tr>
<tr><td>T10261</td><td>GACGGGCCCTGGACCCTGCACGG</td><td></td><td>1000</td><td></td><td></td><td></td><td></td><td></td><td></td><td></td><td></td><td></td></tr>
<tr><td>Oligo fwd</td><td>CACCGACGGGCCCTGGACCCTGCA</td><td></td><td></td><td></td><td></td><td></td><td></td><td></td><td></td><td></td><td></td><td></td></tr>
<tr><td>Oligo rev</td><td>AAACTGCAGGGTCCAGGGCCCGTC</td><td></td><td></td><td></td><td></td><td></td><td></td><td></td><td></td><td></td><td></td><td></td></tr>
<tr><td>Chromosome</td><td>start</td><td></td><td>end</td><td>strand</td><td>MM</td><td>target_seq</td><td>PAM</td><td>alignment</td><td>distance</td><td>position</td><td>gene name</td><td>gene id</td></tr>
<tr><td>GU305914.1</td><td></td><td>135088</td><td>135110</td><td>+</td><td>0</td><td>GACGGGCCCTGGACCCTGCA</td><td>CGG</td><td>||||||||[||||||||||||]PAM</td><td>NA</td><td>Intergenic</td><td>NA</td><td>NA</td></tr>
<tr><td>T10262</td><td>GGGCCCTGGACCCTGCACGGCGG</td><td></td><td>1000</td><td></td><td></td><td></td><td></td><td></td><td></td><td></td><td></td><td></td></tr>
<tr><td>Oligo fwd</td><td>CACCGGGCCCTGGACCCTGCACGG</td><td></td><td></td><td></td><td></td><td></td><td></td><td></td><td></td><td></td><td></td><td></td></tr>
<tr><td>Oligo rev</td><td>AAACCCGTGCAGGGTCCAGGGCCC</td><td></td><td></td><td></td><td></td><td></td><td></td><td></td><td></td><td></td><td></td><td></td></tr>
<tr><td>Chromosome</td><td>start</td><td></td><td>end</td><td>strand</td><td>MM</td><td>target_seq</td><td>PAM</td><td>alignment</td><td>distance</td><td>position</td><td>gene name</td><td>gene id</td></tr>
<tr><td>GU305914.1</td><td></td><td>135091</td><td>135113</td><td>+</td><td>0</td><td>GGGCCCTGGACCCTGCACGG</td><td>CGG</td><td>||||||||[||||||||||||]PAM</td><td>NA</td><td>Intergenic</td><td>NA</td><td>NA</td></tr>
<tr><td>T10263</td><td>CCCTGGACCCTGCACGGCGGCGG</td><td></td><td>1000</td><td></td><td></td><td></td><td></td><td></td><td></td><td></td><td></td><td></td></tr>
<tr><td>Oligo fwd</td><td>CACCCCCTGGACCCTGCACGGCGG</td><td></td><td></td><td></td><td></td><td></td><td></td><td></td><td></td><td></td><td></td><td></td></tr>
<tr><td>Oligo rev</td><td>AAACCCGCCGTGCAGGGTCCAGGG</td><td></td><td></td><td></td><td></td><td></td><td></td><td></td><td></td><td></td><td></td><td></td></tr>
<tr><td>Chromosome</td><td>start</td><td></td><td>end</td><td>strand</td><td>MM</td><td>target_seq</td><td>PAM</td><td>alignment</td><td>distance</td><td>position</td><td>gene name</td><td>gene id</td></tr>
<tr><td>GU305914.1</td><td></td><td>135094</td><td>135116</td><td>+</td><td>0</td><td>CCCTGGACCCTGCACGGCGG</td><td>CGG</td><td>||||||||[||||||||||||]PAM</td><td>NA</td><td>Intergenic</td><td>NA</td><td>NA</td></tr>
<tr><td>T10264</td><td>GGCGACGAGAACTGTCGCTACGG</td><td></td><td>1000</td><td></td><td></td><td></td><td></td><td></td><td></td><td></td><td></td><td></td></tr>
<tr><td>Oligo fwd</td><td>CACCGGCGACGAGAACTGTCGCTA</td><td></td><td></td><td></td><td></td><td></td><td></td><td></td><td></td><td></td><td></td><td></td></tr>
<tr><td>Oligo rev</td><td>AAACTAGCGACAGTTCTCGTCGCC</td><td></td><td></td><td></td><td></td><td></td><td></td><td></td><td></td><td></td><td></td><td></td></tr>
<tr><td>Chromosome</td><td>start</td><td></td><td>end</td><td>strand</td><td>MM</td><td>target_seq</td><td>PAM</td><td>alignment</td><td>distance</td><td>position</td><td>gene name</td><td>gene id</td></tr>
<tr><td>GU305914.1</td><td></td><td>135115</td><td>135137</td><td>+</td><td>0</td><td>GGCGACGAGAACTGTCGCTA</td><td>CGG</td><td>||||||||[||||||||||||]PAM</td><td>NA</td><td>Intergenic</td><td>NA</td><td>NA</td></tr>
<tr><td>T10265</td><td>CTACGGACTCGCCTTCAGCGTGG</td><td></td><td>1000</td><td></td><td></td><td></td><td></td><td></td><td></td><td></td><td></td><td></td></tr>
<tr><td>Oligo fwd</td><td>CACCCTACGGACTCGCCTTCAGCG</td><td></td><td></td><td></td><td></td><td></td><td></td><td></td><td></td><td></td><td></td><td></td></tr>
<tr><td>Oligo rev</td><td>AAACCGCTGAAGGCGAGTCCGTAG</td><td></td><td></td><td></td><td></td><td></td><td></td><td></td><td></td><td></td><td></td><td></td></tr>
<tr><td>Chromosome</td><td>start</td><td></td><td>end</td><td>strand</td><td>MM</td><td>target_seq</td><td>PAM</td><td>alignment</td><td>distance</td><td>position</td><td>gene name</td><td>gene id</td></tr>
<tr><td>GU305914.1</td><td></td><td>135132</td><td>135154</td><td>+</td><td>0</td><td>CTACGGACTCGCCTTCAGCG</td><td>TGG</td><td>||||||||[||||||||||||]PAM</td><td>NA</td><td>Intergenic</td><td>NA</td><td>NA</td></tr>
<tr><td>T10266</td><td>TGCACCTCCGCGACGTCGAGTGG</td><td></td><td>1000</td><td></td><td></td><td></td><td></td><td></td><td></td><td></td><td></td><td></td></tr>
<tr><td>Oligo fwd</td><td>CACCTGCACCTCCGCGACGTCGAG</td><td></td><td></td><td></td><td></td><td></td><td></td><td></td><td></td><td></td><td></td><td></td></tr>
<tr><td>Oligo rev</td><td>AAACCTCGACGTCGCGGAGGTGCA</td><td></td><td></td><td></td><td></td><td></td><td></td><td></td><td></td><td></td><td></td><td></td></tr>
<tr><td>Chromosome</td><td>start</td><td></td><td>end</td><td>strand</td><td>MM</td><td>target_seq</td><td>PAM</td><td>alignment</td><td>distance</td><td>position</td><td>gene name</td><td>gene id</td></tr>
<tr><td>GU305914.1</td><td></td><td>135158</td><td>135180</td><td>+</td><td>0</td><td>TGCACCTCCGCGACGTCGAG</td><td>TGG</td><td>||||||||[||||||||||||]PAM</td><td>NA</td><td>Intergenic</td><td>NA</td><td>NA</td></tr>
<tr><td>T10267</td><td>ACCTCCGCGACGTCGAGTGGTGG</td><td></td><td>1000</td><td></td><td></td><td></td><td></td><td></td><td></td><td></td><td></td><td></td></tr>
<tr><td>Oligo fwd</td><td>CACCACCTCCGCGACGTCGAGTGG</td><td></td><td></td><td></td><td></td><td></td><td></td><td></td><td></td><td></td><td></td><td></td></tr>
<tr><td>Oligo rev</td><td>AAACCCACTCGACGTCGCGGAGGT</td><td></td><td></td><td></td><td></td><td></td><td></td><td></td><td></td><td></td><td></td><td></td></tr>
<tr><td>Chromosome</td><td>start</td><td></td><td>end</td><td>strand</td><td>MM</td><td>target_seq</td><td>PAM</td><td>alignment</td><td>distance</td><td>position</td><td>gene name</td><td>gene id</td></tr>
<tr><td>GU305914.1</td><td></td><td>135161</td><td>135183</td><td>+</td><td>0</td><td>ACCTCCGCGACGTCGAGTGG</td><td>TGG</td><td>||||||||[||||||||||||]PAM</td><td>NA</td><td>Intergenic</td><td>NA</td><td>NA</td></tr>
<tr><td>T10268</td><td>CCGCGACGTCGAGTGGTGGAAGG</td><td></td><td>1000</td><td></td><td></td><td></td><td></td><td></td><td></td><td></td><td></td><td></td></tr>
<tr><td>Oligo fwd</td><td>CACCCCGCGACGTCGAGTGGTGGA</td><td></td><td></td><td></td><td></td><td></td><td></td><td></td><td></td><td></td><td></td><td></td></tr>
<tr><td>Oligo rev</td><td>AAACTCCACCACTCGACGTCGCGG</td><td></td><td></td><td></td><td></td><td></td><td></td><td></td><td></td><td></td><td></td><td></td></tr>
<tr><td>Chromosome</td><td>start</td><td></td><td>end</td><td>strand</td><td>MM</td><td>target_seq</td><td>PAM</td><td>alignment</td><td>distance</td><td>position</td><td>gene name</td><td>gene id</td></tr>
<tr><td>GU305914.1</td><td></td><td>135165</td><td>135187</td><td>+</td><td>0</td><td>CCGCGACGTCGAGTGGTGGA</td><td>AGG</td><td>||||||||[||||||||||||]PAM</td><td>NA</td><td>Intergenic</td><td>NA</td><td>NA</td></tr>
<tr><td>T10269</td><td>GGAAGGACGTGATCAAGTTCAGG</td><td></td><td>1000</td><td></td><td></td><td></td><td></td><td></td><td></td><td></td><td></td><td></td></tr>
<tr><td>Oligo fwd</td><td>CACCGGAAGGACGTGATCAAGTTC</td><td></td><td></td><td></td><td></td><td></td><td></td><td></td><td></td><td></td><td></td><td></td></tr>
<tr><td>Oligo rev</td><td>AAACGAACTTGATCACGTCCTTCC</td><td></td><td></td><td></td><td></td><td></td><td></td><td></td><td></td><td></td><td></td><td></td></tr>
<tr><td>Chromosome</td><td>start</td><td></td><td>end</td><td>strand</td><td>MM</td><td>target_seq</td><td>PAM</td><td>alignment</td><td>distance</td><td>position</td><td>gene name</td><td>gene id</td></tr>
<tr><td>GU305914.1</td><td></td><td>135182</td><td>135204</td><td>+</td><td>0</td><td>GGAAGGACGTGATCAAGTTC</td><td>AGG</td><td>||||||||[||||||||||||]PAM</td><td>NA</td><td>Intergenic</td><td>NA</td><td>NA</td></tr>
<tr><td>T10270</td><td>ATCAAGTTCAGGTTGTTCATCGG</td><td></td><td>1000</td><td></td><td></td><td></td><td></td><td></td><td></td><td></td><td></td><td></td></tr>
<tr><td>Oligo fwd</td><td>CACCATCAAGTTCAGGTTGTTCAT</td><td></td><td></td><td></td><td></td><td></td><td></td><td></td><td></td><td></td><td></td><td></td></tr>
<tr><td>Oligo rev</td><td>AAACATGAACAACCTGAACTTGAT</td><td></td><td></td><td></td><td></td><td></td><td></td><td></td><td></td><td></td><td></td><td></td></tr>
<tr><td>Chromosome</td><td>start</td><td></td><td>end</td><td>strand</td><td>MM</td><td>target_seq</td><td>PAM</td><td>alignment</td><td>distance</td><td>position</td><td>gene name</td><td>gene id</td></tr>
<tr><td>GU305914.1</td><td></td><td>135193</td><td>135215</td><td>+</td><td>0</td><td>ATCAAGTTCAGGTTGTTCAT</td><td>CGG</td><td>||||||||[||||||||||||]PAM</td><td>NA</td><td>Intergenic</td><td>NA</td><td>NA</td></tr>
<tr><td>T10271</td><td>GTTCATCGGAGACGACGAGACGG</td><td></td><td>1000</td><td></td><td></td><td></td><td></td><td></td><td></td><td></td><td></td><td></td></tr>
<tr><td>Oligo fwd</td><td>CACCGTTCATCGGAGACGACGAGA</td><td></td><td></td><td></td><td></td><td></td><td></td><td></td><td></td><td></td><td></td><td></td></tr>
<tr><td>Oligo rev</td><td>AAACTCTCGTCGTCTCCGATGAAC</td><td></td><td></td><td></td><td></td><td></td><td></td><td></td><td></td><td></td><td></td><td></td></tr>
<tr><td>Chromosome</td><td>start</td><td></td><td>end</td><td>strand</td><td>MM</td><td>target_seq</td><td>PAM</td><td>alignment</td><td>distance</td><td>position</td><td>gene name</td><td>gene id</td></tr>
<tr><td>GU305914.1</td><td></td><td>135207</td><td>135229</td><td>+</td><td>0</td><td>GTTCATCGGAGACGACGAGA</td><td>CGG</td><td>||||||||[||||||||||||]PAM</td><td>NA</td><td>Intergenic</td><td>NA</td><td>NA</td></tr>
<tr><td>T10272</td><td>TTCGTCAGACTTCTGCGCGAAGG</td><td></td><td>1000</td><td></td><td></td><td></td><td></td><td></td><td></td><td></td><td></td><td></td></tr>
<tr><td>Oligo fwd</td><td>CACCTTCGTCAGACTTCTGCGCGA</td><td></td><td></td><td></td><td></td><td></td><td></td><td></td><td></td><td></td><td></td><td></td></tr>
<tr><td>Oligo rev</td><td>AAACTCGCGCAGAAGTCTGACGAA</td><td></td><td></td><td></td><td></td><td></td><td></td><td></td><td></td><td></td><td></td><td></td></tr>
<tr><td>Chromosome</td><td>start</td><td></td><td>end</td><td>strand</td><td>MM</td><td>target_seq</td><td>PAM</td><td>alignment</td><td>distance</td><td>position</td><td>gene name</td><td>gene id</td></tr>
<tr><td>GU305914.1</td><td></td><td>135247</td><td>135269</td><td>+</td><td>0</td><td>TTCGTCAGACTTCTGCGCGA</td><td>AGG</td><td>||||||||[||||||||||||]PAM</td><td>NA</td><td>Intergenic</td><td>NA</td><td>NA</td></tr>
<tr><td>T10273</td><td>TCGTCAGACTTCTGCGCGAAGGG</td><td></td><td>1000</td><td></td><td></td><td></td><td></td><td></td><td></td><td></td><td></td><td></td></tr>
<tr><td>Oligo fwd</td><td>CACCTCGTCAGACTTCTGCGCGAA</td><td></td><td></td><td></td><td></td><td></td><td></td><td></td><td></td><td></td><td></td><td></td></tr>
<tr><td>Oligo rev</td><td>AAACTTCGCGCAGAAGTCTGACGA</td><td></td><td></td><td></td><td></td><td></td><td></td><td></td><td></td><td></td><td></td><td></td></tr>
<tr><td>Chromosome</td><td>start</td><td></td><td>end</td><td>strand</td><td>MM</td><td>target_seq</td><td>PAM</td><td>alignment</td><td>distance</td><td>position</td><td>gene name</td><td>gene id</td></tr>
<tr><td>GU305914.1</td><td></td><td>135248</td><td>135270</td><td>+</td><td>0</td><td>TCGTCAGACTTCTGCGCGAA</td><td>GGG</td><td>||||||||[||||||||||||]PAM</td><td>NA</td><td>Intergenic</td><td>NA</td><td>NA</td></tr>
</table>

T10274 CGCGAAGGGCTCCGCTTTTTCGG 1000
Oligo fwd CACCCGCGAAGGGCTCCGCTTTTT
Oligo rev AAACAAAAAGCGGAGCCCTTCGCG
Chromosome start end strand MM target_seq PAM alignment distance position gene name gene id
GU305914.1 135262 135284 + 0 CGCGAAGGGCTCCGCTTTTT CGG ||||||||[||||||||||||]PAM NA Intergenic NA NA

T10275 AGGGCTCCGCTTTTTCGGAGAGG 1000
Oligo fwd CACCAGGGCTCCGCTTTTTCGGAG
Oligo rev AAACCTCCGAAAAAGCGGAGCCCT
Chromosome start end strand MM target_seq PAM alignment distance position gene name gene id
GU305914.1 135267 135289 + 0 AGGGCTCCGCTTTTTCGGAG AGG ||||||||[||||||||||||]PAM NA Intergenic NA NA

T10276 TCCGCTTTTTCGGAGAGGTCAGG 1000
Oligo fwd CACCTCCGCTTTTTCGGAGAGGTC
Oligo rev AAACGACCTCTCCGAAAAAGCGGA
Chromosome start end strand MM target_seq PAM alignment distance position gene name gene id
GU305914.1 135272 135294 + 0 TCCGCTTTTTCGGAGAGGTC AGG ||||||||[||||||||||||]PAM NA Intergenic NA NA

T10277 TCGGAGAGGTCAGGCGTACGCGG 1000
Oligo fwd CACCTCGGAGAGGTCAGGCGTACG
Oligo rev AAACCGTACGCCTGACCTCTCCGA
Chromosome start end strand MM target_seq PAM alignment distance position gene name gene id
GU305914.1 135281 135303 + 0 TCGGAGAGGTCAGGCGTACG CGG ||||||||[||||||||||||]PAM NA Intergenic NA NA

T10278 TCAGGCGTACGCGGCAGCCGCGG 1000
Oligo fwd CACCTCAGGCGTACGCGGCAGCCG
Oligo rev AAACCGGCTGCCGCGTACGCCTGA
Chromosome start end strand MM target_seq PAM alignment distance position gene name gene id
GU305914.1 135290 135312 + 0 TCAGGCGTACGCGGCAGCCG CGG ||||||||[||||||||||||]PAM NA Intergenic NA NA

T10279 CGTACGCGGCAGCCGCGGAGCGG 1000
Oligo fwd CACCCGTACGCGGCAGCCGCGGAG
Oligo rev AAACCTCCGCGGCTGCCGCGTACG
Chromosome start end strand MM target_seq PAM alignment distance position gene name gene id
GU305914.1 135295 135317 + 0 CGTACGCGGCAGCCGCGGAG CGG ||||||||[||||||||||||]PAM NA Intergenic NA NA

T10280 GTACGCGGCAGCCGCGGAGCGGG 1000
Oligo fwd CACCGTACGCGGCAGCCGCGGAGC
Oligo rev AAACGCTCCGCGGCTGCCGCGTAC
Chromosome start end strand MM target_seq PAM alignment distance position gene name gene id
GU305914.1 135296 135318 + 0 GTACGCGGCAGCCGCGGAGC GGG ||||||||[||||||||||||]PAM NA Intergenic NA NA

T10281 GGCCCCGCAGCGCCAGTAGTCGG 1000
Oligo fwd CACCGGCCCCGCAGCGCCAGTAGT
Oligo rev AAACACTACTGGCGCTGCGGGGCC
Chromosome start end strand MM target_seq PAM alignment distance position gene name gene id
GU305914.1 135317 135339 + 0 GGCCCCGCAGCGCCAGTAGT CGG ||||||||[||||||||||||]PAM NA Intergenic NA NA

T10282 ATCATCATCCTCACCATCACCGG 1000
Oligo fwd CACCATCATCATCCTCACCATCAC
Oligo rev AAACGTGATGGTGAGGATGATGAT
Chromosome start end strand MM target_seq PAM alignment distance position gene name gene id
GU305914.1 135347 135369 + 0 ATCATCATCCTCACCATCAC CGG ||||||||[||||||||||||]PAM NA Intergenic NA NA

T10283 TCATCATCCTCACCATCACCGGG 1000
Oligo fwd CACCTCATCATCCTCACCATCACC
Oligo rev AAACGGTGATGGTGAGGATGATGA
Chromosome start end strand MM target_seq PAM alignment distance position gene name gene id
GU305914.1 135348 135370 + 0 TCATCATCCTCACCATCACC GGG ||||||||[||||||||||||]PAM NA Intergenic NA NA

T10284 TCATCCTCACCATCACCGGGCGG 1000
Oligo fwd CACCTCATCCTCACCATCACCGGG
Oligo rev AAACCCCGGTGATGGTGAGGATGA
Chromosome start end strand MM target_seq PAM alignment distance position gene name gene id
GU305914.1 135351 135373 + 0 TCATCCTCACCATCACCGGG CGG ||||||||[||||||||||||]PAM NA Intergenic NA NA

T10285 CATCCTCACCATCACCGGGCGGG 1000
Oligo fwd CACCCATCCTCACCATCACCGGGC
Oligo rev AAACGCCCGGTGATGGTGAGGATG
Chromosome start end strand MM target_seq PAM alignment distance position gene name gene id
GU305914.1 135352 135374 + 0 CATCCTCACCATCACCGGGC GGG ||||||||[||||||||||||]PAM NA Intergenic NA NA

T10286 TCACCATCACCGGGCGGGCGCGG 1000
Oligo fwd CACCTCACCATCACCGGGCGGGCG
Oligo rev AAACCGCCCGCCCGGTGATGGTGA
Chromosome start end strand MM target_seq PAM alignment distance position gene name gene id
GU305914.1 135357 135379 + 0 TCACCATCACCGGGCGGGCG CGG ||||||||[||||||||||||]PAM NA Intergenic NA NA

T10287 CCATCACCGGGCGGGCGCGGCGG 1000
Oligo fwd CACCCCATCACCGGGCGGGCGCGG
Oligo rev AAACCCGCGCCCGCCCGGTGATGG
Chromosome start end strand MM target_seq PAM alignment distance position gene name gene id
GU305914.1 135360 135382 + 0 CCATCACCGGGCGGGCGCGG CGG ||||||||[||||||||||||]PAM NA Intergenic NA NA

T10288 CATCACCGGGCGGGCGCGGCGGG 1000
Oligo fwd CACCCATCACCGGGCGGGCGCGGC
Oligo rev AAACGCCGCGCCCGCCCGGTGATG
Chromosome start end strand MM target_seq PAM alignment distance position gene name gene id

| | | | | | | | | | | | |
|---|---|---|---|---|---|---|---|---|---|---|---|
| GU305914.1 | 135361 | 135383 | + | 0 | CATCACCGGGCGGGCGCGGC | GGG | \|\|\|\|\|\|\|\|[\|\|\|\|\|\|\|\|\|\|\|\|]PAM | NA | Intergenic | NA | NA |
| T10289 | GGCGCGGCGGGTGATGACGACGG | 1000 | | | | | | | | | |
| Oligo fwd | CACCGGCGCGGCGGGTGATGACGA | | | | | | | | | | |
| Oligo rev | AAACTCGTCATCACCCGCCGCGCC | | | | | | | | | | |
| Chromosome | start | end | strand | MM | target_seq | PAM | alignment | distance | position | gene name | gene id |
| GU305914.1 | 135373 | 135395 | + | 0 | GGCGCGGCGGGTGATGACGA | CGG | \|\|\|\|\|\|\|\|[\|\|\|\|\|\|\|\|\|\|\|\|]PAM | NA | Intergenic | NA | NA |
| T10290 | GCGCGGCGGGTGATGACGACGGG | 1000 | | | | | | | | | |
| Oligo fwd | CACCGCGCGGCGGGTGATGACGAC | | | | | | | | | | |
| Oligo rev | AAACGTCGTCATCACCCGCCGCGC | | | | | | | | | | |
| Chromosome | start | end | strand | MM | target_seq | PAM | alignment | distance | position | gene name | gene id |
| GU305914.1 | 135374 | 135396 | + | 0 | GCGCGGCGGGTGATGACGAC | GGG | \|\|\|\|\|\|\|\|[\|\|\|\|\|\|\|\|\|\|\|\|]PAM | NA | Intergenic | NA | NA |
| T10291 | CGCGGCGGGTGATGACGACGGGG | 1000 | | | | | | | | | |
| Oligo fwd | CACCCGCGGCGGGTGATGACGACG | | | | | | | | | | |
| Oligo rev | AAACCGTCGTCATCACCCGCCGCG | | | | | | | | | | |
| Chromosome | start | end | strand | MM | target_seq | PAM | alignment | distance | position | gene name | gene id |
| GU305914.1 | 135375 | 135397 | + | 0 | CGCGGCGGGTGATGACGACG | GGG | \|\|\|\|\|\|\|\|[\|\|\|\|\|\|\|\|\|\|\|\|]PAM | NA | Intergenic | NA | NA |
| T10292 | CGGGGACGTCGATGACGACGAGG | 1000 | | | | | | | | | |
| Oligo fwd | CACCCGGGGACGTCGATGACGACG | | | | | | | | | | |
| Oligo rev | AAACCGTCGTCATCGACGTCCCCG | | | | | | | | | | |
| Chromosome | start | end | strand | MM | target_seq | PAM | alignment | distance | position | gene name | gene id |
| GU305914.1 | 135393 | 135415 | + | 0 | CGGGGACGTCGATGACGACG | AGG | \|\|\|\|\|\|\|\|[\|\|\|\|\|\|\|\|\|\|\|\|]PAM | NA | Intergenic | NA | NA |
| GU305914.1 | 114978 | 115000 | + | 4 | CGACGATGACGATGACGACG | AGG | \|\|--\|\|-\|[-\|\|\|\|\|\|\|\|\|\|\|]PAM | NA | Intergenic | NA | NA |
| T10293 | GGGGACGTCGATGACGACGAGGG | 1000 | | | | | | | | | |
| Oligo fwd | CACCGGGGACGTCGATGACGACGA | | | | | | | | | | |
| Oligo rev | AAACTCGTCGTCATCGACGTCCCC | | | | | | | | | | |
| Chromosome | start | end | strand | MM | target_seq | PAM | alignment | distance | position | gene name | gene id |
| GU305914.1 | 135394 | 135416 | + | 0 | GGGGACGTCGATGACGACGA | GGG | \|\|\|\|\|\|\|\|[\|\|\|\|\|\|\|\|\|\|\|\|]PAM | NA | Intergenic | NA | NA |
| T10294 | TGACGACGAGGGACTGCCCGAGG | 1000 | | | | | | | | | |
| Oligo fwd | CACCTGACGACGAGGGACTGCCCG | | | | | | | | | | |
| Oligo rev | AAACCGGGCAGTCCCTCGTCGTCA | | | | | | | | | | |
| Chromosome | start | end | strand | MM | target_seq | PAM | alignment | distance | position | gene name | gene id |
| GU305914.1 | 135405 | 135427 | + | 0 | TGACGACGAGGGACTGCCCG | AGG | \|\|\|\|\|\|\|\|[\|\|\|\|\|\|\|\|\|\|\|\|]PAM | NA | Intergenic | NA | NA |
| T10295 | GACGACGAGGGACTGCCCGAGGG | 1000 | | | | | | | | | |
| Oligo fwd | CACCGACGACGAGGGACTGCCCGA | | | | | | | | | | |
| Oligo rev | AAACTCGGGCAGTCCCTCGTCGTC | | | | | | | | | | |
| Chromosome | start | end | strand | MM | target_seq | PAM | alignment | distance | position | gene name | gene id |
| GU305914.1 | 135406 | 135428 | + | 0 | GACGACGAGGGACTGCCCGA | GGG | \|\|\|\|\|\|\|\|[\|\|\|\|\|\|\|\|\|\|\|\|]PAM | NA | Intergenic | NA | NA |
| T10296 | GGGACTGCCCGAGGGACTGTTGG | 1000 | | | | | | | | | |
| Oligo fwd | CACCGGGACTGCCCGAGGGACTGT | | | | | | | | | | |
| Oligo rev | AAACACAGTCCCTCGGGCAGTCCC | | | | | | | | | | |
| Chromosome | start | end | strand | MM | target_seq | PAM | alignment | distance | position | gene name | gene id |
| GU305914.1 | 135414 | 135436 | + | 0 | GGGACTGCCCGAGGGACTGT | TGG | \|\|\|\|\|\|\|\|[\|\|\|\|\|\|\|\|\|\|\|\|]PAM | NA | Intergenic | NA | NA |
| T10297 | GGACTGCCCGAGGGACTGTTGGG | 1000 | | | | | | | | | |
| Oligo fwd | CACCGGACTGCCCGAGGGACTGTT | | | | | | | | | | |
| Oligo rev | AAACAACAGTCCCTCGGGCAGTCC | | | | | | | | | | |
| Chromosome | start | end | strand | MM | target_seq | PAM | alignment | distance | position | gene name | gene id |
| GU305914.1 | 135415 | 135437 | + | 0 | GGACTGCCCGAGGGACTGTT | GGG | \|\|\|\|\|\|\|\|[\|\|\|\|\|\|\|\|\|\|\|\|]PAM | NA | Intergenic | NA | NA |
| T10298 | CTGTTGGGCGCCGCCGCCGACGG | 1000 | | | | | | | | | |
| Oligo fwd | CACCCTGTTGGGCGCCGCCGCCGA | | | | | | | | | | |
| Oligo rev | AAACTCGGCGGCGGCGCCCAACAG | | | | | | | | | | |
| Chromosome | start | end | strand | MM | target_seq | PAM | alignment | distance | position | gene name | gene id |
| GU305914.1 | 135430 | 135452 | + | 0 | CTGTTGGGCGCCGCCGCCGA | CGG | \|\|\|\|\|\|\|\|[\|\|\|\|\|\|\|\|\|\|\|\|]PAM | NA | Intergenic | NA | NA |
| T10299 | GGGCGCCGCCGCCGACGGCCCGG | 1000 | | | | | | | | | |
| Oligo fwd | CACCGGGCGCCGCCGCCGACGGCC | | | | | | | | | | |
| Oligo rev | AAACGGCCGTCGGCGGCGGCGCCC | | | | | | | | | | |
| Chromosome | start | end | strand | MM | target_seq | PAM | alignment | distance | position | gene name | gene id |
| GU305914.1 | 135435 | 135457 | + | 0 | GGGCGCCGCCGCCGACGGCC | CGG | \|\|\|\|\|\|\|\|[\|\|\|\|\|\|\|\|\|\|\|\|]PAM | NA | Intergenic | NA | NA |
| GU305914.1 | 97885 | 97907 | + | 4 | GGTCGCTGCCGCCGCCGCCC | GGG | \|\|-\|\|\|-\|[\|\|\|\|\|\|-\|\|-\|\|]PAM | NA | Intergenic | NA | NA |
| T10300 | GGCGCCGCCGCCGACGGCCCGGG | 1000 | | | | | | | | | |
| Oligo fwd | CACCGGCGCCGCCGCCGACGGCCC | | | | | | | | | | |
| Oligo rev | AAACGGGCCGTCGGCGGCGGCGCC | | | | | | | | | | |
| Chromosome | start | end | strand | MM | target_seq | PAM | alignment | distance | position | gene name | gene id |
| GU305914.1 | 135436 | 135458 | + | 0 | GGCGCCGCCGCCGACGGCCC | GGG | \|\|\|\|\|\|\|\|[\|\|\|\|\|\|\|\|\|\|\|\|]PAM | NA | Intergenic | NA | NA |
| T10301 | GGCCCGGGTTTCTTCGATCTCGG | 1000 | | | | | | | | | |
| Oligo fwd | CACCGGCCCGGGTTTCTTCGATCT | | | | | | | | | | |
| Oligo rev | AAACAGATCGAAGAAACCCGGGCC | | | | | | | | | | |
| Chromosome | start | end | strand | MM | target_seq | PAM | alignment | distance | position | gene name | gene id |
| GU305914.1 | 135451 | 135473 | + | 0 | GGCCCGGGTTTCTTCGATCT | CGG | \|\|\|\|\|\|\|\|[\|\|\|\|\|\|\|\|\|\|\|\|]PAM | NA | Intergenic | NA | NA |
| T10302 | CGATAATCGCAAGATGTCCGAGG | 1000 | | | | | | | | | |
| Oligo fwd | CACCCGATAATCGCAAGATGTCCG | | | | | | | | | | |
| Oligo rev | AAACCGGACATCTTGCGATTATCG | | | | | | | | | | |
| Chromosome | start | end | strand | MM | target_seq | PAM | alignment | distance | position | gene name | gene id |
| GU305914.1 | 135474 | 135496 | + | 0 | CGATAATCGCAAGATGTCCG | AGG | \|\|\|\|\|\|\|\|[\|\|\|\|\|\|\|\|\|\|\|\|]PAM | NA | Intergenic | NA | NA |
| T10303 | TCGCAAGATGTCCGAGGAGATGG | 1000 | | | | | | | | | |

<table>
<tr><td>Oligo fwd</td><td>CACCTCGCAAGATGTCCGAGGAGA</td><td></td><td></td><td></td><td></td><td></td><td></td><td></td><td></td><td></td><td></td><td></td></tr>
<tr><td>Oligo rev</td><td>AAACTCTCCTCGGACATCTTGCGA</td><td></td><td></td><td></td><td></td><td></td><td></td><td></td><td></td><td></td><td></td><td></td></tr>
<tr><td>Chromosome</td><td>start</td><td></td><td>end</td><td>strand</td><td>MM</td><td>target_seq</td><td>PAM</td><td>alignment</td><td>distance</td><td>position</td><td>gene name</td><td>gene id</td></tr>
<tr><td>GU305914.1</td><td></td><td>135480</td><td>135502</td><td>+</td><td>0</td><td>TCGCAAGATGTCCGAGGAGA</td><td>TGG</td><td>||||||||[||||||||||||]PAM</td><td>NA</td><td>Intergenic</td><td>NA</td><td>NA</td></tr>
<tr><td></td><td></td><td></td><td></td><td></td><td></td><td></td><td></td><td></td><td></td><td></td><td></td><td></td></tr>
<tr><td>T10304</td><td>CAAGATGTCCGAGGAGATGGAGG</td><td></td><td>1000</td><td></td><td></td><td></td><td></td><td></td><td></td><td></td><td></td><td></td></tr>
<tr><td>Oligo fwd</td><td>CACCCAAGATGTCCGAGGAGATGG</td><td></td><td></td><td></td><td></td><td></td><td></td><td></td><td></td><td></td><td></td><td></td></tr>
<tr><td>Oligo rev</td><td>AAACCCATCTCCTCGGACATCTTG</td><td></td><td></td><td></td><td></td><td></td><td></td><td></td><td></td><td></td><td></td><td></td></tr>
<tr><td>Chromosome</td><td>start</td><td></td><td>end</td><td>strand</td><td>MM</td><td>target_seq</td><td>PAM</td><td>alignment</td><td>distance</td><td>position</td><td>gene name</td><td>gene id</td></tr>
<tr><td>GU305914.1</td><td></td><td>135483</td><td>135505</td><td>+</td><td>0</td><td>CAAGATGTCCGAGGAGATGG</td><td>AGG</td><td>||||||||[||||||||||||]PAM</td><td>NA</td><td>Intergenic</td><td>NA</td><td>NA</td></tr>
<tr><td></td><td></td><td></td><td></td><td></td><td></td><td></td><td></td><td></td><td></td><td></td><td></td><td></td></tr>
<tr><td>T10305</td><td>TGTCCGAGGAGATGGAGGACCGG</td><td></td><td>1000</td><td></td><td></td><td></td><td></td><td></td><td></td><td></td><td></td><td></td></tr>
<tr><td>Oligo fwd</td><td>CACCTGTCCGAGGAGATGGAGGAC</td><td></td><td></td><td></td><td></td><td></td><td></td><td></td><td></td><td></td><td></td><td></td></tr>
<tr><td>Oligo rev</td><td>AAACGTCCTCCATCTCCTCGGACA</td><td></td><td></td><td></td><td></td><td></td><td></td><td></td><td></td><td></td><td></td><td></td></tr>
<tr><td>Chromosome</td><td>start</td><td></td><td>end</td><td>strand</td><td>MM</td><td>target_seq</td><td>PAM</td><td>alignment</td><td>distance</td><td>position</td><td>gene name</td><td>gene id</td></tr>
<tr><td>GU305914.1</td><td></td><td>135488</td><td>135510</td><td>+</td><td>0</td><td>TGTCCGAGGAGATGGAGGAC</td><td>CGG</td><td>||||||||[||||||||||||]PAM</td><td>NA</td><td>Intergenic</td><td>NA</td><td>NA</td></tr>
<tr><td></td><td></td><td></td><td></td><td></td><td></td><td></td><td></td><td></td><td></td><td></td><td></td><td></td></tr>
<tr><td>T10306</td><td>CGAGGAGATGGAGGACCGGCTGG</td><td></td><td>1000</td><td></td><td></td><td></td><td></td><td></td><td></td><td></td><td></td><td></td></tr>
<tr><td>Oligo fwd</td><td>CACCCGAGGAGATGGAGGACCGGC</td><td></td><td></td><td></td><td></td><td></td><td></td><td></td><td></td><td></td><td></td><td></td></tr>
<tr><td>Oligo rev</td><td>AAACGCCGGTCCTCCATCTCCTCG</td><td></td><td></td><td></td><td></td><td></td><td></td><td></td><td></td><td></td><td></td><td></td></tr>
<tr><td>Chromosome</td><td>start</td><td></td><td>end</td><td>strand</td><td>MM</td><td>target_seq</td><td>PAM</td><td>alignment</td><td>distance</td><td>position</td><td>gene name</td><td>gene id</td></tr>
<tr><td>GU305914.1</td><td></td><td>135492</td><td>135514</td><td>+</td><td>0</td><td>CGAGGAGATGGAGGACCGGC</td><td>TGG</td><td>||||||||[||||||||||||]PAM</td><td>NA</td><td>Intergenic</td><td>NA</td><td>NA</td></tr>
<tr><td></td><td></td><td></td><td></td><td></td><td></td><td></td><td></td><td></td><td></td><td></td><td></td><td></td></tr>
<tr><td>T10307</td><td>AGATGGAGGACCGGCTGGCCAGG</td><td></td><td>1000</td><td></td><td></td><td></td><td></td><td></td><td></td><td></td><td></td><td></td></tr>
<tr><td>Oligo fwd</td><td>CACCAGATGGAGGACCGGCTGGCC</td><td></td><td></td><td></td><td></td><td></td><td></td><td></td><td></td><td></td><td></td><td></td></tr>
<tr><td>Oligo rev</td><td>AAACGGCCAGCCGGTCCTCCATCT</td><td></td><td></td><td></td><td></td><td></td><td></td><td></td><td></td><td></td><td></td><td></td></tr>
<tr><td>Chromosome</td><td>start</td><td></td><td>end</td><td>strand</td><td>MM</td><td>target_seq</td><td>PAM</td><td>alignment</td><td>distance</td><td>position</td><td>gene name</td><td>gene id</td></tr>
<tr><td>GU305914.1</td><td></td><td>135497</td><td>135519</td><td>+</td><td>0</td><td>AGATGGAGGACCGGCTGGCC</td><td>AGG</td><td>||||||||[||||||||||||]PAM</td><td>NA</td><td>Intergenic</td><td>NA</td><td>NA</td></tr>
<tr><td></td><td></td><td></td><td></td><td></td><td></td><td></td><td></td><td></td><td></td><td></td><td></td><td></td></tr>
<tr><td>T10308</td><td>CGCGTCTCCGCCGCCGACTTCGG</td><td></td><td>1000</td><td></td><td></td><td></td><td></td><td></td><td></td><td></td><td></td><td></td></tr>
<tr><td>Oligo fwd</td><td>CACCCGCGTCTCCGCCGCCGACTT</td><td></td><td></td><td></td><td></td><td></td><td></td><td></td><td></td><td></td><td></td><td></td></tr>
<tr><td>Oligo rev</td><td>AAACAAGTCGGCGGCGGAGACGCG</td><td></td><td></td><td></td><td></td><td></td><td></td><td></td><td></td><td></td><td></td><td></td></tr>
<tr><td>Chromosome</td><td>start</td><td></td><td>end</td><td>strand</td><td>MM</td><td>target_seq</td><td>PAM</td><td>alignment</td><td>distance</td><td>position</td><td>gene name</td><td>gene id</td></tr>
<tr><td>GU305914.1</td><td></td><td>135532</td><td>135554</td><td>+</td><td>0</td><td>CGCGTCTCCGCCGCCGACTT</td><td>CGG</td><td>||||||||[||||||||||||]PAM</td><td>NA</td><td>Intergenic</td><td>NA</td><td>NA</td></tr>
<tr><td></td><td></td><td></td><td></td><td></td><td></td><td></td><td></td><td></td><td></td><td></td><td></td><td></td></tr>
<tr><td>T10309</td><td>CCCGCTCGAGCCGCCCTCCAAGG</td><td></td><td>1000</td><td></td><td></td><td></td><td></td><td></td><td></td><td></td><td></td><td></td></tr>
<tr><td>Oligo fwd</td><td>CACCCCCGCTCGAGCCGCCCTCCA</td><td></td><td></td><td></td><td></td><td></td><td></td><td></td><td></td><td></td><td></td><td></td></tr>
<tr><td>Oligo rev</td><td>AAACTGGAGGGCGGCTCGAGCGGG</td><td></td><td></td><td></td><td></td><td></td><td></td><td></td><td></td><td></td><td></td><td></td></tr>
<tr><td>Chromosome</td><td>start</td><td></td><td>end</td><td>strand</td><td>MM</td><td>target_seq</td><td>PAM</td><td>alignment</td><td>distance</td><td>position</td><td>gene name</td><td>gene id</td></tr>
<tr><td>GU305914.1</td><td></td><td>135555</td><td>135577</td><td>+</td><td>0</td><td>CCCGCTCGAGCCGCCCTCCA</td><td>AGG</td><td>||||||||[||||||||||||]PAM</td><td>NA</td><td>Intergenic</td><td>NA</td><td>NA</td></tr>
<tr><td></td><td></td><td></td><td></td><td></td><td></td><td></td><td></td><td></td><td></td><td></td><td></td><td></td></tr>
<tr><td>T10310</td><td>TCGAGCCGCCCTCCAAGGTCAGG</td><td></td><td>1000</td><td></td><td></td><td></td><td></td><td></td><td></td><td></td><td></td><td></td></tr>
<tr><td>Oligo fwd</td><td>CACCTCGAGCCGCCCTCCAAGGTC</td><td></td><td></td><td></td><td></td><td></td><td></td><td></td><td></td><td></td><td></td><td></td></tr>
<tr><td>Oligo rev</td><td>AAACGACCTTGGAGGGCGGCTCGA</td><td></td><td></td><td></td><td></td><td></td><td></td><td></td><td></td><td></td><td></td><td></td></tr>
<tr><td>Chromosome</td><td>start</td><td></td><td>end</td><td>strand</td><td>MM</td><td>target_seq</td><td>PAM</td><td>alignment</td><td>distance</td><td>position</td><td>gene name</td><td>gene id</td></tr>
<tr><td>GU305914.1</td><td></td><td>135560</td><td>135582</td><td>+</td><td>0</td><td>TCGAGCCGCCCTCCAAGGTC</td><td>AGG</td><td>||||||||[||||||||||||]PAM</td><td>NA</td><td>Intergenic</td><td>NA</td><td>NA</td></tr>
<tr><td></td><td></td><td></td><td></td><td></td><td></td><td></td><td></td><td></td><td></td><td></td><td></td><td></td></tr>
<tr><td>T10311</td><td>CGAGCCGCCCTCCAAGGTCAGGG</td><td></td><td>1000</td><td></td><td></td><td></td><td></td><td></td><td></td><td></td><td></td><td></td></tr>
<tr><td>Oligo fwd</td><td>CACCCGAGCCGCCCTCCAAGGTCA</td><td></td><td></td><td></td><td></td><td></td><td></td><td></td><td></td><td></td><td></td><td></td></tr>
<tr><td>Oligo rev</td><td>AAACTGACCTTGGAGGGCGGCTCG</td><td></td><td></td><td></td><td></td><td></td><td></td><td></td><td></td><td></td><td></td><td></td></tr>
<tr><td>Chromosome</td><td>start</td><td></td><td>end</td><td>strand</td><td>MM</td><td>target_seq</td><td>PAM</td><td>alignment</td><td>distance</td><td>position</td><td>gene name</td><td>gene id</td></tr>
<tr><td>GU305914.1</td><td></td><td>135561</td><td>135583</td><td>+</td><td>0</td><td>CGAGCCGCCCTCCAAGGTCA</td><td>GGG</td><td>||||||||[||||||||||||]PAM</td><td>NA</td><td>Intergenic</td><td>NA</td><td>NA</td></tr>
<tr><td></td><td></td><td></td><td></td><td></td><td></td><td></td><td></td><td></td><td></td><td></td><td></td><td></td></tr>
<tr><td>T10312</td><td>GAGCCGCCCTCCAAGGTCAGGGG</td><td></td><td>1000</td><td></td><td></td><td></td><td></td><td></td><td></td><td></td><td></td><td></td></tr>
<tr><td>Oligo fwd</td><td>CACCGAGCCGCCCTCCAAGGTCAG</td><td></td><td></td><td></td><td></td><td></td><td></td><td></td><td></td><td></td><td></td><td></td></tr>
<tr><td>Oligo rev</td><td>AAACCTGACCTTGGAGGGCGGCTC</td><td></td><td></td><td></td><td></td><td></td><td></td><td></td><td></td><td></td><td></td><td></td></tr>
<tr><td>Chromosome</td><td>start</td><td></td><td>end</td><td>strand</td><td>MM</td><td>target_seq</td><td>PAM</td><td>alignment</td><td>distance</td><td>position</td><td>gene name</td><td>gene id</td></tr>
<tr><td>GU305914.1</td><td></td><td>135562</td><td>135584</td><td>+</td><td>0</td><td>GAGCCGCCCTCCAAGGTCAG</td><td>GGG</td><td>||||||||[||||||||||||]PAM</td><td>NA</td><td>Intergenic</td><td>NA</td><td>NA</td></tr>
<tr><td></td><td></td><td></td><td></td><td></td><td></td><td></td><td></td><td></td><td></td><td></td><td></td><td></td></tr>
<tr><td>T10313</td><td>GGTCAGGGGCCGCGAGATCAAGG</td><td></td><td>1000</td><td></td><td></td><td></td><td></td><td></td><td></td><td></td><td></td><td></td></tr>
<tr><td>Oligo fwd</td><td>CACCGGTCAGGGGCCGCGAGATCA</td><td></td><td></td><td></td><td></td><td></td><td></td><td></td><td></td><td></td><td></td><td></td></tr>
<tr><td>Oligo rev</td><td>AAACTGATCTCGCGGCCCCTGACC</td><td></td><td></td><td></td><td></td><td></td><td></td><td></td><td></td><td></td><td></td><td></td></tr>
<tr><td>Chromosome</td><td>start</td><td></td><td>end</td><td>strand</td><td>MM</td><td>target_seq</td><td>PAM</td><td>alignment</td><td>distance</td><td>position</td><td>gene name</td><td>gene id</td></tr>
<tr><td>GU305914.1</td><td></td><td>135576</td><td>135598</td><td>+</td><td>0</td><td>GGTCAGGGGCCGCGAGATCA</td><td>AGG</td><td>||||||||[||||||||||||]PAM</td><td>NA</td><td>Intergenic</td><td>NA</td><td>NA</td></tr>
<tr><td></td><td></td><td></td><td></td><td></td><td></td><td></td><td></td><td></td><td></td><td></td><td></td><td></td></tr>
<tr><td>T10314</td><td>CGCGAGATCAAGGCCAGACGCGG</td><td></td><td>1000</td><td></td><td></td><td></td><td></td><td></td><td></td><td></td><td></td><td></td></tr>
<tr><td>Oligo fwd</td><td>CACCCGCGAGATCAAGGCCAGACG</td><td></td><td></td><td></td><td></td><td></td><td></td><td></td><td></td><td></td><td></td><td></td></tr>
<tr><td>Oligo rev</td><td>AAACCGTCTGGCCTTGATCTCGCG</td><td></td><td></td><td></td><td></td><td></td><td></td><td></td><td></td><td></td><td></td><td></td></tr>
<tr><td>Chromosome</td><td>start</td><td></td><td>end</td><td>strand</td><td>MM</td><td>target_seq</td><td>PAM</td><td>alignment</td><td>distance</td><td>position</td><td>gene name</td><td>gene id</td></tr>
<tr><td>GU305914.1</td><td></td><td>135586</td><td>135608</td><td>+</td><td>0</td><td>CGCGAGATCAAGGCCAGACG</td><td>CGG</td><td>||||||||[||||||||||||]PAM</td><td>NA</td><td>Intergenic</td><td>NA</td><td>NA</td></tr>
<tr><td></td><td></td><td></td><td></td><td></td><td></td><td></td><td></td><td></td><td></td><td></td><td></td><td></td></tr>
<tr><td>T10315</td><td>CCAGACGCGGCGAGTTCTTCAGG</td><td></td><td>1000</td><td></td><td></td><td></td><td></td><td></td><td></td><td></td><td></td><td></td></tr>
<tr><td>Oligo fwd</td><td>CACCCCAGACGCGGCGAGTTCTTC</td><td></td><td></td><td></td><td></td><td></td><td></td><td></td><td></td><td></td><td></td><td></td></tr>
<tr><td>Oligo rev</td><td>AAACGAAGAACTCGCCGCGTCTGG</td><td></td><td></td><td></td><td></td><td></td><td></td><td></td><td></td><td></td><td></td><td></td></tr>
<tr><td>Chromosome</td><td>start</td><td></td><td>end</td><td>strand</td><td>MM</td><td>target_seq</td><td>PAM</td><td>alignment</td><td>distance</td><td>position</td><td>gene name</td><td>gene id</td></tr>
<tr><td>GU305914.1</td><td></td><td>135599</td><td>135621</td><td>+</td><td>0</td><td>CCAGACGCGGCGAGTTCTTC</td><td>AGG</td><td>||||||||[||||||||||||]PAM</td><td>NA</td><td>Intergenic</td><td>NA</td><td>NA</td></tr>
<tr><td></td><td></td><td></td><td></td><td></td><td></td><td></td><td></td><td></td><td></td><td></td><td></td><td></td></tr>
<tr><td>T10316</td><td>CAGACGCGGCGAGTTCTTCAGGG</td><td></td><td>1000</td><td></td><td></td><td></td><td></td><td></td><td></td><td></td><td></td><td></td></tr>
<tr><td>Oligo fwd</td><td>CACCCAGACGCGGCGAGTTCTTCA</td><td></td><td></td><td></td><td></td><td></td><td></td><td></td><td></td><td></td><td></td><td></td></tr>
<tr><td>Oligo rev</td><td>AAACTGAAGAACTCGCCGCGTCTG</td><td></td><td></td><td></td><td></td><td></td><td></td><td></td><td></td><td></td><td></td><td></td></tr>
<tr><td>Chromosome</td><td>start</td><td></td><td>end</td><td>strand</td><td>MM</td><td>target_seq</td><td>PAM</td><td>alignment</td><td>distance</td><td>position</td><td>gene name</td><td>gene id</td></tr>
<tr><td>GU305914.1</td><td></td><td>135600</td><td>135622</td><td>+</td><td>0</td><td>CAGACGCGGCGAGTTCTTCA</td><td>GGG</td><td>||||||||[||||||||||||]PAM</td><td>NA</td><td>Intergenic</td><td>NA</td><td>NA</td></tr>
<tr><td>GU305914.1</td><td></td><td>29398</td><td>29420</td><td>-</td><td>4</td><td>CAGACGACGCAAGTTCGTCA</td><td>CGG</td><td>||||||--[||-|||||-|||]PAM</td><td>NA</td><td>Intergenic</td><td>NA</td><td>NA</td></tr>
<tr><td></td><td></td><td></td><td></td><td></td><td></td><td></td><td></td><td></td><td></td><td></td><td></td><td></td></tr>
<tr><td>T10317</td><td>CGCGGCGAGTTCTTCAGGGACGG</td><td></td><td>1000</td><td></td><td></td><td></td><td></td><td></td><td></td><td></td><td></td><td></td></tr>
<tr><td>Oligo fwd</td><td>CACCCGCGGCGAGTTCTTCAGGGA</td><td></td><td></td><td></td><td></td><td></td><td></td><td></td><td></td><td></td><td></td><td></td></tr>
<tr><td>Oligo rev</td><td>AAACTCCCTGAAGAACTCGCCGCG</td><td></td><td></td><td></td><td></td><td></td><td></td><td></td><td></td><td></td><td></td><td></td></tr>
<tr><td>Chromosome</td><td>start</td><td></td><td>end</td><td>strand</td><td>MM</td><td>target_seq</td><td>PAM</td><td>alignment</td><td>distance</td><td>position</td><td>gene name</td><td>gene id</td></tr>
<tr><td>GU305914.1</td><td></td><td>135604</td><td>135626</td><td>+</td><td>0</td><td>CGCGGCGAGTTCTTCAGGGA</td><td>CGG</td><td>||||||||[||||||||||||]PAM</td><td>NA</td><td>Intergenic</td><td>NA</td><td>NA</td></tr>
</table>

T10318 CTTCAGGGACGGTGACGCGCCGG 1000
Oligo fwd CACCCTTCAGGGACGGTGACGCGC
Oligo rev AAACGCGCGTCACCGTCCCTGAAG
Chromosome start end strand MM target_seq PAM alignment distance position gene name gene id
GU305914.1 135615 135637 + 0 CTTCAGGGACGGTGACGCGC CGG ||||||||[||||||||||||]PAM NA Intergenic NA NA

T10319 CGGCCGCGCCCGACCCCGTCAGG 1000
Oligo fwd CACCCGGCCGCGCCCGACCCCGTC
Oligo rev AAACGACGGGGTCGGGCGCGGCCG
Chromosome start end strand MM target_seq PAM alignment distance position gene name gene id
GU305914.1 135635 135657 + 0 CGGCCGCGCCCGACCCCGTC AGG ||||||||[||||||||||||]PAM NA Intergenic NA NA

T10320 CGACCCCGTCAGGAGCCGATCGG 1000
Oligo fwd CACCCGACCCCGTCAGGAGCCGAT
Oligo rev AAACATCGGCTCCTGACGGGGTCG
Chromosome start end strand MM target_seq PAM alignment distance position gene name gene id
GU305914.1 135645 135667 + 0 CGACCCCGTCAGGAGCCGAT CGG ||||||||[||||||||||||]PAM NA Intergenic NA NA

T10321 TCAGGAGCCGATCGGACCGCTGG 1000
Oligo fwd CACCTCAGGAGCCGATCGGACCGC
Oligo rev AAACGCGGTCCGATCGGCTCCTGA
Chromosome start end strand MM target_seq PAM alignment distance position gene name gene id
GU305914.1 135653 135675 + 0 TCAGGAGCCGATCGGACCGC TGG ||||||||[||||||||||||]PAM NA Intergenic NA NA

T10322 AGCCGATCGGACCGCTGGTTCGG 1000
Oligo fwd CACCAGCCGATCGGACCGCTGGTT
Oligo rev AAACAACCAGCGGTCCGATCGGCT
Chromosome start end strand MM target_seq PAM alignment distance position gene name gene id
GU305914.1 135658 135680 + 0 AGCCGATCGGACCGCTGGTT CGG ||||||||[||||||||||||]PAM NA Intergenic NA NA

T10323 CATTCGCATCTCGTACGCCCAGG 1000
Oligo fwd CACCCATTCGCATCTCGTACGCCC
Oligo rev AAACGGGCGTACGAGATGCGAATG
Chromosome start end strand MM target_seq PAM alignment distance position gene name gene id
GU305914.1 135693 135715 + 0 CATTCGCATCTCGTACGCCC AGG ||||||||[||||||||||||]PAM NA Intergenic NA NA

T10324 TCGCATCTCGTACGCCCAGGAGG 1000
Oligo fwd CACCTCGCATCTCGTACGCCCAGG
Oligo rev AAACCCTGGGCGTACGAGATGCGA
Chromosome start end strand MM target_seq PAM alignment distance position gene name gene id
GU305914.1 135696 135718 + 0 TCGCATCTCGTACGCCCAGG AGG ||||||||[||||||||||||]PAM NA Intergenic NA NA

T10325 TCGTACGCCCAGGAGGTCACCGG 1000
Oligo fwd CACCTCGTACGCCCAGGAGGTCAC
Oligo rev AAACGTGACCTCCTGGGCGTACGA
Chromosome start end strand MM target_seq PAM alignment distance position gene name gene id
GU305914.1 135703 135725 + 0 TCGTACGCCCAGGAGGTCAC CGG ||||||||[||||||||||||]PAM NA Intergenic NA NA

T10326 AGGAGGTCACCGGCTCGCGCAGG 1000
Oligo fwd CACCAGGAGGTCACCGGCTCGCGC
Oligo rev AAACGCGCGAGCCGGTGACCTCCT
Chromosome start end strand MM target_seq PAM alignment distance position gene name gene id
GU305914.1 135713 135735 + 0 AGGAGGTCACCGGCTCGCGC AGG ||||||||[||||||||||||]PAM NA Intergenic NA NA

T10327 CGCGCAGGTTCGCCGTGATCTGG 1000
Oligo fwd CACCCGCGCAGGTTCGCCGTGATC
Oligo rev AAACGATCACGGCGAACCTGCGCG
Chromosome start end strand MM target_seq PAM alignment distance position gene name gene id
GU305914.1 135728 135750 + 0 CGCGCAGGTTCGCCGTGATC TGG ||||||||[||||||||||||]PAM NA Intergenic NA NA

T10328 ATCTGGTACTCCGAGTCCATCGG 1000
Oligo fwd CACCATCTGGTACTCCGAGTCCAT
Oligo rev AAACATGGACTCGGAGTACCAGAT
Chromosome start end strand MM target_seq PAM alignment distance position gene name gene id
GU305914.1 135745 135767 + 0 ATCTGGTACTCCGAGTCCAT CGG ||||||||[||||||||||||]PAM NA Intergenic NA NA

T10329 TCGACCGTCGACGTCGACGTCGG 1000
Oligo fwd CACCTCGACCGTCGACGTCGACGT
Oligo rev AAACACGTCGACGTCGACGGTCGA
Chromosome start end strand MM target_seq PAM alignment distance position gene name gene id
GU305914.1 135769 135791 + 0 TCGACCGTCGACGTCGACGT CGG ||||||||[||||||||||||]PAM NA Intergenic NA NA

T10330 CCGTCGACGTCGACGTCGGACGG 1000
Oligo fwd CACCCCGTCGACGTCGACGTCGGA
Oligo rev AAACTCCGACGTCGACGTCGACGG
Chromosome start end strand MM target_seq PAM alignment distance position gene name gene id
GU305914.1 135773 135795 + 0 CCGTCGACGTCGACGTCGGA CGG ||||||||[||||||||||||]PAM NA Intergenic NA NA
GU305914.1 15998 16020 - 4 ACGTCGAGATCGACGTCCGA AGG -||||||-[-||||||||-||]PAM NA Intergenic NA NA
GU305914.1 142265 142287 + 4 ACGTCGTCGTCGACGACGGT GGG -|||||-|[|||||||-|||-]PAM NA Intergenic NA NA

T10331 CGTCGACGTCGACGTCGGACGGG 1000
Oligo fwd CACCCGTCGACGTCGACGTCGGAC
Oligo rev AAACGTCCGACGTCGACGTCGACG
Chromosome start end strand MM target_seq PAM alignment distance position gene name gene id
GU305914.1 135774 135796 + 0 CGTCGACGTCGACGTCGGAC GGG ||||||||[||||||||||||]PAM NA Intergenic NA NA

T10332 CGACGTCGACGTCGGACGGGCGG 1000
Oligo fwd CACCCGACGTCGACGTCGGACGGG

| | | | | | | | | | | | |
|---|---|---|---|---|---|---|---|---|---|---|---|
| Oligo rev | AAACCCCGTCCGACGTCGACGTCG | | | | | | | | | | |
| Chromosome | start | end | strand | MM | target_seq | PAM | alignment | distance | position | gene name | gene id |
| GU305914.1 | 135777 | 135799 | + | 0 | CGACGTCGACGTCGGACGGG | CGG | \|\|\|\|\|\|\|\|[\|\|\|\|\|\|\|\|\|\|\|\|]PAM | NA | Intergenic | NA | NA |
| T10333 | GACGTCGGACGGGCGGACGACGG | 1000 | | | | | | | | | |
| Oligo fwd | CACCGACGTCGGACGGGCGGACGA | | | | | | | | | | |
| Oligo rev | AAACTCGTCCGCCCGTCCGACGTC | | | | | | | | | | |
| Chromosome | start | end | strand | MM | target_seq | PAM | alignment | distance | position | gene name | gene id |
| GU305914.1 | 135784 | 135806 | + | 0 | GACGTCGGACGGGCGGACGA | CGG | \|\|\|\|\|\|\|\|[\|\|\|\|\|\|\|\|\|\|\|\|]PAM | NA | Intergenic | NA | NA |
| GU305914.1 | 64711 | 64733 | - | 4 | CACGGCGGACGCGCGGTCGA | TGG | -\|\|\|-\|\|\|[\|\|\|-\|\|\|\|-\|\|\|]PAM | NA | Intergenic | NA | NA |
| GU305914.1 | 132549 | 132571 | - | 4 | GAACTCGGACGTGCGGAAGA | CGG | \|\|--\|\|\|\|[\|\|\|-\|\|\|\|\|-\|\|]PAM | NA | Intergenic | NA | NA |
| T10334 | ACGTCGGACGGGCGGACGACGGG | 1000 | | | | | | | | | |
| Oligo fwd | CACCACGTCGGACGGGCGGACGAC | | | | | | | | | | |
| Oligo rev | AAACGTCGTCCGCCCGTCCGACGT | | | | | | | | | | |
| Chromosome | start | end | strand | MM | target_seq | PAM | alignment | distance | position | gene name | gene id |
| GU305914.1 | 135785 | 135807 | + | 0 | ACGTCGGACGGGCGGACGAC | GGG | \|\|\|\|\|\|\|\|[\|\|\|\|\|\|\|\|\|\|\|\|]PAM | NA | Intergenic | NA | NA |
| T10335 | CGTCGGACGGGCGGACGACGGGG | 1000 | | | | | | | | | |
| Oligo fwd | CACCCGTCGGACGGGCGGACGACG | | | | | | | | | | |
| Oligo rev | AAACCGTCGTCCGCCCGTCCGACG | | | | | | | | | | |
| Chromosome | start | end | strand | MM | target_seq | PAM | alignment | distance | position | gene name | gene id |
| GU305914.1 | 135786 | 135808 | + | 0 | CGTCGGACGGGCGGACGACG | GGG | \|\|\|\|\|\|\|\|[\|\|\|\|\|\|\|\|\|\|\|\|]PAM | NA | Intergenic | NA | NA |
| T10336 | CGGACGGGCGGACGACGGGGAGG | 1000 | | | | | | | | | |
| Oligo fwd | CACCCGGACGGGCGGACGACGGGG | | | | | | | | | | |
| Oligo rev | AAACCCCCGTCGTCCGCCCGTCCG | | | | | | | | | | |
| Chromosome | start | end | strand | MM | target_seq | PAM | alignment | distance | position | gene name | gene id |
| GU305914.1 | 135789 | 135811 | + | 0 | CGGACGGGCGGACGACGGGG | AGG | \|\|\|\|\|\|\|\|[\|\|\|\|\|\|\|\|\|\|\|\|]PAM | NA | Intergenic | NA | NA |
| T10337 | ACGGGCGGACGACGGGGAGGCGG | 1000 | | | | | | | | | |
| Oligo fwd | CACCACGGGCGGACGACGGGGAGG | | | | | | | | | | |
| Oligo rev | AAACCCTCCCCGTCGTCCGCCCGT | | | | | | | | | | |
| Chromosome | start | end | strand | MM | target_seq | PAM | alignment | distance | position | gene name | gene id |
| GU305914.1 | 135792 | 135814 | + | 0 | ACGGGCGGACGACGGGGAGG | CGG | \|\|\|\|\|\|\|\|[\|\|\|\|\|\|\|\|\|\|\|\|]PAM | NA | Intergenic | NA | NA |
| T10338 | CCTTCGAACACACGCGCCGCTGG | 1000 | | | | | | | | | |
| Oligo fwd | CACCCCTTCGAACACACGCGCCGC | | | | | | | | | | |
| Oligo rev | AAACGCGGCGCGTGTGTTCGAAGG | | | | | | | | | | |
| Chromosome | start | end | strand | MM | target_seq | PAM | alignment | distance | position | gene name | gene id |
| GU305914.1 | 135824 | 135846 | + | 0 | CCTTCGAACACACGCGCCGC | TGG | \|\|\|\|\|\|\|\|[\|\|\|\|\|\|\|\|\|\|\|\|]PAM | NA | Intergenic | NA | NA |
| T10339 | CTTCGAACACACGCGCCGCTGGG | 1000 | | | | | | | | | |
| Oligo fwd | CACCCTTCGAACACACGCGCCGCT | | | | | | | | | | |
| Oligo rev | AAACAGCGGCGCGTGTGTTCGAAG | | | | | | | | | | |
| Chromosome | start | end | strand | MM | target_seq | PAM | alignment | distance | position | gene name | gene id |
| GU305914.1 | 135825 | 135847 | + | 0 | CTTCGAACACACGCGCCGCT | GGG | \|\|\|\|\|\|\|\|[\|\|\|\|\|\|\|\|\|\|\|\|]PAM | NA | Intergenic | NA | NA |
| T10340 | ACACACGCGCCGCTGGGAGACGG | 1000 | | | | | | | | | |
| Oligo fwd | CACCACACACGCGCCGCTGGGAGA | | | | | | | | | | |
| Oligo rev | AAACTCTCCCAGCGGCGCGTGTGT | | | | | | | | | | |
| Chromosome | start | end | strand | MM | target_seq | PAM | alignment | distance | position | gene name | gene id |
| GU305914.1 | 135831 | 135853 | + | 0 | ACACACGCGCCGCTGGGAGA | CGG | \|\|\|\|\|\|\|\|[\|\|\|\|\|\|\|\|\|\|\|\|]PAM | NA | Intergenic | NA | NA |
| T10341 | GCTGGGAGACGGCCGTCACGCGG | 1000 | | | | | | | | | |
| Oligo fwd | CACCGCTGGGAGACGGCCGTCACG | | | | | | | | | | |
| Oligo rev | AAACCGTGACGGCCGTCTCCCAGC | | | | | | | | | | |
| Chromosome | start | end | strand | MM | target_seq | PAM | alignment | distance | position | gene name | gene id |
| GU305914.1 | 135842 | 135864 | + | 0 | GCTGGGAGACGGCCGTCACG | CGG | \|\|\|\|\|\|\|\|[\|\|\|\|\|\|\|\|\|\|\|\|]PAM | NA | Intergenic | NA | NA |
| T10342 | CTGGGAGACGGCCGTCACGCGGG | 1000 | | | | | | | | | |
| Oligo fwd | CACCCTGGGAGACGGCCGTCACGC | | | | | | | | | | |
| Oligo rev | AAACGCGTGACGGCCGTCTCCCAG | | | | | | | | | | |
| Chromosome | start | end | strand | MM | target_seq | PAM | alignment | distance | position | gene name | gene id |
| GU305914.1 | 135843 | 135865 | + | 0 | CTGGGAGACGGCCGTCACGC | GGG | \|\|\|\|\|\|\|\|[\|\|\|\|\|\|\|\|\|\|\|\|]PAM | NA | Intergenic | NA | NA |
| T10343 | ACGGCCGTCACGCGGGTCTACGG | 1000 | | | | | | | | | |
| Oligo fwd | CACCACGGCCGTCACGCGGGTCTA | | | | | | | | | | |
| Oligo rev | AAACTAGACCCGCGTGACGGCCGT | | | | | | | | | | |
| Chromosome | start | end | strand | MM | target_seq | PAM | alignment | distance | position | gene name | gene id |
| GU305914.1 | 135850 | 135872 | + | 0 | ACGGCCGTCACGCGGGTCTA | CGG | \|\|\|\|\|\|\|\|[\|\|\|\|\|\|\|\|\|\|\|\|]PAM | NA | Intergenic | NA | NA |
| T10344 | CACCAGCCGACAGCTCGCCGCGG | 1000 | | | | | | | | | |
| Oligo fwd | CACCCACCAGCCGACAGCTCGCCG | | | | | | | | | | |
| Oligo rev | AAACCGGCGAGCTGTCGGCTGGTG | | | | | | | | | | |
| Chromosome | start | end | strand | MM | target_seq | PAM | alignment | distance | position | gene name | gene id |
| GU305914.1 | 135876 | 135898 | + | 0 | CACCAGCCGACAGCTCGCCG | CGG | \|\|\|\|\|\|\|\|[\|\|\|\|\|\|\|\|\|\|\|\|]PAM | NA | Intergenic | NA | NA |
| T10345 | GCCGCGGCCGTCACGCGCGTCGG | 1000 | | | | | | | | | |
| Oligo fwd | CACCGCCGCGGCCGTCACGCGCGT | | | | | | | | | | |
| Oligo rev | AAACACGCGCGTGACGGCCGCGGC | | | | | | | | | | |
| Chromosome | start | end | strand | MM | target_seq | PAM | alignment | distance | position | gene name | gene id |
| GU305914.1 | 135892 | 135914 | + | 0 | GCCGCGGCCGTCACGCGCGT | CGG | \|\|\|\|\|\|\|\|[\|\|\|\|\|\|\|\|\|\|\|\|]PAM | NA | Intergenic | NA | NA |
| T10346 | CCGCGGCCGTCACGCGCGTCGGG | 1000 | | | | | | | | | |
| Oligo fwd | CACCCCGCGGCCGTCACGCGCGTC | | | | | | | | | | |
| Oligo rev | AAACGACGCGCGTGACGGCCGCGG | | | | | | | | | | |
| Chromosome | start | end | strand | MM | target_seq | PAM | alignment | distance | position | gene name | gene id |
| GU305914.1 | 135893 | 135915 | + | 0 | CCGCGGCCGTCACGCGCGTC | GGG | \|\|\|\|\|\|\|\|[\|\|\|\|\|\|\|\|\|\|\|\|]PAM | NA | Intergenic | NA | NA |

T10347 CGCGGCCGTCACGCGCGTCGGGG 1000
Oligo fwd CACCCGCGGCCGTCACGCGCGTCG
Oligo rev AAACCGACGCGCGTGACGGCCGCG
Chromosome start end strand MM target_seq PAM alignment distance position gene name gene id
GU305914.1 135894 135916 + 0 CGCGGCCGTCACGCGCGTCG GGG ||||||||[||||||||||||]PAM NA Intergenic NA NA

T10348 GCGGCCGTCACGCGCGTCGGGGG 1000
Oligo fwd CACCGCGGCCGTCACGCGCGTCGG
Oligo rev AAACCCGACGCGCGTGACGGCCGC
Chromosome start end strand MM target_seq PAM alignment distance position gene name gene id
GU305914.1 135895 135917 + 0 GCGGCCGTCACGCGCGTCGG GGG ||||||||[||||||||||||]PAM NA Intergenic NA NA

T10349 CGGCCGTCACGCGCGTCGGGGGG 1000
Oligo fwd CACCCGGCCGTCACGCGCGTCGGG
Oligo rev AAACCCCGACGCGCGTGACGGCCG
Chromosome start end strand MM target_seq PAM alignment distance position gene name gene id
GU305914.1 135896 135918 + 0 CGGCCGTCACGCGCGTCGGG GGG ||||||||[||||||||||||]PAM NA Intergenic NA NA

T10350 GCGTCGGGGGGCTCGCGCAGAGG 1000
Oligo fwd CACCGCGTCGGGGGGCTCGCGCAG
Oligo rev AAACCTGCGCGAGCCCCCCGACGC
Chromosome start end strand MM target_seq PAM alignment distance position gene name gene id
GU305914.1 135908 135930 + 0 GCGTCGGGGGGCTCGCGCAG AGG ||||||||[||||||||||||]PAM NA Intergenic NA NA

T10351 TTCTACGTGAATCTCTACGCCGG 1000
Oligo fwd CACCTTCTACGTGAATCTCTACGC
Oligo rev AAACGCGTAGAGATTCACGTAGAA
Chromosome start end strand MM target_seq PAM alignment distance position gene name gene id
GU305914.1 135931 135953 + 0 TTCTACGTGAATCTCTACGC CGG ||||||||[||||||||||||]PAM NA Intergenic NA NA

T10352 TCTACGTGAATCTCTACGCCGGG 1000
Oligo fwd CACCTCTACGTGAATCTCTACGCC
Oligo rev AAACGGCGTAGAGATTCACGTAGA
Chromosome start end strand MM target_seq PAM alignment distance position gene name gene id
GU305914.1 135932 135954 + 0 TCTACGTGAATCTCTACGCC GGG ||||||||[||||||||||||]PAM NA Intergenic NA NA

T10353 TGAATCTCTACGCCGGGCTGCGG 1000
Oligo fwd CACCTGAATCTCTACGCCGGGCTG
Oligo rev AAACCAGCCCGGCGTAGAGATTCA
Chromosome start end strand MM target_seq PAM alignment distance position gene name gene id
GU305914.1 135938 135960 + 0 TGAATCTCTACGCCGGGCTG CGG ||||||||[||||||||||||]PAM NA Intergenic NA NA

T10354 GAATCTCTACGCCGGGCTGCGGG 1000
Oligo fwd CACCGAATCTCTACGCCGGGCTGC
Oligo rev AAACGCAGCCCGGCGTAGAGATTC
Chromosome start end strand MM target_seq PAM alignment distance position gene name gene id
GU305914.1 135939 135961 + 0 GAATCTCTACGCCGGGCTGC GGG ||||||||[||||||||||||]PAM NA Intergenic NA NA

T10355 TCTCTACGCCGGGCTGCGGGCGG 1000
Oligo fwd CACCTCTCTACGCCGGGCTGCGGG
Oligo rev AAACCCCGCAGCCCGGCGTAGAGA
Chromosome start end strand MM target_seq PAM alignment distance position gene name gene id
GU305914.1 135942 135964 + 0 TCTCTACGCCGGGCTGCGGG CGG ||||||||[||||||||||||]PAM NA Intergenic NA NA

T10356 CTCTACGCCGGGCTGCGGGCGGG 1000
Oligo fwd CACCCTCTACGCCGGGCTGCGGGC
Oligo rev AAACGCCCGCAGCCCGGCGTAGAG
Chromosome start end strand MM target_seq PAM alignment distance position gene name gene id
GU305914.1 135943 135965 + 0 CTCTACGCCGGGCTGCGGGC GGG ||||||||[||||||||||||]PAM NA Intergenic NA NA

T10357 TCTACGCCGGGCTGCGGGCGGGG 1000
Oligo fwd CACCTCTACGCCGGGCTGCGGGCG
Oligo rev AAACCGCCCGCAGCCCGGCGTAGA
Chromosome start end strand MM target_seq PAM alignment distance position gene name gene id
GU305914.1 135944 135966 + 0 TCTACGCCGGGCTGCGGGCG GGG ||||||||[||||||||||||]PAM NA Intergenic NA NA

T10358 CTACGCCGGGCTGCGGGCGGGGG 1000
Oligo fwd CACCCTACGCCGGGCTGCGGGCGG
Oligo rev AAACCCGCCCGCAGCCCGGCGTAG
Chromosome start end strand MM target_seq PAM alignment distance position gene name gene id
GU305914.1 135945 135967 + 0 CTACGCCGGGCTGCGGGCGG GGG ||||||||[||||||||||||]PAM NA Intergenic NA NA

T10359 CGGGCTGCGGGCGGGGGATCTGG 1000
Oligo fwd CACCCGGGCTGCGGGCGGGGGATC
Oligo rev AAACGATCCCCCGCCCGCAGCCCG
Chromosome start end strand MM target_seq PAM alignment distance position gene name gene id
GU305914.1 135951 135973 + 0 CGGGCTGCGGGCGGGGGATC TGG ||||||||[||||||||||||]PAM NA Intergenic NA NA

T10360 GCTGCGGGCGGGGGATCTGGTGG 1000
Oligo fwd CACCGCTGCGGGCGGGGGATCTGG
Oligo rev AAACCCAGATCCCCCGCCCGCAGC
Chromosome start end strand MM target_seq PAM alignment distance position gene name gene id
GU305914.1 135954 135976 + 0 GCTGCGGGCGGGGGATCTGG TGG ||||||||[||||||||||||]PAM NA Intergenic NA NA

T10361 TGCTGCTTTCGCGCGACGCCTGG 1000
Oligo fwd CACCTGCTGCTTTCGCGCGACGCC
Oligo rev AAACGGCGTCGCGCGAAAGCAGCA
Chromosome start end strand MM target_seq PAM alignment distance position gene name gene id

GU305914.1 135983 136005 + 0 TGCTGCTTTCGCGCGACGCC TGG ||||||||[||||||||||||]PAM NA Intergenic NA NA

T10362 CGCGACGCCTGGCTCGTGAACGG 1000
Oligo fwd CACCCGCGACGCCTGGCTCGTGAA
Oligo rev AAACTTCACGAGCCAGGCGTCGCG
Chromosome start end strand MM target_seq PAM alignment distance position gene name gene id
GU305914.1 135994 136016 + 0 CGCGACGCCTGGCTCGTGAA CGG ||||||||[||||||||||||]PAM NA Intergenic NA NA

T10363 GCCTGGCTCGTGAACGGCACCGG 1000
Oligo fwd CACCGCCTGGCTCGTGAACGGCAC
Oligo rev AAACGTGCCGTTCACGAGCCAGGC
Chromosome start end strand MM target_seq PAM alignment distance position gene name gene id
GU305914.1 136000 136022 + 0 GCCTGGCTCGTGAACGGCAC CGG ||||||||[||||||||||||]PAM NA Intergenic NA NA

T10364 TGGCTCGTGAACGGCACCGGAGG 1000
Oligo fwd CACCTGGCTCGTGAACGGCACCGG
Oligo rev AAACCCGGTGCCGTTCACGAGCCA
Chromosome start end strand MM target_seq PAM alignment distance position gene name gene id
GU305914.1 136003 136025 + 0 TGGCTCGTGAACGGCACCGG AGG ||||||||[||||||||||||]PAM NA Intergenic NA NA

T10365 GGCTCGTGAACGGCACCGGAGGG 1000
Oligo fwd CACCGGCTCGTGAACGGCACCGGA
Oligo rev AAACTCCGGTGCCGTTCACGAGCC
Chromosome start end strand MM target_seq PAM alignment distance position gene name gene id
GU305914.1 136004 136026 + 0 GGCTCGTGAACGGCACCGGA GGG ||||||||[||||||||||||]PAM NA Intergenic NA NA

T10366 GCTCGTGAACGGCACCGGAGGGG 1000
Oligo fwd CACCGCTCGTGAACGGCACCGGAG
Oligo rev AAACCTCCGGTGCCGTTCACGAGC
Chromosome start end strand MM target_seq PAM alignment distance position gene name gene id
GU305914.1 136005 136027 + 0 GCTCGTGAACGGCACCGGAG GGG ||||||||[||||||||||||]PAM NA Intergenic NA NA

T10367 CTCGTGAACGGCACCGGAGGGGG 1000
Oligo fwd CACCCTCGTGAACGGCACCGGAGG
Oligo rev AAACCCTCCGGTGCCGTTCACGAG
Chromosome start end strand MM target_seq PAM alignment distance position gene name gene id
GU305914.1 136006 136028 + 0 CTCGTGAACGGCACCGGAGG GGG ||||||||[||||||||||||]PAM NA Intergenic NA NA

T10368 CGGAGGGGGCTGCATTATTAAGG 1000
Oligo fwd CACCCGGAGGGGGCTGCATTATTA
Oligo rev AAACTAATAATGCAGCCCCCTCCG
Chromosome start end strand MM target_seq PAM alignment distance position gene name gene id
GU305914.1 136020 136042 + 0 CGGAGGGGGCTGCATTATTA AGG ||||||||[||||||||||||]PAM NA Intergenic NA NA

T10369 CGCTCTCCTCCGATCTTCATCGG 1000
Oligo fwd CACCCGCTCTCCTCCGATCTTCAT
Oligo rev AAACATGAAGATCGGAGGAGAGCG
Chromosome start end strand MM target_seq PAM alignment distance position gene name gene id
GU305914.1 136043 136065 + 0 CGCTCTCCTCCGATCTTCAT CGG ||||||||[||||||||||||]PAM NA Intergenic NA NA

T10370 CCTCCGATCTTCATCGGCGTCGG 1000
Oligo fwd CACCCCTCCGATCTTCATCGGCGT
Oligo rev AAACACGCCGATGAAGATCGGAGG
Chromosome start end strand MM target_seq PAM alignment distance position gene name gene id
GU305914.1 136049 136071 + 0 CCTCCGATCTTCATCGGCGT CGG ||||||||[||||||||||||]PAM NA Intergenic NA NA

T10371 CTTCATCGGCGTCGGCGATCTGG 1000
Oligo fwd CACCCTTCATCGGCGTCGGCGATC
Oligo rev AAACGATCGCCGACGCCGATGAAG
Chromosome start end strand MM target_seq PAM alignment distance position gene name gene id
GU305914.1 136057 136079 + 0 CTTCATCGGCGTCGGCGATC TGG ||||||||[||||||||||||]PAM NA Intergenic NA NA

T10372 CGATCTGGCGACCTCGTCGATGG 1000
Oligo fwd CACCCGATCTGGCGACCTCGTCGA
Oligo rev AAACTCGACGAGGTCGCCAGATCG
Chromosome start end strand MM target_seq PAM alignment distance position gene name gene id
GU305914.1 136072 136094 + 0 CGATCTGGCGACCTCGTCGA TGG ||||||||[||||||||||||]PAM NA Intergenic NA NA

T10373 GATCTGGCGACCTCGTCGATGGG 1000
Oligo fwd CACCGATCTGGCGACCTCGTCGAT
Oligo rev AAACATCGACGAGGTCGCCAGATC
Chromosome start end strand MM target_seq PAM alignment distance position gene name gene id
GU305914.1 136073 136095 + 0 GATCTGGCGACCTCGTCGAT GGG ||||||||[||||||||||||]PAM NA Intergenic NA NA

T10374 CTCGTCGATGGGTGTCGTGTCGG 1000
Oligo fwd CACCCTCGTCGATGGGTGTCGTGT
Oligo rev AAACACACGACACCCATCGACGAG
Chromosome start end strand MM target_seq PAM alignment distance position gene name gene id
GU305914.1 136084 136106 + 0 CTCGTCGATGGGTGTCGTGT CGG ||||||||[||||||||||||]PAM NA Intergenic NA NA

T10375 TCGATGGGTGTCGTGTCGGTTGG 1000
Oligo fwd CACCTCGATGGGTGTCGTGTCGGT
Oligo rev AAACACCGACACGACACCCATCGA
Chromosome start end strand MM target_seq PAM alignment distance position gene name gene id
GU305914.1 136088 136110 + 0 TCGATGGGTGTCGTGTCGGT TGG ||||||||[||||||||||||]PAM NA Intergenic NA NA

T10376 CGATGGGTGTCGTGTCGGTTGGG 1000
Oligo fwd CACCCGATGGGTGTCGTGTCGGTT
Oligo rev AAACAACCGACACGACACCCATCG

| | | | | | | | | | | | |
|---|---|---|---|---|---|---|---|---|---|---|---|
| Chromosome | start | end | strand | MM | target_seq | PAM | alignment | distance | position | gene name | gene id |
| GU305914.1 | 136089 | 136111 | + | 0 | CGATGGGTGTCGTGTCGGTT | GGG | \|\|\|\|\|\|\|\|[\|\|\|\|\|\|\|\|\|\|\|\|]PAM | NA | Intergenic | NA | NA |
| T10377 | TCGGTTGGGCCGATGTGATCTGG | 1000 | | | | | | | | | |
| Oligo fwd | CACCTCGGTTGGGCCGATGTGATC | | | | | | | | | | |
| Oligo rev | AAACGATCACATCGGCCCAACCGA | | | | | | | | | | |
| Chromosome | start | end | strand | MM | target_seq | PAM | alignment | distance | position | gene name | gene id |
| GU305914.1 | 136103 | 136125 | + | 0 | TCGGTTGGGCCGATGTGATC | TGG | \|\|\|\|\|\|\|\|[\|\|\|\|\|\|\|\|\|\|\|\|]PAM | NA | Intergenic | NA | NA |
| GU305914.1 | 131590 | 131612 | + | 4 | TCGATCGGGACGACGTGATC | AGG | \|\|\|-\|-\|\|[\|-\|\|\|-\|\|\|\|\|\|]PAM | NA | Intergenic | NA | NA |
| T10378 | CGATGTGATCTGGTACGCGCAGG | 1000 | | | | | | | | | |
| Oligo fwd | CACCCGATGTGATCTGGTACGCGC | | | | | | | | | | |
| Oligo rev | AAACGCGCGTACCAGATCACATCG | | | | | | | | | | |
| Chromosome | start | end | strand | MM | target_seq | PAM | alignment | distance | position | gene name | gene id |
| GU305914.1 | 136113 | 136135 | + | 0 | CGATGTGATCTGGTACGCGC | AGG | \|\|\|\|\|\|\|\|[\|\|\|\|\|\|\|\|\|\|\|\|]PAM | NA | Intergenic | NA | NA |
| T10379 | GCGCAGGCACGCGTCGCGTCTGG | 1000 | | | | | | | | | |
| Oligo fwd | CACCGCGCAGGCACGCGTCGCGTC | | | | | | | | | | |
| Oligo rev | AAACGACGCGACGCGTGCCTGCGC | | | | | | | | | | |
| Chromosome | start | end | strand | MM | target_seq | PAM | alignment | distance | position | gene name | gene id |
| GU305914.1 | 136129 | 136151 | + | 0 | GCGCAGGCACGCGTCGCGTC | TGG | \|\|\|\|\|\|\|\|[\|\|\|\|\|\|\|\|\|\|\|\|]PAM | NA | Intergenic | NA | NA |
| T10380 | CGCAGGCACGCGTCGCGTCTGGG | 1000 | | | | | | | | | |
| Oligo fwd | CACCCGCAGGCACGCGTCGCGTCT | | | | | | | | | | |
| Oligo rev | AAACAGACGCGACGCGTGCCTGCG | | | | | | | | | | |
| Chromosome | start | end | strand | MM | target_seq | PAM | alignment | distance | position | gene name | gene id |
| GU305914.1 | 136130 | 136152 | + | 0 | CGCAGGCACGCGTCGCGTCT | GGG | \|\|\|\|\|\|\|\|[\|\|\|\|\|\|\|\|\|\|\|\|]PAM | NA | Intergenic | NA | NA |
| T10381 | GGATACCCATAAAGCTCATGAGG | 1000 | | | | | | | | | |
| Oligo fwd | CACCGGATACCCATAAAGCTCATG | | | | | | | | | | |
| Oligo rev | AAACCATGAGCTTTATGGGTATCC | | | | | | | | | | |
| Chromosome | start | end | strand | MM | target_seq | PAM | alignment | distance | position | gene name | gene id |
| GU305914.1 | 136151 | 136173 | + | 0 | GGATACCCATAAAGCTCATG | AGG | \|\|\|\|\|\|\|\|[\|\|\|\|\|\|\|\|\|\|\|\|]PAM | NA | Intergenic | NA | NA |
| T10382 | GATACCCATAAAGCTCATGAGGG | 1000 | | | | | | | | | |
| Oligo fwd | CACCGATACCCATAAAGCTCATGA | | | | | | | | | | |
| Oligo rev | AAACTCATGAGCTTTATGGGTATC | | | | | | | | | | |
| Chromosome | start | end | strand | MM | target_seq | PAM | alignment | distance | position | gene name | gene id |
| GU305914.1 | 136152 | 136174 | + | 0 | GATACCCATAAAGCTCATGA | GGG | \|\|\|\|\|\|\|\|[\|\|\|\|\|\|\|\|\|\|\|\|]PAM | NA | Intergenic | NA | NA |
| T10383 | CCCATAAAGCTCATGAGGGTCGG | 1000 | | | | | | | | | |
| Oligo fwd | CACCCCCATAAAGCTCATGAGGGT | | | | | | | | | | |
| Oligo rev | AAACACCCTCATGAGCTTTATGGG | | | | | | | | | | |
| Chromosome | start | end | strand | MM | target_seq | PAM | alignment | distance | position | gene name | gene id |
| GU305914.1 | 136156 | 136178 | + | 0 | CCCATAAAGCTCATGAGGGT | CGG | \|\|\|\|\|\|\|\|[\|\|\|\|\|\|\|\|\|\|\|\|]PAM | NA | Intergenic | NA | NA |
| T10384 | ATAAAGCTCATGAGGGTCGGTGG | 1000 | | | | | | | | | |
| Oligo fwd | CACCATAAAGCTCATGAGGGTCGG | | | | | | | | | | |
| Oligo rev | AAACCCGACCCTCATGAGCTTTAT | | | | | | | | | | |
| Chromosome | start | end | strand | MM | target_seq | PAM | alignment | distance | position | gene name | gene id |
| GU305914.1 | 136159 | 136181 | + | 0 | ATAAAGCTCATGAGGGTCGG | TGG | \|\|\|\|\|\|\|\|[\|\|\|\|\|\|\|\|\|\|\|\|]PAM | NA | Intergenic | NA | NA |
| T10385 | TAAAGCTCATGAGGGTCGGTGGG | 1000 | | | | | | | | | |
| Oligo fwd | CACCTAAAGCTCATGAGGGTCGGT | | | | | | | | | | |
| Oligo rev | AAACACCGACCCTCATGAGCTTTA | | | | | | | | | | |
| Chromosome | start | end | strand | MM | target_seq | PAM | alignment | distance | position | gene name | gene id |
| GU305914.1 | 136160 | 136182 | + | 0 | TAAAGCTCATGAGGGTCGGT | GGG | \|\|\|\|\|\|\|\|[\|\|\|\|\|\|\|\|\|\|\|\|]PAM | NA | Intergenic | NA | NA |
| T10386 | AAAGCTCATGAGGGTCGGTGGGG | 1000 | | | | | | | | | |
| Oligo fwd | CACCAAAGCTCATGAGGGTCGGTG | | | | | | | | | | |
| Oligo rev | AAACCACCGACCCTCATGAGCTTT | | | | | | | | | | |
| Chromosome | start | end | strand | MM | target_seq | PAM | alignment | distance | position | gene name | gene id |
| GU305914.1 | 136161 | 136183 | + | 0 | AAAGCTCATGAGGGTCGGTG | GGG | \|\|\|\|\|\|\|\|[\|\|\|\|\|\|\|\|\|\|\|\|]PAM | NA | Intergenic | NA | NA |
| T10387 | CTCATGAGGGTCGGTGGGGATGG | 1000 | | | | | | | | | |
| Oligo fwd | CACCCTCATGAGGGTCGGTGGGGA | | | | | | | | | | |
| Oligo rev | AAACTCCCCACCGACCCTCATGAG | | | | | | | | | | |
| Chromosome | start | end | strand | MM | target_seq | PAM | alignment | distance | position | gene name | gene id |
| GU305914.1 | 136165 | 136187 | + | 0 | CTCATGAGGGTCGGTGGGGA | TGG | \|\|\|\|\|\|\|\|[\|\|\|\|\|\|\|\|\|\|\|\|]PAM | NA | Intergenic | NA | NA |
| T10388 | GGGTCGGTGGGGATGGCCTCTGG | 1000 | | | | | | | | | |
| Oligo fwd | CACCGGGTCGGTGGGGATGGCCTC | | | | | | | | | | |
| Oligo rev | AAACGAGGCCATCCCCACCGACCC | | | | | | | | | | |
| Chromosome | start | end | strand | MM | target_seq | PAM | alignment | distance | position | gene name | gene id |
| GU305914.1 | 136172 | 136194 | + | 0 | GGGTCGGTGGGGATGGCCTC | TGG | \|\|\|\|\|\|\|\|[\|\|\|\|\|\|\|\|\|\|\|\|]PAM | NA | Intergenic | NA | NA |
| T10389 | TGGCGATCTTTTCTCGACAGCGG | 1000 | | | | | | | | | |
| Oligo fwd | CACCTGGCGATCTTTTCTCGACAG | | | | | | | | | | |
| Oligo rev | AAACCTGTCGAGAAAAGATCGCCA | | | | | | | | | | |
| Chromosome | start | end | strand | MM | target_seq | PAM | alignment | distance | position | gene name | gene id |
| GU305914.1 | 136192 | 136214 | + | 0 | TGGCGATCTTTTCTCGACAG | CGG | \|\|\|\|\|\|\|\|[\|\|\|\|\|\|\|\|\|\|\|\|]PAM | NA | Intergenic | NA | NA |
| T10390 | TCTCGACAGCGGAAAACTGACGG | 1000 | | | | | | | | | |
| Oligo fwd | CACCTCTCGACAGCGGAAAACTGA | | | | | | | | | | |
| Oligo rev | AAACTCAGTTTTCCGCTGTCGAGA | | | | | | | | | | |
| Chromosome | start | end | strand | MM | target_seq | PAM | alignment | distance | position | gene name | gene id |
| GU305914.1 | 136203 | 136225 | + | 0 | TCTCGACAGCGGAAAACTGA | CGG | \|\|\|\|\|\|\|\|[\|\|\|\|\|\|\|\|\|\|\|\|]PAM | NA | Intergenic | NA | NA |
| T10391 | ACAGCGGAAAACTGACGGACTGG | 1000 | | | | | | | | | |

| | | | | | | | | | | | |
|---|---|---|---|---|---|---|---|---|---|---|---|
| Oligo fwd | CACCACAGCGGAAAACTGACGGAC | | | | | | | | | | |
| Oligo rev | AAACGTCCGTCAGTTTTCCGCTGT | | | | | | | | | | |
| Chromosome | start | end | strand | MM | target_seq | PAM | alignment | distance | position | gene name | gene id |
| GU305914.1 | 136208 | 136230 | + | 0 | ACAGCGGAAAACTGACGGAC | TGG | ||||||||[||||||||||||]PAM | NA | Intergenic | NA | NA |
| T10392 | CAGCGGAAAACTGACGGACTGGG | 1000 | | | | | | | | | |
| Oligo fwd | CACCCAGCGGAAAACTGACGGACT | | | | | | | | | | |
| Oligo rev | AAACAGTCCGTCAGTTTTCCGCTG | | | | | | | | | | |
| Chromosome | start | end | strand | MM | target_seq | PAM | alignment | distance | position | gene name | gene id |
| GU305914.1 | 136209 | 136231 | + | 0 | CAGCGGAAAACTGACGGACT | GGG | ||||||||[||||||||||||]PAM | NA | Intergenic | NA | NA |
| T10393 | CTGTGTCCTCTACGCCTTCGAGG | 1000 | | | | | | | | | |
| Oligo fwd | CACCCTGTGTCCTCTACGCCTTCG | | | | | | | | | | |
| Oligo rev | AAACCGAAGGCGTAGAGGACACAG | | | | | | | | | | |
| Chromosome | start | end | strand | MM | target_seq | PAM | alignment | distance | position | gene name | gene id |
| GU305914.1 | 136248 | 136270 | + | 0 | CTGTGTCCTCTACGCCTTCG | AGG | ||||||||[||||||||||||]PAM | NA | Intergenic | NA | NA |
| T10394 | TCGAGGACCGCAGCCTCTTCTGG | 1000 | | | | | | | | | |
| Oligo fwd | CACCTCGAGGACCGCAGCCTCTTC | | | | | | | | | | |
| Oligo rev | AAACGAAGAGGCTGCGGTCCTCGA | | | | | | | | | | |
| Chromosome | start | end | strand | MM | target_seq | PAM | alignment | distance | position | gene name | gene id |
| GU305914.1 | 136265 | 136287 | + | 0 | TCGAGGACCGCAGCCTCTTC | TGG | ||||||||[||||||||||||]PAM | NA | Intergenic | NA | NA |
| T10395 | CGAGGACCGCAGCCTCTTCTGGG | 1000 | | | | | | | | | |
| Oligo fwd | CACCCGAGGACCGCAGCCTCTTCT | | | | | | | | | | |
| Oligo rev | AAACAGAAGAGGCTGCGGTCCTCG | | | | | | | | | | |
| Chromosome | start | end | strand | MM | target_seq | PAM | alignment | distance | position | gene name | gene id |
| GU305914.1 | 136266 | 136288 | + | 0 | CGAGGACCGCAGCCTCTTCT | GGG | ||||||||[||||||||||||]PAM | NA | Intergenic | NA | NA |
| T10396 | AGCCTCTTCTGGGTGCTTCCCGG | 1000 | | | | | | | | | |
| Oligo fwd | CACCAGCCTCTTCTGGGTGCTTCC | | | | | | | | | | |
| Oligo rev | AAACGGAAGCACCCAGAAGAGGCT | | | | | | | | | | |
| Chromosome | start | end | strand | MM | target_seq | PAM | alignment | distance | position | gene name | gene id |
| GU305914.1 | 136276 | 136298 | + | 0 | AGCCTCTTCTGGGTGCTTCC | CGG | ||||||||[||||||||||||]PAM | NA | Intergenic | NA | NA |
| T10397 | CTCTTCTGGGTGCTTCCCGGCGG | 1000 | | | | | | | | | |
| Oligo fwd | CACCCTCTTCTGGGTGCTTCCCGG | | | | | | | | | | |
| Oligo rev | AAACCCGGGAAGCACCCAGAAGAG | | | | | | | | | | |
| Chromosome | start | end | strand | MM | target_seq | PAM | alignment | distance | position | gene name | gene id |
| GU305914.1 | 136279 | 136301 | + | 0 | CTCTTCTGGGTGCTTCCCGG | CGG | ||||||||[||||||||||||]PAM | NA | Intergenic | NA | NA |
| T10398 | CTGCGCGACCTTCCAGCTGCAGG | 1000 | | | | | | | | | |
| Oligo fwd | CACCCTGCGCGACCTTCCAGCTGC | | | | | | | | | | |
| Oligo rev | AAACGCAGCTGGAAGGTCGCGCAG | | | | | | | | | | |
| Chromosome | start | end | strand | MM | target_seq | PAM | alignment | distance | position | gene name | gene id |
| GU305914.1 | 136305 | 136327 | + | 0 | CTGCGCGACCTTCCAGCTGC | AGG | ||||||||[||||||||||||]PAM | NA | Intergenic | NA | NA |
| T10399 | TTCCAGCTGCAGGTAGACCATGG | 1000 | | | | | | | | | |
| Oligo fwd | CACCTTCCAGCTGCAGGTAGACCA | | | | | | | | | | |
| Oligo rev | AAACTGGTCTACCTGCAGCTGGAA | | | | | | | | | | |
| Chromosome | start | end | strand | MM | target_seq | PAM | alignment | distance | position | gene name | gene id |
| GU305914.1 | 136315 | 136337 | + | 0 | TTCCAGCTGCAGGTAGACCA | TGG | ||||||||[||||||||||||]PAM | NA | Intergenic | NA | NA |
| T10400 | AAAGTTTGAGAAGCATCAGTAGG | 1000 | | | | | | | | | |
| Oligo fwd | CACCAAAGTTTGAGAAGCATCAGT | | | | | | | | | | |
| Oligo rev | AAACACTGATGCTTCTCAAACTTT | | | | | | | | | | |
| Chromosome | start | end | strand | MM | target_seq | PAM | alignment | distance | position | gene name | gene id |
| GU305914.1 | 136359 | 136381 | + | 0 | AAAGTTTGAGAAGCATCAGT | AGG | ||||||||[||||||||||||]PAM | NA | Intergenic | NA | NA |
| T10401 | GCATCAGTAGGTTCGTCCAGTGG | 1000 | | | | | | | | | |
| Oligo fwd | CACCGCATCAGTAGGTTCGTCCAG | | | | | | | | | | |
| Oligo rev | AAACCTGGACGAACCTACTGATGC | | | | | | | | | | |
| Chromosome | start | end | strand | MM | target_seq | PAM | alignment | distance | position | gene name | gene id |
| GU305914.1 | 136371 | 136393 | + | 0 | GCATCAGTAGGTTCGTCCAG | TGG | ||||||||[||||||||||||]PAM | NA | Intergenic | NA | NA |
| T10402 | CATCAGTAGGTTCGTCCAGTGGG | 1000 | | | | | | | | | |
| Oligo fwd | CACCCATCAGTAGGTTCGTCCAGT | | | | | | | | | | |
| Oligo rev | AAACACTGGACGAACCTACTGATG | | | | | | | | | | |
| Chromosome | start | end | strand | MM | target_seq | PAM | alignment | distance | position | gene name | gene id |
| GU305914.1 | 136372 | 136394 | + | 0 | CATCAGTAGGTTCGTCCAGT | GGG | ||||||||[||||||||||||]PAM | NA | Intergenic | NA | NA |
| T10403 | TCGTCCAGTGGGAGTGTTGTTGG | 1000 | | | | | | | | | |
| Oligo fwd | CACCTCGTCCAGTGGGAGTGTTGT | | | | | | | | | | |
| Oligo rev | AAACACAACACTCCCACTGGACGA | | | | | | | | | | |
| Chromosome | start | end | strand | MM | target_seq | PAM | alignment | distance | position | gene name | gene id |
| GU305914.1 | 136383 | 136405 | + | 0 | TCGTCCAGTGGGAGTGTTGT | TGG | ||||||||[||||||||||||]PAM | NA | Intergenic | NA | NA |
| T10404 | CGCCGTCACCAGCCAGTCGCCGG | 1000 | | | | | | | | | |
| Oligo fwd | CACCCGCCGTCACCAGCCAGTCGC | | | | | | | | | | |
| Oligo rev | AAACGCGACTGGCTGGTGACGGCG | | | | | | | | | | |
| Chromosome | start | end | strand | MM | target_seq | PAM | alignment | distance | position | gene name | gene id |
| GU305914.1 | 136444 | 136466 | + | 0 | CGCCGTCACCAGCCAGTCGC | CGG | ||||||||[||||||||||||]PAM | NA | Intergenic | NA | NA |
| T10405 | GCCGTCACCAGCCAGTCGCCGGG | 1000 | | | | | | | | | |
| Oligo fwd | CACCGCCGTCACCAGCCAGTCGCC | | | | | | | | | | |
| Oligo rev | AAACGGCGACTGGCTGGTGACGGC | | | | | | | | | | |
| Chromosome | start | end | strand | MM | target_seq | PAM | alignment | distance | position | gene name | gene id |
| GU305914.1 | 136445 | 136467 | + | 0 | GCCGTCACCAGCCAGTCGCC | GGG | ||||||||[||||||||||||]PAM | NA | Intergenic | NA | NA |

<table>
<tr><td>T10406</td><td>CCGTCACCAGCCAGTCGCCGGGG</td><td>1000</td><td></td><td></td><td></td><td></td><td></td><td></td><td></td><td></td><td></td></tr>
<tr><td>Oligo fwd</td><td>CACCCCGTCACCAGCCAGTCGCCG</td><td></td><td></td><td></td><td></td><td></td><td></td><td></td><td></td><td></td><td></td></tr>
<tr><td>Oligo rev</td><td>AAACCGGCGACTGGCTGGTGACGG</td><td></td><td></td><td></td><td></td><td></td><td></td><td></td><td></td><td></td><td></td></tr>
<tr><td>Chromosome</td><td>start</td><td>end</td><td>strand</td><td>MM</td><td>target_seq</td><td>PAM</td><td>alignment</td><td>distance</td><td>position</td><td>gene name</td><td>gene id</td></tr>
<tr><td>GU305914.1</td><td>136446</td><td>136468</td><td>+</td><td>0</td><td>CCGTCACCAGCCAGTCGCCG</td><td>GGG</td><td>||||||||[||||||||||||]PAM</td><td>NA</td><td>Intergenic</td><td>NA</td><td>NA</td></tr>
<tr><td>T10407</td><td>CACCAGCCAGTCGCCGGGGCTGG</td><td>1000</td><td></td><td></td><td></td><td></td><td></td><td></td><td></td><td></td><td></td></tr>
<tr><td>Oligo fwd</td><td>CACCCACCAGCCAGTCGCCGGGGC</td><td></td><td></td><td></td><td></td><td></td><td></td><td></td><td></td><td></td><td></td></tr>
<tr><td>Oligo rev</td><td>AAACGCCCCGGCGACTGGCTGGTG</td><td></td><td></td><td></td><td></td><td></td><td></td><td></td><td></td><td></td><td></td></tr>
<tr><td>Chromosome</td><td>start</td><td>end</td><td>strand</td><td>MM</td><td>target_seq</td><td>PAM</td><td>alignment</td><td>distance</td><td>position</td><td>gene name</td><td>gene id</td></tr>
<tr><td>GU305914.1</td><td>136450</td><td>136472</td><td>+</td><td>0</td><td>CACCAGCCAGTCGCCGGGGC</td><td>TGG</td><td>||||||||[||||||||||||]PAM</td><td>NA</td><td>Intergenic</td><td>NA</td><td>NA</td></tr>
<tr><td>T10408</td><td>ACCAGCCAGTCGCCGGGGCTGGG</td><td>1000</td><td></td><td></td><td></td><td></td><td></td><td></td><td></td><td></td><td></td></tr>
<tr><td>Oligo fwd</td><td>CACCACCAGCCAGTCGCCGGGGCT</td><td></td><td></td><td></td><td></td><td></td><td></td><td></td><td></td><td></td><td></td></tr>
<tr><td>Oligo rev</td><td>AAACAGCCCCGGCGACTGGCTGGT</td><td></td><td></td><td></td><td></td><td></td><td></td><td></td><td></td><td></td><td></td></tr>
<tr><td>Chromosome</td><td>start</td><td>end</td><td>strand</td><td>MM</td><td>target_seq</td><td>PAM</td><td>alignment</td><td>distance</td><td>position</td><td>gene name</td><td>gene id</td></tr>
<tr><td>GU305914.1</td><td>136451</td><td>136473</td><td>+</td><td>0</td><td>ACCAGCCAGTCGCCGGGGCT</td><td>GGG</td><td>||||||||[||||||||||||]PAM</td><td>NA</td><td>Intergenic</td><td>NA</td><td>NA</td></tr>
<tr><td>T10409</td><td>CCAGCCAGTCGCCGGGGCTGGGG</td><td>1000</td><td></td><td></td><td></td><td></td><td></td><td></td><td></td><td></td><td></td></tr>
<tr><td>Oligo fwd</td><td>CACCCCAGCCAGTCGCCGGGGCTG</td><td></td><td></td><td></td><td></td><td></td><td></td><td></td><td></td><td></td><td></td></tr>
<tr><td>Oligo rev</td><td>AAACCAGCCCCGGCGACTGGCTGG</td><td></td><td></td><td></td><td></td><td></td><td></td><td></td><td></td><td></td><td></td></tr>
<tr><td>Chromosome</td><td>start</td><td>end</td><td>strand</td><td>MM</td><td>target_seq</td><td>PAM</td><td>alignment</td><td>distance</td><td>position</td><td>gene name</td><td>gene id</td></tr>
<tr><td>GU305914.1</td><td>136452</td><td>136474</td><td>+</td><td>0</td><td>CCAGCCAGTCGCCGGGGCTG</td><td>GGG</td><td>||||||||[||||||||||||]PAM</td><td>NA</td><td>Intergenic</td><td>NA</td><td>NA</td></tr>
<tr><td>T10410</td><td>CCAGTCGCCGGGGCTGGGGAAGG</td><td>1000</td><td></td><td></td><td></td><td></td><td></td><td></td><td></td><td></td><td></td></tr>
<tr><td>Oligo fwd</td><td>CACCCCAGTCGCCGGGGCTGGGGA</td><td></td><td></td><td></td><td></td><td></td><td></td><td></td><td></td><td></td><td></td></tr>
<tr><td>Oligo rev</td><td>AAACTCCCCAGCCCCGGCGACTGG</td><td></td><td></td><td></td><td></td><td></td><td></td><td></td><td></td><td></td><td></td></tr>
<tr><td>Chromosome</td><td>start</td><td>end</td><td>strand</td><td>MM</td><td>target_seq</td><td>PAM</td><td>alignment</td><td>distance</td><td>position</td><td>gene name</td><td>gene id</td></tr>
<tr><td>GU305914.1</td><td>136456</td><td>136478</td><td>+</td><td>0</td><td>CCAGTCGCCGGGGCTGGGGA</td><td>AGG</td><td>||||||||[||||||||||||]PAM</td><td>NA</td><td>Intergenic</td><td>NA</td><td>NA</td></tr>
<tr><td>T10411</td><td>AAGGTCTCCTCCACCGACGACGG</td><td>1000</td><td></td><td></td><td></td><td></td><td></td><td></td><td></td><td></td><td></td></tr>
<tr><td>Oligo fwd</td><td>CACCAAGGTCTCCTCCACCGACGA</td><td></td><td></td><td></td><td></td><td></td><td></td><td></td><td></td><td></td><td></td></tr>
<tr><td>Oligo rev</td><td>AAACTCGTCGGTGGAGGAGACCTT</td><td></td><td></td><td></td><td></td><td></td><td></td><td></td><td></td><td></td><td></td></tr>
<tr><td>Chromosome</td><td>start</td><td>end</td><td>strand</td><td>MM</td><td>target_seq</td><td>PAM</td><td>alignment</td><td>distance</td><td>position</td><td>gene name</td><td>gene id</td></tr>
<tr><td>GU305914.1</td><td>136475</td><td>136497</td><td>+</td><td>0</td><td>AAGGTCTCCTCCACCGACGA</td><td>CGG</td><td>||||||||[||||||||||||]PAM</td><td>NA</td><td>Intergenic</td><td>NA</td><td>NA</td></tr>
<tr><td>T10412</td><td>AGGTCTCCTCCACCGACGACGGG</td><td>1000</td><td></td><td></td><td></td><td></td><td></td><td></td><td></td><td></td><td></td></tr>
<tr><td>Oligo fwd</td><td>CACCAGGTCTCCTCCACCGACGAC</td><td></td><td></td><td></td><td></td><td></td><td></td><td></td><td></td><td></td><td></td></tr>
<tr><td>Oligo rev</td><td>AAACGTCGTCGGTGGAGGAGACCT</td><td></td><td></td><td></td><td></td><td></td><td></td><td></td><td></td><td></td><td></td></tr>
<tr><td>Chromosome</td><td>start</td><td>end</td><td>strand</td><td>MM</td><td>target_seq</td><td>PAM</td><td>alignment</td><td>distance</td><td>position</td><td>gene name</td><td>gene id</td></tr>
<tr><td>GU305914.1</td><td>136476</td><td>136498</td><td>+</td><td>0</td><td>AGGTCTCCTCCACCGACGAC</td><td>GGG</td><td>||||||||[||||||||||||]PAM</td><td>NA</td><td>Intergenic</td><td>NA</td><td>NA</td></tr>
<tr><td>T10413</td><td>ATGATGCTTTTCAGACGTGACGG</td><td>1000</td><td></td><td></td><td></td><td></td><td></td><td></td><td></td><td></td><td></td></tr>
<tr><td>Oligo fwd</td><td>CACCATGATGCTTTTCAGACGTGA</td><td></td><td></td><td></td><td></td><td></td><td></td><td></td><td></td><td></td><td></td></tr>
<tr><td>Oligo rev</td><td>AAACTCACGTCTGAAAAGCATCAT</td><td></td><td></td><td></td><td></td><td></td><td></td><td></td><td></td><td></td><td></td></tr>
<tr><td>Chromosome</td><td>start</td><td>end</td><td>strand</td><td>MM</td><td>target_seq</td><td>PAM</td><td>alignment</td><td>distance</td><td>position</td><td>gene name</td><td>gene id</td></tr>
<tr><td>GU305914.1</td><td>136517</td><td>136539</td><td>+</td><td>0</td><td>ATGATGCTTTTCAGACGTGA</td><td>CGG</td><td>||||||||[||||||||||||]PAM</td><td>NA</td><td>Intergenic</td><td>NA</td><td>NA</td></tr>
<tr><td>T10414</td><td>CGGTAATTTCGTCTTCTGTCTGG</td><td>1000</td><td></td><td></td><td></td><td></td><td></td><td></td><td></td><td></td><td></td></tr>
<tr><td>Oligo fwd</td><td>CACCCGGTAATTTCGTCTTCTGTC</td><td></td><td></td><td></td><td></td><td></td><td></td><td></td><td></td><td></td><td></td></tr>
<tr><td>Oligo rev</td><td>AAACGACAGAAGACGAAATTACCG</td><td></td><td></td><td></td><td></td><td></td><td></td><td></td><td></td><td></td><td></td></tr>
<tr><td>Chromosome</td><td>start</td><td>end</td><td>strand</td><td>MM</td><td>target_seq</td><td>PAM</td><td>alignment</td><td>distance</td><td>position</td><td>gene name</td><td>gene id</td></tr>
<tr><td>GU305914.1</td><td>136537</td><td>136559</td><td>+</td><td>0</td><td>CGGTAATTTCGTCTTCTGTC</td><td>TGG</td><td>||||||||[||||||||||||]PAM</td><td>NA</td><td>Intergenic</td><td>NA</td><td>NA</td></tr>
<tr><td>GU305914.1</td><td>14921</td><td>14943</td><td>-</td><td>4</td><td>GTATAATTTCGTCTTCTGTG</td><td>TGG</td><td>---|||||[|||||||||||-]PAM</td><td>NA</td><td>Intergenic</td><td>NA</td><td>NA</td></tr>
<tr><td>T10415</td><td>CTTCTGTCTGGTCGTCAATAAGG</td><td>1000</td><td></td><td></td><td></td><td></td><td></td><td></td><td></td><td></td><td></td></tr>
<tr><td>Oligo fwd</td><td>CACCCTTCTGTCTGGTCGTCAATA</td><td></td><td></td><td></td><td></td><td></td><td></td><td></td><td></td><td></td><td></td></tr>
<tr><td>Oligo rev</td><td>AAACTATTGACGACCAGACAGAAG</td><td></td><td></td><td></td><td></td><td></td><td></td><td></td><td></td><td></td><td></td></tr>
<tr><td>Chromosome</td><td>start</td><td>end</td><td>strand</td><td>MM</td><td>target_seq</td><td>PAM</td><td>alignment</td><td>distance</td><td>position</td><td>gene name</td><td>gene id</td></tr>
<tr><td>GU305914.1</td><td>136549</td><td>136571</td><td>+</td><td>0</td><td>CTTCTGTCTGGTCGTCAATA</td><td>AGG</td><td>||||||||[||||||||||||]PAM</td><td>NA</td><td>Intergenic</td><td>NA</td><td>NA</td></tr>
<tr><td>T10416</td><td>TCTGGTCGTCAATAAGGAGCCGG</td><td>1000</td><td></td><td></td><td></td><td></td><td></td><td></td><td></td><td></td><td></td></tr>
<tr><td>Oligo fwd</td><td>CACCTCTGGTCGTCAATAAGGAGC</td><td></td><td></td><td></td><td></td><td></td><td></td><td></td><td></td><td></td><td></td></tr>
<tr><td>Oligo rev</td><td>AAACGCTCCTTATTGACGACCAGA</td><td></td><td></td><td></td><td></td><td></td><td></td><td></td><td></td><td></td><td></td></tr>
<tr><td>Chromosome</td><td>start</td><td>end</td><td>strand</td><td>MM</td><td>target_seq</td><td>PAM</td><td>alignment</td><td>distance</td><td>position</td><td>gene name</td><td>gene id</td></tr>
<tr><td>GU305914.1</td><td>136555</td><td>136577</td><td>+</td><td>0</td><td>TCTGGTCGTCAATAAGGAGC</td><td>CGG</td><td>||||||||[||||||||||||]PAM</td><td>NA</td><td>Intergenic</td><td>NA</td><td>NA</td></tr>
<tr><td>T10417</td><td>GGTCGTCAATAAGGAGCCGGTGG</td><td>1000</td><td></td><td></td><td></td><td></td><td></td><td></td><td></td><td></td><td></td></tr>
<tr><td>Oligo fwd</td><td>CACCGGTCGTCAATAAGGAGCCGG</td><td></td><td></td><td></td><td></td><td></td><td></td><td></td><td></td><td></td><td></td></tr>
<tr><td>Oligo rev</td><td>AAACCCGGCTCCTTATTGACGACC</td><td></td><td></td><td></td><td></td><td></td><td></td><td></td><td></td><td></td><td></td></tr>
<tr><td>Chromosome</td><td>start</td><td>end</td><td>strand</td><td>MM</td><td>target_seq</td><td>PAM</td><td>alignment</td><td>distance</td><td>position</td><td>gene name</td><td>gene id</td></tr>
<tr><td>GU305914.1</td><td>136558</td><td>136580</td><td>+</td><td>0</td><td>GGTCGTCAATAAGGAGCCGG</td><td>TGG</td><td>||||||||[||||||||||||]PAM</td><td>NA</td><td>Intergenic</td><td>NA</td><td>NA</td></tr>
<tr><td>T10418</td><td>GTCGTCAATAAGGAGCCGGTGGG</td><td>1000</td><td></td><td></td><td></td><td></td><td></td><td></td><td></td><td></td><td></td></tr>
<tr><td>Oligo fwd</td><td>CACCGTCGTCAATAAGGAGCCGGT</td><td></td><td></td><td></td><td></td><td></td><td></td><td></td><td></td><td></td><td></td></tr>
<tr><td>Oligo rev</td><td>AAACACCGGCTCCTTATTGACGAC</td><td></td><td></td><td></td><td></td><td></td><td></td><td></td><td></td><td></td><td></td></tr>
<tr><td>Chromosome</td><td>start</td><td>end</td><td>strand</td><td>MM</td><td>target_seq</td><td>PAM</td><td>alignment</td><td>distance</td><td>position</td><td>gene name</td><td>gene id</td></tr>
<tr><td>GU305914.1</td><td>136559</td><td>136581</td><td>+</td><td>0</td><td>GTCGTCAATAAGGAGCCGGT</td><td>GGG</td><td>||||||||[||||||||||||]PAM</td><td>NA</td><td>Intergenic</td><td>NA</td><td>NA</td></tr>
<tr><td>T10419</td><td>AAGGAGCCGGTGGGTCAGTTCGG</td><td>1000</td><td></td><td></td><td></td><td></td><td></td><td></td><td></td><td></td><td></td></tr>
<tr><td>Oligo fwd</td><td>CACCAAGGAGCCGGTGGGTCAGTT</td><td></td><td></td><td></td><td></td><td></td><td></td><td></td><td></td><td></td><td></td></tr>
<tr><td>Oligo rev</td><td>AAACAACTGACCCACCGGCTCCTT</td><td></td><td></td><td></td><td></td><td></td><td></td><td></td><td></td><td></td><td></td></tr>
<tr><td>Chromosome</td><td>start</td><td>end</td><td>strand</td><td>MM</td><td>target_seq</td><td>PAM</td><td>alignment</td><td>distance</td><td>position</td><td>gene name</td><td>gene id</td></tr>
<tr><td>GU305914.1</td><td>136568</td><td>136590</td><td>+</td><td>0</td><td>AAGGAGCCGGTGGGTCAGTT</td><td>CGG</td><td>||||||||[||||||||||||]PAM</td><td>NA</td><td>Intergenic</td><td>NA</td><td>NA</td></tr>
<tr><td>T10420</td><td>GTGGGTCAGTTCGGCTGCAGTGG</td><td>1000</td><td></td><td></td><td></td><td></td><td></td><td></td><td></td><td></td><td></td></tr>
<tr><td>Oligo fwd</td><td>CACCGTGGGTCAGTTCGGCTGCAG</td><td></td><td></td><td></td><td></td><td></td><td></td><td></td><td></td><td></td><td></td></tr>
<tr><td>Oligo rev</td><td>AAACCTGCAGCCGAACTGACCCAC</td><td></td><td></td><td></td><td></td><td></td><td></td><td></td><td></td><td></td><td></td></tr>
<tr><td>Chromosome</td><td>start</td><td>end</td><td>strand</td><td>MM</td><td>target_seq</td><td>PAM</td><td>alignment</td><td>distance</td><td>position</td><td>gene name</td><td>gene id</td></tr>
</table>

| | | | | | | | | | | | |
|---|---|---|---|---|---|---|---|---|---|---|---|
| GU305914.1 | 136577 | 136599 | + | 0 | GTGGGTCAGTTCGGCTGCAG | TGG | \|\|\|\|\|\|\|\|[\|\|\|\|\|\|\|\|\|\|\|\|]PAM | NA | Intergenic | NA | NA |
| T10421 | AGTTCGGCTGCAGTGGCATGCGG | 1000 | | | | | | | | | |
| Oligo fwd | CACCAGTTCGGCTGCAGTGGCATG | | | | | | | | | | |
| Oligo rev | AAACCATGCCACTGCAGCCGAACT | | | | | | | | | | |
| Chromosome | start | end | strand | MM | target_seq | PAM | alignment | distance | position | gene name | gene id |
| GU305914.1 | 136584 | 136606 | + | 0 | AGTTCGGCTGCAGTGGCATG | CGG | \|\|\|\|\|\|\|\|[\|\|\|\|\|\|\|\|\|\|\|\|]PAM | NA | Intergenic | NA | NA |
| T10422 | TGGCATGCGGCGCGAGAAGATGG | 1000 | | | | | | | | | |
| Oligo fwd | CACCTGGCATGCGGCGCGAGAAGA | | | | | | | | | | |
| Oligo rev | AAACTCTTCTCGCGCCGCATGCCA | | | | | | | | | | |
| Chromosome | start | end | strand | MM | target_seq | PAM | alignment | distance | position | gene name | gene id |
| GU305914.1 | 136597 | 136619 | + | 0 | TGGCATGCGGCGCGAGAAGA | TGG | \|\|\|\|\|\|\|\|[\|\|\|\|\|\|\|\|\|\|\|\|]PAM | NA | Intergenic | NA | NA |
| T10423 | CGCGAGAAGATGGTCATCGATGG | 1000 | | | | | | | | | |
| Oligo fwd | CACCCGCGAGAAGATGGTCATCGA | | | | | | | | | | |
| Oligo rev | AAACTCGATGACCATCTTCTCGCG | | | | | | | | | | |
| Chromosome | start | end | strand | MM | target_seq | PAM | alignment | distance | position | gene name | gene id |
| GU305914.1 | 136607 | 136629 | + | 0 | CGCGAGAAGATGGTCATCGA | TGG | \|\|\|\|\|\|\|\|[\|\|\|\|\|\|\|\|\|\|\|\|]PAM | NA | Intergenic | NA | NA |
| T10424 | GATGGTCATCGATGGACTCCAGG | 1000 | | | | | | | | | |
| Oligo fwd | CACCGATGGTCATCGATGGACTCC | | | | | | | | | | |
| Oligo rev | AAACGGAGTCCATCGATGACCATC | | | | | | | | | | |
| Chromosome | start | end | strand | MM | target_seq | PAM | alignment | distance | position | gene name | gene id |
| GU305914.1 | 136615 | 136637 | + | 0 | GATGGTCATCGATGGACTCC | AGG | \|\|\|\|\|\|\|\|[\|\|\|\|\|\|\|\|\|\|\|\|]PAM | NA | Intergenic | NA | NA |
| T10425 | CGTCTACGTGATGCGTCTCCTGG | 1000 | | | | | | | | | |
| Oligo fwd | CACCCGTCTACGTGATGCGTCTCC | | | | | | | | | | |
| Oligo rev | AAACGGAGACGCATCACGTAGACG | | | | | | | | | | |
| Chromosome | start | end | strand | MM | target_seq | PAM | alignment | distance | position | gene name | gene id |
| GU305914.1 | 136642 | 136664 | + | 0 | CGTCTACGTGATGCGTCTCC | TGG | \|\|\|\|\|\|\|\|[\|\|\|\|\|\|\|\|\|\|\|\|]PAM | NA | Intergenic | NA | NA |
| T10426 | CCCATCATCCCCGTCAAGCTAGG | 1000 | | | | | | | | | |
| Oligo fwd | CACCCCCATCATCCCCGTCAAGCT | | | | | | | | | | |
| Oligo rev | AAACAGCTTGACGGGGATGATGGG | | | | | | | | | | |
| Chromosome | start | end | strand | MM | target_seq | PAM | alignment | distance | position | gene name | gene id |
| GU305914.1 | 136667 | 136689 | + | 0 | CCCATCATCCCCGTCAAGCT | AGG | \|\|\|\|\|\|\|\|[\|\|\|\|\|\|\|\|\|\|\|\|]PAM | NA | Intergenic | NA | NA |
| T10427 | ATGTTGCCGCCTAAGAGCATCGG | 1000 | | | | | | | | | |
| Oligo fwd | CACCATGTTGCCGCCTAAGAGCAT | | | | | | | | | | |
| Oligo rev | AAACATGCTCTTAGGCGGCAACAT | | | | | | | | | | |
| Chromosome | start | end | strand | MM | target_seq | PAM | alignment | distance | position | gene name | gene id |
| GU305914.1 | 136703 | 136725 | + | 0 | ATGTTGCCGCCTAAGAGCAT | CGG | \|\|\|\|\|\|\|\|[\|\|\|\|\|\|\|\|\|\|\|\|]PAM | NA | Intergenic | NA | NA |
| T10428 | TTGCCGCCTAAGAGCATCGGCGG | 1000 | | | | | | | | | |
| Oligo fwd | CACCTTGCCGCCTAAGAGCATCGG | | | | | | | | | | |
| Oligo rev | AAACCCGATGCTCTTAGGCGGCAA | | | | | | | | | | |
| Chromosome | start | end | strand | MM | target_seq | PAM | alignment | distance | position | gene name | gene id |
| GU305914.1 | 136706 | 136728 | + | 0 | TTGCCGCCTAAGAGCATCGG | CGG | \|\|\|\|\|\|\|\|[\|\|\|\|\|\|\|\|\|\|\|\|]PAM | NA | Intergenic | NA | NA |
| T10429 | AAGAGCATCGGCGGCTCCAGCGG | 1000 | | | | | | | | | |
| Oligo fwd | CACCAAGAGCATCGGCGGCTCCAG | | | | | | | | | | |
| Oligo rev | AAACCTGGAGCCGCCGATGCTCTT | | | | | | | | | | |
| Chromosome | start | end | strand | MM | target_seq | PAM | alignment | distance | position | gene name | gene id |
| GU305914.1 | 136715 | 136737 | + | 0 | AAGAGCATCGGCGGCTCCAG | CGG | \|\|\|\|\|\|\|\|[\|\|\|\|\|\|\|\|\|\|\|\|]PAM | NA | Intergenic | NA | NA |
| T10430 | CATCGGCGGCTCCAGCGGTCTGG | 1000 | | | | | | | | | |
| Oligo fwd | CACCCATCGGCGGCTCCAGCGGTC | | | | | | | | | | |
| Oligo rev | AAACGACCGCTGGAGCCGCCGATG | | | | | | | | | | |
| Chromosome | start | end | strand | MM | target_seq | PAM | alignment | distance | position | gene name | gene id |
| GU305914.1 | 136720 | 136742 | + | 0 | CATCGGCGGCTCCAGCGGTC | TGG | \|\|\|\|\|\|\|\|[\|\|\|\|\|\|\|\|\|\|\|\|]PAM | NA | Intergenic | NA | NA |
| T10431 | CATCTACCAGAACGCGAGTGTGG | 1000 | | | | | | | | | |
| Oligo fwd | CACCCATCTACCAGAACGCGAGTG | | | | | | | | | | |
| Oligo rev | AAACCACTCGCGTTCTGGTAGATG | | | | | | | | | | |
| Chromosome | start | end | strand | MM | target_seq | PAM | alignment | distance | position | gene name | gene id |
| GU305914.1 | 136753 | 136775 | + | 0 | CATCTACCAGAACGCGAGTG | TGG | \|\|\|\|\|\|\|\|[\|\|\|\|\|\|\|\|\|\|\|\|]PAM | NA | Intergenic | NA | NA |
| GU305914.1 | 2743 | 2765 | + | 3 | CATCTTTCAGAACGCGAGCG | TGG | \|\|\|\|\|--\|[\|\|\|\|\|\|\|\|\|\|-\|]PAM | NA | Intergenic | NA | NA |
| T10432 | GAGTGTGGTCACGCCCGAAGAGG | 1000 | | | | | | | | | |
| Oligo fwd | CACCGAGTGTGGTCACGCCCGAAG | | | | | | | | | | |
| Oligo rev | AAACCTTCGGGCGTGACCACACTC | | | | | | | | | | |
| Chromosome | start | end | strand | MM | target_seq | PAM | alignment | distance | position | gene name | gene id |
| GU305914.1 | 136768 | 136790 | + | 0 | GAGTGTGGTCACGCCCGAAG | AGG | \|\|\|\|\|\|\|\|[\|\|\|\|\|\|\|\|\|\|\|\|]PAM | NA | Intergenic | NA | NA |
| T10433 | GGCCGCCACCGTCACTATGCAGG | 1000 | | | | | | | | | |
| Oligo fwd | CACCGGCCGCCACCGTCACTATGC | | | | | | | | | | |
| Oligo rev | AAACGCATAGTGACGGTGGCGGCC | | | | | | | | | | |
| Chromosome | start | end | strand | MM | target_seq | PAM | alignment | distance | position | gene name | gene id |
| GU305914.1 | 136789 | 136811 | + | 0 | GGCCGCCACCGTCACTATGC | AGG | \|\|\|\|\|\|\|\|[\|\|\|\|\|\|\|\|\|\|\|\|]PAM | NA | Intergenic | NA | NA |
| T10434 | GCCGCCACCGTCACTATGCAGGG | 1000 | | | | | | | | | |
| Oligo fwd | CACCGCCGCCACCGTCACTATGCA | | | | | | | | | | |
| Oligo rev | AAACTGCATAGTGACGGTGGCGGC | | | | | | | | | | |
| Chromosome | start | end | strand | MM | target_seq | PAM | alignment | distance | position | gene name | gene id |
| GU305914.1 | 136790 | 136812 | + | 0 | GCCGCCACCGTCACTATGCA | GGG | \|\|\|\|\|\|\|\|[\|\|\|\|\|\|\|\|\|\|\|\|]PAM | NA | Intergenic | NA | NA |
| T10435 | ACCGTCACTATGCAGGGTTCCGG | 1000 | | | | | | | | | |
| Oligo fwd | CACCACCGTCACTATGCAGGGTTC | | | | | | | | | | |

<table>
<tr><td>Oligo rev</td><td>AAACGAACCCTGCATAGTGACGGT</td></tr>
<tr><td>Chromosome</td><td>start</td><td>end</td><td>strand</td><td>MM</td><td>target_seq</td><td>PAM</td><td>alignment</td><td>distance</td><td>position</td><td>gene name</td><td>gene id</td></tr>
<tr><td>GU305914.1</td><td>136796</td><td>136818</td><td>+</td><td>0</td><td>ACCGTCACTATGCAGGGTTC</td><td>CGG</td><td>||||||||[||||||||||||]PAM</td><td>NA</td><td>Intergenic</td><td>NA</td><td>NA</td></tr>
<tr><td>T10436</td><td>GGGTTCCGGCATCGTGACCGTGG</td><td>1000</td></tr>
<tr><td>Oligo fwd</td><td>CACCGGGTTCCGGCATCGTGACCG</td></tr>
<tr><td>Oligo rev</td><td>AAACCGGTCACGATGCCGGAACCC</td></tr>
<tr><td>Chromosome</td><td>start</td><td>end</td><td>strand</td><td>MM</td><td>target_seq</td><td>PAM</td><td>alignment</td><td>distance</td><td>position</td><td>gene name</td><td>gene id</td></tr>
<tr><td>GU305914.1</td><td>136810</td><td>136832</td><td>+</td><td>0</td><td>GGGTTCCGGCATCGTGACCG</td><td>TGG</td><td>||||||||[||||||||||||]PAM</td><td>NA</td><td>Intergenic</td><td>NA</td><td>NA</td></tr>
<tr><td>GU305914.1</td><td>195187</td><td>195209</td><td>+</td><td>4</td><td>GGGCTTCGGCATCGAGACCA</td><td>CGG</td><td>|||-|-||[||||||-||||-]PAM</td><td>NA</td><td>Intergenic</td><td>NA</td><td>NA</td></tr>
<tr><td>T10437</td><td>GGTTCCGGCATCGTGACCGTGGG</td><td>1000</td></tr>
<tr><td>Oligo fwd</td><td>CACCGGTTCCGGCATCGTGACCGT</td></tr>
<tr><td>Oligo rev</td><td>AAACACGGTCACGATGCCGGAACC</td></tr>
<tr><td>Chromosome</td><td>start</td><td>end</td><td>strand</td><td>MM</td><td>target_seq</td><td>PAM</td><td>alignment</td><td>distance</td><td>position</td><td>gene name</td><td>gene id</td></tr>
<tr><td>GU305914.1</td><td>136811</td><td>136833</td><td>+</td><td>0</td><td>GGTTCCGGCATCGTGACCGT</td><td>GGG</td><td>||||||||[||||||||||||]PAM</td><td>NA</td><td>Intergenic</td><td>NA</td><td>NA</td></tr>
<tr><td>T10438</td><td>ATCGTGACCGTGGGACTCAGTGG</td><td>1000</td></tr>
<tr><td>Oligo fwd</td><td>CACCATCGTGACCGTGGGACTCAG</td></tr>
<tr><td>Oligo rev</td><td>AAACCTGAGTCCCACGGTCACGAT</td></tr>
<tr><td>Chromosome</td><td>start</td><td>end</td><td>strand</td><td>MM</td><td>target_seq</td><td>PAM</td><td>alignment</td><td>distance</td><td>position</td><td>gene name</td><td>gene id</td></tr>
<tr><td>GU305914.1</td><td>136820</td><td>136842</td><td>+</td><td>0</td><td>ATCGTGACCGTGGGACTCAG</td><td>TGG</td><td>||||||||[||||||||||||]PAM</td><td>NA</td><td>Intergenic</td><td>NA</td><td>NA</td></tr>
<tr><td>T10439</td><td>ACCGTGGGACTCAGTGGCGTTGG</td><td>1000</td></tr>
<tr><td>Oligo fwd</td><td>CACCACCGTGGGACTCAGTGGCGT</td></tr>
<tr><td>Oligo rev</td><td>AAACACGCCACTGAGTCCCACGGT</td></tr>
<tr><td>Chromosome</td><td>start</td><td>end</td><td>strand</td><td>MM</td><td>target_seq</td><td>PAM</td><td>alignment</td><td>distance</td><td>position</td><td>gene name</td><td>gene id</td></tr>
<tr><td>GU305914.1</td><td>136826</td><td>136848</td><td>+</td><td>0</td><td>ACCGTGGGACTCAGTGGCGT</td><td>TGG</td><td>||||||||[||||||||||||]PAM</td><td>NA</td><td>Intergenic</td><td>NA</td><td>NA</td></tr>
<tr><td>T10440</td><td>GACTCAGTGGCGTTGGCTCCTGG</td><td>1000</td></tr>
<tr><td>Oligo fwd</td><td>CACCGACTCAGTGGCGTTGGCTCC</td></tr>
<tr><td>Oligo rev</td><td>AAACGGAGCCAACGCCACTGAGTC</td></tr>
<tr><td>Chromosome</td><td>start</td><td>end</td><td>strand</td><td>MM</td><td>target_seq</td><td>PAM</td><td>alignment</td><td>distance</td><td>position</td><td>gene name</td><td>gene id</td></tr>
<tr><td>GU305914.1</td><td>136833</td><td>136855</td><td>+</td><td>0</td><td>GACTCAGTGGCGTTGGCTCC</td><td>TGG</td><td>||||||||[||||||||||||]PAM</td><td>NA</td><td>Intergenic</td><td>NA</td><td>NA</td></tr>
<tr><td>T10441</td><td>ACTCAGTGGCGTTGGCTCCTGGG</td><td>1000</td></tr>
<tr><td>Oligo fwd</td><td>CACCACTCAGTGGCGTTGGCTCCT</td></tr>
<tr><td>Oligo rev</td><td>AAACAGGAGCCAACGCCACTGAGT</td></tr>
<tr><td>Chromosome</td><td>start</td><td>end</td><td>strand</td><td>MM</td><td>target_seq</td><td>PAM</td><td>alignment</td><td>distance</td><td>position</td><td>gene name</td><td>gene id</td></tr>
<tr><td>GU305914.1</td><td>136834</td><td>136856</td><td>+</td><td>0</td><td>ACTCAGTGGCGTTGGCTCCT</td><td>GGG</td><td>||||||||[||||||||||||]PAM</td><td>NA</td><td>Intergenic</td><td>NA</td><td>NA</td></tr>
<tr><td>T10442</td><td>TGGCTCCTGGGTGCAGATCAAGG</td><td>1000</td></tr>
<tr><td>Oligo fwd</td><td>CACCTGGCTCCTGGGTGCAGATCA</td></tr>
<tr><td>Oligo rev</td><td>AAACTGATCTGCACCCAGGAGCCA</td></tr>
<tr><td>Chromosome</td><td>start</td><td>end</td><td>strand</td><td>MM</td><td>target_seq</td><td>PAM</td><td>alignment</td><td>distance</td><td>position</td><td>gene name</td><td>gene id</td></tr>
<tr><td>GU305914.1</td><td>136846</td><td>136868</td><td>+</td><td>0</td><td>TGGCTCCTGGGTGCAGATCA</td><td>AGG</td><td>||||||||[||||||||||||]PAM</td><td>NA</td><td>Intergenic</td><td>NA</td><td>NA</td></tr>
<tr><td>T10443</td><td>TCCTGGGTGCAGATCAAGGATGG</td><td>1000</td></tr>
<tr><td>Oligo fwd</td><td>CACCTCCTGGGTGCAGATCAAGGA</td></tr>
<tr><td>Oligo rev</td><td>AAACTCCTTGATCTGCACCCAGGA</td></tr>
<tr><td>Chromosome</td><td>start</td><td>end</td><td>strand</td><td>MM</td><td>target_seq</td><td>PAM</td><td>alignment</td><td>distance</td><td>position</td><td>gene name</td><td>gene id</td></tr>
<tr><td>GU305914.1</td><td>136850</td><td>136872</td><td>+</td><td>0</td><td>TCCTGGGTGCAGATCAAGGA</td><td>TGG</td><td>||||||||[||||||||||||]PAM</td><td>NA</td><td>Intergenic</td><td>NA</td><td>NA</td></tr>
<tr><td>T10444</td><td>TGGGTGCAGATCAAGGATGGTGG</td><td>1000</td></tr>
<tr><td>Oligo fwd</td><td>CACCTGGGTGCAGATCAAGGATGG</td></tr>
<tr><td>Oligo rev</td><td>AAACCCATCCTTGATCTGCACCCA</td></tr>
<tr><td>Chromosome</td><td>start</td><td>end</td><td>strand</td><td>MM</td><td>target_seq</td><td>PAM</td><td>alignment</td><td>distance</td><td>position</td><td>gene name</td><td>gene id</td></tr>
<tr><td>GU305914.1</td><td>136853</td><td>136875</td><td>+</td><td>0</td><td>TGGGTGCAGATCAAGGATGG</td><td>TGG</td><td>||||||||[||||||||||||]PAM</td><td>NA</td><td>Intergenic</td><td>NA</td><td>NA</td></tr>
<tr><td>T10445</td><td>GGGTGCAGATCAAGGATGGTGGG</td><td>1000</td></tr>
<tr><td>Oligo fwd</td><td>CACCGGGTGCAGATCAAGGATGGT</td></tr>
<tr><td>Oligo rev</td><td>AAACACCATCCTTGATCTGCACCC</td></tr>
<tr><td>Chromosome</td><td>start</td><td>end</td><td>strand</td><td>MM</td><td>target_seq</td><td>PAM</td><td>alignment</td><td>distance</td><td>position</td><td>gene name</td><td>gene id</td></tr>
<tr><td>GU305914.1</td><td>136854</td><td>136876</td><td>+</td><td>0</td><td>GGGTGCAGATCAAGGATGGT</td><td>GGG</td><td>||||||||[||||||||||||]PAM</td><td>NA</td><td>Intergenic</td><td>NA</td><td>NA</td></tr>
<tr><td>T10446</td><td>AACATGAAGCTCTTCGTCTTCGG</td><td>1000</td></tr>
<tr><td>Oligo fwd</td><td>CACCAACATGAAGCTCTTCGTCTT</td></tr>
<tr><td>Oligo rev</td><td>AAACAAGACGAAGAGCTTCATGTT</td></tr>
<tr><td>Chromosome</td><td>start</td><td>end</td><td>strand</td><td>MM</td><td>target_seq</td><td>PAM</td><td>alignment</td><td>distance</td><td>position</td><td>gene name</td><td>gene id</td></tr>
<tr><td>GU305914.1</td><td>136877</td><td>136899</td><td>+</td><td>0</td><td>AACATGAAGCTCTTCGTCTT</td><td>CGG</td><td>||||||||[||||||||||||]PAM</td><td>NA</td><td>Intergenic</td><td>NA</td><td>NA</td></tr>
<tr><td>GU305914.1</td><td>83851</td><td>83873</td><td>-</td><td>4</td><td>AGCAGGAAGATCTTCGTCGT</td><td>GGG</td><td>|-||-|||[|-||||||||-|]PAM</td><td>NA</td><td>Intergenic</td><td>NA</td><td>NA</td></tr>
<tr><td>T10447</td><td>TCTTTACCGCCTGCTGCGATCGG</td><td>1000</td></tr>
<tr><td>Oligo fwd</td><td>CACCTCTTTACCGCCTGCTGCGAT</td></tr>
<tr><td>Oligo rev</td><td>AAACATCGCAGCAGGCGGTAAAGA</td></tr>
<tr><td>Chromosome</td><td>start</td><td>end</td><td>strand</td><td>MM</td><td>target_seq</td><td>PAM</td><td>alignment</td><td>distance</td><td>position</td><td>gene name</td><td>gene id</td></tr>
<tr><td>GU305914.1</td><td>136914</td><td>136936</td><td>+</td><td>0</td><td>TCTTTACCGCCTGCTGCGAT</td><td>CGG</td><td>||||||||[||||||||||||]PAM</td><td>NA</td><td>Intergenic</td><td>NA</td><td>NA</td></tr>
<tr><td>T10448</td><td>TCGGCTCGCCTTCCCGTCCCTGG</td><td>1000</td></tr>
<tr><td>Oligo fwd</td><td>CACCTCGGCTCGCCTTCCCGTCCC</td></tr>
<tr><td>Oligo rev</td><td>AAACGGGACGGGAAGGCGAGCCGA</td></tr>
<tr><td>Chromosome</td><td>start</td><td>end</td><td>strand</td><td>MM</td><td>target_seq</td><td>PAM</td><td>alignment</td><td>distance</td><td>position</td><td>gene name</td><td>gene id</td></tr>
<tr><td>GU305914.1</td><td>136933</td><td>136955</td><td>+</td><td>0</td><td>TCGGCTCGCCTTCCCGTCCC</td><td>TGG</td><td>||||||||[||||||||||||]PAM</td><td>NA</td><td>Intergenic</td><td>NA</td><td>NA</td></tr>
<tr><td>T10449</td><td>CAGCGAAACTGTGTCCTGCGAGG</td><td>1000</td></tr>
<tr><td>Oligo fwd</td><td>CACCCAGCGAAACTGTGTCCTGCG</td></tr>
<tr><td>Oligo rev</td><td>AAACCGCAGGACACAGTTTCGCTG</td></tr>
<tr><td>Chromosome</td><td>start</td><td>end</td><td>strand</td><td>MM</td><td>target_seq</td><td>PAM</td><td>alignment</td><td>distance</td><td>position</td><td>gene name</td><td>gene id</td></tr>
<tr><td>GU305914.1</td><td>136966</td><td>136988</td><td>+</td><td>0</td><td>CAGCGAAACTGTGTCCTGCG</td><td>AGG</td><td>||||||||[||||||||||||]PAM</td><td>NA</td><td>Intergenic</td><td>NA</td><td>NA</td></tr>
</table>

T10450 TCCTGCGAGGCCGACAAGTGCGG 1000
Oligo fwd CACCTCCTGCGAGGCCGACAAGTG
Oligo rev AAACCACTTGTCGGCCTCGCAGGA
Chromosome start end strand MM target_seq PAM alignment distance position gene name gene id
GU305914.1 136979 137001 + 0 TCCTGCGAGGCCGACAAGTG CGG ||||||||[||||||||||||]PAM NA Intergenic NA NA

T10451 TGCGGATTCTGTCGAGATTCCGG 1000
Oligo fwd CACCTGCGGATTCTGTCGAGATTC
Oligo rev AAACGAATCTCGACAGAATCCGCA
Chromosome start end strand MM target_seq PAM alignment distance position gene name gene id
GU305914.1 136997 137019 + 0 TGCGGATTCTGTCGAGATTC CGG ||||||||[||||||||||||]PAM NA Intergenic NA NA

T10452 GATTCTGTCGAGATTCCGGCAGG 1000
Oligo fwd CACCGATTCTGTCGAGATTCCGGC
Oligo rev AAACGCCGGAATCTCGACAGAATC
Chromosome start end strand MM target_seq PAM alignment distance position gene name gene id
GU305914.1 137001 137023 + 0 GATTCTGTCGAGATTCCGGC AGG ||||||||[||||||||||||]PAM NA Intergenic NA NA

T10453 GGCAGGCACGTCGACCCCACCGG 1000
Oligo fwd CACCGGCAGGCACGTCGACCCCAC
Oligo rev AAACGTGGGGTCGACGTGCCTGCC
Chromosome start end strand MM target_seq PAM alignment distance position gene name gene id
GU305914.1 137018 137040 + 0 GGCAGGCACGTCGACCCCAC CGG ||||||||[||||||||||||]PAM NA Intergenic NA NA

T10454 GACCCCACCGGCCGCTTCGTCGG 1000
Oligo fwd CACCGACCCCACCGGCCGCTTCGT
Oligo rev AAACACGAAGCGGCCGGTGGGGTC
Chromosome start end strand MM target_seq PAM alignment distance position gene name gene id
GU305914.1 137030 137052 + 0 GACCCCACCGGCCGCTTCGT CGG ||||||||[||||||||||||]PAM NA Intergenic NA NA

T10455 GTCGGCTGCGTCCCCGACAGTGG 1000
Oligo fwd CACCGTCGGCTGCGTCCCCGACAG
Oligo rev AAACCTGTCGGGGACGCAGCCGAC
Chromosome start end strand MM target_seq PAM alignment distance position gene name gene id
GU305914.1 137048 137070 + 0 GTCGGCTGCGTCCCCGACAG TGG ||||||||[||||||||||||]PAM NA Intergenic NA NA

T10456 CTCTGTTACTCGCCGTGTCGCGG 1000
Oligo fwd CACCCTCTGTTACTCGCCGTGTCG
Oligo rev AAACCGACACGGCGAGTAACAGAG
Chromosome start end strand MM target_seq PAM alignment distance position gene name gene id
GU305914.1 137078 137100 + 0 CTCTGTTACTCGCCGTGTCG CGG ||||||||[||||||||||||]PAM NA Intergenic NA NA

T10457 TCTGTTACTCGCCGTGTCGCGGG 1000
Oligo fwd CACCTCTGTTACTCGCCGTGTCGC
Oligo rev AAACGCGACACGGCGAGTAACAGA
Chromosome start end strand MM target_seq PAM alignment distance position gene name gene id
GU305914.1 137079 137101 + 0 TCTGTTACTCGCCGTGTCGC GGG ||||||||[||||||||||||]PAM NA Intergenic NA NA

T10458 TTACTCGCCGTGTCGCGGGACGG 1000
Oligo fwd CACCTTACTCGCCGTGTCGCGGGA
Oligo rev AAACTCCCGCGACACGGCGAGTAA
Chromosome start end strand MM target_seq PAM alignment distance position gene name gene id
GU305914.1 137083 137105 + 0 TTACTCGCCGTGTCGCGGGA CGG ||||||||[||||||||||||]PAM NA Intergenic NA NA

T10459 GTGTCGCGGGACGGATGCCGTGG 1000
Oligo fwd CACCGTGTCGCGGGACGGATGCCG
Oligo rev AAACCGGCATCCGTCCCGCGACAC
Chromosome start end strand MM target_seq PAM alignment distance position gene name gene id
GU305914.1 137092 137114 + 0 GTGTCGCGGGACGGATGCCG TGG ||||||||[||||||||||||]PAM NA Intergenic NA NA

T10460 CCGTGGTCAGCGTCAGAAGCTGG 1000
Oligo fwd CACCCCGTGGTCAGCGTCAGAAGC
Oligo rev AAACGCTTCTGACGCTGACCACGG
Chromosome start end strand MM target_seq PAM alignment distance position gene name gene id
GU305914.1 137109 137131 + 0 CCGTGGTCAGCGTCAGAAGC TGG ||||||||[||||||||||||]PAM NA Intergenic NA NA

T10461 CAGAAGCTGGTTACCTTACCTGG 1000
Oligo fwd CACCCAGAAGCTGGTTACCTTACC
Oligo rev AAACGGTAAGGTAACCAGCTTCTG
Chromosome start end strand MM target_seq PAM alignment distance position gene name gene id
GU305914.1 137122 137144 + 0 CAGAAGCTGGTTACCTTACC TGG ||||||||[||||||||||||]PAM NA Intergenic NA NA

T10462 CTGGTTACCTTACCTGGAACTGG 1000
Oligo fwd CACCCTGGTTACCTTACCTGGAAC
Oligo rev AAACGTTCCAGGTAAGGTAACCAG
Chromosome start end strand MM target_seq PAM alignment distance position gene name gene id
GU305914.1 137128 137150 + 0 CTGGTTACCTTACCTGGAAC TGG ||||||||[||||||||||||]PAM NA Intergenic NA NA

T10463 CCTTACCTGGAACTGGAAGACGG 1000
Oligo fwd CACCCCTTACCTGGAACTGGAAGA
Oligo rev AAACTCTTCCAGTTCCAGGTAAGG
Chromosome start end strand MM target_seq PAM alignment distance position gene name gene id
GU305914.1 137135 137157 + 0 CCTTACCTGGAACTGGAAGA CGG ||||||||[||||||||||||]PAM NA Intergenic NA NA

T10464 ACACGCACAGCCTCTTCGTGCGG 1000
Oligo fwd CACCACACGCACAGCCTCTTCGTG
Oligo rev AAACCACGAAGAGGCTGTGCGTGT
Chromosome start end strand MM target_seq PAM alignment distance position gene name gene id

| | | | | | | | | | | | |
|---|---|---|---|---|---|---|---|---|---|---|---|
| GU305914.1 | 137163 | 137185 | + | 0 | ACACGCACAGCCTCTTCGTG | CGG | \|\|\|\|\|\|\|\|[\|\|\|\|\|\|\|\|\|\|\|\|]PAM | NA | Intergenic | NA | NA |
| T10465 | CTCTTCGTGCGGCGCTACGACGG | 1000 | | | | | | | | | |
| Oligo fwd | CACCCTCTTCGTGCGGCGCTACGA | | | | | | | | | | |
| Oligo rev | AAACTCGTAGCGCCGCACGAAGAG | | | | | | | | | | |
| Chromosome | start | end | strand | MM | target_seq | PAM | alignment | distance | position | gene name | gene id |
| GU305914.1 | 137174 | 137196 | + | 0 | CTCTTCGTGCGGCGCTACGA | CGG | \|\|\|\|\|\|\|\|[\|\|\|\|\|\|\|\|\|\|\|\|]PAM | NA | Intergenic | NA | NA |
| T10466 | TCGTGCGGCGCTACGACGGCAGG | 1000 | | | | | | | | | |
| Oligo fwd | CACCTCGTGCGGCGCTACGACGGC | | | | | | | | | | |
| Oligo rev | AAACGCCGTCGTAGCGCCGCACGA | | | | | | | | | | |
| Chromosome | start | end | strand | MM | target_seq | PAM | alignment | distance | position | gene name | gene id |
| GU305914.1 | 137178 | 137200 | + | 0 | TCGTGCGGCGCTACGACGGC | AGG | \|\|\|\|\|\|\|\|[\|\|\|\|\|\|\|\|\|\|\|\|]PAM | NA | Intergenic | NA | NA |
| T10467 | CGGCGCTACGACGGCAGGAAAGG | 1000 | | | | | | | | | |
| Oligo fwd | CACCCGGCGCTACGACGGCAGGAA | | | | | | | | | | |
| Oligo rev | AAACTTCCTGCCGTCGTAGCGCCG | | | | | | | | | | |
| Chromosome | start | end | strand | MM | target_seq | PAM | alignment | distance | position | gene name | gene id |
| GU305914.1 | 137183 | 137205 | + | 0 | CGGCGCTACGACGGCAGGAA | AGG | \|\|\|\|\|\|\|\|[\|\|\|\|\|\|\|\|\|\|\|\|]PAM | NA | Intergenic | NA | NA |
| T10468 | CGACGGCAGGAAAGGACTGCCGG | 1000 | | | | | | | | | |
| Oligo fwd | CACCCGACGGCAGGAAAGGACTGC | | | | | | | | | | |
| Oligo rev | AAACGCAGTCCTTTCCTGCCGTCG | | | | | | | | | | |
| Chromosome | start | end | strand | MM | target_seq | PAM | alignment | distance | position | gene name | gene id |
| GU305914.1 | 137191 | 137213 | + | 0 | CGACGGCAGGAAAGGACTGC | CGG | \|\|\|\|\|\|\|\|[\|\|\|\|\|\|\|\|\|\|\|\|]PAM | NA | Intergenic | NA | NA |
| T10469 | GCCACGATATCTGACTACCTCGG | 1000 | | | | | | | | | |
| Oligo fwd | CACCGCCACGATATCTGACTACCT | | | | | | | | | | |
| Oligo rev | AAACAGGTAGTCAGATATCGTGGC | | | | | | | | | | |
| Chromosome | start | end | strand | MM | target_seq | PAM | alignment | distance | position | gene name | gene id |
| GU305914.1 | 137213 | 137235 | + | 0 | GCCACGATATCTGACTACCT | CGG | \|\|\|\|\|\|\|\|[\|\|\|\|\|\|\|\|\|\|\|\|]PAM | NA | Intergenic | NA | NA |
| T10470 | TATCTGACTACCTCGGCGCCAGG | 1000 | | | | | | | | | |
| Oligo fwd | CACCTATCTGACTACCTCGGCGCC | | | | | | | | | | |
| Oligo rev | AAACGGCGCCGAGGTAGTCAGATA | | | | | | | | | | |
| Chromosome | start | end | strand | MM | target_seq | PAM | alignment | distance | position | gene name | gene id |
| GU305914.1 | 137220 | 137242 | + | 0 | TATCTGACTACCTCGGCGCC | AGG | \|\|\|\|\|\|\|\|[\|\|\|\|\|\|\|\|\|\|\|\|]PAM | NA | Intergenic | NA | NA |
| T10471 | CCTCGGCGCCAGGAACAGCGAGG | 1000 | | | | | | | | | |
| Oligo fwd | CACCCCTCGGCGCCAGGAACAGCG | | | | | | | | | | |
| Oligo rev | AAACCGCTGTTCCTGGCGCCGAGG | | | | | | | | | | |
| Chromosome | start | end | strand | MM | target_seq | PAM | alignment | distance | position | gene name | gene id |
| GU305914.1 | 137230 | 137252 | + | 0 | CCTCGGCGCCAGGAACAGCG | AGG | \|\|\|\|\|\|\|\|[\|\|\|\|\|\|\|\|\|\|\|\|]PAM | NA | Intergenic | NA | NA |
| T10472 | CTCGGCGCCAGGAACAGCGAGGG | 1000 | | | | | | | | | |
| Oligo fwd | CACCCTCGGCGCCAGGAACAGCGA | | | | | | | | | | |
| Oligo rev | AAACTCGCTGTTCCTGGCGCCGAG | | | | | | | | | | |
| Chromosome | start | end | strand | MM | target_seq | PAM | alignment | distance | position | gene name | gene id |
| GU305914.1 | 137231 | 137253 | + | 0 | CTCGGCGCCAGGAACAGCGA | GGG | \|\|\|\|\|\|\|\|[\|\|\|\|\|\|\|\|\|\|\|\|]PAM | NA | Intergenic | NA | NA |
| T10473 | CGCCAGGAACAGCGAGGGCGAGG | 1000 | | | | | | | | | |
| Oligo fwd | CACCCGCCAGGAACAGCGAGGGCG | | | | | | | | | | |
| Oligo rev | AAACCGCCCTCGCTGTTCCTGGCG | | | | | | | | | | |
| Chromosome | start | end | strand | MM | target_seq | PAM | alignment | distance | position | gene name | gene id |
| GU305914.1 | 137236 | 137258 | + | 0 | CGCCAGGAACAGCGAGGGCG | AGG | \|\|\|\|\|\|\|\|[\|\|\|\|\|\|\|\|\|\|\|\|]PAM | NA | Intergenic | NA | NA |
| T10474 | AGGGCGAGGAGATCCCGCTGAGG | 1000 | | | | | | | | | |
| Oligo fwd | CACCAGGGCGAGGAGATCCCGCTG | | | | | | | | | | |
| Oligo rev | AAACCAGCGGGATCTCCTCGCCCT | | | | | | | | | | |
| Chromosome | start | end | strand | MM | target_seq | PAM | alignment | distance | position | gene name | gene id |
| GU305914.1 | 137250 | 137272 | + | 0 | AGGGCGAGGAGATCCCGCTG | AGG | \|\|\|\|\|\|\|\|[\|\|\|\|\|\|\|\|\|\|\|\|]PAM | NA | Intergenic | NA | NA |
| T10475 | TCCCGCTGAGGACCGAGCCCTGG | 1000 | | | | | | | | | |
| Oligo fwd | CACCTCCCGCTGAGGACCGAGCCC | | | | | | | | | | |
| Oligo rev | AAACGGGCTCGGTCCTCAGCGGGA | | | | | | | | | | |
| Chromosome | start | end | strand | MM | target_seq | PAM | alignment | distance | position | gene name | gene id |
| GU305914.1 | 137262 | 137284 | + | 0 | TCCCGCTGAGGACCGAGCCC | TGG | \|\|\|\|\|\|\|\|[\|\|\|\|\|\|\|\|\|\|\|\|]PAM | NA | Intergenic | NA | NA |
| T10476 | GCTGTCCGCCATGATCATCATGG | 1000 | | | | | | | | | |
| Oligo fwd | CACCGCTGTCCGCCATGATCATCA | | | | | | | | | | |
| Oligo rev | AAACTGATGATCATGGCGGACAGC | | | | | | | | | | |
| Chromosome | start | end | strand | MM | target_seq | PAM | alignment | distance | position | gene name | gene id |
| GU305914.1 | 137308 | 137330 | + | 0 | GCTGTCCGCCATGATCATCA | TGG | \|\|\|\|\|\|\|\|[\|\|\|\|\|\|\|\|\|\|\|\|]PAM | NA | Intergenic | NA | NA |
| T10477 | CAAATCAACCAACCCATCCATGG | 1000 | | | | | | | | | |
| Oligo fwd | CACCCAAATCAACCAACCCATCCA | | | | | | | | | | |
| Oligo rev | AAACTGGATGGGTTGGTTGATTTG | | | | | | | | | | |
| Chromosome | start | end | strand | MM | target_seq | PAM | alignment | distance | position | gene name | gene id |
| GU305914.1 | 137388 | 137410 | + | 0 | CAAATCAACCAACCCATCCA | TGG | \|\|\|\|\|\|\|\|[\|\|\|\|\|\|\|\|\|\|\|\|]PAM | NA | Intergenic | NA | NA |
| T10478 | TCCCTCCCGCATATCTTGTCTGG | 1000 | | | | | | | | | |
| Oligo fwd | CACCTCCCTCCCGCATATCTTGTC | | | | | | | | | | |
| Oligo rev | AAACGACAAGATATGCGGGAGGGA | | | | | | | | | | |
| Chromosome | start | end | strand | MM | target_seq | PAM | alignment | distance | position | gene name | gene id |
| GU305914.1 | 137440 | 137462 | + | 0 | TCCCTCCCGCATATCTTGTC | TGG | \|\|\|\|\|\|\|\|[\|\|\|\|\|\|\|\|\|\|\|\|]PAM | NA | Intergenic | NA | NA |
| T10479 | CACGTTAGTGTTGTAGCAGCTGG | 1000 | | | | | | | | | |
| Oligo fwd | CACCCACGTTAGTGTTGTAGCAGC | | | | | | | | | | |
| Oligo rev | AAACGCTGCTACAACACTAACGTG | | | | | | | | | | |

Chromosome start end strand MM target_seq PAM alignment distance position gene name gene id
GU305914.1 137487 137509 + 0 CACGTTAGTGTTGTAGCAGC TGG ||||||||[||||||||||||]PAM NA Intergenic NA NA

T10480 TGTTGTAGCAGCTGGCGAACAGG 1000
Oligo fwd CACCTGTTGTAGCAGCTGGCGAAC
Oligo rev AAACGTTCGCCAGCTGCTACAACA
Chromosome start end strand MM target_seq PAM alignment distance position gene name gene id
GU305914.1 137495 137517 + 0 TGTTGTAGCAGCTGGCGAAC AGG ||||||||[||||||||||||]PAM NA Intergenic NA NA

T10481 GTTGTAGCAGCTGGCGAACAGGG 1000
Oligo fwd CACCGTTGTAGCAGCTGGCGAACA
Oligo rev AAACTGTTCGCCAGCTGCTACAAC
Chromosome start end strand MM target_seq PAM alignment distance position gene name gene id
GU305914.1 137496 137518 + 0 GTTGTAGCAGCTGGCGAACA GGG ||||||||[||||||||||||]PAM NA Intergenic NA NA

T10482 GGCGAACAGGGCCGTGCTCTTGG 1000
Oligo fwd CACCGGCGAACAGGGCCGTGCTCT
Oligo rev AAACAGAGCACGGCCCTGTTCGCC
Chromosome start end strand MM target_seq PAM alignment distance position gene name gene id
GU305914.1 137508 137530 + 0 GGCGAACAGGGCCGTGCTCT TGG ||||||||[||||||||||||]PAM NA Intergenic NA NA

T10483 CCGTGCTCTTGGCGATCTTGTGG 1000
Oligo fwd CACCCCGTGCTCTTGGCGATCTTG
Oligo rev AAACCAAGATCGCCAAGAGCACGG
Chromosome start end strand MM target_seq PAM alignment distance position gene name gene id
GU305914.1 137519 137541 + 0 CCGTGCTCTTGGCGATCTTG TGG ||||||||[||||||||||||]PAM NA Intergenic NA NA

T10484 CCTTGTTCTCGACCATGAACCGG 1000
Oligo fwd CACCCCTTGTTCTCGACCATGAAC
Oligo rev AAACGTTCATGGTCGAGAACAAGG
Chromosome start end strand MM target_seq PAM alignment distance position gene name gene id
GU305914.1 137561 137583 + 0 CCTTGTTCTCGACCATGAAC CGG ||||||||[||||||||||||]PAM NA Intergenic NA NA

T10485 CGACCTCTTTGAGCACGTACTGG 1000
Oligo fwd CACCCGACCTCTTTGAGCACGTAC
Oligo rev AAACGTACGTGCTCAAAGAGGTCG
Chromosome start end strand MM target_seq PAM alignment distance position gene name gene id
GU305914.1 137612 137634 + 0 CGACCTCTTTGAGCACGTAC TGG ||||||||[||||||||||||]PAM NA Intergenic NA NA

T10486 GACCTCTTTGAGCACGTACTGGG 1000
Oligo fwd CACCGACCTCTTTGAGCACGTACT
Oligo rev AAACAGTACGTGCTCAAAGAGGTC
Chromosome start end strand MM target_seq PAM alignment distance position gene name gene id
GU305914.1 137613 137635 + 0 GACCTCTTTGAGCACGTACT GGG ||||||||[||||||||||||]PAM NA Intergenic NA NA

T10487 CGTACTGGGAGACGTGCTTCTGG 1000
Oligo fwd CACCCGTACTGGGAGACGTGCTTC
Oligo rev AAACGAAGCACGTCTCCCAGTACG
Chromosome start end strand MM target_seq PAM alignment distance position gene name gene id
GU305914.1 137627 137649 + 0 CGTACTGGGAGACGTGCTTC TGG ||||||||[||||||||||||]PAM NA Intergenic NA NA

T10488 ACTGGGAGACGTGCTTCTGGTGG 1000
Oligo fwd CACCACTGGGAGACGTGCTTCTGG
Oligo rev AAACCCAGAAGCACGTCTCCCAGT
Chromosome start end strand MM target_seq PAM alignment distance position gene name gene id
GU305914.1 137630 137652 + 0 ACTGGGAGACGTGCTTCTGG TGG ||||||||[||||||||||||]PAM NA Intergenic NA NA

T10489 CTGGGAGACGTGCTTCTGGTGGG 1000
Oligo fwd CACCCTGGGAGACGTGCTTCTGGT
Oligo rev AAACACCAGAAGCACGTCTCCCAG
Chromosome start end strand MM target_seq PAM alignment distance position gene name gene id
GU305914.1 137631 137653 + 0 CTGGGAGACGTGCTTCTGGT GGG ||||||||[||||||||||||]PAM NA Intergenic NA NA

T10490 TCCCGATGATGTTCTTCAGTAGG 1000
Oligo fwd CACCTCCCGATGATGTTCTTCAGT
Oligo rev AAACACTGAAGAACATCATCGGGA
Chromosome start end strand MM target_seq PAM alignment distance position gene name gene id
GU305914.1 137672 137694 + 0 TCCCGATGATGTTCTTCAGT AGG ||||||||[||||||||||||]PAM NA Intergenic NA NA

T10491 CTTCAGTAGGAAGCAGATCACGG 1000
Oligo fwd CACCCTTCAGTAGGAAGCAGATCA
Oligo rev AAACTGATCTGCTTCCTACTGAAG
Chromosome start end strand MM target_seq PAM alignment distance position gene name gene id
GU305914.1 137685 137707 + 0 CTTCAGTAGGAAGCAGATCA CGG ||||||||[||||||||||||]PAM NA Intergenic NA NA

T10492 TTCAGTAGGAAGCAGATCACGGG 1000
Oligo fwd CACCTTCAGTAGGAAGCAGATCAC
Oligo rev AAACGTGATCTGCTTCCTACTGAA
Chromosome start end strand MM target_seq PAM alignment distance position gene name gene id
GU305914.1 137686 137708 + 0 TTCAGTAGGAAGCAGATCAC GGG ||||||||[||||||||||||]PAM NA Intergenic NA NA

T10493 TCAGTAGGAAGCAGATCACGGGG 1000
Oligo fwd CACCTCAGTAGGAAGCAGATCACG
Oligo rev AAACCGTGATCTGCTTCCTACTGA
Chromosome start end strand MM target_seq PAM alignment distance position gene name gene id
GU305914.1 137687 137709 + 0 TCAGTAGGAAGCAGATCACG GGG ||||||||[||||||||||||]PAM NA Intergenic NA NA

T10494 TAGGAAGCAGATCACGGGGATGG 1000
Oligo fwd CACCTAGGAAGCAGATCACGGGGA

| | | | | | | | | | | | |
|---|---|---|---|---|---|---|---|---|---|---|---|
| Oligo rev | AAACTCCCCGTGATCTGCTTCCTA | | | | | | | | | | |
| Chromosome | start | end | strand | MM | target_seq | PAM | alignment | distance | position | gene name | gene id |
| GU305914.1 | 137691 | 137713 | + | 0 | TAGGAAGCAGATCACGGGGA | TGG | ||||||||[||||||||||||]PAM | NA | Intergenic | NA | NA |
| T10495 | GATCACGGGGATGGTGAACCAGG | 1000 | | | | | | | | | |
| Oligo fwd | CACCGATCACGGGGATGGTGAACC | | | | | | | | | | |
| Oligo rev | AAACGGTTCACCATCCCCGTGATC | | | | | | | | | | |
| Chromosome | start | end | strand | MM | target_seq | PAM | alignment | distance | position | gene name | gene id |
| GU305914.1 | 137700 | 137722 | + | 0 | GATCACGGGGATGGTGAACC | AGG | ||||||||[||||||||||||]PAM | NA | Intergenic | NA | NA |
| T10496 | ACCAGGTCTTCCCGTGCCGCCGG | 1000 | | | | | | | | | |
| Oligo fwd | CACCACCAGGTCTTCCCGTGCCGC | | | | | | | | | | |
| Oligo rev | AAACGCGGCACGGGAAGACCTGGT | | | | | | | | | | |
| Chromosome | start | end | strand | MM | target_seq | PAM | alignment | distance | position | gene name | gene id |
| GU305914.1 | 137717 | 137739 | + | 0 | ACCAGGTCTTCCCGTGCCGC | CGG | ||||||||[||||||||||||]PAM | NA | Intergenic | NA | NA |
| T10497 | CCAGGTCTTCCCGTGCCGCCGGG | 1000 | | | | | | | | | |
| Oligo fwd | CACCCCAGGTCTTCCCGTGCCGCC | | | | | | | | | | |
| Oligo rev | AAACGGCGGCACGGGAAGACCTGG | | | | | | | | | | |
| Chromosome | start | end | strand | MM | target_seq | PAM | alignment | distance | position | gene name | gene id |
| GU305914.1 | 137718 | 137740 | + | 0 | CCAGGTCTTCCCGTGCCGCC | GGG | ||||||||[||||||||||||]PAM | NA | Intergenic | NA | NA |
| T10498 | CAGGTCTTCCCGTGCCGCCGGGG | 1000 | | | | | | | | | |
| Oligo fwd | CACCCAGGTCTTCCCGTGCCGCCG | | | | | | | | | | |
| Oligo rev | AAACCGGCGGCACGGGAAGACCTG | | | | | | | | | | |
| Chromosome | start | end | strand | MM | target_seq | PAM | alignment | distance | position | gene name | gene id |
| GU305914.1 | 137719 | 137741 | + | 0 | CAGGTCTTCCCGTGCCGCCG | GGG | ||||||||[||||||||||||]PAM | NA | Intergenic | NA | NA |
| T10499 | TCCCGTGCCGCCGGGGCACGAGG | 1000 | | | | | | | | | |
| Oligo fwd | CACCTCCCGTGCCGCCGGGGCACG | | | | | | | | | | |
| Oligo rev | AAACCGTGCCCCGGCGGCACGGGA | | | | | | | | | | |
| Chromosome | start | end | strand | MM | target_seq | PAM | alignment | distance | position | gene name | gene id |
| GU305914.1 | 137726 | 137748 | + | 0 | TCCCGTGCCGCCGGGGCACG | AGG | ||||||||[||||||||||||]PAM | NA | Intergenic | NA | NA |
| T10500 | CCGGGGCACGAGGAACACCGTGG | 1000 | | | | | | | | | |
| Oligo fwd | CACCCCGGGGCACGAGGAACACCG | | | | | | | | | | |
| Oligo rev | AAACCGGTGTTCCTCGTGCCCCGG | | | | | | | | | | |
| Chromosome | start | end | strand | MM | target_seq | PAM | alignment | distance | position | gene name | gene id |
| GU305914.1 | 137736 | 137758 | + | 0 | CCGGGGCACGAGGAACACCG | TGG | ||||||||[||||||||||||]PAM | NA | Intergenic | NA | NA |
| T10501 | CCGTGGCCTTCTGCTTGAACAGG | 1000 | | | | | | | | | |
| Oligo fwd | CACCCCGTGGCCTTCTGCTTGAAC | | | | | | | | | | |
| Oligo rev | AAACGTTCAAGCAGAAGGCCACGG | | | | | | | | | | |
| Chromosome | start | end | strand | MM | target_seq | PAM | alignment | distance | position | gene name | gene id |
| GU305914.1 | 137753 | 137775 | + | 0 | CCGTGGCCTTCTGCTTGAAC | AGG | ||||||||[||||||||||||]PAM | NA | Intergenic | NA | NA |
| T10502 | GAACAGGTCTATCGACTGCGAGG | 1000 | | | | | | | | | |
| Oligo fwd | CACCGAACAGGTCTATCGACTGCG | | | | | | | | | | |
| Oligo rev | AAACCGCAGTCGATAGACCTGTTC | | | | | | | | | | |
| Chromosome | start | end | strand | MM | target_seq | PAM | alignment | distance | position | gene name | gene id |
| GU305914.1 | 137769 | 137791 | + | 0 | GAACAGGTCTATCGACTGCG | AGG | ||||||||[||||||||||||]PAM | NA | Intergenic | NA | NA |
| T10503 | ACTGCGAGGACAGAAAATGCAGG | 1000 | | | | | | | | | |
| Oligo fwd | CACCACTGCGAGGACAGAAAATGC | | | | | | | | | | |
| Oligo rev | AAACGCATTTTCTGTCCTCGCAGT | | | | | | | | | | |
| Chromosome | start | end | strand | MM | target_seq | PAM | alignment | distance | position | gene name | gene id |
| GU305914.1 | 137783 | 137805 | + | 0 | ACTGCGAGGACAGAAAATGC | AGG | ||||||||[||||||||||||]PAM | NA | Intergenic | NA | NA |
| T10504 | GGACAGAAAATGCAGGTCGAAGG | 1000 | | | | | | | | | |
| Oligo fwd | CACCGGACAGAAAATGCAGGTCGA | | | | | | | | | | |
| Oligo rev | AAACTCGACCTGCATTTTCTGTCC | | | | | | | | | | |
| Chromosome | start | end | strand | MM | target_seq | PAM | alignment | distance | position | gene name | gene id |
| GU305914.1 | 137790 | 137812 | + | 0 | GGACAGAAAATGCAGGTCGA | AGG | ||||||||[||||||||||||]PAM | NA | Intergenic | NA | NA |
| T10505 | GCAGGTCGAAGGCGTGCGTCAGG | 1000 | | | | | | | | | |
| Oligo fwd | CACCGCAGGTCGAAGGCGTGCGTC | | | | | | | | | | |
| Oligo rev | AAACGACGCACGCCTTCGACCTGC | | | | | | | | | | |
| Chromosome | start | end | strand | MM | target_seq | PAM | alignment | distance | position | gene name | gene id |
| GU305914.1 | 137801 | 137823 | + | 0 | GCAGGTCGAAGGCGTGCGTC | AGG | ||||||||[||||||||||||]PAM | NA | Intergenic | NA | NA |
| T10506 | GCGTCAGGTACTCCGTCACGCGG | 1000 | | | | | | | | | |
| Oligo fwd | CACCGCGTCAGGTACTCCGTCACG | | | | | | | | | | |
| Oligo rev | AAACCGTGACGGAGTACCTGACGC | | | | | | | | | | |
| Chromosome | start | end | strand | MM | target_seq | PAM | alignment | distance | position | gene name | gene id |
| GU305914.1 | 137816 | 137838 | + | 0 | GCGTCAGGTACTCCGTCACG | CGG | ||||||||[||||||||||||]PAM | NA | Intergenic | NA | NA |
| T10507 | CAGGTACTCCGTCACGCGGTTGG | 1000 | | | | | | | | | |
| Oligo fwd | CACCCAGGTACTCCGTCACGCGGT | | | | | | | | | | |
| Oligo rev | AAACACCGCGTGACGGAGTACCTG | | | | | | | | | | |
| Chromosome | start | end | strand | MM | target_seq | PAM | alignment | distance | position | gene name | gene id |
| GU305914.1 | 137820 | 137842 | + | 0 | CAGGTACTCCGTCACGCGGT | TGG | ||||||||[||||||||||||]PAM | NA | Intergenic | NA | NA |
| T10508 | CGTCACGCGGTTGGCCAGCGTGG | 1000 | | | | | | | | | |
| Oligo fwd | CACCCGTCACGCGGTTGGCCAGCG | | | | | | | | | | |
| Oligo rev | AAACCGCTGGCCAACCGCGTGACG | | | | | | | | | | |
| Chromosome | start | end | strand | MM | target_seq | PAM | alignment | distance | position | gene name | gene id |
| GU305914.1 | 137829 | 137851 | + | 0 | CGTCACGCGGTTGGCCAGCG | TGG | ||||||||[||||||||||||]PAM | NA | Intergenic | NA | NA |
| T10509 | GTCACGCGGTTGGCCAGCGTGGG | 1000 | | | | | | | | | |

<table>
<tr><td>Oligo fwd</td><td>CACCGTCACGCGGTTGGCCAGCGT</td><td></td><td></td><td></td><td></td><td></td><td></td><td></td><td></td><td></td><td></td></tr>
<tr><td>Oligo rev</td><td>AAACACGCTGGCCAACCGCGTGAC</td><td></td><td></td><td></td><td></td><td></td><td></td><td></td><td></td><td></td><td></td></tr>
<tr><td>Chromosome</td><td>start</td><td>end</td><td>strand</td><td>MM</td><td>target_seq</td><td>PAM</td><td>alignment</td><td>distance</td><td>position</td><td>gene name</td><td>gene id</td></tr>
<tr><td>GU305914.1</td><td>137830</td><td>137852</td><td>+</td><td>0</td><td>GTCACGCGGTTGGCCAGCGT</td><td>GGG</td><td>||||||||[||||||||||||]PAM</td><td>NA</td><td>Intergenic</td><td>NA</td><td>NA</td></tr>
<tr><td>T10510</td><td>TCACGCGGTTGGCCAGCGTGGGG</td><td>1000</td><td></td><td></td><td></td><td></td><td></td><td></td><td></td><td></td><td></td></tr>
<tr><td>Oligo fwd</td><td>CACCTCACGCGGTTGGCCAGCGTG</td><td></td><td></td><td></td><td></td><td></td><td></td><td></td><td></td><td></td><td></td></tr>
<tr><td>Oligo rev</td><td>AAACCACGCTGGCCAACCGCGTGA</td><td></td><td></td><td></td><td></td><td></td><td></td><td></td><td></td><td></td><td></td></tr>
<tr><td>Chromosome</td><td>start</td><td>end</td><td>strand</td><td>MM</td><td>target_seq</td><td>PAM</td><td>alignment</td><td>distance</td><td>position</td><td>gene name</td><td>gene id</td></tr>
<tr><td>GU305914.1</td><td>137831</td><td>137853</td><td>+</td><td>0</td><td>TCACGCGGTTGGCCAGCGTG</td><td>GGG</td><td>||||||||[||||||||||||]PAM</td><td>NA</td><td>Intergenic</td><td>NA</td><td>NA</td></tr>
<tr><td>T10511</td><td>GTTGGCCAGCGTGGGGTTCTTGG</td><td>1000</td><td></td><td></td><td></td><td></td><td></td><td></td><td></td><td></td><td></td></tr>
<tr><td>Oligo fwd</td><td>CACCGTTGGCCAGCGTGGGGTTCT</td><td></td><td></td><td></td><td></td><td></td><td></td><td></td><td></td><td></td><td></td></tr>
<tr><td>Oligo rev</td><td>AAACAGAACCCCACGCTGGCCAAC</td><td></td><td></td><td></td><td></td><td></td><td></td><td></td><td></td><td></td><td></td></tr>
<tr><td>Chromosome</td><td>start</td><td>end</td><td>strand</td><td>MM</td><td>target_seq</td><td>PAM</td><td>alignment</td><td>distance</td><td>position</td><td>gene name</td><td>gene id</td></tr>
<tr><td>GU305914.1</td><td>137838</td><td>137860</td><td>+</td><td>0</td><td>GTTGGCCAGCGTGGGGTTCT</td><td>TGG</td><td>||||||||[||||||||||||]PAM</td><td>NA</td><td>Intergenic</td><td>NA</td><td>NA</td></tr>
<tr><td>T10512</td><td>CAGCGTGGGGTTCTTGGTAGTGG</td><td>1000</td><td></td><td></td><td></td><td></td><td></td><td></td><td></td><td></td><td></td></tr>
<tr><td>Oligo fwd</td><td>CACCCAGCGTGGGGTTCTTGGTAG</td><td></td><td></td><td></td><td></td><td></td><td></td><td></td><td></td><td></td><td></td></tr>
<tr><td>Oligo rev</td><td>AAACCTACCAAGAACCCCACGCTG</td><td></td><td></td><td></td><td></td><td></td><td></td><td></td><td></td><td></td><td></td></tr>
<tr><td>Chromosome</td><td>start</td><td>end</td><td>strand</td><td>MM</td><td>target_seq</td><td>PAM</td><td>alignment</td><td>distance</td><td>position</td><td>gene name</td><td>gene id</td></tr>
<tr><td>GU305914.1</td><td>137844</td><td>137866</td><td>+</td><td>0</td><td>CAGCGTGGGGTTCTTGGTAG</td><td>TGG</td><td>||||||||[||||||||||||]PAM</td><td>NA</td><td>Intergenic</td><td>NA</td><td>NA</td></tr>
<tr><td>T10513</td><td>TGGGGTTCTTGGTAGTGGCGAGG</td><td>1000</td><td></td><td></td><td></td><td></td><td></td><td></td><td></td><td></td><td></td></tr>
<tr><td>Oligo fwd</td><td>CACCTGGGGTTCTTGGTAGTGGCG</td><td></td><td></td><td></td><td></td><td></td><td></td><td></td><td></td><td></td><td></td></tr>
<tr><td>Oligo rev</td><td>AAACCGCCACTACCAAGAACCCCA</td><td></td><td></td><td></td><td></td><td></td><td></td><td></td><td></td><td></td><td></td></tr>
<tr><td>Chromosome</td><td>start</td><td>end</td><td>strand</td><td>MM</td><td>target_seq</td><td>PAM</td><td>alignment</td><td>distance</td><td>position</td><td>gene name</td><td>gene id</td></tr>
<tr><td>GU305914.1</td><td>137849</td><td>137871</td><td>+</td><td>0</td><td>TGGGGTTCTTGGTAGTGGCG</td><td>AGG</td><td>||||||||[||||||||||||]PAM</td><td>NA</td><td>Intergenic</td><td>NA</td><td>NA</td></tr>
<tr><td>T10514</td><td>TGGCGAGGAAGAAGAGCACGTGG</td><td>1000</td><td></td><td></td><td></td><td></td><td></td><td></td><td></td><td></td><td></td></tr>
<tr><td>Oligo fwd</td><td>CACCTGGCGAGGAAGAAGAGCACG</td><td></td><td></td><td></td><td></td><td></td><td></td><td></td><td></td><td></td><td></td></tr>
<tr><td>Oligo rev</td><td>AAACCGTGCTCTTCTTCCTCGCCA</td><td></td><td></td><td></td><td></td><td></td><td></td><td></td><td></td><td></td><td></td></tr>
<tr><td>Chromosome</td><td>start</td><td>end</td><td>strand</td><td>MM</td><td>target_seq</td><td>PAM</td><td>alignment</td><td>distance</td><td>position</td><td>gene name</td><td>gene id</td></tr>
<tr><td>GU305914.1</td><td>137864</td><td>137886</td><td>+</td><td>0</td><td>TGGCGAGGAAGAAGAGCACG</td><td>TGG</td><td>||||||||[||||||||||||]PAM</td><td>NA</td><td>Intergenic</td><td>NA</td><td>NA</td></tr>
<tr><td>T10515</td><td>GGCGAGGAAGAAGAGCACGTGGG</td><td>1000</td><td></td><td></td><td></td><td></td><td></td><td></td><td></td><td></td><td></td></tr>
<tr><td>Oligo fwd</td><td>CACCGGCGAGGAAGAAGAGCACGT</td><td></td><td></td><td></td><td></td><td></td><td></td><td></td><td></td><td></td><td></td></tr>
<tr><td>Oligo rev</td><td>AAACACGTGCTCTTCTTCCTCGCC</td><td></td><td></td><td></td><td></td><td></td><td></td><td></td><td></td><td></td><td></td></tr>
<tr><td>Chromosome</td><td>start</td><td>end</td><td>strand</td><td>MM</td><td>target_seq</td><td>PAM</td><td>alignment</td><td>distance</td><td>position</td><td>gene name</td><td>gene id</td></tr>
<tr><td>GU305914.1</td><td>137865</td><td>137887</td><td>+</td><td>0</td><td>GGCGAGGAAGAAGAGCACGT</td><td>GGG</td><td>||||||||[||||||||||||]PAM</td><td>NA</td><td>Intergenic</td><td>NA</td><td>NA</td></tr>
<tr><td>GU305914.1</td><td>64906</td><td>64928</td><td>-</td><td>3</td><td>GCCGGGGTAGAAGAGCACGT</td><td>TGG</td><td>|-||-||-[||||||||||||]PAM</td><td>NA</td><td>Intergenic</td><td>NA</td><td>NA</td></tr>
<tr><td>T10516</td><td>AGAAGAGCACGTGGGCTAGCAGG</td><td>1000</td><td></td><td></td><td></td><td></td><td></td><td></td><td></td><td></td><td></td></tr>
<tr><td>Oligo fwd</td><td>CACCAGAAGAGCACGTGGGCTAGC</td><td></td><td></td><td></td><td></td><td></td><td></td><td></td><td></td><td></td><td></td></tr>
<tr><td>Oligo rev</td><td>AAACGCTAGCCCACGTGCTCTTCT</td><td></td><td></td><td></td><td></td><td></td><td></td><td></td><td></td><td></td><td></td></tr>
<tr><td>Chromosome</td><td>start</td><td>end</td><td>strand</td><td>MM</td><td>target_seq</td><td>PAM</td><td>alignment</td><td>distance</td><td>position</td><td>gene name</td><td>gene id</td></tr>
<tr><td>GU305914.1</td><td>137873</td><td>137895</td><td>+</td><td>0</td><td>AGAAGAGCACGTGGGCTAGC</td><td>AGG</td><td>||||||||[||||||||||||]PAM</td><td>NA</td><td>Intergenic</td><td>NA</td><td>NA</td></tr>
<tr><td>T10517</td><td>CGTGGGCTAGCAGGTTGCGTTGG</td><td>1000</td><td></td><td></td><td></td><td></td><td></td><td></td><td></td><td></td><td></td></tr>
<tr><td>Oligo fwd</td><td>CACCCGTGGGCTAGCAGGTTGCGT</td><td></td><td></td><td></td><td></td><td></td><td></td><td></td><td></td><td></td><td></td></tr>
<tr><td>Oligo rev</td><td>AAACACGCAACCTGCTAGCCCACG</td><td></td><td></td><td></td><td></td><td></td><td></td><td></td><td></td><td></td><td></td></tr>
<tr><td>Chromosome</td><td>start</td><td>end</td><td>strand</td><td>MM</td><td>target_seq</td><td>PAM</td><td>alignment</td><td>distance</td><td>position</td><td>gene name</td><td>gene id</td></tr>
<tr><td>GU305914.1</td><td>137882</td><td>137904</td><td>+</td><td>0</td><td>CGTGGGCTAGCAGGTTGCGT</td><td>TGG</td><td>||||||||[||||||||||||]PAM</td><td>NA</td><td>Intergenic</td><td>NA</td><td>NA</td></tr>
<tr><td>T10518</td><td>GGCTAGCAGGTTGCGTTGGAAGG</td><td>1000</td><td></td><td></td><td></td><td></td><td></td><td></td><td></td><td></td><td></td></tr>
<tr><td>Oligo fwd</td><td>CACCGGCTAGCAGGTTGCGTTGGA</td><td></td><td></td><td></td><td></td><td></td><td></td><td></td><td></td><td></td><td></td></tr>
<tr><td>Oligo rev</td><td>AAACTCCAACGCAACCTGCTAGCC</td><td></td><td></td><td></td><td></td><td></td><td></td><td></td><td></td><td></td><td></td></tr>
<tr><td>Chromosome</td><td>start</td><td>end</td><td>strand</td><td>MM</td><td>target_seq</td><td>PAM</td><td>alignment</td><td>distance</td><td>position</td><td>gene name</td><td>gene id</td></tr>
<tr><td>GU305914.1</td><td>137886</td><td>137908</td><td>+</td><td>0</td><td>GGCTAGCAGGTTGCGTTGGA</td><td>AGG</td><td>||||||||[||||||||||||]PAM</td><td>NA</td><td>Intergenic</td><td>NA</td><td>NA</td></tr>
<tr><td>T10519</td><td>GCTAGCAGGTTGCGTTGGAAGGG</td><td>1000</td><td></td><td></td><td></td><td></td><td></td><td></td><td></td><td></td><td></td></tr>
<tr><td>Oligo fwd</td><td>CACCGCTAGCAGGTTGCGTTGGAA</td><td></td><td></td><td></td><td></td><td></td><td></td><td></td><td></td><td></td><td></td></tr>
<tr><td>Oligo rev</td><td>AAACTTCCAACGCAACCTGCTAGC</td><td></td><td></td><td></td><td></td><td></td><td></td><td></td><td></td><td></td><td></td></tr>
<tr><td>Chromosome</td><td>start</td><td>end</td><td>strand</td><td>MM</td><td>target_seq</td><td>PAM</td><td>alignment</td><td>distance</td><td>position</td><td>gene name</td><td>gene id</td></tr>
<tr><td>GU305914.1</td><td>137887</td><td>137909</td><td>+</td><td>0</td><td>GCTAGCAGGTTGCGTTGGAA</td><td>GGG</td><td>||||||||[||||||||||||]PAM</td><td>NA</td><td>Intergenic</td><td>NA</td><td>NA</td></tr>
<tr><td>T10520</td><td>GGTTGCGTTGGAAGGGCTCTAGG</td><td>1000</td><td></td><td></td><td></td><td></td><td></td><td></td><td></td><td></td><td></td></tr>
<tr><td>Oligo fwd</td><td>CACCGGTTGCGTTGGAAGGGCTCT</td><td></td><td></td><td></td><td></td><td></td><td></td><td></td><td></td><td></td><td></td></tr>
<tr><td>Oligo rev</td><td>AAACAGAGCCCTTCCAACGCAACC</td><td></td><td></td><td></td><td></td><td></td><td></td><td></td><td></td><td></td><td></td></tr>
<tr><td>Chromosome</td><td>start</td><td>end</td><td>strand</td><td>MM</td><td>target_seq</td><td>PAM</td><td>alignment</td><td>distance</td><td>position</td><td>gene name</td><td>gene id</td></tr>
<tr><td>GU305914.1</td><td>137894</td><td>137916</td><td>+</td><td>0</td><td>GGTTGCGTTGGAAGGGCTCT</td><td>AGG</td><td>||||||||[||||||||||||]PAM</td><td>NA</td><td>Intergenic</td><td>NA</td><td>NA</td></tr>
<tr><td>T10521</td><td>ATCTTGTTCCGTCGTTCGCTCGG</td><td>1000</td><td></td><td></td><td></td><td></td><td></td><td></td><td></td><td></td><td></td></tr>
<tr><td>Oligo fwd</td><td>CACCATCTTGTTCCGTCGTTCGCT</td><td></td><td></td><td></td><td></td><td></td><td></td><td></td><td></td><td></td><td></td></tr>
<tr><td>Oligo rev</td><td>AAACAGCGAACGACGGAACAAGAT</td><td></td><td></td><td></td><td></td><td></td><td></td><td></td><td></td><td></td><td></td></tr>
<tr><td>Chromosome</td><td>start</td><td>end</td><td>strand</td><td>MM</td><td>target_seq</td><td>PAM</td><td>alignment</td><td>distance</td><td>position</td><td>gene name</td><td>gene id</td></tr>
<tr><td>GU305914.1</td><td>137920</td><td>137942</td><td>+</td><td>0</td><td>ATCTTGTTCCGTCGTTCGCT</td><td>CGG</td><td>||||||||[||||||||||||]PAM</td><td>NA</td><td>Intergenic</td><td>NA</td><td>NA</td></tr>
<tr><td>T10522</td><td>TCTTGTTCCGTCGTTCGCTCGGG</td><td>1000</td><td></td><td></td><td></td><td></td><td></td><td></td><td></td><td></td><td></td></tr>
<tr><td>Oligo fwd</td><td>CACCTCTTGTTCCGTCGTTCGCTC</td><td></td><td></td><td></td><td></td><td></td><td></td><td></td><td></td><td></td><td></td></tr>
<tr><td>Oligo rev</td><td>AAACGAGCGAACGACGGAACAAGA</td><td></td><td></td><td></td><td></td><td></td><td></td><td></td><td></td><td></td><td></td></tr>
<tr><td>Chromosome</td><td>start</td><td>end</td><td>strand</td><td>MM</td><td>target_seq</td><td>PAM</td><td>alignment</td><td>distance</td><td>position</td><td>gene name</td><td>gene id</td></tr>
<tr><td>GU305914.1</td><td>137921</td><td>137943</td><td>+</td><td>0</td><td>TCTTGTTCCGTCGTTCGCTC</td><td>GGG</td><td>||||||||[||||||||||||]PAM</td><td>NA</td><td>Intergenic</td><td>NA</td><td>NA</td></tr>
<tr><td>T10523</td><td>CGGGCAGTTCTCGTCACGACCGG</td><td>1000</td><td></td><td></td><td></td><td></td><td></td><td></td><td></td><td></td><td></td></tr>
<tr><td>Oligo fwd</td><td>CACCCGGGCAGTTCTCGTCACGAC</td><td></td><td></td><td></td><td></td><td></td><td></td><td></td><td></td><td></td><td></td></tr>
<tr><td>Oligo rev</td><td>AAACGTCGTGACGAGAACTGCCCG</td><td></td><td></td><td></td><td></td><td></td><td></td><td></td><td></td><td></td><td></td></tr>
<tr><td>Chromosome</td><td>start</td><td>end</td><td>strand</td><td>MM</td><td>target_seq</td><td>PAM</td><td>alignment</td><td>distance</td><td>position</td><td>gene name</td><td>gene id</td></tr>
<tr><td>GU305914.1</td><td>137940</td><td>137962</td><td>+</td><td>0</td><td>CGGGCAGTTCTCGTCACGAC</td><td>CGG</td><td>||||||||[||||||||||||]PAM</td><td>NA</td><td>Intergenic</td><td>NA</td><td>NA</td></tr>
</table>

| | | | | | | | | | | | |
|---|---|---|---|---|---|---|---|---|---|---|---|
| T10524 | TCCAGTTCTCGAAGTCCTGTAGG | 1000 | | | | | | | | | |
| Oligo fwd | CACCTCCAGTTCTCGAAGTCCTGT | | | | | | | | | | |
| Oligo rev | AAACACAGGACTTCGAGAACTGGA | | | | | | | | | | |
| Chromosome | start | end | strand | MM | target_seq | PAM | alignment | distance | position | gene name | gene id |
| GU305914.1 | 137969 | 137991 | + | 0 | TCCAGTTCTCGAAGTCCTGT | AGG | \|\|\|\|\|\|\|\|[\|\|\|\|\|\|\|\|\|\|\|\|]PAM | NA | Intergenic | NA | NA |
| T10525 | TTCACTATCTGCACGAACTCCGG | 1000 | | | | | | | | | |
| Oligo fwd | CACCTTCACTATCTGCACGAACTC | | | | | | | | | | |
| Oligo rev | AAACGAGTTCGTGCAGATAGTGAA | | | | | | | | | | |
| Chromosome | start | end | strand | MM | target_seq | PAM | alignment | distance | position | gene name | gene id |
| GU305914.1 | 137995 | 138017 | + | 0 | TTCACTATCTGCACGAACTC | CGG | \|\|\|\|\|\|\|\|[\|\|\|\|\|\|\|\|\|\|\|\|]PAM | NA | Intergenic | NA | NA |
| T10526 | CTCCGGATCTCTATATAAGTCGG | 1000 | | | | | | | | | |
| Oligo fwd | CACCCTCCGGATCTCTATATAAGT | | | | | | | | | | |
| Oligo rev | AAACACTTATATAGAGATCCGGAG | | | | | | | | | | |
| Chromosome | start | end | strand | MM | target_seq | PAM | alignment | distance | position | gene name | gene id |
| GU305914.1 | 138012 | 138034 | + | 0 | CTCCGGATCTCTATATAAGT | CGG | \|\|\|\|\|\|\|\|[\|\|\|\|\|\|\|\|\|\|\|\|]PAM | NA | Intergenic | NA | NA |
| T10527 | AGAACGCGTTCATCTTGCTGCGG | 1000 | | | | | | | | | |
| Oligo fwd | CACCAGAACGCGTTCATCTTGCTG | | | | | | | | | | |
| Oligo rev | AAACCAGCAAGATGAACGCGTTCT | | | | | | | | | | |
| Chromosome | start | end | strand | MM | target_seq | PAM | alignment | distance | position | gene name | gene id |
| GU305914.1 | 138035 | 138057 | + | 0 | AGAACGCGTTCATCTTGCTG | CGG | \|\|\|\|\|\|\|\|[\|\|\|\|\|\|\|\|\|\|\|\|]PAM | NA | Intergenic | NA | NA |
| T10528 | CTGCGGTACTCCAGCTGTTCCGG | 1000 | | | | | | | | | |
| Oligo fwd | CACCCTGCGGTACTCCAGCTGTTC | | | | | | | | | | |
| Oligo rev | AAACGAACAGCTGGAGTACCGCAG | | | | | | | | | | |
| Chromosome | start | end | strand | MM | target_seq | PAM | alignment | distance | position | gene name | gene id |
| GU305914.1 | 138052 | 138074 | + | 0 | CTGCGGTACTCCAGCTGTTC | CGG | \|\|\|\|\|\|\|\|[\|\|\|\|\|\|\|\|\|\|\|\|]PAM | NA | Intergenic | NA | NA |
| T10529 | TGCGGTACTCCAGCTGTTCCGGG | 1000 | | | | | | | | | |
| Oligo fwd | CACCTGCGGTACTCCAGCTGTTCC | | | | | | | | | | |
| Oligo rev | AAACGGAACAGCTGGAGTACCGCA | | | | | | | | | | |
| Chromosome | start | end | strand | MM | target_seq | PAM | alignment | distance | position | gene name | gene id |
| GU305914.1 | 138053 | 138075 | + | 0 | TGCGGTACTCCAGCTGTTCC | GGG | \|\|\|\|\|\|\|\|[\|\|\|\|\|\|\|\|\|\|\|\|]PAM | NA | Intergenic | NA | NA |
| T10530 | GCGGTACTCCAGCTGTTCCGGGG | 1000 | | | | | | | | | |
| Oligo fwd | CACCGCGGTACTCCAGCTGTTCCG | | | | | | | | | | |
| Oligo rev | AAACCGGAACAGCTGGAGTACCGC | | | | | | | | | | |
| Chromosome | start | end | strand | MM | target_seq | PAM | alignment | distance | position | gene name | gene id |
| GU305914.1 | 138054 | 138076 | + | 0 | GCGGTACTCCAGCTGTTCCG | GGG | \|\|\|\|\|\|\|\|[\|\|\|\|\|\|\|\|\|\|\|\|]PAM | NA | Intergenic | NA | NA |
| T10531 | TCCAGCTGTTCCGGGGTACACGG | 1000 | | | | | | | | | |
| Oligo fwd | CACCTCCAGCTGTTCCGGGGTACA | | | | | | | | | | |
| Oligo rev | AAACTGTACCCCGGAACAGCTGGA | | | | | | | | | | |
| Chromosome | start | end | strand | MM | target_seq | PAM | alignment | distance | position | gene name | gene id |
| GU305914.1 | 138061 | 138083 | + | 0 | TCCAGCTGTTCCGGGGTACA | CGG | \|\|\|\|\|\|\|\|[\|\|\|\|\|\|\|\|\|\|\|\|]PAM | NA | Intergenic | NA | NA |
| T10532 | TCGATCGCCGCCGACAGCGTCGG | 1000 | | | | | | | | | |
| Oligo fwd | CACCTCGATCGCCGCCGACAGCGT | | | | | | | | | | |
| Oligo rev | AAACACGCTGTCGGCGGCGATCGA | | | | | | | | | | |
| Chromosome | start | end | strand | MM | target_seq | PAM | alignment | distance | position | gene name | gene id |
| GU305914.1 | 138100 | 138122 | + | 0 | TCGATCGCCGCCGACAGCGT | CGG | \|\|\|\|\|\|\|\|[\|\|\|\|\|\|\|\|\|\|\|\|]PAM | NA | Intergenic | NA | NA |
| T10533 | CGATCGCCGCCGACAGCGTCGGG | 1000 | | | | | | | | | |
| Oligo fwd | CACCCGATCGCCGCCGACAGCGTC | | | | | | | | | | |
| Oligo rev | AAACGACGCTGTCGGCGGCGATCG | | | | | | | | | | |
| Chromosome | start | end | strand | MM | target_seq | PAM | alignment | distance | position | gene name | gene id |
| GU305914.1 | 138101 | 138123 | + | 0 | CGATCGCCGCCGACAGCGTC | GGG | \|\|\|\|\|\|\|\|[\|\|\|\|\|\|\|\|\|\|\|\|]PAM | NA | Intergenic | NA | NA |
| GU305914.1 | 130591 | 130613 | - | 4 | GGCGCGGCGCCGACAGCGTC | CGG | -\|--\|\|-\|[\|\|\|\|\|\|\|\|\|\|\|\|]PAM | NA | Intergenic | NA | NA |
| GU305914.1 | 5916 | 5938 | + | 4 | CGAGCCTCGCAGACAGCGTC | CGG | \|\|\|-\|--\|[\|\|-\|\|\|\|\|\|\|\|\|]PAM | NA | Intergenic | NA | NA |
| T10534 | GATCGCCGCCGACAGCGTCGGGG | 1000 | | | | | | | | | |
| Oligo fwd | CACCGATCGCCGCCGACAGCGTCG | | | | | | | | | | |
| Oligo rev | AAACCGACGCTGTCGGCGGCGATC | | | | | | | | | | |
| Chromosome | start | end | strand | MM | target_seq | PAM | alignment | distance | position | gene name | gene id |
| GU305914.1 | 138102 | 138124 | + | 0 | GATCGCCGCCGACAGCGTCG | GGG | \|\|\|\|\|\|\|\|[\|\|\|\|\|\|\|\|\|\|\|\|]PAM | NA | Intergenic | NA | NA |
| GU305914.1 | 113163 | 113185 | + | 4 | CATCTCCACCGACAGCGCCG | CGG | -\|\|\|-\|\|-[\|\|\|\|\|\|\|\|\|-\|\|]PAM | NA | Intergenic | NA | NA |
| T10535 | ATCTTCAGCGCGCTCGTCTTCGG | 1000 | | | | | | | | | |
| Oligo fwd | CACCATCTTCAGCGCGCTCGTCTT | | | | | | | | | | |
| Oligo rev | AAACAAGACGAGCGCGCTGAAGAT | | | | | | | | | | |
| Chromosome | start | end | strand | MM | target_seq | PAM | alignment | distance | position | gene name | gene id |
| GU305914.1 | 138127 | 138149 | + | 0 | ATCTTCAGCGCGCTCGTCTT | CGG | \|\|\|\|\|\|\|\|[\|\|\|\|\|\|\|\|\|\|\|\|]PAM | NA | Intergenic | NA | NA |
| T10536 | TCTTCAGCGCGCTCGTCTTCGGG | 1000 | | | | | | | | | |
| Oligo fwd | CACCTCTTCAGCGCGCTCGTCTTC | | | | | | | | | | |
| Oligo rev | AAACGAAGACGAGCGCGCTGAAGA | | | | | | | | | | |
| Chromosome | start | end | strand | MM | target_seq | PAM | alignment | distance | position | gene name | gene id |
| GU305914.1 | 138128 | 138150 | + | 0 | TCTTCAGCGCGCTCGTCTTC | GGG | \|\|\|\|\|\|\|\|[\|\|\|\|\|\|\|\|\|\|\|\|]PAM | NA | Intergenic | NA | NA |
| T10537 | GGCTGCTCTCGTTGTAGTCCCGG | 1000 | | | | | | | | | |
| Oligo fwd | CACCGGCTGCTCTCGTTGTAGTCC | | | | | | | | | | |
| Oligo rev | AAACGGACTACAACGAGAGCAGCC | | | | | | | | | | |
| Chromosome | start | end | strand | MM | target_seq | PAM | alignment | distance | position | gene name | gene id |
| GU305914.1 | 138149 | 138171 | + | 0 | GGCTGCTCTCGTTGTAGTCC | CGG | \|\|\|\|\|\|\|\|[\|\|\|\|\|\|\|\|\|\|\|\|]PAM | NA | Intergenic | NA | NA |
| T10538 | CCGGTATTGCACGCAGAACTTGG | 1000 | | | | | | | | | |

| | | | | | | | | | | | |
|---|---|---|---|---|---|---|---|---|---|---|---|
| Oligo fwd | CACCCCGGTATTGCACGCAGAACT | | | | | | | | | | |
| Oligo rev | AAACAGTTCTGCGTGCAATACCGG | | | | | | | | | | |
| Chromosome | start | end | strand | MM | target_seq | PAM | alignment | distance | position | gene name | gene id |
| GU305914.1 | 138168 | 138190 | + | 0 | CCGGTATTGCACGCAGAACT | TGG | \|\|\|\|\|\|\|\|[\|\|\|\|\|\|\|\|\|\|\|\|]PAM | NA | Intergenic | NA | NA |
| T10539 | ATTGCACGCAGAACTTGGTCAGG | 1000 | | | | | | | | | |
| Oligo fwd | CACCATTGCACGCAGAACTTGGTC | | | | | | | | | | |
| Oligo rev | AAACGACCAAGTTCTGCGTGCAAT | | | | | | | | | | |
| Chromosome | start | end | strand | MM | target_seq | PAM | alignment | distance | position | gene name | gene id |
| GU305914.1 | 138173 | 138195 | + | 0 | ATTGCACGCAGAACTTGGTC | AGG | \|\|\|\|\|\|\|\|[\|\|\|\|\|\|\|\|\|\|\|\|]PAM | NA | Intergenic | NA | NA |
| T10540 | TTGCACGCAGAACTTGGTCAGGG | 1000 | | | | | | | | | |
| Oligo fwd | CACCTTGCACGCAGAACTTGGTCA | | | | | | | | | | |
| Oligo rev | AAACTGACCAAGTTCTGCGTGCAA | | | | | | | | | | |
| Chromosome | start | end | strand | MM | target_seq | PAM | alignment | distance | position | gene name | gene id |
| GU305914.1 | 138174 | 138196 | + | 0 | TTGCACGCAGAACTTGGTCA | GGG | \|\|\|\|\|\|\|\|[\|\|\|\|\|\|\|\|\|\|\|\|]PAM | NA | Intergenic | NA | NA |
| T10541 | TGCACGCAGAACTTGGTCAGGGG | 1000 | | | | | | | | | |
| Oligo fwd | CACCTGCACGCAGAACTTGGTCAG | | | | | | | | | | |
| Oligo rev | AAACCTGACCAAGTTCTGCGTGCA | | | | | | | | | | |
| Chromosome | start | end | strand | MM | target_seq | PAM | alignment | distance | position | gene name | gene id |
| GU305914.1 | 138175 | 138197 | + | 0 | TGCACGCAGAACTTGGTCAG | GGG | \|\|\|\|\|\|\|\|[\|\|\|\|\|\|\|\|\|\|\|\|]PAM | NA | Intergenic | NA | NA |
| T10542 | GCACGCAGAACTTGGTCAGGGGG | 1000 | | | | | | | | | |
| Oligo fwd | CACCGCACGCAGAACTTGGTCAGG | | | | | | | | | | |
| Oligo rev | AAACCCTGACCAAGTTCTGCGTGC | | | | | | | | | | |
| Chromosome | start | end | strand | MM | target_seq | PAM | alignment | distance | position | gene name | gene id |
| GU305914.1 | 138176 | 138198 | + | 0 | GCACGCAGAACTTGGTCAGG | GGG | \|\|\|\|\|\|\|\|[\|\|\|\|\|\|\|\|\|\|\|\|]PAM | NA | Intergenic | NA | NA |
| T10543 | CACGCAGAACTTGGTCAGGGGGG | 1000 | | | | | | | | | |
| Oligo fwd | CACCCACGCAGAACTTGGTCAGGG | | | | | | | | | | |
| Oligo rev | AAACCCCTGACCAAGTTCTGCGTG | | | | | | | | | | |
| Chromosome | start | end | strand | MM | target_seq | PAM | alignment | distance | position | gene name | gene id |
| GU305914.1 | 138177 | 138199 | + | 0 | CACGCAGAACTTGGTCAGGG | GGG | \|\|\|\|\|\|\|\|[\|\|\|\|\|\|\|\|\|\|\|\|]PAM | NA | Intergenic | NA | NA |
| T10544 | TCTCGTTGCACGCGTTCGTCAGG | 1000 | | | | | | | | | |
| Oligo fwd | CACCTCTCGTTGCACGCGTTCGTC | | | | | | | | | | |
| Oligo rev | AAACGACGAACGCGTGCAACGAGA | | | | | | | | | | |
| Chromosome | start | end | strand | MM | target_seq | PAM | alignment | distance | position | gene name | gene id |
| GU305914.1 | 138206 | 138228 | + | 0 | TCTCGTTGCACGCGTTCGTC | AGG | \|\|\|\|\|\|\|\|[\|\|\|\|\|\|\|\|\|\|\|\|]PAM | NA | Intergenic | NA | NA |
| T10545 | GCGTTCGTCAGGCCCAGCTCCGG | 1000 | | | | | | | | | |
| Oligo fwd | CACCGCGTTCGTCAGGCCCAGCTC | | | | | | | | | | |
| Oligo rev | AAACGAGCTGGGCCTGACGAACGC | | | | | | | | | | |
| Chromosome | start | end | strand | MM | target_seq | PAM | alignment | distance | position | gene name | gene id |
| GU305914.1 | 138217 | 138239 | + | 0 | GCGTTCGTCAGGCCCAGCTC | CGG | \|\|\|\|\|\|\|\|[\|\|\|\|\|\|\|\|\|\|\|\|]PAM | NA | Intergenic | NA | NA |
| GU305914.1 | 87710 | 87732 | + | 4 | TCGATCGTCAGCGCCAGCTC | GGG | -\|\|-\|\|\|\|[\|\|\|--\|\|\|\|\|\|\|]PAM | NA | Intergenic | NA | NA |
| T10546 | CGTTCGTCAGGCCCAGCTCCGGG | 1000 | | | | | | | | | |
| Oligo fwd | CACCCGTTCGTCAGGCCCAGCTCC | | | | | | | | | | |
| Oligo rev | AAACGGAGCTGGGCCTGACGAACG | | | | | | | | | | |
| Chromosome | start | end | strand | MM | target_seq | PAM | alignment | distance | position | gene name | gene id |
| GU305914.1 | 138218 | 138240 | + | 0 | CGTTCGTCAGGCCCAGCTCC | GGG | \|\|\|\|\|\|\|\|[\|\|\|\|\|\|\|\|\|\|\|\|]PAM | NA | Intergenic | NA | NA |
| T10547 | CAGGCCCAGCTCCGGGTGTACGG | 1000 | | | | | | | | | |
| Oligo fwd | CACCCAGGCCCAGCTCCGGGTGTA | | | | | | | | | | |
| Oligo rev | AAACTACACCCGGAGCTGGGCCTG | | | | | | | | | | |
| Chromosome | start | end | strand | MM | target_seq | PAM | alignment | distance | position | gene name | gene id |
| GU305914.1 | 138225 | 138247 | + | 0 | CAGGCCCAGCTCCGGGTGTA | CGG | \|\|\|\|\|\|\|\|[\|\|\|\|\|\|\|\|\|\|\|\|]PAM | NA | Intergenic | NA | NA |
| T10548 | GCGACAGAAGATACCCACGTCGG | 1000 | | | | | | | | | |
| Oligo fwd | CACCGCGACAGAAGATACCCACGT | | | | | | | | | | |
| Oligo rev | AAACACGTGGGTATCTTCTGTCGC | | | | | | | | | | |
| Chromosome | start | end | strand | MM | target_seq | PAM | alignment | distance | position | gene name | gene id |
| GU305914.1 | 138261 | 138283 | + | 0 | GCGACAGAAGATACCCACGT | CGG | \|\|\|\|\|\|\|\|[\|\|\|\|\|\|\|\|\|\|\|\|]PAM | NA | Intergenic | NA | NA |
| T10549 | CGACAGAAGATACCCACGTCGGG | 1000 | | | | | | | | | |
| Oligo fwd | CACCCGACAGAAGATACCCACGTC | | | | | | | | | | |
| Oligo rev | AAACGACGTGGGTATCTTCTGTCG | | | | | | | | | | |
| Chromosome | start | end | strand | MM | target_seq | PAM | alignment | distance | position | gene name | gene id |
| GU305914.1 | 138262 | 138284 | + | 0 | CGACAGAAGATACCCACGTC | GGG | \|\|\|\|\|\|\|\|[\|\|\|\|\|\|\|\|\|\|\|\|]PAM | NA | Intergenic | NA | NA |
| T10550 | ACCCACGTCGGGAAACTCCGTGG | 1000 | | | | | | | | | |
| Oligo fwd | CACCACCCACGTCGGGAAACTCCG | | | | | | | | | | |
| Oligo rev | AAACCGGAGTTTCCCGACGTGGGT | | | | | | | | | | |
| Chromosome | start | end | strand | MM | target_seq | PAM | alignment | distance | position | gene name | gene id |
| GU305914.1 | 138273 | 138295 | + | 0 | ACCCACGTCGGGAAACTCCG | TGG | \|\|\|\|\|\|\|\|[\|\|\|\|\|\|\|\|\|\|\|\|]PAM | NA | Intergenic | NA | NA |
| T10551 | GGGAAACTCCGTGGTGATCACGG | 1000 | | | | | | | | | |
| Oligo fwd | CACCGGGAAACTCCGTGGTGATCA | | | | | | | | | | |
| Oligo rev | AAACTGATCACCACGGAGTTTCCC | | | | | | | | | | |
| Chromosome | start | end | strand | MM | target_seq | PAM | alignment | distance | position | gene name | gene id |
| GU305914.1 | 138282 | 138304 | + | 0 | GGGAAACTCCGTGGTGATCA | CGG | \|\|\|\|\|\|\|\|[\|\|\|\|\|\|\|\|\|\|\|\|]PAM | NA | Intergenic | NA | NA |
| T10552 | CTCCGTGGTGATCACGGTCGTGG | 1000 | | | | | | | | | |
| Oligo fwd | CACCCTCCGTGGTGATCACGGTCG | | | | | | | | | | |
| Oligo rev | AAACCGACCGTGATCACCACGGAG | | | | | | | | | | |
| Chromosome | start | end | strand | MM | target_seq | PAM | alignment | distance | position | gene name | gene id |
| GU305914.1 | 138288 | 138310 | + | 0 | CTCCGTGGTGATCACGGTCG | TGG | \|\|\|\|\|\|\|\|[\|\|\|\|\|\|\|\|\|\|\|\|]PAM | NA | Intergenic | NA | NA |

<table>
<tr><td>T10553</td><td>TCCGTGGTGATCACGGTCGTGGG</td><td>1000</td><td></td><td></td><td></td><td></td><td></td><td></td><td></td><td></td><td></td></tr>
<tr><td>Oligo fwd</td><td>CACCTCCGTGGTGATCACGGTCGT</td><td></td><td></td><td></td><td></td><td></td><td></td><td></td><td></td><td></td><td></td></tr>
<tr><td>Oligo rev</td><td>AAACACGACCGTGATCACCACGGA</td><td></td><td></td><td></td><td></td><td></td><td></td><td></td><td></td><td></td><td></td></tr>
<tr><td>Chromosome</td><td>start</td><td>end</td><td>strand</td><td>MM</td><td>target_seq</td><td>PAM</td><td>alignment</td><td>distance</td><td>position</td><td>gene name</td><td>gene id</td></tr>
<tr><td>GU305914.1</td><td>138289</td><td>138311</td><td>+</td><td>0</td><td>TCCGTGGTGATCACGGTCGT</td><td>GGG</td><td>||||||||[||||||||||||]PAM</td><td>NA</td><td>Intergenic</td><td>NA</td><td>NA</td></tr>
<tr><td>T10554</td><td>CAATTTCATCATAGCGACGTCGG</td><td>1000</td><td></td><td></td><td></td><td></td><td></td><td></td><td></td><td></td><td></td></tr>
<tr><td>Oligo fwd</td><td>CACCCAATTTCATCATAGCGACGT</td><td></td><td></td><td></td><td></td><td></td><td></td><td></td><td></td><td></td><td></td></tr>
<tr><td>Oligo rev</td><td>AAACACGTCGCTATGATGAAATTG</td><td></td><td></td><td></td><td></td><td></td><td></td><td></td><td></td><td></td><td></td></tr>
<tr><td>Chromosome</td><td>start</td><td>end</td><td>strand</td><td>MM</td><td>target_seq</td><td>PAM</td><td>alignment</td><td>distance</td><td>position</td><td>gene name</td><td>gene id</td></tr>
<tr><td>GU305914.1</td><td>138329</td><td>138351</td><td>+</td><td>0</td><td>CAATTTCATCATAGCGACGT</td><td>CGG</td><td>||||||||[||||||||||||]PAM</td><td>NA</td><td>Intergenic</td><td>NA</td><td>NA</td></tr>
<tr><td>T10555</td><td>ATAGCGACGTCGGACCGCGACGG</td><td>1000</td><td></td><td></td><td></td><td></td><td></td><td></td><td></td><td></td><td></td></tr>
<tr><td>Oligo fwd</td><td>CACCATAGCGACGTCGGACCGCGA</td><td></td><td></td><td></td><td></td><td></td><td></td><td></td><td></td><td></td><td></td></tr>
<tr><td>Oligo rev</td><td>AAACTCGCGGTCCGACGTCGCTAT</td><td></td><td></td><td></td><td></td><td></td><td></td><td></td><td></td><td></td><td></td></tr>
<tr><td>Chromosome</td><td>start</td><td>end</td><td>strand</td><td>MM</td><td>target_seq</td><td>PAM</td><td>alignment</td><td>distance</td><td>position</td><td>gene name</td><td>gene id</td></tr>
<tr><td>GU305914.1</td><td>138339</td><td>138361</td><td>+</td><td>0</td><td>ATAGCGACGTCGGACCGCGA</td><td>CGG</td><td>||||||||[||||||||||||]PAM</td><td>NA</td><td>Intergenic</td><td>NA</td><td>NA</td></tr>
<tr><td>T10556</td><td>GACCGCGACGGCGATCGCGTCGG</td><td>1000</td><td></td><td></td><td></td><td></td><td></td><td></td><td></td><td></td><td></td></tr>
<tr><td>Oligo fwd</td><td>CACCGACCGCGACGGCGATCGCGT</td><td></td><td></td><td></td><td></td><td></td><td></td><td></td><td></td><td></td><td></td></tr>
<tr><td>Oligo rev</td><td>AAACACGCGATCGCCGTCGCGGTC</td><td></td><td></td><td></td><td></td><td></td><td></td><td></td><td></td><td></td><td></td></tr>
<tr><td>Chromosome</td><td>start</td><td>end</td><td>strand</td><td>MM</td><td>target_seq</td><td>PAM</td><td>alignment</td><td>distance</td><td>position</td><td>gene name</td><td>gene id</td></tr>
<tr><td>GU305914.1</td><td>138351</td><td>138373</td><td>+</td><td>0</td><td>GACCGCGACGGCGATCGCGT</td><td>CGG</td><td>||||||||[||||||||||||]PAM</td><td>NA</td><td>Intergenic</td><td>NA</td><td>NA</td></tr>
<tr><td>T10557</td><td>GGCGATCGCGTCGGTGAGCATGG</td><td>1000</td><td></td><td></td><td></td><td></td><td></td><td></td><td></td><td></td><td></td></tr>
<tr><td>Oligo fwd</td><td>CACCGGCGATCGCGTCGGTGAGCA</td><td></td><td></td><td></td><td></td><td></td><td></td><td></td><td></td><td></td><td></td></tr>
<tr><td>Oligo rev</td><td>AAACTGCTCACCGACGCGATCGCC</td><td></td><td></td><td></td><td></td><td></td><td></td><td></td><td></td><td></td><td></td></tr>
<tr><td>Chromosome</td><td>start</td><td>end</td><td>strand</td><td>MM</td><td>target_seq</td><td>PAM</td><td>alignment</td><td>distance</td><td>position</td><td>gene name</td><td>gene id</td></tr>
<tr><td>GU305914.1</td><td>138360</td><td>138382</td><td>+</td><td>0</td><td>GGCGATCGCGTCGGTGAGCA</td><td>TGG</td><td>||||||||[||||||||||||]PAM</td><td>NA</td><td>Intergenic</td><td>NA</td><td>NA</td></tr>
<tr><td>T10558</td><td>GTCGGTGAGCATGGCTACAGCGG</td><td>1000</td><td></td><td></td><td></td><td></td><td></td><td></td><td></td><td></td><td></td></tr>
<tr><td>Oligo fwd</td><td>CACCGTCGGTGAGCATGGCTACAG</td><td></td><td></td><td></td><td></td><td></td><td></td><td></td><td></td><td></td><td></td></tr>
<tr><td>Oligo rev</td><td>AAACCTGTAGCCATGCTCACCGAC</td><td></td><td></td><td></td><td></td><td></td><td></td><td></td><td></td><td></td><td></td></tr>
<tr><td>Chromosome</td><td>start</td><td>end</td><td>strand</td><td>MM</td><td>target_seq</td><td>PAM</td><td>alignment</td><td>distance</td><td>position</td><td>gene name</td><td>gene id</td></tr>
<tr><td>GU305914.1</td><td>138369</td><td>138391</td><td>+</td><td>0</td><td>GTCGGTGAGCATGGCTACAG</td><td>CGG</td><td>||||||||[||||||||||||]PAM</td><td>NA</td><td>Intergenic</td><td>NA</td><td>NA</td></tr>
<tr><td>T10559</td><td>GTGAGCATGGCTACAGCGGCCGG</td><td>1000</td><td></td><td></td><td></td><td></td><td></td><td></td><td></td><td></td><td></td></tr>
<tr><td>Oligo fwd</td><td>CACCGTGAGCATGGCTACAGCGGC</td><td></td><td></td><td></td><td></td><td></td><td></td><td></td><td></td><td></td><td></td></tr>
<tr><td>Oligo rev</td><td>AAACGCCGCTGTAGCCATGCTCAC</td><td></td><td></td><td></td><td></td><td></td><td></td><td></td><td></td><td></td><td></td></tr>
<tr><td>Chromosome</td><td>start</td><td>end</td><td>strand</td><td>MM</td><td>target_seq</td><td>PAM</td><td>alignment</td><td>distance</td><td>position</td><td>gene name</td><td>gene id</td></tr>
<tr><td>GU305914.1</td><td>138373</td><td>138395</td><td>+</td><td>0</td><td>GTGAGCATGGCTACAGCGGC</td><td>CGG</td><td>||||||||[||||||||||||]PAM</td><td>NA</td><td>Intergenic</td><td>NA</td><td>NA</td></tr>
<tr><td>T10560</td><td>AGCATGGCTACAGCGGCCGGCGG</td><td>1000</td><td></td><td></td><td></td><td></td><td></td><td></td><td></td><td></td><td></td></tr>
<tr><td>Oligo fwd</td><td>CACCAGCATGGCTACAGCGGCCGG</td><td></td><td></td><td></td><td></td><td></td><td></td><td></td><td></td><td></td><td></td></tr>
<tr><td>Oligo rev</td><td>AAACCCGGCCGCTGTAGCCATGCT</td><td></td><td></td><td></td><td></td><td></td><td></td><td></td><td></td><td></td><td></td></tr>
<tr><td>Chromosome</td><td>start</td><td>end</td><td>strand</td><td>MM</td><td>target_seq</td><td>PAM</td><td>alignment</td><td>distance</td><td>position</td><td>gene name</td><td>gene id</td></tr>
<tr><td>GU305914.1</td><td>138376</td><td>138398</td><td>+</td><td>0</td><td>AGCATGGCTACAGCGGCCGG</td><td>CGG</td><td>||||||||[||||||||||||]PAM</td><td>NA</td><td>Intergenic</td><td>NA</td><td>NA</td></tr>
<tr><td>T10561</td><td>GCGGCCGGCGGTGTTTACGCTGG</td><td>1000</td><td></td><td></td><td></td><td></td><td></td><td></td><td></td><td></td><td></td></tr>
<tr><td>Oligo fwd</td><td>CACCGCGGCCGGCGGTGTTTACGC</td><td></td><td></td><td></td><td></td><td></td><td></td><td></td><td></td><td></td><td></td></tr>
<tr><td>Oligo rev</td><td>AAACGCGTAAACACCGCCGGCCGC</td><td></td><td></td><td></td><td></td><td></td><td></td><td></td><td></td><td></td><td></td></tr>
<tr><td>Chromosome</td><td>start</td><td>end</td><td>strand</td><td>MM</td><td>target_seq</td><td>PAM</td><td>alignment</td><td>distance</td><td>position</td><td>gene name</td><td>gene id</td></tr>
<tr><td>GU305914.1</td><td>138388</td><td>138410</td><td>+</td><td>0</td><td>GCGGCCGGCGGTGTTTACGC</td><td>TGG</td><td>||||||||[||||||||||||]PAM</td><td>NA</td><td>Intergenic</td><td>NA</td><td>NA</td></tr>
<tr><td>T10562</td><td>GCCGGCGGTGTTTACGCTGGTGG</td><td>1000</td><td></td><td></td><td></td><td></td><td></td><td></td><td></td><td></td><td></td></tr>
<tr><td>Oligo fwd</td><td>CACCGCCGGCGGTGTTTACGCTGG</td><td></td><td></td><td></td><td></td><td></td><td></td><td></td><td></td><td></td><td></td></tr>
<tr><td>Oligo rev</td><td>AAACCCAGCGTAAACACCGCCGGC</td><td></td><td></td><td></td><td></td><td></td><td></td><td></td><td></td><td></td><td></td></tr>
<tr><td>Chromosome</td><td>start</td><td>end</td><td>strand</td><td>MM</td><td>target_seq</td><td>PAM</td><td>alignment</td><td>distance</td><td>position</td><td>gene name</td><td>gene id</td></tr>
<tr><td>GU305914.1</td><td>138391</td><td>138413</td><td>+</td><td>0</td><td>GCCGGCGGTGTTTACGCTGG</td><td>TGG</td><td>||||||||[||||||||||||]PAM</td><td>NA</td><td>Intergenic</td><td>NA</td><td>NA</td></tr>
<tr><td>T10563</td><td>CGGTGTTTACGCTGGTGGCGAGG</td><td>1000</td><td></td><td></td><td></td><td></td><td></td><td></td><td></td><td></td><td></td></tr>
<tr><td>Oligo fwd</td><td>CACCCGGTGTTTACGCTGGTGGCG</td><td></td><td></td><td></td><td></td><td></td><td></td><td></td><td></td><td></td><td></td></tr>
<tr><td>Oligo rev</td><td>AAACCGCCACCAGCGTAAACACCG</td><td></td><td></td><td></td><td></td><td></td><td></td><td></td><td></td><td></td><td></td></tr>
<tr><td>Chromosome</td><td>start</td><td>end</td><td>strand</td><td>MM</td><td>target_seq</td><td>PAM</td><td>alignment</td><td>distance</td><td>position</td><td>gene name</td><td>gene id</td></tr>
<tr><td>GU305914.1</td><td>138396</td><td>138418</td><td>+</td><td>0</td><td>CGGTGTTTACGCTGGTGGCG</td><td>AGG</td><td>||||||||[||||||||||||]PAM</td><td>NA</td><td>Intergenic</td><td>NA</td><td>NA</td></tr>
<tr><td>T10564</td><td>GCGAGGATCAACAACAACAACGG</td><td>1000</td><td></td><td></td><td></td><td></td><td></td><td></td><td></td><td></td><td></td></tr>
<tr><td>Oligo fwd</td><td>CACCGCGAGGATCAACAACAACAA</td><td></td><td></td><td></td><td></td><td></td><td></td><td></td><td></td><td></td><td></td></tr>
<tr><td>Oligo rev</td><td>AAACTTGTTGTTGTTGATCCTCGC</td><td></td><td></td><td></td><td></td><td></td><td></td><td></td><td></td><td></td><td></td></tr>
<tr><td>Chromosome</td><td>start</td><td>end</td><td>strand</td><td>MM</td><td>target_seq</td><td>PAM</td><td>alignment</td><td>distance</td><td>position</td><td>gene name</td><td>gene id</td></tr>
<tr><td>GU305914.1</td><td>138413</td><td>138435</td><td>+</td><td>0</td><td>GCGAGGATCAACAACAACAA</td><td>CGG</td><td>||||||||[||||||||||||]PAM</td><td>NA</td><td>Intergenic</td><td>NA</td><td>NA</td></tr>
<tr><td>T10565</td><td>ATCAACAACAACAACGGCCACGG</td><td>1000</td><td></td><td></td><td></td><td></td><td></td><td></td><td></td><td></td><td></td></tr>
<tr><td>Oligo fwd</td><td>CACCATCAACAACAACAACGGCCA</td><td></td><td></td><td></td><td></td><td></td><td></td><td></td><td></td><td></td><td></td></tr>
<tr><td>Oligo rev</td><td>AAACTGGCCGTTGTTGTTGTTGAT</td><td></td><td></td><td></td><td></td><td></td><td></td><td></td><td></td><td></td><td></td></tr>
<tr><td>Chromosome</td><td>start</td><td>end</td><td>strand</td><td>MM</td><td>target_seq</td><td>PAM</td><td>alignment</td><td>distance</td><td>position</td><td>gene name</td><td>gene id</td></tr>
<tr><td>GU305914.1</td><td>138419</td><td>138441</td><td>+</td><td>0</td><td>ATCAACAACAACAACGGCCA</td><td>CGG</td><td>||||||||[||||||||||||]PAM</td><td>NA</td><td>Intergenic</td><td>NA</td><td>NA</td></tr>
<tr><td>T10566</td><td>TCAACAACAACAACGGCCACGGG</td><td>1000</td><td></td><td></td><td></td><td></td><td></td><td></td><td></td><td></td><td></td></tr>
<tr><td>Oligo fwd</td><td>CACCTCAACAACAACAACGGCCAC</td><td></td><td></td><td></td><td></td><td></td><td></td><td></td><td></td><td></td><td></td></tr>
<tr><td>Oligo rev</td><td>AAACGTGGCCGTTGTTGTTGTTGA</td><td></td><td></td><td></td><td></td><td></td><td></td><td></td><td></td><td></td><td></td></tr>
<tr><td>Chromosome</td><td>start</td><td>end</td><td>strand</td><td>MM</td><td>target_seq</td><td>PAM</td><td>alignment</td><td>distance</td><td>position</td><td>gene name</td><td>gene id</td></tr>
<tr><td>GU305914.1</td><td>138420</td><td>138442</td><td>+</td><td>0</td><td>TCAACAACAACAACGGCCAC</td><td>GGG</td><td>||||||||[||||||||||||]PAM</td><td>NA</td><td>Intergenic</td><td>NA</td><td>NA</td></tr>
<tr><td>T10567</td><td>GACCGCGCTCTGCGATGTCGAGG</td><td>1000</td><td></td><td></td><td></td><td></td><td></td><td></td><td></td><td></td><td></td></tr>
<tr><td>Oligo fwd</td><td>CACCGACCGCGCTCTGCGATGTCG</td><td></td><td></td><td></td><td></td><td></td><td></td><td></td><td></td><td></td><td></td></tr>
<tr><td>Oligo rev</td><td>AAACCGACATCGCAGAGCGCGGTC</td><td></td><td></td><td></td><td></td><td></td><td></td><td></td><td></td><td></td><td></td></tr>
<tr><td>Chromosome</td><td>start</td><td>end</td><td>strand</td><td>MM</td><td>target_seq</td><td>PAM</td><td>alignment</td><td>distance</td><td>position</td><td>gene name</td><td>gene id</td></tr>
</table>

| | | | | | | | | | | | |
|---|---|---|---|---|---|---|---|---|---|---|---|
| GU305914.1 | 138456 | 138478 | + | 0 | GACCGCGCTCTGCGATGTCG | AGG | \|\|\|\|\|\|\|\|[\|\|\|\|\|\|\|\|\|\|\|\|]PAM | NA | Intergenic | NA | NA |
| T10568 | CGAGGCGCTCGCCGCCGTCGAGG | 1000 | | | | | | | | | |
| Oligo fwd | CACCCGAGGCGCTCGCCGCCGTCG | | | | | | | | | | |
| Oligo rev | AAACCGACGGCGGCGAGCGCCTCG | | | | | | | | | | |
| Chromosome | start | end | strand | MM | target_seq | PAM | alignment | distance | position | gene name | gene id |
| GU305914.1 | 138474 | 138496 | + | 0 | CGAGGCGCTCGCCGCCGTCG | AGG | \|\|\|\|\|\|\|\|[\|\|\|\|\|\|\|\|\|\|\|\|]PAM | NA | Intergenic | NA | NA |
| T10569 | GGCGCTCGCCGCCGTCGAGGAGG | 1000 | | | | | | | | | |
| Oligo fwd | CACCGGCGCTCGCCGCCGTCGAGG | | | | | | | | | | |
| Oligo rev | AAACCCTCGACGGCGGCGAGCGCC | | | | | | | | | | |
| Chromosome | start | end | strand | MM | target_seq | PAM | alignment | distance | position | gene name | gene id |
| GU305914.1 | 138477 | 138499 | + | 0 | GGCGCTCGCCGCCGTCGAGG | AGG | \|\|\|\|\|\|\|\|[\|\|\|\|\|\|\|\|\|\|\|\|]PAM | NA | Intergenic | NA | NA |
| GU305914.1 | 81258 | 81280 | - | 4 | GGCGGCCGCCGCGATCGAGG | AGG | \|\|\|\|--\|\|[\|\|\|\|--\|\|\|\|\|\|]PAM | NA | Intergenic | NA | NA |
| GU305914.1 | 91034 | 91056 | - | 4 | GAAGCTCACCGCCGTCGAGA | GGG | \|--\|\|\|\|-[\|\|\|\|\|\|\|\|\|\|\|-]PAM | NA | Intergenic | NA | NA |
| GU305914.1 | 88512 | 88534 | + | 4 | GGAGCCCGCCGCCGTCGCGT | TGG | \|\|-\|\|-\|\|[\|\|\|\|\|\|\|\|\|-\|-]PAM | NA | Intergenic | NA | NA |
| T10570 | GCGCTCGCCGCCGTCGAGGAGGG | 1000 | | | | | | | | | |
| Oligo fwd | CACCGCGCTCGCCGCCGTCGAGGA | | | | | | | | | | |
| Oligo rev | AAACTCCTCGACGGCGGCGAGCGC | | | | | | | | | | |
| Chromosome | start | end | strand | MM | target_seq | PAM | alignment | distance | position | gene name | gene id |
| GU305914.1 | 138478 | 138500 | + | 0 | GCGCTCGCCGCCGTCGAGGA | GGG | \|\|\|\|\|\|\|\|[\|\|\|\|\|\|\|\|\|\|\|\|]PAM | NA | Intergenic | NA | NA |
| T10571 | TCGCCGCCGTCGAGGAGGGTCGG | 1000 | | | | | | | | | |
| Oligo fwd | CACCTCGCCGCCGTCGAGGAGGGT | | | | | | | | | | |
| Oligo rev | AAACACCCTCCTCGACGGCGGCGA | | | | | | | | | | |
| Chromosome | start | end | strand | MM | target_seq | PAM | alignment | distance | position | gene name | gene id |
| GU305914.1 | 138482 | 138504 | + | 0 | TCGCCGCCGTCGAGGAGGGT | CGG | \|\|\|\|\|\|\|\|[\|\|\|\|\|\|\|\|\|\|\|\|]PAM | NA | Intergenic | NA | NA |
| T10572 | CGCCGCCGTCGAGGAGGGTCGGG | 1000 | | | | | | | | | |
| Oligo fwd | CACCCGCCGCCGTCGAGGAGGGTC | | | | | | | | | | |
| Oligo rev | AAACGACCCTCCTCGACGGCGGCG | | | | | | | | | | |
| Chromosome | start | end | strand | MM | target_seq | PAM | alignment | distance | position | gene name | gene id |
| GU305914.1 | 138483 | 138505 | + | 0 | CGCCGCCGTCGAGGAGGGTC | GGG | \|\|\|\|\|\|\|\|[\|\|\|\|\|\|\|\|\|\|\|\|]PAM | NA | Intergenic | NA | NA |
| T10573 | CGAGGAGGGTCGGGTGTCCGAGG | 1000 | | | | | | | | | |
| Oligo fwd | CACCCGAGGAGGGTCGGGTGTCCG | | | | | | | | | | |
| Oligo rev | AAACCGGACACCCGACCCTCCTCG | | | | | | | | | | |
| Chromosome | start | end | strand | MM | target_seq | PAM | alignment | distance | position | gene name | gene id |
| GU305914.1 | 138492 | 138514 | + | 0 | CGAGGAGGGTCGGGTGTCCG | AGG | \|\|\|\|\|\|\|\|[\|\|\|\|\|\|\|\|\|\|\|\|]PAM | NA | Intergenic | NA | NA |
| T10574 | GGAGGGTCGGGTGTCCGAGGCGG | 1000 | | | | | | | | | |
| Oligo fwd | CACCGGAGGGTCGGGTGTCCGAGG | | | | | | | | | | |
| Oligo rev | AAACCCTCGGACACCCGACCCTCC | | | | | | | | | | |
| Chromosome | start | end | strand | MM | target_seq | PAM | alignment | distance | position | gene name | gene id |
| GU305914.1 | 138495 | 138517 | + | 0 | GGAGGGTCGGGTGTCCGAGG | CGG | \|\|\|\|\|\|\|\|[\|\|\|\|\|\|\|\|\|\|\|\|]PAM | NA | Intergenic | NA | NA |
| T10575 | TGTCCGAGGCGGACGTCAACAGG | 1000 | | | | | | | | | |
| Oligo fwd | CACCTGTCCGAGGCGGACGTCAAC | | | | | | | | | | |
| Oligo rev | AAACGTTGACGTCCGCCTCGGACA | | | | | | | | | | |
| Chromosome | start | end | strand | MM | target_seq | PAM | alignment | distance | position | gene name | gene id |
| GU305914.1 | 138506 | 138528 | + | 0 | TGTCCGAGGCGGACGTCAAC | AGG | \|\|\|\|\|\|\|\|[\|\|\|\|\|\|\|\|\|\|\|\|]PAM | NA | Intergenic | NA | NA |
| T10576 | AGGCGGACGTCAACAGGTACAGG | 1000 | | | | | | | | | |
| Oligo fwd | CACCAGGCGGACGTCAACAGGTAC | | | | | | | | | | |
| Oligo rev | AAACGTACCTGTTGACGTCCGCCT | | | | | | | | | | |
| Chromosome | start | end | strand | MM | target_seq | PAM | alignment | distance | position | gene name | gene id |
| GU305914.1 | 138512 | 138534 | + | 0 | AGGCGGACGTCAACAGGTAC | AGG | \|\|\|\|\|\|\|\|[\|\|\|\|\|\|\|\|\|\|\|\|]PAM | NA | Intergenic | NA | NA |
| T10577 | GGCGGACGTCAACAGGTACAGGG | 1000 | | | | | | | | | |
| Oligo fwd | CACCGGCGGACGTCAACAGGTACA | | | | | | | | | | |
| Oligo rev | AAACTGTACCTGTTGACGTCCGCC | | | | | | | | | | |
| Chromosome | start | end | strand | MM | target_seq | PAM | alignment | distance | position | gene name | gene id |
| GU305914.1 | 138513 | 138535 | + | 0 | GGCGGACGTCAACAGGTACA | GGG | \|\|\|\|\|\|\|\|[\|\|\|\|\|\|\|\|\|\|\|\|]PAM | NA | Intergenic | NA | NA |
| T10578 | GGACGTCAACAGGTACAGGGAGG | 1000 | | | | | | | | | |
| Oligo fwd | CACCGGACGTCAACAGGTACAGGG | | | | | | | | | | |
| Oligo rev | AAACCCCTGTACCTGTTGACGTCC | | | | | | | | | | |
| Chromosome | start | end | strand | MM | target_seq | PAM | alignment | distance | position | gene name | gene id |
| GU305914.1 | 138516 | 138538 | + | 0 | GGACGTCAACAGGTACAGGG | AGG | \|\|\|\|\|\|\|\|[\|\|\|\|\|\|\|\|\|\|\|\|]PAM | NA | Intergenic | NA | NA |
| T10579 | CAACAGGTACAGGGAGGCCGTGG | 1000 | | | | | | | | | |
| Oligo fwd | CACCCAACAGGTACAGGGAGGCCG | | | | | | | | | | |
| Oligo rev | AAACCGGCCTCCCTGTACCTGTTG | | | | | | | | | | |
| Chromosome | start | end | strand | MM | target_seq | PAM | alignment | distance | position | gene name | gene id |
| GU305914.1 | 138522 | 138544 | + | 0 | CAACAGGTACAGGGAGGCCG | TGG | \|\|\|\|\|\|\|\|[\|\|\|\|\|\|\|\|\|\|\|\|]PAM | NA | Intergenic | NA | NA |
| T10580 | GTACAGGGAGGCCGTGGACGCGG | 1000 | | | | | | | | | |
| Oligo fwd | CACCGTACAGGGAGGCCGTGGACG | | | | | | | | | | |
| Oligo rev | AAACCGTCCACGGCCTCCCTGTAC | | | | | | | | | | |
| Chromosome | start | end | strand | MM | target_seq | PAM | alignment | distance | position | gene name | gene id |
| GU305914.1 | 138528 | 138550 | + | 0 | GTACAGGGAGGCCGTGGACG | CGG | \|\|\|\|\|\|\|\|[\|\|\|\|\|\|\|\|\|\|\|\|]PAM | NA | Intergenic | NA | NA |
| T10581 | CGCGGCCCTGATCGCGTGCGAGG | 1000 | | | | | | | | | |
| Oligo fwd | CACCCGCGGCCCTGATCGCGTGCG | | | | | | | | | | |
| Oligo rev | AAACCGCACGCGATCAGGGCCGCG | | | | | | | | | | |
| Chromosome | start | end | strand | MM | target_seq | PAM | alignment | distance | position | gene name | gene id |
| GU305914.1 | 138546 | 138568 | + | 0 | CGCGGCCCTGATCGCGTGCG | AGG | \|\|\|\|\|\|\|\|[\|\|\|\|\|\|\|\|\|\|\|\|]PAM | NA | Intergenic | NA | NA |

T10582 CGTGCGAGGCCTCCTCACCCAGG 1000
Oligo fwd CACCCGTGCGAGGCCTCCTCACCC
Oligo rev AAACGGGTGAGGAGGCCTCGCACG
Chromosome start end strand MM target_seq PAM alignment distance position gene name gene id
GU305914.1 138560 138582 + 0 CGTGCGAGGCCTCCTCACCC AGG ||||||||[||||||||||||]PAM NA Intergenic NA NA

T10583 GTGCGAGGCCTCCTCACCCAGGG 1000
Oligo fwd CACCGTGCGAGGCCTCCTCACCCA
Oligo rev AAACTGGGTGAGGAGGCCTCGCAC
Chromosome start end strand MM target_seq PAM alignment distance position gene name gene id
GU305914.1 138561 138583 + 0 GTGCGAGGCCTCCTCACCCA GGG ||||||||[||||||||||||]PAM NA Intergenic NA NA

T10584 AGGCCTCCTCACCCAGGGACAGG 1000
Oligo fwd CACCAGGCCTCCTCACCCAGGGAC
Oligo rev AAACGTCCCTGGGTGAGGAGGCCT
Chromosome start end strand MM target_seq PAM alignment distance position gene name gene id
GU305914.1 138566 138588 + 0 AGGCCTCCTCACCCAGGGAC AGG ||||||||[||||||||||||]PAM NA Intergenic NA NA

T10585 CAGGTTTCGCCTGATCGAGACGG 1000
Oligo fwd CACCCAGGTTTCGCCTGATCGAGA
Oligo rev AAACTCTCGATCAGGCGAAACCTG
Chromosome start end strand MM target_seq PAM alignment distance position gene name gene id
GU305914.1 138585 138607 + 0 CAGGTTTCGCCTGATCGAGA CGG ||||||||[||||||||||||]PAM NA Intergenic NA NA

T10586 TTTCGCCTGATCGAGACGGCCGG 1000
Oligo fwd CACCTTTCGCCTGATCGAGACGGC
Oligo rev AAACGCCGTCTCGATCAGGCGAAA
Chromosome start end strand MM target_seq PAM alignment distance position gene name gene id
GU305914.1 138589 138611 + 0 TTTCGCCTGATCGAGACGGC CGG ||||||||[||||||||||||]PAM NA Intergenic NA NA

T10587 TTCGCCTGATCGAGACGGCCGGG 1000
Oligo fwd CACCTTCGCCTGATCGAGACGGCC
Oligo rev AAACGGCCGTCTCGATCAGGCGAA
Chromosome start end strand MM target_seq PAM alignment distance position gene name gene id
GU305914.1 138590 138612 + 0 TTCGCCTGATCGAGACGGCC GGG ||||||||[||||||||||||]PAM NA Intergenic NA NA

T10588 TCGCCTGATCGAGACGGCCGGGG 1000
Oligo fwd CACCTCGCCTGATCGAGACGGCCG
Oligo rev AAACCGGCCGTCTCGATCAGGCGA
Chromosome start end strand MM target_seq PAM alignment distance position gene name gene id
GU305914.1 138591 138613 + 0 TCGCCTGATCGAGACGGCCG GGG ||||||||[||||||||||||]PAM NA Intergenic NA NA

T10589 CGCCTGATCGAGACGGCCGGGGG 1000
Oligo fwd CACCCGCCTGATCGAGACGGCCGG
Oligo rev AAACCCGGCCGTCTCGATCAGGCG
Chromosome start end strand MM target_seq PAM alignment distance position gene name gene id
GU305914.1 138592 138614 + 0 CGCCTGATCGAGACGGCCGG GGG ||||||||[||||||||||||]PAM NA Intergenic NA NA

T10590 CGTCACCAACGCGCTGCCCAAGG 1000
Oligo fwd CACCCGTCACCAACGCGCTGCCCA
Oligo rev AAACTGGGCAGCGCGTTGGTGACG
Chromosome start end strand MM target_seq PAM alignment distance position gene name gene id
GU305914.1 138627 138649 + 0 CGTCACCAACGCGCTGCCCA AGG ||||||||[||||||||||||]PAM NA Intergenic NA NA

T10591 CCAACGCGCTGCCCAAGGACCGG 1000
Oligo fwd CACCCCAACGCGCTGCCCAAGGAC
Oligo rev AAACGTCCTTGGGCAGCGCGTTGG
Chromosome start end strand MM target_seq PAM alignment distance position gene name gene id
GU305914.1 138632 138654 + 0 CCAACGCGCTGCCCAAGGAC CGG ||||||||[||||||||||||]PAM NA Intergenic NA NA
GU305914.1 110791 110813 + 3 CCATCACGCTGCCCAAGCAC AGG |||-|-||[|||||||||-||]PAM NA Intergenic NA NA

T10592 CGCGCTGCCCAAGGACCGGACGG 1000
Oligo fwd CACCCGCGCTGCCCAAGGACCGGA
Oligo rev AAACTCCGGTCCTTGGGCAGCGCG
Chromosome start end strand MM target_seq PAM alignment distance position gene name gene id
GU305914.1 138636 138658 + 0 CGCGCTGCCCAAGGACCGGA CGG ||||||||[||||||||||||]PAM NA Intergenic NA NA

T10593 ACAACCGCCGTGCGTCCTCGAGG 1000
Oligo fwd CACCACAACCGCCGTGCGTCCTCG
Oligo rev AAACCGAGGACGCACGGCGGTTGT
Chromosome start end strand MM target_seq PAM alignment distance position gene name gene id
GU305914.1 138663 138685 + 0 ACAACCGCCGTGCGTCCTCG AGG ||||||||[||||||||||||]PAM NA Intergenic NA NA

T10594 CAACCGCCGTGCGTCCTCGAGGG 1000
Oligo fwd CACCCAACCGCCGTGCGTCCTCGA
Oligo rev AAACTCGAGGACGCACGGCGGTTG
Chromosome start end strand MM target_seq PAM alignment distance position gene name gene id
GU305914.1 138664 138686 + 0 CAACCGCCGTGCGTCCTCGA GGG ||||||||[||||||||||||]PAM NA Intergenic NA NA

T10595 AACCGCCGTGCGTCCTCGAGGGG 1000
Oligo fwd CACCAACCGCCGTGCGTCCTCGAG
Oligo rev AAACCTCGAGGACGCACGGCGGTT
Chromosome start end strand MM target_seq PAM alignment distance position gene name gene id
GU305914.1 138665 138687 + 0 AACCGCCGTGCGTCCTCGAG GGG ||||||||[||||||||||||]PAM NA Intergenic NA NA
GU305914.1 41413 41435 + 4 TCCCGCCGCGCGCCCTCGAG CGG --||||||[-|||-|||||||]PAM NA Intergenic NA NA

T10596 GCCGTGCGTCCTCGAGGGGTCGG 1000
Oligo fwd CACCGCCGTGCGTCCTCGAGGGGT
Oligo rev AAACACCCCTCGAGGACGCACGGC

<table>
<tr><td>Chromosome</td><td>start</td><td>end</td><td>strand</td><td>MM</td><td>target_seq</td><td>PAM</td><td>alignment</td><td>distance</td><td>position</td><td>gene name</td><td>gene id</td></tr>
<tr><td>GU305914.1</td><td>138669</td><td>138691</td><td>+</td><td>0</td><td>GCCGTGCGTCCTCGAGGGGT</td><td>CGG</td><td>||||||||[||||||||||||]PAM</td><td>NA</td><td>Intergenic</td><td>NA</td><td>NA</td></tr>
<tr><td>T10597</td><td>CCGTGCGTCCTCGAGGGGTCGGG</td><td>1000</td></tr>
<tr><td>Oligo fwd</td><td>CACCCCGTGCGTCCTCGAGGGGTC</td></tr>
<tr><td>Oligo rev</td><td>AAACGACCCCTCGAGGACGCACGG</td></tr>
<tr><td>Chromosome</td><td>start</td><td>end</td><td>strand</td><td>MM</td><td>target_seq</td><td>PAM</td><td>alignment</td><td>distance</td><td>position</td><td>gene name</td><td>gene id</td></tr>
<tr><td>GU305914.1</td><td>138670</td><td>138692</td><td>+</td><td>0</td><td>CCGTGCGTCCTCGAGGGGTC</td><td>GGG</td><td>||||||||[||||||||||||]PAM</td><td>NA</td><td>Intergenic</td><td>NA</td><td>NA</td></tr>
<tr><td>T10598</td><td>CGTGCGTCCTCGAGGGGTCGGGG</td><td>1000</td></tr>
<tr><td>Oligo fwd</td><td>CACCCGTGCGTCCTCGAGGGGTCG</td></tr>
<tr><td>Oligo rev</td><td>AAACCGACCCCTCGAGGACGCACG</td></tr>
<tr><td>Chromosome</td><td>start</td><td>end</td><td>strand</td><td>MM</td><td>target_seq</td><td>PAM</td><td>alignment</td><td>distance</td><td>position</td><td>gene name</td><td>gene id</td></tr>
<tr><td>GU305914.1</td><td>138671</td><td>138693</td><td>+</td><td>0</td><td>CGTGCGTCCTCGAGGGGTCG</td><td>GGG</td><td>||||||||[||||||||||||]PAM</td><td>NA</td><td>Intergenic</td><td>NA</td><td>NA</td></tr>
<tr><td>T10599</td><td>GCGTCCTCGAGGGGTCGGGGCGG</td><td>1000</td></tr>
<tr><td>Oligo fwd</td><td>CACCGCGTCCTCGAGGGGTCGGGG</td></tr>
<tr><td>Oligo rev</td><td>AAACCCCCGACCCCTCGAGGACGC</td></tr>
<tr><td>Chromosome</td><td>start</td><td>end</td><td>strand</td><td>MM</td><td>target_seq</td><td>PAM</td><td>alignment</td><td>distance</td><td>position</td><td>gene name</td><td>gene id</td></tr>
<tr><td>GU305914.1</td><td>138674</td><td>138696</td><td>+</td><td>0</td><td>GCGTCCTCGAGGGGTCGGGG</td><td>CGG</td><td>||||||||[||||||||||||]PAM</td><td>NA</td><td>Intergenic</td><td>NA</td><td>NA</td></tr>
<tr><td>T10600</td><td>CGTCCTCGAGGGGTCGGGGCGGG</td><td>1000</td></tr>
<tr><td>Oligo fwd</td><td>CACCCGTCCTCGAGGGGTCGGGGC</td></tr>
<tr><td>Oligo rev</td><td>AAACGCCCCGACCCCTCGAGGACG</td></tr>
<tr><td>Chromosome</td><td>start</td><td>end</td><td>strand</td><td>MM</td><td>target_seq</td><td>PAM</td><td>alignment</td><td>distance</td><td>position</td><td>gene name</td><td>gene id</td></tr>
<tr><td>GU305914.1</td><td>138675</td><td>138697</td><td>+</td><td>0</td><td>CGTCCTCGAGGGGTCGGGGC</td><td>GGG</td><td>||||||||[||||||||||||]PAM</td><td>NA</td><td>Intergenic</td><td>NA</td><td>NA</td></tr>
<tr><td>T10601</td><td>AGCAGCAACAACAACTACGAAGG</td><td>1000</td></tr>
<tr><td>Oligo fwd</td><td>CACCAGCAGCAACAACAACTACGA</td></tr>
<tr><td>Oligo rev</td><td>AAACTCGTAGTTGTTGTTGCTGCT</td></tr>
<tr><td>Chromosome</td><td>start</td><td>end</td><td>strand</td><td>MM</td><td>target_seq</td><td>PAM</td><td>alignment</td><td>distance</td><td>position</td><td>gene name</td><td>gene id</td></tr>
<tr><td>GU305914.1</td><td>138700</td><td>138722</td><td>+</td><td>0</td><td>AGCAGCAACAACAACTACGA</td><td>AGG</td><td>||||||||[||||||||||||]PAM</td><td>NA</td><td>Intergenic</td><td>NA</td><td>NA</td></tr>
<tr><td>T10602</td><td>GCAGCAACAACAACTACGAAGGG</td><td>1000</td></tr>
<tr><td>Oligo fwd</td><td>CACCGCAGCAACAACAACTACGAA</td></tr>
<tr><td>Oligo rev</td><td>AAACTTCGTAGTTGTTGTTGCTGC</td></tr>
<tr><td>Chromosome</td><td>start</td><td>end</td><td>strand</td><td>MM</td><td>target_seq</td><td>PAM</td><td>alignment</td><td>distance</td><td>position</td><td>gene name</td><td>gene id</td></tr>
<tr><td>GU305914.1</td><td>138701</td><td>138723</td><td>+</td><td>0</td><td>GCAGCAACAACAACTACGAA</td><td>GGG</td><td>||||||||[||||||||||||]PAM</td><td>NA</td><td>Intergenic</td><td>NA</td><td>NA</td></tr>
<tr><td>T10603</td><td>AACAACAACTACGAAGGGATAGG</td><td>1000</td></tr>
<tr><td>Oligo fwd</td><td>CACCAACAACAACTACGAAGGGAT</td></tr>
<tr><td>Oligo rev</td><td>AAACATCCCTTCGTAGTTGTTGTT</td></tr>
<tr><td>Chromosome</td><td>start</td><td>end</td><td>strand</td><td>MM</td><td>target_seq</td><td>PAM</td><td>alignment</td><td>distance</td><td>position</td><td>gene name</td><td>gene id</td></tr>
<tr><td>GU305914.1</td><td>138706</td><td>138728</td><td>+</td><td>0</td><td>AACAACAACTACGAAGGGAT</td><td>AGG</td><td>||||||||[||||||||||||]PAM</td><td>NA</td><td>Intergenic</td><td>NA</td><td>NA</td></tr>
<tr><td>T10604</td><td>GGGATAGGAACGCCGTCCGCAGG</td><td>1000</td></tr>
<tr><td>Oligo fwd</td><td>CACCGGGATAGGAACGCCGTCCGC</td></tr>
<tr><td>Oligo rev</td><td>AAACGCGGACGGCGTTCCTATCCC</td></tr>
<tr><td>Chromosome</td><td>start</td><td>end</td><td>strand</td><td>MM</td><td>target_seq</td><td>PAM</td><td>alignment</td><td>distance</td><td>position</td><td>gene name</td><td>gene id</td></tr>
<tr><td>GU305914.1</td><td>138721</td><td>138743</td><td>+</td><td>0</td><td>GGGATAGGAACGCCGTCCGC</td><td>AGG</td><td>||||||||[||||||||||||]PAM</td><td>NA</td><td>Intergenic</td><td>NA</td><td>NA</td></tr>
<tr><td>T10605</td><td>GGAACGCCGTCCGCAGGTAGCGG</td><td>1000</td></tr>
<tr><td>Oligo fwd</td><td>CACCGGAACGCCGTCCGCAGGTAG</td></tr>
<tr><td>Oligo rev</td><td>AAACCTACCTGCGGACGGCGTTCC</td></tr>
<tr><td>Chromosome</td><td>start</td><td>end</td><td>strand</td><td>MM</td><td>target_seq</td><td>PAM</td><td>alignment</td><td>distance</td><td>position</td><td>gene name</td><td>gene id</td></tr>
<tr><td>GU305914.1</td><td>138727</td><td>138749</td><td>+</td><td>0</td><td>GGAACGCCGTCCGCAGGTAG</td><td>CGG</td><td>||||||||[||||||||||||]PAM</td><td>NA</td><td>Intergenic</td><td>NA</td><td>NA</td></tr>
<tr><td>T10606</td><td>GGTAGCGGTAACGCGTTCGACGG</td><td>1000</td></tr>
<tr><td>Oligo fwd</td><td>CACCGGTAGCGGTAACGCGTTCGA</td></tr>
<tr><td>Oligo rev</td><td>AAACTCGAACGCGTTACCGCTACC</td></tr>
<tr><td>Chromosome</td><td>start</td><td>end</td><td>strand</td><td>MM</td><td>target_seq</td><td>PAM</td><td>alignment</td><td>distance</td><td>position</td><td>gene name</td><td>gene id</td></tr>
<tr><td>GU305914.1</td><td>138742</td><td>138764</td><td>+</td><td>0</td><td>GGTAGCGGTAACGCGTTCGA</td><td>CGG</td><td>||||||||[||||||||||||]PAM</td><td>NA</td><td>Intergenic</td><td>NA</td><td>NA</td></tr>
<tr><td>T10607</td><td>GTAGCGGTAACGCGTTCGACGGG</td><td>1000</td></tr>
<tr><td>Oligo fwd</td><td>CACCGTAGCGGTAACGCGTTCGAC</td></tr>
<tr><td>Oligo rev</td><td>AAACGTCGAACGCGTTACCGCTAC</td></tr>
<tr><td>Chromosome</td><td>start</td><td>end</td><td>strand</td><td>MM</td><td>target_seq</td><td>PAM</td><td>alignment</td><td>distance</td><td>position</td><td>gene name</td><td>gene id</td></tr>
<tr><td>GU305914.1</td><td>138743</td><td>138765</td><td>+</td><td>0</td><td>GTAGCGGTAACGCGTTCGAC</td><td>GGG</td><td>||||||||[||||||||||||]PAM</td><td>NA</td><td>Intergenic</td><td>NA</td><td>NA</td></tr>
<tr><td>T10608</td><td>TAACGCGTTCGACGGGCTGTTGG</td><td>1000</td></tr>
<tr><td>Oligo fwd</td><td>CACCTAACGCGTTCGACGGGCTGT</td></tr>
<tr><td>Oligo rev</td><td>AAACACAGCCCGTCGAACGCGTTA</td></tr>
<tr><td>Chromosome</td><td>start</td><td>end</td><td>strand</td><td>MM</td><td>target_seq</td><td>PAM</td><td>alignment</td><td>distance</td><td>position</td><td>gene name</td><td>gene id</td></tr>
<tr><td>GU305914.1</td><td>138750</td><td>138772</td><td>+</td><td>0</td><td>TAACGCGTTCGACGGGCTGT</td><td>TGG</td><td>||||||||[||||||||||||]PAM</td><td>NA</td><td>Intergenic</td><td>NA</td><td>NA</td></tr>
<tr><td>T10609</td><td>GGGCTGTTGGCGCTCGAACGCGG</td><td>1000</td></tr>
<tr><td>Oligo fwd</td><td>CACCGGGCTGTTGGCGCTCGAACG</td></tr>
<tr><td>Oligo rev</td><td>AAACCGTTCGAGCGCCAACAGCCC</td></tr>
<tr><td>Chromosome</td><td>start</td><td>end</td><td>strand</td><td>MM</td><td>target_seq</td><td>PAM</td><td>alignment</td><td>distance</td><td>position</td><td>gene name</td><td>gene id</td></tr>
<tr><td>GU305914.1</td><td>138763</td><td>138785</td><td>+</td><td>0</td><td>GGGCTGTTGGCGCTCGAACG</td><td>CGG</td><td>||||||||[||||||||||||]PAM</td><td>NA</td><td>Intergenic</td><td>NA</td><td>NA</td></tr>
<tr><td>T10610</td><td>GCGCTCGAACGCGGCACGAGCGG</td><td>1000</td></tr>
<tr><td>Oligo fwd</td><td>CACCGCGCTCGAACGCGGCACGAG</td></tr>
<tr><td>Oligo rev</td><td>AAACCTCGTGCCGCGTTCGAGCGC</td></tr>
<tr><td>Chromosome</td><td>start</td><td>end</td><td>strand</td><td>MM</td><td>target_seq</td><td>PAM</td><td>alignment</td><td>distance</td><td>position</td><td>gene name</td><td>gene id</td></tr>
<tr><td>GU305914.1</td><td>138772</td><td>138794</td><td>+</td><td>0</td><td>GCGCTCGAACGCGGCACGAG</td><td>CGG</td><td>||||||||[||||||||||||]PAM</td><td>NA</td><td>Intergenic</td><td>NA</td><td>NA</td></tr>
<tr><td>T10611</td><td>CTCGAACGCGGCACGAGCGGCGG</td><td>1000</td></tr>
<tr><td>Oligo fwd</td><td>CACCCTCGAACGCGGCACGAGCGG</td></tr>
</table>

<table>
<tr><td>Oligo rev</td><td>AAACCCGCTCGTGCCGCGTTCGAG</td><td></td><td></td><td></td><td></td><td></td><td></td><td></td><td></td><td></td><td></td></tr>
<tr><td>Chromosome</td><td>start</td><td>end</td><td>strand</td><td>MM</td><td>target_seq</td><td>PAM</td><td>alignment</td><td>distance</td><td>position</td><td>gene name</td><td>gene id</td></tr>
<tr><td>GU305914.1</td><td>138775</td><td>138797</td><td>+</td><td>0</td><td>CTCGAACGCGGCACGAGCGG</td><td>CGG</td><td>||||||||[||||||||||||]PAM</td><td>NA</td><td>Intergenic</td><td>NA</td><td>NA</td></tr>
<tr><td></td><td></td><td></td><td></td><td></td><td></td><td></td><td></td><td></td><td></td><td></td><td></td></tr>
<tr><td>T10612</td><td>GAACGCGGCACGAGCGGCGGCGG</td><td>1000</td><td></td><td></td><td></td><td></td><td></td><td></td><td></td><td></td><td></td></tr>
<tr><td>Oligo fwd</td><td>CACCGAACGCGGCACGAGCGGCGG</td><td></td><td></td><td></td><td></td><td></td><td></td><td></td><td></td><td></td><td></td></tr>
<tr><td>Oligo rev</td><td>AAACCCGCCGCTCGTGCCGCGTTC</td><td></td><td></td><td></td><td></td><td></td><td></td><td></td><td></td><td></td><td></td></tr>
<tr><td>Chromosome</td><td>start</td><td>end</td><td>strand</td><td>MM</td><td>target_seq</td><td>PAM</td><td>alignment</td><td>distance</td><td>position</td><td>gene name</td><td>gene id</td></tr>
<tr><td>GU305914.1</td><td>138778</td><td>138800</td><td>+</td><td>0</td><td>GAACGCGGCACGAGCGGCGG</td><td>CGG</td><td>||||||||[||||||||||||]PAM</td><td>NA</td><td>Intergenic</td><td>NA</td><td>NA</td></tr>
<tr><td></td><td></td><td></td><td></td><td></td><td></td><td></td><td></td><td></td><td></td><td></td><td></td></tr>
<tr><td>T10613</td><td>CACGAGCGGCGGCGGACTGACGG</td><td>1000</td><td></td><td></td><td></td><td></td><td></td><td></td><td></td><td></td><td></td></tr>
<tr><td>Oligo fwd</td><td>CACCCACGAGCGGCGGCGGACTGA</td><td></td><td></td><td></td><td></td><td></td><td></td><td></td><td></td><td></td><td></td></tr>
<tr><td>Oligo rev</td><td>AAACTCAGTCCGCCGCCGCTCGTG</td><td></td><td></td><td></td><td></td><td></td><td></td><td></td><td></td><td></td><td></td></tr>
<tr><td>Chromosome</td><td>start</td><td>end</td><td>strand</td><td>MM</td><td>target_seq</td><td>PAM</td><td>alignment</td><td>distance</td><td>position</td><td>gene name</td><td>gene id</td></tr>
<tr><td>GU305914.1</td><td>138786</td><td>138808</td><td>+</td><td>0</td><td>CACGAGCGGCGGCGGACTGA</td><td>CGG</td><td>||||||||[||||||||||||]PAM</td><td>NA</td><td>Intergenic</td><td>NA</td><td>NA</td></tr>
<tr><td></td><td></td><td></td><td></td><td></td><td></td><td></td><td></td><td></td><td></td><td></td><td></td></tr>
<tr><td>T10614</td><td>CGGCGGCGGACTGACGGCGACGG</td><td>1000</td><td></td><td></td><td></td><td></td><td></td><td></td><td></td><td></td><td></td></tr>
<tr><td>Oligo fwd</td><td>CACCCGGCGGCGGACTGACGGCGA</td><td></td><td></td><td></td><td></td><td></td><td></td><td></td><td></td><td></td><td></td></tr>
<tr><td>Oligo rev</td><td>AAACTCGCCGTCAGTCCGCCGCCG</td><td></td><td></td><td></td><td></td><td></td><td></td><td></td><td></td><td></td><td></td></tr>
<tr><td>Chromosome</td><td>start</td><td>end</td><td>strand</td><td>MM</td><td>target_seq</td><td>PAM</td><td>alignment</td><td>distance</td><td>position</td><td>gene name</td><td>gene id</td></tr>
<tr><td>GU305914.1</td><td>138792</td><td>138814</td><td>+</td><td>0</td><td>CGGCGGCGGACTGACGGCGA</td><td>CGG</td><td>||||||||[||||||||||||]PAM</td><td>NA</td><td>Intergenic</td><td>NA</td><td>NA</td></tr>
<tr><td></td><td></td><td></td><td></td><td></td><td></td><td></td><td></td><td></td><td></td><td></td><td></td></tr>
<tr><td>T10615</td><td>GGCGACGGTGCCGTCCGCTCCGG</td><td>1000</td><td></td><td></td><td></td><td></td><td></td><td></td><td></td><td></td><td></td></tr>
<tr><td>Oligo fwd</td><td>CACCGGCGACGGTGCCGTCCGCTC</td><td></td><td></td><td></td><td></td><td></td><td></td><td></td><td></td><td></td><td></td></tr>
<tr><td>Oligo rev</td><td>AAACGAGCGGACGGCACCGTCGCC</td><td></td><td></td><td></td><td></td><td></td><td></td><td></td><td></td><td></td><td></td></tr>
<tr><td>Chromosome</td><td>start</td><td>end</td><td>strand</td><td>MM</td><td>target_seq</td><td>PAM</td><td>alignment</td><td>distance</td><td>position</td><td>gene name</td><td>gene id</td></tr>
<tr><td>GU305914.1</td><td>138807</td><td>138829</td><td>+</td><td>0</td><td>GGCGACGGTGCCGTCCGCTC</td><td>CGG</td><td>||||||||[||||||||||||]PAM</td><td>NA</td><td>Intergenic</td><td>NA</td><td>NA</td></tr>
<tr><td></td><td></td><td></td><td></td><td></td><td></td><td></td><td></td><td></td><td></td><td></td><td></td></tr>
<tr><td>T10616</td><td>GCGACGGTGCCGTCCGCTCCGGG</td><td>1000</td><td></td><td></td><td></td><td></td><td></td><td></td><td></td><td></td><td></td></tr>
<tr><td>Oligo fwd</td><td>CACCGCGACGGTGCCGTCCGCTCC</td><td></td><td></td><td></td><td></td><td></td><td></td><td></td><td></td><td></td><td></td></tr>
<tr><td>Oligo rev</td><td>AAACGGAGCGGACGGCACCGTCGC</td><td></td><td></td><td></td><td></td><td></td><td></td><td></td><td></td><td></td><td></td></tr>
<tr><td>Chromosome</td><td>start</td><td>end</td><td>strand</td><td>MM</td><td>target_seq</td><td>PAM</td><td>alignment</td><td>distance</td><td>position</td><td>gene name</td><td>gene id</td></tr>
<tr><td>GU305914.1</td><td>138808</td><td>138830</td><td>+</td><td>0</td><td>GCGACGGTGCCGTCCGCTCC</td><td>GGG</td><td>||||||||[||||||||||||]PAM</td><td>NA</td><td>Intergenic</td><td>NA</td><td>NA</td></tr>
<tr><td></td><td></td><td></td><td></td><td></td><td></td><td></td><td></td><td></td><td></td><td></td><td></td></tr>
<tr><td>T10617</td><td>TCCGGGATATGTAGCGAAATCGG</td><td>1000</td><td></td><td></td><td></td><td></td><td></td><td></td><td></td><td></td><td></td></tr>
<tr><td>Oligo fwd</td><td>CACCTCCGGGATATGTAGCGAAAT</td><td></td><td></td><td></td><td></td><td></td><td></td><td></td><td></td><td></td><td></td></tr>
<tr><td>Oligo rev</td><td>AAACATTTCGCTACATATCCCGGA</td><td></td><td></td><td></td><td></td><td></td><td></td><td></td><td></td><td></td><td></td></tr>
<tr><td>Chromosome</td><td>start</td><td>end</td><td>strand</td><td>MM</td><td>target_seq</td><td>PAM</td><td>alignment</td><td>distance</td><td>position</td><td>gene name</td><td>gene id</td></tr>
<tr><td>GU305914.1</td><td>138825</td><td>138847</td><td>+</td><td>0</td><td>TCCGGGATATGTAGCGAAAT</td><td>CGG</td><td>||||||||[||||||||||||]PAM</td><td>NA</td><td>Intergenic</td><td>NA</td><td>NA</td></tr>
<tr><td></td><td></td><td></td><td></td><td></td><td></td><td></td><td></td><td></td><td></td><td></td><td></td></tr>
<tr><td>T10618</td><td>GTGAACACGCTGTCTTACGACGG</td><td>1000</td><td></td><td></td><td></td><td></td><td></td><td></td><td></td><td></td><td></td></tr>
<tr><td>Oligo fwd</td><td>CACCGTGAACACGCTGTCTTACGA</td><td></td><td></td><td></td><td></td><td></td><td></td><td></td><td></td><td></td><td></td></tr>
<tr><td>Oligo rev</td><td>AAACTCGTAAGACAGCGTGTTCAC</td><td></td><td></td><td></td><td></td><td></td><td></td><td></td><td></td><td></td><td></td></tr>
<tr><td>Chromosome</td><td>start</td><td>end</td><td>strand</td><td>MM</td><td>target_seq</td><td>PAM</td><td>alignment</td><td>distance</td><td>position</td><td>gene name</td><td>gene id</td></tr>
<tr><td>GU305914.1</td><td>138847</td><td>138869</td><td>+</td><td>0</td><td>GTGAACACGCTGTCTTACGA</td><td>CGG</td><td>||||||||[||||||||||||]PAM</td><td>NA</td><td>Intergenic</td><td>NA</td><td>NA</td></tr>
<tr><td></td><td></td><td></td><td></td><td></td><td></td><td></td><td></td><td></td><td></td><td></td><td></td></tr>
<tr><td>T10619</td><td>TGAACACGCTGTCTTACGACGGG</td><td>1000</td><td></td><td></td><td></td><td></td><td></td><td></td><td></td><td></td><td></td></tr>
<tr><td>Oligo fwd</td><td>CACCTGAACACGCTGTCTTACGAC</td><td></td><td></td><td></td><td></td><td></td><td></td><td></td><td></td><td></td><td></td></tr>
<tr><td>Oligo rev</td><td>AAACGTCGTAAGACAGCGTGTTCA</td><td></td><td></td><td></td><td></td><td></td><td></td><td></td><td></td><td></td><td></td></tr>
<tr><td>Chromosome</td><td>start</td><td>end</td><td>strand</td><td>MM</td><td>target_seq</td><td>PAM</td><td>alignment</td><td>distance</td><td>position</td><td>gene name</td><td>gene id</td></tr>
<tr><td>GU305914.1</td><td>138848</td><td>138870</td><td>+</td><td>0</td><td>TGAACACGCTGTCTTACGAC</td><td>GGG</td><td>||||||||[||||||||||||]PAM</td><td>NA</td><td>Intergenic</td><td>NA</td><td>NA</td></tr>
<tr><td></td><td></td><td></td><td></td><td></td><td></td><td></td><td></td><td></td><td></td><td></td><td></td></tr>
<tr><td>T10620</td><td>CTCCTACGTGCTTTACACTAAGG</td><td>1000</td><td></td><td></td><td></td><td></td><td></td><td></td><td></td><td></td><td></td></tr>
<tr><td>Oligo fwd</td><td>CACCCTCCTACGTGCTTTACACTA</td><td></td><td></td><td></td><td></td><td></td><td></td><td></td><td></td><td></td><td></td></tr>
<tr><td>Oligo rev</td><td>AAACTAGTGTAAAGCACGTAGGAG</td><td></td><td></td><td></td><td></td><td></td><td></td><td></td><td></td><td></td><td></td></tr>
<tr><td>Chromosome</td><td>start</td><td>end</td><td>strand</td><td>MM</td><td>target_seq</td><td>PAM</td><td>alignment</td><td>distance</td><td>position</td><td>gene name</td><td>gene id</td></tr>
<tr><td>GU305914.1</td><td>138885</td><td>138907</td><td>+</td><td>0</td><td>CTCCTACGTGCTTTACACTA</td><td>AGG</td><td>||||||||[||||||||||||]PAM</td><td>NA</td><td>Intergenic</td><td>NA</td><td>NA</td></tr>
<tr><td></td><td></td><td></td><td></td><td></td><td></td><td></td><td></td><td></td><td></td><td></td><td></td></tr>
<tr><td>T10621</td><td>TTTACACTAAGGAACAGCTGAGG</td><td>1000</td><td></td><td></td><td></td><td></td><td></td><td></td><td></td><td></td><td></td></tr>
<tr><td>Oligo fwd</td><td>CACCTTTACACTAAGGAACAGCTG</td><td></td><td></td><td></td><td></td><td></td><td></td><td></td><td></td><td></td><td></td></tr>
<tr><td>Oligo rev</td><td>AAACCAGCTGTTCCTTAGTGTAAA</td><td></td><td></td><td></td><td></td><td></td><td></td><td></td><td></td><td></td><td></td></tr>
<tr><td>Chromosome</td><td>start</td><td>end</td><td>strand</td><td>MM</td><td>target_seq</td><td>PAM</td><td>alignment</td><td>distance</td><td>position</td><td>gene name</td><td>gene id</td></tr>
<tr><td>GU305914.1</td><td>138896</td><td>138918</td><td>+</td><td>0</td><td>TTTACACTAAGGAACAGCTG</td><td>AGG</td><td>||||||||[||||||||||||]PAM</td><td>NA</td><td>Intergenic</td><td>NA</td><td>NA</td></tr>
<tr><td></td><td></td><td></td><td></td><td></td><td></td><td></td><td></td><td></td><td></td><td></td><td></td></tr>
<tr><td>T10622</td><td>AGTCGCTCTCTCCCGATAAGCGG</td><td>1000</td><td></td><td></td><td></td><td></td><td></td><td></td><td></td><td></td><td></td></tr>
<tr><td>Oligo fwd</td><td>CACCAGTCGCTCTCTCCCGATAAG</td><td></td><td></td><td></td><td></td><td></td><td></td><td></td><td></td><td></td><td></td></tr>
<tr><td>Oligo rev</td><td>AAACCTTATCGGGAGAGAGCGACT</td><td></td><td></td><td></td><td></td><td></td><td></td><td></td><td></td><td></td><td></td></tr>
<tr><td>Chromosome</td><td>start</td><td>end</td><td>strand</td><td>MM</td><td>target_seq</td><td>PAM</td><td>alignment</td><td>distance</td><td>position</td><td>gene name</td><td>gene id</td></tr>
<tr><td>GU305914.1</td><td>138920</td><td>138942</td><td>+</td><td>0</td><td>AGTCGCTCTCTCCCGATAAG</td><td>CGG</td><td>||||||||[||||||||||||]PAM</td><td>NA</td><td>Intergenic</td><td>NA</td><td>NA</td></tr>
<tr><td></td><td></td><td></td><td></td><td></td><td></td><td></td><td></td><td></td><td></td><td></td><td></td></tr>
<tr><td>T10623</td><td>GTCGCTCTCTCCCGATAAGCGGG</td><td>1000</td><td></td><td></td><td></td><td></td><td></td><td></td><td></td><td></td><td></td></tr>
<tr><td>Oligo fwd</td><td>CACCGTCGCTCTCTCCCGATAAGC</td><td></td><td></td><td></td><td></td><td></td><td></td><td></td><td></td><td></td><td></td></tr>
<tr><td>Oligo rev</td><td>AAACGCTTATCGGGAGAGAGCGAC</td><td></td><td></td><td></td><td></td><td></td><td></td><td></td><td></td><td></td><td></td></tr>
<tr><td>Chromosome</td><td>start</td><td>end</td><td>strand</td><td>MM</td><td>target_seq</td><td>PAM</td><td>alignment</td><td>distance</td><td>position</td><td>gene name</td><td>gene id</td></tr>
<tr><td>GU305914.1</td><td>138921</td><td>138943</td><td>+</td><td>0</td><td>GTCGCTCTCTCCCGATAAGC</td><td>GGG</td><td>||||||||[||||||||||||]PAM</td><td>NA</td><td>Intergenic</td><td>NA</td><td>NA</td></tr>
<tr><td></td><td></td><td></td><td></td><td></td><td></td><td></td><td></td><td></td><td></td><td></td><td></td></tr>
<tr><td>T10624</td><td>TCCCGATAAGCGGGCGATCGTGG</td><td>1000</td><td></td><td></td><td></td><td></td><td></td><td></td><td></td><td></td><td></td></tr>
<tr><td>Oligo fwd</td><td>CACCTCCCGATAAGCGGGCGATCG</td><td></td><td></td><td></td><td></td><td></td><td></td><td></td><td></td><td></td><td></td></tr>
<tr><td>Oligo rev</td><td>AAACCGATCGCCCGCTTATCGGGA</td><td></td><td></td><td></td><td></td><td></td><td></td><td></td><td></td><td></td><td></td></tr>
<tr><td>Chromosome</td><td>start</td><td>end</td><td>strand</td><td>MM</td><td>target_seq</td><td>PAM</td><td>alignment</td><td>distance</td><td>position</td><td>gene name</td><td>gene id</td></tr>
<tr><td>GU305914.1</td><td>138930</td><td>138952</td><td>+</td><td>0</td><td>TCCCGATAAGCGGGCGATCG</td><td>TGG</td><td>||||||||[||||||||||||]PAM</td><td>NA</td><td>Intergenic</td><td>NA</td><td>NA</td></tr>
<tr><td></td><td></td><td></td><td></td><td></td><td></td><td></td><td></td><td></td><td></td><td></td><td></td></tr>
<tr><td>T10625</td><td>ATAAGCGGGCGATCGTGGAGCGG</td><td>1000</td><td></td><td></td><td></td><td></td><td></td><td></td><td></td><td></td><td></td></tr>
<tr><td>Oligo fwd</td><td>CACCATAAGCGGGCGATCGTGGAG</td><td></td><td></td><td></td><td></td><td></td><td></td><td></td><td></td><td></td><td></td></tr>
<tr><td>Oligo rev</td><td>AAACCTCCACGATCGCCCGCTTAT</td><td></td><td></td><td></td><td></td><td></td><td></td><td></td><td></td><td></td><td></td></tr>
<tr><td>Chromosome</td><td>start</td><td>end</td><td>strand</td><td>MM</td><td>target_seq</td><td>PAM</td><td>alignment</td><td>distance</td><td>position</td><td>gene name</td><td>gene id</td></tr>
<tr><td>GU305914.1</td><td>138935</td><td>138957</td><td>+</td><td>0</td><td>ATAAGCGGGCGATCGTGGAG</td><td>CGG</td><td>||||||||[||||||||||||]PAM</td><td>NA</td><td>Intergenic</td><td>NA</td><td>NA</td></tr>
<tr><td></td><td></td><td></td><td></td><td></td><td></td><td></td><td></td><td></td><td></td><td></td><td></td></tr>
<tr><td>T10626</td><td>CGATCGTGGAGCGGATCCTCAGG</td><td>1000</td><td></td><td></td><td></td><td></td><td></td><td></td><td></td><td></td><td></td></tr>
</table>

<table>
<tr><td>Oligo fwd</td><td>CACCCGATCGTGGAGCGGATCCTC</td></tr>
<tr><td>Oligo rev</td><td>AAACGAGGATCCGCTCCACGATCG</td></tr>
<tr><td>Chromosome</td><td>start</td><td>end</td><td>strand</td><td>MM</td><td>target_seq</td><td>PAM</td><td>alignment</td><td>distance</td><td>position</td><td>gene name</td><td>gene id</td></tr>
<tr><td>GU305914.1</td><td>138944</td><td>138966</td><td>+</td><td>0</td><td>CGATCGTGGAGCGGATCCTC</td><td>AGG</td><td>||||||||[||||||||||||]PAM</td><td>NA</td><td>Intergenic</td><td>NA</td><td>NA</td></tr>
<tr><td>T10627</td><td>GGAGCGGATCCTCAGGTTCGTGG</td><td>1000</td></tr>
<tr><td>Oligo fwd</td><td>CACCGGAGCGGATCCTCAGGTTCG</td></tr>
<tr><td>Oligo rev</td><td>AAACCGAACCTGAGGATCCGCTCC</td></tr>
<tr><td>Chromosome</td><td>start</td><td>end</td><td>strand</td><td>MM</td><td>target_seq</td><td>PAM</td><td>alignment</td><td>distance</td><td>position</td><td>gene name</td><td>gene id</td></tr>
<tr><td>GU305914.1</td><td>138951</td><td>138973</td><td>+</td><td>0</td><td>GGAGCGGATCCTCAGGTTCG</td><td>TGG</td><td>||||||||[||||||||||||]PAM</td><td>NA</td><td>Intergenic</td><td>NA</td><td>NA</td></tr>
<tr><td>T10628</td><td>CCTCAGGTTCGTGGACACGCCGG</td><td>1000</td></tr>
<tr><td>Oligo fwd</td><td>CACCCCTCAGGTTCGTGGACACGC</td></tr>
<tr><td>Oligo rev</td><td>AAACGCGTGTCCACGAACCTGAGG</td></tr>
<tr><td>Chromosome</td><td>start</td><td>end</td><td>strand</td><td>MM</td><td>target_seq</td><td>PAM</td><td>alignment</td><td>distance</td><td>position</td><td>gene name</td><td>gene id</td></tr>
<tr><td>GU305914.1</td><td>138960</td><td>138982</td><td>+</td><td>0</td><td>CCTCAGGTTCGTGGACACGC</td><td>CGG</td><td>||||||||[||||||||||||]PAM</td><td>NA</td><td>Intergenic</td><td>NA</td><td>NA</td></tr>
<tr><td>T10629</td><td>CTCAGGTTCGTGGACACGCCGGG</td><td>1000</td></tr>
<tr><td>Oligo fwd</td><td>CACCCTCAGGTTCGTGGACACGCC</td></tr>
<tr><td>Oligo rev</td><td>AAACGGCGTGTCCACGAACCTGAG</td></tr>
<tr><td>Chromosome</td><td>start</td><td>end</td><td>strand</td><td>MM</td><td>target_seq</td><td>PAM</td><td>alignment</td><td>distance</td><td>position</td><td>gene name</td><td>gene id</td></tr>
<tr><td>GU305914.1</td><td>138961</td><td>138983</td><td>+</td><td>0</td><td>CTCAGGTTCGTGGACACGCC</td><td>GGG</td><td>||||||||[||||||||||||]PAM</td><td>NA</td><td>Intergenic</td><td>NA</td><td>NA</td></tr>
<tr><td>T10630</td><td>TCAGGTTCGTGGACACGCCGGGG</td><td>1000</td></tr>
<tr><td>Oligo fwd</td><td>CACCTCAGGTTCGTGGACACGCCG</td></tr>
<tr><td>Oligo rev</td><td>AAACCGGCGTGTCCACGAACCTGA</td></tr>
<tr><td>Chromosome</td><td>start</td><td>end</td><td>strand</td><td>MM</td><td>target_seq</td><td>PAM</td><td>alignment</td><td>distance</td><td>position</td><td>gene name</td><td>gene id</td></tr>
<tr><td>GU305914.1</td><td>138962</td><td>138984</td><td>+</td><td>0</td><td>TCAGGTTCGTGGACACGCCG</td><td>GGG</td><td>||||||||[||||||||||||]PAM</td><td>NA</td><td>Intergenic</td><td>NA</td><td>NA</td></tr>
<tr><td>T10631</td><td>CTTAGACCACAATAACGTGCAGG</td><td>1000</td></tr>
<tr><td>Oligo fwd</td><td>CACCCTTAGACCACAATAACGTGC</td></tr>
<tr><td>Oligo rev</td><td>AAACGCACGTTATTGTGGTCTAAG</td></tr>
<tr><td>Chromosome</td><td>start</td><td>end</td><td>strand</td><td>MM</td><td>target_seq</td><td>PAM</td><td>alignment</td><td>distance</td><td>position</td><td>gene name</td><td>gene id</td></tr>
<tr><td>GU305914.1</td><td>138987</td><td>139009</td><td>+</td><td>0</td><td>CTTAGACCACAATAACGTGC</td><td>AGG</td><td>||||||||[||||||||||||]PAM</td><td>NA</td><td>Intergenic</td><td>NA</td><td>NA</td></tr>
<tr><td>T10632</td><td>CAATAACGTGCAGGATGTCGAGG</td><td>1000</td></tr>
<tr><td>Oligo fwd</td><td>CACCCAATAACGTGCAGGATGTCG</td></tr>
<tr><td>Oligo rev</td><td>AAACCGACATCCTGCACGTTATTG</td></tr>
<tr><td>Chromosome</td><td>start</td><td>end</td><td>strand</td><td>MM</td><td>target_seq</td><td>PAM</td><td>alignment</td><td>distance</td><td>position</td><td>gene name</td><td>gene id</td></tr>
<tr><td>GU305914.1</td><td>138996</td><td>139018</td><td>+</td><td>0</td><td>CAATAACGTGCAGGATGTCG</td><td>AGG</td><td>||||||||[||||||||||||]PAM</td><td>NA</td><td>Intergenic</td><td>NA</td><td>NA</td></tr>
<tr><td>T10633</td><td>AGGATGTCGAGGCCGTGCTCTGG</td><td>1000</td></tr>
<tr><td>Oligo fwd</td><td>CACCAGGATGTCGAGGCCGTGCTC</td></tr>
<tr><td>Oligo rev</td><td>AAACGAGCACGGCCTCGACATCCT</td></tr>
<tr><td>Chromosome</td><td>start</td><td>end</td><td>strand</td><td>MM</td><td>target_seq</td><td>PAM</td><td>alignment</td><td>distance</td><td>position</td><td>gene name</td><td>gene id</td></tr>
<tr><td>GU305914.1</td><td>139007</td><td>139029</td><td>+</td><td>0</td><td>AGGATGTCGAGGCCGTGCTC</td><td>TGG</td><td>||||||||[||||||||||||]PAM</td><td>NA</td><td>Intergenic</td><td>NA</td><td>NA</td></tr>
<tr><td>T10634</td><td>GTGCTCTGGCTGCTGTTCTGCGG</td><td>1000</td></tr>
<tr><td>Oligo fwd</td><td>CACCGTGCTCTGGCTGCTGTTCTG</td></tr>
<tr><td>Oligo rev</td><td>AAACCAGAACAGCAGCCAGAGCAC</td></tr>
<tr><td>Chromosome</td><td>start</td><td>end</td><td>strand</td><td>MM</td><td>target_seq</td><td>PAM</td><td>alignment</td><td>distance</td><td>position</td><td>gene name</td><td>gene id</td></tr>
<tr><td>GU305914.1</td><td>139021</td><td>139043</td><td>+</td><td>0</td><td>GTGCTCTGGCTGCTGTTCTG</td><td>CGG</td><td>||||||||[||||||||||||]PAM</td><td>NA</td><td>Intergenic</td><td>NA</td><td>NA</td></tr>
<tr><td>T10635</td><td>TGTCAGAACCCTACCTGTTTCGG</td><td>1000</td></tr>
<tr><td>Oligo fwd</td><td>CACCTGTCAGAACCCTACCTGTTT</td></tr>
<tr><td>Oligo rev</td><td>AAACAAACAGGTAGGGTTCTGACA</td></tr>
<tr><td>Chromosome</td><td>start</td><td>end</td><td>strand</td><td>MM</td><td>target_seq</td><td>PAM</td><td>alignment</td><td>distance</td><td>position</td><td>gene name</td><td>gene id</td></tr>
<tr><td>GU305914.1</td><td>139057</td><td>139079</td><td>+</td><td>0</td><td>TGTCAGAACCCTACCTGTTT</td><td>CGG</td><td>||||||||[||||||||||||]PAM</td><td>NA</td><td>Intergenic</td><td>NA</td><td>NA</td></tr>
<tr><td>T10636</td><td>GTCAGAACCCTACCTGTTTCGGG</td><td>1000</td></tr>
<tr><td>Oligo fwd</td><td>CACCGTCAGAACCCTACCTGTTTC</td></tr>
<tr><td>Oligo rev</td><td>AAACGAAACAGGTAGGGTTCTGAC</td></tr>
<tr><td>Chromosome</td><td>start</td><td>end</td><td>strand</td><td>MM</td><td>target_seq</td><td>PAM</td><td>alignment</td><td>distance</td><td>position</td><td>gene name</td><td>gene id</td></tr>
<tr><td>GU305914.1</td><td>139058</td><td>139080</td><td>+</td><td>0</td><td>GTCAGAACCCTACCTGTTTC</td><td>GGG</td><td>||||||||[||||||||||||]PAM</td><td>NA</td><td>Intergenic</td><td>NA</td><td>NA</td></tr>
<tr><td>T10637</td><td>CGGGCGCGACCGAGAGTGTGAGG</td><td>1000</td></tr>
<tr><td>Oligo fwd</td><td>CACCCGGGCGCGACCGAGAGTGTG</td></tr>
<tr><td>Oligo rev</td><td>AAACCACACTCTCGGTCGCGCCCG</td></tr>
<tr><td>Chromosome</td><td>start</td><td>end</td><td>strand</td><td>MM</td><td>target_seq</td><td>PAM</td><td>alignment</td><td>distance</td><td>position</td><td>gene name</td><td>gene id</td></tr>
<tr><td>GU305914.1</td><td>139077</td><td>139099</td><td>+</td><td>0</td><td>CGGGCGCGACCGAGAGTGTG</td><td>AGG</td><td>||||||||[||||||||||||]PAM</td><td>NA</td><td>Intergenic</td><td>NA</td><td>NA</td></tr>
<tr><td>T10638</td><td>CGACCCCATCACCGACTACTCGG</td><td>1000</td></tr>
<tr><td>Oligo fwd</td><td>CACCCGACCCCATCACCGACTACT</td></tr>
<tr><td>Oligo rev</td><td>AAACAGTAGTCGGTGATGGGGTCG</td></tr>
<tr><td>Chromosome</td><td>start</td><td>end</td><td>strand</td><td>MM</td><td>target_seq</td><td>PAM</td><td>alignment</td><td>distance</td><td>position</td><td>gene name</td><td>gene id</td></tr>
<tr><td>GU305914.1</td><td>139134</td><td>139156</td><td>+</td><td>0</td><td>CGACCCCATCACCGACTACT</td><td>CGG</td><td>||||||||[||||||||||||]PAM</td><td>NA</td><td>Intergenic</td><td>NA</td><td>NA</td></tr>
<tr><td>T10639</td><td>CTACTCGGCCTACATCAACCTGG</td><td>1000</td></tr>
<tr><td>Oligo fwd</td><td>CACCCTACTCGGCCTACATCAACC</td></tr>
<tr><td>Oligo rev</td><td>AAACGGTTGATGTAGGCCGAGTAG</td></tr>
<tr><td>Chromosome</td><td>start</td><td>end</td><td>strand</td><td>MM</td><td>target_seq</td><td>PAM</td><td>alignment</td><td>distance</td><td>position</td><td>gene name</td><td>gene id</td></tr>
<tr><td>GU305914.1</td><td>139149</td><td>139171</td><td>+</td><td>0</td><td>CTACTCGGCCTACATCAACC</td><td>TGG</td><td>||||||||[||||||||||||]PAM</td><td>NA</td><td>Intergenic</td><td>NA</td><td>NA</td></tr>
<tr><td>T10640</td><td>CCGAGCTCTACGTGTACGTGTGG</td><td>1000</td></tr>
<tr><td>Oligo fwd</td><td>CACCCCGAGCTCTACGTGTACGTG</td></tr>
<tr><td>Oligo rev</td><td>AAACCACGTACACGTAGAGCTCGG</td></tr>
<tr><td>Chromosome</td><td>start</td><td>end</td><td>strand</td><td>MM</td><td>target_seq</td><td>PAM</td><td>alignment</td><td>distance</td><td>position</td><td>gene name</td><td>gene id</td></tr>
<tr><td>GU305914.1</td><td>139172</td><td>139194</td><td>+</td><td>0</td><td>CCGAGCTCTACGTGTACGTG</td><td>TGG</td><td>||||||||[||||||||||||]PAM</td><td>NA</td><td>Intergenic</td><td>NA</td><td>NA</td></tr>
</table>

<table>
<tr><td>T10641</td><td>TCTACGTGTACGTGTGGTATAGG</td><td>1000</td><td></td><td></td><td></td><td></td><td></td><td></td><td></td><td></td><td></td></tr>
<tr><td>Oligo fwd</td><td>CACCTCTACGTGTACGTGTGGTAT</td><td></td><td></td><td></td><td></td><td></td><td></td><td></td><td></td><td></td><td></td></tr>
<tr><td>Oligo rev</td><td>AAACATACCACACGTACACGTAGA</td><td></td><td></td><td></td><td></td><td></td><td></td><td></td><td></td><td></td><td></td></tr>
<tr><td>Chromosome</td><td>start</td><td>end</td><td>strand</td><td>MM</td><td>target_seq</td><td>PAM</td><td>alignment</td><td>distance</td><td>position</td><td>gene name</td><td>gene id</td></tr>
<tr><td>GU305914.1</td><td>139178</td><td>139200</td><td>+</td><td>0</td><td>TCTACGTGTACGTGTGGTAT</td><td>AGG</td><td>||||||||[||||||||||||]PAM</td><td>NA</td><td>Intergenic</td><td>NA</td><td>NA</td></tr>
<tr><td>T10642</td><td>ATTTCGATTCCGAGCCCACGCGG</td><td>1000</td><td></td><td></td><td></td><td></td><td></td><td></td><td></td><td></td><td></td></tr>
<tr><td>Oligo fwd</td><td>CACCATTTCGATTCCGAGCCCACG</td><td></td><td></td><td></td><td></td><td></td><td></td><td></td><td></td><td></td><td></td></tr>
<tr><td>Oligo rev</td><td>AAACCGTGGGCTCGGAATCGAAAT</td><td></td><td></td><td></td><td></td><td></td><td></td><td></td><td></td><td></td><td></td></tr>
<tr><td>Chromosome</td><td>start</td><td>end</td><td>strand</td><td>MM</td><td>target_seq</td><td>PAM</td><td>alignment</td><td>distance</td><td>position</td><td>gene name</td><td>gene id</td></tr>
<tr><td>GU305914.1</td><td>139208</td><td>139230</td><td>+</td><td>0</td><td>ATTTCGATTCCGAGCCCACG</td><td>CGG</td><td>||||||||[||||||||||||]PAM</td><td>NA</td><td>Intergenic</td><td>NA</td><td>NA</td></tr>
<tr><td>T10643</td><td>CCCACGCGGTGCTACGAGCTCGG</td><td>1000</td><td></td><td></td><td></td><td></td><td></td><td></td><td></td><td></td><td></td></tr>
<tr><td>Oligo fwd</td><td>CACCCCCACGCGGTGCTACGAGCT</td><td></td><td></td><td></td><td></td><td></td><td></td><td></td><td></td><td></td><td></td></tr>
<tr><td>Oligo rev</td><td>AAACAGCTCGTAGCACCGCGTGGG</td><td></td><td></td><td></td><td></td><td></td><td></td><td></td><td></td><td></td><td></td></tr>
<tr><td>Chromosome</td><td>start</td><td>end</td><td>strand</td><td>MM</td><td>target_seq</td><td>PAM</td><td>alignment</td><td>distance</td><td>position</td><td>gene name</td><td>gene id</td></tr>
<tr><td>GU305914.1</td><td>139222</td><td>139244</td><td>+</td><td>0</td><td>CCCACGCGGTGCTACGAGCT</td><td>CGG</td><td>||||||||[||||||||||||]PAM</td><td>NA</td><td>Intergenic</td><td>NA</td><td>NA</td></tr>
<tr><td>T10644</td><td>CCACGCGGTGCTACGAGCTCGGG</td><td>1000</td><td></td><td></td><td></td><td></td><td></td><td></td><td></td><td></td><td></td></tr>
<tr><td>Oligo fwd</td><td>CACCCCACGCGGTGCTACGAGCTC</td><td></td><td></td><td></td><td></td><td></td><td></td><td></td><td></td><td></td><td></td></tr>
<tr><td>Oligo rev</td><td>AAACGAGCTCGTAGCACCGCGTGG</td><td></td><td></td><td></td><td></td><td></td><td></td><td></td><td></td><td></td><td></td></tr>
<tr><td>Chromosome</td><td>start</td><td>end</td><td>strand</td><td>MM</td><td>target_seq</td><td>PAM</td><td>alignment</td><td>distance</td><td>position</td><td>gene name</td><td>gene id</td></tr>
<tr><td>GU305914.1</td><td>139223</td><td>139245</td><td>+</td><td>0</td><td>CCACGCGGTGCTACGAGCTC</td><td>GGG</td><td>||||||||[||||||||||||]PAM</td><td>NA</td><td>Intergenic</td><td>NA</td><td>NA</td></tr>
<tr><td>T10645</td><td>GCGGTGCTACGAGCTCGGGACGG</td><td>1000</td><td></td><td></td><td></td><td></td><td></td><td></td><td></td><td></td><td></td></tr>
<tr><td>Oligo fwd</td><td>CACCGCGGTGCTACGAGCTCGGGA</td><td></td><td></td><td></td><td></td><td></td><td></td><td></td><td></td><td></td><td></td></tr>
<tr><td>Oligo rev</td><td>AAACTCCCGAGCTCGTAGCACCGC</td><td></td><td></td><td></td><td></td><td></td><td></td><td></td><td></td><td></td><td></td></tr>
<tr><td>Chromosome</td><td>start</td><td>end</td><td>strand</td><td>MM</td><td>target_seq</td><td>PAM</td><td>alignment</td><td>distance</td><td>position</td><td>gene name</td><td>gene id</td></tr>
<tr><td>GU305914.1</td><td>139227</td><td>139249</td><td>+</td><td>0</td><td>GCGGTGCTACGAGCTCGGGA</td><td>CGG</td><td>||||||||[||||||||||||]PAM</td><td>NA</td><td>Intergenic</td><td>NA</td><td>NA</td></tr>
<tr><td>T10646</td><td>GTGCTACGAGCTCGGGACGGTGG</td><td>1000</td><td></td><td></td><td></td><td></td><td></td><td></td><td></td><td></td><td></td></tr>
<tr><td>Oligo fwd</td><td>CACCGTGCTACGAGCTCGGGACGG</td><td></td><td></td><td></td><td></td><td></td><td></td><td></td><td></td><td></td><td></td></tr>
<tr><td>Oligo rev</td><td>AAACCCGTCCCGAGCTCGTAGCAC</td><td></td><td></td><td></td><td></td><td></td><td></td><td></td><td></td><td></td><td></td></tr>
<tr><td>Chromosome</td><td>start</td><td>end</td><td>strand</td><td>MM</td><td>target_seq</td><td>PAM</td><td>alignment</td><td>distance</td><td>position</td><td>gene name</td><td>gene id</td></tr>
<tr><td>GU305914.1</td><td>139230</td><td>139252</td><td>+</td><td>0</td><td>GTGCTACGAGCTCGGGACGG</td><td>TGG</td><td>||||||||[||||||||||||]PAM</td><td>NA</td><td>Intergenic</td><td>NA</td><td>NA</td></tr>
<tr><td>T10647</td><td>CGAGCTCGGGACGGTGGCCATGG</td><td>1000</td><td></td><td></td><td></td><td></td><td></td><td></td><td></td><td></td><td></td></tr>
<tr><td>Oligo fwd</td><td>CACCCGAGCTCGGGACGGTGGCCA</td><td></td><td></td><td></td><td></td><td></td><td></td><td></td><td></td><td></td><td></td></tr>
<tr><td>Oligo rev</td><td>AAACTGGCCACCGTCCCGAGCTCG</td><td></td><td></td><td></td><td></td><td></td><td></td><td></td><td></td><td></td><td></td></tr>
<tr><td>Chromosome</td><td>start</td><td>end</td><td>strand</td><td>MM</td><td>target_seq</td><td>PAM</td><td>alignment</td><td>distance</td><td>position</td><td>gene name</td><td>gene id</td></tr>
<tr><td>GU305914.1</td><td>139236</td><td>139258</td><td>+</td><td>0</td><td>CGAGCTCGGGACGGTGGCCA</td><td>TGG</td><td>||||||||[||||||||||||]PAM</td><td>NA</td><td>Intergenic</td><td>NA</td><td>NA</td></tr>
<tr><td>T10648</td><td>CGCTGCAGTCCGTCCGCCAGAGG</td><td>1000</td><td></td><td></td><td></td><td></td><td></td><td></td><td></td><td></td><td></td></tr>
<tr><td>Oligo fwd</td><td>CACCCGCTGCAGTCCGTCCGCCAG</td><td></td><td></td><td></td><td></td><td></td><td></td><td></td><td></td><td></td><td></td></tr>
<tr><td>Oligo rev</td><td>AAACCTGGCGGACGGACTGCAGCG</td><td></td><td></td><td></td><td></td><td></td><td></td><td></td><td></td><td></td><td></td></tr>
<tr><td>Chromosome</td><td>start</td><td>end</td><td>strand</td><td>MM</td><td>target_seq</td><td>PAM</td><td>alignment</td><td>distance</td><td>position</td><td>gene name</td><td>gene id</td></tr>
<tr><td>GU305914.1</td><td>139274</td><td>139296</td><td>+</td><td>0</td><td>CGCTGCAGTCCGTCCGCCAG</td><td>AGG</td><td>||||||||[||||||||||||]PAM</td><td>NA</td><td>Intergenic</td><td>NA</td><td>NA</td></tr>
<tr><td>T10649</td><td>CCGACCGCAGCGTGCCCGTCTGG</td><td>1000</td><td></td><td></td><td></td><td></td><td></td><td></td><td></td><td></td><td></td></tr>
<tr><td>Oligo fwd</td><td>CACCCCGACCGCAGCGTGCCCGTC</td><td></td><td></td><td></td><td></td><td></td><td></td><td></td><td></td><td></td><td></td></tr>
<tr><td>Oligo rev</td><td>AAACGACGGGCACGCTGCGGTCGG</td><td></td><td></td><td></td><td></td><td></td><td></td><td></td><td></td><td></td><td></td></tr>
<tr><td>Chromosome</td><td>start</td><td>end</td><td>strand</td><td>MM</td><td>target_seq</td><td>PAM</td><td>alignment</td><td>distance</td><td>position</td><td>gene name</td><td>gene id</td></tr>
<tr><td>GU305914.1</td><td>139301</td><td>139323</td><td>+</td><td>0</td><td>CCGACCGCAGCGTGCCCGTC</td><td>TGG</td><td>||||||||[||||||||||||]PAM</td><td>NA</td><td>Intergenic</td><td>NA</td><td>NA</td></tr>
<tr><td>GU305914.1</td><td>29816</td><td>29838</td><td>-</td><td>4</td><td>CCGAACTCCGCATGCCCGTC</td><td>GGG</td><td>||||-|-|[-||-||||||||]PAM</td><td>NA</td><td>Intergenic</td><td>NA</td><td>NA</td></tr>
<tr><td>T10650</td><td>TAATCAGAACCGTGTGTGTCTGG</td><td>1000</td><td></td><td></td><td></td><td></td><td></td><td></td><td></td><td></td><td></td></tr>
<tr><td>Oligo fwd</td><td>CACCTAATCAGAACCGTGTGTGTC</td><td></td><td></td><td></td><td></td><td></td><td></td><td></td><td></td><td></td><td></td></tr>
<tr><td>Oligo rev</td><td>AAACGACACACACGGTTCTGATTA</td><td></td><td></td><td></td><td></td><td></td><td></td><td></td><td></td><td></td><td></td></tr>
<tr><td>Chromosome</td><td>start</td><td>end</td><td>strand</td><td>MM</td><td>target_seq</td><td>PAM</td><td>alignment</td><td>distance</td><td>position</td><td>gene name</td><td>gene id</td></tr>
<tr><td>GU305914.1</td><td>139362</td><td>139384</td><td>+</td><td>0</td><td>TAATCAGAACCGTGTGTGTC</td><td>TGG</td><td>||||||||[||||||||||||]PAM</td><td>NA</td><td>Intergenic</td><td>NA</td><td>NA</td></tr>
<tr><td>T10651</td><td>CTACAGCCCGATCATCATCAAGG</td><td>1000</td><td></td><td></td><td></td><td></td><td></td><td></td><td></td><td></td><td></td></tr>
<tr><td>Oligo fwd</td><td>CACCCTACAGCCCGATCATCATCA</td><td></td><td></td><td></td><td></td><td></td><td></td><td></td><td></td><td></td><td></td></tr>
<tr><td>Oligo rev</td><td>AAACTGATGATGATCGGGCTGTAG</td><td></td><td></td><td></td><td></td><td></td><td></td><td></td><td></td><td></td><td></td></tr>
<tr><td>Chromosome</td><td>start</td><td>end</td><td>strand</td><td>MM</td><td>target_seq</td><td>PAM</td><td>alignment</td><td>distance</td><td>position</td><td>gene name</td><td>gene id</td></tr>
<tr><td>GU305914.1</td><td>139416</td><td>139438</td><td>+</td><td>0</td><td>CTACAGCCCGATCATCATCA</td><td>AGG</td><td>||||||||[||||||||||||]PAM</td><td>NA</td><td>Intergenic</td><td>NA</td><td>NA</td></tr>
<tr><td>T10652</td><td>CGATCATCATCAAGGACTGTCGG</td><td>1000</td><td></td><td></td><td></td><td></td><td></td><td></td><td></td><td></td><td></td></tr>
<tr><td>Oligo fwd</td><td>CACCCGATCATCATCAAGGACTGT</td><td></td><td></td><td></td><td></td><td></td><td></td><td></td><td></td><td></td><td></td></tr>
<tr><td>Oligo rev</td><td>AAACACAGTCCTTGATGATGATCG</td><td></td><td></td><td></td><td></td><td></td><td></td><td></td><td></td><td></td><td></td></tr>
<tr><td>Chromosome</td><td>start</td><td>end</td><td>strand</td><td>MM</td><td>target_seq</td><td>PAM</td><td>alignment</td><td>distance</td><td>position</td><td>gene name</td><td>gene id</td></tr>
<tr><td>GU305914.1</td><td>139424</td><td>139446</td><td>+</td><td>0</td><td>CGATCATCATCAAGGACTGT</td><td>CGG</td><td>||||||||[||||||||||||]PAM</td><td>NA</td><td>Intergenic</td><td>NA</td><td>NA</td></tr>
<tr><td>T10653</td><td>GATCATCATCAAGGACTGTCGGG</td><td>1000</td><td></td><td></td><td></td><td></td><td></td><td></td><td></td><td></td><td></td></tr>
<tr><td>Oligo fwd</td><td>CACCGATCATCATCAAGGACTGTC</td><td></td><td></td><td></td><td></td><td></td><td></td><td></td><td></td><td></td><td></td></tr>
<tr><td>Oligo rev</td><td>AAACGACAGTCCTTGATGATGATC</td><td></td><td></td><td></td><td></td><td></td><td></td><td></td><td></td><td></td><td></td></tr>
<tr><td>Chromosome</td><td>start</td><td>end</td><td>strand</td><td>MM</td><td>target_seq</td><td>PAM</td><td>alignment</td><td>distance</td><td>position</td><td>gene name</td><td>gene id</td></tr>
<tr><td>GU305914.1</td><td>139425</td><td>139447</td><td>+</td><td>0</td><td>GATCATCATCAAGGACTGTC</td><td>GGG</td><td>||||||||[||||||||||||]PAM</td><td>NA</td><td>Intergenic</td><td>NA</td><td>NA</td></tr>
<tr><td>T10654</td><td>CATCAAGGACTGTCGGGATGCGG</td><td>1000</td><td></td><td></td><td></td><td></td><td></td><td></td><td></td><td></td><td></td></tr>
<tr><td>Oligo fwd</td><td>CACCCATCAAGGACTGTCGGGATG</td><td></td><td></td><td></td><td></td><td></td><td></td><td></td><td></td><td></td><td></td></tr>
<tr><td>Oligo rev</td><td>AAACCATCCCGACAGTCCTTGATG</td><td></td><td></td><td></td><td></td><td></td><td></td><td></td><td></td><td></td><td></td></tr>
<tr><td>Chromosome</td><td>start</td><td>end</td><td>strand</td><td>MM</td><td>target_seq</td><td>PAM</td><td>alignment</td><td>distance</td><td>position</td><td>gene name</td><td>gene id</td></tr>
<tr><td>GU305914.1</td><td>139431</td><td>139453</td><td>+</td><td>0</td><td>CATCAAGGACTGTCGGGATG</td><td>CGG</td><td>||||||||[||||||||||||]PAM</td><td>NA</td><td>Intergenic</td><td>NA</td><td>NA</td></tr>
<tr><td>T10655</td><td>GACCCTCAGTCACGTCCTGCCGG</td><td>1000</td><td></td><td></td><td></td><td></td><td></td><td></td><td></td><td></td><td></td></tr>
<tr><td>Oligo fwd</td><td>CACCGACCCTCAGTCACGTCCTGC</td><td></td><td></td><td></td><td></td><td></td><td></td><td></td><td></td><td></td><td></td></tr>
<tr><td>Oligo rev</td><td>AAACGCAGGACGTGACTGAGGGTC</td><td></td><td></td><td></td><td></td><td></td><td></td><td></td><td></td><td></td><td></td></tr>
<tr><td>Chromosome</td><td>start</td><td>end</td><td>strand</td><td>MM</td><td>target_seq</td><td>PAM</td><td>alignment</td><td>distance</td><td>position</td><td>gene name</td><td>gene id</td></tr>
</table>

| | | | | | | | | | | | |
|---|---|---|---|---|---|---|---|---|---|---|---|
| GU305914.1 | | 139464 | 139486 | + | 0 | GACCCTCAGTCACGTCCTGC | CGG | \|\|\|\|\|\|\|\|[\|\|\|\|\|\|\|\|\|\|\|\|]PAM | NA | Intergenic | NA | NA |
| T10656 | ACCCTCAGTCACGTCCTGCCGGG | | 1000 | | | | | | | | | |
| Oligo fwd | CACCACCCTCAGTCACGTCCTGCC | | | | | | | | | | | |
| Oligo rev | AAACGGCAGGACGTGACTGAGGGT | | | | | | | | | | | |
| Chromosome | start | | end | strand | MM | target_seq | PAM | alignment | distance | position | gene name | gene id |
| GU305914.1 | | 139465 | 139487 | + | 0 | ACCCTCAGTCACGTCCTGCC | GGG | \|\|\|\|\|\|\|\|[\|\|\|\|\|\|\|\|\|\|\|\|]PAM | NA | Intergenic | NA | NA |
| T10657 | CTGTTTCCCGTCTACGACATCGG | | 1000 | | | | | | | | | |
| Oligo fwd | CACCCTGTTTCCCGTCTACGACAT | | | | | | | | | | | |
| Oligo rev | AAACATGTCGTAGACGGGAAACAG | | | | | | | | | | | |
| Chromosome | start | | end | strand | MM | target_seq | PAM | alignment | distance | position | gene name | gene id |
| GU305914.1 | | 139504 | 139526 | + | 0 | CTGTTTCCCGTCTACGACAT | CGG | \|\|\|\|\|\|\|\|[\|\|\|\|\|\|\|\|\|\|\|\|]PAM | NA | Intergenic | NA | NA |
| T10658 | TGTTTCCCGTCTACGACATCGGG | | 1000 | | | | | | | | | |
| Oligo fwd | CACCTGTTTCCCGTCTACGACATC | | | | | | | | | | | |
| Oligo rev | AAACGATGTCGTAGACGGGAAACA | | | | | | | | | | | |
| Chromosome | start | | end | strand | MM | target_seq | PAM | alignment | distance | position | gene name | gene id |
| GU305914.1 | | 139505 | 139527 | + | 0 | TGTTTCCCGTCTACGACATC | GGG | \|\|\|\|\|\|\|\|[\|\|\|\|\|\|\|\|\|\|\|\|]PAM | NA | Intergenic | NA | NA |
| T10659 | GCGCTCTGCGACTCCAACGACGG | | 1000 | | | | | | | | | |
| Oligo fwd | CACCGCGCTCTGCGACTCCAACGA | | | | | | | | | | | |
| Oligo rev | AAACTCGTTGGAGTCGCAGAGCGC | | | | | | | | | | | |
| Chromosome | start | | end | strand | MM | target_seq | PAM | alignment | distance | position | gene name | gene id |
| GU305914.1 | | 139540 | 139562 | + | 0 | GCGCTCTGCGACTCCAACGA | CGG | \|\|\|\|\|\|\|\|[\|\|\|\|\|\|\|\|\|\|\|\|]PAM | NA | Intergenic | NA | NA |
| T10660 | CTGCGACTCCAACGACGGCGAGG | | 1000 | | | | | | | | | |
| Oligo fwd | CACCCTGCGACTCCAACGACGGCG | | | | | | | | | | | |
| Oligo rev | AAACCGCCGTCGTTGGAGTCGCAG | | | | | | | | | | | |
| Chromosome | start | | end | strand | MM | target_seq | PAM | alignment | distance | position | gene name | gene id |
| GU305914.1 | | 139545 | 139567 | + | 0 | CTGCGACTCCAACGACGGCG | AGG | \|\|\|\|\|\|\|\|[\|\|\|\|\|\|\|\|\|\|\|\|]PAM | NA | Intergenic | NA | NA |
| T10661 | CCAACGACGGCGAGGAGCGCAGG | | 1000 | | | | | | | | | |
| Oligo fwd | CACCCCAACGACGGCGAGGAGCGC | | | | | | | | | | | |
| Oligo rev | AAACGCGCTCCTCGCCGTCGTTGG | | | | | | | | | | | |
| Chromosome | start | | end | strand | MM | target_seq | PAM | alignment | distance | position | gene name | gene id |
| GU305914.1 | | 139553 | 139575 | + | 0 | CCAACGACGGCGAGGAGCGC | AGG | \|\|\|\|\|\|\|\|[\|\|\|\|\|\|\|\|\|\|\|\|]PAM | NA | Intergenic | NA | NA |
| T10662 | AGGAGCGCAGGAAACGCATGCGG | | 1000 | | | | | | | | | |
| Oligo fwd | CACCAGGAGCGCAGGAAACGCATG | | | | | | | | | | | |
| Oligo rev | AAACCATGCGTTTCCTGCGCTCCT | | | | | | | | | | | |
| Chromosome | start | | end | strand | MM | target_seq | PAM | alignment | distance | position | gene name | gene id |
| GU305914.1 | | 139565 | 139587 | + | 0 | AGGAGCGCAGGAAACGCATG | CGG | \|\|\|\|\|\|\|\|[\|\|\|\|\|\|\|\|\|\|\|\|]PAM | NA | Intergenic | NA | NA |
| T10663 | GGAGCGCAGGAAACGCATGCGGG | | 1000 | | | | | | | | | |
| Oligo fwd | CACCGGAGCGCAGGAAACGCATGC | | | | | | | | | | | |
| Oligo rev | AAACGCATGCGTTTCCTGCGCTCC | | | | | | | | | | | |
| Chromosome | start | | end | strand | MM | target_seq | PAM | alignment | distance | position | gene name | gene id |
| GU305914.1 | | 139566 | 139588 | + | 0 | GGAGCGCAGGAAACGCATGC | GGG | \|\|\|\|\|\|\|\|[\|\|\|\|\|\|\|\|\|\|\|\|]PAM | NA | Intergenic | NA | NA |
| T10664 | CATGCGGGAGACCATCGATTCGG | | 1000 | | | | | | | | | |
| Oligo fwd | CACCCATGCGGGAGACCATCGATT | | | | | | | | | | | |
| Oligo rev | AAACAATCGATGGTCTCCCGCATG | | | | | | | | | | | |
| Chromosome | start | | end | strand | MM | target_seq | PAM | alignment | distance | position | gene name | gene id |
| GU305914.1 | | 139581 | 139603 | + | 0 | CATGCGGGAGACCATCGATT | CGG | \|\|\|\|\|\|\|\|[\|\|\|\|\|\|\|\|\|\|\|\|]PAM | NA | Intergenic | NA | NA |
| T10665 | GCTCTCCACCACCGACGACGCGG | | 1000 | | | | | | | | | |
| Oligo fwd | CACCGCTCTCCACCACCGACGACG | | | | | | | | | | | |
| Oligo rev | AAACCGTCGTCGGTGGTGGAGAGC | | | | | | | | | | | |
| Chromosome | start | | end | strand | MM | target_seq | PAM | alignment | distance | position | gene name | gene id |
| GU305914.1 | | 139605 | 139627 | + | 0 | GCTCTCCACCACCGACGACG | CGG | \|\|\|\|\|\|\|\|[\|\|\|\|\|\|\|\|\|\|\|\|]PAM | NA | Intergenic | NA | NA |
| GU305914.1 | | 96820 | 96842 | - | 4 | GCTCGCGACCGCCGGCGACG | AGG | \|\|\|\|-\|-\|[\|\|-\|\|\|-\|\|\|\|\|]PAM | NA | Intergenic | NA | NA |
| T10666 | CCACCGACGACGCGGTCTGATGG | | 1000 | | | | | | | | | |
| Oligo fwd | CACCCCACCGACGACGCGGTCTGA | | | | | | | | | | | |
| Oligo rev | AAACTCAGACCGCGTCGTCGGTGG | | | | | | | | | | | |
| Chromosome | start | | end | strand | MM | target_seq | PAM | alignment | distance | position | gene name | gene id |
| GU305914.1 | | 139613 | 139635 | + | 0 | CCACCGACGACGCGGTCTGA | TGG | \|\|\|\|\|\|\|\|[\|\|\|\|\|\|\|\|\|\|\|\|]PAM | NA | Intergenic | NA | NA |
| T10667 | CACCGACGACGCGGTCTGATGGG | | 1000 | | | | | | | | | |
| Oligo fwd | CACCCACCGACGACGCGGTCTGAT | | | | | | | | | | | |
| Oligo rev | AAACATCAGACCGCGTCGTCGGTG | | | | | | | | | | | |
| Chromosome | start | | end | strand | MM | target_seq | PAM | alignment | distance | position | gene name | gene id |
| GU305914.1 | | 139614 | 139636 | + | 0 | CACCGACGACGCGGTCTGAT | GGG | \|\|\|\|\|\|\|\|[\|\|\|\|\|\|\|\|\|\|\|\|]PAM | NA | Intergenic | NA | NA |
| T10668 | ACCGACGACGCGGTCTGATGGGG | | 1000 | | | | | | | | | |
| Oligo fwd | CACCACCGACGACGCGGTCTGATG | | | | | | | | | | | |
| Oligo rev | AAACCATCAGACCGCGTCGTCGGT | | | | | | | | | | | |
| Chromosome | start | | end | strand | MM | target_seq | PAM | alignment | distance | position | gene name | gene id |
| GU305914.1 | | 139615 | 139637 | + | 0 | ACCGACGACGCGGTCTGATG | GGG | \|\|\|\|\|\|\|\|[\|\|\|\|\|\|\|\|\|\|\|\|]PAM | NA | Intergenic | NA | NA |
| T10669 | GCGGTCTGATGGGGTCGCGACGG | | 1000 | | | | | | | | | |
| Oligo fwd | CACCGCGGTCTGATGGGGTCGCGA | | | | | | | | | | | |
| Oligo rev | AAACTCGCGACCCCATCAGACCGC | | | | | | | | | | | |
| Chromosome | start | | end | strand | MM | target_seq | PAM | alignment | distance | position | gene name | gene id |
| GU305914.1 | | 139624 | 139646 | + | 0 | GCGGTCTGATGGGGTCGCGA | CGG | \|\|\|\|\|\|\|\|[\|\|\|\|\|\|\|\|\|\|\|\|]PAM | NA | Intergenic | NA | NA |
| T10670 | CAGCGAGAGTGCGCTGCTCAAGG | | 1000 | | | | | | | | | |
| Oligo fwd | CACCCAGCGAGAGTGCGCTGCTCA | | | | | | | | | | | |

<table>
<tr><td>Oligo rev</td><td>AAACTGAGCAGCGCACTCTCGCTG</td><td></td><td></td><td></td><td></td><td></td><td></td><td></td><td></td><td></td><td></td></tr>
<tr><td>Chromosome</td><td>start</td><td>end</td><td>strand</td><td>MM</td><td>target_seq</td><td>PAM</td><td>alignment</td><td>distance</td><td>position</td><td>gene name</td><td>gene id</td></tr>
<tr><td>GU305914.1</td><td>139655</td><td>139677</td><td>+</td><td>0</td><td>CAGCGAGAGTGCGCTGCTCA</td><td>AGG</td><td>||||||||[||||||||||||]PAM</td><td>NA</td><td>Intergenic</td><td>NA</td><td>NA</td></tr>
<tr><td></td><td></td><td></td><td></td><td></td><td></td><td></td><td></td><td></td><td></td><td></td><td></td></tr>
<tr><td>T10671</td><td>CGCTGCTCAAGGACGCTATGCGG</td><td>1000</td><td></td><td></td><td></td><td></td><td></td><td></td><td></td><td></td><td></td></tr>
<tr><td>Oligo fwd</td><td>CACCCGCTGCTCAAGGACGCTATG</td><td></td><td></td><td></td><td></td><td></td><td></td><td></td><td></td><td></td><td></td></tr>
<tr><td>Oligo rev</td><td>AAACCATAGCGTCCTTGAGCAGCG</td><td></td><td></td><td></td><td></td><td></td><td></td><td></td><td></td><td></td><td></td></tr>
<tr><td>Chromosome</td><td>start</td><td>end</td><td>strand</td><td>MM</td><td>target_seq</td><td>PAM</td><td>alignment</td><td>distance</td><td>position</td><td>gene name</td><td>gene id</td></tr>
<tr><td>GU305914.1</td><td>139666</td><td>139688</td><td>+</td><td>0</td><td>CGCTGCTCAAGGACGCTATG</td><td>CGG</td><td>||||||||[||||||||||||]PAM</td><td>NA</td><td>Intergenic</td><td>NA</td><td>NA</td></tr>
<tr><td></td><td></td><td></td><td></td><td></td><td></td><td></td><td></td><td></td><td></td><td></td><td></td></tr>
<tr><td>T10672</td><td>GCTGCTCAAGGACGCTATGCGGG</td><td>1000</td><td></td><td></td><td></td><td></td><td></td><td></td><td></td><td></td><td></td></tr>
<tr><td>Oligo fwd</td><td>CACCGCTGCTCAAGGACGCTATGC</td><td></td><td></td><td></td><td></td><td></td><td></td><td></td><td></td><td></td><td></td></tr>
<tr><td>Oligo rev</td><td>AAACGCATAGCGTCCTTGAGCAGC</td><td></td><td></td><td></td><td></td><td></td><td></td><td></td><td></td><td></td><td></td></tr>
<tr><td>Chromosome</td><td>start</td><td>end</td><td>strand</td><td>MM</td><td>target_seq</td><td>PAM</td><td>alignment</td><td>distance</td><td>position</td><td>gene name</td><td>gene id</td></tr>
<tr><td>GU305914.1</td><td>139667</td><td>139689</td><td>+</td><td>0</td><td>GCTGCTCAAGGACGCTATGC</td><td>GGG</td><td>||||||||[||||||||||||]PAM</td><td>NA</td><td>Intergenic</td><td>NA</td><td>NA</td></tr>
<tr><td></td><td></td><td></td><td></td><td></td><td></td><td></td><td></td><td></td><td></td><td></td><td></td></tr>
<tr><td>T10673</td><td>GCTCAAGGACGCTATGCGGGCGG</td><td>1000</td><td></td><td></td><td></td><td></td><td></td><td></td><td></td><td></td><td></td></tr>
<tr><td>Oligo fwd</td><td>CACCGCTCAAGGACGCTATGCGGG</td><td></td><td></td><td></td><td></td><td></td><td></td><td></td><td></td><td></td><td></td></tr>
<tr><td>Oligo rev</td><td>AAACCCCGCATAGCGTCCTTGAGC</td><td></td><td></td><td></td><td></td><td></td><td></td><td></td><td></td><td></td><td></td></tr>
<tr><td>Chromosome</td><td>start</td><td>end</td><td>strand</td><td>MM</td><td>target_seq</td><td>PAM</td><td>alignment</td><td>distance</td><td>position</td><td>gene name</td><td>gene id</td></tr>
<tr><td>GU305914.1</td><td>139670</td><td>139692</td><td>+</td><td>0</td><td>GCTCAAGGACGCTATGCGGG</td><td>CGG</td><td>||||||||[||||||||||||]PAM</td><td>NA</td><td>Intergenic</td><td>NA</td><td>NA</td></tr>
<tr><td></td><td></td><td></td><td></td><td></td><td></td><td></td><td></td><td></td><td></td><td></td><td></td></tr>
<tr><td>T10674</td><td>GGACGCTATGCGGGCGGACCTGG</td><td>1000</td><td></td><td></td><td></td><td></td><td></td><td></td><td></td><td></td><td></td></tr>
<tr><td>Oligo fwd</td><td>CACCGGACGCTATGCGGGCGGACC</td><td></td><td></td><td></td><td></td><td></td><td></td><td></td><td></td><td></td><td></td></tr>
<tr><td>Oligo rev</td><td>AAACGGTCCGCCCGCATAGCGTCC</td><td></td><td></td><td></td><td></td><td></td><td></td><td></td><td></td><td></td><td></td></tr>
<tr><td>Chromosome</td><td>start</td><td>end</td><td>strand</td><td>MM</td><td>target_seq</td><td>PAM</td><td>alignment</td><td>distance</td><td>position</td><td>gene name</td><td>gene id</td></tr>
<tr><td>GU305914.1</td><td>139676</td><td>139698</td><td>+</td><td>0</td><td>GGACGCTATGCGGGCGGACC</td><td>TGG</td><td>||||||||[||||||||||||]PAM</td><td>NA</td><td>Intergenic</td><td>NA</td><td>NA</td></tr>
<tr><td></td><td></td><td></td><td></td><td></td><td></td><td></td><td></td><td></td><td></td><td></td><td></td></tr>
<tr><td>T10675</td><td>TGCGGGCGGACCTGGAGAAACGG</td><td>1000</td><td></td><td></td><td></td><td></td><td></td><td></td><td></td><td></td><td></td></tr>
<tr><td>Oligo fwd</td><td>CACCTGCGGGCGGACCTGGAGAAA</td><td></td><td></td><td></td><td></td><td></td><td></td><td></td><td></td><td></td><td></td></tr>
<tr><td>Oligo rev</td><td>AAACTTTCTCCAGGTCCGCCCGCA</td><td></td><td></td><td></td><td></td><td></td><td></td><td></td><td></td><td></td><td></td></tr>
<tr><td>Chromosome</td><td>start</td><td>end</td><td>strand</td><td>MM</td><td>target_seq</td><td>PAM</td><td>alignment</td><td>distance</td><td>position</td><td>gene name</td><td>gene id</td></tr>
<tr><td>GU305914.1</td><td>139684</td><td>139706</td><td>+</td><td>0</td><td>TGCGGGCGGACCTGGAGAAA</td><td>CGG</td><td>||||||||[||||||||||||]PAM</td><td>NA</td><td>Intergenic</td><td>NA</td><td>NA</td></tr>
<tr><td></td><td></td><td></td><td></td><td></td><td></td><td></td><td></td><td></td><td></td><td></td><td></td></tr>
<tr><td>T10676</td><td>GCAGAAGCGATTTTTCGTCGAGG</td><td>1000</td><td></td><td></td><td></td><td></td><td></td><td></td><td></td><td></td><td></td></tr>
<tr><td>Oligo fwd</td><td>CACCGCAGAAGCGATTTTTCGTCG</td><td></td><td></td><td></td><td></td><td></td><td></td><td></td><td></td><td></td><td></td></tr>
<tr><td>Oligo rev</td><td>AAACCGACGAAAAATCGCTTCTGC</td><td></td><td></td><td></td><td></td><td></td><td></td><td></td><td></td><td></td><td></td></tr>
<tr><td>Chromosome</td><td>start</td><td>end</td><td>strand</td><td>MM</td><td>target_seq</td><td>PAM</td><td>alignment</td><td>distance</td><td>position</td><td>gene name</td><td>gene id</td></tr>
<tr><td>GU305914.1</td><td>139706</td><td>139728</td><td>+</td><td>0</td><td>GCAGAAGCGATTTTTCGTCG</td><td>AGG</td><td>||||||||[||||||||||||]PAM</td><td>NA</td><td>Intergenic</td><td>NA</td><td>NA</td></tr>
<tr><td></td><td></td><td></td><td></td><td></td><td></td><td></td><td></td><td></td><td></td><td></td><td></td></tr>
<tr><td>T10677</td><td>CGATTTTTCGTCGAGGCGTTCGG</td><td>1000</td><td></td><td></td><td></td><td></td><td></td><td></td><td></td><td></td><td></td></tr>
<tr><td>Oligo fwd</td><td>CACCCGATTTTTCGTCGAGGCGTT</td><td></td><td></td><td></td><td></td><td></td><td></td><td></td><td></td><td></td><td></td></tr>
<tr><td>Oligo rev</td><td>AAACAACGCCTCGACGAAAAATCG</td><td></td><td></td><td></td><td></td><td></td><td></td><td></td><td></td><td></td><td></td></tr>
<tr><td>Chromosome</td><td>start</td><td>end</td><td>strand</td><td>MM</td><td>target_seq</td><td>PAM</td><td>alignment</td><td>distance</td><td>position</td><td>gene name</td><td>gene id</td></tr>
<tr><td>GU305914.1</td><td>139713</td><td>139735</td><td>+</td><td>0</td><td>CGATTTTTCGTCGAGGCGTT</td><td>CGG</td><td>||||||||[||||||||||||]PAM</td><td>NA</td><td>Intergenic</td><td>NA</td><td>NA</td></tr>
<tr><td></td><td></td><td></td><td></td><td></td><td></td><td></td><td></td><td></td><td></td><td></td><td></td></tr>
<tr><td>T10678</td><td>GTTCGGTCCTGATCATCCCCTGG</td><td>1000</td><td></td><td></td><td></td><td></td><td></td><td></td><td></td><td></td><td></td></tr>
<tr><td>Oligo fwd</td><td>CACCGTTCGGTCCTGATCATCCCC</td><td></td><td></td><td></td><td></td><td></td><td></td><td></td><td></td><td></td><td></td></tr>
<tr><td>Oligo rev</td><td>AAACGGGGATGATCAGGACCGAAC</td><td></td><td></td><td></td><td></td><td></td><td></td><td></td><td></td><td></td><td></td></tr>
<tr><td>Chromosome</td><td>start</td><td>end</td><td>strand</td><td>MM</td><td>target_seq</td><td>PAM</td><td>alignment</td><td>distance</td><td>position</td><td>gene name</td><td>gene id</td></tr>
<tr><td>GU305914.1</td><td>139730</td><td>139752</td><td>+</td><td>0</td><td>GTTCGGTCCTGATCATCCCC</td><td>TGG</td><td>||||||||[||||||||||||]PAM</td><td>NA</td><td>Intergenic</td><td>NA</td><td>NA</td></tr>
<tr><td></td><td></td><td></td><td></td><td></td><td></td><td></td><td></td><td></td><td></td><td></td><td></td></tr>
<tr><td>T10679</td><td>TCATCCCCTGGTGCGTATTCAGG</td><td>1000</td><td></td><td></td><td></td><td></td><td></td><td></td><td></td><td></td><td></td></tr>
<tr><td>Oligo fwd</td><td>CACCTCATCCCCTGGTGCGTATTC</td><td></td><td></td><td></td><td></td><td></td><td></td><td></td><td></td><td></td><td></td></tr>
<tr><td>Oligo rev</td><td>AAACGAATACGCACCAGGGGATGA</td><td></td><td></td><td></td><td></td><td></td><td></td><td></td><td></td><td></td><td></td></tr>
<tr><td>Chromosome</td><td>start</td><td>end</td><td>strand</td><td>MM</td><td>target_seq</td><td>PAM</td><td>alignment</td><td>distance</td><td>position</td><td>gene name</td><td>gene id</td></tr>
<tr><td>GU305914.1</td><td>139742</td><td>139764</td><td>+</td><td>0</td><td>TCATCCCCTGGTGCGTATTC</td><td>AGG</td><td>||||||||[||||||||||||]PAM</td><td>NA</td><td>Intergenic</td><td>NA</td><td>NA</td></tr>
<tr><td></td><td></td><td></td><td></td><td></td><td></td><td></td><td></td><td></td><td></td><td></td><td></td></tr>
<tr><td>T10680</td><td>CATCCCCTGGTGCGTATTCAGGG</td><td>1000</td><td></td><td></td><td></td><td></td><td></td><td></td><td></td><td></td><td></td></tr>
<tr><td>Oligo fwd</td><td>CACCCATCCCCTGGTGCGTATTCA</td><td></td><td></td><td></td><td></td><td></td><td></td><td></td><td></td><td></td><td></td></tr>
<tr><td>Oligo rev</td><td>AAACTGAATACGCACCAGGGGATG</td><td></td><td></td><td></td><td></td><td></td><td></td><td></td><td></td><td></td><td></td></tr>
<tr><td>Chromosome</td><td>start</td><td>end</td><td>strand</td><td>MM</td><td>target_seq</td><td>PAM</td><td>alignment</td><td>distance</td><td>position</td><td>gene name</td><td>gene id</td></tr>
<tr><td>GU305914.1</td><td>139743</td><td>139765</td><td>+</td><td>0</td><td>CATCCCCTGGTGCGTATTCA</td><td>GGG</td><td>||||||||[||||||||||||]PAM</td><td>NA</td><td>Intergenic</td><td>NA</td><td>NA</td></tr>
<tr><td></td><td></td><td></td><td></td><td></td><td></td><td></td><td></td><td></td><td></td><td></td><td></td></tr>
<tr><td>T10681</td><td>ATCCCCTGGTGCGTATTCAGGGG</td><td>1000</td><td></td><td></td><td></td><td></td><td></td><td></td><td></td><td></td><td></td></tr>
<tr><td>Oligo fwd</td><td>CACCATCCCCTGGTGCGTATTCAG</td><td></td><td></td><td></td><td></td><td></td><td></td><td></td><td></td><td></td><td></td></tr>
<tr><td>Oligo rev</td><td>AAACCTGAATACGCACCAGGGGAT</td><td></td><td></td><td></td><td></td><td></td><td></td><td></td><td></td><td></td><td></td></tr>
<tr><td>Chromosome</td><td>start</td><td>end</td><td>strand</td><td>MM</td><td>target_seq</td><td>PAM</td><td>alignment</td><td>distance</td><td>position</td><td>gene name</td><td>gene id</td></tr>
<tr><td>GU305914.1</td><td>139744</td><td>139766</td><td>+</td><td>0</td><td>ATCCCCTGGTGCGTATTCAG</td><td>GGG</td><td>||||||||[||||||||||||]PAM</td><td>NA</td><td>Intergenic</td><td>NA</td><td>NA</td></tr>
<tr><td></td><td></td><td></td><td></td><td></td><td></td><td></td><td></td><td></td><td></td><td></td><td></td></tr>
<tr><td>T10682</td><td>TGGTGCGTATTCAGGGGATCAGG</td><td>1000</td><td></td><td></td><td></td><td></td><td></td><td></td><td></td><td></td><td></td></tr>
<tr><td>Oligo fwd</td><td>CACCTGGTGCGTATTCAGGGGATC</td><td></td><td></td><td></td><td></td><td></td><td></td><td></td><td></td><td></td><td></td></tr>
<tr><td>Oligo rev</td><td>AAACGATCCCCTGAATACGCACCA</td><td></td><td></td><td></td><td></td><td></td><td></td><td></td><td></td><td></td><td></td></tr>
<tr><td>Chromosome</td><td>start</td><td>end</td><td>strand</td><td>MM</td><td>target_seq</td><td>PAM</td><td>alignment</td><td>distance</td><td>position</td><td>gene name</td><td>gene id</td></tr>
<tr><td>GU305914.1</td><td>139750</td><td>139772</td><td>+</td><td>0</td><td>TGGTGCGTATTCAGGGGATC</td><td>AGG</td><td>||||||||[||||||||||||]PAM</td><td>NA</td><td>Intergenic</td><td>NA</td><td>NA</td></tr>
<tr><td></td><td></td><td></td><td></td><td></td><td></td><td></td><td></td><td></td><td></td><td></td><td></td></tr>
<tr><td>T10683</td><td>TATTCAGGGGATCAGGACGTCGG</td><td>1000</td><td></td><td></td><td></td><td></td><td></td><td></td><td></td><td></td><td></td></tr>
<tr><td>Oligo fwd</td><td>CACCTATTCAGGGGATCAGGACGT</td><td></td><td></td><td></td><td></td><td></td><td></td><td></td><td></td><td></td><td></td></tr>
<tr><td>Oligo rev</td><td>AAACACGTCCTGATCCCCTGAATA</td><td></td><td></td><td></td><td></td><td></td><td></td><td></td><td></td><td></td><td></td></tr>
<tr><td>Chromosome</td><td>start</td><td>end</td><td>strand</td><td>MM</td><td>target_seq</td><td>PAM</td><td>alignment</td><td>distance</td><td>position</td><td>gene name</td><td>gene id</td></tr>
<tr><td>GU305914.1</td><td>139757</td><td>139779</td><td>+</td><td>0</td><td>TATTCAGGGGATCAGGACGT</td><td>CGG</td><td>||||||||[||||||||||||]PAM</td><td>NA</td><td>Intergenic</td><td>NA</td><td>NA</td></tr>
<tr><td></td><td></td><td></td><td></td><td></td><td></td><td></td><td></td><td></td><td></td><td></td><td></td></tr>
<tr><td>T10684</td><td>GAGAATGCGCGCAGAGCACTCGG</td><td>1000</td><td></td><td></td><td></td><td></td><td></td><td></td><td></td><td></td><td></td></tr>
<tr><td>Oligo fwd</td><td>CACCGAGAATGCGCGCAGAGCACT</td><td></td><td></td><td></td><td></td><td></td><td></td><td></td><td></td><td></td><td></td></tr>
<tr><td>Oligo rev</td><td>AAACAGTGCTCTGCGCGCATTCTC</td><td></td><td></td><td></td><td></td><td></td><td></td><td></td><td></td><td></td><td></td></tr>
<tr><td>Chromosome</td><td>start</td><td>end</td><td>strand</td><td>MM</td><td>target_seq</td><td>PAM</td><td>alignment</td><td>distance</td><td>position</td><td>gene name</td><td>gene id</td></tr>
<tr><td>GU305914.1</td><td>139804</td><td>139826</td><td>+</td><td>0</td><td>GAGAATGCGCGCAGAGCACT</td><td>CGG</td><td>||||||||[||||||||||||]PAM</td><td>NA</td><td>Intergenic</td><td>NA</td><td>NA</td></tr>
<tr><td></td><td></td><td></td><td></td><td></td><td></td><td></td><td></td><td></td><td></td><td></td><td></td></tr>
<tr><td>T10685</td><td>AGAATGCGCGCAGAGCACTCGGG</td><td>1000</td><td></td><td></td><td></td><td></td><td></td><td></td><td></td><td></td><td></td></tr>
</table>

| | | | | | | | | | | | |
|---|---|---|---|---|---|---|---|---|---|---|---|
| Oligo fwd | CACCAGAATGCGCGCAGAGCACTC | | | | | | | | | | |
| Oligo rev | AAACGAGTGCTCTGCGCGCATTCT | | | | | | | | | | |
| Chromosome | start | end | strand | MM | target_seq | PAM | alignment | distance | position | gene name | gene id |
| GU305914.1 | 139805 | 139827 | + | 0 | AGAATGCGCGCAGAGCACTC | GGG | \|\|\|\|\|\|\|\|[\|\|\|\|\|\|\|\|\|\|\|\|]PAM | NA | Intergenic | NA | NA |
| T10686 | ATGCGCGCAGAGCACTCGGGAGG | 1000 | | | | | | | | | |
| Oligo fwd | CACCATGCGCGCAGAGCACTCGGG | | | | | | | | | | |
| Oligo rev | AAACCCCGAGTGCTCTGCGCGCAT | | | | | | | | | | |
| Chromosome | start | end | strand | MM | target_seq | PAM | alignment | distance | position | gene name | gene id |
| GU305914.1 | 139808 | 139830 | + | 0 | ATGCGCGCAGAGCACTCGGG | AGG | \|\|\|\|\|\|\|\|[\|\|\|\|\|\|\|\|\|\|\|\|]PAM | NA | Intergenic | NA | NA |
| T10687 | CGCGCAGAGCACTCGGGAGGTGG | 1000 | | | | | | | | | |
| Oligo fwd | CACCCGCGCAGAGCACTCGGGAGG | | | | | | | | | | |
| Oligo rev | AAACCCTCCCGAGTGCTCTGCGCG | | | | | | | | | | |
| Chromosome | start | end | strand | MM | target_seq | PAM | alignment | distance | position | gene name | gene id |
| GU305914.1 | 139811 | 139833 | + | 0 | CGCGCAGAGCACTCGGGAGG | TGG | \|\|\|\|\|\|\|\|[\|\|\|\|\|\|\|\|\|\|\|\|]PAM | NA | Intergenic | NA | NA |
| T10688 | GAGCACTCGGGAGGTGGCCGCGG | 1000 | | | | | | | | | |
| Oligo fwd | CACCGAGCACTCGGGAGGTGGCCG | | | | | | | | | | |
| Oligo rev | AAACCGGCCACCTCCCGAGTGCTC | | | | | | | | | | |
| Chromosome | start | end | strand | MM | target_seq | PAM | alignment | distance | position | gene name | gene id |
| GU305914.1 | 139817 | 139839 | + | 0 | GAGCACTCGGGAGGTGGCCG | CGG | \|\|\|\|\|\|\|\|[\|\|\|\|\|\|\|\|\|\|\|\|]PAM | NA | Intergenic | NA | NA |
| T10689 | GTGGCCGCGGCGATCGCCGTTGG | 1000 | | | | | | | | | |
| Oligo fwd | CACCGTGGCCGCGGCGATCGCCGT | | | | | | | | | | |
| Oligo rev | AAACACGGCGATCGCCGCGGCCAC | | | | | | | | | | |
| Chromosome | start | end | strand | MM | target_seq | PAM | alignment | distance | position | gene name | gene id |
| GU305914.1 | 139830 | 139852 | + | 0 | GTGGCCGCGGCGATCGCCGT | TGG | \|\|\|\|\|\|\|\|[\|\|\|\|\|\|\|\|\|\|\|\|]PAM | NA | Intergenic | NA | NA |
| GU305914.1 | 22026 | 22048 | - | 4 | GTGCCGGCGGCGATGACCGT | CGG | \|\|\|-\|-\|\|[\|\|\|\|\|\|--\|\|\|\|]PAM | NA | Intergenic | NA | NA |
| T10690 | CGCGGCGATCGCCGTTGGCAAGG | 1000 | | | | | | | | | |
| Oligo fwd | CACCCGCGGCGATCGCCGTTGGCA | | | | | | | | | | |
| Oligo rev | AAACTGCCAACGGCGATCGCCGCG | | | | | | | | | | |
| Chromosome | start | end | strand | MM | target_seq | PAM | alignment | distance | position | gene name | gene id |
| GU305914.1 | 139835 | 139857 | + | 0 | CGCGGCGATCGCCGTTGGCA | AGG | \|\|\|\|\|\|\|\|[\|\|\|\|\|\|\|\|\|\|\|\|]PAM | NA | Intergenic | NA | NA |
| T10691 | CCGTTGGCAAGGCGCAGATGCGG | 1000 | | | | | | | | | |
| Oligo fwd | CACCCCGTTGGCAAGGCGCAGATG | | | | | | | | | | |
| Oligo rev | AAACCATCTGCGCCTTGCCAACGG | | | | | | | | | | |
| Chromosome | start | end | strand | MM | target_seq | PAM | alignment | distance | position | gene name | gene id |
| GU305914.1 | 139846 | 139868 | + | 0 | CCGTTGGCAAGGCGCAGATG | CGG | \|\|\|\|\|\|\|\|[\|\|\|\|\|\|\|\|\|\|\|\|]PAM | NA | Intergenic | NA | NA |
| T10692 | GGCAAGGCGCAGATGCGGAGCGG | 1000 | | | | | | | | | |
| Oligo fwd | CACCGGCAAGGCGCAGATGCGGAG | | | | | | | | | | |
| Oligo rev | AAACCTCCGCATCTGCGCCTTGCC | | | | | | | | | | |
| Chromosome | start | end | strand | MM | target_seq | PAM | alignment | distance | position | gene name | gene id |
| GU305914.1 | 139851 | 139873 | + | 0 | GGCAAGGCGCAGATGCGGAG | CGG | \|\|\|\|\|\|\|\|[\|\|\|\|\|\|\|\|\|\|\|\|]PAM | NA | Intergenic | NA | NA |
| T10693 | AAGGCGCAGATGCGGAGCGGTGG | 1000 | | | | | | | | | |
| Oligo fwd | CACCAAGGCGCAGATGCGGAGCGG | | | | | | | | | | |
| Oligo rev | AAACCCGCTCCGCATCTGCGCCTT | | | | | | | | | | |
| Chromosome | start | end | strand | MM | target_seq | PAM | alignment | distance | position | gene name | gene id |
| GU305914.1 | 139854 | 139876 | + | 0 | AAGGCGCAGATGCGGAGCGG | TGG | \|\|\|\|\|\|\|\|[\|\|\|\|\|\|\|\|\|\|\|\|]PAM | NA | Intergenic | NA | NA |
| T10694 | GAGCGGTGGCGTGTCGACGACGG | 1000 | | | | | | | | | |
| Oligo fwd | CACCGAGCGGTGGCGTGTCGACGA | | | | | | | | | | |
| Oligo rev | AAACTCGTCGACACGCCACCGCTC | | | | | | | | | | |
| Chromosome | start | end | strand | MM | target_seq | PAM | alignment | distance | position | gene name | gene id |
| GU305914.1 | 139868 | 139890 | + | 0 | GAGCGGTGGCGTGTCGACGA | CGG | \|\|\|\|\|\|\|\|[\|\|\|\|\|\|\|\|\|\|\|\|]PAM | NA | Intergenic | NA | NA |
| T10695 | AGCGGTGGCGTGTCGACGACGGG | 1000 | | | | | | | | | |
| Oligo fwd | CACCAGCGGTGGCGTGTCGACGAC | | | | | | | | | | |
| Oligo rev | AAACGTCGTCGACACGCCACCGCT | | | | | | | | | | |
| Chromosome | start | end | strand | MM | target_seq | PAM | alignment | distance | position | gene name | gene id |
| GU305914.1 | 139869 | 139891 | + | 0 | AGCGGTGGCGTGTCGACGAC | GGG | \|\|\|\|\|\|\|\|[\|\|\|\|\|\|\|\|\|\|\|\|]PAM | NA | Intergenic | NA | NA |
| T10696 | GCGGTGGCGTGTCGACGACGGGG | 1000 | | | | | | | | | |
| Oligo fwd | CACCGCGGTGGCGTGTCGACGACG | | | | | | | | | | |
| Oligo rev | AAACCGTCGTCGACACGCCACCGC | | | | | | | | | | |
| Chromosome | start | end | strand | MM | target_seq | PAM | alignment | distance | position | gene name | gene id |
| GU305914.1 | 139870 | 139892 | + | 0 | GCGGTGGCGTGTCGACGACG | GGG | \|\|\|\|\|\|\|\|[\|\|\|\|\|\|\|\|\|\|\|\|]PAM | NA | Intergenic | NA | NA |
| T10697 | TGGCGTGTCGACGACGGGGAAGG | 1000 | | | | | | | | | |
| Oligo fwd | CACCTGGCGTGTCGACGACGGGGA | | | | | | | | | | |
| Oligo rev | AAACTCCCCGTCGTCGACACGCCA | | | | | | | | | | |
| Chromosome | start | end | strand | MM | target_seq | PAM | alignment | distance | position | gene name | gene id |
| GU305914.1 | 139874 | 139896 | + | 0 | TGGCGTGTCGACGACGGGGA | AGG | \|\|\|\|\|\|\|\|[\|\|\|\|\|\|\|\|\|\|\|\|]PAM | NA | Intergenic | NA | NA |
| T10698 | CGGGGAAGGACTCGTCGTCTTGG | 1000 | | | | | | | | | |
| Oligo fwd | CACCCGGGGAAGGACTCGTCGTCT | | | | | | | | | | |
| Oligo rev | AAACAGACGACGAGTCCTTCCCCG | | | | | | | | | | |
| Chromosome | start | end | strand | MM | target_seq | PAM | alignment | distance | position | gene name | gene id |
| GU305914.1 | 139888 | 139910 | + | 0 | CGGGGAAGGACTCGTCGTCT | TGG | \|\|\|\|\|\|\|\|[\|\|\|\|\|\|\|\|\|\|\|\|]PAM | NA | Intergenic | NA | NA |
| T10699 | GGGGAAGGACTCGTCGTCTTGGG | 1000 | | | | | | | | | |
| Oligo fwd | CACCGGGGAAGGACTCGTCGTCTT | | | | | | | | | | |
| Oligo rev | AAACAAGACGACGAGTCCTTCCCC | | | | | | | | | | |
| Chromosome | start | end | strand | MM | target_seq | PAM | alignment | distance | position | gene name | gene id |
| GU305914.1 | 139889 | 139911 | + | 0 | GGGGAAGGACTCGTCGTCTT | GGG | \|\|\|\|\|\|\|\|[\|\|\|\|\|\|\|\|\|\|\|\|]PAM | NA | Intergenic | NA | NA |

| | | | | | | | | | | | |
|---|---|---|---|---|---|---|---|---|---|---|---|
| T10700 | GGATCGCGTTCATCGCGACGCGG | 1000 | | | | | | | | | |
| Oligo fwd | CACCGGATCGCGTTCATCGCGACG | | | | | | | | | | |
| Oligo rev | AAACCGTCGCGATGAACGCGATCC | | | | | | | | | | |
| Chromosome | start | end | strand | MM | target_seq | PAM | alignment | distance | position | gene name | gene id |
| GU305914.1 | 139910 | 139932 | + | 0 | GGATCGCGTTCATCGCGACG | CGG | \|\|\|\|\|\|\|\|[\|\|\|\|\|\|\|\|\|\|\|\|]PAM | NA | Intergenic | NA | NA |

| | | | | | | | | | | | |
|---|---|---|---|---|---|---|---|---|---|---|---|
| T10701 | TCGCGACGCGGACGAGTTCGCGG | 1000 | | | | | | | | | |
| Oligo fwd | CACCTCGCGACGCGGACGAGTTCG | | | | | | | | | | |
| Oligo rev | AAACCGAACTCGTCCGCGTCGCGA | | | | | | | | | | |
| Chromosome | start | end | strand | MM | target_seq | PAM | alignment | distance | position | gene name | gene id |
| GU305914.1 | 139922 | 139944 | + | 0 | TCGCGACGCGGACGAGTTCG | CGG | \|\|\|\|\|\|\|\|[\|\|\|\|\|\|\|\|\|\|\|\|]PAM | NA | Intergenic | NA | NA |
| GU305914.1 | 71620 | 71642 | + | 4 | TCGCGGGACGGACGTGTTCG | GGG | \|\|\|\|\|---[\|\|\|\|\|\|-\|\|\|\|\|]PAM | NA | Intergenic | NA | NA |
| GU305914.1 | 118347 | 118369 | + | 4 | TCGCCGCGCGGACGAGGTTG | CGG | \|\|\|\|--\|\|[\|\|\|\|\|\|\|\|-\|-\|]PAM | NA | Intergenic | NA | NA |

| | | | | | | | | | | | |
|---|---|---|---|---|---|---|---|---|---|---|---|
| T10702 | CGGACGCGCTTGCCGAGATGAGG | 1000 | | | | | | | | | |
| Oligo fwd | CACCCGGACGCGCTTGCCGAGATG | | | | | | | | | | |
| Oligo rev | AAACCATCTCGGCAAGCGCGTCCG | | | | | | | | | | |
| Chromosome | start | end | strand | MM | target_seq | PAM | alignment | distance | position | gene name | gene id |
| GU305914.1 | 139942 | 139964 | + | 0 | CGGACGCGCTTGCCGAGATG | AGG | \|\|\|\|\|\|\|\|[\|\|\|\|\|\|\|\|\|\|\|\|]PAM | NA | Intergenic | NA | NA |

| | | | | | | | | | | | |
|---|---|---|---|---|---|---|---|---|---|---|---|
| T10703 | GGACGCGCTTGCCGAGATGAGGG | 1000 | | | | | | | | | |
| Oligo fwd | CACCGGACGCGCTTGCCGAGATGA | | | | | | | | | | |
| Oligo rev | AAACTCATCTCGGCAAGCGCGTCC | | | | | | | | | | |
| Chromosome | start | end | strand | MM | target_seq | PAM | alignment | distance | position | gene name | gene id |
| GU305914.1 | 139943 | 139965 | + | 0 | GGACGCGCTTGCCGAGATGA | GGG | \|\|\|\|\|\|\|\|[\|\|\|\|\|\|\|\|\|\|\|\|]PAM | NA | Intergenic | NA | NA |

| | | | | | | | | | | | |
|---|---|---|---|---|---|---|---|---|---|---|---|
| T10704 | GCTTGCCGAGATGAGGGACGTGG | 1000 | | | | | | | | | |
| Oligo fwd | CACCGCTTGCCGAGATGAGGGACG | | | | | | | | | | |
| Oligo rev | AAACCGTCCCTCATCTCGGCAAGC | | | | | | | | | | |
| Chromosome | start | end | strand | MM | target_seq | PAM | alignment | distance | position | gene name | gene id |
| GU305914.1 | 139949 | 139971 | + | 0 | GCTTGCCGAGATGAGGGACG | TGG | \|\|\|\|\|\|\|\|[\|\|\|\|\|\|\|\|\|\|\|\|]PAM | NA | Intergenic | NA | NA |

| | | | | | | | | | | | |
|---|---|---|---|---|---|---|---|---|---|---|---|
| T10705 | CGAGATGAGGGACGTGGCCGAGG | 1000 | | | | | | | | | |
| Oligo fwd | CACCCGAGATGAGGGACGTGGCCG | | | | | | | | | | |
| Oligo rev | AAACCGGCCACGTCCCTCATCTCG | | | | | | | | | | |
| Chromosome | start | end | strand | MM | target_seq | PAM | alignment | distance | position | gene name | gene id |
| GU305914.1 | 139955 | 139977 | + | 0 | CGAGATGAGGGACGTGGCCG | AGG | \|\|\|\|\|\|\|\|[\|\|\|\|\|\|\|\|\|\|\|\|]PAM | NA | Intergenic | NA | NA |

| | | | | | | | | | | | |
|---|---|---|---|---|---|---|---|---|---|---|---|
| T10706 | GTTCACGTCATCAGACAGTGAGG | 1000 | | | | | | | | | |
| Oligo fwd | CACCGTTCACGTCATCAGACAGTG | | | | | | | | | | |
| Oligo rev | AAACCACTGTCTGATGACGTGAAC | | | | | | | | | | |
| Chromosome | start | end | strand | MM | target_seq | PAM | alignment | distance | position | gene name | gene id |
| GU305914.1 | 139991 | 140013 | + | 0 | GTTCACGTCATCAGACAGTG | AGG | \|\|\|\|\|\|\|\|[\|\|\|\|\|\|\|\|\|\|\|\|]PAM | NA | Intergenic | NA | NA |

| | | | | | | | | | | | |
|---|---|---|---|---|---|---|---|---|---|---|---|
| T10707 | CACGTCATCAGACAGTGAGGAGG | 1000 | | | | | | | | | |
| Oligo fwd | CACCCACGTCATCAGACAGTGAGG | | | | | | | | | | |
| Oligo rev | AAACCCTCACTGTCTGATGACGTG | | | | | | | | | | |
| Chromosome | start | end | strand | MM | target_seq | PAM | alignment | distance | position | gene name | gene id |
| GU305914.1 | 139994 | 140016 | + | 0 | CACGTCATCAGACAGTGAGG | AGG | \|\|\|\|\|\|\|\|[\|\|\|\|\|\|\|\|\|\|\|\|]PAM | NA | Intergenic | NA | NA |

| | | | | | | | | | | | |
|---|---|---|---|---|---|---|---|---|---|---|---|
| T10708 | CAGTGAGGAGGTGTGACTGACGG | 1000 | | | | | | | | | |
| Oligo fwd | CACCCAGTGAGGAGGTGTGACTGA | | | | | | | | | | |
| Oligo rev | AAACTCAGTCACACCTCCTCACTG | | | | | | | | | | |
| Chromosome | start | end | strand | MM | target_seq | PAM | alignment | distance | position | gene name | gene id |
| GU305914.1 | 140006 | 140028 | + | 0 | CAGTGAGGAGGTGTGACTGA | CGG | \|\|\|\|\|\|\|\|[\|\|\|\|\|\|\|\|\|\|\|\|]PAM | NA | Intergenic | NA | NA |

| | | | | | | | | | | | |
|---|---|---|---|---|---|---|---|---|---|---|---|
| T10709 | CTGCCGCTGTGTTGCGCGCTCGG | 1000 | | | | | | | | | |
| Oligo fwd | CACCCTGCCGCTGTGTTGCGCGCT | | | | | | | | | | |
| Oligo rev | AAACAGCGCGCAACACAGCGGCAG | | | | | | | | | | |
| Chromosome | start | end | strand | MM | target_seq | PAM | alignment | distance | position | gene name | gene id |
| GU305914.1 | 140054 | 140076 | + | 0 | CTGCCGCTGTGTTGCGCGCT | CGG | \|\|\|\|\|\|\|\|[\|\|\|\|\|\|\|\|\|\|\|\|]PAM | NA | Intergenic | NA | NA |

| | | | | | | | | | | | |
|---|---|---|---|---|---|---|---|---|---|---|---|
| T10710 | GTTGCGCGCTCGGATCACTCTGG | 1000 | | | | | | | | | |
| Oligo fwd | CACCGTTGCGCGCTCGGATCACTC | | | | | | | | | | |
| Oligo rev | AAACGAGTGATCCGAGCGCGCAAC | | | | | | | | | | |
| Chromosome | start | end | strand | MM | target_seq | PAM | alignment | distance | position | gene name | gene id |
| GU305914.1 | 140064 | 140086 | + | 0 | GTTGCGCGCTCGGATCACTC | TGG | \|\|\|\|\|\|\|\|[\|\|\|\|\|\|\|\|\|\|\|\|]PAM | NA | Intergenic | NA | NA |

| | | | | | | | | | | | |
|---|---|---|---|---|---|---|---|---|---|---|---|
| T10711 | CTTTTTATTATTAAGTGAGCAGG | 1000 | | | | | | | | | |
| Oligo fwd | CACCCTTTTTATTATTAAGTGAGC | | | | | | | | | | |
| Oligo rev | AAACGCTCACTTAATAATAAAAAG | | | | | | | | | | |
| Chromosome | start | end | strand | MM | target_seq | PAM | alignment | distance | position | gene name | gene id |
| GU305914.1 | 140094 | 140116 | + | 0 | CTTTTTATTATTAAGTGAGC | AGG | \|\|\|\|\|\|\|\|[\|\|\|\|\|\|\|\|\|\|\|\|]PAM | NA | Intergenic | NA | NA |

| | | | | | | | | | | | |
|---|---|---|---|---|---|---|---|---|---|---|---|
| T10712 | TTTTTATTATTAAGTGAGCAGGG | 1000 | | | | | | | | | |
| Oligo fwd | CACCTTTTTATTATTAAGTGAGCA | | | | | | | | | | |
| Oligo rev | AAACTGCTCACTTAATAATAAAAA | | | | | | | | | | |
| Chromosome | start | end | strand | MM | target_seq | PAM | alignment | distance | position | gene name | gene id |
| GU305914.1 | 140095 | 140117 | + | 0 | TTTTTATTATTAAGTGAGCA | GGG | \|\|\|\|\|\|\|\|[\|\|\|\|\|\|\|\|\|\|\|\|]PAM | NA | Intergenic | NA | NA |

| | | | | | | | | | | | |
|---|---|---|---|---|---|---|---|---|---|---|---|
| T10713 | TAAGTGAGCAGGGTCCGTGACGG | 1000 | | | | | | | | | |
| Oligo fwd | CACCTAAGTGAGCAGGGTCCGTGA | | | | | | | | | | |
| Oligo rev | AAACTCACGGACCCTGCTCACTTA | | | | | | | | | | |
| Chromosome | start | end | strand | MM | target_seq | PAM | alignment | distance | position | gene name | gene id |
| GU305914.1 | 140105 | 140127 | + | 0 | TAAGTGAGCAGGGTCCGTGA | CGG | \|\|\|\|\|\|\|\|[\|\|\|\|\|\|\|\|\|\|\|\|]PAM | NA | Intergenic | NA | NA |

| | | |
|---|---|---|
| T10714 | GTGAGCAGGGTCCGTGACGGTGG | 1000 |
| Oligo fwd | CACCGTGAGCAGGGTCCGTGACGG | |

Oligo rev AAACCCGTCACGGACCCTGCTCAC
Chromosome start end strand MM target_seq PAM alignment distance position gene name gene id
GU305914.1 140108 140130 + 0 GTGAGCAGGGTCCGTGACGG TGG ||||||||[||||||||||||]PAM NA Intergenic NA NA

T10715 CAGGGTCCGTGACGGTGGTCCGG 1000
Oligo fwd CACCCAGGGTCCGTGACGGTGGTC
Oligo rev AAACGACCACCGTCACGGACCCTG
Chromosome start end strand MM target_seq PAM alignment distance position gene name gene id
GU305914.1 140113 140135 + 0 CAGGGTCCGTGACGGTGGTC CGG ||||||||[||||||||||||]PAM NA Intergenic NA NA

T10716 AGGGTCCGTGACGGTGGTCCGGG 1000
Oligo fwd CACCAGGGTCCGTGACGGTGGTCC
Oligo rev AAACGGACCACCGTCACGGACCCT
Chromosome start end strand MM target_seq PAM alignment distance position gene name gene id
GU305914.1 140114 140136 + 0 AGGGTCCGTGACGGTGGTCC GGG ||||||||[||||||||||||]PAM NA Intergenic NA NA

T10717 CCGTGACGGTGGTCCGGGTGAGG 1000
Oligo fwd CACCCCGTGACGGTGGTCCGGGTG
Oligo rev AAACCACCCGGACCACCGTCACGG
Chromosome start end strand MM target_seq PAM alignment distance position gene name gene id
GU305914.1 140119 140141 + 0 CCGTGACGGTGGTCCGGGTG AGG ||||||||[||||||||||||]PAM NA Intergenic NA NA

T10718 CGTGACGGTGGTCCGGGTGAGGG 1000
Oligo fwd CACCCGTGACGGTGGTCCGGGTGA
Oligo rev AAACTCACCCGGACCACCGTCACG
Chromosome start end strand MM target_seq PAM alignment distance position gene name gene id
GU305914.1 140120 140142 + 0 CGTGACGGTGGTCCGGGTGA GGG ||||||||[||||||||||||]PAM NA Intergenic NA NA

T10719 GAGGGTGATCGCATCCCCCGAGG 1000
Oligo fwd CACCGAGGGTGATCGCATCCCCCG
Oligo rev AAACCGGGGGATGCGATCACCCTC
Chromosome start end strand MM target_seq PAM alignment distance position gene name gene id
GU305914.1 140138 140160 + 0 GAGGGTGATCGCATCCCCCG AGG ||||||||[||||||||||||]PAM NA Intergenic NA NA

T10720 AGGGTGATCGCATCCCCCGAGGG 1000
Oligo fwd CACCAGGGTGATCGCATCCCCCGA
Oligo rev AAACTCGGGGGATGCGATCACCCT
Chromosome start end strand MM target_seq PAM alignment distance position gene name gene id
GU305914.1 140139 140161 + 0 AGGGTGATCGCATCCCCCGA GGG ||||||||[||||||||||||]PAM NA Intergenic NA NA

T10721 GGGTGATCGCATCCCCCGAGGGG 1000
Oligo fwd CACCGGGTGATCGCATCCCCCGAG
Oligo rev AAACCTCGGGGGATGCGATCACCC
Chromosome start end strand MM target_seq PAM alignment distance position gene name gene id
GU305914.1 140140 140162 + 0 GGGTGATCGCATCCCCCGAG GGG ||||||||[||||||||||||]PAM NA Intergenic NA NA

T10722 GATCGCATCCCCCGAGGGGTCGG 1000
Oligo fwd CACCGATCGCATCCCCCGAGGGGT
Oligo rev AAACACCCCTCGGGGGATGCGATC
Chromosome start end strand MM target_seq PAM alignment distance position gene name gene id
GU305914.1 140144 140166 + 0 GATCGCATCCCCCGAGGGGT CGG ||||||||[||||||||||||]PAM NA Intergenic NA NA

T10723 CCGAGGGGTCGGTCGACGAACGG 1000
Oligo fwd CACCCCGAGGGGTCGGTCGACGAA
Oligo rev AAACTTCGTCGACCGACCCCTCGG
Chromosome start end strand MM target_seq PAM alignment distance position gene name gene id
GU305914.1 140155 140177 + 0 CCGAGGGGTCGGTCGACGAA CGG ||||||||[||||||||||||]PAM NA Intergenic NA NA

T10724 GGGGTCGGTCGACGAACGGACGG 1000
Oligo fwd CACCGGGGTCGGTCGACGAACGGA
Oligo rev AAACTCCGTTCGTCGACCGACCCC
Chromosome start end strand MM target_seq PAM alignment distance position gene name gene id
GU305914.1 140159 140181 + 0 GGGGTCGGTCGACGAACGGA CGG ||||||||[||||||||||||]PAM NA Intergenic NA NA

T10725 GGGTCGGTCGACGAACGGACGGG 1000
Oligo fwd CACCGGGTCGGTCGACGAACGGAC
Oligo rev AAACGTCCGTTCGTCGACCGACCC
Chromosome start end strand MM target_seq PAM alignment distance position gene name gene id
GU305914.1 140160 140182 + 0 GGGTCGGTCGACGAACGGAC GGG ||||||||[||||||||||||]PAM NA Intergenic NA NA

T10726 CCCCGCCGTCGCCATGAGCGTGG 1000
Oligo fwd CACCCCCCGCCGTCGCCATGAGCG
Oligo rev AAACCGCTCATGGCGACGGCGGGG
Chromosome start end strand MM target_seq PAM alignment distance position gene name gene id
GU305914.1 140225 140247 + 0 CCCCGCCGTCGCCATGAGCG TGG ||||||||[||||||||||||]PAM NA Intergenic NA NA

T10727 CTGACGCCGCCGAGAAGCGACGG 1000
Oligo fwd CACCCTGACGCCGCCGAGAAGCGA
Oligo rev AAACTCGCTTCTCGGCGGCGTCAG
Chromosome start end strand MM target_seq PAM alignment distance position gene name gene id
GU305914.1 140250 140272 + 0 CTGACGCCGCCGAGAAGCGA CGG ||||||||[||||||||||||]PAM NA Intergenic NA NA

T10728 CCGAGAAGCGACGGCTCCGTCGG 1000
Oligo fwd CACCCCGAGAAGCGACGGCTCCGT
Oligo rev AAACACGGAGCCGTCGCTTCTCGG
Chromosome start end strand MM target_seq PAM alignment distance position gene name gene id
GU305914.1 140259 140281 + 0 CCGAGAAGCGACGGCTCCGT CGG ||||||||[||||||||||||]PAM NA Intergenic NA NA

T10729 CGAGAAGCGACGGCTCCGTCGGG 1000

<table>
<tr><td>Oligo fwd</td><td>CACCCGAGAAGCGACGGCTCCGTC</td><td></td><td></td><td></td><td></td><td></td><td></td><td></td><td></td><td></td><td></td></tr>
<tr><td>Oligo rev</td><td>AAACGACGGAGCCGTCGCTTCTCG</td><td></td><td></td><td></td><td></td><td></td><td></td><td></td><td></td><td></td><td></td></tr>
<tr><td>Chromosome</td><td>start</td><td>end</td><td>strand</td><td>MM</td><td>target_seq</td><td>PAM</td><td>alignment</td><td>distance</td><td>position</td><td>gene name</td><td>gene id</td></tr>
<tr><td>GU305914.1</td><td>140260</td><td>140282</td><td>+</td><td>0</td><td>CGAGAAGCGACGGCTCCGTC</td><td>GGG</td><td>||||||||[||||||||||||]PAM</td><td>NA</td><td>Intergenic</td><td>NA</td><td>NA</td></tr>
<tr><td></td><td></td><td></td><td></td><td></td><td></td><td></td><td></td><td></td><td></td><td></td><td></td></tr>
<tr><td>T10730</td><td>CGCTCCCGTCGTCGTCCCTCCGG</td><td>1000</td><td></td><td></td><td></td><td></td><td></td><td></td><td></td><td></td><td></td></tr>
<tr><td>Oligo fwd</td><td>CACCCGCTCCCGTCGTCGTCCCTC</td><td></td><td></td><td></td><td></td><td></td><td></td><td></td><td></td><td></td><td></td></tr>
<tr><td>Oligo rev</td><td>AAACGAGGGACGACGACGGGAGCG</td><td></td><td></td><td></td><td></td><td></td><td></td><td></td><td></td><td></td><td></td></tr>
<tr><td>Chromosome</td><td>start</td><td>end</td><td>strand</td><td>MM</td><td>target_seq</td><td>PAM</td><td>alignment</td><td>distance</td><td>position</td><td>gene name</td><td>gene id</td></tr>
<tr><td>GU305914.1</td><td>140285</td><td>140307</td><td>+</td><td>0</td><td>CGCTCCCGTCGTCGTCCCTC</td><td>CGG</td><td>||||||||[||||||||||||]PAM</td><td>NA</td><td>Intergenic</td><td>NA</td><td>NA</td></tr>
<tr><td></td><td></td><td></td><td></td><td></td><td></td><td></td><td></td><td></td><td></td><td></td><td></td></tr>
<tr><td>T10731</td><td>CGGCTCCCAGAAAGCCGTTGAGG</td><td>1000</td><td></td><td></td><td></td><td></td><td></td><td></td><td></td><td></td><td></td></tr>
<tr><td>Oligo fwd</td><td>CACCCGGCTCCCAGAAAGCCGTTG</td><td></td><td></td><td></td><td></td><td></td><td></td><td></td><td></td><td></td><td></td></tr>
<tr><td>Oligo rev</td><td>AAACCAACGGCTTTCTGGGAGCCG</td><td></td><td></td><td></td><td></td><td></td><td></td><td></td><td></td><td></td><td></td></tr>
<tr><td>Chromosome</td><td>start</td><td>end</td><td>strand</td><td>MM</td><td>target_seq</td><td>PAM</td><td>alignment</td><td>distance</td><td>position</td><td>gene name</td><td>gene id</td></tr>
<tr><td>GU305914.1</td><td>140305</td><td>140327</td><td>+</td><td>0</td><td>CGGCTCCCAGAAAGCCGTTG</td><td>AGG</td><td>||||||||[||||||||||||]PAM</td><td>NA</td><td>Intergenic</td><td>NA</td><td>NA</td></tr>
<tr><td></td><td></td><td></td><td></td><td></td><td></td><td></td><td></td><td></td><td></td><td></td><td></td></tr>
<tr><td>T10732</td><td>GCGACGCGCCGTGAGCGATCTGG</td><td>1000</td><td></td><td></td><td></td><td></td><td></td><td></td><td></td><td></td><td></td></tr>
<tr><td>Oligo fwd</td><td>CACCGCGACGCGCCGTGAGCGATC</td><td></td><td></td><td></td><td></td><td></td><td></td><td></td><td></td><td></td><td></td></tr>
<tr><td>Oligo rev</td><td>AAACGATCGCTCACGGCGCGTCGC</td><td></td><td></td><td></td><td></td><td></td><td></td><td></td><td></td><td></td><td></td></tr>
<tr><td>Chromosome</td><td>start</td><td>end</td><td>strand</td><td>MM</td><td>target_seq</td><td>PAM</td><td>alignment</td><td>distance</td><td>position</td><td>gene name</td><td>gene id</td></tr>
<tr><td>GU305914.1</td><td>140327</td><td>140349</td><td>+</td><td>0</td><td>GCGACGCGCCGTGAGCGATC</td><td>TGG</td><td>||||||||[||||||||||||]PAM</td><td>NA</td><td>Intergenic</td><td>NA</td><td>NA</td></tr>
<tr><td></td><td></td><td></td><td></td><td></td><td></td><td></td><td></td><td></td><td></td><td></td><td></td></tr>
<tr><td>T10733</td><td>AAAAACTGTATAAAGTCAAGAGG</td><td>1000</td><td></td><td></td><td></td><td></td><td></td><td></td><td></td><td></td><td></td></tr>
<tr><td>Oligo fwd</td><td>CACCAAAAACTGTATAAAGTCAAG</td><td></td><td></td><td></td><td></td><td></td><td></td><td></td><td></td><td></td><td></td></tr>
<tr><td>Oligo rev</td><td>AAACCTTGACTTTATACAGTTTTT</td><td></td><td></td><td></td><td></td><td></td><td></td><td></td><td></td><td></td><td></td></tr>
<tr><td>Chromosome</td><td>start</td><td>end</td><td>strand</td><td>MM</td><td>target_seq</td><td>PAM</td><td>alignment</td><td>distance</td><td>position</td><td>gene name</td><td>gene id</td></tr>
<tr><td>GU305914.1</td><td>140350</td><td>140372</td><td>+</td><td>0</td><td>AAAAACTGTATAAAGTCAAG</td><td>AGG</td><td>||||||||[||||||||||||]PAM</td><td>NA</td><td>Intergenic</td><td>NA</td><td>NA</td></tr>
<tr><td></td><td></td><td></td><td></td><td></td><td></td><td></td><td></td><td></td><td></td><td></td><td></td></tr>
<tr><td>T10734</td><td>GTATAAAGTCAAGAGGAGATTGG</td><td>1000</td><td></td><td></td><td></td><td></td><td></td><td></td><td></td><td></td><td></td></tr>
<tr><td>Oligo fwd</td><td>CACCGTATAAAGTCAAGAGGAGAT</td><td></td><td></td><td></td><td></td><td></td><td></td><td></td><td></td><td></td><td></td></tr>
<tr><td>Oligo rev</td><td>AAACATCTCCTCTTGACTTTATAC</td><td></td><td></td><td></td><td></td><td></td><td></td><td></td><td></td><td></td><td></td></tr>
<tr><td>Chromosome</td><td>start</td><td>end</td><td>strand</td><td>MM</td><td>target_seq</td><td>PAM</td><td>alignment</td><td>distance</td><td>position</td><td>gene name</td><td>gene id</td></tr>
<tr><td>GU305914.1</td><td>140357</td><td>140379</td><td>+</td><td>0</td><td>GTATAAAGTCAAGAGGAGAT</td><td>TGG</td><td>||||||||[||||||||||||]PAM</td><td>NA</td><td>Intergenic</td><td>NA</td><td>NA</td></tr>
<tr><td></td><td></td><td></td><td></td><td></td><td></td><td></td><td></td><td></td><td></td><td></td><td></td></tr>
<tr><td>T10735</td><td>GTCAAGAGGAGATTGGTGTTCGG</td><td>1000</td><td></td><td></td><td></td><td></td><td></td><td></td><td></td><td></td><td></td></tr>
<tr><td>Oligo fwd</td><td>CACCGTCAAGAGGAGATTGGTGTT</td><td></td><td></td><td></td><td></td><td></td><td></td><td></td><td></td><td></td><td></td></tr>
<tr><td>Oligo rev</td><td>AAACAACACCAATCTCCTCTTGAC</td><td></td><td></td><td></td><td></td><td></td><td></td><td></td><td></td><td></td><td></td></tr>
<tr><td>Chromosome</td><td>start</td><td>end</td><td>strand</td><td>MM</td><td>target_seq</td><td>PAM</td><td>alignment</td><td>distance</td><td>position</td><td>gene name</td><td>gene id</td></tr>
<tr><td>GU305914.1</td><td>140364</td><td>140386</td><td>+</td><td>0</td><td>GTCAAGAGGAGATTGGTGTT</td><td>CGG</td><td>||||||||[||||||||||||]PAM</td><td>NA</td><td>Intergenic</td><td>NA</td><td>NA</td></tr>
<tr><td></td><td></td><td></td><td></td><td></td><td></td><td></td><td></td><td></td><td></td><td></td><td></td></tr>
<tr><td>T10736</td><td>TCAAGAGGAGATTGGTGTTCGGG</td><td>1000</td><td></td><td></td><td></td><td></td><td></td><td></td><td></td><td></td><td></td></tr>
<tr><td>Oligo fwd</td><td>CACCTCAAGAGGAGATTGGTGTTC</td><td></td><td></td><td></td><td></td><td></td><td></td><td></td><td></td><td></td><td></td></tr>
<tr><td>Oligo rev</td><td>AAACGAACACCAATCTCCTCTTGA</td><td></td><td></td><td></td><td></td><td></td><td></td><td></td><td></td><td></td><td></td></tr>
<tr><td>Chromosome</td><td>start</td><td>end</td><td>strand</td><td>MM</td><td>target_seq</td><td>PAM</td><td>alignment</td><td>distance</td><td>position</td><td>gene name</td><td>gene id</td></tr>
<tr><td>GU305914.1</td><td>140365</td><td>140387</td><td>+</td><td>0</td><td>TCAAGAGGAGATTGGTGTTC</td><td>GGG</td><td>||||||||[||||||||||||]PAM</td><td>NA</td><td>Intergenic</td><td>NA</td><td>NA</td></tr>
<tr><td></td><td></td><td></td><td></td><td></td><td></td><td></td><td></td><td></td><td></td><td></td><td></td></tr>
<tr><td>T10737</td><td>CAAGAGGAGATTGGTGTTCGGGG</td><td>1000</td><td></td><td></td><td></td><td></td><td></td><td></td><td></td><td></td><td></td></tr>
<tr><td>Oligo fwd</td><td>CACCCAAGAGGAGATTGGTGTTCG</td><td></td><td></td><td></td><td></td><td></td><td></td><td></td><td></td><td></td><td></td></tr>
<tr><td>Oligo rev</td><td>AAACCGAACACCAATCTCCTCTTG</td><td></td><td></td><td></td><td></td><td></td><td></td><td></td><td></td><td></td><td></td></tr>
<tr><td>Chromosome</td><td>start</td><td>end</td><td>strand</td><td>MM</td><td>target_seq</td><td>PAM</td><td>alignment</td><td>distance</td><td>position</td><td>gene name</td><td>gene id</td></tr>
<tr><td>GU305914.1</td><td>140366</td><td>140388</td><td>+</td><td>0</td><td>CAAGAGGAGATTGGTGTTCG</td><td>GGG</td><td>||||||||[||||||||||||]PAM</td><td>NA</td><td>Intergenic</td><td>NA</td><td>NA</td></tr>
<tr><td></td><td></td><td></td><td></td><td></td><td></td><td></td><td></td><td></td><td></td><td></td><td></td></tr>
<tr><td>T10738</td><td>TTGGTGTTCGGGGCCGACGATGG</td><td>1000</td><td></td><td></td><td></td><td></td><td></td><td></td><td></td><td></td><td></td></tr>
<tr><td>Oligo fwd</td><td>CACCTTGGTGTTCGGGGCCGACGA</td><td></td><td></td><td></td><td></td><td></td><td></td><td></td><td></td><td></td><td></td></tr>
<tr><td>Oligo rev</td><td>AAACTCGTCGGCCCCGAACACCAA</td><td></td><td></td><td></td><td></td><td></td><td></td><td></td><td></td><td></td><td></td></tr>
<tr><td>Chromosome</td><td>start</td><td>end</td><td>strand</td><td>MM</td><td>target_seq</td><td>PAM</td><td>alignment</td><td>distance</td><td>position</td><td>gene name</td><td>gene id</td></tr>
<tr><td>GU305914.1</td><td>140376</td><td>140398</td><td>+</td><td>0</td><td>TTGGTGTTCGGGGCCGACGA</td><td>TGG</td><td>||||||||[||||||||||||]PAM</td><td>NA</td><td>Intergenic</td><td>NA</td><td>NA</td></tr>
<tr><td></td><td></td><td></td><td></td><td></td><td></td><td></td><td></td><td></td><td></td><td></td><td></td></tr>
<tr><td>T10739</td><td>GATACCAGTAACAACAACAGCGG</td><td>1000</td><td></td><td></td><td></td><td></td><td></td><td></td><td></td><td></td><td></td></tr>
<tr><td>Oligo fwd</td><td>CACCGATACCAGTAACAACAACAG</td><td></td><td></td><td></td><td></td><td></td><td></td><td></td><td></td><td></td><td></td></tr>
<tr><td>Oligo rev</td><td>AAACCTGTTGTTGTTACTGGTATC</td><td></td><td></td><td></td><td></td><td></td><td></td><td></td><td></td><td></td><td></td></tr>
<tr><td>Chromosome</td><td>start</td><td>end</td><td>strand</td><td>MM</td><td>target_seq</td><td>PAM</td><td>alignment</td><td>distance</td><td>position</td><td>gene name</td><td>gene id</td></tr>
<tr><td>GU305914.1</td><td>140412</td><td>140434</td><td>+</td><td>0</td><td>GATACCAGTAACAACAACAG</td><td>CGG</td><td>||||||||[||||||||||||]PAM</td><td>NA</td><td>Intergenic</td><td>NA</td><td>NA</td></tr>
<tr><td></td><td></td><td></td><td></td><td></td><td></td><td></td><td></td><td></td><td></td><td></td><td></td></tr>
<tr><td>T10740</td><td>GCAGTAGCAGCACCACCAGCAGG</td><td>1000</td><td></td><td></td><td></td><td></td><td></td><td></td><td></td><td></td><td></td></tr>
<tr><td>Oligo fwd</td><td>CACCGCAGTAGCAGCACCACCAGC</td><td></td><td></td><td></td><td></td><td></td><td></td><td></td><td></td><td></td><td></td></tr>
<tr><td>Oligo rev</td><td>AAACGCTGGTGGTGCTGCTACTGC</td><td></td><td></td><td></td><td></td><td></td><td></td><td></td><td></td><td></td><td></td></tr>
<tr><td>Chromosome</td><td>start</td><td>end</td><td>strand</td><td>MM</td><td>target_seq</td><td>PAM</td><td>alignment</td><td>distance</td><td>position</td><td>gene name</td><td>gene id</td></tr>
<tr><td>GU305914.1</td><td>140434</td><td>140456</td><td>+</td><td>0</td><td>GCAGTAGCAGCACCACCAGC</td><td>AGG</td><td>||||||||[||||||||||||]PAM</td><td>NA</td><td>Intergenic</td><td>NA</td><td>NA</td></tr>
<tr><td></td><td></td><td></td><td></td><td></td><td></td><td></td><td></td><td></td><td></td><td></td><td></td></tr>
<tr><td>T10741</td><td>GCAGCACCACCAGCAGGTCGCGG</td><td>1000</td><td></td><td></td><td></td><td></td><td></td><td></td><td></td><td></td><td></td></tr>
<tr><td>Oligo fwd</td><td>CACCGCAGCACCACCAGCAGGTCG</td><td></td><td></td><td></td><td></td><td></td><td></td><td></td><td></td><td></td><td></td></tr>
<tr><td>Oligo rev</td><td>AAACCGACCTGCTGGTGGTGCTGC</td><td></td><td></td><td></td><td></td><td></td><td></td><td></td><td></td><td></td><td></td></tr>
<tr><td>Chromosome</td><td>start</td><td>end</td><td>strand</td><td>MM</td><td>target_seq</td><td>PAM</td><td>alignment</td><td>distance</td><td>position</td><td>gene name</td><td>gene id</td></tr>
<tr><td>GU305914.1</td><td>140440</td><td>140462</td><td>+</td><td>0</td><td>GCAGCACCACCAGCAGGTCG</td><td>CGG</td><td>||||||||[||||||||||||]PAM</td><td>NA</td><td>Intergenic</td><td>NA</td><td>NA</td></tr>
<tr><td>GU305914.1</td><td>107638</td><td>107660</td><td>+</td><td>4</td><td>TGACCACCACCAGGAGGTCG</td><td>TGG</td><td>--|-||||[|||||-||||||]PAM</td><td>NA</td><td>Intergenic</td><td>NA</td><td>NA</td></tr>
<tr><td></td><td></td><td></td><td></td><td></td><td></td><td></td><td></td><td></td><td></td><td></td><td></td></tr>
<tr><td>T10742</td><td>CAGCAGGTCGCGGCGCAAGACGG</td><td>1000</td><td></td><td></td><td></td><td></td><td></td><td></td><td></td><td></td><td></td></tr>
<tr><td>Oligo fwd</td><td>CACCCAGCAGGTCGCGGCGCAAGA</td><td></td><td></td><td></td><td></td><td></td><td></td><td></td><td></td><td></td><td></td></tr>
<tr><td>Oligo rev</td><td>AAACTCTTGCGCCGCGACCTGCTG</td><td></td><td></td><td></td><td></td><td></td><td></td><td></td><td></td><td></td><td></td></tr>
<tr><td>Chromosome</td><td>start</td><td>end</td><td>strand</td><td>MM</td><td>target_seq</td><td>PAM</td><td>alignment</td><td>distance</td><td>position</td><td>gene name</td><td>gene id</td></tr>
<tr><td>GU305914.1</td><td>140450</td><td>140472</td><td>+</td><td>0</td><td>CAGCAGGTCGCGGCGCAAGA</td><td>CGG</td><td>||||||||[||||||||||||]PAM</td><td>NA</td><td>Intergenic</td><td>NA</td><td>NA</td></tr>
<tr><td></td><td></td><td></td><td></td><td></td><td></td><td></td><td></td><td></td><td></td><td></td><td></td></tr>
<tr><td>T10743</td><td>CACTGATGATTCTTCCACCGCGG</td><td>1000</td><td></td><td></td><td></td><td></td><td></td><td></td><td></td><td></td><td></td></tr>
<tr><td>Oligo fwd</td><td>CACCCACTGATGATTCTTCCACCG</td><td></td><td></td><td></td><td></td><td></td><td></td><td></td><td></td><td></td><td></td></tr>
<tr><td>Oligo rev</td><td>AAACCGGTGGAAGAATCATCAGTG</td><td></td><td></td><td></td><td></td><td></td><td></td><td></td><td></td><td></td><td></td></tr>
<tr><td>Chromosome</td><td>start</td><td>end</td><td>strand</td><td>MM</td><td>target_seq</td><td>PAM</td><td>alignment</td><td>distance</td><td>position</td><td>gene name</td><td>gene id</td></tr>
<tr><td>GU305914.1</td><td>140504</td><td>140526</td><td>+</td><td>0</td><td>CACTGATGATTCTTCCACCG</td><td>CGG</td><td>||||||||[||||||||||||]PAM</td><td>NA</td><td>Intergenic</td><td>NA</td><td>NA</td></tr>
</table>

<table>
<tr><td>T10744</td><td>ACTGATGATTCTTCCACCGCGGG</td><td>1000</td><td></td><td></td><td></td><td></td><td></td><td></td><td></td><td></td><td></td></tr>
<tr><td>Oligo fwd</td><td>CACCACTGATGATTCTTCCACCGC</td><td></td><td></td><td></td><td></td><td></td><td></td><td></td><td></td><td></td><td></td></tr>
<tr><td>Oligo rev</td><td>AAACGCGGTGGAAGAATCATCAGT</td><td></td><td></td><td></td><td></td><td></td><td></td><td></td><td></td><td></td><td></td></tr>
<tr><td>Chromosome</td><td>start</td><td>end</td><td>strand</td><td>MM</td><td>target_seq</td><td>PAM</td><td>alignment</td><td>distance</td><td>position</td><td>gene name</td><td>gene id</td></tr>
<tr><td>GU305914.1</td><td>140505</td><td>140527</td><td>+</td><td>0</td><td>ACTGATGATTCTTCCACCGC</td><td>GGG</td><td>||||||||[||||||||||||]PAM</td><td>NA</td><td>Intergenic</td><td>NA</td><td>NA</td></tr>
<tr><td>T10745</td><td>TGATTCTTCCACCGCGGGCGAGG</td><td>1000</td><td></td><td></td><td></td><td></td><td></td><td></td><td></td><td></td><td></td></tr>
<tr><td>Oligo fwd</td><td>CACCTGATTCTTCCACCGCGGGCG</td><td></td><td></td><td></td><td></td><td></td><td></td><td></td><td></td><td></td><td></td></tr>
<tr><td>Oligo rev</td><td>AAACCGCCCGCGGTGGAAGAATCA</td><td></td><td></td><td></td><td></td><td></td><td></td><td></td><td></td><td></td><td></td></tr>
<tr><td>Chromosome</td><td>start</td><td>end</td><td>strand</td><td>MM</td><td>target_seq</td><td>PAM</td><td>alignment</td><td>distance</td><td>position</td><td>gene name</td><td>gene id</td></tr>
<tr><td>GU305914.1</td><td>140510</td><td>140532</td><td>+</td><td>0</td><td>TGATTCTTCCACCGCGGGCG</td><td>AGG</td><td>||||||||[||||||||||||]PAM</td><td>NA</td><td>Intergenic</td><td>NA</td><td>NA</td></tr>
<tr><td>T10746</td><td>TCTTCCACCGCGGGCGAGGATGG</td><td>1000</td><td></td><td></td><td></td><td></td><td></td><td></td><td></td><td></td><td></td></tr>
<tr><td>Oligo fwd</td><td>CACCTCTTCCACCGCGGGCGAGGA</td><td></td><td></td><td></td><td></td><td></td><td></td><td></td><td></td><td></td><td></td></tr>
<tr><td>Oligo rev</td><td>AAACTCCTCGCCCGCGGTGGAAGA</td><td></td><td></td><td></td><td></td><td></td><td></td><td></td><td></td><td></td><td></td></tr>
<tr><td>Chromosome</td><td>start</td><td>end</td><td>strand</td><td>MM</td><td>target_seq</td><td>PAM</td><td>alignment</td><td>distance</td><td>position</td><td>gene name</td><td>gene id</td></tr>
<tr><td>GU305914.1</td><td>140514</td><td>140536</td><td>+</td><td>0</td><td>TCTTCCACCGCGGGCGAGGA</td><td>TGG</td><td>||||||||[||||||||||||]PAM</td><td>NA</td><td>Intergenic</td><td>NA</td><td>NA</td></tr>
<tr><td>T10747</td><td>CATTCCTGCGCCTGTACCCCCGG</td><td>1000</td><td></td><td></td><td></td><td></td><td></td><td></td><td></td><td></td><td></td></tr>
<tr><td>Oligo fwd</td><td>CACCCATTCCTGCGCCTGTACCCC</td><td></td><td></td><td></td><td></td><td></td><td></td><td></td><td></td><td></td><td></td></tr>
<tr><td>Oligo rev</td><td>AAACGGGGTACAGGCGCAGGAATG</td><td></td><td></td><td></td><td></td><td></td><td></td><td></td><td></td><td></td><td></td></tr>
<tr><td>Chromosome</td><td>start</td><td>end</td><td>strand</td><td>MM</td><td>target_seq</td><td>PAM</td><td>alignment</td><td>distance</td><td>position</td><td>gene name</td><td>gene id</td></tr>
<tr><td>GU305914.1</td><td>140553</td><td>140575</td><td>+</td><td>0</td><td>CATTCCTGCGCCTGTACCCC</td><td>CGG</td><td>||||||||[||||||||||||]PAM</td><td>NA</td><td>Intergenic</td><td>NA</td><td>NA</td></tr>
<tr><td>T10748</td><td>TGCTGCGAGCTCGTCTCCATTGG</td><td>1000</td><td></td><td></td><td></td><td></td><td></td><td></td><td></td><td></td><td></td></tr>
<tr><td>Oligo fwd</td><td>CACCTGCTGCGAGCTCGTCTCCAT</td><td></td><td></td><td></td><td></td><td></td><td></td><td></td><td></td><td></td><td></td></tr>
<tr><td>Oligo rev</td><td>AAACATGGAGACGAGCTCGCAGCA</td><td></td><td></td><td></td><td></td><td></td><td></td><td></td><td></td><td></td><td></td></tr>
<tr><td>Chromosome</td><td>start</td><td>end</td><td>strand</td><td>MM</td><td>target_seq</td><td>PAM</td><td>alignment</td><td>distance</td><td>position</td><td>gene name</td><td>gene id</td></tr>
<tr><td>GU305914.1</td><td>140592</td><td>140614</td><td>+</td><td>0</td><td>TGCTGCGAGCTCGTCTCCAT</td><td>TGG</td><td>||||||||[||||||||||||]PAM</td><td>NA</td><td>Intergenic</td><td>NA</td><td>NA</td></tr>
<tr><td>T10749</td><td>GCTGCGAGCTCGTCTCCATTGGG</td><td>1000</td><td></td><td></td><td></td><td></td><td></td><td></td><td></td><td></td><td></td></tr>
<tr><td>Oligo fwd</td><td>CACCGCTGCGAGCTCGTCTCCATT</td><td></td><td></td><td></td><td></td><td></td><td></td><td></td><td></td><td></td><td></td></tr>
<tr><td>Oligo rev</td><td>AAACAATGGAGACGAGCTCGCAGC</td><td></td><td></td><td></td><td></td><td></td><td></td><td></td><td></td><td></td><td></td></tr>
<tr><td>Chromosome</td><td>start</td><td>end</td><td>strand</td><td>MM</td><td>target_seq</td><td>PAM</td><td>alignment</td><td>distance</td><td>position</td><td>gene name</td><td>gene id</td></tr>
<tr><td>GU305914.1</td><td>140593</td><td>140615</td><td>+</td><td>0</td><td>GCTGCGAGCTCGTCTCCATT</td><td>GGG</td><td>||||||||[||||||||||||]PAM</td><td>NA</td><td>Intergenic</td><td>NA</td><td>NA</td></tr>
<tr><td>T10750</td><td>CTGCGAGCTCGTCTCCATTGGGG</td><td>1000</td><td></td><td></td><td></td><td></td><td></td><td></td><td></td><td></td><td></td></tr>
<tr><td>Oligo fwd</td><td>CACCCTGCGAGCTCGTCTCCATTG</td><td></td><td></td><td></td><td></td><td></td><td></td><td></td><td></td><td></td><td></td></tr>
<tr><td>Oligo rev</td><td>AAACCAATGGAGACGAGCTCGCAG</td><td></td><td></td><td></td><td></td><td></td><td></td><td></td><td></td><td></td><td></td></tr>
<tr><td>Chromosome</td><td>start</td><td>end</td><td>strand</td><td>MM</td><td>target_seq</td><td>PAM</td><td>alignment</td><td>distance</td><td>position</td><td>gene name</td><td>gene id</td></tr>
<tr><td>GU305914.1</td><td>140594</td><td>140616</td><td>+</td><td>0</td><td>CTGCGAGCTCGTCTCCATTG</td><td>GGG</td><td>||||||||[||||||||||||]PAM</td><td>NA</td><td>Intergenic</td><td>NA</td><td>NA</td></tr>
<tr><td>T10751</td><td>CTCCATTGGGGATTCCGTGTCGG</td><td>1000</td><td></td><td></td><td></td><td></td><td></td><td></td><td></td><td></td><td></td></tr>
<tr><td>Oligo fwd</td><td>CACCCTCCATTGGGGATTCCGTGT</td><td></td><td></td><td></td><td></td><td></td><td></td><td></td><td></td><td></td><td></td></tr>
<tr><td>Oligo rev</td><td>AAACACACGGAATCCCCAATGGAG</td><td></td><td></td><td></td><td></td><td></td><td></td><td></td><td></td><td></td><td></td></tr>
<tr><td>Chromosome</td><td>start</td><td>end</td><td>strand</td><td>MM</td><td>target_seq</td><td>PAM</td><td>alignment</td><td>distance</td><td>position</td><td>gene name</td><td>gene id</td></tr>
<tr><td>GU305914.1</td><td>140606</td><td>140628</td><td>+</td><td>0</td><td>CTCCATTGGGGATTCCGTGT</td><td>CGG</td><td>||||||||[||||||||||||]PAM</td><td>NA</td><td>Intergenic</td><td>NA</td><td>NA</td></tr>
<tr><td>T10752</td><td>GGGATTCCGTGTCGGTCGCCCGG</td><td>1000</td><td></td><td></td><td></td><td></td><td></td><td></td><td></td><td></td><td></td></tr>
<tr><td>Oligo fwd</td><td>CACCGGGATTCCGTGTCGGTCGCC</td><td></td><td></td><td></td><td></td><td></td><td></td><td></td><td></td><td></td><td></td></tr>
<tr><td>Oligo rev</td><td>AAACGGCGACCGACACGGAATCCC</td><td></td><td></td><td></td><td></td><td></td><td></td><td></td><td></td><td></td><td></td></tr>
<tr><td>Chromosome</td><td>start</td><td>end</td><td>strand</td><td>MM</td><td>target_seq</td><td>PAM</td><td>alignment</td><td>distance</td><td>position</td><td>gene name</td><td>gene id</td></tr>
<tr><td>GU305914.1</td><td>140614</td><td>140636</td><td>+</td><td>0</td><td>GGGATTCCGTGTCGGTCGCC</td><td>CGG</td><td>||||||||[||||||||||||]PAM</td><td>NA</td><td>Intergenic</td><td>NA</td><td>NA</td></tr>
<tr><td>T10753</td><td>CGGTGTCCCCTGTGTTCTTTAGG</td><td>1000</td><td></td><td></td><td></td><td></td><td></td><td></td><td></td><td></td><td></td></tr>
<tr><td>Oligo fwd</td><td>CACCCGGTGTCCCCTGTGTTCTTT</td><td></td><td></td><td></td><td></td><td></td><td></td><td></td><td></td><td></td><td></td></tr>
<tr><td>Oligo rev</td><td>AAACAAAGAACACAGGGGACACCG</td><td></td><td></td><td></td><td></td><td></td><td></td><td></td><td></td><td></td><td></td></tr>
<tr><td>Chromosome</td><td>start</td><td>end</td><td>strand</td><td>MM</td><td>target_seq</td><td>PAM</td><td>alignment</td><td>distance</td><td>position</td><td>gene name</td><td>gene id</td></tr>
<tr><td>GU305914.1</td><td>140634</td><td>140656</td><td>+</td><td>0</td><td>CGGTGTCCCCTGTGTTCTTT</td><td>AGG</td><td>||||||||[||||||||||||]PAM</td><td>NA</td><td>Intergenic</td><td>NA</td><td>NA</td></tr>
<tr><td>T10754</td><td>AGCACTACGTATCTGAGCCGCGG</td><td>1000</td><td></td><td></td><td></td><td></td><td></td><td></td><td></td><td></td><td></td></tr>
<tr><td>Oligo fwd</td><td>CACCAGCACTACGTATCTGAGCCG</td><td></td><td></td><td></td><td></td><td></td><td></td><td></td><td></td><td></td><td></td></tr>
<tr><td>Oligo rev</td><td>AAACCGGCTCAGATACGTAGTGCT</td><td></td><td></td><td></td><td></td><td></td><td></td><td></td><td></td><td></td><td></td></tr>
<tr><td>Chromosome</td><td>start</td><td>end</td><td>strand</td><td>MM</td><td>target_seq</td><td>PAM</td><td>alignment</td><td>distance</td><td>position</td><td>gene name</td><td>gene id</td></tr>
<tr><td>GU305914.1</td><td>140661</td><td>140683</td><td>+</td><td>0</td><td>AGCACTACGTATCTGAGCCG</td><td>CGG</td><td>||||||||[||||||||||||]PAM</td><td>NA</td><td>Intergenic</td><td>NA</td><td>NA</td></tr>
<tr><td>T10755</td><td>CTGAGCCGCGGCTGTTGCCGCGG</td><td>1000</td><td></td><td></td><td></td><td></td><td></td><td></td><td></td><td></td><td></td></tr>
<tr><td>Oligo fwd</td><td>CACCCTGAGCCGCGGCTGTTGCCG</td><td></td><td></td><td></td><td></td><td></td><td></td><td></td><td></td><td></td><td></td></tr>
<tr><td>Oligo rev</td><td>AAACCGGCAACAGCCGCGGCTCAG</td><td></td><td></td><td></td><td></td><td></td><td></td><td></td><td></td><td></td><td></td></tr>
<tr><td>Chromosome</td><td>start</td><td>end</td><td>strand</td><td>MM</td><td>target_seq</td><td>PAM</td><td>alignment</td><td>distance</td><td>position</td><td>gene name</td><td>gene id</td></tr>
<tr><td>GU305914.1</td><td>140673</td><td>140695</td><td>+</td><td>0</td><td>CTGAGCCGCGGCTGTTGCCG</td><td>CGG</td><td>||||||||[||||||||||||]PAM</td><td>NA</td><td>Intergenic</td><td>NA</td><td>NA</td></tr>
<tr><td>T10756</td><td>CTGTTGCCGCGGCCGAAGTAAGG</td><td>1000</td><td></td><td></td><td></td><td></td><td></td><td></td><td></td><td></td><td></td></tr>
<tr><td>Oligo fwd</td><td>CACCCTGTTGCCGCGGCCGAAGTA</td><td></td><td></td><td></td><td></td><td></td><td></td><td></td><td></td><td></td><td></td></tr>
<tr><td>Oligo rev</td><td>AAACTACTTCGGCCGCGGCAACAG</td><td></td><td></td><td></td><td></td><td></td><td></td><td></td><td></td><td></td><td></td></tr>
<tr><td>Chromosome</td><td>start</td><td>end</td><td>strand</td><td>MM</td><td>target_seq</td><td>PAM</td><td>alignment</td><td>distance</td><td>position</td><td>gene name</td><td>gene id</td></tr>
<tr><td>GU305914.1</td><td>140684</td><td>140706</td><td>+</td><td>0</td><td>CTGTTGCCGCGGCCGAAGTA</td><td>AGG</td><td>||||||||[||||||||||||]PAM</td><td>NA</td><td>Intergenic</td><td>NA</td><td>NA</td></tr>
<tr><td>T10757</td><td>CCGCGGCCGAAGTAAGGTGACGG</td><td>1000</td><td></td><td></td><td></td><td></td><td></td><td></td><td></td><td></td><td></td></tr>
<tr><td>Oligo fwd</td><td>CACCCCGCGGCCGAAGTAAGGTGA</td><td></td><td></td><td></td><td></td><td></td><td></td><td></td><td></td><td></td><td></td></tr>
<tr><td>Oligo rev</td><td>AAACTCACCTTACTTCGGCCGCGG</td><td></td><td></td><td></td><td></td><td></td><td></td><td></td><td></td><td></td><td></td></tr>
<tr><td>Chromosome</td><td>start</td><td>end</td><td>strand</td><td>MM</td><td>target_seq</td><td>PAM</td><td>alignment</td><td>distance</td><td>position</td><td>gene name</td><td>gene id</td></tr>
<tr><td>GU305914.1</td><td>140690</td><td>140712</td><td>+</td><td>0</td><td>CCGCGGCCGAAGTAAGGTGA</td><td>CGG</td><td>||||||||[||||||||||||]PAM</td><td>NA</td><td>Intergenic</td><td>NA</td><td>NA</td></tr>
<tr><td>T10758</td><td>CGCGGCCGAAGTAAGGTGACGGG</td><td>1000</td><td></td><td></td><td></td><td></td><td></td><td></td><td></td><td></td><td></td></tr>
<tr><td>Oligo fwd</td><td>CACCCGCGGCCGAAGTAAGGTGAC</td><td></td><td></td><td></td><td></td><td></td><td></td><td></td><td></td><td></td><td></td></tr>
<tr><td>Oligo rev</td><td>AAACGTCACCTTACTTCGGCCGCG</td><td></td><td></td><td></td><td></td><td></td><td></td><td></td><td></td><td></td><td></td></tr>
<tr><td>Chromosome</td><td>start</td><td>end</td><td>strand</td><td>MM</td><td>target_seq</td><td>PAM</td><td>alignment</td><td>distance</td><td>position</td><td>gene name</td><td>gene id</td></tr>
</table>

| | | | | | | | | | | | |
|---|---|---|---|---|---|---|---|---|---|---|---|
| GU305914.1 | 140691 | 140713 | + | 0 | CGCGGCCGAAGTAAGGTGAC | GGG | \|\|\|\|\|\|\|\|[\|\|\|\|\|\|\|\|\|\|\|\|]PAM | NA | Intergenic | NA | NA |
| T10759 | GCGGCCGAAGTAAGGTGACGGGG | 1000 | | | | | | | | | |
| Oligo fwd | CACCGCGGCCGAAGTAAGGTGACG | | | | | | | | | | |
| Oligo rev | AAACCGTCACCTTACTTCGGCCGC | | | | | | | | | | |
| Chromosome | start | end | strand | MM | target_seq | PAM | alignment | distance | position | gene name | gene id |
| GU305914.1 | 140692 | 140714 | + | 0 | GCGGCCGAAGTAAGGTGACG | GGG | \|\|\|\|\|\|\|\|[\|\|\|\|\|\|\|\|\|\|\|\|]PAM | NA | Intergenic | NA | NA |
| T10760 | CGGCCGAAGTAAGGTGACGGGGG | 1000 | | | | | | | | | |
| Oligo fwd | CACCCGGCCGAAGTAAGGTGACGG | | | | | | | | | | |
| Oligo rev | AAACCCGTCACCTTACTTCGGCCG | | | | | | | | | | |
| Chromosome | start | end | strand | MM | target_seq | PAM | alignment | distance | position | gene name | gene id |
| GU305914.1 | 140693 | 140715 | + | 0 | CGGCCGAAGTAAGGTGACGG | GGG | \|\|\|\|\|\|\|\|[\|\|\|\|\|\|\|\|\|\|\|\|]PAM | NA | Intergenic | NA | NA |
| T10761 | GGCCGAAGTAAGGTGACGGGGGG | 1000 | | | | | | | | | |
| Oligo fwd | CACCGGCCGAAGTAAGGTGACGGG | | | | | | | | | | |
| Oligo rev | AAACCCCGTCACCTTACTTCGGCC | | | | | | | | | | |
| Chromosome | start | end | strand | MM | target_seq | PAM | alignment | distance | position | gene name | gene id |
| GU305914.1 | 140694 | 140716 | + | 0 | GGCCGAAGTAAGGTGACGGG | GGG | \|\|\|\|\|\|\|\|[\|\|\|\|\|\|\|\|\|\|\|\|]PAM | NA | Intergenic | NA | NA |
| T10762 | TAAGGTGACGGGGGGAGATGAGG | 1000 | | | | | | | | | |
| Oligo fwd | CACCTAAGGTGACGGGGGGAGATG | | | | | | | | | | |
| Oligo rev | AAACCATCTCCCCCCGTCACCTTA | | | | | | | | | | |
| Chromosome | start | end | strand | MM | target_seq | PAM | alignment | distance | position | gene name | gene id |
| GU305914.1 | 140702 | 140724 | + | 0 | TAAGGTGACGGGGGGAGATG | AGG | \|\|\|\|\|\|\|\|[\|\|\|\|\|\|\|\|\|\|\|\|]PAM | NA | Intergenic | NA | NA |
| T10763 | GACGGGGGGAGATGAGGACGAGG | 1000 | | | | | | | | | |
| Oligo fwd | CACCGACGGGGGGAGATGAGGACG | | | | | | | | | | |
| Oligo rev | AAACCGTCCTCATCTCCCCCCGTC | | | | | | | | | | |
| Chromosome | start | end | strand | MM | target_seq | PAM | alignment | distance | position | gene name | gene id |
| GU305914.1 | 140708 | 140730 | + | 0 | GACGGGGGGAGATGAGGACG | AGG | \|\|\|\|\|\|\|\|[\|\|\|\|\|\|\|\|\|\|\|\|]PAM | NA | Intergenic | NA | NA |
| T10764 | GGGGGGAGATGAGGACGAGGAGG | 1000 | | | | | | | | | |
| Oligo fwd | CACCGGGGGGAGATGAGGACGAGG | | | | | | | | | | |
| Oligo rev | AAACCCTCGTCCTCATCTCCCCCC | | | | | | | | | | |
| Chromosome | start | end | strand | MM | target_seq | PAM | alignment | distance | position | gene name | gene id |
| GU305914.1 | 140711 | 140733 | + | 0 | GGGGGGAGATGAGGACGAGG | AGG | \|\|\|\|\|\|\|\|[\|\|\|\|\|\|\|\|\|\|\|\|]PAM | NA | Intergenic | NA | NA |
| GU305914.1 | 42858 | 42880 | - | 4 | GGGGGAGGATGAGGAAGAGA | AGG | \|\|\|\|\|--\|[\|\|\|\|\|\|\|-\|\|\|-]PAM | NA | Intergenic | NA | NA |
| T10765 | AGATGAGGACGAGGAGGATGAGG | 1000 | | | | | | | | | |
| Oligo fwd | CACCAGATGAGGACGAGGAGGATG | | | | | | | | | | |
| Oligo rev | AAACCATCCTCCTCGTCCTCATCT | | | | | | | | | | |
| Chromosome | start | end | strand | MM | target_seq | PAM | alignment | distance | position | gene name | gene id |
| GU305914.1 | 140717 | 140739 | + | 0 | AGATGAGGACGAGGAGGATG | AGG | \|\|\|\|\|\|\|\|[\|\|\|\|\|\|\|\|\|\|\|\|]PAM | NA | Intergenic | NA | NA |
| GU305914.1 | 140723 | 140745 | + | 4 | GGACGAGGAGGATGAGGATG | AGG | -\|\|-\|\|\|\|[\|-\|\|-\|\|\|\|\|\|\|]PAM | NA | Intergenic | NA | NA |
| GU305914.1 | 118506 | 118528 | - | 4 | CGATGATGAAGAGGAGGACG | AGG | -\|\|\|\|\|-\|[\|-\|\|\|\|\|\|\|\|-\|]PAM | NA | Intergenic | NA | NA |
| GU305914.1 | 65114 | 65136 | + | 4 | AGAAGACGACGACGAGGAGG | AGG | \|\|\|-\|\|-\|[\|\|\|\|-\|\|\|\|\|-\|]PAM | NA | Intergenic | NA | NA |
| GU305914.1 | 152954 | 152976 | + | 4 | CGATGAGTATGAGGAGGATC | TGG | -\|\|\|\|\|\|-[\|-\|\|\|\|\|\|\|\|\|-]PAM | NA | Intergenic | NA | NA |
| T10766 | GGACGAGGAGGATGAGGATGAGG | 1000 | | | | | | | | | |
| Oligo fwd | CACCGGACGAGGAGGATGAGGATG | | | | | | | | | | |
| Oligo rev | AAACCATCCTCATCCTCCTCGTCC | | | | | | | | | | |
| Chromosome | start | end | strand | MM | target_seq | PAM | alignment | distance | position | gene name | gene id |
| GU305914.1 | 140723 | 140745 | + | 0 | GGACGAGGAGGATGAGGATG | AGG | \|\|\|\|\|\|\|\|[\|\|\|\|\|\|\|\|\|\|\|\|]PAM | NA | Intergenic | NA | NA |
| GU305914.1 | 140717 | 140739 | + | 4 | AGATGAGGACGAGGAGGATG | AGG | -\|\|-\|\|\|\|[\|-\|\|-\|\|\|\|\|\|\|]PAM | NA | Intergenic | NA | NA |
| GU305914.1 | 77037 | 77059 | - | 3 | GGTCGACGAGGATGAGGAGG | AGG | \|\|-\|\|\|-\|[\|\|\|\|\|\|\|\|\|\|-\|]PAM | NA | Intergenic | NA | NA |
| T10767 | CGAGGAGGATGAGGATGAGGAGG | 1000 | | | | | | | | | |
| Oligo fwd | CACCCGAGGAGGATGAGGATGAGG | | | | | | | | | | |
| Oligo rev | AAACCCTCATCCTCATCCTCCTCG | | | | | | | | | | |
| Chromosome | start | end | strand | MM | target_seq | PAM | alignment | distance | position | gene name | gene id |
| GU305914.1 | 140726 | 140748 | + | 0 | CGAGGAGGATGAGGATGAGG | AGG | \|\|\|\|\|\|\|\|[\|\|\|\|\|\|\|\|\|\|\|\|]PAM | NA | Intergenic | NA | NA |
| GU305914.1 | 140759 | 140781 | + | 4 | GGATGAGGATAGGGATGAGG | AGG | -\|\|-\|\|\|\|[\|\|--\|\|\|\|\|\|\|\|]PAM | NA | Intergenic | NA | NA |
| GU305914.1 | 118512 | 118534 | - | 4 | TGAGGACGATGATGAAGAGG | AGG | -\|\|\|\|\|-\|[\|\|\|\|-\|\|-\|\|\|\|]PAM | NA | Intergenic | NA | NA |
| GU305914.1 | 118809 | 118831 | - | 4 | CGACGATGATGACGACGAGG | AGG | \|\|\|-\|\|-\|[\|\|\|\|-\|\|-\|\|\|\|]PAM | NA | Intergenic | NA | NA |
| GU305914.1 | 42858 | 42880 | - | 4 | GGGGGAGGATGAGGAAGAGA | AGG | -\|-\|\|\|\|\|[\|\|\|\|\|\|\|-\|\|\|-]PAM | NA | Intergenic | NA | NA |
| T10768 | GGATGAGGAGGAGAATAGTCAGG | 1000 | | | | | | | | | |
| Oligo fwd | CACCGGATGAGGAGGAGAATAGTC | | | | | | | | | | |
| Oligo rev | AAACGACTATTCTCCTCCTCATCC | | | | | | | | | | |
| Chromosome | start | end | strand | MM | target_seq | PAM | alignment | distance | position | gene name | gene id |
| GU305914.1 | 140738 | 140760 | + | 0 | GGATGAGGAGGAGAATAGTC | AGG | \|\|\|\|\|\|\|\|[\|\|\|\|\|\|\|\|\|\|\|\|]PAM | NA | Intergenic | NA | NA |
| T10769 | GGAGGAGAATAGTCAGGATGAGG | 1000 | | | | | | | | | |
| Oligo fwd | CACCGGAGGAGAATAGTCAGGATG | | | | | | | | | | |
| Oligo rev | AAACCATCCTGACTATTCTCCTCC | | | | | | | | | | |
| Chromosome | start | end | strand | MM | target_seq | PAM | alignment | distance | position | gene name | gene id |
| GU305914.1 | 140744 | 140766 | + | 0 | GGAGGAGAATAGTCAGGATG | AGG | \|\|\|\|\|\|\|\|[\|\|\|\|\|\|\|\|\|\|\|\|]PAM | NA | Intergenic | NA | NA |
| T10770 | AGAATAGTCAGGATGAGGATAGG | 1000 | | | | | | | | | |
| Oligo fwd | CACCAGAATAGTCAGGATGAGGAT | | | | | | | | | | |
| Oligo rev | AAACATCCTCATCCTGACTATTCT | | | | | | | | | | |
| Chromosome | start | end | strand | MM | target_seq | PAM | alignment | distance | position | gene name | gene id |
| GU305914.1 | 140749 | 140771 | + | 0 | AGAATAGTCAGGATGAGGAT | AGG | \|\|\|\|\|\|\|\|[\|\|\|\|\|\|\|\|\|\|\|\|]PAM | NA | Intergenic | NA | NA |
| T10771 | GAATAGTCAGGATGAGGATAGGG | 1000 | | | | | | | | | |
| Oligo fwd | CACCGAATAGTCAGGATGAGGATA | | | | | | | | | | |
| Oligo rev | AAACTATCCTCATCCTGACTATTC | | | | | | | | | | |
| Chromosome | start | end | strand | MM | target_seq | PAM | alignment | distance | position | gene name | gene id |

<table>
<tr><td>GU305914.1</td><td>140750</td><td>140772</td><td>+</td><td>0</td><td>GAATAGTCAGGATGAGGATA</td><td>GGG</td><td>||||||||[||||||||||||]PAM</td><td>NA</td><td>Intergenic</td><td>NA</td><td>NA</td></tr>
<tr><td>T10772</td><td>TCAGGATGAGGATAGGGATGAGG</td><td>1000</td><td></td><td></td><td></td><td></td><td></td><td></td><td></td><td></td><td></td></tr>
<tr><td>Oligo fwd</td><td>CACCTCAGGATGAGGATAGGGATG</td><td></td><td></td><td></td><td></td><td></td><td></td><td></td><td></td><td></td><td></td></tr>
<tr><td>Oligo rev</td><td>AAACCATCCCTATCCTCATCCTGA</td><td></td><td></td><td></td><td></td><td></td><td></td><td></td><td></td><td></td><td></td></tr>
<tr><td>Chromosome</td><td>start</td><td>end</td><td>strand</td><td>MM</td><td>target_seq</td><td>PAM</td><td>alignment</td><td>distance</td><td>position</td><td>gene name</td><td>gene id</td></tr>
<tr><td>GU305914.1</td><td>140756</td><td>140778</td><td>+</td><td>0</td><td>TCAGGATGAGGATAGGGATG</td><td>AGG</td><td>||||||||[||||||||||||]PAM</td><td>NA</td><td>Intergenic</td><td>NA</td><td>NA</td></tr>
<tr><td>T10773</td><td>GGATGAGGATAGGGATGAGGAGG</td><td>1000</td><td></td><td></td><td></td><td></td><td></td><td></td><td></td><td></td><td></td></tr>
<tr><td>Oligo fwd</td><td>CACCGGATGAGGATAGGGATGAGG</td><td></td><td></td><td></td><td></td><td></td><td></td><td></td><td></td><td></td><td></td></tr>
<tr><td>Oligo rev</td><td>AAACCCTCATCCCTATCCTCATCC</td><td></td><td></td><td></td><td></td><td></td><td></td><td></td><td></td><td></td><td></td></tr>
<tr><td>Chromosome</td><td>start</td><td>end</td><td>strand</td><td>MM</td><td>target_seq</td><td>PAM</td><td>alignment</td><td>distance</td><td>position</td><td>gene name</td><td>gene id</td></tr>
<tr><td>GU305914.1</td><td>140759</td><td>140781</td><td>+</td><td>0</td><td>GGATGAGGATAGGGATGAGG</td><td>AGG</td><td>||||||||[||||||||||||]PAM</td><td>NA</td><td>Intergenic</td><td>NA</td><td>NA</td></tr>
<tr><td>GU305914.1</td><td>140726</td><td>140748</td><td>+</td><td>4</td><td>CGAGGAGGATGAGGATGAGG</td><td>AGG</td><td>-||-||||[||--||||||||]PAM</td><td>NA</td><td>Intergenic</td><td>NA</td><td>NA</td></tr>
<tr><td>T10774</td><td>TGAGGATAGGGATGAGGAGGAGG</td><td>1000</td><td></td><td></td><td></td><td></td><td></td><td></td><td></td><td></td><td></td></tr>
<tr><td>Oligo fwd</td><td>CACCTGAGGATAGGGATGAGGAGG</td><td></td><td></td><td></td><td></td><td></td><td></td><td></td><td></td><td></td><td></td></tr>
<tr><td>Oligo rev</td><td>AAACCCTCCTCATCCCTATCCTCA</td><td></td><td></td><td></td><td></td><td></td><td></td><td></td><td></td><td></td><td></td></tr>
<tr><td>Chromosome</td><td>start</td><td>end</td><td>strand</td><td>MM</td><td>target_seq</td><td>PAM</td><td>alignment</td><td>distance</td><td>position</td><td>gene name</td><td>gene id</td></tr>
<tr><td>GU305914.1</td><td>140762</td><td>140784</td><td>+</td><td>0</td><td>TGAGGATAGGGATGAGGAGG</td><td>AGG</td><td>||||||||[||||||||||||]PAM</td><td>NA</td><td>Intergenic</td><td>NA</td><td>NA</td></tr>
<tr><td>T10775</td><td>GGATAGGGATGAGGAGGAGGCGG</td><td>1000</td><td></td><td></td><td></td><td></td><td></td><td></td><td></td><td></td><td></td></tr>
<tr><td>Oligo fwd</td><td>CACCGGATAGGGATGAGGAGGAGG</td><td></td><td></td><td></td><td></td><td></td><td></td><td></td><td></td><td></td><td></td></tr>
<tr><td>Oligo rev</td><td>AAACCCTCCTCCTCATCCCTATCC</td><td></td><td></td><td></td><td></td><td></td><td></td><td></td><td></td><td></td><td></td></tr>
<tr><td>Chromosome</td><td>start</td><td>end</td><td>strand</td><td>MM</td><td>target_seq</td><td>PAM</td><td>alignment</td><td>distance</td><td>position</td><td>gene name</td><td>gene id</td></tr>
<tr><td>GU305914.1</td><td>140765</td><td>140787</td><td>+</td><td>0</td><td>GGATAGGGATGAGGAGGAGG</td><td>CGG</td><td>||||||||[||||||||||||]PAM</td><td>NA</td><td>Intergenic</td><td>NA</td><td>NA</td></tr>
<tr><td>T10776</td><td>GGATGAGGAGGAGGCGGCGTCGG</td><td>1000</td><td></td><td></td><td></td><td></td><td></td><td></td><td></td><td></td><td></td></tr>
<tr><td>Oligo fwd</td><td>CACCGGATGAGGAGGAGGCGGCGT</td><td></td><td></td><td></td><td></td><td></td><td></td><td></td><td></td><td></td><td></td></tr>
<tr><td>Oligo rev</td><td>AAACACGCCGCCTCCTCCTCATCC</td><td></td><td></td><td></td><td></td><td></td><td></td><td></td><td></td><td></td><td></td></tr>
<tr><td>Chromosome</td><td>start</td><td>end</td><td>strand</td><td>MM</td><td>target_seq</td><td>PAM</td><td>alignment</td><td>distance</td><td>position</td><td>gene name</td><td>gene id</td></tr>
<tr><td>GU305914.1</td><td>140771</td><td>140793</td><td>+</td><td>0</td><td>GGATGAGGAGGAGGCGGCGT</td><td>CGG</td><td>||||||||[||||||||||||]PAM</td><td>NA</td><td>Intergenic</td><td>NA</td><td>NA</td></tr>
<tr><td>GU305914.1</td><td>161757</td><td>161779</td><td>-</td><td>4</td><td>GGAGGCCGAGGAGGCGGCGG</td><td>CGG</td><td>|||-|--|[|||||||||||-]PAM</td><td>NA</td><td>Intergenic</td><td>NA</td><td>NA</td></tr>
<tr><td>GU305914.1</td><td>161754</td><td>161776</td><td>-</td><td>4</td><td>GGCCGAGGAGGCGGCGGCGG</td><td>AGG</td><td>||--||||[|||-|||||||-]PAM</td><td>NA</td><td>Intergenic</td><td>NA</td><td>NA</td></tr>
<tr><td>T10777</td><td>GCGGCGTCGGCTTCCTCTAGCGG</td><td>1000</td><td></td><td></td><td></td><td></td><td></td><td></td><td></td><td></td><td></td></tr>
<tr><td>Oligo fwd</td><td>CACCGCGGCGTCGGCTTCCTCTAG</td><td></td><td></td><td></td><td></td><td></td><td></td><td></td><td></td><td></td><td></td></tr>
<tr><td>Oligo rev</td><td>AAACCTAGAGGAAGCCGACGCCGC</td><td></td><td></td><td></td><td></td><td></td><td></td><td></td><td></td><td></td><td></td></tr>
<tr><td>Chromosome</td><td>start</td><td>end</td><td>strand</td><td>MM</td><td>target_seq</td><td>PAM</td><td>alignment</td><td>distance</td><td>position</td><td>gene name</td><td>gene id</td></tr>
<tr><td>GU305914.1</td><td>140784</td><td>140806</td><td>+</td><td>0</td><td>GCGGCGTCGGCTTCCTCTAG</td><td>CGG</td><td>||||||||[||||||||||||]PAM</td><td>NA</td><td>Intergenic</td><td>NA</td><td>NA</td></tr>
<tr><td>T10778</td><td>GCGTCGGCTTCCTCTAGCGGCGG</td><td>1000</td><td></td><td></td><td></td><td></td><td></td><td></td><td></td><td></td><td></td></tr>
<tr><td>Oligo fwd</td><td>CACCGCGTCGGCTTCCTCTAGCGG</td><td></td><td></td><td></td><td></td><td></td><td></td><td></td><td></td><td></td><td></td></tr>
<tr><td>Oligo rev</td><td>AAACCCGCTAGAGGAAGCCGACGC</td><td></td><td></td><td></td><td></td><td></td><td></td><td></td><td></td><td></td><td></td></tr>
<tr><td>Chromosome</td><td>start</td><td>end</td><td>strand</td><td>MM</td><td>target_seq</td><td>PAM</td><td>alignment</td><td>distance</td><td>position</td><td>gene name</td><td>gene id</td></tr>
<tr><td>GU305914.1</td><td>140787</td><td>140809</td><td>+</td><td>0</td><td>GCGTCGGCTTCCTCTAGCGG</td><td>CGG</td><td>||||||||[||||||||||||]PAM</td><td>NA</td><td>Intergenic</td><td>NA</td><td>NA</td></tr>
<tr><td>T10779</td><td>CGTCGGCTTCCTCTAGCGGCGGG</td><td>1000</td><td></td><td></td><td></td><td></td><td></td><td></td><td></td><td></td><td></td></tr>
<tr><td>Oligo fwd</td><td>CACCCGTCGGCTTCCTCTAGCGGC</td><td></td><td></td><td></td><td></td><td></td><td></td><td></td><td></td><td></td><td></td></tr>
<tr><td>Oligo rev</td><td>AAACGCCGCTAGAGGAAGCCGACG</td><td></td><td></td><td></td><td></td><td></td><td></td><td></td><td></td><td></td><td></td></tr>
<tr><td>Chromosome</td><td>start</td><td>end</td><td>strand</td><td>MM</td><td>target_seq</td><td>PAM</td><td>alignment</td><td>distance</td><td>position</td><td>gene name</td><td>gene id</td></tr>
<tr><td>GU305914.1</td><td>140788</td><td>140810</td><td>+</td><td>0</td><td>CGTCGGCTTCCTCTAGCGGC</td><td>GGG</td><td>||||||||[||||||||||||]PAM</td><td>NA</td><td>Intergenic</td><td>NA</td><td>NA</td></tr>
<tr><td>T10780</td><td>GGCTTCCTCTAGCGGCGGGCTGG</td><td>1000</td><td></td><td></td><td></td><td></td><td></td><td></td><td></td><td></td><td></td></tr>
<tr><td>Oligo fwd</td><td>CACCGGCTTCCTCTAGCGGCGGGC</td><td></td><td></td><td></td><td></td><td></td><td></td><td></td><td></td><td></td><td></td></tr>
<tr><td>Oligo rev</td><td>AAACGCCCGCCGCTAGAGGAAGCC</td><td></td><td></td><td></td><td></td><td></td><td></td><td></td><td></td><td></td><td></td></tr>
<tr><td>Chromosome</td><td>start</td><td>end</td><td>strand</td><td>MM</td><td>target_seq</td><td>PAM</td><td>alignment</td><td>distance</td><td>position</td><td>gene name</td><td>gene id</td></tr>
<tr><td>GU305914.1</td><td>140792</td><td>140814</td><td>+</td><td>0</td><td>GGCTTCCTCTAGCGGCGGGC</td><td>TGG</td><td>||||||||[||||||||||||]PAM</td><td>NA</td><td>Intergenic</td><td>NA</td><td>NA</td></tr>
<tr><td>T10781</td><td>CCTCTAGCGGCGGGCTGGAGTGG</td><td>1000</td><td></td><td></td><td></td><td></td><td></td><td></td><td></td><td></td><td></td></tr>
<tr><td>Oligo fwd</td><td>CACCCCTCTAGCGGCGGGCTGGAG</td><td></td><td></td><td></td><td></td><td></td><td></td><td></td><td></td><td></td><td></td></tr>
<tr><td>Oligo rev</td><td>AAACCTCCAGCCCGCCGCTAGAGG</td><td></td><td></td><td></td><td></td><td></td><td></td><td></td><td></td><td></td><td></td></tr>
<tr><td>Chromosome</td><td>start</td><td>end</td><td>strand</td><td>MM</td><td>target_seq</td><td>PAM</td><td>alignment</td><td>distance</td><td>position</td><td>gene name</td><td>gene id</td></tr>
<tr><td>GU305914.1</td><td>140797</td><td>140819</td><td>+</td><td>0</td><td>CCTCTAGCGGCGGGCTGGAG</td><td>TGG</td><td>||||||||[||||||||||||]PAM</td><td>NA</td><td>Intergenic</td><td>NA</td><td>NA</td></tr>
<tr><td>T10782</td><td>ATAGTAATAGCGCGCTCAGTTGG</td><td>1000</td><td></td><td></td><td></td><td></td><td></td><td></td><td></td><td></td><td></td></tr>
<tr><td>Oligo fwd</td><td>CACCATAGTAATAGCGCGCTCAGT</td><td></td><td></td><td></td><td></td><td></td><td></td><td></td><td></td><td></td><td></td></tr>
<tr><td>Oligo rev</td><td>AAACACTGAGCGCGCTATTACTAT</td><td></td><td></td><td></td><td></td><td></td><td></td><td></td><td></td><td></td><td></td></tr>
<tr><td>Chromosome</td><td>start</td><td>end</td><td>strand</td><td>MM</td><td>target_seq</td><td>PAM</td><td>alignment</td><td>distance</td><td>position</td><td>gene name</td><td>gene id</td></tr>
<tr><td>GU305914.1</td><td>140827</td><td>140849</td><td>+</td><td>0</td><td>ATAGTAATAGCGCGCTCAGT</td><td>TGG</td><td>||||||||[||||||||||||]PAM</td><td>NA</td><td>Intergenic</td><td>NA</td><td>NA</td></tr>
<tr><td>T10783</td><td>TAATAGCGCGCTCAGTTGGTCGG</td><td>1000</td><td></td><td></td><td></td><td></td><td></td><td></td><td></td><td></td><td></td></tr>
<tr><td>Oligo fwd</td><td>CACCTAATAGCGCGCTCAGTTGGT</td><td></td><td></td><td></td><td></td><td></td><td></td><td></td><td></td><td></td><td></td></tr>
<tr><td>Oligo rev</td><td>AAACACCAACTGAGCGCGCTATTA</td><td></td><td></td><td></td><td></td><td></td><td></td><td></td><td></td><td></td><td></td></tr>
<tr><td>Chromosome</td><td>start</td><td>end</td><td>strand</td><td>MM</td><td>target_seq</td><td>PAM</td><td>alignment</td><td>distance</td><td>position</td><td>gene name</td><td>gene id</td></tr>
<tr><td>GU305914.1</td><td>140831</td><td>140853</td><td>+</td><td>0</td><td>TAATAGCGCGCTCAGTTGGT</td><td>CGG</td><td>||||||||[||||||||||||]PAM</td><td>NA</td><td>Intergenic</td><td>NA</td><td>NA</td></tr>
<tr><td>T10784</td><td>ATCATAATCTCCCCGTTTCCCGG</td><td>1000</td><td></td><td></td><td></td><td></td><td></td><td></td><td></td><td></td><td></td></tr>
<tr><td>Oligo fwd</td><td>CACCATCATAATCTCCCCGTTTCC</td><td></td><td></td><td></td><td></td><td></td><td></td><td></td><td></td><td></td><td></td></tr>
<tr><td>Oligo rev</td><td>AAACGGAAACGGGGAGATTATGAT</td><td></td><td></td><td></td><td></td><td></td><td></td><td></td><td></td><td></td><td></td></tr>
<tr><td>Chromosome</td><td>start</td><td>end</td><td>strand</td><td>MM</td><td>target_seq</td><td>PAM</td><td>alignment</td><td>distance</td><td>position</td><td>gene name</td><td>gene id</td></tr>
<tr><td>GU305914.1</td><td>140862</td><td>140884</td><td>+</td><td>0</td><td>ATCATAATCTCCCCGTTTCC</td><td>CGG</td><td>||||||||[||||||||||||]PAM</td><td>NA</td><td>Intergenic</td><td>NA</td><td>NA</td></tr>
<tr><td>T10785</td><td>TCATAATCTCCCCGTTTCCCGGG</td><td>1000</td><td></td><td></td><td></td><td></td><td></td><td></td><td></td><td></td><td></td></tr>
<tr><td>Oligo fwd</td><td>CACCTCATAATCTCCCCGTTTCCC</td><td></td><td></td><td></td><td></td><td></td><td></td><td></td><td></td><td></td><td></td></tr>
<tr><td>Oligo rev</td><td>AAACGGGAAACGGGGAGATTATGA</td><td></td><td></td><td></td><td></td><td></td><td></td><td></td><td></td><td></td><td></td></tr>
<tr><td>Chromosome</td><td>start</td><td>end</td><td>strand</td><td>MM</td><td>target_seq</td><td>PAM</td><td>alignment</td><td>distance</td><td>position</td><td>gene name</td><td>gene id</td></tr>
<tr><td>GU305914.1</td><td>140863</td><td>140885</td><td>+</td><td>0</td><td>TCATAATCTCCCCGTTTCCC</td><td>GGG</td><td>||||||||[||||||||||||]PAM</td><td>NA</td><td>Intergenic</td><td>NA</td><td>NA</td></tr>
</table>

| | | | | | | | | | | | |
|---|---|---|---|---|---|---|---|---|---|---|---|
| T10786 | ATGTTATGTCACGACATTCGAGG | 1000 | | | | | | | | | |
| Oligo fwd | CACCATGTTATGTCACGACATTCG | | | | | | | | | | |
| Oligo rev | AAACCGAATGTCGTGACATAACAT | | | | | | | | | | |
| Chromosome | start | end | strand | MM | target_seq | PAM | alignment | distance | position | gene name | gene id |
| GU305914.1 | 140891 | 140913 | + | 0 | ATGTTATGTCACGACATTCG | AGG | \|\|\|\|\|\|\|\|[\|\|\|\|\|\|\|\|\|\|\|\|]PAM | NA | Intergenic | NA | NA |
| T10787 | ACAGCCCGTGCTGCTCGAGACGG | 1000 | | | | | | | | | |
| Oligo fwd | CACCACAGCCCGTGCTGCTCGAGA | | | | | | | | | | |
| Oligo rev | AAACTCTCGAGCAGCACGGGCTGT | | | | | | | | | | |
| Chromosome | start | end | strand | MM | target_seq | PAM | alignment | distance | position | gene name | gene id |
| GU305914.1 | 140921 | 140943 | + | 0 | ACAGCCCGTGCTGCTCGAGA | CGG | \|\|\|\|\|\|\|\|[\|\|\|\|\|\|\|\|\|\|\|\|]PAM | NA | Intergenic | NA | NA |
| T10788 | CAGCCCGTGCTGCTCGAGACGGG | 1000 | | | | | | | | | |
| Oligo fwd | CACCCAGCCCGTGCTGCTCGAGAC | | | | | | | | | | |
| Oligo rev | AAACGTCTCGAGCAGCACGGGCTG | | | | | | | | | | |
| Chromosome | start | end | strand | MM | target_seq | PAM | alignment | distance | position | gene name | gene id |
| GU305914.1 | 140922 | 140944 | + | 0 | CAGCCCGTGCTGCTCGAGAC | GGG | \|\|\|\|\|\|\|\|[\|\|\|\|\|\|\|\|\|\|\|\|]PAM | NA | Intergenic | NA | NA |
| T10789 | CGTATGACGAGAGTTTCTGTCGG | 1000 | | | | | | | | | |
| Oligo fwd | CACCCGTATGACGAGAGTTTCTGT | | | | | | | | | | |
| Oligo rev | AAACACAGAAACTCTCGTCATACG | | | | | | | | | | |
| Chromosome | start | end | strand | MM | target_seq | PAM | alignment | distance | position | gene name | gene id |
| GU305914.1 | 140971 | 140993 | + | 0 | CGTATGACGAGAGTTTCTGT | CGG | \|\|\|\|\|\|\|\|[\|\|\|\|\|\|\|\|\|\|\|\|]PAM | NA | Intergenic | NA | NA |
| T10790 | ACGAGAGTTTCTGTCGGAACAGG | 1000 | | | | | | | | | |
| Oligo fwd | CACCACGAGAGTTTCTGTCGGAAC | | | | | | | | | | |
| Oligo rev | AAACGTTCCGACAGAAACTCTCGT | | | | | | | | | | |
| Chromosome | start | end | strand | MM | target_seq | PAM | alignment | distance | position | gene name | gene id |
| GU305914.1 | 140977 | 140999 | + | 0 | ACGAGAGTTTCTGTCGGAAC | AGG | \|\|\|\|\|\|\|\|[\|\|\|\|\|\|\|\|\|\|\|\|]PAM | NA | Intergenic | NA | NA |
| T10791 | TTTCTGTCGGAACAGGTGTCTGG | 1000 | | | | | | | | | |
| Oligo fwd | CACCTTTCTGTCGGAACAGGTGTC | | | | | | | | | | |
| Oligo rev | AAACGACACCTGTTCCGACAGAAA | | | | | | | | | | |
| Chromosome | start | end | strand | MM | target_seq | PAM | alignment | distance | position | gene name | gene id |
| GU305914.1 | 140984 | 141006 | + | 0 | TTTCTGTCGGAACAGGTGTC | TGG | \|\|\|\|\|\|\|\|[\|\|\|\|\|\|\|\|\|\|\|\|]PAM | NA | Intergenic | NA | NA |
| T10792 | CGGAACAGGTGTCTGGAACGTGG | 1000 | | | | | | | | | |
| Oligo fwd | CACCCGGAACAGGTGTCTGGAACG | | | | | | | | | | |
| Oligo rev | AAACCGTTCCAGACACCTGTTCCG | | | | | | | | | | |
| Chromosome | start | end | strand | MM | target_seq | PAM | alignment | distance | position | gene name | gene id |
| GU305914.1 | 140991 | 141013 | + | 0 | CGGAACAGGTGTCTGGAACG | TGG | \|\|\|\|\|\|\|\|[\|\|\|\|\|\|\|\|\|\|\|\|]PAM | NA | Intergenic | NA | NA |
| T10793 | AACAGGTGTCTGGAACGTGGTGG | 1000 | | | | | | | | | |
| Oligo fwd | CACCAACAGGTGTCTGGAACGTGG | | | | | | | | | | |
| Oligo rev | AAACCCACGTTCCAGACACCTGTT | | | | | | | | | | |
| Chromosome | start | end | strand | MM | target_seq | PAM | alignment | distance | position | gene name | gene id |
| GU305914.1 | 140994 | 141016 | + | 0 | AACAGGTGTCTGGAACGTGG | TGG | \|\|\|\|\|\|\|\|[\|\|\|\|\|\|\|\|\|\|\|\|]PAM | NA | Intergenic | NA | NA |
| T10794 | ACAGGTGTCTGGAACGTGGTGGG | 1000 | | | | | | | | | |
| Oligo fwd | CACCACAGGTGTCTGGAACGTGGT | | | | | | | | | | |
| Oligo rev | AAACACCACGTTCCAGACACCTGT | | | | | | | | | | |
| Chromosome | start | end | strand | MM | target_seq | PAM | alignment | distance | position | gene name | gene id |
| GU305914.1 | 140995 | 141017 | + | 0 | ACAGGTGTCTGGAACGTGGT | GGG | \|\|\|\|\|\|\|\|[\|\|\|\|\|\|\|\|\|\|\|\|]PAM | NA | Intergenic | NA | NA |
| T10795 | CAGGTGTCTGGAACGTGGTGGGG | 1000 | | | | | | | | | |
| Oligo fwd | CACCCAGGTGTCTGGAACGTGGTG | | | | | | | | | | |
| Oligo rev | AAACCACCACGTTCCAGACACCTG | | | | | | | | | | |
| Chromosome | start | end | strand | MM | target_seq | PAM | alignment | distance | position | gene name | gene id |
| GU305914.1 | 140996 | 141018 | + | 0 | CAGGTGTCTGGAACGTGGTG | GGG | \|\|\|\|\|\|\|\|[\|\|\|\|\|\|\|\|\|\|\|\|]PAM | NA | Intergenic | NA | NA |
| T10796 | GAGAGAGACGCGACGCTCATCGG | 1000 | | | | | | | | | |
| Oligo fwd | CACCGAGAGAGACGCGACGCTCAT | | | | | | | | | | |
| Oligo rev | AAACATGAGCGTCGCGTCTCTCTC | | | | | | | | | | |
| Chromosome | start | end | strand | MM | target_seq | PAM | alignment | distance | position | gene name | gene id |
| GU305914.1 | 141030 | 141052 | + | 0 | GAGAGAGACGCGACGCTCAT | CGG | \|\|\|\|\|\|\|\|[\|\|\|\|\|\|\|\|\|\|\|\|]PAM | NA | Intergenic | NA | NA |
| T10797 | AGAGAGACGCGACGCTCATCGGG | 1000 | | | | | | | | | |
| Oligo fwd | CACCAGAGAGACGCGACGCTCATC | | | | | | | | | | |
| Oligo rev | AAACGATGAGCGTCGCGTCTCTCT | | | | | | | | | | |
| Chromosome | start | end | strand | MM | target_seq | PAM | alignment | distance | position | gene name | gene id |
| GU305914.1 | 141031 | 141053 | + | 0 | AGAGAGACGCGACGCTCATC | GGG | \|\|\|\|\|\|\|\|[\|\|\|\|\|\|\|\|\|\|\|\|]PAM | NA | Intergenic | NA | NA |
| GU305914.1 | 44906 | 44928 | - | 4 | AGCGAGAGGCGACGCTGCTC | CGG | \|\|-\|\|\|\|-[\|\|\|\|\|\|\|\|--\|\|]PAM | NA | Intergenic | NA | NA |
| T10798 | AGACGCGACGCTCATCGGGAAGG | 1000 | | | | | | | | | |
| Oligo fwd | CACCAGACGCGACGCTCATCGGGA | | | | | | | | | | |
| Oligo rev | AAACTCCCGATGAGCGTCGCGTCT | | | | | | | | | | |
| Chromosome | start | end | strand | MM | target_seq | PAM | alignment | distance | position | gene name | gene id |
| GU305914.1 | 141035 | 141057 | + | 0 | AGACGCGACGCTCATCGGGA | AGG | \|\|\|\|\|\|\|\|[\|\|\|\|\|\|\|\|\|\|\|\|]PAM | NA | Intergenic | NA | NA |
| T10799 | GACGCGACGCTCATCGGGAAGGG | 1000 | | | | | | | | | |
| Oligo fwd | CACCGACGCGACGCTCATCGGGAA | | | | | | | | | | |
| Oligo rev | AAACTTCCCGATGAGCGTCGCGTC | | | | | | | | | | |
| Chromosome | start | end | strand | MM | target_seq | PAM | alignment | distance | position | gene name | gene id |
| GU305914.1 | 141036 | 141058 | + | 0 | GACGCGACGCTCATCGGGAA | GGG | \|\|\|\|\|\|\|\|[\|\|\|\|\|\|\|\|\|\|\|\|]PAM | NA | Intergenic | NA | NA |
| T10800 | ACGCGACGCTCATCGGGAAGGGG | 1000 | | | | | | | | | |
| Oligo fwd | CACCACGCGACGCTCATCGGGAAG | | | | | | | | | | |
| Oligo rev | AAACCTTCCCGATGAGCGTCGCGT | | | | | | | | | | |
| Chromosome | start | end | strand | MM | target_seq | PAM | alignment | distance | position | gene name | gene id |

```
GU305914.1                    141037  141059  +        0  ACGCGACGCTCATCGGGAAG  GGG  ||||||||[||||||||||||]PAM  NA        Intergenic  NA         NA

T10801       CTCATCGGGAAGGGGTCTTTCGG         1000
Oligo fwd    CACCCTCATCGGGAAGGGGTCTTT
Oligo rev    AAACAAAGACCCCTTCCCGATGAG
Chromosome   start                    end     strand  MM  target_seq            PAM  alignment                  distance  position    gene name  gene id
GU305914.1                    141045  141067  +        0  CTCATCGGGAAGGGGTCTTT  CGG  ||||||||[||||||||||||]PAM  NA        Intergenic  NA         NA

T10802       CGGGAAGGGGTCTTTCGGACAGG         1000
Oligo fwd    CACCCGGGAAGGGGTCTTTCGGAC
Oligo rev    AAACGTCCGAAAGACCCCTTCCCG
Chromosome   start                    end     strand  MM  target_seq            PAM  alignment                  distance  position    gene name  gene id
GU305914.1                    141050  141072  +        0  CGGGAAGGGGTCTTTCGGAC  AGG  ||||||||[||||||||||||]PAM  NA        Intergenic  NA         NA

T10803       AGGGGTCTTTCGGACAGGTGTGG         1000
Oligo fwd    CACCAGGGGTCTTTCGGACAGGTG
Oligo rev    AAACCACCTGTCCGAAAGACCCCT
Chromosome   start                    end     strand  MM  target_seq            PAM  alignment                  distance  position    gene name  gene id
GU305914.1                    141055  141077  +        0  AGGGGTCTTTCGGACAGGTG  TGG  ||||||||[||||||||||||]PAM  NA        Intergenic  NA         NA

T10804       GAGACTCTCCGACAAGAAGACGG         1000
Oligo fwd    CACCGAGACTCTCCGACAAGAAGA
Oligo rev    AAACTCTTCTTGTCGGAGAGTCTC
Chromosome   start                    end     strand  MM  target_seq            PAM  alignment                  distance  position    gene name  gene id
GU305914.1                    141077  141099  +        0  GAGACTCTCCGACAAGAAGA  CGG  ||||||||[||||||||||||]PAM  NA        Intergenic  NA         NA

T10805       CGACAAGAAGACGGCCCTGAAGG         1000
Oligo fwd    CACCCGACAAGAAGACGGCCCTGA
Oligo rev    AAACTCAGGGCCGTCTTCTTGTCG
Chromosome   start                    end     strand  MM  target_seq            PAM  alignment                  distance  position    gene name  gene id
GU305914.1                    141086  141108  +        0  CGACAAGAAGACGGCCCTGA  AGG  ||||||||[||||||||||||]PAM  NA        Intergenic  NA         NA

T10806       ACGAGACCCTGTTGACCGTCTGG         1000
Oligo fwd    CACCACGAGACCCTGTTGACCGTC
Oligo rev    AAACGACGGTCAACAGGGTCTCGT
Chromosome   start                    end     strand  MM  target_seq            PAM  alignment                  distance  position    gene name  gene id
GU305914.1                    141130  141152  +        0  ACGAGACCCTGTTGACCGTC  TGG  ||||||||[||||||||||||]PAM  NA        Intergenic  NA         NA

T10807       CCTGTTGACCGTCTGGATCTCGG         1000
Oligo fwd    CACCCCTGTTGACCGTCTGGATCT
Oligo rev    AAACAGATCCAGACGGTCAACAGG
Chromosome   start                    end     strand  MM  target_seq            PAM  alignment                  distance  position    gene name  gene id
GU305914.1                    141137  141159  +        0  CCTGTTGACCGTCTGGATCT  CGG  ||||||||[||||||||||||]PAM  NA        Intergenic  NA         NA

T10808       CTGTTGACCGTCTGGATCTCGGG         1000
Oligo fwd    CACCCTGTTGACCGTCTGGATCTC
Oligo rev    AAACGAGATCCAGACGGTCAACAG
Chromosome   start                    end     strand  MM  target_seq            PAM  alignment                  distance  position    gene name  gene id
GU305914.1                    141138  141160  +        0  CTGTTGACCGTCTGGATCTC  GGG  ||||||||[||||||||||||]PAM  NA        Intergenic  NA         NA

T10809       CGTAGTACGTTCCCGAGCCCAGG         1000
Oligo fwd    CACCCGTAGTACGTTCCCGAGCCC
Oligo rev    AAACGGGCTCGGGAACGTACTACG
Chromosome   start                    end     strand  MM  target_seq            PAM  alignment                  distance  position    gene name  gene id
GU305914.1                    141161  141183  +        0  CGTAGTACGTTCCCGAGCCC  AGG  ||||||||[||||||||||||]PAM  NA        Intergenic  NA         NA

T10810       CGTTCCCGAGCCCAGGACGCCGG         1000
Oligo fwd    CACCCGTTCCCGAGCCCAGGACGC
Oligo rev    AAACGCGTCCTGGGCTCGGGAACG
Chromosome   start                    end     strand  MM  target_seq            PAM  alignment                  distance  position    gene name  gene id
GU305914.1                    141168  141190  +        0  CGTTCCCGAGCCCAGGACGC  CGG  ||||||||[||||||||||||]PAM  NA        Intergenic  NA         NA

T10811       GAGCCCAGGACGCCGGCTACCGG         1000
Oligo fwd    CACCGAGCCCAGGACGCCGGCTAC
Oligo rev    AAACGTAGCCGGCGTCCTGGGCTC
Chromosome   start                    end     strand  MM  target_seq            PAM  alignment                  distance  position    gene name  gene id
GU305914.1                    141175  141197  +        0  GAGCCCAGGACGCCGGCTAC  CGG  ||||||||[||||||||||||]PAM  NA        Intergenic  NA         NA

T10812       AGCCCAGGACGCCGGCTACCGGG         1000
Oligo fwd    CACCAGCCCAGGACGCCGGCTACC
Oligo rev    AAACGGTAGCCGGCGTCCTGGGCT
Chromosome   start                    end     strand  MM  target_seq            PAM  alignment                  distance  position    gene name  gene id
GU305914.1                    141176  141198  +        0  AGCCCAGGACGCCGGCTACC  GGG  ||||||||[||||||||||||]PAM  NA        Intergenic  NA         NA

T10813       GCCCAGGACGCCGGCTACCGGGG         1000
Oligo fwd    CACCGCCCAGGACGCCGGCTACCG
Oligo rev    AAACCGGTAGCCGGCGTCCTGGGC
Chromosome   start                    end     strand  MM  target_seq            PAM  alignment                  distance  position    gene name  gene id
GU305914.1                    141177  141199  +        0  GCCCAGGACGCCGGCTACCG  GGG  ||||||||[||||||||||||]PAM  NA        Intergenic  NA         NA

T10814       CGCCGGCTACCGGGGAGAACTGG         1000
Oligo fwd    CACCCGCCGGCTACCGGGGAGAAC
Oligo rev    AAACGTTCTCCCCGGTAGCCGGCG
Chromosome   start                    end     strand  MM  target_seq            PAM  alignment                  distance  position    gene name  gene id
GU305914.1                    141185  141207  +        0  CGCCGGCTACCGGGGAGAAC  TGG  ||||||||[||||||||||||]PAM  NA        Intergenic  NA         NA

T10815       TAGCGTCTACTGCAACATCCTGG         1000
Oligo fwd    CACCTAGCGTCTACTGCAACATCC
Oligo rev    AAACGGATGTTGCAGTAGACGCTA
```

<table>
<tr><td>Chromosome</td><td>start</td><td>end</td><td>strand</td><td>MM</td><td>target_seq</td><td>PAM</td><td>alignment</td><td>distance</td><td>position</td><td>gene name</td><td>gene id</td></tr>
<tr><td>GU305914.1</td><td>141212</td><td>141234</td><td>+</td><td>0</td><td>TAGCGTCTACTGCAACATCC</td><td>TGG</td><td>||||||||[||||||||||||]PAM</td><td>NA</td><td>Intergenic</td><td>NA</td><td>NA</td></tr>
<tr><td></td><td></td><td></td><td></td><td></td><td></td><td></td><td></td><td></td><td></td><td></td><td></td></tr>
<tr><td>T10816</td><td>CGTCTACTGCAACATCCTGGTGG</td><td>1000</td><td></td><td></td><td></td><td></td><td></td><td></td><td></td><td></td><td></td></tr>
<tr><td>Oligo fwd</td><td>CACCCGTCTACTGCAACATCCTGG</td><td></td><td></td><td></td><td></td><td></td><td></td><td></td><td></td><td></td><td></td></tr>
<tr><td>Oligo rev</td><td>AAACCCAGGATGTTGCAGTAGACG</td><td></td><td></td><td></td><td></td><td></td><td></td><td></td><td></td><td></td><td></td></tr>
<tr><td>Chromosome</td><td>start</td><td>end</td><td>strand</td><td>MM</td><td>target_seq</td><td>PAM</td><td>alignment</td><td>distance</td><td>position</td><td>gene name</td><td>gene id</td></tr>
<tr><td>GU305914.1</td><td>141215</td><td>141237</td><td>+</td><td>0</td><td>CGTCTACTGCAACATCCTGG</td><td>TGG</td><td>||||||||[||||||||||||]PAM</td><td>NA</td><td>Intergenic</td><td>NA</td><td>NA</td></tr>
<tr><td></td><td></td><td></td><td></td><td></td><td></td><td></td><td></td><td></td><td></td><td></td><td></td></tr>
<tr><td>T10817</td><td>TGCAACATCCTGGTGGCCACCGG</td><td>1000</td><td></td><td></td><td></td><td></td><td></td><td></td><td></td><td></td><td></td></tr>
<tr><td>Oligo fwd</td><td>CACCTGCAACATCCTGGTGGCCAC</td><td></td><td></td><td></td><td></td><td></td><td></td><td></td><td></td><td></td><td></td></tr>
<tr><td>Oligo rev</td><td>AAACGTGGCCACCAGGATGTTGCA</td><td></td><td></td><td></td><td></td><td></td><td></td><td></td><td></td><td></td><td></td></tr>
<tr><td>Chromosome</td><td>start</td><td>end</td><td>strand</td><td>MM</td><td>target_seq</td><td>PAM</td><td>alignment</td><td>distance</td><td>position</td><td>gene name</td><td>gene id</td></tr>
<tr><td>GU305914.1</td><td>141222</td><td>141244</td><td>+</td><td>0</td><td>TGCAACATCCTGGTGGCCAC</td><td>CGG</td><td>||||||||[||||||||||||]PAM</td><td>NA</td><td>Intergenic</td><td>NA</td><td>NA</td></tr>
<tr><td></td><td></td><td></td><td></td><td></td><td></td><td></td><td></td><td></td><td></td><td></td><td></td></tr>
<tr><td>T10818</td><td>TGTCGTTCGCCAGTTTCGATCGG</td><td>1000</td><td></td><td></td><td></td><td></td><td></td><td></td><td></td><td></td><td></td></tr>
<tr><td>Oligo fwd</td><td>CACCTGTCGTTCGCCAGTTTCGAT</td><td></td><td></td><td></td><td></td><td></td><td></td><td></td><td></td><td></td><td></td></tr>
<tr><td>Oligo rev</td><td>AAACATCGAAACTGGCGAACGACA</td><td></td><td></td><td></td><td></td><td></td><td></td><td></td><td></td><td></td><td></td></tr>
<tr><td>Chromosome</td><td>start</td><td>end</td><td>strand</td><td>MM</td><td>target_seq</td><td>PAM</td><td>alignment</td><td>distance</td><td>position</td><td>gene name</td><td>gene id</td></tr>
<tr><td>GU305914.1</td><td>141268</td><td>141290</td><td>+</td><td>0</td><td>TGTCGTTCGCCAGTTTCGAT</td><td>CGG</td><td>||||||||[||||||||||||]PAM</td><td>NA</td><td>Intergenic</td><td>NA</td><td>NA</td></tr>
<tr><td></td><td></td><td></td><td></td><td></td><td></td><td></td><td></td><td></td><td></td><td></td><td></td></tr>
<tr><td>T10819</td><td>GTCGTTCGCCAGTTTCGATCGGG</td><td>1000</td><td></td><td></td><td></td><td></td><td></td><td></td><td></td><td></td><td></td></tr>
<tr><td>Oligo fwd</td><td>CACCGTCGTTCGCCAGTTTCGATC</td><td></td><td></td><td></td><td></td><td></td><td></td><td></td><td></td><td></td><td></td></tr>
<tr><td>Oligo rev</td><td>AAACGATCGAAACTGGCGAACGAC</td><td></td><td></td><td></td><td></td><td></td><td></td><td></td><td></td><td></td><td></td></tr>
<tr><td>Chromosome</td><td>start</td><td>end</td><td>strand</td><td>MM</td><td>target_seq</td><td>PAM</td><td>alignment</td><td>distance</td><td>position</td><td>gene name</td><td>gene id</td></tr>
<tr><td>GU305914.1</td><td>141269</td><td>141291</td><td>+</td><td>0</td><td>GTCGTTCGCCAGTTTCGATC</td><td>GGG</td><td>||||||||[||||||||||||]PAM</td><td>NA</td><td>Intergenic</td><td>NA</td><td>NA</td></tr>
<tr><td></td><td></td><td></td><td></td><td></td><td></td><td></td><td></td><td></td><td></td><td></td><td></td></tr>
<tr><td>T10820</td><td>ATCGGGACCTCTACAACTACCGG</td><td>1000</td><td></td><td></td><td></td><td></td><td></td><td></td><td></td><td></td><td></td></tr>
<tr><td>Oligo fwd</td><td>CACCATCGGGACCTCTACAACTAC</td><td></td><td></td><td></td><td></td><td></td><td></td><td></td><td></td><td></td><td></td></tr>
<tr><td>Oligo rev</td><td>AAACGTAGTTGTAGAGGTCCCGAT</td><td></td><td></td><td></td><td></td><td></td><td></td><td></td><td></td><td></td><td></td></tr>
<tr><td>Chromosome</td><td>start</td><td>end</td><td>strand</td><td>MM</td><td>target_seq</td><td>PAM</td><td>alignment</td><td>distance</td><td>position</td><td>gene name</td><td>gene id</td></tr>
<tr><td>GU305914.1</td><td>141286</td><td>141308</td><td>+</td><td>0</td><td>ATCGGGACCTCTACAACTAC</td><td>CGG</td><td>||||||||[||||||||||||]PAM</td><td>NA</td><td>Intergenic</td><td>NA</td><td>NA</td></tr>
<tr><td></td><td></td><td></td><td></td><td></td><td></td><td></td><td></td><td></td><td></td><td></td><td></td></tr>
<tr><td>T10821</td><td>TCGGGACCTCTACAACTACCGGG</td><td>1000</td><td></td><td></td><td></td><td></td><td></td><td></td><td></td><td></td><td></td></tr>
<tr><td>Oligo fwd</td><td>CACCTCGGGACCTCTACAACTACC</td><td></td><td></td><td></td><td></td><td></td><td></td><td></td><td></td><td></td><td></td></tr>
<tr><td>Oligo rev</td><td>AAACGGTAGTTGTAGAGGTCCCGA</td><td></td><td></td><td></td><td></td><td></td><td></td><td></td><td></td><td></td><td></td></tr>
<tr><td>Chromosome</td><td>start</td><td>end</td><td>strand</td><td>MM</td><td>target_seq</td><td>PAM</td><td>alignment</td><td>distance</td><td>position</td><td>gene name</td><td>gene id</td></tr>
<tr><td>GU305914.1</td><td>141287</td><td>141309</td><td>+</td><td>0</td><td>TCGGGACCTCTACAACTACC</td><td>GGG</td><td>||||||||[||||||||||||]PAM</td><td>NA</td><td>Intergenic</td><td>NA</td><td>NA</td></tr>
<tr><td></td><td></td><td></td><td></td><td></td><td></td><td></td><td></td><td></td><td></td><td></td><td></td></tr>
<tr><td>T10822</td><td>CGGGACCTCTACAACTACCGGGG</td><td>1000</td><td></td><td></td><td></td><td></td><td></td><td></td><td></td><td></td><td></td></tr>
<tr><td>Oligo fwd</td><td>CACCCGGGACCTCTACAACTACCG</td><td></td><td></td><td></td><td></td><td></td><td></td><td></td><td></td><td></td><td></td></tr>
<tr><td>Oligo rev</td><td>AAACCGGTAGTTGTAGAGGTCCCG</td><td></td><td></td><td></td><td></td><td></td><td></td><td></td><td></td><td></td><td></td></tr>
<tr><td>Chromosome</td><td>start</td><td>end</td><td>strand</td><td>MM</td><td>target_seq</td><td>PAM</td><td>alignment</td><td>distance</td><td>position</td><td>gene name</td><td>gene id</td></tr>
<tr><td>GU305914.1</td><td>141288</td><td>141310</td><td>+</td><td>0</td><td>CGGGACCTCTACAACTACCG</td><td>GGG</td><td>||||||||[||||||||||||]PAM</td><td>NA</td><td>Intergenic</td><td>NA</td><td>NA</td></tr>
<tr><td></td><td></td><td></td><td></td><td></td><td></td><td></td><td></td><td></td><td></td><td></td><td></td></tr>
<tr><td>T10823</td><td>ACCTCTACAACTACCGGGGCTGG</td><td>1000</td><td></td><td></td><td></td><td></td><td></td><td></td><td></td><td></td><td></td></tr>
<tr><td>Oligo fwd</td><td>CACCACCTCTACAACTACCGGGGC</td><td></td><td></td><td></td><td></td><td></td><td></td><td></td><td></td><td></td><td></td></tr>
<tr><td>Oligo rev</td><td>AAACGCCCCGGTAGTTGTAGAGGT</td><td></td><td></td><td></td><td></td><td></td><td></td><td></td><td></td><td></td><td></td></tr>
<tr><td>Chromosome</td><td>start</td><td>end</td><td>strand</td><td>MM</td><td>target_seq</td><td>PAM</td><td>alignment</td><td>distance</td><td>position</td><td>gene name</td><td>gene id</td></tr>
<tr><td>GU305914.1</td><td>141292</td><td>141314</td><td>+</td><td>0</td><td>ACCTCTACAACTACCGGGGC</td><td>TGG</td><td>||||||||[||||||||||||]PAM</td><td>NA</td><td>Intergenic</td><td>NA</td><td>NA</td></tr>
<tr><td></td><td></td><td></td><td></td><td></td><td></td><td></td><td></td><td></td><td></td><td></td><td></td></tr>
<tr><td>T10824</td><td>CTACCGGGGCTGGCACTACGCGG</td><td>1000</td><td></td><td></td><td></td><td></td><td></td><td></td><td></td><td></td><td></td></tr>
<tr><td>Oligo fwd</td><td>CACCCTACCGGGGCTGGCACTACG</td><td></td><td></td><td></td><td></td><td></td><td></td><td></td><td></td><td></td><td></td></tr>
<tr><td>Oligo rev</td><td>AAACCGTAGTGCCAGCCCCGGTAG</td><td></td><td></td><td></td><td></td><td></td><td></td><td></td><td></td><td></td><td></td></tr>
<tr><td>Chromosome</td><td>start</td><td>end</td><td>strand</td><td>MM</td><td>target_seq</td><td>PAM</td><td>alignment</td><td>distance</td><td>position</td><td>gene name</td><td>gene id</td></tr>
<tr><td>GU305914.1</td><td>141302</td><td>141324</td><td>+</td><td>0</td><td>CTACCGGGGCTGGCACTACG</td><td>CGG</td><td>||||||||[||||||||||||]PAM</td><td>NA</td><td>Intergenic</td><td>NA</td><td>NA</td></tr>
<tr><td></td><td></td><td></td><td></td><td></td><td></td><td></td><td></td><td></td><td></td><td></td><td></td></tr>
<tr><td>T10825</td><td>TACCGGGGCTGGCACTACGCGGG</td><td>1000</td><td></td><td></td><td></td><td></td><td></td><td></td><td></td><td></td><td></td></tr>
<tr><td>Oligo fwd</td><td>CACCTACCGGGGCTGGCACTACGC</td><td></td><td></td><td></td><td></td><td></td><td></td><td></td><td></td><td></td><td></td></tr>
<tr><td>Oligo rev</td><td>AAACGCGTAGTGCCAGCCCCGGTA</td><td></td><td></td><td></td><td></td><td></td><td></td><td></td><td></td><td></td><td></td></tr>
<tr><td>Chromosome</td><td>start</td><td>end</td><td>strand</td><td>MM</td><td>target_seq</td><td>PAM</td><td>alignment</td><td>distance</td><td>position</td><td>gene name</td><td>gene id</td></tr>
<tr><td>GU305914.1</td><td>141303</td><td>141325</td><td>+</td><td>0</td><td>TACCGGGGCTGGCACTACGC</td><td>GGG</td><td>||||||||[||||||||||||]PAM</td><td>NA</td><td>Intergenic</td><td>NA</td><td>NA</td></tr>
<tr><td></td><td></td><td></td><td></td><td></td><td></td><td></td><td></td><td></td><td></td><td></td><td></td></tr>
<tr><td>T10826</td><td>GGGCTGGCACTACGCGGGCCTGG</td><td>1000</td><td></td><td></td><td></td><td></td><td></td><td></td><td></td><td></td><td></td></tr>
<tr><td>Oligo fwd</td><td>CACCGGGCTGGCACTACGCGGGCC</td><td></td><td></td><td></td><td></td><td></td><td></td><td></td><td></td><td></td><td></td></tr>
<tr><td>Oligo rev</td><td>AAACGGCCCGCGTAGTGCCAGCCC</td><td></td><td></td><td></td><td></td><td></td><td></td><td></td><td></td><td></td><td></td></tr>
<tr><td>Chromosome</td><td>start</td><td>end</td><td>strand</td><td>MM</td><td>target_seq</td><td>PAM</td><td>alignment</td><td>distance</td><td>position</td><td>gene name</td><td>gene id</td></tr>
<tr><td>GU305914.1</td><td>141308</td><td>141330</td><td>+</td><td>0</td><td>GGGCTGGCACTACGCGGGCC</td><td>TGG</td><td>||||||||[||||||||||||]PAM</td><td>NA</td><td>Intergenic</td><td>NA</td><td>NA</td></tr>
<tr><td></td><td></td><td></td><td></td><td></td><td></td><td></td><td></td><td></td><td></td><td></td><td></td></tr>
<tr><td>T10827</td><td>AGCTACCGCCGCGCCTTCAGCGG</td><td>1000</td><td></td><td></td><td></td><td></td><td></td><td></td><td></td><td></td><td></td></tr>
<tr><td>Oligo fwd</td><td>CACCAGCTACCGCCGCGCCTTCAG</td><td></td><td></td><td></td><td></td><td></td><td></td><td></td><td></td><td></td><td></td></tr>
<tr><td>Oligo rev</td><td>AAACCTGAAGGCGCGGCGGTAGCT</td><td></td><td></td><td></td><td></td><td></td><td></td><td></td><td></td><td></td><td></td></tr>
<tr><td>Chromosome</td><td>start</td><td>end</td><td>strand</td><td>MM</td><td>target_seq</td><td>PAM</td><td>alignment</td><td>distance</td><td>position</td><td>gene name</td><td>gene id</td></tr>
<tr><td>GU305914.1</td><td>141333</td><td>141355</td><td>+</td><td>0</td><td>AGCTACCGCCGCGCCTTCAG</td><td>CGG</td><td>||||||||[||||||||||||]PAM</td><td>NA</td><td>Intergenic</td><td>NA</td><td>NA</td></tr>
<tr><td></td><td></td><td></td><td></td><td></td><td></td><td></td><td></td><td></td><td></td><td></td><td></td></tr>
<tr><td>T10828</td><td>GCGGCATCGCCGACGCGCTCAGG</td><td>1000</td><td></td><td></td><td></td><td></td><td></td><td></td><td></td><td></td><td></td></tr>
<tr><td>Oligo fwd</td><td>CACCGCGGCATCGCCGACGCGCTC</td><td></td><td></td><td></td><td></td><td></td><td></td><td></td><td></td><td></td><td></td></tr>
<tr><td>Oligo rev</td><td>AAACGAGCGCGTCGGCGATGCCGC</td><td></td><td></td><td></td><td></td><td></td><td></td><td></td><td></td><td></td><td></td></tr>
<tr><td>Chromosome</td><td>start</td><td>end</td><td>strand</td><td>MM</td><td>target_seq</td><td>PAM</td><td>alignment</td><td>distance</td><td>position</td><td>gene name</td><td>gene id</td></tr>
<tr><td>GU305914.1</td><td>141352</td><td>141374</td><td>+</td><td>0</td><td>GCGGCATCGCCGACGCGCTC</td><td>AGG</td><td>||||||||[||||||||||||]PAM</td><td>NA</td><td>Intergenic</td><td>NA</td><td>NA</td></tr>
<tr><td></td><td></td><td></td><td></td><td></td><td></td><td></td><td></td><td></td><td></td><td></td><td></td></tr>
<tr><td>T10829</td><td>AGGTTCCTCAACCTGCGTTGCGG</td><td>1000</td><td></td><td></td><td></td><td></td><td></td><td></td><td></td><td></td><td></td></tr>
<tr><td>Oligo fwd</td><td>CACCAGGTTCCTCAACCTGCGTTG</td><td></td><td></td><td></td><td></td><td></td><td></td><td></td><td></td><td></td><td></td></tr>
<tr><td>Oligo rev</td><td>AAACCAACGCAGGTTGAGGAACCT</td><td></td><td></td><td></td><td></td><td></td><td></td><td></td><td></td><td></td><td></td></tr>
<tr><td>Chromosome</td><td>start</td><td>end</td><td>strand</td><td>MM</td><td>target_seq</td><td>PAM</td><td>alignment</td><td>distance</td><td>position</td><td>gene name</td><td>gene id</td></tr>
<tr><td>GU305914.1</td><td>141372</td><td>141394</td><td>+</td><td>0</td><td>AGGTTCCTCAACCTGCGTTG</td><td>CGG</td><td>||||||||[||||||||||||]PAM</td><td>NA</td><td>Intergenic</td><td>NA</td><td>NA</td></tr>
<tr><td></td><td></td><td></td><td></td><td></td><td></td><td></td><td></td><td></td><td></td><td></td><td></td></tr>
<tr><td>T10830</td><td>CTCAACCTGCGTTGCGGCGTCGG</td><td>1000</td><td></td><td></td><td></td><td></td><td></td><td></td><td></td><td></td><td></td></tr>
<tr><td>Oligo fwd</td><td>CACCCTCAACCTGCGTTGCGGCGT</td><td></td><td></td><td></td><td></td><td></td><td></td><td></td><td></td><td></td><td></td></tr>
</table>

| | | | | | | | | | | | |
|---|---|---|---|---|---|---|---|---|---|---|---|
| Oligo rev | AAACACGCCGCAACGCAGGTTGAG | | | | | | | | | | |
| Chromosome | start | end | strand | MM | target_seq | PAM | alignment | distance | position | gene name | gene id |
| GU305914.1 | 141378 | 141400 | + | 0 | CTCAACCTGCGTTGCGGCGT | CGG | \|\|\|\|\|\|\|\|[\|\|\|\|\|\|\|\|\|\|\|\|]PAM | NA | Intergenic | NA | NA |
| T10831 | ACTATGACCGCGCCGACGACAGG | 1000 | | | | | | | | | |
| Oligo fwd | CACCACTATGACCGCGCCGACGAC | | | | | | | | | | |
| Oligo rev | AAACGTCGTCGGCGCGGTCATAGT | | | | | | | | | | |
| Chromosome | start | end | strand | MM | target_seq | PAM | alignment | distance | position | gene name | gene id |
| GU305914.1 | 141436 | 141458 | + | 0 | ACTATGACCGCGCCGACGAC | AGG | \|\|\|\|\|\|\|\|[\|\|\|\|\|\|\|\|\|\|\|\|]PAM | NA | Intergenic | NA | NA |
| T10832 | CGACAGGCAGATCGCGCGTGCGG | 1000 | | | | | | | | | |
| Oligo fwd | CACCCGACAGGCAGATCGCGCGTG | | | | | | | | | | |
| Oligo rev | AAACCACGCGCGATCTGCCTGTCG | | | | | | | | | | |
| Chromosome | start | end | strand | MM | target_seq | PAM | alignment | distance | position | gene name | gene id |
| GU305914.1 | 141452 | 141474 | + | 0 | CGACAGGCAGATCGCGCGTG | CGG | \|\|\|\|\|\|\|\|[\|\|\|\|\|\|\|\|\|\|\|\|]PAM | NA | Intergenic | NA | NA |
| T10833 | CCTGTCGCAGTGTCACACCGAGG | 1000 | | | | | | | | | |
| Oligo fwd | CACCCCTGTCGCAGTGTCACACCG | | | | | | | | | | |
| Oligo rev | AAACCGGTGTGACACTGCGACAGG | | | | | | | | | | |
| Chromosome | start | end | strand | MM | target_seq | PAM | alignment | distance | position | gene name | gene id |
| GU305914.1 | 141491 | 141513 | + | 0 | CCTGTCGCAGTGTCACACCG | AGG | \|\|\|\|\|\|\|\|[\|\|\|\|\|\|\|\|\|\|\|\|]PAM | NA | Intergenic | NA | NA |
| T10834 | CTGTCGCAGTGTCACACCGAGGG | 1000 | | | | | | | | | |
| Oligo fwd | CACCCTGTCGCAGTGTCACACCGA | | | | | | | | | | |
| Oligo rev | AAACTCGGTGTGACACTGCGACAG | | | | | | | | | | |
| Chromosome | start | end | strand | MM | target_seq | PAM | alignment | distance | position | gene name | gene id |
| GU305914.1 | 141492 | 141514 | + | 0 | CTGTCGCAGTGTCACACCGA | GGG | \|\|\|\|\|\|\|\|[\|\|\|\|\|\|\|\|\|\|\|\|]PAM | NA | Intergenic | NA | NA |
| T10835 | TGTCACACCGAGGGTACTACCGG | 1000 | | | | | | | | | |
| Oligo fwd | CACCTGTCACACCGAGGGTACTAC | | | | | | | | | | |
| Oligo rev | AAACGTAGTACCCTCGGTGTGACA | | | | | | | | | | |
| Chromosome | start | end | strand | MM | target_seq | PAM | alignment | distance | position | gene name | gene id |
| GU305914.1 | 141501 | 141523 | + | 0 | TGTCACACCGAGGGTACTAC | CGG | \|\|\|\|\|\|\|\|[\|\|\|\|\|\|\|\|\|\|\|\|]PAM | NA | Intergenic | NA | NA |
| T10836 | TACTACCGGACACTGCGTCGTGG | 1000 | | | | | | | | | |
| Oligo fwd | CACCTACTACCGGACACTGCGTCG | | | | | | | | | | |
| Oligo rev | AAACCGACGCAGTGTCCGGTAGTA | | | | | | | | | | |
| Chromosome | start | end | strand | MM | target_seq | PAM | alignment | distance | position | gene name | gene id |
| GU305914.1 | 141515 | 141537 | + | 0 | TACTACCGGACACTGCGTCG | TGG | \|\|\|\|\|\|\|\|[\|\|\|\|\|\|\|\|\|\|\|\|]PAM | NA | Intergenic | NA | NA |
| T10837 | TCCAGCAGACCAAGACCGTCAGG | 1000 | | | | | | | | | |
| Oligo fwd | CACCTCCAGCAGACCAAGACCGTC | | | | | | | | | | |
| Oligo rev | AAACGACGGTCTTGGTCTGCTGGA | | | | | | | | | | |
| Chromosome | start | end | strand | MM | target_seq | PAM | alignment | distance | position | gene name | gene id |
| GU305914.1 | 141541 | 141563 | + | 0 | TCCAGCAGACCAAGACCGTC | AGG | \|\|\|\|\|\|\|\|[\|\|\|\|\|\|\|\|\|\|\|\|]PAM | NA | Intergenic | NA | NA |
| T10838 | CCAGCAGACCAAGACCGTCAGGG | 1000 | | | | | | | | | |
| Oligo fwd | CACCCCAGCAGACCAAGACCGTCA | | | | | | | | | | |
| Oligo rev | AAACTGACGGTCTTGGTCTGCTGG | | | | | | | | | | |
| Chromosome | start | end | strand | MM | target_seq | PAM | alignment | distance | position | gene name | gene id |
| GU305914.1 | 141542 | 141564 | + | 0 | CCAGCAGACCAAGACCGTCA | GGG | \|\|\|\|\|\|\|\|[\|\|\|\|\|\|\|\|\|\|\|\|]PAM | NA | Intergenic | NA | NA |
| T10839 | CAAGAGCGCCTACTACCTCACGG | 1000 | | | | | | | | | |
| Oligo fwd | CACCCAAGAGCGCCTACTACCTCA | | | | | | | | | | |
| Oligo rev | AAACTGAGGTAGTAGGCGCTCTTG | | | | | | | | | | |
| Chromosome | start | end | strand | MM | target_seq | PAM | alignment | distance | position | gene name | gene id |
| GU305914.1 | 141572 | 141594 | + | 0 | CAAGAGCGCCTACTACCTCA | CGG | \|\|\|\|\|\|\|\|[\|\|\|\|\|\|\|\|\|\|\|\|]PAM | NA | Intergenic | NA | NA |
| T10840 | CAAGCAGTTCCCCAAGCCCTCGG | 1000 | | | | | | | | | |
| Oligo fwd | CACCCAAGCAGTTCCCCAAGCCCT | | | | | | | | | | |
| Oligo rev | AAACAGGGCTTGGGGAACTGCTTG | | | | | | | | | | |
| Chromosome | start | end | strand | MM | target_seq | PAM | alignment | distance | position | gene name | gene id |
| GU305914.1 | 141656 | 141678 | + | 0 | CAAGCAGTTCCCCAAGCCCT | CGG | \|\|\|\|\|\|\|\|[\|\|\|\|\|\|\|\|\|\|\|\|]PAM | NA | Intergenic | NA | NA |
| T10841 | GGCGAACCGCTTCTGCGTCTCGG | 1000 | | | | | | | | | |
| Oligo fwd | CACCGGCGAACCGCTTCTGCGTCT | | | | | | | | | | |
| Oligo rev | AAACAGACGCAGAAGCGGTTCGCC | | | | | | | | | | |
| Chromosome | start | end | strand | MM | target_seq | PAM | alignment | distance | position | gene name | gene id |
| GU305914.1 | 141677 | 141699 | + | 0 | GGCGAACCGCTTCTGCGTCT | CGG | \|\|\|\|\|\|\|\|[\|\|\|\|\|\|\|\|\|\|\|\|]PAM | NA | Intergenic | NA | NA |
| T10842 | CGTCTCGGACCTCTGCGCCCTGG | 1000 | | | | | | | | | |
| Oligo fwd | CACCCGTCTCGGACCTCTGCGCCC | | | | | | | | | | |
| Oligo rev | AAACGGGCGCAGAGGTCCGAGACG | | | | | | | | | | |
| Chromosome | start | end | strand | MM | target_seq | PAM | alignment | distance | position | gene name | gene id |
| GU305914.1 | 141692 | 141714 | + | 0 | CGTCTCGGACCTCTGCGCCC | TGG | \|\|\|\|\|\|\|\|[\|\|\|\|\|\|\|\|\|\|\|\|]PAM | NA | Intergenic | NA | NA |
| GU305914.1 | 58778 | 58800 | - | 4 | CACCTCGGACCTCAGCGCCA | AGG | \|--\|\|\|\|\|[\|\|\|\|\|-\|\|\|\|\|-]PAM | NA | Intergenic | NA | NA |
| T10843 | GTCTCGGACCTCTGCGCCCTGGG | 1000 | | | | | | | | | |
| Oligo fwd | CACCGTCTCGGACCTCTGCGCCCT | | | | | | | | | | |
| Oligo rev | AAACAGGGCGCAGAGGTCCGAGAC | | | | | | | | | | |
| Chromosome | start | end | strand | MM | target_seq | PAM | alignment | distance | position | gene name | gene id |
| GU305914.1 | 141693 | 141715 | + | 0 | GTCTCGGACCTCTGCGCCCT | GGG | \|\|\|\|\|\|\|\|[\|\|\|\|\|\|\|\|\|\|\|\|]PAM | NA | Intergenic | NA | NA |
| T10844 | CCTCTGCGCCCTGGGACACGTGG | 1000 | | | | | | | | | |
| Oligo fwd | CACCCCTCTGCGCCCTGGGACACG | | | | | | | | | | |
| Oligo rev | AAACCGTGTCCCAGGGCGCAGAGG | | | | | | | | | | |
| Chromosome | start | end | strand | MM | target_seq | PAM | alignment | distance | position | gene name | gene id |
| GU305914.1 | 141701 | 141723 | + | 0 | CCTCTGCGCCCTGGGACACG | TGG | \|\|\|\|\|\|\|\|[\|\|\|\|\|\|\|\|\|\|\|\|]PAM | NA | Intergenic | NA | NA |

| | | | | | | | | | | | |
|---|---|---|---|---|---|---|---|---|---|---|---|
| T10845 | ACACGTGGCCGCTTTCTGCCTGG | 1000 | | | | | | | | | |
| Oligo fwd | CACCACACGTGGCCGCTTTCTGCC | | | | | | | | | | |
| Oligo rev | AAACGGCAGAAAGCGGCCACGTGT | | | | | | | | | | |
| Chromosome | start | end | strand | MM | target_seq | PAM | alignment | distance | position | gene name | gene id |
| GU305914.1 | 141716 | 141738 | + | 0 | ACACGTGGCCGCTTTCTGCC | TGG | \|\|\|\|\|\|\|\|[\|\|\|\|\|\|\|\|\|\|\|\|]PAM | NA | Intergenic | NA | NA |
| T10846 | TGGCCGCTTTCTGCCTGGTCAGG | 1000 | | | | | | | | | |
| Oligo fwd | CACCTGGCCGCTTTCTGCCTGGTC | | | | | | | | | | |
| Oligo rev | AAACGACCAGGCAGAAAGCGGCCA | | | | | | | | | | |
| Chromosome | start | end | strand | MM | target_seq | PAM | alignment | distance | position | gene name | gene id |
| GU305914.1 | 141721 | 141743 | + | 0 | TGGCCGCTTTCTGCCTGGTC | AGG | \|\|\|\|\|\|\|\|[\|\|\|\|\|\|\|\|\|\|\|\|]PAM | NA | Intergenic | NA | NA |
| T10847 | GGCCGCTTTCTGCCTGGTCAGGG | 1000 | | | | | | | | | |
| Oligo fwd | CACCGGCCGCTTTCTGCCTGGTCA | | | | | | | | | | |
| Oligo rev | AAACTGACCAGGCAGAAAGCGGCC | | | | | | | | | | |
| Chromosome | start | end | strand | MM | target_seq | PAM | alignment | distance | position | gene name | gene id |
| GU305914.1 | 141722 | 141744 | + | 0 | GGCCGCTTTCTGCCTGGTCA | GGG | \|\|\|\|\|\|\|\|[\|\|\|\|\|\|\|\|\|\|\|\|]PAM | NA | Intergenic | NA | NA |
| T10848 | TTTCTGCCTGGTCAGGGTCCTGG | 1000 | | | | | | | | | |
| Oligo fwd | CACCTTTCTGCCTGGTCAGGGTCC | | | | | | | | | | |
| Oligo rev | AAACGGACCCTGACCAGGCAGAAA | | | | | | | | | | |
| Chromosome | start | end | strand | MM | target_seq | PAM | alignment | distance | position | gene name | gene id |
| GU305914.1 | 141728 | 141750 | + | 0 | TTTCTGCCTGGTCAGGGTCC | TGG | \|\|\|\|\|\|\|\|[\|\|\|\|\|\|\|\|\|\|\|\|]PAM | NA | Intergenic | NA | NA |
| T10849 | GTCAGGGTCCTGGACGAGCGCGG | 1000 | | | | | | | | | |
| Oligo fwd | CACCGTCAGGGTCCTGGACGAGCG | | | | | | | | | | |
| Oligo rev | AAACCGCTCGTCCAGGACCCTGAC | | | | | | | | | | |
| Chromosome | start | end | strand | MM | target_seq | PAM | alignment | distance | position | gene name | gene id |
| GU305914.1 | 141738 | 141760 | + | 0 | GTCAGGGTCCTGGACGAGCG | CGG | \|\|\|\|\|\|\|\|[\|\|\|\|\|\|\|\|\|\|\|\|]PAM | NA | Intergenic | NA | NA |
| GU305914.1 | 195326 | 195348 | - | 4 | GCCCTGGTCATGGACGAGCG | CGG | \|-\|--\|\|\|[\|-\|\|\|\|\|\|\|\|\|\|]PAM | NA | Intergenic | NA | NA |
| T10850 | GGACGAGCGCGGACAGCTCAAGG | 1000 | | | | | | | | | |
| Oligo fwd | CACCGGACGAGCGCGGACAGCTCA | | | | | | | | | | |
| Oligo rev | AAACTGAGCTGTCCGCGCTCGTCC | | | | | | | | | | |
| Chromosome | start | end | strand | MM | target_seq | PAM | alignment | distance | position | gene name | gene id |
| GU305914.1 | 141749 | 141771 | + | 0 | GGACGAGCGCGGACAGCTCA | AGG | \|\|\|\|\|\|\|\|[\|\|\|\|\|\|\|\|\|\|\|\|]PAM | NA | Intergenic | NA | NA |
| T10851 | AGCGCGGACAGCTCAAGGTCAGG | 1000 | | | | | | | | | |
| Oligo fwd | CACCAGCGCGGACAGCTCAAGGTC | | | | | | | | | | |
| Oligo rev | AAACGACCTTGAGCTGTCCGCGCT | | | | | | | | | | |
| Chromosome | start | end | strand | MM | target_seq | PAM | alignment | distance | position | gene name | gene id |
| GU305914.1 | 141754 | 141776 | + | 0 | AGCGCGGACAGCTCAAGGTC | AGG | \|\|\|\|\|\|\|\|[\|\|\|\|\|\|\|\|\|\|\|\|]PAM | NA | Intergenic | NA | NA |
| T10852 | CAAGGTCAGGAGCACCTCCGAGG | 1000 | | | | | | | | | |
| Oligo fwd | CACCCAAGGTCAGGAGCACCTCCG | | | | | | | | | | |
| Oligo rev | AAACCGGAGGTGCTCCTGACCTTG | | | | | | | | | | |
| Chromosome | start | end | strand | MM | target_seq | PAM | alignment | distance | position | gene name | gene id |
| GU305914.1 | 141767 | 141789 | + | 0 | CAAGGTCAGGAGCACCTCCG | AGG | \|\|\|\|\|\|\|\|[\|\|\|\|\|\|\|\|\|\|\|\|]PAM | NA | Intergenic | NA | NA |
| T10853 | ACCTCCGAGGACGCGCTCTTCGG | 1000 | | | | | | | | | |
| Oligo fwd | CACCACCTCCGAGGACGCGCTCTT | | | | | | | | | | |
| Oligo rev | AAACAAGAGCGCGTCCTCGGAGGT | | | | | | | | | | |
| Chromosome | start | end | strand | MM | target_seq | PAM | alignment | distance | position | gene name | gene id |
| GU305914.1 | 141780 | 141802 | + | 0 | ACCTCCGAGGACGCGCTCTT | CGG | \|\|\|\|\|\|\|\|[\|\|\|\|\|\|\|\|\|\|\|\|]PAM | NA | Intergenic | NA | NA |
| T10854 | AGACCTGCGACGCCCTCGCCCGG | 1000 | | | | | | | | | |
| Oligo fwd | CACCAGACCTGCGACGCCCTCGCC | | | | | | | | | | |
| Oligo rev | AAACGGCGAGGGCGTCGCAGGTCT | | | | | | | | | | |
| Chromosome | start | end | strand | MM | target_seq | PAM | alignment | distance | position | gene name | gene id |
| GU305914.1 | 141814 | 141836 | + | 0 | AGACCTGCGACGCCCTCGCC | CGG | \|\|\|\|\|\|\|\|[\|\|\|\|\|\|\|\|\|\|\|\|]PAM | NA | Intergenic | NA | NA |
| T10855 | CGACGCCCTCGCCCGGCACTCGG | 1000 | | | | | | | | | |
| Oligo fwd | CACCCGACGCCCTCGCCCGGCACT | | | | | | | | | | |
| Oligo rev | AAACAGTGCCGGGCGAGGGCGTCG | | | | | | | | | | |
| Chromosome | start | end | strand | MM | target_seq | PAM | alignment | distance | position | gene name | gene id |
| GU305914.1 | 141821 | 141843 | + | 0 | CGACGCCCTCGCCCGGCACT | CGG | \|\|\|\|\|\|\|\|[\|\|\|\|\|\|\|\|\|\|\|\|]PAM | NA | Intergenic | NA | NA |
| T10856 | CGCCCTCGCCCGGCACTCGGTGG | 1000 | | | | | | | | | |
| Oligo fwd | CACCCGCCCTCGCCCGGCACTCGG | | | | | | | | | | |
| Oligo rev | AAACCCGAGTGCCGGGCGAGGGCG | | | | | | | | | | |
| Chromosome | start | end | strand | MM | target_seq | PAM | alignment | distance | position | gene name | gene id |
| GU305914.1 | 141824 | 141846 | + | 0 | CGCCCTCGCCCGGCACTCGG | TGG | \|\|\|\|\|\|\|\|[\|\|\|\|\|\|\|\|\|\|\|\|]PAM | NA | Intergenic | NA | NA |
| T10857 | CGCCCGGCACTCGGTGGACGAGG | 1000 | | | | | | | | | |
| Oligo fwd | CACCCGCCCGGCACTCGGTGGACG | | | | | | | | | | |
| Oligo rev | AAACCGTCCACCGAGTGCCGGGCG | | | | | | | | | | |
| Chromosome | start | end | strand | MM | target_seq | PAM | alignment | distance | position | gene name | gene id |
| GU305914.1 | 141830 | 141852 | + | 0 | CGCCCGGCACTCGGTGGACG | AGG | \|\|\|\|\|\|\|\|[\|\|\|\|\|\|\|\|\|\|\|\|]PAM | NA | Intergenic | NA | NA |
| T10858 | CCGGCACTCGGTGGACGAGGTGG | 1000 | | | | | | | | | |
| Oligo fwd | CACCCCGGCACTCGGTGGACGAGG | | | | | | | | | | |
| Oligo rev | AAACCCTCGTCCACCGAGTGCCGG | | | | | | | | | | |
| Chromosome | start | end | strand | MM | target_seq | PAM | alignment | distance | position | gene name | gene id |
| GU305914.1 | 141833 | 141855 | + | 0 | CCGGCACTCGGTGGACGAGG | TGG | \|\|\|\|\|\|\|\|[\|\|\|\|\|\|\|\|\|\|\|\|]PAM | NA | Intergenic | NA | NA |
| T10859 | GCTGCTCATCACGCGCCAGCTGG | 1000 | | | | | | | | | |
| Oligo fwd | CACCGCTGCTCATCACGCGCCAGC | | | | | | | | | | |
| Oligo rev | AAACGCTGGCGCGTGATGAGCAGC | | | | | | | | | | |
| Chromosome | start | end | strand | MM | target_seq | PAM | alignment | distance | position | gene name | gene id |

| | | | | | | | | | | | |
|---|---|---|---|---|---|---|---|---|---|---|---|
| GU305914.1 | 141869 | 141891 | + | 0 | GCTGCTCATCACGCGCCAGC | TGG | ||||||||[||||||||||||]PAM | NA | Intergenic | NA | NA |
| T10860 | GGCCTACACCGCCACCTTGCTGG | 1000 | | | | | | | | | |
| Oligo fwd | CACCGGCCTACACCGCCACCTTGC | | | | | | | | | | |
| Oligo rev | AAACGCAAGGTGGCGGTGTAGGCC | | | | | | | | | | |
| Chromosome | start | end | strand | MM | target_seq | PAM | alignment | distance | position | gene name | gene id |
| GU305914.1 | 141890 | 141912 | + | 0 | GGCCTACACCGCCACCTTGC | TGG | ||||||||[||||||||||||]PAM | NA | Intergenic | NA | NA |
| T10861 | GCCTACACCGCCACCTTGCTGGG | 1000 | | | | | | | | | |
| Oligo fwd | CACCGCCTACACCGCCACCTTGCT | | | | | | | | | | |
| Oligo rev | AAACAGCAAGGTGGCGGTGTAGGC | | | | | | | | | | |
| Chromosome | start | end | strand | MM | target_seq | PAM | alignment | distance | position | gene name | gene id |
| GU305914.1 | 141891 | 141913 | + | 0 | GCCTACACCGCCACCTTGCT | GGG | ||||||||[||||||||||||]PAM | NA | Intergenic | NA | NA |
| T10862 | TGCTGGGCAGCGACGACATGCGG | 1000 | | | | | | | | | |
| Oligo fwd | CACCTGCTGGGCAGCGACGACATG | | | | | | | | | | |
| Oligo rev | AAACCATGTCGTCGCTGCCCAGCA | | | | | | | | | | |
| Chromosome | start | end | strand | MM | target_seq | PAM | alignment | distance | position | gene name | gene id |
| GU305914.1 | 141907 | 141929 | + | 0 | TGCTGGGCAGCGACGACATG | CGG | ||||||||[||||||||||||]PAM | NA | Intergenic | NA | NA |
| T10863 | GCTGGGCAGCGACGACATGCGGG | 1000 | | | | | | | | | |
| Oligo fwd | CACCGCTGGGCAGCGACGACATGC | | | | | | | | | | |
| Oligo rev | AAACGCATGTCGTCGCTGCCCAGC | | | | | | | | | | |
| Chromosome | start | end | strand | MM | target_seq | PAM | alignment | distance | position | gene name | gene id |
| GU305914.1 | 141908 | 141930 | + | 0 | GCTGGGCAGCGACGACATGC | GGG | ||||||||[||||||||||||]PAM | NA | Intergenic | NA | NA |
| T10864 | CGACGACATGCGGGAGCCTATGG | 1000 | | | | | | | | | |
| Oligo fwd | CACCCGACGACATGCGGGAGCCTA | | | | | | | | | | |
| Oligo rev | AAACTAGGCTCCCGCATGTCGTCG | | | | | | | | | | |
| Chromosome | start | end | strand | MM | target_seq | PAM | alignment | distance | position | gene name | gene id |
| GU305914.1 | 141917 | 141939 | + | 0 | CGACGACATGCGGGAGCCTA | TGG | ||||||||[||||||||||||]PAM | NA | Intergenic | NA | NA |
| GU305914.1 | 120088 | 120110 | + | 4 | CGACGAGGTGCGGGAGTCCA | AGG | ||||||--[||||||||-|-|]PAM | NA | Intergenic | NA | NA |
| T10865 | ACATGCGGGAGCCTATGGCGCGG | 1000 | | | | | | | | | |
| Oligo fwd | CACCACATGCGGGAGCCTATGGCG | | | | | | | | | | |
| Oligo rev | AAACCGCCATAGGCTCCCGCATGT | | | | | | | | | | |
| Chromosome | start | end | strand | MM | target_seq | PAM | alignment | distance | position | gene name | gene id |
| GU305914.1 | 141922 | 141944 | + | 0 | ACATGCGGGAGCCTATGGCG | CGG | ||||||||[||||||||||||]PAM | NA | Intergenic | NA | NA |
| T10866 | GTTGTGTGACTATTTCGAAACGG | 1000 | | | | | | | | | |
| Oligo fwd | CACCGTTGTGTGACTATTTCGAAA | | | | | | | | | | |
| Oligo rev | AAACTTTCGAAATAGTCACACAAC | | | | | | | | | | |
| Chromosome | start | end | strand | MM | target_seq | PAM | alignment | distance | position | gene name | gene id |
| GU305914.1 | 141944 | 141966 | + | 0 | GTTGTGTGACTATTTCGAAA | CGG | ||||||||[||||||||||||]PAM | NA | Intergenic | NA | NA |
| T10867 | TTTCGAAACGGTTAGCGACAAGG | 1000 | | | | | | | | | |
| Oligo fwd | CACCTTTCGAAACGGTTAGCGACA | | | | | | | | | | |
| Oligo rev | AAACTGTCGCTAACCGTTTCGAAA | | | | | | | | | | |
| Chromosome | start | end | strand | MM | target_seq | PAM | alignment | distance | position | gene name | gene id |
| GU305914.1 | 141956 | 141978 | + | 0 | TTTCGAAACGGTTAGCGACA | AGG | ||||||||[||||||||||||]PAM | NA | Intergenic | NA | NA |
| T10868 | AGGACGCTCCCGACCGTTTTCGG | 1000 | | | | | | | | | |
| Oligo fwd | CACCAGGACGCTCCCGACCGTTTT | | | | | | | | | | |
| Oligo rev | AAACAAAACGGTCGGGAGCGTCCT | | | | | | | | | | |
| Chromosome | start | end | strand | MM | target_seq | PAM | alignment | distance | position | gene name | gene id |
| GU305914.1 | 141976 | 141998 | + | 0 | AGGACGCTCCCGACCGTTTT | CGG | ||||||||[||||||||||||]PAM | NA | Intergenic | NA | NA |
| T10869 | AGAGCTAGACGCGAGATAGACGG | 1000 | | | | | | | | | |
| Oligo fwd | CACCAGAGCTAGACGCGAGATAGA | | | | | | | | | | |
| Oligo rev | AAACTCTATCTCGCGTCTAGCTCT | | | | | | | | | | |
| Chromosome | start | end | strand | MM | target_seq | PAM | alignment | distance | position | gene name | gene id |
| GU305914.1 | 142011 | 142033 | + | 0 | AGAGCTAGACGCGAGATAGA | CGG | ||||||||[||||||||||||]PAM | NA | Intergenic | NA | NA |
| T10870 | GAGCTAGACGCGAGATAGACGGG | 1000 | | | | | | | | | |
| Oligo fwd | CACCGAGCTAGACGCGAGATAGAC | | | | | | | | | | |
| Oligo rev | AAACGTCTATCTCGCGTCTAGCTC | | | | | | | | | | |
| Chromosome | start | end | strand | MM | target_seq | PAM | alignment | distance | position | gene name | gene id |
| GU305914.1 | 142012 | 142034 | + | 0 | GAGCTAGACGCGAGATAGAC | GGG | ||||||||[||||||||||||]PAM | NA | Intergenic | NA | NA |
| T10871 | CGAGATAGACGGGTCATACATGG | 1000 | | | | | | | | | |
| Oligo fwd | CACCCGAGATAGACGGGTCATACA | | | | | | | | | | |
| Oligo rev | AAACTGTATGACCCGTCTATCTCG | | | | | | | | | | |
| Chromosome | start | end | strand | MM | target_seq | PAM | alignment | distance | position | gene name | gene id |
| GU305914.1 | 142022 | 142044 | + | 0 | CGAGATAGACGGGTCATACA | TGG | ||||||||[||||||||||||]PAM | NA | Intergenic | NA | NA |
| T10872 | TAGACGGGTCATACATGGTCCGG | 1000 | | | | | | | | | |
| Oligo fwd | CACCTAGACGGGTCATACATGGTC | | | | | | | | | | |
| Oligo rev | AAACGACCATGTATGACCCGTCTA | | | | | | | | | | |
| Chromosome | start | end | strand | MM | target_seq | PAM | alignment | distance | position | gene name | gene id |
| GU305914.1 | 142027 | 142049 | + | 0 | TAGACGGGTCATACATGGTC | CGG | ||||||||[||||||||||||]PAM | NA | Intergenic | NA | NA |
| T10873 | CCGGCTCCTGTTAGCCGCGTCGG | 1000 | | | | | | | | | |
| Oligo fwd | CACCCCGGCTCCTGTTAGCCGCGT | | | | | | | | | | |
| Oligo rev | AAACACGCGGCTAACAGGAGCCGG | | | | | | | | | | |
| Chromosome | start | end | strand | MM | target_seq | PAM | alignment | distance | position | gene name | gene id |
| GU305914.1 | 142046 | 142068 | + | 0 | CCGGCTCCTGTTAGCCGCGT | CGG | ||||||||[||||||||||||]PAM | NA | Intergenic | NA | NA |
| T10874 | CCTGTTAGCCGCGTCGGAGACGG | 1000 | | | | | | | | | |
| Oligo fwd | CACCCCTGTTAGCCGCGTCGGAGA | | | | | | | | | | |

<table>
<tr><td>Oligo rev</td><td>AAACTCTCCGACGCGGCTAACAGG</td><td></td><td></td><td></td><td></td><td></td><td></td><td></td><td></td><td></td><td></td><td></td></tr>
<tr><td>Chromosome</td><td>start</td><td></td><td>end</td><td>strand</td><td>MM</td><td>target_seq</td><td>PAM</td><td>alignment</td><td>distance</td><td>position</td><td>gene name</td><td>gene id</td></tr>
<tr><td>GU305914.1</td><td></td><td>142052</td><td>142074</td><td>+</td><td>0</td><td>CCTGTTAGCCGCGTCGGAGA</td><td>CGG</td><td>||||||||[||||||||||||]PAM</td><td>NA</td><td>Intergenic</td><td>NA</td><td>NA</td></tr>
<tr><td>T10875</td><td>GTTAGCCGCGTCGGAGACGGAGG</td><td></td><td>1000</td><td></td><td></td><td></td><td></td><td></td><td></td><td></td><td></td><td></td></tr>
<tr><td>Oligo fwd</td><td>CACCGTTAGCCGCGTCGGAGACGG</td><td></td><td></td><td></td><td></td><td></td><td></td><td></td><td></td><td></td><td></td><td></td></tr>
<tr><td>Oligo rev</td><td>AAACCCGTCTCCGACGCGGCTAAC</td><td></td><td></td><td></td><td></td><td></td><td></td><td></td><td></td><td></td><td></td><td></td></tr>
<tr><td>Chromosome</td><td>start</td><td></td><td>end</td><td>strand</td><td>MM</td><td>target_seq</td><td>PAM</td><td>alignment</td><td>distance</td><td>position</td><td>gene name</td><td>gene id</td></tr>
<tr><td>GU305914.1</td><td></td><td>142055</td><td>142077</td><td>+</td><td>0</td><td>GTTAGCCGCGTCGGAGACGG</td><td>AGG</td><td>||||||||[||||||||||||]PAM</td><td>NA</td><td>Intergenic</td><td>NA</td><td>NA</td></tr>
<tr><td>T10876</td><td>GCCGCGTCGGAGACGGAGGACGG</td><td></td><td>1000</td><td></td><td></td><td></td><td></td><td></td><td></td><td></td><td></td><td></td></tr>
<tr><td>Oligo fwd</td><td>CACCGCCGCGTCGGAGACGGAGGA</td><td></td><td></td><td></td><td></td><td></td><td></td><td></td><td></td><td></td><td></td><td></td></tr>
<tr><td>Oligo rev</td><td>AAACTCCTCCGTCTCCGACGCGGC</td><td></td><td></td><td></td><td></td><td></td><td></td><td></td><td></td><td></td><td></td><td></td></tr>
<tr><td>Chromosome</td><td>start</td><td></td><td>end</td><td>strand</td><td>MM</td><td>target_seq</td><td>PAM</td><td>alignment</td><td>distance</td><td>position</td><td>gene name</td><td>gene id</td></tr>
<tr><td>GU305914.1</td><td></td><td>142059</td><td>142081</td><td>+</td><td>0</td><td>GCCGCGTCGGAGACGGAGGA</td><td>CGG</td><td>||||||||[||||||||||||]PAM</td><td>NA</td><td>Intergenic</td><td>NA</td><td>NA</td></tr>
<tr><td>T10877</td><td>CGTCGGAGACGGAGGACGGTAGG</td><td></td><td>1000</td><td></td><td></td><td></td><td></td><td></td><td></td><td></td><td></td><td></td></tr>
<tr><td>Oligo fwd</td><td>CACCCGTCGGAGACGGAGGACGGT</td><td></td><td></td><td></td><td></td><td></td><td></td><td></td><td></td><td></td><td></td><td></td></tr>
<tr><td>Oligo rev</td><td>AAACACCGTCCTCCGTCTCCGACG</td><td></td><td></td><td></td><td></td><td></td><td></td><td></td><td></td><td></td><td></td><td></td></tr>
<tr><td>Chromosome</td><td>start</td><td></td><td>end</td><td>strand</td><td>MM</td><td>target_seq</td><td>PAM</td><td>alignment</td><td>distance</td><td>position</td><td>gene name</td><td>gene id</td></tr>
<tr><td>GU305914.1</td><td></td><td>142063</td><td>142085</td><td>+</td><td>0</td><td>CGTCGGAGACGGAGGACGGT</td><td>AGG</td><td>||||||||[||||||||||||]PAM</td><td>NA</td><td>Intergenic</td><td>NA</td><td>NA</td></tr>
<tr><td>T10878</td><td>CATACGTGCGACGTGTTTAATGG</td><td></td><td>1000</td><td></td><td></td><td></td><td></td><td></td><td></td><td></td><td></td><td></td></tr>
<tr><td>Oligo fwd</td><td>CACCCATACGTGCGACGTGTTTAA</td><td></td><td></td><td></td><td></td><td></td><td></td><td></td><td></td><td></td><td></td><td></td></tr>
<tr><td>Oligo rev</td><td>AAACTTAAACACGTCGCACGTATG</td><td></td><td></td><td></td><td></td><td></td><td></td><td></td><td></td><td></td><td></td><td></td></tr>
<tr><td>Chromosome</td><td>start</td><td></td><td>end</td><td>strand</td><td>MM</td><td>target_seq</td><td>PAM</td><td>alignment</td><td>distance</td><td>position</td><td>gene name</td><td>gene id</td></tr>
<tr><td>GU305914.1</td><td></td><td>142100</td><td>142122</td><td>+</td><td>0</td><td>CATACGTGCGACGTGTTTAA</td><td>TGG</td><td>||||||||[||||||||||||]PAM</td><td>NA</td><td>Intergenic</td><td>NA</td><td>NA</td></tr>
<tr><td>T10879</td><td>ACGTGCGACGTGTTTAATGGTGG</td><td></td><td>1000</td><td></td><td></td><td></td><td></td><td></td><td></td><td></td><td></td><td></td></tr>
<tr><td>Oligo fwd</td><td>CACCACGTGCGACGTGTTTAATGG</td><td></td><td></td><td></td><td></td><td></td><td></td><td></td><td></td><td></td><td></td><td></td></tr>
<tr><td>Oligo rev</td><td>AAACCCATTAAACACGTCGCACGT</td><td></td><td></td><td></td><td></td><td></td><td></td><td></td><td></td><td></td><td></td><td></td></tr>
<tr><td>Chromosome</td><td>start</td><td></td><td>end</td><td>strand</td><td>MM</td><td>target_seq</td><td>PAM</td><td>alignment</td><td>distance</td><td>position</td><td>gene name</td><td>gene id</td></tr>
<tr><td>GU305914.1</td><td></td><td>142103</td><td>142125</td><td>+</td><td>0</td><td>ACGTGCGACGTGTTTAATGG</td><td>TGG</td><td>||||||||[||||||||||||]PAM</td><td>NA</td><td>Intergenic</td><td>NA</td><td>NA</td></tr>
<tr><td>T10880</td><td>GTGTTTAATGGTGGATAGCGAGG</td><td></td><td>1000</td><td></td><td></td><td></td><td></td><td></td><td></td><td></td><td></td><td></td></tr>
<tr><td>Oligo fwd</td><td>CACCGTGTTTAATGGTGGATAGCG</td><td></td><td></td><td></td><td></td><td></td><td></td><td></td><td></td><td></td><td></td><td></td></tr>
<tr><td>Oligo rev</td><td>AAACCGCTATCCACCATTAAACAC</td><td></td><td></td><td></td><td></td><td></td><td></td><td></td><td></td><td></td><td></td><td></td></tr>
<tr><td>Chromosome</td><td>start</td><td></td><td>end</td><td>strand</td><td>MM</td><td>target_seq</td><td>PAM</td><td>alignment</td><td>distance</td><td>position</td><td>gene name</td><td>gene id</td></tr>
<tr><td>GU305914.1</td><td></td><td>142112</td><td>142134</td><td>+</td><td>0</td><td>GTGTTTAATGGTGGATAGCG</td><td>AGG</td><td>||||||||[||||||||||||]PAM</td><td>NA</td><td>Intergenic</td><td>NA</td><td>NA</td></tr>
<tr><td>T10881</td><td>ATCTTTCCCTGATCGTCCTCCGG</td><td></td><td>1000</td><td></td><td></td><td></td><td></td><td></td><td></td><td></td><td></td><td></td></tr>
<tr><td>Oligo fwd</td><td>CACCATCTTTCCCTGATCGTCCTC</td><td></td><td></td><td></td><td></td><td></td><td></td><td></td><td></td><td></td><td></td><td></td></tr>
<tr><td>Oligo rev</td><td>AAACGAGGACGATCAGGGAAAGAT</td><td></td><td></td><td></td><td></td><td></td><td></td><td></td><td></td><td></td><td></td><td></td></tr>
<tr><td>Chromosome</td><td>start</td><td></td><td>end</td><td>strand</td><td>MM</td><td>target_seq</td><td>PAM</td><td>alignment</td><td>distance</td><td>position</td><td>gene name</td><td>gene id</td></tr>
<tr><td>GU305914.1</td><td></td><td>142158</td><td>142180</td><td>+</td><td>0</td><td>ATCTTTCCCTGATCGTCCTC</td><td>CGG</td><td>||||||||[||||||||||||]PAM</td><td>NA</td><td>Intergenic</td><td>NA</td><td>NA</td></tr>
<tr><td>T10882</td><td>TCTTTCCCTGATCGTCCTCCGGG</td><td></td><td>1000</td><td></td><td></td><td></td><td></td><td></td><td></td><td></td><td></td><td></td></tr>
<tr><td>Oligo fwd</td><td>CACCTCTTTCCCTGATCGTCCTCC</td><td></td><td></td><td></td><td></td><td></td><td></td><td></td><td></td><td></td><td></td><td></td></tr>
<tr><td>Oligo rev</td><td>AAACGGAGGACGATCAGGGAAAGA</td><td></td><td></td><td></td><td></td><td></td><td></td><td></td><td></td><td></td><td></td><td></td></tr>
<tr><td>Chromosome</td><td>start</td><td></td><td>end</td><td>strand</td><td>MM</td><td>target_seq</td><td>PAM</td><td>alignment</td><td>distance</td><td>position</td><td>gene name</td><td>gene id</td></tr>
<tr><td>GU305914.1</td><td></td><td>142159</td><td>142181</td><td>+</td><td>0</td><td>TCTTTCCCTGATCGTCCTCC</td><td>GGG</td><td>||||||||[||||||||||||]PAM</td><td>NA</td><td>Intergenic</td><td>NA</td><td>NA</td></tr>
<tr><td>T10883</td><td>TCCCTGATCGTCCTCCGGGTCGG</td><td></td><td>1000</td><td></td><td></td><td></td><td></td><td></td><td></td><td></td><td></td><td></td></tr>
<tr><td>Oligo fwd</td><td>CACCTCCCTGATCGTCCTCCGGGT</td><td></td><td></td><td></td><td></td><td></td><td></td><td></td><td></td><td></td><td></td><td></td></tr>
<tr><td>Oligo rev</td><td>AAACACCCGGAGGACGATCAGGGA</td><td></td><td></td><td></td><td></td><td></td><td></td><td></td><td></td><td></td><td></td><td></td></tr>
<tr><td>Chromosome</td><td>start</td><td></td><td>end</td><td>strand</td><td>MM</td><td>target_seq</td><td>PAM</td><td>alignment</td><td>distance</td><td>position</td><td>gene name</td><td>gene id</td></tr>
<tr><td>GU305914.1</td><td></td><td>142163</td><td>142185</td><td>+</td><td>0</td><td>TCCCTGATCGTCCTCCGGGT</td><td>CGG</td><td>||||||||[||||||||||||]PAM</td><td>NA</td><td>Intergenic</td><td>NA</td><td>NA</td></tr>
<tr><td>T10884</td><td>GGGTCGGCGAAATCCAGATATGG</td><td></td><td>1000</td><td></td><td></td><td></td><td></td><td></td><td></td><td></td><td></td><td></td></tr>
<tr><td>Oligo fwd</td><td>CACCGGGTCGGCGAAATCCAGATA</td><td></td><td></td><td></td><td></td><td></td><td></td><td></td><td></td><td></td><td></td><td></td></tr>
<tr><td>Oligo rev</td><td>AAACTATCTGGATTTCGCCGACCC</td><td></td><td></td><td></td><td></td><td></td><td></td><td></td><td></td><td></td><td></td><td></td></tr>
<tr><td>Chromosome</td><td>start</td><td></td><td>end</td><td>strand</td><td>MM</td><td>target_seq</td><td>PAM</td><td>alignment</td><td>distance</td><td>position</td><td>gene name</td><td>gene id</td></tr>
<tr><td>GU305914.1</td><td></td><td>142179</td><td>142201</td><td>+</td><td>0</td><td>GGGTCGGCGAAATCCAGATA</td><td>TGG</td><td>||||||||[||||||||||||]PAM</td><td>NA</td><td>Intergenic</td><td>NA</td><td>NA</td></tr>
<tr><td>T10885</td><td>AAATCCAGATATGGCGACGACGG</td><td></td><td>1000</td><td></td><td></td><td></td><td></td><td></td><td></td><td></td><td></td><td></td></tr>
<tr><td>Oligo fwd</td><td>CACCAAATCCAGATATGGCGACGA</td><td></td><td></td><td></td><td></td><td></td><td></td><td></td><td></td><td></td><td></td><td></td></tr>
<tr><td>Oligo rev</td><td>AAACTCGTCGCCATATCTGGATTT</td><td></td><td></td><td></td><td></td><td></td><td></td><td></td><td></td><td></td><td></td><td></td></tr>
<tr><td>Chromosome</td><td>start</td><td></td><td>end</td><td>strand</td><td>MM</td><td>target_seq</td><td>PAM</td><td>alignment</td><td>distance</td><td>position</td><td>gene name</td><td>gene id</td></tr>
<tr><td>GU305914.1</td><td></td><td>142188</td><td>142210</td><td>+</td><td>0</td><td>AAATCCAGATATGGCGACGA</td><td>CGG</td><td>||||||||[||||||||||||]PAM</td><td>NA</td><td>Intergenic</td><td>NA</td><td>NA</td></tr>
<tr><td>T10886</td><td>GGCGACGACGGTCCTCCCTCCGG</td><td></td><td>1000</td><td></td><td></td><td></td><td></td><td></td><td></td><td></td><td></td><td></td></tr>
<tr><td>Oligo fwd</td><td>CACCGGCGACGACGGTCCTCCCTC</td><td></td><td></td><td></td><td></td><td></td><td></td><td></td><td></td><td></td><td></td><td></td></tr>
<tr><td>Oligo rev</td><td>AAACGAGGGAGGACCGTCGTCGCC</td><td></td><td></td><td></td><td></td><td></td><td></td><td></td><td></td><td></td><td></td><td></td></tr>
<tr><td>Chromosome</td><td>start</td><td></td><td>end</td><td>strand</td><td>MM</td><td>target_seq</td><td>PAM</td><td>alignment</td><td>distance</td><td>position</td><td>gene name</td><td>gene id</td></tr>
<tr><td>GU305914.1</td><td></td><td>142200</td><td>142222</td><td>+</td><td>0</td><td>GGCGACGACGGTCCTCCCTC</td><td>CGG</td><td>||||||||[||||||||||||]PAM</td><td>NA</td><td>Intergenic</td><td>NA</td><td>NA</td></tr>
<tr><td>T10887</td><td>GACGACGGTCCTCCCTCCGGAGG</td><td></td><td>1000</td><td></td><td></td><td></td><td></td><td></td><td></td><td></td><td></td><td></td></tr>
<tr><td>Oligo fwd</td><td>CACCGACGACGGTCCTCCCTCCGG</td><td></td><td></td><td></td><td></td><td></td><td></td><td></td><td></td><td></td><td></td><td></td></tr>
<tr><td>Oligo rev</td><td>AAACCCGGAGGGAGGACCGTCGTC</td><td></td><td></td><td></td><td></td><td></td><td></td><td></td><td></td><td></td><td></td><td></td></tr>
<tr><td>Chromosome</td><td>start</td><td></td><td>end</td><td>strand</td><td>MM</td><td>target_seq</td><td>PAM</td><td>alignment</td><td>distance</td><td>position</td><td>gene name</td><td>gene id</td></tr>
<tr><td>GU305914.1</td><td></td><td>142203</td><td>142225</td><td>+</td><td>0</td><td>GACGACGGTCCTCCCTCCGG</td><td>AGG</td><td>||||||||[||||||||||||]PAM</td><td>NA</td><td>Intergenic</td><td>NA</td><td>NA</td></tr>
<tr><td>T10888</td><td>GGTCCTCCCTCCGGAGGCGACGG</td><td></td><td>1000</td><td></td><td></td><td></td><td></td><td></td><td></td><td></td><td></td><td></td></tr>
<tr><td>Oligo fwd</td><td>CACCGGTCCTCCCTCCGGAGGCGA</td><td></td><td></td><td></td><td></td><td></td><td></td><td></td><td></td><td></td><td></td><td></td></tr>
<tr><td>Oligo rev</td><td>AAACTCGCCTCCGGAGGGAGGACC</td><td></td><td></td><td></td><td></td><td></td><td></td><td></td><td></td><td></td><td></td><td></td></tr>
<tr><td>Chromosome</td><td>start</td><td></td><td>end</td><td>strand</td><td>MM</td><td>target_seq</td><td>PAM</td><td>alignment</td><td>distance</td><td>position</td><td>gene name</td><td>gene id</td></tr>
<tr><td>GU305914.1</td><td></td><td>142209</td><td>142231</td><td>+</td><td>0</td><td>GGTCCTCCCTCCGGAGGCGA</td><td>CGG</td><td>||||||||[||||||||||||]PAM</td><td>NA</td><td>Intergenic</td><td>NA</td><td>NA</td></tr>
<tr><td>T10889</td><td>CGGCGATGACGACGATACCGCGG</td><td></td><td>1000</td><td></td><td></td><td></td><td></td><td></td><td></td><td></td><td></td><td></td></tr>
</table>

| | | | | | | | | | | | |
|---|---|---|---|---|---|---|---|---|---|---|---|
| Oligo fwd | CACCCGGCGATGACGACGATACCG | | | | | | | | | | |
| Oligo rev | AAACCGGTATCGTCGTCATCGCCG | | | | | | | | | | |
| Chromosome | start | end | strand | MM | target_seq | PAM | alignment | distance | position | gene name | gene id |
| GU305914.1 | 142229 | 142251 | + | 0 | CGGCGATGACGACGATACCG | CGG | \|\|\|\|\|\|\|\|[\|\|\|\|\|\|\|\|\|\|\|\|]PAM | NA | Intergenic | NA | NA |
| GU305914.1 | 170516 | 170538 | - | 4 | CGGCCACGGCCACGATACCG | GGG | \|\|\|\|-\|-\|[-\|-\|\|\|\|\|\|\|\|\|]PAM | NA | Intergenic | NA | NA |
| T10890 | CGATGACGACGATACCGCGGCGG | 1000 | | | | | | | | | |
| Oligo fwd | CACCCGATGACGACGATACCGCGG | | | | | | | | | | |
| Oligo rev | AAACCCGCGGTATCGTCGTCATCG | | | | | | | | | | |
| Chromosome | start | end | strand | MM | target_seq | PAM | alignment | distance | position | gene name | gene id |
| GU305914.1 | 142232 | 142254 | + | 0 | CGATGACGACGATACCGCGG | CGG | \|\|\|\|\|\|\|\|[\|\|\|\|\|\|\|\|\|\|\|\|]PAM | NA | Intergenic | NA | NA |
| GU305914.1 | 98743 | 98765 | - | 4 | CGACGAGGACGATGCCGCGA | CGG | \|\|\|-\|\|-\|[\|\|\|\|\|-\|\|\|\|\|-]PAM | NA | Intergenic | NA | NA |
| T10891 | GACGACGTCGTCGTCGACGACGG | 1000 | | | | | | | | | |
| Oligo fwd | CACCGACGACGTCGTCGTCGACGA | | | | | | | | | | |
| Oligo rev | AAACTCGTCGACGACGACGTCGTC | | | | | | | | | | |
| Chromosome | start | end | strand | MM | target_seq | PAM | alignment | distance | position | gene name | gene id |
| GU305914.1 | 142261 | 142283 | + | 0 | GACGACGTCGTCGTCGACGA | CGG | \|\|\|\|\|\|\|\|[\|\|\|\|\|\|\|\|\|\|\|\|]PAM | NA | Intergenic | NA | NA |
| GU305914.1 | 142264 | 142286 | + | 3 | GACGTCGTCGTCGACGACGG | TGG | \|\|\|\|-\|\|\|[\|\|\|\|\|-\|\|\|\|\|-]PAM | NA | Intergenic | NA | NA |
| T10892 | GACGTCGTCGTCGACGACGGTGG | 1000 | | | | | | | | | |
| Oligo fwd | CACCGACGTCGTCGTCGACGACGG | | | | | | | | | | |
| Oligo rev | AAACCCGTCGTCGACGACGACGTC | | | | | | | | | | |
| Chromosome | start | end | strand | MM | target_seq | PAM | alignment | distance | position | gene name | gene id |
| GU305914.1 | 142264 | 142286 | + | 0 | GACGTCGTCGTCGACGACGG | TGG | \|\|\|\|\|\|\|\|[\|\|\|\|\|\|\|\|\|\|\|\|]PAM | NA | Intergenic | NA | NA |
| GU305914.1 | 142261 | 142283 | + | 3 | GACGACGTCGTCGTCGACGA | CGG | \|\|\|\|-\|\|\|[\|\|\|\|\|-\|\|\|\|\|-]PAM | NA | Intergenic | NA | NA |
| T10893 | ACGTCGTCGTCGACGACGGTGGG | 1000 | | | | | | | | | |
| Oligo fwd | CACCACGTCGTCGTCGACGACGGT | | | | | | | | | | |
| Oligo rev | AAACACCGTCGTCGACGACGACGT | | | | | | | | | | |
| Chromosome | start | end | strand | MM | target_seq | PAM | alignment | distance | position | gene name | gene id |
| GU305914.1 | 142265 | 142287 | + | 0 | ACGTCGTCGTCGACGACGGT | GGG | \|\|\|\|\|\|\|\|[\|\|\|\|\|\|\|\|\|\|\|\|]PAM | NA | Intergenic | NA | NA |
| GU305914.1 | 135773 | 135795 | + | 4 | CCGTCGACGTCGACGTCGGA | CGG | -\|\|\|\|\|-\|[\|\|\|\|\|\|\|-\|\|\|-]PAM | NA | Intergenic | NA | NA |
| T10894 | CGTCGTCGTCGACGACGGTGGGG | 1000 | | | | | | | | | |
| Oligo fwd | CACCCGTCGTCGTCGACGACGGTG | | | | | | | | | | |
| Oligo rev | AAACCACCGTCGTCGACGACGACG | | | | | | | | | | |
| Chromosome | start | end | strand | MM | target_seq | PAM | alignment | distance | position | gene name | gene id |
| GU305914.1 | 142266 | 142288 | + | 0 | CGTCGTCGTCGACGACGGTG | GGG | \|\|\|\|\|\|\|\|[\|\|\|\|\|\|\|\|\|\|\|\|]PAM | NA | Intergenic | NA | NA |
| GU305914.1 | 89324 | 89346 | + | 3 | CGTAGTCGTCGGTGACGGTG | AGG | \|\|\|-\|\|\|\|[\|\|\|--\|\|\|\|\|\|\|]PAM | NA | Intergenic | NA | NA |
| GU305914.1 | 201033 | 201055 | - | 4 | CGTCGAGGTCCACGACGGGG | CGG | \|\|\|\|\|--\|[\|\|-\|\|\|\|\|\|\|-\|]PAM | NA | Intergenic | NA | NA |
| T10895 | GTCGTCGTCGACGACGGTGGGGG | 1000 | | | | | | | | | |
| Oligo fwd | CACCGTCGTCGTCGACGACGGTGG | | | | | | | | | | |
| Oligo rev | AAACCCACCGTCGTCGACGACGAC | | | | | | | | | | |
| Chromosome | start | end | strand | MM | target_seq | PAM | alignment | distance | position | gene name | gene id |
| GU305914.1 | 142267 | 142289 | + | 0 | GTCGTCGTCGACGACGGTGG | GGG | \|\|\|\|\|\|\|\|[\|\|\|\|\|\|\|\|\|\|\|\|]PAM | NA | Intergenic | NA | NA |
| T10896 | CATAATATCGACAGCGCGTTCGG | 1000 | | | | | | | | | |
| Oligo fwd | CACCCATAATATCGACAGCGCGTT | | | | | | | | | | |
| Oligo rev | AAACAACGCGCTGTCGATATTATG | | | | | | | | | | |
| Chromosome | start | end | strand | MM | target_seq | PAM | alignment | distance | position | gene name | gene id |
| GU305914.1 | 142300 | 142322 | + | 0 | CATAATATCGACAGCGCGTT | CGG | \|\|\|\|\|\|\|\|[\|\|\|\|\|\|\|\|\|\|\|\|]PAM | NA | Intergenic | NA | NA |
| T10897 | AATATCGACAGCGCGTTCGGCGG | 1000 | | | | | | | | | |
| Oligo fwd | CACCAATATCGACAGCGCGTTCGG | | | | | | | | | | |
| Oligo rev | AAACCCGAACGCGCTGTCGATATT | | | | | | | | | | |
| Chromosome | start | end | strand | MM | target_seq | PAM | alignment | distance | position | gene name | gene id |
| GU305914.1 | 142303 | 142325 | + | 0 | AATATCGACAGCGCGTTCGG | CGG | \|\|\|\|\|\|\|\|[\|\|\|\|\|\|\|\|\|\|\|\|]PAM | NA | Intergenic | NA | NA |
| T10898 | ATATCGACAGCGCGTTCGGCGGG | 1000 | | | | | | | | | |
| Oligo fwd | CACCATATCGACAGCGCGTTCGGC | | | | | | | | | | |
| Oligo rev | AAACGCCGAACGCGCTGTCGATAT | | | | | | | | | | |
| Chromosome | start | end | strand | MM | target_seq | PAM | alignment | distance | position | gene name | gene id |
| GU305914.1 | 142304 | 142326 | + | 0 | ATATCGACAGCGCGTTCGGC | GGG | \|\|\|\|\|\|\|\|[\|\|\|\|\|\|\|\|\|\|\|\|]PAM | NA | Intergenic | NA | NA |
| T10899 | GCGCGTTCGGCGGGTCCGAACGG | 1000 | | | | | | | | | |
| Oligo fwd | CACCGCGCGTTCGGCGGGTCCGAA | | | | | | | | | | |
| Oligo rev | AAACTTCGGACCCGCCGAACGCGC | | | | | | | | | | |
| Chromosome | start | end | strand | MM | target_seq | PAM | alignment | distance | position | gene name | gene id |
| GU305914.1 | 142313 | 142335 | + | 0 | GCGCGTTCGGCGGGTCCGAA | CGG | \|\|\|\|\|\|\|\|[\|\|\|\|\|\|\|\|\|\|\|\|]PAM | NA | Intergenic | NA | NA |
| T10900 | AGAGCGAGCAGTGTCTCGCTCGG | 1000 | | | | | | | | | |
| Oligo fwd | CACCAGAGCGAGCAGTGTCTCGCT | | | | | | | | | | |
| Oligo rev | AAACAGCGAGACACTGCTCGCTCT | | | | | | | | | | |
| Chromosome | start | end | strand | MM | target_seq | PAM | alignment | distance | position | gene name | gene id |
| GU305914.1 | 142358 | 142380 | + | 0 | AGAGCGAGCAGTGTCTCGCT | CGG | \|\|\|\|\|\|\|\|[\|\|\|\|\|\|\|\|\|\|\|\|]PAM | NA | Intergenic | NA | NA |
| T10901 | GAGCGAGCAGTGTCTCGCTCGGG | 1000 | | | | | | | | | |
| Oligo fwd | CACCGAGCGAGCAGTGTCTCGCTC | | | | | | | | | | |
| Oligo rev | AAACGAGCGAGACACTGCTCGCTC | | | | | | | | | | |
| Chromosome | start | end | strand | MM | target_seq | PAM | alignment | distance | position | gene name | gene id |
| GU305914.1 | 142359 | 142381 | + | 0 | GAGCGAGCAGTGTCTCGCTC | GGG | \|\|\|\|\|\|\|\|[\|\|\|\|\|\|\|\|\|\|\|\|]PAM | NA | Intergenic | NA | NA |
| T10902 | TGTCTCGCTCGGGTGTTCGACGG | 1000 | | | | | | | | | |
| Oligo fwd | CACCTGTCTCGCTCGGGTGTTCGA | | | | | | | | | | |
| Oligo rev | AAACTCGAACACCCGAGCGAGACA | | | | | | | | | | |
| Chromosome | start | end | strand | MM | target_seq | PAM | alignment | distance | position | gene name | gene id |
| GU305914.1 | 142369 | 142391 | + | 0 | TGTCTCGCTCGGGTGTTCGA | CGG | \|\|\|\|\|\|\|\|[\|\|\|\|\|\|\|\|\|\|\|\|]PAM | NA | Intergenic | NA | NA |

<table>
<tr><td>T10903</td><td>GTCTCGCTCGGGTGTTCGACGGG</td><td>1000</td><td></td><td></td><td></td><td></td><td></td><td></td><td></td><td></td><td></td></tr>
<tr><td>Oligo fwd</td><td>CACCGTCTCGCTCGGGTGTTCGAC</td><td></td><td></td><td></td><td></td><td></td><td></td><td></td><td></td><td></td><td></td></tr>
<tr><td>Oligo rev</td><td>AAACGTCGAACACCCGAGCGAGAC</td><td></td><td></td><td></td><td></td><td></td><td></td><td></td><td></td><td></td><td></td></tr>
<tr><td>Chromosome</td><td>start</td><td>end</td><td>strand</td><td>MM</td><td>target_seq</td><td>PAM</td><td>alignment</td><td>distance</td><td>position</td><td>gene name</td><td>gene id</td></tr>
<tr><td>GU305914.1</td><td>142370</td><td>142392</td><td>+</td><td>0</td><td>GTCTCGCTCGGGTGTTCGAC</td><td>GGG</td><td>||||||||[||||||||||||]PAM</td><td>NA</td><td>Intergenic</td><td>NA</td><td>NA</td></tr>
<tr><td>T10904</td><td>GTATAACCTCCTAGATATCAAGG</td><td>1000</td><td></td><td></td><td></td><td></td><td></td><td></td><td></td><td></td><td></td></tr>
<tr><td>Oligo fwd</td><td>CACCGTATAACCTCCTAGATATCA</td><td></td><td></td><td></td><td></td><td></td><td></td><td></td><td></td><td></td><td></td></tr>
<tr><td>Oligo rev</td><td>AAACTGATATCTAGGAGGTTATAC</td><td></td><td></td><td></td><td></td><td></td><td></td><td></td><td></td><td></td><td></td></tr>
<tr><td>Chromosome</td><td>start</td><td>end</td><td>strand</td><td>MM</td><td>target_seq</td><td>PAM</td><td>alignment</td><td>distance</td><td>position</td><td>gene name</td><td>gene id</td></tr>
<tr><td>GU305914.1</td><td>142425</td><td>142447</td><td>+</td><td>0</td><td>GTATAACCTCCTAGATATCA</td><td>AGG</td><td>||||||||[||||||||||||]PAM</td><td>NA</td><td>Intergenic</td><td>NA</td><td>NA</td></tr>
<tr><td>T10905</td><td>CCCCGTGTGCGCGTTGCGCCTGG</td><td>1000</td><td></td><td></td><td></td><td></td><td></td><td></td><td></td><td></td><td></td></tr>
<tr><td>Oligo fwd</td><td>CACCCCCCGTGTGCGCGTTGCGCC</td><td></td><td></td><td></td><td></td><td></td><td></td><td></td><td></td><td></td><td></td></tr>
<tr><td>Oligo rev</td><td>AAACGGCGCAACGCGCACACGGGG</td><td></td><td></td><td></td><td></td><td></td><td></td><td></td><td></td><td></td><td></td></tr>
<tr><td>Chromosome</td><td>start</td><td>end</td><td>strand</td><td>MM</td><td>target_seq</td><td>PAM</td><td>alignment</td><td>distance</td><td>position</td><td>gene name</td><td>gene id</td></tr>
<tr><td>GU305914.1</td><td>142464</td><td>142486</td><td>+</td><td>0</td><td>CCCCGTGTGCGCGTTGCGCC</td><td>TGG</td><td>||||||||[||||||||||||]PAM</td><td>NA</td><td>Intergenic</td><td>NA</td><td>NA</td></tr>
<tr><td>T10906</td><td>CGTTGCGCCTGGCGTATTTTCGG</td><td>1000</td><td></td><td></td><td></td><td></td><td></td><td></td><td></td><td></td><td></td></tr>
<tr><td>Oligo fwd</td><td>CACCCGTTGCGCCTGGCGTATTTT</td><td></td><td></td><td></td><td></td><td></td><td></td><td></td><td></td><td></td><td></td></tr>
<tr><td>Oligo rev</td><td>AAACAAAATACGCCAGGCGCAACG</td><td></td><td></td><td></td><td></td><td></td><td></td><td></td><td></td><td></td><td></td></tr>
<tr><td>Chromosome</td><td>start</td><td>end</td><td>strand</td><td>MM</td><td>target_seq</td><td>PAM</td><td>alignment</td><td>distance</td><td>position</td><td>gene name</td><td>gene id</td></tr>
<tr><td>GU305914.1</td><td>142475</td><td>142497</td><td>+</td><td>0</td><td>CGTTGCGCCTGGCGTATTTT</td><td>CGG</td><td>||||||||[||||||||||||]PAM</td><td>NA</td><td>Intergenic</td><td>NA</td><td>NA</td></tr>
<tr><td>T10907</td><td>GCGCCTGGCGTATTTTCGGTCGG</td><td>1000</td><td></td><td></td><td></td><td></td><td></td><td></td><td></td><td></td><td></td></tr>
<tr><td>Oligo fwd</td><td>CACCGCGCCTGGCGTATTTTCGGT</td><td></td><td></td><td></td><td></td><td></td><td></td><td></td><td></td><td></td><td></td></tr>
<tr><td>Oligo rev</td><td>AAACACCGAAAATACGCCAGGCGC</td><td></td><td></td><td></td><td></td><td></td><td></td><td></td><td></td><td></td><td></td></tr>
<tr><td>Chromosome</td><td>start</td><td>end</td><td>strand</td><td>MM</td><td>target_seq</td><td>PAM</td><td>alignment</td><td>distance</td><td>position</td><td>gene name</td><td>gene id</td></tr>
<tr><td>GU305914.1</td><td>142479</td><td>142501</td><td>+</td><td>0</td><td>GCGCCTGGCGTATTTTCGGT</td><td>CGG</td><td>||||||||[||||||||||||]PAM</td><td>NA</td><td>Intergenic</td><td>NA</td><td>NA</td></tr>
<tr><td>T10908</td><td>AAAATCAACCTGACACACGATGG</td><td>1000</td><td></td><td></td><td></td><td></td><td></td><td></td><td></td><td></td><td></td></tr>
<tr><td>Oligo fwd</td><td>CACCAAAATCAACCTGACACACGA</td><td></td><td></td><td></td><td></td><td></td><td></td><td></td><td></td><td></td><td></td></tr>
<tr><td>Oligo rev</td><td>AAACTCGTGTGTCAGGTTGATTTT</td><td></td><td></td><td></td><td></td><td></td><td></td><td></td><td></td><td></td><td></td></tr>
<tr><td>Chromosome</td><td>start</td><td>end</td><td>strand</td><td>MM</td><td>target_seq</td><td>PAM</td><td>alignment</td><td>distance</td><td>position</td><td>gene name</td><td>gene id</td></tr>
<tr><td>GU305914.1</td><td>142510</td><td>142532</td><td>+</td><td>0</td><td>AAAATCAACCTGACACACGA</td><td>TGG</td><td>||||||||[||||||||||||]PAM</td><td>NA</td><td>Intergenic</td><td>NA</td><td>NA</td></tr>
<tr><td>T10909</td><td>AAATCAACCTGACACACGATGGG</td><td>1000</td><td></td><td></td><td></td><td></td><td></td><td></td><td></td><td></td><td></td></tr>
<tr><td>Oligo fwd</td><td>CACCAAATCAACCTGACACACGAT</td><td></td><td></td><td></td><td></td><td></td><td></td><td></td><td></td><td></td><td></td></tr>
<tr><td>Oligo rev</td><td>AAACATCGTGTGTCAGGTTGATTT</td><td></td><td></td><td></td><td></td><td></td><td></td><td></td><td></td><td></td><td></td></tr>
<tr><td>Chromosome</td><td>start</td><td>end</td><td>strand</td><td>MM</td><td>target_seq</td><td>PAM</td><td>alignment</td><td>distance</td><td>position</td><td>gene name</td><td>gene id</td></tr>
<tr><td>GU305914.1</td><td>142511</td><td>142533</td><td>+</td><td>0</td><td>AAATCAACCTGACACACGAT</td><td>GGG</td><td>||||||||[||||||||||||]PAM</td><td>NA</td><td>Intergenic</td><td>NA</td><td>NA</td></tr>
<tr><td>T10910</td><td>AACCTGACACACGATGGGCCCGG</td><td>1000</td><td></td><td></td><td></td><td></td><td></td><td></td><td></td><td></td><td></td></tr>
<tr><td>Oligo fwd</td><td>CACCAACCTGACACACGATGGGCC</td><td></td><td></td><td></td><td></td><td></td><td></td><td></td><td></td><td></td><td></td></tr>
<tr><td>Oligo rev</td><td>AAACGGCCCATCGTGTGTCAGGTT</td><td></td><td></td><td></td><td></td><td></td><td></td><td></td><td></td><td></td><td></td></tr>
<tr><td>Chromosome</td><td>start</td><td>end</td><td>strand</td><td>MM</td><td>target_seq</td><td>PAM</td><td>alignment</td><td>distance</td><td>position</td><td>gene name</td><td>gene id</td></tr>
<tr><td>GU305914.1</td><td>142516</td><td>142538</td><td>+</td><td>0</td><td>AACCTGACACACGATGGGCC</td><td>CGG</td><td>||||||||[||||||||||||]PAM</td><td>NA</td><td>Intergenic</td><td>NA</td><td>NA</td></tr>
<tr><td>T10911</td><td>ACCTGACACACGATGGGCCCGGG</td><td>1000</td><td></td><td></td><td></td><td></td><td></td><td></td><td></td><td></td><td></td></tr>
<tr><td>Oligo fwd</td><td>CACCACCTGACACACGATGGGCCC</td><td></td><td></td><td></td><td></td><td></td><td></td><td></td><td></td><td></td><td></td></tr>
<tr><td>Oligo rev</td><td>AAACGGGCCCATCGTGTGTCAGGT</td><td></td><td></td><td></td><td></td><td></td><td></td><td></td><td></td><td></td><td></td></tr>
<tr><td>Chromosome</td><td>start</td><td>end</td><td>strand</td><td>MM</td><td>target_seq</td><td>PAM</td><td>alignment</td><td>distance</td><td>position</td><td>gene name</td><td>gene id</td></tr>
<tr><td>GU305914.1</td><td>142517</td><td>142539</td><td>+</td><td>0</td><td>ACCTGACACACGATGGGCCC</td><td>GGG</td><td>||||||||[||||||||||||]PAM</td><td>NA</td><td>Intergenic</td><td>NA</td><td>NA</td></tr>
<tr><td>T10912</td><td>GACACACGATGGGCCCGGGTTGG</td><td>1000</td><td></td><td></td><td></td><td></td><td></td><td></td><td></td><td></td><td></td></tr>
<tr><td>Oligo fwd</td><td>CACCGACACACGATGGGCCCGGGT</td><td></td><td></td><td></td><td></td><td></td><td></td><td></td><td></td><td></td><td></td></tr>
<tr><td>Oligo rev</td><td>AAACACCCGGGCCCATCGTGTGTC</td><td></td><td></td><td></td><td></td><td></td><td></td><td></td><td></td><td></td><td></td></tr>
<tr><td>Chromosome</td><td>start</td><td>end</td><td>strand</td><td>MM</td><td>target_seq</td><td>PAM</td><td>alignment</td><td>distance</td><td>position</td><td>gene name</td><td>gene id</td></tr>
<tr><td>GU305914.1</td><td>142521</td><td>142543</td><td>+</td><td>0</td><td>GACACACGATGGGCCCGGGT</td><td>TGG</td><td>||||||||[||||||||||||]PAM</td><td>NA</td><td>Intergenic</td><td>NA</td><td>NA</td></tr>
<tr><td>T10913</td><td>CTTGTCGATCGTCGACCGCGAGG</td><td>1000</td><td></td><td></td><td></td><td></td><td></td><td></td><td></td><td></td><td></td></tr>
<tr><td>Oligo fwd</td><td>CACCCTTGTCGATCGTCGACCGCG</td><td></td><td></td><td></td><td></td><td></td><td></td><td></td><td></td><td></td><td></td></tr>
<tr><td>Oligo rev</td><td>AAACCGCGGTCGACGATCGACAAG</td><td></td><td></td><td></td><td></td><td></td><td></td><td></td><td></td><td></td><td></td></tr>
<tr><td>Chromosome</td><td>start</td><td>end</td><td>strand</td><td>MM</td><td>target_seq</td><td>PAM</td><td>alignment</td><td>distance</td><td>position</td><td>gene name</td><td>gene id</td></tr>
<tr><td>GU305914.1</td><td>142554</td><td>142576</td><td>+</td><td>0</td><td>CTTGTCGATCGTCGACCGCG</td><td>AGG</td><td>||||||||[||||||||||||]PAM</td><td>NA</td><td>Intergenic</td><td>NA</td><td>NA</td></tr>
<tr><td>GU305914.1</td><td>76938</td><td>76960</td><td>-</td><td>4</td><td>CGAGTCGATCGTCGTCAGCG</td><td>AGG</td><td>|--|||||[||||||-|-|||]PAM</td><td>NA</td><td>Intergenic</td><td>NA</td><td>NA</td></tr>
<tr><td>T10914</td><td>GACCGCGAGGTCTCGAACGCCGG</td><td>1000</td><td></td><td></td><td></td><td></td><td></td><td></td><td></td><td></td><td></td></tr>
<tr><td>Oligo fwd</td><td>CACCGACCGCGAGGTCTCGAACGC</td><td></td><td></td><td></td><td></td><td></td><td></td><td></td><td></td><td></td><td></td></tr>
<tr><td>Oligo rev</td><td>AAACGCGTTCGAGACCTCGCGGTC</td><td></td><td></td><td></td><td></td><td></td><td></td><td></td><td></td><td></td><td></td></tr>
<tr><td>Chromosome</td><td>start</td><td>end</td><td>strand</td><td>MM</td><td>target_seq</td><td>PAM</td><td>alignment</td><td>distance</td><td>position</td><td>gene name</td><td>gene id</td></tr>
<tr><td>GU305914.1</td><td>142567</td><td>142589</td><td>+</td><td>0</td><td>GACCGCGAGGTCTCGAACGC</td><td>CGG</td><td>||||||||[||||||||||||]PAM</td><td>NA</td><td>Intergenic</td><td>NA</td><td>NA</td></tr>
<tr><td>T10915</td><td>CGCGAGGTCTCGAACGCCGGCGG</td><td>1000</td><td></td><td></td><td></td><td></td><td></td><td></td><td></td><td></td><td></td></tr>
<tr><td>Oligo fwd</td><td>CACCCGCGAGGTCTCGAACGCCGG</td><td></td><td></td><td></td><td></td><td></td><td></td><td></td><td></td><td></td><td></td></tr>
<tr><td>Oligo rev</td><td>AAACCCGGCGTTCGAGACCTCGCG</td><td></td><td></td><td></td><td></td><td></td><td></td><td></td><td></td><td></td><td></td></tr>
<tr><td>Chromosome</td><td>start</td><td>end</td><td>strand</td><td>MM</td><td>target_seq</td><td>PAM</td><td>alignment</td><td>distance</td><td>position</td><td>gene name</td><td>gene id</td></tr>
<tr><td>GU305914.1</td><td>142570</td><td>142592</td><td>+</td><td>0</td><td>CGCGAGGTCTCGAACGCCGG</td><td>CGG</td><td>||||||||[||||||||||||]PAM</td><td>NA</td><td>Intergenic</td><td>NA</td><td>NA</td></tr>
<tr><td>T10916</td><td>CGAGAGCACCGTCAGAGATGCGG</td><td>1000</td><td></td><td></td><td></td><td></td><td></td><td></td><td></td><td></td><td></td></tr>
<tr><td>Oligo fwd</td><td>CACCCGAGAGCACCGTCAGAGATG</td><td></td><td></td><td></td><td></td><td></td><td></td><td></td><td></td><td></td><td></td></tr>
<tr><td>Oligo rev</td><td>AAACCATCTCTGACGGTGCTCTCG</td><td></td><td></td><td></td><td></td><td></td><td></td><td></td><td></td><td></td><td></td></tr>
<tr><td>Chromosome</td><td>start</td><td>end</td><td>strand</td><td>MM</td><td>target_seq</td><td>PAM</td><td>alignment</td><td>distance</td><td>position</td><td>gene name</td><td>gene id</td></tr>
<tr><td>GU305914.1</td><td>142593</td><td>142615</td><td>+</td><td>0</td><td>CGAGAGCACCGTCAGAGATG</td><td>CGG</td><td>||||||||[||||||||||||]PAM</td><td>NA</td><td>Intergenic</td><td>NA</td><td>NA</td></tr>
<tr><td>T10917</td><td>AGCACCGTCAGAGATGCGGTCGG</td><td>1000</td><td></td><td></td><td></td><td></td><td></td><td></td><td></td><td></td><td></td></tr>
<tr><td>Oligo fwd</td><td>CACCAGCACCGTCAGAGATGCGGT</td><td></td><td></td><td></td><td></td><td></td><td></td><td></td><td></td><td></td><td></td></tr>
<tr><td>Oligo rev</td><td>AAACACCGCATCTCTGACGGTGCT</td><td></td><td></td><td></td><td></td><td></td><td></td><td></td><td></td><td></td><td></td></tr>
</table>

| | | | | | | | | | | | |
|---|---|---|---|---|---|---|---|---|---|---|---|
| Chromosome | start | end | strand | MM | target_seq | PAM | alignment | distance | position | gene name | gene id |
| GU305914.1 | 142597 | 142619 | + | 0 | AGCACCGTCAGAGATGCGGT | CGG | \|\|\|\|\|\|\|\|[\|\|\|\|\|\|\|\|\|\|\|\|\|\|]PAM | NA | Intergenic | NA | NA |
| T10918 | GCACCGTCAGAGATGCGGTCGGG | 1000 | | | | | | | | | |
| Oligo fwd | CACCGCACCGTCAGAGATGCGGTC | | | | | | | | | | |
| Oligo rev | AAACGACCGCATCTCTGACGGTGC | | | | | | | | | | |
| Chromosome | start | end | strand | MM | target_seq | PAM | alignment | distance | position | gene name | gene id |
| GU305914.1 | 142598 | 142620 | + | 0 | GCACCGTCAGAGATGCGGTC | GGG | \|\|\|\|\|\|\|\|[\|\|\|\|\|\|\|\|\|\|\|\|\|\|]PAM | NA | Intergenic | NA | NA |
| T10919 | CACCGTCAGAGATGCGGTCGGGG | 1000 | | | | | | | | | |
| Oligo fwd | CACCCACCGTCAGAGATGCGGTCG | | | | | | | | | | |
| Oligo rev | AAACCGACCGCATCTCTGACGGTG | | | | | | | | | | |
| Chromosome | start | end | strand | MM | target_seq | PAM | alignment | distance | position | gene name | gene id |
| GU305914.1 | 142599 | 142621 | + | 0 | CACCGTCAGAGATGCGGTCG | GGG | \|\|\|\|\|\|\|\|[\|\|\|\|\|\|\|\|\|\|\|\|\|\|]PAM | NA | Intergenic | NA | NA |
| T10920 | ACCGTCAGAGATGCGGTCGGGGG | 1000 | | | | | | | | | |
| Oligo fwd | CACCACCGTCAGAGATGCGGTCGG | | | | | | | | | | |
| Oligo rev | AAACCCGACCGCATCTCTGACGGT | | | | | | | | | | |
| Chromosome | start | end | strand | MM | target_seq | PAM | alignment | distance | position | gene name | gene id |
| GU305914.1 | 142600 | 142622 | + | 0 | ACCGTCAGAGATGCGGTCGG | GGG | \|\|\|\|\|\|\|\|[\|\|\|\|\|\|\|\|\|\|\|\|\|\|]PAM | NA | Intergenic | NA | NA |
| T10921 | GATGCGGTCGGGGGTGTCGAAGG | 1000 | | | | | | | | | |
| Oligo fwd | CACCGATGCGGTCGGGGGTGTCGA | | | | | | | | | | |
| Oligo rev | AAACTCGACACCCCCGACCGCATC | | | | | | | | | | |
| Chromosome | start | end | strand | MM | target_seq | PAM | alignment | distance | position | gene name | gene id |
| GU305914.1 | 142609 | 142631 | + | 0 | GATGCGGTCGGGGGTGTCGA | AGG | \|\|\|\|\|\|\|\|[\|\|\|\|\|\|\|\|\|\|\|\|\|\|]PAM | NA | Intergenic | NA | NA |
| T10922 | GCGCGCAGTTCGCGCTCGCGCGG | 1000 | | | | | | | | | |
| Oligo fwd | CACCGCGCGCAGTTCGCGCTCGCG | | | | | | | | | | |
| Oligo rev | AAACCGCGAGCGCGAACTGCGCGC | | | | | | | | | | |
| Chromosome | start | end | strand | MM | target_seq | PAM | alignment | distance | position | gene name | gene id |
| GU305914.1 | 142631 | 142653 | + | 0 | GCGCGCAGTTCGCGCTCGCG | CGG | \|\|\|\|\|\|\|\|[\|\|\|\|\|\|\|\|\|\|\|\|\|\|]PAM | NA | Intergenic | NA | NA |
| T10923 | CGCGCAGTTCGCGCTCGCGCGGG | 1000 | | | | | | | | | |
| Oligo fwd | CACCCGCGCAGTTCGCGCTCGCGC | | | | | | | | | | |
| Oligo rev | AAACGCGCGAGCGCGAACTGCGCG | | | | | | | | | | |
| Chromosome | start | end | strand | MM | target_seq | PAM | alignment | distance | position | gene name | gene id |
| GU305914.1 | 142632 | 142654 | + | 0 | CGCGCAGTTCGCGCTCGCGC | GGG | \|\|\|\|\|\|\|\|[\|\|\|\|\|\|\|\|\|\|\|\|\|\|]PAM | NA | Intergenic | NA | NA |
| T10924 | GCGGGCCATCTTTCGCCTGCCGG | 1000 | | | | | | | | | |
| Oligo fwd | CACCGCGGGCCATCTTTCGCCTGC | | | | | | | | | | |
| Oligo rev | AAACGCAGGCGAAAGATGGCCCGC | | | | | | | | | | |
| Chromosome | start | end | strand | MM | target_seq | PAM | alignment | distance | position | gene name | gene id |
| GU305914.1 | 142650 | 142672 | + | 0 | GCGGGCCATCTTTCGCCTGC | CGG | \|\|\|\|\|\|\|\|[\|\|\|\|\|\|\|\|\|\|\|\|\|\|]PAM | NA | Intergenic | NA | NA |
| T10925 | TCTTTCGCCTGCCGGCGCGCCGG | 1000 | | | | | | | | | |
| Oligo fwd | CACCTCTTTCGCCTGCCGGCGCGC | | | | | | | | | | |
| Oligo rev | AAACGCGCGCCGGCAGGCGAAAGA | | | | | | | | | | |
| Chromosome | start | end | strand | MM | target_seq | PAM | alignment | distance | position | gene name | gene id |
| GU305914.1 | 142658 | 142680 | + | 0 | TCTTTCGCCTGCCGGCGCGC | CGG | \|\|\|\|\|\|\|\|[\|\|\|\|\|\|\|\|\|\|\|\|\|\|]PAM | NA | Intergenic | NA | NA |
| T10926 | TGCCGGCGCGCCGGCTCCTCAGG | 1000 | | | | | | | | | |
| Oligo fwd | CACCTGCCGGCGCGCCGGCTCCTC | | | | | | | | | | |
| Oligo rev | AAACGAGGAGCCGGCGCGCCGGCA | | | | | | | | | | |
| Chromosome | start | end | strand | MM | target_seq | PAM | alignment | distance | position | gene name | gene id |
| GU305914.1 | 142667 | 142689 | + | 0 | TGCCGGCGCGCCGGCTCCTC | AGG | \|\|\|\|\|\|\|\|[\|\|\|\|\|\|\|\|\|\|\|\|\|\|]PAM | NA | Intergenic | NA | NA |
| T10927 | GCCGGCGCGCCGGCTCCTCAGGG | 1000 | | | | | | | | | |
| Oligo fwd | CACCGCCGGCGCGCCGGCTCCTCA | | | | | | | | | | |
| Oligo rev | AAACTGAGGAGCCGGCGCGCCGGC | | | | | | | | | | |
| Chromosome | start | end | strand | MM | target_seq | PAM | alignment | distance | position | gene name | gene id |
| GU305914.1 | 142668 | 142690 | + | 0 | GCCGGCGCGCCGGCTCCTCA | GGG | \|\|\|\|\|\|\|\|[\|\|\|\|\|\|\|\|\|\|\|\|\|\|]PAM | NA | Intergenic | NA | NA |
| T10928 | CCGGCTCCTCAGGGTCATGAAGG | 1000 | | | | | | | | | |
| Oligo fwd | CACCCCGGCTCCTCAGGGTCATGA | | | | | | | | | | |
| Oligo rev | AAACTCATGACCCTGAGGAGCCGG | | | | | | | | | | |
| Chromosome | start | end | strand | MM | target_seq | PAM | alignment | distance | position | gene name | gene id |
| GU305914.1 | 142677 | 142699 | + | 0 | CCGGCTCCTCAGGGTCATGA | AGG | \|\|\|\|\|\|\|\|[\|\|\|\|\|\|\|\|\|\|\|\|\|\|]PAM | NA | Intergenic | NA | NA |
| T10929 | GGGTCATGAAGGCGATCGAGAGG | 1000 | | | | | | | | | |
| Oligo fwd | CACCGGGTCATGAAGGCGATCGAG | | | | | | | | | | |
| Oligo rev | AAACCTCGATCGCCTTCATGACCC | | | | | | | | | | |
| Chromosome | start | end | strand | MM | target_seq | PAM | alignment | distance | position | gene name | gene id |
| GU305914.1 | 142688 | 142710 | + | 0 | GGGTCATGAAGGCGATCGAG | AGG | \|\|\|\|\|\|\|\|[\|\|\|\|\|\|\|\|\|\|\|\|\|\|]PAM | NA | Intergenic | NA | NA |
| T10930 | GGTCATGAAGGCGATCGAGAGGG | 1000 | | | | | | | | | |
| Oligo fwd | CACCGGTCATGAAGGCGATCGAGA | | | | | | | | | | |
| Oligo rev | AAACTCTCGATCGCCTTCATGACC | | | | | | | | | | |
| Chromosome | start | end | strand | MM | target_seq | PAM | alignment | distance | position | gene name | gene id |
| GU305914.1 | 142689 | 142711 | + | 0 | GGTCATGAAGGCGATCGAGA | GGG | \|\|\|\|\|\|\|\|[\|\|\|\|\|\|\|\|\|\|\|\|\|\|]PAM | NA | Intergenic | NA | NA |
| T10931 | AGGCGATCGAGAGGGATTCGCGG | 1000 | | | | | | | | | |
| Oligo fwd | CACCAGGCGATCGAGAGGGATTCG | | | | | | | | | | |
| Oligo rev | AAACCGAATCCCTCTCGATCGCCT | | | | | | | | | | |
| Chromosome | start | end | strand | MM | target_seq | PAM | alignment | distance | position | gene name | gene id |
| GU305914.1 | 142697 | 142719 | + | 0 | AGGCGATCGAGAGGGATTCG | CGG | \|\|\|\|\|\|\|\|[\|\|\|\|\|\|\|\|\|\|\|\|\|\|]PAM | NA | Intergenic | NA | NA |
| T10932 | GGCGATCGAGAGGGATTCGCGGG | 1000 | | | | | | | | | |
| Oligo fwd | CACCGGCGATCGAGAGGGATTCGC | | | | | | | | | | |

| | | | | | | | | | | | | |
|---|---|---|---|---|---|---|---|---|---|---|---|---|
| Oligo rev | AAACGCGAATCCCTCTCGATCGCC | | | | | | | | | | | |
| Chromosome | start | | end | strand | MM | target_seq | PAM | alignment | distance | position | gene name | gene id |
| GU305914.1 | | 142698 | 142720 | + | 0 | GGCGATCGAGAGGGATTCGC | GGG | ||||||||[||||||||||||]PAM | NA | Intergenic | NA | NA |
| T10933 | GCGATCGAGAGGGATTCGCGGGG | | 1000 | | | | | | | | | |
| Oligo fwd | CACCGCGATCGAGAGGGATTCGCG | | | | | | | | | | | |
| Oligo rev | AAACCGCGAATCCCTCTCGATCGC | | | | | | | | | | | |
| Chromosome | start | | end | strand | MM | target_seq | PAM | alignment | distance | position | gene name | gene id |
| GU305914.1 | | 142699 | 142721 | + | 0 | GCGATCGAGAGGGATTCGCG | GGG | ||||||||[||||||||||||]PAM | NA | Intergenic | NA | NA |
| T10934 | CGAGAGGGATTCGCGGGGACAGG | | 1000 | | | | | | | | | |
| Oligo fwd | CACCCGAGAGGGATTCGCGGGGAC | | | | | | | | | | | |
| Oligo rev | AAACGTCCCCGCGAATCCCTCTCG | | | | | | | | | | | |
| Chromosome | start | | end | strand | MM | target_seq | PAM | alignment | distance | position | gene name | gene id |
| GU305914.1 | | 142704 | 142726 | + | 0 | CGAGAGGGATTCGCGGGGAC | AGG | ||||||||[||||||||||||]PAM | NA | Intergenic | NA | NA |
| T10935 | GAGGGATTCGCGGGGACAGGCGG | | 1000 | | | | | | | | | |
| Oligo fwd | CACCGAGGGATTCGCGGGGACAGG | | | | | | | | | | | |
| Oligo rev | AAACCCTGTCCCCGCGAATCCCTC | | | | | | | | | | | |
| Chromosome | start | | end | strand | MM | target_seq | PAM | alignment | distance | position | gene name | gene id |
| GU305914.1 | | 142707 | 142729 | + | 0 | GAGGGATTCGCGGGGACAGG | CGG | ||||||||[||||||||||||]PAM | NA | Intergenic | NA | NA |
| T10936 | GACAGGCGGCTAACCCCGTCTGG | | 1000 | | | | | | | | | |
| Oligo fwd | CACCGACAGGCGGCTAACCCCGTC | | | | | | | | | | | |
| Oligo rev | AAACGACGGGGTTAGCCGCCTGTC | | | | | | | | | | | |
| Chromosome | start | | end | strand | MM | target_seq | PAM | alignment | distance | position | gene name | gene id |
| GU305914.1 | | 142721 | 142743 | + | 0 | GACAGGCGGCTAACCCCGTC | TGG | ||||||||[||||||||||||]PAM | NA | Intergenic | NA | NA |
| T10937 | TCGACACCGTGTCAGCGACGCGG | | 1000 | | | | | | | | | |
| Oligo fwd | CACCTCGACACCGTGTCAGCGACG | | | | | | | | | | | |
| Oligo rev | AAACCGTCGCTGACACGGTGTCGA | | | | | | | | | | | |
| Chromosome | start | | end | strand | MM | target_seq | PAM | alignment | distance | position | gene name | gene id |
| GU305914.1 | | 142757 | 142779 | + | 0 | TCGACACCGTGTCAGCGACG | CGG | ||||||||[||||||||||||]PAM | NA | Intergenic | NA | NA |
| T10938 | CGACGCGTAAGATCGCCTTCAGG | | 1000 | | | | | | | | | |
| Oligo fwd | CACCCGACGCGTAAGATCGCCTTC | | | | | | | | | | | |
| Oligo rev | AAACGAAGGCGATCTTACGCGTCG | | | | | | | | | | | |
| Chromosome | start | | end | strand | MM | target_seq | PAM | alignment | distance | position | gene name | gene id |
| GU305914.1 | | 142796 | 142818 | + | 0 | CGACGCGTAAGATCGCCTTC | AGG | ||||||||[||||||||||||]PAM | NA | Intergenic | NA | NA |
| T10939 | GCGTAAGATCGCCTTCAGGAAGG | | 1000 | | | | | | | | | |
| Oligo fwd | CACCGCGTAAGATCGCCTTCAGGA | | | | | | | | | | | |
| Oligo rev | AAACTCCTGAAGGCGATCTTACGC | | | | | | | | | | | |
| Chromosome | start | | end | strand | MM | target_seq | PAM | alignment | distance | position | gene name | gene id |
| GU305914.1 | | 142800 | 142822 | + | 0 | GCGTAAGATCGCCTTCAGGA | AGG | ||||||||[||||||||||||]PAM | NA | Intergenic | NA | NA |
| T10940 | GATCGCCTTCAGGAAGGATCTGG | | 1000 | | | | | | | | | |
| Oligo fwd | CACCGATCGCCTTCAGGAAGGATC | | | | | | | | | | | |
| Oligo rev | AAACGATCCTTCCTGAAGGCGATC | | | | | | | | | | | |
| Chromosome | start | | end | strand | MM | target_seq | PAM | alignment | distance | position | gene name | gene id |
| GU305914.1 | | 142806 | 142828 | + | 0 | GATCGCCTTCAGGAAGGATC | TGG | ||||||||[||||||||||||]PAM | NA | Intergenic | NA | NA |
| T10941 | TGTTCGCCACAGCAGCGAAGCGG | | 1000 | | | | | | | | | |
| Oligo fwd | CACCTGTTCGCCACAGCAGCGAAG | | | | | | | | | | | |
| Oligo rev | AAACCTTCGCTGCTGTGGCGAACA | | | | | | | | | | | |
| Chromosome | start | | end | strand | MM | target_seq | PAM | alignment | distance | position | gene name | gene id |
| GU305914.1 | | 142830 | 142852 | + | 0 | TGTTCGCCACAGCAGCGAAG | CGG | ||||||||[||||||||||||]PAM | NA | Intergenic | NA | NA |
| T10942 | AGCAGCGAAGCGGTGCGCTTCGG | | 1000 | | | | | | | | | |
| Oligo fwd | CACCAGCAGCGAAGCGGTGCGCTT | | | | | | | | | | | |
| Oligo rev | AAACAAGCGCACCGCTTCGCTGCT | | | | | | | | | | | |
| Chromosome | start | | end | strand | MM | target_seq | PAM | alignment | distance | position | gene name | gene id |
| GU305914.1 | | 142840 | 142862 | + | 0 | AGCAGCGAAGCGGTGCGCTT | CGG | ||||||||[||||||||||||]PAM | NA | Intergenic | NA | NA |
| T10943 | GTGCGAATCCGCGATCGCGCAGG | | 1000 | | | | | | | | | |
| Oligo fwd | CACCGTGCGAATCCGCGATCGCGC | | | | | | | | | | | |
| Oligo rev | AAACGCGCGATCGCGGATTCGCAC | | | | | | | | | | | |
| Chromosome | start | | end | strand | MM | target_seq | PAM | alignment | distance | position | gene name | gene id |
| GU305914.1 | | 142869 | 142891 | + | 0 | GTGCGAATCCGCGATCGCGC | AGG | ||||||||[||||||||||||]PAM | NA | Intergenic | NA | NA |
| T10944 | CCGCGATCGCGCAGGTGCTCAGG | | 1000 | | | | | | | | | |
| Oligo fwd | CACCCCGCGATCGCGCAGGTGCTC | | | | | | | | | | | |
| Oligo rev | AAACGAGCACCTGCGCGATCGCGG | | | | | | | | | | | |
| Chromosome | start | | end | strand | MM | target_seq | PAM | alignment | distance | position | gene name | gene id |
| GU305914.1 | | 142877 | 142899 | + | 0 | CCGCGATCGCGCAGGTGCTC | AGG | ||||||||[||||||||||||]PAM | NA | Intergenic | NA | NA |
| T10945 | CGCGATCGCGCAGGTGCTCAGGG | | 1000 | | | | | | | | | |
| Oligo fwd | CACCCGCGATCGCGCAGGTGCTCA | | | | | | | | | | | |
| Oligo rev | AAACTGAGCACCTGCGCGATCGCG | | | | | | | | | | | |
| Chromosome | start | | end | strand | MM | target_seq | PAM | alignment | distance | position | gene name | gene id |
| GU305914.1 | | 142878 | 142900 | + | 0 | CGCGATCGCGCAGGTGCTCA | GGG | ||||||||[||||||||||||]PAM | NA | Intergenic | NA | NA |
| T10946 | GCAGGTGCTCAGGGAGTTCGTGG | | 1000 | | | | | | | | | |
| Oligo fwd | CACCGCAGGTGCTCAGGGAGTTCG | | | | | | | | | | | |
| Oligo rev | AAACCGAACTCCCTGAGCACCTGC | | | | | | | | | | | |
| Chromosome | start | | end | strand | MM | target_seq | PAM | alignment | distance | position | gene name | gene id |
| GU305914.1 | | 142887 | 142909 | + | 0 | GCAGGTGCTCAGGGAGTTCG | TGG | ||||||||[||||||||||||]PAM | NA | Intergenic | NA | NA |
| T10947 | GCTCAGGGAGTTCGTGGCCGAGG | | 1000 | | | | | | | | | |

<table>
<tr><td>Oligo fwd</td><td>CACCGCTCAGGGAGTTCGTGGCCG</td><td></td><td></td><td></td><td></td><td></td><td></td><td></td><td></td><td></td><td></td></tr>
<tr><td>Oligo rev</td><td>AAACCGGCCACGAACTCCCTGAGC</td><td></td><td></td><td></td><td></td><td></td><td></td><td></td><td></td><td></td><td></td></tr>
<tr><td>Chromosome</td><td>start</td><td>end</td><td>strand</td><td>MM</td><td>target_seq</td><td>PAM</td><td>alignment</td><td>distance</td><td>position</td><td>gene name</td><td>gene id</td></tr>
<tr><td>GU305914.1</td><td>142893</td><td>142915</td><td>+</td><td>0</td><td>GCTCAGGGAGTTCGTGGCCG</td><td>AGG</td><td>||||||||[||||||||||||]PAM</td><td>NA</td><td>Intergenic</td><td>NA</td><td>NA</td></tr>
<tr><td></td><td></td><td></td><td></td><td></td><td></td><td></td><td></td><td></td><td></td><td></td><td></td></tr>
<tr><td>T10948</td><td>CTCAGGGAGTTCGTGGCCGAGGG</td><td>1000</td><td></td><td></td><td></td><td></td><td></td><td></td><td></td><td></td><td></td></tr>
<tr><td>Oligo fwd</td><td>CACCCTCAGGGAGTTCGTGGCCGA</td><td></td><td></td><td></td><td></td><td></td><td></td><td></td><td></td><td></td><td></td></tr>
<tr><td>Oligo rev</td><td>AAACTCGGCCACGAACTCCCTGAG</td><td></td><td></td><td></td><td></td><td></td><td></td><td></td><td></td><td></td><td></td></tr>
<tr><td>Chromosome</td><td>start</td><td>end</td><td>strand</td><td>MM</td><td>target_seq</td><td>PAM</td><td>alignment</td><td>distance</td><td>position</td><td>gene name</td><td>gene id</td></tr>
<tr><td>GU305914.1</td><td>142894</td><td>142916</td><td>+</td><td>0</td><td>CTCAGGGAGTTCGTGGCCGA</td><td>GGG</td><td>||||||||[||||||||||||]PAM</td><td>NA</td><td>Intergenic</td><td>NA</td><td>NA</td></tr>
<tr><td></td><td></td><td></td><td></td><td></td><td></td><td></td><td></td><td></td><td></td><td></td><td></td></tr>
<tr><td>T10949</td><td>TCAGGGAGTTCGTGGCCGAGGGG</td><td>1000</td><td></td><td></td><td></td><td></td><td></td><td></td><td></td><td></td><td></td></tr>
<tr><td>Oligo fwd</td><td>CACCTCAGGGAGTTCGTGGCCGAG</td><td></td><td></td><td></td><td></td><td></td><td></td><td></td><td></td><td></td><td></td></tr>
<tr><td>Oligo rev</td><td>AAACCTCGGCCACGAACTCCCTGA</td><td></td><td></td><td></td><td></td><td></td><td></td><td></td><td></td><td></td><td></td></tr>
<tr><td>Chromosome</td><td>start</td><td>end</td><td>strand</td><td>MM</td><td>target_seq</td><td>PAM</td><td>alignment</td><td>distance</td><td>position</td><td>gene name</td><td>gene id</td></tr>
<tr><td>GU305914.1</td><td>142895</td><td>142917</td><td>+</td><td>0</td><td>TCAGGGAGTTCGTGGCCGAG</td><td>GGG</td><td>||||||||[||||||||||||]PAM</td><td>NA</td><td>Intergenic</td><td>NA</td><td>NA</td></tr>
<tr><td></td><td></td><td></td><td></td><td></td><td></td><td></td><td></td><td></td><td></td><td></td><td></td></tr>
<tr><td>T10950</td><td>GAGTTCGTGGCCGAGGGGCGAGG</td><td>1000</td><td></td><td></td><td></td><td></td><td></td><td></td><td></td><td></td><td></td></tr>
<tr><td>Oligo fwd</td><td>CACCGAGTTCGTGGCCGAGGGGCG</td><td></td><td></td><td></td><td></td><td></td><td></td><td></td><td></td><td></td><td></td></tr>
<tr><td>Oligo rev</td><td>AAACCGCCCCTCGGCCACGAACTC</td><td></td><td></td><td></td><td></td><td></td><td></td><td></td><td></td><td></td><td></td></tr>
<tr><td>Chromosome</td><td>start</td><td>end</td><td>strand</td><td>MM</td><td>target_seq</td><td>PAM</td><td>alignment</td><td>distance</td><td>position</td><td>gene name</td><td>gene id</td></tr>
<tr><td>GU305914.1</td><td>142900</td><td>142922</td><td>+</td><td>0</td><td>GAGTTCGTGGCCGAGGGGCG</td><td>AGG</td><td>||||||||[||||||||||||]PAM</td><td>NA</td><td>Intergenic</td><td>NA</td><td>NA</td></tr>
<tr><td></td><td></td><td></td><td></td><td></td><td></td><td></td><td></td><td></td><td></td><td></td><td></td></tr>
<tr><td>T10951</td><td>AGTTCGTGGCCGAGGGGCGAGGG</td><td>1000</td><td></td><td></td><td></td><td></td><td></td><td></td><td></td><td></td><td></td></tr>
<tr><td>Oligo fwd</td><td>CACCAGTTCGTGGCCGAGGGGCGA</td><td></td><td></td><td></td><td></td><td></td><td></td><td></td><td></td><td></td><td></td></tr>
<tr><td>Oligo rev</td><td>AAACTCGCCCCTCGGCCACGAACT</td><td></td><td></td><td></td><td></td><td></td><td></td><td></td><td></td><td></td><td></td></tr>
<tr><td>Chromosome</td><td>start</td><td>end</td><td>strand</td><td>MM</td><td>target_seq</td><td>PAM</td><td>alignment</td><td>distance</td><td>position</td><td>gene name</td><td>gene id</td></tr>
<tr><td>GU305914.1</td><td>142901</td><td>142923</td><td>+</td><td>0</td><td>AGTTCGTGGCCGAGGGGCGA</td><td>GGG</td><td>||||||||[||||||||||||]PAM</td><td>NA</td><td>Intergenic</td><td>NA</td><td>NA</td></tr>
<tr><td></td><td></td><td></td><td></td><td></td><td></td><td></td><td></td><td></td><td></td><td></td><td></td></tr>
<tr><td>T10952</td><td>GTTCGTGGCCGAGGGGCGAGGGG</td><td>1000</td><td></td><td></td><td></td><td></td><td></td><td></td><td></td><td></td><td></td></tr>
<tr><td>Oligo fwd</td><td>CACCGTTCGTGGCCGAGGGGCGAG</td><td></td><td></td><td></td><td></td><td></td><td></td><td></td><td></td><td></td><td></td></tr>
<tr><td>Oligo rev</td><td>AAACCTCGCCCCTCGGCCACGAAC</td><td></td><td></td><td></td><td></td><td></td><td></td><td></td><td></td><td></td><td></td></tr>
<tr><td>Chromosome</td><td>start</td><td>end</td><td>strand</td><td>MM</td><td>target_seq</td><td>PAM</td><td>alignment</td><td>distance</td><td>position</td><td>gene name</td><td>gene id</td></tr>
<tr><td>GU305914.1</td><td>142902</td><td>142924</td><td>+</td><td>0</td><td>GTTCGTGGCCGAGGGGCGAG</td><td>GGG</td><td>||||||||[||||||||||||]PAM</td><td>NA</td><td>Intergenic</td><td>NA</td><td>NA</td></tr>
<tr><td></td><td></td><td></td><td></td><td></td><td></td><td></td><td></td><td></td><td></td><td></td><td></td></tr>
<tr><td>T10953</td><td>TTCGTGGCCGAGGGGCGAGGGGG</td><td>1000</td><td></td><td></td><td></td><td></td><td></td><td></td><td></td><td></td><td></td></tr>
<tr><td>Oligo fwd</td><td>CACCTTCGTGGCCGAGGGGCGAGG</td><td></td><td></td><td></td><td></td><td></td><td></td><td></td><td></td><td></td><td></td></tr>
<tr><td>Oligo rev</td><td>AAACCCTCGCCCCTCGGCCACGAA</td><td></td><td></td><td></td><td></td><td></td><td></td><td></td><td></td><td></td><td></td></tr>
<tr><td>Chromosome</td><td>start</td><td>end</td><td>strand</td><td>MM</td><td>target_seq</td><td>PAM</td><td>alignment</td><td>distance</td><td>position</td><td>gene name</td><td>gene id</td></tr>
<tr><td>GU305914.1</td><td>142903</td><td>142925</td><td>+</td><td>0</td><td>TTCGTGGCCGAGGGGCGAGG</td><td>GGG</td><td>||||||||[||||||||||||]PAM</td><td>NA</td><td>Intergenic</td><td>NA</td><td>NA</td></tr>
<tr><td></td><td></td><td></td><td></td><td></td><td></td><td></td><td></td><td></td><td></td><td></td><td></td></tr>
<tr><td>T10954</td><td>CGAGGGGGAGTGAGCGATATCGG</td><td>1000</td><td></td><td></td><td></td><td></td><td></td><td></td><td></td><td></td><td></td></tr>
<tr><td>Oligo fwd</td><td>CACCCGAGGGGGAGTGAGCGATAT</td><td></td><td></td><td></td><td></td><td></td><td></td><td></td><td></td><td></td><td></td></tr>
<tr><td>Oligo rev</td><td>AAACATATCGCTCACTCCCCCTCG</td><td></td><td></td><td></td><td></td><td></td><td></td><td></td><td></td><td></td><td></td></tr>
<tr><td>Chromosome</td><td>start</td><td>end</td><td>strand</td><td>MM</td><td>target_seq</td><td>PAM</td><td>alignment</td><td>distance</td><td>position</td><td>gene name</td><td>gene id</td></tr>
<tr><td>GU305914.1</td><td>142918</td><td>142940</td><td>+</td><td>0</td><td>CGAGGGGGAGTGAGCGATAT</td><td>CGG</td><td>||||||||[||||||||||||]PAM</td><td>NA</td><td>Intergenic</td><td>NA</td><td>NA</td></tr>
<tr><td></td><td></td><td></td><td></td><td></td><td></td><td></td><td></td><td></td><td></td><td></td><td></td></tr>
<tr><td>T10955</td><td>GAGGGGGAGTGAGCGATATCGGG</td><td>1000</td><td></td><td></td><td></td><td></td><td></td><td></td><td></td><td></td><td></td></tr>
<tr><td>Oligo fwd</td><td>CACCGAGGGGGAGTGAGCGATATC</td><td></td><td></td><td></td><td></td><td></td><td></td><td></td><td></td><td></td><td></td></tr>
<tr><td>Oligo rev</td><td>AAACGATATCGCTCACTCCCCCTC</td><td></td><td></td><td></td><td></td><td></td><td></td><td></td><td></td><td></td><td></td></tr>
<tr><td>Chromosome</td><td>start</td><td>end</td><td>strand</td><td>MM</td><td>target_seq</td><td>PAM</td><td>alignment</td><td>distance</td><td>position</td><td>gene name</td><td>gene id</td></tr>
<tr><td>GU305914.1</td><td>142919</td><td>142941</td><td>+</td><td>0</td><td>GAGGGGGAGTGAGCGATATC</td><td>GGG</td><td>||||||||[||||||||||||]PAM</td><td>NA</td><td>Intergenic</td><td>NA</td><td>NA</td></tr>
<tr><td></td><td></td><td></td><td></td><td></td><td></td><td></td><td></td><td></td><td></td><td></td><td></td></tr>
<tr><td>T10956</td><td>GAGCGATATCGGGCTGCTCCTGG</td><td>1000</td><td></td><td></td><td></td><td></td><td></td><td></td><td></td><td></td><td></td></tr>
<tr><td>Oligo fwd</td><td>CACCGAGCGATATCGGGCTGCTCC</td><td></td><td></td><td></td><td></td><td></td><td></td><td></td><td></td><td></td><td></td></tr>
<tr><td>Oligo rev</td><td>AAACGGAGCAGCCCGATATCGCTC</td><td></td><td></td><td></td><td></td><td></td><td></td><td></td><td></td><td></td><td></td></tr>
<tr><td>Chromosome</td><td>start</td><td>end</td><td>strand</td><td>MM</td><td>target_seq</td><td>PAM</td><td>alignment</td><td>distance</td><td>position</td><td>gene name</td><td>gene id</td></tr>
<tr><td>GU305914.1</td><td>142929</td><td>142951</td><td>+</td><td>0</td><td>GAGCGATATCGGGCTGCTCC</td><td>TGG</td><td>||||||||[||||||||||||]PAM</td><td>NA</td><td>Intergenic</td><td>NA</td><td>NA</td></tr>
<tr><td></td><td></td><td></td><td></td><td></td><td></td><td></td><td></td><td></td><td></td><td></td><td></td></tr>
<tr><td>T10957</td><td>TATCGGGCTGCTCCTGGACCCGG</td><td>1000</td><td></td><td></td><td></td><td></td><td></td><td></td><td></td><td></td><td></td></tr>
<tr><td>Oligo fwd</td><td>CACCTATCGGGCTGCTCCTGGACC</td><td></td><td></td><td></td><td></td><td></td><td></td><td></td><td></td><td></td><td></td></tr>
<tr><td>Oligo rev</td><td>AAACGGTCCAGGAGCAGCCCGATA</td><td></td><td></td><td></td><td></td><td></td><td></td><td></td><td></td><td></td><td></td></tr>
<tr><td>Chromosome</td><td>start</td><td>end</td><td>strand</td><td>MM</td><td>target_seq</td><td>PAM</td><td>alignment</td><td>distance</td><td>position</td><td>gene name</td><td>gene id</td></tr>
<tr><td>GU305914.1</td><td>142935</td><td>142957</td><td>+</td><td>0</td><td>TATCGGGCTGCTCCTGGACC</td><td>CGG</td><td>||||||||[||||||||||||]PAM</td><td>NA</td><td>Intergenic</td><td>NA</td><td>NA</td></tr>
<tr><td></td><td></td><td></td><td></td><td></td><td></td><td></td><td></td><td></td><td></td><td></td><td></td></tr>
<tr><td>T10958</td><td>GCTGCTCCTGGACCCGGCGTCGG</td><td>1000</td><td></td><td></td><td></td><td></td><td></td><td></td><td></td><td></td><td></td></tr>
<tr><td>Oligo fwd</td><td>CACCGCTGCTCCTGGACCCGGCGT</td><td></td><td></td><td></td><td></td><td></td><td></td><td></td><td></td><td></td><td></td></tr>
<tr><td>Oligo rev</td><td>AAACACGCCGGGTCCAGGAGCAGC</td><td></td><td></td><td></td><td></td><td></td><td></td><td></td><td></td><td></td><td></td></tr>
<tr><td>Chromosome</td><td>start</td><td>end</td><td>strand</td><td>MM</td><td>target_seq</td><td>PAM</td><td>alignment</td><td>distance</td><td>position</td><td>gene name</td><td>gene id</td></tr>
<tr><td>GU305914.1</td><td>142941</td><td>142963</td><td>+</td><td>0</td><td>GCTGCTCCTGGACCCGGCGT</td><td>CGG</td><td>||||||||[||||||||||||]PAM</td><td>NA</td><td>Intergenic</td><td>NA</td><td>NA</td></tr>
<tr><td></td><td></td><td></td><td></td><td></td><td></td><td></td><td></td><td></td><td></td><td></td><td></td></tr>
<tr><td>T10959</td><td>CTGCTCCTGGACCCGGCGTCGGG</td><td>1000</td><td></td><td></td><td></td><td></td><td></td><td></td><td></td><td></td><td></td></tr>
<tr><td>Oligo fwd</td><td>CACCCTGCTCCTGGACCCGGCGTC</td><td></td><td></td><td></td><td></td><td></td><td></td><td></td><td></td><td></td><td></td></tr>
<tr><td>Oligo rev</td><td>AAACGACGCCGGGTCCAGGAGCAG</td><td></td><td></td><td></td><td></td><td></td><td></td><td></td><td></td><td></td><td></td></tr>
<tr><td>Chromosome</td><td>start</td><td>end</td><td>strand</td><td>MM</td><td>target_seq</td><td>PAM</td><td>alignment</td><td>distance</td><td>position</td><td>gene name</td><td>gene id</td></tr>
<tr><td>GU305914.1</td><td>142942</td><td>142964</td><td>+</td><td>0</td><td>CTGCTCCTGGACCCGGCGTC</td><td>GGG</td><td>||||||||[||||||||||||]PAM</td><td>NA</td><td>Intergenic</td><td>NA</td><td>NA</td></tr>
<tr><td></td><td></td><td></td><td></td><td></td><td></td><td></td><td></td><td></td><td></td><td></td><td></td></tr>
<tr><td>T10960</td><td>TGCTCCTGGACCCGGCGTCGGGG</td><td>1000</td><td></td><td></td><td></td><td></td><td></td><td></td><td></td><td></td><td></td></tr>
<tr><td>Oligo fwd</td><td>CACCTGCTCCTGGACCCGGCGTCG</td><td></td><td></td><td></td><td></td><td></td><td></td><td></td><td></td><td></td><td></td></tr>
<tr><td>Oligo rev</td><td>AAACCGACGCCGGGTCCAGGAGCA</td><td></td><td></td><td></td><td></td><td></td><td></td><td></td><td></td><td></td><td></td></tr>
<tr><td>Chromosome</td><td>start</td><td>end</td><td>strand</td><td>MM</td><td>target_seq</td><td>PAM</td><td>alignment</td><td>distance</td><td>position</td><td>gene name</td><td>gene id</td></tr>
<tr><td>GU305914.1</td><td>142943</td><td>142965</td><td>+</td><td>0</td><td>TGCTCCTGGACCCGGCGTCG</td><td>GGG</td><td>||||||||[||||||||||||]PAM</td><td>NA</td><td>Intergenic</td><td>NA</td><td>NA</td></tr>
<tr><td></td><td></td><td></td><td></td><td></td><td></td><td></td><td></td><td></td><td></td><td></td><td></td></tr>
<tr><td>T10961</td><td>GCTCCTGGACCCGGCGTCGGGGG</td><td>1000</td><td></td><td></td><td></td><td></td><td></td><td></td><td></td><td></td><td></td></tr>
<tr><td>Oligo fwd</td><td>CACCGCTCCTGGACCCGGCGTCGG</td><td></td><td></td><td></td><td></td><td></td><td></td><td></td><td></td><td></td><td></td></tr>
<tr><td>Oligo rev</td><td>AAACCCGACGCCGGGTCCAGGAGC</td><td></td><td></td><td></td><td></td><td></td><td></td><td></td><td></td><td></td><td></td></tr>
<tr><td>Chromosome</td><td>start</td><td>end</td><td>strand</td><td>MM</td><td>target_seq</td><td>PAM</td><td>alignment</td><td>distance</td><td>position</td><td>gene name</td><td>gene id</td></tr>
<tr><td>GU305914.1</td><td>142944</td><td>142966</td><td>+</td><td>0</td><td>GCTCCTGGACCCGGCGTCGG</td><td>GGG</td><td>||||||||[||||||||||||]PAM</td><td>NA</td><td>Intergenic</td><td>NA</td><td>NA</td></tr>
</table>

T10962 GGACCCGGCGTCGGGGGTCCTGG 1000
Oligo fwd CACCGGACCCGGCGTCGGGGGTCC
Oligo rev AAACGGACCCCCGACGCCGGGTCC
Chromosome start end strand MM target_seq PAM alignment distance position gene name gene id
GU305914.1 142950 142972 + 0 GGACCCGGCGTCGGGGGTCC TGG ||||||||[||||||||||||]PAM NA Intergenic NA NA

T10963 GACCCGGCGTCGGGGGTCCTGGG 1000
Oligo fwd CACCGACCCGGCGTCGGGGGTCCT
Oligo rev AAACAGGACCCCCGACGCCGGGTC
Chromosome start end strand MM target_seq PAM alignment distance position gene name gene id
GU305914.1 142951 142973 + 0 GACCCGGCGTCGGGGGTCCT GGG ||||||||[||||||||||||]PAM NA Intergenic NA NA

T10964 GCCTCGCTCGACTTCTGTAGCGG 1000
Oligo fwd CACCGCCTCGCTCGACTTCTGTAG
Oligo rev AAACCTACAGAAGTCGAGCGAGGC
Chromosome start end strand MM target_seq PAM alignment distance position gene name gene id
GU305914.1 142975 142997 + 0 GCCTCGCTCGACTTCTGTAG CGG ||||||||[||||||||||||]PAM NA Intergenic NA NA

T10965 GGTCTCTCCCGCGACGACGACGG 1000
Oligo fwd CACCGGTCTCTCCCGCGACGACGA
Oligo rev AAACTCGTCGTCGCGGGAGAGACC
Chromosome start end strand MM target_seq PAM alignment distance position gene name gene id
GU305914.1 142996 143018 + 0 GGTCTCTCCCGCGACGACGA CGG ||||||||[||||||||||||]PAM NA Intergenic NA NA

T10966 CCGCGACGACGACGGCCTGCTGG 1000
Oligo fwd CACCCCGCGACGACGACGGCCTGC
Oligo rev AAACGCAGGCCGTCGTCGTCGCGG
Chromosome start end strand MM target_seq PAM alignment distance position gene name gene id
GU305914.1 143004 143026 + 0 CCGCGACGACGACGGCCTGC TGG ||||||||[||||||||||||]PAM NA Intergenic NA NA

T10967 CGACGACGACGGCCTGCTGGTGG 1000
Oligo fwd CACCCGACGACGACGGCCTGCTGG
Oligo rev AAACCCAGCAGGCCGTCGTCGTCG
Chromosome start end strand MM target_seq PAM alignment distance position gene name gene id
GU305914.1 143007 143029 + 0 CGACGACGACGGCCTGCTGG TGG ||||||||[||||||||||||]PAM NA Intergenic NA NA
GU305914.1 74525 74547 - 4 CGCCGAGGACGGCGTGCTCG AGG ||-|||-|[|||||-||||-|]PAM NA Intergenic NA NA

T10968 CGACGACGGCCTGCTGGTGGTGG 1000
Oligo fwd CACCCGACGACGGCCTGCTGGTGG
Oligo rev AAACCCACCAGCAGGCCGTCGTCG
Chromosome start end strand MM target_seq PAM alignment distance position gene name gene id
GU305914.1 143010 143032 + 0 CGACGACGGCCTGCTGGTGG TGG ||||||||[||||||||||||]PAM NA Intergenic NA NA
GU305914.1 107641 107663 - 3 CGACCACGACCTCCTGGTGG TGG ||||-|||[-|||-|||||||]PAM NA Intergenic NA NA

T10969 CGGCCTGCTGGTGGTGGCGCCGG 1000
Oligo fwd CACCCGGCCTGCTGGTGGTGGCGC
Oligo rev AAACGCGCCACCACCAGCAGGCCG
Chromosome start end strand MM target_seq PAM alignment distance position gene name gene id
GU305914.1 143016 143038 + 0 CGGCCTGCTGGTGGTGGCGC CGG ||||||||[||||||||||||]PAM NA Intergenic NA NA

T10970 GGCCTGCTGGTGGTGGCGCCGGG 1000
Oligo fwd CACCGGCCTGCTGGTGGTGGCGCC
Oligo rev AAACGGCGCCACCACCAGCAGGCC
Chromosome start end strand MM target_seq PAM alignment distance position gene name gene id
GU305914.1 143017 143039 + 0 GGCCTGCTGGTGGTGGCGCC GGG ||||||||[||||||||||||]PAM NA Intergenic NA NA

T10971 GCCTGCTGGTGGTGGCGCCGGGG 1000
Oligo fwd CACCGCCTGCTGGTGGTGGCGCCG
Oligo rev AAACCGGCGCCACCACCAGCAGGC
Chromosome start end strand MM target_seq PAM alignment distance position gene name gene id
GU305914.1 143018 143040 + 0 GCCTGCTGGTGGTGGCGCCG GGG ||||||||[||||||||||||]PAM NA Intergenic NA NA

T10972 CCTGCTGGTGGTGGCGCCGGGGG 1000
Oligo fwd CACCCCTGCTGGTGGTGGCGCCGG
Oligo rev AAACCCGGCGCCACCACCAGCAGG
Chromosome start end strand MM target_seq PAM alignment distance position gene name gene id
GU305914.1 143019 143041 + 0 CCTGCTGGTGGTGGCGCCGG GGG ||||||||[||||||||||||]PAM NA Intergenic NA NA

T10973 GCTGGTGGTGGCGCCGGGGGCGG 1000
Oligo fwd CACCGCTGGTGGTGGCGCCGGGGG
Oligo rev AAACCCCCCGGCGCCACCACCAGC
Chromosome start end strand MM target_seq PAM alignment distance position gene name gene id
GU305914.1 143022 143044 + 0 GCTGGTGGTGGCGCCGGGGG CGG ||||||||[||||||||||||]PAM NA Intergenic NA NA
GU305914.1 62449 62471 + 3 GGGGGTGGTGGCGGCGGGGG TGG |--|||||[|||||-||||||]PAM NA Intergenic NA NA
GU305914.1 58000 58022 - 4 GTGGGTGGAGGGGCCGGGGG AGG |--|||||[-||-||||||||]PAM NA Intergenic NA NA
GU305914.1 62434 62456 + 4 GACGGCGGTGGCGGCGGGGG TGG |--||-||[|||||-||||||]PAM NA Intergenic NA NA
GU305914.1 168893 168915 + 4 GCTGGCGCTGGTGCCGCGGG CGG |||||-|-[|||-||||-|||]PAM NA Intergenic NA NA
GU305914.1 58129 58151 - 4 GCGGGCGGTGGCGGCGGCGG CGG ||-||-||[|||||-|||-||]PAM NA Intergenic NA NA

T10974 AATGCCGCTTCAAGTACCTGCGG 1000
Oligo fwd CACCAATGCCGCTTCAAGTACCTG
Oligo rev AAACCAGGTACTTGAAGCGGCATT
Chromosome start end strand MM target_seq PAM alignment distance position gene name gene id
GU305914.1 143060 143082 + 0 AATGCCGCTTCAAGTACCTG CGG ||||||||[||||||||||||]PAM NA Intergenic NA NA

T10975 GCGGTCGCGCGACGACCGCGCGG 1000
Oligo fwd CACCGCGGTCGCGCGACGACCGCG
Oligo rev AAACCGCGGTCGTCGCGCGACCGC
Chromosome start end strand MM target_seq PAM alignment distance position gene name gene id

GU305914.1 143079 143101 + 0 GCGGTCGCGCGACGACCGCG CGG ||||||||[||||||||||||]PAM NA Intergenic NA NA

T10976 GCGCGACGACCGCGCGGTGCAGG 1000
Oligo fwd CACCGCGCGACGACCGCGCGGTGC
Oligo rev AAACGCACCGCGCGGTCGTCGCGC
Chromosome start end strand MM target_seq PAM alignment distance position gene name gene id
GU305914.1 143085 143107 + 0 GCGCGACGACCGCGCGGTGC AGG ||||||||[||||||||||||]PAM NA Intergenic NA NA

T10977 CGCGACGACCGCGCGGTGCAGGG 1000
Oligo fwd CACCCGCGACGACCGCGCGGTGCA
Oligo rev AAACTGCACCGCGCGGTCGTCGCG
Chromosome start end strand MM target_seq PAM alignment distance position gene name gene id
GU305914.1 143086 143108 + 0 CGCGACGACCGCGCGGTGCA GGG ||||||||[||||||||||||]PAM NA Intergenic NA NA
GU305914.1 126743 126765 - 3 CGGGACGATCGCGCGGCGCA TGG ||-|||||[-|||||||-|||]PAM NA Intergenic NA NA

T10978 GCGACGACCGCGCGGTGCAGGGG 1000
Oligo fwd CACCGCGACGACCGCGCGGTGCAG
Oligo rev AAACCTGCACCGCGCGGTCGTCGC
Chromosome start end strand MM target_seq PAM alignment distance position gene name gene id
GU305914.1 143087 143109 + 0 GCGACGACCGCGCGGTGCAG GGG ||||||||[||||||||||||]PAM NA Intergenic NA NA

T10979 CCGCGCGGTGCAGGGGCTCCTGG 1000
Oligo fwd CACCCCGCGCGGTGCAGGGGCTCC
Oligo rev AAACGGAGCCCCTGCACCGCGCGG
Chromosome start end strand MM target_seq PAM alignment distance position gene name gene id
GU305914.1 143094 143116 + 0 CCGCGCGGTGCAGGGGCTCC TGG ||||||||[||||||||||||]PAM NA Intergenic NA NA

T10980 CAGGGGCTCCTGGACGACCCCGG 1000
Oligo fwd CACCCAGGGGCTCCTGGACGACCC
Oligo rev AAACGGGTCGTCCAGGAGCCCCTG
Chromosome start end strand MM target_seq PAM alignment distance position gene name gene id
GU305914.1 143104 143126 + 0 CAGGGGCTCCTGGACGACCC CGG ||||||||[||||||||||||]PAM NA Intergenic NA NA

T10981 GAGCTTCGCCGATTTCATCCTGG 1000
Oligo fwd CACCGAGCTTCGCCGATTTCATCC
Oligo rev AAACGGATGAAATCGGCGAAGCTC
Chromosome start end strand MM target_seq PAM alignment distance position gene name gene id
GU305914.1 143133 143155 + 0 GAGCTTCGCCGATTTCATCC TGG ||||||||[||||||||||||]PAM NA Intergenic NA NA

T10982 CATCCTGGACCACCACACGCCGG 1000
Oligo fwd CACCCATCCTGGACCACCACACGC
Oligo rev AAACGCGTGTGGTGGTCCAGGATG
Chromosome start end strand MM target_seq PAM alignment distance position gene name gene id
GU305914.1 143148 143170 + 0 CATCCTGGACCACCACACGC CGG ||||||||[||||||||||||]PAM NA Intergenic NA NA

T10983 ACACGCCGGCCGTCGAGTTCCGG 1000
Oligo fwd CACCACACGCCGGCCGTCGAGTTC
Oligo rev AAACGAACTCGACGGCCGGCGTGT
Chromosome start end strand MM target_seq PAM alignment distance position gene name gene id
GU305914.1 143162 143184 + 0 ACACGCCGGCCGTCGAGTTC CGG ||||||||[||||||||||||]PAM NA Intergenic NA NA

T10984 GGCCGTCGAGTTCCGGCACCAGG 1000
Oligo fwd CACCGGCCGTCGAGTTCCGGCACC
Oligo rev AAACGGTGCCGGAACTCGACGGCC
Chromosome start end strand MM target_seq PAM alignment distance position gene name gene id
GU305914.1 143169 143191 + 0 GGCCGTCGAGTTCCGGCACC AGG ||||||||[||||||||||||]PAM NA Intergenic NA NA

T10985 GCCGTCGAGTTCCGGCACCAGGG 1000
Oligo fwd CACCGCCGTCGAGTTCCGGCACCA
Oligo rev AAACTGGTGCCGGAACTCGACGGC
Chromosome start end strand MM target_seq PAM alignment distance position gene name gene id
GU305914.1 143170 143192 + 0 GCCGTCGAGTTCCGGCACCA GGG ||||||||[||||||||||||]PAM NA Intergenic NA NA

T10986 GCCCACCTCGCGCGAGTGTCTGG 1000
Oligo fwd CACCGCCCACCTCGCGCGAGTGTC
Oligo rev AAACGACACTCGCGCGAGGTGGGC
Chromosome start end strand MM target_seq PAM alignment distance position gene name gene id
GU305914.1 143199 143221 + 0 GCCCACCTCGCGCGAGTGTC TGG ||||||||[||||||||||||]PAM NA Intergenic NA NA

T10987 TGTCCTACGACCGCGTCTTTCGG 1000
Oligo fwd CACCTGTCCTACGACCGCGTCTTT
Oligo rev AAACAAAGACGCGGTCGTAGGACA
Chromosome start end strand MM target_seq PAM alignment distance position gene name gene id
GU305914.1 143222 143244 + 0 TGTCCTACGACCGCGTCTTT CGG ||||||||[||||||||||||]PAM NA Intergenic NA NA

T10988 TCGTGCAAACGACGTCGCACCGG 1000
Oligo fwd CACCTCGTGCAAACGACGTCGCAC
Oligo rev AAACGTGCGACGTCGTTTGCACGA
Chromosome start end strand MM target_seq PAM alignment distance position gene name gene id
GU305914.1 143248 143270 + 0 TCGTGCAAACGACGTCGCAC CGG ||||||||[||||||||||||]PAM NA Intergenic NA NA

T10989 ACGTCGCACCGGCGTCGTTTCGG 1000
Oligo fwd CACCACGTCGCACCGGCGTCGTTT
Oligo rev AAACAAACGACGCCGGTGCGACGT
Chromosome start end strand MM target_seq PAM alignment distance position gene name gene id
GU305914.1 143259 143281 + 0 ACGTCGCACCGGCGTCGTTT CGG ||||||||[||||||||||||]PAM NA Intergenic NA NA

T10990 TTTCGGAGTCCCTGCGCCTCTGG 1000
Oligo fwd CACCTTTCGGAGTCCCTGCGCCTC

<table>
<tr><td>Oligo rev</td><td>AAACGAGGCGCAGGGACTCCGAAA</td><td></td><td></td><td></td><td></td><td></td><td></td><td></td><td></td><td></td><td></td></tr>
<tr><td>Chromosome</td><td>start</td><td>end</td><td>strand</td><td>MM</td><td>target_seq</td><td>PAM</td><td>alignment</td><td>distance</td><td>position</td><td>gene name</td><td>gene id</td></tr>
<tr><td>GU305914.1</td><td>143276</td><td>143298</td><td>+</td><td>0</td><td>TTTCGGAGTCCCTGCGCCTC</td><td>TGG</td><td>||||||||[||||||||||||]PAM</td><td>NA</td><td>Intergenic</td><td>NA</td><td>NA</td></tr>
<tr><td>T10991</td><td>TCCCTGCGCCTCTGGATCGACGG</td><td>1000</td><td></td><td></td><td></td><td></td><td></td><td></td><td></td><td></td><td></td></tr>
<tr><td>Oligo fwd</td><td>CACCTCCCTGCGCCTCTGGATCGA</td><td></td><td></td><td></td><td></td><td></td><td></td><td></td><td></td><td></td><td></td></tr>
<tr><td>Oligo rev</td><td>AAACTCGATCCAGAGGCGCAGGGA</td><td></td><td></td><td></td><td></td><td></td><td></td><td></td><td></td><td></td><td></td></tr>
<tr><td>Chromosome</td><td>start</td><td>end</td><td>strand</td><td>MM</td><td>target_seq</td><td>PAM</td><td>alignment</td><td>distance</td><td>position</td><td>gene name</td><td>gene id</td></tr>
<tr><td>GU305914.1</td><td>143284</td><td>143306</td><td>+</td><td>0</td><td>TCCCTGCGCCTCTGGATCGA</td><td>CGG</td><td>||||||||[||||||||||||]PAM</td><td>NA</td><td>Intergenic</td><td>NA</td><td>NA</td></tr>
<tr><td>T10992</td><td>CTGGATCGACGGCCTCATCAAGG</td><td>1000</td><td></td><td></td><td></td><td></td><td></td><td></td><td></td><td></td><td></td></tr>
<tr><td>Oligo fwd</td><td>CACCCTGGATCGACGGCCTCATCA</td><td></td><td></td><td></td><td></td><td></td><td></td><td></td><td></td><td></td><td></td></tr>
<tr><td>Oligo rev</td><td>AAACTGATGAGGCCGTCGATCCAG</td><td></td><td></td><td></td><td></td><td></td><td></td><td></td><td></td><td></td><td></td></tr>
<tr><td>Chromosome</td><td>start</td><td>end</td><td>strand</td><td>MM</td><td>target_seq</td><td>PAM</td><td>alignment</td><td>distance</td><td>position</td><td>gene name</td><td>gene id</td></tr>
<tr><td>GU305914.1</td><td>143295</td><td>143317</td><td>+</td><td>0</td><td>CTGGATCGACGGCCTCATCA</td><td>AGG</td><td>||||||||[||||||||||||]PAM</td><td>NA</td><td>Intergenic</td><td>NA</td><td>NA</td></tr>
<tr><td>T10993</td><td>CGGCCTCATCAAGGAGAACTCGG</td><td>1000</td><td></td><td></td><td></td><td></td><td></td><td></td><td></td><td></td><td></td></tr>
<tr><td>Oligo fwd</td><td>CACCCGGCCTCATCAAGGAGAACT</td><td></td><td></td><td></td><td></td><td></td><td></td><td></td><td></td><td></td><td></td></tr>
<tr><td>Oligo rev</td><td>AAACAGTTCTCCTTGATGAGGCCG</td><td></td><td></td><td></td><td></td><td></td><td></td><td></td><td></td><td></td><td></td></tr>
<tr><td>Chromosome</td><td>start</td><td>end</td><td>strand</td><td>MM</td><td>target_seq</td><td>PAM</td><td>alignment</td><td>distance</td><td>position</td><td>gene name</td><td>gene id</td></tr>
<tr><td>GU305914.1</td><td>143304</td><td>143326</td><td>+</td><td>0</td><td>CGGCCTCATCAAGGAGAACT</td><td>CGG</td><td>||||||||[||||||||||||]PAM</td><td>NA</td><td>Intergenic</td><td>NA</td><td>NA</td></tr>
<tr><td>T10994</td><td>CCTCATCAAGGAGAACTCGGAGG</td><td>1000</td><td></td><td></td><td></td><td></td><td></td><td></td><td></td><td></td><td></td></tr>
<tr><td>Oligo fwd</td><td>CACCCCTCATCAAGGAGAACTCGG</td><td></td><td></td><td></td><td></td><td></td><td></td><td></td><td></td><td></td><td></td></tr>
<tr><td>Oligo rev</td><td>AAACCCGAGTTCTCCTTGATGAGG</td><td></td><td></td><td></td><td></td><td></td><td></td><td></td><td></td><td></td><td></td></tr>
<tr><td>Chromosome</td><td>start</td><td>end</td><td>strand</td><td>MM</td><td>target_seq</td><td>PAM</td><td>alignment</td><td>distance</td><td>position</td><td>gene name</td><td>gene id</td></tr>
<tr><td>GU305914.1</td><td>143307</td><td>143329</td><td>+</td><td>0</td><td>CCTCATCAAGGAGAACTCGG</td><td>AGG</td><td>||||||||[||||||||||||]PAM</td><td>NA</td><td>Intergenic</td><td>NA</td><td>NA</td></tr>
<tr><td>T10995</td><td>TTTCGACGCGCGCGCCGCCGAGG</td><td>1000</td><td></td><td></td><td></td><td></td><td></td><td></td><td></td><td></td><td></td></tr>
<tr><td>Oligo fwd</td><td>CACCTTTCGACGCGCGCGCCGCCG</td><td></td><td></td><td></td><td></td><td></td><td></td><td></td><td></td><td></td><td></td></tr>
<tr><td>Oligo rev</td><td>AAACCGGCGGCGCGCGCGTCGAAA</td><td></td><td></td><td></td><td></td><td></td><td></td><td></td><td></td><td></td><td></td></tr>
<tr><td>Chromosome</td><td>start</td><td>end</td><td>strand</td><td>MM</td><td>target_seq</td><td>PAM</td><td>alignment</td><td>distance</td><td>position</td><td>gene name</td><td>gene id</td></tr>
<tr><td>GU305914.1</td><td>143349</td><td>143371</td><td>+</td><td>0</td><td>TTTCGACGCGCGCGCCGCCG</td><td>AGG</td><td>||||||||[||||||||||||]PAM</td><td>NA</td><td>Intergenic</td><td>NA</td><td>NA</td></tr>
<tr><td>T10996</td><td>TTCGACGCGCGCGCCGCCGAGGG</td><td>1000</td><td></td><td></td><td></td><td></td><td></td><td></td><td></td><td></td><td></td></tr>
<tr><td>Oligo fwd</td><td>CACCTTCGACGCGCGCGCCGCCGA</td><td></td><td></td><td></td><td></td><td></td><td></td><td></td><td></td><td></td><td></td></tr>
<tr><td>Oligo rev</td><td>AAACTCGGCGGCGCGCGCGTCGAA</td><td></td><td></td><td></td><td></td><td></td><td></td><td></td><td></td><td></td><td></td></tr>
<tr><td>Chromosome</td><td>start</td><td>end</td><td>strand</td><td>MM</td><td>target_seq</td><td>PAM</td><td>alignment</td><td>distance</td><td>position</td><td>gene name</td><td>gene id</td></tr>
<tr><td>GU305914.1</td><td>143350</td><td>143372</td><td>+</td><td>0</td><td>TTCGACGCGCGCGCCGCCGA</td><td>GGG</td><td>||||||||[||||||||||||]PAM</td><td>NA</td><td>Intergenic</td><td>NA</td><td>NA</td></tr>
<tr><td>T10997</td><td>CGCGCCGCCGAGGGCGAGACCGG</td><td>1000</td><td></td><td></td><td></td><td></td><td></td><td></td><td></td><td></td><td></td></tr>
<tr><td>Oligo fwd</td><td>CACCCGCGCCGCCGAGGGCGAGAC</td><td></td><td></td><td></td><td></td><td></td><td></td><td></td><td></td><td></td><td></td></tr>
<tr><td>Oligo rev</td><td>AAACGTCTCGCCCTCGGCGGCGCG</td><td></td><td></td><td></td><td></td><td></td><td></td><td></td><td></td><td></td><td></td></tr>
<tr><td>Chromosome</td><td>start</td><td>end</td><td>strand</td><td>MM</td><td>target_seq</td><td>PAM</td><td>alignment</td><td>distance</td><td>position</td><td>gene name</td><td>gene id</td></tr>
<tr><td>GU305914.1</td><td>143359</td><td>143381</td><td>+</td><td>0</td><td>CGCGCCGCCGAGGGCGAGAC</td><td>CGG</td><td>||||||||[||||||||||||]PAM</td><td>NA</td><td>Intergenic</td><td>NA</td><td>NA</td></tr>
<tr><td>T10998</td><td>ACCGGCGCCGCGACCTGCGTCGG</td><td>1000</td><td></td><td></td><td></td><td></td><td></td><td></td><td></td><td></td><td></td></tr>
<tr><td>Oligo fwd</td><td>CACCACCGGCGCCGCGACCTGCGT</td><td></td><td></td><td></td><td></td><td></td><td></td><td></td><td></td><td></td><td></td></tr>
<tr><td>Oligo rev</td><td>AAACACGCAGGTCGCGGCGCCGGT</td><td></td><td></td><td></td><td></td><td></td><td></td><td></td><td></td><td></td><td></td></tr>
<tr><td>Chromosome</td><td>start</td><td>end</td><td>strand</td><td>MM</td><td>target_seq</td><td>PAM</td><td>alignment</td><td>distance</td><td>position</td><td>gene name</td><td>gene id</td></tr>
<tr><td>GU305914.1</td><td>143377</td><td>143399</td><td>+</td><td>0</td><td>ACCGGCGCCGCGACCTGCGT</td><td>CGG</td><td>||||||||[||||||||||||]PAM</td><td>NA</td><td>Intergenic</td><td>NA</td><td>NA</td></tr>
<tr><td>T10999</td><td>CTGCGTCGGTGATGACGACGAGG</td><td>1000</td><td></td><td></td><td></td><td></td><td></td><td></td><td></td><td></td><td></td></tr>
<tr><td>Oligo fwd</td><td>CACCCTGCGTCGGTGATGACGACG</td><td></td><td></td><td></td><td></td><td></td><td></td><td></td><td></td><td></td><td></td></tr>
<tr><td>Oligo rev</td><td>AAACCGTCGTCATCACCGACGCAG</td><td></td><td></td><td></td><td></td><td></td><td></td><td></td><td></td><td></td><td></td></tr>
<tr><td>Chromosome</td><td>start</td><td>end</td><td>strand</td><td>MM</td><td>target_seq</td><td>PAM</td><td>alignment</td><td>distance</td><td>position</td><td>gene name</td><td>gene id</td></tr>
<tr><td>GU305914.1</td><td>143391</td><td>143413</td><td>+</td><td>0</td><td>CTGCGTCGGTGATGACGACG</td><td>AGG</td><td>||||||||[||||||||||||]PAM</td><td>NA</td><td>Intergenic</td><td>NA</td><td>NA</td></tr>
<tr><td>T11000</td><td>ACTTCATCCTGTCCGCCGAGCGG</td><td>1000</td><td></td><td></td><td></td><td></td><td></td><td></td><td></td><td></td><td></td></tr>
<tr><td>Oligo fwd</td><td>CACCACTTCATCCTGTCCGCCGAG</td><td></td><td></td><td></td><td></td><td></td><td></td><td></td><td></td><td></td><td></td></tr>
<tr><td>Oligo rev</td><td>AAACCTCGGCGGACAGGATGAAGT</td><td></td><td></td><td></td><td></td><td></td><td></td><td></td><td></td><td></td><td></td></tr>
<tr><td>Chromosome</td><td>start</td><td>end</td><td>strand</td><td>MM</td><td>target_seq</td><td>PAM</td><td>alignment</td><td>distance</td><td>position</td><td>gene name</td><td>gene id</td></tr>
<tr><td>GU305914.1</td><td>143414</td><td>143436</td><td>+</td><td>0</td><td>ACTTCATCCTGTCCGCCGAG</td><td>CGG</td><td>||||||||[||||||||||||]PAM</td><td>NA</td><td>Intergenic</td><td>NA</td><td>NA</td></tr>
<tr><td>T11001</td><td>CTACCTAGATCTGTTCCTCAAGG</td><td>1000</td><td></td><td></td><td></td><td></td><td></td><td></td><td></td><td></td><td></td></tr>
<tr><td>Oligo fwd</td><td>CACCCTACCTAGATCTGTTCCTCA</td><td></td><td></td><td></td><td></td><td></td><td></td><td></td><td></td><td></td><td></td></tr>
<tr><td>Oligo rev</td><td>AAACTGAGGAACAGATCTAGGTAG</td><td></td><td></td><td></td><td></td><td></td><td></td><td></td><td></td><td></td><td></td></tr>
<tr><td>Chromosome</td><td>start</td><td>end</td><td>strand</td><td>MM</td><td>target_seq</td><td>PAM</td><td>alignment</td><td>distance</td><td>position</td><td>gene name</td><td>gene id</td></tr>
<tr><td>GU305914.1</td><td>143445</td><td>143467</td><td>+</td><td>0</td><td>CTACCTAGATCTGTTCCTCA</td><td>AGG</td><td>||||||||[||||||||||||]PAM</td><td>NA</td><td>Intergenic</td><td>NA</td><td>NA</td></tr>
<tr><td>T11002</td><td>CCCGTACTACTGTCAGACCCTGG</td><td>1000</td><td></td><td></td><td></td><td></td><td></td><td></td><td></td><td></td><td></td></tr>
<tr><td>Oligo fwd</td><td>CACCCCCGTACTACTGTCAGACCC</td><td></td><td></td><td></td><td></td><td></td><td></td><td></td><td></td><td></td><td></td></tr>
<tr><td>Oligo rev</td><td>AAACGGGTCTGACAGTAGTACGGG</td><td></td><td></td><td></td><td></td><td></td><td></td><td></td><td></td><td></td><td></td></tr>
<tr><td>Chromosome</td><td>start</td><td>end</td><td>strand</td><td>MM</td><td>target_seq</td><td>PAM</td><td>alignment</td><td>distance</td><td>position</td><td>gene name</td><td>gene id</td></tr>
<tr><td>GU305914.1</td><td>143505</td><td>143527</td><td>+</td><td>0</td><td>CCCGTACTACTGTCAGACCC</td><td>TGG</td><td>||||||||[||||||||||||]PAM</td><td>NA</td><td>Intergenic</td><td>NA</td><td>NA</td></tr>
<tr><td>T11003</td><td>CGAACTACCCTCCGTGTACCTGG</td><td>1000</td><td></td><td></td><td></td><td></td><td></td><td></td><td></td><td></td><td></td></tr>
<tr><td>Oligo fwd</td><td>CACCCGAACTACCCTCCGTGTACC</td><td></td><td></td><td></td><td></td><td></td><td></td><td></td><td></td><td></td><td></td></tr>
<tr><td>Oligo rev</td><td>AAACGGTACACGGAGGGTAGTTCG</td><td></td><td></td><td></td><td></td><td></td><td></td><td></td><td></td><td></td><td></td></tr>
<tr><td>Chromosome</td><td>start</td><td>end</td><td>strand</td><td>MM</td><td>target_seq</td><td>PAM</td><td>alignment</td><td>distance</td><td>position</td><td>gene name</td><td>gene id</td></tr>
<tr><td>GU305914.1</td><td>143595</td><td>143617</td><td>+</td><td>0</td><td>CGAACTACCCTCCGTGTACC</td><td>TGG</td><td>||||||||[||||||||||||]PAM</td><td>NA</td><td>Intergenic</td><td>NA</td><td>NA</td></tr>
<tr><td>T11004</td><td>ACCCTCCGTGTACCTGGTCTCGG</td><td>1000</td><td></td><td></td><td></td><td></td><td></td><td></td><td></td><td></td><td></td></tr>
<tr><td>Oligo fwd</td><td>CACCACCCTCCGTGTACCTGGTCT</td><td></td><td></td><td></td><td></td><td></td><td></td><td></td><td></td><td></td><td></td></tr>
<tr><td>Oligo rev</td><td>AAACAGACCAGGTACACGGAGGGT</td><td></td><td></td><td></td><td></td><td></td><td></td><td></td><td></td><td></td><td></td></tr>
<tr><td>Chromosome</td><td>start</td><td>end</td><td>strand</td><td>MM</td><td>target_seq</td><td>PAM</td><td>alignment</td><td>distance</td><td>position</td><td>gene name</td><td>gene id</td></tr>
<tr><td>GU305914.1</td><td>143601</td><td>143623</td><td>+</td><td>0</td><td>ACCCTCCGTGTACCTGGTCT</td><td>CGG</td><td>||||||||[||||||||||||]PAM</td><td>NA</td><td>Intergenic</td><td>NA</td><td>NA</td></tr>
<tr><td>T11005</td><td>TCTCGGCCATACTGAGAAAACGG</td><td>1000</td><td></td><td></td><td></td><td></td><td></td><td></td><td></td><td></td><td></td></tr>
</table>

<table>
<tr><td>Oligo fwd</td><td>CACCTCTCGGCCATACTGAGAAAA</td><td></td><td></td><td></td><td></td><td></td><td></td><td></td><td></td><td></td><td></td></tr>
<tr><td>Oligo rev</td><td>AAACTTTTCTCAGTATGGCCGAGA</td><td></td><td></td><td></td><td></td><td></td><td></td><td></td><td></td><td></td><td></td></tr>
<tr><td>Chromosome</td><td>start</td><td>end</td><td>strand</td><td>MM</td><td>target_seq</td><td>PAM</td><td>alignment</td><td>distance</td><td>position</td><td>gene name</td><td>gene id</td></tr>
<tr><td>GU305914.1</td><td>143618</td><td>143640</td><td>+</td><td>0</td><td>TCTCGGCCATACTGAGAAAA</td><td>CGG</td><td>||||||||[||||||||||||]PAM</td><td>NA</td><td>Intergenic</td><td>NA</td><td>NA</td></tr>
<tr><td></td><td></td><td></td><td></td><td></td><td></td><td></td><td></td><td></td><td></td><td></td><td></td></tr>
<tr><td>T11006</td><td>CTCGGCCATACTGAGAAAACGGG</td><td>1000</td><td></td><td></td><td></td><td></td><td></td><td></td><td></td><td></td><td></td></tr>
<tr><td>Oligo fwd</td><td>CACCCTCGGCCATACTGAGAAAAC</td><td></td><td></td><td></td><td></td><td></td><td></td><td></td><td></td><td></td><td></td></tr>
<tr><td>Oligo rev</td><td>AAACGTTTTCTCAGTATGGCCGAG</td><td></td><td></td><td></td><td></td><td></td><td></td><td></td><td></td><td></td><td></td></tr>
<tr><td>Chromosome</td><td>start</td><td>end</td><td>strand</td><td>MM</td><td>target_seq</td><td>PAM</td><td>alignment</td><td>distance</td><td>position</td><td>gene name</td><td>gene id</td></tr>
<tr><td>GU305914.1</td><td>143619</td><td>143641</td><td>+</td><td>0</td><td>CTCGGCCATACTGAGAAAAC</td><td>GGG</td><td>||||||||[||||||||||||]PAM</td><td>NA</td><td>Intergenic</td><td>NA</td><td>NA</td></tr>
<tr><td></td><td></td><td></td><td></td><td></td><td></td><td></td><td></td><td></td><td></td><td></td><td></td></tr>
<tr><td>T11007</td><td>AAACGGGACGAGAGCGAACGCGG</td><td>1000</td><td></td><td></td><td></td><td></td><td></td><td></td><td></td><td></td><td></td></tr>
<tr><td>Oligo fwd</td><td>CACCAAACGGGACGAGAGCGAACG</td><td></td><td></td><td></td><td></td><td></td><td></td><td></td><td></td><td></td><td></td></tr>
<tr><td>Oligo rev</td><td>AAACCGTTCGCTCTCGTCCCGTTT</td><td></td><td></td><td></td><td></td><td></td><td></td><td></td><td></td><td></td><td></td></tr>
<tr><td>Chromosome</td><td>start</td><td>end</td><td>strand</td><td>MM</td><td>target_seq</td><td>PAM</td><td>alignment</td><td>distance</td><td>position</td><td>gene name</td><td>gene id</td></tr>
<tr><td>GU305914.1</td><td>143635</td><td>143657</td><td>+</td><td>0</td><td>AAACGGGACGAGAGCGAACG</td><td>CGG</td><td>||||||||[||||||||||||]PAM</td><td>NA</td><td>Intergenic</td><td>NA</td><td>NA</td></tr>
<tr><td></td><td></td><td></td><td></td><td></td><td></td><td></td><td></td><td></td><td></td><td></td><td></td></tr>
<tr><td>T11008</td><td>AACGGGACGAGAGCGAACGCGGG</td><td>1000</td><td></td><td></td><td></td><td></td><td></td><td></td><td></td><td></td><td></td></tr>
<tr><td>Oligo fwd</td><td>CACCAACGGGACGAGAGCGAACGC</td><td></td><td></td><td></td><td></td><td></td><td></td><td></td><td></td><td></td><td></td></tr>
<tr><td>Oligo rev</td><td>AAACGCGTTCGCTCTCGTCCCGTT</td><td></td><td></td><td></td><td></td><td></td><td></td><td></td><td></td><td></td><td></td></tr>
<tr><td>Chromosome</td><td>start</td><td>end</td><td>strand</td><td>MM</td><td>target_seq</td><td>PAM</td><td>alignment</td><td>distance</td><td>position</td><td>gene name</td><td>gene id</td></tr>
<tr><td>GU305914.1</td><td>143636</td><td>143658</td><td>+</td><td>0</td><td>AACGGGACGAGAGCGAACGC</td><td>GGG</td><td>||||||||[||||||||||||]PAM</td><td>NA</td><td>Intergenic</td><td>NA</td><td>NA</td></tr>
<tr><td></td><td></td><td></td><td></td><td></td><td></td><td></td><td></td><td></td><td></td><td></td><td></td></tr>
<tr><td>T11009</td><td>GGGAGAGTGATCAGAATCAACGG</td><td>1000</td><td></td><td></td><td></td><td></td><td></td><td></td><td></td><td></td><td></td></tr>
<tr><td>Oligo fwd</td><td>CACCGGGAGAGTGATCAGAATCAA</td><td></td><td></td><td></td><td></td><td></td><td></td><td></td><td></td><td></td><td></td></tr>
<tr><td>Oligo rev</td><td>AAACTTGATTCTGATCACTCTCCC</td><td></td><td></td><td></td><td></td><td></td><td></td><td></td><td></td><td></td><td></td></tr>
<tr><td>Chromosome</td><td>start</td><td>end</td><td>strand</td><td>MM</td><td>target_seq</td><td>PAM</td><td>alignment</td><td>distance</td><td>position</td><td>gene name</td><td>gene id</td></tr>
<tr><td>GU305914.1</td><td>143656</td><td>143678</td><td>+</td><td>0</td><td>GGGAGAGTGATCAGAATCAA</td><td>CGG</td><td>||||||||[||||||||||||]PAM</td><td>NA</td><td>Intergenic</td><td>NA</td><td>NA</td></tr>
<tr><td></td><td></td><td></td><td></td><td></td><td></td><td></td><td></td><td></td><td></td><td></td><td></td></tr>
<tr><td>T11010</td><td>GGAGAGTGATCAGAATCAACGGG</td><td>1000</td><td></td><td></td><td></td><td></td><td></td><td></td><td></td><td></td><td></td></tr>
<tr><td>Oligo fwd</td><td>CACCGGAGAGTGATCAGAATCAAC</td><td></td><td></td><td></td><td></td><td></td><td></td><td></td><td></td><td></td><td></td></tr>
<tr><td>Oligo rev</td><td>AAACGTTGATTCTGATCACTCTCC</td><td></td><td></td><td></td><td></td><td></td><td></td><td></td><td></td><td></td><td></td></tr>
<tr><td>Chromosome</td><td>start</td><td>end</td><td>strand</td><td>MM</td><td>target_seq</td><td>PAM</td><td>alignment</td><td>distance</td><td>position</td><td>gene name</td><td>gene id</td></tr>
<tr><td>GU305914.1</td><td>143657</td><td>143679</td><td>+</td><td>0</td><td>GGAGAGTGATCAGAATCAAC</td><td>GGG</td><td>||||||||[||||||||||||]PAM</td><td>NA</td><td>Intergenic</td><td>NA</td><td>NA</td></tr>
<tr><td></td><td></td><td></td><td></td><td></td><td></td><td></td><td></td><td></td><td></td><td></td><td></td></tr>
<tr><td>T11011</td><td>GCACCGTTCCGACTGCGACGAGG</td><td>1000</td><td></td><td></td><td></td><td></td><td></td><td></td><td></td><td></td><td></td></tr>
<tr><td>Oligo fwd</td><td>CACCGCACCGTTCCGACTGCGACG</td><td></td><td></td><td></td><td></td><td></td><td></td><td></td><td></td><td></td><td></td></tr>
<tr><td>Oligo rev</td><td>AAACCGTCGCAGTCGGAACGGTGC</td><td></td><td></td><td></td><td></td><td></td><td></td><td></td><td></td><td></td><td></td></tr>
<tr><td>Chromosome</td><td>start</td><td>end</td><td>strand</td><td>MM</td><td>target_seq</td><td>PAM</td><td>alignment</td><td>distance</td><td>position</td><td>gene name</td><td>gene id</td></tr>
<tr><td>GU305914.1</td><td>143679</td><td>143701</td><td>+</td><td>0</td><td>GCACCGTTCCGACTGCGACG</td><td>AGG</td><td>||||||||[||||||||||||]PAM</td><td>NA</td><td>Intergenic</td><td>NA</td><td>NA</td></tr>
<tr><td></td><td></td><td></td><td></td><td></td><td></td><td></td><td></td><td></td><td></td><td></td><td></td></tr>
<tr><td>T11012</td><td>CGTCGTGACGCCCGTGCGTCTGG</td><td>1000</td><td></td><td></td><td></td><td></td><td></td><td></td><td></td><td></td><td></td></tr>
<tr><td>Oligo fwd</td><td>CACCCGTCGTGACGCCCGTGCGTC</td><td></td><td></td><td></td><td></td><td></td><td></td><td></td><td></td><td></td><td></td></tr>
<tr><td>Oligo rev</td><td>AAACGACGCACGGGCGTCACGACG</td><td></td><td></td><td></td><td></td><td></td><td></td><td></td><td></td><td></td><td></td></tr>
<tr><td>Chromosome</td><td>start</td><td>end</td><td>strand</td><td>MM</td><td>target_seq</td><td>PAM</td><td>alignment</td><td>distance</td><td>position</td><td>gene name</td><td>gene id</td></tr>
<tr><td>GU305914.1</td><td>143715</td><td>143737</td><td>+</td><td>0</td><td>CGTCGTGACGCCCGTGCGTC</td><td>TGG</td><td>||||||||[||||||||||||]PAM</td><td>NA</td><td>Intergenic</td><td>NA</td><td>NA</td></tr>
<tr><td></td><td></td><td></td><td></td><td></td><td></td><td></td><td></td><td></td><td></td><td></td><td></td></tr>
<tr><td>T11013</td><td>GTCTGGATCCCCACTTCGCCCGG</td><td>1000</td><td></td><td></td><td></td><td></td><td></td><td></td><td></td><td></td><td></td></tr>
<tr><td>Oligo fwd</td><td>CACCGTCTGGATCCCCACTTCGCC</td><td></td><td></td><td></td><td></td><td></td><td></td><td></td><td></td><td></td><td></td></tr>
<tr><td>Oligo rev</td><td>AAACGGCGAAGTGGGGATCCAGAC</td><td></td><td></td><td></td><td></td><td></td><td></td><td></td><td></td><td></td><td></td></tr>
<tr><td>Chromosome</td><td>start</td><td>end</td><td>strand</td><td>MM</td><td>target_seq</td><td>PAM</td><td>alignment</td><td>distance</td><td>position</td><td>gene name</td><td>gene id</td></tr>
<tr><td>GU305914.1</td><td>143732</td><td>143754</td><td>+</td><td>0</td><td>GTCTGGATCCCCACTTCGCC</td><td>CGG</td><td>||||||||[||||||||||||]PAM</td><td>NA</td><td>Intergenic</td><td>NA</td><td>NA</td></tr>
<tr><td></td><td></td><td></td><td></td><td></td><td></td><td></td><td></td><td></td><td></td><td></td><td></td></tr>
<tr><td>T11014</td><td>TCTGGATCCCCACTTCGCCCGGG</td><td>1000</td><td></td><td></td><td></td><td></td><td></td><td></td><td></td><td></td><td></td></tr>
<tr><td>Oligo fwd</td><td>CACCTCTGGATCCCCACTTCGCCC</td><td></td><td></td><td></td><td></td><td></td><td></td><td></td><td></td><td></td><td></td></tr>
<tr><td>Oligo rev</td><td>AAACGGGCGAAGTGGGGATCCAGA</td><td></td><td></td><td></td><td></td><td></td><td></td><td></td><td></td><td></td><td></td></tr>
<tr><td>Chromosome</td><td>start</td><td>end</td><td>strand</td><td>MM</td><td>target_seq</td><td>PAM</td><td>alignment</td><td>distance</td><td>position</td><td>gene name</td><td>gene id</td></tr>
<tr><td>GU305914.1</td><td>143733</td><td>143755</td><td>+</td><td>0</td><td>TCTGGATCCCCACTTCGCCC</td><td>GGG</td><td>||||||||[||||||||||||]PAM</td><td>NA</td><td>Intergenic</td><td>NA</td><td>NA</td></tr>
<tr><td></td><td></td><td></td><td></td><td></td><td></td><td></td><td></td><td></td><td></td><td></td><td></td></tr>
<tr><td>T11015</td><td>TGAGCAGCGTCCTCGACGTCTGG</td><td>1000</td><td></td><td></td><td></td><td></td><td></td><td></td><td></td><td></td><td></td></tr>
<tr><td>Oligo fwd</td><td>CACCTGAGCAGCGTCCTCGACGTC</td><td></td><td></td><td></td><td></td><td></td><td></td><td></td><td></td><td></td><td></td></tr>
<tr><td>Oligo rev</td><td>AAACGACGTCGAGGACGCTGCTCA</td><td></td><td></td><td></td><td></td><td></td><td></td><td></td><td></td><td></td><td></td></tr>
<tr><td>Chromosome</td><td>start</td><td>end</td><td>strand</td><td>MM</td><td>target_seq</td><td>PAM</td><td>alignment</td><td>distance</td><td>position</td><td>gene name</td><td>gene id</td></tr>
<tr><td>GU305914.1</td><td>143762</td><td>143784</td><td>+</td><td>0</td><td>TGAGCAGCGTCCTCGACGTC</td><td>TGG</td><td>||||||||[||||||||||||]PAM</td><td>NA</td><td>Intergenic</td><td>NA</td><td>NA</td></tr>
<tr><td></td><td></td><td></td><td></td><td></td><td></td><td></td><td></td><td></td><td></td><td></td><td></td></tr>
<tr><td>T11016</td><td>GAGCAGCGTCCTCGACGTCTGGG</td><td>1000</td><td></td><td></td><td></td><td></td><td></td><td></td><td></td><td></td><td></td></tr>
<tr><td>Oligo fwd</td><td>CACCGAGCAGCGTCCTCGACGTCT</td><td></td><td></td><td></td><td></td><td></td><td></td><td></td><td></td><td></td><td></td></tr>
<tr><td>Oligo rev</td><td>AAACAGACGTCGAGGACGCTGCTC</td><td></td><td></td><td></td><td></td><td></td><td></td><td></td><td></td><td></td><td></td></tr>
<tr><td>Chromosome</td><td>start</td><td>end</td><td>strand</td><td>MM</td><td>target_seq</td><td>PAM</td><td>alignment</td><td>distance</td><td>position</td><td>gene name</td><td>gene id</td></tr>
<tr><td>GU305914.1</td><td>143763</td><td>143785</td><td>+</td><td>0</td><td>GAGCAGCGTCCTCGACGTCT</td><td>GGG</td><td>||||||||[||||||||||||]PAM</td><td>NA</td><td>Intergenic</td><td>NA</td><td>NA</td></tr>
<tr><td></td><td></td><td></td><td></td><td></td><td></td><td></td><td></td><td></td><td></td><td></td><td></td></tr>
<tr><td>T11017</td><td>CAGCGTCCTCGACGTCTGGGAGG</td><td>1000</td><td></td><td></td><td></td><td></td><td></td><td></td><td></td><td></td><td></td></tr>
<tr><td>Oligo fwd</td><td>CACCCAGCGTCCTCGACGTCTGGG</td><td></td><td></td><td></td><td></td><td></td><td></td><td></td><td></td><td></td><td></td></tr>
<tr><td>Oligo rev</td><td>AAACCCCAGACGTCGAGGACGCTG</td><td></td><td></td><td></td><td></td><td></td><td></td><td></td><td></td><td></td><td></td></tr>
<tr><td>Chromosome</td><td>start</td><td>end</td><td>strand</td><td>MM</td><td>target_seq</td><td>PAM</td><td>alignment</td><td>distance</td><td>position</td><td>gene name</td><td>gene id</td></tr>
<tr><td>GU305914.1</td><td>143766</td><td>143788</td><td>+</td><td>0</td><td>CAGCGTCCTCGACGTCTGGG</td><td>AGG</td><td>||||||||[||||||||||||]PAM</td><td>NA</td><td>Intergenic</td><td>NA</td><td>NA</td></tr>
<tr><td></td><td></td><td></td><td></td><td></td><td></td><td></td><td></td><td></td><td></td><td></td><td></td></tr>
<tr><td>T11018</td><td>AGCGTCCTCGACGTCTGGGAGGG</td><td>1000</td><td></td><td></td><td></td><td></td><td></td><td></td><td></td><td></td><td></td></tr>
<tr><td>Oligo fwd</td><td>CACCAGCGTCCTCGACGTCTGGGA</td><td></td><td></td><td></td><td></td><td></td><td></td><td></td><td></td><td></td><td></td></tr>
<tr><td>Oligo rev</td><td>AAACTCCCAGACGTCGAGGACGCT</td><td></td><td></td><td></td><td></td><td></td><td></td><td></td><td></td><td></td><td></td></tr>
<tr><td>Chromosome</td><td>start</td><td>end</td><td>strand</td><td>MM</td><td>target_seq</td><td>PAM</td><td>alignment</td><td>distance</td><td>position</td><td>gene name</td><td>gene id</td></tr>
<tr><td>GU305914.1</td><td>143767</td><td>143789</td><td>+</td><td>0</td><td>AGCGTCCTCGACGTCTGGGA</td><td>GGG</td><td>||||||||[||||||||||||]PAM</td><td>NA</td><td>Intergenic</td><td>NA</td><td>NA</td></tr>
<tr><td></td><td></td><td></td><td></td><td></td><td></td><td></td><td></td><td></td><td></td><td></td><td></td></tr>
<tr><td>T11019</td><td>GCGTCCTCGACGTCTGGGAGGGG</td><td>1000</td><td></td><td></td><td></td><td></td><td></td><td></td><td></td><td></td><td></td></tr>
<tr><td>Oligo fwd</td><td>CACCGCGTCCTCGACGTCTGGGAG</td><td></td><td></td><td></td><td></td><td></td><td></td><td></td><td></td><td></td><td></td></tr>
<tr><td>Oligo rev</td><td>AAACCTCCCAGACGTCGAGGACGC</td><td></td><td></td><td></td><td></td><td></td><td></td><td></td><td></td><td></td><td></td></tr>
<tr><td>Chromosome</td><td>start</td><td>end</td><td>strand</td><td>MM</td><td>target_seq</td><td>PAM</td><td>alignment</td><td>distance</td><td>position</td><td>gene name</td><td>gene id</td></tr>
<tr><td>GU305914.1</td><td>143768</td><td>143790</td><td>+</td><td>0</td><td>GCGTCCTCGACGTCTGGGAG</td><td>GGG</td><td>||||||||[||||||||||||]PAM</td><td>NA</td><td>Intergenic</td><td>NA</td><td>NA</td></tr>
</table>

T11020 CGTCCTCGACGTCTGGGAGGGGG 1000
Oligo fwd CACCCGTCCTCGACGTCTGGGAGG
Oligo rev AAACCCTCCCAGACGTCGAGGACG
Chromosome start end strand MM target_seq PAM alignment distance position gene name gene id
GU305914.1 143769 143791 + 0 CGTCCTCGACGTCTGGGAGG GGG ||||||||[||||||||||||]PAM NA Intergenic NA NA

T11021 GACGTCTGGGAGGGGGACATCGG 1000
Oligo fwd CACCGACGTCTGGGAGGGGGACAT
Oligo rev AAACATGTCCCCCTCCCAGACGTC
Chromosome start end strand MM target_seq PAM alignment distance position gene name gene id
GU305914.1 143776 143798 + 0 GACGTCTGGGAGGGGGACAT CGG ||||||||[||||||||||||]PAM NA Intergenic NA NA

T11022 GGGGGACATCGGTAAGAAGACGG 1000
Oligo fwd CACCGGGGGACATCGGTAAGAAGA
Oligo rev AAACTCTTCTTACCGATGTCCCCC
Chromosome start end strand MM target_seq PAM alignment distance position gene name gene id
GU305914.1 143787 143809 + 0 GGGGGACATCGGTAAGAAGA CGG ||||||||[||||||||||||]PAM NA Intergenic NA NA

T11023 GGGGACATCGGTAAGAAGACGGG 1000
Oligo fwd CACCGGGGACATCGGTAAGAAGAC
Oligo rev AAACGTCTTCTTACCGATGTCCCC
Chromosome start end strand MM target_seq PAM alignment distance position gene name gene id
GU305914.1 143788 143810 + 0 GGGGACATCGGTAAGAAGAC GGG ||||||||[||||||||||||]PAM NA Intergenic NA NA

T11024 AGAAGACGGGTCTCGCGCTATGG 1000
Oligo fwd CACCAGAAGACGGGTCTCGCGCTA
Oligo rev AAACTAGCGCGAGACCCGTCTTCT
Chromosome start end strand MM target_seq PAM alignment distance position gene name gene id
GU305914.1 143801 143823 + 0 AGAAGACGGGTCTCGCGCTA TGG ||||||||[||||||||||||]PAM NA Intergenic NA NA

T11025 GAAGACGGGTCTCGCGCTATGGG 1000
Oligo fwd CACCGAAGACGGGTCTCGCGCTAT
Oligo rev AAACATAGCGCGAGACCCGTCTTC
Chromosome start end strand MM target_seq PAM alignment distance position gene name gene id
GU305914.1 143802 143824 + 0 GAAGACGGGTCTCGCGCTAT GGG ||||||||[||||||||||||]PAM NA Intergenic NA NA

T11026 CAGCCTGCATCCCTACGCCGCGG 1000
Oligo fwd CACCCAGCCTGCATCCCTACGCCG
Oligo rev AAACCGGCGTAGGGATGCAGGCTG
Chromosome start end strand MM target_seq PAM alignment distance position gene name gene id
GU305914.1 143852 143874 + 0 CAGCCTGCATCCCTACGCCG CGG ||||||||[||||||||||||]PAM NA Intergenic NA NA

T11027 CTACGCCGCGGACTCCCTGAAGG 1000
Oligo fwd CACCCTACGCCGCGGACTCCCTGA
Oligo rev AAACTCAGGGAGTCCGCGGCGTAG
Chromosome start end strand MM target_seq PAM alignment distance position gene name gene id
GU305914.1 143864 143886 + 0 CTACGCCGCGGACTCCCTGA AGG ||||||||[||||||||||||]PAM NA Intergenic NA NA

T11028 GGACTCCCTGAAGGATTCCTCGG 1000
Oligo fwd CACCGGACTCCCTGAAGGATTCCT
Oligo rev AAACAGGAATCCTTCAGGGAGTCC
Chromosome start end strand MM target_seq PAM alignment distance position gene name gene id
GU305914.1 143873 143895 + 0 GGACTCCCTGAAGGATTCCT CGG ||||||||[||||||||||||]PAM NA Intergenic NA NA

T11029 GACTCCCTGAAGGATTCCTCGGG 1000
Oligo fwd CACCGACTCCCTGAAGGATTCCTC
Oligo rev AAACGAGGAATCCTTCAGGGAGTC
Chromosome start end strand MM target_seq PAM alignment distance position gene name gene id
GU305914.1 143874 143896 + 0 GACTCCCTGAAGGATTCCTC GGG ||||||||[||||||||||||]PAM NA Intergenic NA NA

T11030 ACTCCCTGAAGGATTCCTCGGGG 1000
Oligo fwd CACCACTCCCTGAAGGATTCCTCG
Oligo rev AAACCGAGGAATCCTTCAGGGAGT
Chromosome start end strand MM target_seq PAM alignment distance position gene name gene id
GU305914.1 143875 143897 + 0 ACTCCCTGAAGGATTCCTCG GGG ||||||||[||||||||||||]PAM NA Intergenic NA NA

T11031 GGATTCCTCGGGGCGACGCGTGG 1000
Oligo fwd CACCGGATTCCTCGGGGCGACGCG
Oligo rev AAACCGCGTCGCCCCGAGGAATCC
Chromosome start end strand MM target_seq PAM alignment distance position gene name gene id
GU305914.1 143885 143907 + 0 GGATTCCTCGGGGCGACGCG TGG ||||||||[||||||||||||]PAM NA Intergenic NA NA

T11032 TCGGGGCGACGCGTGGACCTCGG 1000
Oligo fwd CACCTCGGGGCGACGCGTGGACCT
Oligo rev AAACAGGTCCACGCGTCGCCCCGA
Chromosome start end strand MM target_seq PAM alignment distance position gene name gene id
GU305914.1 143892 143914 + 0 TCGGGGCGACGCGTGGACCT CGG ||||||||[||||||||||||]PAM NA Intergenic NA NA

T11033 CGGGGCGACGCGTGGACCTCGGG 1000
Oligo fwd CACCCGGGGCGACGCGTGGACCTC
Oligo rev AAACGAGGTCCACGCGTCGCCCCG
Chromosome start end strand MM target_seq PAM alignment distance position gene name gene id
GU305914.1 143893 143915 + 0 CGGGGCGACGCGTGGACCTC GGG ||||||||[||||||||||||]PAM NA Intergenic NA NA

T11034 GACCGAATTCTCTGTGCTGACGG 1000
Oligo fwd CACCGACCGAATTCTCTGTGCTGA
Oligo rev AAACTCAGCACAGAGAATTCGGTC
Chromosome start end strand MM target_seq PAM alignment distance position gene name gene id
GU305914.1 143915 143937 + 0 GACCGAATTCTCTGTGCTGA CGG ||||||||[||||||||||||]PAM NA Intergenic NA NA

<table>
<tr><td>T11035</td><td>TACGTCTGACGACGAAGACCAGG</td><td>1000</td><td></td><td></td><td></td><td></td><td></td><td></td><td></td><td></td><td></td></tr>
<tr><td>Oligo fwd</td><td>CACCTACGTCTGACGACGAAGACC</td><td></td><td></td><td></td><td></td><td></td><td></td><td></td><td></td><td></td><td></td></tr>
<tr><td>Oligo rev</td><td>AAACGGTCTTCGTCGTCAGACGTA</td><td></td><td></td><td></td><td></td><td></td><td></td><td></td><td></td><td></td><td></td></tr>
<tr><td>Chromosome</td><td>start</td><td>end</td><td>strand</td><td>MM</td><td>target_seq</td><td>PAM</td><td>alignment</td><td>distance</td><td>position</td><td>gene name</td><td>gene id</td></tr>
<tr><td>GU305914.1</td><td>143939</td><td>143961</td><td>+</td><td>0</td><td>TACGTCTGACGACGAAGACC</td><td>AGG</td><td>||||||||[||||||||||||]PAM</td><td>NA</td><td>Intergenic</td><td>NA</td><td>NA</td></tr>
<tr><td>T11036</td><td>TCTGACGACGAAGACCAGGTCGG</td><td>1000</td><td></td><td></td><td></td><td></td><td></td><td></td><td></td><td></td><td></td></tr>
<tr><td>Oligo fwd</td><td>CACCTCTGACGACGAAGACCAGGT</td><td></td><td></td><td></td><td></td><td></td><td></td><td></td><td></td><td></td><td></td></tr>
<tr><td>Oligo rev</td><td>AAACACCTGGTCTTCGTCGTCAGA</td><td></td><td></td><td></td><td></td><td></td><td></td><td></td><td></td><td></td><td></td></tr>
<tr><td>Chromosome</td><td>start</td><td>end</td><td>strand</td><td>MM</td><td>target_seq</td><td>PAM</td><td>alignment</td><td>distance</td><td>position</td><td>gene name</td><td>gene id</td></tr>
<tr><td>GU305914.1</td><td>143943</td><td>143965</td><td>+</td><td>0</td><td>TCTGACGACGAAGACCAGGT</td><td>CGG</td><td>||||||||[||||||||||||]PAM</td><td>NA</td><td>Intergenic</td><td>NA</td><td>NA</td></tr>
<tr><td>T11037</td><td>GACGAAGACCAGGTCGGCGCCGG</td><td>1000</td><td></td><td></td><td></td><td></td><td></td><td></td><td></td><td></td><td></td></tr>
<tr><td>Oligo fwd</td><td>CACCGACGAAGACCAGGTCGGCGC</td><td></td><td></td><td></td><td></td><td></td><td></td><td></td><td></td><td></td><td></td></tr>
<tr><td>Oligo rev</td><td>AAACGCGCCGACCTGGTCTTCGTC</td><td></td><td></td><td></td><td></td><td></td><td></td><td></td><td></td><td></td><td></td></tr>
<tr><td>Chromosome</td><td>start</td><td>end</td><td>strand</td><td>MM</td><td>target_seq</td><td>PAM</td><td>alignment</td><td>distance</td><td>position</td><td>gene name</td><td>gene id</td></tr>
<tr><td>GU305914.1</td><td>143949</td><td>143971</td><td>+</td><td>0</td><td>GACGAAGACCAGGTCGGCGC</td><td>CGG</td><td>||||||||[||||||||||||]PAM</td><td>NA</td><td>Intergenic</td><td>NA</td><td>NA</td></tr>
<tr><td>T11038</td><td>TAGCGCTATACCCCTGACCGTGG</td><td>1000</td><td></td><td></td><td></td><td></td><td></td><td></td><td></td><td></td><td></td></tr>
<tr><td>Oligo fwd</td><td>CACCTAGCGCTATACCCCTGACCG</td><td></td><td></td><td></td><td></td><td></td><td></td><td></td><td></td><td></td><td></td></tr>
<tr><td>Oligo rev</td><td>AAACCGGTCAGGGGTATAGCGCTA</td><td></td><td></td><td></td><td></td><td></td><td></td><td></td><td></td><td></td><td></td></tr>
<tr><td>Chromosome</td><td>start</td><td>end</td><td>strand</td><td>MM</td><td>target_seq</td><td>PAM</td><td>alignment</td><td>distance</td><td>position</td><td>gene name</td><td>gene id</td></tr>
<tr><td>GU305914.1</td><td>144029</td><td>144051</td><td>+</td><td>0</td><td>TAGCGCTATACCCCTGACCG</td><td>TGG</td><td>||||||||[||||||||||||]PAM</td><td>NA</td><td>Intergenic</td><td>NA</td><td>NA</td></tr>
<tr><td>T11039</td><td>GTGGTCACTCCGATACAGCTCGG</td><td>1000</td><td></td><td></td><td></td><td></td><td></td><td></td><td></td><td></td><td></td></tr>
<tr><td>Oligo fwd</td><td>CACCGTGGTCACTCCGATACAGCT</td><td></td><td></td><td></td><td></td><td></td><td></td><td></td><td></td><td></td><td></td></tr>
<tr><td>Oligo rev</td><td>AAACAGCTGTATCGGAGTGACCAC</td><td></td><td></td><td></td><td></td><td></td><td></td><td></td><td></td><td></td><td></td></tr>
<tr><td>Chromosome</td><td>start</td><td>end</td><td>strand</td><td>MM</td><td>target_seq</td><td>PAM</td><td>alignment</td><td>distance</td><td>position</td><td>gene name</td><td>gene id</td></tr>
<tr><td>GU305914.1</td><td>144048</td><td>144070</td><td>+</td><td>0</td><td>GTGGTCACTCCGATACAGCT</td><td>CGG</td><td>||||||||[||||||||||||]PAM</td><td>NA</td><td>Intergenic</td><td>NA</td><td>NA</td></tr>
<tr><td>T11040</td><td>TGGTCACTCCGATACAGCTCGGG</td><td>1000</td><td></td><td></td><td></td><td></td><td></td><td></td><td></td><td></td><td></td></tr>
<tr><td>Oligo fwd</td><td>CACCTGGTCACTCCGATACAGCTC</td><td></td><td></td><td></td><td></td><td></td><td></td><td></td><td></td><td></td><td></td></tr>
<tr><td>Oligo rev</td><td>AAACGAGCTGTATCGGAGTGACCA</td><td></td><td></td><td></td><td></td><td></td><td></td><td></td><td></td><td></td><td></td></tr>
<tr><td>Chromosome</td><td>start</td><td>end</td><td>strand</td><td>MM</td><td>target_seq</td><td>PAM</td><td>alignment</td><td>distance</td><td>position</td><td>gene name</td><td>gene id</td></tr>
<tr><td>GU305914.1</td><td>144049</td><td>144071</td><td>+</td><td>0</td><td>TGGTCACTCCGATACAGCTC</td><td>GGG</td><td>||||||||[||||||||||||]PAM</td><td>NA</td><td>Intergenic</td><td>NA</td><td>NA</td></tr>
<tr><td>T11041</td><td>GGTCACTCCGATACAGCTCGGGG</td><td>1000</td><td></td><td></td><td></td><td></td><td></td><td></td><td></td><td></td><td></td></tr>
<tr><td>Oligo fwd</td><td>CACCGGTCACTCCGATACAGCTCG</td><td></td><td></td><td></td><td></td><td></td><td></td><td></td><td></td><td></td><td></td></tr>
<tr><td>Oligo rev</td><td>AAACCGAGCTGTATCGGAGTGACC</td><td></td><td></td><td></td><td></td><td></td><td></td><td></td><td></td><td></td><td></td></tr>
<tr><td>Chromosome</td><td>start</td><td>end</td><td>strand</td><td>MM</td><td>target_seq</td><td>PAM</td><td>alignment</td><td>distance</td><td>position</td><td>gene name</td><td>gene id</td></tr>
<tr><td>GU305914.1</td><td>144050</td><td>144072</td><td>+</td><td>0</td><td>GGTCACTCCGATACAGCTCG</td><td>GGG</td><td>||||||||[||||||||||||]PAM</td><td>NA</td><td>Intergenic</td><td>NA</td><td>NA</td></tr>
<tr><td>T11042</td><td>GATACAGCTCGGGGCGAACAAGG</td><td>1000</td><td></td><td></td><td></td><td></td><td></td><td></td><td></td><td></td><td></td></tr>
<tr><td>Oligo fwd</td><td>CACCGATACAGCTCGGGGCGAACA</td><td></td><td></td><td></td><td></td><td></td><td></td><td></td><td></td><td></td><td></td></tr>
<tr><td>Oligo rev</td><td>AAACTGTTCGCCCCGAGCTGTATC</td><td></td><td></td><td></td><td></td><td></td><td></td><td></td><td></td><td></td><td></td></tr>
<tr><td>Chromosome</td><td>start</td><td>end</td><td>strand</td><td>MM</td><td>target_seq</td><td>PAM</td><td>alignment</td><td>distance</td><td>position</td><td>gene name</td><td>gene id</td></tr>
<tr><td>GU305914.1</td><td>144059</td><td>144081</td><td>+</td><td>0</td><td>GATACAGCTCGGGGCGAACA</td><td>AGG</td><td>||||||||[||||||||||||]PAM</td><td>NA</td><td>Intergenic</td><td>NA</td><td>NA</td></tr>
<tr><td>T11043</td><td>ATACAGCTCGGGGCGAACAAGGG</td><td>1000</td><td></td><td></td><td></td><td></td><td></td><td></td><td></td><td></td><td></td></tr>
<tr><td>Oligo fwd</td><td>CACCATACAGCTCGGGGCGAACAA</td><td></td><td></td><td></td><td></td><td></td><td></td><td></td><td></td><td></td><td></td></tr>
<tr><td>Oligo rev</td><td>AAACTTGTTCGCCCCGAGCTGTAT</td><td></td><td></td><td></td><td></td><td></td><td></td><td></td><td></td><td></td><td></td></tr>
<tr><td>Chromosome</td><td>start</td><td>end</td><td>strand</td><td>MM</td><td>target_seq</td><td>PAM</td><td>alignment</td><td>distance</td><td>position</td><td>gene name</td><td>gene id</td></tr>
<tr><td>GU305914.1</td><td>144060</td><td>144082</td><td>+</td><td>0</td><td>ATACAGCTCGGGGCGAACAA</td><td>GGG</td><td>||||||||[||||||||||||]PAM</td><td>NA</td><td>Intergenic</td><td>NA</td><td>NA</td></tr>
<tr><td>T11044</td><td>CAGCTCGGGGCGAACAAGGGAGG</td><td>1000</td><td></td><td></td><td></td><td></td><td></td><td></td><td></td><td></td><td></td></tr>
<tr><td>Oligo fwd</td><td>CACCCAGCTCGGGGCGAACAAGGG</td><td></td><td></td><td></td><td></td><td></td><td></td><td></td><td></td><td></td><td></td></tr>
<tr><td>Oligo rev</td><td>AAACCCCTTGTTCGCCCCGAGCTG</td><td></td><td></td><td></td><td></td><td></td><td></td><td></td><td></td><td></td><td></td></tr>
<tr><td>Chromosome</td><td>start</td><td>end</td><td>strand</td><td>MM</td><td>target_seq</td><td>PAM</td><td>alignment</td><td>distance</td><td>position</td><td>gene name</td><td>gene id</td></tr>
<tr><td>GU305914.1</td><td>144063</td><td>144085</td><td>+</td><td>0</td><td>CAGCTCGGGGCGAACAAGGG</td><td>AGG</td><td>||||||||[||||||||||||]PAM</td><td>NA</td><td>Intergenic</td><td>NA</td><td>NA</td></tr>
<tr><td>T11045</td><td>CTCGGGGCGAACAAGGGAGGCGG</td><td>1000</td><td></td><td></td><td></td><td></td><td></td><td></td><td></td><td></td><td></td></tr>
<tr><td>Oligo fwd</td><td>CACCCTCGGGGCGAACAAGGGAGG</td><td></td><td></td><td></td><td></td><td></td><td></td><td></td><td></td><td></td><td></td></tr>
<tr><td>Oligo rev</td><td>AAACCCTCCCTTGTTCGCCCCGAG</td><td></td><td></td><td></td><td></td><td></td><td></td><td></td><td></td><td></td><td></td></tr>
<tr><td>Chromosome</td><td>start</td><td>end</td><td>strand</td><td>MM</td><td>target_seq</td><td>PAM</td><td>alignment</td><td>distance</td><td>position</td><td>gene name</td><td>gene id</td></tr>
<tr><td>GU305914.1</td><td>144066</td><td>144088</td><td>+</td><td>0</td><td>CTCGGGGCGAACAAGGGAGG</td><td>CGG</td><td>||||||||[||||||||||||]PAM</td><td>NA</td><td>Intergenic</td><td>NA</td><td>NA</td></tr>
<tr><td>T11046</td><td>CGAACAAGGGAGGCGGAAAGCGG</td><td>1000</td><td></td><td></td><td></td><td></td><td></td><td></td><td></td><td></td><td></td></tr>
<tr><td>Oligo fwd</td><td>CACCCGAACAAGGGAGGCGGAAAG</td><td></td><td></td><td></td><td></td><td></td><td></td><td></td><td></td><td></td><td></td></tr>
<tr><td>Oligo rev</td><td>AAACCTTTCCGCCTCCCTTGTTCG</td><td></td><td></td><td></td><td></td><td></td><td></td><td></td><td></td><td></td><td></td></tr>
<tr><td>Chromosome</td><td>start</td><td>end</td><td>strand</td><td>MM</td><td>target_seq</td><td>PAM</td><td>alignment</td><td>distance</td><td>position</td><td>gene name</td><td>gene id</td></tr>
<tr><td>GU305914.1</td><td>144073</td><td>144095</td><td>+</td><td>0</td><td>CGAACAAGGGAGGCGGAAAG</td><td>CGG</td><td>||||||||[||||||||||||]PAM</td><td>NA</td><td>Intergenic</td><td>NA</td><td>NA</td></tr>
<tr><td>T11047</td><td>TCTTTGAAATCCGCCAAGAACGG</td><td>1000</td><td></td><td></td><td></td><td></td><td></td><td></td><td></td><td></td><td></td></tr>
<tr><td>Oligo fwd</td><td>CACCTCTTTGAAATCCGCCAAGAA</td><td></td><td></td><td></td><td></td><td></td><td></td><td></td><td></td><td></td><td></td></tr>
<tr><td>Oligo rev</td><td>AAACTTCTTGGCGGATTTCAAAGA</td><td></td><td></td><td></td><td></td><td></td><td></td><td></td><td></td><td></td><td></td></tr>
<tr><td>Chromosome</td><td>start</td><td>end</td><td>strand</td><td>MM</td><td>target_seq</td><td>PAM</td><td>alignment</td><td>distance</td><td>position</td><td>gene name</td><td>gene id</td></tr>
<tr><td>GU305914.1</td><td>144102</td><td>144124</td><td>+</td><td>0</td><td>TCTTTGAAATCCGCCAAGAA</td><td>CGG</td><td>||||||||[||||||||||||]PAM</td><td>NA</td><td>Intergenic</td><td>NA</td><td>NA</td></tr>
<tr><td>T11048</td><td>AAATCCGCCAAGAACGGCGCCGG</td><td>1000</td><td></td><td></td><td></td><td></td><td></td><td></td><td></td><td></td><td></td></tr>
<tr><td>Oligo fwd</td><td>CACCAAATCCGCCAAGAACGGCGC</td><td></td><td></td><td></td><td></td><td></td><td></td><td></td><td></td><td></td><td></td></tr>
<tr><td>Oligo rev</td><td>AAACGCGCCGTTCTTGGCGGATTT</td><td></td><td></td><td></td><td></td><td></td><td></td><td></td><td></td><td></td><td></td></tr>
<tr><td>Chromosome</td><td>start</td><td>end</td><td>strand</td><td>MM</td><td>target_seq</td><td>PAM</td><td>alignment</td><td>distance</td><td>position</td><td>gene name</td><td>gene id</td></tr>
<tr><td>GU305914.1</td><td>144108</td><td>144130</td><td>+</td><td>0</td><td>AAATCCGCCAAGAACGGCGC</td><td>CGG</td><td>||||||||[||||||||||||]PAM</td><td>NA</td><td>Intergenic</td><td>NA</td><td>NA</td></tr>
<tr><td>T11049</td><td>CCGGCGTGAAGAAAAAAGTCAGG</td><td>1000</td><td></td><td></td><td></td><td></td><td></td><td></td><td></td><td></td><td></td></tr>
<tr><td>Oligo fwd</td><td>CACCCCGGCGTGAAGAAAAAAGTC</td><td></td><td></td><td></td><td></td><td></td><td></td><td></td><td></td><td></td><td></td></tr>
<tr><td>Oligo rev</td><td>AAACGACTTTTTTCTTCACGCCGG</td><td></td><td></td><td></td><td></td><td></td><td></td><td></td><td></td><td></td><td></td></tr>
<tr><td>Chromosome</td><td>start</td><td>end</td><td>strand</td><td>MM</td><td>target_seq</td><td>PAM</td><td>alignment</td><td>distance</td><td>position</td><td>gene name</td><td>gene id</td></tr>
</table>

GU305914.1 144127 144149 + 0 CCGGCGTGAAGAAAAAAGTC AGG ||||||||[||||||||||||]PAM NA Intergenic NA NA

T11050 CGGCGTGAAGAAAAAAGTCAGGG 1000
Oligo fwd CACCGGCGTGAAGAAAAAAGTCA
Oligo rev AAACTGACTTTTTTCTTCACGCCG
Chromosome start end strand MM target_seq PAM alignment distance position gene name gene id
GU305914.1 144128 144150 + 0 CGGCGTGAAGAAAAAAGTCA GGG ||||||||[||||||||||||]PAM NA Intergenic NA NA

T11051 CTATCTATCGATCGACCGTGAGG 1000
Oligo fwd CACCCTATCTATCGATCGACCGTG
Oligo rev AAACCACGGTCGATCGATAGATAG
Chromosome start end strand MM target_seq PAM alignment distance position gene name gene id
GU305914.1 144162 144184 + 0 CTATCTATCGATCGACCGTG AGG ||||||||[||||||||||||]PAM NA Intergenic NA NA

T11052 TATCTATCGATCGACCGTGAGGG 1000
Oligo fwd CACCTATCTATCGATCGACCGTGA
Oligo rev AAACTCACGGTCGATCGATAGATA
Chromosome start end strand MM target_seq PAM alignment distance position gene name gene id
GU305914.1 144163 144185 + 0 TATCTATCGATCGACCGTGA GGG ||||||||[||||||||||||]PAM NA Intergenic NA NA

T11053 ATCTATCGATCGACCGTGAGGGG 1000
Oligo fwd CACCATCTATCGATCGACCGTGAG
Oligo rev AAACCTCACGGTCGATCGATAGAT
Chromosome start end strand MM target_seq PAM alignment distance position gene name gene id
GU305914.1 144164 144186 + 0 ATCTATCGATCGACCGTGAG GGG ||||||||[||||||||||||]PAM NA Intergenic NA NA

T11054 TCTATCGATCGACCGTGAGGGGG 1000
Oligo fwd CACCTCTATCGATCGACCGTGAGG
Oligo rev AAACCCTCACGGTCGATCGATAGA
Chromosome start end strand MM target_seq PAM alignment distance position gene name gene id
GU305914.1 144165 144187 + 0 TCTATCGATCGACCGTGAGG GGG ||||||||[||||||||||||]PAM NA Intergenic NA NA

T11055 CTATCGATCGACCGTGAGGGGGG 1000
Oligo fwd CACCCTATCGATCGACCGTGAGGG
Oligo rev AAACCCCTCACGGTCGATCGATAG
Chromosome start end strand MM target_seq PAM alignment distance position gene name gene id
GU305914.1 144166 144188 + 0 CTATCGATCGACCGTGAGGG GGG ||||||||[||||||||||||]PAM NA Intergenic NA NA

T11056 TATCGATCGACCGTGAGGGGGGG 1000
Oligo fwd CACCTATCGATCGACCGTGAGGGG
Oligo rev AAACCCCCTCACGGTCGATCGATA
Chromosome start end strand MM target_seq PAM alignment distance position gene name gene id
GU305914.1 144167 144189 + 0 TATCGATCGACCGTGAGGGG GGG ||||||||[||||||||||||]PAM NA Intergenic NA NA

T11057 CGACCGTGAGGGGGGGAGAAAGG 1000
Oligo fwd CACCCGACCGTGAGGGGGGGAGAA
Oligo rev AAACTTCTCCCCCCCTCACGGTCG
Chromosome start end strand MM target_seq PAM alignment distance position gene name gene id
GU305914.1 144174 144196 + 0 CGACCGTGAGGGGGGGAGAA AGG ||||||||[||||||||||||]PAM NA Intergenic NA NA

T11058 GACCGTGAGGGGGGGAGAAAGGG 1000
Oligo fwd CACCGACCGTGAGGGGGGGAGAAA
Oligo rev AAACTTTCTCCCCCCCTCACGGTC
Chromosome start end strand MM target_seq PAM alignment distance position gene name gene id
GU305914.1 144175 144197 + 0 GACCGTGAGGGGGGGAGAAA GGG ||||||||[||||||||||||]PAM NA Intergenic NA NA

T11059 GGGGAGAAAGGGAGAAAGAGTGG 1000
Oligo fwd CACCGGGGAGAAAGGGAGAAAGAG
Oligo rev AAACCTCTTTCTCCCTTTCTCCCC
Chromosome start end strand MM target_seq PAM alignment distance position gene name gene id
GU305914.1 144186 144208 + 0 GGGGAGAAAGGGAGAAAGAG TGG ||||||||[||||||||||||]PAM NA Intergenic NA NA

T11060 GAGAAAGGGAGAAAGAGTGGAGG 1000
Oligo fwd CACCGAGAAAGGGAGAAAGAGTGG
Oligo rev AAACCCACTCTTTCTCCCTTTCTC
Chromosome start end strand MM target_seq PAM alignment distance position gene name gene id
GU305914.1 144189 144211 + 0 GAGAAAGGGAGAAAGAGTGG AGG ||||||||[||||||||||||]PAM NA Intergenic NA NA

T11061 ATTTATTAAAAAAAACGACATGG 1000
Oligo fwd CACCATTTATTAAAAAAAACGACA
Oligo rev AAACTGTCGTTTTTTTTAATAAAT
Chromosome start end strand MM target_seq PAM alignment distance position gene name gene id
GU305914.1 144214 144236 + 0 ATTTATTAAAAAAAACGACA TGG ||||||||[||||||||||||]PAM NA Intergenic NA NA

T11062 TGATTGATCTTTATTATAAATGG 1000
Oligo fwd CACCTGATTGATCTTTATTATAAA
Oligo rev AAACTTTATAATAAAGATCAATCA
Chromosome start end strand MM target_seq PAM alignment distance position gene name gene id
GU305914.1 144276 144298 + 0 TGATTGATCTTTATTATAAA TGG ||||||||[||||||||||||]PAM NA Intergenic NA NA

T11063 CTTTATTATAAATGGCATCTAGG 1000
Oligo fwd CACCCTTTATTATAAATGGCATCT
Oligo rev AAACAGATGCCATTTATAATAAAG
Chromosome start end strand MM target_seq PAM alignment distance position gene name gene id
GU305914.1 144284 144306 + 0 CTTTATTATAAATGGCATCT AGG ||||||||[||||||||||||]PAM NA Intergenic NA NA

T11064 TTTATTATAAATGGCATCTAGGG 1000
Oligo fwd CACCTTTATTATAAATGGCATCTA
Oligo rev AAACTAGATGCCATTTATAATAAA

Chromosome start end strand MM target_seq PAM alignment distance position gene name gene id
GU305914.1 144285 144307 + 0 TTTATTATAAATGGCATCTA GGG ||||||||[||||||||||||]PAM NA Intergenic NA NA

T11065 TTATTATAAATGGCATCTAGGGG 1000
Oligo fwd CACCTTATTATAAATGGCATCTAG
Oligo rev AAACCTAGATGCCATTTATAATAA
Chromosome start end strand MM target_seq PAM alignment distance position gene name gene id
GU305914.1 144286 144308 + 0 TTATTATAAATGGCATCTAG GGG ||||||||[||||||||||||]PAM NA Intergenic NA NA

T11066 AAATGGCATCTAGGGGAAAAAGG 1000
Oligo fwd CACCAAATGGCATCTAGGGGAAAA
Oligo rev AAACTTTTCCCCTAGATGCCATTT
Chromosome start end strand MM target_seq PAM alignment distance position gene name gene id
GU305914.1 144293 144315 + 0 AAATGGCATCTAGGGGAAAA AGG ||||||||[||||||||||||]PAM NA Intergenic NA NA

T11067 AATGGCATCTAGGGGAAAAAGGG 1000
Oligo fwd CACCAATGGCATCTAGGGGAAAAA
Oligo rev AAACTTTTTCCCCTAGATGCCATT
Chromosome start end strand MM target_seq PAM alignment distance position gene name gene id
GU305914.1 144294 144316 + 0 AATGGCATCTAGGGGAAAAA GGG ||||||||[||||||||||||]PAM NA Intergenic NA NA

T11068 ATGGCATCTAGGGGAAAAAGGGG 1000
Oligo fwd CACCATGGCATCTAGGGGAAAAAG
Oligo rev AAACCTTTTTCCCCTAGATGCCAT
Chromosome start end strand MM target_seq PAM alignment distance position gene name gene id
GU305914.1 144295 144317 + 0 ATGGCATCTAGGGGAAAAAG GGG ||||||||[||||||||||||]PAM NA Intergenic NA NA

T11069 CTCACTCTGCGTCGCTTCGCAGG 1000
Oligo fwd CACCCTCACTCTGCGTCGCTTCGC
Oligo rev AAACGCGAAGCGACGCAGAGTGAG
Chromosome start end strand MM target_seq PAM alignment distance position gene name gene id
GU305914.1 144392 144414 + 0 CTCACTCTGCGTCGCTTCGC AGG ||||||||[||||||||||||]PAM NA Intergenic NA NA

T11070 TCACTCTGCGTCGCTTCGCAGGG 1000
Oligo fwd CACCTCACTCTGCGTCGCTTCGCA
Oligo rev AAACTGCGAAGCGACGCAGAGTGA
Chromosome start end strand MM target_seq PAM alignment distance position gene name gene id
GU305914.1 144393 144415 + 0 TCACTCTGCGTCGCTTCGCA GGG ||||||||[||||||||||||]PAM NA Intergenic NA NA

T11071 CGCGTATCTCTTCGTTGTCCAGG 1000
Oligo fwd CACCCGCGTATCTCTTCGTTGTCC
Oligo rev AAACGGACAACGAAGAGATACGCG
Chromosome start end strand MM target_seq PAM alignment distance position gene name gene id
GU305914.1 144416 144438 + 0 CGCGTATCTCTTCGTTGTCC AGG ||||||||[||||||||||||]PAM NA Intergenic NA NA

T11072 GCGTATCTCTTCGTTGTCCAGGG 1000
Oligo fwd CACCGCGTATCTCTTCGTTGTCCA
Oligo rev AAACTGGACAACGAAGAGATACGC
Chromosome start end strand MM target_seq PAM alignment distance position gene name gene id
GU305914.1 144417 144439 + 0 GCGTATCTCTTCGTTGTCCA GGG ||||||||[||||||||||||]PAM NA Intergenic NA NA

T11073 TCTCTTCGTTGTCCAGGGCCAGG 1000
Oligo fwd CACCTCTCTTCGTTGTCCAGGGCC
Oligo rev AAACGGCCCTGGACAACGAAGAGA
Chromosome start end strand MM target_seq PAM alignment distance position gene name gene id
GU305914.1 144422 144444 + 0 TCTCTTCGTTGTCCAGGGCC AGG ||||||||[||||||||||||]PAM NA Intergenic NA NA

T11074 CTCTTCGTTGTCCAGGGCCAGGG 1000
Oligo fwd CACCCTCTTCGTTGTCCAGGGCCA
Oligo rev AAACTGGCCCTGGACAACGAAGAG
Chromosome start end strand MM target_seq PAM alignment distance position gene name gene id
GU305914.1 144423 144445 + 0 CTCTTCGTTGTCCAGGGCCA GGG ||||||||[||||||||||||]PAM NA Intergenic NA NA

T11075 GGGCTCGATATCGCTTGTTGCGG 1000
Oligo fwd CACCGGGCTCGATATCGCTTGTTG
Oligo rev AAACCAACAAGCGATATCGAGCCC
Chromosome start end strand MM target_seq PAM alignment distance position gene name gene id
GU305914.1 144443 144465 + 0 GGGCTCGATATCGCTTGTTG CGG ||||||||[||||||||||||]PAM NA Intergenic NA NA

T11076 GCAGCAAGAACCTCACCGTACGG 1000
Oligo fwd CACCGCAGCAAGAACCTCACCGTA
Oligo rev AAACTACGGTGAGGTTCTTGCTGC
Chromosome start end strand MM target_seq PAM alignment distance position gene name gene id
GU305914.1 144467 144489 + 0 GCAGCAAGAACCTCACCGTA CGG ||||||||[||||||||||||]PAM NA Intergenic NA NA

T11077 GATGTCCCTCGCGTAGCTCGAGG 1000
Oligo fwd CACCGATGTCCCTCGCGTAGCTCG
Oligo rev AAACCGAGCTACGCGAGGGACATC
Chromosome start end strand MM target_seq PAM alignment distance position gene name gene id
GU305914.1 144540 144562 + 0 GATGTCCCTCGCGTAGCTCG AGG ||||||||[||||||||||||]PAM NA Intergenic NA NA

T11078 AGGCGTTCAGCGCCTCGTAGCGG 1000
Oligo fwd CACCAGGCGTTCAGCGCCTCGTAG
Oligo rev AAACCTACGAGGCGCTGAACGCCT
Chromosome start end strand MM target_seq PAM alignment distance position gene name gene id
GU305914.1 144560 144582 + 0 AGGCGTTCAGCGCCTCGTAG CGG ||||||||[||||||||||||]PAM NA Intergenic NA NA

T11079 CAGCGCCTCGTAGCGGATGATGG 1000
Oligo fwd CACCCAGCGCCTCGTAGCGGATGA

<table>
<tr><td>Oligo rev</td><td>AAACTCATCCGCTACGAGGCGCTG</td><td></td><td></td><td></td><td></td><td></td><td></td><td></td><td></td><td></td><td></td></tr>
<tr><td>Chromosome</td><td>start</td><td>end</td><td>strand</td><td>MM</td><td>target_seq</td><td>PAM</td><td>alignment</td><td>distance</td><td>position</td><td>gene name</td><td>gene id</td></tr>
<tr><td>GU305914.1</td><td>144567</td><td>144589</td><td>+</td><td>0</td><td>CAGCGCCTCGTAGCGGATGA</td><td>TGG</td><td>||||||||[||||||||||||]PAM</td><td>NA</td><td>Intergenic</td><td>NA</td><td>NA</td></tr>
<tr><td>T11080</td><td>AGCGCCTCGTAGCGGATGATGGG</td><td>1000</td><td></td><td></td><td></td><td></td><td></td><td></td><td></td><td></td><td></td></tr>
<tr><td>Oligo fwd</td><td>CACCAGCGCCTCGTAGCGGATGAT</td><td></td><td></td><td></td><td></td><td></td><td></td><td></td><td></td><td></td><td></td></tr>
<tr><td>Oligo rev</td><td>AAACATCATCCGCTACGAGGCGCT</td><td></td><td></td><td></td><td></td><td></td><td></td><td></td><td></td><td></td><td></td></tr>
<tr><td>Chromosome</td><td>start</td><td>end</td><td>strand</td><td>MM</td><td>target_seq</td><td>PAM</td><td>alignment</td><td>distance</td><td>position</td><td>gene name</td><td>gene id</td></tr>
<tr><td>GU305914.1</td><td>144568</td><td>144590</td><td>+</td><td>0</td><td>AGCGCCTCGTAGCGGATGAT</td><td>GGG</td><td>||||||||[||||||||||||]PAM</td><td>NA</td><td>Intergenic</td><td>NA</td><td>NA</td></tr>
<tr><td>T11081</td><td>GCGCCTCGTAGCGGATGATGGGG</td><td>1000</td><td></td><td></td><td></td><td></td><td></td><td></td><td></td><td></td><td></td></tr>
<tr><td>Oligo fwd</td><td>CACCGCGCCTCGTAGCGGATGATG</td><td></td><td></td><td></td><td></td><td></td><td></td><td></td><td></td><td></td><td></td></tr>
<tr><td>Oligo rev</td><td>AAACCATCATCCGCTACGAGGCGC</td><td></td><td></td><td></td><td></td><td></td><td></td><td></td><td></td><td></td><td></td></tr>
<tr><td>Chromosome</td><td>start</td><td>end</td><td>strand</td><td>MM</td><td>target_seq</td><td>PAM</td><td>alignment</td><td>distance</td><td>position</td><td>gene name</td><td>gene id</td></tr>
<tr><td>GU305914.1</td><td>144569</td><td>144591</td><td>+</td><td>0</td><td>GCGCCTCGTAGCGGATGATG</td><td>GGG</td><td>||||||||[||||||||||||]PAM</td><td>NA</td><td>Intergenic</td><td>NA</td><td>NA</td></tr>
<tr><td>T11082</td><td>GTAGCGGATGATGGGGTACATGG</td><td>1000</td><td></td><td></td><td></td><td></td><td></td><td></td><td></td><td></td><td></td></tr>
<tr><td>Oligo fwd</td><td>CACCGTAGCGGATGATGGGGTACA</td><td></td><td></td><td></td><td></td><td></td><td></td><td></td><td></td><td></td><td></td></tr>
<tr><td>Oligo rev</td><td>AAACTGTACCCCATCATCCGCTAC</td><td></td><td></td><td></td><td></td><td></td><td></td><td></td><td></td><td></td><td></td></tr>
<tr><td>Chromosome</td><td>start</td><td>end</td><td>strand</td><td>MM</td><td>target_seq</td><td>PAM</td><td>alignment</td><td>distance</td><td>position</td><td>gene name</td><td>gene id</td></tr>
<tr><td>GU305914.1</td><td>144576</td><td>144598</td><td>+</td><td>0</td><td>GTAGCGGATGATGGGGTACA</td><td>TGG</td><td>||||||||[||||||||||||]PAM</td><td>NA</td><td>Intergenic</td><td>NA</td><td>NA</td></tr>
<tr><td>T11083</td><td>CGCCCAGTATCGTCCCGATGTGG</td><td>1000</td><td></td><td></td><td></td><td></td><td></td><td></td><td></td><td></td><td></td></tr>
<tr><td>Oligo fwd</td><td>CACCCGCCCAGTATCGTCCCGATG</td><td></td><td></td><td></td><td></td><td></td><td></td><td></td><td></td><td></td><td></td></tr>
<tr><td>Oligo rev</td><td>AAACCATCGGGACGATACTGGGCG</td><td></td><td></td><td></td><td></td><td></td><td></td><td></td><td></td><td></td><td></td></tr>
<tr><td>Chromosome</td><td>start</td><td>end</td><td>strand</td><td>MM</td><td>target_seq</td><td>PAM</td><td>alignment</td><td>distance</td><td>position</td><td>gene name</td><td>gene id</td></tr>
<tr><td>GU305914.1</td><td>144608</td><td>144630</td><td>+</td><td>0</td><td>CGCCCAGTATCGTCCCGATG</td><td>TGG</td><td>||||||||[||||||||||||]PAM</td><td>NA</td><td>Intergenic</td><td>NA</td><td>NA</td></tr>
<tr><td>T11084</td><td>CACGCACGCCACGATCGCGAAGG</td><td>1000</td><td></td><td></td><td></td><td></td><td></td><td></td><td></td><td></td><td></td></tr>
<tr><td>Oligo fwd</td><td>CACCCACGCACGCCACGATCGCGA</td><td></td><td></td><td></td><td></td><td></td><td></td><td></td><td></td><td></td><td></td></tr>
<tr><td>Oligo rev</td><td>AAACTCGCGATCGTGGCGTGCGTG</td><td></td><td></td><td></td><td></td><td></td><td></td><td></td><td></td><td></td><td></td></tr>
<tr><td>Chromosome</td><td>start</td><td>end</td><td>strand</td><td>MM</td><td>target_seq</td><td>PAM</td><td>alignment</td><td>distance</td><td>position</td><td>gene name</td><td>gene id</td></tr>
<tr><td>GU305914.1</td><td>144687</td><td>144709</td><td>+</td><td>0</td><td>CACGCACGCCACGATCGCGA</td><td>AGG</td><td>||||||||[||||||||||||]PAM</td><td>NA</td><td>Intergenic</td><td>NA</td><td>NA</td></tr>
<tr><td>T11085</td><td>CACGATCGCGAAGGCGACCATGG</td><td>1000</td><td></td><td></td><td></td><td></td><td></td><td></td><td></td><td></td><td></td></tr>
<tr><td>Oligo fwd</td><td>CACCCACGATCGCGAAGGCGACCA</td><td></td><td></td><td></td><td></td><td></td><td></td><td></td><td></td><td></td><td></td></tr>
<tr><td>Oligo rev</td><td>AAACTGGTCGCCTTCGCGATCGTG</td><td></td><td></td><td></td><td></td><td></td><td></td><td></td><td></td><td></td><td></td></tr>
<tr><td>Chromosome</td><td>start</td><td>end</td><td>strand</td><td>MM</td><td>target_seq</td><td>PAM</td><td>alignment</td><td>distance</td><td>position</td><td>gene name</td><td>gene id</td></tr>
<tr><td>GU305914.1</td><td>144696</td><td>144718</td><td>+</td><td>0</td><td>CACGATCGCGAAGGCGACCA</td><td>TGG</td><td>||||||||[||||||||||||]PAM</td><td>NA</td><td>Intergenic</td><td>NA</td><td>NA</td></tr>
<tr><td>T11086</td><td>GAAGCCGAGCGCCAGCGACATGG</td><td>1000</td><td></td><td></td><td></td><td></td><td></td><td></td><td></td><td></td><td></td></tr>
<tr><td>Oligo fwd</td><td>CACCGAAGCCGAGCGCCAGCGACA</td><td></td><td></td><td></td><td></td><td></td><td></td><td></td><td></td><td></td><td></td></tr>
<tr><td>Oligo rev</td><td>AAACTGTCGCTGGCGCTCGGCTTC</td><td></td><td></td><td></td><td></td><td></td><td></td><td></td><td></td><td></td><td></td></tr>
<tr><td>Chromosome</td><td>start</td><td>end</td><td>strand</td><td>MM</td><td>target_seq</td><td>PAM</td><td>alignment</td><td>distance</td><td>position</td><td>gene name</td><td>gene id</td></tr>
<tr><td>GU305914.1</td><td>144762</td><td>144784</td><td>+</td><td>0</td><td>GAAGCCGAGCGCCAGCGACA</td><td>TGG</td><td>||||||||[||||||||||||]PAM</td><td>NA</td><td>Intergenic</td><td>NA</td><td>NA</td></tr>
<tr><td>T11087</td><td>CAGCGACATGGCCAGCACCATGG</td><td>1000</td><td></td><td></td><td></td><td></td><td></td><td></td><td></td><td></td><td></td></tr>
<tr><td>Oligo fwd</td><td>CACCCAGCGACATGGCCAGCACCA</td><td></td><td></td><td></td><td></td><td></td><td></td><td></td><td></td><td></td><td></td></tr>
<tr><td>Oligo rev</td><td>AAACTGGTGCTGGCCATGTCGCTG</td><td></td><td></td><td></td><td></td><td></td><td></td><td></td><td></td><td></td><td></td></tr>
<tr><td>Chromosome</td><td>start</td><td>end</td><td>strand</td><td>MM</td><td>target_seq</td><td>PAM</td><td>alignment</td><td>distance</td><td>position</td><td>gene name</td><td>gene id</td></tr>
<tr><td>GU305914.1</td><td>144774</td><td>144796</td><td>+</td><td>0</td><td>CAGCGACATGGCCAGCACCA</td><td>TGG</td><td>||||||||[||||||||||||]PAM</td><td>NA</td><td>Intergenic</td><td>NA</td><td>NA</td></tr>
<tr><td>T11088</td><td>TCGCGAAGCCCAAGATCAGCTGG</td><td>1000</td><td></td><td></td><td></td><td></td><td></td><td></td><td></td><td></td><td></td></tr>
<tr><td>Oligo fwd</td><td>CACCTCGCGAAGCCCAAGATCAGC</td><td></td><td></td><td></td><td></td><td></td><td></td><td></td><td></td><td></td><td></td></tr>
<tr><td>Oligo rev</td><td>AAACGCTGATCTTGGGCTTCGCGA</td><td></td><td></td><td></td><td></td><td></td><td></td><td></td><td></td><td></td><td></td></tr>
<tr><td>Chromosome</td><td>start</td><td>end</td><td>strand</td><td>MM</td><td>target_seq</td><td>PAM</td><td>alignment</td><td>distance</td><td>position</td><td>gene name</td><td>gene id</td></tr>
<tr><td>GU305914.1</td><td>144797</td><td>144819</td><td>+</td><td>0</td><td>TCGCGAAGCCCAAGATCAGC</td><td>TGG</td><td>||||||||[||||||||||||]PAM</td><td>NA</td><td>Intergenic</td><td>NA</td><td>NA</td></tr>
<tr><td>GU305914.1</td><td>122976</td><td>122998</td><td>+</td><td>4</td><td>TCGGGACGCCCAGGATCATC</td><td>TGG</td><td>|||-||-|[||||-|||||-|]PAM</td><td>NA</td><td>Intergenic</td><td>NA</td><td>NA</td></tr>
<tr><td>T11089</td><td>CGCGAAGCCCAAGATCAGCTGGG</td><td>1000</td><td></td><td></td><td></td><td></td><td></td><td></td><td></td><td></td><td></td></tr>
<tr><td>Oligo fwd</td><td>CACCCGCGAAGCCCAAGATCAGCT</td><td></td><td></td><td></td><td></td><td></td><td></td><td></td><td></td><td></td><td></td></tr>
<tr><td>Oligo rev</td><td>AAACAGCTGATCTTGGGCTTCGCG</td><td></td><td></td><td></td><td></td><td></td><td></td><td></td><td></td><td></td><td></td></tr>
<tr><td>Chromosome</td><td>start</td><td>end</td><td>strand</td><td>MM</td><td>target_seq</td><td>PAM</td><td>alignment</td><td>distance</td><td>position</td><td>gene name</td><td>gene id</td></tr>
<tr><td>GU305914.1</td><td>144798</td><td>144820</td><td>+</td><td>0</td><td>CGCGAAGCCCAAGATCAGCT</td><td>GGG</td><td>||||||||[||||||||||||]PAM</td><td>NA</td><td>Intergenic</td><td>NA</td><td>NA</td></tr>
<tr><td>T11090</td><td>AGCCCAAGATCAGCTGGGTCAGG</td><td>1000</td><td></td><td></td><td></td><td></td><td></td><td></td><td></td><td></td><td></td></tr>
<tr><td>Oligo fwd</td><td>CACCAGCCCAAGATCAGCTGGGTC</td><td></td><td></td><td></td><td></td><td></td><td></td><td></td><td></td><td></td><td></td></tr>
<tr><td>Oligo rev</td><td>AAACGACCCAGCTGATCTTGGGCT</td><td></td><td></td><td></td><td></td><td></td><td></td><td></td><td></td><td></td><td></td></tr>
<tr><td>Chromosome</td><td>start</td><td>end</td><td>strand</td><td>MM</td><td>target_seq</td><td>PAM</td><td>alignment</td><td>distance</td><td>position</td><td>gene name</td><td>gene id</td></tr>
<tr><td>GU305914.1</td><td>144803</td><td>144825</td><td>+</td><td>0</td><td>AGCCCAAGATCAGCTGGGTC</td><td>AGG</td><td>||||||||[||||||||||||]PAM</td><td>NA</td><td>Intergenic</td><td>NA</td><td>NA</td></tr>
<tr><td>T11091</td><td>TCACCACCGCCGTCCGATAGCGG</td><td>1000</td><td></td><td></td><td></td><td></td><td></td><td></td><td></td><td></td><td></td></tr>
<tr><td>Oligo fwd</td><td>CACCTCACCACCGCCGTCCGATAG</td><td></td><td></td><td></td><td></td><td></td><td></td><td></td><td></td><td></td><td></td></tr>
<tr><td>Oligo rev</td><td>AAACCTATCGGACGGCGGTGGTGA</td><td></td><td></td><td></td><td></td><td></td><td></td><td></td><td></td><td></td><td></td></tr>
<tr><td>Chromosome</td><td>start</td><td>end</td><td>strand</td><td>MM</td><td>target_seq</td><td>PAM</td><td>alignment</td><td>distance</td><td>position</td><td>gene name</td><td>gene id</td></tr>
<tr><td>GU305914.1</td><td>144827</td><td>144849</td><td>+</td><td>0</td><td>TCACCACCGCCGTCCGATAG</td><td>CGG</td><td>||||||||[||||||||||||]PAM</td><td>NA</td><td>Intergenic</td><td>NA</td><td>NA</td></tr>
<tr><td>T11092</td><td>CACCGCCGTCCGATAGCGGATGG</td><td>1000</td><td></td><td></td><td></td><td></td><td></td><td></td><td></td><td></td><td></td></tr>
<tr><td>Oligo fwd</td><td>CACCCACCGCCGTCCGATAGCGGA</td><td></td><td></td><td></td><td></td><td></td><td></td><td></td><td></td><td></td><td></td></tr>
<tr><td>Oligo rev</td><td>AAACTCCGCTATCGGACGGCGGTG</td><td></td><td></td><td></td><td></td><td></td><td></td><td></td><td></td><td></td><td></td></tr>
<tr><td>Chromosome</td><td>start</td><td>end</td><td>strand</td><td>MM</td><td>target_seq</td><td>PAM</td><td>alignment</td><td>distance</td><td>position</td><td>gene name</td><td>gene id</td></tr>
<tr><td>GU305914.1</td><td>144831</td><td>144853</td><td>+</td><td>0</td><td>CACCGCCGTCCGATAGCGGA</td><td>TGG</td><td>||||||||[||||||||||||]PAM</td><td>NA</td><td>Intergenic</td><td>NA</td><td>NA</td></tr>
<tr><td>T11093</td><td>CGGATGGTCCCCTGCAGCTTCGG</td><td>1000</td><td></td><td></td><td></td><td></td><td></td><td></td><td></td><td></td><td></td></tr>
<tr><td>Oligo fwd</td><td>CACCCGGATGGTCCCCTGCAGCTT</td><td></td><td></td><td></td><td></td><td></td><td></td><td></td><td></td><td></td><td></td></tr>
<tr><td>Oligo rev</td><td>AAACAAGCTGCAGGGGACCATCCG</td><td></td><td></td><td></td><td></td><td></td><td></td><td></td><td></td><td></td><td></td></tr>
<tr><td>Chromosome</td><td>start</td><td>end</td><td>strand</td><td>MM</td><td>target_seq</td><td>PAM</td><td>alignment</td><td>distance</td><td>position</td><td>gene name</td><td>gene id</td></tr>
<tr><td>GU305914.1</td><td>144847</td><td>144869</td><td>+</td><td>0</td><td>CGGATGGTCCCCTGCAGCTT</td><td>CGG</td><td>||||||||[||||||||||||]PAM</td><td>NA</td><td>Intergenic</td><td>NA</td><td>NA</td></tr>
</table>

T11094 GGATGGTCCCCTGCAGCTTCGGG 1000
Oligo fwd CACCGGATGGTCCCCTGCAGCTTC
Oligo rev AAACGAAGCTGCAGGGGACCATCC
Chromosome start end strand MM target_seq PAM alignment distance position gene name gene id
GU305914.1 144848 144870 + 0 GGATGGTCCCCTGCAGCTTC GGG ||||||||[||||||||||||]PAM NA Intergenic NA NA

T11095 TGGTCCCCTGCAGCTTCGGGTGG 1000
Oligo fwd CACCTGGTCCCCTGCAGCTTCGGG
Oligo rev AAACCCCGAAGCTGCAGGGGACCA
Chromosome start end strand MM target_seq PAM alignment distance position gene name gene id
GU305914.1 144851 144873 + 0 TGGTCCCCTGCAGCTTCGGG TGG ||||||||[||||||||||||]PAM NA Intergenic NA NA

T11096 TGCGCTCGCGCGACTCGTAGTGG 1000
Oligo fwd CACCTGCGCTCGCGCGACTCGTAG
Oligo rev AAACCTACGAGTCGCGCGAGCGCA
Chromosome start end strand MM target_seq PAM alignment distance position gene name gene id
GU305914.1 144896 144918 + 0 TGCGCTCGCGCGACTCGTAG TGG ||||||||[||||||||||||]PAM NA Intergenic NA NA

T11097 GCGCTCGCGCGACTCGTAGTGGG 1000
Oligo fwd CACCGCGCTCGCGCGACTCGTAGT
Oligo rev AAACACTACGAGTCGCGCGAGCGC
Chromosome start end strand MM target_seq PAM alignment distance position gene name gene id
GU305914.1 144897 144919 + 0 GCGCTCGCGCGACTCGTAGT GGG ||||||||[||||||||||||]PAM NA Intergenic NA NA

T11098 GGTCACGAGCGTCACCACGAAGG 1000
Oligo fwd CACCGGTCACGAGCGTCACCACGA
Oligo rev AAACTCGTGGTGACGCTCGTGACC
Chromosome start end strand MM target_seq PAM alignment distance position gene name gene id
GU305914.1 144918 144940 + 0 GGTCACGAGCGTCACCACGA AGG ||||||||[||||||||||||]PAM NA Intergenic NA NA
GU305914.1 192132 192154 - 4 AGTCGCGGCCGTCACCACGA CGG -|||-||-[-|||||||||||]PAM NA Intergenic NA NA

T11099 GCGTCACCACGAAGGCCGTCAGG 1000
Oligo fwd CACCGCGTCACCACGAAGGCCGTC
Oligo rev AAACGACGGCCTTCGTGGTGACGC
Chromosome start end strand MM target_seq PAM alignment distance position gene name gene id
GU305914.1 144926 144948 + 0 GCGTCACCACGAAGGCCGTC AGG ||||||||[||||||||||||]PAM NA Intergenic NA NA

T11100 GCACTTGCTGAACGCGACCATGG 1000
Oligo fwd CACCGCACTTGCTGAACGCGACCA
Oligo rev AAACTGGTCGCGTTCAGCAAGTGC
Chromosome start end strand MM target_seq PAM alignment distance position gene name gene id
GU305914.1 144963 144985 + 0 GCACTTGCTGAACGCGACCA TGG ||||||||[||||||||||||]PAM NA Intergenic NA NA

T11101 TTGCTGAACGCGACCATGGACGG 1000
Oligo fwd CACCTTGCTGAACGCGACCATGGA
Oligo rev AAACTCCATGGTCGCGTTCAGCAA
Chromosome start end strand MM target_seq PAM alignment distance position gene name gene id
GU305914.1 144967 144989 + 0 TTGCTGAACGCGACCATGGA CGG ||||||||[||||||||||||]PAM NA Intergenic NA NA

T11102 TGAACGCGACCATGGACGGTAGG 1000
Oligo fwd CACCTGAACGCGACCATGGACGGT
Oligo rev AAACACCGTCCATGGTCGCGTTCA
Chromosome start end strand MM target_seq PAM alignment distance position gene name gene id
GU305914.1 144971 144993 + 0 TGAACGCGACCATGGACGGT AGG ||||||||[||||||||||||]PAM NA Intergenic NA NA

T11103 ACGCGACCATGGACGGTAGGCGG 1000
Oligo fwd CACCACGCGACCATGGACGGTAGG
Oligo rev AAACCCTACCGTCCATGGTCGCGT
Chromosome start end strand MM target_seq PAM alignment distance position gene name gene id
GU305914.1 144974 144996 + 0 ACGCGACCATGGACGGTAGG CGG ||||||||[||||||||||||]PAM NA Intergenic NA NA

T11104 TGGACGGTAGGCGGAACGACAGG 1000
Oligo fwd CACCTGGACGGTAGGCGGAACGAC
Oligo rev AAACGTCGTTCCGCCTACCGTCCA
Chromosome start end strand MM target_seq PAM alignment distance position gene name gene id
GU305914.1 144983 145005 + 0 TGGACGGTAGGCGGAACGAC AGG ||||||||[||||||||||||]PAM NA Intergenic NA NA

T11105 GGCGGAACGACAGGCTCAGCAGG 1000
Oligo fwd CACCGGCGGAACGACAGGCTCAGC
Oligo rev AAACGCTGAGCCTGTCGTTCCGCC
Chromosome start end strand MM target_seq PAM alignment distance position gene name gene id
GU305914.1 144992 145014 + 0 GGCGGAACGACAGGCTCAGC AGG ||||||||[||||||||||||]PAM NA Intergenic NA NA

T11106 GCTCAGCAGGAAGATCTGAAAGG 1000
Oligo fwd CACCGCTCAGCAGGAAGATCTGAA
Oligo rev AAACTTCAGATCTTCCTGCTGAGC
Chromosome start end strand MM target_seq PAM alignment distance position gene name gene id
GU305914.1 145005 145027 + 0 GCTCAGCAGGAAGATCTGAA AGG ||||||||[||||||||||||]PAM NA Intergenic NA NA

T11107 GGTGTCCATCGTCAGCACGAAGG 1000
Oligo fwd CACCGGTGTCCATCGTCAGCACGA
Oligo rev AAACTCGTGCTGACGATGGACACC
Chromosome start end strand MM target_seq PAM alignment distance position gene name gene id
GU305914.1 145026 145048 + 0 GGTGTCCATCGTCAGCACGA AGG ||||||||[||||||||||||]PAM NA Intergenic NA NA

T11108 CGCCCATGAAGTTGATGTCGCGG 1000
Oligo fwd CACCCGCCCATGAAGTTGATGTCG
Oligo rev AAACCGACATCAACTTCATGGGCG
Chromosome start end strand MM target_seq PAM alignment distance position gene name gene id

GU305914.1 145067 145089 + 0 CGCCCATGAAGTTGATGTCG CGG ||||||||[||||||||||||]PAM NA Intergenic NA NA

T11109 GCCCATGAAGTTGATGTCGCGGG 1000
Oligo fwd CACCGCCCATGAAGTTGATGTCGC
Oligo rev AAACGCGACATCAACTTCATGGGC
Chromosome start end strand MM target_seq PAM alignment distance position gene name gene id
GU305914.1 145068 145090 + 0 GCCCATGAAGTTGATGTCGC GGG ||||||||[||||||||||||]PAM NA Intergenic NA NA

T11110 AGTTGATGTCGCGGGTCGACTGG 1000
Oligo fwd CACCAGTTGATGTCGCGGGTCGAC
Oligo rev AAACGTCGACCCGCGACATCAACT
Chromosome start end strand MM target_seq PAM alignment distance position gene name gene id
GU305914.1 145076 145098 + 0 AGTTGATGTCGCGGGTCGAC TGG ||||||||[||||||||||||]PAM NA Intergenic NA NA

T11111 CCTGCGTGCCGTGCTCGCTGCGG 1000
Oligo fwd CACCCCTGCGTGCCGTGCTCGCTG
Oligo rev AAACCAGCGAGCACGGCACGCAGG
Chromosome start end strand MM target_seq PAM alignment distance position gene name gene id
GU305914.1 145103 145125 + 0 CCTGCGTGCCGTGCTCGCTG CGG ||||||||[||||||||||||]PAM NA Intergenic NA NA

T11112 CACGTACACGATGAGCTGCACGG 1000
Oligo fwd CACCCACGTACACGATGAGCTGCA
Oligo rev AAACTGCAGCTCATCGTGTACGTG
Chromosome start end strand MM target_seq PAM alignment distance position gene name gene id
GU305914.1 145194 145216 + 0 CACGTACACGATGAGCTGCA CGG ||||||||[||||||||||||]PAM NA Intergenic NA NA

T11113 ACGTACACGATGAGCTGCACGGG 1000
Oligo fwd CACCACGTACACGATGAGCTGCAC
Oligo rev AAACGTGCAGCTCATCGTGTACGT
Chromosome start end strand MM target_seq PAM alignment distance position gene name gene id
GU305914.1 145195 145217 + 0 ACGTACACGATGAGCTGCAC GGG ||||||||[||||||||||||]PAM NA Intergenic NA NA

T11114 CGTACACGATGAGCTGCACGGGG 1000
Oligo fwd CACCCGTACACGATGAGCTGCACG
Oligo rev AAACCGTGCAGCTCATCGTGTACG
Chromosome start end strand MM target_seq PAM alignment distance position gene name gene id
GU305914.1 145196 145218 + 0 CGTACACGATGAGCTGCACG GGG ||||||||[||||||||||||]PAM NA Intergenic NA NA

T11115 CGATGAGCTGCACGGGGTCCAGG 1000
Oligo fwd CACCCGATGAGCTGCACGGGGTCC
Oligo rev AAACGGACCCCGTGCAGCTCATCG
Chromosome start end strand MM target_seq PAM alignment distance position gene name gene id
GU305914.1 145202 145224 + 0 CGATGAGCTGCACGGGGTCC AGG ||||||||[||||||||||||]PAM NA Intergenic NA NA

T11116 ACGGGGTCCAGGAACAGCTGCGG 1000
Oligo fwd CACCACGGGGTCCAGGAACAGCTG
Oligo rev AAACCAGCTGTTCCTGGACCCCGT
Chromosome start end strand MM target_seq PAM alignment distance position gene name gene id
GU305914.1 145213 145235 + 0 ACGGGGTCCAGGAACAGCTG CGG ||||||||[||||||||||||]PAM NA Intergenic NA NA

T11117 GGAACAGCTGCGGCGTCAGATGG 1000
Oligo fwd CACCGGAACAGCTGCGGCGTCAGA
Oligo rev AAACTCTGACGCCGCAGCTGTTCC
Chromosome start end strand MM target_seq PAM alignment distance position gene name gene id
GU305914.1 145223 145245 + 0 GGAACAGCTGCGGCGTCAGA TGG ||||||||[||||||||||||]PAM NA Intergenic NA NA

T11118 ACAGCTGCGGCGTCAGATGGTGG 1000
Oligo fwd CACCACAGCTGCGGCGTCAGATGG
Oligo rev AAACCCATCTGACGCCGCAGCTGT
Chromosome start end strand MM target_seq PAM alignment distance position gene name gene id
GU305914.1 145226 145248 + 0 ACAGCTGCGGCGTCAGATGG TGG ||||||||[||||||||||||]PAM NA Intergenic NA NA

T11119 GGATCGCGTTGTACATCGTCAGG 1000
Oligo fwd CACCGGATCGCGTTGTACATCGTC
Oligo rev AAACGACGATGTACAACGCGATCC
Chromosome start end strand MM target_seq PAM alignment distance position gene name gene id
GU305914.1 145247 145269 + 0 GGATCGCGTTGTACATCGTC AGG ||||||||[||||||||||||]PAM NA Intergenic NA NA

T11120 TTGTCGAAGTCGATGATGCGCGG 1000
Oligo fwd CACCTTGTCGAAGTCGATGATGCG
Oligo rev AAACCGCATCATCGACTTCGACAA
Chromosome start end strand MM target_seq PAM alignment distance position gene name gene id
GU305914.1 145276 145298 + 0 TTGTCGAAGTCGATGATGCG CGG ||||||||[||||||||||||]PAM NA Intergenic NA NA

T11121 TGTCGAAGTCGATGATGCGCGGG 1000
Oligo fwd CACCTGTCGAAGTCGATGATGCGC
Oligo rev AAACGCGCATCATCGACTTCGACA
Chromosome start end strand MM target_seq PAM alignment distance position gene name gene id
GU305914.1 145277 145299 + 0 TGTCGAAGTCGATGATGCGC GGG ||||||||[||||||||||||]PAM NA Intergenic NA NA

T11122 ATGATGCGCGGGTAATAGCACGG 1000
Oligo fwd CACCATGATGCGCGGGTAATAGCA
Oligo rev AAACTGCTATTACCCGCGCATCAT
Chromosome start end strand MM target_seq PAM alignment distance position gene name gene id
GU305914.1 145288 145310 + 0 ATGATGCGCGGGTAATAGCA CGG ||||||||[||||||||||||]PAM NA Intergenic NA NA

T11123 ATAGCACGGAAAACCGATGCCGG 1000
Oligo fwd CACCATAGCACGGAAAACCGATGC
Oligo rev AAACGCATCGGTTTTCCGTGCTAT

<table>
<tr><td>Chromosome</td><td>start</td><td>end</td><td>strand</td><td>MM</td><td>target_seq</td><td>PAM</td><td>alignment</td><td>distance</td><td>position</td><td>gene name</td><td>gene id</td></tr>
<tr><td>GU305914.1</td><td>145302</td><td>145324</td><td>+</td><td>0</td><td>ATAGCACGGAAAACCGATGC</td><td>CGG</td><td>||||||||[||||||||||||]PAM</td><td>NA</td><td>Intergenic</td><td>NA</td><td>NA</td></tr>
<tr><td></td><td></td><td></td><td></td><td></td><td></td><td></td><td></td><td></td><td></td><td></td><td></td></tr>
<tr><td>T11124</td><td>TAGCACGGAAAACCGATGCCGGG</td><td>1000</td><td></td><td></td><td></td><td></td><td></td><td></td><td></td><td></td><td></td></tr>
<tr><td>Oligo fwd</td><td>CACCTAGCACGGAAAACCGATGCC</td><td></td><td></td><td></td><td></td><td></td><td></td><td></td><td></td><td></td><td></td></tr>
<tr><td>Oligo rev</td><td>AAACGGCATCGGTTTTCCGTGCTA</td><td></td><td></td><td></td><td></td><td></td><td></td><td></td><td></td><td></td><td></td></tr>
<tr><td>Chromosome</td><td>start</td><td>end</td><td>strand</td><td>MM</td><td>target_seq</td><td>PAM</td><td>alignment</td><td>distance</td><td>position</td><td>gene name</td><td>gene id</td></tr>
<tr><td>GU305914.1</td><td>145303</td><td>145325</td><td>+</td><td>0</td><td>TAGCACGGAAAACCGATGCC</td><td>GGG</td><td>||||||||[||||||||||||]PAM</td><td>NA</td><td>Intergenic</td><td>NA</td><td>NA</td></tr>
<tr><td></td><td></td><td></td><td></td><td></td><td></td><td></td><td></td><td></td><td></td><td></td><td></td></tr>
<tr><td>T11125</td><td>AGCACGGAAAACCGATGCCGGGG</td><td>1000</td><td></td><td></td><td></td><td></td><td></td><td></td><td></td><td></td><td></td></tr>
<tr><td>Oligo fwd</td><td>CACCAGCACGGAAAACCGATGCCG</td><td></td><td></td><td></td><td></td><td></td><td></td><td></td><td></td><td></td><td></td></tr>
<tr><td>Oligo rev</td><td>AAACCGGCATCGGTTTTCCGTGCT</td><td></td><td></td><td></td><td></td><td></td><td></td><td></td><td></td><td></td><td></td></tr>
<tr><td>Chromosome</td><td>start</td><td>end</td><td>strand</td><td>MM</td><td>target_seq</td><td>PAM</td><td>alignment</td><td>distance</td><td>position</td><td>gene name</td><td>gene id</td></tr>
<tr><td>GU305914.1</td><td>145304</td><td>145326</td><td>+</td><td>0</td><td>AGCACGGAAAACCGATGCCG</td><td>GGG</td><td>||||||||[||||||||||||]PAM</td><td>NA</td><td>Intergenic</td><td>NA</td><td>NA</td></tr>
<tr><td></td><td></td><td></td><td></td><td></td><td></td><td></td><td></td><td></td><td></td><td></td><td></td></tr>
<tr><td>T11126</td><td>CGATGTTCACGAACGACAGCAGG</td><td>1000</td><td></td><td></td><td></td><td></td><td></td><td></td><td></td><td></td><td></td></tr>
<tr><td>Oligo fwd</td><td>CACCCGATGTTCACGAACGACAGC</td><td></td><td></td><td></td><td></td><td></td><td></td><td></td><td></td><td></td><td></td></tr>
<tr><td>Oligo rev</td><td>AAACGCTGTCGTTCGTGAACATCG</td><td></td><td></td><td></td><td></td><td></td><td></td><td></td><td></td><td></td><td></td></tr>
<tr><td>Chromosome</td><td>start</td><td>end</td><td>strand</td><td>MM</td><td>target_seq</td><td>PAM</td><td>alignment</td><td>distance</td><td>position</td><td>gene name</td><td>gene id</td></tr>
<tr><td>GU305914.1</td><td>145352</td><td>145374</td><td>+</td><td>0</td><td>CGATGTTCACGAACGACAGC</td><td>AGG</td><td>||||||||[||||||||||||]PAM</td><td>NA</td><td>Intergenic</td><td>NA</td><td>NA</td></tr>
<tr><td></td><td></td><td></td><td></td><td></td><td></td><td></td><td></td><td></td><td></td><td></td><td></td></tr>
<tr><td>T11127</td><td>CGACAGCAGGCAGCACGCGATGG</td><td>1000</td><td></td><td></td><td></td><td></td><td></td><td></td><td></td><td></td><td></td></tr>
<tr><td>Oligo fwd</td><td>CACCCGACAGCAGGCAGCACGCGA</td><td></td><td></td><td></td><td></td><td></td><td></td><td></td><td></td><td></td><td></td></tr>
<tr><td>Oligo rev</td><td>AAACTCGCGTGCTGCCTGCTGTCG</td><td></td><td></td><td></td><td></td><td></td><td></td><td></td><td></td><td></td><td></td></tr>
<tr><td>Chromosome</td><td>start</td><td>end</td><td>strand</td><td>MM</td><td>target_seq</td><td>PAM</td><td>alignment</td><td>distance</td><td>position</td><td>gene name</td><td>gene id</td></tr>
<tr><td>GU305914.1</td><td>145365</td><td>145387</td><td>+</td><td>0</td><td>CGACAGCAGGCAGCACGCGA</td><td>TGG</td><td>||||||||[||||||||||||]PAM</td><td>NA</td><td>Intergenic</td><td>NA</td><td>NA</td></tr>
<tr><td></td><td></td><td></td><td></td><td></td><td></td><td></td><td></td><td></td><td></td><td></td><td></td></tr>
<tr><td>T11128</td><td>TGGCCATCGTCCACGTCCGCAGG</td><td>1000</td><td></td><td></td><td></td><td></td><td></td><td></td><td></td><td></td><td></td></tr>
<tr><td>Oligo fwd</td><td>CACCTGGCCATCGTCCACGTCCGC</td><td></td><td></td><td></td><td></td><td></td><td></td><td></td><td></td><td></td><td></td></tr>
<tr><td>Oligo rev</td><td>AAACGCGGACGTGGACGATGGCCA</td><td></td><td></td><td></td><td></td><td></td><td></td><td></td><td></td><td></td><td></td></tr>
<tr><td>Chromosome</td><td>start</td><td>end</td><td>strand</td><td>MM</td><td>target_seq</td><td>PAM</td><td>alignment</td><td>distance</td><td>position</td><td>gene name</td><td>gene id</td></tr>
<tr><td>GU305914.1</td><td>145385</td><td>145407</td><td>+</td><td>0</td><td>TGGCCATCGTCCACGTCCGC</td><td>AGG</td><td>||||||||[||||||||||||]PAM</td><td>NA</td><td>Intergenic</td><td>NA</td><td>NA</td></tr>
<tr><td></td><td></td><td></td><td></td><td></td><td></td><td></td><td></td><td></td><td></td><td></td><td></td></tr>
<tr><td>T11129</td><td>TCCACGTCCGCAGGTTCATGCGG</td><td>1000</td><td></td><td></td><td></td><td></td><td></td><td></td><td></td><td></td><td></td></tr>
<tr><td>Oligo fwd</td><td>CACCTCCACGTCCGCAGGTTCATG</td><td></td><td></td><td></td><td></td><td></td><td></td><td></td><td></td><td></td><td></td></tr>
<tr><td>Oligo rev</td><td>AAACCATGAACCTGCGGACGTGGA</td><td></td><td></td><td></td><td></td><td></td><td></td><td></td><td></td><td></td><td></td></tr>
<tr><td>Chromosome</td><td>start</td><td>end</td><td>strand</td><td>MM</td><td>target_seq</td><td>PAM</td><td>alignment</td><td>distance</td><td>position</td><td>gene name</td><td>gene id</td></tr>
<tr><td>GU305914.1</td><td>145394</td><td>145416</td><td>+</td><td>0</td><td>TCCACGTCCGCAGGTTCATG</td><td>CGG</td><td>||||||||[||||||||||||]PAM</td><td>NA</td><td>Intergenic</td><td>NA</td><td>NA</td></tr>
<tr><td></td><td></td><td></td><td></td><td></td><td></td><td></td><td></td><td></td><td></td><td></td><td></td></tr>
<tr><td>T11130</td><td>GCAGGTTCATGCGGTCCACGTGG</td><td>1000</td><td></td><td></td><td></td><td></td><td></td><td></td><td></td><td></td><td></td></tr>
<tr><td>Oligo fwd</td><td>CACCGCAGGTTCATGCGGTCCACG</td><td></td><td></td><td></td><td></td><td></td><td></td><td></td><td></td><td></td><td></td></tr>
<tr><td>Oligo rev</td><td>AAACCGTGGACCGCATGAACCTGC</td><td></td><td></td><td></td><td></td><td></td><td></td><td></td><td></td><td></td><td></td></tr>
<tr><td>Chromosome</td><td>start</td><td>end</td><td>strand</td><td>MM</td><td>target_seq</td><td>PAM</td><td>alignment</td><td>distance</td><td>position</td><td>gene name</td><td>gene id</td></tr>
<tr><td>GU305914.1</td><td>145403</td><td>145425</td><td>+</td><td>0</td><td>GCAGGTTCATGCGGTCCACG</td><td>TGG</td><td>||||||||[||||||||||||]PAM</td><td>NA</td><td>Intergenic</td><td>NA</td><td>NA</td></tr>
<tr><td></td><td></td><td></td><td></td><td></td><td></td><td></td><td></td><td></td><td></td><td></td><td></td></tr>
<tr><td>T11131</td><td>CAGGTTCATGCGGTCCACGTGGG</td><td>1000</td><td></td><td></td><td></td><td></td><td></td><td></td><td></td><td></td><td></td></tr>
<tr><td>Oligo fwd</td><td>CACCCAGGTTCATGCGGTCCACGT</td><td></td><td></td><td></td><td></td><td></td><td></td><td></td><td></td><td></td><td></td></tr>
<tr><td>Oligo rev</td><td>AAACACGTGGACCGCATGAACCTG</td><td></td><td></td><td></td><td></td><td></td><td></td><td></td><td></td><td></td><td></td></tr>
<tr><td>Chromosome</td><td>start</td><td>end</td><td>strand</td><td>MM</td><td>target_seq</td><td>PAM</td><td>alignment</td><td>distance</td><td>position</td><td>gene name</td><td>gene id</td></tr>
<tr><td>GU305914.1</td><td>145404</td><td>145426</td><td>+</td><td>0</td><td>CAGGTTCATGCGGTCCACGT</td><td>GGG</td><td>||||||||[||||||||||||]PAM</td><td>NA</td><td>Intergenic</td><td>NA</td><td>NA</td></tr>
<tr><td></td><td></td><td></td><td></td><td></td><td></td><td></td><td></td><td></td><td></td><td></td><td></td></tr>
<tr><td>T11132</td><td>GCCCGCTTTCGCCATCGTGCAGG</td><td>1000</td><td></td><td></td><td></td><td></td><td></td><td></td><td></td><td></td><td></td></tr>
<tr><td>Oligo fwd</td><td>CACCGCCCGCTTTCGCCATCGTGC</td><td></td><td></td><td></td><td></td><td></td><td></td><td></td><td></td><td></td><td></td></tr>
<tr><td>Oligo rev</td><td>AAACGCACGATGGCGAAAGCGGGC</td><td></td><td></td><td></td><td></td><td></td><td></td><td></td><td></td><td></td><td></td></tr>
<tr><td>Chromosome</td><td>start</td><td>end</td><td>strand</td><td>MM</td><td>target_seq</td><td>PAM</td><td>alignment</td><td>distance</td><td>position</td><td>gene name</td><td>gene id</td></tr>
<tr><td>GU305914.1</td><td>145440</td><td>145462</td><td>+</td><td>0</td><td>GCCCGCTTTCGCCATCGTGC</td><td>AGG</td><td>||||||||[||||||||||||]PAM</td><td>NA</td><td>Intergenic</td><td>NA</td><td>NA</td></tr>
<tr><td></td><td></td><td></td><td></td><td></td><td></td><td></td><td></td><td></td><td></td><td></td><td></td></tr>
<tr><td>T11133</td><td>GTGCAGGTGTCTCGTGTCCGCGG</td><td>1000</td><td></td><td></td><td></td><td></td><td></td><td></td><td></td><td></td><td></td></tr>
<tr><td>Oligo fwd</td><td>CACCGTGCAGGTGTCTCGTGTCCG</td><td></td><td></td><td></td><td></td><td></td><td></td><td></td><td></td><td></td><td></td></tr>
<tr><td>Oligo rev</td><td>AAACCGGACACGAGACACCTGCAC</td><td></td><td></td><td></td><td></td><td></td><td></td><td></td><td></td><td></td><td></td></tr>
<tr><td>Chromosome</td><td>start</td><td>end</td><td>strand</td><td>MM</td><td>target_seq</td><td>PAM</td><td>alignment</td><td>distance</td><td>position</td><td>gene name</td><td>gene id</td></tr>
<tr><td>GU305914.1</td><td>145456</td><td>145478</td><td>+</td><td>0</td><td>GTGCAGGTGTCTCGTGTCCG</td><td>CGG</td><td>||||||||[||||||||||||]PAM</td><td>NA</td><td>Intergenic</td><td>NA</td><td>NA</td></tr>
<tr><td></td><td></td><td></td><td></td><td></td><td></td><td></td><td></td><td></td><td></td><td></td><td></td></tr>
<tr><td>T11134</td><td>TGTCTCGTGTCCGCGGTCGACGG</td><td>1000</td><td></td><td></td><td></td><td></td><td></td><td></td><td></td><td></td><td></td></tr>
<tr><td>Oligo fwd</td><td>CACCTGTCTCGTGTCCGCGGTCGA</td><td></td><td></td><td></td><td></td><td></td><td></td><td></td><td></td><td></td><td></td></tr>
<tr><td>Oligo rev</td><td>AAACTCGACCGCGGACACGAGACA</td><td></td><td></td><td></td><td></td><td></td><td></td><td></td><td></td><td></td><td></td></tr>
<tr><td>Chromosome</td><td>start</td><td>end</td><td>strand</td><td>MM</td><td>target_seq</td><td>PAM</td><td>alignment</td><td>distance</td><td>position</td><td>gene name</td><td>gene id</td></tr>
<tr><td>GU305914.1</td><td>145463</td><td>145485</td><td>+</td><td>0</td><td>TGTCTCGTGTCCGCGGTCGA</td><td>CGG</td><td>||||||||[||||||||||||]PAM</td><td>NA</td><td>Intergenic</td><td>NA</td><td>NA</td></tr>
<tr><td></td><td></td><td></td><td></td><td></td><td></td><td></td><td></td><td></td><td></td><td></td><td></td></tr>
<tr><td>T11135</td><td>GTCTCGTGTCCGCGGTCGACGGG</td><td>1000</td><td></td><td></td><td></td><td></td><td></td><td></td><td></td><td></td><td></td></tr>
<tr><td>Oligo fwd</td><td>CACCGTCTCGTGTCCGCGGTCGAC</td><td></td><td></td><td></td><td></td><td></td><td></td><td></td><td></td><td></td><td></td></tr>
<tr><td>Oligo rev</td><td>AAACGTCGACCGCGGACACGAGAC</td><td></td><td></td><td></td><td></td><td></td><td></td><td></td><td></td><td></td><td></td></tr>
<tr><td>Chromosome</td><td>start</td><td>end</td><td>strand</td><td>MM</td><td>target_seq</td><td>PAM</td><td>alignment</td><td>distance</td><td>position</td><td>gene name</td><td>gene id</td></tr>
<tr><td>GU305914.1</td><td>145464</td><td>145486</td><td>+</td><td>0</td><td>GTCTCGTGTCCGCGGTCGAC</td><td>GGG</td><td>||||||||[||||||||||||]PAM</td><td>NA</td><td>Intergenic</td><td>NA</td><td>NA</td></tr>
<tr><td></td><td></td><td></td><td></td><td></td><td></td><td></td><td></td><td></td><td></td><td></td><td></td></tr>
<tr><td>T11136</td><td>TCTCGTGTCCGCGGTCGACGGGG</td><td>1000</td><td></td><td></td><td></td><td></td><td></td><td></td><td></td><td></td><td></td></tr>
<tr><td>Oligo fwd</td><td>CACCTCTCGTGTCCGCGGTCGACG</td><td></td><td></td><td></td><td></td><td></td><td></td><td></td><td></td><td></td><td></td></tr>
<tr><td>Oligo rev</td><td>AAACCGTCGACCGCGGACACGAGA</td><td></td><td></td><td></td><td></td><td></td><td></td><td></td><td></td><td></td><td></td></tr>
<tr><td>Chromosome</td><td>start</td><td>end</td><td>strand</td><td>MM</td><td>target_seq</td><td>PAM</td><td>alignment</td><td>distance</td><td>position</td><td>gene name</td><td>gene id</td></tr>
<tr><td>GU305914.1</td><td>145465</td><td>145487</td><td>+</td><td>0</td><td>TCTCGTGTCCGCGGTCGACG</td><td>GGG</td><td>||||||||[||||||||||||]PAM</td><td>NA</td><td>Intergenic</td><td>NA</td><td>NA</td></tr>
<tr><td></td><td></td><td></td><td></td><td></td><td></td><td></td><td></td><td></td><td></td><td></td><td></td></tr>
<tr><td>T11137</td><td>CGTGTCCGCGGTCGACGGGGCGG</td><td>1000</td><td></td><td></td><td></td><td></td><td></td><td></td><td></td><td></td><td></td></tr>
<tr><td>Oligo fwd</td><td>CACCCGTGTCCGCGGTCGACGGGG</td><td></td><td></td><td></td><td></td><td></td><td></td><td></td><td></td><td></td><td></td></tr>
<tr><td>Oligo rev</td><td>AAACCCCCGTCGACCGCGGACACG</td><td></td><td></td><td></td><td></td><td></td><td></td><td></td><td></td><td></td><td></td></tr>
<tr><td>Chromosome</td><td>start</td><td>end</td><td>strand</td><td>MM</td><td>target_seq</td><td>PAM</td><td>alignment</td><td>distance</td><td>position</td><td>gene name</td><td>gene id</td></tr>
<tr><td>GU305914.1</td><td>145468</td><td>145490</td><td>+</td><td>0</td><td>CGTGTCCGCGGTCGACGGGG</td><td>CGG</td><td>||||||||[||||||||||||]PAM</td><td>NA</td><td>Intergenic</td><td>NA</td><td>NA</td></tr>
<tr><td></td><td></td><td></td><td></td><td></td><td></td><td></td><td></td><td></td><td></td><td></td><td></td></tr>
<tr><td>T11138</td><td>GTGTCCGCGGTCGACGGGGCGGG</td><td>1000</td><td></td><td></td><td></td><td></td><td></td><td></td><td></td><td></td><td></td></tr>
<tr><td>Oligo fwd</td><td>CACCGTGTCCGCGGTCGACGGGGC</td><td></td><td></td><td></td><td></td><td></td><td></td><td></td><td></td><td></td><td></td></tr>
</table>

Oligo rev AAACGCCCCGTCGACCGCGGACAC
Chromosome start end strand MM target_seq PAM alignment distance position gene name gene id
GU305914.1 145469 145491 + 0 GTGTCCGCGGTCGACGGGGC GGG ||||||||[||||||||||||]PAM NA Intergenic NA NA

T11139 TCCGAGCCTCGCCGAGACGACGG 1000
Oligo fwd CACCTCCGAGCCTCGCCGAGACGA
Oligo rev AAACTCGTCTCGGCGAGGCTCGGA
Chromosome start end strand MM target_seq PAM alignment distance position gene name gene id
GU305914.1 145523 145545 + 0 TCCGAGCCTCGCCGAGACGA CGG ||||||||[||||||||||||]PAM NA Intergenic NA NA

T11140 CCGAGCCTCGCCGAGACGACGGG 1000
Oligo fwd CACCCCGAGCCTCGCCGAGACGAC
Oligo rev AAACGTCGTCTCGGCGAGGCTCGG
Chromosome start end strand MM target_seq PAM alignment distance position gene name gene id
GU305914.1 145524 145546 + 0 CCGAGCCTCGCCGAGACGAC GGG ||||||||[||||||||||||]PAM NA Intergenic NA NA

T11141 CGAGCCTCGCCGAGACGACGGGG 1000
Oligo fwd CACCCGAGCCTCGCCGAGACGACG
Oligo rev AAACCGTCGTCTCGGCGAGGCTCG
Chromosome start end strand MM target_seq PAM alignment distance position gene name gene id
GU305914.1 145525 145547 + 0 CGAGCCTCGCCGAGACGACG GGG ||||||||[||||||||||||]PAM NA Intergenic NA NA

T11142 ACGGGGTAACGTCGCCGCTTCGG 1000
Oligo fwd CACCACGGGGTAACGTCGCCGCTT
Oligo rev AAACAAGCGGCGACGTTACCCCGT
Chromosome start end strand MM target_seq PAM alignment distance position gene name gene id
GU305914.1 145542 145564 + 0 ACGGGGTAACGTCGCCGCTT CGG ||||||||[||||||||||||]PAM NA Intergenic NA NA

T11143 CGGGGTAACGTCGCCGCTTCGGG 1000
Oligo fwd CACCCGGGGTAACGTCGCCGCTTC
Oligo rev AAACGAAGCGGCGACGTTACCCCG
Chromosome start end strand MM target_seq PAM alignment distance position gene name gene id
GU305914.1 145543 145565 + 0 CGGGGTAACGTCGCCGCTTC GGG ||||||||[||||||||||||]PAM NA Intergenic NA NA

T11144 TGCCGAGCGACACTCGACACAGG 1000
Oligo fwd CACCTGCCGAGCGACACTCGACAC
Oligo rev AAACGTGTCGAGTGTCGCTCGGCA
Chromosome start end strand MM target_seq PAM alignment distance position gene name gene id
GU305914.1 145566 145588 + 0 TGCCGAGCGACACTCGACAC AGG ||||||||[||||||||||||]PAM NA Intergenic NA NA

T11145 GCCGAGCGACACTCGACACAGGG 1000
Oligo fwd CACCGCCGAGCGACACTCGACACA
Oligo rev AAACTGTGTCGAGTGTCGCTCGGC
Chromosome start end strand MM target_seq PAM alignment distance position gene name gene id
GU305914.1 145567 145589 + 0 GCCGAGCGACACTCGACACA GGG ||||||||[||||||||||||]PAM NA Intergenic NA NA

T11146 CGACACAGGGCGCCTCATCTCGG 1000
Oligo fwd CACCCGACACAGGGCGCCTCATCT
Oligo rev AAACAGATGAGGCGCCCTGTGTCG
Chromosome start end strand MM target_seq PAM alignment distance position gene name gene id
GU305914.1 145580 145602 + 0 CGACACAGGGCGCCTCATCT CGG ||||||||[||||||||||||]PAM NA Intergenic NA NA

T11147 CGCCTCATCTCGGCTGCCCCCGG 1000
Oligo fwd CACCCGCCTCATCTCGGCTGCCCC
Oligo rev AAACGGGGCAGCCGAGATGAGGCG
Chromosome start end strand MM target_seq PAM alignment distance position gene name gene id
GU305914.1 145590 145612 + 0 CGCCTCATCTCGGCTGCCCC CGG ||||||||[||||||||||||]PAM NA Intergenic NA NA

T11148 CCCCCGGTACTATAAATCCCAGG 1000
Oligo fwd CACCCCCCCGGTACTATAAATCCC
Oligo rev AAACGGGATTTATAGTACCGGGGG
Chromosome start end strand MM target_seq PAM alignment distance position gene name gene id
GU305914.1 145606 145628 + 0 CCCCCGGTACTATAAATCCC AGG ||||||||[||||||||||||]PAM NA Intergenic NA NA

T11149 CCCCGGTACTATAAATCCCAGGG 1000
Oligo fwd CACCCCCCGGTACTATAAATCCCA
Oligo rev AAACTGGGATTTATAGTACCGGGG
Chromosome start end strand MM target_seq PAM alignment distance position gene name gene id
GU305914.1 145607 145629 + 0 CCCCGGTACTATAAATCCCA GGG ||||||||[||||||||||||]PAM NA Intergenic NA NA

T11150 ATAAATCCCAGGGCTCCTGCTGG 1000
Oligo fwd CACCATAAATCCCAGGGCTCCTGC
Oligo rev AAACGCAGGAGCCCTGGGATTTAT
Chromosome start end strand MM target_seq PAM alignment distance position gene name gene id
GU305914.1 145617 145639 + 0 ATAAATCCCAGGGCTCCTGC TGG ||||||||[||||||||||||]PAM NA Intergenic NA NA

T11151 TAAATCCCAGGGCTCCTGCTGGG 1000
Oligo fwd CACCTAAATCCCAGGGCTCCTGCT
Oligo rev AAACAGCAGGAGCCCTGGGATTTA
Chromosome start end strand MM target_seq PAM alignment distance position gene name gene id
GU305914.1 145618 145640 + 0 TAAATCCCAGGGCTCCTGCT GGG ||||||||[||||||||||||]PAM NA Intergenic NA NA

T11152 CACACATTGTCTCAGACTCTCGG 1000
Oligo fwd CACCCACACATTGTCTCAGACTCT
Oligo rev AAACAGAGTCTGAGACAATGTGTG
Chromosome start end strand MM target_seq PAM alignment distance position gene name gene id
GU305914.1 145648 145670 + 0 CACACATTGTCTCAGACTCT CGG ||||||||[||||||||||||]PAM NA Intergenic NA NA

T11153 ACATTGTCTCAGACTCTCGGCGG 1000

| | | | | | | | | | | | |
|---|---|---|---|---|---|---|---|---|---|---|---|
| Oligo fwd | CACCACATTGTCTCAGACTCTCGG | | | | | | | | | | |
| Oligo rev | AAACCCGAGAGTCTGAGACAATGT | | | | | | | | | | |
| Chromosome | start | end | strand | MM | target_seq | PAM | alignment | distance | position | gene name | gene id |
| GU305914.1 | 145651 | 145673 | + | 0 | ACATTGTCTCAGACTCTCGG | CGG | \|\|\|\|\|\|\|\|[\|\|\|\|\|\|\|\|\|\|\|\|]PAM | NA | Intergenic | NA | NA |
| T11154 | TTGTCTCAGACTCTCGGCGGCGG | 1000 | | | | | | | | | |
| Oligo fwd | CACCTTGTCTCAGACTCTCGGCGG | | | | | | | | | | |
| Oligo rev | AAACCCGCCGAGAGTCTGAGACAA | | | | | | | | | | |
| Chromosome | start | end | strand | MM | target_seq | PAM | alignment | distance | position | gene name | gene id |
| GU305914.1 | 145654 | 145676 | + | 0 | TTGTCTCAGACTCTCGGCGG | CGG | \|\|\|\|\|\|\|\|[\|\|\|\|\|\|\|\|\|\|\|\|]PAM | NA | Intergenic | NA | NA |
| T11155 | GACTCTCGGCGGCGGCGCTTAGG | 1000 | | | | | | | | | |
| Oligo fwd | CACCGACTCTCGGCGGCGGCGCTT | | | | | | | | | | |
| Oligo rev | AAACAAGCGCCGCCGCCGAGAGTC | | | | | | | | | | |
| Chromosome | start | end | strand | MM | target_seq | PAM | alignment | distance | position | gene name | gene id |
| GU305914.1 | 145662 | 145684 | + | 0 | GACTCTCGGCGGCGGCGCTT | AGG | \|\|\|\|\|\|\|\|[\|\|\|\|\|\|\|\|\|\|\|\|]PAM | NA | Intergenic | NA | NA |
| T11156 | ACTCTCGGCGGCGGCGCTTAGGG | 1000 | | | | | | | | | |
| Oligo fwd | CACCACTCTCGGCGGCGGCGCTTA | | | | | | | | | | |
| Oligo rev | AAACTAAGCGCCGCCGCCGAGAGT | | | | | | | | | | |
| Chromosome | start | end | strand | MM | target_seq | PAM | alignment | distance | position | gene name | gene id |
| GU305914.1 | 145663 | 145685 | + | 0 | ACTCTCGGCGGCGGCGCTTA | GGG | \|\|\|\|\|\|\|\|[\|\|\|\|\|\|\|\|\|\|\|\|]PAM | NA | Intergenic | NA | NA |
| T11157 | CGGCGCTTAGGGAATCGCGATGG | 1000 | | | | | | | | | |
| Oligo fwd | CACCCGGCGCTTAGGGAATCGCGA | | | | | | | | | | |
| Oligo rev | AAACTCGCGATTCCCTAAGCGCCG | | | | | | | | | | |
| Chromosome | start | end | strand | MM | target_seq | PAM | alignment | distance | position | gene name | gene id |
| GU305914.1 | 145674 | 145696 | + | 0 | CGGCGCTTAGGGAATCGCGA | TGG | \|\|\|\|\|\|\|\|[\|\|\|\|\|\|\|\|\|\|\|\|]PAM | NA | Intergenic | NA | NA |
| T11158 | CTTAGGGAATCGCGATGGAGAGG | 1000 | | | | | | | | | |
| Oligo fwd | CACCCTTAGGGAATCGCGATGGAG | | | | | | | | | | |
| Oligo rev | AAACCTCCATCGCGATTCCCTAAG | | | | | | | | | | |
| Chromosome | start | end | strand | MM | target_seq | PAM | alignment | distance | position | gene name | gene id |
| GU305914.1 | 145679 | 145701 | + | 0 | CTTAGGGAATCGCGATGGAG | AGG | \|\|\|\|\|\|\|\|[\|\|\|\|\|\|\|\|\|\|\|\|]PAM | NA | Intergenic | NA | NA |
| T11159 | GAATCGCGATGGAGAGGCGATGG | 1000 | | | | | | | | | |
| Oligo fwd | CACCGAATCGCGATGGAGAGGCGA | | | | | | | | | | |
| Oligo rev | AAACTCGCCTCTCCATCGCGATTC | | | | | | | | | | |
| Chromosome | start | end | strand | MM | target_seq | PAM | alignment | distance | position | gene name | gene id |
| GU305914.1 | 145685 | 145707 | + | 0 | GAATCGCGATGGAGAGGCGA | TGG | \|\|\|\|\|\|\|\|[\|\|\|\|\|\|\|\|\|\|\|\|]PAM | NA | Intergenic | NA | NA |
| T11160 | CGATGGAGAGGCGATGGCGTCGG | 1000 | | | | | | | | | |
| Oligo fwd | CACCCGATGGAGAGGCGATGGCGT | | | | | | | | | | |
| Oligo rev | AAACACGCCATCGCCTCTCCATCG | | | | | | | | | | |
| Chromosome | start | end | strand | MM | target_seq | PAM | alignment | distance | position | gene name | gene id |
| GU305914.1 | 145691 | 145713 | + | 0 | CGATGGAGAGGCGATGGCGT | CGG | \|\|\|\|\|\|\|\|[\|\|\|\|\|\|\|\|\|\|\|\|]PAM | NA | Intergenic | NA | NA |
| T11161 | GATGGAGAGGCGATGGCGTCGGG | 1000 | | | | | | | | | |
| Oligo fwd | CACCGATGGAGAGGCGATGGCGTC | | | | | | | | | | |
| Oligo rev | AAACGACGCCATCGCCTCTCCATC | | | | | | | | | | |
| Chromosome | start | end | strand | MM | target_seq | PAM | alignment | distance | position | gene name | gene id |
| GU305914.1 | 145692 | 145714 | + | 0 | GATGGAGAGGCGATGGCGTC | GGG | \|\|\|\|\|\|\|\|[\|\|\|\|\|\|\|\|\|\|\|\|]PAM | NA | Intergenic | NA | NA |
| T11162 | ATGGAGAGGCGATGGCGTCGGGG | 1000 | | | | | | | | | |
| Oligo fwd | CACCATGGAGAGGCGATGGCGTCG | | | | | | | | | | |
| Oligo rev | AAACCGACGCCATCGCCTCTCCAT | | | | | | | | | | |
| Chromosome | start | end | strand | MM | target_seq | PAM | alignment | distance | position | gene name | gene id |
| GU305914.1 | 145693 | 145715 | + | 0 | ATGGAGAGGCGATGGCGTCG | GGG | \|\|\|\|\|\|\|\|[\|\|\|\|\|\|\|\|\|\|\|\|]PAM | NA | Intergenic | NA | NA |
| T11163 | TGGAGAGGCGATGGCGTCGGGGG | 1000 | | | | | | | | | |
| Oligo fwd | CACCTGGAGAGGCGATGGCGTCGG | | | | | | | | | | |
| Oligo rev | AAACCCGACGCCATCGCCTCTCCA | | | | | | | | | | |
| Chromosome | start | end | strand | MM | target_seq | PAM | alignment | distance | position | gene name | gene id |
| GU305914.1 | 145694 | 145716 | + | 0 | TGGAGAGGCGATGGCGTCGG | GGG | \|\|\|\|\|\|\|\|[\|\|\|\|\|\|\|\|\|\|\|\|]PAM | NA | Intergenic | NA | NA |
| GU305914.1 | 176448 | 176470 | + | 4 | TGCTGAGACGATGGCGACGG | CGG | \|\|--\|\|\|-[\|\|\|\|\|\|\|\|-\|\|\|]PAM | NA | Intergenic | NA | NA |
| T11164 | GCGATGGCGTCGGGGGCTGCCGG | 1000 | | | | | | | | | |
| Oligo fwd | CACCGCGATGGCGTCGGGGGCTGC | | | | | | | | | | |
| Oligo rev | AAACGCAGCCCCCGACGCCATCGC | | | | | | | | | | |
| Chromosome | start | end | strand | MM | target_seq | PAM | alignment | distance | position | gene name | gene id |
| GU305914.1 | 145701 | 145723 | + | 0 | GCGATGGCGTCGGGGGCTGC | CGG | \|\|\|\|\|\|\|\|[\|\|\|\|\|\|\|\|\|\|\|\|]PAM | NA | Intergenic | NA | NA |
| T11165 | CATCTATGCTTTCCTGCCCGAGG | 1000 | | | | | | | | | |
| Oligo fwd | CACCCATCTATGCTTTCCTGCCCG | | | | | | | | | | |
| Oligo rev | AAACCGGGCAGGAAAGCATAGATG | | | | | | | | | | |
| Chromosome | start | end | strand | MM | target_seq | PAM | alignment | distance | position | gene name | gene id |
| GU305914.1 | 145734 | 145756 | + | 0 | CATCTATGCTTTCCTGCCCG | AGG | \|\|\|\|\|\|\|\|[\|\|\|\|\|\|\|\|\|\|\|\|]PAM | NA | Intergenic | NA | NA |
| T11166 | TTTCCTGCCCGAGGAGAACCAGG | 1000 | | | | | | | | | |
| Oligo fwd | CACCTTTCCTGCCCGAGGAGAACC | | | | | | | | | | |
| Oligo rev | AAACGGTTCTCCTCGGGCAGGAAA | | | | | | | | | | |
| Chromosome | start | end | strand | MM | target_seq | PAM | alignment | distance | position | gene name | gene id |
| GU305914.1 | 145743 | 145765 | + | 0 | TTTCCTGCCCGAGGAGAACC | AGG | \|\|\|\|\|\|\|\|[\|\|\|\|\|\|\|\|\|\|\|\|]PAM | NA | Intergenic | NA | NA |
| T11167 | GCAGTGTTTCTTCTGCGAAGTGG | 1000 | | | | | | | | | |
| Oligo fwd | CACCGCAGTGTTTCTTCTGCGAAG | | | | | | | | | | |
| Oligo rev | AAACCTTCGCAGAAGAAACACTGC | | | | | | | | | | |
| Chromosome | start | end | strand | MM | target_seq | PAM | alignment | distance | position | gene name | gene id |
| GU305914.1 | 145773 | 145795 | + | 0 | GCAGTGTTTCTTCTGCGAAG | TGG | \|\|\|\|\|\|\|\|[\|\|\|\|\|\|\|\|\|\|\|\|]PAM | NA | Intergenic | NA | NA |

| | | | | | | | | | | | |
|---|---|---|---|---|---|---|---|---|---|---|---|
| T11168 | TTCTTCTGCGAAGTGGCCGACGG | 1000 | | | | | | | | | |
| Oligo fwd | CACCTTCTTCTGCGAAGTGGCCGA | | | | | | | | | | |
| Oligo rev | AAACTCGGCCACTTCGCAGAAGAA | | | | | | | | | | |
| Chromosome | start | end | strand | MM | target_seq | PAM | alignment | distance | position | gene name | gene id |
| GU305914.1 | 145780 | 145802 | + | 0 | TTCTTCTGCGAAGTGGCCGA | CGG | \|\|\|\|\|\|\|\|[\|\|\|\|\|\|\|\|\|\|\|\|]PAM | NA | Intergenic | NA | NA |
| T11169 | GACCAGACTCTTCGTTTTCCCGG | 1000 | | | | | | | | | |
| Oligo fwd | CACCGACCAGACTCTTCGTTTTCC | | | | | | | | | | |
| Oligo rev | AAACGGAAAACGAAGAGTCTGGTC | | | | | | | | | | |
| Chromosome | start | end | strand | MM | target_seq | PAM | alignment | distance | position | gene name | gene id |
| GU305914.1 | 145815 | 145837 | + | 0 | GACCAGACTCTTCGTTTTCC | CGG | \|\|\|\|\|\|\|\|[\|\|\|\|\|\|\|\|\|\|\|\|]PAM | NA | Intergenic | NA | NA |
| T11170 | CCGGTCAATATCCCCGCCGAAGG | 1000 | | | | | | | | | |
| Oligo fwd | CACCCCGGTCAATATCCCCGCCGA | | | | | | | | | | |
| Oligo rev | AAACTCGGCGGGGATATTGACCGG | | | | | | | | | | |
| Chromosome | start | end | strand | MM | target_seq | PAM | alignment | distance | position | gene name | gene id |
| GU305914.1 | 145834 | 145856 | + | 0 | CCGGTCAATATCCCCGCCGA | AGG | \|\|\|\|\|\|\|\|[\|\|\|\|\|\|\|\|\|\|\|\|]PAM | NA | Intergenic | NA | NA |
| T11171 | CGGTCAATATCCCCGCCGAAGGG | 1000 | | | | | | | | | |
| Oligo fwd | CACCCGGTCAATATCCCCGCCGAA | | | | | | | | | | |
| Oligo rev | AAACTTCGGCGGGGATATTGACCG | | | | | | | | | | |
| Chromosome | start | end | strand | MM | target_seq | PAM | alignment | distance | position | gene name | gene id |
| GU305914.1 | 145835 | 145857 | + | 0 | CGGTCAATATCCCCGCCGAA | GGG | \|\|\|\|\|\|\|\|[\|\|\|\|\|\|\|\|\|\|\|\|]PAM | NA | Intergenic | NA | NA |
| T11172 | GGTCAATATCCCCGCCGAAGGGG | 1000 | | | | | | | | | |
| Oligo fwd | CACCGGTCAATATCCCCGCCGAAG | | | | | | | | | | |
| Oligo rev | AAACCTTCGGCGGGGATATTGACC | | | | | | | | | | |
| Chromosome | start | end | strand | MM | target_seq | PAM | alignment | distance | position | gene name | gene id |
| GU305914.1 | 145836 | 145858 | + | 0 | GGTCAATATCCCCGCCGAAG | GGG | \|\|\|\|\|\|\|\|[\|\|\|\|\|\|\|\|\|\|\|\|]PAM | NA | Intergenic | NA | NA |
| T11173 | CCTGCTCGCTAATTACACGTCGG | 1000 | | | | | | | | | |
| Oligo fwd | CACCCCTGCTCGCTAATTACACGT | | | | | | | | | | |
| Oligo rev | AAACACGTGTAATTAGCGAGCAGG | | | | | | | | | | |
| Chromosome | start | end | strand | MM | target_seq | PAM | alignment | distance | position | gene name | gene id |
| GU305914.1 | 145872 | 145894 | + | 0 | CCTGCTCGCTAATTACACGT | CGG | \|\|\|\|\|\|\|\|[\|\|\|\|\|\|\|\|\|\|\|\|]PAM | NA | Intergenic | NA | NA |
| T11174 | TTCTTCCCTCGACACGATCCCGG | 1000 | | | | | | | | | |
| Oligo fwd | CACCTTCTTCCCTCGACACGATCC | | | | | | | | | | |
| Oligo rev | AAACGGATCGTGTCGAGGGAAGAA | | | | | | | | | | |
| Chromosome | start | end | strand | MM | target_seq | PAM | alignment | distance | position | gene name | gene id |
| GU305914.1 | 145905 | 145927 | + | 0 | TTCTTCCCTCGACACGATCC | CGG | \|\|\|\|\|\|\|\|[\|\|\|\|\|\|\|\|\|\|\|\|]PAM | NA | Intergenic | NA | NA |
| T11175 | GATTCGACGACCGCCGCTAACGG | 1000 | | | | | | | | | |
| Oligo fwd | CACCGATTCGACGACCGCCGCTAA | | | | | | | | | | |
| Oligo rev | AAACTTAGCGGCGGTCGTCGAATC | | | | | | | | | | |
| Chromosome | start | end | strand | MM | target_seq | PAM | alignment | distance | position | gene name | gene id |
| GU305914.1 | 145927 | 145949 | + | 0 | GATTCGACGACCGCCGCTAA | CGG | \|\|\|\|\|\|\|\|[\|\|\|\|\|\|\|\|\|\|\|\|]PAM | NA | Intergenic | NA | NA |
| T11176 | ATTCGACGACCGCCGCTAACGGG | 1000 | | | | | | | | | |
| Oligo fwd | CACCATTCGACGACCGCCGCTAAC | | | | | | | | | | |
| Oligo rev | AAACGTTAGCGGCGGTCGTCGAAT | | | | | | | | | | |
| Chromosome | start | end | strand | MM | target_seq | PAM | alignment | distance | position | gene name | gene id |
| GU305914.1 | 145928 | 145950 | + | 0 | ATTCGACGACCGCCGCTAAC | GGG | \|\|\|\|\|\|\|\|[\|\|\|\|\|\|\|\|\|\|\|\|]PAM | NA | Intergenic | NA | NA |
| T11177 | GACGACCGCCGCTAACGGGTCGG | 1000 | | | | | | | | | |
| Oligo fwd | CACCGACGACCGCCGCTAACGGGT | | | | | | | | | | |
| Oligo rev | AAACACCCGTTAGCGGCGGTCGTC | | | | | | | | | | |
| Chromosome | start | end | strand | MM | target_seq | PAM | alignment | distance | position | gene name | gene id |
| GU305914.1 | 145932 | 145954 | + | 0 | GACGACCGCCGCTAACGGGT | CGG | \|\|\|\|\|\|\|\|[\|\|\|\|\|\|\|\|\|\|\|\|]PAM | NA | Intergenic | NA | NA |
| T11178 | ACGACCGCCGCTAACGGGTCGGG | 1000 | | | | | | | | | |
| Oligo fwd | CACCACGACCGCCGCTAACGGGTC | | | | | | | | | | |
| Oligo rev | AAACGACCCGTTAGCGGCGGTCGT | | | | | | | | | | |
| Chromosome | start | end | strand | MM | target_seq | PAM | alignment | distance | position | gene name | gene id |
| GU305914.1 | 145933 | 145955 | + | 0 | ACGACCGCCGCTAACGGGTC | GGG | \|\|\|\|\|\|\|\|[\|\|\|\|\|\|\|\|\|\|\|\|]PAM | NA | Intergenic | NA | NA |
| T11179 | CGACCGCCGCTAACGGGTCGGGG | 1000 | | | | | | | | | |
| Oligo fwd | CACCCGACCGCCGCTAACGGGTCG | | | | | | | | | | |
| Oligo rev | AAACCGACCCGTTAGCGGCGGTCG | | | | | | | | | | |
| Chromosome | start | end | strand | MM | target_seq | PAM | alignment | distance | position | gene name | gene id |
| GU305914.1 | 145934 | 145956 | + | 0 | CGACCGCCGCTAACGGGTCG | GGG | \|\|\|\|\|\|\|\|[\|\|\|\|\|\|\|\|\|\|\|\|]PAM | NA | Intergenic | NA | NA |
| T11180 | GACCGCCGCTAACGGGTCGGGGG | 1000 | | | | | | | | | |
| Oligo fwd | CACCGACCGCCGCTAACGGGTCGG | | | | | | | | | | |
| Oligo rev | AAACCCGACCCGTTAGCGGCGGTC | | | | | | | | | | |
| Chromosome | start | end | strand | MM | target_seq | PAM | alignment | distance | position | gene name | gene id |
| GU305914.1 | 145935 | 145957 | + | 0 | GACCGCCGCTAACGGGTCGG | GGG | \|\|\|\|\|\|\|\|[\|\|\|\|\|\|\|\|\|\|\|\|]PAM | NA | Intergenic | NA | NA |
| T11181 | GCCGCTAACGGGTCGGGGGTAGG | 1000 | | | | | | | | | |
| Oligo fwd | CACCGCCGCTAACGGGTCGGGGGT | | | | | | | | | | |
| Oligo rev | AAACACCCCCGACCCGTTAGCGGC | | | | | | | | | | |
| Chromosome | start | end | strand | MM | target_seq | PAM | alignment | distance | position | gene name | gene id |
| GU305914.1 | 145939 | 145961 | + | 0 | GCCGCTAACGGGTCGGGGGT | AGG | \|\|\|\|\|\|\|\|[\|\|\|\|\|\|\|\|\|\|\|\|]PAM | NA | Intergenic | NA | NA |
| T11182 | GCTAACGGGTCGGGGGTAGGAGG | 1000 | | | | | | | | | |
| Oligo fwd | CACCGCTAACGGGTCGGGGGTAGG | | | | | | | | | | |
| Oligo rev | AAACCCTACCCCCGACCCGTTAGC | | | | | | | | | | |
| Chromosome | start | end | strand | MM | target_seq | PAM | alignment | distance | position | gene name | gene id |

| | | | | | | | | | | | |
|---|---|---|---|---|---|---|---|---|---|---|---|
| GU305914.1 | | 145942 | 145964 | + | 0 | GCTAACGGGTCGGGGGTAGG | AGG | \|\|\|\|\|\|\|\|[\|\|\|\|\|\|\|\|\|\|\|\|]PAM | NA | Intergenic | NA | NA |
| T11183 | CGGGTCGGGGGTAGGAGGAGAGG | 1000 | | | | | | | | | |
| Oligo fwd | CACCGGGTCGGGGGTAGGAGGAG | | | | | | | | | | |
| Oligo rev | AAACCTCCTCCTACCCCCGACCCG | | | | | | | | | | |
| Chromosome | start | end | strand | MM | target_seq | PAM | alignment | distance | position | gene name | gene id |
| GU305914.1 | 145947 | 145969 | + | 0 | CGGGTCGGGGGTAGGAGGAG | AGG | \|\|\|\|\|\|\|\|[\|\|\|\|\|\|\|\|\|\|\|\|]PAM | NA | Intergenic | NA | NA |
| T11184 | TCGGGGGTAGGAGGAGAGGTCGG | 1000 | | | | | | | | | |
| Oligo fwd | CACCTCGGGGGTAGGAGGAGAGGT | | | | | | | | | | |
| Oligo rev | AAACACCTCTCCTCCTACCCCCGA | | | | | | | | | | |
| Chromosome | start | end | strand | MM | target_seq | PAM | alignment | distance | position | gene name | gene id |
| GU305914.1 | 145951 | 145973 | + | 0 | TCGGGGGTAGGAGGAGAGGT | CGG | \|\|\|\|\|\|\|\|[\|\|\|\|\|\|\|\|\|\|\|\|]PAM | NA | Intergenic | NA | NA |
| T11185 | GGGGTAGGAGGAGAGGTCGGTGG | 1000 | | | | | | | | | |
| Oligo fwd | CACCGGGGTAGGAGGAGAGGTCGG | | | | | | | | | | |
| Oligo rev | AAACCCGACCTCTCCTCCTACCCC | | | | | | | | | | |
| Chromosome | start | end | strand | MM | target_seq | PAM | alignment | distance | position | gene name | gene id |
| GU305914.1 | 145954 | 145976 | + | 0 | GGGGTAGGAGGAGAGGTCGG | TGG | \|\|\|\|\|\|\|\|[\|\|\|\|\|\|\|\|\|\|\|\|]PAM | NA | Intergenic | NA | NA |
| T11186 | GTAGGAGGAGAGGTCGGTGGAGG | 1000 | | | | | | | | | |
| Oligo fwd | CACCGTAGGAGGAGAGGTCGGTGG | | | | | | | | | | |
| Oligo rev | AAACCCACCGACCTCTCCTCCTAC | | | | | | | | | | |
| Chromosome | start | end | strand | MM | target_seq | PAM | alignment | distance | position | gene name | gene id |
| GU305914.1 | 145957 | 145979 | + | 0 | GTAGGAGGAGAGGTCGGTGG | AGG | \|\|\|\|\|\|\|\|[\|\|\|\|\|\|\|\|\|\|\|\|]PAM | NA | Intergenic | NA | NA |
| T11187 | GGAGGAGAGGTCGGTGGAGGAGG | 1000 | | | | | | | | | |
| Oligo fwd | CACCGGAGGAGAGGTCGGTGGAGG | | | | | | | | | | |
| Oligo rev | AAACCCTCCACCGACCTCTCCTCC | | | | | | | | | | |
| Chromosome | start | end | strand | MM | target_seq | PAM | alignment | distance | position | gene name | gene id |
| GU305914.1 | 145960 | 145982 | + | 0 | GGAGGAGAGGTCGGTGGAGG | AGG | \|\|\|\|\|\|\|\|[\|\|\|\|\|\|\|\|\|\|\|\|]PAM | NA | Intergenic | NA | NA |
| GU305914.1 | 57997 | 58019 | - | 4 | GGTGGAGGGGCCGGGGGAGG | CGG | \|\|-\|\|\|\|-[\|\|-\|\|\|-\|\|\|\|\|]PAM | NA | Intergenic | NA | NA |
| T11188 | GGAGAGGTCGGTGGAGGAGGCGG | 1000 | | | | | | | | | |
| Oligo fwd | CACCGGAGAGGTCGGTGGAGGAGG | | | | | | | | | | |
| Oligo rev | AAACCCTCCTCCACCGACCTCTCC | | | | | | | | | | |
| Chromosome | start | end | strand | MM | target_seq | PAM | alignment | distance | position | gene name | gene id |
| GU305914.1 | 145963 | 145985 | + | 0 | GGAGAGGTCGGTGGAGGAGG | CGG | \|\|\|\|\|\|\|\|[\|\|\|\|\|\|\|\|\|\|\|\|]PAM | NA | Intergenic | NA | NA |
| GU305914.1 | 77071 | 77093 | + | 4 | GGAGACGATGCTGGAGGAGG | AGG | \|\|\|\|\|-\|-[-\|-\|\|\|\|\|\|\|\|\|]PAM | NA | Intergenic | NA | NA |
| T11189 | GAGAGGTCGGTGGAGGAGGCGGG | 1000 | | | | | | | | | |
| Oligo fwd | CACCGAGAGGTCGGTGGAGGAGGC | | | | | | | | | | |
| Oligo rev | AAACGCCTCCTCCACCGACCTCTC | | | | | | | | | | |
| Chromosome | start | end | strand | MM | target_seq | PAM | alignment | distance | position | gene name | gene id |
| GU305914.1 | 145964 | 145986 | + | 0 | GAGAGGTCGGTGGAGGAGGC | GGG | \|\|\|\|\|\|\|\|[\|\|\|\|\|\|\|\|\|\|\|\|]PAM | NA | Intergenic | NA | NA |
| T11190 | GTCGGTGGAGGAGGCGGGAGCGG | 1000 | | | | | | | | | |
| Oligo fwd | CACCGTCGGTGGAGGAGGCGGGAG | | | | | | | | | | |
| Oligo rev | AAACCTCCCGCCTCCTCCACCGAC | | | | | | | | | | |
| Chromosome | start | end | strand | MM | target_seq | PAM | alignment | distance | position | gene name | gene id |
| GU305914.1 | 145969 | 145991 | + | 0 | GTCGGTGGAGGAGGCGGGAG | CGG | \|\|\|\|\|\|\|\|[\|\|\|\|\|\|\|\|\|\|\|\|]PAM | NA | Intergenic | NA | NA |
| GU305914.1 | 73780 | 73802 | - | 4 | GGCGGCGGAGGAGGAGGAAG | CGG | \|-\|\|\|-\|\|[\|\|\|\|\|\|-\|\|-\|\|]PAM | NA | Intergenic | NA | NA |
| T11191 | GCGCCGATCGCGTCGCGCGCCGG | 1000 | | | | | | | | | |
| Oligo fwd | CACCGCGCCGATCGCGTCGCGCGC | | | | | | | | | | |
| Oligo rev | AAACGCGCGCGACGCGATCGGCGC | | | | | | | | | | |
| Chromosome | start | end | strand | MM | target_seq | PAM | alignment | distance | position | gene name | gene id |
| GU305914.1 | 146000 | 146022 | + | 0 | GCGCCGATCGCGTCGCGCGC | CGG | \|\|\|\|\|\|\|\|[\|\|\|\|\|\|\|\|\|\|\|\|]PAM | NA | Intergenic | NA | NA |
| T11192 | ATCGCGTCGCGCGCCGGTCGCGG | 1000 | | | | | | | | | |
| Oligo fwd | CACCATCGCGTCGCGCGCCGGTCG | | | | | | | | | | |
| Oligo rev | AAACCGACCGGCGCGCGACGCGAT | | | | | | | | | | |
| Chromosome | start | end | strand | MM | target_seq | PAM | alignment | distance | position | gene name | gene id |
| GU305914.1 | 146006 | 146028 | + | 0 | ATCGCGTCGCGCGCCGGTCG | CGG | \|\|\|\|\|\|\|\|[\|\|\|\|\|\|\|\|\|\|\|\|]PAM | NA | Intergenic | NA | NA |
| GU305914.1 | 201079 | 201101 | + | 4 | CTCTCGTCGCGCGGCGGTGG | CGG | -\|\|-\|\|\|\|[\|\|\|\|\|-\|\|\|\|-\|]PAM | NA | Intergenic | NA | NA |
| T11193 | TCGCGTCGCGCGCCGGTCGCGGG | 1000 | | | | | | | | | |
| Oligo fwd | CACCTCGCGTCGCGCGCCGGTCGC | | | | | | | | | | |
| Oligo rev | AAACGCGACCGGCGCGCGACGCGA | | | | | | | | | | |
| Chromosome | start | end | strand | MM | target_seq | PAM | alignment | distance | position | gene name | gene id |
| GU305914.1 | 146007 | 146029 | + | 0 | TCGCGTCGCGCGCCGGTCGC | GGG | \|\|\|\|\|\|\|\|[\|\|\|\|\|\|\|\|\|\|\|\|]PAM | NA | Intergenic | NA | NA |
| GU305914.1 | 201080 | 201102 | + | 3 | TCTCGTCGCGCGGCGGTGGC | GGG | \|\|-\|\|\|\|\|[\|\|\|\|-\|\|\|\|-\|\|]PAM | NA | Intergenic | NA | NA |
| T11194 | GCACAGATCCTCATGCACGTCGG | 1000 | | | | | | | | | |
| Oligo fwd | CACCGCACAGATCCTCATGCACGT | | | | | | | | | | |
| Oligo rev | AAACACGTGCATGAGGATCTGTGC | | | | | | | | | | |
| Chromosome | start | end | strand | MM | target_seq | PAM | alignment | distance | position | gene name | gene id |
| GU305914.1 | 146035 | 146057 | + | 0 | GCACAGATCCTCATGCACGT | CGG | \|\|\|\|\|\|\|\|[\|\|\|\|\|\|\|\|\|\|\|\|]PAM | NA | Intergenic | NA | NA |
| T11195 | CACAGATCCTCATGCACGTCGGG | 1000 | | | | | | | | | |
| Oligo fwd | CACCCACAGATCCTCATGCACGTC | | | | | | | | | | |
| Oligo rev | AAACGACGTGCATGAGGATCTGTG | | | | | | | | | | |
| Chromosome | start | end | strand | MM | target_seq | PAM | alignment | distance | position | gene name | gene id |
| GU305914.1 | 146036 | 146058 | + | 0 | CACAGATCCTCATGCACGTC | GGG | \|\|\|\|\|\|\|\|[\|\|\|\|\|\|\|\|\|\|\|\|]PAM | NA | Intergenic | NA | NA |
| T11196 | GATCCTCATGCACGTCGGGCCGG | 1000 | | | | | | | | | |
| Oligo fwd | CACCGATCCTCATGCACGTCGGGC | | | | | | | | | | |
| Oligo rev | AAACGCCCGACGTGCATGAGGATC | | | | | | | | | | |
| Chromosome | start | end | strand | MM | target_seq | PAM | alignment | distance | position | gene name | gene id |

<table>
<tr><td>GU305914.1</td><td></td><td>146040</td><td>146062</td><td>+</td><td>0</td><td>GATCCTCATGCACGTCGGGC</td><td>CGG</td><td>||||||||[||||||||||||]PAM</td><td>NA</td><td>Intergenic</td><td>NA</td><td>NA</td></tr>
<tr><td>T11197</td><td>CGGACTTCGAGCCCCTGACTAGG</td><td></td><td>1000</td><td></td><td></td><td></td><td></td><td></td><td></td><td></td><td></td><td></td></tr>
<tr><td>Oligo fwd</td><td>CACCCGGACTTCGAGCCCCTGACT</td><td></td><td></td><td></td><td></td><td></td><td></td><td></td><td></td><td></td><td></td><td></td></tr>
<tr><td>Oligo rev</td><td>AAACAGTCAGGGGCTCGAAGTCCG</td><td></td><td></td><td></td><td></td><td></td><td></td><td></td><td></td><td></td><td></td><td></td></tr>
<tr><td>Chromosome</td><td>start</td><td></td><td>end</td><td>strand</td><td>MM</td><td>target_seq</td><td>PAM</td><td>alignment</td><td>distance</td><td>position</td><td>gene name</td><td>gene id</td></tr>
<tr><td>GU305914.1</td><td></td><td>146060</td><td>146082</td><td>+</td><td>0</td><td>CGGACTTCGAGCCCCTGACT</td><td>AGG</td><td>||||||||[||||||||||||]PAM</td><td>NA</td><td>Intergenic</td><td>NA</td><td>NA</td></tr>
<tr><td>T11198</td><td>CCCGTCGAGATCTTCTCCGACGG</td><td></td><td>1000</td><td></td><td></td><td></td><td></td><td></td><td></td><td></td><td></td><td></td></tr>
<tr><td>Oligo fwd</td><td>CACCCCCGTCGAGATCTTCTCCGA</td><td></td><td></td><td></td><td></td><td></td><td></td><td></td><td></td><td></td><td></td><td></td></tr>
<tr><td>Oligo rev</td><td>AAACTCGGAGAAGATCTCGACGGG</td><td></td><td></td><td></td><td></td><td></td><td></td><td></td><td></td><td></td><td></td><td></td></tr>
<tr><td>Chromosome</td><td>start</td><td></td><td>end</td><td>strand</td><td>MM</td><td>target_seq</td><td>PAM</td><td>alignment</td><td>distance</td><td>position</td><td>gene name</td><td>gene id</td></tr>
<tr><td>GU305914.1</td><td></td><td>146092</td><td>146114</td><td>+</td><td>0</td><td>CCCGTCGAGATCTTCTCCGA</td><td>CGG</td><td>||||||||[||||||||||||]PAM</td><td>NA</td><td>Intergenic</td><td>NA</td><td>NA</td></tr>
<tr><td>T11199</td><td>TCCGACGGCGTCGAGATCCTTGG</td><td></td><td>1000</td><td></td><td></td><td></td><td></td><td></td><td></td><td></td><td></td><td></td></tr>
<tr><td>Oligo fwd</td><td>CACCTCCGACGGCGTCGAGATCCT</td><td></td><td></td><td></td><td></td><td></td><td></td><td></td><td></td><td></td><td></td><td></td></tr>
<tr><td>Oligo rev</td><td>AAACAGGATCTCGACGCCGTCGGA</td><td></td><td></td><td></td><td></td><td></td><td></td><td></td><td></td><td></td><td></td><td></td></tr>
<tr><td>Chromosome</td><td>start</td><td></td><td>end</td><td>strand</td><td>MM</td><td>target_seq</td><td>PAM</td><td>alignment</td><td>distance</td><td>position</td><td>gene name</td><td>gene id</td></tr>
<tr><td>GU305914.1</td><td></td><td>146107</td><td>146129</td><td>+</td><td>0</td><td>TCCGACGGCGTCGAGATCCT</td><td>TGG</td><td>||||||||[||||||||||||]PAM</td><td>NA</td><td>Intergenic</td><td>NA</td><td>NA</td></tr>
<tr><td>T11200</td><td>GCGTCGAGATCCTTGGCTCCCGG</td><td></td><td>1000</td><td></td><td></td><td></td><td></td><td></td><td></td><td></td><td></td><td></td></tr>
<tr><td>Oligo fwd</td><td>CACCGCGTCGAGATCCTTGGCTCC</td><td></td><td></td><td></td><td></td><td></td><td></td><td></td><td></td><td></td><td></td><td></td></tr>
<tr><td>Oligo rev</td><td>AAACGGAGCCAAGGATCTCGACGC</td><td></td><td></td><td></td><td></td><td></td><td></td><td></td><td></td><td></td><td></td><td></td></tr>
<tr><td>Chromosome</td><td>start</td><td></td><td>end</td><td>strand</td><td>MM</td><td>target_seq</td><td>PAM</td><td>alignment</td><td>distance</td><td>position</td><td>gene name</td><td>gene id</td></tr>
<tr><td>GU305914.1</td><td></td><td>146114</td><td>146136</td><td>+</td><td>0</td><td>GCGTCGAGATCCTTGGCTCC</td><td>CGG</td><td>||||||||[||||||||||||]PAM</td><td>NA</td><td>Intergenic</td><td>NA</td><td>NA</td></tr>
<tr><td>T11201</td><td>AGATCCTTGGCTCCCGGAGATGG</td><td></td><td>1000</td><td></td><td></td><td></td><td></td><td></td><td></td><td></td><td></td><td></td></tr>
<tr><td>Oligo fwd</td><td>CACCAGATCCTTGGCTCCCGGAGA</td><td></td><td></td><td></td><td></td><td></td><td></td><td></td><td></td><td></td><td></td><td></td></tr>
<tr><td>Oligo rev</td><td>AAACTCTCCGGGAGCCAAGGATCT</td><td></td><td></td><td></td><td></td><td></td><td></td><td></td><td></td><td></td><td></td><td></td></tr>
<tr><td>Chromosome</td><td>start</td><td></td><td>end</td><td>strand</td><td>MM</td><td>target_seq</td><td>PAM</td><td>alignment</td><td>distance</td><td>position</td><td>gene name</td><td>gene id</td></tr>
<tr><td>GU305914.1</td><td></td><td>146120</td><td>146142</td><td>+</td><td>0</td><td>AGATCCTTGGCTCCCGGAGA</td><td>TGG</td><td>||||||||[||||||||||||]PAM</td><td>NA</td><td>Intergenic</td><td>NA</td><td>NA</td></tr>
<tr><td>T11202</td><td>GATCCTTGGCTCCCGGAGATGGG</td><td></td><td>1000</td><td></td><td></td><td></td><td></td><td></td><td></td><td></td><td></td><td></td></tr>
<tr><td>Oligo fwd</td><td>CACCGATCCTTGGCTCCCGGAGAT</td><td></td><td></td><td></td><td></td><td></td><td></td><td></td><td></td><td></td><td></td><td></td></tr>
<tr><td>Oligo rev</td><td>AAACATCTCCGGGAGCCAAGGATC</td><td></td><td></td><td></td><td></td><td></td><td></td><td></td><td></td><td></td><td></td><td></td></tr>
<tr><td>Chromosome</td><td>start</td><td></td><td>end</td><td>strand</td><td>MM</td><td>target_seq</td><td>PAM</td><td>alignment</td><td>distance</td><td>position</td><td>gene name</td><td>gene id</td></tr>
<tr><td>GU305914.1</td><td></td><td>146121</td><td>146143</td><td>+</td><td>0</td><td>GATCCTTGGCTCCCGGAGAT</td><td>GGG</td><td>||||||||[||||||||||||]PAM</td><td>NA</td><td>Intergenic</td><td>NA</td><td>NA</td></tr>
<tr><td>T11203</td><td>CGGAGATGGGAGAATGTGCGCGG</td><td></td><td>1000</td><td></td><td></td><td></td><td></td><td></td><td></td><td></td><td></td><td></td></tr>
<tr><td>Oligo fwd</td><td>CACCCGGAGATGGGAGAATGTGCG</td><td></td><td></td><td></td><td></td><td></td><td></td><td></td><td></td><td></td><td></td><td></td></tr>
<tr><td>Oligo rev</td><td>AAACCGCACATTCTCCCATCTCCG</td><td></td><td></td><td></td><td></td><td></td><td></td><td></td><td></td><td></td><td></td><td></td></tr>
<tr><td>Chromosome</td><td>start</td><td></td><td>end</td><td>strand</td><td>MM</td><td>target_seq</td><td>PAM</td><td>alignment</td><td>distance</td><td>position</td><td>gene name</td><td>gene id</td></tr>
<tr><td>GU305914.1</td><td></td><td>146134</td><td>146156</td><td>+</td><td>0</td><td>CGGAGATGGGAGAATGTGCG</td><td>CGG</td><td>||||||||[||||||||||||]PAM</td><td>NA</td><td>Intergenic</td><td>NA</td><td>NA</td></tr>
<tr><td>T11204</td><td>CAACAGCATCCTGCAGACCCTGG</td><td></td><td>1000</td><td></td><td></td><td></td><td></td><td></td><td></td><td></td><td></td><td></td></tr>
<tr><td>Oligo fwd</td><td>CACCCAACAGCATCCTGCAGACCC</td><td></td><td></td><td></td><td></td><td></td><td></td><td></td><td></td><td></td><td></td><td></td></tr>
<tr><td>Oligo rev</td><td>AAACGGGTCTGCAGGATGCTGTTG</td><td></td><td></td><td></td><td></td><td></td><td></td><td></td><td></td><td></td><td></td><td></td></tr>
<tr><td>Chromosome</td><td>start</td><td></td><td>end</td><td>strand</td><td>MM</td><td>target_seq</td><td>PAM</td><td>alignment</td><td>distance</td><td>position</td><td>gene name</td><td>gene id</td></tr>
<tr><td>GU305914.1</td><td></td><td>146163</td><td>146185</td><td>+</td><td>0</td><td>CAACAGCATCCTGCAGACCC</td><td>TGG</td><td>||||||||[||||||||||||]PAM</td><td>NA</td><td>Intergenic</td><td>NA</td><td>NA</td></tr>
<tr><td>T11205</td><td>GCAGACCCTGGCCGCCCTGACGG</td><td></td><td>1000</td><td></td><td></td><td></td><td></td><td></td><td></td><td></td><td></td><td></td></tr>
<tr><td>Oligo fwd</td><td>CACCGCAGACCCTGGCCGCCCTGA</td><td></td><td></td><td></td><td></td><td></td><td></td><td></td><td></td><td></td><td></td><td></td></tr>
<tr><td>Oligo rev</td><td>AAACTCAGGGCGGCCAGGGTCTGC</td><td></td><td></td><td></td><td></td><td></td><td></td><td></td><td></td><td></td><td></td><td></td></tr>
<tr><td>Chromosome</td><td>start</td><td></td><td>end</td><td>strand</td><td>MM</td><td>target_seq</td><td>PAM</td><td>alignment</td><td>distance</td><td>position</td><td>gene name</td><td>gene id</td></tr>
<tr><td>GU305914.1</td><td></td><td>146175</td><td>146197</td><td>+</td><td>0</td><td>GCAGACCCTGGCCGCCCTGA</td><td>CGG</td><td>||||||||[||||||||||||]PAM</td><td>NA</td><td>Intergenic</td><td>NA</td><td>NA</td></tr>
<tr><td>T11206</td><td>ACCCTGGCCGCCCTGACGGTCGG</td><td></td><td>1000</td><td></td><td></td><td></td><td></td><td></td><td></td><td></td><td></td><td></td></tr>
<tr><td>Oligo fwd</td><td>CACCACCCTGGCCGCCCTGACGGT</td><td></td><td></td><td></td><td></td><td></td><td></td><td></td><td></td><td></td><td></td><td></td></tr>
<tr><td>Oligo rev</td><td>AAACACCGTCAGGGCGGCCAGGGT</td><td></td><td></td><td></td><td></td><td></td><td></td><td></td><td></td><td></td><td></td><td></td></tr>
<tr><td>Chromosome</td><td>start</td><td></td><td>end</td><td>strand</td><td>MM</td><td>target_seq</td><td>PAM</td><td>alignment</td><td>distance</td><td>position</td><td>gene name</td><td>gene id</td></tr>
<tr><td>GU305914.1</td><td></td><td>146179</td><td>146201</td><td>+</td><td>0</td><td>ACCCTGGCCGCCCTGACGGT</td><td>CGG</td><td>||||||||[||||||||||||]PAM</td><td>NA</td><td>Intergenic</td><td>NA</td><td>NA</td></tr>
<tr><td>T11207</td><td>TGGCCGCCCTGACGGTCGGCAGG</td><td></td><td>1000</td><td></td><td></td><td></td><td></td><td></td><td></td><td></td><td></td><td></td></tr>
<tr><td>Oligo fwd</td><td>CACCTGGCCGCCCTGACGGTCGGC</td><td></td><td></td><td></td><td></td><td></td><td></td><td></td><td></td><td></td><td></td><td></td></tr>
<tr><td>Oligo rev</td><td>AAACGCCGACCGTCAGGGCGGCCA</td><td></td><td></td><td></td><td></td><td></td><td></td><td></td><td></td><td></td><td></td><td></td></tr>
<tr><td>Chromosome</td><td>start</td><td></td><td>end</td><td>strand</td><td>MM</td><td>target_seq</td><td>PAM</td><td>alignment</td><td>distance</td><td>position</td><td>gene name</td><td>gene id</td></tr>
<tr><td>GU305914.1</td><td></td><td>146183</td><td>146205</td><td>+</td><td>0</td><td>TGGCCGCCCTGACGGTCGGC</td><td>AGG</td><td>||||||||[||||||||||||]PAM</td><td>NA</td><td>Intergenic</td><td>NA</td><td>NA</td></tr>
<tr><td>T11208</td><td>GTCGGCAGGATCCGAGCCATCGG</td><td></td><td>1000</td><td></td><td></td><td></td><td></td><td></td><td></td><td></td><td></td><td></td></tr>
<tr><td>Oligo fwd</td><td>CACCGTCGGCAGGATCCGAGCCAT</td><td></td><td></td><td></td><td></td><td></td><td></td><td></td><td></td><td></td><td></td><td></td></tr>
<tr><td>Oligo rev</td><td>AAACATGGCTCGGATCCTGCCGAC</td><td></td><td></td><td></td><td></td><td></td><td></td><td></td><td></td><td></td><td></td><td></td></tr>
<tr><td>Chromosome</td><td>start</td><td></td><td>end</td><td>strand</td><td>MM</td><td>target_seq</td><td>PAM</td><td>alignment</td><td>distance</td><td>position</td><td>gene name</td><td>gene id</td></tr>
<tr><td>GU305914.1</td><td></td><td>146197</td><td>146219</td><td>+</td><td>0</td><td>GTCGGCAGGATCCGAGCCAT</td><td>CGG</td><td>||||||||[||||||||||||]PAM</td><td>NA</td><td>Intergenic</td><td>NA</td><td>NA</td></tr>
<tr><td>T11209</td><td>CATCGGCTCCTATCACCAGATGG</td><td></td><td>1000</td><td></td><td></td><td></td><td></td><td></td><td></td><td></td><td></td><td></td></tr>
<tr><td>Oligo fwd</td><td>CACCCATCGGCTCCTATCACCAGA</td><td></td><td></td><td></td><td></td><td></td><td></td><td></td><td></td><td></td><td></td><td></td></tr>
<tr><td>Oligo rev</td><td>AAACTCTGGTGATAGGAGCCGATG</td><td></td><td></td><td></td><td></td><td></td><td></td><td></td><td></td><td></td><td></td><td></td></tr>
<tr><td>Chromosome</td><td>start</td><td></td><td>end</td><td>strand</td><td>MM</td><td>target_seq</td><td>PAM</td><td>alignment</td><td>distance</td><td>position</td><td>gene name</td><td>gene id</td></tr>
<tr><td>GU305914.1</td><td></td><td>146214</td><td>146236</td><td>+</td><td>0</td><td>CATCGGCTCCTATCACCAGA</td><td>TGG</td><td>||||||||[||||||||||||]PAM</td><td>NA</td><td>Intergenic</td><td>NA</td><td>NA</td></tr>
<tr><td>T11210</td><td>CGGCTCCTATCACCAGATGGCGG</td><td></td><td>1000</td><td></td><td></td><td></td><td></td><td></td><td></td><td></td><td></td><td></td></tr>
<tr><td>Oligo fwd</td><td>CACCCGGCTCCTATCACCAGATGG</td><td></td><td></td><td></td><td></td><td></td><td></td><td></td><td></td><td></td><td></td><td></td></tr>
<tr><td>Oligo rev</td><td>AAACCCATCTGGTGATAGGAGCCG</td><td></td><td></td><td></td><td></td><td></td><td></td><td></td><td></td><td></td><td></td><td></td></tr>
<tr><td>Chromosome</td><td>start</td><td></td><td>end</td><td>strand</td><td>MM</td><td>target_seq</td><td>PAM</td><td>alignment</td><td>distance</td><td>position</td><td>gene name</td><td>gene id</td></tr>
<tr><td>GU305914.1</td><td></td><td>146217</td><td>146239</td><td>+</td><td>0</td><td>CGGCTCCTATCACCAGATGG</td><td>CGG</td><td>||||||||[||||||||||||]PAM</td><td>NA</td><td>Intergenic</td><td>NA</td><td>NA</td></tr>
<tr><td>T11211</td><td>CGTCTCGATCGAGACCAGTACGG</td><td></td><td>1000</td><td></td><td></td><td></td><td></td><td></td><td></td><td></td><td></td><td></td></tr>
<tr><td>Oligo fwd</td><td>CACCCGTCTCGATCGAGACCAGTA</td><td></td><td></td><td></td><td></td><td></td><td></td><td></td><td></td><td></td><td></td><td></td></tr>
<tr><td>Oligo rev</td><td>AAACTACTGGTCTCGATCGAGACG</td><td></td><td></td><td></td><td></td><td></td><td></td><td></td><td></td><td></td><td></td><td></td></tr>
</table>

Chromosome start end strand MM target_seq PAM alignment distance position gene name gene id
GU305914.1 146244 146266 + 0 CGTCTCGATCGAGACCAGTA CGG ||||||||[||||||||||||]PAM NA Intergenic NA NA

T11212 ATCCAGTCCATCATCGCCGAAGG 1000
Oligo fwd CACCATCCAGTCCATCATCGCCGA
Oligo rev AAACTCGGCGATGATGGACTGGAT
Chromosome start end strand MM target_seq PAM alignment distance position gene name gene id
GU305914.1 146269 146291 + 0 ATCCAGTCCATCATCGCCGA AGG ||||||||[||||||||||||]PAM NA Intergenic NA NA

T11213 CGCCGAAGGCGCCACGACCGAGG 1000
Oligo fwd CACCCGCCGAAGGCGCCACGACCG
Oligo rev AAACCGGTCGTGGCGCCTTCGGCG
Chromosome start end strand MM target_seq PAM alignment distance position gene name gene id
GU305914.1 146283 146305 + 0 CGCCGAAGGCGCCACGACCG AGG ||||||||[||||||||||||]PAM NA Intergenic NA NA

T11214 CGCCACGACCGAGGATGCCGAGG 1000
Oligo fwd CACCCGCCACGACCGAGGATGCCG
Oligo rev AAACCGGCATCCTCGGTCGTGGCG
Chromosome start end strand MM target_seq PAM alignment distance position gene name gene id
GU305914.1 146292 146314 + 0 CGCCACGACCGAGGATGCCG AGG ||||||||[||||||||||||]PAM NA Intergenic NA NA

T11215 ACCGCAACACGCACGCCTACAGG 1000
Oligo fwd CACCACCGCAACACGCACGCCTAC
Oligo rev AAACGTAGGCGTGCGTGTTGCGGT
Chromosome start end strand MM target_seq PAM alignment distance position gene name gene id
GU305914.1 146327 146349 + 0 ACCGCAACACGCACGCCTAC AGG ||||||||[||||||||||||]PAM NA Intergenic NA NA

T11216 GCCTACAGGAAGCTCCGCCGCGG 1000
Oligo fwd CACCGCCTACAGGAAGCTCCGCCG
Oligo rev AAACCGGCGGAGCTTCCTGTAGGC
Chromosome start end strand MM target_seq PAM alignment distance position gene name gene id
GU305914.1 146341 146363 + 0 GCCTACAGGAAGCTCCGCCG CGG ||||||||[||||||||||||]PAM NA Intergenic NA NA

T11217 ATCACCTTCTCCATCCGCATCGG 1000
Oligo fwd CACCATCACCTTCTCCATCCGCAT
Oligo rev AAACATGCGGATGGAGAAGGTGAT
Chromosome start end strand MM target_seq PAM alignment distance position gene name gene id
GU305914.1 146380 146402 + 0 ATCACCTTCTCCATCCGCAT CGG ||||||||[||||||||||||]PAM NA Intergenic NA NA

T11218 GCTCAGCGAGCCCTCCGCCGTGG 1000
Oligo fwd CACCGCTCAGCGAGCCCTCCGCCG
Oligo rev AAACCGGCGGAGGGCTCGCTGAGC
Chromosome start end strand MM target_seq PAM alignment distance position gene name gene id
GU305914.1 146418 146440 + 0 GCTCAGCGAGCCCTCCGCCG TGG ||||||||[||||||||||||]PAM NA Intergenic NA NA

T11219 CTCAGCGAGCCCTCCGCCGTGGG 1000
Oligo fwd CACCCTCAGCGAGCCCTCCGCCGT
Oligo rev AAACACGGCGGAGGGCTCGCTGAG
Chromosome start end strand MM target_seq PAM alignment distance position gene name gene id
GU305914.1 146419 146441 + 0 CTCAGCGAGCCCTCCGCCGT GGG ||||||||[||||||||||||]PAM NA Intergenic NA NA

T11220 CTCCGACGTCTTCGTGATCCAGG 1000
Oligo fwd CACCCTCCGACGTCTTCGTGATCC
Oligo rev AAACGGATCACGAAGACGTCGGAG
Chromosome start end strand MM target_seq PAM alignment distance position gene name gene id
GU305914.1 146457 146479 + 0 CTCCGACGTCTTCGTGATCC AGG ||||||||[||||||||||||]PAM NA Intergenic NA NA

T11221 TCGTGATCCAGGATGTCAAGTGG 1000
Oligo fwd CACCTCGTGATCCAGGATGTCAAG
Oligo rev AAACCTTGACATCCTGGATCACGA
Chromosome start end strand MM target_seq PAM alignment distance position gene name gene id
GU305914.1 146468 146490 + 0 TCGTGATCCAGGATGTCAAG TGG ||||||||[||||||||||||]PAM NA Intergenic NA NA

T11222 ATCCAGGATGTCAAGTGGCGCGG 1000
Oligo fwd CACCATCCAGGATGTCAAGTGGCG
Oligo rev AAACCGCCACTTGACATCCTGGAT
Chromosome start end strand MM target_seq PAM alignment distance position gene name gene id
GU305914.1 146473 146495 + 0 ATCCAGGATGTCAAGTGGCG CGG ||||||||[||||||||||||]PAM NA Intergenic NA NA

T11223 GTGCTTTCCGCGCGAGTTCCTGG 1000
Oligo fwd CACCGTGCTTTCCGCGCGAGTTCC
Oligo rev AAACGGAACTCGCGCGGAAAGCAC
Chromosome start end strand MM target_seq PAM alignment distance position gene name gene id
GU305914.1 146511 146533 + 0 GTGCTTTCCGCGCGAGTTCC TGG ||||||||[||||||||||||]PAM NA Intergenic NA NA

T11224 GTTCCTGGCCTTCGTCTTCTCGG 1000
Oligo fwd CACCGTTCCTGGCCTTCGTCTTCT
Oligo rev AAACAGAAGACGAAGGCCAGGAAC
Chromosome start end strand MM target_seq PAM alignment distance position gene name gene id
GU305914.1 146526 146548 + 0 GTTCCTGGCCTTCGTCTTCT CGG ||||||||[||||||||||||]PAM NA Intergenic NA NA

T11225 TGCCATGCAGCGCCTCTTCAAGG 1000
Oligo fwd CACCTGCCATGCAGCGCCTCTTCA
Oligo rev AAACTGAAGAGGCGCTGCATGGCA
Chromosome start end strand MM target_seq PAM alignment distance position gene name gene id
GU305914.1 146577 146599 + 0 TGCCATGCAGCGCCTCTTCA AGG ||||||||[||||||||||||]PAM NA Intergenic NA NA

T11226 CATGCAGCGCCTCTTCAAGGAGG 1000
Oligo fwd CACCCATGCAGCGCCTCTTCAAGG

<table>
<tr><td>Oligo rev</td><td>AAACCCTTGAAGAGGCGCTGCATG</td><td></td><td></td><td></td><td></td><td></td><td></td><td></td><td></td><td></td><td></td></tr>
<tr><td>Chromosome</td><td>start</td><td>end</td><td>strand</td><td>MM</td><td>target_seq</td><td>PAM</td><td>alignment</td><td>distance</td><td>position</td><td>gene name</td><td>gene id</td></tr>
<tr><td>GU305914.1</td><td>146580</td><td>146602</td><td>+</td><td>0</td><td>CATGCAGCGCCTCTTCAAGG</td><td>AGG</td><td>||||||||[||||||||||||]PAM</td><td>NA</td><td>Intergenic</td><td>NA</td><td>NA</td></tr>
<tr><td>T11227</td><td>CGCCTCTTCAAGGAGGTCTACGG</td><td>1000</td><td></td><td></td><td></td><td></td><td></td><td></td><td></td><td></td><td></td></tr>
<tr><td>Oligo fwd</td><td>CACCCGCCTCTTCAAGGAGGTCTA</td><td></td><td></td><td></td><td></td><td></td><td></td><td></td><td></td><td></td><td></td></tr>
<tr><td>Oligo rev</td><td>AAACTAGACCTCCTTGAAGAGGCG</td><td></td><td></td><td></td><td></td><td></td><td></td><td></td><td></td><td></td><td></td></tr>
<tr><td>Chromosome</td><td>start</td><td>end</td><td>strand</td><td>MM</td><td>target_seq</td><td>PAM</td><td>alignment</td><td>distance</td><td>position</td><td>gene name</td><td>gene id</td></tr>
<tr><td>GU305914.1</td><td>146587</td><td>146609</td><td>+</td><td>0</td><td>CGCCTCTTCAAGGAGGTCTA</td><td>CGG</td><td>||||||||[||||||||||||]PAM</td><td>NA</td><td>Intergenic</td><td>NA</td><td>NA</td></tr>
<tr><td>T11228</td><td>CTCTTCAAGGAGGTCTACGGCGG</td><td>1000</td><td></td><td></td><td></td><td></td><td></td><td></td><td></td><td></td><td></td></tr>
<tr><td>Oligo fwd</td><td>CACCCTCTTCAAGGAGGTCTACGG</td><td></td><td></td><td></td><td></td><td></td><td></td><td></td><td></td><td></td><td></td></tr>
<tr><td>Oligo rev</td><td>AAACCCGTAGACCTCCTTGAAGAG</td><td></td><td></td><td></td><td></td><td></td><td></td><td></td><td></td><td></td><td></td></tr>
<tr><td>Chromosome</td><td>start</td><td>end</td><td>strand</td><td>MM</td><td>target_seq</td><td>PAM</td><td>alignment</td><td>distance</td><td>position</td><td>gene name</td><td>gene id</td></tr>
<tr><td>GU305914.1</td><td>146590</td><td>146612</td><td>+</td><td>0</td><td>CTCTTCAAGGAGGTCTACGG</td><td>CGG</td><td>||||||||[||||||||||||]PAM</td><td>NA</td><td>Intergenic</td><td>NA</td><td>NA</td></tr>
<tr><td>T11229</td><td>GAGGTCTACGGCGGATTCTCCGG</td><td>1000</td><td></td><td></td><td></td><td></td><td></td><td></td><td></td><td></td><td></td></tr>
<tr><td>Oligo fwd</td><td>CACCGAGGTCTACGGCGGATTCTC</td><td></td><td></td><td></td><td></td><td></td><td></td><td></td><td></td><td></td><td></td></tr>
<tr><td>Oligo rev</td><td>AAACGAGAATCCGCCGTAGACCTC</td><td></td><td></td><td></td><td></td><td></td><td></td><td></td><td></td><td></td><td></td></tr>
<tr><td>Chromosome</td><td>start</td><td>end</td><td>strand</td><td>MM</td><td>target_seq</td><td>PAM</td><td>alignment</td><td>distance</td><td>position</td><td>gene name</td><td>gene id</td></tr>
<tr><td>GU305914.1</td><td>146599</td><td>146621</td><td>+</td><td>0</td><td>GAGGTCTACGGCGGATTCTC</td><td>CGG</td><td>||||||||[||||||||||||]PAM</td><td>NA</td><td>Intergenic</td><td>NA</td><td>NA</td></tr>
<tr><td>T11230</td><td>TACCCCGTCTTCGACTTCTTCGG</td><td>1000</td><td></td><td></td><td></td><td></td><td></td><td></td><td></td><td></td><td></td></tr>
<tr><td>Oligo fwd</td><td>CACCTACCCCGTCTTCGACTTCTT</td><td></td><td></td><td></td><td></td><td></td><td></td><td></td><td></td><td></td><td></td></tr>
<tr><td>Oligo rev</td><td>AAACAAGAAGTCGAAGACGGGGTA</td><td></td><td></td><td></td><td></td><td></td><td></td><td></td><td></td><td></td><td></td></tr>
<tr><td>Chromosome</td><td>start</td><td>end</td><td>strand</td><td>MM</td><td>target_seq</td><td>PAM</td><td>alignment</td><td>distance</td><td>position</td><td>gene name</td><td>gene id</td></tr>
<tr><td>GU305914.1</td><td>146626</td><td>146648</td><td>+</td><td>0</td><td>TACCCCGTCTTCGACTTCTT</td><td>CGG</td><td>||||||||[||||||||||||]PAM</td><td>NA</td><td>Intergenic</td><td>NA</td><td>NA</td></tr>
<tr><td>GU305914.1</td><td>131458</td><td>131480</td><td>-</td><td>4</td><td>GACCCCGACATCGACTTCTG</td><td>CGG</td><td>-||||||-[|-|||||||||-]PAM</td><td>NA</td><td>Intergenic</td><td>NA</td><td>NA</td></tr>
<tr><td>T11231</td><td>ACCCCGTCTTCGACTTCTTCGGG</td><td>1000</td><td></td><td></td><td></td><td></td><td></td><td></td><td></td><td></td><td></td></tr>
<tr><td>Oligo fwd</td><td>CACCACCCCGTCTTCGACTTCTTC</td><td></td><td></td><td></td><td></td><td></td><td></td><td></td><td></td><td></td><td></td></tr>
<tr><td>Oligo rev</td><td>AAACGAAGAAGTCGAAGACGGGGT</td><td></td><td></td><td></td><td></td><td></td><td></td><td></td><td></td><td></td><td></td></tr>
<tr><td>Chromosome</td><td>start</td><td>end</td><td>strand</td><td>MM</td><td>target_seq</td><td>PAM</td><td>alignment</td><td>distance</td><td>position</td><td>gene name</td><td>gene id</td></tr>
<tr><td>GU305914.1</td><td>146627</td><td>146649</td><td>+</td><td>0</td><td>ACCCCGTCTTCGACTTCTTC</td><td>GGG</td><td>||||||||[||||||||||||]PAM</td><td>NA</td><td>Intergenic</td><td>NA</td><td>NA</td></tr>
<tr><td>T11232</td><td>GGGCCCAATATGCTGCGCTCCGG</td><td>1000</td><td></td><td></td><td></td><td></td><td></td><td></td><td></td><td></td><td></td></tr>
<tr><td>Oligo fwd</td><td>CACCGGGCCCAATATGCTGCGCTC</td><td></td><td></td><td></td><td></td><td></td><td></td><td></td><td></td><td></td><td></td></tr>
<tr><td>Oligo rev</td><td>AAACGAGCGCAGCATATTGGGCCC</td><td></td><td></td><td></td><td></td><td></td><td></td><td></td><td></td><td></td><td></td></tr>
<tr><td>Chromosome</td><td>start</td><td>end</td><td>strand</td><td>MM</td><td>target_seq</td><td>PAM</td><td>alignment</td><td>distance</td><td>position</td><td>gene name</td><td>gene id</td></tr>
<tr><td>GU305914.1</td><td>146647</td><td>146669</td><td>+</td><td>0</td><td>GGGCCCAATATGCTGCGCTC</td><td>CGG</td><td>||||||||[||||||||||||]PAM</td><td>NA</td><td>Intergenic</td><td>NA</td><td>NA</td></tr>
<tr><td>T11233</td><td>GGCCCAATATGCTGCGCTCCGGG</td><td>1000</td><td></td><td></td><td></td><td></td><td></td><td></td><td></td><td></td><td></td></tr>
<tr><td>Oligo fwd</td><td>CACCGGCCCAATATGCTGCGCTCC</td><td></td><td></td><td></td><td></td><td></td><td></td><td></td><td></td><td></td><td></td></tr>
<tr><td>Oligo rev</td><td>AAACGGAGCGCAGCATATTGGGCC</td><td></td><td></td><td></td><td></td><td></td><td></td><td></td><td></td><td></td><td></td></tr>
<tr><td>Chromosome</td><td>start</td><td>end</td><td>strand</td><td>MM</td><td>target_seq</td><td>PAM</td><td>alignment</td><td>distance</td><td>position</td><td>gene name</td><td>gene id</td></tr>
<tr><td>GU305914.1</td><td>146648</td><td>146670</td><td>+</td><td>0</td><td>GGCCCAATATGCTGCGCTCC</td><td>GGG</td><td>||||||||[||||||||||||]PAM</td><td>NA</td><td>Intergenic</td><td>NA</td><td>NA</td></tr>
<tr><td>T11234</td><td>GCCCAATATGCTGCGCTCCGGGG</td><td>1000</td><td></td><td></td><td></td><td></td><td></td><td></td><td></td><td></td><td></td></tr>
<tr><td>Oligo fwd</td><td>CACCGCCCAATATGCTGCGCTCCG</td><td></td><td></td><td></td><td></td><td></td><td></td><td></td><td></td><td></td><td></td></tr>
<tr><td>Oligo rev</td><td>AAACCGGAGCGCAGCATATTGGGC</td><td></td><td></td><td></td><td></td><td></td><td></td><td></td><td></td><td></td><td></td></tr>
<tr><td>Chromosome</td><td>start</td><td>end</td><td>strand</td><td>MM</td><td>target_seq</td><td>PAM</td><td>alignment</td><td>distance</td><td>position</td><td>gene name</td><td>gene id</td></tr>
<tr><td>GU305914.1</td><td>146649</td><td>146671</td><td>+</td><td>0</td><td>GCCCAATATGCTGCGCTCCG</td><td>GGG</td><td>||||||||[||||||||||||]PAM</td><td>NA</td><td>Intergenic</td><td>NA</td><td>NA</td></tr>
<tr><td>T11235</td><td>CCCAATATGCTGCGCTCCGGGGG</td><td>1000</td><td></td><td></td><td></td><td></td><td></td><td></td><td></td><td></td><td></td></tr>
<tr><td>Oligo fwd</td><td>CACCCCCAATATGCTGCGCTCCGG</td><td></td><td></td><td></td><td></td><td></td><td></td><td></td><td></td><td></td><td></td></tr>
<tr><td>Oligo rev</td><td>AAACCCGGAGCGCAGCATATTGGG</td><td></td><td></td><td></td><td></td><td></td><td></td><td></td><td></td><td></td><td></td></tr>
<tr><td>Chromosome</td><td>start</td><td>end</td><td>strand</td><td>MM</td><td>target_seq</td><td>PAM</td><td>alignment</td><td>distance</td><td>position</td><td>gene name</td><td>gene id</td></tr>
<tr><td>GU305914.1</td><td>146650</td><td>146672</td><td>+</td><td>0</td><td>CCCAATATGCTGCGCTCCGG</td><td>GGG</td><td>||||||||[||||||||||||]PAM</td><td>NA</td><td>Intergenic</td><td>NA</td><td>NA</td></tr>
<tr><td>T11236</td><td>CCGCGCTCCGTCTTCTTCCCCGG</td><td>1000</td><td></td><td></td><td></td><td></td><td></td><td></td><td></td><td></td><td></td></tr>
<tr><td>Oligo fwd</td><td>CACCCCGCGCTCCGTCTTCTTCCC</td><td></td><td></td><td></td><td></td><td></td><td></td><td></td><td></td><td></td><td></td></tr>
<tr><td>Oligo rev</td><td>AAACGGGAAGAAGACGGAGCGCGG</td><td></td><td></td><td></td><td></td><td></td><td></td><td></td><td></td><td></td><td></td></tr>
<tr><td>Chromosome</td><td>start</td><td>end</td><td>strand</td><td>MM</td><td>target_seq</td><td>PAM</td><td>alignment</td><td>distance</td><td>position</td><td>gene name</td><td>gene id</td></tr>
<tr><td>GU305914.1</td><td>146674</td><td>146696</td><td>+</td><td>0</td><td>CCGCGCTCCGTCTTCTTCCC</td><td>CGG</td><td>||||||||[||||||||||||]PAM</td><td>NA</td><td>Intergenic</td><td>NA</td><td>NA</td></tr>
<tr><td>T11237</td><td>TCGCCGTCTACTCCGTGCCCTGG</td><td>1000</td><td></td><td></td><td></td><td></td><td></td><td></td><td></td><td></td><td></td></tr>
<tr><td>Oligo fwd</td><td>CACCTCGCCGTCTACTCCGTGCCC</td><td></td><td></td><td></td><td></td><td></td><td></td><td></td><td></td><td></td><td></td></tr>
<tr><td>Oligo rev</td><td>AAACGGGCACGGAGTAGACGGCGA</td><td></td><td></td><td></td><td></td><td></td><td></td><td></td><td></td><td></td><td></td></tr>
<tr><td>Chromosome</td><td>start</td><td>end</td><td>strand</td><td>MM</td><td>target_seq</td><td>PAM</td><td>alignment</td><td>distance</td><td>position</td><td>gene name</td><td>gene id</td></tr>
<tr><td>GU305914.1</td><td>146708</td><td>146730</td><td>+</td><td>0</td><td>TCGCCGTCTACTCCGTGCCC</td><td>TGG</td><td>||||||||[||||||||||||]PAM</td><td>NA</td><td>Intergenic</td><td>NA</td><td>NA</td></tr>
<tr><td>GU305914.1</td><td>29453</td><td>29475</td><td>-</td><td>4</td><td>GCACCGTCTTCTCCCTGCCC</td><td>TGG</td><td>-|-|||||[|-||||-|||||]PAM</td><td>NA</td><td>Intergenic</td><td>NA</td><td>NA</td></tr>
<tr><td>T11238</td><td>CTGGCGCTACGACATGAATACGG</td><td>1000</td><td></td><td></td><td></td><td></td><td></td><td></td><td></td><td></td><td></td></tr>
<tr><td>Oligo fwd</td><td>CACCCTGGCGCTACGACATGAATA</td><td></td><td></td><td></td><td></td><td></td><td></td><td></td><td></td><td></td><td></td></tr>
<tr><td>Oligo rev</td><td>AAACTATTCATGTCGTAGCGCCAG</td><td></td><td></td><td></td><td></td><td></td><td></td><td></td><td></td><td></td><td></td></tr>
<tr><td>Chromosome</td><td>start</td><td>end</td><td>strand</td><td>MM</td><td>target_seq</td><td>PAM</td><td>alignment</td><td>distance</td><td>position</td><td>gene name</td><td>gene id</td></tr>
<tr><td>GU305914.1</td><td>146727</td><td>146749</td><td>+</td><td>0</td><td>CTGGCGCTACGACATGAATA</td><td>CGG</td><td>||||||||[||||||||||||]PAM</td><td>NA</td><td>Intergenic</td><td>NA</td><td>NA</td></tr>
<tr><td>T11239</td><td>TACGACATGAATACGGAGAACGG</td><td>1000</td><td></td><td></td><td></td><td></td><td></td><td></td><td></td><td></td><td></td></tr>
<tr><td>Oligo fwd</td><td>CACCTACGACATGAATACGGAGAA</td><td></td><td></td><td></td><td></td><td></td><td></td><td></td><td></td><td></td><td></td></tr>
<tr><td>Oligo rev</td><td>AAACTTCTCCGTATTCATGTCGTA</td><td></td><td></td><td></td><td></td><td></td><td></td><td></td><td></td><td></td><td></td></tr>
<tr><td>Chromosome</td><td>start</td><td>end</td><td>strand</td><td>MM</td><td>target_seq</td><td>PAM</td><td>alignment</td><td>distance</td><td>position</td><td>gene name</td><td>gene id</td></tr>
<tr><td>GU305914.1</td><td>146734</td><td>146756</td><td>+</td><td>0</td><td>TACGACATGAATACGGAGAA</td><td>CGG</td><td>||||||||[||||||||||||]PAM</td><td>NA</td><td>Intergenic</td><td>NA</td><td>NA</td></tr>
<tr><td>T11240</td><td>CCGACGCCATCAACGAGATCAGG</td><td>1000</td><td></td><td></td><td></td><td></td><td></td><td></td><td></td><td></td><td></td></tr>
<tr><td>Oligo fwd</td><td>CACCCCGACGCCATCAACGAGATC</td><td></td><td></td><td></td><td></td><td></td><td></td><td></td><td></td><td></td><td></td></tr>
<tr><td>Oligo rev</td><td>AAACGATCTCGTTGATGGCGTCGG</td><td></td><td></td><td></td><td></td><td></td><td></td><td></td><td></td><td></td><td></td></tr>
<tr><td>Chromosome</td><td>start</td><td>end</td><td>strand</td><td>MM</td><td>target_seq</td><td>PAM</td><td>alignment</td><td>distance</td><td>position</td><td>gene name</td><td>gene id</td></tr>
<tr><td>GU305914.1</td><td>146759</td><td>146781</td><td>+</td><td>0</td><td>CCGACGCCATCAACGAGATC</td><td>AGG</td><td>||||||||[||||||||||||]PAM</td><td>NA</td><td>Intergenic</td><td>NA</td><td>NA</td></tr>
</table>

| | | | | | | | | | | | |
|---|---|---|---|---|---|---|---|---|---|---|---|
| T11241 | CATCAACGAGATCAGGTCCTTGG | 1000 | | | | | | | | | |
| Oligo fwd | CACCCATCAACGAGATCAGGTCCT | | | | | | | | | | |
| Oligo rev | AAACAGGACCTGATCTCGTTGATG | | | | | | | | | | |
| Chromosome | start | end | strand | MM | target_seq | PAM | alignment | distance | position | gene name | gene id |
| GU305914.1 | 146766 | 146788 | + | 0 | CATCAACGAGATCAGGTCCT | TGG | ||||||||[||||||||||||]PAM | NA | Intergenic | NA | NA |

| | | | | | | | | | | | |
|---|---|---|---|---|---|---|---|---|---|---|---|
| T11242 | AACGAGATCAGGTCCTTGGTCGG | 1000 | | | | | | | | | |
| Oligo fwd | CACCAACGAGATCAGGTCCTTGGT | | | | | | | | | | |
| Oligo rev | AAACACCAAGGACCTGATCTCGTT | | | | | | | | | | |
| Chromosome | start | end | strand | MM | target_seq | PAM | alignment | distance | position | gene name | gene id |
| GU305914.1 | 146770 | 146792 | + | 0 | AACGAGATCAGGTCCTTGGT | CGG | ||||||||[||||||||||||]PAM | NA | Intergenic | NA | NA |

| | | | | | | | | | | | |
|---|---|---|---|---|---|---|---|---|---|---|---|
| T11243 | GTCGGCCTGCCCGACATCGTCGG | 1000 | | | | | | | | | |
| Oligo fwd | CACCGTCGGCCTGCCCGACATCGT | | | | | | | | | | |
| Oligo rev | AAACACGATGTCGGGCAGGCCGAC | | | | | | | | | | |
| Chromosome | start | end | strand | MM | target_seq | PAM | alignment | distance | position | gene name | gene id |
| GU305914.1 | 146788 | 146810 | + | 0 | GTCGGCCTGCCCGACATCGT | CGG | ||||||||[||||||||||||]PAM | NA | Intergenic | NA | NA |

| | | | | | | | | | | | |
|---|---|---|---|---|---|---|---|---|---|---|---|
| T11244 | GCCCGACATCGTCGGCGTCGCGG | 1000 | | | | | | | | | |
| Oligo fwd | CACCGCCCGACATCGTCGGCGTCG | | | | | | | | | | |
| Oligo rev | AAACCGACGCCGACGATGTCGGGC | | | | | | | | | | |
| Chromosome | start | end | strand | MM | target_seq | PAM | alignment | distance | position | gene name | gene id |
| GU305914.1 | 146796 | 146818 | + | 0 | GCCCGACATCGTCGGCGTCG | CGG | ||||||||[||||||||||||]PAM | NA | Intergenic | NA | NA |
| GU305914.1 | 61963 | 61985 | + | 3 | GACCGTCATCGTCGGGGTCG | GGG | |-|||-||[|||||||-||||]PAM | NA | Intergenic | NA | NA |

| | | | | | | | | | | | |
|---|---|---|---|---|---|---|---|---|---|---|---|
| T11245 | CCCGACATCGTCGGCGTCGCGGG | 1000 | | | | | | | | | |
| Oligo fwd | CACCCCCGACATCGTCGGCGTCGC | | | | | | | | | | |
| Oligo rev | AAACGCGACGCCGACGATGTCGGG | | | | | | | | | | |
| Chromosome | start | end | strand | MM | target_seq | PAM | alignment | distance | position | gene name | gene id |
| GU305914.1 | 146797 | 146819 | + | 0 | CCCGACATCGTCGGCGTCGC | GGG | ||||||||[||||||||||||]PAM | NA | Intergenic | NA | NA |
| GU305914.1 | 61964 | 61986 | + | 4 | ACCGTCATCGTCGGGGTCGG | GGG | -|||-|||[||||||-||||-]PAM | NA | Intergenic | NA | NA |

| | | | | | | | | | | | |
|---|---|---|---|---|---|---|---|---|---|---|---|
| T11246 | CATCGTCGGCGTCGCGGGTAAGG | 1000 | | | | | | | | | |
| Oligo fwd | CACCCATCGTCGGCGTCGCGGGTA | | | | | | | | | | |
| Oligo rev | AAACTACCCGCGACGCCGACGATG | | | | | | | | | | |
| Chromosome | start | end | strand | MM | target_seq | PAM | alignment | distance | position | gene name | gene id |
| GU305914.1 | 146802 | 146824 | + | 0 | CATCGTCGGCGTCGCGGGTA | AGG | ||||||||[||||||||||||]PAM | NA | Intergenic | NA | NA |

| | | | | | | | | | | | |
|---|---|---|---|---|---|---|---|---|---|---|---|
| T11247 | GGGTAAGGTGCCGCTCGTGCCGG | 1000 | | | | | | | | | |
| Oligo fwd | CACCGGGTAAGGTGCCGCTCGTGC | | | | | | | | | | |
| Oligo rev | AAACGCACGAGCGGCACCTTACCC | | | | | | | | | | |
| Chromosome | start | end | strand | MM | target_seq | PAM | alignment | distance | position | gene name | gene id |
| GU305914.1 | 146817 | 146839 | + | 0 | GGGTAAGGTGCCGCTCGTGC | CGG | ||||||||[||||||||||||]PAM | NA | Intergenic | NA | NA |

| | | | | | | | | | | | |
|---|---|---|---|---|---|---|---|---|---|---|---|
| T11248 | GTGCCGCTCGTGCCGGAGCCCGG | 1000 | | | | | | | | | |
| Oligo fwd | CACCGTGCCGCTCGTGCCGGAGCC | | | | | | | | | | |
| Oligo rev | AAACGGCTCCGGCACGAGCGGCAC | | | | | | | | | | |
| Chromosome | start | end | strand | MM | target_seq | PAM | alignment | distance | position | gene name | gene id |
| GU305914.1 | 146824 | 146846 | + | 0 | GTGCCGCTCGTGCCGGAGCC | CGG | ||||||||[||||||||||||]PAM | NA | Intergenic | NA | NA |

| | | | | | | | | | | | |
|---|---|---|---|---|---|---|---|---|---|---|---|
| T11249 | CGGCAACGCCGTCGACACCCTGG | 1000 | | | | | | | | | |
| Oligo fwd | CACCCGGCAACGCCGTCGACACCC | | | | | | | | | | |
| Oligo rev | AAACGGGTGTCGACGGCGTTGCCG | | | | | | | | | | |
| Chromosome | start | end | strand | MM | target_seq | PAM | alignment | distance | position | gene name | gene id |
| GU305914.1 | 146844 | 146866 | + | 0 | CGGCAACGCCGTCGACACCC | TGG | ||||||||[||||||||||||]PAM | NA | Intergenic | NA | NA |

| | | | | | | | | | | | |
|---|---|---|---|---|---|---|---|---|---|---|---|
| T11250 | TCGACACCCTGGACGCCGTCAGG | 1000 | | | | | | | | | |
| Oligo fwd | CACCTCGACACCCTGGACGCCGTC | | | | | | | | | | |
| Oligo rev | AAACGACGGCGTCCAGGGTGTCGA | | | | | | | | | | |
| Chromosome | start | end | strand | MM | target_seq | PAM | alignment | distance | position | gene name | gene id |
| GU305914.1 | 146855 | 146877 | + | 0 | TCGACACCCTGGACGCCGTC | AGG | ||||||||[||||||||||||]PAM | NA | Intergenic | NA | NA |

| | | | | | | | | | | | |
|---|---|---|---|---|---|---|---|---|---|---|---|
| T11251 | ACCTGCACTACGCCGACATGAGG | 1000 | | | | | | | | | |
| Oligo fwd | CACCACCTGCACTACGCCGACATG | | | | | | | | | | |
| Oligo rev | AAACCATGTCGGCGTAGTGCAGGT | | | | | | | | | | |
| Chromosome | start | end | strand | MM | target_seq | PAM | alignment | distance | position | gene name | gene id |
| GU305914.1 | 146891 | 146913 | + | 0 | ACCTGCACTACGCCGACATG | AGG | ||||||||[||||||||||||]PAM | NA | Intergenic | NA | NA |

| | | | | | | | | | | | |
|---|---|---|---|---|---|---|---|---|---|---|---|
| T11252 | GGCACTTTCGCATCAACGCCAGG | 1000 | | | | | | | | | |
| Oligo fwd | CACCGGCACTTTCGCATCAACGCC | | | | | | | | | | |
| Oligo rev | AAACGGCGTTGATGCGAAAGTGCC | | | | | | | | | | |
| Chromosome | start | end | strand | MM | target_seq | PAM | alignment | distance | position | gene name | gene id |
| GU305914.1 | 146912 | 146934 | + | 0 | GGCACTTTCGCATCAACGCC | AGG | ||||||||[||||||||||||]PAM | NA | Intergenic | NA | NA |

| | | | | | | | | | | | |
|---|---|---|---|---|---|---|---|---|---|---|---|
| T11253 | GCTCTGCGTTACCCACGACATGG | 1000 | | | | | | | | | |
| Oligo fwd | CACCGCTCTGCGTTACCCACGACA | | | | | | | | | | |
| Oligo rev | AAACTGTCGTGGGTAACGCAGAGC | | | | | | | | | | |
| Chromosome | start | end | strand | MM | target_seq | PAM | alignment | distance | position | gene name | gene id |
| GU305914.1 | 146934 | 146956 | + | 0 | GCTCTGCGTTACCCACGACA | TGG | ||||||||[||||||||||||]PAM | NA | Intergenic | NA | NA |

| | | | | | | | | | | | |
|---|---|---|---|---|---|---|---|---|---|---|---|
| T11254 | CTCTGCGTTACCCACGACATGGG | 1000 | | | | | | | | | |
| Oligo fwd | CACCCTCTGCGTTACCCACGACAT | | | | | | | | | | |
| Oligo rev | AAACATGTCGTGGGTAACGCAGAG | | | | | | | | | | |
| Chromosome | start | end | strand | MM | target_seq | PAM | alignment | distance | position | gene name | gene id |
| GU305914.1 | 146935 | 146957 | + | 0 | CTCTGCGTTACCCACGACAT | GGG | ||||||||[||||||||||||]PAM | NA | Intergenic | NA | NA |

| | | |
|---|---|---|
| T11255 | CTCCGTCGCCCACATCTATGTGG | 1000 |
| Oligo fwd | CACCCTCCGTCGCCCACATCTATG | |

| | | | | | | | | | | | |
|---|---|---|---|---|---|---|---|---|---|---|---|
| Oligo rev | AAACCATAGATGTGGGCGACGGAG | | | | | | | | | | |
| Chromosome | start | end | strand | MM | target_seq | PAM | alignment | distance | position | gene name | gene id |
| GU305914.1 | 146982 | 147004 | + | 0 | CTCCGTCGCCCACATCTATG | TGG | ||||||||[||||||||||||]PAM | NA | Intergenic | NA | NA |
| GU305914.1 | 146991 | 147013 | - | 4 | CGCCGACGCCCACATAGATG | TGG | |-|||-||[|||||||--|||]PAM | NA | Intergenic | NA | NA |
| T11256 | TCCGTCGCCCACATCTATGTGGG | 1000 | | | | | | | | | |
| Oligo fwd | CACCTCCGTCGCCCACATCTATGT | | | | | | | | | | |
| Oligo rev | AAACACATAGATGTGGGCGACGGA | | | | | | | | | | |
| Chromosome | start | end | strand | MM | target_seq | PAM | alignment | distance | position | gene name | gene id |
| GU305914.1 | 146983 | 147005 | + | 0 | TCCGTCGCCCACATCTATGT | GGG | ||||||||[||||||||||||]PAM | NA | Intergenic | NA | NA |
| GU305914.1 | 146990 | 147012 | - | 4 | GCCGACGCCCACATAGATGT | GGG | -|||-|||[||||||--||||]PAM | NA | Intergenic | NA | NA |
| T11257 | GCCCACATCTATGTGGGCGTCGG | 1000 | | | | | | | | | |
| Oligo fwd | CACCGCCCACATCTATGTGGGCGT | | | | | | | | | | |
| Oligo rev | AAACACGCCCACATAGATGTGGGC | | | | | | | | | | |
| Chromosome | start | end | strand | MM | target_seq | PAM | alignment | distance | position | gene name | gene id |
| GU305914.1 | 146989 | 147011 | + | 0 | GCCCACATCTATGTGGGCGT | CGG | ||||||||[||||||||||||]PAM | NA | Intergenic | NA | NA |
| T11258 | CACATCTATGTGGGCGTCGGCGG | 1000 | | | | | | | | | |
| Oligo fwd | CACCCACATCTATGTGGGCGTCGG | | | | | | | | | | |
| Oligo rev | AAACCCGACGCCCACATAGATGTG | | | | | | | | | | |
| Chromosome | start | end | strand | MM | target_seq | PAM | alignment | distance | position | gene name | gene id |
| GU305914.1 | 146992 | 147014 | + | 0 | CACATCTATGTGGGCGTCGG | CGG | ||||||||[||||||||||||]PAM | NA | Intergenic | NA | NA |
| T11259 | GCATCTCCATCGTCGATCTCAGG | 1000 | | | | | | | | | |
| Oligo fwd | CACCGCATCTCCATCGTCGATCTC | | | | | | | | | | |
| Oligo rev | AAACGAGATCGACGATGGAGATGC | | | | | | | | | | |
| Chromosome | start | end | strand | MM | target_seq | PAM | alignment | distance | position | gene name | gene id |
| GU305914.1 | 147023 | 147045 | + | 0 | GCATCTCCATCGTCGATCTC | AGG | ||||||||[||||||||||||]PAM | NA | Intergenic | NA | NA |
| T11260 | ATCTCAGGTTCGCCGTCCTGCGG | 1000 | | | | | | | | | |
| Oligo fwd | CACCATCTCAGGTTCGCCGTCCTG | | | | | | | | | | |
| Oligo rev | AAACCAGGACGGCGAACCTGAGAT | | | | | | | | | | |
| Chromosome | start | end | strand | MM | target_seq | PAM | alignment | distance | position | gene name | gene id |
| GU305914.1 | 147038 | 147060 | + | 0 | ATCTCAGGTTCGCCGTCCTG | CGG | ||||||||[||||||||||||]PAM | NA | Intergenic | NA | NA |
| T11261 | GTCCTGCGGATGTGCCTGCCCGG | 1000 | | | | | | | | | |
| Oligo fwd | CACCGTCCTGCGGATGTGCCTGCC | | | | | | | | | | |
| Oligo rev | AAACGGCAGGCACATCCGCAGGAC | | | | | | | | | | |
| Chromosome | start | end | strand | MM | target_seq | PAM | alignment | distance | position | gene name | gene id |
| GU305914.1 | 147052 | 147074 | + | 0 | GTCCTGCGGATGTGCCTGCC | CGG | ||||||||[||||||||||||]PAM | NA | Intergenic | NA | NA |
| T11262 | TCCTGCGGATGTGCCTGCCCGGG | 1000 | | | | | | | | | |
| Oligo fwd | CACCTCCTGCGGATGTGCCTGCCC | | | | | | | | | | |
| Oligo rev | AAACGGGCAGGCACATCCGCAGGA | | | | | | | | | | |
| Chromosome | start | end | strand | MM | target_seq | PAM | alignment | distance | position | gene name | gene id |
| GU305914.1 | 147053 | 147075 | + | 0 | TCCTGCGGATGTGCCTGCCC | GGG | ||||||||[||||||||||||]PAM | NA | Intergenic | NA | NA |
| T11263 | CTACATCTGCAAGCGCGTCATGG | 1000 | | | | | | | | | |
| Oligo fwd | CACCCTACATCTGCAAGCGCGTCA | | | | | | | | | | |
| Oligo rev | AAACTGACGCGCTTGCAGATGTAG | | | | | | | | | | |
| Chromosome | start | end | strand | MM | target_seq | PAM | alignment | distance | position | gene name | gene id |
| GU305914.1 | 147201 | 147223 | + | 0 | CTACATCTGCAAGCGCGTCA | TGG | ||||||||[||||||||||||]PAM | NA | Intergenic | NA | NA |
| T11264 | CATGGACGCCTGCGAAGACGAGG | 1000 | | | | | | | | | |
| Oligo fwd | CACCCATGGACGCCTGCGAAGACG | | | | | | | | | | |
| Oligo rev | AAACCGTCTTCGCAGGCGTCCATG | | | | | | | | | | |
| Chromosome | start | end | strand | MM | target_seq | PAM | alignment | distance | position | gene name | gene id |
| GU305914.1 | 147219 | 147241 | + | 0 | CATGGACGCCTGCGAAGACG | AGG | ||||||||[||||||||||||]PAM | NA | Intergenic | NA | NA |
| T11265 | ATGGACGCCTGCGAAGACGAGGG | 1000 | | | | | | | | | |
| Oligo fwd | CACCATGGACGCCTGCGAAGACGA | | | | | | | | | | |
| Oligo rev | AAACTCGTCTTCGCAGGCGTCCAT | | | | | | | | | | |
| Chromosome | start | end | strand | MM | target_seq | PAM | alignment | distance | position | gene name | gene id |
| GU305914.1 | 147220 | 147242 | + | 0 | ATGGACGCCTGCGAAGACGA | GGG | ||||||||[||||||||||||]PAM | NA | Intergenic | NA | NA |
| T11266 | GCGAAGACGAGGGTTACCCCTGG | 1000 | | | | | | | | | |
| Oligo fwd | CACCGCGAAGACGAGGGTTACCCC | | | | | | | | | | |
| Oligo rev | AAACGGGGTAACCCTCGTCTTCGC | | | | | | | | | | |
| Chromosome | start | end | strand | MM | target_seq | PAM | alignment | distance | position | gene name | gene id |
| GU305914.1 | 147230 | 147252 | + | 0 | GCGAAGACGAGGGTTACCCC | TGG | ||||||||[||||||||||||]PAM | NA | Intergenic | NA | NA |
| T11267 | CGAGGGTTACCCCTGGATCCTGG | 1000 | | | | | | | | | |
| Oligo fwd | CACCCGAGGGTTACCCCTGGATCC | | | | | | | | | | |
| Oligo rev | AAACGGATCCAGGGGTAACCCTCG | | | | | | | | | | |
| Chromosome | start | end | strand | MM | target_seq | PAM | alignment | distance | position | gene name | gene id |
| GU305914.1 | 147237 | 147259 | + | 0 | CGAGGGTTACCCCTGGATCC | TGG | ||||||||[||||||||||||]PAM | NA | Intergenic | NA | NA |
| T11268 | GTTACCCCTGGATCCTGGTCCGG | 1000 | | | | | | | | | |
| Oligo fwd | CACCGTTACCCCTGGATCCTGGTC | | | | | | | | | | |
| Oligo rev | AAACGACCAGGATCCAGGGGTAAC | | | | | | | | | | |
| Chromosome | start | end | strand | MM | target_seq | PAM | alignment | distance | position | gene name | gene id |
| GU305914.1 | 147242 | 147264 | + | 0 | GTTACCCCTGGATCCTGGTC | CGG | ||||||||[||||||||||||]PAM | NA | Intergenic | NA | NA |
| T11269 | TTACCCCTGGATCCTGGTCCGGG | 1000 | | | | | | | | | |
| Oligo fwd | CACCTTACCCCTGGATCCTGGTCC | | | | | | | | | | |
| Oligo rev | AAACGGACCAGGATCCAGGGGTAA | | | | | | | | | | |
| Chromosome | start | end | strand | MM | target_seq | PAM | alignment | distance | position | gene name | gene id |
| GU305914.1 | 147243 | 147265 | + | 0 | TTACCCCTGGATCCTGGTCC | GGG | ||||||||[||||||||||||]PAM | NA | Intergenic | NA | NA |

| | | | | | | | | | | | |
|---|---|---|---|---|---|---|---|---|---|---|---|
| T11270 | ACGCCCCGTCGACTGCGACCAGG | | 1000 | | | | | | | | |
| Oligo fwd | CACCACGCCCCGTCGACTGCGACC | | | | | | | | | | |
| Oligo rev | AAACGGTCGCAGTCGACGGGGCGT | | | | | | | | | | |
| Chromosome | start | | end | strand | MM | target_seq | PAM | alignment | distance | position | gene name | gene id |
| GU305914.1 | | 147288 | 147310 | + | 0 | ACGCCCCGTCGACTGCGACC | AGG | \|\|\|\|\|\|\|\|[\|\|\|\|\|\|\|\|\|\|\|\|]PAM | NA | Intergenic | NA | NA |

| | | | | | | | | | | | | |
|---|---|---|---|---|---|---|---|---|---|---|---|---|
| T11271 | TCAACTTCGACACCATCGTCAGG | | 1000 | | | | | | | | | |
| Oligo fwd | CACCTCAACTTCGACACCATCGTC | | | | | | | | | | | |
| Oligo rev | AAACGACGATGGTGTCGAAGTTGA | | | | | | | | | | | |
| Chromosome | start | | end | strand | MM | target_seq | PAM | alignment | distance | position | gene name | gene id |
| GU305914.1 | | 147311 | 147333 | + | 0 | TCAACTTCGACACCATCGTC | AGG | \|\|\|\|\|\|\|\|[\|\|\|\|\|\|\|\|\|\|\|\|]PAM | NA | Intergenic | NA | NA |

| | | | | | | | | | | | | |
|---|---|---|---|---|---|---|---|---|---|---|---|---|
| T11272 | CAACTTCGACACCATCGTCAGGG | | 1000 | | | | | | | | | |
| Oligo fwd | CACCCAACTTCGACACCATCGTCA | | | | | | | | | | | |
| Oligo rev | AAACTGACGATGGTGTCGAAGTTG | | | | | | | | | | | |
| Chromosome | start | | end | strand | MM | target_seq | PAM | alignment | distance | position | gene name | gene id |
| GU305914.1 | | 147312 | 147334 | + | 0 | CAACTTCGACACCATCGTCA | GGG | \|\|\|\|\|\|\|\|[\|\|\|\|\|\|\|\|\|\|\|\|]PAM | NA | Intergenic | NA | NA |
| GU305914.1 | | 77368 | 77390 | + | 4 | CGCCTTCGACAACATCTTCA | AGG | \|--\|\|\|\|\|[\|\|\|-\|\|\|\|-\|\|\|]PAM | NA | Intergenic | NA | NA |

| | | | | | | | | | | | | |
|---|---|---|---|---|---|---|---|---|---|---|---|---|
| T11273 | CACCATCGTCAGGGCCAACCTGG | | 1000 | | | | | | | | | |
| Oligo fwd | CACCCACCATCGTCAGGGCCAACC | | | | | | | | | | | |
| Oligo rev | AAACGGTTGGCCCTGACGATGGTG | | | | | | | | | | | |
| Chromosome | start | | end | strand | MM | target_seq | PAM | alignment | distance | position | gene name | gene id |
| GU305914.1 | | 147321 | 147343 | + | 0 | CACCATCGTCAGGGCCAACC | TGG | \|\|\|\|\|\|\|\|[\|\|\|\|\|\|\|\|\|\|\|\|]PAM | NA | Intergenic | NA | NA |

| | | | | | | | | | | | | |
|---|---|---|---|---|---|---|---|---|---|---|---|---|
| T11274 | TCGTCAGGGCCAACCTGGCCAGG | | 1000 | | | | | | | | | |
| Oligo fwd | CACCTCGTCAGGGCCAACCTGGCC | | | | | | | | | | | |
| Oligo rev | AAACGGCCAGGTTGGCCCTGACGA | | | | | | | | | | | |
| Chromosome | start | | end | strand | MM | target_seq | PAM | alignment | distance | position | gene name | gene id |
| GU305914.1 | | 147326 | 147348 | + | 0 | TCGTCAGGGCCAACCTGGCC | AGG | \|\|\|\|\|\|\|\|[\|\|\|\|\|\|\|\|\|\|\|\|]PAM | NA | Intergenic | NA | NA |

| | | | | | | | | | | | | |
|---|---|---|---|---|---|---|---|---|---|---|---|---|
| T11275 | CGTCAGGGCCAACCTGGCCAGGG | | 1000 | | | | | | | | | |
| Oligo fwd | CACCCGTCAGGGCCAACCTGGCCA | | | | | | | | | | | |
| Oligo rev | AAACTGGCCAGGTTGGCCCTGACG | | | | | | | | | | | |
| Chromosome | start | | end | strand | MM | target_seq | PAM | alignment | distance | position | gene name | gene id |
| GU305914.1 | | 147327 | 147349 | + | 0 | CGTCAGGGCCAACCTGGCCA | GGG | \|\|\|\|\|\|\|\|[\|\|\|\|\|\|\|\|\|\|\|\|]PAM | NA | Intergenic | NA | NA |

| | | | | | | | | | | | | |
|---|---|---|---|---|---|---|---|---|---|---|---|---|
| T11276 | GGGCCAACCTGGCCAGGGTCTGG | | 1000 | | | | | | | | | |
| Oligo fwd | CACCGGGCCAACCTGGCCAGGGTC | | | | | | | | | | | |
| Oligo rev | AAACGACCCTGGCCAGGTTGGCCC | | | | | | | | | | | |
| Chromosome | start | | end | strand | MM | target_seq | PAM | alignment | distance | position | gene name | gene id |
| GU305914.1 | | 147332 | 147354 | + | 0 | GGGCCAACCTGGCCAGGGTC | TGG | \|\|\|\|\|\|\|\|[\|\|\|\|\|\|\|\|\|\|\|\|]PAM | NA | Intergenic | NA | NA |

| | | | | | | | | | | | | |
|---|---|---|---|---|---|---|---|---|---|---|---|---|
| T11277 | GGCCAACCTGGCCAGGGTCTGGG | | 1000 | | | | | | | | | |
| Oligo fwd | CACCGGCCAACCTGGCCAGGGTCT | | | | | | | | | | | |
| Oligo rev | AAACAGACCCTGGCCAGGTTGGCC | | | | | | | | | | | |
| Chromosome | start | | end | strand | MM | target_seq | PAM | alignment | distance | position | gene name | gene id |
| GU305914.1 | | 147333 | 147355 | + | 0 | GGCCAACCTGGCCAGGGTCT | GGG | \|\|\|\|\|\|\|\|[\|\|\|\|\|\|\|\|\|\|\|\|]PAM | NA | Intergenic | NA | NA |

| | | | | | | | | | | | | |
|---|---|---|---|---|---|---|---|---|---|---|---|---|
| T11278 | AGGGTCTGGGCCGAGCTCTTCGG | | 1000 | | | | | | | | | |
| Oligo fwd | CACCAGGGTCTGGGCCGAGCTCTT | | | | | | | | | | | |
| Oligo rev | AAACAAGAGCTCGGCCCAGACCCT | | | | | | | | | | | |
| Chromosome | start | | end | strand | MM | target_seq | PAM | alignment | distance | position | gene name | gene id |
| GU305914.1 | | 147346 | 147368 | + | 0 | AGGGTCTGGGCCGAGCTCTT | CGG | \|\|\|\|\|\|\|\|[\|\|\|\|\|\|\|\|\|\|\|\|]PAM | NA | Intergenic | NA | NA |

| | | | | | | | | | | | | |
|---|---|---|---|---|---|---|---|---|---|---|---|---|
| T11279 | CCCCGTCTGCCGCATCACCGTGG | | 1000 | | | | | | | | | |
| Oligo fwd | CACCCCCCGTCTGCCGCATCACCG | | | | | | | | | | | |
| Oligo rev | AAACCGGTGATGCGGCAGACGGGG | | | | | | | | | | | |
| Chromosome | start | | end | strand | MM | target_seq | PAM | alignment | distance | position | gene name | gene id |
| GU305914.1 | | 147384 | 147406 | + | 0 | CCCCGTCTGCCGCATCACCG | TGG | \|\|\|\|\|\|\|\|[\|\|\|\|\|\|\|\|\|\|\|\|]PAM | NA | Intergenic | NA | NA |

| | | | | | | | | | | | | |
|---|---|---|---|---|---|---|---|---|---|---|---|---|
| T11280 | CGCATCACCGTGGCCATGAGCGG | | 1000 | | | | | | | | | |
| Oligo fwd | CACCCGCATCACCGTGGCCATGAG | | | | | | | | | | | |
| Oligo rev | AAACCTCATGGCCACGGTGATGCG | | | | | | | | | | | |
| Chromosome | start | | end | strand | MM | target_seq | PAM | alignment | distance | position | gene name | gene id |
| GU305914.1 | | 147394 | 147416 | + | 0 | CGCATCACCGTGGCCATGAG | CGG | \|\|\|\|\|\|\|\|[\|\|\|\|\|\|\|\|\|\|\|\|]PAM | NA | Intergenic | NA | NA |

| | | | | | | | | | | | | |
|---|---|---|---|---|---|---|---|---|---|---|---|---|
| T11281 | AGCGGCGTGCTCTTCGCCACCGG | | 1000 | | | | | | | | | |
| Oligo fwd | CACCAGCGGCGTGCTCTTCGCCAC | | | | | | | | | | | |
| Oligo rev | AAACGTGGCGAAGAGCACGCCGCT | | | | | | | | | | | |
| Chromosome | start | | end | strand | MM | target_seq | PAM | alignment | distance | position | gene name | gene id |
| GU305914.1 | | 147412 | 147434 | + | 0 | AGCGGCGTGCTCTTCGCCAC | CGG | \|\|\|\|\|\|\|\|[\|\|\|\|\|\|\|\|\|\|\|\|]PAM | NA | Intergenic | NA | NA |

| | | | | | | | | | | | | |
|---|---|---|---|---|---|---|---|---|---|---|---|---|
| T11282 | ACCGGCAAGTACCTGCTCTCCGG | | 1000 | | | | | | | | | |
| Oligo fwd | CACCACCGGCAAGTACCTGCTCTC | | | | | | | | | | | |
| Oligo rev | AAACGAGAGCAGGTACTTGCCGGT | | | | | | | | | | | |
| Chromosome | start | | end | strand | MM | target_seq | PAM | alignment | distance | position | gene name | gene id |
| GU305914.1 | | 147430 | 147452 | + | 0 | ACCGGCAAGTACCTGCTCTC | CGG | \|\|\|\|\|\|\|\|[\|\|\|\|\|\|\|\|\|\|\|\|]PAM | NA | Intergenic | NA | NA |

| | | | | | | | | | | | | |
|---|---|---|---|---|---|---|---|---|---|---|---|---|
| T11283 | CAGAAGTACTTCCCCACCCCCGG | | 1000 | | | | | | | | | |
| Oligo fwd | CACCCAGAAGTACTTCCCCACCCC | | | | | | | | | | | |
| Oligo rev | AAACGGGGTGGGGAAGTACTTCTG | | | | | | | | | | | |
| Chromosome | start | | end | strand | MM | target_seq | PAM | alignment | distance | position | gene name | gene id |
| GU305914.1 | | 147466 | 147488 | + | 0 | CAGAAGTACTTCCCCACCCC | CGG | \|\|\|\|\|\|\|\|[\|\|\|\|\|\|\|\|\|\|\|\|]PAM | NA | Intergenic | NA | NA |

| | | | |
|---|---|---|---|
| T11284 | AGTACTTCCCCACCCCCGGATGG | | 1000 |
| Oligo fwd | CACCAGTACTTCCCCACCCCCGGA | | |
| Oligo rev | AAACTCCGGGGGTGGGGAAGTACT | | |

Chromosome start end strand MM target_seq PAM alignment distance position gene name gene id
GU305914.1 147470 147492 + 0 AGTACTTCCCCACCCCCGGA TGG ||||||||[||||||||||||]PAM NA Intergenic NA NA

T11285 GTACTTCCCCACCCCCGGATGGG 1000
Oligo fwd CACCGTACTTCCCCACCCCCGGAT
Oligo rev AAACATCCGGGGGTGGGGAAGTAC
Chromosome start end strand MM target_seq PAM alignment distance position gene name gene id
GU305914.1 147471 147493 + 0 GTACTTCCCCACCCCCGGAT GGG ||||||||[||||||||||||]PAM NA Intergenic NA NA

T11286 TTCCCCACCCCCGGATGGGTCGG 1000
Oligo fwd CACCTTCCCCACCCCCGGATGGGT
Oligo rev AAACACCCATCCGGGGGTGGGGAA
Chromosome start end strand MM target_seq PAM alignment distance position gene name gene id
GU305914.1 147475 147497 + 0 TTCCCCACCCCCGGATGGGT CGG ||||||||[||||||||||||]PAM NA Intergenic NA NA

T11287 CCCGGATGGGTCGGCGCCACCGG 1000
Oligo fwd CACCCCCGGATGGGTCGGCGCCAC
Oligo rev AAACGTGGCGCCGACCCATCCGGG
Chromosome start end strand MM target_seq PAM alignment distance position gene name gene id
GU305914.1 147484 147506 + 0 CCCGGATGGGTCGGCGCCAC CGG ||||||||[||||||||||||]PAM NA Intergenic NA NA

T11288 CACCGGACGCCTGCTCTCCGAGG 1000
Oligo fwd CACCCACCGGACGCCTGCTCTCCG
Oligo rev AAACCGGAGAGCAGGCGTCCGGTG
Chromosome start end strand MM target_seq PAM alignment distance position gene name gene id
GU305914.1 147501 147523 + 0 CACCGGACGCCTGCTCTCCG AGG ||||||||[||||||||||||]PAM NA Intergenic NA NA

T11289 GCGCCTTCCAGTCCCCCGATTGG 1000
Oligo fwd CACCGCGCCTTCCAGTCCCCCGAT
Oligo rev AAACATCGGGGGACTGGAAGGCGC
Chromosome start end strand MM target_seq PAM alignment distance position gene name gene id
GU305914.1 147533 147555 + 0 GCGCCTTCCAGTCCCCCGAT TGG ||||||||[||||||||||||]PAM NA Intergenic NA NA

T11290 CGCCTTCCAGTCCCCCGATTGGG 1000
Oligo fwd CACCCGCCTTCCAGTCCCCCGATT
Oligo rev AAACAATCGGGGGACTGGAAGGCG
Chromosome start end strand MM target_seq PAM alignment distance position gene name gene id
GU305914.1 147534 147556 + 0 CGCCTTCCAGTCCCCCGATT GGG ||||||||[||||||||||||]PAM NA Intergenic NA NA

T11291 GCCTTCCAGTCCCCCGATTGGGG 1000
Oligo fwd CACCGCCTTCCAGTCCCCCGATTG
Oligo rev AAACCAATCGGGGGACTGGAAGGC
Chromosome start end strand MM target_seq PAM alignment distance position gene name gene id
GU305914.1 147535 147557 + 0 GCCTTCCAGTCCCCCGATTG GGG ||||||||[||||||||||||]PAM NA Intergenic NA NA

T11292 AGACAAGATCATCCGCTTCATGG 1000
Oligo fwd CACCAGACAAGATCATCCGCTTCA
Oligo rev AAACTGAAGCGGATGATCTTGTCT
Chromosome start end strand MM target_seq PAM alignment distance position gene name gene id
GU305914.1 147564 147586 + 0 AGACAAGATCATCCGCTTCA TGG ||||||||[||||||||||||]PAM NA Intergenic NA NA

T11293 CACGCACATGCTGCGCCTCTCGG 1000
Oligo fwd CACCCACGCACATGCTGCGCCTCT
Oligo rev AAACAGAGGCGCAGCATGTGCGTG
Chromosome start end strand MM target_seq PAM alignment distance position gene name gene id
GU305914.1 147588 147610 + 0 CACGCACATGCTGCGCCTCT CGG ||||||||[||||||||||||]PAM NA Intergenic NA NA

T11294 ACATGCTGCGCCTCTCGGCCAGG 1000
Oligo fwd CACCACATGCTGCGCCTCTCGGCC
Oligo rev AAACGGCCGAGAGGCGCAGCATGT
Chromosome start end strand MM target_seq PAM alignment distance position gene name gene id
GU305914.1 147593 147615 + 0 ACATGCTGCGCCTCTCGGCC AGG ||||||||[||||||||||||]PAM NA Intergenic NA NA

T11295 TGCTGCGCCTCTCGGCCAGGAGG 1000
Oligo fwd CACCTGCTGCGCCTCTCGGCCAGG
Oligo rev AAACCCTGGCCGAGAGGCGCAGCA
Chromosome start end strand MM target_seq PAM alignment distance position gene name gene id
GU305914.1 147596 147618 + 0 TGCTGCGCCTCTCGGCCAGG AGG ||||||||[||||||||||||]PAM NA Intergenic NA NA

T11296 GGAGGCACGAGACCCGCTTCTGG 1000
Oligo fwd CACCGGAGGCACGAGACCCGCTTC
Oligo rev AAACGAAGCGGGTCTCGTGCCTCC
Chromosome start end strand MM target_seq PAM alignment distance position gene name gene id
GU305914.1 147614 147636 + 0 GGAGGCACGAGACCCGCTTC TGG ||||||||[||||||||||||]PAM NA Intergenic NA NA

T11297 GAGGCACGAGACCCGCTTCTGGG 1000
Oligo fwd CACCGAGGCACGAGACCCGCTTCT
Oligo rev AAACAGAAGCGGGTCTCGTGCCTC
Chromosome start end strand MM target_seq PAM alignment distance position gene name gene id
GU305914.1 147615 147637 + 0 GAGGCACGAGACCCGCTTCT GGG ||||||||[||||||||||||]PAM NA Intergenic NA NA

T11298 AGACCCGCTTCTGGGCGCAGAGG 1000
Oligo fwd CACCAGACCCGCTTCTGGGCGCAG
Oligo rev AAACCTGCGCCCAGAAGCGGGTCT
Chromosome start end strand MM target_seq PAM alignment distance position gene name gene id
GU305914.1 147623 147645 + 0 AGACCCGCTTCTGGGCGCAG AGG ||||||||[||||||||||||]PAM NA Intergenic NA NA

T11299 TGGGCGCAGAGGTTCGCCCCCGG 1000
Oligo fwd CACCTGGGCGCAGAGGTTCGCCCC

| | | | | | | | | | | | |
|---|---|---|---|---|---|---|---|---|---|---|---|
| Oligo rev | AAACGGGGCGAACCTCTGCGCCCA | | | | | | | | | | |
| Chromosome | start | end | strand | MM | target_seq | PAM | alignment | distance | position | gene name | gene id |
| GU305914.1 | 147634 | 147656 | + | 0 | TGGGCGCAGAGGTTCGCCCC | CGG | \|\|\|\|\|\|\|\|[\|\|\|\|\|\|\|\|\|\|\|\|]PAM | NA | Intergenic | NA | NA |
| T11300 | AACCGCGTCCAGCAGCACGACGG | 1000 | | | | | | | | | |
| Oligo fwd | CACCAACCGCGTCCAGCAGCACGA | | | | | | | | | | |
| Oligo rev | AAACTCGTGCTGCTGGACGCGGTT | | | | | | | | | | |
| Chromosome | start | end | strand | MM | target_seq | PAM | alignment | distance | position | gene name | gene id |
| GU305914.1 | 147661 | 147683 | + | 0 | AACCGCGTCCAGCAGCACGA | CGG | \|\|\|\|\|\|\|\|[\|\|\|\|\|\|\|\|\|\|\|\|]PAM | NA | Intergenic | NA | NA |
| T11301 | CCAGCAGCACGACGGCGTCATGG | 1000 | | | | | | | | | |
| Oligo fwd | CACCCCAGCAGCACGACGGCGTCA | | | | | | | | | | |
| Oligo rev | AAACTGACGCCGTCGTGCTGCTGG | | | | | | | | | | |
| Chromosome | start | end | strand | MM | target_seq | PAM | alignment | distance | position | gene name | gene id |
| GU305914.1 | 147669 | 147691 | + | 0 | CCAGCAGCACGACGGCGTCA | TGG | \|\|\|\|\|\|\|\|[\|\|\|\|\|\|\|\|\|\|\|\|]PAM | NA | Intergenic | NA | NA |
| T11302 | ATGGACGCCAACGAGTTCTGCGG | 1000 | | | | | | | | | |
| Oligo fwd | CACCATGGACGCCAACGAGTTCTG | | | | | | | | | | |
| Oligo rev | AAACCAGAACTCGTTGGCGTCCAT | | | | | | | | | | |
| Chromosome | start | end | strand | MM | target_seq | PAM | alignment | distance | position | gene name | gene id |
| GU305914.1 | 147688 | 147710 | + | 0 | ATGGACGCCAACGAGTTCTG | CGG | \|\|\|\|\|\|\|\|[\|\|\|\|\|\|\|\|\|\|\|\|]PAM | NA | Intergenic | NA | NA |
| T11303 | TGGACGCCAACGAGTTCTGCGGG | 1000 | | | | | | | | | |
| Oligo fwd | CACCTGGACGCCAACGAGTTCTGC | | | | | | | | | | |
| Oligo rev | AAACGCAGAACTCGTTGGCGTCCA | | | | | | | | | | |
| Chromosome | start | end | strand | MM | target_seq | PAM | alignment | distance | position | gene name | gene id |
| GU305914.1 | 147689 | 147711 | + | 0 | TGGACGCCAACGAGTTCTGC | GGG | \|\|\|\|\|\|\|\|[\|\|\|\|\|\|\|\|\|\|\|\|]PAM | NA | Intergenic | NA | NA |
| T11304 | TTCTGCGGGATCCACGTCTGCGG | 1000 | | | | | | | | | |
| Oligo fwd | CACCTTCTGCGGGATCCACGTCTG | | | | | | | | | | |
| Oligo rev | AAACCAGACGTGGATCCCGCAGAA | | | | | | | | | | |
| Chromosome | start | end | strand | MM | target_seq | PAM | alignment | distance | position | gene name | gene id |
| GU305914.1 | 147703 | 147725 | + | 0 | TTCTGCGGGATCCACGTCTG | CGG | \|\|\|\|\|\|\|\|[\|\|\|\|\|\|\|\|\|\|\|\|]PAM | NA | Intergenic | NA | NA |
| T11305 | GATCCACGTCTGCGGCCGCATGG | 1000 | | | | | | | | | |
| Oligo fwd | CACCGATCCACGTCTGCGGCCGCA | | | | | | | | | | |
| Oligo rev | AAACTGCGGCCGCAGACGTGGATC | | | | | | | | | | |
| Chromosome | start | end | strand | MM | target_seq | PAM | alignment | distance | position | gene name | gene id |
| GU305914.1 | 147711 | 147733 | + | 0 | GATCCACGTCTGCGGCCGCA | TGG | \|\|\|\|\|\|\|\|[\|\|\|\|\|\|\|\|\|\|\|\|]PAM | NA | Intergenic | NA | NA |
| T11306 | CCACGTCTGCGGCCGCATGGTGG | 1000 | | | | | | | | | |
| Oligo fwd | CACCCCACGTCTGCGGCCGCATGG | | | | | | | | | | |
| Oligo rev | AAACCCATGCGGCCGCAGACGTGG | | | | | | | | | | |
| Chromosome | start | end | strand | MM | target_seq | PAM | alignment | distance | position | gene name | gene id |
| GU305914.1 | 147714 | 147736 | + | 0 | CCACGTCTGCGGCCGCATGG | TGG | \|\|\|\|\|\|\|\|[\|\|\|\|\|\|\|\|\|\|\|\|]PAM | NA | Intergenic | NA | NA |
| T11307 | CATGGTGGCCGTCCAGCCCCTGG | 1000 | | | | | | | | | |
| Oligo fwd | CACCCATGGTGGCCGTCCAGCCCC | | | | | | | | | | |
| Oligo rev | AAACGGGGCTGGACGGCCACCATG | | | | | | | | | | |
| Chromosome | start | end | strand | MM | target_seq | PAM | alignment | distance | position | gene name | gene id |
| GU305914.1 | 147729 | 147751 | + | 0 | CATGGTGGCCGTCCAGCCCC | TGG | \|\|\|\|\|\|\|\|[\|\|\|\|\|\|\|\|\|\|\|\|]PAM | NA | Intergenic | NA | NA |
| T11308 | GTGGCCGTCCAGCCCCTGGACGG | 1000 | | | | | | | | | |
| Oligo fwd | CACCGTGGCCGTCCAGCCCCTGGA | | | | | | | | | | |
| Oligo rev | AAACTCCAGGGGCTGGACGGCCAC | | | | | | | | | | |
| Chromosome | start | end | strand | MM | target_seq | PAM | alignment | distance | position | gene name | gene id |
| GU305914.1 | 147733 | 147755 | + | 0 | GTGGCCGTCCAGCCCCTGGA | CGG | \|\|\|\|\|\|\|\|[\|\|\|\|\|\|\|\|\|\|\|\|]PAM | NA | Intergenic | NA | NA |
| T11309 | TGGCCGTCCAGCCCCTGGACGGG | 1000 | | | | | | | | | |
| Oligo fwd | CACCTGGCCGTCCAGCCCCTGGAC | | | | | | | | | | |
| Oligo rev | AAACGTCCAGGGGCTGGACGGCCA | | | | | | | | | | |
| Chromosome | start | end | strand | MM | target_seq | PAM | alignment | distance | position | gene name | gene id |
| GU305914.1 | 147734 | 147756 | + | 0 | TGGCCGTCCAGCCCCTGGAC | GGG | \|\|\|\|\|\|\|\|[\|\|\|\|\|\|\|\|\|\|\|\|]PAM | NA | Intergenic | NA | NA |
| T11310 | GGCCGTCCAGCCCCTGGACGGGG | 1000 | | | | | | | | | |
| Oligo fwd | CACCGGCCGTCCAGCCCCTGGACG | | | | | | | | | | |
| Oligo rev | AAACCGTCCAGGGGCTGGACGGCC | | | | | | | | | | |
| Chromosome | start | end | strand | MM | target_seq | PAM | alignment | distance | position | gene name | gene id |
| GU305914.1 | 147735 | 147757 | + | 0 | GGCCGTCCAGCCCCTGGACG | GGG | \|\|\|\|\|\|\|\|[\|\|\|\|\|\|\|\|\|\|\|\|]PAM | NA | Intergenic | NA | NA |
| GU305914.1 | 212045 | 212067 | + | 3 | GACCGTCCAGCCGCTCGACG | AGG | \|-\|\|\|\|\|\|[\|\|\|\|-\|\|-\|\|\|\|]PAM | NA | Intergenic | NA | NA |
| T11311 | CTACGTCAAGAAGACCTTCGAGG | 1000 | | | | | | | | | |
| Oligo fwd | CACCCTACGTCAAGAAGACCTTCG | | | | | | | | | | |
| Oligo rev | AAACCGAAGGTCTTCTTGACGTAG | | | | | | | | | | |
| Chromosome | start | end | strand | MM | target_seq | PAM | alignment | distance | position | gene name | gene id |
| GU305914.1 | 147786 | 147808 | + | 0 | CTACGTCAAGAAGACCTTCG | AGG | \|\|\|\|\|\|\|\|[\|\|\|\|\|\|\|\|\|\|\|\|]PAM | NA | Intergenic | NA | NA |
| GU305914.1 | 126929 | 126951 | - | 4 | CCACCTGAAGGAGACCTTCG | AGG | \|-\|\|-\|-\|[\|\|-\|\|\|\|\|\|\|\|\|]PAM | NA | Intergenic | NA | NA |
| T11312 | AGAAGACCTTCGAGGTGCTCAGG | 1000 | | | | | | | | | |
| Oligo fwd | CACCAGAAGACCTTCGAGGTGCTC | | | | | | | | | | |
| Oligo rev | AAACGAGCACCTCGAAGGTCTTCT | | | | | | | | | | |
| Chromosome | start | end | strand | MM | target_seq | PAM | alignment | distance | position | gene name | gene id |
| GU305914.1 | 147794 | 147816 | + | 0 | AGAAGACCTTCGAGGTGCTC | AGG | \|\|\|\|\|\|\|\|[\|\|\|\|\|\|\|\|\|\|\|\|]PAM | NA | Intergenic | NA | NA |
| T11313 | GAAGACCTTCGAGGTGCTCAGGG | 1000 | | | | | | | | | |
| Oligo fwd | CACCGAAGACCTTCGAGGTGCTCA | | | | | | | | | | |
| Oligo rev | AAACTGAGCACCTCGAAGGTCTTC | | | | | | | | | | |
| Chromosome | start | end | strand | MM | target_seq | PAM | alignment | distance | position | gene name | gene id |
| GU305914.1 | 147795 | 147817 | + | 0 | GAAGACCTTCGAGGTGCTCA | GGG | \|\|\|\|\|\|\|\|[\|\|\|\|\|\|\|\|\|\|\|\|]PAM | NA | Intergenic | NA | NA |

T11314 GCTCAGGGTGACCCTCGAGAAGG 1000
Oligo fwd CACCGCTCAGGGTGACCCTCGAGA
Oligo rev AAACTCTCGAGGGTCACCCTGAGC
Chromosome start end strand MM target_seq PAM alignment distance position gene name gene id
GU305914.1 147810 147832 + 0 GCTCAGGGTGACCCTCGAGA AGG ||||||||[||||||||||||]PAM NA Intergenic NA NA

T11315 GGTCATCATCGTCCTCTCGCAGG 1000
Oligo fwd CACCGGTCATCATCGTCCTCTCGC
Oligo rev AAACGCGAGAGGACGATGATGACC
Chromosome start end strand MM target_seq PAM alignment distance position gene name gene id
GU305914.1 147831 147853 + 0 GGTCATCATCGTCCTCTCGC AGG ||||||||[||||||||||||]PAM NA Intergenic NA NA

T11316 GCAGGACCAGACCAACACCGCGG 1000
Oligo fwd CACCGCAGGACCAGACCAACACCG
Oligo rev AAACCGGTGTTGGTCTGGTCCTGC
Chromosome start end strand MM target_seq PAM alignment distance position gene name gene id
GU305914.1 147849 147871 + 0 GCAGGACCAGACCAACACCG CGG ||||||||[||||||||||||]PAM NA Intergenic NA NA

T11317 CGCGGTCGTCAACAACAGCACGG 1000
Oligo fwd CACCCGCGGTCGTCAACAACAGCA
Oligo rev AAACTGCTGTTGTTGACGACCGCG
Chromosome start end strand MM target_seq PAM alignment distance position gene name gene id
GU305914.1 147867 147889 + 0 CGCGGTCGTCAACAACAGCA CGG ||||||||[||||||||||||]PAM NA Intergenic NA NA

T11318 GCGGTCGTCAACAACAGCACGGG 1000
Oligo fwd CACCGCGGTCGTCAACAACAGCAC
Oligo rev AAACGTGCTGTTGTTGACGACCGC
Chromosome start end strand MM target_seq PAM alignment distance position gene name gene id
GU305914.1 147868 147890 + 0 GCGGTCGTCAACAACAGCAC GGG ||||||||[||||||||||||]PAM NA Intergenic NA NA

T11319 CGACTGCGTCGCCATCAAGCAGG 1000
Oligo fwd CACCCGACTGCGTCGCCATCAAGC
Oligo rev AAACGCTTGATGGCGACGCAGTCG
Chromosome start end strand MM target_seq PAM alignment distance position gene name gene id
GU305914.1 147897 147919 + 0 CGACTGCGTCGCCATCAAGC AGG ||||||||[||||||||||||]PAM NA Intergenic NA NA

T11320 CTGCGTCGCCATCAAGCAGGAGG 1000
Oligo fwd CACCCTGCGTCGCCATCAAGCAGG
Oligo rev AAACCCTGCTTGATGGCGACGCAG
Chromosome start end strand MM target_seq PAM alignment distance position gene name gene id
GU305914.1 147900 147922 + 0 CTGCGTCGCCATCAAGCAGG AGG ||||||||[||||||||||||]PAM NA Intergenic NA NA

T11321 GCGATCGTCAGCGACACCACCGG 1000
Oligo fwd CACCGCGATCGTCAGCGACACCAC
Oligo rev AAACGTGGTGTCGCTGACGATCGC
Chromosome start end strand MM target_seq PAM alignment distance position gene name gene id
GU305914.1 147934 147956 + 0 GCGATCGTCAGCGACACCAC CGG ||||||||[||||||||||||]PAM NA Intergenic NA NA

T11322 CGTCAGCGACACCACCGGCTCGG 1000
Oligo fwd CACCCGTCAGCGACACCACCGGCT
Oligo rev AAACAGCCGGTGGTGTCGCTGACG
Chromosome start end strand MM target_seq PAM alignment distance position gene name gene id
GU305914.1 147939 147961 + 0 CGTCAGCGACACCACCGGCT CGG ||||||||[||||||||||||]PAM NA Intergenic NA NA

T11323 CACCACCGGCTCGGATGAGCTGG 1000
Oligo fwd CACCCACCACCGGCTCGGATGAGC
Oligo rev AAACGCTCATCCGAGCCGGTGGTG
Chromosome start end strand MM target_seq PAM alignment distance position gene name gene id
GU305914.1 147948 147970 + 0 CACCACCGGCTCGGATGAGC TGG ||||||||[||||||||||||]PAM NA Intergenic NA NA

T11324 ATGAGCTGGACGCCGCGCTACGG 1000
Oligo fwd CACCATGAGCTGGACGCCGCGCTA
Oligo rev AAACTAGCGCGGCGTCCAGCTCAT
Chromosome start end strand MM target_seq PAM alignment distance position gene name gene id
GU305914.1 147962 147984 + 0 ATGAGCTGGACGCCGCGCTA CGG ||||||||[||||||||||||]PAM NA Intergenic NA NA

T11325 TGAGCTGGACGCCGCGCTACGGG 1000
Oligo fwd CACCTGAGCTGGACGCCGCGCTAC
Oligo rev AAACGTAGCGCGGCGTCCAGCTCA
Chromosome start end strand MM target_seq PAM alignment distance position gene name gene id
GU305914.1 147963 147985 + 0 TGAGCTGGACGCCGCGCTAC GGG ||||||||[||||||||||||]PAM NA Intergenic NA NA

T11326 ACGCCGCGCTACGGGCCCTCAGG 1000
Oligo fwd CACCACGCCGCGCTACGGGCCCTC
Oligo rev AAACGAGGGCCCGTAGCGCGGCGT
Chromosome start end strand MM target_seq PAM alignment distance position gene name gene id
GU305914.1 147971 147993 + 0 ACGCCGCGCTACGGGCCCTC AGG ||||||||[||||||||||||]PAM NA Intergenic NA NA

T11327 CGCCGCGCTACGGGCCCTCAGGG 1000
Oligo fwd CACCCGCCGCGCTACGGGCCCTCA
Oligo rev AAACTGAGGGCCCGTAGCGCGGCG
Chromosome start end strand MM target_seq PAM alignment distance position gene name gene id
GU305914.1 147972 147994 + 0 CGCCGCGCTACGGGCCCTCA GGG ||||||||[||||||||||||]PAM NA Intergenic NA NA

T11328 ACGGGCCCTCAGGGAGACCACGG 1000
Oligo fwd CACCACGGGCCCTCAGGGAGACCA
Oligo rev AAACTGGTCTCCCTGAGGGCCCGT
Chromosome start end strand MM target_seq PAM alignment distance position gene name gene id

| | | | | | | | | | | | |
|---|---|---|---|---|---|---|---|---|---|---|---|
| GU305914.1 | | 147981 | 148003 | + | 0 | ACGGGCCCTCAGGGAGACCA | CGG | \|\|\|\|\|\|\|\|[\|\|\|\|\|\|\|\|\|\|\|\|]PAM | NA | Intergenic | NA | NA |
| T11329 | GAGACCACGGCCAACACCGTCGG | | 1000 | | | | | | | | | |
| Oligo fwd | CACCGAGACCACGGCCAACACCGT | | | | | | | | | | | |
| Oligo rev | AAACACGGTGTTGGCCGTGGTCTC | | | | | | | | | | | |
| Chromosome | start | | end | strand | MM | target_seq | PAM | alignment | distance | position | gene name | gene id |
| GU305914.1 | | 147994 | 148016 | + | 0 | GAGACCACGGCCAACACCGT | CGG | \|\|\|\|\|\|\|\|[\|\|\|\|\|\|\|\|\|\|\|\|]PAM | NA | Intergenic | NA | NA |
| T11330 | CACGGCCAACACCGTCGGCATGG | | 1000 | | | | | | | | | |
| Oligo fwd | CACCCACGGCCAACACCGTCGGCA | | | | | | | | | | | |
| Oligo rev | AAACTGCCGACGGTGTTGGCCGTG | | | | | | | | | | | |
| Chromosome | start | | end | strand | MM | target_seq | PAM | alignment | distance | position | gene name | gene id |
| GU305914.1 | | 147999 | 148021 | + | 0 | CACGGCCAACACCGTCGGCA | TGG | \|\|\|\|\|\|\|\|[\|\|\|\|\|\|\|\|\|\|\|\|]PAM | NA | Intergenic | NA | NA |
| T11331 | GGCCAACACCGTCGGCATGGTGG | | 1000 | | | | | | | | | |
| Oligo fwd | CACCGGCCAACACCGTCGGCATGG | | | | | | | | | | | |
| Oligo rev | AAACCCATGCCGACGGTGTTGGCC | | | | | | | | | | | |
| Chromosome | start | | end | strand | MM | target_seq | PAM | alignment | distance | position | gene name | gene id |
| GU305914.1 | | 148002 | 148024 | + | 0 | GGCCAACACCGTCGGCATGG | TGG | \|\|\|\|\|\|\|\|[\|\|\|\|\|\|\|\|\|\|\|\|]PAM | NA | Intergenic | NA | NA |
| T11332 | CAACACCGTCGGCATGGTGGTGG | | 1000 | | | | | | | | | |
| Oligo fwd | CACCCAACACCGTCGGCATGGTGG | | | | | | | | | | | |
| Oligo rev | AAACCCACCATGCCGACGGTGTTG | | | | | | | | | | | |
| Chromosome | start | | end | strand | MM | target_seq | PAM | alignment | distance | position | gene name | gene id |
| GU305914.1 | | 148005 | 148027 | + | 0 | CAACACCGTCGGCATGGTGG | TGG | \|\|\|\|\|\|\|\|[\|\|\|\|\|\|\|\|\|\|\|\|]PAM | NA | Intergenic | NA | NA |
| GU305914.1 | | 61554 | 61576 | - | 3 | CAAGACCTACGGCATGGTGG | TGG | \|\|\|-\|\|\|-[-\|\|\|\|\|\|\|\|\|\|\|]PAM | NA | Intergenic | NA | NA |
| T11333 | CGATGTCTTTCAACAGACTATGG | | 1000 | | | | | | | | | |
| Oligo fwd | CACCCGATGTCTTTCAACAGACTA | | | | | | | | | | | |
| Oligo rev | AAACTAGTCTGTTGAAAGACATCG | | | | | | | | | | | |
| Chromosome | start | | end | strand | MM | target_seq | PAM | alignment | distance | position | gene name | gene id |
| GU305914.1 | | 148065 | 148087 | + | 0 | CGATGTCTTTCAACAGACTA | TGG | \|\|\|\|\|\|\|\|[\|\|\|\|\|\|\|\|\|\|\|\|]PAM | NA | Intergenic | NA | NA |
| T11334 | ACAGACTATGGACTTTGTCGTGG | | 1000 | | | | | | | | | |
| Oligo fwd | CACCACAGACTATGGACTTTGTCG | | | | | | | | | | | |
| Oligo rev | AAACCGACAAAGTCCATAGTCTGT | | | | | | | | | | | |
| Chromosome | start | | end | strand | MM | target_seq | PAM | alignment | distance | position | gene name | gene id |
| GU305914.1 | | 148077 | 148099 | + | 0 | ACAGACTATGGACTTTGTCG | TGG | \|\|\|\|\|\|\|\|[\|\|\|\|\|\|\|\|\|\|\|\|]PAM | NA | Intergenic | NA | NA |
| T11335 | CTTTGTCGTGGAACGCTACTCGG | | 1000 | | | | | | | | | |
| Oligo fwd | CACCCTTTGTCGTGGAACGCTACT | | | | | | | | | | | |
| Oligo rev | AAACAGTAGCGTTCCACGACAAAG | | | | | | | | | | | |
| Chromosome | start | | end | strand | MM | target_seq | PAM | alignment | distance | position | gene name | gene id |
| GU305914.1 | | 148089 | 148111 | + | 0 | CTTTGTCGTGGAACGCTACT | CGG | \|\|\|\|\|\|\|\|[\|\|\|\|\|\|\|\|\|\|\|\|]PAM | NA | Intergenic | NA | NA |
| T11336 | TTTCTATGAACTGACTGAGTTGG | | 1000 | | | | | | | | | |
| Oligo fwd | CACCTTTCTATGAACTGACTGAGT | | | | | | | | | | | |
| Oligo rev | AAACACTCAGTCAGTTCATAGAAA | | | | | | | | | | | |
| Chromosome | start | | end | strand | MM | target_seq | PAM | alignment | distance | position | gene name | gene id |
| GU305914.1 | | 148118 | 148140 | + | 0 | TTTCTATGAACTGACTGAGT | TGG | \|\|\|\|\|\|\|\|[\|\|\|\|\|\|\|\|\|\|\|\|]PAM | NA | Intergenic | NA | NA |
| T11337 | TTCTATGAACTGACTGAGTTGGG | | 1000 | | | | | | | | | |
| Oligo fwd | CACCTTCTATGAACTGACTGAGTT | | | | | | | | | | | |
| Oligo rev | AAACAACTCAGTCAGTTCATAGAA | | | | | | | | | | | |
| Chromosome | start | | end | strand | MM | target_seq | PAM | alignment | distance | position | gene name | gene id |
| GU305914.1 | | 148119 | 148141 | + | 0 | TTCTATGAACTGACTGAGTT | GGG | \|\|\|\|\|\|\|\|[\|\|\|\|\|\|\|\|\|\|\|\|]PAM | NA | Intergenic | NA | NA |
| T11338 | TCTATGAACTGACTGAGTTGGGG | | 1000 | | | | | | | | | |
| Oligo fwd | CACCTCTATGAACTGACTGAGTTG | | | | | | | | | | | |
| Oligo rev | AAACCAACTCAGTCAGTTCATAGA | | | | | | | | | | | |
| Chromosome | start | | end | strand | MM | target_seq | PAM | alignment | distance | position | gene name | gene id |
| GU305914.1 | | 148120 | 148142 | + | 0 | TCTATGAACTGACTGAGTTG | GGG | \|\|\|\|\|\|\|\|[\|\|\|\|\|\|\|\|\|\|\|\|]PAM | NA | Intergenic | NA | NA |
| T11339 | ACTGACTGAGTTGGGGAATCAGG | | 1000 | | | | | | | | | |
| Oligo fwd | CACCACTGACTGAGTTGGGGAATC | | | | | | | | | | | |
| Oligo rev | AAACGATTCCCCAACTCAGTCAGT | | | | | | | | | | | |
| Chromosome | start | | end | strand | MM | target_seq | PAM | alignment | distance | position | gene name | gene id |
| GU305914.1 | | 148127 | 148149 | + | 0 | ACTGACTGAGTTGGGGAATC | AGG | \|\|\|\|\|\|\|\|[\|\|\|\|\|\|\|\|\|\|\|\|]PAM | NA | Intergenic | NA | NA |
| T11340 | GACTGAGTTGGGGAATCAGGAGG | | 1000 | | | | | | | | | |
| Oligo fwd | CACCGACTGAGTTGGGGAATCAGG | | | | | | | | | | | |
| Oligo rev | AAACCCTGATTCCCCAACTCAGTC | | | | | | | | | | | |
| Chromosome | start | | end | strand | MM | target_seq | PAM | alignment | distance | position | gene name | gene id |
| GU305914.1 | | 148130 | 148152 | + | 0 | GACTGAGTTGGGGAATCAGG | AGG | \|\|\|\|\|\|\|\|[\|\|\|\|\|\|\|\|\|\|\|\|]PAM | NA | Intergenic | NA | NA |
| T11341 | ACTGAGTTGGGGAATCAGGAGGG | | 1000 | | | | | | | | | |
| Oligo fwd | CACCACTGAGTTGGGGAATCAGGA | | | | | | | | | | | |
| Oligo rev | AAACTCCTGATTCCCCAACTCAGT | | | | | | | | | | | |
| Chromosome | start | | end | strand | MM | target_seq | PAM | alignment | distance | position | gene name | gene id |
| GU305914.1 | | 148131 | 148153 | + | 0 | ACTGAGTTGGGGAATCAGGA | GGG | \|\|\|\|\|\|\|\|[\|\|\|\|\|\|\|\|\|\|\|\|]PAM | NA | Intergenic | NA | NA |
| T11342 | TTGGGGAATCAGGAGGGTGATGG | | 1000 | | | | | | | | | |
| Oligo fwd | CACCTTGGGGAATCAGGAGGGTGA | | | | | | | | | | | |
| Oligo rev | AAACTCACCCTCCTGATTCCCCAA | | | | | | | | | | | |
| Chromosome | start | | end | strand | MM | target_seq | PAM | alignment | distance | position | gene name | gene id |
| GU305914.1 | | 148137 | 148159 | + | 0 | TTGGGGAATCAGGAGGGTGA | TGG | \|\|\|\|\|\|\|\|[\|\|\|\|\|\|\|\|\|\|\|\|]PAM | NA | Intergenic | NA | NA |
| T11343 | GAATCAGGAGGGTGATGGACAGG | | 1000 | | | | | | | | | |
| Oligo fwd | CACCGAATCAGGAGGGTGATGGAC | | | | | | | | | | | |

| | | | | | | | | | | | |
|---|---|---|---|---|---|---|---|---|---|---|---|
| Oligo rev | AAACGTCCATCACCCTCCTGATTC | | | | | | | | | | |
| Chromosome | start | end | strand | MM | target_seq | PAM | alignment | distance | position | gene name | gene id |
| GU305914.1 | 148142 | 148164 | + | 0 | GAATCAGGAGGGTGATGGAC | AGG | \|\|\|\|\|\|\|\|[\|\|\|\|\|\|\|\|\|\|\|\|]PAM | NA | Intergenic | NA | NA |
| T11344 | ACACTGCAAAAGACGATCAACGG | 1000 | | | | | | | | | |
| Oligo fwd | CACCACACTGCAAAAGACGATCAA | | | | | | | | | | |
| Oligo rev | AAACTTGATCGTCTTTTGCAGTGT | | | | | | | | | | |
| Chromosome | start | end | strand | MM | target_seq | PAM | alignment | distance | position | gene name | gene id |
| GU305914.1 | 148243 | 148265 | + | 0 | ACACTGCAAAAGACGATCAA | CGG | \|\|\|\|\|\|\|\|[\|\|\|\|\|\|\|\|\|\|\|\|]PAM | NA | Intergenic | NA | NA |
| T11345 | ATATCATTTTTTATTGTTCGAGG | 1000 | | | | | | | | | |
| Oligo fwd | CACCATATCATTTTTTATTGTTCG | | | | | | | | | | |
| Oligo rev | AAACCGAACAATAAAAAATGATAT | | | | | | | | | | |
| Chromosome | start | end | strand | MM | target_seq | PAM | alignment | distance | position | gene name | gene id |
| GU305914.1 | 148276 | 148298 | + | 0 | ATATCATTTTTTATTGTTCG | AGG | \|\|\|\|\|\|\|\|[\|\|\|\|\|\|\|\|\|\|\|\|]PAM | NA | Intergenic | NA | NA |
| T11346 | CGAGGCGCTTTTCTTAGTGTCGG | 1000 | | | | | | | | | |
| Oligo fwd | CACCCGAGGCGCTTTTCTTAGTGT | | | | | | | | | | |
| Oligo rev | AAACACACTAAGAAAAGCGCCTCG | | | | | | | | | | |
| Chromosome | start | end | strand | MM | target_seq | PAM | alignment | distance | position | gene name | gene id |
| GU305914.1 | 148294 | 148316 | + | 0 | CGAGGCGCTTTTCTTAGTGT | CGG | \|\|\|\|\|\|\|\|[\|\|\|\|\|\|\|\|\|\|\|\|]PAM | NA | Intergenic | NA | NA |
| T11347 | GCTTTTCTTAGTGTCGGCGCCGG | 1000 | | | | | | | | | |
| Oligo fwd | CACCGCTTTTCTTAGTGTCGGCGC | | | | | | | | | | |
| Oligo rev | AAACGCGCCGACACTAAGAAAAGC | | | | | | | | | | |
| Chromosome | start | end | strand | MM | target_seq | PAM | alignment | distance | position | gene name | gene id |
| GU305914.1 | 148300 | 148322 | + | 0 | GCTTTTCTTAGTGTCGGCGC | CGG | \|\|\|\|\|\|\|\|[\|\|\|\|\|\|\|\|\|\|\|\|]PAM | NA | Intergenic | NA | NA |
| T11348 | CTTTTCTTAGTGTCGGCGCCGGG | 1000 | | | | | | | | | |
| Oligo fwd | CACCCTTTTCTTAGTGTCGGCGCC | | | | | | | | | | |
| Oligo rev | AAACGGCGCCGACACTAAGAAAAG | | | | | | | | | | |
| Chromosome | start | end | strand | MM | target_seq | PAM | alignment | distance | position | gene name | gene id |
| GU305914.1 | 148301 | 148323 | + | 0 | CTTTTCTTAGTGTCGGCGCC | GGG | \|\|\|\|\|\|\|\|[\|\|\|\|\|\|\|\|\|\|\|\|]PAM | NA | Intergenic | NA | NA |
| T11349 | AGTGTCGGCGCCGGGAGTAGTGG | 1000 | | | | | | | | | |
| Oligo fwd | CACCAGTGTCGGCGCCGGGAGTAG | | | | | | | | | | |
| Oligo rev | AAACCTACTCCCGGCGCCGACACT | | | | | | | | | | |
| Chromosome | start | end | strand | MM | target_seq | PAM | alignment | distance | position | gene name | gene id |
| GU305914.1 | 148309 | 148331 | + | 0 | AGTGTCGGCGCCGGGAGTAG | TGG | \|\|\|\|\|\|\|\|[\|\|\|\|\|\|\|\|\|\|\|\|]PAM | NA | Intergenic | NA | NA |
| T11350 | GTCGGCGCCGGGAGTAGTGGTGG | 1000 | | | | | | | | | |
| Oligo fwd | CACCGTCGGCGCCGGGAGTAGTGG | | | | | | | | | | |
| Oligo rev | AAACCCACTACTCCCGGCGCCGAC | | | | | | | | | | |
| Chromosome | start | end | strand | MM | target_seq | PAM | alignment | distance | position | gene name | gene id |
| GU305914.1 | 148312 | 148334 | + | 0 | GTCGGCGCCGGGAGTAGTGG | TGG | \|\|\|\|\|\|\|\|[\|\|\|\|\|\|\|\|\|\|\|\|]PAM | NA | Intergenic | NA | NA |
| T11351 | GGCGCCGGGAGTAGTGGTGGTGG | 1000 | | | | | | | | | |
| Oligo fwd | CACCGGCGCCGGGAGTAGTGGTGG | | | | | | | | | | |
| Oligo rev | AAACCCACCACTACTCCCGGCGCC | | | | | | | | | | |
| Chromosome | start | end | strand | MM | target_seq | PAM | alignment | distance | position | gene name | gene id |
| GU305914.1 | 148315 | 148337 | + | 0 | GGCGCCGGGAGTAGTGGTGG | TGG | \|\|\|\|\|\|\|\|[\|\|\|\|\|\|\|\|\|\|\|\|]PAM | NA | Intergenic | NA | NA |
| T11352 | AGTCGTCATATTCTTCTTAGTGG | 1000 | | | | | | | | | |
| Oligo fwd | CACCAGTCGTCATATTCTTCTTAG | | | | | | | | | | |
| Oligo rev | AAACCTAAGAAGAATATGACGACT | | | | | | | | | | |
| Chromosome | start | end | strand | MM | target_seq | PAM | alignment | distance | position | gene name | gene id |
| GU305914.1 | 148390 | 148412 | + | 0 | AGTCGTCATATTCTTCTTAG | TGG | \|\|\|\|\|\|\|\|[\|\|\|\|\|\|\|\|\|\|\|\|]PAM | NA | Intergenic | NA | NA |
| T11353 | CGTCATATTCTTCTTAGTGGTGG | 1000 | | | | | | | | | |
| Oligo fwd | CACCCGTCATATTCTTCTTAGTGG | | | | | | | | | | |
| Oligo rev | AAACCCACTAAGAAGAATATGACG | | | | | | | | | | |
| Chromosome | start | end | strand | MM | target_seq | PAM | alignment | distance | position | gene name | gene id |
| GU305914.1 | 148393 | 148415 | + | 0 | CGTCATATTCTTCTTAGTGG | TGG | \|\|\|\|\|\|\|\|[\|\|\|\|\|\|\|\|\|\|\|\|]PAM | NA | Intergenic | NA | NA |
| T11354 | GTCATATTCTTCTTAGTGGTGGG | 1000 | | | | | | | | | |
| Oligo fwd | CACCGTCATATTCTTCTTAGTGGT | | | | | | | | | | |
| Oligo rev | AAACACCACTAAGAAGAATATGAC | | | | | | | | | | |
| Chromosome | start | end | strand | MM | target_seq | PAM | alignment | distance | position | gene name | gene id |
| GU305914.1 | 148394 | 148416 | + | 0 | GTCATATTCTTCTTAGTGGT | GGG | \|\|\|\|\|\|\|\|[\|\|\|\|\|\|\|\|\|\|\|\|]PAM | NA | Intergenic | NA | NA |
| GU305914.1 | 49571 | 49593 | - | 4 | GGCATGTTCTTCTTCGTGTT | CGG | \|-\|\|\|-\|\|[\|\|\|\|\|\|-\|\|\|-\|]PAM | NA | Intergenic | NA | NA |
| T11355 | ATATTCTTCTTAGTGGTGGGAGG | 1000 | | | | | | | | | |
| Oligo fwd | CACCATATTCTTCTTAGTGGTGGG | | | | | | | | | | |
| Oligo rev | AAACCCCACCACTAAGAAGAATAT | | | | | | | | | | |
| Chromosome | start | end | strand | MM | target_seq | PAM | alignment | distance | position | gene name | gene id |
| GU305914.1 | 148397 | 148419 | + | 0 | ATATTCTTCTTAGTGGTGGG | AGG | \|\|\|\|\|\|\|\|[\|\|\|\|\|\|\|\|\|\|\|\|]PAM | NA | Intergenic | NA | NA |
| T11356 | AGTGGTGGGAGGAGTAGTGATGG | 1000 | | | | | | | | | |
| Oligo fwd | CACCAGTGGTGGGAGGAGTAGTGA | | | | | | | | | | |
| Oligo rev | AAACTCACTACTCCTCCCACCACT | | | | | | | | | | |
| Chromosome | start | end | strand | MM | target_seq | PAM | alignment | distance | position | gene name | gene id |
| GU305914.1 | 148408 | 148430 | + | 0 | AGTGGTGGGAGGAGTAGTGA | TGG | \|\|\|\|\|\|\|\|[\|\|\|\|\|\|\|\|\|\|\|\|]PAM | NA | Intergenic | NA | NA |
| T11357 | GGTGGGAGGAGTAGTGATGGTGG | 1000 | | | | | | | | | |
| Oligo fwd | CACCGGTGGGAGGAGTAGTGATGG | | | | | | | | | | |
| Oligo rev | AAACCCATCACTACTCCTCCCACC | | | | | | | | | | |
| Chromosome | start | end | strand | MM | target_seq | PAM | alignment | distance | position | gene name | gene id |
| GU305914.1 | 148411 | 148433 | + | 0 | GGTGGGAGGAGTAGTGATGG | TGG | \|\|\|\|\|\|\|\|[\|\|\|\|\|\|\|\|\|\|\|\|]PAM | NA | Intergenic | NA | NA |

<table>
<tr><td>T11358</td><td>AGGAGTAGTGATGGTGGTATCGG</td><td></td><td>1000</td><td></td><td></td><td></td><td></td><td></td><td></td><td></td><td></td><td></td></tr>
<tr><td>Oligo fwd</td><td>CACCAGGAGTAGTGATGGTGGTAT</td><td></td><td></td><td></td><td></td><td></td><td></td><td></td><td></td><td></td><td></td><td></td></tr>
<tr><td>Oligo rev</td><td>AAACATACCACCATCACTACTCCT</td><td></td><td></td><td></td><td></td><td></td><td></td><td></td><td></td><td></td><td></td><td></td></tr>
<tr><td>Chromosome</td><td>start</td><td></td><td>end</td><td>strand</td><td>MM</td><td>target_seq</td><td>PAM</td><td>alignment</td><td>distance</td><td>position</td><td>gene name</td><td>gene id</td></tr>
<tr><td>GU305914.1</td><td></td><td>148417</td><td>148439</td><td>+</td><td>0</td><td>AGGAGTAGTGATGGTGGTAT</td><td>CGG</td><td>||||||||[||||||||||||]PAM</td><td>NA</td><td>Intergenic</td><td>NA</td><td>NA</td></tr>
<tr><td>T11359</td><td>GTATCGGTAGTAGTAGTGTCTGG</td><td></td><td>1000</td><td></td><td></td><td></td><td></td><td></td><td></td><td></td><td></td><td></td></tr>
<tr><td>Oligo fwd</td><td>CACCGTATCGGTAGTAGTAGTGTC</td><td></td><td></td><td></td><td></td><td></td><td></td><td></td><td></td><td></td><td></td><td></td></tr>
<tr><td>Oligo rev</td><td>AAACGACACTACTACTACCGATAC</td><td></td><td></td><td></td><td></td><td></td><td></td><td></td><td></td><td></td><td></td><td></td></tr>
<tr><td>Chromosome</td><td>start</td><td></td><td>end</td><td>strand</td><td>MM</td><td>target_seq</td><td>PAM</td><td>alignment</td><td>distance</td><td>position</td><td>gene name</td><td>gene id</td></tr>
<tr><td>GU305914.1</td><td></td><td>148433</td><td>148455</td><td>+</td><td>0</td><td>GTATCGGTAGTAGTAGTGTC</td><td>TGG</td><td>||||||||[||||||||||||]PAM</td><td>NA</td><td>Intergenic</td><td>NA</td><td>NA</td></tr>
<tr><td>T11360</td><td>GTCTGGATACATGCGCTGCTTGG</td><td></td><td>1000</td><td></td><td></td><td></td><td></td><td></td><td></td><td></td><td></td><td></td></tr>
<tr><td>Oligo fwd</td><td>CACCGTCTGGATACATGCGCTGCT</td><td></td><td></td><td></td><td></td><td></td><td></td><td></td><td></td><td></td><td></td><td></td></tr>
<tr><td>Oligo rev</td><td>AAACAGCAGCGCATGTATCCAGAC</td><td></td><td></td><td></td><td></td><td></td><td></td><td></td><td></td><td></td><td></td><td></td></tr>
<tr><td>Chromosome</td><td>start</td><td></td><td>end</td><td>strand</td><td>MM</td><td>target_seq</td><td>PAM</td><td>alignment</td><td>distance</td><td>position</td><td>gene name</td><td>gene id</td></tr>
<tr><td>GU305914.1</td><td></td><td>148450</td><td>148472</td><td>+</td><td>0</td><td>GTCTGGATACATGCGCTGCT</td><td>TGG</td><td>||||||||[||||||||||||]PAM</td><td>NA</td><td>Intergenic</td><td>NA</td><td>NA</td></tr>
<tr><td>T11361</td><td>TCTGGATACATGCGCTGCTTGGG</td><td></td><td>1000</td><td></td><td></td><td></td><td></td><td></td><td></td><td></td><td></td><td></td></tr>
<tr><td>Oligo fwd</td><td>CACCTCTGGATACATGCGCTGCTT</td><td></td><td></td><td></td><td></td><td></td><td></td><td></td><td></td><td></td><td></td><td></td></tr>
<tr><td>Oligo rev</td><td>AAACAAGCAGCGCATGTATCCAGA</td><td></td><td></td><td></td><td></td><td></td><td></td><td></td><td></td><td></td><td></td><td></td></tr>
<tr><td>Chromosome</td><td>start</td><td></td><td>end</td><td>strand</td><td>MM</td><td>target_seq</td><td>PAM</td><td>alignment</td><td>distance</td><td>position</td><td>gene name</td><td>gene id</td></tr>
<tr><td>GU305914.1</td><td></td><td>148451</td><td>148473</td><td>+</td><td>0</td><td>TCTGGATACATGCGCTGCTT</td><td>GGG</td><td>||||||||[||||||||||||]PAM</td><td>NA</td><td>Intergenic</td><td>NA</td><td>NA</td></tr>
<tr><td>T11362</td><td>CTGGATACATGCGCTGCTTGGGG</td><td></td><td>1000</td><td></td><td></td><td></td><td></td><td></td><td></td><td></td><td></td><td></td></tr>
<tr><td>Oligo fwd</td><td>CACCCTGGATACATGCGCTGCTTG</td><td></td><td></td><td></td><td></td><td></td><td></td><td></td><td></td><td></td><td></td><td></td></tr>
<tr><td>Oligo rev</td><td>AAACCAAGCAGCGCATGTATCCAG</td><td></td><td></td><td></td><td></td><td></td><td></td><td></td><td></td><td></td><td></td><td></td></tr>
<tr><td>Chromosome</td><td>start</td><td></td><td>end</td><td>strand</td><td>MM</td><td>target_seq</td><td>PAM</td><td>alignment</td><td>distance</td><td>position</td><td>gene name</td><td>gene id</td></tr>
<tr><td>GU305914.1</td><td></td><td>148452</td><td>148474</td><td>+</td><td>0</td><td>CTGGATACATGCGCTGCTTG</td><td>GGG</td><td>||||||||[||||||||||||]PAM</td><td>NA</td><td>Intergenic</td><td>NA</td><td>NA</td></tr>
<tr><td>T11363</td><td>TGTTGTCGTTCGATAACCAGCGG</td><td></td><td>1000</td><td></td><td></td><td></td><td></td><td></td><td></td><td></td><td></td><td></td></tr>
<tr><td>Oligo fwd</td><td>CACCTGTTGTCGTTCGATAACCAG</td><td></td><td></td><td></td><td></td><td></td><td></td><td></td><td></td><td></td><td></td><td></td></tr>
<tr><td>Oligo rev</td><td>AAACCTGGTTATCGAACGACAACA</td><td></td><td></td><td></td><td></td><td></td><td></td><td></td><td></td><td></td><td></td><td></td></tr>
<tr><td>Chromosome</td><td>start</td><td></td><td>end</td><td>strand</td><td>MM</td><td>target_seq</td><td>PAM</td><td>alignment</td><td>distance</td><td>position</td><td>gene name</td><td>gene id</td></tr>
<tr><td>GU305914.1</td><td></td><td>148476</td><td>148498</td><td>+</td><td>0</td><td>TGTTGTCGTTCGATAACCAG</td><td>CGG</td><td>||||||||[||||||||||||]PAM</td><td>NA</td><td>Intergenic</td><td>NA</td><td>NA</td></tr>
<tr><td>T11364</td><td>GTTGTCGTTCGATAACCAGCGGG</td><td></td><td>1000</td><td></td><td></td><td></td><td></td><td></td><td></td><td></td><td></td><td></td></tr>
<tr><td>Oligo fwd</td><td>CACCGTTGTCGTTCGATAACCAGC</td><td></td><td></td><td></td><td></td><td></td><td></td><td></td><td></td><td></td><td></td><td></td></tr>
<tr><td>Oligo rev</td><td>AAACGCTGGTTATCGAACGACAAC</td><td></td><td></td><td></td><td></td><td></td><td></td><td></td><td></td><td></td><td></td><td></td></tr>
<tr><td>Chromosome</td><td>start</td><td></td><td>end</td><td>strand</td><td>MM</td><td>target_seq</td><td>PAM</td><td>alignment</td><td>distance</td><td>position</td><td>gene name</td><td>gene id</td></tr>
<tr><td>GU305914.1</td><td></td><td>148477</td><td>148499</td><td>+</td><td>0</td><td>GTTGTCGTTCGATAACCAGC</td><td>GGG</td><td>||||||||[||||||||||||]PAM</td><td>NA</td><td>Intergenic</td><td>NA</td><td>NA</td></tr>
<tr><td>T11365</td><td>TAACCAGCGGGTCACGACACCGG</td><td></td><td>1000</td><td></td><td></td><td></td><td></td><td></td><td></td><td></td><td></td><td></td></tr>
<tr><td>Oligo fwd</td><td>CACCTAACCAGCGGGTCACGACAC</td><td></td><td></td><td></td><td></td><td></td><td></td><td></td><td></td><td></td><td></td><td></td></tr>
<tr><td>Oligo rev</td><td>AAACGTGTCGTGACCCGCTGGTTA</td><td></td><td></td><td></td><td></td><td></td><td></td><td></td><td></td><td></td><td></td><td></td></tr>
<tr><td>Chromosome</td><td>start</td><td></td><td>end</td><td>strand</td><td>MM</td><td>target_seq</td><td>PAM</td><td>alignment</td><td>distance</td><td>position</td><td>gene name</td><td>gene id</td></tr>
<tr><td>GU305914.1</td><td></td><td>148489</td><td>148511</td><td>+</td><td>0</td><td>TAACCAGCGGGTCACGACAC</td><td>CGG</td><td>||||||||[||||||||||||]PAM</td><td>NA</td><td>Intergenic</td><td>NA</td><td>NA</td></tr>
<tr><td>T11366</td><td>GGTCACGACACCGGAACCGTTGG</td><td></td><td>1000</td><td></td><td></td><td></td><td></td><td></td><td></td><td></td><td></td><td></td></tr>
<tr><td>Oligo fwd</td><td>CACCGGTCACGACACCGGAACCGT</td><td></td><td></td><td></td><td></td><td></td><td></td><td></td><td></td><td></td><td></td><td></td></tr>
<tr><td>Oligo rev</td><td>AAACACGGTTCCGGTGTCGTGACC</td><td></td><td></td><td></td><td></td><td></td><td></td><td></td><td></td><td></td><td></td><td></td></tr>
<tr><td>Chromosome</td><td>start</td><td></td><td>end</td><td>strand</td><td>MM</td><td>target_seq</td><td>PAM</td><td>alignment</td><td>distance</td><td>position</td><td>gene name</td><td>gene id</td></tr>
<tr><td>GU305914.1</td><td></td><td>148498</td><td>148520</td><td>+</td><td>0</td><td>GGTCACGACACCGGAACCGT</td><td>TGG</td><td>||||||||[||||||||||||]PAM</td><td>NA</td><td>Intergenic</td><td>NA</td><td>NA</td></tr>
<tr><td>T11367</td><td>CCGGAACCGTTGGTCCCCAGTGG</td><td></td><td>1000</td><td></td><td></td><td></td><td></td><td></td><td></td><td></td><td></td><td></td></tr>
<tr><td>Oligo fwd</td><td>CACCCCGGAACCGTTGGTCCCCAG</td><td></td><td></td><td></td><td></td><td></td><td></td><td></td><td></td><td></td><td></td><td></td></tr>
<tr><td>Oligo rev</td><td>AAACCTGGGGACCAACGGTTCCGG</td><td></td><td></td><td></td><td></td><td></td><td></td><td></td><td></td><td></td><td></td><td></td></tr>
<tr><td>Chromosome</td><td>start</td><td></td><td>end</td><td>strand</td><td>MM</td><td>target_seq</td><td>PAM</td><td>alignment</td><td>distance</td><td>position</td><td>gene name</td><td>gene id</td></tr>
<tr><td>GU305914.1</td><td></td><td>148508</td><td>148530</td><td>+</td><td>0</td><td>CCGGAACCGTTGGTCCCCAG</td><td>TGG</td><td>||||||||[||||||||||||]PAM</td><td>NA</td><td>Intergenic</td><td>NA</td><td>NA</td></tr>
<tr><td>T11368</td><td>CACGTAAAACACGTTACACACGG</td><td></td><td>1000</td><td></td><td></td><td></td><td></td><td></td><td></td><td></td><td></td><td></td></tr>
<tr><td>Oligo fwd</td><td>CACCCACGTAAAACACGTTACACA</td><td></td><td></td><td></td><td></td><td></td><td></td><td></td><td></td><td></td><td></td><td></td></tr>
<tr><td>Oligo rev</td><td>AAACTGTGTAACGTGTTTTACGTG</td><td></td><td></td><td></td><td></td><td></td><td></td><td></td><td></td><td></td><td></td><td></td></tr>
<tr><td>Chromosome</td><td>start</td><td></td><td>end</td><td>strand</td><td>MM</td><td>target_seq</td><td>PAM</td><td>alignment</td><td>distance</td><td>position</td><td>gene name</td><td>gene id</td></tr>
<tr><td>GU305914.1</td><td></td><td>148531</td><td>148553</td><td>+</td><td>0</td><td>CACGTAAAACACGTTACACA</td><td>CGG</td><td>||||||||[||||||||||||]PAM</td><td>NA</td><td>Intergenic</td><td>NA</td><td>NA</td></tr>
<tr><td>T11369</td><td>ACACACGGTGTTCAGCACCTCGG</td><td></td><td>1000</td><td></td><td></td><td></td><td></td><td></td><td></td><td></td><td></td><td></td></tr>
<tr><td>Oligo fwd</td><td>CACCACACACGGTGTTCAGCACCT</td><td></td><td></td><td></td><td></td><td></td><td></td><td></td><td></td><td></td><td></td><td></td></tr>
<tr><td>Oligo rev</td><td>AAACAGGTGCTGAACACCGTGTGT</td><td></td><td></td><td></td><td></td><td></td><td></td><td></td><td></td><td></td><td></td><td></td></tr>
<tr><td>Chromosome</td><td>start</td><td></td><td>end</td><td>strand</td><td>MM</td><td>target_seq</td><td>PAM</td><td>alignment</td><td>distance</td><td>position</td><td>gene name</td><td>gene id</td></tr>
<tr><td>GU305914.1</td><td></td><td>148546</td><td>148568</td><td>+</td><td>0</td><td>ACACACGGTGTTCAGCACCT</td><td>CGG</td><td>||||||||[||||||||||||]PAM</td><td>NA</td><td>Intergenic</td><td>NA</td><td>NA</td></tr>
<tr><td>T11370</td><td>TGTTCAGCACCTCGGCCGCCAGG</td><td></td><td>1000</td><td></td><td></td><td></td><td></td><td></td><td></td><td></td><td></td><td></td></tr>
<tr><td>Oligo fwd</td><td>CACCTGTTCAGCACCTCGGCCGCC</td><td></td><td></td><td></td><td></td><td></td><td></td><td></td><td></td><td></td><td></td><td></td></tr>
<tr><td>Oligo rev</td><td>AAACGGCGGCCGAGGTGCTGAACA</td><td></td><td></td><td></td><td></td><td></td><td></td><td></td><td></td><td></td><td></td><td></td></tr>
<tr><td>Chromosome</td><td>start</td><td></td><td>end</td><td>strand</td><td>MM</td><td>target_seq</td><td>PAM</td><td>alignment</td><td>distance</td><td>position</td><td>gene name</td><td>gene id</td></tr>
<tr><td>GU305914.1</td><td></td><td>148554</td><td>148576</td><td>+</td><td>0</td><td>TGTTCAGCACCTCGGCCGCC</td><td>AGG</td><td>||||||||[||||||||||||]PAM</td><td>NA</td><td>Intergenic</td><td>NA</td><td>NA</td></tr>
<tr><td>GU305914.1</td><td></td><td>124778</td><td>124800</td><td>+</td><td>4</td><td>GGAACAGCTCCTCGGCCGCC</td><td>TGG</td><td>-|--||||[-|||||||||||]PAM</td><td>NA</td><td>Intergenic</td><td>NA</td><td>NA</td></tr>
<tr><td>T11371</td><td>CTCGGCCGCCAGGTCGCCCACGG</td><td></td><td>1000</td><td></td><td></td><td></td><td></td><td></td><td></td><td></td><td></td><td></td></tr>
<tr><td>Oligo fwd</td><td>CACCCTCGGCCGCCAGGTCGCCCA</td><td></td><td></td><td></td><td></td><td></td><td></td><td></td><td></td><td></td><td></td><td></td></tr>
<tr><td>Oligo rev</td><td>AAACTGGGCGACCTGGCGGCCGAG</td><td></td><td></td><td></td><td></td><td></td><td></td><td></td><td></td><td></td><td></td><td></td></tr>
<tr><td>Chromosome</td><td>start</td><td></td><td>end</td><td>strand</td><td>MM</td><td>target_seq</td><td>PAM</td><td>alignment</td><td>distance</td><td>position</td><td>gene name</td><td>gene id</td></tr>
<tr><td>GU305914.1</td><td></td><td>148564</td><td>148586</td><td>+</td><td>0</td><td>CTCGGCCGCCAGGTCGCCCA</td><td>CGG</td><td>||||||||[||||||||||||]PAM</td><td>NA</td><td>Intergenic</td><td>NA</td><td>NA</td></tr>
<tr><td>T11372</td><td>GCCCACGGTGTCGCCGTCCTTGG</td><td></td><td>1000</td><td></td><td></td><td></td><td></td><td></td><td></td><td></td><td></td><td></td></tr>
<tr><td>Oligo fwd</td><td>CACCGCCCACGGTGTCGCCGTCCT</td><td></td><td></td><td></td><td></td><td></td><td></td><td></td><td></td><td></td><td></td><td></td></tr>
<tr><td>Oligo rev</td><td>AAACAGGACGGCGACACCGTGGGC</td><td></td><td></td><td></td><td></td><td></td><td></td><td></td><td></td><td></td><td></td><td></td></tr>
<tr><td>Chromosome</td><td>start</td><td></td><td>end</td><td>strand</td><td>MM</td><td>target_seq</td><td>PAM</td><td>alignment</td><td>distance</td><td>position</td><td>gene name</td><td>gene id</td></tr>
</table>

| | | | | | | | | | | | |
|---|---|---|---|---|---|---|---|---|---|---|---|
| GU305914.1 | | 148579 | 148601 | + | 0 | GCCCACGGTGTCGCCGTCCT | TGG | \|\|\|\|\|\|\|\|[\|\|\|\|\|\|\|\|\|\|\|\|]PAM | NA | Intergenic | NA | NA |
| T11373 | TGTCGCCGTCCTTGGTCACCCGG | 1000 | | | | | | | | | | |
| Oligo fwd | CACCTGTCGCCGTCCTTGGTCACC | | | | | | | | | | | |
| Oligo rev | AAACGGTGACCAAGGACGGCGACA | | | | | | | | | | | |
| Chromosome | start | | end | strand | MM | target_seq | PAM | alignment | distance | position | gene name | gene id |
| GU305914.1 | | 148587 | 148609 | + | 0 | TGTCGCCGTCCTTGGTCACC | CGG | \|\|\|\|\|\|\|\|[\|\|\|\|\|\|\|\|\|\|\|\|]PAM | NA | Intergenic | NA | NA |
| T11374 | GAGACATGCTGCTGATGTCCTGG | 1000 | | | | | | | | | | |
| Oligo fwd | CACCGAGACATGCTGCTGATGTCC | | | | | | | | | | | |
| Oligo rev | AAACGGACATCAGCAGCATGTCTC | | | | | | | | | | | |
| Chromosome | start | | end | strand | MM | target_seq | PAM | alignment | distance | position | gene name | gene id |
| GU305914.1 | | 148632 | 148654 | + | 0 | GAGACATGCTGCTGATGTCC | TGG | \|\|\|\|\|\|\|\|[\|\|\|\|\|\|\|\|\|\|\|\|]PAM | NA | Intergenic | NA | NA |
| T11375 | CTGGACGCCGAAGTACGTCTCGG | 1000 | | | | | | | | | | |
| Oligo fwd | CACCCTGGACGCCGAAGTACGTCT | | | | | | | | | | | |
| Oligo rev | AAACAGACGTACTTCGGCGTCCAG | | | | | | | | | | | |
| Chromosome | start | | end | strand | MM | target_seq | PAM | alignment | distance | position | gene name | gene id |
| GU305914.1 | | 148651 | 148673 | + | 0 | CTGGACGCCGAAGTACGTCT | CGG | \|\|\|\|\|\|\|\|[\|\|\|\|\|\|\|\|\|\|\|\|]PAM | NA | Intergenic | NA | NA |
| T11376 | GAAGTACGTCTCGGCCAGCATGG | 1000 | | | | | | | | | | |
| Oligo fwd | CACCGAAGTACGTCTCGGCCAGCA | | | | | | | | | | | |
| Oligo rev | AAACTGCTGGCCGAGACGTACTTC | | | | | | | | | | | |
| Chromosome | start | | end | strand | MM | target_seq | PAM | alignment | distance | position | gene name | gene id |
| GU305914.1 | | 148660 | 148682 | + | 0 | GAAGTACGTCTCGGCCAGCA | TGG | \|\|\|\|\|\|\|\|[\|\|\|\|\|\|\|\|\|\|\|\|]PAM | NA | Intergenic | NA | NA |
| T11377 | TCTCGGCCAGCATGGTCGCCTGG | 1000 | | | | | | | | | | |
| Oligo fwd | CACCTCTCGGCCAGCATGGTCGCC | | | | | | | | | | | |
| Oligo rev | AAACGGCGACCATGCTGGCCGAGA | | | | | | | | | | | |
| Chromosome | start | | end | strand | MM | target_seq | PAM | alignment | distance | position | gene name | gene id |
| GU305914.1 | | 148668 | 148690 | + | 0 | TCTCGGCCAGCATGGTCGCC | TGG | \|\|\|\|\|\|\|\|[\|\|\|\|\|\|\|\|\|\|\|\|]PAM | NA | Intergenic | NA | NA |
| GU305914.1 | | 111541 | 111563 | + | 4 | TCTGGGTCACCGTGGTCGCC | TGG | \|\|\|-\|\|-\|[\|-\|-\|\|\|\|\|\|\|\|]PAM | NA | Intergenic | NA | NA |
| T11378 | CTCGGCCAGCATGGTCGCCTGGG | 1000 | | | | | | | | | | |
| Oligo fwd | CACCCTCGGCCAGCATGGTCGCCT | | | | | | | | | | | |
| Oligo rev | AAACAGGCGACCATGCTGGCCGAG | | | | | | | | | | | |
| Chromosome | start | | end | strand | MM | target_seq | PAM | alignment | distance | position | gene name | gene id |
| GU305914.1 | | 148669 | 148691 | + | 0 | CTCGGCCAGCATGGTCGCCT | GGG | \|\|\|\|\|\|\|\|[\|\|\|\|\|\|\|\|\|\|\|\|]PAM | NA | Intergenic | NA | NA |
| T11379 | GCATGGTCGCCTGGGCGCGTTGG | 1000 | | | | | | | | | | |
| Oligo fwd | CACCGCATGGTCGCCTGGGCGCGT | | | | | | | | | | | |
| Oligo rev | AAACACGCGCCCAGGCGACCATGC | | | | | | | | | | | |
| Chromosome | start | | end | strand | MM | target_seq | PAM | alignment | distance | position | gene name | gene id |
| GU305914.1 | | 148677 | 148699 | + | 0 | GCATGGTCGCCTGGGCGCGT | TGG | \|\|\|\|\|\|\|\|[\|\|\|\|\|\|\|\|\|\|\|\|]PAM | NA | Intergenic | NA | NA |
| T11380 | CATGGTCGCCTGGGCGCGTTGGG | 1000 | | | | | | | | | | |
| Oligo fwd | CACCCATGGTCGCCTGGGCGCGTT | | | | | | | | | | | |
| Oligo rev | AAACAACGCGCCCAGGCGACCATG | | | | | | | | | | | |
| Chromosome | start | | end | strand | MM | target_seq | PAM | alignment | distance | position | gene name | gene id |
| GU305914.1 | | 148678 | 148700 | + | 0 | CATGGTCGCCTGGGCGCGTT | GGG | \|\|\|\|\|\|\|\|[\|\|\|\|\|\|\|\|\|\|\|\|]PAM | NA | Intergenic | NA | NA |
| T11381 | CTTGTACAGCATGCAGATCTCGG | 1000 | | | | | | | | | | |
| Oligo fwd | CACCCTTGTACAGCATGCAGATCT | | | | | | | | | | | |
| Oligo rev | AAACAGATCTGCATGCTGTACAAG | | | | | | | | | | | |
| Chromosome | start | | end | strand | MM | target_seq | PAM | alignment | distance | position | gene name | gene id |
| GU305914.1 | | 148705 | 148727 | + | 0 | CTTGTACAGCATGCAGATCT | CGG | \|\|\|\|\|\|\|\|[\|\|\|\|\|\|\|\|\|\|\|\|]PAM | NA | Intergenic | NA | NA |
| T11382 | TTGTACAGCATGCAGATCTCGGG | 1000 | | | | | | | | | | |
| Oligo fwd | CACCTTGTACAGCATGCAGATCTC | | | | | | | | | | | |
| Oligo rev | AAACGAGATCTGCATGCTGTACAA | | | | | | | | | | | |
| Chromosome | start | | end | strand | MM | target_seq | PAM | alignment | distance | position | gene name | gene id |
| GU305914.1 | | 148706 | 148728 | + | 0 | TTGTACAGCATGCAGATCTC | GGG | \|\|\|\|\|\|\|\|[\|\|\|\|\|\|\|\|\|\|\|\|]PAM | NA | Intergenic | NA | NA |
| T11383 | TGTACAGCATGCAGATCTCGGGG | 1000 | | | | | | | | | | |
| Oligo fwd | CACCTGTACAGCATGCAGATCTCG | | | | | | | | | | | |
| Oligo rev | AAACCGAGATCTGCATGCTGTACA | | | | | | | | | | | |
| Chromosome | start | | end | strand | MM | target_seq | PAM | alignment | distance | position | gene name | gene id |
| GU305914.1 | | 148707 | 148729 | + | 0 | TGTACAGCATGCAGATCTCG | GGG | \|\|\|\|\|\|\|\|[\|\|\|\|\|\|\|\|\|\|\|\|]PAM | NA | Intergenic | NA | NA |
| T11384 | GCATGCAGATCTCGGGGACGTGG | 1000 | | | | | | | | | | |
| Oligo fwd | CACCGCATGCAGATCTCGGGGACG | | | | | | | | | | | |
| Oligo rev | AAACCGTCCCCGAGATCTGCATGC | | | | | | | | | | | |
| Chromosome | start | | end | strand | MM | target_seq | PAM | alignment | distance | position | gene name | gene id |
| GU305914.1 | | 148713 | 148735 | + | 0 | GCATGCAGATCTCGGGGACG | TGG | \|\|\|\|\|\|\|\|[\|\|\|\|\|\|\|\|\|\|\|\|]PAM | NA | Intergenic | NA | NA |
| T11385 | GGGACGTGGCGCGTGATCCTCGG | 1000 | | | | | | | | | | |
| Oligo fwd | CACCGGGACGTGGCGCGTGATCCT | | | | | | | | | | | |
| Oligo rev | AAACAGGATCACGCGCCACGTCCC | | | | | | | | | | | |
| Chromosome | start | | end | strand | MM | target_seq | PAM | alignment | distance | position | gene name | gene id |
| GU305914.1 | | 148727 | 148749 | + | 0 | GGGACGTGGCGCGTGATCCT | CGG | \|\|\|\|\|\|\|\|[\|\|\|\|\|\|\|\|\|\|\|\|]PAM | NA | Intergenic | NA | NA |
| T11386 | GGACGTGGCGCGTGATCCTCGGG | 1000 | | | | | | | | | | |
| Oligo fwd | CACCGGACGTGGCGCGTGATCCTC | | | | | | | | | | | |
| Oligo rev | AAACGAGGATCACGCGCCACGTCC | | | | | | | | | | | |
| Chromosome | start | | end | strand | MM | target_seq | PAM | alignment | distance | position | gene name | gene id |
| GU305914.1 | | 148728 | 148750 | + | 0 | GGACGTGGCGCGTGATCCTC | GGG | \|\|\|\|\|\|\|\|[\|\|\|\|\|\|\|\|\|\|\|\|]PAM | NA | Intergenic | NA | NA |
| T11387 | GGTCCGTCTCGTCGAAGAACAGG | 1000 | | | | | | | | | | |
| Oligo fwd | CACCGGTCCGTCTCGTCGAAGAAC | | | | | | | | | | | |

Oligo rev AAACGTTCTTCGACGAGACGGACC
Chromosome start end strand MM target_seq PAM alignment distance position gene name gene id
GU305914.1 148749 148771 + 0 GGTCCGTCTCGTCGAAGAAC AGG ||||||||[||||||||||||]PAM NA Intergenic NA NA

T11388 TCTCGTCGAAGAACAGGTACAGG 1000
Oligo fwd CACCTCTCGTCGAAGAACAGGTAC
Oligo rev AAACGTACCTGTTCTTCGACGAGA
Chromosome start end strand MM target_seq PAM alignment distance position gene name gene id
GU305914.1 148755 148777 + 0 TCTCGTCGAAGAACAGGTAC AGG ||||||||[||||||||||||]PAM NA Intergenic NA NA

T11389 CTCGTCGAAGAACAGGTACAGGG 1000
Oligo fwd CACCCTCGTCGAAGAACAGGTACA
Oligo rev AAACTGTACCTGTTCTTCGACGAG
Chromosome start end strand MM target_seq PAM alignment distance position gene name gene id
GU305914.1 148756 148778 + 0 CTCGTCGAAGAACAGGTACA GGG ||||||||[||||||||||||]PAM NA Intergenic NA NA

T11390 TGCAGCTCTTGCAGATCATCCGG 1000
Oligo fwd CACCTGCAGCTCTTGCAGATCATC
Oligo rev AAACGATGATCTGCAAGAGCTGCA
Chromosome start end strand MM target_seq PAM alignment distance position gene name gene id
GU305914.1 148797 148819 + 0 TGCAGCTCTTGCAGATCATC CGG ||||||||[||||||||||||]PAM NA Intergenic NA NA

T11391 GCAGCTCTTGCAGATCATCCGGG 1000
Oligo fwd CACCGCAGCTCTTGCAGATCATCC
Oligo rev AAACGGATGATCTGCAAGAGCTGC
Chromosome start end strand MM target_seq PAM alignment distance position gene name gene id
GU305914.1 148798 148820 + 0 GCAGCTCTTGCAGATCATCC GGG ||||||||[||||||||||||]PAM NA Intergenic NA NA

T11392 TCATCCGGGTCAGCCCCGTGCGG 1000
Oligo fwd CACCTCATCCGGGTCAGCCCCGTG
Oligo rev AAACCACGGGGCTGACCCGGATGA
Chromosome start end strand MM target_seq PAM alignment distance position gene name gene id
GU305914.1 148812 148834 + 0 TCATCCGGGTCAGCCCCGTG CGG ||||||||[||||||||||||]PAM NA Intergenic NA NA

T11393 CATCCGGGTCAGCCCCGTGCGGG 1000
Oligo fwd CACCCATCCGGGTCAGCCCCGTGC
Oligo rev AAACGCACGGGGCTGACCCGGATG
Chromosome start end strand MM target_seq PAM alignment distance position gene name gene id
GU305914.1 148813 148835 + 0 CATCCGGGTCAGCCCCGTGC GGG ||||||||[||||||||||||]PAM NA Intergenic NA NA

T11394 ATCCGGGTCAGCCCCGTGCGGGG 1000
Oligo fwd CACCATCCGGGTCAGCCCCGTGCG
Oligo rev AAACCGCACGGGGCTGACCCGGAT
Chromosome start end strand MM target_seq PAM alignment distance position gene name gene id
GU305914.1 148814 148836 + 0 ATCCGGGTCAGCCCCGTGCG GGG ||||||||[||||||||||||]PAM NA Intergenic NA NA

T11395 GGTCAGCCCCGTGCGGGGCCTGG 1000
Oligo fwd CACCGGTCAGCCCCGTGCGGGGCC
Oligo rev AAACGGCCCCGCACGGGGCTGACC
Chromosome start end strand MM target_seq PAM alignment distance position gene name gene id
GU305914.1 148819 148841 + 0 GGTCAGCCCCGTGCGGGGCC TGG ||||||||[||||||||||||]PAM NA Intergenic NA NA

T11396 CCCCGTGCGGGGCCTGGTGCCGG 1000
Oligo fwd CACCCCCCGTGCGGGGCCTGGTGC
Oligo rev AAACGCACCAGGCCCCGCACGGGG
Chromosome start end strand MM target_seq PAM alignment distance position gene name gene id
GU305914.1 148825 148847 + 0 CCCCGTGCGGGGCCTGGTGC CGG ||||||||[||||||||||||]PAM NA Intergenic NA NA

T11397 TGCGGGGCCTGGTGCCGGTCAGG 1000
Oligo fwd CACCTGCGGGGCCTGGTGCCGGTC
Oligo rev AAACGACCGGCACCAGGCCCCGCA
Chromosome start end strand MM target_seq PAM alignment distance position gene name gene id
GU305914.1 148830 148852 + 0 TGCGGGGCCTGGTGCCGGTC AGG ||||||||[||||||||||||]PAM NA Intergenic NA NA

T11398 GGGCCTGGTGCCGGTCAGGAAGG 1000
Oligo fwd CACCGGGCCTGGTGCCGGTCAGGA
Oligo rev AAACTCCTGACCGGCACCAGGCCC
Chromosome start end strand MM target_seq PAM alignment distance position gene name gene id
GU305914.1 148834 148856 + 0 GGGCCTGGTGCCGGTCAGGA AGG ||||||||[||||||||||||]PAM NA Intergenic NA NA

T11399 TGCCGGTCAGGAAGGTCACCAGG 1000
Oligo fwd CACCTGCCGGTCAGGAAGGTCACC
Oligo rev AAACGGTGACCTTCCTGACCGGCA
Chromosome start end strand MM target_seq PAM alignment distance position gene name gene id
GU305914.1 148842 148864 + 0 TGCCGGTCAGGAAGGTCACC AGG ||||||||[||||||||||||]PAM NA Intergenic NA NA

T11400 CAGGTGCTCCAGCGTCTGTCTGG 1000
Oligo fwd CACCCAGGTGCTCCAGCGTCTGTC
Oligo rev AAACGACAGACGCTGGAGCACCTG
Chromosome start end strand MM target_seq PAM alignment distance position gene name gene id
GU305914.1 148861 148883 + 0 CAGGTGCTCCAGCGTCTGTC TGG ||||||||[||||||||||||]PAM NA Intergenic NA NA

T11401 TCCAGCGTCTGTCTGGACAGCGG 1000
Oligo fwd CACCTCCAGCGTCTGTCTGGACAG
Oligo rev AAACCTGTCCAGACAGACGCTGGA
Chromosome start end strand MM target_seq PAM alignment distance position gene name gene id
GU305914.1 148868 148890 + 0 TCCAGCGTCTGTCTGGACAG CGG ||||||||[||||||||||||]PAM NA Intergenic NA NA

T11402 GGACAGCGGCAGCGACTCGATGG 1000

<table>
<tr><td>Oligo fwd</td><td>CACCGGACAGCGGCAGCGACTCGA</td><td></td><td></td><td></td><td></td><td></td><td></td><td></td><td></td><td></td><td></td></tr>
<tr><td>Oligo rev</td><td>AAACTCGAGTCGCTGCCGCTGTCC</td><td></td><td></td><td></td><td></td><td></td><td></td><td></td><td></td><td></td><td></td></tr>
<tr><td>Chromosome</td><td>start</td><td>end</td><td>strand</td><td>MM</td><td>target_seq</td><td>PAM</td><td>alignment</td><td>distance</td><td>position</td><td>gene name</td><td>gene id</td></tr>
<tr><td>GU305914.1</td><td>148882</td><td>148904</td><td>+</td><td>0</td><td>GGACAGCGGCAGCGACTCGA</td><td>TGG</td><td>||||||||[||||||||||||]PAM</td><td>NA</td><td>Intergenic</td><td>NA</td><td>NA</td></tr>
<tr><td>T11403</td><td>TGGCGAACAGCAGCATGCACAGG</td><td>1000</td><td></td><td></td><td></td><td></td><td></td><td></td><td></td><td></td><td></td></tr>
<tr><td>Oligo fwd</td><td>CACCTGGCGAACAGCAGCATGCAC</td><td></td><td></td><td></td><td></td><td></td><td></td><td></td><td></td><td></td><td></td></tr>
<tr><td>Oligo rev</td><td>AAACGTGCATGCTGCTGTTCGCCA</td><td></td><td></td><td></td><td></td><td></td><td></td><td></td><td></td><td></td><td></td></tr>
<tr><td>Chromosome</td><td>start</td><td>end</td><td>strand</td><td>MM</td><td>target_seq</td><td>PAM</td><td>alignment</td><td>distance</td><td>position</td><td>gene name</td><td>gene id</td></tr>
<tr><td>GU305914.1</td><td>148902</td><td>148924</td><td>+</td><td>0</td><td>TGGCGAACAGCAGCATGCAC</td><td>AGG</td><td>||||||||[||||||||||||]PAM</td><td>NA</td><td>Intergenic</td><td>NA</td><td>NA</td></tr>
<tr><td>T11404</td><td>GGCGAACAGCAGCATGCACAGGG</td><td>1000</td><td></td><td></td><td></td><td></td><td></td><td></td><td></td><td></td><td></td></tr>
<tr><td>Oligo fwd</td><td>CACCGGCGAACAGCAGCATGCACA</td><td></td><td></td><td></td><td></td><td></td><td></td><td></td><td></td><td></td><td></td></tr>
<tr><td>Oligo rev</td><td>AAACTGTGCATGCTGCTGTTCGCC</td><td></td><td></td><td></td><td></td><td></td><td></td><td></td><td></td><td></td><td></td></tr>
<tr><td>Chromosome</td><td>start</td><td>end</td><td>strand</td><td>MM</td><td>target_seq</td><td>PAM</td><td>alignment</td><td>distance</td><td>position</td><td>gene name</td><td>gene id</td></tr>
<tr><td>GU305914.1</td><td>148903</td><td>148925</td><td>+</td><td>0</td><td>GGCGAACAGCAGCATGCACA</td><td>GGG</td><td>||||||||[||||||||||||]PAM</td><td>NA</td><td>Intergenic</td><td>NA</td><td>NA</td></tr>
<tr><td>T11405</td><td>GCACAGGGTGAAGTCCGTGTTGG</td><td>1000</td><td></td><td></td><td></td><td></td><td></td><td></td><td></td><td></td><td></td></tr>
<tr><td>Oligo fwd</td><td>CACCGCACAGGGTGAAGTCCGTGT</td><td></td><td></td><td></td><td></td><td></td><td></td><td></td><td></td><td></td><td></td></tr>
<tr><td>Oligo rev</td><td>AAACACACGGACTTCACCCTGTGC</td><td></td><td></td><td></td><td></td><td></td><td></td><td></td><td></td><td></td><td></td></tr>
<tr><td>Chromosome</td><td>start</td><td>end</td><td>strand</td><td>MM</td><td>target_seq</td><td>PAM</td><td>alignment</td><td>distance</td><td>position</td><td>gene name</td><td>gene id</td></tr>
<tr><td>GU305914.1</td><td>148918</td><td>148940</td><td>+</td><td>0</td><td>GCACAGGGTGAAGTCCGTGT</td><td>TGG</td><td>||||||||[||||||||||||]PAM</td><td>NA</td><td>Intergenic</td><td>NA</td><td>NA</td></tr>
<tr><td>T11406</td><td>GTGAAGTCCGTGTTGGTCTTCGG</td><td>1000</td><td></td><td></td><td></td><td></td><td></td><td></td><td></td><td></td><td></td></tr>
<tr><td>Oligo fwd</td><td>CACCGTGAAGTCCGTGTTGGTCTT</td><td></td><td></td><td></td><td></td><td></td><td></td><td></td><td></td><td></td><td></td></tr>
<tr><td>Oligo rev</td><td>AAACAAGACCAACACGGACTTCAC</td><td></td><td></td><td></td><td></td><td></td><td></td><td></td><td></td><td></td><td></td></tr>
<tr><td>Chromosome</td><td>start</td><td>end</td><td>strand</td><td>MM</td><td>target_seq</td><td>PAM</td><td>alignment</td><td>distance</td><td>position</td><td>gene name</td><td>gene id</td></tr>
<tr><td>GU305914.1</td><td>148925</td><td>148947</td><td>+</td><td>0</td><td>GTGAAGTCCGTGTTGGTCTT</td><td>CGG</td><td>||||||||[||||||||||||]PAM</td><td>NA</td><td>Intergenic</td><td>NA</td><td>NA</td></tr>
<tr><td>T11407</td><td>TGAAGTCCGTGTTGGTCTTCGGG</td><td>1000</td><td></td><td></td><td></td><td></td><td></td><td></td><td></td><td></td><td></td></tr>
<tr><td>Oligo fwd</td><td>CACCTGAAGTCCGTGTTGGTCTTC</td><td></td><td></td><td></td><td></td><td></td><td></td><td></td><td></td><td></td><td></td></tr>
<tr><td>Oligo rev</td><td>AAACGAAGACCAACACGGACTTCA</td><td></td><td></td><td></td><td></td><td></td><td></td><td></td><td></td><td></td><td></td></tr>
<tr><td>Chromosome</td><td>start</td><td>end</td><td>strand</td><td>MM</td><td>target_seq</td><td>PAM</td><td>alignment</td><td>distance</td><td>position</td><td>gene name</td><td>gene id</td></tr>
<tr><td>GU305914.1</td><td>148926</td><td>148948</td><td>+</td><td>0</td><td>TGAAGTCCGTGTTGGTCTTC</td><td>GGG</td><td>||||||||[||||||||||||]PAM</td><td>NA</td><td>Intergenic</td><td>NA</td><td>NA</td></tr>
<tr><td>T11408</td><td>AGTCCGTGTTGGTCTTCGGGCGG</td><td>1000</td><td></td><td></td><td></td><td></td><td></td><td></td><td></td><td></td><td></td></tr>
<tr><td>Oligo fwd</td><td>CACCAGTCCGTGTTGGTCTTCGGG</td><td></td><td></td><td></td><td></td><td></td><td></td><td></td><td></td><td></td><td></td></tr>
<tr><td>Oligo rev</td><td>AAACCCCGAAGACCAACACGGACT</td><td></td><td></td><td></td><td></td><td></td><td></td><td></td><td></td><td></td><td></td></tr>
<tr><td>Chromosome</td><td>start</td><td>end</td><td>strand</td><td>MM</td><td>target_seq</td><td>PAM</td><td>alignment</td><td>distance</td><td>position</td><td>gene name</td><td>gene id</td></tr>
<tr><td>GU305914.1</td><td>148929</td><td>148951</td><td>+</td><td>0</td><td>AGTCCGTGTTGGTCTTCGGG</td><td>CGG</td><td>||||||||[||||||||||||]PAM</td><td>NA</td><td>Intergenic</td><td>NA</td><td>NA</td></tr>
<tr><td>T11409</td><td>GTTGGTCTTCGGGCGGAACACGG</td><td>1000</td><td></td><td></td><td></td><td></td><td></td><td></td><td></td><td></td><td></td></tr>
<tr><td>Oligo fwd</td><td>CACCGTTGGTCTTCGGGCGGAACA</td><td></td><td></td><td></td><td></td><td></td><td></td><td></td><td></td><td></td><td></td></tr>
<tr><td>Oligo rev</td><td>AAACTGTTCCGCCCGAAGACCAAC</td><td></td><td></td><td></td><td></td><td></td><td></td><td></td><td></td><td></td><td></td></tr>
<tr><td>Chromosome</td><td>start</td><td>end</td><td>strand</td><td>MM</td><td>target_seq</td><td>PAM</td><td>alignment</td><td>distance</td><td>position</td><td>gene name</td><td>gene id</td></tr>
<tr><td>GU305914.1</td><td>148936</td><td>148958</td><td>+</td><td>0</td><td>GTTGGTCTTCGGGCGGAACA</td><td>CGG</td><td>||||||||[||||||||||||]PAM</td><td>NA</td><td>Intergenic</td><td>NA</td><td>NA</td></tr>
<tr><td>T11410</td><td>TCGGGCGGAACACGGCGATCCGG</td><td>1000</td><td></td><td></td><td></td><td></td><td></td><td></td><td></td><td></td><td></td></tr>
<tr><td>Oligo fwd</td><td>CACCTCGGGCGGAACACGGCGATC</td><td></td><td></td><td></td><td></td><td></td><td></td><td></td><td></td><td></td><td></td></tr>
<tr><td>Oligo rev</td><td>AAACGATCGCCGTGTTCCGCCCGA</td><td></td><td></td><td></td><td></td><td></td><td></td><td></td><td></td><td></td><td></td></tr>
<tr><td>Chromosome</td><td>start</td><td>end</td><td>strand</td><td>MM</td><td>target_seq</td><td>PAM</td><td>alignment</td><td>distance</td><td>position</td><td>gene name</td><td>gene id</td></tr>
<tr><td>GU305914.1</td><td>148944</td><td>148966</td><td>+</td><td>0</td><td>TCGGGCGGAACACGGCGATC</td><td>CGG</td><td>||||||||[||||||||||||]PAM</td><td>NA</td><td>Intergenic</td><td>NA</td><td>NA</td></tr>
<tr><td>T11411</td><td>GAACACGGCGATCCGGTGCTTGG</td><td>1000</td><td></td><td></td><td></td><td></td><td></td><td></td><td></td><td></td><td></td></tr>
<tr><td>Oligo fwd</td><td>CACCGAACACGGCGATCCGGTGCT</td><td></td><td></td><td></td><td></td><td></td><td></td><td></td><td></td><td></td><td></td></tr>
<tr><td>Oligo rev</td><td>AAACAGCACCGGATCGCCGTGTTC</td><td></td><td></td><td></td><td></td><td></td><td></td><td></td><td></td><td></td><td></td></tr>
<tr><td>Chromosome</td><td>start</td><td>end</td><td>strand</td><td>MM</td><td>target_seq</td><td>PAM</td><td>alignment</td><td>distance</td><td>position</td><td>gene name</td><td>gene id</td></tr>
<tr><td>GU305914.1</td><td>148951</td><td>148973</td><td>+</td><td>0</td><td>GAACACGGCGATCCGGTGCT</td><td>TGG</td><td>||||||||[||||||||||||]PAM</td><td>NA</td><td>Intergenic</td><td>NA</td><td>NA</td></tr>
<tr><td>T11412</td><td>CGGCGATCCGGTGCTTGGTCCGG</td><td>1000</td><td></td><td></td><td></td><td></td><td></td><td></td><td></td><td></td><td></td></tr>
<tr><td>Oligo fwd</td><td>CACCCGGCGATCCGGTGCTTGGTC</td><td></td><td></td><td></td><td></td><td></td><td></td><td></td><td></td><td></td><td></td></tr>
<tr><td>Oligo rev</td><td>AAACGACCAAGCACCGGATCGCCG</td><td></td><td></td><td></td><td></td><td></td><td></td><td></td><td></td><td></td><td></td></tr>
<tr><td>Chromosome</td><td>start</td><td>end</td><td>strand</td><td>MM</td><td>target_seq</td><td>PAM</td><td>alignment</td><td>distance</td><td>position</td><td>gene name</td><td>gene id</td></tr>
<tr><td>GU305914.1</td><td>148956</td><td>148978</td><td>+</td><td>0</td><td>CGGCGATCCGGTGCTTGGTC</td><td>CGG</td><td>||||||||[||||||||||||]PAM</td><td>NA</td><td>Intergenic</td><td>NA</td><td>NA</td></tr>
<tr><td>T11413</td><td>CCGGTGCTTGGTCCGGATCGAGG</td><td>1000</td><td></td><td></td><td></td><td></td><td></td><td></td><td></td><td></td><td></td></tr>
<tr><td>Oligo fwd</td><td>CACCCCGGTGCTTGGTCCGGATCG</td><td></td><td></td><td></td><td></td><td></td><td></td><td></td><td></td><td></td><td></td></tr>
<tr><td>Oligo rev</td><td>AAACCGATCCGGACCAAGCACCGG</td><td></td><td></td><td></td><td></td><td></td><td></td><td></td><td></td><td></td><td></td></tr>
<tr><td>Chromosome</td><td>start</td><td>end</td><td>strand</td><td>MM</td><td>target_seq</td><td>PAM</td><td>alignment</td><td>distance</td><td>position</td><td>gene name</td><td>gene id</td></tr>
<tr><td>GU305914.1</td><td>148963</td><td>148985</td><td>+</td><td>0</td><td>CCGGTGCTTGGTCCGGATCG</td><td>AGG</td><td>||||||||[||||||||||||]PAM</td><td>NA</td><td>Intergenic</td><td>NA</td><td>NA</td></tr>
<tr><td>T11414</td><td>AGGACAGCCGCAGCTCCGACAGG</td><td>1000</td><td></td><td></td><td></td><td></td><td></td><td></td><td></td><td></td><td></td></tr>
<tr><td>Oligo fwd</td><td>CACCAGGACAGCCGCAGCTCCGAC</td><td></td><td></td><td></td><td></td><td></td><td></td><td></td><td></td><td></td><td></td></tr>
<tr><td>Oligo rev</td><td>AAACGTCGGAGCTGCGGCTGTCCT</td><td></td><td></td><td></td><td></td><td></td><td></td><td></td><td></td><td></td><td></td></tr>
<tr><td>Chromosome</td><td>start</td><td>end</td><td>strand</td><td>MM</td><td>target_seq</td><td>PAM</td><td>alignment</td><td>distance</td><td>position</td><td>gene name</td><td>gene id</td></tr>
<tr><td>GU305914.1</td><td>148983</td><td>149005</td><td>+</td><td>0</td><td>AGGACAGCCGCAGCTCCGAC</td><td>AGG</td><td>||||||||[||||||||||||]PAM</td><td>NA</td><td>Intergenic</td><td>NA</td><td>NA</td></tr>
<tr><td>T11415</td><td>GGACAGCCGCAGCTCCGACAGGG</td><td>1000</td><td></td><td></td><td></td><td></td><td></td><td></td><td></td><td></td><td></td></tr>
<tr><td>Oligo fwd</td><td>CACCGGACAGCCGCAGCTCCGACA</td><td></td><td></td><td></td><td></td><td></td><td></td><td></td><td></td><td></td><td></td></tr>
<tr><td>Oligo rev</td><td>AAACTGTCGGAGCTGCGGCTGTCC</td><td></td><td></td><td></td><td></td><td></td><td></td><td></td><td></td><td></td><td></td></tr>
<tr><td>Chromosome</td><td>start</td><td>end</td><td>strand</td><td>MM</td><td>target_seq</td><td>PAM</td><td>alignment</td><td>distance</td><td>position</td><td>gene name</td><td>gene id</td></tr>
<tr><td>GU305914.1</td><td>148984</td><td>149006</td><td>+</td><td>0</td><td>GGACAGCCGCAGCTCCGACA</td><td>GGG</td><td>||||||||[||||||||||||]PAM</td><td>NA</td><td>Intergenic</td><td>NA</td><td>NA</td></tr>
<tr><td>T11416</td><td>CAGCTCCGACAGGGCGTCCTCGG</td><td>1000</td><td></td><td></td><td></td><td></td><td></td><td></td><td></td><td></td><td></td></tr>
<tr><td>Oligo fwd</td><td>CACCCAGCTCCGACAGGGCGTCCT</td><td></td><td></td><td></td><td></td><td></td><td></td><td></td><td></td><td></td><td></td></tr>
<tr><td>Oligo rev</td><td>AAACAGGACGCCCTGTCGGAGCTG</td><td></td><td></td><td></td><td></td><td></td><td></td><td></td><td></td><td></td><td></td></tr>
<tr><td>Chromosome</td><td>start</td><td>end</td><td>strand</td><td>MM</td><td>target_seq</td><td>PAM</td><td>alignment</td><td>distance</td><td>position</td><td>gene name</td><td>gene id</td></tr>
<tr><td>GU305914.1</td><td>148993</td><td>149015</td><td>+</td><td>0</td><td>CAGCTCCGACAGGGCGTCCT</td><td>CGG</td><td>||||||||[||||||||||||]PAM</td><td>NA</td><td>Intergenic</td><td>NA</td><td>NA</td></tr>
</table>

<table>
<tr><td>T11417</td><td>ACAGGGCGTCCTCGGCGTCCAGG</td><td>1000</td></tr>
<tr><td>Oligo fwd</td><td>CACCACAGGGCGTCCTCGGCGTCC</td></tr>
<tr><td>Oligo rev</td><td>AAACGGACGCCGAGGACGCCCTGT</td></tr>
<tr><td>Chromosome</td><td>start</td><td>end</td><td>strand</td><td>MM</td><td>target_seq</td><td>PAM</td><td>alignment</td><td>distance</td><td>position</td><td>gene name</td><td>gene id</td></tr>
<tr><td>GU305914.1</td><td>149001</td><td>149023</td><td>+</td><td>0</td><td>ACAGGGCGTCCTCGGCGTCC</td><td>AGG</td><td>||||||||[||||||||||||]PAM</td><td>NA</td><td>Intergenic</td><td>NA</td><td>NA</td></tr>
<tr><td></td></tr>
<tr><td>T11418</td><td>CGGCGTCCAGGATGAAGAACAGG</td><td>1000</td></tr>
<tr><td>Oligo fwd</td><td>CACCCGGCGTCCAGGATGAAGAAC</td></tr>
<tr><td>Oligo rev</td><td>AAACGTTCTTCATCCTGGACGCCG</td></tr>
<tr><td>Chromosome</td><td>start</td><td>end</td><td>strand</td><td>MM</td><td>target_seq</td><td>PAM</td><td>alignment</td><td>distance</td><td>position</td><td>gene name</td><td>gene id</td></tr>
<tr><td>GU305914.1</td><td>149013</td><td>149035</td><td>+</td><td>0</td><td>CGGCGTCCAGGATGAAGAAC</td><td>AGG</td><td>||||||||[||||||||||||]PAM</td><td>NA</td><td>Intergenic</td><td>NA</td><td>NA</td></tr>
<tr><td></td></tr>
<tr><td>T11419</td><td>TGAAGAACAGGCAGAATCCCAGG</td><td>1000</td></tr>
<tr><td>Oligo fwd</td><td>CACCTGAAGAACAGGCAGAATCCC</td></tr>
<tr><td>Oligo rev</td><td>AAACGGGATTCTGCCTGTTCTTCA</td></tr>
<tr><td>Chromosome</td><td>start</td><td>end</td><td>strand</td><td>MM</td><td>target_seq</td><td>PAM</td><td>alignment</td><td>distance</td><td>position</td><td>gene name</td><td>gene id</td></tr>
<tr><td>GU305914.1</td><td>149025</td><td>149047</td><td>+</td><td>0</td><td>TGAAGAACAGGCAGAATCCC</td><td>AGG</td><td>||||||||[||||||||||||]PAM</td><td>NA</td><td>Intergenic</td><td>NA</td><td>NA</td></tr>
<tr><td></td></tr>
<tr><td>T11420</td><td>AGAATCCCAGGAACGACGACCGG</td><td>1000</td></tr>
<tr><td>Oligo fwd</td><td>CACCAGAATCCCAGGAACGACGAC</td></tr>
<tr><td>Oligo rev</td><td>AAACGTCGTCGTTCCTGGGATTCT</td></tr>
<tr><td>Chromosome</td><td>start</td><td>end</td><td>strand</td><td>MM</td><td>target_seq</td><td>PAM</td><td>alignment</td><td>distance</td><td>position</td><td>gene name</td><td>gene id</td></tr>
<tr><td>GU305914.1</td><td>149037</td><td>149059</td><td>+</td><td>0</td><td>AGAATCCCAGGAACGACGAC</td><td>CGG</td><td>||||||||[||||||||||||]PAM</td><td>NA</td><td>Intergenic</td><td>NA</td><td>NA</td></tr>
<tr><td></td></tr>
<tr><td>T11421</td><td>ACCGGCAGTCCGAGTCCGCGCGG</td><td>1000</td></tr>
<tr><td>Oligo fwd</td><td>CACCACCGGCAGTCCGAGTCCGCG</td></tr>
<tr><td>Oligo rev</td><td>AAACCGCGGACTCGGACTGCCGGT</td></tr>
<tr><td>Chromosome</td><td>start</td><td>end</td><td>strand</td><td>MM</td><td>target_seq</td><td>PAM</td><td>alignment</td><td>distance</td><td>position</td><td>gene name</td><td>gene id</td></tr>
<tr><td>GU305914.1</td><td>149055</td><td>149077</td><td>+</td><td>0</td><td>ACCGGCAGTCCGAGTCCGCG</td><td>CGG</td><td>||||||||[||||||||||||]PAM</td><td>NA</td><td>Intergenic</td><td>NA</td><td>NA</td></tr>
<tr><td>GU305914.1</td><td>230353</td><td>230375</td><td>+</td><td>4</td><td>CCCGGCCGTCTGAGTGCGCG</td><td>CGG</td><td>-|||||-|[||-||||-||||]PAM</td><td>NA</td><td>Intergenic</td><td>NA</td><td>NA</td></tr>
<tr><td>GU305914.1</td><td>38</td><td>60</td><td>+</td><td>4</td><td>CCCGGCCGTCTGAGTGCGCG</td><td>CGG</td><td>-|||||-|[||-||||-||||]PAM</td><td>NA</td><td>Intergenic</td><td>NA</td><td>NA</td></tr>
<tr><td></td></tr>
<tr><td>T11422</td><td>GTCCGCGCGGTACTGCGCGTTGG</td><td>1000</td></tr>
<tr><td>Oligo fwd</td><td>CACCGTCCGCGCGGTACTGCGCGT</td></tr>
<tr><td>Oligo rev</td><td>AAACACGCGCAGTACCGCGCGGAC</td></tr>
<tr><td>Chromosome</td><td>start</td><td>end</td><td>strand</td><td>MM</td><td>target_seq</td><td>PAM</td><td>alignment</td><td>distance</td><td>position</td><td>gene name</td><td>gene id</td></tr>
<tr><td>GU305914.1</td><td>149068</td><td>149090</td><td>+</td><td>0</td><td>GTCCGCGCGGTACTGCGCGT</td><td>TGG</td><td>||||||||[||||||||||||]PAM</td><td>NA</td><td>Intergenic</td><td>NA</td><td>NA</td></tr>
<tr><td></td></tr>
<tr><td>T11423</td><td>TCCGCGCGGTACTGCGCGTTGGG</td><td>1000</td></tr>
<tr><td>Oligo fwd</td><td>CACCTCCGCGCGGTACTGCGCGTT</td></tr>
<tr><td>Oligo rev</td><td>AAACAACGCGCAGTACCGCGCGGA</td></tr>
<tr><td>Chromosome</td><td>start</td><td>end</td><td>strand</td><td>MM</td><td>target_seq</td><td>PAM</td><td>alignment</td><td>distance</td><td>position</td><td>gene name</td><td>gene id</td></tr>
<tr><td>GU305914.1</td><td>149069</td><td>149091</td><td>+</td><td>0</td><td>TCCGCGCGGTACTGCGCGTT</td><td>GGG</td><td>||||||||[||||||||||||]PAM</td><td>NA</td><td>Intergenic</td><td>NA</td><td>NA</td></tr>
<tr><td></td></tr>
<tr><td>T11424</td><td>CCGCGCGGTACTGCGCGTTGGGG</td><td>1000</td></tr>
<tr><td>Oligo fwd</td><td>CACCCCGCGCGGTACTGCGCGTTG</td></tr>
<tr><td>Oligo rev</td><td>AAACCAACGCGCAGTACCGCGCGG</td></tr>
<tr><td>Chromosome</td><td>start</td><td>end</td><td>strand</td><td>MM</td><td>target_seq</td><td>PAM</td><td>alignment</td><td>distance</td><td>position</td><td>gene name</td><td>gene id</td></tr>
<tr><td>GU305914.1</td><td>149070</td><td>149092</td><td>+</td><td>0</td><td>CCGCGCGGTACTGCGCGTTG</td><td>GGG</td><td>||||||||[||||||||||||]PAM</td><td>NA</td><td>Intergenic</td><td>NA</td><td>NA</td></tr>
<tr><td></td></tr>
<tr><td>T11425</td><td>GGGGATCAGCTCTCCCGACTCGG</td><td>1000</td></tr>
<tr><td>Oligo fwd</td><td>CACCGGGGATCAGCTCTCCCGACT</td></tr>
<tr><td>Oligo rev</td><td>AAACAGTCGGGAGAGCTGATCCCC</td></tr>
<tr><td>Chromosome</td><td>start</td><td>end</td><td>strand</td><td>MM</td><td>target_seq</td><td>PAM</td><td>alignment</td><td>distance</td><td>position</td><td>gene name</td><td>gene id</td></tr>
<tr><td>GU305914.1</td><td>149089</td><td>149111</td><td>+</td><td>0</td><td>GGGGATCAGCTCTCCCGACT</td><td>CGG</td><td>||||||||[||||||||||||]PAM</td><td>NA</td><td>Intergenic</td><td>NA</td><td>NA</td></tr>
<tr><td></td></tr>
<tr><td>T11426</td><td>TCAGCTCTCCCGACTCGGTGTGG</td><td>1000</td></tr>
<tr><td>Oligo fwd</td><td>CACCTCAGCTCTCCCGACTCGGTG</td></tr>
<tr><td>Oligo rev</td><td>AAACCACCGAGTCGGGAGAGCTGA</td></tr>
<tr><td>Chromosome</td><td>start</td><td>end</td><td>strand</td><td>MM</td><td>target_seq</td><td>PAM</td><td>alignment</td><td>distance</td><td>position</td><td>gene name</td><td>gene id</td></tr>
<tr><td>GU305914.1</td><td>149094</td><td>149116</td><td>+</td><td>0</td><td>TCAGCTCTCCCGACTCGGTG</td><td>TGG</td><td>||||||||[||||||||||||]PAM</td><td>NA</td><td>Intergenic</td><td>NA</td><td>NA</td></tr>
<tr><td></td></tr>
<tr><td>T11427</td><td>ACCGTGATGCTCACGATCTCCGG</td><td>1000</td></tr>
<tr><td>Oligo fwd</td><td>CACCACCGTGATGCTCACGATCTC</td></tr>
<tr><td>Oligo rev</td><td>AAACGAGATCGTGAGCATCACGGT</td></tr>
<tr><td>Chromosome</td><td>start</td><td>end</td><td>strand</td><td>MM</td><td>target_seq</td><td>PAM</td><td>alignment</td><td>distance</td><td>position</td><td>gene name</td><td>gene id</td></tr>
<tr><td>GU305914.1</td><td>149138</td><td>149160</td><td>+</td><td>0</td><td>ACCGTGATGCTCACGATCTC</td><td>CGG</td><td>||||||||[||||||||||||]PAM</td><td>NA</td><td>Intergenic</td><td>NA</td><td>NA</td></tr>
<tr><td></td></tr>
<tr><td>T11428</td><td>CGATCTCCGGCGCCATCAGTCGG</td><td>1000</td></tr>
<tr><td>Oligo fwd</td><td>CACCCGATCTCCGGCGCCATCAGT</td></tr>
<tr><td>Oligo rev</td><td>AAACACTGATGGCGCCGGAGATCG</td></tr>
<tr><td>Chromosome</td><td>start</td><td>end</td><td>strand</td><td>MM</td><td>target_seq</td><td>PAM</td><td>alignment</td><td>distance</td><td>position</td><td>gene name</td><td>gene id</td></tr>
<tr><td>GU305914.1</td><td>149151</td><td>149173</td><td>+</td><td>0</td><td>CGATCTCCGGCGCCATCAGT</td><td>CGG</td><td>||||||||[||||||||||||]PAM</td><td>NA</td><td>Intergenic</td><td>NA</td><td>NA</td></tr>
<tr><td></td></tr>
<tr><td>T11429</td><td>CCATCAGTCGGACGTAGTCCCGG</td><td>1000</td></tr>
<tr><td>Oligo fwd</td><td>CACCCCATCAGTCGGACGTAGTCC</td></tr>
<tr><td>Oligo rev</td><td>AAACGGACTACGTCCGACTGATGG</td></tr>
<tr><td>Chromosome</td><td>start</td><td>end</td><td>strand</td><td>MM</td><td>target_seq</td><td>PAM</td><td>alignment</td><td>distance</td><td>position</td><td>gene name</td><td>gene id</td></tr>
<tr><td>GU305914.1</td><td>149163</td><td>149185</td><td>+</td><td>0</td><td>CCATCAGTCGGACGTAGTCC</td><td>CGG</td><td>||||||||[||||||||||||]PAM</td><td>NA</td><td>Intergenic</td><td>NA</td><td>NA</td></tr>
<tr><td></td></tr>
<tr><td>T11430</td><td>CCGGTCGTTGTGCACCTCCATGG</td><td>1000</td></tr>
<tr><td>Oligo fwd</td><td>CACCCCGGTCGTTGTGCACCTCCA</td></tr>
<tr><td>Oligo rev</td><td>AAACTGGAGGTGCACAACGACCGG</td></tr>
<tr><td>Chromosome</td><td>start</td><td>end</td><td>strand</td><td>MM</td><td>target_seq</td><td>PAM</td><td>alignment</td><td>distance</td><td>position</td><td>gene name</td><td>gene id</td></tr>
<tr><td>GU305914.1</td><td>149182</td><td>149204</td><td>+</td><td>0</td><td>CCGGTCGTTGTGCACCTCCA</td><td>TGG</td><td>||||||||[||||||||||||]PAM</td><td>NA</td><td>Intergenic</td><td>NA</td><td>NA</td></tr>
<tr><td></td></tr>
<tr><td>T11431</td><td>CATGGCGCCCCTAACCATTAAGG</td><td>1000</td></tr>
<tr><td>Oligo fwd</td><td>CACCCATGGCGCCCCTAACCATTA</td></tr>
<tr><td>Oligo rev</td><td>AAACTAATGGTTAGGGGCGCCATG</td></tr>
</table>

| | | | | | | | | | | | |
|---|---|---|---|---|---|---|---|---|---|---|---|
| Chromosome | start | end | strand | MM | target_seq | PAM | alignment | distance | position | gene name | gene id |
| GU305914.1 | 149200 | 149222 | + | 0 | CATGGCGCCCCTAACCATTA | AGG | \|\|\|\|\|\|\|\|[\|\|\|\|\|\|\|\|\|\|\|\|]PAM | NA | Intergenic | NA | NA |
| T11432 | GCCCCTAACCATTAAGGTGCTGG | 1000 | | | | | | | | | |
| Oligo fwd | CACCGCCCCTAACCATTAAGGTGC | | | | | | | | | | |
| Oligo rev | AAACGCACCTTAATGGTTAGGGGC | | | | | | | | | | |
| Chromosome | start | end | strand | MM | target_seq | PAM | alignment | distance | position | gene name | gene id |
| GU305914.1 | 149206 | 149228 | + | 0 | GCCCCTAACCATTAAGGTGC | TGG | \|\|\|\|\|\|\|\|[\|\|\|\|\|\|\|\|\|\|\|\|]PAM | NA | Intergenic | NA | NA |
| T11433 | AGCGTCTCGAGCTTCTGCTGCGG | 1000 | | | | | | | | | |
| Oligo fwd | CACCAGCGTCTCGAGCTTCTGCTG | | | | | | | | | | |
| Oligo rev | AAACCAGCAGAAGCTCGAGACGCT | | | | | | | | | | |
| Chromosome | start | end | strand | MM | target_seq | PAM | alignment | distance | position | gene name | gene id |
| GU305914.1 | 149244 | 149266 | + | 0 | AGCGTCTCGAGCTTCTGCTG | CGG | \|\|\|\|\|\|\|\|[\|\|\|\|\|\|\|\|\|\|\|\|]PAM | NA | Intergenic | NA | NA |
| T11434 | CGAGCTTCTGCTGCGGCGATCGG | 1000 | | | | | | | | | |
| Oligo fwd | CACCCGAGCTTCTGCTGCGGCGAT | | | | | | | | | | |
| Oligo rev | AAACATCGCCGCAGCAGAAGCTCG | | | | | | | | | | |
| Chromosome | start | end | strand | MM | target_seq | PAM | alignment | distance | position | gene name | gene id |
| GU305914.1 | 149251 | 149273 | + | 0 | CGAGCTTCTGCTGCGGCGAT | CGG | \|\|\|\|\|\|\|\|[\|\|\|\|\|\|\|\|\|\|\|\|]PAM | NA | Intergenic | NA | NA |
| T11435 | TGCTGCGGCGATCGGAACCTCGG | 1000 | | | | | | | | | |
| Oligo fwd | CACCTGCTGCGGCGATCGGAACCT | | | | | | | | | | |
| Oligo rev | AAACAGGTTCCGATCGCCGCAGCA | | | | | | | | | | |
| Chromosome | start | end | strand | MM | target_seq | PAM | alignment | distance | position | gene name | gene id |
| GU305914.1 | 149259 | 149281 | + | 0 | TGCTGCGGCGATCGGAACCT | CGG | \|\|\|\|\|\|\|\|[\|\|\|\|\|\|\|\|\|\|\|\|]PAM | NA | Intergenic | NA | NA |
| T11436 | GCTGCGGCGATCGGAACCTCGGG | 1000 | | | | | | | | | |
| Oligo fwd | CACCGCTGCGGCGATCGGAACCTC | | | | | | | | | | |
| Oligo rev | AAACGAGGTTCCGATCGCCGCAGC | | | | | | | | | | |
| Chromosome | start | end | strand | MM | target_seq | PAM | alignment | distance | position | gene name | gene id |
| GU305914.1 | 149260 | 149282 | + | 0 | GCTGCGGCGATCGGAACCTC | GGG | \|\|\|\|\|\|\|\|[\|\|\|\|\|\|\|\|\|\|\|\|]PAM | NA | Intergenic | NA | NA |
| T11437 | TCGGAACCTCGGGCTGACCTCGG | 1000 | | | | | | | | | |
| Oligo fwd | CACCTCGGAACCTCGGGCTGACCT | | | | | | | | | | |
| Oligo rev | AAACAGGTCAGCCCGAGGTTCCGA | | | | | | | | | | |
| Chromosome | start | end | strand | MM | target_seq | PAM | alignment | distance | position | gene name | gene id |
| GU305914.1 | 149270 | 149292 | + | 0 | TCGGAACCTCGGGCTGACCT | CGG | \|\|\|\|\|\|\|\|[\|\|\|\|\|\|\|\|\|\|\|\|]PAM | NA | Intergenic | NA | NA |
| T11438 | CGGAACCTCGGGCTGACCTCGGG | 1000 | | | | | | | | | |
| Oligo fwd | CACCCGGAACCTCGGGCTGACCTC | | | | | | | | | | |
| Oligo rev | AAACGAGGTCAGCCCGAGGTTCCG | | | | | | | | | | |
| Chromosome | start | end | strand | MM | target_seq | PAM | alignment | distance | position | gene name | gene id |
| GU305914.1 | 149271 | 149293 | + | 0 | CGGAACCTCGGGCTGACCTC | GGG | \|\|\|\|\|\|\|\|[\|\|\|\|\|\|\|\|\|\|\|\|]PAM | NA | Intergenic | NA | NA |
| T11439 | CCTCGGGCTGACCTCGGGAGCGG | 1000 | | | | | | | | | |
| Oligo fwd | CACCCCTCGGGCTGACCTCGGGAG | | | | | | | | | | |
| Oligo rev | AAACCTCCCGAGGTCAGCCCGAGG | | | | | | | | | | |
| Chromosome | start | end | strand | MM | target_seq | PAM | alignment | distance | position | gene name | gene id |
| GU305914.1 | 149276 | 149298 | + | 0 | CCTCGGGCTGACCTCGGGAG | CGG | \|\|\|\|\|\|\|\|[\|\|\|\|\|\|\|\|\|\|\|\|]PAM | NA | Intergenic | NA | NA |
| T11440 | CTCGGGCTGACCTCGGGAGCGGG | 1000 | | | | | | | | | |
| Oligo fwd | CACCCTCGGGCTGACCTCGGGAGC | | | | | | | | | | |
| Oligo rev | AAACGCTCCCGAGGTCAGCCCGAG | | | | | | | | | | |
| Chromosome | start | end | strand | MM | target_seq | PAM | alignment | distance | position | gene name | gene id |
| GU305914.1 | 149277 | 149299 | + | 0 | CTCGGGCTGACCTCGGGAGC | GGG | \|\|\|\|\|\|\|\|[\|\|\|\|\|\|\|\|\|\|\|\|]PAM | NA | Intergenic | NA | NA |
| T11441 | ACCTCGGGAGCGGGCGACGACGG | 1000 | | | | | | | | | |
| Oligo fwd | CACCACCTCGGGAGCGGGCGACGA | | | | | | | | | | |
| Oligo rev | AAACTCGTCGCCCGCTCCCGAGGT | | | | | | | | | | |
| Chromosome | start | end | strand | MM | target_seq | PAM | alignment | distance | position | gene name | gene id |
| GU305914.1 | 149286 | 149308 | + | 0 | ACCTCGGGAGCGGGCGACGA | CGG | \|\|\|\|\|\|\|\|[\|\|\|\|\|\|\|\|\|\|\|\|]PAM | NA | Intergenic | NA | NA |
| T11442 | TCGGGAGCGGGCGACGACGGCGG | 1000 | | | | | | | | | |
| Oligo fwd | CACCTCGGGAGCGGGCGACGACGG | | | | | | | | | | |
| Oligo rev | AAACCCGTCGTCGCCCGCTCCCGA | | | | | | | | | | |
| Chromosome | start | end | strand | MM | target_seq | PAM | alignment | distance | position | gene name | gene id |
| GU305914.1 | 149289 | 149311 | + | 0 | TCGGGAGCGGGCGACGACGG | CGG | \|\|\|\|\|\|\|\|[\|\|\|\|\|\|\|\|\|\|\|\|]PAM | NA | Intergenic | NA | NA |
| GU305914.1 | 149491 | 149513 | - | 4 | GAGGGAGCAGGCGACGGCGG | CGG | --\|\|\|\|\|\|[-\|\|\|\|\|\|\|-\|\|\|]PAM | NA | Intergenic | NA | NA |
| T11443 | GGAGCGGGCGACGACGGCGGCGG | 1000 | | | | | | | | | |
| Oligo fwd | CACCGGAGCGGGCGACGACGGCGG | | | | | | | | | | |
| Oligo rev | AAACCCGCCGTCGTCGCCCGCTCC | | | | | | | | | | |
| Chromosome | start | end | strand | MM | target_seq | PAM | alignment | distance | position | gene name | gene id |
| GU305914.1 | 149292 | 149314 | + | 0 | GGAGCGGGCGACGACGGCGG | CGG | \|\|\|\|\|\|\|\|[\|\|\|\|\|\|\|\|\|\|\|\|]PAM | NA | Intergenic | NA | NA |
| GU305914.1 | 149488 | 149510 | - | 2 | GGAGCAGGCGACGGCGGCGG | CGG | \|\|\|\|\|-\|\|[\|\|\|\|\|-\|\|\|\|\|\|]PAM | NA | Intergenic | NA | NA |
| GU305914.1 | 30000 | 30022 | + | 4 | GGTGCTGGCGGCGGCGGCGG | CGG | \|\|-\|\|-\|\|[\|\|-\|\|-\|\|\|\|\|\|]PAM | NA | Intergenic | NA | NA |
| GU305914.1 | 163896 | 163918 | + | 4 | GGATCGAGCGGCGGCGGCGG | CGG | \|\|\|-\|\|-\|[\|\|-\|\|-\|\|\|\|\|\|]PAM | NA | Intergenic | NA | NA |
| GU305914.1 | 90262 | 90284 | - | 4 | GAGGCGGGCGACGGCGGAGG | CGG | \|--\|\|\|\|\|[\|\|\|\|\|-\|\|\|-\|\|]PAM | NA | Intergenic | NA | NA |
| T11444 | CGGGCGACGACGGCGGCGGCAGG | 1000 | | | | | | | | | |
| Oligo fwd | CACCCGGGCGACGACGGCGGCGGC | | | | | | | | | | |
| Oligo rev | AAACGCCGCCGCCGTCGTCGCCCG | | | | | | | | | | |
| Chromosome | start | end | strand | MM | target_seq | PAM | alignment | distance | position | gene name | gene id |
| GU305914.1 | 149296 | 149318 | + | 0 | CGGGCGACGACGGCGGCGGC | AGG | \|\|\|\|\|\|\|\|[\|\|\|\|\|\|\|\|\|\|\|\|]PAM | NA | Intergenic | NA | NA |
| GU305914.1 | 67743 | 67765 | + | 4 | GGAGCGGCGGCGGCGGCGGC | GGG | -\|-\|\|\|-\|[\|-\|\|\|\|\|\|\|\|\|\|]PAM | NA | Intergenic | NA | NA |
| T11445 | GACGGCGGCGGCAGGCCGATCGG | 1000 | | | | | | | | | |
| Oligo fwd | CACCGACGGCGGCGGCAGGCCGAT | | | | | | | | | | |

<table>
<tr><td>Oligo rev</td><td>AAACATCGGCCTGCCGCCGCCGTC</td><td></td><td></td><td></td><td></td><td></td><td></td><td></td><td></td><td></td><td></td></tr>
<tr><td>Chromosome</td><td>start</td><td>end</td><td>strand</td><td>MM</td><td>target_seq</td><td>PAM</td><td>alignment</td><td>distance</td><td>position</td><td>gene name</td><td>gene id</td></tr>
<tr><td>GU305914.1</td><td>149304</td><td>149326</td><td>+</td><td>0</td><td>GACGGCGGCGGCAGGCCGAT</td><td>CGG</td><td>||||||||[||||||||||||]PAM</td><td>NA</td><td>Intergenic</td><td>NA</td><td>NA</td></tr>
<tr><td>T11446</td><td>ACGGCGGCGGCAGGCCGATCGGG</td><td>1000</td><td></td><td></td><td></td><td></td><td></td><td></td><td></td><td></td><td></td></tr>
<tr><td>Oligo fwd</td><td>CACCACGGCGGCGGCAGGCCGATC</td><td></td><td></td><td></td><td></td><td></td><td></td><td></td><td></td><td></td><td></td></tr>
<tr><td>Oligo rev</td><td>AAACGATCGGCCTGCCGCCGCCGT</td><td></td><td></td><td></td><td></td><td></td><td></td><td></td><td></td><td></td><td></td></tr>
<tr><td>Chromosome</td><td>start</td><td>end</td><td>strand</td><td>MM</td><td>target_seq</td><td>PAM</td><td>alignment</td><td>distance</td><td>position</td><td>gene name</td><td>gene id</td></tr>
<tr><td>GU305914.1</td><td>149305</td><td>149327</td><td>+</td><td>0</td><td>ACGGCGGCGGCAGGCCGATC</td><td>GGG</td><td>||||||||[||||||||||||]PAM</td><td>NA</td><td>Intergenic</td><td>NA</td><td>NA</td></tr>
<tr><td>GU305914.1</td><td>102644</td><td>102666</td><td>-</td><td>4</td><td>ACGACGACGGCACGCCGTTC</td><td>AGG</td><td>|||-||-|[||||-||||-||]PAM</td><td>NA</td><td>Intergenic</td><td>NA</td><td>NA</td></tr>
<tr><td>T11447</td><td>CGGCGGCGGCAGGCCGATCGGGG</td><td>1000</td><td></td><td></td><td></td><td></td><td></td><td></td><td></td><td></td><td></td></tr>
<tr><td>Oligo fwd</td><td>CACCCGGCGGCGGCAGGCCGATCG</td><td></td><td></td><td></td><td></td><td></td><td></td><td></td><td></td><td></td><td></td></tr>
<tr><td>Oligo rev</td><td>AAACCGATCGGCCTGCCGCCGCCG</td><td></td><td></td><td></td><td></td><td></td><td></td><td></td><td></td><td></td><td></td></tr>
<tr><td>Chromosome</td><td>start</td><td>end</td><td>strand</td><td>MM</td><td>target_seq</td><td>PAM</td><td>alignment</td><td>distance</td><td>position</td><td>gene name</td><td>gene id</td></tr>
<tr><td>GU305914.1</td><td>149306</td><td>149328</td><td>+</td><td>0</td><td>CGGCGGCGGCAGGCCGATCG</td><td>GGG</td><td>||||||||[||||||||||||]PAM</td><td>NA</td><td>Intergenic</td><td>NA</td><td>NA</td></tr>
<tr><td>T11448</td><td>TCGTCACGATCGTCTTCTGCTGG</td><td>1000</td><td></td><td></td><td></td><td></td><td></td><td></td><td></td><td></td><td></td></tr>
<tr><td>Oligo fwd</td><td>CACCTCGTCACGATCGTCTTCTGC</td><td></td><td></td><td></td><td></td><td></td><td></td><td></td><td></td><td></td><td></td></tr>
<tr><td>Oligo rev</td><td>AAACGCAGAAGACGATCGTGACGA</td><td></td><td></td><td></td><td></td><td></td><td></td><td></td><td></td><td></td><td></td></tr>
<tr><td>Chromosome</td><td>start</td><td>end</td><td>strand</td><td>MM</td><td>target_seq</td><td>PAM</td><td>alignment</td><td>distance</td><td>position</td><td>gene name</td><td>gene id</td></tr>
<tr><td>GU305914.1</td><td>149332</td><td>149354</td><td>+</td><td>0</td><td>TCGTCACGATCGTCTTCTGC</td><td>TGG</td><td>||||||||[||||||||||||]PAM</td><td>NA</td><td>Intergenic</td><td>NA</td><td>NA</td></tr>
<tr><td>T11449</td><td>CGTCACGATCGTCTTCTGCTGGG</td><td>1000</td><td></td><td></td><td></td><td></td><td></td><td></td><td></td><td></td><td></td></tr>
<tr><td>Oligo fwd</td><td>CACCCGTCACGATCGTCTTCTGCT</td><td></td><td></td><td></td><td></td><td></td><td></td><td></td><td></td><td></td><td></td></tr>
<tr><td>Oligo rev</td><td>AAACAGCAGAAGACGATCGTGACG</td><td></td><td></td><td></td><td></td><td></td><td></td><td></td><td></td><td></td><td></td></tr>
<tr><td>Chromosome</td><td>start</td><td>end</td><td>strand</td><td>MM</td><td>target_seq</td><td>PAM</td><td>alignment</td><td>distance</td><td>position</td><td>gene name</td><td>gene id</td></tr>
<tr><td>GU305914.1</td><td>149333</td><td>149355</td><td>+</td><td>0</td><td>CGTCACGATCGTCTTCTGCT</td><td>GGG</td><td>||||||||[||||||||||||]PAM</td><td>NA</td><td>Intergenic</td><td>NA</td><td>NA</td></tr>
<tr><td>T11450</td><td>GTCACGATCGTCTTCTGCTGGGG</td><td>1000</td><td></td><td></td><td></td><td></td><td></td><td></td><td></td><td></td><td></td></tr>
<tr><td>Oligo fwd</td><td>CACCGTCACGATCGTCTTCTGCTG</td><td></td><td></td><td></td><td></td><td></td><td></td><td></td><td></td><td></td><td></td></tr>
<tr><td>Oligo rev</td><td>AAACCAGCAGAAGACGATCGTGAC</td><td></td><td></td><td></td><td></td><td></td><td></td><td></td><td></td><td></td><td></td></tr>
<tr><td>Chromosome</td><td>start</td><td>end</td><td>strand</td><td>MM</td><td>target_seq</td><td>PAM</td><td>alignment</td><td>distance</td><td>position</td><td>gene name</td><td>gene id</td></tr>
<tr><td>GU305914.1</td><td>149334</td><td>149356</td><td>+</td><td>0</td><td>GTCACGATCGTCTTCTGCTG</td><td>GGG</td><td>||||||||[||||||||||||]PAM</td><td>NA</td><td>Intergenic</td><td>NA</td><td>NA</td></tr>
<tr><td>T11451</td><td>CGATCGTCTTCTGCTGGGGACGG</td><td>1000</td><td></td><td></td><td></td><td></td><td></td><td></td><td></td><td></td><td></td></tr>
<tr><td>Oligo fwd</td><td>CACCCGATCGTCTTCTGCTGGGGA</td><td></td><td></td><td></td><td></td><td></td><td></td><td></td><td></td><td></td><td></td></tr>
<tr><td>Oligo rev</td><td>AAACTCCCCAGCAGAAGACGATCG</td><td></td><td></td><td></td><td></td><td></td><td></td><td></td><td></td><td></td><td></td></tr>
<tr><td>Chromosome</td><td>start</td><td>end</td><td>strand</td><td>MM</td><td>target_seq</td><td>PAM</td><td>alignment</td><td>distance</td><td>position</td><td>gene name</td><td>gene id</td></tr>
<tr><td>GU305914.1</td><td>149338</td><td>149360</td><td>+</td><td>0</td><td>CGATCGTCTTCTGCTGGGGA</td><td>CGG</td><td>||||||||[||||||||||||]PAM</td><td>NA</td><td>Intergenic</td><td>NA</td><td>NA</td></tr>
<tr><td>T11452</td><td>GGACGGTAATTCGCCGTGCCTGG</td><td>1000</td><td></td><td></td><td></td><td></td><td></td><td></td><td></td><td></td><td></td></tr>
<tr><td>Oligo fwd</td><td>CACCGGACGGTAATTCGCCGTGCC</td><td></td><td></td><td></td><td></td><td></td><td></td><td></td><td></td><td></td><td></td></tr>
<tr><td>Oligo rev</td><td>AAACGGCACGGCGAATTACCGTCC</td><td></td><td></td><td></td><td></td><td></td><td></td><td></td><td></td><td></td><td></td></tr>
<tr><td>Chromosome</td><td>start</td><td>end</td><td>strand</td><td>MM</td><td>target_seq</td><td>PAM</td><td>alignment</td><td>distance</td><td>position</td><td>gene name</td><td>gene id</td></tr>
<tr><td>GU305914.1</td><td>149355</td><td>149377</td><td>+</td><td>0</td><td>GGACGGTAATTCGCCGTGCC</td><td>TGG</td><td>||||||||[||||||||||||]PAM</td><td>NA</td><td>Intergenic</td><td>NA</td><td>NA</td></tr>
<tr><td>T11453</td><td>TCGAGTCGCCTGATCTTCGTCGG</td><td>1000</td><td></td><td></td><td></td><td></td><td></td><td></td><td></td><td></td><td></td></tr>
<tr><td>Oligo fwd</td><td>CACCTCGAGTCGCCTGATCTTCGT</td><td></td><td></td><td></td><td></td><td></td><td></td><td></td><td></td><td></td><td></td></tr>
<tr><td>Oligo rev</td><td>AAACACGAAGATCAGGCGACTCGA</td><td></td><td></td><td></td><td></td><td></td><td></td><td></td><td></td><td></td><td></td></tr>
<tr><td>Chromosome</td><td>start</td><td>end</td><td>strand</td><td>MM</td><td>target_seq</td><td>PAM</td><td>alignment</td><td>distance</td><td>position</td><td>gene name</td><td>gene id</td></tr>
<tr><td>GU305914.1</td><td>149388</td><td>149410</td><td>+</td><td>0</td><td>TCGAGTCGCCTGATCTTCGT</td><td>CGG</td><td>||||||||[||||||||||||]PAM</td><td>NA</td><td>Intergenic</td><td>NA</td><td>NA</td></tr>
<tr><td>T11454</td><td>CGAGTCGCCTGATCTTCGTCGGG</td><td>1000</td><td></td><td></td><td></td><td></td><td></td><td></td><td></td><td></td><td></td></tr>
<tr><td>Oligo fwd</td><td>CACCCGAGTCGCCTGATCTTCGTC</td><td></td><td></td><td></td><td></td><td></td><td></td><td></td><td></td><td></td><td></td></tr>
<tr><td>Oligo rev</td><td>AAACGACGAAGATCAGGCGACTCG</td><td></td><td></td><td></td><td></td><td></td><td></td><td></td><td></td><td></td><td></td></tr>
<tr><td>Chromosome</td><td>start</td><td>end</td><td>strand</td><td>MM</td><td>target_seq</td><td>PAM</td><td>alignment</td><td>distance</td><td>position</td><td>gene name</td><td>gene id</td></tr>
<tr><td>GU305914.1</td><td>149389</td><td>149411</td><td>+</td><td>0</td><td>CGAGTCGCCTGATCTTCGTC</td><td>GGG</td><td>||||||||[||||||||||||]PAM</td><td>NA</td><td>Intergenic</td><td>NA</td><td>NA</td></tr>
<tr><td>T11455</td><td>GAGTCGCCTGATCTTCGTCGGGG</td><td>1000</td><td></td><td></td><td></td><td></td><td></td><td></td><td></td><td></td><td></td></tr>
<tr><td>Oligo fwd</td><td>CACCGAGTCGCCTGATCTTCGTCG</td><td></td><td></td><td></td><td></td><td></td><td></td><td></td><td></td><td></td><td></td></tr>
<tr><td>Oligo rev</td><td>AAACCGACGAAGATCAGGCGACTC</td><td></td><td></td><td></td><td></td><td></td><td></td><td></td><td></td><td></td><td></td></tr>
<tr><td>Chromosome</td><td>start</td><td>end</td><td>strand</td><td>MM</td><td>target_seq</td><td>PAM</td><td>alignment</td><td>distance</td><td>position</td><td>gene name</td><td>gene id</td></tr>
<tr><td>GU305914.1</td><td>149390</td><td>149412</td><td>+</td><td>0</td><td>GAGTCGCCTGATCTTCGTCG</td><td>GGG</td><td>||||||||[||||||||||||]PAM</td><td>NA</td><td>Intergenic</td><td>NA</td><td>NA</td></tr>
<tr><td>T11456</td><td>AGTCGCCTGATCTTCGTCGGGGG</td><td>1000</td><td></td><td></td><td></td><td></td><td></td><td></td><td></td><td></td><td></td></tr>
<tr><td>Oligo fwd</td><td>CACCAGTCGCCTGATCTTCGTCGG</td><td></td><td></td><td></td><td></td><td></td><td></td><td></td><td></td><td></td><td></td></tr>
<tr><td>Oligo rev</td><td>AAACCCGACGAAGATCAGGCGACT</td><td></td><td></td><td></td><td></td><td></td><td></td><td></td><td></td><td></td><td></td></tr>
<tr><td>Chromosome</td><td>start</td><td>end</td><td>strand</td><td>MM</td><td>target_seq</td><td>PAM</td><td>alignment</td><td>distance</td><td>position</td><td>gene name</td><td>gene id</td></tr>
<tr><td>GU305914.1</td><td>149391</td><td>149413</td><td>+</td><td>0</td><td>AGTCGCCTGATCTTCGTCGG</td><td>GGG</td><td>||||||||[||||||||||||]PAM</td><td>NA</td><td>Intergenic</td><td>NA</td><td>NA</td></tr>
<tr><td>T11457</td><td>GTCGCCTGATCTTCGTCGGGGGG</td><td>1000</td><td></td><td></td><td></td><td></td><td></td><td></td><td></td><td></td><td></td></tr>
<tr><td>Oligo fwd</td><td>CACCGTCGCCTGATCTTCGTCGGG</td><td></td><td></td><td></td><td></td><td></td><td></td><td></td><td></td><td></td><td></td></tr>
<tr><td>Oligo rev</td><td>AAACCCCGACGAAGATCAGGCGAC</td><td></td><td></td><td></td><td></td><td></td><td></td><td></td><td></td><td></td><td></td></tr>
<tr><td>Chromosome</td><td>start</td><td>end</td><td>strand</td><td>MM</td><td>target_seq</td><td>PAM</td><td>alignment</td><td>distance</td><td>position</td><td>gene name</td><td>gene id</td></tr>
<tr><td>GU305914.1</td><td>149392</td><td>149414</td><td>+</td><td>0</td><td>GTCGCCTGATCTTCGTCGGG</td><td>GGG</td><td>||||||||[||||||||||||]PAM</td><td>NA</td><td>Intergenic</td><td>NA</td><td>NA</td></tr>
<tr><td>T11458</td><td>CCTGATCTTCGTCGGGGGGCCGG</td><td>1000</td><td></td><td></td><td></td><td></td><td></td><td></td><td></td><td></td><td></td></tr>
<tr><td>Oligo fwd</td><td>CACCCCTGATCTTCGTCGGGGGGC</td><td></td><td></td><td></td><td></td><td></td><td></td><td></td><td></td><td></td><td></td></tr>
<tr><td>Oligo rev</td><td>AAACGCCCCCCGACGAAGATCAGG</td><td></td><td></td><td></td><td></td><td></td><td></td><td></td><td></td><td></td><td></td></tr>
<tr><td>Chromosome</td><td>start</td><td>end</td><td>strand</td><td>MM</td><td>target_seq</td><td>PAM</td><td>alignment</td><td>distance</td><td>position</td><td>gene name</td><td>gene id</td></tr>
<tr><td>GU305914.1</td><td>149396</td><td>149418</td><td>+</td><td>0</td><td>CCTGATCTTCGTCGGGGGGC</td><td>CGG</td><td>||||||||[||||||||||||]PAM</td><td>NA</td><td>Intergenic</td><td>NA</td><td>NA</td></tr>
<tr><td>T11459</td><td>TCGTCGGGGGGCCGGAGACCTGG</td><td>1000</td><td></td><td></td><td></td><td></td><td></td><td></td><td></td><td></td><td></td></tr>
<tr><td>Oligo fwd</td><td>CACCTCGTCGGGGGGCCGGAGACC</td><td></td><td></td><td></td><td></td><td></td><td></td><td></td><td></td><td></td><td></td></tr>
<tr><td>Oligo rev</td><td>AAACGGTCTCCGGCCCCCCGACGA</td><td></td><td></td><td></td><td></td><td></td><td></td><td></td><td></td><td></td><td></td></tr>
<tr><td>Chromosome</td><td>start</td><td>end</td><td>strand</td><td>MM</td><td>target_seq</td><td>PAM</td><td>alignment</td><td>distance</td><td>position</td><td>gene name</td><td>gene id</td></tr>
<tr><td>GU305914.1</td><td>149404</td><td>149426</td><td>+</td><td>0</td><td>TCGTCGGGGGGCCGGAGACC</td><td>TGG</td><td>||||||||[||||||||||||]PAM</td><td>NA</td><td>Intergenic</td><td>NA</td><td>NA</td></tr>
</table>

<table>
<tr><td>T11460</td><td>GGGGGGCCGGAGACCTGGCCTGG</td><td>1000</td><td></td><td></td><td></td><td></td><td></td><td></td><td></td><td></td><td></td></tr>
<tr><td>Oligo fwd</td><td>CACCGGGGGGCCGGAGACCTGGCC</td><td></td><td></td><td></td><td></td><td></td><td></td><td></td><td></td><td></td><td></td></tr>
<tr><td>Oligo rev</td><td>AAACGGCCAGGTCTCCGGCCCCCC</td><td></td><td></td><td></td><td></td><td></td><td></td><td></td><td></td><td></td><td></td></tr>
<tr><td>Chromosome</td><td>start</td><td>end</td><td>strand</td><td>MM</td><td>target_seq</td><td>PAM</td><td>alignment</td><td>distance</td><td>position</td><td>gene name</td><td>gene id</td></tr>
<tr><td>GU305914.1</td><td>149409</td><td>149431</td><td>+</td><td>0</td><td>GGGGGGCCGGAGACCTGGCC</td><td>TGG</td><td>||||||||[||||||||||||]PAM</td><td>NA</td><td>Intergenic</td><td>NA</td><td>NA</td></tr>
<tr><td>GU305914.1</td><td>202665</td><td>202687</td><td>-</td><td>4</td><td>GGGTGACCGGAGATCCGGCC</td><td>TGG</td><td>|||-|-||[|||||-|-||||]PAM</td><td>NA</td><td>Intergenic</td><td>NA</td><td>NA</td></tr>
<tr><td>T11461</td><td>GCTCGTCCGATCCGCTCGCTCGG</td><td>1000</td><td></td><td></td><td></td><td></td><td></td><td></td><td></td><td></td><td></td></tr>
<tr><td>Oligo fwd</td><td>CACCGCTCGTCCGATCCGCTCGCT</td><td></td><td></td><td></td><td></td><td></td><td></td><td></td><td></td><td></td><td></td></tr>
<tr><td>Oligo rev</td><td>AAACAGCGAGCGGATCGGACGAGC</td><td></td><td></td><td></td><td></td><td></td><td></td><td></td><td></td><td></td><td></td></tr>
<tr><td>Chromosome</td><td>start</td><td>end</td><td>strand</td><td>MM</td><td>target_seq</td><td>PAM</td><td>alignment</td><td>distance</td><td>position</td><td>gene name</td><td>gene id</td></tr>
<tr><td>GU305914.1</td><td>149433</td><td>149455</td><td>+</td><td>0</td><td>GCTCGTCCGATCCGCTCGCT</td><td>CGG</td><td>||||||||[||||||||||||]PAM</td><td>NA</td><td>Intergenic</td><td>NA</td><td>NA</td></tr>
<tr><td>T11462</td><td>CGCCTGCTCCCTCTCTAAAAAGG</td><td>1000</td><td></td><td></td><td></td><td></td><td></td><td></td><td></td><td></td><td></td></tr>
<tr><td>Oligo fwd</td><td>CACCCGCCTGCTCCCTCTCTAAAA</td><td></td><td></td><td></td><td></td><td></td><td></td><td></td><td></td><td></td><td></td></tr>
<tr><td>Oligo rev</td><td>AAACTTTTAGAGAGGGAGCAGGCG</td><td></td><td></td><td></td><td></td><td></td><td></td><td></td><td></td><td></td><td></td></tr>
<tr><td>Chromosome</td><td>start</td><td>end</td><td>strand</td><td>MM</td><td>target_seq</td><td>PAM</td><td>alignment</td><td>distance</td><td>position</td><td>gene name</td><td>gene id</td></tr>
<tr><td>GU305914.1</td><td>149501</td><td>149523</td><td>+</td><td>0</td><td>CGCCTGCTCCCTCTCTAAAA</td><td>AGG</td><td>||||||||[||||||||||||]PAM</td><td>NA</td><td>Intergenic</td><td>NA</td><td>NA</td></tr>
<tr><td>T11463</td><td>GCTCCCTCTCTAAAAAGGCTTGG</td><td>1000</td><td></td><td></td><td></td><td></td><td></td><td></td><td></td><td></td><td></td></tr>
<tr><td>Oligo fwd</td><td>CACCGCTCCCTCTCTAAAAAGGCT</td><td></td><td></td><td></td><td></td><td></td><td></td><td></td><td></td><td></td><td></td></tr>
<tr><td>Oligo rev</td><td>AAACAGCCTTTTTAGAGAGGGAGC</td><td></td><td></td><td></td><td></td><td></td><td></td><td></td><td></td><td></td><td></td></tr>
<tr><td>Chromosome</td><td>start</td><td>end</td><td>strand</td><td>MM</td><td>target_seq</td><td>PAM</td><td>alignment</td><td>distance</td><td>position</td><td>gene name</td><td>gene id</td></tr>
<tr><td>GU305914.1</td><td>149506</td><td>149528</td><td>+</td><td>0</td><td>GCTCCCTCTCTAAAAAGGCT</td><td>TGG</td><td>||||||||[||||||||||||]PAM</td><td>NA</td><td>Intergenic</td><td>NA</td><td>NA</td></tr>
<tr><td>T11464</td><td>TAAAAAGGCTTGGACGCGAATGG</td><td>1000</td><td></td><td></td><td></td><td></td><td></td><td></td><td></td><td></td><td></td></tr>
<tr><td>Oligo fwd</td><td>CACCTAAAAAGGCTTGGACGCGAA</td><td></td><td></td><td></td><td></td><td></td><td></td><td></td><td></td><td></td><td></td></tr>
<tr><td>Oligo rev</td><td>AAACTTCGCGTCCAAGCCTTTTTA</td><td></td><td></td><td></td><td></td><td></td><td></td><td></td><td></td><td></td><td></td></tr>
<tr><td>Chromosome</td><td>start</td><td>end</td><td>strand</td><td>MM</td><td>target_seq</td><td>PAM</td><td>alignment</td><td>distance</td><td>position</td><td>gene name</td><td>gene id</td></tr>
<tr><td>GU305914.1</td><td>149516</td><td>149538</td><td>+</td><td>0</td><td>TAAAAAGGCTTGGACGCGAA</td><td>TGG</td><td>||||||||[||||||||||||]PAM</td><td>NA</td><td>Intergenic</td><td>NA</td><td>NA</td></tr>
<tr><td>T11465</td><td>CTTGGACGCGAATGGTCTCACGG</td><td>1000</td><td></td><td></td><td></td><td></td><td></td><td></td><td></td><td></td><td></td></tr>
<tr><td>Oligo fwd</td><td>CACCCTTGGACGCGAATGGTCTCA</td><td></td><td></td><td></td><td></td><td></td><td></td><td></td><td></td><td></td><td></td></tr>
<tr><td>Oligo rev</td><td>AAACTGAGACCATTCGCGTCCAAG</td><td></td><td></td><td></td><td></td><td></td><td></td><td></td><td></td><td></td><td></td></tr>
<tr><td>Chromosome</td><td>start</td><td>end</td><td>strand</td><td>MM</td><td>target_seq</td><td>PAM</td><td>alignment</td><td>distance</td><td>position</td><td>gene name</td><td>gene id</td></tr>
<tr><td>GU305914.1</td><td>149524</td><td>149546</td><td>+</td><td>0</td><td>CTTGGACGCGAATGGTCTCA</td><td>CGG</td><td>||||||||[||||||||||||]PAM</td><td>NA</td><td>Intergenic</td><td>NA</td><td>NA</td></tr>
<tr><td>T11466</td><td>CGCGAATGGTCTCACGGTAGCGG</td><td>1000</td><td></td><td></td><td></td><td></td><td></td><td></td><td></td><td></td><td></td></tr>
<tr><td>Oligo fwd</td><td>CACCCGCGAATGGTCTCACGGTAG</td><td></td><td></td><td></td><td></td><td></td><td></td><td></td><td></td><td></td><td></td></tr>
<tr><td>Oligo rev</td><td>AAACCTACCGTGAGACCATTCGCG</td><td></td><td></td><td></td><td></td><td></td><td></td><td></td><td></td><td></td><td></td></tr>
<tr><td>Chromosome</td><td>start</td><td>end</td><td>strand</td><td>MM</td><td>target_seq</td><td>PAM</td><td>alignment</td><td>distance</td><td>position</td><td>gene name</td><td>gene id</td></tr>
<tr><td>GU305914.1</td><td>149530</td><td>149552</td><td>+</td><td>0</td><td>CGCGAATGGTCTCACGGTAG</td><td>CGG</td><td>||||||||[||||||||||||]PAM</td><td>NA</td><td>Intergenic</td><td>NA</td><td>NA</td></tr>
<tr><td>T11467</td><td>CTCACGGTAGCGGATGTAGTCGG</td><td>1000</td><td></td><td></td><td></td><td></td><td></td><td></td><td></td><td></td><td></td></tr>
<tr><td>Oligo fwd</td><td>CACCCTCACGGTAGCGGATGTAGT</td><td></td><td></td><td></td><td></td><td></td><td></td><td></td><td></td><td></td><td></td></tr>
<tr><td>Oligo rev</td><td>AAACACTACATCCGCTACCGTGAG</td><td></td><td></td><td></td><td></td><td></td><td></td><td></td><td></td><td></td><td></td></tr>
<tr><td>Chromosome</td><td>start</td><td>end</td><td>strand</td><td>MM</td><td>target_seq</td><td>PAM</td><td>alignment</td><td>distance</td><td>position</td><td>gene name</td><td>gene id</td></tr>
<tr><td>GU305914.1</td><td>149540</td><td>149562</td><td>+</td><td>0</td><td>CTCACGGTAGCGGATGTAGT</td><td>CGG</td><td>||||||||[||||||||||||]PAM</td><td>NA</td><td>Intergenic</td><td>NA</td><td>NA</td></tr>
<tr><td>T11468</td><td>GACAGAAATAAATACTCTTCTGG</td><td>1000</td><td></td><td></td><td></td><td></td><td></td><td></td><td></td><td></td><td></td></tr>
<tr><td>Oligo fwd</td><td>CACCGACAGAAATAAATACTCTTC</td><td></td><td></td><td></td><td></td><td></td><td></td><td></td><td></td><td></td><td></td></tr>
<tr><td>Oligo rev</td><td>AAACGAAGAGTATTTATTTCTGTC</td><td></td><td></td><td></td><td></td><td></td><td></td><td></td><td></td><td></td><td></td></tr>
<tr><td>Chromosome</td><td>start</td><td>end</td><td>strand</td><td>MM</td><td>target_seq</td><td>PAM</td><td>alignment</td><td>distance</td><td>position</td><td>gene name</td><td>gene id</td></tr>
<tr><td>GU305914.1</td><td>149616</td><td>149638</td><td>+</td><td>0</td><td>GACAGAAATAAATACTCTTC</td><td>TGG</td><td>||||||||[||||||||||||]PAM</td><td>NA</td><td>Intergenic</td><td>NA</td><td>NA</td></tr>
<tr><td>T11469</td><td>ACAGAAATAAATACTCTTCTGGG</td><td>1000</td><td></td><td></td><td></td><td></td><td></td><td></td><td></td><td></td><td></td></tr>
<tr><td>Oligo fwd</td><td>CACCACAGAAATAAATACTCTTCT</td><td></td><td></td><td></td><td></td><td></td><td></td><td></td><td></td><td></td><td></td></tr>
<tr><td>Oligo rev</td><td>AAACAGAAGAGTATTTATTTCTGT</td><td></td><td></td><td></td><td></td><td></td><td></td><td></td><td></td><td></td><td></td></tr>
<tr><td>Chromosome</td><td>start</td><td>end</td><td>strand</td><td>MM</td><td>target_seq</td><td>PAM</td><td>alignment</td><td>distance</td><td>position</td><td>gene name</td><td>gene id</td></tr>
<tr><td>GU305914.1</td><td>149617</td><td>149639</td><td>+</td><td>0</td><td>ACAGAAATAAATACTCTTCT</td><td>GGG</td><td>||||||||[||||||||||||]PAM</td><td>NA</td><td>Intergenic</td><td>NA</td><td>NA</td></tr>
<tr><td>T11470</td><td>AAATAAATACTCTTCTGGGATGG</td><td>1000</td><td></td><td></td><td></td><td></td><td></td><td></td><td></td><td></td><td></td></tr>
<tr><td>Oligo fwd</td><td>CACCAAATAAATACTCTTCTGGGA</td><td></td><td></td><td></td><td></td><td></td><td></td><td></td><td></td><td></td><td></td></tr>
<tr><td>Oligo rev</td><td>AAACTCCCAGAAGAGTATTTATTT</td><td></td><td></td><td></td><td></td><td></td><td></td><td></td><td></td><td></td><td></td></tr>
<tr><td>Chromosome</td><td>start</td><td>end</td><td>strand</td><td>MM</td><td>target_seq</td><td>PAM</td><td>alignment</td><td>distance</td><td>position</td><td>gene name</td><td>gene id</td></tr>
<tr><td>GU305914.1</td><td>149621</td><td>149643</td><td>+</td><td>0</td><td>AAATAAATACTCTTCTGGGA</td><td>TGG</td><td>||||||||[||||||||||||]PAM</td><td>NA</td><td>Intergenic</td><td>NA</td><td>NA</td></tr>
<tr><td>T11471</td><td>TTCTGGGATGGCATCCAACATGG</td><td>1000</td><td></td><td></td><td></td><td></td><td></td><td></td><td></td><td></td><td></td></tr>
<tr><td>Oligo fwd</td><td>CACCTTCTGGGATGGCATCCAACA</td><td></td><td></td><td></td><td></td><td></td><td></td><td></td><td></td><td></td><td></td></tr>
<tr><td>Oligo rev</td><td>AAACTGTTGGATGCCATCCCAGAA</td><td></td><td></td><td></td><td></td><td></td><td></td><td></td><td></td><td></td><td></td></tr>
<tr><td>Chromosome</td><td>start</td><td>end</td><td>strand</td><td>MM</td><td>target_seq</td><td>PAM</td><td>alignment</td><td>distance</td><td>position</td><td>gene name</td><td>gene id</td></tr>
<tr><td>GU305914.1</td><td>149633</td><td>149655</td><td>+</td><td>0</td><td>TTCTGGGATGGCATCCAACA</td><td>TGG</td><td>||||||||[||||||||||||]PAM</td><td>NA</td><td>Intergenic</td><td>NA</td><td>NA</td></tr>
<tr><td>T11472</td><td>TCTGGGATGGCATCCAACATGGG</td><td>1000</td><td></td><td></td><td></td><td></td><td></td><td></td><td></td><td></td><td></td></tr>
<tr><td>Oligo fwd</td><td>CACCTCTGGGATGGCATCCAACAT</td><td></td><td></td><td></td><td></td><td></td><td></td><td></td><td></td><td></td><td></td></tr>
<tr><td>Oligo rev</td><td>AAACATGTTGGATGCCATCCCAGA</td><td></td><td></td><td></td><td></td><td></td><td></td><td></td><td></td><td></td><td></td></tr>
<tr><td>Chromosome</td><td>start</td><td>end</td><td>strand</td><td>MM</td><td>target_seq</td><td>PAM</td><td>alignment</td><td>distance</td><td>position</td><td>gene name</td><td>gene id</td></tr>
<tr><td>GU305914.1</td><td>149634</td><td>149656</td><td>+</td><td>0</td><td>TCTGGGATGGCATCCAACAT</td><td>GGG</td><td>||||||||[||||||||||||]PAM</td><td>NA</td><td>Intergenic</td><td>NA</td><td>NA</td></tr>
<tr><td>T11473</td><td>CTGGGATGGCATCCAACATGGGG</td><td>1000</td><td></td><td></td><td></td><td></td><td></td><td></td><td></td><td></td><td></td></tr>
<tr><td>Oligo fwd</td><td>CACCCTGGGATGGCATCCAACATG</td><td></td><td></td><td></td><td></td><td></td><td></td><td></td><td></td><td></td><td></td></tr>
<tr><td>Oligo rev</td><td>AAACCATGTTGGATGCCATCCCAG</td><td></td><td></td><td></td><td></td><td></td><td></td><td></td><td></td><td></td><td></td></tr>
<tr><td>Chromosome</td><td>start</td><td>end</td><td>strand</td><td>MM</td><td>target_seq</td><td>PAM</td><td>alignment</td><td>distance</td><td>position</td><td>gene name</td><td>gene id</td></tr>
<tr><td>GU305914.1</td><td>149635</td><td>149657</td><td>+</td><td>0</td><td>CTGGGATGGCATCCAACATG</td><td>GGG</td><td>||||||||[||||||||||||]PAM</td><td>NA</td><td>Intergenic</td><td>NA</td><td>NA</td></tr>
<tr><td>T11474</td><td>GGAACTCGATGACCGCCGTGAGG</td><td>1000</td><td></td><td></td><td></td><td></td><td></td><td></td><td></td><td></td><td></td></tr>
<tr><td>Oligo fwd</td><td>CACCGGAACTCGATGACCGCCGTG</td><td></td><td></td><td></td><td></td><td></td><td></td><td></td><td></td><td></td><td></td></tr>
<tr><td>Oligo rev</td><td>AAACCACGGCGGTCATCGAGTTCC</td><td></td><td></td><td></td><td></td><td></td><td></td><td></td><td></td><td></td><td></td></tr>
<tr><td>Chromosome</td><td>start</td><td>end</td><td>strand</td><td>MM</td><td>target_seq</td><td>PAM</td><td>alignment</td><td>distance</td><td>position</td><td>gene name</td><td>gene id</td></tr>
</table>

<table>
<tr><td>GU305914.1</td><td>149656</td><td>149678</td><td>+</td><td>0</td><td>GGAACTCGATGACCGCCGTG</td><td>AGG</td><td>||||||||[||||||||||||]PAM</td><td>NA</td><td>Intergenic</td><td>NA</td><td>NA</td></tr>
<tr><td>T11475</td><td>TCGATGACCGCCGTGAGGAACGG</td><td>1000</td><td></td><td></td><td></td><td></td><td></td><td></td><td></td><td></td><td></td></tr>
<tr><td>Oligo fwd</td><td>CACCTCGATGACCGCCGTGAGGAA</td><td></td><td></td><td></td><td></td><td></td><td></td><td></td><td></td><td></td><td></td></tr>
<tr><td>Oligo rev</td><td>AAACTTCCTCACGGCGGTCATCGA</td><td></td><td></td><td></td><td></td><td></td><td></td><td></td><td></td><td></td><td></td></tr>
<tr><td>Chromosome</td><td>start</td><td>end</td><td>strand</td><td>MM</td><td>target_seq</td><td>PAM</td><td>alignment</td><td>distance</td><td>position</td><td>gene name</td><td>gene id</td></tr>
<tr><td>GU305914.1</td><td>149661</td><td>149683</td><td>+</td><td>0</td><td>TCGATGACCGCCGTGAGGAA</td><td>CGG</td><td>||||||||[||||||||||||]PAM</td><td>NA</td><td>Intergenic</td><td>NA</td><td>NA</td></tr>
<tr><td>T11476</td><td>CGATGACCGCCGTGAGGAACGGG</td><td>1000</td><td></td><td></td><td></td><td></td><td></td><td></td><td></td><td></td><td></td></tr>
<tr><td>Oligo fwd</td><td>CACCCGATGACCGCCGTGAGGAAC</td><td></td><td></td><td></td><td></td><td></td><td></td><td></td><td></td><td></td><td></td></tr>
<tr><td>Oligo rev</td><td>AAACGTTCCTCACGGCGGTCATCG</td><td></td><td></td><td></td><td></td><td></td><td></td><td></td><td></td><td></td><td></td></tr>
<tr><td>Chromosome</td><td>start</td><td>end</td><td>strand</td><td>MM</td><td>target_seq</td><td>PAM</td><td>alignment</td><td>distance</td><td>position</td><td>gene name</td><td>gene id</td></tr>
<tr><td>GU305914.1</td><td>149662</td><td>149684</td><td>+</td><td>0</td><td>CGATGACCGCCGTGAGGAAC</td><td>GGG</td><td>||||||||[||||||||||||]PAM</td><td>NA</td><td>Intergenic</td><td>NA</td><td>NA</td></tr>
<tr><td>T11477</td><td>CGAGTACGAGATGCTGTCCTCGG</td><td>1000</td><td></td><td></td><td></td><td></td><td></td><td></td><td></td><td></td><td></td></tr>
<tr><td>Oligo fwd</td><td>CACCCGAGTACGAGATGCTGTCCT</td><td></td><td></td><td></td><td></td><td></td><td></td><td></td><td></td><td></td><td></td></tr>
<tr><td>Oligo rev</td><td>AAACAGGACAGCATCTCGTACTCG</td><td></td><td></td><td></td><td></td><td></td><td></td><td></td><td></td><td></td><td></td></tr>
<tr><td>Chromosome</td><td>start</td><td>end</td><td>strand</td><td>MM</td><td>target_seq</td><td>PAM</td><td>alignment</td><td>distance</td><td>position</td><td>gene name</td><td>gene id</td></tr>
<tr><td>GU305914.1</td><td>149711</td><td>149733</td><td>+</td><td>0</td><td>CGAGTACGAGATGCTGTCCT</td><td>CGG</td><td>||||||||[||||||||||||]PAM</td><td>NA</td><td>Intergenic</td><td>NA</td><td>NA</td></tr>
<tr><td>T11478</td><td>AGATGCTGTCCTCGGCGCCCTGG</td><td>1000</td><td></td><td></td><td></td><td></td><td></td><td></td><td></td><td></td><td></td></tr>
<tr><td>Oligo fwd</td><td>CACCAGATGCTGTCCTCGGCGCCC</td><td></td><td></td><td></td><td></td><td></td><td></td><td></td><td></td><td></td><td></td></tr>
<tr><td>Oligo rev</td><td>AAACGGGCGCCGAGGACAGCATCT</td><td></td><td></td><td></td><td></td><td></td><td></td><td></td><td></td><td></td><td></td></tr>
<tr><td>Chromosome</td><td>start</td><td>end</td><td>strand</td><td>MM</td><td>target_seq</td><td>PAM</td><td>alignment</td><td>distance</td><td>position</td><td>gene name</td><td>gene id</td></tr>
<tr><td>GU305914.1</td><td>149719</td><td>149741</td><td>+</td><td>0</td><td>AGATGCTGTCCTCGGCGCCC</td><td>TGG</td><td>||||||||[||||||||||||]PAM</td><td>NA</td><td>Intergenic</td><td>NA</td><td>NA</td></tr>
<tr><td>T11479</td><td>CCTGGTTCGTCACGTTCCGTCGG</td><td>1000</td><td></td><td></td><td></td><td></td><td></td><td></td><td></td><td></td><td></td></tr>
<tr><td>Oligo fwd</td><td>CACCCCTGGTTCGTCACGTTCCGT</td><td></td><td></td><td></td><td></td><td></td><td></td><td></td><td></td><td></td><td></td></tr>
<tr><td>Oligo rev</td><td>AAACACGGAACGTGACGAACCAGG</td><td></td><td></td><td></td><td></td><td></td><td></td><td></td><td></td><td></td><td></td></tr>
<tr><td>Chromosome</td><td>start</td><td>end</td><td>strand</td><td>MM</td><td>target_seq</td><td>PAM</td><td>alignment</td><td>distance</td><td>position</td><td>gene name</td><td>gene id</td></tr>
<tr><td>GU305914.1</td><td>149737</td><td>149759</td><td>+</td><td>0</td><td>CCTGGTTCGTCACGTTCCGT</td><td>CGG</td><td>||||||||[||||||||||||]PAM</td><td>NA</td><td>Intergenic</td><td>NA</td><td>NA</td></tr>
<tr><td>T11480</td><td>CTTGCTCCCAGAGACGCGACAGG</td><td>1000</td><td></td><td></td><td></td><td></td><td></td><td></td><td></td><td></td><td></td></tr>
<tr><td>Oligo fwd</td><td>CACCCTTGCTCCCAGAGACGCGAC</td><td></td><td></td><td></td><td></td><td></td><td></td><td></td><td></td><td></td><td></td></tr>
<tr><td>Oligo rev</td><td>AAACGTCGCGTCTCTGGGAGCAAG</td><td></td><td></td><td></td><td></td><td></td><td></td><td></td><td></td><td></td><td></td></tr>
<tr><td>Chromosome</td><td>start</td><td>end</td><td>strand</td><td>MM</td><td>target_seq</td><td>PAM</td><td>alignment</td><td>distance</td><td>position</td><td>gene name</td><td>gene id</td></tr>
<tr><td>GU305914.1</td><td>149782</td><td>149804</td><td>+</td><td>0</td><td>CTTGCTCCCAGAGACGCGAC</td><td>AGG</td><td>||||||||[||||||||||||]PAM</td><td>NA</td><td>Intergenic</td><td>NA</td><td>NA</td></tr>
<tr><td>T11481</td><td>CTCGTCCGACTCGCGCTCGTCGG</td><td>1000</td><td></td><td></td><td></td><td></td><td></td><td></td><td></td><td></td><td></td></tr>
<tr><td>Oligo fwd</td><td>CACCCTCGTCCGACTCGCGCTCGT</td><td></td><td></td><td></td><td></td><td></td><td></td><td></td><td></td><td></td><td></td></tr>
<tr><td>Oligo rev</td><td>AAACACGAGCGCGAGTCGGACGAG</td><td></td><td></td><td></td><td></td><td></td><td></td><td></td><td></td><td></td><td></td></tr>
<tr><td>Chromosome</td><td>start</td><td>end</td><td>strand</td><td>MM</td><td>target_seq</td><td>PAM</td><td>alignment</td><td>distance</td><td>position</td><td>gene name</td><td>gene id</td></tr>
<tr><td>GU305914.1</td><td>149807</td><td>149829</td><td>+</td><td>0</td><td>CTCGTCCGACTCGCGCTCGT</td><td>CGG</td><td>||||||||[||||||||||||]PAM</td><td>NA</td><td>Intergenic</td><td>NA</td><td>NA</td></tr>
<tr><td>T11482</td><td>TCGTCCGACTCGCGCTCGTCGGG</td><td>1000</td><td></td><td></td><td></td><td></td><td></td><td></td><td></td><td></td><td></td></tr>
<tr><td>Oligo fwd</td><td>CACCTCGTCCGACTCGCGCTCGTC</td><td></td><td></td><td></td><td></td><td></td><td></td><td></td><td></td><td></td><td></td></tr>
<tr><td>Oligo rev</td><td>AAACGACGAGCGCGAGTCGGACGA</td><td></td><td></td><td></td><td></td><td></td><td></td><td></td><td></td><td></td><td></td></tr>
<tr><td>Chromosome</td><td>start</td><td>end</td><td>strand</td><td>MM</td><td>target_seq</td><td>PAM</td><td>alignment</td><td>distance</td><td>position</td><td>gene name</td><td>gene id</td></tr>
<tr><td>GU305914.1</td><td>149808</td><td>149830</td><td>+</td><td>0</td><td>TCGTCCGACTCGCGCTCGTC</td><td>GGG</td><td>||||||||[||||||||||||]PAM</td><td>NA</td><td>Intergenic</td><td>NA</td><td>NA</td></tr>
<tr><td>T11483</td><td>AGAAGCTGTTGCAGACGACCCGG</td><td>1000</td><td></td><td></td><td></td><td></td><td></td><td></td><td></td><td></td><td></td></tr>
<tr><td>Oligo fwd</td><td>CACCAGAAGCTGTTGCAGACGACC</td><td></td><td></td><td></td><td></td><td></td><td></td><td></td><td></td><td></td><td></td></tr>
<tr><td>Oligo rev</td><td>AAACGGTCGTCTGCAACAGCTTCT</td><td></td><td></td><td></td><td></td><td></td><td></td><td></td><td></td><td></td><td></td></tr>
<tr><td>Chromosome</td><td>start</td><td>end</td><td>strand</td><td>MM</td><td>target_seq</td><td>PAM</td><td>alignment</td><td>distance</td><td>position</td><td>gene name</td><td>gene id</td></tr>
<tr><td>GU305914.1</td><td>149845</td><td>149867</td><td>+</td><td>0</td><td>AGAAGCTGTTGCAGACGACC</td><td>CGG</td><td>||||||||[||||||||||||]PAM</td><td>NA</td><td>Intergenic</td><td>NA</td><td>NA</td></tr>
<tr><td>T11484</td><td>CGGTTGTCGTCCACCGTCACCGG</td><td>1000</td><td></td><td></td><td></td><td></td><td></td><td></td><td></td><td></td><td></td></tr>
<tr><td>Oligo fwd</td><td>CACCCGGTTGTCGTCCACCGTCAC</td><td></td><td></td><td></td><td></td><td></td><td></td><td></td><td></td><td></td><td></td></tr>
<tr><td>Oligo rev</td><td>AAACGTGACGGTGGACGACAACCG</td><td></td><td></td><td></td><td></td><td></td><td></td><td></td><td></td><td></td><td></td></tr>
<tr><td>Chromosome</td><td>start</td><td>end</td><td>strand</td><td>MM</td><td>target_seq</td><td>PAM</td><td>alignment</td><td>distance</td><td>position</td><td>gene name</td><td>gene id</td></tr>
<tr><td>GU305914.1</td><td>149865</td><td>149887</td><td>+</td><td>0</td><td>CGGTTGTCGTCCACCGTCAC</td><td>CGG</td><td>||||||||[||||||||||||]PAM</td><td>NA</td><td>Intergenic</td><td>NA</td><td>NA</td></tr>
<tr><td>T11485</td><td>TCCACCGTCACCGGCGCCGTCGG</td><td>1000</td><td></td><td></td><td></td><td></td><td></td><td></td><td></td><td></td><td></td></tr>
<tr><td>Oligo fwd</td><td>CACCTCCACCGTCACCGGCGCCGT</td><td></td><td></td><td></td><td></td><td></td><td></td><td></td><td></td><td></td><td></td></tr>
<tr><td>Oligo rev</td><td>AAACACGGCGCCGGTGACGGTGGA</td><td></td><td></td><td></td><td></td><td></td><td></td><td></td><td></td><td></td><td></td></tr>
<tr><td>Chromosome</td><td>start</td><td>end</td><td>strand</td><td>MM</td><td>target_seq</td><td>PAM</td><td>alignment</td><td>distance</td><td>position</td><td>gene name</td><td>gene id</td></tr>
<tr><td>GU305914.1</td><td>149874</td><td>149896</td><td>+</td><td>0</td><td>TCCACCGTCACCGGCGCCGT</td><td>CGG</td><td>||||||||[||||||||||||]PAM</td><td>NA</td><td>Intergenic</td><td>NA</td><td>NA</td></tr>
<tr><td>GU305914.1</td><td>194927</td><td>194949</td><td>-</td><td>3</td><td>TGGACCGTCATCGGCGCCGT</td><td>GGG</td><td>|--|||||[||-|||||||||]PAM</td><td>NA</td><td>Intergenic</td><td>NA</td><td>NA</td></tr>
<tr><td>T11486</td><td>CCGTCACCGGCGCCGTCGGCAGG</td><td>1000</td><td></td><td></td><td></td><td></td><td></td><td></td><td></td><td></td><td></td></tr>
<tr><td>Oligo fwd</td><td>CACCCCGTCACCGGCGCCGTCGGC</td><td></td><td></td><td></td><td></td><td></td><td></td><td></td><td></td><td></td><td></td></tr>
<tr><td>Oligo rev</td><td>AAACGCCGACGGCGCCGGTGACGG</td><td></td><td></td><td></td><td></td><td></td><td></td><td></td><td></td><td></td><td></td></tr>
<tr><td>Chromosome</td><td>start</td><td>end</td><td>strand</td><td>MM</td><td>target_seq</td><td>PAM</td><td>alignment</td><td>distance</td><td>position</td><td>gene name</td><td>gene id</td></tr>
<tr><td>GU305914.1</td><td>149878</td><td>149900</td><td>+</td><td>0</td><td>CCGTCACCGGCGCCGTCGGC</td><td>AGG</td><td>||||||||[||||||||||||]PAM</td><td>NA</td><td>Intergenic</td><td>NA</td><td>NA</td></tr>
<tr><td>T11487</td><td>GCAGCGCGTTGACCGTGTCCAGG</td><td>1000</td><td></td><td></td><td></td><td></td><td></td><td></td><td></td><td></td><td></td></tr>
<tr><td>Oligo fwd</td><td>CACCGCAGCGCGTTGACCGTGTCC</td><td></td><td></td><td></td><td></td><td></td><td></td><td></td><td></td><td></td><td></td></tr>
<tr><td>Oligo rev</td><td>AAACGGACACGGTCAACGCGCTGC</td><td></td><td></td><td></td><td></td><td></td><td></td><td></td><td></td><td></td><td></td></tr>
<tr><td>Chromosome</td><td>start</td><td>end</td><td>strand</td><td>MM</td><td>target_seq</td><td>PAM</td><td>alignment</td><td>distance</td><td>position</td><td>gene name</td><td>gene id</td></tr>
<tr><td>GU305914.1</td><td>149914</td><td>149936</td><td>+</td><td>0</td><td>GCAGCGCGTTGACCGTGTCC</td><td>AGG</td><td>||||||||[||||||||||||]PAM</td><td>NA</td><td>Intergenic</td><td>NA</td><td>NA</td></tr>
<tr><td>T11488</td><td>CAGCGCGTTGACCGTGTCCAGGG</td><td>1000</td><td></td><td></td><td></td><td></td><td></td><td></td><td></td><td></td><td></td></tr>
<tr><td>Oligo fwd</td><td>CACCCAGCGCGTTGACCGTGTCCA</td><td></td><td></td><td></td><td></td><td></td><td></td><td></td><td></td><td></td><td></td></tr>
<tr><td>Oligo rev</td><td>AAACTGGACACGGTCAACGCGCTG</td><td></td><td></td><td></td><td></td><td></td><td></td><td></td><td></td><td></td><td></td></tr>
<tr><td>Chromosome</td><td>start</td><td>end</td><td>strand</td><td>MM</td><td>target_seq</td><td>PAM</td><td>alignment</td><td>distance</td><td>position</td><td>gene name</td><td>gene id</td></tr>
<tr><td>GU305914.1</td><td>149915</td><td>149937</td><td>+</td><td>0</td><td>CAGCGCGTTGACCGTGTCCA</td><td>GGG</td><td>||||||||[||||||||||||]PAM</td><td>NA</td><td>Intergenic</td><td>NA</td><td>NA</td></tr>
<tr><td>T11489</td><td>TGACCGTGTCCAGGGTCTGCAGG</td><td>1000</td><td></td><td></td><td></td><td></td><td></td><td></td><td></td><td></td><td></td></tr>
<tr><td>Oligo fwd</td><td>CACCTGACCGTGTCCAGGGTCTGC</td><td></td><td></td><td></td><td></td><td></td><td></td><td></td><td></td><td></td><td></td></tr>
</table>

<table>
<tr><td>Oligo rev</td><td>AAACGCAGACCCTGGACACGGTCA</td><td></td><td></td><td></td><td></td><td></td><td></td><td></td><td></td><td></td><td></td></tr>
<tr><td>Chromosome</td><td>start</td><td>end</td><td>strand</td><td>MM</td><td>target_seq</td><td>PAM</td><td>alignment</td><td>distance</td><td>position</td><td>gene name</td><td>gene id</td></tr>
<tr><td>GU305914.1</td><td>149923</td><td>149945</td><td>+</td><td>0</td><td>TGACCGTGTCCAGGGTCTGC</td><td>AGG</td><td>||||||||[||||||||||||]PAM</td><td>NA</td><td>Intergenic</td><td>NA</td><td>NA</td></tr>
<tr><td>T11490</td><td>AGGGTCTGCAGGTCGAACGCCGG</td><td>1000</td><td></td><td></td><td></td><td></td><td></td><td></td><td></td><td></td><td></td></tr>
<tr><td>Oligo fwd</td><td>CACCAGGGTCTGCAGGTCGAACGC</td><td></td><td></td><td></td><td></td><td></td><td></td><td></td><td></td><td></td><td></td></tr>
<tr><td>Oligo rev</td><td>AAACGCGTTCGACCTGCAGACCCT</td><td></td><td></td><td></td><td></td><td></td><td></td><td></td><td></td><td></td><td></td></tr>
<tr><td>Chromosome</td><td>start</td><td>end</td><td>strand</td><td>MM</td><td>target_seq</td><td>PAM</td><td>alignment</td><td>distance</td><td>position</td><td>gene name</td><td>gene id</td></tr>
<tr><td>GU305914.1</td><td>149934</td><td>149956</td><td>+</td><td>0</td><td>AGGGTCTGCAGGTCGAACGC</td><td>CGG</td><td>||||||||[||||||||||||]PAM</td><td>NA</td><td>Intergenic</td><td>NA</td><td>NA</td></tr>
<tr><td>T11491</td><td>GGGTCTGCAGGTCGAACGCCGGG</td><td>1000</td><td></td><td></td><td></td><td></td><td></td><td></td><td></td><td></td><td></td></tr>
<tr><td>Oligo fwd</td><td>CACCGGGTCTGCAGGTCGAACGCC</td><td></td><td></td><td></td><td></td><td></td><td></td><td></td><td></td><td></td><td></td></tr>
<tr><td>Oligo rev</td><td>AAACGGCGTTCGACCTGCAGACCC</td><td></td><td></td><td></td><td></td><td></td><td></td><td></td><td></td><td></td><td></td></tr>
<tr><td>Chromosome</td><td>start</td><td>end</td><td>strand</td><td>MM</td><td>target_seq</td><td>PAM</td><td>alignment</td><td>distance</td><td>position</td><td>gene name</td><td>gene id</td></tr>
<tr><td>GU305914.1</td><td>149935</td><td>149957</td><td>+</td><td>0</td><td>GGGTCTGCAGGTCGAACGCC</td><td>GGG</td><td>||||||||[||||||||||||]PAM</td><td>NA</td><td>Intergenic</td><td>NA</td><td>NA</td></tr>
<tr><td>T11492</td><td>GGTCTGCAGGTCGAACGCCGGGG</td><td>1000</td><td></td><td></td><td></td><td></td><td></td><td></td><td></td><td></td><td></td></tr>
<tr><td>Oligo fwd</td><td>CACCGGTCTGCAGGTCGAACGCCG</td><td></td><td></td><td></td><td></td><td></td><td></td><td></td><td></td><td></td><td></td></tr>
<tr><td>Oligo rev</td><td>AAACCGGCGTTCGACCTGCAGACC</td><td></td><td></td><td></td><td></td><td></td><td></td><td></td><td></td><td></td><td></td></tr>
<tr><td>Chromosome</td><td>start</td><td>end</td><td>strand</td><td>MM</td><td>target_seq</td><td>PAM</td><td>alignment</td><td>distance</td><td>position</td><td>gene name</td><td>gene id</td></tr>
<tr><td>GU305914.1</td><td>149936</td><td>149958</td><td>+</td><td>0</td><td>GGTCTGCAGGTCGAACGCCG</td><td>GGG</td><td>||||||||[||||||||||||]PAM</td><td>NA</td><td>Intergenic</td><td>NA</td><td>NA</td></tr>
<tr><td>T11493</td><td>GTCTGCAGGTCGAACGCCGGGGG</td><td>1000</td><td></td><td></td><td></td><td></td><td></td><td></td><td></td><td></td><td></td></tr>
<tr><td>Oligo fwd</td><td>CACCGTCTGCAGGTCGAACGCCGG</td><td></td><td></td><td></td><td></td><td></td><td></td><td></td><td></td><td></td><td></td></tr>
<tr><td>Oligo rev</td><td>AAACCCGGCGTTCGACCTGCAGAC</td><td></td><td></td><td></td><td></td><td></td><td></td><td></td><td></td><td></td><td></td></tr>
<tr><td>Chromosome</td><td>start</td><td>end</td><td>strand</td><td>MM</td><td>target_seq</td><td>PAM</td><td>alignment</td><td>distance</td><td>position</td><td>gene name</td><td>gene id</td></tr>
<tr><td>GU305914.1</td><td>149937</td><td>149959</td><td>+</td><td>0</td><td>GTCTGCAGGTCGAACGCCGG</td><td>GGG</td><td>||||||||[||||||||||||]PAM</td><td>NA</td><td>Intergenic</td><td>NA</td><td>NA</td></tr>
<tr><td>T11494</td><td>TCTGCAGGTCGAACGCCGGGGGG</td><td>1000</td><td></td><td></td><td></td><td></td><td></td><td></td><td></td><td></td><td></td></tr>
<tr><td>Oligo fwd</td><td>CACCTCTGCAGGTCGAACGCCGGG</td><td></td><td></td><td></td><td></td><td></td><td></td><td></td><td></td><td></td><td></td></tr>
<tr><td>Oligo rev</td><td>AAACCCCGGCGTTCGACCTGCAGA</td><td></td><td></td><td></td><td></td><td></td><td></td><td></td><td></td><td></td><td></td></tr>
<tr><td>Chromosome</td><td>start</td><td>end</td><td>strand</td><td>MM</td><td>target_seq</td><td>PAM</td><td>alignment</td><td>distance</td><td>position</td><td>gene name</td><td>gene id</td></tr>
<tr><td>GU305914.1</td><td>149938</td><td>149960</td><td>+</td><td>0</td><td>TCTGCAGGTCGAACGCCGGG</td><td>GGG</td><td>||||||||[||||||||||||]PAM</td><td>NA</td><td>Intergenic</td><td>NA</td><td>NA</td></tr>
<tr><td>T11495</td><td>GGTCGAACGCCGGGGGGCCGCGG</td><td>1000</td><td></td><td></td><td></td><td></td><td></td><td></td><td></td><td></td><td></td></tr>
<tr><td>Oligo fwd</td><td>CACCGGTCGAACGCCGGGGGGCCG</td><td></td><td></td><td></td><td></td><td></td><td></td><td></td><td></td><td></td><td></td></tr>
<tr><td>Oligo rev</td><td>AAACCGGCCCCCCGGCGTTCGACC</td><td></td><td></td><td></td><td></td><td></td><td></td><td></td><td></td><td></td><td></td></tr>
<tr><td>Chromosome</td><td>start</td><td>end</td><td>strand</td><td>MM</td><td>target_seq</td><td>PAM</td><td>alignment</td><td>distance</td><td>position</td><td>gene name</td><td>gene id</td></tr>
<tr><td>GU305914.1</td><td>149944</td><td>149966</td><td>+</td><td>0</td><td>GGTCGAACGCCGGGGGGCCG</td><td>CGG</td><td>||||||||[||||||||||||]PAM</td><td>NA</td><td>Intergenic</td><td>NA</td><td>NA</td></tr>
<tr><td>T11496</td><td>CCGGGGGGCCGCGGCCGCGCCGG</td><td>1000</td><td></td><td></td><td></td><td></td><td></td><td></td><td></td><td></td><td></td></tr>
<tr><td>Oligo fwd</td><td>CACCCCGGGGGGCCGCGGCCGCGC</td><td></td><td></td><td></td><td></td><td></td><td></td><td></td><td></td><td></td><td></td></tr>
<tr><td>Oligo rev</td><td>AAACGCGCGGCCGCGGCCCCCCGG</td><td></td><td></td><td></td><td></td><td></td><td></td><td></td><td></td><td></td><td></td></tr>
<tr><td>Chromosome</td><td>start</td><td>end</td><td>strand</td><td>MM</td><td>target_seq</td><td>PAM</td><td>alignment</td><td>distance</td><td>position</td><td>gene name</td><td>gene id</td></tr>
<tr><td>GU305914.1</td><td>149953</td><td>149975</td><td>+</td><td>0</td><td>CCGGGGGGCCGCGGCCGCGC</td><td>CGG</td><td>||||||||[||||||||||||]PAM</td><td>NA</td><td>Intergenic</td><td>NA</td><td>NA</td></tr>
<tr><td>T11497</td><td>GCCGCGGCCGCGCCGGCCCTCGG</td><td>1000</td><td></td><td></td><td></td><td></td><td></td><td></td><td></td><td></td><td></td></tr>
<tr><td>Oligo fwd</td><td>CACCGCCGCGGCCGCGCCGGCCCT</td><td></td><td></td><td></td><td></td><td></td><td></td><td></td><td></td><td></td><td></td></tr>
<tr><td>Oligo rev</td><td>AAACAGGGCCGGCGCGGCCGCGGC</td><td></td><td></td><td></td><td></td><td></td><td></td><td></td><td></td><td></td><td></td></tr>
<tr><td>Chromosome</td><td>start</td><td>end</td><td>strand</td><td>MM</td><td>target_seq</td><td>PAM</td><td>alignment</td><td>distance</td><td>position</td><td>gene name</td><td>gene id</td></tr>
<tr><td>GU305914.1</td><td>149960</td><td>149982</td><td>+</td><td>0</td><td>GCCGCGGCCGCGCCGGCCCT</td><td>CGG</td><td>||||||||[||||||||||||]PAM</td><td>NA</td><td>Intergenic</td><td>NA</td><td>NA</td></tr>
<tr><td>T11498</td><td>GCCGGCCCTCGGACTCATAGCGG</td><td>1000</td><td></td><td></td><td></td><td></td><td></td><td></td><td></td><td></td><td></td></tr>
<tr><td>Oligo fwd</td><td>CACCGCCGGCCCTCGGACTCATAG</td><td></td><td></td><td></td><td></td><td></td><td></td><td></td><td></td><td></td><td></td></tr>
<tr><td>Oligo rev</td><td>AAACCTATGAGTCCGAGGGCCGGC</td><td></td><td></td><td></td><td></td><td></td><td></td><td></td><td></td><td></td><td></td></tr>
<tr><td>Chromosome</td><td>start</td><td>end</td><td>strand</td><td>MM</td><td>target_seq</td><td>PAM</td><td>alignment</td><td>distance</td><td>position</td><td>gene name</td><td>gene id</td></tr>
<tr><td>GU305914.1</td><td>149971</td><td>149993</td><td>+</td><td>0</td><td>GCCGGCCCTCGGACTCATAG</td><td>CGG</td><td>||||||||[||||||||||||]PAM</td><td>NA</td><td>Intergenic</td><td>NA</td><td>NA</td></tr>
<tr><td>T11499</td><td>GATTCTCTATCTCCTCGACATGG</td><td>1000</td><td></td><td></td><td></td><td></td><td></td><td></td><td></td><td></td><td></td></tr>
<tr><td>Oligo fwd</td><td>CACCGATTCTCTATCTCCTCGACA</td><td></td><td></td><td></td><td></td><td></td><td></td><td></td><td></td><td></td><td></td></tr>
<tr><td>Oligo rev</td><td>AAACTGTCGAGGAGATAGAGAATC</td><td></td><td></td><td></td><td></td><td></td><td></td><td></td><td></td><td></td><td></td></tr>
<tr><td>Chromosome</td><td>start</td><td>end</td><td>strand</td><td>MM</td><td>target_seq</td><td>PAM</td><td>alignment</td><td>distance</td><td>position</td><td>gene name</td><td>gene id</td></tr>
<tr><td>GU305914.1</td><td>150052</td><td>150074</td><td>+</td><td>0</td><td>GATTCTCTATCTCCTCGACA</td><td>TGG</td><td>||||||||[||||||||||||]PAM</td><td>NA</td><td>Intergenic</td><td>NA</td><td>NA</td></tr>
<tr><td>T11500</td><td>CTATCTCCTCGACATGGTCTTGG</td><td>1000</td><td></td><td></td><td></td><td></td><td></td><td></td><td></td><td></td><td></td></tr>
<tr><td>Oligo fwd</td><td>CACCCTATCTCCTCGACATGGTCT</td><td></td><td></td><td></td><td></td><td></td><td></td><td></td><td></td><td></td><td></td></tr>
<tr><td>Oligo rev</td><td>AAACAGACCATGTCGAGGAGATAG</td><td></td><td></td><td></td><td></td><td></td><td></td><td></td><td></td><td></td><td></td></tr>
<tr><td>Chromosome</td><td>start</td><td>end</td><td>strand</td><td>MM</td><td>target_seq</td><td>PAM</td><td>alignment</td><td>distance</td><td>position</td><td>gene name</td><td>gene id</td></tr>
<tr><td>GU305914.1</td><td>150058</td><td>150080</td><td>+</td><td>0</td><td>CTATCTCCTCGACATGGTCT</td><td>TGG</td><td>||||||||[||||||||||||]PAM</td><td>NA</td><td>Intergenic</td><td>NA</td><td>NA</td></tr>
<tr><td>T11501</td><td>CCTCGACATGGTCTTGGATCTGG</td><td>1000</td><td></td><td></td><td></td><td></td><td></td><td></td><td></td><td></td><td></td></tr>
<tr><td>Oligo fwd</td><td>CACCCCTCGACATGGTCTTGGATC</td><td></td><td></td><td></td><td></td><td></td><td></td><td></td><td></td><td></td><td></td></tr>
<tr><td>Oligo rev</td><td>AAACGATCCAAGACCATGTCGAGG</td><td></td><td></td><td></td><td></td><td></td><td></td><td></td><td></td><td></td><td></td></tr>
<tr><td>Chromosome</td><td>start</td><td>end</td><td>strand</td><td>MM</td><td>target_seq</td><td>PAM</td><td>alignment</td><td>distance</td><td>position</td><td>gene name</td><td>gene id</td></tr>
<tr><td>GU305914.1</td><td>150064</td><td>150086</td><td>+</td><td>0</td><td>CCTCGACATGGTCTTGGATC</td><td>TGG</td><td>||||||||[||||||||||||]PAM</td><td>NA</td><td>Intergenic</td><td>NA</td><td>NA</td></tr>
<tr><td>T11502</td><td>GGTCTTGGATCTGGTCCTCCAGG</td><td>1000</td><td></td><td></td><td></td><td></td><td></td><td></td><td></td><td></td><td></td></tr>
<tr><td>Oligo fwd</td><td>CACCGGTCTTGGATCTGGTCCTCC</td><td></td><td></td><td></td><td></td><td></td><td></td><td></td><td></td><td></td><td></td></tr>
<tr><td>Oligo rev</td><td>AAACGGAGGACCAGATCCAAGACC</td><td></td><td></td><td></td><td></td><td></td><td></td><td></td><td></td><td></td><td></td></tr>
<tr><td>Chromosome</td><td>start</td><td>end</td><td>strand</td><td>MM</td><td>target_seq</td><td>PAM</td><td>alignment</td><td>distance</td><td>position</td><td>gene name</td><td>gene id</td></tr>
<tr><td>GU305914.1</td><td>150073</td><td>150095</td><td>+</td><td>0</td><td>GGTCTTGGATCTGGTCCTCC</td><td>AGG</td><td>||||||||[||||||||||||]PAM</td><td>NA</td><td>Intergenic</td><td>NA</td><td>NA</td></tr>
<tr><td>T11503</td><td>CCTGTCGCCTAAAGAGCTCTCGG</td><td>1000</td><td></td><td></td><td></td><td></td><td></td><td></td><td></td><td></td><td></td></tr>
<tr><td>Oligo fwd</td><td>CACCCCTGTCGCCTAAAGAGCTCT</td><td></td><td></td><td></td><td></td><td></td><td></td><td></td><td></td><td></td><td></td></tr>
<tr><td>Oligo rev</td><td>AAACAGAGCTCTTTAGGCGACAGG</td><td></td><td></td><td></td><td></td><td></td><td></td><td></td><td></td><td></td><td></td></tr>
<tr><td>Chromosome</td><td>start</td><td>end</td><td>strand</td><td>MM</td><td>target_seq</td><td>PAM</td><td>alignment</td><td>distance</td><td>position</td><td>gene name</td><td>gene id</td></tr>
<tr><td>GU305914.1</td><td>150106</td><td>150128</td><td>+</td><td>0</td><td>CCTGTCGCCTAAAGAGCTCT</td><td>CGG</td><td>||||||||[||||||||||||]PAM</td><td>NA</td><td>Intergenic</td><td>NA</td><td>NA</td></tr>
<tr><td>T11504</td><td>GTCCTGCGCGCCCGAACCCGAGG</td><td>1000</td><td></td><td></td><td></td><td></td><td></td><td></td><td></td><td></td><td></td></tr>
</table>

<table>
<tr><td>Oligo fwd</td><td>CACCGTCCTGCGCGCCCGAACCCG</td><td></td><td></td><td></td><td></td><td></td><td></td><td></td><td></td><td></td><td></td></tr>
<tr><td>Oligo rev</td><td>AAACCGGGTTCGGGCGCGCAGGAC</td><td></td><td></td><td></td><td></td><td></td><td></td><td></td><td></td><td></td><td></td></tr>
<tr><td>Chromosome</td><td>start</td><td>end</td><td>strand</td><td>MM</td><td>target_seq</td><td>PAM</td><td>alignment</td><td>distance</td><td>position</td><td>gene name</td><td>gene id</td></tr>
<tr><td>GU305914.1</td><td>150131</td><td>150153</td><td>+</td><td>0</td><td>GTCCTGCGCGCCCGAACCCG</td><td>AGG</td><td>||||||||[||||||||||||]PAM</td><td>NA</td><td>Intergenic</td><td>NA</td><td>NA</td></tr>
<tr><td></td><td></td><td></td><td></td><td></td><td></td><td></td><td></td><td></td><td></td><td></td><td></td></tr>
<tr><td>T11505</td><td>TCCTGCGCGCCCGAACCCGAGGG</td><td>1000</td><td></td><td></td><td></td><td></td><td></td><td></td><td></td><td></td><td></td></tr>
<tr><td>Oligo fwd</td><td>CACCTCCTGCGCGCCCGAACCCGA</td><td></td><td></td><td></td><td></td><td></td><td></td><td></td><td></td><td></td><td></td></tr>
<tr><td>Oligo rev</td><td>AAACTCGGGTTCGGGCGCGCAGGA</td><td></td><td></td><td></td><td></td><td></td><td></td><td></td><td></td><td></td><td></td></tr>
<tr><td>Chromosome</td><td>start</td><td>end</td><td>strand</td><td>MM</td><td>target_seq</td><td>PAM</td><td>alignment</td><td>distance</td><td>position</td><td>gene name</td><td>gene id</td></tr>
<tr><td>GU305914.1</td><td>150132</td><td>150154</td><td>+</td><td>0</td><td>TCCTGCGCGCCCGAACCCGA</td><td>GGG</td><td>||||||||[||||||||||||]PAM</td><td>NA</td><td>Intergenic</td><td>NA</td><td>NA</td></tr>
<tr><td></td><td></td><td></td><td></td><td></td><td></td><td></td><td></td><td></td><td></td><td></td><td></td></tr>
<tr><td>T11506</td><td>GCGCCCGAACCCGAGGGTGCCGG</td><td>1000</td><td></td><td></td><td></td><td></td><td></td><td></td><td></td><td></td><td></td></tr>
<tr><td>Oligo fwd</td><td>CACCGCGCCCGAACCCGAGGGTGC</td><td></td><td></td><td></td><td></td><td></td><td></td><td></td><td></td><td></td><td></td></tr>
<tr><td>Oligo rev</td><td>AAACGCACCCTCGGGTTCGGGCGC</td><td></td><td></td><td></td><td></td><td></td><td></td><td></td><td></td><td></td><td></td></tr>
<tr><td>Chromosome</td><td>start</td><td>end</td><td>strand</td><td>MM</td><td>target_seq</td><td>PAM</td><td>alignment</td><td>distance</td><td>position</td><td>gene name</td><td>gene id</td></tr>
<tr><td>GU305914.1</td><td>150138</td><td>150160</td><td>+</td><td>0</td><td>GCGCCCGAACCCGAGGGTGC</td><td>CGG</td><td>||||||||[||||||||||||]PAM</td><td>NA</td><td>Intergenic</td><td>NA</td><td>NA</td></tr>
<tr><td></td><td></td><td></td><td></td><td></td><td></td><td></td><td></td><td></td><td></td><td></td><td></td></tr>
<tr><td>T11507</td><td>CCGAACCCGAGGGTGCCGGTTGG</td><td>1000</td><td></td><td></td><td></td><td></td><td></td><td></td><td></td><td></td><td></td></tr>
<tr><td>Oligo fwd</td><td>CACCCCGAACCCGAGGGTGCCGGT</td><td></td><td></td><td></td><td></td><td></td><td></td><td></td><td></td><td></td><td></td></tr>
<tr><td>Oligo rev</td><td>AAACACCGGCACCCTCGGGTTCGG</td><td></td><td></td><td></td><td></td><td></td><td></td><td></td><td></td><td></td><td></td></tr>
<tr><td>Chromosome</td><td>start</td><td>end</td><td>strand</td><td>MM</td><td>target_seq</td><td>PAM</td><td>alignment</td><td>distance</td><td>position</td><td>gene name</td><td>gene id</td></tr>
<tr><td>GU305914.1</td><td>150142</td><td>150164</td><td>+</td><td>0</td><td>CCGAACCCGAGGGTGCCGGT</td><td>TGG</td><td>||||||||[||||||||||||]PAM</td><td>NA</td><td>Intergenic</td><td>NA</td><td>NA</td></tr>
<tr><td></td><td></td><td></td><td></td><td></td><td></td><td></td><td></td><td></td><td></td><td></td><td></td></tr>
<tr><td>T11508</td><td>CCGAGGGTGCCGGTTGGATGAGG</td><td>1000</td><td></td><td></td><td></td><td></td><td></td><td></td><td></td><td></td><td></td></tr>
<tr><td>Oligo fwd</td><td>CACCCCGAGGGTGCCGGTTGGATG</td><td></td><td></td><td></td><td></td><td></td><td></td><td></td><td></td><td></td><td></td></tr>
<tr><td>Oligo rev</td><td>AAACCATCCAACCGGCACCCTCGG</td><td></td><td></td><td></td><td></td><td></td><td></td><td></td><td></td><td></td><td></td></tr>
<tr><td>Chromosome</td><td>start</td><td>end</td><td>strand</td><td>MM</td><td>target_seq</td><td>PAM</td><td>alignment</td><td>distance</td><td>position</td><td>gene name</td><td>gene id</td></tr>
<tr><td>GU305914.1</td><td>150148</td><td>150170</td><td>+</td><td>0</td><td>CCGAGGGTGCCGGTTGGATG</td><td>AGG</td><td>||||||||[||||||||||||]PAM</td><td>NA</td><td>Intergenic</td><td>NA</td><td>NA</td></tr>
<tr><td></td><td></td><td></td><td></td><td></td><td></td><td></td><td></td><td></td><td></td><td></td><td></td></tr>
<tr><td>T11509</td><td>CGAGGGTGCCGGTTGGATGAGGG</td><td>1000</td><td></td><td></td><td></td><td></td><td></td><td></td><td></td><td></td><td></td></tr>
<tr><td>Oligo fwd</td><td>CACCCGAGGGTGCCGGTTGGATGA</td><td></td><td></td><td></td><td></td><td></td><td></td><td></td><td></td><td></td><td></td></tr>
<tr><td>Oligo rev</td><td>AAACTCATCCAACCGGCACCCTCG</td><td></td><td></td><td></td><td></td><td></td><td></td><td></td><td></td><td></td><td></td></tr>
<tr><td>Chromosome</td><td>start</td><td>end</td><td>strand</td><td>MM</td><td>target_seq</td><td>PAM</td><td>alignment</td><td>distance</td><td>position</td><td>gene name</td><td>gene id</td></tr>
<tr><td>GU305914.1</td><td>150149</td><td>150171</td><td>+</td><td>0</td><td>CGAGGGTGCCGGTTGGATGA</td><td>GGG</td><td>||||||||[||||||||||||]PAM</td><td>NA</td><td>Intergenic</td><td>NA</td><td>NA</td></tr>
<tr><td></td><td></td><td></td><td></td><td></td><td></td><td></td><td></td><td></td><td></td><td></td><td></td></tr>
<tr><td>T11510</td><td>GAGGGTGCCGGTTGGATGAGGGG</td><td>1000</td><td></td><td></td><td></td><td></td><td></td><td></td><td></td><td></td><td></td></tr>
<tr><td>Oligo fwd</td><td>CACCGAGGGTGCCGGTTGGATGAG</td><td></td><td></td><td></td><td></td><td></td><td></td><td></td><td></td><td></td><td></td></tr>
<tr><td>Oligo rev</td><td>AAACCTCATCCAACCGGCACCCTC</td><td></td><td></td><td></td><td></td><td></td><td></td><td></td><td></td><td></td><td></td></tr>
<tr><td>Chromosome</td><td>start</td><td>end</td><td>strand</td><td>MM</td><td>target_seq</td><td>PAM</td><td>alignment</td><td>distance</td><td>position</td><td>gene name</td><td>gene id</td></tr>
<tr><td>GU305914.1</td><td>150150</td><td>150172</td><td>+</td><td>0</td><td>GAGGGTGCCGGTTGGATGAG</td><td>GGG</td><td>||||||||[||||||||||||]PAM</td><td>NA</td><td>Intergenic</td><td>NA</td><td>NA</td></tr>
<tr><td></td><td></td><td></td><td></td><td></td><td></td><td></td><td></td><td></td><td></td><td></td><td></td></tr>
<tr><td>T11511</td><td>CCGGTTGGATGAGGGGCGCGCGG</td><td>1000</td><td></td><td></td><td></td><td></td><td></td><td></td><td></td><td></td><td></td></tr>
<tr><td>Oligo fwd</td><td>CACCCCGGTTGGATGAGGGGCGCG</td><td></td><td></td><td></td><td></td><td></td><td></td><td></td><td></td><td></td><td></td></tr>
<tr><td>Oligo rev</td><td>AAACCGCGCCCCTCATCCAACCGG</td><td></td><td></td><td></td><td></td><td></td><td></td><td></td><td></td><td></td><td></td></tr>
<tr><td>Chromosome</td><td>start</td><td>end</td><td>strand</td><td>MM</td><td>target_seq</td><td>PAM</td><td>alignment</td><td>distance</td><td>position</td><td>gene name</td><td>gene id</td></tr>
<tr><td>GU305914.1</td><td>150157</td><td>150179</td><td>+</td><td>0</td><td>CCGGTTGGATGAGGGGCGCG</td><td>CGG</td><td>||||||||[||||||||||||]PAM</td><td>NA</td><td>Intergenic</td><td>NA</td><td>NA</td></tr>
<tr><td></td><td></td><td></td><td></td><td></td><td></td><td></td><td></td><td></td><td></td><td></td><td></td></tr>
<tr><td>T11512</td><td>TGGATGAGGGGCGCGCGGCTCGG</td><td>1000</td><td></td><td></td><td></td><td></td><td></td><td></td><td></td><td></td><td></td></tr>
<tr><td>Oligo fwd</td><td>CACCTGGATGAGGGGCGCGCGGCT</td><td></td><td></td><td></td><td></td><td></td><td></td><td></td><td></td><td></td><td></td></tr>
<tr><td>Oligo rev</td><td>AAACAGCCGCGCGCCCCTCATCCA</td><td></td><td></td><td></td><td></td><td></td><td></td><td></td><td></td><td></td><td></td></tr>
<tr><td>Chromosome</td><td>start</td><td>end</td><td>strand</td><td>MM</td><td>target_seq</td><td>PAM</td><td>alignment</td><td>distance</td><td>position</td><td>gene name</td><td>gene id</td></tr>
<tr><td>GU305914.1</td><td>150162</td><td>150184</td><td>+</td><td>0</td><td>TGGATGAGGGGCGCGCGGCT</td><td>CGG</td><td>||||||||[||||||||||||]PAM</td><td>NA</td><td>Intergenic</td><td>NA</td><td>NA</td></tr>
<tr><td></td><td></td><td></td><td></td><td></td><td></td><td></td><td></td><td></td><td></td><td></td><td></td></tr>
<tr><td>T11513</td><td>TGAGGGGCGCGCGGCTCGGCAGG</td><td>1000</td><td></td><td></td><td></td><td></td><td></td><td></td><td></td><td></td><td></td></tr>
<tr><td>Oligo fwd</td><td>CACCTGAGGGGCGCGCGGCTCGGC</td><td></td><td></td><td></td><td></td><td></td><td></td><td></td><td></td><td></td><td></td></tr>
<tr><td>Oligo rev</td><td>AAACGCCGAGCCGCGCGCCCCTCA</td><td></td><td></td><td></td><td></td><td></td><td></td><td></td><td></td><td></td><td></td></tr>
<tr><td>Chromosome</td><td>start</td><td>end</td><td>strand</td><td>MM</td><td>target_seq</td><td>PAM</td><td>alignment</td><td>distance</td><td>position</td><td>gene name</td><td>gene id</td></tr>
<tr><td>GU305914.1</td><td>150166</td><td>150188</td><td>+</td><td>0</td><td>TGAGGGGCGCGCGGCTCGGC</td><td>AGG</td><td>||||||||[||||||||||||]PAM</td><td>NA</td><td>Intergenic</td><td>NA</td><td>NA</td></tr>
<tr><td></td><td></td><td></td><td></td><td></td><td></td><td></td><td></td><td></td><td></td><td></td><td></td></tr>
<tr><td>T11514</td><td>GGGGCGCGCGGCTCGGCAGGAGG</td><td>1000</td><td></td><td></td><td></td><td></td><td></td><td></td><td></td><td></td><td></td></tr>
<tr><td>Oligo fwd</td><td>CACCGGGGCGCGCGGCTCGGCAGG</td><td></td><td></td><td></td><td></td><td></td><td></td><td></td><td></td><td></td><td></td></tr>
<tr><td>Oligo rev</td><td>AAACCCTGCCGAGCCGCGCGCCCC</td><td></td><td></td><td></td><td></td><td></td><td></td><td></td><td></td><td></td><td></td></tr>
<tr><td>Chromosome</td><td>start</td><td>end</td><td>strand</td><td>MM</td><td>target_seq</td><td>PAM</td><td>alignment</td><td>distance</td><td>position</td><td>gene name</td><td>gene id</td></tr>
<tr><td>GU305914.1</td><td>150169</td><td>150191</td><td>+</td><td>0</td><td>GGGGCGCGCGGCTCGGCAGG</td><td>AGG</td><td>||||||||[||||||||||||]PAM</td><td>NA</td><td>Intergenic</td><td>NA</td><td>NA</td></tr>
<tr><td></td><td></td><td></td><td></td><td></td><td></td><td></td><td></td><td></td><td></td><td></td><td></td></tr>
<tr><td>T11515</td><td>GCGCGCGGCTCGGCAGGAGGCGG</td><td>1000</td><td></td><td></td><td></td><td></td><td></td><td></td><td></td><td></td><td></td></tr>
<tr><td>Oligo fwd</td><td>CACCGCGCGCGGCTCGGCAGGAGG</td><td></td><td></td><td></td><td></td><td></td><td></td><td></td><td></td><td></td><td></td></tr>
<tr><td>Oligo rev</td><td>AAACCCTCCTGCCGAGCCGCGCGC</td><td></td><td></td><td></td><td></td><td></td><td></td><td></td><td></td><td></td><td></td></tr>
<tr><td>Chromosome</td><td>start</td><td>end</td><td>strand</td><td>MM</td><td>target_seq</td><td>PAM</td><td>alignment</td><td>distance</td><td>position</td><td>gene name</td><td>gene id</td></tr>
<tr><td>GU305914.1</td><td>150172</td><td>150194</td><td>+</td><td>0</td><td>GCGCGCGGCTCGGCAGGAGG</td><td>CGG</td><td>||||||||[||||||||||||]PAM</td><td>NA</td><td>Intergenic</td><td>NA</td><td>NA</td></tr>
<tr><td></td><td></td><td></td><td></td><td></td><td></td><td></td><td></td><td></td><td></td><td></td><td></td></tr>
<tr><td>T11516</td><td>CGCGCGGCTCGGCAGGAGGCGGG</td><td>1000</td><td></td><td></td><td></td><td></td><td></td><td></td><td></td><td></td><td></td></tr>
<tr><td>Oligo fwd</td><td>CACCCGCGCGGCTCGGCAGGAGGC</td><td></td><td></td><td></td><td></td><td></td><td></td><td></td><td></td><td></td><td></td></tr>
<tr><td>Oligo rev</td><td>AAACGCCTCCTGCCGAGCCGCGCG</td><td></td><td></td><td></td><td></td><td></td><td></td><td></td><td></td><td></td><td></td></tr>
<tr><td>Chromosome</td><td>start</td><td>end</td><td>strand</td><td>MM</td><td>target_seq</td><td>PAM</td><td>alignment</td><td>distance</td><td>position</td><td>gene name</td><td>gene id</td></tr>
<tr><td>GU305914.1</td><td>150173</td><td>150195</td><td>+</td><td>0</td><td>CGCGCGGCTCGGCAGGAGGC</td><td>GGG</td><td>||||||||[||||||||||||]PAM</td><td>NA</td><td>Intergenic</td><td>NA</td><td>NA</td></tr>
<tr><td></td><td></td><td></td><td></td><td></td><td></td><td></td><td></td><td></td><td></td><td></td><td></td></tr>
<tr><td>T11517</td><td>AGGCGGGCTTGATCCACAGACGG</td><td>1000</td><td></td><td></td><td></td><td></td><td></td><td></td><td></td><td></td><td></td></tr>
<tr><td>Oligo fwd</td><td>CACCAGGCGGGCTTGATCCACAGA</td><td></td><td></td><td></td><td></td><td></td><td></td><td></td><td></td><td></td><td></td></tr>
<tr><td>Oligo rev</td><td>AAACTCTGTGGATCAAGCCCGCCT</td><td></td><td></td><td></td><td></td><td></td><td></td><td></td><td></td><td></td><td></td></tr>
<tr><td>Chromosome</td><td>start</td><td>end</td><td>strand</td><td>MM</td><td>target_seq</td><td>PAM</td><td>alignment</td><td>distance</td><td>position</td><td>gene name</td><td>gene id</td></tr>
<tr><td>GU305914.1</td><td>150189</td><td>150211</td><td>+</td><td>0</td><td>AGGCGGGCTTGATCCACAGA</td><td>CGG</td><td>||||||||[||||||||||||]PAM</td><td>NA</td><td>Intergenic</td><td>NA</td><td>NA</td></tr>
<tr><td></td><td></td><td></td><td></td><td></td><td></td><td></td><td></td><td></td><td></td><td></td><td></td></tr>
<tr><td>T11518</td><td>CTTGATCCACAGACGGCGTCAGG</td><td>1000</td><td></td><td></td><td></td><td></td><td></td><td></td><td></td><td></td><td></td></tr>
<tr><td>Oligo fwd</td><td>CACCCTTGATCCACAGACGGCGTC</td><td></td><td></td><td></td><td></td><td></td><td></td><td></td><td></td><td></td><td></td></tr>
<tr><td>Oligo rev</td><td>AAACGACGCCGTCTGTGGATCAAG</td><td></td><td></td><td></td><td></td><td></td><td></td><td></td><td></td><td></td><td></td></tr>
<tr><td>Chromosome</td><td>start</td><td>end</td><td>strand</td><td>MM</td><td>target_seq</td><td>PAM</td><td>alignment</td><td>distance</td><td>position</td><td>gene name</td><td>gene id</td></tr>
<tr><td>GU305914.1</td><td>150196</td><td>150218</td><td>+</td><td>0</td><td>CTTGATCCACAGACGGCGTC</td><td>AGG</td><td>||||||||[||||||||||||]PAM</td><td>NA</td><td>Intergenic</td><td>NA</td><td>NA</td></tr>
</table>

<table>
<tr><td>T11519</td><td>CAGACGGCGTCAGGTCCTGCAGG</td><td>1000</td><td></td><td></td><td></td><td></td><td></td><td></td><td></td><td></td><td></td></tr>
<tr><td>Oligo fwd</td><td>CACCCAGACGGCGTCAGGTCCTGC</td><td></td><td></td><td></td><td></td><td></td><td></td><td></td><td></td><td></td><td></td></tr>
<tr><td>Oligo rev</td><td>AAACGCAGGACCTGACGCCGTCTG</td><td></td><td></td><td></td><td></td><td></td><td></td><td></td><td></td><td></td><td></td></tr>
<tr><td>Chromosome</td><td>start</td><td>end</td><td>strand</td><td>MM</td><td>target_seq</td><td>PAM</td><td>alignment</td><td>distance</td><td>position</td><td>gene name</td><td>gene id</td></tr>
<tr><td>GU305914.1</td><td>150205</td><td>150227</td><td>+</td><td>0</td><td>CAGACGGCGTCAGGTCCTGC</td><td>AGG</td><td>||||||||[||||||||||||]PAM</td><td>NA</td><td>Intergenic</td><td>NA</td><td>NA</td></tr>
<tr><td>T11520</td><td>TCCGATCTTCGACTTGCTGTCGG</td><td>1000</td><td></td><td></td><td></td><td></td><td></td><td></td><td></td><td></td><td></td></tr>
<tr><td>Oligo fwd</td><td>CACCTCCGATCTTCGACTTGCTGT</td><td></td><td></td><td></td><td></td><td></td><td></td><td></td><td></td><td></td><td></td></tr>
<tr><td>Oligo rev</td><td>AAACACAGCAAGTCGAAGATCGGA</td><td></td><td></td><td></td><td></td><td></td><td></td><td></td><td></td><td></td><td></td></tr>
<tr><td>Chromosome</td><td>start</td><td>end</td><td>strand</td><td>MM</td><td>target_seq</td><td>PAM</td><td>alignment</td><td>distance</td><td>position</td><td>gene name</td><td>gene id</td></tr>
<tr><td>GU305914.1</td><td>150251</td><td>150273</td><td>+</td><td>0</td><td>TCCGATCTTCGACTTGCTGT</td><td>CGG</td><td>||||||||[||||||||||||]PAM</td><td>NA</td><td>Intergenic</td><td>NA</td><td>NA</td></tr>
<tr><td>T11521</td><td>TGCTGTCGGACACGTTCAGCAGG</td><td>1000</td><td></td><td></td><td></td><td></td><td></td><td></td><td></td><td></td><td></td></tr>
<tr><td>Oligo fwd</td><td>CACCTGCTGTCGGACACGTTCAGC</td><td></td><td></td><td></td><td></td><td></td><td></td><td></td><td></td><td></td><td></td></tr>
<tr><td>Oligo rev</td><td>AAACGCTGAACGTGTCCGACAGCA</td><td></td><td></td><td></td><td></td><td></td><td></td><td></td><td></td><td></td><td></td></tr>
<tr><td>Chromosome</td><td>start</td><td>end</td><td>strand</td><td>MM</td><td>target_seq</td><td>PAM</td><td>alignment</td><td>distance</td><td>position</td><td>gene name</td><td>gene id</td></tr>
<tr><td>GU305914.1</td><td>150265</td><td>150287</td><td>+</td><td>0</td><td>TGCTGTCGGACACGTTCAGC</td><td>AGG</td><td>||||||||[||||||||||||]PAM</td><td>NA</td><td>Intergenic</td><td>NA</td><td>NA</td></tr>
<tr><td>T11522</td><td>GTTCAGCAGGAACTTGACGATGG</td><td>1000</td><td></td><td></td><td></td><td></td><td></td><td></td><td></td><td></td><td></td></tr>
<tr><td>Oligo fwd</td><td>CACCGTTCAGCAGGAACTTGACGA</td><td></td><td></td><td></td><td></td><td></td><td></td><td></td><td></td><td></td><td></td></tr>
<tr><td>Oligo rev</td><td>AAACTCGTCAAGTTCCTGCTGAAC</td><td></td><td></td><td></td><td></td><td></td><td></td><td></td><td></td><td></td><td></td></tr>
<tr><td>Chromosome</td><td>start</td><td>end</td><td>strand</td><td>MM</td><td>target_seq</td><td>PAM</td><td>alignment</td><td>distance</td><td>position</td><td>gene name</td><td>gene id</td></tr>
<tr><td>GU305914.1</td><td>150278</td><td>150300</td><td>+</td><td>0</td><td>GTTCAGCAGGAACTTGACGA</td><td>TGG</td><td>||||||||[||||||||||||]PAM</td><td>NA</td><td>Intergenic</td><td>NA</td><td>NA</td></tr>
<tr><td>T11523</td><td>GAACTTGACGATGGACGTCTTGG</td><td>1000</td><td></td><td></td><td></td><td></td><td></td><td></td><td></td><td></td><td></td></tr>
<tr><td>Oligo fwd</td><td>CACCGAACTTGACGATGGACGTCT</td><td></td><td></td><td></td><td></td><td></td><td></td><td></td><td></td><td></td><td></td></tr>
<tr><td>Oligo rev</td><td>AAACAGACGTCCATCGTCAAGTTC</td><td></td><td></td><td></td><td></td><td></td><td></td><td></td><td></td><td></td><td></td></tr>
<tr><td>Chromosome</td><td>start</td><td>end</td><td>strand</td><td>MM</td><td>target_seq</td><td>PAM</td><td>alignment</td><td>distance</td><td>position</td><td>gene name</td><td>gene id</td></tr>
<tr><td>GU305914.1</td><td>150287</td><td>150309</td><td>+</td><td>0</td><td>GAACTTGACGATGGACGTCT</td><td>TGG</td><td>||||||||[||||||||||||]PAM</td><td>NA</td><td>Intergenic</td><td>NA</td><td>NA</td></tr>
<tr><td>T11524</td><td>ACGTCTTGGCGTCCACCTGCCGG</td><td>1000</td><td></td><td></td><td></td><td></td><td></td><td></td><td></td><td></td><td></td></tr>
<tr><td>Oligo fwd</td><td>CACCACGTCTTGGCGTCCACCTGC</td><td></td><td></td><td></td><td></td><td></td><td></td><td></td><td></td><td></td><td></td></tr>
<tr><td>Oligo rev</td><td>AAACGCAGGTGGACGCCAAGACGT</td><td></td><td></td><td></td><td></td><td></td><td></td><td></td><td></td><td></td><td></td></tr>
<tr><td>Chromosome</td><td>start</td><td>end</td><td>strand</td><td>MM</td><td>target_seq</td><td>PAM</td><td>alignment</td><td>distance</td><td>position</td><td>gene name</td><td>gene id</td></tr>
<tr><td>GU305914.1</td><td>150301</td><td>150323</td><td>+</td><td>0</td><td>ACGTCTTGGCGTCCACCTGC</td><td>CGG</td><td>||||||||[||||||||||||]PAM</td><td>NA</td><td>Intergenic</td><td>NA</td><td>NA</td></tr>
<tr><td>T11525</td><td>GCTTCGTGTTCATCTCGTGCCGG</td><td>1000</td><td></td><td></td><td></td><td></td><td></td><td></td><td></td><td></td><td></td></tr>
<tr><td>Oligo fwd</td><td>CACCGCTTCGTGTTCATCTCGTGC</td><td></td><td></td><td></td><td></td><td></td><td></td><td></td><td></td><td></td><td></td></tr>
<tr><td>Oligo rev</td><td>AAACGCACGAGATGAACACGAAGC</td><td></td><td></td><td></td><td></td><td></td><td></td><td></td><td></td><td></td><td></td></tr>
<tr><td>Chromosome</td><td>start</td><td>end</td><td>strand</td><td>MM</td><td>target_seq</td><td>PAM</td><td>alignment</td><td>distance</td><td>position</td><td>gene name</td><td>gene id</td></tr>
<tr><td>GU305914.1</td><td>150352</td><td>150374</td><td>+</td><td>0</td><td>GCTTCGTGTTCATCTCGTGC</td><td>CGG</td><td>||||||||[||||||||||||]PAM</td><td>NA</td><td>Intergenic</td><td>NA</td><td>NA</td></tr>
<tr><td>T11526</td><td>GTTCATCTCGTGCCGGCTCGTGG</td><td>1000</td><td></td><td></td><td></td><td></td><td></td><td></td><td></td><td></td><td></td></tr>
<tr><td>Oligo fwd</td><td>CACCGTTCATCTCGTGCCGGCTCG</td><td></td><td></td><td></td><td></td><td></td><td></td><td></td><td></td><td></td><td></td></tr>
<tr><td>Oligo rev</td><td>AAACCGAGCCGGCACGAGATGAAC</td><td></td><td></td><td></td><td></td><td></td><td></td><td></td><td></td><td></td><td></td></tr>
<tr><td>Chromosome</td><td>start</td><td>end</td><td>strand</td><td>MM</td><td>target_seq</td><td>PAM</td><td>alignment</td><td>distance</td><td>position</td><td>gene name</td><td>gene id</td></tr>
<tr><td>GU305914.1</td><td>150359</td><td>150381</td><td>+</td><td>0</td><td>GTTCATCTCGTGCCGGCTCG</td><td>TGG</td><td>||||||||[||||||||||||]PAM</td><td>NA</td><td>Intergenic</td><td>NA</td><td>NA</td></tr>
<tr><td>T11527</td><td>CTCGTGGTCACCACCTTGATCGG</td><td>1000</td><td></td><td></td><td></td><td></td><td></td><td></td><td></td><td></td><td></td></tr>
<tr><td>Oligo fwd</td><td>CACCCTCGTGGTCACCACCTTGAT</td><td></td><td></td><td></td><td></td><td></td><td></td><td></td><td></td><td></td><td></td></tr>
<tr><td>Oligo rev</td><td>AAACATCAAGGTGGTGACCACGAG</td><td></td><td></td><td></td><td></td><td></td><td></td><td></td><td></td><td></td><td></td></tr>
<tr><td>Chromosome</td><td>start</td><td>end</td><td>strand</td><td>MM</td><td>target_seq</td><td>PAM</td><td>alignment</td><td>distance</td><td>position</td><td>gene name</td><td>gene id</td></tr>
<tr><td>GU305914.1</td><td>150375</td><td>150397</td><td>+</td><td>0</td><td>CTCGTGGTCACCACCTTGAT</td><td>CGG</td><td>||||||||[||||||||||||]PAM</td><td>NA</td><td>Intergenic</td><td>NA</td><td>NA</td></tr>
<tr><td>T11528</td><td>TCGTGGTCACCACCTTGATCGGG</td><td>1000</td><td></td><td></td><td></td><td></td><td></td><td></td><td></td><td></td><td></td></tr>
<tr><td>Oligo fwd</td><td>CACCTCGTGGTCACCACCTTGATC</td><td></td><td></td><td></td><td></td><td></td><td></td><td></td><td></td><td></td><td></td></tr>
<tr><td>Oligo rev</td><td>AAACGATCAAGGTGGTGACCACGA</td><td></td><td></td><td></td><td></td><td></td><td></td><td></td><td></td><td></td><td></td></tr>
<tr><td>Chromosome</td><td>start</td><td>end</td><td>strand</td><td>MM</td><td>target_seq</td><td>PAM</td><td>alignment</td><td>distance</td><td>position</td><td>gene name</td><td>gene id</td></tr>
<tr><td>GU305914.1</td><td>150376</td><td>150398</td><td>+</td><td>0</td><td>TCGTGGTCACCACCTTGATC</td><td>GGG</td><td>||||||||[||||||||||||]PAM</td><td>NA</td><td>Intergenic</td><td>NA</td><td>NA</td></tr>
<tr><td>T11529</td><td>GGTCACCACCTTGATCGGGTTGG</td><td>1000</td><td></td><td></td><td></td><td></td><td></td><td></td><td></td><td></td><td></td></tr>
<tr><td>Oligo fwd</td><td>CACCGGTCACCACCTTGATCGGGT</td><td></td><td></td><td></td><td></td><td></td><td></td><td></td><td></td><td></td><td></td></tr>
<tr><td>Oligo rev</td><td>AAACACCCGATCAAGGTGGTGACC</td><td></td><td></td><td></td><td></td><td></td><td></td><td></td><td></td><td></td><td></td></tr>
<tr><td>Chromosome</td><td>start</td><td>end</td><td>strand</td><td>MM</td><td>target_seq</td><td>PAM</td><td>alignment</td><td>distance</td><td>position</td><td>gene name</td><td>gene id</td></tr>
<tr><td>GU305914.1</td><td>150380</td><td>150402</td><td>+</td><td>0</td><td>GGTCACCACCTTGATCGGGT</td><td>TGG</td><td>||||||||[||||||||||||]PAM</td><td>NA</td><td>Intergenic</td><td>NA</td><td>NA</td></tr>
<tr><td>T11530</td><td>TCGGGTTGGTGTTCAGCAGCTGG</td><td>1000</td><td></td><td></td><td></td><td></td><td></td><td></td><td></td><td></td><td></td></tr>
<tr><td>Oligo fwd</td><td>CACCTCGGGTTGGTGTTCAGCAGC</td><td></td><td></td><td></td><td></td><td></td><td></td><td></td><td></td><td></td><td></td></tr>
<tr><td>Oligo rev</td><td>AAACGCTGCTGAACACCAACCCGA</td><td></td><td></td><td></td><td></td><td></td><td></td><td></td><td></td><td></td><td></td></tr>
<tr><td>Chromosome</td><td>start</td><td>end</td><td>strand</td><td>MM</td><td>target_seq</td><td>PAM</td><td>alignment</td><td>distance</td><td>position</td><td>gene name</td><td>gene id</td></tr>
<tr><td>GU305914.1</td><td>150394</td><td>150416</td><td>+</td><td>0</td><td>TCGGGTTGGTGTTCAGCAGC</td><td>TGG</td><td>||||||||[||||||||||||]PAM</td><td>NA</td><td>Intergenic</td><td>NA</td><td>NA</td></tr>
<tr><td>T11531</td><td>GCAGCTGGCAGATCTTCTTGTGG</td><td>1000</td><td></td><td></td><td></td><td></td><td></td><td></td><td></td><td></td><td></td></tr>
<tr><td>Oligo fwd</td><td>CACCGCAGCTGGCAGATCTTCTTG</td><td></td><td></td><td></td><td></td><td></td><td></td><td></td><td></td><td></td><td></td></tr>
<tr><td>Oligo rev</td><td>AAACCAAGAAGATCTGCCAGCTGC</td><td></td><td></td><td></td><td></td><td></td><td></td><td></td><td></td><td></td><td></td></tr>
<tr><td>Chromosome</td><td>start</td><td>end</td><td>strand</td><td>MM</td><td>target_seq</td><td>PAM</td><td>alignment</td><td>distance</td><td>position</td><td>gene name</td><td>gene id</td></tr>
<tr><td>GU305914.1</td><td>150409</td><td>150431</td><td>+</td><td>0</td><td>GCAGCTGGCAGATCTTCTTG</td><td>TGG</td><td>||||||||[||||||||||||]PAM</td><td>NA</td><td>Intergenic</td><td>NA</td><td>NA</td></tr>
<tr><td>T11532</td><td>TGTGTCGTTTGCGCTCGTAGCGG</td><td>1000</td><td></td><td></td><td></td><td></td><td></td><td></td><td></td><td></td><td></td></tr>
<tr><td>Oligo fwd</td><td>CACCTGTGTCGTTTGCGCTCGTAG</td><td></td><td></td><td></td><td></td><td></td><td></td><td></td><td></td><td></td><td></td></tr>
<tr><td>Oligo rev</td><td>AAACCTACGAGCGCAAACGACACA</td><td></td><td></td><td></td><td></td><td></td><td></td><td></td><td></td><td></td><td></td></tr>
<tr><td>Chromosome</td><td>start</td><td>end</td><td>strand</td><td>MM</td><td>target_seq</td><td>PAM</td><td>alignment</td><td>distance</td><td>position</td><td>gene name</td><td>gene id</td></tr>
<tr><td>GU305914.1</td><td>150445</td><td>150467</td><td>+</td><td>0</td><td>TGTGTCGTTTGCGCTCGTAG</td><td>CGG</td><td>||||||||[||||||||||||]PAM</td><td>NA</td><td>Intergenic</td><td>NA</td><td>NA</td></tr>
<tr><td>T11533</td><td>GTTTGCGCTCGTAGCGGATCCGG</td><td>1000</td><td></td><td></td><td></td><td></td><td></td><td></td><td></td><td></td><td></td></tr>
<tr><td>Oligo fwd</td><td>CACCGTTTGCGCTCGTAGCGGATC</td><td></td><td></td><td></td><td></td><td></td><td></td><td></td><td></td><td></td><td></td></tr>
<tr><td>Oligo rev</td><td>AAACGATCCGCTACGAGCGCAAAC</td><td></td><td></td><td></td><td></td><td></td><td></td><td></td><td></td><td></td><td></td></tr>
<tr><td>Chromosome</td><td>start</td><td>end</td><td>strand</td><td>MM</td><td>target_seq</td><td>PAM</td><td>alignment</td><td>distance</td><td>position</td><td>gene name</td><td>gene id</td></tr>
<tr><td>GU305914.1</td><td>150451</td><td>150473</td><td>+</td><td>0</td><td>GTTTGCGCTCGTAGCGGATC</td><td>CGG</td><td>||||||||[||||||||||||]PAM</td><td>NA</td><td>Intergenic</td><td>NA</td><td>NA</td></tr>
</table>

<table>
<tr><td>T11534</td><td>TTTGCGCTCGTAGCGGATCCGGG</td><td>1000</td><td></td><td></td><td></td><td></td><td></td><td></td><td></td><td></td><td></td></tr>
<tr><td>Oligo fwd</td><td>CACCTTTGCGCTCGTAGCGGATCC</td><td></td><td></td><td></td><td></td><td></td><td></td><td></td><td></td><td></td><td></td></tr>
<tr><td>Oligo rev</td><td>AAACGGATCCGCTACGAGCGCAAA</td><td></td><td></td><td></td><td></td><td></td><td></td><td></td><td></td><td></td><td></td></tr>
<tr><td>Chromosome</td><td>start</td><td>end</td><td>strand</td><td>MM</td><td>target_seq</td><td>PAM</td><td>alignment</td><td>distance</td><td>position</td><td>gene name</td><td>gene id</td></tr>
<tr><td>GU305914.1</td><td>150452</td><td>150474</td><td>+</td><td>0</td><td>TTTGCGCTCGTAGCGGATCC</td><td>GGG</td><td>||||||||[||||||||||||]PAM</td><td>NA</td><td>Intergenic</td><td>NA</td><td>NA</td></tr>
<tr><td>T11535</td><td>TCGTAGCGGATCCGGGCCAGCGG</td><td>1000</td><td></td><td></td><td></td><td></td><td></td><td></td><td></td><td></td><td></td></tr>
<tr><td>Oligo fwd</td><td>CACCTCGTAGCGGATCCGGGCCAG</td><td></td><td></td><td></td><td></td><td></td><td></td><td></td><td></td><td></td><td></td></tr>
<tr><td>Oligo rev</td><td>AAACCTGGCCCGGATCCGCTACGA</td><td></td><td></td><td></td><td></td><td></td><td></td><td></td><td></td><td></td><td></td></tr>
<tr><td>Chromosome</td><td>start</td><td>end</td><td>strand</td><td>MM</td><td>target_seq</td><td>PAM</td><td>alignment</td><td>distance</td><td>position</td><td>gene name</td><td>gene id</td></tr>
<tr><td>GU305914.1</td><td>150459</td><td>150481</td><td>+</td><td>0</td><td>TCGTAGCGGATCCGGGCCAG</td><td>CGG</td><td>||||||||[||||||||||||]PAM</td><td>NA</td><td>Intergenic</td><td>NA</td><td>NA</td></tr>
<tr><td>T11536</td><td>TCCGGGCCAGCGGCGTCGTCAGG</td><td>1000</td><td></td><td></td><td></td><td></td><td></td><td></td><td></td><td></td><td></td></tr>
<tr><td>Oligo fwd</td><td>CACCTCCGGGCCAGCGGCGTCGTC</td><td></td><td></td><td></td><td></td><td></td><td></td><td></td><td></td><td></td><td></td></tr>
<tr><td>Oligo rev</td><td>AAACGACGACGCCGCTGGCCCGGA</td><td></td><td></td><td></td><td></td><td></td><td></td><td></td><td></td><td></td><td></td></tr>
<tr><td>Chromosome</td><td>start</td><td>end</td><td>strand</td><td>MM</td><td>target_seq</td><td>PAM</td><td>alignment</td><td>distance</td><td>position</td><td>gene name</td><td>gene id</td></tr>
<tr><td>GU305914.1</td><td>150469</td><td>150491</td><td>+</td><td>0</td><td>TCCGGGCCAGCGGCGTCGTC</td><td>AGG</td><td>||||||||[||||||||||||]PAM</td><td>NA</td><td>Intergenic</td><td>NA</td><td>NA</td></tr>
<tr><td>T11537</td><td>GGATCAGCCGCTCCTCCTCGAGG</td><td>1000</td><td></td><td></td><td></td><td></td><td></td><td></td><td></td><td></td><td></td></tr>
<tr><td>Oligo fwd</td><td>CACCGGATCAGCCGCTCCTCCTCG</td><td></td><td></td><td></td><td></td><td></td><td></td><td></td><td></td><td></td><td></td></tr>
<tr><td>Oligo rev</td><td>AAACCGAGGAGGAGCGGCTGATCC</td><td></td><td></td><td></td><td></td><td></td><td></td><td></td><td></td><td></td><td></td></tr>
<tr><td>Chromosome</td><td>start</td><td>end</td><td>strand</td><td>MM</td><td>target_seq</td><td>PAM</td><td>alignment</td><td>distance</td><td>position</td><td>gene name</td><td>gene id</td></tr>
<tr><td>GU305914.1</td><td>150490</td><td>150512</td><td>+</td><td>0</td><td>GGATCAGCCGCTCCTCCTCG</td><td>AGG</td><td>||||||||[||||||||||||]PAM</td><td>NA</td><td>Intergenic</td><td>NA</td><td>NA</td></tr>
<tr><td>T11538</td><td>CCGCTCCTCCTCGAGGATCGTGG</td><td>1000</td><td></td><td></td><td></td><td></td><td></td><td></td><td></td><td></td><td></td></tr>
<tr><td>Oligo fwd</td><td>CACCCCGCTCCTCCTCGAGGATCG</td><td></td><td></td><td></td><td></td><td></td><td></td><td></td><td></td><td></td><td></td></tr>
<tr><td>Oligo rev</td><td>AAACCGATCCTCGAGGAGGAGCGG</td><td></td><td></td><td></td><td></td><td></td><td></td><td></td><td></td><td></td><td></td></tr>
<tr><td>Chromosome</td><td>start</td><td>end</td><td>strand</td><td>MM</td><td>target_seq</td><td>PAM</td><td>alignment</td><td>distance</td><td>position</td><td>gene name</td><td>gene id</td></tr>
<tr><td>GU305914.1</td><td>150497</td><td>150519</td><td>+</td><td>0</td><td>CCGCTCCTCCTCGAGGATCG</td><td>TGG</td><td>||||||||[||||||||||||]PAM</td><td>NA</td><td>Intergenic</td><td>NA</td><td>NA</td></tr>
<tr><td>T11539</td><td>CGCTCCTCCTCGAGGATCGTGGG</td><td>1000</td><td></td><td></td><td></td><td></td><td></td><td></td><td></td><td></td><td></td></tr>
<tr><td>Oligo fwd</td><td>CACCCGCTCCTCCTCGAGGATCGT</td><td></td><td></td><td></td><td></td><td></td><td></td><td></td><td></td><td></td><td></td></tr>
<tr><td>Oligo rev</td><td>AAACACGATCCTCGAGGAGGAGCG</td><td></td><td></td><td></td><td></td><td></td><td></td><td></td><td></td><td></td><td></td></tr>
<tr><td>Chromosome</td><td>start</td><td>end</td><td>strand</td><td>MM</td><td>target_seq</td><td>PAM</td><td>alignment</td><td>distance</td><td>position</td><td>gene name</td><td>gene id</td></tr>
<tr><td>GU305914.1</td><td>150498</td><td>150520</td><td>+</td><td>0</td><td>CGCTCCTCCTCGAGGATCGT</td><td>GGG</td><td>||||||||[||||||||||||]PAM</td><td>NA</td><td>Intergenic</td><td>NA</td><td>NA</td></tr>
<tr><td>T11540</td><td>TGTGCCCGTTGCTCTTCAGCCGG</td><td>1000</td><td></td><td></td><td></td><td></td><td></td><td></td><td></td><td></td><td></td></tr>
<tr><td>Oligo fwd</td><td>CACCTGTGCCCGTTGCTCTTCAGC</td><td></td><td></td><td></td><td></td><td></td><td></td><td></td><td></td><td></td><td></td></tr>
<tr><td>Oligo rev</td><td>AAACGCTGAAGAGCAACGGGCACA</td><td></td><td></td><td></td><td></td><td></td><td></td><td></td><td></td><td></td><td></td></tr>
<tr><td>Chromosome</td><td>start</td><td>end</td><td>strand</td><td>MM</td><td>target_seq</td><td>PAM</td><td>alignment</td><td>distance</td><td>position</td><td>gene name</td><td>gene id</td></tr>
<tr><td>GU305914.1</td><td>150571</td><td>150593</td><td>+</td><td>0</td><td>TGTGCCCGTTGCTCTTCAGC</td><td>CGG</td><td>||||||||[||||||||||||]PAM</td><td>NA</td><td>Intergenic</td><td>NA</td><td>NA</td></tr>
<tr><td>T11541</td><td>GCCCGTTGCTCTTCAGCCGGTGG</td><td>1000</td><td></td><td></td><td></td><td></td><td></td><td></td><td></td><td></td><td></td></tr>
<tr><td>Oligo fwd</td><td>CACCGCCCGTTGCTCTTCAGCCGG</td><td></td><td></td><td></td><td></td><td></td><td></td><td></td><td></td><td></td><td></td></tr>
<tr><td>Oligo rev</td><td>AAACCCGGCTGAAGAGCAACGGGC</td><td></td><td></td><td></td><td></td><td></td><td></td><td></td><td></td><td></td><td></td></tr>
<tr><td>Chromosome</td><td>start</td><td>end</td><td>strand</td><td>MM</td><td>target_seq</td><td>PAM</td><td>alignment</td><td>distance</td><td>position</td><td>gene name</td><td>gene id</td></tr>
<tr><td>GU305914.1</td><td>150574</td><td>150596</td><td>+</td><td>0</td><td>GCCCGTTGCTCTTCAGCCGG</td><td>TGG</td><td>||||||||[||||||||||||]PAM</td><td>NA</td><td>Intergenic</td><td>NA</td><td>NA</td></tr>
<tr><td>T11542</td><td>TGCTCTTCAGCCGGTGGATGCGG</td><td>1000</td><td></td><td></td><td></td><td></td><td></td><td></td><td></td><td></td><td></td></tr>
<tr><td>Oligo fwd</td><td>CACCTGCTCTTCAGCCGGTGGATG</td><td></td><td></td><td></td><td></td><td></td><td></td><td></td><td></td><td></td><td></td></tr>
<tr><td>Oligo rev</td><td>AAACCATCCACCGGCTGAAGAGCA</td><td></td><td></td><td></td><td></td><td></td><td></td><td></td><td></td><td></td><td></td></tr>
<tr><td>Chromosome</td><td>start</td><td>end</td><td>strand</td><td>MM</td><td>target_seq</td><td>PAM</td><td>alignment</td><td>distance</td><td>position</td><td>gene name</td><td>gene id</td></tr>
<tr><td>GU305914.1</td><td>150580</td><td>150602</td><td>+</td><td>0</td><td>TGCTCTTCAGCCGGTGGATG</td><td>CGG</td><td>||||||||[||||||||||||]PAM</td><td>NA</td><td>Intergenic</td><td>NA</td><td>NA</td></tr>
<tr><td>GU305914.1</td><td>229733</td><td>229755</td><td>-</td><td>4</td><td>GGCTCGTCAGGCGGCGGATG</td><td>TGG</td><td>-||||-||[||-|||-|||||]PAM</td><td>NA</td><td>Intergenic</td><td>NA</td><td>NA</td></tr>
<tr><td>T11543</td><td>CAGCCGGTGGATGCGGATGTCGG</td><td>1000</td><td></td><td></td><td></td><td></td><td></td><td></td><td></td><td></td><td></td></tr>
<tr><td>Oligo fwd</td><td>CACCCAGCCGGTGGATGCGGATGT</td><td></td><td></td><td></td><td></td><td></td><td></td><td></td><td></td><td></td><td></td></tr>
<tr><td>Oligo rev</td><td>AAACACATCCGCATCCACCGGCTG</td><td></td><td></td><td></td><td></td><td></td><td></td><td></td><td></td><td></td><td></td></tr>
<tr><td>Chromosome</td><td>start</td><td>end</td><td>strand</td><td>MM</td><td>target_seq</td><td>PAM</td><td>alignment</td><td>distance</td><td>position</td><td>gene name</td><td>gene id</td></tr>
<tr><td>GU305914.1</td><td>150587</td><td>150609</td><td>+</td><td>0</td><td>CAGCCGGTGGATGCGGATGT</td><td>CGG</td><td>||||||||[||||||||||||]PAM</td><td>NA</td><td>Intergenic</td><td>NA</td><td>NA</td></tr>
<tr><td>T11544</td><td>AGCCGGTGGATGCGGATGTCGGG</td><td>1000</td><td></td><td></td><td></td><td></td><td></td><td></td><td></td><td></td><td></td></tr>
<tr><td>Oligo fwd</td><td>CACCAGCCGGTGGATGCGGATGTC</td><td></td><td></td><td></td><td></td><td></td><td></td><td></td><td></td><td></td><td></td></tr>
<tr><td>Oligo rev</td><td>AAACGACATCCGCATCCACCGGCT</td><td></td><td></td><td></td><td></td><td></td><td></td><td></td><td></td><td></td><td></td></tr>
<tr><td>Chromosome</td><td>start</td><td>end</td><td>strand</td><td>MM</td><td>target_seq</td><td>PAM</td><td>alignment</td><td>distance</td><td>position</td><td>gene name</td><td>gene id</td></tr>
<tr><td>GU305914.1</td><td>150588</td><td>150610</td><td>+</td><td>0</td><td>AGCCGGTGGATGCGGATGTC</td><td>GGG</td><td>||||||||[||||||||||||]PAM</td><td>NA</td><td>Intergenic</td><td>NA</td><td>NA</td></tr>
<tr><td>T11545</td><td>ATGCGGATGTCGGGCACGTTCGG</td><td>1000</td><td></td><td></td><td></td><td></td><td></td><td></td><td></td><td></td><td></td></tr>
<tr><td>Oligo fwd</td><td>CACCATGCGGATGTCGGGCACGTT</td><td></td><td></td><td></td><td></td><td></td><td></td><td></td><td></td><td></td><td></td></tr>
<tr><td>Oligo rev</td><td>AAACAACGTGCCCGACATCCGCAT</td><td></td><td></td><td></td><td></td><td></td><td></td><td></td><td></td><td></td><td></td></tr>
<tr><td>Chromosome</td><td>start</td><td>end</td><td>strand</td><td>MM</td><td>target_seq</td><td>PAM</td><td>alignment</td><td>distance</td><td>position</td><td>gene name</td><td>gene id</td></tr>
<tr><td>GU305914.1</td><td>150597</td><td>150619</td><td>+</td><td>0</td><td>ATGCGGATGTCGGGCACGTT</td><td>CGG</td><td>||||||||[||||||||||||]PAM</td><td>NA</td><td>Intergenic</td><td>NA</td><td>NA</td></tr>
<tr><td>T11546</td><td>CGGGCACGTTCGGCACCTCCCGG</td><td>1000</td><td></td><td></td><td></td><td></td><td></td><td></td><td></td><td></td><td></td></tr>
<tr><td>Oligo fwd</td><td>CACCCGGGCACGTTCGGCACCTCC</td><td></td><td></td><td></td><td></td><td></td><td></td><td></td><td></td><td></td><td></td></tr>
<tr><td>Oligo rev</td><td>AAACGGAGGTGCCGAACGTGCCCG</td><td></td><td></td><td></td><td></td><td></td><td></td><td></td><td></td><td></td><td></td></tr>
<tr><td>Chromosome</td><td>start</td><td>end</td><td>strand</td><td>MM</td><td>target_seq</td><td>PAM</td><td>alignment</td><td>distance</td><td>position</td><td>gene name</td><td>gene id</td></tr>
<tr><td>GU305914.1</td><td>150607</td><td>150629</td><td>+</td><td>0</td><td>CGGGCACGTTCGGCACCTCC</td><td>CGG</td><td>||||||||[||||||||||||]PAM</td><td>NA</td><td>Intergenic</td><td>NA</td><td>NA</td></tr>
<tr><td>T11547</td><td>GGGCACGTTCGGCACCTCCCGGG</td><td>1000</td><td></td><td></td><td></td><td></td><td></td><td></td><td></td><td></td><td></td></tr>
<tr><td>Oligo fwd</td><td>CACCGGGCACGTTCGGCACCTCCC</td><td></td><td></td><td></td><td></td><td></td><td></td><td></td><td></td><td></td><td></td></tr>
<tr><td>Oligo rev</td><td>AAACGGGAGGTGCCGAACGTGCCC</td><td></td><td></td><td></td><td></td><td></td><td></td><td></td><td></td><td></td><td></td></tr>
<tr><td>Chromosome</td><td>start</td><td>end</td><td>strand</td><td>MM</td><td>target_seq</td><td>PAM</td><td>alignment</td><td>distance</td><td>position</td><td>gene name</td><td>gene id</td></tr>
<tr><td>GU305914.1</td><td>150608</td><td>150630</td><td>+</td><td>0</td><td>GGGCACGTTCGGCACCTCCC</td><td>GGG</td><td>||||||||[||||||||||||]PAM</td><td>NA</td><td>Intergenic</td><td>NA</td><td>NA</td></tr>
<tr><td>T11548</td><td>CGCAGCGTCCGCGCGCGCAGCGG</td><td>1000</td><td></td><td></td><td></td><td></td><td></td><td></td><td></td><td></td><td></td></tr>
<tr><td>Oligo fwd</td><td>CACCCGCAGCGTCCGCGCGCGCAG</td><td></td><td></td><td></td><td></td><td></td><td></td><td></td><td></td><td></td><td></td></tr>
<tr><td>Oligo rev</td><td>AAACCTGCGCGCGCGGACGCTGCG</td><td></td><td></td><td></td><td></td><td></td><td></td><td></td><td></td><td></td><td></td></tr>
</table>

<table>
<tr><td>Chromosome</td><td>start</td><td>end</td><td>strand</td><td>MM</td><td>target_seq</td><td>PAM</td><td>alignment</td><td>distance</td><td>position</td><td>gene name</td><td>gene id</td></tr>
<tr><td>GU305914.1</td><td>150654</td><td>150676</td><td>+</td><td>0</td><td>CGCAGCGTCCGCGCGCGCAG</td><td>CGG</td><td>||||||||[||||||||||||]PAM</td><td>NA</td><td>Intergenic</td><td>NA</td><td>NA</td></tr>
<tr><td>T11549</td><td>GCAGCGTCCGCGCGCGCAGCGGG</td><td>1000</td><td></td><td></td><td></td><td></td><td></td><td></td><td></td><td></td><td></td></tr>
<tr><td>Oligo fwd</td><td>CACCGCAGCGTCCGCGCGCGCAGC</td><td></td><td></td><td></td><td></td><td></td><td></td><td></td><td></td><td></td><td></td></tr>
<tr><td>Oligo rev</td><td>AAACGCTGCGCGCGCGGACGCTGC</td><td></td><td></td><td></td><td></td><td></td><td></td><td></td><td></td><td></td><td></td></tr>
<tr><td>Chromosome</td><td>start</td><td>end</td><td>strand</td><td>MM</td><td>target_seq</td><td>PAM</td><td>alignment</td><td>distance</td><td>position</td><td>gene name</td><td>gene id</td></tr>
<tr><td>GU305914.1</td><td>150655</td><td>150677</td><td>+</td><td>0</td><td>GCAGCGTCCGCGCGCGCAGC</td><td>GGG</td><td>||||||||[||||||||||||]PAM</td><td>NA</td><td>Intergenic</td><td>NA</td><td>NA</td></tr>
<tr><td>T11550</td><td>GTCCGCGCGCGCAGCGGGTCCGG</td><td>1000</td><td></td><td></td><td></td><td></td><td></td><td></td><td></td><td></td><td></td></tr>
<tr><td>Oligo fwd</td><td>CACCGTCCGCGCGCGCAGCGGGTC</td><td></td><td></td><td></td><td></td><td></td><td></td><td></td><td></td><td></td><td></td></tr>
<tr><td>Oligo rev</td><td>AAACGACCCGCTGCGCGCGCGGAC</td><td></td><td></td><td></td><td></td><td></td><td></td><td></td><td></td><td></td><td></td></tr>
<tr><td>Chromosome</td><td>start</td><td>end</td><td>strand</td><td>MM</td><td>target_seq</td><td>PAM</td><td>alignment</td><td>distance</td><td>position</td><td>gene name</td><td>gene id</td></tr>
<tr><td>GU305914.1</td><td>150660</td><td>150682</td><td>+</td><td>0</td><td>GTCCGCGCGCGCAGCGGGTC</td><td>CGG</td><td>||||||||[||||||||||||]PAM</td><td>NA</td><td>Intergenic</td><td>NA</td><td>NA</td></tr>
<tr><td>T11551</td><td>GCGGGTCCGGCCTGTCCAACAGG</td><td>1000</td><td></td><td></td><td></td><td></td><td></td><td></td><td></td><td></td><td></td></tr>
<tr><td>Oligo fwd</td><td>CACCGCGGGTCCGGCCTGTCCAAC</td><td></td><td></td><td></td><td></td><td></td><td></td><td></td><td></td><td></td><td></td></tr>
<tr><td>Oligo rev</td><td>AAACGTTGGACAGGCCGGACCCGC</td><td></td><td></td><td></td><td></td><td></td><td></td><td></td><td></td><td></td><td></td></tr>
<tr><td>Chromosome</td><td>start</td><td>end</td><td>strand</td><td>MM</td><td>target_seq</td><td>PAM</td><td>alignment</td><td>distance</td><td>position</td><td>gene name</td><td>gene id</td></tr>
<tr><td>GU305914.1</td><td>150673</td><td>150695</td><td>+</td><td>0</td><td>GCGGGTCCGGCCTGTCCAAC</td><td>AGG</td><td>||||||||[||||||||||||]PAM</td><td>NA</td><td>Intergenic</td><td>NA</td><td>NA</td></tr>
<tr><td>T11552</td><td>CCTGTCCAACAGGTGACACTCGG</td><td>1000</td><td></td><td></td><td></td><td></td><td></td><td></td><td></td><td></td><td></td></tr>
<tr><td>Oligo fwd</td><td>CACCCCTGTCCAACAGGTGACACT</td><td></td><td></td><td></td><td></td><td></td><td></td><td></td><td></td><td></td><td></td></tr>
<tr><td>Oligo rev</td><td>AAACAGTGTCACCTGTTGGACAGG</td><td></td><td></td><td></td><td></td><td></td><td></td><td></td><td></td><td></td><td></td></tr>
<tr><td>Chromosome</td><td>start</td><td>end</td><td>strand</td><td>MM</td><td>target_seq</td><td>PAM</td><td>alignment</td><td>distance</td><td>position</td><td>gene name</td><td>gene id</td></tr>
<tr><td>GU305914.1</td><td>150683</td><td>150705</td><td>+</td><td>0</td><td>CCTGTCCAACAGGTGACACT</td><td>CGG</td><td>||||||||[||||||||||||]PAM</td><td>NA</td><td>Intergenic</td><td>NA</td><td>NA</td></tr>
<tr><td>T11553</td><td>CAGGTGACACTCGGCCACCTCGG</td><td>1000</td><td></td><td></td><td></td><td></td><td></td><td></td><td></td><td></td><td></td></tr>
<tr><td>Oligo fwd</td><td>CACCCAGGTGACACTCGGCCACCT</td><td></td><td></td><td></td><td></td><td></td><td></td><td></td><td></td><td></td><td></td></tr>
<tr><td>Oligo rev</td><td>AAACAGGTGGCCGAGTGTCACCTG</td><td></td><td></td><td></td><td></td><td></td><td></td><td></td><td></td><td></td><td></td></tr>
<tr><td>Chromosome</td><td>start</td><td>end</td><td>strand</td><td>MM</td><td>target_seq</td><td>PAM</td><td>alignment</td><td>distance</td><td>position</td><td>gene name</td><td>gene id</td></tr>
<tr><td>GU305914.1</td><td>150692</td><td>150714</td><td>+</td><td>0</td><td>CAGGTGACACTCGGCCACCT</td><td>CGG</td><td>||||||||[||||||||||||]PAM</td><td>NA</td><td>Intergenic</td><td>NA</td><td>NA</td></tr>
<tr><td>T11554</td><td>GACACTCGGCCACCTCGGCCAGG</td><td>1000</td><td></td><td></td><td></td><td></td><td></td><td></td><td></td><td></td><td></td></tr>
<tr><td>Oligo fwd</td><td>CACCGACACTCGGCCACCTCGGCC</td><td></td><td></td><td></td><td></td><td></td><td></td><td></td><td></td><td></td><td></td></tr>
<tr><td>Oligo rev</td><td>AAACGGCCGAGGTGGCCGAGTGTC</td><td></td><td></td><td></td><td></td><td></td><td></td><td></td><td></td><td></td><td></td></tr>
<tr><td>Chromosome</td><td>start</td><td>end</td><td>strand</td><td>MM</td><td>target_seq</td><td>PAM</td><td>alignment</td><td>distance</td><td>position</td><td>gene name</td><td>gene id</td></tr>
<tr><td>GU305914.1</td><td>150697</td><td>150719</td><td>+</td><td>0</td><td>GACACTCGGCCACCTCGGCC</td><td>AGG</td><td>||||||||[||||||||||||]PAM</td><td>NA</td><td>Intergenic</td><td>NA</td><td>NA</td></tr>
<tr><td>T11555</td><td>ATCGACCGCCTTCTCCCGCTCGG</td><td>1000</td><td></td><td></td><td></td><td></td><td></td><td></td><td></td><td></td><td></td></tr>
<tr><td>Oligo fwd</td><td>CACCATCGACCGCCTTCTCCCGCT</td><td></td><td></td><td></td><td></td><td></td><td></td><td></td><td></td><td></td><td></td></tr>
<tr><td>Oligo rev</td><td>AAACAGCGGGAGAAGGCGGTCGAT</td><td></td><td></td><td></td><td></td><td></td><td></td><td></td><td></td><td></td><td></td></tr>
<tr><td>Chromosome</td><td>start</td><td>end</td><td>strand</td><td>MM</td><td>target_seq</td><td>PAM</td><td>alignment</td><td>distance</td><td>position</td><td>gene name</td><td>gene id</td></tr>
<tr><td>GU305914.1</td><td>150737</td><td>150759</td><td>+</td><td>0</td><td>ATCGACCGCCTTCTCCCGCT</td><td>CGG</td><td>||||||||[||||||||||||]PAM</td><td>NA</td><td>Intergenic</td><td>NA</td><td>NA</td></tr>
<tr><td>T11556</td><td>CCGCTCGGCCGTGCGTACCTCGG</td><td>1000</td><td></td><td></td><td></td><td></td><td></td><td></td><td></td><td></td><td></td></tr>
<tr><td>Oligo fwd</td><td>CACCCCGCTCGGCCGTGCGTACCT</td><td></td><td></td><td></td><td></td><td></td><td></td><td></td><td></td><td></td><td></td></tr>
<tr><td>Oligo rev</td><td>AAACAGGTACGCACGGCCGAGCGG</td><td></td><td></td><td></td><td></td><td></td><td></td><td></td><td></td><td></td><td></td></tr>
<tr><td>Chromosome</td><td>start</td><td>end</td><td>strand</td><td>MM</td><td>target_seq</td><td>PAM</td><td>alignment</td><td>distance</td><td>position</td><td>gene name</td><td>gene id</td></tr>
<tr><td>GU305914.1</td><td>150752</td><td>150774</td><td>+</td><td>0</td><td>CCGCTCGGCCGTGCGTACCT</td><td>CGG</td><td>||||||||[||||||||||||]PAM</td><td>NA</td><td>Intergenic</td><td>NA</td><td>NA</td></tr>
<tr><td>T11557</td><td>GGCCGTGCGTACCTCGGCGATGG</td><td>1000</td><td></td><td></td><td></td><td></td><td></td><td></td><td></td><td></td><td></td></tr>
<tr><td>Oligo fwd</td><td>CACCGGCCGTGCGTACCTCGGCGA</td><td></td><td></td><td></td><td></td><td></td><td></td><td></td><td></td><td></td><td></td></tr>
<tr><td>Oligo rev</td><td>AAACTCGCCGAGGTACGCACGGCC</td><td></td><td></td><td></td><td></td><td></td><td></td><td></td><td></td><td></td><td></td></tr>
<tr><td>Chromosome</td><td>start</td><td>end</td><td>strand</td><td>MM</td><td>target_seq</td><td>PAM</td><td>alignment</td><td>distance</td><td>position</td><td>gene name</td><td>gene id</td></tr>
<tr><td>GU305914.1</td><td>150758</td><td>150780</td><td>+</td><td>0</td><td>GGCCGTGCGTACCTCGGCGA</td><td>TGG</td><td>||||||||[||||||||||||]PAM</td><td>NA</td><td>Intergenic</td><td>NA</td><td>NA</td></tr>
<tr><td>T11558</td><td>GCCGTGCGTACCTCGGCGATGGG</td><td>1000</td><td></td><td></td><td></td><td></td><td></td><td></td><td></td><td></td><td></td></tr>
<tr><td>Oligo fwd</td><td>CACCGCCGTGCGTACCTCGGCGAT</td><td></td><td></td><td></td><td></td><td></td><td></td><td></td><td></td><td></td><td></td></tr>
<tr><td>Oligo rev</td><td>AAACATCGCCGAGGTACGCACGGC</td><td></td><td></td><td></td><td></td><td></td><td></td><td></td><td></td><td></td><td></td></tr>
<tr><td>Chromosome</td><td>start</td><td>end</td><td>strand</td><td>MM</td><td>target_seq</td><td>PAM</td><td>alignment</td><td>distance</td><td>position</td><td>gene name</td><td>gene id</td></tr>
<tr><td>GU305914.1</td><td>150759</td><td>150781</td><td>+</td><td>0</td><td>GCCGTGCGTACCTCGGCGAT</td><td>GGG</td><td>||||||||[||||||||||||]PAM</td><td>NA</td><td>Intergenic</td><td>NA</td><td>NA</td></tr>
<tr><td>T11559</td><td>TATAAGCTTCTCCAGCAGCGCGG</td><td>1000</td><td></td><td></td><td></td><td></td><td></td><td></td><td></td><td></td><td></td></tr>
<tr><td>Oligo fwd</td><td>CACCTATAAGCTTCTCCAGCAGCG</td><td></td><td></td><td></td><td></td><td></td><td></td><td></td><td></td><td></td><td></td></tr>
<tr><td>Oligo rev</td><td>AAACCGCTGCTGGAGAAGCTTATA</td><td></td><td></td><td></td><td></td><td></td><td></td><td></td><td></td><td></td><td></td></tr>
<tr><td>Chromosome</td><td>start</td><td>end</td><td>strand</td><td>MM</td><td>target_seq</td><td>PAM</td><td>alignment</td><td>distance</td><td>position</td><td>gene name</td><td>gene id</td></tr>
<tr><td>GU305914.1</td><td>150848</td><td>150870</td><td>+</td><td>0</td><td>TATAAGCTTCTCCAGCAGCG</td><td>CGG</td><td>||||||||[||||||||||||]PAM</td><td>NA</td><td>Intergenic</td><td>NA</td><td>NA</td></tr>
<tr><td>T11560</td><td>CCGCGCGCACGAACCGCCGCCGG</td><td>1000</td><td></td><td></td><td></td><td></td><td></td><td></td><td></td><td></td><td></td></tr>
<tr><td>Oligo fwd</td><td>CACCCCGCGCGCACGAACCGCCGC</td><td></td><td></td><td></td><td></td><td></td><td></td><td></td><td></td><td></td><td></td></tr>
<tr><td>Oligo rev</td><td>AAACGCGGCGGTTCGTGCGCGCGG</td><td></td><td></td><td></td><td></td><td></td><td></td><td></td><td></td><td></td><td></td></tr>
<tr><td>Chromosome</td><td>start</td><td>end</td><td>strand</td><td>MM</td><td>target_seq</td><td>PAM</td><td>alignment</td><td>distance</td><td>position</td><td>gene name</td><td>gene id</td></tr>
<tr><td>GU305914.1</td><td>150949</td><td>150971</td><td>+</td><td>0</td><td>CCGCGCGCACGAACCGCCGC</td><td>CGG</td><td>||||||||[||||||||||||]PAM</td><td>NA</td><td>Intergenic</td><td>NA</td><td>NA</td></tr>
<tr><td>T11561</td><td>CGCGCACGAACCGCCGCCGGTGG</td><td>1000</td><td></td><td></td><td></td><td></td><td></td><td></td><td></td><td></td><td></td></tr>
<tr><td>Oligo fwd</td><td>CACCCGCGCACGAACCGCCGCCGG</td><td></td><td></td><td></td><td></td><td></td><td></td><td></td><td></td><td></td><td></td></tr>
<tr><td>Oligo rev</td><td>AAACCCGGCGGCGGTTCGTGCGCG</td><td></td><td></td><td></td><td></td><td></td><td></td><td></td><td></td><td></td><td></td></tr>
<tr><td>Chromosome</td><td>start</td><td>end</td><td>strand</td><td>MM</td><td>target_seq</td><td>PAM</td><td>alignment</td><td>distance</td><td>position</td><td>gene name</td><td>gene id</td></tr>
<tr><td>GU305914.1</td><td>150952</td><td>150974</td><td>+</td><td>0</td><td>CGCGCACGAACCGCCGCCGG</td><td>TGG</td><td>||||||||[||||||||||||]PAM</td><td>NA</td><td>Intergenic</td><td>NA</td><td>NA</td></tr>
<tr><td>T11562</td><td>GCGCACGAACCGCCGCCGGTGGG</td><td>1000</td><td></td><td></td><td></td><td></td><td></td><td></td><td></td><td></td><td></td></tr>
<tr><td>Oligo fwd</td><td>CACCGCGCACGAACCGCCGCCGGT</td><td></td><td></td><td></td><td></td><td></td><td></td><td></td><td></td><td></td><td></td></tr>
<tr><td>Oligo rev</td><td>AAACACCGGCGGCGGTTCGTGCGC</td><td></td><td></td><td></td><td></td><td></td><td></td><td></td><td></td><td></td><td></td></tr>
<tr><td>Chromosome</td><td>start</td><td>end</td><td>strand</td><td>MM</td><td>target_seq</td><td>PAM</td><td>alignment</td><td>distance</td><td>position</td><td>gene name</td><td>gene id</td></tr>
<tr><td>GU305914.1</td><td>150953</td><td>150975</td><td>+</td><td>0</td><td>GCGCACGAACCGCCGCCGGT</td><td>GGG</td><td>||||||||[||||||||||||]PAM</td><td>NA</td><td>Intergenic</td><td>NA</td><td>NA</td></tr>
<tr><td>T11563</td><td>CACGAACCGCCGCCGGTGGGAGG</td><td>1000</td><td></td><td></td><td></td><td></td><td></td><td></td><td></td><td></td><td></td></tr>
<tr><td>Oligo fwd</td><td>CACCCACGAACCGCCGCCGGTGGG</td><td></td><td></td><td></td><td></td><td></td><td></td><td></td><td></td><td></td><td></td></tr>
</table>

<table>
<tr><td>Oligo rev</td><td>AAACCCCACCGGCGGCGGTTCGTG</td><td></td><td></td><td></td><td></td><td></td><td></td><td></td><td></td><td></td><td></td></tr>
<tr><td>Chromosome</td><td>start</td><td>end</td><td>strand</td><td>MM</td><td>target_seq</td><td>PAM</td><td>alignment</td><td>distance</td><td>position</td><td>gene name</td><td>gene id</td></tr>
<tr><td>GU305914.1</td><td>150956</td><td>150978</td><td>+</td><td>0</td><td>CACGAACCGCCGCCGGTGGG</td><td>AGG</td><td>||||||||[||||||||||||]PAM</td><td>NA</td><td>Intergenic</td><td>NA</td><td>NA</td></tr>
<tr><td></td><td></td><td></td><td></td><td></td><td></td><td></td><td></td><td></td><td></td><td></td><td></td></tr>
<tr><td>T11564</td><td>CCGCCGCCGGTGGGAGGCCCAGG</td><td>1000</td><td></td><td></td><td></td><td></td><td></td><td></td><td></td><td></td><td></td></tr>
<tr><td>Oligo fwd</td><td>CACCCCGCCGCCGGTGGGAGGCCC</td><td></td><td></td><td></td><td></td><td></td><td></td><td></td><td></td><td></td><td></td></tr>
<tr><td>Oligo rev</td><td>AAACGGGCCTCCCACCGGCGGCGG</td><td></td><td></td><td></td><td></td><td></td><td></td><td></td><td></td><td></td><td></td></tr>
<tr><td>Chromosome</td><td>start</td><td>end</td><td>strand</td><td>MM</td><td>target_seq</td><td>PAM</td><td>alignment</td><td>distance</td><td>position</td><td>gene name</td><td>gene id</td></tr>
<tr><td>GU305914.1</td><td>150962</td><td>150984</td><td>+</td><td>0</td><td>CCGCCGCCGGTGGGAGGCCC</td><td>AGG</td><td>||||||||[||||||||||||]PAM</td><td>NA</td><td>Intergenic</td><td>NA</td><td>NA</td></tr>
<tr><td>GU305914.1</td><td>71507</td><td>71529</td><td>+</td><td>4</td><td>CCGGCTCTGGTGGGTGGCCC</td><td>CGG</td><td>|||-|-|-[||||||-|||||]PAM</td><td>NA</td><td>Intergenic</td><td>NA</td><td>NA</td></tr>
<tr><td></td><td></td><td></td><td></td><td></td><td></td><td></td><td></td><td></td><td></td><td></td><td></td></tr>
<tr><td>T11565</td><td>CCACGCTGATGCCGTTCAGCAGG</td><td>1000</td><td></td><td></td><td></td><td></td><td></td><td></td><td></td><td></td><td></td></tr>
<tr><td>Oligo fwd</td><td>CACCCCACGCTGATGCCGTTCAGC</td><td></td><td></td><td></td><td></td><td></td><td></td><td></td><td></td><td></td><td></td></tr>
<tr><td>Oligo rev</td><td>AAACGCTGAACGGCATCAGCGTGG</td><td></td><td></td><td></td><td></td><td></td><td></td><td></td><td></td><td></td><td></td></tr>
<tr><td>Chromosome</td><td>start</td><td>end</td><td>strand</td><td>MM</td><td>target_seq</td><td>PAM</td><td>alignment</td><td>distance</td><td>position</td><td>gene name</td><td>gene id</td></tr>
<tr><td>GU305914.1</td><td>151000</td><td>151022</td><td>+</td><td>0</td><td>CCACGCTGATGCCGTTCAGC</td><td>AGG</td><td>||||||||[||||||||||||]PAM</td><td>NA</td><td>Intergenic</td><td>NA</td><td>NA</td></tr>
<tr><td></td><td></td><td></td><td></td><td></td><td></td><td></td><td></td><td></td><td></td><td></td><td></td></tr>
<tr><td>T11566</td><td>TCGCGTAGAAGATCTGCTTGCGG</td><td>1000</td><td></td><td></td><td></td><td></td><td></td><td></td><td></td><td></td><td></td></tr>
<tr><td>Oligo fwd</td><td>CACCTCGCGTAGAAGATCTGCTTG</td><td></td><td></td><td></td><td></td><td></td><td></td><td></td><td></td><td></td><td></td></tr>
<tr><td>Oligo rev</td><td>AAACCAAGCAGATCTTCTACGCGA</td><td></td><td></td><td></td><td></td><td></td><td></td><td></td><td></td><td></td><td></td></tr>
<tr><td>Chromosome</td><td>start</td><td>end</td><td>strand</td><td>MM</td><td>target_seq</td><td>PAM</td><td>alignment</td><td>distance</td><td>position</td><td>gene name</td><td>gene id</td></tr>
<tr><td>GU305914.1</td><td>151033</td><td>151055</td><td>+</td><td>0</td><td>TCGCGTAGAAGATCTGCTTG</td><td>CGG</td><td>||||||||[||||||||||||]PAM</td><td>NA</td><td>Intergenic</td><td>NA</td><td>NA</td></tr>
<tr><td></td><td></td><td></td><td></td><td></td><td></td><td></td><td></td><td></td><td></td><td></td><td></td></tr>
<tr><td>T11567</td><td>ATCTGCTTGCGGAACACGCTCGG</td><td>1000</td><td></td><td></td><td></td><td></td><td></td><td></td><td></td><td></td><td></td></tr>
<tr><td>Oligo fwd</td><td>CACCATCTGCTTGCGGAACACGCT</td><td></td><td></td><td></td><td></td><td></td><td></td><td></td><td></td><td></td><td></td></tr>
<tr><td>Oligo rev</td><td>AAACAGCGTGTTCCGCAAGCAGAT</td><td></td><td></td><td></td><td></td><td></td><td></td><td></td><td></td><td></td><td></td></tr>
<tr><td>Chromosome</td><td>start</td><td>end</td><td>strand</td><td>MM</td><td>target_seq</td><td>PAM</td><td>alignment</td><td>distance</td><td>position</td><td>gene name</td><td>gene id</td></tr>
<tr><td>GU305914.1</td><td>151044</td><td>151066</td><td>+</td><td>0</td><td>ATCTGCTTGCGGAACACGCT</td><td>CGG</td><td>||||||||[||||||||||||]PAM</td><td>NA</td><td>Intergenic</td><td>NA</td><td>NA</td></tr>
<tr><td></td><td></td><td></td><td></td><td></td><td></td><td></td><td></td><td></td><td></td><td></td><td></td></tr>
<tr><td>T11568</td><td>TCTGCTTGCGGAACACGCTCGGG</td><td>1000</td><td></td><td></td><td></td><td></td><td></td><td></td><td></td><td></td><td></td></tr>
<tr><td>Oligo fwd</td><td>CACCTCTGCTTGCGGAACACGCTC</td><td></td><td></td><td></td><td></td><td></td><td></td><td></td><td></td><td></td><td></td></tr>
<tr><td>Oligo rev</td><td>AAACGAGCGTGTTCCGCAAGCAGA</td><td></td><td></td><td></td><td></td><td></td><td></td><td></td><td></td><td></td><td></td></tr>
<tr><td>Chromosome</td><td>start</td><td>end</td><td>strand</td><td>MM</td><td>target_seq</td><td>PAM</td><td>alignment</td><td>distance</td><td>position</td><td>gene name</td><td>gene id</td></tr>
<tr><td>GU305914.1</td><td>151045</td><td>151067</td><td>+</td><td>0</td><td>TCTGCTTGCGGAACACGCTC</td><td>GGG</td><td>||||||||[||||||||||||]PAM</td><td>NA</td><td>Intergenic</td><td>NA</td><td>NA</td></tr>
<tr><td></td><td></td><td></td><td></td><td></td><td></td><td></td><td></td><td></td><td></td><td></td><td></td></tr>
<tr><td>T11569</td><td>GGAACACGCTCGGGTCTCGCAGG</td><td>1000</td><td></td><td></td><td></td><td></td><td></td><td></td><td></td><td></td><td></td></tr>
<tr><td>Oligo fwd</td><td>CACCGGAACACGCTCGGGTCTCGC</td><td></td><td></td><td></td><td></td><td></td><td></td><td></td><td></td><td></td><td></td></tr>
<tr><td>Oligo rev</td><td>AAACGCGAGACCCGAGCGTGTTCC</td><td></td><td></td><td></td><td></td><td></td><td></td><td></td><td></td><td></td><td></td></tr>
<tr><td>Chromosome</td><td>start</td><td>end</td><td>strand</td><td>MM</td><td>target_seq</td><td>PAM</td><td>alignment</td><td>distance</td><td>position</td><td>gene name</td><td>gene id</td></tr>
<tr><td>GU305914.1</td><td>151054</td><td>151076</td><td>+</td><td>0</td><td>GGAACACGCTCGGGTCTCGC</td><td>AGG</td><td>||||||||[||||||||||||]PAM</td><td>NA</td><td>Intergenic</td><td>NA</td><td>NA</td></tr>
<tr><td></td><td></td><td></td><td></td><td></td><td></td><td></td><td></td><td></td><td></td><td></td><td></td></tr>
<tr><td>T11570</td><td>CGCTCGGGTCTCGCAGGATCTGG</td><td>1000</td><td></td><td></td><td></td><td></td><td></td><td></td><td></td><td></td><td></td></tr>
<tr><td>Oligo fwd</td><td>CACCCGCTCGGGTCTCGCAGGATC</td><td></td><td></td><td></td><td></td><td></td><td></td><td></td><td></td><td></td><td></td></tr>
<tr><td>Oligo rev</td><td>AAACGATCCTGCGAGACCCGAGCG</td><td></td><td></td><td></td><td></td><td></td><td></td><td></td><td></td><td></td><td></td></tr>
<tr><td>Chromosome</td><td>start</td><td>end</td><td>strand</td><td>MM</td><td>target_seq</td><td>PAM</td><td>alignment</td><td>distance</td><td>position</td><td>gene name</td><td>gene id</td></tr>
<tr><td>GU305914.1</td><td>151060</td><td>151082</td><td>+</td><td>0</td><td>CGCTCGGGTCTCGCAGGATC</td><td>TGG</td><td>||||||||[||||||||||||]PAM</td><td>NA</td><td>Intergenic</td><td>NA</td><td>NA</td></tr>
<tr><td></td><td></td><td></td><td></td><td></td><td></td><td></td><td></td><td></td><td></td><td></td><td></td></tr>
<tr><td>T11571</td><td>TCGGGTCTCGCAGGATCTGGTGG</td><td>1000</td><td></td><td></td><td></td><td></td><td></td><td></td><td></td><td></td><td></td></tr>
<tr><td>Oligo fwd</td><td>CACCTCGGGTCTCGCAGGATCTGG</td><td></td><td></td><td></td><td></td><td></td><td></td><td></td><td></td><td></td><td></td></tr>
<tr><td>Oligo rev</td><td>AAACCCAGATCCTGCGAGACCCGA</td><td></td><td></td><td></td><td></td><td></td><td></td><td></td><td></td><td></td><td></td></tr>
<tr><td>Chromosome</td><td>start</td><td>end</td><td>strand</td><td>MM</td><td>target_seq</td><td>PAM</td><td>alignment</td><td>distance</td><td>position</td><td>gene name</td><td>gene id</td></tr>
<tr><td>GU305914.1</td><td>151063</td><td>151085</td><td>+</td><td>0</td><td>TCGGGTCTCGCAGGATCTGG</td><td>TGG</td><td>||||||||[||||||||||||]PAM</td><td>NA</td><td>Intergenic</td><td>NA</td><td>NA</td></tr>
<tr><td></td><td></td><td></td><td></td><td></td><td></td><td></td><td></td><td></td><td></td><td></td><td></td></tr>
<tr><td>T11572</td><td>GCCCTCCGTGCCGTTGATCATGG</td><td>1000</td><td></td><td></td><td></td><td></td><td></td><td></td><td></td><td></td><td></td></tr>
<tr><td>Oligo fwd</td><td>CACCGCCCTCCGTGCCGTTGATCA</td><td></td><td></td><td></td><td></td><td></td><td></td><td></td><td></td><td></td><td></td></tr>
<tr><td>Oligo rev</td><td>AAACTGATCAACGGCACGGAGGGC</td><td></td><td></td><td></td><td></td><td></td><td></td><td></td><td></td><td></td><td></td></tr>
<tr><td>Chromosome</td><td>start</td><td>end</td><td>strand</td><td>MM</td><td>target_seq</td><td>PAM</td><td>alignment</td><td>distance</td><td>position</td><td>gene name</td><td>gene id</td></tr>
<tr><td>GU305914.1</td><td>151106</td><td>151128</td><td>+</td><td>0</td><td>GCCCTCCGTGCCGTTGATCA</td><td>TGG</td><td>||||||||[||||||||||||]PAM</td><td>NA</td><td>Intergenic</td><td>NA</td><td>NA</td></tr>
<tr><td></td><td></td><td></td><td></td><td></td><td></td><td></td><td></td><td></td><td></td><td></td><td></td></tr>
<tr><td>T11573</td><td>TGCCGTTGATCATGGAGAAGAGG</td><td>1000</td><td></td><td></td><td></td><td></td><td></td><td></td><td></td><td></td><td></td></tr>
<tr><td>Oligo fwd</td><td>CACCTGCCGTTGATCATGGAGAAG</td><td></td><td></td><td></td><td></td><td></td><td></td><td></td><td></td><td></td><td></td></tr>
<tr><td>Oligo rev</td><td>AAACCTTCTCCATGATCAACGGCA</td><td></td><td></td><td></td><td></td><td></td><td></td><td></td><td></td><td></td><td></td></tr>
<tr><td>Chromosome</td><td>start</td><td>end</td><td>strand</td><td>MM</td><td>target_seq</td><td>PAM</td><td>alignment</td><td>distance</td><td>position</td><td>gene name</td><td>gene id</td></tr>
<tr><td>GU305914.1</td><td>151114</td><td>151136</td><td>+</td><td>0</td><td>TGCCGTTGATCATGGAGAAG</td><td>AGG</td><td>||||||||[||||||||||||]PAM</td><td>NA</td><td>Intergenic</td><td>NA</td><td>NA</td></tr>
<tr><td></td><td></td><td></td><td></td><td></td><td></td><td></td><td></td><td></td><td></td><td></td><td></td></tr>
<tr><td>T11574</td><td>AGGTCCAGCGACGAGTCCGTCGG</td><td>1000</td><td></td><td></td><td></td><td></td><td></td><td></td><td></td><td></td><td></td></tr>
<tr><td>Oligo fwd</td><td>CACCAGGTCCAGCGACGAGTCCGT</td><td></td><td></td><td></td><td></td><td></td><td></td><td></td><td></td><td></td><td></td></tr>
<tr><td>Oligo rev</td><td>AAACACGGACTCGTCGCTGGACCT</td><td></td><td></td><td></td><td></td><td></td><td></td><td></td><td></td><td></td><td></td></tr>
<tr><td>Chromosome</td><td>start</td><td>end</td><td>strand</td><td>MM</td><td>target_seq</td><td>PAM</td><td>alignment</td><td>distance</td><td>position</td><td>gene name</td><td>gene id</td></tr>
<tr><td>GU305914.1</td><td>151134</td><td>151156</td><td>+</td><td>0</td><td>AGGTCCAGCGACGAGTCCGT</td><td>CGG</td><td>||||||||[||||||||||||]PAM</td><td>NA</td><td>Intergenic</td><td>NA</td><td>NA</td></tr>
<tr><td></td><td></td><td></td><td></td><td></td><td></td><td></td><td></td><td></td><td></td><td></td><td></td></tr>
<tr><td>T11575</td><td>GGTCCAGCGACGAGTCCGTCGGG</td><td>1000</td><td></td><td></td><td></td><td></td><td></td><td></td><td></td><td></td><td></td></tr>
<tr><td>Oligo fwd</td><td>CACCGGTCCAGCGACGAGTCCGTC</td><td></td><td></td><td></td><td></td><td></td><td></td><td></td><td></td><td></td><td></td></tr>
<tr><td>Oligo rev</td><td>AAACGACGGACTCGTCGCTGGACC</td><td></td><td></td><td></td><td></td><td></td><td></td><td></td><td></td><td></td><td></td></tr>
<tr><td>Chromosome</td><td>start</td><td>end</td><td>strand</td><td>MM</td><td>target_seq</td><td>PAM</td><td>alignment</td><td>distance</td><td>position</td><td>gene name</td><td>gene id</td></tr>
<tr><td>GU305914.1</td><td>151135</td><td>151157</td><td>+</td><td>0</td><td>GGTCCAGCGACGAGTCCGTC</td><td>GGG</td><td>||||||||[||||||||||||]PAM</td><td>NA</td><td>Intergenic</td><td>NA</td><td>NA</td></tr>
<tr><td></td><td></td><td></td><td></td><td></td><td></td><td></td><td></td><td></td><td></td><td></td><td></td></tr>
<tr><td>T11576</td><td>CGACGAGTCCGTCGGGAACAAGG</td><td>1000</td><td></td><td></td><td></td><td></td><td></td><td></td><td></td><td></td><td></td></tr>
<tr><td>Oligo fwd</td><td>CACCCGACGAGTCCGTCGGGAACA</td><td></td><td></td><td></td><td></td><td></td><td></td><td></td><td></td><td></td><td></td></tr>
<tr><td>Oligo rev</td><td>AAACTGTTCCCGACGGACTCGTCG</td><td></td><td></td><td></td><td></td><td></td><td></td><td></td><td></td><td></td><td></td></tr>
<tr><td>Chromosome</td><td>start</td><td>end</td><td>strand</td><td>MM</td><td>target_seq</td><td>PAM</td><td>alignment</td><td>distance</td><td>position</td><td>gene name</td><td>gene id</td></tr>
<tr><td>GU305914.1</td><td>151142</td><td>151164</td><td>+</td><td>0</td><td>CGACGAGTCCGTCGGGAACA</td><td>AGG</td><td>||||||||[||||||||||||]PAM</td><td>NA</td><td>Intergenic</td><td>NA</td><td>NA</td></tr>
<tr><td></td><td></td><td></td><td></td><td></td><td></td><td></td><td></td><td></td><td></td><td></td><td></td></tr>
<tr><td>T11577</td><td>GACGAGTCCGTCGGGAACAAGGG</td><td>1000</td><td></td><td></td><td></td><td></td><td></td><td></td><td></td><td></td><td></td></tr>
<tr><td>Oligo fwd</td><td>CACCGACGAGTCCGTCGGGAACAA</td><td></td><td></td><td></td><td></td><td></td><td></td><td></td><td></td><td></td><td></td></tr>
<tr><td>Oligo rev</td><td>AAACTTGTTCCCGACGGACTCGTC</td><td></td><td></td><td></td><td></td><td></td><td></td><td></td><td></td><td></td><td></td></tr>
<tr><td>Chromosome</td><td>start</td><td>end</td><td>strand</td><td>MM</td><td>target_seq</td><td>PAM</td><td>alignment</td><td>distance</td><td>position</td><td>gene name</td><td>gene id</td></tr>
<tr><td>GU305914.1</td><td>151143</td><td>151165</td><td>+</td><td>0</td><td>GACGAGTCCGTCGGGAACAA</td><td>GGG</td><td>||||||||[||||||||||||]PAM</td><td>NA</td><td>Intergenic</td><td>NA</td><td>NA</td></tr>
</table>

<table>
<tr><td>T11578</td><td>ACGAGTCCGTCGGGAACAAGGGG</td><td>1000</td><td></td><td></td><td></td><td></td><td></td><td></td><td></td><td></td><td></td></tr>
<tr><td>Oligo fwd</td><td>CACCACGAGTCCGTCGGGAACAAG</td><td></td><td></td><td></td><td></td><td></td><td></td><td></td><td></td><td></td><td></td></tr>
<tr><td>Oligo rev</td><td>AAACCTTGTTCCCGACGGACTCGT</td><td></td><td></td><td></td><td></td><td></td><td></td><td></td><td></td><td></td><td></td></tr>
<tr><td>Chromosome</td><td>start</td><td>end</td><td>strand</td><td>MM</td><td>target_seq</td><td>PAM</td><td>alignment</td><td>distance</td><td>position</td><td>gene name</td><td>gene id</td></tr>
<tr><td>GU305914.1</td><td>151144</td><td>151166</td><td>+</td><td>0</td><td>ACGAGTCCGTCGGGAACAAG</td><td>GGG</td><td>||||||||[||||||||||||]PAM</td><td>NA</td><td>Intergenic</td><td>NA</td><td>NA</td></tr>
<tr><td>T11579</td><td>GTCGGGAACAAGGGGTCCGACGG</td><td>1000</td><td></td><td></td><td></td><td></td><td></td><td></td><td></td><td></td><td></td></tr>
<tr><td>Oligo fwd</td><td>CACCGTCGGGAACAAGGGGTCCGA</td><td></td><td></td><td></td><td></td><td></td><td></td><td></td><td></td><td></td><td></td></tr>
<tr><td>Oligo rev</td><td>AAACTCGGACCCCTTGTTCCCGAC</td><td></td><td></td><td></td><td></td><td></td><td></td><td></td><td></td><td></td><td></td></tr>
<tr><td>Chromosome</td><td>start</td><td>end</td><td>strand</td><td>MM</td><td>target_seq</td><td>PAM</td><td>alignment</td><td>distance</td><td>position</td><td>gene name</td><td>gene id</td></tr>
<tr><td>GU305914.1</td><td>151152</td><td>151174</td><td>+</td><td>0</td><td>GTCGGGAACAAGGGGTCCGA</td><td>CGG</td><td>||||||||[||||||||||||]PAM</td><td>NA</td><td>Intergenic</td><td>NA</td><td>NA</td></tr>
<tr><td>T11580</td><td>GGGAACAAGGGGTCCGACGGCGG</td><td>1000</td><td></td><td></td><td></td><td></td><td></td><td></td><td></td><td></td><td></td></tr>
<tr><td>Oligo fwd</td><td>CACCGGGAACAAGGGGTCCGACGG</td><td></td><td></td><td></td><td></td><td></td><td></td><td></td><td></td><td></td><td></td></tr>
<tr><td>Oligo rev</td><td>AAACCCGTCGGACCCCTTGTTCCC</td><td></td><td></td><td></td><td></td><td></td><td></td><td></td><td></td><td></td><td></td></tr>
<tr><td>Chromosome</td><td>start</td><td>end</td><td>strand</td><td>MM</td><td>target_seq</td><td>PAM</td><td>alignment</td><td>distance</td><td>position</td><td>gene name</td><td>gene id</td></tr>
<tr><td>GU305914.1</td><td>151155</td><td>151177</td><td>+</td><td>0</td><td>GGGAACAAGGGGTCCGACGG</td><td>CGG</td><td>||||||||[||||||||||||]PAM</td><td>NA</td><td>Intergenic</td><td>NA</td><td>NA</td></tr>
<tr><td>T11581</td><td>AAGGGGTCCGACGGCGGCTCCGG</td><td>1000</td><td></td><td></td><td></td><td></td><td></td><td></td><td></td><td></td><td></td></tr>
<tr><td>Oligo fwd</td><td>CACCAAGGGGTCCGACGGCGGCTC</td><td></td><td></td><td></td><td></td><td></td><td></td><td></td><td></td><td></td><td></td></tr>
<tr><td>Oligo rev</td><td>AAACGAGCCGCCGTCGGACCCCTT</td><td></td><td></td><td></td><td></td><td></td><td></td><td></td><td></td><td></td><td></td></tr>
<tr><td>Chromosome</td><td>start</td><td>end</td><td>strand</td><td>MM</td><td>target_seq</td><td>PAM</td><td>alignment</td><td>distance</td><td>position</td><td>gene name</td><td>gene id</td></tr>
<tr><td>GU305914.1</td><td>151161</td><td>151183</td><td>+</td><td>0</td><td>AAGGGGTCCGACGGCGGCTC</td><td>CGG</td><td>||||||||[||||||||||||]PAM</td><td>NA</td><td>Intergenic</td><td>NA</td><td>NA</td></tr>
<tr><td>GU305914.1</td><td>98430</td><td>98452</td><td>+</td><td>4</td><td>GAGGGTGCCGACGACGGCTC</td><td>GGG</td><td>-||||--|[|||||-||||||]PAM</td><td>NA</td><td>Intergenic</td><td>NA</td><td>NA</td></tr>
<tr><td>T11582</td><td>GGGTCCGACGGCGGCTCCGGCGG</td><td>1000</td><td></td><td></td><td></td><td></td><td></td><td></td><td></td><td></td><td></td></tr>
<tr><td>Oligo fwd</td><td>CACCGGGTCCGACGGCGGCTCCGG</td><td></td><td></td><td></td><td></td><td></td><td></td><td></td><td></td><td></td><td></td></tr>
<tr><td>Oligo rev</td><td>AAACCCGGAGCCGCCGTCGGACCC</td><td></td><td></td><td></td><td></td><td></td><td></td><td></td><td></td><td></td><td></td></tr>
<tr><td>Chromosome</td><td>start</td><td>end</td><td>strand</td><td>MM</td><td>target_seq</td><td>PAM</td><td>alignment</td><td>distance</td><td>position</td><td>gene name</td><td>gene id</td></tr>
<tr><td>GU305914.1</td><td>151164</td><td>151186</td><td>+</td><td>0</td><td>GGGTCCGACGGCGGCTCCGG</td><td>CGG</td><td>||||||||[||||||||||||]PAM</td><td>NA</td><td>Intergenic</td><td>NA</td><td>NA</td></tr>
<tr><td>GU305914.1</td><td>163743</td><td>163765</td><td>+</td><td>4</td><td>AGCTCCGTCGGCGGCGCCGG</td><td>TGG</td><td>-|-||||-[|||||||-||||]PAM</td><td>NA</td><td>Intergenic</td><td>NA</td><td>NA</td></tr>
<tr><td>T11583</td><td>GGTCCGACGGCGGCTCCGGCGGG</td><td>1000</td><td></td><td></td><td></td><td></td><td></td><td></td><td></td><td></td><td></td></tr>
<tr><td>Oligo fwd</td><td>CACCGGTCCGACGGCGGCTCCGGC</td><td></td><td></td><td></td><td></td><td></td><td></td><td></td><td></td><td></td><td></td></tr>
<tr><td>Oligo rev</td><td>AAACGCCGGAGCCGCCGTCGGACC</td><td></td><td></td><td></td><td></td><td></td><td></td><td></td><td></td><td></td><td></td></tr>
<tr><td>Chromosome</td><td>start</td><td>end</td><td>strand</td><td>MM</td><td>target_seq</td><td>PAM</td><td>alignment</td><td>distance</td><td>position</td><td>gene name</td><td>gene id</td></tr>
<tr><td>GU305914.1</td><td>151165</td><td>151187</td><td>+</td><td>0</td><td>GGTCCGACGGCGGCTCCGGC</td><td>GGG</td><td>||||||||[||||||||||||]PAM</td><td>NA</td><td>Intergenic</td><td>NA</td><td>NA</td></tr>
<tr><td>T11584</td><td>GCGGCTCCGGCGGGTTGTCGAGG</td><td>1000</td><td></td><td></td><td></td><td></td><td></td><td></td><td></td><td></td><td></td></tr>
<tr><td>Oligo fwd</td><td>CACCGCGGCTCCGGCGGGTTGTCG</td><td></td><td></td><td></td><td></td><td></td><td></td><td></td><td></td><td></td><td></td></tr>
<tr><td>Oligo rev</td><td>AAACCGACAACCCGCCGGAGCCGC</td><td></td><td></td><td></td><td></td><td></td><td></td><td></td><td></td><td></td><td></td></tr>
<tr><td>Chromosome</td><td>start</td><td>end</td><td>strand</td><td>MM</td><td>target_seq</td><td>PAM</td><td>alignment</td><td>distance</td><td>position</td><td>gene name</td><td>gene id</td></tr>
<tr><td>GU305914.1</td><td>151174</td><td>151196</td><td>+</td><td>0</td><td>GCGGCTCCGGCGGGTTGTCG</td><td>AGG</td><td>||||||||[||||||||||||]PAM</td><td>NA</td><td>Intergenic</td><td>NA</td><td>NA</td></tr>
<tr><td>T11585</td><td>CGTCCGCCTCGAAGATCGAGCGG</td><td>1000</td><td></td><td></td><td></td><td></td><td></td><td></td><td></td><td></td><td></td></tr>
<tr><td>Oligo fwd</td><td>CACCCGTCCGCCTCGAAGATCGAG</td><td></td><td></td><td></td><td></td><td></td><td></td><td></td><td></td><td></td><td></td></tr>
<tr><td>Oligo rev</td><td>AAACCTCGATCTTCGAGGCGGACG</td><td></td><td></td><td></td><td></td><td></td><td></td><td></td><td></td><td></td><td></td></tr>
<tr><td>Chromosome</td><td>start</td><td>end</td><td>strand</td><td>MM</td><td>target_seq</td><td>PAM</td><td>alignment</td><td>distance</td><td>position</td><td>gene name</td><td>gene id</td></tr>
<tr><td>GU305914.1</td><td>151225</td><td>151247</td><td>+</td><td>0</td><td>CGTCCGCCTCGAAGATCGAG</td><td>CGG</td><td>||||||||[||||||||||||]PAM</td><td>NA</td><td>Intergenic</td><td>NA</td><td>NA</td></tr>
<tr><td>T11586</td><td>CGTTGACCGCGTCAAGTCGCTGG</td><td>1000</td><td></td><td></td><td></td><td></td><td></td><td></td><td></td><td></td><td></td></tr>
<tr><td>Oligo fwd</td><td>CACCCGTTGACCGCGTCAAGTCGC</td><td></td><td></td><td></td><td></td><td></td><td></td><td></td><td></td><td></td><td></td></tr>
<tr><td>Oligo rev</td><td>AAACGCGACTTGACGCGGTCAACG</td><td></td><td></td><td></td><td></td><td></td><td></td><td></td><td></td><td></td><td></td></tr>
<tr><td>Chromosome</td><td>start</td><td>end</td><td>strand</td><td>MM</td><td>target_seq</td><td>PAM</td><td>alignment</td><td>distance</td><td>position</td><td>gene name</td><td>gene id</td></tr>
<tr><td>GU305914.1</td><td>151250</td><td>151272</td><td>+</td><td>0</td><td>CGTTGACCGCGTCAAGTCGC</td><td>TGG</td><td>||||||||[||||||||||||]PAM</td><td>NA</td><td>Intergenic</td><td>NA</td><td>NA</td></tr>
<tr><td>T11587</td><td>TCAAGTCGCTGGCCTTGAAGCGG</td><td>1000</td><td></td><td></td><td></td><td></td><td></td><td></td><td></td><td></td><td></td></tr>
<tr><td>Oligo fwd</td><td>CACCTCAAGTCGCTGGCCTTGAAG</td><td></td><td></td><td></td><td></td><td></td><td></td><td></td><td></td><td></td><td></td></tr>
<tr><td>Oligo rev</td><td>AAACCTTCAAGGCCAGCGACTTGA</td><td></td><td></td><td></td><td></td><td></td><td></td><td></td><td></td><td></td><td></td></tr>
<tr><td>Chromosome</td><td>start</td><td>end</td><td>strand</td><td>MM</td><td>target_seq</td><td>PAM</td><td>alignment</td><td>distance</td><td>position</td><td>gene name</td><td>gene id</td></tr>
<tr><td>GU305914.1</td><td>151261</td><td>151283</td><td>+</td><td>0</td><td>TCAAGTCGCTGGCCTTGAAG</td><td>CGG</td><td>||||||||[||||||||||||]PAM</td><td>NA</td><td>Intergenic</td><td>NA</td><td>NA</td></tr>
<tr><td>T11588</td><td>CTCCCGCGAATCTCTGTACAAGG</td><td>1000</td><td></td><td></td><td></td><td></td><td></td><td></td><td></td><td></td><td></td></tr>
<tr><td>Oligo fwd</td><td>CACCCTCCCGCGAATCTCTGTACA</td><td></td><td></td><td></td><td></td><td></td><td></td><td></td><td></td><td></td><td></td></tr>
<tr><td>Oligo rev</td><td>AAACTGTACAGAGATTCGCGGGAG</td><td></td><td></td><td></td><td></td><td></td><td></td><td></td><td></td><td></td><td></td></tr>
<tr><td>Chromosome</td><td>start</td><td>end</td><td>strand</td><td>MM</td><td>target_seq</td><td>PAM</td><td>alignment</td><td>distance</td><td>position</td><td>gene name</td><td>gene id</td></tr>
<tr><td>GU305914.1</td><td>151286</td><td>151308</td><td>+</td><td>0</td><td>CTCCCGCGAATCTCTGTACA</td><td>AGG</td><td>||||||||[||||||||||||]PAM</td><td>NA</td><td>Intergenic</td><td>NA</td><td>NA</td></tr>
<tr><td>T11589</td><td>GCGAATCTCTGTACAAGGACTGG</td><td>1000</td><td></td><td></td><td></td><td></td><td></td><td></td><td></td><td></td><td></td></tr>
<tr><td>Oligo fwd</td><td>CACCGCGAATCTCTGTACAAGGAC</td><td></td><td></td><td></td><td></td><td></td><td></td><td></td><td></td><td></td><td></td></tr>
<tr><td>Oligo rev</td><td>AAACGTCCTTGTACAGAGATTCGC</td><td></td><td></td><td></td><td></td><td></td><td></td><td></td><td></td><td></td><td></td></tr>
<tr><td>Chromosome</td><td>start</td><td>end</td><td>strand</td><td>MM</td><td>target_seq</td><td>PAM</td><td>alignment</td><td>distance</td><td>position</td><td>gene name</td><td>gene id</td></tr>
<tr><td>GU305914.1</td><td>151291</td><td>151313</td><td>+</td><td>0</td><td>GCGAATCTCTGTACAAGGAC</td><td>TGG</td><td>||||||||[||||||||||||]PAM</td><td>NA</td><td>Intergenic</td><td>NA</td><td>NA</td></tr>
<tr><td>T11590</td><td>GGACTGGTTCCGCCACATGTTGG</td><td>1000</td><td></td><td></td><td></td><td></td><td></td><td></td><td></td><td></td><td></td></tr>
<tr><td>Oligo fwd</td><td>CACCGGACTGGTTCCGCCACATGT</td><td></td><td></td><td></td><td></td><td></td><td></td><td></td><td></td><td></td><td></td></tr>
<tr><td>Oligo rev</td><td>AAACACATGTGGCGGAACCAGTCC</td><td></td><td></td><td></td><td></td><td></td><td></td><td></td><td></td><td></td><td></td></tr>
<tr><td>Chromosome</td><td>start</td><td>end</td><td>strand</td><td>MM</td><td>target_seq</td><td>PAM</td><td>alignment</td><td>distance</td><td>position</td><td>gene name</td><td>gene id</td></tr>
<tr><td>GU305914.1</td><td>151307</td><td>151329</td><td>+</td><td>0</td><td>GGACTGGTTCCGCCACATGT</td><td>TGG</td><td>||||||||[||||||||||||]PAM</td><td>NA</td><td>Intergenic</td><td>NA</td><td>NA</td></tr>
<tr><td>T11591</td><td>CCACATGTTGGATCCCTGCGCGG</td><td>1000</td><td></td><td></td><td></td><td></td><td></td><td></td><td></td><td></td><td></td></tr>
<tr><td>Oligo fwd</td><td>CACCCCACATGTTGGATCCCTGCG</td><td></td><td></td><td></td><td></td><td></td><td></td><td></td><td></td><td></td><td></td></tr>
<tr><td>Oligo rev</td><td>AAACCGCAGGGATCCAACATGTGG</td><td></td><td></td><td></td><td></td><td></td><td></td><td></td><td></td><td></td><td></td></tr>
<tr><td>Chromosome</td><td>start</td><td>end</td><td>strand</td><td>MM</td><td>target_seq</td><td>PAM</td><td>alignment</td><td>distance</td><td>position</td><td>gene name</td><td>gene id</td></tr>
<tr><td>GU305914.1</td><td>151319</td><td>151341</td><td>+</td><td>0</td><td>CCACATGTTGGATCCCTGCG</td><td>CGG</td><td>||||||||[||||||||||||]PAM</td><td>NA</td><td>Intergenic</td><td>NA</td><td>NA</td></tr>
<tr><td>T11592</td><td>CACATGTTGGATCCCTGCGCGGG</td><td>1000</td><td></td><td></td><td></td><td></td><td></td><td></td><td></td><td></td><td></td></tr>
<tr><td>Oligo fwd</td><td>CACCCACATGTTGGATCCCTGCGC</td><td></td><td></td><td></td><td></td><td></td><td></td><td></td><td></td><td></td><td></td></tr>
<tr><td>Oligo rev</td><td>AAACGCGCAGGGATCCAACATGTG</td><td></td><td></td><td></td><td></td><td></td><td></td><td></td><td></td><td></td><td></td></tr>
</table>

| Chromosome | start | end | strand | MM | target_seq | PAM | alignment | distance | position | gene name | gene id |
|---|---|---|---|---|---|---|---|---|---|---|---|
| GU305914.1 | 151320 | 151342 | + | 0 | CACATGTTGGATCCCTGCGC | GGG | \|\|\|\|\|\|\|\|[\|\|\|\|\|\|\|\|\|\|\|\|]PAM | NA | Intergenic | NA | NA |
| T11593 | GTTGGATCCCTGCGCGGGCCTGG | 1000 | | | | | | | | | |
| Oligo fwd | CACCGTTGGATCCCTGCGCGGGCC | | | | | | | | | | |
| Oligo rev | AAACGGCCCGCGCAGGGATCCAAC | | | | | | | | | | |
| Chromosome | start | end | strand | MM | target_seq | PAM | alignment | distance | position | gene name | gene id |
| GU305914.1 | 151325 | 151347 | + | 0 | GTTGGATCCCTGCGCGGGCC | TGG | \|\|\|\|\|\|\|\|[\|\|\|\|\|\|\|\|\|\|\|\|]PAM | NA | Intergenic | NA | NA |
| T11594 | GGATCCCTGCGCGGGCCTGGTGG | 1000 | | | | | | | | | |
| Oligo fwd | CACCGGATCCCTGCGCGGGCCTGG | | | | | | | | | | |
| Oligo rev | AAACCCAGGCCCGCGCAGGGATCC | | | | | | | | | | |
| Chromosome | start | end | strand | MM | target_seq | PAM | alignment | distance | position | gene name | gene id |
| GU305914.1 | 151328 | 151350 | + | 0 | GGATCCCTGCGCGGGCCTGG | TGG | \|\|\|\|\|\|\|\|[\|\|\|\|\|\|\|\|\|\|\|\|]PAM | NA | Intergenic | NA | NA |
| T11595 | GGCCTGGTGGCGCCCGAGCTCGG | 1000 | | | | | | | | | |
| Oligo fwd | CACCGGCCTGGTGGCGCCCGAGCT | | | | | | | | | | |
| Oligo rev | AAACAGCTCGGGCGCCACCAGGCC | | | | | | | | | | |
| Chromosome | start | end | strand | MM | target_seq | PAM | alignment | distance | position | gene name | gene id |
| GU305914.1 | 151341 | 151363 | + | 0 | GGCCTGGTGGCGCCCGAGCT | CGG | \|\|\|\|\|\|\|\|[\|\|\|\|\|\|\|\|\|\|\|\|]PAM | NA | Intergenic | NA | NA |
| T11596 | GCGCCCGAGCTCGGCGACGACGG | 1000 | | | | | | | | | |
| Oligo fwd | CACCGCGCCCGAGCTCGGCGACGA | | | | | | | | | | |
| Oligo rev | AAACTCGTCGCCGAGCTCGGGCGC | | | | | | | | | | |
| Chromosome | start | end | strand | MM | target_seq | PAM | alignment | distance | position | gene name | gene id |
| GU305914.1 | 151350 | 151372 | + | 0 | GCGCCCGAGCTCGGCGACGA | CGG | \|\|\|\|\|\|\|\|[\|\|\|\|\|\|\|\|\|\|\|\|]PAM | NA | Intergenic | NA | NA |
| GU305914.1 | 58644 | 58666 | + | 4 | GCGGATGAGCTCGCCGACGA | CGG | \|\|\|---\|\|[\|\|\|\|\|-\|\|\|\|\|\|]PAM | NA | Intergenic | NA | NA |
| T11597 | GGCGACGACGGCAGCAGCGAAGG | 1000 | | | | | | | | | |
| Oligo fwd | CACCGGCGACGACGGCAGCAGCGA | | | | | | | | | | |
| Oligo rev | AAACTCGCTGCTGCCGTCGTCGCC | | | | | | | | | | |
| Chromosome | start | end | strand | MM | target_seq | PAM | alignment | distance | position | gene name | gene id |
| GU305914.1 | 151362 | 151384 | + | 0 | GGCGACGACGGCAGCAGCGA | AGG | \|\|\|\|\|\|\|\|[\|\|\|\|\|\|\|\|\|\|\|\|]PAM | NA | Intergenic | NA | NA |
| GU305914.1 | 113918 | 113940 | - | 4 | GTCGGCGGCGGCAGCGGCGA | CGG | \|-\|\|-\|\|-[\|\|\|\|\|\|\|-\|\|\|\|]PAM | NA | Intergenic | NA | NA |
| GU305914.1 | 164959 | 164981 | - | 4 | GGCGGCGGCGGCAGCAGCTG | CGG | \|\|\|\|-\|\|-[\|\|\|\|\|\|\|\|\|\|--]PAM | NA | Intergenic | NA | NA |
| T11598 | GACGACGGCAGCAGCGAAGGCGG | 1000 | | | | | | | | | |
| Oligo fwd | CACCGACGACGGCAGCAGCGAAGG | | | | | | | | | | |
| Oligo rev | AAACCCTTCGCTGCTGCCGTCGTC | | | | | | | | | | |
| Chromosome | start | end | strand | MM | target_seq | PAM | alignment | distance | position | gene name | gene id |
| GU305914.1 | 151365 | 151387 | + | 0 | GACGACGGCAGCAGCGAAGG | CGG | \|\|\|\|\|\|\|\|[\|\|\|\|\|\|\|\|\|\|\|\|]PAM | NA | Intergenic | NA | NA |
| T11599 | GACGGCAGCAGCGAAGGCGGCGG | 1000 | | | | | | | | | |
| Oligo fwd | CACCGACGGCAGCAGCGAAGGCGG | | | | | | | | | | |
| Oligo rev | AAACCCGCCTTCGCTGCTGCCGTC | | | | | | | | | | |
| Chromosome | start | end | strand | MM | target_seq | PAM | alignment | distance | position | gene name | gene id |
| GU305914.1 | 151368 | 151390 | + | 0 | GACGGCAGCAGCGAAGGCGG | CGG | \|\|\|\|\|\|\|\|[\|\|\|\|\|\|\|\|\|\|\|\|]PAM | NA | Intergenic | NA | NA |
| GU305914.1 | 73792 | 73814 | - | 4 | GGCGCCAGCAGCGGCGGCGG | AGG | \|-\|\|-\|\|\|[\|\|\|\|\|--\|\|\|\|\|]PAM | NA | Intergenic | NA | NA |
| T11600 | GGCAACGCCGCGATGATCGTCGG | 1000 | | | | | | | | | |
| Oligo fwd | CACCGGCAACGCCGCGATGATCGT | | | | | | | | | | |
| Oligo rev | AAACACGATCATCGCGGCGTTGCC | | | | | | | | | | |
| Chromosome | start | end | strand | MM | target_seq | PAM | alignment | distance | position | gene name | gene id |
| GU305914.1 | 151389 | 151411 | + | 0 | GGCAACGCCGCGATGATCGT | CGG | \|\|\|\|\|\|\|\|[\|\|\|\|\|\|\|\|\|\|\|\|]PAM | NA | Intergenic | NA | NA |
| T11601 | GATCGTCGGCGACCGCGAGCTGG | 1000 | | | | | | | | | |
| Oligo fwd | CACCGATCGTCGGCGACCGCGAGC | | | | | | | | | | |
| Oligo rev | AAACGCTCGCGGTCGCCGACGATC | | | | | | | | | | |
| Chromosome | start | end | strand | MM | target_seq | PAM | alignment | distance | position | gene name | gene id |
| GU305914.1 | 151403 | 151425 | + | 0 | GATCGTCGGCGACCGCGAGC | TGG | \|\|\|\|\|\|\|\|[\|\|\|\|\|\|\|\|\|\|\|\|]PAM | NA | Intergenic | NA | NA |
| T11602 | GCGACCGCGAGCTGGCGCGCCGG | 1000 | | | | | | | | | |
| Oligo fwd | CACCGCGACCGCGAGCTGGCGCGC | | | | | | | | | | |
| Oligo rev | AAACGCGCGCCAGCTCGCGGTCGC | | | | | | | | | | |
| Chromosome | start | end | strand | MM | target_seq | PAM | alignment | distance | position | gene name | gene id |
| GU305914.1 | 151411 | 151433 | + | 0 | GCGACCGCGAGCTGGCGCGC | CGG | \|\|\|\|\|\|\|\|[\|\|\|\|\|\|\|\|\|\|\|\|]PAM | NA | Intergenic | NA | NA |
| T11603 | TTCTCCTGCCTCCTGATCACCGG | 1000 | | | | | | | | | |
| Oligo fwd | CACCTTCTCCTGCCTCCTGATCAC | | | | | | | | | | |
| Oligo rev | AAACGTGATCAGGAGGCAGGAGAA | | | | | | | | | | |
| Chromosome | start | end | strand | MM | target_seq | PAM | alignment | distance | position | gene name | gene id |
| GU305914.1 | 151449 | 151471 | + | 0 | TTCTCCTGCCTCCTGATCAC | CGG | \|\|\|\|\|\|\|\|[\|\|\|\|\|\|\|\|\|\|\|\|]PAM | NA | Intergenic | NA | NA |
| T11604 | CTCCTGATCACCGGTACCGCCGG | 1000 | | | | | | | | | |
| Oligo fwd | CACCCTCCTGATCACCGGTACCGC | | | | | | | | | | |
| Oligo rev | AAACGCGGTACCGGTGATCAGGAG | | | | | | | | | | |
| Chromosome | start | end | strand | MM | target_seq | PAM | alignment | distance | position | gene name | gene id |
| GU305914.1 | 151458 | 151480 | + | 0 | CTCCTGATCACCGGTACCGC | CGG | \|\|\|\|\|\|\|\|[\|\|\|\|\|\|\|\|\|\|\|\|]PAM | NA | Intergenic | NA | NA |
| T11605 | GATCACCGGTACCGCCGGCGCGG | 1000 | | | | | | | | | |
| Oligo fwd | CACCGATCACCGGTACCGCCGGCG | | | | | | | | | | |
| Oligo rev | AAACCGCCGGCGGTACCGGTGATC | | | | | | | | | | |
| Chromosome | start | end | strand | MM | target_seq | PAM | alignment | distance | position | gene name | gene id |
| GU305914.1 | 151463 | 151485 | + | 0 | GATCACCGGTACCGCCGGCG | CGG | \|\|\|\|\|\|\|\|[\|\|\|\|\|\|\|\|\|\|\|\|]PAM | NA | Intergenic | NA | NA |
| T11606 | ATCACCGGTACCGCCGGCGCGGG | 1000 | | | | | | | | | |
| Oligo fwd | CACCATCACCGGTACCGCCGGCGC | | | | | | | | | | |
| Oligo rev | AAACGCGCCGGCGGTACCGGTGAT | | | | | | | | | | |
| Chromosome | start | end | strand | MM | target_seq | PAM | alignment | distance | position | gene name | gene id |

<table>
<tr><td>GU305914.1</td><td></td><td>151464</td><td>151486</td><td>+</td><td>0</td><td>ATCACCGGTACCGCCGGCGC</td><td>GGG</td><td>||||||||[||||||||||||]PAM</td><td>NA</td><td>Intergenic</td><td>NA</td><td>NA</td></tr>
<tr><td>T11607</td><td>TCACCGGTACCGCCGGCGCGGGG</td><td></td><td>1000</td><td></td><td></td><td></td><td></td><td></td><td></td><td></td><td></td><td></td></tr>
<tr><td>Oligo fwd</td><td>CACCTCACCGGTACCGCCGGCGCG</td><td></td><td></td><td></td><td></td><td></td><td></td><td></td><td></td><td></td><td></td><td></td></tr>
<tr><td>Oligo rev</td><td>AAACCGCGCCGGCGGTACCGGTGA</td><td></td><td></td><td></td><td></td><td></td><td></td><td></td><td></td><td></td><td></td><td></td></tr>
<tr><td>Chromosome</td><td>start</td><td></td><td>end</td><td>strand</td><td>MM</td><td>target_seq</td><td>PAM</td><td>alignment</td><td>distance</td><td>position</td><td>gene name</td><td>gene id</td></tr>
<tr><td>GU305914.1</td><td></td><td>151465</td><td>151487</td><td>+</td><td>0</td><td>TCACCGGTACCGCCGGCGCG</td><td>GGG</td><td>||||||||[||||||||||||]PAM</td><td>NA</td><td>Intergenic</td><td>NA</td><td>NA</td></tr>
<tr><td>T11608</td><td>GGGGAAGACCTCGAGCGTCCAGG</td><td></td><td>1000</td><td></td><td></td><td></td><td></td><td></td><td></td><td></td><td></td><td></td></tr>
<tr><td>Oligo fwd</td><td>CACCGGGGAAGACCTCGAGCGTCC</td><td></td><td></td><td></td><td></td><td></td><td></td><td></td><td></td><td></td><td></td><td></td></tr>
<tr><td>Oligo rev</td><td>AAACGGACGCTCGAGGTCTTCCCC</td><td></td><td></td><td></td><td></td><td></td><td></td><td></td><td></td><td></td><td></td><td></td></tr>
<tr><td>Chromosome</td><td>start</td><td></td><td>end</td><td>strand</td><td>MM</td><td>target_seq</td><td>PAM</td><td>alignment</td><td>distance</td><td>position</td><td>gene name</td><td>gene id</td></tr>
<tr><td>GU305914.1</td><td></td><td>151484</td><td>151506</td><td>+</td><td>0</td><td>GGGGAAGACCTCGAGCGTCC</td><td>AGG</td><td>||||||||[||||||||||||]PAM</td><td>NA</td><td>Intergenic</td><td>NA</td><td>NA</td></tr>
<tr><td>T11609</td><td>GACCTCGAGCGTCCAGGTCCTGG</td><td></td><td>1000</td><td></td><td></td><td></td><td></td><td></td><td></td><td></td><td></td><td></td></tr>
<tr><td>Oligo fwd</td><td>CACCGACCTCGAGCGTCCAGGTCC</td><td></td><td></td><td></td><td></td><td></td><td></td><td></td><td></td><td></td><td></td><td></td></tr>
<tr><td>Oligo rev</td><td>AAACGGACCTGGACGCTCGAGGTC</td><td></td><td></td><td></td><td></td><td></td><td></td><td></td><td></td><td></td><td></td><td></td></tr>
<tr><td>Chromosome</td><td>start</td><td></td><td>end</td><td>strand</td><td>MM</td><td>target_seq</td><td>PAM</td><td>alignment</td><td>distance</td><td>position</td><td>gene name</td><td>gene id</td></tr>
<tr><td>GU305914.1</td><td></td><td>151490</td><td>151512</td><td>+</td><td>0</td><td>GACCTCGAGCGTCCAGGTCC</td><td>TGG</td><td>||||||||[||||||||||||]PAM</td><td>NA</td><td>Intergenic</td><td>NA</td><td>NA</td></tr>
<tr><td>T11610</td><td>TAACCTCGACTGCGTGATCACGG</td><td></td><td>1000</td><td></td><td></td><td></td><td></td><td></td><td></td><td></td><td></td><td></td></tr>
<tr><td>Oligo fwd</td><td>CACCTAACCTCGACTGCGTGATCA</td><td></td><td></td><td></td><td></td><td></td><td></td><td></td><td></td><td></td><td></td><td></td></tr>
<tr><td>Oligo rev</td><td>AAACTGATCACGCAGTCGAGGTTA</td><td></td><td></td><td></td><td></td><td></td><td></td><td></td><td></td><td></td><td></td><td></td></tr>
<tr><td>Chromosome</td><td>start</td><td></td><td>end</td><td>strand</td><td>MM</td><td>target_seq</td><td>PAM</td><td>alignment</td><td>distance</td><td>position</td><td>gene name</td><td>gene id</td></tr>
<tr><td>GU305914.1</td><td></td><td>151517</td><td>151539</td><td>+</td><td>0</td><td>TAACCTCGACTGCGTGATCA</td><td>CGG</td><td>||||||||[||||||||||||]PAM</td><td>NA</td><td>Intergenic</td><td>NA</td><td>NA</td></tr>
<tr><td>T11611</td><td>AACCTCGACTGCGTGATCACGGG</td><td></td><td>1000</td><td></td><td></td><td></td><td></td><td></td><td></td><td></td><td></td><td></td></tr>
<tr><td>Oligo fwd</td><td>CACCAACCTCGACTGCGTGATCAC</td><td></td><td></td><td></td><td></td><td></td><td></td><td></td><td></td><td></td><td></td><td></td></tr>
<tr><td>Oligo rev</td><td>AAACGTGATCACGCAGTCGAGGTT</td><td></td><td></td><td></td><td></td><td></td><td></td><td></td><td></td><td></td><td></td><td></td></tr>
<tr><td>Chromosome</td><td>start</td><td></td><td>end</td><td>strand</td><td>MM</td><td>target_seq</td><td>PAM</td><td>alignment</td><td>distance</td><td>position</td><td>gene name</td><td>gene id</td></tr>
<tr><td>GU305914.1</td><td></td><td>151518</td><td>151540</td><td>+</td><td>0</td><td>AACCTCGACTGCGTGATCAC</td><td>GGG</td><td>||||||||[||||||||||||]PAM</td><td>NA</td><td>Intergenic</td><td>NA</td><td>NA</td></tr>
<tr><td>T11612</td><td>CAGCACCGTGATCTCCTCCCAGG</td><td></td><td>1000</td><td></td><td></td><td></td><td></td><td></td><td></td><td></td><td></td><td></td></tr>
<tr><td>Oligo fwd</td><td>CACCCAGCACCGTGATCTCCTCCC</td><td></td><td></td><td></td><td></td><td></td><td></td><td></td><td></td><td></td><td></td><td></td></tr>
<tr><td>Oligo rev</td><td>AAACGGGAGGAGATCACGGTGCTG</td><td></td><td></td><td></td><td></td><td></td><td></td><td></td><td></td><td></td><td></td><td></td></tr>
<tr><td>Chromosome</td><td>start</td><td></td><td>end</td><td>strand</td><td>MM</td><td>target_seq</td><td>PAM</td><td>alignment</td><td>distance</td><td>position</td><td>gene name</td><td>gene id</td></tr>
<tr><td>GU305914.1</td><td></td><td>151541</td><td>151563</td><td>+</td><td>0</td><td>CAGCACCGTGATCTCCTCCC</td><td>AGG</td><td>||||||||[||||||||||||]PAM</td><td>NA</td><td>Intergenic</td><td>NA</td><td>NA</td></tr>
<tr><td>T11613</td><td>CTCAGATCAAGACCATCTTCAGG</td><td></td><td>1000</td><td></td><td></td><td></td><td></td><td></td><td></td><td></td><td></td><td></td></tr>
<tr><td>Oligo fwd</td><td>CACCCTCAGATCAAGACCATCTTC</td><td></td><td></td><td></td><td></td><td></td><td></td><td></td><td></td><td></td><td></td><td></td></tr>
<tr><td>Oligo rev</td><td>AAACGAAGATGGTCTTGATCTGAG</td><td></td><td></td><td></td><td></td><td></td><td></td><td></td><td></td><td></td><td></td><td></td></tr>
<tr><td>Chromosome</td><td>start</td><td></td><td>end</td><td>strand</td><td>MM</td><td>target_seq</td><td>PAM</td><td>alignment</td><td>distance</td><td>position</td><td>gene name</td><td>gene id</td></tr>
<tr><td>GU305914.1</td><td></td><td>151597</td><td>151619</td><td>+</td><td>0</td><td>CTCAGATCAAGACCATCTTC</td><td>AGG</td><td>||||||||[||||||||||||]PAM</td><td>NA</td><td>Intergenic</td><td>NA</td><td>NA</td></tr>
<tr><td>T11614</td><td>AAGACCATCTTCAGGACTTTTGG</td><td></td><td>1000</td><td></td><td></td><td></td><td></td><td></td><td></td><td></td><td></td><td></td></tr>
<tr><td>Oligo fwd</td><td>CACCAAGACCATCTTCAGGACTTT</td><td></td><td></td><td></td><td></td><td></td><td></td><td></td><td></td><td></td><td></td><td></td></tr>
<tr><td>Oligo rev</td><td>AAACAAAGTCCTGAAGATGGTCTT</td><td></td><td></td><td></td><td></td><td></td><td></td><td></td><td></td><td></td><td></td><td></td></tr>
<tr><td>Chromosome</td><td>start</td><td></td><td>end</td><td>strand</td><td>MM</td><td>target_seq</td><td>PAM</td><td>alignment</td><td>distance</td><td>position</td><td>gene name</td><td>gene id</td></tr>
<tr><td>GU305914.1</td><td></td><td>151605</td><td>151627</td><td>+</td><td>0</td><td>AAGACCATCTTCAGGACTTT</td><td>TGG</td><td>||||||||[||||||||||||]PAM</td><td>NA</td><td>Intergenic</td><td>NA</td><td>NA</td></tr>
<tr><td>T11615</td><td>AGACCATCTTCAGGACTTTTGGG</td><td></td><td>1000</td><td></td><td></td><td></td><td></td><td></td><td></td><td></td><td></td><td></td></tr>
<tr><td>Oligo fwd</td><td>CACCAGACCATCTTCAGGACTTTT</td><td></td><td></td><td></td><td></td><td></td><td></td><td></td><td></td><td></td><td></td><td></td></tr>
<tr><td>Oligo rev</td><td>AAACAAAAGTCCTGAAGATGGTCT</td><td></td><td></td><td></td><td></td><td></td><td></td><td></td><td></td><td></td><td></td><td></td></tr>
<tr><td>Chromosome</td><td>start</td><td></td><td>end</td><td>strand</td><td>MM</td><td>target_seq</td><td>PAM</td><td>alignment</td><td>distance</td><td>position</td><td>gene name</td><td>gene id</td></tr>
<tr><td>GU305914.1</td><td></td><td>151606</td><td>151628</td><td>+</td><td>0</td><td>AGACCATCTTCAGGACTTTT</td><td>GGG</td><td>||||||||[||||||||||||]PAM</td><td>NA</td><td>Intergenic</td><td>NA</td><td>NA</td></tr>
<tr><td>T11616</td><td>GGACTTTTGGGTTTAATAGTCGG</td><td></td><td>1000</td><td></td><td></td><td></td><td></td><td></td><td></td><td></td><td></td><td></td></tr>
<tr><td>Oligo fwd</td><td>CACCGGACTTTTGGGTTTAATAGT</td><td></td><td></td><td></td><td></td><td></td><td></td><td></td><td></td><td></td><td></td><td></td></tr>
<tr><td>Oligo rev</td><td>AAACACTATTAAACCCAAAAGTCC</td><td></td><td></td><td></td><td></td><td></td><td></td><td></td><td></td><td></td><td></td><td></td></tr>
<tr><td>Chromosome</td><td>start</td><td></td><td>end</td><td>strand</td><td>MM</td><td>target_seq</td><td>PAM</td><td>alignment</td><td>distance</td><td>position</td><td>gene name</td><td>gene id</td></tr>
<tr><td>GU305914.1</td><td></td><td>151618</td><td>151640</td><td>+</td><td>0</td><td>GGACTTTTGGGTTTAATAGT</td><td>CGG</td><td>||||||||[||||||||||||]PAM</td><td>NA</td><td>Intergenic</td><td>NA</td><td>NA</td></tr>
<tr><td>T11617</td><td>TCGCCGACCGCGTTCACCTCAGG</td><td></td><td>1000</td><td></td><td></td><td></td><td></td><td></td><td></td><td></td><td></td><td></td></tr>
<tr><td>Oligo fwd</td><td>CACCTCGCCGACCGCGTTCACCTC</td><td></td><td></td><td></td><td></td><td></td><td></td><td></td><td></td><td></td><td></td><td></td></tr>
<tr><td>Oligo rev</td><td>AAACGAGGTGAACGCGGTCGGCGA</td><td></td><td></td><td></td><td></td><td></td><td></td><td></td><td></td><td></td><td></td><td></td></tr>
<tr><td>Chromosome</td><td>start</td><td></td><td>end</td><td>strand</td><td>MM</td><td>target_seq</td><td>PAM</td><td>alignment</td><td>distance</td><td>position</td><td>gene name</td><td>gene id</td></tr>
<tr><td>GU305914.1</td><td></td><td>151651</td><td>151673</td><td>+</td><td>0</td><td>TCGCCGACCGCGTTCACCTC</td><td>AGG</td><td>||||||||[||||||||||||]PAM</td><td>NA</td><td>Intergenic</td><td>NA</td><td>NA</td></tr>
<tr><td>T11618</td><td>CCGACCGCGTTCACCTCAGGCGG</td><td></td><td>1000</td><td></td><td></td><td></td><td></td><td></td><td></td><td></td><td></td><td></td></tr>
<tr><td>Oligo fwd</td><td>CACCCCGACCGCGTTCACCTCAGG</td><td></td><td></td><td></td><td></td><td></td><td></td><td></td><td></td><td></td><td></td><td></td></tr>
<tr><td>Oligo rev</td><td>AAACCCTGAGGTGAACGCGGTCGG</td><td></td><td></td><td></td><td></td><td></td><td></td><td></td><td></td><td></td><td></td><td></td></tr>
<tr><td>Chromosome</td><td>start</td><td></td><td>end</td><td>strand</td><td>MM</td><td>target_seq</td><td>PAM</td><td>alignment</td><td>distance</td><td>position</td><td>gene name</td><td>gene id</td></tr>
<tr><td>GU305914.1</td><td></td><td>151654</td><td>151676</td><td>+</td><td>0</td><td>CCGACCGCGTTCACCTCAGG</td><td>CGG</td><td>||||||||[||||||||||||]PAM</td><td>NA</td><td>Intergenic</td><td>NA</td><td>NA</td></tr>
<tr><td>T11619</td><td>CGCGACGACGTCGCGTTCGACGG</td><td></td><td>1000</td><td></td><td></td><td></td><td></td><td></td><td></td><td></td><td></td><td></td></tr>
<tr><td>Oligo fwd</td><td>CACCCGCGACGACGTCGCGTTCGA</td><td></td><td></td><td></td><td></td><td></td><td></td><td></td><td></td><td></td><td></td><td></td></tr>
<tr><td>Oligo rev</td><td>AAACTCGAACGCGACGTCGTCGCG</td><td></td><td></td><td></td><td></td><td></td><td></td><td></td><td></td><td></td><td></td><td></td></tr>
<tr><td>Chromosome</td><td>start</td><td></td><td>end</td><td>strand</td><td>MM</td><td>target_seq</td><td>PAM</td><td>alignment</td><td>distance</td><td>position</td><td>gene name</td><td>gene id</td></tr>
<tr><td>GU305914.1</td><td></td><td>151677</td><td>151699</td><td>+</td><td>0</td><td>CGCGACGACGTCGCGTTCGA</td><td>CGG</td><td>||||||||[||||||||||||]PAM</td><td>NA</td><td>Intergenic</td><td>NA</td><td>NA</td></tr>
<tr><td>GU305914.1</td><td></td><td>159727</td><td>159749</td><td>+</td><td>4</td><td>CGAGACGTCGGCGCGTTCGG</td><td>CGG</td><td>||-||||-[||-||||||||-]PAM</td><td>NA</td><td>Intergenic</td><td>NA</td><td>NA</td></tr>
<tr><td>T11620</td><td>ACCCCATCTGCCAGCAGCAGTGG</td><td></td><td>1000</td><td></td><td></td><td></td><td></td><td></td><td></td><td></td><td></td><td></td></tr>
<tr><td>Oligo fwd</td><td>CACCACCCCATCTGCCAGCAGCAG</td><td></td><td></td><td></td><td></td><td></td><td></td><td></td><td></td><td></td><td></td><td></td></tr>
<tr><td>Oligo rev</td><td>AAACCTGCTGCTGGCAGATGGGGT</td><td></td><td></td><td></td><td></td><td></td><td></td><td></td><td></td><td></td><td></td><td></td></tr>
<tr><td>Chromosome</td><td>start</td><td></td><td>end</td><td>strand</td><td>MM</td><td>target_seq</td><td>PAM</td><td>alignment</td><td>distance</td><td>position</td><td>gene name</td><td>gene id</td></tr>
<tr><td>GU305914.1</td><td></td><td>151708</td><td>151730</td><td>+</td><td>0</td><td>ACCCCATCTGCCAGCAGCAG</td><td>TGG</td><td>||||||||[||||||||||||]PAM</td><td>NA</td><td>Intergenic</td><td>NA</td><td>NA</td></tr>
<tr><td>T11621</td><td>GGCGCGATCTCTCGACCTACTGG</td><td></td><td>1000</td><td></td><td></td><td></td><td></td><td></td><td></td><td></td><td></td><td></td></tr>
<tr><td>Oligo fwd</td><td>CACCGGCGCGATCTCTCGACCTAC</td><td></td><td></td><td></td><td></td><td></td><td></td><td></td><td></td><td></td><td></td><td></td></tr>
</table>

<table>
<tr><td>Oligo rev</td><td>AAACGTAGGTCGAGAGATCGCGCC</td><td></td><td></td><td></td><td></td><td></td><td></td><td></td><td></td><td></td><td></td></tr>
<tr><td>Chromosome</td><td>start</td><td>end</td><td>strand</td><td>MM</td><td>target_seq</td><td>PAM</td><td>alignment</td><td>distance</td><td>position</td><td>gene name</td><td>gene id</td></tr>
<tr><td>GU305914.1</td><td>151729</td><td>151751</td><td>+</td><td>0</td><td>GGCGCGATCTCTCGACCTAC</td><td>TGG</td><td>||||||||[||||||||||||]PAM</td><td>NA</td><td>Intergenic</td><td>NA</td><td>NA</td></tr>
<tr><td>T11622</td><td>CGATCTCTCGACCTACTGGCCGG</td><td>1000</td><td></td><td></td><td></td><td></td><td></td><td></td><td></td><td></td><td></td></tr>
<tr><td>Oligo fwd</td><td>CACCCGATCTCTCGACCTACTGGC</td><td></td><td></td><td></td><td></td><td></td><td></td><td></td><td></td><td></td><td></td></tr>
<tr><td>Oligo rev</td><td>AAACGCCAGTAGGTCGAGAGATCG</td><td></td><td></td><td></td><td></td><td></td><td></td><td></td><td></td><td></td><td></td></tr>
<tr><td>Chromosome</td><td>start</td><td>end</td><td>strand</td><td>MM</td><td>target_seq</td><td>PAM</td><td>alignment</td><td>distance</td><td>position</td><td>gene name</td><td>gene id</td></tr>
<tr><td>GU305914.1</td><td>151733</td><td>151755</td><td>+</td><td>0</td><td>CGATCTCTCGACCTACTGGC</td><td>CGG</td><td>||||||||[||||||||||||]PAM</td><td>NA</td><td>Intergenic</td><td>NA</td><td>NA</td></tr>
<tr><td>T11623</td><td>CGACATCGCCATCCGCGCCCTGG</td><td>1000</td><td></td><td></td><td></td><td></td><td></td><td></td><td></td><td></td><td></td></tr>
<tr><td>Oligo fwd</td><td>CACCCGACATCGCCATCCGCGCCC</td><td></td><td></td><td></td><td></td><td></td><td></td><td></td><td></td><td></td><td></td></tr>
<tr><td>Oligo rev</td><td>AAACGGGCGCGGATGGCGATGTCG</td><td></td><td></td><td></td><td></td><td></td><td></td><td></td><td></td><td></td><td></td></tr>
<tr><td>Chromosome</td><td>start</td><td>end</td><td>strand</td><td>MM</td><td>target_seq</td><td>PAM</td><td>alignment</td><td>distance</td><td>position</td><td>gene name</td><td>gene id</td></tr>
<tr><td>GU305914.1</td><td>151763</td><td>151785</td><td>+</td><td>0</td><td>CGACATCGCCATCCGCGCCC</td><td>TGG</td><td>||||||||[||||||||||||]PAM</td><td>NA</td><td>Intergenic</td><td>NA</td><td>NA</td></tr>
<tr><td>T11624</td><td>ATCGCCATCCGCGCCCTGGACGG</td><td>1000</td><td></td><td></td><td></td><td></td><td></td><td></td><td></td><td></td><td></td></tr>
<tr><td>Oligo fwd</td><td>CACCATCGCCATCCGCGCCCTGGA</td><td></td><td></td><td></td><td></td><td></td><td></td><td></td><td></td><td></td><td></td></tr>
<tr><td>Oligo rev</td><td>AAACTCCAGGGCGCGGATGGCGAT</td><td></td><td></td><td></td><td></td><td></td><td></td><td></td><td></td><td></td><td></td></tr>
<tr><td>Chromosome</td><td>start</td><td>end</td><td>strand</td><td>MM</td><td>target_seq</td><td>PAM</td><td>alignment</td><td>distance</td><td>position</td><td>gene name</td><td>gene id</td></tr>
<tr><td>GU305914.1</td><td>151767</td><td>151789</td><td>+</td><td>0</td><td>ATCGCCATCCGCGCCCTGGA</td><td>CGG</td><td>||||||||[||||||||||||]PAM</td><td>NA</td><td>Intergenic</td><td>NA</td><td>NA</td></tr>
<tr><td>T11625</td><td>GCCATCCGCGCCCTGGACGGCGG</td><td>1000</td><td></td><td></td><td></td><td></td><td></td><td></td><td></td><td></td><td></td></tr>
<tr><td>Oligo fwd</td><td>CACCGCCATCCGCGCCCTGGACGG</td><td></td><td></td><td></td><td></td><td></td><td></td><td></td><td></td><td></td><td></td></tr>
<tr><td>Oligo rev</td><td>AAACCCGTCCAGGGCGCGGATGGC</td><td></td><td></td><td></td><td></td><td></td><td></td><td></td><td></td><td></td><td></td></tr>
<tr><td>Chromosome</td><td>start</td><td>end</td><td>strand</td><td>MM</td><td>target_seq</td><td>PAM</td><td>alignment</td><td>distance</td><td>position</td><td>gene name</td><td>gene id</td></tr>
<tr><td>GU305914.1</td><td>151770</td><td>151792</td><td>+</td><td>0</td><td>GCCATCCGCGCCCTGGACGG</td><td>CGG</td><td>||||||||[||||||||||||]PAM</td><td>NA</td><td>Intergenic</td><td>NA</td><td>NA</td></tr>
<tr><td>T11626</td><td>CCGCGCCCTGGACGGCGGCAAGG</td><td>1000</td><td></td><td></td><td></td><td></td><td></td><td></td><td></td><td></td><td></td></tr>
<tr><td>Oligo fwd</td><td>CACCCCGCGCCCTGGACGGCGGCA</td><td></td><td></td><td></td><td></td><td></td><td></td><td></td><td></td><td></td><td></td></tr>
<tr><td>Oligo rev</td><td>AAACTGCCGCCGTCCAGGGCGCGG</td><td></td><td></td><td></td><td></td><td></td><td></td><td></td><td></td><td></td><td></td></tr>
<tr><td>Chromosome</td><td>start</td><td>end</td><td>strand</td><td>MM</td><td>target_seq</td><td>PAM</td><td>alignment</td><td>distance</td><td>position</td><td>gene name</td><td>gene id</td></tr>
<tr><td>GU305914.1</td><td>151775</td><td>151797</td><td>+</td><td>0</td><td>CCGCGCCCTGGACGGCGGCA</td><td>AGG</td><td>||||||||[||||||||||||]PAM</td><td>NA</td><td>Intergenic</td><td>NA</td><td>NA</td></tr>
<tr><td>T11627</td><td>CGCGCCCTGGACGGCGGCAAGGG</td><td>1000</td><td></td><td></td><td></td><td></td><td></td><td></td><td></td><td></td><td></td></tr>
<tr><td>Oligo fwd</td><td>CACCCGCGCCCTGGACGGCGGCAA</td><td></td><td></td><td></td><td></td><td></td><td></td><td></td><td></td><td></td><td></td></tr>
<tr><td>Oligo rev</td><td>AAACTTGCCGCCGTCCAGGGCGCG</td><td></td><td></td><td></td><td></td><td></td><td></td><td></td><td></td><td></td><td></td></tr>
<tr><td>Chromosome</td><td>start</td><td>end</td><td>strand</td><td>MM</td><td>target_seq</td><td>PAM</td><td>alignment</td><td>distance</td><td>position</td><td>gene name</td><td>gene id</td></tr>
<tr><td>GU305914.1</td><td>151776</td><td>151798</td><td>+</td><td>0</td><td>CGCGCCCTGGACGGCGGCAA</td><td>GGG</td><td>||||||||[||||||||||||]PAM</td><td>NA</td><td>Intergenic</td><td>NA</td><td>NA</td></tr>
<tr><td>T11628</td><td>GGACGGCGGCAAGGGCCGCAAGG</td><td>1000</td><td></td><td></td><td></td><td></td><td></td><td></td><td></td><td></td><td></td></tr>
<tr><td>Oligo fwd</td><td>CACCGGACGGCGGCAAGGGCCGCA</td><td></td><td></td><td></td><td></td><td></td><td></td><td></td><td></td><td></td><td></td></tr>
<tr><td>Oligo rev</td><td>AAACTGCGGCCCTTGCCGCCGTCC</td><td></td><td></td><td></td><td></td><td></td><td></td><td></td><td></td><td></td><td></td></tr>
<tr><td>Chromosome</td><td>start</td><td>end</td><td>strand</td><td>MM</td><td>target_seq</td><td>PAM</td><td>alignment</td><td>distance</td><td>position</td><td>gene name</td><td>gene id</td></tr>
<tr><td>GU305914.1</td><td>151784</td><td>151806</td><td>+</td><td>0</td><td>GGACGGCGGCAAGGGCCGCA</td><td>AGG</td><td>||||||||[||||||||||||]PAM</td><td>NA</td><td>Intergenic</td><td>NA</td><td>NA</td></tr>
<tr><td>T11629</td><td>ATCATCGTCATCGACGAGTGCGG</td><td>1000</td><td></td><td></td><td></td><td></td><td></td><td></td><td></td><td></td><td></td></tr>
<tr><td>Oligo fwd</td><td>CACCATCATCGTCATCGACGAGTG</td><td></td><td></td><td></td><td></td><td></td><td></td><td></td><td></td><td></td><td></td></tr>
<tr><td>Oligo rev</td><td>AAACCACTCGTCGATGACGATGAT</td><td></td><td></td><td></td><td></td><td></td><td></td><td></td><td></td><td></td><td></td></tr>
<tr><td>Chromosome</td><td>start</td><td>end</td><td>strand</td><td>MM</td><td>target_seq</td><td>PAM</td><td>alignment</td><td>distance</td><td>position</td><td>gene name</td><td>gene id</td></tr>
<tr><td>GU305914.1</td><td>151833</td><td>151855</td><td>+</td><td>0</td><td>ATCATCGTCATCGACGAGTG</td><td>CGG</td><td>||||||||[||||||||||||]PAM</td><td>NA</td><td>Intergenic</td><td>NA</td><td>NA</td></tr>
<tr><td>T11630</td><td>CGAGCTCTACCGCCAGCGCGCGG</td><td>1000</td><td></td><td></td><td></td><td></td><td></td><td></td><td></td><td></td><td></td></tr>
<tr><td>Oligo fwd</td><td>CACCCGAGCTCTACCGCCAGCGCG</td><td></td><td></td><td></td><td></td><td></td><td></td><td></td><td></td><td></td><td></td></tr>
<tr><td>Oligo rev</td><td>AAACCGCGCTGGCGGTAGAGCTCG</td><td></td><td></td><td></td><td></td><td></td><td></td><td></td><td></td><td></td><td></td></tr>
<tr><td>Chromosome</td><td>start</td><td>end</td><td>strand</td><td>MM</td><td>target_seq</td><td>PAM</td><td>alignment</td><td>distance</td><td>position</td><td>gene name</td><td>gene id</td></tr>
<tr><td>GU305914.1</td><td>151925</td><td>151947</td><td>+</td><td>0</td><td>CGAGCTCTACCGCCAGCGCG</td><td>CGG</td><td>||||||||[||||||||||||]PAM</td><td>NA</td><td>Intergenic</td><td>NA</td><td>NA</td></tr>
<tr><td>T11631</td><td>GCGCCCTGCATCGTCTGCGTCGG</td><td>1000</td><td></td><td></td><td></td><td></td><td></td><td></td><td></td><td></td><td></td></tr>
<tr><td>Oligo fwd</td><td>CACCGCGCCCTGCATCGTCTGCGT</td><td></td><td></td><td></td><td></td><td></td><td></td><td></td><td></td><td></td><td></td></tr>
<tr><td>Oligo rev</td><td>AAACACGCAGACGATGCAGGGCGC</td><td></td><td></td><td></td><td></td><td></td><td></td><td></td><td></td><td></td><td></td></tr>
<tr><td>Chromosome</td><td>start</td><td>end</td><td>strand</td><td>MM</td><td>target_seq</td><td>PAM</td><td>alignment</td><td>distance</td><td>position</td><td>gene name</td><td>gene id</td></tr>
<tr><td>GU305914.1</td><td>151947</td><td>151969</td><td>+</td><td>0</td><td>GCGCCCTGCATCGTCTGCGT</td><td>CGG</td><td>||||||||[||||||||||||]PAM</td><td>NA</td><td>Intergenic</td><td>NA</td><td>NA</td></tr>
<tr><td>T11632</td><td>CGCCCTGCATCGTCTGCGTCGGG</td><td>1000</td><td></td><td></td><td></td><td></td><td></td><td></td><td></td><td></td><td></td></tr>
<tr><td>Oligo fwd</td><td>CACCCGCCCTGCATCGTCTGCGTC</td><td></td><td></td><td></td><td></td><td></td><td></td><td></td><td></td><td></td><td></td></tr>
<tr><td>Oligo rev</td><td>AAACGACGCAGACGATGCAGGGCG</td><td></td><td></td><td></td><td></td><td></td><td></td><td></td><td></td><td></td><td></td></tr>
<tr><td>Chromosome</td><td>start</td><td>end</td><td>strand</td><td>MM</td><td>target_seq</td><td>PAM</td><td>alignment</td><td>distance</td><td>position</td><td>gene name</td><td>gene id</td></tr>
<tr><td>GU305914.1</td><td>151948</td><td>151970</td><td>+</td><td>0</td><td>CGCCCTGCATCGTCTGCGTC</td><td>GGG</td><td>||||||||[||||||||||||]PAM</td><td>NA</td><td>Intergenic</td><td>NA</td><td>NA</td></tr>
<tr><td>T11633</td><td>CGGGTCGCCCACGCAGTCCGAGG</td><td>1000</td><td></td><td></td><td></td><td></td><td></td><td></td><td></td><td></td><td></td></tr>
<tr><td>Oligo fwd</td><td>CACCCGGGTCGCCCACGCAGTCCG</td><td></td><td></td><td></td><td></td><td></td><td></td><td></td><td></td><td></td><td></td></tr>
<tr><td>Oligo rev</td><td>AAACCGGACTGCGTGGGCGACCCG</td><td></td><td></td><td></td><td></td><td></td><td></td><td></td><td></td><td></td><td></td></tr>
<tr><td>Chromosome</td><td>start</td><td>end</td><td>strand</td><td>MM</td><td>target_seq</td><td>PAM</td><td>alignment</td><td>distance</td><td>position</td><td>gene name</td><td>gene id</td></tr>
<tr><td>GU305914.1</td><td>151967</td><td>151989</td><td>+</td><td>0</td><td>CGGGTCGCCCACGCAGTCCG</td><td>AGG</td><td>||||||||[||||||||||||]PAM</td><td>NA</td><td>Intergenic</td><td>NA</td><td>NA</td></tr>
<tr><td>T11634</td><td>ACGACCACCGCACGCAGAATCGG</td><td>1000</td><td></td><td></td><td></td><td></td><td></td><td></td><td></td><td></td><td></td></tr>
<tr><td>Oligo fwd</td><td>CACCACGACCACCGCACGCAGAAT</td><td></td><td></td><td></td><td></td><td></td><td></td><td></td><td></td><td></td><td></td></tr>
<tr><td>Oligo rev</td><td>AAACATTCTGCGTGCGGTGGTCGT</td><td></td><td></td><td></td><td></td><td></td><td></td><td></td><td></td><td></td><td></td></tr>
<tr><td>Chromosome</td><td>start</td><td>end</td><td>strand</td><td>MM</td><td>target_seq</td><td>PAM</td><td>alignment</td><td>distance</td><td>position</td><td>gene name</td><td>gene id</td></tr>
<tr><td>GU305914.1</td><td>152005</td><td>152027</td><td>+</td><td>0</td><td>ACGACCACCGCACGCAGAAT</td><td>CGG</td><td>||||||||[||||||||||||]PAM</td><td>NA</td><td>Intergenic</td><td>NA</td><td>NA</td></tr>
<tr><td>T11635</td><td>CGACCACCGCACGCAGAATCGGG</td><td>1000</td><td></td><td></td><td></td><td></td><td></td><td></td><td></td><td></td><td></td></tr>
<tr><td>Oligo fwd</td><td>CACCCGACCACCGCACGCAGAATC</td><td></td><td></td><td></td><td></td><td></td><td></td><td></td><td></td><td></td><td></td></tr>
<tr><td>Oligo rev</td><td>AAACGATTCTGCGTGCGGTGGTCG</td><td></td><td></td><td></td><td></td><td></td><td></td><td></td><td></td><td></td><td></td></tr>
<tr><td>Chromosome</td><td>start</td><td>end</td><td>strand</td><td>MM</td><td>target_seq</td><td>PAM</td><td>alignment</td><td>distance</td><td>position</td><td>gene name</td><td>gene id</td></tr>
<tr><td>GU305914.1</td><td>152006</td><td>152028</td><td>+</td><td>0</td><td>CGACCACCGCACGCAGAATC</td><td>GGG</td><td>||||||||[||||||||||||]PAM</td><td>NA</td><td>Intergenic</td><td>NA</td><td>NA</td></tr>
<tr><td>T11636</td><td>CAGAATCGGGACGTGCAGCGCGG</td><td>1000</td><td></td><td></td><td></td><td></td><td></td><td></td><td></td><td></td><td></td></tr>
</table>

<table>
<tr><td>Oligo fwd</td><td>CACCCAGAATCGGGACGTGCAGCG</td><td></td><td></td><td></td><td></td><td></td><td></td><td></td><td></td><td></td><td></td><td></td></tr>
<tr><td>Oligo rev</td><td>AAACCGCTGCACGTCCCGATTCTG</td><td></td><td></td><td></td><td></td><td></td><td></td><td></td><td></td><td></td><td></td><td></td></tr>
<tr><td>Chromosome</td><td>start</td><td></td><td>end</td><td>strand</td><td>MM</td><td>target_seq</td><td>PAM</td><td>alignment</td><td>distance</td><td>position</td><td>gene name</td><td>gene id</td></tr>
<tr><td>GU305914.1</td><td></td><td>152019</td><td>152041</td><td>+</td><td>0</td><td>CAGAATCGGGACGTGCAGCG</td><td>CGG</td><td>||||||||[||||||||||||]PAM</td><td>NA</td><td>Intergenic</td><td>NA</td><td>NA</td></tr>
<tr><td>T11637</td><td>TCGGGACGTGCAGCGCGGCATGG</td><td></td><td>1000</td><td></td><td></td><td></td><td></td><td></td><td></td><td></td><td></td><td></td></tr>
<tr><td>Oligo fwd</td><td>CACCTCGGGACGTGCAGCGCGGCA</td><td></td><td></td><td></td><td></td><td></td><td></td><td></td><td></td><td></td><td></td><td></td></tr>
<tr><td>Oligo rev</td><td>AAACTGCCGCGCTGCACGTCCCGA</td><td></td><td></td><td></td><td></td><td></td><td></td><td></td><td></td><td></td><td></td><td></td></tr>
<tr><td>Chromosome</td><td>start</td><td></td><td>end</td><td>strand</td><td>MM</td><td>target_seq</td><td>PAM</td><td>alignment</td><td>distance</td><td>position</td><td>gene name</td><td>gene id</td></tr>
<tr><td>GU305914.1</td><td></td><td>152024</td><td>152046</td><td>+</td><td>0</td><td>TCGGGACGTGCAGCGCGGCA</td><td>TGG</td><td>||||||||[||||||||||||]PAM</td><td>NA</td><td>Intergenic</td><td>NA</td><td>NA</td></tr>
<tr><td>T11638</td><td>CTCCGCGCTCATCAGCGACCCGG</td><td></td><td>1000</td><td></td><td></td><td></td><td></td><td></td><td></td><td></td><td></td><td></td></tr>
<tr><td>Oligo fwd</td><td>CACCCTCCGCGCTCATCAGCGACC</td><td></td><td></td><td></td><td></td><td></td><td></td><td></td><td></td><td></td><td></td><td></td></tr>
<tr><td>Oligo rev</td><td>AAACGGTCGCTGATGAGCGCGGAG</td><td></td><td></td><td></td><td></td><td></td><td></td><td></td><td></td><td></td><td></td><td></td></tr>
<tr><td>Chromosome</td><td>start</td><td></td><td>end</td><td>strand</td><td>MM</td><td>target_seq</td><td>PAM</td><td>alignment</td><td>distance</td><td>position</td><td>gene name</td><td>gene id</td></tr>
<tr><td>GU305914.1</td><td></td><td>152054</td><td>152076</td><td>+</td><td>0</td><td>CTCCGCGCTCATCAGCGACC</td><td>CGG</td><td>||||||||[||||||||||||]PAM</td><td>NA</td><td>Intergenic</td><td>NA</td><td>NA</td></tr>
<tr><td>GU305914.1</td><td></td><td>152234</td><td>152256</td><td>+</td><td>4</td><td>CGCCGGCCTCATCAGGGACC</td><td>CGG</td><td>|-|||--|[|||||||-||||]PAM</td><td>NA</td><td>Intergenic</td><td>NA</td><td>NA</td></tr>
<tr><td>T11639</td><td>ACTGCGACGTCGCCCACAACTGG</td><td></td><td>1000</td><td></td><td></td><td></td><td></td><td></td><td></td><td></td><td></td><td></td></tr>
<tr><td>Oligo fwd</td><td>CACCACTGCGACGTCGCCCACAAC</td><td></td><td></td><td></td><td></td><td></td><td></td><td></td><td></td><td></td><td></td><td></td></tr>
<tr><td>Oligo rev</td><td>AAACGTTGTGGGCGACGTCGCAGT</td><td></td><td></td><td></td><td></td><td></td><td></td><td></td><td></td><td></td><td></td><td></td></tr>
<tr><td>Chromosome</td><td>start</td><td></td><td>end</td><td>strand</td><td>MM</td><td>target_seq</td><td>PAM</td><td>alignment</td><td>distance</td><td>position</td><td>gene name</td><td>gene id</td></tr>
<tr><td>GU305914.1</td><td></td><td>152089</td><td>152111</td><td>+</td><td>0</td><td>ACTGCGACGTCGCCCACAAC</td><td>TGG</td><td>||||||||[||||||||||||]PAM</td><td>NA</td><td>Intergenic</td><td>NA</td><td>NA</td></tr>
<tr><td>T11640</td><td>CTGCGACGTCGCCCACAACTGGG</td><td></td><td>1000</td><td></td><td></td><td></td><td></td><td></td><td></td><td></td><td></td><td></td></tr>
<tr><td>Oligo fwd</td><td>CACCCTGCGACGTCGCCCACAACT</td><td></td><td></td><td></td><td></td><td></td><td></td><td></td><td></td><td></td><td></td><td></td></tr>
<tr><td>Oligo rev</td><td>AAACAGTTGTGGGCGACGTCGCAG</td><td></td><td></td><td></td><td></td><td></td><td></td><td></td><td></td><td></td><td></td><td></td></tr>
<tr><td>Chromosome</td><td>start</td><td></td><td>end</td><td>strand</td><td>MM</td><td>target_seq</td><td>PAM</td><td>alignment</td><td>distance</td><td>position</td><td>gene name</td><td>gene id</td></tr>
<tr><td>GU305914.1</td><td></td><td>152090</td><td>152112</td><td>+</td><td>0</td><td>CTGCGACGTCGCCCACAACT</td><td>GGG</td><td>||||||||[||||||||||||]PAM</td><td>NA</td><td>Intergenic</td><td>NA</td><td>NA</td></tr>
<tr><td>T11641</td><td>CGCTGCCTCGACCTCGAGTTCGG</td><td></td><td>1000</td><td></td><td></td><td></td><td></td><td></td><td></td><td></td><td></td><td></td></tr>
<tr><td>Oligo fwd</td><td>CACCCGCTGCCTCGACCTCGAGTT</td><td></td><td></td><td></td><td></td><td></td><td></td><td></td><td></td><td></td><td></td><td></td></tr>
<tr><td>Oligo rev</td><td>AAACAACTCGAGGTCGAGGCAGCG</td><td></td><td></td><td></td><td></td><td></td><td></td><td></td><td></td><td></td><td></td><td></td></tr>
<tr><td>Chromosome</td><td>start</td><td></td><td>end</td><td>strand</td><td>MM</td><td>target_seq</td><td>PAM</td><td>alignment</td><td>distance</td><td>position</td><td>gene name</td><td>gene id</td></tr>
<tr><td>GU305914.1</td><td></td><td>152133</td><td>152155</td><td>+</td><td>0</td><td>CGCTGCCTCGACCTCGAGTT</td><td>CGG</td><td>||||||||[||||||||||||]PAM</td><td>NA</td><td>Intergenic</td><td>NA</td><td>NA</td></tr>
<tr><td>T11642</td><td>CTGCTCAAGCACATCGAGTTCGG</td><td></td><td>1000</td><td></td><td></td><td></td><td></td><td></td><td></td><td></td><td></td><td></td></tr>
<tr><td>Oligo fwd</td><td>CACCCTGCTCAAGCACATCGAGTT</td><td></td><td></td><td></td><td></td><td></td><td></td><td></td><td></td><td></td><td></td><td></td></tr>
<tr><td>Oligo rev</td><td>AAACAACTCGATGTGCTTGAGCAG</td><td></td><td></td><td></td><td></td><td></td><td></td><td></td><td></td><td></td><td></td><td></td></tr>
<tr><td>Chromosome</td><td>start</td><td></td><td>end</td><td>strand</td><td>MM</td><td>target_seq</td><td>PAM</td><td>alignment</td><td>distance</td><td>position</td><td>gene name</td><td>gene id</td></tr>
<tr><td>GU305914.1</td><td></td><td>152160</td><td>152182</td><td>+</td><td>0</td><td>CTGCTCAAGCACATCGAGTT</td><td>CGG</td><td>||||||||[||||||||||||]PAM</td><td>NA</td><td>Intergenic</td><td>NA</td><td>NA</td></tr>
<tr><td>T11643</td><td>TGCTCAAGCACATCGAGTTCGGG</td><td></td><td>1000</td><td></td><td></td><td></td><td></td><td></td><td></td><td></td><td></td><td></td></tr>
<tr><td>Oligo fwd</td><td>CACCTGCTCAAGCACATCGAGTTC</td><td></td><td></td><td></td><td></td><td></td><td></td><td></td><td></td><td></td><td></td><td></td></tr>
<tr><td>Oligo rev</td><td>AAACGAACTCGATGTGCTTGAGCA</td><td></td><td></td><td></td><td></td><td></td><td></td><td></td><td></td><td></td><td></td><td></td></tr>
<tr><td>Chromosome</td><td>start</td><td></td><td>end</td><td>strand</td><td>MM</td><td>target_seq</td><td>PAM</td><td>alignment</td><td>distance</td><td>position</td><td>gene name</td><td>gene id</td></tr>
<tr><td>GU305914.1</td><td></td><td>152161</td><td>152183</td><td>+</td><td>0</td><td>TGCTCAAGCACATCGAGTTC</td><td>GGG</td><td>||||||||[||||||||||||]PAM</td><td>NA</td><td>Intergenic</td><td>NA</td><td>NA</td></tr>
<tr><td>T11644</td><td>AGCACGTCGAGTACCTCGACAGG</td><td></td><td>1000</td><td></td><td></td><td></td><td></td><td></td><td></td><td></td><td></td><td></td></tr>
<tr><td>Oligo fwd</td><td>CACCAGCACGTCGAGTACCTCGAC</td><td></td><td></td><td></td><td></td><td></td><td></td><td></td><td></td><td></td><td></td><td></td></tr>
<tr><td>Oligo rev</td><td>AAACGTCGAGGTACTCGACGTGCT</td><td></td><td></td><td></td><td></td><td></td><td></td><td></td><td></td><td></td><td></td><td></td></tr>
<tr><td>Chromosome</td><td>start</td><td></td><td>end</td><td>strand</td><td>MM</td><td>target_seq</td><td>PAM</td><td>alignment</td><td>distance</td><td>position</td><td>gene name</td><td>gene id</td></tr>
<tr><td>GU305914.1</td><td></td><td>152200</td><td>152222</td><td>+</td><td>0</td><td>AGCACGTCGAGTACCTCGAC</td><td>AGG</td><td>||||||||[||||||||||||]PAM</td><td>NA</td><td>Intergenic</td><td>NA</td><td>NA</td></tr>
<tr><td>GU305914.1</td><td></td><td>186641</td><td>186663</td><td>-</td><td>4</td><td>ATCACGCCGAGAACCTCGAG</td><td>GGG</td><td>|-||||-|[|||-|||||||-]PAM</td><td>NA</td><td>Intergenic</td><td>NA</td><td>NA</td></tr>
<tr><td>T11645</td><td>GACAGGTTCGTGCGCCCCGCCGG</td><td></td><td>1000</td><td></td><td></td><td></td><td></td><td></td><td></td><td></td><td></td><td></td></tr>
<tr><td>Oligo fwd</td><td>CACCGACAGGTTCGTGCGCCCCGC</td><td></td><td></td><td></td><td></td><td></td><td></td><td></td><td></td><td></td><td></td><td></td></tr>
<tr><td>Oligo rev</td><td>AAACGCGGGGCGCACGAACCTGTC</td><td></td><td></td><td></td><td></td><td></td><td></td><td></td><td></td><td></td><td></td><td></td></tr>
<tr><td>Chromosome</td><td>start</td><td></td><td>end</td><td>strand</td><td>MM</td><td>target_seq</td><td>PAM</td><td>alignment</td><td>distance</td><td>position</td><td>gene name</td><td>gene id</td></tr>
<tr><td>GU305914.1</td><td></td><td>152217</td><td>152239</td><td>+</td><td>0</td><td>GACAGGTTCGTGCGCCCCGC</td><td>CGG</td><td>||||||||[||||||||||||]PAM</td><td>NA</td><td>Intergenic</td><td>NA</td><td>NA</td></tr>
<tr><td>T11646</td><td>TGCGCCCCGCCGGCCTCATCAGG</td><td></td><td>1000</td><td></td><td></td><td></td><td></td><td></td><td></td><td></td><td></td><td></td></tr>
<tr><td>Oligo fwd</td><td>CACCTGCGCCCCGCCGGCCTCATC</td><td></td><td></td><td></td><td></td><td></td><td></td><td></td><td></td><td></td><td></td><td></td></tr>
<tr><td>Oligo rev</td><td>AAACGATGAGGCCGGCGGGGCGCA</td><td></td><td></td><td></td><td></td><td></td><td></td><td></td><td></td><td></td><td></td><td></td></tr>
<tr><td>Chromosome</td><td>start</td><td></td><td>end</td><td>strand</td><td>MM</td><td>target_seq</td><td>PAM</td><td>alignment</td><td>distance</td><td>position</td><td>gene name</td><td>gene id</td></tr>
<tr><td>GU305914.1</td><td></td><td>152227</td><td>152249</td><td>+</td><td>0</td><td>TGCGCCCCGCCGGCCTCATC</td><td>AGG</td><td>||||||||[||||||||||||]PAM</td><td>NA</td><td>Intergenic</td><td>NA</td><td>NA</td></tr>
<tr><td>T11647</td><td>GCGCCCCGCCGGCCTCATCAGGG</td><td></td><td>1000</td><td></td><td></td><td></td><td></td><td></td><td></td><td></td><td></td><td></td></tr>
<tr><td>Oligo fwd</td><td>CACCGCGCCCCGCCGGCCTCATCA</td><td></td><td></td><td></td><td></td><td></td><td></td><td></td><td></td><td></td><td></td><td></td></tr>
<tr><td>Oligo rev</td><td>AAACTGATGAGGCCGGCGGGGCGC</td><td></td><td></td><td></td><td></td><td></td><td></td><td></td><td></td><td></td><td></td><td></td></tr>
<tr><td>Chromosome</td><td>start</td><td></td><td>end</td><td>strand</td><td>MM</td><td>target_seq</td><td>PAM</td><td>alignment</td><td>distance</td><td>position</td><td>gene name</td><td>gene id</td></tr>
<tr><td>GU305914.1</td><td></td><td>152228</td><td>152250</td><td>+</td><td>0</td><td>GCGCCCCGCCGGCCTCATCA</td><td>GGG</td><td>||||||||[||||||||||||]PAM</td><td>NA</td><td>Intergenic</td><td>NA</td><td>NA</td></tr>
<tr><td>T11648</td><td>CGCCGGCCTCATCAGGGACCCGG</td><td></td><td>1000</td><td></td><td></td><td></td><td></td><td></td><td></td><td></td><td></td><td></td></tr>
<tr><td>Oligo fwd</td><td>CACCCGCCGGCCTCATCAGGGACC</td><td></td><td></td><td></td><td></td><td></td><td></td><td></td><td></td><td></td><td></td><td></td></tr>
<tr><td>Oligo rev</td><td>AAACGGTCCCTGATGAGGCCGGCG</td><td></td><td></td><td></td><td></td><td></td><td></td><td></td><td></td><td></td><td></td><td></td></tr>
<tr><td>Chromosome</td><td>start</td><td></td><td>end</td><td>strand</td><td>MM</td><td>target_seq</td><td>PAM</td><td>alignment</td><td>distance</td><td>position</td><td>gene name</td><td>gene id</td></tr>
<tr><td>GU305914.1</td><td></td><td>152234</td><td>152256</td><td>+</td><td>0</td><td>CGCCGGCCTCATCAGGGACC</td><td>CGG</td><td>||||||||[||||||||||||]PAM</td><td>NA</td><td>Intergenic</td><td>NA</td><td>NA</td></tr>
<tr><td>GU305914.1</td><td></td><td>152054</td><td>152076</td><td>+</td><td>4</td><td>CTCCGCGCTCATCAGCGACC</td><td>CGG</td><td>|-|||--|[|||||||-||||]PAM</td><td>NA</td><td>Intergenic</td><td>NA</td><td>NA</td></tr>
<tr><td>T11649</td><td>CCTCTTCATCTCGCACGCCGAGG</td><td></td><td>1000</td><td></td><td></td><td></td><td></td><td></td><td></td><td></td><td></td><td></td></tr>
<tr><td>Oligo fwd</td><td>CACCCCTCTTCATCTCGCACGCCG</td><td></td><td></td><td></td><td></td><td></td><td></td><td></td><td></td><td></td><td></td><td></td></tr>
<tr><td>Oligo rev</td><td>AAACCGGCGTGCGAGATGAAGAGG</td><td></td><td></td><td></td><td></td><td></td><td></td><td></td><td></td><td></td><td></td><td></td></tr>
<tr><td>Chromosome</td><td>start</td><td></td><td>end</td><td>strand</td><td>MM</td><td>target_seq</td><td>PAM</td><td>alignment</td><td>distance</td><td>position</td><td>gene name</td><td>gene id</td></tr>
<tr><td>GU305914.1</td><td></td><td>152279</td><td>152301</td><td>+</td><td>0</td><td>CCTCTTCATCTCGCACGCCG</td><td>AGG</td><td>||||||||[||||||||||||]PAM</td><td>NA</td><td>Intergenic</td><td>NA</td><td>NA</td></tr>
<tr><td>GU305914.1</td><td></td><td>126029</td><td>126051</td><td>-</td><td>4</td><td>CCAGCGCATCTCGCACGCCG</td><td>AGG</td><td>||----||[||||||||||||]PAM</td><td>NA</td><td>Intergenic</td><td>NA</td><td>NA</td></tr>
<tr><td>T11650</td><td>TCTCGCACGCCGAGGTCAAGAGG</td><td></td><td>1000</td><td></td><td></td><td></td><td></td><td></td><td></td><td></td><td></td><td></td></tr>
<tr><td>Oligo fwd</td><td>CACCTCTCGCACGCCGAGGTCAAG</td><td></td><td></td><td></td><td></td><td></td><td></td><td></td><td></td><td></td><td></td><td></td></tr>
</table>

Oligo rev AAACCTTGACCTCGGCGTGCGAGA
Chromosome start end strand MM target_seq PAM alignment distance position gene name gene id
GU305914.1 152287 152309 + 0 TCTCGCACGCCGAGGTCAAG AGG ||||||||[||||||||||||]PAM NA Intergenic NA NA

T11651 CGAGGTCAAGAGGTACTTCACGG 1000
Oligo fwd CACCCGAGGTCAAGAGGTACTTCA
Oligo rev AAACTGAAGTACCTCTTGACCTCG
Chromosome start end strand MM target_seq PAM alignment distance position gene name gene id
GU305914.1 152297 152319 + 0 CGAGGTCAAGAGGTACTTCA CGG ||||||||[||||||||||||]PAM NA Intergenic NA NA

T11652 CAGCCAGCACCTCATCTTCGAGG 1000
Oligo fwd CACCCAGCCAGCACCTCATCTTCG
Oligo rev AAACCGAAGATGAGGTGCTGGCTG
Chromosome start end strand MM target_seq PAM alignment distance position gene name gene id
GU305914.1 152345 152367 + 0 CAGCCAGCACCTCATCTTCG AGG ||||||||[||||||||||||]PAM NA Intergenic NA NA

T11653 GCACCTCATCTTCGAGGTGCCGG 1000
Oligo fwd CACCGCACCTCATCTTCGAGGTGC
Oligo rev AAACGCACCTCGAAGATGAGGTGC
Chromosome start end strand MM target_seq PAM alignment distance position gene name gene id
GU305914.1 152351 152373 + 0 GCACCTCATCTTCGAGGTGC CGG ||||||||[||||||||||||]PAM NA Intergenic NA NA

T11654 CTACTGCGTCCTCAACAACTCGG 1000
Oligo fwd CACCCTACTGCGTCCTCAACAACT
Oligo rev AAACAGTTGTTGAGGACGCAGTAG
Chromosome start end strand MM target_seq PAM alignment distance position gene name gene id
GU305914.1 152375 152397 + 0 CTACTGCGTCCTCAACAACT CGG ||||||||[||||||||||||]PAM NA Intergenic NA NA

T11655 TACTGTGCCAGCATGTGCACCGG 1000
Oligo fwd CACCTACTGTGCCAGCATGTGCAC
Oligo rev AAACGTGCACATGCTGGCACAGTA
Chromosome start end strand MM target_seq PAM alignment distance position gene name gene id
GU305914.1 152409 152431 + 0 TACTGTGCCAGCATGTGCAC CGG ||||||||[||||||||||||]PAM NA Intergenic NA NA

T11656 ACTGTGCCAGCATGTGCACCGGG 1000
Oligo fwd CACCACTGTGCCAGCATGTGCACC
Oligo rev AAACGGTGCACATGCTGGCACAGT
Chromosome start end strand MM target_seq PAM alignment distance position gene name gene id
GU305914.1 152410 152432 + 0 ACTGTGCCAGCATGTGCACC GGG ||||||||[||||||||||||]PAM NA Intergenic NA NA

T11657 CTGTGCCAGCATGTGCACCGGGG 1000
Oligo fwd CACCCTGTGCCAGCATGTGCACCG
Oligo rev AAACCGGTGCACATGCTGGCACAG
Chromosome start end strand MM target_seq PAM alignment distance position gene name gene id
GU305914.1 152411 152433 + 0 CTGTGCCAGCATGTGCACCG GGG ||||||||[||||||||||||]PAM NA Intergenic NA NA

T11658 CGGGGAGCCCACGCCGCGTCCGG 1000
Oligo fwd CACCCGGGGAGCCCACGCCGCGTC
Oligo rev AAACGACGCGGCGTGGGCTCCCCG
Chromosome start end strand MM target_seq PAM alignment distance position gene name gene id
GU305914.1 152429 152451 + 0 CGGGGAGCCCACGCCGCGTC CGG ||||||||[||||||||||||]PAM NA Intergenic NA NA

T11659 CCACGCCGCGTCCGGAGACCTGG 1000
Oligo fwd CACCCCACGCCGCGTCCGGAGACC
Oligo rev AAACGGTCTCCGGACGCGGCGTGG
Chromosome start end strand MM target_seq PAM alignment distance position gene name gene id
GU305914.1 152437 152459 + 0 CCACGCCGCGTCCGGAGACC TGG ||||||||[||||||||||||]PAM NA Intergenic NA NA

T11660 GACCTGGTTCCGCAAGAACCTGG 1000
Oligo fwd CACCGACCTGGTTCCGCAAGAACC
Oligo rev AAACGGTTCTTGCGGAACCAGGTC
Chromosome start end strand MM target_seq PAM alignment distance position gene name gene id
GU305914.1 152453 152475 + 0 GACCTGGTTCCGCAAGAACC TGG ||||||||[||||||||||||]PAM NA Intergenic NA NA

T11661 GTTCACCGACCATAACCTGTCGG 1000
Oligo fwd CACCGTTCACCGACCATAACCTGT
Oligo rev AAACACAGGTTATGGTCGGTGAAC
Chromosome start end strand MM target_seq PAM alignment distance position gene name gene id
GU305914.1 152498 152520 + 0 GTTCACCGACCATAACCTGT CGG ||||||||[||||||||||||]PAM NA Intergenic NA NA

T11662 CACCGACCATAACCTGTCGGAGG 1000
Oligo fwd CACCCACCGACCATAACCTGTCGG
Oligo rev AAACCCGACAGGTTATGGTCGGTG
Chromosome start end strand MM target_seq PAM alignment distance position gene name gene id
GU305914.1 152501 152523 + 0 CACCGACCATAACCTGTCGG AGG ||||||||[||||||||||||]PAM NA Intergenic NA NA

T11663 TAACCTGTCGGAGGACATCCAGG 1000
Oligo fwd CACCTAACCTGTCGGAGGACATCC
Oligo rev AAACGGATGTCCTCCGACAGGTTA
Chromosome start end strand MM target_seq PAM alignment distance position gene name gene id
GU305914.1 152510 152532 + 0 TAACCTGTCGGAGGACATCC AGG ||||||||[||||||||||||]PAM NA Intergenic NA NA

T11664 GTCGGAGGACATCCAGGTCGAGG 1000
Oligo fwd CACCGTCGGAGGACATCCAGGTCG
Oligo rev AAACCGACCTGGATGTCCTCCGAC
Chromosome start end strand MM target_seq PAM alignment distance position gene name gene id
GU305914.1 152516 152538 + 0 GTCGGAGGACATCCAGGTCG AGG ||||||||[||||||||||||]PAM NA Intergenic NA NA

T11665 GAGGAGCTCGCCCAGTCCTGCGG 1000

<table>
<tr><td>Oligo fwd</td><td>CACCGAGGAGCTCGCCCAGTCCTG</td><td></td><td></td><td></td><td></td><td></td><td></td><td></td><td></td><td></td><td></td></tr>
<tr><td>Oligo rev</td><td>AAACCAGGACTGGGCGAGCTCCTC</td><td></td><td></td><td></td><td></td><td></td><td></td><td></td><td></td><td></td><td></td></tr>
<tr><td>Chromosome</td><td>start</td><td>end</td><td>strand</td><td>MM</td><td>target_seq</td><td>PAM</td><td>alignment</td><td>distance</td><td>position</td><td>gene name</td><td>gene id</td></tr>
<tr><td>GU305914.1</td><td>152535</td><td>152557</td><td>+</td><td>0</td><td>GAGGAGCTCGCCCAGTCCTG</td><td>CGG</td><td>||||||||[||||||||||||]PAM</td><td>NA</td><td>Intergenic</td><td>NA</td><td>NA</td></tr>
<tr><td></td><td></td><td></td><td></td><td></td><td></td><td></td><td></td><td></td><td></td><td></td><td></td></tr>
<tr><td>T11666</td><td>GAGCTCGCCCAGTCCTGCGGCGG</td><td>1000</td><td></td><td></td><td></td><td></td><td></td><td></td><td></td><td></td><td></td></tr>
<tr><td>Oligo fwd</td><td>CACCGAGCTCGCCCAGTCCTGCGG</td><td></td><td></td><td></td><td></td><td></td><td></td><td></td><td></td><td></td><td></td></tr>
<tr><td>Oligo rev</td><td>AAACCCGCAGGACTGGGCGAGCTC</td><td></td><td></td><td></td><td></td><td></td><td></td><td></td><td></td><td></td><td></td></tr>
<tr><td>Chromosome</td><td>start</td><td>end</td><td>strand</td><td>MM</td><td>target_seq</td><td>PAM</td><td>alignment</td><td>distance</td><td>position</td><td>gene name</td><td>gene id</td></tr>
<tr><td>GU305914.1</td><td>152538</td><td>152560</td><td>+</td><td>0</td><td>GAGCTCGCCCAGTCCTGCGG</td><td>CGG</td><td>||||||||[||||||||||||]PAM</td><td>NA</td><td>Intergenic</td><td>NA</td><td>NA</td></tr>
<tr><td></td><td></td><td></td><td></td><td></td><td></td><td></td><td></td><td></td><td></td><td></td><td></td></tr>
<tr><td>T11667</td><td>CTCGCCCAGTCCTGCGGCGGCGG</td><td>1000</td><td></td><td></td><td></td><td></td><td></td><td></td><td></td><td></td><td></td></tr>
<tr><td>Oligo fwd</td><td>CACCCTCGCCCAGTCCTGCGGCGG</td><td></td><td></td><td></td><td></td><td></td><td></td><td></td><td></td><td></td><td></td></tr>
<tr><td>Oligo rev</td><td>AAACCCGCCGCAGGACTGGGCGAG</td><td></td><td></td><td></td><td></td><td></td><td></td><td></td><td></td><td></td><td></td></tr>
<tr><td>Chromosome</td><td>start</td><td>end</td><td>strand</td><td>MM</td><td>target_seq</td><td>PAM</td><td>alignment</td><td>distance</td><td>position</td><td>gene name</td><td>gene id</td></tr>
<tr><td>GU305914.1</td><td>152541</td><td>152563</td><td>+</td><td>0</td><td>CTCGCCCAGTCCTGCGGCGG</td><td>CGG</td><td>||||||||[||||||||||||]PAM</td><td>NA</td><td>Intergenic</td><td>NA</td><td>NA</td></tr>
<tr><td></td><td></td><td></td><td></td><td></td><td></td><td></td><td></td><td></td><td></td><td></td><td></td></tr>
<tr><td>T11668</td><td>GCCCAGTCCTGCGGCGGCGGCGG</td><td>1000</td><td></td><td></td><td></td><td></td><td></td><td></td><td></td><td></td><td></td></tr>
<tr><td>Oligo fwd</td><td>CACCGCCCAGTCCTGCGGCGGCGG</td><td></td><td></td><td></td><td></td><td></td><td></td><td></td><td></td><td></td><td></td></tr>
<tr><td>Oligo rev</td><td>AAACCCGCCGCCGCAGGACTGGGC</td><td></td><td></td><td></td><td></td><td></td><td></td><td></td><td></td><td></td><td></td></tr>
<tr><td>Chromosome</td><td>start</td><td>end</td><td>strand</td><td>MM</td><td>target_seq</td><td>PAM</td><td>alignment</td><td>distance</td><td>position</td><td>gene name</td><td>gene id</td></tr>
<tr><td>GU305914.1</td><td>152544</td><td>152566</td><td>+</td><td>0</td><td>GCCCAGTCCTGCGGCGGCGG</td><td>CGG</td><td>||||||||[||||||||||||]PAM</td><td>NA</td><td>Intergenic</td><td>NA</td><td>NA</td></tr>
<tr><td>GU305914.1</td><td>110035</td><td>110057</td><td>+</td><td>4</td><td>TCGCCGTCCTGCTGCGGCGG</td><td>GGG</td><td>-|-|-|||[||||-|||||||]PAM</td><td>NA</td><td>Intergenic</td><td>NA</td><td>NA</td></tr>
<tr><td></td><td></td><td></td><td></td><td></td><td></td><td></td><td></td><td></td><td></td><td></td><td></td></tr>
<tr><td>T11669</td><td>TCCTGCGGCGGCGGCGGTGCCGG</td><td>1000</td><td></td><td></td><td></td><td></td><td></td><td></td><td></td><td></td><td></td></tr>
<tr><td>Oligo fwd</td><td>CACCTCCTGCGGCGGCGGCGGTGC</td><td></td><td></td><td></td><td></td><td></td><td></td><td></td><td></td><td></td><td></td></tr>
<tr><td>Oligo rev</td><td>AAACGCACCGCCGCCGCCGCAGGA</td><td></td><td></td><td></td><td></td><td></td><td></td><td></td><td></td><td></td><td></td></tr>
<tr><td>Chromosome</td><td>start</td><td>end</td><td>strand</td><td>MM</td><td>target_seq</td><td>PAM</td><td>alignment</td><td>distance</td><td>position</td><td>gene name</td><td>gene id</td></tr>
<tr><td>GU305914.1</td><td>152550</td><td>152572</td><td>+</td><td>0</td><td>TCCTGCGGCGGCGGCGGTGC</td><td>CGG</td><td>||||||||[||||||||||||]PAM</td><td>NA</td><td>Intergenic</td><td>NA</td><td>NA</td></tr>
<tr><td>GU305914.1</td><td>19436</td><td>19458</td><td>+</td><td>4</td><td>GCCGGCGTCGGCGGAGGTGC</td><td>CGG</td><td>-||-|||-[||||||-|||||]PAM</td><td>NA</td><td>Intergenic</td><td>NA</td><td>NA</td></tr>
<tr><td>GU305914.1</td><td>48150</td><td>48172</td><td>+</td><td>3</td><td>ACCTGTGGCGGCGGCGGTGG</td><td>CGG</td><td>-||||-||[|||||||||||-]PAM</td><td>NA</td><td>Intergenic</td><td>NA</td><td>NA</td></tr>
<tr><td>GU305914.1</td><td>163746</td><td>163768</td><td>+</td><td>4</td><td>TCCGTCGGCGGCGCCGGTGG</td><td>CGG</td><td>|||--|||[|||||-|||||-]PAM</td><td>NA</td><td>Intergenic</td><td>NA</td><td>NA</td></tr>
<tr><td></td><td></td><td></td><td></td><td></td><td></td><td></td><td></td><td></td><td></td><td></td><td></td></tr>
<tr><td>T11670</td><td>TGCGGCGGCGGCGGTGCCGGCGG</td><td>1000</td><td></td><td></td><td></td><td></td><td></td><td></td><td></td><td></td><td></td></tr>
<tr><td>Oligo fwd</td><td>CACCTGCGGCGGCGGCGGTGCCGG</td><td></td><td></td><td></td><td></td><td></td><td></td><td></td><td></td><td></td><td></td></tr>
<tr><td>Oligo rev</td><td>AAACCCGGCACCGCCGCCGCCGCA</td><td></td><td></td><td></td><td></td><td></td><td></td><td></td><td></td><td></td><td></td></tr>
<tr><td>Chromosome</td><td>start</td><td>end</td><td>strand</td><td>MM</td><td>target_seq</td><td>PAM</td><td>alignment</td><td>distance</td><td>position</td><td>gene name</td><td>gene id</td></tr>
<tr><td>GU305914.1</td><td>152553</td><td>152575</td><td>+</td><td>0</td><td>TGCGGCGGCGGCGGTGCCGG</td><td>CGG</td><td>||||||||[||||||||||||]PAM</td><td>NA</td><td>Intergenic</td><td>NA</td><td>NA</td></tr>
<tr><td>GU305914.1</td><td>19439</td><td>19461</td><td>+</td><td>3</td><td>GGCGTCGGCGGAGGTGCCGG</td><td>AGG</td><td>-|||-|||[|||-||||||||]PAM</td><td>NA</td><td>Intergenic</td><td>NA</td><td>NA</td></tr>
<tr><td>GU305914.1</td><td>90250</td><td>90272</td><td>-</td><td>3</td><td>GGCGGAGGCGGCGGAGCCGG</td><td>CGG</td><td>-||||-||[||||||-|||||]PAM</td><td>NA</td><td>Intergenic</td><td>NA</td><td>NA</td></tr>
<tr><td>GU305914.1</td><td>48153</td><td>48175</td><td>+</td><td>2</td><td>TGTGGCGGCGGCGGTGGCGG</td><td>CGG</td><td>||-|||||[||||||||-|||]PAM</td><td>NA</td><td>Intergenic</td><td>NA</td><td>NA</td></tr>
<tr><td>GU305914.1</td><td>163749</td><td>163771</td><td>+</td><td>4</td><td>GTCGGCGGCGCCGGTGGCGG</td><td>CGG</td><td>--||||||[||-|||||-|||]PAM</td><td>NA</td><td>Intergenic</td><td>NA</td><td>NA</td></tr>
<tr><td>GU305914.1</td><td>127365</td><td>127387</td><td>+</td><td>3</td><td>CGCGGCGGCGGCGCTACCGG</td><td>CGG</td><td>-|||||||[|||||-|-||||]PAM</td><td>NA</td><td>Intergenic</td><td>NA</td><td>NA</td></tr>
<tr><td>GU305914.1</td><td>110083</td><td>110105</td><td>-</td><td>4</td><td>CGCGGCCGCGGCGACGCCGG</td><td>TGG</td><td>-|||||-|[|||||--|||||]PAM</td><td>NA</td><td>Intergenic</td><td>NA</td><td>NA</td></tr>
<tr><td>GU305914.1</td><td>113168</td><td>113190</td><td>-</td><td>4</td><td>TTCGGCCGCGGCGCTGTCGG</td><td>TGG</td><td>|-||||-|[|||||-||-|||]PAM</td><td>NA</td><td>Intergenic</td><td>NA</td><td>NA</td></tr>
<tr><td>GU305914.1</td><td>154407</td><td>154429</td><td>+</td><td>4</td><td>GGTGGCGGCGGCGGCGGCGG</td><td>CGG</td><td>-|-|||||[||||||-|-|||]PAM</td><td>NA</td><td>Intergenic</td><td>NA</td><td>NA</td></tr>
<tr><td>GU305914.1</td><td>58123</td><td>58145</td><td>-</td><td>4</td><td>GGTGGCGGCGGCGGCGGCGG</td><td>AGG</td><td>-|-|||||[||||||-|-|||]PAM</td><td>NA</td><td>Intergenic</td><td>NA</td><td>NA</td></tr>
<tr><td>GU305914.1</td><td>149482</td><td>149504</td><td>-</td><td>4</td><td>GGCGACGGCGGCGGCGGCGG</td><td>CGG</td><td>-|||-|||[||||||-|-|||]PAM</td><td>NA</td><td>Intergenic</td><td>NA</td><td>NA</td></tr>
<tr><td>GU305914.1</td><td>154404</td><td>154426</td><td>+</td><td>4</td><td>GGCGGTGGCGGCGGCGGCGG</td><td>CGG</td><td>-||||-||[||||||-|-|||]PAM</td><td>NA</td><td>Intergenic</td><td>NA</td><td>NA</td></tr>
<tr><td>GU305914.1</td><td>58126</td><td>58148</td><td>-</td><td>4</td><td>GGCGGTGGCGGCGGCGGCGG</td><td>CGG</td><td>-||||-||[||||||-|-|||]PAM</td><td>NA</td><td>Intergenic</td><td>NA</td><td>NA</td></tr>
<tr><td></td><td></td><td></td><td></td><td></td><td></td><td></td><td></td><td></td><td></td><td></td><td></td></tr>
<tr><td>T11671</td><td>GCGGCGGCGGCGGTGCCGGCGGG</td><td>1000</td><td></td><td></td><td></td><td></td><td></td><td></td><td></td><td></td><td></td></tr>
<tr><td>Oligo fwd</td><td>CACCGCGGCGGCGGCGGTGCCGGC</td><td></td><td></td><td></td><td></td><td></td><td></td><td></td><td></td><td></td><td></td></tr>
<tr><td>Oligo rev</td><td>AAACGCCGGCACCGCCGCCGCCGC</td><td></td><td></td><td></td><td></td><td></td><td></td><td></td><td></td><td></td><td></td></tr>
<tr><td>Chromosome</td><td>start</td><td>end</td><td>strand</td><td>MM</td><td>target_seq</td><td>PAM</td><td>alignment</td><td>distance</td><td>position</td><td>gene name</td><td>gene id</td></tr>
<tr><td>GU305914.1</td><td>152554</td><td>152576</td><td>+</td><td>0</td><td>GCGGCGGCGGCGGTGCCGGC</td><td>GGG</td><td>||||||||[||||||||||||]PAM</td><td>NA</td><td>Intergenic</td><td>NA</td><td>NA</td></tr>
<tr><td>GU305914.1</td><td>62444</td><td>62466</td><td>+</td><td>3</td><td>GCGGCGGGGGTGGTGGCGGC</td><td>GGG</td><td>|||||||-[||-||||-||||]PAM</td><td>NA</td><td>Intergenic</td><td>NA</td><td>NA</td></tr>
<tr><td>GU305914.1</td><td>67743</td><td>67765</td><td>+</td><td>4</td><td>GGAGCGGCGGCGGCGGCGGC</td><td>GGG</td><td>|--|||||[|||||-|-||||]PAM</td><td>NA</td><td>Intergenic</td><td>NA</td><td>NA</td></tr>
<tr><td></td><td></td><td></td><td></td><td></td><td></td><td></td><td></td><td></td><td></td><td></td><td></td></tr>
<tr><td>T11672</td><td>CGGCGGCGGCGGTGCCGGCGGGG</td><td>1000</td><td></td><td></td><td></td><td></td><td></td><td></td><td></td><td></td><td></td></tr>
<tr><td>Oligo fwd</td><td>CACCCGGCGGCGGCGGTGCCGGCG</td><td></td><td></td><td></td><td></td><td></td><td></td><td></td><td></td><td></td><td></td></tr>
<tr><td>Oligo rev</td><td>AAACCGCCGGCACCGCCGCCGCCG</td><td></td><td></td><td></td><td></td><td></td><td></td><td></td><td></td><td></td><td></td></tr>
<tr><td>Chromosome</td><td>start</td><td>end</td><td>strand</td><td>MM</td><td>target_seq</td><td>PAM</td><td>alignment</td><td>distance</td><td>position</td><td>gene name</td><td>gene id</td></tr>
<tr><td>GU305914.1</td><td>152555</td><td>152577</td><td>+</td><td>0</td><td>CGGCGGCGGCGGTGCCGGCG</td><td>GGG</td><td>||||||||[||||||||||||]PAM</td><td>NA</td><td>Intergenic</td><td>NA</td><td>NA</td></tr>
<tr><td>GU305914.1</td><td>62445</td><td>62467</td><td>+</td><td>3</td><td>CGGCGGGGGTGGTGGCGGCG</td><td>GGG</td><td>||||||-|[|-||||-|||||]PAM</td><td>NA</td><td>Intergenic</td><td>NA</td><td>NA</td></tr>
<tr><td>GU305914.1</td><td>35655</td><td>35677</td><td>+</td><td>4</td><td>CCACGGCGGCGGCGGCGGCG</td><td>GGG</td><td>|--|||||[||||-|-|||||]PAM</td><td>NA</td><td>Intergenic</td><td>NA</td><td>NA</td></tr>
<tr><td>GU305914.1</td><td>153169</td><td>153191</td><td>+</td><td>4</td><td>TGTCGGCGGCGCTGCCGCCG</td><td>CGG</td><td>-|-|||||[|||-|||||-||]PAM</td><td>NA</td><td>Intergenic</td><td>NA</td><td>NA</td></tr>
<tr><td>GU305914.1</td><td>35629</td><td>35651</td><td>+</td><td>4</td><td>TGGCGTCGGCGGTGTCGGTG</td><td>CGG</td><td>-||||-||[||||||-|||-|]PAM</td><td>NA</td><td>Intergenic</td><td>NA</td><td>NA</td></tr>
<tr><td></td><td></td><td></td><td></td><td></td><td></td><td></td><td></td><td></td><td></td><td></td><td></td></tr>
<tr><td>T11673</td><td>GGTGCCGGCGGGGACGATGACGG</td><td>1000</td><td></td><td></td><td></td><td></td><td></td><td></td><td></td><td></td><td></td></tr>
<tr><td>Oligo fwd</td><td>CACCGGTGCCGGCGGGGACGATGA</td><td></td><td></td><td></td><td></td><td></td><td></td><td></td><td></td><td></td><td></td></tr>
<tr><td>Oligo rev</td><td>AAACTCATCGTCCCCGCCGGCACC</td><td></td><td></td><td></td><td></td><td></td><td></td><td></td><td></td><td></td><td></td></tr>
<tr><td>Chromosome</td><td>start</td><td>end</td><td>strand</td><td>MM</td><td>target_seq</td><td>PAM</td><td>alignment</td><td>distance</td><td>position</td><td>gene name</td><td>gene id</td></tr>
<tr><td>GU305914.1</td><td>152565</td><td>152587</td><td>+</td><td>0</td><td>GGTGCCGGCGGGGACGATGA</td><td>CGG</td><td>||||||||[||||||||||||]PAM</td><td>NA</td><td>Intergenic</td><td>NA</td><td>NA</td></tr>
<tr><td></td><td></td><td></td><td></td><td></td><td></td><td></td><td></td><td></td><td></td><td></td><td></td></tr>
<tr><td>T11674</td><td>CGATGACGGCTTCGACCTCGAGG</td><td>1000</td><td></td><td></td><td></td><td></td><td></td><td></td><td></td><td></td><td></td></tr>
<tr><td>Oligo fwd</td><td>CACCCGATGACGGCTTCGACCTCG</td><td></td><td></td><td></td><td></td><td></td><td></td><td></td><td></td><td></td><td></td></tr>
<tr><td>Oligo rev</td><td>AAACCGAGGTCGAAGCCGTCATCG</td><td></td><td></td><td></td><td></td><td></td><td></td><td></td><td></td><td></td><td></td></tr>
<tr><td>Chromosome</td><td>start</td><td>end</td><td>strand</td><td>MM</td><td>target_seq</td><td>PAM</td><td>alignment</td><td>distance</td><td>position</td><td>gene name</td><td>gene id</td></tr>
<tr><td>GU305914.1</td><td>152579</td><td>152601</td><td>+</td><td>0</td><td>CGATGACGGCTTCGACCTCG</td><td>AGG</td><td>||||||||[||||||||||||]PAM</td><td>NA</td><td>Intergenic</td><td>NA</td><td>NA</td></tr>
<tr><td>GU305914.1</td><td>194895</td><td>194917</td><td>-</td><td>4</td><td>GGACGACGGCTTCGTCTTCG</td><td>AGG</td><td>-||-||||[||||||-|-|||]PAM</td><td>NA</td><td>Intergenic</td><td>NA</td><td>NA</td></tr>
<tr><td></td><td></td><td></td><td></td><td></td><td></td><td></td><td></td><td></td><td></td><td></td><td></td></tr>
<tr><td>T11675</td><td>TGACGGCTTCGACCTCGAGGAGG</td><td>1000</td><td></td><td></td><td></td><td></td><td></td><td></td><td></td><td></td><td></td></tr>
<tr><td>Oligo fwd</td><td>CACCTGACGGCTTCGACCTCGAGG</td><td></td><td></td><td></td><td></td><td></td><td></td><td></td><td></td><td></td><td></td></tr>
<tr><td>Oligo rev</td><td>AAACCCTCGAGGTCGAAGCCGTCA</td><td></td><td></td><td></td><td></td><td></td><td></td><td></td><td></td><td></td><td></td></tr>
<tr><td>Chromosome</td><td>start</td><td>end</td><td>strand</td><td>MM</td><td>target_seq</td><td>PAM</td><td>alignment</td><td>distance</td><td>position</td><td>gene name</td><td>gene id</td></tr>
<tr><td>GU305914.1</td><td>152582</td><td>152604</td><td>+</td><td>0</td><td>TGACGGCTTCGACCTCGAGG</td><td>AGG</td><td>||||||||[||||||||||||]PAM</td><td>NA</td><td>Intergenic</td><td>NA</td><td>NA</td></tr>
<tr><td></td><td></td><td></td><td></td><td></td><td></td><td></td><td></td><td></td><td></td><td></td><td></td></tr>
<tr><td>T11676</td><td>CCTGCCGCATCACCTTCATCAGG</td><td>1000</td><td></td><td></td><td></td><td></td><td></td><td></td><td></td><td></td><td></td></tr>
</table>

| | | | | | | | | | | | | |
|---|---|---|---|---|---|---|---|---|---|---|---|---|
| Oligo fwd | CACCCCTGCCGCATCACCTTCATC | | | | | | | | | | | |
| Oligo rev | AAACGATGAAGGTGATGCGGCAGG | | | | | | | | | | | |
| Chromosome | start | | end | strand | MM | target_seq | PAM | alignment | distance | position | gene name | gene id |
| GU305914.1 | | 152629 | 152651 | + | 0 | CCTGCCGCATCACCTTCATC | AGG | \|\|\|\|\|\|\|\|[\|\|\|\|\|\|\|\|\|\|\|\|]PAM | NA | Intergenic | NA | NA |
| T11677 | CTGCCGCATCACCTTCATCAGGG | | 1000 | | | | | | | | | |
| Oligo fwd | CACCCTGCCGCATCACCTTCATCA | | | | | | | | | | | |
| Oligo rev | AAACTGATGAAGGTGATGCGGCAG | | | | | | | | | | | |
| Chromosome | start | | end | strand | MM | target_seq | PAM | alignment | distance | position | gene name | gene id |
| GU305914.1 | | 152630 | 152652 | + | 0 | CTGCCGCATCACCTTCATCA | GGG | \|\|\|\|\|\|\|\|[\|\|\|\|\|\|\|\|\|\|\|\|]PAM | NA | Intergenic | NA | NA |
| T11678 | TTCATCAGGGACAGCGCCGTCGG | | 1000 | | | | | | | | | |
| Oligo fwd | CACCTTCATCAGGGACAGCGCCGT | | | | | | | | | | | |
| Oligo rev | AAACACGGCGCTGTCCCTGATGAA | | | | | | | | | | | |
| Chromosome | start | | end | strand | MM | target_seq | PAM | alignment | distance | position | gene name | gene id |
| GU305914.1 | | 152643 | 152665 | + | 0 | TTCATCAGGGACAGCGCCGT | CGG | \|\|\|\|\|\|\|\|[\|\|\|\|\|\|\|\|\|\|\|\|]PAM | NA | Intergenic | NA | NA |
| T11679 | GGACAGCGCCGTCGGCGTCACGG | | 1000 | | | | | | | | | |
| Oligo fwd | CACCGGACAGCGCCGTCGGCGTCA | | | | | | | | | | | |
| Oligo rev | AAACTGACGCCGACGGCGCTGTCC | | | | | | | | | | | |
| Chromosome | start | | end | strand | MM | target_seq | PAM | alignment | distance | position | gene name | gene id |
| GU305914.1 | | 152651 | 152673 | + | 0 | GGACAGCGCCGTCGGCGTCA | CGG | \|\|\|\|\|\|\|\|[\|\|\|\|\|\|\|\|\|\|\|\|]PAM | NA | Intergenic | NA | NA |
| T11680 | CGGCGTCACGGCCAAAACCAAGG | | 1000 | | | | | | | | | |
| Oligo fwd | CACCCGGCGTCACGGCCAAAACCA | | | | | | | | | | | |
| Oligo rev | AAACTGGTTTTGGCCGTGACGCCG | | | | | | | | | | | |
| Chromosome | start | | end | strand | MM | target_seq | PAM | alignment | distance | position | gene name | gene id |
| GU305914.1 | | 152663 | 152685 | + | 0 | CGGCGTCACGGCCAAAACCA | AGG | \|\|\|\|\|\|\|\|[\|\|\|\|\|\|\|\|\|\|\|\|]PAM | NA | Intergenic | NA | NA |
| GU305914.1 | | 53379 | 53401 | + | 4 | CAGCACCACGGACAAAACCA | CGG | \|-\|\|--\|\|[\|\|\|-\|\|\|\|\|\|\|\|]PAM | NA | Intergenic | NA | NA |
| T11681 | AAAACCAAGGCCTGCGTCGTCGG | | 1000 | | | | | | | | | |
| Oligo fwd | CACCAAAACCAAGGCCTGCGTCGT | | | | | | | | | | | |
| Oligo rev | AAACACGACGCAGGCCTTGGTTTT | | | | | | | | | | | |
| Chromosome | start | | end | strand | MM | target_seq | PAM | alignment | distance | position | gene name | gene id |
| GU305914.1 | | 152676 | 152698 | + | 0 | AAAACCAAGGCCTGCGTCGT | CGG | \|\|\|\|\|\|\|\|[\|\|\|\|\|\|\|\|\|\|\|\|]PAM | NA | Intergenic | NA | NA |
| T11682 | AAACCAAGGCCTGCGTCGTCGGG | | 1000 | | | | | | | | | |
| Oligo fwd | CACCAAACCAAGGCCTGCGTCGTC | | | | | | | | | | | |
| Oligo rev | AAACGACGACGCAGGCCTTGGTTT | | | | | | | | | | | |
| Chromosome | start | | end | strand | MM | target_seq | PAM | alignment | distance | position | gene name | gene id |
| GU305914.1 | | 152677 | 152699 | + | 0 | AAACCAAGGCCTGCGTCGTC | GGG | \|\|\|\|\|\|\|\|[\|\|\|\|\|\|\|\|\|\|\|\|]PAM | NA | Intergenic | NA | NA |
| T11683 | GCCTGCGTCGTCGGGTACACCGG | | 1000 | | | | | | | | | |
| Oligo fwd | CACCGCCTGCGTCGTCGGGTACAC | | | | | | | | | | | |
| Oligo rev | AAACGTGTACCCGACGACGCAGGC | | | | | | | | | | | |
| Chromosome | start | | end | strand | MM | target_seq | PAM | alignment | distance | position | gene name | gene id |
| GU305914.1 | | 152685 | 152707 | + | 0 | GCCTGCGTCGTCGGGTACAC | CGG | \|\|\|\|\|\|\|\|[\|\|\|\|\|\|\|\|\|\|\|\|]PAM | NA | Intergenic | NA | NA |
| T11684 | TTTCGCCGAGATCCTGCAAAAGG | | 1000 | | | | | | | | | |
| Oligo fwd | CACCTTTCGCCGAGATCCTGCAAA | | | | | | | | | | | |
| Oligo rev | AAACTTTGCAGGATCTCGGCGAAA | | | | | | | | | | | |
| Chromosome | start | | end | strand | MM | target_seq | PAM | alignment | distance | position | gene name | gene id |
| GU305914.1 | | 152720 | 152742 | + | 0 | TTTCGCCGAGATCCTGCAAA | AGG | \|\|\|\|\|\|\|\|[\|\|\|\|\|\|\|\|\|\|\|\|]PAM | NA | Intergenic | NA | NA |
| T11685 | AAAAGGACCTCTTCATCGAGAGG | | 1000 | | | | | | | | | |
| Oligo fwd | CACCAAAAGGACCTCTTCATCGAG | | | | | | | | | | | |
| Oligo rev | AAACCTCGATGAAGAGGTCCTTTT | | | | | | | | | | | |
| Chromosome | start | | end | strand | MM | target_seq | PAM | alignment | distance | position | gene name | gene id |
| GU305914.1 | | 152737 | 152759 | + | 0 | AAAAGGACCTCTTCATCGAG | AGG | \|\|\|\|\|\|\|\|[\|\|\|\|\|\|\|\|\|\|\|\|]PAM | NA | Intergenic | NA | NA |
| T11686 | CGAGAGGACCCCCTGCGAGCAGG | | 1000 | | | | | | | | | |
| Oligo fwd | CACCCGAGAGGACCCCCTGCGAGC | | | | | | | | | | | |
| Oligo rev | AAACGCTCGCAGGGGGTCCTCTCG | | | | | | | | | | | |
| Chromosome | start | | end | strand | MM | target_seq | PAM | alignment | distance | position | gene name | gene id |
| GU305914.1 | | 152753 | 152775 | + | 0 | CGAGAGGACCCCCTGCGAGC | AGG | \|\|\|\|\|\|\|\|[\|\|\|\|\|\|\|\|\|\|\|\|]PAM | NA | Intergenic | NA | NA |
| T11687 | CTACGCCTATTCCCTCATCTCGG | | 1000 | | | | | | | | | |
| Oligo fwd | CACCCTACGCCTATTCCCTCATCT | | | | | | | | | | | |
| Oligo rev | AAACAGATGAGGGAATAGGCGTAG | | | | | | | | | | | |
| Chromosome | start | | end | strand | MM | target_seq | PAM | alignment | distance | position | gene name | gene id |
| GU305914.1 | | 152780 | 152802 | + | 0 | CTACGCCTATTCCCTCATCT | CGG | \|\|\|\|\|\|\|\|[\|\|\|\|\|\|\|\|\|\|\|\|]PAM | NA | Intergenic | NA | NA |
| T11688 | TACGCCTATTCCCTCATCTCGGG | | 1000 | | | | | | | | | |
| Oligo fwd | CACCTACGCCTATTCCCTCATCTC | | | | | | | | | | | |
| Oligo rev | AAACGAGATGAGGGAATAGGCGTA | | | | | | | | | | | |
| Chromosome | start | | end | strand | MM | target_seq | PAM | alignment | distance | position | gene name | gene id |
| GU305914.1 | | 152781 | 152803 | + | 0 | TACGCCTATTCCCTCATCTC | GGG | \|\|\|\|\|\|\|\|[\|\|\|\|\|\|\|\|\|\|\|\|]PAM | NA | Intergenic | NA | NA |
| T11689 | CAGCTCCCCCCTCACCACCCCGG | | 1000 | | | | | | | | | |
| Oligo fwd | CACCCAGCTCCCCCCTCACCACCC | | | | | | | | | | | |
| Oligo rev | AAACGGGTGGTGAGGGGGGAGCTG | | | | | | | | | | | |
| Chromosome | start | | end | strand | MM | target_seq | PAM | alignment | distance | position | gene name | gene id |
| GU305914.1 | | 152834 | 152856 | + | 0 | CAGCTCCCCCCTCACCACCC | CGG | \|\|\|\|\|\|\|\|[\|\|\|\|\|\|\|\|\|\|\|\|]PAM | NA | Intergenic | NA | NA |
| T11690 | GACATCCCGACGCTCGTCATCGG | | 1000 | | | | | | | | | |
| Oligo fwd | CACCGACATCCCGACGCTCGTCAT | | | | | | | | | | | |
| Oligo rev | AAACATGACGAGCGTCGGGATGTC | | | | | | | | | | | |
| Chromosome | start | | end | strand | MM | target_seq | PAM | alignment | distance | position | gene name | gene id |
| GU305914.1 | | 152892 | 152914 | + | 0 | GACATCCCGACGCTCGTCAT | CGG | \|\|\|\|\|\|\|\|[\|\|\|\|\|\|\|\|\|\|\|\|]PAM | NA | Intergenic | NA | NA |

<table>
<tr><td>T11691</td><td>ACGCTCGTCATCGGAGCCAACGG</td><td>1000</td><td></td><td></td><td></td><td></td><td></td><td></td><td></td><td></td><td></td></tr>
<tr><td>Oligo fwd</td><td>CACCACGCTCGTCATCGGAGCCAA</td><td></td><td></td><td></td><td></td><td></td><td></td><td></td><td></td><td></td><td></td></tr>
<tr><td>Oligo rev</td><td>AAACTTGGCTCCGATGACGAGCGT</td><td></td><td></td><td></td><td></td><td></td><td></td><td></td><td></td><td></td><td></td></tr>
<tr><td>Chromosome</td><td>start</td><td>end</td><td>strand</td><td>MM</td><td>target_seq</td><td>PAM</td><td>alignment</td><td>distance</td><td>position</td><td>gene name</td><td>gene id</td></tr>
<tr><td>GU305914.1</td><td>152901</td><td>152923</td><td>+</td><td>0</td><td>ACGCTCGTCATCGGAGCCAA</td><td>CGG</td><td>||||||||[||||||||||||]PAM</td><td>NA</td><td>Intergenic</td><td>NA</td><td>NA</td></tr>
<tr><td>T11692</td><td>CTCGTCATCGGAGCCAACGGCGG</td><td>1000</td><td></td><td></td><td></td><td></td><td></td><td></td><td></td><td></td><td></td></tr>
<tr><td>Oligo fwd</td><td>CACCCTCGTCATCGGAGCCAACGG</td><td></td><td></td><td></td><td></td><td></td><td></td><td></td><td></td><td></td><td></td></tr>
<tr><td>Oligo rev</td><td>AAACCCGTTGGCTCCGATGACGAG</td><td></td><td></td><td></td><td></td><td></td><td></td><td></td><td></td><td></td><td></td></tr>
<tr><td>Chromosome</td><td>start</td><td>end</td><td>strand</td><td>MM</td><td>target_seq</td><td>PAM</td><td>alignment</td><td>distance</td><td>position</td><td>gene name</td><td>gene id</td></tr>
<tr><td>GU305914.1</td><td>152904</td><td>152926</td><td>+</td><td>0</td><td>CTCGTCATCGGAGCCAACGG</td><td>CGG</td><td>||||||||[||||||||||||]PAM</td><td>NA</td><td>Intergenic</td><td>NA</td><td>NA</td></tr>
<tr><td>T11693</td><td>ATCGGAGCCAACGGCGGCGACGG</td><td>1000</td><td></td><td></td><td></td><td></td><td></td><td></td><td></td><td></td><td></td></tr>
<tr><td>Oligo fwd</td><td>CACCATCGGAGCCAACGGCGGCGA</td><td></td><td></td><td></td><td></td><td></td><td></td><td></td><td></td><td></td><td></td></tr>
<tr><td>Oligo rev</td><td>AAACTCGCCGCCGTTGGCTCCGAT</td><td></td><td></td><td></td><td></td><td></td><td></td><td></td><td></td><td></td><td></td></tr>
<tr><td>Chromosome</td><td>start</td><td>end</td><td>strand</td><td>MM</td><td>target_seq</td><td>PAM</td><td>alignment</td><td>distance</td><td>position</td><td>gene name</td><td>gene id</td></tr>
<tr><td>GU305914.1</td><td>152910</td><td>152932</td><td>+</td><td>0</td><td>ATCGGAGCCAACGGCGGCGA</td><td>CGG</td><td>||||||||[||||||||||||]PAM</td><td>NA</td><td>Intergenic</td><td>NA</td><td>NA</td></tr>
<tr><td>T11694</td><td>CCAACGGCGGCGACGGCGCGAGG</td><td>1000</td><td></td><td></td><td></td><td></td><td></td><td></td><td></td><td></td><td></td></tr>
<tr><td>Oligo fwd</td><td>CACCCCAACGGCGGCGACGGCGCG</td><td></td><td></td><td></td><td></td><td></td><td></td><td></td><td></td><td></td><td></td></tr>
<tr><td>Oligo rev</td><td>AAACCGCGCCGTCGCCGCCGTTGG</td><td></td><td></td><td></td><td></td><td></td><td></td><td></td><td></td><td></td><td></td></tr>
<tr><td>Chromosome</td><td>start</td><td>end</td><td>strand</td><td>MM</td><td>target_seq</td><td>PAM</td><td>alignment</td><td>distance</td><td>position</td><td>gene name</td><td>gene id</td></tr>
<tr><td>GU305914.1</td><td>152917</td><td>152939</td><td>+</td><td>0</td><td>CCAACGGCGGCGACGGCGCG</td><td>AGG</td><td>||||||||[||||||||||||]PAM</td><td>NA</td><td>Intergenic</td><td>NA</td><td>NA</td></tr>
<tr><td>T11695</td><td>CAACGGCGGCGACGGCGCGAGGG</td><td>1000</td><td></td><td></td><td></td><td></td><td></td><td></td><td></td><td></td><td></td></tr>
<tr><td>Oligo fwd</td><td>CACCCAACGGCGGCGACGGCGCGA</td><td></td><td></td><td></td><td></td><td></td><td></td><td></td><td></td><td></td><td></td></tr>
<tr><td>Oligo rev</td><td>AAACTCGCGCCGTCGCCGCCGTTG</td><td></td><td></td><td></td><td></td><td></td><td></td><td></td><td></td><td></td><td></td></tr>
<tr><td>Chromosome</td><td>start</td><td>end</td><td>strand</td><td>MM</td><td>target_seq</td><td>PAM</td><td>alignment</td><td>distance</td><td>position</td><td>gene name</td><td>gene id</td></tr>
<tr><td>GU305914.1</td><td>152918</td><td>152940</td><td>+</td><td>0</td><td>CAACGGCGGCGACGGCGCGA</td><td>GGG</td><td>||||||||[||||||||||||]PAM</td><td>NA</td><td>Intergenic</td><td>NA</td><td>NA</td></tr>
<tr><td>GU305914.1</td><td>76452</td><td>76474</td><td>-</td><td>3</td><td>CCGCGGCGGCGCCGGCGCGA</td><td>TGG</td><td>|--|||||[|||-||||||||]PAM</td><td>NA</td><td>Intergenic</td><td>NA</td><td>NA</td></tr>
<tr><td>T11696</td><td>CGATGATAACGATGAGTATGAGG</td><td>1000</td><td></td><td></td><td></td><td></td><td></td><td></td><td></td><td></td><td></td></tr>
<tr><td>Oligo fwd</td><td>CACCCGATGATAACGATGAGTATG</td><td></td><td></td><td></td><td></td><td></td><td></td><td></td><td></td><td></td><td></td></tr>
<tr><td>Oligo rev</td><td>AAACCATACTCATCGTTATCATCG</td><td></td><td></td><td></td><td></td><td></td><td></td><td></td><td></td><td></td><td></td></tr>
<tr><td>Chromosome</td><td>start</td><td>end</td><td>strand</td><td>MM</td><td>target_seq</td><td>PAM</td><td>alignment</td><td>distance</td><td>position</td><td>gene name</td><td>gene id</td></tr>
<tr><td>GU305914.1</td><td>152945</td><td>152967</td><td>+</td><td>0</td><td>CGATGATAACGATGAGTATG</td><td>AGG</td><td>||||||||[||||||||||||]PAM</td><td>NA</td><td>Intergenic</td><td>NA</td><td>NA</td></tr>
<tr><td>GU305914.1</td><td>168577</td><td>168599</td><td>+</td><td>4</td><td>AGATGATGAAGATGAATATG</td><td>AGG</td><td>-||||||-[|-|||||-||||]PAM</td><td>NA</td><td>Intergenic</td><td>NA</td><td>NA</td></tr>
<tr><td>T11697</td><td>TGATAACGATGAGTATGAGGAGG</td><td>1000</td><td></td><td></td><td></td><td></td><td></td><td></td><td></td><td></td><td></td></tr>
<tr><td>Oligo fwd</td><td>CACCTGATAACGATGAGTATGAGG</td><td></td><td></td><td></td><td></td><td></td><td></td><td></td><td></td><td></td><td></td></tr>
<tr><td>Oligo rev</td><td>AAACCCTCATACTCATCGTTATCA</td><td></td><td></td><td></td><td></td><td></td><td></td><td></td><td></td><td></td><td></td></tr>
<tr><td>Chromosome</td><td>start</td><td>end</td><td>strand</td><td>MM</td><td>target_seq</td><td>PAM</td><td>alignment</td><td>distance</td><td>position</td><td>gene name</td><td>gene id</td></tr>
<tr><td>GU305914.1</td><td>152948</td><td>152970</td><td>+</td><td>0</td><td>TGATAACGATGAGTATGAGG</td><td>AGG</td><td>||||||||[||||||||||||]PAM</td><td>NA</td><td>Intergenic</td><td>NA</td><td>NA</td></tr>
<tr><td>T11698</td><td>CGATGAGTATGAGGAGGATCTGG</td><td>1000</td><td></td><td></td><td></td><td></td><td></td><td></td><td></td><td></td><td></td></tr>
<tr><td>Oligo fwd</td><td>CACCCGATGAGTATGAGGAGGATC</td><td></td><td></td><td></td><td></td><td></td><td></td><td></td><td></td><td></td><td></td></tr>
<tr><td>Oligo rev</td><td>AAACGATCCTCCTCATACTCATCG</td><td></td><td></td><td></td><td></td><td></td><td></td><td></td><td></td><td></td><td></td></tr>
<tr><td>Chromosome</td><td>start</td><td>end</td><td>strand</td><td>MM</td><td>target_seq</td><td>PAM</td><td>alignment</td><td>distance</td><td>position</td><td>gene name</td><td>gene id</td></tr>
<tr><td>GU305914.1</td><td>152954</td><td>152976</td><td>+</td><td>0</td><td>CGATGAGTATGAGGAGGATC</td><td>TGG</td><td>||||||||[||||||||||||]PAM</td><td>NA</td><td>Intergenic</td><td>NA</td><td>NA</td></tr>
<tr><td>GU305914.1</td><td>179354</td><td>179376</td><td>-</td><td>4</td><td>GGAGGAGTATCAGGAGGAGC</td><td>AGG</td><td>-||-||||[||-|||||||-|]PAM</td><td>NA</td><td>Intergenic</td><td>NA</td><td>NA</td></tr>
<tr><td>GU305914.1</td><td>140717</td><td>140739</td><td>+</td><td>4</td><td>AGATGAGGACGAGGAGGATG</td><td>AGG</td><td>-||||||-[|-|||||||||-]PAM</td><td>NA</td><td>Intergenic</td><td>NA</td><td>NA</td></tr>
<tr><td>T11699</td><td>TGAGTATGAGGAGGATCTGGAGG</td><td>1000</td><td></td><td></td><td></td><td></td><td></td><td></td><td></td><td></td><td></td></tr>
<tr><td>Oligo fwd</td><td>CACCTGAGTATGAGGAGGATCTGG</td><td></td><td></td><td></td><td></td><td></td><td></td><td></td><td></td><td></td><td></td></tr>
<tr><td>Oligo rev</td><td>AAACCCAGATCCTCCTCATACTCA</td><td></td><td></td><td></td><td></td><td></td><td></td><td></td><td></td><td></td><td></td></tr>
<tr><td>Chromosome</td><td>start</td><td>end</td><td>strand</td><td>MM</td><td>target_seq</td><td>PAM</td><td>alignment</td><td>distance</td><td>position</td><td>gene name</td><td>gene id</td></tr>
<tr><td>GU305914.1</td><td>152957</td><td>152979</td><td>+</td><td>0</td><td>TGAGTATGAGGAGGATCTGG</td><td>AGG</td><td>||||||||[||||||||||||]PAM</td><td>NA</td><td>Intergenic</td><td>NA</td><td>NA</td></tr>
<tr><td>T11700</td><td>GAGTATGAGGAGGATCTGGAGGG</td><td>1000</td><td></td><td></td><td></td><td></td><td></td><td></td><td></td><td></td><td></td></tr>
<tr><td>Oligo fwd</td><td>CACCGAGTATGAGGAGGATCTGGA</td><td></td><td></td><td></td><td></td><td></td><td></td><td></td><td></td><td></td><td></td></tr>
<tr><td>Oligo rev</td><td>AAACTCCAGATCCTCCTCATACTC</td><td></td><td></td><td></td><td></td><td></td><td></td><td></td><td></td><td></td><td></td></tr>
<tr><td>Chromosome</td><td>start</td><td>end</td><td>strand</td><td>MM</td><td>target_seq</td><td>PAM</td><td>alignment</td><td>distance</td><td>position</td><td>gene name</td><td>gene id</td></tr>
<tr><td>GU305914.1</td><td>152958</td><td>152980</td><td>+</td><td>0</td><td>GAGTATGAGGAGGATCTGGA</td><td>GGG</td><td>||||||||[||||||||||||]PAM</td><td>NA</td><td>Intergenic</td><td>NA</td><td>NA</td></tr>
<tr><td>T11701</td><td>TATGAGGAGGATCTGGAGGGTGG</td><td>1000</td><td></td><td></td><td></td><td></td><td></td><td></td><td></td><td></td><td></td></tr>
<tr><td>Oligo fwd</td><td>CACCTATGAGGAGGATCTGGAGGG</td><td></td><td></td><td></td><td></td><td></td><td></td><td></td><td></td><td></td><td></td></tr>
<tr><td>Oligo rev</td><td>AAACCCCTCCAGATCCTCCTCATA</td><td></td><td></td><td></td><td></td><td></td><td></td><td></td><td></td><td></td><td></td></tr>
<tr><td>Chromosome</td><td>start</td><td>end</td><td>strand</td><td>MM</td><td>target_seq</td><td>PAM</td><td>alignment</td><td>distance</td><td>position</td><td>gene name</td><td>gene id</td></tr>
<tr><td>GU305914.1</td><td>152961</td><td>152983</td><td>+</td><td>0</td><td>TATGAGGAGGATCTGGAGGG</td><td>TGG</td><td>||||||||[||||||||||||]PAM</td><td>NA</td><td>Intergenic</td><td>NA</td><td>NA</td></tr>
<tr><td>T11702</td><td>GAGGAGGATCTGGAGGGTGGCGG</td><td>1000</td><td></td><td></td><td></td><td></td><td></td><td></td><td></td><td></td><td></td></tr>
<tr><td>Oligo fwd</td><td>CACCGAGGAGGATCTGGAGGGTGG</td><td></td><td></td><td></td><td></td><td></td><td></td><td></td><td></td><td></td><td></td></tr>
<tr><td>Oligo rev</td><td>AAACCCACCCTCCAGATCCTCCTC</td><td></td><td></td><td></td><td></td><td></td><td></td><td></td><td></td><td></td><td></td></tr>
<tr><td>Chromosome</td><td>start</td><td>end</td><td>strand</td><td>MM</td><td>target_seq</td><td>PAM</td><td>alignment</td><td>distance</td><td>position</td><td>gene name</td><td>gene id</td></tr>
<tr><td>GU305914.1</td><td>152964</td><td>152986</td><td>+</td><td>0</td><td>GAGGAGGATCTGGAGGGTGG</td><td>CGG</td><td>||||||||[||||||||||||]PAM</td><td>NA</td><td>Intergenic</td><td>NA</td><td>NA</td></tr>
<tr><td>T11703</td><td>AGGAGGATCTGGAGGGTGGCGGG</td><td>1000</td><td></td><td></td><td></td><td></td><td></td><td></td><td></td><td></td><td></td></tr>
<tr><td>Oligo fwd</td><td>CACCAGGAGGATCTGGAGGGTGGC</td><td></td><td></td><td></td><td></td><td></td><td></td><td></td><td></td><td></td><td></td></tr>
<tr><td>Oligo rev</td><td>AAACGCCACCCTCCAGATCCTCCT</td><td></td><td></td><td></td><td></td><td></td><td></td><td></td><td></td><td></td><td></td></tr>
<tr><td>Chromosome</td><td>start</td><td>end</td><td>strand</td><td>MM</td><td>target_seq</td><td>PAM</td><td>alignment</td><td>distance</td><td>position</td><td>gene name</td><td>gene id</td></tr>
<tr><td>GU305914.1</td><td>152965</td><td>152987</td><td>+</td><td>0</td><td>AGGAGGATCTGGAGGGTGGC</td><td>GGG</td><td>||||||||[||||||||||||]PAM</td><td>NA</td><td>Intergenic</td><td>NA</td><td>NA</td></tr>
<tr><td>T11704</td><td>GAGGGTGGCGGGTGCTTTGACGG</td><td>1000</td><td></td><td></td><td></td><td></td><td></td><td></td><td></td><td></td><td></td></tr>
<tr><td>Oligo fwd</td><td>CACCGAGGGTGGCGGGTGCTTTGA</td><td></td><td></td><td></td><td></td><td></td><td></td><td></td><td></td><td></td><td></td></tr>
<tr><td>Oligo rev</td><td>AAACTCAAAGCACCCGCCACCCTC</td><td></td><td></td><td></td><td></td><td></td><td></td><td></td><td></td><td></td><td></td></tr>
<tr><td>Chromosome</td><td>start</td><td>end</td><td>strand</td><td>MM</td><td>target_seq</td><td>PAM</td><td>alignment</td><td>distance</td><td>position</td><td>gene name</td><td>gene id</td></tr>
<tr><td>GU305914.1</td><td>152976</td><td>152998</td><td>+</td><td>0</td><td>GAGGGTGGCGGGTGCTTTGA</td><td>CGG</td><td>||||||||[||||||||||||]PAM</td><td>NA</td><td>Intergenic</td><td>NA</td><td>NA</td></tr>
</table>

T11705 CGGGTGCTTTGACGGTGTTTCGG 1000
Oligo fwd CACCCGGGTGCTTTGACGGTGTTT
Oligo rev AAACAAACACCGTCAAAGCACCCG
Chromosome start end strand MM target_seq PAM alignment distance position gene name gene id
GU305914.1 152984 153006 + 0 CGGGTGCTTTGACGGTGTTT CGG ||||||||[||||||||||||]PAM NA Intergenic NA NA

T11706 GGGTGCTTTGACGGTGTTTCGGG 1000
Oligo fwd CACCGGGTGCTTTGACGGTGTTTC
Oligo rev AAACGAAACACCGTCAAAGCACCC
Chromosome start end strand MM target_seq PAM alignment distance position gene name gene id
GU305914.1 152985 153007 + 0 GGGTGCTTTGACGGTGTTTC GGG ||||||||[||||||||||||]PAM NA Intergenic NA NA

T11707 GGTGCTTTGACGGTGTTTCGGGG 1000
Oligo fwd CACCGGTGCTTTGACGGTGTTTCG
Oligo rev AAACCGAAACACCGTCAAAGCACC
Chromosome start end strand MM target_seq PAM alignment distance position gene name gene id
GU305914.1 152986 153008 + 0 GGTGCTTTGACGGTGTTTCG GGG ||||||||[||||||||||||]PAM NA Intergenic NA NA

T11708 GTGCTTTGACGGTGTTTCGGGGG 1000
Oligo fwd CACCGTGCTTTGACGGTGTTTCGG
Oligo rev AAACCCGAAACACCGTCAAAGCAC
Chromosome start end strand MM target_seq PAM alignment distance position gene name gene id
GU305914.1 152987 153009 + 0 GTGCTTTGACGGTGTTTCGG GGG ||||||||[||||||||||||]PAM NA Intergenic NA NA

T11709 TGCTTTGACGGTGTTTCGGGGGG 1000
Oligo fwd CACCTGCTTTGACGGTGTTTCGGG
Oligo rev AAACCCCGAAACACCGTCAAAGCA
Chromosome start end strand MM target_seq PAM alignment distance position gene name gene id
GU305914.1 152988 153010 + 0 TGCTTTGACGGTGTTTCGGG GGG ||||||||[||||||||||||]PAM NA Intergenic NA NA

T11710 TTTGACGGTGTTTCGGGGGGTGG 1000
Oligo fwd CACCTTTGACGGTGTTTCGGGGGG
Oligo rev AAACCCCCCCGAAACACCGTCAAA
Chromosome start end strand MM target_seq PAM alignment distance position gene name gene id
GU305914.1 152991 153013 + 0 TTTGACGGTGTTTCGGGGGG TGG ||||||||[||||||||||||]PAM NA Intergenic NA NA

T11711 GGTGTTTCGGGGGGTGGTAGCGG 1000
Oligo fwd CACCGGTGTTTCGGGGGGTGGTAG
Oligo rev AAACCTACCACCCCCCGAAACACC
Chromosome start end strand MM target_seq PAM alignment distance position gene name gene id
GU305914.1 152997 153019 + 0 GGTGTTTCGGGGGGTGGTAG CGG ||||||||[||||||||||||]PAM NA Intergenic NA NA

T11712 TCGGGGGGTGGTAGCGGTAACGG 1000
Oligo fwd CACCTCGGGGGGTGGTAGCGGTAA
Oligo rev AAACTTACCGCTACCACCCCCCGA
Chromosome start end strand MM target_seq PAM alignment distance position gene name gene id
GU305914.1 153003 153025 + 0 TCGGGGGGTGGTAGCGGTAA CGG ||||||||[||||||||||||]PAM NA Intergenic NA NA

T11713 GGGGGTGGTAGCGGTAACGGTGG 1000
Oligo fwd CACCGGGGGTGGTAGCGGTAACGG
Oligo rev AAACCCGTTACCGCTACCACCCCC
Chromosome start end strand MM target_seq PAM alignment distance position gene name gene id
GU305914.1 153006 153028 + 0 GGGGGTGGTAGCGGTAACGG TGG ||||||||[||||||||||||]PAM NA Intergenic NA NA

T11714 GGTGGTAGCGGTAACGGTGGCGG 1000
Oligo fwd CACCGGTGGTAGCGGTAACGGTGG
Oligo rev AAACCCACCGTTACCGCTACCACC
Chromosome start end strand MM target_seq PAM alignment distance position gene name gene id
GU305914.1 153009 153031 + 0 GGTGGTAGCGGTAACGGTGG CGG ||||||||[||||||||||||]PAM NA Intergenic NA NA

T11715 TCGACTGACCTCCGACGACGAGG 1000
Oligo fwd CACCTCGACTGACCTCCGACGACG
Oligo rev AAACCGTCGTCGGAGGTCAGTCGA
Chromosome start end strand MM target_seq PAM alignment distance position gene name gene id
GU305914.1 153047 153069 + 0 TCGACTGACCTCCGACGACG AGG ||||||||[||||||||||||]PAM NA Intergenic NA NA

T11716 GACCTGTCCTACGTCGATCGAGG 1000
Oligo fwd CACCGACCTGTCCTACGTCGATCG
Oligo rev AAACCGATCGACGTAGGACAGGTC
Chromosome start end strand MM target_seq PAM alignment distance position gene name gene id
GU305914.1 153081 153103 + 0 GACCTGTCCTACGTCGATCG AGG ||||||||[||||||||||||]PAM NA Intergenic NA NA

T11717 TGTCCTACGTCGATCGAGGTCGG 1000
Oligo fwd CACCTGTCCTACGTCGATCGAGGT
Oligo rev AAACACCTCGATCGACGTAGGACA
Chromosome start end strand MM target_seq PAM alignment distance position gene name gene id
GU305914.1 153085 153107 + 0 TGTCCTACGTCGATCGAGGT CGG ||||||||[||||||||||||]PAM NA Intergenic NA NA

T11718 GCAGCCCCGTCTGACGATGTCGG 1000
Oligo fwd CACCGCAGCCCCGTCTGACGATGT
Oligo rev AAACACATCGTCAGACGGGGCTGC
Chromosome start end strand MM target_seq PAM alignment distance position gene name gene id
GU305914.1 153152 153174 + 0 GCAGCCCCGTCTGACGATGT CGG ||||||||[||||||||||||]PAM NA Intergenic NA NA

T11719 GCCCCGTCTGACGATGTCGGCGG 1000
Oligo fwd CACCGCCCCGTCTGACGATGTCGG
Oligo rev AAACCCGACATCGTCAGACGGGGC
Chromosome start end strand MM target_seq PAM alignment distance position gene name gene id
GU305914.1 153155 153177 + 0 GCCCCGTCTGACGATGTCGG CGG ||||||||[||||||||||||]PAM NA Intergenic NA NA

<table>
<tr><td>T11720</td><td>TGTCGGCGGCGCTGCCGCCGCGG</td><td>1000</td><td></td><td></td><td></td><td></td><td></td><td></td><td></td><td></td><td></td></tr>
<tr><td>Oligo fwd</td><td>CACCTGTCGGCGGCGCTGCCGCCG</td><td></td><td></td><td></td><td></td><td></td><td></td><td></td><td></td><td></td><td></td></tr>
<tr><td>Oligo rev</td><td>AAACCGGCGGCAGCGCCGCCGACA</td><td></td><td></td><td></td><td></td><td></td><td></td><td></td><td></td><td></td><td></td></tr>
<tr><td>Chromosome</td><td>start</td><td>end</td><td>strand</td><td>MM</td><td>target_seq</td><td>PAM</td><td>alignment</td><td>distance</td><td>position</td><td>gene name</td><td>gene id</td></tr>
<tr><td>GU305914.1</td><td>153169</td><td>153191</td><td>+</td><td>0</td><td>TGTCGGCGGCGCTGCCGCCG</td><td>CGG</td><td>||||||||[||||||||||||]PAM</td><td>NA</td><td>Intergenic</td><td>NA</td><td>NA</td></tr>
<tr><td>GU305914.1</td><td>1214</td><td>1236</td><td>+</td><td>3</td><td>TGTCGTCGGCGCCGCTGCCG</td><td>GGG</td><td>|||||-||[||||-||-||||]PAM</td><td>NA</td><td>Intergenic</td><td>NA</td><td>NA</td></tr>
<tr><td>GU305914.1</td><td>152555</td><td>152577</td><td>+</td><td>4</td><td>CGGCGGCGGCGGTGCCGGCG</td><td>GGG</td><td>-|-|||||[|||-|||||-||]PAM</td><td>NA</td><td>Intergenic</td><td>NA</td><td>NA</td></tr>
<tr><td></td><td></td><td></td><td></td><td></td><td></td><td></td><td></td><td></td><td></td><td></td><td></td></tr>
<tr><td>T11721</td><td>GCCGCCGCGGCAAATCGACGAGG</td><td>1000</td><td></td><td></td><td></td><td></td><td></td><td></td><td></td><td></td><td></td></tr>
<tr><td>Oligo fwd</td><td>CACCGCCGCCGCGGCAAATCGACG</td><td></td><td></td><td></td><td></td><td></td><td></td><td></td><td></td><td></td><td></td></tr>
<tr><td>Oligo rev</td><td>AAACCGTCGATTTGCCGCGGCGGC</td><td></td><td></td><td></td><td></td><td></td><td></td><td></td><td></td><td></td><td></td></tr>
<tr><td>Chromosome</td><td>start</td><td>end</td><td>strand</td><td>MM</td><td>target_seq</td><td>PAM</td><td>alignment</td><td>distance</td><td>position</td><td>gene name</td><td>gene id</td></tr>
<tr><td>GU305914.1</td><td>153182</td><td>153204</td><td>+</td><td>0</td><td>GCCGCCGCGGCAAATCGACG</td><td>AGG</td><td>||||||||[||||||||||||]PAM</td><td>NA</td><td>Intergenic</td><td>NA</td><td>NA</td></tr>
<tr><td></td><td></td><td></td><td></td><td></td><td></td><td></td><td></td><td></td><td></td><td></td><td></td></tr>
<tr><td>T11722</td><td>CGACGAGGAGATCTCCGACGTGG</td><td>1000</td><td></td><td></td><td></td><td></td><td></td><td></td><td></td><td></td><td></td></tr>
<tr><td>Oligo fwd</td><td>CACCCGACGAGGAGATCTCCGACG</td><td></td><td></td><td></td><td></td><td></td><td></td><td></td><td></td><td></td><td></td></tr>
<tr><td>Oligo rev</td><td>AAACCGTCGGAGATCTCCTCGTCG</td><td></td><td></td><td></td><td></td><td></td><td></td><td></td><td></td><td></td><td></td></tr>
<tr><td>Chromosome</td><td>start</td><td>end</td><td>strand</td><td>MM</td><td>target_seq</td><td>PAM</td><td>alignment</td><td>distance</td><td>position</td><td>gene name</td><td>gene id</td></tr>
<tr><td>GU305914.1</td><td>153197</td><td>153219</td><td>+</td><td>0</td><td>CGACGAGGAGATCTCCGACG</td><td>TGG</td><td>||||||||[||||||||||||]PAM</td><td>NA</td><td>Intergenic</td><td>NA</td><td>NA</td></tr>
<tr><td>GU305914.1</td><td>27375</td><td>27397</td><td>+</td><td>4</td><td>TTGCGAGGAGATCTGCGACG</td><td>GGG</td><td>---|||||[||||||-|||||]PAM</td><td>NA</td><td>Intergenic</td><td>NA</td><td>NA</td></tr>
<tr><td></td><td></td><td></td><td></td><td></td><td></td><td></td><td></td><td></td><td></td><td></td><td></td></tr>
<tr><td>T11723</td><td>CGTTAGCTCCATCAGTTTCGAGG</td><td>1000</td><td></td><td></td><td></td><td></td><td></td><td></td><td></td><td></td><td></td></tr>
<tr><td>Oligo fwd</td><td>CACCCGTTAGCTCCATCAGTTTCG</td><td></td><td></td><td></td><td></td><td></td><td></td><td></td><td></td><td></td><td></td></tr>
<tr><td>Oligo rev</td><td>AAACCGAAACTGATGGAGCTAACG</td><td></td><td></td><td></td><td></td><td></td><td></td><td></td><td></td><td></td><td></td></tr>
<tr><td>Chromosome</td><td>start</td><td>end</td><td>strand</td><td>MM</td><td>target_seq</td><td>PAM</td><td>alignment</td><td>distance</td><td>position</td><td>gene name</td><td>gene id</td></tr>
<tr><td>GU305914.1</td><td>153284</td><td>153306</td><td>+</td><td>0</td><td>CGTTAGCTCCATCAGTTTCG</td><td>AGG</td><td>||||||||[||||||||||||]PAM</td><td>NA</td><td>Intergenic</td><td>NA</td><td>NA</td></tr>
<tr><td></td><td></td><td></td><td></td><td></td><td></td><td></td><td></td><td></td><td></td><td></td><td></td></tr>
<tr><td>T11724</td><td>TCCGAGACATCTTTCTCGCCAGG</td><td>1000</td><td></td><td></td><td></td><td></td><td></td><td></td><td></td><td></td><td></td></tr>
<tr><td>Oligo fwd</td><td>CACCTCCGAGACATCTTTCTCGCC</td><td></td><td></td><td></td><td></td><td></td><td></td><td></td><td></td><td></td><td></td></tr>
<tr><td>Oligo rev</td><td>AAACGGCGAGAAAGATGTCTCGGA</td><td></td><td></td><td></td><td></td><td></td><td></td><td></td><td></td><td></td><td></td></tr>
<tr><td>Chromosome</td><td>start</td><td>end</td><td>strand</td><td>MM</td><td>target_seq</td><td>PAM</td><td>alignment</td><td>distance</td><td>position</td><td>gene name</td><td>gene id</td></tr>
<tr><td>GU305914.1</td><td>153331</td><td>153353</td><td>+</td><td>0</td><td>TCCGAGACATCTTTCTCGCC</td><td>AGG</td><td>||||||||[||||||||||||]PAM</td><td>NA</td><td>Intergenic</td><td>NA</td><td>NA</td></tr>
<tr><td></td><td></td><td></td><td></td><td></td><td></td><td></td><td></td><td></td><td></td><td></td><td></td></tr>
<tr><td>T11725</td><td>ACATCTTTCTCGCCAGGTATCGG</td><td>1000</td><td></td><td></td><td></td><td></td><td></td><td></td><td></td><td></td><td></td></tr>
<tr><td>Oligo fwd</td><td>CACCACATCTTTCTCGCCAGGTAT</td><td></td><td></td><td></td><td></td><td></td><td></td><td></td><td></td><td></td><td></td></tr>
<tr><td>Oligo rev</td><td>AAACATACCTGGCGAGAAAGATGT</td><td></td><td></td><td></td><td></td><td></td><td></td><td></td><td></td><td></td><td></td></tr>
<tr><td>Chromosome</td><td>start</td><td>end</td><td>strand</td><td>MM</td><td>target_seq</td><td>PAM</td><td>alignment</td><td>distance</td><td>position</td><td>gene name</td><td>gene id</td></tr>
<tr><td>GU305914.1</td><td>153337</td><td>153359</td><td>+</td><td>0</td><td>ACATCTTTCTCGCCAGGTAT</td><td>CGG</td><td>||||||||[||||||||||||]PAM</td><td>NA</td><td>Intergenic</td><td>NA</td><td>NA</td></tr>
<tr><td></td><td></td><td></td><td></td><td></td><td></td><td></td><td></td><td></td><td></td><td></td><td></td></tr>
<tr><td>T11726</td><td>GATCATGCAGAAGCACACCAAGG</td><td>1000</td><td></td><td></td><td></td><td></td><td></td><td></td><td></td><td></td><td></td></tr>
<tr><td>Oligo fwd</td><td>CACCGATCATGCAGAAGCACACCA</td><td></td><td></td><td></td><td></td><td></td><td></td><td></td><td></td><td></td><td></td></tr>
<tr><td>Oligo rev</td><td>AAACTGGTGTGCTTCTGCATGATC</td><td></td><td></td><td></td><td></td><td></td><td></td><td></td><td></td><td></td><td></td></tr>
<tr><td>Chromosome</td><td>start</td><td>end</td><td>strand</td><td>MM</td><td>target_seq</td><td>PAM</td><td>alignment</td><td>distance</td><td>position</td><td>gene name</td><td>gene id</td></tr>
<tr><td>GU305914.1</td><td>153359</td><td>153381</td><td>+</td><td>0</td><td>GATCATGCAGAAGCACACCA</td><td>AGG</td><td>||||||||[||||||||||||]PAM</td><td>NA</td><td>Intergenic</td><td>NA</td><td>NA</td></tr>
<tr><td>GU305914.1</td><td>74562</td><td>74584</td><td>+</td><td>4</td><td>GGTCTTGCAGAAGCACCACA</td><td>CGG</td><td>|-||-|||[||||||||--||]PAM</td><td>NA</td><td>Intergenic</td><td>NA</td><td>NA</td></tr>
<tr><td></td><td></td><td></td><td></td><td></td><td></td><td></td><td></td><td></td><td></td><td></td><td></td></tr>
<tr><td>T11727</td><td>ATCATGCAGAAGCACACCAAGGG</td><td>1000</td><td></td><td></td><td></td><td></td><td></td><td></td><td></td><td></td><td></td></tr>
<tr><td>Oligo fwd</td><td>CACCATCATGCAGAAGCACACCAA</td><td></td><td></td><td></td><td></td><td></td><td></td><td></td><td></td><td></td><td></td></tr>
<tr><td>Oligo rev</td><td>AAACTTGGTGTGCTTCTGCATGAT</td><td></td><td></td><td></td><td></td><td></td><td></td><td></td><td></td><td></td><td></td></tr>
<tr><td>Chromosome</td><td>start</td><td>end</td><td>strand</td><td>MM</td><td>target_seq</td><td>PAM</td><td>alignment</td><td>distance</td><td>position</td><td>gene name</td><td>gene id</td></tr>
<tr><td>GU305914.1</td><td>153360</td><td>153382</td><td>+</td><td>0</td><td>ATCATGCAGAAGCACACCAA</td><td>GGG</td><td>||||||||[||||||||||||]PAM</td><td>NA</td><td>Intergenic</td><td>NA</td><td>NA</td></tr>
<tr><td></td><td></td><td></td><td></td><td></td><td></td><td></td><td></td><td></td><td></td><td></td><td></td></tr>
<tr><td>T11728</td><td>AAGCACACCAAGGGCGCCTTCGG</td><td>1000</td><td></td><td></td><td></td><td></td><td></td><td></td><td></td><td></td><td></td></tr>
<tr><td>Oligo fwd</td><td>CACCAAGCACACCAAGGGCGCCTT</td><td></td><td></td><td></td><td></td><td></td><td></td><td></td><td></td><td></td><td></td></tr>
<tr><td>Oligo rev</td><td>AAACAAGGCGCCCTTGGTGTGCTT</td><td></td><td></td><td></td><td></td><td></td><td></td><td></td><td></td><td></td><td></td></tr>
<tr><td>Chromosome</td><td>start</td><td>end</td><td>strand</td><td>MM</td><td>target_seq</td><td>PAM</td><td>alignment</td><td>distance</td><td>position</td><td>gene name</td><td>gene id</td></tr>
<tr><td>GU305914.1</td><td>153369</td><td>153391</td><td>+</td><td>0</td><td>AAGCACACCAAGGGCGCCTT</td><td>CGG</td><td>||||||||[||||||||||||]PAM</td><td>NA</td><td>Intergenic</td><td>NA</td><td>NA</td></tr>
<tr><td></td><td></td><td></td><td></td><td></td><td></td><td></td><td></td><td></td><td></td><td></td><td></td></tr>
<tr><td>T11729</td><td>AGGGCGCCTTCGGCAAGACCCGG</td><td>1000</td><td></td><td></td><td></td><td></td><td></td><td></td><td></td><td></td><td></td></tr>
<tr><td>Oligo fwd</td><td>CACCAGGGCGCCTTCGGCAAGACC</td><td></td><td></td><td></td><td></td><td></td><td></td><td></td><td></td><td></td><td></td></tr>
<tr><td>Oligo rev</td><td>AAACGGTCTTGCCGAAGGCGCCCT</td><td></td><td></td><td></td><td></td><td></td><td></td><td></td><td></td><td></td><td></td></tr>
<tr><td>Chromosome</td><td>start</td><td>end</td><td>strand</td><td>MM</td><td>target_seq</td><td>PAM</td><td>alignment</td><td>distance</td><td>position</td><td>gene name</td><td>gene id</td></tr>
<tr><td>GU305914.1</td><td>153379</td><td>153401</td><td>+</td><td>0</td><td>AGGGCGCCTTCGGCAAGACC</td><td>CGG</td><td>||||||||[||||||||||||]PAM</td><td>NA</td><td>Intergenic</td><td>NA</td><td>NA</td></tr>
<tr><td></td><td></td><td></td><td></td><td></td><td></td><td></td><td></td><td></td><td></td><td></td><td></td></tr>
<tr><td>T11730</td><td>CCCGGCTCGTCACCTACAACAGG</td><td>1000</td><td></td><td></td><td></td><td></td><td></td><td></td><td></td><td></td><td></td></tr>
<tr><td>Oligo fwd</td><td>CACCCCCGGCTCGTCACCTACAAC</td><td></td><td></td><td></td><td></td><td></td><td></td><td></td><td></td><td></td><td></td></tr>
<tr><td>Oligo rev</td><td>AAACGTTGTAGGTGACGAGCCGGG</td><td></td><td></td><td></td><td></td><td></td><td></td><td></td><td></td><td></td><td></td></tr>
<tr><td>Chromosome</td><td>start</td><td>end</td><td>strand</td><td>MM</td><td>target_seq</td><td>PAM</td><td>alignment</td><td>distance</td><td>position</td><td>gene name</td><td>gene id</td></tr>
<tr><td>GU305914.1</td><td>153397</td><td>153419</td><td>+</td><td>0</td><td>CCCGGCTCGTCACCTACAAC</td><td>AGG</td><td>||||||||[||||||||||||]PAM</td><td>NA</td><td>Intergenic</td><td>NA</td><td>NA</td></tr>
<tr><td></td><td></td><td></td><td></td><td></td><td></td><td></td><td></td><td></td><td></td><td></td><td></td></tr>
<tr><td>T11731</td><td>GGCTCGTCACCTACAACAGGAGG</td><td>1000</td><td></td><td></td><td></td><td></td><td></td><td></td><td></td><td></td><td></td></tr>
<tr><td>Oligo fwd</td><td>CACCGGCTCGTCACCTACAACAGG</td><td></td><td></td><td></td><td></td><td></td><td></td><td></td><td></td><td></td><td></td></tr>
<tr><td>Oligo rev</td><td>AAACCCTGTTGTAGGTGACGAGCC</td><td></td><td></td><td></td><td></td><td></td><td></td><td></td><td></td><td></td><td></td></tr>
<tr><td>Chromosome</td><td>start</td><td>end</td><td>strand</td><td>MM</td><td>target_seq</td><td>PAM</td><td>alignment</td><td>distance</td><td>position</td><td>gene name</td><td>gene id</td></tr>
<tr><td>GU305914.1</td><td>153400</td><td>153422</td><td>+</td><td>0</td><td>GGCTCGTCACCTACAACAGG</td><td>AGG</td><td>||||||||[||||||||||||]PAM</td><td>NA</td><td>Intergenic</td><td>NA</td><td>NA</td></tr>
<tr><td></td><td></td><td></td><td></td><td></td><td></td><td></td><td></td><td></td><td></td><td></td><td></td></tr>
<tr><td>T11732</td><td>CCTACAACAGGAGGAACGTGTGG</td><td>1000</td><td></td><td></td><td></td><td></td><td></td><td></td><td></td><td></td><td></td></tr>
<tr><td>Oligo fwd</td><td>CACCCCTACAACAGGAGGAACGTG</td><td></td><td></td><td></td><td></td><td></td><td></td><td></td><td></td><td></td><td></td></tr>
<tr><td>Oligo rev</td><td>AAACCACGTTCCTCCTGTTGTAGG</td><td></td><td></td><td></td><td></td><td></td><td></td><td></td><td></td><td></td><td></td></tr>
<tr><td>Chromosome</td><td>start</td><td>end</td><td>strand</td><td>MM</td><td>target_seq</td><td>PAM</td><td>alignment</td><td>distance</td><td>position</td><td>gene name</td><td>gene id</td></tr>
<tr><td>GU305914.1</td><td>153409</td><td>153431</td><td>+</td><td>0</td><td>CCTACAACAGGAGGAACGTG</td><td>TGG</td><td>||||||||[||||||||||||]PAM</td><td>NA</td><td>Intergenic</td><td>NA</td><td>NA</td></tr>
<tr><td></td><td></td><td></td><td></td><td></td><td></td><td></td><td></td><td></td><td></td><td></td><td></td></tr>
<tr><td>T11733</td><td>ACAGGAGGAACGTGTGGCGCAGG</td><td>1000</td><td></td><td></td><td></td><td></td><td></td><td></td><td></td><td></td><td></td></tr>
<tr><td>Oligo fwd</td><td>CACCACAGGAGGAACGTGTGGCGC</td><td></td><td></td><td></td><td></td><td></td><td></td><td></td><td></td><td></td><td></td></tr>
<tr><td>Oligo rev</td><td>AAACGCGCCACACGTTCCTCCTGT</td><td></td><td></td><td></td><td></td><td></td><td></td><td></td><td></td><td></td><td></td></tr>
<tr><td>Chromosome</td><td>start</td><td>end</td><td>strand</td><td>MM</td><td>target_seq</td><td>PAM</td><td>alignment</td><td>distance</td><td>position</td><td>gene name</td><td>gene id</td></tr>
<tr><td>GU305914.1</td><td>153415</td><td>153437</td><td>+</td><td>0</td><td>ACAGGAGGAACGTGTGGCGC</td><td>AGG</td><td>||||||||[||||||||||||]PAM</td><td>NA</td><td>Intergenic</td><td>NA</td><td>NA</td></tr>
</table>

<table>
<tr><td>T11734</td><td>CTGCGAGATCGAGTCCCAGACGG</td><td></td><td>1000</td><td></td><td></td><td></td><td></td><td></td><td></td><td></td><td></td><td></td></tr>
<tr><td>Oligo fwd</td><td>CACCCTGCGAGATCGAGTCCCAGA</td><td></td><td></td><td></td><td></td><td></td><td></td><td></td><td></td><td></td><td></td><td></td></tr>
<tr><td>Oligo rev</td><td>AAACTCTGGGACTCGATCTCGCAG</td><td></td><td></td><td></td><td></td><td></td><td></td><td></td><td></td><td></td><td></td><td></td></tr>
<tr><td>Chromosome</td><td>start</td><td></td><td>end</td><td>strand</td><td>MM</td><td>target_seq</td><td>PAM</td><td>alignment</td><td>distance</td><td>position</td><td>gene name</td><td>gene id</td></tr>
<tr><td>GU305914.1</td><td></td><td>153443</td><td>153465</td><td>+</td><td>0</td><td>CTGCGAGATCGAGTCCCAGA</td><td>CGG</td><td>||||||||[||||||||||||]PAM</td><td>NA</td><td>Intergenic</td><td>NA</td><td>NA</td></tr>
<tr><td>T11735</td><td>TGCGAGATCGAGTCCCAGACGGG</td><td></td><td>1000</td><td></td><td></td><td></td><td></td><td></td><td></td><td></td><td></td><td></td></tr>
<tr><td>Oligo fwd</td><td>CACCTGCGAGATCGAGTCCCAGAC</td><td></td><td></td><td></td><td></td><td></td><td></td><td></td><td></td><td></td><td></td><td></td></tr>
<tr><td>Oligo rev</td><td>AAACGTCTGGGACTCGATCTCGCA</td><td></td><td></td><td></td><td></td><td></td><td></td><td></td><td></td><td></td><td></td><td></td></tr>
<tr><td>Chromosome</td><td>start</td><td></td><td>end</td><td>strand</td><td>MM</td><td>target_seq</td><td>PAM</td><td>alignment</td><td>distance</td><td>position</td><td>gene name</td><td>gene id</td></tr>
<tr><td>GU305914.1</td><td></td><td>153444</td><td>153466</td><td>+</td><td>0</td><td>TGCGAGATCGAGTCCCAGAC</td><td>GGG</td><td>||||||||[||||||||||||]PAM</td><td>NA</td><td>Intergenic</td><td>NA</td><td>NA</td></tr>
<tr><td>T11736</td><td>GTCCCAGACGGGATCCTTCGTGG</td><td></td><td>1000</td><td></td><td></td><td></td><td></td><td></td><td></td><td></td><td></td><td></td></tr>
<tr><td>Oligo fwd</td><td>CACCGTCCCAGACGGGATCCTTCG</td><td></td><td></td><td></td><td></td><td></td><td></td><td></td><td></td><td></td><td></td><td></td></tr>
<tr><td>Oligo rev</td><td>AAACCGAAGGATCCCGTCTGGGAC</td><td></td><td></td><td></td><td></td><td></td><td></td><td></td><td></td><td></td><td></td><td></td></tr>
<tr><td>Chromosome</td><td>start</td><td></td><td>end</td><td>strand</td><td>MM</td><td>target_seq</td><td>PAM</td><td>alignment</td><td>distance</td><td>position</td><td>gene name</td><td>gene id</td></tr>
<tr><td>GU305914.1</td><td></td><td>153455</td><td>153477</td><td>+</td><td>0</td><td>GTCCCAGACGGGATCCTTCG</td><td>TGG</td><td>||||||||[||||||||||||]PAM</td><td>NA</td><td>Intergenic</td><td>NA</td><td>NA</td></tr>
<tr><td>T11737</td><td>TCCCAGACGGGATCCTTCGTGGG</td><td></td><td>1000</td><td></td><td></td><td></td><td></td><td></td><td></td><td></td><td></td><td></td></tr>
<tr><td>Oligo fwd</td><td>CACCTCCCAGACGGGATCCTTCGT</td><td></td><td></td><td></td><td></td><td></td><td></td><td></td><td></td><td></td><td></td><td></td></tr>
<tr><td>Oligo rev</td><td>AAACACGAAGGATCCCGTCTGGGA</td><td></td><td></td><td></td><td></td><td></td><td></td><td></td><td></td><td></td><td></td><td></td></tr>
<tr><td>Chromosome</td><td>start</td><td></td><td>end</td><td>strand</td><td>MM</td><td>target_seq</td><td>PAM</td><td>alignment</td><td>distance</td><td>position</td><td>gene name</td><td>gene id</td></tr>
<tr><td>GU305914.1</td><td></td><td>153456</td><td>153478</td><td>+</td><td>0</td><td>TCCCAGACGGGATCCTTCGT</td><td>GGG</td><td>||||||||[||||||||||||]PAM</td><td>NA</td><td>Intergenic</td><td>NA</td><td>NA</td></tr>
<tr><td>T11738</td><td>CCCAGACGGGATCCTTCGTGGGG</td><td></td><td>1000</td><td></td><td></td><td></td><td></td><td></td><td></td><td></td><td></td><td></td></tr>
<tr><td>Oligo fwd</td><td>CACCCCCAGACGGGATCCTTCGTG</td><td></td><td></td><td></td><td></td><td></td><td></td><td></td><td></td><td></td><td></td><td></td></tr>
<tr><td>Oligo rev</td><td>AAACCACGAAGGATCCCGTCTGGG</td><td></td><td></td><td></td><td></td><td></td><td></td><td></td><td></td><td></td><td></td><td></td></tr>
<tr><td>Chromosome</td><td>start</td><td></td><td>end</td><td>strand</td><td>MM</td><td>target_seq</td><td>PAM</td><td>alignment</td><td>distance</td><td>position</td><td>gene name</td><td>gene id</td></tr>
<tr><td>GU305914.1</td><td></td><td>153457</td><td>153479</td><td>+</td><td>0</td><td>CCCAGACGGGATCCTTCGTG</td><td>GGG</td><td>||||||||[||||||||||||]PAM</td><td>NA</td><td>Intergenic</td><td>NA</td><td>NA</td></tr>
<tr><td>T11739</td><td>CCCCAGCAACAACTACGTGCTGG</td><td></td><td>1000</td><td></td><td></td><td></td><td></td><td></td><td></td><td></td><td></td><td></td></tr>
<tr><td>Oligo fwd</td><td>CACCCCCCAGCAACAACTACGTGC</td><td></td><td></td><td></td><td></td><td></td><td></td><td></td><td></td><td></td><td></td><td></td></tr>
<tr><td>Oligo rev</td><td>AAACGCACGTAGTTGTTGCTGGGG</td><td></td><td></td><td></td><td></td><td></td><td></td><td></td><td></td><td></td><td></td><td></td></tr>
<tr><td>Chromosome</td><td>start</td><td></td><td>end</td><td>strand</td><td>MM</td><td>target_seq</td><td>PAM</td><td>alignment</td><td>distance</td><td>position</td><td>gene name</td><td>gene id</td></tr>
<tr><td>GU305914.1</td><td></td><td>153497</td><td>153519</td><td>+</td><td>0</td><td>CCCCAGCAACAACTACGTGC</td><td>TGG</td><td>||||||||[||||||||||||]PAM</td><td>NA</td><td>Intergenic</td><td>NA</td><td>NA</td></tr>
<tr><td>T11740</td><td>CAGCAACAACTACGTGCTGGAGG</td><td></td><td>1000</td><td></td><td></td><td></td><td></td><td></td><td></td><td></td><td></td><td></td></tr>
<tr><td>Oligo fwd</td><td>CACCCAGCAACAACTACGTGCTGG</td><td></td><td></td><td></td><td></td><td></td><td></td><td></td><td></td><td></td><td></td><td></td></tr>
<tr><td>Oligo rev</td><td>AAACCCAGCACGTAGTTGTTGCTG</td><td></td><td></td><td></td><td></td><td></td><td></td><td></td><td></td><td></td><td></td><td></td></tr>
<tr><td>Chromosome</td><td>start</td><td></td><td>end</td><td>strand</td><td>MM</td><td>target_seq</td><td>PAM</td><td>alignment</td><td>distance</td><td>position</td><td>gene name</td><td>gene id</td></tr>
<tr><td>GU305914.1</td><td></td><td>153500</td><td>153522</td><td>+</td><td>0</td><td>CAGCAACAACTACGTGCTGG</td><td>AGG</td><td>||||||||[||||||||||||]PAM</td><td>NA</td><td>Intergenic</td><td>NA</td><td>NA</td></tr>
<tr><td>T11741</td><td>AGCAACAACTACGTGCTGGAGGG</td><td></td><td>1000</td><td></td><td></td><td></td><td></td><td></td><td></td><td></td><td></td><td></td></tr>
<tr><td>Oligo fwd</td><td>CACCAGCAACAACTACGTGCTGGA</td><td></td><td></td><td></td><td></td><td></td><td></td><td></td><td></td><td></td><td></td><td></td></tr>
<tr><td>Oligo rev</td><td>AAACTCCAGCACGTAGTTGTTGCT</td><td></td><td></td><td></td><td></td><td></td><td></td><td></td><td></td><td></td><td></td><td></td></tr>
<tr><td>Chromosome</td><td>start</td><td></td><td>end</td><td>strand</td><td>MM</td><td>target_seq</td><td>PAM</td><td>alignment</td><td>distance</td><td>position</td><td>gene name</td><td>gene id</td></tr>
<tr><td>GU305914.1</td><td></td><td>153501</td><td>153523</td><td>+</td><td>0</td><td>AGCAACAACTACGTGCTGGA</td><td>GGG</td><td>||||||||[||||||||||||]PAM</td><td>NA</td><td>Intergenic</td><td>NA</td><td>NA</td></tr>
<tr><td>T11742</td><td>CAACCACAACGTCTTCATCATGG</td><td></td><td>1000</td><td></td><td></td><td></td><td></td><td></td><td></td><td></td><td></td><td></td></tr>
<tr><td>Oligo fwd</td><td>CACCCAACCACAACGTCTTCATCA</td><td></td><td></td><td></td><td></td><td></td><td></td><td></td><td></td><td></td><td></td><td></td></tr>
<tr><td>Oligo rev</td><td>AAACTGATGAAGACGTTGTGGTTG</td><td></td><td></td><td></td><td></td><td></td><td></td><td></td><td></td><td></td><td></td><td></td></tr>
<tr><td>Chromosome</td><td>start</td><td></td><td>end</td><td>strand</td><td>MM</td><td>target_seq</td><td>PAM</td><td>alignment</td><td>distance</td><td>position</td><td>gene name</td><td>gene id</td></tr>
<tr><td>GU305914.1</td><td></td><td>153530</td><td>153552</td><td>+</td><td>0</td><td>CAACCACAACGTCTTCATCA</td><td>TGG</td><td>||||||||[||||||||||||]PAM</td><td>NA</td><td>Intergenic</td><td>NA</td><td>NA</td></tr>
<tr><td>T11743</td><td>TCTTCATCATGGACGCCGAACGG</td><td></td><td>1000</td><td></td><td></td><td></td><td></td><td></td><td></td><td></td><td></td><td></td></tr>
<tr><td>Oligo fwd</td><td>CACCTCTTCATCATGGACGCCGAA</td><td></td><td></td><td></td><td></td><td></td><td></td><td></td><td></td><td></td><td></td><td></td></tr>
<tr><td>Oligo rev</td><td>AAACTTCGGCGTCCATGATGAAGA</td><td></td><td></td><td></td><td></td><td></td><td></td><td></td><td></td><td></td><td></td><td></td></tr>
<tr><td>Chromosome</td><td>start</td><td></td><td>end</td><td>strand</td><td>MM</td><td>target_seq</td><td>PAM</td><td>alignment</td><td>distance</td><td>position</td><td>gene name</td><td>gene id</td></tr>
<tr><td>GU305914.1</td><td></td><td>153541</td><td>153563</td><td>+</td><td>0</td><td>TCTTCATCATGGACGCCGAA</td><td>CGG</td><td>||||||||[||||||||||||]PAM</td><td>NA</td><td>Intergenic</td><td>NA</td><td>NA</td></tr>
<tr><td>T11744</td><td>CCGAACGGAACCGTATACATCGG</td><td></td><td>1000</td><td></td><td></td><td></td><td></td><td></td><td></td><td></td><td></td><td></td></tr>
<tr><td>Oligo fwd</td><td>CACCCCGAACGGAACCGTATACAT</td><td></td><td></td><td></td><td></td><td></td><td></td><td></td><td></td><td></td><td></td><td></td></tr>
<tr><td>Oligo rev</td><td>AAACATGTATACGGTTCCGTTCGG</td><td></td><td></td><td></td><td></td><td></td><td></td><td></td><td></td><td></td><td></td><td></td></tr>
<tr><td>Chromosome</td><td>start</td><td></td><td>end</td><td>strand</td><td>MM</td><td>target_seq</td><td>PAM</td><td>alignment</td><td>distance</td><td>position</td><td>gene name</td><td>gene id</td></tr>
<tr><td>GU305914.1</td><td></td><td>153556</td><td>153578</td><td>+</td><td>0</td><td>CCGAACGGAACCGTATACAT</td><td>CGG</td><td>||||||||[||||||||||||]PAM</td><td>NA</td><td>Intergenic</td><td>NA</td><td>NA</td></tr>
<tr><td>T11745</td><td>AACGGAACCGTATACATCGGCGG</td><td></td><td>1000</td><td></td><td></td><td></td><td></td><td></td><td></td><td></td><td></td><td></td></tr>
<tr><td>Oligo fwd</td><td>CACCAACGGAACCGTATACATCGG</td><td></td><td></td><td></td><td></td><td></td><td></td><td></td><td></td><td></td><td></td><td></td></tr>
<tr><td>Oligo rev</td><td>AAACCCGATGTATACGGTTCCGTT</td><td></td><td></td><td></td><td></td><td></td><td></td><td></td><td></td><td></td><td></td><td></td></tr>
<tr><td>Chromosome</td><td>start</td><td></td><td>end</td><td>strand</td><td>MM</td><td>target_seq</td><td>PAM</td><td>alignment</td><td>distance</td><td>position</td><td>gene name</td><td>gene id</td></tr>
<tr><td>GU305914.1</td><td></td><td>153559</td><td>153581</td><td>+</td><td>0</td><td>AACGGAACCGTATACATCGG</td><td>CGG</td><td>||||||||[||||||||||||]PAM</td><td>NA</td><td>Intergenic</td><td>NA</td><td>NA</td></tr>
<tr><td>T11746</td><td>CCGTATACATCGGCGGATCCTGG</td><td></td><td>1000</td><td></td><td></td><td></td><td></td><td></td><td></td><td></td><td></td><td></td></tr>
<tr><td>Oligo fwd</td><td>CACCCCGTATACATCGGCGGATCC</td><td></td><td></td><td></td><td></td><td></td><td></td><td></td><td></td><td></td><td></td><td></td></tr>
<tr><td>Oligo rev</td><td>AAACGGATCCGCCGATGTATACGG</td><td></td><td></td><td></td><td></td><td></td><td></td><td></td><td></td><td></td><td></td><td></td></tr>
<tr><td>Chromosome</td><td>start</td><td></td><td>end</td><td>strand</td><td>MM</td><td>target_seq</td><td>PAM</td><td>alignment</td><td>distance</td><td>position</td><td>gene name</td><td>gene id</td></tr>
<tr><td>GU305914.1</td><td></td><td>153566</td><td>153588</td><td>+</td><td>0</td><td>CCGTATACATCGGCGGATCC</td><td>TGG</td><td>||||||||[||||||||||||]PAM</td><td>NA</td><td>Intergenic</td><td>NA</td><td>NA</td></tr>
<tr><td>T11747</td><td>ACATCGGCGGATCCTGGAGAAGG</td><td></td><td>1000</td><td></td><td></td><td></td><td></td><td></td><td></td><td></td><td></td><td></td></tr>
<tr><td>Oligo fwd</td><td>CACCACATCGGCGGATCCTGGAGA</td><td></td><td></td><td></td><td></td><td></td><td></td><td></td><td></td><td></td><td></td><td></td></tr>
<tr><td>Oligo rev</td><td>AAACTCTCCAGGATCCGCCGATGT</td><td></td><td></td><td></td><td></td><td></td><td></td><td></td><td></td><td></td><td></td><td></td></tr>
<tr><td>Chromosome</td><td>start</td><td></td><td>end</td><td>strand</td><td>MM</td><td>target_seq</td><td>PAM</td><td>alignment</td><td>distance</td><td>position</td><td>gene name</td><td>gene id</td></tr>
<tr><td>GU305914.1</td><td></td><td>153572</td><td>153594</td><td>+</td><td>0</td><td>ACATCGGCGGATCCTGGAGA</td><td>AGG</td><td>||||||||[||||||||||||]PAM</td><td>NA</td><td>Intergenic</td><td>NA</td><td>NA</td></tr>
<tr><td>T11748</td><td>CATCGGCGGATCCTGGAGAAGGG</td><td></td><td>1000</td><td></td><td></td><td></td><td></td><td></td><td></td><td></td><td></td><td></td></tr>
<tr><td>Oligo fwd</td><td>CACCCATCGGCGGATCCTGGAGAA</td><td></td><td></td><td></td><td></td><td></td><td></td><td></td><td></td><td></td><td></td><td></td></tr>
<tr><td>Oligo rev</td><td>AAACTTCTCCAGGATCCGCCGATG</td><td></td><td></td><td></td><td></td><td></td><td></td><td></td><td></td><td></td><td></td><td></td></tr>
<tr><td>Chromosome</td><td>start</td><td></td><td>end</td><td>strand</td><td>MM</td><td>target_seq</td><td>PAM</td><td>alignment</td><td>distance</td><td>position</td><td>gene name</td><td>gene id</td></tr>
<tr><td>GU305914.1</td><td></td><td>153573</td><td>153595</td><td>+</td><td>0</td><td>CATCGGCGGATCCTGGAGAA</td><td>GGG</td><td>||||||||[||||||||||||]PAM</td><td>NA</td><td>Intergenic</td><td>NA</td><td>NA</td></tr>
</table>

T11749 TCCTGGAGAAGGGCCTCCCCAGG 1000
Oligo fwd CACCTCCTGGAGAAGGGCCTCCCC
Oligo rev AAACGGGGAGGCCCTTCTCCAGGA
Chromosome start end strand MM target_seq PAM alignment distance position gene name gene id
GU305914.1 153583 153605 + 0 TCCTGGAGAAGGGCCTCCCC AGG ||||||||[||||||||||||]PAM NA Intergenic NA NA

T11750 GCCTCCCCAGGCTGATCGTCAGG 1000
Oligo fwd CACCGCCTCCCCAGGCTGATCGTC
Oligo rev AAACGACGATCAGCCTGGGGAGGC
Chromosome start end strand MM target_seq PAM alignment distance position gene name gene id
GU305914.1 153595 153617 + 0 GCCTCCCCAGGCTGATCGTC AGG ||||||||[||||||||||||]PAM NA Intergenic NA NA

T11751 CCTCCCCAGGCTGATCGTCAGGG 1000
Oligo fwd CACCCCTCCCCAGGCTGATCGTCA
Oligo rev AAACTGACGATCAGCCTGGGGAGG
Chromosome start end strand MM target_seq PAM alignment distance position gene name gene id
GU305914.1 153596 153618 + 0 CCTCCCCAGGCTGATCGTCA GGG ||||||||[||||||||||||]PAM NA Intergenic NA NA

T11752 CTGATCGTCAGGGACGCCTGCGG 1000
Oligo fwd CACCCTGATCGTCAGGGACGCCTG
Oligo rev AAACCAGGCGTCCCTGACGATCAG
Chromosome start end strand MM target_seq PAM alignment distance position gene name gene id
GU305914.1 153606 153628 + 0 CTGATCGTCAGGGACGCCTG CGG ||||||||[||||||||||||]PAM NA Intergenic NA NA

T11753 CTGCGGCTTCCTGCTCATCCTGG 1000
Oligo fwd CACCCTGCGGCTTCCTGCTCATCC
Oligo rev AAACGGATGAGCAGGAAGCCGCAG
Chromosome start end strand MM target_seq PAM alignment distance position gene name gene id
GU305914.1 153623 153645 + 0 CTGCGGCTTCCTGCTCATCC TGG ||||||||[||||||||||||]PAM NA Intergenic NA NA

T11754 AAGTTCTCCGACGTGATAGACGG 1000
Oligo fwd CACCAAGTTCTCCGACGTGATAGA
Oligo rev AAACTCTATCACGTCGGAGAACTT
Chromosome start end strand MM target_seq PAM alignment distance position gene name gene id
GU305914.1 153660 153682 + 0 AAGTTCTCCGACGTGATAGA CGG ||||||||[||||||||||||]PAM NA Intergenic NA NA

T11755 GAGCGTGCACATCTGCACCATGG 1000
Oligo fwd CACCGAGCGTGCACATCTGCACCA
Oligo rev AAACTGGTGCAGATGTGCACGCTC
Chromosome start end strand MM target_seq PAM alignment distance position gene name gene id
GU305914.1 153686 153708 + 0 GAGCGTGCACATCTGCACCA TGG ||||||||[||||||||||||]PAM NA Intergenic NA NA

T11756 CGTGCACATCTGCACCATGGTGG 1000
Oligo fwd CACCCGTGCACATCTGCACCATGG
Oligo rev AAACCCATGGTGCAGATGTGCACG
Chromosome start end strand MM target_seq PAM alignment distance position gene name gene id
GU305914.1 153689 153711 + 0 CGTGCACATCTGCACCATGG TGG ||||||||[||||||||||||]PAM NA Intergenic NA NA

T11757 ATCTGCACCATGGTGGATTACGG 1000
Oligo fwd CACCATCTGCACCATGGTGGATTA
Oligo rev AAACTAATCCACCATGGTGCAGAT
Chromosome start end strand MM target_seq PAM alignment distance position gene name gene id
GU305914.1 153696 153718 + 0 ATCTGCACCATGGTGGATTA CGG ||||||||[||||||||||||]PAM NA Intergenic NA NA

T11758 TCTGCACCATGGTGGATTACGGG 1000
Oligo fwd CACCTCTGCACCATGGTGGATTAC
Oligo rev AAACGTAATCCACCATGGTGCAGA
Chromosome start end strand MM target_seq PAM alignment distance position gene name gene id
GU305914.1 153697 153719 + 0 TCTGCACCATGGTGGATTAC GGG ||||||||[||||||||||||]PAM NA Intergenic NA NA

T11759 CTGCACCATGGTGGATTACGGGG 1000
Oligo fwd CACCCTGCACCATGGTGGATTACG
Oligo rev AAACCGTAATCCACCATGGTGCAG
Chromosome start end strand MM target_seq PAM alignment distance position gene name gene id
GU305914.1 153698 153720 + 0 CTGCACCATGGTGGATTACG GGG ||||||||[||||||||||||]PAM NA Intergenic NA NA

T11760 TTACGGGGTGACCTCGCGCATGG 1000
Oligo fwd CACCTTACGGGGTGACCTCGCGCA
Oligo rev AAACTGCGCGAGGTCACCCCGTAA
Chromosome start end strand MM target_seq PAM alignment distance position gene name gene id
GU305914.1 153713 153735 + 0 TTACGGGGTGACCTCGCGCA TGG ||||||||[||||||||||||]PAM NA Intergenic NA NA

T11761 CATGACCATCGCCAAGAGTCAGG 1000
Oligo fwd CACCCATGACCATCGCCAAGAGTC
Oligo rev AAACGACTCTTGGCGATGGTCATG
Chromosome start end strand MM target_seq PAM alignment distance position gene name gene id
GU305914.1 153737 153759 + 0 CATGACCATCGCCAAGAGTC AGG ||||||||[||||||||||||]PAM NA Intergenic NA NA

T11762 ATGACCATCGCCAAGAGTCAGGG 1000
Oligo fwd CACCATGACCATCGCCAAGAGTCA
Oligo rev AAACTGACTCTTGGCGATGGTCAT
Chromosome start end strand MM target_seq PAM alignment distance position gene name gene id
GU305914.1 153738 153760 + 0 ATGACCATCGCCAAGAGTCA GGG ||||||||[||||||||||||]PAM NA Intergenic NA NA

T11763 ATCGCCAAGAGTCAGGGCATCGG 1000
Oligo fwd CACCATCGCCAAGAGTCAGGGCAT
Oligo rev AAACATGCCCTGACTCTTGGCGAT
Chromosome start end strand MM target_seq PAM alignment distance position gene name gene id

<table>
<tr><td>GU305914.1</td><td></td><td>153744</td><td>153766</td><td>+</td><td>0</td><td>ATCGCCAAGAGTCAGGGCAT</td><td>CGG</td><td>||||||||[||||||||||||]PAM</td><td>NA</td><td>Intergenic</td><td>NA</td><td>NA</td></tr>
<tr><td>T11764</td><td>CAAGAGTCAGGGCATCGGTCTGG</td><td></td><td>1000</td><td></td><td></td><td></td><td></td><td></td><td></td><td></td><td></td><td></td></tr>
<tr><td>Oligo fwd</td><td>CACCCAAGAGTCAGGGCATCGGTC</td><td></td><td></td><td></td><td></td><td></td><td></td><td></td><td></td><td></td><td></td><td></td></tr>
<tr><td>Oligo rev</td><td>AAACGACCGATGCCCTGACTCTTG</td><td></td><td></td><td></td><td></td><td></td><td></td><td></td><td></td><td></td><td></td><td></td></tr>
<tr><td>Chromosome</td><td>start</td><td></td><td>end</td><td>strand</td><td>MM</td><td>target_seq</td><td>PAM</td><td>alignment</td><td>distance</td><td>position</td><td>gene name</td><td>gene id</td></tr>
<tr><td>GU305914.1</td><td></td><td>153749</td><td>153771</td><td>+</td><td>0</td><td>CAAGAGTCAGGGCATCGGTC</td><td>TGG</td><td>||||||||[||||||||||||]PAM</td><td>NA</td><td>Intergenic</td><td>NA</td><td>NA</td></tr>
<tr><td>T11765</td><td>GGGCATCGGTCTGGAGAGCGTGG</td><td></td><td>1000</td><td></td><td></td><td></td><td></td><td></td><td></td><td></td><td></td><td></td></tr>
<tr><td>Oligo fwd</td><td>CACCGGGCATCGGTCTGGAGAGCG</td><td></td><td></td><td></td><td></td><td></td><td></td><td></td><td></td><td></td><td></td><td></td></tr>
<tr><td>Oligo rev</td><td>AAACCGCTCTCCAGACCGATGCCC</td><td></td><td></td><td></td><td></td><td></td><td></td><td></td><td></td><td></td><td></td><td></td></tr>
<tr><td>Chromosome</td><td>start</td><td></td><td>end</td><td>strand</td><td>MM</td><td>target_seq</td><td>PAM</td><td>alignment</td><td>distance</td><td>position</td><td>gene name</td><td>gene id</td></tr>
<tr><td>GU305914.1</td><td></td><td>153758</td><td>153780</td><td>+</td><td>0</td><td>GGGCATCGGTCTGGAGAGCG</td><td>TGG</td><td>||||||||[||||||||||||]PAM</td><td>NA</td><td>Intergenic</td><td>NA</td><td>NA</td></tr>
<tr><td>T11766</td><td>GAGAGCGTGGCCATCGATTTCGG</td><td></td><td>1000</td><td></td><td></td><td></td><td></td><td></td><td></td><td></td><td></td><td></td></tr>
<tr><td>Oligo fwd</td><td>CACCGAGAGCGTGGCCATCGATTT</td><td></td><td></td><td></td><td></td><td></td><td></td><td></td><td></td><td></td><td></td><td></td></tr>
<tr><td>Oligo rev</td><td>AAACAAATCGATGGCCACGCTCTC</td><td></td><td></td><td></td><td></td><td></td><td></td><td></td><td></td><td></td><td></td><td></td></tr>
<tr><td>Chromosome</td><td>start</td><td></td><td>end</td><td>strand</td><td>MM</td><td>target_seq</td><td>PAM</td><td>alignment</td><td>distance</td><td>position</td><td>gene name</td><td>gene id</td></tr>
<tr><td>GU305914.1</td><td></td><td>153771</td><td>153793</td><td>+</td><td>0</td><td>GAGAGCGTGGCCATCGATTT</td><td>CGG</td><td>||||||||[||||||||||||]PAM</td><td>NA</td><td>Intergenic</td><td>NA</td><td>NA</td></tr>
<tr><td>T11767</td><td>AAGATGAGCCAGATCTACGTAGG</td><td></td><td>1000</td><td></td><td></td><td></td><td></td><td></td><td></td><td></td><td></td><td></td></tr>
<tr><td>Oligo fwd</td><td>CACCAAGATGAGCCAGATCTACGT</td><td></td><td></td><td></td><td></td><td></td><td></td><td></td><td></td><td></td><td></td><td></td></tr>
<tr><td>Oligo rev</td><td>AAACACGTAGATCTGGCTCATCTT</td><td></td><td></td><td></td><td></td><td></td><td></td><td></td><td></td><td></td><td></td><td></td></tr>
<tr><td>Chromosome</td><td>start</td><td></td><td>end</td><td>strand</td><td>MM</td><td>target_seq</td><td>PAM</td><td>alignment</td><td>distance</td><td>position</td><td>gene name</td><td>gene id</td></tr>
<tr><td>GU305914.1</td><td></td><td>153813</td><td>153835</td><td>+</td><td>0</td><td>AAGATGAGCCAGATCTACGT</td><td>AGG</td><td>||||||||[||||||||||||]PAM</td><td>NA</td><td>Intergenic</td><td>NA</td><td>NA</td></tr>
<tr><td>T11768</td><td>AGATCTACGTAGGTATCTCCCGG</td><td></td><td>1000</td><td></td><td></td><td></td><td></td><td></td><td></td><td></td><td></td><td></td></tr>
<tr><td>Oligo fwd</td><td>CACCAGATCTACGTAGGTATCTCC</td><td></td><td></td><td></td><td></td><td></td><td></td><td></td><td></td><td></td><td></td><td></td></tr>
<tr><td>Oligo rev</td><td>AAACGGAGATACCTACGTAGATCT</td><td></td><td></td><td></td><td></td><td></td><td></td><td></td><td></td><td></td><td></td><td></td></tr>
<tr><td>Chromosome</td><td>start</td><td></td><td>end</td><td>strand</td><td>MM</td><td>target_seq</td><td>PAM</td><td>alignment</td><td>distance</td><td>position</td><td>gene name</td><td>gene id</td></tr>
<tr><td>GU305914.1</td><td></td><td>153823</td><td>153845</td><td>+</td><td>0</td><td>AGATCTACGTAGGTATCTCC</td><td>CGG</td><td>||||||||[||||||||||||]PAM</td><td>NA</td><td>Intergenic</td><td>NA</td><td>NA</td></tr>
<tr><td>T11769</td><td>GATCTACGTAGGTATCTCCCGGG</td><td></td><td>1000</td><td></td><td></td><td></td><td></td><td></td><td></td><td></td><td></td><td></td></tr>
<tr><td>Oligo fwd</td><td>CACCGATCTACGTAGGTATCTCCC</td><td></td><td></td><td></td><td></td><td></td><td></td><td></td><td></td><td></td><td></td><td></td></tr>
<tr><td>Oligo rev</td><td>AAACGGGAGATACCTACGTAGATC</td><td></td><td></td><td></td><td></td><td></td><td></td><td></td><td></td><td></td><td></td><td></td></tr>
<tr><td>Chromosome</td><td>start</td><td></td><td>end</td><td>strand</td><td>MM</td><td>target_seq</td><td>PAM</td><td>alignment</td><td>distance</td><td>position</td><td>gene name</td><td>gene id</td></tr>
<tr><td>GU305914.1</td><td></td><td>153824</td><td>153846</td><td>+</td><td>0</td><td>GATCTACGTAGGTATCTCCC</td><td>GGG</td><td>||||||||[||||||||||||]PAM</td><td>NA</td><td>Intergenic</td><td>NA</td><td>NA</td></tr>
<tr><td>T11770</td><td>CTACGTAGGTATCTCCCGGGTGG</td><td></td><td>1000</td><td></td><td></td><td></td><td></td><td></td><td></td><td></td><td></td><td></td></tr>
<tr><td>Oligo fwd</td><td>CACCCTACGTAGGTATCTCCCGGG</td><td></td><td></td><td></td><td></td><td></td><td></td><td></td><td></td><td></td><td></td><td></td></tr>
<tr><td>Oligo rev</td><td>AAACCCCGGGAGATACCTACGTAG</td><td></td><td></td><td></td><td></td><td></td><td></td><td></td><td></td><td></td><td></td><td></td></tr>
<tr><td>Chromosome</td><td>start</td><td></td><td>end</td><td>strand</td><td>MM</td><td>target_seq</td><td>PAM</td><td>alignment</td><td>distance</td><td>position</td><td>gene name</td><td>gene id</td></tr>
<tr><td>GU305914.1</td><td></td><td>153827</td><td>153849</td><td>+</td><td>0</td><td>CTACGTAGGTATCTCCCGGG</td><td>TGG</td><td>||||||||[||||||||||||]PAM</td><td>NA</td><td>Intergenic</td><td>NA</td><td>NA</td></tr>
<tr><td>T11771</td><td>CGTAGGTATCTCCCGGGTGGTGG</td><td></td><td>1000</td><td></td><td></td><td></td><td></td><td></td><td></td><td></td><td></td><td></td></tr>
<tr><td>Oligo fwd</td><td>CACCCGTAGGTATCTCCCGGGTGG</td><td></td><td></td><td></td><td></td><td></td><td></td><td></td><td></td><td></td><td></td><td></td></tr>
<tr><td>Oligo rev</td><td>AAACCCACCCGGGAGATACCTACG</td><td></td><td></td><td></td><td></td><td></td><td></td><td></td><td></td><td></td><td></td><td></td></tr>
<tr><td>Chromosome</td><td>start</td><td></td><td>end</td><td>strand</td><td>MM</td><td>target_seq</td><td>PAM</td><td>alignment</td><td>distance</td><td>position</td><td>gene name</td><td>gene id</td></tr>
<tr><td>GU305914.1</td><td></td><td>153830</td><td>153852</td><td>+</td><td>0</td><td>CGTAGGTATCTCCCGGGTGG</td><td>TGG</td><td>||||||||[||||||||||||]PAM</td><td>NA</td><td>Intergenic</td><td>NA</td><td>NA</td></tr>
<tr><td>T11772</td><td>ACAGAATAAAGATACCACTCTGG</td><td></td><td>1000</td><td></td><td></td><td></td><td></td><td></td><td></td><td></td><td></td><td></td></tr>
<tr><td>Oligo fwd</td><td>CACCACAGAATAAAGATACCACTC</td><td></td><td></td><td></td><td></td><td></td><td></td><td></td><td></td><td></td><td></td><td></td></tr>
<tr><td>Oligo rev</td><td>AAACGAGTGGTATCTTTATTCTGT</td><td></td><td></td><td></td><td></td><td></td><td></td><td></td><td></td><td></td><td></td><td></td></tr>
<tr><td>Chromosome</td><td>start</td><td></td><td>end</td><td>strand</td><td>MM</td><td>target_seq</td><td>PAM</td><td>alignment</td><td>distance</td><td>position</td><td>gene name</td><td>gene id</td></tr>
<tr><td>GU305914.1</td><td></td><td>153941</td><td>153963</td><td>+</td><td>0</td><td>ACAGAATAAAGATACCACTC</td><td>TGG</td><td>||||||||[||||||||||||]PAM</td><td>NA</td><td>Intergenic</td><td>NA</td><td>NA</td></tr>
<tr><td>T11773</td><td>TACCACCTCGACCACGATTCTGG</td><td></td><td>1000</td><td></td><td></td><td></td><td></td><td></td><td></td><td></td><td></td><td></td></tr>
<tr><td>Oligo fwd</td><td>CACCTACCACCTCGACCACGATTC</td><td></td><td></td><td></td><td></td><td></td><td></td><td></td><td></td><td></td><td></td><td></td></tr>
<tr><td>Oligo rev</td><td>AAACGAATCGTGGTCGAGGTGGTA</td><td></td><td></td><td></td><td></td><td></td><td></td><td></td><td></td><td></td><td></td><td></td></tr>
<tr><td>Chromosome</td><td>start</td><td></td><td>end</td><td>strand</td><td>MM</td><td>target_seq</td><td>PAM</td><td>alignment</td><td>distance</td><td>position</td><td>gene name</td><td>gene id</td></tr>
<tr><td>GU305914.1</td><td></td><td>154016</td><td>154038</td><td>+</td><td>0</td><td>TACCACCTCGACCACGATTC</td><td>TGG</td><td>||||||||[||||||||||||]PAM</td><td>NA</td><td>Intergenic</td><td>NA</td><td>NA</td></tr>
<tr><td>GU305914.1</td><td></td><td>214143</td><td>214165</td><td>-</td><td>4</td><td>TACGGTCTCGACCACGATGC</td><td>GGG</td><td>|||---||[||||||||||-|]PAM</td><td>NA</td><td>Intergenic</td><td>NA</td><td>NA</td></tr>
<tr><td>T11774</td><td>ACCACCTCGACCACGATTCTGGG</td><td></td><td>1000</td><td></td><td></td><td></td><td></td><td></td><td></td><td></td><td></td><td></td></tr>
<tr><td>Oligo fwd</td><td>CACCACCACCTCGACCACGATTCT</td><td></td><td></td><td></td><td></td><td></td><td></td><td></td><td></td><td></td><td></td><td></td></tr>
<tr><td>Oligo rev</td><td>AAACAGAATCGTGGTCGAGGTGGT</td><td></td><td></td><td></td><td></td><td></td><td></td><td></td><td></td><td></td><td></td><td></td></tr>
<tr><td>Chromosome</td><td>start</td><td></td><td>end</td><td>strand</td><td>MM</td><td>target_seq</td><td>PAM</td><td>alignment</td><td>distance</td><td>position</td><td>gene name</td><td>gene id</td></tr>
<tr><td>GU305914.1</td><td></td><td>154017</td><td>154039</td><td>+</td><td>0</td><td>ACCACCTCGACCACGATTCT</td><td>GGG</td><td>||||||||[||||||||||||]PAM</td><td>NA</td><td>Intergenic</td><td>NA</td><td>NA</td></tr>
<tr><td>T11775</td><td>GACCACGATTCTGGGTATCTCGG</td><td></td><td>1000</td><td></td><td></td><td></td><td></td><td></td><td></td><td></td><td></td><td></td></tr>
<tr><td>Oligo fwd</td><td>CACCGACCACGATTCTGGGTATCT</td><td></td><td></td><td></td><td></td><td></td><td></td><td></td><td></td><td></td><td></td><td></td></tr>
<tr><td>Oligo rev</td><td>AAACAGATACCCAGAATCGTGGTC</td><td></td><td></td><td></td><td></td><td></td><td></td><td></td><td></td><td></td><td></td><td></td></tr>
<tr><td>Chromosome</td><td>start</td><td></td><td>end</td><td>strand</td><td>MM</td><td>target_seq</td><td>PAM</td><td>alignment</td><td>distance</td><td>position</td><td>gene name</td><td>gene id</td></tr>
<tr><td>GU305914.1</td><td></td><td>154025</td><td>154047</td><td>+</td><td>0</td><td>GACCACGATTCTGGGTATCT</td><td>CGG</td><td>||||||||[||||||||||||]PAM</td><td>NA</td><td>Intergenic</td><td>NA</td><td>NA</td></tr>
<tr><td>T11776</td><td>ACCACGATTCTGGGTATCTCGGG</td><td></td><td>1000</td><td></td><td></td><td></td><td></td><td></td><td></td><td></td><td></td><td></td></tr>
<tr><td>Oligo fwd</td><td>CACCACCACGATTCTGGGTATCTC</td><td></td><td></td><td></td><td></td><td></td><td></td><td></td><td></td><td></td><td></td><td></td></tr>
<tr><td>Oligo rev</td><td>AAACGAGATACCCAGAATCGTGGT</td><td></td><td></td><td></td><td></td><td></td><td></td><td></td><td></td><td></td><td></td><td></td></tr>
<tr><td>Chromosome</td><td>start</td><td></td><td>end</td><td>strand</td><td>MM</td><td>target_seq</td><td>PAM</td><td>alignment</td><td>distance</td><td>position</td><td>gene name</td><td>gene id</td></tr>
<tr><td>GU305914.1</td><td></td><td>154026</td><td>154048</td><td>+</td><td>0</td><td>ACCACGATTCTGGGTATCTC</td><td>GGG</td><td>||||||||[||||||||||||]PAM</td><td>NA</td><td>Intergenic</td><td>NA</td><td>NA</td></tr>
<tr><td>T11777</td><td>CCACGATTCTGGGTATCTCGGGG</td><td></td><td>1000</td><td></td><td></td><td></td><td></td><td></td><td></td><td></td><td></td><td></td></tr>
<tr><td>Oligo fwd</td><td>CACCCCACGATTCTGGGTATCTCG</td><td></td><td></td><td></td><td></td><td></td><td></td><td></td><td></td><td></td><td></td><td></td></tr>
<tr><td>Oligo rev</td><td>AAACCGAGATACCCAGAATCGTGG</td><td></td><td></td><td></td><td></td><td></td><td></td><td></td><td></td><td></td><td></td><td></td></tr>
<tr><td>Chromosome</td><td>start</td><td></td><td>end</td><td>strand</td><td>MM</td><td>target_seq</td><td>PAM</td><td>alignment</td><td>distance</td><td>position</td><td>gene name</td><td>gene id</td></tr>
<tr><td>GU305914.1</td><td></td><td>154027</td><td>154049</td><td>+</td><td>0</td><td>CCACGATTCTGGGTATCTCG</td><td>GGG</td><td>||||||||[||||||||||||]PAM</td><td>NA</td><td>Intergenic</td><td>NA</td><td>NA</td></tr>
<tr><td>T11778</td><td>GGGTATCTCGGGGATCGTGTTGG</td><td></td><td>1000</td><td></td><td></td><td></td><td></td><td></td><td></td><td></td><td></td><td></td></tr>
<tr><td>Oligo fwd</td><td>CACCGGGTATCTCGGGGATCGTGT</td><td></td><td></td><td></td><td></td><td></td><td></td><td></td><td></td><td></td><td></td><td></td></tr>
</table>

<table>
<tr><td>Oligo rev</td><td>AAACACACGATCCCCGAGATACCC</td><td></td><td></td><td></td><td></td><td></td><td></td><td></td><td></td><td></td><td></td></tr>
<tr><td>Chromosome</td><td>start</td><td>end</td><td>strand</td><td>MM</td><td>target_seq</td><td>PAM</td><td>alignment</td><td>distance</td><td>position</td><td>gene name</td><td>gene id</td></tr>
<tr><td>GU305914.1</td><td>154037</td><td>154059</td><td>+</td><td>0</td><td>GGGTATCTCGGGGATCGTGT</td><td>TGG</td><td>||||||||[||||||||||||]PAM</td><td>NA</td><td>Intergenic</td><td>NA</td><td>NA</td></tr>
<tr><td>T11779</td><td>TATCTCGGGGATCGTGTTGGTGG</td><td>1000</td><td></td><td></td><td></td><td></td><td></td><td></td><td></td><td></td><td></td></tr>
<tr><td>Oligo fwd</td><td>CACCTATCTCGGGGATCGTGTTGG</td><td></td><td></td><td></td><td></td><td></td><td></td><td></td><td></td><td></td><td></td></tr>
<tr><td>Oligo rev</td><td>AAACCCAACACGATCCCCGAGATA</td><td></td><td></td><td></td><td></td><td></td><td></td><td></td><td></td><td></td><td></td></tr>
<tr><td>Chromosome</td><td>start</td><td>end</td><td>strand</td><td>MM</td><td>target_seq</td><td>PAM</td><td>alignment</td><td>distance</td><td>position</td><td>gene name</td><td>gene id</td></tr>
<tr><td>GU305914.1</td><td>154040</td><td>154062</td><td>+</td><td>0</td><td>TATCTCGGGGATCGTGTTGG</td><td>TGG</td><td>||||||||[||||||||||||]PAM</td><td>NA</td><td>Intergenic</td><td>NA</td><td>NA</td></tr>
<tr><td>T11780</td><td>GGATCGTGTTGGTGGATTGTTGG</td><td>1000</td><td></td><td></td><td></td><td></td><td></td><td></td><td></td><td></td><td></td></tr>
<tr><td>Oligo fwd</td><td>CACCGGATCGTGTTGGTGGATTGT</td><td></td><td></td><td></td><td></td><td></td><td></td><td></td><td></td><td></td><td></td></tr>
<tr><td>Oligo rev</td><td>AAACACAATCCACCAACACGATCC</td><td></td><td></td><td></td><td></td><td></td><td></td><td></td><td></td><td></td><td></td></tr>
<tr><td>Chromosome</td><td>start</td><td>end</td><td>strand</td><td>MM</td><td>target_seq</td><td>PAM</td><td>alignment</td><td>distance</td><td>position</td><td>gene name</td><td>gene id</td></tr>
<tr><td>GU305914.1</td><td>154048</td><td>154070</td><td>+</td><td>0</td><td>GGATCGTGTTGGTGGATTGT</td><td>TGG</td><td>||||||||[||||||||||||]PAM</td><td>NA</td><td>Intergenic</td><td>NA</td><td>NA</td></tr>
<tr><td>T11781</td><td>GTTGGTGGATTGTTGGTCATTGG</td><td>1000</td><td></td><td></td><td></td><td></td><td></td><td></td><td></td><td></td><td></td></tr>
<tr><td>Oligo fwd</td><td>CACCGTTGGTGGATTGTTGGTCAT</td><td></td><td></td><td></td><td></td><td></td><td></td><td></td><td></td><td></td><td></td></tr>
<tr><td>Oligo rev</td><td>AAACATGACCAACAATCCACCAAC</td><td></td><td></td><td></td><td></td><td></td><td></td><td></td><td></td><td></td><td></td></tr>
<tr><td>Chromosome</td><td>start</td><td>end</td><td>strand</td><td>MM</td><td>target_seq</td><td>PAM</td><td>alignment</td><td>distance</td><td>position</td><td>gene name</td><td>gene id</td></tr>
<tr><td>GU305914.1</td><td>154055</td><td>154077</td><td>+</td><td>0</td><td>GTTGGTGGATTGTTGGTCAT</td><td>TGG</td><td>||||||||[||||||||||||]PAM</td><td>NA</td><td>Intergenic</td><td>NA</td><td>NA</td></tr>
<tr><td>T11782</td><td>TGTTGGTCATTGGACTGTAACGG</td><td>1000</td><td></td><td></td><td></td><td></td><td></td><td></td><td></td><td></td><td></td></tr>
<tr><td>Oligo fwd</td><td>CACCTGTTGGTCATTGGACTGTAA</td><td></td><td></td><td></td><td></td><td></td><td></td><td></td><td></td><td></td><td></td></tr>
<tr><td>Oligo rev</td><td>AAACTTACAGTCCAATGACCAACA</td><td></td><td></td><td></td><td></td><td></td><td></td><td></td><td></td><td></td><td></td></tr>
<tr><td>Chromosome</td><td>start</td><td>end</td><td>strand</td><td>MM</td><td>target_seq</td><td>PAM</td><td>alignment</td><td>distance</td><td>position</td><td>gene name</td><td>gene id</td></tr>
<tr><td>GU305914.1</td><td>154065</td><td>154087</td><td>+</td><td>0</td><td>TGTTGGTCATTGGACTGTAA</td><td>CGG</td><td>||||||||[||||||||||||]PAM</td><td>NA</td><td>Intergenic</td><td>NA</td><td>NA</td></tr>
<tr><td>T11783</td><td>GACGCTGATCGCGACGACGACGG</td><td>1000</td><td></td><td></td><td></td><td></td><td></td><td></td><td></td><td></td><td></td></tr>
<tr><td>Oligo fwd</td><td>CACCGACGCTGATCGCGACGACGA</td><td></td><td></td><td></td><td></td><td></td><td></td><td></td><td></td><td></td><td></td></tr>
<tr><td>Oligo rev</td><td>AAACTCGTCGTCGCGATCAGCGTC</td><td></td><td></td><td></td><td></td><td></td><td></td><td></td><td></td><td></td><td></td></tr>
<tr><td>Chromosome</td><td>start</td><td>end</td><td>strand</td><td>MM</td><td>target_seq</td><td>PAM</td><td>alignment</td><td>distance</td><td>position</td><td>gene name</td><td>gene id</td></tr>
<tr><td>GU305914.1</td><td>154114</td><td>154136</td><td>+</td><td>0</td><td>GACGCTGATCGCGACGACGA</td><td>CGG</td><td>||||||||[||||||||||||]PAM</td><td>NA</td><td>Intergenic</td><td>NA</td><td>NA</td></tr>
<tr><td>T11784</td><td>GCTGATCGCGACGACGACGGTGG</td><td>1000</td><td></td><td></td><td></td><td></td><td></td><td></td><td></td><td></td><td></td></tr>
<tr><td>Oligo fwd</td><td>CACCGCTGATCGCGACGACGACGG</td><td></td><td></td><td></td><td></td><td></td><td></td><td></td><td></td><td></td><td></td></tr>
<tr><td>Oligo rev</td><td>AAACCCGTCGTCGTCGCGATCAGC</td><td></td><td></td><td></td><td></td><td></td><td></td><td></td><td></td><td></td><td></td></tr>
<tr><td>Chromosome</td><td>start</td><td>end</td><td>strand</td><td>MM</td><td>target_seq</td><td>PAM</td><td>alignment</td><td>distance</td><td>position</td><td>gene name</td><td>gene id</td></tr>
<tr><td>GU305914.1</td><td>154117</td><td>154139</td><td>+</td><td>0</td><td>GCTGATCGCGACGACGACGG</td><td>TGG</td><td>||||||||[||||||||||||]PAM</td><td>NA</td><td>Intergenic</td><td>NA</td><td>NA</td></tr>
<tr><td>GU305914.1</td><td>58647</td><td>58669</td><td>+</td><td>4</td><td>GATGAGCTCGCCGACGACGG</td><td>CGG</td><td>|-|||-|-[||-|||||||||]PAM</td><td>NA</td><td>Intergenic</td><td>NA</td><td>NA</td></tr>
<tr><td>T11785</td><td>GACGACGACGGTGGTGTTCGTGG</td><td>1000</td><td></td><td></td><td></td><td></td><td></td><td></td><td></td><td></td><td></td></tr>
<tr><td>Oligo fwd</td><td>CACCGACGACGACGGTGGTGTTCG</td><td></td><td></td><td></td><td></td><td></td><td></td><td></td><td></td><td></td><td></td></tr>
<tr><td>Oligo rev</td><td>AAACCGAACACCACCGTCGTCGTC</td><td></td><td></td><td></td><td></td><td></td><td></td><td></td><td></td><td></td><td></td></tr>
<tr><td>Chromosome</td><td>start</td><td>end</td><td>strand</td><td>MM</td><td>target_seq</td><td>PAM</td><td>alignment</td><td>distance</td><td>position</td><td>gene name</td><td>gene id</td></tr>
<tr><td>GU305914.1</td><td>154126</td><td>154148</td><td>+</td><td>0</td><td>GACGACGACGGTGGTGTTCG</td><td>TGG</td><td>||||||||[||||||||||||]PAM</td><td>NA</td><td>Intergenic</td><td>NA</td><td>NA</td></tr>
<tr><td>T11786</td><td>ACGACGACGGTGGTGTTCGTGGG</td><td>1000</td><td></td><td></td><td></td><td></td><td></td><td></td><td></td><td></td><td></td></tr>
<tr><td>Oligo fwd</td><td>CACCACGACGACGGTGGTGTTCGT</td><td></td><td></td><td></td><td></td><td></td><td></td><td></td><td></td><td></td><td></td></tr>
<tr><td>Oligo rev</td><td>AAACACGAACACCACCGTCGTCGT</td><td></td><td></td><td></td><td></td><td></td><td></td><td></td><td></td><td></td><td></td></tr>
<tr><td>Chromosome</td><td>start</td><td>end</td><td>strand</td><td>MM</td><td>target_seq</td><td>PAM</td><td>alignment</td><td>distance</td><td>position</td><td>gene name</td><td>gene id</td></tr>
<tr><td>GU305914.1</td><td>154127</td><td>154149</td><td>+</td><td>0</td><td>ACGACGACGGTGGTGTTCGT</td><td>GGG</td><td>||||||||[||||||||||||]PAM</td><td>NA</td><td>Intergenic</td><td>NA</td><td>NA</td></tr>
<tr><td>T11787</td><td>CGACGACGGTGGTGTTCGTGGGG</td><td>1000</td><td></td><td></td><td></td><td></td><td></td><td></td><td></td><td></td><td></td></tr>
<tr><td>Oligo fwd</td><td>CACCCGACGACGGTGGTGTTCGTG</td><td></td><td></td><td></td><td></td><td></td><td></td><td></td><td></td><td></td><td></td></tr>
<tr><td>Oligo rev</td><td>AAACCACGAACACCACCGTCGTCG</td><td></td><td></td><td></td><td></td><td></td><td></td><td></td><td></td><td></td><td></td></tr>
<tr><td>Chromosome</td><td>start</td><td>end</td><td>strand</td><td>MM</td><td>target_seq</td><td>PAM</td><td>alignment</td><td>distance</td><td>position</td><td>gene name</td><td>gene id</td></tr>
<tr><td>GU305914.1</td><td>154128</td><td>154150</td><td>+</td><td>0</td><td>CGACGACGGTGGTGTTCGTG</td><td>GGG</td><td>||||||||[||||||||||||]PAM</td><td>NA</td><td>Intergenic</td><td>NA</td><td>NA</td></tr>
<tr><td>T11788</td><td>TGGTGTTCGTGGGGACCCGCTGG</td><td>1000</td><td></td><td></td><td></td><td></td><td></td><td></td><td></td><td></td><td></td></tr>
<tr><td>Oligo fwd</td><td>CACCTGGTGTTCGTGGGGACCCGC</td><td></td><td></td><td></td><td></td><td></td><td></td><td></td><td></td><td></td><td></td></tr>
<tr><td>Oligo rev</td><td>AAACGCGGGTCCCCACGAACACCA</td><td></td><td></td><td></td><td></td><td></td><td></td><td></td><td></td><td></td><td></td></tr>
<tr><td>Chromosome</td><td>start</td><td>end</td><td>strand</td><td>MM</td><td>target_seq</td><td>PAM</td><td>alignment</td><td>distance</td><td>position</td><td>gene name</td><td>gene id</td></tr>
<tr><td>GU305914.1</td><td>154137</td><td>154159</td><td>+</td><td>0</td><td>TGGTGTTCGTGGGGACCCGC</td><td>TGG</td><td>||||||||[||||||||||||]PAM</td><td>NA</td><td>Intergenic</td><td>NA</td><td>NA</td></tr>
<tr><td>T11789</td><td>TGTTCGTGGGGACCCGCTGGCGG</td><td>1000</td><td></td><td></td><td></td><td></td><td></td><td></td><td></td><td></td><td></td></tr>
<tr><td>Oligo fwd</td><td>CACCTGTTCGTGGGGACCCGCTGG</td><td></td><td></td><td></td><td></td><td></td><td></td><td></td><td></td><td></td><td></td></tr>
<tr><td>Oligo rev</td><td>AAACCCAGCGGGTCCCCACGAACA</td><td></td><td></td><td></td><td></td><td></td><td></td><td></td><td></td><td></td><td></td></tr>
<tr><td>Chromosome</td><td>start</td><td>end</td><td>strand</td><td>MM</td><td>target_seq</td><td>PAM</td><td>alignment</td><td>distance</td><td>position</td><td>gene name</td><td>gene id</td></tr>
<tr><td>GU305914.1</td><td>154140</td><td>154162</td><td>+</td><td>0</td><td>TGTTCGTGGGGACCCGCTGG</td><td>CGG</td><td>||||||||[||||||||||||]PAM</td><td>NA</td><td>Intergenic</td><td>NA</td><td>NA</td></tr>
<tr><td>T11790</td><td>GTGGGGACCCGCTGGCGGTCTGG</td><td>1000</td><td></td><td></td><td></td><td></td><td></td><td></td><td></td><td></td><td></td></tr>
<tr><td>Oligo fwd</td><td>CACCGTGGGGACCCGCTGGCGGTC</td><td></td><td></td><td></td><td></td><td></td><td></td><td></td><td></td><td></td><td></td></tr>
<tr><td>Oligo rev</td><td>AAACGACCGCCAGCGGGTCCCCAC</td><td></td><td></td><td></td><td></td><td></td><td></td><td></td><td></td><td></td><td></td></tr>
<tr><td>Chromosome</td><td>start</td><td>end</td><td>strand</td><td>MM</td><td>target_seq</td><td>PAM</td><td>alignment</td><td>distance</td><td>position</td><td>gene name</td><td>gene id</td></tr>
<tr><td>GU305914.1</td><td>154145</td><td>154167</td><td>+</td><td>0</td><td>GTGGGGACCCGCTGGCGGTC</td><td>TGG</td><td>||||||||[||||||||||||]PAM</td><td>NA</td><td>Intergenic</td><td>NA</td><td>NA</td></tr>
<tr><td>T11791</td><td>TGGGGACCCGCTGGCGGTCTGGG</td><td>1000</td><td></td><td></td><td></td><td></td><td></td><td></td><td></td><td></td><td></td></tr>
<tr><td>Oligo fwd</td><td>CACCTGGGGACCCGCTGGCGGTCT</td><td></td><td></td><td></td><td></td><td></td><td></td><td></td><td></td><td></td><td></td></tr>
<tr><td>Oligo rev</td><td>AAACAGACCGCCAGCGGGTCCCCA</td><td></td><td></td><td></td><td></td><td></td><td></td><td></td><td></td><td></td><td></td></tr>
<tr><td>Chromosome</td><td>start</td><td>end</td><td>strand</td><td>MM</td><td>target_seq</td><td>PAM</td><td>alignment</td><td>distance</td><td>position</td><td>gene name</td><td>gene id</td></tr>
<tr><td>GU305914.1</td><td>154146</td><td>154168</td><td>+</td><td>0</td><td>TGGGGACCCGCTGGCGGTCT</td><td>GGG</td><td>||||||||[||||||||||||]PAM</td><td>NA</td><td>Intergenic</td><td>NA</td><td>NA</td></tr>
<tr><td>T11792</td><td>CCCGCTGGCGGTCTGGGTATAGG</td><td>1000</td><td></td><td></td><td></td><td></td><td></td><td></td><td></td><td></td><td></td></tr>
<tr><td>Oligo fwd</td><td>CACCCCCGCTGGCGGTCTGGGTAT</td><td></td><td></td><td></td><td></td><td></td><td></td><td></td><td></td><td></td><td></td></tr>
<tr><td>Oligo rev</td><td>AAACATACCCAGACCGCCAGCGGG</td><td></td><td></td><td></td><td></td><td></td><td></td><td></td><td></td><td></td><td></td></tr>
<tr><td>Chromosome</td><td>start</td><td>end</td><td>strand</td><td>MM</td><td>target_seq</td><td>PAM</td><td>alignment</td><td>distance</td><td>position</td><td>gene name</td><td>gene id</td></tr>
<tr><td>GU305914.1</td><td>154152</td><td>154174</td><td>+</td><td>0</td><td>CCCGCTGGCGGTCTGGGTAT</td><td>AGG</td><td>||||||||[||||||||||||]PAM</td><td>NA</td><td>Intergenic</td><td>NA</td><td>NA</td></tr>
</table>

<table>
<tr><td>T11793</td><td>CTGGCGGTCTGGGTATAGGACGG</td><td>1000</td><td></td><td></td><td></td><td></td><td></td><td></td><td></td><td></td><td></td></tr>
<tr><td>Oligo fwd</td><td>CACCCTGGCGGTCTGGGTATAGGA</td><td></td><td></td><td></td><td></td><td></td><td></td><td></td><td></td><td></td><td></td></tr>
<tr><td>Oligo rev</td><td>AAACTCCTATACCCAGACCGCCAG</td><td></td><td></td><td></td><td></td><td></td><td></td><td></td><td></td><td></td><td></td></tr>
<tr><td>Chromosome</td><td>start</td><td>end</td><td>strand</td><td>MM</td><td>target_seq</td><td>PAM</td><td>alignment</td><td>distance</td><td>position</td><td>gene name</td><td>gene id</td></tr>
<tr><td>GU305914.1</td><td>154156</td><td>154178</td><td>+</td><td>0</td><td>CTGGCGGTCTGGGTATAGGA</td><td>CGG</td><td>||||||||[||||||||||||]PAM</td><td>NA</td><td>Intergenic</td><td>NA</td><td>NA</td></tr>
<tr><td>T11794</td><td>TGGCGGTCTGGGTATAGGACGGG</td><td>1000</td><td></td><td></td><td></td><td></td><td></td><td></td><td></td><td></td><td></td></tr>
<tr><td>Oligo fwd</td><td>CACCTGGCGGTCTGGGTATAGGAC</td><td></td><td></td><td></td><td></td><td></td><td></td><td></td><td></td><td></td><td></td></tr>
<tr><td>Oligo rev</td><td>AAACGTCCTATACCCAGACCGCCA</td><td></td><td></td><td></td><td></td><td></td><td></td><td></td><td></td><td></td><td></td></tr>
<tr><td>Chromosome</td><td>start</td><td>end</td><td>strand</td><td>MM</td><td>target_seq</td><td>PAM</td><td>alignment</td><td>distance</td><td>position</td><td>gene name</td><td>gene id</td></tr>
<tr><td>GU305914.1</td><td>154157</td><td>154179</td><td>+</td><td>0</td><td>TGGCGGTCTGGGTATAGGAC</td><td>GGG</td><td>||||||||[||||||||||||]PAM</td><td>NA</td><td>Intergenic</td><td>NA</td><td>NA</td></tr>
<tr><td>T11795</td><td>GGCGGTCTGGGTATAGGACGGGG</td><td>1000</td><td></td><td></td><td></td><td></td><td></td><td></td><td></td><td></td><td></td></tr>
<tr><td>Oligo fwd</td><td>CACCGGCGGTCTGGGTATAGGACG</td><td></td><td></td><td></td><td></td><td></td><td></td><td></td><td></td><td></td><td></td></tr>
<tr><td>Oligo rev</td><td>AAACCGTCCTATACCCAGACCGCC</td><td></td><td></td><td></td><td></td><td></td><td></td><td></td><td></td><td></td><td></td></tr>
<tr><td>Chromosome</td><td>start</td><td>end</td><td>strand</td><td>MM</td><td>target_seq</td><td>PAM</td><td>alignment</td><td>distance</td><td>position</td><td>gene name</td><td>gene id</td></tr>
<tr><td>GU305914.1</td><td>154158</td><td>154180</td><td>+</td><td>0</td><td>GGCGGTCTGGGTATAGGACG</td><td>GGG</td><td>||||||||[||||||||||||]PAM</td><td>NA</td><td>Intergenic</td><td>NA</td><td>NA</td></tr>
<tr><td>T11796</td><td>AGCGAATTCTAGCGTTACCGCGG</td><td>1000</td><td></td><td></td><td></td><td></td><td></td><td></td><td></td><td></td><td></td></tr>
<tr><td>Oligo fwd</td><td>CACCAGCGAATTCTAGCGTTACCG</td><td></td><td></td><td></td><td></td><td></td><td></td><td></td><td></td><td></td><td></td></tr>
<tr><td>Oligo rev</td><td>AAACCGGTAACGCTAGAATTCGCT</td><td></td><td></td><td></td><td></td><td></td><td></td><td></td><td></td><td></td><td></td></tr>
<tr><td>Chromosome</td><td>start</td><td>end</td><td>strand</td><td>MM</td><td>target_seq</td><td>PAM</td><td>alignment</td><td>distance</td><td>position</td><td>gene name</td><td>gene id</td></tr>
<tr><td>GU305914.1</td><td>154186</td><td>154208</td><td>+</td><td>0</td><td>AGCGAATTCTAGCGTTACCG</td><td>CGG</td><td>||||||||[||||||||||||]PAM</td><td>NA</td><td>Intergenic</td><td>NA</td><td>NA</td></tr>
<tr><td>T11797</td><td>GCGACTTGTCTGCGCCGTCGCGG</td><td>1000</td><td></td><td></td><td></td><td></td><td></td><td></td><td></td><td></td><td></td></tr>
<tr><td>Oligo fwd</td><td>CACCGCGACTTGTCTGCGCCGTCG</td><td></td><td></td><td></td><td></td><td></td><td></td><td></td><td></td><td></td><td></td></tr>
<tr><td>Oligo rev</td><td>AAACCGACGGCGCAGACAAGTCGC</td><td></td><td></td><td></td><td></td><td></td><td></td><td></td><td></td><td></td><td></td></tr>
<tr><td>Chromosome</td><td>start</td><td>end</td><td>strand</td><td>MM</td><td>target_seq</td><td>PAM</td><td>alignment</td><td>distance</td><td>position</td><td>gene name</td><td>gene id</td></tr>
<tr><td>GU305914.1</td><td>154214</td><td>154236</td><td>+</td><td>0</td><td>GCGACTTGTCTGCGCCGTCG</td><td>CGG</td><td>||||||||[||||||||||||]PAM</td><td>NA</td><td>Intergenic</td><td>NA</td><td>NA</td></tr>
<tr><td>T11798</td><td>TGTCGTCTCTGTCGTGTTCGCGG</td><td>1000</td><td></td><td></td><td></td><td></td><td></td><td></td><td></td><td></td><td></td></tr>
<tr><td>Oligo fwd</td><td>CACCTGTCGTCTCTGTCGTGTTCG</td><td></td><td></td><td></td><td></td><td></td><td></td><td></td><td></td><td></td><td></td></tr>
<tr><td>Oligo rev</td><td>AAACCGAACACGACAGAGACGACA</td><td></td><td></td><td></td><td></td><td></td><td></td><td></td><td></td><td></td><td></td></tr>
<tr><td>Chromosome</td><td>start</td><td>end</td><td>strand</td><td>MM</td><td>target_seq</td><td>PAM</td><td>alignment</td><td>distance</td><td>position</td><td>gene name</td><td>gene id</td></tr>
<tr><td>GU305914.1</td><td>154250</td><td>154272</td><td>+</td><td>0</td><td>TGTCGTCTCTGTCGTGTTCG</td><td>CGG</td><td>||||||||[||||||||||||]PAM</td><td>NA</td><td>Intergenic</td><td>NA</td><td>NA</td></tr>
<tr><td>T11799</td><td>CGTCTCTGTCGTGTTCGCGGCGG</td><td>1000</td><td></td><td></td><td></td><td></td><td></td><td></td><td></td><td></td><td></td></tr>
<tr><td>Oligo fwd</td><td>CACCCGTCTCTGTCGTGTTCGCGG</td><td></td><td></td><td></td><td></td><td></td><td></td><td></td><td></td><td></td><td></td></tr>
<tr><td>Oligo rev</td><td>AAACCCGCGAACACGACAGAGACG</td><td></td><td></td><td></td><td></td><td></td><td></td><td></td><td></td><td></td><td></td></tr>
<tr><td>Chromosome</td><td>start</td><td>end</td><td>strand</td><td>MM</td><td>target_seq</td><td>PAM</td><td>alignment</td><td>distance</td><td>position</td><td>gene name</td><td>gene id</td></tr>
<tr><td>GU305914.1</td><td>154253</td><td>154275</td><td>+</td><td>0</td><td>CGTCTCTGTCGTGTTCGCGG</td><td>CGG</td><td>||||||||[||||||||||||]PAM</td><td>NA</td><td>Intergenic</td><td>NA</td><td>NA</td></tr>
<tr><td>T11800</td><td>CTCTGTCGTGTTCGCGGCGGCGG</td><td>1000</td><td></td><td></td><td></td><td></td><td></td><td></td><td></td><td></td><td></td></tr>
<tr><td>Oligo fwd</td><td>CACCCTCTGTCGTGTTCGCGGCGG</td><td></td><td></td><td></td><td></td><td></td><td></td><td></td><td></td><td></td><td></td></tr>
<tr><td>Oligo rev</td><td>AAACCCGCCGCGAACACGACAGAG</td><td></td><td></td><td></td><td></td><td></td><td></td><td></td><td></td><td></td><td></td></tr>
<tr><td>Chromosome</td><td>start</td><td>end</td><td>strand</td><td>MM</td><td>target_seq</td><td>PAM</td><td>alignment</td><td>distance</td><td>position</td><td>gene name</td><td>gene id</td></tr>
<tr><td>GU305914.1</td><td>154256</td><td>154278</td><td>+</td><td>0</td><td>CTCTGTCGTGTTCGCGGCGG</td><td>CGG</td><td>||||||||[||||||||||||]PAM</td><td>NA</td><td>Intergenic</td><td>NA</td><td>NA</td></tr>
<tr><td>T11801</td><td>TGTCGTGTTCGCGGCGGCGGCGG</td><td>1000</td><td></td><td></td><td></td><td></td><td></td><td></td><td></td><td></td><td></td></tr>
<tr><td>Oligo fwd</td><td>CACCTGTCGTGTTCGCGGCGGCGG</td><td></td><td></td><td></td><td></td><td></td><td></td><td></td><td></td><td></td><td></td></tr>
<tr><td>Oligo rev</td><td>AAACCCGCCGCCGCGAACACGACA</td><td></td><td></td><td></td><td></td><td></td><td></td><td></td><td></td><td></td><td></td></tr>
<tr><td>Chromosome</td><td>start</td><td>end</td><td>strand</td><td>MM</td><td>target_seq</td><td>PAM</td><td>alignment</td><td>distance</td><td>position</td><td>gene name</td><td>gene id</td></tr>
<tr><td>GU305914.1</td><td>154259</td><td>154281</td><td>+</td><td>0</td><td>TGTCGTGTTCGCGGCGGCGG</td><td>CGG</td><td>||||||||[||||||||||||]PAM</td><td>NA</td><td>Intergenic</td><td>NA</td><td>NA</td></tr>
<tr><td>T11802</td><td>GCGGCGACGTTGTCATCGAGAGG</td><td>1000</td><td></td><td></td><td></td><td></td><td></td><td></td><td></td><td></td><td></td></tr>
<tr><td>Oligo fwd</td><td>CACCGCGGCGACGTTGTCATCGAG</td><td></td><td></td><td></td><td></td><td></td><td></td><td></td><td></td><td></td><td></td></tr>
<tr><td>Oligo rev</td><td>AAACCTCGATGACAACGTCGCCGC</td><td></td><td></td><td></td><td></td><td></td><td></td><td></td><td></td><td></td><td></td></tr>
<tr><td>Chromosome</td><td>start</td><td>end</td><td>strand</td><td>MM</td><td>target_seq</td><td>PAM</td><td>alignment</td><td>distance</td><td>position</td><td>gene name</td><td>gene id</td></tr>
<tr><td>GU305914.1</td><td>154278</td><td>154300</td><td>+</td><td>0</td><td>GCGGCGACGTTGTCATCGAG</td><td>AGG</td><td>||||||||[||||||||||||]PAM</td><td>NA</td><td>Intergenic</td><td>NA</td><td>NA</td></tr>
<tr><td>T11803</td><td>ACGTTGTCATCGAGAGGTAGTGG</td><td>1000</td><td></td><td></td><td></td><td></td><td></td><td></td><td></td><td></td><td></td></tr>
<tr><td>Oligo fwd</td><td>CACCACGTTGTCATCGAGAGGTAG</td><td></td><td></td><td></td><td></td><td></td><td></td><td></td><td></td><td></td><td></td></tr>
<tr><td>Oligo rev</td><td>AAACCTACCTCTCGATGACAACGT</td><td></td><td></td><td></td><td></td><td></td><td></td><td></td><td></td><td></td><td></td></tr>
<tr><td>Chromosome</td><td>start</td><td>end</td><td>strand</td><td>MM</td><td>target_seq</td><td>PAM</td><td>alignment</td><td>distance</td><td>position</td><td>gene name</td><td>gene id</td></tr>
<tr><td>GU305914.1</td><td>154284</td><td>154306</td><td>+</td><td>0</td><td>ACGTTGTCATCGAGAGGTAG</td><td>TGG</td><td>||||||||[||||||||||||]PAM</td><td>NA</td><td>Intergenic</td><td>NA</td><td>NA</td></tr>
<tr><td>T11804</td><td>TTGTCATCGAGAGGTAGTGGAGG</td><td>1000</td><td></td><td></td><td></td><td></td><td></td><td></td><td></td><td></td><td></td></tr>
<tr><td>Oligo fwd</td><td>CACCTTGTCATCGAGAGGTAGTGG</td><td></td><td></td><td></td><td></td><td></td><td></td><td></td><td></td><td></td><td></td></tr>
<tr><td>Oligo rev</td><td>AAACCCACTACCTCTCGATGACAA</td><td></td><td></td><td></td><td></td><td></td><td></td><td></td><td></td><td></td><td></td></tr>
<tr><td>Chromosome</td><td>start</td><td>end</td><td>strand</td><td>MM</td><td>target_seq</td><td>PAM</td><td>alignment</td><td>distance</td><td>position</td><td>gene name</td><td>gene id</td></tr>
<tr><td>GU305914.1</td><td>154287</td><td>154309</td><td>+</td><td>0</td><td>TTGTCATCGAGAGGTAGTGG</td><td>AGG</td><td>||||||||[||||||||||||]PAM</td><td>NA</td><td>Intergenic</td><td>NA</td><td>NA</td></tr>
<tr><td>T11805</td><td>CATCGAGAGGTAGTGGAGGATGG</td><td>1000</td><td></td><td></td><td></td><td></td><td></td><td></td><td></td><td></td><td></td></tr>
<tr><td>Oligo fwd</td><td>CACCCATCGAGAGGTAGTGGAGGA</td><td></td><td></td><td></td><td></td><td></td><td></td><td></td><td></td><td></td><td></td></tr>
<tr><td>Oligo rev</td><td>AAACTCCTCCACTACCTCTCGATG</td><td></td><td></td><td></td><td></td><td></td><td></td><td></td><td></td><td></td><td></td></tr>
<tr><td>Chromosome</td><td>start</td><td>end</td><td>strand</td><td>MM</td><td>target_seq</td><td>PAM</td><td>alignment</td><td>distance</td><td>position</td><td>gene name</td><td>gene id</td></tr>
<tr><td>GU305914.1</td><td>154291</td><td>154313</td><td>+</td><td>0</td><td>CATCGAGAGGTAGTGGAGGA</td><td>TGG</td><td>||||||||[||||||||||||]PAM</td><td>NA</td><td>Intergenic</td><td>NA</td><td>NA</td></tr>
<tr><td>T11806</td><td>CGAGAGGTAGTGGAGGATGGAGG</td><td>1000</td><td></td><td></td><td></td><td></td><td></td><td></td><td></td><td></td><td></td></tr>
<tr><td>Oligo fwd</td><td>CACCCGAGAGGTAGTGGAGGATGG</td><td></td><td></td><td></td><td></td><td></td><td></td><td></td><td></td><td></td><td></td></tr>
<tr><td>Oligo rev</td><td>AAACCCATCCTCCACTACCTCTCG</td><td></td><td></td><td></td><td></td><td></td><td></td><td></td><td></td><td></td><td></td></tr>
<tr><td>Chromosome</td><td>start</td><td>end</td><td>strand</td><td>MM</td><td>target_seq</td><td>PAM</td><td>alignment</td><td>distance</td><td>position</td><td>gene name</td><td>gene id</td></tr>
<tr><td>GU305914.1</td><td>154294</td><td>154316</td><td>+</td><td>0</td><td>CGAGAGGTAGTGGAGGATGG</td><td>AGG</td><td>||||||||[||||||||||||]PAM</td><td>NA</td><td>Intergenic</td><td>NA</td><td>NA</td></tr>
<tr><td>T11807</td><td>GATGGAGGCATCCGTGTCCCTGG</td><td>1000</td><td></td><td></td><td></td><td></td><td></td><td></td><td></td><td></td><td></td></tr>
<tr><td>Oligo fwd</td><td>CACCGATGGAGGCATCCGTGTCCC</td><td></td><td></td><td></td><td></td><td></td><td></td><td></td><td></td><td></td><td></td></tr>
<tr><td>Oligo rev</td><td>AAACGGGACACGGATGCCTCCATC</td><td></td><td></td><td></td><td></td><td></td><td></td><td></td><td></td><td></td><td></td></tr>
<tr><td>Chromosome</td><td>start</td><td>end</td><td>strand</td><td>MM</td><td>target_seq</td><td>PAM</td><td>alignment</td><td>distance</td><td>position</td><td>gene name</td><td>gene id</td></tr>
<tr><td>GU305914.1</td><td>154309</td><td>154331</td><td>+</td><td>0</td><td>GATGGAGGCATCCGTGTCCC</td><td>TGG</td><td>||||||||[||||||||||||]PAM</td><td>NA</td><td>Intergenic</td><td>NA</td><td>NA</td></tr>
</table>

<table>
<tr><td>T11808</td><td>GTGTCCGCCTCCTCGCAGCATGG</td><td>1000</td><td></td><td></td><td></td><td></td><td></td><td></td><td></td><td></td><td></td></tr>
<tr><td>Oligo fwd</td><td>CACCGTGTCCGCCTCCTCGCAGCA</td><td></td><td></td><td></td><td></td><td></td><td></td><td></td><td></td><td></td><td></td></tr>
<tr><td>Oligo rev</td><td>AAACTGCTGCGAGGAGGCGGACAC</td><td></td><td></td><td></td><td></td><td></td><td></td><td></td><td></td><td></td><td></td></tr>
<tr><td>Chromosome</td><td>start</td><td>end</td><td>strand</td><td>MM</td><td>target_seq</td><td>PAM</td><td>alignment</td><td>distance</td><td>position</td><td>gene name</td><td>gene id</td></tr>
<tr><td>GU305914.1</td><td>154349</td><td>154371</td><td>+</td><td>0</td><td>GTGTCCGCCTCCTCGCAGCA</td><td>TGG</td><td>||||||||[||||||||||||]PAM</td><td>NA</td><td>Intergenic</td><td>NA</td><td>NA</td></tr>
<tr><td>T11809</td><td>TGTCCGCCTCCTCGCAGCATGGG</td><td>1000</td><td></td><td></td><td></td><td></td><td></td><td></td><td></td><td></td><td></td></tr>
<tr><td>Oligo fwd</td><td>CACCTGTCCGCCTCCTCGCAGCAT</td><td></td><td></td><td></td><td></td><td></td><td></td><td></td><td></td><td></td><td></td></tr>
<tr><td>Oligo rev</td><td>AAACATGCTGCGAGGAGGCGGACA</td><td></td><td></td><td></td><td></td><td></td><td></td><td></td><td></td><td></td><td></td></tr>
<tr><td>Chromosome</td><td>start</td><td>end</td><td>strand</td><td>MM</td><td>target_seq</td><td>PAM</td><td>alignment</td><td>distance</td><td>position</td><td>gene name</td><td>gene id</td></tr>
<tr><td>GU305914.1</td><td>154350</td><td>154372</td><td>+</td><td>0</td><td>TGTCCGCCTCCTCGCAGCAT</td><td>GGG</td><td>||||||||[||||||||||||]PAM</td><td>NA</td><td>Intergenic</td><td>NA</td><td>NA</td></tr>
<tr><td>T11810</td><td>GTCCGCCTCCTCGCAGCATGGGG</td><td>1000</td><td></td><td></td><td></td><td></td><td></td><td></td><td></td><td></td><td></td></tr>
<tr><td>Oligo fwd</td><td>CACCGTCCGCCTCCTCGCAGCATG</td><td></td><td></td><td></td><td></td><td></td><td></td><td></td><td></td><td></td><td></td></tr>
<tr><td>Oligo rev</td><td>AAACCATGCTGCGAGGAGGCGGAC</td><td></td><td></td><td></td><td></td><td></td><td></td><td></td><td></td><td></td><td></td></tr>
<tr><td>Chromosome</td><td>start</td><td>end</td><td>strand</td><td>MM</td><td>target_seq</td><td>PAM</td><td>alignment</td><td>distance</td><td>position</td><td>gene name</td><td>gene id</td></tr>
<tr><td>GU305914.1</td><td>154351</td><td>154373</td><td>+</td><td>0</td><td>GTCCGCCTCCTCGCAGCATG</td><td>GGG</td><td>||||||||[||||||||||||]PAM</td><td>NA</td><td>Intergenic</td><td>NA</td><td>NA</td></tr>
<tr><td>T11811</td><td>ATGGGGATCGTGTGTCCACATGG</td><td>1000</td><td></td><td></td><td></td><td></td><td></td><td></td><td></td><td></td><td></td></tr>
<tr><td>Oligo fwd</td><td>CACCATGGGGATCGTGTGTCCACA</td><td></td><td></td><td></td><td></td><td></td><td></td><td></td><td></td><td></td><td></td></tr>
<tr><td>Oligo rev</td><td>AAACTGTGGACACACGATCCCCAT</td><td></td><td></td><td></td><td></td><td></td><td></td><td></td><td></td><td></td><td></td></tr>
<tr><td>Chromosome</td><td>start</td><td>end</td><td>strand</td><td>MM</td><td>target_seq</td><td>PAM</td><td>alignment</td><td>distance</td><td>position</td><td>gene name</td><td>gene id</td></tr>
<tr><td>GU305914.1</td><td>154368</td><td>154390</td><td>+</td><td>0</td><td>ATGGGGATCGTGTGTCCACA</td><td>TGG</td><td>||||||||[||||||||||||]PAM</td><td>NA</td><td>Intergenic</td><td>NA</td><td>NA</td></tr>
<tr><td>T11812</td><td>TGGGGATCGTGTGTCCACATGGG</td><td>1000</td><td></td><td></td><td></td><td></td><td></td><td></td><td></td><td></td><td></td></tr>
<tr><td>Oligo fwd</td><td>CACCTGGGGATCGTGTGTCCACAT</td><td></td><td></td><td></td><td></td><td></td><td></td><td></td><td></td><td></td><td></td></tr>
<tr><td>Oligo rev</td><td>AAACATGTGGACACACGATCCCCA</td><td></td><td></td><td></td><td></td><td></td><td></td><td></td><td></td><td></td><td></td></tr>
<tr><td>Chromosome</td><td>start</td><td>end</td><td>strand</td><td>MM</td><td>target_seq</td><td>PAM</td><td>alignment</td><td>distance</td><td>position</td><td>gene name</td><td>gene id</td></tr>
<tr><td>GU305914.1</td><td>154369</td><td>154391</td><td>+</td><td>0</td><td>TGGGGATCGTGTGTCCACAT</td><td>GGG</td><td>||||||||[||||||||||||]PAM</td><td>NA</td><td>Intergenic</td><td>NA</td><td>NA</td></tr>
<tr><td>T11813</td><td>TCCACATGGGACGATCGCCCCGG</td><td>1000</td><td></td><td></td><td></td><td></td><td></td><td></td><td></td><td></td><td></td></tr>
<tr><td>Oligo fwd</td><td>CACCTCCACATGGGACGATCGCCC</td><td></td><td></td><td></td><td></td><td></td><td></td><td></td><td></td><td></td><td></td></tr>
<tr><td>Oligo rev</td><td>AAACGGGCGATCGTCCCATGTGGA</td><td></td><td></td><td></td><td></td><td></td><td></td><td></td><td></td><td></td><td></td></tr>
<tr><td>Chromosome</td><td>start</td><td>end</td><td>strand</td><td>MM</td><td>target_seq</td><td>PAM</td><td>alignment</td><td>distance</td><td>position</td><td>gene name</td><td>gene id</td></tr>
<tr><td>GU305914.1</td><td>154382</td><td>154404</td><td>+</td><td>0</td><td>TCCACATGGGACGATCGCCC</td><td>CGG</td><td>||||||||[||||||||||||]PAM</td><td>NA</td><td>Intergenic</td><td>NA</td><td>NA</td></tr>
<tr><td>T11814</td><td>CCACATGGGACGATCGCCCCGGG</td><td>1000</td><td></td><td></td><td></td><td></td><td></td><td></td><td></td><td></td><td></td></tr>
<tr><td>Oligo fwd</td><td>CACCCCACATGGGACGATCGCCCC</td><td></td><td></td><td></td><td></td><td></td><td></td><td></td><td></td><td></td><td></td></tr>
<tr><td>Oligo rev</td><td>AAACGGGGCGATCGTCCCATGTGG</td><td></td><td></td><td></td><td></td><td></td><td></td><td></td><td></td><td></td><td></td></tr>
<tr><td>Chromosome</td><td>start</td><td>end</td><td>strand</td><td>MM</td><td>target_seq</td><td>PAM</td><td>alignment</td><td>distance</td><td>position</td><td>gene name</td><td>gene id</td></tr>
<tr><td>GU305914.1</td><td>154383</td><td>154405</td><td>+</td><td>0</td><td>CCACATGGGACGATCGCCCC</td><td>GGG</td><td>||||||||[||||||||||||]PAM</td><td>NA</td><td>Intergenic</td><td>NA</td><td>NA</td></tr>
<tr><td>T11815</td><td>CATGGGACGATCGCCCCGGGCGG</td><td>1000</td><td></td><td></td><td></td><td></td><td></td><td></td><td></td><td></td><td></td></tr>
<tr><td>Oligo fwd</td><td>CACCCATGGGACGATCGCCCCGGG</td><td></td><td></td><td></td><td></td><td></td><td></td><td></td><td></td><td></td><td></td></tr>
<tr><td>Oligo rev</td><td>AAACCCCGGGGCGATCGTCCCATG</td><td></td><td></td><td></td><td></td><td></td><td></td><td></td><td></td><td></td><td></td></tr>
<tr><td>Chromosome</td><td>start</td><td>end</td><td>strand</td><td>MM</td><td>target_seq</td><td>PAM</td><td>alignment</td><td>distance</td><td>position</td><td>gene name</td><td>gene id</td></tr>
<tr><td>GU305914.1</td><td>154386</td><td>154408</td><td>+</td><td>0</td><td>CATGGGACGATCGCCCCGGG</td><td>CGG</td><td>||||||||[||||||||||||]PAM</td><td>NA</td><td>Intergenic</td><td>NA</td><td>NA</td></tr>
<tr><td>T11816</td><td>GGGACGATCGCCCCGGGCGGTGG</td><td>1000</td><td></td><td></td><td></td><td></td><td></td><td></td><td></td><td></td><td></td></tr>
<tr><td>Oligo fwd</td><td>CACCGGGACGATCGCCCCGGGCGG</td><td></td><td></td><td></td><td></td><td></td><td></td><td></td><td></td><td></td><td></td></tr>
<tr><td>Oligo rev</td><td>AAACCCGCCCGGGGCGATCGTCCC</td><td></td><td></td><td></td><td></td><td></td><td></td><td></td><td></td><td></td><td></td></tr>
<tr><td>Chromosome</td><td>start</td><td>end</td><td>strand</td><td>MM</td><td>target_seq</td><td>PAM</td><td>alignment</td><td>distance</td><td>position</td><td>gene name</td><td>gene id</td></tr>
<tr><td>GU305914.1</td><td>154389</td><td>154411</td><td>+</td><td>0</td><td>GGGACGATCGCCCCGGGCGG</td><td>TGG</td><td>||||||||[||||||||||||]PAM</td><td>NA</td><td>Intergenic</td><td>NA</td><td>NA</td></tr>
<tr><td>T11817</td><td>ACGATCGCCCCGGGCGGTGGCGG</td><td>1000</td><td></td><td></td><td></td><td></td><td></td><td></td><td></td><td></td><td></td></tr>
<tr><td>Oligo fwd</td><td>CACCACGATCGCCCCGGGCGGTGG</td><td></td><td></td><td></td><td></td><td></td><td></td><td></td><td></td><td></td><td></td></tr>
<tr><td>Oligo rev</td><td>AAACCCACCGCCCGGGGCGATCGT</td><td></td><td></td><td></td><td></td><td></td><td></td><td></td><td></td><td></td><td></td></tr>
<tr><td>Chromosome</td><td>start</td><td>end</td><td>strand</td><td>MM</td><td>target_seq</td><td>PAM</td><td>alignment</td><td>distance</td><td>position</td><td>gene name</td><td>gene id</td></tr>
<tr><td>GU305914.1</td><td>154392</td><td>154414</td><td>+</td><td>0</td><td>ACGATCGCCCCGGGCGGTGG</td><td>CGG</td><td>||||||||[||||||||||||]PAM</td><td>NA</td><td>Intergenic</td><td>NA</td><td>NA</td></tr>
<tr><td>T11818</td><td>ATCGCCCCGGGCGGTGGCGGCGG</td><td>1000</td><td></td><td></td><td></td><td></td><td></td><td></td><td></td><td></td><td></td></tr>
<tr><td>Oligo fwd</td><td>CACCATCGCCCCGGGCGGTGGCGG</td><td></td><td></td><td></td><td></td><td></td><td></td><td></td><td></td><td></td><td></td></tr>
<tr><td>Oligo rev</td><td>AAACCCGCCACCGCCCGGGGCGAT</td><td></td><td></td><td></td><td></td><td></td><td></td><td></td><td></td><td></td><td></td></tr>
<tr><td>Chromosome</td><td>start</td><td>end</td><td>strand</td><td>MM</td><td>target_seq</td><td>PAM</td><td>alignment</td><td>distance</td><td>position</td><td>gene name</td><td>gene id</td></tr>
<tr><td>GU305914.1</td><td>154395</td><td>154417</td><td>+</td><td>0</td><td>ATCGCCCCGGGCGGTGGCGG</td><td>CGG</td><td>||||||||[||||||||||||]PAM</td><td>NA</td><td>Intergenic</td><td>NA</td><td>NA</td></tr>
<tr><td>GU305914.1</td><td>52716</td><td>52738</td><td>+</td><td>4</td><td>GCCGCCGCTGGCGGTGGCGG</td><td>TGG</td><td>--||||-|[-|||||||||||]PAM</td><td>NA</td><td>Intergenic</td><td>NA</td><td>NA</td></tr>
<tr><td>GU305914.1</td><td>22081</td><td>22103</td><td>+</td><td>4</td><td>ATCCCTCCGGTCGGTCGCGG</td><td>CGG</td><td>|||-|-||[||-||||-||||]PAM</td><td>NA</td><td>Intergenic</td><td>NA</td><td>NA</td></tr>
<tr><td>T11819</td><td>GCCCCGGGCGGTGGCGGCGGCGG</td><td>1000</td><td></td><td></td><td></td><td></td><td></td><td></td><td></td><td></td><td></td></tr>
<tr><td>Oligo fwd</td><td>CACCGCCCCGGGCGGTGGCGGCGG</td><td></td><td></td><td></td><td></td><td></td><td></td><td></td><td></td><td></td><td></td></tr>
<tr><td>Oligo rev</td><td>AAACCCGCCGCCACCGCCCGGGGC</td><td></td><td></td><td></td><td></td><td></td><td></td><td></td><td></td><td></td><td></td></tr>
<tr><td>Chromosome</td><td>start</td><td>end</td><td>strand</td><td>MM</td><td>target_seq</td><td>PAM</td><td>alignment</td><td>distance</td><td>position</td><td>gene name</td><td>gene id</td></tr>
<tr><td>GU305914.1</td><td>154398</td><td>154420</td><td>+</td><td>0</td><td>GCCCCGGGCGGTGGCGGCGG</td><td>CGG</td><td>||||||||[||||||||||||]PAM</td><td>NA</td><td>Intergenic</td><td>NA</td><td>NA</td></tr>
<tr><td>GU305914.1</td><td>58132</td><td>58154</td><td>-</td><td>4</td><td>AAGGCGGGCGGTGGCGGCGG</td><td>CGG</td><td>----||||[||||||||||||]PAM</td><td>NA</td><td>Intergenic</td><td>NA</td><td>NA</td></tr>
<tr><td>GU305914.1</td><td>48156</td><td>48178</td><td>+</td><td>4</td><td>GGCGGCGGCGGTGGCGGCGG</td><td>CGG</td><td>|-|---||[||||||||||||]PAM</td><td>NA</td><td>Intergenic</td><td>NA</td><td>NA</td></tr>
<tr><td>GU305914.1</td><td>62446</td><td>62468</td><td>+</td><td>4</td><td>GGCGGGGGTGGTGGCGGCGG</td><td>GGG</td><td>|-|--|||[-|||||||||||]PAM</td><td>NA</td><td>Intergenic</td><td>NA</td><td>NA</td></tr>
<tr><td>GU305914.1</td><td>52719</td><td>52741</td><td>+</td><td>3</td><td>GCCGCTGGCGGTGGCGGTGG</td><td>CGG</td><td>|||-|-||[|||||||||-||]PAM</td><td>NA</td><td>Intergenic</td><td>NA</td><td>NA</td></tr>
<tr><td>T11820</td><td>CCGGGCGGTGGCGGCGGCGGCGG</td><td>1000</td><td></td><td></td><td></td><td></td><td></td><td></td><td></td><td></td><td></td></tr>
<tr><td>Oligo fwd</td><td>CACCCCGGGCGGTGGCGGCGGCGG</td><td></td><td></td><td></td><td></td><td></td><td></td><td></td><td></td><td></td><td></td></tr>
<tr><td>Oligo rev</td><td>AAACCCGCCGCCGCCACCGCCCGG</td><td></td><td></td><td></td><td></td><td></td><td></td><td></td><td></td><td></td><td></td></tr>
<tr><td>Chromosome</td><td>start</td><td>end</td><td>strand</td><td>MM</td><td>target_seq</td><td>PAM</td><td>alignment</td><td>distance</td><td>position</td><td>gene name</td><td>gene id</td></tr>
<tr><td>GU305914.1</td><td>154401</td><td>154423</td><td>+</td><td>0</td><td>CCGGGCGGTGGCGGCGGCGG</td><td>CGG</td><td>||||||||[||||||||||||]PAM</td><td>NA</td><td>Intergenic</td><td>NA</td><td>NA</td></tr>
<tr><td>GU305914.1</td><td>58129</td><td>58151</td><td>-</td><td>1</td><td>GCGGGCGGTGGCGGCGGCGG</td><td>CGG</td><td>-|||||||[||||||||||||]PAM</td><td>NA</td><td>Intergenic</td><td>NA</td><td>NA</td></tr>
<tr><td>GU305914.1</td><td>48159</td><td>48181</td><td>+</td><td>3</td><td>GGCGGCGGTGGCGGCGGCGG</td><td>TGG</td><td>---|||||[||||||||||||]PAM</td><td>NA</td><td>Intergenic</td><td>NA</td><td>NA</td></tr>
<tr><td>GU305914.1</td><td>154407</td><td>154429</td><td>+</td><td>4</td><td>GGTGGCGGCGGCGGCGGCGG</td><td>CGG</td><td>---|||||[-|||||||||||]PAM</td><td>NA</td><td>Intergenic</td><td>NA</td><td>NA</td></tr>
<tr><td>GU305914.1</td><td>58123</td><td>58145</td><td>-</td><td>4</td><td>GGTGGCGGCGGCGGCGGCGG</td><td>AGG</td><td>---|||||[-|||||||||||]PAM</td><td>NA</td><td>Intergenic</td><td>NA</td><td>NA</td></tr>
<tr><td>GU305914.1</td><td>48147</td><td>48169</td><td>+</td><td>4</td><td>CGGACCTGTGGCGGCGGCGG</td><td>TGG</td><td>|-|--|-|[||||||||||||]PAM</td><td>NA</td><td>Intergenic</td><td>NA</td><td>NA</td></tr>
</table>

<table>
<tr><td>GU305914.1</td><td>67742</td><td>67764</td><td>+</td><td>4</td><td>GGGAGCGGCGGCGGCGGCGG</td><td>CGG</td><td>--|-||||[-|||||||||||]PAM</td><td>NA</td><td>Intergenic</td><td>NA</td><td>NA</td></tr>
<tr><td>GU305914.1</td><td>29997</td><td>30019</td><td>+</td><td>4</td><td>ACCGGTGCTGGCGGCGGCGG</td><td>CGG</td><td>-|-||-|-[||||||||||||]PAM</td><td>NA</td><td>Intergenic</td><td>NA</td><td>NA</td></tr>
<tr><td>GU305914.1</td><td>149485</td><td>149507</td><td>-</td><td>4</td><td>GCAGGCGACGGCGGCGGCGG</td><td>CGG</td><td>-|-||||-[-|||||||||||]PAM</td><td>NA</td><td>Intergenic</td><td>NA</td><td>NA</td></tr>
<tr><td>GU305914.1</td><td>67739</td><td>67761</td><td>+</td><td>4</td><td>TCGGGGAGCGGCGGCGGCGG</td><td>CGG</td><td>-||||--|[-|||||||||||]PAM</td><td>NA</td><td>Intergenic</td><td>NA</td><td>NA</td></tr>
<tr><td>GU305914.1</td><td>45255</td><td>45277</td><td>-</td><td>4</td><td>GGTGGCGGTGGCGGCAGCGG</td><td>CGG</td><td>---|||||[|||||||-||||]PAM</td><td>NA</td><td>Intergenic</td><td>NA</td><td>NA</td></tr>
<tr><td>GU305914.1</td><td>75848</td><td>75870</td><td>+</td><td>4</td><td>CCCGGGGGAGGCGGAGGCGG</td><td>CGG</td><td>||-||-||[-|||||-|||||]PAM</td><td>NA</td><td>Intergenic</td><td>NA</td><td>NA</td></tr>
<tr><td>GU305914.1</td><td>90259</td><td>90281</td><td>-</td><td>4</td><td>GCGGGCGACGGCGGAGGCGG</td><td>CGG</td><td>-||||||-[-|||||-|||||]PAM</td><td>NA</td><td>Intergenic</td><td>NA</td><td>NA</td></tr>
<tr><td>GU305914.1</td><td>62434</td><td>62456</td><td>+</td><td>4</td><td>GACGGCGGTGGCGGCGGGGG</td><td>TGG</td><td>---|||||[|||||||||-||]PAM</td><td>NA</td><td>Intergenic</td><td>NA</td><td>NA</td></tr>
<tr><td>GU305914.1</td><td>62449</td><td>62471</td><td>+</td><td>4</td><td>GGGGGTGGTGGCGGCGGGGG</td><td>TGG</td><td>--|||-||[|||||||||-||]PAM</td><td>NA</td><td>Intergenic</td><td>NA</td><td>NA</td></tr>
<tr><td></td><td></td><td></td><td></td><td></td><td></td><td></td><td></td><td></td><td></td><td></td><td></td></tr>
<tr><td>T11821</td><td>GGCGGTGGCGGCGGCGGCGG</td><td>1000</td><td></td><td></td><td></td><td></td><td></td><td></td><td></td><td></td><td></td></tr>
<tr><td>Oligo fwd</td><td>CACCGGCGGTGGCGGCGGCGGCGG</td><td></td><td></td><td></td><td></td><td></td><td></td><td></td><td></td><td></td><td></td></tr>
<tr><td>Oligo rev</td><td>AAACCCGCCGCCGCCGCCACCGCC</td><td></td><td></td><td></td><td></td><td></td><td></td><td></td><td></td><td></td><td></td></tr>
<tr><td>Chromosome</td><td>start</td><td>end</td><td>strand</td><td>MM</td><td>target_seq</td><td>PAM</td><td>alignment</td><td>distance</td><td>position</td><td>gene name</td><td>gene id</td></tr>
<tr><td>GU305914.1</td><td>154404</td><td>154426</td><td>+</td><td>0</td><td>GGCGGTGGCGGCGGCGGCGG</td><td>CGG</td><td>||||||||[||||||||||||]PAM</td><td>NA</td><td>Intergenic</td><td>NA</td><td>NA</td></tr>
<tr><td>GU305914.1</td><td>58126</td><td>58148</td><td>-</td><td>0</td><td>GGCGGTGGCGGCGGCGGCGG</td><td>CGG</td><td>||||||||[||||||||||||]PAM</td><td>NA</td><td>Intergenic</td><td>NA</td><td>NA</td></tr>
<tr><td>GU305914.1</td><td>30000</td><td>30022</td><td>+</td><td>2</td><td>GGTGCTGGCGGCGGCGGCGG</td><td>CGG</td><td>||-|-|||[||||||||||||]PAM</td><td>NA</td><td>Intergenic</td><td>NA</td><td>NA</td></tr>
<tr><td>GU305914.1</td><td>154407</td><td>154429</td><td>+</td><td>2</td><td>GGTGGCGGCGGCGGCGGCGG</td><td>CGG</td><td>||-||-||[||||||||||||]PAM</td><td>NA</td><td>Intergenic</td><td>NA</td><td>NA</td></tr>
<tr><td>GU305914.1</td><td>58123</td><td>58145</td><td>-</td><td>2</td><td>GGTGGCGGCGGCGGCGGCGG</td><td>AGG</td><td>||-||-||[||||||||||||]PAM</td><td>NA</td><td>Intergenic</td><td>NA</td><td>NA</td></tr>
<tr><td>GU305914.1</td><td>149482</td><td>149504</td><td>-</td><td>2</td><td>GGCGACGGCGGCGGCGGCGG</td><td>CGG</td><td>||||--||[||||||||||||]PAM</td><td>NA</td><td>Intergenic</td><td>NA</td><td>NA</td></tr>
<tr><td>GU305914.1</td><td>67742</td><td>67764</td><td>+</td><td>3</td><td>GGGAGCGGCGGCGGCGGCGG</td><td>CGG</td><td>||--|-||[||||||||||||]PAM</td><td>NA</td><td>Intergenic</td><td>NA</td><td>NA</td></tr>
<tr><td>GU305914.1</td><td>168820</td><td>168842</td><td>+</td><td>2</td><td>GGTGGTGGCAGCGGCGGCGG</td><td>CGG</td><td>||-|||||[|-||||||||||]PAM</td><td>NA</td><td>Intergenic</td><td>NA</td><td>NA</td></tr>
<tr><td>GU305914.1</td><td>48159</td><td>48181</td><td>+</td><td>2</td><td>GGCGGCGGTGGCGGCGGCGG</td><td>TGG</td><td>|||||-||[-|||||||||||]PAM</td><td>NA</td><td>Intergenic</td><td>NA</td><td>NA</td></tr>
<tr><td>GU305914.1</td><td>56773</td><td>56795</td><td>+</td><td>4</td><td>CGCCGCAGCGGCGGCGGCGG</td><td>CGG</td><td>-||-|--|[||||||||||||]PAM</td><td>NA</td><td>Intergenic</td><td>NA</td><td>NA</td></tr>
<tr><td>GU305914.1</td><td>161754</td><td>161776</td><td>-</td><td>3</td><td>GGCCGAGGAGGCGGCGGCGG</td><td>AGG</td><td>|||-|-||[-|||||||||||]PAM</td><td>NA</td><td>Intergenic</td><td>NA</td><td>NA</td></tr>
<tr><td>GU305914.1</td><td>149485</td><td>149507</td><td>-</td><td>4</td><td>GCAGGCGACGGCGGCGGCGG</td><td>CGG</td><td>|--||-|-[||||||||||||]PAM</td><td>NA</td><td>Intergenic</td><td>NA</td><td>NA</td></tr>
<tr><td>GU305914.1</td><td>58129</td><td>58151</td><td>-</td><td>4</td><td>GCGGGCGGTGGCGGCGGCGG</td><td>CGG</td><td>|--||-||[-|||||||||||]PAM</td><td>NA</td><td>Intergenic</td><td>NA</td><td>NA</td></tr>
<tr><td>GU305914.1</td><td>176454</td><td>176476</td><td>+</td><td>3</td><td>GACGATGGCGACGGCGGCGG</td><td>AGG</td><td>|-||-|||[||-|||||||||]PAM</td><td>NA</td><td>Intergenic</td><td>NA</td><td>NA</td></tr>
<tr><td>GU305914.1</td><td>48156</td><td>48178</td><td>+</td><td>2</td><td>GGCGGCGGCGGTGGCGGCGG</td><td>CGG</td><td>|||||-||[|||-||||||||]PAM</td><td>NA</td><td>Intergenic</td><td>NA</td><td>NA</td></tr>
<tr><td>GU305914.1</td><td>29997</td><td>30019</td><td>+</td><td>4</td><td>ACCGGTGCTGGCGGCGGCGG</td><td>CGG</td><td>--|||||-[-|||||||||||]PAM</td><td>NA</td><td>Intergenic</td><td>NA</td><td>NA</td></tr>
<tr><td>GU305914.1</td><td>164971</td><td>164993</td><td>-</td><td>4</td><td>GGTGGGACCGGCGGCGGCGG</td><td>CGG</td><td>||-||---[||||||||||||]PAM</td><td>NA</td><td>Intergenic</td><td>NA</td><td>NA</td></tr>
<tr><td>GU305914.1</td><td>114658</td><td>114680</td><td>-</td><td>2</td><td>GGCGGTGGTGGTGGCGGCGG</td><td>CGG</td><td>||||||||[-||-||||||||]PAM</td><td>NA</td><td>Intergenic</td><td>NA</td><td>NA</td></tr>
<tr><td>GU305914.1</td><td>149488</td><td>149510</td><td>-</td><td>4</td><td>GGAGCAGGCGACGGCGGCGG</td><td>CGG</td><td>||-|--||[||-|||||||||]PAM</td><td>NA</td><td>Intergenic</td><td>NA</td><td>NA</td></tr>
<tr><td>GU305914.1</td><td>73792</td><td>73814</td><td>-</td><td>4</td><td>GGCGCCAGCAGCGGCGGCGG</td><td>AGG</td><td>||||---|[|-||||||||||]PAM</td><td>NA</td><td>Intergenic</td><td>NA</td><td>NA</td></tr>
<tr><td>GU305914.1</td><td>168817</td><td>168839</td><td>+</td><td>2</td><td>GGCGGTGGTGGCAGCGGCGG</td><td>CGG</td><td>||||||||[-|||-|||||||]PAM</td><td>NA</td><td>Intergenic</td><td>NA</td><td>NA</td></tr>
<tr><td>GU305914.1</td><td>62446</td><td>62468</td><td>+</td><td>3</td><td>GGCGGGGGTGGTGGCGGCGG</td><td>GGG</td><td>|||||-||[-||-||||||||]PAM</td><td>NA</td><td>Intergenic</td><td>NA</td><td>NA</td></tr>
<tr><td>GU305914.1</td><td>113930</td><td>113952</td><td>-</td><td>4</td><td>CGAGGTGTCGGCGGCGTCGGCGG</td><td>CGG</td><td>-|-||||-[|||||-||||||]PAM</td><td>NA</td><td>Intergenic</td><td>NA</td><td>NA</td></tr>
<tr><td>GU305914.1</td><td>48153</td><td>48175</td><td>+</td><td>4</td><td>TGTGGCGGCGGCGGTGGCGG</td><td>CGG</td><td>-|-||-||[||||||-|||||]PAM</td><td>NA</td><td>Intergenic</td><td>NA</td><td>NA</td></tr>
<tr><td>GU305914.1</td><td>18653</td><td>18675</td><td>-</td><td>4</td><td>GGCAGAGGTGGCGGAGGCGG</td><td>GGG</td><td>|||-|-||[-|||||-|||||]PAM</td><td>NA</td><td>Intergenic</td><td>NA</td><td>NA</td></tr>
<tr><td>GU305914.1</td><td>163749</td><td>163771</td><td>+</td><td>4</td><td>GTCGGCGGCGCCGGTGGCGG</td><td>CGG</td><td>|-|||-||[||-|||-|||||]PAM</td><td>NA</td><td>Intergenic</td><td>NA</td><td>NA</td></tr>
<tr><td>GU305914.1</td><td>163911</td><td>163933</td><td>+</td><td>3</td><td>GGCGGCGGCAGCGGCAGCGG</td><td>AGG</td><td>|||||-||[|-|||||-||||]PAM</td><td>NA</td><td>Intergenic</td><td>NA</td><td>NA</td></tr>
<tr><td>GU305914.1</td><td>45255</td><td>45277</td><td>-</td><td>4</td><td>GGTGGCGGTGGCGGCAGCGG</td><td>CGG</td><td>||-||-||[-||||||-||||]PAM</td><td>NA</td><td>Intergenic</td><td>NA</td><td>NA</td></tr>
<tr><td>GU305914.1</td><td>19936</td><td>19958</td><td>+</td><td>4</td><td>GACGGTAGCGGTGGTGGCGG</td><td>TGG</td><td>|-||||-|[|||-||-|||||]PAM</td><td>NA</td><td>Intergenic</td><td>NA</td><td>NA</td></tr>
<tr><td>GU305914.1</td><td>62155</td><td>62177</td><td>+</td><td>4</td><td>GGCGACGGCAGCGGCAGCGG</td><td>CGG</td><td>||||--||[|-|||||-||||]PAM</td><td>NA</td><td>Intergenic</td><td>NA</td><td>NA</td></tr>
<tr><td>GU305914.1</td><td>48150</td><td>48172</td><td>+</td><td>4</td><td>ACCTGTGGCGGCGGCGGTGG</td><td>CGG</td><td>--|-||||[|||||||||-||]PAM</td><td>NA</td><td>Intergenic</td><td>NA</td><td>NA</td></tr>
<tr><td>GU305914.1</td><td>62449</td><td>62471</td><td>+</td><td>3</td><td>GGGGGTGGTGGCGGCGGGGG</td><td>TGG</td><td>||-|||||[-||||||||-||]PAM</td><td>NA</td><td>Intergenic</td><td>NA</td><td>NA</td></tr>
<tr><td>GU305914.1</td><td>30006</td><td>30028</td><td>+</td><td>2</td><td>GGCGGCGGCGGCGGCGGCAG</td><td>TGG</td><td>|||||-||[||||||||||-|]PAM</td><td>NA</td><td>Intergenic</td><td>NA</td><td>NA</td></tr>
<tr><td>GU305914.1</td><td>149476</td><td>149498</td><td>-</td><td>2</td><td>GGCGGCGGCGGCGGCGGCAG</td><td>CGG</td><td>|||||-||[||||||||||-|]PAM</td><td>NA</td><td>Intergenic</td><td>NA</td><td>NA</td></tr>
<tr><td>GU305914.1</td><td>163902</td><td>163924</td><td>+</td><td>3</td><td>AGCGGCGGCGGCGGCGGCAG</td><td>CGG</td><td>-||||-||[||||||||||-|]PAM</td><td>NA</td><td>Intergenic</td><td>NA</td><td>NA</td></tr>
<tr><td>GU305914.1</td><td>62434</td><td>62456</td><td>+</td><td>4</td><td>GACGGCGGTGGCGGCGGGGG</td><td>TGG</td><td>|-|||-||[-||||||||-||]PAM</td><td>NA</td><td>Intergenic</td><td>NA</td><td>NA</td></tr>
<tr><td>GU305914.1</td><td>52719</td><td>52741</td><td>+</td><td>4</td><td>GCCGCTGGCGGTGGCGGTGG</td><td>CGG</td><td>|-||-|||[|||-|||||-||]PAM</td><td>NA</td><td>Intergenic</td><td>NA</td><td>NA</td></tr>
<tr><td>GU305914.1</td><td>90250</td><td>90272</td><td>-</td><td>3</td><td>GGCGGAGGCGGCGGAGCCGG</td><td>CGG</td><td>|||||-||[||||||-|-|||]PAM</td><td>NA</td><td>Intergenic</td><td>NA</td><td>NA</td></tr>
<tr><td>GU305914.1</td><td>113924</td><td>113946</td><td>-</td><td>4</td><td>GTCGGCGTCGGCGGCGGCAG</td><td>CGG</td><td>|-|||-|-[||||||||||-|]PAM</td><td>NA</td><td>Intergenic</td><td>NA</td><td>NA</td></tr>
<tr><td>GU305914.1</td><td>152553</td><td>152575</td><td>+</td><td>4</td><td>TGCGGCGGCGGCGGTGCCGG</td><td>CGG</td><td>-||||-||[||||||-|-|||]PAM</td><td>NA</td><td>Intergenic</td><td>NA</td><td>NA</td></tr>
<tr><td>GU305914.1</td><td>62437</td><td>62459</td><td>+</td><td>2</td><td>GGCGGTGGCGGCGGGGGTGG</td><td>TGG</td><td>||||||||[||||||-||-||]PAM</td><td>NA</td><td>Intergenic</td><td>NA</td><td>NA</td></tr>
<tr><td>GU305914.1</td><td>45258</td><td>45280</td><td>-</td><td>4</td><td>AACGGTGGCGGTGGCGGCAG</td><td>CGG</td><td>--||||||[|||-||||||-|]PAM</td><td>NA</td><td>Intergenic</td><td>NA</td><td>NA</td></tr>
<tr><td>GU305914.1</td><td>62452</td><td>62474</td><td>+</td><td>3</td><td>GGTGGTGGCGGCGGGGGTGG</td><td>CGG</td><td>||-|||||[||||||-||-||]PAM</td><td>NA</td><td>Intergenic</td><td>NA</td><td>NA</td></tr>
<tr><td>GU305914.1</td><td>163908</td><td>163930</td><td>+</td><td>3</td><td>GGCGGCGGCGGCAGCGGCAG</td><td>CGG</td><td>|||||-||[||||-|||||-|]PAM</td><td>NA</td><td>Intergenic</td><td>NA</td><td>NA</td></tr>
<tr><td>GU305914.1</td><td>113918</td><td>113940</td><td>-</td><td>4</td><td>GTCGGCGGCGGCAGCGGCGA</td><td>CGG</td><td>|-|||-||[||||-||||||-]PAM</td><td>NA</td><td>Intergenic</td><td>NA</td><td>NA</td></tr>
<tr><td>GU305914.1</td><td>113912</td><td>113934</td><td>-</td><td>4</td><td>GGCGGCAGCGGCGACGGCGT</td><td>CGG</td><td>|||||--|[|||||-|||||-]PAM</td><td>NA</td><td>Intergenic</td><td>NA</td><td>NA</td></tr>
<tr><td>GU305914.1</td><td>168829</td><td>168851</td><td>+</td><td>4</td><td>AGCGGCGGCGGCGGCGACAG</td><td>CGG</td><td>-||||-||[||||||||-|-|]PAM</td><td>NA</td><td>Intergenic</td><td>NA</td><td>NA</td></tr>
<tr><td>GU305914.1</td><td>170530</td><td>170552</td><td>-</td><td>4</td><td>GATGGTGGCGGCGGCGGCCA</td><td>CGG</td><td>|--|||||[||||||||||--]PAM</td><td>NA</td><td>Intergenic</td><td>NA</td><td>NA</td></tr>
<tr><td></td><td></td><td></td><td></td><td></td><td></td><td></td><td></td><td></td><td></td><td></td><td></td></tr>
<tr><td>T11822</td><td>GGTGGCGGCGGCGGCGGCGG</td><td>1000</td><td></td><td></td><td></td><td></td><td></td><td></td><td></td><td></td><td></td></tr>
<tr><td>Oligo fwd</td><td>CACCGGTGGCGGCGGCGGCGGCGG</td><td></td><td></td><td></td><td></td><td></td><td></td><td></td><td></td><td></td><td></td></tr>
<tr><td>Oligo rev</td><td>AAACCCGCCGCCGCCGCCGCCACC</td><td></td><td></td><td></td><td></td><td></td><td></td><td></td><td></td><td></td><td></td></tr>
<tr><td>Chromosome</td><td>start</td><td>end</td><td>strand</td><td>MM</td><td>target_seq</td><td>PAM</td><td>alignment</td><td>distance</td><td>position</td><td>gene name</td><td>gene id</td></tr>
<tr><td>GU305914.1</td><td>154407</td><td>154429</td><td>+</td><td>0</td><td>GGTGGCGGCGGCGGCGGCGG</td><td>CGG</td><td>||||||||[||||||||||||]PAM</td><td>NA</td><td>Intergenic</td><td>NA</td><td>NA</td></tr>
<tr><td>GU305914.1</td><td>58123</td><td>58145</td><td>-</td><td>0</td><td>GGTGGCGGCGGCGGCGGCGG</td><td>AGG</td><td>||||||||[||||||||||||]PAM</td><td>NA</td><td>Intergenic</td><td>NA</td><td>NA</td></tr>
<tr><td>GU305914.1</td><td>67742</td><td>67764</td><td>+</td><td>2</td><td>GGGAGCGGCGGCGGCGGCGG</td><td>CGG</td><td>||--||||[||||||||||||]PAM</td><td>NA</td><td>Intergenic</td><td>NA</td><td>NA</td></tr>
<tr><td>GU305914.1</td><td>149482</td><td>149504</td><td>-</td><td>2</td><td>GGCGACGGCGGCGGCGGCGG</td><td>CGG</td><td>||-|-|||[||||||||||||]PAM</td><td>NA</td><td>Intergenic</td><td>NA</td><td>NA</td></tr>
<tr><td>GU305914.1</td><td>154404</td><td>154426</td><td>+</td><td>2</td><td>GGCGGTGGCGGCGGCGGCGG</td><td>CGG</td><td>||-||-||[||||||||||||]PAM</td><td>NA</td><td>Intergenic</td><td>NA</td><td>NA</td></tr>
<tr><td>GU305914.1</td><td>58126</td><td>58148</td><td>-</td><td>2</td><td>GGCGGTGGCGGCGGCGGCGG</td><td>CGG</td><td>||-||-||[||||||||||||]PAM</td><td>NA</td><td>Intergenic</td><td>NA</td><td>NA</td></tr>
<tr><td>GU305914.1</td><td>30000</td><td>30022</td><td>+</td><td>2</td><td>GGTGCTGGCGGCGGCGGCGG</td><td>CGG</td><td>||||--||[||||||||||||]PAM</td><td>NA</td><td>Intergenic</td><td>NA</td><td>NA</td></tr>
<tr><td>GU305914.1</td><td>48159</td><td>48181</td><td>+</td><td>2</td><td>GGCGGCGGTGGCGGCGGCGG</td><td>TGG</td><td>||-|||||[-|||||||||||]PAM</td><td>NA</td><td>Intergenic</td><td>NA</td><td>NA</td></tr>
<tr><td>GU305914.1</td><td>149485</td><td>149507</td><td>-</td><td>3</td><td>GCAGGCGACGGCGGCGGCGG</td><td>CGG</td><td>|--||||-[||||||||||||]PAM</td><td>NA</td><td>Intergenic</td><td>NA</td><td>NA</td></tr>
<tr><td>GU305914.1</td><td>58129</td><td>58151</td><td>-</td><td>3</td><td>GCGGGCGGTGGCGGCGGCGG</td><td>CGG</td><td>|--|||||[-|||||||||||]PAM</td><td>NA</td><td>Intergenic</td><td>NA</td><td>NA</td></tr>
<tr><td>GU305914.1</td><td>56773</td><td>56795</td><td>+</td><td>4</td><td>CGCCGCAGCGGCGGCGGCGG</td><td>CGG</td><td>-|--||-|[||||||||||||]PAM</td><td>NA</td><td>Intergenic</td><td>NA</td><td>NA</td></tr>
<tr><td>GU305914.1</td><td>168820</td><td>168842</td><td>+</td><td>2</td><td>GGTGGTGGCAGCGGCGGCGG</td><td>CGG</td><td>|||||-||[|-||||||||||]PAM</td><td>NA</td><td>Intergenic</td><td>NA</td><td>NA</td></tr>
<tr><td>GU305914.1</td><td>154401</td><td>154423</td><td>+</td><td>4</td><td>CCGGGCGGTGGCGGCGGCGG</td><td>CGG</td><td>---|||||[-|||||||||||]PAM</td><td>NA</td><td>Intergenic</td><td>NA</td><td>NA</td></tr>
<tr><td>GU305914.1</td><td>48156</td><td>48178</td><td>+</td><td>2</td><td>GGCGGCGGCGGTGGCGGCGG</td><td>CGG</td><td>||-|||||[||-|||||||||]PAM</td><td>NA</td><td>Intergenic</td><td>NA</td><td>NA</td></tr>
<tr><td>GU305914.1</td><td>164971</td><td>164993</td><td>-</td><td>3</td><td>GGTGGGACCGGCGGCGGCGG</td><td>CGG</td><td>|||||---[||||||||||||]PAM</td><td>NA</td><td>Intergenic</td><td>NA</td><td>NA</td></tr>
<tr><td>GU305914.1</td><td>81246</td><td>81268</td><td>-</td><td>4</td><td>GATCGAGGAGGCGGCGGCGG</td><td>GGG</td><td>|-|-|-||[-|||||||||||]PAM</td><td>NA</td><td>Intergenic</td><td>NA</td><td>NA</td></tr>
<tr><td>GU305914.1</td><td>161754</td><td>161776</td><td>-</td><td>4</td><td>GGCCGAGGAGGCGGCGGCGG</td><td>AGG</td><td>||--|-||[-|||||||||||]PAM</td><td>NA</td><td>Intergenic</td><td>NA</td><td>NA</td></tr>
<tr><td>GU305914.1</td><td>73792</td><td>73814</td><td>-</td><td>4</td><td>GGCGCCAGCAGCGGCGGCGG</td><td>AGG</td><td>||-|-|-|[|-||||||||||]PAM</td><td>NA</td><td>Intergenic</td><td>NA</td><td>NA</td></tr>
<tr><td>GU305914.1</td><td>149488</td><td>149510</td><td>-</td><td>4</td><td>GGAGCAGGCGACGGCGGCGG</td><td>CGG</td><td>||-|--||[||-|||||||||]PAM</td><td>NA</td><td>Intergenic</td><td>NA</td><td>NA</td></tr>
<tr><td>GU305914.1</td><td>48153</td><td>48175</td><td>+</td><td>2</td><td>TGTGGCGGCGGCGGTGGCGG</td><td>CGG</td><td>-|||||||[|||||-||||||]PAM</td><td>NA</td><td>Intergenic</td><td>NA</td><td>NA</td></tr>
<tr><td>GU305914.1</td><td>62446</td><td>62468</td><td>+</td><td>4</td><td>GGCGGGGGTGGTGGCGGCGG</td><td>GGG</td><td>||-||-||[-||-||||||||]PAM</td><td>NA</td><td>Intergenic</td><td>NA</td><td>NA</td></tr>
<tr><td>GU305914.1</td><td>114658</td><td>114680</td><td>-</td><td>4</td><td>GGCGGTGGTGGTGGCGGCGG</td><td>CGG</td><td>||-||-||[-||-||||||||]PAM</td><td>NA</td><td>Intergenic</td><td>NA</td><td>NA</td></tr>
</table>

| | | | | | | | | | | | |
|---|---|---|---|---|---|---|---|---|---|---|---|
| GU305914.1 | 161757 | 161779 | - | 4 | GGAGGCCGAGGAGGCGGCGG | CGG | ||-|||-|[-||-||||||||]PAM | NA | Intergenic | NA | NA |
| GU305914.1 | 168817 | 168839 | + | 4 | GGCGGTGGTGGCAGCGGCGG | CGG | ||-||-||[-|||-|||||||]PAM | NA | Intergenic | NA | NA |
| GU305914.1 | 73795 | 73817 | - | 4 | AGTGGCGCCAGCAGCGGCGG | CGG | -||||||-[|-||-|||||||]PAM | NA | Intergenic | NA | NA |
| GU305914.1 | 90259 | 90281 | - | 4 | GCGGGCGACGGCGGAGGCGG | CGG | |--||||-[||||||-|||||]PAM | NA | Intergenic | NA | NA |
| GU305914.1 | 62672 | 62694 | - | 4 | GCTGCCGCCGGCGGGGGCGG | CGG | |-||-||-[||||||-|||||]PAM | NA | Intergenic | NA | NA |
| GU305914.1 | 45255 | 45277 | - | 2 | GGTGGCGGTGGCGGCAGCGG | CGG | ||||||||[-||||||-||||]PAM | NA | Intergenic | NA | NA |
| GU305914.1 | 163749 | 163771 | + | 4 | GTCGGCGGCGCCGGTGGCGG | CGG | |--|||||[||-|||-|||||]PAM | NA | Intergenic | NA | NA |
| GU305914.1 | 163911 | 163933 | + | 3 | GGCGGCGGCAGCGGCAGCGG | AGG | ||-|||||[|-|||||-||||]PAM | NA | Intergenic | NA | NA |
| GU305914.1 | 110125 | 110147 | - | 4 | GGACGCGACGGCGGCCGCGG | CGG | ||--|||-[|||||||-||||]PAM | NA | Intergenic | NA | NA |
| GU305914.1 | 62155 | 62177 | + | 4 | GGCGACGGCAGCGGCAGCGG | CGG | ||-|-|||[|-|||||-||||]PAM | NA | Intergenic | NA | NA |
| GU305914.1 | 30006 | 30028 | + | 2 | GGCGGCGGCGGCGGCGGCAG | TGG | ||-|||||[||||||||||-|]PAM | NA | Intergenic | NA | NA |
| GU305914.1 | 149476 | 149498 | - | 2 | GGCGGCGGCGGCGGCGGCAG | CGG | ||-|||||[||||||||||-|]PAM | NA | Intergenic | NA | NA |
| GU305914.1 | 163902 | 163924 | + | 3 | AGCGGCGGCGGCGGCGGCAG | CGG | -|-|||||[||||||||||-|]PAM | NA | Intergenic | NA | NA |
| GU305914.1 | 62434 | 62456 | + | 4 | GACGGCGGTGGCGGCGGGGG | TGG | |--|||||[-||||||||-||]PAM | NA | Intergenic | NA | NA |
| GU305914.1 | 62449 | 62471 | + | 4 | GGGGGTGGTGGCGGCGGGGG | TGG | ||-||-||[-||||||||-||]PAM | NA | Intergenic | NA | NA |
| GU305914.1 | 113924 | 113946 | - | 4 | GTCGGCGTCGGCGGCGGCAG | CGG | |--||||-[||||||||||-|]PAM | NA | Intergenic | NA | NA |
| GU305914.1 | 152553 | 152575 | + | 4 | TGCGGCGGCGGCGGTGCCGG | CGG | -|-|||||[||||||-|-|||]PAM | NA | Intergenic | NA | NA |
| GU305914.1 | 90250 | 90272 | - | 4 | GGCGGAGGCGGCGGAGCCGG | CGG | ||-||-||[||||||-|-|||]PAM | NA | Intergenic | NA | NA |
| GU305914.1 | 74673 | 74695 | + | 4 | GGTGCCGTCGGTGGCGGTGG | CGG | ||||-||-[|||-|||||-||]PAM | NA | Intergenic | NA | NA |
| GU305914.1 | 163908 | 163930 | + | 3 | GGCGGCGGCGGCAGCGGCAG | CGG | ||-|||||[||||-|||||-|]PAM | NA | Intergenic | NA | NA |
| GU305914.1 | 62452 | 62474 | + | 3 | GGTGGTGGCGGCGGGGGTGG | CGG | |||||-||[||||||-||-||]PAM | NA | Intergenic | NA | NA |
| GU305914.1 | 62437 | 62459 | + | 4 | GGCGGTGGCGGCGGGGGTGG | TGG | ||-||-||[||||||-||-||]PAM | NA | Intergenic | NA | NA |
| GU305914.1 | 113918 | 113940 | - | 4 | GTCGGCGGCGGCAGCGGCGA | CGG | |--|||||[||||-||||||-]PAM | NA | Intergenic | NA | NA |
| GU305914.1 | 113912 | 113934 | - | 4 | GGCGGCAGCGGCGACGGCGT | CGG | ||-|||-|[|||||-|||||-]PAM | NA | Intergenic | NA | NA |
| GU305914.1 | 168829 | 168851 | + | 4 | AGCGGCGGCGGCGGCGACAG | CGG | -|-|||||[||||||||-|-|]PAM | NA | Intergenic | NA | NA |
| GU305914.1 | 170530 | 170552 | - | 4 | GATGGTGGCGGCGGCGGCCA | CGG | |-|||-||[||||||||||--]PAM | NA | Intergenic | NA | NA |
| T11823 | GCGGCGGCGGCGGCTGTTGCTGG | 1000 | | | | | | | | | |
| Oligo fwd | CACCGCGGCGGCGGCGGCTGTTGC | | | | | | | | | | |
| Oligo rev | AAACGCAACAGCCGCCGCCGCCGC | | | | | | | | | | |
| Chromosome | start | end | strand | MM | target_seq | PAM | alignment | distance | position | gene name | gene id |
| GU305914.1 | 154417 | 154439 | + | 0 | GCGGCGGCGGCGGCTGTTGC | TGG | ||||||||[||||||||||||]PAM | NA | Intergenic | NA | NA |
| GU305914.1 | 74703 | 74725 | - | 4 | GTGGCGGTGGCGGCGGTTCC | GGG | |-|||||-[||||||-|||-|]PAM | NA | Intergenic | NA | NA |
| T11824 | TCTCGCATCCTGAAAAAGAAAGG | 1000 | | | | | | | | | |
| Oligo fwd | CACCTCTCGCATCCTGAAAAAGAA | | | | | | | | | | |
| Oligo rev | AAACTTCTTTTTCAGGATGCGAGA | | | | | | | | | | |
| Chromosome | start | end | strand | MM | target_seq | PAM | alignment | distance | position | gene name | gene id |
| GU305914.1 | 154453 | 154475 | + | 0 | TCTCGCATCCTGAAAAAGAA | AGG | ||||||||[||||||||||||]PAM | NA | Intergenic | NA | NA |
| T11825 | CTGAAAAAGAAAGGAGAGAACGG | 1000 | | | | | | | | | |
| Oligo fwd | CACCCTGAAAAAGAAAGGAGAGAA | | | | | | | | | | |
| Oligo rev | AAACTTCTCTCCTTTCTTTTTCAG | | | | | | | | | | |
| Chromosome | start | end | strand | MM | target_seq | PAM | alignment | distance | position | gene name | gene id |
| GU305914.1 | 154462 | 154484 | + | 0 | CTGAAAAAGAAAGGAGAGAA | CGG | ||||||||[||||||||||||]PAM | NA | Intergenic | NA | NA |
| T11826 | TGAAAAAGAAAGGAGAGAACGGG | 1000 | | | | | | | | | |
| Oligo fwd | CACCTGAAAAAGAAAGGAGAGAAC | | | | | | | | | | |
| Oligo rev | AAACGTTCTCTCCTTTCTTTTTCA | | | | | | | | | | |
| Chromosome | start | end | strand | MM | target_seq | PAM | alignment | distance | position | gene name | gene id |
| GU305914.1 | 154463 | 154485 | + | 0 | TGAAAAAGAAAGGAGAGAAC | GGG | ||||||||[||||||||||||]PAM | NA | Intergenic | NA | NA |
| T11827 | AAAAGAAAGGAGAGAACGGGTGG | 1000 | | | | | | | | | |
| Oligo fwd | CACCAAAAGAAAGGAGAGAACGGG | | | | | | | | | | |
| Oligo rev | AAACCCCGTTCTCTCCTTTCTTTT | | | | | | | | | | |
| Chromosome | start | end | strand | MM | target_seq | PAM | alignment | distance | position | gene name | gene id |
| GU305914.1 | 154466 | 154488 | + | 0 | AAAAGAAAGGAGAGAACGGG | TGG | ||||||||[||||||||||||]PAM | NA | Intergenic | NA | NA |
| T11828 | AAAGAAAGGAGAGAACGGGTGGG | 1000 | | | | | | | | | |
| Oligo fwd | CACCAAAGAAAGGAGAGAACGGGT | | | | | | | | | | |
| Oligo rev | AAACACCCGTTCTCTCCTTTCTTT | | | | | | | | | | |
| Chromosome | start | end | strand | MM | target_seq | PAM | alignment | distance | position | gene name | gene id |
| GU305914.1 | 154467 | 154489 | + | 0 | AAAGAAAGGAGAGAACGGGT | GGG | ||||||||[||||||||||||]PAM | NA | Intergenic | NA | NA |
| T11829 | GAGAGAACGGGTGGGAGCCGAGG | 1000 | | | | | | | | | |
| Oligo fwd | CACCGAGAGAACGGGTGGGAGCCG | | | | | | | | | | |
| Oligo rev | AAACCGGCTCCCACCCGTTCTCTC | | | | | | | | | | |
| Chromosome | start | end | strand | MM | target_seq | PAM | alignment | distance | position | gene name | gene id |
| GU305914.1 | 154475 | 154497 | + | 0 | GAGAGAACGGGTGGGAGCCG | AGG | ||||||||[||||||||||||]PAM | NA | Intergenic | NA | NA |
| T11830 | CAGAAAATTGCGTACAGATGCGG | 1000 | | | | | | | | | |
| Oligo fwd | CACCCAGAAAATTGCGTACAGATG | | | | | | | | | | |
| Oligo rev | AAACCATCTGTACGCAATTTTCTG | | | | | | | | | | |
| Chromosome | start | end | strand | MM | target_seq | PAM | alignment | distance | position | gene name | gene id |
| GU305914.1 | 154527 | 154549 | + | 0 | CAGAAAATTGCGTACAGATG | CGG | ||||||||[||||||||||||]PAM | NA | Intergenic | NA | NA |
| T11831 | AGAAAATTGCGTACAGATGCGGG | 1000 | | | | | | | | | |
| Oligo fwd | CACCAGAAAATTGCGTACAGATGC | | | | | | | | | | |
| Oligo rev | AAACGCATCTGTACGCAATTTTCT | | | | | | | | | | |
| Chromosome | start | end | strand | MM | target_seq | PAM | alignment | distance | position | gene name | gene id |
| GU305914.1 | 154528 | 154550 | + | 0 | AGAAAATTGCGTACAGATGC | GGG | ||||||||[||||||||||||]PAM | NA | Intergenic | NA | NA |
| T11832 | AAATTGCGTACAGATGCGGGTGG | 1000 | | | | | | | | | |
| Oligo fwd | CACCAAATTGCGTACAGATGCGGG | | | | | | | | | | |
| Oligo rev | AAACCCCGCATCTGTACGCAATTT | | | | | | | | | | |
| Chromosome | start | end | strand | MM | target_seq | PAM | alignment | distance | position | gene name | gene id |
| GU305914.1 | 154531 | 154553 | + | 0 | AAATTGCGTACAGATGCGGG | TGG | ||||||||[||||||||||||]PAM | NA | Intergenic | NA | NA |
| T11833 | CCTTATTTTTGCAGAAAAAATAGG | 1000 | | | | | | | | | |

Oligo fwd CACCCCTTATTTTTGCAGAAAAAT
Oligo rev AAACATTTTTCTGCAAAAATAAGG
Chromosome start end strand MM target_seq PAM alignment distance position gene name gene id
GU305914.1 154569 154591 + 0 CCTTATTTTTGCAGAAAAAT AGG ||||||||[||||||||||||]PAM NA Intergenic NA NA

T11834 TTATCTCAGTTCATACATCTAGG 1000
Oligo fwd CACCTTATCTCAGTTCATACATCT
Oligo rev AAACAGATGTATGAACTGAGATAA
Chromosome start end strand MM target_seq PAM alignment distance position gene name gene id
GU305914.1 154631 154653 + 0 TTATCTCAGTTCATACATCT AGG ||||||||[||||||||||||]PAM NA Intergenic NA NA

T11835 TATCTCAGTTCATACATCTAGGG 1000
Oligo fwd CACCTATCTCAGTTCATACATCTA
Oligo rev AAACTAGATGTATGAACTGAGATA
Chromosome start end strand MM target_seq PAM alignment distance position gene name gene id
GU305914.1 154632 154654 + 0 TATCTCAGTTCATACATCTA GGG ||||||||[||||||||||||]PAM NA Intergenic NA NA

T11836 GGAAGTTTAAGATGATTGATTGG 1000
Oligo fwd CACCGGAAGTTTAAGATGATTGAT
Oligo rev AAACATCAATCATCTTAAACTTCC
Chromosome start end strand MM target_seq PAM alignment distance position gene name gene id
GU305914.1 154653 154675 + 0 GGAAGTTTAAGATGATTGAT TGG ||||||||[||||||||||||]PAM NA Intergenic NA NA

T11837 TCATTCTTGATGAAAATGCAAGG 1000
Oligo fwd CACCTCATTCTTGATGAAAATGCA
Oligo rev AAACTGCATTTTCATCAAGAATGA
Chromosome start end strand MM target_seq PAM alignment distance position gene name gene id
GU305914.1 154677 154699 + 0 TCATTCTTGATGAAAATGCA AGG ||||||||[||||||||||||]PAM NA Intergenic NA NA

T11838 CATTCTTGATGAAAATGCAAGGG 1000
Oligo fwd CACCCATTCTTGATGAAAATGCAA
Oligo rev AAACTTGCATTTTCATCAAGAATG
Chromosome start end strand MM target_seq PAM alignment distance position gene name gene id
GU305914.1 154678 154700 + 0 CATTCTTGATGAAAATGCAA GGG ||||||||[||||||||||||]PAM NA Intergenic NA NA

T11839 GGGATATATGTATATGTTGATGG 1000
Oligo fwd CACCGGGATATATGTATATGTTGA
Oligo rev AAACTCAACATATACATATATCCC
Chromosome start end strand MM target_seq PAM alignment distance position gene name gene id
GU305914.1 154698 154720 + 0 GGGATATATGTATATGTTGA TGG ||||||||[||||||||||||]PAM NA Intergenic NA NA

T11840 TGGTATTTTCGCCGACCCAGAGG 1000
Oligo fwd CACCTGGTATTTTCGCCGACCCAG
Oligo rev AAACCTGGGTCGGCGAAAATACCA
Chromosome start end strand MM target_seq PAM alignment distance position gene name gene id
GU305914.1 154718 154740 + 0 TGGTATTTTCGCCGACCCAG AGG ||||||||[||||||||||||]PAM NA Intergenic NA NA

T11841 TTCGCCGACCCAGAGGCGAATGG 1000
Oligo fwd CACCTTCGCCGACCCAGAGGCGAA
Oligo rev AAACTTCGCCTCTGGGTCGGCGAA
Chromosome start end strand MM target_seq PAM alignment distance position gene name gene id
GU305914.1 154725 154747 + 0 TTCGCCGACCCAGAGGCGAA TGG ||||||||[||||||||||||]PAM NA Intergenic NA NA

T11842 TCGCCGACCCAGAGGCGAATGGG 1000
Oligo fwd CACCTCGCCGACCCAGAGGCGAAT
Oligo rev AAACATTCGCCTCTGGGTCGGCGA
Chromosome start end strand MM target_seq PAM alignment distance position gene name gene id
GU305914.1 154726 154748 + 0 TCGCCGACCCAGAGGCGAAT GGG ||||||||[||||||||||||]PAM NA Intergenic NA NA

T11843 CGCCGACCCAGAGGCGAATGGGG 1000
Oligo fwd CACCCGCCGACCCAGAGGCGAATG
Oligo rev AAACCATTCGCCTCTGGGTCGGCG
Chromosome start end strand MM target_seq PAM alignment distance position gene name gene id
GU305914.1 154727 154749 + 0 CGCCGACCCAGAGGCGAATG GGG ||||||||[||||||||||||]PAM NA Intergenic NA NA

T11844 GCCGACCCAGAGGCGAATGGGGG 1000
Oligo fwd CACCGCCGACCCAGAGGCGAATGG
Oligo rev AAACCCATTCGCCTCTGGGTCGGC
Chromosome start end strand MM target_seq PAM alignment distance position gene name gene id
GU305914.1 154728 154750 + 0 GCCGACCCAGAGGCGAATGG GGG ||||||||[||||||||||||]PAM NA Intergenic NA NA

T11845 CCGACCCAGAGGCGAATGGGGGG 1000
Oligo fwd CACCCCGACCCAGAGGCGAATGGG
Oligo rev AAACCCCATTCGCCTCTGGGTCGG
Chromosome start end strand MM target_seq PAM alignment distance position gene name gene id
GU305914.1 154729 154751 + 0 CCGACCCAGAGGCGAATGGG GGG ||||||||[||||||||||||]PAM NA Intergenic NA NA

T11846 CCAGAGGCGAATGGGGGGTATGG 1000
Oligo fwd CACCCCAGAGGCGAATGGGGGGTA
Oligo rev AAACTACCCCCCATTCGCCTCTGG
Chromosome start end strand MM target_seq PAM alignment distance position gene name gene id
GU305914.1 154734 154756 + 0 CCAGAGGCGAATGGGGGGTA TGG ||||||||[||||||||||||]PAM NA Intergenic NA NA

T11847 TGTTATTTGTAATCTATGTGTGG 1000
Oligo fwd CACCTGTTATTTGTAATCTATGTG
Oligo rev AAACCACATAGATTACAAATAACA
Chromosome start end strand MM target_seq PAM alignment distance position gene name gene id
GU305914.1 154782 154804 + 0 TGTTATTTGTAATCTATGTG TGG ||||||||[||||||||||||]PAM NA Intergenic NA NA

<table>
<tr><td>T11848</td><td>CCCGAGCCGCTTCGTGTTGATGG</td><td>1000</td></tr>
<tr><td>Oligo fwd</td><td>CACCCCCGAGCCGCTTCGTGTTGA</td></tr>
<tr><td>Oligo rev</td><td>AAACTCAACACGAAGCGGCTCGGG</td></tr>
<tr><td>Chromosome</td><td>start</td><td>end</td><td>strand</td><td>MM</td><td>target_seq</td><td>PAM</td><td>alignment</td><td>distance</td><td>position</td><td>gene name</td><td>gene id</td></tr>
<tr><td>GU305914.1</td><td>154810</td><td>154832</td><td>+</td><td>0</td><td>CCCGAGCCGCTTCGTGTTGA</td><td>TGG</td><td>||||||||[||||||||||||]PAM</td><td>NA</td><td>Intergenic</td><td>NA</td><td>NA</td></tr>
<tr><td>T11849</td><td>TAAGATCTACTACAACGCAGCGG</td><td>1000</td></tr>
<tr><td>Oligo fwd</td><td>CACCTAAGATCTACTACAACGCAG</td></tr>
<tr><td>Oligo rev</td><td>AAACCTGCGTTGTAGTAGATCTTA</td></tr>
<tr><td>Chromosome</td><td>start</td><td>end</td><td>strand</td><td>MM</td><td>target_seq</td><td>PAM</td><td>alignment</td><td>distance</td><td>position</td><td>gene name</td><td>gene id</td></tr>
<tr><td>GU305914.1</td><td>154898</td><td>154920</td><td>+</td><td>0</td><td>TAAGATCTACTACAACGCAG</td><td>CGG</td><td>||||||||[||||||||||||]PAM</td><td>NA</td><td>Intergenic</td><td>NA</td><td>NA</td></tr>
<tr><td>T11850</td><td>TCTACTACAACGCAGCGGCATGG</td><td>1000</td></tr>
<tr><td>Oligo fwd</td><td>CACCTCTACTACAACGCAGCGGCA</td></tr>
<tr><td>Oligo rev</td><td>AAACTGCCGCTGCGTTGTAGTAGA</td></tr>
<tr><td>Chromosome</td><td>start</td><td>end</td><td>strand</td><td>MM</td><td>target_seq</td><td>PAM</td><td>alignment</td><td>distance</td><td>position</td><td>gene name</td><td>gene id</td></tr>
<tr><td>GU305914.1</td><td>154903</td><td>154925</td><td>+</td><td>0</td><td>TCTACTACAACGCAGCGGCA</td><td>TGG</td><td>||||||||[||||||||||||]PAM</td><td>NA</td><td>Intergenic</td><td>NA</td><td>NA</td></tr>
<tr><td>T11851</td><td>CTACTACAACGCAGCGGCATGGG</td><td>1000</td></tr>
<tr><td>Oligo fwd</td><td>CACCCTACTACAACGCAGCGGCAT</td></tr>
<tr><td>Oligo rev</td><td>AAACATGCCGCTGCGTTGTAGTAG</td></tr>
<tr><td>Chromosome</td><td>start</td><td>end</td><td>strand</td><td>MM</td><td>target_seq</td><td>PAM</td><td>alignment</td><td>distance</td><td>position</td><td>gene name</td><td>gene id</td></tr>
<tr><td>GU305914.1</td><td>154904</td><td>154926</td><td>+</td><td>0</td><td>CTACTACAACGCAGCGGCAT</td><td>GGG</td><td>||||||||[||||||||||||]PAM</td><td>NA</td><td>Intergenic</td><td>NA</td><td>NA</td></tr>
<tr><td>T11852</td><td>TACTACAACGCAGCGGCATGGGG</td><td>1000</td></tr>
<tr><td>Oligo fwd</td><td>CACCTACTACAACGCAGCGGCATG</td></tr>
<tr><td>Oligo rev</td><td>AAACCATGCCGCTGCGTTGTAGTA</td></tr>
<tr><td>Chromosome</td><td>start</td><td>end</td><td>strand</td><td>MM</td><td>target_seq</td><td>PAM</td><td>alignment</td><td>distance</td><td>position</td><td>gene name</td><td>gene id</td></tr>
<tr><td>GU305914.1</td><td>154905</td><td>154927</td><td>+</td><td>0</td><td>TACTACAACGCAGCGGCATG</td><td>GGG</td><td>||||||||[||||||||||||]PAM</td><td>NA</td><td>Intergenic</td><td>NA</td><td>NA</td></tr>
<tr><td>T11853</td><td>TCTATTTATCTAAAATGATATGG</td><td>1000</td></tr>
<tr><td>Oligo fwd</td><td>CACCTCTATTTATCTAAAATGATA</td></tr>
<tr><td>Oligo rev</td><td>AAACTATCATTTTAGATAAATAGA</td></tr>
<tr><td>Chromosome</td><td>start</td><td>end</td><td>strand</td><td>MM</td><td>target_seq</td><td>PAM</td><td>alignment</td><td>distance</td><td>position</td><td>gene name</td><td>gene id</td></tr>
<tr><td>GU305914.1</td><td>154968</td><td>154990</td><td>+</td><td>0</td><td>TCTATTTATCTAAAATGATA</td><td>TGG</td><td>||||||||[||||||||||||]PAM</td><td>NA</td><td>Intergenic</td><td>NA</td><td>NA</td></tr>
<tr><td>T11854</td><td>TATATCTACCTAAAAAATTATGG</td><td>1000</td></tr>
<tr><td>Oligo fwd</td><td>CACCTATATCTACCTAAAAAATTA</td></tr>
<tr><td>Oligo rev</td><td>AAACTAATTTTTTAGGTAGATATA</td></tr>
<tr><td>Chromosome</td><td>start</td><td>end</td><td>strand</td><td>MM</td><td>target_seq</td><td>PAM</td><td>alignment</td><td>distance</td><td>position</td><td>gene name</td><td>gene id</td></tr>
<tr><td>GU305914.1</td><td>155002</td><td>155024</td><td>+</td><td>0</td><td>TATATCTACCTAAAAAATTA</td><td>TGG</td><td>||||||||[||||||||||||]PAM</td><td>NA</td><td>Intergenic</td><td>NA</td><td>NA</td></tr>
<tr><td>T11855</td><td>CTACCTAAAAAATTATGGAGTGG</td><td>1000</td></tr>
<tr><td>Oligo fwd</td><td>CACCCTACCTAAAAAATTATGGAG</td></tr>
<tr><td>Oligo rev</td><td>AAACCTCCATAATTTTTTAGGTAG</td></tr>
<tr><td>Chromosome</td><td>start</td><td>end</td><td>strand</td><td>MM</td><td>target_seq</td><td>PAM</td><td>alignment</td><td>distance</td><td>position</td><td>gene name</td><td>gene id</td></tr>
<tr><td>GU305914.1</td><td>155007</td><td>155029</td><td>+</td><td>0</td><td>CTACCTAAAAAATTATGGAG</td><td>TGG</td><td>||||||||[||||||||||||]PAM</td><td>NA</td><td>Intergenic</td><td>NA</td><td>NA</td></tr>
<tr><td>T11856</td><td>TATGGAGTGGATATCATTCTCGG</td><td>1000</td></tr>
<tr><td>Oligo fwd</td><td>CACCTATGGAGTGGATATCATTCT</td></tr>
<tr><td>Oligo rev</td><td>AAACAGAATGATATCCACTCCATA</td></tr>
<tr><td>Chromosome</td><td>start</td><td>end</td><td>strand</td><td>MM</td><td>target_seq</td><td>PAM</td><td>alignment</td><td>distance</td><td>position</td><td>gene name</td><td>gene id</td></tr>
<tr><td>GU305914.1</td><td>155020</td><td>155042</td><td>+</td><td>0</td><td>TATGGAGTGGATATCATTCT</td><td>CGG</td><td>||||||||[||||||||||||]PAM</td><td>NA</td><td>Intergenic</td><td>NA</td><td>NA</td></tr>
<tr><td>T11857</td><td>ATGGAGTGGATATCATTCTCGGG</td><td>1000</td></tr>
<tr><td>Oligo fwd</td><td>CACCATGGAGTGGATATCATTCTC</td></tr>
<tr><td>Oligo rev</td><td>AAACGAGAATGATATCCACTCCAT</td></tr>
<tr><td>Chromosome</td><td>start</td><td>end</td><td>strand</td><td>MM</td><td>target_seq</td><td>PAM</td><td>alignment</td><td>distance</td><td>position</td><td>gene name</td><td>gene id</td></tr>
<tr><td>GU305914.1</td><td>155021</td><td>155043</td><td>+</td><td>0</td><td>ATGGAGTGGATATCATTCTC</td><td>GGG</td><td>||||||||[||||||||||||]PAM</td><td>NA</td><td>Intergenic</td><td>NA</td><td>NA</td></tr>
<tr><td>T11858</td><td>ATCATTCTCGGGCGCGAATGCGG</td><td>1000</td></tr>
<tr><td>Oligo fwd</td><td>CACCATCATTCTCGGGCGCGAATG</td></tr>
<tr><td>Oligo rev</td><td>AAACCATTCGCGCCCGAGAATGAT</td></tr>
<tr><td>Chromosome</td><td>start</td><td>end</td><td>strand</td><td>MM</td><td>target_seq</td><td>PAM</td><td>alignment</td><td>distance</td><td>position</td><td>gene name</td><td>gene id</td></tr>
<tr><td>GU305914.1</td><td>155032</td><td>155054</td><td>+</td><td>0</td><td>ATCATTCTCGGGCGCGAATG</td><td>CGG</td><td>||||||||[||||||||||||]PAM</td><td>NA</td><td>Intergenic</td><td>NA</td><td>NA</td></tr>
<tr><td>T11859</td><td>CGGTCGCGTAGACTATACACTGG</td><td>1000</td></tr>
<tr><td>Oligo fwd</td><td>CACCCGGTCGCGTAGACTATACAC</td></tr>
<tr><td>Oligo rev</td><td>AAACGTGTATAGTCTACGCGACCG</td></tr>
<tr><td>Chromosome</td><td>start</td><td>end</td><td>strand</td><td>MM</td><td>target_seq</td><td>PAM</td><td>alignment</td><td>distance</td><td>position</td><td>gene name</td><td>gene id</td></tr>
<tr><td>GU305914.1</td><td>155052</td><td>155074</td><td>+</td><td>0</td><td>CGGTCGCGTAGACTATACAC</td><td>TGG</td><td>||||||||[||||||||||||]PAM</td><td>NA</td><td>Intergenic</td><td>NA</td><td>NA</td></tr>
<tr><td>T11860</td><td>ACTATACACTGGAAATGTGATGG</td><td>1000</td></tr>
<tr><td>Oligo fwd</td><td>CACCACTATACACTGGAAATGTGA</td></tr>
<tr><td>Oligo rev</td><td>AAACTCACATTTCCAGTGTATAGT</td></tr>
<tr><td>Chromosome</td><td>start</td><td>end</td><td>strand</td><td>MM</td><td>target_seq</td><td>PAM</td><td>alignment</td><td>distance</td><td>position</td><td>gene name</td><td>gene id</td></tr>
<tr><td>GU305914.1</td><td>155063</td><td>155085</td><td>+</td><td>0</td><td>ACTATACACTGGAAATGTGA</td><td>TGG</td><td>||||||||[||||||||||||]PAM</td><td>NA</td><td>Intergenic</td><td>NA</td><td>NA</td></tr>
<tr><td>T11861</td><td>ATACACTGGAAATGTGATGGTGG</td><td>1000</td></tr>
<tr><td>Oligo fwd</td><td>CACCATACACTGGAAATGTGATGG</td></tr>
<tr><td>Oligo rev</td><td>AAACCCATCACATTTCCAGTGTAT</td></tr>
<tr><td>Chromosome</td><td>start</td><td>end</td><td>strand</td><td>MM</td><td>target_seq</td><td>PAM</td><td>alignment</td><td>distance</td><td>position</td><td>gene name</td><td>gene id</td></tr>
<tr><td>GU305914.1</td><td>155066</td><td>155088</td><td>+</td><td>0</td><td>ATACACTGGAAATGTGATGG</td><td>TGG</td><td>||||||||[||||||||||||]PAM</td><td>NA</td><td>Intergenic</td><td>NA</td><td>NA</td></tr>
<tr><td>T11862</td><td>AACTAACATTACATTTTATGAGG</td><td>1000</td></tr>
<tr><td>Oligo fwd</td><td>CACCAACTAACATTACATTTTATG</td></tr>
<tr><td>Oligo rev</td><td>AAACCATAAAATGTAATGTTAGTT</td></tr>
<tr><td>Chromosome</td><td>start</td><td>end</td><td>strand</td><td>MM</td><td>target_seq</td><td>PAM</td><td>alignment</td><td>distance</td><td>position</td><td>gene name</td><td>gene id</td></tr>
<tr><td>GU305914.1</td><td>155095</td><td>155117</td><td>+</td><td>0</td><td>AACTAACATTACATTTTATG</td><td>AGG</td><td>||||||||[||||||||||||]PAM</td><td>NA</td><td>Intergenic</td><td>NA</td><td>NA</td></tr>
</table>

<table>
<tr><td>T11863</td><td>ATTGTAGACAACATTATAGACGG</td><td>1000</td><td></td><td></td><td></td><td></td><td></td><td></td><td></td><td></td><td></td></tr>
<tr><td>Oligo fwd</td><td>CACCATTGTAGACAACATTATAGA</td><td></td><td></td><td></td><td></td><td></td><td></td><td></td><td></td><td></td><td></td></tr>
<tr><td>Oligo rev</td><td>AAACTCTATAATGTTGTCTACAAT</td><td></td><td></td><td></td><td></td><td></td><td></td><td></td><td></td><td></td><td></td></tr>
<tr><td>Chromosome</td><td>start</td><td>end</td><td>strand</td><td>MM</td><td>target_seq</td><td>PAM</td><td>alignment</td><td>distance</td><td>position</td><td>gene name</td><td>gene id</td></tr>
<tr><td>GU305914.1</td><td>155125</td><td>155147</td><td>+</td><td>0</td><td>ATTGTAGACAACATTATAGA</td><td>CGG</td><td>||||||||[||||||||||||]PAM</td><td>NA</td><td>Intergenic</td><td>NA</td><td>NA</td></tr>
<tr><td>T11864</td><td>TTGTAGACAACATTATAGACGGG</td><td>1000</td><td></td><td></td><td></td><td></td><td></td><td></td><td></td><td></td><td></td></tr>
<tr><td>Oligo fwd</td><td>CACCTTGTAGACAACATTATAGAC</td><td></td><td></td><td></td><td></td><td></td><td></td><td></td><td></td><td></td><td></td></tr>
<tr><td>Oligo rev</td><td>AAACGTCTATAATGTTGTCTACAA</td><td></td><td></td><td></td><td></td><td></td><td></td><td></td><td></td><td></td><td></td></tr>
<tr><td>Chromosome</td><td>start</td><td>end</td><td>strand</td><td>MM</td><td>target_seq</td><td>PAM</td><td>alignment</td><td>distance</td><td>position</td><td>gene name</td><td>gene id</td></tr>
<tr><td>GU305914.1</td><td>155126</td><td>155148</td><td>+</td><td>0</td><td>TTGTAGACAACATTATAGAC</td><td>GGG</td><td>||||||||[||||||||||||]PAM</td><td>NA</td><td>Intergenic</td><td>NA</td><td>NA</td></tr>
<tr><td>T11865</td><td>TGTAGACAACATTATAGACGGGG</td><td>1000</td><td></td><td></td><td></td><td></td><td></td><td></td><td></td><td></td><td></td></tr>
<tr><td>Oligo fwd</td><td>CACCTGTAGACAACATTATAGACG</td><td></td><td></td><td></td><td></td><td></td><td></td><td></td><td></td><td></td><td></td></tr>
<tr><td>Oligo rev</td><td>AAACCGTCTATAATGTTGTCTACA</td><td></td><td></td><td></td><td></td><td></td><td></td><td></td><td></td><td></td><td></td></tr>
<tr><td>Chromosome</td><td>start</td><td>end</td><td>strand</td><td>MM</td><td>target_seq</td><td>PAM</td><td>alignment</td><td>distance</td><td>position</td><td>gene name</td><td>gene id</td></tr>
<tr><td>GU305914.1</td><td>155127</td><td>155149</td><td>+</td><td>0</td><td>TGTAGACAACATTATAGACG</td><td>GGG</td><td>||||||||[||||||||||||]PAM</td><td>NA</td><td>Intergenic</td><td>NA</td><td>NA</td></tr>
<tr><td>T11866</td><td>GATCGCGCGATCGAAGCCGCCGG</td><td>1000</td><td></td><td></td><td></td><td></td><td></td><td></td><td></td><td></td><td></td></tr>
<tr><td>Oligo fwd</td><td>CACCGATCGCGCGATCGAAGCCGC</td><td></td><td></td><td></td><td></td><td></td><td></td><td></td><td></td><td></td><td></td></tr>
<tr><td>Oligo rev</td><td>AAACGCGGCTTCGATCGCGCGATC</td><td></td><td></td><td></td><td></td><td></td><td></td><td></td><td></td><td></td><td></td></tr>
<tr><td>Chromosome</td><td>start</td><td>end</td><td>strand</td><td>MM</td><td>target_seq</td><td>PAM</td><td>alignment</td><td>distance</td><td>position</td><td>gene name</td><td>gene id</td></tr>
<tr><td>GU305914.1</td><td>155155</td><td>155177</td><td>+</td><td>0</td><td>GATCGCGCGATCGAAGCCGC</td><td>CGG</td><td>||||||||[||||||||||||]PAM</td><td>NA</td><td>Intergenic</td><td>NA</td><td>NA</td></tr>
<tr><td>T11867</td><td>ATGATGAAAATTGTGTATTCTGG</td><td>1000</td><td></td><td></td><td></td><td></td><td></td><td></td><td></td><td></td><td></td></tr>
<tr><td>Oligo fwd</td><td>CACCATGATGAAAATTGTGTATTC</td><td></td><td></td><td></td><td></td><td></td><td></td><td></td><td></td><td></td><td></td></tr>
<tr><td>Oligo rev</td><td>AAACGAATACACAATTTTCATCAT</td><td></td><td></td><td></td><td></td><td></td><td></td><td></td><td></td><td></td><td></td></tr>
<tr><td>Chromosome</td><td>start</td><td>end</td><td>strand</td><td>MM</td><td>target_seq</td><td>PAM</td><td>alignment</td><td>distance</td><td>position</td><td>gene name</td><td>gene id</td></tr>
<tr><td>GU305914.1</td><td>155187</td><td>155209</td><td>+</td><td>0</td><td>ATGATGAAAATTGTGTATTC</td><td>TGG</td><td>||||||||[||||||||||||]PAM</td><td>NA</td><td>Intergenic</td><td>NA</td><td>NA</td></tr>
<tr><td>T11868</td><td>TGAAAATTGTGTATTCTGGCCGG</td><td>1000</td><td></td><td></td><td></td><td></td><td></td><td></td><td></td><td></td><td></td></tr>
<tr><td>Oligo fwd</td><td>CACCTGAAAATTGTGTATTCTGGC</td><td></td><td></td><td></td><td></td><td></td><td></td><td></td><td></td><td></td><td></td></tr>
<tr><td>Oligo rev</td><td>AAACGCCAGAATACACAATTTTCA</td><td></td><td></td><td></td><td></td><td></td><td></td><td></td><td></td><td></td><td></td></tr>
<tr><td>Chromosome</td><td>start</td><td>end</td><td>strand</td><td>MM</td><td>target_seq</td><td>PAM</td><td>alignment</td><td>distance</td><td>position</td><td>gene name</td><td>gene id</td></tr>
<tr><td>GU305914.1</td><td>155191</td><td>155213</td><td>+</td><td>0</td><td>TGAAAATTGTGTATTCTGGC</td><td>CGG</td><td>||||||||[||||||||||||]PAM</td><td>NA</td><td>Intergenic</td><td>NA</td><td>NA</td></tr>
<tr><td>T11869</td><td>ATTCTGGCCGGTGCCTGTGATGG</td><td>1000</td><td></td><td></td><td></td><td></td><td></td><td></td><td></td><td></td><td></td></tr>
<tr><td>Oligo fwd</td><td>CACCATTCTGGCCGGTGCCTGTGA</td><td></td><td></td><td></td><td></td><td></td><td></td><td></td><td></td><td></td><td></td></tr>
<tr><td>Oligo rev</td><td>AAACTCACAGGCACCGGCCAGAAT</td><td></td><td></td><td></td><td></td><td></td><td></td><td></td><td></td><td></td><td></td></tr>
<tr><td>Chromosome</td><td>start</td><td>end</td><td>strand</td><td>MM</td><td>target_seq</td><td>PAM</td><td>alignment</td><td>distance</td><td>position</td><td>gene name</td><td>gene id</td></tr>
<tr><td>GU305914.1</td><td>155203</td><td>155225</td><td>+</td><td>0</td><td>ATTCTGGCCGGTGCCTGTGA</td><td>TGG</td><td>||||||||[||||||||||||]PAM</td><td>NA</td><td>Intergenic</td><td>NA</td><td>NA</td></tr>
<tr><td>T11870</td><td>TTATACATCTGTTCTCACATCGG</td><td>1000</td><td></td><td></td><td></td><td></td><td></td><td></td><td></td><td></td><td></td></tr>
<tr><td>Oligo fwd</td><td>CACCTTATACATCTGTTCTCACAT</td><td></td><td></td><td></td><td></td><td></td><td></td><td></td><td></td><td></td><td></td></tr>
<tr><td>Oligo rev</td><td>AAACATGTGAGAACAGATGTATAA</td><td></td><td></td><td></td><td></td><td></td><td></td><td></td><td></td><td></td><td></td></tr>
<tr><td>Chromosome</td><td>start</td><td>end</td><td>strand</td><td>MM</td><td>target_seq</td><td>PAM</td><td>alignment</td><td>distance</td><td>position</td><td>gene name</td><td>gene id</td></tr>
<tr><td>GU305914.1</td><td>155396</td><td>155418</td><td>+</td><td>0</td><td>TTATACATCTGTTCTCACAT</td><td>CGG</td><td>||||||||[||||||||||||]PAM</td><td>NA</td><td>Intergenic</td><td>NA</td><td>NA</td></tr>
<tr><td>T11871</td><td>ATAAGAACATTACTATTGCCAGG</td><td>1000</td><td></td><td></td><td></td><td></td><td></td><td></td><td></td><td></td><td></td></tr>
<tr><td>Oligo fwd</td><td>CACCATAAGAACATTACTATTGCC</td><td></td><td></td><td></td><td></td><td></td><td></td><td></td><td></td><td></td><td></td></tr>
<tr><td>Oligo rev</td><td>AAACGGCAATAGTAATGTTCTTAT</td><td></td><td></td><td></td><td></td><td></td><td></td><td></td><td></td><td></td><td></td></tr>
<tr><td>Chromosome</td><td>start</td><td>end</td><td>strand</td><td>MM</td><td>target_seq</td><td>PAM</td><td>alignment</td><td>distance</td><td>position</td><td>gene name</td><td>gene id</td></tr>
<tr><td>GU305914.1</td><td>155511</td><td>155533</td><td>+</td><td>0</td><td>ATAAGAACATTACTATTGCC</td><td>AGG</td><td>||||||||[||||||||||||]PAM</td><td>NA</td><td>Intergenic</td><td>NA</td><td>NA</td></tr>
<tr><td>T11872</td><td>ATTCCTATCTGTATTCCTAAAGG</td><td>1000</td><td></td><td></td><td></td><td></td><td></td><td></td><td></td><td></td><td></td></tr>
<tr><td>Oligo fwd</td><td>CACCATTCCTATCTGTATTCCTAA</td><td></td><td></td><td></td><td></td><td></td><td></td><td></td><td></td><td></td><td></td></tr>
<tr><td>Oligo rev</td><td>AAACTTAGGAATACAGATAGGAAT</td><td></td><td></td><td></td><td></td><td></td><td></td><td></td><td></td><td></td><td></td></tr>
<tr><td>Chromosome</td><td>start</td><td>end</td><td>strand</td><td>MM</td><td>target_seq</td><td>PAM</td><td>alignment</td><td>distance</td><td>position</td><td>gene name</td><td>gene id</td></tr>
<tr><td>GU305914.1</td><td>155547</td><td>155569</td><td>+</td><td>0</td><td>ATTCCTATCTGTATTCCTAA</td><td>AGG</td><td>||||||||[||||||||||||]PAM</td><td>NA</td><td>Intergenic</td><td>NA</td><td>NA</td></tr>
<tr><td>T11873</td><td>TCCTAAAGGAGATTGTGATTCGG</td><td>1000</td><td></td><td></td><td></td><td></td><td></td><td></td><td></td><td></td><td></td></tr>
<tr><td>Oligo fwd</td><td>CACCTCCTAAAGGAGATTGTGATT</td><td></td><td></td><td></td><td></td><td></td><td></td><td></td><td></td><td></td><td></td></tr>
<tr><td>Oligo rev</td><td>AAACAATCACAATCTCCTTTAGGA</td><td></td><td></td><td></td><td></td><td></td><td></td><td></td><td></td><td></td><td></td></tr>
<tr><td>Chromosome</td><td>start</td><td>end</td><td>strand</td><td>MM</td><td>target_seq</td><td>PAM</td><td>alignment</td><td>distance</td><td>position</td><td>gene name</td><td>gene id</td></tr>
<tr><td>GU305914.1</td><td>155561</td><td>155583</td><td>+</td><td>0</td><td>TCCTAAAGGAGATTGTGATT</td><td>CGG</td><td>||||||||[||||||||||||]PAM</td><td>NA</td><td>Intergenic</td><td>NA</td><td>NA</td></tr>
<tr><td>T11874</td><td>AATTGCTACTGCGATCATACAGG</td><td>1000</td><td></td><td></td><td></td><td></td><td></td><td></td><td></td><td></td><td></td></tr>
<tr><td>Oligo fwd</td><td>CACCAATTGCTACTGCGATCATAC</td><td></td><td></td><td></td><td></td><td></td><td></td><td></td><td></td><td></td><td></td></tr>
<tr><td>Oligo rev</td><td>AAACGTATGATCGCAGTAGCAATT</td><td></td><td></td><td></td><td></td><td></td><td></td><td></td><td></td><td></td><td></td></tr>
<tr><td>Chromosome</td><td>start</td><td>end</td><td>strand</td><td>MM</td><td>target_seq</td><td>PAM</td><td>alignment</td><td>distance</td><td>position</td><td>gene name</td><td>gene id</td></tr>
<tr><td>GU305914.1</td><td>155642</td><td>155664</td><td>+</td><td>0</td><td>AATTGCTACTGCGATCATAC</td><td>AGG</td><td>||||||||[||||||||||||]PAM</td><td>NA</td><td>Intergenic</td><td>NA</td><td>NA</td></tr>
<tr><td>T11875</td><td>ACAGGCTAACGTTTATACAATGG</td><td>1000</td><td></td><td></td><td></td><td></td><td></td><td></td><td></td><td></td><td></td></tr>
<tr><td>Oligo fwd</td><td>CACCACAGGCTAACGTTTATACAA</td><td></td><td></td><td></td><td></td><td></td><td></td><td></td><td></td><td></td><td></td></tr>
<tr><td>Oligo rev</td><td>AAACTTGTATAAACGTTAGCCTGT</td><td></td><td></td><td></td><td></td><td></td><td></td><td></td><td></td><td></td><td></td></tr>
<tr><td>Chromosome</td><td>start</td><td>end</td><td>strand</td><td>MM</td><td>target_seq</td><td>PAM</td><td>alignment</td><td>distance</td><td>position</td><td>gene name</td><td>gene id</td></tr>
<tr><td>GU305914.1</td><td>155660</td><td>155682</td><td>+</td><td>0</td><td>ACAGGCTAACGTTTATACAA</td><td>TGG</td><td>||||||||[||||||||||||]PAM</td><td>NA</td><td>Intergenic</td><td>NA</td><td>NA</td></tr>
<tr><td>T11876</td><td>CTCTATGACTACTGATATATCGG</td><td>1000</td><td></td><td></td><td></td><td></td><td></td><td></td><td></td><td></td><td></td></tr>
<tr><td>Oligo fwd</td><td>CACCCTCTATGACTACTGATATAT</td><td></td><td></td><td></td><td></td><td></td><td></td><td></td><td></td><td></td><td></td></tr>
<tr><td>Oligo rev</td><td>AAACATATATCAGTAGTCATAGAG</td><td></td><td></td><td></td><td></td><td></td><td></td><td></td><td></td><td></td><td></td></tr>
<tr><td>Chromosome</td><td>start</td><td>end</td><td>strand</td><td>MM</td><td>target_seq</td><td>PAM</td><td>alignment</td><td>distance</td><td>position</td><td>gene name</td><td>gene id</td></tr>
<tr><td>GU305914.1</td><td>155697</td><td>155719</td><td>+</td><td>0</td><td>CTCTATGACTACTGATATAT</td><td>CGG</td><td>||||||||[||||||||||||]PAM</td><td>NA</td><td>Intergenic</td><td>NA</td><td>NA</td></tr>
<tr><td>T11877</td><td>TACAAACCCAAACTGACGAGCGG</td><td>1000</td><td></td><td></td><td></td><td></td><td></td><td></td><td></td><td></td><td></td></tr>
<tr><td>Oligo fwd</td><td>CACCTACAAACCCAAACTGACGAG</td><td></td><td></td><td></td><td></td><td></td><td></td><td></td><td></td><td></td><td></td></tr>
<tr><td>Oligo rev</td><td>AAACCTCGTCAGTTTGGGTTTGTA</td><td></td><td></td><td></td><td></td><td></td><td></td><td></td><td></td><td></td><td></td></tr>
<tr><td>Chromosome</td><td>start</td><td>end</td><td>strand</td><td>MM</td><td>target_seq</td><td>PAM</td><td>alignment</td><td>distance</td><td>position</td><td>gene name</td><td>gene id</td></tr>
</table>

GU305914.1 155876 155898 + 0 TACAAACCCAAACTGACGAG CGG ||||||||[||||||||||||]PAM NA Intergenic NA NA

T11878 ACAAACCCAAACTGACGAGCGGG 1000
Oligo fwd CACCACAAACCCAAACTGACGAGC
Oligo rev AAACGCTCGTCAGTTTGGGTTTGT
Chromosome start end strand MM target_seq PAM alignment distance position gene name gene id
GU305914.1 155877 155899 + 0 ACAAACCCAAACTGACGAGC GGG ||||||||[||||||||||||]PAM NA Intergenic NA NA

T11879 CCCAAACTGACGAGCGGGTTTGG 1000
Oligo fwd CACCCCCAAACTGACGAGCGGGTT
Oligo rev AAACAACCCGCTCGTCAGTTTGGG
Chromosome start end strand MM target_seq PAM alignment distance position gene name gene id
GU305914.1 155882 155904 + 0 CCCAAACTGACGAGCGGGTT TGG ||||||||[||||||||||||]PAM NA Intergenic NA NA

T11880 AACTGACGAGCGGGTTTGGTTGG 1000
Oligo fwd CACCAACTGACGAGCGGGTTTGGT
Oligo rev AAACACCAAACCCGCTCGTCAGTT
Chromosome start end strand MM target_seq PAM alignment distance position gene name gene id
GU305914.1 155886 155908 + 0 AACTGACGAGCGGGTTTGGT TGG ||||||||[||||||||||||]PAM NA Intergenic NA NA

T11881 GAAAGAAAAAATATTAAAAATGG 1000
Oligo fwd CACCGAAAGAAAAAATATTAAAAA
Oligo rev AAACTTTTTAATATTTTTTCTTTC
Chromosome start end strand MM target_seq PAM alignment distance position gene name gene id
GU305914.1 155926 155948 + 0 GAAAGAAAAAATATTAAAAA TGG ||||||||[||||||||||||]PAM NA Intergenic NA NA

T11882 AATATTAAAAATGGTGCTCTTGG 1000
Oligo fwd CACCAATATTAAAAATGGTGCTCT
Oligo rev AAACAGAGCACCATTTTTAATATT
Chromosome start end strand MM target_seq PAM alignment distance position gene name gene id
GU305914.1 155935 155957 + 0 AATATTAAAAATGGTGCTCT TGG ||||||||[||||||||||||]PAM NA Intergenic NA NA

T11883 ATGGTGCTCTTGGTTTCTCGTGG 1000
Oligo fwd CACCATGGTGCTCTTGGTTTCTCG
Oligo rev AAACCGAGAAACCAAGAGCACCAT
Chromosome start end strand MM target_seq PAM alignment distance position gene name gene id
GU305914.1 155945 155967 + 0 ATGGTGCTCTTGGTTTCTCG TGG ||||||||[||||||||||||]PAM NA Intergenic NA NA

T11884 CCCATAGTAGTTTACATAGAAGG 1000
Oligo fwd CACCCCCATAGTAGTTTACATAGA
Oligo rev AAACTCTATGTAAACTACTATGGG
Chromosome start end strand MM target_seq PAM alignment distance position gene name gene id
GU305914.1 156082 156104 + 0 CCCATAGTAGTTTACATAGA AGG ||||||||[||||||||||||]PAM NA Intergenic NA NA

T11885 AGAGATGCTGATAAGAAAGAAGG 1000
Oligo fwd CACCAGAGATGCTGATAAGAAAGA
Oligo rev AAACTCTTTCTTATCAGCATCTCT
Chromosome start end strand MM target_seq PAM alignment distance position gene name gene id
GU305914.1 156143 156165 + 0 AGAGATGCTGATAAGAAAGA AGG ||||||||[||||||||||||]PAM NA Intergenic NA NA

T11886 GTAACCTCTCTATACATGCATGG 1000
Oligo fwd CACCGTAACCTCTCTATACATGCA
Oligo rev AAACTGCATGTATAGAGAGGTTAC
Chromosome start end strand MM target_seq PAM alignment distance position gene name gene id
GU305914.1 156207 156229 + 0 GTAACCTCTCTATACATGCA TGG ||||||||[||||||||||||]PAM NA Intergenic NA NA

T11887 CTCTATACATGCATGGATATTGG 1000
Oligo fwd CACCCTCTATACATGCATGGATAT
Oligo rev AAACATATCCATGCATGTATAGAG
Chromosome start end strand MM target_seq PAM alignment distance position gene name gene id
GU305914.1 156214 156236 + 0 CTCTATACATGCATGGATAT TGG ||||||||[||||||||||||]PAM NA Intergenic NA NA

T11888 ATTGGCTAAATATCATTCTTAGG 1000
Oligo fwd CACCATTGGCTAAATATCATTCTT
Oligo rev AAACAAGAATGATATTTAGCCAAT
Chromosome start end strand MM target_seq PAM alignment distance position gene name gene id
GU305914.1 156232 156254 + 0 ATTGGCTAAATATCATTCTT AGG ||||||||[||||||||||||]PAM NA Intergenic NA NA

T11889 TAACAGTTTCGCACAACTGTAGG 1000
Oligo fwd CACCTAACAGTTTCGCACAACTGT
Oligo rev AAACACAGTTGTGCGAAACTGTTA
Chromosome start end strand MM target_seq PAM alignment distance position gene name gene id
GU305914.1 156301 156323 + 0 TAACAGTTTCGCACAACTGT AGG ||||||||[||||||||||||]PAM NA Intergenic NA NA

T11890 GGTTGATTTTGTATTGTATATGG 1000
Oligo fwd CACCGGTTGATTTTGTATTGTATA
Oligo rev AAACTATACAATACAAAATCAACC
Chromosome start end strand MM target_seq PAM alignment distance position gene name gene id
GU305914.1 156322 156344 + 0 GGTTGATTTTGTATTGTATA TGG ||||||||[||||||||||||]PAM NA Intergenic NA NA

T11891 TGTTTTGTGTATCGTGAGTGAGG 1000
Oligo fwd CACCTGTTTTGTGTATCGTGAGTG
Oligo rev AAACCACTCACGATACACAAAACA
Chromosome start end strand MM target_seq PAM alignment distance position gene name gene id
GU305914.1 156433 156455 + 0 TGTTTTGTGTATCGTGAGTG AGG ||||||||[||||||||||||]PAM NA Intergenic NA NA

T11892 GAGTGAGGATGCCGAATCTGTGG 1000
Oligo fwd CACCGAGTGAGGATGCCGAATCTG
Oligo rev AAACCAGATTCGGCATCCTCACTC

<table>
<tr><td>Chromosome</td><td>start</td><td>end</td><td>strand</td><td>MM</td><td>target_seq</td><td>PAM</td><td>alignment</td><td>distance</td><td>position</td><td>gene name</td><td>gene id</td></tr>
<tr><td>GU305914.1</td><td>156448</td><td>156470</td><td>+</td><td>0</td><td>GAGTGAGGATGCCGAATCTG</td><td>TGG</td><td>||||||||[||||||||||||]PAM</td><td>NA</td><td>Intergenic</td><td>NA</td><td>NA</td></tr>
<tr><td></td><td></td><td></td><td></td><td></td><td></td><td></td><td></td><td></td><td></td><td></td><td></td></tr>
<tr><td>T11893</td><td>TCCCCTCACACGCTTTGTTACGG</td><td>1000</td><td></td><td></td><td></td><td></td><td></td><td></td><td></td><td></td><td></td></tr>
<tr><td>Oligo fwd</td><td>CACCTCCCCTCACACGCTTTGTTA</td><td></td><td></td><td></td><td></td><td></td><td></td><td></td><td></td><td></td><td></td></tr>
<tr><td>Oligo rev</td><td>AAACTAACAAAGCGTGTGAGGGGA</td><td></td><td></td><td></td><td></td><td></td><td></td><td></td><td></td><td></td><td></td></tr>
<tr><td>Chromosome</td><td>start</td><td>end</td><td>strand</td><td>MM</td><td>target_seq</td><td>PAM</td><td>alignment</td><td>distance</td><td>position</td><td>gene name</td><td>gene id</td></tr>
<tr><td>GU305914.1</td><td>156474</td><td>156496</td><td>+</td><td>0</td><td>TCCCCTCACACGCTTTGTTA</td><td>CGG</td><td>||||||||[||||||||||||]PAM</td><td>NA</td><td>Intergenic</td><td>NA</td><td>NA</td></tr>
<tr><td></td><td></td><td></td><td></td><td></td><td></td><td></td><td></td><td></td><td></td><td></td><td></td></tr>
<tr><td>T11894</td><td>TCACACGCTTTGTTACGGTTCGG</td><td>1000</td><td></td><td></td><td></td><td></td><td></td><td></td><td></td><td></td><td></td></tr>
<tr><td>Oligo fwd</td><td>CACCTCACACGCTTTGTTACGGTT</td><td></td><td></td><td></td><td></td><td></td><td></td><td></td><td></td><td></td><td></td></tr>
<tr><td>Oligo rev</td><td>AAACAACCGTAACAAAGCGTGTGA</td><td></td><td></td><td></td><td></td><td></td><td></td><td></td><td></td><td></td><td></td></tr>
<tr><td>Chromosome</td><td>start</td><td>end</td><td>strand</td><td>MM</td><td>target_seq</td><td>PAM</td><td>alignment</td><td>distance</td><td>position</td><td>gene name</td><td>gene id</td></tr>
<tr><td>GU305914.1</td><td>156479</td><td>156501</td><td>+</td><td>0</td><td>TCACACGCTTTGTTACGGTT</td><td>CGG</td><td>||||||||[||||||||||||]PAM</td><td>NA</td><td>Intergenic</td><td>NA</td><td>NA</td></tr>
<tr><td></td><td></td><td></td><td></td><td></td><td></td><td></td><td></td><td></td><td></td><td></td><td></td></tr>
<tr><td>T11895</td><td>TACATGTCGTGCTAGTTAGTAGG</td><td>1000</td><td></td><td></td><td></td><td></td><td></td><td></td><td></td><td></td><td></td></tr>
<tr><td>Oligo fwd</td><td>CACCTACATGTCGTGCTAGTTAGT</td><td></td><td></td><td></td><td></td><td></td><td></td><td></td><td></td><td></td><td></td></tr>
<tr><td>Oligo rev</td><td>AAACACTAACTAGCACGACATGTA</td><td></td><td></td><td></td><td></td><td></td><td></td><td></td><td></td><td></td><td></td></tr>
<tr><td>Chromosome</td><td>start</td><td>end</td><td>strand</td><td>MM</td><td>target_seq</td><td>PAM</td><td>alignment</td><td>distance</td><td>position</td><td>gene name</td><td>gene id</td></tr>
<tr><td>GU305914.1</td><td>156540</td><td>156562</td><td>+</td><td>0</td><td>TACATGTCGTGCTAGTTAGT</td><td>AGG</td><td>||||||||[||||||||||||]PAM</td><td>NA</td><td>Intergenic</td><td>NA</td><td>NA</td></tr>
<tr><td></td><td></td><td></td><td></td><td></td><td></td><td></td><td></td><td></td><td></td><td></td><td></td></tr>
<tr><td>T11896</td><td>AGCTACCTAGTAGTTAGAAGAGG</td><td>1000</td><td></td><td></td><td></td><td></td><td></td><td></td><td></td><td></td><td></td></tr>
<tr><td>Oligo fwd</td><td>CACCAGCTACCTAGTAGTTAGAAG</td><td></td><td></td><td></td><td></td><td></td><td></td><td></td><td></td><td></td><td></td></tr>
<tr><td>Oligo rev</td><td>AAACCTTCTAACTACTAGGTAGCT</td><td></td><td></td><td></td><td></td><td></td><td></td><td></td><td></td><td></td><td></td></tr>
<tr><td>Chromosome</td><td>start</td><td>end</td><td>strand</td><td>MM</td><td>target_seq</td><td>PAM</td><td>alignment</td><td>distance</td><td>position</td><td>gene name</td><td>gene id</td></tr>
<tr><td>GU305914.1</td><td>156851</td><td>156873</td><td>+</td><td>0</td><td>AGCTACCTAGTAGTTAGAAG</td><td>AGG</td><td>||||||||[||||||||||||]PAM</td><td>NA</td><td>Intergenic</td><td>NA</td><td>NA</td></tr>
<tr><td></td><td></td><td></td><td></td><td></td><td></td><td></td><td></td><td></td><td></td><td></td><td></td></tr>
<tr><td>T11897</td><td>GCTACCTAGTAGTTAGAAGAGGG</td><td>1000</td><td></td><td></td><td></td><td></td><td></td><td></td><td></td><td></td><td></td></tr>
<tr><td>Oligo fwd</td><td>CACCGCTACCTAGTAGTTAGAAGA</td><td></td><td></td><td></td><td></td><td></td><td></td><td></td><td></td><td></td><td></td></tr>
<tr><td>Oligo rev</td><td>AAACTCTTCTAACTACTAGGTAGC</td><td></td><td></td><td></td><td></td><td></td><td></td><td></td><td></td><td></td><td></td></tr>
<tr><td>Chromosome</td><td>start</td><td>end</td><td>strand</td><td>MM</td><td>target_seq</td><td>PAM</td><td>alignment</td><td>distance</td><td>position</td><td>gene name</td><td>gene id</td></tr>
<tr><td>GU305914.1</td><td>156852</td><td>156874</td><td>+</td><td>0</td><td>GCTACCTAGTAGTTAGAAGA</td><td>GGG</td><td>||||||||[||||||||||||]PAM</td><td>NA</td><td>Intergenic</td><td>NA</td><td>NA</td></tr>
<tr><td></td><td></td><td></td><td></td><td></td><td></td><td></td><td></td><td></td><td></td><td></td><td></td></tr>
<tr><td>T11898</td><td>AGAACTATCAAAACTGCATATGG</td><td>1000</td><td></td><td></td><td></td><td></td><td></td><td></td><td></td><td></td><td></td></tr>
<tr><td>Oligo fwd</td><td>CACCAGAACTATCAAAACTGCATA</td><td></td><td></td><td></td><td></td><td></td><td></td><td></td><td></td><td></td><td></td></tr>
<tr><td>Oligo rev</td><td>AAACTATGCAGTTTTGATAGTTCT</td><td></td><td></td><td></td><td></td><td></td><td></td><td></td><td></td><td></td><td></td></tr>
<tr><td>Chromosome</td><td>start</td><td>end</td><td>strand</td><td>MM</td><td>target_seq</td><td>PAM</td><td>alignment</td><td>distance</td><td>position</td><td>gene name</td><td>gene id</td></tr>
<tr><td>GU305914.1</td><td>156889</td><td>156911</td><td>+</td><td>0</td><td>AGAACTATCAAAACTGCATA</td><td>TGG</td><td>||||||||[||||||||||||]PAM</td><td>NA</td><td>Intergenic</td><td>NA</td><td>NA</td></tr>
<tr><td></td><td></td><td></td><td></td><td></td><td></td><td></td><td></td><td></td><td></td><td></td><td></td></tr>
<tr><td>T11899</td><td>TATCAAAACTGCATATGGTTCGG</td><td>1000</td><td></td><td></td><td></td><td></td><td></td><td></td><td></td><td></td><td></td></tr>
<tr><td>Oligo fwd</td><td>CACCTATCAAAACTGCATATGGTT</td><td></td><td></td><td></td><td></td><td></td><td></td><td></td><td></td><td></td><td></td></tr>
<tr><td>Oligo rev</td><td>AAACAACCATATGCAGTTTTGATA</td><td></td><td></td><td></td><td></td><td></td><td></td><td></td><td></td><td></td><td></td></tr>
<tr><td>Chromosome</td><td>start</td><td>end</td><td>strand</td><td>MM</td><td>target_seq</td><td>PAM</td><td>alignment</td><td>distance</td><td>position</td><td>gene name</td><td>gene id</td></tr>
<tr><td>GU305914.1</td><td>156894</td><td>156916</td><td>+</td><td>0</td><td>TATCAAAACTGCATATGGTT</td><td>CGG</td><td>||||||||[||||||||||||]PAM</td><td>NA</td><td>Intergenic</td><td>NA</td><td>NA</td></tr>
<tr><td></td><td></td><td></td><td></td><td></td><td></td><td></td><td></td><td></td><td></td><td></td><td></td></tr>
<tr><td>T11900</td><td>TATGGTTCGGTATGAGATTACGG</td><td>1000</td><td></td><td></td><td></td><td></td><td></td><td></td><td></td><td></td><td></td></tr>
<tr><td>Oligo fwd</td><td>CACCTATGGTTCGGTATGAGATTA</td><td></td><td></td><td></td><td></td><td></td><td></td><td></td><td></td><td></td><td></td></tr>
<tr><td>Oligo rev</td><td>AAACTAATCTCATACCGAACCATA</td><td></td><td></td><td></td><td></td><td></td><td></td><td></td><td></td><td></td><td></td></tr>
<tr><td>Chromosome</td><td>start</td><td>end</td><td>strand</td><td>MM</td><td>target_seq</td><td>PAM</td><td>alignment</td><td>distance</td><td>position</td><td>gene name</td><td>gene id</td></tr>
<tr><td>GU305914.1</td><td>156907</td><td>156929</td><td>+</td><td>0</td><td>TATGGTTCGGTATGAGATTA</td><td>CGG</td><td>||||||||[||||||||||||]PAM</td><td>NA</td><td>Intergenic</td><td>NA</td><td>NA</td></tr>
<tr><td></td><td></td><td></td><td></td><td></td><td></td><td></td><td></td><td></td><td></td><td></td><td></td></tr>
<tr><td>T11901</td><td>TGAGATTACGGACGATTTGATGG</td><td>1000</td><td></td><td></td><td></td><td></td><td></td><td></td><td></td><td></td><td></td></tr>
<tr><td>Oligo fwd</td><td>CACCTGAGATTACGGACGATTTGA</td><td></td><td></td><td></td><td></td><td></td><td></td><td></td><td></td><td></td><td></td></tr>
<tr><td>Oligo rev</td><td>AAACTCAAATCGTCCGTAATCTCA</td><td></td><td></td><td></td><td></td><td></td><td></td><td></td><td></td><td></td><td></td></tr>
<tr><td>Chromosome</td><td>start</td><td>end</td><td>strand</td><td>MM</td><td>target_seq</td><td>PAM</td><td>alignment</td><td>distance</td><td>position</td><td>gene name</td><td>gene id</td></tr>
<tr><td>GU305914.1</td><td>156919</td><td>156941</td><td>+</td><td>0</td><td>TGAGATTACGGACGATTTGA</td><td>TGG</td><td>||||||||[||||||||||||]PAM</td><td>NA</td><td>Intergenic</td><td>NA</td><td>NA</td></tr>
<tr><td></td><td></td><td></td><td></td><td></td><td></td><td></td><td></td><td></td><td></td><td></td><td></td></tr>
<tr><td>T11902</td><td>TTGTCTTTCTTTATTTATCGCGG</td><td>1000</td><td></td><td></td><td></td><td></td><td></td><td></td><td></td><td></td><td></td></tr>
<tr><td>Oligo fwd</td><td>CACCTTGTCTTTCTTTATTTATCG</td><td></td><td></td><td></td><td></td><td></td><td></td><td></td><td></td><td></td><td></td></tr>
<tr><td>Oligo rev</td><td>AAACCGATAAATAAAGAAAGACAA</td><td></td><td></td><td></td><td></td><td></td><td></td><td></td><td></td><td></td><td></td></tr>
<tr><td>Chromosome</td><td>start</td><td>end</td><td>strand</td><td>MM</td><td>target_seq</td><td>PAM</td><td>alignment</td><td>distance</td><td>position</td><td>gene name</td><td>gene id</td></tr>
<tr><td>GU305914.1</td><td>156986</td><td>157008</td><td>+</td><td>0</td><td>TTGTCTTTCTTTATTTATCG</td><td>CGG</td><td>||||||||[||||||||||||]PAM</td><td>NA</td><td>Intergenic</td><td>NA</td><td>NA</td></tr>
<tr><td>GU305914.1</td><td>39163</td><td>39185</td><td>-</td><td>4</td><td>GTGTTTTTCTTGATTTATTG</td><td>AGG</td><td>-|||-|||[|||-||||||-|]PAM</td><td>NA</td><td>Intergenic</td><td>NA</td><td>NA</td></tr>
<tr><td></td><td></td><td></td><td></td><td></td><td></td><td></td><td></td><td></td><td></td><td></td><td></td></tr>
<tr><td>T11903</td><td>CTTTCTTTATTTATCGCGGATGG</td><td>1000</td><td></td><td></td><td></td><td></td><td></td><td></td><td></td><td></td><td></td></tr>
<tr><td>Oligo fwd</td><td>CACCCTTTCTTTATTTATCGCGGA</td><td></td><td></td><td></td><td></td><td></td><td></td><td></td><td></td><td></td><td></td></tr>
<tr><td>Oligo rev</td><td>AAACTCCGCGATAAATAAAGAAAG</td><td></td><td></td><td></td><td></td><td></td><td></td><td></td><td></td><td></td><td></td></tr>
<tr><td>Chromosome</td><td>start</td><td>end</td><td>strand</td><td>MM</td><td>target_seq</td><td>PAM</td><td>alignment</td><td>distance</td><td>position</td><td>gene name</td><td>gene id</td></tr>
<tr><td>GU305914.1</td><td>156990</td><td>157012</td><td>+</td><td>0</td><td>CTTTCTTTATTTATCGCGGA</td><td>TGG</td><td>||||||||[||||||||||||]PAM</td><td>NA</td><td>Intergenic</td><td>NA</td><td>NA</td></tr>
<tr><td></td><td></td><td></td><td></td><td></td><td></td><td></td><td></td><td></td><td></td><td></td><td></td></tr>
<tr><td>T11904</td><td>ATGATGATACTTTTTCCGTGTGG</td><td>1000</td><td></td><td></td><td></td><td></td><td></td><td></td><td></td><td></td><td></td></tr>
<tr><td>Oligo fwd</td><td>CACCATGATGATACTTTTTCCGTG</td><td></td><td></td><td></td><td></td><td></td><td></td><td></td><td></td><td></td><td></td></tr>
<tr><td>Oligo rev</td><td>AAACCACGGAAAAAGTATCATCAT</td><td></td><td></td><td></td><td></td><td></td><td></td><td></td><td></td><td></td><td></td></tr>
<tr><td>Chromosome</td><td>start</td><td>end</td><td>strand</td><td>MM</td><td>target_seq</td><td>PAM</td><td>alignment</td><td>distance</td><td>position</td><td>gene name</td><td>gene id</td></tr>
<tr><td>GU305914.1</td><td>157064</td><td>157086</td><td>+</td><td>0</td><td>ATGATGATACTTTTTCCGTG</td><td>TGG</td><td>||||||||[||||||||||||]PAM</td><td>NA</td><td>Intergenic</td><td>NA</td><td>NA</td></tr>
<tr><td>GU305914.1</td><td>218796</td><td>218818</td><td>-</td><td>3</td><td>ATAACGGTACTTTTTCCGTG</td><td>TGG</td><td>||-|-|-|[||||||||||||]PAM</td><td>NA</td><td>Intergenic</td><td>NA</td><td>NA</td></tr>
<tr><td></td><td></td><td></td><td></td><td></td><td></td><td></td><td></td><td></td><td></td><td></td><td></td></tr>
<tr><td>T11905</td><td>GAGAACTATAACTAGTAGCGAGG</td><td>1000</td><td></td><td></td><td></td><td></td><td></td><td></td><td></td><td></td><td></td></tr>
<tr><td>Oligo fwd</td><td>CACCGAGAACTATAACTAGTAGCG</td><td></td><td></td><td></td><td></td><td></td><td></td><td></td><td></td><td></td><td></td></tr>
<tr><td>Oligo rev</td><td>AAACCGCTACTAGTTATAGTTCTC</td><td></td><td></td><td></td><td></td><td></td><td></td><td></td><td></td><td></td><td></td></tr>
<tr><td>Chromosome</td><td>start</td><td>end</td><td>strand</td><td>MM</td><td>target_seq</td><td>PAM</td><td>alignment</td><td>distance</td><td>position</td><td>gene name</td><td>gene id</td></tr>
<tr><td>GU305914.1</td><td>157116</td><td>157138</td><td>+</td><td>0</td><td>GAGAACTATAACTAGTAGCG</td><td>AGG</td><td>||||||||[||||||||||||]PAM</td><td>NA</td><td>Intergenic</td><td>NA</td><td>NA</td></tr>
<tr><td></td><td></td><td></td><td></td><td></td><td></td><td></td><td></td><td></td><td></td><td></td><td></td></tr>
<tr><td>T11906</td><td>CTAGCAGCTCGAAAGTTGAACGG</td><td>1000</td><td></td><td></td><td></td><td></td><td></td><td></td><td></td><td></td><td></td></tr>
<tr><td>Oligo fwd</td><td>CACCCTAGCAGCTCGAAAGTTGAA</td><td></td><td></td><td></td><td></td><td></td><td></td><td></td><td></td><td></td><td></td></tr>
<tr><td>Oligo rev</td><td>AAACTTCAACTTTCGAGCTGCTAG</td><td></td><td></td><td></td><td></td><td></td><td></td><td></td><td></td><td></td><td></td></tr>
<tr><td>Chromosome</td><td>start</td><td>end</td><td>strand</td><td>MM</td><td>target_seq</td><td>PAM</td><td>alignment</td><td>distance</td><td>position</td><td>gene name</td><td>gene id</td></tr>
<tr><td>GU305914.1</td><td>157140</td><td>157162</td><td>+</td><td>0</td><td>CTAGCAGCTCGAAAGTTGAA</td><td>CGG</td><td>||||||||[||||||||||||]PAM</td><td>NA</td><td>Intergenic</td><td>NA</td><td>NA</td></tr>
</table>

| | | | | | | | | | | | |
|---|---|---|---|---|---|---|---|---|---|---|---|
| T11907 | TAGCAGCTCGAAAGTTGAACGGG | 1000 | | | | | | | | | |
| Oligo fwd | CACCTAGCAGCTCGAAAGTTGAAC | | | | | | | | | | |
| Oligo rev | AAACGTTCAACTTTCGAGCTGCTA | | | | | | | | | | |
| Chromosome | start | end | strand | MM | target_seq | PAM | alignment | distance | position | gene name | gene id |
| GU305914.1 | 157141 | 157163 | + | 0 | TAGCAGCTCGAAAGTTGAAC | GGG | \|\|\|\|\|\|\|\|[\|\|\|\|\|\|\|\|\|\|\|\|]PAM | NA | Intergenic | NA | NA |
| T11908 | TAGTACGTAGTGTGAACGAACGG | 1000 | | | | | | | | | |
| Oligo fwd | CACCTAGTACGTAGTGTGAACGAA | | | | | | | | | | |
| Oligo rev | AAACTTCGTTCACACTACGTACTA | | | | | | | | | | |
| Chromosome | start | end | strand | MM | target_seq | PAM | alignment | distance | position | gene name | gene id |
| GU305914.1 | 157186 | 157208 | + | 0 | TAGTACGTAGTGTGAACGAA | CGG | \|\|\|\|\|\|\|\|[\|\|\|\|\|\|\|\|\|\|\|\|]PAM | NA | Intergenic | NA | NA |
| T11909 | AGTACGTAGTGTGAACGAACGGG | 1000 | | | | | | | | | |
| Oligo fwd | CACCAGTACGTAGTGTGAACGAAC | | | | | | | | | | |
| Oligo rev | AAACGTTCGTTCACACTACGTACT | | | | | | | | | | |
| Chromosome | start | end | strand | MM | target_seq | PAM | alignment | distance | position | gene name | gene id |
| GU305914.1 | 157187 | 157209 | + | 0 | AGTACGTAGTGTGAACGAAC | GGG | \|\|\|\|\|\|\|\|[\|\|\|\|\|\|\|\|\|\|\|\|]PAM | NA | Intergenic | NA | NA |
| T11910 | GTGTGAACGAACGGGTAAAACGG | 1000 | | | | | | | | | |
| Oligo fwd | CACCGTGTGAACGAACGGGTAAAA | | | | | | | | | | |
| Oligo rev | AAACTTTTACCCGTTCGTTCACAC | | | | | | | | | | |
| Chromosome | start | end | strand | MM | target_seq | PAM | alignment | distance | position | gene name | gene id |
| GU305914.1 | 157195 | 157217 | + | 0 | GTGTGAACGAACGGGTAAAA | CGG | \|\|\|\|\|\|\|\|[\|\|\|\|\|\|\|\|\|\|\|\|]PAM | NA | Intergenic | NA | NA |
| T11911 | TGTGAACGAACGGGTAAAACGGG | 1000 | | | | | | | | | |
| Oligo fwd | CACCTGTGAACGAACGGGTAAAAC | | | | | | | | | | |
| Oligo rev | AAACGTTTTACCCGTTCGTTCACA | | | | | | | | | | |
| Chromosome | start | end | strand | MM | target_seq | PAM | alignment | distance | position | gene name | gene id |
| GU305914.1 | 157196 | 157218 | + | 0 | TGTGAACGAACGGGTAAAAC | GGG | \|\|\|\|\|\|\|\|[\|\|\|\|\|\|\|\|\|\|\|\|]PAM | NA | Intergenic | NA | NA |
| T11912 | ACGAACGGGTAAAACGGGTAAGG | 1000 | | | | | | | | | |
| Oligo fwd | CACCACGAACGGGTAAAACGGGTA | | | | | | | | | | |
| Oligo rev | AAACTACCCGTTTTACCCGTTCGT | | | | | | | | | | |
| Chromosome | start | end | strand | MM | target_seq | PAM | alignment | distance | position | gene name | gene id |
| GU305914.1 | 157201 | 157223 | + | 0 | ACGAACGGGTAAAACGGGTA | AGG | \|\|\|\|\|\|\|\|[\|\|\|\|\|\|\|\|\|\|\|\|]PAM | NA | Intergenic | NA | NA |
| T11913 | CGAACGGGTAAAACGGGTAAGGG | 1000 | | | | | | | | | |
| Oligo fwd | CACCCGAACGGGTAAAACGGGTAA | | | | | | | | | | |
| Oligo rev | AAACTTACCCGTTTTACCCGTTCG | | | | | | | | | | |
| Chromosome | start | end | strand | MM | target_seq | PAM | alignment | distance | position | gene name | gene id |
| GU305914.1 | 157202 | 157224 | + | 0 | CGAACGGGTAAAACGGGTAA | GGG | \|\|\|\|\|\|\|\|[\|\|\|\|\|\|\|\|\|\|\|\|]PAM | NA | Intergenic | NA | NA |
| T11914 | AAGTAGTAAGTAGTGTGTTTTGG | 1000 | | | | | | | | | |
| Oligo fwd | CACCAAGTAGTAAGTAGTGTGTTT | | | | | | | | | | |
| Oligo rev | AAACAAACACACTACTTACTACTT | | | | | | | | | | |
| Chromosome | start | end | strand | MM | target_seq | PAM | alignment | distance | position | gene name | gene id |
| GU305914.1 | 157227 | 157249 | + | 0 | AAGTAGTAAGTAGTGTGTTT | TGG | \|\|\|\|\|\|\|\|[\|\|\|\|\|\|\|\|\|\|\|\|]PAM | NA | Intergenic | NA | NA |
| T11915 | TAGTAAGTAGTGTGTTTTGGTGG | 1000 | | | | | | | | | |
| Oligo fwd | CACCTAGTAAGTAGTGTGTTTTGG | | | | | | | | | | |
| Oligo rev | AAACCCAAAACACACTACTTACTA | | | | | | | | | | |
| Chromosome | start | end | strand | MM | target_seq | PAM | alignment | distance | position | gene name | gene id |
| GU305914.1 | 157230 | 157252 | + | 0 | TAGTAAGTAGTGTGTTTTGG | TGG | \|\|\|\|\|\|\|\|[\|\|\|\|\|\|\|\|\|\|\|\|]PAM | NA | Intergenic | NA | NA |
| T11916 | GTGTGTTTTGGTGGATGACTCGG | 1000 | | | | | | | | | |
| Oligo fwd | CACCGTGTGTTTTGGTGGATGACT | | | | | | | | | | |
| Oligo rev | AAACAGTCATCCACCAAAACACAC | | | | | | | | | | |
| Chromosome | start | end | strand | MM | target_seq | PAM | alignment | distance | position | gene name | gene id |
| GU305914.1 | 157239 | 157261 | + | 0 | GTGTGTTTTGGTGGATGACT | CGG | \|\|\|\|\|\|\|\|[\|\|\|\|\|\|\|\|\|\|\|\|]PAM | NA | Intergenic | NA | NA |
| T11917 | TTTGGTGGATGACTCGGTTATGG | 1000 | | | | | | | | | |
| Oligo fwd | CACCTTTGGTGGATGACTCGGTTA | | | | | | | | | | |
| Oligo rev | AAACTAACCGAGTCATCCACCAAA | | | | | | | | | | |
| Chromosome | start | end | strand | MM | target_seq | PAM | alignment | distance | position | gene name | gene id |
| GU305914.1 | 157245 | 157267 | + | 0 | TTTGGTGGATGACTCGGTTA | TGG | \|\|\|\|\|\|\|\|[\|\|\|\|\|\|\|\|\|\|\|\|]PAM | NA | Intergenic | NA | NA |
| T11918 | TGATGTTCTTCCCGACGTGATGG | 1000 | | | | | | | | | |
| Oligo fwd | CACCTGATGTTCTTCCCGACGTGA | | | | | | | | | | |
| Oligo rev | AAACTCACGTCGGGAAGAACATCA | | | | | | | | | | |
| Chromosome | start | end | strand | MM | target_seq | PAM | alignment | distance | position | gene name | gene id |
| GU305914.1 | 157277 | 157299 | + | 0 | TGATGTTCTTCCCGACGTGA | TGG | \|\|\|\|\|\|\|\|[\|\|\|\|\|\|\|\|\|\|\|\|]PAM | NA | Intergenic | NA | NA |
| T11919 | GATGTTCTTCCCGACGTGATGGG | 1000 | | | | | | | | | |
| Oligo fwd | CACCGATGTTCTTCCCGACGTGAT | | | | | | | | | | |
| Oligo rev | AAACATCACGTCGGGAAGAACATC | | | | | | | | | | |
| Chromosome | start | end | strand | MM | target_seq | PAM | alignment | distance | position | gene name | gene id |
| GU305914.1 | 157278 | 157300 | + | 0 | GATGTTCTTCCCGACGTGAT | GGG | \|\|\|\|\|\|\|\|[\|\|\|\|\|\|\|\|\|\|\|\|]PAM | NA | Intergenic | NA | NA |
| T11920 | GTGATGGGTTAGAACTGACTCGG | 1000 | | | | | | | | | |
| Oligo fwd | CACCGTGATGGGTTAGAACTGACT | | | | | | | | | | |
| Oligo rev | AAACAGTCAGTTCTAACCCATCAC | | | | | | | | | | |
| Chromosome | start | end | strand | MM | target_seq | PAM | alignment | distance | position | gene name | gene id |
| GU305914.1 | 157293 | 157315 | + | 0 | GTGATGGGTTAGAACTGACT | CGG | \|\|\|\|\|\|\|\|[\|\|\|\|\|\|\|\|\|\|\|\|]PAM | NA | Intergenic | NA | NA |
| T11921 | TGGGTTAGAACTGACTCGGTTGG | 1000 | | | | | | | | | |
| Oligo fwd | CACCTGGGTTAGAACTGACTCGGT | | | | | | | | | | |
| Oligo rev | AAACACCGAGTCAGTTCTAACCCA | | | | | | | | | | |
| Chromosome | start | end | strand | MM | target_seq | PAM | alignment | distance | position | gene name | gene id |
| GU305914.1 | 157297 | 157319 | + | 0 | TGGGTTAGAACTGACTCGGT | TGG | \|\|\|\|\|\|\|\|[\|\|\|\|\|\|\|\|\|\|\|\|]PAM | NA | Intergenic | NA | NA |

<table>
<tr><td>T11922</td><td>CTCGGTTGGTCTACGTGTCGCGG</td><td></td><td>1000</td><td></td><td></td><td></td><td></td><td></td><td></td><td></td><td></td><td></td></tr>
<tr><td>Oligo fwd</td><td>CACCCTCGGTTGGTCTACGTGTCG</td><td></td><td></td><td></td><td></td><td></td><td></td><td></td><td></td><td></td><td></td><td></td></tr>
<tr><td>Oligo rev</td><td>AAACCGACACGTAGACCAACCGAG</td><td></td><td></td><td></td><td></td><td></td><td></td><td></td><td></td><td></td><td></td><td></td></tr>
<tr><td>Chromosome</td><td>start</td><td></td><td>end</td><td>strand</td><td>MM</td><td>target_seq</td><td>PAM</td><td>alignment</td><td>distance</td><td>position</td><td>gene name</td><td>gene id</td></tr>
<tr><td>GU305914.1</td><td></td><td>157311</td><td>157333</td><td>+</td><td>0</td><td>CTCGGTTGGTCTACGTGTCG</td><td>CGG</td><td>||||||||[||||||||||||]PAM</td><td>NA</td><td>Intergenic</td><td>NA</td><td>NA</td></tr>
<tr><td>T11923</td><td>CGAAAAGATCGAGTGATTATCGG</td><td></td><td>1000</td><td></td><td></td><td></td><td></td><td></td><td></td><td></td><td></td><td></td></tr>
<tr><td>Oligo fwd</td><td>CACCCGAAAAGATCGAGTGATTAT</td><td></td><td></td><td></td><td></td><td></td><td></td><td></td><td></td><td></td><td></td><td></td></tr>
<tr><td>Oligo rev</td><td>AAACATAATCACTCGATCTTTTCG</td><td></td><td></td><td></td><td></td><td></td><td></td><td></td><td></td><td></td><td></td><td></td></tr>
<tr><td>Chromosome</td><td>start</td><td></td><td>end</td><td>strand</td><td>MM</td><td>target_seq</td><td>PAM</td><td>alignment</td><td>distance</td><td>position</td><td>gene name</td><td>gene id</td></tr>
<tr><td>GU305914.1</td><td></td><td>157335</td><td>157357</td><td>+</td><td>0</td><td>CGAAAAGATCGAGTGATTAT</td><td>CGG</td><td>||||||||[||||||||||||]PAM</td><td>NA</td><td>Intergenic</td><td>NA</td><td>NA</td></tr>
<tr><td>T11924</td><td>AAGATCGAGTGATTATCGGTTGG</td><td></td><td>1000</td><td></td><td></td><td></td><td></td><td></td><td></td><td></td><td></td><td></td></tr>
<tr><td>Oligo fwd</td><td>CACCAAGATCGAGTGATTATCGGT</td><td></td><td></td><td></td><td></td><td></td><td></td><td></td><td></td><td></td><td></td><td></td></tr>
<tr><td>Oligo rev</td><td>AAACACCGATAATCACTCGATCTT</td><td></td><td></td><td></td><td></td><td></td><td></td><td></td><td></td><td></td><td></td><td></td></tr>
<tr><td>Chromosome</td><td>start</td><td></td><td>end</td><td>strand</td><td>MM</td><td>target_seq</td><td>PAM</td><td>alignment</td><td>distance</td><td>position</td><td>gene name</td><td>gene id</td></tr>
<tr><td>GU305914.1</td><td></td><td>157339</td><td>157361</td><td>+</td><td>0</td><td>AAGATCGAGTGATTATCGGT</td><td>TGG</td><td>||||||||[||||||||||||]PAM</td><td>NA</td><td>Intergenic</td><td>NA</td><td>NA</td></tr>
<tr><td>T11925</td><td>TCGAGTGATTATCGGTTGGCCGG</td><td></td><td>1000</td><td></td><td></td><td></td><td></td><td></td><td></td><td></td><td></td><td></td></tr>
<tr><td>Oligo fwd</td><td>CACCTCGAGTGATTATCGGTTGGC</td><td></td><td></td><td></td><td></td><td></td><td></td><td></td><td></td><td></td><td></td><td></td></tr>
<tr><td>Oligo rev</td><td>AAACGCCAACCGATAATCACTCGA</td><td></td><td></td><td></td><td></td><td></td><td></td><td></td><td></td><td></td><td></td><td></td></tr>
<tr><td>Chromosome</td><td>start</td><td></td><td>end</td><td>strand</td><td>MM</td><td>target_seq</td><td>PAM</td><td>alignment</td><td>distance</td><td>position</td><td>gene name</td><td>gene id</td></tr>
<tr><td>GU305914.1</td><td></td><td>157343</td><td>157365</td><td>+</td><td>0</td><td>TCGAGTGATTATCGGTTGGC</td><td>CGG</td><td>||||||||[||||||||||||]PAM</td><td>NA</td><td>Intergenic</td><td>NA</td><td>NA</td></tr>
<tr><td>T11926</td><td>GATTATCGGTTGGCCGGCCGCGG</td><td></td><td>1000</td><td></td><td></td><td></td><td></td><td></td><td></td><td></td><td></td><td></td></tr>
<tr><td>Oligo fwd</td><td>CACCGATTATCGGTTGGCCGGCCG</td><td></td><td></td><td></td><td></td><td></td><td></td><td></td><td></td><td></td><td></td><td></td></tr>
<tr><td>Oligo rev</td><td>AAACCGGCCGGCCAACCGATAATC</td><td></td><td></td><td></td><td></td><td></td><td></td><td></td><td></td><td></td><td></td><td></td></tr>
<tr><td>Chromosome</td><td>start</td><td></td><td>end</td><td>strand</td><td>MM</td><td>target_seq</td><td>PAM</td><td>alignment</td><td>distance</td><td>position</td><td>gene name</td><td>gene id</td></tr>
<tr><td>GU305914.1</td><td></td><td>157349</td><td>157371</td><td>+</td><td>0</td><td>GATTATCGGTTGGCCGGCCG</td><td>CGG</td><td>||||||||[||||||||||||]PAM</td><td>NA</td><td>Intergenic</td><td>NA</td><td>NA</td></tr>
<tr><td>T11927</td><td>TATCGGTTGGCCGGCCGCGGAGG</td><td></td><td>1000</td><td></td><td></td><td></td><td></td><td></td><td></td><td></td><td></td><td></td></tr>
<tr><td>Oligo fwd</td><td>CACCTATCGGTTGGCCGGCCGCGG</td><td></td><td></td><td></td><td></td><td></td><td></td><td></td><td></td><td></td><td></td><td></td></tr>
<tr><td>Oligo rev</td><td>AAACCCGCGGCCGGCCAACCGATA</td><td></td><td></td><td></td><td></td><td></td><td></td><td></td><td></td><td></td><td></td><td></td></tr>
<tr><td>Chromosome</td><td>start</td><td></td><td>end</td><td>strand</td><td>MM</td><td>target_seq</td><td>PAM</td><td>alignment</td><td>distance</td><td>position</td><td>gene name</td><td>gene id</td></tr>
<tr><td>GU305914.1</td><td></td><td>157352</td><td>157374</td><td>+</td><td>0</td><td>TATCGGTTGGCCGGCCGCGG</td><td>AGG</td><td>||||||||[||||||||||||]PAM</td><td>NA</td><td>Intergenic</td><td>NA</td><td>NA</td></tr>
<tr><td>T11928</td><td>GGAGGCGCGACTCGCGAACGAGG</td><td></td><td>1000</td><td></td><td></td><td></td><td></td><td></td><td></td><td></td><td></td><td></td></tr>
<tr><td>Oligo fwd</td><td>CACCGGAGGCGCGACTCGCGAACG</td><td></td><td></td><td></td><td></td><td></td><td></td><td></td><td></td><td></td><td></td><td></td></tr>
<tr><td>Oligo rev</td><td>AAACCGTTCGCGAGTCGCGCCTCC</td><td></td><td></td><td></td><td></td><td></td><td></td><td></td><td></td><td></td><td></td><td></td></tr>
<tr><td>Chromosome</td><td>start</td><td></td><td>end</td><td>strand</td><td>MM</td><td>target_seq</td><td>PAM</td><td>alignment</td><td>distance</td><td>position</td><td>gene name</td><td>gene id</td></tr>
<tr><td>GU305914.1</td><td></td><td>157370</td><td>157392</td><td>+</td><td>0</td><td>GGAGGCGCGACTCGCGAACG</td><td>AGG</td><td>||||||||[||||||||||||]PAM</td><td>NA</td><td>Intergenic</td><td>NA</td><td>NA</td></tr>
<tr><td>T11929</td><td>AGTCGAGATACATTGATCGATGG</td><td></td><td>1000</td><td></td><td></td><td></td><td></td><td></td><td></td><td></td><td></td><td></td></tr>
<tr><td>Oligo fwd</td><td>CACCAGTCGAGATACATTGATCGA</td><td></td><td></td><td></td><td></td><td></td><td></td><td></td><td></td><td></td><td></td><td></td></tr>
<tr><td>Oligo rev</td><td>AAACTCGATCAATGTATCTCGACT</td><td></td><td></td><td></td><td></td><td></td><td></td><td></td><td></td><td></td><td></td><td></td></tr>
<tr><td>Chromosome</td><td>start</td><td></td><td>end</td><td>strand</td><td>MM</td><td>target_seq</td><td>PAM</td><td>alignment</td><td>distance</td><td>position</td><td>gene name</td><td>gene id</td></tr>
<tr><td>GU305914.1</td><td></td><td>157396</td><td>157418</td><td>+</td><td>0</td><td>AGTCGAGATACATTGATCGA</td><td>TGG</td><td>||||||||[||||||||||||]PAM</td><td>NA</td><td>Intergenic</td><td>NA</td><td>NA</td></tr>
<tr><td>T11930</td><td>AGATACATTGATCGATGGTGAGG</td><td></td><td>1000</td><td></td><td></td><td></td><td></td><td></td><td></td><td></td><td></td><td></td></tr>
<tr><td>Oligo fwd</td><td>CACCAGATACATTGATCGATGGTG</td><td></td><td></td><td></td><td></td><td></td><td></td><td></td><td></td><td></td><td></td><td></td></tr>
<tr><td>Oligo rev</td><td>AAACCACCATCGATCAATGTATCT</td><td></td><td></td><td></td><td></td><td></td><td></td><td></td><td></td><td></td><td></td><td></td></tr>
<tr><td>Chromosome</td><td>start</td><td></td><td>end</td><td>strand</td><td>MM</td><td>target_seq</td><td>PAM</td><td>alignment</td><td>distance</td><td>position</td><td>gene name</td><td>gene id</td></tr>
<tr><td>GU305914.1</td><td></td><td>157401</td><td>157423</td><td>+</td><td>0</td><td>AGATACATTGATCGATGGTG</td><td>AGG</td><td>||||||||[||||||||||||]PAM</td><td>NA</td><td>Intergenic</td><td>NA</td><td>NA</td></tr>
<tr><td>T11931</td><td>TGATCGATGGTGAGGATCTGCGG</td><td></td><td>1000</td><td></td><td></td><td></td><td></td><td></td><td></td><td></td><td></td><td></td></tr>
<tr><td>Oligo fwd</td><td>CACCTGATCGATGGTGAGGATCTG</td><td></td><td></td><td></td><td></td><td></td><td></td><td></td><td></td><td></td><td></td><td></td></tr>
<tr><td>Oligo rev</td><td>AAACCAGATCCTCACCATCGATCA</td><td></td><td></td><td></td><td></td><td></td><td></td><td></td><td></td><td></td><td></td><td></td></tr>
<tr><td>Chromosome</td><td>start</td><td></td><td>end</td><td>strand</td><td>MM</td><td>target_seq</td><td>PAM</td><td>alignment</td><td>distance</td><td>position</td><td>gene name</td><td>gene id</td></tr>
<tr><td>GU305914.1</td><td></td><td>157409</td><td>157431</td><td>+</td><td>0</td><td>TGATCGATGGTGAGGATCTG</td><td>CGG</td><td>||||||||[||||||||||||]PAM</td><td>NA</td><td>Intergenic</td><td>NA</td><td>NA</td></tr>
<tr><td>T11932</td><td>GATCGATGGTGAGGATCTGCGGG</td><td></td><td>1000</td><td></td><td></td><td></td><td></td><td></td><td></td><td></td><td></td><td></td></tr>
<tr><td>Oligo fwd</td><td>CACCGATCGATGGTGAGGATCTGC</td><td></td><td></td><td></td><td></td><td></td><td></td><td></td><td></td><td></td><td></td><td></td></tr>
<tr><td>Oligo rev</td><td>AAACGCAGATCCTCACCATCGATC</td><td></td><td></td><td></td><td></td><td></td><td></td><td></td><td></td><td></td><td></td><td></td></tr>
<tr><td>Chromosome</td><td>start</td><td></td><td>end</td><td>strand</td><td>MM</td><td>target_seq</td><td>PAM</td><td>alignment</td><td>distance</td><td>position</td><td>gene name</td><td>gene id</td></tr>
<tr><td>GU305914.1</td><td></td><td>157410</td><td>157432</td><td>+</td><td>0</td><td>GATCGATGGTGAGGATCTGC</td><td>GGG</td><td>||||||||[||||||||||||]PAM</td><td>NA</td><td>Intergenic</td><td>NA</td><td>NA</td></tr>
<tr><td>T11933</td><td>ATCGATGGTGAGGATCTGCGGGG</td><td></td><td>1000</td><td></td><td></td><td></td><td></td><td></td><td></td><td></td><td></td><td></td></tr>
<tr><td>Oligo fwd</td><td>CACCATCGATGGTGAGGATCTGCG</td><td></td><td></td><td></td><td></td><td></td><td></td><td></td><td></td><td></td><td></td><td></td></tr>
<tr><td>Oligo rev</td><td>AAACCGCAGATCCTCACCATCGAT</td><td></td><td></td><td></td><td></td><td></td><td></td><td></td><td></td><td></td><td></td><td></td></tr>
<tr><td>Chromosome</td><td>start</td><td></td><td>end</td><td>strand</td><td>MM</td><td>target_seq</td><td>PAM</td><td>alignment</td><td>distance</td><td>position</td><td>gene name</td><td>gene id</td></tr>
<tr><td>GU305914.1</td><td></td><td>157411</td><td>157433</td><td>+</td><td>0</td><td>ATCGATGGTGAGGATCTGCG</td><td>GGG</td><td>||||||||[||||||||||||]PAM</td><td>NA</td><td>Intergenic</td><td>NA</td><td>NA</td></tr>
<tr><td>T11934</td><td>GATGGTGAGGATCTGCGGGGCGG</td><td></td><td>1000</td><td></td><td></td><td></td><td></td><td></td><td></td><td></td><td></td><td></td></tr>
<tr><td>Oligo fwd</td><td>CACCGATGGTGAGGATCTGCGGGG</td><td></td><td></td><td></td><td></td><td></td><td></td><td></td><td></td><td></td><td></td><td></td></tr>
<tr><td>Oligo rev</td><td>AAACCCCCGCAGATCCTCACCATC</td><td></td><td></td><td></td><td></td><td></td><td></td><td></td><td></td><td></td><td></td><td></td></tr>
<tr><td>Chromosome</td><td>start</td><td></td><td>end</td><td>strand</td><td>MM</td><td>target_seq</td><td>PAM</td><td>alignment</td><td>distance</td><td>position</td><td>gene name</td><td>gene id</td></tr>
<tr><td>GU305914.1</td><td></td><td>157414</td><td>157436</td><td>+</td><td>0</td><td>GATGGTGAGGATCTGCGGGG</td><td>CGG</td><td>||||||||[||||||||||||]PAM</td><td>NA</td><td>Intergenic</td><td>NA</td><td>NA</td></tr>
<tr><td>T11935</td><td>GATCTGCGGGGCGGTGCGAGTGG</td><td></td><td>1000</td><td></td><td></td><td></td><td></td><td></td><td></td><td></td><td></td><td></td></tr>
<tr><td>Oligo fwd</td><td>CACCGATCTGCGGGGCGGTGCGAG</td><td></td><td></td><td></td><td></td><td></td><td></td><td></td><td></td><td></td><td></td><td></td></tr>
<tr><td>Oligo rev</td><td>AAACCTCGCACCGCCCCGCAGATC</td><td></td><td></td><td></td><td></td><td></td><td></td><td></td><td></td><td></td><td></td><td></td></tr>
<tr><td>Chromosome</td><td>start</td><td></td><td>end</td><td>strand</td><td>MM</td><td>target_seq</td><td>PAM</td><td>alignment</td><td>distance</td><td>position</td><td>gene name</td><td>gene id</td></tr>
<tr><td>GU305914.1</td><td></td><td>157423</td><td>157445</td><td>+</td><td>0</td><td>GATCTGCGGGGCGGTGCGAG</td><td>TGG</td><td>||||||||[||||||||||||]PAM</td><td>NA</td><td>Intergenic</td><td>NA</td><td>NA</td></tr>
<tr><td>T11936</td><td>ATCTGCGGGGCGGTGCGAGTGGG</td><td></td><td>1000</td><td></td><td></td><td></td><td></td><td></td><td></td><td></td><td></td><td></td></tr>
<tr><td>Oligo fwd</td><td>CACCATCTGCGGGGCGGTGCGAGT</td><td></td><td></td><td></td><td></td><td></td><td></td><td></td><td></td><td></td><td></td><td></td></tr>
<tr><td>Oligo rev</td><td>AAACACTCGCACCGCCCCGCAGAT</td><td></td><td></td><td></td><td></td><td></td><td></td><td></td><td></td><td></td><td></td><td></td></tr>
<tr><td>Chromosome</td><td>start</td><td></td><td>end</td><td>strand</td><td>MM</td><td>target_seq</td><td>PAM</td><td>alignment</td><td>distance</td><td>position</td><td>gene name</td><td>gene id</td></tr>
</table>

GU305914.1 157424 157446 + 0 ATCTGCGGGGCGGTGCGAGT GGG ||||||||[||||||||||||]PAM NA Intergenic NA NA

T11937 GGTATAGATAATATCATCGTAGG 1000
Oligo fwd CACCGGTATAGATAATATCATCGT
Oligo rev AAACACGATGATATTATCTATACC
Chromosome start end strand MM target_seq PAM alignment distance position gene name gene id
GU305914.1 157445 157467 + 0 GGTATAGATAATATCATCGT AGG ||||||||[||||||||||||]PAM NA Intergenic NA NA

T11938 GTATCTCTTTTATTCCGTTTCGG 1000
Oligo fwd CACCGTATCTCTTTTATTCCGTTT
Oligo rev AAACAAACGGAATAAAAGAGATAC
Chromosome start end strand MM target_seq PAM alignment distance position gene name gene id
GU305914.1 157467 157489 + 0 GTATCTCTTTTATTCCGTTT CGG ||||||||[||||||||||||]PAM NA Intergenic NA NA

T11939 TCTCTTTTATTCCGTTTCGGTGG 1000
Oligo fwd CACCTCTCTTTTATTCCGTTTCGG
Oligo rev AAACCCGAAACGGAATAAAAGAGA
Chromosome start end strand MM target_seq PAM alignment distance position gene name gene id
GU305914.1 157470 157492 + 0 TCTCTTTTATTCCGTTTCGG TGG ||||||||[||||||||||||]PAM NA Intergenic NA NA

T11940 CTCTTTTATTCCGTTTCGGTGGG 1000
Oligo fwd CACCCTCTTTTATTCCGTTTCGGT
Oligo rev AAACACCGAAACGGAATAAAAGAG
Chromosome start end strand MM target_seq PAM alignment distance position gene name gene id
GU305914.1 157471 157493 + 0 CTCTTTTATTCCGTTTCGGT GGG ||||||||[||||||||||||]PAM NA Intergenic NA NA

T11941 TCTTTTATTCCGTTTCGGTGGGG 1000
Oligo fwd CACCTCTTTTATTCCGTTTCGGTG
Oligo rev AAACCACCGAAACGGAATAAAAGA
Chromosome start end strand MM target_seq PAM alignment distance position gene name gene id
GU305914.1 157472 157494 + 0 TCTTTTATTCCGTTTCGGTG GGG ||||||||[||||||||||||]PAM NA Intergenic NA NA

T11942 CTTTTATTCCGTTTCGGTGGGGG 1000
Oligo fwd CACCCTTTTATTCCGTTTCGGTGG
Oligo rev AAACCCACCGAAACGGAATAAAAG
Chromosome start end strand MM target_seq PAM alignment distance position gene name gene id
GU305914.1 157473 157495 + 0 CTTTTATTCCGTTTCGGTGG GGG ||||||||[||||||||||||]PAM NA Intergenic NA NA

T11943 CTCGATTCCTTGACAAAACTAGG 1000
Oligo fwd CACCCTCGATTCCTTGACAAAACT
Oligo rev AAACAGTTTTGTCAAGGAATCGAG
Chromosome start end strand MM target_seq PAM alignment distance position gene name gene id
GU305914.1 157521 157543 + 0 CTCGATTCCTTGACAAAACT AGG ||||||||[||||||||||||]PAM NA Intergenic NA NA

T11944 AAAACTAGGTATAACATATCTGG 1000
Oligo fwd CACCAAAACTAGGTATAACATATC
Oligo rev AAACGATATGTTATACCTAGTTTT
Chromosome start end strand MM target_seq PAM alignment distance position gene name gene id
GU305914.1 157535 157557 + 0 AAAACTAGGTATAACATATC TGG ||||||||[||||||||||||]PAM NA Intergenic NA NA

T11945 GTATAACATATCTGGATTTCTGG 1000
Oligo fwd CACCGTATAACATATCTGGATTTC
Oligo rev AAACGAAATCCAGATATGTTATAC
Chromosome start end strand MM target_seq PAM alignment distance position gene name gene id
GU305914.1 157543 157565 + 0 GTATAACATATCTGGATTTC TGG ||||||||[||||||||||||]PAM NA Intergenic NA NA

T11946 TATAACATATCTGGATTTCTGGG 1000
Oligo fwd CACCTATAACATATCTGGATTTCT
Oligo rev AAACAGAAATCCAGATATGTTATA
Chromosome start end strand MM target_seq PAM alignment distance position gene name gene id
GU305914.1 157544 157566 + 0 TATAACATATCTGGATTTCT GGG ||||||||[||||||||||||]PAM NA Intergenic NA NA

T11947 TCACATGTCCGTATTCCAAGAGG 1000
Oligo fwd CACCTCACATGTCCGTATTCCAAG
Oligo rev AAACCTTGGAATACGGACATGTGA
Chromosome start end strand MM target_seq PAM alignment distance position gene name gene id
GU305914.1 157627 157649 + 0 TCACATGTCCGTATTCCAAG AGG ||||||||[||||||||||||]PAM NA Intergenic NA NA

T11948 CATTGATTTAATTGATACACAGG 1000
Oligo fwd CACCCATTGATTTAATTGATACAC
Oligo rev AAACGTGTATCAATTAAATCAATG
Chromosome start end strand MM target_seq PAM alignment distance position gene name gene id
GU305914.1 157682 157704 + 0 CATTGATTTAATTGATACAC AGG ||||||||[||||||||||||]PAM NA Intergenic NA NA

T11949 AGTAGTATATTTCGATTAGCTGG 1000
Oligo fwd CACCAGTAGTATATTTCGATTAGC
Oligo rev AAACGCTAATCGAAATATACTACT
Chromosome start end strand MM target_seq PAM alignment distance position gene name gene id
GU305914.1 157760 157782 + 0 AGTAGTATATTTCGATTAGC TGG ||||||||[||||||||||||]PAM NA Intergenic NA NA

T11950 GATTAGCTGGCTTTGATTAAAGG 1000
Oligo fwd CACCGATTAGCTGGCTTTGATTAA
Oligo rev AAACTTAATCAAAGCCAGCTAATC
Chromosome start end strand MM target_seq PAM alignment distance position gene name gene id
GU305914.1 157773 157795 + 0 GATTAGCTGGCTTTGATTAA AGG ||||||||[||||||||||||]PAM NA Intergenic NA NA

T11951 TGGCTTTGATTAAAGGTGTTTGG 1000
Oligo fwd CACCTGGCTTTGATTAAAGGTGTT
Oligo rev AAACAACACCTTTAATCAAAGCCA

Chromosome start end strand MM target_seq PAM alignment distance position gene name gene id
GU305914.1 157780 157802 + 0 TGGCTTTGATTAAAGGTGTT TGG ||||||||[||||||||||||]PAM NA Intergenic NA NA

T11952 GTATTCATCATTAAGATTATCGG 1000
Oligo fwd CACCGTATTCATCATTAAGATTAT
Oligo rev AAACATAATCTTAATGATGAATAC
Chromosome start end strand MM target_seq PAM alignment distance position gene name gene id
GU305914.1 157835 157857 + 0 GTATTCATCATTAAGATTAT CGG ||||||||[||||||||||||]PAM NA Intergenic NA NA

T11953 GGATTTTTTTTTTTTTTACGTGG 1000
Oligo fwd CACCGGATTTTTTTTTTTTTTACG
Oligo rev AAACCGTAAAAAAAAAAAAAATCC
Chromosome start end strand MM target_seq PAM alignment distance position gene name gene id
GU305914.1 157856 157878 + 0 GGATTTTTTTTTTTTTTACG TGG ||||||||[||||||||||||]PAM NA Intergenic NA NA

T11954 GATTTTTTTTTTTTTTACGTGGG 1000
Oligo fwd CACCGATTTTTTTTTTTTTTACGT
Oligo rev AAACACGTAAAAAAAAAAAAAATC
Chromosome start end strand MM target_seq PAM alignment distance position gene name gene id
GU305914.1 157857 157879 + 0 GATTTTTTTTTTTTTTACGT GGG ||||||||[||||||||||||]PAM NA Intergenic NA NA

T11955 AAAACAAACTGCATCCCGTATGG 1000
Oligo fwd CACCAAAACAAACTGCATCCCGTA
Oligo rev AAACTACGGGATGCAGTTTGTTTT
Chromosome start end strand MM target_seq PAM alignment distance position gene name gene id
GU305914.1 157889 157911 + 0 AAAACAAACTGCATCCCGTA TGG ||||||||[||||||||||||]PAM NA Intergenic NA NA

T11956 ATCCCGTATGGTGCATACGAAGG 1000
Oligo fwd CACCATCCCGTATGGTGCATACGA
Oligo rev AAACTCGTATGCACCATACGGGAT
Chromosome start end strand MM target_seq PAM alignment distance position gene name gene id
GU305914.1 157901 157923 + 0 ATCCCGTATGGTGCATACGA AGG ||||||||[||||||||||||]PAM NA Intergenic NA NA

T11957 GGTGCATACGAAGGATAGATAGG 1000
Oligo fwd CACCGGTGCATACGAAGGATAGAT
Oligo rev AAACATCTATCCTTCGTATGCACC
Chromosome start end strand MM target_seq PAM alignment distance position gene name gene id
GU305914.1 157910 157932 + 0 GGTGCATACGAAGGATAGAT AGG ||||||||[||||||||||||]PAM NA Intergenic NA NA

T11958 GTGCATACGAAGGATAGATAGGG 1000
Oligo fwd CACCGTGCATACGAAGGATAGATA
Oligo rev AAACTATCTATCCTTCGTATGCAC
Chromosome start end strand MM target_seq PAM alignment distance position gene name gene id
GU305914.1 157911 157933 + 0 GTGCATACGAAGGATAGATA GGG ||||||||[||||||||||||]PAM NA Intergenic NA NA

T11959 TCATTTCTGTTGTTTTCCGCAGG 1000
Oligo fwd CACCTCATTTCTGTTGTTTTCCGC
Oligo rev AAACGCGGAAAACAACAGAAATGA
Chromosome start end strand MM target_seq PAM alignment distance position gene name gene id
GU305914.1 157944 157966 + 0 TCATTTCTGTTGTTTTCCGC AGG ||||||||[||||||||||||]PAM NA Intergenic NA NA

T11960 TTTCCGCAGGTGATCACCGTCGG 1000
Oligo fwd CACCTTTCCGCAGGTGATCACCGT
Oligo rev AAACACGGTGATCACCTGCGGAAA
Chromosome start end strand MM target_seq PAM alignment distance position gene name gene id
GU305914.1 157957 157979 + 0 TTTCCGCAGGTGATCACCGT CGG ||||||||[||||||||||||]PAM NA Intergenic NA NA

T11961 TTCCGCAGGTGATCACCGTCGGG 1000
Oligo fwd CACCTTCCGCAGGTGATCACCGTC
Oligo rev AAACGACGGTGATCACCTGCGGAA
Chromosome start end strand MM target_seq PAM alignment distance position gene name gene id
GU305914.1 157958 157980 + 0 TTCCGCAGGTGATCACCGTC GGG ||||||||[||||||||||||]PAM NA Intergenic NA NA

T11962 GGTGATCACCGTCGGGACCGCGG 1000
Oligo fwd CACCGGTGATCACCGTCGGGACCG
Oligo rev AAACCGGTCCCGACGGTGATCACC
Chromosome start end strand MM target_seq PAM alignment distance position gene name gene id
GU305914.1 157965 157987 + 0 GGTGATCACCGTCGGGACCG CGG ||||||||[||||||||||||]PAM NA Intergenic NA NA

T11963 CACCGTCGGGACCGCGGTCGCGG 1000
Oligo fwd CACCCACCGTCGGGACCGCGGTCG
Oligo rev AAACCGACCGCGGTCCCGACGGTG
Chromosome start end strand MM target_seq PAM alignment distance position gene name gene id
GU305914.1 157971 157993 + 0 CACCGTCGGGACCGCGGTCG CGG ||||||||[||||||||||||]PAM NA Intergenic NA NA

T11964 GTTGTCGCAAAGTGATGAAATGG 1000
Oligo fwd CACCGTTGTCGCAAAGTGATGAAA
Oligo rev AAACTTTCATCACTTTGCGACAAC
Chromosome start end strand MM target_seq PAM alignment distance position gene name gene id
GU305914.1 158055 158077 + 0 GTTGTCGCAAAGTGATGAAA TGG ||||||||[||||||||||||]PAM NA Intergenic NA NA

T11965 AGTGATGAAATGGTGTGTATAGG 1000
Oligo fwd CACCAGTGATGAAATGGTGTGTAT
Oligo rev AAACATACACACCATTTCATCACT
Chromosome start end strand MM target_seq PAM alignment distance position gene name gene id
GU305914.1 158065 158087 + 0 AGTGATGAAATGGTGTGTAT AGG ||||||||[||||||||||||]PAM NA Intergenic NA NA

T11966 ATGAAATGGTGTGTATAGGATGG 1000
Oligo fwd CACCATGAAATGGTGTGTATAGGA

Oligo rev AAACTCCTATACACACCATTTCAT
Chromosome start end strand MM target_seq PAM alignment distance position gene name gene id
GU305914.1 158069 158091 + 0 ATGAAATGGTGTGTATAGGA TGG ||||||||[||||||||||||]PAM NA Intergenic NA NA

T11967 TGAAATGGTGTGTATAGGATGGG 1000
Oligo fwd CACCTGAAATGGTGTGTATAGGAT
Oligo rev AAACATCCTATACACACCATTTCA
Chromosome start end strand MM target_seq PAM alignment distance position gene name gene id
GU305914.1 158070 158092 + 0 TGAAATGGTGTGTATAGGAT GGG ||||||||[||||||||||||]PAM NA Intergenic NA NA

T11968 GCGATTAACACGTTGAGCACTGG 1000
Oligo fwd CACCGCGATTAACACGTTGAGCAC
Oligo rev AAACGTGCTCAACGTGTTAATCGC
Chromosome start end strand MM target_seq PAM alignment distance position gene name gene id
GU305914.1 158102 158124 + 0 GCGATTAACACGTTGAGCAC TGG ||||||||[||||||||||||]PAM NA Intergenic NA NA

T11969 CGATTAACACGTTGAGCACTGGG 1000
Oligo fwd CACCCGATTAACACGTTGAGCACT
Oligo rev AAACAGTGCTCAACGTGTTAATCG
Chromosome start end strand MM target_seq PAM alignment distance position gene name gene id
GU305914.1 158103 158125 + 0 CGATTAACACGTTGAGCACT GGG ||||||||[||||||||||||]PAM NA Intergenic NA NA

T11970 ACCCCTATTGCCTAAAATTATGG 1000
Oligo fwd CACCACCCCTATTGCCTAAAATTA
Oligo rev AAACTAATTTTAGGCAATAGGGGT
Chromosome start end strand MM target_seq PAM alignment distance position gene name gene id
GU305914.1 158239 158261 + 0 ACCCCTATTGCCTAAAATTA TGG ||||||||[||||||||||||]PAM NA Intergenic NA NA

T11971 AAATTATGGAAAATCGTATATGG 1000
Oligo fwd CACCAAATTATGGAAAATCGTATA
Oligo rev AAACTATACGATTTTCCATAATTT
Chromosome start end strand MM target_seq PAM alignment distance position gene name gene id
GU305914.1 158253 158275 + 0 AAATTATGGAAAATCGTATA TGG ||||||||[||||||||||||]PAM NA Intergenic NA NA

T11972 TGGAAAATCGTATATGGAAAAGG 1000
Oligo fwd CACCTGGAAAATCGTATATGGAAA
Oligo rev AAACTTTCCATATACGATTTTCCA
Chromosome start end strand MM target_seq PAM alignment distance position gene name gene id
GU305914.1 158259 158281 + 0 TGGAAAATCGTATATGGAAA AGG ||||||||[||||||||||||]PAM NA Intergenic NA NA

T11973 GGAAAATCGTATATGGAAAAGGG 1000
Oligo fwd CACCGGAAAATCGTATATGGAAAA
Oligo rev AAACTTTTCCATATACGATTTTCC
Chromosome start end strand MM target_seq PAM alignment distance position gene name gene id
GU305914.1 158260 158282 + 0 GGAAAATCGTATATGGAAAA GGG ||||||||[||||||||||||]PAM NA Intergenic NA NA

T11974 CGATAAGACCACGAATTGAGTGG 1000
Oligo fwd CACCCGATAAGACCACGAATTGAG
Oligo rev AAACCTCAATTCGTGGTCTTATCG
Chromosome start end strand MM target_seq PAM alignment distance position gene name gene id
GU305914.1 158315 158337 + 0 CGATAAGACCACGAATTGAG TGG ||||||||[||||||||||||]PAM NA Intergenic NA NA

T11975 GATAAGACCACGAATTGAGTGGG 1000
Oligo fwd CACCGATAAGACCACGAATTGAGT
Oligo rev AAACACTCAATTCGTGGTCTTATC
Chromosome start end strand MM target_seq PAM alignment distance position gene name gene id
GU305914.1 158316 158338 + 0 GATAAGACCACGAATTGAGT GGG ||||||||[||||||||||||]PAM NA Intergenic NA NA

T11976 GTAATTATTAACTAACTGATTGG 1000
Oligo fwd CACCGTAATTATTAACTAACTGAT
Oligo rev AAACATCAGTTAGTTAATAATTAC
Chromosome start end strand MM target_seq PAM alignment distance position gene name gene id
GU305914.1 158416 158438 + 0 GTAATTATTAACTAACTGAT TGG ||||||||[||||||||||||]PAM NA Intergenic NA NA

T11977 GTAGAACGAAAGCCCTAAGTTGG 1000
Oligo fwd CACCGTAGAACGAAAGCCCTAAGT
Oligo rev AAACACTTAGGGCTTTCGTTCTAC
Chromosome start end strand MM target_seq PAM alignment distance position gene name gene id
GU305914.1 158441 158463 + 0 GTAGAACGAAAGCCCTAAGT TGG ||||||||[||||||||||||]PAM NA Intergenic NA NA

T11978 TTAATCCTTTATGTGCTATATGG 1000
Oligo fwd CACCTTAATCCTTTATGTGCTATA
Oligo rev AAACTATAGCACATAAAGGATTAA
Chromosome start end strand MM target_seq PAM alignment distance position gene name gene id
GU305914.1 158482 158504 + 0 TTAATCCTTTATGTGCTATA TGG ||||||||[||||||||||||]PAM NA Intergenic NA NA

T11979 TCTAACTTATTTACCTACCTTGG 1000
Oligo fwd CACCTCTAACTTATTTACCTACCT
Oligo rev AAACAGGTAGGTAAATAAGTTAGA
Chromosome start end strand MM target_seq PAM alignment distance position gene name gene id
GU305914.1 158548 158570 + 0 TCTAACTTATTTACCTACCT TGG ||||||||[||||||||||||]PAM NA Intergenic NA NA

T11980 ATTGATTCGTATGTGTTCTATGG 1000
Oligo fwd CACCATTGATTCGTATGTGTTCTA
Oligo rev AAACTAGAACACATACGAATCAAT
Chromosome start end strand MM target_seq PAM alignment distance position gene name gene id
GU305914.1 158581 158603 + 0 ATTGATTCGTATGTGTTCTA TGG ||||||||[||||||||||||]PAM NA Intergenic NA NA

T11981 AAATAATAAAAGAAAATCTATGG 1000

| | | | | | | | | | | | |
|---|---|---|---|---|---|---|---|---|---|---|---|
| Oligo fwd | CACCAAATAATAAAAGAAAATCTA | | | | | | | | | | |
| Oligo rev | AAACTAGATTTTCTTTTATTATTT | | | | | | | | | | |
| Chromosome | start | end | strand | MM | target_seq | PAM | alignment | distance | position | gene name | gene id |
| GU305914.1 | 158636 | 158658 | + | 0 | AAATAATAAAAGAAAATCTA | TGG | \|\|\|\|\|\|\|\|[\|\|\|\|\|\|\|\|\|\|\|\|]PAM | NA | Intergenic | NA | NA |
| T11982 | CAAGTTAAAGTTATTTTCTTTGG | 1000 | | | | | | | | | |
| Oligo fwd | CACCCAAGTTAAAGTTATTTTCTT | | | | | | | | | | |
| Oligo rev | AAACAAGAAAATAACTTTAACTTG | | | | | | | | | | |
| Chromosome | start | end | strand | MM | target_seq | PAM | alignment | distance | position | gene name | gene id |
| GU305914.1 | 158738 | 158760 | + | 0 | CAAGTTAAAGTTATTTTCTT | TGG | \|\|\|\|\|\|\|\|[\|\|\|\|\|\|\|\|\|\|\|\|]PAM | NA | Intergenic | NA | NA |
| T11983 | TCGCAGACTGCGAGCGCGACCGG | 1000 | | | | | | | | | |
| Oligo fwd | CACCTCGCAGACTGCGAGCGCGAC | | | | | | | | | | |
| Oligo rev | AAACGTCGCGCTCGCAGTCTGCGA | | | | | | | | | | |
| Chromosome | start | end | strand | MM | target_seq | PAM | alignment | distance | position | gene name | gene id |
| GU305914.1 | 158781 | 158803 | + | 0 | TCGCAGACTGCGAGCGCGAC | CGG | \|\|\|\|\|\|\|\|[\|\|\|\|\|\|\|\|\|\|\|\|]PAM | NA | Intergenic | NA | NA |
| T11984 | TCTCTCGCACGCTCATCGCGAGG | 1000 | | | | | | | | | |
| Oligo fwd | CACCTCTCTCGCACGCTCATCGCG | | | | | | | | | | |
| Oligo rev | AAACCGCGATGAGCGTGCGAGAGA | | | | | | | | | | |
| Chromosome | start | end | strand | MM | target_seq | PAM | alignment | distance | position | gene name | gene id |
| GU305914.1 | 158808 | 158830 | + | 0 | TCTCTCGCACGCTCATCGCG | AGG | \|\|\|\|\|\|\|\|[\|\|\|\|\|\|\|\|\|\|\|\|]PAM | NA | Intergenic | NA | NA |
| T11985 | GAGGCAGAAACTTCGAAAAAAGG | 1000 | | | | | | | | | |
| Oligo fwd | CACCGAGGCAGAAACTTCGAAAAA | | | | | | | | | | |
| Oligo rev | AAACTTTTTCGAAGTTTCTGCCTC | | | | | | | | | | |
| Chromosome | start | end | strand | MM | target_seq | PAM | alignment | distance | position | gene name | gene id |
| GU305914.1 | 158827 | 158849 | + | 0 | GAGGCAGAAACTTCGAAAAA | AGG | \|\|\|\|\|\|\|\|[\|\|\|\|\|\|\|\|\|\|\|\|]PAM | NA | Intergenic | NA | NA |
| T11986 | AGGCAGAAACTTCGAAAAAAGGG | 1000 | | | | | | | | | |
| Oligo fwd | CACCAGGCAGAAACTTCGAAAAAA | | | | | | | | | | |
| Oligo rev | AAACTTTTTTCGAAGTTTCTGCCT | | | | | | | | | | |
| Chromosome | start | end | strand | MM | target_seq | PAM | alignment | distance | position | gene name | gene id |
| GU305914.1 | 158828 | 158850 | + | 0 | AGGCAGAAACTTCGAAAAAA | GGG | \|\|\|\|\|\|\|\|[\|\|\|\|\|\|\|\|\|\|\|\|]PAM | NA | Intergenic | NA | NA |
| T11987 | CAGAAACTTCGAAAAAAGGGAGG | 1000 | | | | | | | | | |
| Oligo fwd | CACCCAGAAACTTCGAAAAAAGGG | | | | | | | | | | |
| Oligo rev | AAACCCCTTTTTTCGAAGTTTCTG | | | | | | | | | | |
| Chromosome | start | end | strand | MM | target_seq | PAM | alignment | distance | position | gene name | gene id |
| GU305914.1 | 158831 | 158853 | + | 0 | CAGAAACTTCGAAAAAAGGG | AGG | \|\|\|\|\|\|\|\|[\|\|\|\|\|\|\|\|\|\|\|\|]PAM | NA | Intergenic | NA | NA |
| T11988 | GAAAAAAGGGAGGTTCTCCATGG | 1000 | | | | | | | | | |
| Oligo fwd | CACCGAAAAAAGGGAGGTTCTCCA | | | | | | | | | | |
| Oligo rev | AAACTGGAGAACCTCCCTTTTTTC | | | | | | | | | | |
| Chromosome | start | end | strand | MM | target_seq | PAM | alignment | distance | position | gene name | gene id |
| GU305914.1 | 158841 | 158863 | + | 0 | GAAAAAAGGGAGGTTCTCCA | TGG | \|\|\|\|\|\|\|\|[\|\|\|\|\|\|\|\|\|\|\|\|]PAM | NA | Intergenic | NA | NA |
| T11989 | AAAAAAGGGAGGTTCTCCATGGG | 1000 | | | | | | | | | |
| Oligo fwd | CACCAAAAAAGGGAGGTTCTCCAT | | | | | | | | | | |
| Oligo rev | AAACATGGAGAACCTCCCTTTTTT | | | | | | | | | | |
| Chromosome | start | end | strand | MM | target_seq | PAM | alignment | distance | position | gene name | gene id |
| GU305914.1 | 158842 | 158864 | + | 0 | AAAAAAGGGAGGTTCTCCAT | GGG | \|\|\|\|\|\|\|\|[\|\|\|\|\|\|\|\|\|\|\|\|]PAM | NA | Intergenic | NA | NA |
| T11990 | TCCATGGGAACCCGTGTGTCTGG | 1000 | | | | | | | | | |
| Oligo fwd | CACCTCCATGGGAACCCGTGTGTC | | | | | | | | | | |
| Oligo rev | AAACGACACACGGGTTCCCATGGA | | | | | | | | | | |
| Chromosome | start | end | strand | MM | target_seq | PAM | alignment | distance | position | gene name | gene id |
| GU305914.1 | 158857 | 158879 | + | 0 | TCCATGGGAACCCGTGTGTC | TGG | \|\|\|\|\|\|\|\|[\|\|\|\|\|\|\|\|\|\|\|\|]PAM | NA | Intergenic | NA | NA |
| T11991 | CCATGGGAACCCGTGTGTCTGGG | 1000 | | | | | | | | | |
| Oligo fwd | CACCCCATGGGAACCCGTGTGTCT | | | | | | | | | | |
| Oligo rev | AAACAGACACACGGGTTCCCATGG | | | | | | | | | | |
| Chromosome | start | end | strand | MM | target_seq | PAM | alignment | distance | position | gene name | gene id |
| GU305914.1 | 158858 | 158880 | + | 0 | CCATGGGAACCCGTGTGTCT | GGG | \|\|\|\|\|\|\|\|[\|\|\|\|\|\|\|\|\|\|\|\|]PAM | NA | Intergenic | NA | NA |
| T11992 | CATGGGAACCCGTGTGTCTGGGG | 1000 | | | | | | | | | |
| Oligo fwd | CACCCATGGGAACCCGTGTGTCTG | | | | | | | | | | |
| Oligo rev | AAACCAGACACACGGGTTCCCATG | | | | | | | | | | |
| Chromosome | start | end | strand | MM | target_seq | PAM | alignment | distance | position | gene name | gene id |
| GU305914.1 | 158859 | 158881 | + | 0 | CATGGGAACCCGTGTGTCTG | GGG | \|\|\|\|\|\|\|\|[\|\|\|\|\|\|\|\|\|\|\|\|]PAM | NA | Intergenic | NA | NA |
| T11993 | AACAACAAAAAAAAAACAAACGG | 1000 | | | | | | | | | |
| Oligo fwd | CACCAACAACAAAAAAAAAACAAA | | | | | | | | | | |
| Oligo rev | AAACTTTGTTTTTTTTTTGTTGTT | | | | | | | | | | |
| Chromosome | start | end | strand | MM | target_seq | PAM | alignment | distance | position | gene name | gene id |
| GU305914.1 | 158901 | 158923 | + | 0 | AACAACAAAAAAAAAACAAA | CGG | \|\|\|\|\|\|\|\|[\|\|\|\|\|\|\|\|\|\|\|\|]PAM | NA | Intergenic | NA | NA |
| T11994 | ACGGTTTTTTTCCACTGAAAAGG | 1000 | | | | | | | | | |
| Oligo fwd | CACCACGGTTTTTTTCCACTGAAA | | | | | | | | | | |
| Oligo rev | AAACTTTCAGTGGAAAAAAACCGT | | | | | | | | | | |
| Chromosome | start | end | strand | MM | target_seq | PAM | alignment | distance | position | gene name | gene id |
| GU305914.1 | 158920 | 158942 | + | 0 | ACGGTTTTTTTCCACTGAAA | AGG | \|\|\|\|\|\|\|\|[\|\|\|\|\|\|\|\|\|\|\|\|]PAM | NA | Intergenic | NA | NA |
| T11995 | AGTGAATGCTTTTGAAAGAATGG | 1000 | | | | | | | | | |
| Oligo fwd | CACCAGTGAATGCTTTTGAAAGAA | | | | | | | | | | |
| Oligo rev | AAACTTCTTTCAAAAGCATTCACT | | | | | | | | | | |
| Chromosome | start | end | strand | MM | target_seq | PAM | alignment | distance | position | gene name | gene id |
| GU305914.1 | 158954 | 158976 | + | 0 | AGTGAATGCTTTTGAAAGAA | TGG | \|\|\|\|\|\|\|\|[\|\|\|\|\|\|\|\|\|\|\|\|]PAM | NA | Intergenic | NA | NA |

<table>
<tr><td>T11996</td><td>CGCCCGACGTGCGACCAGCGCGG</td><td></td><td>1000</td><td></td><td></td><td></td><td></td><td></td><td></td><td></td><td></td><td></td></tr>
<tr><td>Oligo fwd</td><td>CACCCGCCCGACGTGCGACCAGCG</td><td></td><td></td><td></td><td></td><td></td><td></td><td></td><td></td><td></td><td></td><td></td></tr>
<tr><td>Oligo rev</td><td>AAACCGCTGGTCGCACGTCGGGCG</td><td></td><td></td><td></td><td></td><td></td><td></td><td></td><td></td><td></td><td></td><td></td></tr>
<tr><td>Chromosome</td><td>start</td><td></td><td>end</td><td>strand</td><td>MM</td><td>target_seq</td><td>PAM</td><td>alignment</td><td>distance</td><td>position</td><td>gene name</td><td>gene id</td></tr>
<tr><td>GU305914.1</td><td></td><td>158983</td><td>159005</td><td>+</td><td>0</td><td>CGCCCGACGTGCGACCAGCG</td><td>CGG</td><td>||||||||[||||||||||||]PAM</td><td>NA</td><td>Intergenic</td><td>NA</td><td>NA</td></tr>
<tr><td>T11997</td><td>CGTGCGACCAGCGCGGCCATCGG</td><td></td><td>1000</td><td></td><td></td><td></td><td></td><td></td><td></td><td></td><td></td><td></td></tr>
<tr><td>Oligo fwd</td><td>CACCCGTGCGACCAGCGCGGCCAT</td><td></td><td></td><td></td><td></td><td></td><td></td><td></td><td></td><td></td><td></td><td></td></tr>
<tr><td>Oligo rev</td><td>AAACATGGCCGCGCTGGTCGCACG</td><td></td><td></td><td></td><td></td><td></td><td></td><td></td><td></td><td></td><td></td><td></td></tr>
<tr><td>Chromosome</td><td>start</td><td></td><td>end</td><td>strand</td><td>MM</td><td>target_seq</td><td>PAM</td><td>alignment</td><td>distance</td><td>position</td><td>gene name</td><td>gene id</td></tr>
<tr><td>GU305914.1</td><td></td><td>158990</td><td>159012</td><td>+</td><td>0</td><td>CGTGCGACCAGCGCGGCCAT</td><td>CGG</td><td>||||||||[||||||||||||]PAM</td><td>NA</td><td>Intergenic</td><td>NA</td><td>NA</td></tr>
<tr><td>T11998</td><td>GTGCGACCAGCGCGGCCATCGGG</td><td></td><td>1000</td><td></td><td></td><td></td><td></td><td></td><td></td><td></td><td></td><td></td></tr>
<tr><td>Oligo fwd</td><td>CACCGTGCGACCAGCGCGGCCATC</td><td></td><td></td><td></td><td></td><td></td><td></td><td></td><td></td><td></td><td></td><td></td></tr>
<tr><td>Oligo rev</td><td>AAACGATGGCCGCGCTGGTCGCAC</td><td></td><td></td><td></td><td></td><td></td><td></td><td></td><td></td><td></td><td></td><td></td></tr>
<tr><td>Chromosome</td><td>start</td><td></td><td>end</td><td>strand</td><td>MM</td><td>target_seq</td><td>PAM</td><td>alignment</td><td>distance</td><td>position</td><td>gene name</td><td>gene id</td></tr>
<tr><td>GU305914.1</td><td></td><td>158991</td><td>159013</td><td>+</td><td>0</td><td>GTGCGACCAGCGCGGCCATC</td><td>GGG</td><td>||||||||[||||||||||||]PAM</td><td>NA</td><td>Intergenic</td><td>NA</td><td>NA</td></tr>
<tr><td>T11999</td><td>CAGCGCGGCCATCGGGAGCACGG</td><td></td><td>1000</td><td></td><td></td><td></td><td></td><td></td><td></td><td></td><td></td><td></td></tr>
<tr><td>Oligo fwd</td><td>CACCCAGCGCGGCCATCGGGAGCA</td><td></td><td></td><td></td><td></td><td></td><td></td><td></td><td></td><td></td><td></td><td></td></tr>
<tr><td>Oligo rev</td><td>AAACTGCTCCCGATGGCCGCGCTG</td><td></td><td></td><td></td><td></td><td></td><td></td><td></td><td></td><td></td><td></td><td></td></tr>
<tr><td>Chromosome</td><td>start</td><td></td><td>end</td><td>strand</td><td>MM</td><td>target_seq</td><td>PAM</td><td>alignment</td><td>distance</td><td>position</td><td>gene name</td><td>gene id</td></tr>
<tr><td>GU305914.1</td><td></td><td>158998</td><td>159020</td><td>+</td><td>0</td><td>CAGCGCGGCCATCGGGAGCA</td><td>CGG</td><td>||||||||[||||||||||||]PAM</td><td>NA</td><td>Intergenic</td><td>NA</td><td>NA</td></tr>
<tr><td>T12000</td><td>CGGCCATCGGGAGCACGGCACGG</td><td></td><td>1000</td><td></td><td></td><td></td><td></td><td></td><td></td><td></td><td></td><td></td></tr>
<tr><td>Oligo fwd</td><td>CACCCGGCCATCGGGAGCACGGCA</td><td></td><td></td><td></td><td></td><td></td><td></td><td></td><td></td><td></td><td></td><td></td></tr>
<tr><td>Oligo rev</td><td>AAACTGCCGTGCTCCCGATGGCCG</td><td></td><td></td><td></td><td></td><td></td><td></td><td></td><td></td><td></td><td></td><td></td></tr>
<tr><td>Chromosome</td><td>start</td><td></td><td>end</td><td>strand</td><td>MM</td><td>target_seq</td><td>PAM</td><td>alignment</td><td>distance</td><td>position</td><td>gene name</td><td>gene id</td></tr>
<tr><td>GU305914.1</td><td></td><td>159003</td><td>159025</td><td>+</td><td>0</td><td>CGGCCATCGGGAGCACGGCA</td><td>CGG</td><td>||||||||[||||||||||||]PAM</td><td>NA</td><td>Intergenic</td><td>NA</td><td>NA</td></tr>
<tr><td>T12001</td><td>ACACACCTTCCCTTGCTGCGAGG</td><td></td><td>1000</td><td></td><td></td><td></td><td></td><td></td><td></td><td></td><td></td><td></td></tr>
<tr><td>Oligo fwd</td><td>CACCACACACCTTCCCTTGCTGCG</td><td></td><td></td><td></td><td></td><td></td><td></td><td></td><td></td><td></td><td></td><td></td></tr>
<tr><td>Oligo rev</td><td>AAACCGCAGCAAGGGAAGGTGTGT</td><td></td><td></td><td></td><td></td><td></td><td></td><td></td><td></td><td></td><td></td><td></td></tr>
<tr><td>Chromosome</td><td>start</td><td></td><td>end</td><td>strand</td><td>MM</td><td>target_seq</td><td>PAM</td><td>alignment</td><td>distance</td><td>position</td><td>gene name</td><td>gene id</td></tr>
<tr><td>GU305914.1</td><td></td><td>159031</td><td>159053</td><td>+</td><td>0</td><td>ACACACCTTCCCTTGCTGCG</td><td>AGG</td><td>||||||||[||||||||||||]PAM</td><td>NA</td><td>Intergenic</td><td>NA</td><td>NA</td></tr>
<tr><td>T12002</td><td>ACCTTCCCTTGCTGCGAGGAAGG</td><td></td><td>1000</td><td></td><td></td><td></td><td></td><td></td><td></td><td></td><td></td><td></td></tr>
<tr><td>Oligo fwd</td><td>CACCACCTTCCCTTGCTGCGAGGA</td><td></td><td></td><td></td><td></td><td></td><td></td><td></td><td></td><td></td><td></td><td></td></tr>
<tr><td>Oligo rev</td><td>AAACTCCTCGCAGCAAGGGAAGGT</td><td></td><td></td><td></td><td></td><td></td><td></td><td></td><td></td><td></td><td></td><td></td></tr>
<tr><td>Chromosome</td><td>start</td><td></td><td>end</td><td>strand</td><td>MM</td><td>target_seq</td><td>PAM</td><td>alignment</td><td>distance</td><td>position</td><td>gene name</td><td>gene id</td></tr>
<tr><td>GU305914.1</td><td></td><td>159035</td><td>159057</td><td>+</td><td>0</td><td>ACCTTCCCTTGCTGCGAGGA</td><td>AGG</td><td>||||||||[||||||||||||]PAM</td><td>NA</td><td>Intergenic</td><td>NA</td><td>NA</td></tr>
<tr><td>T12003</td><td>CCTTCCCTTGCTGCGAGGAAGGG</td><td></td><td>1000</td><td></td><td></td><td></td><td></td><td></td><td></td><td></td><td></td><td></td></tr>
<tr><td>Oligo fwd</td><td>CACCCCTTCCCTTGCTGCGAGGAA</td><td></td><td></td><td></td><td></td><td></td><td></td><td></td><td></td><td></td><td></td><td></td></tr>
<tr><td>Oligo rev</td><td>AAACTTCCTCGCAGCAAGGGAAGG</td><td></td><td></td><td></td><td></td><td></td><td></td><td></td><td></td><td></td><td></td><td></td></tr>
<tr><td>Chromosome</td><td>start</td><td></td><td>end</td><td>strand</td><td>MM</td><td>target_seq</td><td>PAM</td><td>alignment</td><td>distance</td><td>position</td><td>gene name</td><td>gene id</td></tr>
<tr><td>GU305914.1</td><td></td><td>159036</td><td>159058</td><td>+</td><td>0</td><td>CCTTCCCTTGCTGCGAGGAA</td><td>GGG</td><td>||||||||[||||||||||||]PAM</td><td>NA</td><td>Intergenic</td><td>NA</td><td>NA</td></tr>
<tr><td>T12004</td><td>CTTCCCTTGCTGCGAGGAAGGGG</td><td></td><td>1000</td><td></td><td></td><td></td><td></td><td></td><td></td><td></td><td></td><td></td></tr>
<tr><td>Oligo fwd</td><td>CACCCTTCCCTTGCTGCGAGGAAG</td><td></td><td></td><td></td><td></td><td></td><td></td><td></td><td></td><td></td><td></td><td></td></tr>
<tr><td>Oligo rev</td><td>AAACCTTCCTCGCAGCAAGGGAAG</td><td></td><td></td><td></td><td></td><td></td><td></td><td></td><td></td><td></td><td></td><td></td></tr>
<tr><td>Chromosome</td><td>start</td><td></td><td>end</td><td>strand</td><td>MM</td><td>target_seq</td><td>PAM</td><td>alignment</td><td>distance</td><td>position</td><td>gene name</td><td>gene id</td></tr>
<tr><td>GU305914.1</td><td></td><td>159037</td><td>159059</td><td>+</td><td>0</td><td>CTTCCCTTGCTGCGAGGAAG</td><td>GGG</td><td>||||||||[||||||||||||]PAM</td><td>NA</td><td>Intergenic</td><td>NA</td><td>NA</td></tr>
<tr><td>GU305914.1</td><td></td><td>154353</td><td>154375</td><td>-</td><td>4</td><td>ATCCCCATGCTGCGAGGAGG</td><td>CGG</td><td>-|-|||-|[||||||||||-|]PAM</td><td>NA</td><td>Intergenic</td><td>NA</td><td>NA</td></tr>
<tr><td>T12005</td><td>TTCCCTTGCTGCGAGGAAGGGGG</td><td></td><td>1000</td><td></td><td></td><td></td><td></td><td></td><td></td><td></td><td></td><td></td></tr>
<tr><td>Oligo fwd</td><td>CACCTTCCCTTGCTGCGAGGAAGG</td><td></td><td></td><td></td><td></td><td></td><td></td><td></td><td></td><td></td><td></td><td></td></tr>
<tr><td>Oligo rev</td><td>AAACCCTTCCTCGCAGCAAGGGAA</td><td></td><td></td><td></td><td></td><td></td><td></td><td></td><td></td><td></td><td></td><td></td></tr>
<tr><td>Chromosome</td><td>start</td><td></td><td>end</td><td>strand</td><td>MM</td><td>target_seq</td><td>PAM</td><td>alignment</td><td>distance</td><td>position</td><td>gene name</td><td>gene id</td></tr>
<tr><td>GU305914.1</td><td></td><td>159038</td><td>159060</td><td>+</td><td>0</td><td>TTCCCTTGCTGCGAGGAAGG</td><td>GGG</td><td>||||||||[||||||||||||]PAM</td><td>NA</td><td>Intergenic</td><td>NA</td><td>NA</td></tr>
<tr><td>T12006</td><td>GAGGAAGGGGGTCAAGTTGCTGG</td><td></td><td>1000</td><td></td><td></td><td></td><td></td><td></td><td></td><td></td><td></td><td></td></tr>
<tr><td>Oligo fwd</td><td>CACCGAGGAAGGGGGTCAAGTTGC</td><td></td><td></td><td></td><td></td><td></td><td></td><td></td><td></td><td></td><td></td><td></td></tr>
<tr><td>Oligo rev</td><td>AAACGCAACTTGACCCCCTTCCTC</td><td></td><td></td><td></td><td></td><td></td><td></td><td></td><td></td><td></td><td></td><td></td></tr>
<tr><td>Chromosome</td><td>start</td><td></td><td>end</td><td>strand</td><td>MM</td><td>target_seq</td><td>PAM</td><td>alignment</td><td>distance</td><td>position</td><td>gene name</td><td>gene id</td></tr>
<tr><td>GU305914.1</td><td></td><td>159050</td><td>159072</td><td>+</td><td>0</td><td>GAGGAAGGGGGTCAAGTTGC</td><td>TGG</td><td>||||||||[||||||||||||]PAM</td><td>NA</td><td>Intergenic</td><td>NA</td><td>NA</td></tr>
<tr><td>T12007</td><td>TCGAAACGAAAATCGCGTGCAGG</td><td></td><td>1000</td><td></td><td></td><td></td><td></td><td></td><td></td><td></td><td></td><td></td></tr>
<tr><td>Oligo fwd</td><td>CACCTCGAAACGAAAATCGCGTGC</td><td></td><td></td><td></td><td></td><td></td><td></td><td></td><td></td><td></td><td></td><td></td></tr>
<tr><td>Oligo rev</td><td>AAACGCACGCGATTTTCGTTTCGA</td><td></td><td></td><td></td><td></td><td></td><td></td><td></td><td></td><td></td><td></td><td></td></tr>
<tr><td>Chromosome</td><td>start</td><td></td><td>end</td><td>strand</td><td>MM</td><td>target_seq</td><td>PAM</td><td>alignment</td><td>distance</td><td>position</td><td>gene name</td><td>gene id</td></tr>
<tr><td>GU305914.1</td><td></td><td>159090</td><td>159112</td><td>+</td><td>0</td><td>TCGAAACGAAAATCGCGTGC</td><td>AGG</td><td>||||||||[||||||||||||]PAM</td><td>NA</td><td>Intergenic</td><td>NA</td><td>NA</td></tr>
<tr><td>T12008</td><td>GTGCAGGCTGCTGACCGACGCGG</td><td></td><td>1000</td><td></td><td></td><td></td><td></td><td></td><td></td><td></td><td></td><td></td></tr>
<tr><td>Oligo fwd</td><td>CACCGTGCAGGCTGCTGACCGACG</td><td></td><td></td><td></td><td></td><td></td><td></td><td></td><td></td><td></td><td></td><td></td></tr>
<tr><td>Oligo rev</td><td>AAACCGTCGGTCAGCAGCCTGCAC</td><td></td><td></td><td></td><td></td><td></td><td></td><td></td><td></td><td></td><td></td><td></td></tr>
<tr><td>Chromosome</td><td>start</td><td></td><td>end</td><td>strand</td><td>MM</td><td>target_seq</td><td>PAM</td><td>alignment</td><td>distance</td><td>position</td><td>gene name</td><td>gene id</td></tr>
<tr><td>GU305914.1</td><td></td><td>159106</td><td>159128</td><td>+</td><td>0</td><td>GTGCAGGCTGCTGACCGACG</td><td>CGG</td><td>||||||||[||||||||||||]PAM</td><td>NA</td><td>Intergenic</td><td>NA</td><td>NA</td></tr>
<tr><td>T12009</td><td>GCGGTCACGTTACCGAAATAAGG</td><td></td><td>1000</td><td></td><td></td><td></td><td></td><td></td><td></td><td></td><td></td><td></td></tr>
<tr><td>Oligo fwd</td><td>CACCGCGGTCACGTTACCGAAATA</td><td></td><td></td><td></td><td></td><td></td><td></td><td></td><td></td><td></td><td></td><td></td></tr>
<tr><td>Oligo rev</td><td>AAACTATTTCGGTAACGTGACCGC</td><td></td><td></td><td></td><td></td><td></td><td></td><td></td><td></td><td></td><td></td><td></td></tr>
<tr><td>Chromosome</td><td>start</td><td></td><td>end</td><td>strand</td><td>MM</td><td>target_seq</td><td>PAM</td><td>alignment</td><td>distance</td><td>position</td><td>gene name</td><td>gene id</td></tr>
<tr><td>GU305914.1</td><td></td><td>159125</td><td>159147</td><td>+</td><td>0</td><td>GCGGTCACGTTACCGAAATA</td><td>AGG</td><td>||||||||[||||||||||||]PAM</td><td>NA</td><td>Intergenic</td><td>NA</td><td>NA</td></tr>
<tr><td>T12010</td><td>CGGTCACGTTACCGAAATAAGGG</td><td></td><td>1000</td><td></td><td></td><td></td><td></td><td></td><td></td><td></td><td></td><td></td></tr>
<tr><td>Oligo fwd</td><td>CACCCGGTCACGTTACCGAAATAA</td><td></td><td></td><td></td><td></td><td></td><td></td><td></td><td></td><td></td><td></td><td></td></tr>
<tr><td>Oligo rev</td><td>AAACTTATTTCGGTAACGTGACCG</td><td></td><td></td><td></td><td></td><td></td><td></td><td></td><td></td><td></td><td></td><td></td></tr>
<tr><td>Chromosome</td><td>start</td><td></td><td>end</td><td>strand</td><td>MM</td><td>target_seq</td><td>PAM</td><td>alignment</td><td>distance</td><td>position</td><td>gene name</td><td>gene id</td></tr>
</table>

GU305914.1 159126 159148 + 0 CGGTCACGTTACCGAAATAA GGG ||||||||[||||||||||||]PAM NA Intergenic NA NA

T12011 ATTGCTGATCGTTCTTCTTGTGG 1000
Oligo fwd CACCATTGCTGATCGTTCTTCTTG
Oligo rev AAACCAAGAAGAACGATCAGCAAT
Chromosome start end strand MM target_seq PAM alignment distance position gene name gene id
GU305914.1 159153 159175 + 0 ATTGCTGATCGTTCTTCTTG TGG ||||||||[||||||||||||]PAM NA Intergenic NA NA

T12012 TCTTCTTCGTCGACAGCCGTCGG 1000
Oligo fwd CACCTCTTCTTCGTCGACAGCCGT
Oligo rev AAACACGGCTGTCGACGAAGAAGA
Chromosome start end strand MM target_seq PAM alignment distance position gene name gene id
GU305914.1 159222 159244 + 0 TCTTCTTCGTCGACAGCCGT CGG ||||||||[||||||||||||]PAM NA Intergenic NA NA

T12013 CGTCGACAGCCGTCGGCGAGAGG 1000
Oligo fwd CACCCGTCGACAGCCGTCGGCGAG
Oligo rev AAACCTCGCCGACGGCTGTCGACG
Chromosome start end strand MM target_seq PAM alignment distance position gene name gene id
GU305914.1 159229 159251 + 0 CGTCGACAGCCGTCGGCGAG AGG ||||||||[||||||||||||]PAM NA Intergenic NA NA

T12014 GGCTTCTTCCTTGTCCCGTGAGG 1000
Oligo fwd CACCGGCTTCTTCCTTGTCCCGTG
Oligo rev AAACCACGGGACAAGGAAGAAGCC
Chromosome start end strand MM target_seq PAM alignment distance position gene name gene id
GU305914.1 159250 159272 + 0 GGCTTCTTCCTTGTCCCGTG AGG ||||||||[||||||||||||]PAM NA Intergenic NA NA

T12015 GCTTCTTCCTTGTCCCGTGAGGG 1000
Oligo fwd CACCGCTTCTTCCTTGTCCCGTGA
Oligo rev AAACTCACGGGACAAGGAAGAAGC
Chromosome start end strand MM target_seq PAM alignment distance position gene name gene id
GU305914.1 159251 159273 + 0 GCTTCTTCCTTGTCCCGTGA GGG ||||||||[||||||||||||]PAM NA Intergenic NA NA

T12016 CTTCTTCCTTGTCCCGTGAGGGG 1000
Oligo fwd CACCCTTCTTCCTTGTCCCGTGAG
Oligo rev AAACCTCACGGGACAAGGAAGAAG
Chromosome start end strand MM target_seq PAM alignment distance position gene name gene id
GU305914.1 159252 159274 + 0 CTTCTTCCTTGTCCCGTGAG GGG ||||||||[||||||||||||]PAM NA Intergenic NA NA

T12017 TACCATCCCTCCAGAGACCTCGG 1000
Oligo fwd CACCTACCATCCCTCCAGAGACCT
Oligo rev AAACAGGTCTCTGGAGGGATGGTA
Chromosome start end strand MM target_seq PAM alignment distance position gene name gene id
GU305914.1 159278 159300 + 0 TACCATCCCTCCAGAGACCT CGG ||||||||[||||||||||||]PAM NA Intergenic NA NA

T12018 ACCATCCCTCCAGAGACCTCGGG 1000
Oligo fwd CACCACCATCCCTCCAGAGACCTC
Oligo rev AAACGAGGTCTCTGGAGGGATGGT
Chromosome start end strand MM target_seq PAM alignment distance position gene name gene id
GU305914.1 159279 159301 + 0 ACCATCCCTCCAGAGACCTC GGG ||||||||[||||||||||||]PAM NA Intergenic NA NA

T12019 CCTCCAGAGACCTCGGGTCGCGG 1000
Oligo fwd CACCCCTCCAGAGACCTCGGGTCG
Oligo rev AAACCGACCCGAGGTCTCTGGAGG
Chromosome start end strand MM target_seq PAM alignment distance position gene name gene id
GU305914.1 159285 159307 + 0 CCTCCAGAGACCTCGGGTCG CGG ||||||||[||||||||||||]PAM NA Intergenic NA NA

T12020 TCGGGTCGCGGCGACCCATGTGG 1000
Oligo fwd CACCTCGGGTCGCGGCGACCCATG
Oligo rev AAACCATGGGTCGCCGCGACCCGA
Chromosome start end strand MM target_seq PAM alignment distance position gene name gene id
GU305914.1 159297 159319 + 0 TCGGGTCGCGGCGACCCATG TGG ||||||||[||||||||||||]PAM NA Intergenic NA NA
GU305914.1 147485 147507 - 4 TCCGGTGGCGCCGACCCATC CGG ||-|||-|[||-||||||||-]PAM NA Intergenic NA NA

T12021 ATGTGGTCAGAATCGCGATCTGG 1000
Oligo fwd CACCATGTGGTCAGAATCGCGATC
Oligo rev AAACGATCGCGATTCTGACCACAT
Chromosome start end strand MM target_seq PAM alignment distance position gene name gene id
GU305914.1 159314 159336 + 0 ATGTGGTCAGAATCGCGATC TGG ||||||||[||||||||||||]PAM NA Intergenic NA NA

T12022 CTGGACCGACCGCCGCGACCCGG 1000
Oligo fwd CACCCTGGACCGACCGCCGCGACC
Oligo rev AAACGGTCGCGGCGGTCGGTCCAG
Chromosome start end strand MM target_seq PAM alignment distance position gene name gene id
GU305914.1 159333 159355 + 0 CTGGACCGACCGCCGCGACC CGG ||||||||[||||||||||||]PAM NA Intergenic NA NA

T12023 AAGCGTTGAAGAAAAAGTCCTGG 1000
Oligo fwd CACCAAGCGTTGAAGAAAAAGTCC
Oligo rev AAACGGACTTTTTCTTCAACGCTT
Chromosome start end strand MM target_seq PAM alignment distance position gene name gene id
GU305914.1 159382 159404 + 0 AAGCGTTGAAGAAAAAGTCC TGG ||||||||[||||||||||||]PAM NA Intergenic NA NA

T12024 TCGTATTGTAGAGCCCGTTGCGG 1000
Oligo fwd CACCTCGTATTGTAGAGCCCGTTG
Oligo rev AAACCAACGGGCTCTACAATACGA
Chromosome start end strand MM target_seq PAM alignment distance position gene name gene id
GU305914.1 159440 159462 + 0 TCGTATTGTAGAGCCCGTTG CGG ||||||||[||||||||||||]PAM NA Intergenic NA NA

T12025 ATTGTAGAGCCCGTTGCGGCAGG 1000
Oligo fwd CACCATTGTAGAGCCCGTTGCGGC

<table>
<tr><td>Oligo rev</td><td>AAACGCCGCAACGGGCTCTACAAT</td><td></td><td></td><td></td><td></td><td></td><td></td><td></td><td></td><td></td><td></td><td></td></tr>
<tr><td>Chromosome</td><td>start</td><td></td><td>end</td><td>strand</td><td>MM</td><td>target_seq</td><td>PAM</td><td>alignment</td><td>distance</td><td>position</td><td>gene name</td><td>gene id</td></tr>
<tr><td>GU305914.1</td><td></td><td>159444</td><td>159466</td><td>+</td><td>0</td><td>ATTGTAGAGCCCGTTGCGGC</td><td>AGG</td><td>||||||||[||||||||||||]PAM</td><td>NA</td><td>Intergenic</td><td>NA</td><td>NA</td></tr>
<tr><td></td><td></td><td></td><td></td><td></td><td></td><td></td><td></td><td></td><td></td><td></td><td></td><td></td></tr>
<tr><td>T12026</td><td>GCAGGATGATATATATTGTCTGG</td><td></td><td>1000</td><td></td><td></td><td></td><td></td><td></td><td></td><td></td><td></td><td></td></tr>
<tr><td>Oligo fwd</td><td>CACCGCAGGATGATATATATTGTC</td><td></td><td></td><td></td><td></td><td></td><td></td><td></td><td></td><td></td><td></td><td></td></tr>
<tr><td>Oligo rev</td><td>AAACGACAATATATATCATCCTGC</td><td></td><td></td><td></td><td></td><td></td><td></td><td></td><td></td><td></td><td></td><td></td></tr>
<tr><td>Chromosome</td><td>start</td><td></td><td>end</td><td>strand</td><td>MM</td><td>target_seq</td><td>PAM</td><td>alignment</td><td>distance</td><td>position</td><td>gene name</td><td>gene id</td></tr>
<tr><td>GU305914.1</td><td></td><td>159462</td><td>159484</td><td>+</td><td>0</td><td>GCAGGATGATATATATTGTC</td><td>TGG</td><td>||||||||[||||||||||||]PAM</td><td>NA</td><td>Intergenic</td><td>NA</td><td>NA</td></tr>
<tr><td></td><td></td><td></td><td></td><td></td><td></td><td></td><td></td><td></td><td></td><td></td><td></td><td></td></tr>
<tr><td>T12027</td><td>TAGAGACAAAAATCAGTTGAAGG</td><td></td><td>1000</td><td></td><td></td><td></td><td></td><td></td><td></td><td></td><td></td><td></td></tr>
<tr><td>Oligo fwd</td><td>CACCTAGAGACAAAAATCAGTTGA</td><td></td><td></td><td></td><td></td><td></td><td></td><td></td><td></td><td></td><td></td><td></td></tr>
<tr><td>Oligo rev</td><td>AAACTCAACTGATTTTTGTCTCTA</td><td></td><td></td><td></td><td></td><td></td><td></td><td></td><td></td><td></td><td></td><td></td></tr>
<tr><td>Chromosome</td><td>start</td><td></td><td>end</td><td>strand</td><td>MM</td><td>target_seq</td><td>PAM</td><td>alignment</td><td>distance</td><td>position</td><td>gene name</td><td>gene id</td></tr>
<tr><td>GU305914.1</td><td></td><td>159501</td><td>159523</td><td>+</td><td>0</td><td>TAGAGACAAAAATCAGTTGA</td><td>AGG</td><td>||||||||[||||||||||||]PAM</td><td>NA</td><td>Intergenic</td><td>NA</td><td>NA</td></tr>
<tr><td></td><td></td><td></td><td></td><td></td><td></td><td></td><td></td><td></td><td></td><td></td><td></td><td></td></tr>
<tr><td>T12028</td><td>ATATATAGAGATAGATAGATTGG</td><td></td><td>1000</td><td></td><td></td><td></td><td></td><td></td><td></td><td></td><td></td><td></td></tr>
<tr><td>Oligo fwd</td><td>CACCATATATAGAGATAGATAGAT</td><td></td><td></td><td></td><td></td><td></td><td></td><td></td><td></td><td></td><td></td><td></td></tr>
<tr><td>Oligo rev</td><td>AAACATCTATCTATCTCTATATAT</td><td></td><td></td><td></td><td></td><td></td><td></td><td></td><td></td><td></td><td></td><td></td></tr>
<tr><td>Chromosome</td><td>start</td><td></td><td>end</td><td>strand</td><td>MM</td><td>target_seq</td><td>PAM</td><td>alignment</td><td>distance</td><td>position</td><td>gene name</td><td>gene id</td></tr>
<tr><td>GU305914.1</td><td></td><td>159624</td><td>159646</td><td>+</td><td>0</td><td>ATATATAGAGATAGATAGAT</td><td>TGG</td><td>||||||||[||||||||||||]PAM</td><td>NA</td><td>Intergenic</td><td>NA</td><td>NA</td></tr>
<tr><td></td><td></td><td></td><td></td><td></td><td></td><td></td><td></td><td></td><td></td><td></td><td></td><td></td></tr>
<tr><td>T12029</td><td>AGATAGATAGATTGGTAGATAGG</td><td></td><td>1000</td><td></td><td></td><td></td><td></td><td></td><td></td><td></td><td></td><td></td></tr>
<tr><td>Oligo fwd</td><td>CACCAGATAGATAGATTGGTAGAT</td><td></td><td></td><td></td><td></td><td></td><td></td><td></td><td></td><td></td><td></td><td></td></tr>
<tr><td>Oligo rev</td><td>AAACATCTACCAATCTATCTATCT</td><td></td><td></td><td></td><td></td><td></td><td></td><td></td><td></td><td></td><td></td><td></td></tr>
<tr><td>Chromosome</td><td>start</td><td></td><td>end</td><td>strand</td><td>MM</td><td>target_seq</td><td>PAM</td><td>alignment</td><td>distance</td><td>position</td><td>gene name</td><td>gene id</td></tr>
<tr><td>GU305914.1</td><td></td><td>159632</td><td>159654</td><td>+</td><td>0</td><td>AGATAGATAGATTGGTAGAT</td><td>AGG</td><td>||||||||[||||||||||||]PAM</td><td>NA</td><td>Intergenic</td><td>NA</td><td>NA</td></tr>
<tr><td></td><td></td><td></td><td></td><td></td><td></td><td></td><td></td><td></td><td></td><td></td><td></td><td></td></tr>
<tr><td>T12030</td><td>TAAAAGTCACATATAACCGTCGG</td><td></td><td>1000</td><td></td><td></td><td></td><td></td><td></td><td></td><td></td><td></td><td></td></tr>
<tr><td>Oligo fwd</td><td>CACCTAAAAGTCACATATAACCGT</td><td></td><td></td><td></td><td></td><td></td><td></td><td></td><td></td><td></td><td></td><td></td></tr>
<tr><td>Oligo rev</td><td>AAACACGGTTATATGTGACTTTTA</td><td></td><td></td><td></td><td></td><td></td><td></td><td></td><td></td><td></td><td></td><td></td></tr>
<tr><td>Chromosome</td><td>start</td><td></td><td>end</td><td>strand</td><td>MM</td><td>target_seq</td><td>PAM</td><td>alignment</td><td>distance</td><td>position</td><td>gene name</td><td>gene id</td></tr>
<tr><td>GU305914.1</td><td></td><td>159673</td><td>159695</td><td>+</td><td>0</td><td>TAAAAGTCACATATAACCGT</td><td>CGG</td><td>||||||||[||||||||||||]PAM</td><td>NA</td><td>Intergenic</td><td>NA</td><td>NA</td></tr>
<tr><td></td><td></td><td></td><td></td><td></td><td></td><td></td><td></td><td></td><td></td><td></td><td></td><td></td></tr>
<tr><td>T12031</td><td>CGTCGTTGTTTGCGAGACGTCGG</td><td></td><td>1000</td><td></td><td></td><td></td><td></td><td></td><td></td><td></td><td></td><td></td></tr>
<tr><td>Oligo fwd</td><td>CACCCGTCGTTGTTTGCGAGACGT</td><td></td><td></td><td></td><td></td><td></td><td></td><td></td><td></td><td></td><td></td><td></td></tr>
<tr><td>Oligo rev</td><td>AAACACGTCTCGCAAACAACGACG</td><td></td><td></td><td></td><td></td><td></td><td></td><td></td><td></td><td></td><td></td><td></td></tr>
<tr><td>Chromosome</td><td>start</td><td></td><td>end</td><td>strand</td><td>MM</td><td>target_seq</td><td>PAM</td><td>alignment</td><td>distance</td><td>position</td><td>gene name</td><td>gene id</td></tr>
<tr><td>GU305914.1</td><td></td><td>159715</td><td>159737</td><td>+</td><td>0</td><td>CGTCGTTGTTTGCGAGACGT</td><td>CGG</td><td>||||||||[||||||||||||]PAM</td><td>NA</td><td>Intergenic</td><td>NA</td><td>NA</td></tr>
<tr><td></td><td></td><td></td><td></td><td></td><td></td><td></td><td></td><td></td><td></td><td></td><td></td><td></td></tr>
<tr><td>T12032</td><td>TTGCGAGACGTCGGCGCGTTCGG</td><td></td><td>1000</td><td></td><td></td><td></td><td></td><td></td><td></td><td></td><td></td><td></td></tr>
<tr><td>Oligo fwd</td><td>CACCTTGCGAGACGTCGGCGCGTT</td><td></td><td></td><td></td><td></td><td></td><td></td><td></td><td></td><td></td><td></td><td></td></tr>
<tr><td>Oligo rev</td><td>AAACAACGCGCCGACGTCTCGCAA</td><td></td><td></td><td></td><td></td><td></td><td></td><td></td><td></td><td></td><td></td><td></td></tr>
<tr><td>Chromosome</td><td>start</td><td></td><td>end</td><td>strand</td><td>MM</td><td>target_seq</td><td>PAM</td><td>alignment</td><td>distance</td><td>position</td><td>gene name</td><td>gene id</td></tr>
<tr><td>GU305914.1</td><td></td><td>159724</td><td>159746</td><td>+</td><td>0</td><td>TTGCGAGACGTCGGCGCGTT</td><td>CGG</td><td>||||||||[||||||||||||]PAM</td><td>NA</td><td>Intergenic</td><td>NA</td><td>NA</td></tr>
<tr><td></td><td></td><td></td><td></td><td></td><td></td><td></td><td></td><td></td><td></td><td></td><td></td><td></td></tr>
<tr><td>T12033</td><td>CGAGACGTCGGCGCGTTCGGCGG</td><td></td><td>1000</td><td></td><td></td><td></td><td></td><td></td><td></td><td></td><td></td><td></td></tr>
<tr><td>Oligo fwd</td><td>CACCCGAGACGTCGGCGCGTTCGG</td><td></td><td></td><td></td><td></td><td></td><td></td><td></td><td></td><td></td><td></td><td></td></tr>
<tr><td>Oligo rev</td><td>AAACCCGAACGCGCCGACGTCTCG</td><td></td><td></td><td></td><td></td><td></td><td></td><td></td><td></td><td></td><td></td><td></td></tr>
<tr><td>Chromosome</td><td>start</td><td></td><td>end</td><td>strand</td><td>MM</td><td>target_seq</td><td>PAM</td><td>alignment</td><td>distance</td><td>position</td><td>gene name</td><td>gene id</td></tr>
<tr><td>GU305914.1</td><td></td><td>159727</td><td>159749</td><td>+</td><td>0</td><td>CGAGACGTCGGCGCGTTCGG</td><td>CGG</td><td>||||||||[||||||||||||]PAM</td><td>NA</td><td>Intergenic</td><td>NA</td><td>NA</td></tr>
<tr><td>GU305914.1</td><td></td><td>151677</td><td>151699</td><td>+</td><td>4</td><td>CGCGACGACGTCGCGTTCGA</td><td>CGG</td><td>||-||||-[||-||||||||-]PAM</td><td>NA</td><td>Intergenic</td><td>NA</td><td>NA</td></tr>
<tr><td></td><td></td><td></td><td></td><td></td><td></td><td></td><td></td><td></td><td></td><td></td><td></td><td></td></tr>
<tr><td>T12034</td><td>GTCGGCGCGTTCGGCGGAGACGG</td><td></td><td>1000</td><td></td><td></td><td></td><td></td><td></td><td></td><td></td><td></td><td></td></tr>
<tr><td>Oligo fwd</td><td>CACCGTCGGCGCGTTCGGCGGAGA</td><td></td><td></td><td></td><td></td><td></td><td></td><td></td><td></td><td></td><td></td><td></td></tr>
<tr><td>Oligo rev</td><td>AAACTCTCCGCCGAACGCGCCGAC</td><td></td><td></td><td></td><td></td><td></td><td></td><td></td><td></td><td></td><td></td><td></td></tr>
<tr><td>Chromosome</td><td>start</td><td></td><td>end</td><td>strand</td><td>MM</td><td>target_seq</td><td>PAM</td><td>alignment</td><td>distance</td><td>position</td><td>gene name</td><td>gene id</td></tr>
<tr><td>GU305914.1</td><td></td><td>159733</td><td>159755</td><td>+</td><td>0</td><td>GTCGGCGCGTTCGGCGGAGA</td><td>CGG</td><td>||||||||[||||||||||||]PAM</td><td>NA</td><td>Intergenic</td><td>NA</td><td>NA</td></tr>
<tr><td></td><td></td><td></td><td></td><td></td><td></td><td></td><td></td><td></td><td></td><td></td><td></td><td></td></tr>
<tr><td>T12035</td><td>GCGCGTTCGGCGGAGACGGATGG</td><td></td><td>1000</td><td></td><td></td><td></td><td></td><td></td><td></td><td></td><td></td><td></td></tr>
<tr><td>Oligo fwd</td><td>CACCGCGCGTTCGGCGGAGACGGA</td><td></td><td></td><td></td><td></td><td></td><td></td><td></td><td></td><td></td><td></td><td></td></tr>
<tr><td>Oligo rev</td><td>AAACTCCGTCTCCGCCGAACGCGC</td><td></td><td></td><td></td><td></td><td></td><td></td><td></td><td></td><td></td><td></td><td></td></tr>
<tr><td>Chromosome</td><td>start</td><td></td><td>end</td><td>strand</td><td>MM</td><td>target_seq</td><td>PAM</td><td>alignment</td><td>distance</td><td>position</td><td>gene name</td><td>gene id</td></tr>
<tr><td>GU305914.1</td><td></td><td>159737</td><td>159759</td><td>+</td><td>0</td><td>GCGCGTTCGGCGGAGACGGA</td><td>TGG</td><td>||||||||[||||||||||||]PAM</td><td>NA</td><td>Intergenic</td><td>NA</td><td>NA</td></tr>
<tr><td></td><td></td><td></td><td></td><td></td><td></td><td></td><td></td><td></td><td></td><td></td><td></td><td></td></tr>
<tr><td>T12036</td><td>TTCGGCGGAGACGGATGGCGCGG</td><td></td><td>1000</td><td></td><td></td><td></td><td></td><td></td><td></td><td></td><td></td><td></td></tr>
<tr><td>Oligo fwd</td><td>CACCTTCGGCGGAGACGGATGGCG</td><td></td><td></td><td></td><td></td><td></td><td></td><td></td><td></td><td></td><td></td><td></td></tr>
<tr><td>Oligo rev</td><td>AAACCGCCATCCGTCTCCGCCGAA</td><td></td><td></td><td></td><td></td><td></td><td></td><td></td><td></td><td></td><td></td><td></td></tr>
<tr><td>Chromosome</td><td>start</td><td></td><td>end</td><td>strand</td><td>MM</td><td>target_seq</td><td>PAM</td><td>alignment</td><td>distance</td><td>position</td><td>gene name</td><td>gene id</td></tr>
<tr><td>GU305914.1</td><td></td><td>159742</td><td>159764</td><td>+</td><td>0</td><td>TTCGGCGGAGACGGATGGCG</td><td>CGG</td><td>||||||||[||||||||||||]PAM</td><td>NA</td><td>Intergenic</td><td>NA</td><td>NA</td></tr>
<tr><td></td><td></td><td></td><td></td><td></td><td></td><td></td><td></td><td></td><td></td><td></td><td></td><td></td></tr>
<tr><td>T12037</td><td>ACGGATGGCGCGGATTCATTTGG</td><td></td><td>1000</td><td></td><td></td><td></td><td></td><td></td><td></td><td></td><td></td><td></td></tr>
<tr><td>Oligo fwd</td><td>CACCACGGATGGCGCGGATTCATT</td><td></td><td></td><td></td><td></td><td></td><td></td><td></td><td></td><td></td><td></td><td></td></tr>
<tr><td>Oligo rev</td><td>AAACAATGAATCCGCGCCATCCGT</td><td></td><td></td><td></td><td></td><td></td><td></td><td></td><td></td><td></td><td></td><td></td></tr>
<tr><td>Chromosome</td><td>start</td><td></td><td>end</td><td>strand</td><td>MM</td><td>target_seq</td><td>PAM</td><td>alignment</td><td>distance</td><td>position</td><td>gene name</td><td>gene id</td></tr>
<tr><td>GU305914.1</td><td></td><td>159752</td><td>159774</td><td>+</td><td>0</td><td>ACGGATGGCGCGGATTCATT</td><td>TGG</td><td>||||||||[||||||||||||]PAM</td><td>NA</td><td>Intergenic</td><td>NA</td><td>NA</td></tr>
<tr><td></td><td></td><td></td><td></td><td></td><td></td><td></td><td></td><td></td><td></td><td></td><td></td><td></td></tr>
<tr><td>T12038</td><td>CGGATGGCGCGGATTCATTTGGG</td><td></td><td>1000</td><td></td><td></td><td></td><td></td><td></td><td></td><td></td><td></td><td></td></tr>
<tr><td>Oligo fwd</td><td>CACCCGGATGGCGCGGATTCATTT</td><td></td><td></td><td></td><td></td><td></td><td></td><td></td><td></td><td></td><td></td><td></td></tr>
<tr><td>Oligo rev</td><td>AAACAAATGAATCCGCGCCATCCG</td><td></td><td></td><td></td><td></td><td></td><td></td><td></td><td></td><td></td><td></td><td></td></tr>
<tr><td>Chromosome</td><td>start</td><td></td><td>end</td><td>strand</td><td>MM</td><td>target_seq</td><td>PAM</td><td>alignment</td><td>distance</td><td>position</td><td>gene name</td><td>gene id</td></tr>
<tr><td>GU305914.1</td><td></td><td>159753</td><td>159775</td><td>+</td><td>0</td><td>CGGATGGCGCGGATTCATTT</td><td>GGG</td><td>||||||||[||||||||||||]PAM</td><td>NA</td><td>Intergenic</td><td>NA</td><td>NA</td></tr>
<tr><td></td><td></td><td></td><td></td><td></td><td></td><td></td><td></td><td></td><td></td><td></td><td></td><td></td></tr>
<tr><td>T12039</td><td>GGATGGCGCGGATTCATTTGGGG</td><td></td><td>1000</td><td></td><td></td><td></td><td></td><td></td><td></td><td></td><td></td><td></td></tr>
<tr><td>Oligo fwd</td><td>CACCGGATGGCGCGGATTCATTTG</td><td></td><td></td><td></td><td></td><td></td><td></td><td></td><td></td><td></td><td></td><td></td></tr>
<tr><td>Oligo rev</td><td>AAACCAAATGAATCCGCGCCATCC</td><td></td><td></td><td></td><td></td><td></td><td></td><td></td><td></td><td></td><td></td><td></td></tr>
<tr><td>Chromosome</td><td>start</td><td></td><td>end</td><td>strand</td><td>MM</td><td>target_seq</td><td>PAM</td><td>alignment</td><td>distance</td><td>position</td><td>gene name</td><td>gene id</td></tr>
<tr><td>GU305914.1</td><td></td><td>159754</td><td>159776</td><td>+</td><td>0</td><td>GGATGGCGCGGATTCATTTG</td><td>GGG</td><td>||||||||[||||||||||||]PAM</td><td>NA</td><td>Intergenic</td><td>NA</td><td>NA</td></tr>
</table>

<table>
<tr><td>T12040</td><td>GATGGCGCGGATTCATTTGGGGG</td><td></td><td>1000</td><td></td><td></td><td></td><td></td><td></td><td></td><td></td><td></td><td></td></tr>
<tr><td>Oligo fwd</td><td>CACCGATGGCGCGGATTCATTTGG</td><td></td><td></td><td></td><td></td><td></td><td></td><td></td><td></td><td></td><td></td><td></td></tr>
<tr><td>Oligo rev</td><td>AAACCCAAATGAATCCGCGCCATC</td><td></td><td></td><td></td><td></td><td></td><td></td><td></td><td></td><td></td><td></td><td></td></tr>
<tr><td>Chromosome</td><td>start</td><td></td><td>end</td><td>strand</td><td>MM</td><td>target_seq</td><td>PAM</td><td>alignment</td><td>distance</td><td>position</td><td>gene name</td><td>gene id</td></tr>
<tr><td>GU305914.1</td><td></td><td>159755</td><td>159777</td><td>+</td><td>0</td><td>GATGGCGCGGATTCATTTGG</td><td>GGG</td><td>||||||||[||||||||||||]PAM</td><td>NA</td><td>Intergenic</td><td>NA</td><td>NA</td></tr>
<tr><td>T12041</td><td>ATGGCGCGGATTCATTTGGGGGG</td><td></td><td>1000</td><td></td><td></td><td></td><td></td><td></td><td></td><td></td><td></td><td></td></tr>
<tr><td>Oligo fwd</td><td>CACCATGGCGCGGATTCATTTGGG</td><td></td><td></td><td></td><td></td><td></td><td></td><td></td><td></td><td></td><td></td><td></td></tr>
<tr><td>Oligo rev</td><td>AAACCCCAAATGAATCCGCGCCAT</td><td></td><td></td><td></td><td></td><td></td><td></td><td></td><td></td><td></td><td></td><td></td></tr>
<tr><td>Chromosome</td><td>start</td><td></td><td>end</td><td>strand</td><td>MM</td><td>target_seq</td><td>PAM</td><td>alignment</td><td>distance</td><td>position</td><td>gene name</td><td>gene id</td></tr>
<tr><td>GU305914.1</td><td></td><td>159756</td><td>159778</td><td>+</td><td>0</td><td>ATGGCGCGGATTCATTTGGG</td><td>GGG</td><td>||||||||[||||||||||||]PAM</td><td>NA</td><td>Intergenic</td><td>NA</td><td>NA</td></tr>
<tr><td>T12042</td><td>GCGGATTCATTTGGGGGGTCAGG</td><td></td><td>1000</td><td></td><td></td><td></td><td></td><td></td><td></td><td></td><td></td><td></td></tr>
<tr><td>Oligo fwd</td><td>CACCGCGGATTCATTTGGGGGGTC</td><td></td><td></td><td></td><td></td><td></td><td></td><td></td><td></td><td></td><td></td><td></td></tr>
<tr><td>Oligo rev</td><td>AAACGACCCCCCAAATGAATCCGC</td><td></td><td></td><td></td><td></td><td></td><td></td><td></td><td></td><td></td><td></td><td></td></tr>
<tr><td>Chromosome</td><td>start</td><td></td><td>end</td><td>strand</td><td>MM</td><td>target_seq</td><td>PAM</td><td>alignment</td><td>distance</td><td>position</td><td>gene name</td><td>gene id</td></tr>
<tr><td>GU305914.1</td><td></td><td>159761</td><td>159783</td><td>+</td><td>0</td><td>GCGGATTCATTTGGGGGGTC</td><td>AGG</td><td>||||||||[||||||||||||]PAM</td><td>NA</td><td>Intergenic</td><td>NA</td><td>NA</td></tr>
<tr><td>T12043</td><td>CGGATTCATTTGGGGGGTCAGGG</td><td></td><td>1000</td><td></td><td></td><td></td><td></td><td></td><td></td><td></td><td></td><td></td></tr>
<tr><td>Oligo fwd</td><td>CACCCGGATTCATTTGGGGGGTCA</td><td></td><td></td><td></td><td></td><td></td><td></td><td></td><td></td><td></td><td></td><td></td></tr>
<tr><td>Oligo rev</td><td>AAACTGACCCCCCAAATGAATCCG</td><td></td><td></td><td></td><td></td><td></td><td></td><td></td><td></td><td></td><td></td><td></td></tr>
<tr><td>Chromosome</td><td>start</td><td></td><td>end</td><td>strand</td><td>MM</td><td>target_seq</td><td>PAM</td><td>alignment</td><td>distance</td><td>position</td><td>gene name</td><td>gene id</td></tr>
<tr><td>GU305914.1</td><td></td><td>159762</td><td>159784</td><td>+</td><td>0</td><td>CGGATTCATTTGGGGGGTCA</td><td>GGG</td><td>||||||||[||||||||||||]PAM</td><td>NA</td><td>Intergenic</td><td>NA</td><td>NA</td></tr>
<tr><td>T12044</td><td>GTTGAACACACAAAAACACAAGG</td><td></td><td>1000</td><td></td><td></td><td></td><td></td><td></td><td></td><td></td><td></td><td></td></tr>
<tr><td>Oligo fwd</td><td>CACCGTTGAACACACAAAAACACA</td><td></td><td></td><td></td><td></td><td></td><td></td><td></td><td></td><td></td><td></td><td></td></tr>
<tr><td>Oligo rev</td><td>AAACTGTGTTTTTGTGTGTTCAAC</td><td></td><td></td><td></td><td></td><td></td><td></td><td></td><td></td><td></td><td></td><td></td></tr>
<tr><td>Chromosome</td><td>start</td><td></td><td>end</td><td>strand</td><td>MM</td><td>target_seq</td><td>PAM</td><td>alignment</td><td>distance</td><td>position</td><td>gene name</td><td>gene id</td></tr>
<tr><td>GU305914.1</td><td></td><td>159784</td><td>159806</td><td>+</td><td>0</td><td>GTTGAACACACAAAAACACA</td><td>AGG</td><td>||||||||[||||||||||||]PAM</td><td>NA</td><td>Intergenic</td><td>NA</td><td>NA</td></tr>
<tr><td>T12045</td><td>TTGAACACACAAAAACACAAGGG</td><td></td><td>1000</td><td></td><td></td><td></td><td></td><td></td><td></td><td></td><td></td><td></td></tr>
<tr><td>Oligo fwd</td><td>CACCTTGAACACACAAAAACACAA</td><td></td><td></td><td></td><td></td><td></td><td></td><td></td><td></td><td></td><td></td><td></td></tr>
<tr><td>Oligo rev</td><td>AAACTTGTGTTTTTGTGTGTTCAA</td><td></td><td></td><td></td><td></td><td></td><td></td><td></td><td></td><td></td><td></td><td></td></tr>
<tr><td>Chromosome</td><td>start</td><td></td><td>end</td><td>strand</td><td>MM</td><td>target_seq</td><td>PAM</td><td>alignment</td><td>distance</td><td>position</td><td>gene name</td><td>gene id</td></tr>
<tr><td>GU305914.1</td><td></td><td>159785</td><td>159807</td><td>+</td><td>0</td><td>TTGAACACACAAAAACACAA</td><td>GGG</td><td>||||||||[||||||||||||]PAM</td><td>NA</td><td>Intergenic</td><td>NA</td><td>NA</td></tr>
<tr><td>T12046</td><td>TGAACACACAAAAACACAAGGGG</td><td></td><td>1000</td><td></td><td></td><td></td><td></td><td></td><td></td><td></td><td></td><td></td></tr>
<tr><td>Oligo fwd</td><td>CACCTGAACACACAAAAACACAAG</td><td></td><td></td><td></td><td></td><td></td><td></td><td></td><td></td><td></td><td></td><td></td></tr>
<tr><td>Oligo rev</td><td>AAACCTTGTGTTTTTGTGTGTTCA</td><td></td><td></td><td></td><td></td><td></td><td></td><td></td><td></td><td></td><td></td><td></td></tr>
<tr><td>Chromosome</td><td>start</td><td></td><td>end</td><td>strand</td><td>MM</td><td>target_seq</td><td>PAM</td><td>alignment</td><td>distance</td><td>position</td><td>gene name</td><td>gene id</td></tr>
<tr><td>GU305914.1</td><td></td><td>159786</td><td>159808</td><td>+</td><td>0</td><td>TGAACACACAAAAACACAAG</td><td>GGG</td><td>||||||||[||||||||||||]PAM</td><td>NA</td><td>Intergenic</td><td>NA</td><td>NA</td></tr>
<tr><td>T12047</td><td>GAACACACAAAAACACAAGGGGG</td><td></td><td>1000</td><td></td><td></td><td></td><td></td><td></td><td></td><td></td><td></td><td></td></tr>
<tr><td>Oligo fwd</td><td>CACCGAACACACAAAAACACAAGG</td><td></td><td></td><td></td><td></td><td></td><td></td><td></td><td></td><td></td><td></td><td></td></tr>
<tr><td>Oligo rev</td><td>AAACCCTTGTGTTTTTGTGTGTTC</td><td></td><td></td><td></td><td></td><td></td><td></td><td></td><td></td><td></td><td></td><td></td></tr>
<tr><td>Chromosome</td><td>start</td><td></td><td>end</td><td>strand</td><td>MM</td><td>target_seq</td><td>PAM</td><td>alignment</td><td>distance</td><td>position</td><td>gene name</td><td>gene id</td></tr>
<tr><td>GU305914.1</td><td></td><td>159787</td><td>159809</td><td>+</td><td>0</td><td>GAACACACAAAAACACAAGG</td><td>GGG</td><td>||||||||[||||||||||||]PAM</td><td>NA</td><td>Intergenic</td><td>NA</td><td>NA</td></tr>
<tr><td>T12048</td><td>CAAGGGGGAGTGTATCGTAGCGG</td><td></td><td>1000</td><td></td><td></td><td></td><td></td><td></td><td></td><td></td><td></td><td></td></tr>
<tr><td>Oligo fwd</td><td>CACCCAAGGGGGAGTGTATCGTAG</td><td></td><td></td><td></td><td></td><td></td><td></td><td></td><td></td><td></td><td></td><td></td></tr>
<tr><td>Oligo rev</td><td>AAACCTACGATACACTCCCCCTTG</td><td></td><td></td><td></td><td></td><td></td><td></td><td></td><td></td><td></td><td></td><td></td></tr>
<tr><td>Chromosome</td><td>start</td><td></td><td>end</td><td>strand</td><td>MM</td><td>target_seq</td><td>PAM</td><td>alignment</td><td>distance</td><td>position</td><td>gene name</td><td>gene id</td></tr>
<tr><td>GU305914.1</td><td></td><td>159802</td><td>159824</td><td>+</td><td>0</td><td>CAAGGGGGAGTGTATCGTAG</td><td>CGG</td><td>||||||||[||||||||||||]PAM</td><td>NA</td><td>Intergenic</td><td>NA</td><td>NA</td></tr>
<tr><td>T12049</td><td>GGGGGAGTGTATCGTAGCGGTGG</td><td></td><td>1000</td><td></td><td></td><td></td><td></td><td></td><td></td><td></td><td></td><td></td></tr>
<tr><td>Oligo fwd</td><td>CACCGGGGGAGTGTATCGTAGCGG</td><td></td><td></td><td></td><td></td><td></td><td></td><td></td><td></td><td></td><td></td><td></td></tr>
<tr><td>Oligo rev</td><td>AAACCCGCTACGATACACTCCCCC</td><td></td><td></td><td></td><td></td><td></td><td></td><td></td><td></td><td></td><td></td><td></td></tr>
<tr><td>Chromosome</td><td>start</td><td></td><td>end</td><td>strand</td><td>MM</td><td>target_seq</td><td>PAM</td><td>alignment</td><td>distance</td><td>position</td><td>gene name</td><td>gene id</td></tr>
<tr><td>GU305914.1</td><td></td><td>159805</td><td>159827</td><td>+</td><td>0</td><td>GGGGGAGTGTATCGTAGCGG</td><td>TGG</td><td>||||||||[||||||||||||]PAM</td><td>NA</td><td>Intergenic</td><td>NA</td><td>NA</td></tr>
<tr><td>T12050</td><td>ATCGTAGCGGTGGAAAAACTAGG</td><td></td><td>1000</td><td></td><td></td><td></td><td></td><td></td><td></td><td></td><td></td><td></td></tr>
<tr><td>Oligo fwd</td><td>CACCATCGTAGCGGTGGAAAAACT</td><td></td><td></td><td></td><td></td><td></td><td></td><td></td><td></td><td></td><td></td><td></td></tr>
<tr><td>Oligo rev</td><td>AAACAGTTTTTCCACCGCTACGAT</td><td></td><td></td><td></td><td></td><td></td><td></td><td></td><td></td><td></td><td></td><td></td></tr>
<tr><td>Chromosome</td><td>start</td><td></td><td>end</td><td>strand</td><td>MM</td><td>target_seq</td><td>PAM</td><td>alignment</td><td>distance</td><td>position</td><td>gene name</td><td>gene id</td></tr>
<tr><td>GU305914.1</td><td></td><td>159815</td><td>159837</td><td>+</td><td>0</td><td>ATCGTAGCGGTGGAAAAACT</td><td>AGG</td><td>||||||||[||||||||||||]PAM</td><td>NA</td><td>Intergenic</td><td>NA</td><td>NA</td></tr>
<tr><td>T12051</td><td>CGGTGGAAAAACTAGGAGACCGG</td><td></td><td>1000</td><td></td><td></td><td></td><td></td><td></td><td></td><td></td><td></td><td></td></tr>
<tr><td>Oligo fwd</td><td>CACCCGGTGGAAAAACTAGGAGAC</td><td></td><td></td><td></td><td></td><td></td><td></td><td></td><td></td><td></td><td></td><td></td></tr>
<tr><td>Oligo rev</td><td>AAACGTCTCCTAGTTTTTCCACCG</td><td></td><td></td><td></td><td></td><td></td><td></td><td></td><td></td><td></td><td></td><td></td></tr>
<tr><td>Chromosome</td><td>start</td><td></td><td>end</td><td>strand</td><td>MM</td><td>target_seq</td><td>PAM</td><td>alignment</td><td>distance</td><td>position</td><td>gene name</td><td>gene id</td></tr>
<tr><td>GU305914.1</td><td></td><td>159822</td><td>159844</td><td>+</td><td>0</td><td>CGGTGGAAAAACTAGGAGAC</td><td>CGG</td><td>||||||||[||||||||||||]PAM</td><td>NA</td><td>Intergenic</td><td>NA</td><td>NA</td></tr>
<tr><td>T12052</td><td>GACCGGAGACTGCTTTTTTTTGG</td><td></td><td>1000</td><td></td><td></td><td></td><td></td><td></td><td></td><td></td><td></td><td></td></tr>
<tr><td>Oligo fwd</td><td>CACCGACCGGAGACTGCTTTTTTT</td><td></td><td></td><td></td><td></td><td></td><td></td><td></td><td></td><td></td><td></td><td></td></tr>
<tr><td>Oligo rev</td><td>AAACAAAAAAAGCAGTCTCCGGTC</td><td></td><td></td><td></td><td></td><td></td><td></td><td></td><td></td><td></td><td></td><td></td></tr>
<tr><td>Chromosome</td><td>start</td><td></td><td>end</td><td>strand</td><td>MM</td><td>target_seq</td><td>PAM</td><td>alignment</td><td>distance</td><td>position</td><td>gene name</td><td>gene id</td></tr>
<tr><td>GU305914.1</td><td></td><td>159839</td><td>159861</td><td>+</td><td>0</td><td>GACCGGAGACTGCTTTTTTT</td><td>TGG</td><td>||||||||[||||||||||||]PAM</td><td>NA</td><td>Intergenic</td><td>NA</td><td>NA</td></tr>
<tr><td>T12053</td><td>ACCGGAGACTGCTTTTTTTTGGG</td><td></td><td>1000</td><td></td><td></td><td></td><td></td><td></td><td></td><td></td><td></td><td></td></tr>
<tr><td>Oligo fwd</td><td>CACCACCGGAGACTGCTTTTTTTT</td><td></td><td></td><td></td><td></td><td></td><td></td><td></td><td></td><td></td><td></td><td></td></tr>
<tr><td>Oligo rev</td><td>AAACAAAAAAAAGCAGTCTCCGGT</td><td></td><td></td><td></td><td></td><td></td><td></td><td></td><td></td><td></td><td></td><td></td></tr>
<tr><td>Chromosome</td><td>start</td><td></td><td>end</td><td>strand</td><td>MM</td><td>target_seq</td><td>PAM</td><td>alignment</td><td>distance</td><td>position</td><td>gene name</td><td>gene id</td></tr>
<tr><td>GU305914.1</td><td></td><td>159840</td><td>159862</td><td>+</td><td>0</td><td>ACCGGAGACTGCTTTTTTTT</td><td>GGG</td><td>||||||||[||||||||||||]PAM</td><td>NA</td><td>Intergenic</td><td>NA</td><td>NA</td></tr>
<tr><td>T12054</td><td>TGTCTTGTTGAGTCTTTCTCCGG</td><td></td><td>1000</td><td></td><td></td><td></td><td></td><td></td><td></td><td></td><td></td><td></td></tr>
<tr><td>Oligo fwd</td><td>CACCTGTCTTGTTGAGTCTTTCTC</td><td></td><td></td><td></td><td></td><td></td><td></td><td></td><td></td><td></td><td></td><td></td></tr>
<tr><td>Oligo rev</td><td>AAACGAGAAAGACTCAACAAGACA</td><td></td><td></td><td></td><td></td><td></td><td></td><td></td><td></td><td></td><td></td><td></td></tr>
<tr><td>Chromosome</td><td>start</td><td></td><td>end</td><td>strand</td><td>MM</td><td>target_seq</td><td>PAM</td><td>alignment</td><td>distance</td><td>position</td><td>gene name</td><td>gene id</td></tr>
<tr><td>GU305914.1</td><td></td><td>159876</td><td>159898</td><td>+</td><td>0</td><td>TGTCTTGTTGAGTCTTTCTC</td><td>CGG</td><td>||||||||[||||||||||||]PAM</td><td>NA</td><td>Intergenic</td><td>NA</td><td>NA</td></tr>
</table>

T12055 CCGGAGAAAATTAATGACACAGG 1000
Oligo fwd CACCCCGGAGAAAATTAATGACAC
Oligo rev AAACGTGTCATTAATTTTCTCCGG
Chromosome start end strand MM target_seq PAM alignment distance position gene name gene id
GU305914.1 159895 159917 + 0 CCGGAGAAAATTAATGACAC AGG ||||||||[||||||||||||]PAM NA Intergenic NA NA

T12056 TTAATGACACAGGTACAAAAAGG 1000
Oligo fwd CACCTTAATGACACAGGTACAAAA
Oligo rev AAACTTTTGTACCTGTGTCATTAA
Chromosome start end strand MM target_seq PAM alignment distance position gene name gene id
GU305914.1 159905 159927 + 0 TTAATGACACAGGTACAAAA AGG ||||||||[||||||||||||]PAM NA Intergenic NA NA

T12057 TAATGACACAGGTACAAAAAGGG 1000
Oligo fwd CACCTAATGACACAGGTACAAAAA
Oligo rev AAACTTTTTGTACCTGTGTCATTA
Chromosome start end strand MM target_seq PAM alignment distance position gene name gene id
GU305914.1 159906 159928 + 0 TAATGACACAGGTACAAAAA GGG ||||||||[||||||||||||]PAM NA Intergenic NA NA

T12058 AATGACACAGGTACAAAAAGGGG 1000
Oligo fwd CACCAATGACACAGGTACAAAAAG
Oligo rev AAACCTTTTTGTACCTGTGTCATT
Chromosome start end strand MM target_seq PAM alignment distance position gene name gene id
GU305914.1 159907 159929 + 0 AATGACACAGGTACAAAAAG GGG ||||||||[||||||||||||]PAM NA Intergenic NA NA

T12059 ATGACACAGGTACAAAAAGGGGG 1000
Oligo fwd CACCATGACACAGGTACAAAAAGG
Oligo rev AAACCCTTTTTGTACCTGTGTCAT
Chromosome start end strand MM target_seq PAM alignment distance position gene name gene id
GU305914.1 159908 159930 + 0 ATGACACAGGTACAAAAAGG GGG ||||||||[||||||||||||]PAM NA Intergenic NA NA

T12060 GGGGACATATAGCATACCCGAGG 1000
Oligo fwd CACCGGGGACATATAGCATACCCG
Oligo rev AAACCGGGTATGCTATATGTCCCC
Chromosome start end strand MM target_seq PAM alignment distance position gene name gene id
GU305914.1 159927 159949 + 0 GGGGACATATAGCATACCCG AGG ||||||||[||||||||||||]PAM NA Intergenic NA NA

T12061 CGCCGATAGAACGACGACCACGG 1000
Oligo fwd CACCCGCCGATAGAACGACGACCA
Oligo rev AAACTGGTCGTCGTTCTATCGGCG
Chromosome start end strand MM target_seq PAM alignment distance position gene name gene id
GU305914.1 159969 159991 + 0 CGCCGATAGAACGACGACCA CGG ||||||||[||||||||||||]PAM NA Intergenic NA NA

T12062 ACGACCACGGAGATGATACCAGG 1000
Oligo fwd CACCACGACCACGGAGATGATACC
Oligo rev AAACGGTATCATCTCCGTGGTCGT
Chromosome start end strand MM target_seq PAM alignment distance position gene name gene id
GU305914.1 159982 160004 + 0 ACGACCACGGAGATGATACC AGG ||||||||[||||||||||||]PAM NA Intergenic NA NA

T12063 CGACCACGGAGATGATACCAGGG 1000
Oligo fwd CACCCGACCACGGAGATGATACCA
Oligo rev AAACTGGTATCATCTCCGTGGTCG
Chromosome start end strand MM target_seq PAM alignment distance position gene name gene id
GU305914.1 159983 160005 + 0 CGACCACGGAGATGATACCA GGG ||||||||[||||||||||||]PAM NA Intergenic NA NA

T12064 GACCACGGAGATGATACCAGGGG 1000
Oligo fwd CACCGACCACGGAGATGATACCAG
Oligo rev AAACCTGGTATCATCTCCGTGGTC
Chromosome start end strand MM target_seq PAM alignment distance position gene name gene id
GU305914.1 159984 160006 + 0 GACCACGGAGATGATACCAG GGG ||||||||[||||||||||||]PAM NA Intergenic NA NA

T12065 GATACCAGGGGCCAGTCAGCTGG 1000
Oligo fwd CACCGATACCAGGGGCCAGTCAGC
Oligo rev AAACGCTGACTGGCCCCTGGTATC
Chromosome start end strand MM target_seq PAM alignment distance position gene name gene id
GU305914.1 159996 160018 + 0 GATACCAGGGGCCAGTCAGC TGG ||||||||[||||||||||||]PAM NA Intergenic NA NA

T12066 ATGTGTTGACACACACAGCTTGG 1000
Oligo fwd CACCATGTGTTGACACACACAGCT
Oligo rev AAACAGCTGTGTGTGTCAACACAT
Chromosome start end strand MM target_seq PAM alignment distance position gene name gene id
GU305914.1 160023 160045 + 0 ATGTGTTGACACACACAGCT TGG ||||||||[||||||||||||]PAM NA Intergenic NA NA

T12067 GAGCTCGCGTTCTCACCGAACGG 1000
Oligo fwd CACCGAGCTCGCGTTCTCACCGAA
Oligo rev AAACTTCGGTGAGAACGCGAGCTC
Chromosome start end strand MM target_seq PAM alignment distance position gene name gene id
GU305914.1 160052 160074 + 0 GAGCTCGCGTTCTCACCGAA CGG ||||||||[||||||||||||]PAM NA Intergenic NA NA

T12068 TATAGATCTCTCTCGTCGTGTGG 1000
Oligo fwd CACCTATAGATCTCTCTCGTCGTG
Oligo rev AAACCACGACGAGAGAGATCTATA
Chromosome start end strand MM target_seq PAM alignment distance position gene name gene id
GU305914.1 160115 160137 + 0 TATAGATCTCTCTCGTCGTG TGG ||||||||[||||||||||||]PAM NA Intergenic NA NA

T12069 ATAGATCTCTCTCGTCGTGTGGG 1000
Oligo fwd CACCATAGATCTCTCTCGTCGTGT
Oligo rev AAACACACGACGAGAGAGATCTAT
Chromosome start end strand MM target_seq PAM alignment distance position gene name gene id

<table>
<tr><td>GU305914.1</td><td>160116</td><td>160138</td><td>+</td><td>0</td><td>ATAGATCTCTCTCGTCGTGT</td><td>GGG</td><td>||||||||[||||||||||||]PAM</td><td>NA</td><td>Intergenic</td><td>NA</td><td>NA</td></tr>
<tr><td></td><td></td><td></td><td></td><td></td><td></td><td></td><td></td><td></td><td></td><td></td><td></td></tr>
<tr><td>T12070</td><td>TCGTCGTGTGGGTACGTACGTGG</td><td>1000</td><td></td><td></td><td></td><td></td><td></td><td></td><td></td><td></td><td></td></tr>
<tr><td>Oligo fwd</td><td>CACCTCGTCGTGTGGGTACGTACG</td><td></td><td></td><td></td><td></td><td></td><td></td><td></td><td></td><td></td><td></td></tr>
<tr><td>Oligo rev</td><td>AAACCGTACGTACCCACACGACGA</td><td></td><td></td><td></td><td></td><td></td><td></td><td></td><td></td><td></td><td></td></tr>
<tr><td>Chromosome</td><td>start</td><td>end</td><td>strand</td><td>MM</td><td>target_seq</td><td>PAM</td><td>alignment</td><td>distance</td><td>position</td><td>gene name</td><td>gene id</td></tr>
<tr><td>GU305914.1</td><td>160127</td><td>160149</td><td>+</td><td>0</td><td>TCGTCGTGTGGGTACGTACG</td><td>TGG</td><td>||||||||[||||||||||||]PAM</td><td>NA</td><td>Intergenic</td><td>NA</td><td>NA</td></tr>
<tr><td></td><td></td><td></td><td></td><td></td><td></td><td></td><td></td><td></td><td></td><td></td><td></td></tr>
<tr><td>T12071</td><td>CGTCGTGTGGGTACGTACGTGGG</td><td>1000</td><td></td><td></td><td></td><td></td><td></td><td></td><td></td><td></td><td></td></tr>
<tr><td>Oligo fwd</td><td>CACCCGTCGTGTGGGTACGTACGT</td><td></td><td></td><td></td><td></td><td></td><td></td><td></td><td></td><td></td><td></td></tr>
<tr><td>Oligo rev</td><td>AAACACGTACGTACCCACACGACG</td><td></td><td></td><td></td><td></td><td></td><td></td><td></td><td></td><td></td><td></td></tr>
<tr><td>Chromosome</td><td>start</td><td>end</td><td>strand</td><td>MM</td><td>target_seq</td><td>PAM</td><td>alignment</td><td>distance</td><td>position</td><td>gene name</td><td>gene id</td></tr>
<tr><td>GU305914.1</td><td>160128</td><td>160150</td><td>+</td><td>0</td><td>CGTCGTGTGGGTACGTACGT</td><td>GGG</td><td>||||||||[||||||||||||]PAM</td><td>NA</td><td>Intergenic</td><td>NA</td><td>NA</td></tr>
<tr><td></td><td></td><td></td><td></td><td></td><td></td><td></td><td></td><td></td><td></td><td></td><td></td></tr>
<tr><td>T12072</td><td>CGTAACGTTAGTTCGTTCGTCGG</td><td>1000</td><td></td><td></td><td></td><td></td><td></td><td></td><td></td><td></td><td></td></tr>
<tr><td>Oligo fwd</td><td>CACCCGTAACGTTAGTTCGTTCGT</td><td></td><td></td><td></td><td></td><td></td><td></td><td></td><td></td><td></td><td></td></tr>
<tr><td>Oligo rev</td><td>AAACACGAACGAACTAACGTTACG</td><td></td><td></td><td></td><td></td><td></td><td></td><td></td><td></td><td></td><td></td></tr>
<tr><td>Chromosome</td><td>start</td><td>end</td><td>strand</td><td>MM</td><td>target_seq</td><td>PAM</td><td>alignment</td><td>distance</td><td>position</td><td>gene name</td><td>gene id</td></tr>
<tr><td>GU305914.1</td><td>160152</td><td>160174</td><td>+</td><td>0</td><td>CGTAACGTTAGTTCGTTCGT</td><td>CGG</td><td>||||||||[||||||||||||]PAM</td><td>NA</td><td>Intergenic</td><td>NA</td><td>NA</td></tr>
<tr><td></td><td></td><td></td><td></td><td></td><td></td><td></td><td></td><td></td><td></td><td></td><td></td></tr>
<tr><td>T12073</td><td>AACGTTAGTTCGTTCGTCGGTGG</td><td>1000</td><td></td><td></td><td></td><td></td><td></td><td></td><td></td><td></td><td></td></tr>
<tr><td>Oligo fwd</td><td>CACCAACGTTAGTTCGTTCGTCGG</td><td></td><td></td><td></td><td></td><td></td><td></td><td></td><td></td><td></td><td></td></tr>
<tr><td>Oligo rev</td><td>AAACCCGACGAACGAACTAACGTT</td><td></td><td></td><td></td><td></td><td></td><td></td><td></td><td></td><td></td><td></td></tr>
<tr><td>Chromosome</td><td>start</td><td>end</td><td>strand</td><td>MM</td><td>target_seq</td><td>PAM</td><td>alignment</td><td>distance</td><td>position</td><td>gene name</td><td>gene id</td></tr>
<tr><td>GU305914.1</td><td>160155</td><td>160177</td><td>+</td><td>0</td><td>AACGTTAGTTCGTTCGTCGG</td><td>TGG</td><td>||||||||[||||||||||||]PAM</td><td>NA</td><td>Intergenic</td><td>NA</td><td>NA</td></tr>
<tr><td></td><td></td><td></td><td></td><td></td><td></td><td></td><td></td><td></td><td></td><td></td><td></td></tr>
<tr><td>T12074</td><td>GGCGAGACGTTCATTTGATAAGG</td><td>1000</td><td></td><td></td><td></td><td></td><td></td><td></td><td></td><td></td><td></td></tr>
<tr><td>Oligo fwd</td><td>CACCGGCGAGACGTTCATTTGATA</td><td></td><td></td><td></td><td></td><td></td><td></td><td></td><td></td><td></td><td></td></tr>
<tr><td>Oligo rev</td><td>AAACTATCAAATGAACGTCTCGCC</td><td></td><td></td><td></td><td></td><td></td><td></td><td></td><td></td><td></td><td></td></tr>
<tr><td>Chromosome</td><td>start</td><td>end</td><td>strand</td><td>MM</td><td>target_seq</td><td>PAM</td><td>alignment</td><td>distance</td><td>position</td><td>gene name</td><td>gene id</td></tr>
<tr><td>GU305914.1</td><td>160176</td><td>160198</td><td>+</td><td>0</td><td>GGCGAGACGTTCATTTGATA</td><td>AGG</td><td>||||||||[||||||||||||]PAM</td><td>NA</td><td>Intergenic</td><td>NA</td><td>NA</td></tr>
<tr><td></td><td></td><td></td><td></td><td></td><td></td><td></td><td></td><td></td><td></td><td></td><td></td></tr>
<tr><td>T12075</td><td>GATTTTTAACTGTTCTGCTGCGG</td><td>1000</td><td></td><td></td><td></td><td></td><td></td><td></td><td></td><td></td><td></td></tr>
<tr><td>Oligo fwd</td><td>CACCGATTTTTAACTGTTCTGCTG</td><td></td><td></td><td></td><td></td><td></td><td></td><td></td><td></td><td></td><td></td></tr>
<tr><td>Oligo rev</td><td>AAACCAGCAGAACAGTTAAAAATC</td><td></td><td></td><td></td><td></td><td></td><td></td><td></td><td></td><td></td><td></td></tr>
<tr><td>Chromosome</td><td>start</td><td>end</td><td>strand</td><td>MM</td><td>target_seq</td><td>PAM</td><td>alignment</td><td>distance</td><td>position</td><td>gene name</td><td>gene id</td></tr>
<tr><td>GU305914.1</td><td>160231</td><td>160253</td><td>+</td><td>0</td><td>GATTTTTAACTGTTCTGCTG</td><td>CGG</td><td>||||||||[||||||||||||]PAM</td><td>NA</td><td>Intergenic</td><td>NA</td><td>NA</td></tr>
<tr><td></td><td></td><td></td><td></td><td></td><td></td><td></td><td></td><td></td><td></td><td></td><td></td></tr>
<tr><td>T12076</td><td>GGCCGTGCGAACAGTCACAGAGG</td><td>1000</td><td></td><td></td><td></td><td></td><td></td><td></td><td></td><td></td><td></td></tr>
<tr><td>Oligo fwd</td><td>CACCGGCCGTGCGAACAGTCACAG</td><td></td><td></td><td></td><td></td><td></td><td></td><td></td><td></td><td></td><td></td></tr>
<tr><td>Oligo rev</td><td>AAACCTGTGACTGTTCGCACGGCC</td><td></td><td></td><td></td><td></td><td></td><td></td><td></td><td></td><td></td><td></td></tr>
<tr><td>Chromosome</td><td>start</td><td>end</td><td>strand</td><td>MM</td><td>target_seq</td><td>PAM</td><td>alignment</td><td>distance</td><td>position</td><td>gene name</td><td>gene id</td></tr>
<tr><td>GU305914.1</td><td>160252</td><td>160274</td><td>+</td><td>0</td><td>GGCCGTGCGAACAGTCACAG</td><td>AGG</td><td>||||||||[||||||||||||]PAM</td><td>NA</td><td>Intergenic</td><td>NA</td><td>NA</td></tr>
<tr><td></td><td></td><td></td><td></td><td></td><td></td><td></td><td></td><td></td><td></td><td></td><td></td></tr>
<tr><td>T12077</td><td>GCCGTGCGAACAGTCACAGAGGG</td><td>1000</td><td></td><td></td><td></td><td></td><td></td><td></td><td></td><td></td><td></td></tr>
<tr><td>Oligo fwd</td><td>CACCGCCGTGCGAACAGTCACAGA</td><td></td><td></td><td></td><td></td><td></td><td></td><td></td><td></td><td></td><td></td></tr>
<tr><td>Oligo rev</td><td>AAACTCTGTGACTGTTCGCACGGC</td><td></td><td></td><td></td><td></td><td></td><td></td><td></td><td></td><td></td><td></td></tr>
<tr><td>Chromosome</td><td>start</td><td>end</td><td>strand</td><td>MM</td><td>target_seq</td><td>PAM</td><td>alignment</td><td>distance</td><td>position</td><td>gene name</td><td>gene id</td></tr>
<tr><td>GU305914.1</td><td>160253</td><td>160275</td><td>+</td><td>0</td><td>GCCGTGCGAACAGTCACAGA</td><td>GGG</td><td>||||||||[||||||||||||]PAM</td><td>NA</td><td>Intergenic</td><td>NA</td><td>NA</td></tr>
<tr><td></td><td></td><td></td><td></td><td></td><td></td><td></td><td></td><td></td><td></td><td></td><td></td></tr>
<tr><td>T12078</td><td>GGTCGCAGTTATAGATCGAATGG</td><td>1000</td><td></td><td></td><td></td><td></td><td></td><td></td><td></td><td></td><td></td></tr>
<tr><td>Oligo fwd</td><td>CACCGGTCGCAGTTATAGATCGAA</td><td></td><td></td><td></td><td></td><td></td><td></td><td></td><td></td><td></td><td></td></tr>
<tr><td>Oligo rev</td><td>AAACTTCGATCTATAACTGCGACC</td><td></td><td></td><td></td><td></td><td></td><td></td><td></td><td></td><td></td><td></td></tr>
<tr><td>Chromosome</td><td>start</td><td>end</td><td>strand</td><td>MM</td><td>target_seq</td><td>PAM</td><td>alignment</td><td>distance</td><td>position</td><td>gene name</td><td>gene id</td></tr>
<tr><td>GU305914.1</td><td>160274</td><td>160296</td><td>+</td><td>0</td><td>GGTCGCAGTTATAGATCGAA</td><td>TGG</td><td>||||||||[||||||||||||]PAM</td><td>NA</td><td>Intergenic</td><td>NA</td><td>NA</td></tr>
<tr><td></td><td></td><td></td><td></td><td></td><td></td><td></td><td></td><td></td><td></td><td></td><td></td></tr>
<tr><td>T12079</td><td>TTGTAGTTTTTCTGCTAATTTGG</td><td>1000</td><td></td><td></td><td></td><td></td><td></td><td></td><td></td><td></td><td></td></tr>
<tr><td>Oligo fwd</td><td>CACCTTGTAGTTTTTCTGCTAATT</td><td></td><td></td><td></td><td></td><td></td><td></td><td></td><td></td><td></td><td></td></tr>
<tr><td>Oligo rev</td><td>AAACAATTAGCAGAAAAACTACAA</td><td></td><td></td><td></td><td></td><td></td><td></td><td></td><td></td><td></td><td></td></tr>
<tr><td>Chromosome</td><td>start</td><td>end</td><td>strand</td><td>MM</td><td>target_seq</td><td>PAM</td><td>alignment</td><td>distance</td><td>position</td><td>gene name</td><td>gene id</td></tr>
<tr><td>GU305914.1</td><td>160393</td><td>160415</td><td>+</td><td>0</td><td>TTGTAGTTTTTCTGCTAATT</td><td>TGG</td><td>||||||||[||||||||||||]PAM</td><td>NA</td><td>Intergenic</td><td>NA</td><td>NA</td></tr>
<tr><td></td><td></td><td></td><td></td><td></td><td></td><td></td><td></td><td></td><td></td><td></td><td></td></tr>
<tr><td>T12080</td><td>TTGGTCACTGATGTATTGAATGG</td><td>1000</td><td></td><td></td><td></td><td></td><td></td><td></td><td></td><td></td><td></td></tr>
<tr><td>Oligo fwd</td><td>CACCTTGGTCACTGATGTATTGAA</td><td></td><td></td><td></td><td></td><td></td><td></td><td></td><td></td><td></td><td></td></tr>
<tr><td>Oligo rev</td><td>AAACTTCAATACATCAGTGACCAA</td><td></td><td></td><td></td><td></td><td></td><td></td><td></td><td></td><td></td><td></td></tr>
<tr><td>Chromosome</td><td>start</td><td>end</td><td>strand</td><td>MM</td><td>target_seq</td><td>PAM</td><td>alignment</td><td>distance</td><td>position</td><td>gene name</td><td>gene id</td></tr>
<tr><td>GU305914.1</td><td>160412</td><td>160434</td><td>+</td><td>0</td><td>TTGGTCACTGATGTATTGAA</td><td>TGG</td><td>||||||||[||||||||||||]PAM</td><td>NA</td><td>Intergenic</td><td>NA</td><td>NA</td></tr>
<tr><td></td><td></td><td></td><td></td><td></td><td></td><td></td><td></td><td></td><td></td><td></td><td></td></tr>
<tr><td>T12081</td><td>CATTTGTGATTACATAGATCTGG</td><td>1000</td><td></td><td></td><td></td><td></td><td></td><td></td><td></td><td></td><td></td></tr>
<tr><td>Oligo fwd</td><td>CACCCATTTGTGATTACATAGATC</td><td></td><td></td><td></td><td></td><td></td><td></td><td></td><td></td><td></td><td></td></tr>
<tr><td>Oligo rev</td><td>AAACGATCTATGTAATCACAAATG</td><td></td><td></td><td></td><td></td><td></td><td></td><td></td><td></td><td></td><td></td></tr>
<tr><td>Chromosome</td><td>start</td><td>end</td><td>strand</td><td>MM</td><td>target_seq</td><td>PAM</td><td>alignment</td><td>distance</td><td>position</td><td>gene name</td><td>gene id</td></tr>
<tr><td>GU305914.1</td><td>160463</td><td>160485</td><td>+</td><td>0</td><td>CATTTGTGATTACATAGATC</td><td>TGG</td><td>||||||||[||||||||||||]PAM</td><td>NA</td><td>Intergenic</td><td>NA</td><td>NA</td></tr>
<tr><td></td><td></td><td></td><td></td><td></td><td></td><td></td><td></td><td></td><td></td><td></td><td></td></tr>
<tr><td>T12082</td><td>CATATAAGATATACAGAGAAAGG</td><td>1000</td><td></td><td></td><td></td><td></td><td></td><td></td><td></td><td></td><td></td></tr>
<tr><td>Oligo fwd</td><td>CACCCATATAAGATATACAGAGAA</td><td></td><td></td><td></td><td></td><td></td><td></td><td></td><td></td><td></td><td></td></tr>
<tr><td>Oligo rev</td><td>AAACTTCTCTGTATATCTTATATG</td><td></td><td></td><td></td><td></td><td></td><td></td><td></td><td></td><td></td><td></td></tr>
<tr><td>Chromosome</td><td>start</td><td>end</td><td>strand</td><td>MM</td><td>target_seq</td><td>PAM</td><td>alignment</td><td>distance</td><td>position</td><td>gene name</td><td>gene id</td></tr>
<tr><td>GU305914.1</td><td>160541</td><td>160563</td><td>+</td><td>0</td><td>CATATAAGATATACAGAGAA</td><td>AGG</td><td>||||||||[||||||||||||]PAM</td><td>NA</td><td>Intergenic</td><td>NA</td><td>NA</td></tr>
<tr><td></td><td></td><td></td><td></td><td></td><td></td><td></td><td></td><td></td><td></td><td></td><td></td></tr>
<tr><td>T12083</td><td>ATGTTAGTCGTACACTGTATCGG</td><td>1000</td><td></td><td></td><td></td><td></td><td></td><td></td><td></td><td></td><td></td></tr>
<tr><td>Oligo fwd</td><td>CACCATGTTAGTCGTACACTGTAT</td><td></td><td></td><td></td><td></td><td></td><td></td><td></td><td></td><td></td><td></td></tr>
<tr><td>Oligo rev</td><td>AAACATACAGTGTACGACTAACAT</td><td></td><td></td><td></td><td></td><td></td><td></td><td></td><td></td><td></td><td></td></tr>
<tr><td>Chromosome</td><td>start</td><td>end</td><td>strand</td><td>MM</td><td>target_seq</td><td>PAM</td><td>alignment</td><td>distance</td><td>position</td><td>gene name</td><td>gene id</td></tr>
<tr><td>GU305914.1</td><td>160590</td><td>160612</td><td>+</td><td>0</td><td>ATGTTAGTCGTACACTGTAT</td><td>CGG</td><td>||||||||[||||||||||||]PAM</td><td>NA</td><td>Intergenic</td><td>NA</td><td>NA</td></tr>
<tr><td></td><td></td><td></td><td></td><td></td><td></td><td></td><td></td><td></td><td></td><td></td><td></td></tr>
<tr><td>T12084</td><td>CAGATAGAGTTCTGTCTGTAAGG</td><td>1000</td><td></td><td></td><td></td><td></td><td></td><td></td><td></td><td></td><td></td></tr>
<tr><td>Oligo fwd</td><td>CACCCAGATAGAGTTCTGTCTGTA</td><td></td><td></td><td></td><td></td><td></td><td></td><td></td><td></td><td></td><td></td></tr>
<tr><td>Oligo rev</td><td>AAACTACAGACAGAACTCTATCTG</td><td></td><td></td><td></td><td></td><td></td><td></td><td></td><td></td><td></td><td></td></tr>
</table>

| | | | | | | | | | | | |
|---|---|---|---|---|---|---|---|---|---|---|---|
| Chromosome | start | end | strand | MM | target_seq | PAM | alignment | distance | position | gene name | gene id |
| GU305914.1 | 160618 | 160640 | + | 0 | CAGATAGAGTTCTGTCTGTA | AGG | \|\|\|\|\|\|\|\|[\|\|\|\|\|\|\|\|\|\|\|\|]PAM | NA | Intergenic | NA | NA |
| T12085 | TAAAAAGAAAACCAACGAAGAGG | 1000 | | | | | | | | | |
| Oligo fwd | CACCTAAAAAGAAAACCAACGAAG | | | | | | | | | | |
| Oligo rev | AAACCTTCGTTGGTTTTCTTTTTA | | | | | | | | | | |
| Chromosome | start | end | strand | MM | target_seq | PAM | alignment | distance | position | gene name | gene id |
| GU305914.1 | 160673 | 160695 | + | 0 | TAAAAAGAAAACCAACGAAG | AGG | \|\|\|\|\|\|\|\|[\|\|\|\|\|\|\|\|\|\|\|\|]PAM | NA | Intergenic | NA | NA |
| T12086 | AAAGAAAACCAACGAAGAGGAGG | 1000 | | | | | | | | | |
| Oligo fwd | CACCAAAGAAAACCAACGAAGAGG | | | | | | | | | | |
| Oligo rev | AAACCCTCTTCGTTGGTTTTCTTT | | | | | | | | | | |
| Chromosome | start | end | strand | MM | target_seq | PAM | alignment | distance | position | gene name | gene id |
| GU305914.1 | 160676 | 160698 | + | 0 | AAAGAAAACCAACGAAGAGG | AGG | \|\|\|\|\|\|\|\|[\|\|\|\|\|\|\|\|\|\|\|\|]PAM | NA | Intergenic | NA | NA |
| T12087 | AAACCAACGAAGAGGAGGAGAGG | 1000 | | | | | | | | | |
| Oligo fwd | CACCAAACCAACGAAGAGGAGGAG | | | | | | | | | | |
| Oligo rev | AAACCTCCTCCTCTTCGTTGGTTT | | | | | | | | | | |
| Chromosome | start | end | strand | MM | target_seq | PAM | alignment | distance | position | gene name | gene id |
| GU305914.1 | 160681 | 160703 | + | 0 | AAACCAACGAAGAGGAGGAG | AGG | \|\|\|\|\|\|\|\|[\|\|\|\|\|\|\|\|\|\|\|\|]PAM | NA | Intergenic | NA | NA |
| GU305914.1 | 196441 | 196463 | - | 4 | ACACGAACGAGGAGGAGGAC | TGG | \|-\|\|-\|\|\|[\|\|-\|\|\|\|\|\|\|\|-]PAM | NA | Intergenic | NA | NA |
| T12088 | CCAACGAAGAGGAGGAGAGGAGG | 1000 | | | | | | | | | |
| Oligo fwd | CACCCCAACGAAGAGGAGGAGAGG | | | | | | | | | | |
| Oligo rev | AAACCCTCTCCTCCTCTTCGTTGG | | | | | | | | | | |
| Chromosome | start | end | strand | MM | target_seq | PAM | alignment | distance | position | gene name | gene id |
| GU305914.1 | 160684 | 160706 | + | 0 | CCAACGAAGAGGAGGAGAGG | AGG | \|\|\|\|\|\|\|\|[\|\|\|\|\|\|\|\|\|\|\|\|]PAM | NA | Intergenic | NA | NA |
| T12089 | GACAAGATCTTTGATTATAATGG | 1000 | | | | | | | | | |
| Oligo fwd | CACCGACAAGATCTTTGATTATAA | | | | | | | | | | |
| Oligo rev | AAACTTATAATCAAAGATCTTGTC | | | | | | | | | | |
| Chromosome | start | end | strand | MM | target_seq | PAM | alignment | distance | position | gene name | gene id |
| GU305914.1 | 160715 | 160737 | + | 0 | GACAAGATCTTTGATTATAA | TGG | \|\|\|\|\|\|\|\|[\|\|\|\|\|\|\|\|\|\|\|\|]PAM | NA | Intergenic | NA | NA |
| T12090 | TTTTTATTTTTTTTATTTTTCGG | 1000 | | | | | | | | | |
| Oligo fwd | CACCTTTTTATTTTTTTTATTTTT | | | | | | | | | | |
| Oligo rev | AAACAAAAATAAAAAAAATAAAAA | | | | | | | | | | |
| Chromosome | start | end | strand | MM | target_seq | PAM | alignment | distance | position | gene name | gene id |
| GU305914.1 | 160775 | 160797 | + | 0 | TTTTTATTTTTTTTATTTTT | CGG | \|\|\|\|\|\|\|\|[\|\|\|\|\|\|\|\|\|\|\|\|]PAM | NA | Intergenic | NA | NA |
| T12091 | TTTATTTTTCGGTGAGCAGACGG | 1000 | | | | | | | | | |
| Oligo fwd | CACCTTTATTTTTCGGTGAGCAGA | | | | | | | | | | |
| Oligo rev | AAACTCTGCTCACCGAAAAATAAA | | | | | | | | | | |
| Chromosome | start | end | strand | MM | target_seq | PAM | alignment | distance | position | gene name | gene id |
| GU305914.1 | 160786 | 160808 | + | 0 | TTTATTTTTCGGTGAGCAGA | CGG | \|\|\|\|\|\|\|\|[\|\|\|\|\|\|\|\|\|\|\|\|]PAM | NA | Intergenic | NA | NA |
| T12092 | GTTCTGAAAATACAGAAAACCGG | 1000 | | | | | | | | | |
| Oligo fwd | CACCGTTCTGAAAATACAGAAAAC | | | | | | | | | | |
| Oligo rev | AAACGTTTTCTGTATTTTCAGAAC | | | | | | | | | | |
| Chromosome | start | end | strand | MM | target_seq | PAM | alignment | distance | position | gene name | gene id |
| GU305914.1 | 160808 | 160830 | + | 0 | GTTCTGAAAATACAGAAAAC | CGG | \|\|\|\|\|\|\|\|[\|\|\|\|\|\|\|\|\|\|\|\|]PAM | NA | Intergenic | NA | NA |
| T12093 | TACAGAAAACCGGATCACCCTGG | 1000 | | | | | | | | | |
| Oligo fwd | CACCTACAGAAAACCGGATCACCC | | | | | | | | | | |
| Oligo rev | AAACGGGTGATCCGGTTTTCTGTA | | | | | | | | | | |
| Chromosome | start | end | strand | MM | target_seq | PAM | alignment | distance | position | gene name | gene id |
| GU305914.1 | 160818 | 160840 | + | 0 | TACAGAAAACCGGATCACCC | TGG | \|\|\|\|\|\|\|\|[\|\|\|\|\|\|\|\|\|\|\|\|]PAM | NA | Intergenic | NA | NA |
| T12094 | ACAGAAAACCGGATCACCCTGGG | 1000 | | | | | | | | | |
| Oligo fwd | CACCACAGAAAACCGGATCACCCT | | | | | | | | | | |
| Oligo rev | AAACAGGGTGATCCGGTTTTCTGT | | | | | | | | | | |
| Chromosome | start | end | strand | MM | target_seq | PAM | alignment | distance | position | gene name | gene id |
| GU305914.1 | 160819 | 160841 | + | 0 | ACAGAAAACCGGATCACCCT | GGG | \|\|\|\|\|\|\|\|[\|\|\|\|\|\|\|\|\|\|\|\|]PAM | NA | Intergenic | NA | NA |
| T12095 | AAACCGGATCACCCTGGGAATGG | 1000 | | | | | | | | | |
| Oligo fwd | CACCAAACCGGATCACCCTGGGAA | | | | | | | | | | |
| Oligo rev | AAACTTCCCAGGGTGATCCGGTTT | | | | | | | | | | |
| Chromosome | start | end | strand | MM | target_seq | PAM | alignment | distance | position | gene name | gene id |
| GU305914.1 | 160824 | 160846 | + | 0 | AAACCGGATCACCCTGGGAA | TGG | \|\|\|\|\|\|\|\|[\|\|\|\|\|\|\|\|\|\|\|\|]PAM | NA | Intergenic | NA | NA |
| T12096 | CCGGATCACCCTGGGAATGGTGG | 1000 | | | | | | | | | |
| Oligo fwd | CACCCCGGATCACCCTGGGAATGG | | | | | | | | | | |
| Oligo rev | AAACCCATTCCCAGGGTGATCCGG | | | | | | | | | | |
| Chromosome | start | end | strand | MM | target_seq | PAM | alignment | distance | position | gene name | gene id |
| GU305914.1 | 160827 | 160849 | + | 0 | CCGGATCACCCTGGGAATGG | TGG | \|\|\|\|\|\|\|\|[\|\|\|\|\|\|\|\|\|\|\|\|]PAM | NA | Intergenic | NA | NA |
| T12097 | CGGATCACCCTGGGAATGGTGGG | 1000 | | | | | | | | | |
| Oligo fwd | CACCCGGATCACCCTGGGAATGGT | | | | | | | | | | |
| Oligo rev | AAACACCATTCCCAGGGTGATCCG | | | | | | | | | | |
| Chromosome | start | end | strand | MM | target_seq | PAM | alignment | distance | position | gene name | gene id |
| GU305914.1 | 160828 | 160850 | + | 0 | CGGATCACCCTGGGAATGGT | GGG | \|\|\|\|\|\|\|\|[\|\|\|\|\|\|\|\|\|\|\|\|]PAM | NA | Intergenic | NA | NA |
| T12098 | GGTGGGTAACAACTAAGCTAAGG | 1000 | | | | | | | | | |
| Oligo fwd | CACCGGTGGGTAACAACTAAGCTA | | | | | | | | | | |
| Oligo rev | AAACTAGCTTAGTTGTTACCCACC | | | | | | | | | | |
| Chromosome | start | end | strand | MM | target_seq | PAM | alignment | distance | position | gene name | gene id |
| GU305914.1 | 160845 | 160867 | + | 0 | GGTGGGTAACAACTAAGCTA | AGG | \|\|\|\|\|\|\|\|[\|\|\|\|\|\|\|\|\|\|\|\|]PAM | NA | Intergenic | NA | NA |
| T12099 | GCATTGCATGTAGTTTCTAATGG | 1000 | | | | | | | | | |

<table>
<tr><td>Oligo fwd</td><td>CACCGCATTGCATGTAGTTTCTAA</td><td></td><td></td><td></td><td></td><td></td><td></td><td></td><td></td><td></td><td></td></tr>
<tr><td>Oligo rev</td><td>AAACTTAGAAACTACATGCAATGC</td><td></td><td></td><td></td><td></td><td></td><td></td><td></td><td></td><td></td><td></td></tr>
<tr><td>Chromosome</td><td>start</td><td>end</td><td>strand</td><td>MM</td><td>target_seq</td><td>PAM</td><td>alignment</td><td>distance</td><td>position</td><td>gene name</td><td>gene id</td></tr>
<tr><td>GU305914.1</td><td>160889</td><td>160911</td><td>+</td><td>0</td><td>GCATTGCATGTAGTTTCTAA</td><td>TGG</td><td>||||||||[||||||||||||]PAM</td><td>NA</td><td>Intergenic</td><td>NA</td><td>NA</td></tr>
<tr><td></td><td></td><td></td><td></td><td></td><td></td><td></td><td></td><td></td><td></td><td></td><td></td></tr>
<tr><td>T12100</td><td>TAGTAACTAACAAAAAGTCTCGG</td><td>1000</td><td></td><td></td><td></td><td></td><td></td><td></td><td></td><td></td><td></td></tr>
<tr><td>Oligo fwd</td><td>CACCTAGTAACTAACAAAAAGTCT</td><td></td><td></td><td></td><td></td><td></td><td></td><td></td><td></td><td></td><td></td></tr>
<tr><td>Oligo rev</td><td>AAACAGACTTTTTGTTAGTTACTA</td><td></td><td></td><td></td><td></td><td></td><td></td><td></td><td></td><td></td><td></td></tr>
<tr><td>Chromosome</td><td>start</td><td>end</td><td>strand</td><td>MM</td><td>target_seq</td><td>PAM</td><td>alignment</td><td>distance</td><td>position</td><td>gene name</td><td>gene id</td></tr>
<tr><td>GU305914.1</td><td>160912</td><td>160934</td><td>+</td><td>0</td><td>TAGTAACTAACAAAAAGTCT</td><td>CGG</td><td>||||||||[||||||||||||]PAM</td><td>NA</td><td>Intergenic</td><td>NA</td><td>NA</td></tr>
<tr><td></td><td></td><td></td><td></td><td></td><td></td><td></td><td></td><td></td><td></td><td></td><td></td></tr>
<tr><td>T12101</td><td>TCTCTCTCGAGAACCCAAGACGG</td><td>1000</td><td></td><td></td><td></td><td></td><td></td><td></td><td></td><td></td><td></td></tr>
<tr><td>Oligo fwd</td><td>CACCTCTCTCTCGAGAACCCAAGA</td><td></td><td></td><td></td><td></td><td></td><td></td><td></td><td></td><td></td><td></td></tr>
<tr><td>Oligo rev</td><td>AAACTCTTGGGTTCTCGAGAGAGA</td><td></td><td></td><td></td><td></td><td></td><td></td><td></td><td></td><td></td><td></td></tr>
<tr><td>Chromosome</td><td>start</td><td>end</td><td>strand</td><td>MM</td><td>target_seq</td><td>PAM</td><td>alignment</td><td>distance</td><td>position</td><td>gene name</td><td>gene id</td></tr>
<tr><td>GU305914.1</td><td>160936</td><td>160958</td><td>+</td><td>0</td><td>TCTCTCTCGAGAACCCAAGA</td><td>CGG</td><td>||||||||[||||||||||||]PAM</td><td>NA</td><td>Intergenic</td><td>NA</td><td>NA</td></tr>
<tr><td></td><td></td><td></td><td></td><td></td><td></td><td></td><td></td><td></td><td></td><td></td><td></td></tr>
<tr><td>T12102</td><td>CTCTCTCGAGAACCCAAGACGGG</td><td>1000</td><td></td><td></td><td></td><td></td><td></td><td></td><td></td><td></td><td></td></tr>
<tr><td>Oligo fwd</td><td>CACCCTCTCTCGAGAACCCAAGAC</td><td></td><td></td><td></td><td></td><td></td><td></td><td></td><td></td><td></td><td></td></tr>
<tr><td>Oligo rev</td><td>AAACGTCTTGGGTTCTCGAGAGAG</td><td></td><td></td><td></td><td></td><td></td><td></td><td></td><td></td><td></td><td></td></tr>
<tr><td>Chromosome</td><td>start</td><td>end</td><td>strand</td><td>MM</td><td>target_seq</td><td>PAM</td><td>alignment</td><td>distance</td><td>position</td><td>gene name</td><td>gene id</td></tr>
<tr><td>GU305914.1</td><td>160937</td><td>160959</td><td>+</td><td>0</td><td>CTCTCTCGAGAACCCAAGAC</td><td>GGG</td><td>||||||||[||||||||||||]PAM</td><td>NA</td><td>Intergenic</td><td>NA</td><td>NA</td></tr>
<tr><td></td><td></td><td></td><td></td><td></td><td></td><td></td><td></td><td></td><td></td><td></td><td></td></tr>
<tr><td>T12103</td><td>AACACGATGCTCTGTGTCGTCGG</td><td>1000</td><td></td><td></td><td></td><td></td><td></td><td></td><td></td><td></td><td></td></tr>
<tr><td>Oligo fwd</td><td>CACCAACACGATGCTCTGTGTCGT</td><td></td><td></td><td></td><td></td><td></td><td></td><td></td><td></td><td></td><td></td></tr>
<tr><td>Oligo rev</td><td>AAACACGACACAGAGCATCGTGTT</td><td></td><td></td><td></td><td></td><td></td><td></td><td></td><td></td><td></td><td></td></tr>
<tr><td>Chromosome</td><td>start</td><td>end</td><td>strand</td><td>MM</td><td>target_seq</td><td>PAM</td><td>alignment</td><td>distance</td><td>position</td><td>gene name</td><td>gene id</td></tr>
<tr><td>GU305914.1</td><td>160973</td><td>160995</td><td>+</td><td>0</td><td>AACACGATGCTCTGTGTCGT</td><td>CGG</td><td>||||||||[||||||||||||]PAM</td><td>NA</td><td>Intergenic</td><td>NA</td><td>NA</td></tr>
<tr><td></td><td></td><td></td><td></td><td></td><td></td><td></td><td></td><td></td><td></td><td></td><td></td></tr>
<tr><td>T12104</td><td>CGATGCTCTGTGTCGTCGGTCGG</td><td>1000</td><td></td><td></td><td></td><td></td><td></td><td></td><td></td><td></td><td></td></tr>
<tr><td>Oligo fwd</td><td>CACCCGATGCTCTGTGTCGTCGGT</td><td></td><td></td><td></td><td></td><td></td><td></td><td></td><td></td><td></td><td></td></tr>
<tr><td>Oligo rev</td><td>AAACACCGACGACACAGAGCATCG</td><td></td><td></td><td></td><td></td><td></td><td></td><td></td><td></td><td></td><td></td></tr>
<tr><td>Chromosome</td><td>start</td><td>end</td><td>strand</td><td>MM</td><td>target_seq</td><td>PAM</td><td>alignment</td><td>distance</td><td>position</td><td>gene name</td><td>gene id</td></tr>
<tr><td>GU305914.1</td><td>160977</td><td>160999</td><td>+</td><td>0</td><td>CGATGCTCTGTGTCGTCGGT</td><td>CGG</td><td>||||||||[||||||||||||]PAM</td><td>NA</td><td>Intergenic</td><td>NA</td><td>NA</td></tr>
<tr><td></td><td></td><td></td><td></td><td></td><td></td><td></td><td></td><td></td><td></td><td></td><td></td></tr>
<tr><td>T12105</td><td>GCTCTGTGTCGTCGGTCGGTTGG</td><td>1000</td><td></td><td></td><td></td><td></td><td></td><td></td><td></td><td></td><td></td></tr>
<tr><td>Oligo fwd</td><td>CACCGCTCTGTGTCGTCGGTCGGT</td><td></td><td></td><td></td><td></td><td></td><td></td><td></td><td></td><td></td><td></td></tr>
<tr><td>Oligo rev</td><td>AAACACCGACCGACGACACAGAGC</td><td></td><td></td><td></td><td></td><td></td><td></td><td></td><td></td><td></td><td></td></tr>
<tr><td>Chromosome</td><td>start</td><td>end</td><td>strand</td><td>MM</td><td>target_seq</td><td>PAM</td><td>alignment</td><td>distance</td><td>position</td><td>gene name</td><td>gene id</td></tr>
<tr><td>GU305914.1</td><td>160981</td><td>161003</td><td>+</td><td>0</td><td>GCTCTGTGTCGTCGGTCGGT</td><td>TGG</td><td>||||||||[||||||||||||]PAM</td><td>NA</td><td>Intergenic</td><td>NA</td><td>NA</td></tr>
<tr><td></td><td></td><td></td><td></td><td></td><td></td><td></td><td></td><td></td><td></td><td></td><td></td></tr>
<tr><td>T12106</td><td>CGGTTGGACATCTATCGAGTTGG</td><td>1000</td><td></td><td></td><td></td><td></td><td></td><td></td><td></td><td></td><td></td></tr>
<tr><td>Oligo fwd</td><td>CACCCGGTTGGACATCTATCGAGT</td><td></td><td></td><td></td><td></td><td></td><td></td><td></td><td></td><td></td><td></td></tr>
<tr><td>Oligo rev</td><td>AAACACTCGATAGATGTCCAACCG</td><td></td><td></td><td></td><td></td><td></td><td></td><td></td><td></td><td></td><td></td></tr>
<tr><td>Chromosome</td><td>start</td><td>end</td><td>strand</td><td>MM</td><td>target_seq</td><td>PAM</td><td>alignment</td><td>distance</td><td>position</td><td>gene name</td><td>gene id</td></tr>
<tr><td>GU305914.1</td><td>160997</td><td>161019</td><td>+</td><td>0</td><td>CGGTTGGACATCTATCGAGT</td><td>TGG</td><td>||||||||[||||||||||||]PAM</td><td>NA</td><td>Intergenic</td><td>NA</td><td>NA</td></tr>
<tr><td></td><td></td><td></td><td></td><td></td><td></td><td></td><td></td><td></td><td></td><td></td><td></td></tr>
<tr><td>T12107</td><td>TTGGACATCTATCGAGTTGGCGG</td><td>1000</td><td></td><td></td><td></td><td></td><td></td><td></td><td></td><td></td><td></td></tr>
<tr><td>Oligo fwd</td><td>CACCTTGGACATCTATCGAGTTGG</td><td></td><td></td><td></td><td></td><td></td><td></td><td></td><td></td><td></td><td></td></tr>
<tr><td>Oligo rev</td><td>AAACCCAACTCGATAGATGTCCAA</td><td></td><td></td><td></td><td></td><td></td><td></td><td></td><td></td><td></td><td></td></tr>
<tr><td>Chromosome</td><td>start</td><td>end</td><td>strand</td><td>MM</td><td>target_seq</td><td>PAM</td><td>alignment</td><td>distance</td><td>position</td><td>gene name</td><td>gene id</td></tr>
<tr><td>GU305914.1</td><td>161000</td><td>161022</td><td>+</td><td>0</td><td>TTGGACATCTATCGAGTTGG</td><td>CGG</td><td>||||||||[||||||||||||]PAM</td><td>NA</td><td>Intergenic</td><td>NA</td><td>NA</td></tr>
<tr><td></td><td></td><td></td><td></td><td></td><td></td><td></td><td></td><td></td><td></td><td></td><td></td></tr>
<tr><td>T12108</td><td>GTTGTTCAAAACGCGCTCCCCGG</td><td>1000</td><td></td><td></td><td></td><td></td><td></td><td></td><td></td><td></td><td></td></tr>
<tr><td>Oligo fwd</td><td>CACCGTTGTTCAAAACGCGCTCCC</td><td></td><td></td><td></td><td></td><td></td><td></td><td></td><td></td><td></td><td></td></tr>
<tr><td>Oligo rev</td><td>AAACGGGAGCGCGTTTTGAACAAC</td><td></td><td></td><td></td><td></td><td></td><td></td><td></td><td></td><td></td><td></td></tr>
<tr><td>Chromosome</td><td>start</td><td>end</td><td>strand</td><td>MM</td><td>target_seq</td><td>PAM</td><td>alignment</td><td>distance</td><td>position</td><td>gene name</td><td>gene id</td></tr>
<tr><td>GU305914.1</td><td>161032</td><td>161054</td><td>+</td><td>0</td><td>GTTGTTCAAAACGCGCTCCC</td><td>CGG</td><td>||||||||[||||||||||||]PAM</td><td>NA</td><td>Intergenic</td><td>NA</td><td>NA</td></tr>
<tr><td></td><td></td><td></td><td></td><td></td><td></td><td></td><td></td><td></td><td></td><td></td><td></td></tr>
<tr><td>T12109</td><td>TTGTTGTTTTGTGATGAGATTGG</td><td>1000</td><td></td><td></td><td></td><td></td><td></td><td></td><td></td><td></td><td></td></tr>
<tr><td>Oligo fwd</td><td>CACCTTGTTGTTTTGTGATGAGAT</td><td></td><td></td><td></td><td></td><td></td><td></td><td></td><td></td><td></td><td></td></tr>
<tr><td>Oligo rev</td><td>AAACATCTCATCACAAAACAACAA</td><td></td><td></td><td></td><td></td><td></td><td></td><td></td><td></td><td></td><td></td></tr>
<tr><td>Chromosome</td><td>start</td><td>end</td><td>strand</td><td>MM</td><td>target_seq</td><td>PAM</td><td>alignment</td><td>distance</td><td>position</td><td>gene name</td><td>gene id</td></tr>
<tr><td>GU305914.1</td><td>161061</td><td>161083</td><td>+</td><td>0</td><td>TTGTTGTTTTGTGATGAGAT</td><td>TGG</td><td>||||||||[||||||||||||]PAM</td><td>NA</td><td>Intergenic</td><td>NA</td><td>NA</td></tr>
<tr><td></td><td></td><td></td><td></td><td></td><td></td><td></td><td></td><td></td><td></td><td></td><td></td></tr>
<tr><td>T12110</td><td>TGTTGTTTTGTGATGAGATTGGG</td><td>1000</td><td></td><td></td><td></td><td></td><td></td><td></td><td></td><td></td><td></td></tr>
<tr><td>Oligo fwd</td><td>CACCTGTTGTTTTGTGATGAGATT</td><td></td><td></td><td></td><td></td><td></td><td></td><td></td><td></td><td></td><td></td></tr>
<tr><td>Oligo rev</td><td>AAACAATCTCATCACAAAACAACA</td><td></td><td></td><td></td><td></td><td></td><td></td><td></td><td></td><td></td><td></td></tr>
<tr><td>Chromosome</td><td>start</td><td>end</td><td>strand</td><td>MM</td><td>target_seq</td><td>PAM</td><td>alignment</td><td>distance</td><td>position</td><td>gene name</td><td>gene id</td></tr>
<tr><td>GU305914.1</td><td>161062</td><td>161084</td><td>+</td><td>0</td><td>TGTTGTTTTGTGATGAGATT</td><td>GGG</td><td>||||||||[||||||||||||]PAM</td><td>NA</td><td>Intergenic</td><td>NA</td><td>NA</td></tr>
<tr><td></td><td></td><td></td><td></td><td></td><td></td><td></td><td></td><td></td><td></td><td></td><td></td></tr>
<tr><td>T12111</td><td>TTGTTGATATAGATTTTTAGTGG</td><td>1000</td><td></td><td></td><td></td><td></td><td></td><td></td><td></td><td></td><td></td></tr>
<tr><td>Oligo fwd</td><td>CACCTTGTTGATATAGATTTTTAG</td><td></td><td></td><td></td><td></td><td></td><td></td><td></td><td></td><td></td><td></td></tr>
<tr><td>Oligo rev</td><td>AAACCTAAAAATCTATATCAACAA</td><td></td><td></td><td></td><td></td><td></td><td></td><td></td><td></td><td></td><td></td></tr>
<tr><td>Chromosome</td><td>start</td><td>end</td><td>strand</td><td>MM</td><td>target_seq</td><td>PAM</td><td>alignment</td><td>distance</td><td>position</td><td>gene name</td><td>gene id</td></tr>
<tr><td>GU305914.1</td><td>161142</td><td>161164</td><td>+</td><td>0</td><td>TTGTTGATATAGATTTTTAG</td><td>TGG</td><td>||||||||[||||||||||||]PAM</td><td>NA</td><td>Intergenic</td><td>NA</td><td>NA</td></tr>
<tr><td></td><td></td><td></td><td></td><td></td><td></td><td></td><td></td><td></td><td></td><td></td><td></td></tr>
<tr><td>T12112</td><td>GTGAGTAATCTCGTTGATGTAGG</td><td>1000</td><td></td><td></td><td></td><td></td><td></td><td></td><td></td><td></td><td></td></tr>
<tr><td>Oligo fwd</td><td>CACCGTGAGTAATCTCGTTGATGT</td><td></td><td></td><td></td><td></td><td></td><td></td><td></td><td></td><td></td><td></td></tr>
<tr><td>Oligo rev</td><td>AAACACATCAACGAGATTACTCAC</td><td></td><td></td><td></td><td></td><td></td><td></td><td></td><td></td><td></td><td></td></tr>
<tr><td>Chromosome</td><td>start</td><td>end</td><td>strand</td><td>MM</td><td>target_seq</td><td>PAM</td><td>alignment</td><td>distance</td><td>position</td><td>gene name</td><td>gene id</td></tr>
<tr><td>GU305914.1</td><td>161183</td><td>161205</td><td>+</td><td>0</td><td>GTGAGTAATCTCGTTGATGT</td><td>AGG</td><td>||||||||[||||||||||||]PAM</td><td>NA</td><td>Intergenic</td><td>NA</td><td>NA</td></tr>
<tr><td></td><td></td><td></td><td></td><td></td><td></td><td></td><td></td><td></td><td></td><td></td><td></td></tr>
<tr><td>T12113</td><td>TAATCTCGTTGATGTAGGTGAGG</td><td>1000</td><td></td><td></td><td></td><td></td><td></td><td></td><td></td><td></td><td></td></tr>
<tr><td>Oligo fwd</td><td>CACCTAATCTCGTTGATGTAGGTG</td><td></td><td></td><td></td><td></td><td></td><td></td><td></td><td></td><td></td><td></td></tr>
<tr><td>Oligo rev</td><td>AAACCACCTACATCAACGAGATTA</td><td></td><td></td><td></td><td></td><td></td><td></td><td></td><td></td><td></td><td></td></tr>
<tr><td>Chromosome</td><td>start</td><td>end</td><td>strand</td><td>MM</td><td>target_seq</td><td>PAM</td><td>alignment</td><td>distance</td><td>position</td><td>gene name</td><td>gene id</td></tr>
<tr><td>GU305914.1</td><td>161188</td><td>161210</td><td>+</td><td>0</td><td>TAATCTCGTTGATGTAGGTG</td><td>AGG</td><td>||||||||[||||||||||||]PAM</td><td>NA</td><td>Intergenic</td><td>NA</td><td>NA</td></tr>
</table>

<table>
<tr><td>T12114</td><td>TCTCTAAGTAAAGCAGTACGAGG</td><td>1000</td><td></td><td></td><td></td><td></td><td></td><td></td><td></td><td></td><td></td></tr>
<tr><td>Oligo fwd</td><td>CACCTCTCTAAGTAAAGCAGTACG</td><td></td><td></td><td></td><td></td><td></td><td></td><td></td><td></td><td></td><td></td></tr>
<tr><td>Oligo rev</td><td>AAACCGTACTGCTTTACTTAGAGA</td><td></td><td></td><td></td><td></td><td></td><td></td><td></td><td></td><td></td><td></td></tr>
<tr><td>Chromosome</td><td>start</td><td>end</td><td>strand</td><td>MM</td><td>target_seq</td><td>PAM</td><td>alignment</td><td>distance</td><td>position</td><td>gene name</td><td>gene id</td></tr>
<tr><td>GU305914.1</td><td>161265</td><td>161287</td><td>+</td><td>0</td><td>TCTCTAAGTAAAGCAGTACG</td><td>AGG</td><td>||||||||[||||||||||||]PAM</td><td>NA</td><td>Intergenic</td><td>NA</td><td>NA</td></tr>
<tr><td>T12115</td><td>CTAAGTAAAGCAGTACGAGGCGG</td><td>1000</td><td></td><td></td><td></td><td></td><td></td><td></td><td></td><td></td><td></td></tr>
<tr><td>Oligo fwd</td><td>CACCCTAAGTAAAGCAGTACGAGG</td><td></td><td></td><td></td><td></td><td></td><td></td><td></td><td></td><td></td><td></td></tr>
<tr><td>Oligo rev</td><td>AAACCCTCGTACTGCTTTACTTAG</td><td></td><td></td><td></td><td></td><td></td><td></td><td></td><td></td><td></td><td></td></tr>
<tr><td>Chromosome</td><td>start</td><td>end</td><td>strand</td><td>MM</td><td>target_seq</td><td>PAM</td><td>alignment</td><td>distance</td><td>position</td><td>gene name</td><td>gene id</td></tr>
<tr><td>GU305914.1</td><td>161268</td><td>161290</td><td>+</td><td>0</td><td>CTAAGTAAAGCAGTACGAGG</td><td>CGG</td><td>||||||||[||||||||||||]PAM</td><td>NA</td><td>Intergenic</td><td>NA</td><td>NA</td></tr>
<tr><td>GU305914.1</td><td>3968</td><td>3990</td><td>-</td><td>4</td><td>ACAAGTAAATCAGTACGAGA</td><td>AGG</td><td>--||||||[|-|||||||||-]PAM</td><td>NA</td><td>Intergenic</td><td>NA</td><td>NA</td></tr>
<tr><td>T12116</td><td>ACGAGGCGGCATCGACGTGCAGG</td><td>1000</td><td></td><td></td><td></td><td></td><td></td><td></td><td></td><td></td><td></td></tr>
<tr><td>Oligo fwd</td><td>CACCACGAGGCGGCATCGACGTGC</td><td></td><td></td><td></td><td></td><td></td><td></td><td></td><td></td><td></td><td></td></tr>
<tr><td>Oligo rev</td><td>AAACGCACGTCGATGCCGCCTCGT</td><td></td><td></td><td></td><td></td><td></td><td></td><td></td><td></td><td></td><td></td></tr>
<tr><td>Chromosome</td><td>start</td><td>end</td><td>strand</td><td>MM</td><td>target_seq</td><td>PAM</td><td>alignment</td><td>distance</td><td>position</td><td>gene name</td><td>gene id</td></tr>
<tr><td>GU305914.1</td><td>161282</td><td>161304</td><td>+</td><td>0</td><td>ACGAGGCGGCATCGACGTGC</td><td>AGG</td><td>||||||||[||||||||||||]PAM</td><td>NA</td><td>Intergenic</td><td>NA</td><td>NA</td></tr>
<tr><td>T12117</td><td>CGAGGCGGCATCGACGTGCAGGG</td><td>1000</td><td></td><td></td><td></td><td></td><td></td><td></td><td></td><td></td><td></td></tr>
<tr><td>Oligo fwd</td><td>CACCCGAGGCGGCATCGACGTGCA</td><td></td><td></td><td></td><td></td><td></td><td></td><td></td><td></td><td></td><td></td></tr>
<tr><td>Oligo rev</td><td>AAACTGCACGTCGATGCCGCCTCG</td><td></td><td></td><td></td><td></td><td></td><td></td><td></td><td></td><td></td><td></td></tr>
<tr><td>Chromosome</td><td>start</td><td>end</td><td>strand</td><td>MM</td><td>target_seq</td><td>PAM</td><td>alignment</td><td>distance</td><td>position</td><td>gene name</td><td>gene id</td></tr>
<tr><td>GU305914.1</td><td>161283</td><td>161305</td><td>+</td><td>0</td><td>CGAGGCGGCATCGACGTGCA</td><td>GGG</td><td>||||||||[||||||||||||]PAM</td><td>NA</td><td>Intergenic</td><td>NA</td><td>NA</td></tr>
<tr><td>T12118</td><td>GAGGCGGCATCGACGTGCAGGGG</td><td>1000</td><td></td><td></td><td></td><td></td><td></td><td></td><td></td><td></td><td></td></tr>
<tr><td>Oligo fwd</td><td>CACCGAGGCGGCATCGACGTGCAG</td><td></td><td></td><td></td><td></td><td></td><td></td><td></td><td></td><td></td><td></td></tr>
<tr><td>Oligo rev</td><td>AAACCTGCACGTCGATGCCGCCTC</td><td></td><td></td><td></td><td></td><td></td><td></td><td></td><td></td><td></td><td></td></tr>
<tr><td>Chromosome</td><td>start</td><td>end</td><td>strand</td><td>MM</td><td>target_seq</td><td>PAM</td><td>alignment</td><td>distance</td><td>position</td><td>gene name</td><td>gene id</td></tr>
<tr><td>GU305914.1</td><td>161284</td><td>161306</td><td>+</td><td>0</td><td>GAGGCGGCATCGACGTGCAG</td><td>GGG</td><td>||||||||[||||||||||||]PAM</td><td>NA</td><td>Intergenic</td><td>NA</td><td>NA</td></tr>
<tr><td>T12119</td><td>CGGCATCGACGTGCAGGGGCCGG</td><td>1000</td><td></td><td></td><td></td><td></td><td></td><td></td><td></td><td></td><td></td></tr>
<tr><td>Oligo fwd</td><td>CACCCGGCATCGACGTGCAGGGGC</td><td></td><td></td><td></td><td></td><td></td><td></td><td></td><td></td><td></td><td></td></tr>
<tr><td>Oligo rev</td><td>AAACGCCCCTGCACGTCGATGCCG</td><td></td><td></td><td></td><td></td><td></td><td></td><td></td><td></td><td></td><td></td></tr>
<tr><td>Chromosome</td><td>start</td><td>end</td><td>strand</td><td>MM</td><td>target_seq</td><td>PAM</td><td>alignment</td><td>distance</td><td>position</td><td>gene name</td><td>gene id</td></tr>
<tr><td>GU305914.1</td><td>161288</td><td>161310</td><td>+</td><td>0</td><td>CGGCATCGACGTGCAGGGGC</td><td>CGG</td><td>||||||||[||||||||||||]PAM</td><td>NA</td><td>Intergenic</td><td>NA</td><td>NA</td></tr>
<tr><td>T12120</td><td>GGCCGGTCGACGCTATCTATCGG</td><td>1000</td><td></td><td></td><td></td><td></td><td></td><td></td><td></td><td></td><td></td></tr>
<tr><td>Oligo fwd</td><td>CACCGGCCGGTCGACGCTATCTAT</td><td></td><td></td><td></td><td></td><td></td><td></td><td></td><td></td><td></td><td></td></tr>
<tr><td>Oligo rev</td><td>AAACATAGATAGCGTCGACCGGCC</td><td></td><td></td><td></td><td></td><td></td><td></td><td></td><td></td><td></td><td></td></tr>
<tr><td>Chromosome</td><td>start</td><td>end</td><td>strand</td><td>MM</td><td>target_seq</td><td>PAM</td><td>alignment</td><td>distance</td><td>position</td><td>gene name</td><td>gene id</td></tr>
<tr><td>GU305914.1</td><td>161305</td><td>161327</td><td>+</td><td>0</td><td>GGCCGGTCGACGCTATCTAT</td><td>CGG</td><td>||||||||[||||||||||||]PAM</td><td>NA</td><td>Intergenic</td><td>NA</td><td>NA</td></tr>
<tr><td>T12121</td><td>GAAAGAAGTTTGATGTAGTTTGG</td><td>1000</td><td></td><td></td><td></td><td></td><td></td><td></td><td></td><td></td><td></td></tr>
<tr><td>Oligo fwd</td><td>CACCGAAAGAAGTTTGATGTAGTT</td><td></td><td></td><td></td><td></td><td></td><td></td><td></td><td></td><td></td><td></td></tr>
<tr><td>Oligo rev</td><td>AAACAACTACATCAAACTTCTTTC</td><td></td><td></td><td></td><td></td><td></td><td></td><td></td><td></td><td></td><td></td></tr>
<tr><td>Chromosome</td><td>start</td><td>end</td><td>strand</td><td>MM</td><td>target_seq</td><td>PAM</td><td>alignment</td><td>distance</td><td>position</td><td>gene name</td><td>gene id</td></tr>
<tr><td>GU305914.1</td><td>161443</td><td>161465</td><td>+</td><td>0</td><td>GAAAGAAGTTTGATGTAGTT</td><td>TGG</td><td>||||||||[||||||||||||]PAM</td><td>NA</td><td>Intergenic</td><td>NA</td><td>NA</td></tr>
<tr><td>T12122</td><td>AAGTTTGATGTAGTTTGGTTTGG</td><td>1000</td><td></td><td></td><td></td><td></td><td></td><td></td><td></td><td></td><td></td></tr>
<tr><td>Oligo fwd</td><td>CACCAAGTTTGATGTAGTTTGGTT</td><td></td><td></td><td></td><td></td><td></td><td></td><td></td><td></td><td></td><td></td></tr>
<tr><td>Oligo rev</td><td>AAACAACCAAACTACATCAAACTT</td><td></td><td></td><td></td><td></td><td></td><td></td><td></td><td></td><td></td><td></td></tr>
<tr><td>Chromosome</td><td>start</td><td>end</td><td>strand</td><td>MM</td><td>target_seq</td><td>PAM</td><td>alignment</td><td>distance</td><td>position</td><td>gene name</td><td>gene id</td></tr>
<tr><td>GU305914.1</td><td>161448</td><td>161470</td><td>+</td><td>0</td><td>AAGTTTGATGTAGTTTGGTT</td><td>TGG</td><td>||||||||[||||||||||||]PAM</td><td>NA</td><td>Intergenic</td><td>NA</td><td>NA</td></tr>
<tr><td>T12123</td><td>TTTTCAGACTCTACCTAGCCAGG</td><td>1000</td><td></td><td></td><td></td><td></td><td></td><td></td><td></td><td></td><td></td></tr>
<tr><td>Oligo fwd</td><td>CACCTTTTCAGACTCTACCTAGCC</td><td></td><td></td><td></td><td></td><td></td><td></td><td></td><td></td><td></td><td></td></tr>
<tr><td>Oligo rev</td><td>AAACGGCTAGGTAGAGTCTGAAAA</td><td></td><td></td><td></td><td></td><td></td><td></td><td></td><td></td><td></td><td></td></tr>
<tr><td>Chromosome</td><td>start</td><td>end</td><td>strand</td><td>MM</td><td>target_seq</td><td>PAM</td><td>alignment</td><td>distance</td><td>position</td><td>gene name</td><td>gene id</td></tr>
<tr><td>GU305914.1</td><td>161488</td><td>161510</td><td>+</td><td>0</td><td>TTTTCAGACTCTACCTAGCC</td><td>AGG</td><td>||||||||[||||||||||||]PAM</td><td>NA</td><td>Intergenic</td><td>NA</td><td>NA</td></tr>
<tr><td>T12124</td><td>TCTACCTAGCCAGGATTCCTAGG</td><td>1000</td><td></td><td></td><td></td><td></td><td></td><td></td><td></td><td></td><td></td></tr>
<tr><td>Oligo fwd</td><td>CACCTCTACCTAGCCAGGATTCCT</td><td></td><td></td><td></td><td></td><td></td><td></td><td></td><td></td><td></td><td></td></tr>
<tr><td>Oligo rev</td><td>AAACAGGAATCCTGGCTAGGTAGA</td><td></td><td></td><td></td><td></td><td></td><td></td><td></td><td></td><td></td><td></td></tr>
<tr><td>Chromosome</td><td>start</td><td>end</td><td>strand</td><td>MM</td><td>target_seq</td><td>PAM</td><td>alignment</td><td>distance</td><td>position</td><td>gene name</td><td>gene id</td></tr>
<tr><td>GU305914.1</td><td>161497</td><td>161519</td><td>+</td><td>0</td><td>TCTACCTAGCCAGGATTCCT</td><td>AGG</td><td>||||||||[||||||||||||]PAM</td><td>NA</td><td>Intergenic</td><td>NA</td><td>NA</td></tr>
<tr><td>T12125</td><td>GCCAGGATTCCTAGGCCGATTGG</td><td>1000</td><td></td><td></td><td></td><td></td><td></td><td></td><td></td><td></td><td></td></tr>
<tr><td>Oligo fwd</td><td>CACCGCCAGGATTCCTAGGCCGAT</td><td></td><td></td><td></td><td></td><td></td><td></td><td></td><td></td><td></td><td></td></tr>
<tr><td>Oligo rev</td><td>AAACATCGGCCTAGGAATCCTGGC</td><td></td><td></td><td></td><td></td><td></td><td></td><td></td><td></td><td></td><td></td></tr>
<tr><td>Chromosome</td><td>start</td><td>end</td><td>strand</td><td>MM</td><td>target_seq</td><td>PAM</td><td>alignment</td><td>distance</td><td>position</td><td>gene name</td><td>gene id</td></tr>
<tr><td>GU305914.1</td><td>161505</td><td>161527</td><td>+</td><td>0</td><td>GCCAGGATTCCTAGGCCGAT</td><td>TGG</td><td>||||||||[||||||||||||]PAM</td><td>NA</td><td>Intergenic</td><td>NA</td><td>NA</td></tr>
<tr><td>T12126</td><td>ATGAGATCTACGCTCGACCGAGG</td><td>1000</td><td></td><td></td><td></td><td></td><td></td><td></td><td></td><td></td><td></td></tr>
<tr><td>Oligo fwd</td><td>CACCATGAGATCTACGCTCGACCG</td><td></td><td></td><td></td><td></td><td></td><td></td><td></td><td></td><td></td><td></td></tr>
<tr><td>Oligo rev</td><td>AAACCGGTCGAGCGTAGATCTCAT</td><td></td><td></td><td></td><td></td><td></td><td></td><td></td><td></td><td></td><td></td></tr>
<tr><td>Chromosome</td><td>start</td><td>end</td><td>strand</td><td>MM</td><td>target_seq</td><td>PAM</td><td>alignment</td><td>distance</td><td>position</td><td>gene name</td><td>gene id</td></tr>
<tr><td>GU305914.1</td><td>161535</td><td>161557</td><td>+</td><td>0</td><td>ATGAGATCTACGCTCGACCG</td><td>AGG</td><td>||||||||[||||||||||||]PAM</td><td>NA</td><td>Intergenic</td><td>NA</td><td>NA</td></tr>
<tr><td>T12127</td><td>AAAGTCTTTTTATAGCTTTATGG</td><td>1000</td><td></td><td></td><td></td><td></td><td></td><td></td><td></td><td></td><td></td></tr>
<tr><td>Oligo fwd</td><td>CACCAAAGTCTTTTTATAGCTTTA</td><td></td><td></td><td></td><td></td><td></td><td></td><td></td><td></td><td></td><td></td></tr>
<tr><td>Oligo rev</td><td>AAACTAAAGCTATAAAAAGACTTT</td><td></td><td></td><td></td><td></td><td></td><td></td><td></td><td></td><td></td><td></td></tr>
<tr><td>Chromosome</td><td>start</td><td>end</td><td>strand</td><td>MM</td><td>target_seq</td><td>PAM</td><td>alignment</td><td>distance</td><td>position</td><td>gene name</td><td>gene id</td></tr>
<tr><td>GU305914.1</td><td>161571</td><td>161593</td><td>+</td><td>0</td><td>AAAGTCTTTTTATAGCTTTA</td><td>TGG</td><td>||||||||[||||||||||||]PAM</td><td>NA</td><td>Intergenic</td><td>NA</td><td>NA</td></tr>
<tr><td>T12128</td><td>AAAAAAAGTATTCTGTTTTTCGG</td><td>1000</td><td></td><td></td><td></td><td></td><td></td><td></td><td></td><td></td><td></td></tr>
<tr><td>Oligo fwd</td><td>CACCAAAAAAAGTATTCTGTTTTT</td><td></td><td></td><td></td><td></td><td></td><td></td><td></td><td></td><td></td><td></td></tr>
<tr><td>Oligo rev</td><td>AAACAAAAACAGAATACTTTTTTT</td><td></td><td></td><td></td><td></td><td></td><td></td><td></td><td></td><td></td><td></td></tr>
<tr><td>Chromosome</td><td>start</td><td>end</td><td>strand</td><td>MM</td><td>target_seq</td><td>PAM</td><td>alignment</td><td>distance</td><td>position</td><td>gene name</td><td>gene id</td></tr>
</table>

| | | | | | | | | | | | |
|---|---|---|---|---|---|---|---|---|---|---|---|
| GU305914.1 | 161618 | 161640 | + | 0 | AAAAAAAGTATTCTGTTTTT | CGG | \|\|\|\|\|\|\|\|[\|\|\|\|\|\|\|\|\|\|\|\|]PAM | NA | Intergenic | NA | NA |
| T12129 | AAAAAAGTATTCTGTTTTTCGGG | 1000 | | | | | | | | | |
| Oligo fwd | CACCAAAAAAGTATTCTGTTTTTC | | | | | | | | | | |
| Oligo rev | AAACGAAAAACAGAATACTTTTTT | | | | | | | | | | |
| Chromosome | start | end | strand | MM | target_seq | PAM | alignment | distance | position | gene name | gene id |
| GU305914.1 | 161619 | 161641 | + | 0 | AAAAAAGTATTCTGTTTTTC | GGG | \|\|\|\|\|\|\|\|[\|\|\|\|\|\|\|\|\|\|\|\|]PAM | NA | Intergenic | NA | NA |
| T12130 | TTTTATGAATTACCATCTAAAGG | 1000 | | | | | | | | | |
| Oligo fwd | CACCTTTTATGAATTACCATCTAA | | | | | | | | | | |
| Oligo rev | AAACTTAGATGGTAATTCATAAAA | | | | | | | | | | |
| Chromosome | start | end | strand | MM | target_seq | PAM | alignment | distance | position | gene name | gene id |
| GU305914.1 | 161643 | 161665 | + | 0 | TTTTATGAATTACCATCTAA | AGG | \|\|\|\|\|\|\|\|[\|\|\|\|\|\|\|\|\|\|\|\|]PAM | NA | Intergenic | NA | NA |
| T12131 | ATCTAAAGGAACGATAAAAGAGG | 1000 | | | | | | | | | |
| Oligo fwd | CACCATCTAAAGGAACGATAAAAG | | | | | | | | | | |
| Oligo rev | AAACCTTTTATCGTTCCTTTAGAT | | | | | | | | | | |
| Chromosome | start | end | strand | MM | target_seq | PAM | alignment | distance | position | gene name | gene id |
| GU305914.1 | 161657 | 161679 | + | 0 | ATCTAAAGGAACGATAAAAG | AGG | \|\|\|\|\|\|\|\|[\|\|\|\|\|\|\|\|\|\|\|\|]PAM | NA | Intergenic | NA | NA |
| T12132 | TTGTTCCTACCTGAGAGCTCCGG | 1000 | | | | | | | | | |
| Oligo fwd | CACCTTGTTCCTACCTGAGAGCTC | | | | | | | | | | |
| Oligo rev | AAACGAGCTCTCAGGTAGGAACAA | | | | | | | | | | |
| Chromosome | start | end | strand | MM | target_seq | PAM | alignment | distance | position | gene name | gene id |
| GU305914.1 | 161713 | 161735 | + | 0 | TTGTTCCTACCTGAGAGCTC | CGG | \|\|\|\|\|\|\|\|[\|\|\|\|\|\|\|\|\|\|\|\|]PAM | NA | Intergenic | NA | NA |
| T12133 | TGTTCCTACCTGAGAGCTCCGGG | 1000 | | | | | | | | | |
| Oligo fwd | CACCTGTTCCTACCTGAGAGCTCC | | | | | | | | | | |
| Oligo rev | AAACGGAGCTCTCAGGTAGGAACA | | | | | | | | | | |
| Chromosome | start | end | strand | MM | target_seq | PAM | alignment | distance | position | gene name | gene id |
| GU305914.1 | 161714 | 161736 | + | 0 | TGTTCCTACCTGAGAGCTCC | GGG | \|\|\|\|\|\|\|\|[\|\|\|\|\|\|\|\|\|\|\|\|]PAM | NA | Intergenic | NA | NA |
| T12134 | ACCTGAGAGCTCCGGGCCTTCGG | 1000 | | | | | | | | | |
| Oligo fwd | CACCACCTGAGAGCTCCGGGCCTT | | | | | | | | | | |
| Oligo rev | AAACAAGGCCCGGAGCTCTCAGGT | | | | | | | | | | |
| Chromosome | start | end | strand | MM | target_seq | PAM | alignment | distance | position | gene name | gene id |
| GU305914.1 | 161721 | 161743 | + | 0 | ACCTGAGAGCTCCGGGCCTT | CGG | \|\|\|\|\|\|\|\|[\|\|\|\|\|\|\|\|\|\|\|\|]PAM | NA | Intergenic | NA | NA |
| T12135 | CCTGAGAGCTCCGGGCCTTCGGG | 1000 | | | | | | | | | |
| Oligo fwd | CACCCCTGAGAGCTCCGGGCCTTC | | | | | | | | | | |
| Oligo rev | AAACGAAGGCCCGGAGCTCTCAGG | | | | | | | | | | |
| Chromosome | start | end | strand | MM | target_seq | PAM | alignment | distance | position | gene name | gene id |
| GU305914.1 | 161722 | 161744 | + | 0 | CCTGAGAGCTCCGGGCCTTC | GGG | \|\|\|\|\|\|\|\|[\|\|\|\|\|\|\|\|\|\|\|\|]PAM | NA | Intergenic | NA | NA |
| T12136 | CACCTCCGCCGCCGCCTCCTCGG | 1000 | | | | | | | | | |
| Oligo fwd | CACCCACCTCCGCCGCCGCCTCCT | | | | | | | | | | |
| Oligo rev | AAACAGGAGGCGGCGGCGGAGGTG | | | | | | | | | | |
| Chromosome | start | end | strand | MM | target_seq | PAM | alignment | distance | position | gene name | gene id |
| GU305914.1 | 161752 | 161774 | + | 0 | CACCTCCGCCGCCGCCTCCT | CGG | \|\|\|\|\|\|\|\|[\|\|\|\|\|\|\|\|\|\|\|\|]PAM | NA | Intergenic | NA | NA |
| GU305914.1 | 98350 | 98372 | + | 4 | CGCCGCCGCCACCGCCGCCT | AGG | \|-\|\|-\|\|\|[\|\|-\|\|\|\|\|-\|\|\|]PAM | NA | Intergenic | NA | NA |
| GU305914.1 | 35655 | 35677 | - | 4 | CCCCGCCGCCGCCGCCGCCG | TGG | \|-\|\|-\|\|\|[\|\|\|\|\|\|\|\|-\|\|-]PAM | NA | Intergenic | NA | NA |
| T12137 | GCCGCCGCCTCCTCGGCCTCCGG | 1000 | | | | | | | | | |
| Oligo fwd | CACCGCCGCCGCCTCCTCGGCCTC | | | | | | | | | | |
| Oligo rev | AAACGAGGCCGAGGAGGCGGCGGC | | | | | | | | | | |
| Chromosome | start | end | strand | MM | target_seq | PAM | alignment | distance | position | gene name | gene id |
| GU305914.1 | 161759 | 161781 | + | 0 | GCCGCCGCCTCCTCGGCCTC | CGG | \|\|\|\|\|\|\|\|[\|\|\|\|\|\|\|\|\|\|\|\|]PAM | NA | Intergenic | NA | NA |
| GU305914.1 | 221602 | 221624 | + | 2 | GCCGCCGCCACCACGGCCTC | CGG | \|\|\|\|\|\|\|\|[\|-\|\|-\|\|\|\|\|\|\|]PAM | NA | Intergenic | NA | NA |
| T12138 | CCGCCGCCTCCTCGGCCTCCGGG | 1000 | | | | | | | | | |
| Oligo fwd | CACCCCGCCGCCTCCTCGGCCTCC | | | | | | | | | | |
| Oligo rev | AAACGGAGGCCGAGGAGGCGGCGG | | | | | | | | | | |
| Chromosome | start | end | strand | MM | target_seq | PAM | alignment | distance | position | gene name | gene id |
| GU305914.1 | 161760 | 161782 | + | 0 | CCGCCGCCTCCTCGGCCTCC | GGG | \|\|\|\|\|\|\|\|[\|\|\|\|\|\|\|\|\|\|\|\|]PAM | NA | Intergenic | NA | NA |
| T12139 | CCGCCTCACCTGAGCCTGCTCGG | 1000 | | | | | | | | | |
| Oligo fwd | CACCCCGCCTCACCTGAGCCTGCT | | | | | | | | | | |
| Oligo rev | AAACAGCAGGCTCAGGTGAGGCGG | | | | | | | | | | |
| Chromosome | start | end | strand | MM | target_seq | PAM | alignment | distance | position | gene name | gene id |
| GU305914.1 | 161794 | 161816 | + | 0 | CCGCCTCACCTGAGCCTGCT | CGG | \|\|\|\|\|\|\|\|[\|\|\|\|\|\|\|\|\|\|\|\|]PAM | NA | Intergenic | NA | NA |
| T12140 | GCCTGCTCGGCCGTTCGCTCAGG | 1000 | | | | | | | | | |
| Oligo fwd | CACCGCCTGCTCGGCCGTTCGCTC | | | | | | | | | | |
| Oligo rev | AAACGAGCGAACGGCCGAGCAGGC | | | | | | | | | | |
| Chromosome | start | end | strand | MM | target_seq | PAM | alignment | distance | position | gene name | gene id |
| GU305914.1 | 161807 | 161829 | + | 0 | GCCTGCTCGGCCGTTCGCTC | AGG | \|\|\|\|\|\|\|\|[\|\|\|\|\|\|\|\|\|\|\|\|]PAM | NA | Intergenic | NA | NA |
| T12141 | GTACGTTTATATATTCTAGCCGG | 1000 | | | | | | | | | |
| Oligo fwd | CACCGTACGTTTATATATTCTAGC | | | | | | | | | | |
| Oligo rev | AAACGCTAGAATATATAAACGTAC | | | | | | | | | | |
| Chromosome | start | end | strand | MM | target_seq | PAM | alignment | distance | position | gene name | gene id |
| GU305914.1 | 161857 | 161879 | + | 0 | GTACGTTTATATATTCTAGC | CGG | \|\|\|\|\|\|\|\|[\|\|\|\|\|\|\|\|\|\|\|\|]PAM | NA | Intergenic | NA | NA |
| T12142 | TACGTTTATATATTCTAGCCGGG | 1000 | | | | | | | | | |
| Oligo fwd | CACCTACGTTTATATATTCTAGCC | | | | | | | | | | |
| Oligo rev | AAACGGCTAGAATATATAAACGTA | | | | | | | | | | |
| Chromosome | start | end | strand | MM | target_seq | PAM | alignment | distance | position | gene name | gene id |
| GU305914.1 | 161858 | 161880 | + | 0 | TACGTTTATATATTCTAGCC | GGG | \|\|\|\|\|\|\|\|[\|\|\|\|\|\|\|\|\|\|\|\|]PAM | NA | Intergenic | NA | NA |

<table>
<tr><td>T12143</td><td>TATTCTAGCCGGGATGCGTCAGG</td><td>1000</td><td></td><td></td><td></td><td></td><td></td><td></td><td></td><td></td><td></td></tr>
<tr><td>Oligo fwd</td><td>CACCTATTCTAGCCGGGATGCGTC</td><td></td><td></td><td></td><td></td><td></td><td></td><td></td><td></td><td></td><td></td></tr>
<tr><td>Oligo rev</td><td>AAACGACGCATCCCGGCTAGAATA</td><td></td><td></td><td></td><td></td><td></td><td></td><td></td><td></td><td></td><td></td></tr>
<tr><td>Chromosome</td><td>start</td><td>end</td><td>strand</td><td>MM</td><td>target_seq</td><td>PAM</td><td>alignment</td><td>distance</td><td>position</td><td>gene name</td><td>gene id</td></tr>
<tr><td>GU305914.1</td><td>161868</td><td>161890</td><td>+</td><td>0</td><td>TATTCTAGCCGGGATGCGTC</td><td>AGG</td><td>||||||||[||||||||||||]PAM</td><td>NA</td><td>Intergenic</td><td>NA</td><td>NA</td></tr>
<tr><td>T12144</td><td>ATGCGTCAGGTTGCGTCTGACGG</td><td>1000</td><td></td><td></td><td></td><td></td><td></td><td></td><td></td><td></td><td></td></tr>
<tr><td>Oligo fwd</td><td>CACCATGCGTCAGGTTGCGTCTGA</td><td></td><td></td><td></td><td></td><td></td><td></td><td></td><td></td><td></td><td></td></tr>
<tr><td>Oligo rev</td><td>AAACTCAGACGCAACCTGACGCAT</td><td></td><td></td><td></td><td></td><td></td><td></td><td></td><td></td><td></td><td></td></tr>
<tr><td>Chromosome</td><td>start</td><td>end</td><td>strand</td><td>MM</td><td>target_seq</td><td>PAM</td><td>alignment</td><td>distance</td><td>position</td><td>gene name</td><td>gene id</td></tr>
<tr><td>GU305914.1</td><td>161881</td><td>161903</td><td>+</td><td>0</td><td>ATGCGTCAGGTTGCGTCTGA</td><td>CGG</td><td>||||||||[||||||||||||]PAM</td><td>NA</td><td>Intergenic</td><td>NA</td><td>NA</td></tr>
<tr><td>T12145</td><td>ACGGTGAATACCTACACTCCAGG</td><td>1000</td><td></td><td></td><td></td><td></td><td></td><td></td><td></td><td></td><td></td></tr>
<tr><td>Oligo fwd</td><td>CACCACGGTGAATACCTACACTCC</td><td></td><td></td><td></td><td></td><td></td><td></td><td></td><td></td><td></td><td></td></tr>
<tr><td>Oligo rev</td><td>AAACGGAGTGTAGGTATTCACCGT</td><td></td><td></td><td></td><td></td><td></td><td></td><td></td><td></td><td></td><td></td></tr>
<tr><td>Chromosome</td><td>start</td><td>end</td><td>strand</td><td>MM</td><td>target_seq</td><td>PAM</td><td>alignment</td><td>distance</td><td>position</td><td>gene name</td><td>gene id</td></tr>
<tr><td>GU305914.1</td><td>161900</td><td>161922</td><td>+</td><td>0</td><td>ACGGTGAATACCTACACTCC</td><td>AGG</td><td>||||||||[||||||||||||]PAM</td><td>NA</td><td>Intergenic</td><td>NA</td><td>NA</td></tr>
<tr><td>T12146</td><td>TCCAGGTATCCTACCTAGCGCGG</td><td>1000</td><td></td><td></td><td></td><td></td><td></td><td></td><td></td><td></td><td></td></tr>
<tr><td>Oligo fwd</td><td>CACCTCCAGGTATCCTACCTAGCG</td><td></td><td></td><td></td><td></td><td></td><td></td><td></td><td></td><td></td><td></td></tr>
<tr><td>Oligo rev</td><td>AAACCGCTAGGTAGGATACCTGGA</td><td></td><td></td><td></td><td></td><td></td><td></td><td></td><td></td><td></td><td></td></tr>
<tr><td>Chromosome</td><td>start</td><td>end</td><td>strand</td><td>MM</td><td>target_seq</td><td>PAM</td><td>alignment</td><td>distance</td><td>position</td><td>gene name</td><td>gene id</td></tr>
<tr><td>GU305914.1</td><td>161917</td><td>161939</td><td>+</td><td>0</td><td>TCCAGGTATCCTACCTAGCG</td><td>CGG</td><td>||||||||[||||||||||||]PAM</td><td>NA</td><td>Intergenic</td><td>NA</td><td>NA</td></tr>
<tr><td>T12147</td><td>GACGCCGCAAAGTATAATAGCGG</td><td>1000</td><td></td><td></td><td></td><td></td><td></td><td></td><td></td><td></td><td></td></tr>
<tr><td>Oligo fwd</td><td>CACCGACGCCGCAAAGTATAATAG</td><td></td><td></td><td></td><td></td><td></td><td></td><td></td><td></td><td></td><td></td></tr>
<tr><td>Oligo rev</td><td>AAACCTATTATACTTTGCGGCGTC</td><td></td><td></td><td></td><td></td><td></td><td></td><td></td><td></td><td></td><td></td></tr>
<tr><td>Chromosome</td><td>start</td><td>end</td><td>strand</td><td>MM</td><td>target_seq</td><td>PAM</td><td>alignment</td><td>distance</td><td>position</td><td>gene name</td><td>gene id</td></tr>
<tr><td>GU305914.1</td><td>161948</td><td>161970</td><td>+</td><td>0</td><td>GACGCCGCAAAGTATAATAG</td><td>CGG</td><td>||||||||[||||||||||||]PAM</td><td>NA</td><td>Intergenic</td><td>NA</td><td>NA</td></tr>
<tr><td>T12148</td><td>CAAAGTATAATAGCGGAGTACGG</td><td>1000</td><td></td><td></td><td></td><td></td><td></td><td></td><td></td><td></td><td></td></tr>
<tr><td>Oligo fwd</td><td>CACCCAAAGTATAATAGCGGAGTA</td><td></td><td></td><td></td><td></td><td></td><td></td><td></td><td></td><td></td><td></td></tr>
<tr><td>Oligo rev</td><td>AAACTACTCCGCTATTATACTTTG</td><td></td><td></td><td></td><td></td><td></td><td></td><td></td><td></td><td></td><td></td></tr>
<tr><td>Chromosome</td><td>start</td><td>end</td><td>strand</td><td>MM</td><td>target_seq</td><td>PAM</td><td>alignment</td><td>distance</td><td>position</td><td>gene name</td><td>gene id</td></tr>
<tr><td>GU305914.1</td><td>161955</td><td>161977</td><td>+</td><td>0</td><td>CAAAGTATAATAGCGGAGTA</td><td>CGG</td><td>||||||||[||||||||||||]PAM</td><td>NA</td><td>Intergenic</td><td>NA</td><td>NA</td></tr>
<tr><td>T12149</td><td>TAGCGGAGTACGGAGACACACGG</td><td>1000</td><td></td><td></td><td></td><td></td><td></td><td></td><td></td><td></td><td></td></tr>
<tr><td>Oligo fwd</td><td>CACCTAGCGGAGTACGGAGACACA</td><td></td><td></td><td></td><td></td><td></td><td></td><td></td><td></td><td></td><td></td></tr>
<tr><td>Oligo rev</td><td>AAACTGTGTCTCCGTACTCCGCTA</td><td></td><td></td><td></td><td></td><td></td><td></td><td></td><td></td><td></td><td></td></tr>
<tr><td>Chromosome</td><td>start</td><td>end</td><td>strand</td><td>MM</td><td>target_seq</td><td>PAM</td><td>alignment</td><td>distance</td><td>position</td><td>gene name</td><td>gene id</td></tr>
<tr><td>GU305914.1</td><td>161965</td><td>161987</td><td>+</td><td>0</td><td>TAGCGGAGTACGGAGACACA</td><td>CGG</td><td>||||||||[||||||||||||]PAM</td><td>NA</td><td>Intergenic</td><td>NA</td><td>NA</td></tr>
<tr><td>T12150</td><td>CGGAGTACGGAGACACACGGTGG</td><td>1000</td><td></td><td></td><td></td><td></td><td></td><td></td><td></td><td></td><td></td></tr>
<tr><td>Oligo fwd</td><td>CACCCGGAGTACGGAGACACACGG</td><td></td><td></td><td></td><td></td><td></td><td></td><td></td><td></td><td></td><td></td></tr>
<tr><td>Oligo rev</td><td>AAACCCGTGTGTCTCCGTACTCCG</td><td></td><td></td><td></td><td></td><td></td><td></td><td></td><td></td><td></td><td></td></tr>
<tr><td>Chromosome</td><td>start</td><td>end</td><td>strand</td><td>MM</td><td>target_seq</td><td>PAM</td><td>alignment</td><td>distance</td><td>position</td><td>gene name</td><td>gene id</td></tr>
<tr><td>GU305914.1</td><td>161968</td><td>161990</td><td>+</td><td>0</td><td>CGGAGTACGGAGACACACGG</td><td>TGG</td><td>||||||||[||||||||||||]PAM</td><td>NA</td><td>Intergenic</td><td>NA</td><td>NA</td></tr>
<tr><td>T12151</td><td>ATCATTTTATTTCCCCGTGCCGG</td><td>1000</td><td></td><td></td><td></td><td></td><td></td><td></td><td></td><td></td><td></td></tr>
<tr><td>Oligo fwd</td><td>CACCATCATTTTATTTCCCCGTGC</td><td></td><td></td><td></td><td></td><td></td><td></td><td></td><td></td><td></td><td></td></tr>
<tr><td>Oligo rev</td><td>AAACGCACGGGGAAATAAAATGAT</td><td></td><td></td><td></td><td></td><td></td><td></td><td></td><td></td><td></td><td></td></tr>
<tr><td>Chromosome</td><td>start</td><td>end</td><td>strand</td><td>MM</td><td>target_seq</td><td>PAM</td><td>alignment</td><td>distance</td><td>position</td><td>gene name</td><td>gene id</td></tr>
<tr><td>GU305914.1</td><td>161998</td><td>162020</td><td>+</td><td>0</td><td>ATCATTTTATTTCCCCGTGC</td><td>CGG</td><td>||||||||[||||||||||||]PAM</td><td>NA</td><td>Intergenic</td><td>NA</td><td>NA</td></tr>
<tr><td>T12152</td><td>ATTTTATTTCCCCGTGCCGGTGG</td><td>1000</td><td></td><td></td><td></td><td></td><td></td><td></td><td></td><td></td><td></td></tr>
<tr><td>Oligo fwd</td><td>CACCATTTTATTTCCCCGTGCCGG</td><td></td><td></td><td></td><td></td><td></td><td></td><td></td><td></td><td></td><td></td></tr>
<tr><td>Oligo rev</td><td>AAACCCGGCACGGGGAAATAAAAT</td><td></td><td></td><td></td><td></td><td></td><td></td><td></td><td></td><td></td><td></td></tr>
<tr><td>Chromosome</td><td>start</td><td>end</td><td>strand</td><td>MM</td><td>target_seq</td><td>PAM</td><td>alignment</td><td>distance</td><td>position</td><td>gene name</td><td>gene id</td></tr>
<tr><td>GU305914.1</td><td>162001</td><td>162023</td><td>+</td><td>0</td><td>ATTTTATTTCCCCGTGCCGG</td><td>TGG</td><td>||||||||[||||||||||||]PAM</td><td>NA</td><td>Intergenic</td><td>NA</td><td>NA</td></tr>
<tr><td>T12153</td><td>TTTTATTTCCCCGTGCCGGTGGG</td><td>1000</td><td></td><td></td><td></td><td></td><td></td><td></td><td></td><td></td><td></td></tr>
<tr><td>Oligo fwd</td><td>CACCTTTTATTTCCCCGTGCCGGT</td><td></td><td></td><td></td><td></td><td></td><td></td><td></td><td></td><td></td><td></td></tr>
<tr><td>Oligo rev</td><td>AAACACCGGCACGGGGAAATAAAA</td><td></td><td></td><td></td><td></td><td></td><td></td><td></td><td></td><td></td><td></td></tr>
<tr><td>Chromosome</td><td>start</td><td>end</td><td>strand</td><td>MM</td><td>target_seq</td><td>PAM</td><td>alignment</td><td>distance</td><td>position</td><td>gene name</td><td>gene id</td></tr>
<tr><td>GU305914.1</td><td>162002</td><td>162024</td><td>+</td><td>0</td><td>TTTTATTTCCCCGTGCCGGT</td><td>GGG</td><td>||||||||[||||||||||||]PAM</td><td>NA</td><td>Intergenic</td><td>NA</td><td>NA</td></tr>
<tr><td>T12154</td><td>TTTATTTCCCCGTGCCGGTGGGG</td><td>1000</td><td></td><td></td><td></td><td></td><td></td><td></td><td></td><td></td><td></td></tr>
<tr><td>Oligo fwd</td><td>CACCTTTATTTCCCCGTGCCGGTG</td><td></td><td></td><td></td><td></td><td></td><td></td><td></td><td></td><td></td><td></td></tr>
<tr><td>Oligo rev</td><td>AAACCACCGGCACGGGGAAATAAA</td><td></td><td></td><td></td><td></td><td></td><td></td><td></td><td></td><td></td><td></td></tr>
<tr><td>Chromosome</td><td>start</td><td>end</td><td>strand</td><td>MM</td><td>target_seq</td><td>PAM</td><td>alignment</td><td>distance</td><td>position</td><td>gene name</td><td>gene id</td></tr>
<tr><td>GU305914.1</td><td>162003</td><td>162025</td><td>+</td><td>0</td><td>TTTATTTCCCCGTGCCGGTG</td><td>GGG</td><td>||||||||[||||||||||||]PAM</td><td>NA</td><td>Intergenic</td><td>NA</td><td>NA</td></tr>
<tr><td>T12155</td><td>CCGTGCCGGTGGGGTTCCTTCGG</td><td>1000</td><td></td><td></td><td></td><td></td><td></td><td></td><td></td><td></td><td></td></tr>
<tr><td>Oligo fwd</td><td>CACCCCGTGCCGGTGGGGTTCCTT</td><td></td><td></td><td></td><td></td><td></td><td></td><td></td><td></td><td></td><td></td></tr>
<tr><td>Oligo rev</td><td>AAACAAGGAACCCCACCGGCACGG</td><td></td><td></td><td></td><td></td><td></td><td></td><td></td><td></td><td></td><td></td></tr>
<tr><td>Chromosome</td><td>start</td><td>end</td><td>strand</td><td>MM</td><td>target_seq</td><td>PAM</td><td>alignment</td><td>distance</td><td>position</td><td>gene name</td><td>gene id</td></tr>
<tr><td>GU305914.1</td><td>162012</td><td>162034</td><td>+</td><td>0</td><td>CCGTGCCGGTGGGGTTCCTT</td><td>CGG</td><td>||||||||[||||||||||||]PAM</td><td>NA</td><td>Intergenic</td><td>NA</td><td>NA</td></tr>
<tr><td>T12156</td><td>TGGGGTTCCTTCGGCCGACCCGG</td><td>1000</td><td></td><td></td><td></td><td></td><td></td><td></td><td></td><td></td><td></td></tr>
<tr><td>Oligo fwd</td><td>CACCTGGGGTTCCTTCGGCCGACC</td><td></td><td></td><td></td><td></td><td></td><td></td><td></td><td></td><td></td><td></td></tr>
<tr><td>Oligo rev</td><td>AAACGGTCGGCCGAAGGAACCCCA</td><td></td><td></td><td></td><td></td><td></td><td></td><td></td><td></td><td></td><td></td></tr>
<tr><td>Chromosome</td><td>start</td><td>end</td><td>strand</td><td>MM</td><td>target_seq</td><td>PAM</td><td>alignment</td><td>distance</td><td>position</td><td>gene name</td><td>gene id</td></tr>
<tr><td>GU305914.1</td><td>162021</td><td>162043</td><td>+</td><td>0</td><td>TGGGGTTCCTTCGGCCGACC</td><td>CGG</td><td>||||||||[||||||||||||]PAM</td><td>NA</td><td>Intergenic</td><td>NA</td><td>NA</td></tr>
<tr><td>T12157</td><td>TCCTTCGGCCGACCCGGTCACGG</td><td>1000</td><td></td><td></td><td></td><td></td><td></td><td></td><td></td><td></td><td></td></tr>
<tr><td>Oligo fwd</td><td>CACCTCCTTCGGCCGACCCGGTCA</td><td></td><td></td><td></td><td></td><td></td><td></td><td></td><td></td><td></td><td></td></tr>
<tr><td>Oligo rev</td><td>AAACTGACCGGGTCGGCCGAAGGA</td><td></td><td></td><td></td><td></td><td></td><td></td><td></td><td></td><td></td><td></td></tr>
<tr><td>Chromosome</td><td>start</td><td>end</td><td>strand</td><td>MM</td><td>target_seq</td><td>PAM</td><td>alignment</td><td>distance</td><td>position</td><td>gene name</td><td>gene id</td></tr>
<tr><td>GU305914.1</td><td>162027</td><td>162049</td><td>+</td><td>0</td><td>TCCTTCGGCCGACCCGGTCA</td><td>CGG</td><td>||||||||[||||||||||||]PAM</td><td>NA</td><td>Intergenic</td><td>NA</td><td>NA</td></tr>
</table>

T12158 CACGGACTTTCTTTTTAACCCGG 1000
Oligo fwd CACCCACGGACTTTCTTTTTAACC
Oligo rev AAACGGTTAAAAAGAAAGTCCGTG
Chromosome start end strand MM target_seq PAM alignment distance position gene name gene id
GU305914.1 162045 162067 + 0 CACGGACTTTCTTTTTAACC CGG ||||||||[||||||||||||]PAM NA Intergenic NA NA

T12159 ACGGACTTTCTTTTTAACCCGGG 1000
Oligo fwd CACCACGGACTTTCTTTTTAACCC
Oligo rev AAACGGGTTAAAAAGAAAGTCCGT
Chromosome start end strand MM target_seq PAM alignment distance position gene name gene id
GU305914.1 162046 162068 + 0 ACGGACTTTCTTTTTAACCC GGG ||||||||[||||||||||||]PAM NA Intergenic NA NA

T12160 CGGACTTTCTTTTTAACCCGGGG 1000
Oligo fwd CACCCGGACTTTCTTTTTAACCCG
Oligo rev AAACCGGGTTAAAAAGAAAGTCCG
Chromosome start end strand MM target_seq PAM alignment distance position gene name gene id
GU305914.1 162047 162069 + 0 CGGACTTTCTTTTTAACCCG GGG ||||||||[||||||||||||]PAM NA Intergenic NA NA

T12161 GGACTTTCTTTTTAACCCGGGGG 1000
Oligo fwd CACCGGACTTTCTTTTTAACCCGG
Oligo rev AAACCCGGGTTAAAAAGAAAGTCC
Chromosome start end strand MM target_seq PAM alignment distance position gene name gene id
GU305914.1 162048 162070 + 0 GGACTTTCTTTTTAACCCGG GGG ||||||||[||||||||||||]PAM NA Intergenic NA NA

T12162 TTTCTTTTTAACCCGGGGGCTGG 1000
Oligo fwd CACCTTTCTTTTTAACCCGGGGGC
Oligo rev AAACGCCCCCGGGTTAAAAAGAAA
Chromosome start end strand MM target_seq PAM alignment distance position gene name gene id
GU305914.1 162052 162074 + 0 TTTCTTTTTAACCCGGGGGC TGG ||||||||[||||||||||||]PAM NA Intergenic NA NA

T12163 TTTTTAACCCGGGGGCTGGATGG 1000
Oligo fwd CACCTTTTTAACCCGGGGGCTGGA
Oligo rev AAACTCCAGCCCCCGGGTTAAAAA
Chromosome start end strand MM target_seq PAM alignment distance position gene name gene id
GU305914.1 162056 162078 + 0 TTTTTAACCCGGGGGCTGGA TGG ||||||||[||||||||||||]PAM NA Intergenic NA NA

T12164 TTTTAACCCGGGGGCTGGATGGG 1000
Oligo fwd CACCTTTTAACCCGGGGGCTGGAT
Oligo rev AAACATCCAGCCCCCGGGTTAAAA
Chromosome start end strand MM target_seq PAM alignment distance position gene name gene id
GU305914.1 162057 162079 + 0 TTTTAACCCGGGGGCTGGAT GGG ||||||||[||||||||||||]PAM NA Intergenic NA NA

T12165 CCGGGGGCTGGATGGGCGCATGG 1000
Oligo fwd CACCCCGGGGGCTGGATGGGCGCA
Oligo rev AAACTGCGCCCATCCAGCCCCCGG
Chromosome start end strand MM target_seq PAM alignment distance position gene name gene id
GU305914.1 162064 162086 + 0 CCGGGGGCTGGATGGGCGCA TGG ||||||||[||||||||||||]PAM NA Intergenic NA NA

T12166 GGGCTGGATGGGCGCATGGACGG 1000
Oligo fwd CACCGGGCTGGATGGGCGCATGGA
Oligo rev AAACTCCATGCGCCCATCCAGCCC
Chromosome start end strand MM target_seq PAM alignment distance position gene name gene id
GU305914.1 162068 162090 + 0 GGGCTGGATGGGCGCATGGA CGG ||||||||[||||||||||||]PAM NA Intergenic NA NA

T12167 TGGATGGGCGCATGGACGGACGG 1000
Oligo fwd CACCTGGATGGGCGCATGGACGGA
Oligo rev AAACTCCGTCCATGCGCCCATCCA
Chromosome start end strand MM target_seq PAM alignment distance position gene name gene id
GU305914.1 162072 162094 + 0 TGGATGGGCGCATGGACGGA CGG ||||||||[||||||||||||]PAM NA Intergenic NA NA

T12168 TGGGCGCATGGACGGACGGACGG 1000
Oligo fwd CACCTGGGCGCATGGACGGACGGA
Oligo rev AAACTCCGTCCGTCCATGCGCCCA
Chromosome start end strand MM target_seq PAM alignment distance position gene name gene id
GU305914.1 162076 162098 + 0 TGGGCGCATGGACGGACGGA CGG ||||||||[||||||||||||]PAM NA Intergenic NA NA

T12169 GGGCGCATGGACGGACGGACGGG 1000
Oligo fwd CACCGGGCGCATGGACGGACGGAC
Oligo rev AAACGTCCGTCCGTCCATGCGCCC
Chromosome start end strand MM target_seq PAM alignment distance position gene name gene id
GU305914.1 162077 162099 + 0 GGGCGCATGGACGGACGGAC GGG ||||||||[||||||||||||]PAM NA Intergenic NA NA

T12170 CGCATGGACGGACGGACGGGCGG 1000
Oligo fwd CACCCGCATGGACGGACGGACGGG
Oligo rev AAACCCCGTCCGTCCGTCCATGCG
Chromosome start end strand MM target_seq PAM alignment distance position gene name gene id
GU305914.1 162080 162102 + 0 CGCATGGACGGACGGACGGG CGG ||||||||[||||||||||||]PAM NA Intergenic NA NA
GU305914.1 162084 162106 + 4 TGGACGGACGGACGGGCGGG CGG -|-|-|||[|||||||-||||]PAM NA Intergenic NA NA

T12171 GCATGGACGGACGGACGGGCGGG 1000
Oligo fwd CACCGCATGGACGGACGGACGGGC
Oligo rev AAACGCCCGTCCGTCCGTCCATGC
Chromosome start end strand MM target_seq PAM alignment distance position gene name gene id
GU305914.1 162081 162103 + 0 GCATGGACGGACGGACGGGC GGG ||||||||[||||||||||||]PAM NA Intergenic NA NA

T12172 TGGACGGACGGACGGGCGGGCGG 1000
Oligo fwd CACCTGGACGGACGGACGGGCGGG
Oligo rev AAACCCCGCCCGTCCGTCCGTCCA

<table>
<tr><td>Chromosome</td><td>start</td><td>end</td><td>strand</td><td>MM</td><td>target_seq</td><td>PAM</td><td>alignment</td><td>distance</td><td>position</td><td>gene name</td><td>gene id</td></tr>
<tr><td>GU305914.1</td><td>162084</td><td>162106</td><td>+</td><td>0</td><td>TGGACGGACGGACGGGCGGG</td><td>CGG</td><td>||||||||[||||||||||||]PAM</td><td>NA</td><td>Intergenic</td><td>NA</td><td>NA</td></tr>
<tr><td>GU305914.1</td><td>162080</td><td>162102</td><td>+</td><td>4</td><td>CGCATGGACGGACGGACGGG</td><td>CGG</td><td>-|-|-|||[|||||||-||||]PAM</td><td>NA</td><td>Intergenic</td><td>NA</td><td>NA</td></tr>
<tr><td>GU305914.1</td><td>162088</td><td>162110</td><td>+</td><td>3</td><td>CGGACGGACGGGCGGGCGGA</td><td>CGG</td><td>-|||||||[|||-|||||||-]PAM</td><td>NA</td><td>Intergenic</td><td>NA</td><td>NA</td></tr>
<tr><td></td><td></td><td></td><td></td><td></td><td></td><td></td><td></td><td></td><td></td><td></td><td></td></tr>
<tr><td>T12173</td><td>CGGACGGACGGGCGGGCGGACGG</td><td>1000</td><td></td><td></td><td></td><td></td><td></td><td></td><td></td><td></td><td></td></tr>
<tr><td>Oligo fwd</td><td>CACCCGGACGGACGGGCGGGCGGA</td><td></td><td></td><td></td><td></td><td></td><td></td><td></td><td></td><td></td><td></td></tr>
<tr><td>Oligo rev</td><td>AAACTCCGCCCGCCCGTCCGTCCG</td><td></td><td></td><td></td><td></td><td></td><td></td><td></td><td></td><td></td><td></td></tr>
<tr><td>Chromosome</td><td>start</td><td>end</td><td>strand</td><td>MM</td><td>target_seq</td><td>PAM</td><td>alignment</td><td>distance</td><td>position</td><td>gene name</td><td>gene id</td></tr>
<tr><td>GU305914.1</td><td>162088</td><td>162110</td><td>+</td><td>0</td><td>CGGACGGACGGGCGGGCGGA</td><td>CGG</td><td>||||||||[||||||||||||]PAM</td><td>NA</td><td>Intergenic</td><td>NA</td><td>NA</td></tr>
<tr><td>GU305914.1</td><td>162084</td><td>162106</td><td>+</td><td>3</td><td>TGGACGGACGGACGGGCGGG</td><td>CGG</td><td>-|||||||[|||-|||||||-]PAM</td><td>NA</td><td>Intergenic</td><td>NA</td><td>NA</td></tr>
<tr><td></td><td></td><td></td><td></td><td></td><td></td><td></td><td></td><td></td><td></td><td></td><td></td></tr>
<tr><td>T12174</td><td>GGACGGACGGGCGGGCGGACGGG</td><td>1000</td><td></td><td></td><td></td><td></td><td></td><td></td><td></td><td></td><td></td></tr>
<tr><td>Oligo fwd</td><td>CACCGGACGGACGGGCGGGCGGAC</td><td></td><td></td><td></td><td></td><td></td><td></td><td></td><td></td><td></td><td></td></tr>
<tr><td>Oligo rev</td><td>AAACGTCCGCCCGCCCGTCCGTCC</td><td></td><td></td><td></td><td></td><td></td><td></td><td></td><td></td><td></td><td></td></tr>
<tr><td>Chromosome</td><td>start</td><td>end</td><td>strand</td><td>MM</td><td>target_seq</td><td>PAM</td><td>alignment</td><td>distance</td><td>position</td><td>gene name</td><td>gene id</td></tr>
<tr><td>GU305914.1</td><td>162089</td><td>162111</td><td>+</td><td>0</td><td>GGACGGACGGGCGGGCGGAC</td><td>GGG</td><td>||||||||[||||||||||||]PAM</td><td>NA</td><td>Intergenic</td><td>NA</td><td>NA</td></tr>
<tr><td></td><td></td><td></td><td></td><td></td><td></td><td></td><td></td><td></td><td></td><td></td><td></td></tr>
<tr><td>T12175</td><td>GACGGACGGGCGGGCGGACGGGG</td><td>1000</td><td></td><td></td><td></td><td></td><td></td><td></td><td></td><td></td><td></td></tr>
<tr><td>Oligo fwd</td><td>CACCGACGGACGGGCGGGCGGACG</td><td></td><td></td><td></td><td></td><td></td><td></td><td></td><td></td><td></td><td></td></tr>
<tr><td>Oligo rev</td><td>AAACCGTCCGCCCGCCCGTCCGTC</td><td></td><td></td><td></td><td></td><td></td><td></td><td></td><td></td><td></td><td></td></tr>
<tr><td>Chromosome</td><td>start</td><td>end</td><td>strand</td><td>MM</td><td>target_seq</td><td>PAM</td><td>alignment</td><td>distance</td><td>position</td><td>gene name</td><td>gene id</td></tr>
<tr><td>GU305914.1</td><td>162090</td><td>162112</td><td>+</td><td>0</td><td>GACGGACGGGCGGGCGGACG</td><td>GGG</td><td>||||||||[||||||||||||]PAM</td><td>NA</td><td>Intergenic</td><td>NA</td><td>NA</td></tr>
<tr><td></td><td></td><td></td><td></td><td></td><td></td><td></td><td></td><td></td><td></td><td></td><td></td></tr>
<tr><td>T12176</td><td>GACGGGCGGGCGGACGGGGACGG</td><td>1000</td><td></td><td></td><td></td><td></td><td></td><td></td><td></td><td></td><td></td></tr>
<tr><td>Oligo fwd</td><td>CACCGACGGGCGGGCGGACGGGGA</td><td></td><td></td><td></td><td></td><td></td><td></td><td></td><td></td><td></td><td></td></tr>
<tr><td>Oligo rev</td><td>AAACTCCCCGTCCGCCCGCCCGTC</td><td></td><td></td><td></td><td></td><td></td><td></td><td></td><td></td><td></td><td></td></tr>
<tr><td>Chromosome</td><td>start</td><td>end</td><td>strand</td><td>MM</td><td>target_seq</td><td>PAM</td><td>alignment</td><td>distance</td><td>position</td><td>gene name</td><td>gene id</td></tr>
<tr><td>GU305914.1</td><td>162094</td><td>162116</td><td>+</td><td>0</td><td>GACGGGCGGGCGGACGGGGA</td><td>CGG</td><td>||||||||[||||||||||||]PAM</td><td>NA</td><td>Intergenic</td><td>NA</td><td>NA</td></tr>
<tr><td></td><td></td><td></td><td></td><td></td><td></td><td></td><td></td><td></td><td></td><td></td><td></td></tr>
<tr><td>T12177</td><td>ACGGGCGGGCGGACGGGGACGGG</td><td>1000</td><td></td><td></td><td></td><td></td><td></td><td></td><td></td><td></td><td></td></tr>
<tr><td>Oligo fwd</td><td>CACCACGGGCGGGCGGACGGGGAC</td><td></td><td></td><td></td><td></td><td></td><td></td><td></td><td></td><td></td><td></td></tr>
<tr><td>Oligo rev</td><td>AAACGTCCCCGTCCGCCCGCCCGT</td><td></td><td></td><td></td><td></td><td></td><td></td><td></td><td></td><td></td><td></td></tr>
<tr><td>Chromosome</td><td>start</td><td>end</td><td>strand</td><td>MM</td><td>target_seq</td><td>PAM</td><td>alignment</td><td>distance</td><td>position</td><td>gene name</td><td>gene id</td></tr>
<tr><td>GU305914.1</td><td>162095</td><td>162117</td><td>+</td><td>0</td><td>ACGGGCGGGCGGACGGGGAC</td><td>GGG</td><td>||||||||[||||||||||||]PAM</td><td>NA</td><td>Intergenic</td><td>NA</td><td>NA</td></tr>
<tr><td></td><td></td><td></td><td></td><td></td><td></td><td></td><td></td><td></td><td></td><td></td><td></td></tr>
<tr><td>T12178</td><td>CGGGCGGACGGGGACGGGTTCGG</td><td>1000</td><td></td><td></td><td></td><td></td><td></td><td></td><td></td><td></td><td></td></tr>
<tr><td>Oligo fwd</td><td>CACCCGGGCGGACGGGGACGGGTT</td><td></td><td></td><td></td><td></td><td></td><td></td><td></td><td></td><td></td><td></td></tr>
<tr><td>Oligo rev</td><td>AAACAACCCGTCCCCGTCCGCCCG</td><td></td><td></td><td></td><td></td><td></td><td></td><td></td><td></td><td></td><td></td></tr>
<tr><td>Chromosome</td><td>start</td><td>end</td><td>strand</td><td>MM</td><td>target_seq</td><td>PAM</td><td>alignment</td><td>distance</td><td>position</td><td>gene name</td><td>gene id</td></tr>
<tr><td>GU305914.1</td><td>162100</td><td>162122</td><td>+</td><td>0</td><td>CGGGCGGACGGGGACGGGTT</td><td>CGG</td><td>||||||||[||||||||||||]PAM</td><td>NA</td><td>Intergenic</td><td>NA</td><td>NA</td></tr>
<tr><td></td><td></td><td></td><td></td><td></td><td></td><td></td><td></td><td></td><td></td><td></td><td></td></tr>
<tr><td>T12179</td><td>GCGGACGGGGACGGGTTCGGAGG</td><td>1000</td><td></td><td></td><td></td><td></td><td></td><td></td><td></td><td></td><td></td></tr>
<tr><td>Oligo fwd</td><td>CACCGCGGACGGGGACGGGTTCGG</td><td></td><td></td><td></td><td></td><td></td><td></td><td></td><td></td><td></td><td></td></tr>
<tr><td>Oligo rev</td><td>AAACCCGAACCCGTCCCCGTCCGC</td><td></td><td></td><td></td><td></td><td></td><td></td><td></td><td></td><td></td><td></td></tr>
<tr><td>Chromosome</td><td>start</td><td>end</td><td>strand</td><td>MM</td><td>target_seq</td><td>PAM</td><td>alignment</td><td>distance</td><td>position</td><td>gene name</td><td>gene id</td></tr>
<tr><td>GU305914.1</td><td>162103</td><td>162125</td><td>+</td><td>0</td><td>GCGGACGGGGACGGGTTCGG</td><td>AGG</td><td>||||||||[||||||||||||]PAM</td><td>NA</td><td>Intergenic</td><td>NA</td><td>NA</td></tr>
<tr><td>GU305914.1</td><td>170210</td><td>170232</td><td>+</td><td>4</td><td>GGGTTCGGGGACGGCTCGG</td><td>CGG</td><td>|-|--|||[|||||||-||||]PAM</td><td>NA</td><td>Intergenic</td><td>NA</td><td>NA</td></tr>
<tr><td></td><td></td><td></td><td></td><td></td><td></td><td></td><td></td><td></td><td></td><td></td><td></td></tr>
<tr><td>T12180</td><td>CGGACGGGGACGGGTTCGGAGGG</td><td>1000</td><td></td><td></td><td></td><td></td><td></td><td></td><td></td><td></td><td></td></tr>
<tr><td>Oligo fwd</td><td>CACCCGGACGGGGACGGGTTCGGA</td><td></td><td></td><td></td><td></td><td></td><td></td><td></td><td></td><td></td><td></td></tr>
<tr><td>Oligo rev</td><td>AAACTCCGAACCCGTCCCCGTCCG</td><td></td><td></td><td></td><td></td><td></td><td></td><td></td><td></td><td></td><td></td></tr>
<tr><td>Chromosome</td><td>start</td><td>end</td><td>strand</td><td>MM</td><td>target_seq</td><td>PAM</td><td>alignment</td><td>distance</td><td>position</td><td>gene name</td><td>gene id</td></tr>
<tr><td>GU305914.1</td><td>162104</td><td>162126</td><td>+</td><td>0</td><td>CGGACGGGGACGGGTTCGGA</td><td>GGG</td><td>||||||||[||||||||||||]PAM</td><td>NA</td><td>Intergenic</td><td>NA</td><td>NA</td></tr>
<tr><td></td><td></td><td></td><td></td><td></td><td></td><td></td><td></td><td></td><td></td><td></td><td></td></tr>
<tr><td>T12181</td><td>CGGGTTCGGAGGGAGTACGTAGG</td><td>1000</td><td></td><td></td><td></td><td></td><td></td><td></td><td></td><td></td><td></td></tr>
<tr><td>Oligo fwd</td><td>CACCCGGGTTCGGAGGGAGTACGT</td><td></td><td></td><td></td><td></td><td></td><td></td><td></td><td></td><td></td><td></td></tr>
<tr><td>Oligo rev</td><td>AAACACGTACTCCCTCCGAACCCG</td><td></td><td></td><td></td><td></td><td></td><td></td><td></td><td></td><td></td><td></td></tr>
<tr><td>Chromosome</td><td>start</td><td>end</td><td>strand</td><td>MM</td><td>target_seq</td><td>PAM</td><td>alignment</td><td>distance</td><td>position</td><td>gene name</td><td>gene id</td></tr>
<tr><td>GU305914.1</td><td>162114</td><td>162136</td><td>+</td><td>0</td><td>CGGGTTCGGAGGGAGTACGT</td><td>AGG</td><td>||||||||[||||||||||||]PAM</td><td>NA</td><td>Intergenic</td><td>NA</td><td>NA</td></tr>
<tr><td></td><td></td><td></td><td></td><td></td><td></td><td></td><td></td><td></td><td></td><td></td><td></td></tr>
<tr><td>T12182</td><td>GGGTTCGGAGGGAGTACGTAGGG</td><td>1000</td><td></td><td></td><td></td><td></td><td></td><td></td><td></td><td></td><td></td></tr>
<tr><td>Oligo fwd</td><td>CACCGGGTTCGGAGGGAGTACGTA</td><td></td><td></td><td></td><td></td><td></td><td></td><td></td><td></td><td></td><td></td></tr>
<tr><td>Oligo rev</td><td>AAACTACGTACTCCCTCCGAACCC</td><td></td><td></td><td></td><td></td><td></td><td></td><td></td><td></td><td></td><td></td></tr>
<tr><td>Chromosome</td><td>start</td><td>end</td><td>strand</td><td>MM</td><td>target_seq</td><td>PAM</td><td>alignment</td><td>distance</td><td>position</td><td>gene name</td><td>gene id</td></tr>
<tr><td>GU305914.1</td><td>162115</td><td>162137</td><td>+</td><td>0</td><td>GGGTTCGGAGGGAGTACGTA</td><td>GGG</td><td>||||||||[||||||||||||]PAM</td><td>NA</td><td>Intergenic</td><td>NA</td><td>NA</td></tr>
<tr><td></td><td></td><td></td><td></td><td></td><td></td><td></td><td></td><td></td><td></td><td></td><td></td></tr>
<tr><td>T12183</td><td>TTCGGAGGGAGTACGTAGGGAGG</td><td>1000</td><td></td><td></td><td></td><td></td><td></td><td></td><td></td><td></td><td></td></tr>
<tr><td>Oligo fwd</td><td>CACCTTCGGAGGGAGTACGTAGGG</td><td></td><td></td><td></td><td></td><td></td><td></td><td></td><td></td><td></td><td></td></tr>
<tr><td>Oligo rev</td><td>AAACCCCTACGTACTCCCTCCGAA</td><td></td><td></td><td></td><td></td><td></td><td></td><td></td><td></td><td></td><td></td></tr>
<tr><td>Chromosome</td><td>start</td><td>end</td><td>strand</td><td>MM</td><td>target_seq</td><td>PAM</td><td>alignment</td><td>distance</td><td>position</td><td>gene name</td><td>gene id</td></tr>
<tr><td>GU305914.1</td><td>162118</td><td>162140</td><td>+</td><td>0</td><td>TTCGGAGGGAGTACGTAGGG</td><td>AGG</td><td>||||||||[||||||||||||]PAM</td><td>NA</td><td>Intergenic</td><td>NA</td><td>NA</td></tr>
<tr><td></td><td></td><td></td><td></td><td></td><td></td><td></td><td></td><td></td><td></td><td></td><td></td></tr>
<tr><td>T12184</td><td>TACGTAGGGAGGTGCTCCATCGG</td><td>1000</td><td></td><td></td><td></td><td></td><td></td><td></td><td></td><td></td><td></td></tr>
<tr><td>Oligo fwd</td><td>CACCTACGTAGGGAGGTGCTCCAT</td><td></td><td></td><td></td><td></td><td></td><td></td><td></td><td></td><td></td><td></td></tr>
<tr><td>Oligo rev</td><td>AAACATGGAGCACCTCCCTACGTA</td><td></td><td></td><td></td><td></td><td></td><td></td><td></td><td></td><td></td><td></td></tr>
<tr><td>Chromosome</td><td>start</td><td>end</td><td>strand</td><td>MM</td><td>target_seq</td><td>PAM</td><td>alignment</td><td>distance</td><td>position</td><td>gene name</td><td>gene id</td></tr>
<tr><td>GU305914.1</td><td>162129</td><td>162151</td><td>+</td><td>0</td><td>TACGTAGGGAGGTGCTCCAT</td><td>CGG</td><td>||||||||[||||||||||||]PAM</td><td>NA</td><td>Intergenic</td><td>NA</td><td>NA</td></tr>
<tr><td></td><td></td><td></td><td></td><td></td><td></td><td></td><td></td><td></td><td></td><td></td><td></td></tr>
<tr><td>T12185</td><td>GTAGGGAGGTGCTCCATCGGCGG</td><td>1000</td><td></td><td></td><td></td><td></td><td></td><td></td><td></td><td></td><td></td></tr>
<tr><td>Oligo fwd</td><td>CACCGTAGGGAGGTGCTCCATCGG</td><td></td><td></td><td></td><td></td><td></td><td></td><td></td><td></td><td></td><td></td></tr>
<tr><td>Oligo rev</td><td>AAACCCGATGGAGCACCTCCCTAC</td><td></td><td></td><td></td><td></td><td></td><td></td><td></td><td></td><td></td><td></td></tr>
<tr><td>Chromosome</td><td>start</td><td>end</td><td>strand</td><td>MM</td><td>target_seq</td><td>PAM</td><td>alignment</td><td>distance</td><td>position</td><td>gene name</td><td>gene id</td></tr>
<tr><td>GU305914.1</td><td>162132</td><td>162154</td><td>+</td><td>0</td><td>GTAGGGAGGTGCTCCATCGG</td><td>CGG</td><td>||||||||[||||||||||||]PAM</td><td>NA</td><td>Intergenic</td><td>NA</td><td>NA</td></tr>
<tr><td></td><td></td><td></td><td></td><td></td><td></td><td></td><td></td><td></td><td></td><td></td><td></td></tr>
<tr><td>T12186</td><td>CCTGACCGACGCTCGTCGACCGG</td><td>1000</td><td></td><td></td><td></td><td></td><td></td><td></td><td></td><td></td><td></td></tr>
<tr><td>Oligo fwd</td><td>CACCCCTGACCGACGCTCGTCGAC</td><td></td><td></td><td></td><td></td><td></td><td></td><td></td><td></td><td></td><td></td></tr>
<tr><td>Oligo rev</td><td>AAACGTCGACGAGCGTCGGTCAGG</td><td></td><td></td><td></td><td></td><td></td><td></td><td></td><td></td><td></td><td></td></tr>
<tr><td>Chromosome</td><td>start</td><td>end</td><td>strand</td><td>MM</td><td>target_seq</td><td>PAM</td><td>alignment</td><td>distance</td><td>position</td><td>gene name</td><td>gene id</td></tr>
</table>

<table>
<tr><td>GU305914.1</td><td>162155</td><td>162177</td><td>+</td><td>0</td><td>CCTGACCGACGCTCGTCGAC</td><td>CGG</td><td>||||||||[||||||||||||]PAM</td><td>NA</td><td>Intergenic</td><td>NA</td><td>NA</td></tr>
<tr><td>T12187</td><td>AGTCACCGTCCGTAGCGACTCGG</td><td>1000</td></tr>
<tr><td>Oligo fwd</td><td>CACCAGTCACCGTCCGTAGCGACT</td></tr>
<tr><td>Oligo rev</td><td>AAACAGTCGCTACGGACGGTGACT</td></tr>
<tr><td>Chromosome</td><td>start</td><td>end</td><td>strand</td><td>MM</td><td>target_seq</td><td>PAM</td><td>alignment</td><td>distance</td><td>position</td><td>gene name</td><td>gene id</td></tr>
<tr><td>GU305914.1</td><td>162190</td><td>162212</td><td>+</td><td>0</td><td>AGTCACCGTCCGTAGCGACT</td><td>CGG</td><td>||||||||[||||||||||||]PAM</td><td>NA</td><td>Intergenic</td><td>NA</td><td>NA</td></tr>
<tr><td>T12188</td><td>GACCGCTCGATCGAAGCGAACGG</td><td>1000</td></tr>
<tr><td>Oligo fwd</td><td>CACCGACCGCTCGATCGAAGCGAA</td></tr>
<tr><td>Oligo rev</td><td>AAACTTCGCTTCGATCGAGCGGTC</td></tr>
<tr><td>Chromosome</td><td>start</td><td>end</td><td>strand</td><td>MM</td><td>target_seq</td><td>PAM</td><td>alignment</td><td>distance</td><td>position</td><td>gene name</td><td>gene id</td></tr>
<tr><td>GU305914.1</td><td>162334</td><td>162356</td><td>+</td><td>0</td><td>GACCGCTCGATCGAAGCGAA</td><td>CGG</td><td>||||||||[||||||||||||]PAM</td><td>NA</td><td>Intergenic</td><td>NA</td><td>NA</td></tr>
<tr><td>T12189</td><td>ACCGCTCGATCGAAGCGAACGGG</td><td>1000</td></tr>
<tr><td>Oligo fwd</td><td>CACCACCGCTCGATCGAAGCGAAC</td></tr>
<tr><td>Oligo rev</td><td>AAACGTTCGCTTCGATCGAGCGGT</td></tr>
<tr><td>Chromosome</td><td>start</td><td>end</td><td>strand</td><td>MM</td><td>target_seq</td><td>PAM</td><td>alignment</td><td>distance</td><td>position</td><td>gene name</td><td>gene id</td></tr>
<tr><td>GU305914.1</td><td>162335</td><td>162357</td><td>+</td><td>0</td><td>ACCGCTCGATCGAAGCGAAC</td><td>GGG</td><td>||||||||[||||||||||||]PAM</td><td>NA</td><td>Intergenic</td><td>NA</td><td>NA</td></tr>
<tr><td>T12190</td><td>ATGCGATCAAGTAACCGAAGCGG</td><td>1000</td></tr>
<tr><td>Oligo fwd</td><td>CACCATGCGATCAAGTAACCGAAG</td></tr>
<tr><td>Oligo rev</td><td>AAACCTTCGGTTACTTGATCGCAT</td></tr>
<tr><td>Chromosome</td><td>start</td><td>end</td><td>strand</td><td>MM</td><td>target_seq</td><td>PAM</td><td>alignment</td><td>distance</td><td>position</td><td>gene name</td><td>gene id</td></tr>
<tr><td>GU305914.1</td><td>162383</td><td>162405</td><td>+</td><td>0</td><td>ATGCGATCAAGTAACCGAAG</td><td>CGG</td><td>||||||||[||||||||||||]PAM</td><td>NA</td><td>Intergenic</td><td>NA</td><td>NA</td></tr>
<tr><td>T12191</td><td>AACCGAAGCGGACCTGAAAACGG</td><td>1000</td></tr>
<tr><td>Oligo fwd</td><td>CACCAACCGAAGCGGACCTGAAAA</td></tr>
<tr><td>Oligo rev</td><td>AAACTTTTCAGGTCCGCTTCGGTT</td></tr>
<tr><td>Chromosome</td><td>start</td><td>end</td><td>strand</td><td>MM</td><td>target_seq</td><td>PAM</td><td>alignment</td><td>distance</td><td>position</td><td>gene name</td><td>gene id</td></tr>
<tr><td>GU305914.1</td><td>162395</td><td>162417</td><td>+</td><td>0</td><td>AACCGAAGCGGACCTGAAAA</td><td>CGG</td><td>||||||||[||||||||||||]PAM</td><td>NA</td><td>Intergenic</td><td>NA</td><td>NA</td></tr>
<tr><td>T12192</td><td>ACCGAAGCGGACCTGAAAACGGG</td><td>1000</td></tr>
<tr><td>Oligo fwd</td><td>CACCACCGAAGCGGACCTGAAAAC</td></tr>
<tr><td>Oligo rev</td><td>AAACGTTTTCAGGTCCGCTTCGGT</td></tr>
<tr><td>Chromosome</td><td>start</td><td>end</td><td>strand</td><td>MM</td><td>target_seq</td><td>PAM</td><td>alignment</td><td>distance</td><td>position</td><td>gene name</td><td>gene id</td></tr>
<tr><td>GU305914.1</td><td>162396</td><td>162418</td><td>+</td><td>0</td><td>ACCGAAGCGGACCTGAAAAC</td><td>GGG</td><td>||||||||[||||||||||||]PAM</td><td>NA</td><td>Intergenic</td><td>NA</td><td>NA</td></tr>
<tr><td>T12193</td><td>CCGAAGCGGACCTGAAAACGGGG</td><td>1000</td></tr>
<tr><td>Oligo fwd</td><td>CACCCCGAAGCGGACCTGAAAACG</td></tr>
<tr><td>Oligo rev</td><td>AAACCGTTTTCAGGTCCGCTTCGG</td></tr>
<tr><td>Chromosome</td><td>start</td><td>end</td><td>strand</td><td>MM</td><td>target_seq</td><td>PAM</td><td>alignment</td><td>distance</td><td>position</td><td>gene name</td><td>gene id</td></tr>
<tr><td>GU305914.1</td><td>162397</td><td>162419</td><td>+</td><td>0</td><td>CCGAAGCGGACCTGAAAACG</td><td>GGG</td><td>||||||||[||||||||||||]PAM</td><td>NA</td><td>Intergenic</td><td>NA</td><td>NA</td></tr>
<tr><td>T12194</td><td>AGCGGACCTGAAAACGGGGAAGG</td><td>1000</td></tr>
<tr><td>Oligo fwd</td><td>CACCAGCGGACCTGAAAACGGGGA</td></tr>
<tr><td>Oligo rev</td><td>AAACTCCCCGTTTTCAGGTCCGCT</td></tr>
<tr><td>Chromosome</td><td>start</td><td>end</td><td>strand</td><td>MM</td><td>target_seq</td><td>PAM</td><td>alignment</td><td>distance</td><td>position</td><td>gene name</td><td>gene id</td></tr>
<tr><td>GU305914.1</td><td>162401</td><td>162423</td><td>+</td><td>0</td><td>AGCGGACCTGAAAACGGGGA</td><td>AGG</td><td>||||||||[||||||||||||]PAM</td><td>NA</td><td>Intergenic</td><td>NA</td><td>NA</td></tr>
<tr><td>T12195</td><td>GGGGAAGGTCCGTCAGCCCTCGG</td><td>1000</td></tr>
<tr><td>Oligo fwd</td><td>CACCGGGGAAGGTCCGTCAGCCCT</td></tr>
<tr><td>Oligo rev</td><td>AAACAGGGCTGACGGACCTTCCCC</td></tr>
<tr><td>Chromosome</td><td>start</td><td>end</td><td>strand</td><td>MM</td><td>target_seq</td><td>PAM</td><td>alignment</td><td>distance</td><td>position</td><td>gene name</td><td>gene id</td></tr>
<tr><td>GU305914.1</td><td>162416</td><td>162438</td><td>+</td><td>0</td><td>GGGGAAGGTCCGTCAGCCCT</td><td>CGG</td><td>||||||||[||||||||||||]PAM</td><td>NA</td><td>Intergenic</td><td>NA</td><td>NA</td></tr>
<tr><td>T12196</td><td>GAAGGTCCGTCAGCCCTCGGAGG</td><td>1000</td></tr>
<tr><td>Oligo fwd</td><td>CACCGAAGGTCCGTCAGCCCTCGG</td></tr>
<tr><td>Oligo rev</td><td>AAACCCGAGGGCTGACGGACCTTC</td></tr>
<tr><td>Chromosome</td><td>start</td><td>end</td><td>strand</td><td>MM</td><td>target_seq</td><td>PAM</td><td>alignment</td><td>distance</td><td>position</td><td>gene name</td><td>gene id</td></tr>
<tr><td>GU305914.1</td><td>162419</td><td>162441</td><td>+</td><td>0</td><td>GAAGGTCCGTCAGCCCTCGG</td><td>AGG</td><td>||||||||[||||||||||||]PAM</td><td>NA</td><td>Intergenic</td><td>NA</td><td>NA</td></tr>
<tr><td>T12197</td><td>TCCGTCAGCCCTCGGAGGATCGG</td><td>1000</td></tr>
<tr><td>Oligo fwd</td><td>CACCTCCGTCAGCCCTCGGAGGAT</td></tr>
<tr><td>Oligo rev</td><td>AAACATCCTCCGAGGGCTGACGGA</td></tr>
<tr><td>Chromosome</td><td>start</td><td>end</td><td>strand</td><td>MM</td><td>target_seq</td><td>PAM</td><td>alignment</td><td>distance</td><td>position</td><td>gene name</td><td>gene id</td></tr>
<tr><td>GU305914.1</td><td>162424</td><td>162446</td><td>+</td><td>0</td><td>TCCGTCAGCCCTCGGAGGAT</td><td>CGG</td><td>||||||||[||||||||||||]PAM</td><td>NA</td><td>Intergenic</td><td>NA</td><td>NA</td></tr>
<tr><td>T12198</td><td>GTCAGCCCTCGGAGGATCGGAGG</td><td>1000</td></tr>
<tr><td>Oligo fwd</td><td>CACCGTCAGCCCTCGGAGGATCGG</td></tr>
<tr><td>Oligo rev</td><td>AAACCCGATCCTCCGAGGGCTGAC</td></tr>
<tr><td>Chromosome</td><td>start</td><td>end</td><td>strand</td><td>MM</td><td>target_seq</td><td>PAM</td><td>alignment</td><td>distance</td><td>position</td><td>gene name</td><td>gene id</td></tr>
<tr><td>GU305914.1</td><td>162427</td><td>162449</td><td>+</td><td>0</td><td>GTCAGCCCTCGGAGGATCGG</td><td>AGG</td><td>||||||||[||||||||||||]PAM</td><td>NA</td><td>Intergenic</td><td>NA</td><td>NA</td></tr>
<tr><td>T12199</td><td>CCCTCGGAGGATCGGAGGATCGG</td><td>1000</td></tr>
<tr><td>Oligo fwd</td><td>CACCCCCTCGGAGGATCGGAGGAT</td></tr>
<tr><td>Oligo rev</td><td>AAACATCCTCCGATCCTCCGAGGG</td></tr>
<tr><td>Chromosome</td><td>start</td><td>end</td><td>strand</td><td>MM</td><td>target_seq</td><td>PAM</td><td>alignment</td><td>distance</td><td>position</td><td>gene name</td><td>gene id</td></tr>
<tr><td>GU305914.1</td><td>162432</td><td>162454</td><td>+</td><td>0</td><td>CCCTCGGAGGATCGGAGGAT</td><td>CGG</td><td>||||||||[||||||||||||]PAM</td><td>NA</td><td>Intergenic</td><td>NA</td><td>NA</td></tr>
<tr><td>T12200</td><td>GAGCCACCGTCAGAAACGAACGG</td><td>1000</td></tr>
<tr><td>Oligo fwd</td><td>CACCGAGCCACCGTCAGAAACGAA</td></tr>
<tr><td>Oligo rev</td><td>AAACTTCGTTTCTGACGGTGGCTC</td></tr>
<tr><td>Chromosome</td><td>start</td><td>end</td><td>strand</td><td>MM</td><td>target_seq</td><td>PAM</td><td>alignment</td><td>distance</td><td>position</td><td>gene name</td><td>gene id</td></tr>
<tr><td>GU305914.1</td><td>162470</td><td>162492</td><td>+</td><td>0</td><td>GAGCCACCGTCAGAAACGAA</td><td>CGG</td><td>||||||||[||||||||||||]PAM</td><td>NA</td><td>Intergenic</td><td>NA</td><td>NA</td></tr>
<tr><td>T12201</td><td>ATACTTACACGTGCGTAACGAGG</td><td>1000</td></tr>
<tr><td>Oligo fwd</td><td>CACCATACTTACACGTGCGTAACG</td></tr>
<tr><td>Oligo rev</td><td>AAACCGTTACGCACGTGTAAGTAT</td></tr>
</table>

Chromosome start end strand MM target_seq PAM alignment distance position gene name gene id
GU305914.1 162508 162530 + 0 ATACTTACACGTGCGTAACG AGG ||||||||[||||||||||||]PAM NA Intergenic NA NA

T12202 CTTACACGTGCGTAACGAGGCGG 1000
Oligo fwd CACCCTTACACGTGCGTAACGAGG
Oligo rev AAACCCTCGTTACGCACGTGTAAG
Chromosome start end strand MM target_seq PAM alignment distance position gene name gene id
GU305914.1 162511 162533 + 0 CTTACACGTGCGTAACGAGG CGG ||||||||[||||||||||||]PAM NA Intergenic NA NA

T12203 GAGGCGGAAGACCCACATTTCGG 1000
Oligo fwd CACCGAGGCGGAAGACCCACATTT
Oligo rev AAACAAATGTGGGTCTTCCGCCTC
Chromosome start end strand MM target_seq PAM alignment distance position gene name gene id
GU305914.1 162527 162549 + 0 GAGGCGGAAGACCCACATTT CGG ||||||||[||||||||||||]PAM NA Intergenic NA NA

T12204 AGGCGGAAGACCCACATTTCGGG 1000
Oligo fwd CACCAGGCGGAAGACCCACATTTC
Oligo rev AAACGAAATGTGGGTCTTCCGCCT
Chromosome start end strand MM target_seq PAM alignment distance position gene name gene id
GU305914.1 162528 162550 + 0 AGGCGGAAGACCCACATTTC GGG ||||||||[||||||||||||]PAM NA Intergenic NA NA

T12205 CGGAAGACCCACATTTCGGGCGG 1000
Oligo fwd CACCCGGAAGACCCACATTTCGGG
Oligo rev AAACCCCGAAATGTGGGTCTTCCG
Chromosome start end strand MM target_seq PAM alignment distance position gene name gene id
GU305914.1 162531 162553 + 0 CGGAAGACCCACATTTCGGG CGG ||||||||[||||||||||||]PAM NA Intergenic NA NA

T12206 GGCGGTTGCTCGTGAACGAGCGG 1000
Oligo fwd CACCGGCGGTTGCTCGTGAACGAG
Oligo rev AAACCTCGTTCACGAGCAACCGCC
Chromosome start end strand MM target_seq PAM alignment distance position gene name gene id
GU305914.1 162549 162571 + 0 GGCGGTTGCTCGTGAACGAG CGG ||||||||[||||||||||||]PAM NA Intergenic NA NA

T12207 GAACGAGCGGTCGCTCGTCACGG 1000
Oligo fwd CACCGAACGAGCGGTCGCTCGTCA
Oligo rev AAACTGACGAGCGACCGCTCGTTC
Chromosome start end strand MM target_seq PAM alignment distance position gene name gene id
GU305914.1 162562 162584 + 0 GAACGAGCGGTCGCTCGTCA CGG ||||||||[||||||||||||]PAM NA Intergenic NA NA

T12208 GCTCGTCACGGCGATACCGACGG 1000
Oligo fwd CACCGCTCGTCACGGCGATACCGA
Oligo rev AAACTCGGTATCGCCGTGACGAGC
Chromosome start end strand MM target_seq PAM alignment distance position gene name gene id
GU305914.1 162574 162596 + 0 GCTCGTCACGGCGATACCGA CGG ||||||||[||||||||||||]PAM NA Intergenic NA NA

T12209 GACCCCCTCATCGAGCCGATCGG 1000
Oligo fwd CACCGACCCCCTCATCGAGCCGAT
Oligo rev AAACATCGGCTCGATGAGGGGGTC
Chromosome start end strand MM target_seq PAM alignment distance position gene name gene id
GU305914.1 162602 162624 + 0 GACCCCCTCATCGAGCCGAT CGG ||||||||[||||||||||||]PAM NA Intergenic NA NA

T12210 CCTCATCGAGCCGATCGGACCGG 1000
Oligo fwd CACCCCTCATCGAGCCGATCGGAC
Oligo rev AAACGTCCGATCGGCTCGATGAGG
Chromosome start end strand MM target_seq PAM alignment distance position gene name gene id
GU305914.1 162607 162629 + 0 CCTCATCGAGCCGATCGGAC CGG ||||||||[||||||||||||]PAM NA Intergenic NA NA

T12211 CTCATCGAGCCGATCGGACCGGG 1000
Oligo fwd CACCCTCATCGAGCCGATCGGACC
Oligo rev AAACGGTCCGATCGGCTCGATGAG
Chromosome start end strand MM target_seq PAM alignment distance position gene name gene id
GU305914.1 162608 162630 + 0 CTCATCGAGCCGATCGGACC GGG ||||||||[||||||||||||]PAM NA Intergenic NA NA

T12212 GGACCGGGAAAACCCCTCTTCGG 1000
Oligo fwd CACCGGACCGGGAAAACCCCTCTT
Oligo rev AAACAAGAGGGGTTTTCCCGGTCC
Chromosome start end strand MM target_seq PAM alignment distance position gene name gene id
GU305914.1 162623 162645 + 0 GGACCGGGAAAACCCCTCTT CGG ||||||||[||||||||||||]PAM NA Intergenic NA NA

T12213 GGACACCCGCTCGACGCCACCGG 1000
Oligo fwd CACCGGACACCCGCTCGACGCCAC
Oligo rev AAACGTGGCGTCGAGCGGGTGTCC
Chromosome start end strand MM target_seq PAM alignment distance position gene name gene id
GU305914.1 162644 162666 + 0 GGACACCCGCTCGACGCCAC CGG ||||||||[||||||||||||]PAM NA Intergenic NA NA

T12214 CCGGCATCTCGAACGCCGTACGG 1000
Oligo fwd CACCCCGGCATCTCGAACGCCGTA
Oligo rev AAACTACGGCGTTCGAGATGCCGG
Chromosome start end strand MM target_seq PAM alignment distance position gene name gene id
GU305914.1 162663 162685 + 0 CCGGCATCTCGAACGCCGTA CGG ||||||||[||||||||||||]PAM NA Intergenic NA NA

T12215 TCGAACGCCGTACGGTGCGACGG 1000
Oligo fwd CACCTCGAACGCCGTACGGTGCGA
Oligo rev AAACTCGCACCGTACGGCGTTCGA
Chromosome start end strand MM target_seq PAM alignment distance position gene name gene id
GU305914.1 162671 162693 + 0 TCGAACGCCGTACGGTGCGA CGG ||||||||[||||||||||||]PAM NA Intergenic NA NA

T12216 CGCCGCGCTATCTCGAGCCGAGG 1000
Oligo fwd CACCCGCCGCGCTATCTCGAGCCG

Oligo rev AAACCGGCTCGAGATAGCGCGGCG
Chromosome start end strand MM target_seq PAM alignment distance position gene name gene id
GU305914.1 162695 162717 + 0 CGCCGCGCTATCTCGAGCCG AGG ||||||||[||||||||||||]PAM NA Intergenic NA NA

T12217 GAGCCGAGGCCCCTGCGCTTCGG 1000
Oligo fwd CACCGAGCCGAGGCCCCTGCGCTT
Oligo rev AAACAAGCGCAGGGGCCTCGGCTC
Chromosome start end strand MM target_seq PAM alignment distance position gene name gene id
GU305914.1 162709 162731 + 0 GAGCCGAGGCCCCTGCGCTT CGG ||||||||[||||||||||||]PAM NA Intergenic NA NA

T12218 AGGCCCCTGCGCTTCGGCGAAGG 1000
Oligo fwd CACCAGGCCCCTGCGCTTCGGCGA
Oligo rev AAACTCGCCGAAGCGCAGGGGCCT
Chromosome start end strand MM target_seq PAM alignment distance position gene name gene id
GU305914.1 162715 162737 + 0 AGGCCCCTGCGCTTCGGCGA AGG ||||||||[||||||||||||]PAM NA Intergenic NA NA

T12219 GCTTCGGCGAAGGCCTCGCGCGG 1000
Oligo fwd CACCGCTTCGGCGAAGGCCTCGCG
Oligo rev AAACCGCGAGGCCTTCGCCGAAGC
Chromosome start end strand MM target_seq PAM alignment distance position gene name gene id
GU305914.1 162725 162747 + 0 GCTTCGGCGAAGGCCTCGCG CGG ||||||||[||||||||||||]PAM NA Intergenic NA NA

T12220 CGGCGAAGGCCTCGCGCGGCCGG 1000
Oligo fwd CACCCGGCGAAGGCCTCGCGCGGC
Oligo rev AAACGCCGCGCGAGGCCTTCGCCG
Chromosome start end strand MM target_seq PAM alignment distance position gene name gene id
GU305914.1 162729 162751 + 0 CGGCGAAGGCCTCGCGCGGC CGG ||||||||[||||||||||||]PAM NA Intergenic NA NA

T12221 AAGGCCTCGCGCGGCCGGCCAGG 1000
Oligo fwd CACCAAGGCCTCGCGCGGCCGGCC
Oligo rev AAACGGCCGGCCGCGCGAGGCCTT
Chromosome start end strand MM target_seq PAM alignment distance position gene name gene id
GU305914.1 162734 162756 + 0 AAGGCCTCGCGCGGCCGGCC AGG ||||||||[||||||||||||]PAM NA Intergenic NA NA

T12222 TCGCGCGGCCGGCCAGGACGAGG 1000
Oligo fwd CACCTCGCGCGGCCGGCCAGGACG
Oligo rev AAACCGTCCTGGCCGGCCGCGCGA
Chromosome start end strand MM target_seq PAM alignment distance position gene name gene id
GU305914.1 162740 162762 + 0 TCGCGCGGCCGGCCAGGACG AGG ||||||||[||||||||||||]PAM NA Intergenic NA NA

T12223 CCAGGACGAGGAGCGCCACTAGG 1000
Oligo fwd CACCCCAGGACGAGGAGCGCCACT
Oligo rev AAACAGTGGCGCTCCTCGTCCTGG
Chromosome start end strand MM target_seq PAM alignment distance position gene name gene id
GU305914.1 162752 162774 + 0 CCAGGACGAGGAGCGCCACT AGG ||||||||[||||||||||||]PAM NA Intergenic NA NA

T12224 TGAACATCCGCACGAGCCGCCGG 1000
Oligo fwd CACCTGAACATCCGCACGAGCCGC
Oligo rev AAACGCGGCTCGTGCGGATGTTCA
Chromosome start end strand MM target_seq PAM alignment distance position gene name gene id
GU305914.1 162776 162798 + 0 TGAACATCCGCACGAGCCGC CGG ||||||||[||||||||||||]PAM NA Intergenic NA NA

T12225 GAACATCCGCACGAGCCGCCGGG 1000
Oligo fwd CACCGAACATCCGCACGAGCCGCC
Oligo rev AAACGGCGGCTCGTGCGGATGTTC
Chromosome start end strand MM target_seq PAM alignment distance position gene name gene id
GU305914.1 162777 162799 + 0 GAACATCCGCACGAGCCGCC GGG ||||||||[||||||||||||]PAM NA Intergenic NA NA

T12226 TCCGCACGAGCCGCCGGGCCAGG 1000
Oligo fwd CACCTCCGCACGAGCCGCCGGGCC
Oligo rev AAACGGCCCGGCGGCTCGTGCGGA
Chromosome start end strand MM target_seq PAM alignment distance position gene name gene id
GU305914.1 162782 162804 + 0 TCCGCACGAGCCGCCGGGCC AGG ||||||||[||||||||||||]PAM NA Intergenic NA NA

T12227 CGAGCCGCCGGGCCAGGTCTCGG 1000
Oligo fwd CACCCGAGCCGCCGGGCCAGGTCT
Oligo rev AAACAGACCTGGCCCGGCGGCTCG
Chromosome start end strand MM target_seq PAM alignment distance position gene name gene id
GU305914.1 162788 162810 + 0 CGAGCCGCCGGGCCAGGTCT CGG ||||||||[||||||||||||]PAM NA Intergenic NA NA

T12228 CCGCCGGGCCAGGTCTCGGACGG 1000
Oligo fwd CACCCCGCCGGGCCAGGTCTCGGA
Oligo rev AAACTCCGAGACCTGGCCCGGCGG
Chromosome start end strand MM target_seq PAM alignment distance position gene name gene id
GU305914.1 162792 162814 + 0 CCGCCGGGCCAGGTCTCGGA CGG ||||||||[||||||||||||]PAM NA Intergenic NA NA

T12229 CGCCGGGCCAGGTCTCGGACGGG 1000
Oligo fwd CACCCGCCGGGCCAGGTCTCGGAC
Oligo rev AAACGTCCGAGACCTGGCCCGGCG
Chromosome start end strand MM target_seq PAM alignment distance position gene name gene id
GU305914.1 162793 162815 + 0 CGCCGGGCCAGGTCTCGGAC GGG ||||||||[||||||||||||]PAM NA Intergenic NA NA

T12230 AGACTCGATCTCGTGCATGTCGG 1000
Oligo fwd CACCAGACTCGATCTCGTGCATGT
Oligo rev AAACACATGCACGAGATCGAGTCT
Chromosome start end strand MM target_seq PAM alignment distance position gene name gene id
GU305914.1 162822 162844 + 0 AGACTCGATCTCGTGCATGT CGG ||||||||[||||||||||||]PAM NA Intergenic NA NA

T12231 CTCGATCTCGTGCATGTCGGCGG 1000

| | | | | | | | | | | | |
|---|---|---|---|---|---|---|---|---|---|---|---|
| Oligo fwd | CACCCTCGATCTCGTGCATGTCGG | | | | | | | | | | |
| Oligo rev | AAACCCGACATGCACGAGATCGAG | | | | | | | | | | |
| Chromosome | start | end | strand | MM | target_seq | PAM | alignment | distance | position | gene name | gene id |
| GU305914.1 | 162825 | 162847 | + | 0 | CTCGATCTCGTGCATGTCGG | CGG | \|\|\|\|\|\|\|\|[\|\|\|\|\|\|\|\|\|\|\|\|]PAM | NA | Intergenic | NA | NA |
| GU305914.1 | 64750 | 64772 | - | 3 | CTGGATCTGGTGGATGTCGG | TGG | \|\|-\|\|\|\|\|[-\|\|\|-\|\|\|\|\|\|\|]PAM | NA | Intergenic | NA | NA |

| | | | | | | | | | | | |
|---|---|---|---|---|---|---|---|---|---|---|---|
| T12232 | TCGTGCATGTCGGCGGTCCGCGG | 1000 | | | | | | | | | |
| Oligo fwd | CACCTCGTGCATGTCGGCGGTCCG | | | | | | | | | | |
| Oligo rev | AAACCGGACCGCCGACATGCACGA | | | | | | | | | | |
| Chromosome | start | end | strand | MM | target_seq | PAM | alignment | distance | position | gene name | gene id |
| GU305914.1 | 162832 | 162854 | + | 0 | TCGTGCATGTCGGCGGTCCG | CGG | \|\|\|\|\|\|\|\|[\|\|\|\|\|\|\|\|\|\|\|\|]PAM | NA | Intergenic | NA | NA |

| | | | | | | | | | | | |
|---|---|---|---|---|---|---|---|---|---|---|---|
| T12233 | CATGTCGGCGGTCCGCGGTGAGG | 1000 | | | | | | | | | |
| Oligo fwd | CACCCATGTCGGCGGTCCGCGGTG | | | | | | | | | | |
| Oligo rev | AAACCACCGCGGACCGCCGACATG | | | | | | | | | | |
| Chromosome | start | end | strand | MM | target_seq | PAM | alignment | distance | position | gene name | gene id |
| GU305914.1 | 162837 | 162859 | + | 0 | CATGTCGGCGGTCCGCGGTG | AGG | \|\|\|\|\|\|\|\|[\|\|\|\|\|\|\|\|\|\|\|\|]PAM | NA | Intergenic | NA | NA |

| | | | | | | | | | | | |
|---|---|---|---|---|---|---|---|---|---|---|---|
| T12234 | GAGGTTATAGACCATCTGCTAGG | 1000 | | | | | | | | | |
| Oligo fwd | CACCGAGGTTATAGACCATCTGCT | | | | | | | | | | |
| Oligo rev | AAACAGCAGATGGTCTATAACCTC | | | | | | | | | | |
| Chromosome | start | end | strand | MM | target_seq | PAM | alignment | distance | position | gene name | gene id |
| GU305914.1 | 162856 | 162878 | + | 0 | GAGGTTATAGACCATCTGCT | AGG | \|\|\|\|\|\|\|\|[\|\|\|\|\|\|\|\|\|\|\|\|]PAM | NA | Intergenic | NA | NA |

| | | | | | | | | | | | |
|---|---|---|---|---|---|---|---|---|---|---|---|
| T12235 | GTTATAGACCATCTGCTAGGCGG | 1000 | | | | | | | | | |
| Oligo fwd | CACCGTTATAGACCATCTGCTAGG | | | | | | | | | | |
| Oligo rev | AAACCCTAGCAGATGGTCTATAAC | | | | | | | | | | |
| Chromosome | start | end | strand | MM | target_seq | PAM | alignment | distance | position | gene name | gene id |
| GU305914.1 | 162859 | 162881 | + | 0 | GTTATAGACCATCTGCTAGG | CGG | \|\|\|\|\|\|\|\|[\|\|\|\|\|\|\|\|\|\|\|\|]PAM | NA | Intergenic | NA | NA |

| | | | | | | | | | | | |
|---|---|---|---|---|---|---|---|---|---|---|---|
| T12236 | TTATAGACCATCTGCTAGGCGGG | 1000 | | | | | | | | | |
| Oligo fwd | CACCTTATAGACCATCTGCTAGGC | | | | | | | | | | |
| Oligo rev | AAACGCCTAGCAGATGGTCTATAA | | | | | | | | | | |
| Chromosome | start | end | strand | MM | target_seq | PAM | alignment | distance | position | gene name | gene id |
| GU305914.1 | 162860 | 162882 | + | 0 | TTATAGACCATCTGCTAGGC | GGG | \|\|\|\|\|\|\|\|[\|\|\|\|\|\|\|\|\|\|\|\|]PAM | NA | Intergenic | NA | NA |

| | | | | | | | | | | | |
|---|---|---|---|---|---|---|---|---|---|---|---|
| T12237 | GACCATCTGCTAGGCGGGTCCGG | 1000 | | | | | | | | | |
| Oligo fwd | CACCGACCATCTGCTAGGCGGGTC | | | | | | | | | | |
| Oligo rev | AAACGACCCGCCTAGCAGATGGTC | | | | | | | | | | |
| Chromosome | start | end | strand | MM | target_seq | PAM | alignment | distance | position | gene name | gene id |
| GU305914.1 | 162865 | 162887 | + | 0 | GACCATCTGCTAGGCGGGTC | CGG | \|\|\|\|\|\|\|\|[\|\|\|\|\|\|\|\|\|\|\|\|]PAM | NA | Intergenic | NA | NA |

| | | | | | | | | | | | |
|---|---|---|---|---|---|---|---|---|---|---|---|
| T12238 | ACCATCTGCTAGGCGGGTCCGGG | 1000 | | | | | | | | | |
| Oligo fwd | CACCACCATCTGCTAGGCGGGTCC | | | | | | | | | | |
| Oligo rev | AAACGGACCCGCCTAGCAGATGGT | | | | | | | | | | |
| Chromosome | start | end | strand | MM | target_seq | PAM | alignment | distance | position | gene name | gene id |
| GU305914.1 | 162866 | 162888 | + | 0 | ACCATCTGCTAGGCGGGTCC | GGG | \|\|\|\|\|\|\|\|[\|\|\|\|\|\|\|\|\|\|\|\|]PAM | NA | Intergenic | NA | NA |

| | | | | | | | | | | | |
|---|---|---|---|---|---|---|---|---|---|---|---|
| T12239 | CCATCTGCTAGGCGGGTCCGGGG | 1000 | | | | | | | | | |
| Oligo fwd | CACCCCATCTGCTAGGCGGGTCCG | | | | | | | | | | |
| Oligo rev | AAACCGGACCCGCCTAGCAGATGG | | | | | | | | | | |
| Chromosome | start | end | strand | MM | target_seq | PAM | alignment | distance | position | gene name | gene id |
| GU305914.1 | 162867 | 162889 | + | 0 | CCATCTGCTAGGCGGGTCCG | GGG | \|\|\|\|\|\|\|\|[\|\|\|\|\|\|\|\|\|\|\|\|]PAM | NA | Intergenic | NA | NA |

| | | | | | | | | | | | |
|---|---|---|---|---|---|---|---|---|---|---|---|
| T12240 | CTAGGCGGGTCCGGGGAGACAGG | 1000 | | | | | | | | | |
| Oligo fwd | CACCCTAGGCGGGTCCGGGGAGAC | | | | | | | | | | |
| Oligo rev | AAACGTCTCCCCGGACCCGCCTAG | | | | | | | | | | |
| Chromosome | start | end | strand | MM | target_seq | PAM | alignment | distance | position | gene name | gene id |
| GU305914.1 | 162874 | 162896 | + | 0 | CTAGGCGGGTCCGGGGAGAC | AGG | \|\|\|\|\|\|\|\|[\|\|\|\|\|\|\|\|\|\|\|\|]PAM | NA | Intergenic | NA | NA |

| | | | | | | | | | | | |
|---|---|---|---|---|---|---|---|---|---|---|---|
| T12241 | CGGGGAGACAGGCACATTACTGG | 1000 | | | | | | | | | |
| Oligo fwd | CACCCGGGGAGACAGGCACATTAC | | | | | | | | | | |
| Oligo rev | AAACGTAATGTGCCTGTCTCCCCG | | | | | | | | | | |
| Chromosome | start | end | strand | MM | target_seq | PAM | alignment | distance | position | gene name | gene id |
| GU305914.1 | 162885 | 162907 | + | 0 | CGGGGAGACAGGCACATTAC | TGG | \|\|\|\|\|\|\|\|[\|\|\|\|\|\|\|\|\|\|\|\|]PAM | NA | Intergenic | NA | NA |

| | | | | | | | | | | | |
|---|---|---|---|---|---|---|---|---|---|---|---|
| T12242 | GACAGGCACATTACTGGCCTCGG | 1000 | | | | | | | | | |
| Oligo fwd | CACCGACAGGCACATTACTGGCCT | | | | | | | | | | |
| Oligo rev | AAACAGGCCAGTAATGTGCCTGTC | | | | | | | | | | |
| Chromosome | start | end | strand | MM | target_seq | PAM | alignment | distance | position | gene name | gene id |
| GU305914.1 | 162891 | 162913 | + | 0 | GACAGGCACATTACTGGCCT | CGG | \|\|\|\|\|\|\|\|[\|\|\|\|\|\|\|\|\|\|\|\|]PAM | NA | Intergenic | NA | NA |

| | | | | | | | | | | | |
|---|---|---|---|---|---|---|---|---|---|---|---|
| T12243 | CTGGCCTCGGCGCCCAGCCTAGG | 1000 | | | | | | | | | |
| Oligo fwd | CACCCTGGCCTCGGCGCCCAGCCT | | | | | | | | | | |
| Oligo rev | AAACAGGCTGGGCGCCGAGGCCAG | | | | | | | | | | |
| Chromosome | start | end | strand | MM | target_seq | PAM | alignment | distance | position | gene name | gene id |
| GU305914.1 | 162904 | 162926 | + | 0 | CTGGCCTCGGCGCCCAGCCT | AGG | \|\|\|\|\|\|\|\|[\|\|\|\|\|\|\|\|\|\|\|\|]PAM | NA | Intergenic | NA | NA |

| | | | | | | | | | | | |
|---|---|---|---|---|---|---|---|---|---|---|---|
| T12244 | CTAGAGCTCGACCGCGCGTCCGG | 1000 | | | | | | | | | |
| Oligo fwd | CACCCTAGAGCTCGACCGCGCGTC | | | | | | | | | | |
| Oligo rev | AAACGACGCGCGGTCGAGCTCTAG | | | | | | | | | | |
| Chromosome | start | end | strand | MM | target_seq | PAM | alignment | distance | position | gene name | gene id |
| GU305914.1 | 162932 | 162954 | + | 0 | CTAGAGCTCGACCGCGCGTC | CGG | \|\|\|\|\|\|\|\|[\|\|\|\|\|\|\|\|\|\|\|\|]PAM | NA | Intergenic | NA | NA |

| | | | | | | | | | | | |
|---|---|---|---|---|---|---|---|---|---|---|---|
| T12245 | CGTCCGGAGCGCCATTCGACCGG | 1000 | | | | | | | | | |
| Oligo fwd | CACCCGTCCGGAGCGCCATTCGAC | | | | | | | | | | |
| Oligo rev | AAACGTCGAATGGCGCTCCGGACG | | | | | | | | | | |
| Chromosome | start | end | strand | MM | target_seq | PAM | alignment | distance | position | gene name | gene id |
| GU305914.1 | 162948 | 162970 | + | 0 | CGTCCGGAGCGCCATTCGAC | CGG | \|\|\|\|\|\|\|\|[\|\|\|\|\|\|\|\|\|\|\|\|]PAM | NA | Intergenic | NA | NA |

<table>
<tr><td>T12246</td><td>CCGGAGCGCCATTCGACCGGCGG</td><td>1000</td><td></td><td></td><td></td><td></td><td></td><td></td><td></td><td></td><td></td></tr>
<tr><td>Oligo fwd</td><td>CACCCCGGAGCGCCATTCGACCGG</td><td></td><td></td><td></td><td></td><td></td><td></td><td></td><td></td><td></td><td></td></tr>
<tr><td>Oligo rev</td><td>AAACCCGGTCGAATGGCGCTCCGG</td><td></td><td></td><td></td><td></td><td></td><td></td><td></td><td></td><td></td><td></td></tr>
<tr><td>Chromosome</td><td>start</td><td>end</td><td>strand</td><td>MM</td><td>target_seq</td><td>PAM</td><td>alignment</td><td>distance</td><td>position</td><td>gene name</td><td>gene id</td></tr>
<tr><td>GU305914.1</td><td>162951</td><td>162973</td><td>+</td><td>0</td><td>CCGGAGCGCCATTCGACCGG</td><td>CGG</td><td>||||||||[||||||||||||]PAM</td><td>NA</td><td>Intergenic</td><td>NA</td><td>NA</td></tr>
<tr><td>T12247</td><td>CGGAGCGCCATTCGACCGGCGGG</td><td>1000</td><td></td><td></td><td></td><td></td><td></td><td></td><td></td><td></td><td></td></tr>
<tr><td>Oligo fwd</td><td>CACCCGGAGCGCCATTCGACCGGC</td><td></td><td></td><td></td><td></td><td></td><td></td><td></td><td></td><td></td><td></td></tr>
<tr><td>Oligo rev</td><td>AAACGCCGGTCGAATGGCGCTCCG</td><td></td><td></td><td></td><td></td><td></td><td></td><td></td><td></td><td></td><td></td></tr>
<tr><td>Chromosome</td><td>start</td><td>end</td><td>strand</td><td>MM</td><td>target_seq</td><td>PAM</td><td>alignment</td><td>distance</td><td>position</td><td>gene name</td><td>gene id</td></tr>
<tr><td>GU305914.1</td><td>162952</td><td>162974</td><td>+</td><td>0</td><td>CGGAGCGCCATTCGACCGGC</td><td>GGG</td><td>||||||||[||||||||||||]PAM</td><td>NA</td><td>Intergenic</td><td>NA</td><td>NA</td></tr>
<tr><td>T12248</td><td>GCGGGTAGCGAGAAGAACGCCGG</td><td>1000</td><td></td><td></td><td></td><td></td><td></td><td></td><td></td><td></td><td></td></tr>
<tr><td>Oligo fwd</td><td>CACCGCGGGTAGCGAGAAGAACGC</td><td></td><td></td><td></td><td></td><td></td><td></td><td></td><td></td><td></td><td></td></tr>
<tr><td>Oligo rev</td><td>AAACGCGTTCTTCTCGCTACCCGC</td><td></td><td></td><td></td><td></td><td></td><td></td><td></td><td></td><td></td><td></td></tr>
<tr><td>Chromosome</td><td>start</td><td>end</td><td>strand</td><td>MM</td><td>target_seq</td><td>PAM</td><td>alignment</td><td>distance</td><td>position</td><td>gene name</td><td>gene id</td></tr>
<tr><td>GU305914.1</td><td>162970</td><td>162992</td><td>+</td><td>0</td><td>GCGGGTAGCGAGAAGAACGC</td><td>CGG</td><td>||||||||[||||||||||||]PAM</td><td>NA</td><td>Intergenic</td><td>NA</td><td>NA</td></tr>
<tr><td>T12249</td><td>AGAAGAACGCCGGAGACCGCAGG</td><td>1000</td><td></td><td></td><td></td><td></td><td></td><td></td><td></td><td></td><td></td></tr>
<tr><td>Oligo fwd</td><td>CACCAGAAGAACGCCGGAGACCGC</td><td></td><td></td><td></td><td></td><td></td><td></td><td></td><td></td><td></td><td></td></tr>
<tr><td>Oligo rev</td><td>AAACGCGGTCTCCGGCGTTCTTCT</td><td></td><td></td><td></td><td></td><td></td><td></td><td></td><td></td><td></td><td></td></tr>
<tr><td>Chromosome</td><td>start</td><td>end</td><td>strand</td><td>MM</td><td>target_seq</td><td>PAM</td><td>alignment</td><td>distance</td><td>position</td><td>gene name</td><td>gene id</td></tr>
<tr><td>GU305914.1</td><td>162980</td><td>163002</td><td>+</td><td>0</td><td>AGAAGAACGCCGGAGACCGC</td><td>AGG</td><td>||||||||[||||||||||||]PAM</td><td>NA</td><td>Intergenic</td><td>NA</td><td>NA</td></tr>
<tr><td>T12250</td><td>CGTCATGCATAAATTAAGAATGG</td><td>1000</td><td></td><td></td><td></td><td></td><td></td><td></td><td></td><td></td><td></td></tr>
<tr><td>Oligo fwd</td><td>CACCCGTCATGCATAAATTAAGAA</td><td></td><td></td><td></td><td></td><td></td><td></td><td></td><td></td><td></td><td></td></tr>
<tr><td>Oligo rev</td><td>AAACTTCTTAATTTATGCATGACG</td><td></td><td></td><td></td><td></td><td></td><td></td><td></td><td></td><td></td><td></td></tr>
<tr><td>Chromosome</td><td>start</td><td>end</td><td>strand</td><td>MM</td><td>target_seq</td><td>PAM</td><td>alignment</td><td>distance</td><td>position</td><td>gene name</td><td>gene id</td></tr>
<tr><td>GU305914.1</td><td>163012</td><td>163034</td><td>+</td><td>0</td><td>CGTCATGCATAAATTAAGAA</td><td>TGG</td><td>||||||||[||||||||||||]PAM</td><td>NA</td><td>Intergenic</td><td>NA</td><td>NA</td></tr>
<tr><td>T12251</td><td>GTCATGCATAAATTAAGAATGGG</td><td>1000</td><td></td><td></td><td></td><td></td><td></td><td></td><td></td><td></td><td></td></tr>
<tr><td>Oligo fwd</td><td>CACCGTCATGCATAAATTAAGAAT</td><td></td><td></td><td></td><td></td><td></td><td></td><td></td><td></td><td></td><td></td></tr>
<tr><td>Oligo rev</td><td>AAACATTCTTAATTTATGCATGAC</td><td></td><td></td><td></td><td></td><td></td><td></td><td></td><td></td><td></td><td></td></tr>
<tr><td>Chromosome</td><td>start</td><td>end</td><td>strand</td><td>MM</td><td>target_seq</td><td>PAM</td><td>alignment</td><td>distance</td><td>position</td><td>gene name</td><td>gene id</td></tr>
<tr><td>GU305914.1</td><td>163013</td><td>163035</td><td>+</td><td>0</td><td>GTCATGCATAAATTAAGAAT</td><td>GGG</td><td>||||||||[||||||||||||]PAM</td><td>NA</td><td>Intergenic</td><td>NA</td><td>NA</td></tr>
<tr><td>T12252</td><td>AGACTTTATAAATCGCAAGCCGG</td><td>1000</td><td></td><td></td><td></td><td></td><td></td><td></td><td></td><td></td><td></td></tr>
<tr><td>Oligo fwd</td><td>CACCAGACTTTATAAATCGCAAGC</td><td></td><td></td><td></td><td></td><td></td><td></td><td></td><td></td><td></td><td></td></tr>
<tr><td>Oligo rev</td><td>AAACGCTTGCGATTTATAAAGTCT</td><td></td><td></td><td></td><td></td><td></td><td></td><td></td><td></td><td></td><td></td></tr>
<tr><td>Chromosome</td><td>start</td><td>end</td><td>strand</td><td>MM</td><td>target_seq</td><td>PAM</td><td>alignment</td><td>distance</td><td>position</td><td>gene name</td><td>gene id</td></tr>
<tr><td>GU305914.1</td><td>163044</td><td>163066</td><td>+</td><td>0</td><td>AGACTTTATAAATCGCAAGC</td><td>CGG</td><td>||||||||[||||||||||||]PAM</td><td>NA</td><td>Intergenic</td><td>NA</td><td>NA</td></tr>
<tr><td>T12253</td><td>TTATAAATCGCAAGCCGGAGCGG</td><td>1000</td><td></td><td></td><td></td><td></td><td></td><td></td><td></td><td></td><td></td></tr>
<tr><td>Oligo fwd</td><td>CACCTTATAAATCGCAAGCCGGAG</td><td></td><td></td><td></td><td></td><td></td><td></td><td></td><td></td><td></td><td></td></tr>
<tr><td>Oligo rev</td><td>AAACCTCCGGCTTGCGATTTATAA</td><td></td><td></td><td></td><td></td><td></td><td></td><td></td><td></td><td></td><td></td></tr>
<tr><td>Chromosome</td><td>start</td><td>end</td><td>strand</td><td>MM</td><td>target_seq</td><td>PAM</td><td>alignment</td><td>distance</td><td>position</td><td>gene name</td><td>gene id</td></tr>
<tr><td>GU305914.1</td><td>163049</td><td>163071</td><td>+</td><td>0</td><td>TTATAAATCGCAAGCCGGAG</td><td>CGG</td><td>||||||||[||||||||||||]PAM</td><td>NA</td><td>Intergenic</td><td>NA</td><td>NA</td></tr>
<tr><td>T12254</td><td>CGCAAGCCGGAGCGGATCCTAGG</td><td>1000</td><td></td><td></td><td></td><td></td><td></td><td></td><td></td><td></td><td></td></tr>
<tr><td>Oligo fwd</td><td>CACCCGCAAGCCGGAGCGGATCCT</td><td></td><td></td><td></td><td></td><td></td><td></td><td></td><td></td><td></td><td></td></tr>
<tr><td>Oligo rev</td><td>AAACAGGATCCGCTCCGGCTTGCG</td><td></td><td></td><td></td><td></td><td></td><td></td><td></td><td></td><td></td><td></td></tr>
<tr><td>Chromosome</td><td>start</td><td>end</td><td>strand</td><td>MM</td><td>target_seq</td><td>PAM</td><td>alignment</td><td>distance</td><td>position</td><td>gene name</td><td>gene id</td></tr>
<tr><td>GU305914.1</td><td>163057</td><td>163079</td><td>+</td><td>0</td><td>CGCAAGCCGGAGCGGATCCT</td><td>AGG</td><td>||||||||[||||||||||||]PAM</td><td>NA</td><td>Intergenic</td><td>NA</td><td>NA</td></tr>
<tr><td>T12255</td><td>TGCTGATAGTTCCTGCGTCGCGG</td><td>1000</td><td></td><td></td><td></td><td></td><td></td><td></td><td></td><td></td><td></td></tr>
<tr><td>Oligo fwd</td><td>CACCTGCTGATAGTTCCTGCGTCG</td><td></td><td></td><td></td><td></td><td></td><td></td><td></td><td></td><td></td><td></td></tr>
<tr><td>Oligo rev</td><td>AAACCGACGCAGGAACTATCAGCA</td><td></td><td></td><td></td><td></td><td></td><td></td><td></td><td></td><td></td><td></td></tr>
<tr><td>Chromosome</td><td>start</td><td>end</td><td>strand</td><td>MM</td><td>target_seq</td><td>PAM</td><td>alignment</td><td>distance</td><td>position</td><td>gene name</td><td>gene id</td></tr>
<tr><td>GU305914.1</td><td>163118</td><td>163140</td><td>+</td><td>0</td><td>TGCTGATAGTTCCTGCGTCG</td><td>CGG</td><td>||||||||[||||||||||||]PAM</td><td>NA</td><td>Intergenic</td><td>NA</td><td>NA</td></tr>
<tr><td>T12256</td><td>CCCACTTTTCTCGTCGCGACCGG</td><td>1000</td><td></td><td></td><td></td><td></td><td></td><td></td><td></td><td></td><td></td></tr>
<tr><td>Oligo fwd</td><td>CACCCCCACTTTTCTCGTCGCGAC</td><td></td><td></td><td></td><td></td><td></td><td></td><td></td><td></td><td></td><td></td></tr>
<tr><td>Oligo rev</td><td>AAACGTCGCGACGAGAAAAGTGGG</td><td></td><td></td><td></td><td></td><td></td><td></td><td></td><td></td><td></td><td></td></tr>
<tr><td>Chromosome</td><td>start</td><td>end</td><td>strand</td><td>MM</td><td>target_seq</td><td>PAM</td><td>alignment</td><td>distance</td><td>position</td><td>gene name</td><td>gene id</td></tr>
<tr><td>GU305914.1</td><td>163152</td><td>163174</td><td>+</td><td>0</td><td>CCCACTTTTCTCGTCGCGAC</td><td>CGG</td><td>||||||||[||||||||||||]PAM</td><td>NA</td><td>Intergenic</td><td>NA</td><td>NA</td></tr>
<tr><td>T12257</td><td>CCGGTGAAAAGACCTTCGTTCGG</td><td>1000</td><td></td><td></td><td></td><td></td><td></td><td></td><td></td><td></td><td></td></tr>
<tr><td>Oligo fwd</td><td>CACCCCGGTGAAAAGACCTTCGTT</td><td></td><td></td><td></td><td></td><td></td><td></td><td></td><td></td><td></td><td></td></tr>
<tr><td>Oligo rev</td><td>AAACAACGAAGGTCTTTTCACCGG</td><td></td><td></td><td></td><td></td><td></td><td></td><td></td><td></td><td></td><td></td></tr>
<tr><td>Chromosome</td><td>start</td><td>end</td><td>strand</td><td>MM</td><td>target_seq</td><td>PAM</td><td>alignment</td><td>distance</td><td>position</td><td>gene name</td><td>gene id</td></tr>
<tr><td>GU305914.1</td><td>163171</td><td>163193</td><td>+</td><td>0</td><td>CCGGTGAAAAGACCTTCGTT</td><td>CGG</td><td>||||||||[||||||||||||]PAM</td><td>NA</td><td>Intergenic</td><td>NA</td><td>NA</td></tr>
<tr><td>T12258</td><td>AAAGACCTTCGTTCGGACCATGG</td><td>1000</td><td></td><td></td><td></td><td></td><td></td><td></td><td></td><td></td><td></td></tr>
<tr><td>Oligo fwd</td><td>CACCAAAGACCTTCGTTCGGACCA</td><td></td><td></td><td></td><td></td><td></td><td></td><td></td><td></td><td></td><td></td></tr>
<tr><td>Oligo rev</td><td>AAACTGGTCCGAACGAAGGTCTTT</td><td></td><td></td><td></td><td></td><td></td><td></td><td></td><td></td><td></td><td></td></tr>
<tr><td>Chromosome</td><td>start</td><td>end</td><td>strand</td><td>MM</td><td>target_seq</td><td>PAM</td><td>alignment</td><td>distance</td><td>position</td><td>gene name</td><td>gene id</td></tr>
<tr><td>GU305914.1</td><td>163178</td><td>163200</td><td>+</td><td>0</td><td>AAAGACCTTCGTTCGGACCA</td><td>TGG</td><td>||||||||[||||||||||||]PAM</td><td>NA</td><td>Intergenic</td><td>NA</td><td>NA</td></tr>
<tr><td>T12259</td><td>ATGGCCGCGCCAGATCGACGCGG</td><td>1000</td><td></td><td></td><td></td><td></td><td></td><td></td><td></td><td></td><td></td></tr>
<tr><td>Oligo fwd</td><td>CACCATGGCCGCGCCAGATCGACG</td><td></td><td></td><td></td><td></td><td></td><td></td><td></td><td></td><td></td><td></td></tr>
<tr><td>Oligo rev</td><td>AAACCGTCGATCTGGCGCGGCCAT</td><td></td><td></td><td></td><td></td><td></td><td></td><td></td><td></td><td></td><td></td></tr>
<tr><td>Chromosome</td><td>start</td><td>end</td><td>strand</td><td>MM</td><td>target_seq</td><td>PAM</td><td>alignment</td><td>distance</td><td>position</td><td>gene name</td><td>gene id</td></tr>
<tr><td>GU305914.1</td><td>163197</td><td>163219</td><td>+</td><td>0</td><td>ATGGCCGCGCCAGATCGACG</td><td>CGG</td><td>||||||||[||||||||||||]PAM</td><td>NA</td><td>Intergenic</td><td>NA</td><td>NA</td></tr>
<tr><td>T12260</td><td>GACGACGTTACTTCACCTTCCGG</td><td>1000</td><td></td><td></td><td></td><td></td><td></td><td></td><td></td><td></td><td></td></tr>
<tr><td>Oligo fwd</td><td>CACCGACGACGTTACTTCACCTTC</td><td></td><td></td><td></td><td></td><td></td><td></td><td></td><td></td><td></td><td></td></tr>
<tr><td>Oligo rev</td><td>AAACGAAGGTGAAGTAACGTCGTC</td><td></td><td></td><td></td><td></td><td></td><td></td><td></td><td></td><td></td><td></td></tr>
<tr><td>Chromosome</td><td>start</td><td>end</td><td>strand</td><td>MM</td><td>target_seq</td><td>PAM</td><td>alignment</td><td>distance</td><td>position</td><td>gene name</td><td>gene id</td></tr>
</table>

GU305914.1 163237 163259 + 0 GACGACGTTACTTCACCTTC CGG ||||||||[||||||||||||]PAM NA Intergenic NA NA

T12261 TCACCTTCCGGAACTCGAGTCGG 1000
Oligo fwd CACCTCACCTTCCGGAACTCGAGT
Oligo rev AAACACTCGAGTTCCGGAAGGTGA
Chromosome start end strand MM target_seq PAM alignment distance position gene name gene id
GU305914.1 163249 163271 + 0 TCACCTTCCGGAACTCGAGT CGG ||||||||[||||||||||||]PAM NA Intergenic NA NA

T12262 GCATGTTCGACCTCCGTCAGTGG 1000
Oligo fwd CACCGCATGTTCGACCTCCGTCAG
Oligo rev AAACCTGACGGAGGTCGAACATGC
Chromosome start end strand MM target_seq PAM alignment distance position gene name gene id
GU305914.1 163294 163316 + 0 GCATGTTCGACCTCCGTCAG TGG ||||||||[||||||||||||]PAM NA Intergenic NA NA

T12263 CTACGAGAGCGCCCGCGTTTTGG 1000
Oligo fwd CACCCTACGAGAGCGCCCGCGTTT
Oligo rev AAACAAACGCGGGCGCTCTCGTAG
Chromosome start end strand MM target_seq PAM alignment distance position gene name gene id
GU305914.1 163319 163341 + 0 CTACGAGAGCGCCCGCGTTT TGG ||||||||[||||||||||||]PAM NA Intergenic NA NA

T12264 CGCGTTTTGGACTGCACGAACGG 1000
Oligo fwd CACCCGCGTTTTGGACTGCACGAA
Oligo rev AAACTTCGTGCAGTCCAAAACGCG
Chromosome start end strand MM target_seq PAM alignment distance position gene name gene id
GU305914.1 163332 163354 + 0 CGCGTTTTGGACTGCACGAA CGG ||||||||[||||||||||||]PAM NA Intergenic NA NA

T12265 TTTGGACTGCACGAACGGCGAGG 1000
Oligo fwd CACCTTTGGACTGCACGAACGGCG
Oligo rev AAACCGCCGTTCGTGCAGTCCAAA
Chromosome start end strand MM target_seq PAM alignment distance position gene name gene id
GU305914.1 163337 163359 + 0 TTTGGACTGCACGAACGGCG AGG ||||||||[||||||||||||]PAM NA Intergenic NA NA

T12266 TTGGACTGCACGAACGGCGAGGG 1000
Oligo fwd CACCTTGGACTGCACGAACGGCGA
Oligo rev AAACTCGCCGTTCGTGCAGTCCAA
Chromosome start end strand MM target_seq PAM alignment distance position gene name gene id
GU305914.1 163338 163360 + 0 TTGGACTGCACGAACGGCGA GGG ||||||||[||||||||||||]PAM NA Intergenic NA NA

T12267 GCGAGGGCCGCACCGAGAACTGG 1000
Oligo fwd CACCGCGAGGGCCGCACCGAGAAC
Oligo rev AAACGTTCTCGGTGCGGCCCTCGC
Chromosome start end strand MM target_seq PAM alignment distance position gene name gene id
GU305914.1 163354 163376 + 0 GCGAGGGCCGCACCGAGAAC TGG ||||||||[||||||||||||]PAM NA Intergenic NA NA

T12268 CGAGGGCCGCACCGAGAACTGGG 1000
Oligo fwd CACCCGAGGGCCGCACCGAGAACT
Oligo rev AAACAGTTCTCGGTGCGGCCCTCG
Chromosome start end strand MM target_seq PAM alignment distance position gene name gene id
GU305914.1 163355 163377 + 0 CGAGGGCCGCACCGAGAACT GGG ||||||||[||||||||||||]PAM NA Intergenic NA NA

T12269 GAGGGCCGCACCGAGAACTGGGG 1000
Oligo fwd CACCGAGGGCCGCACCGAGAACTG
Oligo rev AAACCAGTTCTCGGTGCGGCCCTC
Chromosome start end strand MM target_seq PAM alignment distance position gene name gene id
GU305914.1 163356 163378 + 0 GAGGGCCGCACCGAGAACTG GGG ||||||||[||||||||||||]PAM NA Intergenic NA NA

T12270 AGGGCCGCACCGAGAACTGGGGG 1000
Oligo fwd CACCAGGGCCGCACCGAGAACTGG
Oligo rev AAACCCAGTTCTCGGTGCGGCCCT
Chromosome start end strand MM target_seq PAM alignment distance position gene name gene id
GU305914.1 163357 163379 + 0 AGGGCCGCACCGAGAACTGG GGG ||||||||[||||||||||||]PAM NA Intergenic NA NA

T12271 CGCACCGAGAACTGGGGGCCCGG 1000
Oligo fwd CACCCGCACCGAGAACTGGGGGCC
Oligo rev AAACGGCCCCCAGTTCTCGGTGCG
Chromosome start end strand MM target_seq PAM alignment distance position gene name gene id
GU305914.1 163362 163384 + 0 CGCACCGAGAACTGGGGGCC CGG ||||||||[||||||||||||]PAM NA Intergenic NA NA

T12272 CCGAGAACTGGGGGCCCGGCTGG 1000
Oligo fwd CACCCCGAGAACTGGGGGCCCGGC
Oligo rev AAACGCCGGGCCCCCAGTTCTCGG
Chromosome start end strand MM target_seq PAM alignment distance position gene name gene id
GU305914.1 163366 163388 + 0 CCGAGAACTGGGGGCCCGGC TGG ||||||||[||||||||||||]PAM NA Intergenic NA NA

T12273 CGCGACCATCATGCAGTCTCCGG 1000
Oligo fwd CACCCGCGACCATCATGCAGTCTC
Oligo rev AAACGAGACTGCATGATGGTCGCG
Chromosome start end strand MM target_seq PAM alignment distance position gene name gene id
GU305914.1 163394 163416 + 0 CGCGACCATCATGCAGTCTC CGG ||||||||[||||||||||||]PAM NA Intergenic NA NA

T12274 ATCATGCAGTCTCCGGACAGCGG 1000
Oligo fwd CACCATCATGCAGTCTCCGGACAG
Oligo rev AAACCTGTCCGGAGACTGCATGAT
Chromosome start end strand MM target_seq PAM alignment distance position gene name gene id
GU305914.1 163401 163423 + 0 ATCATGCAGTCTCCGGACAG CGG ||||||||[||||||||||||]PAM NA Intergenic NA NA

T12275 CCGGACAGCGGCTCGTCGTCCGG 1000
Oligo fwd CACCCCGGACAGCGGCTCGTCGTC
Oligo rev AAACGACGACGAGCCGCTGTCCGG

| Chromosome | start | end | strand | MM | target_seq | PAM | alignment | distance | position | gene name | gene id |
|---|---|---|---|---|---|---|---|---|---|---|---|
| GU305914.1 | 163413 | 163435 | + | 0 | CCGGACAGCGGCTCGTCGTC | CGG | ||||||||[||||||||||||]PAM | NA | Intergenic | NA | NA |
| T12276 | CAGCGGCTCGTCGTCCGGATCGG | 1000 | | | | | | | | | |
| Oligo fwd | CACCCAGCGGCTCGTCGTCCGGAT | | | | | | | | | | |
| Oligo rev | AAACATCCGGACGACGAGCCGCTG | | | | | | | | | | |
| Chromosome | start | end | strand | MM | target_seq | PAM | alignment | distance | position | gene name | gene id |
| GU305914.1 | 163418 | 163440 | + | 0 | CAGCGGCTCGTCGTCCGGAT | CGG | ||||||||[||||||||||||]PAM | NA | Intergenic | NA | NA |
| T12277 | CGGCTCGTCGTCCGGATCGGCGG | 1000 | | | | | | | | | |
| Oligo fwd | CACCCGGCTCGTCGTCCGGATCGG | | | | | | | | | | |
| Oligo rev | AAACCCGATCCGGACGACGAGCCG | | | | | | | | | | |
| Chromosome | start | end | strand | MM | target_seq | PAM | alignment | distance | position | gene name | gene id |
| GU305914.1 | 163421 | 163443 | + | 0 | CGGCTCGTCGTCCGGATCGG | CGG | ||||||||[||||||||||||]PAM | NA | Intergenic | NA | NA |
| T12278 | GTCGTCCGGATCGGCGGCCCAGG | 1000 | | | | | | | | | |
| Oligo fwd | CACCGTCGTCCGGATCGGCGGCCC | | | | | | | | | | |
| Oligo rev | AAACGGGCCGCCGATCCGGACGAC | | | | | | | | | | |
| Chromosome | start | end | strand | MM | target_seq | PAM | alignment | distance | position | gene name | gene id |
| GU305914.1 | 163427 | 163449 | + | 0 | GTCGTCCGGATCGGCGGCCC | AGG | ||||||||[||||||||||||]PAM | NA | Intergenic | NA | NA |
| T12279 | TCGTCCGGATCGGCGGCCCAGGG | 1000 | | | | | | | | | |
| Oligo fwd | CACCTCGTCCGGATCGGCGGCCCA | | | | | | | | | | |
| Oligo rev | AAACTGGGCCGCCGATCCGGACGA | | | | | | | | | | |
| Chromosome | start | end | strand | MM | target_seq | PAM | alignment | distance | position | gene name | gene id |
| GU305914.1 | 163428 | 163450 | + | 0 | TCGTCCGGATCGGCGGCCCA | GGG | ||||||||[||||||||||||]PAM | NA | Intergenic | NA | NA |
| T12280 | GGCCCAGGGATGCATGTCTCTGG | 1000 | | | | | | | | | |
| Oligo fwd | CACCGGCCCAGGGATGCATGTCTC | | | | | | | | | | |
| Oligo rev | AAACGAGACATGCATCCCTGGGCC | | | | | | | | | | |
| Chromosome | start | end | strand | MM | target_seq | PAM | alignment | distance | position | gene name | gene id |
| GU305914.1 | 163442 | 163464 | + | 0 | GGCCCAGGGATGCATGTCTC | TGG | ||||||||[||||||||||||]PAM | NA | Intergenic | NA | NA |
| T12281 | CATGTCTCTGGACATCACCTCGG | 1000 | | | | | | | | | |
| Oligo fwd | CACCCATGTCTCTGGACATCACCT | | | | | | | | | | |
| Oligo rev | AAACAGGTGATGTCCAGAGACATG | | | | | | | | | | |
| Chromosome | start | end | strand | MM | target_seq | PAM | alignment | distance | position | gene name | gene id |
| GU305914.1 | 163454 | 163476 | + | 0 | CATGTCTCTGGACATCACCT | CGG | ||||||||[||||||||||||]PAM | NA | Intergenic | NA | NA |
| T12282 | CTGGACATCACCTCGGACGACGG | 1000 | | | | | | | | | |
| Oligo fwd | CACCCTGGACATCACCTCGGACGA | | | | | | | | | | |
| Oligo rev | AAACTCGTCCGAGGTGATGTCCAG | | | | | | | | | | |
| Chromosome | start | end | strand | MM | target_seq | PAM | alignment | distance | position | gene name | gene id |
| GU305914.1 | 163461 | 163483 | + | 0 | CTGGACATCACCTCGGACGA | CGG | ||||||||[||||||||||||]PAM | NA | Intergenic | NA | NA |
| T12283 | CATCACCTCGGACGACGGACCGG | 1000 | | | | | | | | | |
| Oligo fwd | CACCCATCACCTCGGACGACGGAC | | | | | | | | | | |
| Oligo rev | AAACGTCCGTCGTCCGAGGTGATG | | | | | | | | | | |
| Chromosome | start | end | strand | MM | target_seq | PAM | alignment | distance | position | gene name | gene id |
| GU305914.1 | 163466 | 163488 | + | 0 | CATCACCTCGGACGACGGAC | CGG | ||||||||[||||||||||||]PAM | NA | Intergenic | NA | NA |
| T12284 | CTCGGACGACGGACCGGAGAAGG | 1000 | | | | | | | | | |
| Oligo fwd | CACCCTCGGACGACGGACCGGAGA | | | | | | | | | | |
| Oligo rev | AAACTCTCCGGTCCGTCGTCCGAG | | | | | | | | | | |
| Chromosome | start | end | strand | MM | target_seq | PAM | alignment | distance | position | gene name | gene id |
| GU305914.1 | 163472 | 163494 | + | 0 | CTCGGACGACGGACCGGAGA | AGG | ||||||||[||||||||||||]PAM | NA | Intergenic | NA | NA |
| T12285 | GTCATGAACATGTTCCACCGCGG | 1000 | | | | | | | | | |
| Oligo fwd | CACCGTCATGAACATGTTCCACCG | | | | | | | | | | |
| Oligo rev | AAACCGGTGGAACATGTTCATGAC | | | | | | | | | | |
| Chromosome | start | end | strand | MM | target_seq | PAM | alignment | distance | position | gene name | gene id |
| GU305914.1 | 163494 | 163516 | + | 0 | GTCATGAACATGTTCCACCG | CGG | ||||||||[||||||||||||]PAM | NA | Intergenic | NA | NA |
| GU305914.1 | 194336 | 194358 | - | 4 | GCCATGGACATGTTCGACGG | CGG | |-||||-|[|||||||-||-|]PAM | NA | Intergenic | NA | NA |
| T12286 | TCATGAACATGTTCCACCGCGGG | 1000 | | | | | | | | | |
| Oligo fwd | CACCTCATGAACATGTTCCACCGC | | | | | | | | | | |
| Oligo rev | AAACGCGGTGGAACATGTTCATGA | | | | | | | | | | |
| Chromosome | start | end | strand | MM | target_seq | PAM | alignment | distance | position | gene name | gene id |
| GU305914.1 | 163495 | 163517 | + | 0 | TCATGAACATGTTCCACCGC | GGG | ||||||||[||||||||||||]PAM | NA | Intergenic | NA | NA |
| T12287 | CTGTAACAAGACCGTCTCTTCGG | 1000 | | | | | | | | | |
| Oligo fwd | CACCCTGTAACAAGACCGTCTCTT | | | | | | | | | | |
| Oligo rev | AAACAAGAGACGGTCTTGTTACAG | | | | | | | | | | |
| Chromosome | start | end | strand | MM | target_seq | PAM | alignment | distance | position | gene name | gene id |
| GU305914.1 | 163526 | 163548 | + | 0 | CTGTAACAAGACCGTCTCTT | CGG | ||||||||[||||||||||||]PAM | NA | Intergenic | NA | NA |
| T12288 | CTCTTCGGTCGCGACGCCGCCGG | 1000 | | | | | | | | | |
| Oligo fwd | CACCCTCTTCGGTCGCGACGCCGC | | | | | | | | | | |
| Oligo rev | AAACGCGGCGTCGCGACCGAAGAG | | | | | | | | | | |
| Chromosome | start | end | strand | MM | target_seq | PAM | alignment | distance | position | gene name | gene id |
| GU305914.1 | 163541 | 163563 | + | 0 | CTCTTCGGTCGCGACGCCGC | CGG | ||||||||[||||||||||||]PAM | NA | Intergenic | NA | NA |
| GU305914.1 | 72022 | 72044 | + | 4 | CTGGTCGGAGGCGACGCCGC | CGG | ||--||||[--||||||||||]PAM | NA | Intergenic | NA | NA |
| T12289 | TCTTCGGTCGCGACGCCGCCGGG | 1000 | | | | | | | | | |
| Oligo fwd | CACCTCTTCGGTCGCGACGCCGCC | | | | | | | | | | |
| Oligo rev | AAACGGCGGCGTCGCGACCGAAGA | | | | | | | | | | |
| Chromosome | start | end | strand | MM | target_seq | PAM | alignment | distance | position | gene name | gene id |
| GU305914.1 | 163542 | 163564 | + | 0 | TCTTCGGTCGCGACGCCGCC | GGG | ||||||||[||||||||||||]PAM | NA | Intergenic | NA | NA |

| | | | | | | | | | | | |
|---|---|---|---|---|---|---|---|---|---|---|---|
| T12290 | CTTCGGTCGCGACGCCGCCGGGG | 1000 | | | | | | | | | |
| Oligo fwd | CACCCTTCGGTCGCGACGCCGCCG | | | | | | | | | | |
| Oligo rev | AAACCGGCGGCGTCGCGACCGAAG | | | | | | | | | | |
| Chromosome | start | end | strand | MM | target_seq | PAM | alignment | distance | position | gene name | gene id |
| GU305914.1 | 163543 | 163565 | + | 0 | CTTCGGTCGCGACGCCGCCG | GGG | \|\|\|\|\|\|\|\|[\|\|\|\|\|\|\|\|\|\|\|\|]PAM | NA | Intergenic | NA | NA |

| | | | | | | | | | | | |
|---|---|---|---|---|---|---|---|---|---|---|---|
| T12291 | GCGACGCCGCCGGGGTCCGACGG | 1000 | | | | | | | | | |
| Oligo fwd | CACCGCGACGCCGCCGGGGTCCGA | | | | | | | | | | |
| Oligo rev | AAACTCGGACCCCGGCGGCGTCGC | | | | | | | | | | |
| Chromosome | start | end | strand | MM | target_seq | PAM | alignment | distance | position | gene name | gene id |
| GU305914.1 | 163551 | 163573 | + | 0 | GCGACGCCGCCGGGGTCCGA | CGG | \|\|\|\|\|\|\|\|[\|\|\|\|\|\|\|\|\|\|\|\|]PAM | NA | Intergenic | NA | NA |

| | | | | | | | | | | | |
|---|---|---|---|---|---|---|---|---|---|---|---|
| T12292 | GCCGCCGGGGTCCGACGGCACGG | 1000 | | | | | | | | | |
| Oligo fwd | CACCGCCGCCGGGGTCCGACGGCA | | | | | | | | | | |
| Oligo rev | AAACTGCCGTCGGACCCCGGCGGC | | | | | | | | | | |
| Chromosome | start | end | strand | MM | target_seq | PAM | alignment | distance | position | gene name | gene id |
| GU305914.1 | 163556 | 163578 | + | 0 | GCCGCCGGGGTCCGACGGCA | CGG | \|\|\|\|\|\|\|\|[\|\|\|\|\|\|\|\|\|\|\|\|]PAM | NA | Intergenic | NA | NA |
| GU305914.1 | 130328 | 130350 | + | 4 | GCCGCGGCGGTCCGGCGTCA | TGG | \|\|\|\|\|-\|-[\|\|\|\|\|\|-\|\|-\|\|]PAM | NA | Intergenic | NA | NA |

| | | | | | | | | | | | |
|---|---|---|---|---|---|---|---|---|---|---|---|
| T12293 | CCGCCGGGGTCCGACGGCACGGG | 1000 | | | | | | | | | |
| Oligo fwd | CACCCCGCCGGGGTCCGACGGCAC | | | | | | | | | | |
| Oligo rev | AAACGTGCCGTCGGACCCCGGCGG | | | | | | | | | | |
| Chromosome | start | end | strand | MM | target_seq | PAM | alignment | distance | position | gene name | gene id |
| GU305914.1 | 163557 | 163579 | + | 0 | CCGCCGGGGTCCGACGGCAC | GGG | \|\|\|\|\|\|\|\|[\|\|\|\|\|\|\|\|\|\|\|\|]PAM | NA | Intergenic | NA | NA |

| | | | | | | | | | | | |
|---|---|---|---|---|---|---|---|---|---|---|---|
| T12294 | GGGCGCCAGCCTCTTGACTCTGG | 1000 | | | | | | | | | |
| Oligo fwd | CACCGGGCGCCAGCCTCTTGACTC | | | | | | | | | | |
| Oligo rev | AAACGAGTCAAGAGGCTGGCGCCC | | | | | | | | | | |
| Chromosome | start | end | strand | MM | target_seq | PAM | alignment | distance | position | gene name | gene id |
| GU305914.1 | 163577 | 163599 | + | 0 | GGGCGCCAGCCTCTTGACTC | TGG | \|\|\|\|\|\|\|\|[\|\|\|\|\|\|\|\|\|\|\|\|]PAM | NA | Intergenic | NA | NA |

| | | | | | | | | | | | |
|---|---|---|---|---|---|---|---|---|---|---|---|
| T12295 | CGCCAGCCTCTTGACTCTGGTGG | 1000 | | | | | | | | | |
| Oligo fwd | CACCCGCCAGCCTCTTGACTCTGG | | | | | | | | | | |
| Oligo rev | AAACCCAGAGTCAAGAGGCTGGCG | | | | | | | | | | |
| Chromosome | start | end | strand | MM | target_seq | PAM | alignment | distance | position | gene name | gene id |
| GU305914.1 | 163580 | 163602 | + | 0 | CGCCAGCCTCTTGACTCTGG | TGG | \|\|\|\|\|\|\|\|[\|\|\|\|\|\|\|\|\|\|\|\|]PAM | NA | Intergenic | NA | NA |

| | | | | | | | | | | | |
|---|---|---|---|---|---|---|---|---|---|---|---|
| T12296 | CAGCCTCTTGACTCTGGTGGCGG | 1000 | | | | | | | | | |
| Oligo fwd | CACCCAGCCTCTTGACTCTGGTGG | | | | | | | | | | |
| Oligo rev | AAACCCACCAGAGTCAAGAGGCTG | | | | | | | | | | |
| Chromosome | start | end | strand | MM | target_seq | PAM | alignment | distance | position | gene name | gene id |
| GU305914.1 | 163583 | 163605 | + | 0 | CAGCCTCTTGACTCTGGTGG | CGG | \|\|\|\|\|\|\|\|[\|\|\|\|\|\|\|\|\|\|\|\|]PAM | NA | Intergenic | NA | NA |

| | | | | | | | | | | | |
|---|---|---|---|---|---|---|---|---|---|---|---|
| T12297 | CTCTTGACTCTGGTGGCGGACGG | 1000 | | | | | | | | | |
| Oligo fwd | CACCCTCTTGACTCTGGTGGCGGA | | | | | | | | | | |
| Oligo rev | AAACTCCGCCACCAGAGTCAAGAG | | | | | | | | | | |
| Chromosome | start | end | strand | MM | target_seq | PAM | alignment | distance | position | gene name | gene id |
| GU305914.1 | 163587 | 163609 | + | 0 | CTCTTGACTCTGGTGGCGGA | CGG | \|\|\|\|\|\|\|\|[\|\|\|\|\|\|\|\|\|\|\|\|]PAM | NA | Intergenic | NA | NA |

| | | | | | | | | | | | |
|---|---|---|---|---|---|---|---|---|---|---|---|
| T12298 | CAGCTTGCTGCAAGTCATCCTGG | 1000 | | | | | | | | | |
| Oligo fwd | CACCCAGCTTGCTGCAAGTCATCC | | | | | | | | | | |
| Oligo rev | AAACGGATGACTTGCAGCAAGCTG | | | | | | | | | | |
| Chromosome | start | end | strand | MM | target_seq | PAM | alignment | distance | position | gene name | gene id |
| GU305914.1 | 163610 | 163632 | + | 0 | CAGCTTGCTGCAAGTCATCC | TGG | \|\|\|\|\|\|\|\|[\|\|\|\|\|\|\|\|\|\|\|\|]PAM | NA | Intergenic | NA | NA |

| | | | | | | | | | | | |
|---|---|---|---|---|---|---|---|---|---|---|---|
| T12299 | CATCCTGGTCGAACACTCGCCGG | 1000 | | | | | | | | | |
| Oligo fwd | CACCCATCCTGGTCGAACACTCGC | | | | | | | | | | |
| Oligo rev | AAACGCGAGTGTTCGACCAGGATG | | | | | | | | | | |
| Chromosome | start | end | strand | MM | target_seq | PAM | alignment | distance | position | gene name | gene id |
| GU305914.1 | 163625 | 163647 | + | 0 | CATCCTGGTCGAACACTCGC | CGG | \|\|\|\|\|\|\|\|[\|\|\|\|\|\|\|\|\|\|\|\|]PAM | NA | Intergenic | NA | NA |

| | | | | | | | | | | | |
|---|---|---|---|---|---|---|---|---|---|---|---|
| T12300 | ATCCTGGTCGAACACTCGCCGGG | 1000 | | | | | | | | | |
| Oligo fwd | CACCATCCTGGTCGAACACTCGCC | | | | | | | | | | |
| Oligo rev | AAACGGCGAGTGTTCGACCAGGAT | | | | | | | | | | |
| Chromosome | start | end | strand | MM | target_seq | PAM | alignment | distance | position | gene name | gene id |
| GU305914.1 | 163626 | 163648 | + | 0 | ATCCTGGTCGAACACTCGCC | GGG | \|\|\|\|\|\|\|\|[\|\|\|\|\|\|\|\|\|\|\|\|]PAM | NA | Intergenic | NA | NA |

| | | | | | | | | | | | |
|---|---|---|---|---|---|---|---|---|---|---|---|
| T12301 | GACGATGTCGCCGAGAGCGTCGG | 1000 | | | | | | | | | |
| Oligo fwd | CACCGACGATGTCGCCGAGAGCGT | | | | | | | | | | |
| Oligo rev | AAACACGCTCTCGGCGACATCGTC | | | | | | | | | | |
| Chromosome | start | end | strand | MM | target_seq | PAM | alignment | distance | position | gene name | gene id |
| GU305914.1 | 163668 | 163690 | + | 0 | GACGATGTCGCCGAGAGCGT | CGG | \|\|\|\|\|\|\|\|[\|\|\|\|\|\|\|\|\|\|\|\|]PAM | NA | Intergenic | NA | NA |

| | | | | | | | | | | | |
|---|---|---|---|---|---|---|---|---|---|---|---|
| T12302 | CGAGAGCGTCGGCAACGCCGCGG | 1000 | | | | | | | | | |
| Oligo fwd | CACCCGAGAGCGTCGGCAACGCCG | | | | | | | | | | |
| Oligo rev | AAACCGGCGTTGCCGACGCTCTCG | | | | | | | | | | |
| Chromosome | start | end | strand | MM | target_seq | PAM | alignment | distance | position | gene name | gene id |
| GU305914.1 | 163679 | 163701 | + | 0 | CGAGAGCGTCGGCAACGCCG | CGG | \|\|\|\|\|\|\|\|[\|\|\|\|\|\|\|\|\|\|\|\|]PAM | NA | Intergenic | NA | NA |

| | | | | | | | | | | | |
|---|---|---|---|---|---|---|---|---|---|---|---|
| T12303 | GAGCGTCGGCAACGCCGCGGTGG | 1000 | | | | | | | | | |
| Oligo fwd | CACCGAGCGTCGGCAACGCCGCGG | | | | | | | | | | |
| Oligo rev | AAACCCGCGGCGTTGCCGACGCTC | | | | | | | | | | |
| Chromosome | start | end | strand | MM | target_seq | PAM | alignment | distance | position | gene name | gene id |
| GU305914.1 | 163682 | 163704 | + | 0 | GAGCGTCGGCAACGCCGCGG | TGG | \|\|\|\|\|\|\|\|[\|\|\|\|\|\|\|\|\|\|\|\|]PAM | NA | Intergenic | NA | NA |

| | | | | | | | | | | | |
|---|---|---|---|---|---|---|---|---|---|---|---|
| T12304 | GGTGGCCGCCGCCGTGCAGTCGG | 1000 | | | | | | | | | |
| Oligo fwd | CACCGGTGGCCGCCGCCGTGCAGT | | | | | | | | | | |
| Oligo rev | AAACACTGCACGGCGGCGGCCACC | | | | | | | | | | |
| Chromosome | start | end | strand | MM | target_seq | PAM | alignment | distance | position | gene name | gene id |

| | | | | | | | | | | | |
|---|---|---|---|---|---|---|---|---|---|---|---|
| GU305914.1 | 163700 | 163722 | + | 0 | GGTGGCCGCCGCCGTGCAGT | CGG | \|\|\|\|\|\|\|\|[\|\|\|\|\|\|\|\|\|\|\|\|]PAM | NA | Intergenic | NA | NA |
| T12305 | CGTGCAGTCGGCCACTTCGATGG | 1000 | | | | | | | | | |
| Oligo fwd | CACCCGTGCAGTCGGCCACTTCGA | | | | | | | | | | |
| Oligo rev | AAACTCGAAGTGGCCGACTGCACG | | | | | | | | | | |
| Chromosome | start | end | strand | MM | target_seq | PAM | alignment | distance | position | gene name | gene id |
| GU305914.1 | 163712 | 163734 | + | 0 | CGTGCAGTCGGCCACTTCGA | TGG | \|\|\|\|\|\|\|\|[\|\|\|\|\|\|\|\|\|\|\|\|]PAM | NA | Intergenic | NA | NA |
| T12306 | GTGCAGTCGGCCACTTCGATGGG | 1000 | | | | | | | | | |
| Oligo fwd | CACCGTGCAGTCGGCCACTTCGAT | | | | | | | | | | |
| Oligo rev | AAACATCGAAGTGGCCGACTGCAC | | | | | | | | | | |
| Chromosome | start | end | strand | MM | target_seq | PAM | alignment | distance | position | gene name | gene id |
| GU305914.1 | 163713 | 163735 | + | 0 | GTGCAGTCGGCCACTTCGAT | GGG | \|\|\|\|\|\|\|\|[\|\|\|\|\|\|\|\|\|\|\|\|]PAM | NA | Intergenic | NA | NA |
| T12307 | ATGGGTAGCTCGAGCTCCGTCGG | 1000 | | | | | | | | | |
| Oligo fwd | CACCATGGGTAGCTCGAGCTCCGT | | | | | | | | | | |
| Oligo rev | AAACACGGAGCTCGAGCTACCCAT | | | | | | | | | | |
| Chromosome | start | end | strand | MM | target_seq | PAM | alignment | distance | position | gene name | gene id |
| GU305914.1 | 163731 | 163753 | + | 0 | ATGGGTAGCTCGAGCTCCGT | CGG | \|\|\|\|\|\|\|\|[\|\|\|\|\|\|\|\|\|\|\|\|]PAM | NA | Intergenic | NA | NA |
| T12308 | GGTAGCTCGAGCTCCGTCGGCGG | 1000 | | | | | | | | | |
| Oligo fwd | CACCGGTAGCTCGAGCTCCGTCGG | | | | | | | | | | |
| Oligo rev | AAACCCGACGGAGCTCGAGCTACC | | | | | | | | | | |
| Chromosome | start | end | strand | MM | target_seq | PAM | alignment | distance | position | gene name | gene id |
| GU305914.1 | 163734 | 163756 | + | 0 | GGTAGCTCGAGCTCCGTCGG | CGG | \|\|\|\|\|\|\|\|[\|\|\|\|\|\|\|\|\|\|\|\|]PAM | NA | Intergenic | NA | NA |
| T12309 | TCGAGCTCCGTCGGCGGCGCCGG | 1000 | | | | | | | | | |
| Oligo fwd | CACCTCGAGCTCCGTCGGCGGCGC | | | | | | | | | | |
| Oligo rev | AAACGCGCCGCCGACGGAGCTCGA | | | | | | | | | | |
| Chromosome | start | end | strand | MM | target_seq | PAM | alignment | distance | position | gene name | gene id |
| GU305914.1 | 163740 | 163762 | + | 0 | TCGAGCTCCGTCGGCGGCGC | CGG | \|\|\|\|\|\|\|\|[\|\|\|\|\|\|\|\|\|\|\|\|]PAM | NA | Intergenic | NA | NA |
| T12310 | AGCTCCGTCGGCGGCGCCGGTGG | 1000 | | | | | | | | | |
| Oligo fwd | CACCAGCTCCGTCGGCGGCGCCGG | | | | | | | | | | |
| Oligo rev | AAACCCGGCGCCGCCGACGGAGCT | | | | | | | | | | |
| Chromosome | start | end | strand | MM | target_seq | PAM | alignment | distance | position | gene name | gene id |
| GU305914.1 | 163743 | 163765 | + | 0 | AGCTCCGTCGGCGGCGCCGG | TGG | \|\|\|\|\|\|\|\|[\|\|\|\|\|\|\|\|\|\|\|\|]PAM | NA | Intergenic | NA | NA |
| GU305914.1 | 151164 | 151186 | + | 4 | GGGTCCGACGGCGGCTCCGG | CGG | -\|-\|\|\|\|-[\|\|\|\|\|\|\|-\|\|\|\|]PAM | NA | Intergenic | NA | NA |
| GU305914.1 | 27311 | 27333 | - | 4 | CGCGTCGTCGGCGGCGCCGC | CGG | -\|\|--\|\|\|[\|\|\|\|\|\|\|\|\|\|\|-]PAM | NA | Intergenic | NA | NA |
| T12311 | TCCGTCGGCGGCGCCGGTGGCGG | 1000 | | | | | | | | | |
| Oligo fwd | CACCTCCGTCGGCGGCGCCGGTGG | | | | | | | | | | |
| Oligo rev | AAACCCACCGGCGCCGCCGACGGA | | | | | | | | | | |
| Chromosome | start | end | strand | MM | target_seq | PAM | alignment | distance | position | gene name | gene id |
| GU305914.1 | 163746 | 163768 | + | 0 | TCCGTCGGCGGCGCCGGTGG | CGG | \|\|\|\|\|\|\|\|[\|\|\|\|\|\|\|\|\|\|\|\|]PAM | NA | Intergenic | NA | NA |
| GU305914.1 | 152550 | 152572 | + | 4 | TCCTGCGGCGGCGGCGGTGC | CGG | \|\|\|--\|\|\|[\|\|\|\|\|-\|\|\|\|\|-]PAM | NA | Intergenic | NA | NA |
| T12312 | GTCGGCGGCGCCGGTGGCGGCGG | 1000 | | | | | | | | | |
| Oligo fwd | CACCGTCGGCGGCGCCGGTGGCGG | | | | | | | | | | |
| Oligo rev | AAACCCGCCACCGGCGCCGCCGAC | | | | | | | | | | |
| Chromosome | start | end | strand | MM | target_seq | PAM | alignment | distance | position | gene name | gene id |
| GU305914.1 | 163749 | 163771 | + | 0 | GTCGGCGGCGCCGGTGGCGG | CGG | \|\|\|\|\|\|\|\|[\|\|\|\|\|\|\|\|\|\|\|\|]PAM | NA | Intergenic | NA | NA |
| GU305914.1 | 48153 | 48175 | + | 4 | TGTGGCGGCGGCGGTGGCGG | CGG | ---\|\|\|\|\|[\|\|-\|\|\|\|\|\|\|\|\|]PAM | NA | Intergenic | NA | NA |
| GU305914.1 | 74670 | 74692 | + | 4 | GTTGGTGCCGTCGGTGGCGG | TGG | \|\|-\|\|-\|-[\|\|-\|\|\|\|\|\|\|\|\|]PAM | NA | Intergenic | NA | NA |
| GU305914.1 | 154407 | 154429 | + | 4 | GGTGGCGGCGGCGGCGGCGG | CGG | \|--\|\|\|\|\|[\|\|-\|\|\|-\|\|\|\|\|]PAM | NA | Intergenic | NA | NA |
| GU305914.1 | 58123 | 58145 | - | 4 | GGTGGCGGCGGCGGCGGCGG | AGG | \|--\|\|\|\|\|[\|\|-\|\|\|-\|\|\|\|\|]PAM | NA | Intergenic | NA | NA |
| GU305914.1 | 149875 | 149897 | - | 3 | GCCGACGGCGCCGGTGACGG | TGG | \|-\|\|-\|\|\|[\|\|\|\|\|\|\|\|-\|\|\|]PAM | NA | Intergenic | NA | NA |
| GU305914.1 | 149482 | 149504 | - | 4 | GGCGACGGCGGCGGCGGCGG | CGG | \|-\|\|-\|\|\|[\|\|-\|\|\|-\|\|\|\|\|]PAM | NA | Intergenic | NA | NA |
| GU305914.1 | 154404 | 154426 | + | 4 | GGCGGTGGCGGCGGCGGCGG | CGG | \|-\|\|\|-\|\|[\|\|-\|\|\|-\|\|\|\|\|]PAM | NA | Intergenic | NA | NA |
| GU305914.1 | 58126 | 58148 | - | 4 | GGCGGTGGCGGCGGCGGCGG | CGG | \|-\|\|\|-\|\|[\|\|-\|\|\|-\|\|\|\|\|]PAM | NA | Intergenic | NA | NA |
| GU305914.1 | 152553 | 152575 | + | 4 | TGCGGCGGCGGCGGTGCCGG | CGG | --\|\|\|\|\|\|[\|\|-\|\|\|\|\|-\|\|\|]PAM | NA | Intergenic | NA | NA |
| GU305914.1 | 174915 | 174937 | + | 4 | GTCGGTTGCGCGGGTGCCGG | AGG | \|\|\|\|\|--\|[\|\|\|-\|\|\|\|-\|\|\|]PAM | NA | Intergenic | NA | NA |
| GU305914.1 | 166410 | 166432 | + | 4 | GACGTCGGCGCCGGCGGCAG | CGG | \|-\|\|-\|\|\|[\|\|\|\|\|\|-\|\|\|-\|]PAM | NA | Intergenic | NA | NA |
| T12313 | TGGCGGCGGCTTCAATTCTTCGG | 1000 | | | | | | | | | |
| Oligo fwd | CACCTGGCGGCGGCTTCAATTCTT | | | | | | | | | | |
| Oligo rev | AAACAAGAATTGAAGCCGCCGCCA | | | | | | | | | | |
| Chromosome | start | end | strand | MM | target_seq | PAM | alignment | distance | position | gene name | gene id |
| GU305914.1 | 163763 | 163785 | + | 0 | TGGCGGCGGCTTCAATTCTT | CGG | \|\|\|\|\|\|\|\|[\|\|\|\|\|\|\|\|\|\|\|\|]PAM | NA | Intergenic | NA | NA |
| T12314 | AAGAGATTCGAACTCTCCAACGG | 1000 | | | | | | | | | |
| Oligo fwd | CACCAAGAGATTCGAACTCTCCAA | | | | | | | | | | |
| Oligo rev | AAACTTGGAGAGTTCGAATCTCTT | | | | | | | | | | |
| Chromosome | start | end | strand | MM | target_seq | PAM | alignment | distance | position | gene name | gene id |
| GU305914.1 | 163839 | 163861 | + | 0 | AAGAGATTCGAACTCTCCAA | CGG | \|\|\|\|\|\|\|\|[\|\|\|\|\|\|\|\|\|\|\|\|]PAM | NA | Intergenic | NA | NA |
| T12315 | TCTCCAACGGTACCCTTCATAGG | 1000 | | | | | | | | | |
| Oligo fwd | CACCTCTCCAACGGTACCCTTCAT | | | | | | | | | | |
| Oligo rev | AAACATGAAGGGTACCGTTGGAGA | | | | | | | | | | |
| Chromosome | start | end | strand | MM | target_seq | PAM | alignment | distance | position | gene name | gene id |
| GU305914.1 | 163852 | 163874 | + | 0 | TCTCCAACGGTACCCTTCAT | AGG | \|\|\|\|\|\|\|\|[\|\|\|\|\|\|\|\|\|\|\|\|]PAM | NA | Intergenic | NA | NA |
| T12316 | CTCCAACGGTACCCTTCATAGGG | 1000 | | | | | | | | | |
| Oligo fwd | CACCCTCCAACGGTACCCTTCATA | | | | | | | | | | |
| Oligo rev | AAACTATGAAGGGTACCGTTGGAG | | | | | | | | | | |
| Chromosome | start | end | strand | MM | target_seq | PAM | alignment | distance | position | gene name | gene id |
| GU305914.1 | 163853 | 163875 | + | 0 | CTCCAACGGTACCCTTCATA | GGG | \|\|\|\|\|\|\|\|[\|\|\|\|\|\|\|\|\|\|\|\|]PAM | NA | Intergenic | NA | NA |
| T12317 | ACGGTACCCTTCATAGGGAGCGG | 1000 | | | | | | | | | |

<table>
<tr><td>Oligo fwd</td><td>CACCACGGTACCCTTCATAGGGAG</td><td></td><td></td><td></td><td></td><td></td><td></td><td></td><td></td><td></td><td></td></tr>
<tr><td>Oligo rev</td><td>AAACCTCCCTATGAAGGGTACCGT</td><td></td><td></td><td></td><td></td><td></td><td></td><td></td><td></td><td></td><td></td></tr>
<tr><td>Chromosome</td><td>start</td><td>end</td><td>strand</td><td>MM</td><td>target_seq</td><td>PAM</td><td>alignment</td><td>distance</td><td>position</td><td>gene name</td><td>gene id</td></tr>
<tr><td>GU305914.1</td><td>163858</td><td>163880</td><td>+</td><td>0</td><td>ACGGTACCCTTCATAGGGAG</td><td>CGG</td><td>||||||||[||||||||||||]PAM</td><td>NA</td><td>Intergenic</td><td>NA</td><td>NA</td></tr>
<tr><td></td><td></td><td></td><td></td><td></td><td></td><td></td><td></td><td></td><td></td><td></td><td></td></tr>
<tr><td>T12318</td><td>CCTTCATAGGGAGCGGACCTCGG</td><td>1000</td><td></td><td></td><td></td><td></td><td></td><td></td><td></td><td></td><td></td></tr>
<tr><td>Oligo fwd</td><td>CACCCCTTCATAGGGAGCGGACCT</td><td></td><td></td><td></td><td></td><td></td><td></td><td></td><td></td><td></td><td></td></tr>
<tr><td>Oligo rev</td><td>AAACAGGTCCGCTCCCTATGAAGG</td><td></td><td></td><td></td><td></td><td></td><td></td><td></td><td></td><td></td><td></td></tr>
<tr><td>Chromosome</td><td>start</td><td>end</td><td>strand</td><td>MM</td><td>target_seq</td><td>PAM</td><td>alignment</td><td>distance</td><td>position</td><td>gene name</td><td>gene id</td></tr>
<tr><td>GU305914.1</td><td>163865</td><td>163887</td><td>+</td><td>0</td><td>CCTTCATAGGGAGCGGACCT</td><td>CGG</td><td>||||||||[||||||||||||]PAM</td><td>NA</td><td>Intergenic</td><td>NA</td><td>NA</td></tr>
<tr><td></td><td></td><td></td><td></td><td></td><td></td><td></td><td></td><td></td><td></td><td></td><td></td></tr>
<tr><td>T12319</td><td>CTTCATAGGGAGCGGACCTCGGG</td><td>1000</td><td></td><td></td><td></td><td></td><td></td><td></td><td></td><td></td><td></td></tr>
<tr><td>Oligo fwd</td><td>CACCCTTCATAGGGAGCGGACCTC</td><td></td><td></td><td></td><td></td><td></td><td></td><td></td><td></td><td></td><td></td></tr>
<tr><td>Oligo rev</td><td>AAACGAGGTCCGCTCCCTATGAAG</td><td></td><td></td><td></td><td></td><td></td><td></td><td></td><td></td><td></td><td></td></tr>
<tr><td>Chromosome</td><td>start</td><td>end</td><td>strand</td><td>MM</td><td>target_seq</td><td>PAM</td><td>alignment</td><td>distance</td><td>position</td><td>gene name</td><td>gene id</td></tr>
<tr><td>GU305914.1</td><td>163866</td><td>163888</td><td>+</td><td>0</td><td>CTTCATAGGGAGCGGACCTC</td><td>GGG</td><td>||||||||[||||||||||||]PAM</td><td>NA</td><td>Intergenic</td><td>NA</td><td>NA</td></tr>
<tr><td></td><td></td><td></td><td></td><td></td><td></td><td></td><td></td><td></td><td></td><td></td><td></td></tr>
<tr><td>T12320</td><td>GAGCGGACCTCGGGTAGCTCCGG</td><td>1000</td><td></td><td></td><td></td><td></td><td></td><td></td><td></td><td></td><td></td></tr>
<tr><td>Oligo fwd</td><td>CACCGAGCGGACCTCGGGTAGCTC</td><td></td><td></td><td></td><td></td><td></td><td></td><td></td><td></td><td></td><td></td></tr>
<tr><td>Oligo rev</td><td>AAACGAGCTACCCGAGGTCCGCTC</td><td></td><td></td><td></td><td></td><td></td><td></td><td></td><td></td><td></td><td></td></tr>
<tr><td>Chromosome</td><td>start</td><td>end</td><td>strand</td><td>MM</td><td>target_seq</td><td>PAM</td><td>alignment</td><td>distance</td><td>position</td><td>gene name</td><td>gene id</td></tr>
<tr><td>GU305914.1</td><td>163875</td><td>163897</td><td>+</td><td>0</td><td>GAGCGGACCTCGGGTAGCTC</td><td>CGG</td><td>||||||||[||||||||||||]PAM</td><td>NA</td><td>Intergenic</td><td>NA</td><td>NA</td></tr>
<tr><td></td><td></td><td></td><td></td><td></td><td></td><td></td><td></td><td></td><td></td><td></td><td></td></tr>
<tr><td>T12321</td><td>TCGGGTAGCTCCGGATCGAGCGG</td><td>1000</td><td></td><td></td><td></td><td></td><td></td><td></td><td></td><td></td><td></td></tr>
<tr><td>Oligo fwd</td><td>CACCTCGGGTAGCTCCGGATCGAG</td><td></td><td></td><td></td><td></td><td></td><td></td><td></td><td></td><td></td><td></td></tr>
<tr><td>Oligo rev</td><td>AAACCTCGATCCGGAGCTACCCGA</td><td></td><td></td><td></td><td></td><td></td><td></td><td></td><td></td><td></td><td></td></tr>
<tr><td>Chromosome</td><td>start</td><td>end</td><td>strand</td><td>MM</td><td>target_seq</td><td>PAM</td><td>alignment</td><td>distance</td><td>position</td><td>gene name</td><td>gene id</td></tr>
<tr><td>GU305914.1</td><td>163884</td><td>163906</td><td>+</td><td>0</td><td>TCGGGTAGCTCCGGATCGAG</td><td>CGG</td><td>||||||||[||||||||||||]PAM</td><td>NA</td><td>Intergenic</td><td>NA</td><td>NA</td></tr>
<tr><td></td><td></td><td></td><td></td><td></td><td></td><td></td><td></td><td></td><td></td><td></td><td></td></tr>
<tr><td>T12322</td><td>GGTAGCTCCGGATCGAGCGGCGG</td><td>1000</td><td></td><td></td><td></td><td></td><td></td><td></td><td></td><td></td><td></td></tr>
<tr><td>Oligo fwd</td><td>CACCGGTAGCTCCGGATCGAGCGG</td><td></td><td></td><td></td><td></td><td></td><td></td><td></td><td></td><td></td><td></td></tr>
<tr><td>Oligo rev</td><td>AAACCCGCTCGATCCGGAGCTACC</td><td></td><td></td><td></td><td></td><td></td><td></td><td></td><td></td><td></td><td></td></tr>
<tr><td>Chromosome</td><td>start</td><td>end</td><td>strand</td><td>MM</td><td>target_seq</td><td>PAM</td><td>alignment</td><td>distance</td><td>position</td><td>gene name</td><td>gene id</td></tr>
<tr><td>GU305914.1</td><td>163887</td><td>163909</td><td>+</td><td>0</td><td>GGTAGCTCCGGATCGAGCGG</td><td>CGG</td><td>||||||||[||||||||||||]PAM</td><td>NA</td><td>Intergenic</td><td>NA</td><td>NA</td></tr>
<tr><td></td><td></td><td></td><td></td><td></td><td></td><td></td><td></td><td></td><td></td><td></td><td></td></tr>
<tr><td>T12323</td><td>AGCTCCGGATCGAGCGGCGGCGG</td><td>1000</td><td></td><td></td><td></td><td></td><td></td><td></td><td></td><td></td><td></td></tr>
<tr><td>Oligo fwd</td><td>CACCAGCTCCGGATCGAGCGGCGG</td><td></td><td></td><td></td><td></td><td></td><td></td><td></td><td></td><td></td><td></td></tr>
<tr><td>Oligo rev</td><td>AAACCCGCCGCTCGATCCGGAGCT</td><td></td><td></td><td></td><td></td><td></td><td></td><td></td><td></td><td></td><td></td></tr>
<tr><td>Chromosome</td><td>start</td><td>end</td><td>strand</td><td>MM</td><td>target_seq</td><td>PAM</td><td>alignment</td><td>distance</td><td>position</td><td>gene name</td><td>gene id</td></tr>
<tr><td>GU305914.1</td><td>163890</td><td>163912</td><td>+</td><td>0</td><td>AGCTCCGGATCGAGCGGCGG</td><td>CGG</td><td>||||||||[||||||||||||]PAM</td><td>NA</td><td>Intergenic</td><td>NA</td><td>NA</td></tr>
<tr><td></td><td></td><td></td><td></td><td></td><td></td><td></td><td></td><td></td><td></td><td></td><td></td></tr>
<tr><td>T12324</td><td>TCCGGATCGAGCGGCGGCGGCGG</td><td>1000</td><td></td><td></td><td></td><td></td><td></td><td></td><td></td><td></td><td></td></tr>
<tr><td>Oligo fwd</td><td>CACCTCCGGATCGAGCGGCGGCGG</td><td></td><td></td><td></td><td></td><td></td><td></td><td></td><td></td><td></td><td></td></tr>
<tr><td>Oligo rev</td><td>AAACCCGCCGCCGCTCGATCCGGA</td><td></td><td></td><td></td><td></td><td></td><td></td><td></td><td></td><td></td><td></td></tr>
<tr><td>Chromosome</td><td>start</td><td>end</td><td>strand</td><td>MM</td><td>target_seq</td><td>PAM</td><td>alignment</td><td>distance</td><td>position</td><td>gene name</td><td>gene id</td></tr>
<tr><td>GU305914.1</td><td>163893</td><td>163915</td><td>+</td><td>0</td><td>TCCGGATCGAGCGGCGGCGG</td><td>CGG</td><td>||||||||[||||||||||||]PAM</td><td>NA</td><td>Intergenic</td><td>NA</td><td>NA</td></tr>
<tr><td></td><td></td><td></td><td></td><td></td><td></td><td></td><td></td><td></td><td></td><td></td><td></td></tr>
<tr><td>T12325</td><td>GGATCGAGCGGCGGCGGCGGCGG</td><td>1000</td><td></td><td></td><td></td><td></td><td></td><td></td><td></td><td></td><td></td></tr>
<tr><td>Oligo fwd</td><td>CACCGGATCGAGCGGCGGCGGCGG</td><td></td><td></td><td></td><td></td><td></td><td></td><td></td><td></td><td></td><td></td></tr>
<tr><td>Oligo rev</td><td>AAACCCGCCGCCGCCGCTCGATCC</td><td></td><td></td><td></td><td></td><td></td><td></td><td></td><td></td><td></td><td></td></tr>
<tr><td>Chromosome</td><td>start</td><td>end</td><td>strand</td><td>MM</td><td>target_seq</td><td>PAM</td><td>alignment</td><td>distance</td><td>position</td><td>gene name</td><td>gene id</td></tr>
<tr><td>GU305914.1</td><td>163896</td><td>163918</td><td>+</td><td>0</td><td>GGATCGAGCGGCGGCGGCGG</td><td>CGG</td><td>||||||||[||||||||||||]PAM</td><td>NA</td><td>Intergenic</td><td>NA</td><td>NA</td></tr>
<tr><td>GU305914.1</td><td>30000</td><td>30022</td><td>+</td><td>4</td><td>GGTGCTGGCGGCGGCGGCGG</td><td>CGG</td><td>||--|--|[||||||||||||]PAM</td><td>NA</td><td>Intergenic</td><td>NA</td><td>NA</td></tr>
<tr><td>GU305914.1</td><td>164971</td><td>164993</td><td>-</td><td>4</td><td>GGTGGGACCGGCGGCGGCGG</td><td>CGG</td><td>||---||-[||||||||||||]PAM</td><td>NA</td><td>Intergenic</td><td>NA</td><td>NA</td></tr>
<tr><td>GU305914.1</td><td>73792</td><td>73814</td><td>-</td><td>4</td><td>GGCGCCAGCAGCGGCGGCGG</td><td>AGG</td><td>||--|-||[|-||||||||||]PAM</td><td>NA</td><td>Intergenic</td><td>NA</td><td>NA</td></tr>
<tr><td>GU305914.1</td><td>128639</td><td>128661</td><td>+</td><td>3</td><td>GGGCCGAGCGGCAGCGGCGG</td><td>CGG</td><td>||--||||[||||-|||||||]PAM</td><td>NA</td><td>Intergenic</td><td>NA</td><td>NA</td></tr>
<tr><td>GU305914.1</td><td>149488</td><td>149510</td><td>-</td><td>4</td><td>GGAGCAGGCGACGGCGGCGG</td><td>CGG</td><td>|||-|--|[||-|||||||||]PAM</td><td>NA</td><td>Intergenic</td><td>NA</td><td>NA</td></tr>
<tr><td>GU305914.1</td><td>149292</td><td>149314</td><td>+</td><td>4</td><td>GGAGCGGGCGACGACGGCGG</td><td>CGG</td><td>|||-||-|[||-||-||||||]PAM</td><td>NA</td><td>Intergenic</td><td>NA</td><td>NA</td></tr>
<tr><td></td><td></td><td></td><td></td><td></td><td></td><td></td><td></td><td></td><td></td><td></td><td></td></tr>
<tr><td>T12326</td><td>AGCGGCGGCGGCGGCGGCAGCGG</td><td>1000</td><td></td><td></td><td></td><td></td><td></td><td></td><td></td><td></td><td></td></tr>
<tr><td>Oligo fwd</td><td>CACCAGCGGCGGCGGCGGCGGCAG</td><td></td><td></td><td></td><td></td><td></td><td></td><td></td><td></td><td></td><td></td></tr>
<tr><td>Oligo rev</td><td>AAACCTGCCGCCGCCGCCGCCGCT</td><td></td><td></td><td></td><td></td><td></td><td></td><td></td><td></td><td></td><td></td></tr>
<tr><td>Chromosome</td><td>start</td><td>end</td><td>strand</td><td>MM</td><td>target_seq</td><td>PAM</td><td>alignment</td><td>distance</td><td>position</td><td>gene name</td><td>gene id</td></tr>
<tr><td>GU305914.1</td><td>163902</td><td>163924</td><td>+</td><td>0</td><td>AGCGGCGGCGGCGGCGGCAG</td><td>CGG</td><td>||||||||[||||||||||||]PAM</td><td>NA</td><td>Intergenic</td><td>NA</td><td>NA</td></tr>
<tr><td>GU305914.1</td><td>30006</td><td>30028</td><td>+</td><td>1</td><td>GGCGGCGGCGGCGGCGGCAG</td><td>TGG</td><td>-|||||||[||||||||||||]PAM</td><td>NA</td><td>Intergenic</td><td>NA</td><td>NA</td></tr>
<tr><td>GU305914.1</td><td>149476</td><td>149498</td><td>-</td><td>1</td><td>GGCGGCGGCGGCGGCGGCAG</td><td>CGG</td><td>-|||||||[||||||||||||]PAM</td><td>NA</td><td>Intergenic</td><td>NA</td><td>NA</td></tr>
<tr><td>GU305914.1</td><td>113924</td><td>113946</td><td>-</td><td>3</td><td>GTCGGCGTCGGCGGCGGCAG</td><td>CGG</td><td>--|||||-[||||||||||||]PAM</td><td>NA</td><td>Intergenic</td><td>NA</td><td>NA</td></tr>
<tr><td>GU305914.1</td><td>163908</td><td>163930</td><td>+</td><td>2</td><td>GGCGGCGGCGGCAGCGGCAG</td><td>CGG</td><td>-|||||||[||||-|||||||]PAM</td><td>NA</td><td>Intergenic</td><td>NA</td><td>NA</td></tr>
<tr><td>GU305914.1</td><td>166410</td><td>166432</td><td>+</td><td>4</td><td>GACGTCGGCGCCGGCGGCAG</td><td>CGG</td><td>--||-|||[||-|||||||||]PAM</td><td>NA</td><td>Intergenic</td><td>NA</td><td>NA</td></tr>
<tr><td>GU305914.1</td><td>45258</td><td>45280</td><td>-</td><td>3</td><td>AACGGTGGCGGTGGCGGCAG</td><td>CGG</td><td>|-|||-||[|||-||||||||]PAM</td><td>NA</td><td>Intergenic</td><td>NA</td><td>NA</td></tr>
<tr><td>GU305914.1</td><td>65721</td><td>65743</td><td>-</td><td>4</td><td>TCCTGCGGCGGCAGCGGCAG</td><td>CGG</td><td>--|-||||[||||-|||||||]PAM</td><td>NA</td><td>Intergenic</td><td>NA</td><td>NA</td></tr>
<tr><td>GU305914.1</td><td>62152</td><td>62174</td><td>+</td><td>3</td><td>AGGGGCGACGGCAGCGGCAG</td><td>CGG</td><td>||-||||-[||||-|||||||]PAM</td><td>NA</td><td>Intergenic</td><td>NA</td><td>NA</td></tr>
<tr><td>GU305914.1</td><td>168829</td><td>168851</td><td>+</td><td>1</td><td>AGCGGCGGCGGCGGCGACAG</td><td>CGG</td><td>||||||||[||||||||-|||]PAM</td><td>NA</td><td>Intergenic</td><td>NA</td><td>NA</td></tr>
<tr><td>GU305914.1</td><td>62722</td><td>62744</td><td>+</td><td>4</td><td>ATTGGCGGCGGAGACGGCAG</td><td>CGG</td><td>|--|||||[|||-|-||||||]PAM</td><td>NA</td><td>Intergenic</td><td>NA</td><td>NA</td></tr>
<tr><td>GU305914.1</td><td>24557</td><td>24579</td><td>+</td><td>4</td><td>GGCGGCAGCGACGACGGCAG</td><td>CGG</td><td>-|||||-|[||-||-||||||]PAM</td><td>NA</td><td>Intergenic</td><td>NA</td><td>NA</td></tr>
<tr><td>GU305914.1</td><td>163920</td><td>163942</td><td>+</td><td>3</td><td>AGCGGCAGCGGAGGCAGCAG</td><td>CGG</td><td>||||||-|[|||-|||-|||||]PAM</td><td>NA</td><td>Intergenic</td><td>NA</td><td>NA</td></tr>
<tr><td>GU305914.1</td><td>128715</td><td>128737</td><td>-</td><td>4</td><td>AGCAGCAGCGGCGCTGGCAG</td><td>TGG</td><td>|||-||-|[|||||--|||||]PAM</td><td>NA</td><td>Intergenic</td><td>NA</td><td>NA</td></tr>
<tr><td>GU305914.1</td><td>154407</td><td>154429</td><td>+</td><td>3</td><td>GGTGGCGGCGGCGGCGGCGG</td><td>CGG</td><td>-|-|||||[||||||||||-|]PAM</td><td>NA</td><td>Intergenic</td><td>NA</td><td>NA</td></tr>
<tr><td>GU305914.1</td><td>58123</td><td>58145</td><td>-</td><td>3</td><td>GGTGGCGGCGGCGGCGGCGG</td><td>AGG</td><td>-|-|||||[||||||||||-|]PAM</td><td>NA</td><td>Intergenic</td><td>NA</td><td>NA</td></tr>
<tr><td>GU305914.1</td><td>149482</td><td>149504</td><td>-</td><td>3</td><td>GGCGACGGCGGCGGCGGCGG</td><td>CGG</td><td>-|||-|||[||||||||||-|]PAM</td><td>NA</td><td>Intergenic</td><td>NA</td><td>NA</td></tr>
<tr><td>GU305914.1</td><td>154404</td><td>154426</td><td>+</td><td>3</td><td>GGCGGTGGCGGCGGCGGCGG</td><td>CGG</td><td>-||||-||[||||||||||-|]PAM</td><td>NA</td><td>Intergenic</td><td>NA</td><td>NA</td></tr>
<tr><td>GU305914.1</td><td>58126</td><td>58148</td><td>-</td><td>3</td><td>GGCGGTGGCGGCGGCGGCGG</td><td>CGG</td><td>-||||-||[||||||||||-|]PAM</td><td>NA</td><td>Intergenic</td><td>NA</td><td>NA</td></tr>
<tr><td>GU305914.1</td><td>67742</td><td>67764</td><td>+</td><td>4</td><td>GGGAGCGGCGGCGGCGGCGG</td><td>CGG</td><td>-|--||||[||||||||||-|]PAM</td><td>NA</td><td>Intergenic</td><td>NA</td><td>NA</td></tr>
<tr><td>GU305914.1</td><td>35653</td><td>35675</td><td>+</td><td>4</td><td>ATCCACGGCGGCGGCGGCGG</td><td>CGG</td><td>|-|--|||[||||||||||-|]PAM</td><td>NA</td><td>Intergenic</td><td>NA</td><td>NA</td></tr>
<tr><td>GU305914.1</td><td>48159</td><td>48181</td><td>+</td><td>3</td><td>GGCGGCGGTGGCGGCGGCGG</td><td>TGG</td><td>-|||||||[-|||||||||-|]PAM</td><td>NA</td><td>Intergenic</td><td>NA</td><td>NA</td></tr>
<tr><td>GU305914.1</td><td>56773</td><td>56795</td><td>+</td><td>4</td><td>CGCCGCAGCGGCGGCGGCGG</td><td>CGG</td><td>-||-||-|[||||||||||-|]PAM</td><td>NA</td><td>Intergenic</td><td>NA</td><td>NA</td></tr>
<tr><td>GU305914.1</td><td>48156</td><td>48178</td><td>+</td><td>3</td><td>GGCGGCGGCGGTGGCGGCGG</td><td>CGG</td><td>-|||||||[|||-||||||-|]PAM</td><td>NA</td><td>Intergenic</td><td>NA</td><td>NA</td></tr>
<tr><td>GU305914.1</td><td>71753</td><td>71775</td><td>+</td><td>3</td><td>AGCAGCAGCGGCGGCGGCAA</td><td>CGG</td><td>|||-||-|[|||||||||||-]PAM</td><td>NA</td><td>Intergenic</td><td>NA</td><td>NA</td></tr>
<tr><td>GU305914.1</td><td>48153</td><td>48175</td><td>+</td><td>4</td><td>TGTGGCGGCGGCGGTGGCGG</td><td>CGG</td><td>-|-|||||[||||||-|||-|]PAM</td><td>NA</td><td>Intergenic</td><td>NA</td><td>NA</td></tr>
</table>

T12327 GGCGGCGGCGGCAGCGGCAGCGG 1000
Oligo fwd CACCGGCGGCGGCGGCAGCGGCAG
Oligo rev AAACCTGCCGCTGCCGCCGCCGCC

<table>
<tr><th>Chromosome</th><th>start</th><th>end</th><th>strand</th><th>MM</th><th>target_seq</th><th>PAM</th><th>alignment</th><th>distance</th><th>position</th><th>gene name</th><th>gene id</th></tr>
<tr><td>GU305914.1</td><td>163908</td><td>163930</td><td>+</td><td>0</td><td>GGCGGCGGCGGCAGCGGCAG</td><td>CGG</td><td>||||||||[||||||||||||]PAM</td><td>NA</td><td>Intergenic</td><td>NA</td><td>NA</td></tr>
<tr><td>GU305914.1</td><td>65721</td><td>65743</td><td>-</td><td>3</td><td>TCCTGCGGCGGCAGCGGCAG</td><td>CGG</td><td>--|-||||[||||||||||||]PAM</td><td>NA</td><td>Intergenic</td><td>NA</td><td>NA</td></tr>
<tr><td>GU305914.1</td><td>62152</td><td>62174</td><td>+</td><td>3</td><td>AGGGGCGACGGCAGCGGCAG</td><td>CGG</td><td>-|-||||-[||||||||||||]PAM</td><td>NA</td><td>Intergenic</td><td>NA</td><td>NA</td></tr>
<tr><td>GU305914.1</td><td>128981</td><td>129003</td><td>+</td><td>4</td><td>GCAGCCGTCGGCAGCGGCAG</td><td>CGG</td><td>|--|-||-[||||||||||||]PAM</td><td>NA</td><td>Intergenic</td><td>NA</td><td>NA</td></tr>
<tr><td>GU305914.1</td><td>30006</td><td>30028</td><td>+</td><td>1</td><td>GGCGGCGGCGGCGGCGGCAG</td><td>TGG</td><td>||||||||[||||-|||||||]PAM</td><td>NA</td><td>Intergenic</td><td>NA</td><td>NA</td></tr>
<tr><td>GU305914.1</td><td>149476</td><td>149498</td><td>-</td><td>1</td><td>GGCGGCGGCGGCGGCGGCAG</td><td>CGG</td><td>||||||||[||||-|||||||]PAM</td><td>NA</td><td>Intergenic</td><td>NA</td><td>NA</td></tr>
<tr><td>GU305914.1</td><td>163902</td><td>163924</td><td>+</td><td>2</td><td>AGCGGCGGCGGCGGCGGCAG</td><td>CGG</td><td>-|||||||[||||-|||||||]PAM</td><td>NA</td><td>Intergenic</td><td>NA</td><td>NA</td></tr>
<tr><td>GU305914.1</td><td>113924</td><td>113946</td><td>-</td><td>3</td><td>GTCGGCGTCGGCGGCGGCAG</td><td>CGG</td><td>|-|||||-[||||-|||||||]PAM</td><td>NA</td><td>Intergenic</td><td>NA</td><td>NA</td></tr>
<tr><td>GU305914.1</td><td>166410</td><td>166432</td><td>+</td><td>4</td><td>GACGTCGGCGCCGGCGGCAG</td><td>CGG</td><td>|-||-|||[||-|-|||||||]PAM</td><td>NA</td><td>Intergenic</td><td>NA</td><td>NA</td></tr>
<tr><td>GU305914.1</td><td>128727</td><td>128749</td><td>-</td><td>3</td><td>GGTGGAGGCGGCAGCAGCAG</td><td>CGG</td><td>||-||-||[|||||||-||||]PAM</td><td>NA</td><td>Intergenic</td><td>NA</td><td>NA</td></tr>
<tr><td>GU305914.1</td><td>27408</td><td>27430</td><td>+</td><td>4</td><td>GGCAGCAGCAGCAGCAGCAG</td><td>CGG</td><td>|||-||-|[|-|||||-||||]PAM</td><td>NA</td><td>Intergenic</td><td>NA</td><td>NA</td></tr>
<tr><td>GU305914.1</td><td>168829</td><td>168851</td><td>+</td><td>3</td><td>AGCGGCGGCGGCGGCGACAG</td><td>CGG</td><td>-|||||||[||||-|||-|||]PAM</td><td>NA</td><td>Intergenic</td><td>NA</td><td>NA</td></tr>
<tr><td>GU305914.1</td><td>168817</td><td>168839</td><td>+</td><td>3</td><td>GGCGGTGGTGGCAGCGGCGG</td><td>CGG</td><td>|||||-||[-|||||||||-|]PAM</td><td>NA</td><td>Intergenic</td><td>NA</td><td>NA</td></tr>
<tr><td>GU305914.1</td><td>71759</td><td>71781</td><td>+</td><td>3</td><td>AGCGGCGGCGGCAACGGGAG</td><td>CGG</td><td>-|||||||[|||||-|||-||]PAM</td><td>NA</td><td>Intergenic</td><td>NA</td><td>NA</td></tr>
<tr><td>GU305914.1</td><td>56692</td><td>56714</td><td>+</td><td>4</td><td>GGATGCGGTGGCAGCGGCCG</td><td>CGG</td><td>||--||||[-|||||||||-|]PAM</td><td>NA</td><td>Intergenic</td><td>NA</td><td>NA</td></tr>
<tr><td>GU305914.1</td><td>154407</td><td>154429</td><td>+</td><td>3</td><td>GGTGGCGGCGGCGGCGGCGG</td><td>CGG</td><td>||-|||||[||||-|||||-|]PAM</td><td>NA</td><td>Intergenic</td><td>NA</td><td>NA</td></tr>
<tr><td>GU305914.1</td><td>58123</td><td>58145</td><td>-</td><td>3</td><td>GGTGGCGGCGGCGGCGGCGG</td><td>AGG</td><td>||-|||||[||||-|||||-|]PAM</td><td>NA</td><td>Intergenic</td><td>NA</td><td>NA</td></tr>
<tr><td>GU305914.1</td><td>149482</td><td>149504</td><td>-</td><td>3</td><td>GGCGACGGCGGCGGCGGCGG</td><td>CGG</td><td>||||-|||[||||-|||||-|]PAM</td><td>NA</td><td>Intergenic</td><td>NA</td><td>NA</td></tr>
<tr><td>GU305914.1</td><td>154404</td><td>154426</td><td>+</td><td>3</td><td>GGCGGTGGCGGCGGCGGCGG</td><td>CGG</td><td>|||||-||[||||-|||||-|]PAM</td><td>NA</td><td>Intergenic</td><td>NA</td><td>NA</td></tr>
<tr><td>GU305914.1</td><td>58126</td><td>58148</td><td>-</td><td>3</td><td>GGCGGTGGCGGCGGCGGCGG</td><td>CGG</td><td>|||||-||[||||-|||||-|]PAM</td><td>NA</td><td>Intergenic</td><td>NA</td><td>NA</td></tr>
<tr><td>GU305914.1</td><td>67742</td><td>67764</td><td>+</td><td>4</td><td>GGGAGCGGCGGCGGCGGCGG</td><td>CGG</td><td>||--||||[||||-|||||-|]PAM</td><td>NA</td><td>Intergenic</td><td>NA</td><td>NA</td></tr>
<tr><td>GU305914.1</td><td>164959</td><td>164981</td><td>-</td><td>2</td><td>GGCGGCGGCGGCAGCAGCTG</td><td>CGG</td><td>||||||||[|||||||-||-|]PAM</td><td>NA</td><td>Intergenic</td><td>NA</td><td>NA</td></tr>
<tr><td>GU305914.1</td><td>113918</td><td>113940</td><td>-</td><td>3</td><td>GTCGGCGGCGGCAGCGGCGA</td><td>CGG</td><td>|-||||||[||||||||||--]PAM</td><td>NA</td><td>Intergenic</td><td>NA</td><td>NA</td></tr>
</table>

T12328 GGCGGCGGCAGCGGCAGCGGAGG 1000
Oligo fwd CACCGGCGGCGGCAGCGGCAGCGG
Oligo rev AAACCCGCTGCCGCTGCCGCCGCC

<table>
<tr><th>Chromosome</th><th>start</th><th>end</th><th>strand</th><th>MM</th><th>target_seq</th><th>PAM</th><th>alignment</th><th>distance</th><th>position</th><th>gene name</th><th>gene id</th></tr>
<tr><td>GU305914.1</td><td>163911</td><td>163933</td><td>+</td><td>0</td><td>GGCGGCGGCAGCGGCAGCGG</td><td>AGG</td><td>||||||||[||||||||||||]PAM</td><td>NA</td><td>Intergenic</td><td>NA</td><td>NA</td></tr>
<tr><td>GU305914.1</td><td>62155</td><td>62177</td><td>+</td><td>1</td><td>GGCGACGGCAGCGGCAGCGG</td><td>CGG</td><td>||||-|||[||||||||||||]PAM</td><td>NA</td><td>Intergenic</td><td>NA</td><td>NA</td></tr>
<tr><td>GU305914.1</td><td>45255</td><td>45277</td><td>-</td><td>3</td><td>GGTGGCGGTGGCGGCAGCGG</td><td>CGG</td><td>||-|||||[--||||||||||]PAM</td><td>NA</td><td>Intergenic</td><td>NA</td><td>NA</td></tr>
<tr><td>GU305914.1</td><td>27411</td><td>27433</td><td>+</td><td>4</td><td>AGCAGCAGCAGCAGCAGCGG</td><td>CGG</td><td>-||-||-|[||||-|||||||]PAM</td><td>NA</td><td>Intergenic</td><td>NA</td><td>NA</td></tr>
<tr><td>GU305914.1</td><td>168820</td><td>168842</td><td>+</td><td>3</td><td>GGTGGTGGCAGCGGCGGCGG</td><td>CGG</td><td>||-||-||[|||||||-||||]PAM</td><td>NA</td><td>Intergenic</td><td>NA</td><td>NA</td></tr>
<tr><td>GU305914.1</td><td>73792</td><td>73814</td><td>-</td><td>3</td><td>GGCGCCAGCAGCGGCGGCGG</td><td>AGG</td><td>||||-|-|[|||||||-||||]PAM</td><td>NA</td><td>Intergenic</td><td>NA</td><td>NA</td></tr>
<tr><td>GU305914.1</td><td>154407</td><td>154429</td><td>+</td><td>3</td><td>GGTGGCGGCGGCGGCGGCGG</td><td>CGG</td><td>||-|||||[|-|||||-||||]PAM</td><td>NA</td><td>Intergenic</td><td>NA</td><td>NA</td></tr>
<tr><td>GU305914.1</td><td>58123</td><td>58145</td><td>-</td><td>3</td><td>GGTGGCGGCGGCGGCGGCGG</td><td>AGG</td><td>||-|||||[|-|||||-||||]PAM</td><td>NA</td><td>Intergenic</td><td>NA</td><td>NA</td></tr>
<tr><td>GU305914.1</td><td>149482</td><td>149504</td><td>-</td><td>3</td><td>GGCGACGGCGGCGGCGGCGG</td><td>CGG</td><td>||||-|||[|-|||||-||||]PAM</td><td>NA</td><td>Intergenic</td><td>NA</td><td>NA</td></tr>
<tr><td>GU305914.1</td><td>154404</td><td>154426</td><td>+</td><td>3</td><td>GGCGGTGGCGGCGGCGGCGG</td><td>CGG</td><td>|||||-||[|-|||||-||||]PAM</td><td>NA</td><td>Intergenic</td><td>NA</td><td>NA</td></tr>
<tr><td>GU305914.1</td><td>58126</td><td>58148</td><td>-</td><td>3</td><td>GGCGGTGGCGGCGGCGGCGG</td><td>CGG</td><td>|||||-||[|-|||||-||||]PAM</td><td>NA</td><td>Intergenic</td><td>NA</td><td>NA</td></tr>
<tr><td>GU305914.1</td><td>67742</td><td>67764</td><td>+</td><td>4</td><td>GGGAGCGGCGGCGGCGGCGG</td><td>CGG</td><td>||--||||[|-|||||-||||]PAM</td><td>NA</td><td>Intergenic</td><td>NA</td><td>NA</td></tr>
<tr><td>GU305914.1</td><td>62542</td><td>62564</td><td>+</td><td>4</td><td>GACGGCAGGAGCGGCCGCGG</td><td>CGG</td><td>|-||||-|[-||||||-||||]PAM</td><td>NA</td><td>Intergenic</td><td>NA</td><td>NA</td></tr>
<tr><td>GU305914.1</td><td>27408</td><td>27430</td><td>+</td><td>4</td><td>GGCAGCAGCAGCAGCAGCAG</td><td>CGG</td><td>|||-||-|[||||-|||||-|]PAM</td><td>NA</td><td>Intergenic</td><td>NA</td><td>NA</td></tr>
</table>

T12329 AGCGGCAGCGGAGGCAGCAGCGG 1000
Oligo fwd CACCAGCGGCAGCGGAGGCAGCAG
Oligo rev AAACCTGCTGCCTCCGCTGCCGCT

<table>
<tr><th>Chromosome</th><th>start</th><th>end</th><th>strand</th><th>MM</th><th>target_seq</th><th>PAM</th><th>alignment</th><th>distance</th><th>position</th><th>gene name</th><th>gene id</th></tr>
<tr><td>GU305914.1</td><td>163920</td><td>163942</td><td>+</td><td>0</td><td>AGCGGCAGCGGAGGCAGCAG</td><td>CGG</td><td>||||||||[||||||||||||]PAM</td><td>NA</td><td>Intergenic</td><td>NA</td><td>NA</td></tr>
<tr><td>GU305914.1</td><td>163902</td><td>163924</td><td>+</td><td>3</td><td>AGCGGCGGCGGCGGCGGCAG</td><td>CGG</td><td>||||||-|[|||-|||-||||]PAM</td><td>NA</td><td>Intergenic</td><td>NA</td><td>NA</td></tr>
<tr><td>GU305914.1</td><td>30006</td><td>30028</td><td>+</td><td>4</td><td>GGCGGCGGCGGCGGCGGCAG</td><td>TGG</td><td>-|||||-|[|||-|||-||||]PAM</td><td>NA</td><td>Intergenic</td><td>NA</td><td>NA</td></tr>
<tr><td>GU305914.1</td><td>149476</td><td>149498</td><td>-</td><td>4</td><td>GGCGGCGGCGGCGGCGGCAG</td><td>CGG</td><td>-|||||-|[|||-|||-||||]PAM</td><td>NA</td><td>Intergenic</td><td>NA</td><td>NA</td></tr>
</table>

T12330 CGGCCTCAGTACAAAGCAGAAGG 1000
Oligo fwd CACCCGGCCTCAGTACAAAGCAGA
Oligo rev AAACTCTGCTTTGTACTGAGGCCG

<table>
<tr><th>Chromosome</th><th>start</th><th>end</th><th>strand</th><th>MM</th><th>target_seq</th><th>PAM</th><th>alignment</th><th>distance</th><th>position</th><th>gene name</th><th>gene id</th></tr>
<tr><td>GU305914.1</td><td>163940</td><td>163962</td><td>+</td><td>0</td><td>CGGCCTCAGTACAAAGCAGA</td><td>AGG</td><td>||||||||[||||||||||||]PAM</td><td>NA</td><td>Intergenic</td><td>NA</td><td>NA</td></tr>
</table>

T12331 CAAAGCAGAAGGAGCGCAGCAGG 1000
Oligo fwd CACCCAAAGCAGAAGGAGCGCAGC
Oligo rev AAACGCTGCGCTCCTTCTGCTTTG

<table>
<tr><th>Chromosome</th><th>start</th><th>end</th><th>strand</th><th>MM</th><th>target_seq</th><th>PAM</th><th>alignment</th><th>distance</th><th>position</th><th>gene name</th><th>gene id</th></tr>
<tr><td>GU305914.1</td><td>163951</td><td>163973</td><td>+</td><td>0</td><td>CAAAGCAGAAGGAGCGCAGC</td><td>AGG</td><td>||||||||[||||||||||||]PAM</td><td>NA</td><td>Intergenic</td><td>NA</td><td>NA</td></tr>
</table>

T12332 AGCAGAAGGAGCGCAGCAGGAGG 1000
Oligo fwd CACCAGCAGAAGGAGCGCAGCAGG
Oligo rev AAACCCTGCTGCGCTCCTTCTGCT

<table>
<tr><th>Chromosome</th><th>start</th><th>end</th><th>strand</th><th>MM</th><th>target_seq</th><th>PAM</th><th>alignment</th><th>distance</th><th>position</th><th>gene name</th><th>gene id</th></tr>
<tr><td>GU305914.1</td><td>163954</td><td>163976</td><td>+</td><td>0</td><td>AGCAGAAGGAGCGCAGCAGG</td><td>AGG</td><td>||||||||[||||||||||||]PAM</td><td>NA</td><td>Intergenic</td><td>NA</td><td>NA</td></tr>
</table>

T12333 GCGCAGCAGGAGGCTCGAAGAGG 1000
Oligo fwd CACCGCGCAGCAGGAGGCTCGAAG
Oligo rev AAACCTTCGAGCCTCCTGCTGCGC

<table>
<tr><th>Chromosome</th><th>start</th><th>end</th><th>strand</th><th>MM</th><th>target_seq</th><th>PAM</th><th>alignment</th><th>distance</th><th>position</th><th>gene name</th><th>gene id</th></tr>
<tr><td>GU305914.1</td><td>163964</td><td>163986</td><td>+</td><td>0</td><td>GCGCAGCAGGAGGCTCGAAG</td><td>AGG</td><td>||||||||[||||||||||||]PAM</td><td>NA</td><td>Intergenic</td><td>NA</td><td>NA</td></tr>
</table>

T12334 GGAATGTAAGTAGTTCTGATCGG 1000
Oligo fwd CACCGGAATGTAAGTAGTTCTGAT
Oligo rev AAACATCAGAACTACTTACATTCC

<table>
<tr><th>Chromosome</th><th>start</th><th>end</th><th>strand</th><th>MM</th><th>target_seq</th><th>PAM</th><th>alignment</th><th>distance</th><th>position</th><th>gene name</th><th>gene id</th></tr>
<tr><td>GU305914.1</td><td>163985</td><td>164007</td><td>+</td><td>0</td><td>GGAATGTAAGTAGTTCTGAT</td><td>CGG</td><td>||||||||[||||||||||||]PAM</td><td>NA</td><td>Intergenic</td><td>NA</td><td>NA</td></tr>
</table>

T12335 GAATGTAAGTAGTTCTGATCGGG 1000
Oligo fwd CACCGAATGTAAGTAGTTCTGATC

Oligo rev AAACGATCAGAACTACTTACATTC
Chromosome start end strand MM target_seq PAM alignment distance position gene name gene id
GU305914.1 163986 164008 + 0 GAATGTAAGTAGTTCTGATC GGG ||||||||[||||||||||||]PAM NA Intergenic NA NA

T12336 AATGTAAGTAGTTCTGATCGGGG 1000
Oligo fwd CACCAATGTAAGTAGTTCTGATCG
Oligo rev AAACCGATCAGAACTACTTACATT
Chromosome start end strand MM target_seq PAM alignment distance position gene name gene id
GU305914.1 163987 164009 + 0 AATGTAAGTAGTTCTGATCG GGG ||||||||[||||||||||||]PAM NA Intergenic NA NA

T12337 AGTTCTGATCGGGGTTTTACTGG 1000
Oligo fwd CACCAGTTCTGATCGGGGTTTTAC
Oligo rev AAACGTAAAACCCCGATCAGAACT
Chromosome start end strand MM target_seq PAM alignment distance position gene name gene id
GU305914.1 163996 164018 + 0 AGTTCTGATCGGGGTTTTAC TGG ||||||||[||||||||||||]PAM NA Intergenic NA NA

T12338 TCGCTTTCCCCCTCGACCTTCGG 1000
Oligo fwd CACCTCGCTTTCCCCCTCGACCTT
Oligo rev AAACAAGGTCGAGGGGGAAAGCGA
Chromosome start end strand MM target_seq PAM alignment distance position gene name gene id
GU305914.1 164019 164041 + 0 TCGCTTTCCCCCTCGACCTT CGG ||||||||[||||||||||||]PAM NA Intergenic NA NA

T12339 CGACCTTCGGTGAACCGTCACGG 1000
Oligo fwd CACCCGACCTTCGGTGAACCGTCA
Oligo rev AAACTGACGGTTCACCGAAGGTCG
Chromosome start end strand MM target_seq PAM alignment distance position gene name gene id
GU305914.1 164032 164054 + 0 CGACCTTCGGTGAACCGTCA CGG ||||||||[||||||||||||]PAM NA Intergenic NA NA

T12340 GACCTTCGGTGAACCGTCACGGG 1000
Oligo fwd CACCGACCTTCGGTGAACCGTCAC
Oligo rev AAACGTGACGGTTCACCGAAGGTC
Chromosome start end strand MM target_seq PAM alignment distance position gene name gene id
GU305914.1 164033 164055 + 0 GACCTTCGGTGAACCGTCAC GGG ||||||||[||||||||||||]PAM NA Intergenic NA NA

T12341 ACTAATGATGTTACTCTGTTCGG 1000
Oligo fwd CACCACTAATGATGTTACTCTGTT
Oligo rev AAACAACAGAGTAACATCATTAGT
Chromosome start end strand MM target_seq PAM alignment distance position gene name gene id
GU305914.1 164056 164078 + 0 ACTAATGATGTTACTCTGTT CGG ||||||||[||||||||||||]PAM NA Intergenic NA NA

T12342 TGATGTTACTCTGTTCGGAAAGG 1000
Oligo fwd CACCTGATGTTACTCTGTTCGGAA
Oligo rev AAACTTCCGAACAGAGTAACATCA
Chromosome start end strand MM target_seq PAM alignment distance position gene name gene id
GU305914.1 164061 164083 + 0 TGATGTTACTCTGTTCGGAA AGG ||||||||[||||||||||||]PAM NA Intergenic NA NA

T12343 CGGAAAGGTTCTCCACGCAGCGG 1000
Oligo fwd CACCCGGAAAGGTTCTCCACGCAG
Oligo rev AAACCTGCGTGGAGAACCTTTCCG
Chromosome start end strand MM target_seq PAM alignment distance position gene name gene id
GU305914.1 164076 164098 + 0 CGGAAAGGTTCTCCACGCAG CGG ||||||||[||||||||||||]PAM NA Intergenic NA NA

T12344 GGAAAGGTTCTCCACGCAGCGGG 1000
Oligo fwd CACCGGAAAGGTTCTCCACGCAGC
Oligo rev AAACGCTGCGTGGAGAACCTTTCC
Chromosome start end strand MM target_seq PAM alignment distance position gene name gene id
GU305914.1 164077 164099 + 0 GGAAAGGTTCTCCACGCAGC GGG ||||||||[||||||||||||]PAM NA Intergenic NA NA

T12345 GAAAGGTTCTCCACGCAGCGGGG 1000
Oligo fwd CACCGAAAGGTTCTCCACGCAGCG
Oligo rev AAACCGCTGCGTGGAGAACCTTTC
Chromosome start end strand MM target_seq PAM alignment distance position gene name gene id
GU305914.1 164078 164100 + 0 GAAAGGTTCTCCACGCAGCG GGG ||||||||[||||||||||||]PAM NA Intergenic NA NA

T12346 AAAGGTTCTCCACGCAGCGGGGG 1000
Oligo fwd CACCAAAGGTTCTCCACGCAGCGG
Oligo rev AAACCCGCTGCGTGGAGAACCTTT
Chromosome start end strand MM target_seq PAM alignment distance position gene name gene id
GU305914.1 164079 164101 + 0 AAAGGTTCTCCACGCAGCGG GGG ||||||||[||||||||||||]PAM NA Intergenic NA NA

T12347 AAGGTTCTCCACGCAGCGGGGGG 1000
Oligo fwd CACCAAGGTTCTCCACGCAGCGGG
Oligo rev AAACCCCGCTGCGTGGAGAACCTT
Chromosome start end strand MM target_seq PAM alignment distance position gene name gene id
GU305914.1 164080 164102 + 0 AAGGTTCTCCACGCAGCGGG GGG ||||||||[||||||||||||]PAM NA Intergenic NA NA

T12348 AGGTTCTCCACGCAGCGGGGGGG 1000
Oligo fwd CACCAGGTTCTCCACGCAGCGGGG
Oligo rev AAACCCCCGCTGCGTGGAGAACCT
Chromosome start end strand MM target_seq PAM alignment distance position gene name gene id
GU305914.1 164081 164103 + 0 AGGTTCTCCACGCAGCGGGG GGG ||||||||[||||||||||||]PAM NA Intergenic NA NA

T12349 GACCCACCACGATCCGAGACCGG 1000
Oligo fwd CACCGACCCACCACGATCCGAGAC
Oligo rev AAACGTCTCGGATCGTGGTGGGTC
Chromosome start end strand MM target_seq PAM alignment distance position gene name gene id
GU305914.1 164120 164142 + 0 GACCCACCACGATCCGAGAC CGG ||||||||[||||||||||||]PAM NA Intergenic NA NA

T12350 ACCACGATCCGAGACCGGAGAGG 1000

<table>
<tr><td>Oligo fwd</td><td>CACCACCACGATCCGAGACCGGAG</td><td></td><td></td><td></td><td></td><td></td><td></td><td></td><td></td><td></td><td></td></tr>
<tr><td>Oligo rev</td><td>AAACCTCCGGTCTCGGATCGTGGT</td><td></td><td></td><td></td><td></td><td></td><td></td><td></td><td></td><td></td><td></td></tr>
<tr><td>Chromosome</td><td>start</td><td>end</td><td>strand</td><td>MM</td><td>target_seq</td><td>PAM</td><td>alignment</td><td>distance</td><td>position</td><td>gene name</td><td>gene id</td></tr>
<tr><td>GU305914.1</td><td>164125</td><td>164147</td><td>+</td><td>0</td><td>ACCACGATCCGAGACCGGAG</td><td>AGG</td><td>||||||||[||||||||||||]PAM</td><td>NA</td><td>Intergenic</td><td>NA</td><td>NA</td></tr>
<tr><td>T12351</td><td>CCACGATCCGAGACCGGAGAGGG</td><td>1000</td><td></td><td></td><td></td><td></td><td></td><td></td><td></td><td></td><td></td></tr>
<tr><td>Oligo fwd</td><td>CACCCCACGATCCGAGACCGGAGA</td><td></td><td></td><td></td><td></td><td></td><td></td><td></td><td></td><td></td><td></td></tr>
<tr><td>Oligo rev</td><td>AAACTCTCCGGTCTCGGATCGTGG</td><td></td><td></td><td></td><td></td><td></td><td></td><td></td><td></td><td></td><td></td></tr>
<tr><td>Chromosome</td><td>start</td><td>end</td><td>strand</td><td>MM</td><td>target_seq</td><td>PAM</td><td>alignment</td><td>distance</td><td>position</td><td>gene name</td><td>gene id</td></tr>
<tr><td>GU305914.1</td><td>164126</td><td>164148</td><td>+</td><td>0</td><td>CCACGATCCGAGACCGGAGA</td><td>GGG</td><td>||||||||[||||||||||||]PAM</td><td>NA</td><td>Intergenic</td><td>NA</td><td>NA</td></tr>
<tr><td>T12352</td><td>TCCGAGACCGGAGAGGGATCTGG</td><td>1000</td><td></td><td></td><td></td><td></td><td></td><td></td><td></td><td></td><td></td></tr>
<tr><td>Oligo fwd</td><td>CACCTCCGAGACCGGAGAGGGATC</td><td></td><td></td><td></td><td></td><td></td><td></td><td></td><td></td><td></td><td></td></tr>
<tr><td>Oligo rev</td><td>AAACGATCCCTCTCCGGTCTCGGA</td><td></td><td></td><td></td><td></td><td></td><td></td><td></td><td></td><td></td><td></td></tr>
<tr><td>Chromosome</td><td>start</td><td>end</td><td>strand</td><td>MM</td><td>target_seq</td><td>PAM</td><td>alignment</td><td>distance</td><td>position</td><td>gene name</td><td>gene id</td></tr>
<tr><td>GU305914.1</td><td>164132</td><td>164154</td><td>+</td><td>0</td><td>TCCGAGACCGGAGAGGGATC</td><td>TGG</td><td>||||||||[||||||||||||]PAM</td><td>NA</td><td>Intergenic</td><td>NA</td><td>NA</td></tr>
<tr><td>T12353</td><td>AGAGGGATCTGGAACCGCCTAGG</td><td>1000</td><td></td><td></td><td></td><td></td><td></td><td></td><td></td><td></td><td></td></tr>
<tr><td>Oligo fwd</td><td>CACCAGAGGGATCTGGAACCGCCT</td><td></td><td></td><td></td><td></td><td></td><td></td><td></td><td></td><td></td><td></td></tr>
<tr><td>Oligo rev</td><td>AAACAGGCGGTTCCAGATCCCTCT</td><td></td><td></td><td></td><td></td><td></td><td></td><td></td><td></td><td></td><td></td></tr>
<tr><td>Chromosome</td><td>start</td><td>end</td><td>strand</td><td>MM</td><td>target_seq</td><td>PAM</td><td>alignment</td><td>distance</td><td>position</td><td>gene name</td><td>gene id</td></tr>
<tr><td>GU305914.1</td><td>164143</td><td>164165</td><td>+</td><td>0</td><td>AGAGGGATCTGGAACCGCCT</td><td>AGG</td><td>||||||||[||||||||||||]PAM</td><td>NA</td><td>Intergenic</td><td>NA</td><td>NA</td></tr>
<tr><td>T12354</td><td>AGGAGCAGTACCACTGTCGATGG</td><td>1000</td><td></td><td></td><td></td><td></td><td></td><td></td><td></td><td></td><td></td></tr>
<tr><td>Oligo fwd</td><td>CACCAGGAGCAGTACCACTGTCGA</td><td></td><td></td><td></td><td></td><td></td><td></td><td></td><td></td><td></td><td></td></tr>
<tr><td>Oligo rev</td><td>AAACTCGACAGTGGTACTGCTCCT</td><td></td><td></td><td></td><td></td><td></td><td></td><td></td><td></td><td></td><td></td></tr>
<tr><td>Chromosome</td><td>start</td><td>end</td><td>strand</td><td>MM</td><td>target_seq</td><td>PAM</td><td>alignment</td><td>distance</td><td>position</td><td>gene name</td><td>gene id</td></tr>
<tr><td>GU305914.1</td><td>164163</td><td>164185</td><td>+</td><td>0</td><td>AGGAGCAGTACCACTGTCGA</td><td>TGG</td><td>||||||||[||||||||||||]PAM</td><td>NA</td><td>Intergenic</td><td>NA</td><td>NA</td></tr>
<tr><td>T12355</td><td>GATGGTAATGCAGACGAAGCTGG</td><td>1000</td><td></td><td></td><td></td><td></td><td></td><td></td><td></td><td></td><td></td></tr>
<tr><td>Oligo fwd</td><td>CACCGATGGTAATGCAGACGAAGC</td><td></td><td></td><td></td><td></td><td></td><td></td><td></td><td></td><td></td><td></td></tr>
<tr><td>Oligo rev</td><td>AAACGCTTCGTCTGCATTACCATC</td><td></td><td></td><td></td><td></td><td></td><td></td><td></td><td></td><td></td><td></td></tr>
<tr><td>Chromosome</td><td>start</td><td>end</td><td>strand</td><td>MM</td><td>target_seq</td><td>PAM</td><td>alignment</td><td>distance</td><td>position</td><td>gene name</td><td>gene id</td></tr>
<tr><td>GU305914.1</td><td>164181</td><td>164203</td><td>+</td><td>0</td><td>GATGGTAATGCAGACGAAGC</td><td>TGG</td><td>||||||||[||||||||||||]PAM</td><td>NA</td><td>Intergenic</td><td>NA</td><td>NA</td></tr>
<tr><td>T12356</td><td>TAATGCAGACGAAGCTGGAGCGG</td><td>1000</td><td></td><td></td><td></td><td></td><td></td><td></td><td></td><td></td><td></td></tr>
<tr><td>Oligo fwd</td><td>CACCTAATGCAGACGAAGCTGGAG</td><td></td><td></td><td></td><td></td><td></td><td></td><td></td><td></td><td></td><td></td></tr>
<tr><td>Oligo rev</td><td>AAACCTCCAGCTTCGTCTGCATTA</td><td></td><td></td><td></td><td></td><td></td><td></td><td></td><td></td><td></td><td></td></tr>
<tr><td>Chromosome</td><td>start</td><td>end</td><td>strand</td><td>MM</td><td>target_seq</td><td>PAM</td><td>alignment</td><td>distance</td><td>position</td><td>gene name</td><td>gene id</td></tr>
<tr><td>GU305914.1</td><td>164186</td><td>164208</td><td>+</td><td>0</td><td>TAATGCAGACGAAGCTGGAG</td><td>CGG</td><td>||||||||[||||||||||||]PAM</td><td>NA</td><td>Intergenic</td><td>NA</td><td>NA</td></tr>
<tr><td>T12357</td><td>TCGCTGTCCGCCGTCAGCGATGG</td><td>1000</td><td></td><td></td><td></td><td></td><td></td><td></td><td></td><td></td><td></td></tr>
<tr><td>Oligo fwd</td><td>CACCTCGCTGTCCGCCGTCAGCGA</td><td></td><td></td><td></td><td></td><td></td><td></td><td></td><td></td><td></td><td></td></tr>
<tr><td>Oligo rev</td><td>AAACTCGCTGACGGCGGACAGCGA</td><td></td><td></td><td></td><td></td><td></td><td></td><td></td><td></td><td></td><td></td></tr>
<tr><td>Chromosome</td><td>start</td><td>end</td><td>strand</td><td>MM</td><td>target_seq</td><td>PAM</td><td>alignment</td><td>distance</td><td>position</td><td>gene name</td><td>gene id</td></tr>
<tr><td>GU305914.1</td><td>164238</td><td>164260</td><td>+</td><td>0</td><td>TCGCTGTCCGCCGTCAGCGA</td><td>TGG</td><td>||||||||[||||||||||||]PAM</td><td>NA</td><td>Intergenic</td><td>NA</td><td>NA</td></tr>
<tr><td>T12358</td><td>TGCGTTACCACCCTCCTCTCCGG</td><td>1000</td><td></td><td></td><td></td><td></td><td></td><td></td><td></td><td></td><td></td></tr>
<tr><td>Oligo fwd</td><td>CACCTGCGTTACCACCCTCCTCTC</td><td></td><td></td><td></td><td></td><td></td><td></td><td></td><td></td><td></td><td></td></tr>
<tr><td>Oligo rev</td><td>AAACGAGAGGAGGGTGGTAACGCA</td><td></td><td></td><td></td><td></td><td></td><td></td><td></td><td></td><td></td><td></td></tr>
<tr><td>Chromosome</td><td>start</td><td>end</td><td>strand</td><td>MM</td><td>target_seq</td><td>PAM</td><td>alignment</td><td>distance</td><td>position</td><td>gene name</td><td>gene id</td></tr>
<tr><td>GU305914.1</td><td>164261</td><td>164283</td><td>+</td><td>0</td><td>TGCGTTACCACCCTCCTCTC</td><td>CGG</td><td>||||||||[||||||||||||]PAM</td><td>NA</td><td>Intergenic</td><td>NA</td><td>NA</td></tr>
<tr><td>T12359</td><td>CCGGTGACTATAGAGACACCAGG</td><td>1000</td><td></td><td></td><td></td><td></td><td></td><td></td><td></td><td></td><td></td></tr>
<tr><td>Oligo fwd</td><td>CACCCCGGTGACTATAGAGACACC</td><td></td><td></td><td></td><td></td><td></td><td></td><td></td><td></td><td></td><td></td></tr>
<tr><td>Oligo rev</td><td>AAACGGTGTCTCTATAGTCACCGG</td><td></td><td></td><td></td><td></td><td></td><td></td><td></td><td></td><td></td><td></td></tr>
<tr><td>Chromosome</td><td>start</td><td>end</td><td>strand</td><td>MM</td><td>target_seq</td><td>PAM</td><td>alignment</td><td>distance</td><td>position</td><td>gene name</td><td>gene id</td></tr>
<tr><td>GU305914.1</td><td>164280</td><td>164302</td><td>+</td><td>0</td><td>CCGGTGACTATAGAGACACC</td><td>AGG</td><td>||||||||[||||||||||||]PAM</td><td>NA</td><td>Intergenic</td><td>NA</td><td>NA</td></tr>
<tr><td>T12360</td><td>CGGTGACTATAGAGACACCAGGG</td><td>1000</td><td></td><td></td><td></td><td></td><td></td><td></td><td></td><td></td><td></td></tr>
<tr><td>Oligo fwd</td><td>CACCCGGTGACTATAGAGACACCA</td><td></td><td></td><td></td><td></td><td></td><td></td><td></td><td></td><td></td><td></td></tr>
<tr><td>Oligo rev</td><td>AAACTGGTGTCTCTATAGTCACCG</td><td></td><td></td><td></td><td></td><td></td><td></td><td></td><td></td><td></td><td></td></tr>
<tr><td>Chromosome</td><td>start</td><td>end</td><td>strand</td><td>MM</td><td>target_seq</td><td>PAM</td><td>alignment</td><td>distance</td><td>position</td><td>gene name</td><td>gene id</td></tr>
<tr><td>GU305914.1</td><td>164281</td><td>164303</td><td>+</td><td>0</td><td>CGGTGACTATAGAGACACCA</td><td>GGG</td><td>||||||||[||||||||||||]PAM</td><td>NA</td><td>Intergenic</td><td>NA</td><td>NA</td></tr>
<tr><td>T12361</td><td>TAATGACTTGATAGAATCCGAGG</td><td>1000</td><td></td><td></td><td></td><td></td><td></td><td></td><td></td><td></td><td></td></tr>
<tr><td>Oligo fwd</td><td>CACCTAATGACTTGATAGAATCCG</td><td></td><td></td><td></td><td></td><td></td><td></td><td></td><td></td><td></td><td></td></tr>
<tr><td>Oligo rev</td><td>AAACCGGATTCTATCAAGTCATTA</td><td></td><td></td><td></td><td></td><td></td><td></td><td></td><td></td><td></td><td></td></tr>
<tr><td>Chromosome</td><td>start</td><td>end</td><td>strand</td><td>MM</td><td>target_seq</td><td>PAM</td><td>alignment</td><td>distance</td><td>position</td><td>gene name</td><td>gene id</td></tr>
<tr><td>GU305914.1</td><td>164306</td><td>164328</td><td>+</td><td>0</td><td>TAATGACTTGATAGAATCCG</td><td>AGG</td><td>||||||||[||||||||||||]PAM</td><td>NA</td><td>Intergenic</td><td>NA</td><td>NA</td></tr>
<tr><td>T12362</td><td>TGACTTGATAGAATCCGAGGAGG</td><td>1000</td><td></td><td></td><td></td><td></td><td></td><td></td><td></td><td></td><td></td></tr>
<tr><td>Oligo fwd</td><td>CACCTGACTTGATAGAATCCGAGG</td><td></td><td></td><td></td><td></td><td></td><td></td><td></td><td></td><td></td><td></td></tr>
<tr><td>Oligo rev</td><td>AAACCCTCGGATTCTATCAAGTCA</td><td></td><td></td><td></td><td></td><td></td><td></td><td></td><td></td><td></td><td></td></tr>
<tr><td>Chromosome</td><td>start</td><td>end</td><td>strand</td><td>MM</td><td>target_seq</td><td>PAM</td><td>alignment</td><td>distance</td><td>position</td><td>gene name</td><td>gene id</td></tr>
<tr><td>GU305914.1</td><td>164309</td><td>164331</td><td>+</td><td>0</td><td>TGACTTGATAGAATCCGAGG</td><td>AGG</td><td>||||||||[||||||||||||]PAM</td><td>NA</td><td>Intergenic</td><td>NA</td><td>NA</td></tr>
<tr><td>T12363</td><td>GAATCCGAGGAGGAAGACGATGG</td><td>1000</td><td></td><td></td><td></td><td></td><td></td><td></td><td></td><td></td><td></td></tr>
<tr><td>Oligo fwd</td><td>CACCGAATCCGAGGAGGAAGACGA</td><td></td><td></td><td></td><td></td><td></td><td></td><td></td><td></td><td></td><td></td></tr>
<tr><td>Oligo rev</td><td>AAACTCGTCTTCCTCCTCGGATTC</td><td></td><td></td><td></td><td></td><td></td><td></td><td></td><td></td><td></td><td></td></tr>
<tr><td>Chromosome</td><td>start</td><td>end</td><td>strand</td><td>MM</td><td>target_seq</td><td>PAM</td><td>alignment</td><td>distance</td><td>position</td><td>gene name</td><td>gene id</td></tr>
<tr><td>GU305914.1</td><td>164319</td><td>164341</td><td>+</td><td>0</td><td>GAATCCGAGGAGGAAGACGA</td><td>TGG</td><td>||||||||[||||||||||||]PAM</td><td>NA</td><td>Intergenic</td><td>NA</td><td>NA</td></tr>
<tr><td>GU305914.1</td><td>226629</td><td>226651</td><td>-</td><td>4</td><td>GAAGCCTCCGAGGAAGACGA</td><td>AGG</td><td>|||-||--[-|||||||||||]PAM</td><td>NA</td><td>Intergenic</td><td>NA</td><td>NA</td></tr>
<tr><td>T12364</td><td>AGGAGGAAGACGATGGAACTCGG</td><td>1000</td><td></td><td></td><td></td><td></td><td></td><td></td><td></td><td></td><td></td></tr>
<tr><td>Oligo fwd</td><td>CACCAGGAGGAAGACGATGGAACT</td><td></td><td></td><td></td><td></td><td></td><td></td><td></td><td></td><td></td><td></td></tr>
<tr><td>Oligo rev</td><td>AAACAGTTCCATCGTCTTCCTCCT</td><td></td><td></td><td></td><td></td><td></td><td></td><td></td><td></td><td></td><td></td></tr>
<tr><td>Chromosome</td><td>start</td><td>end</td><td>strand</td><td>MM</td><td>target_seq</td><td>PAM</td><td>alignment</td><td>distance</td><td>position</td><td>gene name</td><td>gene id</td></tr>
<tr><td>GU305914.1</td><td>164326</td><td>164348</td><td>+</td><td>0</td><td>AGGAGGAAGACGATGGAACT</td><td>CGG</td><td>||||||||[||||||||||||]PAM</td><td>NA</td><td>Intergenic</td><td>NA</td><td>NA</td></tr>
</table>

| | | | | | | | | | | | |
|---|---|---|---|---|---|---|---|---|---|---|---|
| T12365 | ACTCGGTGTTCGTCTGCCGATGG | 1000 | | | | | | | | | |
| Oligo fwd | CACCACTCGGTGTTCGTCTGCCGA | | | | | | | | | | |
| Oligo rev | AAACTCGGCAGACGAACACCGAGT | | | | | | | | | | |
| Chromosome | start | end | strand | MM | target_seq | PAM | alignment | distance | position | gene name | gene id |
| GU305914.1 | 164343 | 164365 | + | 0 | ACTCGGTGTTCGTCTGCCGA | TGG | \|\|\|\|\|\|\|\|[\|\|\|\|\|\|\|\|\|\|\|\|]PAM | NA | Intergenic | NA | NA |
| T12366 | GTGTTCGTCTGCCGATGGTACGG | 1000 | | | | | | | | | |
| Oligo fwd | CACCGTGTTCGTCTGCCGATGGTA | | | | | | | | | | |
| Oligo rev | AAACTACCATCGGCAGACGAACAC | | | | | | | | | | |
| Chromosome | start | end | strand | MM | target_seq | PAM | alignment | distance | position | gene name | gene id |
| GU305914.1 | 164348 | 164370 | + | 0 | GTGTTCGTCTGCCGATGGTA | CGG | \|\|\|\|\|\|\|\|[\|\|\|\|\|\|\|\|\|\|\|\|]PAM | NA | Intergenic | NA | NA |
| T12367 | GTCTGCCGATGGTACGGCGACGG | 1000 | | | | | | | | | |
| Oligo fwd | CACCGTCTGCCGATGGTACGGCGA | | | | | | | | | | |
| Oligo rev | AAACTCGCCGTACCATCGGCAGAC | | | | | | | | | | |
| Chromosome | start | end | strand | MM | target_seq | PAM | alignment | distance | position | gene name | gene id |
| GU305914.1 | 164354 | 164376 | + | 0 | GTCTGCCGATGGTACGGCGA | CGG | \|\|\|\|\|\|\|\|[\|\|\|\|\|\|\|\|\|\|\|\|]PAM | NA | Intergenic | NA | NA |
| T12368 | ACGGCGACGGTAACCGAGACCGG | 1000 | | | | | | | | | |
| Oligo fwd | CACCACGGCGACGGTAACCGAGAC | | | | | | | | | | |
| Oligo rev | AAACGTCTCGGTTACCGTCGCCGT | | | | | | | | | | |
| Chromosome | start | end | strand | MM | target_seq | PAM | alignment | distance | position | gene name | gene id |
| GU305914.1 | 164367 | 164389 | + | 0 | ACGGCGACGGTAACCGAGAC | CGG | \|\|\|\|\|\|\|\|[\|\|\|\|\|\|\|\|\|\|\|\|]PAM | NA | Intergenic | NA | NA |
| T12369 | CGGCGACGGTAACCGAGACCGGG | 1000 | | | | | | | | | |
| Oligo fwd | CACCCGGCGACGGTAACCGAGACC | | | | | | | | | | |
| Oligo rev | AAACGGTCTCGGTTACCGTCGCCG | | | | | | | | | | |
| Chromosome | start | end | strand | MM | target_seq | PAM | alignment | distance | position | gene name | gene id |
| GU305914.1 | 164368 | 164390 | + | 0 | CGGCGACGGTAACCGAGACC | GGG | \|\|\|\|\|\|\|\|[\|\|\|\|\|\|\|\|\|\|\|\|]PAM | NA | Intergenic | NA | NA |
| T12370 | GGCGACGGTAACCGAGACCGGGG | 1000 | | | | | | | | | |
| Oligo fwd | CACCGGCGACGGTAACCGAGACCG | | | | | | | | | | |
| Oligo rev | AAACCGGTCTCGGTTACCGTCGCC | | | | | | | | | | |
| Chromosome | start | end | strand | MM | target_seq | PAM | alignment | distance | position | gene name | gene id |
| GU305914.1 | 164369 | 164391 | + | 0 | GGCGACGGTAACCGAGACCG | GGG | \|\|\|\|\|\|\|\|[\|\|\|\|\|\|\|\|\|\|\|\|]PAM | NA | Intergenic | NA | NA |
| T12371 | ATCGAGCAAACGTTCACCGACGG | 1000 | | | | | | | | | |
| Oligo fwd | CACCATCGAGCAAACGTTCACCGA | | | | | | | | | | |
| Oligo rev | AAACTCGGTGAACGTTTGCTCGAT | | | | | | | | | | |
| Chromosome | start | end | strand | MM | target_seq | PAM | alignment | distance | position | gene name | gene id |
| GU305914.1 | 164403 | 164425 | + | 0 | ATCGAGCAAACGTTCACCGA | CGG | \|\|\|\|\|\|\|\|[\|\|\|\|\|\|\|\|\|\|\|\|]PAM | NA | Intergenic | NA | NA |
| T12372 | GAGCAAACGTTCACCGACGGTGG | 1000 | | | | | | | | | |
| Oligo fwd | CACCGAGCAAACGTTCACCGACGG | | | | | | | | | | |
| Oligo rev | AAACCCGTCGGTGAACGTTTGCTC | | | | | | | | | | |
| Chromosome | start | end | strand | MM | target_seq | PAM | alignment | distance | position | gene name | gene id |
| GU305914.1 | 164406 | 164428 | + | 0 | GAGCAAACGTTCACCGACGG | TGG | \|\|\|\|\|\|\|\|[\|\|\|\|\|\|\|\|\|\|\|\|]PAM | NA | Intergenic | NA | NA |
| T12373 | AGCAAACGTTCACCGACGGTGGG | 1000 | | | | | | | | | |
| Oligo fwd | CACCAGCAAACGTTCACCGACGGT | | | | | | | | | | |
| Oligo rev | AAACACCGTCGGTGAACGTTTGCT | | | | | | | | | | |
| Chromosome | start | end | strand | MM | target_seq | PAM | alignment | distance | position | gene name | gene id |
| GU305914.1 | 164407 | 164429 | + | 0 | AGCAAACGTTCACCGACGGT | GGG | \|\|\|\|\|\|\|\|[\|\|\|\|\|\|\|\|\|\|\|\|]PAM | NA | Intergenic | NA | NA |
| T12374 | ACGGTGGGCTAGCATGCATGAGG | 1000 | | | | | | | | | |
| Oligo fwd | CACCACGGTGGGCTAGCATGCATG | | | | | | | | | | |
| Oligo rev | AAACCATGCATGCTAGCCCACCGT | | | | | | | | | | |
| Chromosome | start | end | strand | MM | target_seq | PAM | alignment | distance | position | gene name | gene id |
| GU305914.1 | 164422 | 164444 | + | 0 | ACGGTGGGCTAGCATGCATG | AGG | \|\|\|\|\|\|\|\|[\|\|\|\|\|\|\|\|\|\|\|\|]PAM | NA | Intergenic | NA | NA |
| T12375 | TGAGGTCCTTAATTGACTCGCGG | 1000 | | | | | | | | | |
| Oligo fwd | CACCTGAGGTCCTTAATTGACTCG | | | | | | | | | | |
| Oligo rev | AAACCGAGTCAATTAAGGACCTCA | | | | | | | | | | |
| Chromosome | start | end | strand | MM | target_seq | PAM | alignment | distance | position | gene name | gene id |
| GU305914.1 | 164440 | 164462 | + | 0 | TGAGGTCCTTAATTGACTCG | CGG | \|\|\|\|\|\|\|\|[\|\|\|\|\|\|\|\|\|\|\|\|]PAM | NA | Intergenic | NA | NA |
| T12376 | GAGGTCCTTAATTGACTCGCGGG | 1000 | | | | | | | | | |
| Oligo fwd | CACCGAGGTCCTTAATTGACTCGC | | | | | | | | | | |
| Oligo rev | AAACGCGAGTCAATTAAGGACCTC | | | | | | | | | | |
| Chromosome | start | end | strand | MM | target_seq | PAM | alignment | distance | position | gene name | gene id |
| GU305914.1 | 164441 | 164463 | + | 0 | GAGGTCCTTAATTGACTCGC | GGG | \|\|\|\|\|\|\|\|[\|\|\|\|\|\|\|\|\|\|\|\|]PAM | NA | Intergenic | NA | NA |
| T12377 | AATTGACTCGCGGGCCCTCATGG | 1000 | | | | | | | | | |
| Oligo fwd | CACCAATTGACTCGCGGGCCCTCA | | | | | | | | | | |
| Oligo rev | AAACTGAGGGCCCGCGAGTCAATT | | | | | | | | | | |
| Chromosome | start | end | strand | MM | target_seq | PAM | alignment | distance | position | gene name | gene id |
| GU305914.1 | 164450 | 164472 | + | 0 | AATTGACTCGCGGGCCCTCA | TGG | \|\|\|\|\|\|\|\|[\|\|\|\|\|\|\|\|\|\|\|\|]PAM | NA | Intergenic | NA | NA |
| T12378 | CGCCTTTACGAGCCGCATGACGG | 1000 | | | | | | | | | |
| Oligo fwd | CACCCGCCTTTACGAGCCGCATGA | | | | | | | | | | |
| Oligo rev | AAACTCATGCGGCTCGTAAAGGCG | | | | | | | | | | |
| Chromosome | start | end | strand | MM | target_seq | PAM | alignment | distance | position | gene name | gene id |
| GU305914.1 | 164477 | 164499 | + | 0 | CGCCTTTACGAGCCGCATGA | CGG | \|\|\|\|\|\|\|\|[\|\|\|\|\|\|\|\|\|\|\|\|]PAM | NA | Intergenic | NA | NA |
| T12379 | CTTTACGAGCCGCATGACGGCGG | 1000 | | | | | | | | | |
| Oligo fwd | CACCCTTTACGAGCCGCATGACGG | | | | | | | | | | |
| Oligo rev | AAACCCGTCATGCGGCTCGTAAAG | | | | | | | | | | |
| Chromosome | start | end | strand | MM | target_seq | PAM | alignment | distance | position | gene name | gene id |

GU305914.1 164480 164502 + 0 CTTTACGAGCCGCATGACGG CGG ||||||||[||||||||||||]PAM NA Intergenic NA NA

T12380 CCGCATGACGGCGGTGAAGTCGG 1000
Oligo fwd CACCCCGCATGACGGCGGTGAAGT
Oligo rev AAACACTTCACCGCCGTCATGCGG
Chromosome start end strand MM target_seq PAM alignment distance position gene name gene id
GU305914.1 164489 164511 + 0 CCGCATGACGGCGGTGAAGT CGG ||||||||[||||||||||||]PAM NA Intergenic NA NA

T12381 CGCATGACGGCGGTGAAGTCGGG 1000
Oligo fwd CACCCGCATGACGGCGGTGAAGTC
Oligo rev AAACGACTTCACCGCCGTCATGCG
Chromosome start end strand MM target_seq PAM alignment distance position gene name gene id
GU305914.1 164490 164512 + 0 CGCATGACGGCGGTGAAGTC GGG ||||||||[||||||||||||]PAM NA Intergenic NA NA

T12382 GGCGGTGAAGTCGGGATCTGAGG 1000
Oligo fwd CACCGGCGGTGAAGTCGGGATCTG
Oligo rev AAACCAGATCCCGACTTCACCGCC
Chromosome start end strand MM target_seq PAM alignment distance position gene name gene id
GU305914.1 164498 164520 + 0 GGCGGTGAAGTCGGGATCTG AGG ||||||||[||||||||||||]PAM NA Intergenic NA NA

T12383 GCGGTGAAGTCGGGATCTGAGGG 1000
Oligo fwd CACCGCGGTGAAGTCGGGATCTGA
Oligo rev AAACTCAGATCCCGACTTCACCGC
Chromosome start end strand MM target_seq PAM alignment distance position gene name gene id
GU305914.1 164499 164521 + 0 GCGGTGAAGTCGGGATCTGA GGG ||||||||[||||||||||||]PAM NA Intergenic NA NA

T12384 GAAGTCGGGATCTGAGGGAGCGG 1000
Oligo fwd CACCGAAGTCGGGATCTGAGGGAG
Oligo rev AAACCTCCCTCAGATCCCGACTTC
Chromosome start end strand MM target_seq PAM alignment distance position gene name gene id
GU305914.1 164504 164526 + 0 GAAGTCGGGATCTGAGGGAG CGG ||||||||[||||||||||||]PAM NA Intergenic NA NA

T12385 AAGTCGGGATCTGAGGGAGCGGG 1000
Oligo fwd CACCAAGTCGGGATCTGAGGGAGC
Oligo rev AAACGCTCCCTCAGATCCCGACTT
Chromosome start end strand MM target_seq PAM alignment distance position gene name gene id
GU305914.1 164505 164527 + 0 AAGTCGGGATCTGAGGGAGC GGG ||||||||[||||||||||||]PAM NA Intergenic NA NA

T12386 GGAGCGGGCAACACCTGTCCTGG 1000
Oligo fwd CACCGGAGCGGGCAACACCTGTCC
Oligo rev AAACGGACAGGTGTTGCCCGCTCC
Chromosome start end strand MM target_seq PAM alignment distance position gene name gene id
GU305914.1 164520 164542 + 0 GGAGCGGGCAACACCTGTCC TGG ||||||||[||||||||||||]PAM NA Intergenic NA NA

T12387 GAGCGGGCAACACCTGTCCTGGG 1000
Oligo fwd CACCGAGCGGGCAACACCTGTCCT
Oligo rev AAACAGGACAGGTGTTGCCCGCTC
Chromosome start end strand MM target_seq PAM alignment distance position gene name gene id
GU305914.1 164521 164543 + 0 GAGCGGGCAACACCTGTCCT GGG ||||||||[||||||||||||]PAM NA Intergenic NA NA

T12388 AGCGGGCAACACCTGTCCTGGGG 1000
Oligo fwd CACCAGCGGGCAACACCTGTCCTG
Oligo rev AAACCAGGACAGGTGTTGCCCGCT
Chromosome start end strand MM target_seq PAM alignment distance position gene name gene id
GU305914.1 164522 164544 + 0 AGCGGGCAACACCTGTCCTG GGG ||||||||[||||||||||||]PAM NA Intergenic NA NA

T12389 ACACCTGTCCTGGGGAACAGAGG 1000
Oligo fwd CACCACACCTGTCCTGGGGAACAG
Oligo rev AAACCTGTTCCCCAGGACAGGTGT
Chromosome start end strand MM target_seq PAM alignment distance position gene name gene id
GU305914.1 164530 164552 + 0 ACACCTGTCCTGGGGAACAG AGG ||||||||[||||||||||||]PAM NA Intergenic NA NA

T12390 CAATCTTCCTCCGAATTCACAGG 1000
Oligo fwd CACCCAATCTTCCTCCGAATTCAC
Oligo rev AAACGTGAATTCGGAGGAAGATTG
Chromosome start end strand MM target_seq PAM alignment distance position gene name gene id
GU305914.1 164564 164586 + 0 CAATCTTCCTCCGAATTCAC AGG ||||||||[||||||||||||]PAM NA Intergenic NA NA

T12391 TCTTCCTCCGAATTCACAGGAGG 1000
Oligo fwd CACCTCTTCCTCCGAATTCACAGG
Oligo rev AAACCCTGTGAATTCGGAGGAAGA
Chromosome start end strand MM target_seq PAM alignment distance position gene name gene id
GU305914.1 164567 164589 + 0 TCTTCCTCCGAATTCACAGG AGG ||||||||[||||||||||||]PAM NA Intergenic NA NA

T12392 GACAACAACCCGCGTTTTGAAGG 1000
Oligo fwd CACCGACAACAACCCGCGTTTTGA
Oligo rev AAACTCAAAACGCGGGTTGTTGTC
Chromosome start end strand MM target_seq PAM alignment distance position gene name gene id
GU305914.1 164589 164611 + 0 GACAACAACCCGCGTTTTGA AGG ||||||||[||||||||||||]PAM NA Intergenic NA NA

T12393 CTCGAGTTCCGCCGCCGTACCGG 1000
Oligo fwd CACCCTCGAGTTCCGCCGCCGTAC
Oligo rev AAACGTACGGCGGCGGAACTCGAG
Chromosome start end strand MM target_seq PAM alignment distance position gene name gene id
GU305914.1 164666 164688 + 0 CTCGAGTTCCGCCGCCGTAC CGG ||||||||[||||||||||||]PAM NA Intergenic NA NA

T12394 TCGAGTTCCGCCGCCGTACCGGG 1000
Oligo fwd CACCTCGAGTTCCGCCGCCGTACC
Oligo rev AAACGGTACGGCGGCGGAACTCGA

Chromosome start end strand MM target_seq PAM alignment distance position gene name gene id
GU305914.1 164667 164689 + 0 TCGAGTTCCGCCGCCGTACC GGG ||||||||[||||||||||||]PAM NA Intergenic NA NA

T12395 GTACCGGGATCCAGTTGTACAGG 1000
Oligo fwd CACCGTACCGGGATCCAGTTGTAC
Oligo rev AAACGTACAACTGGATCCCGGTAC
Chromosome start end strand MM target_seq PAM alignment distance position gene name gene id
GU305914.1 164682 164704 + 0 GTACCGGGATCCAGTTGTAC AGG ||||||||[||||||||||||]PAM NA Intergenic NA NA

T12396 TACAGGCACCACGCAGACAGAGG 1000
Oligo fwd CACCTACAGGCACCACGCAGACAG
Oligo rev AAACCTGTCTGCGTGGTGCCTGTA
Chromosome start end strand MM target_seq PAM alignment distance position gene name gene id
GU305914.1 164699 164721 + 0 TACAGGCACCACGCAGACAG AGG ||||||||[||||||||||||]PAM NA Intergenic NA NA

T12397 ACAGGCACCACGCAGACAGAGGG 1000
Oligo fwd CACCACAGGCACCACGCAGACAGA
Oligo rev AAACTCTGTCTGCGTGGTGCCTGT
Chromosome start end strand MM target_seq PAM alignment distance position gene name gene id
GU305914.1 164700 164722 + 0 ACAGGCACCACGCAGACAGA GGG ||||||||[||||||||||||]PAM NA Intergenic NA NA

T12398 CAGGCACCACGCAGACAGAGGGG 1000
Oligo fwd CACCCAGGCACCACGCAGACAGAG
Oligo rev AAACCTCTGTCTGCGTGGTGCCTG
Chromosome start end strand MM target_seq PAM alignment distance position gene name gene id
GU305914.1 164701 164723 + 0 CAGGCACCACGCAGACAGAG GGG ||||||||[||||||||||||]PAM NA Intergenic NA NA

T12399 CAGAGGGGATGCCTCACCACAGG 1000
Oligo fwd CACCCAGAGGGGATGCCTCACCAC
Oligo rev AAACGTGGTGAGGCATCCCCTCTG
Chromosome start end strand MM target_seq PAM alignment distance position gene name gene id
GU305914.1 164716 164738 + 0 CAGAGGGGATGCCTCACCAC AGG ||||||||[||||||||||||]PAM NA Intergenic NA NA

T12400 AGAGGGGATGCCTCACCACAGGG 1000
Oligo fwd CACCAGAGGGGATGCCTCACCACA
Oligo rev AAACTGTGGTGAGGCATCCCCTCT
Chromosome start end strand MM target_seq PAM alignment distance position gene name gene id
GU305914.1 164717 164739 + 0 AGAGGGGATGCCTCACCACA GGG ||||||||[||||||||||||]PAM NA Intergenic NA NA

T12401 GAGGGGATGCCTCACCACAGGGG 1000
Oligo fwd CACCGAGGGGATGCCTCACCACAG
Oligo rev AAACCTGTGGTGAGGCATCCCCTC
Chromosome start end strand MM target_seq PAM alignment distance position gene name gene id
GU305914.1 164718 164740 + 0 GAGGGGATGCCTCACCACAG GGG ||||||||[||||||||||||]PAM NA Intergenic NA NA

T12402 ACCACAGGGGTATCTTCGACAGG 1000
Oligo fwd CACCACCACAGGGGTATCTTCGAC
Oligo rev AAACGTCGAAGATACCCCTGTGGT
Chromosome start end strand MM target_seq PAM alignment distance position gene name gene id
GU305914.1 164731 164753 + 0 ACCACAGGGGTATCTTCGAC AGG ||||||||[||||||||||||]PAM NA Intergenic NA NA

T12403 CCACAGGGGTATCTTCGACAGGG 1000
Oligo fwd CACCCCACAGGGGTATCTTCGACA
Oligo rev AAACTGTCGAAGATACCCCTGTGG
Chromosome start end strand MM target_seq PAM alignment distance position gene name gene id
GU305914.1 164732 164754 + 0 CCACAGGGGTATCTTCGACA GGG ||||||||[||||||||||||]PAM NA Intergenic NA NA

T12404 AGAAACCTGTATGTGAGATACGG 1000
Oligo fwd CACCAGAAACCTGTATGTGAGATA
Oligo rev AAACTATCTCACATACAGGTTTCT
Chromosome start end strand MM target_seq PAM alignment distance position gene name gene id
GU305914.1 164773 164795 + 0 AGAAACCTGTATGTGAGATA CGG ||||||||[||||||||||||]PAM NA Intergenic NA NA

T12405 CATAGCTCGCTGTGTCGAGCCGG 1000
Oligo fwd CACCCATAGCTCGCTGTGTCGAGC
Oligo rev AAACGCTCGACACAGCGAGCTATG
Chromosome start end strand MM target_seq PAM alignment distance position gene name gene id
GU305914.1 164807 164829 + 0 CATAGCTCGCTGTGTCGAGC CGG ||||||||[||||||||||||]PAM NA Intergenic NA NA

T12406 AGCTCGCTGTGTCGAGCCGGCGG 1000
Oligo fwd CACCAGCTCGCTGTGTCGAGCCGG
Oligo rev AAACCCGGCTCGACACAGCGAGCT
Chromosome start end strand MM target_seq PAM alignment distance position gene name gene id
GU305914.1 164810 164832 + 0 AGCTCGCTGTGTCGAGCCGG CGG ||||||||[||||||||||||]PAM NA Intergenic NA NA

T12407 TCGCTGTGTCGAGCCGGCGGCGG 1000
Oligo fwd CACCTCGCTGTGTCGAGCCGGCGG
Oligo rev AAACCCGCCGGCTCGACACAGCGA
Chromosome start end strand MM target_seq PAM alignment distance position gene name gene id
GU305914.1 164813 164835 + 0 TCGCTGTGTCGAGCCGGCGG CGG ||||||||[||||||||||||]PAM NA Intergenic NA NA

T12408 TTCGCGCTCCAGAGCCACCACGG 1000
Oligo fwd CACCTTCGCGCTCCAGAGCCACCA
Oligo rev AAACTGGTGGCTCTGGAGCGCGAA
Chromosome start end strand MM target_seq PAM alignment distance position gene name gene id
GU305914.1 164846 164868 + 0 TTCGCGCTCCAGAGCCACCA CGG ||||||||[||||||||||||]PAM NA Intergenic NA NA

T12409 CCACCACGGCCGCCGCTCGCAGG 1000
Oligo fwd CACCCCACCACGGCCGCCGCTCGC

| | | | | | | | | | | | | |
|---|---|---|---|---|---|---|---|---|---|---|---|---|
| Oligo rev | AAACGCGAGCGGCGGCCGTGGTGG | | | | | | | | | | | |
| Chromosome | start | | end | strand | MM | target_seq | PAM | alignment | distance | position | gene name | gene id |
| GU305914.1 | | 164860 | 164882 | + | 0 | CCACCACGGCCGCCGCTCGC | AGG | \|\|\|\|\|\|\|\|[\|\|\|\|\|\|\|\|\|\|\|\|]PAM | NA | Intergenic | NA | NA |
| T12410 | CCACGGCCGCCGCTCGCAGGCGG | | 1000 | | | | | | | | | |
| Oligo fwd | CACCCCACGGCCGCCGCTCGCAGG | | | | | | | | | | | |
| Oligo rev | AAACCCTGCGAGCGGCGGCCGTGG | | | | | | | | | | | |
| Chromosome | start | | end | strand | MM | target_seq | PAM | alignment | distance | position | gene name | gene id |
| GU305914.1 | | 164863 | 164885 | + | 0 | CCACGGCCGCCGCTCGCAGG | CGG | \|\|\|\|\|\|\|\|[\|\|\|\|\|\|\|\|\|\|\|\|]PAM | NA | Intergenic | NA | NA |
| T12411 | CACGGCCGCCGCTCGCAGGCGGG | | 1000 | | | | | | | | | |
| Oligo fwd | CACCCACGGCCGCCGCTCGCAGGC | | | | | | | | | | | |
| Oligo rev | AAACGCCTGCGAGCGGCGGCCGTG | | | | | | | | | | | |
| Chromosome | start | | end | strand | MM | target_seq | PAM | alignment | distance | position | gene name | gene id |
| GU305914.1 | | 164864 | 164886 | + | 0 | CACGGCCGCCGCTCGCAGGC | GGG | \|\|\|\|\|\|\|\|[\|\|\|\|\|\|\|\|\|\|\|\|]PAM | NA | Intergenic | NA | NA |
| T12412 | ACGGCCGCCGCTCGCAGGCGGGG | | 1000 | | | | | | | | | |
| Oligo fwd | CACCACGGCCGCCGCTCGCAGGCG | | | | | | | | | | | |
| Oligo rev | AAACCGCCTGCGAGCGGCGGCCGT | | | | | | | | | | | |
| Chromosome | start | | end | strand | MM | target_seq | PAM | alignment | distance | position | gene name | gene id |
| GU305914.1 | | 164865 | 164887 | + | 0 | ACGGCCGCCGCTCGCAGGCG | GGG | \|\|\|\|\|\|\|\|[\|\|\|\|\|\|\|\|\|\|\|\|]PAM | NA | Intergenic | NA | NA |
| T12413 | GCCGCTCGCAGGCGGGGAAGAGG | | 1000 | | | | | | | | | |
| Oligo fwd | CACCGCCGCTCGCAGGCGGGGAAG | | | | | | | | | | | |
| Oligo rev | AAACCTTCCCCGCCTGCGAGCGGC | | | | | | | | | | | |
| Chromosome | start | | end | strand | MM | target_seq | PAM | alignment | distance | position | gene name | gene id |
| GU305914.1 | | 164871 | 164893 | + | 0 | GCCGCTCGCAGGCGGGGAAG | AGG | \|\|\|\|\|\|\|\|[\|\|\|\|\|\|\|\|\|\|\|\|]PAM | NA | Intergenic | NA | NA |
| T12414 | GGGGAAGAGGCTCGCAGCCTCGG | | 1000 | | | | | | | | | |
| Oligo fwd | CACCGGGGAAGAGGCTCGCAGCCT | | | | | | | | | | | |
| Oligo rev | AAACAGGCTGCGAGCCTCTTCCCC | | | | | | | | | | | |
| Chromosome | start | | end | strand | MM | target_seq | PAM | alignment | distance | position | gene name | gene id |
| GU305914.1 | | 164884 | 164906 | + | 0 | GGGGAAGAGGCTCGCAGCCT | CGG | \|\|\|\|\|\|\|\|[\|\|\|\|\|\|\|\|\|\|\|\|]PAM | NA | Intergenic | NA | NA |
| T12415 | GAAGAGGCTCGCAGCCTCGGAGG | | 1000 | | | | | | | | | |
| Oligo fwd | CACCGAAGAGGCTCGCAGCCTCGG | | | | | | | | | | | |
| Oligo rev | AAACCCGAGGCTGCGAGCCTCTTC | | | | | | | | | | | |
| Chromosome | start | | end | strand | MM | target_seq | PAM | alignment | distance | position | gene name | gene id |
| GU305914.1 | | 164887 | 164909 | + | 0 | GAAGAGGCTCGCAGCCTCGG | AGG | \|\|\|\|\|\|\|\|[\|\|\|\|\|\|\|\|\|\|\|\|]PAM | NA | Intergenic | NA | NA |
| T12416 | CGCAGCCTCGGAGGAACAGTCGG | | 1000 | | | | | | | | | |
| Oligo fwd | CACCCGCAGCCTCGGAGGAACAGT | | | | | | | | | | | |
| Oligo rev | AAACACTGTTCCTCCGAGGCTGCG | | | | | | | | | | | |
| Chromosome | start | | end | strand | MM | target_seq | PAM | alignment | distance | position | gene name | gene id |
| GU305914.1 | | 164896 | 164918 | + | 0 | CGCAGCCTCGGAGGAACAGT | CGG | \|\|\|\|\|\|\|\|[\|\|\|\|\|\|\|\|\|\|\|\|]PAM | NA | Intergenic | NA | NA |
| T12417 | GCAGCCTCGGAGGAACAGTCGGG | | 1000 | | | | | | | | | |
| Oligo fwd | CACCGCAGCCTCGGAGGAACAGTC | | | | | | | | | | | |
| Oligo rev | AAACGACTGTTCCTCCGAGGCTGC | | | | | | | | | | | |
| Chromosome | start | | end | strand | MM | target_seq | PAM | alignment | distance | position | gene name | gene id |
| GU305914.1 | | 164897 | 164919 | + | 0 | GCAGCCTCGGAGGAACAGTC | GGG | \|\|\|\|\|\|\|\|[\|\|\|\|\|\|\|\|\|\|\|\|]PAM | NA | Intergenic | NA | NA |
| T12418 | TCGGAGGAACAGTCGGGTCGCGG | | 1000 | | | | | | | | | |
| Oligo fwd | CACCTCGGAGGAACAGTCGGGTCG | | | | | | | | | | | |
| Oligo rev | AAACCGACCCGACTGTTCCTCCGA | | | | | | | | | | | |
| Chromosome | start | | end | strand | MM | target_seq | PAM | alignment | distance | position | gene name | gene id |
| GU305914.1 | | 164903 | 164925 | + | 0 | TCGGAGGAACAGTCGGGTCG | CGG | \|\|\|\|\|\|\|\|[\|\|\|\|\|\|\|\|\|\|\|\|]PAM | NA | Intergenic | NA | NA |
| T12419 | GGTCGCGGCGCGCGTCGCCCAGG | | 1000 | | | | | | | | | |
| Oligo fwd | CACCGGTCGCGGCGCGCGTCGCCC | | | | | | | | | | | |
| Oligo rev | AAACGGGCGACGCGCGCCGCGACC | | | | | | | | | | | |
| Chromosome | start | | end | strand | MM | target_seq | PAM | alignment | distance | position | gene name | gene id |
| GU305914.1 | | 164918 | 164940 | + | 0 | GGTCGCGGCGCGCGTCGCCC | AGG | \|\|\|\|\|\|\|\|[\|\|\|\|\|\|\|\|\|\|\|\|]PAM | NA | Intergenic | NA | NA |
| T12420 | GCTGCTGCCGCCGCCGCCGCCGG | | 1000 | | | | | | | | | |
| Oligo fwd | CACCGCTGCTGCCGCCGCCGCCGC | | | | | | | | | | | |
| Oligo rev | AAACGCGGCGGCGGCGGCAGCAGC | | | | | | | | | | | |
| Chromosome | start | | end | strand | MM | target_seq | PAM | alignment | distance | position | gene name | gene id |
| GU305914.1 | | 164964 | 164986 | + | 0 | GCTGCTGCCGCCGCCGCCGC | CGG | \|\|\|\|\|\|\|\|[\|\|\|\|\|\|\|\|\|\|\|\|]PAM | NA | Intergenic | NA | NA |
| GU305914.1 | | 152551 | 152573 | - | 4 | GCCGGCACCGCCGCCGCCGC | AGG | \|\|-\|---\|[\|\|\|\|\|\|\|\|\|\|\|\|]PAM | NA | Intergenic | NA | NA |
| GU305914.1 | | 65720 | 65742 | + | 3 | CCCGCTGCCGCTGCCGCCGC | AGG | -\|-\|\|\|\|\|[\|\|\|-\|\|\|\|\|\|\|\|]PAM | NA | Intergenic | NA | NA |
| GU305914.1 | | 24620 | 24642 | + | 4 | GTCGCCGCCACCGCCGCCGC | CGG | \|--\|\|-\|\|[\|-\|\|\|\|\|\|\|\|\|\|]PAM | NA | Intergenic | NA | NA |
| GU305914.1 | | 86843 | 86865 | + | 3 | GAAGCTGCCGCCCCCGCCGC | GGG | \|--\|\|\|\|\|[\|\|\|\|-\|\|\|\|\|\|\|]PAM | NA | Intergenic | NA | NA |
| GU305914.1 | | 48151 | 48173 | - | 4 | GCCGCCACCGCCGCCGCCAC | AGG | \|\|-\|\|--\|[\|\|\|\|\|\|\|\|\|\|-\|]PAM | NA | Intergenic | NA | NA |
| T12421 | CCGCCGCCGGTCCCACCACGCGG | | 1000 | | | | | | | | | |
| Oligo fwd | CACCCCGCCGCCGGTCCCACCACG | | | | | | | | | | | |
| Oligo rev | AAACCGTGGTGGGACCGGCGGCGG | | | | | | | | | | | |
| Chromosome | start | | end | strand | MM | target_seq | PAM | alignment | distance | position | gene name | gene id |
| GU305914.1 | | 164977 | 164999 | + | 0 | CCGCCGCCGGTCCCACCACG | CGG | \|\|\|\|\|\|\|\|[\|\|\|\|\|\|\|\|\|\|\|\|]PAM | NA | Intergenic | NA | NA |
| T12422 | CGCCGCCGGTCCCACCACGCGGG | | 1000 | | | | | | | | | |
| Oligo fwd | CACCCGCCGCCGGTCCCACCACGC | | | | | | | | | | | |
| Oligo rev | AAACGCGTGGTGGGACCGGCGGCG | | | | | | | | | | | |
| Chromosome | start | | end | strand | MM | target_seq | PAM | alignment | distance | position | gene name | gene id |
| GU305914.1 | | 164978 | 165000 | + | 0 | CGCCGCCGGTCCCACCACGC | GGG | \|\|\|\|\|\|\|\|[\|\|\|\|\|\|\|\|\|\|\|\|]PAM | NA | Intergenic | NA | NA |
| T12423 | GCCGCCGGTCCCACCACGCGGGG | | 1000 | | | | | | | | | |
| Oligo fwd | CACCGCCGCCGGTCCCACCACGCG | | | | | | | | | | | |

Oligo rev AAACCGCGTGGTGGGACCGGCGGC

| Chromosome | start | end | strand | MM | target_seq | PAM | alignment | distance | position | gene name | gene id |
|---|---|---|---|---|---|---|---|---|---|---|---|
| GU305914.1 | 164979 | 165001 | + | 0 | GCCGCCGGTCCCACCACGCG | GGG | \|\|\|\|\|\|\|\|[\|\|\|\|\|\|\|\|\|\|\|\|]PAM | NA | Intergenic | NA | NA |

T12424 CCGGTCCCACCACGCGGGGAAGG 1000
Oligo fwd CACCCCGGTCCCACCACGCGGGGA
Oligo rev AAACTCCCCGCGTGGTGGGACCGG

| Chromosome | start | end | strand | MM | target_seq | PAM | alignment | distance | position | gene name | gene id |
|---|---|---|---|---|---|---|---|---|---|---|---|
| GU305914.1 | 164983 | 165005 | + | 0 | CCGGTCCCACCACGCGGGGA | AGG | \|\|\|\|\|\|\|\|[\|\|\|\|\|\|\|\|\|\|\|\|]PAM | NA | Intergenic | NA | NA |

T12425 CCACCACGCGGGGAAGGAGCAGG 1000
Oligo fwd CACCCCACCACGCGGGGAAGGAGC
Oligo rev AAACGCTCCTTCCCCGCGTGGTGG

| Chromosome | start | end | strand | MM | target_seq | PAM | alignment | distance | position | gene name | gene id |
|---|---|---|---|---|---|---|---|---|---|---|---|
| GU305914.1 | 164989 | 165011 | + | 0 | CCACCACGCGGGGAAGGAGC | AGG | \|\|\|\|\|\|\|\|[\|\|\|\|\|\|\|\|\|\|\|\|]PAM | NA | Intergenic | NA | NA |

T12426 CGCGGGGAAGGAGCAGGAGTAGG 1000
Oligo fwd CACCCGCGGGGAAGGAGCAGGAGT
Oligo rev AAACACTCCTGCTCCTTCCCCGCG

| Chromosome | start | end | strand | MM | target_seq | PAM | alignment | distance | position | gene name | gene id |
|---|---|---|---|---|---|---|---|---|---|---|---|
| GU305914.1 | 164995 | 165017 | + | 0 | CGCGGGGAAGGAGCAGGAGT | AGG | \|\|\|\|\|\|\|\|[\|\|\|\|\|\|\|\|\|\|\|\|]PAM | NA | Intergenic | NA | NA |
| GU305914.1 | 33852 | 33874 | + | 3 | CGAGGGGAAGGAGAAGGGGT | TGG | \|\|-\|\|\|\|\|[\|\|\|\|\|-\|\|\|-\|\|]PAM | NA | Intergenic | NA | NA |

T12427 GGGGAAGGAGCAGGAGTAGGAGG 1000
Oligo fwd CACCGGGGAAGGAGCAGGAGTAGG
Oligo rev AAACCCTACTCCTGCTCCTTCCCC

| Chromosome | start | end | strand | MM | target_seq | PAM | alignment | distance | position | gene name | gene id |
|---|---|---|---|---|---|---|---|---|---|---|---|
| GU305914.1 | 164998 | 165020 | + | 0 | GGGGAAGGAGCAGGAGTAGG | AGG | \|\|\|\|\|\|\|\|[\|\|\|\|\|\|\|\|\|\|\|\|]PAM | NA | Intergenic | NA | NA |

T12428 GGGAAGGAGCAGGAGTAGGAGGG 1000
Oligo fwd CACCGGGAAGGAGCAGGAGTAGGA
Oligo rev AAACTCCTACTCCTGCTCCTTCCC

| Chromosome | start | end | strand | MM | target_seq | PAM | alignment | distance | position | gene name | gene id |
|---|---|---|---|---|---|---|---|---|---|---|---|
| GU305914.1 | 164999 | 165021 | + | 0 | GGGAAGGAGCAGGAGTAGGA | GGG | \|\|\|\|\|\|\|\|[\|\|\|\|\|\|\|\|\|\|\|\|]PAM | NA | Intergenic | NA | NA |

T12429 GGAAGGAGCAGGAGTAGGAGGGG 1000
Oligo fwd CACCGGAAGGAGCAGGAGTAGGAG
Oligo rev AAACCTCCTACTCCTGCTCCTTCC

| Chromosome | start | end | strand | MM | target_seq | PAM | alignment | distance | position | gene name | gene id |
|---|---|---|---|---|---|---|---|---|---|---|---|
| GU305914.1 | 165000 | 165022 | + | 0 | GGAAGGAGCAGGAGTAGGAG | GGG | \|\|\|\|\|\|\|\|[\|\|\|\|\|\|\|\|\|\|\|\|]PAM | NA | Intergenic | NA | NA |

T12430 AGGAGTAGGAGGGGAAGACGAGG 1000
Oligo fwd CACCAGGAGTAGGAGGGGAAGACG
Oligo rev AAACCGTCTTCCCCTCCTACTCCT

| Chromosome | start | end | strand | MM | target_seq | PAM | alignment | distance | position | gene name | gene id |
|---|---|---|---|---|---|---|---|---|---|---|---|
| GU305914.1 | 165009 | 165031 | + | 0 | AGGAGTAGGAGGGGAAGACG | AGG | \|\|\|\|\|\|\|\|[\|\|\|\|\|\|\|\|\|\|\|\|]PAM | NA | Intergenic | NA | NA |
| GU305914.1 | 86796 | 86818 | - | 4 | CGGGGCAGGAGGGGAAGAGG | GGG | -\|\|-\|-\|\|[\|\|\|\|\|\|\|\|\|\|-\|]PAM | NA | Intergenic | NA | NA |
| GU305914.1 | 45056 | 45078 | + | 3 | AGTAGTAGGAGGAGAAGAGG | AGG | \|\|-\|\|\|\|\|[\|\|\|\|-\|\|\|\|\|-\|]PAM | NA | Intergenic | NA | NA |

T12431 AAGACGAGGCCCAGACCTCACGG 1000
Oligo fwd CACCAAGACGAGGCCCAGACCTCA
Oligo rev AAACTGAGGTCTGGGCCTCGTCTT

| Chromosome | start | end | strand | MM | target_seq | PAM | alignment | distance | position | gene name | gene id |
|---|---|---|---|---|---|---|---|---|---|---|---|
| GU305914.1 | 165023 | 165045 | + | 0 | AAGACGAGGCCCAGACCTCA | CGG | \|\|\|\|\|\|\|\|[\|\|\|\|\|\|\|\|\|\|\|\|]PAM | NA | Intergenic | NA | NA |

T12432 ACGAGGCCCAGACCTCACGGAGG 1000
Oligo fwd CACCACGAGGCCCAGACCTCACGG
Oligo rev AAACCCGTGAGGTCTGGGCCTCGT

| Chromosome | start | end | strand | MM | target_seq | PAM | alignment | distance | position | gene name | gene id |
|---|---|---|---|---|---|---|---|---|---|---|---|
| GU305914.1 | 165026 | 165048 | + | 0 | ACGAGGCCCAGACCTCACGG | AGG | \|\|\|\|\|\|\|\|[\|\|\|\|\|\|\|\|\|\|\|\|]PAM | NA | Intergenic | NA | NA |

T12433 GGCCCAGACCTCACGGAGGACGG 1000
Oligo fwd CACCGGCCCAGACCTCACGGAGGA
Oligo rev AAACTCCTCCGTGAGGTCTGGGCC

| Chromosome | start | end | strand | MM | target_seq | PAM | alignment | distance | position | gene name | gene id |
|---|---|---|---|---|---|---|---|---|---|---|---|
| GU305914.1 | 165030 | 165052 | + | 0 | GGCCCAGACCTCACGGAGGA | CGG | \|\|\|\|\|\|\|\|[\|\|\|\|\|\|\|\|\|\|\|\|]PAM | NA | Intergenic | NA | NA |

T12434 GCCCAGACCTCACGGAGGACGGG 1000
Oligo fwd CACCGCCCAGACCTCACGGAGGAC
Oligo rev AAACGTCCTCCGTGAGGTCTGGGC

| Chromosome | start | end | strand | MM | target_seq | PAM | alignment | distance | position | gene name | gene id |
|---|---|---|---|---|---|---|---|---|---|---|---|
| GU305914.1 | 165031 | 165053 | + | 0 | GCCCAGACCTCACGGAGGAC | GGG | \|\|\|\|\|\|\|\|[\|\|\|\|\|\|\|\|\|\|\|\|]PAM | NA | Intergenic | NA | NA |

T12435 AGACCTCACGGAGGACGGGTTGG 1000
Oligo fwd CACCAGACCTCACGGAGGACGGGT
Oligo rev AAACACCCGTCCTCCGTGAGGTCT

| Chromosome | start | end | strand | MM | target_seq | PAM | alignment | distance | position | gene name | gene id |
|---|---|---|---|---|---|---|---|---|---|---|---|
| GU305914.1 | 165035 | 165057 | + | 0 | AGACCTCACGGAGGACGGGT | TGG | \|\|\|\|\|\|\|\|[\|\|\|\|\|\|\|\|\|\|\|\|]PAM | NA | Intergenic | NA | NA |

T12436 CGGGTTGGAGATTGTCGAGACGG 1000
Oligo fwd CACCCGGGTTGGAGATTGTCGAGA
Oligo rev AAACTCTCGACAATCTCCAACCCG

| Chromosome | start | end | strand | MM | target_seq | PAM | alignment | distance | position | gene name | gene id |
|---|---|---|---|---|---|---|---|---|---|---|---|
| GU305914.1 | 165050 | 165072 | + | 0 | CGGGTTGGAGATTGTCGAGA | CGG | \|\|\|\|\|\|\|\|[\|\|\|\|\|\|\|\|\|\|\|\|]PAM | NA | Intergenic | NA | NA |

T12437 GGGTTGGAGATTGTCGAGACGGG 1000
Oligo fwd CACCGGGTTGGAGATTGTCGAGAC
Oligo rev AAACGTCTCGACAATCTCCAACCC

| Chromosome | start | end | strand | MM | target_seq | PAM | alignment | distance | position | gene name | gene id |
|---|---|---|---|---|---|---|---|---|---|---|---|

| | | | | | | | | | | | |
|---|---|---|---|---|---|---|---|---|---|---|---|
| GU305914.1 | 165051 | 165073 | + | 0 | GGGTTGGAGATTGTCGAGAC | GGG | ||||||||[||||||||||||]PAM | NA | Intergenic | NA | NA |
| T12438 | GTCGAGACGGGCGCCACCGCCGG | 1000 | | | | | | | | | |
| Oligo fwd | CACCGTCGAGACGGGCGCCACCGC | | | | | | | | | | |
| Oligo rev | AAACGCGGTGGCGCCCGTCTCGAC | | | | | | | | | | |
| Chromosome | start | end | strand | MM | target_seq | PAM | alignment | distance | position | gene name | gene id |
| GU305914.1 | 165063 | 165085 | + | 0 | GTCGAGACGGGCGCCACCGC | CGG | ||||||||[||||||||||||]PAM | NA | Intergenic | NA | NA |
| T12439 | GGGCGCCACCGCCGGCACTCTGG | 1000 | | | | | | | | | |
| Oligo fwd | CACCGGGCGCCACCGCCGGCACTC | | | | | | | | | | |
| Oligo rev | AAACGAGTGCCGGCGGTGGCGCCC | | | | | | | | | | |
| Chromosome | start | end | strand | MM | target_seq | PAM | alignment | distance | position | gene name | gene id |
| GU305914.1 | 165071 | 165093 | + | 0 | GGGCGCCACCGCCGGCACTC | TGG | ||||||||[||||||||||||]PAM | NA | Intergenic | NA | NA |
| T12440 | CGGCACTCTGGCCGTTTCCGAGG | 1000 | | | | | | | | | |
| Oligo fwd | CACCCGGCACTCTGGCCGTTTCCG | | | | | | | | | | |
| Oligo rev | AAACCGGAAACGGCCAGAGTGCCG | | | | | | | | | | |
| Chromosome | start | end | strand | MM | target_seq | PAM | alignment | distance | position | gene name | gene id |
| GU305914.1 | 165083 | 165105 | + | 0 | CGGCACTCTGGCCGTTTCCG | AGG | ||||||||[||||||||||||]PAM | NA | Intergenic | NA | NA |
| T12441 | CACTCTGGCCGTTTCCGAGGAGG | 1000 | | | | | | | | | |
| Oligo fwd | CACCCACTCTGGCCGTTTCCGAGG | | | | | | | | | | |
| Oligo rev | AAACCCTCGGAAACGGCCAGAGTG | | | | | | | | | | |
| Chromosome | start | end | strand | MM | target_seq | PAM | alignment | distance | position | gene name | gene id |
| GU305914.1 | 165086 | 165108 | + | 0 | CACTCTGGCCGTTTCCGAGG | AGG | ||||||||[||||||||||||]PAM | NA | Intergenic | NA | NA |
| T12442 | TTCCGAGGAGGAAACCGCGATGG | 1000 | | | | | | | | | |
| Oligo fwd | CACCTTCCGAGGAGGAAACCGCGA | | | | | | | | | | |
| Oligo rev | AAACTCGCGGTTTCCTCCTCGGAA | | | | | | | | | | |
| Chromosome | start | end | strand | MM | target_seq | PAM | alignment | distance | position | gene name | gene id |
| GU305914.1 | 165098 | 165120 | + | 0 | TTCCGAGGAGGAAACCGCGA | TGG | ||||||||[||||||||||||]PAM | NA | Intergenic | NA | NA |
| GU305914.1 | 201602 | 201624 | - | 3 | GTCCGAGGAGGAGACCGCGG | TGG | -|||||||[||||-||||||-]PAM | NA | Intergenic | NA | NA |
| T12443 | GATGGCCGCCGCTATGTTAGAGG | 1000 | | | | | | | | | |
| Oligo fwd | CACCGATGGCCGCCGCTATGTTAG | | | | | | | | | | |
| Oligo rev | AAACCTAACATAGCGGCGGCCATC | | | | | | | | | | |
| Chromosome | start | end | strand | MM | target_seq | PAM | alignment | distance | position | gene name | gene id |
| GU305914.1 | 165116 | 165138 | + | 0 | GATGGCCGCCGCTATGTTAG | AGG | ||||||||[||||||||||||]PAM | NA | Intergenic | NA | NA |
| GU305914.1 | 190207 | 190229 | - | 4 | GTTGGCGGCCGCTATATTGG | TGG | |-||||-|[|||||||-||-|]PAM | NA | Intergenic | NA | NA |
| T12444 | CGCCGCTATGTTAGAGGATATGG | 1000 | | | | | | | | | |
| Oligo fwd | CACCCGCCGCTATGTTAGAGGATA | | | | | | | | | | |
| Oligo rev | AAACTATCCTCTAACATAGCGGCG | | | | | | | | | | |
| Chromosome | start | end | strand | MM | target_seq | PAM | alignment | distance | position | gene name | gene id |
| GU305914.1 | 165122 | 165144 | + | 0 | CGCCGCTATGTTAGAGGATA | TGG | ||||||||[||||||||||||]PAM | NA | Intergenic | NA | NA |
| T12445 | GTTAGAGGATATGGTCGACTTGG | 1000 | | | | | | | | | |
| Oligo fwd | CACCGTTAGAGGATATGGTCGACT | | | | | | | | | | |
| Oligo rev | AAACAGTCGACCATATCCTCTAAC | | | | | | | | | | |
| Chromosome | start | end | strand | MM | target_seq | PAM | alignment | distance | position | gene name | gene id |
| GU305914.1 | 165131 | 165153 | + | 0 | GTTAGAGGATATGGTCGACT | TGG | ||||||||[||||||||||||]PAM | NA | Intergenic | NA | NA |
| T12446 | TTAATTGATCTCATGAAAGATGG | 1000 | | | | | | | | | |
| Oligo fwd | CACCTTAATTGATCTCATGAAAGA | | | | | | | | | | |
| Oligo rev | AAACTCTTTCATGAGATCAATTAA | | | | | | | | | | |
| Chromosome | start | end | strand | MM | target_seq | PAM | alignment | distance | position | gene name | gene id |
| GU305914.1 | 165172 | 165194 | + | 0 | TTAATTGATCTCATGAAAGA | TGG | ||||||||[||||||||||||]PAM | NA | Intergenic | NA | NA |
| T12447 | ATCGATCGCCTCGTCTCGCTCGG | 1000 | | | | | | | | | |
| Oligo fwd | CACCATCGATCGCCTCGTCTCGCT | | | | | | | | | | |
| Oligo rev | AAACAGCGAGACGAGGCGATCGAT | | | | | | | | | | |
| Chromosome | start | end | strand | MM | target_seq | PAM | alignment | distance | position | gene name | gene id |
| GU305914.1 | 165236 | 165258 | + | 0 | ATCGATCGCCTCGTCTCGCT | CGG | ||||||||[||||||||||||]PAM | NA | Intergenic | NA | NA |
| T12448 | TCGATCGCCTCGTCTCGCTCGGG | 1000 | | | | | | | | | |
| Oligo fwd | CACCTCGATCGCCTCGTCTCGCTC | | | | | | | | | | |
| Oligo rev | AAACGAGCGAGACGAGGCGATCGA | | | | | | | | | | |
| Chromosome | start | end | strand | MM | target_seq | PAM | alignment | distance | position | gene name | gene id |
| GU305914.1 | 165237 | 165259 | + | 0 | TCGATCGCCTCGTCTCGCTC | GGG | ||||||||[||||||||||||]PAM | NA | Intergenic | NA | NA |
| T12449 | CTATAGCAGTAGAATATGTATGG | 1000 | | | | | | | | | |
| Oligo fwd | CACCCTATAGCAGTAGAATATGTA | | | | | | | | | | |
| Oligo rev | AAACTACATATTCTACTGCTATAG | | | | | | | | | | |
| Chromosome | start | end | strand | MM | target_seq | PAM | alignment | distance | position | gene name | gene id |
| GU305914.1 | 165307 | 165329 | + | 0 | CTATAGCAGTAGAATATGTA | TGG | ||||||||[||||||||||||]PAM | NA | Intergenic | NA | NA |
| T12450 | CTGTGTGTGTCTCTGTCTACCGG | 1000 | | | | | | | | | |
| Oligo fwd | CACCCTGTGTGTGTCTCTGTCTAC | | | | | | | | | | |
| Oligo rev | AAACGTAGACAGAGACACACACAG | | | | | | | | | | |
| Chromosome | start | end | strand | MM | target_seq | PAM | alignment | distance | position | gene name | gene id |
| GU305914.1 | 165469 | 165491 | + | 0 | CTGTGTGTGTCTCTGTCTAC | CGG | ||||||||[||||||||||||]PAM | NA | Intergenic | NA | NA |
| T12451 | TGTGTGTGTCTCTGTCTACCGGG | 1000 | | | | | | | | | |
| Oligo fwd | CACCTGTGTGTGTCTCTGTCTACC | | | | | | | | | | |
| Oligo rev | AAACGGTAGACAGAGACACACACA | | | | | | | | | | |
| Chromosome | start | end | strand | MM | target_seq | PAM | alignment | distance | position | gene name | gene id |
| GU305914.1 | 165470 | 165492 | + | 0 | TGTGTGTGTCTCTGTCTACC | GGG | ||||||||[||||||||||||]PAM | NA | Intergenic | NA | NA |
| T12452 | GTCTCTGTCTACCGGGCCGATGG | 1000 | | | | | | | | | |

<table>
<tr><td>Oligo fwd</td><td>CACCGTCTCTGTCTACCGGGCCGA</td><td></td><td></td><td></td><td></td><td></td><td></td><td></td><td></td><td></td><td></td></tr>
<tr><td>Oligo rev</td><td>AAACTCGGCCCGGTAGACAGAGAC</td><td></td><td></td><td></td><td></td><td></td><td></td><td></td><td></td><td></td><td></td></tr>
<tr><td>Chromosome</td><td>start</td><td>end</td><td>strand</td><td>MM</td><td>target_seq</td><td>PAM</td><td>alignment</td><td>distance</td><td>position</td><td>gene name</td><td>gene id</td></tr>
<tr><td>GU305914.1</td><td>165477</td><td>165499</td><td>+</td><td>0</td><td>GTCTCTGTCTACCGGGCCGA</td><td>TGG</td><td>||||||||[||||||||||||]PAM</td><td>NA</td><td>Intergenic</td><td>NA</td><td>NA</td></tr>
<tr><td>T12453</td><td>GCCGATGGAGATATTATCCCTGG</td><td>1000</td><td></td><td></td><td></td><td></td><td></td><td></td><td></td><td></td><td></td></tr>
<tr><td>Oligo fwd</td><td>CACCGCCGATGGAGATATTATCCC</td><td></td><td></td><td></td><td></td><td></td><td></td><td></td><td></td><td></td><td></td></tr>
<tr><td>Oligo rev</td><td>AAACGGGATAATATCTCCATCGGC</td><td></td><td></td><td></td><td></td><td></td><td></td><td></td><td></td><td></td><td></td></tr>
<tr><td>Chromosome</td><td>start</td><td>end</td><td>strand</td><td>MM</td><td>target_seq</td><td>PAM</td><td>alignment</td><td>distance</td><td>position</td><td>gene name</td><td>gene id</td></tr>
<tr><td>GU305914.1</td><td>165492</td><td>165514</td><td>+</td><td>0</td><td>GCCGATGGAGATATTATCCC</td><td>TGG</td><td>||||||||[||||||||||||]PAM</td><td>NA</td><td>Intergenic</td><td>NA</td><td>NA</td></tr>
<tr><td>T12454</td><td>TCGCCTTTATTTTTTGTGTTTGG</td><td>1000</td><td></td><td></td><td></td><td></td><td></td><td></td><td></td><td></td><td></td></tr>
<tr><td>Oligo fwd</td><td>CACCTCGCCTTTATTTTTTGTGTT</td><td></td><td></td><td></td><td></td><td></td><td></td><td></td><td></td><td></td><td></td></tr>
<tr><td>Oligo rev</td><td>AAACAACACAAAAAATAAAGGCGA</td><td></td><td></td><td></td><td></td><td></td><td></td><td></td><td></td><td></td><td></td></tr>
<tr><td>Chromosome</td><td>start</td><td>end</td><td>strand</td><td>MM</td><td>target_seq</td><td>PAM</td><td>alignment</td><td>distance</td><td>position</td><td>gene name</td><td>gene id</td></tr>
<tr><td>GU305914.1</td><td>165619</td><td>165641</td><td>+</td><td>0</td><td>TCGCCTTTATTTTTTGTGTT</td><td>TGG</td><td>||||||||[||||||||||||]PAM</td><td>NA</td><td>Intergenic</td><td>NA</td><td>NA</td></tr>
<tr><td>T12455</td><td>CCTTTATTTTTTGTGTTTGGTGG</td><td>1000</td><td></td><td></td><td></td><td></td><td></td><td></td><td></td><td></td><td></td></tr>
<tr><td>Oligo fwd</td><td>CACCCCTTTATTTTTTGTGTTTGG</td><td></td><td></td><td></td><td></td><td></td><td></td><td></td><td></td><td></td><td></td></tr>
<tr><td>Oligo rev</td><td>AAACCCAAACACAAAAAATAAAGG</td><td></td><td></td><td></td><td></td><td></td><td></td><td></td><td></td><td></td><td></td></tr>
<tr><td>Chromosome</td><td>start</td><td>end</td><td>strand</td><td>MM</td><td>target_seq</td><td>PAM</td><td>alignment</td><td>distance</td><td>position</td><td>gene name</td><td>gene id</td></tr>
<tr><td>GU305914.1</td><td>165622</td><td>165644</td><td>+</td><td>0</td><td>CCTTTATTTTTTGTGTTTGG</td><td>TGG</td><td>||||||||[||||||||||||]PAM</td><td>NA</td><td>Intergenic</td><td>NA</td><td>NA</td></tr>
<tr><td>T12456</td><td>TTATTTTTTGTGTTTGGTGGTGG</td><td>1000</td><td></td><td></td><td></td><td></td><td></td><td></td><td></td><td></td><td></td></tr>
<tr><td>Oligo fwd</td><td>CACCTTATTTTTTGTGTTTGGTGG</td><td></td><td></td><td></td><td></td><td></td><td></td><td></td><td></td><td></td><td></td></tr>
<tr><td>Oligo rev</td><td>AAACCCACCAAACACAAAAAATAA</td><td></td><td></td><td></td><td></td><td></td><td></td><td></td><td></td><td></td><td></td></tr>
<tr><td>Chromosome</td><td>start</td><td>end</td><td>strand</td><td>MM</td><td>target_seq</td><td>PAM</td><td>alignment</td><td>distance</td><td>position</td><td>gene name</td><td>gene id</td></tr>
<tr><td>GU305914.1</td><td>165625</td><td>165647</td><td>+</td><td>0</td><td>TTATTTTTTGTGTTTGGTGG</td><td>TGG</td><td>||||||||[||||||||||||]PAM</td><td>NA</td><td>Intergenic</td><td>NA</td><td>NA</td></tr>
<tr><td>T12457</td><td>TTTGTGTTTGGTGGTGGAGTTGG</td><td>1000</td><td></td><td></td><td></td><td></td><td></td><td></td><td></td><td></td><td></td></tr>
<tr><td>Oligo fwd</td><td>CACCTTTGTGTTTGGTGGTGGAGT</td><td></td><td></td><td></td><td></td><td></td><td></td><td></td><td></td><td></td><td></td></tr>
<tr><td>Oligo rev</td><td>AAACACTCCACCACCAAACACAAA</td><td></td><td></td><td></td><td></td><td></td><td></td><td></td><td></td><td></td><td></td></tr>
<tr><td>Chromosome</td><td>start</td><td>end</td><td>strand</td><td>MM</td><td>target_seq</td><td>PAM</td><td>alignment</td><td>distance</td><td>position</td><td>gene name</td><td>gene id</td></tr>
<tr><td>GU305914.1</td><td>165631</td><td>165653</td><td>+</td><td>0</td><td>TTTGTGTTTGGTGGTGGAGT</td><td>TGG</td><td>||||||||[||||||||||||]PAM</td><td>NA</td><td>Intergenic</td><td>NA</td><td>NA</td></tr>
<tr><td>T12458</td><td>GTGTTTGGTGGTGGAGTTGGAGG</td><td>1000</td><td></td><td></td><td></td><td></td><td></td><td></td><td></td><td></td><td></td></tr>
<tr><td>Oligo fwd</td><td>CACCGTGTTTGGTGGTGGAGTTGG</td><td></td><td></td><td></td><td></td><td></td><td></td><td></td><td></td><td></td><td></td></tr>
<tr><td>Oligo rev</td><td>AAACCCAACTCCACCACCAAACAC</td><td></td><td></td><td></td><td></td><td></td><td></td><td></td><td></td><td></td><td></td></tr>
<tr><td>Chromosome</td><td>start</td><td>end</td><td>strand</td><td>MM</td><td>target_seq</td><td>PAM</td><td>alignment</td><td>distance</td><td>position</td><td>gene name</td><td>gene id</td></tr>
<tr><td>GU305914.1</td><td>165634</td><td>165656</td><td>+</td><td>0</td><td>GTGTTTGGTGGTGGAGTTGG</td><td>AGG</td><td>||||||||[||||||||||||]PAM</td><td>NA</td><td>Intergenic</td><td>NA</td><td>NA</td></tr>
<tr><td>T12459</td><td>TTTGGTGGTGGAGTTGGAGGAGG</td><td>1000</td><td></td><td></td><td></td><td></td><td></td><td></td><td></td><td></td><td></td></tr>
<tr><td>Oligo fwd</td><td>CACCTTTGGTGGTGGAGTTGGAGG</td><td></td><td></td><td></td><td></td><td></td><td></td><td></td><td></td><td></td><td></td></tr>
<tr><td>Oligo rev</td><td>AAACCCTCCAACTCCACCACCAAA</td><td></td><td></td><td></td><td></td><td></td><td></td><td></td><td></td><td></td><td></td></tr>
<tr><td>Chromosome</td><td>start</td><td>end</td><td>strand</td><td>MM</td><td>target_seq</td><td>PAM</td><td>alignment</td><td>distance</td><td>position</td><td>gene name</td><td>gene id</td></tr>
<tr><td>GU305914.1</td><td>165637</td><td>165659</td><td>+</td><td>0</td><td>TTTGGTGGTGGAGTTGGAGG</td><td>AGG</td><td>||||||||[||||||||||||]PAM</td><td>NA</td><td>Intergenic</td><td>NA</td><td>NA</td></tr>
<tr><td>T12460</td><td>GGTGGAGTTGGAGGAGGTTGTGG</td><td>1000</td><td></td><td></td><td></td><td></td><td></td><td></td><td></td><td></td><td></td></tr>
<tr><td>Oligo fwd</td><td>CACCGGTGGAGTTGGAGGAGGTTG</td><td></td><td></td><td></td><td></td><td></td><td></td><td></td><td></td><td></td><td></td></tr>
<tr><td>Oligo rev</td><td>AAACCAACCTCCTCCAACTCCACC</td><td></td><td></td><td></td><td></td><td></td><td></td><td></td><td></td><td></td><td></td></tr>
<tr><td>Chromosome</td><td>start</td><td>end</td><td>strand</td><td>MM</td><td>target_seq</td><td>PAM</td><td>alignment</td><td>distance</td><td>position</td><td>gene name</td><td>gene id</td></tr>
<tr><td>GU305914.1</td><td>165643</td><td>165665</td><td>+</td><td>0</td><td>GGTGGAGTTGGAGGAGGTTG</td><td>TGG</td><td>||||||||[||||||||||||]PAM</td><td>NA</td><td>Intergenic</td><td>NA</td><td>NA</td></tr>
<tr><td>T12461</td><td>GTGGAGTTGGAGGAGGTTGTGGG</td><td>1000</td><td></td><td></td><td></td><td></td><td></td><td></td><td></td><td></td><td></td></tr>
<tr><td>Oligo fwd</td><td>CACCGTGGAGTTGGAGGAGGTTGT</td><td></td><td></td><td></td><td></td><td></td><td></td><td></td><td></td><td></td><td></td></tr>
<tr><td>Oligo rev</td><td>AAACACAACCTCCTCCAACTCCAC</td><td></td><td></td><td></td><td></td><td></td><td></td><td></td><td></td><td></td><td></td></tr>
<tr><td>Chromosome</td><td>start</td><td>end</td><td>strand</td><td>MM</td><td>target_seq</td><td>PAM</td><td>alignment</td><td>distance</td><td>position</td><td>gene name</td><td>gene id</td></tr>
<tr><td>GU305914.1</td><td>165644</td><td>165666</td><td>+</td><td>0</td><td>GTGGAGTTGGAGGAGGTTGT</td><td>GGG</td><td>||||||||[||||||||||||]PAM</td><td>NA</td><td>Intergenic</td><td>NA</td><td>NA</td></tr>
<tr><td>T12462</td><td>TGGAGTTGGAGGAGGTTGTGGGG</td><td>1000</td><td></td><td></td><td></td><td></td><td></td><td></td><td></td><td></td><td></td></tr>
<tr><td>Oligo fwd</td><td>CACCTGGAGTTGGAGGAGGTTGTG</td><td></td><td></td><td></td><td></td><td></td><td></td><td></td><td></td><td></td><td></td></tr>
<tr><td>Oligo rev</td><td>AAACCACAACCTCCTCCAACTCCA</td><td></td><td></td><td></td><td></td><td></td><td></td><td></td><td></td><td></td><td></td></tr>
<tr><td>Chromosome</td><td>start</td><td>end</td><td>strand</td><td>MM</td><td>target_seq</td><td>PAM</td><td>alignment</td><td>distance</td><td>position</td><td>gene name</td><td>gene id</td></tr>
<tr><td>GU305914.1</td><td>165645</td><td>165667</td><td>+</td><td>0</td><td>TGGAGTTGGAGGAGGTTGTG</td><td>GGG</td><td>||||||||[||||||||||||]PAM</td><td>NA</td><td>Intergenic</td><td>NA</td><td>NA</td></tr>
<tr><td>T12463</td><td>GGAGTTGGAGGAGGTTGTGGGGG</td><td>1000</td><td></td><td></td><td></td><td></td><td></td><td></td><td></td><td></td><td></td></tr>
<tr><td>Oligo fwd</td><td>CACCGGAGTTGGAGGAGGTTGTGG</td><td></td><td></td><td></td><td></td><td></td><td></td><td></td><td></td><td></td><td></td></tr>
<tr><td>Oligo rev</td><td>AAACCCACAACCTCCTCCAACTCC</td><td></td><td></td><td></td><td></td><td></td><td></td><td></td><td></td><td></td><td></td></tr>
<tr><td>Chromosome</td><td>start</td><td>end</td><td>strand</td><td>MM</td><td>target_seq</td><td>PAM</td><td>alignment</td><td>distance</td><td>position</td><td>gene name</td><td>gene id</td></tr>
<tr><td>GU305914.1</td><td>165646</td><td>165668</td><td>+</td><td>0</td><td>GGAGTTGGAGGAGGTTGTGG</td><td>GGG</td><td>||||||||[||||||||||||]PAM</td><td>NA</td><td>Intergenic</td><td>NA</td><td>NA</td></tr>
<tr><td>T12464</td><td>GATAATAATATTATTCTCATCGG</td><td>1000</td><td></td><td></td><td></td><td></td><td></td><td></td><td></td><td></td><td></td></tr>
<tr><td>Oligo fwd</td><td>CACCGATAATAATATTATTCTCAT</td><td></td><td></td><td></td><td></td><td></td><td></td><td></td><td></td><td></td><td></td></tr>
<tr><td>Oligo rev</td><td>AAACATGAGAATAATATTATTATC</td><td></td><td></td><td></td><td></td><td></td><td></td><td></td><td></td><td></td><td></td></tr>
<tr><td>Chromosome</td><td>start</td><td>end</td><td>strand</td><td>MM</td><td>target_seq</td><td>PAM</td><td>alignment</td><td>distance</td><td>position</td><td>gene name</td><td>gene id</td></tr>
<tr><td>GU305914.1</td><td>165670</td><td>165692</td><td>+</td><td>0</td><td>GATAATAATATTATTCTCAT</td><td>CGG</td><td>||||||||[||||||||||||]PAM</td><td>NA</td><td>Intergenic</td><td>NA</td><td>NA</td></tr>
<tr><td>T12465</td><td>AATAATATTATTCTCATCGGTGG</td><td>1000</td><td></td><td></td><td></td><td></td><td></td><td></td><td></td><td></td><td></td></tr>
<tr><td>Oligo fwd</td><td>CACCAATAATATTATTCTCATCGG</td><td></td><td></td><td></td><td></td><td></td><td></td><td></td><td></td><td></td><td></td></tr>
<tr><td>Oligo rev</td><td>AAACCCGATGAGAATAATATTATT</td><td></td><td></td><td></td><td></td><td></td><td></td><td></td><td></td><td></td><td></td></tr>
<tr><td>Chromosome</td><td>start</td><td>end</td><td>strand</td><td>MM</td><td>target_seq</td><td>PAM</td><td>alignment</td><td>distance</td><td>position</td><td>gene name</td><td>gene id</td></tr>
<tr><td>GU305914.1</td><td>165673</td><td>165695</td><td>+</td><td>0</td><td>AATAATATTATTCTCATCGG</td><td>TGG</td><td>||||||||[||||||||||||]PAM</td><td>NA</td><td>Intergenic</td><td>NA</td><td>NA</td></tr>
<tr><td>T12466</td><td>TCCAGTCGACGCGCTCCCCGCGG</td><td>1000</td><td></td><td></td><td></td><td></td><td></td><td></td><td></td><td></td><td></td></tr>
<tr><td>Oligo fwd</td><td>CACCTCCAGTCGACGCGCTCCCCG</td><td></td><td></td><td></td><td></td><td></td><td></td><td></td><td></td><td></td><td></td></tr>
<tr><td>Oligo rev</td><td>AAACCGGGGAGCGCGTCGACTGGA</td><td></td><td></td><td></td><td></td><td></td><td></td><td></td><td></td><td></td><td></td></tr>
<tr><td>Chromosome</td><td>start</td><td>end</td><td>strand</td><td>MM</td><td>target_seq</td><td>PAM</td><td>alignment</td><td>distance</td><td>position</td><td>gene name</td><td>gene id</td></tr>
<tr><td>GU305914.1</td><td>165716</td><td>165738</td><td>+</td><td>0</td><td>TCCAGTCGACGCGCTCCCCG</td><td>CGG</td><td>||||||||[||||||||||||]PAM</td><td>NA</td><td>Intergenic</td><td>NA</td><td>NA</td></tr>
</table>

<table>
<tr><td>T12467</td><td>AGTCGACGCGCTCCCCGCGGCGG</td><td>1000</td><td></td><td></td><td></td><td></td><td></td><td></td><td></td><td></td><td></td></tr>
<tr><td>Oligo fwd</td><td>CACCAGTCGACGCGCTCCCCGCGG</td><td></td><td></td><td></td><td></td><td></td><td></td><td></td><td></td><td></td><td></td></tr>
<tr><td>Oligo rev</td><td>AAACCCGCGGGGAGCGCGTCGACT</td><td></td><td></td><td></td><td></td><td></td><td></td><td></td><td></td><td></td><td></td></tr>
<tr><td>Chromosome</td><td>start</td><td>end</td><td>strand</td><td>MM</td><td>target_seq</td><td>PAM</td><td>alignment</td><td>distance</td><td>position</td><td>gene name</td><td>gene id</td></tr>
<tr><td>GU305914.1</td><td>165719</td><td>165741</td><td>+</td><td>0</td><td>AGTCGACGCGCTCCCCGCGG</td><td>CGG</td><td>||||||||[||||||||||||]PAM</td><td>NA</td><td>Intergenic</td><td>NA</td><td>NA</td></tr>
<tr><td>GU305914.1</td><td>30615</td><td>30637</td><td>-</td><td>4</td><td>GGACGAGGCGCTCGCCGCGG</td><td>CGG</td><td>-|-|||-|[|||||-||||||]PAM</td><td>NA</td><td>Intergenic</td><td>NA</td><td>NA</td></tr>
<tr><td>T12468</td><td>GGCTCAGATACTCGTTCGCCAGG</td><td>1000</td><td></td><td></td><td></td><td></td><td></td><td></td><td></td><td></td><td></td></tr>
<tr><td>Oligo fwd</td><td>CACCGGCTCAGATACTCGTTCGCC</td><td></td><td></td><td></td><td></td><td></td><td></td><td></td><td></td><td></td><td></td></tr>
<tr><td>Oligo rev</td><td>AAACGGCGAACGAGTATCTGAGCC</td><td></td><td></td><td></td><td></td><td></td><td></td><td></td><td></td><td></td><td></td></tr>
<tr><td>Chromosome</td><td>start</td><td>end</td><td>strand</td><td>MM</td><td>target_seq</td><td>PAM</td><td>alignment</td><td>distance</td><td>position</td><td>gene name</td><td>gene id</td></tr>
<tr><td>GU305914.1</td><td>165740</td><td>165762</td><td>+</td><td>0</td><td>GGCTCAGATACTCGTTCGCC</td><td>AGG</td><td>||||||||[||||||||||||]PAM</td><td>NA</td><td>Intergenic</td><td>NA</td><td>NA</td></tr>
<tr><td>T12469</td><td>ACTCGTTCGCCAGGACGAAGTGG</td><td>1000</td><td></td><td></td><td></td><td></td><td></td><td></td><td></td><td></td><td></td></tr>
<tr><td>Oligo fwd</td><td>CACCACTCGTTCGCCAGGACGAAG</td><td></td><td></td><td></td><td></td><td></td><td></td><td></td><td></td><td></td><td></td></tr>
<tr><td>Oligo rev</td><td>AAACCTTCGTCCTGGCGAACGAGT</td><td></td><td></td><td></td><td></td><td></td><td></td><td></td><td></td><td></td><td></td></tr>
<tr><td>Chromosome</td><td>start</td><td>end</td><td>strand</td><td>MM</td><td>target_seq</td><td>PAM</td><td>alignment</td><td>distance</td><td>position</td><td>gene name</td><td>gene id</td></tr>
<tr><td>GU305914.1</td><td>165749</td><td>165771</td><td>+</td><td>0</td><td>ACTCGTTCGCCAGGACGAAG</td><td>TGG</td><td>||||||||[||||||||||||]PAM</td><td>NA</td><td>Intergenic</td><td>NA</td><td>NA</td></tr>
<tr><td>T12470</td><td>GAAGTGGTTGTTCCCCACGAAGG</td><td>1000</td><td></td><td></td><td></td><td></td><td></td><td></td><td></td><td></td><td></td></tr>
<tr><td>Oligo fwd</td><td>CACCGAAGTGGTTGTTCCCCACGA</td><td></td><td></td><td></td><td></td><td></td><td></td><td></td><td></td><td></td><td></td></tr>
<tr><td>Oligo rev</td><td>AAACTCGTGGGGAACAACCACTTC</td><td></td><td></td><td></td><td></td><td></td><td></td><td></td><td></td><td></td><td></td></tr>
<tr><td>Chromosome</td><td>start</td><td>end</td><td>strand</td><td>MM</td><td>target_seq</td><td>PAM</td><td>alignment</td><td>distance</td><td>position</td><td>gene name</td><td>gene id</td></tr>
<tr><td>GU305914.1</td><td>165765</td><td>165787</td><td>+</td><td>0</td><td>GAAGTGGTTGTTCCCCACGA</td><td>AGG</td><td>||||||||[||||||||||||]PAM</td><td>NA</td><td>Intergenic</td><td>NA</td><td>NA</td></tr>
<tr><td>T12471</td><td>AAGTGGTTGTTCCCCACGAAGGG</td><td>1000</td><td></td><td></td><td></td><td></td><td></td><td></td><td></td><td></td><td></td></tr>
<tr><td>Oligo fwd</td><td>CACCAAGTGGTTGTTCCCCACGAA</td><td></td><td></td><td></td><td></td><td></td><td></td><td></td><td></td><td></td><td></td></tr>
<tr><td>Oligo rev</td><td>AAACTTCGTGGGGAACAACCACTT</td><td></td><td></td><td></td><td></td><td></td><td></td><td></td><td></td><td></td><td></td></tr>
<tr><td>Chromosome</td><td>start</td><td>end</td><td>strand</td><td>MM</td><td>target_seq</td><td>PAM</td><td>alignment</td><td>distance</td><td>position</td><td>gene name</td><td>gene id</td></tr>
<tr><td>GU305914.1</td><td>165766</td><td>165788</td><td>+</td><td>0</td><td>AAGTGGTTGTTCCCCACGAA</td><td>GGG</td><td>||||||||[||||||||||||]PAM</td><td>NA</td><td>Intergenic</td><td>NA</td><td>NA</td></tr>
<tr><td>T12472</td><td>AGTGGTTGTTCCCCACGAAGGGG</td><td>1000</td><td></td><td></td><td></td><td></td><td></td><td></td><td></td><td></td><td></td></tr>
<tr><td>Oligo fwd</td><td>CACCAGTGGTTGTTCCCCACGAAG</td><td></td><td></td><td></td><td></td><td></td><td></td><td></td><td></td><td></td><td></td></tr>
<tr><td>Oligo rev</td><td>AAACCTTCGTGGGGAACAACCACT</td><td></td><td></td><td></td><td></td><td></td><td></td><td></td><td></td><td></td><td></td></tr>
<tr><td>Chromosome</td><td>start</td><td>end</td><td>strand</td><td>MM</td><td>target_seq</td><td>PAM</td><td>alignment</td><td>distance</td><td>position</td><td>gene name</td><td>gene id</td></tr>
<tr><td>GU305914.1</td><td>165767</td><td>165789</td><td>+</td><td>0</td><td>AGTGGTTGTTCCCCACGAAG</td><td>GGG</td><td>||||||||[||||||||||||]PAM</td><td>NA</td><td>Intergenic</td><td>NA</td><td>NA</td></tr>
<tr><td>T12473</td><td>TTGCGTGCTCCTTGAGCGCGTGG</td><td>1000</td><td></td><td></td><td></td><td></td><td></td><td></td><td></td><td></td><td></td></tr>
<tr><td>Oligo fwd</td><td>CACCTTGCGTGCTCCTTGAGCGCG</td><td></td><td></td><td></td><td></td><td></td><td></td><td></td><td></td><td></td><td></td></tr>
<tr><td>Oligo rev</td><td>AAACCGCGCTCAAGGAGCACGCAA</td><td></td><td></td><td></td><td></td><td></td><td></td><td></td><td></td><td></td><td></td></tr>
<tr><td>Chromosome</td><td>start</td><td>end</td><td>strand</td><td>MM</td><td>target_seq</td><td>PAM</td><td>alignment</td><td>distance</td><td>position</td><td>gene name</td><td>gene id</td></tr>
<tr><td>GU305914.1</td><td>165790</td><td>165812</td><td>+</td><td>0</td><td>TTGCGTGCTCCTTGAGCGCG</td><td>TGG</td><td>||||||||[||||||||||||]PAM</td><td>NA</td><td>Intergenic</td><td>NA</td><td>NA</td></tr>
<tr><td>T12474</td><td>TGCGTGCTCCTTGAGCGCGTGGG</td><td>1000</td><td></td><td></td><td></td><td></td><td></td><td></td><td></td><td></td><td></td></tr>
<tr><td>Oligo fwd</td><td>CACCTGCGTGCTCCTTGAGCGCGT</td><td></td><td></td><td></td><td></td><td></td><td></td><td></td><td></td><td></td><td></td></tr>
<tr><td>Oligo rev</td><td>AAACACGCGCTCAAGGAGCACGCA</td><td></td><td></td><td></td><td></td><td></td><td></td><td></td><td></td><td></td><td></td></tr>
<tr><td>Chromosome</td><td>start</td><td>end</td><td>strand</td><td>MM</td><td>target_seq</td><td>PAM</td><td>alignment</td><td>distance</td><td>position</td><td>gene name</td><td>gene id</td></tr>
<tr><td>GU305914.1</td><td>165791</td><td>165813</td><td>+</td><td>0</td><td>TGCGTGCTCCTTGAGCGCGT</td><td>GGG</td><td>||||||||[||||||||||||]PAM</td><td>NA</td><td>Intergenic</td><td>NA</td><td>NA</td></tr>
<tr><td>T12475</td><td>GCGTGCTCCTTGAGCGCGTGGGG</td><td>1000</td><td></td><td></td><td></td><td></td><td></td><td></td><td></td><td></td><td></td></tr>
<tr><td>Oligo fwd</td><td>CACCGCGTGCTCCTTGAGCGCGTG</td><td></td><td></td><td></td><td></td><td></td><td></td><td></td><td></td><td></td><td></td></tr>
<tr><td>Oligo rev</td><td>AAACCACGCGCTCAAGGAGCACGC</td><td></td><td></td><td></td><td></td><td></td><td></td><td></td><td></td><td></td><td></td></tr>
<tr><td>Chromosome</td><td>start</td><td>end</td><td>strand</td><td>MM</td><td>target_seq</td><td>PAM</td><td>alignment</td><td>distance</td><td>position</td><td>gene name</td><td>gene id</td></tr>
<tr><td>GU305914.1</td><td>165792</td><td>165814</td><td>+</td><td>0</td><td>GCGTGCTCCTTGAGCGCGTG</td><td>GGG</td><td>||||||||[||||||||||||]PAM</td><td>NA</td><td>Intergenic</td><td>NA</td><td>NA</td></tr>
<tr><td>T12476</td><td>GCTCCTTGAGCGCGTGGGGACGG</td><td>1000</td><td></td><td></td><td></td><td></td><td></td><td></td><td></td><td></td><td></td></tr>
<tr><td>Oligo fwd</td><td>CACCGCTCCTTGAGCGCGTGGGGA</td><td></td><td></td><td></td><td></td><td></td><td></td><td></td><td></td><td></td><td></td></tr>
<tr><td>Oligo rev</td><td>AAACTCCCCACGCGCTCAAGGAGC</td><td></td><td></td><td></td><td></td><td></td><td></td><td></td><td></td><td></td><td></td></tr>
<tr><td>Chromosome</td><td>start</td><td>end</td><td>strand</td><td>MM</td><td>target_seq</td><td>PAM</td><td>alignment</td><td>distance</td><td>position</td><td>gene name</td><td>gene id</td></tr>
<tr><td>GU305914.1</td><td>165796</td><td>165818</td><td>+</td><td>0</td><td>GCTCCTTGAGCGCGTGGGGA</td><td>CGG</td><td>||||||||[||||||||||||]PAM</td><td>NA</td><td>Intergenic</td><td>NA</td><td>NA</td></tr>
<tr><td>T12477</td><td>CTCCTTGAGCGCGTGGGGACGGG</td><td>1000</td><td></td><td></td><td></td><td></td><td></td><td></td><td></td><td></td><td></td></tr>
<tr><td>Oligo fwd</td><td>CACCCTCCTTGAGCGCGTGGGGAC</td><td></td><td></td><td></td><td></td><td></td><td></td><td></td><td></td><td></td><td></td></tr>
<tr><td>Oligo rev</td><td>AAACGTCCCCACGCGCTCAAGGAG</td><td></td><td></td><td></td><td></td><td></td><td></td><td></td><td></td><td></td><td></td></tr>
<tr><td>Chromosome</td><td>start</td><td>end</td><td>strand</td><td>MM</td><td>target_seq</td><td>PAM</td><td>alignment</td><td>distance</td><td>position</td><td>gene name</td><td>gene id</td></tr>
<tr><td>GU305914.1</td><td>165797</td><td>165819</td><td>+</td><td>0</td><td>CTCCTTGAGCGCGTGGGGAC</td><td>GGG</td><td>||||||||[||||||||||||]PAM</td><td>NA</td><td>Intergenic</td><td>NA</td><td>NA</td></tr>
<tr><td>T12478</td><td>CTTGAGCGCGTGGGGACGGGTGG</td><td>1000</td><td></td><td></td><td></td><td></td><td></td><td></td><td></td><td></td><td></td></tr>
<tr><td>Oligo fwd</td><td>CACCCTTGAGCGCGTGGGGACGGG</td><td></td><td></td><td></td><td></td><td></td><td></td><td></td><td></td><td></td><td></td></tr>
<tr><td>Oligo rev</td><td>AAACCCCGTCCCCACGCGCTCAAG</td><td></td><td></td><td></td><td></td><td></td><td></td><td></td><td></td><td></td><td></td></tr>
<tr><td>Chromosome</td><td>start</td><td>end</td><td>strand</td><td>MM</td><td>target_seq</td><td>PAM</td><td>alignment</td><td>distance</td><td>position</td><td>gene name</td><td>gene id</td></tr>
<tr><td>GU305914.1</td><td>165800</td><td>165822</td><td>+</td><td>0</td><td>CTTGAGCGCGTGGGGACGGG</td><td>TGG</td><td>||||||||[||||||||||||]PAM</td><td>NA</td><td>Intergenic</td><td>NA</td><td>NA</td></tr>
<tr><td>T12479</td><td>AGCGCGTGGGGACGGGTGGCTGG</td><td>1000</td><td></td><td></td><td></td><td></td><td></td><td></td><td></td><td></td><td></td></tr>
<tr><td>Oligo fwd</td><td>CACCAGCGCGTGGGGACGGGTGGC</td><td></td><td></td><td></td><td></td><td></td><td></td><td></td><td></td><td></td><td></td></tr>
<tr><td>Oligo rev</td><td>AAACGCCACCCGTCCCCACGCGCT</td><td></td><td></td><td></td><td></td><td></td><td></td><td></td><td></td><td></td><td></td></tr>
<tr><td>Chromosome</td><td>start</td><td>end</td><td>strand</td><td>MM</td><td>target_seq</td><td>PAM</td><td>alignment</td><td>distance</td><td>position</td><td>gene name</td><td>gene id</td></tr>
<tr><td>GU305914.1</td><td>165804</td><td>165826</td><td>+</td><td>0</td><td>AGCGCGTGGGGACGGGTGGC</td><td>TGG</td><td>||||||||[||||||||||||]PAM</td><td>NA</td><td>Intergenic</td><td>NA</td><td>NA</td></tr>
<tr><td>T12480</td><td>GTGGGGACGGGTGGCTGGATCGG</td><td>1000</td><td></td><td></td><td></td><td></td><td></td><td></td><td></td><td></td><td></td></tr>
<tr><td>Oligo fwd</td><td>CACCGTGGGGACGGGTGGCTGGAT</td><td></td><td></td><td></td><td></td><td></td><td></td><td></td><td></td><td></td><td></td></tr>
<tr><td>Oligo rev</td><td>AAACATCCAGCCACCCGTCCCCAC</td><td></td><td></td><td></td><td></td><td></td><td></td><td></td><td></td><td></td><td></td></tr>
<tr><td>Chromosome</td><td>start</td><td>end</td><td>strand</td><td>MM</td><td>target_seq</td><td>PAM</td><td>alignment</td><td>distance</td><td>position</td><td>gene name</td><td>gene id</td></tr>
<tr><td>GU305914.1</td><td>165809</td><td>165831</td><td>+</td><td>0</td><td>GTGGGGACGGGTGGCTGGAT</td><td>CGG</td><td>||||||||[||||||||||||]PAM</td><td>NA</td><td>Intergenic</td><td>NA</td><td>NA</td></tr>
<tr><td>T12481</td><td>GGGACGGGTGGCTGGATCGGAGG</td><td>1000</td><td></td><td></td><td></td><td></td><td></td><td></td><td></td><td></td><td></td></tr>
<tr><td>Oligo fwd</td><td>CACCGGGACGGGTGGCTGGATCGG</td><td></td><td></td><td></td><td></td><td></td><td></td><td></td><td></td><td></td><td></td></tr>
<tr><td>Oligo rev</td><td>AAACCCGATCCAGCCACCCGTCCC</td><td></td><td></td><td></td><td></td><td></td><td></td><td></td><td></td><td></td><td></td></tr>
<tr><td>Chromosome</td><td>start</td><td>end</td><td>strand</td><td>MM</td><td>target_seq</td><td>PAM</td><td>alignment</td><td>distance</td><td>position</td><td>gene name</td><td>gene id</td></tr>
</table>

| | | | | | | | | | | | |
|---|---|---|---|---|---|---|---|---|---|---|---|
| GU305914.1 | 165812 | 165834 | + | 0 | GGGACGGGTGGCTGGATCGG | AGG | \|\|\|\|\|\|\|\|[\|\|\|\|\|\|\|\|\|\|\|\|]PAM | NA | Intergenic | NA | NA |
| T12482 | CGGAGGATCAGATGTTTCCTCGG | 1000 | | | | | | | | | |
| Oligo fwd | CACCCGGAGGATCAGATGTTTCCT | | | | | | | | | | |
| Oligo rev | AAACAGGAAACATCTGATCCTCCG | | | | | | | | | | |
| Chromosome | start | end | strand | MM | target_seq | PAM | alignment | distance | position | gene name | gene id |
| GU305914.1 | 165829 | 165851 | + | 0 | CGGAGGATCAGATGTTTCCT | CGG | \|\|\|\|\|\|\|\|[\|\|\|\|\|\|\|\|\|\|\|\|]PAM | NA | Intergenic | NA | NA |
| T12483 | GGAGGATCAGATGTTTCCTCGGG | 1000 | | | | | | | | | |
| Oligo fwd | CACCGGAGGATCAGATGTTTCCTC | | | | | | | | | | |
| Oligo rev | AAACGAGGAAACATCTGATCCTCC | | | | | | | | | | |
| Chromosome | start | end | strand | MM | target_seq | PAM | alignment | distance | position | gene name | gene id |
| GU305914.1 | 165830 | 165852 | + | 0 | GGAGGATCAGATGTTTCCTC | GGG | \|\|\|\|\|\|\|\|[\|\|\|\|\|\|\|\|\|\|\|\|]PAM | NA | Intergenic | NA | NA |
| T12484 | GATGTTTCCTCGGGTCGATCAGG | 1000 | | | | | | | | | |
| Oligo fwd | CACCGATGTTTCCTCGGGTCGATC | | | | | | | | | | |
| Oligo rev | AAACGATCGACCCGAGGAAACATC | | | | | | | | | | |
| Chromosome | start | end | strand | MM | target_seq | PAM | alignment | distance | position | gene name | gene id |
| GU305914.1 | 165839 | 165861 | + | 0 | GATGTTTCCTCGGGTCGATC | AGG | \|\|\|\|\|\|\|\|[\|\|\|\|\|\|\|\|\|\|\|\|]PAM | NA | Intergenic | NA | NA |
| T12485 | AACCAGTCCTTCTCTCTGCTCGG | 1000 | | | | | | | | | |
| Oligo fwd | CACCAACCAGTCCTTCTCTCTGCT | | | | | | | | | | |
| Oligo rev | AAACAGCAGAGAGAAGGACTGGTT | | | | | | | | | | |
| Chromosome | start | end | strand | MM | target_seq | PAM | alignment | distance | position | gene name | gene id |
| GU305914.1 | 165900 | 165922 | + | 0 | AACCAGTCCTTCTCTCTGCT | CGG | \|\|\|\|\|\|\|\|[\|\|\|\|\|\|\|\|\|\|\|\|]PAM | NA | Intergenic | NA | NA |
| T12486 | TCTGCTCGGACAGAGCCTGCAGG | 1000 | | | | | | | | | |
| Oligo fwd | CACCTCTGCTCGGACAGAGCCTGC | | | | | | | | | | |
| Oligo rev | AAACGCAGGCTCTGTCCGAGCAGA | | | | | | | | | | |
| Chromosome | start | end | strand | MM | target_seq | PAM | alignment | distance | position | gene name | gene id |
| GU305914.1 | 165914 | 165936 | + | 0 | TCTGCTCGGACAGAGCCTGC | AGG | \|\|\|\|\|\|\|\|[\|\|\|\|\|\|\|\|\|\|\|\|]PAM | NA | Intergenic | NA | NA |
| T12487 | CCTGCAGGACGCGATCCGAGAGG | 1000 | | | | | | | | | |
| Oligo fwd | CACCCCTGCAGGACGCGATCCGAG | | | | | | | | | | |
| Oligo rev | AAACCTCGGATCGCGTCCTGCAGG | | | | | | | | | | |
| Chromosome | start | end | strand | MM | target_seq | PAM | alignment | distance | position | gene name | gene id |
| GU305914.1 | 165929 | 165951 | + | 0 | CCTGCAGGACGCGATCCGAG | AGG | \|\|\|\|\|\|\|\|[\|\|\|\|\|\|\|\|\|\|\|\|]PAM | NA | Intergenic | NA | NA |
| T12488 | CCGAGAGGATCTGCCATCCCAGG | 1000 | | | | | | | | | |
| Oligo fwd | CACCCCGAGAGGATCTGCCATCCC | | | | | | | | | | |
| Oligo rev | AAACGGGATGGCAGATCCTCTCGG | | | | | | | | | | |
| Chromosome | start | end | strand | MM | target_seq | PAM | alignment | distance | position | gene name | gene id |
| GU305914.1 | 165944 | 165966 | + | 0 | CCGAGAGGATCTGCCATCCC | AGG | \|\|\|\|\|\|\|\|[\|\|\|\|\|\|\|\|\|\|\|\|]PAM | NA | Intergenic | NA | NA |
| T12489 | CGAGAGGATCTGCCATCCCAGGG | 1000 | | | | | | | | | |
| Oligo fwd | CACCCGAGAGGATCTGCCATCCCA | | | | | | | | | | |
| Oligo rev | AAACTGGGATGGCAGATCCTCTCG | | | | | | | | | | |
| Chromosome | start | end | strand | MM | target_seq | PAM | alignment | distance | position | gene name | gene id |
| GU305914.1 | 165945 | 165967 | + | 0 | CGAGAGGATCTGCCATCCCA | GGG | \|\|\|\|\|\|\|\|[\|\|\|\|\|\|\|\|\|\|\|\|]PAM | NA | Intergenic | NA | NA |
| T12490 | CCCAGGGCCTCGTGTGAGCCCGG | 1000 | | | | | | | | | |
| Oligo fwd | CACCCCCAGGGCCTCGTGTGAGCC | | | | | | | | | | |
| Oligo rev | AAACGGCTCACACGAGGCCCTGGG | | | | | | | | | | |
| Chromosome | start | end | strand | MM | target_seq | PAM | alignment | distance | position | gene name | gene id |
| GU305914.1 | 165961 | 165983 | + | 0 | CCCAGGGCCTCGTGTGAGCC | CGG | \|\|\|\|\|\|\|\|[\|\|\|\|\|\|\|\|\|\|\|\|]PAM | NA | Intergenic | NA | NA |
| T12491 | GCCCGGCTGTCCCTTGACCACGG | 1000 | | | | | | | | | |
| Oligo fwd | CACCGCCCGGCTGTCCCTTGACCA | | | | | | | | | | |
| Oligo rev | AAACTGGTCAAGGGACAGCCGGGC | | | | | | | | | | |
| Chromosome | start | end | strand | MM | target_seq | PAM | alignment | distance | position | gene name | gene id |
| GU305914.1 | 165978 | 166000 | + | 0 | GCCCGGCTGTCCCTTGACCA | CGG | \|\|\|\|\|\|\|\|[\|\|\|\|\|\|\|\|\|\|\|\|]PAM | NA | Intergenic | NA | NA |
| T12492 | TGATTAGCAGCACACCCTCGCGG | 1000 | | | | | | | | | |
| Oligo fwd | CACCTGATTAGCAGCACACCCTCG | | | | | | | | | | |
| Oligo rev | AAACCGAGGGTGTGCTGCTAATCA | | | | | | | | | | |
| Chromosome | start | end | strand | MM | target_seq | PAM | alignment | distance | position | gene name | gene id |
| GU305914.1 | 166013 | 166035 | + | 0 | TGATTAGCAGCACACCCTCG | CGG | \|\|\|\|\|\|\|\|[\|\|\|\|\|\|\|\|\|\|\|\|]PAM | NA | Intergenic | NA | NA |
| T12493 | CCTCGCGGCACCACGCGTCCAGG | 1000 | | | | | | | | | |
| Oligo fwd | CACCCCTCGCGGCACCACGCGTCC | | | | | | | | | | |
| Oligo rev | AAACGGACGCGTGGTGCCGCGAGG | | | | | | | | | | |
| Chromosome | start | end | strand | MM | target_seq | PAM | alignment | distance | position | gene name | gene id |
| GU305914.1 | 166028 | 166050 | + | 0 | CCTCGCGGCACCACGCGTCC | AGG | \|\|\|\|\|\|\|\|[\|\|\|\|\|\|\|\|\|\|\|\|]PAM | NA | Intergenic | NA | NA |
| T12494 | GCGTCCAGGCACCCGCACTTCGG | 1000 | | | | | | | | | |
| Oligo fwd | CACCGCGTCCAGGCACCCGCACTT | | | | | | | | | | |
| Oligo rev | AAACAAGTGCGGGTGCCTGGACGC | | | | | | | | | | |
| Chromosome | start | end | strand | MM | target_seq | PAM | alignment | distance | position | gene name | gene id |
| GU305914.1 | 166042 | 166064 | + | 0 | GCGTCCAGGCACCCGCACTT | CGG | \|\|\|\|\|\|\|\|[\|\|\|\|\|\|\|\|\|\|\|\|]PAM | NA | Intergenic | NA | NA |
| T12495 | CACTTCGGCATCGAAAACTCCGG | 1000 | | | | | | | | | |
| Oligo fwd | CACCCACTTCGGCATCGAAAACTC | | | | | | | | | | |
| Oligo rev | AAACGAGTTTTCGATGCCGAAGTG | | | | | | | | | | |
| Chromosome | start | end | strand | MM | target_seq | PAM | alignment | distance | position | gene name | gene id |
| GU305914.1 | 166057 | 166079 | + | 0 | CACTTCGGCATCGAAAACTC | CGG | \|\|\|\|\|\|\|\|[\|\|\|\|\|\|\|\|\|\|\|\|]PAM | NA | Intergenic | NA | NA |
| T12496 | ACTTCGGCATCGAAAACTCCGGG | 1000 | | | | | | | | | |
| Oligo fwd | CACCACTTCGGCATCGAAAACTCC | | | | | | | | | | |
| Oligo rev | AAACGGAGTTTTCGATGCCGAAGT | | | | | | | | | | |

<table>
<tr><td>Chromosome</td><td>start</td><td></td><td>end</td><td>strand</td><td>MM</td><td>target_seq</td><td>PAM</td><td>alignment</td><td>distance</td><td>position</td><td>gene name</td><td>gene id</td></tr>
<tr><td>GU305914.1</td><td></td><td>166058</td><td>166080</td><td>+</td><td>0</td><td>ACTTCGGCATCGAAAACTCC</td><td>GGG</td><td>||||||||[||||||||||||]PAM</td><td>NA</td><td>Intergenic</td><td>NA</td><td>NA</td></tr>
<tr><td></td><td></td><td></td><td></td><td></td><td></td><td></td><td></td><td></td><td></td><td></td><td></td><td></td></tr>
<tr><td>T12497</td><td>AAAACTCCGGGATGCTCCGCCGG</td><td></td><td>1000</td><td></td><td></td><td></td><td></td><td></td><td></td><td></td><td></td><td></td></tr>
<tr><td>Oligo fwd</td><td>CACCAAAACTCCGGGATGCTCCGC</td><td></td><td></td><td></td><td></td><td></td><td></td><td></td><td></td><td></td><td></td><td></td></tr>
<tr><td>Oligo rev</td><td>AAACGCGGAGCATCCCGGAGTTTT</td><td></td><td></td><td></td><td></td><td></td><td></td><td></td><td></td><td></td><td></td><td></td></tr>
<tr><td>Chromosome</td><td>start</td><td></td><td>end</td><td>strand</td><td>MM</td><td>target_seq</td><td>PAM</td><td>alignment</td><td>distance</td><td>position</td><td>gene name</td><td>gene id</td></tr>
<tr><td>GU305914.1</td><td></td><td>166070</td><td>166092</td><td>+</td><td>0</td><td>AAAACTCCGGGATGCTCCGC</td><td>CGG</td><td>||||||||[||||||||||||]PAM</td><td>NA</td><td>Intergenic</td><td>NA</td><td>NA</td></tr>
<tr><td></td><td></td><td></td><td></td><td></td><td></td><td></td><td></td><td></td><td></td><td></td><td></td><td></td></tr>
<tr><td>T12498</td><td>GTTCCTTGAAGACCGTCACTAGG</td><td></td><td>1000</td><td></td><td></td><td></td><td></td><td></td><td></td><td></td><td></td><td></td></tr>
<tr><td>Oligo fwd</td><td>CACCGTTCCTTGAAGACCGTCACT</td><td></td><td></td><td></td><td></td><td></td><td></td><td></td><td></td><td></td><td></td><td></td></tr>
<tr><td>Oligo rev</td><td>AAACAGTGACGGTCTTCAAGGAAC</td><td></td><td></td><td></td><td></td><td></td><td></td><td></td><td></td><td></td><td></td><td></td></tr>
<tr><td>Chromosome</td><td>start</td><td></td><td>end</td><td>strand</td><td>MM</td><td>target_seq</td><td>PAM</td><td>alignment</td><td>distance</td><td>position</td><td>gene name</td><td>gene id</td></tr>
<tr><td>GU305914.1</td><td></td><td>166094</td><td>166116</td><td>+</td><td>0</td><td>GTTCCTTGAAGACCGTCACT</td><td>AGG</td><td>||||||||[||||||||||||]PAM</td><td>NA</td><td>Intergenic</td><td>NA</td><td>NA</td></tr>
<tr><td></td><td></td><td></td><td></td><td></td><td></td><td></td><td></td><td></td><td></td><td></td><td></td><td></td></tr>
<tr><td>T12499</td><td>TTCCTTGAAGACCGTCACTAGGG</td><td></td><td>1000</td><td></td><td></td><td></td><td></td><td></td><td></td><td></td><td></td><td></td></tr>
<tr><td>Oligo fwd</td><td>CACCTTCCTTGAAGACCGTCACTA</td><td></td><td></td><td></td><td></td><td></td><td></td><td></td><td></td><td></td><td></td><td></td></tr>
<tr><td>Oligo rev</td><td>AAACTAGTGACGGTCTTCAAGGAA</td><td></td><td></td><td></td><td></td><td></td><td></td><td></td><td></td><td></td><td></td><td></td></tr>
<tr><td>Chromosome</td><td>start</td><td></td><td>end</td><td>strand</td><td>MM</td><td>target_seq</td><td>PAM</td><td>alignment</td><td>distance</td><td>position</td><td>gene name</td><td>gene id</td></tr>
<tr><td>GU305914.1</td><td></td><td>166095</td><td>166117</td><td>+</td><td>0</td><td>TTCCTTGAAGACCGTCACTA</td><td>GGG</td><td>||||||||[||||||||||||]PAM</td><td>NA</td><td>Intergenic</td><td>NA</td><td>NA</td></tr>
<tr><td></td><td></td><td></td><td></td><td></td><td></td><td></td><td></td><td></td><td></td><td></td><td></td><td></td></tr>
<tr><td>T12500</td><td>TTGAAGACCGTCACTAGGGACGG</td><td></td><td>1000</td><td></td><td></td><td></td><td></td><td></td><td></td><td></td><td></td><td></td></tr>
<tr><td>Oligo fwd</td><td>CACCTTGAAGACCGTCACTAGGGA</td><td></td><td></td><td></td><td></td><td></td><td></td><td></td><td></td><td></td><td></td><td></td></tr>
<tr><td>Oligo rev</td><td>AAACTCCCTAGTGACGGTCTTCAA</td><td></td><td></td><td></td><td></td><td></td><td></td><td></td><td></td><td></td><td></td><td></td></tr>
<tr><td>Chromosome</td><td>start</td><td></td><td>end</td><td>strand</td><td>MM</td><td>target_seq</td><td>PAM</td><td>alignment</td><td>distance</td><td>position</td><td>gene name</td><td>gene id</td></tr>
<tr><td>GU305914.1</td><td></td><td>166099</td><td>166121</td><td>+</td><td>0</td><td>TTGAAGACCGTCACTAGGGA</td><td>CGG</td><td>||||||||[||||||||||||]PAM</td><td>NA</td><td>Intergenic</td><td>NA</td><td>NA</td></tr>
<tr><td>GU305914.1</td><td></td><td>64683</td><td>64705</td><td>-</td><td>4</td><td>TGGAAGTCCGTCTCTAGGGC</td><td>GGG</td><td>|-||||-|[||||-||||||-]PAM</td><td>NA</td><td>Intergenic</td><td>NA</td><td>NA</td></tr>
<tr><td></td><td></td><td></td><td></td><td></td><td></td><td></td><td></td><td></td><td></td><td></td><td></td><td></td></tr>
<tr><td>T12501</td><td>AAGACCGTCACTAGGGACGGCGG</td><td></td><td>1000</td><td></td><td></td><td></td><td></td><td></td><td></td><td></td><td></td><td></td></tr>
<tr><td>Oligo fwd</td><td>CACCAAGACCGTCACTAGGGACGG</td><td></td><td></td><td></td><td></td><td></td><td></td><td></td><td></td><td></td><td></td><td></td></tr>
<tr><td>Oligo rev</td><td>AAACCCGTCCCTAGTGACGGTCTT</td><td></td><td></td><td></td><td></td><td></td><td></td><td></td><td></td><td></td><td></td><td></td></tr>
<tr><td>Chromosome</td><td>start</td><td></td><td>end</td><td>strand</td><td>MM</td><td>target_seq</td><td>PAM</td><td>alignment</td><td>distance</td><td>position</td><td>gene name</td><td>gene id</td></tr>
<tr><td>GU305914.1</td><td></td><td>166102</td><td>166124</td><td>+</td><td>0</td><td>AAGACCGTCACTAGGGACGG</td><td>CGG</td><td>||||||||[||||||||||||]PAM</td><td>NA</td><td>Intergenic</td><td>NA</td><td>NA</td></tr>
<tr><td></td><td></td><td></td><td></td><td></td><td></td><td></td><td></td><td></td><td></td><td></td><td></td><td></td></tr>
<tr><td>T12502</td><td>AGACCGTCACTAGGGACGGCGGG</td><td></td><td>1000</td><td></td><td></td><td></td><td></td><td></td><td></td><td></td><td></td><td></td></tr>
<tr><td>Oligo fwd</td><td>CACCAGACCGTCACTAGGGACGGC</td><td></td><td></td><td></td><td></td><td></td><td></td><td></td><td></td><td></td><td></td><td></td></tr>
<tr><td>Oligo rev</td><td>AAACGCCGTCCCTAGTGACGGTCT</td><td></td><td></td><td></td><td></td><td></td><td></td><td></td><td></td><td></td><td></td><td></td></tr>
<tr><td>Chromosome</td><td>start</td><td></td><td>end</td><td>strand</td><td>MM</td><td>target_seq</td><td>PAM</td><td>alignment</td><td>distance</td><td>position</td><td>gene name</td><td>gene id</td></tr>
<tr><td>GU305914.1</td><td></td><td>166103</td><td>166125</td><td>+</td><td>0</td><td>AGACCGTCACTAGGGACGGC</td><td>GGG</td><td>||||||||[||||||||||||]PAM</td><td>NA</td><td>Intergenic</td><td>NA</td><td>NA</td></tr>
<tr><td></td><td></td><td></td><td></td><td></td><td></td><td></td><td></td><td></td><td></td><td></td><td></td><td></td></tr>
<tr><td>T12503</td><td>GACCGTCACTAGGGACGGCGGGG</td><td></td><td>1000</td><td></td><td></td><td></td><td></td><td></td><td></td><td></td><td></td><td></td></tr>
<tr><td>Oligo fwd</td><td>CACCGACCGTCACTAGGGACGGCG</td><td></td><td></td><td></td><td></td><td></td><td></td><td></td><td></td><td></td><td></td><td></td></tr>
<tr><td>Oligo rev</td><td>AAACCGCCGTCCCTAGTGACGGTC</td><td></td><td></td><td></td><td></td><td></td><td></td><td></td><td></td><td></td><td></td><td></td></tr>
<tr><td>Chromosome</td><td>start</td><td></td><td>end</td><td>strand</td><td>MM</td><td>target_seq</td><td>PAM</td><td>alignment</td><td>distance</td><td>position</td><td>gene name</td><td>gene id</td></tr>
<tr><td>GU305914.1</td><td></td><td>166104</td><td>166126</td><td>+</td><td>0</td><td>GACCGTCACTAGGGACGGCG</td><td>GGG</td><td>||||||||[||||||||||||]PAM</td><td>NA</td><td>Intergenic</td><td>NA</td><td>NA</td></tr>
<tr><td></td><td></td><td></td><td></td><td></td><td></td><td></td><td></td><td></td><td></td><td></td><td></td><td></td></tr>
<tr><td>T12504</td><td>GTCACTAGGGACGGCGGGGCCGG</td><td></td><td>1000</td><td></td><td></td><td></td><td></td><td></td><td></td><td></td><td></td><td></td></tr>
<tr><td>Oligo fwd</td><td>CACCGTCACTAGGGACGGCGGGGC</td><td></td><td></td><td></td><td></td><td></td><td></td><td></td><td></td><td></td><td></td><td></td></tr>
<tr><td>Oligo rev</td><td>AAACGCCCCGCCGTCCCTAGTGAC</td><td></td><td></td><td></td><td></td><td></td><td></td><td></td><td></td><td></td><td></td><td></td></tr>
<tr><td>Chromosome</td><td>start</td><td></td><td>end</td><td>strand</td><td>MM</td><td>target_seq</td><td>PAM</td><td>alignment</td><td>distance</td><td>position</td><td>gene name</td><td>gene id</td></tr>
<tr><td>GU305914.1</td><td></td><td>166108</td><td>166130</td><td>+</td><td>0</td><td>GTCACTAGGGACGGCGGGGC</td><td>CGG</td><td>||||||||[||||||||||||]PAM</td><td>NA</td><td>Intergenic</td><td>NA</td><td>NA</td></tr>
<tr><td></td><td></td><td></td><td></td><td></td><td></td><td></td><td></td><td></td><td></td><td></td><td></td><td></td></tr>
<tr><td>T12505</td><td>CTAGGGACGGCGGGGCCGGACGG</td><td></td><td>1000</td><td></td><td></td><td></td><td></td><td></td><td></td><td></td><td></td><td></td></tr>
<tr><td>Oligo fwd</td><td>CACCCTAGGGACGGCGGGGCCGGA</td><td></td><td></td><td></td><td></td><td></td><td></td><td></td><td></td><td></td><td></td><td></td></tr>
<tr><td>Oligo rev</td><td>AAACTCCGGCCCCGCCGTCCCTAG</td><td></td><td></td><td></td><td></td><td></td><td></td><td></td><td></td><td></td><td></td><td></td></tr>
<tr><td>Chromosome</td><td>start</td><td></td><td>end</td><td>strand</td><td>MM</td><td>target_seq</td><td>PAM</td><td>alignment</td><td>distance</td><td>position</td><td>gene name</td><td>gene id</td></tr>
<tr><td>GU305914.1</td><td></td><td>166112</td><td>166134</td><td>+</td><td>0</td><td>CTAGGGACGGCGGGGCCGGA</td><td>CGG</td><td>||||||||[||||||||||||]PAM</td><td>NA</td><td>Intergenic</td><td>NA</td><td>NA</td></tr>
<tr><td></td><td></td><td></td><td></td><td></td><td></td><td></td><td></td><td></td><td></td><td></td><td></td><td></td></tr>
<tr><td>T12506</td><td>CGGGGCCGGACGGTCTCTCACGG</td><td></td><td>1000</td><td></td><td></td><td></td><td></td><td></td><td></td><td></td><td></td><td></td></tr>
<tr><td>Oligo fwd</td><td>CACCCGGGGCCGGACGGTCTCTCA</td><td></td><td></td><td></td><td></td><td></td><td></td><td></td><td></td><td></td><td></td><td></td></tr>
<tr><td>Oligo rev</td><td>AAACTGAGAGACCGTCCGGCCCCG</td><td></td><td></td><td></td><td></td><td></td><td></td><td></td><td></td><td></td><td></td><td></td></tr>
<tr><td>Chromosome</td><td>start</td><td></td><td>end</td><td>strand</td><td>MM</td><td>target_seq</td><td>PAM</td><td>alignment</td><td>distance</td><td>position</td><td>gene name</td><td>gene id</td></tr>
<tr><td>GU305914.1</td><td></td><td>166122</td><td>166144</td><td>+</td><td>0</td><td>CGGGGCCGGACGGTCTCTCA</td><td>CGG</td><td>||||||||[||||||||||||]PAM</td><td>NA</td><td>Intergenic</td><td>NA</td><td>NA</td></tr>
<tr><td></td><td></td><td></td><td></td><td></td><td></td><td></td><td></td><td></td><td></td><td></td><td></td><td></td></tr>
<tr><td>T12507</td><td>ACGGTCTCTCACGGTCCCGAAGG</td><td></td><td>1000</td><td></td><td></td><td></td><td></td><td></td><td></td><td></td><td></td><td></td></tr>
<tr><td>Oligo fwd</td><td>CACCACGGTCTCTCACGGTCCCGA</td><td></td><td></td><td></td><td></td><td></td><td></td><td></td><td></td><td></td><td></td><td></td></tr>
<tr><td>Oligo rev</td><td>AAACTCGGGACCGTGAGAGACCGT</td><td></td><td></td><td></td><td></td><td></td><td></td><td></td><td></td><td></td><td></td><td></td></tr>
<tr><td>Chromosome</td><td>start</td><td></td><td>end</td><td>strand</td><td>MM</td><td>target_seq</td><td>PAM</td><td>alignment</td><td>distance</td><td>position</td><td>gene name</td><td>gene id</td></tr>
<tr><td>GU305914.1</td><td></td><td>166131</td><td>166153</td><td>+</td><td>0</td><td>ACGGTCTCTCACGGTCCCGA</td><td>AGG</td><td>||||||||[||||||||||||]PAM</td><td>NA</td><td>Intergenic</td><td>NA</td><td>NA</td></tr>
<tr><td></td><td></td><td></td><td></td><td></td><td></td><td></td><td></td><td></td><td></td><td></td><td></td><td></td></tr>
<tr><td>T12508</td><td>CTCTCACGGTCCCGAAGGCGAGG</td><td></td><td>1000</td><td></td><td></td><td></td><td></td><td></td><td></td><td></td><td></td><td></td></tr>
<tr><td>Oligo fwd</td><td>CACCCTCTCACGGTCCCGAAGGCG</td><td></td><td></td><td></td><td></td><td></td><td></td><td></td><td></td><td></td><td></td><td></td></tr>
<tr><td>Oligo rev</td><td>AAACCGCCTTCGGGACCGTGAGAG</td><td></td><td></td><td></td><td></td><td></td><td></td><td></td><td></td><td></td><td></td><td></td></tr>
<tr><td>Chromosome</td><td>start</td><td></td><td>end</td><td>strand</td><td>MM</td><td>target_seq</td><td>PAM</td><td>alignment</td><td>distance</td><td>position</td><td>gene name</td><td>gene id</td></tr>
<tr><td>GU305914.1</td><td></td><td>166136</td><td>166158</td><td>+</td><td>0</td><td>CTCTCACGGTCCCGAAGGCG</td><td>AGG</td><td>||||||||[||||||||||||]PAM</td><td>NA</td><td>Intergenic</td><td>NA</td><td>NA</td></tr>
<tr><td></td><td></td><td></td><td></td><td></td><td></td><td></td><td></td><td></td><td></td><td></td><td></td><td></td></tr>
<tr><td>T12509</td><td>GGCCGCACGCGCTGCCGTCGTGG</td><td></td><td>1000</td><td></td><td></td><td></td><td></td><td></td><td></td><td></td><td></td><td></td></tr>
<tr><td>Oligo fwd</td><td>CACCGGCCGCACGCGCTGCCGTCG</td><td></td><td></td><td></td><td></td><td></td><td></td><td></td><td></td><td></td><td></td><td></td></tr>
<tr><td>Oligo rev</td><td>AAACCGACGGCAGCGCGTGCGGCC</td><td></td><td></td><td></td><td></td><td></td><td></td><td></td><td></td><td></td><td></td><td></td></tr>
<tr><td>Chromosome</td><td>start</td><td></td><td>end</td><td>strand</td><td>MM</td><td>target_seq</td><td>PAM</td><td>alignment</td><td>distance</td><td>position</td><td>gene name</td><td>gene id</td></tr>
<tr><td>GU305914.1</td><td></td><td>166157</td><td>166179</td><td>+</td><td>0</td><td>GGCCGCACGCGCTGCCGTCG</td><td>TGG</td><td>||||||||[||||||||||||]PAM</td><td>NA</td><td>Intergenic</td><td>NA</td><td>NA</td></tr>
<tr><td></td><td></td><td></td><td></td><td></td><td></td><td></td><td></td><td></td><td></td><td></td><td></td><td></td></tr>
<tr><td>T12510</td><td>CACGCGCTGCCGTCGTGGTACGG</td><td></td><td>1000</td><td></td><td></td><td></td><td></td><td></td><td></td><td></td><td></td><td></td></tr>
<tr><td>Oligo fwd</td><td>CACCCACGCGCTGCCGTCGTGGTA</td><td></td><td></td><td></td><td></td><td></td><td></td><td></td><td></td><td></td><td></td><td></td></tr>
<tr><td>Oligo rev</td><td>AAACTACCACGACGGCAGCGCGTG</td><td></td><td></td><td></td><td></td><td></td><td></td><td></td><td></td><td></td><td></td><td></td></tr>
<tr><td>Chromosome</td><td>start</td><td></td><td>end</td><td>strand</td><td>MM</td><td>target_seq</td><td>PAM</td><td>alignment</td><td>distance</td><td>position</td><td>gene name</td><td>gene id</td></tr>
<tr><td>GU305914.1</td><td></td><td>166162</td><td>166184</td><td>+</td><td>0</td><td>CACGCGCTGCCGTCGTGGTA</td><td>CGG</td><td>||||||||[||||||||||||]PAM</td><td>NA</td><td>Intergenic</td><td>NA</td><td>NA</td></tr>
<tr><td></td><td></td><td></td><td></td><td></td><td></td><td></td><td></td><td></td><td></td><td></td><td></td><td></td></tr>
<tr><td>T12511</td><td>CGTGGTACGGATCTTGTCCTAGG</td><td></td><td>1000</td><td></td><td></td><td></td><td></td><td></td><td></td><td></td><td></td><td></td></tr>
</table>

Oligo fwd CACCCGTGGTACGGATCTTGTCCT
Oligo rev AAACAGGACAAGATCCGTACCACG
Chromosome start end strand MM target_seq PAM alignment distance position gene name gene id
GU305914.1 166175 166197 + 0 CGTGGTACGGATCTTGTCCT AGG ||||||||[||||||||||||]PAM NA Intergenic NA NA

T12512 ATCACCACTCGCACGTCGTACGG 1000
Oligo fwd CACCATCACCACTCGCACGTCGTA
Oligo rev AAACTACGACGTGCGAGTGGTGAT
Chromosome start end strand MM target_seq PAM alignment distance position gene name gene id
GU305914.1 166198 166220 + 0 ATCACCACTCGCACGTCGTA CGG ||||||||[||||||||||||]PAM NA Intergenic NA NA

T12513 TCACCACTCGCACGTCGTACGGG 1000
Oligo fwd CACCTCACCACTCGCACGTCGTAC
Oligo rev AAACGTACGACGTGCGAGTGGTGA
Chromosome start end strand MM target_seq PAM alignment distance position gene name gene id
GU305914.1 166199 166221 + 0 TCACCACTCGCACGTCGTAC GGG ||||||||[||||||||||||]PAM NA Intergenic NA NA

T12514 GGAAACAGAGTCTGCTCCAACGG 1000
Oligo fwd CACCGGAAACAGAGTCTGCTCCAA
Oligo rev AAACTTGGAGCAGACTCTGTTTCC
Chromosome start end strand MM target_seq PAM alignment distance position gene name gene id
GU305914.1 166220 166242 + 0 GGAAACAGAGTCTGCTCCAA CGG ||||||||[||||||||||||]PAM NA Intergenic NA NA

T12515 AACAGAGTCTGCTCCAACGGTGG 1000
Oligo fwd CACCAACAGAGTCTGCTCCAACGG
Oligo rev AAACCCGTTGGAGCAGACTCTGTT
Chromosome start end strand MM target_seq PAM alignment distance position gene name gene id
GU305914.1 166223 166245 + 0 AACAGAGTCTGCTCCAACGG TGG ||||||||[||||||||||||]PAM NA Intergenic NA NA

T12516 ACGGTGGACGTCGCTCTTCTCGG 1000
Oligo fwd CACCACGGTGGACGTCGCTCTTCT
Oligo rev AAACAGAAGAGCGACGTCCACCGT
Chromosome start end strand MM target_seq PAM alignment distance position gene name gene id
GU305914.1 166239 166261 + 0 ACGGTGGACGTCGCTCTTCT CGG ||||||||[||||||||||||]PAM NA Intergenic NA NA

T12517 CGGTGGACGTCGCTCTTCTCGGG 1000
Oligo fwd CACCCGGTGGACGTCGCTCTTCTC
Oligo rev AAACGAGAAGAGCGACGTCCACCG
Chromosome start end strand MM target_seq PAM alignment distance position gene name gene id
GU305914.1 166240 166262 + 0 CGGTGGACGTCGCTCTTCTC GGG ||||||||[||||||||||||]PAM NA Intergenic NA NA

T12518 GGTGGACGTCGCTCTTCTCGGGG 1000
Oligo fwd CACCGGTGGACGTCGCTCTTCTCG
Oligo rev AAACCGAGAAGAGCGACGTCCACC
Chromosome start end strand MM target_seq PAM alignment distance position gene name gene id
GU305914.1 166241 166263 + 0 GGTGGACGTCGCTCTTCTCG GGG ||||||||[||||||||||||]PAM NA Intergenic NA NA

T12519 GTCGCTCTTCTCGGGGTAGATGG 1000
Oligo fwd CACCGTCGCTCTTCTCGGGGTAGA
Oligo rev AAACTCTACCCCGAGAAGAGCGAC
Chromosome start end strand MM target_seq PAM alignment distance position gene name gene id
GU305914.1 166248 166270 + 0 GTCGCTCTTCTCGGGGTAGA TGG ||||||||[||||||||||||]PAM NA Intergenic NA NA

T12520 TCTCTCCTGATCTACCCTGTCGG 1000
Oligo fwd CACCTCTCTCCTGATCTACCCTGT
Oligo rev AAACACAGGGTAGATCAGGAGAGA
Chromosome start end strand MM target_seq PAM alignment distance position gene name gene id
GU305914.1 166281 166303 + 0 TCTCTCCTGATCTACCCTGT CGG ||||||||[||||||||||||]PAM NA Intergenic NA NA

T12521 GATCTACCCTGTCGGTGATGCGG 1000
Oligo fwd CACCGATCTACCCTGTCGGTGATG
Oligo rev AAACCATCACCGACAGGGTAGATC
Chromosome start end strand MM target_seq PAM alignment distance position gene name gene id
GU305914.1 166289 166311 + 0 GATCTACCCTGTCGGTGATG CGG ||||||||[||||||||||||]PAM NA Intergenic NA NA

T12522 GACCAGCTGTGACGCCTGCGTGG 1000
Oligo fwd CACCGACCAGCTGTGACGCCTGCG
Oligo rev AAACCGCAGGCGTCACAGCTGGTC
Chromosome start end strand MM target_seq PAM alignment distance position gene name gene id
GU305914.1 166311 166333 + 0 GACCAGCTGTGACGCCTGCG TGG ||||||||[||||||||||||]PAM NA Intergenic NA NA

T12523 CAGCTGTGACGCCTGCGTGGCGG 1000
Oligo fwd CACCCAGCTGTGACGCCTGCGTGG
Oligo rev AAACCCACGCAGGCGTCACAGCTG
Chromosome start end strand MM target_seq PAM alignment distance position gene name gene id
GU305914.1 166314 166336 + 0 CAGCTGTGACGCCTGCGTGG CGG ||||||||[||||||||||||]PAM NA Intergenic NA NA

T12524 TAGTTTCAGAAAGCTCATCCAGG 1000
Oligo fwd CACCTAGTTTCAGAAAGCTCATCC
Oligo rev AAACGGATGAGCTTTCTGAAACTA
Chromosome start end strand MM target_seq PAM alignment distance position gene name gene id
GU305914.1 166338 166360 + 0 TAGTTTCAGAAAGCTCATCC AGG ||||||||[||||||||||||]PAM NA Intergenic NA NA

T12525 GAAAGCTCATCCAGGCTTCGTGG 1000
Oligo fwd CACCGAAAGCTCATCCAGGCTTCG
Oligo rev AAACCGAAGCCTGGATGAGCTTTC
Chromosome start end strand MM target_seq PAM alignment distance position gene name gene id
GU305914.1 166346 166368 + 0 GAAAGCTCATCCAGGCTTCG TGG ||||||||[||||||||||||]PAM NA Intergenic NA NA

| | | | | | | | | | | | |
|---|---|---|---|---|---|---|---|---|---|---|---|
| T12526 | CGCCTGTTGATGAATCACGTCGG | 1000 | | | | | | | | | |
| Oligo fwd | CACCCGCCTGTTGATGAATCACGT | | | | | | | | | | |
| Oligo rev | AAACACGTGATTCATCAACAGGCG | | | | | | | | | | |
| Chromosome | start | end | strand | MM | target_seq | PAM | alignment | distance | position | gene name | gene id |
| GU305914.1 | 166383 | 166405 | + | 0 | CGCCTGTTGATGAATCACGT | CGG | \|\|\|\|\|\|\|\|[\|\|\|\|\|\|\|\|\|\|\|\|]PAM | NA | Intergenic | NA | NA |

| | | | | | | | | | | | |
|---|---|---|---|---|---|---|---|---|---|---|---|
| T12527 | AATCACGTCGGCAGAGACGTCGG | 1000 | | | | | | | | | |
| Oligo fwd | CACCAATCACGTCGGCAGAGACGT | | | | | | | | | | |
| Oligo rev | AAACACGTCTCTGCCGACGTGATT | | | | | | | | | | |
| Chromosome | start | end | strand | MM | target_seq | PAM | alignment | distance | position | gene name | gene id |
| GU305914.1 | 166395 | 166417 | + | 0 | AATCACGTCGGCAGAGACGT | CGG | \|\|\|\|\|\|\|\|[\|\|\|\|\|\|\|\|\|\|\|\|]PAM | NA | Intergenic | NA | NA |

| | | | | | | | | | | | |
|---|---|---|---|---|---|---|---|---|---|---|---|
| T12528 | GTCGGCAGAGACGTCGGCGCCGG | 1000 | | | | | | | | | |
| Oligo fwd | CACCGTCGGCAGAGACGTCGGCGC | | | | | | | | | | |
| Oligo rev | AAACGCGCCGACGTCTCTGCCGAC | | | | | | | | | | |
| Chromosome | start | end | strand | MM | target_seq | PAM | alignment | distance | position | gene name | gene id |
| GU305914.1 | 166401 | 166423 | + | 0 | GTCGGCAGAGACGTCGGCGC | CGG | \|\|\|\|\|\|\|\|[\|\|\|\|\|\|\|\|\|\|\|\|]PAM | NA | Intergenic | NA | NA |

| | | | | | | | | | | | |
|---|---|---|---|---|---|---|---|---|---|---|---|
| T12529 | GGCAGAGACGTCGGCGCCGGCGG | 1000 | | | | | | | | | |
| Oligo fwd | CACCGGCAGAGACGTCGGCGCCGG | | | | | | | | | | |
| Oligo rev | AAACCCGGCGCCGACGTCTCTGCC | | | | | | | | | | |
| Chromosome | start | end | strand | MM | target_seq | PAM | alignment | distance | position | gene name | gene id |
| GU305914.1 | 166404 | 166426 | + | 0 | GGCAGAGACGTCGGCGCCGG | CGG | \|\|\|\|\|\|\|\|[\|\|\|\|\|\|\|\|\|\|\|\|]PAM | NA | Intergenic | NA | NA |
| GU305914.1 | 90250 | 90272 | - | 4 | GGCGGAGGCGGCGGAGCCGG | CGG | \|\|\|-\|\|\|-[\|\|-\|\|\|-\|\|\|\|\|]PAM | NA | Intergenic | NA | NA |
| GU305914.1 | 128809 | 128831 | + | 3 | GGCAGCGACGTCGGCGGCGA | AGG | \|\|\|\|\|-\|\|[\|\|\|\|\|\|\|\|-\|\|-]PAM | NA | Intergenic | NA | NA |

| | | | | | | | | | | | |
|---|---|---|---|---|---|---|---|---|---|---|---|
| T12530 | GACGTCGGCGCCGGCGGCAGCGG | 1000 | | | | | | | | | |
| Oligo fwd | CACCGACGTCGGCGCCGGCGGCAG | | | | | | | | | | |
| Oligo rev | AAACCTGCCGCCGGCGCCGACGTC | | | | | | | | | | |
| Chromosome | start | end | strand | MM | target_seq | PAM | alignment | distance | position | gene name | gene id |
| GU305914.1 | 166410 | 166432 | + | 0 | GACGTCGGCGCCGGCGGCAG | CGG | \|\|\|\|\|\|\|\|[\|\|\|\|\|\|\|\|\|\|\|\|]PAM | NA | Intergenic | NA | NA |
| GU305914.1 | 30006 | 30028 | + | 3 | GGCGGCGGCGGCGGCGGCAG | TGG | \|-\|\|-\|\|\|[\|\|-\|\|\|\|\|\|\|\|\|]PAM | NA | Intergenic | NA | NA |
| GU305914.1 | 149476 | 149498 | - | 3 | GGCGGCGGCGGCGGCGGCAG | CGG | \|-\|\|-\|\|\|[\|\|-\|\|\|\|\|\|\|\|\|]PAM | NA | Intergenic | NA | NA |
| GU305914.1 | 163902 | 163924 | + | 4 | AGCGGCGGCGGCGGCGGCAG | CGG | --\|\|-\|\|\|[\|\|-\|\|\|\|\|\|\|\|\|]PAM | NA | Intergenic | NA | NA |
| GU305914.1 | 113924 | 113946 | - | 4 | GTCGGCGTCGGCGGCGGCAG | CGG | \|-\|\|-\|\|-[\|\|-\|\|\|\|\|\|\|\|\|]PAM | NA | Intergenic | NA | NA |
| GU305914.1 | 163908 | 163930 | + | 4 | GGCGGCGGCGGCAGCGGCAG | CGG | \|-\|\|-\|\|\|[\|\|-\|-\|\|\|\|\|\|\|]PAM | NA | Intergenic | NA | NA |
| GU305914.1 | 122483 | 122505 | - | 4 | GGTGTCGGCGCGGGCAGCAG | CGG | \|--\|\|\|\|\|[\|\|\|-\|\|\|-\|\|\|\|]PAM | NA | Intergenic | NA | NA |
| GU305914.1 | 149482 | 149504 | - | 4 | GGCGACGGCGGCGGCGGCGG | CGG | \|-\|\|-\|\|\|[\|\|-\|\|\|\|\|\|\|-\|]PAM | NA | Intergenic | NA | NA |
| GU305914.1 | 176454 | 176476 | + | 4 | GACGATGGCGACGGCGGCGG | AGG | \|\|\|\|--\|\|[\|\|-\|\|\|\|\|\|\|-\|]PAM | NA | Intergenic | NA | NA |
| GU305914.1 | 163749 | 163771 | + | 4 | GTCGGCGGCGCCGGTGGCGG | CGG | \|-\|\|-\|\|\|[\|\|\|\|\|\|-\|\|\|-\|]PAM | NA | Intergenic | NA | NA |

| | | | | | | | | | | | |
|---|---|---|---|---|---|---|---|---|---|---|---|
| T12531 | TGTGTCTGAGCATCCACTGCCGG | 1000 | | | | | | | | | |
| Oligo fwd | CACCTGTGTCTGAGCATCCACTGC | | | | | | | | | | |
| Oligo rev | AAACGCAGTGGATGCTCAGACACA | | | | | | | | | | |
| Chromosome | start | end | strand | MM | target_seq | PAM | alignment | distance | position | gene name | gene id |
| GU305914.1 | 166454 | 166476 | + | 0 | TGTGTCTGAGCATCCACTGC | CGG | \|\|\|\|\|\|\|\|[\|\|\|\|\|\|\|\|\|\|\|\|]PAM | NA | Intergenic | NA | NA |

| | | | | | | | | | | | |
|---|---|---|---|---|---|---|---|---|---|---|---|
| T12532 | CCACTGCCGGAGAGCCATCACGG | 1000 | | | | | | | | | |
| Oligo fwd | CACCCCACTGCCGGAGAGCCATCA | | | | | | | | | | |
| Oligo rev | AAACTGATGGCTCTCCGGCAGTGG | | | | | | | | | | |
| Chromosome | start | end | strand | MM | target_seq | PAM | alignment | distance | position | gene name | gene id |
| GU305914.1 | 166467 | 166489 | + | 0 | CCACTGCCGGAGAGCCATCA | CGG | \|\|\|\|\|\|\|\|[\|\|\|\|\|\|\|\|\|\|\|\|]PAM | NA | Intergenic | NA | NA |

| | | | | | | | | | | | |
|---|---|---|---|---|---|---|---|---|---|---|---|
| T12533 | GGTCTCTTTCGTTGATATTGAGG | 1000 | | | | | | | | | |
| Oligo fwd | CACCGGTCTCTTTCGTTGATATTG | | | | | | | | | | |
| Oligo rev | AAACCAATATCAACGAAAGAGACC | | | | | | | | | | |
| Chromosome | start | end | strand | MM | target_seq | PAM | alignment | distance | position | gene name | gene id |
| GU305914.1 | 166488 | 166510 | + | 0 | GGTCTCTTTCGTTGATATTG | AGG | \|\|\|\|\|\|\|\|[\|\|\|\|\|\|\|\|\|\|\|\|]PAM | NA | Intergenic | NA | NA |

| | | | | | | | | | | | |
|---|---|---|---|---|---|---|---|---|---|---|---|
| T12534 | GTCTCTTTCGTTGATATTGAGGG | 1000 | | | | | | | | | |
| Oligo fwd | CACCGTCTCTTTCGTTGATATTGA | | | | | | | | | | |
| Oligo rev | AAACTCAATATCAACGAAAGAGAC | | | | | | | | | | |
| Chromosome | start | end | strand | MM | target_seq | PAM | alignment | distance | position | gene name | gene id |
| GU305914.1 | 166489 | 166511 | + | 0 | GTCTCTTTCGTTGATATTGA | GGG | \|\|\|\|\|\|\|\|[\|\|\|\|\|\|\|\|\|\|\|\|]PAM | NA | Intergenic | NA | NA |

| | | | | | | | | | | | |
|---|---|---|---|---|---|---|---|---|---|---|---|
| T12535 | ATTGAGGGCGATAGTGTTGTCGG | 1000 | | | | | | | | | |
| Oligo fwd | CACCATTGAGGGCGATAGTGTTGT | | | | | | | | | | |
| Oligo rev | AAACACAACACTATCGCCCTCAAT | | | | | | | | | | |
| Chromosome | start | end | strand | MM | target_seq | PAM | alignment | distance | position | gene name | gene id |
| GU305914.1 | 166504 | 166526 | + | 0 | ATTGAGGGCGATAGTGTTGT | CGG | \|\|\|\|\|\|\|\|[\|\|\|\|\|\|\|\|\|\|\|\|]PAM | NA | Intergenic | NA | NA |

| | | | | | | | | | | | |
|---|---|---|---|---|---|---|---|---|---|---|---|
| T12536 | TTGAGGGCGATAGTGTTGTCGGG | 1000 | | | | | | | | | |
| Oligo fwd | CACCTTGAGGGCGATAGTGTTGTC | | | | | | | | | | |
| Oligo rev | AAACGACAACACTATCGCCCTCAA | | | | | | | | | | |
| Chromosome | start | end | strand | MM | target_seq | PAM | alignment | distance | position | gene name | gene id |
| GU305914.1 | 166505 | 166527 | + | 0 | TTGAGGGCGATAGTGTTGTC | GGG | \|\|\|\|\|\|\|\|[\|\|\|\|\|\|\|\|\|\|\|\|]PAM | NA | Intergenic | NA | NA |

| | | | | | | | | | | | |
|---|---|---|---|---|---|---|---|---|---|---|---|
| T12537 | GTTGTCGGGTGATCTTGATCTGG | 1000 | | | | | | | | | |
| Oligo fwd | CACCGTTGTCGGGTGATCTTGATC | | | | | | | | | | |
| Oligo rev | AAACGATCAAGATCACCCGACAAC | | | | | | | | | | |
| Chromosome | start | end | strand | MM | target_seq | PAM | alignment | distance | position | gene name | gene id |
| GU305914.1 | 166519 | 166541 | + | 0 | GTTGTCGGGTGATCTTGATC | TGG | \|\|\|\|\|\|\|\|[\|\|\|\|\|\|\|\|\|\|\|\|]PAM | NA | Intergenic | NA | NA |

| | | | | | | | | | | | |
|---|---|---|---|---|---|---|---|---|---|---|---|
| T12538 | CTGGTCGCTGACTGAAAACACGG | 1000 | | | | | | | | | |
| Oligo fwd | CACCCTGGTCGCTGACTGAAAACA | | | | | | | | | | |
| Oligo rev | AAACTGTTTTCAGTCAGCGACCAG | | | | | | | | | | |
| Chromosome | start | end | strand | MM | target_seq | PAM | alignment | distance | position | gene name | gene id |
| GU305914.1 | 166538 | 166560 | + | 0 | CTGGTCGCTGACTGAAAACA | CGG | \|\|\|\|\|\|\|\|[\|\|\|\|\|\|\|\|\|\|\|\|]PAM | NA | Intergenic | NA | NA |

<table>
<tr><td>T12539</td><td>GTCGCTGACTGAAAACACGGCGG</td><td>1000</td><td></td><td></td><td></td><td></td><td></td><td></td><td></td><td></td><td></td></tr>
<tr><td>Oligo fwd</td><td>CACCGTCGCTGACTGAAAACACGG</td><td></td><td></td><td></td><td></td><td></td><td></td><td></td><td></td><td></td><td></td></tr>
<tr><td>Oligo rev</td><td>AAACCCGTGTTTTCAGTCAGCGAC</td><td></td><td></td><td></td><td></td><td></td><td></td><td></td><td></td><td></td><td></td></tr>
<tr><td>Chromosome</td><td>start</td><td>end</td><td>strand</td><td>MM</td><td>target_seq</td><td>PAM</td><td>alignment</td><td>distance</td><td>position</td><td>gene name</td><td>gene id</td></tr>
<tr><td>GU305914.1</td><td>166541</td><td>166563</td><td>+</td><td>0</td><td>GTCGCTGACTGAAAACACGG</td><td>CGG</td><td>||||||||[||||||||||||]PAM</td><td>NA</td><td>Intergenic</td><td>NA</td><td>NA</td></tr>
<tr><td>T12540</td><td>TCGCTGACTGAAAACACGGCGGG</td><td>1000</td><td></td><td></td><td></td><td></td><td></td><td></td><td></td><td></td><td></td></tr>
<tr><td>Oligo fwd</td><td>CACCTCGCTGACTGAAAACACGGC</td><td></td><td></td><td></td><td></td><td></td><td></td><td></td><td></td><td></td><td></td></tr>
<tr><td>Oligo rev</td><td>AAACGCCGTGTTTTCAGTCAGCGA</td><td></td><td></td><td></td><td></td><td></td><td></td><td></td><td></td><td></td><td></td></tr>
<tr><td>Chromosome</td><td>start</td><td>end</td><td>strand</td><td>MM</td><td>target_seq</td><td>PAM</td><td>alignment</td><td>distance</td><td>position</td><td>gene name</td><td>gene id</td></tr>
<tr><td>GU305914.1</td><td>166542</td><td>166564</td><td>+</td><td>0</td><td>TCGCTGACTGAAAACACGGC</td><td>GGG</td><td>||||||||[||||||||||||]PAM</td><td>NA</td><td>Intergenic</td><td>NA</td><td>NA</td></tr>
<tr><td>T12541</td><td>CGCTGACTGAAAACACGGCGGGG</td><td>1000</td><td></td><td></td><td></td><td></td><td></td><td></td><td></td><td></td><td></td></tr>
<tr><td>Oligo fwd</td><td>CACCCGCTGACTGAAAACACGGCG</td><td></td><td></td><td></td><td></td><td></td><td></td><td></td><td></td><td></td><td></td></tr>
<tr><td>Oligo rev</td><td>AAACCGCCGTGTTTTCAGTCAGCG</td><td></td><td></td><td></td><td></td><td></td><td></td><td></td><td></td><td></td><td></td></tr>
<tr><td>Chromosome</td><td>start</td><td>end</td><td>strand</td><td>MM</td><td>target_seq</td><td>PAM</td><td>alignment</td><td>distance</td><td>position</td><td>gene name</td><td>gene id</td></tr>
<tr><td>GU305914.1</td><td>166543</td><td>166565</td><td>+</td><td>0</td><td>CGCTGACTGAAAACACGGCG</td><td>GGG</td><td>||||||||[||||||||||||]PAM</td><td>NA</td><td>Intergenic</td><td>NA</td><td>NA</td></tr>
<tr><td>T12542</td><td>GGAAGCGATCCCTGTAACTGCGG</td><td>1000</td><td></td><td></td><td></td><td></td><td></td><td></td><td></td><td></td><td></td></tr>
<tr><td>Oligo fwd</td><td>CACCGGAAGCGATCCCTGTAACTG</td><td></td><td></td><td></td><td></td><td></td><td></td><td></td><td></td><td></td><td></td></tr>
<tr><td>Oligo rev</td><td>AAACCAGTTACAGGGATCGCTTCC</td><td></td><td></td><td></td><td></td><td></td><td></td><td></td><td></td><td></td><td></td></tr>
<tr><td>Chromosome</td><td>start</td><td>end</td><td>strand</td><td>MM</td><td>target_seq</td><td>PAM</td><td>alignment</td><td>distance</td><td>position</td><td>gene name</td><td>gene id</td></tr>
<tr><td>GU305914.1</td><td>166564</td><td>166586</td><td>+</td><td>0</td><td>GGAAGCGATCCCTGTAACTG</td><td>CGG</td><td>||||||||[||||||||||||]PAM</td><td>NA</td><td>Intergenic</td><td>NA</td><td>NA</td></tr>
<tr><td>T12543</td><td>AGCGATCCCTGTAACTGCGGAGG</td><td>1000</td><td></td><td></td><td></td><td></td><td></td><td></td><td></td><td></td><td></td></tr>
<tr><td>Oligo fwd</td><td>CACCAGCGATCCCTGTAACTGCGG</td><td></td><td></td><td></td><td></td><td></td><td></td><td></td><td></td><td></td><td></td></tr>
<tr><td>Oligo rev</td><td>AAACCCGCAGTTACAGGGATCGCT</td><td></td><td></td><td></td><td></td><td></td><td></td><td></td><td></td><td></td><td></td></tr>
<tr><td>Chromosome</td><td>start</td><td>end</td><td>strand</td><td>MM</td><td>target_seq</td><td>PAM</td><td>alignment</td><td>distance</td><td>position</td><td>gene name</td><td>gene id</td></tr>
<tr><td>GU305914.1</td><td>166567</td><td>166589</td><td>+</td><td>0</td><td>AGCGATCCCTGTAACTGCGG</td><td>AGG</td><td>||||||||[||||||||||||]PAM</td><td>NA</td><td>Intergenic</td><td>NA</td><td>NA</td></tr>
<tr><td>T12544</td><td>AACTGCGGAGGACTCCTATCAGG</td><td>1000</td><td></td><td></td><td></td><td></td><td></td><td></td><td></td><td></td><td></td></tr>
<tr><td>Oligo fwd</td><td>CACCAACTGCGGAGGACTCCTATC</td><td></td><td></td><td></td><td></td><td></td><td></td><td></td><td></td><td></td><td></td></tr>
<tr><td>Oligo rev</td><td>AAACGATAGGAGTCCTCCGCAGTT</td><td></td><td></td><td></td><td></td><td></td><td></td><td></td><td></td><td></td><td></td></tr>
<tr><td>Chromosome</td><td>start</td><td>end</td><td>strand</td><td>MM</td><td>target_seq</td><td>PAM</td><td>alignment</td><td>distance</td><td>position</td><td>gene name</td><td>gene id</td></tr>
<tr><td>GU305914.1</td><td>166579</td><td>166601</td><td>+</td><td>0</td><td>AACTGCGGAGGACTCCTATC</td><td>AGG</td><td>||||||||[||||||||||||]PAM</td><td>NA</td><td>Intergenic</td><td>NA</td><td>NA</td></tr>
<tr><td>T12545</td><td>CTCCTATCAGGTCCGTGTCTAGG</td><td>1000</td><td></td><td></td><td></td><td></td><td></td><td></td><td></td><td></td><td></td></tr>
<tr><td>Oligo fwd</td><td>CACCCTCCTATCAGGTCCGTGTCT</td><td></td><td></td><td></td><td></td><td></td><td></td><td></td><td></td><td></td><td></td></tr>
<tr><td>Oligo rev</td><td>AAACAGACACGGACCTGATAGGAG</td><td></td><td></td><td></td><td></td><td></td><td></td><td></td><td></td><td></td><td></td></tr>
<tr><td>Chromosome</td><td>start</td><td>end</td><td>strand</td><td>MM</td><td>target_seq</td><td>PAM</td><td>alignment</td><td>distance</td><td>position</td><td>gene name</td><td>gene id</td></tr>
<tr><td>GU305914.1</td><td>166591</td><td>166613</td><td>+</td><td>0</td><td>CTCCTATCAGGTCCGTGTCT</td><td>AGG</td><td>||||||||[||||||||||||]PAM</td><td>NA</td><td>Intergenic</td><td>NA</td><td>NA</td></tr>
<tr><td>T12546</td><td>TCCTATCAGGTCCGTGTCTAGGG</td><td>1000</td><td></td><td></td><td></td><td></td><td></td><td></td><td></td><td></td><td></td></tr>
<tr><td>Oligo fwd</td><td>CACCTCCTATCAGGTCCGTGTCTA</td><td></td><td></td><td></td><td></td><td></td><td></td><td></td><td></td><td></td><td></td></tr>
<tr><td>Oligo rev</td><td>AAACTAGACACGGACCTGATAGGA</td><td></td><td></td><td></td><td></td><td></td><td></td><td></td><td></td><td></td><td></td></tr>
<tr><td>Chromosome</td><td>start</td><td>end</td><td>strand</td><td>MM</td><td>target_seq</td><td>PAM</td><td>alignment</td><td>distance</td><td>position</td><td>gene name</td><td>gene id</td></tr>
<tr><td>GU305914.1</td><td>166592</td><td>166614</td><td>+</td><td>0</td><td>TCCTATCAGGTCCGTGTCTA</td><td>GGG</td><td>||||||||[||||||||||||]PAM</td><td>NA</td><td>Intergenic</td><td>NA</td><td>NA</td></tr>
<tr><td>T12547</td><td>GGTCCGTGTCTAGGGACATATGG</td><td>1000</td><td></td><td></td><td></td><td></td><td></td><td></td><td></td><td></td><td></td></tr>
<tr><td>Oligo fwd</td><td>CACCGGTCCGTGTCTAGGGACATA</td><td></td><td></td><td></td><td></td><td></td><td></td><td></td><td></td><td></td><td></td></tr>
<tr><td>Oligo rev</td><td>AAACTATGTCCCTAGACACGGACC</td><td></td><td></td><td></td><td></td><td></td><td></td><td></td><td></td><td></td><td></td></tr>
<tr><td>Chromosome</td><td>start</td><td>end</td><td>strand</td><td>MM</td><td>target_seq</td><td>PAM</td><td>alignment</td><td>distance</td><td>position</td><td>gene name</td><td>gene id</td></tr>
<tr><td>GU305914.1</td><td>166600</td><td>166622</td><td>+</td><td>0</td><td>GGTCCGTGTCTAGGGACATA</td><td>TGG</td><td>||||||||[||||||||||||]PAM</td><td>NA</td><td>Intergenic</td><td>NA</td><td>NA</td></tr>
<tr><td>T12548</td><td>TAGGGACATATGGCCTAAGATGG</td><td>1000</td><td></td><td></td><td></td><td></td><td></td><td></td><td></td><td></td><td></td></tr>
<tr><td>Oligo fwd</td><td>CACCTAGGGACATATGGCCTAAGA</td><td></td><td></td><td></td><td></td><td></td><td></td><td></td><td></td><td></td><td></td></tr>
<tr><td>Oligo rev</td><td>AAACTCTTAGGCCATATGTCCCTA</td><td></td><td></td><td></td><td></td><td></td><td></td><td></td><td></td><td></td><td></td></tr>
<tr><td>Chromosome</td><td>start</td><td>end</td><td>strand</td><td>MM</td><td>target_seq</td><td>PAM</td><td>alignment</td><td>distance</td><td>position</td><td>gene name</td><td>gene id</td></tr>
<tr><td>GU305914.1</td><td>166610</td><td>166632</td><td>+</td><td>0</td><td>TAGGGACATATGGCCTAAGA</td><td>TGG</td><td>||||||||[||||||||||||]PAM</td><td>NA</td><td>Intergenic</td><td>NA</td><td>NA</td></tr>
<tr><td>T12549</td><td>ACATATGGCCTAAGATGGTACGG</td><td>1000</td><td></td><td></td><td></td><td></td><td></td><td></td><td></td><td></td><td></td></tr>
<tr><td>Oligo fwd</td><td>CACCACATATGGCCTAAGATGGTA</td><td></td><td></td><td></td><td></td><td></td><td></td><td></td><td></td><td></td><td></td></tr>
<tr><td>Oligo rev</td><td>AAACTACCATCTTAGGCCATATGT</td><td></td><td></td><td></td><td></td><td></td><td></td><td></td><td></td><td></td><td></td></tr>
<tr><td>Chromosome</td><td>start</td><td>end</td><td>strand</td><td>MM</td><td>target_seq</td><td>PAM</td><td>alignment</td><td>distance</td><td>position</td><td>gene name</td><td>gene id</td></tr>
<tr><td>GU305914.1</td><td>166615</td><td>166637</td><td>+</td><td>0</td><td>ACATATGGCCTAAGATGGTA</td><td>CGG</td><td>||||||||[||||||||||||]PAM</td><td>NA</td><td>Intergenic</td><td>NA</td><td>NA</td></tr>
<tr><td>T12550</td><td>TATGGCCTAAGATGGTACGGAGG</td><td>1000</td><td></td><td></td><td></td><td></td><td></td><td></td><td></td><td></td><td></td></tr>
<tr><td>Oligo fwd</td><td>CACCTATGGCCTAAGATGGTACGG</td><td></td><td></td><td></td><td></td><td></td><td></td><td></td><td></td><td></td><td></td></tr>
<tr><td>Oligo rev</td><td>AAACCCGTACCATCTTAGGCCATA</td><td></td><td></td><td></td><td></td><td></td><td></td><td></td><td></td><td></td><td></td></tr>
<tr><td>Chromosome</td><td>start</td><td>end</td><td>strand</td><td>MM</td><td>target_seq</td><td>PAM</td><td>alignment</td><td>distance</td><td>position</td><td>gene name</td><td>gene id</td></tr>
<tr><td>GU305914.1</td><td>166618</td><td>166640</td><td>+</td><td>0</td><td>TATGGCCTAAGATGGTACGG</td><td>AGG</td><td>||||||||[||||||||||||]PAM</td><td>NA</td><td>Intergenic</td><td>NA</td><td>NA</td></tr>
<tr><td>T12551</td><td>GGCCTAAGATGGTACGGAGGAGG</td><td>1000</td><td></td><td></td><td></td><td></td><td></td><td></td><td></td><td></td><td></td></tr>
<tr><td>Oligo fwd</td><td>CACCGGCCTAAGATGGTACGGAGG</td><td></td><td></td><td></td><td></td><td></td><td></td><td></td><td></td><td></td><td></td></tr>
<tr><td>Oligo rev</td><td>AAACCCTCCGTACCATCTTAGGCC</td><td></td><td></td><td></td><td></td><td></td><td></td><td></td><td></td><td></td><td></td></tr>
<tr><td>Chromosome</td><td>start</td><td>end</td><td>strand</td><td>MM</td><td>target_seq</td><td>PAM</td><td>alignment</td><td>distance</td><td>position</td><td>gene name</td><td>gene id</td></tr>
<tr><td>GU305914.1</td><td>166621</td><td>166643</td><td>+</td><td>0</td><td>GGCCTAAGATGGTACGGAGG</td><td>AGG</td><td>||||||||[||||||||||||]PAM</td><td>NA</td><td>Intergenic</td><td>NA</td><td>NA</td></tr>
<tr><td>T12552</td><td>GGTTATATGTGCCGTCGAGCAGG</td><td>1000</td><td></td><td></td><td></td><td></td><td></td><td></td><td></td><td></td><td></td></tr>
<tr><td>Oligo fwd</td><td>CACCGGTTATATGTGCCGTCGAGC</td><td></td><td></td><td></td><td></td><td></td><td></td><td></td><td></td><td></td><td></td></tr>
<tr><td>Oligo rev</td><td>AAACGCTCGACGGCACATATAACC</td><td></td><td></td><td></td><td></td><td></td><td></td><td></td><td></td><td></td><td></td></tr>
<tr><td>Chromosome</td><td>start</td><td>end</td><td>strand</td><td>MM</td><td>target_seq</td><td>PAM</td><td>alignment</td><td>distance</td><td>position</td><td>gene name</td><td>gene id</td></tr>
<tr><td>GU305914.1</td><td>166642</td><td>166664</td><td>+</td><td>0</td><td>GGTTATATGTGCCGTCGAGC</td><td>AGG</td><td>||||||||[||||||||||||]PAM</td><td>NA</td><td>Intergenic</td><td>NA</td><td>NA</td></tr>
<tr><td>T12553</td><td>GTTATATGTGCCGTCGAGCAGGG</td><td>1000</td><td></td><td></td><td></td><td></td><td></td><td></td><td></td><td></td><td></td></tr>
<tr><td>Oligo fwd</td><td>CACCGTTATATGTGCCGTCGAGCA</td><td></td><td></td><td></td><td></td><td></td><td></td><td></td><td></td><td></td><td></td></tr>
<tr><td>Oligo rev</td><td>AAACTGCTCGACGGCACATATAAC</td><td></td><td></td><td></td><td></td><td></td><td></td><td></td><td></td><td></td><td></td></tr>
<tr><td>Chromosome</td><td>start</td><td>end</td><td>strand</td><td>MM</td><td>target_seq</td><td>PAM</td><td>alignment</td><td>distance</td><td>position</td><td>gene name</td><td>gene id</td></tr>
<tr><td>GU305914.1</td><td>166643</td><td>166665</td><td>+</td><td>0</td><td>GTTATATGTGCCGTCGAGCA</td><td>GGG</td><td>||||||||[||||||||||||]PAM</td><td>NA</td><td>Intergenic</td><td>NA</td><td>NA</td></tr>
</table>

<table>
<tr><td>T12554</td><td>GTCGAGCAGGGACATGCTGCTGG</td><td>1000</td><td></td><td></td><td></td><td></td><td></td><td></td><td></td><td></td><td></td></tr>
<tr><td>Oligo fwd</td><td>CACCGTCGAGCAGGGACATGCTGC</td><td></td><td></td><td></td><td></td><td></td><td></td><td></td><td></td><td></td><td></td></tr>
<tr><td>Oligo rev</td><td>AAACGCAGCATGTCCCTGCTCGAC</td><td></td><td></td><td></td><td></td><td></td><td></td><td></td><td></td><td></td><td></td></tr>
<tr><td>Chromosome</td><td>start</td><td>end</td><td>strand</td><td>MM</td><td>target_seq</td><td>PAM</td><td>alignment</td><td>distance</td><td>position</td><td>gene name</td><td>gene id</td></tr>
<tr><td>GU305914.1</td><td>166655</td><td>166677</td><td>+</td><td>0</td><td>GTCGAGCAGGGACATGCTGC</td><td>TGG</td><td>||||||||[||||||||||||]PAM</td><td>NA</td><td>Intergenic</td><td>NA</td><td>NA</td></tr>
<tr><td>T12555</td><td>AGCAGGGACATGCTGCTGGTCGG</td><td>1000</td><td></td><td></td><td></td><td></td><td></td><td></td><td></td><td></td><td></td></tr>
<tr><td>Oligo fwd</td><td>CACCAGCAGGGACATGCTGCTGGT</td><td></td><td></td><td></td><td></td><td></td><td></td><td></td><td></td><td></td><td></td></tr>
<tr><td>Oligo rev</td><td>AAACACCAGCAGCATGTCCCTGCT</td><td></td><td></td><td></td><td></td><td></td><td></td><td></td><td></td><td></td><td></td></tr>
<tr><td>Chromosome</td><td>start</td><td>end</td><td>strand</td><td>MM</td><td>target_seq</td><td>PAM</td><td>alignment</td><td>distance</td><td>position</td><td>gene name</td><td>gene id</td></tr>
<tr><td>GU305914.1</td><td>166659</td><td>166681</td><td>+</td><td>0</td><td>AGCAGGGACATGCTGCTGGT</td><td>CGG</td><td>||||||||[||||||||||||]PAM</td><td>NA</td><td>Intergenic</td><td>NA</td><td>NA</td></tr>
<tr><td>T12556</td><td>GCAGGGACATGCTGCTGGTCGGG</td><td>1000</td><td></td><td></td><td></td><td></td><td></td><td></td><td></td><td></td><td></td></tr>
<tr><td>Oligo fwd</td><td>CACCGCAGGGACATGCTGCTGGTC</td><td></td><td></td><td></td><td></td><td></td><td></td><td></td><td></td><td></td><td></td></tr>
<tr><td>Oligo rev</td><td>AAACGACCAGCAGCATGTCCCTGC</td><td></td><td></td><td></td><td></td><td></td><td></td><td></td><td></td><td></td><td></td></tr>
<tr><td>Chromosome</td><td>start</td><td>end</td><td>strand</td><td>MM</td><td>target_seq</td><td>PAM</td><td>alignment</td><td>distance</td><td>position</td><td>gene name</td><td>gene id</td></tr>
<tr><td>GU305914.1</td><td>166660</td><td>166682</td><td>+</td><td>0</td><td>GCAGGGACATGCTGCTGGTC</td><td>GGG</td><td>||||||||[||||||||||||]PAM</td><td>NA</td><td>Intergenic</td><td>NA</td><td>NA</td></tr>
<tr><td>T12557</td><td>ACATGCTGCTGGTCGGGACTCGG</td><td>1000</td><td></td><td></td><td></td><td></td><td></td><td></td><td></td><td></td><td></td></tr>
<tr><td>Oligo fwd</td><td>CACCACATGCTGCTGGTCGGGACT</td><td></td><td></td><td></td><td></td><td></td><td></td><td></td><td></td><td></td><td></td></tr>
<tr><td>Oligo rev</td><td>AAACAGTCCCGACCAGCAGCATGT</td><td></td><td></td><td></td><td></td><td></td><td></td><td></td><td></td><td></td><td></td></tr>
<tr><td>Chromosome</td><td>start</td><td>end</td><td>strand</td><td>MM</td><td>target_seq</td><td>PAM</td><td>alignment</td><td>distance</td><td>position</td><td>gene name</td><td>gene id</td></tr>
<tr><td>GU305914.1</td><td>166666</td><td>166688</td><td>+</td><td>0</td><td>ACATGCTGCTGGTCGGGACT</td><td>CGG</td><td>||||||||[||||||||||||]PAM</td><td>NA</td><td>Intergenic</td><td>NA</td><td>NA</td></tr>
<tr><td>T12558</td><td>TCGGACCACCGACTCCGCCTTGG</td><td>1000</td><td></td><td></td><td></td><td></td><td></td><td></td><td></td><td></td><td></td></tr>
<tr><td>Oligo fwd</td><td>CACCTCGGACCACCGACTCCGCCT</td><td></td><td></td><td></td><td></td><td></td><td></td><td></td><td></td><td></td><td></td></tr>
<tr><td>Oligo rev</td><td>AAACAGGCGGAGTCGGTGGTCCGA</td><td></td><td></td><td></td><td></td><td></td><td></td><td></td><td></td><td></td><td></td></tr>
<tr><td>Chromosome</td><td>start</td><td>end</td><td>strand</td><td>MM</td><td>target_seq</td><td>PAM</td><td>alignment</td><td>distance</td><td>position</td><td>gene name</td><td>gene id</td></tr>
<tr><td>GU305914.1</td><td>166685</td><td>166707</td><td>+</td><td>0</td><td>TCGGACCACCGACTCCGCCT</td><td>TGG</td><td>||||||||[||||||||||||]PAM</td><td>NA</td><td>Intergenic</td><td>NA</td><td>NA</td></tr>
<tr><td>T12559</td><td>CCACCGACTCCGCCTTGGTGTGG</td><td>1000</td><td></td><td></td><td></td><td></td><td></td><td></td><td></td><td></td><td></td></tr>
<tr><td>Oligo fwd</td><td>CACCCCACCGACTCCGCCTTGGTG</td><td></td><td></td><td></td><td></td><td></td><td></td><td></td><td></td><td></td><td></td></tr>
<tr><td>Oligo rev</td><td>AAACCACCAAGGCGGAGTCGGTGG</td><td></td><td></td><td></td><td></td><td></td><td></td><td></td><td></td><td></td><td></td></tr>
<tr><td>Chromosome</td><td>start</td><td>end</td><td>strand</td><td>MM</td><td>target_seq</td><td>PAM</td><td>alignment</td><td>distance</td><td>position</td><td>gene name</td><td>gene id</td></tr>
<tr><td>GU305914.1</td><td>166690</td><td>166712</td><td>+</td><td>0</td><td>CCACCGACTCCGCCTTGGTG</td><td>TGG</td><td>||||||||[||||||||||||]PAM</td><td>NA</td><td>Intergenic</td><td>NA</td><td>NA</td></tr>
<tr><td>T12560</td><td>GTTACGTAGCAGCACGAGCACGG</td><td>1000</td><td></td><td></td><td></td><td></td><td></td><td></td><td></td><td></td><td></td></tr>
<tr><td>Oligo fwd</td><td>CACCGTTACGTAGCAGCACGAGCA</td><td></td><td></td><td></td><td></td><td></td><td></td><td></td><td></td><td></td><td></td></tr>
<tr><td>Oligo rev</td><td>AAACTGCTCGTGCTGCTACGTAAC</td><td></td><td></td><td></td><td></td><td></td><td></td><td></td><td></td><td></td><td></td></tr>
<tr><td>Chromosome</td><td>start</td><td>end</td><td>strand</td><td>MM</td><td>target_seq</td><td>PAM</td><td>alignment</td><td>distance</td><td>position</td><td>gene name</td><td>gene id</td></tr>
<tr><td>GU305914.1</td><td>166715</td><td>166737</td><td>+</td><td>0</td><td>GTTACGTAGCAGCACGAGCA</td><td>CGG</td><td>||||||||[||||||||||||]PAM</td><td>NA</td><td>Intergenic</td><td>NA</td><td>NA</td></tr>
<tr><td>T12561</td><td>CAGCACGAGCACGGACAGCACGG</td><td>1000</td><td></td><td></td><td></td><td></td><td></td><td></td><td></td><td></td><td></td></tr>
<tr><td>Oligo fwd</td><td>CACCCAGCACGAGCACGGACAGCA</td><td></td><td></td><td></td><td></td><td></td><td></td><td></td><td></td><td></td><td></td></tr>
<tr><td>Oligo rev</td><td>AAACTGCTGTCCGTGCTCGTGCTG</td><td></td><td></td><td></td><td></td><td></td><td></td><td></td><td></td><td></td><td></td></tr>
<tr><td>Chromosome</td><td>start</td><td>end</td><td>strand</td><td>MM</td><td>target_seq</td><td>PAM</td><td>alignment</td><td>distance</td><td>position</td><td>gene name</td><td>gene id</td></tr>
<tr><td>GU305914.1</td><td>166724</td><td>166746</td><td>+</td><td>0</td><td>CAGCACGAGCACGGACAGCA</td><td>CGG</td><td>||||||||[||||||||||||]PAM</td><td>NA</td><td>Intergenic</td><td>NA</td><td>NA</td></tr>
<tr><td>T12562</td><td>AGCACGAGCACGGACAGCACGGG</td><td>1000</td><td></td><td></td><td></td><td></td><td></td><td></td><td></td><td></td><td></td></tr>
<tr><td>Oligo fwd</td><td>CACCAGCACGAGCACGGACAGCAC</td><td></td><td></td><td></td><td></td><td></td><td></td><td></td><td></td><td></td><td></td></tr>
<tr><td>Oligo rev</td><td>AAACGTGCTGTCCGTGCTCGTGCT</td><td></td><td></td><td></td><td></td><td></td><td></td><td></td><td></td><td></td><td></td></tr>
<tr><td>Chromosome</td><td>start</td><td>end</td><td>strand</td><td>MM</td><td>target_seq</td><td>PAM</td><td>alignment</td><td>distance</td><td>position</td><td>gene name</td><td>gene id</td></tr>
<tr><td>GU305914.1</td><td>166725</td><td>166747</td><td>+</td><td>0</td><td>AGCACGAGCACGGACAGCAC</td><td>GGG</td><td>||||||||[||||||||||||]PAM</td><td>NA</td><td>Intergenic</td><td>NA</td><td>NA</td></tr>
<tr><td>T12563</td><td>ACGAGCACGGACAGCACGGGCGG</td><td>1000</td><td></td><td></td><td></td><td></td><td></td><td></td><td></td><td></td><td></td></tr>
<tr><td>Oligo fwd</td><td>CACCACGAGCACGGACAGCACGGG</td><td></td><td></td><td></td><td></td><td></td><td></td><td></td><td></td><td></td><td></td></tr>
<tr><td>Oligo rev</td><td>AAACCCCGTGCTGTCCGTGCTCGT</td><td></td><td></td><td></td><td></td><td></td><td></td><td></td><td></td><td></td><td></td></tr>
<tr><td>Chromosome</td><td>start</td><td>end</td><td>strand</td><td>MM</td><td>target_seq</td><td>PAM</td><td>alignment</td><td>distance</td><td>position</td><td>gene name</td><td>gene id</td></tr>
<tr><td>GU305914.1</td><td>166728</td><td>166750</td><td>+</td><td>0</td><td>ACGAGCACGGACAGCACGGG</td><td>CGG</td><td>||||||||[||||||||||||]PAM</td><td>NA</td><td>Intergenic</td><td>NA</td><td>NA</td></tr>
<tr><td>T12564</td><td>GCACGGGCGGTGACACGTCGAGG</td><td>1000</td><td></td><td></td><td></td><td></td><td></td><td></td><td></td><td></td><td></td></tr>
<tr><td>Oligo fwd</td><td>CACCGCACGGGCGGTGACACGTCG</td><td></td><td></td><td></td><td></td><td></td><td></td><td></td><td></td><td></td><td></td></tr>
<tr><td>Oligo rev</td><td>AAACCGACGTGTCACCGCCCGTGC</td><td></td><td></td><td></td><td></td><td></td><td></td><td></td><td></td><td></td><td></td></tr>
<tr><td>Chromosome</td><td>start</td><td>end</td><td>strand</td><td>MM</td><td>target_seq</td><td>PAM</td><td>alignment</td><td>distance</td><td>position</td><td>gene name</td><td>gene id</td></tr>
<tr><td>GU305914.1</td><td>166741</td><td>166763</td><td>+</td><td>0</td><td>GCACGGGCGGTGACACGTCG</td><td>AGG</td><td>||||||||[||||||||||||]PAM</td><td>NA</td><td>Intergenic</td><td>NA</td><td>NA</td></tr>
<tr><td>T12565</td><td>CCAACACGTTCTCCGTGAACAGG</td><td>1000</td><td></td><td></td><td></td><td></td><td></td><td></td><td></td><td></td><td></td></tr>
<tr><td>Oligo fwd</td><td>CACCCCAACACGTTCTCCGTGAAC</td><td></td><td></td><td></td><td></td><td></td><td></td><td></td><td></td><td></td><td></td></tr>
<tr><td>Oligo rev</td><td>AAACGTTCACGGAGAACGTGTTGG</td><td></td><td></td><td></td><td></td><td></td><td></td><td></td><td></td><td></td><td></td></tr>
<tr><td>Chromosome</td><td>start</td><td>end</td><td>strand</td><td>MM</td><td>target_seq</td><td>PAM</td><td>alignment</td><td>distance</td><td>position</td><td>gene name</td><td>gene id</td></tr>
<tr><td>GU305914.1</td><td>166765</td><td>166787</td><td>+</td><td>0</td><td>CCAACACGTTCTCCGTGAAC</td><td>AGG</td><td>||||||||[||||||||||||]PAM</td><td>NA</td><td>Intergenic</td><td>NA</td><td>NA</td></tr>
<tr><td>T12566</td><td>CACGTTCTCCGTGAACAGGTCGG</td><td>1000</td><td></td><td></td><td></td><td></td><td></td><td></td><td></td><td></td><td></td></tr>
<tr><td>Oligo fwd</td><td>CACCCACGTTCTCCGTGAACAGGT</td><td></td><td></td><td></td><td></td><td></td><td></td><td></td><td></td><td></td><td></td></tr>
<tr><td>Oligo rev</td><td>AAACACCTGTTCACGGAGAACGTG</td><td></td><td></td><td></td><td></td><td></td><td></td><td></td><td></td><td></td><td></td></tr>
<tr><td>Chromosome</td><td>start</td><td>end</td><td>strand</td><td>MM</td><td>target_seq</td><td>PAM</td><td>alignment</td><td>distance</td><td>position</td><td>gene name</td><td>gene id</td></tr>
<tr><td>GU305914.1</td><td>166769</td><td>166791</td><td>+</td><td>0</td><td>CACGTTCTCCGTGAACAGGT</td><td>CGG</td><td>||||||||[||||||||||||]PAM</td><td>NA</td><td>Intergenic</td><td>NA</td><td>NA</td></tr>
<tr><td>T12567</td><td>TCGTGTAGTTCAACCGCATCAGG</td><td>1000</td><td></td><td></td><td></td><td></td><td></td><td></td><td></td><td></td><td></td></tr>
<tr><td>Oligo fwd</td><td>CACCTCGTGTAGTTCAACCGCATC</td><td></td><td></td><td></td><td></td><td></td><td></td><td></td><td></td><td></td><td></td></tr>
<tr><td>Oligo rev</td><td>AAACGATGCGGTTGAACTACACGA</td><td></td><td></td><td></td><td></td><td></td><td></td><td></td><td></td><td></td><td></td></tr>
<tr><td>Chromosome</td><td>start</td><td>end</td><td>strand</td><td>MM</td><td>target_seq</td><td>PAM</td><td>alignment</td><td>distance</td><td>position</td><td>gene name</td><td>gene id</td></tr>
<tr><td>GU305914.1</td><td>166792</td><td>166814</td><td>+</td><td>0</td><td>TCGTGTAGTTCAACCGCATC</td><td>AGG</td><td>||||||||[||||||||||||]PAM</td><td>NA</td><td>Intergenic</td><td>NA</td><td>NA</td></tr>
<tr><td>T12568</td><td>GTAGCACACGCTCTCCACGCAGG</td><td>1000</td><td></td><td></td><td></td><td></td><td></td><td></td><td></td><td></td><td></td></tr>
<tr><td>Oligo fwd</td><td>CACCGTAGCACACGCTCTCCACGC</td><td></td><td></td><td></td><td></td><td></td><td></td><td></td><td></td><td></td><td></td></tr>
<tr><td>Oligo rev</td><td>AAACGCGTGGAGAGCGTGTGCTAC</td><td></td><td></td><td></td><td></td><td></td><td></td><td></td><td></td><td></td><td></td></tr>
<tr><td>Chromosome</td><td>start</td><td>end</td><td>strand</td><td>MM</td><td>target_seq</td><td>PAM</td><td>alignment</td><td>distance</td><td>position</td><td>gene name</td><td>gene id</td></tr>
</table>

GU305914.1 166823 166845 + 0 GTAGCACACGCTCTCCACGC AGG ||||||||[||||||||||||]PAM NA Intergenic NA NA

T12569 CAGGTGAACACCACCGACTTCGG 1000
Oligo fwd CACCCAGGTGAACACCACCGACTT
Oligo rev AAACAAGTCGGTGGTGTTCACCTG
Chromosome start end strand MM target_seq PAM alignment distance position gene name gene id
GU305914.1 166842 166864 + 0 CAGGTGAACACCACCGACTT CGG ||||||||[||||||||||||]PAM NA Intergenic NA NA

T12570 CCGACTTCGGCGCGTCGCACTGG 1000
Oligo fwd CACCCCGACTTCGGCGCGTCGCAC
Oligo rev AAACGTGCGACGCGCCGAAGTCGG
Chromosome start end strand MM target_seq PAM alignment distance position gene name gene id
GU305914.1 166855 166877 + 0 CCGACTTCGGCGCGTCGCAC TGG ||||||||[||||||||||||]PAM NA Intergenic NA NA

T12571 CGACTTCGGCGCGTCGCACTGGG 1000
Oligo fwd CACCCGACTTCGGCGCGTCGCACT
Oligo rev AAACAGTGCGACGCGCCGAAGTCG
Chromosome start end strand MM target_seq PAM alignment distance position gene name gene id
GU305914.1 166856 166878 + 0 CGACTTCGGCGCGTCGCACT GGG ||||||||[||||||||||||]PAM NA Intergenic NA NA

T12572 CAAGAAGCTCATCCAGTCGCTGG 1000
Oligo fwd CACCCAAGAAGCTCATCCAGTCGC
Oligo rev AAACGCGACTGGATGAGCTTCTTG
Chromosome start end strand MM target_seq PAM alignment distance position gene name gene id
GU305914.1 166889 166911 + 0 CAAGAAGCTCATCCAGTCGC TGG ||||||||[||||||||||||]PAM NA Intergenic NA NA

T12573 AGCTCATCCAGTCGCTGGCGCGG 1000
Oligo fwd CACCAGCTCATCCAGTCGCTGGCG
Oligo rev AAACCGCCAGCGACTGGATGAGCT
Chromosome start end strand MM target_seq PAM alignment distance position gene name gene id
GU305914.1 166894 166916 + 0 AGCTCATCCAGTCGCTGGCG CGG ||||||||[||||||||||||]PAM NA Intergenic NA NA

T12574 CATCCAGTCGCTGGCGCGGCTGG 1000
Oligo fwd CACCCATCCAGTCGCTGGCGCGGC
Oligo rev AAACGCCGCGCCAGCGACTGGATG
Chromosome start end strand MM target_seq PAM alignment distance position gene name gene id
GU305914.1 166898 166920 + 0 CATCCAGTCGCTGGCGCGGC TGG ||||||||[||||||||||||]PAM NA Intergenic NA NA

T12575 GCATCAGCGTCAGCAGAACTCGG 1000
Oligo fwd CACCGCATCAGCGTCAGCAGAACT
Oligo rev AAACAGTTCTGCTGACGCTGATGC
Chromosome start end strand MM target_seq PAM alignment distance position gene name gene id
GU305914.1 166924 166946 + 0 GCATCAGCGTCAGCAGAACT CGG ||||||||[||||||||||||]PAM NA Intergenic NA NA

T12576 GCGTCAGCAGAACTCGGAGCTGG 1000
Oligo fwd CACCGCGTCAGCAGAACTCGGAGC
Oligo rev AAACGCTCCGAGTTCTGCTGACGC
Chromosome start end strand MM target_seq PAM alignment distance position gene name gene id
GU305914.1 166930 166952 + 0 GCGTCAGCAGAACTCGGAGC TGG ||||||||[||||||||||||]PAM NA Intergenic NA NA

T12577 GGAGCTGGTCCGTGTTGTTGTGG 1000
Oligo fwd CACCGGAGCTGGTCCGTGTTGTTG
Oligo rev AAACCAACAACACGGACCAGCTCC
Chromosome start end strand MM target_seq PAM alignment distance position gene name gene id
GU305914.1 166945 166967 + 0 GGAGCTGGTCCGTGTTGTTG TGG ||||||||[||||||||||||]PAM NA Intergenic NA NA
GU305914.1 104633 104655 + 4 AGAGACGGTACGTGTTGTTG TGG -|||--||[|-||||||||||]PAM NA Intergenic NA NA

T12578 GGTCCGTGTTGTTGTGGAGCAGG 1000
Oligo fwd CACCGGTCCGTGTTGTTGTGGAGC
Oligo rev AAACGCTCCACAACAACACGGACC
Chromosome start end strand MM target_seq PAM alignment distance position gene name gene id
GU305914.1 166951 166973 + 0 GGTCCGTGTTGTTGTGGAGC AGG ||||||||[||||||||||||]PAM NA Intergenic NA NA

T12579 GTCCGTGTTGTTGTGGAGCAGGG 1000
Oligo fwd CACCGTCCGTGTTGTTGTGGAGCA
Oligo rev AAACTGCTCCACAACAACACGGAC
Chromosome start end strand MM target_seq PAM alignment distance position gene name gene id
GU305914.1 166952 166974 + 0 GTCCGTGTTGTTGTGGAGCA GGG ||||||||[||||||||||||]PAM NA Intergenic NA NA

T12580 GGAGCAGGGACAGCTGCTCCAGG 1000
Oligo fwd CACCGGAGCAGGGACAGCTGCTCC
Oligo rev AAACGGAGCAGCTGTCCCTGCTCC
Chromosome start end strand MM target_seq PAM alignment distance position gene name gene id
GU305914.1 166966 166988 + 0 GGAGCAGGGACAGCTGCTCC AGG ||||||||[||||||||||||]PAM NA Intergenic NA NA

T12581 GAACTCGTCGTTGAGCTCGATGG 1000
Oligo fwd CACCGAACTCGTCGTTGAGCTCGA
Oligo rev AAACTCGAGCTCAACGACGAGTTC
Chromosome start end strand MM target_seq PAM alignment distance position gene name gene id
GU305914.1 166988 167010 + 0 GAACTCGTCGTTGAGCTCGA TGG ||||||||[||||||||||||]PAM NA Intergenic NA NA

T12582 AACTCGTCGTTGAGCTCGATGGG 1000
Oligo fwd CACCAACTCGTCGTTGAGCTCGAT
Oligo rev AAACATCGAGCTCAACGACGAGTT
Chromosome start end strand MM target_seq PAM alignment distance position gene name gene id
GU305914.1 166989 167011 + 0 AACTCGTCGTTGAGCTCGAT GGG ||||||||[||||||||||||]PAM NA Intergenic NA NA

T12583 TCGTTGAGCTCGATGGGCAGCGG 1000
Oligo fwd CACCTCGTTGAGCTCGATGGGCAG

<table>
<tr><td>Oligo rev</td><td>AAACCTGCCCATCGAGCTCAACGA</td><td></td><td></td><td></td><td></td><td></td><td></td><td></td><td></td><td></td><td></td></tr>
<tr><td>Chromosome</td><td>start</td><td>end</td><td>strand</td><td>MM</td><td>target_seq</td><td>PAM</td><td>alignment</td><td>distance</td><td>position</td><td>gene name</td><td>gene id</td></tr>
<tr><td>GU305914.1</td><td>166995</td><td>167017</td><td>+</td><td>0</td><td>TCGTTGAGCTCGATGGGCAG</td><td>CGG</td><td>||||||||[||||||||||||]PAM</td><td>NA</td><td>Intergenic</td><td>NA</td><td>NA</td></tr>
<tr><td>T12584</td><td>CGTTGAGCTCGATGGGCAGCGGG</td><td>1000</td><td></td><td></td><td></td><td></td><td></td><td></td><td></td><td></td><td></td></tr>
<tr><td>Oligo fwd</td><td>CACCCGTTGAGCTCGATGGGCAGC</td><td></td><td></td><td></td><td></td><td></td><td></td><td></td><td></td><td></td><td></td></tr>
<tr><td>Oligo rev</td><td>AAACGCTGCCCATCGAGCTCAACG</td><td></td><td></td><td></td><td></td><td></td><td></td><td></td><td></td><td></td><td></td></tr>
<tr><td>Chromosome</td><td>start</td><td>end</td><td>strand</td><td>MM</td><td>target_seq</td><td>PAM</td><td>alignment</td><td>distance</td><td>position</td><td>gene name</td><td>gene id</td></tr>
<tr><td>GU305914.1</td><td>166996</td><td>167018</td><td>+</td><td>0</td><td>CGTTGAGCTCGATGGGCAGC</td><td>GGG</td><td>||||||||[||||||||||||]PAM</td><td>NA</td><td>Intergenic</td><td>NA</td><td>NA</td></tr>
<tr><td>T12585</td><td>ATCATCGTCGAGAACCTATACGG</td><td>1000</td><td></td><td></td><td></td><td></td><td></td><td></td><td></td><td></td><td></td></tr>
<tr><td>Oligo fwd</td><td>CACCATCATCGTCGAGAACCTATA</td><td></td><td></td><td></td><td></td><td></td><td></td><td></td><td></td><td></td><td></td></tr>
<tr><td>Oligo rev</td><td>AAACTATAGGTTCTCGACGATGAT</td><td></td><td></td><td></td><td></td><td></td><td></td><td></td><td></td><td></td><td></td></tr>
<tr><td>Chromosome</td><td>start</td><td>end</td><td>strand</td><td>MM</td><td>target_seq</td><td>PAM</td><td>alignment</td><td>distance</td><td>position</td><td>gene name</td><td>gene id</td></tr>
<tr><td>GU305914.1</td><td>167046</td><td>167068</td><td>+</td><td>0</td><td>ATCATCGTCGAGAACCTATA</td><td>CGG</td><td>||||||||[||||||||||||]PAM</td><td>NA</td><td>Intergenic</td><td>NA</td><td>NA</td></tr>
<tr><td>T12586</td><td>TCATCGTCGAGAACCTATACGGG</td><td>1000</td><td></td><td></td><td></td><td></td><td></td><td></td><td></td><td></td><td></td></tr>
<tr><td>Oligo fwd</td><td>CACCTCATCGTCGAGAACCTATAC</td><td></td><td></td><td></td><td></td><td></td><td></td><td></td><td></td><td></td><td></td></tr>
<tr><td>Oligo rev</td><td>AAACGTATAGGTTCTCGACGATGA</td><td></td><td></td><td></td><td></td><td></td><td></td><td></td><td></td><td></td><td></td></tr>
<tr><td>Chromosome</td><td>start</td><td>end</td><td>strand</td><td>MM</td><td>target_seq</td><td>PAM</td><td>alignment</td><td>distance</td><td>position</td><td>gene name</td><td>gene id</td></tr>
<tr><td>GU305914.1</td><td>167047</td><td>167069</td><td>+</td><td>0</td><td>TCATCGTCGAGAACCTATAC</td><td>GGG</td><td>||||||||[||||||||||||]PAM</td><td>NA</td><td>Intergenic</td><td>NA</td><td>NA</td></tr>
<tr><td>T12587</td><td>GTCGAGAACCTATACGGGTCCGG</td><td>1000</td><td></td><td></td><td></td><td></td><td></td><td></td><td></td><td></td><td></td></tr>
<tr><td>Oligo fwd</td><td>CACCGTCGAGAACCTATACGGGTC</td><td></td><td></td><td></td><td></td><td></td><td></td><td></td><td></td><td></td><td></td></tr>
<tr><td>Oligo rev</td><td>AAACGACCCGTATAGGTTCTCGAC</td><td></td><td></td><td></td><td></td><td></td><td></td><td></td><td></td><td></td><td></td></tr>
<tr><td>Chromosome</td><td>start</td><td>end</td><td>strand</td><td>MM</td><td>target_seq</td><td>PAM</td><td>alignment</td><td>distance</td><td>position</td><td>gene name</td><td>gene id</td></tr>
<tr><td>GU305914.1</td><td>167052</td><td>167074</td><td>+</td><td>0</td><td>GTCGAGAACCTATACGGGTC</td><td>CGG</td><td>||||||||[||||||||||||]PAM</td><td>NA</td><td>Intergenic</td><td>NA</td><td>NA</td></tr>
<tr><td>T12588</td><td>AGAACCTATACGGGTCCGGACGG</td><td>1000</td><td></td><td></td><td></td><td></td><td></td><td></td><td></td><td></td><td></td></tr>
<tr><td>Oligo fwd</td><td>CACCAGAACCTATACGGGTCCGGA</td><td></td><td></td><td></td><td></td><td></td><td></td><td></td><td></td><td></td><td></td></tr>
<tr><td>Oligo rev</td><td>AAACTCCGGACCCGTATAGGTTCT</td><td></td><td></td><td></td><td></td><td></td><td></td><td></td><td></td><td></td><td></td></tr>
<tr><td>Chromosome</td><td>start</td><td>end</td><td>strand</td><td>MM</td><td>target_seq</td><td>PAM</td><td>alignment</td><td>distance</td><td>position</td><td>gene name</td><td>gene id</td></tr>
<tr><td>GU305914.1</td><td>167056</td><td>167078</td><td>+</td><td>0</td><td>AGAACCTATACGGGTCCGGA</td><td>CGG</td><td>||||||||[||||||||||||]PAM</td><td>NA</td><td>Intergenic</td><td>NA</td><td>NA</td></tr>
<tr><td>T12589</td><td>CCTATACGGGTCCGGACGGATGG</td><td>1000</td><td></td><td></td><td></td><td></td><td></td><td></td><td></td><td></td><td></td></tr>
<tr><td>Oligo fwd</td><td>CACCCCTATACGGGTCCGGACGGA</td><td></td><td></td><td></td><td></td><td></td><td></td><td></td><td></td><td></td><td></td></tr>
<tr><td>Oligo rev</td><td>AAACTCCGTCCGGACCCGTATAGG</td><td></td><td></td><td></td><td></td><td></td><td></td><td></td><td></td><td></td><td></td></tr>
<tr><td>Chromosome</td><td>start</td><td>end</td><td>strand</td><td>MM</td><td>target_seq</td><td>PAM</td><td>alignment</td><td>distance</td><td>position</td><td>gene name</td><td>gene id</td></tr>
<tr><td>GU305914.1</td><td>167060</td><td>167082</td><td>+</td><td>0</td><td>CCTATACGGGTCCGGACGGA</td><td>TGG</td><td>||||||||[||||||||||||]PAM</td><td>NA</td><td>Intergenic</td><td>NA</td><td>NA</td></tr>
<tr><td>T12590</td><td>CTATACGGGTCCGGACGGATGGG</td><td>1000</td><td></td><td></td><td></td><td></td><td></td><td></td><td></td><td></td><td></td></tr>
<tr><td>Oligo fwd</td><td>CACCCTATACGGGTCCGGACGGAT</td><td></td><td></td><td></td><td></td><td></td><td></td><td></td><td></td><td></td><td></td></tr>
<tr><td>Oligo rev</td><td>AAACATCCGTCCGGACCCGTATAG</td><td></td><td></td><td></td><td></td><td></td><td></td><td></td><td></td><td></td><td></td></tr>
<tr><td>Chromosome</td><td>start</td><td>end</td><td>strand</td><td>MM</td><td>target_seq</td><td>PAM</td><td>alignment</td><td>distance</td><td>position</td><td>gene name</td><td>gene id</td></tr>
<tr><td>GU305914.1</td><td>167061</td><td>167083</td><td>+</td><td>0</td><td>CTATACGGGTCCGGACGGAT</td><td>GGG</td><td>||||||||[||||||||||||]PAM</td><td>NA</td><td>Intergenic</td><td>NA</td><td>NA</td></tr>
<tr><td>T12591</td><td>TATACGGGTCCGGACGGATGGGG</td><td>1000</td><td></td><td></td><td></td><td></td><td></td><td></td><td></td><td></td><td></td></tr>
<tr><td>Oligo fwd</td><td>CACCTATACGGGTCCGGACGGATG</td><td></td><td></td><td></td><td></td><td></td><td></td><td></td><td></td><td></td><td></td></tr>
<tr><td>Oligo rev</td><td>AAACCATCCGTCCGGACCCGTATA</td><td></td><td></td><td></td><td></td><td></td><td></td><td></td><td></td><td></td><td></td></tr>
<tr><td>Chromosome</td><td>start</td><td>end</td><td>strand</td><td>MM</td><td>target_seq</td><td>PAM</td><td>alignment</td><td>distance</td><td>position</td><td>gene name</td><td>gene id</td></tr>
<tr><td>GU305914.1</td><td>167062</td><td>167084</td><td>+</td><td>0</td><td>TATACGGGTCCGGACGGATG</td><td>GGG</td><td>||||||||[||||||||||||]PAM</td><td>NA</td><td>Intergenic</td><td>NA</td><td>NA</td></tr>
<tr><td>T12592</td><td>ATGGGGCCGTCATCGTGCAGCGG</td><td>1000</td><td></td><td></td><td></td><td></td><td></td><td></td><td></td><td></td><td></td></tr>
<tr><td>Oligo fwd</td><td>CACCATGGGGCCGTCATCGTGCAG</td><td></td><td></td><td></td><td></td><td></td><td></td><td></td><td></td><td></td><td></td></tr>
<tr><td>Oligo rev</td><td>AAACCTGCACGATGACGGCCCCAT</td><td></td><td></td><td></td><td></td><td></td><td></td><td></td><td></td><td></td><td></td></tr>
<tr><td>Chromosome</td><td>start</td><td>end</td><td>strand</td><td>MM</td><td>target_seq</td><td>PAM</td><td>alignment</td><td>distance</td><td>position</td><td>gene name</td><td>gene id</td></tr>
<tr><td>GU305914.1</td><td>167079</td><td>167101</td><td>+</td><td>0</td><td>ATGGGGCCGTCATCGTGCAG</td><td>CGG</td><td>||||||||[||||||||||||]PAM</td><td>NA</td><td>Intergenic</td><td>NA</td><td>NA</td></tr>
<tr><td>T12593</td><td>GGCCGTCATCGTGCAGCGGTTGG</td><td>1000</td><td></td><td></td><td></td><td></td><td></td><td></td><td></td><td></td><td></td></tr>
<tr><td>Oligo fwd</td><td>CACCGGCCGTCATCGTGCAGCGGT</td><td></td><td></td><td></td><td></td><td></td><td></td><td></td><td></td><td></td><td></td></tr>
<tr><td>Oligo rev</td><td>AAACACCGCTGCACGATGACGGCC</td><td></td><td></td><td></td><td></td><td></td><td></td><td></td><td></td><td></td><td></td></tr>
<tr><td>Chromosome</td><td>start</td><td>end</td><td>strand</td><td>MM</td><td>target_seq</td><td>PAM</td><td>alignment</td><td>distance</td><td>position</td><td>gene name</td><td>gene id</td></tr>
<tr><td>GU305914.1</td><td>167083</td><td>167105</td><td>+</td><td>0</td><td>GGCCGTCATCGTGCAGCGGT</td><td>TGG</td><td>||||||||[||||||||||||]PAM</td><td>NA</td><td>Intergenic</td><td>NA</td><td>NA</td></tr>
<tr><td>T12594</td><td>TCCCGTTGTAGCAGCTCGCCAGG</td><td>1000</td><td></td><td></td><td></td><td></td><td></td><td></td><td></td><td></td><td></td></tr>
<tr><td>Oligo fwd</td><td>CACCTCCCGTTGTAGCAGCTCGCC</td><td></td><td></td><td></td><td></td><td></td><td></td><td></td><td></td><td></td><td></td></tr>
<tr><td>Oligo rev</td><td>AAACGGCGAGCTGCTACAACGGGA</td><td></td><td></td><td></td><td></td><td></td><td></td><td></td><td></td><td></td><td></td></tr>
<tr><td>Chromosome</td><td>start</td><td>end</td><td>strand</td><td>MM</td><td>target_seq</td><td>PAM</td><td>alignment</td><td>distance</td><td>position</td><td>gene name</td><td>gene id</td></tr>
<tr><td>GU305914.1</td><td>167113</td><td>167135</td><td>+</td><td>0</td><td>TCCCGTTGTAGCAGCTCGCC</td><td>AGG</td><td>||||||||[||||||||||||]PAM</td><td>NA</td><td>Intergenic</td><td>NA</td><td>NA</td></tr>
<tr><td>T12595</td><td>CCCGTTGTAGCAGCTCGCCAGGG</td><td>1000</td><td></td><td></td><td></td><td></td><td></td><td></td><td></td><td></td><td></td></tr>
<tr><td>Oligo fwd</td><td>CACCCCCGTTGTAGCAGCTCGCCA</td><td></td><td></td><td></td><td></td><td></td><td></td><td></td><td></td><td></td><td></td></tr>
<tr><td>Oligo rev</td><td>AAACTGGCGAGCTGCTACAACGGG</td><td></td><td></td><td></td><td></td><td></td><td></td><td></td><td></td><td></td><td></td></tr>
<tr><td>Chromosome</td><td>start</td><td>end</td><td>strand</td><td>MM</td><td>target_seq</td><td>PAM</td><td>alignment</td><td>distance</td><td>position</td><td>gene name</td><td>gene id</td></tr>
<tr><td>GU305914.1</td><td>167114</td><td>167136</td><td>+</td><td>0</td><td>CCCGTTGTAGCAGCTCGCCA</td><td>GGG</td><td>||||||||[||||||||||||]PAM</td><td>NA</td><td>Intergenic</td><td>NA</td><td>NA</td></tr>
<tr><td>T12596</td><td>AGCAGCTCGCCAGGGCCTCGCGG</td><td>1000</td><td></td><td></td><td></td><td></td><td></td><td></td><td></td><td></td><td></td></tr>
<tr><td>Oligo fwd</td><td>CACCAGCAGCTCGCCAGGGCCTCG</td><td></td><td></td><td></td><td></td><td></td><td></td><td></td><td></td><td></td><td></td></tr>
<tr><td>Oligo rev</td><td>AAACCGAGGCCCTGGCGAGCTGCT</td><td></td><td></td><td></td><td></td><td></td><td></td><td></td><td></td><td></td><td></td></tr>
<tr><td>Chromosome</td><td>start</td><td>end</td><td>strand</td><td>MM</td><td>target_seq</td><td>PAM</td><td>alignment</td><td>distance</td><td>position</td><td>gene name</td><td>gene id</td></tr>
<tr><td>GU305914.1</td><td>167122</td><td>167144</td><td>+</td><td>0</td><td>AGCAGCTCGCCAGGGCCTCG</td><td>CGG</td><td>||||||||[||||||||||||]PAM</td><td>NA</td><td>Intergenic</td><td>NA</td><td>NA</td></tr>
<tr><td>T12597</td><td>CTCGCGGCACTTGTGTTCGACGG</td><td>1000</td><td></td><td></td><td></td><td></td><td></td><td></td><td></td><td></td><td></td></tr>
<tr><td>Oligo fwd</td><td>CACCCTCGCGGCACTTGTGTTCGA</td><td></td><td></td><td></td><td></td><td></td><td></td><td></td><td></td><td></td><td></td></tr>
<tr><td>Oligo rev</td><td>AAACTCGAACACAAGTGCCGCGAG</td><td></td><td></td><td></td><td></td><td></td><td></td><td></td><td></td><td></td><td></td></tr>
<tr><td>Chromosome</td><td>start</td><td>end</td><td>strand</td><td>MM</td><td>target_seq</td><td>PAM</td><td>alignment</td><td>distance</td><td>position</td><td>gene name</td><td>gene id</td></tr>
<tr><td>GU305914.1</td><td>167138</td><td>167160</td><td>+</td><td>0</td><td>CTCGCGGCACTTGTGTTCGA</td><td>CGG</td><td>||||||||[||||||||||||]PAM</td><td>NA</td><td>Intergenic</td><td>NA</td><td>NA</td></tr>
<tr><td>GU305914.1</td><td>135824</td><td>135846</td><td>-</td><td>4</td><td>CCAGCGGCGCGTGTGTTCGA</td><td>AGG</td><td>|--|||||[-|-|||||||||]PAM</td><td>NA</td><td>Intergenic</td><td>NA</td><td>NA</td></tr>
</table>

<table>
<tr><td>T12598</td><td>GCGGCACTTGTGTTCGACGGTGG</td><td>1000</td><td></td><td></td><td></td><td></td><td></td><td></td><td></td><td></td><td></td></tr>
<tr><td>Oligo fwd</td><td>CACCGCGGCACTTGTGTTCGACGG</td><td></td><td></td><td></td><td></td><td></td><td></td><td></td><td></td><td></td><td></td></tr>
<tr><td>Oligo rev</td><td>AAACCCGTCGAACACAAGTGCCGC</td><td></td><td></td><td></td><td></td><td></td><td></td><td></td><td></td><td></td><td></td></tr>
<tr><td>Chromosome</td><td>start</td><td>end</td><td>strand</td><td>MM</td><td>target_seq</td><td>PAM</td><td>alignment</td><td>distance</td><td>position</td><td>gene name</td><td>gene id</td></tr>
<tr><td>GU305914.1</td><td>167141</td><td>167163</td><td>+</td><td>0</td><td>GCGGCACTTGTGTTCGACGG</td><td>TGG</td><td>||||||||[||||||||||||]PAM</td><td>NA</td><td>Intergenic</td><td>NA</td><td>NA</td></tr>
<tr><td>GU305914.1</td><td>135821</td><td>135843</td><td>-</td><td>3</td><td>GCGGCGCGTGTGTTCGAAGG</td><td>AGG</td><td>|||||-|-[|||||||||-||]PAM</td><td>NA</td><td>Intergenic</td><td>NA</td><td>NA</td></tr>
<tr><td>T12599</td><td>GTGTTCGACGGTGGCGTTGATGG</td><td>1000</td><td></td><td></td><td></td><td></td><td></td><td></td><td></td><td></td><td></td></tr>
<tr><td>Oligo fwd</td><td>CACCGTGTTCGACGGTGGCGTTGA</td><td></td><td></td><td></td><td></td><td></td><td></td><td></td><td></td><td></td><td></td></tr>
<tr><td>Oligo rev</td><td>AAACTCAACGCCACCGTCGAACAC</td><td></td><td></td><td></td><td></td><td></td><td></td><td></td><td></td><td></td><td></td></tr>
<tr><td>Chromosome</td><td>start</td><td>end</td><td>strand</td><td>MM</td><td>target_seq</td><td>PAM</td><td>alignment</td><td>distance</td><td>position</td><td>gene name</td><td>gene id</td></tr>
<tr><td>GU305914.1</td><td>167150</td><td>167172</td><td>+</td><td>0</td><td>GTGTTCGACGGTGGCGTTGA</td><td>TGG</td><td>||||||||[||||||||||||]PAM</td><td>NA</td><td>Intergenic</td><td>NA</td><td>NA</td></tr>
<tr><td>T12600</td><td>TGTTCGACGGTGGCGTTGATGGG</td><td>1000</td><td></td><td></td><td></td><td></td><td></td><td></td><td></td><td></td><td></td></tr>
<tr><td>Oligo fwd</td><td>CACCTGTTCGACGGTGGCGTTGAT</td><td></td><td></td><td></td><td></td><td></td><td></td><td></td><td></td><td></td><td></td></tr>
<tr><td>Oligo rev</td><td>AAACATCAACGCCACCGTCGAACA</td><td></td><td></td><td></td><td></td><td></td><td></td><td></td><td></td><td></td><td></td></tr>
<tr><td>Chromosome</td><td>start</td><td>end</td><td>strand</td><td>MM</td><td>target_seq</td><td>PAM</td><td>alignment</td><td>distance</td><td>position</td><td>gene name</td><td>gene id</td></tr>
<tr><td>GU305914.1</td><td>167151</td><td>167173</td><td>+</td><td>0</td><td>TGTTCGACGGTGGCGTTGAT</td><td>GGG</td><td>||||||||[||||||||||||]PAM</td><td>NA</td><td>Intergenic</td><td>NA</td><td>NA</td></tr>
<tr><td>T12601</td><td>GCGTTGATGGGCTCGCTGCGCGG</td><td>1000</td><td></td><td></td><td></td><td></td><td></td><td></td><td></td><td></td><td></td></tr>
<tr><td>Oligo fwd</td><td>CACCGCGTTGATGGGCTCGCTGCG</td><td></td><td></td><td></td><td></td><td></td><td></td><td></td><td></td><td></td><td></td></tr>
<tr><td>Oligo rev</td><td>AAACCGCAGCGAGCCCATCAACGC</td><td></td><td></td><td></td><td></td><td></td><td></td><td></td><td></td><td></td><td></td></tr>
<tr><td>Chromosome</td><td>start</td><td>end</td><td>strand</td><td>MM</td><td>target_seq</td><td>PAM</td><td>alignment</td><td>distance</td><td>position</td><td>gene name</td><td>gene id</td></tr>
<tr><td>GU305914.1</td><td>167163</td><td>167185</td><td>+</td><td>0</td><td>GCGTTGATGGGCTCGCTGCG</td><td>CGG</td><td>||||||||[||||||||||||]PAM</td><td>NA</td><td>Intergenic</td><td>NA</td><td>NA</td></tr>
<tr><td>T12602</td><td>GCTGCGCGGTAACACGTATACGG</td><td>1000</td><td></td><td></td><td></td><td></td><td></td><td></td><td></td><td></td><td></td></tr>
<tr><td>Oligo fwd</td><td>CACCGCTGCGCGGTAACACGTATA</td><td></td><td></td><td></td><td></td><td></td><td></td><td></td><td></td><td></td><td></td></tr>
<tr><td>Oligo rev</td><td>AAACTATACGTGTTACCGCGCAGC</td><td></td><td></td><td></td><td></td><td></td><td></td><td></td><td></td><td></td><td></td></tr>
<tr><td>Chromosome</td><td>start</td><td>end</td><td>strand</td><td>MM</td><td>target_seq</td><td>PAM</td><td>alignment</td><td>distance</td><td>position</td><td>gene name</td><td>gene id</td></tr>
<tr><td>GU305914.1</td><td>167177</td><td>167199</td><td>+</td><td>0</td><td>GCTGCGCGGTAACACGTATA</td><td>CGG</td><td>||||||||[||||||||||||]PAM</td><td>NA</td><td>Intergenic</td><td>NA</td><td>NA</td></tr>
<tr><td>T12603</td><td>GCGCGGTAACACGTATACGGTGG</td><td>1000</td><td></td><td></td><td></td><td></td><td></td><td></td><td></td><td></td><td></td></tr>
<tr><td>Oligo fwd</td><td>CACCGCGCGGTAACACGTATACGG</td><td></td><td></td><td></td><td></td><td></td><td></td><td></td><td></td><td></td><td></td></tr>
<tr><td>Oligo rev</td><td>AAACCCGTATACGTGTTACCGCGC</td><td></td><td></td><td></td><td></td><td></td><td></td><td></td><td></td><td></td><td></td></tr>
<tr><td>Chromosome</td><td>start</td><td>end</td><td>strand</td><td>MM</td><td>target_seq</td><td>PAM</td><td>alignment</td><td>distance</td><td>position</td><td>gene name</td><td>gene id</td></tr>
<tr><td>GU305914.1</td><td>167180</td><td>167202</td><td>+</td><td>0</td><td>GCGCGGTAACACGTATACGG</td><td>TGG</td><td>||||||||[||||||||||||]PAM</td><td>NA</td><td>Intergenic</td><td>NA</td><td>NA</td></tr>
<tr><td>T12604</td><td>GTAACACGTATACGGTGGTGAGG</td><td>1000</td><td></td><td></td><td></td><td></td><td></td><td></td><td></td><td></td><td></td></tr>
<tr><td>Oligo fwd</td><td>CACCGTAACACGTATACGGTGGTG</td><td></td><td></td><td></td><td></td><td></td><td></td><td></td><td></td><td></td><td></td></tr>
<tr><td>Oligo rev</td><td>AAACCACCACCGTATACGTGTTAC</td><td></td><td></td><td></td><td></td><td></td><td></td><td></td><td></td><td></td><td></td></tr>
<tr><td>Chromosome</td><td>start</td><td>end</td><td>strand</td><td>MM</td><td>target_seq</td><td>PAM</td><td>alignment</td><td>distance</td><td>position</td><td>gene name</td><td>gene id</td></tr>
<tr><td>GU305914.1</td><td>167185</td><td>167207</td><td>+</td><td>0</td><td>GTAACACGTATACGGTGGTG</td><td>AGG</td><td>||||||||[||||||||||||]PAM</td><td>NA</td><td>Intergenic</td><td>NA</td><td>NA</td></tr>
<tr><td>T12605</td><td>GTCCAACGCCCGTCGCTCGCCGG</td><td>1000</td><td></td><td></td><td></td><td></td><td></td><td></td><td></td><td></td><td></td></tr>
<tr><td>Oligo fwd</td><td>CACCGTCCAACGCCCGTCGCTCGC</td><td></td><td></td><td></td><td></td><td></td><td></td><td></td><td></td><td></td><td></td></tr>
<tr><td>Oligo rev</td><td>AAACGCGAGCGACGGGCGTTGGAC</td><td></td><td></td><td></td><td></td><td></td><td></td><td></td><td></td><td></td><td></td></tr>
<tr><td>Chromosome</td><td>start</td><td>end</td><td>strand</td><td>MM</td><td>target_seq</td><td>PAM</td><td>alignment</td><td>distance</td><td>position</td><td>gene name</td><td>gene id</td></tr>
<tr><td>GU305914.1</td><td>167207</td><td>167229</td><td>+</td><td>0</td><td>GTCCAACGCCCGTCGCTCGC</td><td>CGG</td><td>||||||||[||||||||||||]PAM</td><td>NA</td><td>Intergenic</td><td>NA</td><td>NA</td></tr>
<tr><td>T12606</td><td>GCCCGTCGCTCGCCGGCGACCGG</td><td>1000</td><td></td><td></td><td></td><td></td><td></td><td></td><td></td><td></td><td></td></tr>
<tr><td>Oligo fwd</td><td>CACCGCCCGTCGCTCGCCGGCGAC</td><td></td><td></td><td></td><td></td><td></td><td></td><td></td><td></td><td></td><td></td></tr>
<tr><td>Oligo rev</td><td>AAACGTCGCCGGCGAGCGACGGGC</td><td></td><td></td><td></td><td></td><td></td><td></td><td></td><td></td><td></td><td></td></tr>
<tr><td>Chromosome</td><td>start</td><td>end</td><td>strand</td><td>MM</td><td>target_seq</td><td>PAM</td><td>alignment</td><td>distance</td><td>position</td><td>gene name</td><td>gene id</td></tr>
<tr><td>GU305914.1</td><td>167214</td><td>167236</td><td>+</td><td>0</td><td>GCCCGTCGCTCGCCGGCGAC</td><td>CGG</td><td>||||||||[||||||||||||]PAM</td><td>NA</td><td>Intergenic</td><td>NA</td><td>NA</td></tr>
<tr><td>T12607</td><td>GCGACTGCGCCGCCCCCAGCAGG</td><td>1000</td><td></td><td></td><td></td><td></td><td></td><td></td><td></td><td></td><td></td></tr>
<tr><td>Oligo fwd</td><td>CACCGCGACTGCGCCGCCCCCAGC</td><td></td><td></td><td></td><td></td><td></td><td></td><td></td><td></td><td></td><td></td></tr>
<tr><td>Oligo rev</td><td>AAACGCTGGGGGCGGCGCAGTCGC</td><td></td><td></td><td></td><td></td><td></td><td></td><td></td><td></td><td></td><td></td></tr>
<tr><td>Chromosome</td><td>start</td><td>end</td><td>strand</td><td>MM</td><td>target_seq</td><td>PAM</td><td>alignment</td><td>distance</td><td>position</td><td>gene name</td><td>gene id</td></tr>
<tr><td>GU305914.1</td><td>167245</td><td>167267</td><td>+</td><td>0</td><td>GCGACTGCGCCGCCCCCAGC</td><td>AGG</td><td>||||||||[||||||||||||]PAM</td><td>NA</td><td>Intergenic</td><td>NA</td><td>NA</td></tr>
<tr><td>T12608</td><td>CGCCCCCAGCAGGTTACGTGTGG</td><td>1000</td><td></td><td></td><td></td><td></td><td></td><td></td><td></td><td></td><td></td></tr>
<tr><td>Oligo fwd</td><td>CACCCGCCCCCAGCAGGTTACGTG</td><td></td><td></td><td></td><td></td><td></td><td></td><td></td><td></td><td></td><td></td></tr>
<tr><td>Oligo rev</td><td>AAACCACGTAACCTGCTGGGGGCG</td><td></td><td></td><td></td><td></td><td></td><td></td><td></td><td></td><td></td><td></td></tr>
<tr><td>Chromosome</td><td>start</td><td>end</td><td>strand</td><td>MM</td><td>target_seq</td><td>PAM</td><td>alignment</td><td>distance</td><td>position</td><td>gene name</td><td>gene id</td></tr>
<tr><td>GU305914.1</td><td>167255</td><td>167277</td><td>+</td><td>0</td><td>CGCCCCCAGCAGGTTACGTG</td><td>TGG</td><td>||||||||[||||||||||||]PAM</td><td>NA</td><td>Intergenic</td><td>NA</td><td>NA</td></tr>
<tr><td>T12609</td><td>AGCAGTATGAGCAATAATAAAGG</td><td>1000</td><td></td><td></td><td></td><td></td><td></td><td></td><td></td><td></td><td></td></tr>
<tr><td>Oligo fwd</td><td>CACCAGCAGTATGAGCAATAATAA</td><td></td><td></td><td></td><td></td><td></td><td></td><td></td><td></td><td></td><td></td></tr>
<tr><td>Oligo rev</td><td>AAACTTATTATTGCTCATACTGCT</td><td></td><td></td><td></td><td></td><td></td><td></td><td></td><td></td><td></td><td></td></tr>
<tr><td>Chromosome</td><td>start</td><td>end</td><td>strand</td><td>MM</td><td>target_seq</td><td>PAM</td><td>alignment</td><td>distance</td><td>position</td><td>gene name</td><td>gene id</td></tr>
<tr><td>GU305914.1</td><td>167280</td><td>167302</td><td>+</td><td>0</td><td>AGCAGTATGAGCAATAATAA</td><td>AGG</td><td>||||||||[||||||||||||]PAM</td><td>NA</td><td>Intergenic</td><td>NA</td><td>NA</td></tr>
<tr><td>T12610</td><td>AAGGCATCATATGAAGTCTGCGG</td><td>1000</td><td></td><td></td><td></td><td></td><td></td><td></td><td></td><td></td><td></td></tr>
<tr><td>Oligo fwd</td><td>CACCAAGGCATCATATGAAGTCTG</td><td></td><td></td><td></td><td></td><td></td><td></td><td></td><td></td><td></td><td></td></tr>
<tr><td>Oligo rev</td><td>AAACCAGACTTCATATGATGCCTT</td><td></td><td></td><td></td><td></td><td></td><td></td><td></td><td></td><td></td><td></td></tr>
<tr><td>Chromosome</td><td>start</td><td>end</td><td>strand</td><td>MM</td><td>target_seq</td><td>PAM</td><td>alignment</td><td>distance</td><td>position</td><td>gene name</td><td>gene id</td></tr>
<tr><td>GU305914.1</td><td>167299</td><td>167321</td><td>+</td><td>0</td><td>AAGGCATCATATGAAGTCTG</td><td>CGG</td><td>||||||||[||||||||||||]PAM</td><td>NA</td><td>Intergenic</td><td>NA</td><td>NA</td></tr>
<tr><td>T12611</td><td>AGGCATCATATGAAGTCTGCGGG</td><td>1000</td><td></td><td></td><td></td><td></td><td></td><td></td><td></td><td></td><td></td></tr>
<tr><td>Oligo fwd</td><td>CACCAGGCATCATATGAAGTCTGC</td><td></td><td></td><td></td><td></td><td></td><td></td><td></td><td></td><td></td><td></td></tr>
<tr><td>Oligo rev</td><td>AAACGCAGACTTCATATGATGCCT</td><td></td><td></td><td></td><td></td><td></td><td></td><td></td><td></td><td></td><td></td></tr>
<tr><td>Chromosome</td><td>start</td><td>end</td><td>strand</td><td>MM</td><td>target_seq</td><td>PAM</td><td>alignment</td><td>distance</td><td>position</td><td>gene name</td><td>gene id</td></tr>
<tr><td>GU305914.1</td><td>167300</td><td>167322</td><td>+</td><td>0</td><td>AGGCATCATATGAAGTCTGC</td><td>GGG</td><td>||||||||[||||||||||||]PAM</td><td>NA</td><td>Intergenic</td><td>NA</td><td>NA</td></tr>
<tr><td>T12612</td><td>TGCGGGTGTGTACGTGTATTCGG</td><td>1000</td><td></td><td></td><td></td><td></td><td></td><td></td><td></td><td></td><td></td></tr>
<tr><td>Oligo fwd</td><td>CACCTGCGGGTGTGTACGTGTATT</td><td></td><td></td><td></td><td></td><td></td><td></td><td></td><td></td><td></td><td></td></tr>
<tr><td>Oligo rev</td><td>AAACAATACACGTACACACCCGCA</td><td></td><td></td><td></td><td></td><td></td><td></td><td></td><td></td><td></td><td></td></tr>
<tr><td>Chromosome</td><td>start</td><td>end</td><td>strand</td><td>MM</td><td>target_seq</td><td>PAM</td><td>alignment</td><td>distance</td><td>position</td><td>gene name</td><td>gene id</td></tr>
</table>

GU305914.1 167317 167339 + 0 TGCGGGTGTGTACGTGTATT CGG ||||||||[||||||||||||]PAM NA Intergenic NA NA

T12613 TGAGAAACACCGAAGCGCCCAGG 1000
Oligo fwd CACCTGAGAAACACCGAAGCGCCC
Oligo rev AAACGGGCGCTTCGGTGTTTCTCA
Chromosome start end strand MM target_seq PAM alignment distance position gene name gene id
GU305914.1 167397 167419 + 0 TGAGAAACACCGAAGCGCCC AGG ||||||||[||||||||||||]PAM NA Intergenic NA NA

T12614 AACACCGAAGCGCCCAGGCCCGG 1000
Oligo fwd CACCAACACCGAAGCGCCCAGGCC
Oligo rev AAACGGCCTGGGCGCTTCGGTGTT
Chromosome start end strand MM target_seq PAM alignment distance position gene name gene id
GU305914.1 167402 167424 + 0 AACACCGAAGCGCCCAGGCC CGG ||||||||[||||||||||||]PAM NA Intergenic NA NA

T12615 AATGAGCGAGTCGTTAAATATGG 1000
Oligo fwd CACCAATGAGCGAGTCGTTAAATA
Oligo rev AAACTATTTAACGACTCGCTCATT
Chromosome start end strand MM target_seq PAM alignment distance position gene name gene id
GU305914.1 167431 167453 + 0 AATGAGCGAGTCGTTAAATA TGG ||||||||[||||||||||||]PAM NA Intergenic NA NA

T12616 CCGCGCGACTGAAAGTCACTCGG 1000
Oligo fwd CACCCCGCGCGACTGAAAGTCACT
Oligo rev AAACAGTGACTTTCAGTCGCGCGG
Chromosome start end strand MM target_seq PAM alignment distance position gene name gene id
GU305914.1 167472 167494 + 0 CCGCGCGACTGAAAGTCACT CGG ||||||||[||||||||||||]PAM NA Intergenic NA NA

T12617 CGCGCGACTGAAAGTCACTCGGG 1000
Oligo fwd CACCCGCGCGACTGAAAGTCACTC
Oligo rev AAACGAGTGACTTTCAGTCGCGCG
Chromosome start end strand MM target_seq PAM alignment distance position gene name gene id
GU305914.1 167473 167495 + 0 CGCGCGACTGAAAGTCACTC GGG ||||||||[||||||||||||]PAM NA Intergenic NA NA

T12618 ACTGAAAGTCACTCGGGTGAAGG 1000
Oligo fwd CACCACTGAAAGTCACTCGGGTGA
Oligo rev AAACTCACCCGAGTGACTTTCAGT
Chromosome start end strand MM target_seq PAM alignment distance position gene name gene id
GU305914.1 167479 167501 + 0 ACTGAAAGTCACTCGGGTGA AGG ||||||||[||||||||||||]PAM NA Intergenic NA NA

T12619 TCACTCGGGTGAAGGCGCTCAGG 1000
Oligo fwd CACCTCACTCGGGTGAAGGCGCTC
Oligo rev AAACGAGCGCCTTCACCCGAGTGA
Chromosome start end strand MM target_seq PAM alignment distance position gene name gene id
GU305914.1 167487 167509 + 0 TCACTCGGGTGAAGGCGCTC AGG ||||||||[||||||||||||]PAM NA Intergenic NA NA

T12620 GTGAAGGCGCTCAGGACGAACGG 1000
Oligo fwd CACCGTGAAGGCGCTCAGGACGAA
Oligo rev AAACTTCGTCCTGAGCGCCTTCAC
Chromosome start end strand MM target_seq PAM alignment distance position gene name gene id
GU305914.1 167495 167517 + 0 GTGAAGGCGCTCAGGACGAA CGG ||||||||[||||||||||||]PAM NA Intergenic NA NA

T12621 GACGAACGGCCCGTATTTCGCGG 1000
Oligo fwd CACCGACGAACGGCCCGTATTTCG
Oligo rev AAACCGAAATACGGGCCGTTCGTC
Chromosome start end strand MM target_seq PAM alignment distance position gene name gene id
GU305914.1 167509 167531 + 0 GACGAACGGCCCGTATTTCG CGG ||||||||[||||||||||||]PAM NA Intergenic NA NA

T12622 ACGAACGGCCCGTATTTCGCGGG 1000
Oligo fwd CACCACGAACGGCCCGTATTTCGC
Oligo rev AAACGCGAAATACGGGCCGTTCGT
Chromosome start end strand MM target_seq PAM alignment distance position gene name gene id
GU305914.1 167510 167532 + 0 ACGAACGGCCCGTATTTCGC GGG ||||||||[||||||||||||]PAM NA Intergenic NA NA

T12623 CGAACGGCCCGTATTTCGCGGGG 1000
Oligo fwd CACCCGAACGGCCCGTATTTCGCG
Oligo rev AAACCGCGAAATACGGGCCGTTCG
Chromosome start end strand MM target_seq PAM alignment distance position gene name gene id
GU305914.1 167511 167533 + 0 CGAACGGCCCGTATTTCGCG GGG ||||||||[||||||||||||]PAM NA Intergenic NA NA

T12624 GGCAGTACCTGCGAGTGCATAGG 1000
Oligo fwd CACCGGCAGTACCTGCGAGTGCAT
Oligo rev AAACATGCACTCGCAGGTACTGCC
Chromosome start end strand MM target_seq PAM alignment distance position gene name gene id
GU305914.1 167532 167554 + 0 GGCAGTACCTGCGAGTGCAT AGG ||||||||[||||||||||||]PAM NA Intergenic NA NA

T12625 CCTGCGAGTGCATAGGTCTCCGG 1000
Oligo fwd CACCCCTGCGAGTGCATAGGTCTC
Oligo rev AAACGAGACCTATGCACTCGCAGG
Chromosome start end strand MM target_seq PAM alignment distance position gene name gene id
GU305914.1 167539 167561 + 0 CCTGCGAGTGCATAGGTCTC CGG ||||||||[||||||||||||]PAM NA Intergenic NA NA

T12626 CTGCGAGTGCATAGGTCTCCGGG 1000
Oligo fwd CACCCTGCGAGTGCATAGGTCTCC
Oligo rev AAACGGAGACCTATGCACTCGCAG
Chromosome start end strand MM target_seq PAM alignment distance position gene name gene id
GU305914.1 167540 167562 + 0 CTGCGAGTGCATAGGTCTCC GGG ||||||||[||||||||||||]PAM NA Intergenic NA NA

T12627 TGCGAGTGCATAGGTCTCCGGGG 1000
Oligo fwd CACCTGCGAGTGCATAGGTCTCCG
Oligo rev AAACCGGAGACCTATGCACTCGCA

<table>
<tr><th>Chromosome</th><th>start</th><th>end</th><th>strand</th><th>MM</th><th>target_seq</th><th>PAM</th><th>alignment</th><th>distance</th><th>position</th><th>gene name</th><th>gene id</th></tr>
<tr><td>GU305914.1</td><td>167541</td><td>167563</td><td>+</td><td>0</td><td>TGCGAGTGCATAGGTCTCCG</td><td>GGG</td><td>||||||||[||||||||||||]PAM</td><td>NA</td><td>Intergenic</td><td>NA</td><td>NA</td></tr>
<tr><td>T12628</td><td>GCATAGGTCTCCGGGGAATGTGG</td><td>1000</td><td></td><td></td><td></td><td></td><td></td><td></td><td></td><td></td><td></td></tr>
<tr><td>Oligo fwd</td><td>CACCGCATAGGTCTCCGGGGAATG</td><td></td><td></td><td></td><td></td><td></td><td></td><td></td><td></td><td></td><td></td></tr>
<tr><td>Oligo rev</td><td>AAACCATTCCCCGGAGACCTATGC</td><td></td><td></td><td></td><td></td><td></td><td></td><td></td><td></td><td></td><td></td></tr>
<tr><td>Chromosome</td><td>start</td><td>end</td><td>strand</td><td>MM</td><td>target_seq</td><td>PAM</td><td>alignment</td><td>distance</td><td>position</td><td>gene name</td><td>gene id</td></tr>
<tr><td>GU305914.1</td><td>167548</td><td>167570</td><td>+</td><td>0</td><td>GCATAGGTCTCCGGGGAATG</td><td>TGG</td><td>||||||||[||||||||||||]PAM</td><td>NA</td><td>Intergenic</td><td>NA</td><td>NA</td></tr>
<tr><td>T12629</td><td>GTCTCCGGGGAATGTGGTGATGG</td><td>1000</td><td></td><td></td><td></td><td></td><td></td><td></td><td></td><td></td><td></td></tr>
<tr><td>Oligo fwd</td><td>CACCGTCTCCGGGGAATGTGGTGA</td><td></td><td></td><td></td><td></td><td></td><td></td><td></td><td></td><td></td><td></td></tr>
<tr><td>Oligo rev</td><td>AAACTCACCACATTCCCCGGAGAC</td><td></td><td></td><td></td><td></td><td></td><td></td><td></td><td></td><td></td><td></td></tr>
<tr><td>Chromosome</td><td>start</td><td>end</td><td>strand</td><td>MM</td><td>target_seq</td><td>PAM</td><td>alignment</td><td>distance</td><td>position</td><td>gene name</td><td>gene id</td></tr>
<tr><td>GU305914.1</td><td>167554</td><td>167576</td><td>+</td><td>0</td><td>GTCTCCGGGGAATGTGGTGA</td><td>TGG</td><td>||||||||[||||||||||||]PAM</td><td>NA</td><td>Intergenic</td><td>NA</td><td>NA</td></tr>
<tr><td>T12630</td><td>TCCGGGGAATGTGGTGATGGTGG</td><td>1000</td><td></td><td></td><td></td><td></td><td></td><td></td><td></td><td></td><td></td></tr>
<tr><td>Oligo fwd</td><td>CACCTCCGGGGAATGTGGTGATGG</td><td></td><td></td><td></td><td></td><td></td><td></td><td></td><td></td><td></td><td></td></tr>
<tr><td>Oligo rev</td><td>AAACCCATCACCACATTCCCCGGA</td><td></td><td></td><td></td><td></td><td></td><td></td><td></td><td></td><td></td><td></td></tr>
<tr><td>Chromosome</td><td>start</td><td>end</td><td>strand</td><td>MM</td><td>target_seq</td><td>PAM</td><td>alignment</td><td>distance</td><td>position</td><td>gene name</td><td>gene id</td></tr>
<tr><td>GU305914.1</td><td>167557</td><td>167579</td><td>+</td><td>0</td><td>TCCGGGGAATGTGGTGATGG</td><td>TGG</td><td>||||||||[||||||||||||]PAM</td><td>NA</td><td>Intergenic</td><td>NA</td><td>NA</td></tr>
<tr><td>T12631</td><td>CCGGGGAATGTGGTGATGGTGGG</td><td>1000</td><td></td><td></td><td></td><td></td><td></td><td></td><td></td><td></td><td></td></tr>
<tr><td>Oligo fwd</td><td>CACCCCGGGGAATGTGGTGATGGT</td><td></td><td></td><td></td><td></td><td></td><td></td><td></td><td></td><td></td><td></td></tr>
<tr><td>Oligo rev</td><td>AAACACCATCACCACATTCCCCGG</td><td></td><td></td><td></td><td></td><td></td><td></td><td></td><td></td><td></td><td></td></tr>
<tr><td>Chromosome</td><td>start</td><td>end</td><td>strand</td><td>MM</td><td>target_seq</td><td>PAM</td><td>alignment</td><td>distance</td><td>position</td><td>gene name</td><td>gene id</td></tr>
<tr><td>GU305914.1</td><td>167558</td><td>167580</td><td>+</td><td>0</td><td>CCGGGGAATGTGGTGATGGT</td><td>GGG</td><td>||||||||[||||||||||||]PAM</td><td>NA</td><td>Intergenic</td><td>NA</td><td>NA</td></tr>
<tr><td>T12632</td><td>TGTGGTGATGGTGGGATTCGTGG</td><td>1000</td><td></td><td></td><td></td><td></td><td></td><td></td><td></td><td></td><td></td></tr>
<tr><td>Oligo fwd</td><td>CACCTGTGGTGATGGTGGGATTCG</td><td></td><td></td><td></td><td></td><td></td><td></td><td></td><td></td><td></td><td></td></tr>
<tr><td>Oligo rev</td><td>AAACCGAATCCCACCATCACCACA</td><td></td><td></td><td></td><td></td><td></td><td></td><td></td><td></td><td></td><td></td></tr>
<tr><td>Chromosome</td><td>start</td><td>end</td><td>strand</td><td>MM</td><td>target_seq</td><td>PAM</td><td>alignment</td><td>distance</td><td>position</td><td>gene name</td><td>gene id</td></tr>
<tr><td>GU305914.1</td><td>167566</td><td>167588</td><td>+</td><td>0</td><td>TGTGGTGATGGTGGGATTCG</td><td>TGG</td><td>||||||||[||||||||||||]PAM</td><td>NA</td><td>Intergenic</td><td>NA</td><td>NA</td></tr>
<tr><td>T12633</td><td>GATGGTGGGATTCGTGGTAGTGG</td><td>1000</td><td></td><td></td><td></td><td></td><td></td><td></td><td></td><td></td><td></td></tr>
<tr><td>Oligo fwd</td><td>CACCGATGGTGGGATTCGTGGTAG</td><td></td><td></td><td></td><td></td><td></td><td></td><td></td><td></td><td></td><td></td></tr>
<tr><td>Oligo rev</td><td>AAACCTACCACGAATCCCACCATC</td><td></td><td></td><td></td><td></td><td></td><td></td><td></td><td></td><td></td><td></td></tr>
<tr><td>Chromosome</td><td>start</td><td>end</td><td>strand</td><td>MM</td><td>target_seq</td><td>PAM</td><td>alignment</td><td>distance</td><td>position</td><td>gene name</td><td>gene id</td></tr>
<tr><td>GU305914.1</td><td>167572</td><td>167594</td><td>+</td><td>0</td><td>GATGGTGGGATTCGTGGTAG</td><td>TGG</td><td>||||||||[||||||||||||]PAM</td><td>NA</td><td>Intergenic</td><td>NA</td><td>NA</td></tr>
<tr><td>T12634</td><td>GGTGGGATTCGTGGTAGTGGTGG</td><td>1000</td><td></td><td></td><td></td><td></td><td></td><td></td><td></td><td></td><td></td></tr>
<tr><td>Oligo fwd</td><td>CACCGGTGGGATTCGTGGTAGTGG</td><td></td><td></td><td></td><td></td><td></td><td></td><td></td><td></td><td></td><td></td></tr>
<tr><td>Oligo rev</td><td>AAACCCACTACCACGAATCCCACC</td><td></td><td></td><td></td><td></td><td></td><td></td><td></td><td></td><td></td><td></td></tr>
<tr><td>Chromosome</td><td>start</td><td>end</td><td>strand</td><td>MM</td><td>target_seq</td><td>PAM</td><td>alignment</td><td>distance</td><td>position</td><td>gene name</td><td>gene id</td></tr>
<tr><td>GU305914.1</td><td>167575</td><td>167597</td><td>+</td><td>0</td><td>GGTGGGATTCGTGGTAGTGG</td><td>TGG</td><td>||||||||[||||||||||||]PAM</td><td>NA</td><td>Intergenic</td><td>NA</td><td>NA</td></tr>
<tr><td>T12635</td><td>GGGATTCGTGGTAGTGGTGGTGG</td><td>1000</td><td></td><td></td><td></td><td></td><td></td><td></td><td></td><td></td><td></td></tr>
<tr><td>Oligo fwd</td><td>CACCGGGATTCGTGGTAGTGGTGG</td><td></td><td></td><td></td><td></td><td></td><td></td><td></td><td></td><td></td><td></td></tr>
<tr><td>Oligo rev</td><td>AAACCCACCACTACCACGAATCCC</td><td></td><td></td><td></td><td></td><td></td><td></td><td></td><td></td><td></td><td></td></tr>
<tr><td>Chromosome</td><td>start</td><td>end</td><td>strand</td><td>MM</td><td>target_seq</td><td>PAM</td><td>alignment</td><td>distance</td><td>position</td><td>gene name</td><td>gene id</td></tr>
<tr><td>GU305914.1</td><td>167578</td><td>167600</td><td>+</td><td>0</td><td>GGGATTCGTGGTAGTGGTGG</td><td>TGG</td><td>||||||||[||||||||||||]PAM</td><td>NA</td><td>Intergenic</td><td>NA</td><td>NA</td></tr>
<tr><td>T12636</td><td>TGGTGGTGACGACCTGCGTGCGG</td><td>1000</td><td></td><td></td><td></td><td></td><td></td><td></td><td></td><td></td><td></td></tr>
<tr><td>Oligo fwd</td><td>CACCTGGTGGTGACGACCTGCGTG</td><td></td><td></td><td></td><td></td><td></td><td></td><td></td><td></td><td></td><td></td></tr>
<tr><td>Oligo rev</td><td>AAACCACGCAGGTCGTCACCACCA</td><td></td><td></td><td></td><td></td><td></td><td></td><td></td><td></td><td></td><td></td></tr>
<tr><td>Chromosome</td><td>start</td><td>end</td><td>strand</td><td>MM</td><td>target_seq</td><td>PAM</td><td>alignment</td><td>distance</td><td>position</td><td>gene name</td><td>gene id</td></tr>
<tr><td>GU305914.1</td><td>167595</td><td>167617</td><td>+</td><td>0</td><td>TGGTGGTGACGACCTGCGTG</td><td>CGG</td><td>||||||||[||||||||||||]PAM</td><td>NA</td><td>Intergenic</td><td>NA</td><td>NA</td></tr>
<tr><td>T12637</td><td>GACGACCTGCGTGCGGAACGCGG</td><td>1000</td><td></td><td></td><td></td><td></td><td></td><td></td><td></td><td></td><td></td></tr>
<tr><td>Oligo fwd</td><td>CACCGACGACCTGCGTGCGGAACG</td><td></td><td></td><td></td><td></td><td></td><td></td><td></td><td></td><td></td><td></td></tr>
<tr><td>Oligo rev</td><td>AAACCGTTCCGCACGCAGGTCGTC</td><td></td><td></td><td></td><td></td><td></td><td></td><td></td><td></td><td></td><td></td></tr>
<tr><td>Chromosome</td><td>start</td><td>end</td><td>strand</td><td>MM</td><td>target_seq</td><td>PAM</td><td>alignment</td><td>distance</td><td>position</td><td>gene name</td><td>gene id</td></tr>
<tr><td>GU305914.1</td><td>167602</td><td>167624</td><td>+</td><td>0</td><td>GACGACCTGCGTGCGGAACG</td><td>CGG</td><td>||||||||[||||||||||||]PAM</td><td>NA</td><td>Intergenic</td><td>NA</td><td>NA</td></tr>
<tr><td>T12638</td><td>TGCGTGCGGAACGCGGCGCTAGG</td><td>1000</td><td></td><td></td><td></td><td></td><td></td><td></td><td></td><td></td><td></td></tr>
<tr><td>Oligo fwd</td><td>CACCTGCGTGCGGAACGCGGCGCT</td><td></td><td></td><td></td><td></td><td></td><td></td><td></td><td></td><td></td><td></td></tr>
<tr><td>Oligo rev</td><td>AAACAGCGCCGCGTTCCGCACGCA</td><td></td><td></td><td></td><td></td><td></td><td></td><td></td><td></td><td></td><td></td></tr>
<tr><td>Chromosome</td><td>start</td><td>end</td><td>strand</td><td>MM</td><td>target_seq</td><td>PAM</td><td>alignment</td><td>distance</td><td>position</td><td>gene name</td><td>gene id</td></tr>
<tr><td>GU305914.1</td><td>167609</td><td>167631</td><td>+</td><td>0</td><td>TGCGTGCGGAACGCGGCGCT</td><td>AGG</td><td>||||||||[||||||||||||]PAM</td><td>NA</td><td>Intergenic</td><td>NA</td><td>NA</td></tr>
<tr><td>T12639</td><td>GCTAGGAGAAGACACAGACTCGG</td><td>1000</td><td></td><td></td><td></td><td></td><td></td><td></td><td></td><td></td><td></td></tr>
<tr><td>Oligo fwd</td><td>CACCGCTAGGAGAAGACACAGACT</td><td></td><td></td><td></td><td></td><td></td><td></td><td></td><td></td><td></td><td></td></tr>
<tr><td>Oligo rev</td><td>AAACAGTCTGTGTCTTCTCCTAGC</td><td></td><td></td><td></td><td></td><td></td><td></td><td></td><td></td><td></td><td></td></tr>
<tr><td>Chromosome</td><td>start</td><td>end</td><td>strand</td><td>MM</td><td>target_seq</td><td>PAM</td><td>alignment</td><td>distance</td><td>position</td><td>gene name</td><td>gene id</td></tr>
<tr><td>GU305914.1</td><td>167626</td><td>167648</td><td>+</td><td>0</td><td>GCTAGGAGAAGACACAGACT</td><td>CGG</td><td>||||||||[||||||||||||]PAM</td><td>NA</td><td>Intergenic</td><td>NA</td><td>NA</td></tr>
<tr><td>T12640</td><td>GTGTGACACTCAGTTGTGTCTGG</td><td>1000</td><td></td><td></td><td></td><td></td><td></td><td></td><td></td><td></td><td></td></tr>
<tr><td>Oligo fwd</td><td>CACCGTGTGACACTCAGTTGTGTC</td><td></td><td></td><td></td><td></td><td></td><td></td><td></td><td></td><td></td><td></td></tr>
<tr><td>Oligo rev</td><td>AAACGACACAACTGAGTGTCACAC</td><td></td><td></td><td></td><td></td><td></td><td></td><td></td><td></td><td></td><td></td></tr>
<tr><td>Chromosome</td><td>start</td><td>end</td><td>strand</td><td>MM</td><td>target_seq</td><td>PAM</td><td>alignment</td><td>distance</td><td>position</td><td>gene name</td><td>gene id</td></tr>
<tr><td>GU305914.1</td><td>167688</td><td>167710</td><td>+</td><td>0</td><td>GTGTGACACTCAGTTGTGTC</td><td>TGG</td><td>||||||||[||||||||||||]PAM</td><td>NA</td><td>Intergenic</td><td>NA</td><td>NA</td></tr>
<tr><td>T12641</td><td>TGTGACACTCAGTTGTGTCTGGG</td><td>1000</td><td></td><td></td><td></td><td></td><td></td><td></td><td></td><td></td><td></td></tr>
<tr><td>Oligo fwd</td><td>CACCTGTGACACTCAGTTGTGTCT</td><td></td><td></td><td></td><td></td><td></td><td></td><td></td><td></td><td></td><td></td></tr>
<tr><td>Oligo rev</td><td>AAACAGACACAACTGAGTGTCACA</td><td></td><td></td><td></td><td></td><td></td><td></td><td></td><td></td><td></td><td></td></tr>
<tr><td>Chromosome</td><td>start</td><td>end</td><td>strand</td><td>MM</td><td>target_seq</td><td>PAM</td><td>alignment</td><td>distance</td><td>position</td><td>gene name</td><td>gene id</td></tr>
<tr><td>GU305914.1</td><td>167689</td><td>167711</td><td>+</td><td>0</td><td>TGTGACACTCAGTTGTGTCT</td><td>GGG</td><td>||||||||[||||||||||||]PAM</td><td>NA</td><td>Intergenic</td><td>NA</td><td>NA</td></tr>
<tr><td>T12642</td><td>GTGACACTCAGTTGTGTCTGGGG</td><td>1000</td><td></td><td></td><td></td><td></td><td></td><td></td><td></td><td></td><td></td></tr>
<tr><td>Oligo fwd</td><td>CACCGTGACACTCAGTTGTGTCTG</td><td></td><td></td><td></td><td></td><td></td><td></td><td></td><td></td><td></td><td></td></tr>
</table>

<table>
<tr><td>Oligo rev</td><td>AAACCAGACACAACTGAGTGTCAC</td><td></td><td></td><td></td><td></td><td></td><td></td><td></td><td></td><td></td><td></td></tr>
<tr><td>Chromosome</td><td>start</td><td>end</td><td>strand</td><td>MM</td><td>target_seq</td><td>PAM</td><td>alignment</td><td>distance</td><td>position</td><td>gene name</td><td>gene id</td></tr>
<tr><td>GU305914.1</td><td>167690</td><td>167712</td><td>+</td><td>0</td><td>GTGACACTCAGTTGTGTCTG</td><td>GGG</td><td>||||||||[||||||||||||]PAM</td><td>NA</td><td>Intergenic</td><td>NA</td><td>NA</td></tr>
<tr><td>T12643</td><td>AGTTGTGTCTGGGGCTTCGACGG</td><td>1000</td><td></td><td></td><td></td><td></td><td></td><td></td><td></td><td></td><td></td></tr>
<tr><td>Oligo fwd</td><td>CACCAGTTGTGTCTGGGGCTTCGA</td><td></td><td></td><td></td><td></td><td></td><td></td><td></td><td></td><td></td><td></td></tr>
<tr><td>Oligo rev</td><td>AAACTCGAAGCCCCAGACACAACT</td><td></td><td></td><td></td><td></td><td></td><td></td><td></td><td></td><td></td><td></td></tr>
<tr><td>Chromosome</td><td>start</td><td>end</td><td>strand</td><td>MM</td><td>target_seq</td><td>PAM</td><td>alignment</td><td>distance</td><td>position</td><td>gene name</td><td>gene id</td></tr>
<tr><td>GU305914.1</td><td>167699</td><td>167721</td><td>+</td><td>0</td><td>AGTTGTGTCTGGGGCTTCGA</td><td>CGG</td><td>||||||||[||||||||||||]PAM</td><td>NA</td><td>Intergenic</td><td>NA</td><td>NA</td></tr>
<tr><td>T12644</td><td>GTTGTGTCTGGGGCTTCGACGGG</td><td>1000</td><td></td><td></td><td></td><td></td><td></td><td></td><td></td><td></td><td></td></tr>
<tr><td>Oligo fwd</td><td>CACCGTTGTGTCTGGGGCTTCGAC</td><td></td><td></td><td></td><td></td><td></td><td></td><td></td><td></td><td></td><td></td></tr>
<tr><td>Oligo rev</td><td>AAACGTCGAAGCCCCAGACACAAC</td><td></td><td></td><td></td><td></td><td></td><td></td><td></td><td></td><td></td><td></td></tr>
<tr><td>Chromosome</td><td>start</td><td>end</td><td>strand</td><td>MM</td><td>target_seq</td><td>PAM</td><td>alignment</td><td>distance</td><td>position</td><td>gene name</td><td>gene id</td></tr>
<tr><td>GU305914.1</td><td>167700</td><td>167722</td><td>+</td><td>0</td><td>GTTGTGTCTGGGGCTTCGAC</td><td>GGG</td><td>||||||||[||||||||||||]PAM</td><td>NA</td><td>Intergenic</td><td>NA</td><td>NA</td></tr>
<tr><td>T12645</td><td>TGTCTGGGGCTTCGACGGGAAGG</td><td>1000</td><td></td><td></td><td></td><td></td><td></td><td></td><td></td><td></td><td></td></tr>
<tr><td>Oligo fwd</td><td>CACCTGTCTGGGGCTTCGACGGGA</td><td></td><td></td><td></td><td></td><td></td><td></td><td></td><td></td><td></td><td></td></tr>
<tr><td>Oligo rev</td><td>AAACTCCCGTCGAAGCCCCAGACA</td><td></td><td></td><td></td><td></td><td></td><td></td><td></td><td></td><td></td><td></td></tr>
<tr><td>Chromosome</td><td>start</td><td>end</td><td>strand</td><td>MM</td><td>target_seq</td><td>PAM</td><td>alignment</td><td>distance</td><td>position</td><td>gene name</td><td>gene id</td></tr>
<tr><td>GU305914.1</td><td>167704</td><td>167726</td><td>+</td><td>0</td><td>TGTCTGGGGCTTCGACGGGA</td><td>AGG</td><td>||||||||[||||||||||||]PAM</td><td>NA</td><td>Intergenic</td><td>NA</td><td>NA</td></tr>
<tr><td>T12646</td><td>CTGGGGCTTCGACGGGAAGGAGG</td><td>1000</td><td></td><td></td><td></td><td></td><td></td><td></td><td></td><td></td><td></td></tr>
<tr><td>Oligo fwd</td><td>CACCCTGGGGCTTCGACGGGAAGG</td><td></td><td></td><td></td><td></td><td></td><td></td><td></td><td></td><td></td><td></td></tr>
<tr><td>Oligo rev</td><td>AAACCCTTCCCGTCGAAGCCCCAG</td><td></td><td></td><td></td><td></td><td></td><td></td><td></td><td></td><td></td><td></td></tr>
<tr><td>Chromosome</td><td>start</td><td>end</td><td>strand</td><td>MM</td><td>target_seq</td><td>PAM</td><td>alignment</td><td>distance</td><td>position</td><td>gene name</td><td>gene id</td></tr>
<tr><td>GU305914.1</td><td>167707</td><td>167729</td><td>+</td><td>0</td><td>CTGGGGCTTCGACGGGAAGG</td><td>AGG</td><td>||||||||[||||||||||||]PAM</td><td>NA</td><td>Intergenic</td><td>NA</td><td>NA</td></tr>
<tr><td>T12647</td><td>CTTCGACGGGAAGGAGGAGAAGG</td><td>1000</td><td></td><td></td><td></td><td></td><td></td><td></td><td></td><td></td><td></td></tr>
<tr><td>Oligo fwd</td><td>CACCCTTCGACGGGAAGGAGGAGA</td><td></td><td></td><td></td><td></td><td></td><td></td><td></td><td></td><td></td><td></td></tr>
<tr><td>Oligo rev</td><td>AAACTCTCCTCCTTCCCGTCGAAG</td><td></td><td></td><td></td><td></td><td></td><td></td><td></td><td></td><td></td><td></td></tr>
<tr><td>Chromosome</td><td>start</td><td>end</td><td>strand</td><td>MM</td><td>target_seq</td><td>PAM</td><td>alignment</td><td>distance</td><td>position</td><td>gene name</td><td>gene id</td></tr>
<tr><td>GU305914.1</td><td>167713</td><td>167735</td><td>+</td><td>0</td><td>CTTCGACGGGAAGGAGGAGA</td><td>AGG</td><td>||||||||[||||||||||||]PAM</td><td>NA</td><td>Intergenic</td><td>NA</td><td>NA</td></tr>
<tr><td>T12648</td><td>CGGGAAGGAGGAGAAGGATATGG</td><td>1000</td><td></td><td></td><td></td><td></td><td></td><td></td><td></td><td></td><td></td></tr>
<tr><td>Oligo fwd</td><td>CACCCGGGAAGGAGGAGAAGGATA</td><td></td><td></td><td></td><td></td><td></td><td></td><td></td><td></td><td></td><td></td></tr>
<tr><td>Oligo rev</td><td>AAACTATCCTTCTCCTCCTTCCCG</td><td></td><td></td><td></td><td></td><td></td><td></td><td></td><td></td><td></td><td></td></tr>
<tr><td>Chromosome</td><td>start</td><td>end</td><td>strand</td><td>MM</td><td>target_seq</td><td>PAM</td><td>alignment</td><td>distance</td><td>position</td><td>gene name</td><td>gene id</td></tr>
<tr><td>GU305914.1</td><td>167719</td><td>167741</td><td>+</td><td>0</td><td>CGGGAAGGAGGAGAAGGATA</td><td>TGG</td><td>||||||||[||||||||||||]PAM</td><td>NA</td><td>Intergenic</td><td>NA</td><td>NA</td></tr>
<tr><td>T12649</td><td>TGGTACTACTGCTGCTGCTGAGG</td><td>1000</td><td></td><td></td><td></td><td></td><td></td><td></td><td></td><td></td><td></td></tr>
<tr><td>Oligo fwd</td><td>CACCTGGTACTACTGCTGCTGCTG</td><td></td><td></td><td></td><td></td><td></td><td></td><td></td><td></td><td></td><td></td></tr>
<tr><td>Oligo rev</td><td>AAACCAGCAGCAGCAGTAGTACCA</td><td></td><td></td><td></td><td></td><td></td><td></td><td></td><td></td><td></td><td></td></tr>
<tr><td>Chromosome</td><td>start</td><td>end</td><td>strand</td><td>MM</td><td>target_seq</td><td>PAM</td><td>alignment</td><td>distance</td><td>position</td><td>gene name</td><td>gene id</td></tr>
<tr><td>GU305914.1</td><td>167739</td><td>167761</td><td>+</td><td>0</td><td>TGGTACTACTGCTGCTGCTG</td><td>AGG</td><td>||||||||[||||||||||||]PAM</td><td>NA</td><td>Intergenic</td><td>NA</td><td>NA</td></tr>
<tr><td>T12650</td><td>CTACTGCTGCTGCTGAGGAGAGG</td><td>1000</td><td></td><td></td><td></td><td></td><td></td><td></td><td></td><td></td><td></td></tr>
<tr><td>Oligo fwd</td><td>CACCCTACTGCTGCTGCTGAGGAG</td><td></td><td></td><td></td><td></td><td></td><td></td><td></td><td></td><td></td><td></td></tr>
<tr><td>Oligo rev</td><td>AAACCTCCTCAGCAGCAGCAGTAG</td><td></td><td></td><td></td><td></td><td></td><td></td><td></td><td></td><td></td><td></td></tr>
<tr><td>Chromosome</td><td>start</td><td>end</td><td>strand</td><td>MM</td><td>target_seq</td><td>PAM</td><td>alignment</td><td>distance</td><td>position</td><td>gene name</td><td>gene id</td></tr>
<tr><td>GU305914.1</td><td>167744</td><td>167766</td><td>+</td><td>0</td><td>CTACTGCTGCTGCTGAGGAG</td><td>AGG</td><td>||||||||[||||||||||||]PAM</td><td>NA</td><td>Intergenic</td><td>NA</td><td>NA</td></tr>
<tr><td>T12651</td><td>GCTGCTGAGGAGAGGAGTGACGG</td><td>1000</td><td></td><td></td><td></td><td></td><td></td><td></td><td></td><td></td><td></td></tr>
<tr><td>Oligo fwd</td><td>CACCGCTGCTGAGGAGAGGAGTGA</td><td></td><td></td><td></td><td></td><td></td><td></td><td></td><td></td><td></td><td></td></tr>
<tr><td>Oligo rev</td><td>AAACTCACTCCTCTCCTCAGCAGC</td><td></td><td></td><td></td><td></td><td></td><td></td><td></td><td></td><td></td><td></td></tr>
<tr><td>Chromosome</td><td>start</td><td>end</td><td>strand</td><td>MM</td><td>target_seq</td><td>PAM</td><td>alignment</td><td>distance</td><td>position</td><td>gene name</td><td>gene id</td></tr>
<tr><td>GU305914.1</td><td>167752</td><td>167774</td><td>+</td><td>0</td><td>GCTGCTGAGGAGAGGAGTGA</td><td>CGG</td><td>||||||||[||||||||||||]PAM</td><td>NA</td><td>Intergenic</td><td>NA</td><td>NA</td></tr>
<tr><td>T12652</td><td>ACGGTCAGAGATGCTCTCGAAGG</td><td>1000</td><td></td><td></td><td></td><td></td><td></td><td></td><td></td><td></td><td></td></tr>
<tr><td>Oligo fwd</td><td>CACCACGGTCAGAGATGCTCTCGA</td><td></td><td></td><td></td><td></td><td></td><td></td><td></td><td></td><td></td><td></td></tr>
<tr><td>Oligo rev</td><td>AAACTCGAGAGCATCTCTGACCGT</td><td></td><td></td><td></td><td></td><td></td><td></td><td></td><td></td><td></td><td></td></tr>
<tr><td>Chromosome</td><td>start</td><td>end</td><td>strand</td><td>MM</td><td>target_seq</td><td>PAM</td><td>alignment</td><td>distance</td><td>position</td><td>gene name</td><td>gene id</td></tr>
<tr><td>GU305914.1</td><td>167771</td><td>167793</td><td>+</td><td>0</td><td>ACGGTCAGAGATGCTCTCGA</td><td>AGG</td><td>||||||||[||||||||||||]PAM</td><td>NA</td><td>Intergenic</td><td>NA</td><td>NA</td></tr>
<tr><td>T12653</td><td>CAGAGATGCTCTCGAAGGTACGG</td><td>1000</td><td></td><td></td><td></td><td></td><td></td><td></td><td></td><td></td><td></td></tr>
<tr><td>Oligo fwd</td><td>CACCCAGAGATGCTCTCGAAGGTA</td><td></td><td></td><td></td><td></td><td></td><td></td><td></td><td></td><td></td><td></td></tr>
<tr><td>Oligo rev</td><td>AAACTACCTTCGAGAGCATCTCTG</td><td></td><td></td><td></td><td></td><td></td><td></td><td></td><td></td><td></td><td></td></tr>
<tr><td>Chromosome</td><td>start</td><td>end</td><td>strand</td><td>MM</td><td>target_seq</td><td>PAM</td><td>alignment</td><td>distance</td><td>position</td><td>gene name</td><td>gene id</td></tr>
<tr><td>GU305914.1</td><td>167776</td><td>167798</td><td>+</td><td>0</td><td>CAGAGATGCTCTCGAAGGTA</td><td>CGG</td><td>||||||||[||||||||||||]PAM</td><td>NA</td><td>Intergenic</td><td>NA</td><td>NA</td></tr>
<tr><td>T12654</td><td>TGCTCTCGAAGGTACGGCTGTGG</td><td>1000</td><td></td><td></td><td></td><td></td><td></td><td></td><td></td><td></td><td></td></tr>
<tr><td>Oligo fwd</td><td>CACCTGCTCTCGAAGGTACGGCTG</td><td></td><td></td><td></td><td></td><td></td><td></td><td></td><td></td><td></td><td></td></tr>
<tr><td>Oligo rev</td><td>AAACCAGCCGTACCTTCGAGAGCA</td><td></td><td></td><td></td><td></td><td></td><td></td><td></td><td></td><td></td><td></td></tr>
<tr><td>Chromosome</td><td>start</td><td>end</td><td>strand</td><td>MM</td><td>target_seq</td><td>PAM</td><td>alignment</td><td>distance</td><td>position</td><td>gene name</td><td>gene id</td></tr>
<tr><td>GU305914.1</td><td>167782</td><td>167804</td><td>+</td><td>0</td><td>TGCTCTCGAAGGTACGGCTG</td><td>TGG</td><td>||||||||[||||||||||||]PAM</td><td>NA</td><td>Intergenic</td><td>NA</td><td>NA</td></tr>
<tr><td>T12655</td><td>CGAAGGTACGGCTGTGGTAGCGG</td><td>1000</td><td></td><td></td><td></td><td></td><td></td><td></td><td></td><td></td><td></td></tr>
<tr><td>Oligo fwd</td><td>CACCCGAAGGTACGGCTGTGGTAG</td><td></td><td></td><td></td><td></td><td></td><td></td><td></td><td></td><td></td><td></td></tr>
<tr><td>Oligo rev</td><td>AAACCTACCACAGCCGTACCTTCG</td><td></td><td></td><td></td><td></td><td></td><td></td><td></td><td></td><td></td><td></td></tr>
<tr><td>Chromosome</td><td>start</td><td>end</td><td>strand</td><td>MM</td><td>target_seq</td><td>PAM</td><td>alignment</td><td>distance</td><td>position</td><td>gene name</td><td>gene id</td></tr>
<tr><td>GU305914.1</td><td>167788</td><td>167810</td><td>+</td><td>0</td><td>CGAAGGTACGGCTGTGGTAG</td><td>CGG</td><td>||||||||[||||||||||||]PAM</td><td>NA</td><td>Intergenic</td><td>NA</td><td>NA</td></tr>
<tr><td>T12656</td><td>GAAGGTACGGCTGTGGTAGCGGG</td><td>1000</td><td></td><td></td><td></td><td></td><td></td><td></td><td></td><td></td><td></td></tr>
<tr><td>Oligo fwd</td><td>CACCGAAGGTACGGCTGTGGTAGC</td><td></td><td></td><td></td><td></td><td></td><td></td><td></td><td></td><td></td><td></td></tr>
<tr><td>Oligo rev</td><td>AAACGCTACCACAGCCGTACCTTC</td><td></td><td></td><td></td><td></td><td></td><td></td><td></td><td></td><td></td><td></td></tr>
<tr><td>Chromosome</td><td>start</td><td>end</td><td>strand</td><td>MM</td><td>target_seq</td><td>PAM</td><td>alignment</td><td>distance</td><td>position</td><td>gene name</td><td>gene id</td></tr>
<tr><td>GU305914.1</td><td>167789</td><td>167811</td><td>+</td><td>0</td><td>GAAGGTACGGCTGTGGTAGC</td><td>GGG</td><td>||||||||[||||||||||||]PAM</td><td>NA</td><td>Intergenic</td><td>NA</td><td>NA</td></tr>
<tr><td>T12657</td><td>AAGGTACGGCTGTGGTAGCGGGG</td><td>1000</td><td></td><td></td><td></td><td></td><td></td><td></td><td></td><td></td><td></td></tr>
</table>

<table>
<tr><td>Oligo fwd</td><td>CACCAAGGTACGGCTGTGGTAGCG</td><td></td><td></td><td></td><td></td><td></td><td></td><td></td><td></td><td></td><td></td></tr>
<tr><td>Oligo rev</td><td>AAACCGCTACCACAGCCGTACCTT</td><td></td><td></td><td></td><td></td><td></td><td></td><td></td><td></td><td></td><td></td></tr>
<tr><td>Chromosome</td><td>start</td><td>end</td><td>strand</td><td>MM</td><td>target_seq</td><td>PAM</td><td>alignment</td><td>distance</td><td>position</td><td>gene name</td><td>gene id</td></tr>
<tr><td>GU305914.1</td><td>167790</td><td>167812</td><td>+</td><td>0</td><td>AAGGTACGGCTGTGGTAGCG</td><td>GGG</td><td>||||||||[||||||||||||]PAM</td><td>NA</td><td>Intergenic</td><td>NA</td><td>NA</td></tr>
<tr><td></td><td></td><td></td><td></td><td></td><td></td><td></td><td></td><td></td><td></td><td></td><td></td></tr>
<tr><td>T12658</td><td>GGCTGTGGTAGCGGGGTCCTTGG</td><td>1000</td><td></td><td></td><td></td><td></td><td></td><td></td><td></td><td></td><td></td></tr>
<tr><td>Oligo fwd</td><td>CACCGGCTGTGGTAGCGGGGTCCT</td><td></td><td></td><td></td><td></td><td></td><td></td><td></td><td></td><td></td><td></td></tr>
<tr><td>Oligo rev</td><td>AAACAGGACCCCGCTACCACAGCC</td><td></td><td></td><td></td><td></td><td></td><td></td><td></td><td></td><td></td><td></td></tr>
<tr><td>Chromosome</td><td>start</td><td>end</td><td>strand</td><td>MM</td><td>target_seq</td><td>PAM</td><td>alignment</td><td>distance</td><td>position</td><td>gene name</td><td>gene id</td></tr>
<tr><td>GU305914.1</td><td>167797</td><td>167819</td><td>+</td><td>0</td><td>GGCTGTGGTAGCGGGGTCCT</td><td>TGG</td><td>||||||||[||||||||||||]PAM</td><td>NA</td><td>Intergenic</td><td>NA</td><td>NA</td></tr>
<tr><td></td><td></td><td></td><td></td><td></td><td></td><td></td><td></td><td></td><td></td><td></td><td></td></tr>
<tr><td>T12659</td><td>TGTGGTAGCGGGGTCCTTGGTGG</td><td>1000</td><td></td><td></td><td></td><td></td><td></td><td></td><td></td><td></td><td></td></tr>
<tr><td>Oligo fwd</td><td>CACCTGTGGTAGCGGGGTCCTTGG</td><td></td><td></td><td></td><td></td><td></td><td></td><td></td><td></td><td></td><td></td></tr>
<tr><td>Oligo rev</td><td>AAACCCAAGGACCCCGCTACCACA</td><td></td><td></td><td></td><td></td><td></td><td></td><td></td><td></td><td></td><td></td></tr>
<tr><td>Chromosome</td><td>start</td><td>end</td><td>strand</td><td>MM</td><td>target_seq</td><td>PAM</td><td>alignment</td><td>distance</td><td>position</td><td>gene name</td><td>gene id</td></tr>
<tr><td>GU305914.1</td><td>167800</td><td>167822</td><td>+</td><td>0</td><td>TGTGGTAGCGGGGTCCTTGG</td><td>TGG</td><td>||||||||[||||||||||||]PAM</td><td>NA</td><td>Intergenic</td><td>NA</td><td>NA</td></tr>
<tr><td></td><td></td><td></td><td></td><td></td><td></td><td></td><td></td><td></td><td></td><td></td><td></td></tr>
<tr><td>T12660</td><td>AGCGGGGTCCTTGGTGGTGATGG</td><td>1000</td><td></td><td></td><td></td><td></td><td></td><td></td><td></td><td></td><td></td></tr>
<tr><td>Oligo fwd</td><td>CACCAGCGGGGTCCTTGGTGGTGA</td><td></td><td></td><td></td><td></td><td></td><td></td><td></td><td></td><td></td><td></td></tr>
<tr><td>Oligo rev</td><td>AAACTCACCACCAAGGACCCCGCT</td><td></td><td></td><td></td><td></td><td></td><td></td><td></td><td></td><td></td><td></td></tr>
<tr><td>Chromosome</td><td>start</td><td>end</td><td>strand</td><td>MM</td><td>target_seq</td><td>PAM</td><td>alignment</td><td>distance</td><td>position</td><td>gene name</td><td>gene id</td></tr>
<tr><td>GU305914.1</td><td>167806</td><td>167828</td><td>+</td><td>0</td><td>AGCGGGGTCCTTGGTGGTGA</td><td>TGG</td><td>||||||||[||||||||||||]PAM</td><td>NA</td><td>Intergenic</td><td>NA</td><td>NA</td></tr>
<tr><td></td><td></td><td></td><td></td><td></td><td></td><td></td><td></td><td></td><td></td><td></td><td></td></tr>
<tr><td>T12661</td><td>GGGGTCCTTGGTGGTGATGGCGG</td><td>1000</td><td></td><td></td><td></td><td></td><td></td><td></td><td></td><td></td><td></td></tr>
<tr><td>Oligo fwd</td><td>CACCGGGGTCCTTGGTGGTGATGG</td><td></td><td></td><td></td><td></td><td></td><td></td><td></td><td></td><td></td><td></td></tr>
<tr><td>Oligo rev</td><td>AAACCCATCACCACCAAGGACCCC</td><td></td><td></td><td></td><td></td><td></td><td></td><td></td><td></td><td></td><td></td></tr>
<tr><td>Chromosome</td><td>start</td><td>end</td><td>strand</td><td>MM</td><td>target_seq</td><td>PAM</td><td>alignment</td><td>distance</td><td>position</td><td>gene name</td><td>gene id</td></tr>
<tr><td>GU305914.1</td><td>167809</td><td>167831</td><td>+</td><td>0</td><td>GGGGTCCTTGGTGGTGATGG</td><td>CGG</td><td>||||||||[||||||||||||]PAM</td><td>NA</td><td>Intergenic</td><td>NA</td><td>NA</td></tr>
<tr><td>GU305914.1</td><td>44657</td><td>44679</td><td>-</td><td>4</td><td>GGGGATGATGGTGGTGATGG</td><td>TGG</td><td>||||----[||||||||||||]PAM</td><td>NA</td><td>Intergenic</td><td>NA</td><td>NA</td></tr>
<tr><td></td><td></td><td></td><td></td><td></td><td></td><td></td><td></td><td></td><td></td><td></td><td></td></tr>
<tr><td>T12662</td><td>GATGGCGGTGATGATTGTGTTGG</td><td>1000</td><td></td><td></td><td></td><td></td><td></td><td></td><td></td><td></td><td></td></tr>
<tr><td>Oligo fwd</td><td>CACCGATGGCGGTGATGATTGTGT</td><td></td><td></td><td></td><td></td><td></td><td></td><td></td><td></td><td></td><td></td></tr>
<tr><td>Oligo rev</td><td>AAACACACAATCATCACCGCCATC</td><td></td><td></td><td></td><td></td><td></td><td></td><td></td><td></td><td></td><td></td></tr>
<tr><td>Chromosome</td><td>start</td><td>end</td><td>strand</td><td>MM</td><td>target_seq</td><td>PAM</td><td>alignment</td><td>distance</td><td>position</td><td>gene name</td><td>gene id</td></tr>
<tr><td>GU305914.1</td><td>167824</td><td>167846</td><td>+</td><td>0</td><td>GATGGCGGTGATGATTGTGT</td><td>TGG</td><td>||||||||[||||||||||||]PAM</td><td>NA</td><td>Intergenic</td><td>NA</td><td>NA</td></tr>
<tr><td></td><td></td><td></td><td></td><td></td><td></td><td></td><td></td><td></td><td></td><td></td><td></td></tr>
<tr><td>T12663</td><td>GTTGTTGTCGACGTCTGATGTGG</td><td>1000</td><td></td><td></td><td></td><td></td><td></td><td></td><td></td><td></td><td></td></tr>
<tr><td>Oligo fwd</td><td>CACCGTTGTTGTCGACGTCTGATG</td><td></td><td></td><td></td><td></td><td></td><td></td><td></td><td></td><td></td><td></td></tr>
<tr><td>Oligo rev</td><td>AAACCATCAGACGTCGACAACAAC</td><td></td><td></td><td></td><td></td><td></td><td></td><td></td><td></td><td></td><td></td></tr>
<tr><td>Chromosome</td><td>start</td><td>end</td><td>strand</td><td>MM</td><td>target_seq</td><td>PAM</td><td>alignment</td><td>distance</td><td>position</td><td>gene name</td><td>gene id</td></tr>
<tr><td>GU305914.1</td><td>167848</td><td>167870</td><td>+</td><td>0</td><td>GTTGTTGTCGACGTCTGATG</td><td>TGG</td><td>||||||||[||||||||||||]PAM</td><td>NA</td><td>Intergenic</td><td>NA</td><td>NA</td></tr>
<tr><td></td><td></td><td></td><td></td><td></td><td></td><td></td><td></td><td></td><td></td><td></td><td></td></tr>
<tr><td>T12664</td><td>TTGTTGTCGACGTCTGATGTGGG</td><td>1000</td><td></td><td></td><td></td><td></td><td></td><td></td><td></td><td></td><td></td></tr>
<tr><td>Oligo fwd</td><td>CACCTTGTTGTCGACGTCTGATGT</td><td></td><td></td><td></td><td></td><td></td><td></td><td></td><td></td><td></td><td></td></tr>
<tr><td>Oligo rev</td><td>AAACACATCAGACGTCGACAACAA</td><td></td><td></td><td></td><td></td><td></td><td></td><td></td><td></td><td></td><td></td></tr>
<tr><td>Chromosome</td><td>start</td><td>end</td><td>strand</td><td>MM</td><td>target_seq</td><td>PAM</td><td>alignment</td><td>distance</td><td>position</td><td>gene name</td><td>gene id</td></tr>
<tr><td>GU305914.1</td><td>167849</td><td>167871</td><td>+</td><td>0</td><td>TTGTTGTCGACGTCTGATGT</td><td>GGG</td><td>||||||||[||||||||||||]PAM</td><td>NA</td><td>Intergenic</td><td>NA</td><td>NA</td></tr>
<tr><td></td><td></td><td></td><td></td><td></td><td></td><td></td><td></td><td></td><td></td><td></td><td></td></tr>
<tr><td>T12665</td><td>TCGACGTCTGATGTGGGTGCTGG</td><td>1000</td><td></td><td></td><td></td><td></td><td></td><td></td><td></td><td></td><td></td></tr>
<tr><td>Oligo fwd</td><td>CACCTCGACGTCTGATGTGGGTGC</td><td></td><td></td><td></td><td></td><td></td><td></td><td></td><td></td><td></td><td></td></tr>
<tr><td>Oligo rev</td><td>AAACGCACCCACATCAGACGTCGA</td><td></td><td></td><td></td><td></td><td></td><td></td><td></td><td></td><td></td><td></td></tr>
<tr><td>Chromosome</td><td>start</td><td>end</td><td>strand</td><td>MM</td><td>target_seq</td><td>PAM</td><td>alignment</td><td>distance</td><td>position</td><td>gene name</td><td>gene id</td></tr>
<tr><td>GU305914.1</td><td>167855</td><td>167877</td><td>+</td><td>0</td><td>TCGACGTCTGATGTGGGTGC</td><td>TGG</td><td>||||||||[||||||||||||]PAM</td><td>NA</td><td>Intergenic</td><td>NA</td><td>NA</td></tr>
<tr><td></td><td></td><td></td><td></td><td></td><td></td><td></td><td></td><td></td><td></td><td></td><td></td></tr>
<tr><td>T12666</td><td>ACGTCTGATGTGGGTGCTGGCGG</td><td>1000</td><td></td><td></td><td></td><td></td><td></td><td></td><td></td><td></td><td></td></tr>
<tr><td>Oligo fwd</td><td>CACCACGTCTGATGTGGGTGCTGG</td><td></td><td></td><td></td><td></td><td></td><td></td><td></td><td></td><td></td><td></td></tr>
<tr><td>Oligo rev</td><td>AAACCCAGCACCCACATCAGACGT</td><td></td><td></td><td></td><td></td><td></td><td></td><td></td><td></td><td></td><td></td></tr>
<tr><td>Chromosome</td><td>start</td><td>end</td><td>strand</td><td>MM</td><td>target_seq</td><td>PAM</td><td>alignment</td><td>distance</td><td>position</td><td>gene name</td><td>gene id</td></tr>
<tr><td>GU305914.1</td><td>167858</td><td>167880</td><td>+</td><td>0</td><td>ACGTCTGATGTGGGTGCTGG</td><td>CGG</td><td>||||||||[||||||||||||]PAM</td><td>NA</td><td>Intergenic</td><td>NA</td><td>NA</td></tr>
<tr><td></td><td></td><td></td><td></td><td></td><td></td><td></td><td></td><td></td><td></td><td></td><td></td></tr>
<tr><td>T12667</td><td>CGTCTGATGTGGGTGCTGGCGGG</td><td>1000</td><td></td><td></td><td></td><td></td><td></td><td></td><td></td><td></td><td></td></tr>
<tr><td>Oligo fwd</td><td>CACCCGTCTGATGTGGGTGCTGGC</td><td></td><td></td><td></td><td></td><td></td><td></td><td></td><td></td><td></td><td></td></tr>
<tr><td>Oligo rev</td><td>AAACGCCAGCACCCACATCAGACG</td><td></td><td></td><td></td><td></td><td></td><td></td><td></td><td></td><td></td><td></td></tr>
<tr><td>Chromosome</td><td>start</td><td>end</td><td>strand</td><td>MM</td><td>target_seq</td><td>PAM</td><td>alignment</td><td>distance</td><td>position</td><td>gene name</td><td>gene id</td></tr>
<tr><td>GU305914.1</td><td>167859</td><td>167881</td><td>+</td><td>0</td><td>CGTCTGATGTGGGTGCTGGC</td><td>GGG</td><td>||||||||[||||||||||||]PAM</td><td>NA</td><td>Intergenic</td><td>NA</td><td>NA</td></tr>
<tr><td></td><td></td><td></td><td></td><td></td><td></td><td></td><td></td><td></td><td></td><td></td><td></td></tr>
<tr><td>T12668</td><td>GTCTGATGTGGGTGCTGGCGGGG</td><td>1000</td><td></td><td></td><td></td><td></td><td></td><td></td><td></td><td></td><td></td></tr>
<tr><td>Oligo fwd</td><td>CACCGTCTGATGTGGGTGCTGGCG</td><td></td><td></td><td></td><td></td><td></td><td></td><td></td><td></td><td></td><td></td></tr>
<tr><td>Oligo rev</td><td>AAACCGCCAGCACCCACATCAGAC</td><td></td><td></td><td></td><td></td><td></td><td></td><td></td><td></td><td></td><td></td></tr>
<tr><td>Chromosome</td><td>start</td><td>end</td><td>strand</td><td>MM</td><td>target_seq</td><td>PAM</td><td>alignment</td><td>distance</td><td>position</td><td>gene name</td><td>gene id</td></tr>
<tr><td>GU305914.1</td><td>167860</td><td>167882</td><td>+</td><td>0</td><td>GTCTGATGTGGGTGCTGGCG</td><td>GGG</td><td>||||||||[||||||||||||]PAM</td><td>NA</td><td>Intergenic</td><td>NA</td><td>NA</td></tr>
<tr><td></td><td></td><td></td><td></td><td></td><td></td><td></td><td></td><td></td><td></td><td></td><td></td></tr>
<tr><td>T12669</td><td>TGGCGGGGCTGTCGTTAGTGTGG</td><td>1000</td><td></td><td></td><td></td><td></td><td></td><td></td><td></td><td></td><td></td></tr>
<tr><td>Oligo fwd</td><td>CACCTGGCGGGGCTGTCGTTAGTG</td><td></td><td></td><td></td><td></td><td></td><td></td><td></td><td></td><td></td><td></td></tr>
<tr><td>Oligo rev</td><td>AAACCACTAACGACAGCCCCGCCA</td><td></td><td></td><td></td><td></td><td></td><td></td><td></td><td></td><td></td><td></td></tr>
<tr><td>Chromosome</td><td>start</td><td>end</td><td>strand</td><td>MM</td><td>target_seq</td><td>PAM</td><td>alignment</td><td>distance</td><td>position</td><td>gene name</td><td>gene id</td></tr>
<tr><td>GU305914.1</td><td>167875</td><td>167897</td><td>+</td><td>0</td><td>TGGCGGGGCTGTCGTTAGTG</td><td>TGG</td><td>||||||||[||||||||||||]PAM</td><td>NA</td><td>Intergenic</td><td>NA</td><td>NA</td></tr>
<tr><td></td><td></td><td></td><td></td><td></td><td></td><td></td><td></td><td></td><td></td><td></td><td></td></tr>
<tr><td>T12670</td><td>GGGCTGTCGTTAGTGTGGTCAGG</td><td>1000</td><td></td><td></td><td></td><td></td><td></td><td></td><td></td><td></td><td></td></tr>
<tr><td>Oligo fwd</td><td>CACCGGGCTGTCGTTAGTGTGGTC</td><td></td><td></td><td></td><td></td><td></td><td></td><td></td><td></td><td></td><td></td></tr>
<tr><td>Oligo rev</td><td>AAACGACCACACTAACGACAGCCC</td><td></td><td></td><td></td><td></td><td></td><td></td><td></td><td></td><td></td><td></td></tr>
<tr><td>Chromosome</td><td>start</td><td>end</td><td>strand</td><td>MM</td><td>target_seq</td><td>PAM</td><td>alignment</td><td>distance</td><td>position</td><td>gene name</td><td>gene id</td></tr>
<tr><td>GU305914.1</td><td>167880</td><td>167902</td><td>+</td><td>0</td><td>GGGCTGTCGTTAGTGTGGTC</td><td>AGG</td><td>||||||||[||||||||||||]PAM</td><td>NA</td><td>Intergenic</td><td>NA</td><td>NA</td></tr>
<tr><td></td><td></td><td></td><td></td><td></td><td></td><td></td><td></td><td></td><td></td><td></td><td></td></tr>
<tr><td>T12671</td><td>GTCGTTAGTGTGGTCAGGTACGG</td><td>1000</td><td></td><td></td><td></td><td></td><td></td><td></td><td></td><td></td><td></td></tr>
<tr><td>Oligo fwd</td><td>CACCGTCGTTAGTGTGGTCAGGTA</td><td></td><td></td><td></td><td></td><td></td><td></td><td></td><td></td><td></td><td></td></tr>
<tr><td>Oligo rev</td><td>AAACTACCTGACCACACTAACGAC</td><td></td><td></td><td></td><td></td><td></td><td></td><td></td><td></td><td></td><td></td></tr>
<tr><td>Chromosome</td><td>start</td><td>end</td><td>strand</td><td>MM</td><td>target_seq</td><td>PAM</td><td>alignment</td><td>distance</td><td>position</td><td>gene name</td><td>gene id</td></tr>
<tr><td>GU305914.1</td><td>167885</td><td>167907</td><td>+</td><td>0</td><td>GTCGTTAGTGTGGTCAGGTA</td><td>CGG</td><td>||||||||[||||||||||||]PAM</td><td>NA</td><td>Intergenic</td><td>NA</td><td>NA</td></tr>
</table>

| | | | | | | | | | | | |
|---|---|---|---|---|---|---|---|---|---|---|---|
| T12672 | TCGTTAGTGTGGTCAGGTACGGG | 1000 | | | | | | | | | |
| Oligo fwd | CACCTCGTTAGTGTGGTCAGGTAC | | | | | | | | | | |
| Oligo rev | AAACGTACCTGACCACACTAACGA | | | | | | | | | | |
| Chromosome | start | end | strand | MM | target_seq | PAM | alignment | distance | position | gene name | gene id |
| GU305914.1 | 167886 | 167908 | + | 0 | TCGTTAGTGTGGTCAGGTAC | GGG | \|\|\|\|\|\|\|\|[\|\|\|\|\|\|\|\|\|\|\|\|]PAM | NA | Intergenic | NA | NA |
| T12673 | GTCAGGTACGGGATGCTCGTCGG | 1000 | | | | | | | | | |
| Oligo fwd | CACCGTCAGGTACGGGATGCTCGT | | | | | | | | | | |
| Oligo rev | AAACACGAGCATCCCGTACCTGAC | | | | | | | | | | |
| Chromosome | start | end | strand | MM | target_seq | PAM | alignment | distance | position | gene name | gene id |
| GU305914.1 | 167897 | 167919 | + | 0 | GTCAGGTACGGGATGCTCGT | CGG | \|\|\|\|\|\|\|\|[\|\|\|\|\|\|\|\|\|\|\|\|]PAM | NA | Intergenic | NA | NA |
| T12674 | AGGTACGGGATGCTCGTCGGCGG | 1000 | | | | | | | | | |
| Oligo fwd | CACCAGGTACGGGATGCTCGTCGG | | | | | | | | | | |
| Oligo rev | AAACCCGACGAGCATCCCGTACCT | | | | | | | | | | |
| Chromosome | start | end | strand | MM | target_seq | PAM | alignment | distance | position | gene name | gene id |
| GU305914.1 | 167900 | 167922 | + | 0 | AGGTACGGGATGCTCGTCGG | CGG | \|\|\|\|\|\|\|\|[\|\|\|\|\|\|\|\|\|\|\|\|]PAM | NA | Intergenic | NA | NA |
| T12675 | GGTACGGGATGCTCGTCGGCGGG | 1000 | | | | | | | | | |
| Oligo fwd | CACCGGTACGGGATGCTCGTCGGC | | | | | | | | | | |
| Oligo rev | AAACGCCGACGAGCATCCCGTACC | | | | | | | | | | |
| Chromosome | start | end | strand | MM | target_seq | PAM | alignment | distance | position | gene name | gene id |
| GU305914.1 | 167901 | 167923 | + | 0 | GGTACGGGATGCTCGTCGGC | GGG | \|\|\|\|\|\|\|\|[\|\|\|\|\|\|\|\|\|\|\|\|]PAM | NA | Intergenic | NA | NA |
| T12676 | CGGGATGCTCGTCGGCGGGACGG | 1000 | | | | | | | | | |
| Oligo fwd | CACCCGGGATGCTCGTCGGCGGGA | | | | | | | | | | |
| Oligo rev | AAACTCCCGCCGACGAGCATCCCG | | | | | | | | | | |
| Chromosome | start | end | strand | MM | target_seq | PAM | alignment | distance | position | gene name | gene id |
| GU305914.1 | 167905 | 167927 | + | 0 | CGGGATGCTCGTCGGCGGGA | CGG | \|\|\|\|\|\|\|\|[\|\|\|\|\|\|\|\|\|\|\|\|]PAM | NA | Intergenic | NA | NA |
| T12677 | GGGACGGCCGTGACCGTGCTCGG | 1000 | | | | | | | | | |
| Oligo fwd | CACCGGGACGGCCGTGACCGTGCT | | | | | | | | | | |
| Oligo rev | AAACAGCACGGTCACGGCCGTCCC | | | | | | | | | | |
| Chromosome | start | end | strand | MM | target_seq | PAM | alignment | distance | position | gene name | gene id |
| GU305914.1 | 167921 | 167943 | + | 0 | GGGACGGCCGTGACCGTGCT | CGG | \|\|\|\|\|\|\|\|[\|\|\|\|\|\|\|\|\|\|\|\|]PAM | NA | Intergenic | NA | NA |
| GU305914.1 | 161930 | 161952 | - | 4 | GCGTCGGACGTGACCGCGCT | AGG | \|-\|-\|\|\|-[\|\|\|\|\|\|\|\|-\|\|\|]PAM | NA | Intergenic | NA | NA |
| T12678 | CGAGCTCTGCTGCAGTTCTCTGG | 1000 | | | | | | | | | |
| Oligo fwd | CACCCGAGCTCTGCTGCAGTTCTC | | | | | | | | | | |
| Oligo rev | AAACGAGAACTGCAGCAGAGCTCG | | | | | | | | | | |
| Chromosome | start | end | strand | MM | target_seq | PAM | alignment | distance | position | gene name | gene id |
| GU305914.1 | 167977 | 167999 | + | 0 | CGAGCTCTGCTGCAGTTCTC | TGG | \|\|\|\|\|\|\|\|[\|\|\|\|\|\|\|\|\|\|\|\|]PAM | NA | Intergenic | NA | NA |
| T12679 | CTGTCGACCCGCTCTCTTCAAGG | 1000 | | | | | | | | | |
| Oligo fwd | CACCCTGTCGACCCGCTCTCTTCA | | | | | | | | | | |
| Oligo rev | AAACTGAAGAGAGCGGGTCGACAG | | | | | | | | | | |
| Chromosome | start | end | strand | MM | target_seq | PAM | alignment | distance | position | gene name | gene id |
| GU305914.1 | 168001 | 168023 | + | 0 | CTGTCGACCCGCTCTCTTCA | AGG | \|\|\|\|\|\|\|\|[\|\|\|\|\|\|\|\|\|\|\|\|]PAM | NA | Intergenic | NA | NA |
| T12680 | CCGCTCTCTTCAAGGTCGATAGG | 1000 | | | | | | | | | |
| Oligo fwd | CACCCCGCTCTCTTCAAGGTCGAT | | | | | | | | | | |
| Oligo rev | AAACATCGACCTTGAAGAGAGCGG | | | | | | | | | | |
| Chromosome | start | end | strand | MM | target_seq | PAM | alignment | distance | position | gene name | gene id |
| GU305914.1 | 168009 | 168031 | + | 0 | CCGCTCTCTTCAAGGTCGAT | AGG | \|\|\|\|\|\|\|\|[\|\|\|\|\|\|\|\|\|\|\|\|]PAM | NA | Intergenic | NA | NA |
| T12681 | CGCTCTCTTCAAGGTCGATAGGG | 1000 | | | | | | | | | |
| Oligo fwd | CACCCGCTCTCTTCAAGGTCGATA | | | | | | | | | | |
| Oligo rev | AAACTATCGACCTTGAAGAGAGCG | | | | | | | | | | |
| Chromosome | start | end | strand | MM | target_seq | PAM | alignment | distance | position | gene name | gene id |
| GU305914.1 | 168010 | 168032 | + | 0 | CGCTCTCTTCAAGGTCGATA | GGG | \|\|\|\|\|\|\|\|[\|\|\|\|\|\|\|\|\|\|\|\|]PAM | NA | Intergenic | NA | NA |
| T12682 | GATAGGGACGTCGATTTCTTCGG | 1000 | | | | | | | | | |
| Oligo fwd | CACCGATAGGGACGTCGATTTCTT | | | | | | | | | | |
| Oligo rev | AAACAAGAAATCGACGTCCCTATC | | | | | | | | | | |
| Chromosome | start | end | strand | MM | target_seq | PAM | alignment | distance | position | gene name | gene id |
| GU305914.1 | 168026 | 168048 | + | 0 | GATAGGGACGTCGATTTCTT | CGG | \|\|\|\|\|\|\|\|[\|\|\|\|\|\|\|\|\|\|\|\|]PAM | NA | Intergenic | NA | NA |
| T12683 | CGGCGTCTCTTTCTGTAGACCGG | 1000 | | | | | | | | | |
| Oligo fwd | CACCCGGCGTCTCTTTCTGTAGAC | | | | | | | | | | |
| Oligo rev | AAACGTCTACAGAAAGAGACGCCG | | | | | | | | | | |
| Chromosome | start | end | strand | MM | target_seq | PAM | alignment | distance | position | gene name | gene id |
| GU305914.1 | 168046 | 168068 | + | 0 | CGGCGTCTCTTTCTGTAGAC | CGG | \|\|\|\|\|\|\|\|[\|\|\|\|\|\|\|\|\|\|\|\|]PAM | NA | Intergenic | NA | NA |
| T12684 | CGGCGTGACCGCTTTCCGTCTGG | 1000 | | | | | | | | | |
| Oligo fwd | CACCCGGCGTGACCGCTTTCCGTC | | | | | | | | | | |
| Oligo rev | AAACGACGGAAAGCGGTCACGCCG | | | | | | | | | | |
| Chromosome | start | end | strand | MM | target_seq | PAM | alignment | distance | position | gene name | gene id |
| GU305914.1 | 168066 | 168088 | + | 0 | CGGCGTGACCGCTTTCCGTC | TGG | \|\|\|\|\|\|\|\|[\|\|\|\|\|\|\|\|\|\|\|\|]PAM | NA | Intergenic | NA | NA |
| T12685 | ACTCTGTGTTGACCTGTTATCGG | 1000 | | | | | | | | | |
| Oligo fwd | CACCACTCTGTGTTGACCTGTTAT | | | | | | | | | | |
| Oligo rev | AAACATAACAGGTCAACACAGAGT | | | | | | | | | | |
| Chromosome | start | end | strand | MM | target_seq | PAM | alignment | distance | position | gene name | gene id |
| GU305914.1 | 168100 | 168122 | + | 0 | ACTCTGTGTTGACCTGTTAT | CGG | \|\|\|\|\|\|\|\|[\|\|\|\|\|\|\|\|\|\|\|\|]PAM | NA | Intergenic | NA | NA |
| T12686 | GTTGACCTGTTATCGGAGAGCGG | 1000 | | | | | | | | | |
| Oligo fwd | CACCGTTGACCTGTTATCGGAGAG | | | | | | | | | | |
| Oligo rev | AAACCTCTCCGATAACAGGTCAAC | | | | | | | | | | |

<table>
<tr><td>Chromosome</td><td>start</td><td>end</td><td>strand</td><td>MM</td><td>target_seq</td><td>PAM</td><td>alignment</td><td>distance</td><td>position</td><td>gene name</td><td>gene id</td></tr>
<tr><td>GU305914.1</td><td>168107</td><td>168129</td><td>+</td><td>0</td><td>GTTGACCTGTTATCGGAGAG</td><td>CGG</td><td>||||||||[||||||||||||]PAM</td><td>NA</td><td>Intergenic</td><td>NA</td><td>NA</td></tr>
<tr><td></td><td></td><td></td><td></td><td></td><td></td><td></td><td></td><td></td><td></td><td></td><td></td></tr>
<tr><td>T12687</td><td>GGAGAGCGGAGAGTTAGTGATGG</td><td>1000</td><td></td><td></td><td></td><td></td><td></td><td></td><td></td><td></td><td></td></tr>
<tr><td>Oligo fwd</td><td>CACCGGAGAGCGGAGAGTTAGTGA</td><td></td><td></td><td></td><td></td><td></td><td></td><td></td><td></td><td></td><td></td></tr>
<tr><td>Oligo rev</td><td>AAACTCACTAACTCTCCGCTCTCC</td><td></td><td></td><td></td><td></td><td></td><td></td><td></td><td></td><td></td><td></td></tr>
<tr><td>Chromosome</td><td>start</td><td>end</td><td>strand</td><td>MM</td><td>target_seq</td><td>PAM</td><td>alignment</td><td>distance</td><td>position</td><td>gene name</td><td>gene id</td></tr>
<tr><td>GU305914.1</td><td>168121</td><td>168143</td><td>+</td><td>0</td><td>GGAGAGCGGAGAGTTAGTGA</td><td>TGG</td><td>||||||||[||||||||||||]PAM</td><td>NA</td><td>Intergenic</td><td>NA</td><td>NA</td></tr>
<tr><td></td><td></td><td></td><td></td><td></td><td></td><td></td><td></td><td></td><td></td><td></td><td></td></tr>
<tr><td>T12688</td><td>CAACGCATCATCGCCGCCAGTGG</td><td>1000</td><td></td><td></td><td></td><td></td><td></td><td></td><td></td><td></td><td></td></tr>
<tr><td>Oligo fwd</td><td>CACCCAACGCATCATCGCCGCCAG</td><td></td><td></td><td></td><td></td><td></td><td></td><td></td><td></td><td></td><td></td></tr>
<tr><td>Oligo rev</td><td>AAACCTGGCGGCGATGATGCGTTG</td><td></td><td></td><td></td><td></td><td></td><td></td><td></td><td></td><td></td><td></td></tr>
<tr><td>Chromosome</td><td>start</td><td>end</td><td>strand</td><td>MM</td><td>target_seq</td><td>PAM</td><td>alignment</td><td>distance</td><td>position</td><td>gene name</td><td>gene id</td></tr>
<tr><td>GU305914.1</td><td>168163</td><td>168185</td><td>+</td><td>0</td><td>CAACGCATCATCGCCGCCAG</td><td>TGG</td><td>||||||||[||||||||||||]PAM</td><td>NA</td><td>Intergenic</td><td>NA</td><td>NA</td></tr>
<tr><td></td><td></td><td></td><td></td><td></td><td></td><td></td><td></td><td></td><td></td><td></td><td></td></tr>
<tr><td>T12689</td><td>GTGGCGTTAGTTACCTGCTGAGG</td><td>1000</td><td></td><td></td><td></td><td></td><td></td><td></td><td></td><td></td><td></td></tr>
<tr><td>Oligo fwd</td><td>CACCGTGGCGTTAGTTACCTGCTG</td><td></td><td></td><td></td><td></td><td></td><td></td><td></td><td></td><td></td><td></td></tr>
<tr><td>Oligo rev</td><td>AAACCAGCAGGTAACTAACGCCAC</td><td></td><td></td><td></td><td></td><td></td><td></td><td></td><td></td><td></td><td></td></tr>
<tr><td>Chromosome</td><td>start</td><td>end</td><td>strand</td><td>MM</td><td>target_seq</td><td>PAM</td><td>alignment</td><td>distance</td><td>position</td><td>gene name</td><td>gene id</td></tr>
<tr><td>GU305914.1</td><td>168182</td><td>168204</td><td>+</td><td>0</td><td>GTGGCGTTAGTTACCTGCTG</td><td>AGG</td><td>||||||||[||||||||||||]PAM</td><td>NA</td><td>Intergenic</td><td>NA</td><td>NA</td></tr>
<tr><td></td><td></td><td></td><td></td><td></td><td></td><td></td><td></td><td></td><td></td><td></td><td></td></tr>
<tr><td>T12690</td><td>TACCTGCTGAGGAGTAGTCTTGG</td><td>1000</td><td></td><td></td><td></td><td></td><td></td><td></td><td></td><td></td><td></td></tr>
<tr><td>Oligo fwd</td><td>CACCTACCTGCTGAGGAGTAGTCT</td><td></td><td></td><td></td><td></td><td></td><td></td><td></td><td></td><td></td><td></td></tr>
<tr><td>Oligo rev</td><td>AAACAGACTACTCCTCAGCAGGTA</td><td></td><td></td><td></td><td></td><td></td><td></td><td></td><td></td><td></td><td></td></tr>
<tr><td>Chromosome</td><td>start</td><td>end</td><td>strand</td><td>MM</td><td>target_seq</td><td>PAM</td><td>alignment</td><td>distance</td><td>position</td><td>gene name</td><td>gene id</td></tr>
<tr><td>GU305914.1</td><td>168193</td><td>168215</td><td>+</td><td>0</td><td>TACCTGCTGAGGAGTAGTCT</td><td>TGG</td><td>||||||||[||||||||||||]PAM</td><td>NA</td><td>Intergenic</td><td>NA</td><td>NA</td></tr>
<tr><td></td><td></td><td></td><td></td><td></td><td></td><td></td><td></td><td></td><td></td><td></td><td></td></tr>
<tr><td>T12691</td><td>AGGAGTAGTCTTGGCAGCGTCGG</td><td>1000</td><td></td><td></td><td></td><td></td><td></td><td></td><td></td><td></td><td></td></tr>
<tr><td>Oligo fwd</td><td>CACCAGGAGTAGTCTTGGCAGCGT</td><td></td><td></td><td></td><td></td><td></td><td></td><td></td><td></td><td></td><td></td></tr>
<tr><td>Oligo rev</td><td>AAACACGCTGCCAAGACTACTCCT</td><td></td><td></td><td></td><td></td><td></td><td></td><td></td><td></td><td></td><td></td></tr>
<tr><td>Chromosome</td><td>start</td><td>end</td><td>strand</td><td>MM</td><td>target_seq</td><td>PAM</td><td>alignment</td><td>distance</td><td>position</td><td>gene name</td><td>gene id</td></tr>
<tr><td>GU305914.1</td><td>168202</td><td>168224</td><td>+</td><td>0</td><td>AGGAGTAGTCTTGGCAGCGT</td><td>CGG</td><td>||||||||[||||||||||||]PAM</td><td>NA</td><td>Intergenic</td><td>NA</td><td>NA</td></tr>
<tr><td></td><td></td><td></td><td></td><td></td><td></td><td></td><td></td><td></td><td></td><td></td><td></td></tr>
<tr><td>T12692</td><td>GCAGCGTCGGCGAGCTTCGTCGG</td><td>1000</td><td></td><td></td><td></td><td></td><td></td><td></td><td></td><td></td><td></td></tr>
<tr><td>Oligo fwd</td><td>CACCGCAGCGTCGGCGAGCTTCGT</td><td></td><td></td><td></td><td></td><td></td><td></td><td></td><td></td><td></td><td></td></tr>
<tr><td>Oligo rev</td><td>AAACACGAAGCTCGCCGACGCTGC</td><td></td><td></td><td></td><td></td><td></td><td></td><td></td><td></td><td></td><td></td></tr>
<tr><td>Chromosome</td><td>start</td><td>end</td><td>strand</td><td>MM</td><td>target_seq</td><td>PAM</td><td>alignment</td><td>distance</td><td>position</td><td>gene name</td><td>gene id</td></tr>
<tr><td>GU305914.1</td><td>168215</td><td>168237</td><td>+</td><td>0</td><td>GCAGCGTCGGCGAGCTTCGT</td><td>CGG</td><td>||||||||[||||||||||||]PAM</td><td>NA</td><td>Intergenic</td><td>NA</td><td>NA</td></tr>
<tr><td></td><td></td><td></td><td></td><td></td><td></td><td></td><td></td><td></td><td></td><td></td><td></td></tr>
<tr><td>T12693</td><td>CGTCGGCTTGTCCACGATCGAGG</td><td>1000</td><td></td><td></td><td></td><td></td><td></td><td></td><td></td><td></td><td></td></tr>
<tr><td>Oligo fwd</td><td>CACCCGTCGGCTTGTCCACGATCG</td><td></td><td></td><td></td><td></td><td></td><td></td><td></td><td></td><td></td><td></td></tr>
<tr><td>Oligo rev</td><td>AAACCGATCGTGGACAAGCCGACG</td><td></td><td></td><td></td><td></td><td></td><td></td><td></td><td></td><td></td><td></td></tr>
<tr><td>Chromosome</td><td>start</td><td>end</td><td>strand</td><td>MM</td><td>target_seq</td><td>PAM</td><td>alignment</td><td>distance</td><td>position</td><td>gene name</td><td>gene id</td></tr>
<tr><td>GU305914.1</td><td>168232</td><td>168254</td><td>+</td><td>0</td><td>CGTCGGCTTGTCCACGATCG</td><td>AGG</td><td>||||||||[||||||||||||]PAM</td><td>NA</td><td>Intergenic</td><td>NA</td><td>NA</td></tr>
<tr><td></td><td></td><td></td><td></td><td></td><td></td><td></td><td></td><td></td><td></td><td></td><td></td></tr>
<tr><td>T12694</td><td>CGGCTTGTCCACGATCGAGGCGG</td><td>1000</td><td></td><td></td><td></td><td></td><td></td><td></td><td></td><td></td><td></td></tr>
<tr><td>Oligo fwd</td><td>CACCCGGCTTGTCCACGATCGAGG</td><td></td><td></td><td></td><td></td><td></td><td></td><td></td><td></td><td></td><td></td></tr>
<tr><td>Oligo rev</td><td>AAACCCTCGATCGTGGACAAGCCG</td><td></td><td></td><td></td><td></td><td></td><td></td><td></td><td></td><td></td><td></td></tr>
<tr><td>Chromosome</td><td>start</td><td>end</td><td>strand</td><td>MM</td><td>target_seq</td><td>PAM</td><td>alignment</td><td>distance</td><td>position</td><td>gene name</td><td>gene id</td></tr>
<tr><td>GU305914.1</td><td>168235</td><td>168257</td><td>+</td><td>0</td><td>CGGCTTGTCCACGATCGAGG</td><td>CGG</td><td>||||||||[||||||||||||]PAM</td><td>NA</td><td>Intergenic</td><td>NA</td><td>NA</td></tr>
<tr><td>GU305914.1</td><td>123471</td><td>123493</td><td>+</td><td>3</td><td>GGGCTTCTCCACGATCGAGT</td><td>TGG</td><td>-|||||-|[|||||||||||-]PAM</td><td>NA</td><td>Intergenic</td><td>NA</td><td>NA</td></tr>
<tr><td></td><td></td><td></td><td></td><td></td><td></td><td></td><td></td><td></td><td></td><td></td><td></td></tr>
<tr><td>T12695</td><td>ACAGCTCCGTCGACGCGTCTCGG</td><td>1000</td><td></td><td></td><td></td><td></td><td></td><td></td><td></td><td></td><td></td></tr>
<tr><td>Oligo fwd</td><td>CACCACAGCTCCGTCGACGCGTCT</td><td></td><td></td><td></td><td></td><td></td><td></td><td></td><td></td><td></td><td></td></tr>
<tr><td>Oligo rev</td><td>AAACAGACGCGTCGACGGAGCTGT</td><td></td><td></td><td></td><td></td><td></td><td></td><td></td><td></td><td></td><td></td></tr>
<tr><td>Chromosome</td><td>start</td><td>end</td><td>strand</td><td>MM</td><td>target_seq</td><td>PAM</td><td>alignment</td><td>distance</td><td>position</td><td>gene name</td><td>gene id</td></tr>
<tr><td>GU305914.1</td><td>168273</td><td>168295</td><td>+</td><td>0</td><td>ACAGCTCCGTCGACGCGTCT</td><td>CGG</td><td>||||||||[||||||||||||]PAM</td><td>NA</td><td>Intergenic</td><td>NA</td><td>NA</td></tr>
<tr><td></td><td></td><td></td><td></td><td></td><td></td><td></td><td></td><td></td><td></td><td></td><td></td></tr>
<tr><td>T12696</td><td>CAGCTCCGTCGACGCGTCTCGGG</td><td>1000</td><td></td><td></td><td></td><td></td><td></td><td></td><td></td><td></td><td></td></tr>
<tr><td>Oligo fwd</td><td>CACCCAGCTCCGTCGACGCGTCTC</td><td></td><td></td><td></td><td></td><td></td><td></td><td></td><td></td><td></td><td></td></tr>
<tr><td>Oligo rev</td><td>AAACGAGACGCGTCGACGGAGCTG</td><td></td><td></td><td></td><td></td><td></td><td></td><td></td><td></td><td></td><td></td></tr>
<tr><td>Chromosome</td><td>start</td><td>end</td><td>strand</td><td>MM</td><td>target_seq</td><td>PAM</td><td>alignment</td><td>distance</td><td>position</td><td>gene name</td><td>gene id</td></tr>
<tr><td>GU305914.1</td><td>168274</td><td>168296</td><td>+</td><td>0</td><td>CAGCTCCGTCGACGCGTCTC</td><td>GGG</td><td>||||||||[||||||||||||]PAM</td><td>NA</td><td>Intergenic</td><td>NA</td><td>NA</td></tr>
<tr><td></td><td></td><td></td><td></td><td></td><td></td><td></td><td></td><td></td><td></td><td></td><td></td></tr>
<tr><td>T12697</td><td>TCCGTCGACGCGTCTCGGGCCGG</td><td>1000</td><td></td><td></td><td></td><td></td><td></td><td></td><td></td><td></td><td></td></tr>
<tr><td>Oligo fwd</td><td>CACCTCCGTCGACGCGTCTCGGGC</td><td></td><td></td><td></td><td></td><td></td><td></td><td></td><td></td><td></td><td></td></tr>
<tr><td>Oligo rev</td><td>AAACGCCCGAGACGCGTCGACGGA</td><td></td><td></td><td></td><td></td><td></td><td></td><td></td><td></td><td></td><td></td></tr>
<tr><td>Chromosome</td><td>start</td><td>end</td><td>strand</td><td>MM</td><td>target_seq</td><td>PAM</td><td>alignment</td><td>distance</td><td>position</td><td>gene name</td><td>gene id</td></tr>
<tr><td>GU305914.1</td><td>168278</td><td>168300</td><td>+</td><td>0</td><td>TCCGTCGACGCGTCTCGGGC</td><td>CGG</td><td>||||||||[||||||||||||]PAM</td><td>NA</td><td>Intergenic</td><td>NA</td><td>NA</td></tr>
<tr><td></td><td></td><td></td><td></td><td></td><td></td><td></td><td></td><td></td><td></td><td></td><td></td></tr>
<tr><td>T12698</td><td>TCGGGCCGGTTCTGTGTTCGCGG</td><td>1000</td><td></td><td></td><td></td><td></td><td></td><td></td><td></td><td></td><td></td></tr>
<tr><td>Oligo fwd</td><td>CACCTCGGGCCGGTTCTGTGTTCG</td><td></td><td></td><td></td><td></td><td></td><td></td><td></td><td></td><td></td><td></td></tr>
<tr><td>Oligo rev</td><td>AAACCGAACACAGAACCGGCCCGA</td><td></td><td></td><td></td><td></td><td></td><td></td><td></td><td></td><td></td><td></td></tr>
<tr><td>Chromosome</td><td>start</td><td>end</td><td>strand</td><td>MM</td><td>target_seq</td><td>PAM</td><td>alignment</td><td>distance</td><td>position</td><td>gene name</td><td>gene id</td></tr>
<tr><td>GU305914.1</td><td>168292</td><td>168314</td><td>+</td><td>0</td><td>TCGGGCCGGTTCTGTGTTCG</td><td>CGG</td><td>||||||||[||||||||||||]PAM</td><td>NA</td><td>Intergenic</td><td>NA</td><td>NA</td></tr>
<tr><td></td><td></td><td></td><td></td><td></td><td></td><td></td><td></td><td></td><td></td><td></td><td></td></tr>
<tr><td>T12699</td><td>CGGGCCGGTTCTGTGTTCGCGGG</td><td>1000</td><td></td><td></td><td></td><td></td><td></td><td></td><td></td><td></td><td></td></tr>
<tr><td>Oligo fwd</td><td>CACCCGGGCCGGTTCTGTGTTCGC</td><td></td><td></td><td></td><td></td><td></td><td></td><td></td><td></td><td></td><td></td></tr>
<tr><td>Oligo rev</td><td>AAACGCGAACACAGAACCGGCCCG</td><td></td><td></td><td></td><td></td><td></td><td></td><td></td><td></td><td></td><td></td></tr>
<tr><td>Chromosome</td><td>start</td><td>end</td><td>strand</td><td>MM</td><td>target_seq</td><td>PAM</td><td>alignment</td><td>distance</td><td>position</td><td>gene name</td><td>gene id</td></tr>
<tr><td>GU305914.1</td><td>168293</td><td>168315</td><td>+</td><td>0</td><td>CGGGCCGGTTCTGTGTTCGC</td><td>GGG</td><td>||||||||[||||||||||||]PAM</td><td>NA</td><td>Intergenic</td><td>NA</td><td>NA</td></tr>
<tr><td></td><td></td><td></td><td></td><td></td><td></td><td></td><td></td><td></td><td></td><td></td><td></td></tr>
<tr><td>T12700</td><td>GGGCCGGTTCTGTGTTCGCGGGG</td><td>1000</td><td></td><td></td><td></td><td></td><td></td><td></td><td></td><td></td><td></td></tr>
<tr><td>Oligo fwd</td><td>CACCGGGCCGGTTCTGTGTTCGCG</td><td></td><td></td><td></td><td></td><td></td><td></td><td></td><td></td><td></td><td></td></tr>
<tr><td>Oligo rev</td><td>AAACCGCGAACACAGAACCGGCCC</td><td></td><td></td><td></td><td></td><td></td><td></td><td></td><td></td><td></td><td></td></tr>
<tr><td>Chromosome</td><td>start</td><td>end</td><td>strand</td><td>MM</td><td>target_seq</td><td>PAM</td><td>alignment</td><td>distance</td><td>position</td><td>gene name</td><td>gene id</td></tr>
<tr><td>GU305914.1</td><td>168294</td><td>168316</td><td>+</td><td>0</td><td>GGGCCGGTTCTGTGTTCGCG</td><td>GGG</td><td>||||||||[||||||||||||]PAM</td><td>NA</td><td>Intergenic</td><td>NA</td><td>NA</td></tr>
<tr><td></td><td></td><td></td><td></td><td></td><td></td><td></td><td></td><td></td><td></td><td></td><td></td></tr>
<tr><td>T12701</td><td>GGCCGGTTCTGTGTTCGCGGGGG</td><td>1000</td><td></td><td></td><td></td><td></td><td></td><td></td><td></td><td></td><td></td></tr>
</table>

<table>
<tr><td>Oligo fwd</td><td>CACCGGCCGGTTCTGTGTTCGCGG</td><td></td><td></td><td></td><td></td><td></td><td></td><td></td><td></td><td></td><td></td></tr>
<tr><td>Oligo rev</td><td>AAACCCGCGAACACAGAACCGGCC</td><td></td><td></td><td></td><td></td><td></td><td></td><td></td><td></td><td></td><td></td></tr>
<tr><td>Chromosome</td><td>start</td><td>end</td><td>strand</td><td>MM</td><td>target_seq</td><td>PAM</td><td>alignment</td><td>distance</td><td>position</td><td>gene name</td><td>gene id</td></tr>
<tr><td>GU305914.1</td><td>168295</td><td>168317</td><td>+</td><td>0</td><td>GGCCGGTTCTGTGTTCGCGG</td><td>GGG</td><td>||||||||[||||||||||||]PAM</td><td>NA</td><td>Intergenic</td><td>NA</td><td>NA</td></tr>
<tr><td></td><td></td><td></td><td></td><td></td><td></td><td></td><td></td><td></td><td></td><td></td><td></td></tr>
<tr><td>T12702</td><td>TCTGTGTTCGCGGGGGCTGCTGG</td><td>1000</td><td></td><td></td><td></td><td></td><td></td><td></td><td></td><td></td><td></td></tr>
<tr><td>Oligo fwd</td><td>CACCTCTGTGTTCGCGGGGGCTGC</td><td></td><td></td><td></td><td></td><td></td><td></td><td></td><td></td><td></td><td></td></tr>
<tr><td>Oligo rev</td><td>AAACGCAGCCCCCGCGAACACAGA</td><td></td><td></td><td></td><td></td><td></td><td></td><td></td><td></td><td></td><td></td></tr>
<tr><td>Chromosome</td><td>start</td><td>end</td><td>strand</td><td>MM</td><td>target_seq</td><td>PAM</td><td>alignment</td><td>distance</td><td>position</td><td>gene name</td><td>gene id</td></tr>
<tr><td>GU305914.1</td><td>168302</td><td>168324</td><td>+</td><td>0</td><td>TCTGTGTTCGCGGGGGCTGC</td><td>TGG</td><td>||||||||[||||||||||||]PAM</td><td>NA</td><td>Intergenic</td><td>NA</td><td>NA</td></tr>
<tr><td></td><td></td><td></td><td></td><td></td><td></td><td></td><td></td><td></td><td></td><td></td><td></td></tr>
<tr><td>T12703</td><td>GGGGCTGCTGGTGCTGCTCCTGG</td><td>1000</td><td></td><td></td><td></td><td></td><td></td><td></td><td></td><td></td><td></td></tr>
<tr><td>Oligo fwd</td><td>CACCGGGGCTGCTGGTGCTGCTCC</td><td></td><td></td><td></td><td></td><td></td><td></td><td></td><td></td><td></td><td></td></tr>
<tr><td>Oligo rev</td><td>AAACGGAGCAGCACCAGCAGCCCC</td><td></td><td></td><td></td><td></td><td></td><td></td><td></td><td></td><td></td><td></td></tr>
<tr><td>Chromosome</td><td>start</td><td>end</td><td>strand</td><td>MM</td><td>target_seq</td><td>PAM</td><td>alignment</td><td>distance</td><td>position</td><td>gene name</td><td>gene id</td></tr>
<tr><td>GU305914.1</td><td>168314</td><td>168336</td><td>+</td><td>0</td><td>GGGGCTGCTGGTGCTGCTCC</td><td>TGG</td><td>||||||||[||||||||||||]PAM</td><td>NA</td><td>Intergenic</td><td>NA</td><td>NA</td></tr>
<tr><td>GU305914.1</td><td>27406</td><td>27428</td><td>-</td><td>4</td><td>GCTGCTGCTGCTGCTGCTGC</td><td>CGG</td><td>|--|||||[||-|||||||-|]PAM</td><td>NA</td><td>Intergenic</td><td>NA</td><td>NA</td></tr>
<tr><td>GU305914.1</td><td>27021</td><td>27043</td><td>-</td><td>4</td><td>GAGGCTGTTGTTGCTGCTGC</td><td>TGG</td><td>|-|||||-[||-|||||||-|]PAM</td><td>NA</td><td>Intergenic</td><td>NA</td><td>NA</td></tr>
<tr><td></td><td></td><td></td><td></td><td></td><td></td><td></td><td></td><td></td><td></td><td></td><td></td></tr>
<tr><td>T12704</td><td>GCTGGTGCTGCTCCTGGTTGCGG</td><td>1000</td><td></td><td></td><td></td><td></td><td></td><td></td><td></td><td></td><td></td></tr>
<tr><td>Oligo fwd</td><td>CACCGCTGGTGCTGCTCCTGGTTG</td><td></td><td></td><td></td><td></td><td></td><td></td><td></td><td></td><td></td><td></td></tr>
<tr><td>Oligo rev</td><td>AAACCAACCAGGAGCAGCACCAGC</td><td></td><td></td><td></td><td></td><td></td><td></td><td></td><td></td><td></td><td></td></tr>
<tr><td>Chromosome</td><td>start</td><td>end</td><td>strand</td><td>MM</td><td>target_seq</td><td>PAM</td><td>alignment</td><td>distance</td><td>position</td><td>gene name</td><td>gene id</td></tr>
<tr><td>GU305914.1</td><td>168320</td><td>168342</td><td>+</td><td>0</td><td>GCTGGTGCTGCTCCTGGTTG</td><td>CGG</td><td>||||||||[||||||||||||]PAM</td><td>NA</td><td>Intergenic</td><td>NA</td><td>NA</td></tr>
<tr><td>GU305914.1</td><td>18975</td><td>18997</td><td>+</td><td>4</td><td>GTTGCTGCTGCACCTGGGTG</td><td>TGG</td><td>|-||-|||[|||-|||||-||]PAM</td><td>NA</td><td>Intergenic</td><td>NA</td><td>NA</td></tr>
<tr><td>GU305914.1</td><td>110249</td><td>110271</td><td>-</td><td>4</td><td>GGTAGTGCTGCTCCTGGAGG</td><td>AGG</td><td>|-|-||||[|||||||||--|]PAM</td><td>NA</td><td>Intergenic</td><td>NA</td><td>NA</td></tr>
<tr><td></td><td></td><td></td><td></td><td></td><td></td><td></td><td></td><td></td><td></td><td></td><td></td></tr>
<tr><td>T12705</td><td>CGAACTCGCGCGAGTAGATGCGG</td><td>1000</td><td></td><td></td><td></td><td></td><td></td><td></td><td></td><td></td><td></td></tr>
<tr><td>Oligo fwd</td><td>CACCCGAACTCGCGCGAGTAGATG</td><td></td><td></td><td></td><td></td><td></td><td></td><td></td><td></td><td></td><td></td></tr>
<tr><td>Oligo rev</td><td>AAACCATCTACTCGCGCGAGTTCG</td><td></td><td></td><td></td><td></td><td></td><td></td><td></td><td></td><td></td><td></td></tr>
<tr><td>Chromosome</td><td>start</td><td>end</td><td>strand</td><td>MM</td><td>target_seq</td><td>PAM</td><td>alignment</td><td>distance</td><td>position</td><td>gene name</td><td>gene id</td></tr>
<tr><td>GU305914.1</td><td>168343</td><td>168365</td><td>+</td><td>0</td><td>CGAACTCGCGCGAGTAGATG</td><td>CGG</td><td>||||||||[||||||||||||]PAM</td><td>NA</td><td>Intergenic</td><td>NA</td><td>NA</td></tr>
<tr><td></td><td></td><td></td><td></td><td></td><td></td><td></td><td></td><td></td><td></td><td></td><td></td></tr>
<tr><td>T12706</td><td>GAACTCGCGCGAGTAGATGCGGG</td><td>1000</td><td></td><td></td><td></td><td></td><td></td><td></td><td></td><td></td><td></td></tr>
<tr><td>Oligo fwd</td><td>CACCGAACTCGCGCGAGTAGATGC</td><td></td><td></td><td></td><td></td><td></td><td></td><td></td><td></td><td></td><td></td></tr>
<tr><td>Oligo rev</td><td>AAACGCATCTACTCGCGCGAGTTC</td><td></td><td></td><td></td><td></td><td></td><td></td><td></td><td></td><td></td><td></td></tr>
<tr><td>Chromosome</td><td>start</td><td>end</td><td>strand</td><td>MM</td><td>target_seq</td><td>PAM</td><td>alignment</td><td>distance</td><td>position</td><td>gene name</td><td>gene id</td></tr>
<tr><td>GU305914.1</td><td>168344</td><td>168366</td><td>+</td><td>0</td><td>GAACTCGCGCGAGTAGATGC</td><td>GGG</td><td>||||||||[||||||||||||]PAM</td><td>NA</td><td>Intergenic</td><td>NA</td><td>NA</td></tr>
<tr><td></td><td></td><td></td><td></td><td></td><td></td><td></td><td></td><td></td><td></td><td></td><td></td></tr>
<tr><td>T12707</td><td>TTGATCTTTTTCTGAGTCTGAGG</td><td>1000</td><td></td><td></td><td></td><td></td><td></td><td></td><td></td><td></td><td></td></tr>
<tr><td>Oligo fwd</td><td>CACCTTGATCTTTTTCTGAGTCTG</td><td></td><td></td><td></td><td></td><td></td><td></td><td></td><td></td><td></td><td></td></tr>
<tr><td>Oligo rev</td><td>AAACCAGACTCAGAAAAAGATCAA</td><td></td><td></td><td></td><td></td><td></td><td></td><td></td><td></td><td></td><td></td></tr>
<tr><td>Chromosome</td><td>start</td><td>end</td><td>strand</td><td>MM</td><td>target_seq</td><td>PAM</td><td>alignment</td><td>distance</td><td>position</td><td>gene name</td><td>gene id</td></tr>
<tr><td>GU305914.1</td><td>168391</td><td>168413</td><td>+</td><td>0</td><td>TTGATCTTTTTCTGAGTCTG</td><td>AGG</td><td>||||||||[||||||||||||]PAM</td><td>NA</td><td>Intergenic</td><td>NA</td><td>NA</td></tr>
<tr><td></td><td></td><td></td><td></td><td></td><td></td><td></td><td></td><td></td><td></td><td></td><td></td></tr>
<tr><td>T12708</td><td>TGAGGACTCCGATTCCGATTTGG</td><td>1000</td><td></td><td></td><td></td><td></td><td></td><td></td><td></td><td></td><td></td></tr>
<tr><td>Oligo fwd</td><td>CACCTGAGGACTCCGATTCCGATT</td><td></td><td></td><td></td><td></td><td></td><td></td><td></td><td></td><td></td><td></td></tr>
<tr><td>Oligo rev</td><td>AAACAATCGGAATCGGAGTCCTCA</td><td></td><td></td><td></td><td></td><td></td><td></td><td></td><td></td><td></td><td></td></tr>
<tr><td>Chromosome</td><td>start</td><td>end</td><td>strand</td><td>MM</td><td>target_seq</td><td>PAM</td><td>alignment</td><td>distance</td><td>position</td><td>gene name</td><td>gene id</td></tr>
<tr><td>GU305914.1</td><td>168409</td><td>168431</td><td>+</td><td>0</td><td>TGAGGACTCCGATTCCGATT</td><td>TGG</td><td>||||||||[||||||||||||]PAM</td><td>NA</td><td>Intergenic</td><td>NA</td><td>NA</td></tr>
<tr><td></td><td></td><td></td><td></td><td></td><td></td><td></td><td></td><td></td><td></td><td></td><td></td></tr>
<tr><td>T12709</td><td>GAGGACTCCGATTCCGATTTGGG</td><td>1000</td><td></td><td></td><td></td><td></td><td></td><td></td><td></td><td></td><td></td></tr>
<tr><td>Oligo fwd</td><td>CACCGAGGACTCCGATTCCGATTT</td><td></td><td></td><td></td><td></td><td></td><td></td><td></td><td></td><td></td><td></td></tr>
<tr><td>Oligo rev</td><td>AAACAAATCGGAATCGGAGTCCTC</td><td></td><td></td><td></td><td></td><td></td><td></td><td></td><td></td><td></td><td></td></tr>
<tr><td>Chromosome</td><td>start</td><td>end</td><td>strand</td><td>MM</td><td>target_seq</td><td>PAM</td><td>alignment</td><td>distance</td><td>position</td><td>gene name</td><td>gene id</td></tr>
<tr><td>GU305914.1</td><td>168410</td><td>168432</td><td>+</td><td>0</td><td>GAGGACTCCGATTCCGATTT</td><td>GGG</td><td>||||||||[||||||||||||]PAM</td><td>NA</td><td>Intergenic</td><td>NA</td><td>NA</td></tr>
<tr><td></td><td></td><td></td><td></td><td></td><td></td><td></td><td></td><td></td><td></td><td></td><td></td></tr>
<tr><td>T12710</td><td>TTTGGGCGACGTCGAGCCTATGG</td><td>1000</td><td></td><td></td><td></td><td></td><td></td><td></td><td></td><td></td><td></td></tr>
<tr><td>Oligo fwd</td><td>CACCTTTGGGCGACGTCGAGCCTA</td><td></td><td></td><td></td><td></td><td></td><td></td><td></td><td></td><td></td><td></td></tr>
<tr><td>Oligo rev</td><td>AAACTAGGCTCGACGTCGCCCAAA</td><td></td><td></td><td></td><td></td><td></td><td></td><td></td><td></td><td></td><td></td></tr>
<tr><td>Chromosome</td><td>start</td><td>end</td><td>strand</td><td>MM</td><td>target_seq</td><td>PAM</td><td>alignment</td><td>distance</td><td>position</td><td>gene name</td><td>gene id</td></tr>
<tr><td>GU305914.1</td><td>168427</td><td>168449</td><td>+</td><td>0</td><td>TTTGGGCGACGTCGAGCCTA</td><td>TGG</td><td>||||||||[||||||||||||]PAM</td><td>NA</td><td>Intergenic</td><td>NA</td><td>NA</td></tr>
<tr><td></td><td></td><td></td><td></td><td></td><td></td><td></td><td></td><td></td><td></td><td></td><td></td></tr>
<tr><td>T12711</td><td>CGACGTCGAGCCTATGGATTCGG</td><td>1000</td><td></td><td></td><td></td><td></td><td></td><td></td><td></td><td></td><td></td></tr>
<tr><td>Oligo fwd</td><td>CACCCGACGTCGAGCCTATGGATT</td><td></td><td></td><td></td><td></td><td></td><td></td><td></td><td></td><td></td><td></td></tr>
<tr><td>Oligo rev</td><td>AAACAATCCATAGGCTCGACGTCG</td><td></td><td></td><td></td><td></td><td></td><td></td><td></td><td></td><td></td><td></td></tr>
<tr><td>Chromosome</td><td>start</td><td>end</td><td>strand</td><td>MM</td><td>target_seq</td><td>PAM</td><td>alignment</td><td>distance</td><td>position</td><td>gene name</td><td>gene id</td></tr>
<tr><td>GU305914.1</td><td>168433</td><td>168455</td><td>+</td><td>0</td><td>CGACGTCGAGCCTATGGATT</td><td>CGG</td><td>||||||||[||||||||||||]PAM</td><td>NA</td><td>Intergenic</td><td>NA</td><td>NA</td></tr>
<tr><td></td><td></td><td></td><td></td><td></td><td></td><td></td><td></td><td></td><td></td><td></td><td></td></tr>
<tr><td>T12712</td><td>TCGGATCCAGAATCCGAAGATGG</td><td>1000</td><td></td><td></td><td></td><td></td><td></td><td></td><td></td><td></td><td></td></tr>
<tr><td>Oligo fwd</td><td>CACCTCGGATCCAGAATCCGAAGA</td><td></td><td></td><td></td><td></td><td></td><td></td><td></td><td></td><td></td><td></td></tr>
<tr><td>Oligo rev</td><td>AAACTCTTCGGATTCTGGATCCGA</td><td></td><td></td><td></td><td></td><td></td><td></td><td></td><td></td><td></td><td></td></tr>
<tr><td>Chromosome</td><td>start</td><td>end</td><td>strand</td><td>MM</td><td>target_seq</td><td>PAM</td><td>alignment</td><td>distance</td><td>position</td><td>gene name</td><td>gene id</td></tr>
<tr><td>GU305914.1</td><td>168452</td><td>168474</td><td>+</td><td>0</td><td>TCGGATCCAGAATCCGAAGA</td><td>TGG</td><td>||||||||[||||||||||||]PAM</td><td>NA</td><td>Intergenic</td><td>NA</td><td>NA</td></tr>
<tr><td></td><td></td><td></td><td></td><td></td><td></td><td></td><td></td><td></td><td></td><td></td><td></td></tr>
<tr><td>T12713</td><td>CGATGAAGCGCCGTCCGAGCTGG</td><td>1000</td><td></td><td></td><td></td><td></td><td></td><td></td><td></td><td></td><td></td></tr>
<tr><td>Oligo fwd</td><td>CACCCGATGAAGCGCCGTCCGAGC</td><td></td><td></td><td></td><td></td><td></td><td></td><td></td><td></td><td></td><td></td></tr>
<tr><td>Oligo rev</td><td>AAACGCTCGGACGGCGCTTCATCG</td><td></td><td></td><td></td><td></td><td></td><td></td><td></td><td></td><td></td><td></td></tr>
<tr><td>Chromosome</td><td>start</td><td>end</td><td>strand</td><td>MM</td><td>target_seq</td><td>PAM</td><td>alignment</td><td>distance</td><td>position</td><td>gene name</td><td>gene id</td></tr>
<tr><td>GU305914.1</td><td>168487</td><td>168509</td><td>+</td><td>0</td><td>CGATGAAGCGCCGTCCGAGC</td><td>TGG</td><td>||||||||[||||||||||||]PAM</td><td>NA</td><td>Intergenic</td><td>NA</td><td>NA</td></tr>
<tr><td></td><td></td><td></td><td></td><td></td><td></td><td></td><td></td><td></td><td></td><td></td><td></td></tr>
<tr><td>T12714</td><td>GACGATGATGATGATGATGATGG</td><td>1000</td><td></td><td></td><td></td><td></td><td></td><td></td><td></td><td></td><td></td></tr>
<tr><td>Oligo fwd</td><td>CACCGACGATGATGATGATGATGA</td><td></td><td></td><td></td><td></td><td></td><td></td><td></td><td></td><td></td><td></td></tr>
<tr><td>Oligo rev</td><td>AAACTCATCATCATCATCATCGTC</td><td></td><td></td><td></td><td></td><td></td><td></td><td></td><td></td><td></td><td></td></tr>
<tr><td>Chromosome</td><td>start</td><td>end</td><td>strand</td><td>MM</td><td>target_seq</td><td>PAM</td><td>alignment</td><td>distance</td><td>position</td><td>gene name</td><td>gene id</td></tr>
<tr><td>GU305914.1</td><td>168515</td><td>168537</td><td>+</td><td>0</td><td>GACGATGATGATGATGATGA</td><td>TGG</td><td>||||||||[||||||||||||]PAM</td><td>NA</td><td>Intergenic</td><td>NA</td><td>NA</td></tr>
<tr><td>GU305914.1</td><td>173679</td><td>173701</td><td>+</td><td>4</td><td>GTAGAGGATGATGTTGATGA</td><td>TGG</td><td>|--||-||[|||||-||||||]PAM</td><td>NA</td><td>Intergenic</td><td>NA</td><td>NA</td></tr>
<tr><td>GU305914.1</td><td>168590</td><td>168612</td><td>+</td><td>4</td><td>GAATATGAGGATGATGAAGA</td><td>GGG</td><td>||--||||[-||||||||-||]PAM</td><td>NA</td><td>Intergenic</td><td>NA</td><td>NA</td></tr>
</table>

| | | | | | | | | | | | |
|---|---|---|---|---|---|---|---|---|---|---|---|
| T12715 | TGATGGTTCATCCTCTGAGTCGG | 1000 | | | | | | | | | |
| Oligo fwd | CACCTGATGGTTCATCCTCTGAGT | | | | | | | | | | |
| Oligo rev | AAACACTCAGAGGATGAACCATCA | | | | | | | | | | |
| Chromosome | start | end | strand | MM | target_seq | PAM | alignment | distance | position | gene name | gene id |
| GU305914.1 | 168532 | 168554 | + | 0 | TGATGGTTCATCCTCTGAGT | CGG | \|\|\|\|\|\|\|\|[\|\|\|\|\|\|\|\|\|\|\|\|]PAM | NA | Intergenic | NA | NA |
| T12716 | TGGTTCATCCTCTGAGTCGGAGG | 1000 | | | | | | | | | |
| Oligo fwd | CACCTGGTTCATCCTCTGAGTCGG | | | | | | | | | | |
| Oligo rev | AAACCCGACTCAGAGGATGAACCA | | | | | | | | | | |
| Chromosome | start | end | strand | MM | target_seq | PAM | alignment | distance | position | gene name | gene id |
| GU305914.1 | 168535 | 168557 | + | 0 | TGGTTCATCCTCTGAGTCGG | AGG | \|\|\|\|\|\|\|\|[\|\|\|\|\|\|\|\|\|\|\|\|]PAM | NA | Intergenic | NA | NA |
| T12717 | TTCATCCTCTGAGTCGGAGGAGG | 1000 | | | | | | | | | |
| Oligo fwd | CACCTTCATCCTCTGAGTCGGAGG | | | | | | | | | | |
| Oligo rev | AAACCCTCCGACTCAGAGGATGAA | | | | | | | | | | |
| Chromosome | start | end | strand | MM | target_seq | PAM | alignment | distance | position | gene name | gene id |
| GU305914.1 | 168538 | 168560 | + | 0 | TTCATCCTCTGAGTCGGAGG | AGG | \|\|\|\|\|\|\|\|[\|\|\|\|\|\|\|\|\|\|\|\|]PAM | NA | Intergenic | NA | NA |
| T12718 | TCCTCTGAGTCGGAGGAGGATGG | 1000 | | | | | | | | | |
| Oligo fwd | CACCTCCTCTGAGTCGGAGGAGGA | | | | | | | | | | |
| Oligo rev | AAACTCCTCCTCCGACTCAGAGGA | | | | | | | | | | |
| Chromosome | start | end | strand | MM | target_seq | PAM | alignment | distance | position | gene name | gene id |
| GU305914.1 | 168542 | 168564 | + | 0 | TCCTCTGAGTCGGAGGAGGA | TGG | \|\|\|\|\|\|\|\|[\|\|\|\|\|\|\|\|\|\|\|\|]PAM | NA | Intergenic | NA | NA |
| GU305914.1 | 191587 | 191609 | + | 4 | CTCTCTGAGGCGGAGCAGGA | TGG | --\|\|\|\|\|\|[\|-\|\|\|\|\|-\|\|\|\|]PAM | NA | Intergenic | NA | NA |
| T12719 | TGAGTCGGAGGAGGATGGAGAGG | 1000 | | | | | | | | | |
| Oligo fwd | CACCTGAGTCGGAGGAGGATGGAG | | | | | | | | | | |
| Oligo rev | AAACCTCCATCCTCCTCCGACTCA | | | | | | | | | | |
| Chromosome | start | end | strand | MM | target_seq | PAM | alignment | distance | position | gene name | gene id |
| GU305914.1 | 168547 | 168569 | + | 0 | TGAGTCGGAGGAGGATGGAG | AGG | \|\|\|\|\|\|\|\|[\|\|\|\|\|\|\|\|\|\|\|\|]PAM | NA | Intergenic | NA | NA |
| T12720 | AGATGATGAAGATGAATATGAGG | 1000 | | | | | | | | | |
| Oligo fwd | CACCAGATGATGAAGATGAATATG | | | | | | | | | | |
| Oligo rev | AAACCATATTCATCTTCATCATCT | | | | | | | | | | |
| Chromosome | start | end | strand | MM | target_seq | PAM | alignment | distance | position | gene name | gene id |
| GU305914.1 | 168577 | 168599 | + | 0 | AGATGATGAAGATGAATATG | AGG | \|\|\|\|\|\|\|\|[\|\|\|\|\|\|\|\|\|\|\|\|]PAM | NA | Intergenic | NA | NA |
| GU305914.1 | 152945 | 152967 | + | 4 | CGATGATAACGATGAGTATG | AGG | -\|\|\|\|\|\|-[\|-\|\|\|\|\|-\|\|\|\|]PAM | NA | Intergenic | NA | NA |
| T12721 | TGAATATGAGGATGATGAAGAGG | 1000 | | | | | | | | | |
| Oligo fwd | CACCTGAATATGAGGATGATGAAG | | | | | | | | | | |
| Oligo rev | AAACCTTCATCATCCTCATATTCA | | | | | | | | | | |
| Chromosome | start | end | strand | MM | target_seq | PAM | alignment | distance | position | gene name | gene id |
| GU305914.1 | 168589 | 168611 | + | 0 | TGAATATGAGGATGATGAAG | AGG | \|\|\|\|\|\|\|\|[\|\|\|\|\|\|\|\|\|\|\|\|]PAM | NA | Intergenic | NA | NA |
| GU305914.1 | 27345 | 27367 | + | 4 | CGAAGATGATGATGAAGAAG | AGG | -\|\|\|-\|\|\|[\|-\|\|\|\|\|-\|\|\|\|]PAM | NA | Intergenic | NA | NA |
| GU305914.1 | 180051 | 180073 | - | 4 | TGATTCTGAAGATGATGATG | AGG | \|\|\|-\|-\|\|[\|-\|\|\|\|\|\|\|\|-\|]PAM | NA | Intergenic | NA | NA |
| T12722 | GAATATGAGGATGATGAAGAGGG | 1000 | | | | | | | | | |
| Oligo fwd | CACCGAATATGAGGATGATGAAGA | | | | | | | | | | |
| Oligo rev | AAACTCTTCATCATCCTCATATTC | | | | | | | | | | |
| Chromosome | start | end | strand | MM | target_seq | PAM | alignment | distance | position | gene name | gene id |
| GU305914.1 | 168590 | 168612 | + | 0 | GAATATGAGGATGATGAAGA | GGG | \|\|\|\|\|\|\|\|[\|\|\|\|\|\|\|\|\|\|\|\|]PAM | NA | Intergenic | NA | NA |
| GU305914.1 | 168515 | 168537 | + | 4 | GACGATGATGATGATGATGA | TGG | \|\|--\|\|\|\|[-\|\|\|\|\|\|\|\|-\|\|]PAM | NA | Intergenic | NA | NA |
| T12723 | TGAGGATGATGAAGAGGGTTCGG | 1000 | | | | | | | | | |
| Oligo fwd | CACCTGAGGATGATGAAGAGGGTT | | | | | | | | | | |
| Oligo rev | AAACAACCCTCTTCATCATCCTCA | | | | | | | | | | |
| Chromosome | start | end | strand | MM | target_seq | PAM | alignment | distance | position | gene name | gene id |
| GU305914.1 | 168595 | 168617 | + | 0 | TGAGGATGATGAAGAGGGTT | CGG | \|\|\|\|\|\|\|\|[\|\|\|\|\|\|\|\|\|\|\|\|]PAM | NA | Intergenic | NA | NA |
| T12724 | GAGGGTTCGGTCTCCGATCCCGG | 1000 | | | | | | | | | |
| Oligo fwd | CACCGAGGGTTCGGTCTCCGATCC | | | | | | | | | | |
| Oligo rev | AAACGGATCGGAGACCGAACCCTC | | | | | | | | | | |
| Chromosome | start | end | strand | MM | target_seq | PAM | alignment | distance | position | gene name | gene id |
| GU305914.1 | 168608 | 168630 | + | 0 | GAGGGTTCGGTCTCCGATCC | CGG | \|\|\|\|\|\|\|\|[\|\|\|\|\|\|\|\|\|\|\|\|]PAM | NA | Intergenic | NA | NA |
| T12725 | GGTTCGGTCTCCGATCCCGGTGG | 1000 | | | | | | | | | |
| Oligo fwd | CACCGGTTCGGTCTCCGATCCCGG | | | | | | | | | | |
| Oligo rev | AAACCCGGGATCGGAGACCGAACC | | | | | | | | | | |
| Chromosome | start | end | strand | MM | target_seq | PAM | alignment | distance | position | gene name | gene id |
| GU305914.1 | 168611 | 168633 | + | 0 | GGTTCGGTCTCCGATCCCGG | TGG | \|\|\|\|\|\|\|\|[\|\|\|\|\|\|\|\|\|\|\|\|]PAM | NA | Intergenic | NA | NA |
| T12726 | GGTCTCCGATCCCGGTGGTAAGG | 1000 | | | | | | | | | |
| Oligo fwd | CACCGGTCTCCGATCCCGGTGGTA | | | | | | | | | | |
| Oligo rev | AAACTACCACCGGGATCGGAGACC | | | | | | | | | | |
| Chromosome | start | end | strand | MM | target_seq | PAM | alignment | distance | position | gene name | gene id |
| GU305914.1 | 168616 | 168638 | + | 0 | GGTCTCCGATCCCGGTGGTA | AGG | \|\|\|\|\|\|\|\|[\|\|\|\|\|\|\|\|\|\|\|\|]PAM | NA | Intergenic | NA | NA |
| T12727 | GGTGGTAAGGACGAGTTAGACGG | 1000 | | | | | | | | | |
| Oligo fwd | CACCGGTGGTAAGGACGAGTTAGA | | | | | | | | | | |
| Oligo rev | AAACTCTAACTCGTCCTTACCACC | | | | | | | | | | |
| Chromosome | start | end | strand | MM | target_seq | PAM | alignment | distance | position | gene name | gene id |
| GU305914.1 | 168629 | 168651 | + | 0 | GGTGGTAAGGACGAGTTAGA | CGG | \|\|\|\|\|\|\|\|[\|\|\|\|\|\|\|\|\|\|\|\|]PAM | NA | Intergenic | NA | NA |
| T12728 | TTCGCTATCCGAGTCGTCTGAGG | 1000 | | | | | | | | | |
| Oligo fwd | CACCTTCGCTATCCGAGTCGTCTG | | | | | | | | | | |
| Oligo rev | AAACCAGACGACTCGGATAGCGAA | | | | | | | | | | |
| Chromosome | start | end | strand | MM | target_seq | PAM | alignment | distance | position | gene name | gene id |
| GU305914.1 | 168673 | 168695 | + | 0 | TTCGCTATCCGAGTCGTCTG | AGG | \|\|\|\|\|\|\|\|[\|\|\|\|\|\|\|\|\|\|\|\|]PAM | NA | Intergenic | NA | NA |

| | | | | | | | | | | | |
|---|---|---|---|---|---|---|---|---|---|---|---|
| T12729 | CGAAGAGTCCGATGACGAAGAGG | 1000 | | | | | | | | | |
| Oligo fwd | CACCCGAAGAGTCCGATGACGAAG | | | | | | | | | | |
| Oligo rev | AAACCTTCGTCATCGGACTCTTCG | | | | | | | | | | |
| Chromosome | start | end | strand | MM | target_seq | PAM | alignment | distance | position | gene name | gene id |
| GU305914.1 | 168703 | 168725 | + | 0 | CGAAGAGTCCGATGACGAAG | AGG | \|\|\|\|\|\|\|\|[\|\|\|\|\|\|\|\|\|\|\|\|]PAM | NA | Intergenic | NA | NA |
| GU305914.1 | 118515 | 118537 | - | 4 | CGATGAGGACGATGATGAAG | AGG | \|\|\|-\|\|\|-[-\|\|\|\|\|\|-\|\|\|\|]PAM | NA | Intergenic | NA | NA |

| | | | | | | | | | | | |
|---|---|---|---|---|---|---|---|---|---|---|---|
| T12730 | GATTAAGATCTTCGTCCCGTCGG | 1000 | | | | | | | | | |
| Oligo fwd | CACCGATTAAGATCTTCGTCCCGT | | | | | | | | | | |
| Oligo rev | AAACACGGGACGAAGATCTTAATC | | | | | | | | | | |
| Chromosome | start | end | strand | MM | target_seq | PAM | alignment | distance | position | gene name | gene id |
| GU305914.1 | 168730 | 168752 | + | 0 | GATTAAGATCTTCGTCCCGT | CGG | \|\|\|\|\|\|\|\|[\|\|\|\|\|\|\|\|\|\|\|\|]PAM | NA | Intergenic | NA | NA |

| | | | | | | | | | | | |
|---|---|---|---|---|---|---|---|---|---|---|---|
| T12731 | TAAGATCTTCGTCCCGTCGGTGG | 1000 | | | | | | | | | |
| Oligo fwd | CACCTAAGATCTTCGTCCCGTCGG | | | | | | | | | | |
| Oligo rev | AAACCCGACGGGACGAAGATCTTA | | | | | | | | | | |
| Chromosome | start | end | strand | MM | target_seq | PAM | alignment | distance | position | gene name | gene id |
| GU305914.1 | 168733 | 168755 | + | 0 | TAAGATCTTCGTCCCGTCGG | TGG | \|\|\|\|\|\|\|\|[\|\|\|\|\|\|\|\|\|\|\|\|]PAM | NA | Intergenic | NA | NA |

| | | | | | | | | | | | |
|---|---|---|---|---|---|---|---|---|---|---|---|
| T12732 | AAGATCTTCGTCCCGTCGGTGGG | 1000 | | | | | | | | | |
| Oligo fwd | CACCAAGATCTTCGTCCCGTCGGT | | | | | | | | | | |
| Oligo rev | AAACACCGACGGGACGAAGATCTT | | | | | | | | | | |
| Chromosome | start | end | strand | MM | target_seq | PAM | alignment | distance | position | gene name | gene id |
| GU305914.1 | 168734 | 168756 | + | 0 | AAGATCTTCGTCCCGTCGGT | GGG | \|\|\|\|\|\|\|\|[\|\|\|\|\|\|\|\|\|\|\|\|]PAM | NA | Intergenic | NA | NA |

| | | | | | | | | | | | |
|---|---|---|---|---|---|---|---|---|---|---|---|
| T12733 | AGATCTTCGTCCCGTCGGTGGGG | 1000 | | | | | | | | | |
| Oligo fwd | CACCAGATCTTCGTCCCGTCGGTG | | | | | | | | | | |
| Oligo rev | AAACCACCGACGGGACGAAGATCT | | | | | | | | | | |
| Chromosome | start | end | strand | MM | target_seq | PAM | alignment | distance | position | gene name | gene id |
| GU305914.1 | 168735 | 168757 | + | 0 | AGATCTTCGTCCCGTCGGTG | GGG | \|\|\|\|\|\|\|\|[\|\|\|\|\|\|\|\|\|\|\|\|]PAM | NA | Intergenic | NA | NA |

| | | | | | | | | | | | |
|---|---|---|---|---|---|---|---|---|---|---|---|
| T12734 | GATCTTCGTCCCGTCGGTGGGGG | 1000 | | | | | | | | | |
| Oligo fwd | CACCGATCTTCGTCCCGTCGGTGG | | | | | | | | | | |
| Oligo rev | AAACCCACCGACGGGACGAAGATC | | | | | | | | | | |
| Chromosome | start | end | strand | MM | target_seq | PAM | alignment | distance | position | gene name | gene id |
| GU305914.1 | 168736 | 168758 | + | 0 | GATCTTCGTCCCGTCGGTGG | GGG | \|\|\|\|\|\|\|\|[\|\|\|\|\|\|\|\|\|\|\|\|]PAM | NA | Intergenic | NA | NA |

| | | | | | | | | | | | |
|---|---|---|---|---|---|---|---|---|---|---|---|
| T12735 | GTCGCTCGCTCGAGATAGTATGG | 1000 | | | | | | | | | |
| Oligo fwd | CACCGTCGCTCGCTCGAGATAGTA | | | | | | | | | | |
| Oligo rev | AAACTACTATCTCGAGCGAGCGAC | | | | | | | | | | |
| Chromosome | start | end | strand | MM | target_seq | PAM | alignment | distance | position | gene name | gene id |
| GU305914.1 | 168775 | 168797 | + | 0 | GTCGCTCGCTCGAGATAGTA | TGG | \|\|\|\|\|\|\|\|[\|\|\|\|\|\|\|\|\|\|\|\|]PAM | NA | Intergenic | NA | NA |

| | | | | | | | | | | | |
|---|---|---|---|---|---|---|---|---|---|---|---|
| T12736 | GGTCGAAACCCGTTTTTTAGCGG | 1000 | | | | | | | | | |
| Oligo fwd | CACCGGTCGAAACCCGTTTTTTAG | | | | | | | | | | |
| Oligo rev | AAACCTAAAAAACGGGTTTCGACC | | | | | | | | | | |
| Chromosome | start | end | strand | MM | target_seq | PAM | alignment | distance | position | gene name | gene id |
| GU305914.1 | 168796 | 168818 | + | 0 | GGTCGAAACCCGTTTTTTAG | CGG | \|\|\|\|\|\|\|\|[\|\|\|\|\|\|\|\|\|\|\|\|]PAM | NA | Intergenic | NA | NA |

| | | | | | | | | | | | |
|---|---|---|---|---|---|---|---|---|---|---|---|
| T12737 | CGAAACCCGTTTTTTAGCGGCGG | 1000 | | | | | | | | | |
| Oligo fwd | CACCCGAAACCCGTTTTTTAGCGG | | | | | | | | | | |
| Oligo rev | AAACCCGCTAAAAAACGGGTTTCG | | | | | | | | | | |
| Chromosome | start | end | strand | MM | target_seq | PAM | alignment | distance | position | gene name | gene id |
| GU305914.1 | 168799 | 168821 | + | 0 | CGAAACCCGTTTTTTAGCGG | CGG | \|\|\|\|\|\|\|\|[\|\|\|\|\|\|\|\|\|\|\|\|]PAM | NA | Intergenic | NA | NA |

| | | | | | | | | | | | |
|---|---|---|---|---|---|---|---|---|---|---|---|
| T12738 | AACCCGTTTTTTAGCGGCGGTGG | 1000 | | | | | | | | | |
| Oligo fwd | CACCAACCCGTTTTTTAGCGGCGG | | | | | | | | | | |
| Oligo rev | AAACCCGCCGCTAAAAAACGGGTT | | | | | | | | | | |
| Chromosome | start | end | strand | MM | target_seq | PAM | alignment | distance | position | gene name | gene id |
| GU305914.1 | 168802 | 168824 | + | 0 | AACCCGTTTTTTAGCGGCGG | TGG | \|\|\|\|\|\|\|\|[\|\|\|\|\|\|\|\|\|\|\|\|]PAM | NA | Intergenic | NA | NA |

| | | | | | | | | | | | |
|---|---|---|---|---|---|---|---|---|---|---|---|
| T12739 | CCGTTTTTTAGCGGCGGTGGTGG | 1000 | | | | | | | | | |
| Oligo fwd | CACCCCGTTTTTTAGCGGCGGTGG | | | | | | | | | | |
| Oligo rev | AAACCCACCGCCGCTAAAAAACGG | | | | | | | | | | |
| Chromosome | start | end | strand | MM | target_seq | PAM | alignment | distance | position | gene name | gene id |
| GU305914.1 | 168805 | 168827 | + | 0 | CCGTTTTTTAGCGGCGGTGG | TGG | \|\|\|\|\|\|\|\|[\|\|\|\|\|\|\|\|\|\|\|\|]PAM | NA | Intergenic | NA | NA |

| | | | | | | | | | | | |
|---|---|---|---|---|---|---|---|---|---|---|---|
| T12740 | TTTAGCGGCGGTGGTGGCAGCGG | 1000 | | | | | | | | | |
| Oligo fwd | CACCTTTAGCGGCGGTGGTGGCAG | | | | | | | | | | |
| Oligo rev | AAACCTGCCACCACCGCCGCTAAA | | | | | | | | | | |
| Chromosome | start | end | strand | MM | target_seq | PAM | alignment | distance | position | gene name | gene id |
| GU305914.1 | 168811 | 168833 | + | 0 | TTTAGCGGCGGTGGTGGCAG | CGG | \|\|\|\|\|\|\|\|[\|\|\|\|\|\|\|\|\|\|\|\|]PAM | NA | Intergenic | NA | NA |
| GU305914.1 | 48153 | 48175 | + | 4 | TGTGGCGGCGGCGGTGGCGG | CGG | \|-\|-\|\|\|\|[\|\|\|-\|\|\|\|\|\|-\|]PAM | NA | Intergenic | NA | NA |

| | | | | | | | | | | | |
|---|---|---|---|---|---|---|---|---|---|---|---|
| T12741 | AGCGGCGGTGGTGGCAGCGGCGG | 1000 | | | | | | | | | |
| Oligo fwd | CACCAGCGGCGGTGGTGGCAGCGG | | | | | | | | | | |
| Oligo rev | AAACCCGCTGCCACCACCGCCGCT | | | | | | | | | | |
| Chromosome | start | end | strand | MM | target_seq | PAM | alignment | distance | position | gene name | gene id |
| GU305914.1 | 168814 | 168836 | + | 0 | AGCGGCGGTGGTGGCAGCGG | CGG | \|\|\|\|\|\|\|\|[\|\|\|\|\|\|\|\|\|\|\|\|]PAM | NA | Intergenic | NA | NA |
| GU305914.1 | 45255 | 45277 | - | 3 | GGTGGCGGTGGCGGCAGCGG | CGG | -\|-\|\|\|\|\|[\|\|\|-\|\|\|\|\|\|\|\|]PAM | NA | Intergenic | NA | NA |
| GU305914.1 | 62446 | 62468 | + | 3 | GGCGGGGGTGGTGGCGGCGG | GGG | -\|\|\|\|-\|\|[\|\|\|\|\|\|\|-\|\|\|\|]PAM | NA | Intergenic | NA | NA |
| GU305914.1 | 114658 | 114680 | - | 3 | GGCGGTGGTGGTGGCGGCGG | CGG | -\|\|\|\|-\|\|[\|\|\|\|\|\|\|-\|\|\|\|]PAM | NA | Intergenic | NA | NA |
| GU305914.1 | 48156 | 48178 | + | 3 | GGCGGCGGCGGTGGCGGCGG | CGG | -\|\|\|\|\|\|\|[-\|\|\|\|\|\|-\|\|\|\|]PAM | NA | Intergenic | NA | NA |
| GU305914.1 | 48159 | 48181 | + | 3 | GGCGGCGGTGGCGGCGGCGG | TGG | -\|\|\|\|\|\|\|[\|\|\|-\|\|\|-\|\|\|\|]PAM | NA | Intergenic | NA | NA |
| GU305914.1 | 114661 | 114683 | - | 4 | GACGGCGGTGGTGGTGGCGG | CGG | --\|\|\|\|\|\|[\|\|\|\|\|\|--\|\|\|\|]PAM | NA | Intergenic | NA | NA |

| | | |
|---|---|---|
| T12742 | GGCGGTGGTGGCAGCGGCGGCGG | 1000 |
| Oligo fwd | CACCGGCGGTGGTGGCAGCGGCGG | |
| Oligo rev | AAACCCGCCGCTGCCACCACCGCC | |

<table>
<tr><th>Chromosome</th><th>start</th><th>end</th><th>strand</th><th>MM</th><th>target_seq</th><th>PAM</th><th>alignment</th><th>distance</th><th>position</th><th>gene name</th><th>gene id</th></tr>
<tr><td>GU305914.1</td><td>168817</td><td>168839</td><td>+</td><td>0</td><td>GGCGGTGGTGGCAGCGGCGG</td><td>CGG</td><td>||||||||[||||||||||||]PAM</td><td>NA</td><td>Intergenic</td><td>NA</td><td>NA</td></tr>
<tr><td>GU305914.1</td><td>48159</td><td>48181</td><td>+</td><td>2</td><td>GGCGGCGGTGGCGGCGGCGG</td><td>TGG</td><td>|||||-||[||||-|||||||]PAM</td><td>NA</td><td>Intergenic</td><td>NA</td><td>NA</td></tr>
<tr><td>GU305914.1</td><td>154404</td><td>154426</td><td>+</td><td>2</td><td>GGCGGTGGCGGCGGCGGCGG</td><td>CGG</td><td>||||||||[-|||-|||||||]PAM</td><td>NA</td><td>Intergenic</td><td>NA</td><td>NA</td></tr>
<tr><td>GU305914.1</td><td>58126</td><td>58148</td><td>-</td><td>2</td><td>GGCGGTGGCGGCGGCGGCGG</td><td>CGG</td><td>||||||||[-|||-|||||||]PAM</td><td>NA</td><td>Intergenic</td><td>NA</td><td>NA</td></tr>
<tr><td>GU305914.1</td><td>58129</td><td>58151</td><td>-</td><td>4</td><td>GCGGGCGGTGGCGGCGGCGG</td><td>CGG</td><td>|--||-||[||||-|||||||]PAM</td><td>NA</td><td>Intergenic</td><td>NA</td><td>NA</td></tr>
<tr><td>GU305914.1</td><td>29997</td><td>30019</td><td>+</td><td>4</td><td>ACCGGTGCTGGCGGCGGCGG</td><td>CGG</td><td>--|||||-[||||-|||||||]PAM</td><td>NA</td><td>Intergenic</td><td>NA</td><td>NA</td></tr>
<tr><td>GU305914.1</td><td>114658</td><td>114680</td><td>-</td><td>2</td><td>GGCGGTGGTGGTGGCGGCGG</td><td>CGG</td><td>||||||||[|||--|||||||]PAM</td><td>NA</td><td>Intergenic</td><td>NA</td><td>NA</td></tr>
<tr><td>GU305914.1</td><td>30000</td><td>30022</td><td>+</td><td>4</td><td>GGTGCTGGCGGCGGCGGCGG</td><td>CGG</td><td>||-|-|||[-|||-|||||||]PAM</td><td>NA</td><td>Intergenic</td><td>NA</td><td>NA</td></tr>
<tr><td>GU305914.1</td><td>154407</td><td>154429</td><td>+</td><td>4</td><td>GGTGGCGGCGGCGGCGGCGG</td><td>CGG</td><td>||-||-||[-|||-|||||||]PAM</td><td>NA</td><td>Intergenic</td><td>NA</td><td>NA</td></tr>
<tr><td>GU305914.1</td><td>58123</td><td>58145</td><td>-</td><td>4</td><td>GGTGGCGGCGGCGGCGGCGG</td><td>AGG</td><td>||-||-||[-|||-|||||||]PAM</td><td>NA</td><td>Intergenic</td><td>NA</td><td>NA</td></tr>
<tr><td>GU305914.1</td><td>161754</td><td>161776</td><td>-</td><td>4</td><td>GGCCGAGGAGGCGGCGGCGG</td><td>AGG</td><td>|||-|-||[-|||-|||||||]PAM</td><td>NA</td><td>Intergenic</td><td>NA</td><td>NA</td></tr>
<tr><td>GU305914.1</td><td>149482</td><td>149504</td><td>-</td><td>4</td><td>GGCGACGGCGGCGGCGGCGG</td><td>CGG</td><td>||||--||[-|||-|||||||]PAM</td><td>NA</td><td>Intergenic</td><td>NA</td><td>NA</td></tr>
<tr><td>GU305914.1</td><td>62446</td><td>62468</td><td>+</td><td>3</td><td>GGCGGGGGTGGTGGCGGCGG</td><td>GGG</td><td>|||||-||[|||--|||||||]PAM</td><td>NA</td><td>Intergenic</td><td>NA</td><td>NA</td></tr>
<tr><td>GU305914.1</td><td>18653</td><td>18675</td><td>-</td><td>4</td><td>GGCAGAGGTGGCGGAGGCGG</td><td>GGG</td><td>|||-|-||[||||-|-|||||]PAM</td><td>NA</td><td>Intergenic</td><td>NA</td><td>NA</td></tr>
<tr><td>GU305914.1</td><td>45255</td><td>45277</td><td>-</td><td>4</td><td>GGTGGCGGTGGCGGCAGCGG</td><td>CGG</td><td>||-||-||[||||-||-||||]PAM</td><td>NA</td><td>Intergenic</td><td>NA</td><td>NA</td></tr>
<tr><td>GU305914.1</td><td>56692</td><td>56714</td><td>+</td><td>4</td><td>GGATGCGGTGGCAGCGGCCG</td><td>CGG</td><td>||--|-||[||||||||||-|]PAM</td><td>NA</td><td>Intergenic</td><td>NA</td><td>NA</td></tr>
<tr><td>GU305914.1</td><td>62449</td><td>62471</td><td>+</td><td>3</td><td>GGGGGTGGTGGCGGCGGGGG</td><td>TGG</td><td>||-|||||[||||-||||-||]PAM</td><td>NA</td><td>Intergenic</td><td>NA</td><td>NA</td></tr>
<tr><td>GU305914.1</td><td>163908</td><td>163930</td><td>+</td><td>3</td><td>GGCGGCGGCGGCAGCGGCAG</td><td>CGG</td><td>|||||-||[-|||||||||-|]PAM</td><td>NA</td><td>Intergenic</td><td>NA</td><td>NA</td></tr>
<tr><td>GU305914.1</td><td>62434</td><td>62456</td><td>+</td><td>4</td><td>GACGGCGGTGGCGGCGGGGG</td><td>TGG</td><td>|-|||-||[||||-||||-||]PAM</td><td>NA</td><td>Intergenic</td><td>NA</td><td>NA</td></tr>
<tr><td>GU305914.1</td><td>113918</td><td>113940</td><td>-</td><td>4</td><td>GTCGGCGGCGGCAGCGGCGA</td><td>CGG</td><td>|-|||-||[-||||||||||-]PAM</td><td>NA</td><td>Intergenic</td><td>NA</td><td>NA</td></tr>
</table>

T12743 GGTGGTGGCAGCGGCGGCGGCGG 1000
Oligo fwd CACCGGTGGTGGCAGCGGCGGCGG
Oligo rev AAACCCGCCGCCGCTGCCACCACC

<table>
<tr><th>Chromosome</th><th>start</th><th>end</th><th>strand</th><th>MM</th><th>target_seq</th><th>PAM</th><th>alignment</th><th>distance</th><th>position</th><th>gene name</th><th>gene id</th></tr>
<tr><td>GU305914.1</td><td>168820</td><td>168842</td><td>+</td><td>0</td><td>GGTGGTGGCAGCGGCGGCGG</td><td>CGG</td><td>||||||||[||||||||||||]PAM</td><td>NA</td><td>Intergenic</td><td>NA</td><td>NA</td></tr>
<tr><td>GU305914.1</td><td>154404</td><td>154426</td><td>+</td><td>2</td><td>GGCGGTGGCGGCGGCGGCGG</td><td>CGG</td><td>||-|||||[|-||||||||||]PAM</td><td>NA</td><td>Intergenic</td><td>NA</td><td>NA</td></tr>
<tr><td>GU305914.1</td><td>58126</td><td>58148</td><td>-</td><td>2</td><td>GGCGGTGGCGGCGGCGGCGG</td><td>CGG</td><td>||-|||||[|-||||||||||]PAM</td><td>NA</td><td>Intergenic</td><td>NA</td><td>NA</td></tr>
<tr><td>GU305914.1</td><td>30000</td><td>30022</td><td>+</td><td>2</td><td>GGTGCTGGCGGCGGCGGCGG</td><td>CGG</td><td>||||-|||[|-||||||||||]PAM</td><td>NA</td><td>Intergenic</td><td>NA</td><td>NA</td></tr>
<tr><td>GU305914.1</td><td>154407</td><td>154429</td><td>+</td><td>2</td><td>GGTGGCGGCGGCGGCGGCGG</td><td>CGG</td><td>|||||-||[|-||||||||||]PAM</td><td>NA</td><td>Intergenic</td><td>NA</td><td>NA</td></tr>
<tr><td>GU305914.1</td><td>58123</td><td>58145</td><td>-</td><td>2</td><td>GGTGGCGGCGGCGGCGGCGG</td><td>AGG</td><td>|||||-||[|-||||||||||]PAM</td><td>NA</td><td>Intergenic</td><td>NA</td><td>NA</td></tr>
<tr><td>GU305914.1</td><td>73792</td><td>73814</td><td>-</td><td>4</td><td>GGCGCCAGCAGCGGCGGCGG</td><td>AGG</td><td>||-|---|[||||||||||||]PAM</td><td>NA</td><td>Intergenic</td><td>NA</td><td>NA</td></tr>
<tr><td>GU305914.1</td><td>67742</td><td>67764</td><td>+</td><td>4</td><td>GGGAGCGGCGGCGGCGGCGG</td><td>CGG</td><td>||--|-||[|-||||||||||]PAM</td><td>NA</td><td>Intergenic</td><td>NA</td><td>NA</td></tr>
<tr><td>GU305914.1</td><td>149482</td><td>149504</td><td>-</td><td>4</td><td>GGCGACGGCGGCGGCGGCGG</td><td>CGG</td><td>||-|--||[|-||||||||||]PAM</td><td>NA</td><td>Intergenic</td><td>NA</td><td>NA</td></tr>
<tr><td>GU305914.1</td><td>164974</td><td>164996</td><td>-</td><td>3</td><td>CGTGGTGGGACCGGCGGCGG</td><td>CGG</td><td>-|||||||[-|-|||||||||]PAM</td><td>NA</td><td>Intergenic</td><td>NA</td><td>NA</td></tr>
<tr><td>GU305914.1</td><td>48159</td><td>48181</td><td>+</td><td>4</td><td>GGCGGCGGTGGCGGCGGCGG</td><td>TGG</td><td>||-||-||[--||||||||||]PAM</td><td>NA</td><td>Intergenic</td><td>NA</td><td>NA</td></tr>
<tr><td>GU305914.1</td><td>164971</td><td>164993</td><td>-</td><td>4</td><td>GGTGGGACCGGCGGCGGCGG</td><td>CGG</td><td>|||||---[|-||||||||||]PAM</td><td>NA</td><td>Intergenic</td><td>NA</td><td>NA</td></tr>
<tr><td>GU305914.1</td><td>73795</td><td>73817</td><td>-</td><td>4</td><td>AGTGGCGCCAGCAGCGGCGG</td><td>CGG</td><td>-||||-|-[||||-|||||||]PAM</td><td>NA</td><td>Intergenic</td><td>NA</td><td>NA</td></tr>
<tr><td>GU305914.1</td><td>48156</td><td>48178</td><td>+</td><td>4</td><td>GGCGGCGGCGGTGGCGGCGG</td><td>CGG</td><td>||-||-||[|-|-||||||||]PAM</td><td>NA</td><td>Intergenic</td><td>NA</td><td>NA</td></tr>
<tr><td>GU305914.1</td><td>163911</td><td>163933</td><td>+</td><td>3</td><td>GGCGGCGGCAGCGGCAGCGG</td><td>AGG</td><td>||-||-||[|||||||-||||]PAM</td><td>NA</td><td>Intergenic</td><td>NA</td><td>NA</td></tr>
<tr><td>GU305914.1</td><td>62155</td><td>62177</td><td>+</td><td>4</td><td>GGCGACGGCAGCGGCAGCGG</td><td>CGG</td><td>||-|--||[|||||||-||||]PAM</td><td>NA</td><td>Intergenic</td><td>NA</td><td>NA</td></tr>
<tr><td>GU305914.1</td><td>48153</td><td>48175</td><td>+</td><td>4</td><td>TGTGGCGGCGGCGGTGGCGG</td><td>CGG</td><td>-||||-||[|-||||-|||||]PAM</td><td>NA</td><td>Intergenic</td><td>NA</td><td>NA</td></tr>
<tr><td>GU305914.1</td><td>30006</td><td>30028</td><td>+</td><td>4</td><td>GGCGGCGGCGGCGGCGGCAG</td><td>TGG</td><td>||-||-||[|-||||||||-|]PAM</td><td>NA</td><td>Intergenic</td><td>NA</td><td>NA</td></tr>
<tr><td>GU305914.1</td><td>149476</td><td>149498</td><td>-</td><td>4</td><td>GGCGGCGGCGGCGGCGGCAG</td><td>CGG</td><td>||-||-||[|-||||||||-|]PAM</td><td>NA</td><td>Intergenic</td><td>NA</td><td>NA</td></tr>
</table>

T12744 AGCGGCGGCGGCGGCGACAGCGG 1000
Oligo fwd CACCAGCGGCGGCGGCGGCGACAG
Oligo rev AAACCTGTCGCCGCCGCCGCCGCT

<table>
<tr><th>Chromosome</th><th>start</th><th>end</th><th>strand</th><th>MM</th><th>target_seq</th><th>PAM</th><th>alignment</th><th>distance</th><th>position</th><th>gene name</th><th>gene id</th></tr>
<tr><td>GU305914.1</td><td>168829</td><td>168851</td><td>+</td><td>0</td><td>AGCGGCGGCGGCGGCGACAG</td><td>CGG</td><td>||||||||[||||||||||||]PAM</td><td>NA</td><td>Intergenic</td><td>NA</td><td>NA</td></tr>
<tr><td>GU305914.1</td><td>163902</td><td>163924</td><td>+</td><td>1</td><td>AGCGGCGGCGGCGGCGGCAG</td><td>CGG</td><td>||||||||[||||||||-|||]PAM</td><td>NA</td><td>Intergenic</td><td>NA</td><td>NA</td></tr>
<tr><td>GU305914.1</td><td>30006</td><td>30028</td><td>+</td><td>2</td><td>GGCGGCGGCGGCGGCGGCAG</td><td>TGG</td><td>-|||||||[||||||||-|||]PAM</td><td>NA</td><td>Intergenic</td><td>NA</td><td>NA</td></tr>
<tr><td>GU305914.1</td><td>149476</td><td>149498</td><td>-</td><td>2</td><td>GGCGGCGGCGGCGGCGGCAG</td><td>CGG</td><td>-|||||||[||||||||-|||]PAM</td><td>NA</td><td>Intergenic</td><td>NA</td><td>NA</td></tr>
<tr><td>GU305914.1</td><td>113924</td><td>113946</td><td>-</td><td>4</td><td>GTCGGCGTCGGCGGCGGCAG</td><td>CGG</td><td>--|||||-[||||||||-|||]PAM</td><td>NA</td><td>Intergenic</td><td>NA</td><td>NA</td></tr>
<tr><td>GU305914.1</td><td>163908</td><td>163930</td><td>+</td><td>3</td><td>GGCGGCGGCGGCAGCGGCAG</td><td>CGG</td><td>-|||||||[||||-|||-|||]PAM</td><td>NA</td><td>Intergenic</td><td>NA</td><td>NA</td></tr>
<tr><td>GU305914.1</td><td>45258</td><td>45280</td><td>-</td><td>4</td><td>AACGGTGGCGGTGGCGGCAG</td><td>CGG</td><td>|-|||-||[|||-||||-|||]PAM</td><td>NA</td><td>Intergenic</td><td>NA</td><td>NA</td></tr>
<tr><td>GU305914.1</td><td>62152</td><td>62174</td><td>+</td><td>4</td><td>AGGGGCGACGGCAGCGGCAG</td><td>CGG</td><td>||-||||-[||||-|||-|||]PAM</td><td>NA</td><td>Intergenic</td><td>NA</td><td>NA</td></tr>
<tr><td>GU305914.1</td><td>154407</td><td>154429</td><td>+</td><td>4</td><td>GGTGGCGGCGGCGGCGGCGG</td><td>CGG</td><td>-|-|||||[||||||||-|-|]PAM</td><td>NA</td><td>Intergenic</td><td>NA</td><td>NA</td></tr>
<tr><td>GU305914.1</td><td>58123</td><td>58145</td><td>-</td><td>4</td><td>GGTGGCGGCGGCGGCGGCGG</td><td>AGG</td><td>-|-|||||[||||||||-|-|]PAM</td><td>NA</td><td>Intergenic</td><td>NA</td><td>NA</td></tr>
<tr><td>GU305914.1</td><td>149482</td><td>149504</td><td>-</td><td>4</td><td>GGCGACGGCGGCGGCGGCGG</td><td>CGG</td><td>-|||-|||[||||||||-|-|]PAM</td><td>NA</td><td>Intergenic</td><td>NA</td><td>NA</td></tr>
<tr><td>GU305914.1</td><td>154404</td><td>154426</td><td>+</td><td>4</td><td>GGCGGTGGCGGCGGCGGCGG</td><td>CGG</td><td>-||||-||[||||||||-|-|]PAM</td><td>NA</td><td>Intergenic</td><td>NA</td><td>NA</td></tr>
<tr><td>GU305914.1</td><td>58126</td><td>58148</td><td>-</td><td>4</td><td>GGCGGTGGCGGCGGCGGCGG</td><td>CGG</td><td>-||||-||[||||||||-|-|]PAM</td><td>NA</td><td>Intergenic</td><td>NA</td><td>NA</td></tr>
<tr><td>GU305914.1</td><td>71753</td><td>71775</td><td>+</td><td>4</td><td>AGCAGCAGCGGCGGCGGCAA</td><td>CGG</td><td>|||-||-|[||||||||-||-]PAM</td><td>NA</td><td>Intergenic</td><td>NA</td><td>NA</td></tr>
</table>

T12745 GCGGCGGCGGCGGCGACAGCGGG 1000
Oligo fwd CACCGCGGCGGCGGCGGCGACAGC
Oligo rev AAACGCTGTCGCCGCCGCCGCCGC

<table>
<tr><th>Chromosome</th><th>start</th><th>end</th><th>strand</th><th>MM</th><th>target_seq</th><th>PAM</th><th>alignment</th><th>distance</th><th>position</th><th>gene name</th><th>gene id</th></tr>
<tr><td>GU305914.1</td><td>168830</td><td>168852</td><td>+</td><td>0</td><td>GCGGCGGCGGCGGCGACAGC</td><td>GGG</td><td>||||||||[||||||||||||]PAM</td><td>NA</td><td>Intergenic</td><td>NA</td><td>NA</td></tr>
<tr><td>GU305914.1</td><td>149475</td><td>149497</td><td>-</td><td>1</td><td>GCGGCGGCGGCGGCGGCAGC</td><td>GGG</td><td>||||||||[|||||||-||||]PAM</td><td>NA</td><td>Intergenic</td><td>NA</td><td>NA</td></tr>
<tr><td>GU305914.1</td><td>128610</td><td>128632</td><td>+</td><td>4</td><td>GGGGCTTCGGCGGCGAGAGC</td><td>CGG</td><td>|-|||--|[||||||||-|||]PAM</td><td>NA</td><td>Intergenic</td><td>NA</td><td>NA</td></tr>
<tr><td>GU305914.1</td><td>65720</td><td>65742</td><td>-</td><td>4</td><td>CCTGCGGCGGCAGCGGCAGC</td><td>GGG</td><td>-|-|||||[|||-|||-||||]PAM</td><td>NA</td><td>Intergenic</td><td>NA</td><td>NA</td></tr>
<tr><td>GU305914.1</td><td>225977</td><td>225999</td><td>-</td><td>3</td><td>GCGACGGCGGCGGCGGGAGC</td><td>CGG</td><td>|||-||||[|||||||--|||]PAM</td><td>NA</td><td>Intergenic</td><td>NA</td><td>NA</td></tr>
<tr><td>GU305914.1</td><td>67743</td><td>67765</td><td>+</td><td>4</td><td>GGAGCGGCGGCGGCGGCGGC</td><td>GGG</td><td>|--|||||[|||||||-|-||]PAM</td><td>NA</td><td>Intergenic</td><td>NA</td><td>NA</td></tr>
<tr><td>GU305914.1</td><td>71754</td><td>71776</td><td>+</td><td>4</td><td>GCAGCAGCGGCGGCGGCAAC</td><td>GGG</td><td>||-||-||[|||||||-||-|]PAM</td><td>NA</td><td>Intergenic</td><td>NA</td><td>NA</td></tr>
</table>

T12746 GGCGGCGACAGCGGGCGTCTCGG 1000
Oligo fwd CACCGGCGGCGACAGCGGGCGTCT
Oligo rev AAACAGACGCCCGCTGTCGCCGCC

<table>
<tr><th>Chromosome</th><th>start</th><th>end</th><th>strand</th><th>MM</th><th>target_seq</th><th>PAM</th><th>alignment</th><th>distance</th><th>position</th><th>gene name</th><th>gene id</th></tr>
<tr><td>GU305914.1</td><td>168838</td><td>168860</td><td>+</td><td>0</td><td>GGCGGCGACAGCGGGCGTCT</td><td>CGG</td><td>||||||||[||||||||||||]PAM</td><td>NA</td><td>Intergenic</td><td>NA</td><td>NA</td></tr>
</table>

T12747 CTCGGCCCGAGTGCTAGCTGCGG 1000
Oligo fwd CACCCTCGGCCCGAGTGCTAGCTG
Oligo rev AAACCAGCTAGCACTCGGGCCGAG

<table>
<tr><th>Chromosome</th><th>start</th><th>end</th><th>strand</th><th>MM</th><th>target_seq</th><th>PAM</th><th>alignment</th><th>distance</th><th>position</th><th>gene name</th><th>gene id</th></tr>
<tr><td>GU305914.1</td><td>168856</td><td>168878</td><td>+</td><td>0</td><td>CTCGGCCCGAGTGCTAGCTG</td><td>CGG</td><td>||||||||[||||||||||||]PAM</td><td>NA</td><td>Intergenic</td><td>NA</td><td>NA</td></tr>
</table>

<table>
<tr><td>T12748</td><td>GGCCCGAGTGCTAGCTGCGGAGG</td><td>1000</td><td></td><td></td><td></td><td></td><td></td><td></td><td></td><td></td><td></td></tr>
<tr><td>Oligo fwd</td><td>CACCGGCCCGAGTGCTAGCTGCGG</td><td></td><td></td><td></td><td></td><td></td><td></td><td></td><td></td><td></td><td></td></tr>
<tr><td>Oligo rev</td><td>AAACCCGCAGCTAGCACTCGGGCC</td><td></td><td></td><td></td><td></td><td></td><td></td><td></td><td></td><td></td><td></td></tr>
<tr><td>Chromosome</td><td>start</td><td>end</td><td>strand</td><td>MM</td><td>target_seq</td><td>PAM</td><td>alignment</td><td>distance</td><td>position</td><td>gene name</td><td>gene id</td></tr>
<tr><td>GU305914.1</td><td>168859</td><td>168881</td><td>+</td><td>0</td><td>GGCCCGAGTGCTAGCTGCGG</td><td>AGG</td><td>||||||||[||||||||||||]PAM</td><td>NA</td><td>Intergenic</td><td>NA</td><td>NA</td></tr>
<tr><td>T12749</td><td>GCTAGCTGCGGAGGCAGAAGCGG</td><td>1000</td><td></td><td></td><td></td><td></td><td></td><td></td><td></td><td></td><td></td></tr>
<tr><td>Oligo fwd</td><td>CACCGCTAGCTGCGGAGGCAGAAG</td><td></td><td></td><td></td><td></td><td></td><td></td><td></td><td></td><td></td><td></td></tr>
<tr><td>Oligo rev</td><td>AAACCTTCTGCCTCCGCAGCTAGC</td><td></td><td></td><td></td><td></td><td></td><td></td><td></td><td></td><td></td><td></td></tr>
<tr><td>Chromosome</td><td>start</td><td>end</td><td>strand</td><td>MM</td><td>target_seq</td><td>PAM</td><td>alignment</td><td>distance</td><td>position</td><td>gene name</td><td>gene id</td></tr>
<tr><td>GU305914.1</td><td>168868</td><td>168890</td><td>+</td><td>0</td><td>GCTAGCTGCGGAGGCAGAAG</td><td>CGG</td><td>||||||||[||||||||||||]PAM</td><td>NA</td><td>Intergenic</td><td>NA</td><td>NA</td></tr>
<tr><td>T12750</td><td>GCGGAGGCAGAAGCGGCTGCTGG</td><td>1000</td><td></td><td></td><td></td><td></td><td></td><td></td><td></td><td></td><td></td></tr>
<tr><td>Oligo fwd</td><td>CACCGCGGAGGCAGAAGCGGCTGC</td><td></td><td></td><td></td><td></td><td></td><td></td><td></td><td></td><td></td><td></td></tr>
<tr><td>Oligo rev</td><td>AAACGCAGCCGCTTCTGCCTCCGC</td><td></td><td></td><td></td><td></td><td></td><td></td><td></td><td></td><td></td><td></td></tr>
<tr><td>Chromosome</td><td>start</td><td>end</td><td>strand</td><td>MM</td><td>target_seq</td><td>PAM</td><td>alignment</td><td>distance</td><td>position</td><td>gene name</td><td>gene id</td></tr>
<tr><td>GU305914.1</td><td>168875</td><td>168897</td><td>+</td><td>0</td><td>GCGGAGGCAGAAGCGGCTGC</td><td>TGG</td><td>||||||||[||||||||||||]PAM</td><td>NA</td><td>Intergenic</td><td>NA</td><td>NA</td></tr>
<tr><td>T12751</td><td>GCAGAAGCGGCTGCTGGCGCTGG</td><td>1000</td><td></td><td></td><td></td><td></td><td></td><td></td><td></td><td></td><td></td></tr>
<tr><td>Oligo fwd</td><td>CACCGCAGAAGCGGCTGCTGGCGC</td><td></td><td></td><td></td><td></td><td></td><td></td><td></td><td></td><td></td><td></td></tr>
<tr><td>Oligo rev</td><td>AAACGCGCCAGCAGCCGCTTCTGC</td><td></td><td></td><td></td><td></td><td></td><td></td><td></td><td></td><td></td><td></td></tr>
<tr><td>Chromosome</td><td>start</td><td>end</td><td>strand</td><td>MM</td><td>target_seq</td><td>PAM</td><td>alignment</td><td>distance</td><td>position</td><td>gene name</td><td>gene id</td></tr>
<tr><td>GU305914.1</td><td>168881</td><td>168903</td><td>+</td><td>0</td><td>GCAGAAGCGGCTGCTGGCGC</td><td>TGG</td><td>||||||||[||||||||||||]PAM</td><td>NA</td><td>Intergenic</td><td>NA</td><td>NA</td></tr>
<tr><td>GU305914.1</td><td>62684</td><td>62706</td><td>-</td><td>4</td><td>GCTGCCGCGGCTGCTGCCGC</td><td>CGG</td><td>||-|--||[||||||||-|||]PAM</td><td>NA</td><td>Intergenic</td><td>NA</td><td>NA</td></tr>
<tr><td>T12752</td><td>GGCTGCTGGCGCTGGTGCCGCGG</td><td>1000</td><td></td><td></td><td></td><td></td><td></td><td></td><td></td><td></td><td></td></tr>
<tr><td>Oligo fwd</td><td>CACCGGCTGCTGGCGCTGGTGCCG</td><td></td><td></td><td></td><td></td><td></td><td></td><td></td><td></td><td></td><td></td></tr>
<tr><td>Oligo rev</td><td>AAACCGGCACCAGCGCCAGCAGCC</td><td></td><td></td><td></td><td></td><td></td><td></td><td></td><td></td><td></td><td></td></tr>
<tr><td>Chromosome</td><td>start</td><td>end</td><td>strand</td><td>MM</td><td>target_seq</td><td>PAM</td><td>alignment</td><td>distance</td><td>position</td><td>gene name</td><td>gene id</td></tr>
<tr><td>GU305914.1</td><td>168889</td><td>168911</td><td>+</td><td>0</td><td>GGCTGCTGGCGCTGGTGCCG</td><td>CGG</td><td>||||||||[||||||||||||]PAM</td><td>NA</td><td>Intergenic</td><td>NA</td><td>NA</td></tr>
<tr><td>GU305914.1</td><td>202491</td><td>202513</td><td>-</td><td>4</td><td>GGTGGCTGGCCGTGGTGCCG</td><td>CGG</td><td>||--||||[||--||||||||]PAM</td><td>NA</td><td>Intergenic</td><td>NA</td><td>NA</td></tr>
<tr><td>T12753</td><td>GCTGCTGGCGCTGGTGCCGCGGG</td><td>1000</td><td></td><td></td><td></td><td></td><td></td><td></td><td></td><td></td><td></td></tr>
<tr><td>Oligo fwd</td><td>CACCGCTGCTGGCGCTGGTGCCGC</td><td></td><td></td><td></td><td></td><td></td><td></td><td></td><td></td><td></td><td></td></tr>
<tr><td>Oligo rev</td><td>AAACGCGGCACCAGCGCCAGCAGC</td><td></td><td></td><td></td><td></td><td></td><td></td><td></td><td></td><td></td><td></td></tr>
<tr><td>Chromosome</td><td>start</td><td>end</td><td>strand</td><td>MM</td><td>target_seq</td><td>PAM</td><td>alignment</td><td>distance</td><td>position</td><td>gene name</td><td>gene id</td></tr>
<tr><td>GU305914.1</td><td>168890</td><td>168912</td><td>+</td><td>0</td><td>GCTGCTGGCGCTGGTGCCGC</td><td>GGG</td><td>||||||||[||||||||||||]PAM</td><td>NA</td><td>Intergenic</td><td>NA</td><td>NA</td></tr>
<tr><td>GU305914.1</td><td>202490</td><td>202512</td><td>-</td><td>4</td><td>GTGGCTGGCCGTGGTGCCGC</td><td>GGG</td><td>|--|||||[|--|||||||||]PAM</td><td>NA</td><td>Intergenic</td><td>NA</td><td>NA</td></tr>
<tr><td>GU305914.1</td><td>62684</td><td>62706</td><td>-</td><td>4</td><td>GCTGCCGCGGCTGCTGCCGC</td><td>CGG</td><td>|||||-|-[-||||-||||||]PAM</td><td>NA</td><td>Intergenic</td><td>NA</td><td>NA</td></tr>
<tr><td>T12754</td><td>GCTGGCGCTGGTGCCGCGGGCGG</td><td>1000</td><td></td><td></td><td></td><td></td><td></td><td></td><td></td><td></td><td></td></tr>
<tr><td>Oligo fwd</td><td>CACCGCTGGCGCTGGTGCCGCGGG</td><td></td><td></td><td></td><td></td><td></td><td></td><td></td><td></td><td></td><td></td></tr>
<tr><td>Oligo rev</td><td>AAACCCCGCGGCACCAGCGCCAGC</td><td></td><td></td><td></td><td></td><td></td><td></td><td></td><td></td><td></td><td></td></tr>
<tr><td>Chromosome</td><td>start</td><td>end</td><td>strand</td><td>MM</td><td>target_seq</td><td>PAM</td><td>alignment</td><td>distance</td><td>position</td><td>gene name</td><td>gene id</td></tr>
<tr><td>GU305914.1</td><td>168893</td><td>168915</td><td>+</td><td>0</td><td>GCTGGCGCTGGTGCCGCGGG</td><td>CGG</td><td>||||||||[||||||||||||]PAM</td><td>NA</td><td>Intergenic</td><td>NA</td><td>NA</td></tr>
<tr><td>GU305914.1</td><td>143022</td><td>143044</td><td>+</td><td>4</td><td>GCTGGTGGTGGCGCCGGGGG</td><td>CGG</td><td>|||||-|-[|||-||||-|||]PAM</td><td>NA</td><td>Intergenic</td><td>NA</td><td>NA</td></tr>
<tr><td>T12755</td><td>GCTGGTGCCGCGGGCGGAGCCGG</td><td>1000</td><td></td><td></td><td></td><td></td><td></td><td></td><td></td><td></td><td></td></tr>
<tr><td>Oligo fwd</td><td>CACCGCTGGTGCCGCGGGCGGAGC</td><td></td><td></td><td></td><td></td><td></td><td></td><td></td><td></td><td></td><td></td></tr>
<tr><td>Oligo rev</td><td>AAACGCTCCGCCCGCGGCACCAGC</td><td></td><td></td><td></td><td></td><td></td><td></td><td></td><td></td><td></td><td></td></tr>
<tr><td>Chromosome</td><td>start</td><td>end</td><td>strand</td><td>MM</td><td>target_seq</td><td>PAM</td><td>alignment</td><td>distance</td><td>position</td><td>gene name</td><td>gene id</td></tr>
<tr><td>GU305914.1</td><td>168899</td><td>168921</td><td>+</td><td>0</td><td>GCTGGTGCCGCGGGCGGAGC</td><td>CGG</td><td>||||||||[||||||||||||]PAM</td><td>NA</td><td>Intergenic</td><td>NA</td><td>NA</td></tr>
<tr><td>T12756</td><td>TGCCGCGGGCGGAGCCGGCGCGG</td><td>1000</td><td></td><td></td><td></td><td></td><td></td><td></td><td></td><td></td><td></td></tr>
<tr><td>Oligo fwd</td><td>CACCTGCCGCGGGCGGAGCCGGCG</td><td></td><td></td><td></td><td></td><td></td><td></td><td></td><td></td><td></td><td></td></tr>
<tr><td>Oligo rev</td><td>AAACCGCCGGCTCCGCCCGCGGCA</td><td></td><td></td><td></td><td></td><td></td><td></td><td></td><td></td><td></td><td></td></tr>
<tr><td>Chromosome</td><td>start</td><td>end</td><td>strand</td><td>MM</td><td>target_seq</td><td>PAM</td><td>alignment</td><td>distance</td><td>position</td><td>gene name</td><td>gene id</td></tr>
<tr><td>GU305914.1</td><td>168904</td><td>168926</td><td>+</td><td>0</td><td>TGCCGCGGGCGGAGCCGGCG</td><td>CGG</td><td>||||||||[||||||||||||]PAM</td><td>NA</td><td>Intergenic</td><td>NA</td><td>NA</td></tr>
<tr><td>T12757</td><td>GCCGTTCTTGCCCGCTTGTGCGG</td><td>1000</td><td></td><td></td><td></td><td></td><td></td><td></td><td></td><td></td><td></td></tr>
<tr><td>Oligo fwd</td><td>CACCGCCGTTCTTGCCCGCTTGTG</td><td></td><td></td><td></td><td></td><td></td><td></td><td></td><td></td><td></td><td></td></tr>
<tr><td>Oligo rev</td><td>AAACCACAAGCGGGCAAGAACGGC</td><td></td><td></td><td></td><td></td><td></td><td></td><td></td><td></td><td></td><td></td></tr>
<tr><td>Chromosome</td><td>start</td><td>end</td><td>strand</td><td>MM</td><td>target_seq</td><td>PAM</td><td>alignment</td><td>distance</td><td>position</td><td>gene name</td><td>gene id</td></tr>
<tr><td>GU305914.1</td><td>168943</td><td>168965</td><td>+</td><td>0</td><td>GCCGTTCTTGCCCGCTTGTG</td><td>CGG</td><td>||||||||[||||||||||||]PAM</td><td>NA</td><td>Intergenic</td><td>NA</td><td>NA</td></tr>
<tr><td>T12758</td><td>CCCGCTTGTGCGGTTTGTGCAGG</td><td>1000</td><td></td><td></td><td></td><td></td><td></td><td></td><td></td><td></td><td></td></tr>
<tr><td>Oligo fwd</td><td>CACCCCCGCTTGTGCGGTTTGTGC</td><td></td><td></td><td></td><td></td><td></td><td></td><td></td><td></td><td></td><td></td></tr>
<tr><td>Oligo rev</td><td>AAACGCACAAACCGCACAAGCGGG</td><td></td><td></td><td></td><td></td><td></td><td></td><td></td><td></td><td></td><td></td></tr>
<tr><td>Chromosome</td><td>start</td><td>end</td><td>strand</td><td>MM</td><td>target_seq</td><td>PAM</td><td>alignment</td><td>distance</td><td>position</td><td>gene name</td><td>gene id</td></tr>
<tr><td>GU305914.1</td><td>168953</td><td>168975</td><td>+</td><td>0</td><td>CCCGCTTGTGCGGTTTGTGC</td><td>AGG</td><td>||||||||[||||||||||||]PAM</td><td>NA</td><td>Intergenic</td><td>NA</td><td>NA</td></tr>
<tr><td>T12759</td><td>CCGCTTGTGCGGTTTGTGCAGGG</td><td>1000</td><td></td><td></td><td></td><td></td><td></td><td></td><td></td><td></td><td></td></tr>
<tr><td>Oligo fwd</td><td>CACCCCGCTTGTGCGGTTTGTGCA</td><td></td><td></td><td></td><td></td><td></td><td></td><td></td><td></td><td></td><td></td></tr>
<tr><td>Oligo rev</td><td>AAACTGCACAAACCGCACAAGCGG</td><td></td><td></td><td></td><td></td><td></td><td></td><td></td><td></td><td></td><td></td></tr>
<tr><td>Chromosome</td><td>start</td><td>end</td><td>strand</td><td>MM</td><td>target_seq</td><td>PAM</td><td>alignment</td><td>distance</td><td>position</td><td>gene name</td><td>gene id</td></tr>
<tr><td>GU305914.1</td><td>168954</td><td>168976</td><td>+</td><td>0</td><td>CCGCTTGTGCGGTTTGTGCA</td><td>GGG</td><td>||||||||[||||||||||||]PAM</td><td>NA</td><td>Intergenic</td><td>NA</td><td>NA</td></tr>
<tr><td>T12760</td><td>TGTGCGGTTTGTGCAGGGTTCGG</td><td>1000</td><td></td><td></td><td></td><td></td><td></td><td></td><td></td><td></td><td></td></tr>
<tr><td>Oligo fwd</td><td>CACCTGTGCGGTTTGTGCAGGGTT</td><td></td><td></td><td></td><td></td><td></td><td></td><td></td><td></td><td></td><td></td></tr>
<tr><td>Oligo rev</td><td>AAACAACCCTGCACAAACCGCACA</td><td></td><td></td><td></td><td></td><td></td><td></td><td></td><td></td><td></td><td></td></tr>
<tr><td>Chromosome</td><td>start</td><td>end</td><td>strand</td><td>MM</td><td>target_seq</td><td>PAM</td><td>alignment</td><td>distance</td><td>position</td><td>gene name</td><td>gene id</td></tr>
<tr><td>GU305914.1</td><td>168959</td><td>168981</td><td>+</td><td>0</td><td>TGTGCGGTTTGTGCAGGGTT</td><td>CGG</td><td>||||||||[||||||||||||]PAM</td><td>NA</td><td>Intergenic</td><td>NA</td><td>NA</td></tr>
<tr><td>T12761</td><td>CGACGACATCGTGCTCTTTTCGG</td><td>1000</td><td></td><td></td><td></td><td></td><td></td><td></td><td></td><td></td><td></td></tr>
<tr><td>Oligo fwd</td><td>CACCCGACGACATCGTGCTCTTTT</td><td></td><td></td><td></td><td></td><td></td><td></td><td></td><td></td><td></td><td></td></tr>
<tr><td>Oligo rev</td><td>AAACAAAAGAGCACGATGTCGTCG</td><td></td><td></td><td></td><td></td><td></td><td></td><td></td><td></td><td></td><td></td></tr>
<tr><td>Chromosome</td><td>start</td><td>end</td><td>strand</td><td>MM</td><td>target_seq</td><td>PAM</td><td>alignment</td><td>distance</td><td>position</td><td>gene name</td><td>gene id</td></tr>
<tr><td>GU305914.1</td><td>168982</td><td>169004</td><td>+</td><td>0</td><td>CGACGACATCGTGCTCTTTT</td><td>CGG</td><td>||||||||[||||||||||||]PAM</td><td>NA</td><td>Intergenic</td><td>NA</td><td>NA</td></tr>
</table>

T12762 GACGACATCGTGCTCTTTTCGGG 1000
Oligo fwd CACCGACGACATCGTGCTCTTTTC
Oligo rev AAACGAAAAGAGCACGATGTCGTC
Chromosome start end strand MM target_seq PAM alignment distance position gene name gene id
GU305914.1 168983 169005 + 0 GACGACATCGTGCTCTTTTC GGG ||||||||[||||||||||||]PAM NA Intergenic NA NA
GU305914.1 71360 71382 - 4 GCCGCCGTCGTGCTCTCTTC AGG |-||-|-|[||||||||-|||]PAM NA Intergenic NA NA

T12763 GGGCGCTGCGTCTGAAGAATTGG 1000
Oligo fwd CACCGGGCGCTGCGTCTGAAGAAT
Oligo rev AAACATTCTTCAGACGCAGCGCCC
Chromosome start end strand MM target_seq PAM alignment distance position gene name gene id
GU305914.1 169003 169025 + 0 GGGCGCTGCGTCTGAAGAAT TGG ||||||||[||||||||||||]PAM NA Intergenic NA NA

T12764 GTCTGAAGAATTGGCGTTCGCGG 1000
Oligo fwd CACCGTCTGAAGAATTGGCGTTCG
Oligo rev AAACCGAACGCCAATTCTTCAGAC
Chromosome start end strand MM target_seq PAM alignment distance position gene name gene id
GU305914.1 169012 169034 + 0 GTCTGAAGAATTGGCGTTCG CGG ||||||||[||||||||||||]PAM NA Intergenic NA NA

T12765 GAAGAATTGGCGTTCGCGGCCGG 1000
Oligo fwd CACCGAAGAATTGGCGTTCGCGGC
Oligo rev AAACGCCGCGAACGCCAATTCTTC
Chromosome start end strand MM target_seq PAM alignment distance position gene name gene id
GU305914.1 169016 169038 + 0 GAAGAATTGGCGTTCGCGGC CGG ||||||||[||||||||||||]PAM NA Intergenic NA NA

T12766 GGCGTTCGCGGCCGGTTGAGAGG 1000
Oligo fwd CACCGGCGTTCGCGGCCGGTTGAG
Oligo rev AAACCTCAACCGGCCGCGAACGCC
Chromosome start end strand MM target_seq PAM alignment distance position gene name gene id
GU305914.1 169024 169046 + 0 GGCGTTCGCGGCCGGTTGAG AGG ||||||||[||||||||||||]PAM NA Intergenic NA NA

T12767 GCGGCCGGTTGAGAGGTGCCTGG 1000
Oligo fwd CACCGCGGCCGGTTGAGAGGTGCC
Oligo rev AAACGGCACCTCTCAACCGGCCGC
Chromosome start end strand MM target_seq PAM alignment distance position gene name gene id
GU305914.1 169031 169053 + 0 GCGGCCGGTTGAGAGGTGCC TGG ||||||||[||||||||||||]PAM NA Intergenic NA NA

T12768 TCTCAGTCCTCTATCCAGCTTGG 1000
Oligo fwd CACCTCTCAGTCCTCTATCCAGCT
Oligo rev AAACAGCTGGATAGAGGACTGAGA
Chromosome start end strand MM target_seq PAM alignment distance position gene name gene id
GU305914.1 169069 169091 + 0 TCTCAGTCCTCTATCCAGCT TGG ||||||||[||||||||||||]PAM NA Intergenic NA NA

T12769 CTCAGTCCTCTATCCAGCTTGGG 1000
Oligo fwd CACCCTCAGTCCTCTATCCAGCTT
Oligo rev AAACAAGCTGGATAGAGGACTGAG
Chromosome start end strand MM target_seq PAM alignment distance position gene name gene id
GU305914.1 169070 169092 + 0 CTCAGTCCTCTATCCAGCTT GGG ||||||||[||||||||||||]PAM NA Intergenic NA NA

T12770 TCTATCCAGCTTGGGCTGTTTGG 1000
Oligo fwd CACCTCTATCCAGCTTGGGCTGTT
Oligo rev AAACAACAGCCCAAGCTGGATAGA
Chromosome start end strand MM target_seq PAM alignment distance position gene name gene id
GU305914.1 169078 169100 + 0 TCTATCCAGCTTGGGCTGTT TGG ||||||||[||||||||||||]PAM NA Intergenic NA NA

T12771 CTATCCAGCTTGGGCTGTTTGGG 1000
Oligo fwd CACCCTATCCAGCTTGGGCTGTTT
Oligo rev AAACAAACAGCCCAAGCTGGATAG
Chromosome start end strand MM target_seq PAM alignment distance position gene name gene id
GU305914.1 169079 169101 + 0 CTATCCAGCTTGGGCTGTTT GGG ||||||||[||||||||||||]PAM NA Intergenic NA NA

T12772 TTGGGCTGTTTGGGTATGCTCGG 1000
Oligo fwd CACCTTGGGCTGTTTGGGTATGCT
Oligo rev AAACAGCATACCCAAACAGCCCAA
Chromosome start end strand MM target_seq PAM alignment distance position gene name gene id
GU305914.1 169088 169110 + 0 TTGGGCTGTTTGGGTATGCT CGG ||||||||[||||||||||||]PAM NA Intergenic NA NA

T12773 TTTGGGTATGCTCGGCTGCTTGG 1000
Oligo fwd CACCTTTGGGTATGCTCGGCTGCT
Oligo rev AAACAGCAGCCGAGCATACCCAAA
Chromosome start end strand MM target_seq PAM alignment distance position gene name gene id
GU305914.1 169096 169118 + 0 TTTGGGTATGCTCGGCTGCT TGG ||||||||[||||||||||||]PAM NA Intergenic NA NA

T12774 GGGTATGCTCGGCTGCTTGGAGG 1000
Oligo fwd CACCGGGTATGCTCGGCTGCTTGG
Oligo rev AAACCCAAGCAGCCGAGCATACCC
Chromosome start end strand MM target_seq PAM alignment distance position gene name gene id
GU305914.1 169099 169121 + 0 GGGTATGCTCGGCTGCTTGG AGG ||||||||[||||||||||||]PAM NA Intergenic NA NA

T12775 GCTCGGCTGCTTGGAGGTCACGG 1000
Oligo fwd CACCGCTCGGCTGCTTGGAGGTCA
Oligo rev AAACTGACCTCCAAGCAGCCGAGC
Chromosome start end strand MM target_seq PAM alignment distance position gene name gene id
GU305914.1 169105 169127 + 0 GCTCGGCTGCTTGGAGGTCA CGG ||||||||[||||||||||||]PAM NA Intergenic NA NA

T12776 TTTTGAGCTCGCCCTCCACCAGG 1000
Oligo fwd CACCTTTTGAGCTCGCCCTCCACC
Oligo rev AAACGGTGGAGGGCGAGCTCAAAA
Chromosome start end strand MM target_seq PAM alignment distance position gene name gene id

GU305914.1 169130 169152 + 0 TTTTGAGCTCGCCCTCCACC AGG ||||||||[||||||||||||]PAM NA Intergenic NA NA

T12777 CTCCACCAGGCGAATGAGTCTGG 1000
Oligo fwd CACCCTCCACCAGGCGAATGAGTC
Oligo rev AAACGACTCATTCGCCTGGTGGAG
Chromosome start end strand MM target_seq PAM alignment distance position gene name gene id
GU305914.1 169143 169165 + 0 CTCCACCAGGCGAATGAGTC TGG ||||||||[||||||||||||]PAM NA Intergenic NA NA

T12778 CACCAGGCGAATGAGTCTGGAGG 1000
Oligo fwd CACCCACCAGGCGAATGAGTCTGG
Oligo rev AAACCCAGACTCATTCGCCTGGTG
Chromosome start end strand MM target_seq PAM alignment distance position gene name gene id
GU305914.1 169146 169168 + 0 CACCAGGCGAATGAGTCTGG AGG ||||||||[||||||||||||]PAM NA Intergenic NA NA

T12779 GTCTGGAGGTGCTTGACATCCGG 1000
Oligo fwd CACCGTCTGGAGGTGCTTGACATC
Oligo rev AAACGATGTCAAGCACCTCCAGAC
Chromosome start end strand MM target_seq PAM alignment distance position gene name gene id
GU305914.1 169160 169182 + 0 GTCTGGAGGTGCTTGACATC CGG ||||||||[||||||||||||]PAM NA Intergenic NA NA

T12780 GACATCCGGTGTTATGTATACGG 1000
Oligo fwd CACCGACATCCGGTGTTATGTATA
Oligo rev AAACTATACATAACACCGGATGTC
Chromosome start end strand MM target_seq PAM alignment distance position gene name gene id
GU305914.1 169174 169196 + 0 GACATCCGGTGTTATGTATA CGG ||||||||[||||||||||||]PAM NA Intergenic NA NA

T12781 TGTTATGTATACGGCCGTCACGG 1000
Oligo fwd CACCTGTTATGTATACGGCCGTCA
Oligo rev AAACTGACGGCCGTATACATAACA
Chromosome start end strand MM target_seq PAM alignment distance position gene name gene id
GU305914.1 169183 169205 + 0 TGTTATGTATACGGCCGTCA CGG ||||||||[||||||||||||]PAM NA Intergenic NA NA

T12782 AACATGATGTCGTGCCCTCACGG 1000
Oligo fwd CACCAACATGATGTCGTGCCCTCA
Oligo rev AAACTGAGGGCACGACATCATGTT
Chromosome start end strand MM target_seq PAM alignment distance position gene name gene id
GU305914.1 169238 169260 + 0 AACATGATGTCGTGCCCTCA CGG ||||||||[||||||||||||]PAM NA Intergenic NA NA

T12783 TCACGGCGAGCGTGTCCGAGAGG 1000
Oligo fwd CACCTCACGGCGAGCGTGTCCGAG
Oligo rev AAACCTCGGACACGCTCGCCGTGA
Chromosome start end strand MM target_seq PAM alignment distance position gene name gene id
GU305914.1 169255 169277 + 0 TCACGGCGAGCGTGTCCGAG AGG ||||||||[||||||||||||]PAM NA Intergenic NA NA

T12784 CCGAGAGGTTAAATAGTTGATGG 1000
Oligo fwd CACCCCGAGAGGTTAAATAGTTGA
Oligo rev AAACTCAACTATTTAACCTCTCGG
Chromosome start end strand MM target_seq PAM alignment distance position gene name gene id
GU305914.1 169270 169292 + 0 CCGAGAGGTTAAATAGTTGA TGG ||||||||[||||||||||||]PAM NA Intergenic NA NA

T12785 CGAGAGGTTAAATAGTTGATGGG 1000
Oligo fwd CACCCGAGAGGTTAAATAGTTGAT
Oligo rev AAACATCAACTATTTAACCTCTCG
Chromosome start end strand MM target_seq PAM alignment distance position gene name gene id
GU305914.1 169271 169293 + 0 CGAGAGGTTAAATAGTTGAT GGG ||||||||[||||||||||||]PAM NA Intergenic NA NA

T12786 GAGAGGTTAAATAGTTGATGGGG 1000
Oligo fwd CACCGAGAGGTTAAATAGTTGATG
Oligo rev AAACCATCAACTATTTAACCTCTC
Chromosome start end strand MM target_seq PAM alignment distance position gene name gene id
GU305914.1 169272 169294 + 0 GAGAGGTTAAATAGTTGATG GGG ||||||||[||||||||||||]PAM NA Intergenic NA NA

T12787 AGAGGTTAAATAGTTGATGGGGG 1000
Oligo fwd CACCAGAGGTTAAATAGTTGATGG
Oligo rev AAACCCATCAACTATTTAACCTCT
Chromosome start end strand MM target_seq PAM alignment distance position gene name gene id
GU305914.1 169273 169295 + 0 AGAGGTTAAATAGTTGATGG GGG ||||||||[||||||||||||]PAM NA Intergenic NA NA

T12788 TAAATAGTTGATGGGGGTGACGG 1000
Oligo fwd CACCTAAATAGTTGATGGGGGTGA
Oligo rev AAACTCACCCCCATCAACTATTTA
Chromosome start end strand MM target_seq PAM alignment distance position gene name gene id
GU305914.1 169279 169301 + 0 TAAATAGTTGATGGGGGTGA CGG ||||||||[||||||||||||]PAM NA Intergenic NA NA

T12789 AAATAGTTGATGGGGGTGACGGG 1000
Oligo fwd CACCAAATAGTTGATGGGGGTGAC
Oligo rev AAACGTCACCCCCATCAACTATTT
Chromosome start end strand MM target_seq PAM alignment distance position gene name gene id
GU305914.1 169280 169302 + 0 AAATAGTTGATGGGGGTGAC GGG ||||||||[||||||||||||]PAM NA Intergenic NA NA

T12790 AATAGTTGATGGGGGTGACGGGG 1000
Oligo fwd CACCAATAGTTGATGGGGGTGACG
Oligo rev AAACCGTCACCCCCATCAACTATT
Chromosome start end strand MM target_seq PAM alignment distance position gene name gene id
GU305914.1 169281 169303 + 0 AATAGTTGATGGGGGTGACG GGG ||||||||[||||||||||||]PAM NA Intergenic NA NA

T12791 GCATTTGATACCCGTCTTCTTGG 1000
Oligo fwd CACCGCATTTGATACCCGTCTTCT
Oligo rev AAACAGAAGACGGGTATCAAATGC

Chromosome start end strand MM target_seq PAM alignment distance position gene name gene id
GU305914.1 169352 169374 + 0 GCATTTGATACCCGTCTTCT TGG ||||||||[||||||||||||]PAM NA Intergenic NA NA

T12792 TGGCCACGTCCTGACAGACGCGG 1000
Oligo fwd CACCTGGCCACGTCCTGACAGACG
Oligo rev AAACCGTCTGTCAGGACGTGGCCA
Chromosome start end strand MM target_seq PAM alignment distance position gene name gene id
GU305914.1 169372 169394 + 0 TGGCCACGTCCTGACAGACG CGG ||||||||[||||||||||||]PAM NA Intergenic NA NA

T12793 GGCCACGTCCTGACAGACGCGGG 1000
Oligo fwd CACCGGCCACGTCCTGACAGACGC
Oligo rev AAACGCGTCTGTCAGGACGTGGCC
Chromosome start end strand MM target_seq PAM alignment distance position gene name gene id
GU305914.1 169373 169395 + 0 GGCCACGTCCTGACAGACGC GGG ||||||||[||||||||||||]PAM NA Intergenic NA NA

T12794 GTCCTGACAGACGCGGGCCGCGG 1000
Oligo fwd CACCGTCCTGACAGACGCGGGCCG
Oligo rev AAACCGGCCCGCGTCTGTCAGGAC
Chromosome start end strand MM target_seq PAM alignment distance position gene name gene id
GU305914.1 169379 169401 + 0 GTCCTGACAGACGCGGGCCG CGG ||||||||[||||||||||||]PAM NA Intergenic NA NA

T12795 GAGCGCACGATCACTTCGTACGG 1000
Oligo fwd CACCGAGCGCACGATCACTTCGTA
Oligo rev AAACTACGAAGTGATCGTGCGCTC
Chromosome start end strand MM target_seq PAM alignment distance position gene name gene id
GU305914.1 169425 169447 + 0 GAGCGCACGATCACTTCGTA CGG ||||||||[||||||||||||]PAM NA Intergenic NA NA

T12796 AGCGCACGATCACTTCGTACGGG 1000
Oligo fwd CACCAGCGCACGATCACTTCGTAC
Oligo rev AAACGTACGAAGTGATCGTGCGCT
Chromosome start end strand MM target_seq PAM alignment distance position gene name gene id
GU305914.1 169426 169448 + 0 AGCGCACGATCACTTCGTAC GGG ||||||||[||||||||||||]PAM NA Intergenic NA NA

T12797 CACGATCACTTCGTACGGGCCGG 1000
Oligo fwd CACCCACGATCACTTCGTACGGGC
Oligo rev AAACGCCCGTACGAAGTGATCGTG
Chromosome start end strand MM target_seq PAM alignment distance position gene name gene id
GU305914.1 169430 169452 + 0 CACGATCACTTCGTACGGGC CGG ||||||||[||||||||||||]PAM NA Intergenic NA NA

T12798 TGAGTGTTCGCCGCCGTCCTCGG 1000
Oligo fwd CACCTGAGTGTTCGCCGCCGTCCT
Oligo rev AAACAGGACGGCGGCGAACACTCA
Chromosome start end strand MM target_seq PAM alignment distance position gene name gene id
GU305914.1 169461 169483 + 0 TGAGTGTTCGCCGCCGTCCT CGG ||||||||[||||||||||||]PAM NA Intergenic NA NA

T12799 TGTTCGCCGCCGTCCTCGGTTGG 1000
Oligo fwd CACCTGTTCGCCGCCGTCCTCGGT
Oligo rev AAACACCGAGGACGGCGGCGAACA
Chromosome start end strand MM target_seq PAM alignment distance position gene name gene id
GU305914.1 169465 169487 + 0 TGTTCGCCGCCGTCCTCGGT TGG ||||||||[||||||||||||]PAM NA Intergenic NA NA

T12800 GTAGTTGATCTCGTCGAACTTGG 1000
Oligo fwd CACCGTAGTTGATCTCGTCGAACT
Oligo rev AAACAGTTCGACGAGATCAACTAC
Chromosome start end strand MM target_seq PAM alignment distance position gene name gene id
GU305914.1 169487 169509 + 0 GTAGTTGATCTCGTCGAACT TGG ||||||||[||||||||||||]PAM NA Intergenic NA NA

T12801 TAGTTGATCTCGTCGAACTTGGG 1000
Oligo fwd CACCTAGTTGATCTCGTCGAACTT
Oligo rev AAACAAGTTCGACGAGATCAACTA
Chromosome start end strand MM target_seq PAM alignment distance position gene name gene id
GU305914.1 169488 169510 + 0 TAGTTGATCTCGTCGAACTT GGG ||||||||[||||||||||||]PAM NA Intergenic NA NA

T12802 ACTTGGGTTTCTCGTGACAGCGG 1000
Oligo fwd CACCACTTGGGTTTCTCGTGACAG
Oligo rev AAACCTGTCACGAGAAACCCAAGT
Chromosome start end strand MM target_seq PAM alignment distance position gene name gene id
GU305914.1 169504 169526 + 0 ACTTGGGTTTCTCGTGACAG CGG ||||||||[||||||||||||]PAM NA Intergenic NA NA

T12803 GCGCGCCCATCACGATCTTCAGG 1000
Oligo fwd CACCGCGCGCCCATCACGATCTTC
Oligo rev AAACGAAGATCGTGATGGGCGCGC
Chromosome start end strand MM target_seq PAM alignment distance position gene name gene id
GU305914.1 169546 169568 + 0 GCGCGCCCATCACGATCTTC AGG ||||||||[||||||||||||]PAM NA Intergenic NA NA

T12804 GCCCATCACGATCTTCAGGTTGG 1000
Oligo fwd CACCGCCCATCACGATCTTCAGGT
Oligo rev AAACACCTGAAGATCGTGATGGGC
Chromosome start end strand MM target_seq PAM alignment distance position gene name gene id
GU305914.1 169550 169572 + 0 GCCCATCACGATCTTCAGGT TGG ||||||||[||||||||||||]PAM NA Intergenic NA NA

T12805 CCCATCACGATCTTCAGGTTGGG 1000
Oligo fwd CACCCCCATCACGATCTTCAGGTT
Oligo rev AAACAACCTGAAGATCGTGATGGG
Chromosome start end strand MM target_seq PAM alignment distance position gene name gene id
GU305914.1 169551 169573 + 0 CCCATCACGATCTTCAGGTT GGG ||||||||[||||||||||||]PAM NA Intergenic NA NA

T12806 CCATCACGATCTTCAGGTTGGGG 1000
Oligo fwd CACCCCATCACGATCTTCAGGTTG

| | | | | | | | | | | | |
|---|---|---|---|---|---|---|---|---|---|---|---|
| Oligo rev | AAACCAACCTGAAGATCGTGATGG | | | | | | | | | | |
| Chromosome | start | end | strand | MM | target_seq | PAM | alignment | distance | position | gene name | gene id |
| GU305914.1 | 169552 | 169574 | + | 0 | CCATCACGATCTTCAGGTTG | GGG | \|\|\|\|\|\|\|\|[\|\|\|\|\|\|\|\|\|\|\|\|]PAM | NA | Intergenic | NA | NA |
| T12807 | GAAACTTCTGACACGAGAAGTGG | 1000 | | | | | | | | | |
| Oligo fwd | CACCGAAACTTCTGACACGAGAAG | | | | | | | | | | |
| Oligo rev | AAACCTTCTCGTGTCAGAAGTTTC | | | | | | | | | | |
| Chromosome | start | end | strand | MM | target_seq | PAM | alignment | distance | position | gene name | gene id |
| GU305914.1 | 169612 | 169634 | + | 0 | GAAACTTCTGACACGAGAAG | TGG | \|\|\|\|\|\|\|\|[\|\|\|\|\|\|\|\|\|\|\|\|]PAM | NA | Intergenic | NA | NA |
| T12808 | GATCCGTATGTCCGAGACGTCGG | 1000 | | | | | | | | | |
| Oligo fwd | CACCGATCCGTATGTCCGAGACGT | | | | | | | | | | |
| Oligo rev | AAACACGTCTCGGACATACGGATC | | | | | | | | | | |
| Chromosome | start | end | strand | MM | target_seq | PAM | alignment | distance | position | gene name | gene id |
| GU305914.1 | 169634 | 169656 | + | 0 | GATCCGTATGTCCGAGACGT | CGG | \|\|\|\|\|\|\|\|[\|\|\|\|\|\|\|\|\|\|\|\|]PAM | NA | Intergenic | NA | NA |
| T12809 | CGGTATGCCTGTCGCTGCACAGG | 1000 | | | | | | | | | |
| Oligo fwd | CACCCGGTATGCCTGTCGCTGCAC | | | | | | | | | | |
| Oligo rev | AAACGTGCAGCGACAGGCATACCG | | | | | | | | | | |
| Chromosome | start | end | strand | MM | target_seq | PAM | alignment | distance | position | gene name | gene id |
| GU305914.1 | 169654 | 169676 | + | 0 | CGGTATGCCTGTCGCTGCAC | AGG | \|\|\|\|\|\|\|\|[\|\|\|\|\|\|\|\|\|\|\|\|]PAM | NA | Intergenic | NA | NA |
| T12810 | TCGTTTCCGAACTCTTCGTCCGG | 1000 | | | | | | | | | |
| Oligo fwd | CACCTCGTTTCCGAACTCTTCGTC | | | | | | | | | | |
| Oligo rev | AAACGACGAAGAGTTCGGAAACGA | | | | | | | | | | |
| Chromosome | start | end | strand | MM | target_seq | PAM | alignment | distance | position | gene name | gene id |
| GU305914.1 | 169698 | 169720 | + | 0 | TCGTTTCCGAACTCTTCGTC | CGG | \|\|\|\|\|\|\|\|[\|\|\|\|\|\|\|\|\|\|\|\|]PAM | NA | Intergenic | NA | NA |
| T12811 | CGTTTCCGAACTCTTCGTCCGGG | 1000 | | | | | | | | | |
| Oligo fwd | CACCCGTTTCCGAACTCTTCGTCC | | | | | | | | | | |
| Oligo rev | AAACGGACGAAGAGTTCGGAAACG | | | | | | | | | | |
| Chromosome | start | end | strand | MM | target_seq | PAM | alignment | distance | position | gene name | gene id |
| GU305914.1 | 169699 | 169721 | + | 0 | CGTTTCCGAACTCTTCGTCC | GGG | \|\|\|\|\|\|\|\|[\|\|\|\|\|\|\|\|\|\|\|\|]PAM | NA | Intergenic | NA | NA |
| T12812 | GTTTCCGAACTCTTCGTCCGGGG | 1000 | | | | | | | | | |
| Oligo fwd | CACCGTTTCCGAACTCTTCGTCCG | | | | | | | | | | |
| Oligo rev | AAACCGGACGAAGAGTTCGGAAAC | | | | | | | | | | |
| Chromosome | start | end | strand | MM | target_seq | PAM | alignment | distance | position | gene name | gene id |
| GU305914.1 | 169700 | 169722 | + | 0 | GTTTCCGAACTCTTCGTCCG | GGG | \|\|\|\|\|\|\|\|[\|\|\|\|\|\|\|\|\|\|\|\|]PAM | NA | Intergenic | NA | NA |
| T12813 | TTCGTCCGGGGCGCCTTCCGCGG | 1000 | | | | | | | | | |
| Oligo fwd | CACCTTCGTCCGGGGCGCCTTCCG | | | | | | | | | | |
| Oligo rev | AAACCGGAAGGCGCCCCGGACGAA | | | | | | | | | | |
| Chromosome | start | end | strand | MM | target_seq | PAM | alignment | distance | position | gene name | gene id |
| GU305914.1 | 169712 | 169734 | + | 0 | TTCGTCCGGGGCGCCTTCCG | CGG | \|\|\|\|\|\|\|\|[\|\|\|\|\|\|\|\|\|\|\|\|]PAM | NA | Intergenic | NA | NA |
| GU305914.1 | 72995 | 73017 | - | 4 | GTCGCCGGGGGCGCCTTCAG | CGG | -\|\|\|-\|-\|[\|\|\|\|\|\|\|\|\|\|-\|]PAM | NA | Intergenic | NA | NA |
| T12814 | CTTTCTCTTCGCCGTGATCGTGG | 1000 | | | | | | | | | |
| Oligo fwd | CACCCTTTCTCTTCGCCGTGATCG | | | | | | | | | | |
| Oligo rev | AAACCGATCACGGCGAAGAGAAAG | | | | | | | | | | |
| Chromosome | start | end | strand | MM | target_seq | PAM | alignment | distance | position | gene name | gene id |
| GU305914.1 | 169736 | 169758 | + | 0 | CTTTCTCTTCGCCGTGATCG | TGG | \|\|\|\|\|\|\|\|[\|\|\|\|\|\|\|\|\|\|\|\|]PAM | NA | Intergenic | NA | NA |
| T12815 | TCTCTTCGCCGTGATCGTGGCGG | 1000 | | | | | | | | | |
| Oligo fwd | CACCTCTCTTCGCCGTGATCGTGG | | | | | | | | | | |
| Oligo rev | AAACCCACGATCACGGCGAAGAGA | | | | | | | | | | |
| Chromosome | start | end | strand | MM | target_seq | PAM | alignment | distance | position | gene name | gene id |
| GU305914.1 | 169739 | 169761 | + | 0 | TCTCTTCGCCGTGATCGTGG | CGG | \|\|\|\|\|\|\|\|[\|\|\|\|\|\|\|\|\|\|\|\|]PAM | NA | Intergenic | NA | NA |
| T12816 | CTCTTCGCCGTGATCGTGGCGGG | 1000 | | | | | | | | | |
| Oligo fwd | CACCCTCTTCGCCGTGATCGTGGC | | | | | | | | | | |
| Oligo rev | AAACGCCACGATCACGGCGAAGAG | | | | | | | | | | |
| Chromosome | start | end | strand | MM | target_seq | PAM | alignment | distance | position | gene name | gene id |
| GU305914.1 | 169740 | 169762 | + | 0 | CTCTTCGCCGTGATCGTGGC | GGG | \|\|\|\|\|\|\|\|[\|\|\|\|\|\|\|\|\|\|\|\|]PAM | NA | Intergenic | NA | NA |
| T12817 | GTGGCGGGTACCACGATCTTCGG | 1000 | | | | | | | | | |
| Oligo fwd | CACCGTGGCGGGTACCACGATCTT | | | | | | | | | | |
| Oligo rev | AAACAAGATCGTGGTACCCGCCAC | | | | | | | | | | |
| Chromosome | start | end | strand | MM | target_seq | PAM | alignment | distance | position | gene name | gene id |
| GU305914.1 | 169755 | 169777 | + | 0 | GTGGCGGGTACCACGATCTT | CGG | \|\|\|\|\|\|\|\|[\|\|\|\|\|\|\|\|\|\|\|\|]PAM | NA | Intergenic | NA | NA |
| T12818 | GGTATCTTCTTGTGTTTCTTTGG | 1000 | | | | | | | | | |
| Oligo fwd | CACCGGTATCTTCTTGTGTTTCTT | | | | | | | | | | |
| Oligo rev | AAACAAGAAACACAAGAAGATACC | | | | | | | | | | |
| Chromosome | start | end | strand | MM | target_seq | PAM | alignment | distance | position | gene name | gene id |
| GU305914.1 | 169776 | 169798 | + | 0 | GGTATCTTCTTGTGTTTCTT | TGG | \|\|\|\|\|\|\|\|[\|\|\|\|\|\|\|\|\|\|\|\|]PAM | NA | Intergenic | NA | NA |
| T12819 | AGAAGACGCCGCCGCCGTCTCGG | 1000 | | | | | | | | | |
| Oligo fwd | CACCAGAAGACGCCGCCGCCGTCT | | | | | | | | | | |
| Oligo rev | AAACAGACGGCGGCGGCGTCTTCT | | | | | | | | | | |
| Chromosome | start | end | strand | MM | target_seq | PAM | alignment | distance | position | gene name | gene id |
| GU305914.1 | 169811 | 169833 | + | 0 | AGAAGACGCCGCCGCCGTCT | CGG | \|\|\|\|\|\|\|\|[\|\|\|\|\|\|\|\|\|\|\|\|]PAM | NA | Intergenic | NA | NA |
| T12820 | GCGCGCTGTTGCCGCTGTCGCGG | 1000 | | | | | | | | | |
| Oligo fwd | CACCGCGCGCTGTTGCCGCTGTCG | | | | | | | | | | |
| Oligo rev | AAACCGACAGCGGCAACAGCGCGC | | | | | | | | | | |
| Chromosome | start | end | strand | MM | target_seq | PAM | alignment | distance | position | gene name | gene id |
| GU305914.1 | 169839 | 169861 | + | 0 | GCGCGCTGTTGCCGCTGTCG | CGG | \|\|\|\|\|\|\|\|[\|\|\|\|\|\|\|\|\|\|\|\|]PAM | NA | Intergenic | NA | NA |

T12821 TGTTGCCGCTGTCGCGGCTCCGG 1000
Oligo fwd CACCTGTTGCCGCTGTCGCGGCTC
Oligo rev AAACGAGCCGCGACAGCGGCAACA
Chromosome start end strand MM target_seq PAM alignment distance position gene name gene id
GU305914.1 169845 169867 + 0 TGTTGCCGCTGTCGCGGCTC CGG ||||||||[||||||||||||]PAM NA Intergenic NA NA

T12822 GCCGCTGTCGCGGCTCCGGCCGG 1000
Oligo fwd CACCGCCGCTGTCGCGGCTCCGGC
Oligo rev AAACGCCGGAGCCGCGACAGCGGC
Chromosome start end strand MM target_seq PAM alignment distance position gene name gene id
GU305914.1 169849 169871 + 0 GCCGCTGTCGCGGCTCCGGC CGG ||||||||[||||||||||||]PAM NA Intergenic NA NA
GU305914.1 45582 45604 - 4 GCCGCCGCCGCGGCGCCGGG TGG |||||-|-[||||||-||||-]PAM NA Intergenic NA NA

T12823 TCGCGGCTCCGGCCGGTTCGCGG 1000
Oligo fwd CACCTCGCGGCTCCGGCCGGTTCG
Oligo rev AAACCGAACCGGCCGGAGCCGCGA
Chromosome start end strand MM target_seq PAM alignment distance position gene name gene id
GU305914.1 169856 169878 + 0 TCGCGGCTCCGGCCGGTTCG CGG ||||||||[||||||||||||]PAM NA Intergenic NA NA

T12824 CGCGGCTCCGGCCGGTTCGCGGG 1000
Oligo fwd CACCCGCGGCTCCGGCCGGTTCGC
Oligo rev AAACGCGAACCGGCCGGAGCCGCG
Chromosome start end strand MM target_seq PAM alignment distance position gene name gene id
GU305914.1 169857 169879 + 0 CGCGGCTCCGGCCGGTTCGC GGG ||||||||[||||||||||||]PAM NA Intergenic NA NA
GU305914.1 78745 78767 - 4 CGCCGCGCCGGCCGCTGCGC TGG |||-||-|[||||||-|-|||]PAM NA Intergenic NA NA

T12825 GCGGCTCCGGCCGGTTCGCGGGG 1000
Oligo fwd CACCGCGGCTCCGGCCGGTTCGCG
Oligo rev AAACCGCGAACCGGCCGGAGCCGC
Chromosome start end strand MM target_seq PAM alignment distance position gene name gene id
GU305914.1 169858 169880 + 0 GCGGCTCCGGCCGGTTCGCG GGG ||||||||[||||||||||||]PAM NA Intergenic NA NA

T12826 CGGCTCCGGCCGGTTCGCGGGGG 1000
Oligo fwd CACCCGGCTCCGGCCGGTTCGCGG
Oligo rev AAACCCGCGAACCGGCCGGAGCCG
Chromosome start end strand MM target_seq PAM alignment distance position gene name gene id
GU305914.1 169859 169881 + 0 CGGCTCCGGCCGGTTCGCGG GGG ||||||||[||||||||||||]PAM NA Intergenic NA NA

T12827 GTCACGTCGCTTAACCTCGTCGG 1000
Oligo fwd CACCGTCACGTCGCTTAACCTCGT
Oligo rev AAACACGAGGTTAAGCGACGTGAC
Chromosome start end strand MM target_seq PAM alignment distance position gene name gene id
GU305914.1 169884 169906 + 0 GTCACGTCGCTTAACCTCGT CGG ||||||||[||||||||||||]PAM NA Intergenic NA NA

T12828 TCGCTTAACCTCGTCGGAACCGG 1000
Oligo fwd CACCTCGCTTAACCTCGTCGGAAC
Oligo rev AAACGTTCCGACGAGGTTAAGCGA
Chromosome start end strand MM target_seq PAM alignment distance position gene name gene id
GU305914.1 169890 169912 + 0 TCGCTTAACCTCGTCGGAAC CGG ||||||||[||||||||||||]PAM NA Intergenic NA NA

T12829 GGAACCGGCCTCACCACGATCGG 1000
Oligo fwd CACCGGAACCGGCCTCACCACGAT
Oligo rev AAACATCGTGGTGAGGCCGGTTCC
Chromosome start end strand MM target_seq PAM alignment distance position gene name gene id
GU305914.1 169905 169927 + 0 GGAACCGGCCTCACCACGAT CGG ||||||||[||||||||||||]PAM NA Intergenic NA NA

T12830 CTCACCACGATCGGTGCGCGTGG 1000
Oligo fwd CACCCTCACCACGATCGGTGCGCG
Oligo rev AAACCGCGCACCGATCGTGGTGAG
Chromosome start end strand MM target_seq PAM alignment distance position gene name gene id
GU305914.1 169914 169936 + 0 CTCACCACGATCGGTGCGCG TGG ||||||||[||||||||||||]PAM NA Intergenic NA NA

T12831 ACCTTCTTGATGATGAGTTTCGG 1000
Oligo fwd CACCACCTTCTTGATGATGAGTTT
Oligo rev AAACAAACTCATCATCAAGAAGGT
Chromosome start end strand MM target_seq PAM alignment distance position gene name gene id
GU305914.1 169938 169960 + 0 ACCTTCTTGATGATGAGTTT CGG ||||||||[||||||||||||]PAM NA Intergenic NA NA

T12832 GATGATGAGTTTCGGTACCTTGG 1000
Oligo fwd CACCGATGATGAGTTTCGGTACCT
Oligo rev AAACAGGTACCGAAACTCATCATC
Chromosome start end strand MM target_seq PAM alignment distance position gene name gene id
GU305914.1 169946 169968 + 0 GATGATGAGTTTCGGTACCT TGG ||||||||[||||||||||||]PAM NA Intergenic NA NA

T12833 ATGATGAGTTTCGGTACCTTGGG 1000
Oligo fwd CACCATGATGAGTTTCGGTACCTT
Oligo rev AAACAAGGTACCGAAACTCATCAT
Chromosome start end strand MM target_seq PAM alignment distance position gene name gene id
GU305914.1 169947 169969 + 0 ATGATGAGTTTCGGTACCTT GGG ||||||||[||||||||||||]PAM NA Intergenic NA NA

T12834 CGGTACCTTGGGCACTTTCGCGG 1000
Oligo fwd CACCCGGTACCTTGGGCACTTTCG
Oligo rev AAACCGAAAGTGCCCAAGGTACCG
Chromosome start end strand MM target_seq PAM alignment distance position gene name gene id
GU305914.1 169958 169980 + 0 CGGTACCTTGGGCACTTTCG CGG ||||||||[||||||||||||]PAM NA Intergenic NA NA

T12835 GGTACCTTGGGCACTTTCGCGGG 1000
Oligo fwd CACCGGTACCTTGGGCACTTTCGC
Oligo rev AAACGCGAAAGTGCCCAAGGTACC

| | | | | | | | | | | | |
|---|---|---|---|---|---|---|---|---|---|---|---|
| Chromosome | start | end | strand | MM | target_seq | PAM | alignment | distance | position | gene name | gene id |
| GU305914.1 | 169959 | 169981 | + | 0 | GGTACCTTGGGCACTTTCGC | GGG | \|\|\|\|\|\|\|\|[\|\|\|\|\|\|\|\|\|\|\|\|]PAM | NA | Intergenic | NA | NA |
| T12836 | TTCGCGGGCCTAGCGTCCGTCGG | 1000 | | | | | | | | | |
| Oligo fwd | CACCTTCGCGGGCCTAGCGTCCGT | | | | | | | | | | |
| Oligo rev | AAACACGGACGCTAGGCCCGCGAA | | | | | | | | | | |
| Chromosome | start | end | strand | MM | target_seq | PAM | alignment | distance | position | gene name | gene id |
| GU305914.1 | 169974 | 169996 | + | 0 | TTCGCGGGCCTAGCGTCCGT | CGG | \|\|\|\|\|\|\|\|[\|\|\|\|\|\|\|\|\|\|\|\|]PAM | NA | Intergenic | NA | NA |
| T12837 | GGCCTAGCGTCCGTCGGCTGCGG | 1000 | | | | | | | | | |
| Oligo fwd | CACCGGCCTAGCGTCCGTCGGCTG | | | | | | | | | | |
| Oligo rev | AAACCAGCCGACGGACGCTAGGCC | | | | | | | | | | |
| Chromosome | start | end | strand | MM | target_seq | PAM | alignment | distance | position | gene name | gene id |
| GU305914.1 | 169980 | 170002 | + | 0 | GGCCTAGCGTCCGTCGGCTG | CGG | \|\|\|\|\|\|\|\|[\|\|\|\|\|\|\|\|\|\|\|\|]PAM | NA | Intergenic | NA | NA |
| T12838 | GCCTAGCGTCCGTCGGCTGCGGG | 1000 | | | | | | | | | |
| Oligo fwd | CACCGCCTAGCGTCCGTCGGCTGC | | | | | | | | | | |
| Oligo rev | AAACGCAGCCGACGGACGCTAGGC | | | | | | | | | | |
| Chromosome | start | end | strand | MM | target_seq | PAM | alignment | distance | position | gene name | gene id |
| GU305914.1 | 169981 | 170003 | + | 0 | GCCTAGCGTCCGTCGGCTGC | GGG | \|\|\|\|\|\|\|\|[\|\|\|\|\|\|\|\|\|\|\|\|]PAM | NA | Intergenic | NA | NA |
| T12839 | AGCGTCCGTCGGCTGCGGGACGG | 1000 | | | | | | | | | |
| Oligo fwd | CACCAGCGTCCGTCGGCTGCGGGA | | | | | | | | | | |
| Oligo rev | AAACTCCCGCAGCCGACGGACGCT | | | | | | | | | | |
| Chromosome | start | end | strand | MM | target_seq | PAM | alignment | distance | position | gene name | gene id |
| GU305914.1 | 169985 | 170007 | + | 0 | AGCGTCCGTCGGCTGCGGGA | CGG | \|\|\|\|\|\|\|\|[\|\|\|\|\|\|\|\|\|\|\|\|]PAM | NA | Intergenic | NA | NA |
| T12840 | GCGTCCGTCGGCTGCGGGACGGG | 1000 | | | | | | | | | |
| Oligo fwd | CACCGCGTCCGTCGGCTGCGGGAC | | | | | | | | | | |
| Oligo rev | AAACGTCCCGCAGCCGACGGACGC | | | | | | | | | | |
| Chromosome | start | end | strand | MM | target_seq | PAM | alignment | distance | position | gene name | gene id |
| GU305914.1 | 169986 | 170008 | + | 0 | GCGTCCGTCGGCTGCGGGAC | GGG | \|\|\|\|\|\|\|\|[\|\|\|\|\|\|\|\|\|\|\|\|]PAM | NA | Intergenic | NA | NA |
| T12841 | CCTTTTTAGTATGATGAGCTCGG | 1000 | | | | | | | | | |
| Oligo fwd | CACCCCTTTTTAGTATGATGAGCT | | | | | | | | | | |
| Oligo rev | AAACAGCTCATCATACTAAAAAGG | | | | | | | | | | |
| Chromosome | start | end | strand | MM | target_seq | PAM | alignment | distance | position | gene name | gene id |
| GU305914.1 | 170021 | 170043 | + | 0 | CCTTTTTAGTATGATGAGCT | CGG | \|\|\|\|\|\|\|\|[\|\|\|\|\|\|\|\|\|\|\|\|]PAM | NA | Intergenic | NA | NA |
| T12842 | GATGAGCTCGGAATCCGCGACGG | 1000 | | | | | | | | | |
| Oligo fwd | CACCGATGAGCTCGGAATCCGCGA | | | | | | | | | | |
| Oligo rev | AAACTCGCGGATTCCGAGCTCATC | | | | | | | | | | |
| Chromosome | start | end | strand | MM | target_seq | PAM | alignment | distance | position | gene name | gene id |
| GU305914.1 | 170033 | 170055 | + | 0 | GATGAGCTCGGAATCCGCGA | CGG | \|\|\|\|\|\|\|\|[\|\|\|\|\|\|\|\|\|\|\|\|]PAM | NA | Intergenic | NA | NA |
| T12843 | ATGAGCTCGGAATCCGCGACGGG | 1000 | | | | | | | | | |
| Oligo fwd | CACCATGAGCTCGGAATCCGCGAC | | | | | | | | | | |
| Oligo rev | AAACGTCGCGGATTCCGAGCTCAT | | | | | | | | | | |
| Chromosome | start | end | strand | MM | target_seq | PAM | alignment | distance | position | gene name | gene id |
| GU305914.1 | 170034 | 170056 | + | 0 | ATGAGCTCGGAATCCGCGAC | GGG | \|\|\|\|\|\|\|\|[\|\|\|\|\|\|\|\|\|\|\|\|]PAM | NA | Intergenic | NA | NA |
| GU305914.1 | 170112 | 170134 | + | 4 | ACGGGCTCGGGAACCGCGAC | GGG | \|-\|-\|\|\|\|[\|\|-\|-\|\|\|\|\|\|\|]PAM | NA | Intergenic | NA | NA |
| T12844 | CTCGGAATCCGCGACGGGTTTGG | 1000 | | | | | | | | | |
| Oligo fwd | CACCCTCGGAATCCGCGACGGGTT | | | | | | | | | | |
| Oligo rev | AAACAACCCGTCGCGGATTCCGAG | | | | | | | | | | |
| Chromosome | start | end | strand | MM | target_seq | PAM | alignment | distance | position | gene name | gene id |
| GU305914.1 | 170039 | 170061 | + | 0 | CTCGGAATCCGCGACGGGTT | TGG | \|\|\|\|\|\|\|\|[\|\|\|\|\|\|\|\|\|\|\|\|]PAM | NA | Intergenic | NA | NA |
| GU305914.1 | 170117 | 170139 | + | 2 | CTCGGGAACCGCGACGGGTT | CGG | \|\|\|\|\|-\|-[\|\|\|\|\|\|\|\|\|\|\|\|]PAM | NA | Intergenic | NA | NA |
| T12845 | TCGGAATCCGCGACGGGTTTGGG | 1000 | | | | | | | | | |
| Oligo fwd | CACCTCGGAATCCGCGACGGGTTT | | | | | | | | | | |
| Oligo rev | AAACAAACCCGTCGCGGATTCCGA | | | | | | | | | | |
| Chromosome | start | end | strand | MM | target_seq | PAM | alignment | distance | position | gene name | gene id |
| GU305914.1 | 170040 | 170062 | + | 0 | TCGGAATCCGCGACGGGTTT | GGG | \|\|\|\|\|\|\|\|[\|\|\|\|\|\|\|\|\|\|\|\|]PAM | NA | Intergenic | NA | NA |
| GU305914.1 | 170118 | 170140 | + | 3 | TCGGGAACCGCGACGGGTTC | GGG | \|\|\|\|-\|-\|[\|\|\|\|\|\|\|\|\|\|\|-]PAM | NA | Intergenic | NA | NA |
| T12846 | ATCCGCGACGGGTTTGGGAGCGG | 1000 | | | | | | | | | |
| Oligo fwd | CACCATCCGCGACGGGTTTGGGAG | | | | | | | | | | |
| Oligo rev | AAACCTCCCAAACCCGTCGCGGAT | | | | | | | | | | |
| Chromosome | start | end | strand | MM | target_seq | PAM | alignment | distance | position | gene name | gene id |
| GU305914.1 | 170045 | 170067 | + | 0 | ATCCGCGACGGGTTTGGGAG | CGG | \|\|\|\|\|\|\|\|[\|\|\|\|\|\|\|\|\|\|\|\|]PAM | NA | Intergenic | NA | NA |
| T12847 | TCCGCGACGGGTTTGGGAGCGGG | 1000 | | | | | | | | | |
| Oligo fwd | CACCTCCGCGACGGGTTTGGGAGC | | | | | | | | | | |
| Oligo rev | AAACGCTCCCAAACCCGTCGCGGA | | | | | | | | | | |
| Chromosome | start | end | strand | MM | target_seq | PAM | alignment | distance | position | gene name | gene id |
| GU305914.1 | 170046 | 170068 | + | 0 | TCCGCGACGGGTTTGGGAGC | GGG | \|\|\|\|\|\|\|\|[\|\|\|\|\|\|\|\|\|\|\|\|]PAM | NA | Intergenic | NA | NA |
| T12848 | GACGGGTTTGGGAGCGGGAACGG | 1000 | | | | | | | | | |
| Oligo fwd | CACCGACGGGTTTGGGAGCGGGAA | | | | | | | | | | |
| Oligo rev | AAACTTCCCGCTCCCAAACCCGTC | | | | | | | | | | |
| Chromosome | start | end | strand | MM | target_seq | PAM | alignment | distance | position | gene name | gene id |
| GU305914.1 | 170051 | 170073 | + | 0 | GACGGGTTTGGGAGCGGGAA | CGG | \|\|\|\|\|\|\|\|[\|\|\|\|\|\|\|\|\|\|\|\|]PAM | NA | Intergenic | NA | NA |
| T12849 | ACGGGTTTGGGAGCGGGAACGGG | 1000 | | | | | | | | | |
| Oligo fwd | CACCACGGGTTTGGGAGCGGGAAC | | | | | | | | | | |
| Oligo rev | AAACGTTCCCGCTCCCAAACCCGT | | | | | | | | | | |
| Chromosome | start | end | strand | MM | target_seq | PAM | alignment | distance | position | gene name | gene id |
| GU305914.1 | 170052 | 170074 | + | 0 | ACGGGTTTGGGAGCGGGAAC | GGG | \|\|\|\|\|\|\|\|[\|\|\|\|\|\|\|\|\|\|\|\|]PAM | NA | Intergenic | NA | NA |

T12850 TTTGGGAGCGGGAACGGGAACGG 1000
Oligo fwd CACCTTTGGGAGCGGGAACGGGAA
Oligo rev AAACTTCCCGTTCCCGCTCCCAAA
Chromosome start end strand MM target_seq PAM alignment distance position gene name gene id
GU305914.1 170057 170079 + 0 TTTGGGAGCGGGAACGGGAA CGG ||||||||[||||||||||||]PAM NA Intergenic NA NA

T12851 TTGGGAGCGGGAACGGGAACGGG 1000
Oligo fwd CACCTTGGGAGCGGGAACGGGAAC
Oligo rev AAACGTTCCCGTTCCCGCTCCCAA
Chromosome start end strand MM target_seq PAM alignment distance position gene name gene id
GU305914.1 170058 170080 + 0 TTGGGAGCGGGAACGGGAAC GGG ||||||||[||||||||||||]PAM NA Intergenic NA NA

T12852 GCGGGAACGGGAACGGGCTCTGG 1000
Oligo fwd CACCGCGGGAACGGGAACGGGCTC
Oligo rev AAACGAGCCCGTTCCCGTTCCCGC
Chromosome start end strand MM target_seq PAM alignment distance position gene name gene id
GU305914.1 170064 170086 + 0 GCGGGAACGGGAACGGGCTC TGG ||||||||[||||||||||||]PAM NA Intergenic NA NA

T12853 CGGGAACGGGAACGGGCTCTGGG 1000
Oligo fwd CACCCGGGAACGGGAACGGGCTCT
Oligo rev AAACAGAGCCCGTTCCCGTTCCCG
Chromosome start end strand MM target_seq PAM alignment distance position gene name gene id
GU305914.1 170065 170087 + 0 CGGGAACGGGAACGGGCTCT GGG ||||||||[||||||||||||]PAM NA Intergenic NA NA

T12854 AACGGGCTCTGGGAGCTTGACGG 1000
Oligo fwd CACCAACGGGCTCTGGGAGCTTGA
Oligo rev AAACTCAAGCTCCCAGAGCCCGTT
Chromosome start end strand MM target_seq PAM alignment distance position gene name gene id
GU305914.1 170075 170097 + 0 AACGGGCTCTGGGAGCTTGA CGG ||||||||[||||||||||||]PAM NA Intergenic NA NA
GU305914.1 170093 170115 + 3 GACGGGCTCTGGGATCTTAA CGG -|||||||[||||||-|||-|]PAM NA Intergenic NA NA

T12855 ACGGGCTCTGGGAGCTTGACGGG 1000
Oligo fwd CACCACGGGCTCTGGGAGCTTGAC
Oligo rev AAACGTCAAGCTCCCAGAGCCCGT
Chromosome start end strand MM target_seq PAM alignment distance position gene name gene id
GU305914.1 170076 170098 + 0 ACGGGCTCTGGGAGCTTGAC GGG ||||||||[||||||||||||]PAM NA Intergenic NA NA
GU305914.1 170094 170116 + 2 ACGGGCTCTGGGATCTTAAC GGG ||||||||[|||||-|||-||]PAM NA Intergenic NA NA

T12856 TCTGGGAGCTTGACGGGCTCTGG 1000
Oligo fwd CACCTCTGGGAGCTTGACGGGCTC
Oligo rev AAACGAGCCCGTCAAGCTCCCAGA
Chromosome start end strand MM target_seq PAM alignment distance position gene name gene id
GU305914.1 170082 170104 + 0 TCTGGGAGCTTGACGGGCTC TGG ||||||||[||||||||||||]PAM NA Intergenic NA NA
GU305914.1 170100 170122 + 2 TCTGGGATCTTAACGGGCTC GGG |||||||-[|||-||||||||]PAM NA Intergenic NA NA
GU305914.1 98247 98269 + 4 TCTAGCAGCTTGATGGGGTC GGG |||-|-||[|||||-|||-||]PAM NA Intergenic NA NA

T12857 CTGGGAGCTTGACGGGCTCTGGG 1000
Oligo fwd CACCCTGGGAGCTTGACGGGCTCT
Oligo rev AAACAGAGCCCGTCAAGCTCCCAG
Chromosome start end strand MM target_seq PAM alignment distance position gene name gene id
GU305914.1 170083 170105 + 0 CTGGGAGCTTGACGGGCTCT GGG ||||||||[||||||||||||]PAM NA Intergenic NA NA

T12858 GACGGGCTCTGGGATCTTAACGG 1000
Oligo fwd CACCGACGGGCTCTGGGATCTTAA
Oligo rev AAACTTAAGATCCCAGAGCCCGTC
Chromosome start end strand MM target_seq PAM alignment distance position gene name gene id
GU305914.1 170093 170115 + 0 GACGGGCTCTGGGATCTTAA CGG ||||||||[||||||||||||]PAM NA Intergenic NA NA
GU305914.1 170075 170097 + 3 AACGGGCTCTGGGAGCTTGA CGG -|||||||[||||||-|||-|]PAM NA Intergenic NA NA

T12859 ACGGGCTCTGGGATCTTAACGGG 1000
Oligo fwd CACCACGGGCTCTGGGATCTTAAC
Oligo rev AAACGTTAAGATCCCAGAGCCCGT
Chromosome start end strand MM target_seq PAM alignment distance position gene name gene id
GU305914.1 170094 170116 + 0 ACGGGCTCTGGGATCTTAAC GGG ||||||||[||||||||||||]PAM NA Intergenic NA NA
GU305914.1 170076 170098 + 2 ACGGGCTCTGGGAGCTTGAC GGG ||||||||[|||||-|||-||]PAM NA Intergenic NA NA

T12860 CTCTGGGATCTTAACGGGCTCGG 1000
Oligo fwd CACCCTCTGGGATCTTAACGGGCT
Oligo rev AAACAGCCCGTTAAGATCCCAGAG
Chromosome start end strand MM target_seq PAM alignment distance position gene name gene id
GU305914.1 170099 170121 + 0 CTCTGGGATCTTAACGGGCT CGG ||||||||[||||||||||||]PAM NA Intergenic NA NA

T12861 TCTGGGATCTTAACGGGCTCGGG 1000
Oligo fwd CACCTCTGGGATCTTAACGGGCTC
Oligo rev AAACGAGCCCGTTAAGATCCCAGA
Chromosome start end strand MM target_seq PAM alignment distance position gene name gene id
GU305914.1 170100 170122 + 0 TCTGGGATCTTAACGGGCTC GGG ||||||||[||||||||||||]PAM NA Intergenic NA NA
GU305914.1 170082 170104 + 2 TCTGGGAGCTTGACGGGCTC TGG |||||||-[|||-||||||||]PAM NA Intergenic NA NA

T12862 AACGGGCTCGGGAACCGCGACGG 1000
Oligo fwd CACCAACGGGCTCGGGAACCGCGA
Oligo rev AAACTCGCGGTTCCCGAGCCCGTT
Chromosome start end strand MM target_seq PAM alignment distance position gene name gene id
GU305914.1 170111 170133 + 0 AACGGGCTCGGGAACCGCGA CGG ||||||||[||||||||||||]PAM NA Intergenic NA NA

T12863 ACGGGCTCGGGAACCGCGACGGG 1000
Oligo fwd CACCACGGGCTCGGGAACCGCGAC
Oligo rev AAACGTCGCGGTTCCCGAGCCCGT

| | | | | | | | | | | | |
|---|---|---|---|---|---|---|---|---|---|---|---|
| Chromosome | start | end | strand | MM | target_seq | PAM | alignment | distance | position | gene name | gene id |
| GU305914.1 | 170112 | 170134 | + | 0 | ACGGGCTCGGGAACCGCGAC | GGG | \|\|\|\|\|\|\|\|[\|\|\|\|\|\|\|\|\|\|\|\|]PAM | NA | Intergenic | NA | NA |
| GU305914.1 | 170034 | 170056 | + | 4 | ATGAGCTCGGAATCCGCGAC | GGG | \|-\|-\|\|\|\|[\|\|-\|-\|\|\|\|\|\|\|]PAM | NA | Intergenic | NA | NA |
| T12864 | CTCGGGAACCGCGACGGGTTCGG | 1000 | | | | | | | | | |
| Oligo fwd | CACCCTCGGGAACCGCGACGGGTT | | | | | | | | | | |
| Oligo rev | AAACAACCCGTCGCGGTTCCCGAG | | | | | | | | | | |
| Chromosome | start | end | strand | MM | target_seq | PAM | alignment | distance | position | gene name | gene id |
| GU305914.1 | 170117 | 170139 | + | 0 | CTCGGGAACCGCGACGGGTT | CGG | \|\|\|\|\|\|\|\|[\|\|\|\|\|\|\|\|\|\|\|\|]PAM | NA | Intergenic | NA | NA |
| GU305914.1 | 170039 | 170061 | + | 2 | CTCGGAATCCGCGACGGGTT | TGG | \|\|\|\|\|-\|-[\|\|\|\|\|\|\|\|\|\|\|\|]PAM | NA | Intergenic | NA | NA |
| T12865 | TCGGGAACCGCGACGGGTTCGGG | 1000 | | | | | | | | | |
| Oligo fwd | CACCTCGGGAACCGCGACGGGTTC | | | | | | | | | | |
| Oligo rev | AAACGAACCCGTCGCGGTTCCCGA | | | | | | | | | | |
| Chromosome | start | end | strand | MM | target_seq | PAM | alignment | distance | position | gene name | gene id |
| GU305914.1 | 170118 | 170140 | + | 0 | TCGGGAACCGCGACGGGTTC | GGG | \|\|\|\|\|\|\|\|[\|\|\|\|\|\|\|\|\|\|\|\|]PAM | NA | Intergenic | NA | NA |
| GU305914.1 | 170421 | 170443 | + | 4 | TCTGGAAACGCGTCGGGGTC | GGG | \|\|-\|\|\|\|-[\|\|\|\|-\|\|\|\|-\|\|]PAM | NA | Intergenic | NA | NA |
| GU305914.1 | 170040 | 170062 | + | 3 | TCGGAATCCGCGACGGGTTT | GGG | \|\|\|\|-\|-\|[\|\|\|\|\|\|\|\|\|\|\|-]PAM | NA | Intergenic | NA | NA |
| T12866 | CGGGAACCGCGACGGGTTCGGGG | 1000 | | | | | | | | | |
| Oligo fwd | CACCCGGGAACCGCGACGGGTTCG | | | | | | | | | | |
| Oligo rev | AAACCGAACCCGTCGCGGTTCCCG | | | | | | | | | | |
| Chromosome | start | end | strand | MM | target_seq | PAM | alignment | distance | position | gene name | gene id |
| GU305914.1 | 170119 | 170141 | + | 0 | CGGGAACCGCGACGGGTTCG | GGG | \|\|\|\|\|\|\|\|[\|\|\|\|\|\|\|\|\|\|\|\|]PAM | NA | Intergenic | NA | NA |
| GU305914.1 | 170260 | 170282 | + | 4 | CGGGGACGGTGACGGGCTCG | AGG | \|\|\|\|-\|\|-[\|-\|\|\|\|\|\|-\|\|\|]PAM | NA | Intergenic | NA | NA |
| T12867 | AACCGCGACGGGTTCGGGGACGG | 1000 | | | | | | | | | |
| Oligo fwd | CACCAACCGCGACGGGTTCGGGGA | | | | | | | | | | |
| Oligo rev | AAACTCCCCGAACCCGTCGCGGTT | | | | | | | | | | |
| Chromosome | start | end | strand | MM | target_seq | PAM | alignment | distance | position | gene name | gene id |
| GU305914.1 | 170123 | 170145 | + | 0 | AACCGCGACGGGTTCGGGGA | CGG | \|\|\|\|\|\|\|\|[\|\|\|\|\|\|\|\|\|\|\|\|]PAM | NA | Intergenic | NA | NA |
| T12868 | GACGGGTTCGGGGACGGTGACGG | 1000 | | | | | | | | | |
| Oligo fwd | CACCGACGGGTTCGGGGACGGTGA | | | | | | | | | | |
| Oligo rev | AAACTCACCGTCCCCGAACCCGTC | | | | | | | | | | |
| Chromosome | start | end | strand | MM | target_seq | PAM | alignment | distance | position | gene name | gene id |
| GU305914.1 | 170129 | 170151 | + | 0 | GACGGGTTCGGGGACGGTGA | CGG | \|\|\|\|\|\|\|\|[\|\|\|\|\|\|\|\|\|\|\|\|]PAM | NA | Intergenic | NA | NA |
| GU305914.1 | 170252 | 170274 | + | 3 | TTCGGATTCGGGGACGGTGA | CGG | --\|\|\|-\|\|[\|\|\|\|\|\|\|\|\|\|\|\|]PAM | NA | Intergenic | NA | NA |
| T12869 | ACGGGTTCGGGGACGGTGACGGG | 1000 | | | | | | | | | |
| Oligo fwd | CACCACGGGTTCGGGGACGGTGAC | | | | | | | | | | |
| Oligo rev | AAACGTCACCGTCCCCGAACCCGT | | | | | | | | | | |
| Chromosome | start | end | strand | MM | target_seq | PAM | alignment | distance | position | gene name | gene id |
| GU305914.1 | 170130 | 170152 | + | 0 | ACGGGTTCGGGGACGGTGAC | GGG | \|\|\|\|\|\|\|\|[\|\|\|\|\|\|\|\|\|\|\|\|]PAM | NA | Intergenic | NA | NA |
| GU305914.1 | 170253 | 170275 | + | 2 | TCGGATTCGGGGACGGTGAC | GGG | -\|\|\|-\|\|\|[\|\|\|\|\|\|\|\|\|\|\|\|]PAM | NA | Intergenic | NA | NA |
| T12870 | TTCGGGGACGGTGACGGGCTCGG | 1000 | | | | | | | | | |
| Oligo fwd | CACCTTCGGGGACGGTGACGGGCT | | | | | | | | | | |
| Oligo rev | AAACAGCCCGTCACCGTCCCCGAA | | | | | | | | | | |
| Chromosome | start | end | strand | MM | target_seq | PAM | alignment | distance | position | gene name | gene id |
| GU305914.1 | 170135 | 170157 | + | 0 | TTCGGGGACGGTGACGGGCT | CGG | \|\|\|\|\|\|\|\|[\|\|\|\|\|\|\|\|\|\|\|\|]PAM | NA | Intergenic | NA | NA |
| GU305914.1 | 127644 | 127666 | - | 4 | CTCGTCGACGGTGACGGCCT | GGG | -\|\|\|--\|\|[\|\|\|\|\|\|\|\|\|-\|\|]PAM | NA | Intergenic | NA | NA |
| T12871 | ACGGTGACGGGCTCGGAATCAGG | 1000 | | | | | | | | | |
| Oligo fwd | CACCACGGTGACGGGCTCGGAATC | | | | | | | | | | |
| Oligo rev | AAACGATTCCGAGCCCGTCACCGT | | | | | | | | | | |
| Chromosome | start | end | strand | MM | target_seq | PAM | alignment | distance | position | gene name | gene id |
| GU305914.1 | 170142 | 170164 | + | 0 | ACGGTGACGGGCTCGGAATC | AGG | \|\|\|\|\|\|\|\|[\|\|\|\|\|\|\|\|\|\|\|\|]PAM | NA | Intergenic | NA | NA |
| T12872 | ACGGGCTCGGAATCAGGCTCAGG | 1000 | | | | | | | | | |
| Oligo fwd | CACCACGGGCTCGGAATCAGGCTC | | | | | | | | | | |
| Oligo rev | AAACGAGCCTGATTCCGAGCCCGT | | | | | | | | | | |
| Chromosome | start | end | strand | MM | target_seq | PAM | alignment | distance | position | gene name | gene id |
| GU305914.1 | 170148 | 170170 | + | 0 | ACGGGCTCGGAATCAGGCTC | AGG | \|\|\|\|\|\|\|\|[\|\|\|\|\|\|\|\|\|\|\|\|]PAM | NA | Intergenic | NA | NA |
| T12873 | TCGGAATCAGGCTCAGGCTCAGG | 1000 | | | | | | | | | |
| Oligo fwd | CACCTCGGAATCAGGCTCAGGCTC | | | | | | | | | | |
| Oligo rev | AAACGAGCCTGAGCCTGATTCCGA | | | | | | | | | | |
| Chromosome | start | end | strand | MM | target_seq | PAM | alignment | distance | position | gene name | gene id |
| GU305914.1 | 170154 | 170176 | + | 0 | TCGGAATCAGGCTCAGGCTC | AGG | \|\|\|\|\|\|\|\|[\|\|\|\|\|\|\|\|\|\|\|\|]PAM | NA | Intergenic | NA | NA |
| GU305914.1 | 170160 | 170182 | + | 3 | TCAGGCTCAGGCTCAGGCTC | AGG | \|\|-\|--\|\|[\|\|\|\|\|\|\|\|\|\|\|\|]PAM | NA | Intergenic | NA | NA |
| GU305914.1 | 170166 | 170188 | + | 3 | TCAGGCTCAGGCTCAGGCTC | GGG | \|\|-\|--\|\|[\|\|\|\|\|\|\|\|\|\|\|\|]PAM | NA | Intergenic | NA | NA |
| T12874 | TCAGGCTCAGGCTCAGGCTCAGG | 1000 | | | | | | | | | |
| Oligo fwd | CACCTCAGGCTCAGGCTCAGGCTC | | | | | | | | | | |
| Oligo rev | AAACGAGCCTGAGCCTGAGCCTGA | | | | | | | | | | |
| Chromosome | start | end | strand | MM | target_seq | PAM | alignment | distance | position | gene name | gene id |
| GU305914.1 | 170160 | 170182 | + | 0 | TCAGGCTCAGGCTCAGGCTC | AGG | \|\|\|\|\|\|\|\|[\|\|\|\|\|\|\|\|\|\|\|\|]PAM | NA | Intergenic | NA | NA |
| GU305914.1 | 170166 | 170188 | + | 0 | TCAGGCTCAGGCTCAGGCTC | GGG | \|\|\|\|\|\|\|\|[\|\|\|\|\|\|\|\|\|\|\|\|]PAM | NA | Intergenic | NA | NA |
| GU305914.1 | 170154 | 170176 | + | 3 | TCGGAATCAGGCTCAGGCTC | AGG | \|\|-\|--\|\|[\|\|\|\|\|\|\|\|\|\|\|\|]PAM | NA | Intergenic | NA | NA |
| T12875 | CTCAGGCTCAGGCTCAGGCTCGG | 1000 | | | | | | | | | |
| Oligo fwd | CACCCTCAGGCTCAGGCTCAGGCT | | | | | | | | | | |
| Oligo rev | AAACAGCCTGAGCCTGAGCCTGAG | | | | | | | | | | |
| Chromosome | start | end | strand | MM | target_seq | PAM | alignment | distance | position | gene name | gene id |
| GU305914.1 | 170165 | 170187 | + | 0 | CTCAGGCTCAGGCTCAGGCT | CGG | \|\|\|\|\|\|\|\|[\|\|\|\|\|\|\|\|\|\|\|\|]PAM | NA | Intergenic | NA | NA |
| T12876 | TCAGGCTCAGGCTCAGGCTCGGG | 1000 | | | | | | | | | |
| Oligo fwd | CACCTCAGGCTCAGGCTCAGGCTC | | | | | | | | | | |

<table>
<tr><td>Oligo rev</td><td>AAACGAGCCTGAGCCTGAGCCTGA</td><td></td><td></td><td></td><td></td><td></td><td></td><td></td><td></td><td></td><td></td></tr>
<tr><td>Chromosome</td><td>start</td><td>end</td><td>strand</td><td>MM</td><td>target_seq</td><td>PAM</td><td>alignment</td><td>distance</td><td>position</td><td>gene name</td><td>gene id</td></tr>
<tr><td>GU305914.1</td><td>170160</td><td>170182</td><td>+</td><td>0</td><td>TCAGGCTCAGGCTCAGGCTC</td><td>AGG</td><td>||||||||[||||||||||||]PAM</td><td>NA</td><td>Intergenic</td><td>NA</td><td>NA</td></tr>
<tr><td>GU305914.1</td><td>170166</td><td>170188</td><td>+</td><td>0</td><td>TCAGGCTCAGGCTCAGGCTC</td><td>GGG</td><td>||||||||[||||||||||||]PAM</td><td>NA</td><td>Intergenic</td><td>NA</td><td>NA</td></tr>
<tr><td>GU305914.1</td><td>170154</td><td>170176</td><td>+</td><td>3</td><td>TCGGAATCAGGCTCAGGCTC</td><td>AGG</td><td>||-|--||[||||||||||||]PAM</td><td>NA</td><td>Intergenic</td><td>NA</td><td>NA</td></tr>
<tr><td></td><td></td><td></td><td></td><td></td><td></td><td></td><td></td><td></td><td></td><td></td><td></td></tr>
<tr><td>T12877</td><td>CAGGCTCAGGCTCAGGCTCGGGG</td><td>1000</td><td></td><td></td><td></td><td></td><td></td><td></td><td></td><td></td><td></td></tr>
<tr><td>Oligo fwd</td><td>CACCCAGGCTCAGGCTCAGGCTCG</td><td></td><td></td><td></td><td></td><td></td><td></td><td></td><td></td><td></td><td></td></tr>
<tr><td>Oligo rev</td><td>AAACCGAGCCTGAGCCTGAGCCTG</td><td></td><td></td><td></td><td></td><td></td><td></td><td></td><td></td><td></td><td></td></tr>
<tr><td>Chromosome</td><td>start</td><td>end</td><td>strand</td><td>MM</td><td>target_seq</td><td>PAM</td><td>alignment</td><td>distance</td><td>position</td><td>gene name</td><td>gene id</td></tr>
<tr><td>GU305914.1</td><td>170167</td><td>170189</td><td>+</td><td>0</td><td>CAGGCTCAGGCTCAGGCTCG</td><td>GGG</td><td>||||||||[||||||||||||]PAM</td><td>NA</td><td>Intergenic</td><td>NA</td><td>NA</td></tr>
<tr><td></td><td></td><td></td><td></td><td></td><td></td><td></td><td></td><td></td><td></td><td></td><td></td></tr>
<tr><td>T12878</td><td>CTCAGGCTCAGGCTCGGGGACGG</td><td>1000</td><td></td><td></td><td></td><td></td><td></td><td></td><td></td><td></td><td></td></tr>
<tr><td>Oligo fwd</td><td>CACCCTCAGGCTCAGGCTCGGGGA</td><td></td><td></td><td></td><td></td><td></td><td></td><td></td><td></td><td></td><td></td></tr>
<tr><td>Oligo rev</td><td>AAACTCCCCGAGCCTGAGCCTGAG</td><td></td><td></td><td></td><td></td><td></td><td></td><td></td><td></td><td></td><td></td></tr>
<tr><td>Chromosome</td><td>start</td><td>end</td><td>strand</td><td>MM</td><td>target_seq</td><td>PAM</td><td>alignment</td><td>distance</td><td>position</td><td>gene name</td><td>gene id</td></tr>
<tr><td>GU305914.1</td><td>170171</td><td>170193</td><td>+</td><td>0</td><td>CTCAGGCTCAGGCTCGGGGA</td><td>CGG</td><td>||||||||[||||||||||||]PAM</td><td>NA</td><td>Intergenic</td><td>NA</td><td>NA</td></tr>
<tr><td></td><td></td><td></td><td></td><td></td><td></td><td></td><td></td><td></td><td></td><td></td><td></td></tr>
<tr><td>T12879</td><td>TCAGGCTCAGGCTCGGGGACGGG</td><td>1000</td><td></td><td></td><td></td><td></td><td></td><td></td><td></td><td></td><td></td></tr>
<tr><td>Oligo fwd</td><td>CACCTCAGGCTCAGGCTCGGGGAC</td><td></td><td></td><td></td><td></td><td></td><td></td><td></td><td></td><td></td><td></td></tr>
<tr><td>Oligo rev</td><td>AAACGTCCCCGAGCCTGAGCCTGA</td><td></td><td></td><td></td><td></td><td></td><td></td><td></td><td></td><td></td><td></td></tr>
<tr><td>Chromosome</td><td>start</td><td>end</td><td>strand</td><td>MM</td><td>target_seq</td><td>PAM</td><td>alignment</td><td>distance</td><td>position</td><td>gene name</td><td>gene id</td></tr>
<tr><td>GU305914.1</td><td>170172</td><td>170194</td><td>+</td><td>0</td><td>TCAGGCTCAGGCTCGGGGAC</td><td>GGG</td><td>||||||||[||||||||||||]PAM</td><td>NA</td><td>Intergenic</td><td>NA</td><td>NA</td></tr>
<tr><td></td><td></td><td></td><td></td><td></td><td></td><td></td><td></td><td></td><td></td><td></td><td></td></tr>
<tr><td>T12880</td><td>GCTCAGGCTCGGGGACGGGTTGG</td><td>1000</td><td></td><td></td><td></td><td></td><td></td><td></td><td></td><td></td><td></td></tr>
<tr><td>Oligo fwd</td><td>CACCGCTCAGGCTCGGGGACGGGT</td><td></td><td></td><td></td><td></td><td></td><td></td><td></td><td></td><td></td><td></td></tr>
<tr><td>Oligo rev</td><td>AAACACCCGTCCCCGAGCCTGAGC</td><td></td><td></td><td></td><td></td><td></td><td></td><td></td><td></td><td></td><td></td></tr>
<tr><td>Chromosome</td><td>start</td><td>end</td><td>strand</td><td>MM</td><td>target_seq</td><td>PAM</td><td>alignment</td><td>distance</td><td>position</td><td>gene name</td><td>gene id</td></tr>
<tr><td>GU305914.1</td><td>170176</td><td>170198</td><td>+</td><td>0</td><td>GCTCAGGCTCGGGGACGGGT</td><td>TGG</td><td>||||||||[||||||||||||]PAM</td><td>NA</td><td>Intergenic</td><td>NA</td><td>NA</td></tr>
<tr><td>GU305914.1</td><td>116369</td><td>116391</td><td>+</td><td>4</td><td>GCGGCGGCTCGGGGCCGGGT</td><td>CGG</td><td>||---|||[||||||-|||||]PAM</td><td>NA</td><td>Intergenic</td><td>NA</td><td>NA</td></tr>
<tr><td></td><td></td><td></td><td></td><td></td><td></td><td></td><td></td><td></td><td></td><td></td><td></td></tr>
<tr><td>T12881</td><td>CTCAGGCTCGGGGACGGGTTGGG</td><td>1000</td><td></td><td></td><td></td><td></td><td></td><td></td><td></td><td></td><td></td></tr>
<tr><td>Oligo fwd</td><td>CACCCTCAGGCTCGGGGACGGGTT</td><td></td><td></td><td></td><td></td><td></td><td></td><td></td><td></td><td></td><td></td></tr>
<tr><td>Oligo rev</td><td>AAACAACCCGTCCCCGAGCCTGAG</td><td></td><td></td><td></td><td></td><td></td><td></td><td></td><td></td><td></td><td></td></tr>
<tr><td>Chromosome</td><td>start</td><td>end</td><td>strand</td><td>MM</td><td>target_seq</td><td>PAM</td><td>alignment</td><td>distance</td><td>position</td><td>gene name</td><td>gene id</td></tr>
<tr><td>GU305914.1</td><td>170177</td><td>170199</td><td>+</td><td>0</td><td>CTCAGGCTCGGGGACGGGTT</td><td>GGG</td><td>||||||||[||||||||||||]PAM</td><td>NA</td><td>Intergenic</td><td>NA</td><td>NA</td></tr>
<tr><td></td><td></td><td></td><td></td><td></td><td></td><td></td><td></td><td></td><td></td><td></td><td></td></tr>
<tr><td>T12882</td><td>TCAGGCTCGGGGACGGGTTGGGG</td><td>1000</td><td></td><td></td><td></td><td></td><td></td><td></td><td></td><td></td><td></td></tr>
<tr><td>Oligo fwd</td><td>CACCTCAGGCTCGGGGACGGGTTG</td><td></td><td></td><td></td><td></td><td></td><td></td><td></td><td></td><td></td><td></td></tr>
<tr><td>Oligo rev</td><td>AAACCAACCCGTCCCCGAGCCTGA</td><td></td><td></td><td></td><td></td><td></td><td></td><td></td><td></td><td></td><td></td></tr>
<tr><td>Chromosome</td><td>start</td><td>end</td><td>strand</td><td>MM</td><td>target_seq</td><td>PAM</td><td>alignment</td><td>distance</td><td>position</td><td>gene name</td><td>gene id</td></tr>
<tr><td>GU305914.1</td><td>170178</td><td>170200</td><td>+</td><td>0</td><td>TCAGGCTCGGGGACGGGTTG</td><td>GGG</td><td>||||||||[||||||||||||]PAM</td><td>NA</td><td>Intergenic</td><td>NA</td><td>NA</td></tr>
<tr><td></td><td></td><td></td><td></td><td></td><td></td><td></td><td></td><td></td><td></td><td></td><td></td></tr>
<tr><td>T12883</td><td>CAGGCTCGGGGACGGGTTGGGGG</td><td>1000</td><td></td><td></td><td></td><td></td><td></td><td></td><td></td><td></td><td></td></tr>
<tr><td>Oligo fwd</td><td>CACCCAGGCTCGGGGACGGGTTGG</td><td></td><td></td><td></td><td></td><td></td><td></td><td></td><td></td><td></td><td></td></tr>
<tr><td>Oligo rev</td><td>AAACCCAACCCGTCCCCGAGCCTG</td><td></td><td></td><td></td><td></td><td></td><td></td><td></td><td></td><td></td><td></td></tr>
<tr><td>Chromosome</td><td>start</td><td>end</td><td>strand</td><td>MM</td><td>target_seq</td><td>PAM</td><td>alignment</td><td>distance</td><td>position</td><td>gene name</td><td>gene id</td></tr>
<tr><td>GU305914.1</td><td>170179</td><td>170201</td><td>+</td><td>0</td><td>CAGGCTCGGGGACGGGTTGG</td><td>GGG</td><td>||||||||[||||||||||||]PAM</td><td>NA</td><td>Intergenic</td><td>NA</td><td>NA</td></tr>
<tr><td></td><td></td><td></td><td></td><td></td><td></td><td></td><td></td><td></td><td></td><td></td><td></td></tr>
<tr><td>T12884</td><td>GACGGGTTGGGGGATGATAACGG</td><td>1000</td><td></td><td></td><td></td><td></td><td></td><td></td><td></td><td></td><td></td></tr>
<tr><td>Oligo fwd</td><td>CACCGACGGGTTGGGGGATGATAA</td><td></td><td></td><td></td><td></td><td></td><td></td><td></td><td></td><td></td><td></td></tr>
<tr><td>Oligo rev</td><td>AAACTTATCATCCCCCAACCCGTC</td><td></td><td></td><td></td><td></td><td></td><td></td><td></td><td></td><td></td><td></td></tr>
<tr><td>Chromosome</td><td>start</td><td>end</td><td>strand</td><td>MM</td><td>target_seq</td><td>PAM</td><td>alignment</td><td>distance</td><td>position</td><td>gene name</td><td>gene id</td></tr>
<tr><td>GU305914.1</td><td>170189</td><td>170211</td><td>+</td><td>0</td><td>GACGGGTTGGGGGATGATAA</td><td>CGG</td><td>||||||||[||||||||||||]PAM</td><td>NA</td><td>Intergenic</td><td>NA</td><td>NA</td></tr>
<tr><td></td><td></td><td></td><td></td><td></td><td></td><td></td><td></td><td></td><td></td><td></td><td></td></tr>
<tr><td>T12885</td><td>ACGGGTTGGGGGATGATAACGGG</td><td>1000</td><td></td><td></td><td></td><td></td><td></td><td></td><td></td><td></td><td></td></tr>
<tr><td>Oligo fwd</td><td>CACCACGGGTTGGGGGATGATAAC</td><td></td><td></td><td></td><td></td><td></td><td></td><td></td><td></td><td></td><td></td></tr>
<tr><td>Oligo rev</td><td>AAACGTTATCATCCCCCAACCCGT</td><td></td><td></td><td></td><td></td><td></td><td></td><td></td><td></td><td></td><td></td></tr>
<tr><td>Chromosome</td><td>start</td><td>end</td><td>strand</td><td>MM</td><td>target_seq</td><td>PAM</td><td>alignment</td><td>distance</td><td>position</td><td>gene name</td><td>gene id</td></tr>
<tr><td>GU305914.1</td><td>170190</td><td>170212</td><td>+</td><td>0</td><td>ACGGGTTGGGGGATGATAAC</td><td>GGG</td><td>||||||||[||||||||||||]PAM</td><td>NA</td><td>Intergenic</td><td>NA</td><td>NA</td></tr>
<tr><td></td><td></td><td></td><td></td><td></td><td></td><td></td><td></td><td></td><td></td><td></td><td></td></tr>
<tr><td>T12886</td><td>TTGGGGGATGATAACGGGTTCGG</td><td>1000</td><td></td><td></td><td></td><td></td><td></td><td></td><td></td><td></td><td></td></tr>
<tr><td>Oligo fwd</td><td>CACCTTGGGGGATGATAACGGGTT</td><td></td><td></td><td></td><td></td><td></td><td></td><td></td><td></td><td></td><td></td></tr>
<tr><td>Oligo rev</td><td>AAACAACCCGTTATCATCCCCCAA</td><td></td><td></td><td></td><td></td><td></td><td></td><td></td><td></td><td></td><td></td></tr>
<tr><td>Chromosome</td><td>start</td><td>end</td><td>strand</td><td>MM</td><td>target_seq</td><td>PAM</td><td>alignment</td><td>distance</td><td>position</td><td>gene name</td><td>gene id</td></tr>
<tr><td>GU305914.1</td><td>170195</td><td>170217</td><td>+</td><td>0</td><td>TTGGGGGATGATAACGGGTT</td><td>CGG</td><td>||||||||[||||||||||||]PAM</td><td>NA</td><td>Intergenic</td><td>NA</td><td>NA</td></tr>
<tr><td></td><td></td><td></td><td></td><td></td><td></td><td></td><td></td><td></td><td></td><td></td><td></td></tr>
<tr><td>T12887</td><td>TGGGGGATGATAACGGGTTCGGG</td><td>1000</td><td></td><td></td><td></td><td></td><td></td><td></td><td></td><td></td><td></td></tr>
<tr><td>Oligo fwd</td><td>CACCTGGGGGATGATAACGGGTTC</td><td></td><td></td><td></td><td></td><td></td><td></td><td></td><td></td><td></td><td></td></tr>
<tr><td>Oligo rev</td><td>AAACGAACCCGTTATCATCCCCCA</td><td></td><td></td><td></td><td></td><td></td><td></td><td></td><td></td><td></td><td></td></tr>
<tr><td>Chromosome</td><td>start</td><td>end</td><td>strand</td><td>MM</td><td>target_seq</td><td>PAM</td><td>alignment</td><td>distance</td><td>position</td><td>gene name</td><td>gene id</td></tr>
<tr><td>GU305914.1</td><td>170196</td><td>170218</td><td>+</td><td>0</td><td>TGGGGGATGATAACGGGTTC</td><td>GGG</td><td>||||||||[||||||||||||]PAM</td><td>NA</td><td>Intergenic</td><td>NA</td><td>NA</td></tr>
<tr><td></td><td></td><td></td><td></td><td></td><td></td><td></td><td></td><td></td><td></td><td></td><td></td></tr>
<tr><td>T12888</td><td>GGGGGATGATAACGGGTTCGGGG</td><td>1000</td><td></td><td></td><td></td><td></td><td></td><td></td><td></td><td></td><td></td></tr>
<tr><td>Oligo fwd</td><td>CACCGGGGGATGATAACGGGTTCG</td><td></td><td></td><td></td><td></td><td></td><td></td><td></td><td></td><td></td><td></td></tr>
<tr><td>Oligo rev</td><td>AAACCGAACCCGTTATCATCCCCC</td><td></td><td></td><td></td><td></td><td></td><td></td><td></td><td></td><td></td><td></td></tr>
<tr><td>Chromosome</td><td>start</td><td>end</td><td>strand</td><td>MM</td><td>target_seq</td><td>PAM</td><td>alignment</td><td>distance</td><td>position</td><td>gene name</td><td>gene id</td></tr>
<tr><td>GU305914.1</td><td>170197</td><td>170219</td><td>+</td><td>0</td><td>GGGGGATGATAACGGGTTCG</td><td>GGG</td><td>||||||||[||||||||||||]PAM</td><td>NA</td><td>Intergenic</td><td>NA</td><td>NA</td></tr>
<tr><td></td><td></td><td></td><td></td><td></td><td></td><td></td><td></td><td></td><td></td><td></td><td></td></tr>
<tr><td>T12889</td><td>GATGATAACGGGTTCGGGGACGG</td><td>1000</td><td></td><td></td><td></td><td></td><td></td><td></td><td></td><td></td><td></td></tr>
<tr><td>Oligo fwd</td><td>CACCGATGATAACGGGTTCGGGGA</td><td></td><td></td><td></td><td></td><td></td><td></td><td></td><td></td><td></td><td></td></tr>
<tr><td>Oligo rev</td><td>AAACTCCCCGAACCCGTTATCATC</td><td></td><td></td><td></td><td></td><td></td><td></td><td></td><td></td><td></td><td></td></tr>
<tr><td>Chromosome</td><td>start</td><td>end</td><td>strand</td><td>MM</td><td>target_seq</td><td>PAM</td><td>alignment</td><td>distance</td><td>position</td><td>gene name</td><td>gene id</td></tr>
<tr><td>GU305914.1</td><td>170201</td><td>170223</td><td>+</td><td>0</td><td>GATGATAACGGGTTCGGGGA</td><td>CGG</td><td>||||||||[||||||||||||]PAM</td><td>NA</td><td>Intergenic</td><td>NA</td><td>NA</td></tr>
<tr><td></td><td></td><td></td><td></td><td></td><td></td><td></td><td></td><td></td><td></td><td></td><td></td></tr>
<tr><td>T12890</td><td>ATGATAACGGGTTCGGGGACGGG</td><td>1000</td><td></td><td></td><td></td><td></td><td></td><td></td><td></td><td></td><td></td></tr>
<tr><td>Oligo fwd</td><td>CACCATGATAACGGGTTCGGGGAC</td><td></td><td></td><td></td><td></td><td></td><td></td><td></td><td></td><td></td><td></td></tr>
<tr><td>Oligo rev</td><td>AAACGTCCCCGAACCCGTTATCAT</td><td></td><td></td><td></td><td></td><td></td><td></td><td></td><td></td><td></td><td></td></tr>
<tr><td>Chromosome</td><td>start</td><td>end</td><td>strand</td><td>MM</td><td>target_seq</td><td>PAM</td><td>alignment</td><td>distance</td><td>position</td><td>gene name</td><td>gene id</td></tr>
</table>

| | | | | | | | | | | | |
|---|---|---|---|---|---|---|---|---|---|---|---|
| GU305914.1 | 170202 | 170224 | + | 0 | ATGATAACGGGTTCGGGGAC | GGG | \|\|\|\|\|\|\|\|[\|\|\|\|\|\|\|\|\|\|\|\|]PAM | NA | Intergenic | NA | NA |
| T12891 | AACGGGTTCGGGGACGGGCTCGG | 1000 | | | | | | | | | |
| Oligo fwd | CACCAACGGGTTCGGGGACGGGCT | | | | | | | | | | |
| Oligo rev | AAACAGCCCGTCCCCGAACCCGTT | | | | | | | | | | |
| Chromosome | start | end | strand | MM | target_seq | PAM | alignment | distance | position | gene name | gene id |
| GU305914.1 | 170207 | 170229 | + | 0 | AACGGGTTCGGGGACGGGCT | CGG | \|\|\|\|\|\|\|\|[\|\|\|\|\|\|\|\|\|\|\|\|]PAM | NA | Intergenic | NA | NA |
| T12892 | GGGTTCGGGGACGGGCTCGGCGG | 1000 | | | | | | | | | |
| Oligo fwd | CACCGGGTTCGGGGACGGGCTCGG | | | | | | | | | | |
| Oligo rev | AAACCCGAGCCCGTCCCCGAACCC | | | | | | | | | | |
| Chromosome | start | end | strand | MM | target_seq | PAM | alignment | distance | position | gene name | gene id |
| GU305914.1 | 170210 | 170232 | + | 0 | GGGTTCGGGGACGGGCTCGG | CGG | \|\|\|\|\|\|\|\|[\|\|\|\|\|\|\|\|\|\|\|\|]PAM | NA | Intergenic | NA | NA |
| GU305914.1 | 162103 | 162125 | + | 4 | GCGGACGGGGACGGGTTCGG | AGG | \|-\|--\|\|\|[\|\|\|\|\|\|\|-\|\|\|\|]PAM | NA | Intergenic | NA | NA |
| T12893 | GGTTCGGGGACGGGCTCGGCGGG | 1000 | | | | | | | | | |
| Oligo fwd | CACCGGTTCGGGGACGGGCTCGGC | | | | | | | | | | |
| Oligo rev | AAACGCCGAGCCCGTCCCCGAACC | | | | | | | | | | |
| Chromosome | start | end | strand | MM | target_seq | PAM | alignment | distance | position | gene name | gene id |
| GU305914.1 | 170211 | 170233 | + | 0 | GGTTCGGGGACGGGCTCGGC | GGG | \|\|\|\|\|\|\|\|[\|\|\|\|\|\|\|\|\|\|\|\|]PAM | NA | Intergenic | NA | NA |
| T12894 | GGCGGGAGAAGAAGTACCGACGG | 1000 | | | | | | | | | |
| Oligo fwd | CACCGGCGGGAGAAGAAGTACCGA | | | | | | | | | | |
| Oligo rev | AAACTCGGTACTTCTTCTCCCGCC | | | | | | | | | | |
| Chromosome | start | end | strand | MM | target_seq | PAM | alignment | distance | position | gene name | gene id |
| GU305914.1 | 170228 | 170250 | + | 0 | GGCGGGAGAAGAAGTACCGA | CGG | \|\|\|\|\|\|\|\|[\|\|\|\|\|\|\|\|\|\|\|\|]PAM | NA | Intergenic | NA | NA |
| T12895 | GCGGGAGAAGAAGTACCGACGGG | 1000 | | | | | | | | | |
| Oligo fwd | CACCGCGGGAGAAGAAGTACCGAC | | | | | | | | | | |
| Oligo rev | AAACGTCGGTACTTCTTCTCCCGC | | | | | | | | | | |
| Chromosome | start | end | strand | MM | target_seq | PAM | alignment | distance | position | gene name | gene id |
| GU305914.1 | 170229 | 170251 | + | 0 | GCGGGAGAAGAAGTACCGAC | GGG | \|\|\|\|\|\|\|\|[\|\|\|\|\|\|\|\|\|\|\|\|]PAM | NA | Intergenic | NA | NA |
| T12896 | AGAAGAAGTACCGACGGGTTCGG | 1000 | | | | | | | | | |
| Oligo fwd | CACCAGAAGAAGTACCGACGGGTT | | | | | | | | | | |
| Oligo rev | AAACAACCCGTCGGTACTTCTTCT | | | | | | | | | | |
| Chromosome | start | end | strand | MM | target_seq | PAM | alignment | distance | position | gene name | gene id |
| GU305914.1 | 170234 | 170256 | + | 0 | AGAAGAAGTACCGACGGGTT | CGG | \|\|\|\|\|\|\|\|[\|\|\|\|\|\|\|\|\|\|\|\|]PAM | NA | Intergenic | NA | NA |
| T12897 | AGTACCGACGGGTTCGGATTCGG | 1000 | | | | | | | | | |
| Oligo fwd | CACCAGTACCGACGGGTTCGGATT | | | | | | | | | | |
| Oligo rev | AAACAATCCGAACCCGTCGGTACT | | | | | | | | | | |
| Chromosome | start | end | strand | MM | target_seq | PAM | alignment | distance | position | gene name | gene id |
| GU305914.1 | 170240 | 170262 | + | 0 | AGTACCGACGGGTTCGGATT | CGG | \|\|\|\|\|\|\|\|[\|\|\|\|\|\|\|\|\|\|\|\|]PAM | NA | Intergenic | NA | NA |
| T12898 | GTACCGACGGGTTCGGATTCGGG | 1000 | | | | | | | | | |
| Oligo fwd | CACCGTACCGACGGGTTCGGATTC | | | | | | | | | | |
| Oligo rev | AAACGAATCCGAACCCGTCGGTAC | | | | | | | | | | |
| Chromosome | start | end | strand | MM | target_seq | PAM | alignment | distance | position | gene name | gene id |
| GU305914.1 | 170241 | 170263 | + | 0 | GTACCGACGGGTTCGGATTC | GGG | \|\|\|\|\|\|\|\|[\|\|\|\|\|\|\|\|\|\|\|\|]PAM | NA | Intergenic | NA | NA |
| T12899 | TACCGACGGGTTCGGATTCGGGG | 1000 | | | | | | | | | |
| Oligo fwd | CACCTACCGACGGGTTCGGATTCG | | | | | | | | | | |
| Oligo rev | AAACCGAATCCGAACCCGTCGGTA | | | | | | | | | | |
| Chromosome | start | end | strand | MM | target_seq | PAM | alignment | distance | position | gene name | gene id |
| GU305914.1 | 170242 | 170264 | + | 0 | TACCGACGGGTTCGGATTCG | GGG | \|\|\|\|\|\|\|\|[\|\|\|\|\|\|\|\|\|\|\|\|]PAM | NA | Intergenic | NA | NA |
| T12900 | GACGGGTTCGGATTCGGGGACGG | 1000 | | | | | | | | | |
| Oligo fwd | CACCGACGGGTTCGGATTCGGGGA | | | | | | | | | | |
| Oligo rev | AAACTCCCCGAATCCGAACCCGTC | | | | | | | | | | |
| Chromosome | start | end | strand | MM | target_seq | PAM | alignment | distance | position | gene name | gene id |
| GU305914.1 | 170246 | 170268 | + | 0 | GACGGGTTCGGATTCGGGGA | CGG | \|\|\|\|\|\|\|\|[\|\|\|\|\|\|\|\|\|\|\|\|]PAM | NA | Intergenic | NA | NA |
| GU305914.1 | 26115 | 26137 | - | 4 | GCAGGGTTCGGATGCGGAGA | TGG | \|--\|\|\|\|\|[\|\|\|\|\|-\|\|\|-\|\|]PAM | NA | Intergenic | NA | NA |
| T12901 | TTCGGATTCGGGGACGGTGACGG | 1000 | | | | | | | | | |
| Oligo fwd | CACCTTCGGATTCGGGGACGGTGA | | | | | | | | | | |
| Oligo rev | AAACTCACCGTCCCCGAATCCGAA | | | | | | | | | | |
| Chromosome | start | end | strand | MM | target_seq | PAM | alignment | distance | position | gene name | gene id |
| GU305914.1 | 170252 | 170274 | + | 0 | TTCGGATTCGGGGACGGTGA | CGG | \|\|\|\|\|\|\|\|[\|\|\|\|\|\|\|\|\|\|\|\|]PAM | NA | Intergenic | NA | NA |
| GU305914.1 | 170129 | 170151 | + | 3 | GACGGGTTCGGGGACGGTGA | CGG | --\|\|\|-\|\|[\|\|\|\|\|\|\|\|\|\|\|\|]PAM | NA | Intergenic | NA | NA |
| GU305914.1 | 31158 | 31180 | - | 4 | GTCGGAGACGGAGACGGTGA | CGG | -\|\|\|\|\|--[\|\|\|-\|\|\|\|\|\|\|\|]PAM | NA | Intergenic | NA | NA |
| T12902 | TCGGATTCGGGGACGGTGACGGG | 1000 | | | | | | | | | |
| Oligo fwd | CACCTCGGATTCGGGGACGGTGAC | | | | | | | | | | |
| Oligo rev | AAACGTCACCGTCCCCGAATCCGA | | | | | | | | | | |
| Chromosome | start | end | strand | MM | target_seq | PAM | alignment | distance | position | gene name | gene id |
| GU305914.1 | 170253 | 170275 | + | 0 | TCGGATTCGGGGACGGTGAC | GGG | \|\|\|\|\|\|\|\|[\|\|\|\|\|\|\|\|\|\|\|\|]PAM | NA | Intergenic | NA | NA |
| GU305914.1 | 170130 | 170152 | + | 2 | ACGGGTTCGGGGACGGTGAC | GGG | -\|\|\|-\|\|\|[\|\|\|\|\|\|\|\|\|\|\|\|]PAM | NA | Intergenic | NA | NA |
| T12903 | CGGGGACGGTGACGGGCTCGAGG | 1000 | | | | | | | | | |
| Oligo fwd | CACCCGGGGACGGTGACGGGCTCG | | | | | | | | | | |
| Oligo rev | AAACCGAGCCCGTCACCGTCCCCG | | | | | | | | | | |
| Chromosome | start | end | strand | MM | target_seq | PAM | alignment | distance | position | gene name | gene id |
| GU305914.1 | 170260 | 170282 | + | 0 | CGGGGACGGTGACGGGCTCG | AGG | \|\|\|\|\|\|\|\|[\|\|\|\|\|\|\|\|\|\|\|\|]PAM | NA | Intergenic | NA | NA |
| GU305914.1 | 170119 | 170141 | + | 4 | CGGGAACCGCGACGGGTTCG | GGG | \|\|\|\|-\|\|-[\|-\|\|\|\|\|\|-\|\|\|]PAM | NA | Intergenic | NA | NA |
| T12904 | GGGCTCGAGGATGAGCTCTATGG | 1000 | | | | | | | | | |
| Oligo fwd | CACCGGGCTCGAGGATGAGCTCTA | | | | | | | | | | |
| Oligo rev | AAACTAGAGCTCATCCTCGAGCCC | | | | | | | | | | |

<table>
<tr><td>Chromosome</td><td>start</td><td>end</td><td>strand</td><td>MM</td><td>target_seq</td><td>PAM</td><td>alignment</td><td>distance</td><td>position</td><td>gene name</td><td>gene id</td></tr>
<tr><td>GU305914.1</td><td>170273</td><td>170295</td><td>+</td><td>0</td><td>GGGCTCGAGGATGAGCTCTA</td><td>TGG</td><td>||||||||[||||||||||||]PAM</td><td>NA</td><td>Intergenic</td><td>NA</td><td>NA</td></tr>
<tr><td></td><td></td><td></td><td></td><td></td><td></td><td></td><td></td><td></td><td></td><td></td><td></td></tr>
<tr><td>T12905</td><td>GAGGATGAGCTCTATGGTATCGG</td><td>1000</td><td></td><td></td><td></td><td></td><td></td><td></td><td></td><td></td><td></td></tr>
<tr><td>Oligo fwd</td><td>CACCGAGGATGAGCTCTATGGTAT</td><td></td><td></td><td></td><td></td><td></td><td></td><td></td><td></td><td></td><td></td></tr>
<tr><td>Oligo rev</td><td>AAACATACCATAGAGCTCATCCTC</td><td></td><td></td><td></td><td></td><td></td><td></td><td></td><td></td><td></td><td></td></tr>
<tr><td>Chromosome</td><td>start</td><td>end</td><td>strand</td><td>MM</td><td>target_seq</td><td>PAM</td><td>alignment</td><td>distance</td><td>position</td><td>gene name</td><td>gene id</td></tr>
<tr><td>GU305914.1</td><td>170279</td><td>170301</td><td>+</td><td>0</td><td>GAGGATGAGCTCTATGGTAT</td><td>CGG</td><td>||||||||[||||||||||||]PAM</td><td>NA</td><td>Intergenic</td><td>NA</td><td>NA</td></tr>
<tr><td></td><td></td><td></td><td></td><td></td><td></td><td></td><td></td><td></td><td></td><td></td><td></td></tr>
<tr><td>T12906</td><td>AGGATGAGCTCTATGGTATCGGG</td><td>1000</td><td></td><td></td><td></td><td></td><td></td><td></td><td></td><td></td><td></td></tr>
<tr><td>Oligo fwd</td><td>CACCAGGATGAGCTCTATGGTATC</td><td></td><td></td><td></td><td></td><td></td><td></td><td></td><td></td><td></td><td></td></tr>
<tr><td>Oligo rev</td><td>AAACGATACCATAGAGCTCATCCT</td><td></td><td></td><td></td><td></td><td></td><td></td><td></td><td></td><td></td><td></td></tr>
<tr><td>Chromosome</td><td>start</td><td>end</td><td>strand</td><td>MM</td><td>target_seq</td><td>PAM</td><td>alignment</td><td>distance</td><td>position</td><td>gene name</td><td>gene id</td></tr>
<tr><td>GU305914.1</td><td>170280</td><td>170302</td><td>+</td><td>0</td><td>AGGATGAGCTCTATGGTATC</td><td>GGG</td><td>||||||||[||||||||||||]PAM</td><td>NA</td><td>Intergenic</td><td>NA</td><td>NA</td></tr>
<tr><td></td><td></td><td></td><td></td><td></td><td></td><td></td><td></td><td></td><td></td><td></td><td></td></tr>
<tr><td>T12907</td><td>GAGCTCTATGGTATCGGGATCGG</td><td>1000</td><td></td><td></td><td></td><td></td><td></td><td></td><td></td><td></td><td></td></tr>
<tr><td>Oligo fwd</td><td>CACCGAGCTCTATGGTATCGGGAT</td><td></td><td></td><td></td><td></td><td></td><td></td><td></td><td></td><td></td><td></td></tr>
<tr><td>Oligo rev</td><td>AAACATCCCGATACCATAGAGCTC</td><td></td><td></td><td></td><td></td><td></td><td></td><td></td><td></td><td></td><td></td></tr>
<tr><td>Chromosome</td><td>start</td><td>end</td><td>strand</td><td>MM</td><td>target_seq</td><td>PAM</td><td>alignment</td><td>distance</td><td>position</td><td>gene name</td><td>gene id</td></tr>
<tr><td>GU305914.1</td><td>170285</td><td>170307</td><td>+</td><td>0</td><td>GAGCTCTATGGTATCGGGAT</td><td>CGG</td><td>||||||||[||||||||||||]PAM</td><td>NA</td><td>Intergenic</td><td>NA</td><td>NA</td></tr>
<tr><td></td><td></td><td></td><td></td><td></td><td></td><td></td><td></td><td></td><td></td><td></td><td></td></tr>
<tr><td>T12908</td><td>GTATCGGGATCGGTAGTAGTAGG</td><td>1000</td><td></td><td></td><td></td><td></td><td></td><td></td><td></td><td></td><td></td></tr>
<tr><td>Oligo fwd</td><td>CACCGTATCGGGATCGGTAGTAGT</td><td></td><td></td><td></td><td></td><td></td><td></td><td></td><td></td><td></td><td></td></tr>
<tr><td>Oligo rev</td><td>AAACACTACTACCGATCCCGATAC</td><td></td><td></td><td></td><td></td><td></td><td></td><td></td><td></td><td></td><td></td></tr>
<tr><td>Chromosome</td><td>start</td><td>end</td><td>strand</td><td>MM</td><td>target_seq</td><td>PAM</td><td>alignment</td><td>distance</td><td>position</td><td>gene name</td><td>gene id</td></tr>
<tr><td>GU305914.1</td><td>170295</td><td>170317</td><td>+</td><td>0</td><td>GTATCGGGATCGGTAGTAGT</td><td>AGG</td><td>||||||||[||||||||||||]PAM</td><td>NA</td><td>Intergenic</td><td>NA</td><td>NA</td></tr>
<tr><td></td><td></td><td></td><td></td><td></td><td></td><td></td><td></td><td></td><td></td><td></td><td></td></tr>
<tr><td>T12909</td><td>GGGATCGGTAGTAGTAGGTTTGG</td><td>1000</td><td></td><td></td><td></td><td></td><td></td><td></td><td></td><td></td><td></td></tr>
<tr><td>Oligo fwd</td><td>CACCGGGATCGGTAGTAGTAGGTT</td><td></td><td></td><td></td><td></td><td></td><td></td><td></td><td></td><td></td><td></td></tr>
<tr><td>Oligo rev</td><td>AAACAACCTACTACTACCGATCCC</td><td></td><td></td><td></td><td></td><td></td><td></td><td></td><td></td><td></td><td></td></tr>
<tr><td>Chromosome</td><td>start</td><td>end</td><td>strand</td><td>MM</td><td>target_seq</td><td>PAM</td><td>alignment</td><td>distance</td><td>position</td><td>gene name</td><td>gene id</td></tr>
<tr><td>GU305914.1</td><td>170300</td><td>170322</td><td>+</td><td>0</td><td>GGGATCGGTAGTAGTAGGTT</td><td>TGG</td><td>||||||||[||||||||||||]PAM</td><td>NA</td><td>Intergenic</td><td>NA</td><td>NA</td></tr>
<tr><td></td><td></td><td></td><td></td><td></td><td></td><td></td><td></td><td></td><td></td><td></td><td></td></tr>
<tr><td>T12910</td><td>ATCGGTAGTAGTAGGTTTGGTGG</td><td>1000</td><td></td><td></td><td></td><td></td><td></td><td></td><td></td><td></td><td></td></tr>
<tr><td>Oligo fwd</td><td>CACCATCGGTAGTAGTAGGTTTGG</td><td></td><td></td><td></td><td></td><td></td><td></td><td></td><td></td><td></td><td></td></tr>
<tr><td>Oligo rev</td><td>AAACCCAAACCTACTACTACCGAT</td><td></td><td></td><td></td><td></td><td></td><td></td><td></td><td></td><td></td><td></td></tr>
<tr><td>Chromosome</td><td>start</td><td>end</td><td>strand</td><td>MM</td><td>target_seq</td><td>PAM</td><td>alignment</td><td>distance</td><td>position</td><td>gene name</td><td>gene id</td></tr>
<tr><td>GU305914.1</td><td>170303</td><td>170325</td><td>+</td><td>0</td><td>ATCGGTAGTAGTAGGTTTGG</td><td>TGG</td><td>||||||||[||||||||||||]PAM</td><td>NA</td><td>Intergenic</td><td>NA</td><td>NA</td></tr>
<tr><td></td><td></td><td></td><td></td><td></td><td></td><td></td><td></td><td></td><td></td><td></td><td></td></tr>
<tr><td>T12911</td><td>TCGGTAGTAGTAGGTTTGGTGGG</td><td>1000</td><td></td><td></td><td></td><td></td><td></td><td></td><td></td><td></td><td></td></tr>
<tr><td>Oligo fwd</td><td>CACCTCGGTAGTAGTAGGTTTGGT</td><td></td><td></td><td></td><td></td><td></td><td></td><td></td><td></td><td></td><td></td></tr>
<tr><td>Oligo rev</td><td>AAACACCAAACCTACTACTACCGA</td><td></td><td></td><td></td><td></td><td></td><td></td><td></td><td></td><td></td><td></td></tr>
<tr><td>Chromosome</td><td>start</td><td>end</td><td>strand</td><td>MM</td><td>target_seq</td><td>PAM</td><td>alignment</td><td>distance</td><td>position</td><td>gene name</td><td>gene id</td></tr>
<tr><td>GU305914.1</td><td>170304</td><td>170326</td><td>+</td><td>0</td><td>TCGGTAGTAGTAGGTTTGGT</td><td>GGG</td><td>||||||||[||||||||||||]PAM</td><td>NA</td><td>Intergenic</td><td>NA</td><td>NA</td></tr>
<tr><td></td><td></td><td></td><td></td><td></td><td></td><td></td><td></td><td></td><td></td><td></td><td></td></tr>
<tr><td>T12912</td><td>GTCTGTAAAGACGATGAGTTCGG</td><td>1000</td><td></td><td></td><td></td><td></td><td></td><td></td><td></td><td></td><td></td></tr>
<tr><td>Oligo fwd</td><td>CACCGTCTGTAAAGACGATGAGTT</td><td></td><td></td><td></td><td></td><td></td><td></td><td></td><td></td><td></td><td></td></tr>
<tr><td>Oligo rev</td><td>AAACAACTCATCGTCTTTACAGAC</td><td></td><td></td><td></td><td></td><td></td><td></td><td></td><td></td><td></td><td></td></tr>
<tr><td>Chromosome</td><td>start</td><td>end</td><td>strand</td><td>MM</td><td>target_seq</td><td>PAM</td><td>alignment</td><td>distance</td><td>position</td><td>gene name</td><td>gene id</td></tr>
<tr><td>GU305914.1</td><td>170330</td><td>170352</td><td>+</td><td>0</td><td>GTCTGTAAAGACGATGAGTT</td><td>CGG</td><td>||||||||[||||||||||||]PAM</td><td>NA</td><td>Intergenic</td><td>NA</td><td>NA</td></tr>
<tr><td></td><td></td><td></td><td></td><td></td><td></td><td></td><td></td><td></td><td></td><td></td><td></td></tr>
<tr><td>T12913</td><td>TCTGTAAAGACGATGAGTTCGGG</td><td>1000</td><td></td><td></td><td></td><td></td><td></td><td></td><td></td><td></td><td></td></tr>
<tr><td>Oligo fwd</td><td>CACCTCTGTAAAGACGATGAGTTC</td><td></td><td></td><td></td><td></td><td></td><td></td><td></td><td></td><td></td><td></td></tr>
<tr><td>Oligo rev</td><td>AAACGAACTCATCGTCTTTACAGA</td><td></td><td></td><td></td><td></td><td></td><td></td><td></td><td></td><td></td><td></td></tr>
<tr><td>Chromosome</td><td>start</td><td>end</td><td>strand</td><td>MM</td><td>target_seq</td><td>PAM</td><td>alignment</td><td>distance</td><td>position</td><td>gene name</td><td>gene id</td></tr>
<tr><td>GU305914.1</td><td>170331</td><td>170353</td><td>+</td><td>0</td><td>TCTGTAAAGACGATGAGTTC</td><td>GGG</td><td>||||||||[||||||||||||]PAM</td><td>NA</td><td>Intergenic</td><td>NA</td><td>NA</td></tr>
<tr><td></td><td></td><td></td><td></td><td></td><td></td><td></td><td></td><td></td><td></td><td></td><td></td></tr>
<tr><td>T12914</td><td>GTAAAGACGATGAGTTCGGGTGG</td><td>1000</td><td></td><td></td><td></td><td></td><td></td><td></td><td></td><td></td><td></td></tr>
<tr><td>Oligo fwd</td><td>CACCGTAAAGACGATGAGTTCGGG</td><td></td><td></td><td></td><td></td><td></td><td></td><td></td><td></td><td></td><td></td></tr>
<tr><td>Oligo rev</td><td>AAACCCCGAACTCATCGTCTTTAC</td><td></td><td></td><td></td><td></td><td></td><td></td><td></td><td></td><td></td><td></td></tr>
<tr><td>Chromosome</td><td>start</td><td>end</td><td>strand</td><td>MM</td><td>target_seq</td><td>PAM</td><td>alignment</td><td>distance</td><td>position</td><td>gene name</td><td>gene id</td></tr>
<tr><td>GU305914.1</td><td>170334</td><td>170356</td><td>+</td><td>0</td><td>GTAAAGACGATGAGTTCGGG</td><td>TGG</td><td>||||||||[||||||||||||]PAM</td><td>NA</td><td>Intergenic</td><td>NA</td><td>NA</td></tr>
<tr><td></td><td></td><td></td><td></td><td></td><td></td><td></td><td></td><td></td><td></td><td></td><td></td></tr>
<tr><td>T12915</td><td>GATGAGTTCGGGTGGTGTGATGG</td><td>1000</td><td></td><td></td><td></td><td></td><td></td><td></td><td></td><td></td><td></td></tr>
<tr><td>Oligo fwd</td><td>CACCGATGAGTTCGGGTGGTGTGA</td><td></td><td></td><td></td><td></td><td></td><td></td><td></td><td></td><td></td><td></td></tr>
<tr><td>Oligo rev</td><td>AAACTCACACCACCCGAACTCATC</td><td></td><td></td><td></td><td></td><td></td><td></td><td></td><td></td><td></td><td></td></tr>
<tr><td>Chromosome</td><td>start</td><td>end</td><td>strand</td><td>MM</td><td>target_seq</td><td>PAM</td><td>alignment</td><td>distance</td><td>position</td><td>gene name</td><td>gene id</td></tr>
<tr><td>GU305914.1</td><td>170342</td><td>170364</td><td>+</td><td>0</td><td>GATGAGTTCGGGTGGTGTGA</td><td>TGG</td><td>||||||||[||||||||||||]PAM</td><td>NA</td><td>Intergenic</td><td>NA</td><td>NA</td></tr>
<tr><td></td><td></td><td></td><td></td><td></td><td></td><td></td><td></td><td></td><td></td><td></td><td></td></tr>
<tr><td>T12916</td><td>ATGAGTTCGGGTGGTGTGATGGG</td><td>1000</td><td></td><td></td><td></td><td></td><td></td><td></td><td></td><td></td><td></td></tr>
<tr><td>Oligo fwd</td><td>CACCATGAGTTCGGGTGGTGTGAT</td><td></td><td></td><td></td><td></td><td></td><td></td><td></td><td></td><td></td><td></td></tr>
<tr><td>Oligo rev</td><td>AAACATCACACCACCCGAACTCAT</td><td></td><td></td><td></td><td></td><td></td><td></td><td></td><td></td><td></td><td></td></tr>
<tr><td>Chromosome</td><td>start</td><td>end</td><td>strand</td><td>MM</td><td>target_seq</td><td>PAM</td><td>alignment</td><td>distance</td><td>position</td><td>gene name</td><td>gene id</td></tr>
<tr><td>GU305914.1</td><td>170343</td><td>170365</td><td>+</td><td>0</td><td>ATGAGTTCGGGTGGTGTGAT</td><td>GGG</td><td>||||||||[||||||||||||]PAM</td><td>NA</td><td>Intergenic</td><td>NA</td><td>NA</td></tr>
<tr><td></td><td></td><td></td><td></td><td></td><td></td><td></td><td></td><td></td><td></td><td></td><td></td></tr>
<tr><td>T12917</td><td>TCGGGTGGTGTGATGGGCTCCGG</td><td>1000</td><td></td><td></td><td></td><td></td><td></td><td></td><td></td><td></td><td></td></tr>
<tr><td>Oligo fwd</td><td>CACCTCGGGTGGTGTGATGGGCTC</td><td></td><td></td><td></td><td></td><td></td><td></td><td></td><td></td><td></td><td></td></tr>
<tr><td>Oligo rev</td><td>AAACGAGCCCATCACACCACCCGA</td><td></td><td></td><td></td><td></td><td></td><td></td><td></td><td></td><td></td><td></td></tr>
<tr><td>Chromosome</td><td>start</td><td>end</td><td>strand</td><td>MM</td><td>target_seq</td><td>PAM</td><td>alignment</td><td>distance</td><td>position</td><td>gene name</td><td>gene id</td></tr>
<tr><td>GU305914.1</td><td>170349</td><td>170371</td><td>+</td><td>0</td><td>TCGGGTGGTGTGATGGGCTC</td><td>CGG</td><td>||||||||[||||||||||||]PAM</td><td>NA</td><td>Intergenic</td><td>NA</td><td>NA</td></tr>
<tr><td>GU305914.1</td><td>170403</td><td>170425</td><td>+</td><td>4</td><td>TCGGGTATCGAGATGGGCTC</td><td>TGG</td><td>||||||--[-|-|||||||||]PAM</td><td>NA</td><td>Intergenic</td><td>NA</td><td>NA</td></tr>
<tr><td></td><td></td><td></td><td></td><td></td><td></td><td></td><td></td><td></td><td></td><td></td><td></td></tr>
<tr><td>T12918</td><td>CGGGTGGTGTGATGGGCTCCGGG</td><td>1000</td><td></td><td></td><td></td><td></td><td></td><td></td><td></td><td></td><td></td></tr>
<tr><td>Oligo fwd</td><td>CACCCGGGTGGTGTGATGGGCTCC</td><td></td><td></td><td></td><td></td><td></td><td></td><td></td><td></td><td></td><td></td></tr>
<tr><td>Oligo rev</td><td>AAACGGAGCCCATCACACCACCCG</td><td></td><td></td><td></td><td></td><td></td><td></td><td></td><td></td><td></td><td></td></tr>
<tr><td>Chromosome</td><td>start</td><td>end</td><td>strand</td><td>MM</td><td>target_seq</td><td>PAM</td><td>alignment</td><td>distance</td><td>position</td><td>gene name</td><td>gene id</td></tr>
<tr><td>GU305914.1</td><td>170350</td><td>170372</td><td>+</td><td>0</td><td>CGGGTGGTGTGATGGGCTCC</td><td>GGG</td><td>||||||||[||||||||||||]PAM</td><td>NA</td><td>Intergenic</td><td>NA</td><td>NA</td></tr>
<tr><td></td><td></td><td></td><td></td><td></td><td></td><td></td><td></td><td></td><td></td><td></td><td></td></tr>
<tr><td>T12919</td><td>GATGGGCTCCGGGAACGCGATGG</td><td>1000</td><td></td><td></td><td></td><td></td><td></td><td></td><td></td><td></td><td></td></tr>
</table>

| | | | | | | | | | | | |
|---|---|---|---|---|---|---|---|---|---|---|---|
| Oligo fwd | CACCGATGGGCTCCGGGAACGCGA | | | | | | | | | | |
| Oligo rev | AAACTCGCGTTCCCGGAGCCCATC | | | | | | | | | | |
| Chromosome | start | end | strand | MM | target_seq | PAM | alignment | distance | position | gene name | gene id |
| GU305914.1 | 170360 | 170382 | + | 0 | GATGGGCTCCGGGAACGCGA | TGG | \|\|\|\|\|\|\|\|[\|\|\|\|\|\|\|\|\|\|\|\|]PAM | NA | Intergenic | NA | NA |
| T12920 | ATGGGCTCCGGGAACGCGATGGG | 1000 | | | | | | | | | |
| Oligo fwd | CACCATGGGCTCCGGGAACGCGAT | | | | | | | | | | |
| Oligo rev | AAACATCGCGTTCCCGGAGCCCAT | | | | | | | | | | |
| Chromosome | start | end | strand | MM | target_seq | PAM | alignment | distance | position | gene name | gene id |
| GU305914.1 | 170361 | 170383 | + | 0 | ATGGGCTCCGGGAACGCGAT | GGG | \|\|\|\|\|\|\|\|[\|\|\|\|\|\|\|\|\|\|\|\|]PAM | NA | Intergenic | NA | NA |
| T12921 | CTCCGGGAACGCGATGGGCTCGG | 1000 | | | | | | | | | |
| Oligo fwd | CACCCTCCGGGAACGCGATGGGCT | | | | | | | | | | |
| Oligo rev | AAACAGCCCATCGCGTTCCCGGAG | | | | | | | | | | |
| Chromosome | start | end | strand | MM | target_seq | PAM | alignment | distance | position | gene name | gene id |
| GU305914.1 | 170366 | 170388 | + | 0 | CTCCGGGAACGCGATGGGCT | CGG | \|\|\|\|\|\|\|\|[\|\|\|\|\|\|\|\|\|\|\|\|]PAM | NA | Intergenic | NA | NA |
| GU305914.1 | 196693 | 196715 | + | 4 | CGCCCCGAACGCGATGGCCT | TGG | \|-\|\|--\|\|[\|\|\|\|\|\|\|\|\|-\|\|]PAM | NA | Intergenic | NA | NA |
| T12922 | GATGGGCTCGGTGAACCTGCCGG | 1000 | | | | | | | | | |
| Oligo fwd | CACCGATGGGCTCGGTGAACCTGC | | | | | | | | | | |
| Oligo rev | AAACGCAGGTTCACCGAGCCCATC | | | | | | | | | | |
| Chromosome | start | end | strand | MM | target_seq | PAM | alignment | distance | position | gene name | gene id |
| GU305914.1 | 170378 | 170400 | + | 0 | GATGGGCTCGGTGAACCTGC | CGG | \|\|\|\|\|\|\|\|[\|\|\|\|\|\|\|\|\|\|\|\|]PAM | NA | Intergenic | NA | NA |
| T12923 | ATGGGCTCGGTGAACCTGCCGGG | 1000 | | | | | | | | | |
| Oligo fwd | CACCATGGGCTCGGTGAACCTGCC | | | | | | | | | | |
| Oligo rev | AAACGGCAGGTTCACCGAGCCCAT | | | | | | | | | | |
| Chromosome | start | end | strand | MM | target_seq | PAM | alignment | distance | position | gene name | gene id |
| GU305914.1 | 170379 | 170401 | + | 0 | ATGGGCTCGGTGAACCTGCC | GGG | \|\|\|\|\|\|\|\|[\|\|\|\|\|\|\|\|\|\|\|\|]PAM | NA | Intergenic | NA | NA |
| T12924 | CTCGGTGAACCTGCCGGGATCGG | 1000 | | | | | | | | | |
| Oligo fwd | CACCCTCGGTGAACCTGCCGGGAT | | | | | | | | | | |
| Oligo rev | AAACATCCCGGCAGGTTCACCGAG | | | | | | | | | | |
| Chromosome | start | end | strand | MM | target_seq | PAM | alignment | distance | position | gene name | gene id |
| GU305914.1 | 170384 | 170406 | + | 0 | CTCGGTGAACCTGCCGGGAT | CGG | \|\|\|\|\|\|\|\|[\|\|\|\|\|\|\|\|\|\|\|\|]PAM | NA | Intergenic | NA | NA |
| T12925 | TCGGTGAACCTGCCGGGATCGGG | 1000 | | | | | | | | | |
| Oligo fwd | CACCTCGGTGAACCTGCCGGGATC | | | | | | | | | | |
| Oligo rev | AAACGATCCCGGCAGGTTCACCGA | | | | | | | | | | |
| Chromosome | start | end | strand | MM | target_seq | PAM | alignment | distance | position | gene name | gene id |
| GU305914.1 | 170385 | 170407 | + | 0 | TCGGTGAACCTGCCGGGATC | GGG | \|\|\|\|\|\|\|\|[\|\|\|\|\|\|\|\|\|\|\|\|]PAM | NA | Intergenic | NA | NA |
| T12926 | GCCGGGATCGGGTATCGAGATGG | 1000 | | | | | | | | | |
| Oligo fwd | CACCGCCGGGATCGGGTATCGAGA | | | | | | | | | | |
| Oligo rev | AAACTCTCGATACCCGATCCCGGC | | | | | | | | | | |
| Chromosome | start | end | strand | MM | target_seq | PAM | alignment | distance | position | gene name | gene id |
| GU305914.1 | 170396 | 170418 | + | 0 | GCCGGGATCGGGTATCGAGA | TGG | \|\|\|\|\|\|\|\|[\|\|\|\|\|\|\|\|\|\|\|\|]PAM | NA | Intergenic | NA | NA |
| T12927 | CCGGGATCGGGTATCGAGATGGG | 1000 | | | | | | | | | |
| Oligo fwd | CACCCCGGGATCGGGTATCGAGAT | | | | | | | | | | |
| Oligo rev | AAACATCTCGATACCCGATCCCGG | | | | | | | | | | |
| Chromosome | start | end | strand | MM | target_seq | PAM | alignment | distance | position | gene name | gene id |
| GU305914.1 | 170397 | 170419 | + | 0 | CCGGGATCGGGTATCGAGAT | GGG | \|\|\|\|\|\|\|\|[\|\|\|\|\|\|\|\|\|\|\|\|]PAM | NA | Intergenic | NA | NA |
| GU305914.1 | 62288 | 62310 | + | 4 | CGGGGAGCGGGAATCGGGAT | AGG | \|-\|\|\|\|-\|[\|\|\|-\|\|\|\|-\|\|\|]PAM | NA | Intergenic | NA | NA |
| T12928 | TCGGGTATCGAGATGGGCTCTGG | 1000 | | | | | | | | | |
| Oligo fwd | CACCTCGGGTATCGAGATGGGCTC | | | | | | | | | | |
| Oligo rev | AAACGAGCCCATCTCGATACCCGA | | | | | | | | | | |
| Chromosome | start | end | strand | MM | target_seq | PAM | alignment | distance | position | gene name | gene id |
| GU305914.1 | 170403 | 170425 | + | 0 | TCGGGTATCGAGATGGGCTC | TGG | \|\|\|\|\|\|\|\|[\|\|\|\|\|\|\|\|\|\|\|\|]PAM | NA | Intergenic | NA | NA |
| GU305914.1 | 170349 | 170371 | + | 4 | TCGGGTGGTGTGATGGGCTC | CGG | \|\|\|\|\|\|--[-\|-\|\|\|\|\|\|\|\|\|]PAM | NA | Intergenic | NA | NA |
| T12929 | GATGGGCTCTGGAAACGCGTCGG | 1000 | | | | | | | | | |
| Oligo fwd | CACCGATGGGCTCTGGAAACGCGT | | | | | | | | | | |
| Oligo rev | AAACACGCGTTTCCAGAGCCCATC | | | | | | | | | | |
| Chromosome | start | end | strand | MM | target_seq | PAM | alignment | distance | position | gene name | gene id |
| GU305914.1 | 170414 | 170436 | + | 0 | GATGGGCTCTGGAAACGCGT | CGG | \|\|\|\|\|\|\|\|[\|\|\|\|\|\|\|\|\|\|\|\|]PAM | NA | Intergenic | NA | NA |
| T12930 | ATGGGCTCTGGAAACGCGTCGGG | 1000 | | | | | | | | | |
| Oligo fwd | CACCATGGGCTCTGGAAACGCGTC | | | | | | | | | | |
| Oligo rev | AAACGACGCGTTTCCAGAGCCCAT | | | | | | | | | | |
| Chromosome | start | end | strand | MM | target_seq | PAM | alignment | distance | position | gene name | gene id |
| GU305914.1 | 170415 | 170437 | + | 0 | ATGGGCTCTGGAAACGCGTC | GGG | \|\|\|\|\|\|\|\|[\|\|\|\|\|\|\|\|\|\|\|\|]PAM | NA | Intergenic | NA | NA |
| T12931 | TGGGCTCTGGAAACGCGTCGGGG | 1000 | | | | | | | | | |
| Oligo fwd | CACCTGGGCTCTGGAAACGCGTCG | | | | | | | | | | |
| Oligo rev | AAACCGACGCGTTTCCAGAGCCCA | | | | | | | | | | |
| Chromosome | start | end | strand | MM | target_seq | PAM | alignment | distance | position | gene name | gene id |
| GU305914.1 | 170416 | 170438 | + | 0 | TGGGCTCTGGAAACGCGTCG | GGG | \|\|\|\|\|\|\|\|[\|\|\|\|\|\|\|\|\|\|\|\|]PAM | NA | Intergenic | NA | NA |
| T12932 | CTCTGGAAACGCGTCGGGGTCGG | 1000 | | | | | | | | | |
| Oligo fwd | CACCCTCTGGAAACGCGTCGGGGT | | | | | | | | | | |
| Oligo rev | AAACACCCCGACGCGTTTCCAGAG | | | | | | | | | | |
| Chromosome | start | end | strand | MM | target_seq | PAM | alignment | distance | position | gene name | gene id |
| GU305914.1 | 170420 | 170442 | + | 0 | CTCTGGAAACGCGTCGGGGT | CGG | \|\|\|\|\|\|\|\|[\|\|\|\|\|\|\|\|\|\|\|\|]PAM | NA | Intergenic | NA | NA |
| T12933 | TCTGGAAACGCGTCGGGGTCGGG | 1000 | | | | | | | | | |
| Oligo fwd | CACCTCTGGAAACGCGTCGGGGTC | | | | | | | | | | |
| Oligo rev | AAACGACCCCGACGCGTTTCCAGA | | | | | | | | | | |

<table>
<tr><td>Chromosome</td><td>start</td><td>end</td><td>strand</td><td>MM</td><td>target_seq</td><td>PAM</td><td>alignment</td><td>distance</td><td>position</td><td>gene name</td><td>gene id</td></tr>
<tr><td>GU305914.1</td><td>170421</td><td>170443</td><td>+</td><td>0</td><td>TCTGGAAACGCGTCGGGTC</td><td>GGG</td><td>||||||||[||||||||||||]PAM</td><td>NA</td><td>Intergenic</td><td>NA</td><td>NA</td></tr>
<tr><td>GU305914.1</td><td>170118</td><td>170140</td><td>+</td><td>4</td><td>TCGGGAACCGCGACGGGTTC</td><td>GGG</td><td>||-||||-[||||-||||-||]PAM</td><td>NA</td><td>Intergenic</td><td>NA</td><td>NA</td></tr>
<tr><td></td><td></td><td></td><td></td><td></td><td></td><td></td><td></td><td></td><td></td><td></td><td></td></tr>
<tr><td>T12934</td><td>CGTCGGGGTCGGGAAATGTGAGG</td><td>1000</td><td></td><td></td><td></td><td></td><td></td><td></td><td></td><td></td><td></td></tr>
<tr><td>Oligo fwd</td><td>CACCCGTCGGGGTCGGGAAATGTG</td><td></td><td></td><td></td><td></td><td></td><td></td><td></td><td></td><td></td><td></td></tr>
<tr><td>Oligo rev</td><td>AAACCACATTTCCCGACCCCGACG</td><td></td><td></td><td></td><td></td><td></td><td></td><td></td><td></td><td></td><td></td></tr>
<tr><td>Chromosome</td><td>start</td><td>end</td><td>strand</td><td>MM</td><td>target_seq</td><td>PAM</td><td>alignment</td><td>distance</td><td>position</td><td>gene name</td><td>gene id</td></tr>
<tr><td>GU305914.1</td><td>170431</td><td>170453</td><td>+</td><td>0</td><td>CGTCGGGGTCGGGAAATGTG</td><td>AGG</td><td>||||||||[||||||||||||]PAM</td><td>NA</td><td>Intergenic</td><td>NA</td><td>NA</td></tr>
<tr><td></td><td></td><td></td><td></td><td></td><td></td><td></td><td></td><td></td><td></td><td></td><td></td></tr>
<tr><td>T12935</td><td>GTCGGGGTCGGGAAATGTGAGGG</td><td>1000</td><td></td><td></td><td></td><td></td><td></td><td></td><td></td><td></td><td></td></tr>
<tr><td>Oligo fwd</td><td>CACCGTCGGGGTCGGGAAATGTGA</td><td></td><td></td><td></td><td></td><td></td><td></td><td></td><td></td><td></td><td></td></tr>
<tr><td>Oligo rev</td><td>AAACTCACATTTCCCGACCCCGAC</td><td></td><td></td><td></td><td></td><td></td><td></td><td></td><td></td><td></td><td></td></tr>
<tr><td>Chromosome</td><td>start</td><td>end</td><td>strand</td><td>MM</td><td>target_seq</td><td>PAM</td><td>alignment</td><td>distance</td><td>position</td><td>gene name</td><td>gene id</td></tr>
<tr><td>GU305914.1</td><td>170432</td><td>170454</td><td>+</td><td>0</td><td>GTCGGGGTCGGGAAATGTGA</td><td>GGG</td><td>||||||||[||||||||||||]PAM</td><td>NA</td><td>Intergenic</td><td>NA</td><td>NA</td></tr>
<tr><td></td><td></td><td></td><td></td><td></td><td></td><td></td><td></td><td></td><td></td><td></td><td></td></tr>
<tr><td>T12936</td><td>TCGGGGTCGGGAAATGTGAGGGG</td><td>1000</td><td></td><td></td><td></td><td></td><td></td><td></td><td></td><td></td><td></td></tr>
<tr><td>Oligo fwd</td><td>CACCTCGGGGTCGGGAAATGTGAG</td><td></td><td></td><td></td><td></td><td></td><td></td><td></td><td></td><td></td><td></td></tr>
<tr><td>Oligo rev</td><td>AAACCTCACATTTCCCGACCCCGA</td><td></td><td></td><td></td><td></td><td></td><td></td><td></td><td></td><td></td><td></td></tr>
<tr><td>Chromosome</td><td>start</td><td>end</td><td>strand</td><td>MM</td><td>target_seq</td><td>PAM</td><td>alignment</td><td>distance</td><td>position</td><td>gene name</td><td>gene id</td></tr>
<tr><td>GU305914.1</td><td>170433</td><td>170455</td><td>+</td><td>0</td><td>TCGGGGTCGGGAAATGTGAG</td><td>GGG</td><td>||||||||[||||||||||||]PAM</td><td>NA</td><td>Intergenic</td><td>NA</td><td>NA</td></tr>
<tr><td></td><td></td><td></td><td></td><td></td><td></td><td></td><td></td><td></td><td></td><td></td><td></td></tr>
<tr><td>T12937</td><td>GTCGGGAAATGTGAGGGGATCGG</td><td>1000</td><td></td><td></td><td></td><td></td><td></td><td></td><td></td><td></td><td></td></tr>
<tr><td>Oligo fwd</td><td>CACCGTCGGGAAATGTGAGGGGAT</td><td></td><td></td><td></td><td></td><td></td><td></td><td></td><td></td><td></td><td></td></tr>
<tr><td>Oligo rev</td><td>AAACATCCCCTCACATTTCCCGAC</td><td></td><td></td><td></td><td></td><td></td><td></td><td></td><td></td><td></td><td></td></tr>
<tr><td>Chromosome</td><td>start</td><td>end</td><td>strand</td><td>MM</td><td>target_seq</td><td>PAM</td><td>alignment</td><td>distance</td><td>position</td><td>gene name</td><td>gene id</td></tr>
<tr><td>GU305914.1</td><td>170438</td><td>170460</td><td>+</td><td>0</td><td>GTCGGGAAATGTGAGGGGAT</td><td>CGG</td><td>||||||||[||||||||||||]PAM</td><td>NA</td><td>Intergenic</td><td>NA</td><td>NA</td></tr>
<tr><td></td><td></td><td></td><td></td><td></td><td></td><td></td><td></td><td></td><td></td><td></td><td></td></tr>
<tr><td>T12938</td><td>GGGAAATGTGAGGGGATCGGAGG</td><td>1000</td><td></td><td></td><td></td><td></td><td></td><td></td><td></td><td></td><td></td></tr>
<tr><td>Oligo fwd</td><td>CACCGGGAAATGTGAGGGGATCGG</td><td></td><td></td><td></td><td></td><td></td><td></td><td></td><td></td><td></td><td></td></tr>
<tr><td>Oligo rev</td><td>AAACCCGATCCCCTCACATTTCCC</td><td></td><td></td><td></td><td></td><td></td><td></td><td></td><td></td><td></td><td></td></tr>
<tr><td>Chromosome</td><td>start</td><td>end</td><td>strand</td><td>MM</td><td>target_seq</td><td>PAM</td><td>alignment</td><td>distance</td><td>position</td><td>gene name</td><td>gene id</td></tr>
<tr><td>GU305914.1</td><td>170441</td><td>170463</td><td>+</td><td>0</td><td>GGGAAATGTGAGGGGATCGG</td><td>AGG</td><td>||||||||[||||||||||||]PAM</td><td>NA</td><td>Intergenic</td><td>NA</td><td>NA</td></tr>
<tr><td>GU305914.1</td><td>93737</td><td>93759</td><td>-</td><td>4</td><td>GGGAAGGGTGCGGGGATCCG</td><td>AGG</td><td>|||||--|[||-|||||||-|]PAM</td><td>NA</td><td>Intergenic</td><td>NA</td><td>NA</td></tr>
<tr><td></td><td></td><td></td><td></td><td></td><td></td><td></td><td></td><td></td><td></td><td></td><td></td></tr>
<tr><td>T12939</td><td>TGTGAGGGGATCGGAGGAGATGG</td><td>1000</td><td></td><td></td><td></td><td></td><td></td><td></td><td></td><td></td><td></td></tr>
<tr><td>Oligo fwd</td><td>CACCTGTGAGGGGATCGGAGGAGA</td><td></td><td></td><td></td><td></td><td></td><td></td><td></td><td></td><td></td><td></td></tr>
<tr><td>Oligo rev</td><td>AAACTCTCCTCCGATCCCCTCACA</td><td></td><td></td><td></td><td></td><td></td><td></td><td></td><td></td><td></td><td></td></tr>
<tr><td>Chromosome</td><td>start</td><td>end</td><td>strand</td><td>MM</td><td>target_seq</td><td>PAM</td><td>alignment</td><td>distance</td><td>position</td><td>gene name</td><td>gene id</td></tr>
<tr><td>GU305914.1</td><td>170447</td><td>170469</td><td>+</td><td>0</td><td>TGTGAGGGGATCGGAGGAGA</td><td>TGG</td><td>||||||||[||||||||||||]PAM</td><td>NA</td><td>Intergenic</td><td>NA</td><td>NA</td></tr>
<tr><td></td><td></td><td></td><td></td><td></td><td></td><td></td><td></td><td></td><td></td><td></td><td></td></tr>
<tr><td>T12940</td><td>GTGAGGGGATCGGAGGAGATGGG</td><td>1000</td><td></td><td></td><td></td><td></td><td></td><td></td><td></td><td></td><td></td></tr>
<tr><td>Oligo fwd</td><td>CACCGTGAGGGGATCGGAGGAGAT</td><td></td><td></td><td></td><td></td><td></td><td></td><td></td><td></td><td></td><td></td></tr>
<tr><td>Oligo rev</td><td>AAACATCTCCTCCGATCCCCTCAC</td><td></td><td></td><td></td><td></td><td></td><td></td><td></td><td></td><td></td><td></td></tr>
<tr><td>Chromosome</td><td>start</td><td>end</td><td>strand</td><td>MM</td><td>target_seq</td><td>PAM</td><td>alignment</td><td>distance</td><td>position</td><td>gene name</td><td>gene id</td></tr>
<tr><td>GU305914.1</td><td>170448</td><td>170470</td><td>+</td><td>0</td><td>GTGAGGGGATCGGAGGAGAT</td><td>GGG</td><td>||||||||[||||||||||||]PAM</td><td>NA</td><td>Intergenic</td><td>NA</td><td>NA</td></tr>
<tr><td></td><td></td><td></td><td></td><td></td><td></td><td></td><td></td><td></td><td></td><td></td><td></td></tr>
<tr><td>T12941</td><td>TGAGGGGATCGGAGGAGATGGGG</td><td>1000</td><td></td><td></td><td></td><td></td><td></td><td></td><td></td><td></td><td></td></tr>
<tr><td>Oligo fwd</td><td>CACCTGAGGGGATCGGAGGAGATG</td><td></td><td></td><td></td><td></td><td></td><td></td><td></td><td></td><td></td><td></td></tr>
<tr><td>Oligo rev</td><td>AAACCATCTCCTCCGATCCCCTCA</td><td></td><td></td><td></td><td></td><td></td><td></td><td></td><td></td><td></td><td></td></tr>
<tr><td>Chromosome</td><td>start</td><td>end</td><td>strand</td><td>MM</td><td>target_seq</td><td>PAM</td><td>alignment</td><td>distance</td><td>position</td><td>gene name</td><td>gene id</td></tr>
<tr><td>GU305914.1</td><td>170449</td><td>170471</td><td>+</td><td>0</td><td>TGAGGGGATCGGAGGAGATG</td><td>GGG</td><td>||||||||[||||||||||||]PAM</td><td>NA</td><td>Intergenic</td><td>NA</td><td>NA</td></tr>
<tr><td></td><td></td><td></td><td></td><td></td><td></td><td></td><td></td><td></td><td></td><td></td><td></td></tr>
<tr><td>T12942</td><td>GGATCGGAGGAGATGGGGATCGG</td><td>1000</td><td></td><td></td><td></td><td></td><td></td><td></td><td></td><td></td><td></td></tr>
<tr><td>Oligo fwd</td><td>CACCGGATCGGAGGAGATGGGGAT</td><td></td><td></td><td></td><td></td><td></td><td></td><td></td><td></td><td></td><td></td></tr>
<tr><td>Oligo rev</td><td>AAACATCCCCATCTCCTCCGATCC</td><td></td><td></td><td></td><td></td><td></td><td></td><td></td><td></td><td></td><td></td></tr>
<tr><td>Chromosome</td><td>start</td><td>end</td><td>strand</td><td>MM</td><td>target_seq</td><td>PAM</td><td>alignment</td><td>distance</td><td>position</td><td>gene name</td><td>gene id</td></tr>
<tr><td>GU305914.1</td><td>170454</td><td>170476</td><td>+</td><td>0</td><td>GGATCGGAGGAGATGGGGAT</td><td>CGG</td><td>||||||||[||||||||||||]PAM</td><td>NA</td><td>Intergenic</td><td>NA</td><td>NA</td></tr>
<tr><td></td><td></td><td></td><td></td><td></td><td></td><td></td><td></td><td></td><td></td><td></td><td></td></tr>
<tr><td>T12943</td><td>GGGATCGGTTCCTCTATCATCGG</td><td>1000</td><td></td><td></td><td></td><td></td><td></td><td></td><td></td><td></td><td></td></tr>
<tr><td>Oligo fwd</td><td>CACCGGGATCGGTTCCTCTATCAT</td><td></td><td></td><td></td><td></td><td></td><td></td><td></td><td></td><td></td><td></td></tr>
<tr><td>Oligo rev</td><td>AAACATGATAGAGGAACCGATCCC</td><td></td><td></td><td></td><td></td><td></td><td></td><td></td><td></td><td></td><td></td></tr>
<tr><td>Chromosome</td><td>start</td><td>end</td><td>strand</td><td>MM</td><td>target_seq</td><td>PAM</td><td>alignment</td><td>distance</td><td>position</td><td>gene name</td><td>gene id</td></tr>
<tr><td>GU305914.1</td><td>170469</td><td>170491</td><td>+</td><td>0</td><td>GGGATCGGTTCCTCTATCAT</td><td>CGG</td><td>||||||||[||||||||||||]PAM</td><td>NA</td><td>Intergenic</td><td>NA</td><td>NA</td></tr>
<tr><td></td><td></td><td></td><td></td><td></td><td></td><td></td><td></td><td></td><td></td><td></td><td></td></tr>
<tr><td>T12944</td><td>GGATCGGTTCCTCTATCATCGGG</td><td>1000</td><td></td><td></td><td></td><td></td><td></td><td></td><td></td><td></td><td></td></tr>
<tr><td>Oligo fwd</td><td>CACCGGATCGGTTCCTCTATCATC</td><td></td><td></td><td></td><td></td><td></td><td></td><td></td><td></td><td></td><td></td></tr>
<tr><td>Oligo rev</td><td>AAACGATGATAGAGGAACCGATCC</td><td></td><td></td><td></td><td></td><td></td><td></td><td></td><td></td><td></td><td></td></tr>
<tr><td>Chromosome</td><td>start</td><td>end</td><td>strand</td><td>MM</td><td>target_seq</td><td>PAM</td><td>alignment</td><td>distance</td><td>position</td><td>gene name</td><td>gene id</td></tr>
<tr><td>GU305914.1</td><td>170470</td><td>170492</td><td>+</td><td>0</td><td>GGATCGGTTCCTCTATCATC</td><td>GGG</td><td>||||||||[||||||||||||]PAM</td><td>NA</td><td>Intergenic</td><td>NA</td><td>NA</td></tr>
<tr><td></td><td></td><td></td><td></td><td></td><td></td><td></td><td></td><td></td><td></td><td></td><td></td></tr>
<tr><td>T12945</td><td>TTCCTCTATCATCGGGAGTTTGG</td><td>1000</td><td></td><td></td><td></td><td></td><td></td><td></td><td></td><td></td><td></td></tr>
<tr><td>Oligo fwd</td><td>CACCTTCCTCTATCATCGGGAGTT</td><td></td><td></td><td></td><td></td><td></td><td></td><td></td><td></td><td></td><td></td></tr>
<tr><td>Oligo rev</td><td>AAACAACTCCCGATGATAGAGGAA</td><td></td><td></td><td></td><td></td><td></td><td></td><td></td><td></td><td></td><td></td></tr>
<tr><td>Chromosome</td><td>start</td><td>end</td><td>strand</td><td>MM</td><td>target_seq</td><td>PAM</td><td>alignment</td><td>distance</td><td>position</td><td>gene name</td><td>gene id</td></tr>
<tr><td>GU305914.1</td><td>170477</td><td>170499</td><td>+</td><td>0</td><td>TTCCTCTATCATCGGGAGTT</td><td>TGG</td><td>||||||||[||||||||||||]PAM</td><td>NA</td><td>Intergenic</td><td>NA</td><td>NA</td></tr>
<tr><td></td><td></td><td></td><td></td><td></td><td></td><td></td><td></td><td></td><td></td><td></td><td></td></tr>
<tr><td>T12946</td><td>TCCTCTATCATCGGGAGTTTGGG</td><td>1000</td><td></td><td></td><td></td><td></td><td></td><td></td><td></td><td></td><td></td></tr>
<tr><td>Oligo fwd</td><td>CACCTCCTCTATCATCGGGAGTTT</td><td></td><td></td><td></td><td></td><td></td><td></td><td></td><td></td><td></td><td></td></tr>
<tr><td>Oligo rev</td><td>AAACAAACTCCCGATGATAGAGGA</td><td></td><td></td><td></td><td></td><td></td><td></td><td></td><td></td><td></td><td></td></tr>
<tr><td>Chromosome</td><td>start</td><td>end</td><td>strand</td><td>MM</td><td>target_seq</td><td>PAM</td><td>alignment</td><td>distance</td><td>position</td><td>gene name</td><td>gene id</td></tr>
<tr><td>GU305914.1</td><td>170478</td><td>170500</td><td>+</td><td>0</td><td>TCCTCTATCATCGGGAGTTT</td><td>GGG</td><td>||||||||[||||||||||||]PAM</td><td>NA</td><td>Intergenic</td><td>NA</td><td>NA</td></tr>
<tr><td></td><td></td><td></td><td></td><td></td><td></td><td></td><td></td><td></td><td></td><td></td><td></td></tr>
<tr><td>T12947</td><td>ATCGGGAGTTTGGGTAGTATTGG</td><td>1000</td><td></td><td></td><td></td><td></td><td></td><td></td><td></td><td></td><td></td></tr>
<tr><td>Oligo fwd</td><td>CACCATCGGGAGTTTGGGTAGTAT</td><td></td><td></td><td></td><td></td><td></td><td></td><td></td><td></td><td></td><td></td></tr>
<tr><td>Oligo rev</td><td>AAACATACTACCCAAACTCCCGAT</td><td></td><td></td><td></td><td></td><td></td><td></td><td></td><td></td><td></td><td></td></tr>
<tr><td>Chromosome</td><td>start</td><td>end</td><td>strand</td><td>MM</td><td>target_seq</td><td>PAM</td><td>alignment</td><td>distance</td><td>position</td><td>gene name</td><td>gene id</td></tr>
<tr><td>GU305914.1</td><td>170487</td><td>170509</td><td>+</td><td>0</td><td>ATCGGGAGTTTGGGTAGTAT</td><td>TGG</td><td>||||||||[||||||||||||]PAM</td><td>NA</td><td>Intergenic</td><td>NA</td><td>NA</td></tr>
</table>

<table>
<tr><td>T12948</td><td>GGTAGTATTGGTCGTTGCCCCGG</td><td>1000</td><td></td><td></td><td></td><td></td><td></td><td></td><td></td><td></td><td></td></tr>
<tr><td>Oligo fwd</td><td>CACCGGTAGTATTGGTCGTTGCCC</td><td></td><td></td><td></td><td></td><td></td><td></td><td></td><td></td><td></td><td></td></tr>
<tr><td>Oligo rev</td><td>AAACGGGCAACGACCAATACTACC</td><td></td><td></td><td></td><td></td><td></td><td></td><td></td><td></td><td></td><td></td></tr>
<tr><td>Chromosome</td><td>start</td><td>end</td><td>strand</td><td>MM</td><td>target_seq</td><td>PAM</td><td>alignment</td><td>distance</td><td>position</td><td>gene name</td><td>gene id</td></tr>
<tr><td>GU305914.1</td><td>170499</td><td>170521</td><td>+</td><td>0</td><td>GGTAGTATTGGTCGTTGCCC</td><td>CGG</td><td>||||||||[||||||||||||]PAM</td><td>NA</td><td>Intergenic</td><td>NA</td><td>NA</td></tr>
<tr><td>T12949</td><td>TGGTCGTTGCCCCGGTATCGTGG</td><td>1000</td><td></td><td></td><td></td><td></td><td></td><td></td><td></td><td></td><td></td></tr>
<tr><td>Oligo fwd</td><td>CACCTGGTCGTTGCCCCGGTATCG</td><td></td><td></td><td></td><td></td><td></td><td></td><td></td><td></td><td></td><td></td></tr>
<tr><td>Oligo rev</td><td>AAACCGATACCGGGGCAACGACCA</td><td></td><td></td><td></td><td></td><td></td><td></td><td></td><td></td><td></td><td></td></tr>
<tr><td>Chromosome</td><td>start</td><td>end</td><td>strand</td><td>MM</td><td>target_seq</td><td>PAM</td><td>alignment</td><td>distance</td><td>position</td><td>gene name</td><td>gene id</td></tr>
<tr><td>GU305914.1</td><td>170507</td><td>170529</td><td>+</td><td>0</td><td>TGGTCGTTGCCCCGGTATCG</td><td>TGG</td><td>||||||||[||||||||||||]PAM</td><td>NA</td><td>Intergenic</td><td>NA</td><td>NA</td></tr>
<tr><td>T12950</td><td>TTGCCCCGGTATCGTGGCCGTGG</td><td>1000</td><td></td><td></td><td></td><td></td><td></td><td></td><td></td><td></td><td></td></tr>
<tr><td>Oligo fwd</td><td>CACCTTGCCCCGGTATCGTGGCCG</td><td></td><td></td><td></td><td></td><td></td><td></td><td></td><td></td><td></td><td></td></tr>
<tr><td>Oligo rev</td><td>AAACCGGCCACGATACCGGGGCAA</td><td></td><td></td><td></td><td></td><td></td><td></td><td></td><td></td><td></td><td></td></tr>
<tr><td>Chromosome</td><td>start</td><td>end</td><td>strand</td><td>MM</td><td>target_seq</td><td>PAM</td><td>alignment</td><td>distance</td><td>position</td><td>gene name</td><td>gene id</td></tr>
<tr><td>GU305914.1</td><td>170513</td><td>170535</td><td>+</td><td>0</td><td>TTGCCCCGGTATCGTGGCCG</td><td>TGG</td><td>||||||||[||||||||||||]PAM</td><td>NA</td><td>Intergenic</td><td>NA</td><td>NA</td></tr>
<tr><td>T12951</td><td>GTGGCCGCCGCCGCCACCATCGG</td><td>1000</td><td></td><td></td><td></td><td></td><td></td><td></td><td></td><td></td><td></td></tr>
<tr><td>Oligo fwd</td><td>CACCGTGGCCGCCGCCGCCACCAT</td><td></td><td></td><td></td><td></td><td></td><td></td><td></td><td></td><td></td><td></td></tr>
<tr><td>Oligo rev</td><td>AAACATGGTGGCGGCGGCGGCCAC</td><td></td><td></td><td></td><td></td><td></td><td></td><td></td><td></td><td></td><td></td></tr>
<tr><td>Chromosome</td><td>start</td><td>end</td><td>strand</td><td>MM</td><td>target_seq</td><td>PAM</td><td>alignment</td><td>distance</td><td>position</td><td>gene name</td><td>gene id</td></tr>
<tr><td>GU305914.1</td><td>170532</td><td>170554</td><td>+</td><td>0</td><td>GTGGCCGCCGCCGCCACCAT</td><td>CGG</td><td>||||||||[||||||||||||]PAM</td><td>NA</td><td>Intergenic</td><td>NA</td><td>NA</td></tr>
<tr><td>GU305914.1</td><td>24876</td><td>24898</td><td>-</td><td>4</td><td>GCCGCCGTCGCCGCCAACAT</td><td>CGG</td><td>|--||||-[||||||||-|||]PAM</td><td>NA</td><td>Intergenic</td><td>NA</td><td>NA</td></tr>
<tr><td>GU305914.1</td><td>24546</td><td>24568</td><td>-</td><td>4</td><td>GTCGCTGCCGCCGCCATCTT</td><td>GGG</td><td>||-||-||[||||||||-|-|]PAM</td><td>NA</td><td>Intergenic</td><td>NA</td><td>NA</td></tr>
<tr><td>GU305914.1</td><td>29998</td><td>30020</td><td>-</td><td>4</td><td>GCCGCCGCCGCCGCCAGCAC</td><td>CGG</td><td>|--|||||[||||||||-||-]PAM</td><td>NA</td><td>Intergenic</td><td>NA</td><td>NA</td></tr>
<tr><td>T12952</td><td>CGCCGCCACCATCGGCCACGCGG</td><td>1000</td><td></td><td></td><td></td><td></td><td></td><td></td><td></td><td></td><td></td></tr>
<tr><td>Oligo fwd</td><td>CACCCGCCGCCACCATCGGCCACG</td><td></td><td></td><td></td><td></td><td></td><td></td><td></td><td></td><td></td><td></td></tr>
<tr><td>Oligo rev</td><td>AAACCGTGGCCGATGGTGGCGGCG</td><td></td><td></td><td></td><td></td><td></td><td></td><td></td><td></td><td></td><td></td></tr>
<tr><td>Chromosome</td><td>start</td><td>end</td><td>strand</td><td>MM</td><td>target_seq</td><td>PAM</td><td>alignment</td><td>distance</td><td>position</td><td>gene name</td><td>gene id</td></tr>
<tr><td>GU305914.1</td><td>170540</td><td>170562</td><td>+</td><td>0</td><td>CGCCGCCACCATCGGCCACG</td><td>CGG</td><td>||||||||[||||||||||||]PAM</td><td>NA</td><td>Intergenic</td><td>NA</td><td>NA</td></tr>
<tr><td>T12953</td><td>GCCACCATCGGCCACGCGGTCGG</td><td>1000</td><td></td><td></td><td></td><td></td><td></td><td></td><td></td><td></td><td></td></tr>
<tr><td>Oligo fwd</td><td>CACCGCCACCATCGGCCACGCGGT</td><td></td><td></td><td></td><td></td><td></td><td></td><td></td><td></td><td></td><td></td></tr>
<tr><td>Oligo rev</td><td>AAACACCGCGTGGCCGATGGTGGC</td><td></td><td></td><td></td><td></td><td></td><td></td><td></td><td></td><td></td><td></td></tr>
<tr><td>Chromosome</td><td>start</td><td>end</td><td>strand</td><td>MM</td><td>target_seq</td><td>PAM</td><td>alignment</td><td>distance</td><td>position</td><td>gene name</td><td>gene id</td></tr>
<tr><td>GU305914.1</td><td>170544</td><td>170566</td><td>+</td><td>0</td><td>GCCACCATCGGCCACGCGGT</td><td>CGG</td><td>||||||||[||||||||||||]PAM</td><td>NA</td><td>Intergenic</td><td>NA</td><td>NA</td></tr>
<tr><td>T12954</td><td>GCGGTCGGCACCATCGCCGCCGG</td><td>1000</td><td></td><td></td><td></td><td></td><td></td><td></td><td></td><td></td><td></td></tr>
<tr><td>Oligo fwd</td><td>CACCGCGGTCGGCACCATCGCCGC</td><td></td><td></td><td></td><td></td><td></td><td></td><td></td><td></td><td></td><td></td></tr>
<tr><td>Oligo rev</td><td>AAACGCGGCGATGGTGCCGACCGC</td><td></td><td></td><td></td><td></td><td></td><td></td><td></td><td></td><td></td><td></td></tr>
<tr><td>Chromosome</td><td>start</td><td>end</td><td>strand</td><td>MM</td><td>target_seq</td><td>PAM</td><td>alignment</td><td>distance</td><td>position</td><td>gene name</td><td>gene id</td></tr>
<tr><td>GU305914.1</td><td>170559</td><td>170581</td><td>+</td><td>0</td><td>GCGGTCGGCACCATCGCCGC</td><td>CGG</td><td>||||||||[||||||||||||]PAM</td><td>NA</td><td>Intergenic</td><td>NA</td><td>NA</td></tr>
<tr><td>T12955</td><td>CGGTCGGCACCATCGCCGCCGGG</td><td>1000</td><td></td><td></td><td></td><td></td><td></td><td></td><td></td><td></td><td></td></tr>
<tr><td>Oligo fwd</td><td>CACCCGGTCGGCACCATCGCCGCC</td><td></td><td></td><td></td><td></td><td></td><td></td><td></td><td></td><td></td><td></td></tr>
<tr><td>Oligo rev</td><td>AAACGGCGGCGATGGTGCCGACCG</td><td></td><td></td><td></td><td></td><td></td><td></td><td></td><td></td><td></td><td></td></tr>
<tr><td>Chromosome</td><td>start</td><td>end</td><td>strand</td><td>MM</td><td>target_seq</td><td>PAM</td><td>alignment</td><td>distance</td><td>position</td><td>gene name</td><td>gene id</td></tr>
<tr><td>GU305914.1</td><td>170560</td><td>170582</td><td>+</td><td>0</td><td>CGGTCGGCACCATCGCCGCC</td><td>GGG</td><td>||||||||[||||||||||||]PAM</td><td>NA</td><td>Intergenic</td><td>NA</td><td>NA</td></tr>
<tr><td>GU305914.1</td><td>115379</td><td>115401</td><td>+</td><td>4</td><td>CCGGCGGCACCGTCGTCGCC</td><td>AGG</td><td>|-|-||||[|||-|||-||||]PAM</td><td>NA</td><td>Intergenic</td><td>NA</td><td>NA</td></tr>
<tr><td>T12956</td><td>CGGCACCATCGCCGCCGGGTCGG</td><td>1000</td><td></td><td></td><td></td><td></td><td></td><td></td><td></td><td></td><td></td></tr>
<tr><td>Oligo fwd</td><td>CACCCGGCACCATCGCCGCCGGGT</td><td></td><td></td><td></td><td></td><td></td><td></td><td></td><td></td><td></td><td></td></tr>
<tr><td>Oligo rev</td><td>AAACACCCGGCGGCGATGGTGCCG</td><td></td><td></td><td></td><td></td><td></td><td></td><td></td><td></td><td></td><td></td></tr>
<tr><td>Chromosome</td><td>start</td><td>end</td><td>strand</td><td>MM</td><td>target_seq</td><td>PAM</td><td>alignment</td><td>distance</td><td>position</td><td>gene name</td><td>gene id</td></tr>
<tr><td>GU305914.1</td><td>170564</td><td>170586</td><td>+</td><td>0</td><td>CGGCACCATCGCCGCCGGGT</td><td>CGG</td><td>||||||||[||||||||||||]PAM</td><td>NA</td><td>Intergenic</td><td>NA</td><td>NA</td></tr>
<tr><td>T12957</td><td>GCCGCCGGGTCGGTAAACATCGG</td><td>1000</td><td></td><td></td><td></td><td></td><td></td><td></td><td></td><td></td><td></td></tr>
<tr><td>Oligo fwd</td><td>CACCGCCGCCGGGTCGGTAAACAT</td><td></td><td></td><td></td><td></td><td></td><td></td><td></td><td></td><td></td><td></td></tr>
<tr><td>Oligo rev</td><td>AAACATGTTTACCGACCCGGCGGC</td><td></td><td></td><td></td><td></td><td></td><td></td><td></td><td></td><td></td><td></td></tr>
<tr><td>Chromosome</td><td>start</td><td>end</td><td>strand</td><td>MM</td><td>target_seq</td><td>PAM</td><td>alignment</td><td>distance</td><td>position</td><td>gene name</td><td>gene id</td></tr>
<tr><td>GU305914.1</td><td>170574</td><td>170596</td><td>+</td><td>0</td><td>GCCGCCGGGTCGGTAAACAT</td><td>CGG</td><td>||||||||[||||||||||||]PAM</td><td>NA</td><td>Intergenic</td><td>NA</td><td>NA</td></tr>
<tr><td>T12958</td><td>CGATGCCGCTGCCGCGAACGTGG</td><td>1000</td><td></td><td></td><td></td><td></td><td></td><td></td><td></td><td></td><td></td></tr>
<tr><td>Oligo fwd</td><td>CACCCGATGCCGCTGCCGCGAACG</td><td></td><td></td><td></td><td></td><td></td><td></td><td></td><td></td><td></td><td></td></tr>
<tr><td>Oligo rev</td><td>AAACCGTTCGCGGCAGCGGCATCG</td><td></td><td></td><td></td><td></td><td></td><td></td><td></td><td></td><td></td><td></td></tr>
<tr><td>Chromosome</td><td>start</td><td>end</td><td>strand</td><td>MM</td><td>target_seq</td><td>PAM</td><td>alignment</td><td>distance</td><td>position</td><td>gene name</td><td>gene id</td></tr>
<tr><td>GU305914.1</td><td>170603</td><td>170625</td><td>+</td><td>0</td><td>CGATGCCGCTGCCGCGAACG</td><td>TGG</td><td>||||||||[||||||||||||]PAM</td><td>NA</td><td>Intergenic</td><td>NA</td><td>NA</td></tr>
<tr><td>GU305914.1</td><td>81261</td><td>81283</td><td>-</td><td>4</td><td>CGAGGCGGCCGCCGCGATCG</td><td>AGG</td><td>|||-||-|[|-|||||||-||]PAM</td><td>NA</td><td>Intergenic</td><td>NA</td><td>NA</td></tr>
<tr><td>T12959</td><td>GCCGCTGCCGCGAACGTGGTCGG</td><td>1000</td><td></td><td></td><td></td><td></td><td></td><td></td><td></td><td></td><td></td></tr>
<tr><td>Oligo fwd</td><td>CACCGCCGCTGCCGCGAACGTGGT</td><td></td><td></td><td></td><td></td><td></td><td></td><td></td><td></td><td></td><td></td></tr>
<tr><td>Oligo rev</td><td>AAACACCACGTTCGCGGCAGCGGC</td><td></td><td></td><td></td><td></td><td></td><td></td><td></td><td></td><td></td><td></td></tr>
<tr><td>Chromosome</td><td>start</td><td>end</td><td>strand</td><td>MM</td><td>target_seq</td><td>PAM</td><td>alignment</td><td>distance</td><td>position</td><td>gene name</td><td>gene id</td></tr>
<tr><td>GU305914.1</td><td>170607</td><td>170629</td><td>+</td><td>0</td><td>GCCGCTGCCGCGAACGTGGT</td><td>CGG</td><td>||||||||[||||||||||||]PAM</td><td>NA</td><td>Intergenic</td><td>NA</td><td>NA</td></tr>
<tr><td>T12960</td><td>CCGCTGCCGCGAACGTGGTCGGG</td><td>1000</td><td></td><td></td><td></td><td></td><td></td><td></td><td></td><td></td><td></td></tr>
<tr><td>Oligo fwd</td><td>CACCCCGCTGCCGCGAACGTGGTC</td><td></td><td></td><td></td><td></td><td></td><td></td><td></td><td></td><td></td><td></td></tr>
<tr><td>Oligo rev</td><td>AAACGACCACGTTCGCGGCAGCGG</td><td></td><td></td><td></td><td></td><td></td><td></td><td></td><td></td><td></td><td></td></tr>
<tr><td>Chromosome</td><td>start</td><td>end</td><td>strand</td><td>MM</td><td>target_seq</td><td>PAM</td><td>alignment</td><td>distance</td><td>position</td><td>gene name</td><td>gene id</td></tr>
<tr><td>GU305914.1</td><td>170608</td><td>170630</td><td>+</td><td>0</td><td>CCGCTGCCGCGAACGTGGTC</td><td>GGG</td><td>||||||||[||||||||||||]PAM</td><td>NA</td><td>Intergenic</td><td>NA</td><td>NA</td></tr>
<tr><td>T12961</td><td>CGTGGTCGGGATGTCGACCGTGG</td><td>1000</td><td></td><td></td><td></td><td></td><td></td><td></td><td></td><td></td><td></td></tr>
<tr><td>Oligo fwd</td><td>CACCCGTGGTCGGGATGTCGACCG</td><td></td><td></td><td></td><td></td><td></td><td></td><td></td><td></td><td></td><td></td></tr>
<tr><td>Oligo rev</td><td>AAACCGGTCGACATCCCGACCACG</td><td></td><td></td><td></td><td></td><td></td><td></td><td></td><td></td><td></td><td></td></tr>
<tr><td>Chromosome</td><td>start</td><td>end</td><td>strand</td><td>MM</td><td>target_seq</td><td>PAM</td><td>alignment</td><td>distance</td><td>position</td><td>gene name</td><td>gene id</td></tr>
<tr><td>GU305914.1</td><td>170621</td><td>170643</td><td>+</td><td>0</td><td>CGTGGTCGGGATGTCGACCG</td><td>TGG</td><td>||||||||[||||||||||||]PAM</td><td>NA</td><td>Intergenic</td><td>NA</td><td>NA</td></tr>
</table>

T12962 GTGGTCGGGATGTCGACCGTGGG 1000
Oligo fwd CACCGTGGTCGGGATGTCGACCGT
Oligo rev AAACACGGTCGACATCCCGACCAC
Chromosome start end strand MM target_seq PAM alignment distance position gene name gene id
GU305914.1 170622 170644 + 0 GTGGTCGGGATGTCGACCGT GGG ||||||||[||||||||||||]PAM NA Intergenic NA NA

T12963 ATGTCGACCGTGGGCATGAAAGG 1000
Oligo fwd CACCATGTCGACCGTGGGCATGAA
Oligo rev AAACTTCATGCCCACGGTCGACAT
Chromosome start end strand MM target_seq PAM alignment distance position gene name gene id
GU305914.1 170631 170653 + 0 ATGTCGACCGTGGGCATGAA AGG ||||||||[||||||||||||]PAM NA Intergenic NA NA

T12964 ACCGTGGGCATGAAAGGCAGAGG 1000
Oligo fwd CACCACCGTGGGCATGAAAGGCAG
Oligo rev AAACCTGCCTTTCATGCCCACGGT
Chromosome start end strand MM target_seq PAM alignment distance position gene name gene id
GU305914.1 170637 170659 + 0 ACCGTGGGCATGAAAGGCAG AGG ||||||||[||||||||||||]PAM NA Intergenic NA NA

T12965 GGCATGAAAGGCAGAGGTAGCGG 1000
Oligo fwd CACCGGCATGAAAGGCAGAGGTAG
Oligo rev AAACCTACCTCTGCCTTTCATGCC
Chromosome start end strand MM target_seq PAM alignment distance position gene name gene id
GU305914.1 170643 170665 + 0 GGCATGAAAGGCAGAGGTAG CGG ||||||||[||||||||||||]PAM NA Intergenic NA NA

T12966 ATGAAAGGCAGAGGTAGCGGAGG 1000
Oligo fwd CACCATGAAAGGCAGAGGTAGCGG
Oligo rev AAACCCGCTACCTCTGCCTTTCAT
Chromosome start end strand MM target_seq PAM alignment distance position gene name gene id
GU305914.1 170646 170668 + 0 ATGAAAGGCAGAGGTAGCGG AGG ||||||||[||||||||||||]PAM NA Intergenic NA NA

T12967 AAAGGCAGAGGTAGCGGAGGCGG 1000
Oligo fwd CACCAAAGGCAGAGGTAGCGGAGG
Oligo rev AAACCCTCCGCTACCTCTGCCTTT
Chromosome start end strand MM target_seq PAM alignment distance position gene name gene id
GU305914.1 170649 170671 + 0 AAAGGCAGAGGTAGCGGAGG CGG ||||||||[||||||||||||]PAM NA Intergenic NA NA
GU305914.1 18656 18678 - 3 GACGGCAGAGGTGGCGGAGG CGG -|-|||||[||||-|||||||]PAM NA Intergenic NA NA

T12968 GGAGGCGGATTCTGCAACAACGG 1000
Oligo fwd CACCGGAGGCGGATTCTGCAACAA
Oligo rev AAACTTGTTGCAGAATCCGCCTCC
Chromosome start end strand MM target_seq PAM alignment distance position gene name gene id
GU305914.1 170664 170686 + 0 GGAGGCGGATTCTGCAACAA CGG ||||||||[||||||||||||]PAM NA Intergenic NA NA

T12969 AACGGCAAGAAGTGTAGCTGCGG 1000
Oligo fwd CACCAACGGCAAGAAGTGTAGCTG
Oligo rev AAACCAGCTACACTTCTTGCCGTT
Chromosome start end strand MM target_seq PAM alignment distance position gene name gene id
GU305914.1 170682 170704 + 0 AACGGCAAGAAGTGTAGCTG CGG ||||||||[||||||||||||]PAM NA Intergenic NA NA

T12970 GGCAAGAAGTGTAGCTGCGGCGG 1000
Oligo fwd CACCGGCAAGAAGTGTAGCTGCGG
Oligo rev AAACCCGCAGCTACACTTCTTGCC
Chromosome start end strand MM target_seq PAM alignment distance position gene name gene id
GU305914.1 170685 170707 + 0 GGCAAGAAGTGTAGCTGCGG CGG ||||||||[||||||||||||]PAM NA Intergenic NA NA

T12971 TAGCTGCGGCGGATCGACGAAGG 1000
Oligo fwd CACCTAGCTGCGGCGGATCGACGA
Oligo rev AAACTCGTCGATCCGCCGCAGCTA
Chromosome start end strand MM target_seq PAM alignment distance position gene name gene id
GU305914.1 170696 170718 + 0 TAGCTGCGGCGGATCGACGA AGG ||||||||[||||||||||||]PAM NA Intergenic NA NA

T12972 GGATCGACGAAGGACGCCGCCGG 1000
Oligo fwd CACCGGATCGACGAAGGACGCCGC
Oligo rev AAACGCGGCGTCCTTCGTCGATCC
Chromosome start end strand MM target_seq PAM alignment distance position gene name gene id
GU305914.1 170706 170728 + 0 GGATCGACGAAGGACGCCGC CGG ||||||||[||||||||||||]PAM NA Intergenic NA NA

T12973 TCGACGAAGGACGCCGCCGGAGG 1000
Oligo fwd CACCTCGACGAAGGACGCCGCCGG
Oligo rev AAACCCGGCGGCGTCCTTCGTCGA
Chromosome start end strand MM target_seq PAM alignment distance position gene name gene id
GU305914.1 170709 170731 + 0 TCGACGAAGGACGCCGCCGG AGG ||||||||[||||||||||||]PAM NA Intergenic NA NA

T12974 CGCTGCCGCCGCAGCCCGCTCGG 1000
Oligo fwd CACCCGCTGCCGCCGCAGCCCGCT
Oligo rev AAACAGCGGGCTGCGGCGGCAGCG
Chromosome start end strand MM target_seq PAM alignment distance position gene name gene id
GU305914.1 170738 170760 + 0 CGCTGCCGCCGCAGCCCGCT CGG ||||||||[||||||||||||]PAM NA Intergenic NA NA

T12975 GAGCTCTTCGTCTGTCACGTCGG 1000
Oligo fwd CACCGAGCTCTTCGTCTGTCACGT
Oligo rev AAACACGTGACAGACGAAGAGCTC
Chromosome start end strand MM target_seq PAM alignment distance position gene name gene id
GU305914.1 170762 170784 + 0 GAGCTCTTCGTCTGTCACGT CGG ||||||||[||||||||||||]PAM NA Intergenic NA NA

T12976 GATACGAACTCGAGTCTCCCAGG 1000
Oligo fwd CACCGATACGAACTCGAGTCTCCC
Oligo rev AAACGGGAGACTCGAGTTCGTATC
Chromosome start end strand MM target_seq PAM alignment distance position gene name gene id

GU305914.1 170792 170814 + 0 GATACGAACTCGAGTCTCCC AGG ||||||||[||||||||||||]PAM NA Intergenic NA NA

T12977 ATACGAACTCGAGTCTCCCAGGG 1000
Oligo fwd CACCATACGAACTCGAGTCTCCCA
Oligo rev AAACTGGGAGACTCGAGTTCGTAT
Chromosome start end strand MM target_seq PAM alignment distance position gene name gene id
GU305914.1 170793 170815 + 0 ATACGAACTCGAGTCTCCCA GGG ||||||||[||||||||||||]PAM NA Intergenic NA NA

T12978 GTCTCCCAGGGAGATTCTCCCGG 1000
Oligo fwd CACCGTCTCCCAGGGAGATTCTCC
Oligo rev AAACGGAGAATCTCCCTGGGAGAC
Chromosome start end strand MM target_seq PAM alignment distance position gene name gene id
GU305914.1 170805 170827 + 0 GTCTCCCAGGGAGATTCTCC CGG ||||||||[||||||||||||]PAM NA Intergenic NA NA

T12979 TCCCAGGGAGATTCTCCCGGAGG 1000
Oligo fwd CACCTCCCAGGGAGATTCTCCCGG
Oligo rev AAACCCGGGAGAATCTCCCTGGGA
Chromosome start end strand MM target_seq PAM alignment distance position gene name gene id
GU305914.1 170808 170830 + 0 TCCCAGGGAGATTCTCCCGG AGG ||||||||[||||||||||||]PAM NA Intergenic NA NA

T12980 AGATTCTCCCGGAGGTATCCTGG 1000
Oligo fwd CACCAGATTCTCCCGGAGGTATCC
Oligo rev AAACGGATACCTCCGGGAGAATCT
Chromosome start end strand MM target_seq PAM alignment distance position gene name gene id
GU305914.1 170816 170838 + 0 AGATTCTCCCGGAGGTATCC TGG ||||||||[||||||||||||]PAM NA Intergenic NA NA

T12981 CGGAGGTATCCTGGCGAACGTGG 1000
Oligo fwd CACCCGGAGGTATCCTGGCGAACG
Oligo rev AAACCGTTCGCCAGGATACCTCCG
Chromosome start end strand MM target_seq PAM alignment distance position gene name gene id
GU305914.1 170825 170847 + 0 CGGAGGTATCCTGGCGAACG TGG ||||||||[||||||||||||]PAM NA Intergenic NA NA

T12982 CGAACGTGGTCTCGTTCGCGCGG 1000
Oligo fwd CACCCGAACGTGGTCTCGTTCGCG
Oligo rev AAACCGCGAACGAGACCACGTTCG
Chromosome start end strand MM target_seq PAM alignment distance position gene name gene id
GU305914.1 170839 170861 + 0 CGAACGTGGTCTCGTTCGCG CGG ||||||||[||||||||||||]PAM NA Intergenic NA NA
GU305914.1 77637 77659 + 3 CGACCGAGGGCTCGTTCGCG CGG |||-||-|[|-||||||||||]PAM NA Intergenic NA NA

T12983 GATCACGACGACTTTGTCGTCGG 1000
Oligo fwd CACCGATCACGACGACTTTGTCGT
Oligo rev AAACACGACAAAGTCGTCGTGATC
Chromosome start end strand MM target_seq PAM alignment distance position gene name gene id
GU305914.1 170861 170883 + 0 GATCACGACGACTTTGTCGT CGG ||||||||[||||||||||||]PAM NA Intergenic NA NA

T12984 GACTTTGTCGTCGGAGACATTGG 1000
Oligo fwd CACCGACTTTGTCGTCGGAGACAT
Oligo rev AAACATGTCTCCGACGACAAAGTC
Chromosome start end strand MM target_seq PAM alignment distance position gene name gene id
GU305914.1 170870 170892 + 0 GACTTTGTCGTCGGAGACAT TGG ||||||||[||||||||||||]PAM NA Intergenic NA NA

T12985 TTGTCGTCGGAGACATTGGCCGG 1000
Oligo fwd CACCTTGTCGTCGGAGACATTGGC
Oligo rev AAACGCCAATGTCTCCGACGACAA
Chromosome start end strand MM target_seq PAM alignment distance position gene name gene id
GU305914.1 170874 170896 + 0 TTGTCGTCGGAGACATTGGC CGG ||||||||[||||||||||||]PAM NA Intergenic NA NA

T12986 CGGAGACATTGGCCGGCGTCAGG 1000
Oligo fwd CACCCGGAGACATTGGCCGGCGTC
Oligo rev AAACGACGCCGGCCAATGTCTCCG
Chromosome start end strand MM target_seq PAM alignment distance position gene name gene id
GU305914.1 170881 170903 + 0 CGGAGACATTGGCCGGCGTC AGG ||||||||[||||||||||||]PAM NA Intergenic NA NA

T12987 GGAGACATTGGCCGGCGTCAGGG 1000
Oligo fwd CACCGGAGACATTGGCCGGCGTCA
Oligo rev AAACTGACGCCGGCCAATGTCTCC
Chromosome start end strand MM target_seq PAM alignment distance position gene name gene id
GU305914.1 170882 170904 + 0 GGAGACATTGGCCGGCGTCA GGG ||||||||[||||||||||||]PAM NA Intergenic NA NA

T12988 GGCGTCAGGGACGTGACCAGAGG 1000
Oligo fwd CACCGGCGTCAGGGACGTGACCAG
Oligo rev AAACCTGGTCACGTCCCTGACGCC
Chromosome start end strand MM target_seq PAM alignment distance position gene name gene id
GU305914.1 170895 170917 + 0 GGCGTCAGGGACGTGACCAG AGG ||||||||[||||||||||||]PAM NA Intergenic NA NA

T12989 AGACACCTTAGCGTACGTGATGG 1000
Oligo fwd CACCAGACACCTTAGCGTACGTGA
Oligo rev AAACTCACGTACGCTAAGGTGTCT
Chromosome start end strand MM target_seq PAM alignment distance position gene name gene id
GU305914.1 170921 170943 + 0 AGACACCTTAGCGTACGTGA TGG ||||||||[||||||||||||]PAM NA Intergenic NA NA

T12990 GTGATGGTCGACCACGCACTCGG 1000
Oligo fwd CACCGTGATGGTCGACCACGCACT
Oligo rev AAACAGTGCGTGGTCGACCATCAC
Chromosome start end strand MM target_seq PAM alignment distance position gene name gene id
GU305914.1 170937 170959 + 0 GTGATGGTCGACCACGCACT CGG ||||||||[||||||||||||]PAM NA Intergenic NA NA

T12991 GTCGACCACGCACTCGGAGCTGG 1000
Oligo fwd CACCGTCGACCACGCACTCGGAGC

| | | | | | | | | | | | |
|---|---|---|---|---|---|---|---|---|---|---|---|
| Oligo rev | AAACGCTCCGAGTGCGTGGTCGAC | | | | | | | | | | |
| Chromosome | start | end | strand | MM | target_seq | PAM | alignment | distance | position | gene name | gene id |
| GU305914.1 | 170943 | 170965 | + | 0 | GTCGACCACGCACTCGGAGC | TGG | \|\|\|\|\|\|\|\|[\|\|\|\|\|\|\|\|\|\|\|\|]PAM | NA | Intergenic | NA | NA |
| T12992 | TGGAAATCGCACGCCGCCCGCGG | 1000 | | | | | | | | | |
| Oligo fwd | CACCTGGAAATCGCACGCCGCCCG | | | | | | | | | | |
| Oligo rev | AAACCGGGCGGCGTGCGATTTCCA | | | | | | | | | | |
| Chromosome | start | end | strand | MM | target_seq | PAM | alignment | distance | position | gene name | gene id |
| GU305914.1 | 170963 | 170985 | + | 0 | TGGAAATCGCACGCCGCCCG | CGG | \|\|\|\|\|\|\|\|[\|\|\|\|\|\|\|\|\|\|\|\|]PAM | NA | Intergenic | NA | NA |
| T12993 | AAACGCTAAGCAGCACGACGCGG | 1000 | | | | | | | | | |
| Oligo fwd | CACCAAACGCTAAGCAGCACGACG | | | | | | | | | | |
| Oligo rev | AAACCGTCGTGCTGCTTAGCGTTT | | | | | | | | | | |
| Chromosome | start | end | strand | MM | target_seq | PAM | alignment | distance | position | gene name | gene id |
| GU305914.1 | 171026 | 171048 | + | 0 | AAACGCTAAGCAGCACGACG | CGG | \|\|\|\|\|\|\|\|[\|\|\|\|\|\|\|\|\|\|\|\|]PAM | NA | Intergenic | NA | NA |
| T12994 | CGACGCGGCTAAGTAAGCGAAGG | 1000 | | | | | | | | | |
| Oligo fwd | CACCCGACGCGGCTAAGTAAGCGA | | | | | | | | | | |
| Oligo rev | AAACTCGCTTACTTAGCCGCGTCG | | | | | | | | | | |
| Chromosome | start | end | strand | MM | target_seq | PAM | alignment | distance | position | gene name | gene id |
| GU305914.1 | 171041 | 171063 | + | 0 | CGACGCGGCTAAGTAAGCGA | AGG | \|\|\|\|\|\|\|\|[\|\|\|\|\|\|\|\|\|\|\|\|]PAM | NA | Intergenic | NA | NA |
| T12995 | CACACTTTATTCGAAATCTCTGG | 1000 | | | | | | | | | |
| Oligo fwd | CACCCACACTTTATTCGAAATCTC | | | | | | | | | | |
| Oligo rev | AAACGAGATTTCGAATAAAGTGTG | | | | | | | | | | |
| Chromosome | start | end | strand | MM | target_seq | PAM | alignment | distance | position | gene name | gene id |
| GU305914.1 | 171074 | 171096 | + | 0 | CACACTTTATTCGAAATCTC | TGG | \|\|\|\|\|\|\|\|[\|\|\|\|\|\|\|\|\|\|\|\|]PAM | NA | Intergenic | NA | NA |
| T12996 | ATTCGAAATCTCTGGTGTAGCGG | 1000 | | | | | | | | | |
| Oligo fwd | CACCATTCGAAATCTCTGGTGTAG | | | | | | | | | | |
| Oligo rev | AAACCTACACCAGAGATTTCGAAT | | | | | | | | | | |
| Chromosome | start | end | strand | MM | target_seq | PAM | alignment | distance | position | gene name | gene id |
| GU305914.1 | 171082 | 171104 | + | 0 | ATTCGAAATCTCTGGTGTAG | CGG | \|\|\|\|\|\|\|\|[\|\|\|\|\|\|\|\|\|\|\|\|]PAM | NA | Intergenic | NA | NA |
| T12997 | CTCTGGTGTAGCGGAAAGCGAGG | 1000 | | | | | | | | | |
| Oligo fwd | CACCCTCTGGTGTAGCGGAAAGCG | | | | | | | | | | |
| Oligo rev | AAACCGCTTTCCGCTACACCAGAG | | | | | | | | | | |
| Chromosome | start | end | strand | MM | target_seq | PAM | alignment | distance | position | gene name | gene id |
| GU305914.1 | 171091 | 171113 | + | 0 | CTCTGGTGTAGCGGAAAGCG | AGG | \|\|\|\|\|\|\|\|[\|\|\|\|\|\|\|\|\|\|\|\|]PAM | NA | Intergenic | NA | NA |
| T12998 | TGGTGTAGCGGAAAGCGAGGCGG | 1000 | | | | | | | | | |
| Oligo fwd | CACCTGGTGTAGCGGAAAGCGAGG | | | | | | | | | | |
| Oligo rev | AAACCCTCGCTTTCCGCTACACCA | | | | | | | | | | |
| Chromosome | start | end | strand | MM | target_seq | PAM | alignment | distance | position | gene name | gene id |
| GU305914.1 | 171094 | 171116 | + | 0 | TGGTGTAGCGGAAAGCGAGG | CGG | \|\|\|\|\|\|\|\|[\|\|\|\|\|\|\|\|\|\|\|\|]PAM | NA | Intergenic | NA | NA |
| T12999 | GGTGTAGCGGAAAGCGAGGCGGG | 1000 | | | | | | | | | |
| Oligo fwd | CACCGGTGTAGCGGAAAGCGAGGC | | | | | | | | | | |
| Oligo rev | AAACGCCTCGCTTTCCGCTACACC | | | | | | | | | | |
| Chromosome | start | end | strand | MM | target_seq | PAM | alignment | distance | position | gene name | gene id |
| GU305914.1 | 171095 | 171117 | + | 0 | GGTGTAGCGGAAAGCGAGGC | GGG | \|\|\|\|\|\|\|\|[\|\|\|\|\|\|\|\|\|\|\|\|]PAM | NA | Intergenic | NA | NA |
| T13000 | GTACGCGATGATCACAAGAGCGG | 1000 | | | | | | | | | |
| Oligo fwd | CACCGTACGCGATGATCACAAGAG | | | | | | | | | | |
| Oligo rev | AAACCTCTTGTGATCATCGCGTAC | | | | | | | | | | |
| Chromosome | start | end | strand | MM | target_seq | PAM | alignment | distance | position | gene name | gene id |
| GU305914.1 | 171149 | 171171 | + | 0 | GTACGCGATGATCACAAGAG | CGG | \|\|\|\|\|\|\|\|[\|\|\|\|\|\|\|\|\|\|\|\|]PAM | NA | Intergenic | NA | NA |
| T13001 | CAAGAGCGGCCAAGACCAGCAGG | 1000 | | | | | | | | | |
| Oligo fwd | CACCCAAGAGCGGCCAAGACCAGC | | | | | | | | | | |
| Oligo rev | AAACGCTGGTCTTGGCCGCTCTTG | | | | | | | | | | |
| Chromosome | start | end | strand | MM | target_seq | PAM | alignment | distance | position | gene name | gene id |
| GU305914.1 | 171163 | 171185 | + | 0 | CAAGAGCGGCCAAGACCAGC | AGG | \|\|\|\|\|\|\|\|[\|\|\|\|\|\|\|\|\|\|\|\|]PAM | NA | Intergenic | NA | NA |
| T13002 | AGCGGCCAAGACCAGCAGGACGG | 1000 | | | | | | | | | |
| Oligo fwd | CACCAGCGGCCAAGACCAGCAGGA | | | | | | | | | | |
| Oligo rev | AAACTCCTGCTGGTCTTGGCCGCT | | | | | | | | | | |
| Chromosome | start | end | strand | MM | target_seq | PAM | alignment | distance | position | gene name | gene id |
| GU305914.1 | 171167 | 171189 | + | 0 | AGCGGCCAAGACCAGCAGGA | CGG | \|\|\|\|\|\|\|\|[\|\|\|\|\|\|\|\|\|\|\|\|]PAM | NA | Intergenic | NA | NA |
| T13003 | CCAAGACCAGCAGGACGGTCAGG | 1000 | | | | | | | | | |
| Oligo fwd | CACCCCAAGACCAGCAGGACGGTC | | | | | | | | | | |
| Oligo rev | AAACGACCGTCCTGCTGGTCTTGG | | | | | | | | | | |
| Chromosome | start | end | strand | MM | target_seq | PAM | alignment | distance | position | gene name | gene id |
| GU305914.1 | 171172 | 171194 | + | 0 | CCAAGACCAGCAGGACGGTC | AGG | \|\|\|\|\|\|\|\|[\|\|\|\|\|\|\|\|\|\|\|\|]PAM | NA | Intergenic | NA | NA |
| T13004 | CAAGACCAGCAGGACGGTCAGGG | 1000 | | | | | | | | | |
| Oligo fwd | CACCCAAGACCAGCAGGACGGTCA | | | | | | | | | | |
| Oligo rev | AAACTGACCGTCCTGCTGGTCTTG | | | | | | | | | | |
| Chromosome | start | end | strand | MM | target_seq | PAM | alignment | distance | position | gene name | gene id |
| GU305914.1 | 171173 | 171195 | + | 0 | CAAGACCAGCAGGACGGTCA | GGG | \|\|\|\|\|\|\|\|[\|\|\|\|\|\|\|\|\|\|\|\|]PAM | NA | Intergenic | NA | NA |
| T13005 | GACCAGCAGGACGGTCAGGGCGG | 1000 | | | | | | | | | |
| Oligo fwd | CACCGACCAGCAGGACGGTCAGGG | | | | | | | | | | |
| Oligo rev | AAACCCCTGACCGTCCTGCTGGTC | | | | | | | | | | |
| Chromosome | start | end | strand | MM | target_seq | PAM | alignment | distance | position | gene name | gene id |
| GU305914.1 | 171176 | 171198 | + | 0 | GACCAGCAGGACGGTCAGGG | CGG | \|\|\|\|\|\|\|\|[\|\|\|\|\|\|\|\|\|\|\|\|]PAM | NA | Intergenic | NA | NA |
| GU305914.1 | 130502 | 130524 | + | 4 | GATCAGAAGGAAGATCAGGG | TGG | \|\|-\|\|\|-\|[\|\|\|-\|-\|\|\|\|\|\|]PAM | NA | Intergenic | NA | NA |

<table>
<tr><td>T13006</td><td>CGGTCAGGGCGGCATAGTATCGG</td><td>1000</td><td></td><td></td><td></td><td></td><td></td><td></td><td></td><td></td><td></td></tr>
<tr><td>Oligo fwd</td><td>CACCCGGTCAGGGCGGCATAGTAT</td><td></td><td></td><td></td><td></td><td></td><td></td><td></td><td></td><td></td><td></td></tr>
<tr><td>Oligo rev</td><td>AAACATACTATGCCGCCCTGACCG</td><td></td><td></td><td></td><td></td><td></td><td></td><td></td><td></td><td></td><td></td></tr>
<tr><td>Chromosome</td><td>start</td><td>end</td><td>strand</td><td>MM</td><td>target_seq</td><td>PAM</td><td>alignment</td><td>distance</td><td>position</td><td>gene name</td><td>gene id</td></tr>
<tr><td>GU305914.1</td><td>171187</td><td>171209</td><td>+</td><td>0</td><td>CGGTCAGGGCGGCATAGTAT</td><td>CGG</td><td>||||||||[||||||||||||]PAM</td><td>NA</td><td>Intergenic</td><td>NA</td><td>NA</td></tr>
<tr><td>T13007</td><td>GGTCAGGGCGGCATAGTATCGGG</td><td>1000</td><td></td><td></td><td></td><td></td><td></td><td></td><td></td><td></td><td></td></tr>
<tr><td>Oligo fwd</td><td>CACCGGTCAGGGCGGCATAGTATC</td><td></td><td></td><td></td><td></td><td></td><td></td><td></td><td></td><td></td><td></td></tr>
<tr><td>Oligo rev</td><td>AAACGATACTATGCCGCCCTGACC</td><td></td><td></td><td></td><td></td><td></td><td></td><td></td><td></td><td></td><td></td></tr>
<tr><td>Chromosome</td><td>start</td><td>end</td><td>strand</td><td>MM</td><td>target_seq</td><td>PAM</td><td>alignment</td><td>distance</td><td>position</td><td>gene name</td><td>gene id</td></tr>
<tr><td>GU305914.1</td><td>171188</td><td>171210</td><td>+</td><td>0</td><td>GGTCAGGGCGGCATAGTATC</td><td>GGG</td><td>||||||||[||||||||||||]PAM</td><td>NA</td><td>Intergenic</td><td>NA</td><td>NA</td></tr>
<tr><td>T13008</td><td>AGTATCGGGTCGAGTGTCTGCGG</td><td>1000</td><td></td><td></td><td></td><td></td><td></td><td></td><td></td><td></td><td></td></tr>
<tr><td>Oligo fwd</td><td>CACCAGTATCGGGTCGAGTGTCTG</td><td></td><td></td><td></td><td></td><td></td><td></td><td></td><td></td><td></td><td></td></tr>
<tr><td>Oligo rev</td><td>AAACCAGACACTCGACCCGATACT</td><td></td><td></td><td></td><td></td><td></td><td></td><td></td><td></td><td></td><td></td></tr>
<tr><td>Chromosome</td><td>start</td><td>end</td><td>strand</td><td>MM</td><td>target_seq</td><td>PAM</td><td>alignment</td><td>distance</td><td>position</td><td>gene name</td><td>gene id</td></tr>
<tr><td>GU305914.1</td><td>171202</td><td>171224</td><td>+</td><td>0</td><td>AGTATCGGGTCGAGTGTCTG</td><td>CGG</td><td>||||||||[||||||||||||]PAM</td><td>NA</td><td>Intergenic</td><td>NA</td><td>NA</td></tr>
<tr><td>T13009</td><td>AGAAGAGTTTCTTGTAGTTATGG</td><td>1000</td><td></td><td></td><td></td><td></td><td></td><td></td><td></td><td></td><td></td></tr>
<tr><td>Oligo fwd</td><td>CACCAGAAGAGTTTCTTGTAGTTA</td><td></td><td></td><td></td><td></td><td></td><td></td><td></td><td></td><td></td><td></td></tr>
<tr><td>Oligo rev</td><td>AAACTAACTACAAGAAACTCTTCT</td><td></td><td></td><td></td><td></td><td></td><td></td><td></td><td></td><td></td><td></td></tr>
<tr><td>Chromosome</td><td>start</td><td>end</td><td>strand</td><td>MM</td><td>target_seq</td><td>PAM</td><td>alignment</td><td>distance</td><td>position</td><td>gene name</td><td>gene id</td></tr>
<tr><td>GU305914.1</td><td>171229</td><td>171251</td><td>+</td><td>0</td><td>AGAAGAGTTTCTTGTAGTTA</td><td>TGG</td><td>||||||||[||||||||||||]PAM</td><td>NA</td><td>Intergenic</td><td>NA</td><td>NA</td></tr>
<tr><td>T13010</td><td>GTTGTCGACCATGACGTCTACGG</td><td>1000</td><td></td><td></td><td></td><td></td><td></td><td></td><td></td><td></td><td></td></tr>
<tr><td>Oligo fwd</td><td>CACCGTTGTCGACCATGACGTCTA</td><td></td><td></td><td></td><td></td><td></td><td></td><td></td><td></td><td></td><td></td></tr>
<tr><td>Oligo rev</td><td>AAACTAGACGTCATGGTCGACAAC</td><td></td><td></td><td></td><td></td><td></td><td></td><td></td><td></td><td></td><td></td></tr>
<tr><td>Chromosome</td><td>start</td><td>end</td><td>strand</td><td>MM</td><td>target_seq</td><td>PAM</td><td>alignment</td><td>distance</td><td>position</td><td>gene name</td><td>gene id</td></tr>
<tr><td>GU305914.1</td><td>171251</td><td>171273</td><td>+</td><td>0</td><td>GTTGTCGACCATGACGTCTA</td><td>CGG</td><td>||||||||[||||||||||||]PAM</td><td>NA</td><td>Intergenic</td><td>NA</td><td>NA</td></tr>
<tr><td>T13011</td><td>TTCCCCTGACTGTCTGCCCACGG</td><td>1000</td><td></td><td></td><td></td><td></td><td></td><td></td><td></td><td></td><td></td></tr>
<tr><td>Oligo fwd</td><td>CACCTTCCCCTGACTGTCTGCCCA</td><td></td><td></td><td></td><td></td><td></td><td></td><td></td><td></td><td></td><td></td></tr>
<tr><td>Oligo rev</td><td>AAACTGGGCAGACAGTCAGGGGAA</td><td></td><td></td><td></td><td></td><td></td><td></td><td></td><td></td><td></td><td></td></tr>
<tr><td>Chromosome</td><td>start</td><td>end</td><td>strand</td><td>MM</td><td>target_seq</td><td>PAM</td><td>alignment</td><td>distance</td><td>position</td><td>gene name</td><td>gene id</td></tr>
<tr><td>GU305914.1</td><td>171279</td><td>171301</td><td>+</td><td>0</td><td>TTCCCCTGACTGTCTGCCCA</td><td>CGG</td><td>||||||||[||||||||||||]PAM</td><td>NA</td><td>Intergenic</td><td>NA</td><td>NA</td></tr>
<tr><td>T13012</td><td>TCCCCTGACTGTCTGCCCACGGG</td><td>1000</td><td></td><td></td><td></td><td></td><td></td><td></td><td></td><td></td><td></td></tr>
<tr><td>Oligo fwd</td><td>CACCTCCCCTGACTGTCTGCCCAC</td><td></td><td></td><td></td><td></td><td></td><td></td><td></td><td></td><td></td><td></td></tr>
<tr><td>Oligo rev</td><td>AAACGTGGGCAGACAGTCAGGGGA</td><td></td><td></td><td></td><td></td><td></td><td></td><td></td><td></td><td></td><td></td></tr>
<tr><td>Chromosome</td><td>start</td><td>end</td><td>strand</td><td>MM</td><td>target_seq</td><td>PAM</td><td>alignment</td><td>distance</td><td>position</td><td>gene name</td><td>gene id</td></tr>
<tr><td>GU305914.1</td><td>171280</td><td>171302</td><td>+</td><td>0</td><td>TCCCCTGACTGTCTGCCCAC</td><td>GGG</td><td>||||||||[||||||||||||]PAM</td><td>NA</td><td>Intergenic</td><td>NA</td><td>NA</td></tr>
<tr><td>T13013</td><td>GACACTGAACGTCATCAGATAGG</td><td>1000</td><td></td><td></td><td></td><td></td><td></td><td></td><td></td><td></td><td></td></tr>
<tr><td>Oligo fwd</td><td>CACCGACACTGAACGTCATCAGAT</td><td></td><td></td><td></td><td></td><td></td><td></td><td></td><td></td><td></td><td></td></tr>
<tr><td>Oligo rev</td><td>AAACATCTGATGACGTTCAGTGTC</td><td></td><td></td><td></td><td></td><td></td><td></td><td></td><td></td><td></td><td></td></tr>
<tr><td>Chromosome</td><td>start</td><td>end</td><td>strand</td><td>MM</td><td>target_seq</td><td>PAM</td><td>alignment</td><td>distance</td><td>position</td><td>gene name</td><td>gene id</td></tr>
<tr><td>GU305914.1</td><td>171302</td><td>171324</td><td>+</td><td>0</td><td>GACACTGAACGTCATCAGAT</td><td>AGG</td><td>||||||||[||||||||||||]PAM</td><td>NA</td><td>Intergenic</td><td>NA</td><td>NA</td></tr>
<tr><td>T13014</td><td>GAACGTCATCAGATAGGTGATGG</td><td>1000</td><td></td><td></td><td></td><td></td><td></td><td></td><td></td><td></td><td></td></tr>
<tr><td>Oligo fwd</td><td>CACCGAACGTCATCAGATAGGTGA</td><td></td><td></td><td></td><td></td><td></td><td></td><td></td><td></td><td></td><td></td></tr>
<tr><td>Oligo rev</td><td>AAACTCACCTATCTGATGACGTTC</td><td></td><td></td><td></td><td></td><td></td><td></td><td></td><td></td><td></td><td></td></tr>
<tr><td>Chromosome</td><td>start</td><td>end</td><td>strand</td><td>MM</td><td>target_seq</td><td>PAM</td><td>alignment</td><td>distance</td><td>position</td><td>gene name</td><td>gene id</td></tr>
<tr><td>GU305914.1</td><td>171308</td><td>171330</td><td>+</td><td>0</td><td>GAACGTCATCAGATAGGTGA</td><td>TGG</td><td>||||||||[||||||||||||]PAM</td><td>NA</td><td>Intergenic</td><td>NA</td><td>NA</td></tr>
<tr><td>T13015</td><td>TGATGGTTCCGTTCTCGTTCTGG</td><td>1000</td><td></td><td></td><td></td><td></td><td></td><td></td><td></td><td></td><td></td></tr>
<tr><td>Oligo fwd</td><td>CACCTGATGGTTCCGTTCTCGTTC</td><td></td><td></td><td></td><td></td><td></td><td></td><td></td><td></td><td></td><td></td></tr>
<tr><td>Oligo rev</td><td>AAACGAACGAGAACGGAACCATCA</td><td></td><td></td><td></td><td></td><td></td><td></td><td></td><td></td><td></td><td></td></tr>
<tr><td>Chromosome</td><td>start</td><td>end</td><td>strand</td><td>MM</td><td>target_seq</td><td>PAM</td><td>alignment</td><td>distance</td><td>position</td><td>gene name</td><td>gene id</td></tr>
<tr><td>GU305914.1</td><td>171325</td><td>171347</td><td>+</td><td>0</td><td>TGATGGTTCCGTTCTCGTTC</td><td>TGG</td><td>||||||||[||||||||||||]PAM</td><td>NA</td><td>Intergenic</td><td>NA</td><td>NA</td></tr>
<tr><td>T13016</td><td>CTCGTTCTGGATCGACGTGTTGG</td><td>1000</td><td></td><td></td><td></td><td></td><td></td><td></td><td></td><td></td><td></td></tr>
<tr><td>Oligo fwd</td><td>CACCCTCGTTCTGGATCGACGTGT</td><td></td><td></td><td></td><td></td><td></td><td></td><td></td><td></td><td></td><td></td></tr>
<tr><td>Oligo rev</td><td>AAACACACGTCGATCCAGAACGAG</td><td></td><td></td><td></td><td></td><td></td><td></td><td></td><td></td><td></td><td></td></tr>
<tr><td>Chromosome</td><td>start</td><td>end</td><td>strand</td><td>MM</td><td>target_seq</td><td>PAM</td><td>alignment</td><td>distance</td><td>position</td><td>gene name</td><td>gene id</td></tr>
<tr><td>GU305914.1</td><td>171338</td><td>171360</td><td>+</td><td>0</td><td>CTCGTTCTGGATCGACGTGT</td><td>TGG</td><td>||||||||[||||||||||||]PAM</td><td>NA</td><td>Intergenic</td><td>NA</td><td>NA</td></tr>
<tr><td>T13017</td><td>GTTCTGGATCGACGTGTTGGAGG</td><td>1000</td><td></td><td></td><td></td><td></td><td></td><td></td><td></td><td></td><td></td></tr>
<tr><td>Oligo fwd</td><td>CACCGTTCTGGATCGACGTGTTGG</td><td></td><td></td><td></td><td></td><td></td><td></td><td></td><td></td><td></td><td></td></tr>
<tr><td>Oligo rev</td><td>AAACCCAACACGTCGATCCAGAAC</td><td></td><td></td><td></td><td></td><td></td><td></td><td></td><td></td><td></td><td></td></tr>
<tr><td>Chromosome</td><td>start</td><td>end</td><td>strand</td><td>MM</td><td>target_seq</td><td>PAM</td><td>alignment</td><td>distance</td><td>position</td><td>gene name</td><td>gene id</td></tr>
<tr><td>GU305914.1</td><td>171341</td><td>171363</td><td>+</td><td>0</td><td>GTTCTGGATCGACGTGTTGG</td><td>AGG</td><td>||||||||[||||||||||||]PAM</td><td>NA</td><td>Intergenic</td><td>NA</td><td>NA</td></tr>
<tr><td>T13018</td><td>TGGATCGACGTGTTGGAGGTCGG</td><td>1000</td><td></td><td></td><td></td><td></td><td></td><td></td><td></td><td></td><td></td></tr>
<tr><td>Oligo fwd</td><td>CACCTGGATCGACGTGTTGGAGGT</td><td></td><td></td><td></td><td></td><td></td><td></td><td></td><td></td><td></td><td></td></tr>
<tr><td>Oligo rev</td><td>AAACACCTCCAACACGTCGATCCA</td><td></td><td></td><td></td><td></td><td></td><td></td><td></td><td></td><td></td><td></td></tr>
<tr><td>Chromosome</td><td>start</td><td>end</td><td>strand</td><td>MM</td><td>target_seq</td><td>PAM</td><td>alignment</td><td>distance</td><td>position</td><td>gene name</td><td>gene id</td></tr>
<tr><td>GU305914.1</td><td>171345</td><td>171367</td><td>+</td><td>0</td><td>TGGATCGACGTGTTGGAGGT</td><td>CGG</td><td>||||||||[||||||||||||]PAM</td><td>NA</td><td>Intergenic</td><td>NA</td><td>NA</td></tr>
<tr><td>T13019</td><td>GTGTTGGAGGTCGGCGTGAATGG</td><td>1000</td><td></td><td></td><td></td><td></td><td></td><td></td><td></td><td></td><td></td></tr>
<tr><td>Oligo fwd</td><td>CACCGTGTTGGAGGTCGGCGTGAA</td><td></td><td></td><td></td><td></td><td></td><td></td><td></td><td></td><td></td><td></td></tr>
<tr><td>Oligo rev</td><td>AAACTTCACGCCGACCTCCAACAC</td><td></td><td></td><td></td><td></td><td></td><td></td><td></td><td></td><td></td><td></td></tr>
<tr><td>Chromosome</td><td>start</td><td>end</td><td>strand</td><td>MM</td><td>target_seq</td><td>PAM</td><td>alignment</td><td>distance</td><td>position</td><td>gene name</td><td>gene id</td></tr>
<tr><td>GU305914.1</td><td>171354</td><td>171376</td><td>+</td><td>0</td><td>GTGTTGGAGGTCGGCGTGAA</td><td>TGG</td><td>||||||||[||||||||||||]PAM</td><td>NA</td><td>Intergenic</td><td>NA</td><td>NA</td></tr>
<tr><td>T13020</td><td>TGAATGGACTGTTCTTCTCGTGG</td><td>1000</td><td></td><td></td><td></td><td></td><td></td><td></td><td></td><td></td><td></td></tr>
<tr><td>Oligo fwd</td><td>CACCTGAATGGACTGTTCTTCTCG</td><td></td><td></td><td></td><td></td><td></td><td></td><td></td><td></td><td></td><td></td></tr>
<tr><td>Oligo rev</td><td>AAACCGAGAAGAACAGTCCATTCA</td><td></td><td></td><td></td><td></td><td></td><td></td><td></td><td></td><td></td><td></td></tr>
<tr><td>Chromosome</td><td>start</td><td>end</td><td>strand</td><td>MM</td><td>target_seq</td><td>PAM</td><td>alignment</td><td>distance</td><td>position</td><td>gene name</td><td>gene id</td></tr>
<tr><td>GU305914.1</td><td>171370</td><td>171392</td><td>+</td><td>0</td><td>TGAATGGACTGTTCTTCTCG</td><td>TGG</td><td>||||||||[||||||||||||]PAM</td><td>NA</td><td>Intergenic</td><td>NA</td><td>NA</td></tr>
</table>

<table>
<tr><td>T13021</td><td>GTGGTACACCGAGATCGACATGG</td><td>1000</td><td></td><td></td><td></td><td></td><td></td><td></td><td></td><td></td><td></td></tr>
<tr><td>Oligo fwd</td><td>CACCGTGGTACACCGAGATCGACA</td><td></td><td></td><td></td><td></td><td></td><td></td><td></td><td></td><td></td><td></td></tr>
<tr><td>Oligo rev</td><td>AAACTGTCGATCTCGGTGTACCAC</td><td></td><td></td><td></td><td></td><td></td><td></td><td></td><td></td><td></td><td></td></tr>
<tr><td>Chromosome</td><td>start</td><td>end</td><td>strand</td><td>MM</td><td>target_seq</td><td>PAM</td><td>alignment</td><td>distance</td><td>position</td><td>gene name</td><td>gene id</td></tr>
<tr><td>GU305914.1</td><td>171389</td><td>171411</td><td>+</td><td>0</td><td>GTGGTACACCGAGATCGACA</td><td>TGG</td><td>||||||||[||||||||||||]PAM</td><td>NA</td><td>Intergenic</td><td>NA</td><td>NA</td></tr>
<tr><td>T13022</td><td>TGGTACACCGAGATCGACATGGG</td><td>1000</td><td></td><td></td><td></td><td></td><td></td><td></td><td></td><td></td><td></td></tr>
<tr><td>Oligo fwd</td><td>CACCTGGTACACCGAGATCGACAT</td><td></td><td></td><td></td><td></td><td></td><td></td><td></td><td></td><td></td><td></td></tr>
<tr><td>Oligo rev</td><td>AAACATGTCGATCTCGGTGTACCA</td><td></td><td></td><td></td><td></td><td></td><td></td><td></td><td></td><td></td><td></td></tr>
<tr><td>Chromosome</td><td>start</td><td>end</td><td>strand</td><td>MM</td><td>target_seq</td><td>PAM</td><td>alignment</td><td>distance</td><td>position</td><td>gene name</td><td>gene id</td></tr>
<tr><td>GU305914.1</td><td>171390</td><td>171412</td><td>+</td><td>0</td><td>TGGTACACCGAGATCGACAT</td><td>GGG</td><td>||||||||[||||||||||||]PAM</td><td>NA</td><td>Intergenic</td><td>NA</td><td>NA</td></tr>
<tr><td>T13023</td><td>GGTACACCGAGATCGACATGGGG</td><td>1000</td><td></td><td></td><td></td><td></td><td></td><td></td><td></td><td></td><td></td></tr>
<tr><td>Oligo fwd</td><td>CACCGGTACACCGAGATCGACATG</td><td></td><td></td><td></td><td></td><td></td><td></td><td></td><td></td><td></td><td></td></tr>
<tr><td>Oligo rev</td><td>AAACCATGTCGATCTCGGTGTACC</td><td></td><td></td><td></td><td></td><td></td><td></td><td></td><td></td><td></td><td></td></tr>
<tr><td>Chromosome</td><td>start</td><td>end</td><td>strand</td><td>MM</td><td>target_seq</td><td>PAM</td><td>alignment</td><td>distance</td><td>position</td><td>gene name</td><td>gene id</td></tr>
<tr><td>GU305914.1</td><td>171391</td><td>171413</td><td>+</td><td>0</td><td>GGTACACCGAGATCGACATG</td><td>GGG</td><td>||||||||[||||||||||||]PAM</td><td>NA</td><td>Intergenic</td><td>NA</td><td>NA</td></tr>
<tr><td>T13024</td><td>ACCGAGATCGACATGGGGTTCGG</td><td>1000</td><td></td><td></td><td></td><td></td><td></td><td></td><td></td><td></td><td></td></tr>
<tr><td>Oligo fwd</td><td>CACCACCGAGATCGACATGGGGTT</td><td></td><td></td><td></td><td></td><td></td><td></td><td></td><td></td><td></td><td></td></tr>
<tr><td>Oligo rev</td><td>AAACAACCCCATGTCGATCTCGGT</td><td></td><td></td><td></td><td></td><td></td><td></td><td></td><td></td><td></td><td></td></tr>
<tr><td>Chromosome</td><td>start</td><td>end</td><td>strand</td><td>MM</td><td>target_seq</td><td>PAM</td><td>alignment</td><td>distance</td><td>position</td><td>gene name</td><td>gene id</td></tr>
<tr><td>GU305914.1</td><td>171396</td><td>171418</td><td>+</td><td>0</td><td>ACCGAGATCGACATGGGGTT</td><td>CGG</td><td>||||||||[||||||||||||]PAM</td><td>NA</td><td>Intergenic</td><td>NA</td><td>NA</td></tr>
<tr><td>T13025</td><td>ATGGGGTTCGGAGATTTTTTTGG</td><td>1000</td><td></td><td></td><td></td><td></td><td></td><td></td><td></td><td></td><td></td></tr>
<tr><td>Oligo fwd</td><td>CACCATGGGGTTCGGAGATTTTTT</td><td></td><td></td><td></td><td></td><td></td><td></td><td></td><td></td><td></td><td></td></tr>
<tr><td>Oligo rev</td><td>AAACAAAAAATCTCCGAACCCCAT</td><td></td><td></td><td></td><td></td><td></td><td></td><td></td><td></td><td></td><td></td></tr>
<tr><td>Chromosome</td><td>start</td><td>end</td><td>strand</td><td>MM</td><td>target_seq</td><td>PAM</td><td>alignment</td><td>distance</td><td>position</td><td>gene name</td><td>gene id</td></tr>
<tr><td>GU305914.1</td><td>171408</td><td>171430</td><td>+</td><td>0</td><td>ATGGGGTTCGGAGATTTTTT</td><td>TGG</td><td>||||||||[||||||||||||]PAM</td><td>NA</td><td>Intergenic</td><td>NA</td><td>NA</td></tr>
<tr><td>T13026</td><td>GTTCGGAGATTTTTTTGGTTTGG</td><td>1000</td><td></td><td></td><td></td><td></td><td></td><td></td><td></td><td></td><td></td></tr>
<tr><td>Oligo fwd</td><td>CACCGTTCGGAGATTTTTTTGGTT</td><td></td><td></td><td></td><td></td><td></td><td></td><td></td><td></td><td></td><td></td></tr>
<tr><td>Oligo rev</td><td>AAACAACCAAAAAAATCTCCGAAC</td><td></td><td></td><td></td><td></td><td></td><td></td><td></td><td></td><td></td><td></td></tr>
<tr><td>Chromosome</td><td>start</td><td>end</td><td>strand</td><td>MM</td><td>target_seq</td><td>PAM</td><td>alignment</td><td>distance</td><td>position</td><td>gene name</td><td>gene id</td></tr>
<tr><td>GU305914.1</td><td>171413</td><td>171435</td><td>+</td><td>0</td><td>GTTCGGAGATTTTTTTGGTT</td><td>TGG</td><td>||||||||[||||||||||||]PAM</td><td>NA</td><td>Intergenic</td><td>NA</td><td>NA</td></tr>
<tr><td>T13027</td><td>TTCGGAGATTTTTTTGGTTTGGG</td><td>1000</td><td></td><td></td><td></td><td></td><td></td><td></td><td></td><td></td><td></td></tr>
<tr><td>Oligo fwd</td><td>CACCTTCGGAGATTTTTTTGGTTT</td><td></td><td></td><td></td><td></td><td></td><td></td><td></td><td></td><td></td><td></td></tr>
<tr><td>Oligo rev</td><td>AAACAAACCAAAAAAATCTCCGAA</td><td></td><td></td><td></td><td></td><td></td><td></td><td></td><td></td><td></td><td></td></tr>
<tr><td>Chromosome</td><td>start</td><td>end</td><td>strand</td><td>MM</td><td>target_seq</td><td>PAM</td><td>alignment</td><td>distance</td><td>position</td><td>gene name</td><td>gene id</td></tr>
<tr><td>GU305914.1</td><td>171414</td><td>171436</td><td>+</td><td>0</td><td>TTCGGAGATTTTTTTGGTTT</td><td>GGG</td><td>||||||||[||||||||||||]PAM</td><td>NA</td><td>Intergenic</td><td>NA</td><td>NA</td></tr>
<tr><td>T13028</td><td>CAGATCCTTGACCTGAAACCCGG</td><td>1000</td><td></td><td></td><td></td><td></td><td></td><td></td><td></td><td></td><td></td></tr>
<tr><td>Oligo fwd</td><td>CACCCAGATCCTTGACCTGAAACC</td><td></td><td></td><td></td><td></td><td></td><td></td><td></td><td></td><td></td><td></td></tr>
<tr><td>Oligo rev</td><td>AAACGGTTTCAGGTCAAGGATCTG</td><td></td><td></td><td></td><td></td><td></td><td></td><td></td><td></td><td></td><td></td></tr>
<tr><td>Chromosome</td><td>start</td><td>end</td><td>strand</td><td>MM</td><td>target_seq</td><td>PAM</td><td>alignment</td><td>distance</td><td>position</td><td>gene name</td><td>gene id</td></tr>
<tr><td>GU305914.1</td><td>171476</td><td>171498</td><td>+</td><td>0</td><td>CAGATCCTTGACCTGAAACC</td><td>CGG</td><td>||||||||[||||||||||||]PAM</td><td>NA</td><td>Intergenic</td><td>NA</td><td>NA</td></tr>
<tr><td>T13029</td><td>CCGGTCGTCAGAGAGATGAGCGG</td><td>1000</td><td></td><td></td><td></td><td></td><td></td><td></td><td></td><td></td><td></td></tr>
<tr><td>Oligo fwd</td><td>CACCCCGGTCGTCAGAGAGATGAG</td><td></td><td></td><td></td><td></td><td></td><td></td><td></td><td></td><td></td><td></td></tr>
<tr><td>Oligo rev</td><td>AAACCTCATCTCTCTGACGACCGG</td><td></td><td></td><td></td><td></td><td></td><td></td><td></td><td></td><td></td><td></td></tr>
<tr><td>Chromosome</td><td>start</td><td>end</td><td>strand</td><td>MM</td><td>target_seq</td><td>PAM</td><td>alignment</td><td>distance</td><td>position</td><td>gene name</td><td>gene id</td></tr>
<tr><td>GU305914.1</td><td>171495</td><td>171517</td><td>+</td><td>0</td><td>CCGGTCGTCAGAGAGATGAG</td><td>CGG</td><td>||||||||[||||||||||||]PAM</td><td>NA</td><td>Intergenic</td><td>NA</td><td>NA</td></tr>
<tr><td>T13030</td><td>CGGTCGTCAGAGAGATGAGCGGG</td><td>1000</td><td></td><td></td><td></td><td></td><td></td><td></td><td></td><td></td><td></td></tr>
<tr><td>Oligo fwd</td><td>CACCCGGTCGTCAGAGAGATGAGC</td><td></td><td></td><td></td><td></td><td></td><td></td><td></td><td></td><td></td><td></td></tr>
<tr><td>Oligo rev</td><td>AAACGCTCATCTCTCTGACGACCG</td><td></td><td></td><td></td><td></td><td></td><td></td><td></td><td></td><td></td><td></td></tr>
<tr><td>Chromosome</td><td>start</td><td>end</td><td>strand</td><td>MM</td><td>target_seq</td><td>PAM</td><td>alignment</td><td>distance</td><td>position</td><td>gene name</td><td>gene id</td></tr>
<tr><td>GU305914.1</td><td>171496</td><td>171518</td><td>+</td><td>0</td><td>CGGTCGTCAGAGAGATGAGC</td><td>GGG</td><td>||||||||[||||||||||||]PAM</td><td>NA</td><td>Intergenic</td><td>NA</td><td>NA</td></tr>
<tr><td>T13031</td><td>CCACGTCGCCTCTTTCCAATCGG</td><td>1000</td><td></td><td></td><td></td><td></td><td></td><td></td><td></td><td></td><td></td></tr>
<tr><td>Oligo fwd</td><td>CACCCCACGTCGCCTCTTTCCAAT</td><td></td><td></td><td></td><td></td><td></td><td></td><td></td><td></td><td></td><td></td></tr>
<tr><td>Oligo rev</td><td>AAACATTGGAAAGAGGCGACGTGG</td><td></td><td></td><td></td><td></td><td></td><td></td><td></td><td></td><td></td><td></td></tr>
<tr><td>Chromosome</td><td>start</td><td>end</td><td>strand</td><td>MM</td><td>target_seq</td><td>PAM</td><td>alignment</td><td>distance</td><td>position</td><td>gene name</td><td>gene id</td></tr>
<tr><td>GU305914.1</td><td>171533</td><td>171555</td><td>+</td><td>0</td><td>CCACGTCGCCTCTTTCCAAT</td><td>CGG</td><td>||||||||[||||||||||||]PAM</td><td>NA</td><td>Intergenic</td><td>NA</td><td>NA</td></tr>
<tr><td>T13032</td><td>CACGTCGCCTCTTTCCAATCGGG</td><td>1000</td><td></td><td></td><td></td><td></td><td></td><td></td><td></td><td></td><td></td></tr>
<tr><td>Oligo fwd</td><td>CACCCACGTCGCCTCTTTCCAATC</td><td></td><td></td><td></td><td></td><td></td><td></td><td></td><td></td><td></td><td></td></tr>
<tr><td>Oligo rev</td><td>AAACGATTGGAAAGAGGCGACGTG</td><td></td><td></td><td></td><td></td><td></td><td></td><td></td><td></td><td></td><td></td></tr>
<tr><td>Chromosome</td><td>start</td><td>end</td><td>strand</td><td>MM</td><td>target_seq</td><td>PAM</td><td>alignment</td><td>distance</td><td>position</td><td>gene name</td><td>gene id</td></tr>
<tr><td>GU305914.1</td><td>171534</td><td>171556</td><td>+</td><td>0</td><td>CACGTCGCCTCTTTCCAATC</td><td>GGG</td><td>||||||||[||||||||||||]PAM</td><td>NA</td><td>Intergenic</td><td>NA</td><td>NA</td></tr>
<tr><td>T13033</td><td>GGGCCCGCCGTCTCTAGTCCTGG</td><td>1000</td><td></td><td></td><td></td><td></td><td></td><td></td><td></td><td></td><td></td></tr>
<tr><td>Oligo fwd</td><td>CACCGGGCCCGCCGTCTCTAGTCC</td><td></td><td></td><td></td><td></td><td></td><td></td><td></td><td></td><td></td><td></td></tr>
<tr><td>Oligo rev</td><td>AAACGGACTAGAGACGGCGGGCCC</td><td></td><td></td><td></td><td></td><td></td><td></td><td></td><td></td><td></td><td></td></tr>
<tr><td>Chromosome</td><td>start</td><td>end</td><td>strand</td><td>MM</td><td>target_seq</td><td>PAM</td><td>alignment</td><td>distance</td><td>position</td><td>gene name</td><td>gene id</td></tr>
<tr><td>GU305914.1</td><td>171554</td><td>171576</td><td>+</td><td>0</td><td>GGGCCCGCCGTCTCTAGTCC</td><td>TGG</td><td>||||||||[||||||||||||]PAM</td><td>NA</td><td>Intergenic</td><td>NA</td><td>NA</td></tr>
<tr><td>T13034</td><td>CGTCTCTAGTCCTGGCGATGTGG</td><td>1000</td><td></td><td></td><td></td><td></td><td></td><td></td><td></td><td></td><td></td></tr>
<tr><td>Oligo fwd</td><td>CACCCGTCTCTAGTCCTGGCGATG</td><td></td><td></td><td></td><td></td><td></td><td></td><td></td><td></td><td></td><td></td></tr>
<tr><td>Oligo rev</td><td>AAACCATCGCCAGGACTAGAGACG</td><td></td><td></td><td></td><td></td><td></td><td></td><td></td><td></td><td></td><td></td></tr>
<tr><td>Chromosome</td><td>start</td><td>end</td><td>strand</td><td>MM</td><td>target_seq</td><td>PAM</td><td>alignment</td><td>distance</td><td>position</td><td>gene name</td><td>gene id</td></tr>
<tr><td>GU305914.1</td><td>171562</td><td>171584</td><td>+</td><td>0</td><td>CGTCTCTAGTCCTGGCGATG</td><td>TGG</td><td>||||||||[||||||||||||]PAM</td><td>NA</td><td>Intergenic</td><td>NA</td><td>NA</td></tr>
<tr><td>T13035</td><td>CACTTGAAGAACGTGACACCCGG</td><td>1000</td><td></td><td></td><td></td><td></td><td></td><td></td><td></td><td></td><td></td></tr>
<tr><td>Oligo fwd</td><td>CACCCACTTGAAGAACGTGACACC</td><td></td><td></td><td></td><td></td><td></td><td></td><td></td><td></td><td></td><td></td></tr>
<tr><td>Oligo rev</td><td>AAACGGTGTCACGTTCTTCAAGTG</td><td></td><td></td><td></td><td></td><td></td><td></td><td></td><td></td><td></td><td></td></tr>
<tr><td>Chromosome</td><td>start</td><td>end</td><td>strand</td><td>MM</td><td>target_seq</td><td>PAM</td><td>alignment</td><td>distance</td><td>position</td><td>gene name</td><td>gene id</td></tr>
</table>

<table>
<tr><td>GU305914.1</td><td></td><td>171594</td><td>171616</td><td>+</td><td>0</td><td>CACTTGAAGAACGTGACACC</td><td>CGG</td><td>||||||||[||||||||||||]PAM</td><td>NA</td><td>Intergenic</td><td>NA</td><td>NA</td></tr>
<tr><td>T13036</td><td>GAACGTGACACCCGGAGCGACGG</td><td></td><td>1000</td><td></td><td></td><td></td><td></td><td></td><td></td><td></td><td></td><td></td></tr>
<tr><td>Oligo fwd</td><td>CACCGAACGTGACACCCGGAGCGA</td><td></td><td></td><td></td><td></td><td></td><td></td><td></td><td></td><td></td><td></td><td></td></tr>
<tr><td>Oligo rev</td><td>AAACTCGCTCCGGGTGTCACGTTC</td><td></td><td></td><td></td><td></td><td></td><td></td><td></td><td></td><td></td><td></td><td></td></tr>
<tr><td>Chromosome</td><td>start</td><td></td><td>end</td><td>strand</td><td>MM</td><td>target_seq</td><td>PAM</td><td>alignment</td><td>distance</td><td>position</td><td>gene name</td><td>gene id</td></tr>
<tr><td>GU305914.1</td><td></td><td>171602</td><td>171624</td><td>+</td><td>0</td><td>GAACGTGACACCCGGAGCGA</td><td>CGG</td><td>||||||||[||||||||||||]PAM</td><td>NA</td><td>Intergenic</td><td>NA</td><td>NA</td></tr>
<tr><td>T13037</td><td>TGTGACTAGCGTTCAATATATGG</td><td></td><td>1000</td><td></td><td></td><td></td><td></td><td></td><td></td><td></td><td></td><td></td></tr>
<tr><td>Oligo fwd</td><td>CACCTGTGACTAGCGTTCAATATA</td><td></td><td></td><td></td><td></td><td></td><td></td><td></td><td></td><td></td><td></td><td></td></tr>
<tr><td>Oligo rev</td><td>AAACTATATTGAACGCTAGTCACA</td><td></td><td></td><td></td><td></td><td></td><td></td><td></td><td></td><td></td><td></td><td></td></tr>
<tr><td>Chromosome</td><td>start</td><td></td><td>end</td><td>strand</td><td>MM</td><td>target_seq</td><td>PAM</td><td>alignment</td><td>distance</td><td>position</td><td>gene name</td><td>gene id</td></tr>
<tr><td>GU305914.1</td><td></td><td>171628</td><td>171650</td><td>+</td><td>0</td><td>TGTGACTAGCGTTCAATATA</td><td>TGG</td><td>||||||||[||||||||||||]PAM</td><td>NA</td><td>Intergenic</td><td>NA</td><td>NA</td></tr>
<tr><td>T13038</td><td>GTCATAACTGAGTTTCAGCGAGG</td><td></td><td>1000</td><td></td><td></td><td></td><td></td><td></td><td></td><td></td><td></td><td></td></tr>
<tr><td>Oligo fwd</td><td>CACCGTCATAACTGAGTTTCAGCG</td><td></td><td></td><td></td><td></td><td></td><td></td><td></td><td></td><td></td><td></td><td></td></tr>
<tr><td>Oligo rev</td><td>AAACCGCTGAAACTCAGTTATGAC</td><td></td><td></td><td></td><td></td><td></td><td></td><td></td><td></td><td></td><td></td><td></td></tr>
<tr><td>Chromosome</td><td>start</td><td></td><td>end</td><td>strand</td><td>MM</td><td>target_seq</td><td>PAM</td><td>alignment</td><td>distance</td><td>position</td><td>gene name</td><td>gene id</td></tr>
<tr><td>GU305914.1</td><td></td><td>171659</td><td>171681</td><td>+</td><td>0</td><td>GTCATAACTGAGTTTCAGCG</td><td>AGG</td><td>||||||||[||||||||||||]PAM</td><td>NA</td><td>Intergenic</td><td>NA</td><td>NA</td></tr>
<tr><td>T13039</td><td>ACTGAGTTTCAGCGAGGCTGTGG</td><td></td><td>1000</td><td></td><td></td><td></td><td></td><td></td><td></td><td></td><td></td><td></td></tr>
<tr><td>Oligo fwd</td><td>CACCACTGAGTTTCAGCGAGGCTG</td><td></td><td></td><td></td><td></td><td></td><td></td><td></td><td></td><td></td><td></td><td></td></tr>
<tr><td>Oligo rev</td><td>AAACCAGCCTCGCTGAAACTCAGT</td><td></td><td></td><td></td><td></td><td></td><td></td><td></td><td></td><td></td><td></td><td></td></tr>
<tr><td>Chromosome</td><td>start</td><td></td><td>end</td><td>strand</td><td>MM</td><td>target_seq</td><td>PAM</td><td>alignment</td><td>distance</td><td>position</td><td>gene name</td><td>gene id</td></tr>
<tr><td>GU305914.1</td><td></td><td>171665</td><td>171687</td><td>+</td><td>0</td><td>ACTGAGTTTCAGCGAGGCTG</td><td>TGG</td><td>||||||||[||||||||||||]PAM</td><td>NA</td><td>Intergenic</td><td>NA</td><td>NA</td></tr>
<tr><td>T13040</td><td>CTGAGTTTCAGCGAGGCTGTGGG</td><td></td><td>1000</td><td></td><td></td><td></td><td></td><td></td><td></td><td></td><td></td><td></td></tr>
<tr><td>Oligo fwd</td><td>CACCCTGAGTTTCAGCGAGGCTGT</td><td></td><td></td><td></td><td></td><td></td><td></td><td></td><td></td><td></td><td></td><td></td></tr>
<tr><td>Oligo rev</td><td>AAACACAGCCTCGCTGAAACTCAG</td><td></td><td></td><td></td><td></td><td></td><td></td><td></td><td></td><td></td><td></td><td></td></tr>
<tr><td>Chromosome</td><td>start</td><td></td><td>end</td><td>strand</td><td>MM</td><td>target_seq</td><td>PAM</td><td>alignment</td><td>distance</td><td>position</td><td>gene name</td><td>gene id</td></tr>
<tr><td>GU305914.1</td><td></td><td>171666</td><td>171688</td><td>+</td><td>0</td><td>CTGAGTTTCAGCGAGGCTGT</td><td>GGG</td><td>||||||||[||||||||||||]PAM</td><td>NA</td><td>Intergenic</td><td>NA</td><td>NA</td></tr>
<tr><td>T13041</td><td>AAAGACAAAGACAGTCGATTAGG</td><td></td><td>1000</td><td></td><td></td><td></td><td></td><td></td><td></td><td></td><td></td><td></td></tr>
<tr><td>Oligo fwd</td><td>CACCAAAGACAAAGACAGTCGATT</td><td></td><td></td><td></td><td></td><td></td><td></td><td></td><td></td><td></td><td></td><td></td></tr>
<tr><td>Oligo rev</td><td>AAACAATCGACTGTCTTTGTCTTT</td><td></td><td></td><td></td><td></td><td></td><td></td><td></td><td></td><td></td><td></td><td></td></tr>
<tr><td>Chromosome</td><td>start</td><td></td><td>end</td><td>strand</td><td>MM</td><td>target_seq</td><td>PAM</td><td>alignment</td><td>distance</td><td>position</td><td>gene name</td><td>gene id</td></tr>
<tr><td>GU305914.1</td><td></td><td>171693</td><td>171715</td><td>+</td><td>0</td><td>AAAGACAAAGACAGTCGATT</td><td>AGG</td><td>||||||||[||||||||||||]PAM</td><td>NA</td><td>Intergenic</td><td>NA</td><td>NA</td></tr>
<tr><td>T13042</td><td>AAGACAAAGACAGTCGATTAGGG</td><td></td><td>1000</td><td></td><td></td><td></td><td></td><td></td><td></td><td></td><td></td><td></td></tr>
<tr><td>Oligo fwd</td><td>CACCAAGACAAAGACAGTCGATTA</td><td></td><td></td><td></td><td></td><td></td><td></td><td></td><td></td><td></td><td></td><td></td></tr>
<tr><td>Oligo rev</td><td>AAACTAATCGACTGTCTTTGTCTT</td><td></td><td></td><td></td><td></td><td></td><td></td><td></td><td></td><td></td><td></td><td></td></tr>
<tr><td>Chromosome</td><td>start</td><td></td><td>end</td><td>strand</td><td>MM</td><td>target_seq</td><td>PAM</td><td>alignment</td><td>distance</td><td>position</td><td>gene name</td><td>gene id</td></tr>
<tr><td>GU305914.1</td><td></td><td>171694</td><td>171716</td><td>+</td><td>0</td><td>AAGACAAAGACAGTCGATTA</td><td>GGG</td><td>||||||||[||||||||||||]PAM</td><td>NA</td><td>Intergenic</td><td>NA</td><td>NA</td></tr>
<tr><td>T13043</td><td>TTAGGGTAGAACCCCGTACGCGG</td><td></td><td>1000</td><td></td><td></td><td></td><td></td><td></td><td></td><td></td><td></td><td></td></tr>
<tr><td>Oligo fwd</td><td>CACCTTAGGGTAGAACCCCGTACG</td><td></td><td></td><td></td><td></td><td></td><td></td><td></td><td></td><td></td><td></td><td></td></tr>
<tr><td>Oligo rev</td><td>AAACCGTACGGGGTTCTACCCTAA</td><td></td><td></td><td></td><td></td><td></td><td></td><td></td><td></td><td></td><td></td><td></td></tr>
<tr><td>Chromosome</td><td>start</td><td></td><td>end</td><td>strand</td><td>MM</td><td>target_seq</td><td>PAM</td><td>alignment</td><td>distance</td><td>position</td><td>gene name</td><td>gene id</td></tr>
<tr><td>GU305914.1</td><td></td><td>171711</td><td>171733</td><td>+</td><td>0</td><td>TTAGGGTAGAACCCCGTACG</td><td>CGG</td><td>||||||||[||||||||||||]PAM</td><td>NA</td><td>Intergenic</td><td>NA</td><td>NA</td></tr>
<tr><td>T13044</td><td>GTACGCGGTTAGCTTAACTACGG</td><td></td><td>1000</td><td></td><td></td><td></td><td></td><td></td><td></td><td></td><td></td><td></td></tr>
<tr><td>Oligo fwd</td><td>CACCGTACGCGGTTAGCTTAACTA</td><td></td><td></td><td></td><td></td><td></td><td></td><td></td><td></td><td></td><td></td><td></td></tr>
<tr><td>Oligo rev</td><td>AAACTAGTTAAGCTAACCGCGTAC</td><td></td><td></td><td></td><td></td><td></td><td></td><td></td><td></td><td></td><td></td><td></td></tr>
<tr><td>Chromosome</td><td>start</td><td></td><td>end</td><td>strand</td><td>MM</td><td>target_seq</td><td>PAM</td><td>alignment</td><td>distance</td><td>position</td><td>gene name</td><td>gene id</td></tr>
<tr><td>GU305914.1</td><td></td><td>171726</td><td>171748</td><td>+</td><td>0</td><td>GTACGCGGTTAGCTTAACTA</td><td>CGG</td><td>||||||||[||||||||||||]PAM</td><td>NA</td><td>Intergenic</td><td>NA</td><td>NA</td></tr>
<tr><td>T13045</td><td>GCGGTTAGCTTAACTACGGACGG</td><td></td><td>1000</td><td></td><td></td><td></td><td></td><td></td><td></td><td></td><td></td><td></td></tr>
<tr><td>Oligo fwd</td><td>CACCGCGGTTAGCTTAACTACGGA</td><td></td><td></td><td></td><td></td><td></td><td></td><td></td><td></td><td></td><td></td><td></td></tr>
<tr><td>Oligo rev</td><td>AAACTCCGTAGTTAAGCTAACCGC</td><td></td><td></td><td></td><td></td><td></td><td></td><td></td><td></td><td></td><td></td><td></td></tr>
<tr><td>Chromosome</td><td>start</td><td></td><td>end</td><td>strand</td><td>MM</td><td>target_seq</td><td>PAM</td><td>alignment</td><td>distance</td><td>position</td><td>gene name</td><td>gene id</td></tr>
<tr><td>GU305914.1</td><td></td><td>171730</td><td>171752</td><td>+</td><td>0</td><td>GCGGTTAGCTTAACTACGGA</td><td>CGG</td><td>||||||||[||||||||||||]PAM</td><td>NA</td><td>Intergenic</td><td>NA</td><td>NA</td></tr>
<tr><td>T13046</td><td>TTAGCTTAACTACGGACGGACGG</td><td></td><td>1000</td><td></td><td></td><td></td><td></td><td></td><td></td><td></td><td></td><td></td></tr>
<tr><td>Oligo fwd</td><td>CACCTTAGCTTAACTACGGACGGA</td><td></td><td></td><td></td><td></td><td></td><td></td><td></td><td></td><td></td><td></td><td></td></tr>
<tr><td>Oligo rev</td><td>AAACTCCGTCCGTAGTTAAGCTAA</td><td></td><td></td><td></td><td></td><td></td><td></td><td></td><td></td><td></td><td></td><td></td></tr>
<tr><td>Chromosome</td><td>start</td><td></td><td>end</td><td>strand</td><td>MM</td><td>target_seq</td><td>PAM</td><td>alignment</td><td>distance</td><td>position</td><td>gene name</td><td>gene id</td></tr>
<tr><td>GU305914.1</td><td></td><td>171734</td><td>171756</td><td>+</td><td>0</td><td>TTAGCTTAACTACGGACGGA</td><td>CGG</td><td>||||||||[||||||||||||]PAM</td><td>NA</td><td>Intergenic</td><td>NA</td><td>NA</td></tr>
<tr><td>T13047</td><td>CTTAACTACGGACGGACGGACGG</td><td></td><td>1000</td><td></td><td></td><td></td><td></td><td></td><td></td><td></td><td></td><td></td></tr>
<tr><td>Oligo fwd</td><td>CACCCTTAACTACGGACGGACGGA</td><td></td><td></td><td></td><td></td><td></td><td></td><td></td><td></td><td></td><td></td><td></td></tr>
<tr><td>Oligo rev</td><td>AAACTCCGTCCGTCCGTAGTTAAG</td><td></td><td></td><td></td><td></td><td></td><td></td><td></td><td></td><td></td><td></td><td></td></tr>
<tr><td>Chromosome</td><td>start</td><td></td><td>end</td><td>strand</td><td>MM</td><td>target_seq</td><td>PAM</td><td>alignment</td><td>distance</td><td>position</td><td>gene name</td><td>gene id</td></tr>
<tr><td>GU305914.1</td><td></td><td>171738</td><td>171760</td><td>+</td><td>0</td><td>CTTAACTACGGACGGACGGA</td><td>CGG</td><td>||||||||[||||||||||||]PAM</td><td>NA</td><td>Intergenic</td><td>NA</td><td>NA</td></tr>
<tr><td>T13048</td><td>GCACTTACTGTTGTAGTACTTGG</td><td></td><td>1000</td><td></td><td></td><td></td><td></td><td></td><td></td><td></td><td></td><td></td></tr>
<tr><td>Oligo fwd</td><td>CACCGCACTTACTGTTGTAGTACT</td><td></td><td></td><td></td><td></td><td></td><td></td><td></td><td></td><td></td><td></td><td></td></tr>
<tr><td>Oligo rev</td><td>AAACAGTACTACAACAGTAAGTGC</td><td></td><td></td><td></td><td></td><td></td><td></td><td></td><td></td><td></td><td></td><td></td></tr>
<tr><td>Chromosome</td><td>start</td><td></td><td>end</td><td>strand</td><td>MM</td><td>target_seq</td><td>PAM</td><td>alignment</td><td>distance</td><td>position</td><td>gene name</td><td>gene id</td></tr>
<tr><td>GU305914.1</td><td></td><td>171776</td><td>171798</td><td>+</td><td>0</td><td>GCACTTACTGTTGTAGTACT</td><td>TGG</td><td>||||||||[||||||||||||]PAM</td><td>NA</td><td>Intergenic</td><td>NA</td><td>NA</td></tr>
<tr><td>T13049</td><td>TGTTGTAGTACTTGGATCTGCGG</td><td></td><td>1000</td><td></td><td></td><td></td><td></td><td></td><td></td><td></td><td></td><td></td></tr>
<tr><td>Oligo fwd</td><td>CACCTGTTGTAGTACTTGGATCTG</td><td></td><td></td><td></td><td></td><td></td><td></td><td></td><td></td><td></td><td></td><td></td></tr>
<tr><td>Oligo rev</td><td>AAACCAGATCCAAGTACTACAACA</td><td></td><td></td><td></td><td></td><td></td><td></td><td></td><td></td><td></td><td></td><td></td></tr>
<tr><td>Chromosome</td><td>start</td><td></td><td>end</td><td>strand</td><td>MM</td><td>target_seq</td><td>PAM</td><td>alignment</td><td>distance</td><td>position</td><td>gene name</td><td>gene id</td></tr>
<tr><td>GU305914.1</td><td></td><td>171784</td><td>171806</td><td>+</td><td>0</td><td>TGTTGTAGTACTTGGATCTG</td><td>CGG</td><td>||||||||[||||||||||||]PAM</td><td>NA</td><td>Intergenic</td><td>NA</td><td>NA</td></tr>
<tr><td>T13050</td><td>GTAGTACTTGGATCTGCGGTCGG</td><td></td><td>1000</td><td></td><td></td><td></td><td></td><td></td><td></td><td></td><td></td><td></td></tr>
<tr><td>Oligo fwd</td><td>CACCGTAGTACTTGGATCTGCGGT</td><td></td><td></td><td></td><td></td><td></td><td></td><td></td><td></td><td></td><td></td><td></td></tr>
<tr><td>Oligo rev</td><td>AAACACCGCAGATCCAAGTACTAC</td><td></td><td></td><td></td><td></td><td></td><td></td><td></td><td></td><td></td><td></td><td></td></tr>
</table>

Chromosome start end strand MM target_seq PAM alignment distance position gene name gene id
GU305914.1 171788 171810 + 0 GTAGTACTTGGATCTGCGGT CGG ||||||||[||||||||||||]PAM NA Intergenic NA NA

T13051 CTCGAAATGAGTCCATCCGTAGG 1000
Oligo fwd CACCCTCGAAATGAGTCCATCCGT
Oligo rev AAACACGGATGGACTCATTTCGAG
Chromosome start end strand MM target_seq PAM alignment distance position gene name gene id
GU305914.1 171812 171834 + 0 CTCGAAATGAGTCCATCCGT AGG ||||||||[||||||||||||]PAM NA Intergenic NA NA

T13052 AGCCTCGAGATCTTGTGCCACGG 1000
Oligo fwd CACCAGCCTCGAGATCTTGTGCCA
Oligo rev AAACTGGCACAAGATCTCGAGGCT
Chromosome start end strand MM target_seq PAM alignment distance position gene name gene id
GU305914.1 171840 171862 + 0 AGCCTCGAGATCTTGTGCCA CGG ||||||||[||||||||||||]PAM NA Intergenic NA NA

T13053 AGCTTTCCAGAAACAATCGACGG 1000
Oligo fwd CACCAGCTTTCCAGAAACAATCGA
Oligo rev AAACTCGATTGTTTCTGGAAAGCT
Chromosome start end strand MM target_seq PAM alignment distance position gene name gene id
GU305914.1 171875 171897 + 0 AGCTTTCCAGAAACAATCGA CGG ||||||||[||||||||||||]PAM NA Intergenic NA NA

T13054 GCGAACATGCCGCCGTTCTTAGG 1000
Oligo fwd CACCGCGAACATGCCGCCGTTCTT
Oligo rev AAACAAGAACGGCGGCATGTTCGC
Chromosome start end strand MM target_seq PAM alignment distance position gene name gene id
GU305914.1 171897 171919 + 0 GCGAACATGCCGCCGTTCTT AGG ||||||||[||||||||||||]PAM NA Intergenic NA NA

T13055 CGAACATGCCGCCGTTCTTAGGG 1000
Oligo fwd CACCCGAACATGCCGCCGTTCTTA
Oligo rev AAACTAAGAACGGCGGCATGTTCG
Chromosome start end strand MM target_seq PAM alignment distance position gene name gene id
GU305914.1 171898 171920 + 0 CGAACATGCCGCCGTTCTTA GGG ||||||||[||||||||||||]PAM NA Intergenic NA NA

T13056 CGTCACGCTACAATTCACGTTGG 1000
Oligo fwd CACCCGTCACGCTACAATTCACGT
Oligo rev AAACACGTGAATTGTAGCGTGACG
Chromosome start end strand MM target_seq PAM alignment distance position gene name gene id
GU305914.1 171926 171948 + 0 CGTCACGCTACAATTCACGT TGG ||||||||[||||||||||||]PAM NA Intergenic NA NA

T13057 CTCCGCATCGTGACACGACACGG 1000
Oligo fwd CACCCTCCGCATCGTGACACGACA
Oligo rev AAACTGTCGTGTCACGATGCGGAG
Chromosome start end strand MM target_seq PAM alignment distance position gene name gene id
GU305914.1 171968 171990 + 0 CTCCGCATCGTGACACGACA CGG ||||||||[||||||||||||]PAM NA Intergenic NA NA

T13058 CCCGTGCGCTGTGTGAAATGCGG 1000
Oligo fwd CACCCCCGTGCGCTGTGTGAAATG
Oligo rev AAACCATTTCACACAGCGCACGGG
Chromosome start end strand MM target_seq PAM alignment distance position gene name gene id
GU305914.1 171998 172020 + 0 CCCGTGCGCTGTGTGAAATG CGG ||||||||[||||||||||||]PAM NA Intergenic NA NA

T13059 TAGCCTTCTGAAACACGAGACGG 1000
Oligo fwd CACCTAGCCTTCTGAAACACGAGA
Oligo rev AAACTCTCGTGTTTCAGAAGGCTA
Chromosome start end strand MM target_seq PAM alignment distance position gene name gene id
GU305914.1 172058 172080 + 0 TAGCCTTCTGAAACACGAGA CGG ||||||||[||||||||||||]PAM NA Intergenic NA NA

T13060 AGCCTTCTGAAACACGAGACGGG 1000
Oligo fwd CACCAGCCTTCTGAAACACGAGAC
Oligo rev AAACGTCTCGTGTTTCAGAAGGCT
Chromosome start end strand MM target_seq PAM alignment distance position gene name gene id
GU305914.1 172059 172081 + 0 AGCCTTCTGAAACACGAGAC GGG ||||||||[||||||||||||]PAM NA Intergenic NA NA

T13061 AAACACGAGACGGGCGCACGCGG 1000
Oligo fwd CACCAAACACGAGACGGGCGCACG
Oligo rev AAACCGTGCGCCCGTCTCGTGTTT
Chromosome start end strand MM target_seq PAM alignment distance position gene name gene id
GU305914.1 172068 172090 + 0 AAACACGAGACGGGCGCACG CGG ||||||||[||||||||||||]PAM NA Intergenic NA NA

T13062 TTTAAATAAGTTTATTGCTACGG 1000
Oligo fwd CACCTTTAAATAAGTTTATTGCTA
Oligo rev AAACTAGCAATAAACTTATTTAAA
Chromosome start end strand MM target_seq PAM alignment distance position gene name gene id
GU305914.1 172098 172120 + 0 TTTAAATAAGTTTATTGCTA CGG ||||||||[||||||||||||]PAM NA Intergenic NA NA

T13063 AAGTTTATTGCTACGGCTGTAGG 1000
Oligo fwd CACCAAGTTTATTGCTACGGCTGT
Oligo rev AAACACAGCCGTAGCAATAAACTT
Chromosome start end strand MM target_seq PAM alignment distance position gene name gene id
GU305914.1 172105 172127 + 0 AAGTTTATTGCTACGGCTGT AGG ||||||||[||||||||||||]PAM NA Intergenic NA NA

T13064 TTTATTGCTACGGCTGTAGGCGG 1000
Oligo fwd CACCTTTATTGCTACGGCTGTAGG
Oligo rev AAACCCTACAGCCGTAGCAATAAA
Chromosome start end strand MM target_seq PAM alignment distance position gene name gene id
GU305914.1 172108 172130 + 0 TTTATTGCTACGGCTGTAGG CGG ||||||||[||||||||||||]PAM NA Intergenic NA NA

T13065 GGCTGTAGGCGGTAACATCTCGG 1000
Oligo fwd CACCGGCTGTAGGCGGTAACATCT

Oligo rev AAACAGATGTTACCGCCTACAGCC
Chromosome start end strand MM target_seq PAM alignment distance position gene name gene id
GU305914.1 172119 172141 + 0 GGCTGTAGGCGGTAACATCT CGG ||||||||[||||||||||||]PAM NA Intergenic NA NA

T13066 TATCATCAGAGTCGCACGACAGG 1000
Oligo fwd CACCTATCATCAGAGTCGCACGAC
Oligo rev AAACGTCGTGCGACTCTGATGATA
Chromosome start end strand MM target_seq PAM alignment distance position gene name gene id
GU305914.1 172151 172173 + 0 TATCATCAGAGTCGCACGAC AGG ||||||||[||||||||||||]PAM NA Intergenic NA NA

T13067 ATCATCAGAGTCGCACGACAGGG 1000
Oligo fwd CACCATCATCAGAGTCGCACGACA
Oligo rev AAACTGTCGTGCGACTCTGATGAT
Chromosome start end strand MM target_seq PAM alignment distance position gene name gene id
GU305914.1 172152 172174 + 0 ATCATCAGAGTCGCACGACA GGG ||||||||[||||||||||||]PAM NA Intergenic NA NA

T13068 ATCAGAGTCGCACGACAGGGCGG 1000
Oligo fwd CACCATCAGAGTCGCACGACAGGG
Oligo rev AAACCCCTGTCGTGCGACTCTGAT
Chromosome start end strand MM target_seq PAM alignment distance position gene name gene id
GU305914.1 172155 172177 + 0 ATCAGAGTCGCACGACAGGG CGG ||||||||[||||||||||||]PAM NA Intergenic NA NA

T13069 CGACAGGGCGGCAGCCTTCTCGG 1000
Oligo fwd CACCCGACAGGGCGGCAGCCTTCT
Oligo rev AAACAGAAGGCTGCCGCCCTGTCG
Chromosome start end strand MM target_seq PAM alignment distance position gene name gene id
GU305914.1 172167 172189 + 0 CGACAGGGCGGCAGCCTTCT CGG ||||||||[||||||||||||]PAM NA Intergenic NA NA

T13070 CAGGGCGGCAGCCTTCTCGGAGG 1000
Oligo fwd CACCCAGGGCGGCAGCCTTCTCGG
Oligo rev AAACCCGAGAAGGCTGCCGCCCTG
Chromosome start end strand MM target_seq PAM alignment distance position gene name gene id
GU305914.1 172170 172192 + 0 CAGGGCGGCAGCCTTCTCGG AGG ||||||||[||||||||||||]PAM NA Intergenic NA NA

T13071 CACCGAGATCGATACGATCGCGG 1000
Oligo fwd CACCCACCGAGATCGATACGATCG
Oligo rev AAACCGATCGTATCGATCTCGGTG
Chromosome start end strand MM target_seq PAM alignment distance position gene name gene id
GU305914.1 172196 172218 + 0 CACCGAGATCGATACGATCG CGG ||||||||[||||||||||||]PAM NA Intergenic NA NA

T13072 GAGATCGATACGATCGCGGACGG 1000
Oligo fwd CACCGAGATCGATACGATCGCGGA
Oligo rev AAACTCCGCGATCGTATCGATCTC
Chromosome start end strand MM target_seq PAM alignment distance position gene name gene id
GU305914.1 172200 172222 + 0 GAGATCGATACGATCGCGGA CGG ||||||||[||||||||||||]PAM NA Intergenic NA NA

T13073 TCCACATGCAGATCGTCGCGAGG 1000
Oligo fwd CACCTCCACATGCAGATCGTCGCG
Oligo rev AAACCGCGACGATCTGCATGTGGA
Chromosome start end strand MM target_seq PAM alignment distance position gene name gene id
GU305914.1 172251 172273 + 0 TCCACATGCAGATCGTCGCG AGG ||||||||[||||||||||||]PAM NA Intergenic NA NA

T13074 TCGCGAGGATCACGCACGAGAGG 1000
Oligo fwd CACCTCGCGAGGATCACGCACGAG
Oligo rev AAACCTCGTGCGTGATCCTCGCGA
Chromosome start end strand MM target_seq PAM alignment distance position gene name gene id
GU305914.1 172266 172288 + 0 TCGCGAGGATCACGCACGAG AGG ||||||||[||||||||||||]PAM NA Intergenic NA NA

T13075 CGAGAGGAAGATCATTCCGATGG 1000
Oligo fwd CACCCGAGAGGAAGATCATTCCGA
Oligo rev AAACTCGGAATGATCTTCCTCTCG
Chromosome start end strand MM target_seq PAM alignment distance position gene name gene id
GU305914.1 172282 172304 + 0 CGAGAGGAAGATCATTCCGA TGG ||||||||[||||||||||||]PAM NA Intergenic NA NA

T13076 ATCGTTCAGAGTCCTCGTCGCGG 1000
Oligo fwd CACCATCGTTCAGAGTCCTCGTCG
Oligo rev AAACCGACGAGGACTCTGAACGAT
Chromosome start end strand MM target_seq PAM alignment distance position gene name gene id
GU305914.1 172320 172342 + 0 ATCGTTCAGAGTCCTCGTCG CGG ||||||||[||||||||||||]PAM NA Intergenic NA NA

T13077 GTCCTCGTCGCGGTCCTCCGTGG 1000
Oligo fwd CACCGTCCTCGTCGCGGTCCTCCG
Oligo rev AAACCGGAGGACCGCGACGAGGAC
Chromosome start end strand MM target_seq PAM alignment distance position gene name gene id
GU305914.1 172330 172352 + 0 GTCCTCGTCGCGGTCCTCCG TGG ||||||||[||||||||||||]PAM NA Intergenic NA NA

T13078 GCGGTCCTCCGTGGCCGCCGCGG 1000
Oligo fwd CACCGCGGTCCTCCGTGGCCGCCG
Oligo rev AAACCGGCGGCCACGGAGGACCGC
Chromosome start end strand MM target_seq PAM alignment distance position gene name gene id
GU305914.1 172339 172361 + 0 GCGGTCCTCCGTGGCCGCCG CGG ||||||||[||||||||||||]PAM NA Intergenic NA NA

T13079 CGTGGCCGCCGCGGTTCGAGCGG 1000
Oligo fwd CACCCGTGGCCGCCGCGGTTCGAG
Oligo rev AAACCTCGAACCGCGGCGGCCACG
Chromosome start end strand MM target_seq PAM alignment distance position gene name gene id
GU305914.1 172348 172370 + 0 CGTGGCCGCCGCGGTTCGAG CGG ||||||||[||||||||||||]PAM NA Intergenic NA NA

T13080 GTGGCCGCCGCGGTTCGAGCGGG 1000

<table>
<tr><td>Oligo fwd</td><td>CACCGTGGCCGCCGCGGTTCGAGC</td><td></td><td></td><td></td><td></td><td></td><td></td><td></td><td></td><td></td><td></td></tr>
<tr><td>Oligo rev</td><td>AAACGCTCGAACCGCGGCGGCCAC</td><td></td><td></td><td></td><td></td><td></td><td></td><td></td><td></td><td></td><td></td></tr>
<tr><td>Chromosome</td><td>start</td><td>end</td><td>strand</td><td>MM</td><td>target_seq</td><td>PAM</td><td>alignment</td><td>distance</td><td>position</td><td>gene name</td><td>gene id</td></tr>
<tr><td>GU305914.1</td><td>172349</td><td>172371</td><td>+</td><td>0</td><td>GTGGCCGCCGCGGTTCGAGC</td><td>GGG</td><td>||||||||[||||||||||||]PAM</td><td>NA</td><td>Intergenic</td><td>NA</td><td>NA</td></tr>
<tr><td>T13081</td><td>TGGCCGCCGCGGTTCGAGCGGGG</td><td>1000</td><td></td><td></td><td></td><td></td><td></td><td></td><td></td><td></td><td></td></tr>
<tr><td>Oligo fwd</td><td>CACCTGGCCGCCGCGGTTCGAGCG</td><td></td><td></td><td></td><td></td><td></td><td></td><td></td><td></td><td></td><td></td></tr>
<tr><td>Oligo rev</td><td>AAACCGCTCGAACCGCGGCGGCCA</td><td></td><td></td><td></td><td></td><td></td><td></td><td></td><td></td><td></td><td></td></tr>
<tr><td>Chromosome</td><td>start</td><td>end</td><td>strand</td><td>MM</td><td>target_seq</td><td>PAM</td><td>alignment</td><td>distance</td><td>position</td><td>gene name</td><td>gene id</td></tr>
<tr><td>GU305914.1</td><td>172350</td><td>172372</td><td>+</td><td>0</td><td>TGGCCGCCGCGGTTCGAGCG</td><td>GGG</td><td>||||||||[||||||||||||]PAM</td><td>NA</td><td>Intergenic</td><td>NA</td><td>NA</td></tr>
<tr><td>T13082</td><td>GGCCGCCGCGGTTCGAGCGGGGG</td><td>1000</td><td></td><td></td><td></td><td></td><td></td><td></td><td></td><td></td><td></td></tr>
<tr><td>Oligo fwd</td><td>CACCGGCCGCCGCGGTTCGAGCGG</td><td></td><td></td><td></td><td></td><td></td><td></td><td></td><td></td><td></td><td></td></tr>
<tr><td>Oligo rev</td><td>AAACCCGCTCGAACCGCGGCGGCC</td><td></td><td></td><td></td><td></td><td></td><td></td><td></td><td></td><td></td><td></td></tr>
<tr><td>Chromosome</td><td>start</td><td>end</td><td>strand</td><td>MM</td><td>target_seq</td><td>PAM</td><td>alignment</td><td>distance</td><td>position</td><td>gene name</td><td>gene id</td></tr>
<tr><td>GU305914.1</td><td>172351</td><td>172373</td><td>+</td><td>0</td><td>GGCCGCCGCGGTTCGAGCGG</td><td>GGG</td><td>||||||||[||||||||||||]PAM</td><td>NA</td><td>Intergenic</td><td>NA</td><td>NA</td></tr>
<tr><td>T13083</td><td>TTCGAGCGGGGGACGTAGCGCGG</td><td>1000</td><td></td><td></td><td></td><td></td><td></td><td></td><td></td><td></td><td></td></tr>
<tr><td>Oligo fwd</td><td>CACCTTCGAGCGGGGGACGTAGCG</td><td></td><td></td><td></td><td></td><td></td><td></td><td></td><td></td><td></td><td></td></tr>
<tr><td>Oligo rev</td><td>AAACCGCTACGTCCCCCGCTCGAA</td><td></td><td></td><td></td><td></td><td></td><td></td><td></td><td></td><td></td><td></td></tr>
<tr><td>Chromosome</td><td>start</td><td>end</td><td>strand</td><td>MM</td><td>target_seq</td><td>PAM</td><td>alignment</td><td>distance</td><td>position</td><td>gene name</td><td>gene id</td></tr>
<tr><td>GU305914.1</td><td>172362</td><td>172384</td><td>+</td><td>0</td><td>TTCGAGCGGGGGACGTAGCG</td><td>CGG</td><td>||||||||[||||||||||||]PAM</td><td>NA</td><td>Intergenic</td><td>NA</td><td>NA</td></tr>
<tr><td>T13084</td><td>CCGTCCGCTTTTCAGAAGTCTGG</td><td>1000</td><td></td><td></td><td></td><td></td><td></td><td></td><td></td><td></td><td></td></tr>
<tr><td>Oligo fwd</td><td>CACCCCGTCCGCTTTTCAGAAGTC</td><td></td><td></td><td></td><td></td><td></td><td></td><td></td><td></td><td></td><td></td></tr>
<tr><td>Oligo rev</td><td>AAACGACTTCTGAAAAGCGGACGG</td><td></td><td></td><td></td><td></td><td></td><td></td><td></td><td></td><td></td><td></td></tr>
<tr><td>Chromosome</td><td>start</td><td>end</td><td>strand</td><td>MM</td><td>target_seq</td><td>PAM</td><td>alignment</td><td>distance</td><td>position</td><td>gene name</td><td>gene id</td></tr>
<tr><td>GU305914.1</td><td>172432</td><td>172454</td><td>+</td><td>0</td><td>CCGTCCGCTTTTCAGAAGTC</td><td>TGG</td><td>||||||||[||||||||||||]PAM</td><td>NA</td><td>Intergenic</td><td>NA</td><td>NA</td></tr>
<tr><td>T13085</td><td>CGTCCGCTTTTCAGAAGTCTGGG</td><td>1000</td><td></td><td></td><td></td><td></td><td></td><td></td><td></td><td></td><td></td></tr>
<tr><td>Oligo fwd</td><td>CACCCGTCCGCTTTTCAGAAGTCT</td><td></td><td></td><td></td><td></td><td></td><td></td><td></td><td></td><td></td><td></td></tr>
<tr><td>Oligo rev</td><td>AAACAGACTTCTGAAAAGCGGACG</td><td></td><td></td><td></td><td></td><td></td><td></td><td></td><td></td><td></td><td></td></tr>
<tr><td>Chromosome</td><td>start</td><td>end</td><td>strand</td><td>MM</td><td>target_seq</td><td>PAM</td><td>alignment</td><td>distance</td><td>position</td><td>gene name</td><td>gene id</td></tr>
<tr><td>GU305914.1</td><td>172433</td><td>172455</td><td>+</td><td>0</td><td>CGTCCGCTTTTCAGAAGTCT</td><td>GGG</td><td>||||||||[||||||||||||]PAM</td><td>NA</td><td>Intergenic</td><td>NA</td><td>NA</td></tr>
<tr><td>T13086</td><td>AAGTCTGGGCGAACCGTCGTCGG</td><td>1000</td><td></td><td></td><td></td><td></td><td></td><td></td><td></td><td></td><td></td></tr>
<tr><td>Oligo fwd</td><td>CACCAAGTCTGGGCGAACCGTCGT</td><td></td><td></td><td></td><td></td><td></td><td></td><td></td><td></td><td></td><td></td></tr>
<tr><td>Oligo rev</td><td>AAACACGACGGTTCGCCCAGACTT</td><td></td><td></td><td></td><td></td><td></td><td></td><td></td><td></td><td></td><td></td></tr>
<tr><td>Chromosome</td><td>start</td><td>end</td><td>strand</td><td>MM</td><td>target_seq</td><td>PAM</td><td>alignment</td><td>distance</td><td>position</td><td>gene name</td><td>gene id</td></tr>
<tr><td>GU305914.1</td><td>172447</td><td>172469</td><td>+</td><td>0</td><td>AAGTCTGGGCGAACCGTCGT</td><td>CGG</td><td>||||||||[||||||||||||]PAM</td><td>NA</td><td>Intergenic</td><td>NA</td><td>NA</td></tr>
<tr><td>T13087</td><td>GAACCGTCGTCGGTCGTCTCCGG</td><td>1000</td><td></td><td></td><td></td><td></td><td></td><td></td><td></td><td></td><td></td></tr>
<tr><td>Oligo fwd</td><td>CACCGAACCGTCGTCGGTCGTCTC</td><td></td><td></td><td></td><td></td><td></td><td></td><td></td><td></td><td></td><td></td></tr>
<tr><td>Oligo rev</td><td>AAACGAGACGACCGACGACGGTTC</td><td></td><td></td><td></td><td></td><td></td><td></td><td></td><td></td><td></td><td></td></tr>
<tr><td>Chromosome</td><td>start</td><td>end</td><td>strand</td><td>MM</td><td>target_seq</td><td>PAM</td><td>alignment</td><td>distance</td><td>position</td><td>gene name</td><td>gene id</td></tr>
<tr><td>GU305914.1</td><td>172457</td><td>172479</td><td>+</td><td>0</td><td>GAACCGTCGTCGGTCGTCTC</td><td>CGG</td><td>||||||||[||||||||||||]PAM</td><td>NA</td><td>Intergenic</td><td>NA</td><td>NA</td></tr>
<tr><td>T13088</td><td>CCGTCGTCGGTCGTCTCCGGAGG</td><td>1000</td><td></td><td></td><td></td><td></td><td></td><td></td><td></td><td></td><td></td></tr>
<tr><td>Oligo fwd</td><td>CACCCCGTCGTCGGTCGTCTCCGG</td><td></td><td></td><td></td><td></td><td></td><td></td><td></td><td></td><td></td><td></td></tr>
<tr><td>Oligo rev</td><td>AAACCCGGAGACGACCGACGACGG</td><td></td><td></td><td></td><td></td><td></td><td></td><td></td><td></td><td></td><td></td></tr>
<tr><td>Chromosome</td><td>start</td><td>end</td><td>strand</td><td>MM</td><td>target_seq</td><td>PAM</td><td>alignment</td><td>distance</td><td>position</td><td>gene name</td><td>gene id</td></tr>
<tr><td>GU305914.1</td><td>172460</td><td>172482</td><td>+</td><td>0</td><td>CCGTCGTCGGTCGTCTCCGG</td><td>AGG</td><td>||||||||[||||||||||||]PAM</td><td>NA</td><td>Intergenic</td><td>NA</td><td>NA</td></tr>
<tr><td>T13089</td><td>CGTCGTCGGTCGTCTCCGGAGGG</td><td>1000</td><td></td><td></td><td></td><td></td><td></td><td></td><td></td><td></td><td></td></tr>
<tr><td>Oligo fwd</td><td>CACCCGTCGTCGGTCGTCTCCGGA</td><td></td><td></td><td></td><td></td><td></td><td></td><td></td><td></td><td></td><td></td></tr>
<tr><td>Oligo rev</td><td>AAACTCCGGAGACGACCGACGACG</td><td></td><td></td><td></td><td></td><td></td><td></td><td></td><td></td><td></td><td></td></tr>
<tr><td>Chromosome</td><td>start</td><td>end</td><td>strand</td><td>MM</td><td>target_seq</td><td>PAM</td><td>alignment</td><td>distance</td><td>position</td><td>gene name</td><td>gene id</td></tr>
<tr><td>GU305914.1</td><td>172461</td><td>172483</td><td>+</td><td>0</td><td>CGTCGTCGGTCGTCTCCGGA</td><td>GGG</td><td>||||||||[||||||||||||]PAM</td><td>NA</td><td>Intergenic</td><td>NA</td><td>NA</td></tr>
<tr><td>T13090</td><td>GCTGATCTCGTCGTCCCCGTCGG</td><td>1000</td><td></td><td></td><td></td><td></td><td></td><td></td><td></td><td></td><td></td></tr>
<tr><td>Oligo fwd</td><td>CACCGCTGATCTCGTCGTCCCCGT</td><td></td><td></td><td></td><td></td><td></td><td></td><td></td><td></td><td></td><td></td></tr>
<tr><td>Oligo rev</td><td>AAACACGGGGACGACGAGATCAGC</td><td></td><td></td><td></td><td></td><td></td><td></td><td></td><td></td><td></td><td></td></tr>
<tr><td>Chromosome</td><td>start</td><td>end</td><td>strand</td><td>MM</td><td>target_seq</td><td>PAM</td><td>alignment</td><td>distance</td><td>position</td><td>gene name</td><td>gene id</td></tr>
<tr><td>GU305914.1</td><td>172486</td><td>172508</td><td>+</td><td>0</td><td>GCTGATCTCGTCGTCCCCGT</td><td>CGG</td><td>||||||||[||||||||||||]PAM</td><td>NA</td><td>Intergenic</td><td>NA</td><td>NA</td></tr>
<tr><td>T13091</td><td>CTGATCTCGTCGTCCCCGTCGGG</td><td>1000</td><td></td><td></td><td></td><td></td><td></td><td></td><td></td><td></td><td></td></tr>
<tr><td>Oligo fwd</td><td>CACCCTGATCTCGTCGTCCCCGTC</td><td></td><td></td><td></td><td></td><td></td><td></td><td></td><td></td><td></td><td></td></tr>
<tr><td>Oligo rev</td><td>AAACGACGGGGACGACGAGATCAG</td><td></td><td></td><td></td><td></td><td></td><td></td><td></td><td></td><td></td><td></td></tr>
<tr><td>Chromosome</td><td>start</td><td>end</td><td>strand</td><td>MM</td><td>target_seq</td><td>PAM</td><td>alignment</td><td>distance</td><td>position</td><td>gene name</td><td>gene id</td></tr>
<tr><td>GU305914.1</td><td>172487</td><td>172509</td><td>+</td><td>0</td><td>CTGATCTCGTCGTCCCCGTC</td><td>GGG</td><td>||||||||[||||||||||||]PAM</td><td>NA</td><td>Intergenic</td><td>NA</td><td>NA</td></tr>
<tr><td>T13092</td><td>TCGGGTCCTCCCGTCACGAACGG</td><td>1000</td><td></td><td></td><td></td><td></td><td></td><td></td><td></td><td></td><td></td></tr>
<tr><td>Oligo fwd</td><td>CACCTCGGGTCCTCCCGTCACGAA</td><td></td><td></td><td></td><td></td><td></td><td></td><td></td><td></td><td></td><td></td></tr>
<tr><td>Oligo rev</td><td>AAACTTCGTGACGGGAGGACCCGA</td><td></td><td></td><td></td><td></td><td></td><td></td><td></td><td></td><td></td><td></td></tr>
<tr><td>Chromosome</td><td>start</td><td>end</td><td>strand</td><td>MM</td><td>target_seq</td><td>PAM</td><td>alignment</td><td>distance</td><td>position</td><td>gene name</td><td>gene id</td></tr>
<tr><td>GU305914.1</td><td>172505</td><td>172527</td><td>+</td><td>0</td><td>TCGGGTCCTCCCGTCACGAA</td><td>CGG</td><td>||||||||[||||||||||||]PAM</td><td>NA</td><td>Intergenic</td><td>NA</td><td>NA</td></tr>
<tr><td>T13093</td><td>GTCACGAACGGCCCCTCGTAAGG</td><td>1000</td><td></td><td></td><td></td><td></td><td></td><td></td><td></td><td></td><td></td></tr>
<tr><td>Oligo fwd</td><td>CACCGTCACGAACGGCCCCTCGTA</td><td></td><td></td><td></td><td></td><td></td><td></td><td></td><td></td><td></td><td></td></tr>
<tr><td>Oligo rev</td><td>AAACTACGAGGGGCCGTTCGTGAC</td><td></td><td></td><td></td><td></td><td></td><td></td><td></td><td></td><td></td><td></td></tr>
<tr><td>Chromosome</td><td>start</td><td>end</td><td>strand</td><td>MM</td><td>target_seq</td><td>PAM</td><td>alignment</td><td>distance</td><td>position</td><td>gene name</td><td>gene id</td></tr>
<tr><td>GU305914.1</td><td>172517</td><td>172539</td><td>+</td><td>0</td><td>GTCACGAACGGCCCCTCGTA</td><td>AGG</td><td>||||||||[||||||||||||]PAM</td><td>NA</td><td>Intergenic</td><td>NA</td><td>NA</td></tr>
<tr><td>T13094</td><td>ACGAACGGCCCCTCGTAAGGCGG</td><td>1000</td><td></td><td></td><td></td><td></td><td></td><td></td><td></td><td></td><td></td></tr>
<tr><td>Oligo fwd</td><td>CACCACGAACGGCCCCTCGTAAGG</td><td></td><td></td><td></td><td></td><td></td><td></td><td></td><td></td><td></td><td></td></tr>
<tr><td>Oligo rev</td><td>AAACCCTTACGAGGGGCCGTTCGT</td><td></td><td></td><td></td><td></td><td></td><td></td><td></td><td></td><td></td><td></td></tr>
<tr><td>Chromosome</td><td>start</td><td>end</td><td>strand</td><td>MM</td><td>target_seq</td><td>PAM</td><td>alignment</td><td>distance</td><td>position</td><td>gene name</td><td>gene id</td></tr>
<tr><td>GU305914.1</td><td>172520</td><td>172542</td><td>+</td><td>0</td><td>ACGAACGGCCCCTCGTAAGG</td><td>CGG</td><td>||||||||[||||||||||||]PAM</td><td>NA</td><td>Intergenic</td><td>NA</td><td>NA</td></tr>
</table>

<table>
<tr><td>T13095</td><td>CCCCTCGTAAGGCGGCCACCTGG</td><td>1000</td><td></td><td></td><td></td><td></td><td></td><td></td><td></td><td></td><td></td></tr>
<tr><td>Oligo fwd</td><td>CACCCCCCTCGTAAGGCGGCCACC</td><td></td><td></td><td></td><td></td><td></td><td></td><td></td><td></td><td></td><td></td></tr>
<tr><td>Oligo rev</td><td>AAACGGTGGCCGCCTTACGAGGGG</td><td></td><td></td><td></td><td></td><td></td><td></td><td></td><td></td><td></td><td></td></tr>
<tr><td>Chromosome</td><td>start</td><td>end</td><td>strand</td><td>MM</td><td>target_seq</td><td>PAM</td><td>alignment</td><td>distance</td><td>position</td><td>gene name</td><td>gene id</td></tr>
<tr><td>GU305914.1</td><td>172528</td><td>172550</td><td>+</td><td>0</td><td>CCCCTCGTAAGGCGGCCACC</td><td>TGG</td><td>||||||||[||||||||||||]PAM</td><td>NA</td><td>Intergenic</td><td>NA</td><td>NA</td></tr>
</table>

<table>
<tr><td>T13096</td><td>GCCGTTACAGACAAAGTACACGG</td><td>1000</td><td></td><td></td><td></td><td></td><td></td><td></td><td></td><td></td><td></td></tr>
<tr><td>Oligo fwd</td><td>CACCGCCGTTACAGACAAAGTACA</td><td></td><td></td><td></td><td></td><td></td><td></td><td></td><td></td><td></td><td></td></tr>
<tr><td>Oligo rev</td><td>AAACTGTACTTTGTCTGTAACGGC</td><td></td><td></td><td></td><td></td><td></td><td></td><td></td><td></td><td></td><td></td></tr>
<tr><td>Chromosome</td><td>start</td><td>end</td><td>strand</td><td>MM</td><td>target_seq</td><td>PAM</td><td>alignment</td><td>distance</td><td>position</td><td>gene name</td><td>gene id</td></tr>
<tr><td>GU305914.1</td><td>172558</td><td>172580</td><td>+</td><td>0</td><td>GCCGTTACAGACAAAGTACA</td><td>CGG</td><td>||||||||[||||||||||||]PAM</td><td>NA</td><td>Intergenic</td><td>NA</td><td>NA</td></tr>
</table>

<table>
<tr><td>T13097</td><td>AGTACACGGTCGCGTTGACTTGG</td><td>1000</td><td></td><td></td><td></td><td></td><td></td><td></td><td></td><td></td><td></td></tr>
<tr><td>Oligo fwd</td><td>CACCAGTACACGGTCGCGTTGACT</td><td></td><td></td><td></td><td></td><td></td><td></td><td></td><td></td><td></td><td></td></tr>
<tr><td>Oligo rev</td><td>AAACAGTCAACGCGACCGTGTACT</td><td></td><td></td><td></td><td></td><td></td><td></td><td></td><td></td><td></td><td></td></tr>
<tr><td>Chromosome</td><td>start</td><td>end</td><td>strand</td><td>MM</td><td>target_seq</td><td>PAM</td><td>alignment</td><td>distance</td><td>position</td><td>gene name</td><td>gene id</td></tr>
<tr><td>GU305914.1</td><td>172572</td><td>172594</td><td>+</td><td>0</td><td>AGTACACGGTCGCGTTGACT</td><td>TGG</td><td>||||||||[||||||||||||]PAM</td><td>NA</td><td>Intergenic</td><td>NA</td><td>NA</td></tr>
</table>

<table>
<tr><td>T13098</td><td>CGCGTTGACTTGGATGACGACGG</td><td>1000</td><td></td><td></td><td></td><td></td><td></td><td></td><td></td><td></td><td></td></tr>
<tr><td>Oligo fwd</td><td>CACCCGCGTTGACTTGGATGACGA</td><td></td><td></td><td></td><td></td><td></td><td></td><td></td><td></td><td></td><td></td></tr>
<tr><td>Oligo rev</td><td>AAACTCGTCATCCAAGTCAACGCG</td><td></td><td></td><td></td><td></td><td></td><td></td><td></td><td></td><td></td><td></td></tr>
<tr><td>Chromosome</td><td>start</td><td>end</td><td>strand</td><td>MM</td><td>target_seq</td><td>PAM</td><td>alignment</td><td>distance</td><td>position</td><td>gene name</td><td>gene id</td></tr>
<tr><td>GU305914.1</td><td>172582</td><td>172604</td><td>+</td><td>0</td><td>CGCGTTGACTTGGATGACGA</td><td>CGG</td><td>||||||||[||||||||||||]PAM</td><td>NA</td><td>Intergenic</td><td>NA</td><td>NA</td></tr>
</table>

<table>
<tr><td>T13099</td><td>GACGACGGTGCAGATCAGCGCGG</td><td>1000</td><td></td><td></td><td></td><td></td><td></td><td></td><td></td><td></td><td></td></tr>
<tr><td>Oligo fwd</td><td>CACCGACGACGGTGCAGATCAGCG</td><td></td><td></td><td></td><td></td><td></td><td></td><td></td><td></td><td></td><td></td></tr>
<tr><td>Oligo rev</td><td>AAACCGCTGATCTGCACCGTCGTC</td><td></td><td></td><td></td><td></td><td></td><td></td><td></td><td></td><td></td><td></td></tr>
<tr><td>Chromosome</td><td>start</td><td>end</td><td>strand</td><td>MM</td><td>target_seq</td><td>PAM</td><td>alignment</td><td>distance</td><td>position</td><td>gene name</td><td>gene id</td></tr>
<tr><td>GU305914.1</td><td>172597</td><td>172619</td><td>+</td><td>0</td><td>GACGACGGTGCAGATCAGCG</td><td>CGG</td><td>||||||||[||||||||||||]PAM</td><td>NA</td><td>Intergenic</td><td>NA</td><td>NA</td></tr>
</table>

<table>
<tr><td>T13100</td><td>GCGCGGCCCTCACCTGACAGTGG</td><td>1000</td><td></td><td></td><td></td><td></td><td></td><td></td><td></td><td></td><td></td></tr>
<tr><td>Oligo fwd</td><td>CACCGCGCGGCCCTCACCTGACAG</td><td></td><td></td><td></td><td></td><td></td><td></td><td></td><td></td><td></td><td></td></tr>
<tr><td>Oligo rev</td><td>AAACCTGTCAGGTGAGGGCCGCGC</td><td></td><td></td><td></td><td></td><td></td><td></td><td></td><td></td><td></td><td></td></tr>
<tr><td>Chromosome</td><td>start</td><td>end</td><td>strand</td><td>MM</td><td>target_seq</td><td>PAM</td><td>alignment</td><td>distance</td><td>position</td><td>gene name</td><td>gene id</td></tr>
<tr><td>GU305914.1</td><td>172614</td><td>172636</td><td>+</td><td>0</td><td>GCGCGGCCCTCACCTGACAG</td><td>TGG</td><td>||||||||[||||||||||||]PAM</td><td>NA</td><td>Intergenic</td><td>NA</td><td>NA</td></tr>
</table>

<table>
<tr><td>T13101</td><td>CTCTCCCCAGAACGAACCCGTGG</td><td>1000</td><td></td><td></td><td></td><td></td><td></td><td></td><td></td><td></td><td></td></tr>
<tr><td>Oligo fwd</td><td>CACCCTCTCCCCAGAACGAACCCG</td><td></td><td></td><td></td><td></td><td></td><td></td><td></td><td></td><td></td><td></td></tr>
<tr><td>Oligo rev</td><td>AAACCGGGTTCGTTCTGGGGAGAG</td><td></td><td></td><td></td><td></td><td></td><td></td><td></td><td></td><td></td><td></td></tr>
<tr><td>Chromosome</td><td>start</td><td>end</td><td>strand</td><td>MM</td><td>target_seq</td><td>PAM</td><td>alignment</td><td>distance</td><td>position</td><td>gene name</td><td>gene id</td></tr>
<tr><td>GU305914.1</td><td>172642</td><td>172664</td><td>+</td><td>0</td><td>CTCTCCCCAGAACGAACCCG</td><td>TGG</td><td>||||||||[||||||||||||]PAM</td><td>NA</td><td>Intergenic</td><td>NA</td><td>NA</td></tr>
</table>

<table>
<tr><td>T13102</td><td>TCCCCAGAACGAACCCGTGGCGG</td><td>1000</td><td></td><td></td><td></td><td></td><td></td><td></td><td></td><td></td><td></td></tr>
<tr><td>Oligo fwd</td><td>CACCTCCCCAGAACGAACCCGTGG</td><td></td><td></td><td></td><td></td><td></td><td></td><td></td><td></td><td></td><td></td></tr>
<tr><td>Oligo rev</td><td>AAACCCACGGGTTCGTTCTGGGGA</td><td></td><td></td><td></td><td></td><td></td><td></td><td></td><td></td><td></td><td></td></tr>
<tr><td>Chromosome</td><td>start</td><td>end</td><td>strand</td><td>MM</td><td>target_seq</td><td>PAM</td><td>alignment</td><td>distance</td><td>position</td><td>gene name</td><td>gene id</td></tr>
<tr><td>GU305914.1</td><td>172645</td><td>172667</td><td>+</td><td>0</td><td>TCCCCAGAACGAACCCGTGG</td><td>CGG</td><td>||||||||[||||||||||||]PAM</td><td>NA</td><td>Intergenic</td><td>NA</td><td>NA</td></tr>
</table>

<table>
<tr><td>T13103</td><td>TAACTCGACTCCTTGTGCCTCGG</td><td>1000</td><td></td><td></td><td></td><td></td><td></td><td></td><td></td><td></td><td></td></tr>
<tr><td>Oligo fwd</td><td>CACCTAACTCGACTCCTTGTGCCT</td><td></td><td></td><td></td><td></td><td></td><td></td><td></td><td></td><td></td><td></td></tr>
<tr><td>Oligo rev</td><td>AAACAGGCACAAGGAGTCGAGTTA</td><td></td><td></td><td></td><td></td><td></td><td></td><td></td><td></td><td></td><td></td></tr>
<tr><td>Chromosome</td><td>start</td><td>end</td><td>strand</td><td>MM</td><td>target_seq</td><td>PAM</td><td>alignment</td><td>distance</td><td>position</td><td>gene name</td><td>gene id</td></tr>
<tr><td>GU305914.1</td><td>172676</td><td>172698</td><td>+</td><td>0</td><td>TAACTCGACTCCTTGTGCCT</td><td>CGG</td><td>||||||||[||||||||||||]PAM</td><td>NA</td><td>Intergenic</td><td>NA</td><td>NA</td></tr>
</table>

<table>
<tr><td>T13104</td><td>TCCTTGTGCCTCGGCAGCCTCGG</td><td>1000</td><td></td><td></td><td></td><td></td><td></td><td></td><td></td><td></td><td></td></tr>
<tr><td>Oligo fwd</td><td>CACCTCCTTGTGCCTCGGCAGCCT</td><td></td><td></td><td></td><td></td><td></td><td></td><td></td><td></td><td></td><td></td></tr>
<tr><td>Oligo rev</td><td>AAACAGGCTGCCGAGGCACAAGGA</td><td></td><td></td><td></td><td></td><td></td><td></td><td></td><td></td><td></td><td></td></tr>
<tr><td>Chromosome</td><td>start</td><td>end</td><td>strand</td><td>MM</td><td>target_seq</td><td>PAM</td><td>alignment</td><td>distance</td><td>position</td><td>gene name</td><td>gene id</td></tr>
<tr><td>GU305914.1</td><td>172685</td><td>172707</td><td>+</td><td>0</td><td>TCCTTGTGCCTCGGCAGCCT</td><td>CGG</td><td>||||||||[||||||||||||]PAM</td><td>NA</td><td>Intergenic</td><td>NA</td><td>NA</td></tr>
</table>

<table>
<tr><td>T13105</td><td>CCTTGTGCCTCGGCAGCCTCGGG</td><td>1000</td><td></td><td></td><td></td><td></td><td></td><td></td><td></td><td></td><td></td></tr>
<tr><td>Oligo fwd</td><td>CACCCCTTGTGCCTCGGCAGCCTC</td><td></td><td></td><td></td><td></td><td></td><td></td><td></td><td></td><td></td><td></td></tr>
<tr><td>Oligo rev</td><td>AAACGAGGCTGCCGAGGCACAAGG</td><td></td><td></td><td></td><td></td><td></td><td></td><td></td><td></td><td></td><td></td></tr>
<tr><td>Chromosome</td><td>start</td><td>end</td><td>strand</td><td>MM</td><td>target_seq</td><td>PAM</td><td>alignment</td><td>distance</td><td>position</td><td>gene name</td><td>gene id</td></tr>
<tr><td>GU305914.1</td><td>172686</td><td>172708</td><td>+</td><td>0</td><td>CCTTGTGCCTCGGCAGCCTC</td><td>GGG</td><td>||||||||[||||||||||||]PAM</td><td>NA</td><td>Intergenic</td><td>NA</td><td>NA</td></tr>
</table>

<table>
<tr><td>T13106</td><td>AAAGTGTATCTCAGAGAGCTCGG</td><td>1000</td><td></td><td></td><td></td><td></td><td></td><td></td><td></td><td></td><td></td></tr>
<tr><td>Oligo fwd</td><td>CACCAAAGTGTATCTCAGAGAGCT</td><td></td><td></td><td></td><td></td><td></td><td></td><td></td><td></td><td></td><td></td></tr>
<tr><td>Oligo rev</td><td>AAACAGCTCTCTGAGATACACTTT</td><td></td><td></td><td></td><td></td><td></td><td></td><td></td><td></td><td></td><td></td></tr>
<tr><td>Chromosome</td><td>start</td><td>end</td><td>strand</td><td>MM</td><td>target_seq</td><td>PAM</td><td>alignment</td><td>distance</td><td>position</td><td>gene name</td><td>gene id</td></tr>
<tr><td>GU305914.1</td><td>172747</td><td>172769</td><td>+</td><td>0</td><td>AAAGTGTATCTCAGAGAGCT</td><td>CGG</td><td>||||||||[||||||||||||]PAM</td><td>NA</td><td>Intergenic</td><td>NA</td><td>NA</td></tr>
</table>

<table>
<tr><td>T13107</td><td>AAGTGTATCTCAGAGAGCTCGGG</td><td>1000</td><td></td><td></td><td></td><td></td><td></td><td></td><td></td><td></td><td></td></tr>
<tr><td>Oligo fwd</td><td>CACCAAGTGTATCTCAGAGAGCTC</td><td></td><td></td><td></td><td></td><td></td><td></td><td></td><td></td><td></td><td></td></tr>
<tr><td>Oligo rev</td><td>AAACGAGCTCTCTGAGATACACTT</td><td></td><td></td><td></td><td></td><td></td><td></td><td></td><td></td><td></td><td></td></tr>
<tr><td>Chromosome</td><td>start</td><td>end</td><td>strand</td><td>MM</td><td>target_seq</td><td>PAM</td><td>alignment</td><td>distance</td><td>position</td><td>gene name</td><td>gene id</td></tr>
<tr><td>GU305914.1</td><td>172748</td><td>172770</td><td>+</td><td>0</td><td>AAGTGTATCTCAGAGAGCTC</td><td>GGG</td><td>||||||||[||||||||||||]PAM</td><td>NA</td><td>Intergenic</td><td>NA</td><td>NA</td></tr>
</table>

<table>
<tr><td>T13108</td><td>AGAGCTCGGGCACCTCGCACCGG</td><td>1000</td><td></td><td></td><td></td><td></td><td></td><td></td><td></td><td></td><td></td></tr>
<tr><td>Oligo fwd</td><td>CACCAGAGCTCGGGCACCTCGCAC</td><td></td><td></td><td></td><td></td><td></td><td></td><td></td><td></td><td></td><td></td></tr>
<tr><td>Oligo rev</td><td>AAACGTGCGAGGTGCCCGAGCTCT</td><td></td><td></td><td></td><td></td><td></td><td></td><td></td><td></td><td></td><td></td></tr>
<tr><td>Chromosome</td><td>start</td><td>end</td><td>strand</td><td>MM</td><td>target_seq</td><td>PAM</td><td>alignment</td><td>distance</td><td>position</td><td>gene name</td><td>gene id</td></tr>
<tr><td>GU305914.1</td><td>172761</td><td>172783</td><td>+</td><td>0</td><td>AGAGCTCGGGCACCTCGCAC</td><td>CGG</td><td>||||||||[||||||||||||]PAM</td><td>NA</td><td>Intergenic</td><td>NA</td><td>NA</td></tr>
</table>

<table>
<tr><td>T13109</td><td>CTCGGGCACCTCGCACCGGACGG</td><td>1000</td><td></td><td></td><td></td><td></td><td></td><td></td><td></td><td></td><td></td></tr>
<tr><td>Oligo fwd</td><td>CACCCTCGGGCACCTCGCACCGGA</td><td></td><td></td><td></td><td></td><td></td><td></td><td></td><td></td><td></td><td></td></tr>
<tr><td>Oligo rev</td><td>AAACTCCGGTGCGAGGTGCCCGAG</td><td></td><td></td><td></td><td></td><td></td><td></td><td></td><td></td><td></td><td></td></tr>
<tr><td>Chromosome</td><td>start</td><td>end</td><td>strand</td><td>MM</td><td>target_seq</td><td>PAM</td><td>alignment</td><td>distance</td><td>position</td><td>gene name</td><td>gene id</td></tr>
<tr><td>GU305914.1</td><td>172765</td><td>172787</td><td>+</td><td>0</td><td>CTCGGGCACCTCGCACCGGA</td><td>CGG</td><td>||||||||[||||||||||||]PAM</td><td>NA</td><td>Intergenic</td><td>NA</td><td>NA</td></tr>
</table>

| | | | | | | | | | | | |
|---|---|---|---|---|---|---|---|---|---|---|---|
| T13110 | TACGACAACGACACGTTACACGG | 1000 | | | | | | | | | |
| Oligo fwd | CACCTACGACAACGACACGTTACA | | | | | | | | | | |
| Oligo rev | AAACTGTAACGTGTCGTTGTCGTA | | | | | | | | | | |
| Chromosome | start | end | strand | MM | target_seq | PAM | alignment | distance | position | gene name | gene id |
| GU305914.1 | 172823 | 172845 | + | 0 | TACGACAACGACACGTTACA | CGG | ||||||||[||||||||||||]PAM | NA | Intergenic | NA | NA |
| T13111 | ACGACAACGACACGTTACACGGG | 1000 | | | | | | | | | |
| Oligo fwd | CACCACGACAACGACACGTTACAC | | | | | | | | | | |
| Oligo rev | AAACGTGTAACGTGTCGTTGTCGT | | | | | | | | | | |
| Chromosome | start | end | strand | MM | target_seq | PAM | alignment | distance | position | gene name | gene id |
| GU305914.1 | 172824 | 172846 | + | 0 | ACGACAACGACACGTTACAC | GGG | ||||||||[||||||||||||]PAM | NA | Intergenic | NA | NA |
| T13112 | AACGACACGTTACACGGGATCGG | 1000 | | | | | | | | | |
| Oligo fwd | CACCAACGACACGTTACACGGGAT | | | | | | | | | | |
| Oligo rev | AAACATCCCGTGTAACGTGTCGTT | | | | | | | | | | |
| Chromosome | start | end | strand | MM | target_seq | PAM | alignment | distance | position | gene name | gene id |
| GU305914.1 | 172829 | 172851 | + | 0 | AACGACACGTTACACGGGAT | CGG | ||||||||[||||||||||||]PAM | NA | Intergenic | NA | NA |
| T13113 | AAGAGACACAGACCACGAGATGG | 1000 | | | | | | | | | |
| Oligo fwd | CACCAAGAGACACAGACCACGAGA | | | | | | | | | | |
| Oligo rev | AAACTCTCGTGGTCTGTGTCTCTT | | | | | | | | | | |
| Chromosome | start | end | strand | MM | target_seq | PAM | alignment | distance | position | gene name | gene id |
| GU305914.1 | 172869 | 172891 | + | 0 | AAGAGACACAGACCACGAGA | TGG | ||||||||[||||||||||||]PAM | NA | Intergenic | NA | NA |
| T13114 | ATTTTTAATGACGATGTCCGTGG | 1000 | | | | | | | | | |
| Oligo fwd | CACCATTTTTAATGACGATGTCCG | | | | | | | | | | |
| Oligo rev | AAACCGGACATCGTCATTAAAAAT | | | | | | | | | | |
| Chromosome | start | end | strand | MM | target_seq | PAM | alignment | distance | position | gene name | gene id |
| GU305914.1 | 172910 | 172932 | + | 0 | ATTTTTAATGACGATGTCCG | TGG | ||||||||[||||||||||||]PAM | NA | Intergenic | NA | NA |
| T13115 | GTTATTTATGCGATGATGTGAGG | 1000 | | | | | | | | | |
| Oligo fwd | CACCGTTATTTATGCGATGATGTG | | | | | | | | | | |
| Oligo rev | AAACCACATCATCGCATAAATAAC | | | | | | | | | | |
| Chromosome | start | end | strand | MM | target_seq | PAM | alignment | distance | position | gene name | gene id |
| GU305914.1 | 172978 | 173000 | + | 0 | GTTATTTATGCGATGATGTG | AGG | ||||||||[||||||||||||]PAM | NA | Intergenic | NA | NA |
| T13116 | TGAGGCGTATGCGAATGAGCTGG | 1000 | | | | | | | | | |
| Oligo fwd | CACCTGAGGCGTATGCGAATGAGC | | | | | | | | | | |
| Oligo rev | AAACGCTCATTCGCATACGCCTCA | | | | | | | | | | |
| Chromosome | start | end | strand | MM | target_seq | PAM | alignment | distance | position | gene name | gene id |
| GU305914.1 | 172996 | 173018 | + | 0 | TGAGGCGTATGCGAATGAGC | TGG | ||||||||[||||||||||||]PAM | NA | Intergenic | NA | NA |
| T13117 | GAGGCGTATGCGAATGAGCTGGG | 1000 | | | | | | | | | |
| Oligo fwd | CACCGAGGCGTATGCGAATGAGCT | | | | | | | | | | |
| Oligo rev | AAACAGCTCATTCGCATACGCCTC | | | | | | | | | | |
| Chromosome | start | end | strand | MM | target_seq | PAM | alignment | distance | position | gene name | gene id |
| GU305914.1 | 172997 | 173019 | + | 0 | GAGGCGTATGCGAATGAGCT | GGG | ||||||||[||||||||||||]PAM | NA | Intergenic | NA | NA |
| T13118 | GCGTATGCGAATGAGCTGGGCGG | 1000 | | | | | | | | | |
| Oligo fwd | CACCGCGTATGCGAATGAGCTGGG | | | | | | | | | | |
| Oligo rev | AAACCCCAGCTCATTCGCATACGC | | | | | | | | | | |
| Chromosome | start | end | strand | MM | target_seq | PAM | alignment | distance | position | gene name | gene id |
| GU305914.1 | 173000 | 173022 | + | 0 | GCGTATGCGAATGAGCTGGG | CGG | ||||||||[||||||||||||]PAM | NA | Intergenic | NA | NA |
| T13119 | CGTATGCGAATGAGCTGGGCGGG | 1000 | | | | | | | | | |
| Oligo fwd | CACCCGTATGCGAATGAGCTGGGC | | | | | | | | | | |
| Oligo rev | AAACGCCCAGCTCATTCGCATACG | | | | | | | | | | |
| Chromosome | start | end | strand | MM | target_seq | PAM | alignment | distance | position | gene name | gene id |
| GU305914.1 | 173001 | 173023 | + | 0 | CGTATGCGAATGAGCTGGGC | GGG | ||||||||[||||||||||||]PAM | NA | Intergenic | NA | NA |
| T13120 | TGCGAATGAGCTGGGCGGGTAGG | 1000 | | | | | | | | | |
| Oligo fwd | CACCTGCGAATGAGCTGGGCGGGT | | | | | | | | | | |
| Oligo rev | AAACACCCGCCCAGCTCATTCGCA | | | | | | | | | | |
| Chromosome | start | end | strand | MM | target_seq | PAM | alignment | distance | position | gene name | gene id |
| GU305914.1 | 173005 | 173027 | + | 0 | TGCGAATGAGCTGGGCGGGT | AGG | ||||||||[||||||||||||]PAM | NA | Intergenic | NA | NA |
| T13121 | AATGAGCTGGGCGGGTAGGAAGG | 1000 | | | | | | | | | |
| Oligo fwd | CACCAATGAGCTGGGCGGGTAGGA | | | | | | | | | | |
| Oligo rev | AAACTCCTACCCGCCCAGCTCATT | | | | | | | | | | |
| Chromosome | start | end | strand | MM | target_seq | PAM | alignment | distance | position | gene name | gene id |
| GU305914.1 | 173009 | 173031 | + | 0 | AATGAGCTGGGCGGGTAGGA | AGG | ||||||||[||||||||||||]PAM | NA | Intergenic | NA | NA |
| T13122 | GGGTAGGAAGGATGCTTAATCGG | 1000 | | | | | | | | | |
| Oligo fwd | CACCGGGTAGGAAGGATGCTTAAT | | | | | | | | | | |
| Oligo rev | AAACATTAAGCATCCTTCCTACCC | | | | | | | | | | |
| Chromosome | start | end | strand | MM | target_seq | PAM | alignment | distance | position | gene name | gene id |
| GU305914.1 | 173021 | 173043 | + | 0 | GGGTAGGAAGGATGCTTAAT | CGG | ||||||||[||||||||||||]PAM | NA | Intergenic | NA | NA |
| T13123 | CTTAATCGGTTTATAGTGTGTGG | 1000 | | | | | | | | | |
| Oligo fwd | CACCCTTAATCGGTTTATAGTGTG | | | | | | | | | | |
| Oligo rev | AAACCACACTATAAACCGATTAAG | | | | | | | | | | |
| Chromosome | start | end | strand | MM | target_seq | PAM | alignment | distance | position | gene name | gene id |
| GU305914.1 | 173035 | 173057 | + | 0 | CTTAATCGGTTTATAGTGTG | TGG | ||||||||[||||||||||||]PAM | NA | Intergenic | NA | NA |
| T13124 | CGTTAAGCCAGTAGACTCAGTGG | 1000 | | | | | | | | | |
| Oligo fwd | CACCCGTTAAGCCAGTAGACTCAG | | | | | | | | | | |
| Oligo rev | AAACCTGAGTCTACTGGCTTAACG | | | | | | | | | | |
| Chromosome | start | end | strand | MM | target_seq | PAM | alignment | distance | position | gene name | gene id |

<table>
<tr><td>GU305914.1</td><td>173093</td><td>173115</td><td>+</td><td>0</td><td>CGTTAAGCCAGTAGACTCAG</td><td>TGG</td><td>||||||||[||||||||||||]PAM</td><td>NA</td><td>Intergenic</td><td>NA</td><td>NA</td></tr>
<tr><td>T13125</td><td>GTTAAGCCAGTAGACTCAGTGGG</td><td>1000</td><td></td><td></td><td></td><td></td><td></td><td></td><td></td><td></td><td></td></tr>
<tr><td>Oligo fwd</td><td>CACCGTTAAGCCAGTAGACTCAGT</td><td></td><td></td><td></td><td></td><td></td><td></td><td></td><td></td><td></td><td></td></tr>
<tr><td>Oligo rev</td><td>AAACACTGAGTCTACTGGCTTAAC</td><td></td><td></td><td></td><td></td><td></td><td></td><td></td><td></td><td></td><td></td></tr>
<tr><td>Chromosome</td><td>start</td><td>end</td><td>strand</td><td>MM</td><td>target_seq</td><td>PAM</td><td>alignment</td><td>distance</td><td>position</td><td>gene name</td><td>gene id</td></tr>
<tr><td>GU305914.1</td><td>173094</td><td>173116</td><td>+</td><td>0</td><td>GTTAAGCCAGTAGACTCAGT</td><td>GGG</td><td>||||||||[||||||||||||]PAM</td><td>NA</td><td>Intergenic</td><td>NA</td><td>NA</td></tr>
<tr><td>T13126</td><td>AGTGGGTTCCGTCACTTCGTTGG</td><td>1000</td><td></td><td></td><td></td><td></td><td></td><td></td><td></td><td></td><td></td></tr>
<tr><td>Oligo fwd</td><td>CACCAGTGGGTTCCGTCACTTCGT</td><td></td><td></td><td></td><td></td><td></td><td></td><td></td><td></td><td></td><td></td></tr>
<tr><td>Oligo rev</td><td>AAACACGAAGTGACGGAACCCACT</td><td></td><td></td><td></td><td></td><td></td><td></td><td></td><td></td><td></td><td></td></tr>
<tr><td>Chromosome</td><td>start</td><td>end</td><td>strand</td><td>MM</td><td>target_seq</td><td>PAM</td><td>alignment</td><td>distance</td><td>position</td><td>gene name</td><td>gene id</td></tr>
<tr><td>GU305914.1</td><td>173111</td><td>173133</td><td>+</td><td>0</td><td>AGTGGGTTCCGTCACTTCGT</td><td>TGG</td><td>||||||||[||||||||||||]PAM</td><td>NA</td><td>Intergenic</td><td>NA</td><td>NA</td></tr>
<tr><td>T13127</td><td>GTGGGTTCCGTCACTTCGTTGGG</td><td>1000</td><td></td><td></td><td></td><td></td><td></td><td></td><td></td><td></td><td></td></tr>
<tr><td>Oligo fwd</td><td>CACCGTGGGTTCCGTCACTTCGTT</td><td></td><td></td><td></td><td></td><td></td><td></td><td></td><td></td><td></td><td></td></tr>
<tr><td>Oligo rev</td><td>AAACAACGAAGTGACGGAACCCAC</td><td></td><td></td><td></td><td></td><td></td><td></td><td></td><td></td><td></td><td></td></tr>
<tr><td>Chromosome</td><td>start</td><td>end</td><td>strand</td><td>MM</td><td>target_seq</td><td>PAM</td><td>alignment</td><td>distance</td><td>position</td><td>gene name</td><td>gene id</td></tr>
<tr><td>GU305914.1</td><td>173112</td><td>173134</td><td>+</td><td>0</td><td>GTGGGTTCCGTCACTTCGTT</td><td>GGG</td><td>||||||||[||||||||||||]PAM</td><td>NA</td><td>Intergenic</td><td>NA</td><td>NA</td></tr>
<tr><td>T13128</td><td>GATGTACATGAAGTCATCCTCGG</td><td>1000</td><td></td><td></td><td></td><td></td><td></td><td></td><td></td><td></td><td></td></tr>
<tr><td>Oligo fwd</td><td>CACCGATGTACATGAAGTCATCCT</td><td></td><td></td><td></td><td></td><td></td><td></td><td></td><td></td><td></td><td></td></tr>
<tr><td>Oligo rev</td><td>AAACAGGATGACTTCATGTACATC</td><td></td><td></td><td></td><td></td><td></td><td></td><td></td><td></td><td></td><td></td></tr>
<tr><td>Chromosome</td><td>start</td><td>end</td><td>strand</td><td>MM</td><td>target_seq</td><td>PAM</td><td>alignment</td><td>distance</td><td>position</td><td>gene name</td><td>gene id</td></tr>
<tr><td>GU305914.1</td><td>173134</td><td>173156</td><td>+</td><td>0</td><td>GATGTACATGAAGTCATCCT</td><td>CGG</td><td>||||||||[||||||||||||]PAM</td><td>NA</td><td>Intergenic</td><td>NA</td><td>NA</td></tr>
<tr><td>T13129</td><td>TTATTCTTCTTTTTTTCTTCAGG</td><td>1000</td><td></td><td></td><td></td><td></td><td></td><td></td><td></td><td></td><td></td></tr>
<tr><td>Oligo fwd</td><td>CACCTTATTCTTCTTTTTTTCTTC</td><td></td><td></td><td></td><td></td><td></td><td></td><td></td><td></td><td></td><td></td></tr>
<tr><td>Oligo rev</td><td>AAACGAAGAAAAAAAGAAGAATAA</td><td></td><td></td><td></td><td></td><td></td><td></td><td></td><td></td><td></td><td></td></tr>
<tr><td>Chromosome</td><td>start</td><td>end</td><td>strand</td><td>MM</td><td>target_seq</td><td>PAM</td><td>alignment</td><td>distance</td><td>position</td><td>gene name</td><td>gene id</td></tr>
<tr><td>GU305914.1</td><td>173192</td><td>173214</td><td>+</td><td>0</td><td>TTATTCTTCTTTTTTTCTTC</td><td>AGG</td><td>||||||||[||||||||||||]PAM</td><td>NA</td><td>Intergenic</td><td>NA</td><td>NA</td></tr>
<tr><td>T13130</td><td>TTTTTTTCTTCAGGTGTCTGCGG</td><td>1000</td><td></td><td></td><td></td><td></td><td></td><td></td><td></td><td></td><td></td></tr>
<tr><td>Oligo fwd</td><td>CACCTTTTTTTCTTCAGGTGTCTG</td><td></td><td></td><td></td><td></td><td></td><td></td><td></td><td></td><td></td><td></td></tr>
<tr><td>Oligo rev</td><td>AAACCAGACACCTGAAGAAAAAAA</td><td></td><td></td><td></td><td></td><td></td><td></td><td></td><td></td><td></td><td></td></tr>
<tr><td>Chromosome</td><td>start</td><td>end</td><td>strand</td><td>MM</td><td>target_seq</td><td>PAM</td><td>alignment</td><td>distance</td><td>position</td><td>gene name</td><td>gene id</td></tr>
<tr><td>GU305914.1</td><td>173201</td><td>173223</td><td>+</td><td>0</td><td>TTTTTTTCTTCAGGTGTCTG</td><td>CGG</td><td>||||||||[||||||||||||]PAM</td><td>NA</td><td>Intergenic</td><td>NA</td><td>NA</td></tr>
<tr><td>T13131</td><td>TATAGTATAGAGCTGATAGCTGG</td><td>1000</td><td></td><td></td><td></td><td></td><td></td><td></td><td></td><td></td><td></td></tr>
<tr><td>Oligo fwd</td><td>CACCTATAGTATAGAGCTGATAGC</td><td></td><td></td><td></td><td></td><td></td><td></td><td></td><td></td><td></td><td></td></tr>
<tr><td>Oligo rev</td><td>AAACGCTATCAGCTCTATACTATA</td><td></td><td></td><td></td><td></td><td></td><td></td><td></td><td></td><td></td><td></td></tr>
<tr><td>Chromosome</td><td>start</td><td>end</td><td>strand</td><td>MM</td><td>target_seq</td><td>PAM</td><td>alignment</td><td>distance</td><td>position</td><td>gene name</td><td>gene id</td></tr>
<tr><td>GU305914.1</td><td>173236</td><td>173258</td><td>+</td><td>0</td><td>TATAGTATAGAGCTGATAGC</td><td>TGG</td><td>||||||||[||||||||||||]PAM</td><td>NA</td><td>Intergenic</td><td>NA</td><td>NA</td></tr>
<tr><td>T13132</td><td>GAAGCTTTTTTAGCTTGTGCAGG</td><td>1000</td><td></td><td></td><td></td><td></td><td></td><td></td><td></td><td></td><td></td></tr>
<tr><td>Oligo fwd</td><td>CACCGAAGCTTTTTTAGCTTGTGC</td><td></td><td></td><td></td><td></td><td></td><td></td><td></td><td></td><td></td><td></td></tr>
<tr><td>Oligo rev</td><td>AAACGCACAAGCTAAAAAAGCTTC</td><td></td><td></td><td></td><td></td><td></td><td></td><td></td><td></td><td></td><td></td></tr>
<tr><td>Chromosome</td><td>start</td><td>end</td><td>strand</td><td>MM</td><td>target_seq</td><td>PAM</td><td>alignment</td><td>distance</td><td>position</td><td>gene name</td><td>gene id</td></tr>
<tr><td>GU305914.1</td><td>173268</td><td>173290</td><td>+</td><td>0</td><td>GAAGCTTTTTTAGCTTGTGC</td><td>AGG</td><td>||||||||[||||||||||||]PAM</td><td>NA</td><td>Intergenic</td><td>NA</td><td>NA</td></tr>
<tr><td>T13133</td><td>TTTTTAGCTTGTGCAGGAATAGG</td><td>1000</td><td></td><td></td><td></td><td></td><td></td><td></td><td></td><td></td><td></td></tr>
<tr><td>Oligo fwd</td><td>CACCTTTTTAGCTTGTGCAGGAAT</td><td></td><td></td><td></td><td></td><td></td><td></td><td></td><td></td><td></td><td></td></tr>
<tr><td>Oligo rev</td><td>AAACATTCCTGCACAAGCTAAAAA</td><td></td><td></td><td></td><td></td><td></td><td></td><td></td><td></td><td></td><td></td></tr>
<tr><td>Chromosome</td><td>start</td><td>end</td><td>strand</td><td>MM</td><td>target_seq</td><td>PAM</td><td>alignment</td><td>distance</td><td>position</td><td>gene name</td><td>gene id</td></tr>
<tr><td>GU305914.1</td><td>173274</td><td>173296</td><td>+</td><td>0</td><td>TTTTTAGCTTGTGCAGGAAT</td><td>AGG</td><td>||||||||[||||||||||||]PAM</td><td>NA</td><td>Intergenic</td><td>NA</td><td>NA</td></tr>
<tr><td>T13134</td><td>AGCTTGTGCAGGAATAGGTTCGG</td><td>1000</td><td></td><td></td><td></td><td></td><td></td><td></td><td></td><td></td><td></td></tr>
<tr><td>Oligo fwd</td><td>CACCAGCTTGTGCAGGAATAGGTT</td><td></td><td></td><td></td><td></td><td></td><td></td><td></td><td></td><td></td><td></td></tr>
<tr><td>Oligo rev</td><td>AAACAACCTATTCCTGCACAAGCT</td><td></td><td></td><td></td><td></td><td></td><td></td><td></td><td></td><td></td><td></td></tr>
<tr><td>Chromosome</td><td>start</td><td>end</td><td>strand</td><td>MM</td><td>target_seq</td><td>PAM</td><td>alignment</td><td>distance</td><td>position</td><td>gene name</td><td>gene id</td></tr>
<tr><td>GU305914.1</td><td>173279</td><td>173301</td><td>+</td><td>0</td><td>AGCTTGTGCAGGAATAGGTT</td><td>CGG</td><td>||||||||[||||||||||||]PAM</td><td>NA</td><td>Intergenic</td><td>NA</td><td>NA</td></tr>
<tr><td>T13135</td><td>TGCAGGAATAGGTTCGGATGCGG</td><td>1000</td><td></td><td></td><td></td><td></td><td></td><td></td><td></td><td></td><td></td></tr>
<tr><td>Oligo fwd</td><td>CACCTGCAGGAATAGGTTCGGATG</td><td></td><td></td><td></td><td></td><td></td><td></td><td></td><td></td><td></td><td></td></tr>
<tr><td>Oligo rev</td><td>AAACCATCCGAACCTATTCCTGCA</td><td></td><td></td><td></td><td></td><td></td><td></td><td></td><td></td><td></td><td></td></tr>
<tr><td>Chromosome</td><td>start</td><td>end</td><td>strand</td><td>MM</td><td>target_seq</td><td>PAM</td><td>alignment</td><td>distance</td><td>position</td><td>gene name</td><td>gene id</td></tr>
<tr><td>GU305914.1</td><td>173285</td><td>173307</td><td>+</td><td>0</td><td>TGCAGGAATAGGTTCGGATG</td><td>CGG</td><td>||||||||[||||||||||||]PAM</td><td>NA</td><td>Intergenic</td><td>NA</td><td>NA</td></tr>
<tr><td>T13136</td><td>GTTCGGATGCGGTGCTCTCATGG</td><td>1000</td><td></td><td></td><td></td><td></td><td></td><td></td><td></td><td></td><td></td></tr>
<tr><td>Oligo fwd</td><td>CACCGTTCGGATGCGGTGCTCTCA</td><td></td><td></td><td></td><td></td><td></td><td></td><td></td><td></td><td></td><td></td></tr>
<tr><td>Oligo rev</td><td>AAACTGAGAGCACCGCATCCGAAC</td><td></td><td></td><td></td><td></td><td></td><td></td><td></td><td></td><td></td><td></td></tr>
<tr><td>Chromosome</td><td>start</td><td>end</td><td>strand</td><td>MM</td><td>target_seq</td><td>PAM</td><td>alignment</td><td>distance</td><td>position</td><td>gene name</td><td>gene id</td></tr>
<tr><td>GU305914.1</td><td>173296</td><td>173318</td><td>+</td><td>0</td><td>GTTCGGATGCGGTGCTCTCA</td><td>TGG</td><td>||||||||[||||||||||||]PAM</td><td>NA</td><td>Intergenic</td><td>NA</td><td>NA</td></tr>
<tr><td>T13137</td><td>CGGATGCGGTGCTCTCATGGCGG</td><td>1000</td><td></td><td></td><td></td><td></td><td></td><td></td><td></td><td></td><td></td></tr>
<tr><td>Oligo fwd</td><td>CACCCGGATGCGGTGCTCTCATGG</td><td></td><td></td><td></td><td></td><td></td><td></td><td></td><td></td><td></td><td></td></tr>
<tr><td>Oligo rev</td><td>AAACCCATGAGAGCACCGCATCCG</td><td></td><td></td><td></td><td></td><td></td><td></td><td></td><td></td><td></td><td></td></tr>
<tr><td>Chromosome</td><td>start</td><td>end</td><td>strand</td><td>MM</td><td>target_seq</td><td>PAM</td><td>alignment</td><td>distance</td><td>position</td><td>gene name</td><td>gene id</td></tr>
<tr><td>GU305914.1</td><td>173299</td><td>173321</td><td>+</td><td>0</td><td>CGGATGCGGTGCTCTCATGG</td><td>CGG</td><td>||||||||[||||||||||||]PAM</td><td>NA</td><td>Intergenic</td><td>NA</td><td>NA</td></tr>
<tr><td>T13138</td><td>CGGTGCTCTCATGGCGGCCACGG</td><td>1000</td><td></td><td></td><td></td><td></td><td></td><td></td><td></td><td></td><td></td></tr>
<tr><td>Oligo fwd</td><td>CACCCGGTGCTCTCATGGCGGCCA</td><td></td><td></td><td></td><td></td><td></td><td></td><td></td><td></td><td></td><td></td></tr>
<tr><td>Oligo rev</td><td>AAACTGGCCGCCATGAGAGCACCG</td><td></td><td></td><td></td><td></td><td></td><td></td><td></td><td></td><td></td><td></td></tr>
<tr><td>Chromosome</td><td>start</td><td>end</td><td>strand</td><td>MM</td><td>target_seq</td><td>PAM</td><td>alignment</td><td>distance</td><td>position</td><td>gene name</td><td>gene id</td></tr>
<tr><td>GU305914.1</td><td>173305</td><td>173327</td><td>+</td><td>0</td><td>CGGTGCTCTCATGGCGGCCA</td><td>CGG</td><td>||||||||[||||||||||||]PAM</td><td>NA</td><td>Intergenic</td><td>NA</td><td>NA</td></tr>
<tr><td>T13139</td><td>TCATGGCGGCCACGGCCATCAGG</td><td>1000</td><td></td><td></td><td></td><td></td><td></td><td></td><td></td><td></td><td></td></tr>
<tr><td>Oligo fwd</td><td>CACCTCATGGCGGCCACGGCCATC</td><td></td><td></td><td></td><td></td><td></td><td></td><td></td><td></td><td></td><td></td></tr>
<tr><td>Oligo rev</td><td>AAACGATGGCCGTGGCCGCCATGA</td><td></td><td></td><td></td><td></td><td></td><td></td><td></td><td></td><td></td><td></td></tr>
</table>

Chromosome start end strand MM target_seq PAM alignment distance position gene name gene id
GU305914.1 173313 173335 + 0 TCATGGCGGCCACGGCCATC AGG ||||||||[||||||||||||]PAM NA Intergenic NA NA

T13140 GGCCACGGCCATCAGGATCCCGG 1000
Oligo fwd CACCGGCCACGGCCATCAGGATCC
Oligo rev AAACGGATCCTGATGGCCGTGGCC
Chromosome start end strand MM target_seq PAM alignment distance position gene name gene id
GU305914.1 173320 173342 + 0 GGCCACGGCCATCAGGATCC CGG ||||||||[||||||||||||]PAM NA Intergenic NA NA

T13141 CGGCCATCAGGATCCCGGAGAGG 1000
Oligo fwd CACCCGGCCATCAGGATCCCGGAG
Oligo rev AAACCTCCGGGATCCTGATGGCCG
Chromosome start end strand MM target_seq PAM alignment distance position gene name gene id
GU305914.1 173325 173347 + 0 CGGCCATCAGGATCCCGGAG AGG ||||||||[||||||||||||]PAM NA Intergenic NA NA

T13142 CTAGCAGTATGCACGAGATGAGG 1000
Oligo fwd CACCCTAGCAGTATGCACGAGATG
Oligo rev AAACCATCTCGTGCATACTGCTAG
Chromosome start end strand MM target_seq PAM alignment distance position gene name gene id
GU305914.1 173358 173380 + 0 CTAGCAGTATGCACGAGATG AGG ||||||||[||||||||||||]PAM NA Intergenic NA NA

T13143 ACGAGATGAGGCCGTACGTGAGG 1000
Oligo fwd CACCACGAGATGAGGCCGTACGTG
Oligo rev AAACCACGTACGGCCTCATCTCGT
Chromosome start end strand MM target_seq PAM alignment distance position gene name gene id
GU305914.1 173370 173392 + 0 ACGAGATGAGGCCGTACGTG AGG ||||||||[||||||||||||]PAM NA Intergenic NA NA

T13144 GAGGCCGTACGTGAGGAGATCGG 1000
Oligo fwd CACCGAGGCCGTACGTGAGGAGAT
Oligo rev AAACATCTCCTCACGTACGGCCTC
Chromosome start end strand MM target_seq PAM alignment distance position gene name gene id
GU305914.1 173377 173399 + 0 GAGGCCGTACGTGAGGAGAT CGG ||||||||[||||||||||||]PAM NA Intergenic NA NA

T13145 TAGACGTCAGTTCCATGACGAGG 1000
Oligo fwd CACCTAGACGTCAGTTCCATGACG
Oligo rev AAACCGTCATGGAACTGACGTCTA
Chromosome start end strand MM target_seq PAM alignment distance position gene name gene id
GU305914.1 173400 173422 + 0 TAGACGTCAGTTCCATGACG AGG ||||||||[||||||||||||]PAM NA Intergenic NA NA

T13146 TTGAAAATAGCGATTGAGCTCGG 1000
Oligo fwd CACCTTGAAAATAGCGATTGAGCT
Oligo rev AAACAGCTCAATCGCTATTTTCAA
Chromosome start end strand MM target_seq PAM alignment distance position gene name gene id
GU305914.1 173423 173445 + 0 TTGAAAATAGCGATTGAGCT CGG ||||||||[||||||||||||]PAM NA Intergenic NA NA

T13147 AAATAGCGATTGAGCTCGGTTGG 1000
Oligo fwd CACCAAATAGCGATTGAGCTCGGT
Oligo rev AAACACCGAGCTCAATCGCTATTT
Chromosome start end strand MM target_seq PAM alignment distance position gene name gene id
GU305914.1 173427 173449 + 0 AAATAGCGATTGAGCTCGGT TGG ||||||||[||||||||||||]PAM NA Intergenic NA NA

T13148 AATAGCGATTGAGCTCGGTTGGG 1000
Oligo fwd CACCAATAGCGATTGAGCTCGGTT
Oligo rev AAACAACCGAGCTCAATCGCTATT
Chromosome start end strand MM target_seq PAM alignment distance position gene name gene id
GU305914.1 173428 173450 + 0 AATAGCGATTGAGCTCGGTT GGG ||||||||[||||||||||||]PAM NA Intergenic NA NA

T13149 ATAGCGATTGAGCTCGGTTGGGG 1000
Oligo fwd CACCATAGCGATTGAGCTCGGTTG
Oligo rev AAACCAACCGAGCTCAATCGCTAT
Chromosome start end strand MM target_seq PAM alignment distance position gene name gene id
GU305914.1 173429 173451 + 0 ATAGCGATTGAGCTCGGTTG GGG ||||||||[||||||||||||]PAM NA Intergenic NA NA

T13150 TAGCGATTGAGCTCGGTTGGGGG 1000
Oligo fwd CACCTAGCGATTGAGCTCGGTTGG
Oligo rev AAACCCAACCGAGCTCAATCGCTA
Chromosome start end strand MM target_seq PAM alignment distance position gene name gene id
GU305914.1 173430 173452 + 0 TAGCGATTGAGCTCGGTTGG GGG ||||||||[||||||||||||]PAM NA Intergenic NA NA

T13151 AGCGATTGAGCTCGGTTGGGGGG 1000
Oligo fwd CACCAGCGATTGAGCTCGGTTGGG
Oligo rev AAACCCCAACCGAGCTCAATCGCT
Chromosome start end strand MM target_seq PAM alignment distance position gene name gene id
GU305914.1 173431 173453 + 0 AGCGATTGAGCTCGGTTGGG GGG ||||||||[||||||||||||]PAM NA Intergenic NA NA

T13152 TTTTCAAAGATGATTCCATGAGG 1000
Oligo fwd CACCTTTTCAAAGATGATTCCATG
Oligo rev AAACCATGGAATCATCTTTGAAAA
Chromosome start end strand MM target_seq PAM alignment distance position gene name gene id
GU305914.1 173472 173494 + 0 TTTTCAAAGATGATTCCATG AGG ||||||||[||||||||||||]PAM NA Intergenic NA NA

T13153 CTCTTCTTTTTCACACGAATCGG 1000
Oligo fwd CACCCTCTTCTTTTTCACACGAAT
Oligo rev AAACATTCGTGTGAAAAAGAAGAG
Chromosome start end strand MM target_seq PAM alignment distance position gene name gene id
GU305914.1 173500 173522 + 0 CTCTTCTTTTTCACACGAAT CGG ||||||||[||||||||||||]PAM NA Intergenic NA NA

T13154 TTCACACGAATCGGTTAGCTAGG 1000
Oligo fwd CACCTTCACACGAATCGGTTAGCT

| | | | | | | | | | | | |
|---|---|---|---|---|---|---|---|---|---|---|---|
| Oligo rev | AAACAGCTAACCGATTCGTGTGAA | | | | | | | | | | |
| Chromosome | start | end | strand | MM | target_seq | PAM | alignment | distance | position | gene name | gene id |
| GU305914.1 | 173509 | 173531 | + | 0 | TTCACACGAATCGGTTAGCT | AGG | \|\|\|\|\|\|\|\|[\|\|\|\|\|\|\|\|\|\|\|\|]PAM | NA | Intergenic | NA | NA |
| T13155 | CACGAATCGGTTAGCTAGGACGG | 1000 | | | | | | | | | |
| Oligo fwd | CACCCACGAATCGGTTAGCTAGGA | | | | | | | | | | |
| Oligo rev | AAACTCCTAGCTAACCGATTCGTG | | | | | | | | | | |
| Chromosome | start | end | strand | MM | target_seq | PAM | alignment | distance | position | gene name | gene id |
| GU305914.1 | 173513 | 173535 | + | 0 | CACGAATCGGTTAGCTAGGA | CGG | \|\|\|\|\|\|\|\|[\|\|\|\|\|\|\|\|\|\|\|\|]PAM | NA | Intergenic | NA | NA |
| T13156 | TCGGTTAGCTAGGACGGCGATGG | 1000 | | | | | | | | | |
| Oligo fwd | CACCTCGGTTAGCTAGGACGGCGA | | | | | | | | | | |
| Oligo rev | AAACTCGCCGTCCTAGCTAACCGA | | | | | | | | | | |
| Chromosome | start | end | strand | MM | target_seq | PAM | alignment | distance | position | gene name | gene id |
| GU305914.1 | 173519 | 173541 | + | 0 | TCGGTTAGCTAGGACGGCGA | TGG | \|\|\|\|\|\|\|\|[\|\|\|\|\|\|\|\|\|\|\|\|]PAM | NA | Intergenic | NA | NA |
| T13157 | TAGGACGGCGATGGCCGTCGAGG | 1000 | | | | | | | | | |
| Oligo fwd | CACCTAGGACGGCGATGGCCGTCG | | | | | | | | | | |
| Oligo rev | AAACCGACGGCCATCGCCGTCCTA | | | | | | | | | | |
| Chromosome | start | end | strand | MM | target_seq | PAM | alignment | distance | position | gene name | gene id |
| GU305914.1 | 173528 | 173550 | + | 0 | TAGGACGGCGATGGCCGTCG | AGG | \|\|\|\|\|\|\|\|[\|\|\|\|\|\|\|\|\|\|\|\|]PAM | NA | Intergenic | NA | NA |
| T13158 | CCGTCGAGGAGAACGCTAGTCGG | 1000 | | | | | | | | | |
| Oligo fwd | CACCCCGTCGAGGAGAACGCTAGT | | | | | | | | | | |
| Oligo rev | AAACACTAGCGTTCTCCTCGACGG | | | | | | | | | | |
| Chromosome | start | end | strand | MM | target_seq | PAM | alignment | distance | position | gene name | gene id |
| GU305914.1 | 173542 | 173564 | + | 0 | CCGTCGAGGAGAACGCTAGT | CGG | \|\|\|\|\|\|\|\|[\|\|\|\|\|\|\|\|\|\|\|\|]PAM | NA | Intergenic | NA | NA |
| T13159 | AGGAGAACGCTAGTCGGAACAGG | 1000 | | | | | | | | | |
| Oligo fwd | CACCAGGAGAACGCTAGTCGGAAC | | | | | | | | | | |
| Oligo rev | AAACGTTCCGACTAGCGTTCTCCT | | | | | | | | | | |
| Chromosome | start | end | strand | MM | target_seq | PAM | alignment | distance | position | gene name | gene id |
| GU305914.1 | 173548 | 173570 | + | 0 | AGGAGAACGCTAGTCGGAAC | AGG | \|\|\|\|\|\|\|\|[\|\|\|\|\|\|\|\|\|\|\|\|]PAM | NA | Intergenic | NA | NA |
| T13160 | AGAACGCTAGTCGGAACAGGAGG | 1000 | | | | | | | | | |
| Oligo fwd | CACCAGAACGCTAGTCGGAACAGG | | | | | | | | | | |
| Oligo rev | AAACCCTGTTCCGACTAGCGTTCT | | | | | | | | | | |
| Chromosome | start | end | strand | MM | target_seq | PAM | alignment | distance | position | gene name | gene id |
| GU305914.1 | 173551 | 173573 | + | 0 | AGAACGCTAGTCGGAACAGG | AGG | \|\|\|\|\|\|\|\|[\|\|\|\|\|\|\|\|\|\|\|\|]PAM | NA | Intergenic | NA | NA |
| T13161 | GAACAGGAGGTTCTTCTTCGAGG | 1000 | | | | | | | | | |
| Oligo fwd | CACCGAACAGGAGGTTCTTCTTCG | | | | | | | | | | |
| Oligo rev | AAACCGAAGAAGAACCTCCTGTTC | | | | | | | | | | |
| Chromosome | start | end | strand | MM | target_seq | PAM | alignment | distance | position | gene name | gene id |
| GU305914.1 | 173564 | 173586 | + | 0 | GAACAGGAGGTTCTTCTTCG | AGG | \|\|\|\|\|\|\|\|[\|\|\|\|\|\|\|\|\|\|\|\|]PAM | NA | Intergenic | NA | NA |
| T13162 | GAGGTTCTTCTTCGAGGTGTCGG | 1000 | | | | | | | | | |
| Oligo fwd | CACCGAGGTTCTTCTTCGAGGTGT | | | | | | | | | | |
| Oligo rev | AAACACACCTCGAAGAAGAACCTC | | | | | | | | | | |
| Chromosome | start | end | strand | MM | target_seq | PAM | alignment | distance | position | gene name | gene id |
| GU305914.1 | 173570 | 173592 | + | 0 | GAGGTTCTTCTTCGAGGTGT | CGG | \|\|\|\|\|\|\|\|[\|\|\|\|\|\|\|\|\|\|\|\|]PAM | NA | Intergenic | NA | NA |
| T13163 | TTCTTCTTCGAGGTGTCGGTCGG | 1000 | | | | | | | | | |
| Oligo fwd | CACCTTCTTCTTCGAGGTGTCGGT | | | | | | | | | | |
| Oligo rev | AAACACCGACACCTCGAAGAAGAA | | | | | | | | | | |
| Chromosome | start | end | strand | MM | target_seq | PAM | alignment | distance | position | gene name | gene id |
| GU305914.1 | 173574 | 173596 | + | 0 | TTCTTCTTCGAGGTGTCGGT | CGG | \|\|\|\|\|\|\|\|[\|\|\|\|\|\|\|\|\|\|\|\|]PAM | NA | Intergenic | NA | NA |
| T13164 | CGAGGTGTCGGTCGGCGACGAGG | 1000 | | | | | | | | | |
| Oligo fwd | CACCCGAGGTGTCGGTCGGCGACG | | | | | | | | | | |
| Oligo rev | AAACCGTCGCCGACCGACACCTCG | | | | | | | | | | |
| Chromosome | start | end | strand | MM | target_seq | PAM | alignment | distance | position | gene name | gene id |
| GU305914.1 | 173582 | 173604 | + | 0 | CGAGGTGTCGGTCGGCGACG | AGG | \|\|\|\|\|\|\|\|[\|\|\|\|\|\|\|\|\|\|\|\|]PAM | NA | Intergenic | NA | NA |
| GU305914.1 | 75860 | 75882 | + | 4 | GGAGGCGGCGGTCGGCGAGG | GGG | -\|\|\|\|-\|-[\|\|\|\|\|\|\|\|\|\|-\|]PAM | NA | Intergenic | NA | NA |
| GU305914.1 | 213268 | 213290 | + | 4 | GGAGGTGGCGGTCGGAGAGG | CGG | -\|\|\|\|\|\|-[\|\|\|\|\|\|\|-\|\|-\|]PAM | NA | Intergenic | NA | NA |
| T13165 | GAGGTGTCGGTCGGCGACGAGGG | 1000 | | | | | | | | | |
| Oligo fwd | CACCGAGGTGTCGGTCGGCGACGA | | | | | | | | | | |
| Oligo rev | AAACTCGTCGCCGACCGACACCTC | | | | | | | | | | |
| Chromosome | start | end | strand | MM | target_seq | PAM | alignment | distance | position | gene name | gene id |
| GU305914.1 | 173583 | 173605 | + | 0 | GAGGTGTCGGTCGGCGACGA | GGG | \|\|\|\|\|\|\|\|[\|\|\|\|\|\|\|\|\|\|\|\|]PAM | NA | Intergenic | NA | NA |
| T13166 | GTCGGTCGGCGACGAGGGTGAGG | 1000 | | | | | | | | | |
| Oligo fwd | CACCGTCGGTCGGCGACGAGGGTG | | | | | | | | | | |
| Oligo rev | AAACCACCCTCGTCGCCGACCGAC | | | | | | | | | | |
| Chromosome | start | end | strand | MM | target_seq | PAM | alignment | distance | position | gene name | gene id |
| GU305914.1 | 173588 | 173610 | + | 0 | GTCGGTCGGCGACGAGGGTG | AGG | \|\|\|\|\|\|\|\|[\|\|\|\|\|\|\|\|\|\|\|\|]PAM | NA | Intergenic | NA | NA |
| T13167 | GGTCGGCGACGAGGGTGAGGTGG | 1000 | | | | | | | | | |
| Oligo fwd | CACCGGTCGGCGACGAGGGTGAGG | | | | | | | | | | |
| Oligo rev | AAACCCTCACCCTCGTCGCCGACC | | | | | | | | | | |
| Chromosome | start | end | strand | MM | target_seq | PAM | alignment | distance | position | gene name | gene id |
| GU305914.1 | 173591 | 173613 | + | 0 | GGTCGGCGACGAGGGTGAGG | TGG | \|\|\|\|\|\|\|\|[\|\|\|\|\|\|\|\|\|\|\|\|]PAM | NA | Intergenic | NA | NA |
| T13168 | GGCAGAGTTCTTAGTAGTAGTGG | 1000 | | | | | | | | | |
| Oligo fwd | CACCGGCAGAGTTCTTAGTAGTAG | | | | | | | | | | |
| Oligo rev | AAACCTACTACTAAGAACTCTGCC | | | | | | | | | | |
| Chromosome | start | end | strand | MM | target_seq | PAM | alignment | distance | position | gene name | gene id |
| GU305914.1 | 173612 | 173634 | + | 0 | GGCAGAGTTCTTAGTAGTAG | TGG | \|\|\|\|\|\|\|\|[\|\|\|\|\|\|\|\|\|\|\|\|]PAM | NA | Intergenic | NA | NA |

<table>
<tr><td>T13169</td><td>TAGTGGAGTTGCTGTAACTTTGG</td><td>1000</td><td></td><td></td><td></td><td></td><td></td><td></td><td></td><td></td><td></td></tr>
<tr><td>Oligo fwd</td><td>CACCTAGTGGAGTTGCTGTAACTT</td><td></td><td></td><td></td><td></td><td></td><td></td><td></td><td></td><td></td><td></td></tr>
<tr><td>Oligo rev</td><td>AAACAAGTTACAGCAACTCCACTA</td><td></td><td></td><td></td><td></td><td></td><td></td><td></td><td></td><td></td><td></td></tr>
<tr><td>Chromosome</td><td>start</td><td>end</td><td>strand</td><td>MM</td><td>target_seq</td><td>PAM</td><td>alignment</td><td>distance</td><td>position</td><td>gene name</td><td>gene id</td></tr>
<tr><td>GU305914.1</td><td>173629</td><td>173651</td><td>+</td><td>0</td><td>TAGTGGAGTTGCTGTAACTT</td><td>TGG</td><td>||||||||[||||||||||||]PAM</td><td>NA</td><td>Intergenic</td><td>NA</td><td>NA</td></tr>
<tr><td>T13170</td><td>TTTGGTGACTGTTTATAGTCTGG</td><td>1000</td><td></td><td></td><td></td><td></td><td></td><td></td><td></td><td></td><td></td></tr>
<tr><td>Oligo fwd</td><td>CACCTTTGGTGACTGTTTATAGTC</td><td></td><td></td><td></td><td></td><td></td><td></td><td></td><td></td><td></td><td></td></tr>
<tr><td>Oligo rev</td><td>AAACGACTATAAACAGTCACCAAA</td><td></td><td></td><td></td><td></td><td></td><td></td><td></td><td></td><td></td><td></td></tr>
<tr><td>Chromosome</td><td>start</td><td>end</td><td>strand</td><td>MM</td><td>target_seq</td><td>PAM</td><td>alignment</td><td>distance</td><td>position</td><td>gene name</td><td>gene id</td></tr>
<tr><td>GU305914.1</td><td>173647</td><td>173669</td><td>+</td><td>0</td><td>TTTGGTGACTGTTTATAGTC</td><td>TGG</td><td>||||||||[||||||||||||]PAM</td><td>NA</td><td>Intergenic</td><td>NA</td><td>NA</td></tr>
<tr><td>T13171</td><td>AGTCTGGTTCGAAGCAGTAGAGG</td><td>1000</td><td></td><td></td><td></td><td></td><td></td><td></td><td></td><td></td><td></td></tr>
<tr><td>Oligo fwd</td><td>CACCAGTCTGGTTCGAAGCAGTAG</td><td></td><td></td><td></td><td></td><td></td><td></td><td></td><td></td><td></td><td></td></tr>
<tr><td>Oligo rev</td><td>AAACCTACTGCTTCGAACCAGACT</td><td></td><td></td><td></td><td></td><td></td><td></td><td></td><td></td><td></td><td></td></tr>
<tr><td>Chromosome</td><td>start</td><td>end</td><td>strand</td><td>MM</td><td>target_seq</td><td>PAM</td><td>alignment</td><td>distance</td><td>position</td><td>gene name</td><td>gene id</td></tr>
<tr><td>GU305914.1</td><td>173663</td><td>173685</td><td>+</td><td>0</td><td>AGTCTGGTTCGAAGCAGTAG</td><td>AGG</td><td>||||||||[||||||||||||]PAM</td><td>NA</td><td>Intergenic</td><td>NA</td><td>NA</td></tr>
<tr><td>T13172</td><td>GTAGAGGATGATGTTGATGATGG</td><td>1000</td><td></td><td></td><td></td><td></td><td></td><td></td><td></td><td></td><td></td></tr>
<tr><td>Oligo fwd</td><td>CACCGTAGAGGATGATGTTGATGA</td><td></td><td></td><td></td><td></td><td></td><td></td><td></td><td></td><td></td><td></td></tr>
<tr><td>Oligo rev</td><td>AAACTCATCAACATCATCCTCTAC</td><td></td><td></td><td></td><td></td><td></td><td></td><td></td><td></td><td></td><td></td></tr>
<tr><td>Chromosome</td><td>start</td><td>end</td><td>strand</td><td>MM</td><td>target_seq</td><td>PAM</td><td>alignment</td><td>distance</td><td>position</td><td>gene name</td><td>gene id</td></tr>
<tr><td>GU305914.1</td><td>173679</td><td>173701</td><td>+</td><td>0</td><td>GTAGAGGATGATGTTGATGA</td><td>TGG</td><td>||||||||[||||||||||||]PAM</td><td>NA</td><td>Intergenic</td><td>NA</td><td>NA</td></tr>
<tr><td>GU305914.1</td><td>168515</td><td>168537</td><td>+</td><td>4</td><td>GACGATGATGATGATGATGA</td><td>TGG</td><td>|--||-||[|||||-||||||]PAM</td><td>NA</td><td>Intergenic</td><td>NA</td><td>NA</td></tr>
<tr><td>T13173</td><td>TAGAGGATGATGTTGATGATGGG</td><td>1000</td><td></td><td></td><td></td><td></td><td></td><td></td><td></td><td></td><td></td></tr>
<tr><td>Oligo fwd</td><td>CACCTAGAGGATGATGTTGATGAT</td><td></td><td></td><td></td><td></td><td></td><td></td><td></td><td></td><td></td><td></td></tr>
<tr><td>Oligo rev</td><td>AAACATCATCAACATCATCCTCTA</td><td></td><td></td><td></td><td></td><td></td><td></td><td></td><td></td><td></td><td></td></tr>
<tr><td>Chromosome</td><td>start</td><td>end</td><td>strand</td><td>MM</td><td>target_seq</td><td>PAM</td><td>alignment</td><td>distance</td><td>position</td><td>gene name</td><td>gene id</td></tr>
<tr><td>GU305914.1</td><td>173680</td><td>173702</td><td>+</td><td>0</td><td>TAGAGGATGATGTTGATGAT</td><td>GGG</td><td>||||||||[||||||||||||]PAM</td><td>NA</td><td>Intergenic</td><td>NA</td><td>NA</td></tr>
<tr><td>T13174</td><td>GGATGATGTTGATGATGGGCTGG</td><td>1000</td><td></td><td></td><td></td><td></td><td></td><td></td><td></td><td></td><td></td></tr>
<tr><td>Oligo fwd</td><td>CACCGGATGATGTTGATGATGGGC</td><td></td><td></td><td></td><td></td><td></td><td></td><td></td><td></td><td></td><td></td></tr>
<tr><td>Oligo rev</td><td>AAACGCCCATCATCAACATCATCC</td><td></td><td></td><td></td><td></td><td></td><td></td><td></td><td></td><td></td><td></td></tr>
<tr><td>Chromosome</td><td>start</td><td>end</td><td>strand</td><td>MM</td><td>target_seq</td><td>PAM</td><td>alignment</td><td>distance</td><td>position</td><td>gene name</td><td>gene id</td></tr>
<tr><td>GU305914.1</td><td>173684</td><td>173706</td><td>+</td><td>0</td><td>GGATGATGTTGATGATGGGC</td><td>TGG</td><td>||||||||[||||||||||||]PAM</td><td>NA</td><td>Intergenic</td><td>NA</td><td>NA</td></tr>
<tr><td>T13175</td><td>GATGATGTTGATGATGGGCTGGG</td><td>1000</td><td></td><td></td><td></td><td></td><td></td><td></td><td></td><td></td><td></td></tr>
<tr><td>Oligo fwd</td><td>CACCGATGATGTTGATGATGGGCT</td><td></td><td></td><td></td><td></td><td></td><td></td><td></td><td></td><td></td><td></td></tr>
<tr><td>Oligo rev</td><td>AAACAGCCCATCATCAACATCATC</td><td></td><td></td><td></td><td></td><td></td><td></td><td></td><td></td><td></td><td></td></tr>
<tr><td>Chromosome</td><td>start</td><td>end</td><td>strand</td><td>MM</td><td>target_seq</td><td>PAM</td><td>alignment</td><td>distance</td><td>position</td><td>gene name</td><td>gene id</td></tr>
<tr><td>GU305914.1</td><td>173685</td><td>173707</td><td>+</td><td>0</td><td>GATGATGTTGATGATGGGCT</td><td>GGG</td><td>||||||||[||||||||||||]PAM</td><td>NA</td><td>Intergenic</td><td>NA</td><td>NA</td></tr>
<tr><td>GU305914.1</td><td>58866</td><td>58888</td><td>+</td><td>4</td><td>GATGGAGTTGTTGATGGTCT</td><td>TGG</td><td>||||--||[||-|||||||-||]PAM</td><td>NA</td><td>Intergenic</td><td>NA</td><td>NA</td></tr>
<tr><td>T13176</td><td>GTTGATGATGGGCTGGGCTTAGG</td><td>1000</td><td></td><td></td><td></td><td></td><td></td><td></td><td></td><td></td><td></td></tr>
<tr><td>Oligo fwd</td><td>CACCGTTGATGATGGGCTGGGCTT</td><td></td><td></td><td></td><td></td><td></td><td></td><td></td><td></td><td></td><td></td></tr>
<tr><td>Oligo rev</td><td>AAACAAGCCCAGCCCATCATCAAC</td><td></td><td></td><td></td><td></td><td></td><td></td><td></td><td></td><td></td><td></td></tr>
<tr><td>Chromosome</td><td>start</td><td>end</td><td>strand</td><td>MM</td><td>target_seq</td><td>PAM</td><td>alignment</td><td>distance</td><td>position</td><td>gene name</td><td>gene id</td></tr>
<tr><td>GU305914.1</td><td>173691</td><td>173713</td><td>+</td><td>0</td><td>GTTGATGATGGGCTGGGCTT</td><td>AGG</td><td>||||||||[||||||||||||]PAM</td><td>NA</td><td>Intergenic</td><td>NA</td><td>NA</td></tr>
<tr><td>T13177</td><td>TTGATGATGGGCTGGGCTTAGGG</td><td>1000</td><td></td><td></td><td></td><td></td><td></td><td></td><td></td><td></td><td></td></tr>
<tr><td>Oligo fwd</td><td>CACCTTGATGATGGGCTGGGCTTA</td><td></td><td></td><td></td><td></td><td></td><td></td><td></td><td></td><td></td><td></td></tr>
<tr><td>Oligo rev</td><td>AAACTAAGCCCAGCCCATCATCAA</td><td></td><td></td><td></td><td></td><td></td><td></td><td></td><td></td><td></td><td></td></tr>
<tr><td>Chromosome</td><td>start</td><td>end</td><td>strand</td><td>MM</td><td>target_seq</td><td>PAM</td><td>alignment</td><td>distance</td><td>position</td><td>gene name</td><td>gene id</td></tr>
<tr><td>GU305914.1</td><td>173692</td><td>173714</td><td>+</td><td>0</td><td>TTGATGATGGGCTGGGCTTA</td><td>GGG</td><td>||||||||[||||||||||||]PAM</td><td>NA</td><td>Intergenic</td><td>NA</td><td>NA</td></tr>
<tr><td>T13178</td><td>TGATGATGGGCTGGGCTTAGGGG</td><td>1000</td><td></td><td></td><td></td><td></td><td></td><td></td><td></td><td></td><td></td></tr>
<tr><td>Oligo fwd</td><td>CACCTGATGATGGGCTGGGCTTAG</td><td></td><td></td><td></td><td></td><td></td><td></td><td></td><td></td><td></td><td></td></tr>
<tr><td>Oligo rev</td><td>AAACCTAAGCCCAGCCCATCATCA</td><td></td><td></td><td></td><td></td><td></td><td></td><td></td><td></td><td></td><td></td></tr>
<tr><td>Chromosome</td><td>start</td><td>end</td><td>strand</td><td>MM</td><td>target_seq</td><td>PAM</td><td>alignment</td><td>distance</td><td>position</td><td>gene name</td><td>gene id</td></tr>
<tr><td>GU305914.1</td><td>173693</td><td>173715</td><td>+</td><td>0</td><td>TGATGATGGGCTGGGCTTAG</td><td>GGG</td><td>||||||||[||||||||||||]PAM</td><td>NA</td><td>Intergenic</td><td>NA</td><td>NA</td></tr>
<tr><td>T13179</td><td>AGGGGTGACGTTCGACGTACCGG</td><td>1000</td><td></td><td></td><td></td><td></td><td></td><td></td><td></td><td></td><td></td></tr>
<tr><td>Oligo fwd</td><td>CACCAGGGGTGACGTTCGACGTAC</td><td></td><td></td><td></td><td></td><td></td><td></td><td></td><td></td><td></td><td></td></tr>
<tr><td>Oligo rev</td><td>AAACGTACGTCGAACGTCACCCCT</td><td></td><td></td><td></td><td></td><td></td><td></td><td></td><td></td><td></td><td></td></tr>
<tr><td>Chromosome</td><td>start</td><td>end</td><td>strand</td><td>MM</td><td>target_seq</td><td>PAM</td><td>alignment</td><td>distance</td><td>position</td><td>gene name</td><td>gene id</td></tr>
<tr><td>GU305914.1</td><td>173711</td><td>173733</td><td>+</td><td>0</td><td>AGGGGTGACGTTCGACGTAC</td><td>CGG</td><td>||||||||[||||||||||||]PAM</td><td>NA</td><td>Intergenic</td><td>NA</td><td>NA</td></tr>
<tr><td>T13180</td><td>GGTGACGTTCGACGTACCGGTGG</td><td>1000</td><td></td><td></td><td></td><td></td><td></td><td></td><td></td><td></td><td></td></tr>
<tr><td>Oligo fwd</td><td>CACCGGTGACGTTCGACGTACCGG</td><td></td><td></td><td></td><td></td><td></td><td></td><td></td><td></td><td></td><td></td></tr>
<tr><td>Oligo rev</td><td>AAACCCGGTACGTCGAACGTCACC</td><td></td><td></td><td></td><td></td><td></td><td></td><td></td><td></td><td></td><td></td></tr>
<tr><td>Chromosome</td><td>start</td><td>end</td><td>strand</td><td>MM</td><td>target_seq</td><td>PAM</td><td>alignment</td><td>distance</td><td>position</td><td>gene name</td><td>gene id</td></tr>
<tr><td>GU305914.1</td><td>173714</td><td>173736</td><td>+</td><td>0</td><td>GGTGACGTTCGACGTACCGG</td><td>TGG</td><td>||||||||[||||||||||||]PAM</td><td>NA</td><td>Intergenic</td><td>NA</td><td>NA</td></tr>
<tr><td>T13181</td><td>GACGTTCGACGTACCGGTGGCGG</td><td>1000</td><td></td><td></td><td></td><td></td><td></td><td></td><td></td><td></td><td></td></tr>
<tr><td>Oligo fwd</td><td>CACCGACGTTCGACGTACCGGTGG</td><td></td><td></td><td></td><td></td><td></td><td></td><td></td><td></td><td></td><td></td></tr>
<tr><td>Oligo rev</td><td>AAACCCACCGGTACGTCGAACGTC</td><td></td><td></td><td></td><td></td><td></td><td></td><td></td><td></td><td></td><td></td></tr>
<tr><td>Chromosome</td><td>start</td><td>end</td><td>strand</td><td>MM</td><td>target_seq</td><td>PAM</td><td>alignment</td><td>distance</td><td>position</td><td>gene name</td><td>gene id</td></tr>
<tr><td>GU305914.1</td><td>173717</td><td>173739</td><td>+</td><td>0</td><td>GACGTTCGACGTACCGGTGG</td><td>CGG</td><td>||||||||[||||||||||||]PAM</td><td>NA</td><td>Intergenic</td><td>NA</td><td>NA</td></tr>
<tr><td>T13182</td><td>GCTCGACGCCAGCACCGAAGCGG</td><td>1000</td><td></td><td></td><td></td><td></td><td></td><td></td><td></td><td></td><td></td></tr>
<tr><td>Oligo fwd</td><td>CACCGCTCGACGCCAGCACCGAAG</td><td></td><td></td><td></td><td></td><td></td><td></td><td></td><td></td><td></td><td></td></tr>
<tr><td>Oligo rev</td><td>AAACCTTCGGTGCTGGCGTCGAGC</td><td></td><td></td><td></td><td></td><td></td><td></td><td></td><td></td><td></td><td></td></tr>
<tr><td>Chromosome</td><td>start</td><td>end</td><td>strand</td><td>MM</td><td>target_seq</td><td>PAM</td><td>alignment</td><td>distance</td><td>position</td><td>gene name</td><td>gene id</td></tr>
<tr><td>GU305914.1</td><td>173741</td><td>173763</td><td>+</td><td>0</td><td>GCTCGACGCCAGCACCGAAG</td><td>CGG</td><td>||||||||[||||||||||||]PAM</td><td>NA</td><td>Intergenic</td><td>NA</td><td>NA</td></tr>
<tr><td>T13183</td><td>CTCGACGCCAGCACCGAAGCGGG</td><td>1000</td><td></td><td></td><td></td><td></td><td></td><td></td><td></td><td></td><td></td></tr>
<tr><td>Oligo fwd</td><td>CACCCTCGACGCCAGCACCGAAGC</td><td></td><td></td><td></td><td></td><td></td><td></td><td></td><td></td><td></td><td></td></tr>
</table>

<table>
<tr><td>Oligo rev</td><td>AAACGCTTCGGTGCTGGCGTCGAG</td><td></td><td></td><td></td><td></td><td></td><td></td><td></td><td></td><td></td><td></td></tr>
<tr><td>Chromosome</td><td>start</td><td>end</td><td>strand</td><td>MM</td><td>target_seq</td><td>PAM</td><td>alignment</td><td>distance</td><td>position</td><td>gene name</td><td>gene id</td></tr>
<tr><td>GU305914.1</td><td>173742</td><td>173764</td><td>+</td><td>0</td><td>CTCGACGCCAGCACCGAAGC</td><td>GGG</td><td>||||||||[||||||||||||]PAM</td><td>NA</td><td>Intergenic</td><td>NA</td><td>NA</td></tr>
<tr><td>GU305914.1</td><td>143390</td><td>143412</td><td>-</td><td>4</td><td>CTCGTCGTCATCACCGACGC</td><td>AGG</td><td>||||-||-[||-||||||-||]PAM</td><td>NA</td><td>Intergenic</td><td>NA</td><td>NA</td></tr>
<tr><td></td><td></td><td></td><td></td><td></td><td></td><td></td><td></td><td></td><td></td><td></td><td></td></tr>
<tr><td>T13184</td><td>CCAGCACCGAAGCGGGATGCAGG</td><td>1000</td><td></td><td></td><td></td><td></td><td></td><td></td><td></td><td></td><td></td></tr>
<tr><td>Oligo fwd</td><td>CACCCCAGCACCGAAGCGGGATGC</td><td></td><td></td><td></td><td></td><td></td><td></td><td></td><td></td><td></td><td></td></tr>
<tr><td>Oligo rev</td><td>AAACGCATCCCGCTTCGGTGCTGG</td><td></td><td></td><td></td><td></td><td></td><td></td><td></td><td></td><td></td><td></td></tr>
<tr><td>Chromosome</td><td>start</td><td>end</td><td>strand</td><td>MM</td><td>target_seq</td><td>PAM</td><td>alignment</td><td>distance</td><td>position</td><td>gene name</td><td>gene id</td></tr>
<tr><td>GU305914.1</td><td>173749</td><td>173771</td><td>+</td><td>0</td><td>CCAGCACCGAAGCGGGATGC</td><td>AGG</td><td>||||||||[||||||||||||]PAM</td><td>NA</td><td>Intergenic</td><td>NA</td><td>NA</td></tr>
<tr><td></td><td></td><td></td><td></td><td></td><td></td><td></td><td></td><td></td><td></td><td></td><td></td></tr>
<tr><td>T13185</td><td>GCACCGAAGCGGGATGCAGGAGG</td><td>1000</td><td></td><td></td><td></td><td></td><td></td><td></td><td></td><td></td><td></td></tr>
<tr><td>Oligo fwd</td><td>CACCGCACCGAAGCGGGATGCAGG</td><td></td><td></td><td></td><td></td><td></td><td></td><td></td><td></td><td></td><td></td></tr>
<tr><td>Oligo rev</td><td>AAACCCTGCATCCCGCTTCGGTGC</td><td></td><td></td><td></td><td></td><td></td><td></td><td></td><td></td><td></td><td></td></tr>
<tr><td>Chromosome</td><td>start</td><td>end</td><td>strand</td><td>MM</td><td>target_seq</td><td>PAM</td><td>alignment</td><td>distance</td><td>position</td><td>gene name</td><td>gene id</td></tr>
<tr><td>GU305914.1</td><td>173752</td><td>173774</td><td>+</td><td>0</td><td>GCACCGAAGCGGGATGCAGG</td><td>AGG</td><td>||||||||[||||||||||||]PAM</td><td>NA</td><td>Intergenic</td><td>NA</td><td>NA</td></tr>
<tr><td></td><td></td><td></td><td></td><td></td><td></td><td></td><td></td><td></td><td></td><td></td><td></td></tr>
<tr><td>T13186</td><td>CACACCATACAGAAGACCAACGG</td><td>1000</td><td></td><td></td><td></td><td></td><td></td><td></td><td></td><td></td><td></td></tr>
<tr><td>Oligo fwd</td><td>CACCCACACCATACAGAAGACCAA</td><td></td><td></td><td></td><td></td><td></td><td></td><td></td><td></td><td></td><td></td></tr>
<tr><td>Oligo rev</td><td>AAACTTGGTCTTCTGTATGGTGTG</td><td></td><td></td><td></td><td></td><td></td><td></td><td></td><td></td><td></td><td></td></tr>
<tr><td>Chromosome</td><td>start</td><td>end</td><td>strand</td><td>MM</td><td>target_seq</td><td>PAM</td><td>alignment</td><td>distance</td><td>position</td><td>gene name</td><td>gene id</td></tr>
<tr><td>GU305914.1</td><td>173784</td><td>173806</td><td>+</td><td>0</td><td>CACACCATACAGAAGACCAA</td><td>CGG</td><td>||||||||[||||||||||||]PAM</td><td>NA</td><td>Intergenic</td><td>NA</td><td>NA</td></tr>
<tr><td></td><td></td><td></td><td></td><td></td><td></td><td></td><td></td><td></td><td></td><td></td><td></td></tr>
<tr><td>T13187</td><td>GACCCGCATCACCGTATCTTCGG</td><td>1000</td><td></td><td></td><td></td><td></td><td></td><td></td><td></td><td></td><td></td></tr>
<tr><td>Oligo fwd</td><td>CACCGACCCGCATCACCGTATCTT</td><td></td><td></td><td></td><td></td><td></td><td></td><td></td><td></td><td></td><td></td></tr>
<tr><td>Oligo rev</td><td>AAACAAGATACGGTGATGCGGGTC</td><td></td><td></td><td></td><td></td><td></td><td></td><td></td><td></td><td></td><td></td></tr>
<tr><td>Chromosome</td><td>start</td><td>end</td><td>strand</td><td>MM</td><td>target_seq</td><td>PAM</td><td>alignment</td><td>distance</td><td>position</td><td>gene name</td><td>gene id</td></tr>
<tr><td>GU305914.1</td><td>173813</td><td>173835</td><td>+</td><td>0</td><td>GACCCGCATCACCGTATCTT</td><td>CGG</td><td>||||||||[||||||||||||]PAM</td><td>NA</td><td>Intergenic</td><td>NA</td><td>NA</td></tr>
<tr><td></td><td></td><td></td><td></td><td></td><td></td><td></td><td></td><td></td><td></td><td></td><td></td></tr>
<tr><td>T13188</td><td>CCGCATCACCGTATCTTCGGCGG</td><td>1000</td><td></td><td></td><td></td><td></td><td></td><td></td><td></td><td></td><td></td></tr>
<tr><td>Oligo fwd</td><td>CACCCCGCATCACCGTATCTTCGG</td><td></td><td></td><td></td><td></td><td></td><td></td><td></td><td></td><td></td><td></td></tr>
<tr><td>Oligo rev</td><td>AAACCCGAAGATACGGTGATGCGG</td><td></td><td></td><td></td><td></td><td></td><td></td><td></td><td></td><td></td><td></td></tr>
<tr><td>Chromosome</td><td>start</td><td>end</td><td>strand</td><td>MM</td><td>target_seq</td><td>PAM</td><td>alignment</td><td>distance</td><td>position</td><td>gene name</td><td>gene id</td></tr>
<tr><td>GU305914.1</td><td>173816</td><td>173838</td><td>+</td><td>0</td><td>CCGCATCACCGTATCTTCGG</td><td>CGG</td><td>||||||||[||||||||||||]PAM</td><td>NA</td><td>Intergenic</td><td>NA</td><td>NA</td></tr>
<tr><td></td><td></td><td></td><td></td><td></td><td></td><td></td><td></td><td></td><td></td><td></td><td></td></tr>
<tr><td>T13189</td><td>CGCATCACCGTATCTTCGGCGGG</td><td>1000</td><td></td><td></td><td></td><td></td><td></td><td></td><td></td><td></td><td></td></tr>
<tr><td>Oligo fwd</td><td>CACCCGCATCACCGTATCTTCGGC</td><td></td><td></td><td></td><td></td><td></td><td></td><td></td><td></td><td></td><td></td></tr>
<tr><td>Oligo rev</td><td>AAACGCCGAAGATACGGTGATGCG</td><td></td><td></td><td></td><td></td><td></td><td></td><td></td><td></td><td></td><td></td></tr>
<tr><td>Chromosome</td><td>start</td><td>end</td><td>strand</td><td>MM</td><td>target_seq</td><td>PAM</td><td>alignment</td><td>distance</td><td>position</td><td>gene name</td><td>gene id</td></tr>
<tr><td>GU305914.1</td><td>173817</td><td>173839</td><td>+</td><td>0</td><td>CGCATCACCGTATCTTCGGC</td><td>GGG</td><td>||||||||[||||||||||||]PAM</td><td>NA</td><td>Intergenic</td><td>NA</td><td>NA</td></tr>
<tr><td></td><td></td><td></td><td></td><td></td><td></td><td></td><td></td><td></td><td></td><td></td><td></td></tr>
<tr><td>T13190</td><td>GCATCACCGTATCTTCGGCGGGG</td><td>1000</td><td></td><td></td><td></td><td></td><td></td><td></td><td></td><td></td><td></td></tr>
<tr><td>Oligo fwd</td><td>CACCGCATCACCGTATCTTCGGCG</td><td></td><td></td><td></td><td></td><td></td><td></td><td></td><td></td><td></td><td></td></tr>
<tr><td>Oligo rev</td><td>AAACCGCCGAAGATACGGTGATGC</td><td></td><td></td><td></td><td></td><td></td><td></td><td></td><td></td><td></td><td></td></tr>
<tr><td>Chromosome</td><td>start</td><td>end</td><td>strand</td><td>MM</td><td>target_seq</td><td>PAM</td><td>alignment</td><td>distance</td><td>position</td><td>gene name</td><td>gene id</td></tr>
<tr><td>GU305914.1</td><td>173818</td><td>173840</td><td>+</td><td>0</td><td>GCATCACCGTATCTTCGGCG</td><td>GGG</td><td>||||||||[||||||||||||]PAM</td><td>NA</td><td>Intergenic</td><td>NA</td><td>NA</td></tr>
<tr><td></td><td></td><td></td><td></td><td></td><td></td><td></td><td></td><td></td><td></td><td></td><td></td></tr>
<tr><td>T13191</td><td>TCACCGTATCTTCGGCGGGGAGG</td><td>1000</td><td></td><td></td><td></td><td></td><td></td><td></td><td></td><td></td><td></td></tr>
<tr><td>Oligo fwd</td><td>CACCTCACCGTATCTTCGGCGGGG</td><td></td><td></td><td></td><td></td><td></td><td></td><td></td><td></td><td></td><td></td></tr>
<tr><td>Oligo rev</td><td>AAACCCCCGCCGAAGATACGGTGA</td><td></td><td></td><td></td><td></td><td></td><td></td><td></td><td></td><td></td><td></td></tr>
<tr><td>Chromosome</td><td>start</td><td>end</td><td>strand</td><td>MM</td><td>target_seq</td><td>PAM</td><td>alignment</td><td>distance</td><td>position</td><td>gene name</td><td>gene id</td></tr>
<tr><td>GU305914.1</td><td>173821</td><td>173843</td><td>+</td><td>0</td><td>TCACCGTATCTTCGGCGGGG</td><td>AGG</td><td>||||||||[||||||||||||]PAM</td><td>NA</td><td>Intergenic</td><td>NA</td><td>NA</td></tr>
<tr><td></td><td></td><td></td><td></td><td></td><td></td><td></td><td></td><td></td><td></td><td></td><td></td></tr>
<tr><td>T13192</td><td>CGTATCTTCGGCGGGGAGGTCGG</td><td>1000</td><td></td><td></td><td></td><td></td><td></td><td></td><td></td><td></td><td></td></tr>
<tr><td>Oligo fwd</td><td>CACCCGTATCTTCGGCGGGGAGGT</td><td></td><td></td><td></td><td></td><td></td><td></td><td></td><td></td><td></td><td></td></tr>
<tr><td>Oligo rev</td><td>AAACACCTCCCCGCCGAAGATACG</td><td></td><td></td><td></td><td></td><td></td><td></td><td></td><td></td><td></td><td></td></tr>
<tr><td>Chromosome</td><td>start</td><td>end</td><td>strand</td><td>MM</td><td>target_seq</td><td>PAM</td><td>alignment</td><td>distance</td><td>position</td><td>gene name</td><td>gene id</td></tr>
<tr><td>GU305914.1</td><td>173825</td><td>173847</td><td>+</td><td>0</td><td>CGTATCTTCGGCGGGGAGGT</td><td>CGG</td><td>||||||||[||||||||||||]PAM</td><td>NA</td><td>Intergenic</td><td>NA</td><td>NA</td></tr>
<tr><td></td><td></td><td></td><td></td><td></td><td></td><td></td><td></td><td></td><td></td><td></td><td></td></tr>
<tr><td>T13193</td><td>ACGTCGTTTCGTTAAATAGATGG</td><td>1000</td><td></td><td></td><td></td><td></td><td></td><td></td><td></td><td></td><td></td></tr>
<tr><td>Oligo fwd</td><td>CACCACGTCGTTTCGTTAAATAGA</td><td></td><td></td><td></td><td></td><td></td><td></td><td></td><td></td><td></td><td></td></tr>
<tr><td>Oligo rev</td><td>AAACTCTATTTAACGAAACGACGT</td><td></td><td></td><td></td><td></td><td></td><td></td><td></td><td></td><td></td><td></td></tr>
<tr><td>Chromosome</td><td>start</td><td>end</td><td>strand</td><td>MM</td><td>target_seq</td><td>PAM</td><td>alignment</td><td>distance</td><td>position</td><td>gene name</td><td>gene id</td></tr>
<tr><td>GU305914.1</td><td>173854</td><td>173876</td><td>+</td><td>0</td><td>ACGTCGTTTCGTTAAATAGA</td><td>TGG</td><td>||||||||[||||||||||||]PAM</td><td>NA</td><td>Intergenic</td><td>NA</td><td>NA</td></tr>
<tr><td></td><td></td><td></td><td></td><td></td><td></td><td></td><td></td><td></td><td></td><td></td><td></td></tr>
<tr><td>T13194</td><td>CGCGTCCGACCGAATTGTGTCGG</td><td>1000</td><td></td><td></td><td></td><td></td><td></td><td></td><td></td><td></td><td></td></tr>
<tr><td>Oligo fwd</td><td>CACCCGCGTCCGACCGAATTGTGT</td><td></td><td></td><td></td><td></td><td></td><td></td><td></td><td></td><td></td><td></td></tr>
<tr><td>Oligo rev</td><td>AAACACACAATTCGGTCGGACGCG</td><td></td><td></td><td></td><td></td><td></td><td></td><td></td><td></td><td></td><td></td></tr>
<tr><td>Chromosome</td><td>start</td><td>end</td><td>strand</td><td>MM</td><td>target_seq</td><td>PAM</td><td>alignment</td><td>distance</td><td>position</td><td>gene name</td><td>gene id</td></tr>
<tr><td>GU305914.1</td><td>173912</td><td>173934</td><td>+</td><td>0</td><td>CGCGTCCGACCGAATTGTGT</td><td>CGG</td><td>||||||||[||||||||||||]PAM</td><td>NA</td><td>Intergenic</td><td>NA</td><td>NA</td></tr>
<tr><td></td><td></td><td></td><td></td><td></td><td></td><td></td><td></td><td></td><td></td><td></td><td></td></tr>
<tr><td>T13195</td><td>TCCGACCGAATTGTGTCGGATGG</td><td>1000</td><td></td><td></td><td></td><td></td><td></td><td></td><td></td><td></td><td></td></tr>
<tr><td>Oligo fwd</td><td>CACCTCCGACCGAATTGTGTCGGA</td><td></td><td></td><td></td><td></td><td></td><td></td><td></td><td></td><td></td><td></td></tr>
<tr><td>Oligo rev</td><td>AAACTCCGACACAATTCGGTCGGA</td><td></td><td></td><td></td><td></td><td></td><td></td><td></td><td></td><td></td><td></td></tr>
<tr><td>Chromosome</td><td>start</td><td>end</td><td>strand</td><td>MM</td><td>target_seq</td><td>PAM</td><td>alignment</td><td>distance</td><td>position</td><td>gene name</td><td>gene id</td></tr>
<tr><td>GU305914.1</td><td>173916</td><td>173938</td><td>+</td><td>0</td><td>TCCGACCGAATTGTGTCGGA</td><td>TGG</td><td>||||||||[||||||||||||]PAM</td><td>NA</td><td>Intergenic</td><td>NA</td><td>NA</td></tr>
<tr><td></td><td></td><td></td><td></td><td></td><td></td><td></td><td></td><td></td><td></td><td></td><td></td></tr>
<tr><td>T13196</td><td>GTCGGATGGTCTCATCTCGACGG</td><td>1000</td><td></td><td></td><td></td><td></td><td></td><td></td><td></td><td></td><td></td></tr>
<tr><td>Oligo fwd</td><td>CACCGTCGGATGGTCTCATCTCGA</td><td></td><td></td><td></td><td></td><td></td><td></td><td></td><td></td><td></td><td></td></tr>
<tr><td>Oligo rev</td><td>AAACTCGAGATGAGACCATCCGAC</td><td></td><td></td><td></td><td></td><td></td><td></td><td></td><td></td><td></td><td></td></tr>
<tr><td>Chromosome</td><td>start</td><td>end</td><td>strand</td><td>MM</td><td>target_seq</td><td>PAM</td><td>alignment</td><td>distance</td><td>position</td><td>gene name</td><td>gene id</td></tr>
<tr><td>GU305914.1</td><td>173930</td><td>173952</td><td>+</td><td>0</td><td>GTCGGATGGTCTCATCTCGA</td><td>CGG</td><td>||||||||[||||||||||||]PAM</td><td>NA</td><td>Intergenic</td><td>NA</td><td>NA</td></tr>
<tr><td></td><td></td><td></td><td></td><td></td><td></td><td></td><td></td><td></td><td></td><td></td><td></td></tr>
<tr><td>T13197</td><td>TCTCATCTCGACGGAAGAGATGG</td><td>1000</td><td></td><td></td><td></td><td></td><td></td><td></td><td></td><td></td><td></td></tr>
<tr><td>Oligo fwd</td><td>CACCTCTCATCTCGACGGAAGAGA</td><td></td><td></td><td></td><td></td><td></td><td></td><td></td><td></td><td></td><td></td></tr>
<tr><td>Oligo rev</td><td>AAACTCTCTTCCGTCGAGATGAGA</td><td></td><td></td><td></td><td></td><td></td><td></td><td></td><td></td><td></td><td></td></tr>
<tr><td>Chromosome</td><td>start</td><td>end</td><td>strand</td><td>MM</td><td>target_seq</td><td>PAM</td><td>alignment</td><td>distance</td><td>position</td><td>gene name</td><td>gene id</td></tr>
<tr><td>GU305914.1</td><td>173939</td><td>173961</td><td>+</td><td>0</td><td>TCTCATCTCGACGGAAGAGA</td><td>TGG</td><td>||||||||[||||||||||||]PAM</td><td>NA</td><td>Intergenic</td><td>NA</td><td>NA</td></tr>
</table>

<table>
<tr><td>T13198</td><td>TCGACGGAAGAGATGGTCTCCGG</td><td>1000</td><td></td><td></td><td></td><td></td><td></td><td></td><td></td><td></td><td></td></tr>
<tr><td>Oligo fwd</td><td>CACCTCGACGGAAGAGATGGTCTC</td><td></td><td></td><td></td><td></td><td></td><td></td><td></td><td></td><td></td><td></td></tr>
<tr><td>Oligo rev</td><td>AAACGAGACCATCTCTTCCGTCGA</td><td></td><td></td><td></td><td></td><td></td><td></td><td></td><td></td><td></td><td></td></tr>
<tr><td>Chromosome</td><td>start</td><td>end</td><td>strand</td><td>MM</td><td>target_seq</td><td>PAM</td><td>alignment</td><td>distance</td><td>position</td><td>gene name</td><td>gene id</td></tr>
<tr><td>GU305914.1</td><td>173946</td><td>173968</td><td>+</td><td>0</td><td>TCGACGGAAGAGATGGTCTC</td><td>CGG</td><td>||||||||[||||||||||||]PAM</td><td>NA</td><td>Intergenic</td><td>NA</td><td>NA</td></tr>
<tr><td>T13199</td><td>GACATAATCCTGTCGACGAGTGG</td><td>1000</td><td></td><td></td><td></td><td></td><td></td><td></td><td></td><td></td><td></td></tr>
<tr><td>Oligo fwd</td><td>CACCGACATAATCCTGTCGACGAG</td><td></td><td></td><td></td><td></td><td></td><td></td><td></td><td></td><td></td><td></td></tr>
<tr><td>Oligo rev</td><td>AAACCTCGTCGACAGGATTATGTC</td><td></td><td></td><td></td><td></td><td></td><td></td><td></td><td></td><td></td><td></td></tr>
<tr><td>Chromosome</td><td>start</td><td>end</td><td>strand</td><td>MM</td><td>target_seq</td><td>PAM</td><td>alignment</td><td>distance</td><td>position</td><td>gene name</td><td>gene id</td></tr>
<tr><td>GU305914.1</td><td>173988</td><td>174010</td><td>+</td><td>0</td><td>GACATAATCCTGTCGACGAG</td><td>TGG</td><td>||||||||[||||||||||||]PAM</td><td>NA</td><td>Intergenic</td><td>NA</td><td>NA</td></tr>
<tr><td>T13200</td><td>ACATAATCCTGTCGACGAGTGGG</td><td>1000</td><td></td><td></td><td></td><td></td><td></td><td></td><td></td><td></td><td></td></tr>
<tr><td>Oligo fwd</td><td>CACCACATAATCCTGTCGACGAGT</td><td></td><td></td><td></td><td></td><td></td><td></td><td></td><td></td><td></td><td></td></tr>
<tr><td>Oligo rev</td><td>AAACACTCGTCGACAGGATTATGT</td><td></td><td></td><td></td><td></td><td></td><td></td><td></td><td></td><td></td><td></td></tr>
<tr><td>Chromosome</td><td>start</td><td>end</td><td>strand</td><td>MM</td><td>target_seq</td><td>PAM</td><td>alignment</td><td>distance</td><td>position</td><td>gene name</td><td>gene id</td></tr>
<tr><td>GU305914.1</td><td>173989</td><td>174011</td><td>+</td><td>0</td><td>ACATAATCCTGTCGACGAGT</td><td>GGG</td><td>||||||||[||||||||||||]PAM</td><td>NA</td><td>Intergenic</td><td>NA</td><td>NA</td></tr>
<tr><td>T13201</td><td>CATAATCCTGTCGACGAGTGGGG</td><td>1000</td><td></td><td></td><td></td><td></td><td></td><td></td><td></td><td></td><td></td></tr>
<tr><td>Oligo fwd</td><td>CACCCATAATCCTGTCGACGAGTG</td><td></td><td></td><td></td><td></td><td></td><td></td><td></td><td></td><td></td><td></td></tr>
<tr><td>Oligo rev</td><td>AAACCACTCGTCGACAGGATTATG</td><td></td><td></td><td></td><td></td><td></td><td></td><td></td><td></td><td></td><td></td></tr>
<tr><td>Chromosome</td><td>start</td><td>end</td><td>strand</td><td>MM</td><td>target_seq</td><td>PAM</td><td>alignment</td><td>distance</td><td>position</td><td>gene name</td><td>gene id</td></tr>
<tr><td>GU305914.1</td><td>173990</td><td>174012</td><td>+</td><td>0</td><td>CATAATCCTGTCGACGAGTG</td><td>GGG</td><td>||||||||[||||||||||||]PAM</td><td>NA</td><td>Intergenic</td><td>NA</td><td>NA</td></tr>
<tr><td>T13202</td><td>ATAATCCTGTCGACGAGTGGGGG</td><td>1000</td><td></td><td></td><td></td><td></td><td></td><td></td><td></td><td></td><td></td></tr>
<tr><td>Oligo fwd</td><td>CACCATAATCCTGTCGACGAGTGG</td><td></td><td></td><td></td><td></td><td></td><td></td><td></td><td></td><td></td><td></td></tr>
<tr><td>Oligo rev</td><td>AAACCCACTCGTCGACAGGATTAT</td><td></td><td></td><td></td><td></td><td></td><td></td><td></td><td></td><td></td><td></td></tr>
<tr><td>Chromosome</td><td>start</td><td>end</td><td>strand</td><td>MM</td><td>target_seq</td><td>PAM</td><td>alignment</td><td>distance</td><td>position</td><td>gene name</td><td>gene id</td></tr>
<tr><td>GU305914.1</td><td>173991</td><td>174013</td><td>+</td><td>0</td><td>ATAATCCTGTCGACGAGTGG</td><td>GGG</td><td>||||||||[||||||||||||]PAM</td><td>NA</td><td>Intergenic</td><td>NA</td><td>NA</td></tr>
<tr><td>T13203</td><td>TAACCCTTTCGTCCGAATGACGG</td><td>1000</td><td></td><td></td><td></td><td></td><td></td><td></td><td></td><td></td><td></td></tr>
<tr><td>Oligo fwd</td><td>CACCTAACCCTTTCGTCCGAATGA</td><td></td><td></td><td></td><td></td><td></td><td></td><td></td><td></td><td></td><td></td></tr>
<tr><td>Oligo rev</td><td>AAACTCATTCGGACGAAAGGGTTA</td><td></td><td></td><td></td><td></td><td></td><td></td><td></td><td></td><td></td><td></td></tr>
<tr><td>Chromosome</td><td>start</td><td>end</td><td>strand</td><td>MM</td><td>target_seq</td><td>PAM</td><td>alignment</td><td>distance</td><td>position</td><td>gene name</td><td>gene id</td></tr>
<tr><td>GU305914.1</td><td>174014</td><td>174036</td><td>+</td><td>0</td><td>TAACCCTTTCGTCCGAATGA</td><td>CGG</td><td>||||||||[||||||||||||]PAM</td><td>NA</td><td>Intergenic</td><td>NA</td><td>NA</td></tr>
<tr><td>T13204</td><td>AGCCTAGTTTTTCGACGATCCGG</td><td>1000</td><td></td><td></td><td></td><td></td><td></td><td></td><td></td><td></td><td></td></tr>
<tr><td>Oligo fwd</td><td>CACCAGCCTAGTTTTTCGACGATC</td><td></td><td></td><td></td><td></td><td></td><td></td><td></td><td></td><td></td><td></td></tr>
<tr><td>Oligo rev</td><td>AAACGATCGTCGAAAAACTAGGCT</td><td></td><td></td><td></td><td></td><td></td><td></td><td></td><td></td><td></td><td></td></tr>
<tr><td>Chromosome</td><td>start</td><td>end</td><td>strand</td><td>MM</td><td>target_seq</td><td>PAM</td><td>alignment</td><td>distance</td><td>position</td><td>gene name</td><td>gene id</td></tr>
<tr><td>GU305914.1</td><td>174047</td><td>174069</td><td>+</td><td>0</td><td>AGCCTAGTTTTTCGACGATC</td><td>CGG</td><td>||||||||[||||||||||||]PAM</td><td>NA</td><td>Intergenic</td><td>NA</td><td>NA</td></tr>
<tr><td>T13205</td><td>TTCGACGATCCGGTCGACCGAGG</td><td>1000</td><td></td><td></td><td></td><td></td><td></td><td></td><td></td><td></td><td></td></tr>
<tr><td>Oligo fwd</td><td>CACCTTCGACGATCCGGTCGACCG</td><td></td><td></td><td></td><td></td><td></td><td></td><td></td><td></td><td></td><td></td></tr>
<tr><td>Oligo rev</td><td>AAACCGGTCGACCGGATCGTCGAA</td><td></td><td></td><td></td><td></td><td></td><td></td><td></td><td></td><td></td><td></td></tr>
<tr><td>Chromosome</td><td>start</td><td>end</td><td>strand</td><td>MM</td><td>target_seq</td><td>PAM</td><td>alignment</td><td>distance</td><td>position</td><td>gene name</td><td>gene id</td></tr>
<tr><td>GU305914.1</td><td>174057</td><td>174079</td><td>+</td><td>0</td><td>TTCGACGATCCGGTCGACCG</td><td>AGG</td><td>||||||||[||||||||||||]PAM</td><td>NA</td><td>Intergenic</td><td>NA</td><td>NA</td></tr>
<tr><td>T13206</td><td>TCGACGATCCGGTCGACCGAGGG</td><td>1000</td><td></td><td></td><td></td><td></td><td></td><td></td><td></td><td></td><td></td></tr>
<tr><td>Oligo fwd</td><td>CACCTCGACGATCCGGTCGACCGA</td><td></td><td></td><td></td><td></td><td></td><td></td><td></td><td></td><td></td><td></td></tr>
<tr><td>Oligo rev</td><td>AAACTCGGTCGACCGGATCGTCGA</td><td></td><td></td><td></td><td></td><td></td><td></td><td></td><td></td><td></td><td></td></tr>
<tr><td>Chromosome</td><td>start</td><td>end</td><td>strand</td><td>MM</td><td>target_seq</td><td>PAM</td><td>alignment</td><td>distance</td><td>position</td><td>gene name</td><td>gene id</td></tr>
<tr><td>GU305914.1</td><td>174058</td><td>174080</td><td>+</td><td>0</td><td>TCGACGATCCGGTCGACCGA</td><td>GGG</td><td>||||||||[||||||||||||]PAM</td><td>NA</td><td>Intergenic</td><td>NA</td><td>NA</td></tr>
<tr><td>T13207</td><td>TCGACCGAGGGAAACTGCCTAGG</td><td>1000</td><td></td><td></td><td></td><td></td><td></td><td></td><td></td><td></td><td></td></tr>
<tr><td>Oligo fwd</td><td>CACCTCGACCGAGGGAAACTGCCT</td><td></td><td></td><td></td><td></td><td></td><td></td><td></td><td></td><td></td><td></td></tr>
<tr><td>Oligo rev</td><td>AAACAGGCAGTTTCCCTCGGTCGA</td><td></td><td></td><td></td><td></td><td></td><td></td><td></td><td></td><td></td><td></td></tr>
<tr><td>Chromosome</td><td>start</td><td>end</td><td>strand</td><td>MM</td><td>target_seq</td><td>PAM</td><td>alignment</td><td>distance</td><td>position</td><td>gene name</td><td>gene id</td></tr>
<tr><td>GU305914.1</td><td>174070</td><td>174092</td><td>+</td><td>0</td><td>TCGACCGAGGGAAACTGCCT</td><td>AGG</td><td>||||||||[||||||||||||]PAM</td><td>NA</td><td>Intergenic</td><td>NA</td><td>NA</td></tr>
<tr><td>T13208</td><td>GGAAACTGCCTAGGATCCTATGG</td><td>1000</td><td></td><td></td><td></td><td></td><td></td><td></td><td></td><td></td><td></td></tr>
<tr><td>Oligo fwd</td><td>CACCGGAAACTGCCTAGGATCCTA</td><td></td><td></td><td></td><td></td><td></td><td></td><td></td><td></td><td></td><td></td></tr>
<tr><td>Oligo rev</td><td>AAACTAGGATCCTAGGCAGTTTCC</td><td></td><td></td><td></td><td></td><td></td><td></td><td></td><td></td><td></td><td></td></tr>
<tr><td>Chromosome</td><td>start</td><td>end</td><td>strand</td><td>MM</td><td>target_seq</td><td>PAM</td><td>alignment</td><td>distance</td><td>position</td><td>gene name</td><td>gene id</td></tr>
<tr><td>GU305914.1</td><td>174079</td><td>174101</td><td>+</td><td>0</td><td>GGAAACTGCCTAGGATCCTA</td><td>TGG</td><td>||||||||[||||||||||||]PAM</td><td>NA</td><td>Intergenic</td><td>NA</td><td>NA</td></tr>
<tr><td>T13209</td><td>GAAACTGCCTAGGATCCTATGGG</td><td>1000</td><td></td><td></td><td></td><td></td><td></td><td></td><td></td><td></td><td></td></tr>
<tr><td>Oligo fwd</td><td>CACCGAAACTGCCTAGGATCCTAT</td><td></td><td></td><td></td><td></td><td></td><td></td><td></td><td></td><td></td><td></td></tr>
<tr><td>Oligo rev</td><td>AAACATAGGATCCTAGGCAGTTTC</td><td></td><td></td><td></td><td></td><td></td><td></td><td></td><td></td><td></td><td></td></tr>
<tr><td>Chromosome</td><td>start</td><td>end</td><td>strand</td><td>MM</td><td>target_seq</td><td>PAM</td><td>alignment</td><td>distance</td><td>position</td><td>gene name</td><td>gene id</td></tr>
<tr><td>GU305914.1</td><td>174080</td><td>174102</td><td>+</td><td>0</td><td>GAAACTGCCTAGGATCCTAT</td><td>GGG</td><td>||||||||[||||||||||||]PAM</td><td>NA</td><td>Intergenic</td><td>NA</td><td>NA</td></tr>
<tr><td>T13210</td><td>CTAGGATCCTATGGGTTTCTCGG</td><td>1000</td><td></td><td></td><td></td><td></td><td></td><td></td><td></td><td></td><td></td></tr>
<tr><td>Oligo fwd</td><td>CACCCTAGGATCCTATGGGTTTCT</td><td></td><td></td><td></td><td></td><td></td><td></td><td></td><td></td><td></td><td></td></tr>
<tr><td>Oligo rev</td><td>AAACAGAAACCCATAGGATCCTAG</td><td></td><td></td><td></td><td></td><td></td><td></td><td></td><td></td><td></td><td></td></tr>
<tr><td>Chromosome</td><td>start</td><td>end</td><td>strand</td><td>MM</td><td>target_seq</td><td>PAM</td><td>alignment</td><td>distance</td><td>position</td><td>gene name</td><td>gene id</td></tr>
<tr><td>GU305914.1</td><td>174088</td><td>174110</td><td>+</td><td>0</td><td>CTAGGATCCTATGGGTTTCT</td><td>CGG</td><td>||||||||[||||||||||||]PAM</td><td>NA</td><td>Intergenic</td><td>NA</td><td>NA</td></tr>
<tr><td>T13211</td><td>CCTATGGGTTTCTCGGAAATCGG</td><td>1000</td><td></td><td></td><td></td><td></td><td></td><td></td><td></td><td></td><td></td></tr>
<tr><td>Oligo fwd</td><td>CACCCCTATGGGTTTCTCGGAAAT</td><td></td><td></td><td></td><td></td><td></td><td></td><td></td><td></td><td></td><td></td></tr>
<tr><td>Oligo rev</td><td>AAACATTTCCGAGAAACCCATAGG</td><td></td><td></td><td></td><td></td><td></td><td></td><td></td><td></td><td></td><td></td></tr>
<tr><td>Chromosome</td><td>start</td><td>end</td><td>strand</td><td>MM</td><td>target_seq</td><td>PAM</td><td>alignment</td><td>distance</td><td>position</td><td>gene name</td><td>gene id</td></tr>
<tr><td>GU305914.1</td><td>174095</td><td>174117</td><td>+</td><td>0</td><td>CCTATGGGTTTCTCGGAAAT</td><td>CGG</td><td>||||||||[||||||||||||]PAM</td><td>NA</td><td>Intergenic</td><td>NA</td><td>NA</td></tr>
<tr><td>T13212</td><td>GGAGACGTTTCTCGAAAACACGG</td><td>1000</td><td></td><td></td><td></td><td></td><td></td><td></td><td></td><td></td><td></td></tr>
<tr><td>Oligo fwd</td><td>CACCGGAGACGTTTCTCGAAAACA</td><td></td><td></td><td></td><td></td><td></td><td></td><td></td><td></td><td></td><td></td></tr>
<tr><td>Oligo rev</td><td>AAACTGTTTTCGAGAAACGTCTCC</td><td></td><td></td><td></td><td></td><td></td><td></td><td></td><td></td><td></td><td></td></tr>
<tr><td>Chromosome</td><td>start</td><td>end</td><td>strand</td><td>MM</td><td>target_seq</td><td>PAM</td><td>alignment</td><td>distance</td><td>position</td><td>gene name</td><td>gene id</td></tr>
<tr><td>GU305914.1</td><td>174116</td><td>174138</td><td>+</td><td>0</td><td>GGAGACGTTTCTCGAAAACA</td><td>CGG</td><td>||||||||[||||||||||||]PAM</td><td>NA</td><td>Intergenic</td><td>NA</td><td>NA</td></tr>
</table>

| | | | | | | | | | | | |
|---|---|---|---|---|---|---|---|---|---|---|---|
| T13213 | CTCGAAAACACGGTAAGATGAGG | 1000 | | | | | | | | | |
| Oligo fwd | CACCCTCGAAAACACGGTAAGATG | | | | | | | | | | |
| Oligo rev | AAACCATCTTACCGTGTTTTCGAG | | | | | | | | | | |
| Chromosome | start | end | strand | MM | target_seq | PAM | alignment | distance | position | gene name | gene id |
| GU305914.1 | 174126 | 174148 | + | 0 | CTCGAAAACACGGTAAGATG | AGG | \|\|\|\|\|\|\|\|[\|\|\|\|\|\|\|\|\|\|\|\|]PAM | NA | Intergenic | NA | NA |
| T13214 | GCGCCGTCACGCCGTGTGTGTGG | 1000 | | | | | | | | | |
| Oligo fwd | CACCGCGCCGTCACGCCGTGTGTG | | | | | | | | | | |
| Oligo rev | AAACCACACACGGCGTGACGGCGC | | | | | | | | | | |
| Chromosome | start | end | strand | MM | target_seq | PAM | alignment | distance | position | gene name | gene id |
| GU305914.1 | 174148 | 174170 | + | 0 | GCGCCGTCACGCCGTGTGTG | TGG | \|\|\|\|\|\|\|\|[\|\|\|\|\|\|\|\|\|\|\|\|]PAM | NA | Intergenic | NA | NA |
| T13215 | CGCCGTCACGCCGTGTGTGTGGG | 1000 | | | | | | | | | |
| Oligo fwd | CACCCGCCGTCACGCCGTGTGTGT | | | | | | | | | | |
| Oligo rev | AAACACACACACGGCGTGACGGCG | | | | | | | | | | |
| Chromosome | start | end | strand | MM | target_seq | PAM | alignment | distance | position | gene name | gene id |
| GU305914.1 | 174149 | 174171 | + | 0 | CGCCGTCACGCCGTGTGTGT | GGG | \|\|\|\|\|\|\|\|[\|\|\|\|\|\|\|\|\|\|\|\|]PAM | NA | Intergenic | NA | NA |
| T13216 | GCCGCCGCCGCCTCAACACTCGG | 1000 | | | | | | | | | |
| Oligo fwd | CACCGCCGCCGCCGCCTCAACACT | | | | | | | | | | |
| Oligo rev | AAACAGTGTTGAGGCGGCGGCGGC | | | | | | | | | | |
| Chromosome | start | end | strand | MM | target_seq | PAM | alignment | distance | position | gene name | gene id |
| GU305914.1 | 174171 | 174193 | + | 0 | GCCGCCGCCGCCTCAACACT | CGG | \|\|\|\|\|\|\|\|[\|\|\|\|\|\|\|\|\|\|\|\|]PAM | NA | Intergenic | NA | NA |
| T13217 | GGTAGATCCTATACACCGCGCGG | 1000 | | | | | | | | | |
| Oligo fwd | CACCGGTAGATCCTATACACCGCG | | | | | | | | | | |
| Oligo rev | AAACCGCGGTGTATAGGATCTACC | | | | | | | | | | |
| Chromosome | start | end | strand | MM | target_seq | PAM | alignment | distance | position | gene name | gene id |
| GU305914.1 | 174192 | 174214 | + | 0 | GGTAGATCCTATACACCGCG | CGG | \|\|\|\|\|\|\|\|[\|\|\|\|\|\|\|\|\|\|\|\|]PAM | NA | Intergenic | NA | NA |
| T13218 | CGCCAACGCGATGATCGCGACGG | 1000 | | | | | | | | | |
| Oligo fwd | CACCCGCCAACGCGATGATCGCGA | | | | | | | | | | |
| Oligo rev | AAACTCGCGATCATCGCGTTGGCG | | | | | | | | | | |
| Chromosome | start | end | strand | MM | target_seq | PAM | alignment | distance | position | gene name | gene id |
| GU305914.1 | 174244 | 174266 | + | 0 | CGCCAACGCGATGATCGCGA | CGG | \|\|\|\|\|\|\|\|[\|\|\|\|\|\|\|\|\|\|\|\|]PAM | NA | Intergenic | NA | NA |
| GU305914.1 | 174246 | 174268 | - | 4 | CGCCGTCGCGATCATCGCGT | TGG | \|\|\|\|--\|\|[\|\|\|\|-\|\|\|\|\|\|-]PAM | NA | Intergenic | NA | NA |
| T13219 | CGACGGCGACGATGACGAACAGG | 1000 | | | | | | | | | |
| Oligo fwd | CACCCGACGGCGACGATGACGAAC | | | | | | | | | | |
| Oligo rev | AAACGTTCGTCATCGTCGCCGTCG | | | | | | | | | | |
| Chromosome | start | end | strand | MM | target_seq | PAM | alignment | distance | position | gene name | gene id |
| GU305914.1 | 174261 | 174283 | + | 0 | CGACGGCGACGATGACGAAC | AGG | \|\|\|\|\|\|\|\|[\|\|\|\|\|\|\|\|\|\|\|\|]PAM | NA | Intergenic | NA | NA |
| GU305914.1 | 68313 | 68335 | + | 4 | CGGCGGTGACGAGGACGGAC | CGG | \|\|-\|\|\|-\|[\|\|\|\|-\|\|\|\|-\|\|]PAM | NA | Intergenic | NA | NA |
| GU305914.1 | 96811 | 96833 | - | 3 | CGCCGGCGACGAGGACGAAG | GGG | \|\|-\|\|\|\|\|[\|\|\|\|-\|\|\|\|\|\|-]PAM | NA | Intergenic | NA | NA |
| GU305914.1 | 114978 | 115000 | + | 4 | CGACGATGACGATGACGACG | AGG | \|\|\|\|\|--\|[\|\|\|\|\|\|\|\|\|\|--]PAM | NA | Intergenic | NA | NA |
| T13220 | ACGATGACGAACAGGAGCTCCGG | 1000 | | | | | | | | | |
| Oligo fwd | CACCACGATGACGAACAGGAGCTC | | | | | | | | | | |
| Oligo rev | AAACGAGCTCCTGTTCGTCATCGT | | | | | | | | | | |
| Chromosome | start | end | strand | MM | target_seq | PAM | alignment | distance | position | gene name | gene id |
| GU305914.1 | 174269 | 174291 | + | 0 | ACGATGACGAACAGGAGCTC | CGG | \|\|\|\|\|\|\|\|[\|\|\|\|\|\|\|\|\|\|\|\|]PAM | NA | Intergenic | NA | NA |
| T13221 | CCGGCAGCATGACGACGAACCGG | 1000 | | | | | | | | | |
| Oligo fwd | CACCCCGGCAGCATGACGACGAAC | | | | | | | | | | |
| Oligo rev | AAACGTTCGTCGTCATGCTGCCGG | | | | | | | | | | |
| Chromosome | start | end | strand | MM | target_seq | PAM | alignment | distance | position | gene name | gene id |
| GU305914.1 | 174288 | 174310 | + | 0 | CCGGCAGCATGACGACGAAC | CGG | \|\|\|\|\|\|\|\|[\|\|\|\|\|\|\|\|\|\|\|\|]PAM | NA | Intergenic | NA | NA |
| T13222 | CATGACGACGAACCGGACGATGG | 1000 | | | | | | | | | |
| Oligo fwd | CACCCATGACGACGAACCGGACGA | | | | | | | | | | |
| Oligo rev | AAACTCGTCCGGTTCGTCGTCATG | | | | | | | | | | |
| Chromosome | start | end | strand | MM | target_seq | PAM | alignment | distance | position | gene name | gene id |
| GU305914.1 | 174295 | 174317 | + | 0 | CATGACGACGAACCGGACGA | TGG | \|\|\|\|\|\|\|\|[\|\|\|\|\|\|\|\|\|\|\|\|]PAM | NA | Intergenic | NA | NA |
| T13223 | ATGACGACGAACCGGACGATGGG | 1000 | | | | | | | | | |
| Oligo fwd | CACCATGACGACGAACCGGACGAT | | | | | | | | | | |
| Oligo rev | AAACATCGTCCGGTTCGTCGTCAT | | | | | | | | | | |
| Chromosome | start | end | strand | MM | target_seq | PAM | alignment | distance | position | gene name | gene id |
| GU305914.1 | 174296 | 174318 | + | 0 | ATGACGACGAACCGGACGAT | GGG | \|\|\|\|\|\|\|\|[\|\|\|\|\|\|\|\|\|\|\|\|]PAM | NA | Intergenic | NA | NA |
| T13224 | ACCGGACGATGGGTGACCAGAGG | 1000 | | | | | | | | | |
| Oligo fwd | CACCACCGGACGATGGGTGACCAG | | | | | | | | | | |
| Oligo rev | AAACCTGGTCACCCATCGTCCGGT | | | | | | | | | | |
| Chromosome | start | end | strand | MM | target_seq | PAM | alignment | distance | position | gene name | gene id |
| GU305914.1 | 174306 | 174328 | + | 0 | ACCGGACGATGGGTGACCAG | AGG | \|\|\|\|\|\|\|\|[\|\|\|\|\|\|\|\|\|\|\|\|]PAM | NA | Intergenic | NA | NA |
| T13225 | CCGGACGATGGGTGACCAGAGGG | 1000 | | | | | | | | | |
| Oligo fwd | CACCCCGGACGATGGGTGACCAGA | | | | | | | | | | |
| Oligo rev | AAACTCTGGTCACCCATCGTCCGG | | | | | | | | | | |
| Chromosome | start | end | strand | MM | target_seq | PAM | alignment | distance | position | gene name | gene id |
| GU305914.1 | 174307 | 174329 | + | 0 | CCGGACGATGGGTGACCAGA | GGG | \|\|\|\|\|\|\|\|[\|\|\|\|\|\|\|\|\|\|\|\|]PAM | NA | Intergenic | NA | NA |
| T13226 | ATGTATCCCGAGACCATGTGTGG | 1000 | | | | | | | | | |
| Oligo fwd | CACCATGTATCCCGAGACCATGTG | | | | | | | | | | |
| Oligo rev | AAACCACATGGTCTCGGGATACAT | | | | | | | | | | |
| Chromosome | start | end | strand | MM | target_seq | PAM | alignment | distance | position | gene name | gene id |
| GU305914.1 | 174338 | 174360 | + | 0 | ATGTATCCCGAGACCATGTG | TGG | \|\|\|\|\|\|\|\|[\|\|\|\|\|\|\|\|\|\|\|\|]PAM | NA | Intergenic | NA | NA |

| | | | | | | | | | | | |
|---|---|---|---|---|---|---|---|---|---|---|---|
| T13227 | ACCATGTGTGGTAGAAACGTCGG | 1000 | | | | | | | | | |
| Oligo fwd | CACCACCATGTGTGGTAGAAACGT | | | | | | | | | | |
| Oligo rev | AAACACGTTTCTACCACACATGGT | | | | | | | | | | |
| Chromosome | start | end | strand | MM | target_seq | PAM | alignment | distance | position | gene name | gene id |
| GU305914.1 | 174350 | 174372 | + | 0 | ACCATGTGTGGTAGAAACGT | CGG | \|\|\|\|\|\|\|\|[\|\|\|\|\|\|\|\|\|\|\|\|]PAM | NA | Intergenic | NA | NA |
| T13228 | TGGTAGAAACGTCGGAAAGTCGG | 1000 | | | | | | | | | |
| Oligo fwd | CACCTGGTAGAAACGTCGGAAAGT | | | | | | | | | | |
| Oligo rev | AAACACTTTCCGACGTTTCTACCA | | | | | | | | | | |
| Chromosome | start | end | strand | MM | target_seq | PAM | alignment | distance | position | gene name | gene id |
| GU305914.1 | 174358 | 174380 | + | 0 | TGGTAGAAACGTCGGAAAGT | CGG | \|\|\|\|\|\|\|\|[\|\|\|\|\|\|\|\|\|\|\|\|]PAM | NA | Intergenic | NA | NA |
| T13229 | GGTAGAAACGTCGGAAAGTCGGG | 1000 | | | | | | | | | |
| Oligo fwd | CACCGGTAGAAACGTCGGAAAGTC | | | | | | | | | | |
| Oligo rev | AAACGACTTTCCGACGTTTCTACC | | | | | | | | | | |
| Chromosome | start | end | strand | MM | target_seq | PAM | alignment | distance | position | gene name | gene id |
| GU305914.1 | 174359 | 174381 | + | 0 | GGTAGAAACGTCGGAAAGTC | GGG | \|\|\|\|\|\|\|\|[\|\|\|\|\|\|\|\|\|\|\|\|]PAM | NA | Intergenic | NA | NA |
| T13230 | AAACGTCGGAAAGTCGGGCTCGG | 1000 | | | | | | | | | |
| Oligo fwd | CACCAAACGTCGGAAAGTCGGGCT | | | | | | | | | | |
| Oligo rev | AAACAGCCCGACTTTCCGACGTTT | | | | | | | | | | |
| Chromosome | start | end | strand | MM | target_seq | PAM | alignment | distance | position | gene name | gene id |
| GU305914.1 | 174364 | 174386 | + | 0 | AAACGTCGGAAAGTCGGGCT | CGG | \|\|\|\|\|\|\|\|[\|\|\|\|\|\|\|\|\|\|\|\|]PAM | NA | Intergenic | NA | NA |
| T13231 | TCGGTTCGATTGCTCCTTAGCGG | 1000 | | | | | | | | | |
| Oligo fwd | CACCTCGGTTCGATTGCTCCTTAG | | | | | | | | | | |
| Oligo rev | AAACCTAAGGAGCAATCGAACCGA | | | | | | | | | | |
| Chromosome | start | end | strand | MM | target_seq | PAM | alignment | distance | position | gene name | gene id |
| GU305914.1 | 174383 | 174405 | + | 0 | TCGGTTCGATTGCTCCTTAG | CGG | \|\|\|\|\|\|\|\|[\|\|\|\|\|\|\|\|\|\|\|\|]PAM | NA | Intergenic | NA | NA |
| T13232 | CGGTTCGATTGCTCCTTAGCGGG | 1000 | | | | | | | | | |
| Oligo fwd | CACCCGGTTCGATTGCTCCTTAGC | | | | | | | | | | |
| Oligo rev | AAACGCTAAGGAGCAATCGAACCG | | | | | | | | | | |
| Chromosome | start | end | strand | MM | target_seq | PAM | alignment | distance | position | gene name | gene id |
| GU305914.1 | 174384 | 174406 | + | 0 | CGGTTCGATTGCTCCTTAGC | GGG | \|\|\|\|\|\|\|\|[\|\|\|\|\|\|\|\|\|\|\|\|]PAM | NA | Intergenic | NA | NA |
| T13233 | GGTTCGATTGCTCCTTAGCGGGG | 1000 | | | | | | | | | |
| Oligo fwd | CACCGGTTCGATTGCTCCTTAGCG | | | | | | | | | | |
| Oligo rev | AAACCGCTAAGGAGCAATCGAACC | | | | | | | | | | |
| Chromosome | start | end | strand | MM | target_seq | PAM | alignment | distance | position | gene name | gene id |
| GU305914.1 | 174385 | 174407 | + | 0 | GGTTCGATTGCTCCTTAGCG | GGG | \|\|\|\|\|\|\|\|[\|\|\|\|\|\|\|\|\|\|\|\|]PAM | NA | Intergenic | NA | NA |
| T13234 | ATTGCTCCTTAGCGGGGCAGCGG | 1000 | | | | | | | | | |
| Oligo fwd | CACCATTGCTCCTTAGCGGGGCAG | | | | | | | | | | |
| Oligo rev | AAACCTGCCCCGCTAAGGAGCAAT | | | | | | | | | | |
| Chromosome | start | end | strand | MM | target_seq | PAM | alignment | distance | position | gene name | gene id |
| GU305914.1 | 174391 | 174413 | + | 0 | ATTGCTCCTTAGCGGGGCAG | CGG | \|\|\|\|\|\|\|\|[\|\|\|\|\|\|\|\|\|\|\|\|]PAM | NA | Intergenic | NA | NA |
| T13235 | TTGCTCCTTAGCGGGGCAGCGGG | 1000 | | | | | | | | | |
| Oligo fwd | CACCTTGCTCCTTAGCGGGGCAGC | | | | | | | | | | |
| Oligo rev | AAACGCTGCCCCGCTAAGGAGCAA | | | | | | | | | | |
| Chromosome | start | end | strand | MM | target_seq | PAM | alignment | distance | position | gene name | gene id |
| GU305914.1 | 174392 | 174414 | + | 0 | TTGCTCCTTAGCGGGGCAGC | GGG | \|\|\|\|\|\|\|\|[\|\|\|\|\|\|\|\|\|\|\|\|]PAM | NA | Intergenic | NA | NA |
| T13236 | TGCTCCTTAGCGGGGCAGCGGGG | 1000 | | | | | | | | | |
| Oligo fwd | CACCTGCTCCTTAGCGGGGCAGCG | | | | | | | | | | |
| Oligo rev | AAACCGCTGCCCCGCTAAGGAGCA | | | | | | | | | | |
| Chromosome | start | end | strand | MM | target_seq | PAM | alignment | distance | position | gene name | gene id |
| GU305914.1 | 174393 | 174415 | + | 0 | TGCTCCTTAGCGGGGCAGCG | GGG | \|\|\|\|\|\|\|\|[\|\|\|\|\|\|\|\|\|\|\|\|]PAM | NA | Intergenic | NA | NA |
| T13237 | AGCGGGGCAGCGGGGTCTTGTGG | 1000 | | | | | | | | | |
| Oligo fwd | CACCAGCGGGGCAGCGGGGTCTTG | | | | | | | | | | |
| Oligo rev | AAACCAAGACCCCGCTGCCCCGCT | | | | | | | | | | |
| Chromosome | start | end | strand | MM | target_seq | PAM | alignment | distance | position | gene name | gene id |
| GU305914.1 | 174401 | 174423 | + | 0 | AGCGGGGCAGCGGGGTCTTG | TGG | \|\|\|\|\|\|\|\|[\|\|\|\|\|\|\|\|\|\|\|\|]PAM | NA | Intergenic | NA | NA |
| T13238 | GCGGGGTCTTGTGGAAATTCTGG | 1000 | | | | | | | | | |
| Oligo fwd | CACCGCGGGGTCTTGTGGAAATTC | | | | | | | | | | |
| Oligo rev | AAACGAATTTCCACAAGACCCCGC | | | | | | | | | | |
| Chromosome | start | end | strand | MM | target_seq | PAM | alignment | distance | position | gene name | gene id |
| GU305914.1 | 174410 | 174432 | + | 0 | GCGGGGTCTTGTGGAAATTC | TGG | \|\|\|\|\|\|\|\|[\|\|\|\|\|\|\|\|\|\|\|\|]PAM | NA | Intergenic | NA | NA |
| T13239 | GTGGAAATTCTGGCATGATACGG | 1000 | | | | | | | | | |
| Oligo fwd | CACCGTGGAAATTCTGGCATGATA | | | | | | | | | | |
| Oligo rev | AAACTATCATGCCAGAATTTCCAC | | | | | | | | | | |
| Chromosome | start | end | strand | MM | target_seq | PAM | alignment | distance | position | gene name | gene id |
| GU305914.1 | 174420 | 174442 | + | 0 | GTGGAAATTCTGGCATGATA | CGG | \|\|\|\|\|\|\|\|[\|\|\|\|\|\|\|\|\|\|\|\|]PAM | NA | Intergenic | NA | NA |
| T13240 | AATTCTGGCATGATACGGCGAGG | 1000 | | | | | | | | | |
| Oligo fwd | CACCAATTCTGGCATGATACGGCG | | | | | | | | | | |
| Oligo rev | AAACCGCCGTATCATGCCAGAATT | | | | | | | | | | |
| Chromosome | start | end | strand | MM | target_seq | PAM | alignment | distance | position | gene name | gene id |
| GU305914.1 | 174425 | 174447 | + | 0 | AATTCTGGCATGATACGGCG | AGG | \|\|\|\|\|\|\|\|[\|\|\|\|\|\|\|\|\|\|\|\|]PAM | NA | Intergenic | NA | NA |
| T13241 | ATTCTGGCATGATACGGCGAGGG | 1000 | | | | | | | | | |
| Oligo fwd | CACCATTCTGGCATGATACGGCGA | | | | | | | | | | |
| Oligo rev | AAACTCGCCGTATCATGCCAGAAT | | | | | | | | | | |
| Chromosome | start | end | strand | MM | target_seq | PAM | alignment | distance | position | gene name | gene id |
| GU305914.1 | 174426 | 174448 | + | 0 | ATTCTGGCATGATACGGCGA | GGG | \|\|\|\|\|\|\|\|[\|\|\|\|\|\|\|\|\|\|\|\|]PAM | NA | Intergenic | NA | NA |

T13242 TTCTGGCATGATACGGCGAGGGG 1000
Oligo fwd CACCTTCTGGCATGATACGGCGAG
Oligo rev AAACCTCGCCGTATCATGCCAGAA
Chromosome start end strand MM target_seq PAM alignment distance position gene name gene id
GU305914.1 174427 174449 + 0 TTCTGGCATGATACGGCGAG GGG ||||||||[||||||||||||]PAM NA Intergenic NA NA

T13243 GGGATAGATGTGTCCCCTCGAGG 1000
Oligo fwd CACCGGGATAGATGTGTCCCCTCG
Oligo rev AAACCGAGGGGACACATCTATCCC
Chromosome start end strand MM target_seq PAM alignment distance position gene name gene id
GU305914.1 174447 174469 + 0 GGGATAGATGTGTCCCCTCG AGG ||||||||[||||||||||||]PAM NA Intergenic NA NA

T13244 GGATAGATGTGTCCCCTCGAGGG 1000
Oligo fwd CACCGGATAGATGTGTCCCCTCGA
Oligo rev AAACTCGAGGGGACACATCTATCC
Chromosome start end strand MM target_seq PAM alignment distance position gene name gene id
GU305914.1 174448 174470 + 0 GGATAGATGTGTCCCCTCGA GGG ||||||||[||||||||||||]PAM NA Intergenic NA NA

T13245 GATAGATGTGTCCCCTCGAGGGG 1000
Oligo fwd CACCGATAGATGTGTCCCCTCGAG
Oligo rev AAACCTCGAGGGGACACATCTATC
Chromosome start end strand MM target_seq PAM alignment distance position gene name gene id
GU305914.1 174449 174471 + 0 GATAGATGTGTCCCCTCGAG GGG ||||||||[||||||||||||]PAM NA Intergenic NA NA

T13246 GTCCCCTCGAGGGGAACACGCGG 1000
Oligo fwd CACCGTCCCCTCGAGGGGAACACG
Oligo rev AAACCGTGTTCCCCTCGAGGGGAC
Chromosome start end strand MM target_seq PAM alignment distance position gene name gene id
GU305914.1 174458 174480 + 0 GTCCCCTCGAGGGGAACACG CGG ||||||||[||||||||||||]PAM NA Intergenic NA NA

T13247 TCCCCTCGAGGGGAACACGCGGG 1000
Oligo fwd CACCTCCCCTCGAGGGGAACACGC
Oligo rev AAACGCGTGTTCCCCTCGAGGGGA
Chromosome start end strand MM target_seq PAM alignment distance position gene name gene id
GU305914.1 174459 174481 + 0 TCCCCTCGAGGGGAACACGC GGG ||||||||[||||||||||||]PAM NA Intergenic NA NA

T13248 GAGGGGAACACGCGGGACGATGG 1000
Oligo fwd CACCGAGGGGAACACGCGGGACGA
Oligo rev AAACTCGTCCCGCGTGTTCCCCTC
Chromosome start end strand MM target_seq PAM alignment distance position gene name gene id
GU305914.1 174466 174488 + 0 GAGGGGAACACGCGGGACGA TGG ||||||||[||||||||||||]PAM NA Intergenic NA NA

T13249 ACACGCGGGACGATGGTCTGTGG 1000
Oligo fwd CACCACACGCGGGACGATGGTCTG
Oligo rev AAACCAGACCATCGTCCCGCGTGT
Chromosome start end strand MM target_seq PAM alignment distance position gene name gene id
GU305914.1 174473 174495 + 0 ACACGCGGGACGATGGTCTG TGG ||||||||[||||||||||||]PAM NA Intergenic NA NA

T13250 GCGGGACGATGGTCTGTGGAAGG 1000
Oligo fwd CACCGCGGGACGATGGTCTGTGGA
Oligo rev AAACTCCACAGACCATCGTCCCGC
Chromosome start end strand MM target_seq PAM alignment distance position gene name gene id
GU305914.1 174477 174499 + 0 GCGGGACGATGGTCTGTGGA AGG ||||||||[||||||||||||]PAM NA Intergenic NA NA

T13251 CGGGACGATGGTCTGTGGAAGGG 1000
Oligo fwd CACCCGGGACGATGGTCTGTGGAA
Oligo rev AAACTTCCACAGACCATCGTCCCG
Chromosome start end strand MM target_seq PAM alignment distance position gene name gene id
GU305914.1 174478 174500 + 0 CGGGACGATGGTCTGTGGAA GGG ||||||||[||||||||||||]PAM NA Intergenic NA NA

T13252 ACGATGGTCTGTGGAAGGGACGG 1000
Oligo fwd CACCACGATGGTCTGTGGAAGGGA
Oligo rev AAACTCCCTTCCACAGACCATCGT
Chromosome start end strand MM target_seq PAM alignment distance position gene name gene id
GU305914.1 174482 174504 + 0 ACGATGGTCTGTGGAAGGGA CGG ||||||||[||||||||||||]PAM NA Intergenic NA NA

T13253 TGATCATCTACCAGCTTGTGTGG 1000
Oligo fwd CACCTGATCATCTACCAGCTTGTG
Oligo rev AAACCACAAGCTGGTAGATGATCA
Chromosome start end strand MM target_seq PAM alignment distance position gene name gene id
GU305914.1 174505 174527 + 0 TGATCATCTACCAGCTTGTG TGG ||||||||[||||||||||||]PAM NA Intergenic NA NA

T13254 GTGTGGTAACGAATTCCTCTTGG 1000
Oligo fwd CACCGTGTGGTAACGAATTCCTCT
Oligo rev AAACAGAGGAATTCGTTACCACAC
Chromosome start end strand MM target_seq PAM alignment distance position gene name gene id
GU305914.1 174522 174544 + 0 GTGTGGTAACGAATTCCTCT TGG ||||||||[||||||||||||]PAM NA Intergenic NA NA

T13255 CGCGACAGCGAACACTAACGCGG 1000
Oligo fwd CACCCGCGACAGCGAACACTAACG
Oligo rev AAACCGTTAGTGTTCGCTGTCGCG
Chromosome start end strand MM target_seq PAM alignment distance position gene name gene id
GU305914.1 174555 174577 + 0 CGCGACAGCGAACACTAACG CGG ||||||||[||||||||||||]PAM NA Intergenic NA NA

T13256 CGAACACTAACGCGGCCAGTAGG 1000
Oligo fwd CACCCGAACACTAACGCGGCCAGT
Oligo rev AAACACTGGCCGCGTTAGTGTTCG
Chromosome start end strand MM target_seq PAM alignment distance position gene name gene id

GU305914.1 174563 174585 + 0 CGAACACTAACGCGGCCAGT AGG ||||||||[||||||||||||]PAM NA Intergenic NA NA

T13257 ACGCGGCCAGTAGGATGATCAGG 1000
Oligo fwd CACCACGCGGCCAGTAGGATGATC
Oligo rev AAACGATCATCCTACTGGCCGCGT
Chromosome start end strand MM target_seq PAM alignment distance position gene name gene id
GU305914.1 174572 174594 + 0 ACGCGGCCAGTAGGATGATC AGG ||||||||[||||||||||||]PAM NA Intergenic NA NA

T13258 CGCGGCCAGTAGGATGATCAGGG 1000
Oligo fwd CACCCGCGGCCAGTAGGATGATCA
Oligo rev AAACTGATCATCCTACTGGCCGCG
Chromosome start end strand MM target_seq PAM alignment distance position gene name gene id
GU305914.1 174573 174595 + 0 CGCGGCCAGTAGGATGATCA GGG ||||||||[||||||||||||]PAM NA Intergenic NA NA

T13259 GGGTCCCCGCGAATGCGATCAGG 1000
Oligo fwd CACCGGGTCCCCGCGAATGCGATC
Oligo rev AAACGATCGCATTCGCGGGGACCC
Chromosome start end strand MM target_seq PAM alignment distance position gene name gene id
GU305914.1 174593 174615 + 0 GGGTCCCCGCGAATGCGATC AGG ||||||||[||||||||||||]PAM NA Intergenic NA NA

T13260 GGTCCCCGCGAATGCGATCAGGG 1000
Oligo fwd CACCGGTCCCCGCGAATGCGATCA
Oligo rev AAACTGATCGCATTCGCGGGGACC
Chromosome start end strand MM target_seq PAM alignment distance position gene name gene id
GU305914.1 174594 174616 + 0 GGTCCCCGCGAATGCGATCA GGG ||||||||[||||||||||||]PAM NA Intergenic NA NA

T13261 ATCAGGGAGATAACGTAGTCAGG 1000
Oligo fwd CACCATCAGGGAGATAACGTAGTC
Oligo rev AAACGACTACGTTATCTCCCTGAT
Chromosome start end strand MM target_seq PAM alignment distance position gene name gene id
GU305914.1 174610 174632 + 0 ATCAGGGAGATAACGTAGTC AGG ||||||||[||||||||||||]PAM NA Intergenic NA NA

T13262 GATAACGTAGTCAGGTCCGCCGG 1000
Oligo fwd CACCGATAACGTAGTCAGGTCCGC
Oligo rev AAACGCGGACCTGACTACGTTATC
Chromosome start end strand MM target_seq PAM alignment distance position gene name gene id
GU305914.1 174618 174640 + 0 GATAACGTAGTCAGGTCCGC CGG ||||||||[||||||||||||]PAM NA Intergenic NA NA

T13263 AACGTAGTCAGGTCCGCCGGAGG 1000
Oligo fwd CACCAACGTAGTCAGGTCCGCCGG
Oligo rev AAACCCGGCGGACCTGACTACGTT
Chromosome start end strand MM target_seq PAM alignment distance position gene name gene id
GU305914.1 174621 174643 + 0 AACGTAGTCAGGTCCGCCGG AGG ||||||||[||||||||||||]PAM NA Intergenic NA NA

T13264 GTCAGGTCCGCCGGAGGAGATGG 1000
Oligo fwd CACCGTCAGGTCCGCCGGAGGAGA
Oligo rev AAACTCTCCTCCGGCGGACCTGAC
Chromosome start end strand MM target_seq PAM alignment distance position gene name gene id
GU305914.1 174627 174649 + 0 GTCAGGTCCGCCGGAGGAGA TGG ||||||||[||||||||||||]PAM NA Intergenic NA NA

T13265 AGGTCCGCCGGAGGAGATGGTGG 1000
Oligo fwd CACCAGGTCCGCCGGAGGAGATGG
Oligo rev AAACCCATCTCCTCCGGCGGACCT
Chromosome start end strand MM target_seq PAM alignment distance position gene name gene id
GU305914.1 174630 174652 + 0 AGGTCCGCCGGAGGAGATGG TGG ||||||||[||||||||||||]PAM NA Intergenic NA NA

T13266 GCCGGAGGAGATGGTGGTAGTGG 1000
Oligo fwd CACCGCCGGAGGAGATGGTGGTAG
Oligo rev AAACCTACCACCATCTCCTCCGGC
Chromosome start end strand MM target_seq PAM alignment distance position gene name gene id
GU305914.1 174636 174658 + 0 GCCGGAGGAGATGGTGGTAG TGG ||||||||[||||||||||||]PAM NA Intergenic NA NA
GU305914.1 102688 102710 - 3 GGCAGAGGAGATGGTAGTAG TGG |-|-||||[|||||||-||||]PAM NA Intergenic NA NA

T13267 GGAGATGGTGGTAGTGGTGACGG 1000
Oligo fwd CACCGGAGATGGTGGTAGTGGTGA
Oligo rev AAACTCACCACTACCACCATCTCC
Chromosome start end strand MM target_seq PAM alignment distance position gene name gene id
GU305914.1 174642 174664 + 0 GGAGATGGTGGTAGTGGTGA CGG ||||||||[||||||||||||]PAM NA Intergenic NA NA
GU305914.1 213586 213608 + 4 GGTGGTGGTGGTAGTGGTAG TGG ||-|-|||[||||||||||--]PAM NA Intergenic NA NA

T13268 AGTGGTGACGGTGTCGTTCATGG 1000
Oligo fwd CACCAGTGGTGACGGTGTCGTTCA
Oligo rev AAACTGAACGACACCGTCACCACT
Chromosome start end strand MM target_seq PAM alignment distance position gene name gene id
GU305914.1 174654 174676 + 0 AGTGGTGACGGTGTCGTTCA TGG ||||||||[||||||||||||]PAM NA Intergenic NA NA

T13269 GTGACGGTGTCGTTCATGGTCGG 1000
Oligo fwd CACCGTGACGGTGTCGTTCATGGT
Oligo rev AAACACCATGAACGACACCGTCAC
Chromosome start end strand MM target_seq PAM alignment distance position gene name gene id
GU305914.1 174658 174680 + 0 GTGACGGTGTCGTTCATGGT CGG ||||||||[||||||||||||]PAM NA Intergenic NA NA

T13270 TGACGGTGTCGTTCATGGTCGGG 1000
Oligo fwd CACCTGACGGTGTCGTTCATGGTC
Oligo rev AAACGACCATGAACGACACCGTCA
Chromosome start end strand MM target_seq PAM alignment distance position gene name gene id
GU305914.1 174659 174681 + 0 TGACGGTGTCGTTCATGGTC GGG ||||||||[||||||||||||]PAM NA Intergenic NA NA

T13271 GTCGGGTAGACGAGAAAGAGCGG 1000

<table>
<tr><td>Oligo fwd</td><td>CACCGTCGGGTAGACGAGAAAGAG</td><td></td><td></td><td></td><td></td><td></td><td></td><td></td><td></td><td></td><td></td></tr>
<tr><td>Oligo rev</td><td>AAACCTCTTTCTCGTCTACCCGAC</td><td></td><td></td><td></td><td></td><td></td><td></td><td></td><td></td><td></td><td></td></tr>
<tr><td>Chromosome</td><td>start</td><td>end</td><td>strand</td><td>MM</td><td>target_seq</td><td>PAM</td><td>alignment</td><td>distance</td><td>position</td><td>gene name</td><td>gene id</td></tr>
<tr><td>GU305914.1</td><td>174676</td><td>174698</td><td>+</td><td>0</td><td>GTCGGGTAGACGAGAAAGAG</td><td>CGG</td><td>||||||||[||||||||||||]PAM</td><td>NA</td><td>Intergenic</td><td>NA</td><td>NA</td></tr>
<tr><td></td><td></td><td></td><td></td><td></td><td></td><td></td><td></td><td></td><td></td><td></td><td></td></tr>
<tr><td>T13272</td><td>TCGGGTAGACGAGAAAGAGCGGG</td><td>1000</td><td></td><td></td><td></td><td></td><td></td><td></td><td></td><td></td><td></td></tr>
<tr><td>Oligo fwd</td><td>CACCTCGGGTAGACGAGAAAGAGC</td><td></td><td></td><td></td><td></td><td></td><td></td><td></td><td></td><td></td><td></td></tr>
<tr><td>Oligo rev</td><td>AAACGCTCTTTCTCGTCTACCCGA</td><td></td><td></td><td></td><td></td><td></td><td></td><td></td><td></td><td></td><td></td></tr>
<tr><td>Chromosome</td><td>start</td><td>end</td><td>strand</td><td>MM</td><td>target_seq</td><td>PAM</td><td>alignment</td><td>distance</td><td>position</td><td>gene name</td><td>gene id</td></tr>
<tr><td>GU305914.1</td><td>174677</td><td>174699</td><td>+</td><td>0</td><td>TCGGGTAGACGAGAAAGAGC</td><td>GGG</td><td>||||||||[||||||||||||]PAM</td><td>NA</td><td>Intergenic</td><td>NA</td><td>NA</td></tr>
<tr><td></td><td></td><td></td><td></td><td></td><td></td><td></td><td></td><td></td><td></td><td></td><td></td></tr>
<tr><td>T13273</td><td>GGTAGACGAGAAAGAGCGGGAGG</td><td>1000</td><td></td><td></td><td></td><td></td><td></td><td></td><td></td><td></td><td></td></tr>
<tr><td>Oligo fwd</td><td>CACCGGTAGACGAGAAAGAGCGGG</td><td></td><td></td><td></td><td></td><td></td><td></td><td></td><td></td><td></td><td></td></tr>
<tr><td>Oligo rev</td><td>AAACCCCGCTCTTTCTCGTCTACC</td><td></td><td></td><td></td><td></td><td></td><td></td><td></td><td></td><td></td><td></td></tr>
<tr><td>Chromosome</td><td>start</td><td>end</td><td>strand</td><td>MM</td><td>target_seq</td><td>PAM</td><td>alignment</td><td>distance</td><td>position</td><td>gene name</td><td>gene id</td></tr>
<tr><td>GU305914.1</td><td>174680</td><td>174702</td><td>+</td><td>0</td><td>GGTAGACGAGAAAGAGCGGG</td><td>AGG</td><td>||||||||[||||||||||||]PAM</td><td>NA</td><td>Intergenic</td><td>NA</td><td>NA</td></tr>
<tr><td></td><td></td><td></td><td></td><td></td><td></td><td></td><td></td><td></td><td></td><td></td><td></td></tr>
<tr><td>T13274</td><td>GTAGACGAGAAAGAGCGGGAGGG</td><td>1000</td><td></td><td></td><td></td><td></td><td></td><td></td><td></td><td></td><td></td></tr>
<tr><td>Oligo fwd</td><td>CACCGTAGACGAGAAAGAGCGGGA</td><td></td><td></td><td></td><td></td><td></td><td></td><td></td><td></td><td></td><td></td></tr>
<tr><td>Oligo rev</td><td>AAACTCCCGCTCTTTCTCGTCTAC</td><td></td><td></td><td></td><td></td><td></td><td></td><td></td><td></td><td></td><td></td></tr>
<tr><td>Chromosome</td><td>start</td><td>end</td><td>strand</td><td>MM</td><td>target_seq</td><td>PAM</td><td>alignment</td><td>distance</td><td>position</td><td>gene name</td><td>gene id</td></tr>
<tr><td>GU305914.1</td><td>174681</td><td>174703</td><td>+</td><td>0</td><td>GTAGACGAGAAAGAGCGGGA</td><td>GGG</td><td>||||||||[||||||||||||]PAM</td><td>NA</td><td>Intergenic</td><td>NA</td><td>NA</td></tr>
<tr><td></td><td></td><td></td><td></td><td></td><td></td><td></td><td></td><td></td><td></td><td></td><td></td></tr>
<tr><td>T13275</td><td>GAGAAAGAGCGGGAGGGACGCGG</td><td>1000</td><td></td><td></td><td></td><td></td><td></td><td></td><td></td><td></td><td></td></tr>
<tr><td>Oligo fwd</td><td>CACCGAGAAAGAGCGGGAGGGACG</td><td></td><td></td><td></td><td></td><td></td><td></td><td></td><td></td><td></td><td></td></tr>
<tr><td>Oligo rev</td><td>AAACCGTCCCTCCCGCTCTTTCTC</td><td></td><td></td><td></td><td></td><td></td><td></td><td></td><td></td><td></td><td></td></tr>
<tr><td>Chromosome</td><td>start</td><td>end</td><td>strand</td><td>MM</td><td>target_seq</td><td>PAM</td><td>alignment</td><td>distance</td><td>position</td><td>gene name</td><td>gene id</td></tr>
<tr><td>GU305914.1</td><td>174687</td><td>174709</td><td>+</td><td>0</td><td>GAGAAAGAGCGGGAGGGACG</td><td>CGG</td><td>||||||||[||||||||||||]PAM</td><td>NA</td><td>Intergenic</td><td>NA</td><td>NA</td></tr>
<tr><td></td><td></td><td></td><td></td><td></td><td></td><td></td><td></td><td></td><td></td><td></td><td></td></tr>
<tr><td>T13276</td><td>GAGCGGGAGGGACGCGGTCACGG</td><td>1000</td><td></td><td></td><td></td><td></td><td></td><td></td><td></td><td></td><td></td></tr>
<tr><td>Oligo fwd</td><td>CACCGAGCGGGAGGGACGCGGTCA</td><td></td><td></td><td></td><td></td><td></td><td></td><td></td><td></td><td></td><td></td></tr>
<tr><td>Oligo rev</td><td>AAACTGACCGCGTCCCTCCCGCTC</td><td></td><td></td><td></td><td></td><td></td><td></td><td></td><td></td><td></td><td></td></tr>
<tr><td>Chromosome</td><td>start</td><td>end</td><td>strand</td><td>MM</td><td>target_seq</td><td>PAM</td><td>alignment</td><td>distance</td><td>position</td><td>gene name</td><td>gene id</td></tr>
<tr><td>GU305914.1</td><td>174693</td><td>174715</td><td>+</td><td>0</td><td>GAGCGGGAGGGACGCGGTCA</td><td>CGG</td><td>||||||||[||||||||||||]PAM</td><td>NA</td><td>Intergenic</td><td>NA</td><td>NA</td></tr>
<tr><td></td><td></td><td></td><td></td><td></td><td></td><td></td><td></td><td></td><td></td><td></td><td></td></tr>
<tr><td>T13277</td><td>ACGGACTCGTACGTGTCTTTTGG</td><td>1000</td><td></td><td></td><td></td><td></td><td></td><td></td><td></td><td></td><td></td></tr>
<tr><td>Oligo fwd</td><td>CACCACGGACTCGTACGTGTCTTT</td><td></td><td></td><td></td><td></td><td></td><td></td><td></td><td></td><td></td><td></td></tr>
<tr><td>Oligo rev</td><td>AAACAAAGACACGTACGAGTCCGT</td><td></td><td></td><td></td><td></td><td></td><td></td><td></td><td></td><td></td><td></td></tr>
<tr><td>Chromosome</td><td>start</td><td>end</td><td>strand</td><td>MM</td><td>target_seq</td><td>PAM</td><td>alignment</td><td>distance</td><td>position</td><td>gene name</td><td>gene id</td></tr>
<tr><td>GU305914.1</td><td>174712</td><td>174734</td><td>+</td><td>0</td><td>ACGGACTCGTACGTGTCTTT</td><td>TGG</td><td>||||||||[||||||||||||]PAM</td><td>NA</td><td>Intergenic</td><td>NA</td><td>NA</td></tr>
<tr><td></td><td></td><td></td><td></td><td></td><td></td><td></td><td></td><td></td><td></td><td></td><td></td></tr>
<tr><td>T13278</td><td>CGGACTCGTACGTGTCTTTTGGG</td><td>1000</td><td></td><td></td><td></td><td></td><td></td><td></td><td></td><td></td><td></td></tr>
<tr><td>Oligo fwd</td><td>CACCCGGACTCGTACGTGTCTTTT</td><td></td><td></td><td></td><td></td><td></td><td></td><td></td><td></td><td></td><td></td></tr>
<tr><td>Oligo rev</td><td>AAACAAAAGACACGTACGAGTCCG</td><td></td><td></td><td></td><td></td><td></td><td></td><td></td><td></td><td></td><td></td></tr>
<tr><td>Chromosome</td><td>start</td><td>end</td><td>strand</td><td>MM</td><td>target_seq</td><td>PAM</td><td>alignment</td><td>distance</td><td>position</td><td>gene name</td><td>gene id</td></tr>
<tr><td>GU305914.1</td><td>174713</td><td>174735</td><td>+</td><td>0</td><td>CGGACTCGTACGTGTCTTTT</td><td>GGG</td><td>||||||||[||||||||||||]PAM</td><td>NA</td><td>Intergenic</td><td>NA</td><td>NA</td></tr>
<tr><td></td><td></td><td></td><td></td><td></td><td></td><td></td><td></td><td></td><td></td><td></td><td></td></tr>
<tr><td>T13279</td><td>TGTCTTTTGGGATCACAACTCGG</td><td>1000</td><td></td><td></td><td></td><td></td><td></td><td></td><td></td><td></td><td></td></tr>
<tr><td>Oligo fwd</td><td>CACCTGTCTTTTGGGATCACAACT</td><td></td><td></td><td></td><td></td><td></td><td></td><td></td><td></td><td></td><td></td></tr>
<tr><td>Oligo rev</td><td>AAACAGTTGTGATCCCAAAAGACA</td><td></td><td></td><td></td><td></td><td></td><td></td><td></td><td></td><td></td><td></td></tr>
<tr><td>Chromosome</td><td>start</td><td>end</td><td>strand</td><td>MM</td><td>target_seq</td><td>PAM</td><td>alignment</td><td>distance</td><td>position</td><td>gene name</td><td>gene id</td></tr>
<tr><td>GU305914.1</td><td>174725</td><td>174747</td><td>+</td><td>0</td><td>TGTCTTTTGGGATCACAACT</td><td>CGG</td><td>||||||||[||||||||||||]PAM</td><td>NA</td><td>Intergenic</td><td>NA</td><td>NA</td></tr>
<tr><td></td><td></td><td></td><td></td><td></td><td></td><td></td><td></td><td></td><td></td><td></td><td></td></tr>
<tr><td>T13280</td><td>GGATCACAACTCGGTGATCTCGG</td><td>1000</td><td></td><td></td><td></td><td></td><td></td><td></td><td></td><td></td><td></td></tr>
<tr><td>Oligo fwd</td><td>CACCGGATCACAACTCGGTGATCT</td><td></td><td></td><td></td><td></td><td></td><td></td><td></td><td></td><td></td><td></td></tr>
<tr><td>Oligo rev</td><td>AAACAGATCACCGAGTTGTGATCC</td><td></td><td></td><td></td><td></td><td></td><td></td><td></td><td></td><td></td><td></td></tr>
<tr><td>Chromosome</td><td>start</td><td>end</td><td>strand</td><td>MM</td><td>target_seq</td><td>PAM</td><td>alignment</td><td>distance</td><td>position</td><td>gene name</td><td>gene id</td></tr>
<tr><td>GU305914.1</td><td>174734</td><td>174756</td><td>+</td><td>0</td><td>GGATCACAACTCGGTGATCT</td><td>CGG</td><td>||||||||[||||||||||||]PAM</td><td>NA</td><td>Intergenic</td><td>NA</td><td>NA</td></tr>
<tr><td></td><td></td><td></td><td></td><td></td><td></td><td></td><td></td><td></td><td></td><td></td><td></td></tr>
<tr><td>T13281</td><td>TCACAACTCGGTGATCTCGGCGG</td><td>1000</td><td></td><td></td><td></td><td></td><td></td><td></td><td></td><td></td><td></td></tr>
<tr><td>Oligo fwd</td><td>CACCTCACAACTCGGTGATCTCGG</td><td></td><td></td><td></td><td></td><td></td><td></td><td></td><td></td><td></td><td></td></tr>
<tr><td>Oligo rev</td><td>AAACCCGAGATCACCGAGTTGTGA</td><td></td><td></td><td></td><td></td><td></td><td></td><td></td><td></td><td></td><td></td></tr>
<tr><td>Chromosome</td><td>start</td><td>end</td><td>strand</td><td>MM</td><td>target_seq</td><td>PAM</td><td>alignment</td><td>distance</td><td>position</td><td>gene name</td><td>gene id</td></tr>
<tr><td>GU305914.1</td><td>174737</td><td>174759</td><td>+</td><td>0</td><td>TCACAACTCGGTGATCTCGG</td><td>CGG</td><td>||||||||[||||||||||||]PAM</td><td>NA</td><td>Intergenic</td><td>NA</td><td>NA</td></tr>
<tr><td></td><td></td><td></td><td></td><td></td><td></td><td></td><td></td><td></td><td></td><td></td><td></td></tr>
<tr><td>T13282</td><td>CTCGGCGGACCTTTGACTCTCGG</td><td>1000</td><td></td><td></td><td></td><td></td><td></td><td></td><td></td><td></td><td></td></tr>
<tr><td>Oligo fwd</td><td>CACCCTCGGCGGACCTTTGACTCT</td><td></td><td></td><td></td><td></td><td></td><td></td><td></td><td></td><td></td><td></td></tr>
<tr><td>Oligo rev</td><td>AAACAGAGTCAAAGGTCCGCCGAG</td><td></td><td></td><td></td><td></td><td></td><td></td><td></td><td></td><td></td><td></td></tr>
<tr><td>Chromosome</td><td>start</td><td>end</td><td>strand</td><td>MM</td><td>target_seq</td><td>PAM</td><td>alignment</td><td>distance</td><td>position</td><td>gene name</td><td>gene id</td></tr>
<tr><td>GU305914.1</td><td>174752</td><td>174774</td><td>+</td><td>0</td><td>CTCGGCGGACCTTTGACTCT</td><td>CGG</td><td>||||||||[||||||||||||]PAM</td><td>NA</td><td>Intergenic</td><td>NA</td><td>NA</td></tr>
<tr><td></td><td></td><td></td><td></td><td></td><td></td><td></td><td></td><td></td><td></td><td></td><td></td></tr>
<tr><td>T13283</td><td>TGACTCTCGGTCCTGTTGCACGG</td><td>1000</td><td></td><td></td><td></td><td></td><td></td><td></td><td></td><td></td><td></td></tr>
<tr><td>Oligo fwd</td><td>CACCTGACTCTCGGTCCTGTTGCA</td><td></td><td></td><td></td><td></td><td></td><td></td><td></td><td></td><td></td><td></td></tr>
<tr><td>Oligo rev</td><td>AAACTGCAACAGGACCGAGAGTCA</td><td></td><td></td><td></td><td></td><td></td><td></td><td></td><td></td><td></td><td></td></tr>
<tr><td>Chromosome</td><td>start</td><td>end</td><td>strand</td><td>MM</td><td>target_seq</td><td>PAM</td><td>alignment</td><td>distance</td><td>position</td><td>gene name</td><td>gene id</td></tr>
<tr><td>GU305914.1</td><td>174765</td><td>174787</td><td>+</td><td>0</td><td>TGACTCTCGGTCCTGTTGCA</td><td>CGG</td><td>||||||||[||||||||||||]PAM</td><td>NA</td><td>Intergenic</td><td>NA</td><td>NA</td></tr>
<tr><td></td><td></td><td></td><td></td><td></td><td></td><td></td><td></td><td></td><td></td><td></td><td></td></tr>
<tr><td>T13284</td><td>CGGTTTATATACCTTCATATCGG</td><td>1000</td><td></td><td></td><td></td><td></td><td></td><td></td><td></td><td></td><td></td></tr>
<tr><td>Oligo fwd</td><td>CACCCGGTTTATATACCTTCATAT</td><td></td><td></td><td></td><td></td><td></td><td></td><td></td><td></td><td></td><td></td></tr>
<tr><td>Oligo rev</td><td>AAACATATGAAGGTATATAAACCG</td><td></td><td></td><td></td><td></td><td></td><td></td><td></td><td></td><td></td><td></td></tr>
<tr><td>Chromosome</td><td>start</td><td>end</td><td>strand</td><td>MM</td><td>target_seq</td><td>PAM</td><td>alignment</td><td>distance</td><td>position</td><td>gene name</td><td>gene id</td></tr>
<tr><td>GU305914.1</td><td>174785</td><td>174807</td><td>+</td><td>0</td><td>CGGTTTATATACCTTCATAT</td><td>CGG</td><td>||||||||[||||||||||||]PAM</td><td>NA</td><td>Intergenic</td><td>NA</td><td>NA</td></tr>
<tr><td></td><td></td><td></td><td></td><td></td><td></td><td></td><td></td><td></td><td></td><td></td><td></td></tr>
<tr><td>T13285</td><td>TTATATACCTTCATATCGGAAGG</td><td>1000</td><td></td><td></td><td></td><td></td><td></td><td></td><td></td><td></td><td></td></tr>
<tr><td>Oligo fwd</td><td>CACCTTATATACCTTCATATCGGA</td><td></td><td></td><td></td><td></td><td></td><td></td><td></td><td></td><td></td><td></td></tr>
<tr><td>Oligo rev</td><td>AAACTCCGATATGAAGGTATATAA</td><td></td><td></td><td></td><td></td><td></td><td></td><td></td><td></td><td></td><td></td></tr>
<tr><td>Chromosome</td><td>start</td><td>end</td><td>strand</td><td>MM</td><td>target_seq</td><td>PAM</td><td>alignment</td><td>distance</td><td>position</td><td>gene name</td><td>gene id</td></tr>
<tr><td>GU305914.1</td><td>174789</td><td>174811</td><td>+</td><td>0</td><td>TTATATACCTTCATATCGGA</td><td>AGG</td><td>||||||||[||||||||||||]PAM</td><td>NA</td><td>Intergenic</td><td>NA</td><td>NA</td></tr>
</table>

T13286 TATCGGAAGGACCGCGAGATAGG 1000
Oligo fwd CACCTATCGGAAGGACCGCGAGAT
Oligo rev AAACATCTCGCGGTCCTTCCGATA
Chromosome start end strand MM target_seq PAM alignment distance position gene name gene id
GU305914.1 174802 174824 + 0 TATCGGAAGGACCGCGAGAT AGG ||||||||[||||||||||||]PAM NA Intergenic NA NA

T13287 GGCGCGCGTGTTTTCCTTTGTGG 1000
Oligo fwd CACCGGCGCGCGTGTTTTCCTTTG
Oligo rev AAACCAAAGGAAAACACGCGCGCC
Chromosome start end strand MM target_seq PAM alignment distance position gene name gene id
GU305914.1 174823 174845 + 0 GGCGCGCGTGTTTTCCTTTG TGG ||||||||[||||||||||||]PAM NA Intergenic NA NA

T13288 GTGGTGAATCGTGAATATCGCGG 1000
Oligo fwd CACCGTGGTGAATCGTGAATATCG
Oligo rev AAACCGATATTCACGATTCACCAC
Chromosome start end strand MM target_seq PAM alignment distance position gene name gene id
GU305914.1 174842 174864 + 0 GTGGTGAATCGTGAATATCG CGG ||||||||[||||||||||||]PAM NA Intergenic NA NA

T13289 GTGAATCGTGAATATCGCGGCGG 1000
Oligo fwd CACCGTGAATCGTGAATATCGCGG
Oligo rev AAACCCGCGATATTCACGATTCAC
Chromosome start end strand MM target_seq PAM alignment distance position gene name gene id
GU305914.1 174845 174867 + 0 GTGAATCGTGAATATCGCGG CGG ||||||||[||||||||||||]PAM NA Intergenic NA NA

T13290 TCGCGGCGGCGACCCCCGCTAGG 1000
Oligo fwd CACCTCGCGGCGGCGACCCCCGCT
Oligo rev AAACAGCGGGGGTCGCCGCCGCGA
Chromosome start end strand MM target_seq PAM alignment distance position gene name gene id
GU305914.1 174859 174881 + 0 TCGCGGCGGCGACCCCCGCT AGG ||||||||[||||||||||||]PAM NA Intergenic NA NA

T13291 ACGACGAGAAACACTCCCGTCGG 1000
Oligo fwd CACCACGACGAGAAACACTCCCGT
Oligo rev AAACACGGGAGTGTTTCTCGTCGT
Chromosome start end strand MM target_seq PAM alignment distance position gene name gene id
GU305914.1 174897 174919 + 0 ACGACGAGAAACACTCCCGT CGG ||||||||[||||||||||||]PAM NA Intergenic NA NA

T13292 AAACACTCCCGTCGGTTGCGCGG 1000
Oligo fwd CACCAAACACTCCCGTCGGTTGCG
Oligo rev AAACCGCAACCGACGGGAGTGTTT
Chromosome start end strand MM target_seq PAM alignment distance position gene name gene id
GU305914.1 174905 174927 + 0 AAACACTCCCGTCGGTTGCG CGG ||||||||[||||||||||||]PAM NA Intergenic NA NA

T13293 AACACTCCCGTCGGTTGCGCGGG 1000
Oligo fwd CACCAACACTCCCGTCGGTTGCGC
Oligo rev AAACGCGCAACCGACGGGAGTGTT
Chromosome start end strand MM target_seq PAM alignment distance position gene name gene id
GU305914.1 174906 174928 + 0 AACACTCCCGTCGGTTGCGC GGG ||||||||[||||||||||||]PAM NA Intergenic NA NA

T13294 CCCGTCGGTTGCGCGGGTGCCGG 1000
Oligo fwd CACCCCCGTCGGTTGCGCGGGTGC
Oligo rev AAACGCACCCGCGCAACCGACGGG
Chromosome start end strand MM target_seq PAM alignment distance position gene name gene id
GU305914.1 174912 174934 + 0 CCCGTCGGTTGCGCGGGTGC CGG ||||||||[||||||||||||]PAM NA Intergenic NA NA

T13295 GTCGGTTGCGCGGGTGCCGGAGG 1000
Oligo fwd CACCGTCGGTTGCGCGGGTGCCGG
Oligo rev AAACCCGGCACCCGCGCAACCGAC
Chromosome start end strand MM target_seq PAM alignment distance position gene name gene id
GU305914.1 174915 174937 + 0 GTCGGTTGCGCGGGTGCCGG AGG ||||||||[||||||||||||]PAM NA Intergenic NA NA
GU305914.1 163749 163771 + 4 GTCGGCGGCGCCGGTGGCGG CGG |||||--|[|||-||||-|||]PAM NA Intergenic NA NA

T13296 TTGCGCGGGTGCCGGAGGACTGG 1000
Oligo fwd CACCTTGCGCGGGTGCCGGAGGAC
Oligo rev AAACGTCCTCCGGCACCCGCGCAA
Chromosome start end strand MM target_seq PAM alignment distance position gene name gene id
GU305914.1 174920 174942 + 0 TTGCGCGGGTGCCGGAGGAC TGG ||||||||[||||||||||||]PAM NA Intergenic NA NA

T13297 TGCGCGGGTGCCGGAGGACTGGG 1000
Oligo fwd CACCTGCGCGGGTGCCGGAGGACT
Oligo rev AAACAGTCCTCCGGCACCCGCGCA
Chromosome start end strand MM target_seq PAM alignment distance position gene name gene id
GU305914.1 174921 174943 + 0 TGCGCGGGTGCCGGAGGACT GGG ||||||||[||||||||||||]PAM NA Intergenic NA NA

T13298 GCGCGGGTGCCGGAGGACTGGGG 1000
Oligo fwd CACCGCGCGGGTGCCGGAGGACTG
Oligo rev AAACCAGTCCTCCGGCACCCGCGC
Chromosome start end strand MM target_seq PAM alignment distance position gene name gene id
GU305914.1 174922 174944 + 0 GCGCGGGTGCCGGAGGACTG GGG ||||||||[||||||||||||]PAM NA Intergenic NA NA

T13299 CGGAGGACTGGGGTCGTCGCCGG 1000
Oligo fwd CACCCGGAGGACTGGGGTCGTCGC
Oligo rev AAACGCGACGACCCCAGTCCTCCG
Chromosome start end strand MM target_seq PAM alignment distance position gene name gene id
GU305914.1 174932 174954 + 0 CGGAGGACTGGGGTCGTCGC CGG ||||||||[||||||||||||]PAM NA Intergenic NA NA

T13300 GGAGGACTGGGGTCGTCGCCGGG 1000
Oligo fwd CACCGGAGGACTGGGGTCGTCGCC
Oligo rev AAACGGCGACGACCCCAGTCCTCC
Chromosome start end strand MM target_seq PAM alignment distance position gene name gene id

GU305914.1 174933 174955 + 0 GGAGGACTGGGGTCGTCGCC GGG ||||||||[||||||||||||]PAM NA Intergenic NA NA

T13301 GCCGGGACCTCGTTCTCCGTCGG 1000
Oligo fwd CACCGCCGGGACCTCGTTCTCCGT
Oligo rev AAACACGGAGAACGAGGTCCCGGC
Chromosome start end strand MM target_seq PAM alignment distance position gene name gene id
GU305914.1 174950 174972 + 0 GCCGGGACCTCGTTCTCCGT CGG ||||||||[||||||||||||]PAM NA Intergenic NA NA

T13302 CCGGGACCTCGTTCTCCGTCGGG 1000
Oligo fwd CACCCCGGGACCTCGTTCTCCGTC
Oligo rev AAACGACGGAGAACGAGGTCCCGG
Chromosome start end strand MM target_seq PAM alignment distance position gene name gene id
GU305914.1 174951 174973 + 0 CCGGGACCTCGTTCTCCGTC GGG ||||||||[||||||||||||]PAM NA Intergenic NA NA

T13303 CCGTGTCCGTTTCCGCCGCTCGG 1000
Oligo fwd CACCCCGTGTCCGTTTCCGCCGCT
Oligo rev AAACAGCGGCGGAAACGGACACGG
Chromosome start end strand MM target_seq PAM alignment distance position gene name gene id
GU305914.1 174975 174997 + 0 CCGTGTCCGTTTCCGCCGCT CGG ||||||||[||||||||||||]PAM NA Intergenic NA NA
GU305914.1 119818 119840 + 3 CCGAGTCTGTTTCCGCGGCT GGG |||-|||-[||||||||-|||]PAM NA Intergenic NA NA

T13304 GCCGCTCGGTCTCTCGCCCGTGG 1000
Oligo fwd CACCGCCGCTCGGTCTCTCGCCCG
Oligo rev AAACCGGGCGAGAGACCGAGCGGC
Chromosome start end strand MM target_seq PAM alignment distance position gene name gene id
GU305914.1 174989 175011 + 0 GCCGCTCGGTCTCTCGCCCG TGG ||||||||[||||||||||||]PAM NA Intergenic NA NA

T13305 AAAAAGATCGTTTCAAACCCTGG 1000
Oligo fwd CACCAAAAAGATCGTTTCAAACCC
Oligo rev AAACGGGTTTGAAACGATCTTTTT
Chromosome start end strand MM target_seq PAM alignment distance position gene name gene id
GU305914.1 175032 175054 + 0 AAAAAGATCGTTTCAAACCC TGG ||||||||[||||||||||||]PAM NA Intergenic NA NA

T13306 TTTCATCCCCGTCGTGCATCTGG 1000
Oligo fwd CACCTTTCATCCCCGTCGTGCATC
Oligo rev AAACGATGCACGACGGGGATGAAA
Chromosome start end strand MM target_seq PAM alignment distance position gene name gene id
GU305914.1 175085 175107 + 0 TTTCATCCCCGTCGTGCATC TGG ||||||||[||||||||||||]PAM NA Intergenic NA NA

T13307 CATCCCCGTCGTGCATCTGGTGG 1000
Oligo fwd CACCCATCCCCGTCGTGCATCTGG
Oligo rev AAACCCAGATGCACGACGGGGATG
Chromosome start end strand MM target_seq PAM alignment distance position gene name gene id
GU305914.1 175088 175110 + 0 CATCCCCGTCGTGCATCTGG TGG ||||||||[||||||||||||]PAM NA Intergenic NA NA

T13308 ACTATGCATATTAGACAAGCCGG 1000
Oligo fwd CACCACTATGCATATTAGACAAGC
Oligo rev AAACGCTTGTCTAATATGCATAGT
Chromosome start end strand MM target_seq PAM alignment distance position gene name gene id
GU305914.1 175137 175159 + 0 ACTATGCATATTAGACAAGC CGG ||||||||[||||||||||||]PAM NA Intergenic NA NA

T13309 CATATTAGACAAGCCGGTTGAGG 1000
Oligo fwd CACCCATATTAGACAAGCCGGTTG
Oligo rev AAACCAACCGGCTTGTCTAATATG
Chromosome start end strand MM target_seq PAM alignment distance position gene name gene id
GU305914.1 175143 175165 + 0 CATATTAGACAAGCCGGTTG AGG ||||||||[||||||||||||]PAM NA Intergenic NA NA

T13310 ATTAGACAAGCCGGTTGAGGTGG 1000
Oligo fwd CACCATTAGACAAGCCGGTTGAGG
Oligo rev AAACCCTCAACCGGCTTGTCTAAT
Chromosome start end strand MM target_seq PAM alignment distance position gene name gene id
GU305914.1 175146 175168 + 0 ATTAGACAAGCCGGTTGAGG TGG ||||||||[||||||||||||]PAM NA Intergenic NA NA

T13311 GCCGGTTGAGGTGGTCTCCTCGG 1000
Oligo fwd CACCGCCGGTTGAGGTGGTCTCCT
Oligo rev AAACAGGAGACCACCTCAACCGGC
Chromosome start end strand MM target_seq PAM alignment distance position gene name gene id
GU305914.1 175155 175177 + 0 GCCGGTTGAGGTGGTCTCCT CGG ||||||||[||||||||||||]PAM NA Intergenic NA NA

T13312 CAGATCTGTACCGAAAAATCCGG 1000
Oligo fwd CACCCAGATCTGTACCGAAAAATC
Oligo rev AAACGATTTTTCGGTACAGATCTG
Chromosome start end strand MM target_seq PAM alignment distance position gene name gene id
GU305914.1 175214 175236 + 0 CAGATCTGTACCGAAAAATC CGG ||||||||[||||||||||||]PAM NA Intergenic NA NA

T13313 CGAGTTTAGCATCCATAAGATGG 1000
Oligo fwd CACCCGAGTTTAGCATCCATAAGA
Oligo rev AAACTCTTATGGATGCTAAACTCG
Chromosome start end strand MM target_seq PAM alignment distance position gene name gene id
GU305914.1 175301 175323 + 0 CGAGTTTAGCATCCATAAGA TGG ||||||||[||||||||||||]PAM NA Intergenic NA NA

T13314 AGCATCCATAAGATGGTGTATGG 1000
Oligo fwd CACCAGCATCCATAAGATGGTGTA
Oligo rev AAACTACACCATCTTATGGATGCT
Chromosome start end strand MM target_seq PAM alignment distance position gene name gene id
GU305914.1 175308 175330 + 0 AGCATCCATAAGATGGTGTA TGG ||||||||[||||||||||||]PAM NA Intergenic NA NA

T13315 ATCCATAAGATGGTGTATGGTGG 1000
Oligo fwd CACCATCCATAAGATGGTGTATGG

| | | | | | | | | | | | |
|---|---|---|---|---|---|---|---|---|---|---|---|
| Oligo rev | AAACCCATACACCATCTTATGGAT | | | | | | | | | | |
| Chromosome | start | end | strand | MM | target_seq | PAM | alignment | distance | position | gene name | gene id |
| GU305914.1 | 175311 | 175333 | + | 0 | ATCCATAAGATGGTGTATGG | TGG | \|\|\|\|\|\|\|\|[\|\|\|\|\|\|\|\|\|\|\|\|]PAM | NA | Intergenic | NA | NA |
| T13316 | AGATGGTGTATGGTGGAGTTTGG | 1000 | | | | | | | | | |
| Oligo fwd | CACCAGATGGTGTATGGTGGAGTT | | | | | | | | | | |
| Oligo rev | AAACAACTCCACCATACACCATCT | | | | | | | | | | |
| Chromosome | start | end | strand | MM | target_seq | PAM | alignment | distance | position | gene name | gene id |
| GU305914.1 | 175318 | 175340 | + | 0 | AGATGGTGTATGGTGGAGTT | TGG | \|\|\|\|\|\|\|\|[\|\|\|\|\|\|\|\|\|\|\|\|]PAM | NA | Intergenic | NA | NA |
| T13317 | TATGGTGGAGTTTGGTTATGTGG | 1000 | | | | | | | | | |
| Oligo fwd | CACCTATGGTGGAGTTTGGTTATG | | | | | | | | | | |
| Oligo rev | AAACCATAACCAAACTCCACCATA | | | | | | | | | | |
| Chromosome | start | end | strand | MM | target_seq | PAM | alignment | distance | position | gene name | gene id |
| GU305914.1 | 175326 | 175348 | + | 0 | TATGGTGGAGTTTGGTTATG | TGG | \|\|\|\|\|\|\|\|[\|\|\|\|\|\|\|\|\|\|\|\|]PAM | NA | Intergenic | NA | NA |
| T13318 | GTCGTTGAACTCACTGTCAGAGG | 1000 | | | | | | | | | |
| Oligo fwd | CACCGTCGTTGAACTCACTGTCAG | | | | | | | | | | |
| Oligo rev | AAACCTGACAGTGAGTTCAACGAC | | | | | | | | | | |
| Chromosome | start | end | strand | MM | target_seq | PAM | alignment | distance | position | gene name | gene id |
| GU305914.1 | 175380 | 175402 | + | 0 | GTCGTTGAACTCACTGTCAG | AGG | \|\|\|\|\|\|\|\|[\|\|\|\|\|\|\|\|\|\|\|\|]PAM | NA | Intergenic | NA | NA |
| T13319 | CTTGTAACCCCCGTGTCGTGTGG | 1000 | | | | | | | | | |
| Oligo fwd | CACCCTTGTAACCCCCGTGTCGTG | | | | | | | | | | |
| Oligo rev | AAACCACGACACGGGGGTTACAAG | | | | | | | | | | |
| Chromosome | start | end | strand | MM | target_seq | PAM | alignment | distance | position | gene name | gene id |
| GU305914.1 | 175445 | 175467 | + | 0 | CTTGTAACCCCCGTGTCGTG | TGG | \|\|\|\|\|\|\|\|[\|\|\|\|\|\|\|\|\|\|\|\|]PAM | NA | Intergenic | NA | NA |
| T13320 | TGTGGTGTTCATAGTTAACGTGG | 1000 | | | | | | | | | |
| Oligo fwd | CACCTGTGGTGTTCATAGTTAACG | | | | | | | | | | |
| Oligo rev | AAACCGTTAACTATGAACACCACA | | | | | | | | | | |
| Chromosome | start | end | strand | MM | target_seq | PAM | alignment | distance | position | gene name | gene id |
| GU305914.1 | 175463 | 175485 | + | 0 | TGTGGTGTTCATAGTTAACG | TGG | \|\|\|\|\|\|\|\|[\|\|\|\|\|\|\|\|\|\|\|\|]PAM | NA | Intergenic | NA | NA |
| T13321 | TATCCTCCGTCGAGACCGTAAGG | 1000 | | | | | | | | | |
| Oligo fwd | CACCTATCCTCCGTCGAGACCGTA | | | | | | | | | | |
| Oligo rev | AAACTACGGTCTCGACGGAGGATA | | | | | | | | | | |
| Chromosome | start | end | strand | MM | target_seq | PAM | alignment | distance | position | gene name | gene id |
| GU305914.1 | 175527 | 175549 | + | 0 | TATCCTCCGTCGAGACCGTA | AGG | \|\|\|\|\|\|\|\|[\|\|\|\|\|\|\|\|\|\|\|\|]PAM | NA | Intergenic | NA | NA |
| T13322 | ACCGTAAGGTTCCATGACGACGG | 1000 | | | | | | | | | |
| Oligo fwd | CACCACCGTAAGGTTCCATGACGA | | | | | | | | | | |
| Oligo rev | AAACTCGTCATGGAACCTTACGGT | | | | | | | | | | |
| Chromosome | start | end | strand | MM | target_seq | PAM | alignment | distance | position | gene name | gene id |
| GU305914.1 | 175541 | 175563 | + | 0 | ACCGTAAGGTTCCATGACGA | CGG | \|\|\|\|\|\|\|\|[\|\|\|\|\|\|\|\|\|\|\|\|]PAM | NA | Intergenic | NA | NA |
| T13323 | ACCACGATCCTGTATCACCGTGG | 1000 | | | | | | | | | |
| Oligo fwd | CACCACCACGATCCTGTATCACCG | | | | | | | | | | |
| Oligo rev | AAACCGGTGATACAGGATCGTGGT | | | | | | | | | | |
| Chromosome | start | end | strand | MM | target_seq | PAM | alignment | distance | position | gene name | gene id |
| GU305914.1 | 175615 | 175637 | + | 0 | ACCACGATCCTGTATCACCG | TGG | \|\|\|\|\|\|\|\|[\|\|\|\|\|\|\|\|\|\|\|\|]PAM | NA | Intergenic | NA | NA |
| T13324 | TGTATCACCGTGGCCTATACAGG | 1000 | | | | | | | | | |
| Oligo fwd | CACCTGTATCACCGTGGCCTATAC | | | | | | | | | | |
| Oligo rev | AAACGTATAGGCCACGGTGATACA | | | | | | | | | | |
| Chromosome | start | end | strand | MM | target_seq | PAM | alignment | distance | position | gene name | gene id |
| GU305914.1 | 175625 | 175647 | + | 0 | TGTATCACCGTGGCCTATAC | AGG | \|\|\|\|\|\|\|\|[\|\|\|\|\|\|\|\|\|\|\|\|]PAM | NA | Intergenic | NA | NA |
| T13325 | GGTCATGTTAGTCGTTTTCCAGG | 1000 | | | | | | | | | |
| Oligo fwd | CACCGGTCATGTTAGTCGTTTTCC | | | | | | | | | | |
| Oligo rev | AAACGGAAAACGACTAACATGACC | | | | | | | | | | |
| Chromosome | start | end | strand | MM | target_seq | PAM | alignment | distance | position | gene name | gene id |
| GU305914.1 | 175646 | 175668 | + | 0 | GGTCATGTTAGTCGTTTTCC | AGG | \|\|\|\|\|\|\|\|[\|\|\|\|\|\|\|\|\|\|\|\|]PAM | NA | Intergenic | NA | NA |
| T13326 | GTTAGTCGTTTTCCAGGTCGAGG | 1000 | | | | | | | | | |
| Oligo fwd | CACCGTTAGTCGTTTTCCAGGTCG | | | | | | | | | | |
| Oligo rev | AAACCGACCTGGAAAACGACTAAC | | | | | | | | | | |
| Chromosome | start | end | strand | MM | target_seq | PAM | alignment | distance | position | gene name | gene id |
| GU305914.1 | 175652 | 175674 | + | 0 | GTTAGTCGTTTTCCAGGTCG | AGG | \|\|\|\|\|\|\|\|[\|\|\|\|\|\|\|\|\|\|\|\|]PAM | NA | Intergenic | NA | NA |
| T13327 | TCTTACATTGCACGCGACTCAGG | 1000 | | | | | | | | | |
| Oligo fwd | CACCTCTTACATTGCACGCGACTC | | | | | | | | | | |
| Oligo rev | AAACGAGTCGCGTGCAATGTAAGA | | | | | | | | | | |
| Chromosome | start | end | strand | MM | target_seq | PAM | alignment | distance | position | gene name | gene id |
| GU305914.1 | 175690 | 175712 | + | 0 | TCTTACATTGCACGCGACTC | AGG | \|\|\|\|\|\|\|\|[\|\|\|\|\|\|\|\|\|\|\|\|]PAM | NA | Intergenic | NA | NA |
| T13328 | GTAGATTACGCACAAGATCACGG | 1000 | | | | | | | | | |
| Oligo fwd | CACCGTAGATTACGCACAAGATCA | | | | | | | | | | |
| Oligo rev | AAACTGATCTTGTGCGTAATCTAC | | | | | | | | | | |
| Chromosome | start | end | strand | MM | target_seq | PAM | alignment | distance | position | gene name | gene id |
| GU305914.1 | 175712 | 175734 | + | 0 | GTAGATTACGCACAAGATCA | CGG | \|\|\|\|\|\|\|\|[\|\|\|\|\|\|\|\|\|\|\|\|]PAM | NA | Intergenic | NA | NA |
| T13329 | ACAAGATCACGGCGAAGCACAGG | 1000 | | | | | | | | | |
| Oligo fwd | CACCACAAGATCACGGCGAAGCAC | | | | | | | | | | |
| Oligo rev | AAACGTGCTTCGCCGTGATCTTGT | | | | | | | | | | |
| Chromosome | start | end | strand | MM | target_seq | PAM | alignment | distance | position | gene name | gene id |
| GU305914.1 | 175723 | 175745 | + | 0 | ACAAGATCACGGCGAAGCAC | AGG | \|\|\|\|\|\|\|\|[\|\|\|\|\|\|\|\|\|\|\|\|]PAM | NA | Intergenic | NA | NA |
| T13330 | AGATCACGGCGAAGCACAGGAGG | 1000 | | | | | | | | | |

<table>
<tr><td>Oligo fwd</td><td>CACCAGATCACGGCGAAGCACAGG</td></tr>
<tr><td>Oligo rev</td><td>AAACCCTGTGCTTCGCCGTGATCT</td></tr>
<tr><td>Chromosome</td><td>start</td><td>end</td><td>strand</td><td>MM</td><td>target_seq</td><td>PAM</td><td>alignment</td><td>distance</td><td>position</td><td>gene name</td><td>gene id</td></tr>
<tr><td>GU305914.1</td><td>175726</td><td>175748</td><td>+</td><td>0</td><td>AGATCACGGCGAAGCACAGG</td><td>AGG</td><td>||||||||[||||||||||||]PAM</td><td>NA</td><td>Intergenic</td><td>NA</td><td>NA</td></tr>
<tr><td></td></tr>
<tr><td>T13331</td><td>CGAGCACGCCCCATCGTCCACGG</td><td>1000</td></tr>
<tr><td>Oligo fwd</td><td>CACCCGAGCACGCCCCATCGTCCA</td></tr>
<tr><td>Oligo rev</td><td>AAACTGGACGATGGGGCGTGCTCG</td></tr>
<tr><td>Chromosome</td><td>start</td><td>end</td><td>strand</td><td>MM</td><td>target_seq</td><td>PAM</td><td>alignment</td><td>distance</td><td>position</td><td>gene name</td><td>gene id</td></tr>
<tr><td>GU305914.1</td><td>175752</td><td>175774</td><td>+</td><td>0</td><td>CGAGCACGCCCCATCGTCCA</td><td>CGG</td><td>||||||||[||||||||||||]PAM</td><td>NA</td><td>Intergenic</td><td>NA</td><td>NA</td></tr>
<tr><td></td></tr>
<tr><td>T13332</td><td>CCAACAACACCGCACCTGCGAGG</td><td>1000</td></tr>
<tr><td>Oligo fwd</td><td>CACCCCAACAACACCGCACCTGCG</td></tr>
<tr><td>Oligo rev</td><td>AAACCGCAGGTGCGGTGTTGTTGG</td></tr>
<tr><td>Chromosome</td><td>start</td><td>end</td><td>strand</td><td>MM</td><td>target_seq</td><td>PAM</td><td>alignment</td><td>distance</td><td>position</td><td>gene name</td><td>gene id</td></tr>
<tr><td>GU305914.1</td><td>175838</td><td>175860</td><td>+</td><td>0</td><td>CCAACAACACCGCACCTGCG</td><td>AGG</td><td>||||||||[||||||||||||]PAM</td><td>NA</td><td>Intergenic</td><td>NA</td><td>NA</td></tr>
<tr><td></td></tr>
<tr><td>T13333</td><td>CGAGGCTAATTAATAGCACCTGG</td><td>1000</td></tr>
<tr><td>Oligo fwd</td><td>CACCCGAGGCTAATTAATAGCACC</td></tr>
<tr><td>Oligo rev</td><td>AAACGGTGCTATTAATTAGCCTCG</td></tr>
<tr><td>Chromosome</td><td>start</td><td>end</td><td>strand</td><td>MM</td><td>target_seq</td><td>PAM</td><td>alignment</td><td>distance</td><td>position</td><td>gene name</td><td>gene id</td></tr>
<tr><td>GU305914.1</td><td>175856</td><td>175878</td><td>+</td><td>0</td><td>CGAGGCTAATTAATAGCACC</td><td>TGG</td><td>||||||||[||||||||||||]PAM</td><td>NA</td><td>Intergenic</td><td>NA</td><td>NA</td></tr>
<tr><td></td></tr>
<tr><td>T13334</td><td>CTAATTAATAGCACCTGGAGTGG</td><td>1000</td></tr>
<tr><td>Oligo fwd</td><td>CACCCTAATTAATAGCACCTGGAG</td></tr>
<tr><td>Oligo rev</td><td>AAACCTCCAGGTGCTATTAATTAG</td></tr>
<tr><td>Chromosome</td><td>start</td><td>end</td><td>strand</td><td>MM</td><td>target_seq</td><td>PAM</td><td>alignment</td><td>distance</td><td>position</td><td>gene name</td><td>gene id</td></tr>
<tr><td>GU305914.1</td><td>175861</td><td>175883</td><td>+</td><td>0</td><td>CTAATTAATAGCACCTGGAG</td><td>TGG</td><td>||||||||[||||||||||||]PAM</td><td>NA</td><td>Intergenic</td><td>NA</td><td>NA</td></tr>
<tr><td></td></tr>
<tr><td>T13335</td><td>TAATTAATAGCACCTGGAGTGGG</td><td>1000</td></tr>
<tr><td>Oligo fwd</td><td>CACCTAATTAATAGCACCTGGAGT</td></tr>
<tr><td>Oligo rev</td><td>AAACACTCCAGGTGCTATTAATTA</td></tr>
<tr><td>Chromosome</td><td>start</td><td>end</td><td>strand</td><td>MM</td><td>target_seq</td><td>PAM</td><td>alignment</td><td>distance</td><td>position</td><td>gene name</td><td>gene id</td></tr>
<tr><td>GU305914.1</td><td>175862</td><td>175884</td><td>+</td><td>0</td><td>TAATTAATAGCACCTGGAGT</td><td>GGG</td><td>||||||||[||||||||||||]PAM</td><td>NA</td><td>Intergenic</td><td>NA</td><td>NA</td></tr>
<tr><td></td></tr>
<tr><td>T13336</td><td>ACAAACCACGTCCCCACCTGAGG</td><td>1000</td></tr>
<tr><td>Oligo fwd</td><td>CACCACAAACCACGTCCCCACCTG</td></tr>
<tr><td>Oligo rev</td><td>AAACCAGGTGGGGACGTGGTTTGT</td></tr>
<tr><td>Chromosome</td><td>start</td><td>end</td><td>strand</td><td>MM</td><td>target_seq</td><td>PAM</td><td>alignment</td><td>distance</td><td>position</td><td>gene name</td><td>gene id</td></tr>
<tr><td>GU305914.1</td><td>175891</td><td>175913</td><td>+</td><td>0</td><td>ACAAACCACGTCCCCACCTG</td><td>AGG</td><td>||||||||[||||||||||||]PAM</td><td>NA</td><td>Intergenic</td><td>NA</td><td>NA</td></tr>
<tr><td></td></tr>
<tr><td>T13337</td><td>AGGCGTTACTACCTGTCCGTCGG</td><td>1000</td></tr>
<tr><td>Oligo fwd</td><td>CACCAGGCGTTACTACCTGTCCGT</td></tr>
<tr><td>Oligo rev</td><td>AAACACGGACAGGTAGTAACGCCT</td></tr>
<tr><td>Chromosome</td><td>start</td><td>end</td><td>strand</td><td>MM</td><td>target_seq</td><td>PAM</td><td>alignment</td><td>distance</td><td>position</td><td>gene name</td><td>gene id</td></tr>
<tr><td>GU305914.1</td><td>175911</td><td>175933</td><td>+</td><td>0</td><td>AGGCGTTACTACCTGTCCGT</td><td>CGG</td><td>||||||||[||||||||||||]PAM</td><td>NA</td><td>Intergenic</td><td>NA</td><td>NA</td></tr>
<tr><td></td></tr>
<tr><td>T13338</td><td>TGTCCGTCGGTGACAGACAACGG</td><td>1000</td></tr>
<tr><td>Oligo fwd</td><td>CACCTGTCCGTCGGTGACAGACAA</td></tr>
<tr><td>Oligo rev</td><td>AAACTTGTCTGTCACCGACGGACA</td></tr>
<tr><td>Chromosome</td><td>start</td><td>end</td><td>strand</td><td>MM</td><td>target_seq</td><td>PAM</td><td>alignment</td><td>distance</td><td>position</td><td>gene name</td><td>gene id</td></tr>
<tr><td>GU305914.1</td><td>175924</td><td>175946</td><td>+</td><td>0</td><td>TGTCCGTCGGTGACAGACAA</td><td>CGG</td><td>||||||||[||||||||||||]PAM</td><td>NA</td><td>Intergenic</td><td>NA</td><td>NA</td></tr>
<tr><td></td></tr>
<tr><td>T13339</td><td>GTCCGTCGGTGACAGACAACGGG</td><td>1000</td></tr>
<tr><td>Oligo fwd</td><td>CACCGTCCGTCGGTGACAGACAAC</td></tr>
<tr><td>Oligo rev</td><td>AAACGTTGTCTGTCACCGACGGAC</td></tr>
<tr><td>Chromosome</td><td>start</td><td>end</td><td>strand</td><td>MM</td><td>target_seq</td><td>PAM</td><td>alignment</td><td>distance</td><td>position</td><td>gene name</td><td>gene id</td></tr>
<tr><td>GU305914.1</td><td>175925</td><td>175947</td><td>+</td><td>0</td><td>GTCCGTCGGTGACAGACAAC</td><td>GGG</td><td>||||||||[||||||||||||]PAM</td><td>NA</td><td>Intergenic</td><td>NA</td><td>NA</td></tr>
<tr><td></td></tr>
<tr><td>T13340</td><td>TAATCATGCGTATCGATTTCCGG</td><td>1000</td></tr>
<tr><td>Oligo fwd</td><td>CACCTAATCATGCGTATCGATTTC</td></tr>
<tr><td>Oligo rev</td><td>AAACGAAATCGATACGCATGATTA</td></tr>
<tr><td>Chromosome</td><td>start</td><td>end</td><td>strand</td><td>MM</td><td>target_seq</td><td>PAM</td><td>alignment</td><td>distance</td><td>position</td><td>gene name</td><td>gene id</td></tr>
<tr><td>GU305914.1</td><td>175948</td><td>175970</td><td>+</td><td>0</td><td>TAATCATGCGTATCGATTTC</td><td>CGG</td><td>||||||||[||||||||||||]PAM</td><td>NA</td><td>Intergenic</td><td>NA</td><td>NA</td></tr>
<tr><td></td></tr>
<tr><td>T13341</td><td>TTCCGGTTCACCTGAGACCACGG</td><td>1000</td></tr>
<tr><td>Oligo fwd</td><td>CACCTTCCGGTTCACCTGAGACCA</td></tr>
<tr><td>Oligo rev</td><td>AAACTGGTCTCAGGTGAACCGGAA</td></tr>
<tr><td>Chromosome</td><td>start</td><td>end</td><td>strand</td><td>MM</td><td>target_seq</td><td>PAM</td><td>alignment</td><td>distance</td><td>position</td><td>gene name</td><td>gene id</td></tr>
<tr><td>GU305914.1</td><td>175965</td><td>175987</td><td>+</td><td>0</td><td>TTCCGGTTCACCTGAGACCA</td><td>CGG</td><td>||||||||[||||||||||||]PAM</td><td>NA</td><td>Intergenic</td><td>NA</td><td>NA</td></tr>
<tr><td></td></tr>
<tr><td>T13342</td><td>TCCGGTTCACCTGAGACCACGGG</td><td>1000</td></tr>
<tr><td>Oligo fwd</td><td>CACCTCCGGTTCACCTGAGACCAC</td></tr>
<tr><td>Oligo rev</td><td>AAACGTGGTCTCAGGTGAACCGGA</td></tr>
<tr><td>Chromosome</td><td>start</td><td>end</td><td>strand</td><td>MM</td><td>target_seq</td><td>PAM</td><td>alignment</td><td>distance</td><td>position</td><td>gene name</td><td>gene id</td></tr>
<tr><td>GU305914.1</td><td>175966</td><td>175988</td><td>+</td><td>0</td><td>TCCGGTTCACCTGAGACCAC</td><td>GGG</td><td>||||||||[||||||||||||]PAM</td><td>NA</td><td>Intergenic</td><td>NA</td><td>NA</td></tr>
<tr><td></td></tr>
<tr><td>T13343</td><td>CCGGTTCACCTGAGACCACGGGG</td><td>1000</td></tr>
<tr><td>Oligo fwd</td><td>CACCCCGGTTCACCTGAGACCACG</td></tr>
<tr><td>Oligo rev</td><td>AAACCGTGGTCTCAGGTGAACCGG</td></tr>
<tr><td>Chromosome</td><td>start</td><td>end</td><td>strand</td><td>MM</td><td>target_seq</td><td>PAM</td><td>alignment</td><td>distance</td><td>position</td><td>gene name</td><td>gene id</td></tr>
<tr><td>GU305914.1</td><td>175967</td><td>175989</td><td>+</td><td>0</td><td>CCGGTTCACCTGAGACCACG</td><td>GGG</td><td>||||||||[||||||||||||]PAM</td><td>NA</td><td>Intergenic</td><td>NA</td><td>NA</td></tr>
<tr><td></td></tr>
<tr><td>T13344</td><td>CACCTGAGACCACGGGGAGAAGG</td><td>1000</td></tr>
<tr><td>Oligo fwd</td><td>CACCCACCTGAGACCACGGGGAGA</td></tr>
<tr><td>Oligo rev</td><td>AAACTCTCCCCGTGGTCTCAGGTG</td></tr>
<tr><td>Chromosome</td><td>start</td><td>end</td><td>strand</td><td>MM</td><td>target_seq</td><td>PAM</td><td>alignment</td><td>distance</td><td>position</td><td>gene name</td><td>gene id</td></tr>
<tr><td>GU305914.1</td><td>175973</td><td>175995</td><td>+</td><td>0</td><td>CACCTGAGACCACGGGGAGA</td><td>AGG</td><td>||||||||[||||||||||||]PAM</td><td>NA</td><td>Intergenic</td><td>NA</td><td>NA</td></tr>
</table>

<table>
<tr><td>T13345</td><td>ATAGTCTGCGTGCGAGTAGCAGG</td><td>1000</td><td></td><td></td><td></td><td></td><td></td><td></td><td></td><td></td></tr>
<tr><td>Oligo fwd</td><td>CACCATAGTCTGCGTGCGAGTAGC</td><td></td><td></td><td></td><td></td><td></td><td></td><td></td><td></td><td></td></tr>
<tr><td>Oligo rev</td><td>AAACGCTACTCGCACGCAGACTAT</td><td></td><td></td><td></td><td></td><td></td><td></td><td></td><td></td><td></td></tr>
<tr><td>Chromosome</td><td>start</td><td>end</td><td>strand</td><td>MM</td><td>target_seq</td><td>PAM</td><td>alignment</td><td>distance</td><td>position</td><td>gene name</td><td>gene id</td></tr>
<tr><td>GU305914.1</td><td>176008</td><td>176030</td><td>+</td><td>0</td><td>ATAGTCTGCGTGCGAGTAGC</td><td>AGG</td><td>||||||||[||||||||||||]PAM</td><td>NA</td><td>Intergenic</td><td>NA</td><td>NA</td></tr>
<tr><td>T13346</td><td>CGAGTAGCAGGACGCCCTATAGG</td><td>1000</td><td></td><td></td><td></td><td></td><td></td><td></td><td></td><td></td><td></td></tr>
<tr><td>Oligo fwd</td><td>CACCCGAGTAGCAGGACGCCCTAT</td><td></td><td></td><td></td><td></td><td></td><td></td><td></td><td></td><td></td><td></td></tr>
<tr><td>Oligo rev</td><td>AAACATAGGGCGTCCTGCTACTCG</td><td></td><td></td><td></td><td></td><td></td><td></td><td></td><td></td><td></td><td></td></tr>
<tr><td>Chromosome</td><td>start</td><td>end</td><td>strand</td><td>MM</td><td>target_seq</td><td>PAM</td><td>alignment</td><td>distance</td><td>position</td><td>gene name</td><td>gene id</td></tr>
<tr><td>GU305914.1</td><td>176020</td><td>176042</td><td>+</td><td>0</td><td>CGAGTAGCAGGACGCCCTAT</td><td>AGG</td><td>||||||||[||||||||||||]PAM</td><td>NA</td><td>Intergenic</td><td>NA</td><td>NA</td></tr>
<tr><td>T13347</td><td>TATAGGTGTGTTCGAGTGAGAGG</td><td>1000</td><td></td><td></td><td></td><td></td><td></td><td></td><td></td><td></td><td></td></tr>
<tr><td>Oligo fwd</td><td>CACCTATAGGTGTGTTCGAGTGAG</td><td></td><td></td><td></td><td></td><td></td><td></td><td></td><td></td><td></td><td></td></tr>
<tr><td>Oligo rev</td><td>AAACCTCACTCGAACACACCTATA</td><td></td><td></td><td></td><td></td><td></td><td></td><td></td><td></td><td></td><td></td></tr>
<tr><td>Chromosome</td><td>start</td><td>end</td><td>strand</td><td>MM</td><td>target_seq</td><td>PAM</td><td>alignment</td><td>distance</td><td>position</td><td>gene name</td><td>gene id</td></tr>
<tr><td>GU305914.1</td><td>176037</td><td>176059</td><td>+</td><td>0</td><td>TATAGGTGTGTTCGAGTGAG</td><td>AGG</td><td>||||||||[||||||||||||]PAM</td><td>NA</td><td>Intergenic</td><td>NA</td><td>NA</td></tr>
<tr><td>T13348</td><td>GTTCGAGTGAGAGGCCTGAACGG</td><td>1000</td><td></td><td></td><td></td><td></td><td></td><td></td><td></td><td></td><td></td></tr>
<tr><td>Oligo fwd</td><td>CACCGTTCGAGTGAGAGGCCTGAA</td><td></td><td></td><td></td><td></td><td></td><td></td><td></td><td></td><td></td><td></td></tr>
<tr><td>Oligo rev</td><td>AAACTTCAGGCCTCTCACTCGAAC</td><td></td><td></td><td></td><td></td><td></td><td></td><td></td><td></td><td></td><td></td></tr>
<tr><td>Chromosome</td><td>start</td><td>end</td><td>strand</td><td>MM</td><td>target_seq</td><td>PAM</td><td>alignment</td><td>distance</td><td>position</td><td>gene name</td><td>gene id</td></tr>
<tr><td>GU305914.1</td><td>176046</td><td>176068</td><td>+</td><td>0</td><td>GTTCGAGTGAGAGGCCTGAA</td><td>CGG</td><td>||||||||[||||||||||||]PAM</td><td>NA</td><td>Intergenic</td><td>NA</td><td>NA</td></tr>
<tr><td>T13349</td><td>ACGGTCGCCGAATGAGACCTCGG</td><td>1000</td><td></td><td></td><td></td><td></td><td></td><td></td><td></td><td></td><td></td></tr>
<tr><td>Oligo fwd</td><td>CACCACGGTCGCCGAATGAGACCT</td><td></td><td></td><td></td><td></td><td></td><td></td><td></td><td></td><td></td><td></td></tr>
<tr><td>Oligo rev</td><td>AAACAGGTCTCATTCGGCGACCGT</td><td></td><td></td><td></td><td></td><td></td><td></td><td></td><td></td><td></td><td></td></tr>
<tr><td>Chromosome</td><td>start</td><td>end</td><td>strand</td><td>MM</td><td>target_seq</td><td>PAM</td><td>alignment</td><td>distance</td><td>position</td><td>gene name</td><td>gene id</td></tr>
<tr><td>GU305914.1</td><td>176065</td><td>176087</td><td>+</td><td>0</td><td>ACGGTCGCCGAATGAGACCT</td><td>CGG</td><td>||||||||[||||||||||||]PAM</td><td>NA</td><td>Intergenic</td><td>NA</td><td>NA</td></tr>
<tr><td>T13350</td><td>GTCGCCGAATGAGACCTCGGTGG</td><td>1000</td><td></td><td></td><td></td><td></td><td></td><td></td><td></td><td></td><td></td></tr>
<tr><td>Oligo fwd</td><td>CACCGTCGCCGAATGAGACCTCGG</td><td></td><td></td><td></td><td></td><td></td><td></td><td></td><td></td><td></td><td></td></tr>
<tr><td>Oligo rev</td><td>AAACCCGAGGTCTCATTCGGCGAC</td><td></td><td></td><td></td><td></td><td></td><td></td><td></td><td></td><td></td><td></td></tr>
<tr><td>Chromosome</td><td>start</td><td>end</td><td>strand</td><td>MM</td><td>target_seq</td><td>PAM</td><td>alignment</td><td>distance</td><td>position</td><td>gene name</td><td>gene id</td></tr>
<tr><td>GU305914.1</td><td>176068</td><td>176090</td><td>+</td><td>0</td><td>GTCGCCGAATGAGACCTCGG</td><td>TGG</td><td>||||||||[||||||||||||]PAM</td><td>NA</td><td>Intergenic</td><td>NA</td><td>NA</td></tr>
<tr><td>T13351</td><td>CGAATGAGACCTCGGTGGCGAGG</td><td>1000</td><td></td><td></td><td></td><td></td><td></td><td></td><td></td><td></td><td></td></tr>
<tr><td>Oligo fwd</td><td>CACCCGAATGAGACCTCGGTGGCG</td><td></td><td></td><td></td><td></td><td></td><td></td><td></td><td></td><td></td><td></td></tr>
<tr><td>Oligo rev</td><td>AAACCGCCACCGAGGTCTCATTCG</td><td></td><td></td><td></td><td></td><td></td><td></td><td></td><td></td><td></td><td></td></tr>
<tr><td>Chromosome</td><td>start</td><td>end</td><td>strand</td><td>MM</td><td>target_seq</td><td>PAM</td><td>alignment</td><td>distance</td><td>position</td><td>gene name</td><td>gene id</td></tr>
<tr><td>GU305914.1</td><td>176073</td><td>176095</td><td>+</td><td>0</td><td>CGAATGAGACCTCGGTGGCG</td><td>AGG</td><td>||||||||[||||||||||||]PAM</td><td>NA</td><td>Intergenic</td><td>NA</td><td>NA</td></tr>
<tr><td>T13352</td><td>TACCGCATCGCTCTTTCTGCTGG</td><td>1000</td><td></td><td></td><td></td><td></td><td></td><td></td><td></td><td></td><td></td></tr>
<tr><td>Oligo fwd</td><td>CACCTACCGCATCGCTCTTTCTGC</td><td></td><td></td><td></td><td></td><td></td><td></td><td></td><td></td><td></td><td></td></tr>
<tr><td>Oligo rev</td><td>AAACGCAGAAAGAGCGATGCGGTA</td><td></td><td></td><td></td><td></td><td></td><td></td><td></td><td></td><td></td><td></td></tr>
<tr><td>Chromosome</td><td>start</td><td>end</td><td>strand</td><td>MM</td><td>target_seq</td><td>PAM</td><td>alignment</td><td>distance</td><td>position</td><td>gene name</td><td>gene id</td></tr>
<tr><td>GU305914.1</td><td>176115</td><td>176137</td><td>+</td><td>0</td><td>TACCGCATCGCTCTTTCTGC</td><td>TGG</td><td>||||||||[||||||||||||]PAM</td><td>NA</td><td>Intergenic</td><td>NA</td><td>NA</td></tr>
<tr><td>T13353</td><td>TTCCTTCGAGCTCACTGTTCCGG</td><td>1000</td><td></td><td></td><td></td><td></td><td></td><td></td><td></td><td></td><td></td></tr>
<tr><td>Oligo fwd</td><td>CACCTTCCTTCGAGCTCACTGTTC</td><td></td><td></td><td></td><td></td><td></td><td></td><td></td><td></td><td></td><td></td></tr>
<tr><td>Oligo rev</td><td>AAACGAACAGTGAGCTCGAAGGAA</td><td></td><td></td><td></td><td></td><td></td><td></td><td></td><td></td><td></td><td></td></tr>
<tr><td>Chromosome</td><td>start</td><td>end</td><td>strand</td><td>MM</td><td>target_seq</td><td>PAM</td><td>alignment</td><td>distance</td><td>position</td><td>gene name</td><td>gene id</td></tr>
<tr><td>GU305914.1</td><td>176139</td><td>176161</td><td>+</td><td>0</td><td>TTCCTTCGAGCTCACTGTTC</td><td>CGG</td><td>||||||||[||||||||||||]PAM</td><td>NA</td><td>Intergenic</td><td>NA</td><td>NA</td></tr>
<tr><td>T13354</td><td>CGAGCTCACTGTTCCGGATGCGG</td><td>1000</td><td></td><td></td><td></td><td></td><td></td><td></td><td></td><td></td><td></td></tr>
<tr><td>Oligo fwd</td><td>CACCCGAGCTCACTGTTCCGGATG</td><td></td><td></td><td></td><td></td><td></td><td></td><td></td><td></td><td></td><td></td></tr>
<tr><td>Oligo rev</td><td>AAACCATCCGGAACAGTGAGCTCG</td><td></td><td></td><td></td><td></td><td></td><td></td><td></td><td></td><td></td><td></td></tr>
<tr><td>Chromosome</td><td>start</td><td>end</td><td>strand</td><td>MM</td><td>target_seq</td><td>PAM</td><td>alignment</td><td>distance</td><td>position</td><td>gene name</td><td>gene id</td></tr>
<tr><td>GU305914.1</td><td>176145</td><td>176167</td><td>+</td><td>0</td><td>CGAGCTCACTGTTCCGGATG</td><td>CGG</td><td>||||||||[||||||||||||]PAM</td><td>NA</td><td>Intergenic</td><td>NA</td><td>NA</td></tr>
<tr><td>T13355</td><td>CACTGTTCCGGATGCGGTTGTGG</td><td>1000</td><td></td><td></td><td></td><td></td><td></td><td></td><td></td><td></td><td></td></tr>
<tr><td>Oligo fwd</td><td>CACCCACTGTTCCGGATGCGGTTG</td><td></td><td></td><td></td><td></td><td></td><td></td><td></td><td></td><td></td><td></td></tr>
<tr><td>Oligo rev</td><td>AAACCAACCGCATCCGGAACAGTG</td><td></td><td></td><td></td><td></td><td></td><td></td><td></td><td></td><td></td><td></td></tr>
<tr><td>Chromosome</td><td>start</td><td>end</td><td>strand</td><td>MM</td><td>target_seq</td><td>PAM</td><td>alignment</td><td>distance</td><td>position</td><td>gene name</td><td>gene id</td></tr>
<tr><td>GU305914.1</td><td>176151</td><td>176173</td><td>+</td><td>0</td><td>CACTGTTCCGGATGCGGTTG</td><td>TGG</td><td>||||||||[||||||||||||]PAM</td><td>NA</td><td>Intergenic</td><td>NA</td><td>NA</td></tr>
<tr><td>T13356</td><td>GGTTGTGGCACCTGTTCCATTGG</td><td>1000</td><td></td><td></td><td></td><td></td><td></td><td></td><td></td><td></td><td></td></tr>
<tr><td>Oligo fwd</td><td>CACCGGTTGTGGCACCTGTTCCAT</td><td></td><td></td><td></td><td></td><td></td><td></td><td></td><td></td><td></td><td></td></tr>
<tr><td>Oligo rev</td><td>AAACATGGAACAGGTGCCACAACC</td><td></td><td></td><td></td><td></td><td></td><td></td><td></td><td></td><td></td><td></td></tr>
<tr><td>Chromosome</td><td>start</td><td>end</td><td>strand</td><td>MM</td><td>target_seq</td><td>PAM</td><td>alignment</td><td>distance</td><td>position</td><td>gene name</td><td>gene id</td></tr>
<tr><td>GU305914.1</td><td>176166</td><td>176188</td><td>+</td><td>0</td><td>GGTTGTGGCACCTGTTCCAT</td><td>TGG</td><td>||||||||[||||||||||||]PAM</td><td>NA</td><td>Intergenic</td><td>NA</td><td>NA</td></tr>
<tr><td>T13357</td><td>TGCTGAACTGTTAACACTGCCGG</td><td>1000</td><td></td><td></td><td></td><td></td><td></td><td></td><td></td><td></td><td></td></tr>
<tr><td>Oligo fwd</td><td>CACCTGCTGAACTGTTAACACTGC</td><td></td><td></td><td></td><td></td><td></td><td></td><td></td><td></td><td></td><td></td></tr>
<tr><td>Oligo rev</td><td>AAACGCAGTGTTAACAGTTCAGCA</td><td></td><td></td><td></td><td></td><td></td><td></td><td></td><td></td><td></td><td></td></tr>
<tr><td>Chromosome</td><td>start</td><td>end</td><td>strand</td><td>MM</td><td>target_seq</td><td>PAM</td><td>alignment</td><td>distance</td><td>position</td><td>gene name</td><td>gene id</td></tr>
<tr><td>GU305914.1</td><td>176196</td><td>176218</td><td>+</td><td>0</td><td>TGCTGAACTGTTAACACTGC</td><td>CGG</td><td>||||||||[||||||||||||]PAM</td><td>NA</td><td>Intergenic</td><td>NA</td><td>NA</td></tr>
<tr><td>T13358</td><td>GCCGGTTGCGACGACCCGTACGG</td><td>1000</td><td></td><td></td><td></td><td></td><td></td><td></td><td></td><td></td><td></td></tr>
<tr><td>Oligo fwd</td><td>CACCGCCGGTTGCGACGACCCGTA</td><td></td><td></td><td></td><td></td><td></td><td></td><td></td><td></td><td></td><td></td></tr>
<tr><td>Oligo rev</td><td>AAACTACGGGTCGTCGCAACCGGC</td><td></td><td></td><td></td><td></td><td></td><td></td><td></td><td></td><td></td><td></td></tr>
<tr><td>Chromosome</td><td>start</td><td>end</td><td>strand</td><td>MM</td><td>target_seq</td><td>PAM</td><td>alignment</td><td>distance</td><td>position</td><td>gene name</td><td>gene id</td></tr>
<tr><td>GU305914.1</td><td>176214</td><td>176236</td><td>+</td><td>0</td><td>GCCGGTTGCGACGACCCGTA</td><td>CGG</td><td>||||||||[||||||||||||]PAM</td><td>NA</td><td>Intergenic</td><td>NA</td><td>NA</td></tr>
<tr><td>T13359</td><td>GGTTGCGACGACCCGTACGGAGG</td><td>1000</td><td></td><td></td><td></td><td></td><td></td><td></td><td></td><td></td><td></td></tr>
<tr><td>Oligo fwd</td><td>CACCGGTTGCGACGACCCGTACGG</td><td></td><td></td><td></td><td></td><td></td><td></td><td></td><td></td><td></td><td></td></tr>
<tr><td>Oligo rev</td><td>AAACCCGTACGGGTCGTCGCAACC</td><td></td><td></td><td></td><td></td><td></td><td></td><td></td><td></td><td></td><td></td></tr>
<tr><td>Chromosome</td><td>start</td><td>end</td><td>strand</td><td>MM</td><td>target_seq</td><td>PAM</td><td>alignment</td><td>distance</td><td>position</td><td>gene name</td><td>gene id</td></tr>
<tr><td>GU305914.1</td><td>176217</td><td>176239</td><td>+</td><td>0</td><td>GGTTGCGACGACCCGTACGG</td><td>AGG</td><td>||||||||[||||||||||||]PAM</td><td>NA</td><td>Intergenic</td><td>NA</td><td>NA</td></tr>
</table>

<table>
<tr><td>GU305914.1</td><td>147482</td><td>147504</td><td>-</td><td>4</td><td>GGTGGCGCCGACCCATCCGG</td><td>GGG</td><td>|||-|||-[||||||-|-|||]PAM</td><td>NA</td><td>Intergenic</td><td>NA</td><td>NA</td></tr>
<tr><td>T13360</td><td>AGATTGTCTTTCCTGCACGCTGG</td><td>1000</td><td></td><td></td><td></td><td></td><td></td><td></td><td></td><td></td><td></td></tr>
<tr><td>Oligo fwd</td><td>CACCAGATTGTCTTTCCTGCACGC</td><td></td><td></td><td></td><td></td><td></td><td></td><td></td><td></td><td></td><td></td></tr>
<tr><td>Oligo rev</td><td>AAACGCGTGCAGGAAAGACAATCT</td><td></td><td></td><td></td><td></td><td></td><td></td><td></td><td></td><td></td><td></td></tr>
<tr><td>Chromosome</td><td>start</td><td>end</td><td>strand</td><td>MM</td><td>target_seq</td><td>PAM</td><td>alignment</td><td>distance</td><td>position</td><td>gene name</td><td>gene id</td></tr>
<tr><td>GU305914.1</td><td>176241</td><td>176263</td><td>+</td><td>0</td><td>AGATTGTCTTTCCTGCACGC</td><td>TGG</td><td>||||||||[||||||||||||]PAM</td><td>NA</td><td>Intergenic</td><td>NA</td><td>NA</td></tr>
<tr><td>T13361</td><td>TGTCTTTCCTGCACGCTGGTAGG</td><td>1000</td><td></td><td></td><td></td><td></td><td></td><td></td><td></td><td></td><td></td></tr>
<tr><td>Oligo fwd</td><td>CACCTGTCTTTCCTGCACGCTGGT</td><td></td><td></td><td></td><td></td><td></td><td></td><td></td><td></td><td></td><td></td></tr>
<tr><td>Oligo rev</td><td>AAACACCAGCGTGCAGGAAAGACA</td><td></td><td></td><td></td><td></td><td></td><td></td><td></td><td></td><td></td><td></td></tr>
<tr><td>Chromosome</td><td>start</td><td>end</td><td>strand</td><td>MM</td><td>target_seq</td><td>PAM</td><td>alignment</td><td>distance</td><td>position</td><td>gene name</td><td>gene id</td></tr>
<tr><td>GU305914.1</td><td>176245</td><td>176267</td><td>+</td><td>0</td><td>TGTCTTTCCTGCACGCTGGT</td><td>AGG</td><td>||||||||[||||||||||||]PAM</td><td>NA</td><td>Intergenic</td><td>NA</td><td>NA</td></tr>
<tr><td>T13362</td><td>GTCTTTCCTGCACGCTGGTAGGG</td><td>1000</td><td></td><td></td><td></td><td></td><td></td><td></td><td></td><td></td><td></td></tr>
<tr><td>Oligo fwd</td><td>CACCGTCTTTCCTGCACGCTGGTA</td><td></td><td></td><td></td><td></td><td></td><td></td><td></td><td></td><td></td><td></td></tr>
<tr><td>Oligo rev</td><td>AAACTACCAGCGTGCAGGAAAGAC</td><td></td><td></td><td></td><td></td><td></td><td></td><td></td><td></td><td></td><td></td></tr>
<tr><td>Chromosome</td><td>start</td><td>end</td><td>strand</td><td>MM</td><td>target_seq</td><td>PAM</td><td>alignment</td><td>distance</td><td>position</td><td>gene name</td><td>gene id</td></tr>
<tr><td>GU305914.1</td><td>176246</td><td>176268</td><td>+</td><td>0</td><td>GTCTTTCCTGCACGCTGGTA</td><td>GGG</td><td>||||||||[||||||||||||]PAM</td><td>NA</td><td>Intergenic</td><td>NA</td><td>NA</td></tr>
<tr><td>T13363</td><td>TCTTTCCTGCACGCTGGTAGGGG</td><td>1000</td><td></td><td></td><td></td><td></td><td></td><td></td><td></td><td></td><td></td></tr>
<tr><td>Oligo fwd</td><td>CACCTCTTTCCTGCACGCTGGTAG</td><td></td><td></td><td></td><td></td><td></td><td></td><td></td><td></td><td></td><td></td></tr>
<tr><td>Oligo rev</td><td>AAACCTACCAGCGTGCAGGAAAGA</td><td></td><td></td><td></td><td></td><td></td><td></td><td></td><td></td><td></td><td></td></tr>
<tr><td>Chromosome</td><td>start</td><td>end</td><td>strand</td><td>MM</td><td>target_seq</td><td>PAM</td><td>alignment</td><td>distance</td><td>position</td><td>gene name</td><td>gene id</td></tr>
<tr><td>GU305914.1</td><td>176247</td><td>176269</td><td>+</td><td>0</td><td>TCTTTCCTGCACGCTGGTAG</td><td>GGG</td><td>||||||||[||||||||||||]PAM</td><td>NA</td><td>Intergenic</td><td>NA</td><td>NA</td></tr>
<tr><td>T13364</td><td>TGCACGCTGGTAGGGGTTGATGG</td><td>1000</td><td></td><td></td><td></td><td></td><td></td><td></td><td></td><td></td><td></td></tr>
<tr><td>Oligo fwd</td><td>CACCTGCACGCTGGTAGGGGTTGA</td><td></td><td></td><td></td><td></td><td></td><td></td><td></td><td></td><td></td><td></td></tr>
<tr><td>Oligo rev</td><td>AAACTCAACCCCTACCAGCGTGCA</td><td></td><td></td><td></td><td></td><td></td><td></td><td></td><td></td><td></td><td></td></tr>
<tr><td>Chromosome</td><td>start</td><td>end</td><td>strand</td><td>MM</td><td>target_seq</td><td>PAM</td><td>alignment</td><td>distance</td><td>position</td><td>gene name</td><td>gene id</td></tr>
<tr><td>GU305914.1</td><td>176254</td><td>176276</td><td>+</td><td>0</td><td>TGCACGCTGGTAGGGGTTGA</td><td>TGG</td><td>||||||||[||||||||||||]PAM</td><td>NA</td><td>Intergenic</td><td>NA</td><td>NA</td></tr>
<tr><td>T13365</td><td>GCACGCTGGTAGGGGTTGATGGG</td><td>1000</td><td></td><td></td><td></td><td></td><td></td><td></td><td></td><td></td><td></td></tr>
<tr><td>Oligo fwd</td><td>CACCGCACGCTGGTAGGGGTTGAT</td><td></td><td></td><td></td><td></td><td></td><td></td><td></td><td></td><td></td><td></td></tr>
<tr><td>Oligo rev</td><td>AAACATCAACCCCTACCAGCGTGC</td><td></td><td></td><td></td><td></td><td></td><td></td><td></td><td></td><td></td><td></td></tr>
<tr><td>Chromosome</td><td>start</td><td>end</td><td>strand</td><td>MM</td><td>target_seq</td><td>PAM</td><td>alignment</td><td>distance</td><td>position</td><td>gene name</td><td>gene id</td></tr>
<tr><td>GU305914.1</td><td>176255</td><td>176277</td><td>+</td><td>0</td><td>GCACGCTGGTAGGGGTTGAT</td><td>GGG</td><td>||||||||[||||||||||||]PAM</td><td>NA</td><td>Intergenic</td><td>NA</td><td>NA</td></tr>
<tr><td>T13366</td><td>CACGCTGGTAGGGGTTGATGGGG</td><td>1000</td><td></td><td></td><td></td><td></td><td></td><td></td><td></td><td></td><td></td></tr>
<tr><td>Oligo fwd</td><td>CACCCACGCTGGTAGGGGTTGATG</td><td></td><td></td><td></td><td></td><td></td><td></td><td></td><td></td><td></td><td></td></tr>
<tr><td>Oligo rev</td><td>AAACCATCAACCCCTACCAGCGTG</td><td></td><td></td><td></td><td></td><td></td><td></td><td></td><td></td><td></td><td></td></tr>
<tr><td>Chromosome</td><td>start</td><td>end</td><td>strand</td><td>MM</td><td>target_seq</td><td>PAM</td><td>alignment</td><td>distance</td><td>position</td><td>gene name</td><td>gene id</td></tr>
<tr><td>GU305914.1</td><td>176256</td><td>176278</td><td>+</td><td>0</td><td>CACGCTGGTAGGGGTTGATG</td><td>GGG</td><td>||||||||[||||||||||||]PAM</td><td>NA</td><td>Intergenic</td><td>NA</td><td>NA</td></tr>
<tr><td>T13367</td><td>TTCTTTCGACGTAACTGTCGCGG</td><td>1000</td><td></td><td></td><td></td><td></td><td></td><td></td><td></td><td></td><td></td></tr>
<tr><td>Oligo fwd</td><td>CACCTTCTTTCGACGTAACTGTCG</td><td></td><td></td><td></td><td></td><td></td><td></td><td></td><td></td><td></td><td></td></tr>
<tr><td>Oligo rev</td><td>AAACCGACAGTTACGTCGAAAGAA</td><td></td><td></td><td></td><td></td><td></td><td></td><td></td><td></td><td></td><td></td></tr>
<tr><td>Chromosome</td><td>start</td><td>end</td><td>strand</td><td>MM</td><td>target_seq</td><td>PAM</td><td>alignment</td><td>distance</td><td>position</td><td>gene name</td><td>gene id</td></tr>
<tr><td>GU305914.1</td><td>176313</td><td>176335</td><td>+</td><td>0</td><td>TTCTTTCGACGTAACTGTCG</td><td>CGG</td><td>||||||||[||||||||||||]PAM</td><td>NA</td><td>Intergenic</td><td>NA</td><td>NA</td></tr>
<tr><td>T13368</td><td>TCTTTCGACGTAACTGTCGCGGG</td><td>1000</td><td></td><td></td><td></td><td></td><td></td><td></td><td></td><td></td><td></td></tr>
<tr><td>Oligo fwd</td><td>CACCTCTTTCGACGTAACTGTCGC</td><td></td><td></td><td></td><td></td><td></td><td></td><td></td><td></td><td></td><td></td></tr>
<tr><td>Oligo rev</td><td>AAACGCGACAGTTACGTCGAAAGA</td><td></td><td></td><td></td><td></td><td></td><td></td><td></td><td></td><td></td><td></td></tr>
<tr><td>Chromosome</td><td>start</td><td>end</td><td>strand</td><td>MM</td><td>target_seq</td><td>PAM</td><td>alignment</td><td>distance</td><td>position</td><td>gene name</td><td>gene id</td></tr>
<tr><td>GU305914.1</td><td>176314</td><td>176336</td><td>+</td><td>0</td><td>TCTTTCGACGTAACTGTCGC</td><td>GGG</td><td>||||||||[||||||||||||]PAM</td><td>NA</td><td>Intergenic</td><td>NA</td><td>NA</td></tr>
<tr><td>T13369</td><td>CGACGTAACTGTCGCGGGCCCGG</td><td>1000</td><td></td><td></td><td></td><td></td><td></td><td></td><td></td><td></td><td></td></tr>
<tr><td>Oligo fwd</td><td>CACCCGACGTAACTGTCGCGGGCC</td><td></td><td></td><td></td><td></td><td></td><td></td><td></td><td></td><td></td><td></td></tr>
<tr><td>Oligo rev</td><td>AAACGGCCCGCGACAGTTACGTCG</td><td></td><td></td><td></td><td></td><td></td><td></td><td></td><td></td><td></td><td></td></tr>
<tr><td>Chromosome</td><td>start</td><td>end</td><td>strand</td><td>MM</td><td>target_seq</td><td>PAM</td><td>alignment</td><td>distance</td><td>position</td><td>gene name</td><td>gene id</td></tr>
<tr><td>GU305914.1</td><td>176319</td><td>176341</td><td>+</td><td>0</td><td>CGACGTAACTGTCGCGGGCC</td><td>CGG</td><td>||||||||[||||||||||||]PAM</td><td>NA</td><td>Intergenic</td><td>NA</td><td>NA</td></tr>
<tr><td>T13370</td><td>CGTAACTGTCGCGGGCCCGGTGG</td><td>1000</td><td></td><td></td><td></td><td></td><td></td><td></td><td></td><td></td><td></td></tr>
<tr><td>Oligo fwd</td><td>CACCCGTAACTGTCGCGGGCCCGG</td><td></td><td></td><td></td><td></td><td></td><td></td><td></td><td></td><td></td><td></td></tr>
<tr><td>Oligo rev</td><td>AAACCCGGGCCCGCGACAGTTACG</td><td></td><td></td><td></td><td></td><td></td><td></td><td></td><td></td><td></td><td></td></tr>
<tr><td>Chromosome</td><td>start</td><td>end</td><td>strand</td><td>MM</td><td>target_seq</td><td>PAM</td><td>alignment</td><td>distance</td><td>position</td><td>gene name</td><td>gene id</td></tr>
<tr><td>GU305914.1</td><td>176322</td><td>176344</td><td>+</td><td>0</td><td>CGTAACTGTCGCGGGCCCGG</td><td>TGG</td><td>||||||||[||||||||||||]PAM</td><td>NA</td><td>Intergenic</td><td>NA</td><td>NA</td></tr>
<tr><td>T13371</td><td>CCTCGAGAAGCCGCCGCAAGCGG</td><td>1000</td><td></td><td></td><td></td><td></td><td></td><td></td><td></td><td></td><td></td></tr>
<tr><td>Oligo fwd</td><td>CACCCCTCGAGAAGCCGCCGCAAG</td><td></td><td></td><td></td><td></td><td></td><td></td><td></td><td></td><td></td><td></td></tr>
<tr><td>Oligo rev</td><td>AAACCTTGCGGCGGCTTCTCGAGG</td><td></td><td></td><td></td><td></td><td></td><td></td><td></td><td></td><td></td><td></td></tr>
<tr><td>Chromosome</td><td>start</td><td>end</td><td>strand</td><td>MM</td><td>target_seq</td><td>PAM</td><td>alignment</td><td>distance</td><td>position</td><td>gene name</td><td>gene id</td></tr>
<tr><td>GU305914.1</td><td>176369</td><td>176391</td><td>+</td><td>0</td><td>CCTCGAGAAGCCGCCGCAAG</td><td>CGG</td><td>||||||||[||||||||||||]PAM</td><td>NA</td><td>Intergenic</td><td>NA</td><td>NA</td></tr>
<tr><td>T13372</td><td>CGCCGCAAGCGGTATCCTGACGG</td><td>1000</td><td></td><td></td><td></td><td></td><td></td><td></td><td></td><td></td><td></td></tr>
<tr><td>Oligo fwd</td><td>CACCCGCCGCAAGCGGTATCCTGA</td><td></td><td></td><td></td><td></td><td></td><td></td><td></td><td></td><td></td><td></td></tr>
<tr><td>Oligo rev</td><td>AAACTCAGGATACCGCTTGCGGCG</td><td></td><td></td><td></td><td></td><td></td><td></td><td></td><td></td><td></td><td></td></tr>
<tr><td>Chromosome</td><td>start</td><td>end</td><td>strand</td><td>MM</td><td>target_seq</td><td>PAM</td><td>alignment</td><td>distance</td><td>position</td><td>gene name</td><td>gene id</td></tr>
<tr><td>GU305914.1</td><td>176380</td><td>176402</td><td>+</td><td>0</td><td>CGCCGCAAGCGGTATCCTGA</td><td>CGG</td><td>||||||||[||||||||||||]PAM</td><td>NA</td><td>Intergenic</td><td>NA</td><td>NA</td></tr>
<tr><td>T13373</td><td>GCCGCAAGCGGTATCCTGACGGG</td><td>1000</td><td></td><td></td><td></td><td></td><td></td><td></td><td></td><td></td><td></td></tr>
<tr><td>Oligo fwd</td><td>CACCGCCGCAAGCGGTATCCTGAC</td><td></td><td></td><td></td><td></td><td></td><td></td><td></td><td></td><td></td><td></td></tr>
<tr><td>Oligo rev</td><td>AAACGTCAGGATACCGCTTGCGGC</td><td></td><td></td><td></td><td></td><td></td><td></td><td></td><td></td><td></td><td></td></tr>
<tr><td>Chromosome</td><td>start</td><td>end</td><td>strand</td><td>MM</td><td>target_seq</td><td>PAM</td><td>alignment</td><td>distance</td><td>position</td><td>gene name</td><td>gene id</td></tr>
<tr><td>GU305914.1</td><td>176381</td><td>176403</td><td>+</td><td>0</td><td>GCCGCAAGCGGTATCCTGAC</td><td>GGG</td><td>||||||||[||||||||||||]PAM</td><td>NA</td><td>Intergenic</td><td>NA</td><td>NA</td></tr>
<tr><td>T13374</td><td>CCTCGCGTCGTCTCCTTCTTCGG</td><td>1000</td><td></td><td></td><td></td><td></td><td></td><td></td><td></td><td></td><td></td></tr>
<tr><td>Oligo fwd</td><td>CACCCCTCGCGTCGTCTCCTTCTT</td><td></td><td></td><td></td><td></td><td></td><td></td><td></td><td></td><td></td><td></td></tr>
<tr><td>Oligo rev</td><td>AAACAAGAAGGAGACGACGCGAGG</td><td></td><td></td><td></td><td></td><td></td><td></td><td></td><td></td><td></td><td></td></tr>
</table>

| Chromosome | start | end | strand | MM | target_seq | PAM | alignment | distance | position | gene name | gene id |
|---|---|---|---|---|---|---|---|---|---|---|---|
| GU305914.1 | 176408 | 176430 | + | 0 | CCTCGCGTCGTCTCCTTCTT | CGG | \|\|\|\|\|\|\|\|[\|\|\|\|\|\|\|\|\|\|\|\|]PAM | NA | Intergenic | NA | NA |
| T13375 | TCGTCTCCTTCTTCGGACAGAGG | 1000 | | | | | | | | | |
| Oligo fwd | CACCTCGTCTCCTTCTTCGGACAG | | | | | | | | | | |
| Oligo rev | AAACCTGTCCGAAGAAGGAGACGA | | | | | | | | | | |
| Chromosome | start | end | strand | MM | target_seq | PAM | alignment | distance | position | gene name | gene id |
| GU305914.1 | 176415 | 176437 | + | 0 | TCGTCTCCTTCTTCGGACAG | AGG | \|\|\|\|\|\|\|\|[\|\|\|\|\|\|\|\|\|\|\|\|]PAM | NA | Intergenic | NA | NA |
| T13376 | GCGTCCGTCTGCTGAGACGATGG | 1000 | | | | | | | | | |
| Oligo fwd | CACCGCGTCCGTCTGCTGAGACGA | | | | | | | | | | |
| Oligo rev | AAACTCGTCTCAGCAGACGGACGC | | | | | | | | | | |
| Chromosome | start | end | strand | MM | target_seq | PAM | alignment | distance | position | gene name | gene id |
| GU305914.1 | 176439 | 176461 | + | 0 | GCGTCCGTCTGCTGAGACGA | TGG | \|\|\|\|\|\|\|\|[\|\|\|\|\|\|\|\|\|\|\|\|]PAM | NA | Intergenic | NA | NA |
| T13377 | GTCTGCTGAGACGATGGCGACGG | 1000 | | | | | | | | | |
| Oligo fwd | CACCGTCTGCTGAGACGATGGCGA | | | | | | | | | | |
| Oligo rev | AAACTCGCCATCGTCTCAGCAGAC | | | | | | | | | | |
| Chromosome | start | end | strand | MM | target_seq | PAM | alignment | distance | position | gene name | gene id |
| GU305914.1 | 176445 | 176467 | + | 0 | GTCTGCTGAGACGATGGCGA | CGG | \|\|\|\|\|\|\|\|[\|\|\|\|\|\|\|\|\|\|\|\|]PAM | NA | Intergenic | NA | NA |
| T13378 | TGCTGAGACGATGGCGACGGCGG | 1000 | | | | | | | | | |
| Oligo fwd | CACCTGCTGAGACGATGGCGACGG | | | | | | | | | | |
| Oligo rev | AAACCCGTCGCCATCGTCTCAGCA | | | | | | | | | | |
| Chromosome | start | end | strand | MM | target_seq | PAM | alignment | distance | position | gene name | gene id |
| GU305914.1 | 176448 | 176470 | + | 0 | TGCTGAGACGATGGCGACGG | CGG | \|\|\|\|\|\|\|\|[\|\|\|\|\|\|\|\|\|\|\|\|]PAM | NA | Intergenic | NA | NA |
| GU305914.1 | 145694 | 145716 | + | 4 | TGGAGAGGCGATGGCGTCGG | GGG | \|\|--\|\|\|-[\|\|\|\|\|\|\|\|-\|\|\|]PAM | NA | Intergenic | NA | NA |
| T13379 | TGAGACGATGGCGACGGCGGCGG | 1000 | | | | | | | | | |
| Oligo fwd | CACCTGAGACGATGGCGACGGCGG | | | | | | | | | | |
| Oligo rev | AAACCCGCCGTCGCCATCGTCTCA | | | | | | | | | | |
| Chromosome | start | end | strand | MM | target_seq | PAM | alignment | distance | position | gene name | gene id |
| GU305914.1 | 176451 | 176473 | + | 0 | TGAGACGATGGCGACGGCGG | CGG | \|\|\|\|\|\|\|\|[\|\|\|\|\|\|\|\|\|\|\|\|]PAM | NA | Intergenic | NA | NA |
| T13380 | GACGATGGCGACGGCGGCGGAGG | 1000 | | | | | | | | | |
| Oligo fwd | CACCGACGATGGCGACGGCGGCGG | | | | | | | | | | |
| Oligo rev | AAACCCGCCGCCGTCGCCATCGTC | | | | | | | | | | |
| Chromosome | start | end | strand | MM | target_seq | PAM | alignment | distance | position | gene name | gene id |
| GU305914.1 | 176454 | 176476 | + | 0 | GACGATGGCGACGGCGGCGG | AGG | \|\|\|\|\|\|\|\|[\|\|\|\|\|\|\|\|\|\|\|\|]PAM | NA | Intergenic | NA | NA |
| GU305914.1 | 149488 | 149510 | - | 4 | GGAGCAGGCGACGGCGGCGG | CGG | \|--\|--\|\|[\|\|\|\|\|\|\|\|\|\|\|\|]PAM | NA | Intergenic | NA | NA |
| GU305914.1 | 225984 | 226006 | - | 4 | GTGGAGCGCGACGGCGGCGG | CGG | \|--\|\|--\|[\|\|\|\|\|\|\|\|\|\|\|\|]PAM | NA | Intergenic | NA | NA |
| GU305914.1 | 154404 | 154426 | + | 3 | GGCGGTGGCGGCGGCGGCGG | CGG | \|-\|\|-\|\|\|[\|\|-\|\|\|\|\|\|\|\|\|]PAM | NA | Intergenic | NA | NA |
| GU305914.1 | 58126 | 58148 | - | 3 | GGCGGTGGCGGCGGCGGCGG | CGG | \|-\|\|-\|\|\|[\|\|-\|\|\|\|\|\|\|\|\|]PAM | NA | Intergenic | NA | NA |
| GU305914.1 | 149482 | 149504 | - | 3 | GGCGACGGCGGCGGCGGCGG | CGG | \|-\|\|\|-\|\|[\|\|-\|\|\|\|\|\|\|\|\|]PAM | NA | Intergenic | NA | NA |
| GU305914.1 | 30000 | 30022 | + | 4 | GGTGCTGGCGGCGGCGGCGG | CGG | \|--\|-\|\|\|[\|\|-\|\|\|\|\|\|\|\|\|]PAM | NA | Intergenic | NA | NA |
| GU305914.1 | 96945 | 96967 | + | 4 | GAGCAGGGCGACGGCGGAGG | CGG | \|\|--\|-\|\|[\|\|\|\|\|\|\|\|\|-\|\|]PAM | NA | Intergenic | NA | NA |
| GU305914.1 | 90262 | 90284 | - | 4 | GAGGCGGGCGACGGCGGAGG | CGG | \|\|-\|--\|\|[\|\|\|\|\|\|\|\|\|-\|\|]PAM | NA | Intergenic | NA | NA |
| GU305914.1 | 166410 | 166432 | + | 4 | GACGTCGGCGCCGGCGGCAG | CGG | \|\|\|\|--\|\|[\|\|-\|\|\|\|\|\|\|-\|]PAM | NA | Intergenic | NA | NA |
| T13381 | GCAGCTGTTTCAGAGTCTGACGG | 1000 | | | | | | | | | |
| Oligo fwd | CACCGCAGCTGTTTCAGAGTCTGA | | | | | | | | | | |
| Oligo rev | AAACTCAGACTCTGAAACAGCTGC | | | | | | | | | | |
| Chromosome | start | end | strand | MM | target_seq | PAM | alignment | distance | position | gene name | gene id |
| GU305914.1 | 176476 | 176498 | + | 0 | GCAGCTGTTTCAGAGTCTGA | CGG | \|\|\|\|\|\|\|\|[\|\|\|\|\|\|\|\|\|\|\|\|]PAM | NA | Intergenic | NA | NA |
| T13382 | TTCAGAGTCTGACGGTTCACCGG | 1000 | | | | | | | | | |
| Oligo fwd | CACCTTCAGAGTCTGACGGTTCAC | | | | | | | | | | |
| Oligo rev | AAACGTGAACCGTCAGACTCTGAA | | | | | | | | | | |
| Chromosome | start | end | strand | MM | target_seq | PAM | alignment | distance | position | gene name | gene id |
| GU305914.1 | 176484 | 176506 | + | 0 | TTCAGAGTCTGACGGTTCAC | CGG | \|\|\|\|\|\|\|\|[\|\|\|\|\|\|\|\|\|\|\|\|]PAM | NA | Intergenic | NA | NA |
| T13383 | TCTGACGGTTCACCGGCTCTCGG | 1000 | | | | | | | | | |
| Oligo fwd | CACCTCTGACGGTTCACCGGCTCT | | | | | | | | | | |
| Oligo rev | AAACAGAGCCGGTGAACCGTCAGA | | | | | | | | | | |
| Chromosome | start | end | strand | MM | target_seq | PAM | alignment | distance | position | gene name | gene id |
| GU305914.1 | 176491 | 176513 | + | 0 | TCTGACGGTTCACCGGCTCT | CGG | \|\|\|\|\|\|\|\|[\|\|\|\|\|\|\|\|\|\|\|\|]PAM | NA | Intergenic | NA | NA |
| T13384 | GACGGTTCACCGGCTCTCGGCGG | 1000 | | | | | | | | | |
| Oligo fwd | CACCGACGGTTCACCGGCTCTCGG | | | | | | | | | | |
| Oligo rev | AAACCCGAGAGCCGGTGAACCGTC | | | | | | | | | | |
| Chromosome | start | end | strand | MM | target_seq | PAM | alignment | distance | position | gene name | gene id |
| GU305914.1 | 176494 | 176516 | + | 0 | GACGGTTCACCGGCTCTCGG | CGG | \|\|\|\|\|\|\|\|[\|\|\|\|\|\|\|\|\|\|\|\|]PAM | NA | Intergenic | NA | NA |
| T13385 | GGCGGACCGCGCAGACCTGCTGG | 1000 | | | | | | | | | |
| Oligo fwd | CACCGGCGGACCGCGCAGACCTGC | | | | | | | | | | |
| Oligo rev | AAACGCAGGTCTGCGCGGTCCGCC | | | | | | | | | | |
| Chromosome | start | end | strand | MM | target_seq | PAM | alignment | distance | position | gene name | gene id |
| GU305914.1 | 176512 | 176534 | + | 0 | GGCGGACCGCGCAGACCTGC | TGG | \|\|\|\|\|\|\|\|[\|\|\|\|\|\|\|\|\|\|\|\|]PAM | NA | Intergenic | NA | NA |
| T13386 | GCGGACCGCGCAGACCTGCTGGG | 1000 | | | | | | | | | |
| Oligo fwd | CACCGCGGACCGCGCAGACCTGCT | | | | | | | | | | |
| Oligo rev | AAACAGCAGGTCTGCGCGGTCCGC | | | | | | | | | | |
| Chromosome | start | end | strand | MM | target_seq | PAM | alignment | distance | position | gene name | gene id |
| GU305914.1 | 176513 | 176535 | + | 0 | GCGGACCGCGCAGACCTGCT | GGG | \|\|\|\|\|\|\|\|[\|\|\|\|\|\|\|\|\|\|\|\|]PAM | NA | Intergenic | NA | NA |
| T13387 | GCGCAGACCTGCTGGGCTCGCGG | 1000 | | | | | | | | | |
| Oligo fwd | CACCGCGCAGACCTGCTGGGCTCG | | | | | | | | | | |
| Oligo rev | AAACCGAGCCCAGCAGGTCTGCGC | | | | | | | | | | |
| Chromosome | start | end | strand | MM | target_seq | PAM | alignment | distance | position | gene name | gene id |

GU305914.1 176520 176542 + 0 GCGCAGACCTGCTGGGCTCG CGG ||||||||[||||||||||||]PAM NA Intergenic NA NA

T13388 CTTGCATGCGCTGCTCTAGTTGG 1000
Oligo fwd CACCCTTGCATGCGCTGCTCTAGT
Oligo rev AAACACTAGAGCAGCGCATGCAAG
Chromosome start end strand MM target_seq PAM alignment distance position gene name gene id
GU305914.1 176560 176582 + 0 CTTGCATGCGCTGCTCTAGT TGG ||||||||[||||||||||||]PAM NA Intergenic NA NA

T13389 CGCTGCTCTAGTTGGTACTGAGG 1000
Oligo fwd CACCCGCTGCTCTAGTTGGTACTG
Oligo rev AAACCAGTACCAACTAGAGCAGCG
Chromosome start end strand MM target_seq PAM alignment distance position gene name gene id
GU305914.1 176568 176590 + 0 CGCTGCTCTAGTTGGTACTG AGG ||||||||[||||||||||||]PAM NA Intergenic NA NA

T13390 CTCTAGTTGGTACTGAGGATTGG 1000
Oligo fwd CACCCTCTAGTTGGTACTGAGGAT
Oligo rev AAACATCCTCAGTACCAACTAGAG
Chromosome start end strand MM target_seq PAM alignment distance position gene name gene id
GU305914.1 176573 176595 + 0 CTCTAGTTGGTACTGAGGAT TGG ||||||||[||||||||||||]PAM NA Intergenic NA NA

T13391 CTTGCAAAATCTGATGATACAGG 1000
Oligo fwd CACCCTTGCAAAATCTGATGATAC
Oligo rev AAACGTATCATCAGATTTTGCAAG
Chromosome start end strand MM target_seq PAM alignment distance position gene name gene id
GU305914.1 176614 176636 + 0 CTTGCAAAATCTGATGATAC AGG ||||||||[||||||||||||]PAM NA Intergenic NA NA

T13392 TTGCAAAATCTGATGATACAGGG 1000
Oligo fwd CACCTTGCAAAATCTGATGATACA
Oligo rev AAACTGTATCATCAGATTTTGCAA
Chromosome start end strand MM target_seq PAM alignment distance position gene name gene id
GU305914.1 176615 176637 + 0 TTGCAAAATCTGATGATACA GGG ||||||||[||||||||||||]PAM NA Intergenic NA NA

T13393 TCTGATGATACAGGGAGAGATGG 1000
Oligo fwd CACCTCTGATGATACAGGGAGAGA
Oligo rev AAACTCTCTCCCTGTATCATCAGA
Chromosome start end strand MM target_seq PAM alignment distance position gene name gene id
GU305914.1 176623 176645 + 0 TCTGATGATACAGGGAGAGA TGG ||||||||[||||||||||||]PAM NA Intergenic NA NA

T13394 AGGGAGAGATGGTGTCCTTGCGG 1000
Oligo fwd CACCAGGGAGAGATGGTGTCCTTG
Oligo rev AAACCAAGGACACCATCTCTCCCT
Chromosome start end strand MM target_seq PAM alignment distance position gene name gene id
GU305914.1 176634 176656 + 0 AGGGAGAGATGGTGTCCTTG CGG ||||||||[||||||||||||]PAM NA Intergenic NA NA

T13395 AGATGGTGTCCTTGCGGACGAGG 1000
Oligo fwd CACCAGATGGTGTCCTTGCGGACG
Oligo rev AAACCGTCCGCAAGGACACCATCT
Chromosome start end strand MM target_seq PAM alignment distance position gene name gene id
GU305914.1 176640 176662 + 0 AGATGGTGTCCTTGCGGACG AGG ||||||||[||||||||||||]PAM NA Intergenic NA NA

T13396 GCCACCATCTGCTCGCACCACGG 1000
Oligo fwd CACCGCCACCATCTGCTCGCACCA
Oligo rev AAACTGGTGCGAGCAGATGGTGGC
Chromosome start end strand MM target_seq PAM alignment distance position gene name gene id
GU305914.1 176664 176686 + 0 GCCACCATCTGCTCGCACCA CGG ||||||||[||||||||||||]PAM NA Intergenic NA NA

T13397 CTGCTCGCACCACGGATGATAGG 1000
Oligo fwd CACCCTGCTCGCACCACGGATGAT
Oligo rev AAACATCATCCGTGGTGCGAGCAG
Chromosome start end strand MM target_seq PAM alignment distance position gene name gene id
GU305914.1 176672 176694 + 0 CTGCTCGCACCACGGATGAT AGG ||||||||[||||||||||||]PAM NA Intergenic NA NA

T13398 TGCTCGCACCACGGATGATAGGG 1000
Oligo fwd CACCTGCTCGCACCACGGATGATA
Oligo rev AAACTATCATCCGTGGTGCGAGCA
Chromosome start end strand MM target_seq PAM alignment distance position gene name gene id
GU305914.1 176673 176695 + 0 TGCTCGCACCACGGATGATA GGG ||||||||[||||||||||||]PAM NA Intergenic NA NA

T13399 GGATGATAGGGCGCCGCGCCCGG 1000
Oligo fwd CACCGGATGATAGGGCGCCGCGCC
Oligo rev AAACGGCGCGGCGCCCTATCATCC
Chromosome start end strand MM target_seq PAM alignment distance position gene name gene id
GU305914.1 176685 176707 + 0 GGATGATAGGGCGCCGCGCC CGG ||||||||[||||||||||||]PAM NA Intergenic NA NA

T13400 CGCCGCGCCCGGTGCCATTCTGG 1000
Oligo fwd CACCCGCCGCGCCCGGTGCCATTC
Oligo rev AAACGAATGGCACCGGGCGCGGCG
Chromosome start end strand MM target_seq PAM alignment distance position gene name gene id
GU305914.1 176696 176718 + 0 CGCCGCGCCCGGTGCCATTC TGG ||||||||[||||||||||||]PAM NA Intergenic NA NA

T13401 GCCGCGCCCGGTGCCATTCTGGG 1000
Oligo fwd CACCGCCGCGCCCGGTGCCATTCT
Oligo rev AAACAGAATGGCACCGGGCGCGGC
Chromosome start end strand MM target_seq PAM alignment distance position gene name gene id
GU305914.1 176697 176719 + 0 GCCGCGCCCGGTGCCATTCT GGG ||||||||[||||||||||||]PAM NA Intergenic NA NA

T13402 CCGCGCCCGGTGCCATTCTGGGG 1000
Oligo fwd CACCCCGCGCCCGGTGCCATTCTG
Oligo rev AAACCAGAATGGCACCGGGCGCGG

<table>
<tr><td>Chromosome</td><td>start</td><td>end</td><td>strand</td><td>MM</td><td>target_seq</td><td>PAM</td><td>alignment</td><td>distance</td><td>position</td><td>gene name</td><td>gene id</td></tr>
<tr><td>GU305914.1</td><td>176698</td><td>176720</td><td>+</td><td>0</td><td>CCGCGCCCGGTGCCATTCTG</td><td>GGG</td><td>||||||||[||||||||||||]PAM</td><td>NA</td><td>Intergenic</td><td>NA</td><td>NA</td></tr>
<tr><td>T13403</td><td>ACATGTCTCTGACTAAGATCTGG</td><td>1000</td><td></td><td></td><td></td><td></td><td></td><td></td><td></td><td></td><td></td></tr>
<tr><td>Oligo fwd</td><td>CACCACATGTCTCTGACTAAGATC</td><td></td><td></td><td></td><td></td><td></td><td></td><td></td><td></td><td></td><td></td></tr>
<tr><td>Oligo rev</td><td>AAACGATCTTAGTCAGAGACATGT</td><td></td><td></td><td></td><td></td><td></td><td></td><td></td><td></td><td></td><td></td></tr>
<tr><td>Chromosome</td><td>start</td><td>end</td><td>strand</td><td>MM</td><td>target_seq</td><td>PAM</td><td>alignment</td><td>distance</td><td>position</td><td>gene name</td><td>gene id</td></tr>
<tr><td>GU305914.1</td><td>176728</td><td>176750</td><td>+</td><td>0</td><td>ACATGTCTCTGACTAAGATC</td><td>TGG</td><td>||||||||[||||||||||||]PAM</td><td>NA</td><td>Intergenic</td><td>NA</td><td>NA</td></tr>
<tr><td>T13404</td><td>CATGTCTCTGACTAAGATCTGGG</td><td>1000</td><td></td><td></td><td></td><td></td><td></td><td></td><td></td><td></td><td></td></tr>
<tr><td>Oligo fwd</td><td>CACCCATGTCTCTGACTAAGATCT</td><td></td><td></td><td></td><td></td><td></td><td></td><td></td><td></td><td></td><td></td></tr>
<tr><td>Oligo rev</td><td>AAACAGATCTTAGTCAGAGACATG</td><td></td><td></td><td></td><td></td><td></td><td></td><td></td><td></td><td></td><td></td></tr>
<tr><td>Chromosome</td><td>start</td><td>end</td><td>strand</td><td>MM</td><td>target_seq</td><td>PAM</td><td>alignment</td><td>distance</td><td>position</td><td>gene name</td><td>gene id</td></tr>
<tr><td>GU305914.1</td><td>176729</td><td>176751</td><td>+</td><td>0</td><td>CATGTCTCTGACTAAGATCT</td><td>GGG</td><td>||||||||[||||||||||||]PAM</td><td>NA</td><td>Intergenic</td><td>NA</td><td>NA</td></tr>
<tr><td>T13405</td><td>ATGTCTCTGACTAAGATCTGGGG</td><td>1000</td><td></td><td></td><td></td><td></td><td></td><td></td><td></td><td></td><td></td></tr>
<tr><td>Oligo fwd</td><td>CACCATGTCTCTGACTAAGATCTG</td><td></td><td></td><td></td><td></td><td></td><td></td><td></td><td></td><td></td><td></td></tr>
<tr><td>Oligo rev</td><td>AAACCAGATCTTAGTCAGAGACAT</td><td></td><td></td><td></td><td></td><td></td><td></td><td></td><td></td><td></td><td></td></tr>
<tr><td>Chromosome</td><td>start</td><td>end</td><td>strand</td><td>MM</td><td>target_seq</td><td>PAM</td><td>alignment</td><td>distance</td><td>position</td><td>gene name</td><td>gene id</td></tr>
<tr><td>GU305914.1</td><td>176730</td><td>176752</td><td>+</td><td>0</td><td>ATGTCTCTGACTAAGATCTG</td><td>GGG</td><td>||||||||[||||||||||||]PAM</td><td>NA</td><td>Intergenic</td><td>NA</td><td>NA</td></tr>
<tr><td>T13406</td><td>TGACTAAGATCTGGGGTCCCAGG</td><td>1000</td><td></td><td></td><td></td><td></td><td></td><td></td><td></td><td></td><td></td></tr>
<tr><td>Oligo fwd</td><td>CACCTGACTAAGATCTGGGGTCCC</td><td></td><td></td><td></td><td></td><td></td><td></td><td></td><td></td><td></td><td></td></tr>
<tr><td>Oligo rev</td><td>AAACGGGACCCCAGATCTTAGTCA</td><td></td><td></td><td></td><td></td><td></td><td></td><td></td><td></td><td></td><td></td></tr>
<tr><td>Chromosome</td><td>start</td><td>end</td><td>strand</td><td>MM</td><td>target_seq</td><td>PAM</td><td>alignment</td><td>distance</td><td>position</td><td>gene name</td><td>gene id</td></tr>
<tr><td>GU305914.1</td><td>176737</td><td>176759</td><td>+</td><td>0</td><td>TGACTAAGATCTGGGGTCCC</td><td>AGG</td><td>||||||||[||||||||||||]PAM</td><td>NA</td><td>Intergenic</td><td>NA</td><td>NA</td></tr>
<tr><td>T13407</td><td>TCTGGGGTCCCAGGAACCTTAGG</td><td>1000</td><td></td><td></td><td></td><td></td><td></td><td></td><td></td><td></td><td></td></tr>
<tr><td>Oligo fwd</td><td>CACCTCTGGGGTCCCAGGAACCTT</td><td></td><td></td><td></td><td></td><td></td><td></td><td></td><td></td><td></td><td></td></tr>
<tr><td>Oligo rev</td><td>AAACAAGGTTCCTGGGACCCCAGA</td><td></td><td></td><td></td><td></td><td></td><td></td><td></td><td></td><td></td><td></td></tr>
<tr><td>Chromosome</td><td>start</td><td>end</td><td>strand</td><td>MM</td><td>target_seq</td><td>PAM</td><td>alignment</td><td>distance</td><td>position</td><td>gene name</td><td>gene id</td></tr>
<tr><td>GU305914.1</td><td>176746</td><td>176768</td><td>+</td><td>0</td><td>TCTGGGGTCCCAGGAACCTT</td><td>AGG</td><td>||||||||[||||||||||||]PAM</td><td>NA</td><td>Intergenic</td><td>NA</td><td>NA</td></tr>
<tr><td>T13408</td><td>CCCAGGAACCTTAGGATCTGAGG</td><td>1000</td><td></td><td></td><td></td><td></td><td></td><td></td><td></td><td></td><td></td></tr>
<tr><td>Oligo fwd</td><td>CACCCCCAGGAACCTTAGGATCTG</td><td></td><td></td><td></td><td></td><td></td><td></td><td></td><td></td><td></td><td></td></tr>
<tr><td>Oligo rev</td><td>AAACCAGATCCTAAGGTTCCTGGG</td><td></td><td></td><td></td><td></td><td></td><td></td><td></td><td></td><td></td><td></td></tr>
<tr><td>Chromosome</td><td>start</td><td>end</td><td>strand</td><td>MM</td><td>target_seq</td><td>PAM</td><td>alignment</td><td>distance</td><td>position</td><td>gene name</td><td>gene id</td></tr>
<tr><td>GU305914.1</td><td>176754</td><td>176776</td><td>+</td><td>0</td><td>CCCAGGAACCTTAGGATCTG</td><td>AGG</td><td>||||||||[||||||||||||]PAM</td><td>NA</td><td>Intergenic</td><td>NA</td><td>NA</td></tr>
<tr><td>T13409</td><td>ATCTGAGGTGTTGTTGTTGTTGG</td><td>1000</td><td></td><td></td><td></td><td></td><td></td><td></td><td></td><td></td><td></td></tr>
<tr><td>Oligo fwd</td><td>CACCATCTGAGGTGTTGTTGTTGT</td><td></td><td></td><td></td><td></td><td></td><td></td><td></td><td></td><td></td><td></td></tr>
<tr><td>Oligo rev</td><td>AAACACAACAACAACACCTCAGAT</td><td></td><td></td><td></td><td></td><td></td><td></td><td></td><td></td><td></td><td></td></tr>
<tr><td>Chromosome</td><td>start</td><td>end</td><td>strand</td><td>MM</td><td>target_seq</td><td>PAM</td><td>alignment</td><td>distance</td><td>position</td><td>gene name</td><td>gene id</td></tr>
<tr><td>GU305914.1</td><td>176769</td><td>176791</td><td>+</td><td>0</td><td>ATCTGAGGTGTTGTTGTTGT</td><td>TGG</td><td>||||||||[||||||||||||]PAM</td><td>NA</td><td>Intergenic</td><td>NA</td><td>NA</td></tr>
<tr><td>T13410</td><td>TCTGAGGTGTTGTTGTTGTTGGG</td><td>1000</td><td></td><td></td><td></td><td></td><td></td><td></td><td></td><td></td><td></td></tr>
<tr><td>Oligo fwd</td><td>CACCTCTGAGGTGTTGTTGTTGTT</td><td></td><td></td><td></td><td></td><td></td><td></td><td></td><td></td><td></td><td></td></tr>
<tr><td>Oligo rev</td><td>AAACAACAACAACAACACCTCAGA</td><td></td><td></td><td></td><td></td><td></td><td></td><td></td><td></td><td></td><td></td></tr>
<tr><td>Chromosome</td><td>start</td><td>end</td><td>strand</td><td>MM</td><td>target_seq</td><td>PAM</td><td>alignment</td><td>distance</td><td>position</td><td>gene name</td><td>gene id</td></tr>
<tr><td>GU305914.1</td><td>176770</td><td>176792</td><td>+</td><td>0</td><td>TCTGAGGTGTTGTTGTTGTT</td><td>GGG</td><td>||||||||[||||||||||||]PAM</td><td>NA</td><td>Intergenic</td><td>NA</td><td>NA</td></tr>
<tr><td>T13411</td><td>AGGTGTTGTTGTTGTTGGGAAGG</td><td>1000</td><td></td><td></td><td></td><td></td><td></td><td></td><td></td><td></td><td></td></tr>
<tr><td>Oligo fwd</td><td>CACCAGGTGTTGTTGTTGTTGGGA</td><td></td><td></td><td></td><td></td><td></td><td></td><td></td><td></td><td></td><td></td></tr>
<tr><td>Oligo rev</td><td>AAACTCCCAACAACAACAACACCT</td><td></td><td></td><td></td><td></td><td></td><td></td><td></td><td></td><td></td><td></td></tr>
<tr><td>Chromosome</td><td>start</td><td>end</td><td>strand</td><td>MM</td><td>target_seq</td><td>PAM</td><td>alignment</td><td>distance</td><td>position</td><td>gene name</td><td>gene id</td></tr>
<tr><td>GU305914.1</td><td>176774</td><td>176796</td><td>+</td><td>0</td><td>AGGTGTTGTTGTTGTTGGGA</td><td>AGG</td><td>||||||||[||||||||||||]PAM</td><td>NA</td><td>Intergenic</td><td>NA</td><td>NA</td></tr>
<tr><td>T13412</td><td>GGTGTTGTTGTTGTTGGGAAGGG</td><td>1000</td><td></td><td></td><td></td><td></td><td></td><td></td><td></td><td></td><td></td></tr>
<tr><td>Oligo fwd</td><td>CACCGGTGTTGTTGTTGTTGGGAA</td><td></td><td></td><td></td><td></td><td></td><td></td><td></td><td></td><td></td><td></td></tr>
<tr><td>Oligo rev</td><td>AAACTTCCCAACAACAACAACACC</td><td></td><td></td><td></td><td></td><td></td><td></td><td></td><td></td><td></td><td></td></tr>
<tr><td>Chromosome</td><td>start</td><td>end</td><td>strand</td><td>MM</td><td>target_seq</td><td>PAM</td><td>alignment</td><td>distance</td><td>position</td><td>gene name</td><td>gene id</td></tr>
<tr><td>GU305914.1</td><td>176775</td><td>176797</td><td>+</td><td>0</td><td>GGTGTTGTTGTTGTTGGGAA</td><td>GGG</td><td>||||||||[||||||||||||]PAM</td><td>NA</td><td>Intergenic</td><td>NA</td><td>NA</td></tr>
<tr><td>T13413</td><td>GTGTTGTTGTTGTTGGGAAGGGG</td><td>1000</td><td></td><td></td><td></td><td></td><td></td><td></td><td></td><td></td><td></td></tr>
<tr><td>Oligo fwd</td><td>CACCGTGTTGTTGTTGTTGGGAAG</td><td></td><td></td><td></td><td></td><td></td><td></td><td></td><td></td><td></td><td></td></tr>
<tr><td>Oligo rev</td><td>AAACCTTCCCAACAACAACAACAC</td><td></td><td></td><td></td><td></td><td></td><td></td><td></td><td></td><td></td><td></td></tr>
<tr><td>Chromosome</td><td>start</td><td>end</td><td>strand</td><td>MM</td><td>target_seq</td><td>PAM</td><td>alignment</td><td>distance</td><td>position</td><td>gene name</td><td>gene id</td></tr>
<tr><td>GU305914.1</td><td>176776</td><td>176798</td><td>+</td><td>0</td><td>GTGTTGTTGTTGTTGGGAAG</td><td>GGG</td><td>||||||||[||||||||||||]PAM</td><td>NA</td><td>Intergenic</td><td>NA</td><td>NA</td></tr>
<tr><td>T13414</td><td>GGGCTCAGCGTGAAGATTGTCGG</td><td>1000</td><td></td><td></td><td></td><td></td><td></td><td></td><td></td><td></td><td></td></tr>
<tr><td>Oligo fwd</td><td>CACCGGGCTCAGCGTGAAGATTGT</td><td></td><td></td><td></td><td></td><td></td><td></td><td></td><td></td><td></td><td></td></tr>
<tr><td>Oligo rev</td><td>AAACACAATCTTCACGCTGAGCCC</td><td></td><td></td><td></td><td></td><td></td><td></td><td></td><td></td><td></td><td></td></tr>
<tr><td>Chromosome</td><td>start</td><td>end</td><td>strand</td><td>MM</td><td>target_seq</td><td>PAM</td><td>alignment</td><td>distance</td><td>position</td><td>gene name</td><td>gene id</td></tr>
<tr><td>GU305914.1</td><td>176796</td><td>176818</td><td>+</td><td>0</td><td>GGGCTCAGCGTGAAGATTGT</td><td>CGG</td><td>||||||||[||||||||||||]PAM</td><td>NA</td><td>Intergenic</td><td>NA</td><td>NA</td></tr>
<tr><td>T13415</td><td>GGCTCAGCGTGAAGATTGTCGGG</td><td>1000</td><td></td><td></td><td></td><td></td><td></td><td></td><td></td><td></td><td></td></tr>
<tr><td>Oligo fwd</td><td>CACCGGCTCAGCGTGAAGATTGTC</td><td></td><td></td><td></td><td></td><td></td><td></td><td></td><td></td><td></td><td></td></tr>
<tr><td>Oligo rev</td><td>AAACGACAATCTTCACGCTGAGCC</td><td></td><td></td><td></td><td></td><td></td><td></td><td></td><td></td><td></td><td></td></tr>
<tr><td>Chromosome</td><td>start</td><td>end</td><td>strand</td><td>MM</td><td>target_seq</td><td>PAM</td><td>alignment</td><td>distance</td><td>position</td><td>gene name</td><td>gene id</td></tr>
<tr><td>GU305914.1</td><td>176797</td><td>176819</td><td>+</td><td>0</td><td>GGCTCAGCGTGAAGATTGTC</td><td>GGG</td><td>||||||||[||||||||||||]PAM</td><td>NA</td><td>Intergenic</td><td>NA</td><td>NA</td></tr>
<tr><td>T13416</td><td>GTCGGGTGAACGCCCGAGAACGG</td><td>1000</td><td></td><td></td><td></td><td></td><td></td><td></td><td></td><td></td><td></td></tr>
<tr><td>Oligo fwd</td><td>CACCGTCGGGTGAACGCCCGAGAA</td><td></td><td></td><td></td><td></td><td></td><td></td><td></td><td></td><td></td><td></td></tr>
<tr><td>Oligo rev</td><td>AAACTTCTCGGGCGTTCACCCGAC</td><td></td><td></td><td></td><td></td><td></td><td></td><td></td><td></td><td></td><td></td></tr>
<tr><td>Chromosome</td><td>start</td><td>end</td><td>strand</td><td>MM</td><td>target_seq</td><td>PAM</td><td>alignment</td><td>distance</td><td>position</td><td>gene name</td><td>gene id</td></tr>
<tr><td>GU305914.1</td><td>176814</td><td>176836</td><td>+</td><td>0</td><td>GTCGGGTGAACGCCCGAGAA</td><td>CGG</td><td>||||||||[||||||||||||]PAM</td><td>NA</td><td>Intergenic</td><td>NA</td><td>NA</td></tr>
<tr><td>T13417</td><td>GGGTGAACGCCCGAGAACGGAGG</td><td>1000</td><td></td><td></td><td></td><td></td><td></td><td></td><td></td><td></td><td></td></tr>
<tr><td>Oligo fwd</td><td>CACCGGGTGAACGCCCGAGAACGG</td><td></td><td></td><td></td><td></td><td></td><td></td><td></td><td></td><td></td><td></td></tr>
</table>

<table>
<tr><td>Oligo rev</td><td>AAACCCGTTCTCGGGCGTTCACCC</td><td></td><td></td><td></td><td></td><td></td><td></td><td></td><td></td><td></td><td></td></tr>
<tr><td>Chromosome</td><td>start</td><td>end</td><td>strand</td><td>MM</td><td>target_seq</td><td>PAM</td><td>alignment</td><td>distance</td><td>position</td><td>gene name</td><td>gene id</td></tr>
<tr><td>GU305914.1</td><td>176817</td><td>176839</td><td>+</td><td>0</td><td>GGGTGAACGCCCGAGAACGG</td><td>AGG</td><td>||||||||[||||||||||||]PAM</td><td>NA</td><td>Intergenic</td><td>NA</td><td>NA</td></tr>
<tr><td>T13418</td><td>GAACGCCCGAGAACGGAGGCTGG</td><td>1000</td><td></td><td></td><td></td><td></td><td></td><td></td><td></td><td></td><td></td></tr>
<tr><td>Oligo fwd</td><td>CACCGAACGCCCGAGAACGGAGGC</td><td></td><td></td><td></td><td></td><td></td><td></td><td></td><td></td><td></td><td></td></tr>
<tr><td>Oligo rev</td><td>AAACGCCTCCGTTCTCGGGCGTTC</td><td></td><td></td><td></td><td></td><td></td><td></td><td></td><td></td><td></td><td></td></tr>
<tr><td>Chromosome</td><td>start</td><td>end</td><td>strand</td><td>MM</td><td>target_seq</td><td>PAM</td><td>alignment</td><td>distance</td><td>position</td><td>gene name</td><td>gene id</td></tr>
<tr><td>GU305914.1</td><td>176821</td><td>176843</td><td>+</td><td>0</td><td>GAACGCCCGAGAACGGAGGC</td><td>TGG</td><td>||||||||[||||||||||||]PAM</td><td>NA</td><td>Intergenic</td><td>NA</td><td>NA</td></tr>
<tr><td>T13419</td><td>GTGAGCCATCATTCCCACGCTGG</td><td>1000</td><td></td><td></td><td></td><td></td><td></td><td></td><td></td><td></td><td></td></tr>
<tr><td>Oligo fwd</td><td>CACCGTGAGCCATCATTCCCACGC</td><td></td><td></td><td></td><td></td><td></td><td></td><td></td><td></td><td></td><td></td></tr>
<tr><td>Oligo rev</td><td>AAACGCGTGGGAATGATGGCTCAC</td><td></td><td></td><td></td><td></td><td></td><td></td><td></td><td></td><td></td><td></td></tr>
<tr><td>Chromosome</td><td>start</td><td>end</td><td>strand</td><td>MM</td><td>target_seq</td><td>PAM</td><td>alignment</td><td>distance</td><td>position</td><td>gene name</td><td>gene id</td></tr>
<tr><td>GU305914.1</td><td>176843</td><td>176865</td><td>+</td><td>0</td><td>GTGAGCCATCATTCCCACGC</td><td>TGG</td><td>||||||||[||||||||||||]PAM</td><td>NA</td><td>Intergenic</td><td>NA</td><td>NA</td></tr>
<tr><td>T13420</td><td>TGAGCCATCATTCCCACGCTGGG</td><td>1000</td><td></td><td></td><td></td><td></td><td></td><td></td><td></td><td></td><td></td></tr>
<tr><td>Oligo fwd</td><td>CACCTGAGCCATCATTCCCACGCT</td><td></td><td></td><td></td><td></td><td></td><td></td><td></td><td></td><td></td><td></td></tr>
<tr><td>Oligo rev</td><td>AAACAGCGTGGGAATGATGGCTCA</td><td></td><td></td><td></td><td></td><td></td><td></td><td></td><td></td><td></td><td></td></tr>
<tr><td>Chromosome</td><td>start</td><td>end</td><td>strand</td><td>MM</td><td>target_seq</td><td>PAM</td><td>alignment</td><td>distance</td><td>position</td><td>gene name</td><td>gene id</td></tr>
<tr><td>GU305914.1</td><td>176844</td><td>176866</td><td>+</td><td>0</td><td>TGAGCCATCATTCCCACGCT</td><td>GGG</td><td>||||||||[||||||||||||]PAM</td><td>NA</td><td>Intergenic</td><td>NA</td><td>NA</td></tr>
<tr><td>T13421</td><td>TCATTCCCACGCTGGGATAGCGG</td><td>1000</td><td></td><td></td><td></td><td></td><td></td><td></td><td></td><td></td><td></td></tr>
<tr><td>Oligo fwd</td><td>CACCTCATTCCCACGCTGGGATAG</td><td></td><td></td><td></td><td></td><td></td><td></td><td></td><td></td><td></td><td></td></tr>
<tr><td>Oligo rev</td><td>AAACCTATCCCAGCGTGGGAATGA</td><td></td><td></td><td></td><td></td><td></td><td></td><td></td><td></td><td></td><td></td></tr>
<tr><td>Chromosome</td><td>start</td><td>end</td><td>strand</td><td>MM</td><td>target_seq</td><td>PAM</td><td>alignment</td><td>distance</td><td>position</td><td>gene name</td><td>gene id</td></tr>
<tr><td>GU305914.1</td><td>176851</td><td>176873</td><td>+</td><td>0</td><td>TCATTCCCACGCTGGGATAG</td><td>CGG</td><td>||||||||[||||||||||||]PAM</td><td>NA</td><td>Intergenic</td><td>NA</td><td>NA</td></tr>
<tr><td>T13422</td><td>ACGCTGGGATAGCGGAAGTGTGG</td><td>1000</td><td></td><td></td><td></td><td></td><td></td><td></td><td></td><td></td><td></td></tr>
<tr><td>Oligo fwd</td><td>CACCACGCTGGGATAGCGGAAGTG</td><td></td><td></td><td></td><td></td><td></td><td></td><td></td><td></td><td></td><td></td></tr>
<tr><td>Oligo rev</td><td>AAACCACTTCCGCTATCCCAGCGT</td><td></td><td></td><td></td><td></td><td></td><td></td><td></td><td></td><td></td><td></td></tr>
<tr><td>Chromosome</td><td>start</td><td>end</td><td>strand</td><td>MM</td><td>target_seq</td><td>PAM</td><td>alignment</td><td>distance</td><td>position</td><td>gene name</td><td>gene id</td></tr>
<tr><td>GU305914.1</td><td>176859</td><td>176881</td><td>+</td><td>0</td><td>ACGCTGGGATAGCGGAAGTG</td><td>TGG</td><td>||||||||[||||||||||||]PAM</td><td>NA</td><td>Intergenic</td><td>NA</td><td>NA</td></tr>
<tr><td>T13423</td><td>TAGCGGAAGTGTGGAATTAGAGG</td><td>1000</td><td></td><td></td><td></td><td></td><td></td><td></td><td></td><td></td><td></td></tr>
<tr><td>Oligo fwd</td><td>CACCTAGCGGAAGTGTGGAATTAG</td><td></td><td></td><td></td><td></td><td></td><td></td><td></td><td></td><td></td><td></td></tr>
<tr><td>Oligo rev</td><td>AAACCTAATTCCACACTTCCGCTA</td><td></td><td></td><td></td><td></td><td></td><td></td><td></td><td></td><td></td><td></td></tr>
<tr><td>Chromosome</td><td>start</td><td>end</td><td>strand</td><td>MM</td><td>target_seq</td><td>PAM</td><td>alignment</td><td>distance</td><td>position</td><td>gene name</td><td>gene id</td></tr>
<tr><td>GU305914.1</td><td>176868</td><td>176890</td><td>+</td><td>0</td><td>TAGCGGAAGTGTGGAATTAG</td><td>AGG</td><td>||||||||[||||||||||||]PAM</td><td>NA</td><td>Intergenic</td><td>NA</td><td>NA</td></tr>
<tr><td>T13424</td><td>TGTGGAATTAGAGGCAGACGAGG</td><td>1000</td><td></td><td></td><td></td><td></td><td></td><td></td><td></td><td></td><td></td></tr>
<tr><td>Oligo fwd</td><td>CACCTGTGGAATTAGAGGCAGACG</td><td></td><td></td><td></td><td></td><td></td><td></td><td></td><td></td><td></td><td></td></tr>
<tr><td>Oligo rev</td><td>AAACCGTCTGCCTCTAATTCCACA</td><td></td><td></td><td></td><td></td><td></td><td></td><td></td><td></td><td></td><td></td></tr>
<tr><td>Chromosome</td><td>start</td><td>end</td><td>strand</td><td>MM</td><td>target_seq</td><td>PAM</td><td>alignment</td><td>distance</td><td>position</td><td>gene name</td><td>gene id</td></tr>
<tr><td>GU305914.1</td><td>176877</td><td>176899</td><td>+</td><td>0</td><td>TGTGGAATTAGAGGCAGACG</td><td>AGG</td><td>||||||||[||||||||||||]PAM</td><td>NA</td><td>Intergenic</td><td>NA</td><td>NA</td></tr>
<tr><td>T13425</td><td>GAGGCAGACGAGGATGATAGCGG</td><td>1000</td><td></td><td></td><td></td><td></td><td></td><td></td><td></td><td></td><td></td></tr>
<tr><td>Oligo fwd</td><td>CACCGAGGCAGACGAGGATGATAG</td><td></td><td></td><td></td><td></td><td></td><td></td><td></td><td></td><td></td><td></td></tr>
<tr><td>Oligo rev</td><td>AAACCTATCATCCTCGTCTGCCTC</td><td></td><td></td><td></td><td></td><td></td><td></td><td></td><td></td><td></td><td></td></tr>
<tr><td>Chromosome</td><td>start</td><td>end</td><td>strand</td><td>MM</td><td>target_seq</td><td>PAM</td><td>alignment</td><td>distance</td><td>position</td><td>gene name</td><td>gene id</td></tr>
<tr><td>GU305914.1</td><td>176887</td><td>176909</td><td>+</td><td>0</td><td>GAGGCAGACGAGGATGATAG</td><td>CGG</td><td>||||||||[||||||||||||]PAM</td><td>NA</td><td>Intergenic</td><td>NA</td><td>NA</td></tr>
<tr><td>T13426</td><td>AGGCAGACGAGGATGATAGCGGG</td><td>1000</td><td></td><td></td><td></td><td></td><td></td><td></td><td></td><td></td><td></td></tr>
<tr><td>Oligo fwd</td><td>CACCAGGCAGACGAGGATGATAGC</td><td></td><td></td><td></td><td></td><td></td><td></td><td></td><td></td><td></td><td></td></tr>
<tr><td>Oligo rev</td><td>AAACGCTATCATCCTCGTCTGCCT</td><td></td><td></td><td></td><td></td><td></td><td></td><td></td><td></td><td></td><td></td></tr>
<tr><td>Chromosome</td><td>start</td><td>end</td><td>strand</td><td>MM</td><td>target_seq</td><td>PAM</td><td>alignment</td><td>distance</td><td>position</td><td>gene name</td><td>gene id</td></tr>
<tr><td>GU305914.1</td><td>176888</td><td>176910</td><td>+</td><td>0</td><td>AGGCAGACGAGGATGATAGC</td><td>GGG</td><td>||||||||[||||||||||||]PAM</td><td>NA</td><td>Intergenic</td><td>NA</td><td>NA</td></tr>
<tr><td>T13427</td><td>CGAGGATGATAGCGGGCGCCCGG</td><td>1000</td><td></td><td></td><td></td><td></td><td></td><td></td><td></td><td></td><td></td></tr>
<tr><td>Oligo fwd</td><td>CACCCGAGGATGATAGCGGGCGCC</td><td></td><td></td><td></td><td></td><td></td><td></td><td></td><td></td><td></td><td></td></tr>
<tr><td>Oligo rev</td><td>AAACGGCGCCCGCTATCATCCTCG</td><td></td><td></td><td></td><td></td><td></td><td></td><td></td><td></td><td></td><td></td></tr>
<tr><td>Chromosome</td><td>start</td><td>end</td><td>strand</td><td>MM</td><td>target_seq</td><td>PAM</td><td>alignment</td><td>distance</td><td>position</td><td>gene name</td><td>gene id</td></tr>
<tr><td>GU305914.1</td><td>176895</td><td>176917</td><td>+</td><td>0</td><td>CGAGGATGATAGCGGGCGCC</td><td>CGG</td><td>||||||||[||||||||||||]PAM</td><td>NA</td><td>Intergenic</td><td>NA</td><td>NA</td></tr>
<tr><td>T13428</td><td>CGATCTCTGTCTGTAGATCAAGG</td><td>1000</td><td></td><td></td><td></td><td></td><td></td><td></td><td></td><td></td><td></td></tr>
<tr><td>Oligo fwd</td><td>CACCCGATCTCTGTCTGTAGATCA</td><td></td><td></td><td></td><td></td><td></td><td></td><td></td><td></td><td></td><td></td></tr>
<tr><td>Oligo rev</td><td>AAACTGATCTACAGACAGAGATCG</td><td></td><td></td><td></td><td></td><td></td><td></td><td></td><td></td><td></td><td></td></tr>
<tr><td>Chromosome</td><td>start</td><td>end</td><td>strand</td><td>MM</td><td>target_seq</td><td>PAM</td><td>alignment</td><td>distance</td><td>position</td><td>gene name</td><td>gene id</td></tr>
<tr><td>GU305914.1</td><td>176930</td><td>176952</td><td>+</td><td>0</td><td>CGATCTCTGTCTGTAGATCA</td><td>AGG</td><td>||||||||[||||||||||||]PAM</td><td>NA</td><td>Intergenic</td><td>NA</td><td>NA</td></tr>
<tr><td>T13429</td><td>CTCTGTCTGTAGATCAAGGTCGG</td><td>1000</td><td></td><td></td><td></td><td></td><td></td><td></td><td></td><td></td><td></td></tr>
<tr><td>Oligo fwd</td><td>CACCCTCTGTCTGTAGATCAAGGT</td><td></td><td></td><td></td><td></td><td></td><td></td><td></td><td></td><td></td><td></td></tr>
<tr><td>Oligo rev</td><td>AAACACCTTGATCTACAGACAGAG</td><td></td><td></td><td></td><td></td><td></td><td></td><td></td><td></td><td></td><td></td></tr>
<tr><td>Chromosome</td><td>start</td><td>end</td><td>strand</td><td>MM</td><td>target_seq</td><td>PAM</td><td>alignment</td><td>distance</td><td>position</td><td>gene name</td><td>gene id</td></tr>
<tr><td>GU305914.1</td><td>176934</td><td>176956</td><td>+</td><td>0</td><td>CTCTGTCTGTAGATCAAGGT</td><td>CGG</td><td>||||||||[||||||||||||]PAM</td><td>NA</td><td>Intergenic</td><td>NA</td><td>NA</td></tr>
<tr><td>T13430</td><td>TGTCTGTAGATCAAGGTCGGCGG</td><td>1000</td><td></td><td></td><td></td><td></td><td></td><td></td><td></td><td></td><td></td></tr>
<tr><td>Oligo fwd</td><td>CACCTGTCTGTAGATCAAGGTCGG</td><td></td><td></td><td></td><td></td><td></td><td></td><td></td><td></td><td></td><td></td></tr>
<tr><td>Oligo rev</td><td>AAACCCGACCTTGATCTACAGACA</td><td></td><td></td><td></td><td></td><td></td><td></td><td></td><td></td><td></td><td></td></tr>
<tr><td>Chromosome</td><td>start</td><td>end</td><td>strand</td><td>MM</td><td>target_seq</td><td>PAM</td><td>alignment</td><td>distance</td><td>position</td><td>gene name</td><td>gene id</td></tr>
<tr><td>GU305914.1</td><td>176937</td><td>176959</td><td>+</td><td>0</td><td>TGTCTGTAGATCAAGGTCGG</td><td>CGG</td><td>||||||||[||||||||||||]PAM</td><td>NA</td><td>Intergenic</td><td>NA</td><td>NA</td></tr>
<tr><td>T13431</td><td>GTCTGTAGATCAAGGTCGGCGGG</td><td>1000</td><td></td><td></td><td></td><td></td><td></td><td></td><td></td><td></td><td></td></tr>
<tr><td>Oligo fwd</td><td>CACCGTCTGTAGATCAAGGTCGGC</td><td></td><td></td><td></td><td></td><td></td><td></td><td></td><td></td><td></td><td></td></tr>
<tr><td>Oligo rev</td><td>AAACGCCGACCTTGATCTACAGAC</td><td></td><td></td><td></td><td></td><td></td><td></td><td></td><td></td><td></td><td></td></tr>
<tr><td>Chromosome</td><td>start</td><td>end</td><td>strand</td><td>MM</td><td>target_seq</td><td>PAM</td><td>alignment</td><td>distance</td><td>position</td><td>gene name</td><td>gene id</td></tr>
<tr><td>GU305914.1</td><td>176938</td><td>176960</td><td>+</td><td>0</td><td>GTCTGTAGATCAAGGTCGGC</td><td>GGG</td><td>||||||||[||||||||||||]PAM</td><td>NA</td><td>Intergenic</td><td>NA</td><td>NA</td></tr>
<tr><td>T13432</td><td>GGCGGGTATTGATACCTCACCGG</td><td>1000</td><td></td><td></td><td></td><td></td><td></td><td></td><td></td><td></td><td></td></tr>
</table>

Oligo fwd CACCGGCGGGTATTGATACCTCAC
Oligo rev AAACGTGAGGTATCAATACCCGCC
Chromosome start end strand MM target_seq PAM alignment distance position gene name gene id
GU305914.1 176955 176977 + 0 GGCGGGTATTGATACCTCAC CGG ||||||||[||||||||||||]PAM NA Intergenic NA NA

T13433 CTCACCGGTCCCGCGCGTAGCGG 1000
Oligo fwd CACCCTCACCGGTCCCGCGCGTAG
Oligo rev AAACCTACGCGCGGGACCGGTGAG
Chromosome start end strand MM target_seq PAM alignment distance position gene name gene id
GU305914.1 176970 176992 + 0 CTCACCGGTCCCGCGCGTAG CGG ||||||||[||||||||||||]PAM NA Intergenic NA NA

T13434 TCACCGGTCCCGCGCGTAGCGGG 1000
Oligo fwd CACCTCACCGGTCCCGCGCGTAGC
Oligo rev AAACGCTACGCGCGGGACCGGTGA
Chromosome start end strand MM target_seq PAM alignment distance position gene name gene id
GU305914.1 176971 176993 + 0 TCACCGGTCCCGCGCGTAGC GGG ||||||||[||||||||||||]PAM NA Intergenic NA NA

T13435 CGGTCCCGCGCGTAGCGGGTTGG 1000
Oligo fwd CACCCGGTCCCGCGCGTAGCGGGT
Oligo rev AAACACCCGCTACGCGCGGGACCG
Chromosome start end strand MM target_seq PAM alignment distance position gene name gene id
GU305914.1 176975 176997 + 0 CGGTCCCGCGCGTAGCGGGT TGG ||||||||[||||||||||||]PAM NA Intergenic NA NA

T13436 TGGAGAACGTCATCGTGTGTTGG 1000
Oligo fwd CACCTGGAGAACGTCATCGTGTGT
Oligo rev AAACACACACGATGACGTTCTCCA
Chromosome start end strand MM target_seq PAM alignment distance position gene name gene id
GU305914.1 176995 177017 + 0 TGGAGAACGTCATCGTGTGT TGG ||||||||[||||||||||||]PAM NA Intergenic NA NA

T13437 CGTCATCGTGTGTTGGACCGCGG 1000
Oligo fwd CACCCGTCATCGTGTGTTGGACCG
Oligo rev AAACCGGTCCAACACACGATGACG
Chromosome start end strand MM target_seq PAM alignment distance position gene name gene id
GU305914.1 177002 177024 + 0 CGTCATCGTGTGTTGGACCG CGG ||||||||[||||||||||||]PAM NA Intergenic NA NA

T13438 TGTGTTGGACCGCGGTGTTGTGG 1000
Oligo fwd CACCTGTGTTGGACCGCGGTGTTG
Oligo rev AAACCAACACCGCGGTCCAACACA
Chromosome start end strand MM target_seq PAM alignment distance position gene name gene id
GU305914.1 177010 177032 + 0 TGTGTTGGACCGCGGTGTTG TGG ||||||||[||||||||||||]PAM NA Intergenic NA NA

T13439 GTTGTGGCGAATCAGTGACGCGG 1000
Oligo fwd CACCGTTGTGGCGAATCAGTGACG
Oligo rev AAACCGTCACTGATTCGCCACAAC
Chromosome start end strand MM target_seq PAM alignment distance position gene name gene id
GU305914.1 177026 177048 + 0 GTTGTGGCGAATCAGTGACG CGG ||||||||[||||||||||||]PAM NA Intergenic NA NA

T13440 TTGTGGCGAATCAGTGACGCGGG 1000
Oligo fwd CACCTTGTGGCGAATCAGTGACGC
Oligo rev AAACGCGTCACTGATTCGCCACAA
Chromosome start end strand MM target_seq PAM alignment distance position gene name gene id
GU305914.1 177027 177049 + 0 TTGTGGCGAATCAGTGACGC GGG ||||||||[||||||||||||]PAM NA Intergenic NA NA

T13441 GGGAGACTCCTCGCGCCCCAAGG 1000
Oligo fwd CACCGGGAGACTCCTCGCGCCCCA
Oligo rev AAACTGGGGCGCGAGGAGTCTCCC
Chromosome start end strand MM target_seq PAM alignment distance position gene name gene id
GU305914.1 177047 177069 + 0 GGGAGACTCCTCGCGCCCCA AGG ||||||||[||||||||||||]PAM NA Intergenic NA NA

T13442 TCCTCGCGCCCCAAGGTTTGAGG 1000
Oligo fwd CACCTCCTCGCGCCCCAAGGTTTG
Oligo rev AAACCAAACCTTGGGGCGCGAGGA
Chromosome start end strand MM target_seq PAM alignment distance position gene name gene id
GU305914.1 177054 177076 + 0 TCCTCGCGCCCCAAGGTTTG AGG ||||||||[||||||||||||]PAM NA Intergenic NA NA

T13443 GCGCCCCAAGGTTTGAGGCGAGG 1000
Oligo fwd CACCGCGCCCCAAGGTTTGAGGCG
Oligo rev AAACCGCCTCAAACCTTGGGGCGC
Chromosome start end strand MM target_seq PAM alignment distance position gene name gene id
GU305914.1 177059 177081 + 0 GCGCCCCAAGGTTTGAGGCG AGG ||||||||[||||||||||||]PAM NA Intergenic NA NA

T13444 GCGTGCGCCAGACCGATGAAAGG 1000
Oligo fwd CACCGCGTGCGCCAGACCGATGAA
Oligo rev AAACTTCATCGGTCTGGCGCACGC
Chromosome start end strand MM target_seq PAM alignment distance position gene name gene id
GU305914.1 177083 177105 + 0 GCGTGCGCCAGACCGATGAA AGG ||||||||[||||||||||||]PAM NA Intergenic NA NA

T13445 CGTGCGCCAGACCGATGAAAGGG 1000
Oligo fwd CACCCGTGCGCCAGACCGATGAAA
Oligo rev AAACTTTCATCGGTCTGGCGCACG
Chromosome start end strand MM target_seq PAM alignment distance position gene name gene id
GU305914.1 177084 177106 + 0 CGTGCGCCAGACCGATGAAA GGG ||||||||[||||||||||||]PAM NA Intergenic NA NA

T13446 GGGAGAATTCTTGTTCTTTTTGG 1000
Oligo fwd CACCGGGAGAATTCTTGTTCTTTT
Oligo rev AAACAAAAGAACAAGAATTCTCCC
Chromosome start end strand MM target_seq PAM alignment distance position gene name gene id
GU305914.1 177104 177126 + 0 GGGAGAATTCTTGTTCTTTT TGG ||||||||[||||||||||||]PAM NA Intergenic NA NA

<table>
<tr><td>T13447</td><td>TTTGGACTTTTTCTTGTTCTTGG</td><td>1000</td><td></td><td></td><td></td><td></td><td></td><td></td><td></td><td></td><td></td></tr>
<tr><td>Oligo fwd</td><td>CACCTTTGGACTTTTTCTTGTTCT</td><td></td><td></td><td></td><td></td><td></td><td></td><td></td><td></td><td></td><td></td></tr>
<tr><td>Oligo rev</td><td>AAACAGAACAAGAAAAAGTCCAAA</td><td></td><td></td><td></td><td></td><td></td><td></td><td></td><td></td><td></td><td></td></tr>
<tr><td>Chromosome</td><td>start</td><td>end</td><td>strand</td><td>MM</td><td>target_seq</td><td>PAM</td><td>alignment</td><td>distance</td><td>position</td><td>gene name</td><td>gene id</td></tr>
<tr><td>GU305914.1</td><td>177122</td><td>177144</td><td>+</td><td>0</td><td>TTTGGACTTTTTCTTGTTCT</td><td>TGG</td><td>||||||||[||||||||||||]PAM</td><td>NA</td><td>Intergenic</td><td>NA</td><td>NA</td></tr>
<tr><td>T13448</td><td>TTGGACTTTTTCTTGTTCTTGGG</td><td>1000</td><td></td><td></td><td></td><td></td><td></td><td></td><td></td><td></td><td></td></tr>
<tr><td>Oligo fwd</td><td>CACCTTGGACTTTTTCTTGTTCTT</td><td></td><td></td><td></td><td></td><td></td><td></td><td></td><td></td><td></td><td></td></tr>
<tr><td>Oligo rev</td><td>AAACAAGAACAAGAAAAAGTCCAA</td><td></td><td></td><td></td><td></td><td></td><td></td><td></td><td></td><td></td><td></td></tr>
<tr><td>Chromosome</td><td>start</td><td>end</td><td>strand</td><td>MM</td><td>target_seq</td><td>PAM</td><td>alignment</td><td>distance</td><td>position</td><td>gene name</td><td>gene id</td></tr>
<tr><td>GU305914.1</td><td>177123</td><td>177145</td><td>+</td><td>0</td><td>TTGGACTTTTTCTTGTTCTT</td><td>GGG</td><td>||||||||[||||||||||||]PAM</td><td>NA</td><td>Intergenic</td><td>NA</td><td>NA</td></tr>
<tr><td>T13449</td><td>ACTTTTTCTTGTTCTTGGGTCGG</td><td>1000</td><td></td><td></td><td></td><td></td><td></td><td></td><td></td><td></td><td></td></tr>
<tr><td>Oligo fwd</td><td>CACCACTTTTTCTTGTTCTTGGGT</td><td></td><td></td><td></td><td></td><td></td><td></td><td></td><td></td><td></td><td></td></tr>
<tr><td>Oligo rev</td><td>AAACACCCAAGAACAAGAAAAAGT</td><td></td><td></td><td></td><td></td><td></td><td></td><td></td><td></td><td></td><td></td></tr>
<tr><td>Chromosome</td><td>start</td><td>end</td><td>strand</td><td>MM</td><td>target_seq</td><td>PAM</td><td>alignment</td><td>distance</td><td>position</td><td>gene name</td><td>gene id</td></tr>
<tr><td>GU305914.1</td><td>177127</td><td>177149</td><td>+</td><td>0</td><td>ACTTTTTCTTGTTCTTGGGT</td><td>CGG</td><td>||||||||[||||||||||||]PAM</td><td>NA</td><td>Intergenic</td><td>NA</td><td>NA</td></tr>
<tr><td>T13450</td><td>CTTTTTCTTGTTCTTGGGTCGGG</td><td>1000</td><td></td><td></td><td></td><td></td><td></td><td></td><td></td><td></td><td></td></tr>
<tr><td>Oligo fwd</td><td>CACCCTTTTTCTTGTTCTTGGGTC</td><td></td><td></td><td></td><td></td><td></td><td></td><td></td><td></td><td></td><td></td></tr>
<tr><td>Oligo rev</td><td>AAACGACCCAAGAACAAGAAAAAG</td><td></td><td></td><td></td><td></td><td></td><td></td><td></td><td></td><td></td><td></td></tr>
<tr><td>Chromosome</td><td>start</td><td>end</td><td>strand</td><td>MM</td><td>target_seq</td><td>PAM</td><td>alignment</td><td>distance</td><td>position</td><td>gene name</td><td>gene id</td></tr>
<tr><td>GU305914.1</td><td>177128</td><td>177150</td><td>+</td><td>0</td><td>CTTTTTCTTGTTCTTGGGTC</td><td>GGG</td><td>||||||||[||||||||||||]PAM</td><td>NA</td><td>Intergenic</td><td>NA</td><td>NA</td></tr>
<tr><td>T13451</td><td>TTTCTTGTTCTTGGGTCGGGCGG</td><td>1000</td><td></td><td></td><td></td><td></td><td></td><td></td><td></td><td></td><td></td></tr>
<tr><td>Oligo fwd</td><td>CACCTTTCTTGTTCTTGGGTCGGG</td><td></td><td></td><td></td><td></td><td></td><td></td><td></td><td></td><td></td><td></td></tr>
<tr><td>Oligo rev</td><td>AAACCCCGACCCAAGAACAAGAAA</td><td></td><td></td><td></td><td></td><td></td><td></td><td></td><td></td><td></td><td></td></tr>
<tr><td>Chromosome</td><td>start</td><td>end</td><td>strand</td><td>MM</td><td>target_seq</td><td>PAM</td><td>alignment</td><td>distance</td><td>position</td><td>gene name</td><td>gene id</td></tr>
<tr><td>GU305914.1</td><td>177131</td><td>177153</td><td>+</td><td>0</td><td>TTTCTTGTTCTTGGGTCGGG</td><td>CGG</td><td>||||||||[||||||||||||]PAM</td><td>NA</td><td>Intergenic</td><td>NA</td><td>NA</td></tr>
<tr><td>T13452</td><td>TTGTTCTTGGGTCGGGCGGACGG</td><td>1000</td><td></td><td></td><td></td><td></td><td></td><td></td><td></td><td></td><td></td></tr>
<tr><td>Oligo fwd</td><td>CACCTTGTTCTTGGGTCGGGCGGA</td><td></td><td></td><td></td><td></td><td></td><td></td><td></td><td></td><td></td><td></td></tr>
<tr><td>Oligo rev</td><td>AAACTCCGCCCGACCCAAGAACAA</td><td></td><td></td><td></td><td></td><td></td><td></td><td></td><td></td><td></td><td></td></tr>
<tr><td>Chromosome</td><td>start</td><td>end</td><td>strand</td><td>MM</td><td>target_seq</td><td>PAM</td><td>alignment</td><td>distance</td><td>position</td><td>gene name</td><td>gene id</td></tr>
<tr><td>GU305914.1</td><td>177135</td><td>177157</td><td>+</td><td>0</td><td>TTGTTCTTGGGTCGGGCGGA</td><td>CGG</td><td>||||||||[||||||||||||]PAM</td><td>NA</td><td>Intergenic</td><td>NA</td><td>NA</td></tr>
<tr><td>T13453</td><td>CGGGCGGACGGCTCCCGTCGAGG</td><td>1000</td><td></td><td></td><td></td><td></td><td></td><td></td><td></td><td></td><td></td></tr>
<tr><td>Oligo fwd</td><td>CACCCGGGCGGACGGCTCCCGTCG</td><td></td><td></td><td></td><td></td><td></td><td></td><td></td><td></td><td></td><td></td></tr>
<tr><td>Oligo rev</td><td>AAACCGACGGGAGCCGTCCGCCCG</td><td></td><td></td><td></td><td></td><td></td><td></td><td></td><td></td><td></td><td></td></tr>
<tr><td>Chromosome</td><td>start</td><td>end</td><td>strand</td><td>MM</td><td>target_seq</td><td>PAM</td><td>alignment</td><td>distance</td><td>position</td><td>gene name</td><td>gene id</td></tr>
<tr><td>GU305914.1</td><td>177147</td><td>177169</td><td>+</td><td>0</td><td>CGGGCGGACGGCTCCCGTCG</td><td>AGG</td><td>||||||||[||||||||||||]PAM</td><td>NA</td><td>Intergenic</td><td>NA</td><td>NA</td></tr>
<tr><td>T13454</td><td>GACGGCTCCCGTCGAGGCTCTGG</td><td>1000</td><td></td><td></td><td></td><td></td><td></td><td></td><td></td><td></td><td></td></tr>
<tr><td>Oligo fwd</td><td>CACCGACGGCTCCCGTCGAGGCTC</td><td></td><td></td><td></td><td></td><td></td><td></td><td></td><td></td><td></td><td></td></tr>
<tr><td>Oligo rev</td><td>AAACGAGCCTCGACGGGAGCCGTC</td><td></td><td></td><td></td><td></td><td></td><td></td><td></td><td></td><td></td><td></td></tr>
<tr><td>Chromosome</td><td>start</td><td>end</td><td>strand</td><td>MM</td><td>target_seq</td><td>PAM</td><td>alignment</td><td>distance</td><td>position</td><td>gene name</td><td>gene id</td></tr>
<tr><td>GU305914.1</td><td>177153</td><td>177175</td><td>+</td><td>0</td><td>GACGGCTCCCGTCGAGGCTC</td><td>TGG</td><td>||||||||[||||||||||||]PAM</td><td>NA</td><td>Intergenic</td><td>NA</td><td>NA</td></tr>
<tr><td>T13455</td><td>ACGGCTCCCGTCGAGGCTCTGGG</td><td>1000</td><td></td><td></td><td></td><td></td><td></td><td></td><td></td><td></td><td></td></tr>
<tr><td>Oligo fwd</td><td>CACCACGGCTCCCGTCGAGGCTCT</td><td></td><td></td><td></td><td></td><td></td><td></td><td></td><td></td><td></td><td></td></tr>
<tr><td>Oligo rev</td><td>AAACAGAGCCTCGACGGGAGCCGT</td><td></td><td></td><td></td><td></td><td></td><td></td><td></td><td></td><td></td><td></td></tr>
<tr><td>Chromosome</td><td>start</td><td>end</td><td>strand</td><td>MM</td><td>target_seq</td><td>PAM</td><td>alignment</td><td>distance</td><td>position</td><td>gene name</td><td>gene id</td></tr>
<tr><td>GU305914.1</td><td>177154</td><td>177176</td><td>+</td><td>0</td><td>ACGGCTCCCGTCGAGGCTCT</td><td>GGG</td><td>||||||||[||||||||||||]PAM</td><td>NA</td><td>Intergenic</td><td>NA</td><td>NA</td></tr>
<tr><td>T13456</td><td>CTCCCGTCGAGGCTCTGGGAAGG</td><td>1000</td><td></td><td></td><td></td><td></td><td></td><td></td><td></td><td></td><td></td></tr>
<tr><td>Oligo fwd</td><td>CACCCTCCCGTCGAGGCTCTGGGA</td><td></td><td></td><td></td><td></td><td></td><td></td><td></td><td></td><td></td><td></td></tr>
<tr><td>Oligo rev</td><td>AAACTCCCAGAGCCTCGACGGGAG</td><td></td><td></td><td></td><td></td><td></td><td></td><td></td><td></td><td></td><td></td></tr>
<tr><td>Chromosome</td><td>start</td><td>end</td><td>strand</td><td>MM</td><td>target_seq</td><td>PAM</td><td>alignment</td><td>distance</td><td>position</td><td>gene name</td><td>gene id</td></tr>
<tr><td>GU305914.1</td><td>177158</td><td>177180</td><td>+</td><td>0</td><td>CTCCCGTCGAGGCTCTGGGA</td><td>AGG</td><td>||||||||[||||||||||||]PAM</td><td>NA</td><td>Intergenic</td><td>NA</td><td>NA</td></tr>
<tr><td>T13457</td><td>TCCCGTCGAGGCTCTGGGAAGGG</td><td>1000</td><td></td><td></td><td></td><td></td><td></td><td></td><td></td><td></td><td></td></tr>
<tr><td>Oligo fwd</td><td>CACCTCCCGTCGAGGCTCTGGGAA</td><td></td><td></td><td></td><td></td><td></td><td></td><td></td><td></td><td></td><td></td></tr>
<tr><td>Oligo rev</td><td>AAACTTCCCAGAGCCTCGACGGGA</td><td></td><td></td><td></td><td></td><td></td><td></td><td></td><td></td><td></td><td></td></tr>
<tr><td>Chromosome</td><td>start</td><td>end</td><td>strand</td><td>MM</td><td>target_seq</td><td>PAM</td><td>alignment</td><td>distance</td><td>position</td><td>gene name</td><td>gene id</td></tr>
<tr><td>GU305914.1</td><td>177159</td><td>177181</td><td>+</td><td>0</td><td>TCCCGTCGAGGCTCTGGGAA</td><td>GGG</td><td>||||||||[||||||||||||]PAM</td><td>NA</td><td>Intergenic</td><td>NA</td><td>NA</td></tr>
<tr><td>T13458</td><td>GTCATCTCTCTCTGCTCCTGAGG</td><td>1000</td><td></td><td></td><td></td><td></td><td></td><td></td><td></td><td></td><td></td></tr>
<tr><td>Oligo fwd</td><td>CACCGTCATCTCTCTCTGCTCCTG</td><td></td><td></td><td></td><td></td><td></td><td></td><td></td><td></td><td></td><td></td></tr>
<tr><td>Oligo rev</td><td>AAACCAGGAGCAGAGAGAGATGAC</td><td></td><td></td><td></td><td></td><td></td><td></td><td></td><td></td><td></td><td></td></tr>
<tr><td>Chromosome</td><td>start</td><td>end</td><td>strand</td><td>MM</td><td>target_seq</td><td>PAM</td><td>alignment</td><td>distance</td><td>position</td><td>gene name</td><td>gene id</td></tr>
<tr><td>GU305914.1</td><td>177189</td><td>177211</td><td>+</td><td>0</td><td>GTCATCTCTCTCTGCTCCTG</td><td>AGG</td><td>||||||||[||||||||||||]PAM</td><td>NA</td><td>Intergenic</td><td>NA</td><td>NA</td></tr>
<tr><td>T13459</td><td>TCTCTCTCTGCTCCTGAGGTTGG</td><td>1000</td><td></td><td></td><td></td><td></td><td></td><td></td><td></td><td></td><td></td></tr>
<tr><td>Oligo fwd</td><td>CACCTCTCTCTCTGCTCCTGAGGT</td><td></td><td></td><td></td><td></td><td></td><td></td><td></td><td></td><td></td><td></td></tr>
<tr><td>Oligo rev</td><td>AAACACCTCAGGAGCAGAGAGAGA</td><td></td><td></td><td></td><td></td><td></td><td></td><td></td><td></td><td></td><td></td></tr>
<tr><td>Chromosome</td><td>start</td><td>end</td><td>strand</td><td>MM</td><td>target_seq</td><td>PAM</td><td>alignment</td><td>distance</td><td>position</td><td>gene name</td><td>gene id</td></tr>
<tr><td>GU305914.1</td><td>177193</td><td>177215</td><td>+</td><td>0</td><td>TCTCTCTCTGCTCCTGAGGT</td><td>TGG</td><td>||||||||[||||||||||||]PAM</td><td>NA</td><td>Intergenic</td><td>NA</td><td>NA</td></tr>
<tr><td>T13460</td><td>GAGGTTGGTGCGCCATAGCGTGG</td><td>1000</td><td></td><td></td><td></td><td></td><td></td><td></td><td></td><td></td><td></td></tr>
<tr><td>Oligo fwd</td><td>CACCGAGGTTGGTGCGCCATAGCG</td><td></td><td></td><td></td><td></td><td></td><td></td><td></td><td></td><td></td><td></td></tr>
<tr><td>Oligo rev</td><td>AAACCGCTATGGCGCACCAACCTC</td><td></td><td></td><td></td><td></td><td></td><td></td><td></td><td></td><td></td><td></td></tr>
<tr><td>Chromosome</td><td>start</td><td>end</td><td>strand</td><td>MM</td><td>target_seq</td><td>PAM</td><td>alignment</td><td>distance</td><td>position</td><td>gene name</td><td>gene id</td></tr>
<tr><td>GU305914.1</td><td>177208</td><td>177230</td><td>+</td><td>0</td><td>GAGGTTGGTGCGCCATAGCG</td><td>TGG</td><td>||||||||[||||||||||||]PAM</td><td>NA</td><td>Intergenic</td><td>NA</td><td>NA</td></tr>
<tr><td>T13461</td><td>GGTGCGCCATAGCGTGGCGTCGG</td><td>1000</td><td></td><td></td><td></td><td></td><td></td><td></td><td></td><td></td><td></td></tr>
<tr><td>Oligo fwd</td><td>CACCGGTGCGCCATAGCGTGGCGT</td><td></td><td></td><td></td><td></td><td></td><td></td><td></td><td></td><td></td><td></td></tr>
<tr><td>Oligo rev</td><td>AAACACGCCACGCTATGGCGCACC</td><td></td><td></td><td></td><td></td><td></td><td></td><td></td><td></td><td></td><td></td></tr>
<tr><td>Chromosome</td><td>start</td><td>end</td><td>strand</td><td>MM</td><td>target_seq</td><td>PAM</td><td>alignment</td><td>distance</td><td>position</td><td>gene name</td><td>gene id</td></tr>
<tr><td>GU305914.1</td><td>177214</td><td>177236</td><td>+</td><td>0</td><td>GGTGCGCCATAGCGTGGCGT</td><td>CGG</td><td>||||||||[||||||||||||]PAM</td><td>NA</td><td>Intergenic</td><td>NA</td><td>NA</td></tr>
</table>

T13462 CCCGACCAGCGACGCTCCTCAGG 1000
Oligo fwd CACCCCCGACCAGCGACGCTCCTC
Oligo rev AAACGAGGAGCGTCGCTGGTCGGG
Chromosome start end strand MM target_seq PAM alignment distance position gene name gene id
GU305914.1 177257 177279 + 0 CCCGACCAGCGACGCTCCTC AGG ||||||||[||||||||||||]PAM NA Intergenic NA NA

T13463 CCGACCAGCGACGCTCCTCAGGG 1000
Oligo fwd CACCCCGACCAGCGACGCTCCTCA
Oligo rev AAACTGAGGAGCGTCGCTGGTCGG
Chromosome start end strand MM target_seq PAM alignment distance position gene name gene id
GU305914.1 177258 177280 + 0 CCGACCAGCGACGCTCCTCA GGG ||||||||[||||||||||||]PAM NA Intergenic NA NA

T13464 CGCGTGTCAGCGAACTCCTCCGG 1000
Oligo fwd CACCCGCGTGTCAGCGAACTCCTC
Oligo rev AAACGAGGAGTTCGCTGACACGCG
Chromosome start end strand MM target_seq PAM alignment distance position gene name gene id
GU305914.1 177295 177317 + 0 CGCGTGTCAGCGAACTCCTC CGG ||||||||[||||||||||||]PAM NA Intergenic NA NA

T13465 CAGCGAACTCCTCCGGATCGAGG 1000
Oligo fwd CACCCAGCGAACTCCTCCGGATCG
Oligo rev AAACCGATCCGGAGGAGTTCGCTG
Chromosome start end strand MM target_seq PAM alignment distance position gene name gene id
GU305914.1 177302 177324 + 0 CAGCGAACTCCTCCGGATCG AGG ||||||||[||||||||||||]PAM NA Intergenic NA NA

T13466 CAGCAGAGTGTCAGACTCTGAGG 1000
Oligo fwd CACCCAGCAGAGTGTCAGACTCTG
Oligo rev AAACCAGAGTCTGACACTCTGCTG
Chromosome start end strand MM target_seq PAM alignment distance position gene name gene id
GU305914.1 177339 177361 + 0 CAGCAGAGTGTCAGACTCTG AGG ||||||||[||||||||||||]PAM NA Intergenic NA NA

T13467 TCTGAGGTGCAGAGCGCGCACGG 1000
Oligo fwd CACCTCTGAGGTGCAGAGCGCGCA
Oligo rev AAACTGCGCGCTCTGCACCTCAGA
Chromosome start end strand MM target_seq PAM alignment distance position gene name gene id
GU305914.1 177355 177377 + 0 TCTGAGGTGCAGAGCGCGCA CGG ||||||||[||||||||||||]PAM NA Intergenic NA NA

T13468 CTGAGGTGCAGAGCGCGCACGGG 1000
Oligo fwd CACCCTGAGGTGCAGAGCGCGCAC
Oligo rev AAACGTGCGCGCTCTGCACCTCAG
Chromosome start end strand MM target_seq PAM alignment distance position gene name gene id
GU305914.1 177356 177378 + 0 CTGAGGTGCAGAGCGCGCAC GGG ||||||||[||||||||||||]PAM NA Intergenic NA NA

T13469 TGAGGTGCAGAGCGCGCACGGGG 1000
Oligo fwd CACCTGAGGTGCAGAGCGCGCACG
Oligo rev AAACCGTGCGCGCTCTGCACCTCA
Chromosome start end strand MM target_seq PAM alignment distance position gene name gene id
GU305914.1 177357 177379 + 0 TGAGGTGCAGAGCGCGCACG GGG ||||||||[||||||||||||]PAM NA Intergenic NA NA

T13470 GGTGCAGAGCGCGCACGGGGTGG 1000
Oligo fwd CACCGGTGCAGAGCGCGCACGGGG
Oligo rev AAACCCCCGTGCGCGCTCTGCACC
Chromosome start end strand MM target_seq PAM alignment distance position gene name gene id
GU305914.1 177360 177382 + 0 GGTGCAGAGCGCGCACGGGG TGG ||||||||[||||||||||||]PAM NA Intergenic NA NA

T13471 CGCGCACGGGGTGGCACCCTCGG 1000
Oligo fwd CACCCGCGCACGGGGTGGCACCCT
Oligo rev AAACAGGGTGCCACCCCGTGCGCG
Chromosome start end strand MM target_seq PAM alignment distance position gene name gene id
GU305914.1 177369 177391 + 0 CGCGCACGGGGTGGCACCCT CGG ||||||||[||||||||||||]PAM NA Intergenic NA NA

T13472 CTGTAAGAAGAAGATGATCCTGG 1000
Oligo fwd CACCCTGTAAGAAGAAGATGATCC
Oligo rev AAACGGATCATCTTCTTCTTACAG
Chromosome start end strand MM target_seq PAM alignment distance position gene name gene id
GU305914.1 177399 177421 + 0 CTGTAAGAAGAAGATGATCC TGG ||||||||[||||||||||||]PAM NA Intergenic NA NA

T13473 TGTAAGAAGAAGATGATCCTGGG 1000
Oligo fwd CACCTGTAAGAAGAAGATGATCCT
Oligo rev AAACAGGATCATCTTCTTCTTACA
Chromosome start end strand MM target_seq PAM alignment distance position gene name gene id
GU305914.1 177400 177422 + 0 TGTAAGAAGAAGATGATCCT GGG ||||||||[||||||||||||]PAM NA Intergenic NA NA

T13474 GTAAGAAGAAGATGATCCTGGGG 1000
Oligo fwd CACCGTAAGAAGAAGATGATCCTG
Oligo rev AAACCAGGATCATCTTCTTCTTAC
Chromosome start end strand MM target_seq PAM alignment distance position gene name gene id
GU305914.1 177401 177423 + 0 GTAAGAAGAAGATGATCCTG GGG ||||||||[||||||||||||]PAM NA Intergenic NA NA

T13475 AGAAGAAGATGATCCTGGGGAGG 1000
Oligo fwd CACCAGAAGAAGATGATCCTGGGG
Oligo rev AAACCCCCAGGATCATCTTCTTCT
Chromosome start end strand MM target_seq PAM alignment distance position gene name gene id
GU305914.1 177404 177426 + 0 AGAAGAAGATGATCCTGGGG AGG ||||||||[||||||||||||]PAM NA Intergenic NA NA

T13476 GGGAGGAACTGCGCCTCGATCGG 1000
Oligo fwd CACCGGGAGGAACTGCGCCTCGAT
Oligo rev AAACATCGAGGCGCAGTTCCTCCC
Chromosome start end strand MM target_seq PAM alignment distance position gene name gene id

<table>
<tr><td>GU305914.1</td><td>177421</td><td>177443</td><td>+</td><td>0</td><td>GGGAGGAACTGCGCCTCGAT</td><td>CGG</td><td>||||||||[||||||||||||]PAM</td><td>NA</td><td>Intergenic</td><td>NA</td><td>NA</td></tr>
<tr><td>T13477</td><td>AGGAACTGCGCCTCGATCGGAGG</td><td>1000</td><td></td><td></td><td></td><td></td><td></td><td></td><td></td><td></td><td></td></tr>
<tr><td>Oligo fwd</td><td>CACCAGGAACTGCGCCTCGATCGG</td><td></td><td></td><td></td><td></td><td></td><td></td><td></td><td></td><td></td><td></td></tr>
<tr><td>Oligo rev</td><td>AAACCCGATCGAGGCGCAGTTCCT</td><td></td><td></td><td></td><td></td><td></td><td></td><td></td><td></td><td></td><td></td></tr>
<tr><td>Chromosome</td><td>start</td><td>end</td><td>strand</td><td>MM</td><td>target_seq</td><td>PAM</td><td>alignment</td><td>distance</td><td>position</td><td>gene name</td><td>gene id</td></tr>
<tr><td>GU305914.1</td><td>177424</td><td>177446</td><td>+</td><td>0</td><td>AGGAACTGCGCCTCGATCGG</td><td>AGG</td><td>||||||||[||||||||||||]PAM</td><td>NA</td><td>Intergenic</td><td>NA</td><td>NA</td></tr>
<tr><td>T13478</td><td>CCTCGATCGGAGGAGCGTCCTGG</td><td>1000</td><td></td><td></td><td></td><td></td><td></td><td></td><td></td><td></td><td></td></tr>
<tr><td>Oligo fwd</td><td>CACCCCTCGATCGGAGGAGCGTCC</td><td></td><td></td><td></td><td></td><td></td><td></td><td></td><td></td><td></td><td></td></tr>
<tr><td>Oligo rev</td><td>AAACGGACGCTCCTCCGATCGAGG</td><td></td><td></td><td></td><td></td><td></td><td></td><td></td><td></td><td></td><td></td></tr>
<tr><td>Chromosome</td><td>start</td><td>end</td><td>strand</td><td>MM</td><td>target_seq</td><td>PAM</td><td>alignment</td><td>distance</td><td>position</td><td>gene name</td><td>gene id</td></tr>
<tr><td>GU305914.1</td><td>177434</td><td>177456</td><td>+</td><td>0</td><td>CCTCGATCGGAGGAGCGTCC</td><td>TGG</td><td>||||||||[||||||||||||]PAM</td><td>NA</td><td>Intergenic</td><td>NA</td><td>NA</td></tr>
<tr><td>T13479</td><td>GAACCACTCGAGCGTGCTGACGG</td><td>1000</td><td></td><td></td><td></td><td></td><td></td><td></td><td></td><td></td><td></td></tr>
<tr><td>Oligo fwd</td><td>CACCGAACCACTCGAGCGTGCTGA</td><td></td><td></td><td></td><td></td><td></td><td></td><td></td><td></td><td></td><td></td></tr>
<tr><td>Oligo rev</td><td>AAACTCAGCACGCTCGAGTGGTTC</td><td></td><td></td><td></td><td></td><td></td><td></td><td></td><td></td><td></td><td></td></tr>
<tr><td>Chromosome</td><td>start</td><td>end</td><td>strand</td><td>MM</td><td>target_seq</td><td>PAM</td><td>alignment</td><td>distance</td><td>position</td><td>gene name</td><td>gene id</td></tr>
<tr><td>GU305914.1</td><td>177471</td><td>177493</td><td>+</td><td>0</td><td>GAACCACTCGAGCGTGCTGA</td><td>CGG</td><td>||||||||[||||||||||||]PAM</td><td>NA</td><td>Intergenic</td><td>NA</td><td>NA</td></tr>
<tr><td>T13480</td><td>GAAGAGAGCCTCTCCCTGACTGG</td><td>1000</td><td></td><td></td><td></td><td></td><td></td><td></td><td></td><td></td><td></td></tr>
<tr><td>Oligo fwd</td><td>CACCGAAGAGAGCCTCTCCCTGAC</td><td></td><td></td><td></td><td></td><td></td><td></td><td></td><td></td><td></td><td></td></tr>
<tr><td>Oligo rev</td><td>AAACGTCAGGGAGAGGCTCTCTTC</td><td></td><td></td><td></td><td></td><td></td><td></td><td></td><td></td><td></td><td></td></tr>
<tr><td>Chromosome</td><td>start</td><td>end</td><td>strand</td><td>MM</td><td>target_seq</td><td>PAM</td><td>alignment</td><td>distance</td><td>position</td><td>gene name</td><td>gene id</td></tr>
<tr><td>GU305914.1</td><td>177510</td><td>177532</td><td>+</td><td>0</td><td>GAAGAGAGCCTCTCCCTGAC</td><td>TGG</td><td>||||||||[||||||||||||]PAM</td><td>NA</td><td>Intergenic</td><td>NA</td><td>NA</td></tr>
<tr><td>T13481</td><td>AAGAGAGCCTCTCCCTGACTGGG</td><td>1000</td><td></td><td></td><td></td><td></td><td></td><td></td><td></td><td></td><td></td></tr>
<tr><td>Oligo fwd</td><td>CACCAAGAGAGCCTCTCCCTGACT</td><td></td><td></td><td></td><td></td><td></td><td></td><td></td><td></td><td></td><td></td></tr>
<tr><td>Oligo rev</td><td>AAACAGTCAGGGAGAGGCTCTCTT</td><td></td><td></td><td></td><td></td><td></td><td></td><td></td><td></td><td></td><td></td></tr>
<tr><td>Chromosome</td><td>start</td><td>end</td><td>strand</td><td>MM</td><td>target_seq</td><td>PAM</td><td>alignment</td><td>distance</td><td>position</td><td>gene name</td><td>gene id</td></tr>
<tr><td>GU305914.1</td><td>177511</td><td>177533</td><td>+</td><td>0</td><td>AAGAGAGCCTCTCCCTGACT</td><td>GGG</td><td>||||||||[||||||||||||]PAM</td><td>NA</td><td>Intergenic</td><td>NA</td><td>NA</td></tr>
<tr><td>T13482</td><td>AGAGAGCCTCTCCCTGACTGGGG</td><td>1000</td><td></td><td></td><td></td><td></td><td></td><td></td><td></td><td></td><td></td></tr>
<tr><td>Oligo fwd</td><td>CACCAGAGAGCCTCTCCCTGACTG</td><td></td><td></td><td></td><td></td><td></td><td></td><td></td><td></td><td></td><td></td></tr>
<tr><td>Oligo rev</td><td>AAACCAGTCAGGGAGAGGCTCTCT</td><td></td><td></td><td></td><td></td><td></td><td></td><td></td><td></td><td></td><td></td></tr>
<tr><td>Chromosome</td><td>start</td><td>end</td><td>strand</td><td>MM</td><td>target_seq</td><td>PAM</td><td>alignment</td><td>distance</td><td>position</td><td>gene name</td><td>gene id</td></tr>
<tr><td>GU305914.1</td><td>177512</td><td>177534</td><td>+</td><td>0</td><td>AGAGAGCCTCTCCCTGACTG</td><td>GGG</td><td>||||||||[||||||||||||]PAM</td><td>NA</td><td>Intergenic</td><td>NA</td><td>NA</td></tr>
<tr><td>T13483</td><td>GACTGGGGAATAACATACGTAGG</td><td>1000</td><td></td><td></td><td></td><td></td><td></td><td></td><td></td><td></td><td></td></tr>
<tr><td>Oligo fwd</td><td>CACCGACTGGGGAATAACATACGT</td><td></td><td></td><td></td><td></td><td></td><td></td><td></td><td></td><td></td><td></td></tr>
<tr><td>Oligo rev</td><td>AAACACGTATGTTATTCCCCAGTC</td><td></td><td></td><td></td><td></td><td></td><td></td><td></td><td></td><td></td><td></td></tr>
<tr><td>Chromosome</td><td>start</td><td>end</td><td>strand</td><td>MM</td><td>target_seq</td><td>PAM</td><td>alignment</td><td>distance</td><td>position</td><td>gene name</td><td>gene id</td></tr>
<tr><td>GU305914.1</td><td>177527</td><td>177549</td><td>+</td><td>0</td><td>GACTGGGGAATAACATACGT</td><td>AGG</td><td>||||||||[||||||||||||]PAM</td><td>NA</td><td>Intergenic</td><td>NA</td><td>NA</td></tr>
<tr><td>T13484</td><td>GGGGAATAACATACGTAGGACGG</td><td>1000</td><td></td><td></td><td></td><td></td><td></td><td></td><td></td><td></td><td></td></tr>
<tr><td>Oligo fwd</td><td>CACCGGGGAATAACATACGTAGGA</td><td></td><td></td><td></td><td></td><td></td><td></td><td></td><td></td><td></td><td></td></tr>
<tr><td>Oligo rev</td><td>AAACTCCTACGTATGTTATTCCCC</td><td></td><td></td><td></td><td></td><td></td><td></td><td></td><td></td><td></td><td></td></tr>
<tr><td>Chromosome</td><td>start</td><td>end</td><td>strand</td><td>MM</td><td>target_seq</td><td>PAM</td><td>alignment</td><td>distance</td><td>position</td><td>gene name</td><td>gene id</td></tr>
<tr><td>GU305914.1</td><td>177531</td><td>177553</td><td>+</td><td>0</td><td>GGGGAATAACATACGTAGGA</td><td>CGG</td><td>||||||||[||||||||||||]PAM</td><td>NA</td><td>Intergenic</td><td>NA</td><td>NA</td></tr>
<tr><td>T13485</td><td>ACGGTGATATTCACAACGTGAGG</td><td>1000</td><td></td><td></td><td></td><td></td><td></td><td></td><td></td><td></td><td></td></tr>
<tr><td>Oligo fwd</td><td>CACCACGGTGATATTCACAACGTG</td><td></td><td></td><td></td><td></td><td></td><td></td><td></td><td></td><td></td><td></td></tr>
<tr><td>Oligo rev</td><td>AAACCACGTTGTGAATATCACCGT</td><td></td><td></td><td></td><td></td><td></td><td></td><td></td><td></td><td></td><td></td></tr>
<tr><td>Chromosome</td><td>start</td><td>end</td><td>strand</td><td>MM</td><td>target_seq</td><td>PAM</td><td>alignment</td><td>distance</td><td>position</td><td>gene name</td><td>gene id</td></tr>
<tr><td>GU305914.1</td><td>177550</td><td>177572</td><td>+</td><td>0</td><td>ACGGTGATATTCACAACGTG</td><td>AGG</td><td>||||||||[||||||||||||]PAM</td><td>NA</td><td>Intergenic</td><td>NA</td><td>NA</td></tr>
<tr><td>T13486</td><td>GATATTCACAACGTGAGGCCCGG</td><td>1000</td><td></td><td></td><td></td><td></td><td></td><td></td><td></td><td></td><td></td></tr>
<tr><td>Oligo fwd</td><td>CACCGATATTCACAACGTGAGGCC</td><td></td><td></td><td></td><td></td><td></td><td></td><td></td><td></td><td></td><td></td></tr>
<tr><td>Oligo rev</td><td>AAACGGCCTCACGTTGTGAATATC</td><td></td><td></td><td></td><td></td><td></td><td></td><td></td><td></td><td></td><td></td></tr>
<tr><td>Chromosome</td><td>start</td><td>end</td><td>strand</td><td>MM</td><td>target_seq</td><td>PAM</td><td>alignment</td><td>distance</td><td>position</td><td>gene name</td><td>gene id</td></tr>
<tr><td>GU305914.1</td><td>177555</td><td>177577</td><td>+</td><td>0</td><td>GATATTCACAACGTGAGGCC</td><td>CGG</td><td>||||||||[||||||||||||]PAM</td><td>NA</td><td>Intergenic</td><td>NA</td><td>NA</td></tr>
<tr><td>T13487</td><td>CGTTCTCATAGTTGACGATGAGG</td><td>1000</td><td></td><td></td><td></td><td></td><td></td><td></td><td></td><td></td><td></td></tr>
<tr><td>Oligo fwd</td><td>CACCCGTTCTCATAGTTGACGATG</td><td></td><td></td><td></td><td></td><td></td><td></td><td></td><td></td><td></td><td></td></tr>
<tr><td>Oligo rev</td><td>AAACCATCGTCAACTATGAGAACG</td><td></td><td></td><td></td><td></td><td></td><td></td><td></td><td></td><td></td><td></td></tr>
<tr><td>Chromosome</td><td>start</td><td>end</td><td>strand</td><td>MM</td><td>target_seq</td><td>PAM</td><td>alignment</td><td>distance</td><td>position</td><td>gene name</td><td>gene id</td></tr>
<tr><td>GU305914.1</td><td>177596</td><td>177618</td><td>+</td><td>0</td><td>CGTTCTCATAGTTGACGATG</td><td>AGG</td><td>||||||||[||||||||||||]PAM</td><td>NA</td><td>Intergenic</td><td>NA</td><td>NA</td></tr>
<tr><td>GU305914.1</td><td>136744</td><td>136766</td><td>-</td><td>4</td><td>CGTTCTGGTAGATGACGCTG</td><td>GGG</td><td>||||||--[|||-|||||-||]PAM</td><td>NA</td><td>Intergenic</td><td>NA</td><td>NA</td></tr>
<tr><td>T13488</td><td>GACGATGAGGTCTCCGTTCTCGG</td><td>1000</td><td></td><td></td><td></td><td></td><td></td><td></td><td></td><td></td><td></td></tr>
<tr><td>Oligo fwd</td><td>CACCGACGATGAGGTCTCCGTTCT</td><td></td><td></td><td></td><td></td><td></td><td></td><td></td><td></td><td></td><td></td></tr>
<tr><td>Oligo rev</td><td>AAACAGAACGGAGACCTCATCGTC</td><td></td><td></td><td></td><td></td><td></td><td></td><td></td><td></td><td></td><td></td></tr>
<tr><td>Chromosome</td><td>start</td><td>end</td><td>strand</td><td>MM</td><td>target_seq</td><td>PAM</td><td>alignment</td><td>distance</td><td>position</td><td>gene name</td><td>gene id</td></tr>
<tr><td>GU305914.1</td><td>177609</td><td>177631</td><td>+</td><td>0</td><td>GACGATGAGGTCTCCGTTCT</td><td>CGG</td><td>||||||||[||||||||||||]PAM</td><td>NA</td><td>Intergenic</td><td>NA</td><td>NA</td></tr>
<tr><td>T13489</td><td>GATGAGGTCTCCGTTCTCGGCGG</td><td>1000</td><td></td><td></td><td></td><td></td><td></td><td></td><td></td><td></td><td></td></tr>
<tr><td>Oligo fwd</td><td>CACCGATGAGGTCTCCGTTCTCGG</td><td></td><td></td><td></td><td></td><td></td><td></td><td></td><td></td><td></td><td></td></tr>
<tr><td>Oligo rev</td><td>AAACCCGAGAACGGAGACCTCATC</td><td></td><td></td><td></td><td></td><td></td><td></td><td></td><td></td><td></td><td></td></tr>
<tr><td>Chromosome</td><td>start</td><td>end</td><td>strand</td><td>MM</td><td>target_seq</td><td>PAM</td><td>alignment</td><td>distance</td><td>position</td><td>gene name</td><td>gene id</td></tr>
<tr><td>GU305914.1</td><td>177612</td><td>177634</td><td>+</td><td>0</td><td>GATGAGGTCTCCGTTCTCGG</td><td>CGG</td><td>||||||||[||||||||||||]PAM</td><td>NA</td><td>Intergenic</td><td>NA</td><td>NA</td></tr>
<tr><td>T13490</td><td>GCACGTCACGTTGCTAGCGTCGG</td><td>1000</td><td></td><td></td><td></td><td></td><td></td><td></td><td></td><td></td><td></td></tr>
<tr><td>Oligo fwd</td><td>CACCGCACGTCACGTTGCTAGCGT</td><td></td><td></td><td></td><td></td><td></td><td></td><td></td><td></td><td></td><td></td></tr>
<tr><td>Oligo rev</td><td>AAACACGCTAGCAACGTGACGTGC</td><td></td><td></td><td></td><td></td><td></td><td></td><td></td><td></td><td></td><td></td></tr>
<tr><td>Chromosome</td><td>start</td><td>end</td><td>strand</td><td>MM</td><td>target_seq</td><td>PAM</td><td>alignment</td><td>distance</td><td>position</td><td>gene name</td><td>gene id</td></tr>
<tr><td>GU305914.1</td><td>177639</td><td>177661</td><td>+</td><td>0</td><td>GCACGTCACGTTGCTAGCGT</td><td>CGG</td><td>||||||||[||||||||||||]PAM</td><td>NA</td><td>Intergenic</td><td>NA</td><td>NA</td></tr>
<tr><td>T13491</td><td>GCTAGCGTCGGATGTGCAGCCGG</td><td>1000</td><td></td><td></td><td></td><td></td><td></td><td></td><td></td><td></td><td></td></tr>
<tr><td>Oligo fwd</td><td>CACCGCTAGCGTCGGATGTGCAGC</td><td></td><td></td><td></td><td></td><td></td><td></td><td></td><td></td><td></td><td></td></tr>
</table>

<table>
<tr><td>Oligo rev</td><td>AAACGCTGCACATCCGACGCTAGC</td><td></td><td></td><td></td><td></td><td></td><td></td><td></td><td></td><td></td><td></td></tr>
<tr><td>Chromosome</td><td>start</td><td>end</td><td>strand</td><td>MM</td><td>target_seq</td><td>PAM</td><td>alignment</td><td>distance</td><td>position</td><td>gene name</td><td>gene id</td></tr>
<tr><td>GU305914.1</td><td>177651</td><td>177673</td><td>+</td><td>0</td><td>GCTAGCGTCGGATGTGCAGC</td><td>CGG</td><td>||||||||[||||||||||||]PAM</td><td>NA</td><td>Intergenic</td><td>NA</td><td>NA</td></tr>
<tr><td>T13492</td><td>GCTGACTGATATATCACGTCTGG</td><td>1000</td><td></td><td></td><td></td><td></td><td></td><td></td><td></td><td></td><td></td></tr>
<tr><td>Oligo fwd</td><td>CACCGCTGACTGATATATCACGTC</td><td></td><td></td><td></td><td></td><td></td><td></td><td></td><td></td><td></td><td></td></tr>
<tr><td>Oligo rev</td><td>AAACGACGTGATATATCAGTCAGC</td><td></td><td></td><td></td><td></td><td></td><td></td><td></td><td></td><td></td><td></td></tr>
<tr><td>Chromosome</td><td>start</td><td>end</td><td>strand</td><td>MM</td><td>target_seq</td><td>PAM</td><td>alignment</td><td>distance</td><td>position</td><td>gene name</td><td>gene id</td></tr>
<tr><td>GU305914.1</td><td>177684</td><td>177706</td><td>+</td><td>0</td><td>GCTGACTGATATATCACGTC</td><td>TGG</td><td>||||||||[||||||||||||]PAM</td><td>NA</td><td>Intergenic</td><td>NA</td><td>NA</td></tr>
<tr><td>T13493</td><td>TGCCTCCTTCTTCAATGAAGAGG</td><td>1000</td><td></td><td></td><td></td><td></td><td></td><td></td><td></td><td></td><td></td></tr>
<tr><td>Oligo fwd</td><td>CACCTGCCTCCTTCTTCAATGAAG</td><td></td><td></td><td></td><td></td><td></td><td></td><td></td><td></td><td></td><td></td></tr>
<tr><td>Oligo rev</td><td>AAACCTTCATTGAAGAAGGAGGCA</td><td></td><td></td><td></td><td></td><td></td><td></td><td></td><td></td><td></td><td></td></tr>
<tr><td>Chromosome</td><td>start</td><td>end</td><td>strand</td><td>MM</td><td>target_seq</td><td>PAM</td><td>alignment</td><td>distance</td><td>position</td><td>gene name</td><td>gene id</td></tr>
<tr><td>GU305914.1</td><td>177743</td><td>177765</td><td>+</td><td>0</td><td>TGCCTCCTTCTTCAATGAAG</td><td>AGG</td><td>||||||||[||||||||||||]PAM</td><td>NA</td><td>Intergenic</td><td>NA</td><td>NA</td></tr>
<tr><td>T13494</td><td>TTCTTCAATGAAGAGGCACCAGG</td><td>1000</td><td></td><td></td><td></td><td></td><td></td><td></td><td></td><td></td><td></td></tr>
<tr><td>Oligo fwd</td><td>CACCTTCTTCAATGAAGAGGCACC</td><td></td><td></td><td></td><td></td><td></td><td></td><td></td><td></td><td></td><td></td></tr>
<tr><td>Oligo rev</td><td>AAACGGTGCCTCTTCATTGAAGAA</td><td></td><td></td><td></td><td></td><td></td><td></td><td></td><td></td><td></td><td></td></tr>
<tr><td>Chromosome</td><td>start</td><td>end</td><td>strand</td><td>MM</td><td>target_seq</td><td>PAM</td><td>alignment</td><td>distance</td><td>position</td><td>gene name</td><td>gene id</td></tr>
<tr><td>GU305914.1</td><td>177750</td><td>177772</td><td>+</td><td>0</td><td>TTCTTCAATGAAGAGGCACC</td><td>AGG</td><td>||||||||[||||||||||||]PAM</td><td>NA</td><td>Intergenic</td><td>NA</td><td>NA</td></tr>
<tr><td>T13495</td><td>CCAGGTGTTAGTCACTTGAGCGG</td><td>1000</td><td></td><td></td><td></td><td></td><td></td><td></td><td></td><td></td><td></td></tr>
<tr><td>Oligo fwd</td><td>CACCCCAGGTGTTAGTCACTTGAG</td><td></td><td></td><td></td><td></td><td></td><td></td><td></td><td></td><td></td><td></td></tr>
<tr><td>Oligo rev</td><td>AAACCTCAAGTGACTAACACCTGG</td><td></td><td></td><td></td><td></td><td></td><td></td><td></td><td></td><td></td><td></td></tr>
<tr><td>Chromosome</td><td>start</td><td>end</td><td>strand</td><td>MM</td><td>target_seq</td><td>PAM</td><td>alignment</td><td>distance</td><td>position</td><td>gene name</td><td>gene id</td></tr>
<tr><td>GU305914.1</td><td>177768</td><td>177790</td><td>+</td><td>0</td><td>CCAGGTGTTAGTCACTTGAG</td><td>CGG</td><td>||||||||[||||||||||||]PAM</td><td>NA</td><td>Intergenic</td><td>NA</td><td>NA</td></tr>
<tr><td>T13496</td><td>CCCAACACCCCACCGCCAAAAGG</td><td>1000</td><td></td><td></td><td></td><td></td><td></td><td></td><td></td><td></td><td></td></tr>
<tr><td>Oligo fwd</td><td>CACCCCCAACACCCCACCGCCAAA</td><td></td><td></td><td></td><td></td><td></td><td></td><td></td><td></td><td></td><td></td></tr>
<tr><td>Oligo rev</td><td>AAACTTTGGCGGTGGGGTGTTGGG</td><td></td><td></td><td></td><td></td><td></td><td></td><td></td><td></td><td></td><td></td></tr>
<tr><td>Chromosome</td><td>start</td><td>end</td><td>strand</td><td>MM</td><td>target_seq</td><td>PAM</td><td>alignment</td><td>distance</td><td>position</td><td>gene name</td><td>gene id</td></tr>
<tr><td>GU305914.1</td><td>177827</td><td>177849</td><td>+</td><td>0</td><td>CCCAACACCCCACCGCCAAA</td><td>AGG</td><td>||||||||[||||||||||||]PAM</td><td>NA</td><td>Intergenic</td><td>NA</td><td>NA</td></tr>
<tr><td>T13497</td><td>AACACCCCACCGCCAAAAGGAGG</td><td>1000</td><td></td><td></td><td></td><td></td><td></td><td></td><td></td><td></td><td></td></tr>
<tr><td>Oligo fwd</td><td>CACCAACACCCCACCGCCAAAAGG</td><td></td><td></td><td></td><td></td><td></td><td></td><td></td><td></td><td></td><td></td></tr>
<tr><td>Oligo rev</td><td>AAACCCTTTTGGCGGTGGGGTGTT</td><td></td><td></td><td></td><td></td><td></td><td></td><td></td><td></td><td></td><td></td></tr>
<tr><td>Chromosome</td><td>start</td><td>end</td><td>strand</td><td>MM</td><td>target_seq</td><td>PAM</td><td>alignment</td><td>distance</td><td>position</td><td>gene name</td><td>gene id</td></tr>
<tr><td>GU305914.1</td><td>177830</td><td>177852</td><td>+</td><td>0</td><td>AACACCCCACCGCCAAAAGG</td><td>AGG</td><td>||||||||[||||||||||||]PAM</td><td>NA</td><td>Intergenic</td><td>NA</td><td>NA</td></tr>
<tr><td>T13498</td><td>CCCCACCGCCAAAAGGAGGACGG</td><td>1000</td><td></td><td></td><td></td><td></td><td></td><td></td><td></td><td></td><td></td></tr>
<tr><td>Oligo fwd</td><td>CACCCCCCACCGCCAAAAGGAGGA</td><td></td><td></td><td></td><td></td><td></td><td></td><td></td><td></td><td></td><td></td></tr>
<tr><td>Oligo rev</td><td>AAACTCCTCCTTTTGGCGGTGGGG</td><td></td><td></td><td></td><td></td><td></td><td></td><td></td><td></td><td></td><td></td></tr>
<tr><td>Chromosome</td><td>start</td><td>end</td><td>strand</td><td>MM</td><td>target_seq</td><td>PAM</td><td>alignment</td><td>distance</td><td>position</td><td>gene name</td><td>gene id</td></tr>
<tr><td>GU305914.1</td><td>177834</td><td>177856</td><td>+</td><td>0</td><td>CCCCACCGCCAAAAGGAGGA</td><td>CGG</td><td>||||||||[||||||||||||]PAM</td><td>NA</td><td>Intergenic</td><td>NA</td><td>NA</td></tr>
<tr><td>T13499</td><td>AAAAGGAGGACGGTGTCTACTGG</td><td>1000</td><td></td><td></td><td></td><td></td><td></td><td></td><td></td><td></td><td></td></tr>
<tr><td>Oligo fwd</td><td>CACCAAAAGGAGGACGGTGTCTAC</td><td></td><td></td><td></td><td></td><td></td><td></td><td></td><td></td><td></td><td></td></tr>
<tr><td>Oligo rev</td><td>AAACGTAGACACCGTCCTCCTTTT</td><td></td><td></td><td></td><td></td><td></td><td></td><td></td><td></td><td></td><td></td></tr>
<tr><td>Chromosome</td><td>start</td><td>end</td><td>strand</td><td>MM</td><td>target_seq</td><td>PAM</td><td>alignment</td><td>distance</td><td>position</td><td>gene name</td><td>gene id</td></tr>
<tr><td>GU305914.1</td><td>177844</td><td>177866</td><td>+</td><td>0</td><td>AAAAGGAGGACGGTGTCTAC</td><td>TGG</td><td>||||||||[||||||||||||]PAM</td><td>NA</td><td>Intergenic</td><td>NA</td><td>NA</td></tr>
<tr><td>T13500</td><td>ATGATGTCTCAGAGACAAAGAGG</td><td>1000</td><td></td><td></td><td></td><td></td><td></td><td></td><td></td><td></td><td></td></tr>
<tr><td>Oligo fwd</td><td>CACCATGATGTCTCAGAGACAAAG</td><td></td><td></td><td></td><td></td><td></td><td></td><td></td><td></td><td></td><td></td></tr>
<tr><td>Oligo rev</td><td>AAACCTTTGTCTCTGAGACATCAT</td><td></td><td></td><td></td><td></td><td></td><td></td><td></td><td></td><td></td><td></td></tr>
<tr><td>Chromosome</td><td>start</td><td>end</td><td>strand</td><td>MM</td><td>target_seq</td><td>PAM</td><td>alignment</td><td>distance</td><td>position</td><td>gene name</td><td>gene id</td></tr>
<tr><td>GU305914.1</td><td>177877</td><td>177899</td><td>+</td><td>0</td><td>ATGATGTCTCAGAGACAAAG</td><td>AGG</td><td>||||||||[||||||||||||]PAM</td><td>NA</td><td>Intergenic</td><td>NA</td><td>NA</td></tr>
<tr><td>T13501</td><td>TGATGTCTCAGAGACAAAGAGGG</td><td>1000</td><td></td><td></td><td></td><td></td><td></td><td></td><td></td><td></td><td></td></tr>
<tr><td>Oligo fwd</td><td>CACCTGATGTCTCAGAGACAAAGA</td><td></td><td></td><td></td><td></td><td></td><td></td><td></td><td></td><td></td><td></td></tr>
<tr><td>Oligo rev</td><td>AAACTCTTTGTCTCTGAGACATCA</td><td></td><td></td><td></td><td></td><td></td><td></td><td></td><td></td><td></td><td></td></tr>
<tr><td>Chromosome</td><td>start</td><td>end</td><td>strand</td><td>MM</td><td>target_seq</td><td>PAM</td><td>alignment</td><td>distance</td><td>position</td><td>gene name</td><td>gene id</td></tr>
<tr><td>GU305914.1</td><td>177878</td><td>177900</td><td>+</td><td>0</td><td>TGATGTCTCAGAGACAAAGA</td><td>GGG</td><td>||||||||[||||||||||||]PAM</td><td>NA</td><td>Intergenic</td><td>NA</td><td>NA</td></tr>
<tr><td>T13502</td><td>GTGCTCAATATATATCAAAAAGG</td><td>1000</td><td></td><td></td><td></td><td></td><td></td><td></td><td></td><td></td><td></td></tr>
<tr><td>Oligo fwd</td><td>CACCGTGCTCAATATATATCAAAA</td><td></td><td></td><td></td><td></td><td></td><td></td><td></td><td></td><td></td><td></td></tr>
<tr><td>Oligo rev</td><td>AAACTTTTGATATATATTGAGCAC</td><td></td><td></td><td></td><td></td><td></td><td></td><td></td><td></td><td></td><td></td></tr>
<tr><td>Chromosome</td><td>start</td><td>end</td><td>strand</td><td>MM</td><td>target_seq</td><td>PAM</td><td>alignment</td><td>distance</td><td>position</td><td>gene name</td><td>gene id</td></tr>
<tr><td>GU305914.1</td><td>177912</td><td>177934</td><td>+</td><td>0</td><td>GTGCTCAATATATATCAAAA</td><td>AGG</td><td>||||||||[||||||||||||]PAM</td><td>NA</td><td>Intergenic</td><td>NA</td><td>NA</td></tr>
<tr><td>T13503</td><td>TCAAAAAGGATTTATTTAATAGG</td><td>1000</td><td></td><td></td><td></td><td></td><td></td><td></td><td></td><td></td><td></td></tr>
<tr><td>Oligo fwd</td><td>CACCTCAAAAAGGATTTATTTAAT</td><td></td><td></td><td></td><td></td><td></td><td></td><td></td><td></td><td></td><td></td></tr>
<tr><td>Oligo rev</td><td>AAACATTAAATAAATCCTTTTTGA</td><td></td><td></td><td></td><td></td><td></td><td></td><td></td><td></td><td></td><td></td></tr>
<tr><td>Chromosome</td><td>start</td><td>end</td><td>strand</td><td>MM</td><td>target_seq</td><td>PAM</td><td>alignment</td><td>distance</td><td>position</td><td>gene name</td><td>gene id</td></tr>
<tr><td>GU305914.1</td><td>177926</td><td>177948</td><td>+</td><td>0</td><td>TCAAAAAGGATTTATTTAAT</td><td>AGG</td><td>||||||||[||||||||||||]PAM</td><td>NA</td><td>Intergenic</td><td>NA</td><td>NA</td></tr>
<tr><td>T13504</td><td>CAAAAAGGATTTATTTAATAGGG</td><td>1000</td><td></td><td></td><td></td><td></td><td></td><td></td><td></td><td></td><td></td></tr>
<tr><td>Oligo fwd</td><td>CACCCAAAAAGGATTTATTTAATA</td><td></td><td></td><td></td><td></td><td></td><td></td><td></td><td></td><td></td><td></td></tr>
<tr><td>Oligo rev</td><td>AAACTATTAAATAAATCCTTTTTG</td><td></td><td></td><td></td><td></td><td></td><td></td><td></td><td></td><td></td><td></td></tr>
<tr><td>Chromosome</td><td>start</td><td>end</td><td>strand</td><td>MM</td><td>target_seq</td><td>PAM</td><td>alignment</td><td>distance</td><td>position</td><td>gene name</td><td>gene id</td></tr>
<tr><td>GU305914.1</td><td>177927</td><td>177949</td><td>+</td><td>0</td><td>CAAAAAGGATTTATTTAATA</td><td>GGG</td><td>||||||||[||||||||||||]PAM</td><td>NA</td><td>Intergenic</td><td>NA</td><td>NA</td></tr>
<tr><td>T13505</td><td>GGTTCGACATGAGTTAAGATTGG</td><td>1000</td><td></td><td></td><td></td><td></td><td></td><td></td><td></td><td></td><td></td></tr>
<tr><td>Oligo fwd</td><td>CACCGGTTCGACATGAGTTAAGAT</td><td></td><td></td><td></td><td></td><td></td><td></td><td></td><td></td><td></td><td></td></tr>
<tr><td>Oligo rev</td><td>AAACATCTTAACTCATGTCGAACC</td><td></td><td></td><td></td><td></td><td></td><td></td><td></td><td></td><td></td><td></td></tr>
<tr><td>Chromosome</td><td>start</td><td>end</td><td>strand</td><td>MM</td><td>target_seq</td><td>PAM</td><td>alignment</td><td>distance</td><td>position</td><td>gene name</td><td>gene id</td></tr>
<tr><td>GU305914.1</td><td>177948</td><td>177970</td><td>+</td><td>0</td><td>GGTTCGACATGAGTTAAGAT</td><td>TGG</td><td>||||||||[||||||||||||]PAM</td><td>NA</td><td>Intergenic</td><td>NA</td><td>NA</td></tr>
<tr><td>T13506</td><td>GTTCGACATGAGTTAAGATTGGG</td><td>1000</td><td></td><td></td><td></td><td></td><td></td><td></td><td></td><td></td><td></td></tr>
</table>

<table>
<tr><td>Oligo fwd</td><td>CACCGTTCGACATGAGTTAAGATT</td></tr>
<tr><td>Oligo rev</td><td>AAACAATCTTAACTCATGTCGAAC</td></tr>
<tr><td>Chromosome</td><td>start</td><td>end</td><td>strand</td><td>MM</td><td>target_seq</td><td>PAM</td><td>alignment</td><td>distance</td><td>position</td><td>gene name</td><td>gene id</td></tr>
<tr><td>GU305914.1</td><td>177949</td><td>177971</td><td>+</td><td>0</td><td>GTTCGACATGAGTTAAGATT</td><td>GGG</td><td>||||||||[||||||||||||]PAM</td><td>NA</td><td>Intergenic</td><td>NA</td><td>NA</td></tr>
<tr><td>T13507</td><td>TTCGACATGAGTTAAGATTGGGG</td><td>1000</td></tr>
<tr><td>Oligo fwd</td><td>CACCTTCGACATGAGTTAAGATTG</td></tr>
<tr><td>Oligo rev</td><td>AAACCAATCTTAACTCATGTCGAA</td></tr>
<tr><td>Chromosome</td><td>start</td><td>end</td><td>strand</td><td>MM</td><td>target_seq</td><td>PAM</td><td>alignment</td><td>distance</td><td>position</td><td>gene name</td><td>gene id</td></tr>
<tr><td>GU305914.1</td><td>177950</td><td>177972</td><td>+</td><td>0</td><td>TTCGACATGAGTTAAGATTG</td><td>GGG</td><td>||||||||[||||||||||||]PAM</td><td>NA</td><td>Intergenic</td><td>NA</td><td>NA</td></tr>
<tr><td>T13508</td><td>TCGACATGAGTTAAGATTGGGGG</td><td>1000</td></tr>
<tr><td>Oligo fwd</td><td>CACCTCGACATGAGTTAAGATTGG</td></tr>
<tr><td>Oligo rev</td><td>AAACCCAATCTTAACTCATGTCGA</td></tr>
<tr><td>Chromosome</td><td>start</td><td>end</td><td>strand</td><td>MM</td><td>target_seq</td><td>PAM</td><td>alignment</td><td>distance</td><td>position</td><td>gene name</td><td>gene id</td></tr>
<tr><td>GU305914.1</td><td>177951</td><td>177973</td><td>+</td><td>0</td><td>TCGACATGAGTTAAGATTGG</td><td>GGG</td><td>||||||||[||||||||||||]PAM</td><td>NA</td><td>Intergenic</td><td>NA</td><td>NA</td></tr>
<tr><td>T13509</td><td>GAGTTAAGATTGGGGGTCTCAGG</td><td>1000</td></tr>
<tr><td>Oligo fwd</td><td>CACCGAGTTAAGATTGGGGGTCTC</td></tr>
<tr><td>Oligo rev</td><td>AAACGAGACCCCCAATCTTAACTC</td></tr>
<tr><td>Chromosome</td><td>start</td><td>end</td><td>strand</td><td>MM</td><td>target_seq</td><td>PAM</td><td>alignment</td><td>distance</td><td>position</td><td>gene name</td><td>gene id</td></tr>
<tr><td>GU305914.1</td><td>177958</td><td>177980</td><td>+</td><td>0</td><td>GAGTTAAGATTGGGGGTCTC</td><td>AGG</td><td>||||||||[||||||||||||]PAM</td><td>NA</td><td>Intergenic</td><td>NA</td><td>NA</td></tr>
<tr><td>T13510</td><td>ATCTACATATTTTACAAAACAGG</td><td>1000</td></tr>
<tr><td>Oligo fwd</td><td>CACCATCTACATATTTTACAAAAC</td></tr>
<tr><td>Oligo rev</td><td>AAACGTTTTGTAAAATATGTAGAT</td></tr>
<tr><td>Chromosome</td><td>start</td><td>end</td><td>strand</td><td>MM</td><td>target_seq</td><td>PAM</td><td>alignment</td><td>distance</td><td>position</td><td>gene name</td><td>gene id</td></tr>
<tr><td>GU305914.1</td><td>178025</td><td>178047</td><td>+</td><td>0</td><td>ATCTACATATTTTACAAAAC</td><td>AGG</td><td>||||||||[||||||||||||]PAM</td><td>NA</td><td>Intergenic</td><td>NA</td><td>NA</td></tr>
<tr><td>T13511</td><td>TCTACATATTTTACAAAACAGGG</td><td>1000</td></tr>
<tr><td>Oligo fwd</td><td>CACCTCTACATATTTTACAAAACA</td></tr>
<tr><td>Oligo rev</td><td>AAACTGTTTTGTAAAATATGTAGA</td></tr>
<tr><td>Chromosome</td><td>start</td><td>end</td><td>strand</td><td>MM</td><td>target_seq</td><td>PAM</td><td>alignment</td><td>distance</td><td>position</td><td>gene name</td><td>gene id</td></tr>
<tr><td>GU305914.1</td><td>178026</td><td>178048</td><td>+</td><td>0</td><td>TCTACATATTTTACAAAACA</td><td>GGG</td><td>||||||||[||||||||||||]PAM</td><td>NA</td><td>Intergenic</td><td>NA</td><td>NA</td></tr>
<tr><td>T13512</td><td>CCCCAGACTCATCCTCATGATGG</td><td>1000</td></tr>
<tr><td>Oligo fwd</td><td>CACCCCCCAGACTCATCCTCATGA</td></tr>
<tr><td>Oligo rev</td><td>AAACTCATGAGGATGAGTCTGGGG</td></tr>
<tr><td>Chromosome</td><td>start</td><td>end</td><td>strand</td><td>MM</td><td>target_seq</td><td>PAM</td><td>alignment</td><td>distance</td><td>position</td><td>gene name</td><td>gene id</td></tr>
<tr><td>GU305914.1</td><td>178103</td><td>178125</td><td>+</td><td>0</td><td>CCCCAGACTCATCCTCATGA</td><td>TGG</td><td>||||||||[||||||||||||]PAM</td><td>NA</td><td>Intergenic</td><td>NA</td><td>NA</td></tr>
<tr><td>T13513</td><td>TTATATCATCTGCCTCAAACTGG</td><td>1000</td></tr>
<tr><td>Oligo fwd</td><td>CACCTTATATCATCTGCCTCAAAC</td></tr>
<tr><td>Oligo rev</td><td>AAACGTTTGAGGCAGATGATATAA</td></tr>
<tr><td>Chromosome</td><td>start</td><td>end</td><td>strand</td><td>MM</td><td>target_seq</td><td>PAM</td><td>alignment</td><td>distance</td><td>position</td><td>gene name</td><td>gene id</td></tr>
<tr><td>GU305914.1</td><td>178127</td><td>178149</td><td>+</td><td>0</td><td>TTATATCATCTGCCTCAAAC</td><td>TGG</td><td>||||||||[||||||||||||]PAM</td><td>NA</td><td>Intergenic</td><td>NA</td><td>NA</td></tr>
<tr><td>T13514</td><td>TATATCATCTGCCTCAAACTGGG</td><td>1000</td></tr>
<tr><td>Oligo fwd</td><td>CACCTATATCATCTGCCTCAAACT</td></tr>
<tr><td>Oligo rev</td><td>AAACAGTTTGAGGCAGATGATATA</td></tr>
<tr><td>Chromosome</td><td>start</td><td>end</td><td>strand</td><td>MM</td><td>target_seq</td><td>PAM</td><td>alignment</td><td>distance</td><td>position</td><td>gene name</td><td>gene id</td></tr>
<tr><td>GU305914.1</td><td>178128</td><td>178150</td><td>+</td><td>0</td><td>TATATCATCTGCCTCAAACT</td><td>GGG</td><td>||||||||[||||||||||||]PAM</td><td>NA</td><td>Intergenic</td><td>NA</td><td>NA</td></tr>
<tr><td>T13515</td><td>GCTGACAGTCTCCTCATAGATGG</td><td>1000</td></tr>
<tr><td>Oligo fwd</td><td>CACCGCTGACAGTCTCCTCATAGA</td></tr>
<tr><td>Oligo rev</td><td>AAACTCTATGAGGAGACTGTCAGC</td></tr>
<tr><td>Chromosome</td><td>start</td><td>end</td><td>strand</td><td>MM</td><td>target_seq</td><td>PAM</td><td>alignment</td><td>distance</td><td>position</td><td>gene name</td><td>gene id</td></tr>
<tr><td>GU305914.1</td><td>178164</td><td>178186</td><td>+</td><td>0</td><td>GCTGACAGTCTCCTCATAGA</td><td>TGG</td><td>||||||||[||||||||||||]PAM</td><td>NA</td><td>Intergenic</td><td>NA</td><td>NA</td></tr>
<tr><td>T13516</td><td>CTGACAGTCTCCTCATAGATGGG</td><td>1000</td></tr>
<tr><td>Oligo fwd</td><td>CACCCTGACAGTCTCCTCATAGAT</td></tr>
<tr><td>Oligo rev</td><td>AAACATCTATGAGGAGACTGTCAG</td></tr>
<tr><td>Chromosome</td><td>start</td><td>end</td><td>strand</td><td>MM</td><td>target_seq</td><td>PAM</td><td>alignment</td><td>distance</td><td>position</td><td>gene name</td><td>gene id</td></tr>
<tr><td>GU305914.1</td><td>178165</td><td>178187</td><td>+</td><td>0</td><td>CTGACAGTCTCCTCATAGAT</td><td>GGG</td><td>||||||||[||||||||||||]PAM</td><td>NA</td><td>Intergenic</td><td>NA</td><td>NA</td></tr>
<tr><td>T13517</td><td>GTCTCCTCATAGATGGGAAGAGG</td><td>1000</td></tr>
<tr><td>Oligo fwd</td><td>CACCGTCTCCTCATAGATGGGAAG</td></tr>
<tr><td>Oligo rev</td><td>AAACCTTCCCATCTATGAGGAGAC</td></tr>
<tr><td>Chromosome</td><td>start</td><td>end</td><td>strand</td><td>MM</td><td>target_seq</td><td>PAM</td><td>alignment</td><td>distance</td><td>position</td><td>gene name</td><td>gene id</td></tr>
<tr><td>GU305914.1</td><td>178171</td><td>178193</td><td>+</td><td>0</td><td>GTCTCCTCATAGATGGGAAG</td><td>AGG</td><td>||||||||[||||||||||||]PAM</td><td>NA</td><td>Intergenic</td><td>NA</td><td>NA</td></tr>
<tr><td>T13518</td><td>GAGCCTCATGCACACCTGCTTGG</td><td>1000</td></tr>
<tr><td>Oligo fwd</td><td>CACCGAGCCTCATGCACACCTGCT</td></tr>
<tr><td>Oligo rev</td><td>AAACAGCAGGTGTGCATGAGGCTC</td></tr>
<tr><td>Chromosome</td><td>start</td><td>end</td><td>strand</td><td>MM</td><td>target_seq</td><td>PAM</td><td>alignment</td><td>distance</td><td>position</td><td>gene name</td><td>gene id</td></tr>
<tr><td>GU305914.1</td><td>178218</td><td>178240</td><td>+</td><td>0</td><td>GAGCCTCATGCACACCTGCT</td><td>TGG</td><td>||||||||[||||||||||||]PAM</td><td>NA</td><td>Intergenic</td><td>NA</td><td>NA</td></tr>
<tr><td>T13519</td><td>AGCCTCATGCACACCTGCTTGGG</td><td>1000</td></tr>
<tr><td>Oligo fwd</td><td>CACCAGCCTCATGCACACCTGCTT</td></tr>
<tr><td>Oligo rev</td><td>AAACAAGCAGGTGTGCATGAGGCT</td></tr>
<tr><td>Chromosome</td><td>start</td><td>end</td><td>strand</td><td>MM</td><td>target_seq</td><td>PAM</td><td>alignment</td><td>distance</td><td>position</td><td>gene name</td><td>gene id</td></tr>
<tr><td>GU305914.1</td><td>178219</td><td>178241</td><td>+</td><td>0</td><td>AGCCTCATGCACACCTGCTT</td><td>GGG</td><td>||||||||[||||||||||||]PAM</td><td>NA</td><td>Intergenic</td><td>NA</td><td>NA</td></tr>
<tr><td>T13520</td><td>GCCTCATGCACACCTGCTTGGGG</td><td>1000</td></tr>
<tr><td>Oligo fwd</td><td>CACCGCCTCATGCACACCTGCTTG</td></tr>
<tr><td>Oligo rev</td><td>AAACCAAGCAGGTGTGCATGAGGC</td></tr>
<tr><td>Chromosome</td><td>start</td><td>end</td><td>strand</td><td>MM</td><td>target_seq</td><td>PAM</td><td>alignment</td><td>distance</td><td>position</td><td>gene name</td><td>gene id</td></tr>
<tr><td>GU305914.1</td><td>178220</td><td>178242</td><td>+</td><td>0</td><td>GCCTCATGCACACCTGCTTG</td><td>GGG</td><td>||||||||[||||||||||||]PAM</td><td>NA</td><td>Intergenic</td><td>NA</td><td>NA</td></tr>
</table>

<table>
<tr><td>T13521</td><td>GACCTGACTCAAATGCAGACAGG</td><td>1000</td><td></td><td></td><td></td><td></td><td></td><td></td><td></td><td></td><td></td></tr>
<tr><td>Oligo fwd</td><td>CACCGACCTGACTCAAATGCAGAC</td><td></td><td></td><td></td><td></td><td></td><td></td><td></td><td></td><td></td><td></td></tr>
<tr><td>Oligo rev</td><td>AAACGTCTGCATTTGAGTCAGGTC</td><td></td><td></td><td></td><td></td><td></td><td></td><td></td><td></td><td></td><td></td></tr>
<tr><td>Chromosome</td><td>start</td><td>end</td><td>strand</td><td>MM</td><td>target_seq</td><td>PAM</td><td>alignment</td><td>distance</td><td>position</td><td>gene name</td><td>gene id</td></tr>
<tr><td>GU305914.1</td><td>178245</td><td>178267</td><td>+</td><td>0</td><td>GACCTGACTCAAATGCAGAC</td><td>AGG</td><td>||||||||[||||||||||||]PAM</td><td>NA</td><td>Intergenic</td><td>NA</td><td>NA</td></tr>
<tr><td></td><td></td><td></td><td></td><td></td><td></td><td></td><td></td><td></td><td></td><td></td><td></td></tr>
<tr><td>T13522</td><td>TTCACCGCTCCCAAGAAGTCTGG</td><td>1000</td><td></td><td></td><td></td><td></td><td></td><td></td><td></td><td></td><td></td></tr>
<tr><td>Oligo fwd</td><td>CACCTTCACCGCTCCCAAGAAGTC</td><td></td><td></td><td></td><td></td><td></td><td></td><td></td><td></td><td></td><td></td></tr>
<tr><td>Oligo rev</td><td>AAACGACTTCTTGGGAGCGGTGAA</td><td></td><td></td><td></td><td></td><td></td><td></td><td></td><td></td><td></td><td></td></tr>
<tr><td>Chromosome</td><td>start</td><td>end</td><td>strand</td><td>MM</td><td>target_seq</td><td>PAM</td><td>alignment</td><td>distance</td><td>position</td><td>gene name</td><td>gene id</td></tr>
<tr><td>GU305914.1</td><td>178270</td><td>178292</td><td>+</td><td>0</td><td>TTCACCGCTCCCAAGAAGTC</td><td>TGG</td><td>||||||||[||||||||||||]PAM</td><td>NA</td><td>Intergenic</td><td>NA</td><td>NA</td></tr>
<tr><td>GU305914.1</td><td>122876</td><td>122898</td><td>+</td><td>4</td><td>GTCACCTCTCCCGAGAAGAC</td><td>GGG</td><td>-|||||-|[||||-|||||-|]PAM</td><td>NA</td><td>Intergenic</td><td>NA</td><td>NA</td></tr>
<tr><td></td><td></td><td></td><td></td><td></td><td></td><td></td><td></td><td></td><td></td><td></td><td></td></tr>
<tr><td>T13523</td><td>ACCGCTCCCAAGAAGTCTGGAGG</td><td>1000</td><td></td><td></td><td></td><td></td><td></td><td></td><td></td><td></td><td></td></tr>
<tr><td>Oligo fwd</td><td>CACCACCGCTCCCAAGAAGTCTGG</td><td></td><td></td><td></td><td></td><td></td><td></td><td></td><td></td><td></td><td></td></tr>
<tr><td>Oligo rev</td><td>AAACCCAGACTTCTTGGGAGCGGT</td><td></td><td></td><td></td><td></td><td></td><td></td><td></td><td></td><td></td><td></td></tr>
<tr><td>Chromosome</td><td>start</td><td>end</td><td>strand</td><td>MM</td><td>target_seq</td><td>PAM</td><td>alignment</td><td>distance</td><td>position</td><td>gene name</td><td>gene id</td></tr>
<tr><td>GU305914.1</td><td>178273</td><td>178295</td><td>+</td><td>0</td><td>ACCGCTCCCAAGAAGTCTGG</td><td>AGG</td><td>||||||||[||||||||||||]PAM</td><td>NA</td><td>Intergenic</td><td>NA</td><td>NA</td></tr>
<tr><td></td><td></td><td></td><td></td><td></td><td></td><td></td><td></td><td></td><td></td><td></td><td></td></tr>
<tr><td>T13524</td><td>CCGCTCCCAAGAAGTCTGGAGGG</td><td>1000</td><td></td><td></td><td></td><td></td><td></td><td></td><td></td><td></td><td></td></tr>
<tr><td>Oligo fwd</td><td>CACCCCGCTCCCAAGAAGTCTGGA</td><td></td><td></td><td></td><td></td><td></td><td></td><td></td><td></td><td></td><td></td></tr>
<tr><td>Oligo rev</td><td>AAACTCCAGACTTCTTGGGAGCGG</td><td></td><td></td><td></td><td></td><td></td><td></td><td></td><td></td><td></td><td></td></tr>
<tr><td>Chromosome</td><td>start</td><td>end</td><td>strand</td><td>MM</td><td>target_seq</td><td>PAM</td><td>alignment</td><td>distance</td><td>position</td><td>gene name</td><td>gene id</td></tr>
<tr><td>GU305914.1</td><td>178274</td><td>178296</td><td>+</td><td>0</td><td>CCGCTCCCAAGAAGTCTGGA</td><td>GGG</td><td>||||||||[||||||||||||]PAM</td><td>NA</td><td>Intergenic</td><td>NA</td><td>NA</td></tr>
<tr><td></td><td></td><td></td><td></td><td></td><td></td><td></td><td></td><td></td><td></td><td></td><td></td></tr>
<tr><td>T13525</td><td>TCCCAAGAAGTCTGGAGGGTTGG</td><td>1000</td><td></td><td></td><td></td><td></td><td></td><td></td><td></td><td></td><td></td></tr>
<tr><td>Oligo fwd</td><td>CACCTCCCAAGAAGTCTGGAGGGT</td><td></td><td></td><td></td><td></td><td></td><td></td><td></td><td></td><td></td><td></td></tr>
<tr><td>Oligo rev</td><td>AAACACCCTCCAGACTTCTTGGGA</td><td></td><td></td><td></td><td></td><td></td><td></td><td></td><td></td><td></td><td></td></tr>
<tr><td>Chromosome</td><td>start</td><td>end</td><td>strand</td><td>MM</td><td>target_seq</td><td>PAM</td><td>alignment</td><td>distance</td><td>position</td><td>gene name</td><td>gene id</td></tr>
<tr><td>GU305914.1</td><td>178278</td><td>178300</td><td>+</td><td>0</td><td>TCCCAAGAAGTCTGGAGGGT</td><td>TGG</td><td>||||||||[||||||||||||]PAM</td><td>NA</td><td>Intergenic</td><td>NA</td><td>NA</td></tr>
<tr><td></td><td></td><td></td><td></td><td></td><td></td><td></td><td></td><td></td><td></td><td></td><td></td></tr>
<tr><td>T13526</td><td>TATCATGCTCCTGTAGTCTGAGG</td><td>1000</td><td></td><td></td><td></td><td></td><td></td><td></td><td></td><td></td><td></td></tr>
<tr><td>Oligo fwd</td><td>CACCTATCATGCTCCTGTAGTCTG</td><td></td><td></td><td></td><td></td><td></td><td></td><td></td><td></td><td></td><td></td></tr>
<tr><td>Oligo rev</td><td>AAACCAGACTACAGGAGCATGATA</td><td></td><td></td><td></td><td></td><td></td><td></td><td></td><td></td><td></td><td></td></tr>
<tr><td>Chromosome</td><td>start</td><td>end</td><td>strand</td><td>MM</td><td>target_seq</td><td>PAM</td><td>alignment</td><td>distance</td><td>position</td><td>gene name</td><td>gene id</td></tr>
<tr><td>GU305914.1</td><td>178311</td><td>178333</td><td>+</td><td>0</td><td>TATCATGCTCCTGTAGTCTG</td><td>AGG</td><td>||||||||[||||||||||||]PAM</td><td>NA</td><td>Intergenic</td><td>NA</td><td>NA</td></tr>
<tr><td></td><td></td><td></td><td></td><td></td><td></td><td></td><td></td><td></td><td></td><td></td><td></td></tr>
<tr><td>T13527</td><td>GCTCCTGTAGTCTGAGGTCCTGG</td><td>1000</td><td></td><td></td><td></td><td></td><td></td><td></td><td></td><td></td><td></td></tr>
<tr><td>Oligo fwd</td><td>CACCGCTCCTGTAGTCTGAGGTCC</td><td></td><td></td><td></td><td></td><td></td><td></td><td></td><td></td><td></td><td></td></tr>
<tr><td>Oligo rev</td><td>AAACGGACCTCAGACTACAGGAGC</td><td></td><td></td><td></td><td></td><td></td><td></td><td></td><td></td><td></td><td></td></tr>
<tr><td>Chromosome</td><td>start</td><td>end</td><td>strand</td><td>MM</td><td>target_seq</td><td>PAM</td><td>alignment</td><td>distance</td><td>position</td><td>gene name</td><td>gene id</td></tr>
<tr><td>GU305914.1</td><td>178317</td><td>178339</td><td>+</td><td>0</td><td>GCTCCTGTAGTCTGAGGTCC</td><td>TGG</td><td>||||||||[||||||||||||]PAM</td><td>NA</td><td>Intergenic</td><td>NA</td><td>NA</td></tr>
<tr><td></td><td></td><td></td><td></td><td></td><td></td><td></td><td></td><td></td><td></td><td></td><td></td></tr>
<tr><td>T13528</td><td>CTCCTGTAGTCTGAGGTCCTGGG</td><td>1000</td><td></td><td></td><td></td><td></td><td></td><td></td><td></td><td></td><td></td></tr>
<tr><td>Oligo fwd</td><td>CACCCTCCTGTAGTCTGAGGTCCT</td><td></td><td></td><td></td><td></td><td></td><td></td><td></td><td></td><td></td><td></td></tr>
<tr><td>Oligo rev</td><td>AAACAGGACCTCAGACTACAGGAG</td><td></td><td></td><td></td><td></td><td></td><td></td><td></td><td></td><td></td><td></td></tr>
<tr><td>Chromosome</td><td>start</td><td>end</td><td>strand</td><td>MM</td><td>target_seq</td><td>PAM</td><td>alignment</td><td>distance</td><td>position</td><td>gene name</td><td>gene id</td></tr>
<tr><td>GU305914.1</td><td>178318</td><td>178340</td><td>+</td><td>0</td><td>CTCCTGTAGTCTGAGGTCCT</td><td>GGG</td><td>||||||||[||||||||||||]PAM</td><td>NA</td><td>Intergenic</td><td>NA</td><td>NA</td></tr>
<tr><td></td><td></td><td></td><td></td><td></td><td></td><td></td><td></td><td></td><td></td><td></td><td></td></tr>
<tr><td>T13529</td><td>CCACACTGCCTGAGTACCTGCGG</td><td>1000</td><td></td><td></td><td></td><td></td><td></td><td></td><td></td><td></td><td></td></tr>
<tr><td>Oligo fwd</td><td>CACCCCACACTGCCTGAGTACCTG</td><td></td><td></td><td></td><td></td><td></td><td></td><td></td><td></td><td></td><td></td></tr>
<tr><td>Oligo rev</td><td>AAACCAGGTACTCAGGCAGTGTGG</td><td></td><td></td><td></td><td></td><td></td><td></td><td></td><td></td><td></td><td></td></tr>
<tr><td>Chromosome</td><td>start</td><td>end</td><td>strand</td><td>MM</td><td>target_seq</td><td>PAM</td><td>alignment</td><td>distance</td><td>position</td><td>gene name</td><td>gene id</td></tr>
<tr><td>GU305914.1</td><td>178374</td><td>178396</td><td>+</td><td>0</td><td>CCACACTGCCTGAGTACCTG</td><td>CGG</td><td>||||||||[||||||||||||]PAM</td><td>NA</td><td>Intergenic</td><td>NA</td><td>NA</td></tr>
<tr><td></td><td></td><td></td><td></td><td></td><td></td><td></td><td></td><td></td><td></td><td></td><td></td></tr>
<tr><td>T13530</td><td>TGCCTGAGTACCTGCGGTGCAGG</td><td>1000</td><td></td><td></td><td></td><td></td><td></td><td></td><td></td><td></td><td></td></tr>
<tr><td>Oligo fwd</td><td>CACCTGCCTGAGTACCTGCGGTGC</td><td></td><td></td><td></td><td></td><td></td><td></td><td></td><td></td><td></td><td></td></tr>
<tr><td>Oligo rev</td><td>AAACGCACCGCAGGTACTCAGGCA</td><td></td><td></td><td></td><td></td><td></td><td></td><td></td><td></td><td></td><td></td></tr>
<tr><td>Chromosome</td><td>start</td><td>end</td><td>strand</td><td>MM</td><td>target_seq</td><td>PAM</td><td>alignment</td><td>distance</td><td>position</td><td>gene name</td><td>gene id</td></tr>
<tr><td>GU305914.1</td><td>178380</td><td>178402</td><td>+</td><td>0</td><td>TGCCTGAGTACCTGCGGTGC</td><td>AGG</td><td>||||||||[||||||||||||]PAM</td><td>NA</td><td>Intergenic</td><td>NA</td><td>NA</td></tr>
<tr><td></td><td></td><td></td><td></td><td></td><td></td><td></td><td></td><td></td><td></td><td></td><td></td></tr>
<tr><td>T13531</td><td>ACCTGCGGTGCAGGCCGCTGAGG</td><td>1000</td><td></td><td></td><td></td><td></td><td></td><td></td><td></td><td></td><td></td></tr>
<tr><td>Oligo fwd</td><td>CACCACCTGCGGTGCAGGCCGCTG</td><td></td><td></td><td></td><td></td><td></td><td></td><td></td><td></td><td></td><td></td></tr>
<tr><td>Oligo rev</td><td>AAACCAGCGGCCTGCACCGCAGGT</td><td></td><td></td><td></td><td></td><td></td><td></td><td></td><td></td><td></td><td></td></tr>
<tr><td>Chromosome</td><td>start</td><td>end</td><td>strand</td><td>MM</td><td>target_seq</td><td>PAM</td><td>alignment</td><td>distance</td><td>position</td><td>gene name</td><td>gene id</td></tr>
<tr><td>GU305914.1</td><td>178389</td><td>178411</td><td>+</td><td>0</td><td>ACCTGCGGTGCAGGCCGCTG</td><td>AGG</td><td>||||||||[||||||||||||]PAM</td><td>NA</td><td>Intergenic</td><td>NA</td><td>NA</td></tr>
<tr><td></td><td></td><td></td><td></td><td></td><td></td><td></td><td></td><td></td><td></td><td></td><td></td></tr>
<tr><td>T13532</td><td>GGCCGCTGAGGTTCTATCTATGG</td><td>1000</td><td></td><td></td><td></td><td></td><td></td><td></td><td></td><td></td><td></td></tr>
<tr><td>Oligo fwd</td><td>CACCGGCCGCTGAGGTTCTATCTA</td><td></td><td></td><td></td><td></td><td></td><td></td><td></td><td></td><td></td><td></td></tr>
<tr><td>Oligo rev</td><td>AAACTAGATAGAACCTCAGCGGCC</td><td></td><td></td><td></td><td></td><td></td><td></td><td></td><td></td><td></td><td></td></tr>
<tr><td>Chromosome</td><td>start</td><td>end</td><td>strand</td><td>MM</td><td>target_seq</td><td>PAM</td><td>alignment</td><td>distance</td><td>position</td><td>gene name</td><td>gene id</td></tr>
<tr><td>GU305914.1</td><td>178401</td><td>178423</td><td>+</td><td>0</td><td>GGCCGCTGAGGTTCTATCTA</td><td>TGG</td><td>||||||||[||||||||||||]PAM</td><td>NA</td><td>Intergenic</td><td>NA</td><td>NA</td></tr>
<tr><td></td><td></td><td></td><td></td><td></td><td></td><td></td><td></td><td></td><td></td><td></td><td></td></tr>
<tr><td>T13533</td><td>TATCTATGGTCTCCCTGCTGTGG</td><td>1000</td><td></td><td></td><td></td><td></td><td></td><td></td><td></td><td></td><td></td></tr>
<tr><td>Oligo fwd</td><td>CACCTATCTATGGTCTCCCTGCTG</td><td></td><td></td><td></td><td></td><td></td><td></td><td></td><td></td><td></td><td></td></tr>
<tr><td>Oligo rev</td><td>AAACCAGCAGGGAGACCATAGATA</td><td></td><td></td><td></td><td></td><td></td><td></td><td></td><td></td><td></td><td></td></tr>
<tr><td>Chromosome</td><td>start</td><td>end</td><td>strand</td><td>MM</td><td>target_seq</td><td>PAM</td><td>alignment</td><td>distance</td><td>position</td><td>gene name</td><td>gene id</td></tr>
<tr><td>GU305914.1</td><td>178415</td><td>178437</td><td>+</td><td>0</td><td>TATCTATGGTCTCCCTGCTG</td><td>TGG</td><td>||||||||[||||||||||||]PAM</td><td>NA</td><td>Intergenic</td><td>NA</td><td>NA</td></tr>
<tr><td></td><td></td><td></td><td></td><td></td><td></td><td></td><td></td><td></td><td></td><td></td><td></td></tr>
<tr><td>T13534</td><td>GGTCTCCCTGCTGTGGTTAATGG</td><td>1000</td><td></td><td></td><td></td><td></td><td></td><td></td><td></td><td></td><td></td></tr>
<tr><td>Oligo fwd</td><td>CACCGGTCTCCCTGCTGTGGTTAA</td><td></td><td></td><td></td><td></td><td></td><td></td><td></td><td></td><td></td><td></td></tr>
<tr><td>Oligo rev</td><td>AAACTTAACCACAGCAGGGAGACC</td><td></td><td></td><td></td><td></td><td></td><td></td><td></td><td></td><td></td><td></td></tr>
<tr><td>Chromosome</td><td>start</td><td>end</td><td>strand</td><td>MM</td><td>target_seq</td><td>PAM</td><td>alignment</td><td>distance</td><td>position</td><td>gene name</td><td>gene id</td></tr>
<tr><td>GU305914.1</td><td>178422</td><td>178444</td><td>+</td><td>0</td><td>GGTCTCCCTGCTGTGGTTAA</td><td>TGG</td><td>||||||||[||||||||||||]PAM</td><td>NA</td><td>Intergenic</td><td>NA</td><td>NA</td></tr>
<tr><td></td><td></td><td></td><td></td><td></td><td></td><td></td><td></td><td></td><td></td><td></td><td></td></tr>
<tr><td>T13535</td><td>GTCTCCCTGCTGTGGTTAATGGG</td><td>1000</td><td></td><td></td><td></td><td></td><td></td><td></td><td></td><td></td><td></td></tr>
<tr><td>Oligo fwd</td><td>CACCGTCTCCCTGCTGTGGTTAAT</td><td></td><td></td><td></td><td></td><td></td><td></td><td></td><td></td><td></td><td></td></tr>
<tr><td>Oligo rev</td><td>AAACATTAACCACAGCAGGGAGAC</td><td></td><td></td><td></td><td></td><td></td><td></td><td></td><td></td><td></td><td></td></tr>
<tr><td>Chromosome</td><td>start</td><td>end</td><td>strand</td><td>MM</td><td>target_seq</td><td>PAM</td><td>alignment</td><td>distance</td><td>position</td><td>gene name</td><td>gene id</td></tr>
</table>

<table>
<tr><td>GU305914.1</td><td>178423</td><td>178445</td><td>+</td><td>0</td><td>GTCTCCCTGCTGTGGTTAAT</td><td>GGG</td><td>||||||||[||||||||||||]PAM</td><td>NA</td><td>Intergenic</td><td>NA</td><td>NA</td></tr>
<tr><td>T13536</td><td>CCTGCTGTGGTTAATGGGCTTGG</td><td>1000</td><td></td><td></td><td></td><td></td><td></td><td></td><td></td><td></td><td></td></tr>
<tr><td>Oligo fwd</td><td>CACCCCTGCTGTGGTTAATGGGCT</td><td></td><td></td><td></td><td></td><td></td><td></td><td></td><td></td><td></td><td></td></tr>
<tr><td>Oligo rev</td><td>AAACAGCCCATTAACCACAGCAGG</td><td></td><td></td><td></td><td></td><td></td><td></td><td></td><td></td><td></td><td></td></tr>
<tr><td>Chromosome</td><td>start</td><td>end</td><td>strand</td><td>MM</td><td>target_seq</td><td>PAM</td><td>alignment</td><td>distance</td><td>position</td><td>gene name</td><td>gene id</td></tr>
<tr><td>GU305914.1</td><td>178428</td><td>178450</td><td>+</td><td>0</td><td>CCTGCTGTGGTTAATGGGCT</td><td>TGG</td><td>||||||||[||||||||||||]PAM</td><td>NA</td><td>Intergenic</td><td>NA</td><td>NA</td></tr>
<tr><td>T13537</td><td>CTTGGTGTGCTCTGTCATGAAGG</td><td>1000</td><td></td><td></td><td></td><td></td><td></td><td></td><td></td><td></td><td></td></tr>
<tr><td>Oligo fwd</td><td>CACCCTTGGTGTGCTCTGTCATGA</td><td></td><td></td><td></td><td></td><td></td><td></td><td></td><td></td><td></td><td></td></tr>
<tr><td>Oligo rev</td><td>AAACTCATGACAGAGCACACCAAG</td><td></td><td></td><td></td><td></td><td></td><td></td><td></td><td></td><td></td><td></td></tr>
<tr><td>Chromosome</td><td>start</td><td>end</td><td>strand</td><td>MM</td><td>target_seq</td><td>PAM</td><td>alignment</td><td>distance</td><td>position</td><td>gene name</td><td>gene id</td></tr>
<tr><td>GU305914.1</td><td>178446</td><td>178468</td><td>+</td><td>0</td><td>CTTGGTGTGCTCTGTCATGA</td><td>AGG</td><td>||||||||[||||||||||||]PAM</td><td>NA</td><td>Intergenic</td><td>NA</td><td>NA</td></tr>
<tr><td>T13538</td><td>TTGGTGTGCTCTGTCATGAAGGG</td><td>1000</td><td></td><td></td><td></td><td></td><td></td><td></td><td></td><td></td><td></td></tr>
<tr><td>Oligo fwd</td><td>CACCTTGGTGTGCTCTGTCATGAA</td><td></td><td></td><td></td><td></td><td></td><td></td><td></td><td></td><td></td><td></td></tr>
<tr><td>Oligo rev</td><td>AAACTTCATGACAGAGCACACCAA</td><td></td><td></td><td></td><td></td><td></td><td></td><td></td><td></td><td></td><td></td></tr>
<tr><td>Chromosome</td><td>start</td><td>end</td><td>strand</td><td>MM</td><td>target_seq</td><td>PAM</td><td>alignment</td><td>distance</td><td>position</td><td>gene name</td><td>gene id</td></tr>
<tr><td>GU305914.1</td><td>178447</td><td>178469</td><td>+</td><td>0</td><td>TTGGTGTGCTCTGTCATGAA</td><td>GGG</td><td>||||||||[||||||||||||]PAM</td><td>NA</td><td>Intergenic</td><td>NA</td><td>NA</td></tr>
<tr><td>T13539</td><td>TGGTGTGCTCTGTCATGAAGGGG</td><td>1000</td><td></td><td></td><td></td><td></td><td></td><td></td><td></td><td></td><td></td></tr>
<tr><td>Oligo fwd</td><td>CACCTGGTGTGCTCTGTCATGAAG</td><td></td><td></td><td></td><td></td><td></td><td></td><td></td><td></td><td></td><td></td></tr>
<tr><td>Oligo rev</td><td>AAACCTTCATGACAGAGCACACCA</td><td></td><td></td><td></td><td></td><td></td><td></td><td></td><td></td><td></td><td></td></tr>
<tr><td>Chromosome</td><td>start</td><td>end</td><td>strand</td><td>MM</td><td>target_seq</td><td>PAM</td><td>alignment</td><td>distance</td><td>position</td><td>gene name</td><td>gene id</td></tr>
<tr><td>GU305914.1</td><td>178448</td><td>178470</td><td>+</td><td>0</td><td>TGGTGTGCTCTGTCATGAAG</td><td>GGG</td><td>||||||||[||||||||||||]PAM</td><td>NA</td><td>Intergenic</td><td>NA</td><td>NA</td></tr>
<tr><td>T13540</td><td>GTGCTCTGTCATGAAGGGGCAGG</td><td>1000</td><td></td><td></td><td></td><td></td><td></td><td></td><td></td><td></td><td></td></tr>
<tr><td>Oligo fwd</td><td>CACCGTGCTCTGTCATGAAGGGGC</td><td></td><td></td><td></td><td></td><td></td><td></td><td></td><td></td><td></td><td></td></tr>
<tr><td>Oligo rev</td><td>AAACGCCCCTTCATGACAGAGCAC</td><td></td><td></td><td></td><td></td><td></td><td></td><td></td><td></td><td></td><td></td></tr>
<tr><td>Chromosome</td><td>start</td><td>end</td><td>strand</td><td>MM</td><td>target_seq</td><td>PAM</td><td>alignment</td><td>distance</td><td>position</td><td>gene name</td><td>gene id</td></tr>
<tr><td>GU305914.1</td><td>178452</td><td>178474</td><td>+</td><td>0</td><td>GTGCTCTGTCATGAAGGGGC</td><td>AGG</td><td>||||||||[||||||||||||]PAM</td><td>NA</td><td>Intergenic</td><td>NA</td><td>NA</td></tr>
<tr><td>T13541</td><td>TGTCATGAAGGGGCAGGAGATGG</td><td>1000</td><td></td><td></td><td></td><td></td><td></td><td></td><td></td><td></td><td></td></tr>
<tr><td>Oligo fwd</td><td>CACCTGTCATGAAGGGGCAGGAGA</td><td></td><td></td><td></td><td></td><td></td><td></td><td></td><td></td><td></td><td></td></tr>
<tr><td>Oligo rev</td><td>AAACTCTCCTGCCCCTTCATGACA</td><td></td><td></td><td></td><td></td><td></td><td></td><td></td><td></td><td></td><td></td></tr>
<tr><td>Chromosome</td><td>start</td><td>end</td><td>strand</td><td>MM</td><td>target_seq</td><td>PAM</td><td>alignment</td><td>distance</td><td>position</td><td>gene name</td><td>gene id</td></tr>
<tr><td>GU305914.1</td><td>178458</td><td>178480</td><td>+</td><td>0</td><td>TGTCATGAAGGGGCAGGAGA</td><td>TGG</td><td>||||||||[||||||||||||]PAM</td><td>NA</td><td>Intergenic</td><td>NA</td><td>NA</td></tr>
<tr><td>T13542</td><td>TGAAGGGGCAGGAGATGGACAGG</td><td>1000</td><td></td><td></td><td></td><td></td><td></td><td></td><td></td><td></td><td></td></tr>
<tr><td>Oligo fwd</td><td>CACCTGAAGGGGCAGGAGATGGAC</td><td></td><td></td><td></td><td></td><td></td><td></td><td></td><td></td><td></td><td></td></tr>
<tr><td>Oligo rev</td><td>AAACGTCCATCTCCTGCCCCTTCA</td><td></td><td></td><td></td><td></td><td></td><td></td><td></td><td></td><td></td><td></td></tr>
<tr><td>Chromosome</td><td>start</td><td>end</td><td>strand</td><td>MM</td><td>target_seq</td><td>PAM</td><td>alignment</td><td>distance</td><td>position</td><td>gene name</td><td>gene id</td></tr>
<tr><td>GU305914.1</td><td>178463</td><td>178485</td><td>+</td><td>0</td><td>TGAAGGGGCAGGAGATGGAC</td><td>AGG</td><td>||||||||[||||||||||||]PAM</td><td>NA</td><td>Intergenic</td><td>NA</td><td>NA</td></tr>
<tr><td>T13543</td><td>GGGGCAGGAGATGGACAGGTTGG</td><td>1000</td><td></td><td></td><td></td><td></td><td></td><td></td><td></td><td></td><td></td></tr>
<tr><td>Oligo fwd</td><td>CACCGGGGCAGGAGATGGACAGGT</td><td></td><td></td><td></td><td></td><td></td><td></td><td></td><td></td><td></td><td></td></tr>
<tr><td>Oligo rev</td><td>AAACACCTGTCCATCTCCTGCCCC</td><td></td><td></td><td></td><td></td><td></td><td></td><td></td><td></td><td></td><td></td></tr>
<tr><td>Chromosome</td><td>start</td><td>end</td><td>strand</td><td>MM</td><td>target_seq</td><td>PAM</td><td>alignment</td><td>distance</td><td>position</td><td>gene name</td><td>gene id</td></tr>
<tr><td>GU305914.1</td><td>178467</td><td>178489</td><td>+</td><td>0</td><td>GGGGCAGGAGATGGACAGGT</td><td>TGG</td><td>||||||||[||||||||||||]PAM</td><td>NA</td><td>Intergenic</td><td>NA</td><td>NA</td></tr>
<tr><td>T13544</td><td>GGGCAGGAGATGGACAGGTTGGG</td><td>1000</td><td></td><td></td><td></td><td></td><td></td><td></td><td></td><td></td><td></td></tr>
<tr><td>Oligo fwd</td><td>CACCGGGCAGGAGATGGACAGGTT</td><td></td><td></td><td></td><td></td><td></td><td></td><td></td><td></td><td></td><td></td></tr>
<tr><td>Oligo rev</td><td>AAACAACCTGTCCATCTCCTGCCC</td><td></td><td></td><td></td><td></td><td></td><td></td><td></td><td></td><td></td><td></td></tr>
<tr><td>Chromosome</td><td>start</td><td>end</td><td>strand</td><td>MM</td><td>target_seq</td><td>PAM</td><td>alignment</td><td>distance</td><td>position</td><td>gene name</td><td>gene id</td></tr>
<tr><td>GU305914.1</td><td>178468</td><td>178490</td><td>+</td><td>0</td><td>GGGCAGGAGATGGACAGGTT</td><td>GGG</td><td>||||||||[||||||||||||]PAM</td><td>NA</td><td>Intergenic</td><td>NA</td><td>NA</td></tr>
<tr><td>T13545</td><td>GGCAGGAGATGGACAGGTTGGGG</td><td>1000</td><td></td><td></td><td></td><td></td><td></td><td></td><td></td><td></td><td></td></tr>
<tr><td>Oligo fwd</td><td>CACCGGCAGGAGATGGACAGGTTG</td><td></td><td></td><td></td><td></td><td></td><td></td><td></td><td></td><td></td><td></td></tr>
<tr><td>Oligo rev</td><td>AAACCAACCTGTCCATCTCCTGCC</td><td></td><td></td><td></td><td></td><td></td><td></td><td></td><td></td><td></td><td></td></tr>
<tr><td>Chromosome</td><td>start</td><td>end</td><td>strand</td><td>MM</td><td>target_seq</td><td>PAM</td><td>alignment</td><td>distance</td><td>position</td><td>gene name</td><td>gene id</td></tr>
<tr><td>GU305914.1</td><td>178469</td><td>178491</td><td>+</td><td>0</td><td>GGCAGGAGATGGACAGGTTG</td><td>GGG</td><td>||||||||[||||||||||||]PAM</td><td>NA</td><td>Intergenic</td><td>NA</td><td>NA</td></tr>
<tr><td>T13546</td><td>GGTTGGGGCGCATCCCCATGCGG</td><td>1000</td><td></td><td></td><td></td><td></td><td></td><td></td><td></td><td></td><td></td></tr>
<tr><td>Oligo fwd</td><td>CACCGGTTGGGGCGCATCCCCATG</td><td></td><td></td><td></td><td></td><td></td><td></td><td></td><td></td><td></td><td></td></tr>
<tr><td>Oligo rev</td><td>AAACCATGGGGATGCGCCCCAACC</td><td></td><td></td><td></td><td></td><td></td><td></td><td></td><td></td><td></td><td></td></tr>
<tr><td>Chromosome</td><td>start</td><td>end</td><td>strand</td><td>MM</td><td>target_seq</td><td>PAM</td><td>alignment</td><td>distance</td><td>position</td><td>gene name</td><td>gene id</td></tr>
<tr><td>GU305914.1</td><td>178484</td><td>178506</td><td>+</td><td>0</td><td>GGTTGGGGCGCATCCCCATG</td><td>CGG</td><td>||||||||[||||||||||||]PAM</td><td>NA</td><td>Intergenic</td><td>NA</td><td>NA</td></tr>
<tr><td>T13547</td><td>GTTGGGGCGCATCCCCATGCGGG</td><td>1000</td><td></td><td></td><td></td><td></td><td></td><td></td><td></td><td></td><td></td></tr>
<tr><td>Oligo fwd</td><td>CACCGTTGGGGCGCATCCCCATGC</td><td></td><td></td><td></td><td></td><td></td><td></td><td></td><td></td><td></td><td></td></tr>
<tr><td>Oligo rev</td><td>AAACGCATGGGGATGCGCCCCAAC</td><td></td><td></td><td></td><td></td><td></td><td></td><td></td><td></td><td></td><td></td></tr>
<tr><td>Chromosome</td><td>start</td><td>end</td><td>strand</td><td>MM</td><td>target_seq</td><td>PAM</td><td>alignment</td><td>distance</td><td>position</td><td>gene name</td><td>gene id</td></tr>
<tr><td>GU305914.1</td><td>178485</td><td>178507</td><td>+</td><td>0</td><td>GTTGGGGCGCATCCCCATGC</td><td>GGG</td><td>||||||||[||||||||||||]PAM</td><td>NA</td><td>Intergenic</td><td>NA</td><td>NA</td></tr>
<tr><td>T13548</td><td>CTTCCTCACATCCTGACTCCTGG</td><td>1000</td><td></td><td></td><td></td><td></td><td></td><td></td><td></td><td></td><td></td></tr>
<tr><td>Oligo fwd</td><td>CACCCTTCCTCACATCCTGACTCC</td><td></td><td></td><td></td><td></td><td></td><td></td><td></td><td></td><td></td><td></td></tr>
<tr><td>Oligo rev</td><td>AAACGGAGTCAGGATGTGAGGAAG</td><td></td><td></td><td></td><td></td><td></td><td></td><td></td><td></td><td></td><td></td></tr>
<tr><td>Chromosome</td><td>start</td><td>end</td><td>strand</td><td>MM</td><td>target_seq</td><td>PAM</td><td>alignment</td><td>distance</td><td>position</td><td>gene name</td><td>gene id</td></tr>
<tr><td>GU305914.1</td><td>178527</td><td>178549</td><td>+</td><td>0</td><td>CTTCCTCACATCCTGACTCC</td><td>TGG</td><td>||||||||[||||||||||||]PAM</td><td>NA</td><td>Intergenic</td><td>NA</td><td>NA</td></tr>
<tr><td>T13549</td><td>CATGATGAACTTGTCATGCACGG</td><td>1000</td><td></td><td></td><td></td><td></td><td></td><td></td><td></td><td></td><td></td></tr>
<tr><td>Oligo fwd</td><td>CACCCATGATGAACTTGTCATGCA</td><td></td><td></td><td></td><td></td><td></td><td></td><td></td><td></td><td></td><td></td></tr>
<tr><td>Oligo rev</td><td>AAACTGCATGACAAGTTCATCATG</td><td></td><td></td><td></td><td></td><td></td><td></td><td></td><td></td><td></td><td></td></tr>
<tr><td>Chromosome</td><td>start</td><td>end</td><td>strand</td><td>MM</td><td>target_seq</td><td>PAM</td><td>alignment</td><td>distance</td><td>position</td><td>gene name</td><td>gene id</td></tr>
<tr><td>GU305914.1</td><td>178560</td><td>178582</td><td>+</td><td>0</td><td>CATGATGAACTTGTCATGCA</td><td>CGG</td><td>||||||||[||||||||||||]PAM</td><td>NA</td><td>Intergenic</td><td>NA</td><td>NA</td></tr>
<tr><td>T13550</td><td>CGGTCTCCTCCTTACAGAACTGG</td><td>1000</td><td></td><td></td><td></td><td></td><td></td><td></td><td></td><td></td><td></td></tr>
<tr><td>Oligo fwd</td><td>CACCCGGTCTCCTCCTTACAGAAC</td><td></td><td></td><td></td><td></td><td></td><td></td><td></td><td></td><td></td><td></td></tr>
<tr><td>Oligo rev</td><td>AAACGTTCTGTAAGGAGGAGACCG</td><td></td><td></td><td></td><td></td><td></td><td></td><td></td><td></td><td></td><td></td></tr>
</table>

<table>
<tr><th>Chromosome</th><th>start</th><th>end</th><th>strand</th><th>MM</th><th>target_seq</th><th>PAM</th><th>alignment</th><th>distance</th><th>position</th><th>gene name</th><th>gene id</th></tr>
<tr><td>GU305914.1</td><td>178580</td><td>178602</td><td>+</td><td>0</td><td>CGGTCTCCTCCTTACAGAAC</td><td>TGG</td><td>||||||||[||||||||||||]PAM</td><td>NA</td><td>Intergenic</td><td>NA</td><td>NA</td></tr>
<tr><td>T13551</td><td>CTCCTCCTTACAGAACTGGATGG</td><td>1000</td><td></td><td></td><td></td><td></td><td></td><td></td><td></td><td></td><td></td></tr>
<tr><td>Oligo fwd</td><td>CACCCTCCTCCTTACAGAACTGGA</td><td></td><td></td><td></td><td></td><td></td><td></td><td></td><td></td><td></td><td></td></tr>
<tr><td>Oligo rev</td><td>AAACTCCAGTTCTGTAAGGAGGAG</td><td></td><td></td><td></td><td></td><td></td><td></td><td></td><td></td><td></td><td></td></tr>
<tr><td>Chromosome</td><td>start</td><td>end</td><td>strand</td><td>MM</td><td>target_seq</td><td>PAM</td><td>alignment</td><td>distance</td><td>position</td><td>gene name</td><td>gene id</td></tr>
<tr><td>GU305914.1</td><td>178584</td><td>178606</td><td>+</td><td>0</td><td>CTCCTCCTTACAGAACTGGA</td><td>TGG</td><td>||||||||[||||||||||||]PAM</td><td>NA</td><td>Intergenic</td><td>NA</td><td>NA</td></tr>
<tr><td>T13552</td><td>GCCTTACCCAGTAGATACTGTGG</td><td>1000</td><td></td><td></td><td></td><td></td><td></td><td></td><td></td><td></td><td></td></tr>
<tr><td>Oligo fwd</td><td>CACCGCCTTACCCAGTAGATACTG</td><td></td><td></td><td></td><td></td><td></td><td></td><td></td><td></td><td></td><td></td></tr>
<tr><td>Oligo rev</td><td>AAACCAGTATCTACTGGGTAAGGC</td><td></td><td></td><td></td><td></td><td></td><td></td><td></td><td></td><td></td><td></td></tr>
<tr><td>Chromosome</td><td>start</td><td>end</td><td>strand</td><td>MM</td><td>target_seq</td><td>PAM</td><td>alignment</td><td>distance</td><td>position</td><td>gene name</td><td>gene id</td></tr>
<tr><td>GU305914.1</td><td>178606</td><td>178628</td><td>+</td><td>0</td><td>GCCTTACCCAGTAGATACTG</td><td>TGG</td><td>||||||||[||||||||||||]PAM</td><td>NA</td><td>Intergenic</td><td>NA</td><td>NA</td></tr>
<tr><td>T13553</td><td>CCTTACCCAGTAGATACTGTGGG</td><td>1000</td><td></td><td></td><td></td><td></td><td></td><td></td><td></td><td></td><td></td></tr>
<tr><td>Oligo fwd</td><td>CACCCCTTACCCAGTAGATACTGT</td><td></td><td></td><td></td><td></td><td></td><td></td><td></td><td></td><td></td><td></td></tr>
<tr><td>Oligo rev</td><td>AAACACAGTATCTACTGGGTAAGG</td><td></td><td></td><td></td><td></td><td></td><td></td><td></td><td></td><td></td><td></td></tr>
<tr><td>Chromosome</td><td>start</td><td>end</td><td>strand</td><td>MM</td><td>target_seq</td><td>PAM</td><td>alignment</td><td>distance</td><td>position</td><td>gene name</td><td>gene id</td></tr>
<tr><td>GU305914.1</td><td>178607</td><td>178629</td><td>+</td><td>0</td><td>CCTTACCCAGTAGATACTGT</td><td>GGG</td><td>||||||||[||||||||||||]PAM</td><td>NA</td><td>Intergenic</td><td>NA</td><td>NA</td></tr>
<tr><td>T13554</td><td>CTTACCCAGTAGATACTGTGGGG</td><td>1000</td><td></td><td></td><td></td><td></td><td></td><td></td><td></td><td></td><td></td></tr>
<tr><td>Oligo fwd</td><td>CACCCTTACCCAGTAGATACTGTG</td><td></td><td></td><td></td><td></td><td></td><td></td><td></td><td></td><td></td><td></td></tr>
<tr><td>Oligo rev</td><td>AAACCACAGTATCTACTGGGTAAG</td><td></td><td></td><td></td><td></td><td></td><td></td><td></td><td></td><td></td><td></td></tr>
<tr><td>Chromosome</td><td>start</td><td>end</td><td>strand</td><td>MM</td><td>target_seq</td><td>PAM</td><td>alignment</td><td>distance</td><td>position</td><td>gene name</td><td>gene id</td></tr>
<tr><td>GU305914.1</td><td>178608</td><td>178630</td><td>+</td><td>0</td><td>CTTACCCAGTAGATACTGTG</td><td>GGG</td><td>||||||||[||||||||||||]PAM</td><td>NA</td><td>Intergenic</td><td>NA</td><td>NA</td></tr>
<tr><td>T13555</td><td>AGTAGATACTGTGGGGACTGAGG</td><td>1000</td><td></td><td></td><td></td><td></td><td></td><td></td><td></td><td></td><td></td></tr>
<tr><td>Oligo fwd</td><td>CACCAGTAGATACTGTGGGGACTG</td><td></td><td></td><td></td><td></td><td></td><td></td><td></td><td></td><td></td><td></td></tr>
<tr><td>Oligo rev</td><td>AAACCAGTCCCCACAGTATCTACT</td><td></td><td></td><td></td><td></td><td></td><td></td><td></td><td></td><td></td><td></td></tr>
<tr><td>Chromosome</td><td>start</td><td>end</td><td>strand</td><td>MM</td><td>target_seq</td><td>PAM</td><td>alignment</td><td>distance</td><td>position</td><td>gene name</td><td>gene id</td></tr>
<tr><td>GU305914.1</td><td>178615</td><td>178637</td><td>+</td><td>0</td><td>AGTAGATACTGTGGGGACTG</td><td>AGG</td><td>||||||||[||||||||||||]PAM</td><td>NA</td><td>Intergenic</td><td>NA</td><td>NA</td></tr>
<tr><td>T13556</td><td>CTGTGGGGACTGAGGTCTGAAGG</td><td>1000</td><td></td><td></td><td></td><td></td><td></td><td></td><td></td><td></td><td></td></tr>
<tr><td>Oligo fwd</td><td>CACCCTGTGGGGACTGAGGTCTGA</td><td></td><td></td><td></td><td></td><td></td><td></td><td></td><td></td><td></td><td></td></tr>
<tr><td>Oligo rev</td><td>AAACTCAGACCTCAGTCCCCACAG</td><td></td><td></td><td></td><td></td><td></td><td></td><td></td><td></td><td></td><td></td></tr>
<tr><td>Chromosome</td><td>start</td><td>end</td><td>strand</td><td>MM</td><td>target_seq</td><td>PAM</td><td>alignment</td><td>distance</td><td>position</td><td>gene name</td><td>gene id</td></tr>
<tr><td>GU305914.1</td><td>178623</td><td>178645</td><td>+</td><td>0</td><td>CTGTGGGGACTGAGGTCTGA</td><td>AGG</td><td>||||||||[||||||||||||]PAM</td><td>NA</td><td>Intergenic</td><td>NA</td><td>NA</td></tr>
<tr><td>T13557</td><td>TGTGGGGACTGAGGTCTGAAGGG</td><td>1000</td><td></td><td></td><td></td><td></td><td></td><td></td><td></td><td></td><td></td></tr>
<tr><td>Oligo fwd</td><td>CACCTGTGGGGACTGAGGTCTGAA</td><td></td><td></td><td></td><td></td><td></td><td></td><td></td><td></td><td></td><td></td></tr>
<tr><td>Oligo rev</td><td>AAACTTCAGACCTCAGTCCCCACA</td><td></td><td></td><td></td><td></td><td></td><td></td><td></td><td></td><td></td><td></td></tr>
<tr><td>Chromosome</td><td>start</td><td>end</td><td>strand</td><td>MM</td><td>target_seq</td><td>PAM</td><td>alignment</td><td>distance</td><td>position</td><td>gene name</td><td>gene id</td></tr>
<tr><td>GU305914.1</td><td>178624</td><td>178646</td><td>+</td><td>0</td><td>TGTGGGGACTGAGGTCTGAA</td><td>GGG</td><td>||||||||[||||||||||||]PAM</td><td>NA</td><td>Intergenic</td><td>NA</td><td>NA</td></tr>
<tr><td>T13558</td><td>GGGACTGAGGTCTGAAGGGCAGG</td><td>1000</td><td></td><td></td><td></td><td></td><td></td><td></td><td></td><td></td><td></td></tr>
<tr><td>Oligo fwd</td><td>CACCGGGACTGAGGTCTGAAGGGC</td><td></td><td></td><td></td><td></td><td></td><td></td><td></td><td></td><td></td><td></td></tr>
<tr><td>Oligo rev</td><td>AAACGCCCTTCAGACCTCAGTCCC</td><td></td><td></td><td></td><td></td><td></td><td></td><td></td><td></td><td></td><td></td></tr>
<tr><td>Chromosome</td><td>start</td><td>end</td><td>strand</td><td>MM</td><td>target_seq</td><td>PAM</td><td>alignment</td><td>distance</td><td>position</td><td>gene name</td><td>gene id</td></tr>
<tr><td>GU305914.1</td><td>178628</td><td>178650</td><td>+</td><td>0</td><td>GGGACTGAGGTCTGAAGGGC</td><td>AGG</td><td>||||||||[||||||||||||]PAM</td><td>NA</td><td>Intergenic</td><td>NA</td><td>NA</td></tr>
<tr><td>T13559</td><td>GGGCAGGTTCTTGTACTCCAAGG</td><td>1000</td><td></td><td></td><td></td><td></td><td></td><td></td><td></td><td></td><td></td></tr>
<tr><td>Oligo fwd</td><td>CACCGGGCAGGTTCTTGTACTCCA</td><td></td><td></td><td></td><td></td><td></td><td></td><td></td><td></td><td></td><td></td></tr>
<tr><td>Oligo rev</td><td>AAACTGGAGTACAAGAACCTGCCC</td><td></td><td></td><td></td><td></td><td></td><td></td><td></td><td></td><td></td><td></td></tr>
<tr><td>Chromosome</td><td>start</td><td>end</td><td>strand</td><td>MM</td><td>target_seq</td><td>PAM</td><td>alignment</td><td>distance</td><td>position</td><td>gene name</td><td>gene id</td></tr>
<tr><td>GU305914.1</td><td>178644</td><td>178666</td><td>+</td><td>0</td><td>GGGCAGGTTCTTGTACTCCA</td><td>AGG</td><td>||||||||[||||||||||||]PAM</td><td>NA</td><td>Intergenic</td><td>NA</td><td>NA</td></tr>
<tr><td>T13560</td><td>GTTCTTGTACTCCAAGGCCCTGG</td><td>1000</td><td></td><td></td><td></td><td></td><td></td><td></td><td></td><td></td><td></td></tr>
<tr><td>Oligo fwd</td><td>CACCGTTCTTGTACTCCAAGGCCC</td><td></td><td></td><td></td><td></td><td></td><td></td><td></td><td></td><td></td><td></td></tr>
<tr><td>Oligo rev</td><td>AAACGGGCCTTGGAGTACAAGAAC</td><td></td><td></td><td></td><td></td><td></td><td></td><td></td><td></td><td></td><td></td></tr>
<tr><td>Chromosome</td><td>start</td><td>end</td><td>strand</td><td>MM</td><td>target_seq</td><td>PAM</td><td>alignment</td><td>distance</td><td>position</td><td>gene name</td><td>gene id</td></tr>
<tr><td>GU305914.1</td><td>178650</td><td>178672</td><td>+</td><td>0</td><td>GTTCTTGTACTCCAAGGCCC</td><td>TGG</td><td>||||||||[||||||||||||]PAM</td><td>NA</td><td>Intergenic</td><td>NA</td><td>NA</td></tr>
<tr><td>T13561</td><td>TTGTACTCCAAGGCCCTGGCTGG</td><td>1000</td><td></td><td></td><td></td><td></td><td></td><td></td><td></td><td></td><td></td></tr>
<tr><td>Oligo fwd</td><td>CACCTTGTACTCCAAGGCCCTGGC</td><td></td><td></td><td></td><td></td><td></td><td></td><td></td><td></td><td></td><td></td></tr>
<tr><td>Oligo rev</td><td>AAACGCCAGGGCCTTGGAGTACAA</td><td></td><td></td><td></td><td></td><td></td><td></td><td></td><td></td><td></td><td></td></tr>
<tr><td>Chromosome</td><td>start</td><td>end</td><td>strand</td><td>MM</td><td>target_seq</td><td>PAM</td><td>alignment</td><td>distance</td><td>position</td><td>gene name</td><td>gene id</td></tr>
<tr><td>GU305914.1</td><td>178654</td><td>178676</td><td>+</td><td>0</td><td>TTGTACTCCAAGGCCCTGGC</td><td>TGG</td><td>||||||||[||||||||||||]PAM</td><td>NA</td><td>Intergenic</td><td>NA</td><td>NA</td></tr>
<tr><td>T13562</td><td>TGTACTCCAAGGCCCTGGCTGGG</td><td>1000</td><td></td><td></td><td></td><td></td><td></td><td></td><td></td><td></td><td></td></tr>
<tr><td>Oligo fwd</td><td>CACCTGTACTCCAAGGCCCTGGCT</td><td></td><td></td><td></td><td></td><td></td><td></td><td></td><td></td><td></td><td></td></tr>
<tr><td>Oligo rev</td><td>AAACAGCCAGGGCCTTGGAGTACA</td><td></td><td></td><td></td><td></td><td></td><td></td><td></td><td></td><td></td><td></td></tr>
<tr><td>Chromosome</td><td>start</td><td>end</td><td>strand</td><td>MM</td><td>target_seq</td><td>PAM</td><td>alignment</td><td>distance</td><td>position</td><td>gene name</td><td>gene id</td></tr>
<tr><td>GU305914.1</td><td>178655</td><td>178677</td><td>+</td><td>0</td><td>TGTACTCCAAGGCCCTGGCT</td><td>GGG</td><td>||||||||[||||||||||||]PAM</td><td>NA</td><td>Intergenic</td><td>NA</td><td>NA</td></tr>
<tr><td>T13563</td><td>GGCCCTGGCTGGGTACTTCCTGG</td><td>1000</td><td></td><td></td><td></td><td></td><td></td><td></td><td></td><td></td><td></td></tr>
<tr><td>Oligo fwd</td><td>CACCGGCCCTGGCTGGGTACTTCC</td><td></td><td></td><td></td><td></td><td></td><td></td><td></td><td></td><td></td><td></td></tr>
<tr><td>Oligo rev</td><td>AAACGGAAGTACCCAGCCAGGGCC</td><td></td><td></td><td></td><td></td><td></td><td></td><td></td><td></td><td></td><td></td></tr>
<tr><td>Chromosome</td><td>start</td><td>end</td><td>strand</td><td>MM</td><td>target_seq</td><td>PAM</td><td>alignment</td><td>distance</td><td>position</td><td>gene name</td><td>gene id</td></tr>
<tr><td>GU305914.1</td><td>178665</td><td>178687</td><td>+</td><td>0</td><td>GGCCCTGGCTGGGTACTTCC</td><td>TGG</td><td>||||||||[||||||||||||]PAM</td><td>NA</td><td>Intergenic</td><td>NA</td><td>NA</td></tr>
<tr><td>T13564</td><td>TATCTTCCCTGCACTTCTTGTGG</td><td>1000</td><td></td><td></td><td></td><td></td><td></td><td></td><td></td><td></td><td></td></tr>
<tr><td>Oligo fwd</td><td>CACCTATCTTCCCTGCACTTCTTG</td><td></td><td></td><td></td><td></td><td></td><td></td><td></td><td></td><td></td><td></td></tr>
<tr><td>Oligo rev</td><td>AAACCAAGAAGTGCAGGGAAGATA</td><td></td><td></td><td></td><td></td><td></td><td></td><td></td><td></td><td></td><td></td></tr>
<tr><td>Chromosome</td><td>start</td><td>end</td><td>strand</td><td>MM</td><td>target_seq</td><td>PAM</td><td>alignment</td><td>distance</td><td>position</td><td>gene name</td><td>gene id</td></tr>
<tr><td>GU305914.1</td><td>178691</td><td>178713</td><td>+</td><td>0</td><td>TATCTTCCCTGCACTTCTTG</td><td>TGG</td><td>||||||||[||||||||||||]PAM</td><td>NA</td><td>Intergenic</td><td>NA</td><td>NA</td></tr>
<tr><td>T13565</td><td>ATCTTCCCTGCACTTCTTGTGGG</td><td>1000</td><td></td><td></td><td></td><td></td><td></td><td></td><td></td><td></td><td></td></tr>
<tr><td>Oligo fwd</td><td>CACCATCTTCCCTGCACTTCTTGT</td><td></td><td></td><td></td><td></td><td></td><td></td><td></td><td></td><td></td><td></td></tr>
</table>

Oligo rev AAACACAAGAAGTGCAGGGAAGAT
Chromosome start end strand MM target_seq PAM alignment distance position gene name gene id
GU305914.1 178692 178714 + 0 ATCTTCCCTGCACTTCTTGT GGG ||||||||[||||||||||||]PAM NA Intergenic NA NA

T13566 CCTGCACTTCTTGTGGGCGTTGG 1000
Oligo fwd CACCCCTGCACTTCTTGTGGGCGT
Oligo rev AAACACGCCCACAAGAAGTGCAGG
Chromosome start end strand MM target_seq PAM alignment distance position gene name gene id
GU305914.1 178698 178720 + 0 CCTGCACTTCTTGTGGGCGT TGG ||||||||[||||||||||||]PAM NA Intergenic NA NA

T13567 CTGCACTTCTTGTGGGCGTTGGG 1000
Oligo fwd CACCCTGCACTTCTTGTGGGCGTT
Oligo rev AAACAACGCCCACAAGAAGTGCAG
Chromosome start end strand MM target_seq PAM alignment distance position gene name gene id
GU305914.1 178699 178721 + 0 CTGCACTTCTTGTGGGCGTT GGG ||||||||[||||||||||||]PAM NA Intergenic NA NA

T13568 TGCACTTCTTGTGGGCGTTGGGG 1000
Oligo fwd CACCTGCACTTCTTGTGGGCGTTG
Oligo rev AAACCAACGCCCACAAGAAGTGCA
Chromosome start end strand MM target_seq PAM alignment distance position gene name gene id
GU305914.1 178700 178722 + 0 TGCACTTCTTGTGGGCGTTG GGG ||||||||[||||||||||||]PAM NA Intergenic NA NA

T13569 GCACTTCTTGTGGGCGTTGGGGG 1000
Oligo fwd CACCGCACTTCTTGTGGGCGTTGG
Oligo rev AAACCCAACGCCCACAAGAAGTGC
Chromosome start end strand MM target_seq PAM alignment distance position gene name gene id
GU305914.1 178701 178723 + 0 GCACTTCTTGTGGGCGTTGG GGG ||||||||[||||||||||||]PAM NA Intergenic NA NA

T13570 GTTGGGGGCCACGTAGCCTCCGG 1000
Oligo fwd CACCGTTGGGGGCCACGTAGCCTC
Oligo rev AAACGAGGCTACGTGGCCCCCAAC
Chromosome start end strand MM target_seq PAM alignment distance position gene name gene id
GU305914.1 178716 178738 + 0 GTTGGGGGCCACGTAGCCTC CGG ||||||||[||||||||||||]PAM NA Intergenic NA NA

T13571 TCACTCCCACTACTCATCACAGG 1000
Oligo fwd CACCTCACTCCCACTACTCATCAC
Oligo rev AAACGTGATGAGTAGTGGGAGTGA
Chromosome start end strand MM target_seq PAM alignment distance position gene name gene id
GU305914.1 178768 178790 + 0 TCACTCCCACTACTCATCAC AGG ||||||||[||||||||||||]PAM NA Intergenic NA NA

T13572 CTCATCACAGGCTTCTTGCTAGG 1000
Oligo fwd CACCCTCATCACAGGCTTCTTGCT
Oligo rev AAACAGCAAGAAGCCTGTGATGAG
Chromosome start end strand MM target_seq PAM alignment distance position gene name gene id
GU305914.1 178780 178802 + 0 CTCATCACAGGCTTCTTGCT AGG ||||||||[||||||||||||]PAM NA Intergenic NA NA

T13573 TCATCACAGGCTTCTTGCTAGGG 1000
Oligo fwd CACCTCATCACAGGCTTCTTGCTA
Oligo rev AAACTAGCAAGAAGCCTGTGATGA
Chromosome start end strand MM target_seq PAM alignment distance position gene name gene id
GU305914.1 178781 178803 + 0 TCATCACAGGCTTCTTGCTA GGG ||||||||[||||||||||||]PAM NA Intergenic NA NA

T13574 CATCACAGGCTTCTTGCTAGGGG 1000
Oligo fwd CACCCATCACAGGCTTCTTGCTAG
Oligo rev AAACCTAGCAAGAAGCCTGTGATG
Chromosome start end strand MM target_seq PAM alignment distance position gene name gene id
GU305914.1 178782 178804 + 0 CATCACAGGCTTCTTGCTAG GGG ||||||||[||||||||||||]PAM NA Intergenic NA NA

T13575 GCTGCTGCTGTTGCTTCTTGTGG 1000
Oligo fwd CACCGCTGCTGCTGTTGCTTCTTG
Oligo rev AAACCAAGAAGCAACAGCAGCAGC
Chromosome start end strand MM target_seq PAM alignment distance position gene name gene id
GU305914.1 178808 178830 + 0 GCTGCTGCTGTTGCTTCTTG TGG ||||||||[||||||||||||]PAM NA Intergenic NA NA

T13576 AGAGCTGCTGCTGTGTTTGTTGG 1000
Oligo fwd CACCAGAGCTGCTGCTGTGTTTGT
Oligo rev AAACACAAACACAGCAGCAGCTCT
Chromosome start end strand MM target_seq PAM alignment distance position gene name gene id
GU305914.1 178851 178873 + 0 AGAGCTGCTGCTGTGTTTGT TGG ||||||||[||||||||||||]PAM NA Intergenic NA NA

T13577 GCTGCTGTGTTTGTTGGAGTTGG 1000
Oligo fwd CACCGCTGCTGTGTTTGTTGGAGT
Oligo rev AAACACTCCAACAAACACAGCAGC
Chromosome start end strand MM target_seq PAM alignment distance position gene name gene id
GU305914.1 178857 178879 + 0 GCTGCTGTGTTTGTTGGAGT TGG ||||||||[||||||||||||]PAM NA Intergenic NA NA

T13578 TTGGAGTTGGAACCAGAGCTAGG 1000
Oligo fwd CACCTTGGAGTTGGAACCAGAGCT
Oligo rev AAACAGCTCTGGTTCCAACTCCAA
Chromosome start end strand MM target_seq PAM alignment distance position gene name gene id
GU305914.1 178870 178892 + 0 TTGGAGTTGGAACCAGAGCT AGG ||||||||[||||||||||||]PAM NA Intergenic NA NA

T13579 TGGAGTTGGAACCAGAGCTAGGG 1000
Oligo fwd CACCTGGAGTTGGAACCAGAGCTA
Oligo rev AAACTAGCTCTGGTTCCAACTCCA
Chromosome start end strand MM target_seq PAM alignment distance position gene name gene id
GU305914.1 178871 178893 + 0 TGGAGTTGGAACCAGAGCTA GGG ||||||||[||||||||||||]PAM NA Intergenic NA NA

T13580 GGAGTTGGAACCAGAGCTAGGGG 1000

<table>
<tr><td>Oligo fwd</td><td>CACCGGAGTTGGAACCAGAGCTAG</td><td></td><td></td><td></td><td></td><td></td><td></td><td></td><td></td><td></td><td></td></tr>
<tr><td>Oligo rev</td><td>AAACCTAGCTCTGGTTCCAACTCC</td><td></td><td></td><td></td><td></td><td></td><td></td><td></td><td></td><td></td><td></td></tr>
<tr><td>Chromosome</td><td>start</td><td>end</td><td>strand</td><td>MM</td><td>target_seq</td><td>PAM</td><td>alignment</td><td>distance</td><td>position</td><td>gene name</td><td>gene id</td></tr>
<tr><td>GU305914.1</td><td>178872</td><td>178894</td><td>+</td><td>0</td><td>GGAGTTGGAACCAGAGCTAG</td><td>GGG</td><td>||||||||[||||||||||||]PAM</td><td>NA</td><td>Intergenic</td><td>NA</td><td>NA</td></tr>
<tr><td></td><td></td><td></td><td></td><td></td><td></td><td></td><td></td><td></td><td></td><td></td><td></td></tr>
<tr><td>T13581</td><td>GAGTTGGAACCAGAGCTAGGGGG</td><td>1000</td><td></td><td></td><td></td><td></td><td></td><td></td><td></td><td></td><td></td></tr>
<tr><td>Oligo fwd</td><td>CACCGAGTTGGAACCAGAGCTAGG</td><td></td><td></td><td></td><td></td><td></td><td></td><td></td><td></td><td></td><td></td></tr>
<tr><td>Oligo rev</td><td>AAACCCTAGCTCTGGTTCCAACTC</td><td></td><td></td><td></td><td></td><td></td><td></td><td></td><td></td><td></td><td></td></tr>
<tr><td>Chromosome</td><td>start</td><td>end</td><td>strand</td><td>MM</td><td>target_seq</td><td>PAM</td><td>alignment</td><td>distance</td><td>position</td><td>gene name</td><td>gene id</td></tr>
<tr><td>GU305914.1</td><td>178873</td><td>178895</td><td>+</td><td>0</td><td>GAGTTGGAACCAGAGCTAGG</td><td>GGG</td><td>||||||||[||||||||||||]PAM</td><td>NA</td><td>Intergenic</td><td>NA</td><td>NA</td></tr>
<tr><td></td><td></td><td></td><td></td><td></td><td></td><td></td><td></td><td></td><td></td><td></td><td></td></tr>
<tr><td>T13582</td><td>ACCAGAGCTAGGGGGCCTGAAGG</td><td>1000</td><td></td><td></td><td></td><td></td><td></td><td></td><td></td><td></td><td></td></tr>
<tr><td>Oligo fwd</td><td>CACCACCAGAGCTAGGGGGCCTGA</td><td></td><td></td><td></td><td></td><td></td><td></td><td></td><td></td><td></td><td></td></tr>
<tr><td>Oligo rev</td><td>AAACTCAGGCCCCCTAGCTCTGGT</td><td></td><td></td><td></td><td></td><td></td><td></td><td></td><td></td><td></td><td></td></tr>
<tr><td>Chromosome</td><td>start</td><td>end</td><td>strand</td><td>MM</td><td>target_seq</td><td>PAM</td><td>alignment</td><td>distance</td><td>position</td><td>gene name</td><td>gene id</td></tr>
<tr><td>GU305914.1</td><td>178881</td><td>178903</td><td>+</td><td>0</td><td>ACCAGAGCTAGGGGGCCTGA</td><td>AGG</td><td>||||||||[||||||||||||]PAM</td><td>NA</td><td>Intergenic</td><td>NA</td><td>NA</td></tr>
<tr><td></td><td></td><td></td><td></td><td></td><td></td><td></td><td></td><td></td><td></td><td></td><td></td></tr>
<tr><td>T13583</td><td>GCCTGAAGGACTTGATCTTCAGG</td><td>1000</td><td></td><td></td><td></td><td></td><td></td><td></td><td></td><td></td><td></td></tr>
<tr><td>Oligo fwd</td><td>CACCGCCTGAAGGACTTGATCTTC</td><td></td><td></td><td></td><td></td><td></td><td></td><td></td><td></td><td></td><td></td></tr>
<tr><td>Oligo rev</td><td>AAACGAAGATCAAGTCCTTCAGGC</td><td></td><td></td><td></td><td></td><td></td><td></td><td></td><td></td><td></td><td></td></tr>
<tr><td>Chromosome</td><td>start</td><td>end</td><td>strand</td><td>MM</td><td>target_seq</td><td>PAM</td><td>alignment</td><td>distance</td><td>position</td><td>gene name</td><td>gene id</td></tr>
<tr><td>GU305914.1</td><td>178895</td><td>178917</td><td>+</td><td>0</td><td>GCCTGAAGGACTTGATCTTC</td><td>AGG</td><td>||||||||[||||||||||||]PAM</td><td>NA</td><td>Intergenic</td><td>NA</td><td>NA</td></tr>
<tr><td></td><td></td><td></td><td></td><td></td><td></td><td></td><td></td><td></td><td></td><td></td><td></td></tr>
<tr><td>T13584</td><td>CCTGAAGGACTTGATCTTCAGGG</td><td>1000</td><td></td><td></td><td></td><td></td><td></td><td></td><td></td><td></td><td></td></tr>
<tr><td>Oligo fwd</td><td>CACCCCTGAAGGACTTGATCTTCA</td><td></td><td></td><td></td><td></td><td></td><td></td><td></td><td></td><td></td><td></td></tr>
<tr><td>Oligo rev</td><td>AAACTGAAGATCAAGTCCTTCAGG</td><td></td><td></td><td></td><td></td><td></td><td></td><td></td><td></td><td></td><td></td></tr>
<tr><td>Chromosome</td><td>start</td><td>end</td><td>strand</td><td>MM</td><td>target_seq</td><td>PAM</td><td>alignment</td><td>distance</td><td>position</td><td>gene name</td><td>gene id</td></tr>
<tr><td>GU305914.1</td><td>178896</td><td>178918</td><td>+</td><td>0</td><td>CCTGAAGGACTTGATCTTCA</td><td>GGG</td><td>||||||||[||||||||||||]PAM</td><td>NA</td><td>Intergenic</td><td>NA</td><td>NA</td></tr>
<tr><td></td><td></td><td></td><td></td><td></td><td></td><td></td><td></td><td></td><td></td><td></td><td></td></tr>
<tr><td>T13585</td><td>CAGGGTCCCCCTCTGTACCATGG</td><td>1000</td><td></td><td></td><td></td><td></td><td></td><td></td><td></td><td></td><td></td></tr>
<tr><td>Oligo fwd</td><td>CACCCAGGGTCCCCCTCTGTACCA</td><td></td><td></td><td></td><td></td><td></td><td></td><td></td><td></td><td></td><td></td></tr>
<tr><td>Oligo rev</td><td>AAACTGGTACAGAGGGGGACCCTG</td><td></td><td></td><td></td><td></td><td></td><td></td><td></td><td></td><td></td><td></td></tr>
<tr><td>Chromosome</td><td>start</td><td>end</td><td>strand</td><td>MM</td><td>target_seq</td><td>PAM</td><td>alignment</td><td>distance</td><td>position</td><td>gene name</td><td>gene id</td></tr>
<tr><td>GU305914.1</td><td>178914</td><td>178936</td><td>+</td><td>0</td><td>CAGGGTCCCCCTCTGTACCA</td><td>TGG</td><td>||||||||[||||||||||||]PAM</td><td>NA</td><td>Intergenic</td><td>NA</td><td>NA</td></tr>
<tr><td></td><td></td><td></td><td></td><td></td><td></td><td></td><td></td><td></td><td></td><td></td><td></td></tr>
<tr><td>T13586</td><td>GTCCCCCTCTGTACCATGGTAGG</td><td>1000</td><td></td><td></td><td></td><td></td><td></td><td></td><td></td><td></td><td></td></tr>
<tr><td>Oligo fwd</td><td>CACCGTCCCCCTCTGTACCATGGT</td><td></td><td></td><td></td><td></td><td></td><td></td><td></td><td></td><td></td><td></td></tr>
<tr><td>Oligo rev</td><td>AAACACCATGGTACAGAGGGGGAC</td><td></td><td></td><td></td><td></td><td></td><td></td><td></td><td></td><td></td><td></td></tr>
<tr><td>Chromosome</td><td>start</td><td>end</td><td>strand</td><td>MM</td><td>target_seq</td><td>PAM</td><td>alignment</td><td>distance</td><td>position</td><td>gene name</td><td>gene id</td></tr>
<tr><td>GU305914.1</td><td>178918</td><td>178940</td><td>+</td><td>0</td><td>GTCCCCCTCTGTACCATGGT</td><td>AGG</td><td>||||||||[||||||||||||]PAM</td><td>NA</td><td>Intergenic</td><td>NA</td><td>NA</td></tr>
<tr><td></td><td></td><td></td><td></td><td></td><td></td><td></td><td></td><td></td><td></td><td></td><td></td></tr>
<tr><td>T13587</td><td>TCCCCCTCTGTACCATGGTAGGG</td><td>1000</td><td></td><td></td><td></td><td></td><td></td><td></td><td></td><td></td><td></td></tr>
<tr><td>Oligo fwd</td><td>CACCTCCCCCTCTGTACCATGGTA</td><td></td><td></td><td></td><td></td><td></td><td></td><td></td><td></td><td></td><td></td></tr>
<tr><td>Oligo rev</td><td>AAACTACCATGGTACAGAGGGGGA</td><td></td><td></td><td></td><td></td><td></td><td></td><td></td><td></td><td></td><td></td></tr>
<tr><td>Chromosome</td><td>start</td><td>end</td><td>strand</td><td>MM</td><td>target_seq</td><td>PAM</td><td>alignment</td><td>distance</td><td>position</td><td>gene name</td><td>gene id</td></tr>
<tr><td>GU305914.1</td><td>178919</td><td>178941</td><td>+</td><td>0</td><td>TCCCCCTCTGTACCATGGTA</td><td>GGG</td><td>||||||||[||||||||||||]PAM</td><td>NA</td><td>Intergenic</td><td>NA</td><td>NA</td></tr>
<tr><td></td><td></td><td></td><td></td><td></td><td></td><td></td><td></td><td></td><td></td><td></td><td></td></tr>
<tr><td>T13588</td><td>CCCCCTCTGTACCATGGTAGGGG</td><td>1000</td><td></td><td></td><td></td><td></td><td></td><td></td><td></td><td></td><td></td></tr>
<tr><td>Oligo fwd</td><td>CACCCCCCCTCTGTACCATGGTAG</td><td></td><td></td><td></td><td></td><td></td><td></td><td></td><td></td><td></td><td></td></tr>
<tr><td>Oligo rev</td><td>AAACCTACCATGGTACAGAGGGGG</td><td></td><td></td><td></td><td></td><td></td><td></td><td></td><td></td><td></td><td></td></tr>
<tr><td>Chromosome</td><td>start</td><td>end</td><td>strand</td><td>MM</td><td>target_seq</td><td>PAM</td><td>alignment</td><td>distance</td><td>position</td><td>gene name</td><td>gene id</td></tr>
<tr><td>GU305914.1</td><td>178920</td><td>178942</td><td>+</td><td>0</td><td>CCCCCTCTGTACCATGGTAG</td><td>GGG</td><td>||||||||[||||||||||||]PAM</td><td>NA</td><td>Intergenic</td><td>NA</td><td>NA</td></tr>
<tr><td></td><td></td><td></td><td></td><td></td><td></td><td></td><td></td><td></td><td></td><td></td><td></td></tr>
<tr><td>T13589</td><td>AGGGGCATCATCACTGTCTGAGG</td><td>1000</td><td></td><td></td><td></td><td></td><td></td><td></td><td></td><td></td><td></td></tr>
<tr><td>Oligo fwd</td><td>CACCAGGGGCATCATCACTGTCTG</td><td></td><td></td><td></td><td></td><td></td><td></td><td></td><td></td><td></td><td></td></tr>
<tr><td>Oligo rev</td><td>AAACCAGACAGTGATGATGCCCCT</td><td></td><td></td><td></td><td></td><td></td><td></td><td></td><td></td><td></td><td></td></tr>
<tr><td>Chromosome</td><td>start</td><td>end</td><td>strand</td><td>MM</td><td>target_seq</td><td>PAM</td><td>alignment</td><td>distance</td><td>position</td><td>gene name</td><td>gene id</td></tr>
<tr><td>GU305914.1</td><td>178938</td><td>178960</td><td>+</td><td>0</td><td>AGGGGCATCATCACTGTCTG</td><td>AGG</td><td>||||||||[||||||||||||]PAM</td><td>NA</td><td>Intergenic</td><td>NA</td><td>NA</td></tr>
<tr><td></td><td></td><td></td><td></td><td></td><td></td><td></td><td></td><td></td><td></td><td></td><td></td></tr>
<tr><td>T13590</td><td>TCTGAGGAATCCACACGCACAGG</td><td>1000</td><td></td><td></td><td></td><td></td><td></td><td></td><td></td><td></td><td></td></tr>
<tr><td>Oligo fwd</td><td>CACCTCTGAGGAATCCACACGCAC</td><td></td><td></td><td></td><td></td><td></td><td></td><td></td><td></td><td></td><td></td></tr>
<tr><td>Oligo rev</td><td>AAACGTGCGTGTGGATTCCTCAGA</td><td></td><td></td><td></td><td></td><td></td><td></td><td></td><td></td><td></td><td></td></tr>
<tr><td>Chromosome</td><td>start</td><td>end</td><td>strand</td><td>MM</td><td>target_seq</td><td>PAM</td><td>alignment</td><td>distance</td><td>position</td><td>gene name</td><td>gene id</td></tr>
<tr><td>GU305914.1</td><td>178954</td><td>178976</td><td>+</td><td>0</td><td>TCTGAGGAATCCACACGCAC</td><td>AGG</td><td>||||||||[||||||||||||]PAM</td><td>NA</td><td>Intergenic</td><td>NA</td><td>NA</td></tr>
<tr><td></td><td></td><td></td><td></td><td></td><td></td><td></td><td></td><td></td><td></td><td></td><td></td></tr>
<tr><td>T13591</td><td>AGGACACTCATATCTGCTGCTGG</td><td>1000</td><td></td><td></td><td></td><td></td><td></td><td></td><td></td><td></td><td></td></tr>
<tr><td>Oligo fwd</td><td>CACCAGGACACTCATATCTGCTGC</td><td></td><td></td><td></td><td></td><td></td><td></td><td></td><td></td><td></td><td></td></tr>
<tr><td>Oligo rev</td><td>AAACGCAGCAGATATGAGTGTCCT</td><td></td><td></td><td></td><td></td><td></td><td></td><td></td><td></td><td></td><td></td></tr>
<tr><td>Chromosome</td><td>start</td><td>end</td><td>strand</td><td>MM</td><td>target_seq</td><td>PAM</td><td>alignment</td><td>distance</td><td>position</td><td>gene name</td><td>gene id</td></tr>
<tr><td>GU305914.1</td><td>178974</td><td>178996</td><td>+</td><td>0</td><td>AGGACACTCATATCTGCTGC</td><td>TGG</td><td>||||||||[||||||||||||]PAM</td><td>NA</td><td>Intergenic</td><td>NA</td><td>NA</td></tr>
<tr><td></td><td></td><td></td><td></td><td></td><td></td><td></td><td></td><td></td><td></td><td></td><td></td></tr>
<tr><td>T13592</td><td>GGACACTCATATCTGCTGCTGGG</td><td>1000</td><td></td><td></td><td></td><td></td><td></td><td></td><td></td><td></td><td></td></tr>
<tr><td>Oligo fwd</td><td>CACCGGACACTCATATCTGCTGCT</td><td></td><td></td><td></td><td></td><td></td><td></td><td></td><td></td><td></td><td></td></tr>
<tr><td>Oligo rev</td><td>AAACAGCAGCAGATATGAGTGTCC</td><td></td><td></td><td></td><td></td><td></td><td></td><td></td><td></td><td></td><td></td></tr>
<tr><td>Chromosome</td><td>start</td><td>end</td><td>strand</td><td>MM</td><td>target_seq</td><td>PAM</td><td>alignment</td><td>distance</td><td>position</td><td>gene name</td><td>gene id</td></tr>
<tr><td>GU305914.1</td><td>178975</td><td>178997</td><td>+</td><td>0</td><td>GGACACTCATATCTGCTGCT</td><td>GGG</td><td>||||||||[||||||||||||]PAM</td><td>NA</td><td>Intergenic</td><td>NA</td><td>NA</td></tr>
<tr><td></td><td></td><td></td><td></td><td></td><td></td><td></td><td></td><td></td><td></td><td></td><td></td></tr>
<tr><td>T13593</td><td>GACACTCATATCTGCTGCTGGGG</td><td>1000</td><td></td><td></td><td></td><td></td><td></td><td></td><td></td><td></td><td></td></tr>
<tr><td>Oligo fwd</td><td>CACCGACACTCATATCTGCTGCTG</td><td></td><td></td><td></td><td></td><td></td><td></td><td></td><td></td><td></td><td></td></tr>
<tr><td>Oligo rev</td><td>AAACCAGCAGCAGATATGAGTGTC</td><td></td><td></td><td></td><td></td><td></td><td></td><td></td><td></td><td></td><td></td></tr>
<tr><td>Chromosome</td><td>start</td><td>end</td><td>strand</td><td>MM</td><td>target_seq</td><td>PAM</td><td>alignment</td><td>distance</td><td>position</td><td>gene name</td><td>gene id</td></tr>
<tr><td>GU305914.1</td><td>178976</td><td>178998</td><td>+</td><td>0</td><td>GACACTCATATCTGCTGCTG</td><td>GGG</td><td>||||||||[||||||||||||]PAM</td><td>NA</td><td>Intergenic</td><td>NA</td><td>NA</td></tr>
<tr><td></td><td></td><td></td><td></td><td></td><td></td><td></td><td></td><td></td><td></td><td></td><td></td></tr>
<tr><td>T13594</td><td>ACACTCATATCTGCTGCTGGGGG</td><td>1000</td><td></td><td></td><td></td><td></td><td></td><td></td><td></td><td></td><td></td></tr>
<tr><td>Oligo fwd</td><td>CACCACACTCATATCTGCTGCTGG</td><td></td><td></td><td></td><td></td><td></td><td></td><td></td><td></td><td></td><td></td></tr>
<tr><td>Oligo rev</td><td>AAACCCAGCAGCAGATATGAGTGT</td><td></td><td></td><td></td><td></td><td></td><td></td><td></td><td></td><td></td><td></td></tr>
<tr><td>Chromosome</td><td>start</td><td>end</td><td>strand</td><td>MM</td><td>target_seq</td><td>PAM</td><td>alignment</td><td>distance</td><td>position</td><td>gene name</td><td>gene id</td></tr>
<tr><td>GU305914.1</td><td>178977</td><td>178999</td><td>+</td><td>0</td><td>ACACTCATATCTGCTGCTGG</td><td>GGG</td><td>||||||||[||||||||||||]PAM</td><td>NA</td><td>Intergenic</td><td>NA</td><td>NA</td></tr>
</table>

T13595 ATATCTGCTGCTGGGGGCAGCGG 1000
Oligo fwd CACCATATCTGCTGCTGGGGGCAG
Oligo rev AAACCTGCCCCCAGCAGCAGATAT
Chromosome start end strand MM target_seq PAM alignment distance position gene name gene id
GU305914.1 178983 179005 + 0 ATATCTGCTGCTGGGGGCAG CGG ||||||||[||||||||||||]PAM NA Intergenic NA NA

T13596 TCTGCTGCTGGGGGCAGCGGCGG 1000
Oligo fwd CACCTCTGCTGCTGGGGGCAGCGG
Oligo rev AAACCCGCTGCCCCCAGCAGCAGA
Chromosome start end strand MM target_seq PAM alignment distance position gene name gene id
GU305914.1 178986 179008 + 0 TCTGCTGCTGGGGGCAGCGG CGG ||||||||[||||||||||||]PAM NA Intergenic NA NA

T13597 GGCGGCATACCTCTTCGCAGAGG 1000
Oligo fwd CACCGGCGGCATACCTCTTCGCAG
Oligo rev AAACCTGCGAAGAGGTATGCCGCC
Chromosome start end strand MM target_seq PAM alignment distance position gene name gene id
GU305914.1 179004 179026 + 0 GGCGGCATACCTCTTCGCAG AGG ||||||||[||||||||||||]PAM NA Intergenic NA NA

T13598 GCGGCATACCTCTTCGCAGAGGG 1000
Oligo fwd CACCGCGGCATACCTCTTCGCAGA
Oligo rev AAACTCTGCGAAGAGGTATGCCGC
Chromosome start end strand MM target_seq PAM alignment distance position gene name gene id
GU305914.1 179005 179027 + 0 GCGGCATACCTCTTCGCAGA GGG ||||||||[||||||||||||]PAM NA Intergenic NA NA

T13599 GCATACCTCTTCGCAGAGGGAGG 1000
Oligo fwd CACCGCATACCTCTTCGCAGAGGG
Oligo rev AAACCCCTCTGCGAAGAGGTATGC
Chromosome start end strand MM target_seq PAM alignment distance position gene name gene id
GU305914.1 179008 179030 + 0 GCATACCTCTTCGCAGAGGG AGG ||||||||[||||||||||||]PAM NA Intergenic NA NA

T13600 TTCGCAGAGGGAGGTGTGAAAGG 1000
Oligo fwd CACCTTCGCAGAGGGAGGTGTGAA
Oligo rev AAACTTCACACCTCCCTCTGCGAA
Chromosome start end strand MM target_seq PAM alignment distance position gene name gene id
GU305914.1 179017 179039 + 0 TTCGCAGAGGGAGGTGTGAA AGG ||||||||[||||||||||||]PAM NA Intergenic NA NA

T13601 CAGAGGGAGGTGTGAAAGGATGG 1000
Oligo fwd CACCCAGAGGGAGGTGTGAAAGGA
Oligo rev AAACTCCTTTCACACCTCCCTCTG
Chromosome start end strand MM target_seq PAM alignment distance position gene name gene id
GU305914.1 179021 179043 + 0 CAGAGGGAGGTGTGAAAGGA TGG ||||||||[||||||||||||]PAM NA Intergenic NA NA

T13602 AGGGAGGTGTGAAAGGATGGTGG 1000
Oligo fwd CACCAGGGAGGTGTGAAAGGATGG
Oligo rev AAACCCATCCTTTCACACCTCCCT
Chromosome start end strand MM target_seq PAM alignment distance position gene name gene id
GU305914.1 179024 179046 + 0 AGGGAGGTGTGAAAGGATGG TGG ||||||||[||||||||||||]PAM NA Intergenic NA NA

T13603 GTGAAAGGATGGTGGCGTTCAGG 1000
Oligo fwd CACCGTGAAAGGATGGTGGCGTTC
Oligo rev AAACGAACGCCACCATCCTTTCAC
Chromosome start end strand MM target_seq PAM alignment distance position gene name gene id
GU305914.1 179032 179054 + 0 GTGAAAGGATGGTGGCGTTC AGG ||||||||[||||||||||||]PAM NA Intergenic NA NA

T13604 CAGGTACCGCCTTCCTCTTGTGG 1000
Oligo fwd CACCCAGGTACCGCCTTCCTCTTG
Oligo rev AAACCAAGAGGAAGGCGGTACCTG
Chromosome start end strand MM target_seq PAM alignment distance position gene name gene id
GU305914.1 179051 179073 + 0 CAGGTACCGCCTTCCTCTTG TGG ||||||||[||||||||||||]PAM NA Intergenic NA NA

T13605 GTACCGCCTTCCTCTTGTGGTGG 1000
Oligo fwd CACCGTACCGCCTTCCTCTTGTGG
Oligo rev AAACCCACAAGAGGAAGGCGGTAC
Chromosome start end strand MM target_seq PAM alignment distance position gene name gene id
GU305914.1 179054 179076 + 0 GTACCGCCTTCCTCTTGTGG TGG ||||||||[||||||||||||]PAM NA Intergenic NA NA

T13606 CCTCTTGTGGTGGTGTCTAGAGG 1000
Oligo fwd CACCCCTCTTGTGGTGGTGTCTAG
Oligo rev AAACCTAGACACCACCACAAGAGG
Chromosome start end strand MM target_seq PAM alignment distance position gene name gene id
GU305914.1 179064 179086 + 0 CCTCTTGTGGTGGTGTCTAG AGG ||||||||[||||||||||||]PAM NA Intergenic NA NA

T13607 CCCGCTCCCTGATGATTTAATGG 1000
Oligo fwd CACCCCCGCTCCCTGATGATTTAA
Oligo rev AAACTTAAATCATCAGGGAGCGGG
Chromosome start end strand MM target_seq PAM alignment distance position gene name gene id
GU305914.1 179088 179110 + 0 CCCGCTCCCTGATGATTTAA TGG ||||||||[||||||||||||]PAM NA Intergenic NA NA

T13608 GCTCCCTGATGATTTAATGGAGG 1000
Oligo fwd CACCGCTCCCTGATGATTTAATGG
Oligo rev AAACCCATTAAATCATCAGGGAGC
Chromosome start end strand MM target_seq PAM alignment distance position gene name gene id
GU305914.1 179091 179113 + 0 GCTCCCTGATGATTTAATGG AGG ||||||||[||||||||||||]PAM NA Intergenic NA NA

T13609 TAATGGAGGACCTTGATGAGTGG 1000
Oligo fwd CACCTAATGGAGGACCTTGATGAG
Oligo rev AAACCTCATCAAGGTCCTCCATTA
Chromosome start end strand MM target_seq PAM alignment distance position gene name gene id
GU305914.1 179105 179127 + 0 TAATGGAGGACCTTGATGAG TGG ||||||||[||||||||||||]PAM NA Intergenic NA NA

T13610 AATGGAGGACCTTGATGAGTGGG 1000
Oligo fwd CACCAATGGAGGACCTTGATGAGT
Oligo rev AAACACTCATCAAGGTCCTCCATT
Chromosome start end strand MM target_seq PAM alignment distance position gene name gene id
GU305914.1 179106 179128 + 0 AATGGAGGACCTTGATGAGT GGG ||||||||[||||||||||||]PAM NA Intergenic NA NA

T13611 GATGAGTGGGAGCCTCCGTGTGG 1000
Oligo fwd CACCGATGAGTGGGAGCCTCCGTG
Oligo rev AAACCACGGAGGCTCCCACTCATC
Chromosome start end strand MM target_seq PAM alignment distance position gene name gene id
GU305914.1 179119 179141 + 0 GATGAGTGGGAGCCTCCGTG TGG ||||||||[||||||||||||]PAM NA Intergenic NA NA

T13612 ACTGTGCCCACCATGATCTGAGG 1000
Oligo fwd CACCACTGTGCCCACCATGATCTG
Oligo rev AAACCAGATCATGGTGGGCACAGT
Chromosome start end strand MM target_seq PAM alignment distance position gene name gene id
GU305914.1 179271 179293 + 0 ACTGTGCCCACCATGATCTG AGG ||||||||[||||||||||||]PAM NA Intergenic NA NA

T13613 ATGATCTGAGGAAGACCTGCTGG 1000
Oligo fwd CACCATGATCTGAGGAAGACCTGC
Oligo rev AAACGCAGGTCTTCCTCAGATCAT
Chromosome start end strand MM target_seq PAM alignment distance position gene name gene id
GU305914.1 179283 179305 + 0 ATGATCTGAGGAAGACCTGC TGG ||||||||[||||||||||||]PAM NA Intergenic NA NA

T13614 TGATCTGAGGAAGACCTGCTGGG 1000
Oligo fwd CACCTGATCTGAGGAAGACCTGCT
Oligo rev AAACAGCAGGTCTTCCTCAGATCA
Chromosome start end strand MM target_seq PAM alignment distance position gene name gene id
GU305914.1 179284 179306 + 0 TGATCTGAGGAAGACCTGCT GGG ||||||||[||||||||||||]PAM NA Intergenic NA NA
GU305914.1 198065 198087 - 4 CGATCCGATGAAGACCTACT GGG -||||-||[-|||||||||-||]PAM NA Intergenic NA NA

T13615 GATCTGAGGAAGACCTGCTGGGG 1000
Oligo fwd CACCGATCTGAGGAAGACCTGCTG
Oligo rev AAACCAGCAGGTCTTCCTCAGATC
Chromosome start end strand MM target_seq PAM alignment distance position gene name gene id
GU305914.1 179285 179307 + 0 GATCTGAGGAAGACCTGCTG GGG ||||||||[||||||||||||]PAM NA Intergenic NA NA

T13616 CTCCTCTCATCAAAGTGTTGAGG 1000
Oligo fwd CACCCTCCTCTCATCAAAGTGTTG
Oligo rev AAACCAACACTTTGATGAGAGGAG
Chromosome start end strand MM target_seq PAM alignment distance position gene name gene id
GU305914.1 179308 179330 + 0 CTCCTCTCATCAAAGTGTTG AGG ||||||||[||||||||||||]PAM NA Intergenic NA NA

T13617 CTCATCAAAGTGTTGAGGAGAGG 1000
Oligo fwd CACCCTCATCAAAGTGTTGAGGAG
Oligo rev AAACCTCCTCAACACTTTGATGAG
Chromosome start end strand MM target_seq PAM alignment distance position gene name gene id
GU305914.1 179313 179335 + 0 CTCATCAAAGTGTTGAGGAG AGG ||||||||[||||||||||||]PAM NA Intergenic NA NA

T13618 ATCAAAGTGTTGAGGAGAGGAGG 1000
Oligo fwd CACCATCAAAGTGTTGAGGAGAGG
Oligo rev AAACCCTCTCCTCAACACTTTGAT
Chromosome start end strand MM target_seq PAM alignment distance position gene name gene id
GU305914.1 179316 179338 + 0 ATCAAAGTGTTGAGGAGAGG AGG ||||||||[||||||||||||]PAM NA Intergenic NA NA

T13619 GAGGAGAGGAGGAGATCACCTGG 1000
Oligo fwd CACCGAGGAGAGGAGGAGATCACC
Oligo rev AAACGGTGATCTCCTCCTCTCCTC
Chromosome start end strand MM target_seq PAM alignment distance position gene name gene id
GU305914.1 179327 179349 + 0 GAGGAGAGGAGGAGATCACC TGG ||||||||[||||||||||||]PAM NA Intergenic NA NA

T13620 AGGAGAGGAGGAGATCACCTGGG 1000
Oligo fwd CACCAGGAGAGGAGGAGATCACCT
Oligo rev AAACAGGTGATCTCCTCCTCTCCT
Chromosome start end strand MM target_seq PAM alignment distance position gene name gene id
GU305914.1 179328 179350 + 0 AGGAGAGGAGGAGATCACCT GGG ||||||||[||||||||||||]PAM NA Intergenic NA NA
GU305914.1 10580 10602 - 4 AGGTGACGGGGAAATCACCT AGG |||-||-|[-|||-|||||||]PAM NA Intergenic NA NA

T13621 GGAGAGGAGGAGATCACCTGGGG 1000
Oligo fwd CACCGGAGAGGAGGAGATCACCTG
Oligo rev AAACCAGGTGATCTCCTCCTCTCC
Chromosome start end strand MM target_seq PAM alignment distance position gene name gene id
GU305914.1 179329 179351 + 0 GGAGAGGAGGAGATCACCTG GGG ||||||||[||||||||||||]PAM NA Intergenic NA NA

T13622 TCCTGATACTCCTCCTCCTGAGG 1000
Oligo fwd CACCTCCTGATACTCCTCCTCCTG
Oligo rev AAACCAGGAGGAGGAGTATCAGGA
Chromosome start end strand MM target_seq PAM alignment distance position gene name gene id
GU305914.1 179362 179384 + 0 TCCTGATACTCCTCCTCCTG AGG ||||||||[||||||||||||]PAM NA Intergenic NA NA

T13623 CTCATCTGCCTCACTCTCAGAGG 1000
Oligo fwd CACCCTCATCTGCCTCACTCTCAG
Oligo rev AAACCTGAGAGTGAGGCAGATGAG
Chromosome start end strand MM target_seq PAM alignment distance position gene name gene id
GU305914.1 179403 179425 + 0 CTCATCTGCCTCACTCTCAG AGG ||||||||[||||||||||||]PAM NA Intergenic NA NA

T13624 TCATCTGCCTCACTCTCAGAGGG 1000
Oligo fwd CACCTCATCTGCCTCACTCTCAGA

Oligo rev AAACTCTGAGAGTGAGGCAGATGA
Chromosome start end strand MM target_seq PAM alignment distance position gene name gene id
GU305914.1 179404 179426 + 0 TCATCTGCCTCACTCTCAGA GGG ||||||||[||||||||||||]PAM NA Intergenic NA NA

T13625 CAACTACTTCATCAAGTCTCTGG 1000
Oligo fwd CACCCAACTACTTCATCAAGTCTC
Oligo rev AAACGAGACTTGATGAAGTAGTTG
Chromosome start end strand MM target_seq PAM alignment distance position gene name gene id
GU305914.1 179441 179463 + 0 CAACTACTTCATCAAGTCTC TGG ||||||||[||||||||||||]PAM NA Intergenic NA NA

T13626 TCTGAGTGTCTTCTGAAGTCTGG 1000
Oligo fwd CACCTCTGAGTGTCTTCTGAAGTC
Oligo rev AAACGACTTCAGAAGACACTCAGA
Chromosome start end strand MM target_seq PAM alignment distance position gene name gene id
GU305914.1 179497 179519 + 0 TCTGAGTGTCTTCTGAAGTC TGG ||||||||[||||||||||||]PAM NA Intergenic NA NA

T13627 CTGAGTGTCTTCTGAAGTCTGGG 1000
Oligo fwd CACCCTGAGTGTCTTCTGAAGTCT
Oligo rev AAACAGACTTCAGAAGACACTCAG
Chromosome start end strand MM target_seq PAM alignment distance position gene name gene id
GU305914.1 179498 179520 + 0 CTGAGTGTCTTCTGAAGTCT GGG ||||||||[||||||||||||]PAM NA Intergenic NA NA

T13628 TGAGTGTCTTCTGAAGTCTGGGG 1000
Oligo fwd CACCTGAGTGTCTTCTGAAGTCTG
Oligo rev AAACCAGACTTCAGAAGACACTCA
Chromosome start end strand MM target_seq PAM alignment distance position gene name gene id
GU305914.1 179499 179521 + 0 TGAGTGTCTTCTGAAGTCTG GGG ||||||||[||||||||||||]PAM NA Intergenic NA NA

T13629 CTTCTGAAGTCTGGGGTGTTTGG 1000
Oligo fwd CACCCTTCTGAAGTCTGGGGTGTT
Oligo rev AAACAACACCCCAGACTTCAGAAG
Chromosome start end strand MM target_seq PAM alignment distance position gene name gene id
GU305914.1 179506 179528 + 0 CTTCTGAAGTCTGGGGTGTT TGG ||||||||[||||||||||||]PAM NA Intergenic NA NA

T13630 GGGGTGTTTGGTCTTCTGATTGG 1000
Oligo fwd CACCGGGGTGTTTGGTCTTCTGAT
Oligo rev AAACATCAGAAGACCAAACACCCC
Chromosome start end strand MM target_seq PAM alignment distance position gene name gene id
GU305914.1 179518 179540 + 0 GGGGTGTTTGGTCTTCTGAT TGG ||||||||[||||||||||||]PAM NA Intergenic NA NA

T13631 TGTTTGGTCTTCTGATTGGCAGG 1000
Oligo fwd CACCTGTTTGGTCTTCTGATTGGC
Oligo rev AAACGCCAATCAGAAGACCAAACA
Chromosome start end strand MM target_seq PAM alignment distance position gene name gene id
GU305914.1 179522 179544 + 0 TGTTTGGTCTTCTGATTGGC AGG ||||||||[||||||||||||]PAM NA Intergenic NA NA

T13632 GTTTGGTCTTCTGATTGGCAGGG 1000
Oligo fwd CACCGTTTGGTCTTCTGATTGGCA
Oligo rev AAACTGCCAATCAGAAGACCAAAC
Chromosome start end strand MM target_seq PAM alignment distance position gene name gene id
GU305914.1 179523 179545 + 0 GTTTGGTCTTCTGATTGGCA GGG ||||||||[||||||||||||]PAM NA Intergenic NA NA

T13633 GATTGGCAGGGCGCTGTGTGAGG 1000
Oligo fwd CACCGATTGGCAGGGCGCTGTGTG
Oligo rev AAACCACACAGCGCCCTGCCAATC
Chromosome start end strand MM target_seq PAM alignment distance position gene name gene id
GU305914.1 179535 179557 + 0 GATTGGCAGGGCGCTGTGTG AGG ||||||||[||||||||||||]PAM NA Intergenic NA NA

T13634 GGCAGTAGTTATACCTGACATGG 1000
Oligo fwd CACCGGCAGTAGTTATACCTGACA
Oligo rev AAACTGTCAGGTATAACTACTGCC
Chromosome start end strand MM target_seq PAM alignment distance position gene name gene id
GU305914.1 179556 179578 + 0 GGCAGTAGTTATACCTGACA TGG ||||||||[||||||||||||]PAM NA Intergenic NA NA

T13635 AGTAGTTATACCTGACATGGAGG 1000
Oligo fwd CACCAGTAGTTATACCTGACATGG
Oligo rev AAACCCATGTCAGGTATAACTACT
Chromosome start end strand MM target_seq PAM alignment distance position gene name gene id
GU305914.1 179559 179581 + 0 AGTAGTTATACCTGACATGG AGG ||||||||[||||||||||||]PAM NA Intergenic NA NA

T13636 AACAATATCTATGTTCATCTCGG 1000
Oligo fwd CACCAACAATATCTATGTTCATCT
Oligo rev AAACAGATGAACATAGATATTGTT
Chromosome start end strand MM target_seq PAM alignment distance position gene name gene id
GU305914.1 179592 179614 + 0 AACAATATCTATGTTCATCT CGG ||||||||[||||||||||||]PAM NA Intergenic NA NA

T13637 ACAATATCTATGTTCATCTCGGG 1000
Oligo fwd CACCACAATATCTATGTTCATCTC
Oligo rev AAACGAGATGAACATAGATATTGT
Chromosome start end strand MM target_seq PAM alignment distance position gene name gene id
GU305914.1 179593 179615 + 0 ACAATATCTATGTTCATCTC GGG ||||||||[||||||||||||]PAM NA Intergenic NA NA

T13638 AGTTAAAGAAGTCTTTAATGTGG 1000
Oligo fwd CACCAGTTAAAGAAGTCTTTAATG
Oligo rev AAACCATTAAAGACTTCTTTAACT
Chromosome start end strand MM target_seq PAM alignment distance position gene name gene id
GU305914.1 179633 179655 + 0 AGTTAAAGAAGTCTTTAATG TGG ||||||||[||||||||||||]PAM NA Intergenic NA NA

T13639 GTTAAAGAAGTCTTTAATGTGGG 1000

| | | | | | | | | | | | |
|---|---|---|---|---|---|---|---|---|---|---|---|
| Oligo fwd | CACCGTTAAAGAAGTCTTTAATGT | | | | | | | | | | |
| Oligo rev | AAACACATTAAAGACTTCTTTAAC | | | | | | | | | | |
| Chromosome | start | end | strand | MM | target_seq | PAM | alignment | distance | position | gene name | gene id |
| GU305914.1 | 179634 | 179656 | + | 0 | GTTAAAGAAGTCTTTAATGT | GGG | \|\|\|\|\|\|\|\|[\|\|\|\|\|\|\|\|\|\|\|\|]PAM | NA | Intergenic | NA | NA |
| T13640 | GTCTTTAATGTGGGTACAATAGG | 1000 | | | | | | | | | |
| Oligo fwd | CACCGTCTTTAATGTGGGTACAAT | | | | | | | | | | |
| Oligo rev | AAACATTGTACCCACATTAAAGAC | | | | | | | | | | |
| Chromosome | start | end | strand | MM | target_seq | PAM | alignment | distance | position | gene name | gene id |
| GU305914.1 | 179643 | 179665 | + | 0 | GTCTTTAATGTGGGTACAAT | AGG | \|\|\|\|\|\|\|\|[\|\|\|\|\|\|\|\|\|\|\|\|]PAM | NA | Intergenic | NA | NA |
| T13641 | TTAAGTACAGAGAATAAAAGAGG | 1000 | | | | | | | | | |
| Oligo fwd | CACCTTAAGTACAGAGAATAAAAG | | | | | | | | | | |
| Oligo rev | AAACCTTTTATTCTCTGTACTTAA | | | | | | | | | | |
| Chromosome | start | end | strand | MM | target_seq | PAM | alignment | distance | position | gene name | gene id |
| GU305914.1 | 179686 | 179708 | + | 0 | TTAAGTACAGAGAATAAAAG | AGG | \|\|\|\|\|\|\|\|[\|\|\|\|\|\|\|\|\|\|\|\|]PAM | NA | Intergenic | NA | NA |
| T13642 | TAAGTACAGAGAATAAAAGAGGG | 1000 | | | | | | | | | |
| Oligo fwd | CACCTAAGTACAGAGAATAAAAGA | | | | | | | | | | |
| Oligo rev | AAACTCTTTTATTCTCTGTACTTA | | | | | | | | | | |
| Chromosome | start | end | strand | MM | target_seq | PAM | alignment | distance | position | gene name | gene id |
| GU305914.1 | 179687 | 179709 | + | 0 | TAAGTACAGAGAATAAAAGA | GGG | \|\|\|\|\|\|\|\|[\|\|\|\|\|\|\|\|\|\|\|\|]PAM | NA | Intergenic | NA | NA |
| T13643 | AAGTACAGAGAATAAAAGAGGGG | 1000 | | | | | | | | | |
| Oligo fwd | CACCAAGTACAGAGAATAAAAGAG | | | | | | | | | | |
| Oligo rev | AAACCTCTTTTATTCTCTGTACTT | | | | | | | | | | |
| Chromosome | start | end | strand | MM | target_seq | PAM | alignment | distance | position | gene name | gene id |
| GU305914.1 | 179688 | 179710 | + | 0 | AAGTACAGAGAATAAAAGAG | GGG | \|\|\|\|\|\|\|\|[\|\|\|\|\|\|\|\|\|\|\|\|]PAM | NA | Intergenic | NA | NA |
| T13644 | AGTACAGAGAATAAAAGAGGGGG | 1000 | | | | | | | | | |
| Oligo fwd | CACCAGTACAGAGAATAAAAGAGG | | | | | | | | | | |
| Oligo rev | AAACCCTCTTTTATTCTCTGTACT | | | | | | | | | | |
| Chromosome | start | end | strand | MM | target_seq | PAM | alignment | distance | position | gene name | gene id |
| GU305914.1 | 179689 | 179711 | + | 0 | AGTACAGAGAATAAAAGAGG | GGG | \|\|\|\|\|\|\|\|[\|\|\|\|\|\|\|\|\|\|\|\|]PAM | NA | Intergenic | NA | NA |
| T13645 | AGAGAATAAAAGAGGGGGTGTGG | 1000 | | | | | | | | | |
| Oligo fwd | CACCAGAGAATAAAAGAGGGGGTG | | | | | | | | | | |
| Oligo rev | AAACCACCCCCTCTTTTATTCTCT | | | | | | | | | | |
| Chromosome | start | end | strand | MM | target_seq | PAM | alignment | distance | position | gene name | gene id |
| GU305914.1 | 179694 | 179716 | + | 0 | AGAGAATAAAAGAGGGGGTG | TGG | \|\|\|\|\|\|\|\|[\|\|\|\|\|\|\|\|\|\|\|\|]PAM | NA | Intergenic | NA | NA |
| T13646 | TGTTAGTAATTGTGTCTAAGTGG | 1000 | | | | | | | | | |
| Oligo fwd | CACCTGTTAGTAATTGTGTCTAAG | | | | | | | | | | |
| Oligo rev | AAACCTTAGACACAATTACTAACA | | | | | | | | | | |
| Chromosome | start | end | strand | MM | target_seq | PAM | alignment | distance | position | gene name | gene id |
| GU305914.1 | 179717 | 179739 | + | 0 | TGTTAGTAATTGTGTCTAAG | TGG | \|\|\|\|\|\|\|\|[\|\|\|\|\|\|\|\|\|\|\|\|]PAM | NA | Intergenic | NA | NA |
| T13647 | GTTAGTAATTGTGTCTAAGTGGG | 1000 | | | | | | | | | |
| Oligo fwd | CACCGTTAGTAATTGTGTCTAAGT | | | | | | | | | | |
| Oligo rev | AAACACTTAGACACAATTACTAAC | | | | | | | | | | |
| Chromosome | start | end | strand | MM | target_seq | PAM | alignment | distance | position | gene name | gene id |
| GU305914.1 | 179718 | 179740 | + | 0 | GTTAGTAATTGTGTCTAAGT | GGG | \|\|\|\|\|\|\|\|[\|\|\|\|\|\|\|\|\|\|\|\|]PAM | NA | Intergenic | NA | NA |
| T13648 | GTAATTGTGTCTAAGTGGGCAGG | 1000 | | | | | | | | | |
| Oligo fwd | CACCGTAATTGTGTCTAAGTGGGC | | | | | | | | | | |
| Oligo rev | AAACGCCCACTTAGACACAATTAC | | | | | | | | | | |
| Chromosome | start | end | strand | MM | target_seq | PAM | alignment | distance | position | gene name | gene id |
| GU305914.1 | 179722 | 179744 | + | 0 | GTAATTGTGTCTAAGTGGGC | AGG | \|\|\|\|\|\|\|\|[\|\|\|\|\|\|\|\|\|\|\|\|]PAM | NA | Intergenic | NA | NA |
| T13649 | TGTGTCTAAGTGGGCAGGTGAGG | 1000 | | | | | | | | | |
| Oligo fwd | CACCTGTGTCTAAGTGGGCAGGTG | | | | | | | | | | |
| Oligo rev | AAACCACCTGCCCACTTAGACACA | | | | | | | | | | |
| Chromosome | start | end | strand | MM | target_seq | PAM | alignment | distance | position | gene name | gene id |
| GU305914.1 | 179727 | 179749 | + | 0 | TGTGTCTAAGTGGGCAGGTG | AGG | \|\|\|\|\|\|\|\|[\|\|\|\|\|\|\|\|\|\|\|\|]PAM | NA | Intergenic | NA | NA |
| T13650 | TAGTAAAAAAACATTGTCCATGG | 1000 | | | | | | | | | |
| Oligo fwd | CACCTAGTAAAAAAACATTGTCCA | | | | | | | | | | |
| Oligo rev | AAACTGGACAATGTTTTTTTACTA | | | | | | | | | | |
| Chromosome | start | end | strand | MM | target_seq | PAM | alignment | distance | position | gene name | gene id |
| GU305914.1 | 179809 | 179831 | + | 0 | TAGTAAAAAAACATTGTCCA | TGG | \|\|\|\|\|\|\|\|[\|\|\|\|\|\|\|\|\|\|\|\|]PAM | NA | Intergenic | NA | NA |
| T13651 | AAAAACATTGTCCATGGTGATGG | 1000 | | | | | | | | | |
| Oligo fwd | CACCAAAAACATTGTCCATGGTGA | | | | | | | | | | |
| Oligo rev | AAACTCACCATGGACAATGTTTTT | | | | | | | | | | |
| Chromosome | start | end | strand | MM | target_seq | PAM | alignment | distance | position | gene name | gene id |
| GU305914.1 | 179815 | 179837 | + | 0 | AAAAACATTGTCCATGGTGA | TGG | \|\|\|\|\|\|\|\|[\|\|\|\|\|\|\|\|\|\|\|\|]PAM | NA | Intergenic | NA | NA |
| T13652 | AAAACATTGTCCATGGTGATGGG | 1000 | | | | | | | | | |
| Oligo fwd | CACCAAAACATTGTCCATGGTGAT | | | | | | | | | | |
| Oligo rev | AAACATCACCATGGACAATGTTTT | | | | | | | | | | |
| Chromosome | start | end | strand | MM | target_seq | PAM | alignment | distance | position | gene name | gene id |
| GU305914.1 | 179816 | 179838 | + | 0 | AAAACATTGTCCATGGTGAT | GGG | \|\|\|\|\|\|\|\|[\|\|\|\|\|\|\|\|\|\|\|\|]PAM | NA | Intergenic | NA | NA |
| T13653 | ACATTGTCCATGGTGATGGGTGG | 1000 | | | | | | | | | |
| Oligo fwd | CACCACATTGTCCATGGTGATGGG | | | | | | | | | | |
| Oligo rev | AAACCCCATCACCATGGACAATGT | | | | | | | | | | |
| Chromosome | start | end | strand | MM | target_seq | PAM | alignment | distance | position | gene name | gene id |
| GU305914.1 | 179819 | 179841 | + | 0 | ACATTGTCCATGGTGATGGG | TGG | \|\|\|\|\|\|\|\|[\|\|\|\|\|\|\|\|\|\|\|\|]PAM | NA | Intergenic | NA | NA |

<table>
<tr><td>T13654</td><td>CATTGTCCATGGTGATGGGTGGG</td><td>1000</td></tr>
<tr><td>Oligo fwd</td><td>CACCCATTGTCCATGGTGATGGGT</td></tr>
<tr><td>Oligo rev</td><td>AAACACCCATCACCATGGACAATG</td></tr>
<tr><td>Chromosome</td><td>start</td><td>end</td><td>strand</td><td>MM</td><td>target_seq</td><td>PAM</td><td>alignment</td><td>distance</td><td>position</td><td>gene name</td><td>gene id</td></tr>
<tr><td>GU305914.1</td><td>179820</td><td>179842</td><td>+</td><td>0</td><td>CATTGTCCATGGTGATGGGT</td><td>GGG</td><td>||||||||[||||||||||||]PAM</td><td>NA</td><td>Intergenic</td><td>NA</td><td>NA</td></tr>
<tr><td>T13655</td><td>ATTGTCCATGGTGATGGGTGGGG</td><td>1000</td></tr>
<tr><td>Oligo fwd</td><td>CACCATTGTCCATGGTGATGGGTG</td></tr>
<tr><td>Oligo rev</td><td>AAACCACCCATCACCATGGACAAT</td></tr>
<tr><td>Chromosome</td><td>start</td><td>end</td><td>strand</td><td>MM</td><td>target_seq</td><td>PAM</td><td>alignment</td><td>distance</td><td>position</td><td>gene name</td><td>gene id</td></tr>
<tr><td>GU305914.1</td><td>179821</td><td>179843</td><td>+</td><td>0</td><td>ATTGTCCATGGTGATGGGTG</td><td>GGG</td><td>||||||||[||||||||||||]PAM</td><td>NA</td><td>Intergenic</td><td>NA</td><td>NA</td></tr>
<tr><td>T13656</td><td>CATGGTGATGGGTGGGGAGCTGG</td><td>1000</td></tr>
<tr><td>Oligo fwd</td><td>CACCCATGGTGATGGGTGGGGAGC</td></tr>
<tr><td>Oligo rev</td><td>AAACGCTCCCCACCCATCACCATG</td></tr>
<tr><td>Chromosome</td><td>start</td><td>end</td><td>strand</td><td>MM</td><td>target_seq</td><td>PAM</td><td>alignment</td><td>distance</td><td>position</td><td>gene name</td><td>gene id</td></tr>
<tr><td>GU305914.1</td><td>179827</td><td>179849</td><td>+</td><td>0</td><td>CATGGTGATGGGTGGGGAGC</td><td>TGG</td><td>||||||||[||||||||||||]PAM</td><td>NA</td><td>Intergenic</td><td>NA</td><td>NA</td></tr>
<tr><td>T13657</td><td>ATGGTGATGGGTGGGGAGCTGGG</td><td>1000</td></tr>
<tr><td>Oligo fwd</td><td>CACCATGGTGATGGGTGGGGAGCT</td></tr>
<tr><td>Oligo rev</td><td>AAACAGCTCCCCACCCATCACCAT</td></tr>
<tr><td>Chromosome</td><td>start</td><td>end</td><td>strand</td><td>MM</td><td>target_seq</td><td>PAM</td><td>alignment</td><td>distance</td><td>position</td><td>gene name</td><td>gene id</td></tr>
<tr><td>GU305914.1</td><td>179828</td><td>179850</td><td>+</td><td>0</td><td>ATGGTGATGGGTGGGGAGCT</td><td>GGG</td><td>||||||||[||||||||||||]PAM</td><td>NA</td><td>Intergenic</td><td>NA</td><td>NA</td></tr>
<tr><td>T13658</td><td>TGGGTGGGGAGCTGGGCTTGTGG</td><td>1000</td></tr>
<tr><td>Oligo fwd</td><td>CACCTGGGTGGGGAGCTGGGCTTG</td></tr>
<tr><td>Oligo rev</td><td>AAACCAAGCCCAGCTCCCCACCCA</td></tr>
<tr><td>Chromosome</td><td>start</td><td>end</td><td>strand</td><td>MM</td><td>target_seq</td><td>PAM</td><td>alignment</td><td>distance</td><td>position</td><td>gene name</td><td>gene id</td></tr>
<tr><td>GU305914.1</td><td>179835</td><td>179857</td><td>+</td><td>0</td><td>TGGGTGGGGAGCTGGGCTTG</td><td>TGG</td><td>||||||||[||||||||||||]PAM</td><td>NA</td><td>Intergenic</td><td>NA</td><td>NA</td></tr>
<tr><td>T13659</td><td>TCACTTCTTGCTCTTCTTCTTGG</td><td>1000</td></tr>
<tr><td>Oligo fwd</td><td>CACCTCACTTCTTGCTCTTCTTCT</td></tr>
<tr><td>Oligo rev</td><td>AAACAGAAGAAGAGCAAGAAGTGA</td></tr>
<tr><td>Chromosome</td><td>start</td><td>end</td><td>strand</td><td>MM</td><td>target_seq</td><td>PAM</td><td>alignment</td><td>distance</td><td>position</td><td>gene name</td><td>gene id</td></tr>
<tr><td>GU305914.1</td><td>179860</td><td>179882</td><td>+</td><td>0</td><td>TCACTTCTTGCTCTTCTTCT</td><td>TGG</td><td>||||||||[||||||||||||]PAM</td><td>NA</td><td>Intergenic</td><td>NA</td><td>NA</td></tr>
<tr><td>T13660</td><td>CACTTCTTGCTCTTCTTCTTGGG</td><td>1000</td></tr>
<tr><td>Oligo fwd</td><td>CACCCACTTCTTGCTCTTCTTCTT</td></tr>
<tr><td>Oligo rev</td><td>AAACAAGAAGAAGAGCAAGAAGTG</td></tr>
<tr><td>Chromosome</td><td>start</td><td>end</td><td>strand</td><td>MM</td><td>target_seq</td><td>PAM</td><td>alignment</td><td>distance</td><td>position</td><td>gene name</td><td>gene id</td></tr>
<tr><td>GU305914.1</td><td>179861</td><td>179883</td><td>+</td><td>0</td><td>CACTTCTTGCTCTTCTTCTT</td><td>GGG</td><td>||||||||[||||||||||||]PAM</td><td>NA</td><td>Intergenic</td><td>NA</td><td>NA</td></tr>
<tr><td>T13661</td><td>CTGTTGATTCTCAGAAGACTTGG</td><td>1000</td></tr>
<tr><td>Oligo fwd</td><td>CACCCTGTTGATTCTCAGAAGACT</td></tr>
<tr><td>Oligo rev</td><td>AAACAGTCTTCTGAGAATCAACAG</td></tr>
<tr><td>Chromosome</td><td>start</td><td>end</td><td>strand</td><td>MM</td><td>target_seq</td><td>PAM</td><td>alignment</td><td>distance</td><td>position</td><td>gene name</td><td>gene id</td></tr>
<tr><td>GU305914.1</td><td>179887</td><td>179909</td><td>+</td><td>0</td><td>CTGTTGATTCTCAGAAGACT</td><td>TGG</td><td>||||||||[||||||||||||]PAM</td><td>NA</td><td>Intergenic</td><td>NA</td><td>NA</td></tr>
<tr><td>T13662</td><td>ATTCTCAGAAGACTTGGATCTGG</td><td>1000</td></tr>
<tr><td>Oligo fwd</td><td>CACCATTCTCAGAAGACTTGGATC</td></tr>
<tr><td>Oligo rev</td><td>AAACGATCCAAGTCTTCTGAGAAT</td></tr>
<tr><td>Chromosome</td><td>start</td><td>end</td><td>strand</td><td>MM</td><td>target_seq</td><td>PAM</td><td>alignment</td><td>distance</td><td>position</td><td>gene name</td><td>gene id</td></tr>
<tr><td>GU305914.1</td><td>179893</td><td>179915</td><td>+</td><td>0</td><td>ATTCTCAGAAGACTTGGATC</td><td>TGG</td><td>||||||||[||||||||||||]PAM</td><td>NA</td><td>Intergenic</td><td>NA</td><td>NA</td></tr>
<tr><td>T13663</td><td>CTTGGATCTGGTATGCACCATGG</td><td>1000</td></tr>
<tr><td>Oligo fwd</td><td>CACCCTTGGATCTGGTATGCACCA</td></tr>
<tr><td>Oligo rev</td><td>AAACTGGTGCATACCAGATCCAAG</td></tr>
<tr><td>Chromosome</td><td>start</td><td>end</td><td>strand</td><td>MM</td><td>target_seq</td><td>PAM</td><td>alignment</td><td>distance</td><td>position</td><td>gene name</td><td>gene id</td></tr>
<tr><td>GU305914.1</td><td>179905</td><td>179927</td><td>+</td><td>0</td><td>CTTGGATCTGGTATGCACCA</td><td>TGG</td><td>||||||||[||||||||||||]PAM</td><td>NA</td><td>Intergenic</td><td>NA</td><td>NA</td></tr>
<tr><td>T13664</td><td>GCGTACTGCCTCTTGAGAGCTGG</td><td>1000</td></tr>
<tr><td>Oligo fwd</td><td>CACCGCGTACTGCCTCTTGAGAGC</td></tr>
<tr><td>Oligo rev</td><td>AAACGCTCTCAAGAGGCAGTACGC</td></tr>
<tr><td>Chromosome</td><td>start</td><td>end</td><td>strand</td><td>MM</td><td>target_seq</td><td>PAM</td><td>alignment</td><td>distance</td><td>position</td><td>gene name</td><td>gene id</td></tr>
<tr><td>GU305914.1</td><td>179927</td><td>179949</td><td>+</td><td>0</td><td>GCGTACTGCCTCTTGAGAGC</td><td>TGG</td><td>||||||||[||||||||||||]PAM</td><td>NA</td><td>Intergenic</td><td>NA</td><td>NA</td></tr>
<tr><td>T13665</td><td>CGTACTGCCTCTTGAGAGCTGGG</td><td>1000</td></tr>
<tr><td>Oligo fwd</td><td>CACCCGTACTGCCTCTTGAGAGCT</td></tr>
<tr><td>Oligo rev</td><td>AAACAGCTCTCAAGAGGCAGTACG</td></tr>
<tr><td>Chromosome</td><td>start</td><td>end</td><td>strand</td><td>MM</td><td>target_seq</td><td>PAM</td><td>alignment</td><td>distance</td><td>position</td><td>gene name</td><td>gene id</td></tr>
<tr><td>GU305914.1</td><td>179928</td><td>179950</td><td>+</td><td>0</td><td>CGTACTGCCTCTTGAGAGCT</td><td>GGG</td><td>||||||||[||||||||||||]PAM</td><td>NA</td><td>Intergenic</td><td>NA</td><td>NA</td></tr>
<tr><td>T13666</td><td>TGCCTCTTGAGAGCTGGGCGAGG</td><td>1000</td></tr>
<tr><td>Oligo fwd</td><td>CACCTGCCTCTTGAGAGCTGGGCG</td></tr>
<tr><td>Oligo rev</td><td>AAACCGCCCAGCTCTCAAGAGGCA</td></tr>
<tr><td>Chromosome</td><td>start</td><td>end</td><td>strand</td><td>MM</td><td>target_seq</td><td>PAM</td><td>alignment</td><td>distance</td><td>position</td><td>gene name</td><td>gene id</td></tr>
<tr><td>GU305914.1</td><td>179933</td><td>179955</td><td>+</td><td>0</td><td>TGCCTCTTGAGAGCTGGGCG</td><td>AGG</td><td>||||||||[||||||||||||]PAM</td><td>NA</td><td>Intergenic</td><td>NA</td><td>NA</td></tr>
<tr><td>T13667</td><td>AGCTGCCGCTGTTGCAGATCTGG</td><td>1000</td></tr>
<tr><td>Oligo fwd</td><td>CACCAGCTGCCGCTGTTGCAGATC</td></tr>
<tr><td>Oligo rev</td><td>AAACGATCTGCAACAGCGGCAGCT</td></tr>
<tr><td>Chromosome</td><td>start</td><td>end</td><td>strand</td><td>MM</td><td>target_seq</td><td>PAM</td><td>alignment</td><td>distance</td><td>position</td><td>gene name</td><td>gene id</td></tr>
<tr><td>GU305914.1</td><td>179971</td><td>179993</td><td>+</td><td>0</td><td>AGCTGCCGCTGTTGCAGATC</td><td>TGG</td><td>||||||||[||||||||||||]PAM</td><td>NA</td><td>Intergenic</td><td>NA</td><td>NA</td></tr>
<tr><td>T13668</td><td>GCTGTTGCAGATCTGGTCGTAGG</td><td>1000</td></tr>
<tr><td>Oligo fwd</td><td>CACCGCTGTTGCAGATCTGGTCGT</td></tr>
<tr><td>Oligo rev</td><td>AAACACGACCAGATCTGCAACAGC</td></tr>
<tr><td>Chromosome</td><td>start</td><td>end</td><td>strand</td><td>MM</td><td>target_seq</td><td>PAM</td><td>alignment</td><td>distance</td><td>position</td><td>gene name</td><td>gene id</td></tr>
<tr><td>GU305914.1</td><td>179978</td><td>180000</td><td>+</td><td>0</td><td>GCTGTTGCAGATCTGGTCGT</td><td>AGG</td><td>||||||||[||||||||||||]PAM</td><td>NA</td><td>Intergenic</td><td>NA</td><td>NA</td></tr>
</table>

<table>
<tr><td>T13669</td><td>CAGATCTGGTCGTAGGTTGCAGG</td><td>1000</td><td></td><td></td><td></td><td></td><td></td><td></td><td></td><td></td><td></td></tr>
<tr><td>Oligo fwd</td><td>CACCCAGATCTGGTCGTAGGTTGC</td><td></td><td></td><td></td><td></td><td></td><td></td><td></td><td></td><td></td><td></td></tr>
<tr><td>Oligo rev</td><td>AAACGCAACCTACGACCAGATCTG</td><td></td><td></td><td></td><td></td><td></td><td></td><td></td><td></td><td></td><td></td></tr>
<tr><td>Chromosome</td><td>start</td><td>end</td><td>strand</td><td>MM</td><td>target_seq</td><td>PAM</td><td>alignment</td><td>distance</td><td>position</td><td>gene name</td><td>gene id</td></tr>
<tr><td>GU305914.1</td><td>179985</td><td>180007</td><td>+</td><td>0</td><td>CAGATCTGGTCGTAGGTTGC</td><td>AGG</td><td>||||||||[||||||||||||]PAM</td><td>NA</td><td>Intergenic</td><td>NA</td><td>NA</td></tr>
<tr><td>T13670</td><td>CTGGTCGTAGGTTGCAGGACAGG</td><td>1000</td><td></td><td></td><td></td><td></td><td></td><td></td><td></td><td></td><td></td></tr>
<tr><td>Oligo fwd</td><td>CACCCTGGTCGTAGGTTGCAGGAC</td><td></td><td></td><td></td><td></td><td></td><td></td><td></td><td></td><td></td><td></td></tr>
<tr><td>Oligo rev</td><td>AAACGTCCTGCAACCTACGACCAG</td><td></td><td></td><td></td><td></td><td></td><td></td><td></td><td></td><td></td><td></td></tr>
<tr><td>Chromosome</td><td>start</td><td>end</td><td>strand</td><td>MM</td><td>target_seq</td><td>PAM</td><td>alignment</td><td>distance</td><td>position</td><td>gene name</td><td>gene id</td></tr>
<tr><td>GU305914.1</td><td>179990</td><td>180012</td><td>+</td><td>0</td><td>CTGGTCGTAGGTTGCAGGAC</td><td>AGG</td><td>||||||||[||||||||||||]PAM</td><td>NA</td><td>Intergenic</td><td>NA</td><td>NA</td></tr>
<tr><td>T13671</td><td>GTCATGAGACTGAATCCCACTGG</td><td>1000</td><td></td><td></td><td></td><td></td><td></td><td></td><td></td><td></td><td></td></tr>
<tr><td>Oligo fwd</td><td>CACCGTCATGAGACTGAATCCCAC</td><td></td><td></td><td></td><td></td><td></td><td></td><td></td><td></td><td></td><td></td></tr>
<tr><td>Oligo rev</td><td>AAACGTGGGATTCAGTCTCATGAC</td><td></td><td></td><td></td><td></td><td></td><td></td><td></td><td></td><td></td><td></td></tr>
<tr><td>Chromosome</td><td>start</td><td>end</td><td>strand</td><td>MM</td><td>target_seq</td><td>PAM</td><td>alignment</td><td>distance</td><td>position</td><td>gene name</td><td>gene id</td></tr>
<tr><td>GU305914.1</td><td>180017</td><td>180039</td><td>+</td><td>0</td><td>GTCATGAGACTGAATCCCAC</td><td>TGG</td><td>||||||||[||||||||||||]PAM</td><td>NA</td><td>Intergenic</td><td>NA</td><td>NA</td></tr>
<tr><td>T13672</td><td>TCATGAGACTGAATCCCACTGGG</td><td>1000</td><td></td><td></td><td></td><td></td><td></td><td></td><td></td><td></td><td></td></tr>
<tr><td>Oligo fwd</td><td>CACCTCATGAGACTGAATCCCACT</td><td></td><td></td><td></td><td></td><td></td><td></td><td></td><td></td><td></td><td></td></tr>
<tr><td>Oligo rev</td><td>AAACAGTGGGATTCAGTCTCATGA</td><td></td><td></td><td></td><td></td><td></td><td></td><td></td><td></td><td></td><td></td></tr>
<tr><td>Chromosome</td><td>start</td><td>end</td><td>strand</td><td>MM</td><td>target_seq</td><td>PAM</td><td>alignment</td><td>distance</td><td>position</td><td>gene name</td><td>gene id</td></tr>
<tr><td>GU305914.1</td><td>180018</td><td>180040</td><td>+</td><td>0</td><td>TCATGAGACTGAATCCCACT</td><td>GGG</td><td>||||||||[||||||||||||]PAM</td><td>NA</td><td>Intergenic</td><td>NA</td><td>NA</td></tr>
<tr><td>T13673</td><td>GACTGAATCCCACTGGGATGCGG</td><td>1000</td><td></td><td></td><td></td><td></td><td></td><td></td><td></td><td></td><td></td></tr>
<tr><td>Oligo fwd</td><td>CACCGACTGAATCCCACTGGGATG</td><td></td><td></td><td></td><td></td><td></td><td></td><td></td><td></td><td></td><td></td></tr>
<tr><td>Oligo rev</td><td>AAACCATCCCAGTGGGATTCAGTC</td><td></td><td></td><td></td><td></td><td></td><td></td><td></td><td></td><td></td><td></td></tr>
<tr><td>Chromosome</td><td>start</td><td>end</td><td>strand</td><td>MM</td><td>target_seq</td><td>PAM</td><td>alignment</td><td>distance</td><td>position</td><td>gene name</td><td>gene id</td></tr>
<tr><td>GU305914.1</td><td>180024</td><td>180046</td><td>+</td><td>0</td><td>GACTGAATCCCACTGGGATG</td><td>CGG</td><td>||||||||[||||||||||||]PAM</td><td>NA</td><td>Intergenic</td><td>NA</td><td>NA</td></tr>
<tr><td>T13674</td><td>ACTGAATCCCACTGGGATGCGGG</td><td>1000</td><td></td><td></td><td></td><td></td><td></td><td></td><td></td><td></td><td></td></tr>
<tr><td>Oligo fwd</td><td>CACCACTGAATCCCACTGGGATGC</td><td></td><td></td><td></td><td></td><td></td><td></td><td></td><td></td><td></td><td></td></tr>
<tr><td>Oligo rev</td><td>AAACGCATCCCAGTGGGATTCAGT</td><td></td><td></td><td></td><td></td><td></td><td></td><td></td><td></td><td></td><td></td></tr>
<tr><td>Chromosome</td><td>start</td><td>end</td><td>strand</td><td>MM</td><td>target_seq</td><td>PAM</td><td>alignment</td><td>distance</td><td>position</td><td>gene name</td><td>gene id</td></tr>
<tr><td>GU305914.1</td><td>180025</td><td>180047</td><td>+</td><td>0</td><td>ACTGAATCCCACTGGGATGC</td><td>GGG</td><td>||||||||[||||||||||||]PAM</td><td>NA</td><td>Intergenic</td><td>NA</td><td>NA</td></tr>
<tr><td>T13675</td><td>TCAGTTTCGCTGTCAGTCAGAGG</td><td>1000</td><td></td><td></td><td></td><td></td><td></td><td></td><td></td><td></td><td></td></tr>
<tr><td>Oligo fwd</td><td>CACCTCAGTTTCGCTGTCAGTCAG</td><td></td><td></td><td></td><td></td><td></td><td></td><td></td><td></td><td></td><td></td></tr>
<tr><td>Oligo rev</td><td>AAACCTGACTGACAGCGAAACTGA</td><td></td><td></td><td></td><td></td><td></td><td></td><td></td><td></td><td></td><td></td></tr>
<tr><td>Chromosome</td><td>start</td><td>end</td><td>strand</td><td>MM</td><td>target_seq</td><td>PAM</td><td>alignment</td><td>distance</td><td>position</td><td>gene name</td><td>gene id</td></tr>
<tr><td>GU305914.1</td><td>180077</td><td>180099</td><td>+</td><td>0</td><td>TCAGTTTCGCTGTCAGTCAG</td><td>AGG</td><td>||||||||[||||||||||||]PAM</td><td>NA</td><td>Intergenic</td><td>NA</td><td>NA</td></tr>
<tr><td>T13676</td><td>CAGTTTCGCTGTCAGTCAGAGGG</td><td>1000</td><td></td><td></td><td></td><td></td><td></td><td></td><td></td><td></td><td></td></tr>
<tr><td>Oligo fwd</td><td>CACCCAGTTTCGCTGTCAGTCAGA</td><td></td><td></td><td></td><td></td><td></td><td></td><td></td><td></td><td></td><td></td></tr>
<tr><td>Oligo rev</td><td>AAACTCTGACTGACAGCGAAACTG</td><td></td><td></td><td></td><td></td><td></td><td></td><td></td><td></td><td></td><td></td></tr>
<tr><td>Chromosome</td><td>start</td><td>end</td><td>strand</td><td>MM</td><td>target_seq</td><td>PAM</td><td>alignment</td><td>distance</td><td>position</td><td>gene name</td><td>gene id</td></tr>
<tr><td>GU305914.1</td><td>180078</td><td>180100</td><td>+</td><td>0</td><td>CAGTTTCGCTGTCAGTCAGA</td><td>GGG</td><td>||||||||[||||||||||||]PAM</td><td>NA</td><td>Intergenic</td><td>NA</td><td>NA</td></tr>
<tr><td>T13677</td><td>AGTTTCGCTGTCAGTCAGAGGGG</td><td>1000</td><td></td><td></td><td></td><td></td><td></td><td></td><td></td><td></td><td></td></tr>
<tr><td>Oligo fwd</td><td>CACCAGTTTCGCTGTCAGTCAGAG</td><td></td><td></td><td></td><td></td><td></td><td></td><td></td><td></td><td></td><td></td></tr>
<tr><td>Oligo rev</td><td>AAACCTCTGACTGACAGCGAAACT</td><td></td><td></td><td></td><td></td><td></td><td></td><td></td><td></td><td></td><td></td></tr>
<tr><td>Chromosome</td><td>start</td><td>end</td><td>strand</td><td>MM</td><td>target_seq</td><td>PAM</td><td>alignment</td><td>distance</td><td>position</td><td>gene name</td><td>gene id</td></tr>
<tr><td>GU305914.1</td><td>180079</td><td>180101</td><td>+</td><td>0</td><td>AGTTTCGCTGTCAGTCAGAG</td><td>GGG</td><td>||||||||[||||||||||||]PAM</td><td>NA</td><td>Intergenic</td><td>NA</td><td>NA</td></tr>
<tr><td>T13678</td><td>CTGTCAGTCAGAGGGGCAGCAGG</td><td>1000</td><td></td><td></td><td></td><td></td><td></td><td></td><td></td><td></td><td></td></tr>
<tr><td>Oligo fwd</td><td>CACCCTGTCAGTCAGAGGGGCAGC</td><td></td><td></td><td></td><td></td><td></td><td></td><td></td><td></td><td></td><td></td></tr>
<tr><td>Oligo rev</td><td>AAACGCTGCCCCTCTGACTGACAG</td><td></td><td></td><td></td><td></td><td></td><td></td><td></td><td></td><td></td><td></td></tr>
<tr><td>Chromosome</td><td>start</td><td>end</td><td>strand</td><td>MM</td><td>target_seq</td><td>PAM</td><td>alignment</td><td>distance</td><td>position</td><td>gene name</td><td>gene id</td></tr>
<tr><td>GU305914.1</td><td>180086</td><td>180108</td><td>+</td><td>0</td><td>CTGTCAGTCAGAGGGGCAGC</td><td>AGG</td><td>||||||||[||||||||||||]PAM</td><td>NA</td><td>Intergenic</td><td>NA</td><td>NA</td></tr>
<tr><td>T13679</td><td>AGTCAGAGGGGCAGCAGGTCTGG</td><td>1000</td><td></td><td></td><td></td><td></td><td></td><td></td><td></td><td></td><td></td></tr>
<tr><td>Oligo fwd</td><td>CACCAGTCAGAGGGGCAGCAGGTC</td><td></td><td></td><td></td><td></td><td></td><td></td><td></td><td></td><td></td><td></td></tr>
<tr><td>Oligo rev</td><td>AAACGACCTGCTGCCCCTCTGACT</td><td></td><td></td><td></td><td></td><td></td><td></td><td></td><td></td><td></td><td></td></tr>
<tr><td>Chromosome</td><td>start</td><td>end</td><td>strand</td><td>MM</td><td>target_seq</td><td>PAM</td><td>alignment</td><td>distance</td><td>position</td><td>gene name</td><td>gene id</td></tr>
<tr><td>GU305914.1</td><td>180091</td><td>180113</td><td>+</td><td>0</td><td>AGTCAGAGGGGCAGCAGGTC</td><td>TGG</td><td>||||||||[||||||||||||]PAM</td><td>NA</td><td>Intergenic</td><td>NA</td><td>NA</td></tr>
<tr><td>T13680</td><td>GAGGGGCAGCAGGTCTGGATAGG</td><td>1000</td><td></td><td></td><td></td><td></td><td></td><td></td><td></td><td></td><td></td></tr>
<tr><td>Oligo fwd</td><td>CACCGAGGGGCAGCAGGTCTGGAT</td><td></td><td></td><td></td><td></td><td></td><td></td><td></td><td></td><td></td><td></td></tr>
<tr><td>Oligo rev</td><td>AAACATCCAGACCTGCTGCCCCTC</td><td></td><td></td><td></td><td></td><td></td><td></td><td></td><td></td><td></td><td></td></tr>
<tr><td>Chromosome</td><td>start</td><td>end</td><td>strand</td><td>MM</td><td>target_seq</td><td>PAM</td><td>alignment</td><td>distance</td><td>position</td><td>gene name</td><td>gene id</td></tr>
<tr><td>GU305914.1</td><td>180096</td><td>180118</td><td>+</td><td>0</td><td>GAGGGGCAGCAGGTCTGGAT</td><td>AGG</td><td>||||||||[||||||||||||]PAM</td><td>NA</td><td>Intergenic</td><td>NA</td><td>NA</td></tr>
<tr><td>T13681</td><td>ATTCTCCTTCATCAGCACTTTGG</td><td>1000</td><td></td><td></td><td></td><td></td><td></td><td></td><td></td><td></td><td></td></tr>
<tr><td>Oligo fwd</td><td>CACCATTCTCCTTCATCAGCACTT</td><td></td><td></td><td></td><td></td><td></td><td></td><td></td><td></td><td></td><td></td></tr>
<tr><td>Oligo rev</td><td>AAACAAGTGCTGATGAAGGAGAAT</td><td></td><td></td><td></td><td></td><td></td><td></td><td></td><td></td><td></td><td></td></tr>
<tr><td>Chromosome</td><td>start</td><td>end</td><td>strand</td><td>MM</td><td>target_seq</td><td>PAM</td><td>alignment</td><td>distance</td><td>position</td><td>gene name</td><td>gene id</td></tr>
<tr><td>GU305914.1</td><td>180150</td><td>180172</td><td>+</td><td>0</td><td>ATTCTCCTTCATCAGCACTT</td><td>TGG</td><td>||||||||[||||||||||||]PAM</td><td>NA</td><td>Intergenic</td><td>NA</td><td>NA</td></tr>
<tr><td>T13682</td><td>TTCTCCTTCATCAGCACTTTGGG</td><td>1000</td><td></td><td></td><td></td><td></td><td></td><td></td><td></td><td></td><td></td></tr>
<tr><td>Oligo fwd</td><td>CACCTTCTCCTTCATCAGCACTTT</td><td></td><td></td><td></td><td></td><td></td><td></td><td></td><td></td><td></td><td></td></tr>
<tr><td>Oligo rev</td><td>AAACAAAGTGCTGATGAAGGAGAA</td><td></td><td></td><td></td><td></td><td></td><td></td><td></td><td></td><td></td><td></td></tr>
<tr><td>Chromosome</td><td>start</td><td>end</td><td>strand</td><td>MM</td><td>target_seq</td><td>PAM</td><td>alignment</td><td>distance</td><td>position</td><td>gene name</td><td>gene id</td></tr>
<tr><td>GU305914.1</td><td>180151</td><td>180173</td><td>+</td><td>0</td><td>TTCTCCTTCATCAGCACTTT</td><td>GGG</td><td>||||||||[||||||||||||]PAM</td><td>NA</td><td>Intergenic</td><td>NA</td><td>NA</td></tr>
<tr><td>T13683</td><td>TGGGTCTCCTCATCCTCAGCAGG</td><td>1000</td><td></td><td></td><td></td><td></td><td></td><td></td><td></td><td></td><td></td></tr>
<tr><td>Oligo fwd</td><td>CACCTGGGTCTCCTCATCCTCAGC</td><td></td><td></td><td></td><td></td><td></td><td></td><td></td><td></td><td></td><td></td></tr>
<tr><td>Oligo rev</td><td>AAACGCTGAGGATGAGGAGACCCA</td><td></td><td></td><td></td><td></td><td></td><td></td><td></td><td></td><td></td><td></td></tr>
<tr><td>Chromosome</td><td>start</td><td>end</td><td>strand</td><td>MM</td><td>target_seq</td><td>PAM</td><td>alignment</td><td>distance</td><td>position</td><td>gene name</td><td>gene id</td></tr>
</table>

<table>
<tr><td>GU305914.1</td><td></td><td>180170</td><td>180192</td><td>+</td><td>0</td><td>TGGGTCTCCTCATCCTCAGC</td><td>AGG</td><td>||||||||[||||||||||||]PAM</td><td>NA</td><td>Intergenic</td><td>NA</td><td>NA</td></tr>
<tr><td>T13684</td><td>TCCTCATCCTCAGCAGGTGTTGG</td><td></td><td>1000</td><td></td><td></td><td></td><td></td><td></td><td></td><td></td><td></td><td></td></tr>
<tr><td>Oligo fwd</td><td>CACCTCCTCATCCTCAGCAGGTGT</td><td></td><td></td><td></td><td></td><td></td><td></td><td></td><td></td><td></td><td></td><td></td></tr>
<tr><td>Oligo rev</td><td>AAACACACCTGCTGAGGATGAGGA</td><td></td><td></td><td></td><td></td><td></td><td></td><td></td><td></td><td></td><td></td><td></td></tr>
<tr><td>Chromosome</td><td>start</td><td></td><td>end</td><td>strand</td><td>MM</td><td>target_seq</td><td>PAM</td><td>alignment</td><td>distance</td><td>position</td><td>gene name</td><td>gene id</td></tr>
<tr><td>GU305914.1</td><td></td><td>180176</td><td>180198</td><td>+</td><td>0</td><td>TCCTCATCCTCAGCAGGTGT</td><td>TGG</td><td>||||||||[||||||||||||]PAM</td><td>NA</td><td>Intergenic</td><td>NA</td><td>NA</td></tr>
<tr><td>T13685</td><td>GTTGGTGTCTCTGAAGACTCAGG</td><td></td><td>1000</td><td></td><td></td><td></td><td></td><td></td><td></td><td></td><td></td><td></td></tr>
<tr><td>Oligo fwd</td><td>CACCGTTGGTGTCTCTGAAGACTC</td><td></td><td></td><td></td><td></td><td></td><td></td><td></td><td></td><td></td><td></td><td></td></tr>
<tr><td>Oligo rev</td><td>AAACGAGTCTTCAGAGACACCAAC</td><td></td><td></td><td></td><td></td><td></td><td></td><td></td><td></td><td></td><td></td><td></td></tr>
<tr><td>Chromosome</td><td>start</td><td></td><td>end</td><td>strand</td><td>MM</td><td>target_seq</td><td>PAM</td><td>alignment</td><td>distance</td><td>position</td><td>gene name</td><td>gene id</td></tr>
<tr><td>GU305914.1</td><td></td><td>180194</td><td>180216</td><td>+</td><td>0</td><td>GTTGGTGTCTCTGAAGACTC</td><td>AGG</td><td>||||||||[||||||||||||]PAM</td><td>NA</td><td>Intergenic</td><td>NA</td><td>NA</td></tr>
<tr><td>T13686</td><td>CTGAAGACTCAGGTACCACCTGG</td><td></td><td>1000</td><td></td><td></td><td></td><td></td><td></td><td></td><td></td><td></td><td></td></tr>
<tr><td>Oligo fwd</td><td>CACCCTGAAGACTCAGGTACCACC</td><td></td><td></td><td></td><td></td><td></td><td></td><td></td><td></td><td></td><td></td><td></td></tr>
<tr><td>Oligo rev</td><td>AAACGGTGGTACCTGAGTCTTCAG</td><td></td><td></td><td></td><td></td><td></td><td></td><td></td><td></td><td></td><td></td><td></td></tr>
<tr><td>Chromosome</td><td>start</td><td></td><td>end</td><td>strand</td><td>MM</td><td>target_seq</td><td>PAM</td><td>alignment</td><td>distance</td><td>position</td><td>gene name</td><td>gene id</td></tr>
<tr><td>GU305914.1</td><td></td><td>180204</td><td>180226</td><td>+</td><td>0</td><td>CTGAAGACTCAGGTACCACC</td><td>TGG</td><td>||||||||[||||||||||||]PAM</td><td>NA</td><td>Intergenic</td><td>NA</td><td>NA</td></tr>
<tr><td>T13687</td><td>GGTACCACCTGGTTGTCACTTGG</td><td></td><td>1000</td><td></td><td></td><td></td><td></td><td></td><td></td><td></td><td></td><td></td></tr>
<tr><td>Oligo fwd</td><td>CACCGGTACCACCTGGTTGTCACT</td><td></td><td></td><td></td><td></td><td></td><td></td><td></td><td></td><td></td><td></td><td></td></tr>
<tr><td>Oligo rev</td><td>AAACAGTGACAACCAGGTGGTACC</td><td></td><td></td><td></td><td></td><td></td><td></td><td></td><td></td><td></td><td></td><td></td></tr>
<tr><td>Chromosome</td><td>start</td><td></td><td>end</td><td>strand</td><td>MM</td><td>target_seq</td><td>PAM</td><td>alignment</td><td>distance</td><td>position</td><td>gene name</td><td>gene id</td></tr>
<tr><td>GU305914.1</td><td></td><td>180215</td><td>180237</td><td>+</td><td>0</td><td>GGTACCACCTGGTTGTCACT</td><td>TGG</td><td>||||||||[||||||||||||]PAM</td><td>NA</td><td>Intergenic</td><td>NA</td><td>NA</td></tr>
<tr><td>T13688</td><td>ACCTGGTTGTCACTTGGTCCAGG</td><td></td><td>1000</td><td></td><td></td><td></td><td></td><td></td><td></td><td></td><td></td><td></td></tr>
<tr><td>Oligo fwd</td><td>CACCACCTGGTTGTCACTTGGTCC</td><td></td><td></td><td></td><td></td><td></td><td></td><td></td><td></td><td></td><td></td><td></td></tr>
<tr><td>Oligo rev</td><td>AAACGGACCAAGTGACAACCAGGT</td><td></td><td></td><td></td><td></td><td></td><td></td><td></td><td></td><td></td><td></td><td></td></tr>
<tr><td>Chromosome</td><td>start</td><td></td><td>end</td><td>strand</td><td>MM</td><td>target_seq</td><td>PAM</td><td>alignment</td><td>distance</td><td>position</td><td>gene name</td><td>gene id</td></tr>
<tr><td>GU305914.1</td><td></td><td>180221</td><td>180243</td><td>+</td><td>0</td><td>ACCTGGTTGTCACTTGGTCC</td><td>AGG</td><td>||||||||[||||||||||||]PAM</td><td>NA</td><td>Intergenic</td><td>NA</td><td>NA</td></tr>
<tr><td>T13689</td><td>GGTTGTCACTTGGTCCAGGCTGG</td><td></td><td>1000</td><td></td><td></td><td></td><td></td><td></td><td></td><td></td><td></td><td></td></tr>
<tr><td>Oligo fwd</td><td>CACCGGTTGTCACTTGGTCCAGGC</td><td></td><td></td><td></td><td></td><td></td><td></td><td></td><td></td><td></td><td></td><td></td></tr>
<tr><td>Oligo rev</td><td>AAACGCCTGGACCAAGTGACAACC</td><td></td><td></td><td></td><td></td><td></td><td></td><td></td><td></td><td></td><td></td><td></td></tr>
<tr><td>Chromosome</td><td>start</td><td></td><td>end</td><td>strand</td><td>MM</td><td>target_seq</td><td>PAM</td><td>alignment</td><td>distance</td><td>position</td><td>gene name</td><td>gene id</td></tr>
<tr><td>GU305914.1</td><td></td><td>180225</td><td>180247</td><td>+</td><td>0</td><td>GGTTGTCACTTGGTCCAGGC</td><td>TGG</td><td>||||||||[||||||||||||]PAM</td><td>NA</td><td>Intergenic</td><td>NA</td><td>NA</td></tr>
<tr><td>T13690</td><td>GTTGTCACTTGGTCCAGGCTGGG</td><td></td><td>1000</td><td></td><td></td><td></td><td></td><td></td><td></td><td></td><td></td><td></td></tr>
<tr><td>Oligo fwd</td><td>CACCGTTGTCACTTGGTCCAGGCT</td><td></td><td></td><td></td><td></td><td></td><td></td><td></td><td></td><td></td><td></td><td></td></tr>
<tr><td>Oligo rev</td><td>AAACAGCCTGGACCAAGTGACAAC</td><td></td><td></td><td></td><td></td><td></td><td></td><td></td><td></td><td></td><td></td><td></td></tr>
<tr><td>Chromosome</td><td>start</td><td></td><td>end</td><td>strand</td><td>MM</td><td>target_seq</td><td>PAM</td><td>alignment</td><td>distance</td><td>position</td><td>gene name</td><td>gene id</td></tr>
<tr><td>GU305914.1</td><td></td><td>180226</td><td>180248</td><td>+</td><td>0</td><td>GTTGTCACTTGGTCCAGGCT</td><td>GGG</td><td>||||||||[||||||||||||]PAM</td><td>NA</td><td>Intergenic</td><td>NA</td><td>NA</td></tr>
<tr><td>T13691</td><td>ATCTTCTTCAGCTTCCTCAGTGG</td><td></td><td>1000</td><td></td><td></td><td></td><td></td><td></td><td></td><td></td><td></td><td></td></tr>
<tr><td>Oligo fwd</td><td>CACCATCTTCTTCAGCTTCCTCAG</td><td></td><td></td><td></td><td></td><td></td><td></td><td></td><td></td><td></td><td></td><td></td></tr>
<tr><td>Oligo rev</td><td>AAACCTGAGGAAGCTGAAGAAGAT</td><td></td><td></td><td></td><td></td><td></td><td></td><td></td><td></td><td></td><td></td><td></td></tr>
<tr><td>Chromosome</td><td>start</td><td></td><td>end</td><td>strand</td><td>MM</td><td>target_seq</td><td>PAM</td><td>alignment</td><td>distance</td><td>position</td><td>gene name</td><td>gene id</td></tr>
<tr><td>GU305914.1</td><td></td><td>180253</td><td>180275</td><td>+</td><td>0</td><td>ATCTTCTTCAGCTTCCTCAG</td><td>TGG</td><td>||||||||[||||||||||||]PAM</td><td>NA</td><td>Intergenic</td><td>NA</td><td>NA</td></tr>
<tr><td>T13692</td><td>TCCTCAGTGGCTTCACCAGCAGG</td><td></td><td>1000</td><td></td><td></td><td></td><td></td><td></td><td></td><td></td><td></td><td></td></tr>
<tr><td>Oligo fwd</td><td>CACCTCCTCAGTGGCTTCACCAGC</td><td></td><td></td><td></td><td></td><td></td><td></td><td></td><td></td><td></td><td></td><td></td></tr>
<tr><td>Oligo rev</td><td>AAACGCTGGTGAAGCCACTGAGGA</td><td></td><td></td><td></td><td></td><td></td><td></td><td></td><td></td><td></td><td></td><td></td></tr>
<tr><td>Chromosome</td><td>start</td><td></td><td>end</td><td>strand</td><td>MM</td><td>target_seq</td><td>PAM</td><td>alignment</td><td>distance</td><td>position</td><td>gene name</td><td>gene id</td></tr>
<tr><td>GU305914.1</td><td></td><td>180266</td><td>180288</td><td>+</td><td>0</td><td>TCCTCAGTGGCTTCACCAGC</td><td>AGG</td><td>||||||||[||||||||||||]PAM</td><td>NA</td><td>Intergenic</td><td>NA</td><td>NA</td></tr>
<tr><td>T13693</td><td>CCTCCTTCCTCCTCATCTGCAGG</td><td></td><td>1000</td><td></td><td></td><td></td><td></td><td></td><td></td><td></td><td></td><td></td></tr>
<tr><td>Oligo fwd</td><td>CACCCCTCCTTCCTCCTCATCTGC</td><td></td><td></td><td></td><td></td><td></td><td></td><td></td><td></td><td></td><td></td><td></td></tr>
<tr><td>Oligo rev</td><td>AAACGCAGATGAGGAGGAAGGAGG</td><td></td><td></td><td></td><td></td><td></td><td></td><td></td><td></td><td></td><td></td><td></td></tr>
<tr><td>Chromosome</td><td>start</td><td></td><td>end</td><td>strand</td><td>MM</td><td>target_seq</td><td>PAM</td><td>alignment</td><td>distance</td><td>position</td><td>gene name</td><td>gene id</td></tr>
<tr><td>GU305914.1</td><td></td><td>180317</td><td>180339</td><td>+</td><td>0</td><td>CCTCCTTCCTCCTCATCTGC</td><td>AGG</td><td>||||||||[||||||||||||]PAM</td><td>NA</td><td>Intergenic</td><td>NA</td><td>NA</td></tr>
<tr><td>T13694</td><td>CTCCTTCCTCCTCATCTGCAGGG</td><td></td><td>1000</td><td></td><td></td><td></td><td></td><td></td><td></td><td></td><td></td><td></td></tr>
<tr><td>Oligo fwd</td><td>CACCCTCCTTCCTCCTCATCTGCA</td><td></td><td></td><td></td><td></td><td></td><td></td><td></td><td></td><td></td><td></td><td></td></tr>
<tr><td>Oligo rev</td><td>AAACTGCAGATGAGGAGGAAGGAG</td><td></td><td></td><td></td><td></td><td></td><td></td><td></td><td></td><td></td><td></td><td></td></tr>
<tr><td>Chromosome</td><td>start</td><td></td><td>end</td><td>strand</td><td>MM</td><td>target_seq</td><td>PAM</td><td>alignment</td><td>distance</td><td>position</td><td>gene name</td><td>gene id</td></tr>
<tr><td>GU305914.1</td><td></td><td>180318</td><td>180340</td><td>+</td><td>0</td><td>CTCCTTCCTCCTCATCTGCA</td><td>GGG</td><td>||||||||[||||||||||||]PAM</td><td>NA</td><td>Intergenic</td><td>NA</td><td>NA</td></tr>
<tr><td>T13695</td><td>TCCTTCCTCCTCATCTGCAGGGG</td><td></td><td>1000</td><td></td><td></td><td></td><td></td><td></td><td></td><td></td><td></td><td></td></tr>
<tr><td>Oligo fwd</td><td>CACCTCCTTCCTCCTCATCTGCAG</td><td></td><td></td><td></td><td></td><td></td><td></td><td></td><td></td><td></td><td></td><td></td></tr>
<tr><td>Oligo rev</td><td>AAACCTGCAGATGAGGAGGAAGGA</td><td></td><td></td><td></td><td></td><td></td><td></td><td></td><td></td><td></td><td></td><td></td></tr>
<tr><td>Chromosome</td><td>start</td><td></td><td>end</td><td>strand</td><td>MM</td><td>target_seq</td><td>PAM</td><td>alignment</td><td>distance</td><td>position</td><td>gene name</td><td>gene id</td></tr>
<tr><td>GU305914.1</td><td></td><td>180319</td><td>180341</td><td>+</td><td>0</td><td>TCCTTCCTCCTCATCTGCAG</td><td>GGG</td><td>||||||||[||||||||||||]PAM</td><td>NA</td><td>Intergenic</td><td>NA</td><td>NA</td></tr>
<tr><td>T13696</td><td>CCTTCCTCCTCATCTGCAGGGGG</td><td></td><td>1000</td><td></td><td></td><td></td><td></td><td></td><td></td><td></td><td></td><td></td></tr>
<tr><td>Oligo fwd</td><td>CACCCCTTCCTCCTCATCTGCAGG</td><td></td><td></td><td></td><td></td><td></td><td></td><td></td><td></td><td></td><td></td><td></td></tr>
<tr><td>Oligo rev</td><td>AAACCCTGCAGATGAGGAGGAAGG</td><td></td><td></td><td></td><td></td><td></td><td></td><td></td><td></td><td></td><td></td><td></td></tr>
<tr><td>Chromosome</td><td>start</td><td></td><td>end</td><td>strand</td><td>MM</td><td>target_seq</td><td>PAM</td><td>alignment</td><td>distance</td><td>position</td><td>gene name</td><td>gene id</td></tr>
<tr><td>GU305914.1</td><td></td><td>180320</td><td>180342</td><td>+</td><td>0</td><td>CCTTCCTCCTCATCTGCAGG</td><td>GGG</td><td>||||||||[||||||||||||]PAM</td><td>NA</td><td>Intergenic</td><td>NA</td><td>NA</td></tr>
<tr><td>T13697</td><td>TGCTCTTCCTCTCTTTCCAGAGG</td><td></td><td>1000</td><td></td><td></td><td></td><td></td><td></td><td></td><td></td><td></td><td></td></tr>
<tr><td>Oligo fwd</td><td>CACCTGCTCTTCCTCTCTTTCCAG</td><td></td><td></td><td></td><td></td><td></td><td></td><td></td><td></td><td></td><td></td><td></td></tr>
<tr><td>Oligo rev</td><td>AAACCTGGAAAGAGAGGAAGAGCA</td><td></td><td></td><td></td><td></td><td></td><td></td><td></td><td></td><td></td><td></td><td></td></tr>
<tr><td>Chromosome</td><td>start</td><td></td><td>end</td><td>strand</td><td>MM</td><td>target_seq</td><td>PAM</td><td>alignment</td><td>distance</td><td>position</td><td>gene name</td><td>gene id</td></tr>
<tr><td>GU305914.1</td><td></td><td>180368</td><td>180390</td><td>+</td><td>0</td><td>TGCTCTTCCTCTCTTTCCAG</td><td>AGG</td><td>||||||||[||||||||||||]PAM</td><td>NA</td><td>Intergenic</td><td>NA</td><td>NA</td></tr>
<tr><td>T13698</td><td>GCTCTTCCTCTCTTTCCAGAGGG</td><td></td><td>1000</td><td></td><td></td><td></td><td></td><td></td><td></td><td></td><td></td><td></td></tr>
<tr><td>Oligo fwd</td><td>CACCGCTCTTCCTCTCTTTCCAGA</td><td></td><td></td><td></td><td></td><td></td><td></td><td></td><td></td><td></td><td></td><td></td></tr>
<tr><td>Oligo rev</td><td>AAACTCTGGAAAGAGAGGAAGAGC</td><td></td><td></td><td></td><td></td><td></td><td></td><td></td><td></td><td></td><td></td><td></td></tr>
</table>

<table>
<tr><td>Chromosome</td><td>start</td><td>end</td><td>strand</td><td>MM</td><td>target_seq</td><td>PAM</td><td>alignment</td><td>distance</td><td>position</td><td>gene name</td><td>gene id</td></tr>
<tr><td>GU305914.1</td><td>180369</td><td>180391</td><td>+</td><td>0</td><td>GCTCTTCCTCTCTTTCCAGA</td><td>GGG</td><td>||||||||[||||||||||||]PAM</td><td>NA</td><td>Intergenic</td><td>NA</td><td>NA</td></tr>
<tr><td>T13699</td><td>CTCTTCCTCTCTTTCCAGAGGGG</td><td>1000</td><td></td><td></td><td></td><td></td><td></td><td></td><td></td><td></td><td></td></tr>
<tr><td>Oligo fwd</td><td>CACCCTCTTCCTCTCTTTCCAGAG</td><td></td><td></td><td></td><td></td><td></td><td></td><td></td><td></td><td></td><td></td></tr>
<tr><td>Oligo rev</td><td>AAACCTCTGGAAAGAGAGGAAGAG</td><td></td><td></td><td></td><td></td><td></td><td></td><td></td><td></td><td></td><td></td></tr>
<tr><td>Chromosome</td><td>start</td><td>end</td><td>strand</td><td>MM</td><td>target_seq</td><td>PAM</td><td>alignment</td><td>distance</td><td>position</td><td>gene name</td><td>gene id</td></tr>
<tr><td>GU305914.1</td><td>180370</td><td>180392</td><td>+</td><td>0</td><td>CTCTTCCTCTCTTTCCAGAG</td><td>GGG</td><td>||||||||[||||||||||||]PAM</td><td>NA</td><td>Intergenic</td><td>NA</td><td>NA</td></tr>
<tr><td>T13700</td><td>TCTTCCTCTCTTTCCAGAGGGGG</td><td>1000</td><td></td><td></td><td></td><td></td><td></td><td></td><td></td><td></td><td></td></tr>
<tr><td>Oligo fwd</td><td>CACCTCTTCCTCTCTTTCCAGAGG</td><td></td><td></td><td></td><td></td><td></td><td></td><td></td><td></td><td></td><td></td></tr>
<tr><td>Oligo rev</td><td>AAACCCTCTGGAAAGAGAGGAAGA</td><td></td><td></td><td></td><td></td><td></td><td></td><td></td><td></td><td></td><td></td></tr>
<tr><td>Chromosome</td><td>start</td><td>end</td><td>strand</td><td>MM</td><td>target_seq</td><td>PAM</td><td>alignment</td><td>distance</td><td>position</td><td>gene name</td><td>gene id</td></tr>
<tr><td>GU305914.1</td><td>180371</td><td>180393</td><td>+</td><td>0</td><td>TCTTCCTCTCTTTCCAGAGG</td><td>GGG</td><td>||||||||[||||||||||||]PAM</td><td>NA</td><td>Intergenic</td><td>NA</td><td>NA</td></tr>
<tr><td>T13701</td><td>CTTTCCAGAGGGGGCATCTCAGG</td><td>1000</td><td></td><td></td><td></td><td></td><td></td><td></td><td></td><td></td><td></td></tr>
<tr><td>Oligo fwd</td><td>CACCCTTTCCAGAGGGGGCATCTC</td><td></td><td></td><td></td><td></td><td></td><td></td><td></td><td></td><td></td><td></td></tr>
<tr><td>Oligo rev</td><td>AAACGAGATGCCCCCTCTGGAAAG</td><td></td><td></td><td></td><td></td><td></td><td></td><td></td><td></td><td></td><td></td></tr>
<tr><td>Chromosome</td><td>start</td><td>end</td><td>strand</td><td>MM</td><td>target_seq</td><td>PAM</td><td>alignment</td><td>distance</td><td>position</td><td>gene name</td><td>gene id</td></tr>
<tr><td>GU305914.1</td><td>180380</td><td>180402</td><td>+</td><td>0</td><td>CTTTCCAGAGGGGGCATCTC</td><td>AGG</td><td>||||||||[||||||||||||]PAM</td><td>NA</td><td>Intergenic</td><td>NA</td><td>NA</td></tr>
<tr><td>T13702</td><td>GCTGTTCTTCTATCACTCCCTGG</td><td>1000</td><td></td><td></td><td></td><td></td><td></td><td></td><td></td><td></td><td></td></tr>
<tr><td>Oligo fwd</td><td>CACCGCTGTTCTTCTATCACTCCC</td><td></td><td></td><td></td><td></td><td></td><td></td><td></td><td></td><td></td><td></td></tr>
<tr><td>Oligo rev</td><td>AAACGGGAGTGATAGAAGAACAGC</td><td></td><td></td><td></td><td></td><td></td><td></td><td></td><td></td><td></td><td></td></tr>
<tr><td>Chromosome</td><td>start</td><td>end</td><td>strand</td><td>MM</td><td>target_seq</td><td>PAM</td><td>alignment</td><td>distance</td><td>position</td><td>gene name</td><td>gene id</td></tr>
<tr><td>GU305914.1</td><td>180453</td><td>180475</td><td>+</td><td>0</td><td>GCTGTTCTTCTATCACTCCC</td><td>TGG</td><td>||||||||[||||||||||||]PAM</td><td>NA</td><td>Intergenic</td><td>NA</td><td>NA</td></tr>
<tr><td>T13703</td><td>CTGTTCTTCTATCACTCCCTGGG</td><td>1000</td><td></td><td></td><td></td><td></td><td></td><td></td><td></td><td></td><td></td></tr>
<tr><td>Oligo fwd</td><td>CACCCTGTTCTTCTATCACTCCCT</td><td></td><td></td><td></td><td></td><td></td><td></td><td></td><td></td><td></td><td></td></tr>
<tr><td>Oligo rev</td><td>AAACAGGGAGTGATAGAAGAACAG</td><td></td><td></td><td></td><td></td><td></td><td></td><td></td><td></td><td></td><td></td></tr>
<tr><td>Chromosome</td><td>start</td><td>end</td><td>strand</td><td>MM</td><td>target_seq</td><td>PAM</td><td>alignment</td><td>distance</td><td>position</td><td>gene name</td><td>gene id</td></tr>
<tr><td>GU305914.1</td><td>180454</td><td>180476</td><td>+</td><td>0</td><td>CTGTTCTTCTATCACTCCCT</td><td>GGG</td><td>||||||||[||||||||||||]PAM</td><td>NA</td><td>Intergenic</td><td>NA</td><td>NA</td></tr>
<tr><td>T13704</td><td>TGTTCTTCTATCACTCCCTGGGG</td><td>1000</td><td></td><td></td><td></td><td></td><td></td><td></td><td></td><td></td><td></td></tr>
<tr><td>Oligo fwd</td><td>CACCTGTTCTTCTATCACTCCCTG</td><td></td><td></td><td></td><td></td><td></td><td></td><td></td><td></td><td></td><td></td></tr>
<tr><td>Oligo rev</td><td>AAACCAGGGAGTGATAGAAGAACA</td><td></td><td></td><td></td><td></td><td></td><td></td><td></td><td></td><td></td><td></td></tr>
<tr><td>Chromosome</td><td>start</td><td>end</td><td>strand</td><td>MM</td><td>target_seq</td><td>PAM</td><td>alignment</td><td>distance</td><td>position</td><td>gene name</td><td>gene id</td></tr>
<tr><td>GU305914.1</td><td>180455</td><td>180477</td><td>+</td><td>0</td><td>TGTTCTTCTATCACTCCCTG</td><td>GGG</td><td>||||||||[||||||||||||]PAM</td><td>NA</td><td>Intergenic</td><td>NA</td><td>NA</td></tr>
<tr><td>T13705</td><td>TCTATCACTCCCTGGGGTGATGG</td><td>1000</td><td></td><td></td><td></td><td></td><td></td><td></td><td></td><td></td><td></td></tr>
<tr><td>Oligo fwd</td><td>CACCTCTATCACTCCCTGGGGTGA</td><td></td><td></td><td></td><td></td><td></td><td></td><td></td><td></td><td></td><td></td></tr>
<tr><td>Oligo rev</td><td>AAACTCACCCCAGGGAGTGATAGA</td><td></td><td></td><td></td><td></td><td></td><td></td><td></td><td></td><td></td><td></td></tr>
<tr><td>Chromosome</td><td>start</td><td>end</td><td>strand</td><td>MM</td><td>target_seq</td><td>PAM</td><td>alignment</td><td>distance</td><td>position</td><td>gene name</td><td>gene id</td></tr>
<tr><td>GU305914.1</td><td>180461</td><td>180483</td><td>+</td><td>0</td><td>TCTATCACTCCCTGGGGTGA</td><td>TGG</td><td>||||||||[||||||||||||]PAM</td><td>NA</td><td>Intergenic</td><td>NA</td><td>NA</td></tr>
<tr><td>T13706</td><td>GTGATGGTGCACTCCTGTTATGG</td><td>1000</td><td></td><td></td><td></td><td></td><td></td><td></td><td></td><td></td><td></td></tr>
<tr><td>Oligo fwd</td><td>CACCGTGATGGTGCACTCCTGTTA</td><td></td><td></td><td></td><td></td><td></td><td></td><td></td><td></td><td></td><td></td></tr>
<tr><td>Oligo rev</td><td>AAACTAACAGGAGTGCACCATCAC</td><td></td><td></td><td></td><td></td><td></td><td></td><td></td><td></td><td></td><td></td></tr>
<tr><td>Chromosome</td><td>start</td><td>end</td><td>strand</td><td>MM</td><td>target_seq</td><td>PAM</td><td>alignment</td><td>distance</td><td>position</td><td>gene name</td><td>gene id</td></tr>
<tr><td>GU305914.1</td><td>180477</td><td>180499</td><td>+</td><td>0</td><td>GTGATGGTGCACTCCTGTTA</td><td>TGG</td><td>||||||||[||||||||||||]PAM</td><td>NA</td><td>Intergenic</td><td>NA</td><td>NA</td></tr>
<tr><td>T13707</td><td>ATCAGCTAGCTCTCTGAACTTGG</td><td>1000</td><td></td><td></td><td></td><td></td><td></td><td></td><td></td><td></td><td></td></tr>
<tr><td>Oligo fwd</td><td>CACCATCAGCTAGCTCTCTGAACT</td><td></td><td></td><td></td><td></td><td></td><td></td><td></td><td></td><td></td><td></td></tr>
<tr><td>Oligo rev</td><td>AAACAGTTCAGAGAGCTAGCTGAT</td><td></td><td></td><td></td><td></td><td></td><td></td><td></td><td></td><td></td><td></td></tr>
<tr><td>Chromosome</td><td>start</td><td>end</td><td>strand</td><td>MM</td><td>target_seq</td><td>PAM</td><td>alignment</td><td>distance</td><td>position</td><td>gene name</td><td>gene id</td></tr>
<tr><td>GU305914.1</td><td>180505</td><td>180527</td><td>+</td><td>0</td><td>ATCAGCTAGCTCTCTGAACT</td><td>TGG</td><td>||||||||[||||||||||||]PAM</td><td>NA</td><td>Intergenic</td><td>NA</td><td>NA</td></tr>
<tr><td>T13708</td><td>TCTGAACTTGGCCAGAGAGATGG</td><td>1000</td><td></td><td></td><td></td><td></td><td></td><td></td><td></td><td></td><td></td></tr>
<tr><td>Oligo fwd</td><td>CACCTCTGAACTTGGCCAGAGAGA</td><td></td><td></td><td></td><td></td><td></td><td></td><td></td><td></td><td></td><td></td></tr>
<tr><td>Oligo rev</td><td>AAACTCTCTCTGGCCAAGTTCAGA</td><td></td><td></td><td></td><td></td><td></td><td></td><td></td><td></td><td></td><td></td></tr>
<tr><td>Chromosome</td><td>start</td><td>end</td><td>strand</td><td>MM</td><td>target_seq</td><td>PAM</td><td>alignment</td><td>distance</td><td>position</td><td>gene name</td><td>gene id</td></tr>
<tr><td>GU305914.1</td><td>180517</td><td>180539</td><td>+</td><td>0</td><td>TCTGAACTTGGCCAGAGAGA</td><td>TGG</td><td>||||||||[||||||||||||]PAM</td><td>NA</td><td>Intergenic</td><td>NA</td><td>NA</td></tr>
<tr><td>T13709</td><td>TTCATAGCACCTCTGATCCTAGG</td><td>1000</td><td></td><td></td><td></td><td></td><td></td><td></td><td></td><td></td><td></td></tr>
<tr><td>Oligo fwd</td><td>CACCTTCATAGCACCTCTGATCCT</td><td></td><td></td><td></td><td></td><td></td><td></td><td></td><td></td><td></td><td></td></tr>
<tr><td>Oligo rev</td><td>AAACAGGATCAGAGGTGCTATGAA</td><td></td><td></td><td></td><td></td><td></td><td></td><td></td><td></td><td></td><td></td></tr>
<tr><td>Chromosome</td><td>start</td><td>end</td><td>strand</td><td>MM</td><td>target_seq</td><td>PAM</td><td>alignment</td><td>distance</td><td>position</td><td>gene name</td><td>gene id</td></tr>
<tr><td>GU305914.1</td><td>180572</td><td>180594</td><td>+</td><td>0</td><td>TTCATAGCACCTCTGATCCT</td><td>AGG</td><td>||||||||[||||||||||||]PAM</td><td>NA</td><td>Intergenic</td><td>NA</td><td>NA</td></tr>
<tr><td>T13710</td><td>AGCACCTCTGATCCTAGGCATGG</td><td>1000</td><td></td><td></td><td></td><td></td><td></td><td></td><td></td><td></td><td></td></tr>
<tr><td>Oligo fwd</td><td>CACCAGCACCTCTGATCCTAGGCA</td><td></td><td></td><td></td><td></td><td></td><td></td><td></td><td></td><td></td><td></td></tr>
<tr><td>Oligo rev</td><td>AAACTGCCTAGGATCAGAGGTGCT</td><td></td><td></td><td></td><td></td><td></td><td></td><td></td><td></td><td></td><td></td></tr>
<tr><td>Chromosome</td><td>start</td><td>end</td><td>strand</td><td>MM</td><td>target_seq</td><td>PAM</td><td>alignment</td><td>distance</td><td>position</td><td>gene name</td><td>gene id</td></tr>
<tr><td>GU305914.1</td><td>180577</td><td>180599</td><td>+</td><td>0</td><td>AGCACCTCTGATCCTAGGCA</td><td>TGG</td><td>||||||||[||||||||||||]PAM</td><td>NA</td><td>Intergenic</td><td>NA</td><td>NA</td></tr>
<tr><td>T13711</td><td>AAAAAGTGCTAGATTGTATCTGG</td><td>1000</td><td></td><td></td><td></td><td></td><td></td><td></td><td></td><td></td><td></td></tr>
<tr><td>Oligo fwd</td><td>CACCAAAAAGTGCTAGATTGTATC</td><td></td><td></td><td></td><td></td><td></td><td></td><td></td><td></td><td></td><td></td></tr>
<tr><td>Oligo rev</td><td>AAACGATACAATCTAGCACTTTTT</td><td></td><td></td><td></td><td></td><td></td><td></td><td></td><td></td><td></td><td></td></tr>
<tr><td>Chromosome</td><td>start</td><td>end</td><td>strand</td><td>MM</td><td>target_seq</td><td>PAM</td><td>alignment</td><td>distance</td><td>position</td><td>gene name</td><td>gene id</td></tr>
<tr><td>GU305914.1</td><td>180634</td><td>180656</td><td>+</td><td>0</td><td>AAAAAGTGCTAGATTGTATC</td><td>TGG</td><td>||||||||[||||||||||||]PAM</td><td>NA</td><td>Intergenic</td><td>NA</td><td>NA</td></tr>
<tr><td>T13712</td><td>GGTGCTCCTCAGATCAGCTAAGG</td><td>1000</td><td></td><td></td><td></td><td></td><td></td><td></td><td></td><td></td><td></td></tr>
<tr><td>Oligo fwd</td><td>CACCGGTGCTCCTCAGATCAGCTA</td><td></td><td></td><td></td><td></td><td></td><td></td><td></td><td></td><td></td><td></td></tr>
<tr><td>Oligo rev</td><td>AAACTAGCTGATCTGAGGAGCACC</td><td></td><td></td><td></td><td></td><td></td><td></td><td></td><td></td><td></td><td></td></tr>
<tr><td>Chromosome</td><td>start</td><td>end</td><td>strand</td><td>MM</td><td>target_seq</td><td>PAM</td><td>alignment</td><td>distance</td><td>position</td><td>gene name</td><td>gene id</td></tr>
<tr><td>GU305914.1</td><td>180655</td><td>180677</td><td>+</td><td>0</td><td>GGTGCTCCTCAGATCAGCTA</td><td>AGG</td><td>||||||||[||||||||||||]PAM</td><td>NA</td><td>Intergenic</td><td>NA</td><td>NA</td></tr>
<tr><td>T13713</td><td>CAGATCAGCTAAGGTAGCCAAGG</td><td>1000</td><td></td><td></td><td></td><td></td><td></td><td></td><td></td><td></td><td></td></tr>
<tr><td>Oligo fwd</td><td>CACCCAGATCAGCTAAGGTAGCCA</td><td></td><td></td><td></td><td></td><td></td><td></td><td></td><td></td><td></td><td></td></tr>
</table>

| | | | | | | | | | | | | |
|---|---|---|---|---|---|---|---|---|---|---|---|---|
| Oligo rev | AAACTGGCTACCTTAGCTGATCTG | | | | | | | | | | | |
| Chromosome | start | | end | strand | MM | target_seq | PAM | alignment | distance | position | gene name | gene id |
| GU305914.1 | | 180664 | 180686 | + | 0 | CAGATCAGCTAAGGTAGCCA | AGG | ||||||||[||||||||||||]PAM | NA | Intergenic | NA | NA |
| T13714 | GGTAGCCAAGGTTCTGCCTCTGG | | 1000 | | | | | | | | | |
| Oligo fwd | CACCGGTAGCCAAGGTTCTGCCTC | | | | | | | | | | | |
| Oligo rev | AAACGAGGCAGAACCTTGGCTACC | | | | | | | | | | | |
| Chromosome | start | | end | strand | MM | target_seq | PAM | alignment | distance | position | gene name | gene id |
| GU305914.1 | | 180676 | 180698 | + | 0 | GGTAGCCAAGGTTCTGCCTC | TGG | ||||||||[||||||||||||]PAM | NA | Intergenic | NA | NA |
| T13715 | TCTGCCTCTGGCCAGCATACAGG | | 1000 | | | | | | | | | |
| Oligo fwd | CACCTCTGCCTCTGGCCAGCATAC | | | | | | | | | | | |
| Oligo rev | AAACGTATGCTGGCCAGAGGCAGA | | | | | | | | | | | |
| Chromosome | start | | end | strand | MM | target_seq | PAM | alignment | distance | position | gene name | gene id |
| GU305914.1 | | 180688 | 180710 | + | 0 | TCTGCCTCTGGCCAGCATAC | AGG | ||||||||[||||||||||||]PAM | NA | Intergenic | NA | NA |
| T13716 | GCCTCTGGCCAGCATACAGGTGG | | 1000 | | | | | | | | | |
| Oligo fwd | CACCGCCTCTGGCCAGCATACAGG | | | | | | | | | | | |
| Oligo rev | AAACCCTGTATGCTGGCCAGAGGC | | | | | | | | | | | |
| Chromosome | start | | end | strand | MM | target_seq | PAM | alignment | distance | position | gene name | gene id |
| GU305914.1 | | 180691 | 180713 | + | 0 | GCCTCTGGCCAGCATACAGG | TGG | ||||||||[||||||||||||]PAM | NA | Intergenic | NA | NA |
| T13717 | GCATACAGGTGGTCTCTAGATGG | | 1000 | | | | | | | | | |
| Oligo fwd | CACCGCATACAGGTGGTCTCTAGA | | | | | | | | | | | |
| Oligo rev | AAACTCTAGAGACCACCTGTATGC | | | | | | | | | | | |
| Chromosome | start | | end | strand | MM | target_seq | PAM | alignment | distance | position | gene name | gene id |
| GU305914.1 | | 180702 | 180724 | + | 0 | GCATACAGGTGGTCTCTAGA | TGG | ||||||||[||||||||||||]PAM | NA | Intergenic | NA | NA |
| T13718 | TCTGACACACCCTGTGATATTGG | | 1000 | | | | | | | | | |
| Oligo fwd | CACCTCTGACACACCCTGTGATAT | | | | | | | | | | | |
| Oligo rev | AAACATATCACAGGGTGTGTCAGA | | | | | | | | | | | |
| Chromosome | start | | end | strand | MM | target_seq | PAM | alignment | distance | position | gene name | gene id |
| GU305914.1 | | 180747 | 180769 | + | 0 | TCTGACACACCCTGTGATAT | TGG | ||||||||[||||||||||||]PAM | NA | Intergenic | NA | NA |
| T13719 | GTGATATTGGTGTCTCTCATAGG | | 1000 | | | | | | | | | |
| Oligo fwd | CACCGTGATATTGGTGTCTCTCAT | | | | | | | | | | | |
| Oligo rev | AAACATGAGAGACACCAATATCAC | | | | | | | | | | | |
| Chromosome | start | | end | strand | MM | target_seq | PAM | alignment | distance | position | gene name | gene id |
| GU305914.1 | | 180760 | 180782 | + | 0 | GTGATATTGGTGTCTCTCAT | AGG | ||||||||[||||||||||||]PAM | NA | Intergenic | NA | NA |
| T13720 | TTCCACCACTACCACATGTGTGG | | 1000 | | | | | | | | | |
| Oligo fwd | CACCTTCCACCACTACCACATGTG | | | | | | | | | | | |
| Oligo rev | AAACCACATGTGGTAGTGGTGGAA | | | | | | | | | | | |
| Chromosome | start | | end | strand | MM | target_seq | PAM | alignment | distance | position | gene name | gene id |
| GU305914.1 | | 180832 | 180854 | + | 0 | TTCCACCACTACCACATGTG | TGG | ||||||||[||||||||||||]PAM | NA | Intergenic | NA | NA |
| T13721 | CCATACTGCCAGCTGAGACCTGG | | 1000 | | | | | | | | | |
| Oligo fwd | CACCCCATACTGCCAGCTGAGACC | | | | | | | | | | | |
| Oligo rev | AAACGGTCTCAGCTGGCAGTATGG | | | | | | | | | | | |
| Chromosome | start | | end | strand | MM | target_seq | PAM | alignment | distance | position | gene name | gene id |
| GU305914.1 | | 180889 | 180911 | + | 0 | CCATACTGCCAGCTGAGACC | TGG | ||||||||[||||||||||||]PAM | NA | Intergenic | NA | NA |
| T13722 | CCAGCTGAGACCTGGCATCCAGG | | 1000 | | | | | | | | | |
| Oligo fwd | CACCCCAGCTGAGACCTGGCATCC | | | | | | | | | | | |
| Oligo rev | AAACGGATGCCAGGTCTCAGCTGG | | | | | | | | | | | |
| Chromosome | start | | end | strand | MM | target_seq | PAM | alignment | distance | position | gene name | gene id |
| GU305914.1 | | 180897 | 180919 | + | 0 | CCAGCTGAGACCTGGCATCC | AGG | ||||||||[||||||||||||]PAM | NA | Intergenic | NA | NA |
| T13723 | TGAGACCTGGCATCCAGGAAAGG | | 1000 | | | | | | | | | |
| Oligo fwd | CACCTGAGACCTGGCATCCAGGAA | | | | | | | | | | | |
| Oligo rev | AAACTTCCTGGATGCCAGGTCTCA | | | | | | | | | | | |
| Chromosome | start | | end | strand | MM | target_seq | PAM | alignment | distance | position | gene name | gene id |
| GU305914.1 | | 180902 | 180924 | + | 0 | TGAGACCTGGCATCCAGGAA | AGG | ||||||||[||||||||||||]PAM | NA | Intergenic | NA | NA |
| T13724 | CCTGGCATCCAGGAAAGGCTTGG | | 1000 | | | | | | | | | |
| Oligo fwd | CACCCCTGGCATCCAGGAAAGGCT | | | | | | | | | | | |
| Oligo rev | AAACAGCCTTTCCTGGATGCCAGG | | | | | | | | | | | |
| Chromosome | start | | end | strand | MM | target_seq | PAM | alignment | distance | position | gene name | gene id |
| GU305914.1 | | 180907 | 180929 | + | 0 | CCTGGCATCCAGGAAAGGCT | TGG | ||||||||[||||||||||||]PAM | NA | Intergenic | NA | NA |
| T13725 | CAGGAAAGGCTTGGTGTAACAGG | | 1000 | | | | | | | | | |
| Oligo fwd | CACCCAGGAAAGGCTTGGTGTAAC | | | | | | | | | | | |
| Oligo rev | AAACGTTACACCAAGCCTTTCCTG | | | | | | | | | | | |
| Chromosome | start | | end | strand | MM | target_seq | PAM | alignment | distance | position | gene name | gene id |
| GU305914.1 | | 180916 | 180938 | + | 0 | CAGGAAAGGCTTGGTGTAAC | AGG | ||||||||[||||||||||||]PAM | NA | Intergenic | NA | NA |
| T13726 | AGGAAAGGCTTGGTGTAACAGGG | | 1000 | | | | | | | | | |
| Oligo fwd | CACCAGGAAAGGCTTGGTGTAACA | | | | | | | | | | | |
| Oligo rev | AAACTGTTACACCAAGCCTTTCCT | | | | | | | | | | | |
| Chromosome | start | | end | strand | MM | target_seq | PAM | alignment | distance | position | gene name | gene id |
| GU305914.1 | | 180917 | 180939 | + | 0 | AGGAAAGGCTTGGTGTAACA | GGG | ||||||||[||||||||||||]PAM | NA | Intergenic | NA | NA |
| T13727 | AAAGGCTTGGTGTAACAGGGTGG | | 1000 | | | | | | | | | |
| Oligo fwd | CACCAAAGGCTTGGTGTAACAGGG | | | | | | | | | | | |
| Oligo rev | AAACCCCTGTTACACCAAGCCTTT | | | | | | | | | | | |
| Chromosome | start | | end | strand | MM | target_seq | PAM | alignment | distance | position | gene name | gene id |
| GU305914.1 | | 180920 | 180942 | + | 0 | AAAGGCTTGGTGTAACAGGG | TGG | ||||||||[||||||||||||]PAM | NA | Intergenic | NA | NA |
| T13728 | TGGATCATGAAGCCTCCTGATGG | | 1000 | | | | | | | | | |

<table>
<tr><td>Oligo fwd</td><td>CACCTGGATCATGAAGCCTCCTGA</td><td></td><td></td><td></td><td></td><td></td><td></td><td></td><td></td><td></td><td></td></tr>
<tr><td>Oligo rev</td><td>AAACTCAGGAGGCTTCATGATCCA</td><td></td><td></td><td></td><td></td><td></td><td></td><td></td><td></td><td></td><td></td></tr>
<tr><td>Chromosome</td><td>start</td><td>end</td><td>strand</td><td>MM</td><td>target_seq</td><td>PAM</td><td>alignment</td><td>distance</td><td>position</td><td>gene name</td><td>gene id</td></tr>
<tr><td>GU305914.1</td><td>180940</td><td>180962</td><td>+</td><td>0</td><td>TGGATCATGAAGCCTCCTGA</td><td>TGG</td><td>||||||||[||||||||||||]PAM</td><td>NA</td><td>Intergenic</td><td>NA</td><td>NA</td></tr>
<tr><td>T13729</td><td>GCCTCCTGATGGCCCCAACCAGG</td><td>1000</td><td></td><td></td><td></td><td></td><td></td><td></td><td></td><td></td><td></td></tr>
<tr><td>Oligo fwd</td><td>CACCGCCTCCTGATGGCCCCAACC</td><td></td><td></td><td></td><td></td><td></td><td></td><td></td><td></td><td></td><td></td></tr>
<tr><td>Oligo rev</td><td>AAACGGTTGGGGCCATCAGGAGGC</td><td></td><td></td><td></td><td></td><td></td><td></td><td></td><td></td><td></td><td></td></tr>
<tr><td>Chromosome</td><td>start</td><td>end</td><td>strand</td><td>MM</td><td>target_seq</td><td>PAM</td><td>alignment</td><td>distance</td><td>position</td><td>gene name</td><td>gene id</td></tr>
<tr><td>GU305914.1</td><td>180951</td><td>180973</td><td>+</td><td>0</td><td>GCCTCCTGATGGCCCCAACC</td><td>AGG</td><td>||||||||[||||||||||||]PAM</td><td>NA</td><td>Intergenic</td><td>NA</td><td>NA</td></tr>
<tr><td>T13730</td><td>ATCTCTGACTTAAACTCCCCAGG</td><td>1000</td><td></td><td></td><td></td><td></td><td></td><td></td><td></td><td></td><td></td></tr>
<tr><td>Oligo fwd</td><td>CACCATCTCTGACTTAAACTCCCC</td><td></td><td></td><td></td><td></td><td></td><td></td><td></td><td></td><td></td><td></td></tr>
<tr><td>Oligo rev</td><td>AAACGGGGAGTTTAAGTCAGAGAT</td><td></td><td></td><td></td><td></td><td></td><td></td><td></td><td></td><td></td><td></td></tr>
<tr><td>Chromosome</td><td>start</td><td>end</td><td>strand</td><td>MM</td><td>target_seq</td><td>PAM</td><td>alignment</td><td>distance</td><td>position</td><td>gene name</td><td>gene id</td></tr>
<tr><td>GU305914.1</td><td>180980</td><td>181002</td><td>+</td><td>0</td><td>ATCTCTGACTTAAACTCCCC</td><td>AGG</td><td>||||||||[||||||||||||]PAM</td><td>NA</td><td>Intergenic</td><td>NA</td><td>NA</td></tr>
<tr><td>T13731</td><td>AGGCAATGAACAATCAGTCTTGG</td><td>1000</td><td></td><td></td><td></td><td></td><td></td><td></td><td></td><td></td><td></td></tr>
<tr><td>Oligo fwd</td><td>CACCAGGCAATGAACAATCAGTCT</td><td></td><td></td><td></td><td></td><td></td><td></td><td></td><td></td><td></td><td></td></tr>
<tr><td>Oligo rev</td><td>AAACAGACTGATTGTTCATTGCCT</td><td></td><td></td><td></td><td></td><td></td><td></td><td></td><td></td><td></td><td></td></tr>
<tr><td>Chromosome</td><td>start</td><td>end</td><td>strand</td><td>MM</td><td>target_seq</td><td>PAM</td><td>alignment</td><td>distance</td><td>position</td><td>gene name</td><td>gene id</td></tr>
<tr><td>GU305914.1</td><td>181000</td><td>181022</td><td>+</td><td>0</td><td>AGGCAATGAACAATCAGTCT</td><td>TGG</td><td>||||||||[||||||||||||]PAM</td><td>NA</td><td>Intergenic</td><td>NA</td><td>NA</td></tr>
<tr><td>T13732</td><td>ACAATCAGTCTTGGCCCATGCGG</td><td>1000</td><td></td><td></td><td></td><td></td><td></td><td></td><td></td><td></td><td></td></tr>
<tr><td>Oligo fwd</td><td>CACCACAATCAGTCTTGGCCCATG</td><td></td><td></td><td></td><td></td><td></td><td></td><td></td><td></td><td></td><td></td></tr>
<tr><td>Oligo rev</td><td>AAACCATGGGCCAAGACTGATTGT</td><td></td><td></td><td></td><td></td><td></td><td></td><td></td><td></td><td></td><td></td></tr>
<tr><td>Chromosome</td><td>start</td><td>end</td><td>strand</td><td>MM</td><td>target_seq</td><td>PAM</td><td>alignment</td><td>distance</td><td>position</td><td>gene name</td><td>gene id</td></tr>
<tr><td>GU305914.1</td><td>181009</td><td>181031</td><td>+</td><td>0</td><td>ACAATCAGTCTTGGCCCATG</td><td>CGG</td><td>||||||||[||||||||||||]PAM</td><td>NA</td><td>Intergenic</td><td>NA</td><td>NA</td></tr>
<tr><td>T13733</td><td>CGGCACGCTCATCTAGTGCGTGG</td><td>1000</td><td></td><td></td><td></td><td></td><td></td><td></td><td></td><td></td><td></td></tr>
<tr><td>Oligo fwd</td><td>CACCCGGCACGCTCATCTAGTGCG</td><td></td><td></td><td></td><td></td><td></td><td></td><td></td><td></td><td></td><td></td></tr>
<tr><td>Oligo rev</td><td>AAACCGCACTAGATGAGCGTGCCG</td><td></td><td></td><td></td><td></td><td></td><td></td><td></td><td></td><td></td><td></td></tr>
<tr><td>Chromosome</td><td>start</td><td>end</td><td>strand</td><td>MM</td><td>target_seq</td><td>PAM</td><td>alignment</td><td>distance</td><td>position</td><td>gene name</td><td>gene id</td></tr>
<tr><td>GU305914.1</td><td>181029</td><td>181051</td><td>+</td><td>0</td><td>CGGCACGCTCATCTAGTGCG</td><td>TGG</td><td>||||||||[||||||||||||]PAM</td><td>NA</td><td>Intergenic</td><td>NA</td><td>NA</td></tr>
<tr><td>T13734</td><td>ATCTAGTGCGTGGTCAATGTTGG</td><td>1000</td><td></td><td></td><td></td><td></td><td></td><td></td><td></td><td></td><td></td></tr>
<tr><td>Oligo fwd</td><td>CACCATCTAGTGCGTGGTCAATGT</td><td></td><td></td><td></td><td></td><td></td><td></td><td></td><td></td><td></td><td></td></tr>
<tr><td>Oligo rev</td><td>AAACACATTGACCACGCACTAGAT</td><td></td><td></td><td></td><td></td><td></td><td></td><td></td><td></td><td></td><td></td></tr>
<tr><td>Chromosome</td><td>start</td><td>end</td><td>strand</td><td>MM</td><td>target_seq</td><td>PAM</td><td>alignment</td><td>distance</td><td>position</td><td>gene name</td><td>gene id</td></tr>
<tr><td>GU305914.1</td><td>181039</td><td>181061</td><td>+</td><td>0</td><td>ATCTAGTGCGTGGTCAATGT</td><td>TGG</td><td>||||||||[||||||||||||]PAM</td><td>NA</td><td>Intergenic</td><td>NA</td><td>NA</td></tr>
<tr><td>T13735</td><td>TAGTGCGTGGTCAATGTTGGTGG</td><td>1000</td><td></td><td></td><td></td><td></td><td></td><td></td><td></td><td></td><td></td></tr>
<tr><td>Oligo fwd</td><td>CACCTAGTGCGTGGTCAATGTTGG</td><td></td><td></td><td></td><td></td><td></td><td></td><td></td><td></td><td></td><td></td></tr>
<tr><td>Oligo rev</td><td>AAACCCAACATTGACCACGCACTA</td><td></td><td></td><td></td><td></td><td></td><td></td><td></td><td></td><td></td><td></td></tr>
<tr><td>Chromosome</td><td>start</td><td>end</td><td>strand</td><td>MM</td><td>target_seq</td><td>PAM</td><td>alignment</td><td>distance</td><td>position</td><td>gene name</td><td>gene id</td></tr>
<tr><td>GU305914.1</td><td>181042</td><td>181064</td><td>+</td><td>0</td><td>TAGTGCGTGGTCAATGTTGG</td><td>TGG</td><td>||||||||[||||||||||||]PAM</td><td>NA</td><td>Intergenic</td><td>NA</td><td>NA</td></tr>
<tr><td>T13736</td><td>AATGTTGGTGGCTGCCACTATGG</td><td>1000</td><td></td><td></td><td></td><td></td><td></td><td></td><td></td><td></td><td></td></tr>
<tr><td>Oligo fwd</td><td>CACCAATGTTGGTGGCTGCCACTA</td><td></td><td></td><td></td><td></td><td></td><td></td><td></td><td></td><td></td><td></td></tr>
<tr><td>Oligo rev</td><td>AAACTAGTGGCAGCCACCAACATT</td><td></td><td></td><td></td><td></td><td></td><td></td><td></td><td></td><td></td><td></td></tr>
<tr><td>Chromosome</td><td>start</td><td>end</td><td>strand</td><td>MM</td><td>target_seq</td><td>PAM</td><td>alignment</td><td>distance</td><td>position</td><td>gene name</td><td>gene id</td></tr>
<tr><td>GU305914.1</td><td>181054</td><td>181076</td><td>+</td><td>0</td><td>AATGTTGGTGGCTGCCACTA</td><td>TGG</td><td>||||||||[||||||||||||]PAM</td><td>NA</td><td>Intergenic</td><td>NA</td><td>NA</td></tr>
<tr><td>T13737</td><td>CATCCAGACTCTCTTTTCTGAGG</td><td>1000</td><td></td><td></td><td></td><td></td><td></td><td></td><td></td><td></td><td></td></tr>
<tr><td>Oligo fwd</td><td>CACCCATCCAGACTCTCTTTTCTG</td><td></td><td></td><td></td><td></td><td></td><td></td><td></td><td></td><td></td><td></td></tr>
<tr><td>Oligo rev</td><td>AAACCAGAAAAGAGAGTCTGGATG</td><td></td><td></td><td></td><td></td><td></td><td></td><td></td><td></td><td></td><td></td></tr>
<tr><td>Chromosome</td><td>start</td><td>end</td><td>strand</td><td>MM</td><td>target_seq</td><td>PAM</td><td>alignment</td><td>distance</td><td>position</td><td>gene name</td><td>gene id</td></tr>
<tr><td>GU305914.1</td><td>181093</td><td>181115</td><td>+</td><td>0</td><td>CATCCAGACTCTCTTTTCTG</td><td>AGG</td><td>||||||||[||||||||||||]PAM</td><td>NA</td><td>Intergenic</td><td>NA</td><td>NA</td></tr>
<tr><td>T13738</td><td>ATCCAGACTCTCTTTTCTGAGGG</td><td>1000</td><td></td><td></td><td></td><td></td><td></td><td></td><td></td><td></td><td></td></tr>
<tr><td>Oligo fwd</td><td>CACCATCCAGACTCTCTTTTCTGA</td><td></td><td></td><td></td><td></td><td></td><td></td><td></td><td></td><td></td><td></td></tr>
<tr><td>Oligo rev</td><td>AAACTCAGAAAAGAGAGTCTGGAT</td><td></td><td></td><td></td><td></td><td></td><td></td><td></td><td></td><td></td><td></td></tr>
<tr><td>Chromosome</td><td>start</td><td>end</td><td>strand</td><td>MM</td><td>target_seq</td><td>PAM</td><td>alignment</td><td>distance</td><td>position</td><td>gene name</td><td>gene id</td></tr>
<tr><td>GU305914.1</td><td>181094</td><td>181116</td><td>+</td><td>0</td><td>ATCCAGACTCTCTTTTCTGA</td><td>GGG</td><td>||||||||[||||||||||||]PAM</td><td>NA</td><td>Intergenic</td><td>NA</td><td>NA</td></tr>
<tr><td>T13739</td><td>TTCTGAGGGCCCTAGATTAGTGG</td><td>1000</td><td></td><td></td><td></td><td></td><td></td><td></td><td></td><td></td><td></td></tr>
<tr><td>Oligo fwd</td><td>CACCTTCTGAGGGCCCTAGATTAG</td><td></td><td></td><td></td><td></td><td></td><td></td><td></td><td></td><td></td><td></td></tr>
<tr><td>Oligo rev</td><td>AAACCTAATCTAGGGCCCTCAGAA</td><td></td><td></td><td></td><td></td><td></td><td></td><td></td><td></td><td></td><td></td></tr>
<tr><td>Chromosome</td><td>start</td><td>end</td><td>strand</td><td>MM</td><td>target_seq</td><td>PAM</td><td>alignment</td><td>distance</td><td>position</td><td>gene name</td><td>gene id</td></tr>
<tr><td>GU305914.1</td><td>181108</td><td>181130</td><td>+</td><td>0</td><td>TTCTGAGGGCCCTAGATTAG</td><td>TGG</td><td>||||||||[||||||||||||]PAM</td><td>NA</td><td>Intergenic</td><td>NA</td><td>NA</td></tr>
<tr><td>T13740</td><td>TCTGAGGGCCCTAGATTAGTGGG</td><td>1000</td><td></td><td></td><td></td><td></td><td></td><td></td><td></td><td></td><td></td></tr>
<tr><td>Oligo fwd</td><td>CACCTCTGAGGGCCCTAGATTAGT</td><td></td><td></td><td></td><td></td><td></td><td></td><td></td><td></td><td></td><td></td></tr>
<tr><td>Oligo rev</td><td>AAACACTAATCTAGGGCCCTCAGA</td><td></td><td></td><td></td><td></td><td></td><td></td><td></td><td></td><td></td><td></td></tr>
<tr><td>Chromosome</td><td>start</td><td>end</td><td>strand</td><td>MM</td><td>target_seq</td><td>PAM</td><td>alignment</td><td>distance</td><td>position</td><td>gene name</td><td>gene id</td></tr>
<tr><td>GU305914.1</td><td>181109</td><td>181131</td><td>+</td><td>0</td><td>TCTGAGGGCCCTAGATTAGT</td><td>GGG</td><td>||||||||[||||||||||||]PAM</td><td>NA</td><td>Intergenic</td><td>NA</td><td>NA</td></tr>
<tr><td>T13741</td><td>AGATTAGTGGGCATGAAGTGTGG</td><td>1000</td><td></td><td></td><td></td><td></td><td></td><td></td><td></td><td></td><td></td></tr>
<tr><td>Oligo fwd</td><td>CACCAGATTAGTGGGCATGAAGTG</td><td></td><td></td><td></td><td></td><td></td><td></td><td></td><td></td><td></td><td></td></tr>
<tr><td>Oligo rev</td><td>AAACCACTTCATGCCCACTAATCT</td><td></td><td></td><td></td><td></td><td></td><td></td><td></td><td></td><td></td><td></td></tr>
<tr><td>Chromosome</td><td>start</td><td>end</td><td>strand</td><td>MM</td><td>target_seq</td><td>PAM</td><td>alignment</td><td>distance</td><td>position</td><td>gene name</td><td>gene id</td></tr>
<tr><td>GU305914.1</td><td>181121</td><td>181143</td><td>+</td><td>0</td><td>AGATTAGTGGGCATGAAGTG</td><td>TGG</td><td>||||||||[||||||||||||]PAM</td><td>NA</td><td>Intergenic</td><td>NA</td><td>NA</td></tr>
<tr><td>T13742</td><td>GATTAGTGGGCATGAAGTGTGGG</td><td>1000</td><td></td><td></td><td></td><td></td><td></td><td></td><td></td><td></td><td></td></tr>
<tr><td>Oligo fwd</td><td>CACCGATTAGTGGGCATGAAGTGT</td><td></td><td></td><td></td><td></td><td></td><td></td><td></td><td></td><td></td><td></td></tr>
<tr><td>Oligo rev</td><td>AAACACACTTCATGCCCACTAATC</td><td></td><td></td><td></td><td></td><td></td><td></td><td></td><td></td><td></td><td></td></tr>
<tr><td>Chromosome</td><td>start</td><td>end</td><td>strand</td><td>MM</td><td>target_seq</td><td>PAM</td><td>alignment</td><td>distance</td><td>position</td><td>gene name</td><td>gene id</td></tr>
<tr><td>GU305914.1</td><td>181122</td><td>181144</td><td>+</td><td>0</td><td>GATTAGTGGGCATGAAGTGT</td><td>GGG</td><td>||||||||[||||||||||||]PAM</td><td>NA</td><td>Intergenic</td><td>NA</td><td>NA</td></tr>
</table>

| | | | | | | | | | | | | |
|---|---|---|---|---|---|---|---|---|---|---|---|---|
| T13743 | CATACATGAGCCTCCCTAGCTGG | | 1000 | | | | | | | | | |
| Oligo fwd | CACCCATACATGAGCCTCCCTAGC | | | | | | | | | | | |
| Oligo rev | AAACGCTAGGGAGGCTCATGTATG | | | | | | | | | | | |
| Chromosome | start | | end | strand | MM | target_seq | PAM | alignment | distance | position | gene name | gene id |
| GU305914.1 | | 181152 | 181174 | + | 0 | CATACATGAGCCTCCCTAGC | TGG | \|\|\|\|\|\|\|\|[\|\|\|\|\|\|\|\|\|\|\|\|]PAM | NA | Intergenic | NA | NA |

| | | | | | | | | | | | | |
|---|---|---|---|---|---|---|---|---|---|---|---|---|
| T13744 | GATGCGCTCGAAGATATCATTGG | | 1000 | | | | | | | | | |
| Oligo fwd | CACCGATGCGCTCGAAGATATCAT | | | | | | | | | | | |
| Oligo rev | AAACATGATATCTTCGAGCGCATC | | | | | | | | | | | |
| Chromosome | start | | end | strand | MM | target_seq | PAM | alignment | distance | position | gene name | gene id |
| GU305914.1 | | 181228 | 181250 | + | 0 | GATGCGCTCGAAGATATCAT | TGG | \|\|\|\|\|\|\|\|[\|\|\|\|\|\|\|\|\|\|\|\|]PAM | NA | Intergenic | NA | NA |

| | | | | | | | | | | | | |
|---|---|---|---|---|---|---|---|---|---|---|---|---|
| T13745 | GGCTAGCTGACAGAGAATGTTGG | | 1000 | | | | | | | | | |
| Oligo fwd | CACCGGCTAGCTGACAGAGAATGT | | | | | | | | | | | |
| Oligo rev | AAACACATTCTCTGTCAGCTAGCC | | | | | | | | | | | |
| Chromosome | start | | end | strand | MM | target_seq | PAM | alignment | distance | position | gene name | gene id |
| GU305914.1 | | 181249 | 181271 | + | 0 | GGCTAGCTGACAGAGAATGT | TGG | \|\|\|\|\|\|\|\|[\|\|\|\|\|\|\|\|\|\|\|\|]PAM | NA | Intergenic | NA | NA |

| | | | | | | | | | | | | |
|---|---|---|---|---|---|---|---|---|---|---|---|---|
| T13746 | AATGTTGGTAGCAGAGTTGATGG | | 1000 | | | | | | | | | |
| Oligo fwd | CACCAATGTTGGTAGCAGAGTTGA | | | | | | | | | | | |
| Oligo rev | AAACTCAACTCTGCTACCAACATT | | | | | | | | | | | |
| Chromosome | start | | end | strand | MM | target_seq | PAM | alignment | distance | position | gene name | gene id |
| GU305914.1 | | 181264 | 181286 | + | 0 | AATGTTGGTAGCAGAGTTGA | TGG | \|\|\|\|\|\|\|\|[\|\|\|\|\|\|\|\|\|\|\|\|]PAM | NA | Intergenic | NA | NA |

| | | | | | | | | | | | | |
|---|---|---|---|---|---|---|---|---|---|---|---|---|
| T13747 | GATAGTTCTGTTTTATCATGAGG | | 1000 | | | | | | | | | |
| Oligo fwd | CACCGATAGTTCTGTTTTATCATG | | | | | | | | | | | |
| Oligo rev | AAACCATGATAAAACAGAACTATC | | | | | | | | | | | |
| Chromosome | start | | end | strand | MM | target_seq | PAM | alignment | distance | position | gene name | gene id |
| GU305914.1 | | 181293 | 181315 | + | 0 | GATAGTTCTGTTTTATCATG | AGG | \|\|\|\|\|\|\|\|[\|\|\|\|\|\|\|\|\|\|\|\|]PAM | NA | Intergenic | NA | NA |

| | | | | | | | | | | | | |
|---|---|---|---|---|---|---|---|---|---|---|---|---|
| T13748 | CATGAGGTGTGCAATCTTACAGG | | 1000 | | | | | | | | | |
| Oligo fwd | CACCCATGAGGTGTGCAATCTTAC | | | | | | | | | | | |
| Oligo rev | AAACGTAAGATTGCACACCTCATG | | | | | | | | | | | |
| Chromosome | start | | end | strand | MM | target_seq | PAM | alignment | distance | position | gene name | gene id |
| GU305914.1 | | 181309 | 181331 | + | 0 | CATGAGGTGTGCAATCTTAC | AGG | \|\|\|\|\|\|\|\|[\|\|\|\|\|\|\|\|\|\|\|\|]PAM | NA | Intergenic | NA | NA |

| | | | | | | | | | | | | |
|---|---|---|---|---|---|---|---|---|---|---|---|---|
| T13749 | GACAACAGAACGCTCCTCACTGG | | 1000 | | | | | | | | | |
| Oligo fwd | CACCGACAACAGAACGCTCCTCAC | | | | | | | | | | | |
| Oligo rev | AAACGTGAGGAGCGTTCTGTTGTC | | | | | | | | | | | |
| Chromosome | start | | end | strand | MM | target_seq | PAM | alignment | distance | position | gene name | gene id |
| GU305914.1 | | 181331 | 181353 | + | 0 | GACAACAGAACGCTCCTCAC | TGG | \|\|\|\|\|\|\|\|[\|\|\|\|\|\|\|\|\|\|\|\|]PAM | NA | Intergenic | NA | NA |

| | | | | | | | | | | | | |
|---|---|---|---|---|---|---|---|---|---|---|---|---|
| T13750 | TCCTCACTGGAATATAAGAGTGG | | 1000 | | | | | | | | | |
| Oligo fwd | CACCTCCTCACTGGAATATAAGAG | | | | | | | | | | | |
| Oligo rev | AAACCTCTTATATTCCAGTGAGGA | | | | | | | | | | | |
| Chromosome | start | | end | strand | MM | target_seq | PAM | alignment | distance | position | gene name | gene id |
| GU305914.1 | | 181344 | 181366 | + | 0 | TCCTCACTGGAATATAAGAG | TGG | \|\|\|\|\|\|\|\|[\|\|\|\|\|\|\|\|\|\|\|\|]PAM | NA | Intergenic | NA | NA |

| | | | | | | | | | | | | |
|---|---|---|---|---|---|---|---|---|---|---|---|---|
| T13751 | TATAACTCAAGTGTTCTGTGTGG | | 1000 | | | | | | | | | |
| Oligo fwd | CACCTATAACTCAAGTGTTCTGTG | | | | | | | | | | | |
| Oligo rev | AAACCACAGAACACTTGAGTTATA | | | | | | | | | | | |
| Chromosome | start | | end | strand | MM | target_seq | PAM | alignment | distance | position | gene name | gene id |
| GU305914.1 | | 181371 | 181393 | + | 0 | TATAACTCAAGTGTTCTGTG | TGG | \|\|\|\|\|\|\|\|[\|\|\|\|\|\|\|\|\|\|\|\|]PAM | NA | Intergenic | NA | NA |

| | | | | | | | | | | | | |
|---|---|---|---|---|---|---|---|---|---|---|---|---|
| T13752 | ATAACTCAAGTGTTCTGTGTGGG | | 1000 | | | | | | | | | |
| Oligo fwd | CACCATAACTCAAGTGTTCTGTGT | | | | | | | | | | | |
| Oligo rev | AAACACACAGAACACTTGAGTTAT | | | | | | | | | | | |
| Chromosome | start | | end | strand | MM | target_seq | PAM | alignment | distance | position | gene name | gene id |
| GU305914.1 | | 181372 | 181394 | + | 0 | ATAACTCAAGTGTTCTGTGT | GGG | \|\|\|\|\|\|\|\|[\|\|\|\|\|\|\|\|\|\|\|\|]PAM | NA | Intergenic | NA | NA |

| | | | | | | | | | | | | |
|---|---|---|---|---|---|---|---|---|---|---|---|---|
| T13753 | TGAATATGAAAGACAGAGACAGG | | 1000 | | | | | | | | | |
| Oligo fwd | CACCTGAATATGAAAGACAGAGAC | | | | | | | | | | | |
| Oligo rev | AAACGTCTCTGTCTTTCATATTCA | | | | | | | | | | | |
| Chromosome | start | | end | strand | MM | target_seq | PAM | alignment | distance | position | gene name | gene id |
| GU305914.1 | | 181395 | 181417 | + | 0 | TGAATATGAAAGACAGAGAC | AGG | \|\|\|\|\|\|\|\|[\|\|\|\|\|\|\|\|\|\|\|\|]PAM | NA | Intergenic | NA | NA |

| | | | | | | | | | | | | |
|---|---|---|---|---|---|---|---|---|---|---|---|---|
| T13754 | TGAAAGACAGAGACAGGTAGAGG | | 1000 | | | | | | | | | |
| Oligo fwd | CACCTGAAAGACAGAGACAGGTAG | | | | | | | | | | | |
| Oligo rev | AAACCTACCTGTCTCTGTCTTTCA | | | | | | | | | | | |
| Chromosome | start | | end | strand | MM | target_seq | PAM | alignment | distance | position | gene name | gene id |
| GU305914.1 | | 181401 | 181423 | + | 0 | TGAAAGACAGAGACAGGTAG | AGG | \|\|\|\|\|\|\|\|[\|\|\|\|\|\|\|\|\|\|\|\|]PAM | NA | Intergenic | NA | NA |

| | | | | | | | | | | | | |
|---|---|---|---|---|---|---|---|---|---|---|---|---|
| T13755 | CTCACCTGCAGCATGCTTGATGG | | 1000 | | | | | | | | | |
| Oligo fwd | CACCCTCACCTGCAGCATGCTTGA | | | | | | | | | | | |
| Oligo rev | AAACTCAAGCATGCTGCAGGTGAG | | | | | | | | | | | |
| Chromosome | start | | end | strand | MM | target_seq | PAM | alignment | distance | position | gene name | gene id |
| GU305914.1 | | 181467 | 181489 | + | 0 | CTCACCTGCAGCATGCTTGA | TGG | \|\|\|\|\|\|\|\|[\|\|\|\|\|\|\|\|\|\|\|\|]PAM | NA | Intergenic | NA | NA |

| | | | | | | | | | | | | |
|---|---|---|---|---|---|---|---|---|---|---|---|---|
| T13756 | CGCTGTTATCATTCCCCACGTGG | | 1000 | | | | | | | | | |
| Oligo fwd | CACCCGCTGTTATCATTCCCCACG | | | | | | | | | | | |
| Oligo rev | AAACCGTGGGGAATGATAACAGCG | | | | | | | | | | | |
| Chromosome | start | | end | strand | MM | target_seq | PAM | alignment | distance | position | gene name | gene id |
| GU305914.1 | | 181496 | 181518 | + | 0 | CGCTGTTATCATTCCCCACG | TGG | \|\|\|\|\|\|\|\|[\|\|\|\|\|\|\|\|\|\|\|\|]PAM | NA | Intergenic | NA | NA |

| | | | | | | | | | | | | |
|---|---|---|---|---|---|---|---|---|---|---|---|---|
| T13757 | TCCCCACGTGGTCAGCGATGCGG | | 1000 | | | | | | | | | |
| Oligo fwd | CACCTCCCCACGTGGTCAGCGATG | | | | | | | | | | | |
| Oligo rev | AAACCATCGCTGACCACGTGGGGA | | | | | | | | | | | |
| Chromosome | start | | end | strand | MM | target_seq | PAM | alignment | distance | position | gene name | gene id |
| GU305914.1 | | 181508 | 181530 | + | 0 | TCCCCACGTGGTCAGCGATG | CGG | \|\|\|\|\|\|\|\|[\|\|\|\|\|\|\|\|\|\|\|\|]PAM | NA | Intergenic | NA | NA |

T13758 GGTCAGCGATGCGGTCCATGAGG 1000
Oligo fwd CACCGGTCAGCGATGCGGTCCATG
Oligo rev AAACCATGGACCGCATCGCTGACC
Chromosome start end strand MM target_seq PAM alignment distance position gene name gene id
GU305914.1 181517 181539 + 0 GGTCAGCGATGCGGTCCATG AGG ||||||||[||||||||||||]PAM NA Intergenic NA NA

T13759 GAGGATCTTCTCTTCCTCAGTGG 1000
Oligo fwd CACCGAGGATCTTCTCTTCCTCAG
Oligo rev AAACCTGAGGAAGAGAAGATCCTC
Chromosome start end strand MM target_seq PAM alignment distance position gene name gene id
GU305914.1 181536 181558 + 0 GAGGATCTTCTCTTCCTCAG TGG ||||||||[||||||||||||]PAM NA Intergenic NA NA

T13760 AGGATCTTCTCTTCCTCAGTGGG 1000
Oligo fwd CACCAGGATCTTCTCTTCCTCAGT
Oligo rev AAACACTGAGGAAGAGAAGATCCT
Chromosome start end strand MM target_seq PAM alignment distance position gene name gene id
GU305914.1 181537 181559 + 0 AGGATCTTCTCTTCCTCAGT GGG ||||||||[||||||||||||]PAM NA Intergenic NA NA

T13761 CCAACATGTCCTCCAGAGTCAGG 1000
Oligo fwd CACCCCAACATGTCCTCCAGAGTC
Oligo rev AAACGACTCTGGAGGACATGTTGG
Chromosome start end strand MM target_seq PAM alignment distance position gene name gene id
GU305914.1 181580 181602 + 0 CCAACATGTCCTCCAGAGTC AGG ||||||||[||||||||||||]PAM NA Intergenic NA NA

T13762 CCTCCAGAGTCAGGTCACAGCGG 1000
Oligo fwd CACCCCTCCAGAGTCAGGTCACAG
Oligo rev AAACCTGTGACCTGACTCTGGAGG
Chromosome start end strand MM target_seq PAM alignment distance position gene name gene id
GU305914.1 181589 181611 + 0 CCTCCAGAGTCAGGTCACAG CGG ||||||||[||||||||||||]PAM NA Intergenic NA NA

T13763 CAGAGTCAGGTCACAGCGGACGG 1000
Oligo fwd CACCCAGAGTCAGGTCACAGCGGA
Oligo rev AAACTCCGCTGTGACCTGACTCTG
Chromosome start end strand MM target_seq PAM alignment distance position gene name gene id
GU305914.1 181593 181615 + 0 CAGAGTCAGGTCACAGCGGA CGG ||||||||[||||||||||||]PAM NA Intergenic NA NA

T13764 CAGGTCACAGCGGACGGTTTCGG 1000
Oligo fwd CACCCAGGTCACAGCGGACGGTTT
Oligo rev AAACAAACCGTCCGCTGTGACCTG
Chromosome start end strand MM target_seq PAM alignment distance position gene name gene id
GU305914.1 181599 181621 + 0 CAGGTCACAGCGGACGGTTT CGG ||||||||[||||||||||||]PAM NA Intergenic NA NA

T13765 CGGTTTCGGTTCCAGACTCGAGG 1000
Oligo fwd CACCCGGTTTCGGTTCCAGACTCG
Oligo rev AAACCGAGTCTGGAACCGAAACCG
Chromosome start end strand MM target_seq PAM alignment distance position gene name gene id
GU305914.1 181613 181635 + 0 CGGTTTCGGTTCCAGACTCG AGG ||||||||[||||||||||||]PAM NA Intergenic NA NA

T13766 TCCAGACTCGAGGATCTTGTTGG 1000
Oligo fwd CACCTCCAGACTCGAGGATCTTGT
Oligo rev AAACACAAGATCCTCGAGTCTGGA
Chromosome start end strand MM target_seq PAM alignment distance position gene name gene id
GU305914.1 181623 181645 + 0 TCCAGACTCGAGGATCTTGT TGG ||||||||[||||||||||||]PAM NA Intergenic NA NA

T13767 ATCTTGTTGGCGACATGAGCTGG 1000
Oligo fwd CACCATCTTGTTGGCGACATGAGC
Oligo rev AAACGCTCATGTCGCCAACAAGAT
Chromosome start end strand MM target_seq PAM alignment distance position gene name gene id
GU305914.1 181636 181658 + 0 ATCTTGTTGGCGACATGAGC TGG ||||||||[||||||||||||]PAM NA Intergenic NA NA

T13768 GCACCCTGAGAGAACAAGAGTGG 1000
Oligo fwd CACCGCACCCTGAGAGAACAAGAG
Oligo rev AAACCTCTTGTTCTCTCAGGGTGC
Chromosome start end strand MM target_seq PAM alignment distance position gene name gene id
GU305914.1 181658 181680 + 0 GCACCCTGAGAGAACAAGAG TGG ||||||||[||||||||||||]PAM NA Intergenic NA NA

T13769 CACCCTGAGAGAACAAGAGTGGG 1000
Oligo fwd CACCCACCCTGAGAGAACAAGAGT
Oligo rev AAACACTCTTGTTCTCTCAGGGTG
Chromosome start end strand MM target_seq PAM alignment distance position gene name gene id
GU305914.1 181659 181681 + 0 CACCCTGAGAGAACAAGAGT GGG ||||||||[||||||||||||]PAM NA Intergenic NA NA

T13770 GAGTGGGAAGACCATGAGTCAGG 1000
Oligo fwd CACCGAGTGGGAAGACCATGAGTC
Oligo rev AAACGACTCATGGTCTTCCCACTC
Chromosome start end strand MM target_seq PAM alignment distance position gene name gene id
GU305914.1 181675 181697 + 0 GAGTGGGAAGACCATGAGTC AGG ||||||||[||||||||||||]PAM NA Intergenic NA NA

T13771 AGTGGGAAGACCATGAGTCAGGG 1000
Oligo fwd CACCAGTGGGAAGACCATGAGTCA
Oligo rev AAACTGACTCATGGTCTTCCCACT
Chromosome start end strand MM target_seq PAM alignment distance position gene name gene id
GU305914.1 181676 181698 + 0 AGTGGGAAGACCATGAGTCA GGG ||||||||[||||||||||||]PAM NA Intergenic NA NA

T13772 GTGGGAAGACCATGAGTCAGGGG 1000
Oligo fwd CACCGTGGGAAGACCATGAGTCAG
Oligo rev AAACCTGACTCATGGTCTTCCCAC
Chromosome start end strand MM target_seq PAM alignment distance position gene name gene id

GU305914.1 181677 181699 + 0 GTGGGAAGACCATGAGTCAG GGG ||||||||[||||||||||||]PAM NA Intergenic NA NA

T13773 CATGAAATGTGTCACAAGAGAGG 1000
Oligo fwd CACCCATGAAATGTGTCACAAGAG
Oligo rev AAACCTCTTGTGACACATTTCATG
Chromosome start end strand MM target_seq PAM alignment distance position gene name gene id
GU305914.1 181717 181739 + 0 CATGAAATGTGTCACAAGAG AGG ||||||||[||||||||||||]PAM NA Intergenic NA NA

T13774 ATGTGTCACAAGAGAGGAAATGG 1000
Oligo fwd CACCATGTGTCACAAGAGAGGAAA
Oligo rev AAACTTTCCTCTCTTGTGACACAT
Chromosome start end strand MM target_seq PAM alignment distance position gene name gene id
GU305914.1 181723 181745 + 0 ATGTGTCACAAGAGAGGAAA TGG ||||||||[||||||||||||]PAM NA Intergenic NA NA

T13775 ACAGAAACTTACTTGTCTGATGG 1000
Oligo fwd CACCACAGAAACTTACTTGTCTGA
Oligo rev AAACTCAGACAAGTAAGTTTCTGT
Chromosome start end strand MM target_seq PAM alignment distance position gene name gene id
GU305914.1 181748 181770 + 0 ACAGAAACTTACTTGTCTGA TGG ||||||||[||||||||||||]PAM NA Intergenic NA NA

T13776 CAGAAACTTACTTGTCTGATGGG 1000
Oligo fwd CACCCAGAAACTTACTTGTCTGAT
Oligo rev AAACATCAGACAAGTAAGTTTCTG
Chromosome start end strand MM target_seq PAM alignment distance position gene name gene id
GU305914.1 181749 181771 + 0 CAGAAACTTACTTGTCTGAT GGG ||||||||[||||||||||||]PAM NA Intergenic NA NA

T13777 ACTTGTCTGATGGGTAGACACGG 1000
Oligo fwd CACCACTTGTCTGATGGGTAGACA
Oligo rev AAACTGTCTACCCATCAGACAAGT
Chromosome start end strand MM target_seq PAM alignment distance position gene name gene id
GU305914.1 181758 181780 + 0 ACTTGTCTGATGGGTAGACA CGG ||||||||[||||||||||||]PAM NA Intergenic NA NA

T13778 CACGGTTCTCATCCAAGTGACGG 1000
Oligo fwd CACCCACGGTTCTCATCCAAGTGA
Oligo rev AAACTCACTTGGATGAGAACCGTG
Chromosome start end strand MM target_seq PAM alignment distance position gene name gene id
GU305914.1 181776 181798 + 0 CACGGTTCTCATCCAAGTGA CGG ||||||||[||||||||||||]PAM NA Intergenic NA NA

T13779 TCTCATCCAAGTGACGGCCGTGG 1000
Oligo fwd CACCTCTCATCCAAGTGACGGCCG
Oligo rev AAACCGGCCGTCACTTGGATGAGA
Chromosome start end strand MM target_seq PAM alignment distance position gene name gene id
GU305914.1 181782 181804 + 0 TCTCATCCAAGTGACGGCCG TGG ||||||||[||||||||||||]PAM NA Intergenic NA NA

T13780 CTCATCCAAGTGACGGCCGTGGG 1000
Oligo fwd CACCCTCATCCAAGTGACGGCCGT
Oligo rev AAACACGGCCGTCACTTGGATGAG
Chromosome start end strand MM target_seq PAM alignment distance position gene name gene id
GU305914.1 181783 181805 + 0 CTCATCCAAGTGACGGCCGT GGG ||||||||[||||||||||||]PAM NA Intergenic NA NA

T13781 GTGACGGCCGTGGGCGTTGACGG 1000
Oligo fwd CACCGTGACGGCCGTGGGCGTTGA
Oligo rev AAACTCAACGCCCACGGCCGTCAC
Chromosome start end strand MM target_seq PAM alignment distance position gene name gene id
GU305914.1 181792 181814 + 0 GTGACGGCCGTGGGCGTTGA CGG ||||||||[||||||||||||]PAM NA Intergenic NA NA

T13782 GGCGTTGACGGATGCCTGATCGG 1000
Oligo fwd CACCGGCGTTGACGGATGCCTGAT
Oligo rev AAACATCAGGCATCCGTCAACGCC
Chromosome start end strand MM target_seq PAM alignment distance position gene name gene id
GU305914.1 181804 181826 + 0 GGCGTTGACGGATGCCTGAT CGG ||||||||[||||||||||||]PAM NA Intergenic NA NA

T13783 GATCATGATCATGTTGCAACTGG 1000
Oligo fwd CACCGATCATGATCATGTTGCAAC
Oligo rev AAACGTTGCAACATGATCATGATC
Chromosome start end strand MM target_seq PAM alignment distance position gene name gene id
GU305914.1 181828 181850 + 0 GATCATGATCATGTTGCAAC TGG ||||||||[||||||||||||]PAM NA Intergenic NA NA

T13784 ATCATGATCATGTTGCAACTGGG 1000
Oligo fwd CACCATCATGATCATGTTGCAACT
Oligo rev AAACAGTTGCAACATGATCATGAT
Chromosome start end strand MM target_seq PAM alignment distance position gene name gene id
GU305914.1 181829 181851 + 0 ATCATGATCATGTTGCAACT GGG ||||||||[||||||||||||]PAM NA Intergenic NA NA

T13785 GATCATGTTGCAACTGGGTGCGG 1000
Oligo fwd CACCGATCATGTTGCAACTGGGTG
Oligo rev AAACCACCCAGTTGCAACATGATC
Chromosome start end strand MM target_seq PAM alignment distance position gene name gene id
GU305914.1 181834 181856 + 0 GATCATGTTGCAACTGGGTG CGG ||||||||[||||||||||||]PAM NA Intergenic NA NA

T13786 CATGTTGCAACTGGGTGCGGCGG 1000
Oligo fwd CACCCATGTTGCAACTGGGTGCGG
Oligo rev AAACCCGCACCCAGTTGCAACATG
Chromosome start end strand MM target_seq PAM alignment distance position gene name gene id
GU305914.1 181837 181859 + 0 CATGTTGCAACTGGGTGCGG CGG ||||||||[||||||||||||]PAM NA Intergenic NA NA

T13787 ATGTTGCAACTGGGTGCGGCGGG 1000
Oligo fwd CACCATGTTGCAACTGGGTGCGGC
Oligo rev AAACGCCGCACCCAGTTGCAACAT

Chromosome start end strand MM target_seq PAM alignment distance position gene name gene id
GU305914.1 181838 181860 + 0 ATGTTGCAACTGGGTGCGGC GGG ||||||||[||||||||||||]PAM NA Intergenic NA NA

T13788 AGTTGCTGATAAAAAACAACTGG 1000
Oligo fwd CACCAGTTGCTGATAAAAAACAAC
Oligo rev AAACGTTGTTTTTTATCAGCAACT
Chromosome start end strand MM target_seq PAM alignment distance position gene name gene id
GU305914.1 181982 182004 + 0 AGTTGCTGATAAAAAACAAC TGG ||||||||[||||||||||||]PAM NA Intergenic NA NA

T13789 TGCTGATAAAAAACAACTGGTGG 1000
Oligo fwd CACCTGCTGATAAAAAACAACTGG
Oligo rev AAACCCAGTTGTTTTTTATCAGCA
Chromosome start end strand MM target_seq PAM alignment distance position gene name gene id
GU305914.1 181985 182007 + 0 TGCTGATAAAAAACAACTGG TGG ||||||||[||||||||||||]PAM NA Intergenic NA NA

T13790 CAACTGGTGGAAGAGATCTCAGG 1000
Oligo fwd CACCCAACTGGTGGAAGAGATCTC
Oligo rev AAACGAGATCTCTTCCACCAGTTG
Chromosome start end strand MM target_seq PAM alignment distance position gene name gene id
GU305914.1 181998 182020 + 0 CAACTGGTGGAAGAGATCTC AGG ||||||||[||||||||||||]PAM NA Intergenic NA NA

T13791 TCTCAGGTAAAGAAACAACATGG 1000
Oligo fwd CACCTCTCAGGTAAAGAAACAACA
Oligo rev AAACTGTTGTTTCTTTACCTGAGA
Chromosome start end strand MM target_seq PAM alignment distance position gene name gene id
GU305914.1 182014 182036 + 0 TCTCAGGTAAAGAAACAACA TGG ||||||||[||||||||||||]PAM NA Intergenic NA NA

T13792 AACTCTCCCTTTTTTCTGTCAGG 1000
Oligo fwd CACCAACTCTCCCTTTTTTCTGTC
Oligo rev AAACGACAGAAAAAAGGGAGAGTT
Chromosome start end strand MM target_seq PAM alignment distance position gene name gene id
GU305914.1 182078 182100 + 0 AACTCTCCCTTTTTTCTGTC AGG ||||||||[||||||||||||]PAM NA Intergenic NA NA

T13793 ACTCTCCCTTTTTTCTGTCAGGG 1000
Oligo fwd CACCACTCTCCCTTTTTTCTGTCA
Oligo rev AAACTGACAGAAAAAAGGGAGAGT
Chromosome start end strand MM target_seq PAM alignment distance position gene name gene id
GU305914.1 182079 182101 + 0 ACTCTCCCTTTTTTCTGTCA GGG ||||||||[||||||||||||]PAM NA Intergenic NA NA

T13794 CCTTTTTTCTGTCAGGGCCTAGG 1000
Oligo fwd CACCCCTTTTTTCTGTCAGGGCCT
Oligo rev AAACAGGCCCTGACAGAAAAAAGG
Chromosome start end strand MM target_seq PAM alignment distance position gene name gene id
GU305914.1 182085 182107 + 0 CCTTTTTTCTGTCAGGGCCT AGG ||||||||[||||||||||||]PAM NA Intergenic NA NA

T13795 CTGTCAGGGCCTAGGTTTTCAGG 1000
Oligo fwd CACCCTGTCAGGGCCTAGGTTTTC
Oligo rev AAACGAAAACCTAGGCCCTGACAG
Chromosome start end strand MM target_seq PAM alignment distance position gene name gene id
GU305914.1 182093 182115 + 0 CTGTCAGGGCCTAGGTTTTC AGG ||||||||[||||||||||||]PAM NA Intergenic NA NA

T13796 CAGGGCCTAGGTTTTCAGGACGG 1000
Oligo fwd CACCCAGGGCCTAGGTTTTCAGGA
Oligo rev AAACTCCTGAAAACCTAGGCCCTG
Chromosome start end strand MM target_seq PAM alignment distance position gene name gene id
GU305914.1 182097 182119 + 0 CAGGGCCTAGGTTTTCAGGA CGG ||||||||[||||||||||||]PAM NA Intergenic NA NA

T13797 AGGGCCTAGGTTTTCAGGACGGG 1000
Oligo fwd CACCAGGGCCTAGGTTTTCAGGAC
Oligo rev AAACGTCCTGAAAACCTAGGCCCT
Chromosome start end strand MM target_seq PAM alignment distance position gene name gene id
GU305914.1 182098 182120 + 0 AGGGCCTAGGTTTTCAGGAC GGG ||||||||[||||||||||||]PAM NA Intergenic NA NA

T13798 GGGCCTAGGTTTTCAGGACGGGG 1000
Oligo fwd CACCGGGCCTAGGTTTTCAGGACG
Oligo rev AAACCGTCCTGAAAACCTAGGCCC
Chromosome start end strand MM target_seq PAM alignment distance position gene name gene id
GU305914.1 182099 182121 + 0 GGGCCTAGGTTTTCAGGACG GGG ||||||||[||||||||||||]PAM NA Intergenic NA NA

T13799 CAGTACGAGCAACAACACGCAGG 1000
Oligo fwd CACCCAGTACGAGCAACAACACGC
Oligo rev AAACGCGTGTTGTTGCTCGTACTG
Chromosome start end strand MM target_seq PAM alignment distance position gene name gene id
GU305914.1 182137 182159 + 0 CAGTACGAGCAACAACACGC AGG ||||||||[||||||||||||]PAM NA Intergenic NA NA

T13800 CAACAACACGCAGGAACTGACGG 1000
Oligo fwd CACCCAACAACACGCAGGAACTGA
Oligo rev AAACTCAGTTCCTGCGTGTTGTTG
Chromosome start end strand MM target_seq PAM alignment distance position gene name gene id
GU305914.1 182146 182168 + 0 CAACAACACGCAGGAACTGA CGG ||||||||[||||||||||||]PAM NA Intergenic NA NA

T13801 GGAACTGACGGAGTCCTCCACGG 1000
Oligo fwd CACCGGAACTGACGGAGTCCTCCA
Oligo rev AAACTGGAGGACTCCGTCAGTTCC
Chromosome start end strand MM target_seq PAM alignment distance position gene name gene id
GU305914.1 182158 182180 + 0 GGAACTGACGGAGTCCTCCA CGG ||||||||[||||||||||||]PAM NA Intergenic NA NA

T13802 GAACTGACGGAGTCCTCCACGGG 1000
Oligo fwd CACCGAACTGACGGAGTCCTCCAC

<table>
<tr><td>Oligo rev</td><td>AAACGTGGAGGACTCCGTCAGTTC</td><td></td><td></td><td></td><td></td><td></td><td></td><td></td><td></td><td></td><td></td></tr>
<tr><td>Chromosome</td><td>start</td><td>end</td><td>strand</td><td>MM</td><td>target_seq</td><td>PAM</td><td>alignment</td><td>distance</td><td>position</td><td>gene name</td><td>gene id</td></tr>
<tr><td>GU305914.1</td><td>182159</td><td>182181</td><td>+</td><td>0</td><td>GAACTGACGGAGTCCTCCAC</td><td>GGG</td><td>||||||||[||||||||||||]PAM</td><td>NA</td><td>Intergenic</td><td>NA</td><td>NA</td></tr>
<tr><td>T13803</td><td>CGGGAAAAGAGCACCACAGCAGG</td><td>1000</td><td></td><td></td><td></td><td></td><td></td><td></td><td></td><td></td><td></td></tr>
<tr><td>Oligo fwd</td><td>CACCCGGGAAAAGAGCACCACAGC</td><td></td><td></td><td></td><td></td><td></td><td></td><td></td><td></td><td></td><td></td></tr>
<tr><td>Oligo rev</td><td>AAACGCTGTGGTGCTCTTTTCCCG</td><td></td><td></td><td></td><td></td><td></td><td></td><td></td><td></td><td></td><td></td></tr>
<tr><td>Chromosome</td><td>start</td><td>end</td><td>strand</td><td>MM</td><td>target_seq</td><td>PAM</td><td>alignment</td><td>distance</td><td>position</td><td>gene name</td><td>gene id</td></tr>
<tr><td>GU305914.1</td><td>182178</td><td>182200</td><td>+</td><td>0</td><td>CGGGAAAAGAGCACCACAGC</td><td>AGG</td><td>||||||||[||||||||||||]PAM</td><td>NA</td><td>Intergenic</td><td>NA</td><td>NA</td></tr>
<tr><td>T13804</td><td>ACCACAGCAGGCCACCAAGATGG</td><td>1000</td><td></td><td></td><td></td><td></td><td></td><td></td><td></td><td></td><td></td></tr>
<tr><td>Oligo fwd</td><td>CACCACCACAGCAGGCCACCAAGA</td><td></td><td></td><td></td><td></td><td></td><td></td><td></td><td></td><td></td><td></td></tr>
<tr><td>Oligo rev</td><td>AAACTCTTGGTGGCCTGCTGTGGT</td><td></td><td></td><td></td><td></td><td></td><td></td><td></td><td></td><td></td><td></td></tr>
<tr><td>Chromosome</td><td>start</td><td>end</td><td>strand</td><td>MM</td><td>target_seq</td><td>PAM</td><td>alignment</td><td>distance</td><td>position</td><td>gene name</td><td>gene id</td></tr>
<tr><td>GU305914.1</td><td>182190</td><td>182212</td><td>+</td><td>0</td><td>ACCACAGCAGGCCACCAAGA</td><td>TGG</td><td>||||||||[||||||||||||]PAM</td><td>NA</td><td>Intergenic</td><td>NA</td><td>NA</td></tr>
<tr><td>T13805</td><td>TTCTGTATTCTGCTCATTTTTGG</td><td>1000</td><td></td><td></td><td></td><td></td><td></td><td></td><td></td><td></td><td></td></tr>
<tr><td>Oligo fwd</td><td>CACCTTCTGTATTCTGCTCATTTT</td><td></td><td></td><td></td><td></td><td></td><td></td><td></td><td></td><td></td><td></td></tr>
<tr><td>Oligo rev</td><td>AAACAAAATGAGCAGAATACAGAA</td><td></td><td></td><td></td><td></td><td></td><td></td><td></td><td></td><td></td><td></td></tr>
<tr><td>Chromosome</td><td>start</td><td>end</td><td>strand</td><td>MM</td><td>target_seq</td><td>PAM</td><td>alignment</td><td>distance</td><td>position</td><td>gene name</td><td>gene id</td></tr>
<tr><td>GU305914.1</td><td>182234</td><td>182256</td><td>+</td><td>0</td><td>TTCTGTATTCTGCTCATTTT</td><td>TGG</td><td>||||||||[||||||||||||]PAM</td><td>NA</td><td>Intergenic</td><td>NA</td><td>NA</td></tr>
<tr><td>T13806</td><td>GCTCATTTTTGGACTTATCATGG</td><td>1000</td><td></td><td></td><td></td><td></td><td></td><td></td><td></td><td></td><td></td></tr>
<tr><td>Oligo fwd</td><td>CACCGCTCATTTTTGGACTTATCA</td><td></td><td></td><td></td><td></td><td></td><td></td><td></td><td></td><td></td><td></td></tr>
<tr><td>Oligo rev</td><td>AAACTGATAAGTCCAAAAATGAGC</td><td></td><td></td><td></td><td></td><td></td><td></td><td></td><td></td><td></td><td></td></tr>
<tr><td>Chromosome</td><td>start</td><td>end</td><td>strand</td><td>MM</td><td>target_seq</td><td>PAM</td><td>alignment</td><td>distance</td><td>position</td><td>gene name</td><td>gene id</td></tr>
<tr><td>GU305914.1</td><td>182245</td><td>182267</td><td>+</td><td>0</td><td>GCTCATTTTTGGACTTATCA</td><td>TGG</td><td>||||||||[||||||||||||]PAM</td><td>NA</td><td>Intergenic</td><td>NA</td><td>NA</td></tr>
<tr><td>T13807</td><td>CATTTTTGGACTTATCATGGCGG</td><td>1000</td><td></td><td></td><td></td><td></td><td></td><td></td><td></td><td></td><td></td></tr>
<tr><td>Oligo fwd</td><td>CACCCATTTTTGGACTTATCATGG</td><td></td><td></td><td></td><td></td><td></td><td></td><td></td><td></td><td></td><td></td></tr>
<tr><td>Oligo rev</td><td>AAACCCATGATAAGTCCAAAAATG</td><td></td><td></td><td></td><td></td><td></td><td></td><td></td><td></td><td></td><td></td></tr>
<tr><td>Chromosome</td><td>start</td><td>end</td><td>strand</td><td>MM</td><td>target_seq</td><td>PAM</td><td>alignment</td><td>distance</td><td>position</td><td>gene name</td><td>gene id</td></tr>
<tr><td>GU305914.1</td><td>182248</td><td>182270</td><td>+</td><td>0</td><td>CATTTTTGGACTTATCATGG</td><td>CGG</td><td>||||||||[||||||||||||]PAM</td><td>NA</td><td>Intergenic</td><td>NA</td><td>NA</td></tr>
<tr><td>T13808</td><td>CGGCCGTCGCTTGTAATATCTGG</td><td>1000</td><td></td><td></td><td></td><td></td><td></td><td></td><td></td><td></td><td></td></tr>
<tr><td>Oligo fwd</td><td>CACCCGGCCGTCGCTTGTAATATC</td><td></td><td></td><td></td><td></td><td></td><td></td><td></td><td></td><td></td><td></td></tr>
<tr><td>Oligo rev</td><td>AAACGATATTACAAGCGACGGCCG</td><td></td><td></td><td></td><td></td><td></td><td></td><td></td><td></td><td></td><td></td></tr>
<tr><td>Chromosome</td><td>start</td><td>end</td><td>strand</td><td>MM</td><td>target_seq</td><td>PAM</td><td>alignment</td><td>distance</td><td>position</td><td>gene name</td><td>gene id</td></tr>
<tr><td>GU305914.1</td><td>182268</td><td>182290</td><td>+</td><td>0</td><td>CGGCCGTCGCTTGTAATATC</td><td>TGG</td><td>||||||||[||||||||||||]PAM</td><td>NA</td><td>Intergenic</td><td>NA</td><td>NA</td></tr>
<tr><td>T13809</td><td>AATATCTGGTGTGTGAACACAGG</td><td>1000</td><td></td><td></td><td></td><td></td><td></td><td></td><td></td><td></td><td></td></tr>
<tr><td>Oligo fwd</td><td>CACCAATATCTGGTGTGTGAACAC</td><td></td><td></td><td></td><td></td><td></td><td></td><td></td><td></td><td></td><td></td></tr>
<tr><td>Oligo rev</td><td>AAACGTGTTCACACACCAGATATT</td><td></td><td></td><td></td><td></td><td></td><td></td><td></td><td></td><td></td><td></td></tr>
<tr><td>Chromosome</td><td>start</td><td>end</td><td>strand</td><td>MM</td><td>target_seq</td><td>PAM</td><td>alignment</td><td>distance</td><td>position</td><td>gene name</td><td>gene id</td></tr>
<tr><td>GU305914.1</td><td>182282</td><td>182304</td><td>+</td><td>0</td><td>AATATCTGGTGTGTGAACAC</td><td>AGG</td><td>||||||||[||||||||||||]PAM</td><td>NA</td><td>Intergenic</td><td>NA</td><td>NA</td></tr>
<tr><td>T13810</td><td>GGAGACGATCTGAGCAGTCATGG</td><td>1000</td><td></td><td></td><td></td><td></td><td></td><td></td><td></td><td></td><td></td></tr>
<tr><td>Oligo fwd</td><td>CACCGGAGACGATCTGAGCAGTCA</td><td></td><td></td><td></td><td></td><td></td><td></td><td></td><td></td><td></td><td></td></tr>
<tr><td>Oligo rev</td><td>AAACTGACTGCTCAGATCGTCTCC</td><td></td><td></td><td></td><td></td><td></td><td></td><td></td><td></td><td></td><td></td></tr>
<tr><td>Chromosome</td><td>start</td><td>end</td><td>strand</td><td>MM</td><td>target_seq</td><td>PAM</td><td>alignment</td><td>distance</td><td>position</td><td>gene name</td><td>gene id</td></tr>
<tr><td>GU305914.1</td><td>182303</td><td>182325</td><td>+</td><td>0</td><td>GGAGACGATCTGAGCAGTCA</td><td>TGG</td><td>||||||||[||||||||||||]PAM</td><td>NA</td><td>Intergenic</td><td>NA</td><td>NA</td></tr>
<tr><td>T13811</td><td>CAGTCATGGAGACTCTATCTTGG</td><td>1000</td><td></td><td></td><td></td><td></td><td></td><td></td><td></td><td></td><td></td></tr>
<tr><td>Oligo fwd</td><td>CACCCAGTCATGGAGACTCTATCT</td><td></td><td></td><td></td><td></td><td></td><td></td><td></td><td></td><td></td><td></td></tr>
<tr><td>Oligo rev</td><td>AAACAGATAGAGTCTCCATGACTG</td><td></td><td></td><td></td><td></td><td></td><td></td><td></td><td></td><td></td><td></td></tr>
<tr><td>Chromosome</td><td>start</td><td>end</td><td>strand</td><td>MM</td><td>target_seq</td><td>PAM</td><td>alignment</td><td>distance</td><td>position</td><td>gene name</td><td>gene id</td></tr>
<tr><td>GU305914.1</td><td>182317</td><td>182339</td><td>+</td><td>0</td><td>CAGTCATGGAGACTCTATCT</td><td>TGG</td><td>||||||||[||||||||||||]PAM</td><td>NA</td><td>Intergenic</td><td>NA</td><td>NA</td></tr>
<tr><td>T13812</td><td>ATCGAGCTCGCAAGCTCCGCCGG</td><td>1000</td><td></td><td></td><td></td><td></td><td></td><td></td><td></td><td></td><td></td></tr>
<tr><td>Oligo fwd</td><td>CACCATCGAGCTCGCAAGCTCCGC</td><td></td><td></td><td></td><td></td><td></td><td></td><td></td><td></td><td></td><td></td></tr>
<tr><td>Oligo rev</td><td>AAACGCGGAGCTTGCGAGCTCGAT</td><td></td><td></td><td></td><td></td><td></td><td></td><td></td><td></td><td></td><td></td></tr>
<tr><td>Chromosome</td><td>start</td><td>end</td><td>strand</td><td>MM</td><td>target_seq</td><td>PAM</td><td>alignment</td><td>distance</td><td>position</td><td>gene name</td><td>gene id</td></tr>
<tr><td>GU305914.1</td><td>182342</td><td>182364</td><td>+</td><td>0</td><td>ATCGAGCTCGCAAGCTCCGC</td><td>CGG</td><td>||||||||[||||||||||||]PAM</td><td>NA</td><td>Intergenic</td><td>NA</td><td>NA</td></tr>
<tr><td>T13813</td><td>GCTACAGACACATAGCATCCTGG</td><td>1000</td><td></td><td></td><td></td><td></td><td></td><td></td><td></td><td></td><td></td></tr>
<tr><td>Oligo fwd</td><td>CACCGCTACAGACACATAGCATCC</td><td></td><td></td><td></td><td></td><td></td><td></td><td></td><td></td><td></td><td></td></tr>
<tr><td>Oligo rev</td><td>AAACGGATGCTATGTGTCTGTAGC</td><td></td><td></td><td></td><td></td><td></td><td></td><td></td><td></td><td></td><td></td></tr>
<tr><td>Chromosome</td><td>start</td><td>end</td><td>strand</td><td>MM</td><td>target_seq</td><td>PAM</td><td>alignment</td><td>distance</td><td>position</td><td>gene name</td><td>gene id</td></tr>
<tr><td>GU305914.1</td><td>182366</td><td>182388</td><td>+</td><td>0</td><td>GCTACAGACACATAGCATCC</td><td>TGG</td><td>||||||||[||||||||||||]PAM</td><td>NA</td><td>Intergenic</td><td>NA</td><td>NA</td></tr>
<tr><td>T13814</td><td>CTACAGACACATAGCATCCTGGG</td><td>1000</td><td></td><td></td><td></td><td></td><td></td><td></td><td></td><td></td><td></td></tr>
<tr><td>Oligo fwd</td><td>CACCCTACAGACACATAGCATCCT</td><td></td><td></td><td></td><td></td><td></td><td></td><td></td><td></td><td></td><td></td></tr>
<tr><td>Oligo rev</td><td>AAACAGGATGCTATGTGTCTGTAG</td><td></td><td></td><td></td><td></td><td></td><td></td><td></td><td></td><td></td><td></td></tr>
<tr><td>Chromosome</td><td>start</td><td>end</td><td>strand</td><td>MM</td><td>target_seq</td><td>PAM</td><td>alignment</td><td>distance</td><td>position</td><td>gene name</td><td>gene id</td></tr>
<tr><td>GU305914.1</td><td>182367</td><td>182389</td><td>+</td><td>0</td><td>CTACAGACACATAGCATCCT</td><td>GGG</td><td>||||||||[||||||||||||]PAM</td><td>NA</td><td>Intergenic</td><td>NA</td><td>NA</td></tr>
<tr><td>T13815</td><td>TACAGACACATAGCATCCTGGGG</td><td>1000</td><td></td><td></td><td></td><td></td><td></td><td></td><td></td><td></td><td></td></tr>
<tr><td>Oligo fwd</td><td>CACCTACAGACACATAGCATCCTG</td><td></td><td></td><td></td><td></td><td></td><td></td><td></td><td></td><td></td><td></td></tr>
<tr><td>Oligo rev</td><td>AAACCAGGATGCTATGTGTCTGTA</td><td></td><td></td><td></td><td></td><td></td><td></td><td></td><td></td><td></td><td></td></tr>
<tr><td>Chromosome</td><td>start</td><td>end</td><td>strand</td><td>MM</td><td>target_seq</td><td>PAM</td><td>alignment</td><td>distance</td><td>position</td><td>gene name</td><td>gene id</td></tr>
<tr><td>GU305914.1</td><td>182368</td><td>182390</td><td>+</td><td>0</td><td>TACAGACACATAGCATCCTG</td><td>GGG</td><td>||||||||[||||||||||||]PAM</td><td>NA</td><td>Intergenic</td><td>NA</td><td>NA</td></tr>
<tr><td>T13816</td><td>GGGAGAGACGCGTACGTGTCAGG</td><td>1000</td><td></td><td></td><td></td><td></td><td></td><td></td><td></td><td></td><td></td></tr>
<tr><td>Oligo fwd</td><td>CACCGGGAGAGACGCGTACGTGTC</td><td></td><td></td><td></td><td></td><td></td><td></td><td></td><td></td><td></td><td></td></tr>
<tr><td>Oligo rev</td><td>AAACGACACGTACGCGTCTCTCCC</td><td></td><td></td><td></td><td></td><td></td><td></td><td></td><td></td><td></td><td></td></tr>
<tr><td>Chromosome</td><td>start</td><td>end</td><td>strand</td><td>MM</td><td>target_seq</td><td>PAM</td><td>alignment</td><td>distance</td><td>position</td><td>gene name</td><td>gene id</td></tr>
<tr><td>GU305914.1</td><td>182388</td><td>182410</td><td>+</td><td>0</td><td>GGGAGAGACGCGTACGTGTC</td><td>AGG</td><td>||||||||[||||||||||||]PAM</td><td>NA</td><td>Intergenic</td><td>NA</td><td>NA</td></tr>
<tr><td>T13817</td><td>CCTTTTTCTGTACCACGTTACGG</td><td>1000</td><td></td><td></td><td></td><td></td><td></td><td></td><td></td><td></td><td></td></tr>
</table>

<table>
<tr><td>Oligo fwd</td><td>CACCCCTTTTTCTGTACCACGTTA</td></tr>
<tr><td>Oligo rev</td><td>AAACTAACGTGGTACAGAAAAAGG</td></tr>
<tr><td>Chromosome</td><td>start</td><td>end</td><td>strand</td><td>MM</td><td>target_seq</td><td>PAM</td><td>alignment</td><td>distance</td><td>position</td><td>gene name</td><td>gene id</td></tr>
<tr><td>GU305914.1</td><td>182411</td><td>182433</td><td>+</td><td>0</td><td>CCTTTTTCTGTACCACGTTA</td><td>CGG</td><td>||||||||[||||||||||||]PAM</td><td>NA</td><td>Intergenic</td><td>NA</td><td>NA</td></tr>
<tr><td>T13818</td><td>CTGTACCACGTTACGGTACCCGG</td><td>1000</td></tr>
<tr><td>Oligo fwd</td><td>CACCCTGTACCACGTTACGGTACC</td></tr>
<tr><td>Oligo rev</td><td>AAACGGTACCGTAACGTGGTACAG</td></tr>
<tr><td>Chromosome</td><td>start</td><td>end</td><td>strand</td><td>MM</td><td>target_seq</td><td>PAM</td><td>alignment</td><td>distance</td><td>position</td><td>gene name</td><td>gene id</td></tr>
<tr><td>GU305914.1</td><td>182418</td><td>182440</td><td>+</td><td>0</td><td>CTGTACCACGTTACGGTACC</td><td>CGG</td><td>||||||||[||||||||||||]PAM</td><td>NA</td><td>Intergenic</td><td>NA</td><td>NA</td></tr>
<tr><td>T13819</td><td>CCCGGCACCGAAGCACACCGCGG</td><td>1000</td></tr>
<tr><td>Oligo fwd</td><td>CACCCCCGGCACCGAAGCACACCG</td></tr>
<tr><td>Oligo rev</td><td>AAACCGGTGTGCTTCGGTGCCGGG</td></tr>
<tr><td>Chromosome</td><td>start</td><td>end</td><td>strand</td><td>MM</td><td>target_seq</td><td>PAM</td><td>alignment</td><td>distance</td><td>position</td><td>gene name</td><td>gene id</td></tr>
<tr><td>GU305914.1</td><td>182436</td><td>182458</td><td>+</td><td>0</td><td>CCCGGCACCGAAGCACACCG</td><td>CGG</td><td>||||||||[||||||||||||]PAM</td><td>NA</td><td>Intergenic</td><td>NA</td><td>NA</td></tr>
<tr><td>T13820</td><td>ACCGAAGCACACCGCGGCCATGG</td><td>1000</td></tr>
<tr><td>Oligo fwd</td><td>CACCACCGAAGCACACCGCGGCCA</td></tr>
<tr><td>Oligo rev</td><td>AAACTGGCCGCGGTGTGCTTCGGT</td></tr>
<tr><td>Chromosome</td><td>start</td><td>end</td><td>strand</td><td>MM</td><td>target_seq</td><td>PAM</td><td>alignment</td><td>distance</td><td>position</td><td>gene name</td><td>gene id</td></tr>
<tr><td>GU305914.1</td><td>182442</td><td>182464</td><td>+</td><td>0</td><td>ACCGAAGCACACCGCGGCCA</td><td>TGG</td><td>||||||||[||||||||||||]PAM</td><td>NA</td><td>Intergenic</td><td>NA</td><td>NA</td></tr>
<tr><td>T13821</td><td>GCACACCGCGGCCATGGAAGAGG</td><td>1000</td></tr>
<tr><td>Oligo fwd</td><td>CACCGCACACCGCGGCCATGGAAG</td></tr>
<tr><td>Oligo rev</td><td>AAACCTTCCATGGCCGCGGTGTGC</td></tr>
<tr><td>Chromosome</td><td>start</td><td>end</td><td>strand</td><td>MM</td><td>target_seq</td><td>PAM</td><td>alignment</td><td>distance</td><td>position</td><td>gene name</td><td>gene id</td></tr>
<tr><td>GU305914.1</td><td>182448</td><td>182470</td><td>+</td><td>0</td><td>GCACACCGCGGCCATGGAAG</td><td>AGG</td><td>||||||||[||||||||||||]PAM</td><td>NA</td><td>Intergenic</td><td>NA</td><td>NA</td></tr>
<tr><td>T13822</td><td>CACACCGCGGCCATGGAAGAGGG</td><td>1000</td></tr>
<tr><td>Oligo fwd</td><td>CACCCACACCGCGGCCATGGAAGA</td></tr>
<tr><td>Oligo rev</td><td>AAACTCTTCCATGGCCGCGGTGTG</td></tr>
<tr><td>Chromosome</td><td>start</td><td>end</td><td>strand</td><td>MM</td><td>target_seq</td><td>PAM</td><td>alignment</td><td>distance</td><td>position</td><td>gene name</td><td>gene id</td></tr>
<tr><td>GU305914.1</td><td>182449</td><td>182471</td><td>+</td><td>0</td><td>CACACCGCGGCCATGGAAGA</td><td>GGG</td><td>||||||||[||||||||||||]PAM</td><td>NA</td><td>Intergenic</td><td>NA</td><td>NA</td></tr>
<tr><td>T13823</td><td>CAGCATATATATATAAGAAATGG</td><td>1000</td></tr>
<tr><td>Oligo fwd</td><td>CACCCAGCATATATATATAAGAAA</td></tr>
<tr><td>Oligo rev</td><td>AAACTTTCTTATATATATATGCTG</td></tr>
<tr><td>Chromosome</td><td>start</td><td>end</td><td>strand</td><td>MM</td><td>target_seq</td><td>PAM</td><td>alignment</td><td>distance</td><td>position</td><td>gene name</td><td>gene id</td></tr>
<tr><td>GU305914.1</td><td>182480</td><td>182502</td><td>+</td><td>0</td><td>CAGCATATATATATAAGAAA</td><td>TGG</td><td>||||||||[||||||||||||]PAM</td><td>NA</td><td>Intergenic</td><td>NA</td><td>NA</td></tr>
<tr><td>T13824</td><td>ATATATATAAGAAATGGAGAAGG</td><td>1000</td></tr>
<tr><td>Oligo fwd</td><td>CACCATATATATAAGAAATGGAGA</td></tr>
<tr><td>Oligo rev</td><td>AAACTCTCCATTTCTTATATATAT</td></tr>
<tr><td>Chromosome</td><td>start</td><td>end</td><td>strand</td><td>MM</td><td>target_seq</td><td>PAM</td><td>alignment</td><td>distance</td><td>position</td><td>gene name</td><td>gene id</td></tr>
<tr><td>GU305914.1</td><td>182486</td><td>182508</td><td>+</td><td>0</td><td>ATATATATAAGAAATGGAGA</td><td>AGG</td><td>||||||||[||||||||||||]PAM</td><td>NA</td><td>Intergenic</td><td>NA</td><td>NA</td></tr>
<tr><td>T13825</td><td>TATATAAGAAATGGAGAAGGAGG</td><td>1000</td></tr>
<tr><td>Oligo fwd</td><td>CACCTATATAAGAAATGGAGAAGG</td></tr>
<tr><td>Oligo rev</td><td>AAACCCTTCTCCATTTCTTATATA</td></tr>
<tr><td>Chromosome</td><td>start</td><td>end</td><td>strand</td><td>MM</td><td>target_seq</td><td>PAM</td><td>alignment</td><td>distance</td><td>position</td><td>gene name</td><td>gene id</td></tr>
<tr><td>GU305914.1</td><td>182489</td><td>182511</td><td>+</td><td>0</td><td>TATATAAGAAATGGAGAAGG</td><td>AGG</td><td>||||||||[||||||||||||]PAM</td><td>NA</td><td>Intergenic</td><td>NA</td><td>NA</td></tr>
<tr><td>T13826</td><td>AACAAACACCGTACGCAGTATGG</td><td>1000</td></tr>
<tr><td>Oligo fwd</td><td>CACCAACAAACACCGTACGCAGTA</td></tr>
<tr><td>Oligo rev</td><td>AAACTACTGCGTACGGTGTTTGTT</td></tr>
<tr><td>Chromosome</td><td>start</td><td>end</td><td>strand</td><td>MM</td><td>target_seq</td><td>PAM</td><td>alignment</td><td>distance</td><td>position</td><td>gene name</td><td>gene id</td></tr>
<tr><td>GU305914.1</td><td>182582</td><td>182604</td><td>+</td><td>0</td><td>AACAAACACCGTACGCAGTA</td><td>TGG</td><td>||||||||[||||||||||||]PAM</td><td>NA</td><td>Intergenic</td><td>NA</td><td>NA</td></tr>
<tr><td>T13827</td><td>CAGTATGGAGCTAGAGAGCCAGG</td><td>1000</td></tr>
<tr><td>Oligo fwd</td><td>CACCCAGTATGGAGCTAGAGAGCC</td></tr>
<tr><td>Oligo rev</td><td>AAACGGCTCTCTAGCTCCATACTG</td></tr>
<tr><td>Chromosome</td><td>start</td><td>end</td><td>strand</td><td>MM</td><td>target_seq</td><td>PAM</td><td>alignment</td><td>distance</td><td>position</td><td>gene name</td><td>gene id</td></tr>
<tr><td>GU305914.1</td><td>182597</td><td>182619</td><td>+</td><td>0</td><td>CAGTATGGAGCTAGAGAGCC</td><td>AGG</td><td>||||||||[||||||||||||]PAM</td><td>NA</td><td>Intergenic</td><td>NA</td><td>NA</td></tr>
<tr><td>T13828</td><td>TATGGAGCTAGAGAGCCAGGTGG</td><td>1000</td></tr>
<tr><td>Oligo fwd</td><td>CACCTATGGAGCTAGAGAGCCAGG</td></tr>
<tr><td>Oligo rev</td><td>AAACCCTGGCTCTCTAGCTCCATA</td></tr>
<tr><td>Chromosome</td><td>start</td><td>end</td><td>strand</td><td>MM</td><td>target_seq</td><td>PAM</td><td>alignment</td><td>distance</td><td>position</td><td>gene name</td><td>gene id</td></tr>
<tr><td>GU305914.1</td><td>182600</td><td>182622</td><td>+</td><td>0</td><td>TATGGAGCTAGAGAGCCAGG</td><td>TGG</td><td>||||||||[||||||||||||]PAM</td><td>NA</td><td>Intergenic</td><td>NA</td><td>NA</td></tr>
<tr><td>T13829</td><td>CAGGTGGATAGTAGATAGATAGG</td><td>1000</td></tr>
<tr><td>Oligo fwd</td><td>CACCCAGGTGGATAGTAGATAGAT</td></tr>
<tr><td>Oligo rev</td><td>AAACATCTATCTACTATCCACCTG</td></tr>
<tr><td>Chromosome</td><td>start</td><td>end</td><td>strand</td><td>MM</td><td>target_seq</td><td>PAM</td><td>alignment</td><td>distance</td><td>position</td><td>gene name</td><td>gene id</td></tr>
<tr><td>GU305914.1</td><td>182616</td><td>182638</td><td>+</td><td>0</td><td>CAGGTGGATAGTAGATAGAT</td><td>AGG</td><td>||||||||[||||||||||||]PAM</td><td>NA</td><td>Intergenic</td><td>NA</td><td>NA</td></tr>
<tr><td>T13830</td><td>AGGTGGATAGTAGATAGATAGGG</td><td>1000</td></tr>
<tr><td>Oligo fwd</td><td>CACCAGGTGGATAGTAGATAGATA</td></tr>
<tr><td>Oligo rev</td><td>AAACTATCTATCTACTATCCACCT</td></tr>
<tr><td>Chromosome</td><td>start</td><td>end</td><td>strand</td><td>MM</td><td>target_seq</td><td>PAM</td><td>alignment</td><td>distance</td><td>position</td><td>gene name</td><td>gene id</td></tr>
<tr><td>GU305914.1</td><td>182617</td><td>182639</td><td>+</td><td>0</td><td>AGGTGGATAGTAGATAGATA</td><td>GGG</td><td>||||||||[||||||||||||]PAM</td><td>NA</td><td>Intergenic</td><td>NA</td><td>NA</td></tr>
<tr><td>T13831</td><td>GTAGATAGATAGGGATAGACAGG</td><td>1000</td></tr>
<tr><td>Oligo fwd</td><td>CACCGTAGATAGATAGGGATAGAC</td></tr>
<tr><td>Oligo rev</td><td>AAACGTCTATCCCTATCTATCTAC</td></tr>
<tr><td>Chromosome</td><td>start</td><td>end</td><td>strand</td><td>MM</td><td>target_seq</td><td>PAM</td><td>alignment</td><td>distance</td><td>position</td><td>gene name</td><td>gene id</td></tr>
<tr><td>GU305914.1</td><td>182626</td><td>182648</td><td>+</td><td>0</td><td>GTAGATAGATAGGGATAGAC</td><td>AGG</td><td>||||||||[||||||||||||]PAM</td><td>NA</td><td>Intergenic</td><td>NA</td><td>NA</td></tr>
</table>

<table>
<tr><td>T13832</td><td>TTAACACCGCAGCCAATAAAAGG</td><td>1000</td><td></td><td></td><td></td><td></td><td></td><td></td><td></td><td></td><td></td></tr>
<tr><td>Oligo fwd</td><td>CACCTTAACACCGCAGCCAATAAA</td><td></td><td></td><td></td><td></td><td></td><td></td><td></td><td></td><td></td><td></td></tr>
<tr><td>Oligo rev</td><td>AAACTTTATTGGCTGCGGTGTTAA</td><td></td><td></td><td></td><td></td><td></td><td></td><td></td><td></td><td></td><td></td></tr>
<tr><td>Chromosome</td><td>start</td><td>end</td><td>strand</td><td>MM</td><td>target_seq</td><td>PAM</td><td>alignment</td><td>distance</td><td>position</td><td>gene name</td><td>gene id</td></tr>
<tr><td>GU305914.1</td><td>182658</td><td>182680</td><td>+</td><td>0</td><td>TTAACACCGCAGCCAATAAA</td><td>AGG</td><td>||||||||[||||||||||||]PAM</td><td>NA</td><td>Intergenic</td><td>NA</td><td>NA</td></tr>
<tr><td>T13833</td><td>CCGCAGCCAATAAAAGGCCTTGG</td><td>1000</td><td></td><td></td><td></td><td></td><td></td><td></td><td></td><td></td><td></td></tr>
<tr><td>Oligo fwd</td><td>CACCCCGCAGCCAATAAAAGGCCT</td><td></td><td></td><td></td><td></td><td></td><td></td><td></td><td></td><td></td><td></td></tr>
<tr><td>Oligo rev</td><td>AAACAGGCCTTTTATTGGCTGCGG</td><td></td><td></td><td></td><td></td><td></td><td></td><td></td><td></td><td></td><td></td></tr>
<tr><td>Chromosome</td><td>start</td><td>end</td><td>strand</td><td>MM</td><td>target_seq</td><td>PAM</td><td>alignment</td><td>distance</td><td>position</td><td>gene name</td><td>gene id</td></tr>
<tr><td>GU305914.1</td><td>182664</td><td>182686</td><td>+</td><td>0</td><td>CCGCAGCCAATAAAAGGCCT</td><td>TGG</td><td>||||||||[||||||||||||]PAM</td><td>NA</td><td>Intergenic</td><td>NA</td><td>NA</td></tr>
<tr><td>T13834</td><td>ACTTACATTGTGTGTCCTCGAGG</td><td>1000</td><td></td><td></td><td></td><td></td><td></td><td></td><td></td><td></td><td></td></tr>
<tr><td>Oligo fwd</td><td>CACCACTTACATTGTGTGTCCTCG</td><td></td><td></td><td></td><td></td><td></td><td></td><td></td><td></td><td></td><td></td></tr>
<tr><td>Oligo rev</td><td>AAACCGAGGACACACAATGTAAGT</td><td></td><td></td><td></td><td></td><td></td><td></td><td></td><td></td><td></td><td></td></tr>
<tr><td>Chromosome</td><td>start</td><td>end</td><td>strand</td><td>MM</td><td>target_seq</td><td>PAM</td><td>alignment</td><td>distance</td><td>position</td><td>gene name</td><td>gene id</td></tr>
<tr><td>GU305914.1</td><td>182690</td><td>182712</td><td>+</td><td>0</td><td>ACTTACATTGTGTGTCCTCG</td><td>AGG</td><td>||||||||[||||||||||||]PAM</td><td>NA</td><td>Intergenic</td><td>NA</td><td>NA</td></tr>
<tr><td>T13835</td><td>CTTACATTGTGTGTCCTCGAGGG</td><td>1000</td><td></td><td></td><td></td><td></td><td></td><td></td><td></td><td></td><td></td></tr>
<tr><td>Oligo fwd</td><td>CACCCTTACATTGTGTGTCCTCGA</td><td></td><td></td><td></td><td></td><td></td><td></td><td></td><td></td><td></td><td></td></tr>
<tr><td>Oligo rev</td><td>AAACTCGAGGACACACAATGTAAG</td><td></td><td></td><td></td><td></td><td></td><td></td><td></td><td></td><td></td><td></td></tr>
<tr><td>Chromosome</td><td>start</td><td>end</td><td>strand</td><td>MM</td><td>target_seq</td><td>PAM</td><td>alignment</td><td>distance</td><td>position</td><td>gene name</td><td>gene id</td></tr>
<tr><td>GU305914.1</td><td>182691</td><td>182713</td><td>+</td><td>0</td><td>CTTACATTGTGTGTCCTCGA</td><td>GGG</td><td>||||||||[||||||||||||]PAM</td><td>NA</td><td>Intergenic</td><td>NA</td><td>NA</td></tr>
<tr><td>T13836</td><td>TTACATTGTGTGTCCTCGAGGGG</td><td>1000</td><td></td><td></td><td></td><td></td><td></td><td></td><td></td><td></td><td></td></tr>
<tr><td>Oligo fwd</td><td>CACCTTACATTGTGTGTCCTCGAG</td><td></td><td></td><td></td><td></td><td></td><td></td><td></td><td></td><td></td><td></td></tr>
<tr><td>Oligo rev</td><td>AAACCTCGAGGACACACAATGTAA</td><td></td><td></td><td></td><td></td><td></td><td></td><td></td><td></td><td></td><td></td></tr>
<tr><td>Chromosome</td><td>start</td><td>end</td><td>strand</td><td>MM</td><td>target_seq</td><td>PAM</td><td>alignment</td><td>distance</td><td>position</td><td>gene name</td><td>gene id</td></tr>
<tr><td>GU305914.1</td><td>182692</td><td>182714</td><td>+</td><td>0</td><td>TTACATTGTGTGTCCTCGAG</td><td>GGG</td><td>||||||||[||||||||||||]PAM</td><td>NA</td><td>Intergenic</td><td>NA</td><td>NA</td></tr>
<tr><td>T13837</td><td>TACATTGTGTGTCCTCGAGGGGG</td><td>1000</td><td></td><td></td><td></td><td></td><td></td><td></td><td></td><td></td><td></td></tr>
<tr><td>Oligo fwd</td><td>CACCTACATTGTGTGTCCTCGAGG</td><td></td><td></td><td></td><td></td><td></td><td></td><td></td><td></td><td></td><td></td></tr>
<tr><td>Oligo rev</td><td>AAACCCTCGAGGACACACAATGTA</td><td></td><td></td><td></td><td></td><td></td><td></td><td></td><td></td><td></td><td></td></tr>
<tr><td>Chromosome</td><td>start</td><td>end</td><td>strand</td><td>MM</td><td>target_seq</td><td>PAM</td><td>alignment</td><td>distance</td><td>position</td><td>gene name</td><td>gene id</td></tr>
<tr><td>GU305914.1</td><td>182693</td><td>182715</td><td>+</td><td>0</td><td>TACATTGTGTGTCCTCGAGG</td><td>GGG</td><td>||||||||[||||||||||||]PAM</td><td>NA</td><td>Intergenic</td><td>NA</td><td>NA</td></tr>
<tr><td>T13838</td><td>GTGTGTCCTCGAGGGGGCGCTGG</td><td>1000</td><td></td><td></td><td></td><td></td><td></td><td></td><td></td><td></td><td></td></tr>
<tr><td>Oligo fwd</td><td>CACCGTGTGTCCTCGAGGGGGCGC</td><td></td><td></td><td></td><td></td><td></td><td></td><td></td><td></td><td></td><td></td></tr>
<tr><td>Oligo rev</td><td>AAACGCGCCCCCTCGAGGACACAC</td><td></td><td></td><td></td><td></td><td></td><td></td><td></td><td></td><td></td><td></td></tr>
<tr><td>Chromosome</td><td>start</td><td>end</td><td>strand</td><td>MM</td><td>target_seq</td><td>PAM</td><td>alignment</td><td>distance</td><td>position</td><td>gene name</td><td>gene id</td></tr>
<tr><td>GU305914.1</td><td>182699</td><td>182721</td><td>+</td><td>0</td><td>GTGTGTCCTCGAGGGGGCGC</td><td>TGG</td><td>||||||||[||||||||||||]PAM</td><td>NA</td><td>Intergenic</td><td>NA</td><td>NA</td></tr>
<tr><td>T13839</td><td>TGTGTCCTCGAGGGGGCGCTGGG</td><td>1000</td><td></td><td></td><td></td><td></td><td></td><td></td><td></td><td></td><td></td></tr>
<tr><td>Oligo fwd</td><td>CACCTGTGTCCTCGAGGGGGCGCT</td><td></td><td></td><td></td><td></td><td></td><td></td><td></td><td></td><td></td><td></td></tr>
<tr><td>Oligo rev</td><td>AAACAGCGCCCCCTCGAGGACACA</td><td></td><td></td><td></td><td></td><td></td><td></td><td></td><td></td><td></td><td></td></tr>
<tr><td>Chromosome</td><td>start</td><td>end</td><td>strand</td><td>MM</td><td>target_seq</td><td>PAM</td><td>alignment</td><td>distance</td><td>position</td><td>gene name</td><td>gene id</td></tr>
<tr><td>GU305914.1</td><td>182700</td><td>182722</td><td>+</td><td>0</td><td>TGTGTCCTCGAGGGGGCGCT</td><td>GGG</td><td>||||||||[||||||||||||]PAM</td><td>NA</td><td>Intergenic</td><td>NA</td><td>NA</td></tr>
<tr><td>T13840</td><td>TCGAGGGGGCGCTGGGCAGCCGG</td><td>1000</td><td></td><td></td><td></td><td></td><td></td><td></td><td></td><td></td><td></td></tr>
<tr><td>Oligo fwd</td><td>CACCTCGAGGGGGCGCTGGGCAGC</td><td></td><td></td><td></td><td></td><td></td><td></td><td></td><td></td><td></td><td></td></tr>
<tr><td>Oligo rev</td><td>AAACGCTGCCCAGCGCCCCCTCGA</td><td></td><td></td><td></td><td></td><td></td><td></td><td></td><td></td><td></td><td></td></tr>
<tr><td>Chromosome</td><td>start</td><td>end</td><td>strand</td><td>MM</td><td>target_seq</td><td>PAM</td><td>alignment</td><td>distance</td><td>position</td><td>gene name</td><td>gene id</td></tr>
<tr><td>GU305914.1</td><td>182707</td><td>182729</td><td>+</td><td>0</td><td>TCGAGGGGGCGCTGGGCAGC</td><td>CGG</td><td>||||||||[||||||||||||]PAM</td><td>NA</td><td>Intergenic</td><td>NA</td><td>NA</td></tr>
<tr><td>T13841</td><td>CGAGGGGGCGCTGGGCAGCCGGG</td><td>1000</td><td></td><td></td><td></td><td></td><td></td><td></td><td></td><td></td><td></td></tr>
<tr><td>Oligo fwd</td><td>CACCCGAGGGGGCGCTGGGCAGCC</td><td></td><td></td><td></td><td></td><td></td><td></td><td></td><td></td><td></td><td></td></tr>
<tr><td>Oligo rev</td><td>AAACGGCTGCCCAGCGCCCCCTCG</td><td></td><td></td><td></td><td></td><td></td><td></td><td></td><td></td><td></td><td></td></tr>
<tr><td>Chromosome</td><td>start</td><td>end</td><td>strand</td><td>MM</td><td>target_seq</td><td>PAM</td><td>alignment</td><td>distance</td><td>position</td><td>gene name</td><td>gene id</td></tr>
<tr><td>GU305914.1</td><td>182708</td><td>182730</td><td>+</td><td>0</td><td>CGAGGGGGCGCTGGGCAGCC</td><td>GGG</td><td>||||||||[||||||||||||]PAM</td><td>NA</td><td>Intergenic</td><td>NA</td><td>NA</td></tr>
<tr><td>T13842</td><td>GAGGGGGCGCTGGGCAGCCGGGG</td><td>1000</td><td></td><td></td><td></td><td></td><td></td><td></td><td></td><td></td><td></td></tr>
<tr><td>Oligo fwd</td><td>CACCGAGGGGGCGCTGGGCAGCCG</td><td></td><td></td><td></td><td></td><td></td><td></td><td></td><td></td><td></td><td></td></tr>
<tr><td>Oligo rev</td><td>AAACCGGCTGCCCAGCGCCCCCTC</td><td></td><td></td><td></td><td></td><td></td><td></td><td></td><td></td><td></td><td></td></tr>
<tr><td>Chromosome</td><td>start</td><td>end</td><td>strand</td><td>MM</td><td>target_seq</td><td>PAM</td><td>alignment</td><td>distance</td><td>position</td><td>gene name</td><td>gene id</td></tr>
<tr><td>GU305914.1</td><td>182709</td><td>182731</td><td>+</td><td>0</td><td>GAGGGGGCGCTGGGCAGCCG</td><td>GGG</td><td>||||||||[||||||||||||]PAM</td><td>NA</td><td>Intergenic</td><td>NA</td><td>NA</td></tr>
<tr><td>T13843</td><td>AGGGGGCGCTGGGCAGCCGGGGG</td><td>1000</td><td></td><td></td><td></td><td></td><td></td><td></td><td></td><td></td><td></td></tr>
<tr><td>Oligo fwd</td><td>CACCAGGGGGCGCTGGGCAGCCGG</td><td></td><td></td><td></td><td></td><td></td><td></td><td></td><td></td><td></td><td></td></tr>
<tr><td>Oligo rev</td><td>AAACCCGGCTGCCCAGCGCCCCCT</td><td></td><td></td><td></td><td></td><td></td><td></td><td></td><td></td><td></td><td></td></tr>
<tr><td>Chromosome</td><td>start</td><td>end</td><td>strand</td><td>MM</td><td>target_seq</td><td>PAM</td><td>alignment</td><td>distance</td><td>position</td><td>gene name</td><td>gene id</td></tr>
<tr><td>GU305914.1</td><td>182710</td><td>182732</td><td>+</td><td>0</td><td>AGGGGGCGCTGGGCAGCCGG</td><td>GGG</td><td>||||||||[||||||||||||]PAM</td><td>NA</td><td>Intergenic</td><td>NA</td><td>NA</td></tr>
<tr><td>T13844</td><td>CGCTGGGCAGCCGGGGGTCCTGG</td><td>1000</td><td></td><td></td><td></td><td></td><td></td><td></td><td></td><td></td><td></td></tr>
<tr><td>Oligo fwd</td><td>CACCCGCTGGGCAGCCGGGGGTCC</td><td></td><td></td><td></td><td></td><td></td><td></td><td></td><td></td><td></td><td></td></tr>
<tr><td>Oligo rev</td><td>AAACGGACCCCCGGCTGCCCAGCG</td><td></td><td></td><td></td><td></td><td></td><td></td><td></td><td></td><td></td><td></td></tr>
<tr><td>Chromosome</td><td>start</td><td>end</td><td>strand</td><td>MM</td><td>target_seq</td><td>PAM</td><td>alignment</td><td>distance</td><td>position</td><td>gene name</td><td>gene id</td></tr>
<tr><td>GU305914.1</td><td>182716</td><td>182738</td><td>+</td><td>0</td><td>CGCTGGGCAGCCGGGGGTCC</td><td>TGG</td><td>||||||||[||||||||||||]PAM</td><td>NA</td><td>Intergenic</td><td>NA</td><td>NA</td></tr>
<tr><td>T13845</td><td>GGTCCTGGTGACGTAGCCTCTGG</td><td>1000</td><td></td><td></td><td></td><td></td><td></td><td></td><td></td><td></td><td></td></tr>
<tr><td>Oligo fwd</td><td>CACCGGTCCTGGTGACGTAGCCTC</td><td></td><td></td><td></td><td></td><td></td><td></td><td></td><td></td><td></td><td></td></tr>
<tr><td>Oligo rev</td><td>AAACGAGGCTACGTCACCAGGACC</td><td></td><td></td><td></td><td></td><td></td><td></td><td></td><td></td><td></td><td></td></tr>
<tr><td>Chromosome</td><td>start</td><td>end</td><td>strand</td><td>MM</td><td>target_seq</td><td>PAM</td><td>alignment</td><td>distance</td><td>position</td><td>gene name</td><td>gene id</td></tr>
<tr><td>GU305914.1</td><td>182731</td><td>182753</td><td>+</td><td>0</td><td>GGTCCTGGTGACGTAGCCTC</td><td>TGG</td><td>||||||||[||||||||||||]PAM</td><td>NA</td><td>Intergenic</td><td>NA</td><td>NA</td></tr>
<tr><td>T13846</td><td>GTCCTGGTGACGTAGCCTCTGGG</td><td>1000</td><td></td><td></td><td></td><td></td><td></td><td></td><td></td><td></td><td></td></tr>
<tr><td>Oligo fwd</td><td>CACCGTCCTGGTGACGTAGCCTCT</td><td></td><td></td><td></td><td></td><td></td><td></td><td></td><td></td><td></td><td></td></tr>
<tr><td>Oligo rev</td><td>AAACAGAGGCTACGTCACCAGGAC</td><td></td><td></td><td></td><td></td><td></td><td></td><td></td><td></td><td></td><td></td></tr>
<tr><td>Chromosome</td><td>start</td><td>end</td><td>strand</td><td>MM</td><td>target_seq</td><td>PAM</td><td>alignment</td><td>distance</td><td>position</td><td>gene name</td><td>gene id</td></tr>
<tr><td>GU305914.1</td><td>182732</td><td>182754</td><td>+</td><td>0</td><td>GTCCTGGTGACGTAGCCTCT</td><td>GGG</td><td>||||||||[||||||||||||]PAM</td><td>NA</td><td>Intergenic</td><td>NA</td><td>NA</td></tr>
</table>

<table>
<tr><td>T13847</td><td>GGTGACGTAGCCTCTGGGCCTGG</td><td>1000</td><td></td><td></td><td></td><td></td><td></td><td></td><td></td><td></td><td></td></tr>
<tr><td>Oligo fwd</td><td>CACCGGTGACGTAGCCTCTGGGCC</td><td></td><td></td><td></td><td></td><td></td><td></td><td></td><td></td><td></td><td></td></tr>
<tr><td>Oligo rev</td><td>AAACGGCCCAGAGGCTACGTCACC</td><td></td><td></td><td></td><td></td><td></td><td></td><td></td><td></td><td></td><td></td></tr>
<tr><td>Chromosome</td><td>start</td><td>end</td><td>strand</td><td>MM</td><td>target_seq</td><td>PAM</td><td>alignment</td><td>distance</td><td>position</td><td>gene name</td><td>gene id</td></tr>
<tr><td>GU305914.1</td><td>182737</td><td>182759</td><td>+</td><td>0</td><td>GGTGACGTAGCCTCTGGGCC</td><td>TGG</td><td>||||||||[||||||||||||]PAM</td><td>NA</td><td>Intergenic</td><td>NA</td><td>NA</td></tr>
<tr><td>T13848</td><td>GTGACGTAGCCTCTGGGCCTGGG</td><td>1000</td><td></td><td></td><td></td><td></td><td></td><td></td><td></td><td></td><td></td></tr>
<tr><td>Oligo fwd</td><td>CACCGTGACGTAGCCTCTGGGCCT</td><td></td><td></td><td></td><td></td><td></td><td></td><td></td><td></td><td></td><td></td></tr>
<tr><td>Oligo rev</td><td>AAACAGGCCCAGAGGCTACGTCAC</td><td></td><td></td><td></td><td></td><td></td><td></td><td></td><td></td><td></td><td></td></tr>
<tr><td>Chromosome</td><td>start</td><td>end</td><td>strand</td><td>MM</td><td>target_seq</td><td>PAM</td><td>alignment</td><td>distance</td><td>position</td><td>gene name</td><td>gene id</td></tr>
<tr><td>GU305914.1</td><td>182738</td><td>182760</td><td>+</td><td>0</td><td>GTGACGTAGCCTCTGGGCCT</td><td>GGG</td><td>||||||||[||||||||||||]PAM</td><td>NA</td><td>Intergenic</td><td>NA</td><td>NA</td></tr>
<tr><td>T13849</td><td>AGCCTCTGGGCCTGGGTCGCTGG</td><td>1000</td><td></td><td></td><td></td><td></td><td></td><td></td><td></td><td></td><td></td></tr>
<tr><td>Oligo fwd</td><td>CACCAGCCTCTGGGCCTGGGTCGC</td><td></td><td></td><td></td><td></td><td></td><td></td><td></td><td></td><td></td><td></td></tr>
<tr><td>Oligo rev</td><td>AAACGCGACCCAGGCCCAGAGGCT</td><td></td><td></td><td></td><td></td><td></td><td></td><td></td><td></td><td></td><td></td></tr>
<tr><td>Chromosome</td><td>start</td><td>end</td><td>strand</td><td>MM</td><td>target_seq</td><td>PAM</td><td>alignment</td><td>distance</td><td>position</td><td>gene name</td><td>gene id</td></tr>
<tr><td>GU305914.1</td><td>182745</td><td>182767</td><td>+</td><td>0</td><td>AGCCTCTGGGCCTGGGTCGC</td><td>TGG</td><td>||||||||[||||||||||||]PAM</td><td>NA</td><td>Intergenic</td><td>NA</td><td>NA</td></tr>
<tr><td>T13850</td><td>GCCTCTGGGCCTGGGTCGCTGGG</td><td>1000</td><td></td><td></td><td></td><td></td><td></td><td></td><td></td><td></td><td></td></tr>
<tr><td>Oligo fwd</td><td>CACCGCCTCTGGGCCTGGGTCGCT</td><td></td><td></td><td></td><td></td><td></td><td></td><td></td><td></td><td></td><td></td></tr>
<tr><td>Oligo rev</td><td>AAACAGCGACCCAGGCCCAGAGGC</td><td></td><td></td><td></td><td></td><td></td><td></td><td></td><td></td><td></td><td></td></tr>
<tr><td>Chromosome</td><td>start</td><td>end</td><td>strand</td><td>MM</td><td>target_seq</td><td>PAM</td><td>alignment</td><td>distance</td><td>position</td><td>gene name</td><td>gene id</td></tr>
<tr><td>GU305914.1</td><td>182746</td><td>182768</td><td>+</td><td>0</td><td>GCCTCTGGGCCTGGGTCGCT</td><td>GGG</td><td>||||||||[||||||||||||]PAM</td><td>NA</td><td>Intergenic</td><td>NA</td><td>NA</td></tr>
<tr><td>T13851</td><td>TCTGGGCCTGGGTCGCTGGGTGG</td><td>1000</td><td></td><td></td><td></td><td></td><td></td><td></td><td></td><td></td><td></td></tr>
<tr><td>Oligo fwd</td><td>CACCTCTGGGCCTGGGTCGCTGGG</td><td></td><td></td><td></td><td></td><td></td><td></td><td></td><td></td><td></td><td></td></tr>
<tr><td>Oligo rev</td><td>AAACCCCAGCGACCCAGGCCCAGA</td><td></td><td></td><td></td><td></td><td></td><td></td><td></td><td></td><td></td><td></td></tr>
<tr><td>Chromosome</td><td>start</td><td>end</td><td>strand</td><td>MM</td><td>target_seq</td><td>PAM</td><td>alignment</td><td>distance</td><td>position</td><td>gene name</td><td>gene id</td></tr>
<tr><td>GU305914.1</td><td>182749</td><td>182771</td><td>+</td><td>0</td><td>TCTGGGCCTGGGTCGCTGGG</td><td>TGG</td><td>||||||||[||||||||||||]PAM</td><td>NA</td><td>Intergenic</td><td>NA</td><td>NA</td></tr>
<tr><td>T13852</td><td>CTGGGCCTGGGTCGCTGGGTGGG</td><td>1000</td><td></td><td></td><td></td><td></td><td></td><td></td><td></td><td></td><td></td></tr>
<tr><td>Oligo fwd</td><td>CACCCTGGGCCTGGGTCGCTGGGT</td><td></td><td></td><td></td><td></td><td></td><td></td><td></td><td></td><td></td><td></td></tr>
<tr><td>Oligo rev</td><td>AAACACCCAGCGACCCAGGCCCAG</td><td></td><td></td><td></td><td></td><td></td><td></td><td></td><td></td><td></td><td></td></tr>
<tr><td>Chromosome</td><td>start</td><td>end</td><td>strand</td><td>MM</td><td>target_seq</td><td>PAM</td><td>alignment</td><td>distance</td><td>position</td><td>gene name</td><td>gene id</td></tr>
<tr><td>GU305914.1</td><td>182750</td><td>182772</td><td>+</td><td>0</td><td>CTGGGCCTGGGTCGCTGGGT</td><td>GGG</td><td>||||||||[||||||||||||]PAM</td><td>NA</td><td>Intergenic</td><td>NA</td><td>NA</td></tr>
<tr><td>T13853</td><td>GGCCTGGGTCGCTGGGTGGGCGG</td><td>1000</td><td></td><td></td><td></td><td></td><td></td><td></td><td></td><td></td><td></td></tr>
<tr><td>Oligo fwd</td><td>CACCGGCCTGGGTCGCTGGGTGGG</td><td></td><td></td><td></td><td></td><td></td><td></td><td></td><td></td><td></td><td></td></tr>
<tr><td>Oligo rev</td><td>AAACCCCACCCAGCGACCCAGGCC</td><td></td><td></td><td></td><td></td><td></td><td></td><td></td><td></td><td></td><td></td></tr>
<tr><td>Chromosome</td><td>start</td><td>end</td><td>strand</td><td>MM</td><td>target_seq</td><td>PAM</td><td>alignment</td><td>distance</td><td>position</td><td>gene name</td><td>gene id</td></tr>
<tr><td>GU305914.1</td><td>182753</td><td>182775</td><td>+</td><td>0</td><td>GGCCTGGGTCGCTGGGTGGG</td><td>CGG</td><td>||||||||[||||||||||||]PAM</td><td>NA</td><td>Intergenic</td><td>NA</td><td>NA</td></tr>
<tr><td>T13854</td><td>CTGGGTCGCTGGGTGGGCGGTGG</td><td>1000</td><td></td><td></td><td></td><td></td><td></td><td></td><td></td><td></td><td></td></tr>
<tr><td>Oligo fwd</td><td>CACCCTGGGTCGCTGGGTGGGCGG</td><td></td><td></td><td></td><td></td><td></td><td></td><td></td><td></td><td></td><td></td></tr>
<tr><td>Oligo rev</td><td>AAACCCGCCCACCCAGCGACCCAG</td><td></td><td></td><td></td><td></td><td></td><td></td><td></td><td></td><td></td><td></td></tr>
<tr><td>Chromosome</td><td>start</td><td>end</td><td>strand</td><td>MM</td><td>target_seq</td><td>PAM</td><td>alignment</td><td>distance</td><td>position</td><td>gene name</td><td>gene id</td></tr>
<tr><td>GU305914.1</td><td>182756</td><td>182778</td><td>+</td><td>0</td><td>CTGGGTCGCTGGGTGGGCGG</td><td>TGG</td><td>||||||||[||||||||||||]PAM</td><td>NA</td><td>Intergenic</td><td>NA</td><td>NA</td></tr>
<tr><td>T13855</td><td>GGGTGGGCGGTGGTCCCAATTGG</td><td>1000</td><td></td><td></td><td></td><td></td><td></td><td></td><td></td><td></td><td></td></tr>
<tr><td>Oligo fwd</td><td>CACCGGGTGGGCGGTGGTCCCAAT</td><td></td><td></td><td></td><td></td><td></td><td></td><td></td><td></td><td></td><td></td></tr>
<tr><td>Oligo rev</td><td>AAACATTGGGACCACCGCCCACCC</td><td></td><td></td><td></td><td></td><td></td><td></td><td></td><td></td><td></td><td></td></tr>
<tr><td>Chromosome</td><td>start</td><td>end</td><td>strand</td><td>MM</td><td>target_seq</td><td>PAM</td><td>alignment</td><td>distance</td><td>position</td><td>gene name</td><td>gene id</td></tr>
<tr><td>GU305914.1</td><td>182766</td><td>182788</td><td>+</td><td>0</td><td>GGGTGGGCGGTGGTCCCAAT</td><td>TGG</td><td>||||||||[||||||||||||]PAM</td><td>NA</td><td>Intergenic</td><td>NA</td><td>NA</td></tr>
<tr><td>GU305914.1</td><td>182783</td><td>182805</td><td>+</td><td>3</td><td>AATTGGGCGGTGGTCCCAAT</td><td>AGG</td><td>---|||||[||||||||||||]PAM</td><td>NA</td><td>Intergenic</td><td>NA</td><td>NA</td></tr>
<tr><td>T13856</td><td>GGTGGGCGGTGGTCCCAATTGGG</td><td>1000</td><td></td><td></td><td></td><td></td><td></td><td></td><td></td><td></td><td></td></tr>
<tr><td>Oligo fwd</td><td>CACCGGTGGGCGGTGGTCCCAATT</td><td></td><td></td><td></td><td></td><td></td><td></td><td></td><td></td><td></td><td></td></tr>
<tr><td>Oligo rev</td><td>AAACAATTGGGACCACCGCCCACC</td><td></td><td></td><td></td><td></td><td></td><td></td><td></td><td></td><td></td><td></td></tr>
<tr><td>Chromosome</td><td>start</td><td>end</td><td>strand</td><td>MM</td><td>target_seq</td><td>PAM</td><td>alignment</td><td>distance</td><td>position</td><td>gene name</td><td>gene id</td></tr>
<tr><td>GU305914.1</td><td>182767</td><td>182789</td><td>+</td><td>0</td><td>GGTGGGCGGTGGTCCCAATT</td><td>GGG</td><td>||||||||[||||||||||||]PAM</td><td>NA</td><td>Intergenic</td><td>NA</td><td>NA</td></tr>
<tr><td>GU305914.1</td><td>182784</td><td>182806</td><td>+</td><td>3</td><td>ATTGGGCGGTGGTCCCAATA</td><td>GGG</td><td>--||||||[|||||||||||-]PAM</td><td>NA</td><td>Intergenic</td><td>NA</td><td>NA</td></tr>
<tr><td>T13857</td><td>GGGCGGTGGTCCCAATTGGGCGG</td><td>1000</td><td></td><td></td><td></td><td></td><td></td><td></td><td></td><td></td><td></td></tr>
<tr><td>Oligo fwd</td><td>CACCGGGCGGTGGTCCCAATTGGG</td><td></td><td></td><td></td><td></td><td></td><td></td><td></td><td></td><td></td><td></td></tr>
<tr><td>Oligo rev</td><td>AAACCCCAATTGGGACCACCGCCC</td><td></td><td></td><td></td><td></td><td></td><td></td><td></td><td></td><td></td><td></td></tr>
<tr><td>Chromosome</td><td>start</td><td>end</td><td>strand</td><td>MM</td><td>target_seq</td><td>PAM</td><td>alignment</td><td>distance</td><td>position</td><td>gene name</td><td>gene id</td></tr>
<tr><td>GU305914.1</td><td>182770</td><td>182792</td><td>+</td><td>0</td><td>GGGCGGTGGTCCCAATTGGG</td><td>CGG</td><td>||||||||[||||||||||||]PAM</td><td>NA</td><td>Intergenic</td><td>NA</td><td>NA</td></tr>
<tr><td>T13858</td><td>CGGTGGTCCCAATTGGGCGGTGG</td><td>1000</td><td></td><td></td><td></td><td></td><td></td><td></td><td></td><td></td><td></td></tr>
<tr><td>Oligo fwd</td><td>CACCCGGTGGTCCCAATTGGGCGG</td><td></td><td></td><td></td><td></td><td></td><td></td><td></td><td></td><td></td><td></td></tr>
<tr><td>Oligo rev</td><td>AAACCCGCCCAATTGGGACCACCG</td><td></td><td></td><td></td><td></td><td></td><td></td><td></td><td></td><td></td><td></td></tr>
<tr><td>Chromosome</td><td>start</td><td>end</td><td>strand</td><td>MM</td><td>target_seq</td><td>PAM</td><td>alignment</td><td>distance</td><td>position</td><td>gene name</td><td>gene id</td></tr>
<tr><td>GU305914.1</td><td>182773</td><td>182795</td><td>+</td><td>0</td><td>CGGTGGTCCCAATTGGGCGG</td><td>TGG</td><td>||||||||[||||||||||||]PAM</td><td>NA</td><td>Intergenic</td><td>NA</td><td>NA</td></tr>
<tr><td>T13859</td><td>AATTGGGCGGTGGTCCCAATAGG</td><td>1000</td><td></td><td></td><td></td><td></td><td></td><td></td><td></td><td></td><td></td></tr>
<tr><td>Oligo fwd</td><td>CACCAATTGGGCGGTGGTCCCAAT</td><td></td><td></td><td></td><td></td><td></td><td></td><td></td><td></td><td></td><td></td></tr>
<tr><td>Oligo rev</td><td>AAACATTGGGACCACCGCCCAATT</td><td></td><td></td><td></td><td></td><td></td><td></td><td></td><td></td><td></td><td></td></tr>
<tr><td>Chromosome</td><td>start</td><td>end</td><td>strand</td><td>MM</td><td>target_seq</td><td>PAM</td><td>alignment</td><td>distance</td><td>position</td><td>gene name</td><td>gene id</td></tr>
<tr><td>GU305914.1</td><td>182783</td><td>182805</td><td>+</td><td>0</td><td>AATTGGGCGGTGGTCCCAAT</td><td>AGG</td><td>||||||||[||||||||||||]PAM</td><td>NA</td><td>Intergenic</td><td>NA</td><td>NA</td></tr>
<tr><td>GU305914.1</td><td>182766</td><td>182788</td><td>+</td><td>3</td><td>GGGTGGGCGGTGGTCCCAAT</td><td>TGG</td><td>---|||||[||||||||||||]PAM</td><td>NA</td><td>Intergenic</td><td>NA</td><td>NA</td></tr>
<tr><td>T13860</td><td>ATTGGGCGGTGGTCCCAATAGGG</td><td>1000</td><td></td><td></td><td></td><td></td><td></td><td></td><td></td><td></td><td></td></tr>
<tr><td>Oligo fwd</td><td>CACCATTGGGCGGTGGTCCCAATA</td><td></td><td></td><td></td><td></td><td></td><td></td><td></td><td></td><td></td><td></td></tr>
<tr><td>Oligo rev</td><td>AAACTATTGGGACCACCGCCCAAT</td><td></td><td></td><td></td><td></td><td></td><td></td><td></td><td></td><td></td><td></td></tr>
<tr><td>Chromosome</td><td>start</td><td>end</td><td>strand</td><td>MM</td><td>target_seq</td><td>PAM</td><td>alignment</td><td>distance</td><td>position</td><td>gene name</td><td>gene id</td></tr>
<tr><td>GU305914.1</td><td>182784</td><td>182806</td><td>+</td><td>0</td><td>ATTGGGCGGTGGTCCCAATA</td><td>GGG</td><td>||||||||[||||||||||||]PAM</td><td>NA</td><td>Intergenic</td><td>NA</td><td>NA</td></tr>
<tr><td>GU305914.1</td><td>182767</td><td>182789</td><td>+</td><td>3</td><td>GGTGGGCGGTGGTCCCAATT</td><td>GGG</td><td>--||||||[|||||||||||-]PAM</td><td>NA</td><td>Intergenic</td><td>NA</td><td>NA</td></tr>
</table>

<table>
<tr><td>T13861</td><td>TTGGGCGGTGGTCCCAATAGGGG</td><td>1000</td><td></td><td></td><td></td><td></td><td></td><td></td><td></td><td></td><td></td></tr>
<tr><td>Oligo fwd</td><td>CACCTTGGGCGGTGGTCCCAATAG</td><td></td><td></td><td></td><td></td><td></td><td></td><td></td><td></td><td></td><td></td></tr>
<tr><td>Oligo rev</td><td>AAACCTATTGGGACCACCGCCCAA</td><td></td><td></td><td></td><td></td><td></td><td></td><td></td><td></td><td></td><td></td></tr>
<tr><td>Chromosome</td><td>start</td><td>end</td><td>strand</td><td>MM</td><td>target_seq</td><td>PAM</td><td>alignment</td><td>distance</td><td>position</td><td>gene name</td><td>gene id</td></tr>
<tr><td>GU305914.1</td><td>182785</td><td>182807</td><td>+</td><td>0</td><td>TTGGGCGGTGGTCCCAATAG</td><td>GGG</td><td>||||||||[||||||||||||]PAM</td><td>NA</td><td>Intergenic</td><td>NA</td><td>NA</td></tr>
</table>

<table>
<tr><td>T13862</td><td>GGCGGTGGTCCCAATAGGGGCGG</td><td>1000</td><td></td><td></td><td></td><td></td><td></td><td></td><td></td><td></td><td></td></tr>
<tr><td>Oligo fwd</td><td>CACCGGCGGTGGTCCCAATAGGGG</td><td></td><td></td><td></td><td></td><td></td><td></td><td></td><td></td><td></td><td></td></tr>
<tr><td>Oligo rev</td><td>AAACCCCCTATTGGGACCACCGCC</td><td></td><td></td><td></td><td></td><td></td><td></td><td></td><td></td><td></td><td></td></tr>
<tr><td>Chromosome</td><td>start</td><td>end</td><td>strand</td><td>MM</td><td>target_seq</td><td>PAM</td><td>alignment</td><td>distance</td><td>position</td><td>gene name</td><td>gene id</td></tr>
<tr><td>GU305914.1</td><td>182788</td><td>182810</td><td>+</td><td>0</td><td>GGCGGTGGTCCCAATAGGGG</td><td>CGG</td><td>||||||||[||||||||||||]PAM</td><td>NA</td><td>Intergenic</td><td>NA</td><td>NA</td></tr>
</table>

<table>
<tr><td>T13863</td><td>GGTGGTCCCAATAGGGGCGGTGG</td><td>1000</td><td></td><td></td><td></td><td></td><td></td><td></td><td></td><td></td><td></td></tr>
<tr><td>Oligo fwd</td><td>CACCGGTGGTCCCAATAGGGGCGG</td><td></td><td></td><td></td><td></td><td></td><td></td><td></td><td></td><td></td><td></td></tr>
<tr><td>Oligo rev</td><td>AAACCCGCCCCTATTGGGACCACC</td><td></td><td></td><td></td><td></td><td></td><td></td><td></td><td></td><td></td><td></td></tr>
<tr><td>Chromosome</td><td>start</td><td>end</td><td>strand</td><td>MM</td><td>target_seq</td><td>PAM</td><td>alignment</td><td>distance</td><td>position</td><td>gene name</td><td>gene id</td></tr>
<tr><td>GU305914.1</td><td>182791</td><td>182813</td><td>+</td><td>0</td><td>GGTGGTCCCAATAGGGGCGG</td><td>TGG</td><td>||||||||[||||||||||||]PAM</td><td>NA</td><td>Intergenic</td><td>NA</td><td>NA</td></tr>
</table>

<table>
<tr><td>T13864</td><td>GGCGGTGGTCCCAATCAAGATGG</td><td>1000</td><td></td><td></td><td></td><td></td><td></td><td></td><td></td><td></td><td></td></tr>
<tr><td>Oligo fwd</td><td>CACCGGCGGTGGTCCCAATCAAGA</td><td></td><td></td><td></td><td></td><td></td><td></td><td></td><td></td><td></td><td></td></tr>
<tr><td>Oligo rev</td><td>AAACTCTTGATTGGGACCACCGCC</td><td></td><td></td><td></td><td></td><td></td><td></td><td></td><td></td><td></td><td></td></tr>
<tr><td>Chromosome</td><td>start</td><td>end</td><td>strand</td><td>MM</td><td>target_seq</td><td>PAM</td><td>alignment</td><td>distance</td><td>position</td><td>gene name</td><td>gene id</td></tr>
<tr><td>GU305914.1</td><td>182806</td><td>182828</td><td>+</td><td>0</td><td>GGCGGTGGTCCCAATCAAGA</td><td>TGG</td><td>||||||||[||||||||||||]PAM</td><td>NA</td><td>Intergenic</td><td>NA</td><td>NA</td></tr>
</table>

<table>
<tr><td>T13865</td><td>GGTGGTCCCAATCAAGATGGCGG</td><td>1000</td><td></td><td></td><td></td><td></td><td></td><td></td><td></td><td></td><td></td></tr>
<tr><td>Oligo fwd</td><td>CACCGGTGGTCCCAATCAAGATGG</td><td></td><td></td><td></td><td></td><td></td><td></td><td></td><td></td><td></td><td></td></tr>
<tr><td>Oligo rev</td><td>AAACCCATCTTGATTGGGACCACC</td><td></td><td></td><td></td><td></td><td></td><td></td><td></td><td></td><td></td><td></td></tr>
<tr><td>Chromosome</td><td>start</td><td>end</td><td>strand</td><td>MM</td><td>target_seq</td><td>PAM</td><td>alignment</td><td>distance</td><td>position</td><td>gene name</td><td>gene id</td></tr>
<tr><td>GU305914.1</td><td>182809</td><td>182831</td><td>+</td><td>0</td><td>GGTGGTCCCAATCAAGATGG</td><td>CGG</td><td>||||||||[||||||||||||]PAM</td><td>NA</td><td>Intergenic</td><td>NA</td><td>NA</td></tr>
</table>

<table>
<tr><td>T13866</td><td>AGATGGCGGCGTTACCAATATGG</td><td>1000</td><td></td><td></td><td></td><td></td><td></td><td></td><td></td><td></td><td></td></tr>
<tr><td>Oligo fwd</td><td>CACCAGATGGCGGCGTTACCAATA</td><td></td><td></td><td></td><td></td><td></td><td></td><td></td><td></td><td></td><td></td></tr>
<tr><td>Oligo rev</td><td>AAACTATTGGTAACGCCGCCATCT</td><td></td><td></td><td></td><td></td><td></td><td></td><td></td><td></td><td></td><td></td></tr>
<tr><td>Chromosome</td><td>start</td><td>end</td><td>strand</td><td>MM</td><td>target_seq</td><td>PAM</td><td>alignment</td><td>distance</td><td>position</td><td>gene name</td><td>gene id</td></tr>
<tr><td>GU305914.1</td><td>182823</td><td>182845</td><td>+</td><td>0</td><td>AGATGGCGGCGTTACCAATA</td><td>TGG</td><td>||||||||[||||||||||||]PAM</td><td>NA</td><td>Intergenic</td><td>NA</td><td>NA</td></tr>
</table>

<table>
<tr><td>T13867</td><td>TGGCGGCGTTACCAATATGGCGG</td><td>1000</td><td></td><td></td><td></td><td></td><td></td><td></td><td></td><td></td><td></td></tr>
<tr><td>Oligo fwd</td><td>CACCTGGCGGCGTTACCAATATGG</td><td></td><td></td><td></td><td></td><td></td><td></td><td></td><td></td><td></td><td></td></tr>
<tr><td>Oligo rev</td><td>AAACCCATATTGGTAACGCCGCCA</td><td></td><td></td><td></td><td></td><td></td><td></td><td></td><td></td><td></td><td></td></tr>
<tr><td>Chromosome</td><td>start</td><td>end</td><td>strand</td><td>MM</td><td>target_seq</td><td>PAM</td><td>alignment</td><td>distance</td><td>position</td><td>gene name</td><td>gene id</td></tr>
<tr><td>GU305914.1</td><td>182826</td><td>182848</td><td>+</td><td>0</td><td>TGGCGGCGTTACCAATATGG</td><td>CGG</td><td>||||||||[||||||||||||]PAM</td><td>NA</td><td>Intergenic</td><td>NA</td><td>NA</td></tr>
<tr><td>GU305914.1</td><td>182843</td><td>182865</td><td>+</td><td>3</td><td>TGGCGGCTGTTCCAATATGG</td><td>CGG</td><td>|||||||-[-|-|||||||||]PAM</td><td>NA</td><td>Intergenic</td><td>NA</td><td>NA</td></tr>
</table>

<table>
<tr><td>T13868</td><td>ATATGGCGGCTGTTCCAATATGG</td><td>1000</td><td></td><td></td><td></td><td></td><td></td><td></td><td></td><td></td><td></td></tr>
<tr><td>Oligo fwd</td><td>CACCATATGGCGGCTGTTCCAATA</td><td></td><td></td><td></td><td></td><td></td><td></td><td></td><td></td><td></td><td></td></tr>
<tr><td>Oligo rev</td><td>AAACTATTGGAACAGCCGCCATAT</td><td></td><td></td><td></td><td></td><td></td><td></td><td></td><td></td><td></td><td></td></tr>
<tr><td>Chromosome</td><td>start</td><td>end</td><td>strand</td><td>MM</td><td>target_seq</td><td>PAM</td><td>alignment</td><td>distance</td><td>position</td><td>gene name</td><td>gene id</td></tr>
<tr><td>GU305914.1</td><td>182840</td><td>182862</td><td>+</td><td>0</td><td>ATATGGCGGCTGTTCCAATA</td><td>TGG</td><td>||||||||[||||||||||||]PAM</td><td>NA</td><td>Intergenic</td><td>NA</td><td>NA</td></tr>
</table>

<table>
<tr><td>T13869</td><td>TGGCGGCTGTTCCAATATGGCGG</td><td>1000</td><td></td><td></td><td></td><td></td><td></td><td></td><td></td><td></td><td></td></tr>
<tr><td>Oligo fwd</td><td>CACCTGGCGGCTGTTCCAATATGG</td><td></td><td></td><td></td><td></td><td></td><td></td><td></td><td></td><td></td><td></td></tr>
<tr><td>Oligo rev</td><td>AAACCCATATTGGAACAGCCGCCA</td><td></td><td></td><td></td><td></td><td></td><td></td><td></td><td></td><td></td><td></td></tr>
<tr><td>Chromosome</td><td>start</td><td>end</td><td>strand</td><td>MM</td><td>target_seq</td><td>PAM</td><td>alignment</td><td>distance</td><td>position</td><td>gene name</td><td>gene id</td></tr>
<tr><td>GU305914.1</td><td>182843</td><td>182865</td><td>+</td><td>0</td><td>TGGCGGCTGTTCCAATATGG</td><td>CGG</td><td>||||||||[||||||||||||]PAM</td><td>NA</td><td>Intergenic</td><td>NA</td><td>NA</td></tr>
<tr><td>GU305914.1</td><td>182826</td><td>182848</td><td>+</td><td>3</td><td>TGGCGGCGTTACCAATATGG</td><td>CGG</td><td>|||||||-[-|-|||||||||]PAM</td><td>NA</td><td>Intergenic</td><td>NA</td><td>NA</td></tr>
</table>

<table>
<tr><td>T13870</td><td>GGCTGTTCCAATATGGCGGATGG</td><td>1000</td><td></td><td></td><td></td><td></td><td></td><td></td><td></td><td></td><td></td></tr>
<tr><td>Oligo fwd</td><td>CACCGGCTGTTCCAATATGGCGGA</td><td></td><td></td><td></td><td></td><td></td><td></td><td></td><td></td><td></td><td></td></tr>
<tr><td>Oligo rev</td><td>AAACTCCGCCATATTGGAACAGCC</td><td></td><td></td><td></td><td></td><td></td><td></td><td></td><td></td><td></td><td></td></tr>
<tr><td>Chromosome</td><td>start</td><td>end</td><td>strand</td><td>MM</td><td>target_seq</td><td>PAM</td><td>alignment</td><td>distance</td><td>position</td><td>gene name</td><td>gene id</td></tr>
<tr><td>GU305914.1</td><td>182847</td><td>182869</td><td>+</td><td>0</td><td>GGCTGTTCCAATATGGCGGA</td><td>TGG</td><td>||||||||[||||||||||||]PAM</td><td>NA</td><td>Intergenic</td><td>NA</td><td>NA</td></tr>
</table>

<table>
<tr><td>T13871</td><td>GCTGTTCCAATATGGCGGATGGG</td><td>1000</td><td></td><td></td><td></td><td></td><td></td><td></td><td></td><td></td><td></td></tr>
<tr><td>Oligo fwd</td><td>CACCGCTGTTCCAATATGGCGGAT</td><td></td><td></td><td></td><td></td><td></td><td></td><td></td><td></td><td></td><td></td></tr>
<tr><td>Oligo rev</td><td>AAACATCCGCCATATTGGAACAGC</td><td></td><td></td><td></td><td></td><td></td><td></td><td></td><td></td><td></td><td></td></tr>
<tr><td>Chromosome</td><td>start</td><td>end</td><td>strand</td><td>MM</td><td>target_seq</td><td>PAM</td><td>alignment</td><td>distance</td><td>position</td><td>gene name</td><td>gene id</td></tr>
<tr><td>GU305914.1</td><td>182848</td><td>182870</td><td>+</td><td>0</td><td>GCTGTTCCAATATGGCGGAT</td><td>GGG</td><td>||||||||[||||||||||||]PAM</td><td>NA</td><td>Intergenic</td><td>NA</td><td>NA</td></tr>
</table>

<table>
<tr><td>T13872</td><td>TCCAATATGGCGGATGGGCGTGG</td><td>1000</td><td></td><td></td><td></td><td></td><td></td><td></td><td></td><td></td><td></td></tr>
<tr><td>Oligo fwd</td><td>CACCTCCAATATGGCGGATGGGCG</td><td></td><td></td><td></td><td></td><td></td><td></td><td></td><td></td><td></td><td></td></tr>
<tr><td>Oligo rev</td><td>AAACCGCCCATCCGCCATATTGGA</td><td></td><td></td><td></td><td></td><td></td><td></td><td></td><td></td><td></td><td></td></tr>
<tr><td>Chromosome</td><td>start</td><td>end</td><td>strand</td><td>MM</td><td>target_seq</td><td>PAM</td><td>alignment</td><td>distance</td><td>position</td><td>gene name</td><td>gene id</td></tr>
<tr><td>GU305914.1</td><td>182853</td><td>182875</td><td>+</td><td>0</td><td>TCCAATATGGCGGATGGGCG</td><td>TGG</td><td>||||||||[||||||||||||]PAM</td><td>NA</td><td>Intergenic</td><td>NA</td><td>NA</td></tr>
</table>

<table>
<tr><td>T13873</td><td>GTGGTCCCAATAGAACCGAGCGG</td><td>1000</td><td></td><td></td><td></td><td></td><td></td><td></td><td></td><td></td><td></td></tr>
<tr><td>Oligo fwd</td><td>CACCGTGGTCCCAATAGAACCGAG</td><td></td><td></td><td></td><td></td><td></td><td></td><td></td><td></td><td></td><td></td></tr>
<tr><td>Oligo rev</td><td>AAACCTCGGTTCTATTGGGACCAC</td><td></td><td></td><td></td><td></td><td></td><td></td><td></td><td></td><td></td><td></td></tr>
<tr><td>Chromosome</td><td>start</td><td>end</td><td>strand</td><td>MM</td><td>target_seq</td><td>PAM</td><td>alignment</td><td>distance</td><td>position</td><td>gene name</td><td>gene id</td></tr>
<tr><td>GU305914.1</td><td>182872</td><td>182894</td><td>+</td><td>0</td><td>GTGGTCCCAATAGAACCGAG</td><td>CGG</td><td>||||||||[||||||||||||]PAM</td><td>NA</td><td>Intergenic</td><td>NA</td><td>NA</td></tr>
</table>

<table>
<tr><td>T13874</td><td>TGGTCCCAATAGAACCGAGCGGG</td><td>1000</td><td></td><td></td><td></td><td></td><td></td><td></td><td></td><td></td><td></td></tr>
<tr><td>Oligo fwd</td><td>CACCTGGTCCCAATAGAACCGAGC</td><td></td><td></td><td></td><td></td><td></td><td></td><td></td><td></td><td></td><td></td></tr>
<tr><td>Oligo rev</td><td>AAACGCTCGGTTCTATTGGGACCA</td><td></td><td></td><td></td><td></td><td></td><td></td><td></td><td></td><td></td><td></td></tr>
<tr><td>Chromosome</td><td>start</td><td>end</td><td>strand</td><td>MM</td><td>target_seq</td><td>PAM</td><td>alignment</td><td>distance</td><td>position</td><td>gene name</td><td>gene id</td></tr>
<tr><td>GU305914.1</td><td>182873</td><td>182895</td><td>+</td><td>0</td><td>TGGTCCCAATAGAACCGAGC</td><td>GGG</td><td>||||||||[||||||||||||]PAM</td><td>NA</td><td>Intergenic</td><td>NA</td><td>NA</td></tr>
</table>

<table>
<tr><td>T13875</td><td>GTCCGATGCGCTTCAGTAGCCGG</td><td>1000</td></tr>
<tr><td>Oligo fwd</td><td>CACCGTCCGATGCGCTTCAGTAGC</td><td></td></tr>
<tr><td>Oligo rev</td><td>AAACGCTACTGAAGCGCATCGGAC</td><td></td></tr>
</table>

<table>
<tr><td>Chromosome</td><td>start</td><td>end</td><td>strand</td><td>MM</td><td>target_seq</td><td>PAM</td><td>alignment</td><td>distance</td><td>position</td><td>gene name</td><td>gene id</td></tr>
<tr><td>GU305914.1</td><td>182895</td><td>182917</td><td>+</td><td>0</td><td>GTCCGATGCGCTTCAGTAGC</td><td>CGG</td><td>||||||||[||||||||||||]PAM</td><td>NA</td><td>Intergenic</td><td>NA</td><td>NA</td></tr>
<tr><td>T13876</td><td>TCCGATGCGCTTCAGTAGCCGGG</td><td>1000</td><td></td><td></td><td></td><td></td><td></td><td></td><td></td><td></td><td></td></tr>
<tr><td>Oligo fwd</td><td>CACCTCCGATGCGCTTCAGTAGCC</td><td></td><td></td><td></td><td></td><td></td><td></td><td></td><td></td><td></td><td></td></tr>
<tr><td>Oligo rev</td><td>AAACGGCTACTGAAGCGCATCGGA</td><td></td><td></td><td></td><td></td><td></td><td></td><td></td><td></td><td></td><td></td></tr>
<tr><td>Chromosome</td><td>start</td><td>end</td><td>strand</td><td>MM</td><td>target_seq</td><td>PAM</td><td>alignment</td><td>distance</td><td>position</td><td>gene name</td><td>gene id</td></tr>
<tr><td>GU305914.1</td><td>182896</td><td>182918</td><td>+</td><td>0</td><td>TCCGATGCGCTTCAGTAGCC</td><td>GGG</td><td>||||||||[||||||||||||]PAM</td><td>NA</td><td>Intergenic</td><td>NA</td><td>NA</td></tr>
<tr><td>T13877</td><td>CCGATGCGCTTCAGTAGCCGGGG</td><td>1000</td><td></td><td></td><td></td><td></td><td></td><td></td><td></td><td></td><td></td></tr>
<tr><td>Oligo fwd</td><td>CACCCCGATGCGCTTCAGTAGCCG</td><td></td><td></td><td></td><td></td><td></td><td></td><td></td><td></td><td></td><td></td></tr>
<tr><td>Oligo rev</td><td>AAACCGGCTACTGAAGCGCATCGG</td><td></td><td></td><td></td><td></td><td></td><td></td><td></td><td></td><td></td><td></td></tr>
<tr><td>Chromosome</td><td>start</td><td>end</td><td>strand</td><td>MM</td><td>target_seq</td><td>PAM</td><td>alignment</td><td>distance</td><td>position</td><td>gene name</td><td>gene id</td></tr>
<tr><td>GU305914.1</td><td>182897</td><td>182919</td><td>+</td><td>0</td><td>CCGATGCGCTTCAGTAGCCG</td><td>GGG</td><td>||||||||[||||||||||||]PAM</td><td>NA</td><td>Intergenic</td><td>NA</td><td>NA</td></tr>
<tr><td>T13878</td><td>TGCGCTTCAGTAGCCGGGGAAGG</td><td>1000</td><td></td><td></td><td></td><td></td><td></td><td></td><td></td><td></td><td></td></tr>
<tr><td>Oligo fwd</td><td>CACCTGCGCTTCAGTAGCCGGGGA</td><td></td><td></td><td></td><td></td><td></td><td></td><td></td><td></td><td></td><td></td></tr>
<tr><td>Oligo rev</td><td>AAACTCCCCGGCTACTGAAGCGCA</td><td></td><td></td><td></td><td></td><td></td><td></td><td></td><td></td><td></td><td></td></tr>
<tr><td>Chromosome</td><td>start</td><td>end</td><td>strand</td><td>MM</td><td>target_seq</td><td>PAM</td><td>alignment</td><td>distance</td><td>position</td><td>gene name</td><td>gene id</td></tr>
<tr><td>GU305914.1</td><td>182901</td><td>182923</td><td>+</td><td>0</td><td>TGCGCTTCAGTAGCCGGGGA</td><td>AGG</td><td>||||||||[||||||||||||]PAM</td><td>NA</td><td>Intergenic</td><td>NA</td><td>NA</td></tr>
<tr><td>T13879</td><td>CGGGGAAGGTCGCTCTTTCCTGG</td><td>1000</td><td></td><td></td><td></td><td></td><td></td><td></td><td></td><td></td><td></td></tr>
<tr><td>Oligo fwd</td><td>CACCCGGGGAAGGTCGCTCTTTCC</td><td></td><td></td><td></td><td></td><td></td><td></td><td></td><td></td><td></td><td></td></tr>
<tr><td>Oligo rev</td><td>AAACGGAAAGAGCGACCTTCCCCG</td><td></td><td></td><td></td><td></td><td></td><td></td><td></td><td></td><td></td><td></td></tr>
<tr><td>Chromosome</td><td>start</td><td>end</td><td>strand</td><td>MM</td><td>target_seq</td><td>PAM</td><td>alignment</td><td>distance</td><td>position</td><td>gene name</td><td>gene id</td></tr>
<tr><td>GU305914.1</td><td>182915</td><td>182937</td><td>+</td><td>0</td><td>CGGGGAAGGTCGCTCTTTCC</td><td>TGG</td><td>||||||||[||||||||||||]PAM</td><td>NA</td><td>Intergenic</td><td>NA</td><td>NA</td></tr>
<tr><td>T13880</td><td>GGGGAAGGTCGCTCTTTCCTGGG</td><td>1000</td><td></td><td></td><td></td><td></td><td></td><td></td><td></td><td></td><td></td></tr>
<tr><td>Oligo fwd</td><td>CACCGGGGAAGGTCGCTCTTTCCT</td><td></td><td></td><td></td><td></td><td></td><td></td><td></td><td></td><td></td><td></td></tr>
<tr><td>Oligo rev</td><td>AAACAGGAAAGAGCGACCTTCCCC</td><td></td><td></td><td></td><td></td><td></td><td></td><td></td><td></td><td></td><td></td></tr>
<tr><td>Chromosome</td><td>start</td><td>end</td><td>strand</td><td>MM</td><td>target_seq</td><td>PAM</td><td>alignment</td><td>distance</td><td>position</td><td>gene name</td><td>gene id</td></tr>
<tr><td>GU305914.1</td><td>182916</td><td>182938</td><td>+</td><td>0</td><td>GGGGAAGGTCGCTCTTTCCT</td><td>GGG</td><td>||||||||[||||||||||||]PAM</td><td>NA</td><td>Intergenic</td><td>NA</td><td>NA</td></tr>
<tr><td>T13881</td><td>AAGGTCGCTCTTTCCTGGGCAGG</td><td>1000</td><td></td><td></td><td></td><td></td><td></td><td></td><td></td><td></td><td></td></tr>
<tr><td>Oligo fwd</td><td>CACCAAGGTCGCTCTTTCCTGGGC</td><td></td><td></td><td></td><td></td><td></td><td></td><td></td><td></td><td></td><td></td></tr>
<tr><td>Oligo rev</td><td>AAACGCCCAGGAAAGAGCGACCTT</td><td></td><td></td><td></td><td></td><td></td><td></td><td></td><td></td><td></td><td></td></tr>
<tr><td>Chromosome</td><td>start</td><td>end</td><td>strand</td><td>MM</td><td>target_seq</td><td>PAM</td><td>alignment</td><td>distance</td><td>position</td><td>gene name</td><td>gene id</td></tr>
<tr><td>GU305914.1</td><td>182920</td><td>182942</td><td>+</td><td>0</td><td>AAGGTCGCTCTTTCCTGGGC</td><td>AGG</td><td>||||||||[||||||||||||]PAM</td><td>NA</td><td>Intergenic</td><td>NA</td><td>NA</td></tr>
<tr><td>T13882</td><td>CTGGGCAGGATACTGCAGCGAGG</td><td>1000</td><td></td><td></td><td></td><td></td><td></td><td></td><td></td><td></td><td></td></tr>
<tr><td>Oligo fwd</td><td>CACCCTGGGCAGGATACTGCAGCG</td><td></td><td></td><td></td><td></td><td></td><td></td><td></td><td></td><td></td><td></td></tr>
<tr><td>Oligo rev</td><td>AAACCGCTGCAGTATCCTGCCCAG</td><td></td><td></td><td></td><td></td><td></td><td></td><td></td><td></td><td></td><td></td></tr>
<tr><td>Chromosome</td><td>start</td><td>end</td><td>strand</td><td>MM</td><td>target_seq</td><td>PAM</td><td>alignment</td><td>distance</td><td>position</td><td>gene name</td><td>gene id</td></tr>
<tr><td>GU305914.1</td><td>182934</td><td>182956</td><td>+</td><td>0</td><td>CTGGGCAGGATACTGCAGCG</td><td>AGG</td><td>||||||||[||||||||||||]PAM</td><td>NA</td><td>Intergenic</td><td>NA</td><td>NA</td></tr>
<tr><td>T13883</td><td>GCGAGGAGCTCTGCGTTCTACGG</td><td>1000</td><td></td><td></td><td></td><td></td><td></td><td></td><td></td><td></td><td></td></tr>
<tr><td>Oligo fwd</td><td>CACCGCGAGGAGCTCTGCGTTCTA</td><td></td><td></td><td></td><td></td><td></td><td></td><td></td><td></td><td></td><td></td></tr>
<tr><td>Oligo rev</td><td>AAACTAGAACGCAGAGCTCCTCGC</td><td></td><td></td><td></td><td></td><td></td><td></td><td></td><td></td><td></td><td></td></tr>
<tr><td>Chromosome</td><td>start</td><td>end</td><td>strand</td><td>MM</td><td>target_seq</td><td>PAM</td><td>alignment</td><td>distance</td><td>position</td><td>gene name</td><td>gene id</td></tr>
<tr><td>GU305914.1</td><td>182951</td><td>182973</td><td>+</td><td>0</td><td>GCGAGGAGCTCTGCGTTCTA</td><td>CGG</td><td>||||||||[||||||||||||]PAM</td><td>NA</td><td>Intergenic</td><td>NA</td><td>NA</td></tr>
<tr><td>T13884</td><td>AGGAGCTCTGCGTTCTACGGTGG</td><td>1000</td><td></td><td></td><td></td><td></td><td></td><td></td><td></td><td></td><td></td></tr>
<tr><td>Oligo fwd</td><td>CACCAGGAGCTCTGCGTTCTACGG</td><td></td><td></td><td></td><td></td><td></td><td></td><td></td><td></td><td></td><td></td></tr>
<tr><td>Oligo rev</td><td>AAACCCGTAGAACGCAGAGCTCCT</td><td></td><td></td><td></td><td></td><td></td><td></td><td></td><td></td><td></td><td></td></tr>
<tr><td>Chromosome</td><td>start</td><td>end</td><td>strand</td><td>MM</td><td>target_seq</td><td>PAM</td><td>alignment</td><td>distance</td><td>position</td><td>gene name</td><td>gene id</td></tr>
<tr><td>GU305914.1</td><td>182954</td><td>182976</td><td>+</td><td>0</td><td>AGGAGCTCTGCGTTCTACGG</td><td>TGG</td><td>||||||||[||||||||||||]PAM</td><td>NA</td><td>Intergenic</td><td>NA</td><td>NA</td></tr>
<tr><td>T13885</td><td>GGTCAGACCGAAGACTGCGACGG</td><td>1000</td><td></td><td></td><td></td><td></td><td></td><td></td><td></td><td></td><td></td></tr>
<tr><td>Oligo fwd</td><td>CACCGGTCAGACCGAAGACTGCGA</td><td></td><td></td><td></td><td></td><td></td><td></td><td></td><td></td><td></td><td></td></tr>
<tr><td>Oligo rev</td><td>AAACTCGCAGTCTTCGGTCTGACC</td><td></td><td></td><td></td><td></td><td></td><td></td><td></td><td></td><td></td><td></td></tr>
<tr><td>Chromosome</td><td>start</td><td>end</td><td>strand</td><td>MM</td><td>target_seq</td><td>PAM</td><td>alignment</td><td>distance</td><td>position</td><td>gene name</td><td>gene id</td></tr>
<tr><td>GU305914.1</td><td>182975</td><td>182997</td><td>+</td><td>0</td><td>GGTCAGACCGAAGACTGCGA</td><td>CGG</td><td>||||||||[||||||||||||]PAM</td><td>NA</td><td>Intergenic</td><td>NA</td><td>NA</td></tr>
<tr><td>GU305914.1</td><td>111181</td><td>111203</td><td>-</td><td>4</td><td>GATCAGGCCGACGACGGCGA</td><td>AGG</td><td>|-||||-|[|||-|||-||||]PAM</td><td>NA</td><td>Intergenic</td><td>NA</td><td>NA</td></tr>
<tr><td>T13886</td><td>GACTGCGACGGTACCGACGCTGG</td><td>1000</td><td></td><td></td><td></td><td></td><td></td><td></td><td></td><td></td><td></td></tr>
<tr><td>Oligo fwd</td><td>CACCGACTGCGACGGTACCGACGC</td><td></td><td></td><td></td><td></td><td></td><td></td><td></td><td></td><td></td><td></td></tr>
<tr><td>Oligo rev</td><td>AAACGCGTCGGTACCGTCGCAGTC</td><td></td><td></td><td></td><td></td><td></td><td></td><td></td><td></td><td></td><td></td></tr>
<tr><td>Chromosome</td><td>start</td><td>end</td><td>strand</td><td>MM</td><td>target_seq</td><td>PAM</td><td>alignment</td><td>distance</td><td>position</td><td>gene name</td><td>gene id</td></tr>
<tr><td>GU305914.1</td><td>182987</td><td>183009</td><td>+</td><td>0</td><td>GACTGCGACGGTACCGACGC</td><td>TGG</td><td>||||||||[||||||||||||]PAM</td><td>NA</td><td>Intergenic</td><td>NA</td><td>NA</td></tr>
<tr><td>T13887</td><td>CAATAGAATGCGTGCCAATATGG</td><td>1000</td><td></td><td></td><td></td><td></td><td></td><td></td><td></td><td></td><td></td></tr>
<tr><td>Oligo fwd</td><td>CACCCAATAGAATGCGTGCCAATA</td><td></td><td></td><td></td><td></td><td></td><td></td><td></td><td></td><td></td><td></td></tr>
<tr><td>Oligo rev</td><td>AAACTATTGGCACGCATTCTATTG</td><td></td><td></td><td></td><td></td><td></td><td></td><td></td><td></td><td></td><td></td></tr>
<tr><td>Chromosome</td><td>start</td><td>end</td><td>strand</td><td>MM</td><td>target_seq</td><td>PAM</td><td>alignment</td><td>distance</td><td>position</td><td>gene name</td><td>gene id</td></tr>
<tr><td>GU305914.1</td><td>183046</td><td>183068</td><td>+</td><td>0</td><td>CAATAGAATGCGTGCCAATA</td><td>TGG</td><td>||||||||[||||||||||||]PAM</td><td>NA</td><td>Intergenic</td><td>NA</td><td>NA</td></tr>
<tr><td>T13888</td><td>CGTGCCAATATGGAATTTCCAGG</td><td>1000</td><td></td><td></td><td></td><td></td><td></td><td></td><td></td><td></td><td></td></tr>
<tr><td>Oligo fwd</td><td>CACCCGTGCCAATATGGAATTTCC</td><td></td><td></td><td></td><td></td><td></td><td></td><td></td><td></td><td></td><td></td></tr>
<tr><td>Oligo rev</td><td>AAACGGAAATTCCATATTGGCACG</td><td></td><td></td><td></td><td></td><td></td><td></td><td></td><td></td><td></td><td></td></tr>
<tr><td>Chromosome</td><td>start</td><td>end</td><td>strand</td><td>MM</td><td>target_seq</td><td>PAM</td><td>alignment</td><td>distance</td><td>position</td><td>gene name</td><td>gene id</td></tr>
<tr><td>GU305914.1</td><td>183056</td><td>183078</td><td>+</td><td>0</td><td>CGTGCCAATATGGAATTTCC</td><td>AGG</td><td>||||||||[||||||||||||]PAM</td><td>NA</td><td>Intergenic</td><td>NA</td><td>NA</td></tr>
<tr><td>T13889</td><td>GTGCCAATATGGAATTTCCAGGG</td><td>1000</td><td></td><td></td><td></td><td></td><td></td><td></td><td></td><td></td><td></td></tr>
<tr><td>Oligo fwd</td><td>CACCGTGCCAATATGGAATTTCCA</td><td></td><td></td><td></td><td></td><td></td><td></td><td></td><td></td><td></td><td></td></tr>
<tr><td>Oligo rev</td><td>AAACTGGAAATTCCATATTGGCAC</td><td></td><td></td><td></td><td></td><td></td><td></td><td></td><td></td><td></td><td></td></tr>
<tr><td>Chromosome</td><td>start</td><td>end</td><td>strand</td><td>MM</td><td>target_seq</td><td>PAM</td><td>alignment</td><td>distance</td><td>position</td><td>gene name</td><td>gene id</td></tr>
<tr><td>GU305914.1</td><td>183057</td><td>183079</td><td>+</td><td>0</td><td>GTGCCAATATGGAATTTCCA</td><td>GGG</td><td>||||||||[||||||||||||]PAM</td><td>NA</td><td>Intergenic</td><td>NA</td><td>NA</td></tr>
<tr><td>T13890</td><td>TGCCAATATGGAATTTCCAGGGG</td><td>1000</td><td></td><td></td><td></td><td></td><td></td><td></td><td></td><td></td><td></td></tr>
</table>

<table>
<tr><td>Oligo fwd</td><td>CACCTGCCAATATGGAATTTCCAG</td><td></td><td></td><td></td><td></td><td></td><td></td><td></td><td></td><td></td><td></td></tr>
<tr><td>Oligo rev</td><td>AAACCTGGAAATTCCATATTGGCA</td><td></td><td></td><td></td><td></td><td></td><td></td><td></td><td></td><td></td><td></td></tr>
<tr><td>Chromosome</td><td>start</td><td>end</td><td>strand</td><td>MM</td><td>target_seq</td><td>PAM</td><td>alignment</td><td>distance</td><td>position</td><td>gene name</td><td>gene id</td></tr>
<tr><td>GU305914.1</td><td>183058</td><td>183080</td><td>+</td><td>0</td><td>TGCCAATATGGAATTTCCAG</td><td>GGG</td><td>||||||||[||||||||||||]PAM</td><td>NA</td><td>Intergenic</td><td>NA</td><td>NA</td></tr>
<tr><td>T13891</td><td>TGGAATTTCCAGGGGAAAACCGG</td><td>1000</td><td></td><td></td><td></td><td></td><td></td><td></td><td></td><td></td><td></td></tr>
<tr><td>Oligo fwd</td><td>CACCTGGAATTTCCAGGGGAAAAC</td><td></td><td></td><td></td><td></td><td></td><td></td><td></td><td></td><td></td><td></td></tr>
<tr><td>Oligo rev</td><td>AAACGTTTTCCCCTGGAAATTCCA</td><td></td><td></td><td></td><td></td><td></td><td></td><td></td><td></td><td></td><td></td></tr>
<tr><td>Chromosome</td><td>start</td><td>end</td><td>strand</td><td>MM</td><td>target_seq</td><td>PAM</td><td>alignment</td><td>distance</td><td>position</td><td>gene name</td><td>gene id</td></tr>
<tr><td>GU305914.1</td><td>183066</td><td>183088</td><td>+</td><td>0</td><td>TGGAATTTCCAGGGGAAAAC</td><td>CGG</td><td>||||||||[||||||||||||]PAM</td><td>NA</td><td>Intergenic</td><td>NA</td><td>NA</td></tr>
<tr><td>T13892</td><td>GGAATTTCCAGGGGAAAACCGGG</td><td>1000</td><td></td><td></td><td></td><td></td><td></td><td></td><td></td><td></td><td></td></tr>
<tr><td>Oligo fwd</td><td>CACCGGAATTTCCAGGGGAAAACC</td><td></td><td></td><td></td><td></td><td></td><td></td><td></td><td></td><td></td><td></td></tr>
<tr><td>Oligo rev</td><td>AAACGGTTTTCCCCTGGAAATTCC</td><td></td><td></td><td></td><td></td><td></td><td></td><td></td><td></td><td></td><td></td></tr>
<tr><td>Chromosome</td><td>start</td><td>end</td><td>strand</td><td>MM</td><td>target_seq</td><td>PAM</td><td>alignment</td><td>distance</td><td>position</td><td>gene name</td><td>gene id</td></tr>
<tr><td>GU305914.1</td><td>183067</td><td>183089</td><td>+</td><td>0</td><td>GGAATTTCCAGGGGAAAACC</td><td>GGG</td><td>||||||||[||||||||||||]PAM</td><td>NA</td><td>Intergenic</td><td>NA</td><td>NA</td></tr>
<tr><td>T13893</td><td>GAATTTCCAGGGGAAAACCGGGG</td><td>1000</td><td></td><td></td><td></td><td></td><td></td><td></td><td></td><td></td><td></td></tr>
<tr><td>Oligo fwd</td><td>CACCGAATTTCCAGGGGAAAACCG</td><td></td><td></td><td></td><td></td><td></td><td></td><td></td><td></td><td></td><td></td></tr>
<tr><td>Oligo rev</td><td>AAACCGGTTTTCCCCTGGAAATTC</td><td></td><td></td><td></td><td></td><td></td><td></td><td></td><td></td><td></td><td></td></tr>
<tr><td>Chromosome</td><td>start</td><td>end</td><td>strand</td><td>MM</td><td>target_seq</td><td>PAM</td><td>alignment</td><td>distance</td><td>position</td><td>gene name</td><td>gene id</td></tr>
<tr><td>GU305914.1</td><td>183068</td><td>183090</td><td>+</td><td>0</td><td>GAATTTCCAGGGGAAAACCG</td><td>GGG</td><td>||||||||[||||||||||||]PAM</td><td>NA</td><td>Intergenic</td><td>NA</td><td>NA</td></tr>
<tr><td>T13894</td><td>TTTCCAGGGGAAAACCGGGGCGG</td><td>1000</td><td></td><td></td><td></td><td></td><td></td><td></td><td></td><td></td><td></td></tr>
<tr><td>Oligo fwd</td><td>CACCTTTCCAGGGGAAAACCGGGG</td><td></td><td></td><td></td><td></td><td></td><td></td><td></td><td></td><td></td><td></td></tr>
<tr><td>Oligo rev</td><td>AAACCCCCGGTTTTCCCCTGGAAA</td><td></td><td></td><td></td><td></td><td></td><td></td><td></td><td></td><td></td><td></td></tr>
<tr><td>Chromosome</td><td>start</td><td>end</td><td>strand</td><td>MM</td><td>target_seq</td><td>PAM</td><td>alignment</td><td>distance</td><td>position</td><td>gene name</td><td>gene id</td></tr>
<tr><td>GU305914.1</td><td>183071</td><td>183093</td><td>+</td><td>0</td><td>TTTCCAGGGGAAAACCGGGG</td><td>CGG</td><td>||||||||[||||||||||||]PAM</td><td>NA</td><td>Intergenic</td><td>NA</td><td>NA</td></tr>
<tr><td>T13895</td><td>ACCGGGGCGGTGTTACGTTTTGG</td><td>1000</td><td></td><td></td><td></td><td></td><td></td><td></td><td></td><td></td><td></td></tr>
<tr><td>Oligo fwd</td><td>CACCACCGGGGCGGTGTTACGTTT</td><td></td><td></td><td></td><td></td><td></td><td></td><td></td><td></td><td></td><td></td></tr>
<tr><td>Oligo rev</td><td>AAACAAACGTAACACCGCCCCGGT</td><td></td><td></td><td></td><td></td><td></td><td></td><td></td><td></td><td></td><td></td></tr>
<tr><td>Chromosome</td><td>start</td><td>end</td><td>strand</td><td>MM</td><td>target_seq</td><td>PAM</td><td>alignment</td><td>distance</td><td>position</td><td>gene name</td><td>gene id</td></tr>
<tr><td>GU305914.1</td><td>183084</td><td>183106</td><td>+</td><td>0</td><td>ACCGGGGCGGTGTTACGTTT</td><td>TGG</td><td>||||||||[||||||||||||]PAM</td><td>NA</td><td>Intergenic</td><td>NA</td><td>NA</td></tr>
<tr><td>T13896</td><td>CACTTCCCATTGACGTGTATTGG</td><td>1000</td><td></td><td></td><td></td><td></td><td></td><td></td><td></td><td></td><td></td></tr>
<tr><td>Oligo fwd</td><td>CACCCACTTCCCATTGACGTGTAT</td><td></td><td></td><td></td><td></td><td></td><td></td><td></td><td></td><td></td><td></td></tr>
<tr><td>Oligo rev</td><td>AAACATACACGTCAATGGGAAGTG</td><td></td><td></td><td></td><td></td><td></td><td></td><td></td><td></td><td></td><td></td></tr>
<tr><td>Chromosome</td><td>start</td><td>end</td><td>strand</td><td>MM</td><td>target_seq</td><td>PAM</td><td>alignment</td><td>distance</td><td>position</td><td>gene name</td><td>gene id</td></tr>
<tr><td>GU305914.1</td><td>183116</td><td>183138</td><td>+</td><td>0</td><td>CACTTCCCATTGACGTGTAT</td><td>TGG</td><td>||||||||[||||||||||||]PAM</td><td>NA</td><td>Intergenic</td><td>NA</td><td>NA</td></tr>
<tr><td>T13897</td><td>GACGTGTATTGGCTCGAGAACGG</td><td>1000</td><td></td><td></td><td></td><td></td><td></td><td></td><td></td><td></td><td></td></tr>
<tr><td>Oligo fwd</td><td>CACCGACGTGTATTGGCTCGAGAA</td><td></td><td></td><td></td><td></td><td></td><td></td><td></td><td></td><td></td><td></td></tr>
<tr><td>Oligo rev</td><td>AAACTTCTCGAGCCAATACACGTC</td><td></td><td></td><td></td><td></td><td></td><td></td><td></td><td></td><td></td><td></td></tr>
<tr><td>Chromosome</td><td>start</td><td>end</td><td>strand</td><td>MM</td><td>target_seq</td><td>PAM</td><td>alignment</td><td>distance</td><td>position</td><td>gene name</td><td>gene id</td></tr>
<tr><td>GU305914.1</td><td>183127</td><td>183149</td><td>+</td><td>0</td><td>GACGTGTATTGGCTCGAGAA</td><td>CGG</td><td>||||||||[||||||||||||]PAM</td><td>NA</td><td>Intergenic</td><td>NA</td><td>NA</td></tr>
<tr><td>T13898</td><td>ACTTTCCCATTAATCAGCTATGG</td><td>1000</td><td></td><td></td><td></td><td></td><td></td><td></td><td></td><td></td><td></td></tr>
<tr><td>Oligo fwd</td><td>CACCACTTTCCCATTAATCAGCTA</td><td></td><td></td><td></td><td></td><td></td><td></td><td></td><td></td><td></td><td></td></tr>
<tr><td>Oligo rev</td><td>AAACTAGCTGATTAATGGGAAAGT</td><td></td><td></td><td></td><td></td><td></td><td></td><td></td><td></td><td></td><td></td></tr>
<tr><td>Chromosome</td><td>start</td><td>end</td><td>strand</td><td>MM</td><td>target_seq</td><td>PAM</td><td>alignment</td><td>distance</td><td>position</td><td>gene name</td><td>gene id</td></tr>
<tr><td>GU305914.1</td><td>183151</td><td>183173</td><td>+</td><td>0</td><td>ACTTTCCCATTAATCAGCTA</td><td>TGG</td><td>||||||||[||||||||||||]PAM</td><td>NA</td><td>Intergenic</td><td>NA</td><td>NA</td></tr>
<tr><td>T13899</td><td>CTTTCCCATTAATCAGCTATGGG</td><td>1000</td><td></td><td></td><td></td><td></td><td></td><td></td><td></td><td></td><td></td></tr>
<tr><td>Oligo fwd</td><td>CACCCTTTCCCATTAATCAGCTAT</td><td></td><td></td><td></td><td></td><td></td><td></td><td></td><td></td><td></td><td></td></tr>
<tr><td>Oligo rev</td><td>AAACATAGCTGATTAATGGGAAAG</td><td></td><td></td><td></td><td></td><td></td><td></td><td></td><td></td><td></td><td></td></tr>
<tr><td>Chromosome</td><td>start</td><td>end</td><td>strand</td><td>MM</td><td>target_seq</td><td>PAM</td><td>alignment</td><td>distance</td><td>position</td><td>gene name</td><td>gene id</td></tr>
<tr><td>GU305914.1</td><td>183152</td><td>183174</td><td>+</td><td>0</td><td>CTTTCCCATTAATCAGCTAT</td><td>GGG</td><td>||||||||[||||||||||||]PAM</td><td>NA</td><td>Intergenic</td><td>NA</td><td>NA</td></tr>
<tr><td>T13900</td><td>TATGGGAAAGTACCGTTTAAAGG</td><td>1000</td><td></td><td></td><td></td><td></td><td></td><td></td><td></td><td></td><td></td></tr>
<tr><td>Oligo fwd</td><td>CACCTATGGGAAAGTACCGTTTAA</td><td></td><td></td><td></td><td></td><td></td><td></td><td></td><td></td><td></td><td></td></tr>
<tr><td>Oligo rev</td><td>AAACTTAAACGGTACTTTCCCATA</td><td></td><td></td><td></td><td></td><td></td><td></td><td></td><td></td><td></td><td></td></tr>
<tr><td>Chromosome</td><td>start</td><td>end</td><td>strand</td><td>MM</td><td>target_seq</td><td>PAM</td><td>alignment</td><td>distance</td><td>position</td><td>gene name</td><td>gene id</td></tr>
<tr><td>GU305914.1</td><td>183169</td><td>183191</td><td>+</td><td>0</td><td>TATGGGAAAGTACCGTTTAA</td><td>AGG</td><td>||||||||[||||||||||||]PAM</td><td>NA</td><td>Intergenic</td><td>NA</td><td>NA</td></tr>
<tr><td>T13901</td><td>CAATAGCCCATTGACGTCAATGG</td><td>1000</td><td></td><td></td><td></td><td></td><td></td><td></td><td></td><td></td><td></td></tr>
<tr><td>Oligo fwd</td><td>CACCCAATAGCCCATTGACGTCAA</td><td></td><td></td><td></td><td></td><td></td><td></td><td></td><td></td><td></td><td></td></tr>
<tr><td>Oligo rev</td><td>AAACTTGACGTCAATGGGCTATTG</td><td></td><td></td><td></td><td></td><td></td><td></td><td></td><td></td><td></td><td></td></tr>
<tr><td>Chromosome</td><td>start</td><td>end</td><td>strand</td><td>MM</td><td>target_seq</td><td>PAM</td><td>alignment</td><td>distance</td><td>position</td><td>gene name</td><td>gene id</td></tr>
<tr><td>GU305914.1</td><td>183209</td><td>183231</td><td>+</td><td>0</td><td>CAATAGCCCATTGACGTCAA</td><td>TGG</td><td>||||||||[||||||||||||]PAM</td><td>NA</td><td>Intergenic</td><td>NA</td><td>NA</td></tr>
<tr><td>T13902</td><td>TAGCCCATTGACGTCAATGGTGG</td><td>1000</td><td></td><td></td><td></td><td></td><td></td><td></td><td></td><td></td><td></td></tr>
<tr><td>Oligo fwd</td><td>CACCTAGCCCATTGACGTCAATGG</td><td></td><td></td><td></td><td></td><td></td><td></td><td></td><td></td><td></td><td></td></tr>
<tr><td>Oligo rev</td><td>AAACCCATTGACGTCAATGGGCTA</td><td></td><td></td><td></td><td></td><td></td><td></td><td></td><td></td><td></td><td></td></tr>
<tr><td>Chromosome</td><td>start</td><td>end</td><td>strand</td><td>MM</td><td>target_seq</td><td>PAM</td><td>alignment</td><td>distance</td><td>position</td><td>gene name</td><td>gene id</td></tr>
<tr><td>GU305914.1</td><td>183212</td><td>183234</td><td>+</td><td>0</td><td>TAGCCCATTGACGTCAATGG</td><td>TGG</td><td>||||||||[||||||||||||]PAM</td><td>NA</td><td>Intergenic</td><td>NA</td><td>NA</td></tr>
<tr><td>T13903</td><td>AGCCCATTGACGTCAATGGTGGG</td><td>1000</td><td></td><td></td><td></td><td></td><td></td><td></td><td></td><td></td><td></td></tr>
<tr><td>Oligo fwd</td><td>CACCAGCCCATTGACGTCAATGGT</td><td></td><td></td><td></td><td></td><td></td><td></td><td></td><td></td><td></td><td></td></tr>
<tr><td>Oligo rev</td><td>AAACACCATTGACGTCAATGGGCT</td><td></td><td></td><td></td><td></td><td></td><td></td><td></td><td></td><td></td><td></td></tr>
<tr><td>Chromosome</td><td>start</td><td>end</td><td>strand</td><td>MM</td><td>target_seq</td><td>PAM</td><td>alignment</td><td>distance</td><td>position</td><td>gene name</td><td>gene id</td></tr>
<tr><td>GU305914.1</td><td>183213</td><td>183235</td><td>+</td><td>0</td><td>AGCCCATTGACGTCAATGGT</td><td>GGG</td><td>||||||||[||||||||||||]PAM</td><td>NA</td><td>Intergenic</td><td>NA</td><td>NA</td></tr>
<tr><td>T13904</td><td>GTCAATGGTGGGAAAGTACATGG</td><td>1000</td><td></td><td></td><td></td><td></td><td></td><td></td><td></td><td></td><td></td></tr>
<tr><td>Oligo fwd</td><td>CACCGTCAATGGTGGGAAAGTACA</td><td></td><td></td><td></td><td></td><td></td><td></td><td></td><td></td><td></td><td></td></tr>
<tr><td>Oligo rev</td><td>AAACTGTACTTTCCCACCATTGAC</td><td></td><td></td><td></td><td></td><td></td><td></td><td></td><td></td><td></td><td></td></tr>
<tr><td>Chromosome</td><td>start</td><td>end</td><td>strand</td><td>MM</td><td>target_seq</td><td>PAM</td><td>alignment</td><td>distance</td><td>position</td><td>gene name</td><td>gene id</td></tr>
<tr><td>GU305914.1</td><td>183224</td><td>183246</td><td>+</td><td>0</td><td>GTCAATGGTGGGAAAGTACA</td><td>TGG</td><td>||||||||[||||||||||||]PAM</td><td>NA</td><td>Intergenic</td><td>NA</td><td>NA</td></tr>
</table>

| | | | | | | | | | | | |
|---|---|---|---|---|---|---|---|---|---|---|---|
| T13905 | ACATGGCGTTTTAATTAAATTGG | 1000 | | | | | | | | | |
| Oligo fwd | CACCACATGGCGTTTTAATTAAAT | | | | | | | | | | |
| Oligo rev | AAACATTTAATTAAAACGCCATGT | | | | | | | | | | |
| Chromosome | start | end | strand | MM | target_seq | PAM | alignment | distance | position | gene name | gene id |
| GU305914.1 | 183241 | 183263 | + | 0 | ACATGGCGTTTTAATTAAAT | TGG | \|\|\|\|\|\|\|\|[\|\|\|\|\|\|\|\|\|\|\|\|]PAM | NA | Intergenic | NA | NA |

| | | | | | | | | | | | |
|---|---|---|---|---|---|---|---|---|---|---|---|
| T13906 | GGCGTTTTAATTAAATTGGCTGG | 1000 | | | | | | | | | |
| Oligo fwd | CACCGGCGTTTTAATTAAATTGGC | | | | | | | | | | |
| Oligo rev | AAACGCCAATTTAATTAAAACGCC | | | | | | | | | | |
| Chromosome | start | end | strand | MM | target_seq | PAM | alignment | distance | position | gene name | gene id |
| GU305914.1 | 183245 | 183267 | + | 0 | GGCGTTTTAATTAAATTGGC | TGG | \|\|\|\|\|\|\|\|[\|\|\|\|\|\|\|\|\|\|\|\|]PAM | NA | Intergenic | NA | NA |

| | | | | | | | | | | | |
|---|---|---|---|---|---|---|---|---|---|---|---|
| T13907 | CCTTATGTACGTGCCAATAATGG | 1000 | | | | | | | | | |
| Oligo fwd | CACCCCTTATGTACGTGCCAATAA | | | | | | | | | | |
| Oligo rev | AAACTTATTGGCACGTACATAAGG | | | | | | | | | | |
| Chromosome | start | end | strand | MM | target_seq | PAM | alignment | distance | position | gene name | gene id |
| GU305914.1 | 183295 | 183317 | + | 0 | CCTTATGTACGTGCCAATAA | TGG | \|\|\|\|\|\|\|\|[\|\|\|\|\|\|\|\|\|\|\|\|]PAM | NA | Intergenic | NA | NA |

| | | | | | | | | | | | |
|---|---|---|---|---|---|---|---|---|---|---|---|
| T13908 | CTTATGTACGTGCCAATAATGGG | 1000 | | | | | | | | | |
| Oligo fwd | CACCCTTATGTACGTGCCAATAAT | | | | | | | | | | |
| Oligo rev | AAACATTATTGGCACGTACATAAG | | | | | | | | | | |
| Chromosome | start | end | strand | MM | target_seq | PAM | alignment | distance | position | gene name | gene id |
| GU305914.1 | 183296 | 183318 | + | 0 | CTTATGTACGTGCCAATAAT | GGG | \|\|\|\|\|\|\|\|[\|\|\|\|\|\|\|\|\|\|\|\|]PAM | NA | Intergenic | NA | NA |

| | | | | | | | | | | | |
|---|---|---|---|---|---|---|---|---|---|---|---|
| T13909 | CCCCTATTGACCTTTTGTACTGG | 1000 | | | | | | | | | |
| Oligo fwd | CACCCCCCTATTGACCTTTTGTAC | | | | | | | | | | |
| Oligo rev | AAACGTACAAAAGGTCAATAGGGG | | | | | | | | | | |
| Chromosome | start | end | strand | MM | target_seq | PAM | alignment | distance | position | gene name | gene id |
| GU305914.1 | 183337 | 183359 | + | 0 | CCCCTATTGACCTTTTGTAC | TGG | \|\|\|\|\|\|\|\|[\|\|\|\|\|\|\|\|\|\|\|\|]PAM | NA | Intergenic | NA | NA |
| GU305914.1 | 183708 | 183730 | + | 1 | CCCCTATTGACCTTATGTAC | TGG | \|\|\|\|\|\|\|\|[\|\|\|\|\|\|-\|\|\|\|\|]PAM | NA | Intergenic | NA | NA |
| GU305914.1 | 183615 | 183637 | + | 1 | CCCCTATTGACCTTATGTAC | TGG | \|\|\|\|\|\|\|\|[\|\|\|\|\|\|-\|\|\|\|\|]PAM | NA | Intergenic | NA | NA |
| GU305914.1 | 183429 | 183451 | + | 1 | CCCCTATTGACCTTATGTAC | TGG | \|\|\|\|\|\|\|\|[\|\|\|\|\|\|-\|\|\|\|\|]PAM | NA | Intergenic | NA | NA |
| GU305914.1 | 183801 | 183823 | + | 1 | CCCCTATTGACCTTATGTAC | TGG | \|\|\|\|\|\|\|\|[\|\|\|\|\|\|-\|\|\|\|\|]PAM | NA | Intergenic | NA | NA |
| GU305914.1 | 183522 | 183544 | + | 3 | CTCCCATTGACCTTATGTAC | TGG | \|-\|\|-\|\|\|[\|\|\|\|\|\|-\|\|\|\|\|]PAM | NA | Intergenic | NA | NA |

| | | | | | | | | | | | |
|---|---|---|---|---|---|---|---|---|---|---|---|
| T13910 | CCCTATTGACCTTTTGTACTGGG | 1000 | | | | | | | | | |
| Oligo fwd | CACCCCCTATTGACCTTTTGTACT | | | | | | | | | | |
| Oligo rev | AAACAGTACAAAAGGTCAATAGGG | | | | | | | | | | |
| Chromosome | start | end | strand | MM | target_seq | PAM | alignment | distance | position | gene name | gene id |
| GU305914.1 | 183338 | 183360 | + | 0 | CCCTATTGACCTTTTGTACT | GGG | \|\|\|\|\|\|\|\|[\|\|\|\|\|\|\|\|\|\|\|\|]PAM | NA | Intergenic | NA | NA |
| GU305914.1 | 183709 | 183731 | + | 1 | CCCTATTGACCTTATGTACT | GGG | \|\|\|\|\|\|\|\|[\|\|\|\|\|-\|\|\|\|\|\|]PAM | NA | Intergenic | NA | NA |
| GU305914.1 | 183616 | 183638 | + | 1 | CCCTATTGACCTTATGTACT | GGG | \|\|\|\|\|\|\|\|[\|\|\|\|\|-\|\|\|\|\|\|]PAM | NA | Intergenic | NA | NA |
| GU305914.1 | 183430 | 183452 | + | 1 | CCCTATTGACCTTATGTACT | GGG | \|\|\|\|\|\|\|\|[\|\|\|\|\|-\|\|\|\|\|\|]PAM | NA | Intergenic | NA | NA |
| GU305914.1 | 183802 | 183824 | + | 1 | CCCTATTGACCTTATGTACT | GGG | \|\|\|\|\|\|\|\|[\|\|\|\|\|-\|\|\|\|\|\|]PAM | NA | Intergenic | NA | NA |
| GU305914.1 | 183523 | 183545 | + | 3 | TCCCATTGACCTTATGTACT | GGG | -\|\|-\|\|\|\|[\|\|\|\|\|-\|\|\|\|\|\|]PAM | NA | Intergenic | NA | NA |

| | | | | | | | | | | | |
|---|---|---|---|---|---|---|---|---|---|---|---|
| T13911 | TTGTACTGGGCAAAACCCAATGG | 1000 | | | | | | | | | |
| Oligo fwd | CACCTTGTACTGGGCAAAACCCAA | | | | | | | | | | |
| Oligo rev | AAACTTGGGTTTTGCCCAGTACAA | | | | | | | | | | |
| Chromosome | start | end | strand | MM | target_seq | PAM | alignment | distance | position | gene name | gene id |
| GU305914.1 | 183351 | 183373 | + | 0 | TTGTACTGGGCAAAACCCAA | TGG | \|\|\|\|\|\|\|\|[\|\|\|\|\|\|\|\|\|\|\|\|]PAM | NA | Intergenic | NA | NA |
| GU305914.1 | 183629 | 183651 | + | 1 | ATGTACTGGGCAAAACCCAA | TGG | -\|\|\|\|\|\|\|[\|\|\|\|\|\|\|\|\|\|\|\|]PAM | NA | Intergenic | NA | NA |
| GU305914.1 | 183722 | 183744 | + | 1 | ATGTACTGGGCAAAACCCAA | TGG | -\|\|\|\|\|\|\|[\|\|\|\|\|\|\|\|\|\|\|\|]PAM | NA | Intergenic | NA | NA |
| GU305914.1 | 183443 | 183465 | + | 2 | ATGTACTGGGCAAAACCCAT | TGG | -\|\|\|\|\|\|\|[\|\|\|\|\|\|\|\|\|\|\|-]PAM | NA | Intergenic | NA | NA |
| GU305914.1 | 183536 | 183558 | + | 2 | ATGTACTGGGCAAAACCCAT | TGG | -\|\|\|\|\|\|\|[\|\|\|\|\|\|\|\|\|\|\|-]PAM | NA | Intergenic | NA | NA |

| | | | | | | | | | | | |
|---|---|---|---|---|---|---|---|---|---|---|---|
| T13912 | CCCTATTGACTCAGTGTACTTGG | 1000 | | | | | | | | | |
| Oligo fwd | CACCCCCTATTGACTCAGTGTACT | | | | | | | | | | |
| Oligo rev | AAACAGTACACTGAGTCAATAGGG | | | | | | | | | | |
| Chromosome | start | end | strand | MM | target_seq | PAM | alignment | distance | position | gene name | gene id |
| GU305914.1 | 183379 | 183401 | + | 0 | CCCTATTGACTCAGTGTACT | TGG | \|\|\|\|\|\|\|\|[\|\|\|\|\|\|\|\|\|\|\|\|]PAM | NA | Intergenic | NA | NA |
| GU305914.1 | 183907 | 183929 | + | 1 | CCCTATTGAGTCAGTGTACT | TGG | \|\|\|\|\|\|\|\|[\|-\|\|\|\|\|\|\|\|\|\|]PAM | NA | Intergenic | NA | NA |
| GU305914.1 | 183657 | 183679 | + | 1 | CCCTATTGAGTCAGTGTACT | TGG | \|\|\|\|\|\|\|\|[\|-\|\|\|\|\|\|\|\|\|\|]PAM | NA | Intergenic | NA | NA |
| GU305914.1 | 183750 | 183772 | + | 1 | CCCTATTGACGCAGTGTACT | TGG | \|\|\|\|\|\|\|\|[\|\|-\|\|\|\|\|\|\|\|\|]PAM | NA | Intergenic | NA | NA |
| GU305914.1 | 183564 | 183586 | + | 1 | CCCTATTGACTCAATGTACT | TGG | \|\|\|\|\|\|\|\|[\|\|\|\|\|-\|\|\|\|\|\|]PAM | NA | Intergenic | NA | NA |
| GU305914.1 | 184053 | 184075 | + | 4 | TCCCATTGACCCAATGTACT | TGG | -\|\|-\|\|\|\|[\|\|-\|\|-\|\|\|\|\|\|]PAM | NA | Intergenic | NA | NA |
| GU305914.1 | 183959 | 183981 | + | 4 | TCCCATTGACCCAATGTACT | TGG | -\|\|-\|\|\|\|[\|\|-\|\|-\|\|\|\|\|\|]PAM | NA | Intergenic | NA | NA |

| | | | | | | | | | | | |
|---|---|---|---|---|---|---|---|---|---|---|---|
| T13913 | CTCAGTGTACTTGGCTCCAATGG | 1000 | | | | | | | | | |
| Oligo fwd | CACCCTCAGTGTACTTGGCTCCAA | | | | | | | | | | |
| Oligo rev | AAACTTGGAGCCAAGTACACTGAG | | | | | | | | | | |
| Chromosome | start | end | strand | MM | target_seq | PAM | alignment | distance | position | gene name | gene id |
| GU305914.1 | 183388 | 183410 | + | 0 | CTCAGTGTACTTGGCTCCAA | TGG | \|\|\|\|\|\|\|\|[\|\|\|\|\|\|\|\|\|\|\|\|]PAM | NA | Intergenic | NA | NA |
| GU305914.1 | 183666 | 183688 | + | 1 | GTCAGTGTACTTGGCTCCAA | TGG | -\|\|\|\|\|\|\|[\|\|\|\|\|\|\|\|\|\|\|\|]PAM | NA | Intergenic | NA | NA |
| GU305914.1 | 183759 | 183781 | + | 1 | CGCAGTGTACTTGGCTCCAA | TGG | \|-\|\|\|\|\|\|[\|\|\|\|\|\|\|\|\|\|\|\|]PAM | NA | Intergenic | NA | NA |
| GU305914.1 | 183573 | 183595 | + | 1 | CTCAATGTACTTGGCTCCAA | TGG | \|\|\|\|-\|\|\|[\|\|\|\|\|\|\|\|\|\|\|\|]PAM | NA | Intergenic | NA | NA |
| GU305914.1 | 183916 | 183938 | + | 3 | GTCAGTGTACTTGGCAGCAA | TGG | -\|\|\|\|\|\|\|[\|\|\|\|\|\|\|--\|\|\|]PAM | NA | Intergenic | NA | NA |

| | | | | | | | | | | | |
|---|---|---|---|---|---|---|---|---|---|---|---|
| T13914 | TCAGTGTACTTGGCTCCAATGGG | 1000 | | | | | | | | | |
| Oligo fwd | CACCTCAGTGTACTTGGCTCCAAT | | | | | | | | | | |
| Oligo rev | AAACATTGGAGCCAAGTACACTGA | | | | | | | | | | |
| Chromosome | start | end | strand | MM | target_seq | PAM | alignment | distance | position | gene name | gene id |
| GU305914.1 | 183389 | 183411 | + | 0 | TCAGTGTACTTGGCTCCAAT | GGG | \|\|\|\|\|\|\|\|[\|\|\|\|\|\|\|\|\|\|\|\|]PAM | NA | Intergenic | NA | NA |
| GU305914.1 | 183667 | 183689 | + | 0 | TCAGTGTACTTGGCTCCAAT | GGG | \|\|\|\|\|\|\|\|[\|\|\|\|\|\|\|\|\|\|\|\|]PAM | NA | Intergenic | NA | NA |
| GU305914.1 | 183760 | 183782 | + | 1 | GCAGTGTACTTGGCTCCAAT | GGG | -\|\|\|\|\|\|\|[\|\|\|\|\|\|\|\|\|\|\|\|]PAM | NA | Intergenic | NA | NA |
| GU305914.1 | 183574 | 183596 | + | 1 | TCAATGTACTTGGCTCCAAT | GGG | \|\|\|-\|\|\|\|[\|\|\|\|\|\|\|\|\|\|\|\|]PAM | NA | Intergenic | NA | NA |
| GU305914.1 | 183917 | 183939 | + | 2 | TCAGTGTACTTGGCAGCAAT | GGG | \|\|\|\|\|\|\|\|[\|\|\|\|\|\|--\|\|\|\|]PAM | NA | Intergenic | NA | NA |

| | | |
|---|---|---|
| T13915 | CCCCTATTGACCTTATGTACTGG | 1000 |

<table>
<tr><td>Oligo fwd</td><td>CACCCCCCTATTGACCTTATGTAC</td><td></td><td></td><td></td><td></td><td></td><td></td><td></td><td></td><td></td><td></td></tr>
<tr><td>Oligo rev</td><td>AAACGTACATAAGGTCAATAGGGG</td><td></td><td></td><td></td><td></td><td></td><td></td><td></td><td></td><td></td><td></td></tr>
<tr><td>Chromosome</td><td>start</td><td>end</td><td>strand</td><td>MM</td><td>target_seq</td><td>PAM</td><td>alignment</td><td>distance</td><td>position</td><td>gene name</td><td>gene id</td></tr>
<tr><td>GU305914.1</td><td>183801</td><td>183823</td><td>+</td><td>0</td><td>CCCCTATTGACCTTATGTAC</td><td>TGG</td><td>||||||||[||||||||||||]PAM</td><td>NA</td><td>Intergenic</td><td>NA</td><td>NA</td></tr>
<tr><td>GU305914.1</td><td>183708</td><td>183730</td><td>+</td><td>0</td><td>CCCCTATTGACCTTATGTAC</td><td>TGG</td><td>||||||||[||||||||||||]PAM</td><td>NA</td><td>Intergenic</td><td>NA</td><td>NA</td></tr>
<tr><td>GU305914.1</td><td>183615</td><td>183637</td><td>+</td><td>0</td><td>CCCCTATTGACCTTATGTAC</td><td>TGG</td><td>||||||||[||||||||||||]PAM</td><td>NA</td><td>Intergenic</td><td>NA</td><td>NA</td></tr>
<tr><td>GU305914.1</td><td>183429</td><td>183451</td><td>+</td><td>0</td><td>CCCCTATTGACCTTATGTAC</td><td>TGG</td><td>||||||||[||||||||||||]PAM</td><td>NA</td><td>Intergenic</td><td>NA</td><td>NA</td></tr>
<tr><td>GU305914.1</td><td>183522</td><td>183544</td><td>+</td><td>2</td><td>CTCCCATTGACCTTATGTAC</td><td>TGG</td><td>|-||-|||[||||||||||||]PAM</td><td>NA</td><td>Intergenic</td><td>NA</td><td>NA</td></tr>
<tr><td>GU305914.1</td><td>183337</td><td>183359</td><td>+</td><td>1</td><td>CCCCTATTGACCTTTTGTAC</td><td>TGG</td><td>||||||||[||||||-|||||]PAM</td><td>NA</td><td>Intergenic</td><td>NA</td><td>NA</td></tr>
<tr><td>T13916</td><td>CCCTATTGACCTTATGTACTGGG</td><td>1000</td><td></td><td></td><td></td><td></td><td></td><td></td><td></td><td></td><td></td></tr>
<tr><td>Oligo fwd</td><td>CACCCCCTATTGACCTTATGTACT</td><td></td><td></td><td></td><td></td><td></td><td></td><td></td><td></td><td></td><td></td></tr>
<tr><td>Oligo rev</td><td>AAACAGTACATAAGGTCAATAGGG</td><td></td><td></td><td></td><td></td><td></td><td></td><td></td><td></td><td></td><td></td></tr>
<tr><td>Chromosome</td><td>start</td><td>end</td><td>strand</td><td>MM</td><td>target_seq</td><td>PAM</td><td>alignment</td><td>distance</td><td>position</td><td>gene name</td><td>gene id</td></tr>
<tr><td>GU305914.1</td><td>183616</td><td>183638</td><td>+</td><td>0</td><td>CCCTATTGACCTTATGTACT</td><td>GGG</td><td>||||||||[||||||||||||]PAM</td><td>NA</td><td>Intergenic</td><td>NA</td><td>NA</td></tr>
<tr><td>GU305914.1</td><td>183430</td><td>183452</td><td>+</td><td>0</td><td>CCCTATTGACCTTATGTACT</td><td>GGG</td><td>||||||||[||||||||||||]PAM</td><td>NA</td><td>Intergenic</td><td>NA</td><td>NA</td></tr>
<tr><td>GU305914.1</td><td>183802</td><td>183824</td><td>+</td><td>0</td><td>CCCTATTGACCTTATGTACT</td><td>GGG</td><td>||||||||[||||||||||||]PAM</td><td>NA</td><td>Intergenic</td><td>NA</td><td>NA</td></tr>
<tr><td>GU305914.1</td><td>183709</td><td>183731</td><td>+</td><td>0</td><td>CCCTATTGACCTTATGTACT</td><td>GGG</td><td>||||||||[||||||||||||]PAM</td><td>NA</td><td>Intergenic</td><td>NA</td><td>NA</td></tr>
<tr><td>GU305914.1</td><td>183523</td><td>183545</td><td>+</td><td>2</td><td>TCCCATTGACCTTATGTACT</td><td>GGG</td><td>-||-||||[||||||||||||]PAM</td><td>NA</td><td>Intergenic</td><td>NA</td><td>NA</td></tr>
<tr><td>GU305914.1</td><td>183338</td><td>183360</td><td>+</td><td>1</td><td>CCCTATTGACCTTTTGTACT</td><td>GGG</td><td>||||||||[|||||-||||||]PAM</td><td>NA</td><td>Intergenic</td><td>NA</td><td>NA</td></tr>
<tr><td>GU305914.1</td><td>183959</td><td>183981</td><td>+</td><td>4</td><td>TCCCATTGACCCAATGTACT</td><td>TGG</td><td>-||-||||[|||--|||||||]PAM</td><td>NA</td><td>Intergenic</td><td>NA</td><td>NA</td></tr>
<tr><td>GU305914.1</td><td>184053</td><td>184075</td><td>+</td><td>4</td><td>TCCCATTGACCCAATGTACT</td><td>TGG</td><td>-||-||||[|||--|||||||]PAM</td><td>NA</td><td>Intergenic</td><td>NA</td><td>NA</td></tr>
<tr><td>T13917</td><td>ATGTACTGGGCAAAACCCATTGG</td><td>1000</td><td></td><td></td><td></td><td></td><td></td><td></td><td></td><td></td><td></td></tr>
<tr><td>Oligo fwd</td><td>CACCATGTACTGGGCAAAACCCAT</td><td></td><td></td><td></td><td></td><td></td><td></td><td></td><td></td><td></td><td></td></tr>
<tr><td>Oligo rev</td><td>AAACATGGGTTTTGCCCAGTACAT</td><td></td><td></td><td></td><td></td><td></td><td></td><td></td><td></td><td></td><td></td></tr>
<tr><td>Chromosome</td><td>start</td><td>end</td><td>strand</td><td>MM</td><td>target_seq</td><td>PAM</td><td>alignment</td><td>distance</td><td>position</td><td>gene name</td><td>gene id</td></tr>
<tr><td>GU305914.1</td><td>183536</td><td>183558</td><td>+</td><td>0</td><td>ATGTACTGGGCAAAACCCAT</td><td>TGG</td><td>||||||||[||||||||||||]PAM</td><td>NA</td><td>Intergenic</td><td>NA</td><td>NA</td></tr>
<tr><td>GU305914.1</td><td>183443</td><td>183465</td><td>+</td><td>0</td><td>ATGTACTGGGCAAAACCCAT</td><td>TGG</td><td>||||||||[||||||||||||]PAM</td><td>NA</td><td>Intergenic</td><td>NA</td><td>NA</td></tr>
<tr><td>GU305914.1</td><td>183494</td><td>183516</td><td>+</td><td>4</td><td>CAGTAATGGGAAAAACCCAT</td><td>TGG</td><td>--|||-||[||-|||||||||]PAM</td><td>NA</td><td>Intergenic</td><td>NA</td><td>NA</td></tr>
<tr><td>GU305914.1</td><td>184024</td><td>184046</td><td>+</td><td>4</td><td>CAGTAATGGGAAAAACCCAT</td><td>TGG</td><td>--|||-||[||-|||||||||]PAM</td><td>NA</td><td>Intergenic</td><td>NA</td><td>NA</td></tr>
<tr><td>GU305914.1</td><td>183722</td><td>183744</td><td>+</td><td>1</td><td>ATGTACTGGGCAAAACCCAA</td><td>TGG</td><td>||||||||[|||||||||||-]PAM</td><td>NA</td><td>Intergenic</td><td>NA</td><td>NA</td></tr>
<tr><td>GU305914.1</td><td>183629</td><td>183651</td><td>+</td><td>1</td><td>ATGTACTGGGCAAAACCCAA</td><td>TGG</td><td>||||||||[|||||||||||-]PAM</td><td>NA</td><td>Intergenic</td><td>NA</td><td>NA</td></tr>
<tr><td>GU305914.1</td><td>183351</td><td>183373</td><td>+</td><td>2</td><td>TTGTACTGGGCAAAACCCAA</td><td>TGG</td><td>-|||||||[|||||||||||-]PAM</td><td>NA</td><td>Intergenic</td><td>NA</td><td>NA</td></tr>
<tr><td>T13918</td><td>CTCAGTATACGTGCCAGTAATGG</td><td>1000</td><td></td><td></td><td></td><td></td><td></td><td></td><td></td><td></td><td></td></tr>
<tr><td>Oligo fwd</td><td>CACCCTCAGTATACGTGCCAGTAA</td><td></td><td></td><td></td><td></td><td></td><td></td><td></td><td></td><td></td><td></td></tr>
<tr><td>Oligo rev</td><td>AAACTTACTGGCACGTATACTGAG</td><td></td><td></td><td></td><td></td><td></td><td></td><td></td><td></td><td></td><td></td></tr>
<tr><td>Chromosome</td><td>start</td><td>end</td><td>strand</td><td>MM</td><td>target_seq</td><td>PAM</td><td>alignment</td><td>distance</td><td>position</td><td>gene name</td><td>gene id</td></tr>
<tr><td>GU305914.1</td><td>183480</td><td>183502</td><td>+</td><td>0</td><td>CTCAGTATACGTGCCAGTAA</td><td>TGG</td><td>||||||||[||||||||||||]PAM</td><td>NA</td><td>Intergenic</td><td>NA</td><td>NA</td></tr>
<tr><td>GU305914.1</td><td>184010</td><td>184032</td><td>+</td><td>3</td><td>CTCAGTGTGCTTGCCAGTAA</td><td>TGG</td><td>||||||-|[-|-|||||||||]PAM</td><td>NA</td><td>Intergenic</td><td>NA</td><td>NA</td></tr>
<tr><td>T13919</td><td>TCAGTATACGTGCCAGTAATGGG</td><td>1000</td><td></td><td></td><td></td><td></td><td></td><td></td><td></td><td></td><td></td></tr>
<tr><td>Oligo fwd</td><td>CACCTCAGTATACGTGCCAGTAAT</td><td></td><td></td><td></td><td></td><td></td><td></td><td></td><td></td><td></td><td></td></tr>
<tr><td>Oligo rev</td><td>AAACATTACTGGCACGTATACTGA</td><td></td><td></td><td></td><td></td><td></td><td></td><td></td><td></td><td></td><td></td></tr>
<tr><td>Chromosome</td><td>start</td><td>end</td><td>strand</td><td>MM</td><td>target_seq</td><td>PAM</td><td>alignment</td><td>distance</td><td>position</td><td>gene name</td><td>gene id</td></tr>
<tr><td>GU305914.1</td><td>183481</td><td>183503</td><td>+</td><td>0</td><td>TCAGTATACGTGCCAGTAAT</td><td>GGG</td><td>||||||||[||||||||||||]PAM</td><td>NA</td><td>Intergenic</td><td>NA</td><td>NA</td></tr>
<tr><td>GU305914.1</td><td>184011</td><td>184033</td><td>+</td><td>3</td><td>TCAGTGTGCTTGCCAGTAAT</td><td>GGG</td><td>|||||-|-[|-||||||||||]PAM</td><td>NA</td><td>Intergenic</td><td>NA</td><td>NA</td></tr>
<tr><td>T13920</td><td>CAGTAATGGGAAAAACCCATTGG</td><td>1000</td><td></td><td></td><td></td><td></td><td></td><td></td><td></td><td></td><td></td></tr>
<tr><td>Oligo fwd</td><td>CACCCAGTAATGGGAAAAACCCAT</td><td></td><td></td><td></td><td></td><td></td><td></td><td></td><td></td><td></td><td></td></tr>
<tr><td>Oligo rev</td><td>AAACATGGGTTTTTCCCATTACTG</td><td></td><td></td><td></td><td></td><td></td><td></td><td></td><td></td><td></td><td></td></tr>
<tr><td>Chromosome</td><td>start</td><td>end</td><td>strand</td><td>MM</td><td>target_seq</td><td>PAM</td><td>alignment</td><td>distance</td><td>position</td><td>gene name</td><td>gene id</td></tr>
<tr><td>GU305914.1</td><td>184024</td><td>184046</td><td>+</td><td>0</td><td>CAGTAATGGGAAAAACCCAT</td><td>TGG</td><td>||||||||[||||||||||||]PAM</td><td>NA</td><td>Intergenic</td><td>NA</td><td>NA</td></tr>
<tr><td>GU305914.1</td><td>183494</td><td>183516</td><td>+</td><td>0</td><td>CAGTAATGGGAAAAACCCAT</td><td>TGG</td><td>||||||||[||||||||||||]PAM</td><td>NA</td><td>Intergenic</td><td>NA</td><td>NA</td></tr>
<tr><td>GU305914.1</td><td>183682</td><td>183704</td><td>-</td><td>4</td><td>AGTCAATGGGAAAAACCCAT</td><td>TGG</td><td>----||||[||||||||||||]PAM</td><td>NA</td><td>Intergenic</td><td>NA</td><td>NA</td></tr>
<tr><td>GU305914.1</td><td>183443</td><td>183465</td><td>+</td><td>4</td><td>ATGTACTGGGCAAAACCCAT</td><td>TGG</td><td>--|||-||[||-|||||||||]PAM</td><td>NA</td><td>Intergenic</td><td>NA</td><td>NA</td></tr>
<tr><td>GU305914.1</td><td>183536</td><td>183558</td><td>+</td><td>4</td><td>ATGTACTGGGCAAAACCCAT</td><td>TGG</td><td>--|||-||[||-|||||||||]PAM</td><td>NA</td><td>Intergenic</td><td>NA</td><td>NA</td></tr>
<tr><td>T13921</td><td>CTCCCATTGACCTTATGTACTGG</td><td>1000</td><td></td><td></td><td></td><td></td><td></td><td></td><td></td><td></td><td></td></tr>
<tr><td>Oligo fwd</td><td>CACCCTCCCATTGACCTTATGTAC</td><td></td><td></td><td></td><td></td><td></td><td></td><td></td><td></td><td></td><td></td></tr>
<tr><td>Oligo rev</td><td>AAACGTACATAAGGTCAATGGGAG</td><td></td><td></td><td></td><td></td><td></td><td></td><td></td><td></td><td></td><td></td></tr>
<tr><td>Chromosome</td><td>start</td><td>end</td><td>strand</td><td>MM</td><td>target_seq</td><td>PAM</td><td>alignment</td><td>distance</td><td>position</td><td>gene name</td><td>gene id</td></tr>
<tr><td>GU305914.1</td><td>183522</td><td>183544</td><td>+</td><td>0</td><td>CTCCCATTGACCTTATGTAC</td><td>TGG</td><td>||||||||[||||||||||||]PAM</td><td>NA</td><td>Intergenic</td><td>NA</td><td>NA</td></tr>
<tr><td>GU305914.1</td><td>183708</td><td>183730</td><td>+</td><td>2</td><td>CCCCTATTGACCTTATGTAC</td><td>TGG</td><td>|-||-|||[||||||||||||]PAM</td><td>NA</td><td>Intergenic</td><td>NA</td><td>NA</td></tr>
<tr><td>GU305914.1</td><td>183615</td><td>183637</td><td>+</td><td>2</td><td>CCCCTATTGACCTTATGTAC</td><td>TGG</td><td>|-||-|||[||||||||||||]PAM</td><td>NA</td><td>Intergenic</td><td>NA</td><td>NA</td></tr>
<tr><td>GU305914.1</td><td>183429</td><td>183451</td><td>+</td><td>2</td><td>CCCCTATTGACCTTATGTAC</td><td>TGG</td><td>|-||-|||[||||||||||||]PAM</td><td>NA</td><td>Intergenic</td><td>NA</td><td>NA</td></tr>
<tr><td>GU305914.1</td><td>183801</td><td>183823</td><td>+</td><td>2</td><td>CCCCTATTGACCTTATGTAC</td><td>TGG</td><td>|-||-|||[||||||||||||]PAM</td><td>NA</td><td>Intergenic</td><td>NA</td><td>NA</td></tr>
<tr><td>GU305914.1</td><td>183337</td><td>183359</td><td>+</td><td>3</td><td>CCCCTATTGACCTTTTGTAC</td><td>TGG</td><td>|-||-|||[||||||-|||||]PAM</td><td>NA</td><td>Intergenic</td><td>NA</td><td>NA</td></tr>
<tr><td>T13922</td><td>TCCCATTGACCTTATGTACTGGG</td><td>1000</td><td></td><td></td><td></td><td></td><td></td><td></td><td></td><td></td><td></td></tr>
<tr><td>Oligo fwd</td><td>CACCTCCCATTGACCTTATGTACT</td><td></td><td></td><td></td><td></td><td></td><td></td><td></td><td></td><td></td><td></td></tr>
<tr><td>Oligo rev</td><td>AAACAGTACATAAGGTCAATGGGA</td><td></td><td></td><td></td><td></td><td></td><td></td><td></td><td></td><td></td><td></td></tr>
<tr><td>Chromosome</td><td>start</td><td>end</td><td>strand</td><td>MM</td><td>target_seq</td><td>PAM</td><td>alignment</td><td>distance</td><td>position</td><td>gene name</td><td>gene id</td></tr>
<tr><td>GU305914.1</td><td>183523</td><td>183545</td><td>+</td><td>0</td><td>TCCCATTGACCTTATGTACT</td><td>GGG</td><td>||||||||[||||||||||||]PAM</td><td>NA</td><td>Intergenic</td><td>NA</td><td>NA</td></tr>
<tr><td>GU305914.1</td><td>183430</td><td>183452</td><td>+</td><td>2</td><td>CCCTATTGACCTTATGTACT</td><td>GGG</td><td>-||-||||[||||||||||||]PAM</td><td>NA</td><td>Intergenic</td><td>NA</td><td>NA</td></tr>
<tr><td>GU305914.1</td><td>183802</td><td>183824</td><td>+</td><td>2</td><td>CCCTATTGACCTTATGTACT</td><td>GGG</td><td>-||-||||[||||||||||||]PAM</td><td>NA</td><td>Intergenic</td><td>NA</td><td>NA</td></tr>
<tr><td>GU305914.1</td><td>183709</td><td>183731</td><td>+</td><td>2</td><td>CCCTATTGACCTTATGTACT</td><td>GGG</td><td>-||-||||[||||||||||||]PAM</td><td>NA</td><td>Intergenic</td><td>NA</td><td>NA</td></tr>
<tr><td>GU305914.1</td><td>183616</td><td>183638</td><td>+</td><td>2</td><td>CCCTATTGACCTTATGTACT</td><td>GGG</td><td>-||-||||[||||||||||||]PAM</td><td>NA</td><td>Intergenic</td><td>NA</td><td>NA</td></tr>
<tr><td>GU305914.1</td><td>183338</td><td>183360</td><td>+</td><td>3</td><td>CCCTATTGACCTTTTGTACT</td><td>GGG</td><td>-||-||||[|||||-||||||]PAM</td><td>NA</td><td>Intergenic</td><td>NA</td><td>NA</td></tr>
<tr><td>GU305914.1</td><td>183959</td><td>183981</td><td>+</td><td>2</td><td>TCCCATTGACCCAATGTACT</td><td>TGG</td><td>||||||||[|||--|||||||]PAM</td><td>NA</td><td>Intergenic</td><td>NA</td><td>NA</td></tr>
<tr><td>GU305914.1</td><td>184053</td><td>184075</td><td>+</td><td>2</td><td>TCCCATTGACCCAATGTACT</td><td>TGG</td><td>||||||||[|||--|||||||]PAM</td><td>NA</td><td>Intergenic</td><td>NA</td><td>NA</td></tr>
<tr><td>T13923</td><td>ATGTACTGGGCAAAACCCATTGG</td><td>1000</td><td></td><td></td><td></td><td></td><td></td><td></td><td></td><td></td><td></td></tr>
<tr><td>Oligo fwd</td><td>CACCATGTACTGGGCAAAACCCAT</td><td></td><td></td><td></td><td></td><td></td><td></td><td></td><td></td><td></td><td></td></tr>
<tr><td>Oligo rev</td><td>AAACATGGGTTTTGCCCAGTACAT</td><td></td><td></td><td></td><td></td><td></td><td></td><td></td><td></td><td></td><td></td></tr>
<tr><td>Chromosome</td><td>start</td><td>end</td><td>strand</td><td>MM</td><td>target_seq</td><td>PAM</td><td>alignment</td><td>distance</td><td>position</td><td>gene name</td><td>gene id</td></tr>
<tr><td>GU305914.1</td><td>183536</td><td>183558</td><td>+</td><td>0</td><td>ATGTACTGGGCAAAACCCAT</td><td>TGG</td><td>||||||||[||||||||||||]PAM</td><td>NA</td><td>Intergenic</td><td>NA</td><td>NA</td></tr>
<tr><td>GU305914.1</td><td>183443</td><td>183465</td><td>+</td><td>0</td><td>ATGTACTGGGCAAAACCCAT</td><td>TGG</td><td>||||||||[||||||||||||]PAM</td><td>NA</td><td>Intergenic</td><td>NA</td><td>NA</td></tr>
</table>

<table>
<tr><td>GU305914.1</td><td>183494</td><td>183516</td><td>+</td><td>4</td><td>CAGTAATGGGAAAAACCCAT</td><td>TGG</td><td>--|||-||[||-|||||||||]PAM</td><td>NA</td><td>Intergenic</td><td>NA</td><td>NA</td></tr>
<tr><td>GU305914.1</td><td>184024</td><td>184046</td><td>+</td><td>4</td><td>CAGTAATGGGAAAAACCCAT</td><td>TGG</td><td>--|||-||[||-|||||||||]PAM</td><td>NA</td><td>Intergenic</td><td>NA</td><td>NA</td></tr>
<tr><td>GU305914.1</td><td>183722</td><td>183744</td><td>+</td><td>1</td><td>ATGTACTGGGCAAAACCCAA</td><td>TGG</td><td>||||||||[|||||||||||-]PAM</td><td>NA</td><td>Intergenic</td><td>NA</td><td>NA</td></tr>
<tr><td>GU305914.1</td><td>183629</td><td>183651</td><td>+</td><td>1</td><td>ATGTACTGGGCAAAACCCAA</td><td>TGG</td><td>||||||||[|||||||||||-]PAM</td><td>NA</td><td>Intergenic</td><td>NA</td><td>NA</td></tr>
<tr><td>GU305914.1</td><td>183351</td><td>183373</td><td>+</td><td>2</td><td>TTGTACTGGGCAAAACCCAA</td><td>TGG</td><td>-|||||||[|||||||||||-]PAM</td><td>NA</td><td>Intergenic</td><td>NA</td><td>NA</td></tr>
<tr><td>T13924</td><td>CCCTATTGACTCAATGTACTTGG</td><td>1000</td><td></td><td></td><td></td><td></td><td></td><td></td><td></td><td></td><td></td></tr>
<tr><td>Oligo fwd</td><td>CACCCCCTATTGACTCAATGTACT</td><td></td><td></td><td></td><td></td><td></td><td></td><td></td><td></td><td></td><td></td></tr>
<tr><td>Oligo rev</td><td>AAACAGTACATTGAGTCAATAGGG</td><td></td><td></td><td></td><td></td><td></td><td></td><td></td><td></td><td></td><td></td></tr>
<tr><td>Chromosome</td><td>start</td><td>end</td><td>strand</td><td>MM</td><td>target_seq</td><td>PAM</td><td>alignment</td><td>distance</td><td>position</td><td>gene name</td><td>gene id</td></tr>
<tr><td>GU305914.1</td><td>183564</td><td>183586</td><td>+</td><td>0</td><td>CCCTATTGACTCAATGTACT</td><td>TGG</td><td>||||||||[||||||||||||]PAM</td><td>NA</td><td>Intergenic</td><td>NA</td><td>NA</td></tr>
<tr><td>GU305914.1</td><td>183959</td><td>183981</td><td>+</td><td>3</td><td>TCCCATTGACCCAATGTACT</td><td>TGG</td><td>-||-||||[||-|||||||||]PAM</td><td>NA</td><td>Intergenic</td><td>NA</td><td>NA</td></tr>
<tr><td>GU305914.1</td><td>184053</td><td>184075</td><td>+</td><td>3</td><td>TCCCATTGACCCAATGTACT</td><td>TGG</td><td>-||-||||[||-|||||||||]PAM</td><td>NA</td><td>Intergenic</td><td>NA</td><td>NA</td></tr>
<tr><td>GU305914.1</td><td>183379</td><td>183401</td><td>+</td><td>1</td><td>CCCTATTGACTCAGTGTACT</td><td>TGG</td><td>||||||||[|||||-||||||]PAM</td><td>NA</td><td>Intergenic</td><td>NA</td><td>NA</td></tr>
<tr><td>GU305914.1</td><td>183907</td><td>183929</td><td>+</td><td>2</td><td>CCCTATTGAGTCAGTGTACT</td><td>TGG</td><td>||||||||[|-|||-||||||]PAM</td><td>NA</td><td>Intergenic</td><td>NA</td><td>NA</td></tr>
<tr><td>GU305914.1</td><td>183657</td><td>183679</td><td>+</td><td>2</td><td>CCCTATTGAGTCAGTGTACT</td><td>TGG</td><td>||||||||[|-|||-||||||]PAM</td><td>NA</td><td>Intergenic</td><td>NA</td><td>NA</td></tr>
<tr><td>GU305914.1</td><td>183750</td><td>183772</td><td>+</td><td>2</td><td>CCCTATTGACGCAGTGTACT</td><td>TGG</td><td>||||||||[||-||-||||||]PAM</td><td>NA</td><td>Intergenic</td><td>NA</td><td>NA</td></tr>
<tr><td>T13925</td><td>CTCAATGTACTTGGCTCCAATGG</td><td>1000</td><td></td><td></td><td></td><td></td><td></td><td></td><td></td><td></td><td></td></tr>
<tr><td>Oligo fwd</td><td>CACCCTCAATGTACTTGGCTCCAA</td><td></td><td></td><td></td><td></td><td></td><td></td><td></td><td></td><td></td><td></td></tr>
<tr><td>Oligo rev</td><td>AAACTTGGAGCCAAGTACATTGAG</td><td></td><td></td><td></td><td></td><td></td><td></td><td></td><td></td><td></td><td></td></tr>
<tr><td>Chromosome</td><td>start</td><td>end</td><td>strand</td><td>MM</td><td>target_seq</td><td>PAM</td><td>alignment</td><td>distance</td><td>position</td><td>gene name</td><td>gene id</td></tr>
<tr><td>GU305914.1</td><td>183573</td><td>183595</td><td>+</td><td>0</td><td>CTCAATGTACTTGGCTCCAA</td><td>TGG</td><td>||||||||[||||||||||||]PAM</td><td>NA</td><td>Intergenic</td><td>NA</td><td>NA</td></tr>
<tr><td>GU305914.1</td><td>183388</td><td>183410</td><td>+</td><td>1</td><td>CTCAGTGTACTTGGCTCCAA</td><td>TGG</td><td>||||-|||[||||||||||||]PAM</td><td>NA</td><td>Intergenic</td><td>NA</td><td>NA</td></tr>
<tr><td>GU305914.1</td><td>183666</td><td>183688</td><td>+</td><td>2</td><td>GTCAGTGTACTTGGCTCCAA</td><td>TGG</td><td>-|||-|||[||||||||||||]PAM</td><td>NA</td><td>Intergenic</td><td>NA</td><td>NA</td></tr>
<tr><td>GU305914.1</td><td>183759</td><td>183781</td><td>+</td><td>2</td><td>CGCAGTGTACTTGGCTCCAA</td><td>TGG</td><td>|-||-|||[||||||||||||]PAM</td><td>NA</td><td>Intergenic</td><td>NA</td><td>NA</td></tr>
<tr><td>GU305914.1</td><td>183916</td><td>183938</td><td>+</td><td>4</td><td>GTCAGTGTACTTGGCAGCAA</td><td>TGG</td><td>-|||-|||[|||||||--|||]PAM</td><td>NA</td><td>Intergenic</td><td>NA</td><td>NA</td></tr>
<tr><td>T13926</td><td>TCAATGTACTTGGCTCCAATGGG</td><td>1000</td><td></td><td></td><td></td><td></td><td></td><td></td><td></td><td></td><td></td></tr>
<tr><td>Oligo fwd</td><td>CACCTCAATGTACTTGGCTCCAAT</td><td></td><td></td><td></td><td></td><td></td><td></td><td></td><td></td><td></td><td></td></tr>
<tr><td>Oligo rev</td><td>AAACATTGGAGCCAAGTACATTGA</td><td></td><td></td><td></td><td></td><td></td><td></td><td></td><td></td><td></td><td></td></tr>
<tr><td>Chromosome</td><td>start</td><td>end</td><td>strand</td><td>MM</td><td>target_seq</td><td>PAM</td><td>alignment</td><td>distance</td><td>position</td><td>gene name</td><td>gene id</td></tr>
<tr><td>GU305914.1</td><td>183574</td><td>183596</td><td>+</td><td>0</td><td>TCAATGTACTTGGCTCCAAT</td><td>GGG</td><td>||||||||[||||||||||||]PAM</td><td>NA</td><td>Intergenic</td><td>NA</td><td>NA</td></tr>
<tr><td>GU305914.1</td><td>183389</td><td>183411</td><td>+</td><td>1</td><td>TCAGTGTACTTGGCTCCAAT</td><td>GGG</td><td>|||-||||[||||||||||||]PAM</td><td>NA</td><td>Intergenic</td><td>NA</td><td>NA</td></tr>
<tr><td>GU305914.1</td><td>183667</td><td>183689</td><td>+</td><td>1</td><td>TCAGTGTACTTGGCTCCAAT</td><td>GGG</td><td>|||-||||[||||||||||||]PAM</td><td>NA</td><td>Intergenic</td><td>NA</td><td>NA</td></tr>
<tr><td>GU305914.1</td><td>183760</td><td>183782</td><td>+</td><td>2</td><td>GCAGTGTACTTGGCTCCAAT</td><td>GGG</td><td>-||-||||[||||||||||||]PAM</td><td>NA</td><td>Intergenic</td><td>NA</td><td>NA</td></tr>
<tr><td>GU305914.1</td><td>183917</td><td>183939</td><td>+</td><td>3</td><td>TCAGTGTACTTGGCAGCAAT</td><td>GGG</td><td>|||-||||[||||||--||||]PAM</td><td>NA</td><td>Intergenic</td><td>NA</td><td>NA</td></tr>
<tr><td>T13927</td><td>CCCCTATTGACCTTATGTACTGG</td><td>1000</td><td></td><td></td><td></td><td></td><td></td><td></td><td></td><td></td><td></td></tr>
<tr><td>Oligo fwd</td><td>CACCCCCCTATTGACCTTATGTAC</td><td></td><td></td><td></td><td></td><td></td><td></td><td></td><td></td><td></td><td></td></tr>
<tr><td>Oligo rev</td><td>AAACGTACATAAGGTCAATAGGGG</td><td></td><td></td><td></td><td></td><td></td><td></td><td></td><td></td><td></td><td></td></tr>
<tr><td>Chromosome</td><td>start</td><td>end</td><td>strand</td><td>MM</td><td>target_seq</td><td>PAM</td><td>alignment</td><td>distance</td><td>position</td><td>gene name</td><td>gene id</td></tr>
<tr><td>GU305914.1</td><td>183801</td><td>183823</td><td>+</td><td>0</td><td>CCCCTATTGACCTTATGTAC</td><td>TGG</td><td>||||||||[||||||||||||]PAM</td><td>NA</td><td>Intergenic</td><td>NA</td><td>NA</td></tr>
<tr><td>GU305914.1</td><td>183708</td><td>183730</td><td>+</td><td>0</td><td>CCCCTATTGACCTTATGTAC</td><td>TGG</td><td>||||||||[||||||||||||]PAM</td><td>NA</td><td>Intergenic</td><td>NA</td><td>NA</td></tr>
<tr><td>GU305914.1</td><td>183615</td><td>183637</td><td>+</td><td>0</td><td>CCCCTATTGACCTTATGTAC</td><td>TGG</td><td>||||||||[||||||||||||]PAM</td><td>NA</td><td>Intergenic</td><td>NA</td><td>NA</td></tr>
<tr><td>GU305914.1</td><td>183429</td><td>183451</td><td>+</td><td>0</td><td>CCCCTATTGACCTTATGTAC</td><td>TGG</td><td>||||||||[||||||||||||]PAM</td><td>NA</td><td>Intergenic</td><td>NA</td><td>NA</td></tr>
<tr><td>GU305914.1</td><td>183522</td><td>183544</td><td>+</td><td>2</td><td>CTCCCATTGACCTTATGTAC</td><td>TGG</td><td>|-||-|||[||||||||||||]PAM</td><td>NA</td><td>Intergenic</td><td>NA</td><td>NA</td></tr>
<tr><td>GU305914.1</td><td>183337</td><td>183359</td><td>+</td><td>1</td><td>CCCCTATTGACCTTTTGTAC</td><td>TGG</td><td>||||||||[||||||-|||||]PAM</td><td>NA</td><td>Intergenic</td><td>NA</td><td>NA</td></tr>
<tr><td>T13928</td><td>CCCTATTGACCTTATGTACTGGG</td><td>1000</td><td></td><td></td><td></td><td></td><td></td><td></td><td></td><td></td><td></td></tr>
<tr><td>Oligo fwd</td><td>CACCCCCTATTGACCTTATGTACT</td><td></td><td></td><td></td><td></td><td></td><td></td><td></td><td></td><td></td><td></td></tr>
<tr><td>Oligo rev</td><td>AAACAGTACATAAGGTCAATAGGG</td><td></td><td></td><td></td><td></td><td></td><td></td><td></td><td></td><td></td><td></td></tr>
<tr><td>Chromosome</td><td>start</td><td>end</td><td>strand</td><td>MM</td><td>target_seq</td><td>PAM</td><td>alignment</td><td>distance</td><td>position</td><td>gene name</td><td>gene id</td></tr>
<tr><td>GU305914.1</td><td>183616</td><td>183638</td><td>+</td><td>0</td><td>CCCTATTGACCTTATGTACT</td><td>GGG</td><td>||||||||[||||||||||||]PAM</td><td>NA</td><td>Intergenic</td><td>NA</td><td>NA</td></tr>
<tr><td>GU305914.1</td><td>183430</td><td>183452</td><td>+</td><td>0</td><td>CCCTATTGACCTTATGTACT</td><td>GGG</td><td>||||||||[||||||||||||]PAM</td><td>NA</td><td>Intergenic</td><td>NA</td><td>NA</td></tr>
<tr><td>GU305914.1</td><td>183802</td><td>183824</td><td>+</td><td>0</td><td>CCCTATTGACCTTATGTACT</td><td>GGG</td><td>||||||||[||||||||||||]PAM</td><td>NA</td><td>Intergenic</td><td>NA</td><td>NA</td></tr>
<tr><td>GU305914.1</td><td>183709</td><td>183731</td><td>+</td><td>0</td><td>CCCTATTGACCTTATGTACT</td><td>GGG</td><td>||||||||[||||||||||||]PAM</td><td>NA</td><td>Intergenic</td><td>NA</td><td>NA</td></tr>
<tr><td>GU305914.1</td><td>183523</td><td>183545</td><td>+</td><td>2</td><td>TCCCATTGACCTTATGTACT</td><td>GGG</td><td>-||-||||[||||||||||||]PAM</td><td>NA</td><td>Intergenic</td><td>NA</td><td>NA</td></tr>
<tr><td>GU305914.1</td><td>183338</td><td>183360</td><td>+</td><td>1</td><td>CCCTATTGACCTTTTGTACT</td><td>GGG</td><td>||||||||[|||||-||||||]PAM</td><td>NA</td><td>Intergenic</td><td>NA</td><td>NA</td></tr>
<tr><td>GU305914.1</td><td>183959</td><td>183981</td><td>+</td><td>4</td><td>TCCCATTGACCCAATGTACT</td><td>TGG</td><td>-||-||||[|||--|||||||]PAM</td><td>NA</td><td>Intergenic</td><td>NA</td><td>NA</td></tr>
<tr><td>GU305914.1</td><td>184053</td><td>184075</td><td>+</td><td>4</td><td>TCCCATTGACCCAATGTACT</td><td>TGG</td><td>-||-||||[|||--|||||||]PAM</td><td>NA</td><td>Intergenic</td><td>NA</td><td>NA</td></tr>
<tr><td>T13929</td><td>ATGTACTGGGCAAAACCCAATGG</td><td>1000</td><td></td><td></td><td></td><td></td><td></td><td></td><td></td><td></td><td></td></tr>
<tr><td>Oligo fwd</td><td>CACCATGTACTGGGCAAAACCCAA</td><td></td><td></td><td></td><td></td><td></td><td></td><td></td><td></td><td></td><td></td></tr>
<tr><td>Oligo rev</td><td>AAACTTGGGTTTTGCCCAGTACAT</td><td></td><td></td><td></td><td></td><td></td><td></td><td></td><td></td><td></td><td></td></tr>
<tr><td>Chromosome</td><td>start</td><td>end</td><td>strand</td><td>MM</td><td>target_seq</td><td>PAM</td><td>alignment</td><td>distance</td><td>position</td><td>gene name</td><td>gene id</td></tr>
<tr><td>GU305914.1</td><td>183722</td><td>183744</td><td>+</td><td>0</td><td>ATGTACTGGGCAAAACCCAA</td><td>TGG</td><td>||||||||[||||||||||||]PAM</td><td>NA</td><td>Intergenic</td><td>NA</td><td>NA</td></tr>
<tr><td>GU305914.1</td><td>183629</td><td>183651</td><td>+</td><td>0</td><td>ATGTACTGGGCAAAACCCAA</td><td>TGG</td><td>||||||||[||||||||||||]PAM</td><td>NA</td><td>Intergenic</td><td>NA</td><td>NA</td></tr>
<tr><td>GU305914.1</td><td>183351</td><td>183373</td><td>+</td><td>1</td><td>TTGTACTGGGCAAAACCCAA</td><td>TGG</td><td>-|||||||[||||||||||||]PAM</td><td>NA</td><td>Intergenic</td><td>NA</td><td>NA</td></tr>
<tr><td>GU305914.1</td><td>183536</td><td>183558</td><td>+</td><td>1</td><td>ATGTACTGGGCAAAACCCAT</td><td>TGG</td><td>||||||||[|||||||||||-]PAM</td><td>NA</td><td>Intergenic</td><td>NA</td><td>NA</td></tr>
<tr><td>GU305914.1</td><td>183443</td><td>183465</td><td>+</td><td>1</td><td>ATGTACTGGGCAAAACCCAT</td><td>TGG</td><td>||||||||[|||||||||||-]PAM</td><td>NA</td><td>Intergenic</td><td>NA</td><td>NA</td></tr>
<tr><td>T13930</td><td>CCCTATTGAGTCAGTGTACTTGG</td><td>1000</td><td></td><td></td><td></td><td></td><td></td><td></td><td></td><td></td><td></td></tr>
<tr><td>Oligo fwd</td><td>CACCCCCTATTGAGTCAGTGTACT</td><td></td><td></td><td></td><td></td><td></td><td></td><td></td><td></td><td></td><td></td></tr>
<tr><td>Oligo rev</td><td>AAACAGTACACTGACTCAATAGGG</td><td></td><td></td><td></td><td></td><td></td><td></td><td></td><td></td><td></td><td></td></tr>
<tr><td>Chromosome</td><td>start</td><td>end</td><td>strand</td><td>MM</td><td>target_seq</td><td>PAM</td><td>alignment</td><td>distance</td><td>position</td><td>gene name</td><td>gene id</td></tr>
<tr><td>GU305914.1</td><td>183657</td><td>183679</td><td>+</td><td>0</td><td>CCCTATTGAGTCAGTGTACT</td><td>TGG</td><td>||||||||[||||||||||||]PAM</td><td>NA</td><td>Intergenic</td><td>NA</td><td>NA</td></tr>
<tr><td>GU305914.1</td><td>183907</td><td>183929</td><td>+</td><td>0</td><td>CCCTATTGAGTCAGTGTACT</td><td>TGG</td><td>||||||||[||||||||||||]PAM</td><td>NA</td><td>Intergenic</td><td>NA</td><td>NA</td></tr>
<tr><td>GU305914.1</td><td>183379</td><td>183401</td><td>+</td><td>1</td><td>CCCTATTGACTCAGTGTACT</td><td>TGG</td><td>||||||||[|-||||||||||]PAM</td><td>NA</td><td>Intergenic</td><td>NA</td><td>NA</td></tr>
<tr><td>GU305914.1</td><td>183750</td><td>183772</td><td>+</td><td>2</td><td>CCCTATTGACGCAGTGTACT</td><td>TGG</td><td>||||||||[|--|||||||||]PAM</td><td>NA</td><td>Intergenic</td><td>NA</td><td>NA</td></tr>
<tr><td>GU305914.1</td><td>183564</td><td>183586</td><td>+</td><td>2</td><td>CCCTATTGACTCAATGTACT</td><td>TGG</td><td>||||||||[|-|||-||||||]PAM</td><td>NA</td><td>Intergenic</td><td>NA</td><td>NA</td></tr>
<tr><td>T13931</td><td>GTCAGTGTACTTGGCTCCAATGG</td><td>1000</td><td></td><td></td><td></td><td></td><td></td><td></td><td></td><td></td><td></td></tr>
<tr><td>Oligo fwd</td><td>CACCGTCAGTGTACTTGGCTCCAA</td><td></td><td></td><td></td><td></td><td></td><td></td><td></td><td></td><td></td><td></td></tr>
<tr><td>Oligo rev</td><td>AAACTTGGAGCCAAGTACACTGAC</td><td></td><td></td><td></td><td></td><td></td><td></td><td></td><td></td><td></td><td></td></tr>
<tr><td>Chromosome</td><td>start</td><td>end</td><td>strand</td><td>MM</td><td>target_seq</td><td>PAM</td><td>alignment</td><td>distance</td><td>position</td><td>gene name</td><td>gene id</td></tr>
<tr><td>GU305914.1</td><td>183666</td><td>183688</td><td>+</td><td>0</td><td>GTCAGTGTACTTGGCTCCAA</td><td>TGG</td><td>||||||||[||||||||||||]PAM</td><td>NA</td><td>Intergenic</td><td>NA</td><td>NA</td></tr>
<tr><td>GU305914.1</td><td>183388</td><td>183410</td><td>+</td><td>1</td><td>CTCAGTGTACTTGGCTCCAA</td><td>TGG</td><td>-|||||||[||||||||||||]PAM</td><td>NA</td><td>Intergenic</td><td>NA</td><td>NA</td></tr>
<tr><td>GU305914.1</td><td>183759</td><td>183781</td><td>+</td><td>2</td><td>CGCAGTGTACTTGGCTCCAA</td><td>TGG</td><td>--||||||[||||||||||||]PAM</td><td>NA</td><td>Intergenic</td><td>NA</td><td>NA</td></tr>
</table>

<table>
<tr><td>GU305914.1</td><td>183573</td><td>183595</td><td>+</td><td>2</td><td>CTCAATGTACTTGGCTCCAA</td><td>TGG</td><td>-|||-|||[||||||||||||]PAM</td><td>NA</td><td>Intergenic</td><td>NA</td><td>NA</td></tr>
<tr><td>GU305914.1</td><td>183916</td><td>183938</td><td>+</td><td>2</td><td>GTCAGTGTACTTGGCAGCAA</td><td>TGG</td><td>||||||||[|||||||--|||]PAM</td><td>NA</td><td>Intergenic</td><td>NA</td><td>NA</td></tr>
<tr><td></td><td></td><td></td><td></td><td></td><td></td><td></td><td></td><td></td><td></td><td></td><td></td></tr>
<tr><td>T13932</td><td>TCAGTGTACTTGGCTCCAATGGG</td><td>1000</td><td></td><td></td><td></td><td></td><td></td><td></td><td></td><td></td><td></td></tr>
<tr><td>Oligo fwd</td><td>CACCTCAGTGTACTTGGCTCCAAT</td><td></td><td></td><td></td><td></td><td></td><td></td><td></td><td></td><td></td><td></td></tr>
<tr><td>Oligo rev</td><td>AAACATTGGAGCCAAGTACACTGA</td><td></td><td></td><td></td><td></td><td></td><td></td><td></td><td></td><td></td><td></td></tr>
<tr><td>Chromosome</td><td>start</td><td>end</td><td>strand</td><td>MM</td><td>target_seq</td><td>PAM</td><td>alignment</td><td>distance</td><td>position</td><td>gene name</td><td>gene id</td></tr>
<tr><td>GU305914.1</td><td>183389</td><td>183411</td><td>+</td><td>0</td><td>TCAGTGTACTTGGCTCCAAT</td><td>GGG</td><td>||||||||[||||||||||||]PAM</td><td>NA</td><td>Intergenic</td><td>NA</td><td>NA</td></tr>
<tr><td>GU305914.1</td><td>183667</td><td>183689</td><td>+</td><td>0</td><td>TCAGTGTACTTGGCTCCAAT</td><td>GGG</td><td>||||||||[||||||||||||]PAM</td><td>NA</td><td>Intergenic</td><td>NA</td><td>NA</td></tr>
<tr><td>GU305914.1</td><td>183760</td><td>183782</td><td>+</td><td>1</td><td>GCAGTGTACTTGGCTCCAAT</td><td>GGG</td><td>-|||||||[||||||||||||]PAM</td><td>NA</td><td>Intergenic</td><td>NA</td><td>NA</td></tr>
<tr><td>GU305914.1</td><td>183574</td><td>183596</td><td>+</td><td>1</td><td>TCAATGTACTTGGCTCCAAT</td><td>GGG</td><td>|||-||||[||||||||||||]PAM</td><td>NA</td><td>Intergenic</td><td>NA</td><td>NA</td></tr>
<tr><td>GU305914.1</td><td>183917</td><td>183939</td><td>+</td><td>2</td><td>TCAGTGTACTTGGCAGCAAT</td><td>GGG</td><td>||||||||[||||||--||||]PAM</td><td>NA</td><td>Intergenic</td><td>NA</td><td>NA</td></tr>
<tr><td></td><td></td><td></td><td></td><td></td><td></td><td></td><td></td><td></td><td></td><td></td><td></td></tr>
<tr><td>T13933</td><td>CCCCTATTGACCTTATGTACTGG</td><td>1000</td><td></td><td></td><td></td><td></td><td></td><td></td><td></td><td></td><td></td></tr>
<tr><td>Oligo fwd</td><td>CACCCCCCTATTGACCTTATGTAC</td><td></td><td></td><td></td><td></td><td></td><td></td><td></td><td></td><td></td><td></td></tr>
<tr><td>Oligo rev</td><td>AAACGTACATAAGGTCAATAGGGG</td><td></td><td></td><td></td><td></td><td></td><td></td><td></td><td></td><td></td><td></td></tr>
<tr><td>Chromosome</td><td>start</td><td>end</td><td>strand</td><td>MM</td><td>target_seq</td><td>PAM</td><td>alignment</td><td>distance</td><td>position</td><td>gene name</td><td>gene id</td></tr>
<tr><td>GU305914.1</td><td>183801</td><td>183823</td><td>+</td><td>0</td><td>CCCCTATTGACCTTATGTAC</td><td>TGG</td><td>||||||||[||||||||||||]PAM</td><td>NA</td><td>Intergenic</td><td>NA</td><td>NA</td></tr>
<tr><td>GU305914.1</td><td>183708</td><td>183730</td><td>+</td><td>0</td><td>CCCCTATTGACCTTATGTAC</td><td>TGG</td><td>||||||||[||||||||||||]PAM</td><td>NA</td><td>Intergenic</td><td>NA</td><td>NA</td></tr>
<tr><td>GU305914.1</td><td>183615</td><td>183637</td><td>+</td><td>0</td><td>CCCCTATTGACCTTATGTAC</td><td>TGG</td><td>||||||||[||||||||||||]PAM</td><td>NA</td><td>Intergenic</td><td>NA</td><td>NA</td></tr>
<tr><td>GU305914.1</td><td>183429</td><td>183451</td><td>+</td><td>0</td><td>CCCCTATTGACCTTATGTAC</td><td>TGG</td><td>||||||||[||||||||||||]PAM</td><td>NA</td><td>Intergenic</td><td>NA</td><td>NA</td></tr>
<tr><td>GU305914.1</td><td>183522</td><td>183544</td><td>+</td><td>2</td><td>CTCCCATTGACCTTATGTAC</td><td>TGG</td><td>|-||-|||[||||||||||||]PAM</td><td>NA</td><td>Intergenic</td><td>NA</td><td>NA</td></tr>
<tr><td>GU305914.1</td><td>183337</td><td>183359</td><td>+</td><td>1</td><td>CCCCTATTGACCTTTTGTAC</td><td>TGG</td><td>||||||||[||||||-|||||]PAM</td><td>NA</td><td>Intergenic</td><td>NA</td><td>NA</td></tr>
<tr><td></td><td></td><td></td><td></td><td></td><td></td><td></td><td></td><td></td><td></td><td></td><td></td></tr>
<tr><td>T13934</td><td>CCCTATTGACCTTATGTACTGGG</td><td>1000</td><td></td><td></td><td></td><td></td><td></td><td></td><td></td><td></td><td></td></tr>
<tr><td>Oligo fwd</td><td>CACCCCCTATTGACCTTATGTACT</td><td></td><td></td><td></td><td></td><td></td><td></td><td></td><td></td><td></td><td></td></tr>
<tr><td>Oligo rev</td><td>AAACAGTACATAAGGTCAATAGGG</td><td></td><td></td><td></td><td></td><td></td><td></td><td></td><td></td><td></td><td></td></tr>
<tr><td>Chromosome</td><td>start</td><td>end</td><td>strand</td><td>MM</td><td>target_seq</td><td>PAM</td><td>alignment</td><td>distance</td><td>position</td><td>gene name</td><td>gene id</td></tr>
<tr><td>GU305914.1</td><td>183616</td><td>183638</td><td>+</td><td>0</td><td>CCCTATTGACCTTATGTACT</td><td>GGG</td><td>||||||||[||||||||||||]PAM</td><td>NA</td><td>Intergenic</td><td>NA</td><td>NA</td></tr>
<tr><td>GU305914.1</td><td>183430</td><td>183452</td><td>+</td><td>0</td><td>CCCTATTGACCTTATGTACT</td><td>GGG</td><td>||||||||[||||||||||||]PAM</td><td>NA</td><td>Intergenic</td><td>NA</td><td>NA</td></tr>
<tr><td>GU305914.1</td><td>183802</td><td>183824</td><td>+</td><td>0</td><td>CCCTATTGACCTTATGTACT</td><td>GGG</td><td>||||||||[||||||||||||]PAM</td><td>NA</td><td>Intergenic</td><td>NA</td><td>NA</td></tr>
<tr><td>GU305914.1</td><td>183709</td><td>183731</td><td>+</td><td>0</td><td>CCCTATTGACCTTATGTACT</td><td>GGG</td><td>||||||||[||||||||||||]PAM</td><td>NA</td><td>Intergenic</td><td>NA</td><td>NA</td></tr>
<tr><td>GU305914.1</td><td>183523</td><td>183545</td><td>+</td><td>2</td><td>TCCCATTGACCTTATGTACT</td><td>GGG</td><td>-||-||||[||||||||||||]PAM</td><td>NA</td><td>Intergenic</td><td>NA</td><td>NA</td></tr>
<tr><td>GU305914.1</td><td>183338</td><td>183360</td><td>+</td><td>1</td><td>CCCTATTGACCTTTTGTACT</td><td>GGG</td><td>||||||||[|||||-||||||]PAM</td><td>NA</td><td>Intergenic</td><td>NA</td><td>NA</td></tr>
<tr><td>GU305914.1</td><td>183959</td><td>183981</td><td>+</td><td>4</td><td>TCCCATTGACCCAATGTACT</td><td>TGG</td><td>-||-||||[|||--|||||||]PAM</td><td>NA</td><td>Intergenic</td><td>NA</td><td>NA</td></tr>
<tr><td>GU305914.1</td><td>184053</td><td>184075</td><td>+</td><td>4</td><td>TCCCATTGACCCAATGTACT</td><td>TGG</td><td>-||-||||[|||--|||||||]PAM</td><td>NA</td><td>Intergenic</td><td>NA</td><td>NA</td></tr>
<tr><td></td><td></td><td></td><td></td><td></td><td></td><td></td><td></td><td></td><td></td><td></td><td></td></tr>
<tr><td>T13935</td><td>ATGTACTGGGCAAAACCCAATGG</td><td>1000</td><td></td><td></td><td></td><td></td><td></td><td></td><td></td><td></td><td></td></tr>
<tr><td>Oligo fwd</td><td>CACCATGTACTGGGCAAAACCCAA</td><td></td><td></td><td></td><td></td><td></td><td></td><td></td><td></td><td></td><td></td></tr>
<tr><td>Oligo rev</td><td>AAACTTGGGTTTTGCCCAGTACAT</td><td></td><td></td><td></td><td></td><td></td><td></td><td></td><td></td><td></td><td></td></tr>
<tr><td>Chromosome</td><td>start</td><td>end</td><td>strand</td><td>MM</td><td>target_seq</td><td>PAM</td><td>alignment</td><td>distance</td><td>position</td><td>gene name</td><td>gene id</td></tr>
<tr><td>GU305914.1</td><td>183722</td><td>183744</td><td>+</td><td>0</td><td>ATGTACTGGGCAAAACCCAA</td><td>TGG</td><td>||||||||[||||||||||||]PAM</td><td>NA</td><td>Intergenic</td><td>NA</td><td>NA</td></tr>
<tr><td>GU305914.1</td><td>183629</td><td>183651</td><td>+</td><td>0</td><td>ATGTACTGGGCAAAACCCAA</td><td>TGG</td><td>||||||||[||||||||||||]PAM</td><td>NA</td><td>Intergenic</td><td>NA</td><td>NA</td></tr>
<tr><td>GU305914.1</td><td>183351</td><td>183373</td><td>+</td><td>1</td><td>TTGTACTGGGCAAAACCCAA</td><td>TGG</td><td>-|||||||[||||||||||||]PAM</td><td>NA</td><td>Intergenic</td><td>NA</td><td>NA</td></tr>
<tr><td>GU305914.1</td><td>183536</td><td>183558</td><td>+</td><td>1</td><td>ATGTACTGGGCAAAACCCAT</td><td>TGG</td><td>||||||||[|||||||||||-]PAM</td><td>NA</td><td>Intergenic</td><td>NA</td><td>NA</td></tr>
<tr><td>GU305914.1</td><td>183443</td><td>183465</td><td>+</td><td>1</td><td>ATGTACTGGGCAAAACCCAT</td><td>TGG</td><td>||||||||[|||||||||||-]PAM</td><td>NA</td><td>Intergenic</td><td>NA</td><td>NA</td></tr>
<tr><td></td><td></td><td></td><td></td><td></td><td></td><td></td><td></td><td></td><td></td><td></td><td></td></tr>
<tr><td>T13936</td><td>CCCTATTGACGCAGTGTACTTGG</td><td>1000</td><td></td><td></td><td></td><td></td><td></td><td></td><td></td><td></td><td></td></tr>
<tr><td>Oligo fwd</td><td>CACCCCCTATTGACGCAGTGTACT</td><td></td><td></td><td></td><td></td><td></td><td></td><td></td><td></td><td></td><td></td></tr>
<tr><td>Oligo rev</td><td>AAACAGTACACTGCGTCAATAGGG</td><td></td><td></td><td></td><td></td><td></td><td></td><td></td><td></td><td></td><td></td></tr>
<tr><td>Chromosome</td><td>start</td><td>end</td><td>strand</td><td>MM</td><td>target_seq</td><td>PAM</td><td>alignment</td><td>distance</td><td>position</td><td>gene name</td><td>gene id</td></tr>
<tr><td>GU305914.1</td><td>183750</td><td>183772</td><td>+</td><td>0</td><td>CCCTATTGACGCAGTGTACT</td><td>TGG</td><td>||||||||[||||||||||||]PAM</td><td>NA</td><td>Intergenic</td><td>NA</td><td>NA</td></tr>
<tr><td>GU305914.1</td><td>183379</td><td>183401</td><td>+</td><td>1</td><td>CCCTATTGACTCAGTGTACT</td><td>TGG</td><td>||||||||[||-|||||||||]PAM</td><td>NA</td><td>Intergenic</td><td>NA</td><td>NA</td></tr>
<tr><td>GU305914.1</td><td>183657</td><td>183679</td><td>+</td><td>2</td><td>CCCTATTGAGTCAGTGTACT</td><td>TGG</td><td>||||||||[|--|||||||||]PAM</td><td>NA</td><td>Intergenic</td><td>NA</td><td>NA</td></tr>
<tr><td>GU305914.1</td><td>183907</td><td>183929</td><td>+</td><td>2</td><td>CCCTATTGAGTCAGTGTACT</td><td>TGG</td><td>||||||||[|--|||||||||]PAM</td><td>NA</td><td>Intergenic</td><td>NA</td><td>NA</td></tr>
<tr><td>GU305914.1</td><td>183564</td><td>183586</td><td>+</td><td>2</td><td>CCCTATTGACTCAATGTACT</td><td>TGG</td><td>||||||||[||-||-||||||]PAM</td><td>NA</td><td>Intergenic</td><td>NA</td><td>NA</td></tr>
<tr><td>GU305914.1</td><td>184053</td><td>184075</td><td>+</td><td>4</td><td>TCCCATTGACCCAATGTACT</td><td>TGG</td><td>-||-||||[||-||-||||||]PAM</td><td>NA</td><td>Intergenic</td><td>NA</td><td>NA</td></tr>
<tr><td>GU305914.1</td><td>183959</td><td>183981</td><td>+</td><td>4</td><td>TCCCATTGACCCAATGTACT</td><td>TGG</td><td>-||-||||[||-||-||||||]PAM</td><td>NA</td><td>Intergenic</td><td>NA</td><td>NA</td></tr>
<tr><td></td><td></td><td></td><td></td><td></td><td></td><td></td><td></td><td></td><td></td><td></td><td></td></tr>
<tr><td>T13937</td><td>CGCAGTGTACTTGGCTCCAATGG</td><td>1000</td><td></td><td></td><td></td><td></td><td></td><td></td><td></td><td></td><td></td></tr>
<tr><td>Oligo fwd</td><td>CACCGCAGTGTACTTGGCTCCAA</td><td></td><td></td><td></td><td></td><td></td><td></td><td></td><td></td><td></td><td></td></tr>
<tr><td>Oligo rev</td><td>AAACTTGGAGCCAAGTACACTGCG</td><td></td><td></td><td></td><td></td><td></td><td></td><td></td><td></td><td></td><td></td></tr>
<tr><td>Chromosome</td><td>start</td><td>end</td><td>strand</td><td>MM</td><td>target_seq</td><td>PAM</td><td>alignment</td><td>distance</td><td>position</td><td>gene name</td><td>gene id</td></tr>
<tr><td>GU305914.1</td><td>183759</td><td>183781</td><td>+</td><td>0</td><td>CGCAGTGTACTTGGCTCCAA</td><td>TGG</td><td>||||||||[||||||||||||]PAM</td><td>NA</td><td>Intergenic</td><td>NA</td><td>NA</td></tr>
<tr><td>GU305914.1</td><td>183388</td><td>183410</td><td>+</td><td>1</td><td>CTCAGTGTACTTGGCTCCAA</td><td>TGG</td><td>|-||||||[||||||||||||]PAM</td><td>NA</td><td>Intergenic</td><td>NA</td><td>NA</td></tr>
<tr><td>GU305914.1</td><td>183666</td><td>183688</td><td>+</td><td>2</td><td>GTCAGTGTACTTGGCTCCAA</td><td>TGG</td><td>--||||||[||||||||||||]PAM</td><td>NA</td><td>Intergenic</td><td>NA</td><td>NA</td></tr>
<tr><td>GU305914.1</td><td>183573</td><td>183595</td><td>+</td><td>2</td><td>CTCAATGTACTTGGCTCCAA</td><td>TGG</td><td>|-||-|||[||||||||||||]PAM</td><td>NA</td><td>Intergenic</td><td>NA</td><td>NA</td></tr>
<tr><td>GU305914.1</td><td>183916</td><td>183938</td><td>+</td><td>4</td><td>GTCAGTGTACTTGGCAGCAA</td><td>TGG</td><td>--||||||[|||||||--|||]PAM</td><td>NA</td><td>Intergenic</td><td>NA</td><td>NA</td></tr>
<tr><td></td><td></td><td></td><td></td><td></td><td></td><td></td><td></td><td></td><td></td><td></td><td></td></tr>
<tr><td>T13938</td><td>GCAGTGTACTTGGCTCCAATGGG</td><td>1000</td><td></td><td></td><td></td><td></td><td></td><td></td><td></td><td></td><td></td></tr>
<tr><td>Oligo fwd</td><td>CACCGCAGTGTACTTGGCTCCAAT</td><td></td><td></td><td></td><td></td><td></td><td></td><td></td><td></td><td></td><td></td></tr>
<tr><td>Oligo rev</td><td>AAACATTGGAGCCAAGTACACTGC</td><td></td><td></td><td></td><td></td><td></td><td></td><td></td><td></td><td></td><td></td></tr>
<tr><td>Chromosome</td><td>start</td><td>end</td><td>strand</td><td>MM</td><td>target_seq</td><td>PAM</td><td>alignment</td><td>distance</td><td>position</td><td>gene name</td><td>gene id</td></tr>
<tr><td>GU305914.1</td><td>183760</td><td>183782</td><td>+</td><td>0</td><td>GCAGTGTACTTGGCTCCAAT</td><td>GGG</td><td>||||||||[||||||||||||]PAM</td><td>NA</td><td>Intergenic</td><td>NA</td><td>NA</td></tr>
<tr><td>GU305914.1</td><td>183667</td><td>183689</td><td>+</td><td>1</td><td>TCAGTGTACTTGGCTCCAAT</td><td>GGG</td><td>-|||||||[||||||||||||]PAM</td><td>NA</td><td>Intergenic</td><td>NA</td><td>NA</td></tr>
<tr><td>GU305914.1</td><td>183389</td><td>183411</td><td>+</td><td>1</td><td>TCAGTGTACTTGGCTCCAAT</td><td>GGG</td><td>-|||||||[||||||||||||]PAM</td><td>NA</td><td>Intergenic</td><td>NA</td><td>NA</td></tr>
<tr><td>GU305914.1</td><td>183574</td><td>183596</td><td>+</td><td>2</td><td>TCAATGTACTTGGCTCCAAT</td><td>GGG</td><td>-||-||||[||||||||||||]PAM</td><td>NA</td><td>Intergenic</td><td>NA</td><td>NA</td></tr>
<tr><td>GU305914.1</td><td>183917</td><td>183939</td><td>+</td><td>3</td><td>TCAGTGTACTTGGCAGCAAT</td><td>GGG</td><td>-|||||||[||||||--||||]PAM</td><td>NA</td><td>Intergenic</td><td>NA</td><td>NA</td></tr>
<tr><td></td><td></td><td></td><td></td><td></td><td></td><td></td><td></td><td></td><td></td><td></td><td></td></tr>
<tr><td>T13939</td><td>CCCCTATTGACCTTATGTACTGG</td><td>1000</td><td></td><td></td><td></td><td></td><td></td><td></td><td></td><td></td><td></td></tr>
<tr><td>Oligo fwd</td><td>CACCCCCCTATTGACCTTATGTAC</td><td></td><td></td><td></td><td></td><td></td><td></td><td></td><td></td><td></td><td></td></tr>
<tr><td>Oligo rev</td><td>AAACGTACATAAGGTCAATAGGGG</td><td></td><td></td><td></td><td></td><td></td><td></td><td></td><td></td><td></td><td></td></tr>
<tr><td>Chromosome</td><td>start</td><td>end</td><td>strand</td><td>MM</td><td>target_seq</td><td>PAM</td><td>alignment</td><td>distance</td><td>position</td><td>gene name</td><td>gene id</td></tr>
<tr><td>GU305914.1</td><td>183801</td><td>183823</td><td>+</td><td>0</td><td>CCCCTATTGACCTTATGTAC</td><td>TGG</td><td>||||||||[||||||||||||]PAM</td><td>NA</td><td>Intergenic</td><td>NA</td><td>NA</td></tr>
<tr><td>GU305914.1</td><td>183708</td><td>183730</td><td>+</td><td>0</td><td>CCCCTATTGACCTTATGTAC</td><td>TGG</td><td>||||||||[||||||||||||]PAM</td><td>NA</td><td>Intergenic</td><td>NA</td><td>NA</td></tr>
<tr><td>GU305914.1</td><td>183615</td><td>183637</td><td>+</td><td>0</td><td>CCCCTATTGACCTTATGTAC</td><td>TGG</td><td>||||||||[||||||||||||]PAM</td><td>NA</td><td>Intergenic</td><td>NA</td><td>NA</td></tr>
<tr><td>GU305914.1</td><td>183429</td><td>183451</td><td>+</td><td>0</td><td>CCCCTATTGACCTTATGTAC</td><td>TGG</td><td>||||||||[||||||||||||]PAM</td><td>NA</td><td>Intergenic</td><td>NA</td><td>NA</td></tr>
<tr><td>GU305914.1</td><td>183522</td><td>183544</td><td>+</td><td>2</td><td>CTCCCATTGACCTTATGTAC</td><td>TGG</td><td>|-||-|||[||||||||||||]PAM</td><td>NA</td><td>Intergenic</td><td>NA</td><td>NA</td></tr>
<tr><td>GU305914.1</td><td>183337</td><td>183359</td><td>+</td><td>1</td><td>CCCCTATTGACCTTTTGTAC</td><td>TGG</td><td>||||||||[||||||-|||||]PAM</td><td>NA</td><td>Intergenic</td><td>NA</td><td>NA</td></tr>
</table>

T13940 CCCTATTGACCTTATGTACTGGG 1000
Oligo fwd CACCCCCTATTGACCTTATGTACT
Oligo rev AAACAGTACATAAGGTCAATAGGG

| Chromosome | start | end | strand | MM | target_seq | PAM | alignment | distance | position | gene name | gene id |
|---|---|---|---|---|---|---|---|---|---|---|---|
| GU305914.1 | 183616 | 183638 | + | 0 | CCCTATTGACCTTATGTACT | GGG | \|\|\|\|\|\|\|\|[\|\|\|\|\|\|\|\|\|\|\|\|]PAM | NA | Intergenic | NA | NA |
| GU305914.1 | 183430 | 183452 | + | 0 | CCCTATTGACCTTATGTACT | GGG | \|\|\|\|\|\|\|\|[\|\|\|\|\|\|\|\|\|\|\|\|]PAM | NA | Intergenic | NA | NA |
| GU305914.1 | 183802 | 183824 | + | 0 | CCCTATTGACCTTATGTACT | GGG | \|\|\|\|\|\|\|\|[\|\|\|\|\|\|\|\|\|\|\|\|]PAM | NA | Intergenic | NA | NA |
| GU305914.1 | 183709 | 183731 | + | 0 | CCCTATTGACCTTATGTACT | GGG | \|\|\|\|\|\|\|\|[\|\|\|\|\|\|\|\|\|\|\|\|]PAM | NA | Intergenic | NA | NA |
| GU305914.1 | 183523 | 183545 | + | 2 | TCCCATTGACCTTATGTACT | GGG | -\|\|-\|\|\|\|[\|\|\|\|\|\|\|\|\|\|\|\|]PAM | NA | Intergenic | NA | NA |
| GU305914.1 | 183338 | 183360 | + | 1 | CCCTATTGACCTTTTGTACT | GGG | \|\|\|\|\|\|\|\|[\|\|\|\|\|-\|\|\|\|\|\|]PAM | NA | Intergenic | NA | NA |
| GU305914.1 | 183959 | 183981 | + | 4 | TCCCATTGACCCAATGTACT | TGG | -\|\|-\|\|\|\|[\|\|\|--\|\|\|\|\|\|\|]PAM | NA | Intergenic | NA | NA |
| GU305914.1 | 184053 | 184075 | + | 4 | TCCCATTGACCCAATGTACT | TGG | -\|\|-\|\|\|\|[\|\|\|--\|\|\|\|\|\|\|]PAM | NA | Intergenic | NA | NA |

T13941 ATGTACTGGGCAGAATACAATGG 1000
Oligo fwd CACCATGTACTGGGCAGAATACAA
Oligo rev AAACTTGTATTCTGCCCAGTACAT

| Chromosome | start | end | strand | MM | target_seq | PAM | alignment | distance | position | gene name | gene id |
|---|---|---|---|---|---|---|---|---|---|---|---|
| GU305914.1 | 183815 | 183837 | + | 0 | ATGTACTGGGCAGAATACAA | TGG | \|\|\|\|\|\|\|\|[\|\|\|\|\|\|\|\|\|\|\|\|]PAM | NA | Intergenic | NA | NA |

T13942 CCTTATATGCTTGCCAACAATGG 1000
Oligo fwd CACCCCTTATATGCTTGCCAACAA
Oligo rev AAACTTGTTGGCAAGCATATAAGG

| Chromosome | start | end | strand | MM | target_seq | PAM | alignment | distance | position | gene name | gene id |
|---|---|---|---|---|---|---|---|---|---|---|---|
| GU305914.1 | 183866 | 183888 | + | 0 | CCTTATATGCTTGCCAACAA | TGG | \|\|\|\|\|\|\|\|[\|\|\|\|\|\|\|\|\|\|\|\|]PAM | NA | Intergenic | NA | NA |

T13943 CCAACAATGGAAAAACCCATTGG 1000
Oligo fwd CACCCCAACAATGGAAAAACCCAT
Oligo rev AAACATGGGTTTTTCCATTGTTGG

| Chromosome | start | end | strand | MM | target_seq | PAM | alignment | distance | position | gene name | gene id |
|---|---|---|---|---|---|---|---|---|---|---|---|
| GU305914.1 | 183879 | 183901 | + | 0 | CCAACAATGGAAAAACCCAT | TGG | \|\|\|\|\|\|\|\|[\|\|\|\|\|\|\|\|\|\|\|\|]PAM | NA | Intergenic | NA | NA |

T13944 CCCTATTGAGTCAGTGTACTTGG 1000
Oligo fwd CACCCCCTATTGAGTCAGTGTACT
Oligo rev AAACAGTACACTGACTCAATAGGG

| Chromosome | start | end | strand | MM | target_seq | PAM | alignment | distance | position | gene name | gene id |
|---|---|---|---|---|---|---|---|---|---|---|---|
| GU305914.1 | 183657 | 183679 | + | 0 | CCCTATTGAGTCAGTGTACT | TGG | \|\|\|\|\|\|\|\|[\|\|\|\|\|\|\|\|\|\|\|\|]PAM | NA | Intergenic | NA | NA |
| GU305914.1 | 183907 | 183929 | + | 0 | CCCTATTGAGTCAGTGTACT | TGG | \|\|\|\|\|\|\|\|[\|\|\|\|\|\|\|\|\|\|\|\|]PAM | NA | Intergenic | NA | NA |
| GU305914.1 | 183379 | 183401 | + | 1 | CCCTATTGACTCAGTGTACT | TGG | \|\|\|\|\|\|\|\|[\|-\|\|\|\|\|\|\|\|\|\|]PAM | NA | Intergenic | NA | NA |
| GU305914.1 | 183750 | 183772 | + | 2 | CCCTATTGACGCAGTGTACT | TGG | \|\|\|\|\|\|\|\|[\|--\|\|\|\|\|\|\|\|\|]PAM | NA | Intergenic | NA | NA |
| GU305914.1 | 183564 | 183586 | + | 2 | CCCTATTGACTCAATGTACT | TGG | \|\|\|\|\|\|\|\|[\|-\|\|\|-\|\|\|\|\|\|]PAM | NA | Intergenic | NA | NA |

T13945 GTCAGTGTACTTGGCAGCAATGG 1000
Oligo fwd CACCGTCAGTGTACTTGGCAGCAA
Oligo rev AAACTTGCTGCCAAGTACACTGAC

| Chromosome | start | end | strand | MM | target_seq | PAM | alignment | distance | position | gene name | gene id |
|---|---|---|---|---|---|---|---|---|---|---|---|
| GU305914.1 | 183916 | 183938 | + | 0 | GTCAGTGTACTTGGCAGCAA | TGG | \|\|\|\|\|\|\|\|[\|\|\|\|\|\|\|\|\|\|\|\|]PAM | NA | Intergenic | NA | NA |
| GU305914.1 | 183666 | 183688 | + | 2 | GTCAGTGTACTTGGCTCCAA | TGG | \|\|\|\|\|\|\|\|[\|\|\|\|\|\|\|--\|\|\|]PAM | NA | Intergenic | NA | NA |
| GU305914.1 | 183388 | 183410 | + | 3 | CTCAGTGTACTTGGCTCCAA | TGG | -\|\|\|\|\|\|\|[\|\|\|\|\|\|\|--\|\|\|]PAM | NA | Intergenic | NA | NA |
| GU305914.1 | 183759 | 183781 | + | 4 | CGCAGTGTACTTGGCTCCAA | TGG | --\|\|\|\|\|\|[\|\|\|\|\|\|\|--\|\|\|]PAM | NA | Intergenic | NA | NA |
| GU305914.1 | 183573 | 183595 | + | 4 | CTCAATGTACTTGGCTCCAA | TGG | -\|\|\|-\|\|\|[\|\|\|\|\|\|\|--\|\|\|]PAM | NA | Intergenic | NA | NA |

T13946 TCAGTGTACTTGGCAGCAATGGG 1000
Oligo fwd CACCTCAGTGTACTTGGCAGCAAT
Oligo rev AAACATTGCTGCCAAGTACACTGA

| Chromosome | start | end | strand | MM | target_seq | PAM | alignment | distance | position | gene name | gene id |
|---|---|---|---|---|---|---|---|---|---|---|---|
| GU305914.1 | 183917 | 183939 | + | 0 | TCAGTGTACTTGGCAGCAAT | GGG | \|\|\|\|\|\|\|\|[\|\|\|\|\|\|\|\|\|\|\|\|]PAM | NA | Intergenic | NA | NA |
| GU305914.1 | 183667 | 183689 | + | 2 | TCAGTGTACTTGGCTCCAAT | GGG | \|\|\|\|\|\|\|\|[\|\|\|\|\|\|--\|\|\|\|]PAM | NA | Intergenic | NA | NA |
| GU305914.1 | 183389 | 183411 | + | 2 | TCAGTGTACTTGGCTCCAAT | GGG | \|\|\|\|\|\|\|\|[\|\|\|\|\|\|--\|\|\|\|]PAM | NA | Intergenic | NA | NA |
| GU305914.1 | 183760 | 183782 | + | 3 | GCAGTGTACTTGGCTCCAAT | GGG | -\|\|\|\|\|\|\|[\|\|\|\|\|\|--\|\|\|\|]PAM | NA | Intergenic | NA | NA |
| GU305914.1 | 183574 | 183596 | + | 3 | TCAATGTACTTGGCTCCAAT | GGG | \|\|\|-\|\|\|\|[\|\|\|\|\|\|--\|\|\|\|]PAM | NA | Intergenic | NA | NA |
| GU305914.1 | 184011 | 184033 | + | 3 | TCAGTGTGCTTGCCAGTAAT | GGG | \|\|\|\|\|\|\|-[\|\|\|\|-\|\|\|-\|\|\|]PAM | NA | Intergenic | NA | NA |

T13947 CAGCAATGGGTTTTTCCCATTGG 1000
Oligo fwd CACCCAGCAATGGGTTTTTCCCAT
Oligo rev AAACATGGGAAAAACCCATTGCTG

| Chromosome | start | end | strand | MM | target_seq | PAM | alignment | distance | position | gene name | gene id |
|---|---|---|---|---|---|---|---|---|---|---|---|
| GU305914.1 | 183930 | 183952 | + | 0 | CAGCAATGGGTTTTTCCCAT | TGG | \|\|\|\|\|\|\|\|[\|\|\|\|\|\|\|\|\|\|\|\|]PAM | NA | Intergenic | NA | NA |

T13948 TCCCATTGACCCAATGTACTTGG 1000
Oligo fwd CACCTCCCATTGACCCAATGTACT
Oligo rev AAACAGTACATTGGGTCAATGGGA

| Chromosome | start | end | strand | MM | target_seq | PAM | alignment | distance | position | gene name | gene id |
|---|---|---|---|---|---|---|---|---|---|---|---|
| GU305914.1 | 183959 | 183981 | + | 0 | TCCCATTGACCCAATGTACT | TGG | \|\|\|\|\|\|\|\|[\|\|\|\|\|\|\|\|\|\|\|\|]PAM | NA | Intergenic | NA | NA |
| GU305914.1 | 184053 | 184075 | + | 0 | TCCCATTGACCCAATGTACT | TGG | \|\|\|\|\|\|\|\|[\|\|\|\|\|\|\|\|\|\|\|\|]PAM | NA | Intergenic | NA | NA |
| GU305914.1 | 183564 | 183586 | + | 3 | CCCTATTGACTCAATGTACT | TGG | -\|\|-\|\|\|\|[\|\|-\|\|\|\|\|\|\|\|\|]PAM | NA | Intergenic | NA | NA |
| GU305914.1 | 183523 | 183545 | + | 2 | TCCCATTGACCTTATGTACT | GGG | \|\|\|\|\|\|\|\|[\|\|\|--\|\|\|\|\|\|\|]PAM | NA | Intergenic | NA | NA |
| GU305914.1 | 183430 | 183452 | + | 4 | CCCTATTGACCTTATGTACT | GGG | -\|\|-\|\|\|\|[\|\|\|--\|\|\|\|\|\|\|]PAM | NA | Intergenic | NA | NA |
| GU305914.1 | 183802 | 183824 | + | 4 | CCCTATTGACCTTATGTACT | GGG | -\|\|-\|\|\|\|[\|\|\|--\|\|\|\|\|\|\|]PAM | NA | Intergenic | NA | NA |
| GU305914.1 | 183709 | 183731 | + | 4 | CCCTATTGACCTTATGTACT | GGG | -\|\|-\|\|\|\|[\|\|\|--\|\|\|\|\|\|\|]PAM | NA | Intergenic | NA | NA |
| GU305914.1 | 183616 | 183638 | + | 4 | CCCTATTGACCTTATGTACT | GGG | -\|\|-\|\|\|\|[\|\|\|--\|\|\|\|\|\|\|]PAM | NA | Intergenic | NA | NA |
| GU305914.1 | 183379 | 183401 | + | 4 | CCCTATTGACTCAGTGTACT | TGG | -\|\|-\|\|\|\|[\|\|-\|\|-\|\|\|\|\|\|]PAM | NA | Intergenic | NA | NA |
| GU305914.1 | 183750 | 183772 | + | 4 | CCCTATTGACGCAGTGTACT | TGG | -\|\|-\|\|\|\|[\|\|-\|\|-\|\|\|\|\|\|]PAM | NA | Intergenic | NA | NA |

T13949 CCCATTGACCCAATGTACTTGGG 1000
Oligo fwd CACCCCCATTGACCCAATGTACTT
Oligo rev AAACAAGTACATTGGGTCAATGGG

| Chromosome | start | end | strand | MM | target_seq | PAM | alignment | distance | position | gene name | gene id |
|---|---|---|---|---|---|---|---|---|---|---|---|
| GU305914.1 | 183960 | 183982 | + | 0 | CCCATTGACCCAATGTACTT | GGG | \|\|\|\|\|\|\|\|[\|\|\|\|\|\|\|\|\|\|\|\|]PAM | NA | Intergenic | NA | NA |

GU305914.1 184054 184076 + 0 CCCATTGACCCAATGTACTT GGG ||||||||[||||||||||||]PAM NA Intergenic NA NA

T13950 TGTACTTGGGCAAAACCCAGTGG 1000
Oligo fwd CACCTGTACTTGGGCAAAACCCAG
Oligo rev AAACCTGGGTTTTGCCCAAGTACA
Chromosome start end strand MM target_seq PAM alignment distance position gene name gene id
GU305914.1 183973 183995 + 0 TGTACTTGGGCAAAACCCAG TGG ||||||||[||||||||||||]PAM NA Intergenic NA NA

T13951 CTCAGTGTGCTTGCCAGTAATGG 1000
Oligo fwd CACCCTCAGTGTGCTTGCCAGTAA
Oligo rev AAACTTACTGGCAAGCACACTGAG
Chromosome start end strand MM target_seq PAM alignment distance position gene name gene id
GU305914.1 184010 184032 + 0 CTCAGTGTGCTTGCCAGTAA TGG ||||||||[||||||||||||]PAM NA Intergenic NA NA
GU305914.1 183480 183502 + 3 CTCAGTATACGTGCCAGTAA TGG ||||||-|[-|-|||||||||]PAM NA Intergenic NA NA

T13952 TCAGTGTGCTTGCCAGTAATGGG 1000
Oligo fwd CACCTCAGTGTGCTTGCCAGTAAT
Oligo rev AAACATTACTGGCAAGCACACTGA
Chromosome start end strand MM target_seq PAM alignment distance position gene name gene id
GU305914.1 184011 184033 + 0 TCAGTGTGCTTGCCAGTAAT GGG ||||||||[||||||||||||]PAM NA Intergenic NA NA
GU305914.1 183481 183503 + 3 TCAGTATACGTGCCAGTAAT GGG |||||-|-[|-||||||||||]PAM NA Intergenic NA NA
GU305914.1 183917 183939 + 3 TCAGTGTACTTGGCAGCAAT GGG |||||||-[||||-|||-|||]PAM NA Intergenic NA NA

T13953 CAGTAATGGGAAAAACCCATTGG 1000
Oligo fwd CACCCAGTAATGGGAAAAACCCAT
Oligo rev AAACATGGGTTTTTCCCATTACTG
Chromosome start end strand MM target_seq PAM alignment distance position gene name gene id
GU305914.1 184024 184046 + 0 CAGTAATGGGAAAAACCCAT TGG ||||||||[||||||||||||]PAM NA Intergenic NA NA
GU305914.1 183494 183516 + 0 CAGTAATGGGAAAAACCCAT TGG ||||||||[||||||||||||]PAM NA Intergenic NA NA
GU305914.1 183682 183704 - 4 AGTCAATGGGAAAAACCCAT TGG ----||||[||||||||||||]PAM NA Intergenic NA NA
GU305914.1 183443 183465 + 4 ATGTACTGGGCAAAACCCAT TGG --|||-||[||-|||||||||]PAM NA Intergenic NA NA
GU305914.1 183536 183558 + 4 ATGTACTGGGCAAAACCCAT TGG --|||-||[||-|||||||||]PAM NA Intergenic NA NA

T13954 TCCCATTGACCCAATGTACTTGG 1000
Oligo fwd CACCTCCCATTGACCCAATGTACT
Oligo rev AAACAGTACATTGGGTCAATGGGA
Chromosome start end strand MM target_seq PAM alignment distance position gene name gene id
GU305914.1 183959 183981 + 0 TCCCATTGACCCAATGTACT TGG ||||||||[||||||||||||]PAM NA Intergenic NA NA
GU305914.1 184053 184075 + 0 TCCCATTGACCCAATGTACT TGG ||||||||[||||||||||||]PAM NA Intergenic NA NA
GU305914.1 183564 183586 + 3 CCCTATTGACTCAATGTACT TGG -||-||||[||-|||||||||]PAM NA Intergenic NA NA
GU305914.1 183523 183545 + 2 TCCCATTGACCTTATGTACT GGG ||||||||[|||--|||||||]PAM NA Intergenic NA NA
GU305914.1 183430 183452 + 4 CCCTATTGACCTTATGTACT GGG -||-||||[|||--|||||||]PAM NA Intergenic NA NA
GU305914.1 183802 183824 + 4 CCCTATTGACCTTATGTACT GGG -||-||||[|||--|||||||]PAM NA Intergenic NA NA
GU305914.1 183709 183731 + 4 CCCTATTGACCTTATGTACT GGG -||-||||[|||--|||||||]PAM NA Intergenic NA NA
GU305914.1 183616 183638 + 4 CCCTATTGACCTTATGTACT GGG -||-||||[|||--|||||||]PAM NA Intergenic NA NA
GU305914.1 183379 183401 + 4 CCCTATTGACTCAGTGTACT TGG -||-||||[||-||-||||||]PAM NA Intergenic NA NA
GU305914.1 183750 183772 + 4 CCCTATTGACGCAGTGTACT TGG -||-||||[||-||-||||||]PAM NA Intergenic NA NA

T13955 CCCATTGACCCAATGTACTTGGG 1000
Oligo fwd CACCCCCATTGACCCAATGTACTT
Oligo rev AAACAAGTACATTGGGTCAATGGG
Chromosome start end strand MM target_seq PAM alignment distance position gene name gene id
GU305914.1 183960 183982 + 0 CCCATTGACCCAATGTACTT GGG ||||||||[||||||||||||]PAM NA Intergenic NA NA
GU305914.1 184054 184076 + 0 CCCATTGACCCAATGTACTT GGG ||||||||[||||||||||||]PAM NA Intergenic NA NA

T13956 GACCCAATGTACTTGGGTAAAGG 1000
Oligo fwd CACCGACCCAATGTACTTGGGTAA
Oligo rev AAACTTACCCAAGTACATTGGGTC
Chromosome start end strand MM target_seq PAM alignment distance position gene name gene id
GU305914.1 184060 184082 + 0 GACCCAATGTACTTGGGTAA AGG ||||||||[||||||||||||]PAM NA Intergenic NA NA

T13957 TTGAGTCACCACCCCTATGCTGG 1000
Oligo fwd CACCTTGAGTCACCACCCCTATGC
Oligo rev AAACGCATAGGGGTGGTGACTCAA
Chromosome start end strand MM target_seq PAM alignment distance position gene name gene id
GU305914.1 184086 184108 + 0 TTGAGTCACCACCCCTATGC TGG ||||||||[||||||||||||]PAM NA Intergenic NA NA

T13958 TGAGTCACCACCCCTATGCTGGG 1000
Oligo fwd CACCTGAGTCACCACCCCTATGCT
Oligo rev AAACAGCATAGGGGTGGTGACTCA
Chromosome start end strand MM target_seq PAM alignment distance position gene name gene id
GU305914.1 184087 184109 + 0 TGAGTCACCACCCCTATGCT GGG ||||||||[||||||||||||]PAM NA Intergenic NA NA

T13959 ACCACCCCTATGCTGGGAAATGG 1000
Oligo fwd CACCACCACCCCTATGCTGGGAAA
Oligo rev AAACTTTCCCAGCATAGGGGTGGT
Chromosome start end strand MM target_seq PAM alignment distance position gene name gene id
GU305914.1 184093 184115 + 0 ACCACCCCTATGCTGGGAAA TGG ||||||||[||||||||||||]PAM NA Intergenic NA NA

T13960 AATGGTGAACGCCCCCTATGTGG 1000
Oligo fwd CACCAATGGTGAACGCCCCCTATG
Oligo rev AAACCATAGGGGGCGTTCACCATT
Chromosome start end strand MM target_seq PAM alignment distance position gene name gene id
GU305914.1 184111 184133 + 0 AATGGTGAACGCCCCCTATG TGG ||||||||[||||||||||||]PAM NA Intergenic NA NA

T13961 CCCTATGTGGAAAGTCCCTATGG 1000
Oligo fwd CACCCCCTATGTGGAAAGTCCCTA
Oligo rev AAACTAGGGACTTTCCACATAGGG
Chromosome start end strand MM target_seq PAM alignment distance position gene name gene id

GU305914.1 184124 184146 + 0 CCCTATGTGGAAAGTCCCTA TGG ||||||||[||||||||||||]PAM NA Intergenic NA NA

T13962 CCTATGTGGAAAGTCCCTATGGG 1000
Oligo fwd CACCCCTATGTGGAAAGTCCCTAT
Oligo rev AAACATAGGGACTTTCCACATAGG
Chromosome start end strand MM target_seq PAM alignment distance position gene name gene id
GU305914.1 184125 184147 + 0 CCTATGTGGAAAGTCCCTAT GGG ||||||||[||||||||||||]PAM NA Intergenic NA NA

T13963 ATATGCCATATGAGTGTATTAGG 1000
Oligo fwd CACCATATGCCATATGAGTGTATT
Oligo rev AAACAATACACTCATATGGCATAT
Chromosome start end strand MM target_seq PAM alignment distance position gene name gene id
GU305914.1 184179 184201 + 0 ATATGCCATATGAGTGTATT AGG ||||||||[||||||||||||]PAM NA Intergenic NA NA

T13964 TATGCCATATGAGTGTATTAGGG 1000
Oligo fwd CACCTATGCCATATGAGTGTATTA
Oligo rev AAACTAATACACTCATATGGCATA
Chromosome start end strand MM target_seq PAM alignment distance position gene name gene id
GU305914.1 184180 184202 + 0 TATGCCATATGAGTGTATTA GGG ||||||||[||||||||||||]PAM NA Intergenic NA NA

T13965 ATGCCATATGAGTGTATTAGGGG 1000
Oligo fwd CACCATGCCATATGAGTGTATTAG
Oligo rev AAACCTAATACACTCATATGGCAT
Chromosome start end strand MM target_seq PAM alignment distance position gene name gene id
GU305914.1 184181 184203 + 0 ATGCCATATGAGTGTATTAG GGG ||||||||[||||||||||||]PAM NA Intergenic NA NA

T13966 TGCCATATGAGTGTATTAGGGGG 1000
Oligo fwd CACCTGCCATATGAGTGTATTAGG
Oligo rev AAACCCTAATACACTCATATGGCA
Chromosome start end strand MM target_seq PAM alignment distance position gene name gene id
GU305914.1 184182 184204 + 0 TGCCATATGAGTGTATTAGG GGG ||||||||[||||||||||||]PAM NA Intergenic NA NA

T13967 GTATTAGGGGGCTTTCCGCTTGG 1000
Oligo fwd CACCGTATTAGGGGGCTTTCCGCT
Oligo rev AAACAGCGGAAAGCCCCCTAATAC
Chromosome start end strand MM target_seq PAM alignment distance position gene name gene id
GU305914.1 184194 184216 + 0 GTATTAGGGGGCTTTCCGCT TGG ||||||||[||||||||||||]PAM NA Intergenic NA NA

T13968 TATTAGGGGGCTTTCCGCTTGGG 1000
Oligo fwd CACCTATTAGGGGGCTTTCCGCTT
Oligo rev AAACAAGCGGAAAGCCCCCTAATA
Chromosome start end strand MM target_seq PAM alignment distance position gene name gene id
GU305914.1 184195 184217 + 0 TATTAGGGGGCTTTCCGCTT GGG ||||||||[||||||||||||]PAM NA Intergenic NA NA

T13969 GGGCTTTCCGCTTGGGAAATTGG 1000
Oligo fwd CACCGGGCTTTCCGCTTGGGAAAT
Oligo rev AAACATTTCCCAAGCGGAAAGCCC
Chromosome start end strand MM target_seq PAM alignment distance position gene name gene id
GU305914.1 184202 184224 + 0 GGGCTTTCCGCTTGGGAAAT TGG ||||||||[||||||||||||]PAM NA Intergenic NA NA

T13970 GGCTTTCCGCTTGGGAAATTGGG 1000
Oligo fwd CACCGGCTTTCCGCTTGGGAAATT
Oligo rev AAACAATTTCCCAAGCGGAAAGCC
Chromosome start end strand MM target_seq PAM alignment distance position gene name gene id
GU305914.1 184203 184225 + 0 GGCTTTCCGCTTGGGAAATT GGG ||||||||[||||||||||||]PAM NA Intergenic NA NA

T13971 AGTCCCCGTATTACTCACATAGG 1000
Oligo fwd CACCAGTCCCCGTATTACTCACAT
Oligo rev AAACATGTGAGTAATACGGGGACT
Chromosome start end strand MM target_seq PAM alignment distance position gene name gene id
GU305914.1 184231 184253 + 0 AGTCCCCGTATTACTCACAT AGG ||||||||[||||||||||||]PAM NA Intergenic NA NA

T13972 GTCCCCGTATTACTCACATAGGG 1000
Oligo fwd CACCGTCCCCGTATTACTCACATA
Oligo rev AAACTATGTGAGTAATACGGGGAC
Chromosome start end strand MM target_seq PAM alignment distance position gene name gene id
GU305914.1 184232 184254 + 0 GTCCCCGTATTACTCACATA GGG ||||||||[||||||||||||]PAM NA Intergenic NA NA

T13973 TCCCCGTATTACTCACATAGGGG 1000
Oligo fwd CACCTCCCCGTATTACTCACATAG
Oligo rev AAACCTATGTGAGTAATACGGGGA
Chromosome start end strand MM target_seq PAM alignment distance position gene name gene id
GU305914.1 184233 184255 + 0 TCCCCGTATTACTCACATAG GGG ||||||||[||||||||||||]PAM NA Intergenic NA NA

T13974 CCCCGTATTACTCACATAGGGGG 1000
Oligo fwd CACCCCCCGTATTACTCACATAGG
Oligo rev AAACCCTATGTGAGTAATACGGGG
Chromosome start end strand MM target_seq PAM alignment distance position gene name gene id
GU305914.1 184234 184256 + 0 CCCCGTATTACTCACATAGG GGG ||||||||[||||||||||||]PAM NA Intergenic NA NA

T13975 TTACTCACATAGGGGGCGTTTGG 1000
Oligo fwd CACCTTACTCACATAGGGGGCGTT
Oligo rev AAACAACGCCCCCTATGTGAGTAA
Chromosome start end strand MM target_seq PAM alignment distance position gene name gene id
GU305914.1 184241 184263 + 0 TTACTCACATAGGGGGCGTT TGG ||||||||[||||||||||||]PAM NA Intergenic NA NA

T13976 GGCGTTTGGCTTTGCAAATTAGG 1000
Oligo fwd CACCGGCGTTTGGCTTTGCAAATT
Oligo rev AAACAATTTGCAAAGCCAAACGCC

<table>
<tr><td>Chromosome</td><td>start</td><td>end</td><td>strand</td><td>MM</td><td>target_seq</td><td>PAM</td><td>alignment</td><td>distance</td><td>position</td><td>gene name</td><td>gene id</td></tr>
<tr><td>GU305914.1</td><td>184255</td><td>184277</td><td>+</td><td>0</td><td>GGCGTTTGGCTTTGCAAATT</td><td>AGG</td><td>||||||||[||||||||||||]PAM</td><td>NA</td><td>Intergenic</td><td>NA</td><td>NA</td></tr>
<tr><td>T13977</td><td>GCGTTTGGCTTTGCAAATTAGGG</td><td>1000</td><td></td><td></td><td></td><td></td><td></td><td></td><td></td><td></td><td></td></tr>
<tr><td>Oligo fwd</td><td>CACCGCGTTTGGCTTTGCAAATTA</td><td></td><td></td><td></td><td></td><td></td><td></td><td></td><td></td><td></td><td></td></tr>
<tr><td>Oligo rev</td><td>AAACTAATTTGCAAAGCCAAACGC</td><td></td><td></td><td></td><td></td><td></td><td></td><td></td><td></td><td></td><td></td></tr>
<tr><td>Chromosome</td><td>start</td><td>end</td><td>strand</td><td>MM</td><td>target_seq</td><td>PAM</td><td>alignment</td><td>distance</td><td>position</td><td>gene name</td><td>gene id</td></tr>
<tr><td>GU305914.1</td><td>184256</td><td>184278</td><td>+</td><td>0</td><td>GCGTTTGGCTTTGCAAATTA</td><td>GGG</td><td>||||||||[||||||||||||]PAM</td><td>NA</td><td>Intergenic</td><td>NA</td><td>NA</td></tr>
<tr><td>T13978</td><td>CGTTTGGCTTTGCAAATTAGGGG</td><td>1000</td><td></td><td></td><td></td><td></td><td></td><td></td><td></td><td></td><td></td></tr>
<tr><td>Oligo fwd</td><td>CACCCGTTTGGCTTTGCAAATTAG</td><td></td><td></td><td></td><td></td><td></td><td></td><td></td><td></td><td></td><td></td></tr>
<tr><td>Oligo rev</td><td>AAACCTAATTTGCAAAGCCAAACG</td><td></td><td></td><td></td><td></td><td></td><td></td><td></td><td></td><td></td><td></td></tr>
<tr><td>Chromosome</td><td>start</td><td>end</td><td>strand</td><td>MM</td><td>target_seq</td><td>PAM</td><td>alignment</td><td>distance</td><td>position</td><td>gene name</td><td>gene id</td></tr>
<tr><td>GU305914.1</td><td>184257</td><td>184279</td><td>+</td><td>0</td><td>CGTTTGGCTTTGCAAATTAG</td><td>GGG</td><td>||||||||[||||||||||||]PAM</td><td>NA</td><td>Intergenic</td><td>NA</td><td>NA</td></tr>
<tr><td>T13979</td><td>TTAGGGGATTTCAGTGCATTTGG</td><td>1000</td><td></td><td></td><td></td><td></td><td></td><td></td><td></td><td></td><td></td></tr>
<tr><td>Oligo fwd</td><td>CACCTTAGGGGATTTCAGTGCATT</td><td></td><td></td><td></td><td></td><td></td><td></td><td></td><td></td><td></td><td></td></tr>
<tr><td>Oligo rev</td><td>AAACAATGCACTGAAATCCCCTAA</td><td></td><td></td><td></td><td></td><td></td><td></td><td></td><td></td><td></td><td></td></tr>
<tr><td>Chromosome</td><td>start</td><td>end</td><td>strand</td><td>MM</td><td>target_seq</td><td>PAM</td><td>alignment</td><td>distance</td><td>position</td><td>gene name</td><td>gene id</td></tr>
<tr><td>GU305914.1</td><td>184273</td><td>184295</td><td>+</td><td>0</td><td>TTAGGGGATTTCAGTGCATT</td><td>TGG</td><td>||||||||[||||||||||||]PAM</td><td>NA</td><td>Intergenic</td><td>NA</td><td>NA</td></tr>
<tr><td>T13980</td><td>CATTTGGCATTAAAAACTATTGG</td><td>1000</td><td></td><td></td><td></td><td></td><td></td><td></td><td></td><td></td><td></td></tr>
<tr><td>Oligo fwd</td><td>CACCCATTTGGCATTAAAAACTAT</td><td></td><td></td><td></td><td></td><td></td><td></td><td></td><td></td><td></td><td></td></tr>
<tr><td>Oligo rev</td><td>AAACATAGTTTTTAATGCCAAATG</td><td></td><td></td><td></td><td></td><td></td><td></td><td></td><td></td><td></td><td></td></tr>
<tr><td>Chromosome</td><td>start</td><td>end</td><td>strand</td><td>MM</td><td>target_seq</td><td>PAM</td><td>alignment</td><td>distance</td><td>position</td><td>gene name</td><td>gene id</td></tr>
<tr><td>GU305914.1</td><td>184289</td><td>184311</td><td>+</td><td>0</td><td>CATTTGGCATTAAAAACTAT</td><td>TGG</td><td>||||||||[||||||||||||]PAM</td><td>NA</td><td>Intergenic</td><td>NA</td><td>NA</td></tr>
<tr><td>T13981</td><td>TATTGGTTCTAGTCATAAAACGG</td><td>1000</td><td></td><td></td><td></td><td></td><td></td><td></td><td></td><td></td><td></td></tr>
<tr><td>Oligo fwd</td><td>CACCTATTGGTTCTAGTCATAAAA</td><td></td><td></td><td></td><td></td><td></td><td></td><td></td><td></td><td></td><td></td></tr>
<tr><td>Oligo rev</td><td>AAACTTTTATGACTAGAACCAATA</td><td></td><td></td><td></td><td></td><td></td><td></td><td></td><td></td><td></td><td></td></tr>
<tr><td>Chromosome</td><td>start</td><td>end</td><td>strand</td><td>MM</td><td>target_seq</td><td>PAM</td><td>alignment</td><td>distance</td><td>position</td><td>gene name</td><td>gene id</td></tr>
<tr><td>GU305914.1</td><td>184306</td><td>184328</td><td>+</td><td>0</td><td>TATTGGTTCTAGTCATAAAA</td><td>CGG</td><td>||||||||[||||||||||||]PAM</td><td>NA</td><td>Intergenic</td><td>NA</td><td>NA</td></tr>
<tr><td>T13982</td><td>ATTGGTTCTAGTCATAAAACGGG</td><td>1000</td><td></td><td></td><td></td><td></td><td></td><td></td><td></td><td></td><td></td></tr>
<tr><td>Oligo fwd</td><td>CACCATTGGTTCTAGTCATAAAAC</td><td></td><td></td><td></td><td></td><td></td><td></td><td></td><td></td><td></td><td></td></tr>
<tr><td>Oligo rev</td><td>AAACGTTTTATGACTAGAACCAAT</td><td></td><td></td><td></td><td></td><td></td><td></td><td></td><td></td><td></td><td></td></tr>
<tr><td>Chromosome</td><td>start</td><td>end</td><td>strand</td><td>MM</td><td>target_seq</td><td>PAM</td><td>alignment</td><td>distance</td><td>position</td><td>gene name</td><td>gene id</td></tr>
<tr><td>GU305914.1</td><td>184307</td><td>184329</td><td>+</td><td>0</td><td>ATTGGTTCTAGTCATAAAAC</td><td>GGG</td><td>||||||||[||||||||||||]PAM</td><td>NA</td><td>Intergenic</td><td>NA</td><td>NA</td></tr>
<tr><td>T13983</td><td>GGTTCTAGTCATAAAACGGGCGG</td><td>1000</td><td></td><td></td><td></td><td></td><td></td><td></td><td></td><td></td><td></td></tr>
<tr><td>Oligo fwd</td><td>CACCGGTTCTAGTCATAAAACGGG</td><td></td><td></td><td></td><td></td><td></td><td></td><td></td><td></td><td></td><td></td></tr>
<tr><td>Oligo rev</td><td>AAACCCCGTTTTATGACTAGAACC</td><td></td><td></td><td></td><td></td><td></td><td></td><td></td><td></td><td></td><td></td></tr>
<tr><td>Chromosome</td><td>start</td><td>end</td><td>strand</td><td>MM</td><td>target_seq</td><td>PAM</td><td>alignment</td><td>distance</td><td>position</td><td>gene name</td><td>gene id</td></tr>
<tr><td>GU305914.1</td><td>184310</td><td>184332</td><td>+</td><td>0</td><td>GGTTCTAGTCATAAAACGGG</td><td>CGG</td><td>||||||||[||||||||||||]PAM</td><td>NA</td><td>Intergenic</td><td>NA</td><td>NA</td></tr>
<tr><td>T13984</td><td>ATGCCATTCGAGCCCAGAGCAGG</td><td>1000</td><td></td><td></td><td></td><td></td><td></td><td></td><td></td><td></td><td></td></tr>
<tr><td>Oligo fwd</td><td>CACCATGCCATTCGAGCCCAGAGC</td><td></td><td></td><td></td><td></td><td></td><td></td><td></td><td></td><td></td><td></td></tr>
<tr><td>Oligo rev</td><td>AAACGCTCTGGGCTCGAATGGCAT</td><td></td><td></td><td></td><td></td><td></td><td></td><td></td><td></td><td></td><td></td></tr>
<tr><td>Chromosome</td><td>start</td><td>end</td><td>strand</td><td>MM</td><td>target_seq</td><td>PAM</td><td>alignment</td><td>distance</td><td>position</td><td>gene name</td><td>gene id</td></tr>
<tr><td>GU305914.1</td><td>184359</td><td>184381</td><td>+</td><td>0</td><td>ATGCCATTCGAGCCCAGAGC</td><td>AGG</td><td>||||||||[||||||||||||]PAM</td><td>NA</td><td>Intergenic</td><td>NA</td><td>NA</td></tr>
<tr><td>T13985</td><td>CATTCGAGCCCAGAGCAGGACGG</td><td>1000</td><td></td><td></td><td></td><td></td><td></td><td></td><td></td><td></td><td></td></tr>
<tr><td>Oligo fwd</td><td>CACCCATTCGAGCCCAGAGCAGGA</td><td></td><td></td><td></td><td></td><td></td><td></td><td></td><td></td><td></td><td></td></tr>
<tr><td>Oligo rev</td><td>AAACTCCTGCTCTGGGCTCGAATG</td><td></td><td></td><td></td><td></td><td></td><td></td><td></td><td></td><td></td><td></td></tr>
<tr><td>Chromosome</td><td>start</td><td>end</td><td>strand</td><td>MM</td><td>target_seq</td><td>PAM</td><td>alignment</td><td>distance</td><td>position</td><td>gene name</td><td>gene id</td></tr>
<tr><td>GU305914.1</td><td>184363</td><td>184385</td><td>+</td><td>0</td><td>CATTCGAGCCCAGAGCAGGA</td><td>CGG</td><td>||||||||[||||||||||||]PAM</td><td>NA</td><td>Intergenic</td><td>NA</td><td>NA</td></tr>
<tr><td>T13986</td><td>GCCCAGAGCAGGACGGACCGCGG</td><td>1000</td><td></td><td></td><td></td><td></td><td></td><td></td><td></td><td></td><td></td></tr>
<tr><td>Oligo fwd</td><td>CACCGCCCAGAGCAGGACGGACCG</td><td></td><td></td><td></td><td></td><td></td><td></td><td></td><td></td><td></td><td></td></tr>
<tr><td>Oligo rev</td><td>AAACCGGTCCGTCCTGCTCTGGGC</td><td></td><td></td><td></td><td></td><td></td><td></td><td></td><td></td><td></td><td></td></tr>
<tr><td>Chromosome</td><td>start</td><td>end</td><td>strand</td><td>MM</td><td>target_seq</td><td>PAM</td><td>alignment</td><td>distance</td><td>position</td><td>gene name</td><td>gene id</td></tr>
<tr><td>GU305914.1</td><td>184370</td><td>184392</td><td>+</td><td>0</td><td>GCCCAGAGCAGGACGGACCG</td><td>CGG</td><td>||||||||[||||||||||||]PAM</td><td>NA</td><td>Intergenic</td><td>NA</td><td>NA</td></tr>
<tr><td>T13987</td><td>ACCCTATCTACGTTAACGAGCGG</td><td>1000</td><td></td><td></td><td></td><td></td><td></td><td></td><td></td><td></td><td></td></tr>
<tr><td>Oligo fwd</td><td>CACCACCCTATCTACGTTAACGAG</td><td></td><td></td><td></td><td></td><td></td><td></td><td></td><td></td><td></td><td></td></tr>
<tr><td>Oligo rev</td><td>AAACCTCGTTAACGTAGATAGGGT</td><td></td><td></td><td></td><td></td><td></td><td></td><td></td><td></td><td></td><td></td></tr>
<tr><td>Chromosome</td><td>start</td><td>end</td><td>strand</td><td>MM</td><td>target_seq</td><td>PAM</td><td>alignment</td><td>distance</td><td>position</td><td>gene name</td><td>gene id</td></tr>
<tr><td>GU305914.1</td><td>184402</td><td>184424</td><td>+</td><td>0</td><td>ACCCTATCTACGTTAACGAG</td><td>CGG</td><td>||||||||[||||||||||||]PAM</td><td>NA</td><td>Intergenic</td><td>NA</td><td>NA</td></tr>
<tr><td>T13988</td><td>CCCTATCTACGTTAACGAGCGGG</td><td>1000</td><td></td><td></td><td></td><td></td><td></td><td></td><td></td><td></td><td></td></tr>
<tr><td>Oligo fwd</td><td>CACCCCCTATCTACGTTAACGAGC</td><td></td><td></td><td></td><td></td><td></td><td></td><td></td><td></td><td></td><td></td></tr>
<tr><td>Oligo rev</td><td>AAACGCTCGTTAACGTAGATAGGG</td><td></td><td></td><td></td><td></td><td></td><td></td><td></td><td></td><td></td><td></td></tr>
<tr><td>Chromosome</td><td>start</td><td>end</td><td>strand</td><td>MM</td><td>target_seq</td><td>PAM</td><td>alignment</td><td>distance</td><td>position</td><td>gene name</td><td>gene id</td></tr>
<tr><td>GU305914.1</td><td>184403</td><td>184425</td><td>+</td><td>0</td><td>CCCTATCTACGTTAACGAGC</td><td>GGG</td><td>||||||||[||||||||||||]PAM</td><td>NA</td><td>Intergenic</td><td>NA</td><td>NA</td></tr>
<tr><td>T13989</td><td>TACGTTAACGAGCGGGTGAACGG</td><td>1000</td><td></td><td></td><td></td><td></td><td></td><td></td><td></td><td></td><td></td></tr>
<tr><td>Oligo fwd</td><td>CACCTACGTTAACGAGCGGGTGAA</td><td></td><td></td><td></td><td></td><td></td><td></td><td></td><td></td><td></td><td></td></tr>
<tr><td>Oligo rev</td><td>AAACTTCACCCGCTCGTTAACGTA</td><td></td><td></td><td></td><td></td><td></td><td></td><td></td><td></td><td></td><td></td></tr>
<tr><td>Chromosome</td><td>start</td><td>end</td><td>strand</td><td>MM</td><td>target_seq</td><td>PAM</td><td>alignment</td><td>distance</td><td>position</td><td>gene name</td><td>gene id</td></tr>
<tr><td>GU305914.1</td><td>184410</td><td>184432</td><td>+</td><td>0</td><td>TACGTTAACGAGCGGGTGAA</td><td>CGG</td><td>||||||||[||||||||||||]PAM</td><td>NA</td><td>Intergenic</td><td>NA</td><td>NA</td></tr>
<tr><td>T13990</td><td>GCGGGTGAACGGAGCCCCCCAGG</td><td>1000</td><td></td><td></td><td></td><td></td><td></td><td></td><td></td><td></td><td></td></tr>
<tr><td>Oligo fwd</td><td>CACCGCGGGTGAACGGAGCCCCCC</td><td></td><td></td><td></td><td></td><td></td><td></td><td></td><td></td><td></td><td></td></tr>
<tr><td>Oligo rev</td><td>AAACGGGGGGCTCCGTTCACCCGC</td><td></td><td></td><td></td><td></td><td></td><td></td><td></td><td></td><td></td><td></td></tr>
<tr><td>Chromosome</td><td>start</td><td>end</td><td>strand</td><td>MM</td><td>target_seq</td><td>PAM</td><td>alignment</td><td>distance</td><td>position</td><td>gene name</td><td>gene id</td></tr>
<tr><td>GU305914.1</td><td>184421</td><td>184443</td><td>+</td><td>0</td><td>GCGGGTGAACGGAGCCCCCC</td><td>AGG</td><td>||||||||[||||||||||||]PAM</td><td>NA</td><td>Intergenic</td><td>NA</td><td>NA</td></tr>
<tr><td>T13991</td><td>GTGAACGGAGCCCCCCAGGCCGG</td><td>1000</td><td></td><td></td><td></td><td></td><td></td><td></td><td></td><td></td><td></td></tr>
<tr><td>Oligo fwd</td><td>CACCGTGAACGGAGCCCCCCAGGC</td><td></td><td></td><td></td><td></td><td></td><td></td><td></td><td></td><td></td><td></td></tr>
</table>

Oligo rev AAACGCCTGGGGGGCTCCGTTCAC
Chromosome start end strand MM target_seq PAM alignment distance position gene name gene id
GU305914.1 184425 184447 + 0 GTGAACGGAGCCCCCCAGGC CGG ||||||||[||||||||||||]PAM NA Intergenic NA NA

T13992 TGAACGGAGCCCCCCAGGCCGGG 1000
Oligo fwd CACCTGAACGGAGCCCCCCAGGCC
Oligo rev AAACGGCCTGGGGGGCTCCGTTCA
Chromosome start end strand MM target_seq PAM alignment distance position gene name gene id
GU305914.1 184426 184448 + 0 TGAACGGAGCCCCCCAGGCC GGG ||||||||[||||||||||||]PAM NA Intergenic NA NA

T13993 GGAGCCCCCCAGGCCGGGTAAGG 1000
Oligo fwd CACCGGAGCCCCCCAGGCCGGGTA
Oligo rev AAACTACCCGGCCTGGGGGGCTCC
Chromosome start end strand MM target_seq PAM alignment distance position gene name gene id
GU305914.1 184431 184453 + 0 GGAGCCCCCCAGGCCGGGTA AGG ||||||||[||||||||||||]PAM NA Intergenic NA NA

T13994 CAGGCCGGGTAAGGTTGCATTGG 1000
Oligo fwd CACCCAGGCCGGGTAAGGTTGCAT
Oligo rev AAACATGCAACCTTACCCGGCCTG
Chromosome start end strand MM target_seq PAM alignment distance position gene name gene id
GU305914.1 184440 184462 + 0 CAGGCCGGGTAAGGTTGCAT TGG ||||||||[||||||||||||]PAM NA Intergenic NA NA

T13995 AGGCCGGGTAAGGTTGCATTGGG 1000
Oligo fwd CACCAGGCCGGGTAAGGTTGCATT
Oligo rev AAACAATGCAACCTTACCCGGCCT
Chromosome start end strand MM target_seq PAM alignment distance position gene name gene id
GU305914.1 184441 184463 + 0 AGGCCGGGTAAGGTTGCATT GGG ||||||||[||||||||||||]PAM NA Intergenic NA NA

T13996 GGGTAAGGTTGCATTGGGCTCGG 1000
Oligo fwd CACCGGGTAAGGTTGCATTGGGCT
Oligo rev AAACAGCCCAATGCAACCTTACCC
Chromosome start end strand MM target_seq PAM alignment distance position gene name gene id
GU305914.1 184446 184468 + 0 GGGTAAGGTTGCATTGGGCT CGG ||||||||[||||||||||||]PAM NA Intergenic NA NA

T13997 GGTTGCATTGGGCTCGGCTATGG 1000
Oligo fwd CACCGGTTGCATTGGGCTCGGCTA
Oligo rev AAACTAGCCGAGCCCAATGCAACC
Chromosome start end strand MM target_seq PAM alignment distance position gene name gene id
GU305914.1 184452 184474 + 0 GGTTGCATTGGGCTCGGCTA TGG ||||||||[||||||||||||]PAM NA Intergenic NA NA

T13998 GTTGCATTGGGCTCGGCTATGGG 1000
Oligo fwd CACCGTTGCATTGGGCTCGGCTAT
Oligo rev AAACATAGCCGAGCCCAATGCAAC
Chromosome start end strand MM target_seq PAM alignment distance position gene name gene id
GU305914.1 184453 184475 + 0 GTTGCATTGGGCTCGGCTAT GGG ||||||||[||||||||||||]PAM NA Intergenic NA NA

T13999 CATTGGGCTCGGCTATGGGACGG 1000
Oligo fwd CACCCATTGGGCTCGGCTATGGGA
Oligo rev AAACTCCCATAGCCGAGCCCAATG
Chromosome start end strand MM target_seq PAM alignment distance position gene name gene id
GU305914.1 184457 184479 + 0 CATTGGGCTCGGCTATGGGA CGG ||||||||[||||||||||||]PAM NA Intergenic NA NA

T14000 ATTGGGCTCGGCTATGGGACGGG 1000
Oligo fwd CACCATTGGGCTCGGCTATGGGAC
Oligo rev AAACGTCCCATAGCCGAGCCCAAT
Chromosome start end strand MM target_seq PAM alignment distance position gene name gene id
GU305914.1 184458 184480 + 0 ATTGGGCTCGGCTATGGGAC GGG ||||||||[||||||||||||]PAM NA Intergenic NA NA

T14001 TTGGGCTCGGCTATGGGACGGGG 1000
Oligo fwd CACCTTGGGCTCGGCTATGGGACG
Oligo rev AAACCGTCCCATAGCCGAGCCCAA
Chromosome start end strand MM target_seq PAM alignment distance position gene name gene id
GU305914.1 184459 184481 + 0 TTGGGCTCGGCTATGGGACG GGG ||||||||[||||||||||||]PAM NA Intergenic NA NA

T14002 TCGGCTATGGGACGGGGATTTGG 1000
Oligo fwd CACCTCGGCTATGGGACGGGGATT
Oligo rev AAACAATCCCCGTCCCATAGCCGA
Chromosome start end strand MM target_seq PAM alignment distance position gene name gene id
GU305914.1 184465 184487 + 0 TCGGCTATGGGACGGGGATT TGG ||||||||[||||||||||||]PAM NA Intergenic NA NA

T14003 TATGGGACGGGGATTTGGCCCGG 1000
Oligo fwd CACCTATGGGACGGGGATTTGGCC
Oligo rev AAACGGCCAAATCCCCGTCCCATA
Chromosome start end strand MM target_seq PAM alignment distance position gene name gene id
GU305914.1 184470 184492 + 0 TATGGGACGGGGATTTGGCC CGG ||||||||[||||||||||||]PAM NA Intergenic NA NA

T14004 TTTGGCCCGGTACTTTAATATGG 1000
Oligo fwd CACCTTTGGCCCGGTACTTTAATA
Oligo rev AAACTATTAAAGTACCGGGCCAAA
Chromosome start end strand MM target_seq PAM alignment distance position gene name gene id
GU305914.1 184483 184505 + 0 TTTGGCCCGGTACTTTAATA TGG ||||||||[||||||||||||]PAM NA Intergenic NA NA

T14005 TTGGCCCGGTACTTTAATATGGG 1000
Oligo fwd CACCTTGGCCCGGTACTTTAATAT
Oligo rev AAACATATTAAAGTACCGGGCCAA
Chromosome start end strand MM target_seq PAM alignment distance position gene name gene id
GU305914.1 184484 184506 + 0 TTGGCCCGGTACTTTAATAT GGG ||||||||[||||||||||||]PAM NA Intergenic NA NA

T14006 GCCCGGTACTTTAATATGGGTGG 1000

<table>
<tr><td>Oligo fwd</td><td>CACCGCCCGGTACTTTAATATGGG</td><td></td><td></td><td></td><td></td><td></td><td></td><td></td><td></td><td></td><td></td></tr>
<tr><td>Oligo rev</td><td>AAACCCCATATTAAAGTACCGGGC</td><td></td><td></td><td></td><td></td><td></td><td></td><td></td><td></td><td></td><td></td></tr>
<tr><td>Chromosome</td><td>start</td><td>end</td><td>strand</td><td>MM</td><td>target_seq</td><td>PAM</td><td>alignment</td><td>distance</td><td>position</td><td>gene name</td><td>gene id</td></tr>
<tr><td>GU305914.1</td><td>184487</td><td>184509</td><td>+</td><td>0</td><td>GCCCGGTACTTTAATATGGG</td><td>TGG</td><td>||||||||[||||||||||||]PAM</td><td>NA</td><td>Intergenic</td><td>NA</td><td>NA</td></tr>
<tr><td>T14007</td><td>CCCGGTACTTTAATATGGGTGGG</td><td>1000</td><td></td><td></td><td></td><td></td><td></td><td></td><td></td><td></td><td></td></tr>
<tr><td>Oligo fwd</td><td>CACCCCCGGTACTTTAATATGGGT</td><td></td><td></td><td></td><td></td><td></td><td></td><td></td><td></td><td></td><td></td></tr>
<tr><td>Oligo rev</td><td>AAACACCCATATTAAAGTACCGGG</td><td></td><td></td><td></td><td></td><td></td><td></td><td></td><td></td><td></td><td></td></tr>
<tr><td>Chromosome</td><td>start</td><td>end</td><td>strand</td><td>MM</td><td>target_seq</td><td>PAM</td><td>alignment</td><td>distance</td><td>position</td><td>gene name</td><td>gene id</td></tr>
<tr><td>GU305914.1</td><td>184488</td><td>184510</td><td>+</td><td>0</td><td>CCCGGTACTTTAATATGGGT</td><td>GGG</td><td>||||||||[||||||||||||]PAM</td><td>NA</td><td>Intergenic</td><td>NA</td><td>NA</td></tr>
<tr><td>T14008</td><td>CCGGTACTTTAATATGGGTGGGG</td><td>1000</td><td></td><td></td><td></td><td></td><td></td><td></td><td></td><td></td><td></td></tr>
<tr><td>Oligo fwd</td><td>CACCCCGGTACTTTAATATGGGTG</td><td></td><td></td><td></td><td></td><td></td><td></td><td></td><td></td><td></td><td></td></tr>
<tr><td>Oligo rev</td><td>AAACCACCCATATTAAAGTACCGG</td><td></td><td></td><td></td><td></td><td></td><td></td><td></td><td></td><td></td><td></td></tr>
<tr><td>Chromosome</td><td>start</td><td>end</td><td>strand</td><td>MM</td><td>target_seq</td><td>PAM</td><td>alignment</td><td>distance</td><td>position</td><td>gene name</td><td>gene id</td></tr>
<tr><td>GU305914.1</td><td>184489</td><td>184511</td><td>+</td><td>0</td><td>CCGGTACTTTAATATGGGTG</td><td>GGG</td><td>||||||||[||||||||||||]PAM</td><td>NA</td><td>Intergenic</td><td>NA</td><td>NA</td></tr>
<tr><td>T14009</td><td>TTAATATGGGTGGGGTCTTATGG</td><td>1000</td><td></td><td></td><td></td><td></td><td></td><td></td><td></td><td></td><td></td></tr>
<tr><td>Oligo fwd</td><td>CACCTTAATATGGGTGGGGTCTTA</td><td></td><td></td><td></td><td></td><td></td><td></td><td></td><td></td><td></td><td></td></tr>
<tr><td>Oligo rev</td><td>AAACTAAGACCCCACCCATATTAA</td><td></td><td></td><td></td><td></td><td></td><td></td><td></td><td></td><td></td><td></td></tr>
<tr><td>Chromosome</td><td>start</td><td>end</td><td>strand</td><td>MM</td><td>target_seq</td><td>PAM</td><td>alignment</td><td>distance</td><td>position</td><td>gene name</td><td>gene id</td></tr>
<tr><td>GU305914.1</td><td>184497</td><td>184519</td><td>+</td><td>0</td><td>TTAATATGGGTGGGGTCTTA</td><td>TGG</td><td>||||||||[||||||||||||]PAM</td><td>NA</td><td>Intergenic</td><td>NA</td><td>NA</td></tr>
<tr><td>T14010</td><td>TAATATGGGTGGGGTCTTATGGG</td><td>1000</td><td></td><td></td><td></td><td></td><td></td><td></td><td></td><td></td><td></td></tr>
<tr><td>Oligo fwd</td><td>CACCTAATATGGGTGGGGTCTTAT</td><td></td><td></td><td></td><td></td><td></td><td></td><td></td><td></td><td></td><td></td></tr>
<tr><td>Oligo rev</td><td>AAACATAAGACCCCACCCATATTA</td><td></td><td></td><td></td><td></td><td></td><td></td><td></td><td></td><td></td><td></td></tr>
<tr><td>Chromosome</td><td>start</td><td>end</td><td>strand</td><td>MM</td><td>target_seq</td><td>PAM</td><td>alignment</td><td>distance</td><td>position</td><td>gene name</td><td>gene id</td></tr>
<tr><td>GU305914.1</td><td>184498</td><td>184520</td><td>+</td><td>0</td><td>TAATATGGGTGGGGTCTTAT</td><td>GGG</td><td>||||||||[||||||||||||]PAM</td><td>NA</td><td>Intergenic</td><td>NA</td><td>NA</td></tr>
<tr><td>T14011</td><td>ATGGGTGGGGTCTTATGGGTAGG</td><td>1000</td><td></td><td></td><td></td><td></td><td></td><td></td><td></td><td></td><td></td></tr>
<tr><td>Oligo fwd</td><td>CACCATGGGTGGGGTCTTATGGGT</td><td></td><td></td><td></td><td></td><td></td><td></td><td></td><td></td><td></td><td></td></tr>
<tr><td>Oligo rev</td><td>AAACACCCATAAGACCCCACCCAT</td><td></td><td></td><td></td><td></td><td></td><td></td><td></td><td></td><td></td><td></td></tr>
<tr><td>Chromosome</td><td>start</td><td>end</td><td>strand</td><td>MM</td><td>target_seq</td><td>PAM</td><td>alignment</td><td>distance</td><td>position</td><td>gene name</td><td>gene id</td></tr>
<tr><td>GU305914.1</td><td>184502</td><td>184524</td><td>+</td><td>0</td><td>ATGGGTGGGGTCTTATGGGT</td><td>AGG</td><td>||||||||[||||||||||||]PAM</td><td>NA</td><td>Intergenic</td><td>NA</td><td>NA</td></tr>
<tr><td>T14012</td><td>TGGGTGGGGTCTTATGGGTAGGG</td><td>1000</td><td></td><td></td><td></td><td></td><td></td><td></td><td></td><td></td><td></td></tr>
<tr><td>Oligo fwd</td><td>CACCTGGGTGGGGTCTTATGGGTA</td><td></td><td></td><td></td><td></td><td></td><td></td><td></td><td></td><td></td><td></td></tr>
<tr><td>Oligo rev</td><td>AAACTACCCATAAGACCCCACCCA</td><td></td><td></td><td></td><td></td><td></td><td></td><td></td><td></td><td></td><td></td></tr>
<tr><td>Chromosome</td><td>start</td><td>end</td><td>strand</td><td>MM</td><td>target_seq</td><td>PAM</td><td>alignment</td><td>distance</td><td>position</td><td>gene name</td><td>gene id</td></tr>
<tr><td>GU305914.1</td><td>184503</td><td>184525</td><td>+</td><td>0</td><td>TGGGTGGGGTCTTATGGGTA</td><td>GGG</td><td>||||||||[||||||||||||]PAM</td><td>NA</td><td>Intergenic</td><td>NA</td><td>NA</td></tr>
<tr><td>T14013</td><td>GGGTGGGGTCTTATGGGTAGGGG</td><td>1000</td><td></td><td></td><td></td><td></td><td></td><td></td><td></td><td></td><td></td></tr>
<tr><td>Oligo fwd</td><td>CACCGGGTGGGGTCTTATGGGTAG</td><td></td><td></td><td></td><td></td><td></td><td></td><td></td><td></td><td></td><td></td></tr>
<tr><td>Oligo rev</td><td>AAACCTACCCATAAGACCCCACCC</td><td></td><td></td><td></td><td></td><td></td><td></td><td></td><td></td><td></td><td></td></tr>
<tr><td>Chromosome</td><td>start</td><td>end</td><td>strand</td><td>MM</td><td>target_seq</td><td>PAM</td><td>alignment</td><td>distance</td><td>position</td><td>gene name</td><td>gene id</td></tr>
<tr><td>GU305914.1</td><td>184504</td><td>184526</td><td>+</td><td>0</td><td>GGGTGGGGTCTTATGGGTAG</td><td>GGG</td><td>||||||||[||||||||||||]PAM</td><td>NA</td><td>Intergenic</td><td>NA</td><td>NA</td></tr>
<tr><td>T14014</td><td>GGTGGGGTCTTATGGGTAGGGGG</td><td>1000</td><td></td><td></td><td></td><td></td><td></td><td></td><td></td><td></td><td></td></tr>
<tr><td>Oligo fwd</td><td>CACCGGTGGGGTCTTATGGGTAGG</td><td></td><td></td><td></td><td></td><td></td><td></td><td></td><td></td><td></td><td></td></tr>
<tr><td>Oligo rev</td><td>AAACCCTACCCATAAGACCCCACC</td><td></td><td></td><td></td><td></td><td></td><td></td><td></td><td></td><td></td><td></td></tr>
<tr><td>Chromosome</td><td>start</td><td>end</td><td>strand</td><td>MM</td><td>target_seq</td><td>PAM</td><td>alignment</td><td>distance</td><td>position</td><td>gene name</td><td>gene id</td></tr>
<tr><td>GU305914.1</td><td>184505</td><td>184527</td><td>+</td><td>0</td><td>GGTGGGGTCTTATGGGTAGG</td><td>GGG</td><td>||||||||[||||||||||||]PAM</td><td>NA</td><td>Intergenic</td><td>NA</td><td>NA</td></tr>
<tr><td>T14015</td><td>GAAGCCCCCTACCACGCCATTGG</td><td>1000</td><td></td><td></td><td></td><td></td><td></td><td></td><td></td><td></td><td></td></tr>
<tr><td>Oligo fwd</td><td>CACCGAAGCCCCCTACCACGCCAT</td><td></td><td></td><td></td><td></td><td></td><td></td><td></td><td></td><td></td><td></td></tr>
<tr><td>Oligo rev</td><td>AAACATGGCGTGGTAGGGGGCTTC</td><td></td><td></td><td></td><td></td><td></td><td></td><td></td><td></td><td></td><td></td></tr>
<tr><td>Chromosome</td><td>start</td><td>end</td><td>strand</td><td>MM</td><td>target_seq</td><td>PAM</td><td>alignment</td><td>distance</td><td>position</td><td>gene name</td><td>gene id</td></tr>
<tr><td>GU305914.1</td><td>184576</td><td>184598</td><td>+</td><td>0</td><td>GAAGCCCCCTACCACGCCAT</td><td>TGG</td><td>||||||||[||||||||||||]PAM</td><td>NA</td><td>Intergenic</td><td>NA</td><td>NA</td></tr>
<tr><td>T14016</td><td>CCATTGGCGCCCGCCCATAGCGG</td><td>1000</td><td></td><td></td><td></td><td></td><td></td><td></td><td></td><td></td><td></td></tr>
<tr><td>Oligo fwd</td><td>CACCCCATTGGCGCCCGCCCATAG</td><td></td><td></td><td></td><td></td><td></td><td></td><td></td><td></td><td></td><td></td></tr>
<tr><td>Oligo rev</td><td>AAACCTATGGGCGGGCGCCAATGG</td><td></td><td></td><td></td><td></td><td></td><td></td><td></td><td></td><td></td><td></td></tr>
<tr><td>Chromosome</td><td>start</td><td>end</td><td>strand</td><td>MM</td><td>target_seq</td><td>PAM</td><td>alignment</td><td>distance</td><td>position</td><td>gene name</td><td>gene id</td></tr>
<tr><td>GU305914.1</td><td>184592</td><td>184614</td><td>+</td><td>0</td><td>CCATTGGCGCCCGCCCATAG</td><td>CGG</td><td>||||||||[||||||||||||]PAM</td><td>NA</td><td>Intergenic</td><td>NA</td><td>NA</td></tr>
<tr><td>T14017</td><td>CCCGCCCATAGCGGTACTTTCGG</td><td>1000</td><td></td><td></td><td></td><td></td><td></td><td></td><td></td><td></td><td></td></tr>
<tr><td>Oligo fwd</td><td>CACCCCCGCCCATAGCGGTACTTT</td><td></td><td></td><td></td><td></td><td></td><td></td><td></td><td></td><td></td><td></td></tr>
<tr><td>Oligo rev</td><td>AAACAAAGTACCGCTATGGGCGGG</td><td></td><td></td><td></td><td></td><td></td><td></td><td></td><td></td><td></td><td></td></tr>
<tr><td>Chromosome</td><td>start</td><td>end</td><td>strand</td><td>MM</td><td>target_seq</td><td>PAM</td><td>alignment</td><td>distance</td><td>position</td><td>gene name</td><td>gene id</td></tr>
<tr><td>GU305914.1</td><td>184601</td><td>184623</td><td>+</td><td>0</td><td>CCCGCCCATAGCGGTACTTT</td><td>CGG</td><td>||||||||[||||||||||||]PAM</td><td>NA</td><td>Intergenic</td><td>NA</td><td>NA</td></tr>
<tr><td>T14018</td><td>ACTTTCGGTTCTATACAAGATGG</td><td>1000</td><td></td><td></td><td></td><td></td><td></td><td></td><td></td><td></td><td></td></tr>
<tr><td>Oligo fwd</td><td>CACCACTTTCGGTTCTATACAAGA</td><td></td><td></td><td></td><td></td><td></td><td></td><td></td><td></td><td></td><td></td></tr>
<tr><td>Oligo rev</td><td>AAACTCTTGTATAGAACCGAAAGT</td><td></td><td></td><td></td><td></td><td></td><td></td><td></td><td></td><td></td><td></td></tr>
<tr><td>Chromosome</td><td>start</td><td>end</td><td>strand</td><td>MM</td><td>target_seq</td><td>PAM</td><td>alignment</td><td>distance</td><td>position</td><td>gene name</td><td>gene id</td></tr>
<tr><td>GU305914.1</td><td>184616</td><td>184638</td><td>+</td><td>0</td><td>ACTTTCGGTTCTATACAAGA</td><td>TGG</td><td>||||||||[||||||||||||]PAM</td><td>NA</td><td>Intergenic</td><td>NA</td><td>NA</td></tr>
<tr><td>T14019</td><td>TCTATACAAGATGGCTACCGCGG</td><td>1000</td><td></td><td></td><td></td><td></td><td></td><td></td><td></td><td></td><td></td></tr>
<tr><td>Oligo fwd</td><td>CACCTCTATACAAGATGGCTACCG</td><td></td><td></td><td></td><td></td><td></td><td></td><td></td><td></td><td></td><td></td></tr>
<tr><td>Oligo rev</td><td>AAACCGGTAGCCATCTTGTATAGA</td><td></td><td></td><td></td><td></td><td></td><td></td><td></td><td></td><td></td><td></td></tr>
<tr><td>Chromosome</td><td>start</td><td>end</td><td>strand</td><td>MM</td><td>target_seq</td><td>PAM</td><td>alignment</td><td>distance</td><td>position</td><td>gene name</td><td>gene id</td></tr>
<tr><td>GU305914.1</td><td>184625</td><td>184647</td><td>+</td><td>0</td><td>TCTATACAAGATGGCTACCG</td><td>CGG</td><td>||||||||[||||||||||||]PAM</td><td>NA</td><td>Intergenic</td><td>NA</td><td>NA</td></tr>
<tr><td>T14020</td><td>CCGCGGTTTTCGATCGCTCCCGG</td><td>1000</td><td></td><td></td><td></td><td></td><td></td><td></td><td></td><td></td><td></td></tr>
<tr><td>Oligo fwd</td><td>CACCCCGCGGTTTTCGATCGCTCC</td><td></td><td></td><td></td><td></td><td></td><td></td><td></td><td></td><td></td><td></td></tr>
<tr><td>Oligo rev</td><td>AAACGGAGCGATCGAAAACCGCGG</td><td></td><td></td><td></td><td></td><td></td><td></td><td></td><td></td><td></td><td></td></tr>
<tr><td>Chromosome</td><td>start</td><td>end</td><td>strand</td><td>MM</td><td>target_seq</td><td>PAM</td><td>alignment</td><td>distance</td><td>position</td><td>gene name</td><td>gene id</td></tr>
<tr><td>GU305914.1</td><td>184642</td><td>184664</td><td>+</td><td>0</td><td>CCGCGGTTTTCGATCGCTCC</td><td>CGG</td><td>||||||||[||||||||||||]PAM</td><td>NA</td><td>Intergenic</td><td>NA</td><td>NA</td></tr>
</table>

<table>
<tr><td>T14021</td><td>CGCGGTTTTCGATCGCTCCCGGG</td><td>1000</td><td></td><td></td><td></td><td></td><td></td><td></td><td></td><td></td><td></td></tr>
<tr><td>Oligo fwd</td><td>CACCCGCGGTTTTCGATCGCTCCC</td><td></td><td></td><td></td><td></td><td></td><td></td><td></td><td></td><td></td><td></td></tr>
<tr><td>Oligo rev</td><td>AAACGGGAGCGATCGAAAACCGCG</td><td></td><td></td><td></td><td></td><td></td><td></td><td></td><td></td><td></td><td></td></tr>
<tr><td>Chromosome</td><td>start</td><td>end</td><td>strand</td><td>MM</td><td>target_seq</td><td>PAM</td><td>alignment</td><td>distance</td><td>position</td><td>gene name</td><td>gene id</td></tr>
<tr><td>GU305914.1</td><td>184643</td><td>184665</td><td>+</td><td>0</td><td>CGCGGTTTTCGATCGCTCCC</td><td>GGG</td><td>||||||||[||||||||||||]PAM</td><td>NA</td><td>Intergenic</td><td>NA</td><td>NA</td></tr>
<tr><td>T14022</td><td>GCGGTTTTCGATCGCTCCCGGGG</td><td>1000</td><td></td><td></td><td></td><td></td><td></td><td></td><td></td><td></td><td></td></tr>
<tr><td>Oligo fwd</td><td>CACCGCGGTTTTCGATCGCTCCCG</td><td></td><td></td><td></td><td></td><td></td><td></td><td></td><td></td><td></td><td></td></tr>
<tr><td>Oligo rev</td><td>AAACCGGGAGCGATCGAAAACCGC</td><td></td><td></td><td></td><td></td><td></td><td></td><td></td><td></td><td></td><td></td></tr>
<tr><td>Chromosome</td><td>start</td><td>end</td><td>strand</td><td>MM</td><td>target_seq</td><td>PAM</td><td>alignment</td><td>distance</td><td>position</td><td>gene name</td><td>gene id</td></tr>
<tr><td>GU305914.1</td><td>184644</td><td>184666</td><td>+</td><td>0</td><td>GCGGTTTTCGATCGCTCCCG</td><td>GGG</td><td>||||||||[||||||||||||]PAM</td><td>NA</td><td>Intergenic</td><td>NA</td><td>NA</td></tr>
<tr><td>T14023</td><td>CGGTTTTCGATCGCTCCCGGGGG</td><td>1000</td><td></td><td></td><td></td><td></td><td></td><td></td><td></td><td></td><td></td></tr>
<tr><td>Oligo fwd</td><td>CACCCGGTTTTCGATCGCTCCCGG</td><td></td><td></td><td></td><td></td><td></td><td></td><td></td><td></td><td></td><td></td></tr>
<tr><td>Oligo rev</td><td>AAACCCGGGAGCGATCGAAAACCG</td><td></td><td></td><td></td><td></td><td></td><td></td><td></td><td></td><td></td><td></td></tr>
<tr><td>Chromosome</td><td>start</td><td>end</td><td>strand</td><td>MM</td><td>target_seq</td><td>PAM</td><td>alignment</td><td>distance</td><td>position</td><td>gene name</td><td>gene id</td></tr>
<tr><td>GU305914.1</td><td>184645</td><td>184667</td><td>+</td><td>0</td><td>CGGTTTTCGATCGCTCCCGG</td><td>GGG</td><td>||||||||[||||||||||||]PAM</td><td>NA</td><td>Intergenic</td><td>NA</td><td>NA</td></tr>
<tr><td>T14024</td><td>GGGGGTCTCATTTCTATAGCTGG</td><td>1000</td><td></td><td></td><td></td><td></td><td></td><td></td><td></td><td></td><td></td></tr>
<tr><td>Oligo fwd</td><td>CACCGGGGGTCTCATTTCTATAGC</td><td></td><td></td><td></td><td></td><td></td><td></td><td></td><td></td><td></td><td></td></tr>
<tr><td>Oligo rev</td><td>AAACGCTATAGAAATGAGACCCCC</td><td></td><td></td><td></td><td></td><td></td><td></td><td></td><td></td><td></td><td></td></tr>
<tr><td>Chromosome</td><td>start</td><td>end</td><td>strand</td><td>MM</td><td>target_seq</td><td>PAM</td><td>alignment</td><td>distance</td><td>position</td><td>gene name</td><td>gene id</td></tr>
<tr><td>GU305914.1</td><td>184663</td><td>184685</td><td>+</td><td>0</td><td>GGGGGTCTCATTTCTATAGC</td><td>TGG</td><td>||||||||[||||||||||||]PAM</td><td>NA</td><td>Intergenic</td><td>NA</td><td>NA</td></tr>
<tr><td>T14025</td><td>TGATCTGTGCCTTAGACCCCAGG</td><td>1000</td><td></td><td></td><td></td><td></td><td></td><td></td><td></td><td></td><td></td></tr>
<tr><td>Oligo fwd</td><td>CACCTGATCTGTGCCTTAGACCCC</td><td></td><td></td><td></td><td></td><td></td><td></td><td></td><td></td><td></td><td></td></tr>
<tr><td>Oligo rev</td><td>AAACGGGGTCTAAGGCACAGATCA</td><td></td><td></td><td></td><td></td><td></td><td></td><td></td><td></td><td></td><td></td></tr>
<tr><td>Chromosome</td><td>start</td><td>end</td><td>strand</td><td>MM</td><td>target_seq</td><td>PAM</td><td>alignment</td><td>distance</td><td>position</td><td>gene name</td><td>gene id</td></tr>
<tr><td>GU305914.1</td><td>184703</td><td>184725</td><td>+</td><td>0</td><td>TGATCTGTGCCTTAGACCCC</td><td>AGG</td><td>||||||||[||||||||||||]PAM</td><td>NA</td><td>Intergenic</td><td>NA</td><td>NA</td></tr>
<tr><td>T14026</td><td>GATCTGTGCCTTAGACCCCAGGG</td><td>1000</td><td></td><td></td><td></td><td></td><td></td><td></td><td></td><td></td><td></td></tr>
<tr><td>Oligo fwd</td><td>CACCGATCTGTGCCTTAGACCCCA</td><td></td><td></td><td></td><td></td><td></td><td></td><td></td><td></td><td></td><td></td></tr>
<tr><td>Oligo rev</td><td>AAACTGGGGTCTAAGGCACAGATC</td><td></td><td></td><td></td><td></td><td></td><td></td><td></td><td></td><td></td><td></td></tr>
<tr><td>Chromosome</td><td>start</td><td>end</td><td>strand</td><td>MM</td><td>target_seq</td><td>PAM</td><td>alignment</td><td>distance</td><td>position</td><td>gene name</td><td>gene id</td></tr>
<tr><td>GU305914.1</td><td>184704</td><td>184726</td><td>+</td><td>0</td><td>GATCTGTGCCTTAGACCCCA</td><td>GGG</td><td>||||||||[||||||||||||]PAM</td><td>NA</td><td>Intergenic</td><td>NA</td><td>NA</td></tr>
<tr><td>T14027</td><td>TAGACCCCAGGGTACGTGCCTGG</td><td>1000</td><td></td><td></td><td></td><td></td><td></td><td></td><td></td><td></td><td></td></tr>
<tr><td>Oligo fwd</td><td>CACCTAGACCCCAGGGTACGTGCC</td><td></td><td></td><td></td><td></td><td></td><td></td><td></td><td></td><td></td><td></td></tr>
<tr><td>Oligo rev</td><td>AAACGGCACGTACCCTGGGGTCTA</td><td></td><td></td><td></td><td></td><td></td><td></td><td></td><td></td><td></td><td></td></tr>
<tr><td>Chromosome</td><td>start</td><td>end</td><td>strand</td><td>MM</td><td>target_seq</td><td>PAM</td><td>alignment</td><td>distance</td><td>position</td><td>gene name</td><td>gene id</td></tr>
<tr><td>GU305914.1</td><td>184715</td><td>184737</td><td>+</td><td>0</td><td>TAGACCCCAGGGTACGTGCC</td><td>TGG</td><td>||||||||[||||||||||||]PAM</td><td>NA</td><td>Intergenic</td><td>NA</td><td>NA</td></tr>
<tr><td>T14028</td><td>CCCAGGGTACGTGCCTGGATAGG</td><td>1000</td><td></td><td></td><td></td><td></td><td></td><td></td><td></td><td></td><td></td></tr>
<tr><td>Oligo fwd</td><td>CACCCCCAGGGTACGTGCCTGGAT</td><td></td><td></td><td></td><td></td><td></td><td></td><td></td><td></td><td></td><td></td></tr>
<tr><td>Oligo rev</td><td>AAACATCCAGGCACGTACCCTGGG</td><td></td><td></td><td></td><td></td><td></td><td></td><td></td><td></td><td></td><td></td></tr>
<tr><td>Chromosome</td><td>start</td><td>end</td><td>strand</td><td>MM</td><td>target_seq</td><td>PAM</td><td>alignment</td><td>distance</td><td>position</td><td>gene name</td><td>gene id</td></tr>
<tr><td>GU305914.1</td><td>184720</td><td>184742</td><td>+</td><td>0</td><td>CCCAGGGTACGTGCCTGGAT</td><td>AGG</td><td>||||||||[||||||||||||]PAM</td><td>NA</td><td>Intergenic</td><td>NA</td><td>NA</td></tr>
<tr><td>T14029</td><td>AGGGTACGTGCCTGGATAGGTGG</td><td>1000</td><td></td><td></td><td></td><td></td><td></td><td></td><td></td><td></td><td></td></tr>
<tr><td>Oligo fwd</td><td>CACCAGGGTACGTGCCTGGATAGG</td><td></td><td></td><td></td><td></td><td></td><td></td><td></td><td></td><td></td><td></td></tr>
<tr><td>Oligo rev</td><td>AAACCCTATCCAGGCACGTACCCT</td><td></td><td></td><td></td><td></td><td></td><td></td><td></td><td></td><td></td><td></td></tr>
<tr><td>Chromosome</td><td>start</td><td>end</td><td>strand</td><td>MM</td><td>target_seq</td><td>PAM</td><td>alignment</td><td>distance</td><td>position</td><td>gene name</td><td>gene id</td></tr>
<tr><td>GU305914.1</td><td>184723</td><td>184745</td><td>+</td><td>0</td><td>AGGGTACGTGCCTGGATAGG</td><td>TGG</td><td>||||||||[||||||||||||]PAM</td><td>NA</td><td>Intergenic</td><td>NA</td><td>NA</td></tr>
<tr><td>T14030</td><td>GGGTACGTGCCTGGATAGGTGGG</td><td>1000</td><td></td><td></td><td></td><td></td><td></td><td></td><td></td><td></td><td></td></tr>
<tr><td>Oligo fwd</td><td>CACCGGGTACGTGCCTGGATAGGT</td><td></td><td></td><td></td><td></td><td></td><td></td><td></td><td></td><td></td><td></td></tr>
<tr><td>Oligo rev</td><td>AAACACCTATCCAGGCACGTACCC</td><td></td><td></td><td></td><td></td><td></td><td></td><td></td><td></td><td></td><td></td></tr>
<tr><td>Chromosome</td><td>start</td><td>end</td><td>strand</td><td>MM</td><td>target_seq</td><td>PAM</td><td>alignment</td><td>distance</td><td>position</td><td>gene name</td><td>gene id</td></tr>
<tr><td>GU305914.1</td><td>184724</td><td>184746</td><td>+</td><td>0</td><td>GGGTACGTGCCTGGATAGGT</td><td>GGG</td><td>||||||||[||||||||||||]PAM</td><td>NA</td><td>Intergenic</td><td>NA</td><td>NA</td></tr>
<tr><td>T14031</td><td>GATAGGTGGGCGACCCCTAGCGG</td><td>1000</td><td></td><td></td><td></td><td></td><td></td><td></td><td></td><td></td><td></td></tr>
<tr><td>Oligo fwd</td><td>CACCGATAGGTGGGCGACCCCTAG</td><td></td><td></td><td></td><td></td><td></td><td></td><td></td><td></td><td></td><td></td></tr>
<tr><td>Oligo rev</td><td>AAACCTAGGGGTCGCCCACCTATC</td><td></td><td></td><td></td><td></td><td></td><td></td><td></td><td></td><td></td><td></td></tr>
<tr><td>Chromosome</td><td>start</td><td>end</td><td>strand</td><td>MM</td><td>target_seq</td><td>PAM</td><td>alignment</td><td>distance</td><td>position</td><td>gene name</td><td>gene id</td></tr>
<tr><td>GU305914.1</td><td>184737</td><td>184759</td><td>+</td><td>0</td><td>GATAGGTGGGCGACCCCTAG</td><td>CGG</td><td>||||||||[||||||||||||]PAM</td><td>NA</td><td>Intergenic</td><td>NA</td><td>NA</td></tr>
<tr><td>T14032</td><td>AGCCGCCCGTAACAGCTGTCAGG</td><td>1000</td><td></td><td></td><td></td><td></td><td></td><td></td><td></td><td></td><td></td></tr>
<tr><td>Oligo fwd</td><td>CACCAGCCGCCCGTAACAGCTGTC</td><td></td><td></td><td></td><td></td><td></td><td></td><td></td><td></td><td></td><td></td></tr>
<tr><td>Oligo rev</td><td>AAACGACAGCTGTTACGGGCGGCT</td><td></td><td></td><td></td><td></td><td></td><td></td><td></td><td></td><td></td><td></td></tr>
<tr><td>Chromosome</td><td>start</td><td>end</td><td>strand</td><td>MM</td><td>target_seq</td><td>PAM</td><td>alignment</td><td>distance</td><td>position</td><td>gene name</td><td>gene id</td></tr>
<tr><td>GU305914.1</td><td>184795</td><td>184817</td><td>+</td><td>0</td><td>AGCCGCCCGTAACAGCTGTC</td><td>AGG</td><td>||||||||[||||||||||||]PAM</td><td>NA</td><td>Intergenic</td><td>NA</td><td>NA</td></tr>
<tr><td>T14033</td><td>GCCGCCCGTAACAGCTGTCAGGG</td><td>1000</td><td></td><td></td><td></td><td></td><td></td><td></td><td></td><td></td><td></td></tr>
<tr><td>Oligo fwd</td><td>CACCGCCGCCCGTAACAGCTGTCA</td><td></td><td></td><td></td><td></td><td></td><td></td><td></td><td></td><td></td><td></td></tr>
<tr><td>Oligo rev</td><td>AAACTGACAGCTGTTACGGGCGGC</td><td></td><td></td><td></td><td></td><td></td><td></td><td></td><td></td><td></td><td></td></tr>
<tr><td>Chromosome</td><td>start</td><td>end</td><td>strand</td><td>MM</td><td>target_seq</td><td>PAM</td><td>alignment</td><td>distance</td><td>position</td><td>gene name</td><td>gene id</td></tr>
<tr><td>GU305914.1</td><td>184796</td><td>184818</td><td>+</td><td>0</td><td>GCCGCCCGTAACAGCTGTCA</td><td>GGG</td><td>||||||||[||||||||||||]PAM</td><td>NA</td><td>Intergenic</td><td>NA</td><td>NA</td></tr>
<tr><td>T14034</td><td>GCTGTCAGGGCTATACAAGATGG</td><td>1000</td><td></td><td></td><td></td><td></td><td></td><td></td><td></td><td></td><td></td></tr>
<tr><td>Oligo fwd</td><td>CACCGCTGTCAGGGCTATACAAGA</td><td></td><td></td><td></td><td></td><td></td><td></td><td></td><td></td><td></td><td></td></tr>
<tr><td>Oligo rev</td><td>AAACTCTTGTATAGCCCTGACAGC</td><td></td><td></td><td></td><td></td><td></td><td></td><td></td><td></td><td></td><td></td></tr>
<tr><td>Chromosome</td><td>start</td><td>end</td><td>strand</td><td>MM</td><td>target_seq</td><td>PAM</td><td>alignment</td><td>distance</td><td>position</td><td>gene name</td><td>gene id</td></tr>
<tr><td>GU305914.1</td><td>184809</td><td>184831</td><td>+</td><td>0</td><td>GCTGTCAGGGCTATACAAGA</td><td>TGG</td><td>||||||||[||||||||||||]PAM</td><td>NA</td><td>Intergenic</td><td>NA</td><td>NA</td></tr>
<tr><td>T14035</td><td>TCCGAAACGCAACCCACCCCCGG</td><td>1000</td><td></td><td></td><td></td><td></td><td></td><td></td><td></td><td></td><td></td></tr>
<tr><td>Oligo fwd</td><td>CACCTCCGAAACGCAACCCACCCC</td><td></td><td></td><td></td><td></td><td></td><td></td><td></td><td></td><td></td><td></td></tr>
<tr><td>Oligo rev</td><td>AAACGGGGTGGGTTGCGTTTCGGA</td><td></td><td></td><td></td><td></td><td></td><td></td><td></td><td></td><td></td><td></td></tr>
<tr><td>Chromosome</td><td>start</td><td>end</td><td>strand</td><td>MM</td><td>target_seq</td><td>PAM</td><td>alignment</td><td>distance</td><td>position</td><td>gene name</td><td>gene id</td></tr>
<tr><td>GU305914.1</td><td>184874</td><td>184896</td><td>+</td><td>0</td><td>TCCGAAACGCAACCCACCCC</td><td>CGG</td><td>||||||||[||||||||||||]PAM</td><td>NA</td><td>Intergenic</td><td>NA</td><td>NA</td></tr>
</table>

T14036 CCGAAACGCAACCCACCCCCGGG 1000
Oligo fwd CACCCCGAAACGCAACCCACCCCC
Oligo rev AAACGGGGGTGGGTTGCGTTTCGG
Chromosome start end strand MM target_seq PAM alignment distance position gene name gene id
GU305914.1 184875 184897 + 0 CCGAAACGCAACCCACCCCC GGG ||||||||[||||||||||||]PAM NA Intergenic NA NA

T14037 CGAAACGCAACCCACCCCCGGGG 1000
Oligo fwd CACCCGAAACGCAACCCACCCCCG
Oligo rev AAACCGGGGGTGGGTTGCGTTTCG
Chromosome start end strand MM target_seq PAM alignment distance position gene name gene id
GU305914.1 184876 184898 + 0 CGAAACGCAACCCACCCCCG GGG ||||||||[||||||||||||]PAM NA Intergenic NA NA

T14038 AGACCCTATAGATGCCCCCCAGG 1000
Oligo fwd CACCAGACCCTATAGATGCCCCCC
Oligo rev AAACGGGGGGCATCTATAGGGTCT
Chromosome start end strand MM target_seq PAM alignment distance position gene name gene id
GU305914.1 184909 184931 + 0 AGACCCTATAGATGCCCCCC AGG ||||||||[||||||||||||]PAM NA Intergenic NA NA

T14039 ATAGATGCCCCCCAGGACGCAGG 1000
Oligo fwd CACCATAGATGCCCCCCAGGACGC
Oligo rev AAACGCGTCCTGGGGGGCATCTAT
Chromosome start end strand MM target_seq PAM alignment distance position gene name gene id
GU305914.1 184916 184938 + 0 ATAGATGCCCCCCAGGACGC AGG ||||||||[||||||||||||]PAM NA Intergenic NA NA

T14040 ATTCGAGCGATCGAAATCCGCGG 1000
Oligo fwd CACCATTCGAGCGATCGAAATCCG
Oligo rev AAACCGGATTTCGATCGCTCGAAT
Chromosome start end strand MM target_seq PAM alignment distance position gene name gene id
GU305914.1 184967 184989 + 0 ATTCGAGCGATCGAAATCCG CGG ||||||||[||||||||||||]PAM NA Intergenic NA NA

T14041 ATCGAAATCCGCGGACCGCCCGG 1000
Oligo fwd CACCATCGAAATCCGCGGACCGCC
Oligo rev AAACGGCGGTCCGCGGATTTCGAT
Chromosome start end strand MM target_seq PAM alignment distance position gene name gene id
GU305914.1 184976 184998 + 0 ATCGAAATCCGCGGACCGCC CGG ||||||||[||||||||||||]PAM NA Intergenic NA NA

T14042 TCGAAATCCGCGGACCGCCCGGG 1000
Oligo fwd CACCTCGAAATCCGCGGACCGCCC
Oligo rev AAACGGGCGGTCCGCGGATTTCGA
Chromosome start end strand MM target_seq PAM alignment distance position gene name gene id
GU305914.1 184977 184999 + 0 TCGAAATCCGCGGACCGCCC GGG ||||||||[||||||||||||]PAM NA Intergenic NA NA

T14043 GGTTTTTACTTTCGTTTTCTCGG 1000
Oligo fwd CACCGGTTTTTACTTTCGTTTTCT
Oligo rev AAACAGAAAACGAAAGTAAAAACC
Chromosome start end strand MM target_seq PAM alignment distance position gene name gene id
GU305914.1 184998 185020 + 0 GGTTTTTACTTTCGTTTTCT CGG ||||||||[||||||||||||]PAM NA Intergenic NA NA

T14044 TCGTTTTCTCGGTCGCCGCTCGG 1000
Oligo fwd CACCTCGTTTTCTCGGTCGCCGCT
Oligo rev AAACAGCGGCGACCGAGAAAACGA
Chromosome start end strand MM target_seq PAM alignment distance position gene name gene id
GU305914.1 185009 185031 + 0 TCGTTTTCTCGGTCGCCGCT CGG ||||||||[||||||||||||]PAM NA Intergenic NA NA

T14045 TTCTCGGTCGCCGCTCGGCTCGG 1000
Oligo fwd CACCTTCTCGGTCGCCGCTCGGCT
Oligo rev AAACAGCCGAGCGGCGACCGAGAA
Chromosome start end strand MM target_seq PAM alignment distance position gene name gene id
GU305914.1 185014 185036 + 0 TTCTCGGTCGCCGCTCGGCT CGG ||||||||[||||||||||||]PAM NA Intergenic NA NA

T14046 CTCGGATTCCGAGCAGAAAAAGG 1000
Oligo fwd CACCCTCGGATTCCGAGCAGAAAA
Oligo rev AAACTTTTCTGCTCGGAATCCGAG
Chromosome start end strand MM target_seq PAM alignment distance position gene name gene id
GU305914.1 185032 185054 + 0 CTCGGATTCCGAGCAGAAAA AGG ||||||||[||||||||||||]PAM NA Intergenic NA NA

T14047 TCGGATTCCGAGCAGAAAAAGGG 1000
Oligo fwd CACCTCGGATTCCGAGCAGAAAAA
Oligo rev AAACTTTTTCTGCTCGGAATCCGA
Chromosome start end strand MM target_seq PAM alignment distance position gene name gene id
GU305914.1 185033 185055 + 0 TCGGATTCCGAGCAGAAAAA GGG ||||||||[||||||||||||]PAM NA Intergenic NA NA

T14048 ACCTGCCGAAACGCGTTGTTCGG 1000
Oligo fwd CACCACCTGCCGAAACGCGTTGTT
Oligo rev AAACAACAACGCGTTTCGGCAGGT
Chromosome start end strand MM target_seq PAM alignment distance position gene name gene id
GU305914.1 185072 185094 + 0 ACCTGCCGAAACGCGTTGTT CGG ||||||||[||||||||||||]PAM NA Intergenic NA NA

T14049 ACGCGTTGTTCGGATCATCCCGG 1000
Oligo fwd CACCACGCGTTGTTCGGATCATCC
Oligo rev AAACGGATGATCCGAACAACGCGT
Chromosome start end strand MM target_seq PAM alignment distance position gene name gene id
GU305914.1 185082 185104 + 0 ACGCGTTGTTCGGATCATCC CGG ||||||||[||||||||||||]PAM NA Intergenic NA NA

T14050 CGCGTTGTTCGGATCATCCCGGG 1000
Oligo fwd CACCCGCGTTGTTCGGATCATCCC
Oligo rev AAACGGGATGATCCGAACAACGCG
Chromosome start end strand MM target_seq PAM alignment distance position gene name gene id

GU305914.1 185083 185105 + 0 CGCGTTGTTCGGATCATCCC GGG ||||||||[||||||||||||]PAM NA Intergenic NA NA

T14051 GTTGTTCGGATCATCCCGGGAGG 1000
Oligo fwd CACCGTTGTTCGGATCATCCCGGG
Oligo rev AAACCCCGGGATGATCCGAACAAC
Chromosome start end strand MM target_seq PAM alignment distance position gene name gene id
GU305914.1 185086 185108 + 0 GTTGTTCGGATCATCCCGGG AGG ||||||||[||||||||||||]PAM NA Intergenic NA NA

T14052 TTGTTCGGATCATCCCGGGAGGG 1000
Oligo fwd CACCTTGTTCGGATCATCCCGGGA
Oligo rev AAACTCCCGGGATGATCCGAACAA
Chromosome start end strand MM target_seq PAM alignment distance position gene name gene id
GU305914.1 185087 185109 + 0 TTGTTCGGATCATCCCGGGA GGG ||||||||[||||||||||||]PAM NA Intergenic NA NA

T14053 GCCAGCGTAGTCTCCGTTGCTGG 1000
Oligo fwd CACCGCCAGCGTAGTCTCCGTTGC
Oligo rev AAACGCAACGGAGACTACGCTGGC
Chromosome start end strand MM target_seq PAM alignment distance position gene name gene id
GU305914.1 185109 185131 + 0 GCCAGCGTAGTCTCCGTTGC TGG ||||||||[||||||||||||]PAM NA Intergenic NA NA

T14054 CCAGCGTAGTCTCCGTTGCTGGG 1000
Oligo fwd CACCCCAGCGTAGTCTCCGTTGCT
Oligo rev AAACAGCAACGGAGACTACGCTGG
Chromosome start end strand MM target_seq PAM alignment distance position gene name gene id
GU305914.1 185110 185132 + 0 CCAGCGTAGTCTCCGTTGCT GGG ||||||||[||||||||||||]PAM NA Intergenic NA NA

T14055 GTAGTCTCCGTTGCTGGGTTCGG 1000
Oligo fwd CACCGTAGTCTCCGTTGCTGGGTT
Oligo rev AAACAACCCAGCAACGGAGACTAC
Chromosome start end strand MM target_seq PAM alignment distance position gene name gene id
GU305914.1 185115 185137 + 0 GTAGTCTCCGTTGCTGGGTT CGG ||||||||[||||||||||||]PAM NA Intergenic NA NA

T14056 TCCGTTGCTGGGTTCGGCCGAGG 1000
Oligo fwd CACCTCCGTTGCTGGGTTCGGCCG
Oligo rev AAACCGGCCGAACCCAGCAACGGA
Chromosome start end strand MM target_seq PAM alignment distance position gene name gene id
GU305914.1 185121 185143 + 0 TCCGTTGCTGGGTTCGGCCG AGG ||||||||[||||||||||||]PAM NA Intergenic NA NA

T14057 CCGTTGCTGGGTTCGGCCGAGGG 1000
Oligo fwd CACCCCGTTGCTGGGTTCGGCCGA
Oligo rev AAACTCGGCCGAACCCAGCAACGG
Chromosome start end strand MM target_seq PAM alignment distance position gene name gene id
GU305914.1 185122 185144 + 0 CCGTTGCTGGGTTCGGCCGA GGG ||||||||[||||||||||||]PAM NA Intergenic NA NA

T14058 GGCCGAGGGATCGAGTCGAGCGG 1000
Oligo fwd CACCGGCCGAGGGATCGAGTCGAG
Oligo rev AAACCTCGACTCGATCCCTCGGCC
Chromosome start end strand MM target_seq PAM alignment distance position gene name gene id
GU305914.1 185136 185158 + 0 GGCCGAGGGATCGAGTCGAG CGG ||||||||[||||||||||||]PAM NA Intergenic NA NA

T14059 GCCGAGGGATCGAGTCGAGCGGG 1000
Oligo fwd CACCGCCGAGGGATCGAGTCGAGC
Oligo rev AAACGCTCGACTCGATCCCTCGGC
Chromosome start end strand MM target_seq PAM alignment distance position gene name gene id
GU305914.1 185137 185159 + 0 GCCGAGGGATCGAGTCGAGC GGG ||||||||[||||||||||||]PAM NA Intergenic NA NA

T14060 GAGGGATCGAGTCGAGCGGGCGG 1000
Oligo fwd CACCGAGGGATCGAGTCGAGCGGG
Oligo rev AAACCCCGCTCGACTCGATCCCTC
Chromosome start end strand MM target_seq PAM alignment distance position gene name gene id
GU305914.1 185140 185162 + 0 GAGGGATCGAGTCGAGCGGG CGG ||||||||[||||||||||||]PAM NA Intergenic NA NA

T14061 GGATCGAGTCGAGCGGGCGGCGG 1000
Oligo fwd CACCGGATCGAGTCGAGCGGGCGG
Oligo rev AAACCCGCCCGCTCGACTCGATCC
Chromosome start end strand MM target_seq PAM alignment distance position gene name gene id
GU305914.1 185143 185165 + 0 GGATCGAGTCGAGCGGGCGG CGG ||||||||[||||||||||||]PAM NA Intergenic NA NA

T14062 GATCGAGTCGAGCGGGCGGCGGG 1000
Oligo fwd CACCGATCGAGTCGAGCGGGCGGC
Oligo rev AAACGCCGCCCGCTCGACTCGATC
Chromosome start end strand MM target_seq PAM alignment distance position gene name gene id
GU305914.1 185144 185166 + 0 GATCGAGTCGAGCGGGCGGC GGG ||||||||[||||||||||||]PAM NA Intergenic NA NA

T14063 TCGAGCGGGCGGCGGGACGCCGG 1000
Oligo fwd CACCTCGAGCGGGCGGCGGGACGC
Oligo rev AAACGCGTCCCGCCGCCCGCTCGA
Chromosome start end strand MM target_seq PAM alignment distance position gene name gene id
GU305914.1 185151 185173 + 0 TCGAGCGGGCGGCGGGACGC CGG ||||||||[||||||||||||]PAM NA Intergenic NA NA

T14064 CGAGCGGGCGGCGGGACGCCGGG 1000
Oligo fwd CACCCGAGCGGGCGGCGGGACGCC
Oligo rev AAACGGCGTCCCGCCGCCCGCTCG
Chromosome start end strand MM target_seq PAM alignment distance position gene name gene id
GU305914.1 185152 185174 + 0 CGAGCGGGCGGCGGGACGCC GGG ||||||||[||||||||||||]PAM NA Intergenic NA NA

T14065 CGGGACGCCGGGAACGAGAGAGG 1000
Oligo fwd CACCCGGGACGCCGGGAACGAGAG
Oligo rev AAACCTCTCGTTCCCGGCGTCCCG

<table>
<tr><td>Chromosome</td><td>start</td><td>end</td><td>strand</td><td>MM</td><td>target_seq</td><td>PAM</td><td>alignment</td><td>distance</td><td>position</td><td>gene name</td><td>gene id</td></tr>
<tr><td>GU305914.1</td><td>185163</td><td>185185</td><td>+</td><td>0</td><td>CGGGACGCCGGGAACGAGAG</td><td>AGG</td><td>||||||||[||||||||||||]PAM</td><td>NA</td><td>Intergenic</td><td>NA</td><td>NA</td></tr>
<tr><td>T14066</td><td>CGCCGGGAACGAGAGAGGATCGG</td><td>1000</td><td></td><td></td><td></td><td></td><td></td><td></td><td></td><td></td><td></td></tr>
<tr><td>Oligo fwd</td><td>CACCCGCCGGGAACGAGAGAGGAT</td><td></td><td></td><td></td><td></td><td></td><td></td><td></td><td></td><td></td><td></td></tr>
<tr><td>Oligo rev</td><td>AAACATCCTCTCTCGTTCCCGGCG</td><td></td><td></td><td></td><td></td><td></td><td></td><td></td><td></td><td></td><td></td></tr>
<tr><td>Chromosome</td><td>start</td><td>end</td><td>strand</td><td>MM</td><td>target_seq</td><td>PAM</td><td>alignment</td><td>distance</td><td>position</td><td>gene name</td><td>gene id</td></tr>
<tr><td>GU305914.1</td><td>185168</td><td>185190</td><td>+</td><td>0</td><td>CGCCGGGAACGAGAGAGGAT</td><td>CGG</td><td>||||||||[||||||||||||]PAM</td><td>NA</td><td>Intergenic</td><td>NA</td><td>NA</td></tr>
<tr><td>T14067</td><td>GCCGGGAACGAGAGAGGATCGGG</td><td>1000</td><td></td><td></td><td></td><td></td><td></td><td></td><td></td><td></td><td></td></tr>
<tr><td>Oligo fwd</td><td>CACCGCCGGGAACGAGAGAGGATC</td><td></td><td></td><td></td><td></td><td></td><td></td><td></td><td></td><td></td><td></td></tr>
<tr><td>Oligo rev</td><td>AAACGATCCTCTCTCGTTCCCGGC</td><td></td><td></td><td></td><td></td><td></td><td></td><td></td><td></td><td></td><td></td></tr>
<tr><td>Chromosome</td><td>start</td><td>end</td><td>strand</td><td>MM</td><td>target_seq</td><td>PAM</td><td>alignment</td><td>distance</td><td>position</td><td>gene name</td><td>gene id</td></tr>
<tr><td>GU305914.1</td><td>185169</td><td>185191</td><td>+</td><td>0</td><td>GCCGGGAACGAGAGAGGATC</td><td>GGG</td><td>||||||||[||||||||||||]PAM</td><td>NA</td><td>Intergenic</td><td>NA</td><td>NA</td></tr>
<tr><td>T14068</td><td>GAACGAGAGAGGATCGGGAGCGG</td><td>1000</td><td></td><td></td><td></td><td></td><td></td><td></td><td></td><td></td><td></td></tr>
<tr><td>Oligo fwd</td><td>CACCGAACGAGAGAGGATCGGGAG</td><td></td><td></td><td></td><td></td><td></td><td></td><td></td><td></td><td></td><td></td></tr>
<tr><td>Oligo rev</td><td>AAACCTCCCGATCCTCTCTCGTTC</td><td></td><td></td><td></td><td></td><td></td><td></td><td></td><td></td><td></td><td></td></tr>
<tr><td>Chromosome</td><td>start</td><td>end</td><td>strand</td><td>MM</td><td>target_seq</td><td>PAM</td><td>alignment</td><td>distance</td><td>position</td><td>gene name</td><td>gene id</td></tr>
<tr><td>GU305914.1</td><td>185174</td><td>185196</td><td>+</td><td>0</td><td>GAACGAGAGAGGATCGGGAG</td><td>CGG</td><td>||||||||[||||||||||||]PAM</td><td>NA</td><td>Intergenic</td><td>NA</td><td>NA</td></tr>
<tr><td>T14069</td><td>AACGAGAGAGGATCGGGAGCGGG</td><td>1000</td><td></td><td></td><td></td><td></td><td></td><td></td><td></td><td></td><td></td></tr>
<tr><td>Oligo fwd</td><td>CACCAACGAGAGAGGATCGGGAGC</td><td></td><td></td><td></td><td></td><td></td><td></td><td></td><td></td><td></td><td></td></tr>
<tr><td>Oligo rev</td><td>AAACGCTCCCGATCCTCTCTCGTT</td><td></td><td></td><td></td><td></td><td></td><td></td><td></td><td></td><td></td><td></td></tr>
<tr><td>Chromosome</td><td>start</td><td>end</td><td>strand</td><td>MM</td><td>target_seq</td><td>PAM</td><td>alignment</td><td>distance</td><td>position</td><td>gene name</td><td>gene id</td></tr>
<tr><td>GU305914.1</td><td>185175</td><td>185197</td><td>+</td><td>0</td><td>AACGAGAGAGGATCGGGAGC</td><td>GGG</td><td>||||||||[||||||||||||]PAM</td><td>NA</td><td>Intergenic</td><td>NA</td><td>NA</td></tr>
<tr><td>T14070</td><td>GAGGATCGGGAGCGGGCGCCCGG</td><td>1000</td><td></td><td></td><td></td><td></td><td></td><td></td><td></td><td></td><td></td></tr>
<tr><td>Oligo fwd</td><td>CACCGAGGATCGGGAGCGGGCGCC</td><td></td><td></td><td></td><td></td><td></td><td></td><td></td><td></td><td></td><td></td></tr>
<tr><td>Oligo rev</td><td>AAACGGCGCCCGCTCCCGATCCTC</td><td></td><td></td><td></td><td></td><td></td><td></td><td></td><td></td><td></td><td></td></tr>
<tr><td>Chromosome</td><td>start</td><td>end</td><td>strand</td><td>MM</td><td>target_seq</td><td>PAM</td><td>alignment</td><td>distance</td><td>position</td><td>gene name</td><td>gene id</td></tr>
<tr><td>GU305914.1</td><td>185182</td><td>185204</td><td>+</td><td>0</td><td>GAGGATCGGGAGCGGGCGCC</td><td>CGG</td><td>||||||||[||||||||||||]PAM</td><td>NA</td><td>Intergenic</td><td>NA</td><td>NA</td></tr>
<tr><td>T14071</td><td>AGGATCGGGAGCGGGCGCCCGGG</td><td>1000</td><td></td><td></td><td></td><td></td><td></td><td></td><td></td><td></td><td></td></tr>
<tr><td>Oligo fwd</td><td>CACCAGGATCGGGAGCGGGCGCCC</td><td></td><td></td><td></td><td></td><td></td><td></td><td></td><td></td><td></td><td></td></tr>
<tr><td>Oligo rev</td><td>AAACGGGCGCCCGCTCCCGATCCT</td><td></td><td></td><td></td><td></td><td></td><td></td><td></td><td></td><td></td><td></td></tr>
<tr><td>Chromosome</td><td>start</td><td>end</td><td>strand</td><td>MM</td><td>target_seq</td><td>PAM</td><td>alignment</td><td>distance</td><td>position</td><td>gene name</td><td>gene id</td></tr>
<tr><td>GU305914.1</td><td>185183</td><td>185205</td><td>+</td><td>0</td><td>AGGATCGGGAGCGGGCGCCC</td><td>GGG</td><td>||||||||[||||||||||||]PAM</td><td>NA</td><td>Intergenic</td><td>NA</td><td>NA</td></tr>
<tr><td>T14072</td><td>ATCGGGAGCGGGCGCCCGGGTGG</td><td>1000</td><td></td><td></td><td></td><td></td><td></td><td></td><td></td><td></td><td></td></tr>
<tr><td>Oligo fwd</td><td>CACCATCGGGAGCGGGCGCCCGGG</td><td></td><td></td><td></td><td></td><td></td><td></td><td></td><td></td><td></td><td></td></tr>
<tr><td>Oligo rev</td><td>AAACCCCGGGCGCCCGCTCCCGAT</td><td></td><td></td><td></td><td></td><td></td><td></td><td></td><td></td><td></td><td></td></tr>
<tr><td>Chromosome</td><td>start</td><td>end</td><td>strand</td><td>MM</td><td>target_seq</td><td>PAM</td><td>alignment</td><td>distance</td><td>position</td><td>gene name</td><td>gene id</td></tr>
<tr><td>GU305914.1</td><td>185186</td><td>185208</td><td>+</td><td>0</td><td>ATCGGGAGCGGGCGCCCGGG</td><td>TGG</td><td>||||||||[||||||||||||]PAM</td><td>NA</td><td>Intergenic</td><td>NA</td><td>NA</td></tr>
<tr><td>T14073</td><td>GAGCGGGCGCCCGGGTGGCTCGG</td><td>1000</td><td></td><td></td><td></td><td></td><td></td><td></td><td></td><td></td><td></td></tr>
<tr><td>Oligo fwd</td><td>CACCGAGCGGGCGCCCGGGTGGCT</td><td></td><td></td><td></td><td></td><td></td><td></td><td></td><td></td><td></td><td></td></tr>
<tr><td>Oligo rev</td><td>AAACAGCCACCCGGGCGCCCGCTC</td><td></td><td></td><td></td><td></td><td></td><td></td><td></td><td></td><td></td><td></td></tr>
<tr><td>Chromosome</td><td>start</td><td>end</td><td>strand</td><td>MM</td><td>target_seq</td><td>PAM</td><td>alignment</td><td>distance</td><td>position</td><td>gene name</td><td>gene id</td></tr>
<tr><td>GU305914.1</td><td>185191</td><td>185213</td><td>+</td><td>0</td><td>GAGCGGGCGCCCGGGTGGCT</td><td>CGG</td><td>||||||||[||||||||||||]PAM</td><td>NA</td><td>Intergenic</td><td>NA</td><td>NA</td></tr>
<tr><td>T14074</td><td>AGCGGGCGCCCGGGTGGCTCGGG</td><td>1000</td><td></td><td></td><td></td><td></td><td></td><td></td><td></td><td></td><td></td></tr>
<tr><td>Oligo fwd</td><td>CACCAGCGGGCGCCCGGGTGGCTC</td><td></td><td></td><td></td><td></td><td></td><td></td><td></td><td></td><td></td><td></td></tr>
<tr><td>Oligo rev</td><td>AAACGAGCCACCCGGGCGCCCGCT</td><td></td><td></td><td></td><td></td><td></td><td></td><td></td><td></td><td></td><td></td></tr>
<tr><td>Chromosome</td><td>start</td><td>end</td><td>strand</td><td>MM</td><td>target_seq</td><td>PAM</td><td>alignment</td><td>distance</td><td>position</td><td>gene name</td><td>gene id</td></tr>
<tr><td>GU305914.1</td><td>185192</td><td>185214</td><td>+</td><td>0</td><td>AGCGGGCGCCCGGGTGGCTC</td><td>GGG</td><td>||||||||[||||||||||||]PAM</td><td>NA</td><td>Intergenic</td><td>NA</td><td>NA</td></tr>
<tr><td>GU305914.1</td><td>164854</td><td>164876</td><td>-</td><td>4</td><td>AGCGGCGGCCGTGGTGGCTC</td><td>TGG</td><td>|||||--|[||--||||||||]PAM</td><td>NA</td><td>Intergenic</td><td>NA</td><td>NA</td></tr>
<tr><td>T14075</td><td>GCGCCCGGGTGGCTCGGGAGCGG</td><td>1000</td><td></td><td></td><td></td><td></td><td></td><td></td><td></td><td></td><td></td></tr>
<tr><td>Oligo fwd</td><td>CACCGCGCCCGGGTGGCTCGGGAG</td><td></td><td></td><td></td><td></td><td></td><td></td><td></td><td></td><td></td><td></td></tr>
<tr><td>Oligo rev</td><td>AAACCTCCCGAGCCACCCGGGCGC</td><td></td><td></td><td></td><td></td><td></td><td></td><td></td><td></td><td></td><td></td></tr>
<tr><td>Chromosome</td><td>start</td><td>end</td><td>strand</td><td>MM</td><td>target_seq</td><td>PAM</td><td>alignment</td><td>distance</td><td>position</td><td>gene name</td><td>gene id</td></tr>
<tr><td>GU305914.1</td><td>185197</td><td>185219</td><td>+</td><td>0</td><td>GCGCCCGGGTGGCTCGGGAG</td><td>CGG</td><td>||||||||[||||||||||||]PAM</td><td>NA</td><td>Intergenic</td><td>NA</td><td>NA</td></tr>
<tr><td>T14076</td><td>CCGGGTGGCTCGGGAGCGGCCGG</td><td>1000</td><td></td><td></td><td></td><td></td><td></td><td></td><td></td><td></td><td></td></tr>
<tr><td>Oligo fwd</td><td>CACCCCGGGTGGCTCGGGAGCGGC</td><td></td><td></td><td></td><td></td><td></td><td></td><td></td><td></td><td></td><td></td></tr>
<tr><td>Oligo rev</td><td>AAACGCCGCTCCCGAGCCACCCGG</td><td></td><td></td><td></td><td></td><td></td><td></td><td></td><td></td><td></td><td></td></tr>
<tr><td>Chromosome</td><td>start</td><td>end</td><td>strand</td><td>MM</td><td>target_seq</td><td>PAM</td><td>alignment</td><td>distance</td><td>position</td><td>gene name</td><td>gene id</td></tr>
<tr><td>GU305914.1</td><td>185201</td><td>185223</td><td>+</td><td>0</td><td>CCGGGTGGCTCGGGAGCGGC</td><td>CGG</td><td>||||||||[||||||||||||]PAM</td><td>NA</td><td>Intergenic</td><td>NA</td><td>NA</td></tr>
<tr><td>T14077</td><td>CGGGTGGCTCGGGAGCGGCCGGG</td><td>1000</td><td></td><td></td><td></td><td></td><td></td><td></td><td></td><td></td><td></td></tr>
<tr><td>Oligo fwd</td><td>CACCCGGGTGGCTCGGGAGCGGCC</td><td></td><td></td><td></td><td></td><td></td><td></td><td></td><td></td><td></td><td></td></tr>
<tr><td>Oligo rev</td><td>AAACGGCCGCTCCCGAGCCACCCG</td><td></td><td></td><td></td><td></td><td></td><td></td><td></td><td></td><td></td><td></td></tr>
<tr><td>Chromosome</td><td>start</td><td>end</td><td>strand</td><td>MM</td><td>target_seq</td><td>PAM</td><td>alignment</td><td>distance</td><td>position</td><td>gene name</td><td>gene id</td></tr>
<tr><td>GU305914.1</td><td>185202</td><td>185224</td><td>+</td><td>0</td><td>CGGGTGGCTCGGGAGCGGCC</td><td>GGG</td><td>||||||||[||||||||||||]PAM</td><td>NA</td><td>Intergenic</td><td>NA</td><td>NA</td></tr>
<tr><td>T14078</td><td>GCTCGGGAGCGGCCGGGAGACGG</td><td>1000</td><td></td><td></td><td></td><td></td><td></td><td></td><td></td><td></td><td></td></tr>
<tr><td>Oligo fwd</td><td>CACCGCTCGGGAGCGGCCGGGAGA</td><td></td><td></td><td></td><td></td><td></td><td></td><td></td><td></td><td></td><td></td></tr>
<tr><td>Oligo rev</td><td>AAACTCTCCCGGCCGCTCCCGAGC</td><td></td><td></td><td></td><td></td><td></td><td></td><td></td><td></td><td></td><td></td></tr>
<tr><td>Chromosome</td><td>start</td><td>end</td><td>strand</td><td>MM</td><td>target_seq</td><td>PAM</td><td>alignment</td><td>distance</td><td>position</td><td>gene name</td><td>gene id</td></tr>
<tr><td>GU305914.1</td><td>185208</td><td>185230</td><td>+</td><td>0</td><td>GCTCGGGAGCGGCCGGGAGA</td><td>CGG</td><td>||||||||[||||||||||||]PAM</td><td>NA</td><td>Intergenic</td><td>NA</td><td>NA</td></tr>
<tr><td>T14079</td><td>GTGTCCTGACACTGAACAGTTGG</td><td>1000</td><td></td><td></td><td></td><td></td><td></td><td></td><td></td><td></td><td></td></tr>
<tr><td>Oligo fwd</td><td>CACCGTGTCCTGACACTGAACAGT</td><td></td><td></td><td></td><td></td><td></td><td></td><td></td><td></td><td></td><td></td></tr>
<tr><td>Oligo rev</td><td>AAACACTGTTCAGTGTCAGGACAC</td><td></td><td></td><td></td><td></td><td></td><td></td><td></td><td></td><td></td><td></td></tr>
<tr><td>Chromosome</td><td>start</td><td>end</td><td>strand</td><td>MM</td><td>target_seq</td><td>PAM</td><td>alignment</td><td>distance</td><td>position</td><td>gene name</td><td>gene id</td></tr>
<tr><td>GU305914.1</td><td>185252</td><td>185274</td><td>+</td><td>0</td><td>GTGTCCTGACACTGAACAGT</td><td>TGG</td><td>||||||||[||||||||||||]PAM</td><td>NA</td><td>Intergenic</td><td>NA</td><td>NA</td></tr>
<tr><td>T14080</td><td>TGTCCTGACACTGAACAGTTGGG</td><td>1000</td><td></td><td></td><td></td><td></td><td></td><td></td><td></td><td></td><td></td></tr>
</table>

<table>
<tr><td>Oligo fwd</td><td>CACCTGTCCTGACACTGAACAGTT</td><td></td><td></td><td></td><td></td><td></td><td></td><td></td><td></td><td></td><td></td><td></td></tr>
<tr><td>Oligo rev</td><td>AAACAACTGTTCAGTGTCAGGACA</td><td></td><td></td><td></td><td></td><td></td><td></td><td></td><td></td><td></td><td></td><td></td></tr>
<tr><td>Chromosome</td><td>start</td><td></td><td>end</td><td>strand</td><td>MM</td><td>target_seq</td><td>PAM</td><td>alignment</td><td>distance</td><td>position</td><td>gene name</td><td>gene id</td></tr>
<tr><td>GU305914.1</td><td></td><td>185253</td><td>185275</td><td>+</td><td>0</td><td>TGTCCTGACACTGAACAGTT</td><td>GGG</td><td>||||||||[||||||||||||]PAM</td><td>NA</td><td>Intergenic</td><td>NA</td><td>NA</td></tr>
<tr><td>T14081</td><td>TGAACAGTTGGGAAAAATACTGG</td><td></td><td>1000</td><td></td><td></td><td></td><td></td><td></td><td></td><td></td><td></td><td></td></tr>
<tr><td>Oligo fwd</td><td>CACCTGAACAGTTGGGAAAAATAC</td><td></td><td></td><td></td><td></td><td></td><td></td><td></td><td></td><td></td><td></td><td></td></tr>
<tr><td>Oligo rev</td><td>AAACGTATTTTTCCCAACTGTTCA</td><td></td><td></td><td></td><td></td><td></td><td></td><td></td><td></td><td></td><td></td><td></td></tr>
<tr><td>Chromosome</td><td>start</td><td></td><td>end</td><td>strand</td><td>MM</td><td>target_seq</td><td>PAM</td><td>alignment</td><td>distance</td><td>position</td><td>gene name</td><td>gene id</td></tr>
<tr><td>GU305914.1</td><td></td><td>185264</td><td>185286</td><td>+</td><td>0</td><td>TGAACAGTTGGGAAAAATAC</td><td>TGG</td><td>||||||||[||||||||||||]PAM</td><td>NA</td><td>Intergenic</td><td>NA</td><td>NA</td></tr>
<tr><td>T14082</td><td>ACTGGATCTTATAATGACCCAGG</td><td></td><td>1000</td><td></td><td></td><td></td><td></td><td></td><td></td><td></td><td></td><td></td></tr>
<tr><td>Oligo fwd</td><td>CACCACTGGATCTTATAATGACCC</td><td></td><td></td><td></td><td></td><td></td><td></td><td></td><td></td><td></td><td></td><td></td></tr>
<tr><td>Oligo rev</td><td>AAACGGGTCATTATAAGATCCAGT</td><td></td><td></td><td></td><td></td><td></td><td></td><td></td><td></td><td></td><td></td><td></td></tr>
<tr><td>Chromosome</td><td>start</td><td></td><td>end</td><td>strand</td><td>MM</td><td>target_seq</td><td>PAM</td><td>alignment</td><td>distance</td><td>position</td><td>gene name</td><td>gene id</td></tr>
<tr><td>GU305914.1</td><td></td><td>185282</td><td>185304</td><td>+</td><td>0</td><td>ACTGGATCTTATAATGACCC</td><td>AGG</td><td>||||||||[||||||||||||]PAM</td><td>NA</td><td>Intergenic</td><td>NA</td><td>NA</td></tr>
<tr><td>T14083</td><td>CTGGATCTTATAATGACCCAGGG</td><td></td><td>1000</td><td></td><td></td><td></td><td></td><td></td><td></td><td></td><td></td><td></td></tr>
<tr><td>Oligo fwd</td><td>CACCCTGGATCTTATAATGACCCA</td><td></td><td></td><td></td><td></td><td></td><td></td><td></td><td></td><td></td><td></td><td></td></tr>
<tr><td>Oligo rev</td><td>AAACTGGGTCATTATAAGATCCAG</td><td></td><td></td><td></td><td></td><td></td><td></td><td></td><td></td><td></td><td></td><td></td></tr>
<tr><td>Chromosome</td><td>start</td><td></td><td>end</td><td>strand</td><td>MM</td><td>target_seq</td><td>PAM</td><td>alignment</td><td>distance</td><td>position</td><td>gene name</td><td>gene id</td></tr>
<tr><td>GU305914.1</td><td></td><td>185283</td><td>185305</td><td>+</td><td>0</td><td>CTGGATCTTATAATGACCCA</td><td>GGG</td><td>||||||||[||||||||||||]PAM</td><td>NA</td><td>Intergenic</td><td>NA</td><td>NA</td></tr>
<tr><td>T14084</td><td>TGGATCTTATAATGACCCAGGGG</td><td></td><td>1000</td><td></td><td></td><td></td><td></td><td></td><td></td><td></td><td></td><td></td></tr>
<tr><td>Oligo fwd</td><td>CACCTGGATCTTATAATGACCCAG</td><td></td><td></td><td></td><td></td><td></td><td></td><td></td><td></td><td></td><td></td><td></td></tr>
<tr><td>Oligo rev</td><td>AAACCTGGGTCATTATAAGATCCA</td><td></td><td></td><td></td><td></td><td></td><td></td><td></td><td></td><td></td><td></td><td></td></tr>
<tr><td>Chromosome</td><td>start</td><td></td><td>end</td><td>strand</td><td>MM</td><td>target_seq</td><td>PAM</td><td>alignment</td><td>distance</td><td>position</td><td>gene name</td><td>gene id</td></tr>
<tr><td>GU305914.1</td><td></td><td>185284</td><td>185306</td><td>+</td><td>0</td><td>TGGATCTTATAATGACCCAG</td><td>GGG</td><td>||||||||[||||||||||||]PAM</td><td>NA</td><td>Intergenic</td><td>NA</td><td>NA</td></tr>
<tr><td>T14085</td><td>TCTTATAATGACCCAGGGGAAGG</td><td></td><td>1000</td><td></td><td></td><td></td><td></td><td></td><td></td><td></td><td></td><td></td></tr>
<tr><td>Oligo fwd</td><td>CACCTCTTATAATGACCCAGGGGA</td><td></td><td></td><td></td><td></td><td></td><td></td><td></td><td></td><td></td><td></td><td></td></tr>
<tr><td>Oligo rev</td><td>AAACTCCCCTGGGTCATTATAAGA</td><td></td><td></td><td></td><td></td><td></td><td></td><td></td><td></td><td></td><td></td><td></td></tr>
<tr><td>Chromosome</td><td>start</td><td></td><td>end</td><td>strand</td><td>MM</td><td>target_seq</td><td>PAM</td><td>alignment</td><td>distance</td><td>position</td><td>gene name</td><td>gene id</td></tr>
<tr><td>GU305914.1</td><td></td><td>185288</td><td>185310</td><td>+</td><td>0</td><td>TCTTATAATGACCCAGGGGA</td><td>AGG</td><td>||||||||[||||||||||||]PAM</td><td>NA</td><td>Intergenic</td><td>NA</td><td>NA</td></tr>
<tr><td>T14086</td><td>CTTATAATGACCCAGGGGAAGGG</td><td></td><td>1000</td><td></td><td></td><td></td><td></td><td></td><td></td><td></td><td></td><td></td></tr>
<tr><td>Oligo fwd</td><td>CACCCTTATAATGACCCAGGGGAA</td><td></td><td></td><td></td><td></td><td></td><td></td><td></td><td></td><td></td><td></td><td></td></tr>
<tr><td>Oligo rev</td><td>AAACTTCCCCTGGGTCATTATAAG</td><td></td><td></td><td></td><td></td><td></td><td></td><td></td><td></td><td></td><td></td><td></td></tr>
<tr><td>Chromosome</td><td>start</td><td></td><td>end</td><td>strand</td><td>MM</td><td>target_seq</td><td>PAM</td><td>alignment</td><td>distance</td><td>position</td><td>gene name</td><td>gene id</td></tr>
<tr><td>GU305914.1</td><td></td><td>185289</td><td>185311</td><td>+</td><td>0</td><td>CTTATAATGACCCAGGGGAA</td><td>GGG</td><td>||||||||[||||||||||||]PAM</td><td>NA</td><td>Intergenic</td><td>NA</td><td>NA</td></tr>
<tr><td>T14087</td><td>TTATAATGACCCAGGGGAAGGGG</td><td></td><td>1000</td><td></td><td></td><td></td><td></td><td></td><td></td><td></td><td></td><td></td></tr>
<tr><td>Oligo fwd</td><td>CACCTTATAATGACCCAGGGGAAG</td><td></td><td></td><td></td><td></td><td></td><td></td><td></td><td></td><td></td><td></td><td></td></tr>
<tr><td>Oligo rev</td><td>AAACCTTCCCCTGGGTCATTATAA</td><td></td><td></td><td></td><td></td><td></td><td></td><td></td><td></td><td></td><td></td><td></td></tr>
<tr><td>Chromosome</td><td>start</td><td></td><td>end</td><td>strand</td><td>MM</td><td>target_seq</td><td>PAM</td><td>alignment</td><td>distance</td><td>position</td><td>gene name</td><td>gene id</td></tr>
<tr><td>GU305914.1</td><td></td><td>185290</td><td>185312</td><td>+</td><td>0</td><td>TTATAATGACCCAGGGGAAG</td><td>GGG</td><td>||||||||[||||||||||||]PAM</td><td>NA</td><td>Intergenic</td><td>NA</td><td>NA</td></tr>
<tr><td>T14088</td><td>TTGATAGCAGATGTCCTCTTTGG</td><td></td><td>1000</td><td></td><td></td><td></td><td></td><td></td><td></td><td></td><td></td><td></td></tr>
<tr><td>Oligo fwd</td><td>CACCTTGATAGCAGATGTCCTCTT</td><td></td><td></td><td></td><td></td><td></td><td></td><td></td><td></td><td></td><td></td><td></td></tr>
<tr><td>Oligo rev</td><td>AAACAAGAGGACATCTGCTATCAA</td><td></td><td></td><td></td><td></td><td></td><td></td><td></td><td></td><td></td><td></td><td></td></tr>
<tr><td>Chromosome</td><td>start</td><td></td><td>end</td><td>strand</td><td>MM</td><td>target_seq</td><td>PAM</td><td>alignment</td><td>distance</td><td>position</td><td>gene name</td><td>gene id</td></tr>
<tr><td>GU305914.1</td><td></td><td>185425</td><td>185447</td><td>+</td><td>0</td><td>TTGATAGCAGATGTCCTCTT</td><td>TGG</td><td>||||||||[||||||||||||]PAM</td><td>NA</td><td>Intergenic</td><td>NA</td><td>NA</td></tr>
<tr><td>T14089</td><td>TAGCAGATGTCCTCTTTGGAAGG</td><td></td><td>1000</td><td></td><td></td><td></td><td></td><td></td><td></td><td></td><td></td><td></td></tr>
<tr><td>Oligo fwd</td><td>CACCTAGCAGATGTCCTCTTTGGA</td><td></td><td></td><td></td><td></td><td></td><td></td><td></td><td></td><td></td><td></td><td></td></tr>
<tr><td>Oligo rev</td><td>AAACTCCAAAGAGGACATCTGCTA</td><td></td><td></td><td></td><td></td><td></td><td></td><td></td><td></td><td></td><td></td><td></td></tr>
<tr><td>Chromosome</td><td>start</td><td></td><td>end</td><td>strand</td><td>MM</td><td>target_seq</td><td>PAM</td><td>alignment</td><td>distance</td><td>position</td><td>gene name</td><td>gene id</td></tr>
<tr><td>GU305914.1</td><td></td><td>185429</td><td>185451</td><td>+</td><td>0</td><td>TAGCAGATGTCCTCTTTGGA</td><td>AGG</td><td>||||||||[||||||||||||]PAM</td><td>NA</td><td>Intergenic</td><td>NA</td><td>NA</td></tr>
<tr><td>T14090</td><td>ACAAACACCGTTCCTCACTGAGG</td><td></td><td>1000</td><td></td><td></td><td></td><td></td><td></td><td></td><td></td><td></td><td></td></tr>
<tr><td>Oligo fwd</td><td>CACCACAAACACCGTTCCTCACTG</td><td></td><td></td><td></td><td></td><td></td><td></td><td></td><td></td><td></td><td></td><td></td></tr>
<tr><td>Oligo rev</td><td>AAACCAGTGAGGAACGGTGTTTGT</td><td></td><td></td><td></td><td></td><td></td><td></td><td></td><td></td><td></td><td></td><td></td></tr>
<tr><td>Chromosome</td><td>start</td><td></td><td>end</td><td>strand</td><td>MM</td><td>target_seq</td><td>PAM</td><td>alignment</td><td>distance</td><td>position</td><td>gene name</td><td>gene id</td></tr>
<tr><td>GU305914.1</td><td></td><td>185464</td><td>185486</td><td>+</td><td>0</td><td>ACAAACACCGTTCCTCACTG</td><td>AGG</td><td>||||||||[||||||||||||]PAM</td><td>NA</td><td>Intergenic</td><td>NA</td><td>NA</td></tr>
<tr><td>T14091</td><td>CCTCACTGAGGTGCATTGTCTGG</td><td></td><td>1000</td><td></td><td></td><td></td><td></td><td></td><td></td><td></td><td></td><td></td></tr>
<tr><td>Oligo fwd</td><td>CACCCCTCACTGAGGTGCATTGTC</td><td></td><td></td><td></td><td></td><td></td><td></td><td></td><td></td><td></td><td></td><td></td></tr>
<tr><td>Oligo rev</td><td>AAACGACAATGCACCTCAGTGAGG</td><td></td><td></td><td></td><td></td><td></td><td></td><td></td><td></td><td></td><td></td><td></td></tr>
<tr><td>Chromosome</td><td>start</td><td></td><td>end</td><td>strand</td><td>MM</td><td>target_seq</td><td>PAM</td><td>alignment</td><td>distance</td><td>position</td><td>gene name</td><td>gene id</td></tr>
<tr><td>GU305914.1</td><td></td><td>185476</td><td>185498</td><td>+</td><td>0</td><td>CCTCACTGAGGTGCATTGTC</td><td>TGG</td><td>||||||||[||||||||||||]PAM</td><td>NA</td><td>Intergenic</td><td>NA</td><td>NA</td></tr>
<tr><td>T14092</td><td>CTCACTGAGGTGCATTGTCTGGG</td><td></td><td>1000</td><td></td><td></td><td></td><td></td><td></td><td></td><td></td><td></td><td></td></tr>
<tr><td>Oligo fwd</td><td>CACCCTCACTGAGGTGCATTGTCT</td><td></td><td></td><td></td><td></td><td></td><td></td><td></td><td></td><td></td><td></td><td></td></tr>
<tr><td>Oligo rev</td><td>AAACAGACAATGCACCTCAGTGAG</td><td></td><td></td><td></td><td></td><td></td><td></td><td></td><td></td><td></td><td></td><td></td></tr>
<tr><td>Chromosome</td><td>start</td><td></td><td>end</td><td>strand</td><td>MM</td><td>target_seq</td><td>PAM</td><td>alignment</td><td>distance</td><td>position</td><td>gene name</td><td>gene id</td></tr>
<tr><td>GU305914.1</td><td></td><td>185477</td><td>185499</td><td>+</td><td>0</td><td>CTCACTGAGGTGCATTGTCT</td><td>GGG</td><td>||||||||[||||||||||||]PAM</td><td>NA</td><td>Intergenic</td><td>NA</td><td>NA</td></tr>
<tr><td>T14093</td><td>ACTGAGGTGCATTGTCTGGGAGG</td><td></td><td>1000</td><td></td><td></td><td></td><td></td><td></td><td></td><td></td><td></td><td></td></tr>
<tr><td>Oligo fwd</td><td>CACCACTGAGGTGCATTGTCTGGG</td><td></td><td></td><td></td><td></td><td></td><td></td><td></td><td></td><td></td><td></td><td></td></tr>
<tr><td>Oligo rev</td><td>AAACCCCAGACAATGCACCTCAGT</td><td></td><td></td><td></td><td></td><td></td><td></td><td></td><td></td><td></td><td></td><td></td></tr>
<tr><td>Chromosome</td><td>start</td><td></td><td>end</td><td>strand</td><td>MM</td><td>target_seq</td><td>PAM</td><td>alignment</td><td>distance</td><td>position</td><td>gene name</td><td>gene id</td></tr>
<tr><td>GU305914.1</td><td></td><td>185480</td><td>185502</td><td>+</td><td>0</td><td>ACTGAGGTGCATTGTCTGGG</td><td>AGG</td><td>||||||||[||||||||||||]PAM</td><td>NA</td><td>Intergenic</td><td>NA</td><td>NA</td></tr>
<tr><td>T14094</td><td>CTGAGGTGCATTGTCTGGGAGGG</td><td></td><td>1000</td><td></td><td></td><td></td><td></td><td></td><td></td><td></td><td></td><td></td></tr>
<tr><td>Oligo fwd</td><td>CACCCTGAGGTGCATTGTCTGGGA</td><td></td><td></td><td></td><td></td><td></td><td></td><td></td><td></td><td></td><td></td><td></td></tr>
<tr><td>Oligo rev</td><td>AAACTCCCAGACAATGCACCTCAG</td><td></td><td></td><td></td><td></td><td></td><td></td><td></td><td></td><td></td><td></td><td></td></tr>
<tr><td>Chromosome</td><td>start</td><td></td><td>end</td><td>strand</td><td>MM</td><td>target_seq</td><td>PAM</td><td>alignment</td><td>distance</td><td>position</td><td>gene name</td><td>gene id</td></tr>
<tr><td>GU305914.1</td><td></td><td>185481</td><td>185503</td><td>+</td><td>0</td><td>CTGAGGTGCATTGTCTGGGA</td><td>GGG</td><td>||||||||[||||||||||||]PAM</td><td>NA</td><td>Intergenic</td><td>NA</td><td>NA</td></tr>
</table>

<table>
<tr><td>T14095</td><td>GGTGCATTGTCTGGGAGGGAAGG</td><td>1000</td><td></td><td></td><td></td><td></td><td></td><td></td><td></td><td></td><td></td></tr>
<tr><td>Oligo fwd</td><td>CACCGGTGCATTGTCTGGGAGGGA</td><td></td><td></td><td></td><td></td><td></td><td></td><td></td><td></td><td></td><td></td></tr>
<tr><td>Oligo rev</td><td>AAACTCCCTCCCAGACAATGCACC</td><td></td><td></td><td></td><td></td><td></td><td></td><td></td><td></td><td></td><td></td></tr>
<tr><td>Chromosome</td><td>start</td><td>end</td><td>strand</td><td>MM</td><td>target_seq</td><td>PAM</td><td>alignment</td><td>distance</td><td>position</td><td>gene name</td><td>gene id</td></tr>
<tr><td>GU305914.1</td><td>185485</td><td>185507</td><td>+</td><td>0</td><td>GGTGCATTGTCTGGGAGGGA</td><td>AGG</td><td>||||||||[||||||||||||]PAM</td><td>NA</td><td>Intergenic</td><td>NA</td><td>NA</td></tr>
<tr><td>T14096</td><td>GGTCACCTTCTTGTTCATTGTGG</td><td>1000</td><td></td><td></td><td></td><td></td><td></td><td></td><td></td><td></td><td></td></tr>
<tr><td>Oligo fwd</td><td>CACCGGTCACCTTCTTGTTCATTG</td><td></td><td></td><td></td><td></td><td></td><td></td><td></td><td></td><td></td><td></td></tr>
<tr><td>Oligo rev</td><td>AAACCAATGAACAAGAAGGTGACC</td><td></td><td></td><td></td><td></td><td></td><td></td><td></td><td></td><td></td><td></td></tr>
<tr><td>Chromosome</td><td>start</td><td>end</td><td>strand</td><td>MM</td><td>target_seq</td><td>PAM</td><td>alignment</td><td>distance</td><td>position</td><td>gene name</td><td>gene id</td></tr>
<tr><td>GU305914.1</td><td>185506</td><td>185528</td><td>+</td><td>0</td><td>GGTCACCTTCTTGTTCATTG</td><td>TGG</td><td>||||||||[||||||||||||]PAM</td><td>NA</td><td>Intergenic</td><td>NA</td><td>NA</td></tr>
<tr><td>T14097</td><td>ACCTTCTTGTTCATTGTGGACGG</td><td>1000</td><td></td><td></td><td></td><td></td><td></td><td></td><td></td><td></td><td></td></tr>
<tr><td>Oligo fwd</td><td>CACCACCTTCTTGTTCATTGTGGA</td><td></td><td></td><td></td><td></td><td></td><td></td><td></td><td></td><td></td><td></td></tr>
<tr><td>Oligo rev</td><td>AAACTCCACAATGAACAAGAAGGT</td><td></td><td></td><td></td><td></td><td></td><td></td><td></td><td></td><td></td><td></td></tr>
<tr><td>Chromosome</td><td>start</td><td>end</td><td>strand</td><td>MM</td><td>target_seq</td><td>PAM</td><td>alignment</td><td>distance</td><td>position</td><td>gene name</td><td>gene id</td></tr>
<tr><td>GU305914.1</td><td>185510</td><td>185532</td><td>+</td><td>0</td><td>ACCTTCTTGTTCATTGTGGA</td><td>CGG</td><td>||||||||[||||||||||||]PAM</td><td>NA</td><td>Intergenic</td><td>NA</td><td>NA</td></tr>
<tr><td>T14098</td><td>TGTTCATTGTGGACGGTGTGCGG</td><td>1000</td><td></td><td></td><td></td><td></td><td></td><td></td><td></td><td></td><td></td></tr>
<tr><td>Oligo fwd</td><td>CACCTGTTCATTGTGGACGGTGTG</td><td></td><td></td><td></td><td></td><td></td><td></td><td></td><td></td><td></td><td></td></tr>
<tr><td>Oligo rev</td><td>AAACCACACCGTCCACAATGAACA</td><td></td><td></td><td></td><td></td><td></td><td></td><td></td><td></td><td></td><td></td></tr>
<tr><td>Chromosome</td><td>start</td><td>end</td><td>strand</td><td>MM</td><td>target_seq</td><td>PAM</td><td>alignment</td><td>distance</td><td>position</td><td>gene name</td><td>gene id</td></tr>
<tr><td>GU305914.1</td><td>185517</td><td>185539</td><td>+</td><td>0</td><td>TGTTCATTGTGGACGGTGTG</td><td>CGG</td><td>||||||||[||||||||||||]PAM</td><td>NA</td><td>Intergenic</td><td>NA</td><td>NA</td></tr>
<tr><td>T14099</td><td>TTGTGGACGGTGTGCGGCCCAGG</td><td>1000</td><td></td><td></td><td></td><td></td><td></td><td></td><td></td><td></td><td></td></tr>
<tr><td>Oligo fwd</td><td>CACCTTGTGGACGGTGTGCGGCCC</td><td></td><td></td><td></td><td></td><td></td><td></td><td></td><td></td><td></td><td></td></tr>
<tr><td>Oligo rev</td><td>AAACGGGCCGCACACCGTCCACAA</td><td></td><td></td><td></td><td></td><td></td><td></td><td></td><td></td><td></td><td></td></tr>
<tr><td>Chromosome</td><td>start</td><td>end</td><td>strand</td><td>MM</td><td>target_seq</td><td>PAM</td><td>alignment</td><td>distance</td><td>position</td><td>gene name</td><td>gene id</td></tr>
<tr><td>GU305914.1</td><td>185523</td><td>185545</td><td>+</td><td>0</td><td>TTGTGGACGGTGTGCGGCCC</td><td>AGG</td><td>||||||||[||||||||||||]PAM</td><td>NA</td><td>Intergenic</td><td>NA</td><td>NA</td></tr>
<tr><td>T14100</td><td>ACGGTGTGCGGCCCAGGCCTCGG</td><td>1000</td><td></td><td></td><td></td><td></td><td></td><td></td><td></td><td></td><td></td></tr>
<tr><td>Oligo fwd</td><td>CACCACGGTGTGCGGCCCAGGCCT</td><td></td><td></td><td></td><td></td><td></td><td></td><td></td><td></td><td></td><td></td></tr>
<tr><td>Oligo rev</td><td>AAACAGGCCTGGGCCGCACACCGT</td><td></td><td></td><td></td><td></td><td></td><td></td><td></td><td></td><td></td><td></td></tr>
<tr><td>Chromosome</td><td>start</td><td>end</td><td>strand</td><td>MM</td><td>target_seq</td><td>PAM</td><td>alignment</td><td>distance</td><td>position</td><td>gene name</td><td>gene id</td></tr>
<tr><td>GU305914.1</td><td>185529</td><td>185551</td><td>+</td><td>0</td><td>ACGGTGTGCGGCCCAGGCCT</td><td>CGG</td><td>||||||||[||||||||||||]PAM</td><td>NA</td><td>Intergenic</td><td>NA</td><td>NA</td></tr>
<tr><td>T14101</td><td>GCCCAGGCCTCGGACGACCCTGG</td><td>1000</td><td></td><td></td><td></td><td></td><td></td><td></td><td></td><td></td><td></td></tr>
<tr><td>Oligo fwd</td><td>CACCGCCCAGGCCTCGGACGACCC</td><td></td><td></td><td></td><td></td><td></td><td></td><td></td><td></td><td></td><td></td></tr>
<tr><td>Oligo rev</td><td>AAACGGGTCGTCCGAGGCCTGGGC</td><td></td><td></td><td></td><td></td><td></td><td></td><td></td><td></td><td></td><td></td></tr>
<tr><td>Chromosome</td><td>start</td><td>end</td><td>strand</td><td>MM</td><td>target_seq</td><td>PAM</td><td>alignment</td><td>distance</td><td>position</td><td>gene name</td><td>gene id</td></tr>
<tr><td>GU305914.1</td><td>185539</td><td>185561</td><td>+</td><td>0</td><td>GCCCAGGCCTCGGACGACCC</td><td>TGG</td><td>||||||||[||||||||||||]PAM</td><td>NA</td><td>Intergenic</td><td>NA</td><td>NA</td></tr>
<tr><td>T14102</td><td>GGCCTCGGACGACCCTGGCCCGG</td><td>1000</td><td></td><td></td><td></td><td></td><td></td><td></td><td></td><td></td><td></td></tr>
<tr><td>Oligo fwd</td><td>CACCGGCCTCGGACGACCCTGGCC</td><td></td><td></td><td></td><td></td><td></td><td></td><td></td><td></td><td></td><td></td></tr>
<tr><td>Oligo rev</td><td>AAACGGCCAGGGTCGTCCGAGGCC</td><td></td><td></td><td></td><td></td><td></td><td></td><td></td><td></td><td></td><td></td></tr>
<tr><td>Chromosome</td><td>start</td><td>end</td><td>strand</td><td>MM</td><td>target_seq</td><td>PAM</td><td>alignment</td><td>distance</td><td>position</td><td>gene name</td><td>gene id</td></tr>
<tr><td>GU305914.1</td><td>185544</td><td>185566</td><td>+</td><td>0</td><td>GGCCTCGGACGACCCTGGCC</td><td>CGG</td><td>||||||||[||||||||||||]PAM</td><td>NA</td><td>Intergenic</td><td>NA</td><td>NA</td></tr>
<tr><td>T14103</td><td>GCCTCGGACGACCCTGGCCCGGG</td><td>1000</td><td></td><td></td><td></td><td></td><td></td><td></td><td></td><td></td><td></td></tr>
<tr><td>Oligo fwd</td><td>CACCGCCTCGGACGACCCTGGCCC</td><td></td><td></td><td></td><td></td><td></td><td></td><td></td><td></td><td></td><td></td></tr>
<tr><td>Oligo rev</td><td>AAACGGGCCAGGGTCGTCCGAGGC</td><td></td><td></td><td></td><td></td><td></td><td></td><td></td><td></td><td></td><td></td></tr>
<tr><td>Chromosome</td><td>start</td><td>end</td><td>strand</td><td>MM</td><td>target_seq</td><td>PAM</td><td>alignment</td><td>distance</td><td>position</td><td>gene name</td><td>gene id</td></tr>
<tr><td>GU305914.1</td><td>185545</td><td>185567</td><td>+</td><td>0</td><td>GCCTCGGACGACCCTGGCCC</td><td>GGG</td><td>||||||||[||||||||||||]PAM</td><td>NA</td><td>Intergenic</td><td>NA</td><td>NA</td></tr>
<tr><td>T14104</td><td>GGCTTCCTTCTTCCTGATCCAGG</td><td>1000</td><td></td><td></td><td></td><td></td><td></td><td></td><td></td><td></td><td></td></tr>
<tr><td>Oligo fwd</td><td>CACCGGCTTCCTTCTTCCTGATCC</td><td></td><td></td><td></td><td></td><td></td><td></td><td></td><td></td><td></td><td></td></tr>
<tr><td>Oligo rev</td><td>AAACGGATCAGGAAGAAGGAAGCC</td><td></td><td></td><td></td><td></td><td></td><td></td><td></td><td></td><td></td><td></td></tr>
<tr><td>Chromosome</td><td>start</td><td>end</td><td>strand</td><td>MM</td><td>target_seq</td><td>PAM</td><td>alignment</td><td>distance</td><td>position</td><td>gene name</td><td>gene id</td></tr>
<tr><td>GU305914.1</td><td>185566</td><td>185588</td><td>+</td><td>0</td><td>GGCTTCCTTCTTCCTGATCC</td><td>AGG</td><td>||||||||[||||||||||||]PAM</td><td>NA</td><td>Intergenic</td><td>NA</td><td>NA</td></tr>
<tr><td>T14105</td><td>GATCCAGGTAGATTGACGTGCGG</td><td>1000</td><td></td><td></td><td></td><td></td><td></td><td></td><td></td><td></td><td></td></tr>
<tr><td>Oligo fwd</td><td>CACCGATCCAGGTAGATTGACGTG</td><td></td><td></td><td></td><td></td><td></td><td></td><td></td><td></td><td></td><td></td></tr>
<tr><td>Oligo rev</td><td>AAACCACGTCAATCTACCTGGATC</td><td></td><td></td><td></td><td></td><td></td><td></td><td></td><td></td><td></td><td></td></tr>
<tr><td>Chromosome</td><td>start</td><td>end</td><td>strand</td><td>MM</td><td>target_seq</td><td>PAM</td><td>alignment</td><td>distance</td><td>position</td><td>gene name</td><td>gene id</td></tr>
<tr><td>GU305914.1</td><td>185581</td><td>185603</td><td>+</td><td>0</td><td>GATCCAGGTAGATTGACGTG</td><td>CGG</td><td>||||||||[||||||||||||]PAM</td><td>NA</td><td>Intergenic</td><td>NA</td><td>NA</td></tr>
<tr><td>T14106</td><td>TGATCTCTCTTGTTCTCCATAGG</td><td>1000</td><td></td><td></td><td></td><td></td><td></td><td></td><td></td><td></td><td></td></tr>
<tr><td>Oligo fwd</td><td>CACCTGATCTCTCTTGTTCTCCAT</td><td></td><td></td><td></td><td></td><td></td><td></td><td></td><td></td><td></td><td></td></tr>
<tr><td>Oligo rev</td><td>AAACATGGAGAACAAGAGAGATCA</td><td></td><td></td><td></td><td></td><td></td><td></td><td></td><td></td><td></td><td></td></tr>
<tr><td>Chromosome</td><td>start</td><td>end</td><td>strand</td><td>MM</td><td>target_seq</td><td>PAM</td><td>alignment</td><td>distance</td><td>position</td><td>gene name</td><td>gene id</td></tr>
<tr><td>GU305914.1</td><td>185672</td><td>185694</td><td>+</td><td>0</td><td>TGATCTCTCTTGTTCTCCAT</td><td>AGG</td><td>||||||||[||||||||||||]PAM</td><td>NA</td><td>Intergenic</td><td>NA</td><td>NA</td></tr>
<tr><td>T14107</td><td>GGTCAGCTACGCCCCTGCTCCGG</td><td>1000</td><td></td><td></td><td></td><td></td><td></td><td></td><td></td><td></td><td></td></tr>
<tr><td>Oligo fwd</td><td>CACCGGTCAGCTACGCCCCTGCTC</td><td></td><td></td><td></td><td></td><td></td><td></td><td></td><td></td><td></td><td></td></tr>
<tr><td>Oligo rev</td><td>AAACGAGCAGGGGCGTAGCTGACC</td><td></td><td></td><td></td><td></td><td></td><td></td><td></td><td></td><td></td><td></td></tr>
<tr><td>Chromosome</td><td>start</td><td>end</td><td>strand</td><td>MM</td><td>target_seq</td><td>PAM</td><td>alignment</td><td>distance</td><td>position</td><td>gene name</td><td>gene id</td></tr>
<tr><td>GU305914.1</td><td>185693</td><td>185715</td><td>+</td><td>0</td><td>GGTCAGCTACGCCCCTGCTC</td><td>CGG</td><td>||||||||[||||||||||||]PAM</td><td>NA</td><td>Intergenic</td><td>NA</td><td>NA</td></tr>
<tr><td>T14108</td><td>CTTCAGACTGACGCCGCTAGAGG</td><td>1000</td><td></td><td></td><td></td><td></td><td></td><td></td><td></td><td></td><td></td></tr>
<tr><td>Oligo fwd</td><td>CACCCTTCAGACTGACGCCGCTAG</td><td></td><td></td><td></td><td></td><td></td><td></td><td></td><td></td><td></td><td></td></tr>
<tr><td>Oligo rev</td><td>AAACCTAGCGGCGTCAGTCTGAAG</td><td></td><td></td><td></td><td></td><td></td><td></td><td></td><td></td><td></td><td></td></tr>
<tr><td>Chromosome</td><td>start</td><td>end</td><td>strand</td><td>MM</td><td>target_seq</td><td>PAM</td><td>alignment</td><td>distance</td><td>position</td><td>gene name</td><td>gene id</td></tr>
<tr><td>GU305914.1</td><td>185740</td><td>185762</td><td>+</td><td>0</td><td>CTTCAGACTGACGCCGCTAG</td><td>AGG</td><td>||||||||[||||||||||||]PAM</td><td>NA</td><td>Intergenic</td><td>NA</td><td>NA</td></tr>
<tr><td>T14109</td><td>ACGCCGCTAGAGGACTTCCTCGG</td><td>1000</td><td></td><td></td><td></td><td></td><td></td><td></td><td></td><td></td><td></td></tr>
<tr><td>Oligo fwd</td><td>CACCACGCCGCTAGAGGACTTCCT</td><td></td><td></td><td></td><td></td><td></td><td></td><td></td><td></td><td></td><td></td></tr>
<tr><td>Oligo rev</td><td>AAACAGGAAGTCCTCTAGCGGCGT</td><td></td><td></td><td></td><td></td><td></td><td></td><td></td><td></td><td></td><td></td></tr>
<tr><td>Chromosome</td><td>start</td><td>end</td><td>strand</td><td>MM</td><td>target_seq</td><td>PAM</td><td>alignment</td><td>distance</td><td>position</td><td>gene name</td><td>gene id</td></tr>
<tr><td>GU305914.1</td><td>185750</td><td>185772</td><td>+</td><td>0</td><td>ACGCCGCTAGAGGACTTCCT</td><td>CGG</td><td>||||||||[||||||||||||]PAM</td><td>NA</td><td>Intergenic</td><td>NA</td><td>NA</td></tr>
</table>

<table>
<tr><td>T14110</td><td>CGCCGCTAGAGGACTTCCTCGGG</td><td>1000</td><td></td><td></td><td></td><td></td><td></td><td></td><td></td><td></td><td></td></tr>
<tr><td>Oligo fwd</td><td>CACCCGCCGCTAGAGGACTTCCTC</td><td></td><td></td><td></td><td></td><td></td><td></td><td></td><td></td><td></td><td></td></tr>
<tr><td>Oligo rev</td><td>AAACGAGGAAGTCCTCTAGCGGCG</td><td></td><td></td><td></td><td></td><td></td><td></td><td></td><td></td><td></td><td></td></tr>
<tr><td>Chromosome</td><td>start</td><td>end</td><td>strand</td><td>MM</td><td>target_seq</td><td>PAM</td><td>alignment</td><td>distance</td><td>position</td><td>gene name</td><td>gene id</td></tr>
<tr><td>GU305914.1</td><td>185751</td><td>185773</td><td>+</td><td>0</td><td>CGCCGCTAGAGGACTTCCTC</td><td>GGG</td><td>||||||||[||||||||||||]PAM</td><td>NA</td><td>Intergenic</td><td>NA</td><td>NA</td></tr>
<tr><td>T14111</td><td>GCCGCTAGAGGACTTCCTCGGGG</td><td>1000</td><td></td><td></td><td></td><td></td><td></td><td></td><td></td><td></td><td></td></tr>
<tr><td>Oligo fwd</td><td>CACCGCCGCTAGAGGACTTCCTCG</td><td></td><td></td><td></td><td></td><td></td><td></td><td></td><td></td><td></td><td></td></tr>
<tr><td>Oligo rev</td><td>AAACCGAGGAAGTCCTCTAGCGGC</td><td></td><td></td><td></td><td></td><td></td><td></td><td></td><td></td><td></td><td></td></tr>
<tr><td>Chromosome</td><td>start</td><td>end</td><td>strand</td><td>MM</td><td>target_seq</td><td>PAM</td><td>alignment</td><td>distance</td><td>position</td><td>gene name</td><td>gene id</td></tr>
<tr><td>GU305914.1</td><td>185752</td><td>185774</td><td>+</td><td>0</td><td>GCCGCTAGAGGACTTCCTCG</td><td>GGG</td><td>||||||||[||||||||||||]PAM</td><td>NA</td><td>Intergenic</td><td>NA</td><td>NA</td></tr>
<tr><td>T14112</td><td>AGGACTTCCTCGGGGAGATGAGG</td><td>1000</td><td></td><td></td><td></td><td></td><td></td><td></td><td></td><td></td><td></td></tr>
<tr><td>Oligo fwd</td><td>CACCAGGACTTCCTCGGGGAGATG</td><td></td><td></td><td></td><td></td><td></td><td></td><td></td><td></td><td></td><td></td></tr>
<tr><td>Oligo rev</td><td>AAACCATCTCCCCGAGGAAGTCCT</td><td></td><td></td><td></td><td></td><td></td><td></td><td></td><td></td><td></td><td></td></tr>
<tr><td>Chromosome</td><td>start</td><td>end</td><td>strand</td><td>MM</td><td>target_seq</td><td>PAM</td><td>alignment</td><td>distance</td><td>position</td><td>gene name</td><td>gene id</td></tr>
<tr><td>GU305914.1</td><td>185760</td><td>185782</td><td>+</td><td>0</td><td>AGGACTTCCTCGGGGAGATG</td><td>AGG</td><td>||||||||[||||||||||||]PAM</td><td>NA</td><td>Intergenic</td><td>NA</td><td>NA</td></tr>
<tr><td>T14113</td><td>CCTCGGGGAGATGAGGACACCGG</td><td>1000</td><td></td><td></td><td></td><td></td><td></td><td></td><td></td><td></td><td></td></tr>
<tr><td>Oligo fwd</td><td>CACCCCTCGGGGAGATGAGGACAC</td><td></td><td></td><td></td><td></td><td></td><td></td><td></td><td></td><td></td><td></td></tr>
<tr><td>Oligo rev</td><td>AAACGTGTCCTCATCTCCCCGAGG</td><td></td><td></td><td></td><td></td><td></td><td></td><td></td><td></td><td></td><td></td></tr>
<tr><td>Chromosome</td><td>start</td><td>end</td><td>strand</td><td>MM</td><td>target_seq</td><td>PAM</td><td>alignment</td><td>distance</td><td>position</td><td>gene name</td><td>gene id</td></tr>
<tr><td>GU305914.1</td><td>185767</td><td>185789</td><td>+</td><td>0</td><td>CCTCGGGGAGATGAGGACAC</td><td>CGG</td><td>||||||||[||||||||||||]PAM</td><td>NA</td><td>Intergenic</td><td>NA</td><td>NA</td></tr>
<tr><td>T14114</td><td>CTCGGGGAGATGAGGACACCGGG</td><td>1000</td><td></td><td></td><td></td><td></td><td></td><td></td><td></td><td></td><td></td></tr>
<tr><td>Oligo fwd</td><td>CACCCTCGGGGAGATGAGGACACC</td><td></td><td></td><td></td><td></td><td></td><td></td><td></td><td></td><td></td><td></td></tr>
<tr><td>Oligo rev</td><td>AAACGGTGTCCTCATCTCCCCGAG</td><td></td><td></td><td></td><td></td><td></td><td></td><td></td><td></td><td></td><td></td></tr>
<tr><td>Chromosome</td><td>start</td><td>end</td><td>strand</td><td>MM</td><td>target_seq</td><td>PAM</td><td>alignment</td><td>distance</td><td>position</td><td>gene name</td><td>gene id</td></tr>
<tr><td>GU305914.1</td><td>185768</td><td>185790</td><td>+</td><td>0</td><td>CTCGGGGAGATGAGGACACC</td><td>GGG</td><td>||||||||[||||||||||||]PAM</td><td>NA</td><td>Intergenic</td><td>NA</td><td>NA</td></tr>
<tr><td>T14115</td><td>GGAGATGAGGACACCGGGACAGG</td><td>1000</td><td></td><td></td><td></td><td></td><td></td><td></td><td></td><td></td><td></td></tr>
<tr><td>Oligo fwd</td><td>CACCGGAGATGAGGACACCGGGAC</td><td></td><td></td><td></td><td></td><td></td><td></td><td></td><td></td><td></td><td></td></tr>
<tr><td>Oligo rev</td><td>AAACGTCCCGGTGTCCTCATCTCC</td><td></td><td></td><td></td><td></td><td></td><td></td><td></td><td></td><td></td><td></td></tr>
<tr><td>Chromosome</td><td>start</td><td>end</td><td>strand</td><td>MM</td><td>target_seq</td><td>PAM</td><td>alignment</td><td>distance</td><td>position</td><td>gene name</td><td>gene id</td></tr>
<tr><td>GU305914.1</td><td>185773</td><td>185795</td><td>+</td><td>0</td><td>GGAGATGAGGACACCGGGAC</td><td>AGG</td><td>||||||||[||||||||||||]PAM</td><td>NA</td><td>Intergenic</td><td>NA</td><td>NA</td></tr>
<tr><td>T14116</td><td>GTTGTTACTAGTGAGTACAGTGG</td><td>1000</td><td></td><td></td><td></td><td></td><td></td><td></td><td></td><td></td><td></td></tr>
<tr><td>Oligo fwd</td><td>CACCGTTGTTACTAGTGAGTACAG</td><td></td><td></td><td></td><td></td><td></td><td></td><td></td><td></td><td></td><td></td></tr>
<tr><td>Oligo rev</td><td>AAACCTGTACTCACTAGTAACAAC</td><td></td><td></td><td></td><td></td><td></td><td></td><td></td><td></td><td></td><td></td></tr>
<tr><td>Chromosome</td><td>start</td><td>end</td><td>strand</td><td>MM</td><td>target_seq</td><td>PAM</td><td>alignment</td><td>distance</td><td>position</td><td>gene name</td><td>gene id</td></tr>
<tr><td>GU305914.1</td><td>185808</td><td>185830</td><td>+</td><td>0</td><td>GTTGTTACTAGTGAGTACAG</td><td>TGG</td><td>||||||||[||||||||||||]PAM</td><td>NA</td><td>Intergenic</td><td>NA</td><td>NA</td></tr>
<tr><td>T14117</td><td>TTACTAGTGAGTACAGTGGCTGG</td><td>1000</td><td></td><td></td><td></td><td></td><td></td><td></td><td></td><td></td><td></td></tr>
<tr><td>Oligo fwd</td><td>CACCTTACTAGTGAGTACAGTGGC</td><td></td><td></td><td></td><td></td><td></td><td></td><td></td><td></td><td></td><td></td></tr>
<tr><td>Oligo rev</td><td>AAACGCCACTGTACTCACTAGTAA</td><td></td><td></td><td></td><td></td><td></td><td></td><td></td><td></td><td></td><td></td></tr>
<tr><td>Chromosome</td><td>start</td><td>end</td><td>strand</td><td>MM</td><td>target_seq</td><td>PAM</td><td>alignment</td><td>distance</td><td>position</td><td>gene name</td><td>gene id</td></tr>
<tr><td>GU305914.1</td><td>185812</td><td>185834</td><td>+</td><td>0</td><td>TTACTAGTGAGTACAGTGGC</td><td>TGG</td><td>||||||||[||||||||||||]PAM</td><td>NA</td><td>Intergenic</td><td>NA</td><td>NA</td></tr>
<tr><td>T14118</td><td>GGCTGGCTGACGTGTCCCTGCGG</td><td>1000</td><td></td><td></td><td></td><td></td><td></td><td></td><td></td><td></td><td></td></tr>
<tr><td>Oligo fwd</td><td>CACCGGCTGGCTGACGTGTCCCTG</td><td></td><td></td><td></td><td></td><td></td><td></td><td></td><td></td><td></td><td></td></tr>
<tr><td>Oligo rev</td><td>AAACCAGGGACACGTCAGCCAGCC</td><td></td><td></td><td></td><td></td><td></td><td></td><td></td><td></td><td></td><td></td></tr>
<tr><td>Chromosome</td><td>start</td><td>end</td><td>strand</td><td>MM</td><td>target_seq</td><td>PAM</td><td>alignment</td><td>distance</td><td>position</td><td>gene name</td><td>gene id</td></tr>
<tr><td>GU305914.1</td><td>185829</td><td>185851</td><td>+</td><td>0</td><td>GGCTGGCTGACGTGTCCCTG</td><td>CGG</td><td>||||||||[||||||||||||]PAM</td><td>NA</td><td>Intergenic</td><td>NA</td><td>NA</td></tr>
<tr><td>T14119</td><td>GTCCTGACAGAATAGCATTGTGG</td><td>1000</td><td></td><td></td><td></td><td></td><td></td><td></td><td></td><td></td><td></td></tr>
<tr><td>Oligo fwd</td><td>CACCGTCCTGACAGAATAGCATTG</td><td></td><td></td><td></td><td></td><td></td><td></td><td></td><td></td><td></td><td></td></tr>
<tr><td>Oligo rev</td><td>AAACCAATGCTATTCTGTCAGGAC</td><td></td><td></td><td></td><td></td><td></td><td></td><td></td><td></td><td></td><td></td></tr>
<tr><td>Chromosome</td><td>start</td><td>end</td><td>strand</td><td>MM</td><td>target_seq</td><td>PAM</td><td>alignment</td><td>distance</td><td>position</td><td>gene name</td><td>gene id</td></tr>
<tr><td>GU305914.1</td><td>185873</td><td>185895</td><td>+</td><td>0</td><td>GTCCTGACAGAATAGCATTG</td><td>TGG</td><td>||||||||[||||||||||||]PAM</td><td>NA</td><td>Intergenic</td><td>NA</td><td>NA</td></tr>
<tr><td>T14120</td><td>TCCTGACAGAATAGCATTGTGGG</td><td>1000</td><td></td><td></td><td></td><td></td><td></td><td></td><td></td><td></td><td></td></tr>
<tr><td>Oligo fwd</td><td>CACCTCCTGACAGAATAGCATTGT</td><td></td><td></td><td></td><td></td><td></td><td></td><td></td><td></td><td></td><td></td></tr>
<tr><td>Oligo rev</td><td>AAACACAATGCTATTCTGTCAGGA</td><td></td><td></td><td></td><td></td><td></td><td></td><td></td><td></td><td></td><td></td></tr>
<tr><td>Chromosome</td><td>start</td><td>end</td><td>strand</td><td>MM</td><td>target_seq</td><td>PAM</td><td>alignment</td><td>distance</td><td>position</td><td>gene name</td><td>gene id</td></tr>
<tr><td>GU305914.1</td><td>185874</td><td>185896</td><td>+</td><td>0</td><td>TCCTGACAGAATAGCATTGT</td><td>GGG</td><td>||||||||[||||||||||||]PAM</td><td>NA</td><td>Intergenic</td><td>NA</td><td>NA</td></tr>
<tr><td>T14121</td><td>AGCATTGTGGGAGATTTGCAAGG</td><td>1000</td><td></td><td></td><td></td><td></td><td></td><td></td><td></td><td></td><td></td></tr>
<tr><td>Oligo fwd</td><td>CACCAGCATTGTGGGAGATTTGCA</td><td></td><td></td><td></td><td></td><td></td><td></td><td></td><td></td><td></td><td></td></tr>
<tr><td>Oligo rev</td><td>AAACTGCAAATCTCCCACAATGCT</td><td></td><td></td><td></td><td></td><td></td><td></td><td></td><td></td><td></td><td></td></tr>
<tr><td>Chromosome</td><td>start</td><td>end</td><td>strand</td><td>MM</td><td>target_seq</td><td>PAM</td><td>alignment</td><td>distance</td><td>position</td><td>gene name</td><td>gene id</td></tr>
<tr><td>GU305914.1</td><td>185886</td><td>185908</td><td>+</td><td>0</td><td>AGCATTGTGGGAGATTTGCA</td><td>AGG</td><td>||||||||[||||||||||||]PAM</td><td>NA</td><td>Intergenic</td><td>NA</td><td>NA</td></tr>
<tr><td>T14122</td><td>GGAGATTTGCAAGGTAACCTAGG</td><td>1000</td><td></td><td></td><td></td><td></td><td></td><td></td><td></td><td></td><td></td></tr>
<tr><td>Oligo fwd</td><td>CACCGGAGATTTGCAAGGTAACCT</td><td></td><td></td><td></td><td></td><td></td><td></td><td></td><td></td><td></td><td></td></tr>
<tr><td>Oligo rev</td><td>AAACAGGTTACCTTGCAAATCTCC</td><td></td><td></td><td></td><td></td><td></td><td></td><td></td><td></td><td></td><td></td></tr>
<tr><td>Chromosome</td><td>start</td><td>end</td><td>strand</td><td>MM</td><td>target_seq</td><td>PAM</td><td>alignment</td><td>distance</td><td>position</td><td>gene name</td><td>gene id</td></tr>
<tr><td>GU305914.1</td><td>185895</td><td>185917</td><td>+</td><td>0</td><td>GGAGATTTGCAAGGTAACCT</td><td>AGG</td><td>||||||||[||||||||||||]PAM</td><td>NA</td><td>Intergenic</td><td>NA</td><td>NA</td></tr>
<tr><td>T14123</td><td>CAAGGTAACCTAGGCTGCACAGG</td><td>1000</td><td></td><td></td><td></td><td></td><td></td><td></td><td></td><td></td><td></td></tr>
<tr><td>Oligo fwd</td><td>CACCCAAGGTAACCTAGGCTGCAC</td><td></td><td></td><td></td><td></td><td></td><td></td><td></td><td></td><td></td><td></td></tr>
<tr><td>Oligo rev</td><td>AAACGTGCAGCCTAGGTTACCTTG</td><td></td><td></td><td></td><td></td><td></td><td></td><td></td><td></td><td></td><td></td></tr>
<tr><td>Chromosome</td><td>start</td><td>end</td><td>strand</td><td>MM</td><td>target_seq</td><td>PAM</td><td>alignment</td><td>distance</td><td>position</td><td>gene name</td><td>gene id</td></tr>
<tr><td>GU305914.1</td><td>185904</td><td>185926</td><td>+</td><td>0</td><td>CAAGGTAACCTAGGCTGCAC</td><td>AGG</td><td>||||||||[||||||||||||]PAM</td><td>NA</td><td>Intergenic</td><td>NA</td><td>NA</td></tr>
<tr><td>T14124</td><td>CAGAGCAGAGCGTCCGAGCACGG</td><td>1000</td><td></td><td></td><td></td><td></td><td></td><td></td><td></td><td></td><td></td></tr>
<tr><td>Oligo fwd</td><td>CACCCAGAGCAGAGCGTCCGAGCA</td><td></td><td></td><td></td><td></td><td></td><td></td><td></td><td></td><td></td><td></td></tr>
<tr><td>Oligo rev</td><td>AAACTGCTCGGACGCTCTGCTCTG</td><td></td><td></td><td></td><td></td><td></td><td></td><td></td><td></td><td></td><td></td></tr>
<tr><td>Chromosome</td><td>start</td><td>end</td><td>strand</td><td>MM</td><td>target_seq</td><td>PAM</td><td>alignment</td><td>distance</td><td>position</td><td>gene name</td><td>gene id</td></tr>
</table>

<table>
<tr><td>GU305914.1</td><td>185939</td><td>185961</td><td>+</td><td>0</td><td>CAGAGCAGAGCGTCCGAGCA</td><td>CGG</td><td>||||||||[||||||||||||]PAM</td><td>NA</td><td>Intergenic</td><td>NA</td><td>NA</td></tr>
<tr><td></td><td></td><td></td><td></td><td></td><td></td><td></td><td></td><td></td><td></td><td></td><td></td></tr>
<tr><td>T14125</td><td>AGAGCGTCCGAGCACGGAGCAGG</td><td>1000</td><td></td><td></td><td></td><td></td><td></td><td></td><td></td><td></td><td></td></tr>
<tr><td>Oligo fwd</td><td>CACCAGAGCGTCCGAGCACGGAGC</td><td></td><td></td><td></td><td></td><td></td><td></td><td></td><td></td><td></td><td></td></tr>
<tr><td>Oligo rev</td><td>AAACGCTCCGTGCTCGGACGCTCT</td><td></td><td></td><td></td><td></td><td></td><td></td><td></td><td></td><td></td><td></td></tr>
<tr><td>Chromosome</td><td>start</td><td>end</td><td>strand</td><td>MM</td><td>target_seq</td><td>PAM</td><td>alignment</td><td>distance</td><td>position</td><td>gene name</td><td>gene id</td></tr>
<tr><td>GU305914.1</td><td>185945</td><td>185967</td><td>+</td><td>0</td><td>AGAGCGTCCGAGCACGGAGC</td><td>AGG</td><td>||||||||[||||||||||||]PAM</td><td>NA</td><td>Intergenic</td><td>NA</td><td>NA</td></tr>
<tr><td></td><td></td><td></td><td></td><td></td><td></td><td></td><td></td><td></td><td></td><td></td><td></td></tr>
<tr><td>T14126</td><td>CGAGCACGGAGCAGGTAAGACGG</td><td>1000</td><td></td><td></td><td></td><td></td><td></td><td></td><td></td><td></td><td></td></tr>
<tr><td>Oligo fwd</td><td>CACCCGAGCACGGAGCAGGTAAGA</td><td></td><td></td><td></td><td></td><td></td><td></td><td></td><td></td><td></td><td></td></tr>
<tr><td>Oligo rev</td><td>AAACTCTTACCTGCTCCGTGCTCG</td><td></td><td></td><td></td><td></td><td></td><td></td><td></td><td></td><td></td><td></td></tr>
<tr><td>Chromosome</td><td>start</td><td>end</td><td>strand</td><td>MM</td><td>target_seq</td><td>PAM</td><td>alignment</td><td>distance</td><td>position</td><td>gene name</td><td>gene id</td></tr>
<tr><td>GU305914.1</td><td>185953</td><td>185975</td><td>+</td><td>0</td><td>CGAGCACGGAGCAGGTAAGA</td><td>CGG</td><td>||||||||[||||||||||||]PAM</td><td>NA</td><td>Intergenic</td><td>NA</td><td>NA</td></tr>
<tr><td></td><td></td><td></td><td></td><td></td><td></td><td></td><td></td><td></td><td></td><td></td><td></td></tr>
<tr><td>T14127</td><td>GCACGGAGCAGGTAAGACGGCGG</td><td>1000</td><td></td><td></td><td></td><td></td><td></td><td></td><td></td><td></td><td></td></tr>
<tr><td>Oligo fwd</td><td>CACCGCACGGAGCAGGTAAGACGG</td><td></td><td></td><td></td><td></td><td></td><td></td><td></td><td></td><td></td><td></td></tr>
<tr><td>Oligo rev</td><td>AAACCCGTCTTACCTGCTCCGTGC</td><td></td><td></td><td></td><td></td><td></td><td></td><td></td><td></td><td></td><td></td></tr>
<tr><td>Chromosome</td><td>start</td><td>end</td><td>strand</td><td>MM</td><td>target_seq</td><td>PAM</td><td>alignment</td><td>distance</td><td>position</td><td>gene name</td><td>gene id</td></tr>
<tr><td>GU305914.1</td><td>185956</td><td>185978</td><td>+</td><td>0</td><td>GCACGGAGCAGGTAAGACGG</td><td>CGG</td><td>||||||||[||||||||||||]PAM</td><td>NA</td><td>Intergenic</td><td>NA</td><td>NA</td></tr>
<tr><td></td><td></td><td></td><td></td><td></td><td></td><td></td><td></td><td></td><td></td><td></td><td></td></tr>
<tr><td>T14128</td><td>AGACGGCGGTGTTCTCTGCACGG</td><td>1000</td><td></td><td></td><td></td><td></td><td></td><td></td><td></td><td></td><td></td></tr>
<tr><td>Oligo fwd</td><td>CACCAGACGGCGGTGTTCTCTGCA</td><td></td><td></td><td></td><td></td><td></td><td></td><td></td><td></td><td></td><td></td></tr>
<tr><td>Oligo rev</td><td>AAACTGCAGAGAACACCGCCGTCT</td><td></td><td></td><td></td><td></td><td></td><td></td><td></td><td></td><td></td><td></td></tr>
<tr><td>Chromosome</td><td>start</td><td>end</td><td>strand</td><td>MM</td><td>target_seq</td><td>PAM</td><td>alignment</td><td>distance</td><td>position</td><td>gene name</td><td>gene id</td></tr>
<tr><td>GU305914.1</td><td>185970</td><td>185992</td><td>+</td><td>0</td><td>AGACGGCGGTGTTCTCTGCA</td><td>CGG</td><td>||||||||[||||||||||||]PAM</td><td>NA</td><td>Intergenic</td><td>NA</td><td>NA</td></tr>
<tr><td></td><td></td><td></td><td></td><td></td><td></td><td></td><td></td><td></td><td></td><td></td><td></td></tr>
<tr><td>T14129</td><td>GGTGTTCTCTGCACGGATGCTGG</td><td>1000</td><td></td><td></td><td></td><td></td><td></td><td></td><td></td><td></td><td></td></tr>
<tr><td>Oligo fwd</td><td>CACCGGTGTTCTCTGCACGGATGC</td><td></td><td></td><td></td><td></td><td></td><td></td><td></td><td></td><td></td><td></td></tr>
<tr><td>Oligo rev</td><td>AAACGCATCCGTGCAGAGAACACC</td><td></td><td></td><td></td><td></td><td></td><td></td><td></td><td></td><td></td><td></td></tr>
<tr><td>Chromosome</td><td>start</td><td>end</td><td>strand</td><td>MM</td><td>target_seq</td><td>PAM</td><td>alignment</td><td>distance</td><td>position</td><td>gene name</td><td>gene id</td></tr>
<tr><td>GU305914.1</td><td>185977</td><td>185999</td><td>+</td><td>0</td><td>GGTGTTCTCTGCACGGATGC</td><td>TGG</td><td>||||||||[||||||||||||]PAM</td><td>NA</td><td>Intergenic</td><td>NA</td><td>NA</td></tr>
<tr><td></td><td></td><td></td><td></td><td></td><td></td><td></td><td></td><td></td><td></td><td></td><td></td></tr>
<tr><td>T14130</td><td>TCTGCACGGATGCTGGTGACCGG</td><td>1000</td><td></td><td></td><td></td><td></td><td></td><td></td><td></td><td></td><td></td></tr>
<tr><td>Oligo fwd</td><td>CACCTCTGCACGGATGCTGGTGAC</td><td></td><td></td><td></td><td></td><td></td><td></td><td></td><td></td><td></td><td></td></tr>
<tr><td>Oligo rev</td><td>AAACGTCACCAGCATCCGTGCAGA</td><td></td><td></td><td></td><td></td><td></td><td></td><td></td><td></td><td></td><td></td></tr>
<tr><td>Chromosome</td><td>start</td><td>end</td><td>strand</td><td>MM</td><td>target_seq</td><td>PAM</td><td>alignment</td><td>distance</td><td>position</td><td>gene name</td><td>gene id</td></tr>
<tr><td>GU305914.1</td><td>185984</td><td>186006</td><td>+</td><td>0</td><td>TCTGCACGGATGCTGGTGAC</td><td>CGG</td><td>||||||||[||||||||||||]PAM</td><td>NA</td><td>Intergenic</td><td>NA</td><td>NA</td></tr>
<tr><td></td><td></td><td></td><td></td><td></td><td></td><td></td><td></td><td></td><td></td><td></td><td></td></tr>
<tr><td>T14131</td><td>GCTGGTGACCGGTCATTTCTTGG</td><td>1000</td><td></td><td></td><td></td><td></td><td></td><td></td><td></td><td></td><td></td></tr>
<tr><td>Oligo fwd</td><td>CACCGCTGGTGACCGGTCATTTCT</td><td></td><td></td><td></td><td></td><td></td><td></td><td></td><td></td><td></td><td></td></tr>
<tr><td>Oligo rev</td><td>AAACAGAAATGACCGGTCACCAGC</td><td></td><td></td><td></td><td></td><td></td><td></td><td></td><td></td><td></td><td></td></tr>
<tr><td>Chromosome</td><td>start</td><td>end</td><td>strand</td><td>MM</td><td>target_seq</td><td>PAM</td><td>alignment</td><td>distance</td><td>position</td><td>gene name</td><td>gene id</td></tr>
<tr><td>GU305914.1</td><td>185995</td><td>186017</td><td>+</td><td>0</td><td>GCTGGTGACCGGTCATTTCT</td><td>TGG</td><td>||||||||[||||||||||||]PAM</td><td>NA</td><td>Intergenic</td><td>NA</td><td>NA</td></tr>
<tr><td></td><td></td><td></td><td></td><td></td><td></td><td></td><td></td><td></td><td></td><td></td><td></td></tr>
<tr><td>T14132</td><td>GACCGGTCATTTCTTGGTGAAGG</td><td>1000</td><td></td><td></td><td></td><td></td><td></td><td></td><td></td><td></td><td></td></tr>
<tr><td>Oligo fwd</td><td>CACCGACCGGTCATTTCTTGGTGA</td><td></td><td></td><td></td><td></td><td></td><td></td><td></td><td></td><td></td><td></td></tr>
<tr><td>Oligo rev</td><td>AAACTCACCAAGAAATGACCGGTC</td><td></td><td></td><td></td><td></td><td></td><td></td><td></td><td></td><td></td><td></td></tr>
<tr><td>Chromosome</td><td>start</td><td>end</td><td>strand</td><td>MM</td><td>target_seq</td><td>PAM</td><td>alignment</td><td>distance</td><td>position</td><td>gene name</td><td>gene id</td></tr>
<tr><td>GU305914.1</td><td>186001</td><td>186023</td><td>+</td><td>0</td><td>GACCGGTCATTTCTTGGTGA</td><td>AGG</td><td>||||||||[||||||||||||]PAM</td><td>NA</td><td>Intergenic</td><td>NA</td><td>NA</td></tr>
<tr><td></td><td></td><td></td><td></td><td></td><td></td><td></td><td></td><td></td><td></td><td></td><td></td></tr>
<tr><td>T14133</td><td>ACCGGTCATTTCTTGGTGAAGGG</td><td>1000</td><td></td><td></td><td></td><td></td><td></td><td></td><td></td><td></td><td></td></tr>
<tr><td>Oligo fwd</td><td>CACCACCGGTCATTTCTTGGTGAA</td><td></td><td></td><td></td><td></td><td></td><td></td><td></td><td></td><td></td><td></td></tr>
<tr><td>Oligo rev</td><td>AAACTTCACCAAGAAATGACCGGT</td><td></td><td></td><td></td><td></td><td></td><td></td><td></td><td></td><td></td><td></td></tr>
<tr><td>Chromosome</td><td>start</td><td>end</td><td>strand</td><td>MM</td><td>target_seq</td><td>PAM</td><td>alignment</td><td>distance</td><td>position</td><td>gene name</td><td>gene id</td></tr>
<tr><td>GU305914.1</td><td>186002</td><td>186024</td><td>+</td><td>0</td><td>ACCGGTCATTTCTTGGTGAA</td><td>GGG</td><td>||||||||[||||||||||||]PAM</td><td>NA</td><td>Intergenic</td><td>NA</td><td>NA</td></tr>
<tr><td></td><td></td><td></td><td></td><td></td><td></td><td></td><td></td><td></td><td></td><td></td><td></td></tr>
<tr><td>T14134</td><td>CTTGCTCTCGCCCGCCCGCCCGG</td><td>1000</td><td></td><td></td><td></td><td></td><td></td><td></td><td></td><td></td><td></td></tr>
<tr><td>Oligo fwd</td><td>CACCCTTGCTCTCGCCCGCCCGCC</td><td></td><td></td><td></td><td></td><td></td><td></td><td></td><td></td><td></td><td></td></tr>
<tr><td>Oligo rev</td><td>AAACGGCGGGCGGGCGAGAGCAAG</td><td></td><td></td><td></td><td></td><td></td><td></td><td></td><td></td><td></td><td></td></tr>
<tr><td>Chromosome</td><td>start</td><td>end</td><td>strand</td><td>MM</td><td>target_seq</td><td>PAM</td><td>alignment</td><td>distance</td><td>position</td><td>gene name</td><td>gene id</td></tr>
<tr><td>GU305914.1</td><td>186028</td><td>186050</td><td>+</td><td>0</td><td>CTTGCTCTCGCCCGCCCGCC</td><td>CGG</td><td>||||||||[||||||||||||]PAM</td><td>NA</td><td>Intergenic</td><td>NA</td><td>NA</td></tr>
<tr><td></td><td></td><td></td><td></td><td></td><td></td><td></td><td></td><td></td><td></td><td></td><td></td></tr>
<tr><td>T14135</td><td>TCTTCTTTCTGATTGCAGCACGG</td><td>1000</td><td></td><td></td><td></td><td></td><td></td><td></td><td></td><td></td><td></td></tr>
<tr><td>Oligo fwd</td><td>CACCTCTTCTTTCTGATTGCAGCA</td><td></td><td></td><td></td><td></td><td></td><td></td><td></td><td></td><td></td><td></td></tr>
<tr><td>Oligo rev</td><td>AAACTGCTGCAATCAGAAAGAAGA</td><td></td><td></td><td></td><td></td><td></td><td></td><td></td><td></td><td></td><td></td></tr>
<tr><td>Chromosome</td><td>start</td><td>end</td><td>strand</td><td>MM</td><td>target_seq</td><td>PAM</td><td>alignment</td><td>distance</td><td>position</td><td>gene name</td><td>gene id</td></tr>
<tr><td>GU305914.1</td><td>186076</td><td>186098</td><td>+</td><td>0</td><td>TCTTCTTTCTGATTGCAGCA</td><td>CGG</td><td>||||||||[||||||||||||]PAM</td><td>NA</td><td>Intergenic</td><td>NA</td><td>NA</td></tr>
<tr><td></td><td></td><td></td><td></td><td></td><td></td><td></td><td></td><td></td><td></td><td></td><td></td></tr>
<tr><td>T14136</td><td>TCTTTCTGATTGCAGCACGGAGG</td><td>1000</td><td></td><td></td><td></td><td></td><td></td><td></td><td></td><td></td><td></td></tr>
<tr><td>Oligo fwd</td><td>CACCTCTTTCTGATTGCAGCACGG</td><td></td><td></td><td></td><td></td><td></td><td></td><td></td><td></td><td></td><td></td></tr>
<tr><td>Oligo rev</td><td>AAACCCGTGCTGCAATCAGAAAGA</td><td></td><td></td><td></td><td></td><td></td><td></td><td></td><td></td><td></td><td></td></tr>
<tr><td>Chromosome</td><td>start</td><td>end</td><td>strand</td><td>MM</td><td>target_seq</td><td>PAM</td><td>alignment</td><td>distance</td><td>position</td><td>gene name</td><td>gene id</td></tr>
<tr><td>GU305914.1</td><td>186079</td><td>186101</td><td>+</td><td>0</td><td>TCTTTCTGATTGCAGCACGG</td><td>AGG</td><td>||||||||[||||||||||||]PAM</td><td>NA</td><td>Intergenic</td><td>NA</td><td>NA</td></tr>
<tr><td></td><td></td><td></td><td></td><td></td><td></td><td></td><td></td><td></td><td></td><td></td><td></td></tr>
<tr><td>T14137</td><td>TCTGATTGCAGCACGGAGGTCGG</td><td>1000</td><td></td><td></td><td></td><td></td><td></td><td></td><td></td><td></td><td></td></tr>
<tr><td>Oligo fwd</td><td>CACCTCTGATTGCAGCACGGAGGT</td><td></td><td></td><td></td><td></td><td></td><td></td><td></td><td></td><td></td><td></td></tr>
<tr><td>Oligo rev</td><td>AAACACCTCCGTGCTGCAATCAGA</td><td></td><td></td><td></td><td></td><td></td><td></td><td></td><td></td><td></td><td></td></tr>
<tr><td>Chromosome</td><td>start</td><td>end</td><td>strand</td><td>MM</td><td>target_seq</td><td>PAM</td><td>alignment</td><td>distance</td><td>position</td><td>gene name</td><td>gene id</td></tr>
<tr><td>GU305914.1</td><td>186083</td><td>186105</td><td>+</td><td>0</td><td>TCTGATTGCAGCACGGAGGT</td><td>CGG</td><td>||||||||[||||||||||||]PAM</td><td>NA</td><td>Intergenic</td><td>NA</td><td>NA</td></tr>
<tr><td></td><td></td><td></td><td></td><td></td><td></td><td></td><td></td><td></td><td></td><td></td><td></td></tr>
<tr><td>T14138</td><td>CTGATTGCAGCACGGAGGTCGGG</td><td>1000</td><td></td><td></td><td></td><td></td><td></td><td></td><td></td><td></td><td></td></tr>
<tr><td>Oligo fwd</td><td>CACCCTGATTGCAGCACGGAGGTC</td><td></td><td></td><td></td><td></td><td></td><td></td><td></td><td></td><td></td><td></td></tr>
<tr><td>Oligo rev</td><td>AAACGACCTCCGTGCTGCAATCAG</td><td></td><td></td><td></td><td></td><td></td><td></td><td></td><td></td><td></td><td></td></tr>
<tr><td>Chromosome</td><td>start</td><td>end</td><td>strand</td><td>MM</td><td>target_seq</td><td>PAM</td><td>alignment</td><td>distance</td><td>position</td><td>gene name</td><td>gene id</td></tr>
<tr><td>GU305914.1</td><td>186084</td><td>186106</td><td>+</td><td>0</td><td>CTGATTGCAGCACGGAGGTC</td><td>GGG</td><td>||||||||[||||||||||||]PAM</td><td>NA</td><td>Intergenic</td><td>NA</td><td>NA</td></tr>
<tr><td></td><td></td><td></td><td></td><td></td><td></td><td></td><td></td><td></td><td></td><td></td><td></td></tr>
<tr><td>T14139</td><td>TGATTGCAGCACGGAGGTCGGGG</td><td>1000</td><td></td><td></td><td></td><td></td><td></td><td></td><td></td><td></td><td></td></tr>
<tr><td>Oligo fwd</td><td>CACCTGATTGCAGCACGGAGGTCG</td><td></td><td></td><td></td><td></td><td></td><td></td><td></td><td></td><td></td><td></td></tr>
<tr><td>Oligo rev</td><td>AAACCGACCTCCGTGCTGCAATCA</td><td></td><td></td><td></td><td></td><td></td><td></td><td></td><td></td><td></td><td></td></tr>
</table>

<table>
<tr><td>Chromosome</td><td>start</td><td>end</td><td>strand</td><td>MM</td><td>target_seq</td><td>PAM</td><td>alignment</td><td>distance</td><td>position</td><td>gene name</td><td>gene id</td></tr>
<tr><td>GU305914.1</td><td>186085</td><td>186107</td><td>+</td><td>0</td><td>TGATTGCAGCACGGAGGTCG</td><td>GGG</td><td>||||||||[||||||||||||]PAM</td><td>NA</td><td>Intergenic</td><td>NA</td><td>NA</td></tr>
<tr><td></td><td></td><td></td><td></td><td></td><td></td><td></td><td></td><td></td><td></td><td></td><td></td></tr>
<tr><td>T14140</td><td>CAGCTGTAGTGAGAGTCTCTCGG</td><td>1000</td><td></td><td></td><td></td><td></td><td></td><td></td><td></td><td></td><td></td></tr>
<tr><td>Oligo fwd</td><td>CACCCAGCTGTAGTGAGAGTCTCT</td><td></td><td></td><td></td><td></td><td></td><td></td><td></td><td></td><td></td><td></td></tr>
<tr><td>Oligo rev</td><td>AAACAGAGACTCTCACTACAGCTG</td><td></td><td></td><td></td><td></td><td></td><td></td><td></td><td></td><td></td><td></td></tr>
<tr><td>Chromosome</td><td>start</td><td>end</td><td>strand</td><td>MM</td><td>target_seq</td><td>PAM</td><td>alignment</td><td>distance</td><td>position</td><td>gene name</td><td>gene id</td></tr>
<tr><td>GU305914.1</td><td>186111</td><td>186133</td><td>+</td><td>0</td><td>CAGCTGTAGTGAGAGTCTCT</td><td>CGG</td><td>||||||||[||||||||||||]PAM</td><td>NA</td><td>Intergenic</td><td>NA</td><td>NA</td></tr>
<tr><td></td><td></td><td></td><td></td><td></td><td></td><td></td><td></td><td></td><td></td><td></td><td></td></tr>
<tr><td>T14141</td><td>AGCTGTAGTGAGAGTCTCTCGGG</td><td>1000</td><td></td><td></td><td></td><td></td><td></td><td></td><td></td><td></td><td></td></tr>
<tr><td>Oligo fwd</td><td>CACCAGCTGTAGTGAGAGTCTCTC</td><td></td><td></td><td></td><td></td><td></td><td></td><td></td><td></td><td></td><td></td></tr>
<tr><td>Oligo rev</td><td>AAACGAGAGACTCTCACTACAGCT</td><td></td><td></td><td></td><td></td><td></td><td></td><td></td><td></td><td></td><td></td></tr>
<tr><td>Chromosome</td><td>start</td><td>end</td><td>strand</td><td>MM</td><td>target_seq</td><td>PAM</td><td>alignment</td><td>distance</td><td>position</td><td>gene name</td><td>gene id</td></tr>
<tr><td>GU305914.1</td><td>186112</td><td>186134</td><td>+</td><td>0</td><td>AGCTGTAGTGAGAGTCTCTC</td><td>GGG</td><td>||||||||[||||||||||||]PAM</td><td>NA</td><td>Intergenic</td><td>NA</td><td>NA</td></tr>
<tr><td></td><td></td><td></td><td></td><td></td><td></td><td></td><td></td><td></td><td></td><td></td><td></td></tr>
<tr><td>T14142</td><td>TAGTGAGAGTCTCTCGGGTCCGG</td><td>1000</td><td></td><td></td><td></td><td></td><td></td><td></td><td></td><td></td><td></td></tr>
<tr><td>Oligo fwd</td><td>CACCTAGTGAGAGTCTCTCGGGTC</td><td></td><td></td><td></td><td></td><td></td><td></td><td></td><td></td><td></td><td></td></tr>
<tr><td>Oligo rev</td><td>AAACGACCCGAGAGACTCTCACTA</td><td></td><td></td><td></td><td></td><td></td><td></td><td></td><td></td><td></td><td></td></tr>
<tr><td>Chromosome</td><td>start</td><td>end</td><td>strand</td><td>MM</td><td>target_seq</td><td>PAM</td><td>alignment</td><td>distance</td><td>position</td><td>gene name</td><td>gene id</td></tr>
<tr><td>GU305914.1</td><td>186117</td><td>186139</td><td>+</td><td>0</td><td>TAGTGAGAGTCTCTCGGGTC</td><td>CGG</td><td>||||||||[||||||||||||]PAM</td><td>NA</td><td>Intergenic</td><td>NA</td><td>NA</td></tr>
<tr><td></td><td></td><td></td><td></td><td></td><td></td><td></td><td></td><td></td><td></td><td></td><td></td></tr>
<tr><td>T14143</td><td>AGTGAGAGTCTCTCGGGTCCGGG</td><td>1000</td><td></td><td></td><td></td><td></td><td></td><td></td><td></td><td></td><td></td></tr>
<tr><td>Oligo fwd</td><td>CACCAGTGAGAGTCTCTCGGGTCC</td><td></td><td></td><td></td><td></td><td></td><td></td><td></td><td></td><td></td><td></td></tr>
<tr><td>Oligo rev</td><td>AAACGGACCCGAGAGACTCTCACT</td><td></td><td></td><td></td><td></td><td></td><td></td><td></td><td></td><td></td><td></td></tr>
<tr><td>Chromosome</td><td>start</td><td>end</td><td>strand</td><td>MM</td><td>target_seq</td><td>PAM</td><td>alignment</td><td>distance</td><td>position</td><td>gene name</td><td>gene id</td></tr>
<tr><td>GU305914.1</td><td>186118</td><td>186140</td><td>+</td><td>0</td><td>AGTGAGAGTCTCTCGGGTCC</td><td>GGG</td><td>||||||||[||||||||||||]PAM</td><td>NA</td><td>Intergenic</td><td>NA</td><td>NA</td></tr>
<tr><td></td><td></td><td></td><td></td><td></td><td></td><td></td><td></td><td></td><td></td><td></td><td></td></tr>
<tr><td>T14144</td><td>GTGAGAGTCTCTCGGGTCCGGGG</td><td>1000</td><td></td><td></td><td></td><td></td><td></td><td></td><td></td><td></td><td></td></tr>
<tr><td>Oligo fwd</td><td>CACCGTGAGAGTCTCTCGGGTCCG</td><td></td><td></td><td></td><td></td><td></td><td></td><td></td><td></td><td></td><td></td></tr>
<tr><td>Oligo rev</td><td>AAACCGGACCCGAGAGACTCTCAC</td><td></td><td></td><td></td><td></td><td></td><td></td><td></td><td></td><td></td><td></td></tr>
<tr><td>Chromosome</td><td>start</td><td>end</td><td>strand</td><td>MM</td><td>target_seq</td><td>PAM</td><td>alignment</td><td>distance</td><td>position</td><td>gene name</td><td>gene id</td></tr>
<tr><td>GU305914.1</td><td>186119</td><td>186141</td><td>+</td><td>0</td><td>GTGAGAGTCTCTCGGGTCCG</td><td>GGG</td><td>||||||||[||||||||||||]PAM</td><td>NA</td><td>Intergenic</td><td>NA</td><td>NA</td></tr>
<tr><td></td><td></td><td></td><td></td><td></td><td></td><td></td><td></td><td></td><td></td><td></td><td></td></tr>
<tr><td>T14145</td><td>TGAGAGTCTCTCGGGTCCGGGGG</td><td>1000</td><td></td><td></td><td></td><td></td><td></td><td></td><td></td><td></td><td></td></tr>
<tr><td>Oligo fwd</td><td>CACCTGAGAGTCTCTCGGGTCCGG</td><td></td><td></td><td></td><td></td><td></td><td></td><td></td><td></td><td></td><td></td></tr>
<tr><td>Oligo rev</td><td>AAACCCGGACCCGAGAGACTCTCA</td><td></td><td></td><td></td><td></td><td></td><td></td><td></td><td></td><td></td><td></td></tr>
<tr><td>Chromosome</td><td>start</td><td>end</td><td>strand</td><td>MM</td><td>target_seq</td><td>PAM</td><td>alignment</td><td>distance</td><td>position</td><td>gene name</td><td>gene id</td></tr>
<tr><td>GU305914.1</td><td>186120</td><td>186142</td><td>+</td><td>0</td><td>TGAGAGTCTCTCGGGTCCGG</td><td>GGG</td><td>||||||||[||||||||||||]PAM</td><td>NA</td><td>Intergenic</td><td>NA</td><td>NA</td></tr>
<tr><td></td><td></td><td></td><td></td><td></td><td></td><td></td><td></td><td></td><td></td><td></td><td></td></tr>
<tr><td>T14146</td><td>CTCTCGGGTCCGGGGGCGTGAGG</td><td>1000</td><td></td><td></td><td></td><td></td><td></td><td></td><td></td><td></td><td></td></tr>
<tr><td>Oligo fwd</td><td>CACCCTCTCGGGTCCGGGGGCGTG</td><td></td><td></td><td></td><td></td><td></td><td></td><td></td><td></td><td></td><td></td></tr>
<tr><td>Oligo rev</td><td>AAACCACGCCCCCGGACCCGAGAG</td><td></td><td></td><td></td><td></td><td></td><td></td><td></td><td></td><td></td><td></td></tr>
<tr><td>Chromosome</td><td>start</td><td>end</td><td>strand</td><td>MM</td><td>target_seq</td><td>PAM</td><td>alignment</td><td>distance</td><td>position</td><td>gene name</td><td>gene id</td></tr>
<tr><td>GU305914.1</td><td>186127</td><td>186149</td><td>+</td><td>0</td><td>CTCTCGGGTCCGGGGGCGTG</td><td>AGG</td><td>||||||||[||||||||||||]PAM</td><td>NA</td><td>Intergenic</td><td>NA</td><td>NA</td></tr>
<tr><td></td><td></td><td></td><td></td><td></td><td></td><td></td><td></td><td></td><td></td><td></td><td></td></tr>
<tr><td>T14147</td><td>TCTCGGGTCCGGGGGCGTGAGGG</td><td>1000</td><td></td><td></td><td></td><td></td><td></td><td></td><td></td><td></td><td></td></tr>
<tr><td>Oligo fwd</td><td>CACCTCTCGGGTCCGGGGGCGTGA</td><td></td><td></td><td></td><td></td><td></td><td></td><td></td><td></td><td></td><td></td></tr>
<tr><td>Oligo rev</td><td>AAACTCACGCCCCCGGACCCGAGA</td><td></td><td></td><td></td><td></td><td></td><td></td><td></td><td></td><td></td><td></td></tr>
<tr><td>Chromosome</td><td>start</td><td>end</td><td>strand</td><td>MM</td><td>target_seq</td><td>PAM</td><td>alignment</td><td>distance</td><td>position</td><td>gene name</td><td>gene id</td></tr>
<tr><td>GU305914.1</td><td>186128</td><td>186150</td><td>+</td><td>0</td><td>TCTCGGGTCCGGGGGCGTGA</td><td>GGG</td><td>||||||||[||||||||||||]PAM</td><td>NA</td><td>Intergenic</td><td>NA</td><td>NA</td></tr>
<tr><td></td><td></td><td></td><td></td><td></td><td></td><td></td><td></td><td></td><td></td><td></td><td></td></tr>
<tr><td>T14148</td><td>GTGAGGGACGCGACTGCTCACGG</td><td>1000</td><td></td><td></td><td></td><td></td><td></td><td></td><td></td><td></td><td></td></tr>
<tr><td>Oligo fwd</td><td>CACCGTGAGGGACGCGACTGCTCA</td><td></td><td></td><td></td><td></td><td></td><td></td><td></td><td></td><td></td><td></td></tr>
<tr><td>Oligo rev</td><td>AAACTGAGCAGTCGCGTCCCTCAC</td><td></td><td></td><td></td><td></td><td></td><td></td><td></td><td></td><td></td><td></td></tr>
<tr><td>Chromosome</td><td>start</td><td>end</td><td>strand</td><td>MM</td><td>target_seq</td><td>PAM</td><td>alignment</td><td>distance</td><td>position</td><td>gene name</td><td>gene id</td></tr>
<tr><td>GU305914.1</td><td>186144</td><td>186166</td><td>+</td><td>0</td><td>GTGAGGGACGCGACTGCTCA</td><td>CGG</td><td>||||||||[||||||||||||]PAM</td><td>NA</td><td>Intergenic</td><td>NA</td><td>NA</td></tr>
<tr><td></td><td></td><td></td><td></td><td></td><td></td><td></td><td></td><td></td><td></td><td></td><td></td></tr>
<tr><td>T14149</td><td>GTTCTGTTTGTCTGTAGATATGG</td><td>1000</td><td></td><td></td><td></td><td></td><td></td><td></td><td></td><td></td><td></td></tr>
<tr><td>Oligo fwd</td><td>CACCGTTCTGTTTGTCTGTAGATA</td><td></td><td></td><td></td><td></td><td></td><td></td><td></td><td></td><td></td><td></td></tr>
<tr><td>Oligo rev</td><td>AAACTATCTACAGACAAACAGAAC</td><td></td><td></td><td></td><td></td><td></td><td></td><td></td><td></td><td></td><td></td></tr>
<tr><td>Chromosome</td><td>start</td><td>end</td><td>strand</td><td>MM</td><td>target_seq</td><td>PAM</td><td>alignment</td><td>distance</td><td>position</td><td>gene name</td><td>gene id</td></tr>
<tr><td>GU305914.1</td><td>186166</td><td>186188</td><td>+</td><td>0</td><td>GTTCTGTTTGTCTGTAGATA</td><td>TGG</td><td>||||||||[||||||||||||]PAM</td><td>NA</td><td>Intergenic</td><td>NA</td><td>NA</td></tr>
<tr><td></td><td></td><td></td><td></td><td></td><td></td><td></td><td></td><td></td><td></td><td></td><td></td></tr>
<tr><td>T14150</td><td>CTGTAGATATGGAGCGTGTTCGG</td><td>1000</td><td></td><td></td><td></td><td></td><td></td><td></td><td></td><td></td><td></td></tr>
<tr><td>Oligo fwd</td><td>CACCCTGTAGATATGGAGCGTGTT</td><td></td><td></td><td></td><td></td><td></td><td></td><td></td><td></td><td></td><td></td></tr>
<tr><td>Oligo rev</td><td>AAACAACACGCTCCATATCTACAG</td><td></td><td></td><td></td><td></td><td></td><td></td><td></td><td></td><td></td><td></td></tr>
<tr><td>Chromosome</td><td>start</td><td>end</td><td>strand</td><td>MM</td><td>target_seq</td><td>PAM</td><td>alignment</td><td>distance</td><td>position</td><td>gene name</td><td>gene id</td></tr>
<tr><td>GU305914.1</td><td>186177</td><td>186199</td><td>+</td><td>0</td><td>CTGTAGATATGGAGCGTGTT</td><td>CGG</td><td>||||||||[||||||||||||]PAM</td><td>NA</td><td>Intergenic</td><td>NA</td><td>NA</td></tr>
<tr><td></td><td></td><td></td><td></td><td></td><td></td><td></td><td></td><td></td><td></td><td></td><td></td></tr>
<tr><td>T14151</td><td>TGTAGATATGGAGCGTGTTCGGG</td><td>1000</td><td></td><td></td><td></td><td></td><td></td><td></td><td></td><td></td><td></td></tr>
<tr><td>Oligo fwd</td><td>CACCTGTAGATATGGAGCGTGTTC</td><td></td><td></td><td></td><td></td><td></td><td></td><td></td><td></td><td></td><td></td></tr>
<tr><td>Oligo rev</td><td>AAACGAACACGCTCCATATCTACA</td><td></td><td></td><td></td><td></td><td></td><td></td><td></td><td></td><td></td><td></td></tr>
<tr><td>Chromosome</td><td>start</td><td>end</td><td>strand</td><td>MM</td><td>target_seq</td><td>PAM</td><td>alignment</td><td>distance</td><td>position</td><td>gene name</td><td>gene id</td></tr>
<tr><td>GU305914.1</td><td>186178</td><td>186200</td><td>+</td><td>0</td><td>TGTAGATATGGAGCGTGTTC</td><td>GGG</td><td>||||||||[||||||||||||]PAM</td><td>NA</td><td>Intergenic</td><td>NA</td><td>NA</td></tr>
<tr><td></td><td></td><td></td><td></td><td></td><td></td><td></td><td></td><td></td><td></td><td></td><td></td></tr>
<tr><td>T14152</td><td>GTAGATATGGAGCGTGTTCGGGG</td><td>1000</td><td></td><td></td><td></td><td></td><td></td><td></td><td></td><td></td><td></td></tr>
<tr><td>Oligo fwd</td><td>CACCGTAGATATGGAGCGTGTTCG</td><td></td><td></td><td></td><td></td><td></td><td></td><td></td><td></td><td></td><td></td></tr>
<tr><td>Oligo rev</td><td>AAACCGAACACGCTCCATATCTAC</td><td></td><td></td><td></td><td></td><td></td><td></td><td></td><td></td><td></td><td></td></tr>
<tr><td>Chromosome</td><td>start</td><td>end</td><td>strand</td><td>MM</td><td>target_seq</td><td>PAM</td><td>alignment</td><td>distance</td><td>position</td><td>gene name</td><td>gene id</td></tr>
<tr><td>GU305914.1</td><td>186179</td><td>186201</td><td>+</td><td>0</td><td>GTAGATATGGAGCGTGTTCG</td><td>GGG</td><td>||||||||[||||||||||||]PAM</td><td>NA</td><td>Intergenic</td><td>NA</td><td>NA</td></tr>
<tr><td></td><td></td><td></td><td></td><td></td><td></td><td></td><td></td><td></td><td></td><td></td><td></td></tr>
<tr><td>T14153</td><td>GGGAGCTTCCTCCTCAGACTCGG</td><td>1000</td><td></td><td></td><td></td><td></td><td></td><td></td><td></td><td></td><td></td></tr>
<tr><td>Oligo fwd</td><td>CACCGGGAGCTTCCTCCTCAGACT</td><td></td><td></td><td></td><td></td><td></td><td></td><td></td><td></td><td></td><td></td></tr>
<tr><td>Oligo rev</td><td>AAACAGTCTGAGGAGGAAGCTCCC</td><td></td><td></td><td></td><td></td><td></td><td></td><td></td><td></td><td></td><td></td></tr>
<tr><td>Chromosome</td><td>start</td><td>end</td><td>strand</td><td>MM</td><td>target_seq</td><td>PAM</td><td>alignment</td><td>distance</td><td>position</td><td>gene name</td><td>gene id</td></tr>
<tr><td>GU305914.1</td><td>186199</td><td>186221</td><td>+</td><td>0</td><td>GGGAGCTTCCTCCTCAGACT</td><td>CGG</td><td>||||||||[||||||||||||]PAM</td><td>NA</td><td>Intergenic</td><td>NA</td><td>NA</td></tr>
<tr><td></td><td></td><td></td><td></td><td></td><td></td><td></td><td></td><td></td><td></td><td></td><td></td></tr>
<tr><td>T14154</td><td>AGCTTCCTCCTCAGACTCGGTGG</td><td>1000</td><td></td><td></td><td></td><td></td><td></td><td></td><td></td><td></td><td></td></tr>
<tr><td>Oligo fwd</td><td>CACCAGCTTCCTCCTCAGACTCGG</td><td></td><td></td><td></td><td></td><td></td><td></td><td></td><td></td><td></td><td></td></tr>
</table>

Oligo rev AAACCCGAGTCTGAGGAGGAAGCT
Chromosome start end strand MM target_seq PAM alignment distance position gene name gene id
GU305914.1 186202 186224 + 0 AGCTTCCTCCTCAGACTCGG TGG ||||||||[||||||||||||]PAM NA Intergenic NA NA

T14155 GACTCGGTGGCGACCCTCCACGG 1000
Oligo fwd CACCGACTCGGTGGCGACCCTCCA
Oligo rev AAACTGGAGGGTCGCCACCGAGTC
Chromosome start end strand MM target_seq PAM alignment distance position gene name gene id
GU305914.1 186215 186237 + 0 GACTCGGTGGCGACCCTCCA CGG ||||||||[||||||||||||]PAM NA Intergenic NA NA

T14156 CGATCACAACTACCACTGCATGG 1000
Oligo fwd CACCCGATCACAACTACCACTGCA
Oligo rev AAACTGCAGTGGTAGTTGTGATCG
Chromosome start end strand MM target_seq PAM alignment distance position gene name gene id
GU305914.1 186238 186260 + 0 CGATCACAACTACCACTGCA TGG ||||||||[||||||||||||]PAM NA Intergenic NA NA

T14157 CTGCATGGCGACCTTCAAGCTGG 1000
Oligo fwd CACCCTGCATGGCGACCTTCAAGC
Oligo rev AAACGCTTGAAGGTCGCCATGCAG
Chromosome start end strand MM target_seq PAM alignment distance position gene name gene id
GU305914.1 186253 186275 + 0 CTGCATGGCGACCTTCAAGC TGG ||||||||[||||||||||||]PAM NA Intergenic NA NA

T14158 CTTCAAGCTGGATTGCACTTCGG 1000
Oligo fwd CACCCTTCAAGCTGGATTGCACTT
Oligo rev AAACAAGTGCAATCCAGCTTGAAG
Chromosome start end strand MM target_seq PAM alignment distance position gene name gene id
GU305914.1 186265 186287 + 0 CTTCAAGCTGGATTGCACTT CGG ||||||||[||||||||||||]PAM NA Intergenic NA NA

T14159 AGCTGGATTGCACTTCGGCTCGG 1000
Oligo fwd CACCAGCTGGATTGCACTTCGGCT
Oligo rev AAACAGCCGAAGTGCAATCCAGCT
Chromosome start end strand MM target_seq PAM alignment distance position gene name gene id
GU305914.1 186270 186292 + 0 AGCTGGATTGCACTTCGGCT CGG ||||||||[||||||||||||]PAM NA Intergenic NA NA

T14160 ATTGCACTTCGGCTCGGACCCGG 1000
Oligo fwd CACCATTGCACTTCGGCTCGGACC
Oligo rev AAACGGTCCGAGCCGAAGTGCAAT
Chromosome start end strand MM target_seq PAM alignment distance position gene name gene id
GU305914.1 186276 186298 + 0 ATTGCACTTCGGCTCGGACC CGG ||||||||[||||||||||||]PAM NA Intergenic NA NA

T14161 TTGCACTTCGGCTCGGACCCGGG 1000
Oligo fwd CACCTTGCACTTCGGCTCGGACCC
Oligo rev AAACGGGTCCGAGCCGAAGTGCAA
Chromosome start end strand MM target_seq PAM alignment distance position gene name gene id
GU305914.1 186277 186299 + 0 TTGCACTTCGGCTCGGACCC GGG ||||||||[||||||||||||]PAM NA Intergenic NA NA

T14162 TGCACTTCGGCTCGGACCCGGGG 1000
Oligo fwd CACCTGCACTTCGGCTCGGACCCG
Oligo rev AAACCGGGTCCGAGCCGAAGTGCA
Chromosome start end strand MM target_seq PAM alignment distance position gene name gene id
GU305914.1 186278 186300 + 0 TGCACTTCGGCTCGGACCCG GGG ||||||||[||||||||||||]PAM NA Intergenic NA NA

T14163 GCACTTCGGCTCGGACCCGGGGG 1000
Oligo fwd CACCGCACTTCGGCTCGGACCCGG
Oligo rev AAACCCGGGTCCGAGCCGAAGTGC
Chromosome start end strand MM target_seq PAM alignment distance position gene name gene id
GU305914.1 186279 186301 + 0 GCACTTCGGCTCGGACCCGG GGG ||||||||[||||||||||||]PAM NA Intergenic NA NA

T14164 CACTTCGGCTCGGACCCGGGGGG 1000
Oligo fwd CACCCACTTCGGCTCGGACCCGGG
Oligo rev AAACCCCGGGTCCGAGCCGAAGTG
Chromosome start end strand MM target_seq PAM alignment distance position gene name gene id
GU305914.1 186280 186302 + 0 CACTTCGGCTCGGACCCGGG GGG ||||||||[||||||||||||]PAM NA Intergenic NA NA

T14165 CCGGGGGGTTGTAAGTATCCTGG 1000
Oligo fwd CACCCCGGGGGGTTGTAAGTATCC
Oligo rev AAACGGATACTTACAACCCCCCGG
Chromosome start end strand MM target_seq PAM alignment distance position gene name gene id
GU305914.1 186295 186317 + 0 CCGGGGGGTTGTAAGTATCC TGG ||||||||[||||||||||||]PAM NA Intergenic NA NA

T14166 TTGTAAGTATCCTGGAGCACAGG 1000
Oligo fwd CACCTTGTAAGTATCCTGGAGCAC
Oligo rev AAACGTGCTCCAGGATACTTACAA
Chromosome start end strand MM target_seq PAM alignment distance position gene name gene id
GU305914.1 186303 186325 + 0 TTGTAAGTATCCTGGAGCAC AGG ||||||||[||||||||||||]PAM NA Intergenic NA NA

T14167 TGTAAGTATCCTGGAGCACAGGG 1000
Oligo fwd CACCTGTAAGTATCCTGGAGCACA
Oligo rev AAACTGTGCTCCAGGATACTTACA
Chromosome start end strand MM target_seq PAM alignment distance position gene name gene id
GU305914.1 186304 186326 + 0 TGTAAGTATCCTGGAGCACA GGG ||||||||[||||||||||||]PAM NA Intergenic NA NA

T14168 GCACAGGGACCCCGCGACCGTGG 1000
Oligo fwd CACCGCACAGGGACCCCGCGACCG
Oligo rev AAACCGGTCGCGGGGTCCCTGTGC
Chromosome start end strand MM target_seq PAM alignment distance position gene name gene id
GU305914.1 186319 186341 + 0 GCACAGGGACCCCGCGACCG TGG ||||||||[||||||||||||]PAM NA Intergenic NA NA

T14169 TGGACGATATCTTTATGCAGCGG 1000

<table>
<tr><td>Oligo fwd</td><td>CACCTGGACGATATCTTTATGCAG</td></tr>
<tr><td>Oligo rev</td><td>AAACCTGCATAAAGATATCGTCCA</td></tr>
<tr><td>Chromosome</td><td>start</td><td>end</td><td>strand</td><td>MM</td><td>target_seq</td><td>PAM</td><td>alignment</td><td>distance</td><td>position</td><td>gene name</td><td>gene id</td></tr>
<tr><td>GU305914.1</td><td>186339</td><td>186361</td><td>+</td><td>0</td><td>TGGACGATATCTTTATGCAG</td><td>CGG</td><td>||||||||[||||||||||||]PAM</td><td>NA</td><td>Intergenic</td><td>NA</td><td>NA</td></tr>
<tr><td></td></tr>
<tr><td>T14170</td><td>GGACGATATCTTTATGCAGCGGG</td><td>1000</td></tr>
<tr><td>Oligo fwd</td><td>CACCGGACGATATCTTTATGCAGC</td></tr>
<tr><td>Oligo rev</td><td>AAACGCTGCATAAAGATATCGTCC</td></tr>
<tr><td>Chromosome</td><td>start</td><td>end</td><td>strand</td><td>MM</td><td>target_seq</td><td>PAM</td><td>alignment</td><td>distance</td><td>position</td><td>gene name</td><td>gene id</td></tr>
<tr><td>GU305914.1</td><td>186340</td><td>186362</td><td>+</td><td>0</td><td>GGACGATATCTTTATGCAGC</td><td>GGG</td><td>||||||||[||||||||||||]PAM</td><td>NA</td><td>Intergenic</td><td>NA</td><td>NA</td></tr>
<tr><td></td></tr>
<tr><td>T14171</td><td>ACGACATCAACGACGTGCTGCGG</td><td>1000</td></tr>
<tr><td>Oligo fwd</td><td>CACCACGACATCAACGACGTGCTG</td></tr>
<tr><td>Oligo rev</td><td>AAACCAGCACGTCGTTGATGTCGT</td></tr>
<tr><td>Chromosome</td><td>start</td><td>end</td><td>strand</td><td>MM</td><td>target_seq</td><td>PAM</td><td>alignment</td><td>distance</td><td>position</td><td>gene name</td><td>gene id</td></tr>
<tr><td>GU305914.1</td><td>186369</td><td>186391</td><td>+</td><td>0</td><td>ACGACATCAACGACGTGCTG</td><td>CGG</td><td>||||||||[||||||||||||]PAM</td><td>NA</td><td>Intergenic</td><td>NA</td><td>NA</td></tr>
<tr><td></td></tr>
<tr><td>T14172</td><td>CATCAACGACGTGCTGCGGCCGG</td><td>1000</td></tr>
<tr><td>Oligo fwd</td><td>CACCCATCAACGACGTGCTGCGGC</td></tr>
<tr><td>Oligo rev</td><td>AAACGCCGCAGCACGTCGTTGATG</td></tr>
<tr><td>Chromosome</td><td>start</td><td>end</td><td>strand</td><td>MM</td><td>target_seq</td><td>PAM</td><td>alignment</td><td>distance</td><td>position</td><td>gene name</td><td>gene id</td></tr>
<tr><td>GU305914.1</td><td>186373</td><td>186395</td><td>+</td><td>0</td><td>CATCAACGACGTGCTGCGGC</td><td>CGG</td><td>||||||||[||||||||||||]PAM</td><td>NA</td><td>Intergenic</td><td>NA</td><td>NA</td></tr>
<tr><td></td></tr>
<tr><td>T14173</td><td>CCGAATACAGAAGAACCCTGAGG</td><td>1000</td></tr>
<tr><td>Oligo fwd</td><td>CACCCCGAATACAGAAGAACCCTG</td></tr>
<tr><td>Oligo rev</td><td>AAACCAGGGTTCTTCTGTATTCGG</td></tr>
<tr><td>Chromosome</td><td>start</td><td>end</td><td>strand</td><td>MM</td><td>target_seq</td><td>PAM</td><td>alignment</td><td>distance</td><td>position</td><td>gene name</td><td>gene id</td></tr>
<tr><td>GU305914.1</td><td>186417</td><td>186439</td><td>+</td><td>0</td><td>CCGAATACAGAAGAACCCTG</td><td>AGG</td><td>||||||||[||||||||||||]PAM</td><td>NA</td><td>Intergenic</td><td>NA</td><td>NA</td></tr>
<tr><td></td></tr>
<tr><td>T14174</td><td>CGAATACAGAAGAACCCTGAGGG</td><td>1000</td></tr>
<tr><td>Oligo fwd</td><td>CACCCGAATACAGAAGAACCCTGA</td></tr>
<tr><td>Oligo rev</td><td>AAACTCAGGGTTCTTCTGTATTCG</td></tr>
<tr><td>Chromosome</td><td>start</td><td>end</td><td>strand</td><td>MM</td><td>target_seq</td><td>PAM</td><td>alignment</td><td>distance</td><td>position</td><td>gene name</td><td>gene id</td></tr>
<tr><td>GU305914.1</td><td>186418</td><td>186440</td><td>+</td><td>0</td><td>CGAATACAGAAGAACCCTGA</td><td>GGG</td><td>||||||||[||||||||||||]PAM</td><td>NA</td><td>Intergenic</td><td>NA</td><td>NA</td></tr>
<tr><td></td></tr>
<tr><td>T14175</td><td>ATACAGAAGAACCCTGAGGGTGG</td><td>1000</td></tr>
<tr><td>Oligo fwd</td><td>CACCATACAGAAGAACCCTGAGGG</td></tr>
<tr><td>Oligo rev</td><td>AAACCCCTCAGGGTTCTTCTGTAT</td></tr>
<tr><td>Chromosome</td><td>start</td><td>end</td><td>strand</td><td>MM</td><td>target_seq</td><td>PAM</td><td>alignment</td><td>distance</td><td>position</td><td>gene name</td><td>gene id</td></tr>
<tr><td>GU305914.1</td><td>186421</td><td>186443</td><td>+</td><td>0</td><td>ATACAGAAGAACCCTGAGGG</td><td>TGG</td><td>||||||||[||||||||||||]PAM</td><td>NA</td><td>Intergenic</td><td>NA</td><td>NA</td></tr>
<tr><td></td></tr>
<tr><td>T14176</td><td>AAGAACCCTGAGGGTGGCCGTGG</td><td>1000</td></tr>
<tr><td>Oligo fwd</td><td>CACCAAGAACCCTGAGGGTGGCCG</td></tr>
<tr><td>Oligo rev</td><td>AAACCGGCCACCCTCAGGGTTCTT</td></tr>
<tr><td>Chromosome</td><td>start</td><td>end</td><td>strand</td><td>MM</td><td>target_seq</td><td>PAM</td><td>alignment</td><td>distance</td><td>position</td><td>gene name</td><td>gene id</td></tr>
<tr><td>GU305914.1</td><td>186427</td><td>186449</td><td>+</td><td>0</td><td>AAGAACCCTGAGGGTGGCCG</td><td>TGG</td><td>||||||||[||||||||||||]PAM</td><td>NA</td><td>Intergenic</td><td>NA</td><td>NA</td></tr>
<tr><td></td></tr>
<tr><td>T14177</td><td>AACCCTGAGGGTGGCCGTGGAGG</td><td>1000</td></tr>
<tr><td>Oligo fwd</td><td>CACCAACCCTGAGGGTGGCCGTGG</td></tr>
<tr><td>Oligo rev</td><td>AAACCCACGGCCACCCTCAGGGTT</td></tr>
<tr><td>Chromosome</td><td>start</td><td>end</td><td>strand</td><td>MM</td><td>target_seq</td><td>PAM</td><td>alignment</td><td>distance</td><td>position</td><td>gene name</td><td>gene id</td></tr>
<tr><td>GU305914.1</td><td>186430</td><td>186452</td><td>+</td><td>0</td><td>AACCCTGAGGGTGGCCGTGG</td><td>AGG</td><td>||||||||[||||||||||||]PAM</td><td>NA</td><td>Intergenic</td><td>NA</td><td>NA</td></tr>
<tr><td></td></tr>
<tr><td>T14178</td><td>GAGGCCGCCACCGAACTCCACGG</td><td>1000</td></tr>
<tr><td>Oligo fwd</td><td>CACCGAGGCCGCCACCGAACTCCA</td></tr>
<tr><td>Oligo rev</td><td>AAACTGGAGTTCGGTGGCGGCCTC</td></tr>
<tr><td>Chromosome</td><td>start</td><td>end</td><td>strand</td><td>MM</td><td>target_seq</td><td>PAM</td><td>alignment</td><td>distance</td><td>position</td><td>gene name</td><td>gene id</td></tr>
<tr><td>GU305914.1</td><td>186449</td><td>186471</td><td>+</td><td>0</td><td>GAGGCCGCCACCGAACTCCA</td><td>CGG</td><td>||||||||[||||||||||||]PAM</td><td>NA</td><td>Intergenic</td><td>NA</td><td>NA</td></tr>
<tr><td></td></tr>
<tr><td>T14179</td><td>CACGGCGTCTCCGCTCTGTCCGG</td><td>1000</td></tr>
<tr><td>Oligo fwd</td><td>CACCCACGGCGTCTCCGCTCTGTC</td></tr>
<tr><td>Oligo rev</td><td>AAACGACAGAGCGGAGACGCCGTG</td></tr>
<tr><td>Chromosome</td><td>start</td><td>end</td><td>strand</td><td>MM</td><td>target_seq</td><td>PAM</td><td>alignment</td><td>distance</td><td>position</td><td>gene name</td><td>gene id</td></tr>
<tr><td>GU305914.1</td><td>186467</td><td>186489</td><td>+</td><td>0</td><td>CACGGCGTCTCCGCTCTGTC</td><td>CGG</td><td>||||||||[||||||||||||]PAM</td><td>NA</td><td>Intergenic</td><td>NA</td><td>NA</td></tr>
<tr><td></td></tr>
<tr><td>T14180</td><td>CGGCCAGCTGACCAAGTTCTTGG</td><td>1000</td></tr>
<tr><td>Oligo fwd</td><td>CACCCGGCCAGCTGACCAAGTTCT</td></tr>
<tr><td>Oligo rev</td><td>AAACAGAACTTGGTCAGCTGGCCG</td></tr>
<tr><td>Chromosome</td><td>start</td><td>end</td><td>strand</td><td>MM</td><td>target_seq</td><td>PAM</td><td>alignment</td><td>distance</td><td>position</td><td>gene name</td><td>gene id</td></tr>
<tr><td>GU305914.1</td><td>186487</td><td>186509</td><td>+</td><td>0</td><td>CGGCCAGCTGACCAAGTTCT</td><td>TGG</td><td>||||||||[||||||||||||]PAM</td><td>NA</td><td>Intergenic</td><td>NA</td><td>NA</td></tr>
<tr><td></td></tr>
<tr><td>T14181</td><td>CCAGCTGACCAAGTTCTTGGAGG</td><td>1000</td></tr>
<tr><td>Oligo fwd</td><td>CACCCCAGCTGACCAAGTTCTTGG</td></tr>
<tr><td>Oligo rev</td><td>AAACCCAAGAACTTGGTCAGCTGG</td></tr>
<tr><td>Chromosome</td><td>start</td><td>end</td><td>strand</td><td>MM</td><td>target_seq</td><td>PAM</td><td>alignment</td><td>distance</td><td>position</td><td>gene name</td><td>gene id</td></tr>
<tr><td>GU305914.1</td><td>186490</td><td>186512</td><td>+</td><td>0</td><td>CCAGCTGACCAAGTTCTTGG</td><td>AGG</td><td>||||||||[||||||||||||]PAM</td><td>NA</td><td>Intergenic</td><td>NA</td><td>NA</td></tr>
<tr><td></td></tr>
<tr><td>T14182</td><td>AAGTTCTTGGAGGACCACGATGG</td><td>1000</td></tr>
<tr><td>Oligo fwd</td><td>CACCAAGTTCTTGGAGGACCACGA</td></tr>
<tr><td>Oligo rev</td><td>AAACTCGTGGTCCTCCAAGAACTT</td></tr>
<tr><td>Chromosome</td><td>start</td><td>end</td><td>strand</td><td>MM</td><td>target_seq</td><td>PAM</td><td>alignment</td><td>distance</td><td>position</td><td>gene name</td><td>gene id</td></tr>
<tr><td>GU305914.1</td><td>186500</td><td>186522</td><td>+</td><td>0</td><td>AAGTTCTTGGAGGACCACGA</td><td>TGG</td><td>||||||||[||||||||||||]PAM</td><td>NA</td><td>Intergenic</td><td>NA</td><td>NA</td></tr>
<tr><td></td></tr>
<tr><td>T14183</td><td>CGATGGAGCGATGCTGCCCCTGG</td><td>1000</td></tr>
<tr><td>Oligo fwd</td><td>CACCCGATGGAGCGATGCTGCCCC</td></tr>
<tr><td>Oligo rev</td><td>AAACGGGGCAGCATCGCTCCATCG</td></tr>
<tr><td>Chromosome</td><td>start</td><td>end</td><td>strand</td><td>MM</td><td>target_seq</td><td>PAM</td><td>alignment</td><td>distance</td><td>position</td><td>gene name</td><td>gene id</td></tr>
<tr><td>GU305914.1</td><td>186517</td><td>186539</td><td>+</td><td>0</td><td>CGATGGAGCGATGCTGCCCC</td><td>TGG</td><td>||||||||[||||||||||||]PAM</td><td>NA</td><td>Intergenic</td><td>NA</td><td>NA</td></tr>
</table>

| | | | | | | | | | | | |
|---|---|---|---|---|---|---|---|---|---|---|---|
| T14184 | GAGCGATGCTGCCCCTGGCCTGG | 1000 | | | | | | | | | |
| Oligo fwd | CACCGAGCGATGCTGCCCCTGGCC | | | | | | | | | | |
| Oligo rev | AAACGGCCAGGGGCAGCATCGCTC | | | | | | | | | | |
| Chromosome | start | end | strand | MM | target_seq | PAM | alignment | distance | position | gene name | gene id |
| GU305914.1 | 186522 | 186544 | + | 0 | GAGCGATGCTGCCCCTGGCC | TGG | \|\|\|\|\|\|\|\|[\|\|\|\|\|\|\|\|\|\|\|\|]PAM | NA | Intergenic | NA | NA |
| T14185 | CCCTGGCCTGGCCCCCGAACAGG | 1000 | | | | | | | | | |
| Oligo fwd | CACCCCCTGGCCTGGCCCCCGAAC | | | | | | | | | | |
| Oligo rev | AAACGTTCGGGGGCCAGGCCAGGG | | | | | | | | | | |
| Chromosome | start | end | strand | MM | target_seq | PAM | alignment | distance | position | gene name | gene id |
| GU305914.1 | 186534 | 186556 | + | 0 | CCCTGGCCTGGCCCCCGAAC | AGG | \|\|\|\|\|\|\|\|[\|\|\|\|\|\|\|\|\|\|\|\|]PAM | NA | Intergenic | NA | NA |
| T14186 | CAGGTACCTTCGCCTGACGTCGG | 1000 | | | | | | | | | |
| Oligo fwd | CACCCAGGTACCTTCGCCTGACGT | | | | | | | | | | |
| Oligo rev | AAACACGTCAGGCGAAGGTACCTG | | | | | | | | | | |
| Chromosome | start | end | strand | MM | target_seq | PAM | alignment | distance | position | gene name | gene id |
| GU305914.1 | 186553 | 186575 | + | 0 | CAGGTACCTTCGCCTGACGT | CGG | \|\|\|\|\|\|\|\|[\|\|\|\|\|\|\|\|\|\|\|\|]PAM | NA | Intergenic | NA | NA |
| T14187 | AGGTACCTTCGCCTGACGTCGGG | 1000 | | | | | | | | | |
| Oligo fwd | CACCAGGTACCTTCGCCTGACGTC | | | | | | | | | | |
| Oligo rev | AAACGACGTCAGGCGAAGGTACCT | | | | | | | | | | |
| Chromosome | start | end | strand | MM | target_seq | PAM | alignment | distance | position | gene name | gene id |
| GU305914.1 | 186554 | 186576 | + | 0 | AGGTACCTTCGCCTGACGTC | GGG | \|\|\|\|\|\|\|\|[\|\|\|\|\|\|\|\|\|\|\|\|]PAM | NA | Intergenic | NA | NA |
| T14188 | CCTGACGTCGGGCAGAAAGCTGG | 1000 | | | | | | | | | |
| Oligo fwd | CACCCCTGACGTCGGGCAGAAAGC | | | | | | | | | | |
| Oligo rev | AAACGCTTTCTGCCCGACGTCAGG | | | | | | | | | | |
| Chromosome | start | end | strand | MM | target_seq | PAM | alignment | distance | position | gene name | gene id |
| GU305914.1 | 186565 | 186587 | + | 0 | CCTGACGTCGGGCAGAAAGC | TGG | \|\|\|\|\|\|\|\|[\|\|\|\|\|\|\|\|\|\|\|\|]PAM | NA | Intergenic | NA | NA |
| T14189 | CTGACGTCGGGCAGAAAGCTGGG | 1000 | | | | | | | | | |
| Oligo fwd | CACCCTGACGTCGGGCAGAAAGCT | | | | | | | | | | |
| Oligo rev | AAACAGCTTTCTGCCCGACGTCAG | | | | | | | | | | |
| Chromosome | start | end | strand | MM | target_seq | PAM | alignment | distance | position | gene name | gene id |
| GU305914.1 | 186566 | 186588 | + | 0 | CTGACGTCGGGCAGAAAGCT | GGG | \|\|\|\|\|\|\|\|[\|\|\|\|\|\|\|\|\|\|\|\|]PAM | NA | Intergenic | NA | NA |
| T14190 | AAAGCTGGGTTATCTCGACGTGG | 1000 | | | | | | | | | |
| Oligo fwd | CACCAAAGCTGGGTTATCTCGACG | | | | | | | | | | |
| Oligo rev | AAACCGTCGAGATAACCCAGCTTT | | | | | | | | | | |
| Chromosome | start | end | strand | MM | target_seq | PAM | alignment | distance | position | gene name | gene id |
| GU305914.1 | 186580 | 186602 | + | 0 | AAAGCTGGGTTATCTCGACG | TGG | \|\|\|\|\|\|\|\|[\|\|\|\|\|\|\|\|\|\|\|\|]PAM | NA | Intergenic | NA | NA |
| T14191 | GCTGGGTTATCTCGACGTGGCGG | 1000 | | | | | | | | | |
| Oligo fwd | CACCGCTGGGTTATCTCGACGTGG | | | | | | | | | | |
| Oligo rev | AAACCCACGTCGAGATAACCCAGC | | | | | | | | | | |
| Chromosome | start | end | strand | MM | target_seq | PAM | alignment | distance | position | gene name | gene id |
| GU305914.1 | 186583 | 186605 | + | 0 | GCTGGGTTATCTCGACGTGG | CGG | \|\|\|\|\|\|\|\|[\|\|\|\|\|\|\|\|\|\|\|\|]PAM | NA | Intergenic | NA | NA |
| T14192 | CGTGGCGGAGAAGCTGCGCAAGG | 1000 | | | | | | | | | |
| Oligo fwd | CACCCGTGGCGGAGAAGCTGCGCA | | | | | | | | | | |
| Oligo rev | AAACTGCGCAGCTTCTCCGCCACG | | | | | | | | | | |
| Chromosome | start | end | strand | MM | target_seq | PAM | alignment | distance | position | gene name | gene id |
| GU305914.1 | 186598 | 186620 | + | 0 | CGTGGCGGAGAAGCTGCGCA | AGG | \|\|\|\|\|\|\|\|[\|\|\|\|\|\|\|\|\|\|\|\|]PAM | NA | Intergenic | NA | NA |
| T14193 | AAGGAATACCCCATGTGCAACGG | 1000 | | | | | | | | | |
| Oligo fwd | CACCAAGGAATACCCCATGTGCAA | | | | | | | | | | |
| Oligo rev | AAACTTGCACATGGGGTATTCCTT | | | | | | | | | | |
| Chromosome | start | end | strand | MM | target_seq | PAM | alignment | distance | position | gene name | gene id |
| GU305914.1 | 186617 | 186639 | + | 0 | AAGGAATACCCCATGTGCAA | CGG | \|\|\|\|\|\|\|\|[\|\|\|\|\|\|\|\|\|\|\|\|]PAM | NA | Intergenic | NA | NA |
| T14194 | CATGTGCAACGGTCCCCTCGAGG | 1000 | | | | | | | | | |
| Oligo fwd | CACCCATGTGCAACGGTCCCCTCG | | | | | | | | | | |
| Oligo rev | AAACCGAGGGGACCGTTGCACATG | | | | | | | | | | |
| Chromosome | start | end | strand | MM | target_seq | PAM | alignment | distance | position | gene name | gene id |
| GU305914.1 | 186628 | 186650 | + | 0 | CATGTGCAACGGTCCCCTCG | AGG | \|\|\|\|\|\|\|\|[\|\|\|\|\|\|\|\|\|\|\|\|]PAM | NA | Intergenic | NA | NA |
| T14195 | AACGGTCCCCTCGAGGTTCTCGG | 1000 | | | | | | | | | |
| Oligo fwd | CACCAACGGTCCCCTCGAGGTTCT | | | | | | | | | | |
| Oligo rev | AAACAGAACCTCGAGGGGACCGTT | | | | | | | | | | |
| Chromosome | start | end | strand | MM | target_seq | PAM | alignment | distance | position | gene name | gene id |
| GU305914.1 | 186635 | 186657 | + | 0 | AACGGTCCCCTCGAGGTTCT | CGG | \|\|\|\|\|\|\|\|[\|\|\|\|\|\|\|\|\|\|\|\|]PAM | NA | Intergenic | NA | NA |
| T14196 | GAGGTTCTCGGCGTGATCCTCGG | 1000 | | | | | | | | | |
| Oligo fwd | CACCGAGGTTCTCGGCGTGATCCT | | | | | | | | | | |
| Oligo rev | AAACAGGATCACGCCGAGAACCTC | | | | | | | | | | |
| Chromosome | start | end | strand | MM | target_seq | PAM | alignment | distance | position | gene name | gene id |
| GU305914.1 | 186647 | 186669 | + | 0 | GAGGTTCTCGGCGTGATCCT | CGG | \|\|\|\|\|\|\|\|[\|\|\|\|\|\|\|\|\|\|\|\|]PAM | NA | Intergenic | NA | NA |
| T14197 | CTCGGCGTGATCCTCGGACCCGG | 1000 | | | | | | | | | |
| Oligo fwd | CACCCTCGGCGTGATCCTCGGACC | | | | | | | | | | |
| Oligo rev | AAACGGTCCGAGGATCACGCCGAG | | | | | | | | | | |
| Chromosome | start | end | strand | MM | target_seq | PAM | alignment | distance | position | gene name | gene id |
| GU305914.1 | 186653 | 186675 | + | 0 | CTCGGCGTGATCCTCGGACC | CGG | \|\|\|\|\|\|\|\|[\|\|\|\|\|\|\|\|\|\|\|\|]PAM | NA | Intergenic | NA | NA |
| T14198 | CCAGCACCCCGTCCTCTTGATGG | 1000 | | | | | | | | | |
| Oligo fwd | CACCCCAGCACCCCGTCCTCTTGA | | | | | | | | | | |
| Oligo rev | AAACTCAAGAGGACGGGGTGCTGG | | | | | | | | | | |
| Chromosome | start | end | strand | MM | target_seq | PAM | alignment | distance | position | gene name | gene id |
| GU305914.1 | 186694 | 186716 | + | 0 | CCAGCACCCCGTCCTCTTGA | TGG | \|\|\|\|\|\|\|\|[\|\|\|\|\|\|\|\|\|\|\|\|]PAM | NA | Intergenic | NA | NA |

<table>
<tr><td>T14199</td><td>CAGCACCCCGTCCTCTTGATGGG</td><td>1000</td></tr>
<tr><td>Oligo fwd</td><td>CACCCAGCACCCCGTCCTCTTGAT</td></tr>
<tr><td>Oligo rev</td><td>AAACATCAAGAGGACGGGGTGCTG</td></tr>
<tr><td>Chromosome</td><td>start</td><td>end</td><td>strand</td><td>MM</td><td>target_seq</td><td>PAM</td><td>alignment</td><td>distance</td><td>position</td><td>gene name</td><td>gene id</td></tr>
<tr><td>GU305914.1</td><td>186695</td><td>186717</td><td>+</td><td>0</td><td>CAGCACCCCGTCCTCTTGAT</td><td>GGG</td><td>||||||||[||||||||||||]PAM</td><td>NA</td><td>Intergenic</td><td>NA</td><td>NA</td></tr>
<tr><td>T14200</td><td>GTCCTCTTGATGGGCCTCACCGG</td><td>1000</td></tr>
<tr><td>Oligo fwd</td><td>CACCGTCCTCTTGATGGGCCTCAC</td></tr>
<tr><td>Oligo rev</td><td>AAACGTGAGGCCCATCAAGAGGAC</td></tr>
<tr><td>Chromosome</td><td>start</td><td>end</td><td>strand</td><td>MM</td><td>target_seq</td><td>PAM</td><td>alignment</td><td>distance</td><td>position</td><td>gene name</td><td>gene id</td></tr>
<tr><td>GU305914.1</td><td>186704</td><td>186726</td><td>+</td><td>0</td><td>GTCCTCTTGATGGGCCTCAC</td><td>CGG</td><td>||||||||[||||||||||||]PAM</td><td>NA</td><td>Intergenic</td><td>NA</td><td>NA</td></tr>
<tr><td>T14201</td><td>TCCTCTTGATGGGCCTCACCGGG</td><td>1000</td></tr>
<tr><td>Oligo fwd</td><td>CACCTCCTCTTGATGGGCCTCACC</td></tr>
<tr><td>Oligo rev</td><td>AAACGGTGAGGCCCATCAAGAGGA</td></tr>
<tr><td>Chromosome</td><td>start</td><td>end</td><td>strand</td><td>MM</td><td>target_seq</td><td>PAM</td><td>alignment</td><td>distance</td><td>position</td><td>gene name</td><td>gene id</td></tr>
<tr><td>GU305914.1</td><td>186705</td><td>186727</td><td>+</td><td>0</td><td>TCCTCTTGATGGGCCTCACC</td><td>GGG</td><td>||||||||[||||||||||||]PAM</td><td>NA</td><td>Intergenic</td><td>NA</td><td>NA</td></tr>
<tr><td>T14202</td><td>ACCATCTTCCTCCACGCCAGAGG</td><td>1000</td></tr>
<tr><td>Oligo fwd</td><td>CACCACCATCTTCCTCCACGCCAG</td></tr>
<tr><td>Oligo rev</td><td>AAACCTGGCGTGGAGGAAGATGGT</td></tr>
<tr><td>Chromosome</td><td>start</td><td>end</td><td>strand</td><td>MM</td><td>target_seq</td><td>PAM</td><td>alignment</td><td>distance</td><td>position</td><td>gene name</td><td>gene id</td></tr>
<tr><td>GU305914.1</td><td>186728</td><td>186750</td><td>+</td><td>0</td><td>ACCATCTTCCTCCACGCCAG</td><td>AGG</td><td>||||||||[||||||||||||]PAM</td><td>NA</td><td>Intergenic</td><td>NA</td><td>NA</td></tr>
<tr><td>T14203</td><td>TCTTCCTCCACGCCAGAGGCAGG</td><td>1000</td></tr>
<tr><td>Oligo fwd</td><td>CACCTCTTCCTCCACGCCAGAGGC</td></tr>
<tr><td>Oligo rev</td><td>AAACGCCTCTGGCGTGGAGGAAGA</td></tr>
<tr><td>Chromosome</td><td>start</td><td>end</td><td>strand</td><td>MM</td><td>target_seq</td><td>PAM</td><td>alignment</td><td>distance</td><td>position</td><td>gene name</td><td>gene id</td></tr>
<tr><td>GU305914.1</td><td>186732</td><td>186754</td><td>+</td><td>0</td><td>TCTTCCTCCACGCCAGAGGC</td><td>AGG</td><td>||||||||[||||||||||||]PAM</td><td>NA</td><td>Intergenic</td><td>NA</td><td>NA</td></tr>
<tr><td>T14204</td><td>ACGCCAGAGGCAGGCCCGTCTGG</td><td>1000</td></tr>
<tr><td>Oligo fwd</td><td>CACCACGCCAGAGGCAGGCCCGTC</td></tr>
<tr><td>Oligo rev</td><td>AAACGACGGGCCTGCCTCTGGCGT</td></tr>
<tr><td>Chromosome</td><td>start</td><td>end</td><td>strand</td><td>MM</td><td>target_seq</td><td>PAM</td><td>alignment</td><td>distance</td><td>position</td><td>gene name</td><td>gene id</td></tr>
<tr><td>GU305914.1</td><td>186741</td><td>186763</td><td>+</td><td>0</td><td>ACGCCAGAGGCAGGCCCGTC</td><td>TGG</td><td>||||||||[||||||||||||]PAM</td><td>NA</td><td>Intergenic</td><td>NA</td><td>NA</td></tr>
<tr><td>T14205</td><td>CTGGAGTCCCGACTACGATCCGG</td><td>1000</td></tr>
<tr><td>Oligo fwd</td><td>CACCCTGGAGTCCCGACTACGATC</td></tr>
<tr><td>Oligo rev</td><td>AAACGATCGTAGTCGGGACTCCAG</td></tr>
<tr><td>Chromosome</td><td>start</td><td>end</td><td>strand</td><td>MM</td><td>target_seq</td><td>PAM</td><td>alignment</td><td>distance</td><td>position</td><td>gene name</td><td>gene id</td></tr>
<tr><td>GU305914.1</td><td>186760</td><td>186782</td><td>+</td><td>0</td><td>CTGGAGTCCCGACTACGATC</td><td>CGG</td><td>||||||||[||||||||||||]PAM</td><td>NA</td><td>Intergenic</td><td>NA</td><td>NA</td></tr>
<tr><td>T14206</td><td>GTCCCGACTACGATCCGGAGCGG</td><td>1000</td></tr>
<tr><td>Oligo fwd</td><td>CACCGTCCCGACTACGATCCGGAG</td></tr>
<tr><td>Oligo rev</td><td>AAACCTCCGGATCGTAGTCGGGAC</td></tr>
<tr><td>Chromosome</td><td>start</td><td>end</td><td>strand</td><td>MM</td><td>target_seq</td><td>PAM</td><td>alignment</td><td>distance</td><td>position</td><td>gene name</td><td>gene id</td></tr>
<tr><td>GU305914.1</td><td>186765</td><td>186787</td><td>+</td><td>0</td><td>GTCCCGACTACGATCCGGAG</td><td>CGG</td><td>||||||||[||||||||||||]PAM</td><td>NA</td><td>Intergenic</td><td>NA</td><td>NA</td></tr>
<tr><td>T14207</td><td>TCCCGACTACGATCCGGAGCGGG</td><td>1000</td></tr>
<tr><td>Oligo fwd</td><td>CACCTCCCGACTACGATCCGGAGC</td></tr>
<tr><td>Oligo rev</td><td>AAACGCTCCGGATCGTAGTCGGGA</td></tr>
<tr><td>Chromosome</td><td>start</td><td>end</td><td>strand</td><td>MM</td><td>target_seq</td><td>PAM</td><td>alignment</td><td>distance</td><td>position</td><td>gene name</td><td>gene id</td></tr>
<tr><td>GU305914.1</td><td>186766</td><td>186788</td><td>+</td><td>0</td><td>TCCCGACTACGATCCGGAGC</td><td>GGG</td><td>||||||||[||||||||||||]PAM</td><td>NA</td><td>Intergenic</td><td>NA</td><td>NA</td></tr>
<tr><td>T14208</td><td>GGACGCCGATCGACTCTTCCTGG</td><td>1000</td></tr>
<tr><td>Oligo fwd</td><td>CACCGGACGCCGATCGACTCTTCC</td></tr>
<tr><td>Oligo rev</td><td>AAACGGAAGAGTCGATCGGCGTCC</td></tr>
<tr><td>Chromosome</td><td>start</td><td>end</td><td>strand</td><td>MM</td><td>target_seq</td><td>PAM</td><td>alignment</td><td>distance</td><td>position</td><td>gene name</td><td>gene id</td></tr>
<tr><td>GU305914.1</td><td>186787</td><td>186809</td><td>+</td><td>0</td><td>GGACGCCGATCGACTCTTCC</td><td>TGG</td><td>||||||||[||||||||||||]PAM</td><td>NA</td><td>Intergenic</td><td>NA</td><td>NA</td></tr>
<tr><td>T14209</td><td>GCCGAGTCCCTGCAGTCCTTCGG</td><td>1000</td></tr>
<tr><td>Oligo fwd</td><td>CACCGCCGAGTCCCTGCAGTCCTT</td></tr>
<tr><td>Oligo rev</td><td>AAACAAGGACTGCAGGGACTCGGC</td></tr>
<tr><td>Chromosome</td><td>start</td><td>end</td><td>strand</td><td>MM</td><td>target_seq</td><td>PAM</td><td>alignment</td><td>distance</td><td>position</td><td>gene name</td><td>gene id</td></tr>
<tr><td>GU305914.1</td><td>186812</td><td>186834</td><td>+</td><td>0</td><td>GCCGAGTCCCTGCAGTCCTT</td><td>CGG</td><td>||||||||[||||||||||||]PAM</td><td>NA</td><td>Intergenic</td><td>NA</td><td>NA</td></tr>
<tr><td>T14210</td><td>AGTCCCTGCAGTCCTTCGGTAGG</td><td>1000</td></tr>
<tr><td>Oligo fwd</td><td>CACCAGTCCCTGCAGTCCTTCGGT</td></tr>
<tr><td>Oligo rev</td><td>AAACACCGAAGGACTGCAGGGACT</td></tr>
<tr><td>Chromosome</td><td>start</td><td>end</td><td>strand</td><td>MM</td><td>target_seq</td><td>PAM</td><td>alignment</td><td>distance</td><td>position</td><td>gene name</td><td>gene id</td></tr>
<tr><td>GU305914.1</td><td>186816</td><td>186838</td><td>+</td><td>0</td><td>AGTCCCTGCAGTCCTTCGGT</td><td>AGG</td><td>||||||||[||||||||||||]PAM</td><td>NA</td><td>Intergenic</td><td>NA</td><td>NA</td></tr>
<tr><td>T14211</td><td>GTCCCTGCAGTCCTTCGGTAGGG</td><td>1000</td></tr>
<tr><td>Oligo fwd</td><td>CACCGTCCCTGCAGTCCTTCGGTA</td></tr>
<tr><td>Oligo rev</td><td>AAACTACCGAAGGACTGCAGGGAC</td></tr>
<tr><td>Chromosome</td><td>start</td><td>end</td><td>strand</td><td>MM</td><td>target_seq</td><td>PAM</td><td>alignment</td><td>distance</td><td>position</td><td>gene name</td><td>gene id</td></tr>
<tr><td>GU305914.1</td><td>186817</td><td>186839</td><td>+</td><td>0</td><td>GTCCCTGCAGTCCTTCGGTA</td><td>GGG</td><td>||||||||[||||||||||||]PAM</td><td>NA</td><td>Intergenic</td><td>NA</td><td>NA</td></tr>
<tr><td>T14212</td><td>CTGCAGTCCTTCGGTAGGGAAGG</td><td>1000</td></tr>
<tr><td>Oligo fwd</td><td>CACCCTGCAGTCCTTCGGTAGGGA</td></tr>
<tr><td>Oligo rev</td><td>AAACTCCCTACCGAAGGACTGCAG</td></tr>
<tr><td>Chromosome</td><td>start</td><td>end</td><td>strand</td><td>MM</td><td>target_seq</td><td>PAM</td><td>alignment</td><td>distance</td><td>position</td><td>gene name</td><td>gene id</td></tr>
<tr><td>GU305914.1</td><td>186821</td><td>186843</td><td>+</td><td>0</td><td>CTGCAGTCCTTCGGTAGGGA</td><td>AGG</td><td>||||||||[||||||||||||]PAM</td><td>NA</td><td>Intergenic</td><td>NA</td><td>NA</td></tr>
<tr><td>T14213</td><td>TCGGTAGGGAAGGACTCTGCAGG</td><td>1000</td></tr>
<tr><td>Oligo fwd</td><td>CACCTCGGTAGGGAAGGACTCTGC</td></tr>
<tr><td>Oligo rev</td><td>AAACGCAGAGTCCTTCCCTACCGA</td></tr>
<tr><td>Chromosome</td><td>start</td><td>end</td><td>strand</td><td>MM</td><td>target_seq</td><td>PAM</td><td>alignment</td><td>distance</td><td>position</td><td>gene name</td><td>gene id</td></tr>
</table>

GU305914.1 186831 186853 + 0 TCGGTAGGGAAGGACTCTGC AGG ||||||||[||||||||||||]PAM NA Intergenic NA NA

T14214 GTGCGACAACGTCTACACCGAGG 1000
Oligo fwd CACCGTGCGACAACGTCTACACCG
Oligo rev AAACCGGTGTAGACGTTGTCGCAC
Chromosome start end strand MM target_seq PAM alignment distance position gene name gene id
GU305914.1 186853 186875 + 0 GTGCGACAACGTCTACACCG AGG ||||||||[||||||||||||]PAM NA Intergenic NA NA

T14215 GACAACGTCTACACCGAGGACGG 1000
Oligo fwd CACCGACAACGTCTACACCGAGGA
Oligo rev AAACTCCTCGGTGTAGACGTTGTC
Chromosome start end strand MM target_seq PAM alignment distance position gene name gene id
GU305914.1 186857 186879 + 0 GACAACGTCTACACCGAGGA CGG ||||||||[||||||||||||]PAM NA Intergenic NA NA

T14216 AACGTCTACACCGAGGACGGCGG 1000
Oligo fwd CACCAACGTCTACACCGAGGACGG
Oligo rev AAACCCGTCCTCGGTGTAGACGTT
Chromosome start end strand MM target_seq PAM alignment distance position gene name gene id
GU305914.1 186860 186882 + 0 AACGTCTACACCGAGGACGG CGG ||||||||[||||||||||||]PAM NA Intergenic NA NA

T14217 CGCTCCGTACGCTACTCCCGAGG 1000
Oligo fwd CACCCGCTCCGTACGCTACTCCCG
Oligo rev AAACCGGGAGTAGCGTACGGAGCG
Chromosome start end strand MM target_seq PAM alignment distance position gene name gene id
GU305914.1 186883 186905 + 0 CGCTCCGTACGCTACTCCCG AGG ||||||||[||||||||||||]PAM NA Intergenic NA NA

T14218 ATCGTGTTCACGCCCCATCTCGG 1000
Oligo fwd CACCATCGTGTTCACGCCCCATCT
Oligo rev AAACAGATGGGGCGTGAACACGAT
Chromosome start end strand MM target_seq PAM alignment distance position gene name gene id
GU305914.1 186923 186945 + 0 ATCGTGTTCACGCCCCATCT CGG ||||||||[||||||||||||]PAM NA Intergenic NA NA

T14219 GTGTTCACGCCCCATCTCGGCGG 1000
Oligo fwd CACCGTGTTCACGCCCCATCTCGG
Oligo rev AAACCCGAGATGGGGCGTGAACAC
Chromosome start end strand MM target_seq PAM alignment distance position gene name gene id
GU305914.1 186926 186948 + 0 GTGTTCACGCCCCATCTCGG CGG ||||||||[||||||||||||]PAM NA Intergenic NA NA

T14220 TGTTCACGCCCCATCTCGGCGGG 1000
Oligo fwd CACCTGTTCACGCCCCATCTCGGC
Oligo rev AAACGCCGAGATGGGGCGTGAACA
Chromosome start end strand MM target_seq PAM alignment distance position gene name gene id
GU305914.1 186927 186949 + 0 TGTTCACGCCCCATCTCGGC GGG ||||||||[||||||||||||]PAM NA Intergenic NA NA

T14221 CACGCCCCATCTCGGCGGGAAGG 1000
Oligo fwd CACCCACGCCCCATCTCGGCGGGA
Oligo rev AAACTCCCGCCGAGATGGGGCGTG
Chromosome start end strand MM target_seq PAM alignment distance position gene name gene id
GU305914.1 186931 186953 + 0 CACGCCCCATCTCGGCGGGA AGG ||||||||[||||||||||||]PAM NA Intergenic NA NA

T14222 CCAGCAGATCTGTAAGATCAAGG 1000
Oligo fwd CACCCCAGCAGATCTGTAAGATCA
Oligo rev AAACTGATCTTACAGATCTGCTGG
Chromosome start end strand MM target_seq PAM alignment distance position gene name gene id
GU305914.1 186961 186983 + 0 CCAGCAGATCTGTAAGATCA AGG ||||||||[||||||||||||]PAM NA Intergenic NA NA

T14223 CAGCAGATCTGTAAGATCAAGGG 1000
Oligo fwd CACCCAGCAGATCTGTAAGATCAA
Oligo rev AAACTTGATCTTACAGATCTGCTG
Chromosome start end strand MM target_seq PAM alignment distance position gene name gene id
GU305914.1 186962 186984 + 0 CAGCAGATCTGTAAGATCAA GGG ||||||||[||||||||||||]PAM NA Intergenic NA NA

T14224 GTAAGATCAAGGGACACACCTGG 1000
Oligo fwd CACCGTAAGATCAAGGGACACACC
Oligo rev AAACGGTGTGTCCCTTGATCTTAC
Chromosome start end strand MM target_seq PAM alignment distance position gene name gene id
GU305914.1 186972 186994 + 0 GTAAGATCAAGGGACACACC TGG ||||||||[||||||||||||]PAM NA Intergenic NA NA

T14225 GGACACACCTGGTACCTGAACGG 1000
Oligo fwd CACCGGACACACCTGGTACCTGAA
Oligo rev AAACTTCAGGTACCAGGTGTGTCC
Chromosome start end strand MM target_seq PAM alignment distance position gene name gene id
GU305914.1 186983 187005 + 0 GGACACACCTGGTACCTGAA CGG ||||||||[||||||||||||]PAM NA Intergenic NA NA
GU305914.1 67136 67158 + 4 CTACACACCTGTTACCTGAT GGG --||||||[|||-|||||||-]PAM NA Intergenic NA NA

T14226 TGGTACCTGAACGGTTGCCCCGG 1000
Oligo fwd CACCTGGTACCTGAACGGTTGCCC
Oligo rev AAACGGGCAACCGTTCAGGTACCA
Chromosome start end strand MM target_seq PAM alignment distance position gene name gene id
GU305914.1 186992 187014 + 0 TGGTACCTGAACGGTTGCCC CGG ||||||||[||||||||||||]PAM NA Intergenic NA NA

T14227 CGGTTGCCCCGGCATGCTGAAGG 1000
Oligo fwd CACCCGGTTGCCCCGGCATGCTGA
Oligo rev AAACTCAGCATGCCGGGGCAACCG
Chromosome start end strand MM target_seq PAM alignment distance position gene name gene id
GU305914.1 187003 187025 + 0 CGGTTGCCCCGGCATGCTGA AGG ||||||||[||||||||||||]PAM NA Intergenic NA NA

T14228 GCCCCGGCATGCTGAAGGACAGG 1000
Oligo fwd CACCGCCCCGGCATGCTGAAGGAC

<table>
<tr><td>Oligo rev</td><td>AAACGTCCTTCAGCATGCCGGGGC</td><td></td><td></td><td></td><td></td><td></td><td></td><td></td><td></td><td></td><td></td></tr>
<tr><td>Chromosome</td><td>start</td><td>end</td><td>strand</td><td>MM</td><td>target_seq</td><td>PAM</td><td>alignment</td><td>distance</td><td>position</td><td>gene name</td><td>gene id</td></tr>
<tr><td>GU305914.1</td><td>187008</td><td>187030</td><td>+</td><td>0</td><td>GCCCCGGCATGCTGAAGGAC</td><td>AGG</td><td>||||||||[||||||||||||]PAM</td><td>NA</td><td>Intergenic</td><td>NA</td><td>NA</td></tr>
<tr><td></td><td></td><td></td><td></td><td></td><td></td><td></td><td></td><td></td><td></td><td></td><td></td></tr>
<tr><td>T14229</td><td>CCCCGGCATGCTGAAGGACAGGG</td><td>1000</td><td></td><td></td><td></td><td></td><td></td><td></td><td></td><td></td><td></td></tr>
<tr><td>Oligo fwd</td><td>CACCCCCCGGCATGCTGAAGGACA</td><td></td><td></td><td></td><td></td><td></td><td></td><td></td><td></td><td></td><td></td></tr>
<tr><td>Oligo rev</td><td>AAACTGTCCTTCAGCATGCCGGGG</td><td></td><td></td><td></td><td></td><td></td><td></td><td></td><td></td><td></td><td></td></tr>
<tr><td>Chromosome</td><td>start</td><td>end</td><td>strand</td><td>MM</td><td>target_seq</td><td>PAM</td><td>alignment</td><td>distance</td><td>position</td><td>gene name</td><td>gene id</td></tr>
<tr><td>GU305914.1</td><td>187009</td><td>187031</td><td>+</td><td>0</td><td>CCCCGGCATGCTGAAGGACA</td><td>GGG</td><td>||||||||[||||||||||||]PAM</td><td>NA</td><td>Intergenic</td><td>NA</td><td>NA</td></tr>
<tr><td></td><td></td><td></td><td></td><td></td><td></td><td></td><td></td><td></td><td></td><td></td><td></td></tr>
<tr><td>T14230</td><td>CAGGGTGTTCGTCGCCACGCCGG</td><td>1000</td><td></td><td></td><td></td><td></td><td></td><td></td><td></td><td></td><td></td></tr>
<tr><td>Oligo fwd</td><td>CACCCAGGGTGTTCGTCGCCACGC</td><td></td><td></td><td></td><td></td><td></td><td></td><td></td><td></td><td></td><td></td></tr>
<tr><td>Oligo rev</td><td>AAACGCGTGGCGACGAACACCCTG</td><td></td><td></td><td></td><td></td><td></td><td></td><td></td><td></td><td></td><td></td></tr>
<tr><td>Chromosome</td><td>start</td><td>end</td><td>strand</td><td>MM</td><td>target_seq</td><td>PAM</td><td>alignment</td><td>distance</td><td>position</td><td>gene name</td><td>gene id</td></tr>
<tr><td>GU305914.1</td><td>187027</td><td>187049</td><td>+</td><td>0</td><td>CAGGGTGTTCGTCGCCACGC</td><td>CGG</td><td>||||||||[||||||||||||]PAM</td><td>NA</td><td>Intergenic</td><td>NA</td><td>NA</td></tr>
<tr><td></td><td></td><td></td><td></td><td></td><td></td><td></td><td></td><td></td><td></td><td></td><td></td></tr>
<tr><td>T14231</td><td>CTTCGTCCGCCACGTGAACCTGG</td><td>1000</td><td></td><td></td><td></td><td></td><td></td><td></td><td></td><td></td><td></td></tr>
<tr><td>Oligo fwd</td><td>CACCCTTCGTCCGCCACGTGAACC</td><td></td><td></td><td></td><td></td><td></td><td></td><td></td><td></td><td></td><td></td></tr>
<tr><td>Oligo rev</td><td>AAACGGTTCACGTGGCGGACGAAG</td><td></td><td></td><td></td><td></td><td></td><td></td><td></td><td></td><td></td><td></td></tr>
<tr><td>Chromosome</td><td>start</td><td>end</td><td>strand</td><td>MM</td><td>target_seq</td><td>PAM</td><td>alignment</td><td>distance</td><td>position</td><td>gene name</td><td>gene id</td></tr>
<tr><td>GU305914.1</td><td>187060</td><td>187082</td><td>+</td><td>0</td><td>CTTCGTCCGCCACGTGAACC</td><td>TGG</td><td>||||||||[||||||||||||]PAM</td><td>NA</td><td>Intergenic</td><td>NA</td><td>NA</td></tr>
<tr><td></td><td></td><td></td><td></td><td></td><td></td><td></td><td></td><td></td><td></td><td></td><td></td></tr>
<tr><td>T14232</td><td>CACGTGAACCTGGAACTCTTTGG</td><td>1000</td><td></td><td></td><td></td><td></td><td></td><td></td><td></td><td></td><td></td></tr>
<tr><td>Oligo fwd</td><td>CACCCACGTGAACCTGGAACTCTT</td><td></td><td></td><td></td><td></td><td></td><td></td><td></td><td></td><td></td><td></td></tr>
<tr><td>Oligo rev</td><td>AAACAAGAGTTCCAGGTTCACGTG</td><td></td><td></td><td></td><td></td><td></td><td></td><td></td><td></td><td></td><td></td></tr>
<tr><td>Chromosome</td><td>start</td><td>end</td><td>strand</td><td>MM</td><td>target_seq</td><td>PAM</td><td>alignment</td><td>distance</td><td>position</td><td>gene name</td><td>gene id</td></tr>
<tr><td>GU305914.1</td><td>187070</td><td>187092</td><td>+</td><td>0</td><td>CACGTGAACCTGGAACTCTT</td><td>TGG</td><td>||||||||[||||||||||||]PAM</td><td>NA</td><td>Intergenic</td><td>NA</td><td>NA</td></tr>
<tr><td></td><td></td><td></td><td></td><td></td><td></td><td></td><td></td><td></td><td></td><td></td><td></td></tr>
<tr><td>T14233</td><td>ACCTGGAACTCTTTGGACATAGG</td><td>1000</td><td></td><td></td><td></td><td></td><td></td><td></td><td></td><td></td><td></td></tr>
<tr><td>Oligo fwd</td><td>CACCACCTGGAACTCTTTGGACAT</td><td></td><td></td><td></td><td></td><td></td><td></td><td></td><td></td><td></td><td></td></tr>
<tr><td>Oligo rev</td><td>AAACATGTCCAAAGAGTTCCAGGT</td><td></td><td></td><td></td><td></td><td></td><td></td><td></td><td></td><td></td><td></td></tr>
<tr><td>Chromosome</td><td>start</td><td>end</td><td>strand</td><td>MM</td><td>target_seq</td><td>PAM</td><td>alignment</td><td>distance</td><td>position</td><td>gene name</td><td>gene id</td></tr>
<tr><td>GU305914.1</td><td>187077</td><td>187099</td><td>+</td><td>0</td><td>ACCTGGAACTCTTTGGACAT</td><td>AGG</td><td>||||||||[||||||||||||]PAM</td><td>NA</td><td>Intergenic</td><td>NA</td><td>NA</td></tr>
<tr><td></td><td></td><td></td><td></td><td></td><td></td><td></td><td></td><td></td><td></td><td></td><td></td></tr>
<tr><td>T14234</td><td>GGACATAGGTTTCTACCCATCGG</td><td>1000</td><td></td><td></td><td></td><td></td><td></td><td></td><td></td><td></td><td></td></tr>
<tr><td>Oligo fwd</td><td>CACCGGACATAGGTTTCTACCCAT</td><td></td><td></td><td></td><td></td><td></td><td></td><td></td><td></td><td></td><td></td></tr>
<tr><td>Oligo rev</td><td>AAACATGGGTAGAAACCTATGTCC</td><td></td><td></td><td></td><td></td><td></td><td></td><td></td><td></td><td></td><td></td></tr>
<tr><td>Chromosome</td><td>start</td><td>end</td><td>strand</td><td>MM</td><td>target_seq</td><td>PAM</td><td>alignment</td><td>distance</td><td>position</td><td>gene name</td><td>gene id</td></tr>
<tr><td>GU305914.1</td><td>187091</td><td>187113</td><td>+</td><td>0</td><td>GGACATAGGTTTCTACCCAT</td><td>CGG</td><td>||||||||[||||||||||||]PAM</td><td>NA</td><td>Intergenic</td><td>NA</td><td>NA</td></tr>
<tr><td></td><td></td><td></td><td></td><td></td><td></td><td></td><td></td><td></td><td></td><td></td><td></td></tr>
<tr><td>T14235</td><td>ATAGGTTTCTACCCATCGGTAGG</td><td>1000</td><td></td><td></td><td></td><td></td><td></td><td></td><td></td><td></td><td></td></tr>
<tr><td>Oligo fwd</td><td>CACCATAGGTTTCTACCCATCGGT</td><td></td><td></td><td></td><td></td><td></td><td></td><td></td><td></td><td></td><td></td></tr>
<tr><td>Oligo rev</td><td>AAACACCGATGGGTAGAAACCTAT</td><td></td><td></td><td></td><td></td><td></td><td></td><td></td><td></td><td></td><td></td></tr>
<tr><td>Chromosome</td><td>start</td><td>end</td><td>strand</td><td>MM</td><td>target_seq</td><td>PAM</td><td>alignment</td><td>distance</td><td>position</td><td>gene name</td><td>gene id</td></tr>
<tr><td>GU305914.1</td><td>187095</td><td>187117</td><td>+</td><td>0</td><td>ATAGGTTTCTACCCATCGGT</td><td>AGG</td><td>||||||||[||||||||||||]PAM</td><td>NA</td><td>Intergenic</td><td>NA</td><td>NA</td></tr>
<tr><td></td><td></td><td></td><td></td><td></td><td></td><td></td><td></td><td></td><td></td><td></td><td></td></tr>
<tr><td>T14236</td><td>TAGGTTTCTACCCATCGGTAGGG</td><td>1000</td><td></td><td></td><td></td><td></td><td></td><td></td><td></td><td></td><td></td></tr>
<tr><td>Oligo fwd</td><td>CACCTAGGTTTCTACCCATCGGTA</td><td></td><td></td><td></td><td></td><td></td><td></td><td></td><td></td><td></td><td></td></tr>
<tr><td>Oligo rev</td><td>AAACTACCGATGGGTAGAAACCTA</td><td></td><td></td><td></td><td></td><td></td><td></td><td></td><td></td><td></td><td></td></tr>
<tr><td>Chromosome</td><td>start</td><td>end</td><td>strand</td><td>MM</td><td>target_seq</td><td>PAM</td><td>alignment</td><td>distance</td><td>position</td><td>gene name</td><td>gene id</td></tr>
<tr><td>GU305914.1</td><td>187096</td><td>187118</td><td>+</td><td>0</td><td>TAGGTTTCTACCCATCGGTA</td><td>GGG</td><td>||||||||[||||||||||||]PAM</td><td>NA</td><td>Intergenic</td><td>NA</td><td>NA</td></tr>
<tr><td></td><td></td><td></td><td></td><td></td><td></td><td></td><td></td><td></td><td></td><td></td><td></td></tr>
<tr><td>T14237</td><td>TAGGGTGACCCGTTCCCCCGAGG</td><td>1000</td><td></td><td></td><td></td><td></td><td></td><td></td><td></td><td></td><td></td></tr>
<tr><td>Oligo fwd</td><td>CACCTAGGGTGACCCGTTCCCCCG</td><td></td><td></td><td></td><td></td><td></td><td></td><td></td><td></td><td></td><td></td></tr>
<tr><td>Oligo rev</td><td>AAACCGGGGGAACGGGTCACCCTA</td><td></td><td></td><td></td><td></td><td></td><td></td><td></td><td></td><td></td><td></td></tr>
<tr><td>Chromosome</td><td>start</td><td>end</td><td>strand</td><td>MM</td><td>target_seq</td><td>PAM</td><td>alignment</td><td>distance</td><td>position</td><td>gene name</td><td>gene id</td></tr>
<tr><td>GU305914.1</td><td>187114</td><td>187136</td><td>+</td><td>0</td><td>TAGGGTGACCCGTTCCCCCG</td><td>AGG</td><td>||||||||[||||||||||||]PAM</td><td>NA</td><td>Intergenic</td><td>NA</td><td>NA</td></tr>
<tr><td></td><td></td><td></td><td></td><td></td><td></td><td></td><td></td><td></td><td></td><td></td><td></td></tr>
<tr><td>T14238</td><td>CGAGTGTGAGATGTTTATCATGG</td><td>1000</td><td></td><td></td><td></td><td></td><td></td><td></td><td></td><td></td><td></td></tr>
<tr><td>Oligo fwd</td><td>CACCCGAGTGTGAGATGTTTATCA</td><td></td><td></td><td></td><td></td><td></td><td></td><td></td><td></td><td></td><td></td></tr>
<tr><td>Oligo rev</td><td>AAACTGATAAACATCTCACACTCG</td><td></td><td></td><td></td><td></td><td></td><td></td><td></td><td></td><td></td><td></td></tr>
<tr><td>Chromosome</td><td>start</td><td>end</td><td>strand</td><td>MM</td><td>target_seq</td><td>PAM</td><td>alignment</td><td>distance</td><td>position</td><td>gene name</td><td>gene id</td></tr>
<tr><td>GU305914.1</td><td>187141</td><td>187163</td><td>+</td><td>0</td><td>CGAGTGTGAGATGTTTATCA</td><td>TGG</td><td>||||||||[||||||||||||]PAM</td><td>NA</td><td>Intergenic</td><td>NA</td><td>NA</td></tr>
<tr><td></td><td></td><td></td><td></td><td></td><td></td><td></td><td></td><td></td><td></td><td></td><td></td></tr>
<tr><td>T14239</td><td>GTGTGAGATGTTTATCATGGTGG</td><td>1000</td><td></td><td></td><td></td><td></td><td></td><td></td><td></td><td></td><td></td></tr>
<tr><td>Oligo fwd</td><td>CACCGTGTGAGATGTTTATCATGG</td><td></td><td></td><td></td><td></td><td></td><td></td><td></td><td></td><td></td><td></td></tr>
<tr><td>Oligo rev</td><td>AAACCCATGATAAACATCTCACAC</td><td></td><td></td><td></td><td></td><td></td><td></td><td></td><td></td><td></td><td></td></tr>
<tr><td>Chromosome</td><td>start</td><td>end</td><td>strand</td><td>MM</td><td>target_seq</td><td>PAM</td><td>alignment</td><td>distance</td><td>position</td><td>gene name</td><td>gene id</td></tr>
<tr><td>GU305914.1</td><td>187144</td><td>187166</td><td>+</td><td>0</td><td>GTGTGAGATGTTTATCATGG</td><td>TGG</td><td>||||||||[||||||||||||]PAM</td><td>NA</td><td>Intergenic</td><td>NA</td><td>NA</td></tr>
<tr><td></td><td></td><td></td><td></td><td></td><td></td><td></td><td></td><td></td><td></td><td></td><td></td></tr>
<tr><td>T14240</td><td>ATGTTTATCATGGTGGACGCCGG</td><td>1000</td><td></td><td></td><td></td><td></td><td></td><td></td><td></td><td></td><td></td></tr>
<tr><td>Oligo fwd</td><td>CACCATGTTTATCATGGTGGACGC</td><td></td><td></td><td></td><td></td><td></td><td></td><td></td><td></td><td></td><td></td></tr>
<tr><td>Oligo rev</td><td>AAACGCGTCCACCATGATAAACAT</td><td></td><td></td><td></td><td></td><td></td><td></td><td></td><td></td><td></td><td></td></tr>
<tr><td>Chromosome</td><td>start</td><td>end</td><td>strand</td><td>MM</td><td>target_seq</td><td>PAM</td><td>alignment</td><td>distance</td><td>position</td><td>gene name</td><td>gene id</td></tr>
<tr><td>GU305914.1</td><td>187151</td><td>187173</td><td>+</td><td>0</td><td>ATGTTTATCATGGTGGACGC</td><td>CGG</td><td>||||||||[||||||||||||]PAM</td><td>NA</td><td>Intergenic</td><td>NA</td><td>NA</td></tr>
<tr><td></td><td></td><td></td><td></td><td></td><td></td><td></td><td></td><td></td><td></td><td></td><td></td></tr>
<tr><td>T14241</td><td>TGTTTATCATGGTGGACGCCGGG</td><td>1000</td><td></td><td></td><td></td><td></td><td></td><td></td><td></td><td></td><td></td></tr>
<tr><td>Oligo fwd</td><td>CACCTGTTTATCATGGTGGACGCC</td><td></td><td></td><td></td><td></td><td></td><td></td><td></td><td></td><td></td><td></td></tr>
<tr><td>Oligo rev</td><td>AAACGGCGTCCACCATGATAAACA</td><td></td><td></td><td></td><td></td><td></td><td></td><td></td><td></td><td></td><td></td></tr>
<tr><td>Chromosome</td><td>start</td><td>end</td><td>strand</td><td>MM</td><td>target_seq</td><td>PAM</td><td>alignment</td><td>distance</td><td>position</td><td>gene name</td><td>gene id</td></tr>
<tr><td>GU305914.1</td><td>187152</td><td>187174</td><td>+</td><td>0</td><td>TGTTTATCATGGTGGACGCC</td><td>GGG</td><td>||||||||[||||||||||||]PAM</td><td>NA</td><td>Intergenic</td><td>NA</td><td>NA</td></tr>
<tr><td></td><td></td><td></td><td></td><td></td><td></td><td></td><td></td><td></td><td></td><td></td><td></td></tr>
<tr><td>T14242</td><td>GTTTATCATGGTGGACGCCGGGG</td><td>1000</td><td></td><td></td><td></td><td></td><td></td><td></td><td></td><td></td><td></td></tr>
<tr><td>Oligo fwd</td><td>CACCGTTTATCATGGTGGACGCCG</td><td></td><td></td><td></td><td></td><td></td><td></td><td></td><td></td><td></td><td></td></tr>
<tr><td>Oligo rev</td><td>AAACCGGCGTCCACCATGATAAAC</td><td></td><td></td><td></td><td></td><td></td><td></td><td></td><td></td><td></td><td></td></tr>
<tr><td>Chromosome</td><td>start</td><td>end</td><td>strand</td><td>MM</td><td>target_seq</td><td>PAM</td><td>alignment</td><td>distance</td><td>position</td><td>gene name</td><td>gene id</td></tr>
<tr><td>GU305914.1</td><td>187153</td><td>187175</td><td>+</td><td>0</td><td>GTTTATCATGGTGGACGCCG</td><td>GGG</td><td>||||||||[||||||||||||]PAM</td><td>NA</td><td>Intergenic</td><td>NA</td><td>NA</td></tr>
<tr><td>GU305914.1</td><td>149010</td><td>149032</td><td>-</td><td>4</td><td>GTTCTTCATCCTGGACGCCG</td><td>AGG</td><td>|||--|||[|--|||||||||]PAM</td><td>NA</td><td>Intergenic</td><td>NA</td><td>NA</td></tr>
</table>

T14243 TTTATCATGGTGGACGCCGGGGG 1000
Oligo fwd CACCTTTATCATGGTGGACGCCGG
Oligo rev AAACCCGGCGTCCACCATGATAAA
Chromosome start end strand MM target_seq PAM alignment distance position gene name gene id
GU305914.1 187154 187176 + 0 TTTATCATGGTGGACGCCGG GGG ||||||||[||||||||||||]PAM NA Intergenic NA NA

T14244 TTATCATGGTGGACGCCGGGGGG 1000
Oligo fwd CACCTTATCATGGTGGACGCCGGG
Oligo rev AAACCCCGGCGTCCACCATGATAA
Chromosome start end strand MM target_seq PAM alignment distance position gene name gene id
GU305914.1 187155 187177 + 0 TTATCATGGTGGACGCCGGG GGG ||||||||[||||||||||||]PAM NA Intergenic NA NA

T14245 TATCATGGTGGACGCCGGGGGGG 1000
Oligo fwd CACCTATCATGGTGGACGCCGGGG
Oligo rev AAACCCCCGGCGTCCACCATGATA
Chromosome start end strand MM target_seq PAM alignment distance position gene name gene id
GU305914.1 187156 187178 + 0 TATCATGGTGGACGCCGGGG GGG ||||||||[||||||||||||]PAM NA Intergenic NA NA

T14246 GACGCCGGGGGGGCCATCTACGG 1000
Oligo fwd CACCGACGCCGGGGGGGCCATCTA
Oligo rev AAACTAGATGGCCCCCCCGGCGTC
Chromosome start end strand MM target_seq PAM alignment distance position gene name gene id
GU305914.1 187166 187188 + 0 GACGCCGGGGGGGCCATCTA CGG ||||||||[||||||||||||]PAM NA Intergenic NA NA

T14247 ACGCCGGGGGGGCCATCTACGGG 1000
Oligo fwd CACCACGCCGGGGGGGCCATCTAC
Oligo rev AAACGTAGATGGCCCCCCCGGCGT
Chromosome start end strand MM target_seq PAM alignment distance position gene name gene id
GU305914.1 187167 187189 + 0 ACGCCGGGGGGGCCATCTAC GGG ||||||||[||||||||||||]PAM NA Intergenic NA NA

T14248 TACGGGCACATGTTAGATAGCGG 1000
Oligo fwd CACCTACGGGCACATGTTAGATAG
Oligo rev AAACCTATCTAACATGTGCCCGTA
Chromosome start end strand MM target_seq PAM alignment distance position gene name gene id
GU305914.1 187184 187206 + 0 TACGGGCACATGTTAGATAG CGG ||||||||[||||||||||||]PAM NA Intergenic NA NA

T14249 ACGGGCACATGTTAGATAGCGGG 1000
Oligo fwd CACCACGGGCACATGTTAGATAGC
Oligo rev AAACGCTATCTAACATGTGCCCGT
Chromosome start end strand MM target_seq PAM alignment distance position gene name gene id
GU305914.1 187185 187207 + 0 ACGGGCACATGTTAGATAGC GGG ||||||||[||||||||||||]PAM NA Intergenic NA NA

T14250 GCACATGTTAGATAGCGGGAAGG 1000
Oligo fwd CACCGCACATGTTAGATAGCGGGA
Oligo rev AAACTCCCGCTATCTAACATGTGC
Chromosome start end strand MM target_seq PAM alignment distance position gene name gene id
GU305914.1 187189 187211 + 0 GCACATGTTAGATAGCGGGA AGG ||||||||[||||||||||||]PAM NA Intergenic NA NA

T14251 TGTTAGATAGCGGGAAGGTGAGG 1000
Oligo fwd CACCTGTTAGATAGCGGGAAGGTG
Oligo rev AAACCACCTTCCCGCTATCTAACA
Chromosome start end strand MM target_seq PAM alignment distance position gene name gene id
GU305914.1 187194 187216 + 0 TGTTAGATAGCGGGAAGGTG AGG ||||||||[||||||||||||]PAM NA Intergenic NA NA

T14252 TAGATAGCGGGAAGGTGAGGAGG 1000
Oligo fwd CACCTAGATAGCGGGAAGGTGAGG
Oligo rev AAACCCTCACCTTCCCGCTATCTA
Chromosome start end strand MM target_seq PAM alignment distance position gene name gene id
GU305914.1 187197 187219 + 0 TAGATAGCGGGAAGGTGAGG AGG ||||||||[||||||||||||]PAM NA Intergenic NA NA

T14253 ATAACTTTGACCAGTTCATGCGG 1000
Oligo fwd CACCATAACTTTGACCAGTTCATG
Oligo rev AAACCATGAACTGGTCAAAGTTAT
Chromosome start end strand MM target_seq PAM alignment distance position gene name gene id
GU305914.1 187227 187249 + 0 ATAACTTTGACCAGTTCATG CGG ||||||||[||||||||||||]PAM NA Intergenic NA NA

T14254 CTTTGACCAGTTCATGCGGATGG 1000
Oligo fwd CACCCTTTGACCAGTTCATGCGGA
Oligo rev AAACTCCGCATGAACTGGTCAAAG
Chromosome start end strand MM target_seq PAM alignment distance position gene name gene id
GU305914.1 187231 187253 + 0 CTTTGACCAGTTCATGCGGA TGG ||||||||[||||||||||||]PAM NA Intergenic NA NA

T14255 TTTGACCAGTTCATGCGGATGGG 1000
Oligo fwd CACCTTTGACCAGTTCATGCGGAT
Oligo rev AAACATCCGCATGAACTGGTCAAA
Chromosome start end strand MM target_seq PAM alignment distance position gene name gene id
GU305914.1 187232 187254 + 0 TTTGACCAGTTCATGCGGAT GGG ||||||||[||||||||||||]PAM NA Intergenic NA NA

T14256 AGTTCATGCGGATGGGCACCCGG 1000
Oligo fwd CACCAGTTCATGCGGATGGGCACC
Oligo rev AAACGGTGCCCATCCGCATGAACT
Chromosome start end strand MM target_seq PAM alignment distance position gene name gene id
GU305914.1 187239 187261 + 0 AGTTCATGCGGATGGGCACC CGG ||||||||[||||||||||||]PAM NA Intergenic NA NA

T14257 TCATGCGGATGGGCACCCGGAGG 1000
Oligo fwd CACCTCATGCGGATGGGCACCCGG
Oligo rev AAACCCGGGTGCCCATCCGCATGA
Chromosome start end strand MM target_seq PAM alignment distance position gene name gene id
GU305914.1 187242 187264 + 0 TCATGCGGATGGGCACCCGG AGG ||||||||[||||||||||||]PAM NA Intergenic NA NA

T14258 CATGCGGATGGCACCCGGAGGG 1000
Oligo fwd CACCCATGCGGATGGCACCCGGA
Oligo rev AAACTCCGGGTGCCCATCCGCATG
Chromosome start end strand MM target_seq PAM alignment distance position gene name gene id
GU305914.1 187243 187265 + 0 CATGCGGATGGCACCCGGA GGG ||||||||[||||||||||||]PAM NA Intergenic NA NA

T14259 GGCTTATTTCAATTTCCAGATGG 1000
Oligo fwd CACCGGCTTATTTCAATTTCCAGA
Oligo rev AAACTCTGGAAATTGAAATAAGCC
Chromosome start end strand MM target_seq PAM alignment distance position gene name gene id
GU305914.1 187264 187286 + 0 GGCTTATTTCAATTTCCAGA TGG ||||||||[||||||||||||]PAM NA Intergenic NA NA

T14260 TTTCAATTTCCAGATGGTTAAGG 1000
Oligo fwd CACCTTTCAATTTCCAGATGGTTA
Oligo rev AAACTAACCATCTGGAAATTGAAA
Chromosome start end strand MM target_seq PAM alignment distance position gene name gene id
GU305914.1 187270 187292 + 0 TTTCAATTTCCAGATGGTTA AGG ||||||||[||||||||||||]PAM NA Intergenic NA NA

T14261 TTCAATTTCCAGATGGTTAAGGG 1000
Oligo fwd CACCTTCAATTTCCAGATGGTTAA
Oligo rev AAACTTAACCATCTGGAAATTGAA
Chromosome start end strand MM target_seq PAM alignment distance position gene name gene id
GU305914.1 187271 187293 + 0 TTCAATTTCCAGATGGTTAA GGG ||||||||[||||||||||||]PAM NA Intergenic NA NA

T14262 TAGTCTCAACACCGAGTACGAGG 1000
Oligo fwd CACCTAGTCTCAACACCGAGTACG
Oligo rev AAACCGTACTCGGTGTTGAGACTA
Chromosome start end strand MM target_seq PAM alignment distance position gene name gene id
GU305914.1 187297 187319 + 0 TAGTCTCAACACCGAGTACG AGG ||||||||[||||||||||||]PAM NA Intergenic NA NA

T14263 GAGTACGAGGAGACCTCGTTTGG 1000
Oligo fwd CACCGAGTACGAGGAGACCTCGTT
Oligo rev AAACAACGAGGTCTCCTCGTACTC
Chromosome start end strand MM target_seq PAM alignment distance position gene name gene id
GU305914.1 187310 187332 + 0 GAGTACGAGGAGACCTCGTT TGG ||||||||[||||||||||||]PAM NA Intergenic NA NA

T14264 AGTACGAGGAGACCTCGTTTGGG 1000
Oligo fwd CACCAGTACGAGGAGACCTCGTTT
Oligo rev AAACAAACGAGGTCTCCTCGTACT
Chromosome start end strand MM target_seq PAM alignment distance position gene name gene id
GU305914.1 187311 187333 + 0 AGTACGAGGAGACCTCGTTT GGG ||||||||[||||||||||||]PAM NA Intergenic NA NA

T14265 TCGTTTGGGACCAAGAACCACGG 1000
Oligo fwd CACCTCGTTTGGGACCAAGAACCA
Oligo rev AAACTGGTTCTTGGTCCCAAACGA
Chromosome start end strand MM target_seq PAM alignment distance position gene name gene id
GU305914.1 187325 187347 + 0 TCGTTTGGGACCAAGAACCA CGG ||||||||[||||||||||||]PAM NA Intergenic NA NA

T14266 CGTTTGGGACCAAGAACCACGGG 1000
Oligo fwd CACCCGTTTGGGACCAAGAACCAC
Oligo rev AAACGTGGTTCTTGGTCCCAAACG
Chromosome start end strand MM target_seq PAM alignment distance position gene name gene id
GU305914.1 187326 187348 + 0 CGTTTGGGACCAAGAACCAC GGG ||||||||[||||||||||||]PAM NA Intergenic NA NA

T14267 CTTTCTCTTGACCAGAGACCTGG 1000
Oligo fwd CACCCTTTCTCTTGACCAGAGACC
Oligo rev AAACGGTCTCTGGTCAAGAGAAAG
Chromosome start end strand MM target_seq PAM alignment distance position gene name gene id
GU305914.1 187351 187373 + 0 CTTTCTCTTGACCAGAGACC TGG ||||||||[||||||||||||]PAM NA Intergenic NA NA

T14268 CCAGAGACCTGGTGACCGTCAGG 1000
Oligo fwd CACCCCAGAGACCTGGTGACCGTC
Oligo rev AAACGACGGTCACCAGGTCTCTGG
Chromosome start end strand MM target_seq PAM alignment distance position gene name gene id
GU305914.1 187362 187384 + 0 CCAGAGACCTGGTGACCGTC AGG ||||||||[||||||||||||]PAM NA Intergenic NA NA

T14269 CAGGAAGAAGATTCAGTGAGAGG 1000
Oligo fwd CACCCAGGAAGAAGATTCAGTGAG
Oligo rev AAACCTCACTGAATCTTCTTCCTG
Chromosome start end strand MM target_seq PAM alignment distance position gene name gene id
GU305914.1 187381 187403 + 0 CAGGAAGAAGATTCAGTGAG AGG ||||||||[||||||||||||]PAM NA Intergenic NA NA

T14270 AGATTCAGTGAGAGGACACGAGG 1000
Oligo fwd CACCAGATTCAGTGAGAGGACACG
Oligo rev AAACCGTGTCCTCTCACTGAATCT
Chromosome start end strand MM target_seq PAM alignment distance position gene name gene id
GU305914.1 187389 187411 + 0 AGATTCAGTGAGAGGACACG AGG ||||||||[||||||||||||]PAM NA Intergenic NA NA

T14271 AGAGGACACGAGGTATGTCATGG 1000
Oligo fwd CACCAGAGGACACGAGGTATGTCA
Oligo rev AAACTGACATACCTCGTGTCCTCT
Chromosome start end strand MM target_seq PAM alignment distance position gene name gene id
GU305914.1 187399 187421 + 0 AGAGGACACGAGGTATGTCA TGG ||||||||[||||||||||||]PAM NA Intergenic NA NA

T14272 TTTTTAATCAATAAATAAAGAGG 1000
Oligo fwd CACCTTTTTAATCAATAAATAAAG
Oligo rev AAACCTTTATTTATTGATTAAAAA
Chromosome start end strand MM target_seq PAM alignment distance position gene name gene id

<table>
<tr><td>GU305914.1</td><td></td><td>187422</td><td>187444</td><td>+</td><td>0</td><td>TTTTTAATCAATAAATAAAG</td><td>AGG</td><td>||||||||[||||||||||||]PAM</td><td>NA</td><td>Intergenic</td><td>NA</td><td>NA</td></tr>
<tr><td>T14273</td><td>AATAAAGAGGTTTTATTCATCGG</td><td></td><td>1000</td><td></td><td></td><td></td><td></td><td></td><td></td><td></td><td></td><td></td></tr>
<tr><td>Oligo fwd</td><td>CACCAATAAAGAGGTTTTATTCAT</td><td></td><td></td><td></td><td></td><td></td><td></td><td></td><td></td><td></td><td></td><td></td></tr>
<tr><td>Oligo rev</td><td>AAACATGAATAAAACCTCTTTATT</td><td></td><td></td><td></td><td></td><td></td><td></td><td></td><td></td><td></td><td></td><td></td></tr>
<tr><td>Chromosome</td><td>start</td><td></td><td>end</td><td>strand</td><td>MM</td><td>target_seq</td><td>PAM</td><td>alignment</td><td>distance</td><td>position</td><td>gene name</td><td>gene id</td></tr>
<tr><td>GU305914.1</td><td></td><td>187435</td><td>187457</td><td>+</td><td>0</td><td>AATAAAGAGGTTTTATTCAT</td><td>CGG</td><td>||||||||[||||||||||||]PAM</td><td>NA</td><td>Intergenic</td><td>NA</td><td>NA</td></tr>
<tr><td>T14274</td><td>AGCCTGTAAAAGACTCGCCCCGG</td><td></td><td>1000</td><td></td><td></td><td></td><td></td><td></td><td></td><td></td><td></td><td></td></tr>
<tr><td>Oligo fwd</td><td>CACCAGCCTGTAAAAGACTCGCCC</td><td></td><td></td><td></td><td></td><td></td><td></td><td></td><td></td><td></td><td></td><td></td></tr>
<tr><td>Oligo rev</td><td>AAACGGGCGAGTCTTTTACAGGCT</td><td></td><td></td><td></td><td></td><td></td><td></td><td></td><td></td><td></td><td></td><td></td></tr>
<tr><td>Chromosome</td><td>start</td><td></td><td>end</td><td>strand</td><td>MM</td><td>target_seq</td><td>PAM</td><td>alignment</td><td>distance</td><td>position</td><td>gene name</td><td>gene id</td></tr>
<tr><td>GU305914.1</td><td></td><td>187469</td><td>187491</td><td>+</td><td>0</td><td>AGCCTGTAAAAGACTCGCCC</td><td>CGG</td><td>||||||||[||||||||||||]PAM</td><td>NA</td><td>Intergenic</td><td>NA</td><td>NA</td></tr>
<tr><td>T14275</td><td>CTGTAAAAGACTCGCCCCGGAGG</td><td></td><td>1000</td><td></td><td></td><td></td><td></td><td></td><td></td><td></td><td></td><td></td></tr>
<tr><td>Oligo fwd</td><td>CACCCTGTAAAAGACTCGCCCCGG</td><td></td><td></td><td></td><td></td><td></td><td></td><td></td><td></td><td></td><td></td><td></td></tr>
<tr><td>Oligo rev</td><td>AAACCCGGGGCGAGTCTTTTACAG</td><td></td><td></td><td></td><td></td><td></td><td></td><td></td><td></td><td></td><td></td><td></td></tr>
<tr><td>Chromosome</td><td>start</td><td></td><td>end</td><td>strand</td><td>MM</td><td>target_seq</td><td>PAM</td><td>alignment</td><td>distance</td><td>position</td><td>gene name</td><td>gene id</td></tr>
<tr><td>GU305914.1</td><td></td><td>187472</td><td>187494</td><td>+</td><td>0</td><td>CTGTAAAAGACTCGCCCCGG</td><td>AGG</td><td>||||||||[||||||||||||]PAM</td><td>NA</td><td>Intergenic</td><td>NA</td><td>NA</td></tr>
<tr><td>T14276</td><td>TGTAAAAGACTCGCCCCGGAGGG</td><td></td><td>1000</td><td></td><td></td><td></td><td></td><td></td><td></td><td></td><td></td><td></td></tr>
<tr><td>Oligo fwd</td><td>CACCTGTAAAAGACTCGCCCCGGA</td><td></td><td></td><td></td><td></td><td></td><td></td><td></td><td></td><td></td><td></td><td></td></tr>
<tr><td>Oligo rev</td><td>AAACTCCGGGGCGAGTCTTTTACA</td><td></td><td></td><td></td><td></td><td></td><td></td><td></td><td></td><td></td><td></td><td></td></tr>
<tr><td>Chromosome</td><td>start</td><td></td><td>end</td><td>strand</td><td>MM</td><td>target_seq</td><td>PAM</td><td>alignment</td><td>distance</td><td>position</td><td>gene name</td><td>gene id</td></tr>
<tr><td>GU305914.1</td><td></td><td>187473</td><td>187495</td><td>+</td><td>0</td><td>TGTAAAAGACTCGCCCCGGA</td><td>GGG</td><td>||||||||[||||||||||||]PAM</td><td>NA</td><td>Intergenic</td><td>NA</td><td>NA</td></tr>
<tr><td>T14277</td><td>GTAAAAGACTCGCCCCGGAGGGG</td><td></td><td>1000</td><td></td><td></td><td></td><td></td><td></td><td></td><td></td><td></td><td></td></tr>
<tr><td>Oligo fwd</td><td>CACCGTAAAAGACTCGCCCCGGAG</td><td></td><td></td><td></td><td></td><td></td><td></td><td></td><td></td><td></td><td></td><td></td></tr>
<tr><td>Oligo rev</td><td>AAACCTCCGGGGCGAGTCTTTTAC</td><td></td><td></td><td></td><td></td><td></td><td></td><td></td><td></td><td></td><td></td><td></td></tr>
<tr><td>Chromosome</td><td>start</td><td></td><td>end</td><td>strand</td><td>MM</td><td>target_seq</td><td>PAM</td><td>alignment</td><td>distance</td><td>position</td><td>gene name</td><td>gene id</td></tr>
<tr><td>GU305914.1</td><td></td><td>187474</td><td>187496</td><td>+</td><td>0</td><td>GTAAAAGACTCGCCCCGGAG</td><td>GGG</td><td>||||||||[||||||||||||]PAM</td><td>NA</td><td>Intergenic</td><td>NA</td><td>NA</td></tr>
<tr><td>T14278</td><td>TAAAAGACTCGCCCCGGAGGGGG</td><td></td><td>1000</td><td></td><td></td><td></td><td></td><td></td><td></td><td></td><td></td><td></td></tr>
<tr><td>Oligo fwd</td><td>CACCTAAAAGACTCGCCCCGGAGG</td><td></td><td></td><td></td><td></td><td></td><td></td><td></td><td></td><td></td><td></td><td></td></tr>
<tr><td>Oligo rev</td><td>AAACCCTCCGGGGCGAGTCTTTTA</td><td></td><td></td><td></td><td></td><td></td><td></td><td></td><td></td><td></td><td></td><td></td></tr>
<tr><td>Chromosome</td><td>start</td><td></td><td>end</td><td>strand</td><td>MM</td><td>target_seq</td><td>PAM</td><td>alignment</td><td>distance</td><td>position</td><td>gene name</td><td>gene id</td></tr>
<tr><td>GU305914.1</td><td></td><td>187475</td><td>187497</td><td>+</td><td>0</td><td>TAAAAGACTCGCCCCGGAGG</td><td>GGG</td><td>||||||||[||||||||||||]PAM</td><td>NA</td><td>Intergenic</td><td>NA</td><td>NA</td></tr>
<tr><td>T14279</td><td>GAGGGGGTTCCCCCGATGTGAGG</td><td></td><td>1000</td><td></td><td></td><td></td><td></td><td></td><td></td><td></td><td></td><td></td></tr>
<tr><td>Oligo fwd</td><td>CACCGAGGGGGTTCCCCCGATGTG</td><td></td><td></td><td></td><td></td><td></td><td></td><td></td><td></td><td></td><td></td><td></td></tr>
<tr><td>Oligo rev</td><td>AAACCACATCGGGGGAACCCCCTC</td><td></td><td></td><td></td><td></td><td></td><td></td><td></td><td></td><td></td><td></td><td></td></tr>
<tr><td>Chromosome</td><td>start</td><td></td><td>end</td><td>strand</td><td>MM</td><td>target_seq</td><td>PAM</td><td>alignment</td><td>distance</td><td>position</td><td>gene name</td><td>gene id</td></tr>
<tr><td>GU305914.1</td><td></td><td>187491</td><td>187513</td><td>+</td><td>0</td><td>GAGGGGGTTCCCCCGATGTG</td><td>AGG</td><td>||||||||[||||||||||||]PAM</td><td>NA</td><td>Intergenic</td><td>NA</td><td>NA</td></tr>
<tr><td>T14280</td><td>AGGGGGTTCCCCCGATGTGAGGG</td><td></td><td>1000</td><td></td><td></td><td></td><td></td><td></td><td></td><td></td><td></td><td></td></tr>
<tr><td>Oligo fwd</td><td>CACCAGGGGGTTCCCCCGATGTGA</td><td></td><td></td><td></td><td></td><td></td><td></td><td></td><td></td><td></td><td></td><td></td></tr>
<tr><td>Oligo rev</td><td>AAACTCACATCGGGGGAACCCCCT</td><td></td><td></td><td></td><td></td><td></td><td></td><td></td><td></td><td></td><td></td><td></td></tr>
<tr><td>Chromosome</td><td>start</td><td></td><td>end</td><td>strand</td><td>MM</td><td>target_seq</td><td>PAM</td><td>alignment</td><td>distance</td><td>position</td><td>gene name</td><td>gene id</td></tr>
<tr><td>GU305914.1</td><td></td><td>187492</td><td>187514</td><td>+</td><td>0</td><td>AGGGGGTTCCCCCGATGTGA</td><td>GGG</td><td>||||||||[||||||||||||]PAM</td><td>NA</td><td>Intergenic</td><td>NA</td><td>NA</td></tr>
<tr><td>T14281</td><td>GGGGGTTCCCCCGATGTGAGGGG</td><td></td><td>1000</td><td></td><td></td><td></td><td></td><td></td><td></td><td></td><td></td><td></td></tr>
<tr><td>Oligo fwd</td><td>CACCGGGGGTTCCCCCGATGTGAG</td><td></td><td></td><td></td><td></td><td></td><td></td><td></td><td></td><td></td><td></td><td></td></tr>
<tr><td>Oligo rev</td><td>AAACCTCACATCGGGGGAACCCCC</td><td></td><td></td><td></td><td></td><td></td><td></td><td></td><td></td><td></td><td></td><td></td></tr>
<tr><td>Chromosome</td><td>start</td><td></td><td>end</td><td>strand</td><td>MM</td><td>target_seq</td><td>PAM</td><td>alignment</td><td>distance</td><td>position</td><td>gene name</td><td>gene id</td></tr>
<tr><td>GU305914.1</td><td></td><td>187493</td><td>187515</td><td>+</td><td>0</td><td>GGGGGTTCCCCCGATGTGAG</td><td>GGG</td><td>||||||||[||||||||||||]PAM</td><td>NA</td><td>Intergenic</td><td>NA</td><td>NA</td></tr>
<tr><td>T14282</td><td>GTGAGGGGCATGCAGTAGTATGG</td><td></td><td>1000</td><td></td><td></td><td></td><td></td><td></td><td></td><td></td><td></td><td></td></tr>
<tr><td>Oligo fwd</td><td>CACCGTGAGGGGCATGCAGTAGTA</td><td></td><td></td><td></td><td></td><td></td><td></td><td></td><td></td><td></td><td></td><td></td></tr>
<tr><td>Oligo rev</td><td>AAACTACTACTGCATGCCCCTCAC</td><td></td><td></td><td></td><td></td><td></td><td></td><td></td><td></td><td></td><td></td><td></td></tr>
<tr><td>Chromosome</td><td>start</td><td></td><td>end</td><td>strand</td><td>MM</td><td>target_seq</td><td>PAM</td><td>alignment</td><td>distance</td><td>position</td><td>gene name</td><td>gene id</td></tr>
<tr><td>GU305914.1</td><td></td><td>187508</td><td>187530</td><td>+</td><td>0</td><td>GTGAGGGGCATGCAGTAGTA</td><td>TGG</td><td>||||||||[||||||||||||]PAM</td><td>NA</td><td>Intergenic</td><td>NA</td><td>NA</td></tr>
<tr><td>T14283</td><td>GTATGGTGTCCTGAGTGTCTCGG</td><td></td><td>1000</td><td></td><td></td><td></td><td></td><td></td><td></td><td></td><td></td><td></td></tr>
<tr><td>Oligo fwd</td><td>CACCGTATGGTGTCCTGAGTGTCT</td><td></td><td></td><td></td><td></td><td></td><td></td><td></td><td></td><td></td><td></td><td></td></tr>
<tr><td>Oligo rev</td><td>AAACAGACACTCAGGACACCATAC</td><td></td><td></td><td></td><td></td><td></td><td></td><td></td><td></td><td></td><td></td><td></td></tr>
<tr><td>Chromosome</td><td>start</td><td></td><td>end</td><td>strand</td><td>MM</td><td>target_seq</td><td>PAM</td><td>alignment</td><td>distance</td><td>position</td><td>gene name</td><td>gene id</td></tr>
<tr><td>GU305914.1</td><td></td><td>187525</td><td>187547</td><td>+</td><td>0</td><td>GTATGGTGTCCTGAGTGTCT</td><td>CGG</td><td>||||||||[||||||||||||]PAM</td><td>NA</td><td>Intergenic</td><td>NA</td><td>NA</td></tr>
<tr><td>T14284</td><td>GAACTCGCACTCTACCGCCGTGG</td><td></td><td>1000</td><td></td><td></td><td></td><td></td><td></td><td></td><td></td><td></td><td></td></tr>
<tr><td>Oligo fwd</td><td>CACCGAACTCGCACTCTACCGCCG</td><td></td><td></td><td></td><td></td><td></td><td></td><td></td><td></td><td></td><td></td><td></td></tr>
<tr><td>Oligo rev</td><td>AAACCGGCGGTAGAGTGCGAGTTC</td><td></td><td></td><td></td><td></td><td></td><td></td><td></td><td></td><td></td><td></td><td></td></tr>
<tr><td>Chromosome</td><td>start</td><td></td><td>end</td><td>strand</td><td>MM</td><td>target_seq</td><td>PAM</td><td>alignment</td><td>distance</td><td>position</td><td>gene name</td><td>gene id</td></tr>
<tr><td>GU305914.1</td><td></td><td>187558</td><td>187580</td><td>+</td><td>0</td><td>GAACTCGCACTCTACCGCCG</td><td>TGG</td><td>||||||||[||||||||||||]PAM</td><td>NA</td><td>Intergenic</td><td>NA</td><td>NA</td></tr>
<tr><td>T14285</td><td>AACTCGCACTCTACCGCCGTGGG</td><td></td><td>1000</td><td></td><td></td><td></td><td></td><td></td><td></td><td></td><td></td><td></td></tr>
<tr><td>Oligo fwd</td><td>CACCAACTCGCACTCTACCGCCGT</td><td></td><td></td><td></td><td></td><td></td><td></td><td></td><td></td><td></td><td></td><td></td></tr>
<tr><td>Oligo rev</td><td>AAACACGGCGGTAGAGTGCGAGTT</td><td></td><td></td><td></td><td></td><td></td><td></td><td></td><td></td><td></td><td></td><td></td></tr>
<tr><td>Chromosome</td><td>start</td><td></td><td>end</td><td>strand</td><td>MM</td><td>target_seq</td><td>PAM</td><td>alignment</td><td>distance</td><td>position</td><td>gene name</td><td>gene id</td></tr>
<tr><td>GU305914.1</td><td></td><td>187559</td><td>187581</td><td>+</td><td>0</td><td>AACTCGCACTCTACCGCCGT</td><td>GGG</td><td>||||||||[||||||||||||]PAM</td><td>NA</td><td>Intergenic</td><td>NA</td><td>NA</td></tr>
<tr><td>T14286</td><td>ACTCGCACTCTACCGCCGTGGGG</td><td></td><td>1000</td><td></td><td></td><td></td><td></td><td></td><td></td><td></td><td></td><td></td></tr>
<tr><td>Oligo fwd</td><td>CACCACTCGCACTCTACCGCCGTG</td><td></td><td></td><td></td><td></td><td></td><td></td><td></td><td></td><td></td><td></td><td></td></tr>
<tr><td>Oligo rev</td><td>AAACCACGGCGGTAGAGTGCGAGT</td><td></td><td></td><td></td><td></td><td></td><td></td><td></td><td></td><td></td><td></td><td></td></tr>
<tr><td>Chromosome</td><td>start</td><td></td><td>end</td><td>strand</td><td>MM</td><td>target_seq</td><td>PAM</td><td>alignment</td><td>distance</td><td>position</td><td>gene name</td><td>gene id</td></tr>
<tr><td>GU305914.1</td><td></td><td>187560</td><td>187582</td><td>+</td><td>0</td><td>ACTCGCACTCTACCGCCGTG</td><td>GGG</td><td>||||||||[||||||||||||]PAM</td><td>NA</td><td>Intergenic</td><td>NA</td><td>NA</td></tr>
<tr><td>T14287</td><td>CTCGCACTCTACCGCCGTGGGGG</td><td></td><td>1000</td><td></td><td></td><td></td><td></td><td></td><td></td><td></td><td></td><td></td></tr>
<tr><td>Oligo fwd</td><td>CACCCTCGCACTCTACCGCCGTGG</td><td></td><td></td><td></td><td></td><td></td><td></td><td></td><td></td><td></td><td></td><td></td></tr>
<tr><td>Oligo rev</td><td>AAACCCACGGCGGTAGAGTGCGAG</td><td></td><td></td><td></td><td></td><td></td><td></td><td></td><td></td><td></td><td></td><td></td></tr>
</table>

Chromosome start end strand MM target_seq PAM alignment distance position gene name gene id
GU305914.1 187561 187583 + 0 CTCGCACTCTACCGCCGTGG GGG ||||||||[||||||||||||]PAM NA Intergenic NA NA

T14288 TAATAAAGTTTTTGCTGCGCCGG 1000
Oligo fwd CACCTAATAAAGTTTTTGCTGCGC
Oligo rev AAACGCGCAGCAAAAACTTTATTA
Chromosome start end strand MM target_seq PAM alignment distance position gene name gene id
GU305914.1 187585 187607 + 0 TAATAAAGTTTTTGCTGCGC CGG ||||||||[||||||||||||]PAM NA Intergenic NA NA

T14289 AAAGTTTTTGCTGCGCCGGTAGG 1000
Oligo fwd CACCAAAGTTTTTGCTGCGCCGGT
Oligo rev AAACACCGGCGCAGCAAAAACTTT
Chromosome start end strand MM target_seq PAM alignment distance position gene name gene id
GU305914.1 187589 187611 + 0 AAAGTTTTTGCTGCGCCGGT AGG ||||||||[||||||||||||]PAM NA Intergenic NA NA

T14290 AAGTTTTTGCTGCGCCGGTAGGG 1000
Oligo fwd CACCAAGTTTTTGCTGCGCCGGTA
Oligo rev AAACTACCGGCGCAGCAAAAACTT
Chromosome start end strand MM target_seq PAM alignment distance position gene name gene id
GU305914.1 187590 187612 + 0 AAGTTTTTGCTGCGCCGGTA GGG ||||||||[||||||||||||]PAM NA Intergenic NA NA

T14291 AGTTTTTGCTGCGCCGGTAGGGG 1000
Oligo fwd CACCAGTTTTTGCTGCGCCGGTAG
Oligo rev AAACCTACCGGCGCAGCAAAAACT
Chromosome start end strand MM target_seq PAM alignment distance position gene name gene id
GU305914.1 187591 187613 + 0 AGTTTTTGCTGCGCCGGTAG GGG ||||||||[||||||||||||]PAM NA Intergenic NA NA

T14292 GTTTTTGCTGCGCCGGTAGGGGG 1000
Oligo fwd CACCGTTTTTGCTGCGCCGGTAGG
Oligo rev AAACCCTACCGGCGCAGCAAAAAC
Chromosome start end strand MM target_seq PAM alignment distance position gene name gene id
GU305914.1 187592 187614 + 0 GTTTTTGCTGCGCCGGTAGG GGG ||||||||[||||||||||||]PAM NA Intergenic NA NA

T14293 TTTTTGCTGCGCCGGTAGGGGGG 1000
Oligo fwd CACCTTTTTGCTGCGCCGGTAGGG
Oligo rev AAACCCCTACCGGCGCAGCAAAAA
Chromosome start end strand MM target_seq PAM alignment distance position gene name gene id
GU305914.1 187593 187615 + 0 TTTTTGCTGCGCCGGTAGGG GGG ||||||||[||||||||||||]PAM NA Intergenic NA NA

T14294 TTTTGCTGCGCCGGTAGGGGGGG 1000
Oligo fwd CACCTTTTGCTGCGCCGGTAGGGG
Oligo rev AAACCCCCTACCGGCGCAGCAAAA
Chromosome start end strand MM target_seq PAM alignment distance position gene name gene id
GU305914.1 187594 187616 + 0 TTTTGCTGCGCCGGTAGGGG GGG ||||||||[||||||||||||]PAM NA Intergenic NA NA

T14295 TGCTGCGCCGGTAGGGGGGGAGG 1000
Oligo fwd CACCTGCTGCGCCGGTAGGGGGGG
Oligo rev AAACCCCCCCCTACCGGCGCAGCA
Chromosome start end strand MM target_seq PAM alignment distance position gene name gene id
GU305914.1 187597 187619 + 0 TGCTGCGCCGGTAGGGGGGG AGG ||||||||[||||||||||||]PAM NA Intergenic NA NA

T14296 GCCGGTAGGGGGGGAGGCCGAGG 1000
Oligo fwd CACCGCCGGTAGGGGGGGAGGCCG
Oligo rev AAACCGGCCTCCCCCCCTACCGGC
Chromosome start end strand MM target_seq PAM alignment distance position gene name gene id
GU305914.1 187603 187625 + 0 GCCGGTAGGGGGGGAGGCCG AGG ||||||||[||||||||||||]PAM NA Intergenic NA NA

T14297 ATAATAAAGTTGCTACGTACTGG 1000
Oligo fwd CACCATAATAAAGTTGCTACGTAC
Oligo rev AAACGTACGTAGCAACTTTATTAT
Chromosome start end strand MM target_seq PAM alignment distance position gene name gene id
GU305914.1 187626 187648 + 0 ATAATAAAGTTGCTACGTAC TGG ||||||||[||||||||||||]PAM NA Intergenic NA NA

T14298 GTTGAAGTCTAACAATCTCTCGG 1000
Oligo fwd CACCGTTGAAGTCTAACAATCTCT
Oligo rev AAACAGAGATTGTTAGACTTCAAC
Chromosome start end strand MM target_seq PAM alignment distance position gene name gene id
GU305914.1 187648 187670 + 0 GTTGAAGTCTAACAATCTCT CGG ||||||||[||||||||||||]PAM NA Intergenic NA NA

T14299 TTGAAGTCTAACAATCTCTCGGG 1000
Oligo fwd CACCTTGAAGTCTAACAATCTCTC
Oligo rev AAACGAGAGATTGTTAGACTTCAA
Chromosome start end strand MM target_seq PAM alignment distance position gene name gene id
GU305914.1 187649 187671 + 0 TTGAAGTCTAACAATCTCTC GGG ||||||||[||||||||||||]PAM NA Intergenic NA NA

T14300 TGAAGTCTAACAATCTCTCGGGG 1000
Oligo fwd CACCTGAAGTCTAACAATCTCTCG
Oligo rev AAACCGAGAGATTGTTAGACTTCA
Chromosome start end strand MM target_seq PAM alignment distance position gene name gene id
GU305914.1 187650 187672 + 0 TGAAGTCTAACAATCTCTCG GGG ||||||||[||||||||||||]PAM NA Intergenic NA NA

T14301 GAAGTCTAACAATCTCTCGGGGG 1000
Oligo fwd CACCGAAGTCTAACAATCTCTCGG
Oligo rev AAACCCGAGAGATTGTTAGACTTC
Chromosome start end strand MM target_seq PAM alignment distance position gene name gene id
GU305914.1 187651 187673 + 0 GAAGTCTAACAATCTCTCGG GGG ||||||||[||||||||||||]PAM NA Intergenic NA NA

T14302 AAGTCTAACAATCTCTCGGGGGG 1000
Oligo fwd CACCAAGTCTAACAATCTCTCGGG

<table>
<tr><td>Oligo rev</td><td>AAACCCCGAGAGATTGTTAGACTT</td><td></td><td></td><td></td><td></td><td></td><td></td><td></td><td></td><td></td><td></td></tr>
<tr><td>Chromosome</td><td>start</td><td>end</td><td>strand</td><td>MM</td><td>target_seq</td><td>PAM</td><td>alignment</td><td>distance</td><td>position</td><td>gene name</td><td>gene id</td></tr>
<tr><td>GU305914.1</td><td>187652</td><td>187674</td><td>+</td><td>0</td><td>AAGTCTAACAATCTCTCGGG</td><td>GGG</td><td>||||||||[||||||||||||]PAM</td><td>NA</td><td>Intergenic</td><td>NA</td><td>NA</td></tr>
<tr><td>T14303</td><td>CAATCTCTCGGGGGGATCCACGG</td><td>1000</td><td></td><td></td><td></td><td></td><td></td><td></td><td></td><td></td><td></td></tr>
<tr><td>Oligo fwd</td><td>CACCCAATCTCTCGGGGGGATCCA</td><td></td><td></td><td></td><td></td><td></td><td></td><td></td><td></td><td></td><td></td></tr>
<tr><td>Oligo rev</td><td>AAACTGGATCCCCCCGAGAGATTG</td><td></td><td></td><td></td><td></td><td></td><td></td><td></td><td></td><td></td><td></td></tr>
<tr><td>Chromosome</td><td>start</td><td>end</td><td>strand</td><td>MM</td><td>target_seq</td><td>PAM</td><td>alignment</td><td>distance</td><td>position</td><td>gene name</td><td>gene id</td></tr>
<tr><td>GU305914.1</td><td>187660</td><td>187682</td><td>+</td><td>0</td><td>CAATCTCTCGGGGGGATCCA</td><td>CGG</td><td>||||||||[||||||||||||]PAM</td><td>NA</td><td>Intergenic</td><td>NA</td><td>NA</td></tr>
<tr><td>T14304</td><td>ATCCACGGTCCACCGTCTCTCGG</td><td>1000</td><td></td><td></td><td></td><td></td><td></td><td></td><td></td><td></td><td></td></tr>
<tr><td>Oligo fwd</td><td>CACCATCCACGGTCCACCGTCTCT</td><td></td><td></td><td></td><td></td><td></td><td></td><td></td><td></td><td></td><td></td></tr>
<tr><td>Oligo rev</td><td>AAACAGAGACGGTGGACCGTGGAT</td><td></td><td></td><td></td><td></td><td></td><td></td><td></td><td></td><td></td><td></td></tr>
<tr><td>Chromosome</td><td>start</td><td>end</td><td>strand</td><td>MM</td><td>target_seq</td><td>PAM</td><td>alignment</td><td>distance</td><td>position</td><td>gene name</td><td>gene id</td></tr>
<tr><td>GU305914.1</td><td>187675</td><td>187697</td><td>+</td><td>0</td><td>ATCCACGGTCCACCGTCTCT</td><td>CGG</td><td>||||||||[||||||||||||]PAM</td><td>NA</td><td>Intergenic</td><td>NA</td><td>NA</td></tr>
<tr><td>T14305</td><td>CGAGCTGTTGATGCAGCGTGAGG</td><td>1000</td><td></td><td></td><td></td><td></td><td></td><td></td><td></td><td></td><td></td></tr>
<tr><td>Oligo fwd</td><td>CACCCGAGCTGTTGATGCAGCGTG</td><td></td><td></td><td></td><td></td><td></td><td></td><td></td><td></td><td></td><td></td></tr>
<tr><td>Oligo rev</td><td>AAACCACGCTGCATCAACAGCTCG</td><td></td><td></td><td></td><td></td><td></td><td></td><td></td><td></td><td></td><td></td></tr>
<tr><td>Chromosome</td><td>start</td><td>end</td><td>strand</td><td>MM</td><td>target_seq</td><td>PAM</td><td>alignment</td><td>distance</td><td>position</td><td>gene name</td><td>gene id</td></tr>
<tr><td>GU305914.1</td><td>187705</td><td>187727</td><td>+</td><td>0</td><td>CGAGCTGTTGATGCAGCGTG</td><td>AGG</td><td>||||||||[||||||||||||]PAM</td><td>NA</td><td>Intergenic</td><td>NA</td><td>NA</td></tr>
<tr><td>T14306</td><td>GCTGTTGATGCAGCGTGAGGAGG</td><td>1000</td><td></td><td></td><td></td><td></td><td></td><td></td><td></td><td></td><td></td></tr>
<tr><td>Oligo fwd</td><td>CACCGCTGTTGATGCAGCGTGAGG</td><td></td><td></td><td></td><td></td><td></td><td></td><td></td><td></td><td></td><td></td></tr>
<tr><td>Oligo rev</td><td>AAACCCTCACGCTGCATCAACAGC</td><td></td><td></td><td></td><td></td><td></td><td></td><td></td><td></td><td></td><td></td></tr>
<tr><td>Chromosome</td><td>start</td><td>end</td><td>strand</td><td>MM</td><td>target_seq</td><td>PAM</td><td>alignment</td><td>distance</td><td>position</td><td>gene name</td><td>gene id</td></tr>
<tr><td>GU305914.1</td><td>187708</td><td>187730</td><td>+</td><td>0</td><td>GCTGTTGATGCAGCGTGAGG</td><td>AGG</td><td>||||||||[||||||||||||]PAM</td><td>NA</td><td>Intergenic</td><td>NA</td><td>NA</td></tr>
<tr><td>T14307</td><td>CTGTTGATGCAGCGTGAGGAGGG</td><td>1000</td><td></td><td></td><td></td><td></td><td></td><td></td><td></td><td></td><td></td></tr>
<tr><td>Oligo fwd</td><td>CACCCTGTTGATGCAGCGTGAGGA</td><td></td><td></td><td></td><td></td><td></td><td></td><td></td><td></td><td></td><td></td></tr>
<tr><td>Oligo rev</td><td>AAACTCCTCACGCTGCATCAACAG</td><td></td><td></td><td></td><td></td><td></td><td></td><td></td><td></td><td></td><td></td></tr>
<tr><td>Chromosome</td><td>start</td><td>end</td><td>strand</td><td>MM</td><td>target_seq</td><td>PAM</td><td>alignment</td><td>distance</td><td>position</td><td>gene name</td><td>gene id</td></tr>
<tr><td>GU305914.1</td><td>187709</td><td>187731</td><td>+</td><td>0</td><td>CTGTTGATGCAGCGTGAGGA</td><td>GGG</td><td>||||||||[||||||||||||]PAM</td><td>NA</td><td>Intergenic</td><td>NA</td><td>NA</td></tr>
<tr><td>T14308</td><td>TGTTGATGCAGCGTGAGGAGGGG</td><td>1000</td><td></td><td></td><td></td><td></td><td></td><td></td><td></td><td></td><td></td></tr>
<tr><td>Oligo fwd</td><td>CACCTGTTGATGCAGCGTGAGGAG</td><td></td><td></td><td></td><td></td><td></td><td></td><td></td><td></td><td></td><td></td></tr>
<tr><td>Oligo rev</td><td>AAACCTCCTCACGCTGCATCAACA</td><td></td><td></td><td></td><td></td><td></td><td></td><td></td><td></td><td></td><td></td></tr>
<tr><td>Chromosome</td><td>start</td><td>end</td><td>strand</td><td>MM</td><td>target_seq</td><td>PAM</td><td>alignment</td><td>distance</td><td>position</td><td>gene name</td><td>gene id</td></tr>
<tr><td>GU305914.1</td><td>187710</td><td>187732</td><td>+</td><td>0</td><td>TGTTGATGCAGCGTGAGGAG</td><td>GGG</td><td>||||||||[||||||||||||]PAM</td><td>NA</td><td>Intergenic</td><td>NA</td><td>NA</td></tr>
<tr><td>T14309</td><td>GGAGGGGTCGTTATCCTTGCCGG</td><td>1000</td><td></td><td></td><td></td><td></td><td></td><td></td><td></td><td></td><td></td></tr>
<tr><td>Oligo fwd</td><td>CACCGGAGGGGTCGTTATCCTTGC</td><td></td><td></td><td></td><td></td><td></td><td></td><td></td><td></td><td></td><td></td></tr>
<tr><td>Oligo rev</td><td>AAACGCAAGGATAACGACCCCTCC</td><td></td><td></td><td></td><td></td><td></td><td></td><td></td><td></td><td></td><td></td></tr>
<tr><td>Chromosome</td><td>start</td><td>end</td><td>strand</td><td>MM</td><td>target_seq</td><td>PAM</td><td>alignment</td><td>distance</td><td>position</td><td>gene name</td><td>gene id</td></tr>
<tr><td>GU305914.1</td><td>187726</td><td>187748</td><td>+</td><td>0</td><td>GGAGGGGTCGTTATCCTTGC</td><td>CGG</td><td>||||||||[||||||||||||]PAM</td><td>NA</td><td>Intergenic</td><td>NA</td><td>NA</td></tr>
<tr><td>T14310</td><td>CGTGTACTGTATCCACACCGTGG</td><td>1000</td><td></td><td></td><td></td><td></td><td></td><td></td><td></td><td></td><td></td></tr>
<tr><td>Oligo fwd</td><td>CACCCGTGTACTGTATCCACACCG</td><td></td><td></td><td></td><td></td><td></td><td></td><td></td><td></td><td></td><td></td></tr>
<tr><td>Oligo rev</td><td>AAACCGGTGTGGATACAGTACACG</td><td></td><td></td><td></td><td></td><td></td><td></td><td></td><td></td><td></td><td></td></tr>
<tr><td>Chromosome</td><td>start</td><td>end</td><td>strand</td><td>MM</td><td>target_seq</td><td>PAM</td><td>alignment</td><td>distance</td><td>position</td><td>gene name</td><td>gene id</td></tr>
<tr><td>GU305914.1</td><td>187765</td><td>187787</td><td>+</td><td>0</td><td>CGTGTACTGTATCCACACCG</td><td>TGG</td><td>||||||||[||||||||||||]PAM</td><td>NA</td><td>Intergenic</td><td>NA</td><td>NA</td></tr>
<tr><td>T14311</td><td>GTGTACTGTATCCACACCGTGGG</td><td>1000</td><td></td><td></td><td></td><td></td><td></td><td></td><td></td><td></td><td></td></tr>
<tr><td>Oligo fwd</td><td>CACCGTGTACTGTATCCACACCGT</td><td></td><td></td><td></td><td></td><td></td><td></td><td></td><td></td><td></td><td></td></tr>
<tr><td>Oligo rev</td><td>AAACACGGTGTGGATACAGTACAC</td><td></td><td></td><td></td><td></td><td></td><td></td><td></td><td></td><td></td><td></td></tr>
<tr><td>Chromosome</td><td>start</td><td>end</td><td>strand</td><td>MM</td><td>target_seq</td><td>PAM</td><td>alignment</td><td>distance</td><td>position</td><td>gene name</td><td>gene id</td></tr>
<tr><td>GU305914.1</td><td>187766</td><td>187788</td><td>+</td><td>0</td><td>GTGTACTGTATCCACACCGT</td><td>GGG</td><td>||||||||[||||||||||||]PAM</td><td>NA</td><td>Intergenic</td><td>NA</td><td>NA</td></tr>
<tr><td>T14312</td><td>ATCCACACCGTGGGTTTGAACGG</td><td>1000</td><td></td><td></td><td></td><td></td><td></td><td></td><td></td><td></td><td></td></tr>
<tr><td>Oligo fwd</td><td>CACCATCCACACCGTGGGTTTGAA</td><td></td><td></td><td></td><td></td><td></td><td></td><td></td><td></td><td></td><td></td></tr>
<tr><td>Oligo rev</td><td>AAACTTCAAACCCACGGTGTGGAT</td><td></td><td></td><td></td><td></td><td></td><td></td><td></td><td></td><td></td><td></td></tr>
<tr><td>Chromosome</td><td>start</td><td>end</td><td>strand</td><td>MM</td><td>target_seq</td><td>PAM</td><td>alignment</td><td>distance</td><td>position</td><td>gene name</td><td>gene id</td></tr>
<tr><td>GU305914.1</td><td>187775</td><td>187797</td><td>+</td><td>0</td><td>ATCCACACCGTGGGTTTGAA</td><td>CGG</td><td>||||||||[||||||||||||]PAM</td><td>NA</td><td>Intergenic</td><td>NA</td><td>NA</td></tr>
<tr><td>T14313</td><td>TTGAACGGATGTATGCCGAGCGG</td><td>1000</td><td></td><td></td><td></td><td></td><td></td><td></td><td></td><td></td><td></td></tr>
<tr><td>Oligo fwd</td><td>CACCTTGAACGGATGTATGCCGAG</td><td></td><td></td><td></td><td></td><td></td><td></td><td></td><td></td><td></td><td></td></tr>
<tr><td>Oligo rev</td><td>AAACCTCGGCATACATCCGTTCAA</td><td></td><td></td><td></td><td></td><td></td><td></td><td></td><td></td><td></td><td></td></tr>
<tr><td>Chromosome</td><td>start</td><td>end</td><td>strand</td><td>MM</td><td>target_seq</td><td>PAM</td><td>alignment</td><td>distance</td><td>position</td><td>gene name</td><td>gene id</td></tr>
<tr><td>GU305914.1</td><td>187790</td><td>187812</td><td>+</td><td>0</td><td>TTGAACGGATGTATGCCGAG</td><td>CGG</td><td>||||||||[||||||||||||]PAM</td><td>NA</td><td>Intergenic</td><td>NA</td><td>NA</td></tr>
<tr><td>T14314</td><td>GTTCGTACCGTTCCCGACCCAGG</td><td>1000</td><td></td><td></td><td></td><td></td><td></td><td></td><td></td><td></td><td></td></tr>
<tr><td>Oligo fwd</td><td>CACCGTTCGTACCGTTCCCGACCC</td><td></td><td></td><td></td><td></td><td></td><td></td><td></td><td></td><td></td><td></td></tr>
<tr><td>Oligo rev</td><td>AAACGGGTCGGGAACGGTACGAAC</td><td></td><td></td><td></td><td></td><td></td><td></td><td></td><td></td><td></td><td></td></tr>
<tr><td>Chromosome</td><td>start</td><td>end</td><td>strand</td><td>MM</td><td>target_seq</td><td>PAM</td><td>alignment</td><td>distance</td><td>position</td><td>gene name</td><td>gene id</td></tr>
<tr><td>GU305914.1</td><td>187816</td><td>187838</td><td>+</td><td>0</td><td>GTTCGTACCGTTCCCGACCC</td><td>AGG</td><td>||||||||[||||||||||||]PAM</td><td>NA</td><td>Intergenic</td><td>NA</td><td>NA</td></tr>
<tr><td>T14315</td><td>TTCACCGTCCTTCACCTCGTAGG</td><td>1000</td><td></td><td></td><td></td><td></td><td></td><td></td><td></td><td></td><td></td></tr>
<tr><td>Oligo fwd</td><td>CACCTTCACCGTCCTTCACCTCGT</td><td></td><td></td><td></td><td></td><td></td><td></td><td></td><td></td><td></td><td></td></tr>
<tr><td>Oligo rev</td><td>AAACACGAGGTGAAGGACGGTGAA</td><td></td><td></td><td></td><td></td><td></td><td></td><td></td><td></td><td></td><td></td></tr>
<tr><td>Chromosome</td><td>start</td><td>end</td><td>strand</td><td>MM</td><td>target_seq</td><td>PAM</td><td>alignment</td><td>distance</td><td>position</td><td>gene name</td><td>gene id</td></tr>
<tr><td>GU305914.1</td><td>187867</td><td>187889</td><td>+</td><td>0</td><td>TTCACCGTCCTTCACCTCGT</td><td>AGG</td><td>||||||||[||||||||||||]PAM</td><td>NA</td><td>Intergenic</td><td>NA</td><td>NA</td></tr>
<tr><td>T14316</td><td>TCACCGTCCTTCACCTCGTAGGG</td><td>1000</td><td></td><td></td><td></td><td></td><td></td><td></td><td></td><td></td><td></td></tr>
<tr><td>Oligo fwd</td><td>CACCTCACCGTCCTTCACCTCGTA</td><td></td><td></td><td></td><td></td><td></td><td></td><td></td><td></td><td></td><td></td></tr>
<tr><td>Oligo rev</td><td>AAACTACGAGGTGAAGGACGGTGA</td><td></td><td></td><td></td><td></td><td></td><td></td><td></td><td></td><td></td><td></td></tr>
<tr><td>Chromosome</td><td>start</td><td>end</td><td>strand</td><td>MM</td><td>target_seq</td><td>PAM</td><td>alignment</td><td>distance</td><td>position</td><td>gene name</td><td>gene id</td></tr>
<tr><td>GU305914.1</td><td>187868</td><td>187890</td><td>+</td><td>0</td><td>TCACCGTCCTTCACCTCGTA</td><td>GGG</td><td>||||||||[||||||||||||]PAM</td><td>NA</td><td>Intergenic</td><td>NA</td><td>NA</td></tr>
<tr><td>T14317</td><td>CACCGTCCTTCACCTCGTAGGGG</td><td>1000</td><td></td><td></td><td></td><td></td><td></td><td></td><td></td><td></td><td></td></tr>
</table>

| | | | | | | | | | | | |
|---|---|---|---|---|---|---|---|---|---|---|---|
| Oligo fwd | CACCCACCGTCCTTCACCTCGTAG | | | | | | | | | | |
| Oligo rev | AAACCTACGAGGTGAAGGACGGTG | | | | | | | | | | |
| Chromosome | start | end | strand | MM | target_seq | PAM | alignment | distance | position | gene name | gene id |
| GU305914.1 | 187869 | 187891 | + | 0 | CACCGTCCTTCACCTCGTAG | GGG | \|\|\|\|\|\|\|\|[\|\|\|\|\|\|\|\|\|\|\|\|]PAM | NA | Intergenic | NA | NA |
| T14318 | TACTTGCCCACCGTGATGCGTGG | 1000 | | | | | | | | | |
| Oligo fwd | CACCTACTTGCCCACCGTGATGCG | | | | | | | | | | |
| Oligo rev | AAACCGCATCACGGTGGGCAAGTA | | | | | | | | | | |
| Chromosome | start | end | strand | MM | target_seq | PAM | alignment | distance | position | gene name | gene id |
| GU305914.1 | 187892 | 187914 | + | 0 | TACTTGCCCACCGTGATGCG | TGG | \|\|\|\|\|\|\|\|[\|\|\|\|\|\|\|\|\|\|\|\|]PAM | NA | Intergenic | NA | NA |
| T14319 | TGGCGATTCCGTCATCACATCGG | 1000 | | | | | | | | | |
| Oligo fwd | CACCTGGCGATTCCGTCATCACAT | | | | | | | | | | |
| Oligo rev | AAACATGTGATGACGGAATCGCCA | | | | | | | | | | |
| Chromosome | start | end | strand | MM | target_seq | PAM | alignment | distance | position | gene name | gene id |
| GU305914.1 | 187912 | 187934 | + | 0 | TGGCGATTCCGTCATCACAT | CGG | \|\|\|\|\|\|\|\|[\|\|\|\|\|\|\|\|\|\|\|\|]PAM | NA | Intergenic | NA | NA |
| T14320 | ATCGGCCACGTACATCATGTCGG | 1000 | | | | | | | | | |
| Oligo fwd | CACCATCGGCCACGTACATCATGT | | | | | | | | | | |
| Oligo rev | AAACACATGATGTACGTGGCCGAT | | | | | | | | | | |
| Chromosome | start | end | strand | MM | target_seq | PAM | alignment | distance | position | gene name | gene id |
| GU305914.1 | 187930 | 187952 | + | 0 | ATCGGCCACGTACATCATGT | CGG | \|\|\|\|\|\|\|\|[\|\|\|\|\|\|\|\|\|\|\|\|]PAM | NA | Intergenic | NA | NA |
| T14321 | GGCCACGTACATCATGTCGGCGG | 1000 | | | | | | | | | |
| Oligo fwd | CACCGGCCACGTACATCATGTCGG | | | | | | | | | | |
| Oligo rev | AAACCCGACATGATGTACGTGGCC | | | | | | | | | | |
| Chromosome | start | end | strand | MM | target_seq | PAM | alignment | distance | position | gene name | gene id |
| GU305914.1 | 187933 | 187955 | + | 0 | GGCCACGTACATCATGTCGG | CGG | \|\|\|\|\|\|\|\|[\|\|\|\|\|\|\|\|\|\|\|\|]PAM | NA | Intergenic | NA | NA |
| T14322 | GCCACGTACATCATGTCGGCGGG | 1000 | | | | | | | | | |
| Oligo fwd | CACCGCCACGTACATCATGTCGGC | | | | | | | | | | |
| Oligo rev | AAACGCCGACATGATGTACGTGGC | | | | | | | | | | |
| Chromosome | start | end | strand | MM | target_seq | PAM | alignment | distance | position | gene name | gene id |
| GU305914.1 | 187934 | 187956 | + | 0 | GCCACGTACATCATGTCGGC | GGG | \|\|\|\|\|\|\|\|[\|\|\|\|\|\|\|\|\|\|\|\|]PAM | NA | Intergenic | NA | NA |
| T14323 | CCACGTACATCATGTCGGCGGGG | 1000 | | | | | | | | | |
| Oligo fwd | CACCCCACGTACATCATGTCGGCG | | | | | | | | | | |
| Oligo rev | AAACCGCCGACATGATGTACGTGG | | | | | | | | | | |
| Chromosome | start | end | strand | MM | target_seq | PAM | alignment | distance | position | gene name | gene id |
| GU305914.1 | 187935 | 187957 | + | 0 | CCACGTACATCATGTCGGCG | GGG | \|\|\|\|\|\|\|\|[\|\|\|\|\|\|\|\|\|\|\|\|]PAM | NA | Intergenic | NA | NA |
| T14324 | CATCATGTCGGCGGGGTCCACGG | 1000 | | | | | | | | | |
| Oligo fwd | CACCCATCATGTCGGCGGGGTCCA | | | | | | | | | | |
| Oligo rev | AAACTGGACCCCGCCGACATGATG | | | | | | | | | | |
| Chromosome | start | end | strand | MM | target_seq | PAM | alignment | distance | position | gene name | gene id |
| GU305914.1 | 187942 | 187964 | + | 0 | CATCATGTCGGCGGGGTCCA | CGG | \|\|\|\|\|\|\|\|[\|\|\|\|\|\|\|\|\|\|\|\|]PAM | NA | Intergenic | NA | NA |
| T14325 | ATCATGTCGGCGGGGTCCACGGG | 1000 | | | | | | | | | |
| Oligo fwd | CACCATCATGTCGGCGGGGTCCAC | | | | | | | | | | |
| Oligo rev | AAACGTGGACCCCGCCGACATGAT | | | | | | | | | | |
| Chromosome | start | end | strand | MM | target_seq | PAM | alignment | distance | position | gene name | gene id |
| GU305914.1 | 187943 | 187965 | + | 0 | ATCATGTCGGCGGGGTCCAC | GGG | \|\|\|\|\|\|\|\|[\|\|\|\|\|\|\|\|\|\|\|\|]PAM | NA | Intergenic | NA | NA |
| T14326 | GTCCACGGGCACGAGCCTGATGG | 1000 | | | | | | | | | |
| Oligo fwd | CACCGTCCACGGGCACGAGCCTGA | | | | | | | | | | |
| Oligo rev | AAACTCAGGCTCGTGCCCGTGGAC | | | | | | | | | | |
| Chromosome | start | end | strand | MM | target_seq | PAM | alignment | distance | position | gene name | gene id |
| GU305914.1 | 187957 | 187979 | + | 0 | GTCCACGGGCACGAGCCTGA | TGG | \|\|\|\|\|\|\|\|[\|\|\|\|\|\|\|\|\|\|\|\|]PAM | NA | Intergenic | NA | NA |
| GU305914.1 | 111337 | 111359 | - | 4 | GACGACGCGCACGAGCCAGA | TGG | \|-\|-\|\|\|-[\|\|\|\|\|\|\|\|\|-\|\|]PAM | NA | Intergenic | NA | NA |
| T14327 | CACGGGCACGAGCCTGATGGTGG | 1000 | | | | | | | | | |
| Oligo fwd | CACCCACGGGCACGAGCCTGATGG | | | | | | | | | | |
| Oligo rev | AAACCCATCAGGCTCGTGCCCGTG | | | | | | | | | | |
| Chromosome | start | end | strand | MM | target_seq | PAM | alignment | distance | position | gene name | gene id |
| GU305914.1 | 187960 | 187982 | + | 0 | CACGGGCACGAGCCTGATGG | TGG | \|\|\|\|\|\|\|\|[\|\|\|\|\|\|\|\|\|\|\|\|]PAM | NA | Intergenic | NA | NA |
| GU305914.1 | 89043 | 89065 | + | 3 | CACGGGCTCGCGCGTGATGG | TGG | \|\|\|\|\|\|\|-[\|\|-\|\|-\|\|\|\|\|\|]PAM | NA | Intergenic | NA | NA |
| T14328 | GTAGCGCACCAAAGATAAGCAGG | 1000 | | | | | | | | | |
| Oligo fwd | CACCGTAGCGCACCAAAGATAAGC | | | | | | | | | | |
| Oligo rev | AAACGCTTATCTTTGGTGCGCTAC | | | | | | | | | | |
| Chromosome | start | end | strand | MM | target_seq | PAM | alignment | distance | position | gene name | gene id |
| GU305914.1 | 187998 | 188020 | + | 0 | GTAGCGCACCAAAGATAAGC | AGG | \|\|\|\|\|\|\|\|[\|\|\|\|\|\|\|\|\|\|\|\|]PAM | NA | Intergenic | NA | NA |
| T14329 | GATAAGCAGGACACGCAGTCTGG | 1000 | | | | | | | | | |
| Oligo fwd | CACCGATAAGCAGGACACGCAGTC | | | | | | | | | | |
| Oligo rev | AAACGACTGCGTGTCCTGCTTATC | | | | | | | | | | |
| Chromosome | start | end | strand | MM | target_seq | PAM | alignment | distance | position | gene name | gene id |
| GU305914.1 | 188011 | 188033 | + | 0 | GATAAGCAGGACACGCAGTC | TGG | \|\|\|\|\|\|\|\|[\|\|\|\|\|\|\|\|\|\|\|\|]PAM | NA | Intergenic | NA | NA |
| T14330 | CAAGAGTCTGTCCGACCACCAGG | 1000 | | | | | | | | | |
| Oligo fwd | CACCCAAGAGTCTGTCCGACCACC | | | | | | | | | | |
| Oligo rev | AAACGGTGGTCGGACAGACTCTTG | | | | | | | | | | |
| Chromosome | start | end | strand | MM | target_seq | PAM | alignment | distance | position | gene name | gene id |
| GU305914.1 | 188141 | 188163 | + | 0 | CAAGAGTCTGTCCGACCACC | AGG | \|\|\|\|\|\|\|\|[\|\|\|\|\|\|\|\|\|\|\|\|]PAM | NA | Intergenic | NA | NA |
| T14331 | GCACACCCTCATCCCAGAGAAGG | 1000 | | | | | | | | | |
| Oligo fwd | CACCGCACACCCTCATCCCAGAGA | | | | | | | | | | |
| Oligo rev | AAACTCTCTGGGATGAGGGTGTGC | | | | | | | | | | |
| Chromosome | start | end | strand | MM | target_seq | PAM | alignment | distance | position | gene name | gene id |

<table>
<tr><td>GU305914.1</td><td></td><td>188174</td><td>188196</td><td>+</td><td>0</td><td>GCACACCCTCATCCCAGAGA</td><td>AGG</td><td>||||||||[||||||||||||]PAM</td><td>NA</td><td>Intergenic</td><td>NA</td><td>NA</td></tr>
<tr><td>T14332</td><td>GCACGAAGTGTTAGTGTATTCGG</td><td></td><td>1000</td><td></td><td></td><td></td><td></td><td></td><td></td><td></td><td></td><td></td></tr>
<tr><td>Oligo fwd</td><td>CACCGCACGAAGTGTTAGTGTATT</td><td></td><td></td><td></td><td></td><td></td><td></td><td></td><td></td><td></td><td></td><td></td></tr>
<tr><td>Oligo rev</td><td>AAACAATACACTAACACTTCGTGC</td><td></td><td></td><td></td><td></td><td></td><td></td><td></td><td></td><td></td><td></td><td></td></tr>
<tr><td>Chromosome</td><td>start</td><td></td><td>end</td><td>strand</td><td>MM</td><td>target_seq</td><td>PAM</td><td>alignment</td><td>distance</td><td>position</td><td>gene name</td><td>gene id</td></tr>
<tr><td>GU305914.1</td><td></td><td>188219</td><td>188241</td><td>+</td><td>0</td><td>GCACGAAGTGTTAGTGTATT</td><td>CGG</td><td>||||||||[||||||||||||]PAM</td><td>NA</td><td>Intergenic</td><td>NA</td><td>NA</td></tr>
<tr><td>T14333</td><td>GCGCTCTGTACCGCGTACAGCGG</td><td></td><td>1000</td><td></td><td></td><td></td><td></td><td></td><td></td><td></td><td></td><td></td></tr>
<tr><td>Oligo fwd</td><td>CACCGCGCTCTGTACCGCGTACAG</td><td></td><td></td><td></td><td></td><td></td><td></td><td></td><td></td><td></td><td></td><td></td></tr>
<tr><td>Oligo rev</td><td>AAACCTGTACGCGGTACAGAGCGC</td><td></td><td></td><td></td><td></td><td></td><td></td><td></td><td></td><td></td><td></td><td></td></tr>
<tr><td>Chromosome</td><td>start</td><td></td><td>end</td><td>strand</td><td>MM</td><td>target_seq</td><td>PAM</td><td>alignment</td><td>distance</td><td>position</td><td>gene name</td><td>gene id</td></tr>
<tr><td>GU305914.1</td><td></td><td>188241</td><td>188263</td><td>+</td><td>0</td><td>GCGCTCTGTACCGCGTACAG</td><td>CGG</td><td>||||||||[||||||||||||]PAM</td><td>NA</td><td>Intergenic</td><td>NA</td><td>NA</td></tr>
<tr><td>T14334</td><td>CGCTCTGTACCGCGTACAGCGGG</td><td></td><td>1000</td><td></td><td></td><td></td><td></td><td></td><td></td><td></td><td></td><td></td></tr>
<tr><td>Oligo fwd</td><td>CACCCGCTCTGTACCGCGTACAGC</td><td></td><td></td><td></td><td></td><td></td><td></td><td></td><td></td><td></td><td></td><td></td></tr>
<tr><td>Oligo rev</td><td>AAACGCTGTACGCGGTACAGAGCG</td><td></td><td></td><td></td><td></td><td></td><td></td><td></td><td></td><td></td><td></td><td></td></tr>
<tr><td>Chromosome</td><td>start</td><td></td><td>end</td><td>strand</td><td>MM</td><td>target_seq</td><td>PAM</td><td>alignment</td><td>distance</td><td>position</td><td>gene name</td><td>gene id</td></tr>
<tr><td>GU305914.1</td><td></td><td>188242</td><td>188264</td><td>+</td><td>0</td><td>CGCTCTGTACCGCGTACAGC</td><td>GGG</td><td>||||||||[||||||||||||]PAM</td><td>NA</td><td>Intergenic</td><td>NA</td><td>NA</td></tr>
<tr><td>T14335</td><td>TGTACCGCGTACAGCGGGAGAGG</td><td></td><td>1000</td><td></td><td></td><td></td><td></td><td></td><td></td><td></td><td></td><td></td></tr>
<tr><td>Oligo fwd</td><td>CACCTGTACCGCGTACAGCGGGAG</td><td></td><td></td><td></td><td></td><td></td><td></td><td></td><td></td><td></td><td></td><td></td></tr>
<tr><td>Oligo rev</td><td>AAACCTCCCGCTGTACGCGGTACA</td><td></td><td></td><td></td><td></td><td></td><td></td><td></td><td></td><td></td><td></td><td></td></tr>
<tr><td>Chromosome</td><td>start</td><td></td><td>end</td><td>strand</td><td>MM</td><td>target_seq</td><td>PAM</td><td>alignment</td><td>distance</td><td>position</td><td>gene name</td><td>gene id</td></tr>
<tr><td>GU305914.1</td><td></td><td>188247</td><td>188269</td><td>+</td><td>0</td><td>TGTACCGCGTACAGCGGGAG</td><td>AGG</td><td>||||||||[||||||||||||]PAM</td><td>NA</td><td>Intergenic</td><td>NA</td><td>NA</td></tr>
<tr><td>T14336</td><td>GTACCGCGTACAGCGGGAGAGGG</td><td></td><td>1000</td><td></td><td></td><td></td><td></td><td></td><td></td><td></td><td></td><td></td></tr>
<tr><td>Oligo fwd</td><td>CACCGTACCGCGTACAGCGGGAGA</td><td></td><td></td><td></td><td></td><td></td><td></td><td></td><td></td><td></td><td></td><td></td></tr>
<tr><td>Oligo rev</td><td>AAACTCTCCCGCTGTACGCGGTAC</td><td></td><td></td><td></td><td></td><td></td><td></td><td></td><td></td><td></td><td></td><td></td></tr>
<tr><td>Chromosome</td><td>start</td><td></td><td>end</td><td>strand</td><td>MM</td><td>target_seq</td><td>PAM</td><td>alignment</td><td>distance</td><td>position</td><td>gene name</td><td>gene id</td></tr>
<tr><td>GU305914.1</td><td></td><td>188248</td><td>188270</td><td>+</td><td>0</td><td>GTACCGCGTACAGCGGGAGA</td><td>GGG</td><td>||||||||[||||||||||||]PAM</td><td>NA</td><td>Intergenic</td><td>NA</td><td>NA</td></tr>
<tr><td>T14337</td><td>AGCGGGAGAGGGTGCAGCTGCGG</td><td></td><td>1000</td><td></td><td></td><td></td><td></td><td></td><td></td><td></td><td></td><td></td></tr>
<tr><td>Oligo fwd</td><td>CACCAGCGGGAGAGGGTGCAGCTG</td><td></td><td></td><td></td><td></td><td></td><td></td><td></td><td></td><td></td><td></td><td></td></tr>
<tr><td>Oligo rev</td><td>AAACCAGCTGCACCCTCTCCCGCT</td><td></td><td></td><td></td><td></td><td></td><td></td><td></td><td></td><td></td><td></td><td></td></tr>
<tr><td>Chromosome</td><td>start</td><td></td><td>end</td><td>strand</td><td>MM</td><td>target_seq</td><td>PAM</td><td>alignment</td><td>distance</td><td>position</td><td>gene name</td><td>gene id</td></tr>
<tr><td>GU305914.1</td><td></td><td>188259</td><td>188281</td><td>+</td><td>0</td><td>AGCGGGAGAGGGTGCAGCTG</td><td>CGG</td><td>||||||||[||||||||||||]PAM</td><td>NA</td><td>Intergenic</td><td>NA</td><td>NA</td></tr>
<tr><td>T14338</td><td>AGCTGCGGCCGCGCGCAACATGG</td><td></td><td>1000</td><td></td><td></td><td></td><td></td><td></td><td></td><td></td><td></td><td></td></tr>
<tr><td>Oligo fwd</td><td>CACCAGCTGCGGCCGCGCGCAACA</td><td></td><td></td><td></td><td></td><td></td><td></td><td></td><td></td><td></td><td></td><td></td></tr>
<tr><td>Oligo rev</td><td>AAACTGTTGCGCGCGGCCGCAGCT</td><td></td><td></td><td></td><td></td><td></td><td></td><td></td><td></td><td></td><td></td><td></td></tr>
<tr><td>Chromosome</td><td>start</td><td></td><td>end</td><td>strand</td><td>MM</td><td>target_seq</td><td>PAM</td><td>alignment</td><td>distance</td><td>position</td><td>gene name</td><td>gene id</td></tr>
<tr><td>GU305914.1</td><td></td><td>188274</td><td>188296</td><td>+</td><td>0</td><td>AGCTGCGGCCGCGCGCAACA</td><td>TGG</td><td>||||||||[||||||||||||]PAM</td><td>NA</td><td>Intergenic</td><td>NA</td><td>NA</td></tr>
<tr><td>T14339</td><td>GCCGCGCGCAACATGGCTCGCGG</td><td></td><td>1000</td><td></td><td></td><td></td><td></td><td></td><td></td><td></td><td></td><td></td></tr>
<tr><td>Oligo fwd</td><td>CACCGCCGCGCGCAACATGGCTCG</td><td></td><td></td><td></td><td></td><td></td><td></td><td></td><td></td><td></td><td></td><td></td></tr>
<tr><td>Oligo rev</td><td>AAACCGAGCCATGTTGCGCGCGGC</td><td></td><td></td><td></td><td></td><td></td><td></td><td></td><td></td><td></td><td></td><td></td></tr>
<tr><td>Chromosome</td><td>start</td><td></td><td>end</td><td>strand</td><td>MM</td><td>target_seq</td><td>PAM</td><td>alignment</td><td>distance</td><td>position</td><td>gene name</td><td>gene id</td></tr>
<tr><td>GU305914.1</td><td></td><td>188281</td><td>188303</td><td>+</td><td>0</td><td>GCCGCGCGCAACATGGCTCG</td><td>CGG</td><td>||||||||[||||||||||||]PAM</td><td>NA</td><td>Intergenic</td><td>NA</td><td>NA</td></tr>
<tr><td>T14340</td><td>GCGCGCAACATGGCTCGCGGAGG</td><td></td><td>1000</td><td></td><td></td><td></td><td></td><td></td><td></td><td></td><td></td><td></td></tr>
<tr><td>Oligo fwd</td><td>CACCGCGCGCAACATGGCTCGCGG</td><td></td><td></td><td></td><td></td><td></td><td></td><td></td><td></td><td></td><td></td><td></td></tr>
<tr><td>Oligo rev</td><td>AAACCCGCGAGCCATGTTGCGCGC</td><td></td><td></td><td></td><td></td><td></td><td></td><td></td><td></td><td></td><td></td><td></td></tr>
<tr><td>Chromosome</td><td>start</td><td></td><td>end</td><td>strand</td><td>MM</td><td>target_seq</td><td>PAM</td><td>alignment</td><td>distance</td><td>position</td><td>gene name</td><td>gene id</td></tr>
<tr><td>GU305914.1</td><td></td><td>188284</td><td>188306</td><td>+</td><td>0</td><td>GCGCGCAACATGGCTCGCGG</td><td>AGG</td><td>||||||||[||||||||||||]PAM</td><td>NA</td><td>Intergenic</td><td>NA</td><td>NA</td></tr>
<tr><td>T14341</td><td>TGGCTCGCGGAGGTCCGCGACGG</td><td></td><td>1000</td><td></td><td></td><td></td><td></td><td></td><td></td><td></td><td></td><td></td></tr>
<tr><td>Oligo fwd</td><td>CACCTGGCTCGCGGAGGTCCGCGA</td><td></td><td></td><td></td><td></td><td></td><td></td><td></td><td></td><td></td><td></td><td></td></tr>
<tr><td>Oligo rev</td><td>AAACTCGCGGACCTCCGCGAGCCA</td><td></td><td></td><td></td><td></td><td></td><td></td><td></td><td></td><td></td><td></td><td></td></tr>
<tr><td>Chromosome</td><td>start</td><td></td><td>end</td><td>strand</td><td>MM</td><td>target_seq</td><td>PAM</td><td>alignment</td><td>distance</td><td>position</td><td>gene name</td><td>gene id</td></tr>
<tr><td>GU305914.1</td><td></td><td>188294</td><td>188316</td><td>+</td><td>0</td><td>TGGCTCGCGGAGGTCCGCGA</td><td>CGG</td><td>||||||||[||||||||||||]PAM</td><td>NA</td><td>Intergenic</td><td>NA</td><td>NA</td></tr>
<tr><td>T14342</td><td>CCAACACGACGCAACAGACGAGG</td><td></td><td>1000</td><td></td><td></td><td></td><td></td><td></td><td></td><td></td><td></td><td></td></tr>
<tr><td>Oligo fwd</td><td>CACCCCAACACGACGCAACAGACG</td><td></td><td></td><td></td><td></td><td></td><td></td><td></td><td></td><td></td><td></td><td></td></tr>
<tr><td>Oligo rev</td><td>AAACCGTCTGTTGCGTCGTGTTGG</td><td></td><td></td><td></td><td></td><td></td><td></td><td></td><td></td><td></td><td></td><td></td></tr>
<tr><td>Chromosome</td><td>start</td><td></td><td>end</td><td>strand</td><td>MM</td><td>target_seq</td><td>PAM</td><td>alignment</td><td>distance</td><td>position</td><td>gene name</td><td>gene id</td></tr>
<tr><td>GU305914.1</td><td></td><td>188326</td><td>188348</td><td>+</td><td>0</td><td>CCAACACGACGCAACAGACG</td><td>AGG</td><td>||||||||[||||||||||||]PAM</td><td>NA</td><td>Intergenic</td><td>NA</td><td>NA</td></tr>
<tr><td>T14343</td><td>AACAGACGAGGCAGCACACGAGG</td><td></td><td>1000</td><td></td><td></td><td></td><td></td><td></td><td></td><td></td><td></td><td></td></tr>
<tr><td>Oligo fwd</td><td>CACCAACAGACGAGGCAGCACACG</td><td></td><td></td><td></td><td></td><td></td><td></td><td></td><td></td><td></td><td></td><td></td></tr>
<tr><td>Oligo rev</td><td>AAACCGTGTGCTGCCTCGTCTGTT</td><td></td><td></td><td></td><td></td><td></td><td></td><td></td><td></td><td></td><td></td><td></td></tr>
<tr><td>Chromosome</td><td>start</td><td></td><td>end</td><td>strand</td><td>MM</td><td>target_seq</td><td>PAM</td><td>alignment</td><td>distance</td><td>position</td><td>gene name</td><td>gene id</td></tr>
<tr><td>GU305914.1</td><td></td><td>188338</td><td>188360</td><td>+</td><td>0</td><td>AACAGACGAGGCAGCACACG</td><td>AGG</td><td>||||||||[||||||||||||]PAM</td><td>NA</td><td>Intergenic</td><td>NA</td><td>NA</td></tr>
<tr><td>GU305914.1</td><td></td><td>97146</td><td>97168</td><td>-</td><td>4</td><td>AGCAGGCGCGGCAGCAGACG</td><td>GGG</td><td>|-|||-||[-|||||||-|||]PAM</td><td>NA</td><td>Intergenic</td><td>NA</td><td>NA</td></tr>
<tr><td>T14344</td><td>GCAAAAACATAAATATACATAGG</td><td></td><td>1000</td><td></td><td></td><td></td><td></td><td></td><td></td><td></td><td></td><td></td></tr>
<tr><td>Oligo fwd</td><td>CACCGCAAAAACATAAATATACAT</td><td></td><td></td><td></td><td></td><td></td><td></td><td></td><td></td><td></td><td></td><td></td></tr>
<tr><td>Oligo rev</td><td>AAACATGTATATTTATGTTTTTGC</td><td></td><td></td><td></td><td></td><td></td><td></td><td></td><td></td><td></td><td></td><td></td></tr>
<tr><td>Chromosome</td><td>start</td><td></td><td>end</td><td>strand</td><td>MM</td><td>target_seq</td><td>PAM</td><td>alignment</td><td>distance</td><td>position</td><td>gene name</td><td>gene id</td></tr>
<tr><td>GU305914.1</td><td></td><td>188410</td><td>188432</td><td>+</td><td>0</td><td>GCAAAAACATAAATATACAT</td><td>AGG</td><td>||||||||[||||||||||||]PAM</td><td>NA</td><td>Intergenic</td><td>NA</td><td>NA</td></tr>
<tr><td>T14345</td><td>ACATAAATATACATAGGCCGTGG</td><td></td><td>1000</td><td></td><td></td><td></td><td></td><td></td><td></td><td></td><td></td><td></td></tr>
<tr><td>Oligo fwd</td><td>CACCACATAAATATACATAGGCCG</td><td></td><td></td><td></td><td></td><td></td><td></td><td></td><td></td><td></td><td></td><td></td></tr>
<tr><td>Oligo rev</td><td>AAACCGGCCTATGTATATTTATGT</td><td></td><td></td><td></td><td></td><td></td><td></td><td></td><td></td><td></td><td></td><td></td></tr>
<tr><td>Chromosome</td><td>start</td><td></td><td>end</td><td>strand</td><td>MM</td><td>target_seq</td><td>PAM</td><td>alignment</td><td>distance</td><td>position</td><td>gene name</td><td>gene id</td></tr>
<tr><td>GU305914.1</td><td></td><td>188416</td><td>188438</td><td>+</td><td>0</td><td>ACATAAATATACATAGGCCG</td><td>TGG</td><td>||||||||[||||||||||||]PAM</td><td>NA</td><td>Intergenic</td><td>NA</td><td>NA</td></tr>
<tr><td>T14346</td><td>CATAAATATACATAGGCCGTGGG</td><td></td><td>1000</td><td></td><td></td><td></td><td></td><td></td><td></td><td></td><td></td><td></td></tr>
<tr><td>Oligo fwd</td><td>CACCCATAAATATACATAGGCCGT</td><td></td><td></td><td></td><td></td><td></td><td></td><td></td><td></td><td></td><td></td><td></td></tr>
</table>

| | | | | | | | | | | | | |
|---|---|---|---|---|---|---|---|---|---|---|---|---|
| Oligo rev | AAACACGGCCTATGTATATTTATG | | | | | | | | | | | |
| Chromosome | start | | end | strand | MM | target_seq | PAM | alignment | distance | position | gene name | gene id |
| GU305914.1 | | 188417 | 188439 | + | 0 | CATAAATATACATAGGCCGT | GGG | \|\|\|\|\|\|\|\|[\|\|\|\|\|\|\|\|\|\|\|\|]PAM | NA | Intergenic | NA | NA |
| T14347 | ATAAATATACATAGGCCGTGGGG | | 1000 | | | | | | | | | |
| Oligo fwd | CACCATAAATATACATAGGCCGTG | | | | | | | | | | | |
| Oligo rev | AAACCACGGCCTATGTATATTTAT | | | | | | | | | | | |
| Chromosome | start | | end | strand | MM | target_seq | PAM | alignment | distance | position | gene name | gene id |
| GU305914.1 | | 188418 | 188440 | + | 0 | ATAAATATACATAGGCCGTG | GGG | \|\|\|\|\|\|\|\|[\|\|\|\|\|\|\|\|\|\|\|\|]PAM | NA | Intergenic | NA | NA |
| T14348 | TGGGGTCTCCAGCTTCCCATAGG | | 1000 | | | | | | | | | |
| Oligo fwd | CACCTGGGGTCTCCAGCTTCCCAT | | | | | | | | | | | |
| Oligo rev | AAACATGGGAAGCTGGAGACCCCA | | | | | | | | | | | |
| Chromosome | start | | end | strand | MM | target_seq | PAM | alignment | distance | position | gene name | gene id |
| GU305914.1 | | 188436 | 188458 | + | 0 | TGGGGTCTCCAGCTTCCCAT | AGG | \|\|\|\|\|\|\|\|[\|\|\|\|\|\|\|\|\|\|\|\|]PAM | NA | Intergenic | NA | NA |
| T14349 | AGCATTCAACACCTTTTCCCGG | | 1000 | | | | | | | | | |
| Oligo fwd | CACCAGCATTCAACACCTTTTTCC | | | | | | | | | | | |
| Oligo rev | AAACGGAAAAAGGTGTTGAATGCT | | | | | | | | | | | |
| Chromosome | start | | end | strand | MM | target_seq | PAM | alignment | distance | position | gene name | gene id |
| GU305914.1 | | 188512 | 188534 | + | 0 | AGCATTCAACACCTTTTTCC | CGG | \|\|\|\|\|\|\|\|[\|\|\|\|\|\|\|\|\|\|\|\|]PAM | NA | Intergenic | NA | NA |
| T14350 | GCATTCAACACCTTTTTCCCGGG | | 1000 | | | | | | | | | |
| Oligo fwd | CACCGCATTCAACACCTTTTTCCC | | | | | | | | | | | |
| Oligo rev | AAACGGGAAAAAGGTGTTGAATGC | | | | | | | | | | | |
| Chromosome | start | | end | strand | MM | target_seq | PAM | alignment | distance | position | gene name | gene id |
| GU305914.1 | | 188513 | 188535 | + | 0 | GCATTCAACACCTTTTTCCC | GGG | \|\|\|\|\|\|\|\|[\|\|\|\|\|\|\|\|\|\|\|\|]PAM | NA | Intergenic | NA | NA |
| T14351 | CTTTTTCCCGGGTTGTGTCGAGG | | 1000 | | | | | | | | | |
| Oligo fwd | CACCCTTTTTCCCGGGTTGTGTCG | | | | | | | | | | | |
| Oligo rev | AAACCGACACAACCCGGGAAAAAG | | | | | | | | | | | |
| Chromosome | start | | end | strand | MM | target_seq | PAM | alignment | distance | position | gene name | gene id |
| GU305914.1 | | 188524 | 188546 | + | 0 | CTTTTTCCCGGGTTGTGTCG | AGG | \|\|\|\|\|\|\|\|[\|\|\|\|\|\|\|\|\|\|\|\|]PAM | NA | Intergenic | NA | NA |
| T14352 | TTTCCCGGGTTGTGTCGAGGCGG | | 1000 | | | | | | | | | |
| Oligo fwd | CACCTTTCCCGGGTTGTGTCGAGG | | | | | | | | | | | |
| Oligo rev | AAACCCTCGACACAACCCGGGAAA | | | | | | | | | | | |
| Chromosome | start | | end | strand | MM | target_seq | PAM | alignment | distance | position | gene name | gene id |
| GU305914.1 | | 188527 | 188549 | + | 0 | TTTCCCGGGTTGTGTCGAGG | CGG | \|\|\|\|\|\|\|\|[\|\|\|\|\|\|\|\|\|\|\|\|]PAM | NA | Intergenic | NA | NA |
| T14353 | CGGGTTGTGTCGAGGCGGACAGG | | 1000 | | | | | | | | | |
| Oligo fwd | CACCCGGGTTGTGTCGAGGCGGAC | | | | | | | | | | | |
| Oligo rev | AAACGTCCGCCTCGACACAACCCG | | | | | | | | | | | |
| Chromosome | start | | end | strand | MM | target_seq | PAM | alignment | distance | position | gene name | gene id |
| GU305914.1 | | 188532 | 188554 | + | 0 | CGGGTTGTGTCGAGGCGGAC | AGG | \|\|\|\|\|\|\|\|[\|\|\|\|\|\|\|\|\|\|\|\|]PAM | NA | Intergenic | NA | NA |
| T14354 | TTGTGTCGAGGCGGACAGGTCGG | | 1000 | | | | | | | | | |
| Oligo fwd | CACCTTGTGTCGAGGCGGACAGGT | | | | | | | | | | | |
| Oligo rev | AAACACCTGTCCGCCTCGACACAA | | | | | | | | | | | |
| Chromosome | start | | end | strand | MM | target_seq | PAM | alignment | distance | position | gene name | gene id |
| GU305914.1 | | 188536 | 188558 | + | 0 | TTGTGTCGAGGCGGACAGGT | CGG | \|\|\|\|\|\|\|\|[\|\|\|\|\|\|\|\|\|\|\|\|]PAM | NA | Intergenic | NA | NA |
| T14355 | TGTCGAGGCGGACAGGTCGGCGG | | 1000 | | | | | | | | | |
| Oligo fwd | CACCTGTCGAGGCGGACAGGTCGG | | | | | | | | | | | |
| Oligo rev | AAACCCGACCTGTCCGCCTCGACA | | | | | | | | | | | |
| Chromosome | start | | end | strand | MM | target_seq | PAM | alignment | distance | position | gene name | gene id |
| GU305914.1 | | 188539 | 188561 | + | 0 | TGTCGAGGCGGACAGGTCGG | CGG | \|\|\|\|\|\|\|\|[\|\|\|\|\|\|\|\|\|\|\|\|]PAM | NA | Intergenic | NA | NA |
| T14356 | GGACAGGTCGGCGGTCATCTTGG | | 1000 | | | | | | | | | |
| Oligo fwd | CACCGGACAGGTCGGCGGTCATCT | | | | | | | | | | | |
| Oligo rev | AAACAGATGACCGCCGACCTGTCC | | | | | | | | | | | |
| Chromosome | start | | end | strand | MM | target_seq | PAM | alignment | distance | position | gene name | gene id |
| GU305914.1 | | 188548 | 188570 | + | 0 | GGACAGGTCGGCGGTCATCT | TGG | \|\|\|\|\|\|\|\|[\|\|\|\|\|\|\|\|\|\|\|\|]PAM | NA | Intergenic | NA | NA |
| T14357 | CGCGTTGTAACTTCCCAGCACGG | | 1000 | | | | | | | | | |
| Oligo fwd | CACCCGCGTTGTAACTTCCCAGCA | | | | | | | | | | | |
| Oligo rev | AAACTGCTGGGAAGTTACAACGCG | | | | | | | | | | | |
| Chromosome | start | | end | strand | MM | target_seq | PAM | alignment | distance | position | gene name | gene id |
| GU305914.1 | | 188578 | 188600 | + | 0 | CGCGTTGTAACTTCCCAGCA | CGG | \|\|\|\|\|\|\|\|[\|\|\|\|\|\|\|\|\|\|\|\|]PAM | NA | Intergenic | NA | NA |
| T14358 | GCGTTGTAACTTCCCAGCACGGG | | 1000 | | | | | | | | | |
| Oligo fwd | CACCGCGTTGTAACTTCCCAGCAC | | | | | | | | | | | |
| Oligo rev | AAACGTGCTGGGAAGTTACAACGC | | | | | | | | | | | |
| Chromosome | start | | end | strand | MM | target_seq | PAM | alignment | distance | position | gene name | gene id |
| GU305914.1 | | 188579 | 188601 | + | 0 | GCGTTGTAACTTCCCAGCAC | GGG | \|\|\|\|\|\|\|\|[\|\|\|\|\|\|\|\|\|\|\|\|]PAM | NA | Intergenic | NA | NA |
| T14359 | CGTTGTAACTTCCCAGCACGGGG | | 1000 | | | | | | | | | |
| Oligo fwd | CACCCGTTGTAACTTCCCAGCACG | | | | | | | | | | | |
| Oligo rev | AAACCGTGCTGGGAAGTTACAACG | | | | | | | | | | | |
| Chromosome | start | | end | strand | MM | target_seq | PAM | alignment | distance | position | gene name | gene id |
| GU305914.1 | | 188580 | 188602 | + | 0 | CGTTGTAACTTCCCAGCACG | GGG | \|\|\|\|\|\|\|\|[\|\|\|\|\|\|\|\|\|\|\|\|]PAM | NA | Intergenic | NA | NA |
| T14360 | TCCCAGCACGGGGCGTACGACGG | | 1000 | | | | | | | | | |
| Oligo fwd | CACCTCCCAGCACGGGGCGTACGA | | | | | | | | | | | |
| Oligo rev | AAACTCGTACGCCCCGTGCTGGGA | | | | | | | | | | | |
| Chromosome | start | | end | strand | MM | target_seq | PAM | alignment | distance | position | gene name | gene id |
| GU305914.1 | | 188590 | 188612 | + | 0 | TCCCAGCACGGGGCGTACGA | CGG | \|\|\|\|\|\|\|\|[\|\|\|\|\|\|\|\|\|\|\|\|]PAM | NA | Intergenic | NA | NA |
| T14361 | TCCAGCCTGAAAGACAAAAACGG | | 1000 | | | | | | | | | |

<table>
<tr><td>Oligo fwd</td><td>CACCTCCAGCCTGAAAGACAAAAA</td><td></td><td></td><td></td><td></td><td></td><td></td><td></td><td></td><td></td><td></td></tr>
<tr><td>Oligo rev</td><td>AAACTTTTTGTCTTTCAGGCTGGA</td><td></td><td></td><td></td><td></td><td></td><td></td><td></td><td></td><td></td><td></td></tr>
<tr><td>Chromosome</td><td>start</td><td>end</td><td>strand</td><td>MM</td><td>target_seq</td><td>PAM</td><td>alignment</td><td>distance</td><td>position</td><td>gene name</td><td>gene id</td></tr>
<tr><td>GU305914.1</td><td>188696</td><td>188718</td><td>+</td><td>0</td><td>TCCAGCCTGAAAGACAAAAA</td><td>CGG</td><td>||||||||[||||||||||||]PAM</td><td>NA</td><td>Intergenic</td><td>NA</td><td>NA</td></tr>
<tr><td></td><td></td><td></td><td></td><td></td><td></td><td></td><td></td><td></td><td></td><td></td><td></td></tr>
<tr><td>T14362</td><td>CCAGCCTGAAAGACAAAAACGGG</td><td>1000</td><td></td><td></td><td></td><td></td><td></td><td></td><td></td><td></td><td></td></tr>
<tr><td>Oligo fwd</td><td>CACCCCAGCCTGAAAGACAAAAAC</td><td></td><td></td><td></td><td></td><td></td><td></td><td></td><td></td><td></td><td></td></tr>
<tr><td>Oligo rev</td><td>AAACGTTTTTGTCTTTCAGGCTGG</td><td></td><td></td><td></td><td></td><td></td><td></td><td></td><td></td><td></td><td></td></tr>
<tr><td>Chromosome</td><td>start</td><td>end</td><td>strand</td><td>MM</td><td>target_seq</td><td>PAM</td><td>alignment</td><td>distance</td><td>position</td><td>gene name</td><td>gene id</td></tr>
<tr><td>GU305914.1</td><td>188697</td><td>188719</td><td>+</td><td>0</td><td>CCAGCCTGAAAGACAAAAAC</td><td>GGG</td><td>||||||||[||||||||||||]PAM</td><td>NA</td><td>Intergenic</td><td>NA</td><td>NA</td></tr>
<tr><td></td><td></td><td></td><td></td><td></td><td></td><td></td><td></td><td></td><td></td><td></td><td></td></tr>
<tr><td>T14363</td><td>AAACGGGCACACGTCAGAGCCGG</td><td>1000</td><td></td><td></td><td></td><td></td><td></td><td></td><td></td><td></td><td></td></tr>
<tr><td>Oligo fwd</td><td>CACCAAACGGGCACACGTCAGAGC</td><td></td><td></td><td></td><td></td><td></td><td></td><td></td><td></td><td></td><td></td></tr>
<tr><td>Oligo rev</td><td>AAACGCTCTGACGTGTGCCCGTTT</td><td></td><td></td><td></td><td></td><td></td><td></td><td></td><td></td><td></td><td></td></tr>
<tr><td>Chromosome</td><td>start</td><td>end</td><td>strand</td><td>MM</td><td>target_seq</td><td>PAM</td><td>alignment</td><td>distance</td><td>position</td><td>gene name</td><td>gene id</td></tr>
<tr><td>GU305914.1</td><td>188713</td><td>188735</td><td>+</td><td>0</td><td>AAACGGGCACACGTCAGAGC</td><td>CGG</td><td>||||||||[||||||||||||]PAM</td><td>NA</td><td>Intergenic</td><td>NA</td><td>NA</td></tr>
<tr><td></td><td></td><td></td><td></td><td></td><td></td><td></td><td></td><td></td><td></td><td></td><td></td></tr>
<tr><td>T14364</td><td>AACGGGCACACGTCAGAGCCGGG</td><td>1000</td><td></td><td></td><td></td><td></td><td></td><td></td><td></td><td></td><td></td></tr>
<tr><td>Oligo fwd</td><td>CACCAACGGGCACACGTCAGAGCC</td><td></td><td></td><td></td><td></td><td></td><td></td><td></td><td></td><td></td><td></td></tr>
<tr><td>Oligo rev</td><td>AAACGGCTCTGACGTGTGCCCGTT</td><td></td><td></td><td></td><td></td><td></td><td></td><td></td><td></td><td></td><td></td></tr>
<tr><td>Chromosome</td><td>start</td><td>end</td><td>strand</td><td>MM</td><td>target_seq</td><td>PAM</td><td>alignment</td><td>distance</td><td>position</td><td>gene name</td><td>gene id</td></tr>
<tr><td>GU305914.1</td><td>188714</td><td>188736</td><td>+</td><td>0</td><td>AACGGGCACACGTCAGAGCC</td><td>GGG</td><td>||||||||[||||||||||||]PAM</td><td>NA</td><td>Intergenic</td><td>NA</td><td>NA</td></tr>
<tr><td></td><td></td><td></td><td></td><td></td><td></td><td></td><td></td><td></td><td></td><td></td><td></td></tr>
<tr><td>T14365</td><td>ACGGGCACACGTCAGAGCCGGGG</td><td>1000</td><td></td><td></td><td></td><td></td><td></td><td></td><td></td><td></td><td></td></tr>
<tr><td>Oligo fwd</td><td>CACCACGGGCACACGTCAGAGCCG</td><td></td><td></td><td></td><td></td><td></td><td></td><td></td><td></td><td></td><td></td></tr>
<tr><td>Oligo rev</td><td>AAACCGGCTCTGACGTGTGCCCGT</td><td></td><td></td><td></td><td></td><td></td><td></td><td></td><td></td><td></td><td></td></tr>
<tr><td>Chromosome</td><td>start</td><td>end</td><td>strand</td><td>MM</td><td>target_seq</td><td>PAM</td><td>alignment</td><td>distance</td><td>position</td><td>gene name</td><td>gene id</td></tr>
<tr><td>GU305914.1</td><td>188715</td><td>188737</td><td>+</td><td>0</td><td>ACGGGCACACGTCAGAGCCG</td><td>GGG</td><td>||||||||[||||||||||||]PAM</td><td>NA</td><td>Intergenic</td><td>NA</td><td>NA</td></tr>
<tr><td></td><td></td><td></td><td></td><td></td><td></td><td></td><td></td><td></td><td></td><td></td><td></td></tr>
<tr><td>T14366</td><td>CGGGCACACGTCAGAGCCGGGGG</td><td>1000</td><td></td><td></td><td></td><td></td><td></td><td></td><td></td><td></td><td></td></tr>
<tr><td>Oligo fwd</td><td>CACCCGGGCACACGTCAGAGCCGG</td><td></td><td></td><td></td><td></td><td></td><td></td><td></td><td></td><td></td><td></td></tr>
<tr><td>Oligo rev</td><td>AAACCCGGCTCTGACGTGTGCCCG</td><td></td><td></td><td></td><td></td><td></td><td></td><td></td><td></td><td></td><td></td></tr>
<tr><td>Chromosome</td><td>start</td><td>end</td><td>strand</td><td>MM</td><td>target_seq</td><td>PAM</td><td>alignment</td><td>distance</td><td>position</td><td>gene name</td><td>gene id</td></tr>
<tr><td>GU305914.1</td><td>188716</td><td>188738</td><td>+</td><td>0</td><td>CGGGCACACGTCAGAGCCGG</td><td>GGG</td><td>||||||||[||||||||||||]PAM</td><td>NA</td><td>Intergenic</td><td>NA</td><td>NA</td></tr>
<tr><td></td><td></td><td></td><td></td><td></td><td></td><td></td><td></td><td></td><td></td><td></td><td></td></tr>
<tr><td>T14367</td><td>GGGCACACGTCAGAGCCGGGGGG</td><td>1000</td><td></td><td></td><td></td><td></td><td></td><td></td><td></td><td></td><td></td></tr>
<tr><td>Oligo fwd</td><td>CACCGGGCACACGTCAGAGCCGGG</td><td></td><td></td><td></td><td></td><td></td><td></td><td></td><td></td><td></td><td></td></tr>
<tr><td>Oligo rev</td><td>AAACCCCGGCTCTGACGTGTGCCC</td><td></td><td></td><td></td><td></td><td></td><td></td><td></td><td></td><td></td><td></td></tr>
<tr><td>Chromosome</td><td>start</td><td>end</td><td>strand</td><td>MM</td><td>target_seq</td><td>PAM</td><td>alignment</td><td>distance</td><td>position</td><td>gene name</td><td>gene id</td></tr>
<tr><td>GU305914.1</td><td>188717</td><td>188739</td><td>+</td><td>0</td><td>GGGCACACGTCAGAGCCGGG</td><td>GGG</td><td>||||||||[||||||||||||]PAM</td><td>NA</td><td>Intergenic</td><td>NA</td><td>NA</td></tr>
<tr><td></td><td></td><td></td><td></td><td></td><td></td><td></td><td></td><td></td><td></td><td></td><td></td></tr>
<tr><td>T14368</td><td>GTCAGAGAGAGAGCCCTTCTCGG</td><td>1000</td><td></td><td></td><td></td><td></td><td></td><td></td><td></td><td></td><td></td></tr>
<tr><td>Oligo fwd</td><td>CACCGTCAGAGAGAGAGCCCTTCT</td><td></td><td></td><td></td><td></td><td></td><td></td><td></td><td></td><td></td><td></td></tr>
<tr><td>Oligo rev</td><td>AAACAGAAGGGCTCTCTCTCTGAC</td><td></td><td></td><td></td><td></td><td></td><td></td><td></td><td></td><td></td><td></td></tr>
<tr><td>Chromosome</td><td>start</td><td>end</td><td>strand</td><td>MM</td><td>target_seq</td><td>PAM</td><td>alignment</td><td>distance</td><td>position</td><td>gene name</td><td>gene id</td></tr>
<tr><td>GU305914.1</td><td>188746</td><td>188768</td><td>+</td><td>0</td><td>GTCAGAGAGAGAGCCCTTCT</td><td>CGG</td><td>||||||||[||||||||||||]PAM</td><td>NA</td><td>Intergenic</td><td>NA</td><td>NA</td></tr>
<tr><td></td><td></td><td></td><td></td><td></td><td></td><td></td><td></td><td></td><td></td><td></td><td></td></tr>
<tr><td>T14369</td><td>CTCGGAACCCGACCGTTTCTCGG</td><td>1000</td><td></td><td></td><td></td><td></td><td></td><td></td><td></td><td></td><td></td></tr>
<tr><td>Oligo fwd</td><td>CACCCTCGGAACCCGACCGTTTCT</td><td></td><td></td><td></td><td></td><td></td><td></td><td></td><td></td><td></td><td></td></tr>
<tr><td>Oligo rev</td><td>AAACAGAAACGGTCGGGTTCCGAG</td><td></td><td></td><td></td><td></td><td></td><td></td><td></td><td></td><td></td><td></td></tr>
<tr><td>Chromosome</td><td>start</td><td>end</td><td>strand</td><td>MM</td><td>target_seq</td><td>PAM</td><td>alignment</td><td>distance</td><td>position</td><td>gene name</td><td>gene id</td></tr>
<tr><td>GU305914.1</td><td>188764</td><td>188786</td><td>+</td><td>0</td><td>CTCGGAACCCGACCGTTTCT</td><td>CGG</td><td>||||||||[||||||||||||]PAM</td><td>NA</td><td>Intergenic</td><td>NA</td><td>NA</td></tr>
<tr><td></td><td></td><td></td><td></td><td></td><td></td><td></td><td></td><td></td><td></td><td></td><td></td></tr>
<tr><td>T14370</td><td>TCTCCAGCGCCAGTTTCATTTGG</td><td>1000</td><td></td><td></td><td></td><td></td><td></td><td></td><td></td><td></td><td></td></tr>
<tr><td>Oligo fwd</td><td>CACCTCTCCAGCGCCAGTTTCATT</td><td></td><td></td><td></td><td></td><td></td><td></td><td></td><td></td><td></td><td></td></tr>
<tr><td>Oligo rev</td><td>AAACAATGAAACTGGCGCTGGAGA</td><td></td><td></td><td></td><td></td><td></td><td></td><td></td><td></td><td></td><td></td></tr>
<tr><td>Chromosome</td><td>start</td><td>end</td><td>strand</td><td>MM</td><td>target_seq</td><td>PAM</td><td>alignment</td><td>distance</td><td>position</td><td>gene name</td><td>gene id</td></tr>
<tr><td>GU305914.1</td><td>188797</td><td>188819</td><td>+</td><td>0</td><td>TCTCCAGCGCCAGTTTCATT</td><td>TGG</td><td>||||||||[||||||||||||]PAM</td><td>NA</td><td>Intergenic</td><td>NA</td><td>NA</td></tr>
<tr><td></td><td></td><td></td><td></td><td></td><td></td><td></td><td></td><td></td><td></td><td></td><td></td></tr>
<tr><td>T14371</td><td>CTCCAGCGCCAGTTTCATTTGGG</td><td>1000</td><td></td><td></td><td></td><td></td><td></td><td></td><td></td><td></td><td></td></tr>
<tr><td>Oligo fwd</td><td>CACCCTCCAGCGCCAGTTTCATTT</td><td></td><td></td><td></td><td></td><td></td><td></td><td></td><td></td><td></td><td></td></tr>
<tr><td>Oligo rev</td><td>AAACAAATGAAACTGGCGCTGGAG</td><td></td><td></td><td></td><td></td><td></td><td></td><td></td><td></td><td></td><td></td></tr>
<tr><td>Chromosome</td><td>start</td><td>end</td><td>strand</td><td>MM</td><td>target_seq</td><td>PAM</td><td>alignment</td><td>distance</td><td>position</td><td>gene name</td><td>gene id</td></tr>
<tr><td>GU305914.1</td><td>188798</td><td>188820</td><td>+</td><td>0</td><td>CTCCAGCGCCAGTTTCATTT</td><td>GGG</td><td>||||||||[||||||||||||]PAM</td><td>NA</td><td>Intergenic</td><td>NA</td><td>NA</td></tr>
<tr><td></td><td></td><td></td><td></td><td></td><td></td><td></td><td></td><td></td><td></td><td></td><td></td></tr>
<tr><td>T14372</td><td>GTTTCATTTGGGACGCGTTCAGG</td><td>1000</td><td></td><td></td><td></td><td></td><td></td><td></td><td></td><td></td><td></td></tr>
<tr><td>Oligo fwd</td><td>CACCGTTTCATTTGGGACGCGTTC</td><td></td><td></td><td></td><td></td><td></td><td></td><td></td><td></td><td></td><td></td></tr>
<tr><td>Oligo rev</td><td>AAACGAACGCGTCCCAAATGAAAC</td><td></td><td></td><td></td><td></td><td></td><td></td><td></td><td></td><td></td><td></td></tr>
<tr><td>Chromosome</td><td>start</td><td>end</td><td>strand</td><td>MM</td><td>target_seq</td><td>PAM</td><td>alignment</td><td>distance</td><td>position</td><td>gene name</td><td>gene id</td></tr>
<tr><td>GU305914.1</td><td>188809</td><td>188831</td><td>+</td><td>0</td><td>GTTTCATTTGGGACGCGTTC</td><td>AGG</td><td>||||||||[||||||||||||]PAM</td><td>NA</td><td>Intergenic</td><td>NA</td><td>NA</td></tr>
<tr><td></td><td></td><td></td><td></td><td></td><td></td><td></td><td></td><td></td><td></td><td></td><td></td></tr>
<tr><td>T14373</td><td>TTTCATTTGGGACGCGTTCAGGG</td><td>1000</td><td></td><td></td><td></td><td></td><td></td><td></td><td></td><td></td><td></td></tr>
<tr><td>Oligo fwd</td><td>CACCTTTCATTTGGGACGCGTTCA</td><td></td><td></td><td></td><td></td><td></td><td></td><td></td><td></td><td></td><td></td></tr>
<tr><td>Oligo rev</td><td>AAACTGAACGCGTCCCAAATGAAA</td><td></td><td></td><td></td><td></td><td></td><td></td><td></td><td></td><td></td><td></td></tr>
<tr><td>Chromosome</td><td>start</td><td>end</td><td>strand</td><td>MM</td><td>target_seq</td><td>PAM</td><td>alignment</td><td>distance</td><td>position</td><td>gene name</td><td>gene id</td></tr>
<tr><td>GU305914.1</td><td>188810</td><td>188832</td><td>+</td><td>0</td><td>TTTCATTTGGGACGCGTTCA</td><td>GGG</td><td>||||||||[||||||||||||]PAM</td><td>NA</td><td>Intergenic</td><td>NA</td><td>NA</td></tr>
<tr><td></td><td></td><td></td><td></td><td></td><td></td><td></td><td></td><td></td><td></td><td></td><td></td></tr>
<tr><td>T14374</td><td>GGGACGCGTTCAGGGATCGATGG</td><td>1000</td><td></td><td></td><td></td><td></td><td></td><td></td><td></td><td></td><td></td></tr>
<tr><td>Oligo fwd</td><td>CACCGGGACGCGTTCAGGGATCGA</td><td></td><td></td><td></td><td></td><td></td><td></td><td></td><td></td><td></td><td></td></tr>
<tr><td>Oligo rev</td><td>AAACTCGATCCCTGAACGCGTCCC</td><td></td><td></td><td></td><td></td><td></td><td></td><td></td><td></td><td></td><td></td></tr>
<tr><td>Chromosome</td><td>start</td><td>end</td><td>strand</td><td>MM</td><td>target_seq</td><td>PAM</td><td>alignment</td><td>distance</td><td>position</td><td>gene name</td><td>gene id</td></tr>
<tr><td>GU305914.1</td><td>188818</td><td>188840</td><td>+</td><td>0</td><td>GGGACGCGTTCAGGGATCGA</td><td>TGG</td><td>||||||||[||||||||||||]PAM</td><td>NA</td><td>Intergenic</td><td>NA</td><td>NA</td></tr>
<tr><td></td><td></td><td></td><td></td><td></td><td></td><td></td><td></td><td></td><td></td><td></td><td></td></tr>
<tr><td>T14375</td><td>GTTGAACCCCAGCTCGAGCCCGG</td><td>1000</td><td></td><td></td><td></td><td></td><td></td><td></td><td></td><td></td><td></td></tr>
<tr><td>Oligo fwd</td><td>CACCGTTGAACCCCAGCTCGAGCC</td><td></td><td></td><td></td><td></td><td></td><td></td><td></td><td></td><td></td><td></td></tr>
<tr><td>Oligo rev</td><td>AAACGGCTCGAGCTGGGGTTCAAC</td><td></td><td></td><td></td><td></td><td></td><td></td><td></td><td></td><td></td><td></td></tr>
<tr><td>Chromosome</td><td>start</td><td>end</td><td>strand</td><td>MM</td><td>target_seq</td><td>PAM</td><td>alignment</td><td>distance</td><td>position</td><td>gene name</td><td>gene id</td></tr>
<tr><td>GU305914.1</td><td>188840</td><td>188862</td><td>+</td><td>0</td><td>GTTGAACCCCAGCTCGAGCC</td><td>CGG</td><td>||||||||[||||||||||||]PAM</td><td>NA</td><td>Intergenic</td><td>NA</td><td>NA</td></tr>
</table>

<table>
<tr><td>T14376</td><td>CCAGCTCGAGCCCGGAGAGCAGG</td><td></td><td>1000</td></tr>
<tr><td>Oligo fwd</td><td>CACCCCAGCTCGAGCCCGGAGAGC</td></tr>
<tr><td>Oligo rev</td><td>AAACGCTCTCCGGGCTCGAGCTGG</td></tr>
<tr><td>Chromosome</td><td>start</td><td></td><td>end</td><td>strand</td><td>MM</td><td>target_seq</td><td>PAM</td><td>alignment</td><td>distance</td><td>position</td><td>gene name</td><td>gene id</td></tr>
<tr><td>GU305914.1</td><td></td><td>188848</td><td>188870</td><td>+</td><td>0</td><td>CCAGCTCGAGCCCGGAGAGC</td><td>AGG</td><td>||||||||[||||||||||||]PAM</td><td>NA</td><td>Intergenic</td><td>NA</td><td>NA</td></tr>
<tr><td>T14377</td><td>GGAGAGCAGGACGCTCTCCACGG</td><td></td><td>1000</td></tr>
<tr><td>Oligo fwd</td><td>CACCGGAGAGCAGGACGCTCTCCA</td></tr>
<tr><td>Oligo rev</td><td>AAACTGGAGAGCGTCCTGCTCTCC</td></tr>
<tr><td>Chromosome</td><td>start</td><td></td><td>end</td><td>strand</td><td>MM</td><td>target_seq</td><td>PAM</td><td>alignment</td><td>distance</td><td>position</td><td>gene name</td><td>gene id</td></tr>
<tr><td>GU305914.1</td><td></td><td>188861</td><td>188883</td><td>+</td><td>0</td><td>GGAGAGCAGGACGCTCTCCA</td><td>CGG</td><td>||||||||[||||||||||||]PAM</td><td>NA</td><td>Intergenic</td><td>NA</td><td>NA</td></tr>
<tr><td>T14378</td><td>CCACGGCCAGCACCCCGTCTCGG</td><td></td><td>1000</td></tr>
<tr><td>Oligo fwd</td><td>CACCCCACGGCCAGCACCCCGTCT</td></tr>
<tr><td>Oligo rev</td><td>AAACAGACGGGGTGCTGGCCGTGG</td></tr>
<tr><td>Chromosome</td><td>start</td><td></td><td>end</td><td>strand</td><td>MM</td><td>target_seq</td><td>PAM</td><td>alignment</td><td>distance</td><td>position</td><td>gene name</td><td>gene id</td></tr>
<tr><td>GU305914.1</td><td></td><td>188878</td><td>188900</td><td>+</td><td>0</td><td>CCACGGCCAGCACCCCGTCT</td><td>CGG</td><td>||||||||[||||||||||||]PAM</td><td>NA</td><td>Intergenic</td><td>NA</td><td>NA</td></tr>
<tr><td>T14379</td><td>CACCCCGTCTCGGTCCGTCACGG</td><td></td><td>1000</td></tr>
<tr><td>Oligo fwd</td><td>CACCCACCCCGTCTCGGTCCGTCA</td></tr>
<tr><td>Oligo rev</td><td>AAACTGACGGACCGAGACGGGGTG</td></tr>
<tr><td>Chromosome</td><td>start</td><td></td><td>end</td><td>strand</td><td>MM</td><td>target_seq</td><td>PAM</td><td>alignment</td><td>distance</td><td>position</td><td>gene name</td><td>gene id</td></tr>
<tr><td>GU305914.1</td><td></td><td>188888</td><td>188910</td><td>+</td><td>0</td><td>CACCCCGTCTCGGTCCGTCA</td><td>CGG</td><td>||||||||[||||||||||||]PAM</td><td>NA</td><td>Intergenic</td><td>NA</td><td>NA</td></tr>
<tr><td>T14380</td><td>CGTCACGGCGACCGTGTCGCCGG</td><td></td><td>1000</td></tr>
<tr><td>Oligo fwd</td><td>CACCCGTCACGGCGACCGTGTCGC</td></tr>
<tr><td>Oligo rev</td><td>AAACGCGACACGGTCGCCGTGACG</td></tr>
<tr><td>Chromosome</td><td>start</td><td></td><td>end</td><td>strand</td><td>MM</td><td>target_seq</td><td>PAM</td><td>alignment</td><td>distance</td><td>position</td><td>gene name</td><td>gene id</td></tr>
<tr><td>GU305914.1</td><td></td><td>188903</td><td>188925</td><td>+</td><td>0</td><td>CGTCACGGCGACCGTGTCGC</td><td>CGG</td><td>||||||||[||||||||||||]PAM</td><td>NA</td><td>Intergenic</td><td>NA</td><td>NA</td></tr>
<tr><td>T14381</td><td>CACGGCGACCGTGTCGCCGGCGG</td><td></td><td>1000</td></tr>
<tr><td>Oligo fwd</td><td>CACCCACGGCGACCGTGTCGCCGG</td></tr>
<tr><td>Oligo rev</td><td>AAACCCGGCGACACGGTCGCCGTG</td></tr>
<tr><td>Chromosome</td><td>start</td><td></td><td>end</td><td>strand</td><td>MM</td><td>target_seq</td><td>PAM</td><td>alignment</td><td>distance</td><td>position</td><td>gene name</td><td>gene id</td></tr>
<tr><td>GU305914.1</td><td></td><td>188906</td><td>188928</td><td>+</td><td>0</td><td>CACGGCGACCGTGTCGCCGG</td><td>CGG</td><td>||||||||[||||||||||||]PAM</td><td>NA</td><td>Intergenic</td><td>NA</td><td>NA</td></tr>
<tr><td>T14382</td><td>ACGTACCACACCACGACCAGCGG</td><td></td><td>1000</td></tr>
<tr><td>Oligo fwd</td><td>CACCACGTACCACACCACGACCAG</td></tr>
<tr><td>Oligo rev</td><td>AAACCTGGTCGTGGTGTGGTACGT</td></tr>
<tr><td>Chromosome</td><td>start</td><td></td><td>end</td><td>strand</td><td>MM</td><td>target_seq</td><td>PAM</td><td>alignment</td><td>distance</td><td>position</td><td>gene name</td><td>gene id</td></tr>
<tr><td>GU305914.1</td><td></td><td>188934</td><td>188956</td><td>+</td><td>0</td><td>ACGTACCACACCACGACCAG</td><td>CGG</td><td>||||||||[||||||||||||]PAM</td><td>NA</td><td>Intergenic</td><td>NA</td><td>NA</td></tr>
<tr><td>T14383</td><td>CACACCACGACCAGCGGTAACGG</td><td></td><td>1000</td></tr>
<tr><td>Oligo fwd</td><td>CACCCACACCACGACCAGCGGTAA</td></tr>
<tr><td>Oligo rev</td><td>AAACTTACCGCTGGTCGTGGTGTG</td></tr>
<tr><td>Chromosome</td><td>start</td><td></td><td>end</td><td>strand</td><td>MM</td><td>target_seq</td><td>PAM</td><td>alignment</td><td>distance</td><td>position</td><td>gene name</td><td>gene id</td></tr>
<tr><td>GU305914.1</td><td></td><td>188940</td><td>188962</td><td>+</td><td>0</td><td>CACACCACGACCAGCGGTAA</td><td>CGG</td><td>||||||||[||||||||||||]PAM</td><td>NA</td><td>Intergenic</td><td>NA</td><td>NA</td></tr>
<tr><td>T14384</td><td>ACGACCAGCGGTAACGGTAACGG</td><td></td><td>1000</td></tr>
<tr><td>Oligo fwd</td><td>CACCACGACCAGCGGTAACGGTAA</td></tr>
<tr><td>Oligo rev</td><td>AAACTTACCGTTACCGCTGGTCGT</td></tr>
<tr><td>Chromosome</td><td>start</td><td></td><td>end</td><td>strand</td><td>MM</td><td>target_seq</td><td>PAM</td><td>alignment</td><td>distance</td><td>position</td><td>gene name</td><td>gene id</td></tr>
<tr><td>GU305914.1</td><td></td><td>188946</td><td>188968</td><td>+</td><td>0</td><td>ACGACCAGCGGTAACGGTAA</td><td>CGG</td><td>||||||||[||||||||||||]PAM</td><td>NA</td><td>Intergenic</td><td>NA</td><td>NA</td></tr>
<tr><td>T14385</td><td>CGAACATTTTTTTCACCAGCCGG</td><td></td><td>1000</td></tr>
<tr><td>Oligo fwd</td><td>CACCCGAACATTTTTTTCACCAGC</td></tr>
<tr><td>Oligo rev</td><td>AAACGCTGGTGAAAAAAATGTTCG</td></tr>
<tr><td>Chromosome</td><td>start</td><td></td><td>end</td><td>strand</td><td>MM</td><td>target_seq</td><td>PAM</td><td>alignment</td><td>distance</td><td>position</td><td>gene name</td><td>gene id</td></tr>
<tr><td>GU305914.1</td><td></td><td>188983</td><td>189005</td><td>+</td><td>0</td><td>CGAACATTTTTTTCACCAGC</td><td>CGG</td><td>||||||||[||||||||||||]PAM</td><td>NA</td><td>Intergenic</td><td>NA</td><td>NA</td></tr>
<tr><td>T14386</td><td>GAACATTTTTTTCACCAGCCGGG</td><td></td><td>1000</td></tr>
<tr><td>Oligo fwd</td><td>CACCGAACATTTTTTTCACCAGCC</td></tr>
<tr><td>Oligo rev</td><td>AAACGGCTGGTGAAAAAAATGTTC</td></tr>
<tr><td>Chromosome</td><td>start</td><td></td><td>end</td><td>strand</td><td>MM</td><td>target_seq</td><td>PAM</td><td>alignment</td><td>distance</td><td>position</td><td>gene name</td><td>gene id</td></tr>
<tr><td>GU305914.1</td><td></td><td>188984</td><td>189006</td><td>+</td><td>0</td><td>GAACATTTTTTTCACCAGCC</td><td>GGG</td><td>||||||||[||||||||||||]PAM</td><td>NA</td><td>Intergenic</td><td>NA</td><td>NA</td></tr>
<tr><td>T14387</td><td>CCAGCCGGGCGACGCTGTCCAGG</td><td></td><td>1000</td></tr>
<tr><td>Oligo fwd</td><td>CACCCCAGCCGGGCGACGCTGTCC</td></tr>
<tr><td>Oligo rev</td><td>AAACGGACAGCGTCGCCCGGCTGG</td></tr>
<tr><td>Chromosome</td><td>start</td><td></td><td>end</td><td>strand</td><td>MM</td><td>target_seq</td><td>PAM</td><td>alignment</td><td>distance</td><td>position</td><td>gene name</td><td>gene id</td></tr>
<tr><td>GU305914.1</td><td></td><td>188998</td><td>189020</td><td>+</td><td>0</td><td>CCAGCCGGGCGACGCTGTCC</td><td>AGG</td><td>||||||||[||||||||||||]PAM</td><td>NA</td><td>Intergenic</td><td>NA</td><td>NA</td></tr>
<tr><td>T14388</td><td>CGCTGTCCAGGCTTAATAGCAGG</td><td></td><td>1000</td></tr>
<tr><td>Oligo fwd</td><td>CACCCGCTGTCCAGGCTTAATAGC</td></tr>
<tr><td>Oligo rev</td><td>AAACGCTATTAAGCCTGGACAGCG</td></tr>
<tr><td>Chromosome</td><td>start</td><td></td><td>end</td><td>strand</td><td>MM</td><td>target_seq</td><td>PAM</td><td>alignment</td><td>distance</td><td>position</td><td>gene name</td><td>gene id</td></tr>
<tr><td>GU305914.1</td><td></td><td>189010</td><td>189032</td><td>+</td><td>0</td><td>CGCTGTCCAGGCTTAATAGC</td><td>AGG</td><td>||||||||[||||||||||||]PAM</td><td>NA</td><td>Intergenic</td><td>NA</td><td>NA</td></tr>
<tr><td>T14389</td><td>CAGGCTTAATAGCAGGCGCACGG</td><td></td><td>1000</td></tr>
<tr><td>Oligo fwd</td><td>CACCCAGGCTTAATAGCAGGCGCA</td></tr>
<tr><td>Oligo rev</td><td>AAACTGCGCCTGCTATTAAGCCTG</td></tr>
<tr><td>Chromosome</td><td>start</td><td></td><td>end</td><td>strand</td><td>MM</td><td>target_seq</td><td>PAM</td><td>alignment</td><td>distance</td><td>position</td><td>gene name</td><td>gene id</td></tr>
<tr><td>GU305914.1</td><td></td><td>189017</td><td>189039</td><td>+</td><td>0</td><td>CAGGCTTAATAGCAGGCGCA</td><td>CGG</td><td>||||||||[||||||||||||]PAM</td><td>NA</td><td>Intergenic</td><td>NA</td><td>NA</td></tr>
<tr><td>T14390</td><td>GCTTAATAGCAGGCGCACGGCGG</td><td></td><td>1000</td></tr>
<tr><td>Oligo fwd</td><td>CACCGCTTAATAGCAGGCGCACGG</td></tr>
<tr><td>Oligo rev</td><td>AAACCCGTGCGCCTGCTATTAAGC</td></tr>
<tr><td>Chromosome</td><td>start</td><td></td><td>end</td><td>strand</td><td>MM</td><td>target_seq</td><td>PAM</td><td>alignment</td><td>distance</td><td>position</td><td>gene name</td><td>gene id</td></tr>
<tr><td>GU305914.1</td><td></td><td>189020</td><td>189042</td><td>+</td><td>0</td><td>GCTTAATAGCAGGCGCACGG</td><td>CGG</td><td>||||||||[||||||||||||]PAM</td><td>NA</td><td>Intergenic</td><td>NA</td><td>NA</td></tr>
</table>

<table>
<tr><td>T14391</td><td>GCACGGCGGCGAAGCCCACCAGG</td><td>1000</td><td></td><td></td><td></td><td></td><td></td><td></td><td></td><td></td><td></td></tr>
<tr><td>Oligo fwd</td><td>CACCGCACGGCGGCGAAGCCCACC</td><td></td><td></td><td></td><td></td><td></td><td></td><td></td><td></td><td></td><td></td></tr>
<tr><td>Oligo rev</td><td>AAACGGTGGGCTTCGCCGCCGTGC</td><td></td><td></td><td></td><td></td><td></td><td></td><td></td><td></td><td></td><td></td></tr>
<tr><td>Chromosome</td><td>start</td><td>end</td><td>strand</td><td>MM</td><td>target_seq</td><td>PAM</td><td>alignment</td><td>distance</td><td>position</td><td>gene name</td><td>gene id</td></tr>
<tr><td>GU305914.1</td><td>189034</td><td>189056</td><td>+</td><td>0</td><td>GCACGGCGGCGAAGCCCACC</td><td>AGG</td><td>||||||||[||||||||||||]PAM</td><td>NA</td><td>Intergenic</td><td>NA</td><td>NA</td></tr>
<tr><td>GU305914.1</td><td>192672</td><td>192694</td><td>-</td><td>4</td><td>TCACCGTGACGAAGCCCACC</td><td>AGG</td><td>-|||-|-|[-|||||||||||]PAM</td><td>NA</td><td>Intergenic</td><td>NA</td><td>NA</td></tr>
<tr><td>T14392</td><td>CACGGCGGCGAAGCCCACCAGGG</td><td>1000</td><td></td><td></td><td></td><td></td><td></td><td></td><td></td><td></td><td></td></tr>
<tr><td>Oligo fwd</td><td>CACCCACGGCGGCGAAGCCCACCA</td><td></td><td></td><td></td><td></td><td></td><td></td><td></td><td></td><td></td><td></td></tr>
<tr><td>Oligo rev</td><td>AAACTGGTGGGCTTCGCCGCCGTG</td><td></td><td></td><td></td><td></td><td></td><td></td><td></td><td></td><td></td><td></td></tr>
<tr><td>Chromosome</td><td>start</td><td>end</td><td>strand</td><td>MM</td><td>target_seq</td><td>PAM</td><td>alignment</td><td>distance</td><td>position</td><td>gene name</td><td>gene id</td></tr>
<tr><td>GU305914.1</td><td>189035</td><td>189057</td><td>+</td><td>0</td><td>CACGGCGGCGAAGCCCACCA</td><td>GGG</td><td>||||||||[||||||||||||]PAM</td><td>NA</td><td>Intergenic</td><td>NA</td><td>NA</td></tr>
<tr><td>T14393</td><td>CGTGCACGATGCGCACGTTTCGG</td><td>1000</td><td></td><td></td><td></td><td></td><td></td><td></td><td></td><td></td><td></td></tr>
<tr><td>Oligo fwd</td><td>CACCCGTGCACGATGCGCACGTTT</td><td></td><td></td><td></td><td></td><td></td><td></td><td></td><td></td><td></td><td></td></tr>
<tr><td>Oligo rev</td><td>AAACAAACGTGCGCATCGTGCACG</td><td></td><td></td><td></td><td></td><td></td><td></td><td></td><td></td><td></td><td></td></tr>
<tr><td>Chromosome</td><td>start</td><td>end</td><td>strand</td><td>MM</td><td>target_seq</td><td>PAM</td><td>alignment</td><td>distance</td><td>position</td><td>gene name</td><td>gene id</td></tr>
<tr><td>GU305914.1</td><td>189091</td><td>189113</td><td>+</td><td>0</td><td>CGTGCACGATGCGCACGTTT</td><td>CGG</td><td>||||||||[||||||||||||]PAM</td><td>NA</td><td>Intergenic</td><td>NA</td><td>NA</td></tr>
<tr><td>T14394</td><td>GCACGTTTCGGATGTATTCCTGG</td><td>1000</td><td></td><td></td><td></td><td></td><td></td><td></td><td></td><td></td><td></td></tr>
<tr><td>Oligo fwd</td><td>CACCGCACGTTTCGGATGTATTCC</td><td></td><td></td><td></td><td></td><td></td><td></td><td></td><td></td><td></td><td></td></tr>
<tr><td>Oligo rev</td><td>AAACGGAATACATCCGAAACGTGC</td><td></td><td></td><td></td><td></td><td></td><td></td><td></td><td></td><td></td><td></td></tr>
<tr><td>Chromosome</td><td>start</td><td>end</td><td>strand</td><td>MM</td><td>target_seq</td><td>PAM</td><td>alignment</td><td>distance</td><td>position</td><td>gene name</td><td>gene id</td></tr>
<tr><td>GU305914.1</td><td>189103</td><td>189125</td><td>+</td><td>0</td><td>GCACGTTTCGGATGTATTCC</td><td>TGG</td><td>||||||||[||||||||||||]PAM</td><td>NA</td><td>Intergenic</td><td>NA</td><td>NA</td></tr>
<tr><td>T14395</td><td>CCTGGTGCCGCTTGTCCGTCAGG</td><td>1000</td><td></td><td></td><td></td><td></td><td></td><td></td><td></td><td></td><td></td></tr>
<tr><td>Oligo fwd</td><td>CACCCCTGGTGCCGCTTGTCCGTC</td><td></td><td></td><td></td><td></td><td></td><td></td><td></td><td></td><td></td><td></td></tr>
<tr><td>Oligo rev</td><td>AAACGACGGACAAGCGGCACCAGG</td><td></td><td></td><td></td><td></td><td></td><td></td><td></td><td></td><td></td><td></td></tr>
<tr><td>Chromosome</td><td>start</td><td>end</td><td>strand</td><td>MM</td><td>target_seq</td><td>PAM</td><td>alignment</td><td>distance</td><td>position</td><td>gene name</td><td>gene id</td></tr>
<tr><td>GU305914.1</td><td>189121</td><td>189143</td><td>+</td><td>0</td><td>CCTGGTGCCGCTTGTCCGTC</td><td>AGG</td><td>||||||||[||||||||||||]PAM</td><td>NA</td><td>Intergenic</td><td>NA</td><td>NA</td></tr>
<tr><td>T14396</td><td>TGCCGCTTGTCCGTCAGGCATGG</td><td>1000</td><td></td><td></td><td></td><td></td><td></td><td></td><td></td><td></td><td></td></tr>
<tr><td>Oligo fwd</td><td>CACCTGCCGCTTGTCCGTCAGGCA</td><td></td><td></td><td></td><td></td><td></td><td></td><td></td><td></td><td></td><td></td></tr>
<tr><td>Oligo rev</td><td>AAACTGCCTGACGGACAAGCGGCA</td><td></td><td></td><td></td><td></td><td></td><td></td><td></td><td></td><td></td><td></td></tr>
<tr><td>Chromosome</td><td>start</td><td>end</td><td>strand</td><td>MM</td><td>target_seq</td><td>PAM</td><td>alignment</td><td>distance</td><td>position</td><td>gene name</td><td>gene id</td></tr>
<tr><td>GU305914.1</td><td>189126</td><td>189148</td><td>+</td><td>0</td><td>TGCCGCTTGTCCGTCAGGCA</td><td>TGG</td><td>||||||||[||||||||||||]PAM</td><td>NA</td><td>Intergenic</td><td>NA</td><td>NA</td></tr>
<tr><td>T14397</td><td>ACTTGTTGTCCGTCTTCAGCAGG</td><td>1000</td><td></td><td></td><td></td><td></td><td></td><td></td><td></td><td></td><td></td></tr>
<tr><td>Oligo fwd</td><td>CACCACTTGTTGTCCGTCTTCAGC</td><td></td><td></td><td></td><td></td><td></td><td></td><td></td><td></td><td></td><td></td></tr>
<tr><td>Oligo rev</td><td>AAACGCTGAAGACGGACAACAAGT</td><td></td><td></td><td></td><td></td><td></td><td></td><td></td><td></td><td></td><td></td></tr>
<tr><td>Chromosome</td><td>start</td><td>end</td><td>strand</td><td>MM</td><td>target_seq</td><td>PAM</td><td>alignment</td><td>distance</td><td>position</td><td>gene name</td><td>gene id</td></tr>
<tr><td>GU305914.1</td><td>189166</td><td>189188</td><td>+</td><td>0</td><td>ACTTGTTGTCCGTCTTCAGC</td><td>AGG</td><td>||||||||[||||||||||||]PAM</td><td>NA</td><td>Intergenic</td><td>NA</td><td>NA</td></tr>
<tr><td>T14398</td><td>CCGTCTTCAGCAGGCATACGCGG</td><td>1000</td><td></td><td></td><td></td><td></td><td></td><td></td><td></td><td></td><td></td></tr>
<tr><td>Oligo fwd</td><td>CACCCCGTCTTCAGCAGGCATACG</td><td></td><td></td><td></td><td></td><td></td><td></td><td></td><td></td><td></td><td></td></tr>
<tr><td>Oligo rev</td><td>AAACCGTATGCCTGCTGAAGACGG</td><td></td><td></td><td></td><td></td><td></td><td></td><td></td><td></td><td></td><td></td></tr>
<tr><td>Chromosome</td><td>start</td><td>end</td><td>strand</td><td>MM</td><td>target_seq</td><td>PAM</td><td>alignment</td><td>distance</td><td>position</td><td>gene name</td><td>gene id</td></tr>
<tr><td>GU305914.1</td><td>189175</td><td>189197</td><td>+</td><td>0</td><td>CCGTCTTCAGCAGGCATACG</td><td>CGG</td><td>||||||||[||||||||||||]PAM</td><td>NA</td><td>Intergenic</td><td>NA</td><td>NA</td></tr>
<tr><td>T14399</td><td>CTTCAGCAGGCATACGCGGCAGG</td><td>1000</td><td></td><td></td><td></td><td></td><td></td><td></td><td></td><td></td><td></td></tr>
<tr><td>Oligo fwd</td><td>CACCCTTCAGCAGGCATACGCGGC</td><td></td><td></td><td></td><td></td><td></td><td></td><td></td><td></td><td></td><td></td></tr>
<tr><td>Oligo rev</td><td>AAACGCCGCGTATGCCTGCTGAAG</td><td></td><td></td><td></td><td></td><td></td><td></td><td></td><td></td><td></td><td></td></tr>
<tr><td>Chromosome</td><td>start</td><td>end</td><td>strand</td><td>MM</td><td>target_seq</td><td>PAM</td><td>alignment</td><td>distance</td><td>position</td><td>gene name</td><td>gene id</td></tr>
<tr><td>GU305914.1</td><td>189179</td><td>189201</td><td>+</td><td>0</td><td>CTTCAGCAGGCATACGCGGC</td><td>AGG</td><td>||||||||[||||||||||||]PAM</td><td>NA</td><td>Intergenic</td><td>NA</td><td>NA</td></tr>
<tr><td>T14400</td><td>AGGCATACGCGGCAGGTGCGCGG</td><td>1000</td><td></td><td></td><td></td><td></td><td></td><td></td><td></td><td></td><td></td></tr>
<tr><td>Oligo fwd</td><td>CACCAGGCATACGCGGCAGGTGCG</td><td></td><td></td><td></td><td></td><td></td><td></td><td></td><td></td><td></td><td></td></tr>
<tr><td>Oligo rev</td><td>AAACCGCACCTGCCGCGTATGCCT</td><td></td><td></td><td></td><td></td><td></td><td></td><td></td><td></td><td></td><td></td></tr>
<tr><td>Chromosome</td><td>start</td><td>end</td><td>strand</td><td>MM</td><td>target_seq</td><td>PAM</td><td>alignment</td><td>distance</td><td>position</td><td>gene name</td><td>gene id</td></tr>
<tr><td>GU305914.1</td><td>189186</td><td>189208</td><td>+</td><td>0</td><td>AGGCATACGCGGCAGGTGCG</td><td>CGG</td><td>||||||||[||||||||||||]PAM</td><td>NA</td><td>Intergenic</td><td>NA</td><td>NA</td></tr>
<tr><td>T14401</td><td>GGCATACGCGGCAGGTGCGCGGG</td><td>1000</td><td></td><td></td><td></td><td></td><td></td><td></td><td></td><td></td><td></td></tr>
<tr><td>Oligo fwd</td><td>CACCGGCATACGCGGCAGGTGCGC</td><td></td><td></td><td></td><td></td><td></td><td></td><td></td><td></td><td></td><td></td></tr>
<tr><td>Oligo rev</td><td>AAACGCGCACCTGCCGCGTATGCC</td><td></td><td></td><td></td><td></td><td></td><td></td><td></td><td></td><td></td><td></td></tr>
<tr><td>Chromosome</td><td>start</td><td>end</td><td>strand</td><td>MM</td><td>target_seq</td><td>PAM</td><td>alignment</td><td>distance</td><td>position</td><td>gene name</td><td>gene id</td></tr>
<tr><td>GU305914.1</td><td>189187</td><td>189209</td><td>+</td><td>0</td><td>GGCATACGCGGCAGGTGCGC</td><td>GGG</td><td>||||||||[||||||||||||]PAM</td><td>NA</td><td>Intergenic</td><td>NA</td><td>NA</td></tr>
<tr><td>T14402</td><td>GCGGCAGGTGCGCGGGTCTCCGG</td><td>1000</td><td></td><td></td><td></td><td></td><td></td><td></td><td></td><td></td><td></td></tr>
<tr><td>Oligo fwd</td><td>CACCGCGGCAGGTGCGCGGGTCTC</td><td></td><td></td><td></td><td></td><td></td><td></td><td></td><td></td><td></td><td></td></tr>
<tr><td>Oligo rev</td><td>AAACGAGACCCGCGCACCTGCCGC</td><td></td><td></td><td></td><td></td><td></td><td></td><td></td><td></td><td></td><td></td></tr>
<tr><td>Chromosome</td><td>start</td><td>end</td><td>strand</td><td>MM</td><td>target_seq</td><td>PAM</td><td>alignment</td><td>distance</td><td>position</td><td>gene name</td><td>gene id</td></tr>
<tr><td>GU305914.1</td><td>189194</td><td>189216</td><td>+</td><td>0</td><td>GCGGCAGGTGCGCGGGTCTC</td><td>CGG</td><td>||||||||[||||||||||||]PAM</td><td>NA</td><td>Intergenic</td><td>NA</td><td>NA</td></tr>
<tr><td>T14403</td><td>CGGCAGGTGCGCGGGTCTCCGGG</td><td>1000</td><td></td><td></td><td></td><td></td><td></td><td></td><td></td><td></td><td></td></tr>
<tr><td>Oligo fwd</td><td>CACCCGGCAGGTGCGCGGGTCTCC</td><td></td><td></td><td></td><td></td><td></td><td></td><td></td><td></td><td></td><td></td></tr>
<tr><td>Oligo rev</td><td>AAACGGAGACCCGCGCACCTGCCG</td><td></td><td></td><td></td><td></td><td></td><td></td><td></td><td></td><td></td><td></td></tr>
<tr><td>Chromosome</td><td>start</td><td>end</td><td>strand</td><td>MM</td><td>target_seq</td><td>PAM</td><td>alignment</td><td>distance</td><td>position</td><td>gene name</td><td>gene id</td></tr>
<tr><td>GU305914.1</td><td>189195</td><td>189217</td><td>+</td><td>0</td><td>CGGCAGGTGCGCGGGTCTCC</td><td>GGG</td><td>||||||||[||||||||||||]PAM</td><td>NA</td><td>Intergenic</td><td>NA</td><td>NA</td></tr>
<tr><td>T14404</td><td>CGGGTCTCCGGGTTTCACCTTGG</td><td>1000</td><td></td><td></td><td></td><td></td><td></td><td></td><td></td><td></td><td></td></tr>
<tr><td>Oligo fwd</td><td>CACCCGGGTCTCCGGGTTTCACCT</td><td></td><td></td><td></td><td></td><td></td><td></td><td></td><td></td><td></td><td></td></tr>
<tr><td>Oligo rev</td><td>AAACAGGTGAAACCCGGAGACCCG</td><td></td><td></td><td></td><td></td><td></td><td></td><td></td><td></td><td></td><td></td></tr>
<tr><td>Chromosome</td><td>start</td><td>end</td><td>strand</td><td>MM</td><td>target_seq</td><td>PAM</td><td>alignment</td><td>distance</td><td>position</td><td>gene name</td><td>gene id</td></tr>
<tr><td>GU305914.1</td><td>189206</td><td>189228</td><td>+</td><td>0</td><td>CGGGTCTCCGGGTTTCACCT</td><td>TGG</td><td>||||||||[||||||||||||]PAM</td><td>NA</td><td>Intergenic</td><td>NA</td><td>NA</td></tr>
<tr><td>T14405</td><td>GGGTTTCACCTTGGCGCTCATGG</td><td>1000</td><td></td><td></td><td></td><td></td><td></td><td></td><td></td><td></td><td></td></tr>
<tr><td>Oligo fwd</td><td>CACCGGGTTTCACCTTGGCGCTCA</td><td></td><td></td><td></td><td></td><td></td><td></td><td></td><td></td><td></td><td></td></tr>
<tr><td>Oligo rev</td><td>AAACTGAGCGCCAAGGTGAAACCC</td><td></td><td></td><td></td><td></td><td></td><td></td><td></td><td></td><td></td><td></td></tr>
</table>

Chromosome start end strand MM target_seq PAM alignment distance position gene name gene id
GU305914.1 189215 189237 + 0 GGGTTTCACCTTGGCGCTCA TGG ||||||||[||||||||||||]PAM NA Intergenic NA NA

T14406 CACGTATCTTCCCTCGCGTATGG 1000
Oligo fwd CACCCACGTATCTTCCCTCGCGTA
Oligo rev AAACTACGCGAGGGAAGATACGTG
Chromosome start end strand MM target_seq PAM alignment distance position gene name gene id
GU305914.1 189239 189261 + 0 CACGTATCTTCCCTCGCGTA TGG ||||||||[||||||||||||]PAM NA Intergenic NA NA

T14407 GCGTATGGATTTCTTGTAGCTGG 1000
Oligo fwd CACCGCGTATGGATTTCTTGTAGC
Oligo rev AAACGCTACAAGAAATCCATACGC
Chromosome start end strand MM target_seq PAM alignment distance position gene name gene id
GU305914.1 189254 189276 + 0 GCGTATGGATTTCTTGTAGC TGG ||||||||[||||||||||||]PAM NA Intergenic NA NA

T14408 TTTCTTGTAGCTGGCCGTTTTGG 1000
Oligo fwd CACCTTTCTTGTAGCTGGCCGTTT
Oligo rev AAACAAACGGCCAGCTACAAGAAA
Chromosome start end strand MM target_seq PAM alignment distance position gene name gene id
GU305914.1 189263 189285 + 0 TTTCTTGTAGCTGGCCGTTT TGG ||||||||[||||||||||||]PAM NA Intergenic NA NA

T14409 GTAGCTGGCCGTTTTGGCGCAGG 1000
Oligo fwd CACCGTAGCTGGCCGTTTTGGCGC
Oligo rev AAACGCGCCAAAACGGCCAGCTAC
Chromosome start end strand MM target_seq PAM alignment distance position gene name gene id
GU305914.1 189269 189291 + 0 GTAGCTGGCCGTTTTGGCGC AGG ||||||||[||||||||||||]PAM NA Intergenic NA NA

T14410 GTTTTGGCGCAGGCCTCGATCGG 1000
Oligo fwd CACCGTTTTGGCGCAGGCCTCGAT
Oligo rev AAACATCGAGGCCTGCGCCAAAAC
Chromosome start end strand MM target_seq PAM alignment distance position gene name gene id
GU305914.1 189279 189301 + 0 GTTTTGGCGCAGGCCTCGAT CGG ||||||||[||||||||||||]PAM NA Intergenic NA NA

T14411 CTCGATCGGATCTTTCGTTTTGG 1000
Oligo fwd CACCCTCGATCGGATCTTTCGTTT
Oligo rev AAACAAACGAAAGATCCGATCGAG
Chromosome start end strand MM target_seq PAM alignment distance position gene name gene id
GU305914.1 189293 189315 + 0 CTCGATCGGATCTTTCGTTT TGG ||||||||[||||||||||||]PAM NA Intergenic NA NA

T14412 ATCTTTCGTTTTGGCGCCGAAGG 1000
Oligo fwd CACCATCTTTCGTTTTGGCGCCGA
Oligo rev AAACTCGGCGCCAAAACGAAAGAT
Chromosome start end strand MM target_seq PAM alignment distance position gene name gene id
GU305914.1 189302 189324 + 0 ATCTTTCGTTTTGGCGCCGA AGG ||||||||[||||||||||||]PAM NA Intergenic NA NA

T14413 ACCACGCCGTCTCGATCTTCTGG 1000
Oligo fwd CACCACCACGCCGTCTCGATCTTC
Oligo rev AAACGAAGATCGAGACGGCGTGGT
Chromosome start end strand MM target_seq PAM alignment distance position gene name gene id
GU305914.1 189340 189362 + 0 ACCACGCCGTCTCGATCTTC TGG ||||||||[||||||||||||]PAM NA Intergenic NA NA

T14414 CCACGCCGTCTCGATCTTCTGGG 1000
Oligo fwd CACCCCACGCCGTCTCGATCTTCT
Oligo rev AAACAGAAGATCGAGACGGCGTGG
Chromosome start end strand MM target_seq PAM alignment distance position gene name gene id
GU305914.1 189341 189363 + 0 CCACGCCGTCTCGATCTTCT GGG ||||||||[||||||||||||]PAM NA Intergenic NA NA

T14415 GTCTCGATCTTCTGGGTGTCCGG 1000
Oligo fwd CACCGTCTCGATCTTCTGGGTGTC
Oligo rev AAACGACACCCAGAAGATCGAGAC
Chromosome start end strand MM target_seq PAM alignment distance position gene name gene id
GU305914.1 189348 189370 + 0 GTCTCGATCTTCTGGGTGTC CGG ||||||||[||||||||||||]PAM NA Intergenic NA NA

T14416 TGTCCGGAGTTACGCGAAACCGG 1000
Oligo fwd CACCTGTCCGGAGTTACGCGAAAC
Oligo rev AAACGTTTCGCGTAACTCCGGACA
Chromosome start end strand MM target_seq PAM alignment distance position gene name gene id
GU305914.1 189364 189386 + 0 TGTCCGGAGTTACGCGAAAC CGG ||||||||[||||||||||||]PAM NA Intergenic NA NA

T14417 CGAAACCGGCACGTGAAATAAGG 1000
Oligo fwd CACCCGAAACCGGCACGTGAAATA
Oligo rev AAACTATTTCACGTGCCGGTTTCG
Chromosome start end strand MM target_seq PAM alignment distance position gene name gene id
GU305914.1 189378 189400 + 0 CGAAACCGGCACGTGAAATA AGG ||||||||[||||||||||||]PAM NA Intergenic NA NA

T14418 ACCGGCACGTGAAATAAGGCCGG 1000
Oligo fwd CACCACCGGCACGTGAAATAAGGC
Oligo rev AAACGCCTTATTTCACGTGCCGGT
Chromosome start end strand MM target_seq PAM alignment distance position gene name gene id
GU305914.1 189382 189404 + 0 ACCGGCACGTGAAATAAGGC CGG ||||||||[||||||||||||]PAM NA Intergenic NA NA

T14419 GAAATAAGGCCGGTGATTGACGG 1000
Oligo fwd CACCGAAATAAGGCCGGTGATTGA
Oligo rev AAACTCAATCACCGGCCTTATTTC
Chromosome start end strand MM target_seq PAM alignment distance position gene name gene id
GU305914.1 189392 189414 + 0 GAAATAAGGCCGGTGATTGA CGG ||||||||[||||||||||||]PAM NA Intergenic NA NA

T14420 GTGATTGACGGTCTCCACCGTGG 1000
Oligo fwd CACCGTGATTGACGGTCTCCACCG

| | | | | | | | | | | | |
|---|---|---|---|---|---|---|---|---|---|---|---|
| Oligo rev | AAACCGGTGGAGACCGTCAATCAC | | | | | | | | | | |
| Chromosome | start | end | strand | MM | target_seq | PAM | alignment | distance | position | gene name | gene id |
| GU305914.1 | 189404 | 189426 | + | 0 | GTGATTGACGGTCTCCACCG | TGG | \|\|\|\|\|\|\|\|[\|\|\|\|\|\|\|\|\|\|\|\|]PAM | NA | Intergenic | NA | NA |
| T14421 | TGACGGTCTCCACCGTGGCCCGG | 1000 | | | | | | | | | |
| Oligo fwd | CACCTGACGGTCTCCACCGTGGCC | | | | | | | | | | |
| Oligo rev | AAACGGCCACGGTGGAGACCGTCA | | | | | | | | | | |
| Chromosome | start | end | strand | MM | target_seq | PAM | alignment | distance | position | gene name | gene id |
| GU305914.1 | 189409 | 189431 | + | 0 | TGACGGTCTCCACCGTGGCC | CGG | \|\|\|\|\|\|\|\|[\|\|\|\|\|\|\|\|\|\|\|\|]PAM | NA | Intergenic | NA | NA |
| T14422 | ACCGTGGCCCGGACCAACATAGG | 1000 | | | | | | | | | |
| Oligo fwd | CACCACCGTGGCCCGGACCAACAT | | | | | | | | | | |
| Oligo rev | AAACATGTTGGTCCGGGCCACGGT | | | | | | | | | | |
| Chromosome | start | end | strand | MM | target_seq | PAM | alignment | distance | position | gene name | gene id |
| GU305914.1 | 189420 | 189442 | + | 0 | ACCGTGGCCCGGACCAACAT | AGG | \|\|\|\|\|\|\|\|[\|\|\|\|\|\|\|\|\|\|\|\|]PAM | NA | Intergenic | NA | NA |
| T14423 | AGGCGAAAAATCGCGCCGTTTGG | 1000 | | | | | | | | | |
| Oligo fwd | CACCAGGCGAAAAATCGCGCCGTT | | | | | | | | | | |
| Oligo rev | AAACAACGGCGCGATTTTTCGCCT | | | | | | | | | | |
| Chromosome | start | end | strand | MM | target_seq | PAM | alignment | distance | position | gene name | gene id |
| GU305914.1 | 189440 | 189462 | + | 0 | AGGCGAAAAATCGCGCCGTT | TGG | \|\|\|\|\|\|\|\|[\|\|\|\|\|\|\|\|\|\|\|\|]PAM | NA | Intergenic | NA | NA |
| T14424 | GGCGAAAAATCGCGCCGTTTGGG | 1000 | | | | | | | | | |
| Oligo fwd | CACCGGCGAAAAATCGCGCCGTTT | | | | | | | | | | |
| Oligo rev | AAACAAACGGCGCGATTTTTCGCC | | | | | | | | | | |
| Chromosome | start | end | strand | MM | target_seq | PAM | alignment | distance | position | gene name | gene id |
| GU305914.1 | 189441 | 189463 | + | 0 | GGCGAAAAATCGCGCCGTTT | GGG | \|\|\|\|\|\|\|\|[\|\|\|\|\|\|\|\|\|\|\|\|]PAM | NA | Intergenic | NA | NA |
| T14425 | TCCATATCCATCGAATCGCCCGG | 1000 | | | | | | | | | |
| Oligo fwd | CACCTCCATATCCATCGAATCGCC | | | | | | | | | | |
| Oligo rev | AAACGGCGATTCGATGGATATGGA | | | | | | | | | | |
| Chromosome | start | end | strand | MM | target_seq | PAM | alignment | distance | position | gene name | gene id |
| GU305914.1 | 189522 | 189544 | + | 0 | TCCATATCCATCGAATCGCC | CGG | \|\|\|\|\|\|\|\|[\|\|\|\|\|\|\|\|\|\|\|\|]PAM | NA | Intergenic | NA | NA |
| T14426 | GGTCTCTCGCAGTAAAGACACGG | 1000 | | | | | | | | | |
| Oligo fwd | CACCGGTCTCTCGCAGTAAAGACA | | | | | | | | | | |
| Oligo rev | AAACTGTCTTTACTGCGAGAGACC | | | | | | | | | | |
| Chromosome | start | end | strand | MM | target_seq | PAM | alignment | distance | position | gene name | gene id |
| GU305914.1 | 189543 | 189565 | + | 0 | GGTCTCTCGCAGTAAAGACA | CGG | \|\|\|\|\|\|\|\|[\|\|\|\|\|\|\|\|\|\|\|\|]PAM | NA | Intergenic | NA | NA |
| T14427 | GTCTCTCGCAGTAAAGACACGGG | 1000 | | | | | | | | | |
| Oligo fwd | CACCGTCTCTCGCAGTAAAGACAC | | | | | | | | | | |
| Oligo rev | AAACGTGTCTTTACTGCGAGAGAC | | | | | | | | | | |
| Chromosome | start | end | strand | MM | target_seq | PAM | alignment | distance | position | gene name | gene id |
| GU305914.1 | 189544 | 189566 | + | 0 | GTCTCTCGCAGTAAAGACAC | GGG | \|\|\|\|\|\|\|\|[\|\|\|\|\|\|\|\|\|\|\|\|]PAM | NA | Intergenic | NA | NA |
| T14428 | GGGTGACGTTTCTCAAACTCCGG | 1000 | | | | | | | | | |
| Oligo fwd | CACCGGGTGACGTTTCTCAAACTC | | | | | | | | | | |
| Oligo rev | AAACGAGTTTGAGAAACGTCACCC | | | | | | | | | | |
| Chromosome | start | end | strand | MM | target_seq | PAM | alignment | distance | position | gene name | gene id |
| GU305914.1 | 189564 | 189586 | + | 0 | GGGTGACGTTTCTCAAACTC | CGG | \|\|\|\|\|\|\|\|[\|\|\|\|\|\|\|\|\|\|\|\|]PAM | NA | Intergenic | NA | NA |
| T14429 | GGTGACGTTTCTCAAACTCCGGG | 1000 | | | | | | | | | |
| Oligo fwd | CACCGGTGACGTTTCTCAAACTCC | | | | | | | | | | |
| Oligo rev | AAACGGAGTTTGAGAAACGTCACC | | | | | | | | | | |
| Chromosome | start | end | strand | MM | target_seq | PAM | alignment | distance | position | gene name | gene id |
| GU305914.1 | 189565 | 189587 | + | 0 | GGTGACGTTTCTCAAACTCC | GGG | \|\|\|\|\|\|\|\|[\|\|\|\|\|\|\|\|\|\|\|\|]PAM | NA | Intergenic | NA | NA |
| T14430 | CCGGGTCTCTGAACGACAGCCGG | 1000 | | | | | | | | | |
| Oligo fwd | CACCCCGGGTCTCTGAACGACAGC | | | | | | | | | | |
| Oligo rev | AAACGCTGTCGTTCAGAGACCCGG | | | | | | | | | | |
| Chromosome | start | end | strand | MM | target_seq | PAM | alignment | distance | position | gene name | gene id |
| GU305914.1 | 189583 | 189605 | + | 0 | CCGGGTCTCTGAACGACAGC | CGG | \|\|\|\|\|\|\|\|[\|\|\|\|\|\|\|\|\|\|\|\|]PAM | NA | Intergenic | NA | NA |
| T14431 | CAGCCGGCGATGCCGCTTGCCGG | 1000 | | | | | | | | | |
| Oligo fwd | CACCCAGCCGGCGATGCCGCTTGC | | | | | | | | | | |
| Oligo rev | AAACGCAAGCGGCATCGCCGGCTG | | | | | | | | | | |
| Chromosome | start | end | strand | MM | target_seq | PAM | alignment | distance | position | gene name | gene id |
| GU305914.1 | 189599 | 189621 | + | 0 | CAGCCGGCGATGCCGCTTGC | CGG | \|\|\|\|\|\|\|\|[\|\|\|\|\|\|\|\|\|\|\|\|]PAM | NA | Intergenic | NA | NA |
| T14432 | CGAACTATCCTCCACCAACTTGG | 1000 | | | | | | | | | |
| Oligo fwd | CACCCGAACTATCCTCCACCAACT | | | | | | | | | | |
| Oligo rev | AAACAGTTGGTGGAGGATAGTTCG | | | | | | | | | | |
| Chromosome | start | end | strand | MM | target_seq | PAM | alignment | distance | position | gene name | gene id |
| GU305914.1 | 189623 | 189645 | + | 0 | CGAACTATCCTCCACCAACT | TGG | \|\|\|\|\|\|\|\|[\|\|\|\|\|\|\|\|\|\|\|\|]PAM | NA | Intergenic | NA | NA |
| T14433 | ACTATCCTCCACCAACTTGGTGG | 1000 | | | | | | | | | |
| Oligo fwd | CACCACTATCCTCCACCAACTTGG | | | | | | | | | | |
| Oligo rev | AAACCCAAGTTGGTGGAGGATAGT | | | | | | | | | | |
| Chromosome | start | end | strand | MM | target_seq | PAM | alignment | distance | position | gene name | gene id |
| GU305914.1 | 189626 | 189648 | + | 0 | ACTATCCTCCACCAACTTGG | TGG | \|\|\|\|\|\|\|\|[\|\|\|\|\|\|\|\|\|\|\|\|]PAM | NA | Intergenic | NA | NA |
| T14434 | CCTCCACCAACTTGGTGGTCAGG | 1000 | | | | | | | | | |
| Oligo fwd | CACCCCTCCACCAACTTGGTGGTC | | | | | | | | | | |
| Oligo rev | AAACGACCACCAAGTTGGTGGAGG | | | | | | | | | | |
| Chromosome | start | end | strand | MM | target_seq | PAM | alignment | distance | position | gene name | gene id |
| GU305914.1 | 189631 | 189653 | + | 0 | CCTCCACCAACTTGGTGGTC | AGG | \|\|\|\|\|\|\|\|[\|\|\|\|\|\|\|\|\|\|\|\|]PAM | NA | Intergenic | NA | NA |
| T14435 | TGAAGCAAGATCGTGCCGTTCGG | 1000 | | | | | | | | | |

<table>
<tr><td>Oligo fwd</td><td>CACCTGAAGCAAGATCGTGCCGTT</td><td></td><td></td><td></td><td></td><td></td><td></td><td></td><td></td><td></td><td></td></tr>
<tr><td>Oligo rev</td><td>AAACAACGGCACGATCTTGCTTCA</td><td></td><td></td><td></td><td></td><td></td><td></td><td></td><td></td><td></td><td></td></tr>
<tr><td>Chromosome</td><td>start</td><td>end</td><td>strand</td><td>MM</td><td>target_seq</td><td>PAM</td><td>alignment</td><td>distance</td><td>position</td><td>gene name</td><td>gene id</td></tr>
<tr><td>GU305914.1</td><td>189663</td><td>189685</td><td>+</td><td>0</td><td>TGAAGCAAGATCGTGCCGTT</td><td>CGG</td><td>||||||||[||||||||||||]PAM</td><td>NA</td><td>Intergenic</td><td>NA</td><td>NA</td></tr>
<tr><td></td><td></td><td></td><td></td><td></td><td></td><td></td><td></td><td></td><td></td><td></td><td></td></tr>
<tr><td>T14436</td><td>GTTCGGCGTCACCGTCGCCGAGG</td><td>1000</td><td></td><td></td><td></td><td></td><td></td><td></td><td></td><td></td><td></td></tr>
<tr><td>Oligo fwd</td><td>CACCGTTCGGCGTCACCGTCGCCG</td><td></td><td></td><td></td><td></td><td></td><td></td><td></td><td></td><td></td><td></td></tr>
<tr><td>Oligo rev</td><td>AAACCGGCGACGGTGACGCCGAAC</td><td></td><td></td><td></td><td></td><td></td><td></td><td></td><td></td><td></td><td></td></tr>
<tr><td>Chromosome</td><td>start</td><td>end</td><td>strand</td><td>MM</td><td>target_seq</td><td>PAM</td><td>alignment</td><td>distance</td><td>position</td><td>gene name</td><td>gene id</td></tr>
<tr><td>GU305914.1</td><td>189680</td><td>189702</td><td>+</td><td>0</td><td>GTTCGGCGTCACCGTCGCCG</td><td>AGG</td><td>||||||||[||||||||||||]PAM</td><td>NA</td><td>Intergenic</td><td>NA</td><td>NA</td></tr>
<tr><td></td><td></td><td></td><td></td><td></td><td></td><td></td><td></td><td></td><td></td><td></td><td></td></tr>
<tr><td>T14437</td><td>CGCCGAGGATTTGATTTTCTCGG</td><td>1000</td><td></td><td></td><td></td><td></td><td></td><td></td><td></td><td></td><td></td></tr>
<tr><td>Oligo fwd</td><td>CACCCGCCGAGGATTTGATTTTCT</td><td></td><td></td><td></td><td></td><td></td><td></td><td></td><td></td><td></td><td></td></tr>
<tr><td>Oligo rev</td><td>AAACAGAAAATCAAATCCTCGGCG</td><td></td><td></td><td></td><td></td><td></td><td></td><td></td><td></td><td></td><td></td></tr>
<tr><td>Chromosome</td><td>start</td><td>end</td><td>strand</td><td>MM</td><td>target_seq</td><td>PAM</td><td>alignment</td><td>distance</td><td>position</td><td>gene name</td><td>gene id</td></tr>
<tr><td>GU305914.1</td><td>189695</td><td>189717</td><td>+</td><td>0</td><td>CGCCGAGGATTTGATTTTCT</td><td>CGG</td><td>||||||||[||||||||||||]PAM</td><td>NA</td><td>Intergenic</td><td>NA</td><td>NA</td></tr>
<tr><td></td><td></td><td></td><td></td><td></td><td></td><td></td><td></td><td></td><td></td><td></td><td></td></tr>
<tr><td>T14438</td><td>GCCGAGGATTTGATTTTCTCGGG</td><td>1000</td><td></td><td></td><td></td><td></td><td></td><td></td><td></td><td></td><td></td></tr>
<tr><td>Oligo fwd</td><td>CACCGCCGAGGATTTGATTTTCTC</td><td></td><td></td><td></td><td></td><td></td><td></td><td></td><td></td><td></td><td></td></tr>
<tr><td>Oligo rev</td><td>AAACGAGAAAATCAAATCCTCGGC</td><td></td><td></td><td></td><td></td><td></td><td></td><td></td><td></td><td></td><td></td></tr>
<tr><td>Chromosome</td><td>start</td><td>end</td><td>strand</td><td>MM</td><td>target_seq</td><td>PAM</td><td>alignment</td><td>distance</td><td>position</td><td>gene name</td><td>gene id</td></tr>
<tr><td>GU305914.1</td><td>189696</td><td>189718</td><td>+</td><td>0</td><td>GCCGAGGATTTGATTTTCTC</td><td>GGG</td><td>||||||||[||||||||||||]PAM</td><td>NA</td><td>Intergenic</td><td>NA</td><td>NA</td></tr>
<tr><td></td><td></td><td></td><td></td><td></td><td></td><td></td><td></td><td></td><td></td><td></td><td></td></tr>
<tr><td>T14439</td><td>AGGATTTGATTTTCTCGGGAAGG</td><td>1000</td><td></td><td></td><td></td><td></td><td></td><td></td><td></td><td></td><td></td></tr>
<tr><td>Oligo fwd</td><td>CACCAGGATTTGATTTTCTCGGGA</td><td></td><td></td><td></td><td></td><td></td><td></td><td></td><td></td><td></td><td></td></tr>
<tr><td>Oligo rev</td><td>AAACTCCCGAGAAAATCAAATCCT</td><td></td><td></td><td></td><td></td><td></td><td></td><td></td><td></td><td></td><td></td></tr>
<tr><td>Chromosome</td><td>start</td><td>end</td><td>strand</td><td>MM</td><td>target_seq</td><td>PAM</td><td>alignment</td><td>distance</td><td>position</td><td>gene name</td><td>gene id</td></tr>
<tr><td>GU305914.1</td><td>189700</td><td>189722</td><td>+</td><td>0</td><td>AGGATTTGATTTTCTCGGGA</td><td>AGG</td><td>||||||||[||||||||||||]PAM</td><td>NA</td><td>Intergenic</td><td>NA</td><td>NA</td></tr>
<tr><td></td><td></td><td></td><td></td><td></td><td></td><td></td><td></td><td></td><td></td><td></td><td></td></tr>
<tr><td>T14440</td><td>TTTCTCGGGAAGGTAGCAGTTGG</td><td>1000</td><td></td><td></td><td></td><td></td><td></td><td></td><td></td><td></td><td></td></tr>
<tr><td>Oligo fwd</td><td>CACCTTTCTCGGGAAGGTAGCAGT</td><td></td><td></td><td></td><td></td><td></td><td></td><td></td><td></td><td></td><td></td></tr>
<tr><td>Oligo rev</td><td>AAACACTGCTACCTTCCCGAGAAA</td><td></td><td></td><td></td><td></td><td></td><td></td><td></td><td></td><td></td><td></td></tr>
<tr><td>Chromosome</td><td>start</td><td>end</td><td>strand</td><td>MM</td><td>target_seq</td><td>PAM</td><td>alignment</td><td>distance</td><td>position</td><td>gene name</td><td>gene id</td></tr>
<tr><td>GU305914.1</td><td>189710</td><td>189732</td><td>+</td><td>0</td><td>TTTCTCGGGAAGGTAGCAGT</td><td>TGG</td><td>||||||||[||||||||||||]PAM</td><td>NA</td><td>Intergenic</td><td>NA</td><td>NA</td></tr>
<tr><td></td><td></td><td></td><td></td><td></td><td></td><td></td><td></td><td></td><td></td><td></td><td></td></tr>
<tr><td>T14441</td><td>GGAAGGTAGCAGTTGGTAAACGG</td><td>1000</td><td></td><td></td><td></td><td></td><td></td><td></td><td></td><td></td><td></td></tr>
<tr><td>Oligo fwd</td><td>CACCGGAAGGTAGCAGTTGGTAAA</td><td></td><td></td><td></td><td></td><td></td><td></td><td></td><td></td><td></td><td></td></tr>
<tr><td>Oligo rev</td><td>AAACTTTACCAACTGCTACCTTCC</td><td></td><td></td><td></td><td></td><td></td><td></td><td></td><td></td><td></td><td></td></tr>
<tr><td>Chromosome</td><td>start</td><td>end</td><td>strand</td><td>MM</td><td>target_seq</td><td>PAM</td><td>alignment</td><td>distance</td><td>position</td><td>gene name</td><td>gene id</td></tr>
<tr><td>GU305914.1</td><td>189717</td><td>189739</td><td>+</td><td>0</td><td>GGAAGGTAGCAGTTGGTAAA</td><td>CGG</td><td>||||||||[||||||||||||]PAM</td><td>NA</td><td>Intergenic</td><td>NA</td><td>NA</td></tr>
<tr><td></td><td></td><td></td><td></td><td></td><td></td><td></td><td></td><td></td><td></td><td></td><td></td></tr>
<tr><td>T14442</td><td>TAGCAGTTGGTAAACGGCAGCGG</td><td>1000</td><td></td><td></td><td></td><td></td><td></td><td></td><td></td><td></td><td></td></tr>
<tr><td>Oligo fwd</td><td>CACCTAGCAGTTGGTAAACGGCAG</td><td></td><td></td><td></td><td></td><td></td><td></td><td></td><td></td><td></td><td></td></tr>
<tr><td>Oligo rev</td><td>AAACCTGCCGTTTACCAACTGCTA</td><td></td><td></td><td></td><td></td><td></td><td></td><td></td><td></td><td></td><td></td></tr>
<tr><td>Chromosome</td><td>start</td><td>end</td><td>strand</td><td>MM</td><td>target_seq</td><td>PAM</td><td>alignment</td><td>distance</td><td>position</td><td>gene name</td><td>gene id</td></tr>
<tr><td>GU305914.1</td><td>189723</td><td>189745</td><td>+</td><td>0</td><td>TAGCAGTTGGTAAACGGCAG</td><td>CGG</td><td>||||||||[||||||||||||]PAM</td><td>NA</td><td>Intergenic</td><td>NA</td><td>NA</td></tr>
<tr><td></td><td></td><td></td><td></td><td></td><td></td><td></td><td></td><td></td><td></td><td></td><td></td></tr>
<tr><td>T14443</td><td>AAACGGCAGCGGCATGTTGTTGG</td><td>1000</td><td></td><td></td><td></td><td></td><td></td><td></td><td></td><td></td><td></td></tr>
<tr><td>Oligo fwd</td><td>CACCAAACGGCAGCGGCATGTTGT</td><td></td><td></td><td></td><td></td><td></td><td></td><td></td><td></td><td></td><td></td></tr>
<tr><td>Oligo rev</td><td>AAACACAACATGCCGCTGCCGTTT</td><td></td><td></td><td></td><td></td><td></td><td></td><td></td><td></td><td></td><td></td></tr>
<tr><td>Chromosome</td><td>start</td><td>end</td><td>strand</td><td>MM</td><td>target_seq</td><td>PAM</td><td>alignment</td><td>distance</td><td>position</td><td>gene name</td><td>gene id</td></tr>
<tr><td>GU305914.1</td><td>189734</td><td>189756</td><td>+</td><td>0</td><td>AAACGGCAGCGGCATGTTGT</td><td>TGG</td><td>||||||||[||||||||||||]PAM</td><td>NA</td><td>Intergenic</td><td>NA</td><td>NA</td></tr>
<tr><td></td><td></td><td></td><td></td><td></td><td></td><td></td><td></td><td></td><td></td><td></td><td></td></tr>
<tr><td>T14444</td><td>AACGGCAGCGGCATGTTGTTGGG</td><td>1000</td><td></td><td></td><td></td><td></td><td></td><td></td><td></td><td></td><td></td></tr>
<tr><td>Oligo fwd</td><td>CACCAACGGCAGCGGCATGTTGTT</td><td></td><td></td><td></td><td></td><td></td><td></td><td></td><td></td><td></td><td></td></tr>
<tr><td>Oligo rev</td><td>AAACAACAACATGCCGCTGCCGTT</td><td></td><td></td><td></td><td></td><td></td><td></td><td></td><td></td><td></td><td></td></tr>
<tr><td>Chromosome</td><td>start</td><td>end</td><td>strand</td><td>MM</td><td>target_seq</td><td>PAM</td><td>alignment</td><td>distance</td><td>position</td><td>gene name</td><td>gene id</td></tr>
<tr><td>GU305914.1</td><td>189735</td><td>189757</td><td>+</td><td>0</td><td>AACGGCAGCGGCATGTTGTT</td><td>GGG</td><td>||||||||[||||||||||||]PAM</td><td>NA</td><td>Intergenic</td><td>NA</td><td>NA</td></tr>
<tr><td></td><td></td><td></td><td></td><td></td><td></td><td></td><td></td><td></td><td></td><td></td><td></td></tr>
<tr><td>T14445</td><td>AGCGGCATGTTGTTGGGTCTCGG</td><td>1000</td><td></td><td></td><td></td><td></td><td></td><td></td><td></td><td></td><td></td></tr>
<tr><td>Oligo fwd</td><td>CACCAGCGGCATGTTGTTGGGTCT</td><td></td><td></td><td></td><td></td><td></td><td></td><td></td><td></td><td></td><td></td></tr>
<tr><td>Oligo rev</td><td>AAACAGACCCAACAACATGCCGCT</td><td></td><td></td><td></td><td></td><td></td><td></td><td></td><td></td><td></td><td></td></tr>
<tr><td>Chromosome</td><td>start</td><td>end</td><td>strand</td><td>MM</td><td>target_seq</td><td>PAM</td><td>alignment</td><td>distance</td><td>position</td><td>gene name</td><td>gene id</td></tr>
<tr><td>GU305914.1</td><td>189741</td><td>189763</td><td>+</td><td>0</td><td>AGCGGCATGTTGTTGGGTCT</td><td>CGG</td><td>||||||||[||||||||||||]PAM</td><td>NA</td><td>Intergenic</td><td>NA</td><td>NA</td></tr>
<tr><td></td><td></td><td></td><td></td><td></td><td></td><td></td><td></td><td></td><td></td><td></td><td></td></tr>
<tr><td>T14446</td><td>CAATACGAATTCACAGCTGTAGG</td><td>1000</td><td></td><td></td><td></td><td></td><td></td><td></td><td></td><td></td><td></td></tr>
<tr><td>Oligo fwd</td><td>CACCCAATACGAATTCACAGCTGT</td><td></td><td></td><td></td><td></td><td></td><td></td><td></td><td></td><td></td><td></td></tr>
<tr><td>Oligo rev</td><td>AAACACAGCTGTGAATTCGTATTG</td><td></td><td></td><td></td><td></td><td></td><td></td><td></td><td></td><td></td><td></td></tr>
<tr><td>Chromosome</td><td>start</td><td>end</td><td>strand</td><td>MM</td><td>target_seq</td><td>PAM</td><td>alignment</td><td>distance</td><td>position</td><td>gene name</td><td>gene id</td></tr>
<tr><td>GU305914.1</td><td>189764</td><td>189786</td><td>+</td><td>0</td><td>CAATACGAATTCACAGCTGT</td><td>AGG</td><td>||||||||[||||||||||||]PAM</td><td>NA</td><td>Intergenic</td><td>NA</td><td>NA</td></tr>
<tr><td></td><td></td><td></td><td></td><td></td><td></td><td></td><td></td><td></td><td></td><td></td><td></td></tr>
<tr><td>T14447</td><td>GTAGGTTCCGTTTTCATAATCGG</td><td>1000</td><td></td><td></td><td></td><td></td><td></td><td></td><td></td><td></td><td></td></tr>
<tr><td>Oligo fwd</td><td>CACCGTAGGTTCCGTTTTCATAAT</td><td></td><td></td><td></td><td></td><td></td><td></td><td></td><td></td><td></td><td></td></tr>
<tr><td>Oligo rev</td><td>AAACATTATGAAAACGGAACCTAC</td><td></td><td></td><td></td><td></td><td></td><td></td><td></td><td></td><td></td><td></td></tr>
<tr><td>Chromosome</td><td>start</td><td>end</td><td>strand</td><td>MM</td><td>target_seq</td><td>PAM</td><td>alignment</td><td>distance</td><td>position</td><td>gene name</td><td>gene id</td></tr>
<tr><td>GU305914.1</td><td>189782</td><td>189804</td><td>+</td><td>0</td><td>GTAGGTTCCGTTTTCATAAT</td><td>CGG</td><td>||||||||[||||||||||||]PAM</td><td>NA</td><td>Intergenic</td><td>NA</td><td>NA</td></tr>
<tr><td></td><td></td><td></td><td></td><td></td><td></td><td></td><td></td><td></td><td></td><td></td><td></td></tr>
<tr><td>T14448</td><td>AAATTGCTATAGCCCGCGAGCGG</td><td>1000</td><td></td><td></td><td></td><td></td><td></td><td></td><td></td><td></td><td></td></tr>
<tr><td>Oligo fwd</td><td>CACCAAATTGCTATAGCCCGCGAG</td><td></td><td></td><td></td><td></td><td></td><td></td><td></td><td></td><td></td><td></td></tr>
<tr><td>Oligo rev</td><td>AAACCTCGCGGGCTATAGCAATTT</td><td></td><td></td><td></td><td></td><td></td><td></td><td></td><td></td><td></td><td></td></tr>
<tr><td>Chromosome</td><td>start</td><td>end</td><td>strand</td><td>MM</td><td>target_seq</td><td>PAM</td><td>alignment</td><td>distance</td><td>position</td><td>gene name</td><td>gene id</td></tr>
<tr><td>GU305914.1</td><td>189819</td><td>189841</td><td>+</td><td>0</td><td>AAATTGCTATAGCCCGCGAG</td><td>CGG</td><td>||||||||[||||||||||||]PAM</td><td>NA</td><td>Intergenic</td><td>NA</td><td>NA</td></tr>
<tr><td></td><td></td><td></td><td></td><td></td><td></td><td></td><td></td><td></td><td></td><td></td><td></td></tr>
<tr><td>T14449</td><td>GCTATAGCCCGCGAGCGGCCTGG</td><td>1000</td><td></td><td></td><td></td><td></td><td></td><td></td><td></td><td></td><td></td></tr>
<tr><td>Oligo fwd</td><td>CACCGCTATAGCCCGCGAGCGGCC</td><td></td><td></td><td></td><td></td><td></td><td></td><td></td><td></td><td></td><td></td></tr>
<tr><td>Oligo rev</td><td>AAACGGCCGCTCGCGGGCTATAGC</td><td></td><td></td><td></td><td></td><td></td><td></td><td></td><td></td><td></td><td></td></tr>
<tr><td>Chromosome</td><td>start</td><td>end</td><td>strand</td><td>MM</td><td>target_seq</td><td>PAM</td><td>alignment</td><td>distance</td><td>position</td><td>gene name</td><td>gene id</td></tr>
<tr><td>GU305914.1</td><td>189824</td><td>189846</td><td>+</td><td>0</td><td>GCTATAGCCCGCGAGCGGCC</td><td>TGG</td><td>||||||||[||||||||||||]PAM</td><td>NA</td><td>Intergenic</td><td>NA</td><td>NA</td></tr>
</table>

T14450 AGAGAAATCGTGAACAGCAGCGG 1000
Oligo fwd CACCAGAGAAATCGTGAACAGCAG
Oligo rev AAACCTGCTGTTCACGATTTCTCT
Chromosome start end strand MM target_seq PAM alignment distance position gene name gene id
GU305914.1 189864 189886 + 0 AGAGAAATCGTGAACAGCAG CGG ||||||||[||||||||||||]PAM NA Intergenic NA NA

T14451 GAAATCGTGAACAGCAGCGGTGG 1000
Oligo fwd CACCGAAATCGTGAACAGCAGCGG
Oligo rev AAACCCGCTGCTGTTCACGATTTC
Chromosome start end strand MM target_seq PAM alignment distance position gene name gene id
GU305914.1 189867 189889 + 0 GAAATCGTGAACAGCAGCGG TGG ||||||||[||||||||||||]PAM NA Intergenic NA NA

T14452 GGTGGTGACATTCTTCCTGTCGG 1000
Oligo fwd CACCGGTGGTGACATTCTTCCTGT
Oligo rev AAACACAGGAAGAATGTCACCACC
Chromosome start end strand MM target_seq PAM alignment distance position gene name gene id
GU305914.1 189885 189907 + 0 GGTGGTGACATTCTTCCTGT CGG ||||||||[||||||||||||]PAM NA Intergenic NA NA

T14453 GTGGTGACATTCTTCCTGTCGGG 1000
Oligo fwd CACCGTGGTGACATTCTTCCTGTC
Oligo rev AAACGACAGGAAGAATGTCACCAC
Chromosome start end strand MM target_seq PAM alignment distance position gene name gene id
GU305914.1 189886 189908 + 0 GTGGTGACATTCTTCCTGTC GGG ||||||||[||||||||||||]PAM NA Intergenic NA NA

T14454 GGTGCGAAAGTGTGTAGATGCGG 1000
Oligo fwd CACCGGTGCGAAAGTGTGTAGATG
Oligo rev AAACCATCTACACACTTTCGCACC
Chromosome start end strand MM target_seq PAM alignment distance position gene name gene id
GU305914.1 189907 189929 + 0 GGTGCGAAAGTGTGTAGATG CGG ||||||||[||||||||||||]PAM NA Intergenic NA NA

T14455 GTGCGAAAGTGTGTAGATGCGGG 1000
Oligo fwd CACCGTGCGAAAGTGTGTAGATGC
Oligo rev AAACGCATCTACACACTTTCGCAC
Chromosome start end strand MM target_seq PAM alignment distance position gene name gene id
GU305914.1 189908 189930 + 0 GTGCGAAAGTGTGTAGATGC GGG ||||||||[||||||||||||]PAM NA Intergenic NA NA

T14456 CGAAAGTGTGTAGATGCGGGAGG 1000
Oligo fwd CACCCGAAAGTGTGTAGATGCGGG
Oligo rev AAACCCCGCATCTACACACTTTCG
Chromosome start end strand MM target_seq PAM alignment distance position gene name gene id
GU305914.1 189911 189933 + 0 CGAAAGTGTGTAGATGCGGG AGG ||||||||[||||||||||||]PAM NA Intergenic NA NA

T14457 GATATCTACTCTTTATACACCGG 1000
Oligo fwd CACCGATATCTACTCTTTATACAC
Oligo rev AAACGTGTATAAAGAGTAGATATC
Chromosome start end strand MM target_seq PAM alignment distance position gene name gene id
GU305914.1 189936 189958 + 0 GATATCTACTCTTTATACAC CGG ||||||||[||||||||||||]PAM NA Intergenic NA NA

T14458 ATATCTACTCTTTATACACCGGG 1000
Oligo fwd CACCATATCTACTCTTTATACACC
Oligo rev AAACGGTGTATAAAGAGTAGATAT
Chromosome start end strand MM target_seq PAM alignment distance position gene name gene id
GU305914.1 189937 189959 + 0 ATATCTACTCTTTATACACC GGG ||||||||[||||||||||||]PAM NA Intergenic NA NA

T14459 TACTCTTTATACACCGGGTTTGG 1000
Oligo fwd CACCTACTCTTTATACACCGGGTT
Oligo rev AAACAACCCGGTGTATAAAGAGTA
Chromosome start end strand MM target_seq PAM alignment distance position gene name gene id
GU305914.1 189942 189964 + 0 TACTCTTTATACACCGGGTT TGG ||||||||[||||||||||||]PAM NA Intergenic NA NA

T14460 ACTCTTTATACACCGGGTTTGGG 1000
Oligo fwd CACCACTCTTTATACACCGGGTTT
Oligo rev AAACAAACCCGGTGTATAAAGAGT
Chromosome start end strand MM target_seq PAM alignment distance position gene name gene id
GU305914.1 189943 189965 + 0 ACTCTTTATACACCGGGTTT GGG ||||||||[||||||||||||]PAM NA Intergenic NA NA

T14461 CTCTTTATACACCGGGTTTGGGG 1000
Oligo fwd CACCCTCTTTATACACCGGGTTTG
Oligo rev AAACCAAACCCGGTGTATAAAGAG
Chromosome start end strand MM target_seq PAM alignment distance position gene name gene id
GU305914.1 189944 189966 + 0 CTCTTTATACACCGGGTTTG GGG ||||||||[||||||||||||]PAM NA Intergenic NA NA

T14462 TCTTTATACACCGGGTTTGGGGG 1000
Oligo fwd CACCTCTTTATACACCGGGTTTGG
Oligo rev AAACCCAAACCCGGTGTATAAAGA
Chromosome start end strand MM target_seq PAM alignment distance position gene name gene id
GU305914.1 189945 189967 + 0 TCTTTATACACCGGGTTTGG GGG ||||||||[||||||||||||]PAM NA Intergenic NA NA

T14463 TATACACCGGGTTTGGGGGATGG 1000
Oligo fwd CACCTATACACCGGGTTTGGGGGA
Oligo rev AAACTCCCCCAAACCCGGTGTATA
Chromosome start end strand MM target_seq PAM alignment distance position gene name gene id
GU305914.1 189949 189971 + 0 TATACACCGGGTTTGGGGGA TGG ||||||||[||||||||||||]PAM NA Intergenic NA NA

T14464 CTCTCGTAAATCTCGTCCAGCGG 1000
Oligo fwd CACCCTCTCGTAAATCTCGTCCAG
Oligo rev AAACCTGGACGAGATTTACGAGAG
Chromosome start end strand MM target_seq PAM alignment distance position gene name gene id
GU305914.1 190005 190027 + 0 CTCTCGTAAATCTCGTCCAG CGG ||||||||[||||||||||||]PAM NA Intergenic NA NA

| | | | | | | | | | | | |
|---|---|---|---|---|---|---|---|---|---|---|---|
| T14465 | TCTCGTAAATCTCGTCCAGCGGG | 1000 | | | | | | | | | |
| Oligo fwd | CACCTCTCGTAAATCTCGTCCAGC | | | | | | | | | | |
| Oligo rev | AAACGCTGGACGAGATTTACGAGA | | | | | | | | | | |
| Chromosome | start | end | strand | MM | target_seq | PAM | alignment | distance | position | gene name | gene id |
| GU305914.1 | 190006 | 190028 | + | 0 | TCTCGTAAATCTCGTCCAGC | GGG | \|\|\|\|\|\|\|\|[\|\|\|\|\|\|\|\|\|\|\|\|]PAM | NA | Intergenic | NA | NA |
| GU305914.1 | 83915 | 83937 | + | 4 | TCTCGTTGTTCTCGCCCAGC | TGG | \|\|\|\|\|\|--[-\|\|\|\|\|-\|\|\|\|\|]PAM | NA | Intergenic | NA | NA |

| | | | | | | | | | | | |
|---|---|---|---|---|---|---|---|---|---|---|---|
| T14466 | GTAAATCTCGTCCAGCGGGAAGG | 1000 | | | | | | | | | |
| Oligo fwd | CACCGTAAATCTCGTCCAGCGGGA | | | | | | | | | | |
| Oligo rev | AAACTCCCGCTGGACGAGATTTAC | | | | | | | | | | |
| Chromosome | start | end | strand | MM | target_seq | PAM | alignment | distance | position | gene name | gene id |
| GU305914.1 | 190010 | 190032 | + | 0 | GTAAATCTCGTCCAGCGGGA | AGG | \|\|\|\|\|\|\|\|[\|\|\|\|\|\|\|\|\|\|\|\|]PAM | NA | Intergenic | NA | NA |

| | | | | | | | | | | | |
|---|---|---|---|---|---|---|---|---|---|---|---|
| T14467 | TCGTCCAGCGGGAAGGAGTGCGG | 1000 | | | | | | | | | |
| Oligo fwd | CACCTCGTCCAGCGGGAAGGAGTG | | | | | | | | | | |
| Oligo rev | AAACCACTCCTTCCCGCTGGACGA | | | | | | | | | | |
| Chromosome | start | end | strand | MM | target_seq | PAM | alignment | distance | position | gene name | gene id |
| GU305914.1 | 190017 | 190039 | + | 0 | TCGTCCAGCGGGAAGGAGTG | CGG | \|\|\|\|\|\|\|\|[\|\|\|\|\|\|\|\|\|\|\|\|]PAM | NA | Intergenic | NA | NA |

| | | | | | | | | | | | |
|---|---|---|---|---|---|---|---|---|---|---|---|
| T14468 | CGTCCAGCGGGAAGGAGTGCGGG | 1000 | | | | | | | | | |
| Oligo fwd | CACCCGTCCAGCGGGAAGGAGTGC | | | | | | | | | | |
| Oligo rev | AAACGCACTCCTTCCCGCTGGACG | | | | | | | | | | |
| Chromosome | start | end | strand | MM | target_seq | PAM | alignment | distance | position | gene name | gene id |
| GU305914.1 | 190018 | 190040 | + | 0 | CGTCCAGCGGGAAGGAGTGC | GGG | \|\|\|\|\|\|\|\|[\|\|\|\|\|\|\|\|\|\|\|\|]PAM | NA | Intergenic | NA | NA |

| | | | | | | | | | | | |
|---|---|---|---|---|---|---|---|---|---|---|---|
| T14469 | CGGGCTCATGTCTTCGTACCCGG | 1000 | | | | | | | | | |
| Oligo fwd | CACCCGGGCTCATGTCTTCGTACC | | | | | | | | | | |
| Oligo rev | AAACGGTACGAAGACATGAGCCCG | | | | | | | | | | |
| Chromosome | start | end | strand | MM | target_seq | PAM | alignment | distance | position | gene name | gene id |
| GU305914.1 | 190037 | 190059 | + | 0 | CGGGCTCATGTCTTCGTACC | CGG | \|\|\|\|\|\|\|\|[\|\|\|\|\|\|\|\|\|\|\|\|]PAM | NA | Intergenic | NA | NA |

| | | | | | | | | | | | |
|---|---|---|---|---|---|---|---|---|---|---|---|
| T14470 | GCTCATGTCTTCGTACCCGGCGG | 1000 | | | | | | | | | |
| Oligo fwd | CACCGCTCATGTCTTCGTACCCGG | | | | | | | | | | |
| Oligo rev | AAACCCGGGTACGAAGACATGAGC | | | | | | | | | | |
| Chromosome | start | end | strand | MM | target_seq | PAM | alignment | distance | position | gene name | gene id |
| GU305914.1 | 190040 | 190062 | + | 0 | GCTCATGTCTTCGTACCCGG | CGG | \|\|\|\|\|\|\|\|[\|\|\|\|\|\|\|\|\|\|\|\|]PAM | NA | Intergenic | NA | NA |

| | | | | | | | | | | | |
|---|---|---|---|---|---|---|---|---|---|---|---|
| T14471 | CTCATGTCTTCGTACCCGGCGGG | 1000 | | | | | | | | | |
| Oligo fwd | CACCCTCATGTCTTCGTACCCGGC | | | | | | | | | | |
| Oligo rev | AAACGCCGGGTACGAAGACATGAG | | | | | | | | | | |
| Chromosome | start | end | strand | MM | target_seq | PAM | alignment | distance | position | gene name | gene id |
| GU305914.1 | 190041 | 190063 | + | 0 | CTCATGTCTTCGTACCCGGC | GGG | \|\|\|\|\|\|\|\|[\|\|\|\|\|\|\|\|\|\|\|\|]PAM | NA | Intergenic | NA | NA |

| | | | | | | | | | | | |
|---|---|---|---|---|---|---|---|---|---|---|---|
| T14472 | TCATGTCTTCGTACCCGGCGGGG | 1000 | | | | | | | | | |
| Oligo fwd | CACCTCATGTCTTCGTACCCGGCG | | | | | | | | | | |
| Oligo rev | AAACCGCCGGGTACGAAGACATGA | | | | | | | | | | |
| Chromosome | start | end | strand | MM | target_seq | PAM | alignment | distance | position | gene name | gene id |
| GU305914.1 | 190042 | 190064 | + | 0 | TCATGTCTTCGTACCCGGCG | GGG | \|\|\|\|\|\|\|\|[\|\|\|\|\|\|\|\|\|\|\|\|]PAM | NA | Intergenic | NA | NA |

| | | | | | | | | | | | |
|---|---|---|---|---|---|---|---|---|---|---|---|
| T14473 | GCGGGGCGCCCCAGAAACCCCGG | 1000 | | | | | | | | | |
| Oligo fwd | CACCGCGGGGCGCCCCAGAAACCC | | | | | | | | | | |
| Oligo rev | AAACGGGTTTCTGGGGCGCCCCGC | | | | | | | | | | |
| Chromosome | start | end | strand | MM | target_seq | PAM | alignment | distance | position | gene name | gene id |
| GU305914.1 | 190059 | 190081 | + | 0 | GCGGGGCGCCCCAGAAACCC | CGG | \|\|\|\|\|\|\|\|[\|\|\|\|\|\|\|\|\|\|\|\|]PAM | NA | Intergenic | NA | NA |

| | | | | | | | | | | | |
|---|---|---|---|---|---|---|---|---|---|---|---|
| T14474 | CGGGGCGCCCCAGAAACCCCGGG | 1000 | | | | | | | | | |
| Oligo fwd | CACCCGGGGCGCCCCAGAAACCCC | | | | | | | | | | |
| Oligo rev | AAACGGGGTTTCTGGGGCGCCCCG | | | | | | | | | | |
| Chromosome | start | end | strand | MM | target_seq | PAM | alignment | distance | position | gene name | gene id |
| GU305914.1 | 190060 | 190082 | + | 0 | CGGGGCGCCCCAGAAACCCC | GGG | \|\|\|\|\|\|\|\|[\|\|\|\|\|\|\|\|\|\|\|\|]PAM | NA | Intergenic | NA | NA |

| | | | | | | | | | | | |
|---|---|---|---|---|---|---|---|---|---|---|---|
| T14475 | GGGGCGCCCCAGAAACCCCGGGG | 1000 | | | | | | | | | |
| Oligo fwd | CACCGGGGCGCCCCAGAAACCCCG | | | | | | | | | | |
| Oligo rev | AAACCGGGGTTTCTGGGGCGCCCC | | | | | | | | | | |
| Chromosome | start | end | strand | MM | target_seq | PAM | alignment | distance | position | gene name | gene id |
| GU305914.1 | 190061 | 190083 | + | 0 | GGGGCGCCCCAGAAACCCCG | GGG | \|\|\|\|\|\|\|\|[\|\|\|\|\|\|\|\|\|\|\|\|]PAM | NA | Intergenic | NA | NA |

| | | | | | | | | | | | |
|---|---|---|---|---|---|---|---|---|---|---|---|
| T14476 | GGGCGCCCCAGAAACCCCGGGGG | 1000 | | | | | | | | | |
| Oligo fwd | CACCGGGCGCCCCAGAAACCCCGG | | | | | | | | | | |
| Oligo rev | AAACCCGGGGTTTCTGGGGCGCCC | | | | | | | | | | |
| Chromosome | start | end | strand | MM | target_seq | PAM | alignment | distance | position | gene name | gene id |
| GU305914.1 | 190062 | 190084 | + | 0 | GGGCGCCCCAGAAACCCCGG | GGG | \|\|\|\|\|\|\|\|[\|\|\|\|\|\|\|\|\|\|\|\|]PAM | NA | Intergenic | NA | NA |

| | | | | | | | | | | | |
|---|---|---|---|---|---|---|---|---|---|---|---|
| T14477 | GGCGCCCCAGAAACCCCGGGGGG | 1000 | | | | | | | | | |
| Oligo fwd | CACCGGCGCCCCAGAAACCCCGGG | | | | | | | | | | |
| Oligo rev | AAACCCCGGGGTTTCTGGGGCGCC | | | | | | | | | | |
| Chromosome | start | end | strand | MM | target_seq | PAM | alignment | distance | position | gene name | gene id |
| GU305914.1 | 190063 | 190085 | + | 0 | GGCGCCCCAGAAACCCCGGG | GGG | \|\|\|\|\|\|\|\|[\|\|\|\|\|\|\|\|\|\|\|\|]PAM | NA | Intergenic | NA | NA |

| | | | | | | | | | | | |
|---|---|---|---|---|---|---|---|---|---|---|---|
| T14478 | AAATAGACGCTACATTCGATCGG | 1000 | | | | | | | | | |
| Oligo fwd | CACCAAATAGACGCTACATTCGAT | | | | | | | | | | |
| Oligo rev | AAACATCGAATGTAGCGTCTATTT | | | | | | | | | | |
| Chromosome | start | end | strand | MM | target_seq | PAM | alignment | distance | position | gene name | gene id |
| GU305914.1 | 190119 | 190141 | + | 0 | AAATAGACGCTACATTCGAT | CGG | \|\|\|\|\|\|\|\|[\|\|\|\|\|\|\|\|\|\|\|\|]PAM | NA | Intergenic | NA | NA |

| | | |
|---|---|---|
| T14479 | AATAGACGCTACATTCGATCGGG | 1000 |
| Oligo fwd | CACCAATAGACGCTACATTCGATC | |
| Oligo rev | AAACGATCGAATGTAGCGTCTATT | |

| Chromosome | start | end | strand | MM | target_seq | PAM | alignment | distance | position | gene name | gene id |
|---|---|---|---|---|---|---|---|---|---|---|---|
| GU305914.1 | 190120 | 190142 | + | 0 | AATAGACGCTACATTCGATC | GGG | \|\|\|\|\|\|\|\|[\|\|\|\|\|\|\|\|\|\|\|\|]PAM | NA | Intergenic | NA | NA |
| T14480 | GGGACCTGCTCGTCGCCGCGAGG | 1000 | | | | | | | | | |
| Oligo fwd | CACCGGGACCTGCTCGTCGCCGCG | | | | | | | | | | |
| Oligo rev | AAACCGCGGCGACGAGCAGGTCCC | | | | | | | | | | |
| Chromosome | start | end | strand | MM | target_seq | PAM | alignment | distance | position | gene name | gene id |
| GU305914.1 | 190140 | 190162 | + | 0 | GGGACCTGCTCGTCGCCGCG | AGG | \|\|\|\|\|\|\|\|[\|\|\|\|\|\|\|\|\|\|\|\|]PAM | NA | Intergenic | NA | NA |
| GU305914.1 | 70168 | 70190 | - | 4 | CGGTGCTGCTCGTCGCCTCG | GGG | -\|\|--\|\|\|[\|\|\|\|\|\|\|\|\|-\|\|]PAM | NA | Intergenic | NA | NA |
| T14481 | GGACCTGCTCGTCGCCGCGAGGG | 1000 | | | | | | | | | |
| Oligo fwd | CACCGGACCTGCTCGTCGCCGCGA | | | | | | | | | | |
| Oligo rev | AAACTCGCGGCGACGAGCAGGTCC | | | | | | | | | | |
| Chromosome | start | end | strand | MM | target_seq | PAM | alignment | distance | position | gene name | gene id |
| GU305914.1 | 190141 | 190163 | + | 0 | GGACCTGCTCGTCGCCGCGA | GGG | \|\|\|\|\|\|\|\|[\|\|\|\|\|\|\|\|\|\|\|\|]PAM | NA | Intergenic | NA | NA |
| T14482 | GACCTGCTCGTCGCCGCGAGGGG | 1000 | | | | | | | | | |
| Oligo fwd | CACCGACCTGCTCGTCGCCGCGAG | | | | | | | | | | |
| Oligo rev | AAACCTCGCGGCGACGAGCAGGTC | | | | | | | | | | |
| Chromosome | start | end | strand | MM | target_seq | PAM | alignment | distance | position | gene name | gene id |
| GU305914.1 | 190142 | 190164 | + | 0 | GACCTGCTCGTCGCCGCGAG | GGG | \|\|\|\|\|\|\|\|[\|\|\|\|\|\|\|\|\|\|\|\|]PAM | NA | Intergenic | NA | NA |
| T14483 | CGCGAGGGGTTTGCTGCCGTCGG | 1000 | | | | | | | | | |
| Oligo fwd | CACCCGCGAGGGGTTTGCTGCCGT | | | | | | | | | | |
| Oligo rev | AAACACGGCAGCAAACCCCTCGCG | | | | | | | | | | |
| Chromosome | start | end | strand | MM | target_seq | PAM | alignment | distance | position | gene name | gene id |
| GU305914.1 | 190156 | 190178 | + | 0 | CGCGAGGGGTTTGCTGCCGT | CGG | \|\|\|\|\|\|\|\|[\|\|\|\|\|\|\|\|\|\|\|\|]PAM | NA | Intergenic | NA | NA |
| T14484 | TCGGCAGCGTCTCACGAGCGCGG | 1000 | | | | | | | | | |
| Oligo fwd | CACCTCGGCAGCGTCTCACGAGCG | | | | | | | | | | |
| Oligo rev | AAACCGCTCGTGAGACGCTGCCGA | | | | | | | | | | |
| Chromosome | start | end | strand | MM | target_seq | PAM | alignment | distance | position | gene name | gene id |
| GU305914.1 | 190175 | 190197 | + | 0 | TCGGCAGCGTCTCACGAGCG | CGG | \|\|\|\|\|\|\|\|[\|\|\|\|\|\|\|\|\|\|\|\|]PAM | NA | Intergenic | NA | NA |
| T14485 | GATCGCCACCACCAATATAGCGG | 1000 | | | | | | | | | |
| Oligo fwd | CACCGATCGCCACCACCAATATAG | | | | | | | | | | |
| Oligo rev | AAACCTATATTGGTGGTGGCGATC | | | | | | | | | | |
| Chromosome | start | end | strand | MM | target_seq | PAM | alignment | distance | position | gene name | gene id |
| GU305914.1 | 190199 | 190221 | + | 0 | GATCGCCACCACCAATATAG | CGG | \|\|\|\|\|\|\|\|[\|\|\|\|\|\|\|\|\|\|\|\|]PAM | NA | Intergenic | NA | NA |
| T14486 | CCGCCAACAACACCGAAGTCAGG | 1000 | | | | | | | | | |
| Oligo fwd | CACCCCGCCAACAACACCGAAGTC | | | | | | | | | | |
| Oligo rev | AAACGACTTCGGTGTTGTTGGCGG | | | | | | | | | | |
| Chromosome | start | end | strand | MM | target_seq | PAM | alignment | distance | position | gene name | gene id |
| GU305914.1 | 190222 | 190244 | + | 0 | CCGCCAACAACACCGAAGTC | AGG | \|\|\|\|\|\|\|\|[\|\|\|\|\|\|\|\|\|\|\|\|]PAM | NA | Intergenic | NA | NA |
| T14487 | CGCCAACAACACCGAAGTCAGGG | 1000 | | | | | | | | | |
| Oligo fwd | CACCCGCCAACAACACCGAAGTCA | | | | | | | | | | |
| Oligo rev | AAACTGACTTCGGTGTTGTTGGCG | | | | | | | | | | |
| Chromosome | start | end | strand | MM | target_seq | PAM | alignment | distance | position | gene name | gene id |
| GU305914.1 | 190223 | 190245 | + | 0 | CGCCAACAACACCGAAGTCA | GGG | \|\|\|\|\|\|\|\|[\|\|\|\|\|\|\|\|\|\|\|\|]PAM | NA | Intergenic | NA | NA |
| T14488 | CAACAACACCGAAGTCAGGGCGG | 1000 | | | | | | | | | |
| Oligo fwd | CACCCAACAACACCGAAGTCAGGG | | | | | | | | | | |
| Oligo rev | AAACCCCTGACTTCGGTGTTGTTG | | | | | | | | | | |
| Chromosome | start | end | strand | MM | target_seq | PAM | alignment | distance | position | gene name | gene id |
| GU305914.1 | 190226 | 190248 | + | 0 | CAACAACACCGAAGTCAGGG | CGG | \|\|\|\|\|\|\|\|[\|\|\|\|\|\|\|\|\|\|\|\|]PAM | NA | Intergenic | NA | NA |
| T14489 | CACCGAAGTCAGGGCGGCGCAGG | 1000 | | | | | | | | | |
| Oligo fwd | CACCCACCGAAGTCAGGGCGGCGC | | | | | | | | | | |
| Oligo rev | AAACGCGCCGCCCTGACTTCGGTG | | | | | | | | | | |
| Chromosome | start | end | strand | MM | target_seq | PAM | alignment | distance | position | gene name | gene id |
| GU305914.1 | 190232 | 190254 | + | 0 | CACCGAAGTCAGGGCGGCGC | AGG | \|\|\|\|\|\|\|\|[\|\|\|\|\|\|\|\|\|\|\|\|]PAM | NA | Intergenic | NA | NA |
| T14490 | GGCGGCGCAGGTCAGCACGATGG | 1000 | | | | | | | | | |
| Oligo fwd | CACCGGCGGCGCAGGTCAGCACGA | | | | | | | | | | |
| Oligo rev | AAACTCGTGCTGACCTGCGCCGCC | | | | | | | | | | |
| Chromosome | start | end | strand | MM | target_seq | PAM | alignment | distance | position | gene name | gene id |
| GU305914.1 | 190244 | 190266 | + | 0 | GGCGGCGCAGGTCAGCACGA | TGG | \|\|\|\|\|\|\|\|[\|\|\|\|\|\|\|\|\|\|\|\|]PAM | NA | Intergenic | NA | NA |
| T14491 | GGTCAGCACGATGGTCTGAGCGG | 1000 | | | | | | | | | |
| Oligo fwd | CACCGGTCAGCACGATGGTCTGAG | | | | | | | | | | |
| Oligo rev | AAACCTCAGACCATCGTGCTGACC | | | | | | | | | | |
| Chromosome | start | end | strand | MM | target_seq | PAM | alignment | distance | position | gene name | gene id |
| GU305914.1 | 190253 | 190275 | + | 0 | GGTCAGCACGATGGTCTGAG | CGG | \|\|\|\|\|\|\|\|[\|\|\|\|\|\|\|\|\|\|\|\|]PAM | NA | Intergenic | NA | NA |
| T14492 | CACGATGGTCTGAGCGGACACGG | 1000 | | | | | | | | | |
| Oligo fwd | CACCCACGATGGTCTGAGCGGACA | | | | | | | | | | |
| Oligo rev | AAACTGTCCGCTCAGACCATCGTG | | | | | | | | | | |
| Chromosome | start | end | strand | MM | target_seq | PAM | alignment | distance | position | gene name | gene id |
| GU305914.1 | 190259 | 190281 | + | 0 | CACGATGGTCTGAGCGGACA | CGG | \|\|\|\|\|\|\|\|[\|\|\|\|\|\|\|\|\|\|\|\|]PAM | NA | Intergenic | NA | NA |
| T14493 | CTGAGCGGACACGGTCTCAGAGG | 1000 | | | | | | | | | |
| Oligo fwd | CACCCTGAGCGGACACGGTCTCAG | | | | | | | | | | |
| Oligo rev | AAACCTGAGACCGTGTCCGCTCAG | | | | | | | | | | |
| Chromosome | start | end | strand | MM | target_seq | PAM | alignment | distance | position | gene name | gene id |
| GU305914.1 | 190268 | 190290 | + | 0 | CTGAGCGGACACGGTCTCAG | AGG | \|\|\|\|\|\|\|\|[\|\|\|\|\|\|\|\|\|\|\|\|]PAM | NA | Intergenic | NA | NA |
| T14494 | TGAGCGGACACGGTCTCAGAGGG | 1000 | | | | | | | | | |

Oligo fwd CACCTGAGCGGACACGGTCTCAGA
Oligo rev AAACTCTGAGACCGTGTCCGCTCA
Chromosome start end strand MM target_seq PAM alignment distance position gene name gene id
GU305914.1 190269 190291 + 0 TGAGCGGACACGGTCTCAGA GGG ||||||||[||||||||||||]PAM NA Intergenic NA NA

T14495 GGACACGGTCTCAGAGGGAGAGG 1000
Oligo fwd CACCGGACACGGTCTCAGAGGGAG
Oligo rev AAACCTCCCTCTGAGACCGTGTCC
Chromosome start end strand MM target_seq PAM alignment distance position gene name gene id
GU305914.1 190274 190296 + 0 GGACACGGTCTCAGAGGGAG AGG ||||||||[||||||||||||]PAM NA Intergenic NA NA

T14496 GACACGGTCTCAGAGGGAGAGGG 1000
Oligo fwd CACCGACACGGTCTCAGAGGGAGA
Oligo rev AAACTCTCCCTCTGAGACCGTGTC
Chromosome start end strand MM target_seq PAM alignment distance position gene name gene id
GU305914.1 190275 190297 + 0 GACACGGTCTCAGAGGGAGA GGG ||||||||[||||||||||||]PAM NA Intergenic NA NA

T14497 AGAGGGAGAGGGACATTCTGTGG 1000
Oligo fwd CACCAGAGGGAGAGGGACATTCTG
Oligo rev AAACCAGAATGTCCCTCTCCCTCT
Chromosome start end strand MM target_seq PAM alignment distance position gene name gene id
GU305914.1 190286 190308 + 0 AGAGGGAGAGGGACATTCTG TGG ||||||||[||||||||||||]PAM NA Intergenic NA NA

T14498 TTCTGTCCGACAATGAGCCGCGG 1000
Oligo fwd CACCTTCTGTCCGACAATGAGCCG
Oligo rev AAACCGGCTCATTGTCGGACAGAA
Chromosome start end strand MM target_seq PAM alignment distance position gene name gene id
GU305914.1 190366 190388 + 0 TTCTGTCCGACAATGAGCCG CGG ||||||||[||||||||||||]PAM NA Intergenic NA NA

T14499 CCGACAATGAGCCGCGGCACCGG 1000
Oligo fwd CACCCCGACAATGAGCCGCGGCAC
Oligo rev AAACGTGCCGCGGCTCATTGTCGG
Chromosome start end strand MM target_seq PAM alignment distance position gene name gene id
GU305914.1 190372 190394 + 0 CCGACAATGAGCCGCGGCAC CGG ||||||||[||||||||||||]PAM NA Intergenic NA NA

T14500 ACAATGAGCCGCGGCACCGGAGG 1000
Oligo fwd CACCACAATGAGCCGCGGCACCGG
Oligo rev AAACCCGGTGCCGCGGCTCATTGT
Chromosome start end strand MM target_seq PAM alignment distance position gene name gene id
GU305914.1 190375 190397 + 0 ACAATGAGCCGCGGCACCGG AGG ||||||||[||||||||||||]PAM NA Intergenic NA NA

T14501 CGCGAAGTTTTTATTCCCCCCGG 1000
Oligo fwd CACCCGCGAAGTTTTTATTCCCCC
Oligo rev AAACGGGGGAATAAAAACTTCGCG
Chromosome start end strand MM target_seq PAM alignment distance position gene name gene id
GU305914.1 190400 190422 + 0 CGCGAAGTTTTTATTCCCCC CGG ||||||||[||||||||||||]PAM NA Intergenic NA NA

T14502 TTTTATTCCCCCCGGCGCTCCGG 1000
Oligo fwd CACCTTTTATTCCCCCCGGCGCTC
Oligo rev AAACGAGCGCCGGGGGGAATAAAA
Chromosome start end strand MM target_seq PAM alignment distance position gene name gene id
GU305914.1 190408 190430 + 0 TTTTATTCCCCCCGGCGCTC CGG ||||||||[||||||||||||]PAM NA Intergenic NA NA

T14503 CTCTGATACAGCTTGTAACGCGG 1000
Oligo fwd CACCCTCTGATACAGCTTGTAACG
Oligo rev AAACCGTTACAAGCTGTATCAGAG
Chromosome start end strand MM target_seq PAM alignment distance position gene name gene id
GU305914.1 190441 190463 + 0 CTCTGATACAGCTTGTAACG CGG ||||||||[||||||||||||]PAM NA Intergenic NA NA

T14504 TCTGATACAGCTTGTAACGCGGG 1000
Oligo fwd CACCTCTGATACAGCTTGTAACGC
Oligo rev AAACGCGTTACAAGCTGTATCAGA
Chromosome start end strand MM target_seq PAM alignment distance position gene name gene id
GU305914.1 190442 190464 + 0 TCTGATACAGCTTGTAACGC GGG ||||||||[||||||||||||]PAM NA Intergenic NA NA

T14505 ATGATCAGATCGATCTCCCTCGG 1000
Oligo fwd CACCATGATCAGATCGATCTCCCT
Oligo rev AAACAGGGAGATCGATCTGATCAT
Chromosome start end strand MM target_seq PAM alignment distance position gene name gene id
GU305914.1 190474 190496 + 0 ATGATCAGATCGATCTCCCT CGG ||||||||[||||||||||||]PAM NA Intergenic NA NA

T14506 ATCAGATCGATCTCCCTCGGTGG 1000
Oligo fwd CACCATCAGATCGATCTCCCTCGG
Oligo rev AAACCCGAGGGAGATCGATCTGAT
Chromosome start end strand MM target_seq PAM alignment distance position gene name gene id
GU305914.1 190477 190499 + 0 ATCAGATCGATCTCCCTCGG TGG ||||||||[||||||||||||]PAM NA Intergenic NA NA

T14507 TCGATCTCCCTCGGTGGACACGG 1000
Oligo fwd CACCTCGATCTCCCTCGGTGGACA
Oligo rev AAACTGTCCACCGAGGGAGATCGA
Chromosome start end strand MM target_seq PAM alignment distance position gene name gene id
GU305914.1 190483 190505 + 0 TCGATCTCCCTCGGTGGACA CGG ||||||||[||||||||||||]PAM NA Intergenic NA NA

T14508 AGCACTGTGACAGTGACGAGCGG 1000
Oligo fwd CACCAGCACTGTGACAGTGACGAG
Oligo rev AAACCTCGTCACTGTCACAGTGCT
Chromosome start end strand MM target_seq PAM alignment distance position gene name gene id
GU305914.1 190510 190532 + 0 AGCACTGTGACAGTGACGAG CGG ||||||||[||||||||||||]PAM NA Intergenic NA NA

T14509 GACAGTGACGAGCGGCCCGCCGG 1000
Oligo fwd CACCGACAGTGACGAGCGGCCCGC
Oligo rev AAACGCGGGCCGCTCGTCACTGTC
Chromosome start end strand MM target_seq PAM alignment distance position gene name gene id
GU305914.1 190518 190540 + 0 GACAGTGACGAGCGGCCCGC CGG ||||||||[||||||||||||]PAM NA Intergenic NA NA

T14510 ACAGTGACGAGCGGCCCGCCGGG 1000
Oligo fwd CACCACAGTGACGAGCGGCCCGCC
Oligo rev AAACGGCGGGCCGCTCGTCACTGT
Chromosome start end strand MM target_seq PAM alignment distance position gene name gene id
GU305914.1 190519 190541 + 0 ACAGTGACGAGCGGCCCGCC GGG ||||||||[||||||||||||]PAM NA Intergenic NA NA

T14511 CGGGTCCTAAGTAACCCGATAGG 1000
Oligo fwd CACCCGGGTCCTAAGTAACCCGAT
Oligo rev AAACATCGGGTTACTTAGGACCCG
Chromosome start end strand MM target_seq PAM alignment distance position gene name gene id
GU305914.1 190538 190560 + 0 CGGGTCCTAAGTAACCCGAT AGG ||||||||[||||||||||||]PAM NA Intergenic NA NA

T14512 TACTCGATATACGCGCCCCCTGG 1000
Oligo fwd CACCTACTCGATATACGCGCCCCC
Oligo rev AAACGGGGGCGCGTATATCGAGTA
Chromosome start end strand MM target_seq PAM alignment distance position gene name gene id
GU305914.1 190576 190598 + 0 TACTCGATATACGCGCCCCC TGG ||||||||[||||||||||||]PAM NA Intergenic NA NA

T14513 ATATACGCGCCCCCTGGCTCTGG 1000
Oligo fwd CACCATATACGCGCCCCCTGGCTC
Oligo rev AAACGAGCCAGGGGGCGCGTATAT
Chromosome start end strand MM target_seq PAM alignment distance position gene name gene id
GU305914.1 190582 190604 + 0 ATATACGCGCCCCCTGGCTC TGG ||||||||[||||||||||||]PAM NA Intergenic NA NA

T14514 ACGCGCCCCCTGGCTCTGGTAGG 1000
Oligo fwd CACCACGCGCCCCCTGGCTCTGGT
Oligo rev AAACACCAGAGCCAGGGGGCGCGT
Chromosome start end strand MM target_seq PAM alignment distance position gene name gene id
GU305914.1 190586 190608 + 0 ACGCGCCCCCTGGCTCTGGT AGG ||||||||[||||||||||||]PAM NA Intergenic NA NA

T14515 CGCCCCCTGGCTCTGGTAGGTGG 1000
Oligo fwd CACCCGCCCCCTGGCTCTGGTAGG
Oligo rev AAACCCTACCAGAGCCAGGGGGCG
Chromosome start end strand MM target_seq PAM alignment distance position gene name gene id
GU305914.1 190589 190611 + 0 CGCCCCCTGGCTCTGGTAGG TGG ||||||||[||||||||||||]PAM NA Intergenic NA NA

T14516 TTCGCTTCCATTCACACGAACGG 1000
Oligo fwd CACCTTCGCTTCCATTCACACGAA
Oligo rev AAACTTCGTGTGAATGGAAGCGAA
Chromosome start end strand MM target_seq PAM alignment distance position gene name gene id
GU305914.1 190620 190642 + 0 TTCGCTTCCATTCACACGAA CGG ||||||||[||||||||||||]PAM NA Intergenic NA NA

T14517 TCCATTCACACGAACGGTCGCGG 1000
Oligo fwd CACCTCCATTCACACGAACGGTCG
Oligo rev AAACCGACCGTTCGTGTGAATGGA
Chromosome start end strand MM target_seq PAM alignment distance position gene name gene id
GU305914.1 190626 190648 + 0 TCCATTCACACGAACGGTCG CGG ||||||||[||||||||||||]PAM NA Intergenic NA NA

T14518 GGTCGCGGTCAAGTATCCTTTGG 1000
Oligo fwd CACCGGTCGCGGTCAAGTATCCTT
Oligo rev AAACAAGGATACTTGACCGCGACC
Chromosome start end strand MM target_seq PAM alignment distance position gene name gene id
GU305914.1 190641 190663 + 0 GGTCGCGGTCAAGTATCCTT TGG ||||||||[||||||||||||]PAM NA Intergenic NA NA

T14519 GTCAGCCACCCGTAGCCTTCAGG 1000
Oligo fwd CACCGTCAGCCACCCGTAGCCTTC
Oligo rev AAACGAAGGCTACGGGTGGCTGAC
Chromosome start end strand MM target_seq PAM alignment distance position gene name gene id
GU305914.1 190663 190685 + 0 GTCAGCCACCCGTAGCCTTC AGG ||||||||[||||||||||||]PAM NA Intergenic NA NA

T14520 TTCAGGCTCGCAGCTGTATCTGG 1000
Oligo fwd CACCTTCAGGCTCGCAGCTGTATC
Oligo rev AAACGATACAGCTGCGAGCCTGAA
Chromosome start end strand MM target_seq PAM alignment distance position gene name gene id
GU305914.1 190680 190702 + 0 TTCAGGCTCGCAGCTGTATC TGG ||||||||[||||||||||||]PAM NA Intergenic NA NA

T14521 ACCTGATTGTGTGTTGATGTAGG 1000
Oligo fwd CACCACCTGATTGTGTGTTGATGT
Oligo rev AAACACATCAACACACAATCAGGT
Chromosome start end strand MM target_seq PAM alignment distance position gene name gene id
GU305914.1 190776 190798 + 0 ACCTGATTGTGTGTTGATGT AGG ||||||||[||||||||||||]PAM NA Intergenic NA NA

T14522 CGAATGACCCCGTTGAATCGTGG 1000
Oligo fwd CACCCGAATGACCCCGTTGAATCG
Oligo rev AAACCGATTCAACGGGGTCATTCG
Chromosome start end strand MM target_seq PAM alignment distance position gene name gene id
GU305914.1 190802 190824 + 0 CGAATGACCCCGTTGAATCG TGG ||||||||[||||||||||||]PAM NA Intergenic NA NA

T14523 CCCCGTTGAATCGTGGTTTATGG 1000
Oligo fwd CACCCCCCGTTGAATCGTGGTTTA
Oligo rev AAACTAAACCACGATTCAACGGGG
Chromosome start end strand MM target_seq PAM alignment distance position gene name gene id
GU305914.1 190809 190831 + 0 CCCCGTTGAATCGTGGTTTA TGG ||||||||[||||||||||||]PAM NA Intergenic NA NA

| | | | | | | | | | | | |
|---|---|---|---|---|---|---|---|---|---|---|---|
| T14524 | TCTTCCATCTTTTTGTTGTCAGG | 1000 | | | | | | | | | |
| Oligo fwd | CACCTCTTCCATCTTTTTGTTGTC | | | | | | | | | | |
| Oligo rev | AAACGACAACAAAAAGATGGAAGA | | | | | | | | | | |
| Chromosome | start | end | strand | MM | target_seq | PAM | alignment | distance | position | gene name | gene id |
| GU305914.1 | 190855 | 190877 | + | 0 | TCTTCCATCTTTTTGTTGTC | AGG | \|\|\|\|\|\|\|\|[\|\|\|\|\|\|\|\|\|\|\|\|]PAM | NA | Intergenic | NA | NA |
| T14525 | AAATCTGAGTATCTGTATCATGG | 1000 | | | | | | | | | |
| Oligo fwd | CACCAAATCTGAGTATCTGTATCA | | | | | | | | | | |
| Oligo rev | AAACTGATACAGATACTCAGATTT | | | | | | | | | | |
| Chromosome | start | end | strand | MM | target_seq | PAM | alignment | distance | position | gene name | gene id |
| GU305914.1 | 190881 | 190903 | + | 0 | AAATCTGAGTATCTGTATCA | TGG | \|\|\|\|\|\|\|\|[\|\|\|\|\|\|\|\|\|\|\|\|]PAM | NA | Intergenic | NA | NA |
| T14526 | TATCTGTATCATGGCGTCCTCGG | 1000 | | | | | | | | | |
| Oligo fwd | CACCTATCTGTATCATGGCGTCCT | | | | | | | | | | |
| Oligo rev | AAACAGGACGCCATGATACAGATA | | | | | | | | | | |
| Chromosome | start | end | strand | MM | target_seq | PAM | alignment | distance | position | gene name | gene id |
| GU305914.1 | 190890 | 190912 | + | 0 | TATCTGTATCATGGCGTCCT | CGG | \|\|\|\|\|\|\|\|[\|\|\|\|\|\|\|\|\|\|\|\|]PAM | NA | Intergenic | NA | NA |
| T14527 | TGTCGTGAATGTAGTCCACGAGG | 1000 | | | | | | | | | |
| Oligo fwd | CACCTGTCGTGAATGTAGTCCACG | | | | | | | | | | |
| Oligo rev | AAACCGTGGACTACATTCACGACA | | | | | | | | | | |
| Chromosome | start | end | strand | MM | target_seq | PAM | alignment | distance | position | gene name | gene id |
| GU305914.1 | 190944 | 190966 | + | 0 | TGTCGTGAATGTAGTCCACG | AGG | \|\|\|\|\|\|\|\|[\|\|\|\|\|\|\|\|\|\|\|\|]PAM | NA | Intergenic | NA | NA |
| T14528 | TAGTCCACGAGGTCTCGAAGAGG | 1000 | | | | | | | | | |
| Oligo fwd | CACCTAGTCCACGAGGTCTCGAAG | | | | | | | | | | |
| Oligo rev | AAACCTTCGAGACCTCGTGGACTA | | | | | | | | | | |
| Chromosome | start | end | strand | MM | target_seq | PAM | alignment | distance | position | gene name | gene id |
| GU305914.1 | 190955 | 190977 | + | 0 | TAGTCCACGAGGTCTCGAAG | AGG | \|\|\|\|\|\|\|\|[\|\|\|\|\|\|\|\|\|\|\|\|]PAM | NA | Intergenic | NA | NA |
| T14529 | TCCTGTTCCACGTTACGTTTAGG | 1000 | | | | | | | | | |
| Oligo fwd | CACCTCCTGTTCCACGTTACGTTT | | | | | | | | | | |
| Oligo rev | AAACAAACGTAACGTGGAACAGGA | | | | | | | | | | |
| Chromosome | start | end | strand | MM | target_seq | PAM | alignment | distance | position | gene name | gene id |
| GU305914.1 | 190997 | 191019 | + | 0 | TCCTGTTCCACGTTACGTTT | AGG | \|\|\|\|\|\|\|\|[\|\|\|\|\|\|\|\|\|\|\|\|]PAM | NA | Intergenic | NA | NA |
| T14530 | CCTGTTCCACGTTACGTTTAGGG | 1000 | | | | | | | | | |
| Oligo fwd | CACCCCTGTTCCACGTTACGTTTA | | | | | | | | | | |
| Oligo rev | AAACTAAACGTAACGTGGAACAGG | | | | | | | | | | |
| Chromosome | start | end | strand | MM | target_seq | PAM | alignment | distance | position | gene name | gene id |
| GU305914.1 | 190998 | 191020 | + | 0 | CCTGTTCCACGTTACGTTTA | GGG | \|\|\|\|\|\|\|\|[\|\|\|\|\|\|\|\|\|\|\|\|]PAM | NA | Intergenic | NA | NA |
| T14531 | ACGAAAGCACCACTCGATGTAGG | 1000 | | | | | | | | | |
| Oligo fwd | CACCACGAAAGCACCACTCGATGT | | | | | | | | | | |
| Oligo rev | AAACACATCGAGTGGTGCTTTCGT | | | | | | | | | | |
| Chromosome | start | end | strand | MM | target_seq | PAM | alignment | distance | position | gene name | gene id |
| GU305914.1 | 191040 | 191062 | + | 0 | ACGAAAGCACCACTCGATGT | AGG | \|\|\|\|\|\|\|\|[\|\|\|\|\|\|\|\|\|\|\|\|]PAM | NA | Intergenic | NA | NA |
| T14532 | GATGTAGGTTTTGTTACGTGCGG | 1000 | | | | | | | | | |
| Oligo fwd | CACCGATGTAGGTTTTGTTACGTG | | | | | | | | | | |
| Oligo rev | AAACCACGTAACAAAACCTACATC | | | | | | | | | | |
| Chromosome | start | end | strand | MM | target_seq | PAM | alignment | distance | position | gene name | gene id |
| GU305914.1 | 191055 | 191077 | + | 0 | GATGTAGGTTTTGTTACGTG | CGG | \|\|\|\|\|\|\|\|[\|\|\|\|\|\|\|\|\|\|\|\|]PAM | NA | Intergenic | NA | NA |
| T14533 | GTAACCTCGCGTCTGCGAGACGG | 1000 | | | | | | | | | |
| Oligo fwd | CACCGTAACCTCGCGTCTGCGAGA | | | | | | | | | | |
| Oligo rev | AAACTCTCGCAGACGCGAGGTTAC | | | | | | | | | | |
| Chromosome | start | end | strand | MM | target_seq | PAM | alignment | distance | position | gene name | gene id |
| GU305914.1 | 191082 | 191104 | + | 0 | GTAACCTCGCGTCTGCGAGA | CGG | \|\|\|\|\|\|\|\|[\|\|\|\|\|\|\|\|\|\|\|\|]PAM | NA | Intergenic | NA | NA |
| T14534 | TCGCGTCTGCGAGACGGTAGTGG | 1000 | | | | | | | | | |
| Oligo fwd | CACCTCGCGTCTGCGAGACGGTAG | | | | | | | | | | |
| Oligo rev | AAACCTACCGTCTCGCAGACGCGA | | | | | | | | | | |
| Chromosome | start | end | strand | MM | target_seq | PAM | alignment | distance | position | gene name | gene id |
| GU305914.1 | 191088 | 191110 | + | 0 | TCGCGTCTGCGAGACGGTAG | TGG | \|\|\|\|\|\|\|\|[\|\|\|\|\|\|\|\|\|\|\|\|]PAM | NA | Intergenic | NA | NA |
| T14535 | AATAACCACGCCGCCGCCGTCGG | 1000 | | | | | | | | | |
| Oligo fwd | CACCAATAACCACGCCGCCGCCGT | | | | | | | | | | |
| Oligo rev | AAACACGGCGGCGGCGTGGTTATT | | | | | | | | | | |
| Chromosome | start | end | strand | MM | target_seq | PAM | alignment | distance | position | gene name | gene id |
| GU305914.1 | 191140 | 191162 | + | 0 | AATAACCACGCCGCCGCCGT | CGG | \|\|\|\|\|\|\|\|[\|\|\|\|\|\|\|\|\|\|\|\|]PAM | NA | Intergenic | NA | NA |
| T14536 | ACAGTTCTTGTAAATCTTCACGG | 1000 | | | | | | | | | |
| Oligo fwd | CACCACAGTTCTTGTAAATCTTCA | | | | | | | | | | |
| Oligo rev | AAACTGAAGATTTACAAGAACTGT | | | | | | | | | | |
| Chromosome | start | end | strand | MM | target_seq | PAM | alignment | distance | position | gene name | gene id |
| GU305914.1 | 191190 | 191212 | + | 0 | ACAGTTCTTGTAAATCTTCA | CGG | \|\|\|\|\|\|\|\|[\|\|\|\|\|\|\|\|\|\|\|\|]PAM | NA | Intergenic | NA | NA |
| T14537 | GTAAATCTTCACGGTCTATCAGG | 1000 | | | | | | | | | |
| Oligo fwd | CACCGTAAATCTTCACGGTCTATC | | | | | | | | | | |
| Oligo rev | AAACGATAGACCGTGAAGATTTAC | | | | | | | | | | |
| Chromosome | start | end | strand | MM | target_seq | PAM | alignment | distance | position | gene name | gene id |
| GU305914.1 | 191199 | 191221 | + | 0 | GTAAATCTTCACGGTCTATC | AGG | \|\|\|\|\|\|\|\|[\|\|\|\|\|\|\|\|\|\|\|\|]PAM | NA | Intergenic | NA | NA |
| T14538 | CTTCACGGTCTATCAGGCCTCGG | 1000 | | | | | | | | | |
| Oligo fwd | CACCCTTCACGGTCTATCAGGCCT | | | | | | | | | | |
| Oligo rev | AAACAGGCCTGATAGACCGTGAAG | | | | | | | | | | |
| Chromosome | start | end | strand | MM | target_seq | PAM | alignment | distance | position | gene name | gene id |

<table>
<tr><td>GU305914.1</td><td>191205</td><td>191227</td><td>+</td><td>0</td><td>CTTCACGGTCTATCAGGCCT</td><td>CGG</td><td>||||||||[||||||||||||]PAM</td><td>NA</td><td>Intergenic</td><td>NA</td><td>NA</td></tr>
<tr><td>T14539</td><td>TCCGATGAACATCGCGCACCAGG</td><td>1000</td><td></td><td></td><td></td><td></td><td></td><td></td><td></td><td></td><td></td></tr>
<tr><td>Oligo fwd</td><td>CACCTCCGATGAACATCGCGCACC</td><td></td><td></td><td></td><td></td><td></td><td></td><td></td><td></td><td></td><td></td></tr>
<tr><td>Oligo rev</td><td>AAACGGTGCGCGATGTTCATCGGA</td><td></td><td></td><td></td><td></td><td></td><td></td><td></td><td></td><td></td><td></td></tr>
<tr><td>Chromosome</td><td>start</td><td>end</td><td>strand</td><td>MM</td><td>target_seq</td><td>PAM</td><td>alignment</td><td>distance</td><td>position</td><td>gene name</td><td>gene id</td></tr>
<tr><td>GU305914.1</td><td>191236</td><td>191258</td><td>+</td><td>0</td><td>TCCGATGAACATCGCGCACC</td><td>AGG</td><td>||||||||[||||||||||||]PAM</td><td>NA</td><td>Intergenic</td><td>NA</td><td>NA</td></tr>
<tr><td>T14540</td><td>ATGAATGCCTTCTCCGTGTGCGG</td><td>1000</td><td></td><td></td><td></td><td></td><td></td><td></td><td></td><td></td><td></td></tr>
<tr><td>Oligo fwd</td><td>CACCATGAATGCCTTCTCCGTGTG</td><td></td><td></td><td></td><td></td><td></td><td></td><td></td><td></td><td></td><td></td></tr>
<tr><td>Oligo rev</td><td>AAACCACACGGAGAAGGCATTCAT</td><td></td><td></td><td></td><td></td><td></td><td></td><td></td><td></td><td></td><td></td></tr>
<tr><td>Chromosome</td><td>start</td><td>end</td><td>strand</td><td>MM</td><td>target_seq</td><td>PAM</td><td>alignment</td><td>distance</td><td>position</td><td>gene name</td><td>gene id</td></tr>
<tr><td>GU305914.1</td><td>191261</td><td>191283</td><td>+</td><td>0</td><td>ATGAATGCCTTCTCCGTGTG</td><td>CGG</td><td>||||||||[||||||||||||]PAM</td><td>NA</td><td>Intergenic</td><td>NA</td><td>NA</td></tr>
<tr><td>T14541</td><td>TGAATGCCTTCTCCGTGTGCGGG</td><td>1000</td><td></td><td></td><td></td><td></td><td></td><td></td><td></td><td></td><td></td></tr>
<tr><td>Oligo fwd</td><td>CACCTGAATGCCTTCTCCGTGTGC</td><td></td><td></td><td></td><td></td><td></td><td></td><td></td><td></td><td></td><td></td></tr>
<tr><td>Oligo rev</td><td>AAACGCACACGGAGAAGGCATTCA</td><td></td><td></td><td></td><td></td><td></td><td></td><td></td><td></td><td></td><td></td></tr>
<tr><td>Chromosome</td><td>start</td><td>end</td><td>strand</td><td>MM</td><td>target_seq</td><td>PAM</td><td>alignment</td><td>distance</td><td>position</td><td>gene name</td><td>gene id</td></tr>
<tr><td>GU305914.1</td><td>191262</td><td>191284</td><td>+</td><td>0</td><td>TGAATGCCTTCTCCGTGTGC</td><td>GGG</td><td>||||||||[||||||||||||]PAM</td><td>NA</td><td>Intergenic</td><td>NA</td><td>NA</td></tr>
<tr><td>T14542</td><td>GAATGCCTTCTCCGTGTGCGGGG</td><td>1000</td><td></td><td></td><td></td><td></td><td></td><td></td><td></td><td></td><td></td></tr>
<tr><td>Oligo fwd</td><td>CACCGAATGCCTTCTCCGTGTGCG</td><td></td><td></td><td></td><td></td><td></td><td></td><td></td><td></td><td></td><td></td></tr>
<tr><td>Oligo rev</td><td>AAACCGCACACGGAGAAGGCATTC</td><td></td><td></td><td></td><td></td><td></td><td></td><td></td><td></td><td></td><td></td></tr>
<tr><td>Chromosome</td><td>start</td><td>end</td><td>strand</td><td>MM</td><td>target_seq</td><td>PAM</td><td>alignment</td><td>distance</td><td>position</td><td>gene name</td><td>gene id</td></tr>
<tr><td>GU305914.1</td><td>191263</td><td>191285</td><td>+</td><td>0</td><td>GAATGCCTTCTCCGTGTGCG</td><td>GGG</td><td>||||||||[||||||||||||]PAM</td><td>NA</td><td>Intergenic</td><td>NA</td><td>NA</td></tr>
<tr><td>T14543</td><td>CGAAGTACTGTTCGCCCAGCAGG</td><td>1000</td><td></td><td></td><td></td><td></td><td></td><td></td><td></td><td></td><td></td></tr>
<tr><td>Oligo fwd</td><td>CACCCGAAGTACTGTTCGCCCAGC</td><td></td><td></td><td></td><td></td><td></td><td></td><td></td><td></td><td></td><td></td></tr>
<tr><td>Oligo rev</td><td>AAACGCTGGGCGAACAGTACTTCG</td><td></td><td></td><td></td><td></td><td></td><td></td><td></td><td></td><td></td><td></td></tr>
<tr><td>Chromosome</td><td>start</td><td>end</td><td>strand</td><td>MM</td><td>target_seq</td><td>PAM</td><td>alignment</td><td>distance</td><td>position</td><td>gene name</td><td>gene id</td></tr>
<tr><td>GU305914.1</td><td>191289</td><td>191311</td><td>+</td><td>0</td><td>CGAAGTACTGTTCGCCCAGC</td><td>AGG</td><td>||||||||[||||||||||||]PAM</td><td>NA</td><td>Intergenic</td><td>NA</td><td>NA</td></tr>
<tr><td>T14544</td><td>GAAGTACTGTTCGCCCAGCAGGG</td><td>1000</td><td></td><td></td><td></td><td></td><td></td><td></td><td></td><td></td><td></td></tr>
<tr><td>Oligo fwd</td><td>CACCGAAGTACTGTTCGCCCAGCA</td><td></td><td></td><td></td><td></td><td></td><td></td><td></td><td></td><td></td><td></td></tr>
<tr><td>Oligo rev</td><td>AAACTGCTGGGCGAACAGTACTTC</td><td></td><td></td><td></td><td></td><td></td><td></td><td></td><td></td><td></td><td></td></tr>
<tr><td>Chromosome</td><td>start</td><td>end</td><td>strand</td><td>MM</td><td>target_seq</td><td>PAM</td><td>alignment</td><td>distance</td><td>position</td><td>gene name</td><td>gene id</td></tr>
<tr><td>GU305914.1</td><td>191290</td><td>191312</td><td>+</td><td>0</td><td>GAAGTACTGTTCGCCCAGCA</td><td>GGG</td><td>||||||||[||||||||||||]PAM</td><td>NA</td><td>Intergenic</td><td>NA</td><td>NA</td></tr>
<tr><td>T14545</td><td>GCCCAGCAGGGCCAGTAGCACGG</td><td>1000</td><td></td><td></td><td></td><td></td><td></td><td></td><td></td><td></td><td></td></tr>
<tr><td>Oligo fwd</td><td>CACCGCCCAGCAGGGCCAGTAGCA</td><td></td><td></td><td></td><td></td><td></td><td></td><td></td><td></td><td></td><td></td></tr>
<tr><td>Oligo rev</td><td>AAACTGCTACTGGCCCTGCTGGGC</td><td></td><td></td><td></td><td></td><td></td><td></td><td></td><td></td><td></td><td></td></tr>
<tr><td>Chromosome</td><td>start</td><td>end</td><td>strand</td><td>MM</td><td>target_seq</td><td>PAM</td><td>alignment</td><td>distance</td><td>position</td><td>gene name</td><td>gene id</td></tr>
<tr><td>GU305914.1</td><td>191302</td><td>191324</td><td>+</td><td>0</td><td>GCCCAGCAGGGCCAGTAGCA</td><td>CGG</td><td>||||||||[||||||||||||]PAM</td><td>NA</td><td>Intergenic</td><td>NA</td><td>NA</td></tr>
<tr><td>T14546</td><td>TAGCACGGTCGCGACGATCGTGG</td><td>1000</td><td></td><td></td><td></td><td></td><td></td><td></td><td></td><td></td><td></td></tr>
<tr><td>Oligo fwd</td><td>CACCTAGCACGGTCGCGACGATCG</td><td></td><td></td><td></td><td></td><td></td><td></td><td></td><td></td><td></td><td></td></tr>
<tr><td>Oligo rev</td><td>AAACCGATCGTCGCGACCGTGCTA</td><td></td><td></td><td></td><td></td><td></td><td></td><td></td><td></td><td></td><td></td></tr>
<tr><td>Chromosome</td><td>start</td><td>end</td><td>strand</td><td>MM</td><td>target_seq</td><td>PAM</td><td>alignment</td><td>distance</td><td>position</td><td>gene name</td><td>gene id</td></tr>
<tr><td>GU305914.1</td><td>191317</td><td>191339</td><td>+</td><td>0</td><td>TAGCACGGTCGCGACGATCG</td><td>TGG</td><td>||||||||[||||||||||||]PAM</td><td>NA</td><td>Intergenic</td><td>NA</td><td>NA</td></tr>
<tr><td>T14547</td><td>TTCCACCAGAACCCCCGCGCCGG</td><td>1000</td><td></td><td></td><td></td><td></td><td></td><td></td><td></td><td></td><td></td></tr>
<tr><td>Oligo fwd</td><td>CACCTTCCACCAGAACCCCCGCGC</td><td></td><td></td><td></td><td></td><td></td><td></td><td></td><td></td><td></td><td></td></tr>
<tr><td>Oligo rev</td><td>AAACGCGCGGGGGTTCTGGTGGAA</td><td></td><td></td><td></td><td></td><td></td><td></td><td></td><td></td><td></td><td></td></tr>
<tr><td>Chromosome</td><td>start</td><td>end</td><td>strand</td><td>MM</td><td>target_seq</td><td>PAM</td><td>alignment</td><td>distance</td><td>position</td><td>gene name</td><td>gene id</td></tr>
<tr><td>GU305914.1</td><td>191354</td><td>191376</td><td>+</td><td>0</td><td>TTCCACCAGAACCCCCGCGC</td><td>CGG</td><td>||||||||[||||||||||||]PAM</td><td>NA</td><td>Intergenic</td><td>NA</td><td>NA</td></tr>
<tr><td>T14548</td><td>ATGCTGAAACCCAGATCGAACGG</td><td>1000</td><td></td><td></td><td></td><td></td><td></td><td></td><td></td><td></td><td></td></tr>
<tr><td>Oligo fwd</td><td>CACCATGCTGAAACCCAGATCGAA</td><td></td><td></td><td></td><td></td><td></td><td></td><td></td><td></td><td></td><td></td></tr>
<tr><td>Oligo rev</td><td>AAACTTCGATCTGGGTTTCAGCAT</td><td></td><td></td><td></td><td></td><td></td><td></td><td></td><td></td><td></td><td></td></tr>
<tr><td>Chromosome</td><td>start</td><td>end</td><td>strand</td><td>MM</td><td>target_seq</td><td>PAM</td><td>alignment</td><td>distance</td><td>position</td><td>gene name</td><td>gene id</td></tr>
<tr><td>GU305914.1</td><td>191435</td><td>191457</td><td>+</td><td>0</td><td>ATGCTGAAACCCAGATCGAA</td><td>CGG</td><td>||||||||[||||||||||||]PAM</td><td>NA</td><td>Intergenic</td><td>NA</td><td>NA</td></tr>
<tr><td>T14549</td><td>TGCTGAAACCCAGATCGAACGGG</td><td>1000</td><td></td><td></td><td></td><td></td><td></td><td></td><td></td><td></td><td></td></tr>
<tr><td>Oligo fwd</td><td>CACCTGCTGAAACCCAGATCGAAC</td><td></td><td></td><td></td><td></td><td></td><td></td><td></td><td></td><td></td><td></td></tr>
<tr><td>Oligo rev</td><td>AAACGTTCGATCTGGGTTTCAGCA</td><td></td><td></td><td></td><td></td><td></td><td></td><td></td><td></td><td></td><td></td></tr>
<tr><td>Chromosome</td><td>start</td><td>end</td><td>strand</td><td>MM</td><td>target_seq</td><td>PAM</td><td>alignment</td><td>distance</td><td>position</td><td>gene name</td><td>gene id</td></tr>
<tr><td>GU305914.1</td><td>191436</td><td>191458</td><td>+</td><td>0</td><td>TGCTGAAACCCAGATCGAAC</td><td>GGG</td><td>||||||||[||||||||||||]PAM</td><td>NA</td><td>Intergenic</td><td>NA</td><td>NA</td></tr>
<tr><td>T14550</td><td>CAGATCGAACGGGTCGATCTTGG</td><td>1000</td><td></td><td></td><td></td><td></td><td></td><td></td><td></td><td></td><td></td></tr>
<tr><td>Oligo fwd</td><td>CACCCAGATCGAACGGGTCGATCT</td><td></td><td></td><td></td><td></td><td></td><td></td><td></td><td></td><td></td><td></td></tr>
<tr><td>Oligo rev</td><td>AAACAGATCGACCCGTTCGATCTG</td><td></td><td></td><td></td><td></td><td></td><td></td><td></td><td></td><td></td><td></td></tr>
<tr><td>Chromosome</td><td>start</td><td>end</td><td>strand</td><td>MM</td><td>target_seq</td><td>PAM</td><td>alignment</td><td>distance</td><td>position</td><td>gene name</td><td>gene id</td></tr>
<tr><td>GU305914.1</td><td>191446</td><td>191468</td><td>+</td><td>0</td><td>CAGATCGAACGGGTCGATCT</td><td>TGG</td><td>||||||||[||||||||||||]PAM</td><td>NA</td><td>Intergenic</td><td>NA</td><td>NA</td></tr>
<tr><td>T14551</td><td>CGTTTTCGAGAAGTTCAGCCCGG</td><td>1000</td><td></td><td></td><td></td><td></td><td></td><td></td><td></td><td></td><td></td></tr>
<tr><td>Oligo fwd</td><td>CACCCGTTTTCGAGAAGTTCAGCC</td><td></td><td></td><td></td><td></td><td></td><td></td><td></td><td></td><td></td><td></td></tr>
<tr><td>Oligo rev</td><td>AAACGGCTGAACTTCTCGAAAACG</td><td></td><td></td><td></td><td></td><td></td><td></td><td></td><td></td><td></td><td></td></tr>
<tr><td>Chromosome</td><td>start</td><td>end</td><td>strand</td><td>MM</td><td>target_seq</td><td>PAM</td><td>alignment</td><td>distance</td><td>position</td><td>gene name</td><td>gene id</td></tr>
<tr><td>GU305914.1</td><td>191506</td><td>191528</td><td>+</td><td>0</td><td>CGTTTTCGAGAAGTTCAGCC</td><td>CGG</td><td>||||||||[||||||||||||]PAM</td><td>NA</td><td>Intergenic</td><td>NA</td><td>NA</td></tr>
<tr><td>T14552</td><td>GTTTTCGAGAAGTTCAGCCCGGG</td><td>1000</td><td></td><td></td><td></td><td></td><td></td><td></td><td></td><td></td><td></td></tr>
<tr><td>Oligo fwd</td><td>CACCGTTTTCGAGAAGTTCAGCCC</td><td></td><td></td><td></td><td></td><td></td><td></td><td></td><td></td><td></td><td></td></tr>
<tr><td>Oligo rev</td><td>AAACGGGCTGAACTTCTCGAAAAC</td><td></td><td></td><td></td><td></td><td></td><td></td><td></td><td></td><td></td><td></td></tr>
<tr><td>Chromosome</td><td>start</td><td>end</td><td>strand</td><td>MM</td><td>target_seq</td><td>PAM</td><td>alignment</td><td>distance</td><td>position</td><td>gene name</td><td>gene id</td></tr>
<tr><td>GU305914.1</td><td>191507</td><td>191529</td><td>+</td><td>0</td><td>GTTTTCGAGAAGTTCAGCCC</td><td>GGG</td><td>||||||||[||||||||||||]PAM</td><td>NA</td><td>Intergenic</td><td>NA</td><td>NA</td></tr>
<tr><td>T14553</td><td>TTTTCGAGAAGTTCAGCCCGGGG</td><td>1000</td><td></td><td></td><td></td><td></td><td></td><td></td><td></td><td></td><td></td></tr>
<tr><td>Oligo fwd</td><td>CACCTTTTCGAGAAGTTCAGCCCG</td><td></td><td></td><td></td><td></td><td></td><td></td><td></td><td></td><td></td><td></td></tr>
<tr><td>Oligo rev</td><td>AAACCGGGCTGAACTTCTCGAAAA</td><td></td><td></td><td></td><td></td><td></td><td></td><td></td><td></td><td></td><td></td></tr>
</table>

<table>
<tr><td>Chromosome</td><td>start</td><td>end</td><td>strand</td><td>MM</td><td>target_seq</td><td>PAM</td><td>alignment</td><td>distance</td><td>position</td><td>gene name</td><td>gene id</td></tr>
<tr><td>GU305914.1</td><td>191508</td><td>191530</td><td>+</td><td>0</td><td>TTTTCGAGAAGTTCAGCCCG</td><td>GGG</td><td>||||||||[||||||||||||]PAM</td><td>NA</td><td>Intergenic</td><td>NA</td><td>NA</td></tr>
<tr><td>T14554</td><td>CCTCGTCTCGAAGTGCACTTCGG</td><td>1000</td><td></td><td></td><td></td><td></td><td></td><td></td><td></td><td></td><td></td></tr>
<tr><td>Oligo fwd</td><td>CACCCCTCGTCTCGAAGTGCACTT</td><td></td><td></td><td></td><td></td><td></td><td></td><td></td><td></td><td></td><td></td></tr>
<tr><td>Oligo rev</td><td>AAACAAGTGCACTTCGAGACGAGG</td><td></td><td></td><td></td><td></td><td></td><td></td><td></td><td></td><td></td><td></td></tr>
<tr><td>Chromosome</td><td>start</td><td>end</td><td>strand</td><td>MM</td><td>target_seq</td><td>PAM</td><td>alignment</td><td>distance</td><td>position</td><td>gene name</td><td>gene id</td></tr>
<tr><td>GU305914.1</td><td>191560</td><td>191582</td><td>+</td><td>0</td><td>CCTCGTCTCGAAGTGCACTT</td><td>CGG</td><td>||||||||[||||||||||||]PAM</td><td>NA</td><td>Intergenic</td><td>NA</td><td>NA</td></tr>
<tr><td>T14555</td><td>CTCGTCTCGAAGTGCACTTCGGG</td><td>1000</td><td></td><td></td><td></td><td></td><td></td><td></td><td></td><td></td><td></td></tr>
<tr><td>Oligo fwd</td><td>CACCCTCGTCTCGAAGTGCACTTC</td><td></td><td></td><td></td><td></td><td></td><td></td><td></td><td></td><td></td><td></td></tr>
<tr><td>Oligo rev</td><td>AAACGAAGTGCACTTCGAGACGAG</td><td></td><td></td><td></td><td></td><td></td><td></td><td></td><td></td><td></td><td></td></tr>
<tr><td>Chromosome</td><td>start</td><td>end</td><td>strand</td><td>MM</td><td>target_seq</td><td>PAM</td><td>alignment</td><td>distance</td><td>position</td><td>gene name</td><td>gene id</td></tr>
<tr><td>GU305914.1</td><td>191561</td><td>191583</td><td>+</td><td>0</td><td>CTCGTCTCGAAGTGCACTTC</td><td>GGG</td><td>||||||||[||||||||||||]PAM</td><td>NA</td><td>Intergenic</td><td>NA</td><td>NA</td></tr>
<tr><td>T14556</td><td>TCGTCTCGAAGTGCACTTCGGGG</td><td>1000</td><td></td><td></td><td></td><td></td><td></td><td></td><td></td><td></td><td></td></tr>
<tr><td>Oligo fwd</td><td>CACCTCGTCTCGAAGTGCACTTCG</td><td></td><td></td><td></td><td></td><td></td><td></td><td></td><td></td><td></td><td></td></tr>
<tr><td>Oligo rev</td><td>AAACCGAAGTGCACTTCGAGACGA</td><td></td><td></td><td></td><td></td><td></td><td></td><td></td><td></td><td></td><td></td></tr>
<tr><td>Chromosome</td><td>start</td><td>end</td><td>strand</td><td>MM</td><td>target_seq</td><td>PAM</td><td>alignment</td><td>distance</td><td>position</td><td>gene name</td><td>gene id</td></tr>
<tr><td>GU305914.1</td><td>191562</td><td>191584</td><td>+</td><td>0</td><td>TCGTCTCGAAGTGCACTTCG</td><td>GGG</td><td>||||||||[||||||||||||]PAM</td><td>NA</td><td>Intergenic</td><td>NA</td><td>NA</td></tr>
<tr><td>T14557</td><td>GCACTTCGGGGCCCTCTCTGAGG</td><td>1000</td><td></td><td></td><td></td><td></td><td></td><td></td><td></td><td></td><td></td></tr>
<tr><td>Oligo fwd</td><td>CACCGCACTTCGGGGCCCTCTCTG</td><td></td><td></td><td></td><td></td><td></td><td></td><td></td><td></td><td></td><td></td></tr>
<tr><td>Oligo rev</td><td>AAACCAGAGAGGGCCCCGAAGTGC</td><td></td><td></td><td></td><td></td><td></td><td></td><td></td><td></td><td></td><td></td></tr>
<tr><td>Chromosome</td><td>start</td><td>end</td><td>strand</td><td>MM</td><td>target_seq</td><td>PAM</td><td>alignment</td><td>distance</td><td>position</td><td>gene name</td><td>gene id</td></tr>
<tr><td>GU305914.1</td><td>191574</td><td>191596</td><td>+</td><td>0</td><td>GCACTTCGGGGCCCTCTCTG</td><td>AGG</td><td>||||||||[||||||||||||]PAM</td><td>NA</td><td>Intergenic</td><td>NA</td><td>NA</td></tr>
<tr><td>GU305914.1</td><td>135307</td><td>135329</td><td>-</td><td>4</td><td>GCGCTGCGGGGCCCGCTCCG</td><td>CGG</td><td>||-||-||[||||||-|||-|]PAM</td><td>NA</td><td>Intergenic</td><td>NA</td><td>NA</td></tr>
<tr><td>T14558</td><td>CTTCGGGGCCCTCTCTGAGGCGG</td><td>1000</td><td></td><td></td><td></td><td></td><td></td><td></td><td></td><td></td><td></td></tr>
<tr><td>Oligo fwd</td><td>CACCCTTCGGGGCCCTCTCTGAGG</td><td></td><td></td><td></td><td></td><td></td><td></td><td></td><td></td><td></td><td></td></tr>
<tr><td>Oligo rev</td><td>AAACCCTCAGAGAGGGCCCCGAAG</td><td></td><td></td><td></td><td></td><td></td><td></td><td></td><td></td><td></td><td></td></tr>
<tr><td>Chromosome</td><td>start</td><td>end</td><td>strand</td><td>MM</td><td>target_seq</td><td>PAM</td><td>alignment</td><td>distance</td><td>position</td><td>gene name</td><td>gene id</td></tr>
<tr><td>GU305914.1</td><td>191577</td><td>191599</td><td>+</td><td>0</td><td>CTTCGGGGCCCTCTCTGAGG</td><td>CGG</td><td>||||||||[||||||||||||]PAM</td><td>NA</td><td>Intergenic</td><td>NA</td><td>NA</td></tr>
<tr><td>T14559</td><td>GGCCCTCTCTGAGGCGGAGCAGG</td><td>1000</td><td></td><td></td><td></td><td></td><td></td><td></td><td></td><td></td><td></td></tr>
<tr><td>Oligo fwd</td><td>CACCGGCCCTCTCTGAGGCGGAGC</td><td></td><td></td><td></td><td></td><td></td><td></td><td></td><td></td><td></td><td></td></tr>
<tr><td>Oligo rev</td><td>AAACGCTCCGCCTCAGAGAGGGCC</td><td></td><td></td><td></td><td></td><td></td><td></td><td></td><td></td><td></td><td></td></tr>
<tr><td>Chromosome</td><td>start</td><td>end</td><td>strand</td><td>MM</td><td>target_seq</td><td>PAM</td><td>alignment</td><td>distance</td><td>position</td><td>gene name</td><td>gene id</td></tr>
<tr><td>GU305914.1</td><td>191583</td><td>191605</td><td>+</td><td>0</td><td>GGCCCTCTCTGAGGCGGAGC</td><td>AGG</td><td>||||||||[||||||||||||]PAM</td><td>NA</td><td>Intergenic</td><td>NA</td><td>NA</td></tr>
<tr><td>T14560</td><td>CTCTCTGAGGCGGAGCAGGATGG</td><td>1000</td><td></td><td></td><td></td><td></td><td></td><td></td><td></td><td></td><td></td></tr>
<tr><td>Oligo fwd</td><td>CACCCTCTCTGAGGCGGAGCAGGA</td><td></td><td></td><td></td><td></td><td></td><td></td><td></td><td></td><td></td><td></td></tr>
<tr><td>Oligo rev</td><td>AAACTCCTGCTCCGCCTCAGAGAG</td><td></td><td></td><td></td><td></td><td></td><td></td><td></td><td></td><td></td><td></td></tr>
<tr><td>Chromosome</td><td>start</td><td>end</td><td>strand</td><td>MM</td><td>target_seq</td><td>PAM</td><td>alignment</td><td>distance</td><td>position</td><td>gene name</td><td>gene id</td></tr>
<tr><td>GU305914.1</td><td>191587</td><td>191609</td><td>+</td><td>0</td><td>CTCTCTGAGGCGGAGCAGGA</td><td>TGG</td><td>||||||||[||||||||||||]PAM</td><td>NA</td><td>Intergenic</td><td>NA</td><td>NA</td></tr>
<tr><td>GU305914.1</td><td>168542</td><td>168564</td><td>+</td><td>4</td><td>TCCTCTGAGTCGGAGGAGGA</td><td>TGG</td><td>--||||||[|-|||||-||||]PAM</td><td>NA</td><td>Intergenic</td><td>NA</td><td>NA</td></tr>
<tr><td>T14561</td><td>GAGGCGGAGCAGGATGGTGCCGG</td><td>1000</td><td></td><td></td><td></td><td></td><td></td><td></td><td></td><td></td><td></td></tr>
<tr><td>Oligo fwd</td><td>CACCGAGGCGGAGCAGGATGGTGC</td><td></td><td></td><td></td><td></td><td></td><td></td><td></td><td></td><td></td><td></td></tr>
<tr><td>Oligo rev</td><td>AAACGCACCATCCTGCTCCGCCTC</td><td></td><td></td><td></td><td></td><td></td><td></td><td></td><td></td><td></td><td></td></tr>
<tr><td>Chromosome</td><td>start</td><td>end</td><td>strand</td><td>MM</td><td>target_seq</td><td>PAM</td><td>alignment</td><td>distance</td><td>position</td><td>gene name</td><td>gene id</td></tr>
<tr><td>GU305914.1</td><td>191593</td><td>191615</td><td>+</td><td>0</td><td>GAGGCGGAGCAGGATGGTGC</td><td>CGG</td><td>||||||||[||||||||||||]PAM</td><td>NA</td><td>Intergenic</td><td>NA</td><td>NA</td></tr>
<tr><td>T14562</td><td>ACACCATCGCGCTCGACATGTGG</td><td>1000</td><td></td><td></td><td></td><td></td><td></td><td></td><td></td><td></td><td></td></tr>
<tr><td>Oligo fwd</td><td>CACCACACCATCGCGCTCGACATG</td><td></td><td></td><td></td><td></td><td></td><td></td><td></td><td></td><td></td><td></td></tr>
<tr><td>Oligo rev</td><td>AAACCATGTCGAGCGCGATGGTGT</td><td></td><td></td><td></td><td></td><td></td><td></td><td></td><td></td><td></td><td></td></tr>
<tr><td>Chromosome</td><td>start</td><td>end</td><td>strand</td><td>MM</td><td>target_seq</td><td>PAM</td><td>alignment</td><td>distance</td><td>position</td><td>gene name</td><td>gene id</td></tr>
<tr><td>GU305914.1</td><td>191625</td><td>191647</td><td>+</td><td>0</td><td>ACACCATCGCGCTCGACATG</td><td>TGG</td><td>||||||||[||||||||||||]PAM</td><td>NA</td><td>Intergenic</td><td>NA</td><td>NA</td></tr>
<tr><td>T14563</td><td>CGCGCTCGACATGTGGAATTTGG</td><td>1000</td><td></td><td></td><td></td><td></td><td></td><td></td><td></td><td></td><td></td></tr>
<tr><td>Oligo fwd</td><td>CACCCGCGCTCGACATGTGGAATT</td><td></td><td></td><td></td><td></td><td></td><td></td><td></td><td></td><td></td><td></td></tr>
<tr><td>Oligo rev</td><td>AAACAATTCCACATGTCGAGCGCG</td><td></td><td></td><td></td><td></td><td></td><td></td><td></td><td></td><td></td><td></td></tr>
<tr><td>Chromosome</td><td>start</td><td>end</td><td>strand</td><td>MM</td><td>target_seq</td><td>PAM</td><td>alignment</td><td>distance</td><td>position</td><td>gene name</td><td>gene id</td></tr>
<tr><td>GU305914.1</td><td>191632</td><td>191654</td><td>+</td><td>0</td><td>CGCGCTCGACATGTGGAATT</td><td>TGG</td><td>||||||||[||||||||||||]PAM</td><td>NA</td><td>Intergenic</td><td>NA</td><td>NA</td></tr>
<tr><td>T14564</td><td>ACATGTGGAATTTGGCCGACAGG</td><td>1000</td><td></td><td></td><td></td><td></td><td></td><td></td><td></td><td></td><td></td></tr>
<tr><td>Oligo fwd</td><td>CACCACATGTGGAATTTGGCCGAC</td><td></td><td></td><td></td><td></td><td></td><td></td><td></td><td></td><td></td><td></td></tr>
<tr><td>Oligo rev</td><td>AAACGTCGGCCAAATTCCACATGT</td><td></td><td></td><td></td><td></td><td></td><td></td><td></td><td></td><td></td><td></td></tr>
<tr><td>Chromosome</td><td>start</td><td>end</td><td>strand</td><td>MM</td><td>target_seq</td><td>PAM</td><td>alignment</td><td>distance</td><td>position</td><td>gene name</td><td>gene id</td></tr>
<tr><td>GU305914.1</td><td>191640</td><td>191662</td><td>+</td><td>0</td><td>ACATGTGGAATTTGGCCGAC</td><td>AGG</td><td>||||||||[||||||||||||]PAM</td><td>NA</td><td>Intergenic</td><td>NA</td><td>NA</td></tr>
<tr><td>T14565</td><td>GTGGAATTTGGCCGACAGGATGG</td><td>1000</td><td></td><td></td><td></td><td></td><td></td><td></td><td></td><td></td><td></td></tr>
<tr><td>Oligo fwd</td><td>CACCGTGGAATTTGGCCGACAGGA</td><td></td><td></td><td></td><td></td><td></td><td></td><td></td><td></td><td></td><td></td></tr>
<tr><td>Oligo rev</td><td>AAACTCCTGTCGGCCAAATTCCAC</td><td></td><td></td><td></td><td></td><td></td><td></td><td></td><td></td><td></td><td></td></tr>
<tr><td>Chromosome</td><td>start</td><td>end</td><td>strand</td><td>MM</td><td>target_seq</td><td>PAM</td><td>alignment</td><td>distance</td><td>position</td><td>gene name</td><td>gene id</td></tr>
<tr><td>GU305914.1</td><td>191644</td><td>191666</td><td>+</td><td>0</td><td>GTGGAATTTGGCCGACAGGA</td><td>TGG</td><td>||||||||[||||||||||||]PAM</td><td>NA</td><td>Intergenic</td><td>NA</td><td>NA</td></tr>
<tr><td>T14566</td><td>AGTCTTTCACGACGAAGACTCGG</td><td>1000</td><td></td><td></td><td></td><td></td><td></td><td></td><td></td><td></td><td></td></tr>
<tr><td>Oligo fwd</td><td>CACCAGTCTTTCACGACGAAGACT</td><td></td><td></td><td></td><td></td><td></td><td></td><td></td><td></td><td></td><td></td></tr>
<tr><td>Oligo rev</td><td>AAACAGTCTTCGTCGTGAAAGACT</td><td></td><td></td><td></td><td></td><td></td><td></td><td></td><td></td><td></td><td></td></tr>
<tr><td>Chromosome</td><td>start</td><td>end</td><td>strand</td><td>MM</td><td>target_seq</td><td>PAM</td><td>alignment</td><td>distance</td><td>position</td><td>gene name</td><td>gene id</td></tr>
<tr><td>GU305914.1</td><td>191667</td><td>191689</td><td>+</td><td>0</td><td>AGTCTTTCACGACGAAGACT</td><td>CGG</td><td>||||||||[||||||||||||]PAM</td><td>NA</td><td>Intergenic</td><td>NA</td><td>NA</td></tr>
<tr><td>T14567</td><td>CACCCTCGCCGTGAGTATGAAGG</td><td>1000</td><td></td><td></td><td></td><td></td><td></td><td></td><td></td><td></td><td></td></tr>
<tr><td>Oligo fwd</td><td>CACCCACCCTCGCCGTGAGTATGA</td><td></td><td></td><td></td><td></td><td></td><td></td><td></td><td></td><td></td><td></td></tr>
<tr><td>Oligo rev</td><td>AAACTCATACTCACGGCGAGGGTG</td><td></td><td></td><td></td><td></td><td></td><td></td><td></td><td></td><td></td><td></td></tr>
<tr><td>Chromosome</td><td>start</td><td>end</td><td>strand</td><td>MM</td><td>target_seq</td><td>PAM</td><td>alignment</td><td>distance</td><td>position</td><td>gene name</td><td>gene id</td></tr>
<tr><td>GU305914.1</td><td>191692</td><td>191714</td><td>+</td><td>0</td><td>CACCCTCGCCGTGAGTATGA</td><td>AGG</td><td>||||||||[||||||||||||]PAM</td><td>NA</td><td>Intergenic</td><td>NA</td><td>NA</td></tr>
</table>

<table>
<tr><td>T14568</td><td>GGTCGCGCGCGAGATGTGCTCGG</td><td>1000</td><td></td><td></td><td></td><td></td><td></td><td></td><td></td><td></td><td></td></tr>
<tr><td>Oligo fwd</td><td>CACCGGTCGCGCGCGAGATGTGCT</td><td></td><td></td><td></td><td></td><td></td><td></td><td></td><td></td><td></td><td></td></tr>
<tr><td>Oligo rev</td><td>AAACAGCACATCTCGCGCGCGACC</td><td></td><td></td><td></td><td></td><td></td><td></td><td></td><td></td><td></td><td></td></tr>
<tr><td>Chromosome</td><td>start</td><td>end</td><td>strand</td><td>MM</td><td>target_seq</td><td>PAM</td><td>alignment</td><td>distance</td><td>position</td><td>gene name</td><td>gene id</td></tr>
<tr><td>GU305914.1</td><td>191713</td><td>191735</td><td>+</td><td>0</td><td>GGTCGCGCGCGAGATGTGCT</td><td>CGG</td><td>||||||||[||||||||||||]PAM</td><td>NA</td><td>Intergenic</td><td>NA</td><td>NA</td></tr>
<tr><td></td><td></td><td></td><td></td><td></td><td></td><td></td><td></td><td></td><td></td><td></td><td></td></tr>
<tr><td>T14569</td><td>GATGTGCTCGGTGACGACCGAGG</td><td>1000</td><td></td><td></td><td></td><td></td><td></td><td></td><td></td><td></td><td></td></tr>
<tr><td>Oligo fwd</td><td>CACCGATGTGCTCGGTGACGACCG</td><td></td><td></td><td></td><td></td><td></td><td></td><td></td><td></td><td></td><td></td></tr>
<tr><td>Oligo rev</td><td>AAACCGGTCGTCACCGAGCACATC</td><td></td><td></td><td></td><td></td><td></td><td></td><td></td><td></td><td></td><td></td></tr>
<tr><td>Chromosome</td><td>start</td><td>end</td><td>strand</td><td>MM</td><td>target_seq</td><td>PAM</td><td>alignment</td><td>distance</td><td>position</td><td>gene name</td><td>gene id</td></tr>
<tr><td>GU305914.1</td><td>191725</td><td>191747</td><td>+</td><td>0</td><td>GATGTGCTCGGTGACGACCG</td><td>AGG</td><td>||||||||[||||||||||||]PAM</td><td>NA</td><td>Intergenic</td><td>NA</td><td>NA</td></tr>
<tr><td>GU305914.1</td><td>124349</td><td>124371</td><td>-</td><td>4</td><td>GATGTCCCCGGTGTCGATCG</td><td>CGG</td><td>|||||-|-[|||||-|||-||]PAM</td><td>NA</td><td>Intergenic</td><td>NA</td><td>NA</td></tr>
<tr><td></td><td></td><td></td><td></td><td></td><td></td><td></td><td></td><td></td><td></td><td></td><td></td></tr>
<tr><td>T14570</td><td>CCGAGGAGTCCTCGACGCCGAGG</td><td>1000</td><td></td><td></td><td></td><td></td><td></td><td></td><td></td><td></td><td></td></tr>
<tr><td>Oligo fwd</td><td>CACCCCGAGGAGTCCTCGACGCCG</td><td></td><td></td><td></td><td></td><td></td><td></td><td></td><td></td><td></td><td></td></tr>
<tr><td>Oligo rev</td><td>AAACCGGCGTCGAGGACTCCTCGG</td><td></td><td></td><td></td><td></td><td></td><td></td><td></td><td></td><td></td><td></td></tr>
<tr><td>Chromosome</td><td>start</td><td>end</td><td>strand</td><td>MM</td><td>target_seq</td><td>PAM</td><td>alignment</td><td>distance</td><td>position</td><td>gene name</td><td>gene id</td></tr>
<tr><td>GU305914.1</td><td>191742</td><td>191764</td><td>+</td><td>0</td><td>CCGAGGAGTCCTCGACGCCG</td><td>AGG</td><td>||||||||[||||||||||||]PAM</td><td>NA</td><td>Intergenic</td><td>NA</td><td>NA</td></tr>
<tr><td></td><td></td><td></td><td></td><td></td><td></td><td></td><td></td><td></td><td></td><td></td><td></td></tr>
<tr><td>T14571</td><td>GAGTCCTCGACGCCGAGGTGCGG</td><td>1000</td><td></td><td></td><td></td><td></td><td></td><td></td><td></td><td></td><td></td></tr>
<tr><td>Oligo fwd</td><td>CACCGAGTCCTCGACGCCGAGGTG</td><td></td><td></td><td></td><td></td><td></td><td></td><td></td><td></td><td></td><td></td></tr>
<tr><td>Oligo rev</td><td>AAACCACCTCGGCGTCGAGGACTC</td><td></td><td></td><td></td><td></td><td></td><td></td><td></td><td></td><td></td><td></td></tr>
<tr><td>Chromosome</td><td>start</td><td>end</td><td>strand</td><td>MM</td><td>target_seq</td><td>PAM</td><td>alignment</td><td>distance</td><td>position</td><td>gene name</td><td>gene id</td></tr>
<tr><td>GU305914.1</td><td>191747</td><td>191769</td><td>+</td><td>0</td><td>GAGTCCTCGACGCCGAGGTG</td><td>CGG</td><td>||||||||[||||||||||||]PAM</td><td>NA</td><td>Intergenic</td><td>NA</td><td>NA</td></tr>
<tr><td>GU305914.1</td><td>166054</td><td>166076</td><td>-</td><td>4</td><td>GAGTTTTCGATGCCGAAGTG</td><td>CGG</td><td>||||--||[||-|||||-|||]PAM</td><td>NA</td><td>Intergenic</td><td>NA</td><td>NA</td></tr>
<tr><td></td><td></td><td></td><td></td><td></td><td></td><td></td><td></td><td></td><td></td><td></td><td></td></tr>
<tr><td>T14572</td><td>CCACGCACGCGTGTCCCCCCAGG</td><td>1000</td><td></td><td></td><td></td><td></td><td></td><td></td><td></td><td></td><td></td></tr>
<tr><td>Oligo fwd</td><td>CACCCCACGCACGCGTGTCCCCCC</td><td></td><td></td><td></td><td></td><td></td><td></td><td></td><td></td><td></td><td></td></tr>
<tr><td>Oligo rev</td><td>AAACGGGGGGACACGCGTGCGTGG</td><td></td><td></td><td></td><td></td><td></td><td></td><td></td><td></td><td></td><td></td></tr>
<tr><td>Chromosome</td><td>start</td><td>end</td><td>strand</td><td>MM</td><td>target_seq</td><td>PAM</td><td>alignment</td><td>distance</td><td>position</td><td>gene name</td><td>gene id</td></tr>
<tr><td>GU305914.1</td><td>191775</td><td>191797</td><td>+</td><td>0</td><td>CCACGCACGCGTGTCCCCCC</td><td>AGG</td><td>||||||||[||||||||||||]PAM</td><td>NA</td><td>Intergenic</td><td>NA</td><td>NA</td></tr>
<tr><td>GU305914.1</td><td>182384</td><td>182406</td><td>-</td><td>4</td><td>ACACGTACGCGTCTCTCCCC</td><td>AGG</td><td>-||||-||[||||-||-||||]PAM</td><td>NA</td><td>Intergenic</td><td>NA</td><td>NA</td></tr>
<tr><td></td><td></td><td></td><td></td><td></td><td></td><td></td><td></td><td></td><td></td><td></td><td></td></tr>
<tr><td>T14573</td><td>CCCCCCAGGTTCTCGTGCCTCGG</td><td>1000</td><td></td><td></td><td></td><td></td><td></td><td></td><td></td><td></td><td></td></tr>
<tr><td>Oligo fwd</td><td>CACCCCCCCCAGGTTCTCGTGCCT</td><td></td><td></td><td></td><td></td><td></td><td></td><td></td><td></td><td></td><td></td></tr>
<tr><td>Oligo rev</td><td>AAACAGGCACGAGAACCTGGGGGG</td><td></td><td></td><td></td><td></td><td></td><td></td><td></td><td></td><td></td><td></td></tr>
<tr><td>Chromosome</td><td>start</td><td>end</td><td>strand</td><td>MM</td><td>target_seq</td><td>PAM</td><td>alignment</td><td>distance</td><td>position</td><td>gene name</td><td>gene id</td></tr>
<tr><td>GU305914.1</td><td>191789</td><td>191811</td><td>+</td><td>0</td><td>CCCCCCAGGTTCTCGTGCCT</td><td>CGG</td><td>||||||||[||||||||||||]PAM</td><td>NA</td><td>Intergenic</td><td>NA</td><td>NA</td></tr>
<tr><td></td><td></td><td></td><td></td><td></td><td></td><td></td><td></td><td></td><td></td><td></td><td></td></tr>
<tr><td>T14574</td><td>TCGGCTCCAGCTCGATCCCGAGG</td><td>1000</td><td></td><td></td><td></td><td></td><td></td><td></td><td></td><td></td><td></td></tr>
<tr><td>Oligo fwd</td><td>CACCTCGGCTCCAGCTCGATCCCG</td><td></td><td></td><td></td><td></td><td></td><td></td><td></td><td></td><td></td><td></td></tr>
<tr><td>Oligo rev</td><td>AAACCGGGATCGAGCTGGAGCCGA</td><td></td><td></td><td></td><td></td><td></td><td></td><td></td><td></td><td></td><td></td></tr>
<tr><td>Chromosome</td><td>start</td><td>end</td><td>strand</td><td>MM</td><td>target_seq</td><td>PAM</td><td>alignment</td><td>distance</td><td>position</td><td>gene name</td><td>gene id</td></tr>
<tr><td>GU305914.1</td><td>191808</td><td>191830</td><td>+</td><td>0</td><td>TCGGCTCCAGCTCGATCCCG</td><td>AGG</td><td>||||||||[||||||||||||]PAM</td><td>NA</td><td>Intergenic</td><td>NA</td><td>NA</td></tr>
<tr><td></td><td></td><td></td><td></td><td></td><td></td><td></td><td></td><td></td><td></td><td></td><td></td></tr>
<tr><td>T14575</td><td>CCCGAGGTCCACGTCCAGCAAGG</td><td>1000</td><td></td><td></td><td></td><td></td><td></td><td></td><td></td><td></td><td></td></tr>
<tr><td>Oligo fwd</td><td>CACCCCCGAGGTCCACGTCCAGCA</td><td></td><td></td><td></td><td></td><td></td><td></td><td></td><td></td><td></td><td></td></tr>
<tr><td>Oligo rev</td><td>AAACTGCTGGACGTGGACCTCGGG</td><td></td><td></td><td></td><td></td><td></td><td></td><td></td><td></td><td></td><td></td></tr>
<tr><td>Chromosome</td><td>start</td><td>end</td><td>strand</td><td>MM</td><td>target_seq</td><td>PAM</td><td>alignment</td><td>distance</td><td>position</td><td>gene name</td><td>gene id</td></tr>
<tr><td>GU305914.1</td><td>191824</td><td>191846</td><td>+</td><td>0</td><td>CCCGAGGTCCACGTCCAGCA</td><td>AGG</td><td>||||||||[||||||||||||]PAM</td><td>NA</td><td>Intergenic</td><td>NA</td><td>NA</td></tr>
<tr><td></td><td></td><td></td><td></td><td></td><td></td><td></td><td></td><td></td><td></td><td></td><td></td></tr>
<tr><td>T14576</td><td>CCGAGGTCCACGTCCAGCAAGGG</td><td>1000</td><td></td><td></td><td></td><td></td><td></td><td></td><td></td><td></td><td></td></tr>
<tr><td>Oligo fwd</td><td>CACCCCGAGGTCCACGTCCAGCAA</td><td></td><td></td><td></td><td></td><td></td><td></td><td></td><td></td><td></td><td></td></tr>
<tr><td>Oligo rev</td><td>AAACTTGCTGGACGTGGACCTCGG</td><td></td><td></td><td></td><td></td><td></td><td></td><td></td><td></td><td></td><td></td></tr>
<tr><td>Chromosome</td><td>start</td><td>end</td><td>strand</td><td>MM</td><td>target_seq</td><td>PAM</td><td>alignment</td><td>distance</td><td>position</td><td>gene name</td><td>gene id</td></tr>
<tr><td>GU305914.1</td><td>191825</td><td>191847</td><td>+</td><td>0</td><td>CCGAGGTCCACGTCCAGCAA</td><td>GGG</td><td>||||||||[||||||||||||]PAM</td><td>NA</td><td>Intergenic</td><td>NA</td><td>NA</td></tr>
<tr><td></td><td></td><td></td><td></td><td></td><td></td><td></td><td></td><td></td><td></td><td></td><td></td></tr>
<tr><td>T14577</td><td>AGGTCCACGTCCAGCAAGGGCGG</td><td>1000</td><td></td><td></td><td></td><td></td><td></td><td></td><td></td><td></td><td></td></tr>
<tr><td>Oligo fwd</td><td>CACCAGGTCCACGTCCAGCAAGGG</td><td></td><td></td><td></td><td></td><td></td><td></td><td></td><td></td><td></td><td></td></tr>
<tr><td>Oligo rev</td><td>AAACCCCTTGCTGGACGTGGACCT</td><td></td><td></td><td></td><td></td><td></td><td></td><td></td><td></td><td></td><td></td></tr>
<tr><td>Chromosome</td><td>start</td><td>end</td><td>strand</td><td>MM</td><td>target_seq</td><td>PAM</td><td>alignment</td><td>distance</td><td>position</td><td>gene name</td><td>gene id</td></tr>
<tr><td>GU305914.1</td><td>191828</td><td>191850</td><td>+</td><td>0</td><td>AGGTCCACGTCCAGCAAGGG</td><td>CGG</td><td>||||||||[||||||||||||]PAM</td><td>NA</td><td>Intergenic</td><td>NA</td><td>NA</td></tr>
<tr><td></td><td></td><td></td><td></td><td></td><td></td><td></td><td></td><td></td><td></td><td></td><td></td></tr>
<tr><td>T14578</td><td>CTCACTCTTCCGCACGCGTGCGG</td><td>1000</td><td></td><td></td><td></td><td></td><td></td><td></td><td></td><td></td><td></td></tr>
<tr><td>Oligo fwd</td><td>CACCCTCACTCTTCCGCACGCGTG</td><td></td><td></td><td></td><td></td><td></td><td></td><td></td><td></td><td></td><td></td></tr>
<tr><td>Oligo rev</td><td>AAACCACGCGTGCGGAAGAGTGAG</td><td></td><td></td><td></td><td></td><td></td><td></td><td></td><td></td><td></td><td></td></tr>
<tr><td>Chromosome</td><td>start</td><td>end</td><td>strand</td><td>MM</td><td>target_seq</td><td>PAM</td><td>alignment</td><td>distance</td><td>position</td><td>gene name</td><td>gene id</td></tr>
<tr><td>GU305914.1</td><td>191851</td><td>191873</td><td>+</td><td>0</td><td>CTCACTCTTCCGCACGCGTG</td><td>CGG</td><td>||||||||[||||||||||||]PAM</td><td>NA</td><td>Intergenic</td><td>NA</td><td>NA</td></tr>
<tr><td></td><td></td><td></td><td></td><td></td><td></td><td></td><td></td><td></td><td></td><td></td><td></td></tr>
<tr><td>T14579</td><td>TTCCGCACGCGTGCGGTGATCGG</td><td>1000</td><td></td><td></td><td></td><td></td><td></td><td></td><td></td><td></td><td></td></tr>
<tr><td>Oligo fwd</td><td>CACCTTCCGCACGCGTGCGGTGAT</td><td></td><td></td><td></td><td></td><td></td><td></td><td></td><td></td><td></td><td></td></tr>
<tr><td>Oligo rev</td><td>AAACATCACCGCACGCGTGCGGAA</td><td></td><td></td><td></td><td></td><td></td><td></td><td></td><td></td><td></td><td></td></tr>
<tr><td>Chromosome</td><td>start</td><td>end</td><td>strand</td><td>MM</td><td>target_seq</td><td>PAM</td><td>alignment</td><td>distance</td><td>position</td><td>gene name</td><td>gene id</td></tr>
<tr><td>GU305914.1</td><td>191858</td><td>191880</td><td>+</td><td>0</td><td>TTCCGCACGCGTGCGGTGAT</td><td>CGG</td><td>||||||||[||||||||||||]PAM</td><td>NA</td><td>Intergenic</td><td>NA</td><td>NA</td></tr>
<tr><td></td><td></td><td></td><td></td><td></td><td></td><td></td><td></td><td></td><td></td><td></td><td></td></tr>
<tr><td>T14580</td><td>CATCGAACGACATCTCTTCTTGG</td><td>1000</td><td></td><td></td><td></td><td></td><td></td><td></td><td></td><td></td><td></td></tr>
<tr><td>Oligo fwd</td><td>CACCCATCGAACGACATCTCTTCT</td><td></td><td></td><td></td><td></td><td></td><td></td><td></td><td></td><td></td><td></td></tr>
<tr><td>Oligo rev</td><td>AAACAGAAGAGATGTCGTTCGATG</td><td></td><td></td><td></td><td></td><td></td><td></td><td></td><td></td><td></td><td></td></tr>
<tr><td>Chromosome</td><td>start</td><td>end</td><td>strand</td><td>MM</td><td>target_seq</td><td>PAM</td><td>alignment</td><td>distance</td><td>position</td><td>gene name</td><td>gene id</td></tr>
<tr><td>GU305914.1</td><td>191947</td><td>191969</td><td>+</td><td>0</td><td>CATCGAACGACATCTCTTCT</td><td>TGG</td><td>||||||||[||||||||||||]PAM</td><td>NA</td><td>Intergenic</td><td>NA</td><td>NA</td></tr>
<tr><td></td><td></td><td></td><td></td><td></td><td></td><td></td><td></td><td></td><td></td><td></td><td></td></tr>
<tr><td>T14581</td><td>ATCGAACGACATCTCTTCTTGGG</td><td>1000</td><td></td><td></td><td></td><td></td><td></td><td></td><td></td><td></td><td></td></tr>
<tr><td>Oligo fwd</td><td>CACCATCGAACGACATCTCTTCTT</td><td></td><td></td><td></td><td></td><td></td><td></td><td></td><td></td><td></td><td></td></tr>
<tr><td>Oligo rev</td><td>AAACAAGAAGAGATGTCGTTCGAT</td><td></td><td></td><td></td><td></td><td></td><td></td><td></td><td></td><td></td><td></td></tr>
<tr><td>Chromosome</td><td>start</td><td>end</td><td>strand</td><td>MM</td><td>target_seq</td><td>PAM</td><td>alignment</td><td>distance</td><td>position</td><td>gene name</td><td>gene id</td></tr>
<tr><td>GU305914.1</td><td>191948</td><td>191970</td><td>+</td><td>0</td><td>ATCGAACGACATCTCTTCTT</td><td>GGG</td><td>||||||||[||||||||||||]PAM</td><td>NA</td><td>Intergenic</td><td>NA</td><td>NA</td></tr>
<tr><td></td><td></td><td></td><td></td><td></td><td></td><td></td><td></td><td></td><td></td><td></td><td></td></tr>
<tr><td>T14582</td><td>TGACCCAGCCCACCACCCCGCGG</td><td>1000</td><td></td><td></td><td></td><td></td><td></td><td></td><td></td><td></td><td></td></tr>
<tr><td>Oligo fwd</td><td>CACCTGACCCAGCCCACCACCCCG</td><td></td><td></td><td></td><td></td><td></td><td></td><td></td><td></td><td></td><td></td></tr>
</table>

Oligo rev   AAACCGGGGTGGTGGGCTGGGTCA
Chromosome   start   end   strand   MM   target_seq   PAM   alignment   distance   position   gene name   gene id
GU305914.1   191988   192010   +   0   TGACCCAGCCCACCACCCCG   CGG   ||||||||[||||||||||||]PAM   NA   Intergenic   NA   NA

T14583   GACCCAGCCCACCACCCCGCGGG   1000
Oligo fwd   CACCGACCCAGCCCACCACCCCGC
Oligo rev   AAACGCGGGGTGGTGGGCTGGGTC
Chromosome   start   end   strand   MM   target_seq   PAM   alignment   distance   position   gene name   gene id
GU305914.1   191989   192011   +   0   GACCCAGCCCACCACCCCGC   GGG   ||||||||[||||||||||||]PAM   NA   Intergenic   NA   NA

T14584   CCACCACCCCGCGGGAGAATCGG   1000
Oligo fwd   CACCCCACCACCCCGCGGGAGAAT
Oligo rev   AAACATTCTCCCGCGGGGTGGTGG
Chromosome   start   end   strand   MM   target_seq   PAM   alignment   distance   position   gene name   gene id
GU305914.1   191997   192019   +   0   CCACCACCCCGCGGGAGAAT   CGG   ||||||||[||||||||||||]PAM   NA   Intergenic   NA   NA

T14585   CACCACCCCGCGGGAGAATCGGG   1000
Oligo fwd   CACCCACCACCCCGCGGGAGAATC
Oligo rev   AAACGATTCTCCCGCGGGGTGGTG
Chromosome   start   end   strand   MM   target_seq   PAM   alignment   distance   position   gene name   gene id
GU305914.1   191998   192020   +   0   CACCACCCCGCGGGAGAATC   GGG   ||||||||[||||||||||||]PAM   NA   Intergenic   NA   NA

T14586   CGTCAGAGAACAGTTCAACACGG   1000
Oligo fwd   CACCCGTCAGAGAACAGTTCAACA
Oligo rev   AAACTGTTGAACTGTTCTCTGACG
Chromosome   start   end   strand   MM   target_seq   PAM   alignment   distance   position   gene name   gene id
GU305914.1   192022   192044   +   0   CGTCAGAGAACAGTTCAACA   CGG   ||||||||[||||||||||||]PAM   NA   Intergenic   NA   NA

T14587   CAGAGAACAGTTCAACACGGTGG   1000
Oligo fwd   CACCCAGAGAACAGTTCAACACGG
Oligo rev   AAACCCGTGTTGAACTGTTCTCTG
Chromosome   start   end   strand   MM   target_seq   PAM   alignment   distance   position   gene name   gene id
GU305914.1   192025   192047   +   0   CAGAGAACAGTTCAACACGG   TGG   ||||||||[||||||||||||]PAM   NA   Intergenic   NA   NA

T14588   AGAGAACAGTTCAACACGGTGGG   1000
Oligo fwd   CACCAGAGAACAGTTCAACACGGT
Oligo rev   AAACACCGTGTTGAACTGTTCTCT
Chromosome   start   end   strand   MM   target_seq   PAM   alignment   distance   position   gene name   gene id
GU305914.1   192026   192048   +   0   AGAGAACAGTTCAACACGGT   GGG   ||||||||[||||||||||||]PAM   NA   Intergenic   NA   NA

T14589   AACAGTTCAACACGGTGGGCTGG   1000
Oligo fwd   CACCAACAGTTCAACACGGTGGGC
Oligo rev   AAACGCCCACCGTGTTGAACTGTT
Chromosome   start   end   strand   MM   target_seq   PAM   alignment   distance   position   gene name   gene id
GU305914.1   192030   192052   +   0   AACAGTTCAACACGGTGGGC   TGG   ||||||||[||||||||||||]PAM   NA   Intergenic   NA   NA

T14590   ACAGTTCAACACGGTGGGCTGGG   1000
Oligo fwd   CACCACAGTTCAACACGGTGGGCT
Oligo rev   AAACAGCCCACCGTGTTGAACTGT
Chromosome   start   end   strand   MM   target_seq   PAM   alignment   distance   position   gene name   gene id
GU305914.1   192031   192053   +   0   ACAGTTCAACACGGTGGGCT   GGG   ||||||||[||||||||||||]PAM   NA   Intergenic   NA   NA

T14591   GTACATGTTCCTCACGAACTCGG   1000
Oligo fwd   CACCGTACATGTTCCTCACGAACT
Oligo rev   AAACAGTTCGTGAGGAACATGTAC
Chromosome   start   end   strand   MM   target_seq   PAM   alignment   distance   position   gene name   gene id
GU305914.1   192058   192080   +   0   GTACATGTTCCTCACGAACT   CGG   ||||||||[||||||||||||]PAM   NA   Intergenic   NA   NA

T14592   TACATGTTCCTCACGAACTCGGG   1000
Oligo fwd   CACCTACATGTTCCTCACGAACTC
Oligo rev   AAACGAGTTCGTGAGGAACATGTA
Chromosome   start   end   strand   MM   target_seq   PAM   alignment   distance   position   gene name   gene id
GU305914.1   192059   192081   +   0   TACATGTTCCTCACGAACTC   GGG   ||||||||[||||||||||||]PAM   NA   Intergenic   NA   NA

T14593   GGGCTGTTCGAACGACAAGCCGG   1000
Oligo fwd   CACCGGGCTGTTCGAACGACAAGC
Oligo rev   AAACGCTTGTCGTTCGAACAGCCC
Chromosome   start   end   strand   MM   target_seq   PAM   alignment   distance   position   gene name   gene id
GU305914.1   192079   192101   +   0   GGGCTGTTCGAACGACAAGC   CGG   ||||||||[||||||||||||]PAM   NA   Intergenic   NA   NA

T14594   CTGTTCGAACGACAAGCCGGAGG   1000
Oligo fwd   CACCCTGTTCGAACGACAAGCCGG
Oligo rev   AAACCCGGCTTGTCGTTCGAACAG
Chromosome   start   end   strand   MM   target_seq   PAM   alignment   distance   position   gene name   gene id
GU305914.1   192082   192104   +   0   CTGTTCGAACGACAAGCCGG   AGG   ||||||||[||||||||||||]PAM   NA   Intergenic   NA   NA

T14595   TGTTCGAACGACAAGCCGGAGGG   1000
Oligo fwd   CACCTGTTCGAACGACAAGCCGGA
Oligo rev   AAACTCCGGCTTGTCGTTCGAACA
Chromosome   start   end   strand   MM   target_seq   PAM   alignment   distance   position   gene name   gene id
GU305914.1   192083   192105   +   0   TGTTCGAACGACAAGCCGGA   GGG   ||||||||[||||||||||||]PAM   NA   Intergenic   NA   NA

T14596   CCGGAGGGATCCCACCCCCCCGG   1000
Oligo fwd   CACCCCGGAGGGATCCCACCCCCC
Oligo rev   AAACGGGGGGTGGGATCCCTCCGG
Chromosome   start   end   strand   MM   target_seq   PAM   alignment   distance   position   gene name   gene id
GU305914.1   192098   192120   +   0   CCGGAGGGATCCCACCCCCC   CGG   ||||||||[||||||||||||]PAM   NA   Intergenic   NA   NA

T14597   CGGAGGGATCCCACCCCCCCGGG   1000

Oligo fwd CACCCGGAGGGATCCCACCCCCCC
Oligo rev AAACGGGGGGGTGGGATCCCTCCG
Chromosome start end strand MM target_seq PAM alignment distance position gene name gene id
GU305914.1 192099 192121 + 0 CGGAGGGATCCCACCCCCCC GGG ||||||||[||||||||||||]PAM NA Intergenic NA NA

T14598 GGAGGGATCCCACCCCCCCGGGG 1000
Oligo fwd CACCGGAGGGATCCCACCCCCCCG
Oligo rev AAACCGGGGGGGTGGGATCCCTCC
Chromosome start end strand MM target_seq PAM alignment distance position gene name gene id
GU305914.1 192100 192122 + 0 GGAGGGATCCCACCCCCCCG GGG ||||||||[||||||||||||]PAM NA Intergenic NA NA

T14599 CGGGGCTATCAGTGCCGTCGTGG 1000
Oligo fwd CACCCGGGGCTATCAGTGCCGTCG
Oligo rev AAACCGACGGCACTGATAGCCCCG
Chromosome start end strand MM target_seq PAM alignment distance position gene name gene id
GU305914.1 192118 192140 + 0 CGGGGCTATCAGTGCCGTCG TGG ||||||||[||||||||||||]PAM NA Intergenic NA NA

T14600 TATCAGTGCCGTCGTGGTGACGG 1000
Oligo fwd CACCTATCAGTGCCGTCGTGGTGA
Oligo rev AAACTCACCACGACGGCACTGATA
Chromosome start end strand MM target_seq PAM alignment distance position gene name gene id
GU305914.1 192124 192146 + 0 TATCAGTGCCGTCGTGGTGA CGG ||||||||[||||||||||||]PAM NA Intergenic NA NA

T14601 GGCCGCGACTATCGCCGCGATGG 1000
Oligo fwd CACCGGCCGCGACTATCGCCGCGA
Oligo rev AAACTCGCGGCGATAGTCGCGGCC
Chromosome start end strand MM target_seq PAM alignment distance position gene name gene id
GU305914.1 192145 192167 + 0 GGCCGCGACTATCGCCGCGA TGG ||||||||[||||||||||||]PAM NA Intergenic NA NA

T14602 CCGATACCAGTATCGCCACATGG 1000
Oligo fwd CACCCCGATACCAGTATCGCCACA
Oligo rev AAACTGTGGCGATACTGGTATCGG
Chromosome start end strand MM target_seq PAM alignment distance position gene name gene id
GU305914.1 192172 192194 + 0 CCGATACCAGTATCGCCACA TGG ||||||||[||||||||||||]PAM NA Intergenic NA NA

T14603 GTATCGCCACATGGAGAGAGAGG 1000
Oligo fwd CACCGTATCGCCACATGGAGAGAG
Oligo rev AAACCTCTCTCCATGTGGCGATAC
Chromosome start end strand MM target_seq PAM alignment distance position gene name gene id
GU305914.1 192181 192203 + 0 GTATCGCCACATGGAGAGAG AGG ||||||||[||||||||||||]PAM NA Intergenic NA NA

T14604 GCAAGAGAGCCGTGACCCGTCGG 1000
Oligo fwd CACCGCAAGAGAGCCGTGACCCGT
Oligo rev AAACACGGGTCACGGCTCTCTTGC
Chromosome start end strand MM target_seq PAM alignment distance position gene name gene id
GU305914.1 192203 192225 + 0 GCAAGAGAGCCGTGACCCGT CGG ||||||||[||||||||||||]PAM NA Intergenic NA NA

T14605 CGAAAATAACAGAGTAATGAAGG 1000
Oligo fwd CACCCGAAAATAACAGAGTAATGA
Oligo rev AAACTCATTACTCTGTTATTTTCG
Chromosome start end strand MM target_seq PAM alignment distance position gene name gene id
GU305914.1 192230 192252 + 0 CGAAAATAACAGAGTAATGA AGG ||||||||[||||||||||||]PAM NA Intergenic NA NA

T14606 GAGCTTTTATTTATATATGCTGG 1000
Oligo fwd CACCGAGCTTTTATTTATATATGC
Oligo rev AAACGCATATATAAATAAAAGCTC
Chromosome start end strand MM target_seq PAM alignment distance position gene name gene id
GU305914.1 192270 192292 + 0 GAGCTTTTATTTATATATGC TGG ||||||||[||||||||||||]PAM NA Intergenic NA NA

T14607 AGCTTTTATTTATATATGCTGGG 1000
Oligo fwd CACCAGCTTTTATTTATATATGCT
Oligo rev AAACAGCATATATAAATAAAAGCT
Chromosome start end strand MM target_seq PAM alignment distance position gene name gene id
GU305914.1 192271 192293 + 0 AGCTTTTATTTATATATGCT GGG ||||||||[||||||||||||]PAM NA Intergenic NA NA

T14608 GCTTTTATTTATATATGCTGGGG 1000
Oligo fwd CACCGCTTTTATTTATATATGCTG
Oligo rev AAACCAGCATATATAAATAAAAGC
Chromosome start end strand MM target_seq PAM alignment distance position gene name gene id
GU305914.1 192272 192294 + 0 GCTTTTATTTATATATGCTG GGG ||||||||[||||||||||||]PAM NA Intergenic NA NA

T14609 CTTTTATTTATATATGCTGGGGG 1000
Oligo fwd CACCCTTTTATTTATATATGCTGG
Oligo rev AAACCCAGCATATATAAATAAAAG
Chromosome start end strand MM target_seq PAM alignment distance position gene name gene id
GU305914.1 192273 192295 + 0 CTTTTATTTATATATGCTGG GGG ||||||||[||||||||||||]PAM NA Intergenic NA NA

T14610 TACGTGTGACGTACGCAACCCGG 1000
Oligo fwd CACCTACGTGTGACGTACGCAACC
Oligo rev AAACGGTTGCGTACGTCACACGTA
Chromosome start end strand MM target_seq PAM alignment distance position gene name gene id
GU305914.1 192334 192356 + 0 TACGTGTGACGTACGCAACC CGG ||||||||[||||||||||||]PAM NA Intergenic NA NA

T14611 TGTGACGTACGCAACCCGGCAGG 1000
Oligo fwd CACCTGTGACGTACGCAACCCGGC
Oligo rev AAACGCCGGGTTGCGTACGTCACA
Chromosome start end strand MM target_seq PAM alignment distance position gene name gene id
GU305914.1 192338 192360 + 0 TGTGACGTACGCAACCCGGC AGG ||||||||[||||||||||||]PAM NA Intergenic NA NA

<table>
<tr><td>T14612</td><td>ACGCAACCCGGCAGGTTACACGG</td><td>1000</td><td></td><td></td><td></td><td></td><td></td><td></td><td></td><td></td><td></td></tr>
<tr><td>Oligo fwd</td><td>CACCACGCAACCCGGCAGGTTACA</td><td></td><td></td><td></td><td></td><td></td><td></td><td></td><td></td><td></td><td></td></tr>
<tr><td>Oligo rev</td><td>AAACTGTAACCTGCCGGGTTGCGT</td><td></td><td></td><td></td><td></td><td></td><td></td><td></td><td></td><td></td><td></td></tr>
<tr><td>Chromosome</td><td>start</td><td>end</td><td>strand</td><td>MM</td><td>target_seq</td><td>PAM</td><td>alignment</td><td>distance</td><td>position</td><td>gene name</td><td>gene id</td></tr>
<tr><td>GU305914.1</td><td>192346</td><td>192368</td><td>+</td><td>0</td><td>ACGCAACCCGGCAGGTTACA</td><td>CGG</td><td>||||||||[||||||||||||]PAM</td><td>NA</td><td>Intergenic</td><td>NA</td><td>NA</td></tr>
<tr><td></td><td></td><td></td><td></td><td></td><td></td><td></td><td></td><td></td><td></td><td></td><td></td></tr>
<tr><td>T14613</td><td>CAACCCGGCAGGTTACACGGCGG</td><td>1000</td><td></td><td></td><td></td><td></td><td></td><td></td><td></td><td></td><td></td></tr>
<tr><td>Oligo fwd</td><td>CACCCAACCCGGCAGGTTACACGG</td><td></td><td></td><td></td><td></td><td></td><td></td><td></td><td></td><td></td><td></td></tr>
<tr><td>Oligo rev</td><td>AAACCCGTGTAACCTGCCGGGTTG</td><td></td><td></td><td></td><td></td><td></td><td></td><td></td><td></td><td></td><td></td></tr>
<tr><td>Chromosome</td><td>start</td><td>end</td><td>strand</td><td>MM</td><td>target_seq</td><td>PAM</td><td>alignment</td><td>distance</td><td>position</td><td>gene name</td><td>gene id</td></tr>
<tr><td>GU305914.1</td><td>192349</td><td>192371</td><td>+</td><td>0</td><td>CAACCCGGCAGGTTACACGG</td><td>CGG</td><td>||||||||[||||||||||||]PAM</td><td>NA</td><td>Intergenic</td><td>NA</td><td>NA</td></tr>
<tr><td></td><td></td><td></td><td></td><td></td><td></td><td></td><td></td><td></td><td></td><td></td><td></td></tr>
<tr><td>T14614</td><td>ACGAGATCCAGAACGAGATGTGG</td><td>1000</td><td></td><td></td><td></td><td></td><td></td><td></td><td></td><td></td><td></td></tr>
<tr><td>Oligo fwd</td><td>CACCACGAGATCCAGAACGAGATG</td><td></td><td></td><td></td><td></td><td></td><td></td><td></td><td></td><td></td><td></td></tr>
<tr><td>Oligo rev</td><td>AAACCATCTCGTTCTGGATCTCGT</td><td></td><td></td><td></td><td></td><td></td><td></td><td></td><td></td><td></td><td></td></tr>
<tr><td>Chromosome</td><td>start</td><td>end</td><td>strand</td><td>MM</td><td>target_seq</td><td>PAM</td><td>alignment</td><td>distance</td><td>position</td><td>gene name</td><td>gene id</td></tr>
<tr><td>GU305914.1</td><td>192374</td><td>192396</td><td>+</td><td>0</td><td>ACGAGATCCAGAACGAGATG</td><td>TGG</td><td>||||||||[||||||||||||]PAM</td><td>NA</td><td>Intergenic</td><td>NA</td><td>NA</td></tr>
<tr><td></td><td></td><td></td><td></td><td></td><td></td><td></td><td></td><td></td><td></td><td></td><td></td></tr>
<tr><td>T14615</td><td>ACGAGATGTGGACCAACACGAGG</td><td>1000</td><td></td><td></td><td></td><td></td><td></td><td></td><td></td><td></td><td></td></tr>
<tr><td>Oligo fwd</td><td>CACCACGAGATGTGGACCAACACG</td><td></td><td></td><td></td><td></td><td></td><td></td><td></td><td></td><td></td><td></td></tr>
<tr><td>Oligo rev</td><td>AAACCGTGTTGGTCCACATCTCGT</td><td></td><td></td><td></td><td></td><td></td><td></td><td></td><td></td><td></td><td></td></tr>
<tr><td>Chromosome</td><td>start</td><td>end</td><td>strand</td><td>MM</td><td>target_seq</td><td>PAM</td><td>alignment</td><td>distance</td><td>position</td><td>gene name</td><td>gene id</td></tr>
<tr><td>GU305914.1</td><td>192386</td><td>192408</td><td>+</td><td>0</td><td>ACGAGATGTGGACCAACACG</td><td>AGG</td><td>||||||||[||||||||||||]PAM</td><td>NA</td><td>Intergenic</td><td>NA</td><td>NA</td></tr>
<tr><td></td><td></td><td></td><td></td><td></td><td></td><td></td><td></td><td></td><td></td><td></td><td></td></tr>
<tr><td>T14616</td><td>GTGGACCAACACGAGGATCGCGG</td><td>1000</td><td></td><td></td><td></td><td></td><td></td><td></td><td></td><td></td><td></td></tr>
<tr><td>Oligo fwd</td><td>CACCGTGGACCAACACGAGGATCG</td><td></td><td></td><td></td><td></td><td></td><td></td><td></td><td></td><td></td><td></td></tr>
<tr><td>Oligo rev</td><td>AAACCGATCCTCGTGTTGGTCCAC</td><td></td><td></td><td></td><td></td><td></td><td></td><td></td><td></td><td></td><td></td></tr>
<tr><td>Chromosome</td><td>start</td><td>end</td><td>strand</td><td>MM</td><td>target_seq</td><td>PAM</td><td>alignment</td><td>distance</td><td>position</td><td>gene name</td><td>gene id</td></tr>
<tr><td>GU305914.1</td><td>192393</td><td>192415</td><td>+</td><td>0</td><td>GTGGACCAACACGAGGATCG</td><td>CGG</td><td>||||||||[||||||||||||]PAM</td><td>NA</td><td>Intergenic</td><td>NA</td><td>NA</td></tr>
<tr><td></td><td></td><td></td><td></td><td></td><td></td><td></td><td></td><td></td><td></td><td></td><td></td></tr>
<tr><td>T14617</td><td>CCAACACGAGGATCGCGGCTAGG</td><td>1000</td><td></td><td></td><td></td><td></td><td></td><td></td><td></td><td></td><td></td></tr>
<tr><td>Oligo fwd</td><td>CACCCCAACACGAGGATCGCGGCT</td><td></td><td></td><td></td><td></td><td></td><td></td><td></td><td></td><td></td><td></td></tr>
<tr><td>Oligo rev</td><td>AAACAGCCGCGATCCTCGTGTTGG</td><td></td><td></td><td></td><td></td><td></td><td></td><td></td><td></td><td></td><td></td></tr>
<tr><td>Chromosome</td><td>start</td><td>end</td><td>strand</td><td>MM</td><td>target_seq</td><td>PAM</td><td>alignment</td><td>distance</td><td>position</td><td>gene name</td><td>gene id</td></tr>
<tr><td>GU305914.1</td><td>192398</td><td>192420</td><td>+</td><td>0</td><td>CCAACACGAGGATCGCGGCT</td><td>AGG</td><td>||||||||[||||||||||||]PAM</td><td>NA</td><td>Intergenic</td><td>NA</td><td>NA</td></tr>
<tr><td></td><td></td><td></td><td></td><td></td><td></td><td></td><td></td><td></td><td></td><td></td><td></td></tr>
<tr><td>T14618</td><td>GGCTAGGAAGAAACAAGCGACGG</td><td>1000</td><td></td><td></td><td></td><td></td><td></td><td></td><td></td><td></td><td></td></tr>
<tr><td>Oligo fwd</td><td>CACCGGCTAGGAAGAAACAAGCGA</td><td></td><td></td><td></td><td></td><td></td><td></td><td></td><td></td><td></td><td></td></tr>
<tr><td>Oligo rev</td><td>AAACTCGCTTGTTTCTTCCTAGCC</td><td></td><td></td><td></td><td></td><td></td><td></td><td></td><td></td><td></td><td></td></tr>
<tr><td>Chromosome</td><td>start</td><td>end</td><td>strand</td><td>MM</td><td>target_seq</td><td>PAM</td><td>alignment</td><td>distance</td><td>position</td><td>gene name</td><td>gene id</td></tr>
<tr><td>GU305914.1</td><td>192414</td><td>192436</td><td>+</td><td>0</td><td>GGCTAGGAAGAAACAAGCGA</td><td>CGG</td><td>||||||||[||||||||||||]PAM</td><td>NA</td><td>Intergenic</td><td>NA</td><td>NA</td></tr>
<tr><td></td><td></td><td></td><td></td><td></td><td></td><td></td><td></td><td></td><td></td><td></td><td></td></tr>
<tr><td>T14619</td><td>AACAAGCGACGGCCGCCAGACGG</td><td>1000</td><td></td><td></td><td></td><td></td><td></td><td></td><td></td><td></td><td></td></tr>
<tr><td>Oligo fwd</td><td>CACCAACAAGCGACGGCCGCCAGA</td><td></td><td></td><td></td><td></td><td></td><td></td><td></td><td></td><td></td><td></td></tr>
<tr><td>Oligo rev</td><td>AAACTCTGGCGGCCGTCGCTTGTT</td><td></td><td></td><td></td><td></td><td></td><td></td><td></td><td></td><td></td><td></td></tr>
<tr><td>Chromosome</td><td>start</td><td>end</td><td>strand</td><td>MM</td><td>target_seq</td><td>PAM</td><td>alignment</td><td>distance</td><td>position</td><td>gene name</td><td>gene id</td></tr>
<tr><td>GU305914.1</td><td>192425</td><td>192447</td><td>+</td><td>0</td><td>AACAAGCGACGGCCGCCAGA</td><td>CGG</td><td>||||||||[||||||||||||]PAM</td><td>NA</td><td>Intergenic</td><td>NA</td><td>NA</td></tr>
<tr><td></td><td></td><td></td><td></td><td></td><td></td><td></td><td></td><td></td><td></td><td></td><td></td></tr>
<tr><td>T14620</td><td>GCGCTGACGCGAGCGAATAGAGG</td><td>1000</td><td></td><td></td><td></td><td></td><td></td><td></td><td></td><td></td><td></td></tr>
<tr><td>Oligo fwd</td><td>CACCGCGCTGACGCGAGCGAATAG</td><td></td><td></td><td></td><td></td><td></td><td></td><td></td><td></td><td></td><td></td></tr>
<tr><td>Oligo rev</td><td>AAACCTATTCGCTCGCGTCAGCGC</td><td></td><td></td><td></td><td></td><td></td><td></td><td></td><td></td><td></td><td></td></tr>
<tr><td>Chromosome</td><td>start</td><td>end</td><td>strand</td><td>MM</td><td>target_seq</td><td>PAM</td><td>alignment</td><td>distance</td><td>position</td><td>gene name</td><td>gene id</td></tr>
<tr><td>GU305914.1</td><td>192454</td><td>192476</td><td>+</td><td>0</td><td>GCGCTGACGCGAGCGAATAG</td><td>AGG</td><td>||||||||[||||||||||||]PAM</td><td>NA</td><td>Intergenic</td><td>NA</td><td>NA</td></tr>
<tr><td></td><td></td><td></td><td></td><td></td><td></td><td></td><td></td><td></td><td></td><td></td><td></td></tr>
<tr><td>T14621</td><td>GCGAGCGAATAGAGGATAGTAGG</td><td>1000</td><td></td><td></td><td></td><td></td><td></td><td></td><td></td><td></td><td></td></tr>
<tr><td>Oligo fwd</td><td>CACCGCGAGCGAATAGAGGATAGT</td><td></td><td></td><td></td><td></td><td></td><td></td><td></td><td></td><td></td><td></td></tr>
<tr><td>Oligo rev</td><td>AAACACTATCCTCTATTCGCTCGC</td><td></td><td></td><td></td><td></td><td></td><td></td><td></td><td></td><td></td><td></td></tr>
<tr><td>Chromosome</td><td>start</td><td>end</td><td>strand</td><td>MM</td><td>target_seq</td><td>PAM</td><td>alignment</td><td>distance</td><td>position</td><td>gene name</td><td>gene id</td></tr>
<tr><td>GU305914.1</td><td>192462</td><td>192484</td><td>+</td><td>0</td><td>GCGAGCGAATAGAGGATAGT</td><td>AGG</td><td>||||||||[||||||||||||]PAM</td><td>NA</td><td>Intergenic</td><td>NA</td><td>NA</td></tr>
<tr><td></td><td></td><td></td><td></td><td></td><td></td><td></td><td></td><td></td><td></td><td></td><td></td></tr>
<tr><td>T14622</td><td>CGAGCGAATAGAGGATAGTAGGG</td><td>1000</td><td></td><td></td><td></td><td></td><td></td><td></td><td></td><td></td><td></td></tr>
<tr><td>Oligo fwd</td><td>CACCCGAGCGAATAGAGGATAGTA</td><td></td><td></td><td></td><td></td><td></td><td></td><td></td><td></td><td></td><td></td></tr>
<tr><td>Oligo rev</td><td>AAACTACTATCCTCTATTCGCTCG</td><td></td><td></td><td></td><td></td><td></td><td></td><td></td><td></td><td></td><td></td></tr>
<tr><td>Chromosome</td><td>start</td><td>end</td><td>strand</td><td>MM</td><td>target_seq</td><td>PAM</td><td>alignment</td><td>distance</td><td>position</td><td>gene name</td><td>gene id</td></tr>
<tr><td>GU305914.1</td><td>192463</td><td>192485</td><td>+</td><td>0</td><td>CGAGCGAATAGAGGATAGTA</td><td>GGG</td><td>||||||||[||||||||||||]PAM</td><td>NA</td><td>Intergenic</td><td>NA</td><td>NA</td></tr>
<tr><td></td><td></td><td></td><td></td><td></td><td></td><td></td><td></td><td></td><td></td><td></td><td></td></tr>
<tr><td>T14623</td><td>GAGCGAATAGAGGATAGTAGGGG</td><td>1000</td><td></td><td></td><td></td><td></td><td></td><td></td><td></td><td></td><td></td></tr>
<tr><td>Oligo fwd</td><td>CACCGAGCGAATAGAGGATAGTAG</td><td></td><td></td><td></td><td></td><td></td><td></td><td></td><td></td><td></td><td></td></tr>
<tr><td>Oligo rev</td><td>AAACCTACTATCCTCTATTCGCTC</td><td></td><td></td><td></td><td></td><td></td><td></td><td></td><td></td><td></td><td></td></tr>
<tr><td>Chromosome</td><td>start</td><td>end</td><td>strand</td><td>MM</td><td>target_seq</td><td>PAM</td><td>alignment</td><td>distance</td><td>position</td><td>gene name</td><td>gene id</td></tr>
<tr><td>GU305914.1</td><td>192464</td><td>192486</td><td>+</td><td>0</td><td>GAGCGAATAGAGGATAGTAG</td><td>GGG</td><td>||||||||[||||||||||||]PAM</td><td>NA</td><td>Intergenic</td><td>NA</td><td>NA</td></tr>
<tr><td></td><td></td><td></td><td></td><td></td><td></td><td></td><td></td><td></td><td></td><td></td><td></td></tr>
<tr><td>T14624</td><td>TAGTAGGGGTAGCTTTTGAGCGG</td><td>1000</td><td></td><td></td><td></td><td></td><td></td><td></td><td></td><td></td><td></td></tr>
<tr><td>Oligo fwd</td><td>CACCTAGTAGGGGTAGCTTTTGAG</td><td></td><td></td><td></td><td></td><td></td><td></td><td></td><td></td><td></td><td></td></tr>
<tr><td>Oligo rev</td><td>AAACCTCAAAAGCTACCCCTACTA</td><td></td><td></td><td></td><td></td><td></td><td></td><td></td><td></td><td></td><td></td></tr>
<tr><td>Chromosome</td><td>start</td><td>end</td><td>strand</td><td>MM</td><td>target_seq</td><td>PAM</td><td>alignment</td><td>distance</td><td>position</td><td>gene name</td><td>gene id</td></tr>
<tr><td>GU305914.1</td><td>192478</td><td>192500</td><td>+</td><td>0</td><td>TAGTAGGGGTAGCTTTTGAG</td><td>CGG</td><td>||||||||[||||||||||||]PAM</td><td>NA</td><td>Intergenic</td><td>NA</td><td>NA</td></tr>
<tr><td></td><td></td><td></td><td></td><td></td><td></td><td></td><td></td><td></td><td></td><td></td><td></td></tr>
<tr><td>T14625</td><td>CAACAGAAAGTCATCGTCGTCGG</td><td>1000</td><td></td><td></td><td></td><td></td><td></td><td></td><td></td><td></td><td></td></tr>
<tr><td>Oligo fwd</td><td>CACCCAACAGAAAGTCATCGTCGT</td><td></td><td></td><td></td><td></td><td></td><td></td><td></td><td></td><td></td><td></td></tr>
<tr><td>Oligo rev</td><td>AAACACGACGATGACTTTCTGTTG</td><td></td><td></td><td></td><td></td><td></td><td></td><td></td><td></td><td></td><td></td></tr>
<tr><td>Chromosome</td><td>start</td><td>end</td><td>strand</td><td>MM</td><td>target_seq</td><td>PAM</td><td>alignment</td><td>distance</td><td>position</td><td>gene name</td><td>gene id</td></tr>
<tr><td>GU305914.1</td><td>192513</td><td>192535</td><td>+</td><td>0</td><td>CAACAGAAAGTCATCGTCGT</td><td>CGG</td><td>||||||||[||||||||||||]PAM</td><td>NA</td><td>Intergenic</td><td>NA</td><td>NA</td></tr>
<tr><td></td><td></td><td></td><td></td><td></td><td></td><td></td><td></td><td></td><td></td><td></td><td></td></tr>
<tr><td>T14626</td><td>TCTGATCTCTTCTAGTTTCTTGG</td><td>1000</td><td></td><td></td><td></td><td></td><td></td><td></td><td></td><td></td><td></td></tr>
<tr><td>Oligo fwd</td><td>CACCTCTGATCTCTTCTAGTTTCT</td><td></td><td></td><td></td><td></td><td></td><td></td><td></td><td></td><td></td><td></td></tr>
<tr><td>Oligo rev</td><td>AAACAGAAACTAGAAGAGATCAGA</td><td></td><td></td><td></td><td></td><td></td><td></td><td></td><td></td><td></td><td></td></tr>
<tr><td>Chromosome</td><td>start</td><td>end</td><td>strand</td><td>MM</td><td>target_seq</td><td>PAM</td><td>alignment</td><td>distance</td><td>position</td><td>gene name</td><td>gene id</td></tr>
<tr><td>GU305914.1</td><td>192552</td><td>192574</td><td>+</td><td>0</td><td>TCTGATCTCTTCTAGTTTCT</td><td>TGG</td><td>||||||||[||||||||||||]PAM</td><td>NA</td><td>Intergenic</td><td>NA</td><td>NA</td></tr>
</table>

| | | | | | | | | | | | | |
|---|---|---|---|---|---|---|---|---|---|---|---|---|
| T14627 | GATCTCTTCTAGTTTCTTGGCGG | | 1000 | | | | | | | | | |
| Oligo fwd | CACCGATCTCTTCTAGTTTCTTGG | | | | | | | | | | | |
| Oligo rev | AAACCCAAGAAACTAGAAGAGATC | | | | | | | | | | | |
| Chromosome | start | | end | strand | MM | target_seq | PAM | alignment | distance | position | gene name | gene id |
| GU305914.1 | | 192555 | 192577 | + | 0 | GATCTCTTCTAGTTTCTTGG | CGG | \|\|\|\|\|\|\|\|[\|\|\|\|\|\|\|\|\|\|\|\|]PAM | NA | Intergenic | NA | NA |
| T14628 | ATCTCTTCTAGTTTCTTGGCGGG | | 1000 | | | | | | | | | |
| Oligo fwd | CACCATCTCTTCTAGTTTCTTGGC | | | | | | | | | | | |
| Oligo rev | AAACGCCAAGAAACTAGAAGAGAT | | | | | | | | | | | |
| Chromosome | start | | end | strand | MM | target_seq | PAM | alignment | distance | position | gene name | gene id |
| GU305914.1 | | 192556 | 192578 | + | 0 | ATCTCTTCTAGTTTCTTGGC | GGG | \|\|\|\|\|\|\|\|[\|\|\|\|\|\|\|\|\|\|\|\|]PAM | NA | Intergenic | NA | NA |
| T14629 | TTCTAGTTTCTTGGCGGGTGAGG | | 1000 | | | | | | | | | |
| Oligo fwd | CACCTTCTAGTTTCTTGGCGGGTG | | | | | | | | | | | |
| Oligo rev | AAACCACCCGCCAAGAAACTAGAA | | | | | | | | | | | |
| Chromosome | start | | end | strand | MM | target_seq | PAM | alignment | distance | position | gene name | gene id |
| GU305914.1 | | 192561 | 192583 | + | 0 | TTCTAGTTTCTTGGCGGGTG | AGG | \|\|\|\|\|\|\|\|[\|\|\|\|\|\|\|\|\|\|\|\|]PAM | NA | Intergenic | NA | NA |
| T14630 | TCTAGTTTCTTGGCGGGTGAGGG | | 1000 | | | | | | | | | |
| Oligo fwd | CACCTCTAGTTTCTTGGCGGGTGA | | | | | | | | | | | |
| Oligo rev | AAACTCACCCGCCAAGAAACTAGA | | | | | | | | | | | |
| Chromosome | start | | end | strand | MM | target_seq | PAM | alignment | distance | position | gene name | gene id |
| GU305914.1 | | 192562 | 192584 | + | 0 | TCTAGTTTCTTGGCGGGTGA | GGG | \|\|\|\|\|\|\|\|[\|\|\|\|\|\|\|\|\|\|\|\|]PAM | NA | Intergenic | NA | NA |
| T14631 | CTAGTTTCTTGGCGGGTGAGGGG | | 1000 | | | | | | | | | |
| Oligo fwd | CACCCTAGTTTCTTGGCGGGTGAG | | | | | | | | | | | |
| Oligo rev | AAACCTCACCCGCCAAGAAACTAG | | | | | | | | | | | |
| Chromosome | start | | end | strand | MM | target_seq | PAM | alignment | distance | position | gene name | gene id |
| GU305914.1 | | 192563 | 192585 | + | 0 | CTAGTTTCTTGGCGGGTGAG | GGG | \|\|\|\|\|\|\|\|[\|\|\|\|\|\|\|\|\|\|\|\|]PAM | NA | Intergenic | NA | NA |
| T14632 | TAGTTTCTTGGCGGGTGAGGGGG | | 1000 | | | | | | | | | |
| Oligo fwd | CACCTAGTTTCTTGGCGGGTGAGG | | | | | | | | | | | |
| Oligo rev | AAACCCTCACCCGCCAAGAAACTA | | | | | | | | | | | |
| Chromosome | start | | end | strand | MM | target_seq | PAM | alignment | distance | position | gene name | gene id |
| GU305914.1 | | 192564 | 192586 | + | 0 | TAGTTTCTTGGCGGGTGAGG | GGG | \|\|\|\|\|\|\|\|[\|\|\|\|\|\|\|\|\|\|\|\|]PAM | NA | Intergenic | NA | NA |
| T14633 | AGTTTCTTGGCGGGTGAGGGGGG | | 1000 | | | | | | | | | |
| Oligo fwd | CACCAGTTTCTTGGCGGGTGAGGG | | | | | | | | | | | |
| Oligo rev | AAACCCCTCACCCGCCAAGAAACT | | | | | | | | | | | |
| Chromosome | start | | end | strand | MM | target_seq | PAM | alignment | distance | position | gene name | gene id |
| GU305914.1 | | 192565 | 192587 | + | 0 | AGTTTCTTGGCGGGTGAGGG | GGG | \|\|\|\|\|\|\|\|[\|\|\|\|\|\|\|\|\|\|\|\|]PAM | NA | Intergenic | NA | NA |
| T14634 | TTGGCGGGTGAGGGGGGTGATGG | | 1000 | | | | | | | | | |
| Oligo fwd | CACCTTGGCGGGTGAGGGGGGTGA | | | | | | | | | | | |
| Oligo rev | AAACTCACCCCCCTCACCCGCCAA | | | | | | | | | | | |
| Chromosome | start | | end | strand | MM | target_seq | PAM | alignment | distance | position | gene name | gene id |
| GU305914.1 | | 192571 | 192593 | + | 0 | TTGGCGGGTGAGGGGGGTGA | TGG | \|\|\|\|\|\|\|\|[\|\|\|\|\|\|\|\|\|\|\|\|]PAM | NA | Intergenic | NA | NA |
| T14635 | TGAGGGGGGTGATGGTGTTGTGG | | 1000 | | | | | | | | | |
| Oligo fwd | CACCTGAGGGGGGTGATGGTGTTG | | | | | | | | | | | |
| Oligo rev | AAACCAACACCATCACCCCCCTCA | | | | | | | | | | | |
| Chromosome | start | | end | strand | MM | target_seq | PAM | alignment | distance | position | gene name | gene id |
| GU305914.1 | | 192579 | 192601 | + | 0 | TGAGGGGGGTGATGGTGTTG | TGG | \|\|\|\|\|\|\|\|[\|\|\|\|\|\|\|\|\|\|\|\|]PAM | NA | Intergenic | NA | NA |
| GU305914.1 | | 44676 | 44698 | - | 3 | TGAGGCGAGTGATGGTGGTG | GGG | \|\|\|\|\|-\|-[\|\|\|\|\|\|\|\|\|-\|\|]PAM | NA | Intergenic | NA | NA |
| T14636 | TGTGGTCGTCGTAGTAGTAGTGG | | 1000 | | | | | | | | | |
| Oligo fwd | CACCTGTGGTCGTCGTAGTAGTAG | | | | | | | | | | | |
| Oligo rev | AAACCTACTACTACGACGACCACA | | | | | | | | | | | |
| Chromosome | start | | end | strand | MM | target_seq | PAM | alignment | distance | position | gene name | gene id |
| GU305914.1 | | 192597 | 192619 | + | 0 | TGTGGTCGTCGTAGTAGTAG | TGG | \|\|\|\|\|\|\|\|[\|\|\|\|\|\|\|\|\|\|\|\|]PAM | NA | Intergenic | NA | NA |
| GU305914.1 | | 213586 | 213608 | + | 4 | GGTGGTGGTGGTAGTGGTAG | TGG | -\|\|\|\|\|-\|[\|-\|\|\|\|\|-\|\|\|\|]PAM | NA | Intergenic | NA | NA |
| GU305914.1 | | 192600 | 192622 | + | 4 | GGTCGTCGTAGTAGTAGTGG | TGG | -\|\|-\|\|\|\|[\|-\|\|\|\|\|\|\|\|-\|]PAM | NA | Intergenic | NA | NA |
| T14637 | GGTCGTCGTAGTAGTAGTGGTGG | | 1000 | | | | | | | | | |
| Oligo fwd | CACCGGTCGTCGTAGTAGTAGTGG | | | | | | | | | | | |
| Oligo rev | AAACCCACTACTACTACGACGACC | | | | | | | | | | | |
| Chromosome | start | | end | strand | MM | target_seq | PAM | alignment | distance | position | gene name | gene id |
| GU305914.1 | | 192600 | 192622 | + | 0 | GGTCGTCGTAGTAGTAGTGG | TGG | \|\|\|\|\|\|\|\|[\|\|\|\|\|\|\|\|\|\|\|\|]PAM | NA | Intergenic | NA | NA |
| GU305914.1 | | 192603 | 192625 | + | 3 | CGTCGTAGTAGTAGTGGTGG | TGG | -\|\|\|\|\|-\|[\|\|\|\|\|\|\|-\|\|\|\|]PAM | NA | Intergenic | NA | NA |
| GU305914.1 | | 192597 | 192619 | + | 4 | TGTGGTCGTCGTAGTAGTAG | TGG | -\|\|-\|\|\|\|[\|-\|\|\|\|\|\|\|\|-\|]PAM | NA | Intergenic | NA | NA |
| T14638 | CGTCGTAGTAGTAGTGGTGGTGG | | 1000 | | | | | | | | | |
| Oligo fwd | CACCCGTCGTAGTAGTAGTGGTGG | | | | | | | | | | | |
| Oligo rev | AAACCCACCACTACTACTACGACG | | | | | | | | | | | |
| Chromosome | start | | end | strand | MM | target_seq | PAM | alignment | distance | position | gene name | gene id |
| GU305914.1 | | 192603 | 192625 | + | 0 | CGTCGTAGTAGTAGTGGTGG | TGG | \|\|\|\|\|\|\|\|[\|\|\|\|\|\|\|\|\|\|\|\|]PAM | NA | Intergenic | NA | NA |
| GU305914.1 | | 192600 | 192622 | + | 3 | GGTCGTCGTAGTAGTAGTGG | TGG | -\|\|\|\|\|-\|[\|\|\|\|\|\|\|-\|\|\|\|]PAM | NA | Intergenic | NA | NA |
| GU305914.1 | | 213640 | 213662 | + | 4 | CGCAGTAGTAGTGGCGGTGG | TGG | \|\|--\|\|\|\|[\|\|\|\|-\|-\|\|\|\|\|]PAM | NA | Intergenic | NA | NA |
| T14639 | GTAGTAGTGGTGGTGGTCTGAGG | | 1000 | | | | | | | | | |
| Oligo fwd | CACCGTAGTAGTGGTGGTGGTCTG | | | | | | | | | | | |
| Oligo rev | AAACCAGACCACCACCACTACTAC | | | | | | | | | | | |
| Chromosome | start | | end | strand | MM | target_seq | PAM | alignment | distance | position | gene name | gene id |
| GU305914.1 | | 192610 | 192632 | + | 0 | GTAGTAGTGGTGGTGGTCTG | AGG | \|\|\|\|\|\|\|\|[\|\|\|\|\|\|\|\|\|\|\|\|]PAM | NA | Intergenic | NA | NA |
| T14640 | TAGTAGTGGTGGTGGTCTGAGGG | | 1000 | | | | | | | | | |
| Oligo fwd | CACCTAGTAGTGGTGGTGGTCTGA | | | | | | | | | | | |
| Oligo rev | AAACTCAGACCACCACCACTACTA | | | | | | | | | | | |

<table>
<tr><td>Chromosome</td><td>start</td><td>end</td><td>strand</td><td>MM</td><td>target_seq</td><td>PAM</td><td>alignment</td><td>distance</td><td>position</td><td>gene name</td><td>gene id</td></tr>
<tr><td>GU305914.1</td><td>192611</td><td>192633</td><td>+</td><td>0</td><td>TAGTAGTGGTGGTGGTCTGA</td><td>GGG</td><td>||||||||[||||||||||||]PAM</td><td>NA</td><td>Intergenic</td><td>NA</td><td>NA</td></tr>
<tr><td>T14641</td><td>AGTAGTGGTGGTGGTCTGAGGGG</td><td>1000</td><td></td><td></td><td></td><td></td><td></td><td></td><td></td><td></td><td></td></tr>
<tr><td>Oligo fwd</td><td>CACCAGTAGTGGTGGTGGTCTGAG</td><td></td><td></td><td></td><td></td><td></td><td></td><td></td><td></td><td></td><td></td></tr>
<tr><td>Oligo rev</td><td>AAACCTCAGACCACCACCACTACT</td><td></td><td></td><td></td><td></td><td></td><td></td><td></td><td></td><td></td><td></td></tr>
<tr><td>Chromosome</td><td>start</td><td>end</td><td>strand</td><td>MM</td><td>target_seq</td><td>PAM</td><td>alignment</td><td>distance</td><td>position</td><td>gene name</td><td>gene id</td></tr>
<tr><td>GU305914.1</td><td>192612</td><td>192634</td><td>+</td><td>0</td><td>AGTAGTGGTGGTGGTCTGAG</td><td>GGG</td><td>||||||||[||||||||||||]PAM</td><td>NA</td><td>Intergenic</td><td>NA</td><td>NA</td></tr>
<tr><td>T14642</td><td>GGTGGTCTGAGGGGACGTGACGG</td><td>1000</td><td></td><td></td><td></td><td></td><td></td><td></td><td></td><td></td><td></td></tr>
<tr><td>Oligo fwd</td><td>CACCGGTGGTCTGAGGGGACGTGA</td><td></td><td></td><td></td><td></td><td></td><td></td><td></td><td></td><td></td><td></td></tr>
<tr><td>Oligo rev</td><td>AAACTCACGTCCCCTCAGACCACC</td><td></td><td></td><td></td><td></td><td></td><td></td><td></td><td></td><td></td><td></td></tr>
<tr><td>Chromosome</td><td>start</td><td>end</td><td>strand</td><td>MM</td><td>target_seq</td><td>PAM</td><td>alignment</td><td>distance</td><td>position</td><td>gene name</td><td>gene id</td></tr>
<tr><td>GU305914.1</td><td>192621</td><td>192643</td><td>+</td><td>0</td><td>GGTGGTCTGAGGGGACGTGA</td><td>CGG</td><td>||||||||[||||||||||||]PAM</td><td>NA</td><td>Intergenic</td><td>NA</td><td>NA</td></tr>
<tr><td>T14643</td><td>GTGGTCTGAGGGGACGTGACGGG</td><td>1000</td><td></td><td></td><td></td><td></td><td></td><td></td><td></td><td></td><td></td></tr>
<tr><td>Oligo fwd</td><td>CACCGTGGTCTGAGGGGACGTGAC</td><td></td><td></td><td></td><td></td><td></td><td></td><td></td><td></td><td></td><td></td></tr>
<tr><td>Oligo rev</td><td>AAACGTCACGTCCCCTCAGACCAC</td><td></td><td></td><td></td><td></td><td></td><td></td><td></td><td></td><td></td><td></td></tr>
<tr><td>Chromosome</td><td>start</td><td>end</td><td>strand</td><td>MM</td><td>target_seq</td><td>PAM</td><td>alignment</td><td>distance</td><td>position</td><td>gene name</td><td>gene id</td></tr>
<tr><td>GU305914.1</td><td>192622</td><td>192644</td><td>+</td><td>0</td><td>GTGGTCTGAGGGGACGTGAC</td><td>GGG</td><td>||||||||[||||||||||||]PAM</td><td>NA</td><td>Intergenic</td><td>NA</td><td>NA</td></tr>
<tr><td>T14644</td><td>TGACGGGCTCCTCTGTCACGAGG</td><td>1000</td><td></td><td></td><td></td><td></td><td></td><td></td><td></td><td></td><td></td></tr>
<tr><td>Oligo fwd</td><td>CACCTGACGGGCTCCTCTGTCACG</td><td></td><td></td><td></td><td></td><td></td><td></td><td></td><td></td><td></td><td></td></tr>
<tr><td>Oligo rev</td><td>AAACCGTGACAGAGGAGCCCGTCA</td><td></td><td></td><td></td><td></td><td></td><td></td><td></td><td></td><td></td><td></td></tr>
<tr><td>Chromosome</td><td>start</td><td>end</td><td>strand</td><td>MM</td><td>target_seq</td><td>PAM</td><td>alignment</td><td>distance</td><td>position</td><td>gene name</td><td>gene id</td></tr>
<tr><td>GU305914.1</td><td>192638</td><td>192660</td><td>+</td><td>0</td><td>TGACGGGCTCCTCTGTCACG</td><td>AGG</td><td>||||||||[||||||||||||]PAM</td><td>NA</td><td>Intergenic</td><td>NA</td><td>NA</td></tr>
<tr><td>T14645</td><td>GACGGGCTCCTCTGTCACGAGGG</td><td>1000</td><td></td><td></td><td></td><td></td><td></td><td></td><td></td><td></td><td></td></tr>
<tr><td>Oligo fwd</td><td>CACCGACGGGCTCCTCTGTCACGA</td><td></td><td></td><td></td><td></td><td></td><td></td><td></td><td></td><td></td><td></td></tr>
<tr><td>Oligo rev</td><td>AAACTCGTGACAGAGGAGCCCGTC</td><td></td><td></td><td></td><td></td><td></td><td></td><td></td><td></td><td></td><td></td></tr>
<tr><td>Chromosome</td><td>start</td><td>end</td><td>strand</td><td>MM</td><td>target_seq</td><td>PAM</td><td>alignment</td><td>distance</td><td>position</td><td>gene name</td><td>gene id</td></tr>
<tr><td>GU305914.1</td><td>192639</td><td>192661</td><td>+</td><td>0</td><td>GACGGGCTCCTCTGTCACGA</td><td>GGG</td><td>||||||||[||||||||||||]PAM</td><td>NA</td><td>Intergenic</td><td>NA</td><td>NA</td></tr>
<tr><td>T14646</td><td>CACGAGGGCGTATTCCATCCTGG</td><td>1000</td><td></td><td></td><td></td><td></td><td></td><td></td><td></td><td></td><td></td></tr>
<tr><td>Oligo fwd</td><td>CACCCACGAGGGCGTATTCCATCC</td><td></td><td></td><td></td><td></td><td></td><td></td><td></td><td></td><td></td><td></td></tr>
<tr><td>Oligo rev</td><td>AAACGGATGGAATACGCCCTCGTG</td><td></td><td></td><td></td><td></td><td></td><td></td><td></td><td></td><td></td><td></td></tr>
<tr><td>Chromosome</td><td>start</td><td>end</td><td>strand</td><td>MM</td><td>target_seq</td><td>PAM</td><td>alignment</td><td>distance</td><td>position</td><td>gene name</td><td>gene id</td></tr>
<tr><td>GU305914.1</td><td>192654</td><td>192676</td><td>+</td><td>0</td><td>CACGAGGGCGTATTCCATCC</td><td>TGG</td><td>||||||||[||||||||||||]PAM</td><td>NA</td><td>Intergenic</td><td>NA</td><td>NA</td></tr>
<tr><td>T14647</td><td>GAGGGCGTATTCCATCCTGGTGG</td><td>1000</td><td></td><td></td><td></td><td></td><td></td><td></td><td></td><td></td><td></td></tr>
<tr><td>Oligo fwd</td><td>CACCGAGGGCGTATTCCATCCTGG</td><td></td><td></td><td></td><td></td><td></td><td></td><td></td><td></td><td></td><td></td></tr>
<tr><td>Oligo rev</td><td>AAACCCAGGATGGAATACGCCCTC</td><td></td><td></td><td></td><td></td><td></td><td></td><td></td><td></td><td></td><td></td></tr>
<tr><td>Chromosome</td><td>start</td><td>end</td><td>strand</td><td>MM</td><td>target_seq</td><td>PAM</td><td>alignment</td><td>distance</td><td>position</td><td>gene name</td><td>gene id</td></tr>
<tr><td>GU305914.1</td><td>192657</td><td>192679</td><td>+</td><td>0</td><td>GAGGGCGTATTCCATCCTGG</td><td>TGG</td><td>||||||||[||||||||||||]PAM</td><td>NA</td><td>Intergenic</td><td>NA</td><td>NA</td></tr>
<tr><td>T14648</td><td>AGGGCGTATTCCATCCTGGTGGG</td><td>1000</td><td></td><td></td><td></td><td></td><td></td><td></td><td></td><td></td><td></td></tr>
<tr><td>Oligo fwd</td><td>CACCAGGGCGTATTCCATCCTGGT</td><td></td><td></td><td></td><td></td><td></td><td></td><td></td><td></td><td></td><td></td></tr>
<tr><td>Oligo rev</td><td>AAACACCAGGATGGAATACGCCCT</td><td></td><td></td><td></td><td></td><td></td><td></td><td></td><td></td><td></td><td></td></tr>
<tr><td>Chromosome</td><td>start</td><td>end</td><td>strand</td><td>MM</td><td>target_seq</td><td>PAM</td><td>alignment</td><td>distance</td><td>position</td><td>gene name</td><td>gene id</td></tr>
<tr><td>GU305914.1</td><td>192658</td><td>192680</td><td>+</td><td>0</td><td>AGGGCGTATTCCATCCTGGT</td><td>GGG</td><td>||||||||[||||||||||||]PAM</td><td>NA</td><td>Intergenic</td><td>NA</td><td>NA</td></tr>
<tr><td>T14649</td><td>CATCCTGGTGGGCTTCGTCACGG</td><td>1000</td><td></td><td></td><td></td><td></td><td></td><td></td><td></td><td></td><td></td></tr>
<tr><td>Oligo fwd</td><td>CACCCATCCTGGTGGGCTTCGTCA</td><td></td><td></td><td></td><td></td><td></td><td></td><td></td><td></td><td></td><td></td></tr>
<tr><td>Oligo rev</td><td>AAACTGACGAAGCCCACCAGGATG</td><td></td><td></td><td></td><td></td><td></td><td></td><td></td><td></td><td></td><td></td></tr>
<tr><td>Chromosome</td><td>start</td><td>end</td><td>strand</td><td>MM</td><td>target_seq</td><td>PAM</td><td>alignment</td><td>distance</td><td>position</td><td>gene name</td><td>gene id</td></tr>
<tr><td>GU305914.1</td><td>192669</td><td>192691</td><td>+</td><td>0</td><td>CATCCTGGTGGGCTTCGTCA</td><td>CGG</td><td>||||||||[||||||||||||]PAM</td><td>NA</td><td>Intergenic</td><td>NA</td><td>NA</td></tr>
<tr><td>T14650</td><td>GCTTCGTCACGGTGATCGCTCGG</td><td>1000</td><td></td><td></td><td></td><td></td><td></td><td></td><td></td><td></td><td></td></tr>
<tr><td>Oligo fwd</td><td>CACCGCTTCGTCACGGTGATCGCT</td><td></td><td></td><td></td><td></td><td></td><td></td><td></td><td></td><td></td><td></td></tr>
<tr><td>Oligo rev</td><td>AAACAGCGATCACCGTGACGAAGC</td><td></td><td></td><td></td><td></td><td></td><td></td><td></td><td></td><td></td><td></td></tr>
<tr><td>Chromosome</td><td>start</td><td>end</td><td>strand</td><td>MM</td><td>target_seq</td><td>PAM</td><td>alignment</td><td>distance</td><td>position</td><td>gene name</td><td>gene id</td></tr>
<tr><td>GU305914.1</td><td>192680</td><td>192702</td><td>+</td><td>0</td><td>GCTTCGTCACGGTGATCGCT</td><td>CGG</td><td>||||||||[||||||||||||]PAM</td><td>NA</td><td>Intergenic</td><td>NA</td><td>NA</td></tr>
<tr><td>T14651</td><td>CTTCGTCACGGTGATCGCTCGGG</td><td>1000</td><td></td><td></td><td></td><td></td><td></td><td></td><td></td><td></td><td></td></tr>
<tr><td>Oligo fwd</td><td>CACCCTTCGTCACGGTGATCGCTC</td><td></td><td></td><td></td><td></td><td></td><td></td><td></td><td></td><td></td><td></td></tr>
<tr><td>Oligo rev</td><td>AAACGAGCGATCACCGTGACGAAG</td><td></td><td></td><td></td><td></td><td></td><td></td><td></td><td></td><td></td><td></td></tr>
<tr><td>Chromosome</td><td>start</td><td>end</td><td>strand</td><td>MM</td><td>target_seq</td><td>PAM</td><td>alignment</td><td>distance</td><td>position</td><td>gene name</td><td>gene id</td></tr>
<tr><td>GU305914.1</td><td>192681</td><td>192703</td><td>+</td><td>0</td><td>CTTCGTCACGGTGATCGCTC</td><td>GGG</td><td>||||||||[||||||||||||]PAM</td><td>NA</td><td>Intergenic</td><td>NA</td><td>NA</td></tr>
<tr><td>T14652</td><td>CCAGATTCAAGCATCCCGAGCGG</td><td>1000</td><td></td><td></td><td></td><td></td><td></td><td></td><td></td><td></td><td></td></tr>
<tr><td>Oligo fwd</td><td>CACCCCAGATTCAAGCATCCCGAG</td><td></td><td></td><td></td><td></td><td></td><td></td><td></td><td></td><td></td><td></td></tr>
<tr><td>Oligo rev</td><td>AAACCTCGGGATGCTTGAATCTGG</td><td></td><td></td><td></td><td></td><td></td><td></td><td></td><td></td><td></td><td></td></tr>
<tr><td>Chromosome</td><td>start</td><td>end</td><td>strand</td><td>MM</td><td>target_seq</td><td>PAM</td><td>alignment</td><td>distance</td><td>position</td><td>gene name</td><td>gene id</td></tr>
<tr><td>GU305914.1</td><td>192707</td><td>192729</td><td>+</td><td>0</td><td>CCAGATTCAAGCATCCCGAG</td><td>CGG</td><td>||||||||[||||||||||||]PAM</td><td>NA</td><td>Intergenic</td><td>NA</td><td>NA</td></tr>
<tr><td>T14653</td><td>TCAAGCATCCCGAGCGGACGCGG</td><td>1000</td><td></td><td></td><td></td><td></td><td></td><td></td><td></td><td></td><td></td></tr>
<tr><td>Oligo fwd</td><td>CACCTCAAGCATCCCGAGCGGACG</td><td></td><td></td><td></td><td></td><td></td><td></td><td></td><td></td><td></td><td></td></tr>
<tr><td>Oligo rev</td><td>AAACCGTCCGCTCGGGATGCTTGA</td><td></td><td></td><td></td><td></td><td></td><td></td><td></td><td></td><td></td><td></td></tr>
<tr><td>Chromosome</td><td>start</td><td>end</td><td>strand</td><td>MM</td><td>target_seq</td><td>PAM</td><td>alignment</td><td>distance</td><td>position</td><td>gene name</td><td>gene id</td></tr>
<tr><td>GU305914.1</td><td>192713</td><td>192735</td><td>+</td><td>0</td><td>TCAAGCATCCCGAGCGGACG</td><td>CGG</td><td>||||||||[||||||||||||]PAM</td><td>NA</td><td>Intergenic</td><td>NA</td><td>NA</td></tr>
<tr><td>T14654</td><td>AGCGGACGCGGCACCTAAACCGG</td><td>1000</td><td></td><td></td><td></td><td></td><td></td><td></td><td></td><td></td><td></td></tr>
<tr><td>Oligo fwd</td><td>CACCAGCGGACGCGGCACCTAAAC</td><td></td><td></td><td></td><td></td><td></td><td></td><td></td><td></td><td></td><td></td></tr>
<tr><td>Oligo rev</td><td>AAACGTTTAGGTGCCGCGTCCGCT</td><td></td><td></td><td></td><td></td><td></td><td></td><td></td><td></td><td></td><td></td></tr>
<tr><td>Chromosome</td><td>start</td><td>end</td><td>strand</td><td>MM</td><td>target_seq</td><td>PAM</td><td>alignment</td><td>distance</td><td>position</td><td>gene name</td><td>gene id</td></tr>
<tr><td>GU305914.1</td><td>192725</td><td>192747</td><td>+</td><td>0</td><td>AGCGGACGCGGCACCTAAAC</td><td>CGG</td><td>||||||||[||||||||||||]PAM</td><td>NA</td><td>Intergenic</td><td>NA</td><td>NA</td></tr>
<tr><td>T14655</td><td>TCACGCCATCGTCTTCTCTCAGG</td><td>1000</td><td></td><td></td><td></td><td></td><td></td><td></td><td></td><td></td><td></td></tr>
<tr><td>Oligo fwd</td><td>CACCTCACGCCATCGTCTTCTCTC</td><td></td><td></td><td></td><td></td><td></td><td></td><td></td><td></td><td></td><td></td></tr>
</table>

Oligo rev AAACGAGAGAAGACGATGGCGTGA
Chromosome start end strand MM target_seq PAM alignment distance position gene name gene id
GU305914.1 192752 192774 + 0 TCACGCCATCGTCTTCTCTC AGG ||||||||[||||||||||||]PAM NA Intergenic NA NA

T14656 ATCGTCTTCTCTCAGGTACACGG 1000
Oligo fwd CACCATCGTCTTCTCTCAGGTACA
Oligo rev AAACTGTACCTGAGAGAAGACGAT
Chromosome start end strand MM target_seq PAM alignment distance position gene name gene id
GU305914.1 192759 192781 + 0 ATCGTCTTCTCTCAGGTACA CGG ||||||||[||||||||||||]PAM NA Intergenic NA NA

T14657 GGACACCGAGATCTTGCCCTCGG 1000
Oligo fwd CACCGGACACCGAGATCTTGCCCT
Oligo rev AAACAGGGCAAGATCTCGGTGTCC
Chromosome start end strand MM target_seq PAM alignment distance position gene name gene id
GU305914.1 192780 192802 + 0 GGACACCGAGATCTTGCCCT CGG ||||||||[||||||||||||]PAM NA Intergenic NA NA

T14658 CCGCGAGCCGTTGATCGAGATGG 1000
Oligo fwd CACCCCGCGAGCCGTTGATCGAGA
Oligo rev AAACTCTCGATCAACGGCTCGCGG
Chromosome start end strand MM target_seq PAM alignment distance position gene name gene id
GU305914.1 192807 192829 + 0 CCGCGAGCCGTTGATCGAGA TGG ||||||||[||||||||||||]PAM NA Intergenic NA NA

T14659 GTTGATCGAGATGGAGTCGTCGG 1000
Oligo fwd CACCGTTGATCGAGATGGAGTCGT
Oligo rev AAACACGACTCCATCTCGATCAAC
Chromosome start end strand MM target_seq PAM alignment distance position gene name gene id
GU305914.1 192816 192838 + 0 GTTGATCGAGATGGAGTCGT CGG ||||||||[||||||||||||]PAM NA Intergenic NA NA

T14660 GATGGAGTCGTCGGAGACGAAGG 1000
Oligo fwd CACCGATGGAGTCGTCGGAGACGA
Oligo rev AAACTCGTCTCCGACGACTCCATC
Chromosome start end strand MM target_seq PAM alignment distance position gene name gene id
GU305914.1 192825 192847 + 0 GATGGAGTCGTCGGAGACGA AGG ||||||||[||||||||||||]PAM NA Intergenic NA NA
GU305914.1 60607 60629 + 4 GACGAAGCCGCCGGAGACGA CGG ||-|-||-[||-|||||||||]PAM NA Intergenic NA NA

T14661 AGGACTCGTTAGTGCCGACACGG 1000
Oligo fwd CACCAGGACTCGTTAGTGCCGACA
Oligo rev AAACTGTCGGCACTAACGAGTCCT
Chromosome start end strand MM target_seq PAM alignment distance position gene name gene id
GU305914.1 192845 192867 + 0 AGGACTCGTTAGTGCCGACA CGG ||||||||[||||||||||||]PAM NA Intergenic NA NA

T14662 CCGACACGGCTCTGATACACCGG 1000
Oligo fwd CACCCCGACACGGCTCTGATACAC
Oligo rev AAACGTGTATCAGAGCCGTGTCGG
Chromosome start end strand MM target_seq PAM alignment distance position gene name gene id
GU305914.1 192859 192881 + 0 CCGACACGGCTCTGATACAC CGG ||||||||[||||||||||||]PAM NA Intergenic NA NA

T14663 CTCCAGCGTGAACGCGCCGTCGG 1000
Oligo fwd CACCCTCCAGCGTGAACGCGCCGT
Oligo rev AAACACGGCGCGTTCACGCTGGAG
Chromosome start end strand MM target_seq PAM alignment distance position gene name gene id
GU305914.1 192900 192922 + 0 CTCCAGCGTGAACGCGCCGT CGG ||||||||[||||||||||||]PAM NA Intergenic NA NA

T14664 GCCGTCGGACACCGCGTCGAAGG 1000
Oligo fwd CACCGCCGTCGGACACCGCGTCGA
Oligo rev AAACTCGACGCGGTGTCCGACGGC
Chromosome start end strand MM target_seq PAM alignment distance position gene name gene id
GU305914.1 192915 192937 + 0 GCCGTCGGACACCGCGTCGA AGG ||||||||[||||||||||||]PAM NA Intergenic NA NA

T14665 CGTCGAAGGTACACGTTATTTGG 1000
Oligo fwd CACCCGTCGAAGGTACACGTTATT
Oligo rev AAACAATAACGTGTACCTTCGACG
Chromosome start end strand MM target_seq PAM alignment distance position gene name gene id
GU305914.1 192929 192951 + 0 CGTCGAAGGTACACGTTATT TGG ||||||||[||||||||||||]PAM NA Intergenic NA NA

T14666 TGCCGTGTACCATCAGCGACCGG 1000
Oligo fwd CACCTGCCGTGTACCATCAGCGAC
Oligo rev AAACGTCGCTGATGGTACACGGCA
Chromosome start end strand MM target_seq PAM alignment distance position gene name gene id
GU305914.1 193004 193026 + 0 TGCCGTGTACCATCAGCGAC CGG ||||||||[||||||||||||]PAM NA Intergenic NA NA

T14667 TCTCTCTCTCGAAGATACACTGG 1000
Oligo fwd CACCTCTCTCTCTCGAAGATACAC
Oligo rev AAACGTGTATCTTCGAGAGAGAGA
Chromosome start end strand MM target_seq PAM alignment distance position gene name gene id
GU305914.1 193043 193065 + 0 TCTCTCTCTCGAAGATACAC TGG ||||||||[||||||||||||]PAM NA Intergenic NA NA

T14668 AAGATACACTGGTACTCACCCGG 1000
Oligo fwd CACCAAGATACACTGGTACTCACC
Oligo rev AAACGGTGAGTACCAGTGTATCTT
Chromosome start end strand MM target_seq PAM alignment distance position gene name gene id
GU305914.1 193054 193076 + 0 AAGATACACTGGTACTCACC CGG ||||||||[||||||||||||]PAM NA Intergenic NA NA

T14669 AGATACACTGGTACTCACCCGGG 1000
Oligo fwd CACCAGATACACTGGTACTCACCC
Oligo rev AAACGGGTGAGTACCAGTGTATCT
Chromosome start end strand MM target_seq PAM alignment distance position gene name gene id
GU305914.1 193055 193077 + 0 AGATACACTGGTACTCACCC GGG ||||||||[||||||||||||]PAM NA Intergenic NA NA

T14670 GATACACTGGTACTCACCCGGGG 1000
Oligo fwd CACCGATACACTGGTACTCACCCG
Oligo rev AAACCGGGTGAGTACCAGTGTATC
Chromosome start end strand MM target_seq PAM alignment distance position gene name gene id
GU305914.1 193056 193078 + 0 GATACACTGGTACTCACCCG GGG ||||||||[||||||||||||]PAM NA Intergenic NA NA

T14671 ATACACTGGTACTCACCCGGGGG 1000
Oligo fwd CACCATACACTGGTACTCACCCGG
Oligo rev AAACCCGGGTGAGTACCAGTGTAT
Chromosome start end strand MM target_seq PAM alignment distance position gene name gene id
GU305914.1 193057 193079 + 0 ATACACTGGTACTCACCCGG GGG ||||||||[||||||||||||]PAM NA Intergenic NA NA

T14672 TACACTGGTACTCACCCGGGGGG 1000
Oligo fwd CACCTACACTGGTACTCACCCGGG
Oligo rev AAACCCCGGGTGAGTACCAGTGTA
Chromosome start end strand MM target_seq PAM alignment distance position gene name gene id
GU305914.1 193058 193080 + 0 TACACTGGTACTCACCCGGG GGG ||||||||[||||||||||||]PAM NA Intergenic NA NA

T14673 ACTGGTACTCACCCGGGGGGAGG 1000
Oligo fwd CACCACTGGTACTCACCCGGGGGG
Oligo rev AAACCCCCCCGGGTGAGTACCAGT
Chromosome start end strand MM target_seq PAM alignment distance position gene name gene id
GU305914.1 193061 193083 + 0 ACTGGTACTCACCCGGGGGG AGG ||||||||[||||||||||||]PAM NA Intergenic NA NA

T14674 TGTTTGTATGCATACGCCACTGG 1000
Oligo fwd CACCTGTTTGTATGCATACGCCAC
Oligo rev AAACGTGGCGTATGCATACAAACA
Chromosome start end strand MM target_seq PAM alignment distance position gene name gene id
GU305914.1 193104 193126 + 0 TGTTTGTATGCATACGCCAC TGG ||||||||[||||||||||||]PAM NA Intergenic NA NA

T14675 TTGTATGCATACGCCACTGGAGG 1000
Oligo fwd CACCTTGTATGCATACGCCACTGG
Oligo rev AAACCCAGTGGCGTATGCATACAA
Chromosome start end strand MM target_seq PAM alignment distance position gene name gene id
GU305914.1 193107 193129 + 0 TTGTATGCATACGCCACTGG AGG ||||||||[||||||||||||]PAM NA Intergenic NA NA

T14676 AGCATCAAGACCGATTCCAGCGG 1000
Oligo fwd CACCAGCATCAAGACCGATTCCAG
Oligo rev AAACCTGGAATCGGTCTTGATGCT
Chromosome start end strand MM target_seq PAM alignment distance position gene name gene id
GU305914.1 193162 193184 + 0 AGCATCAAGACCGATTCCAG CGG ||||||||[||||||||||||]PAM NA Intergenic NA NA

T14677 TTCCAGCGGCTTTTCCACCGCGG 1000
Oligo fwd CACCTTCCAGCGGCTTTTCCACCG
Oligo rev AAACCGGTGGAAAAGCCGCTGGAA
Chromosome start end strand MM target_seq PAM alignment distance position gene name gene id
GU305914.1 193176 193198 + 0 TTCCAGCGGCTTTTCCACCG CGG ||||||||[||||||||||||]PAM NA Intergenic NA NA

T14678 GCGGCTATGTAGAACATGTACGG 1000
Oligo fwd CACCGCGGCTATGTAGAACATGTA
Oligo rev AAACTACATGTTCTACATAGCCGC
Chromosome start end strand MM target_seq PAM alignment distance position gene name gene id
GU305914.1 193195 193217 + 0 GCGGCTATGTAGAACATGTA CGG ||||||||[||||||||||||]PAM NA Intergenic NA NA

T14679 GCTATGTAGAACATGTACGGCGG 1000
Oligo fwd CACCGCTATGTAGAACATGTACGG
Oligo rev AAACCCGTACATGTTCTACATAGC
Chromosome start end strand MM target_seq PAM alignment distance position gene name gene id
GU305914.1 193198 193220 + 0 GCTATGTAGAACATGTACGG CGG ||||||||[||||||||||||]PAM NA Intergenic NA NA

T14680 CTATGTAGAACATGTACGGCGGG 1000
Oligo fwd CACCCTATGTAGAACATGTACGGC
Oligo rev AAACGCCGTACATGTTCTACATAG
Chromosome start end strand MM target_seq PAM alignment distance position gene name gene id
GU305914.1 193199 193221 + 0 CTATGTAGAACATGTACGGC GGG ||||||||[||||||||||||]PAM NA Intergenic NA NA

T14681 CATGTACGGCGGGATGTCGAAGG 1000
Oligo fwd CACCCATGTACGGCGGGATGTCGA
Oligo rev AAACTCGACATCCCGCCGTACATG
Chromosome start end strand MM target_seq PAM alignment distance position gene name gene id
GU305914.1 193209 193231 + 0 CATGTACGGCGGGATGTCGA AGG ||||||||[||||||||||||]PAM NA Intergenic NA NA

T14682 TGTCGAAGGCAAACAAGAACTGG 1000
Oligo fwd CACCTGTCGAAGGCAAACAAGAAC
Oligo rev AAACGTTCTTGTTTGCCTTCGACA
Chromosome start end strand MM target_seq PAM alignment distance position gene name gene id
GU305914.1 193223 193245 + 0 TGTCGAAGGCAAACAAGAAC TGG ||||||||[||||||||||||]PAM NA Intergenic NA NA

T14683 GTCGAAGGCAAACAAGAACTGGG 1000
Oligo fwd CACCGTCGAAGGCAAACAAGAACT
Oligo rev AAACAGTTCTTGTTTGCCTTCGAC
Chromosome start end strand MM target_seq PAM alignment distance position gene name gene id
GU305914.1 193224 193246 + 0 GTCGAAGGCAAACAAGAACT GGG ||||||||[||||||||||||]PAM NA Intergenic NA NA

T14684 TCGAAGGCAAACAAGAACTGGGG 1000
Oligo fwd CACCTCGAAGGCAAACAAGAACTG
Oligo rev AAACCAGTTCTTGTTTGCCTTCGA
Chromosome start end strand MM target_seq PAM alignment distance position gene name gene id
GU305914.1 193225 193247 + 0 TCGAAGGCAAACAAGAACTG GGG ||||||||[||||||||||||]PAM NA Intergenic NA NA

<table>
<tr><td>T14685</td><td>AGAACTGGGGTTCTGCGCACTGG</td><td>1000</td></tr>
<tr><td>Oligo fwd</td><td>CACCAGAACTGGGGTTCTGCGCAC</td></tr>
<tr><td>Oligo rev</td><td>AAACGTGCGCAGAACCCCAGTTCT</td></tr>
<tr><td>Chromosome</td><td>start</td><td>end</td><td>strand</td><td>MM</td><td>target_seq</td><td>PAM</td><td>alignment</td><td>distance</td><td>position</td><td>gene name</td><td>gene id</td></tr>
<tr><td>GU305914.1</td><td>193238</td><td>193260</td><td>+</td><td>0</td><td>AGAACTGGGGTTCTGCGCAC</td><td>TGG</td><td>||||||||[||||||||||||]PAM</td><td>NA</td><td>Intergenic</td><td>NA</td><td>NA</td></tr>
<tr><td>T14686</td><td>CTGGGGTTCTGCGCACTGGACGG</td><td>1000</td></tr>
<tr><td>Oligo fwd</td><td>CACCCTGGGGTTCTGCGCACTGGA</td></tr>
<tr><td>Oligo rev</td><td>AAACTCCAGTGCGCAGAACCCCAG</td></tr>
<tr><td>Chromosome</td><td>start</td><td>end</td><td>strand</td><td>MM</td><td>target_seq</td><td>PAM</td><td>alignment</td><td>distance</td><td>position</td><td>gene name</td><td>gene id</td></tr>
<tr><td>GU305914.1</td><td>193242</td><td>193264</td><td>+</td><td>0</td><td>CTGGGGTTCTGCGCACTGGA</td><td>CGG</td><td>||||||||[||||||||||||]PAM</td><td>NA</td><td>Intergenic</td><td>NA</td><td>NA</td></tr>
<tr><td>T14687</td><td>TGCGCACTGGACGGTCGTTCTGG</td><td>1000</td></tr>
<tr><td>Oligo fwd</td><td>CACCTGCGCACTGGACGGTCGTTC</td></tr>
<tr><td>Oligo rev</td><td>AAACGAACGACCGTCCAGTGCGCA</td></tr>
<tr><td>Chromosome</td><td>start</td><td>end</td><td>strand</td><td>MM</td><td>target_seq</td><td>PAM</td><td>alignment</td><td>distance</td><td>position</td><td>gene name</td><td>gene id</td></tr>
<tr><td>GU305914.1</td><td>193251</td><td>193273</td><td>+</td><td>0</td><td>TGCGCACTGGACGGTCGTTC</td><td>TGG</td><td>||||||||[||||||||||||]PAM</td><td>NA</td><td>Intergenic</td><td>NA</td><td>NA</td></tr>
<tr><td>T14688</td><td>CTGGACGGTCGTTCTGGTCACGG</td><td>1000</td></tr>
<tr><td>Oligo fwd</td><td>CACCCTGGACGGTCGTTCTGGTCA</td></tr>
<tr><td>Oligo rev</td><td>AAACTGACCAGAACGACCGTCCAG</td></tr>
<tr><td>Chromosome</td><td>start</td><td>end</td><td>strand</td><td>MM</td><td>target_seq</td><td>PAM</td><td>alignment</td><td>distance</td><td>position</td><td>gene name</td><td>gene id</td></tr>
<tr><td>GU305914.1</td><td>193257</td><td>193279</td><td>+</td><td>0</td><td>CTGGACGGTCGTTCTGGTCA</td><td>CGG</td><td>||||||||[||||||||||||]PAM</td><td>NA</td><td>Intergenic</td><td>NA</td><td>NA</td></tr>
<tr><td>T14689</td><td>TCTGGTCACGGTGACGCGCTTGG</td><td>1000</td></tr>
<tr><td>Oligo fwd</td><td>CACCTCTGGTCACGGTGACGCGCT</td></tr>
<tr><td>Oligo rev</td><td>AAACAGCGCGTCACCGTGACCAGA</td></tr>
<tr><td>Chromosome</td><td>start</td><td>end</td><td>strand</td><td>MM</td><td>target_seq</td><td>PAM</td><td>alignment</td><td>distance</td><td>position</td><td>gene name</td><td>gene id</td></tr>
<tr><td>GU305914.1</td><td>193269</td><td>193291</td><td>+</td><td>0</td><td>TCTGGTCACGGTGACGCGCT</td><td>TGG</td><td>||||||||[||||||||||||]PAM</td><td>NA</td><td>Intergenic</td><td>NA</td><td>NA</td></tr>
<tr><td>T14690</td><td>GCTTGGCGAAATCCCAGTTATGG</td><td>1000</td></tr>
<tr><td>Oligo fwd</td><td>CACCGCTTGGCGAAATCCCAGTTA</td></tr>
<tr><td>Oligo rev</td><td>AAACTAACTGGGATTTCGCCAAGC</td></tr>
<tr><td>Chromosome</td><td>start</td><td>end</td><td>strand</td><td>MM</td><td>target_seq</td><td>PAM</td><td>alignment</td><td>distance</td><td>position</td><td>gene name</td><td>gene id</td></tr>
<tr><td>GU305914.1</td><td>193286</td><td>193308</td><td>+</td><td>0</td><td>GCTTGGCGAAATCCCAGTTA</td><td>TGG</td><td>||||||||[||||||||||||]PAM</td><td>NA</td><td>Intergenic</td><td>NA</td><td>NA</td></tr>
<tr><td>T14691</td><td>GAGCTTACAGCAGACTCTGTTGG</td><td>1000</td></tr>
<tr><td>Oligo fwd</td><td>CACCGAGCTTACAGCAGACTCTGT</td></tr>
<tr><td>Oligo rev</td><td>AAACACAGAGTCTGCTGTAAGCTC</td></tr>
<tr><td>Chromosome</td><td>start</td><td>end</td><td>strand</td><td>MM</td><td>target_seq</td><td>PAM</td><td>alignment</td><td>distance</td><td>position</td><td>gene name</td><td>gene id</td></tr>
<tr><td>GU305914.1</td><td>193317</td><td>193339</td><td>+</td><td>0</td><td>GAGCTTACAGCAGACTCTGT</td><td>TGG</td><td>||||||||[||||||||||||]PAM</td><td>NA</td><td>Intergenic</td><td>NA</td><td>NA</td></tr>
<tr><td>T14692</td><td>TCTGTTGGCGACTTCAGTCCCGG</td><td>1000</td></tr>
<tr><td>Oligo fwd</td><td>CACCTCTGTTGGCGACTTCAGTCC</td></tr>
<tr><td>Oligo rev</td><td>AAACGGACTGAAGTCGCCAACAGA</td></tr>
<tr><td>Chromosome</td><td>start</td><td>end</td><td>strand</td><td>MM</td><td>target_seq</td><td>PAM</td><td>alignment</td><td>distance</td><td>position</td><td>gene name</td><td>gene id</td></tr>
<tr><td>GU305914.1</td><td>193332</td><td>193354</td><td>+</td><td>0</td><td>TCTGTTGGCGACTTCAGTCC</td><td>CGG</td><td>||||||||[||||||||||||]PAM</td><td>NA</td><td>Intergenic</td><td>NA</td><td>NA</td></tr>
<tr><td>T14693</td><td>GTTGGCGACTTCAGTCCCGGTGG</td><td>1000</td></tr>
<tr><td>Oligo fwd</td><td>CACCGTTGGCGACTTCAGTCCCGG</td></tr>
<tr><td>Oligo rev</td><td>AAACCCGGGACTGAAGTCGCCAAC</td></tr>
<tr><td>Chromosome</td><td>start</td><td>end</td><td>strand</td><td>MM</td><td>target_seq</td><td>PAM</td><td>alignment</td><td>distance</td><td>position</td><td>gene name</td><td>gene id</td></tr>
<tr><td>GU305914.1</td><td>193335</td><td>193357</td><td>+</td><td>0</td><td>GTTGGCGACTTCAGTCCCGG</td><td>TGG</td><td>||||||||[||||||||||||]PAM</td><td>NA</td><td>Intergenic</td><td>NA</td><td>NA</td></tr>
<tr><td>T14694</td><td>TTCAGTCCCGGTGGAGTCCTTGG</td><td>1000</td></tr>
<tr><td>Oligo fwd</td><td>CACCTTCAGTCCCGGTGGAGTCCT</td></tr>
<tr><td>Oligo rev</td><td>AAACAGGACTCCACCGGGACTGAA</td></tr>
<tr><td>Chromosome</td><td>start</td><td>end</td><td>strand</td><td>MM</td><td>target_seq</td><td>PAM</td><td>alignment</td><td>distance</td><td>position</td><td>gene name</td><td>gene id</td></tr>
<tr><td>GU305914.1</td><td>193344</td><td>193366</td><td>+</td><td>0</td><td>TTCAGTCCCGGTGGAGTCCT</td><td>TGG</td><td>||||||||[||||||||||||]PAM</td><td>NA</td><td>Intergenic</td><td>NA</td><td>NA</td></tr>
<tr><td>T14695</td><td>GTGTACCTTTTCACTCCCGATGG</td><td>1000</td></tr>
<tr><td>Oligo fwd</td><td>CACCGTGTACCTTTTCACTCCCGA</td></tr>
<tr><td>Oligo rev</td><td>AAACTCGGGAGTGAAAAGGTACAC</td></tr>
<tr><td>Chromosome</td><td>start</td><td>end</td><td>strand</td><td>MM</td><td>target_seq</td><td>PAM</td><td>alignment</td><td>distance</td><td>position</td><td>gene name</td><td>gene id</td></tr>
<tr><td>GU305914.1</td><td>193366</td><td>193388</td><td>+</td><td>0</td><td>GTGTACCTTTTCACTCCCGA</td><td>TGG</td><td>||||||||[||||||||||||]PAM</td><td>NA</td><td>Intergenic</td><td>NA</td><td>NA</td></tr>
<tr><td>T14696</td><td>CCTTTTCACTCCCGATGGCATGG</td><td>1000</td></tr>
<tr><td>Oligo fwd</td><td>CACCCCTTTTCACTCCCGATGGCA</td></tr>
<tr><td>Oligo rev</td><td>AAACTGCCATCGGGAGTGAAAAGG</td></tr>
<tr><td>Chromosome</td><td>start</td><td>end</td><td>strand</td><td>MM</td><td>target_seq</td><td>PAM</td><td>alignment</td><td>distance</td><td>position</td><td>gene name</td><td>gene id</td></tr>
<tr><td>GU305914.1</td><td>193371</td><td>193393</td><td>+</td><td>0</td><td>CCTTTTCACTCCCGATGGCA</td><td>TGG</td><td>||||||||[||||||||||||]PAM</td><td>NA</td><td>Intergenic</td><td>NA</td><td>NA</td></tr>
<tr><td>T14697</td><td>TGTCTGTATTCCAACATCAGAGG</td><td>1000</td></tr>
<tr><td>Oligo fwd</td><td>CACCTGTCTGTATTCCAACATCAG</td></tr>
<tr><td>Oligo rev</td><td>AAACCTGATGTTGGAATACAGACA</td></tr>
<tr><td>Chromosome</td><td>start</td><td>end</td><td>strand</td><td>MM</td><td>target_seq</td><td>PAM</td><td>alignment</td><td>distance</td><td>position</td><td>gene name</td><td>gene id</td></tr>
<tr><td>GU305914.1</td><td>193396</td><td>193418</td><td>+</td><td>0</td><td>TGTCTGTATTCCAACATCAG</td><td>AGG</td><td>||||||||[||||||||||||]PAM</td><td>NA</td><td>Intergenic</td><td>NA</td><td>NA</td></tr>
<tr><td>T14698</td><td>AGAGGCTTGCTGACCGCAAAAGG</td><td>1000</td></tr>
<tr><td>Oligo fwd</td><td>CACCAGAGGCTTGCTGACCGCAAA</td></tr>
<tr><td>Oligo rev</td><td>AAACTTTGCGGTCAGCAAGCCTCT</td></tr>
<tr><td>Chromosome</td><td>start</td><td>end</td><td>strand</td><td>MM</td><td>target_seq</td><td>PAM</td><td>alignment</td><td>distance</td><td>position</td><td>gene name</td><td>gene id</td></tr>
<tr><td>GU305914.1</td><td>193414</td><td>193436</td><td>+</td><td>0</td><td>AGAGGCTTGCTGACCGCAAA</td><td>AGG</td><td>||||||||[||||||||||||]PAM</td><td>NA</td><td>Intergenic</td><td>NA</td><td>NA</td></tr>
<tr><td>T14699</td><td>GCAAAAGGAACGCACGAGAGAGG</td><td>1000</td></tr>
<tr><td>Oligo fwd</td><td>CACCGCAAAAGGAACGCACGAGAG</td></tr>
<tr><td>Oligo rev</td><td>AAACCTCTCGTGCGTTCCTTTTGC</td></tr>
<tr><td>Chromosome</td><td>start</td><td>end</td><td>strand</td><td>MM</td><td>target_seq</td><td>PAM</td><td>alignment</td><td>distance</td><td>position</td><td>gene name</td><td>gene id</td></tr>
</table>

| | | | | | | | | | | | |
|---|---|---|---|---|---|---|---|---|---|---|---|
| GU305914.1 | 193429 | 193451 | + | 0 | GCAAAAGGAACGCACGAGAG | AGG | ||||||||[||||||||||||]PAM | NA | Intergenic | NA | NA |
| T14700 | CTGATATTTCACGTGCGTCATGG | 1000 | | | | | | | | | |
| Oligo fwd | CACCCTGATATTTCACGTGCGTCA | | | | | | | | | | |
| Oligo rev | AAACTGACGCACGTGAAATATCAG | | | | | | | | | | |
| Chromosome | start | end | strand | MM | target_seq | PAM | alignment | distance | position | gene name | gene id |
| GU305914.1 | 193464 | 193486 | + | 0 | CTGATATTTCACGTGCGTCA | TGG | ||||||||[||||||||||||]PAM | NA | Intergenic | NA | NA |
| T14701 | TGATATTTCACGTGCGTCATGGG | 1000 | | | | | | | | | |
| Oligo fwd | CACCTGATATTTCACGTGCGTCAT | | | | | | | | | | |
| Oligo rev | AAACATGACGCACGTGAAATATCA | | | | | | | | | | |
| Chromosome | start | end | strand | MM | target_seq | PAM | alignment | distance | position | gene name | gene id |
| GU305914.1 | 193465 | 193487 | + | 0 | TGATATTTCACGTGCGTCAT | GGG | ||||||||[||||||||||||]PAM | NA | Intergenic | NA | NA |
| T14702 | GATCGAGACGAGAAATCGCATGG | 1000 | | | | | | | | | |
| Oligo fwd | CACCGATCGAGACGAGAAATCGCA | | | | | | | | | | |
| Oligo rev | AAACTGCGATTTCTCGTCTCGATC | | | | | | | | | | |
| Chromosome | start | end | strand | MM | target_seq | PAM | alignment | distance | position | gene name | gene id |
| GU305914.1 | 193497 | 193519 | + | 0 | GATCGAGACGAGAAATCGCA | TGG | ||||||||[||||||||||||]PAM | NA | Intergenic | NA | NA |
| T14703 | TGGTACCGTTGCGATGTGAGAGG | 1000 | | | | | | | | | |
| Oligo fwd | CACCTGGTACCGTTGCGATGTGAG | | | | | | | | | | |
| Oligo rev | AAACCTCACATCGCAACGGTACCA | | | | | | | | | | |
| Chromosome | start | end | strand | MM | target_seq | PAM | alignment | distance | position | gene name | gene id |
| GU305914.1 | 193517 | 193539 | + | 0 | TGGTACCGTTGCGATGTGAG | AGG | ||||||||[||||||||||||]PAM | NA | Intergenic | NA | NA |
| T14704 | TTTTTTCACTGCATATCGTCCGG | 1000 | | | | | | | | | |
| Oligo fwd | CACCTTTTTTCACTGCATATCGTC | | | | | | | | | | |
| Oligo rev | AAACGACGATATGCAGTGAAAAAA | | | | | | | | | | |
| Chromosome | start | end | strand | MM | target_seq | PAM | alignment | distance | position | gene name | gene id |
| GU305914.1 | 193557 | 193579 | + | 0 | TTTTTTCACTGCATATCGTC | CGG | ||||||||[||||||||||||]PAM | NA | Intergenic | NA | NA |
| T14705 | TTTTTCACTGCATATCGTCCGGG | 1000 | | | | | | | | | |
| Oligo fwd | CACCTTTTTCACTGCATATCGTCC | | | | | | | | | | |
| Oligo rev | AAACGGACGATATGCAGTGAAAAA | | | | | | | | | | |
| Chromosome | start | end | strand | MM | target_seq | PAM | alignment | distance | position | gene name | gene id |
| GU305914.1 | 193558 | 193580 | + | 0 | TTTTTCACTGCATATCGTCC | GGG | ||||||||[||||||||||||]PAM | NA | Intergenic | NA | NA |
| T14706 | CTTCGAGAAGATGCCCGTCACGG | 1000 | | | | | | | | | |
| Oligo fwd | CACCCTTCGAGAAGATGCCCGTCA | | | | | | | | | | |
| Oligo rev | AAACTGACGGGCATCTTCTCGAAG | | | | | | | | | | |
| Chromosome | start | end | strand | MM | target_seq | PAM | alignment | distance | position | gene name | gene id |
| GU305914.1 | 193638 | 193660 | + | 0 | CTTCGAGAAGATGCCCGTCA | CGG | ||||||||[||||||||||||]PAM | NA | Intergenic | NA | NA |
| T14707 | AGATGCCCGTCACGGTGCAGAGG | 1000 | | | | | | | | | |
| Oligo fwd | CACCAGATGCCCGTCACGGTGCAG | | | | | | | | | | |
| Oligo rev | AAACCTGCACCGTGACGGGCATCT | | | | | | | | | | |
| Chromosome | start | end | strand | MM | target_seq | PAM | alignment | distance | position | gene name | gene id |
| GU305914.1 | 193646 | 193668 | + | 0 | AGATGCCCGTCACGGTGCAG | AGG | ||||||||[||||||||||||]PAM | NA | Intergenic | NA | NA |
| T14708 | CTCATGCCCCAGATCGTCATCGG | 1000 | | | | | | | | | |
| Oligo fwd | CACCCTCATGCCCCAGATCGTCAT | | | | | | | | | | |
| Oligo rev | AAACATGACGATCTGGGGCATGAG | | | | | | | | | | |
| Chromosome | start | end | strand | MM | target_seq | PAM | alignment | distance | position | gene name | gene id |
| GU305914.1 | 193696 | 193718 | + | 0 | CTCATGCCCCAGATCGTCAT | CGG | ||||||||[||||||||||||]PAM | NA | Intergenic | NA | NA |
| T14709 | ATGCCCCAGATCGTCATCGGTGG | 1000 | | | | | | | | | |
| Oligo fwd | CACCATGCCCCAGATCGTCATCGG | | | | | | | | | | |
| Oligo rev | AAACCCGATGACGATCTGGGGCAT | | | | | | | | | | |
| Chromosome | start | end | strand | MM | target_seq | PAM | alignment | distance | position | gene name | gene id |
| GU305914.1 | 193699 | 193721 | + | 0 | ATGCCCCAGATCGTCATCGG | TGG | ||||||||[||||||||||||]PAM | NA | Intergenic | NA | NA |
| GU305914.1 | 228173 | 228195 | + | 4 | ATTCCGCAGATCGTCGTCTG | GGG | ||-||-||[|||||||-||-|]PAM | NA | Intergenic | NA | NA |
| T14710 | TGCCCCAGATCGTCATCGGTGGG | 1000 | | | | | | | | | |
| Oligo fwd | CACCTGCCCCAGATCGTCATCGGT | | | | | | | | | | |
| Oligo rev | AAACACCGATGACGATCTGGGGCA | | | | | | | | | | |
| Chromosome | start | end | strand | MM | target_seq | PAM | alignment | distance | position | gene name | gene id |
| GU305914.1 | 193700 | 193722 | + | 0 | TGCCCCAGATCGTCATCGGT | GGG | ||||||||[||||||||||||]PAM | NA | Intergenic | NA | NA |
| GU305914.1 | 202977 | 202999 | + | 4 | GGAACCAGATGGTCATCGGT | AGG | -|--||||[||-|||||||||]PAM | NA | Intergenic | NA | NA |
| T14711 | GCCCCAGATCGTCATCGGTGGGG | 1000 | | | | | | | | | |
| Oligo fwd | CACCGCCCCAGATCGTCATCGGTG | | | | | | | | | | |
| Oligo rev | AAACCACCGATGACGATCTGGGGC | | | | | | | | | | |
| Chromosome | start | end | strand | MM | target_seq | PAM | alignment | distance | position | gene name | gene id |
| GU305914.1 | 193701 | 193723 | + | 0 | GCCCCAGATCGTCATCGGTG | GGG | ||||||||[||||||||||||]PAM | NA | Intergenic | NA | NA |
| T14712 | GTGGGGTGATCAGTTGCGTCAGG | 1000 | | | | | | | | | |
| Oligo fwd | CACCGTGGGGTGATCAGTTGCGTC | | | | | | | | | | |
| Oligo rev | AAACGACGCAACTGATCACCCCAC | | | | | | | | | | |
| Chromosome | start | end | strand | MM | target_seq | PAM | alignment | distance | position | gene name | gene id |
| GU305914.1 | 193718 | 193740 | + | 0 | GTGGGGTGATCAGTTGCGTC | AGG | ||||||||[||||||||||||]PAM | NA | Intergenic | NA | NA |
| T14713 | TGGGGTGATCAGTTGCGTCAGGG | 1000 | | | | | | | | | |
| Oligo fwd | CACCTGGGGTGATCAGTTGCGTCA | | | | | | | | | | |
| Oligo rev | AAACTGACGCAACTGATCACCCCA | | | | | | | | | | |
| Chromosome | start | end | strand | MM | target_seq | PAM | alignment | distance | position | gene name | gene id |
| GU305914.1 | 193719 | 193741 | + | 0 | TGGGGTGATCAGTTGCGTCA | GGG | ||||||||[||||||||||||]PAM | NA | Intergenic | NA | NA |
| T14714 | GGGGTGATCAGTTGCGTCAGGGG | 1000 | | | | | | | | | |

Oligo fwd CACCGGGGTGATCAGTTGCGTCAG
Oligo rev AAACCTGACGCAACTGATCACCCC
Chromosome start end strand MM target_seq PAM alignment distance position gene name gene id
GU305914.1 193720 193742 + 0 GGGGTGATCAGTTGCGTCAG GGG ||||||||[||||||||||||]PAM NA Intergenic NA NA

T14715 GTTGCGTCAGGGGAGTCGTCAGG 1000
Oligo fwd CACCGTTGCGTCAGGGGAGTCGTC
Oligo rev AAACGACGACTCCCCTGACGCAAC
Chromosome start end strand MM target_seq PAM alignment distance position gene name gene id
GU305914.1 193730 193752 + 0 GTTGCGTCAGGGGAGTCGTC AGG ||||||||[||||||||||||]PAM NA Intergenic NA NA

T14716 GGGAGTCGTCAGGTTACGCTCGG 1000
Oligo fwd CACCGGGAGTCGTCAGGTTACGCT
Oligo rev AAACAGCGTAACCTGACGACTCCC
Chromosome start end strand MM target_seq PAM alignment distance position gene name gene id
GU305914.1 193740 193762 + 0 GGGAGTCGTCAGGTTACGCT CGG ||||||||[||||||||||||]PAM NA Intergenic NA NA

T14717 GACGTAGTAGACCCCGATGTCGG 1000
Oligo fwd CACCGACGTAGTAGACCCCGATGT
Oligo rev AAACACATCGGGGTCTACTACGTC
Chromosome start end strand MM target_seq PAM alignment distance position gene name gene id
GU305914.1 193770 193792 + 0 GACGTAGTAGACCCCGATGT CGG ||||||||[||||||||||||]PAM NA Intergenic NA NA

T14718 GTAGTAGACCCCGATGTCGGCGG 1000
Oligo fwd CACCGTAGTAGACCCCGATGTCGG
Oligo rev AAACCCGACATCGGGGTCTACTAC
Chromosome start end strand MM target_seq PAM alignment distance position gene name gene id
GU305914.1 193773 193795 + 0 GTAGTAGACCCCGATGTCGG CGG ||||||||[||||||||||||]PAM NA Intergenic NA NA

T14719 GGCGGTCGAGTAGTTCCACCTGG 1000
Oligo fwd CACCGGCGGTCGAGTAGTTCCACC
Oligo rev AAACGGTGGAACTACTCGACCGCC
Chromosome start end strand MM target_seq PAM alignment distance position gene name gene id
GU305914.1 193791 193813 + 0 GGCGGTCGAGTAGTTCCACC TGG ||||||||[||||||||||||]PAM NA Intergenic NA NA

T14720 GGTCGAGTAGTTCCACCTGGCGG 1000
Oligo fwd CACCGGTCGAGTAGTTCCACCTGG
Oligo rev AAACCCAGGTGGAACTACTCGACC
Chromosome start end strand MM target_seq PAM alignment distance position gene name gene id
GU305914.1 193794 193816 + 0 GGTCGAGTAGTTCCACCTGG CGG ||||||||[||||||||||||]PAM NA Intergenic NA NA

T14721 AATGTAATTTGTCTCCGCGCCGG 1000
Oligo fwd CACCAATGTAATTTGTCTCCGCGC
Oligo rev AAACGCGCGGAGACAAATTACATT
Chromosome start end strand MM target_seq PAM alignment distance position gene name gene id
GU305914.1 193820 193842 + 0 AATGTAATTTGTCTCCGCGC CGG ||||||||[||||||||||||]PAM NA Intergenic NA NA

T14722 ATGTAATTTGTCTCCGCGCCGGG 1000
Oligo fwd CACCATGTAATTTGTCTCCGCGCC
Oligo rev AAACGGCGCGGAGACAAATTACAT
Chromosome start end strand MM target_seq PAM alignment distance position gene name gene id
GU305914.1 193821 193843 + 0 ATGTAATTTGTCTCCGCGCC GGG ||||||||[||||||||||||]PAM NA Intergenic NA NA

T14723 GGCCATCCCGATCCATCCCTCGG 1000
Oligo fwd CACCGGCCATCCCGATCCATCCCT
Oligo rev AAACAGGGATGGATCGGGATGGCC
Chromosome start end strand MM target_seq PAM alignment distance position gene name gene id
GU305914.1 193842 193864 + 0 GGCCATCCCGATCCATCCCT CGG ||||||||[||||||||||||]PAM NA Intergenic NA NA

T14724 TAGCATTGCCTTCGCCGCAGAGG 1000
Oligo fwd CACCTAGCATTGCCTTCGCCGCAG
Oligo rev AAACCTGCGGCGAAGGCAATGCTA
Chromosome start end strand MM target_seq PAM alignment distance position gene name gene id
GU305914.1 193865 193887 + 0 TAGCATTGCCTTCGCCGCAG AGG ||||||||[||||||||||||]PAM NA Intergenic NA NA

T14725 GTCTCGCCAGACCTTGACCGTGG 1000
Oligo fwd CACCGTCTCGCCAGACCTTGACCG
Oligo rev AAACCGGTCAAGGTCTGGCGAGAC
Chromosome start end strand MM target_seq PAM alignment distance position gene name gene id
GU305914.1 193887 193909 + 0 GTCTCGCCAGACCTTGACCG TGG ||||||||[||||||||||||]PAM NA Intergenic NA NA

T14726 TGACCGTGGAGCCGTTCACCAGG 1000
Oligo fwd CACCTGACCGTGGAGCCGTTCACC
Oligo rev AAACGGTGAACGGCTCCACGGTCA
Chromosome start end strand MM target_seq PAM alignment distance position gene name gene id
GU305914.1 193901 193923 + 0 TGACCGTGGAGCCGTTCACC AGG ||||||||[||||||||||||]PAM NA Intergenic NA NA

T14727 GGAGCCGTTCACCAGGTCGCAGG 1000
Oligo fwd CACCGGAGCCGTTCACCAGGTCGC
Oligo rev AAACGCGACCTGGTGAACGGCTCC
Chromosome start end strand MM target_seq PAM alignment distance position gene name gene id
GU305914.1 193908 193930 + 0 GGAGCCGTTCACCAGGTCGC AGG ||||||||[||||||||||||]PAM NA Intergenic NA NA

T14728 AGGTCGCAGGAGATATTGAACGG 1000
Oligo fwd CACCAGGTCGCAGGAGATATTGAA
Oligo rev AAACTTCAATATCTCCTGCGACCT
Chromosome start end strand MM target_seq PAM alignment distance position gene name gene id
GU305914.1 193921 193943 + 0 AGGTCGCAGGAGATATTGAA CGG ||||||||[||||||||||||]PAM NA Intergenic NA NA

<table>
<tr><td>T14729</td><td>GGTCGCAGGAGATATTGAACGGG</td><td>1000</td><td></td><td></td><td></td><td></td><td></td><td></td><td></td><td></td><td></td></tr>
<tr><td>Oligo fwd</td><td>CACCGGTCGCAGGAGATATTGAAC</td><td></td><td></td><td></td><td></td><td></td><td></td><td></td><td></td><td></td><td></td></tr>
<tr><td>Oligo rev</td><td>AAACGTTCAATATCTCCTGCGACC</td><td></td><td></td><td></td><td></td><td></td><td></td><td></td><td></td><td></td><td></td></tr>
<tr><td>Chromosome</td><td>start</td><td>end</td><td>strand</td><td>MM</td><td>target_seq</td><td>PAM</td><td>alignment</td><td>distance</td><td>position</td><td>gene name</td><td>gene id</td></tr>
<tr><td>GU305914.1</td><td>193922</td><td>193944</td><td>+</td><td>0</td><td>GGTCGCAGGAGATATTGAAC</td><td>GGG</td><td>||||||||[||||||||||||]PAM</td><td>NA</td><td>Intergenic</td><td>NA</td><td>NA</td></tr>
<tr><td>T14730</td><td>GCAGGAGATATTGAACGGGACGG</td><td>1000</td><td></td><td></td><td></td><td></td><td></td><td></td><td></td><td></td><td></td></tr>
<tr><td>Oligo fwd</td><td>CACCGCAGGAGATATTGAACGGGA</td><td></td><td></td><td></td><td></td><td></td><td></td><td></td><td></td><td></td><td></td></tr>
<tr><td>Oligo rev</td><td>AAACTCCCGTTCAATATCTCCTGC</td><td></td><td></td><td></td><td></td><td></td><td></td><td></td><td></td><td></td><td></td></tr>
<tr><td>Chromosome</td><td>start</td><td>end</td><td>strand</td><td>MM</td><td>target_seq</td><td>PAM</td><td>alignment</td><td>distance</td><td>position</td><td>gene name</td><td>gene id</td></tr>
<tr><td>GU305914.1</td><td>193926</td><td>193948</td><td>+</td><td>0</td><td>GCAGGAGATATTGAACGGGA</td><td>CGG</td><td>||||||||[||||||||||||]PAM</td><td>NA</td><td>Intergenic</td><td>NA</td><td>NA</td></tr>
<tr><td>T14731</td><td>GGAGATATTGAACGGGACGGAGG</td><td>1000</td><td></td><td></td><td></td><td></td><td></td><td></td><td></td><td></td><td></td></tr>
<tr><td>Oligo fwd</td><td>CACCGGAGATATTGAACGGGACGG</td><td></td><td></td><td></td><td></td><td></td><td></td><td></td><td></td><td></td><td></td></tr>
<tr><td>Oligo rev</td><td>AAACCCGTCCCGTTCAATATCTCC</td><td></td><td></td><td></td><td></td><td></td><td></td><td></td><td></td><td></td><td></td></tr>
<tr><td>Chromosome</td><td>start</td><td>end</td><td>strand</td><td>MM</td><td>target_seq</td><td>PAM</td><td>alignment</td><td>distance</td><td>position</td><td>gene name</td><td>gene id</td></tr>
<tr><td>GU305914.1</td><td>193929</td><td>193951</td><td>+</td><td>0</td><td>GGAGATATTGAACGGGACGG</td><td>AGG</td><td>||||||||[||||||||||||]PAM</td><td>NA</td><td>Intergenic</td><td>NA</td><td>NA</td></tr>
<tr><td>T14732</td><td>GAACGGGACGGAGGTCTCTGTGG</td><td>1000</td><td></td><td></td><td></td><td></td><td></td><td></td><td></td><td></td><td></td></tr>
<tr><td>Oligo fwd</td><td>CACCGAACGGGACGGAGGTCTCTG</td><td></td><td></td><td></td><td></td><td></td><td></td><td></td><td></td><td></td><td></td></tr>
<tr><td>Oligo rev</td><td>AAACCAGAGACCTCCGTCCCGTTC</td><td></td><td></td><td></td><td></td><td></td><td></td><td></td><td></td><td></td><td></td></tr>
<tr><td>Chromosome</td><td>start</td><td>end</td><td>strand</td><td>MM</td><td>target_seq</td><td>PAM</td><td>alignment</td><td>distance</td><td>position</td><td>gene name</td><td>gene id</td></tr>
<tr><td>GU305914.1</td><td>193938</td><td>193960</td><td>+</td><td>0</td><td>GAACGGGACGGAGGTCTCTG</td><td>TGG</td><td>||||||||[||||||||||||]PAM</td><td>NA</td><td>Intergenic</td><td>NA</td><td>NA</td></tr>
<tr><td>T14733</td><td>AACGGGACGGAGGTCTCTGTGGG</td><td>1000</td><td></td><td></td><td></td><td></td><td></td><td></td><td></td><td></td><td></td></tr>
<tr><td>Oligo fwd</td><td>CACCAACGGGACGGAGGTCTCTGT</td><td></td><td></td><td></td><td></td><td></td><td></td><td></td><td></td><td></td><td></td></tr>
<tr><td>Oligo rev</td><td>AAACACAGAGACCTCCGTCCCGTT</td><td></td><td></td><td></td><td></td><td></td><td></td><td></td><td></td><td></td><td></td></tr>
<tr><td>Chromosome</td><td>start</td><td>end</td><td>strand</td><td>MM</td><td>target_seq</td><td>PAM</td><td>alignment</td><td>distance</td><td>position</td><td>gene name</td><td>gene id</td></tr>
<tr><td>GU305914.1</td><td>193939</td><td>193961</td><td>+</td><td>0</td><td>AACGGGACGGAGGTCTCTGT</td><td>GGG</td><td>||||||||[||||||||||||]PAM</td><td>NA</td><td>Intergenic</td><td>NA</td><td>NA</td></tr>
<tr><td>T14734</td><td>TCTCTGTGGGCAAGCGCCACTGG</td><td>1000</td><td></td><td></td><td></td><td></td><td></td><td></td><td></td><td></td><td></td></tr>
<tr><td>Oligo fwd</td><td>CACCTCTCTGTGGGCAAGCGCCAC</td><td></td><td></td><td></td><td></td><td></td><td></td><td></td><td></td><td></td><td></td></tr>
<tr><td>Oligo rev</td><td>AAACGTGGCGCTTGCCCACAGAGA</td><td></td><td></td><td></td><td></td><td></td><td></td><td></td><td></td><td></td><td></td></tr>
<tr><td>Chromosome</td><td>start</td><td>end</td><td>strand</td><td>MM</td><td>target_seq</td><td>PAM</td><td>alignment</td><td>distance</td><td>position</td><td>gene name</td><td>gene id</td></tr>
<tr><td>GU305914.1</td><td>193952</td><td>193974</td><td>+</td><td>0</td><td>TCTCTGTGGGCAAGCGCCAC</td><td>TGG</td><td>||||||||[||||||||||||]PAM</td><td>NA</td><td>Intergenic</td><td>NA</td><td>NA</td></tr>
<tr><td>T14735</td><td>GGCAAGCGCCACTGGCACGCAGG</td><td>1000</td><td></td><td></td><td></td><td></td><td></td><td></td><td></td><td></td><td></td></tr>
<tr><td>Oligo fwd</td><td>CACCGGCAAGCGCCACTGGCACGC</td><td></td><td></td><td></td><td></td><td></td><td></td><td></td><td></td><td></td><td></td></tr>
<tr><td>Oligo rev</td><td>AAACGCGTGCCAGTGGCGCTTGCC</td><td></td><td></td><td></td><td></td><td></td><td></td><td></td><td></td><td></td><td></td></tr>
<tr><td>Chromosome</td><td>start</td><td>end</td><td>strand</td><td>MM</td><td>target_seq</td><td>PAM</td><td>alignment</td><td>distance</td><td>position</td><td>gene name</td><td>gene id</td></tr>
<tr><td>GU305914.1</td><td>193960</td><td>193982</td><td>+</td><td>0</td><td>GGCAAGCGCCACTGGCACGC</td><td>AGG</td><td>||||||||[||||||||||||]PAM</td><td>NA</td><td>Intergenic</td><td>NA</td><td>NA</td></tr>
<tr><td>T14736</td><td>AGGTAATTGATGCGCCGTCACGG</td><td>1000</td><td></td><td></td><td></td><td></td><td></td><td></td><td></td><td></td><td></td></tr>
<tr><td>Oligo fwd</td><td>CACCAGGTAATTGATGCGCCGTCA</td><td></td><td></td><td></td><td></td><td></td><td></td><td></td><td></td><td></td><td></td></tr>
<tr><td>Oligo rev</td><td>AAACTGACGGCGCATCAATTACCT</td><td></td><td></td><td></td><td></td><td></td><td></td><td></td><td></td><td></td><td></td></tr>
<tr><td>Chromosome</td><td>start</td><td>end</td><td>strand</td><td>MM</td><td>target_seq</td><td>PAM</td><td>alignment</td><td>distance</td><td>position</td><td>gene name</td><td>gene id</td></tr>
<tr><td>GU305914.1</td><td>193980</td><td>194002</td><td>+</td><td>0</td><td>AGGTAATTGATGCGCCGTCA</td><td>CGG</td><td>||||||||[||||||||||||]PAM</td><td>NA</td><td>Intergenic</td><td>NA</td><td>NA</td></tr>
<tr><td>T14737</td><td>GATGCGCCGTCACGGCCGCGAGG</td><td>1000</td><td></td><td></td><td></td><td></td><td></td><td></td><td></td><td></td><td></td></tr>
<tr><td>Oligo fwd</td><td>CACCGATGCGCCGTCACGGCCGCG</td><td></td><td></td><td></td><td></td><td></td><td></td><td></td><td></td><td></td><td></td></tr>
<tr><td>Oligo rev</td><td>AAACCGCGGCCGTGACGGCGCATC</td><td></td><td></td><td></td><td></td><td></td><td></td><td></td><td></td><td></td><td></td></tr>
<tr><td>Chromosome</td><td>start</td><td>end</td><td>strand</td><td>MM</td><td>target_seq</td><td>PAM</td><td>alignment</td><td>distance</td><td>position</td><td>gene name</td><td>gene id</td></tr>
<tr><td>GU305914.1</td><td>193988</td><td>194010</td><td>+</td><td>0</td><td>GATGCGCCGTCACGGCCGCG</td><td>AGG</td><td>||||||||[||||||||||||]PAM</td><td>NA</td><td>Intergenic</td><td>NA</td><td>NA</td></tr>
<tr><td>T14738</td><td>GCGCCGTCACGGCCGCGAGGAGG</td><td>1000</td><td></td><td></td><td></td><td></td><td></td><td></td><td></td><td></td><td></td></tr>
<tr><td>Oligo fwd</td><td>CACCGCGCCGTCACGGCCGCGAGG</td><td></td><td></td><td></td><td></td><td></td><td></td><td></td><td></td><td></td><td></td></tr>
<tr><td>Oligo rev</td><td>AAACCCTCGCGGCCGTGACGGCGC</td><td></td><td></td><td></td><td></td><td></td><td></td><td></td><td></td><td></td><td></td></tr>
<tr><td>Chromosome</td><td>start</td><td>end</td><td>strand</td><td>MM</td><td>target_seq</td><td>PAM</td><td>alignment</td><td>distance</td><td>position</td><td>gene name</td><td>gene id</td></tr>
<tr><td>GU305914.1</td><td>193991</td><td>194013</td><td>+</td><td>0</td><td>GCGCCGTCACGGCCGCGAGG</td><td>AGG</td><td>||||||||[||||||||||||]PAM</td><td>NA</td><td>Intergenic</td><td>NA</td><td>NA</td></tr>
<tr><td>T14739</td><td>CAGAGTGCGATCAGCGTCGAAGG</td><td>1000</td><td></td><td></td><td></td><td></td><td></td><td></td><td></td><td></td><td></td></tr>
<tr><td>Oligo fwd</td><td>CACCCAGAGTGCGATCAGCGTCGA</td><td></td><td></td><td></td><td></td><td></td><td></td><td></td><td></td><td></td><td></td></tr>
<tr><td>Oligo rev</td><td>AAACTCGACGCTGATCGCACTCTG</td><td></td><td></td><td></td><td></td><td></td><td></td><td></td><td></td><td></td><td></td></tr>
<tr><td>Chromosome</td><td>start</td><td>end</td><td>strand</td><td>MM</td><td>target_seq</td><td>PAM</td><td>alignment</td><td>distance</td><td>position</td><td>gene name</td><td>gene id</td></tr>
<tr><td>GU305914.1</td><td>194014</td><td>194036</td><td>+</td><td>0</td><td>CAGAGTGCGATCAGCGTCGA</td><td>AGG</td><td>||||||||[||||||||||||]PAM</td><td>NA</td><td>Intergenic</td><td>NA</td><td>NA</td></tr>
<tr><td>T14740</td><td>CGTCGAAGGCGCCATCGCCTCGG</td><td>1000</td><td></td><td></td><td></td><td></td><td></td><td></td><td></td><td></td><td></td></tr>
<tr><td>Oligo fwd</td><td>CACCCGTCGAAGGCGCCATCGCCT</td><td></td><td></td><td></td><td></td><td></td><td></td><td></td><td></td><td></td><td></td></tr>
<tr><td>Oligo rev</td><td>AAACAGGCGATGGCGCCTTCGACG</td><td></td><td></td><td></td><td></td><td></td><td></td><td></td><td></td><td></td><td></td></tr>
<tr><td>Chromosome</td><td>start</td><td>end</td><td>strand</td><td>MM</td><td>target_seq</td><td>PAM</td><td>alignment</td><td>distance</td><td>position</td><td>gene name</td><td>gene id</td></tr>
<tr><td>GU305914.1</td><td>194028</td><td>194050</td><td>+</td><td>0</td><td>CGTCGAAGGCGCCATCGCCT</td><td>CGG</td><td>||||||||[||||||||||||]PAM</td><td>NA</td><td>Intergenic</td><td>NA</td><td>NA</td></tr>
<tr><td>T14741</td><td>CTCTGCCGACCGAGAGATGATGG</td><td>1000</td><td></td><td></td><td></td><td></td><td></td><td></td><td></td><td></td><td></td></tr>
<tr><td>Oligo fwd</td><td>CACCCTCTGCCGACCGAGAGATGA</td><td></td><td></td><td></td><td></td><td></td><td></td><td></td><td></td><td></td><td></td></tr>
<tr><td>Oligo rev</td><td>AAACTCATCTCTCGGTCGGCAGAG</td><td></td><td></td><td></td><td></td><td></td><td></td><td></td><td></td><td></td><td></td></tr>
<tr><td>Chromosome</td><td>start</td><td>end</td><td>strand</td><td>MM</td><td>target_seq</td><td>PAM</td><td>alignment</td><td>distance</td><td>position</td><td>gene name</td><td>gene id</td></tr>
<tr><td>GU305914.1</td><td>194051</td><td>194073</td><td>+</td><td>0</td><td>CTCTGCCGACCGAGAGATGA</td><td>TGG</td><td>||||||||[||||||||||||]PAM</td><td>NA</td><td>Intergenic</td><td>NA</td><td>NA</td></tr>
<tr><td>GU305914.1</td><td>196154</td><td>196176</td><td>-</td><td>3</td><td>CTCTGCAGACCGTGAGTTGA</td><td>CGG</td><td>||||||-|[||||-|||-|||]PAM</td><td>NA</td><td>Intergenic</td><td>NA</td><td>NA</td></tr>
<tr><td>T14742</td><td>GTGACAGAGTTTCTGTGATGCGG</td><td>1000</td><td></td><td></td><td></td><td></td><td></td><td></td><td></td><td></td><td></td></tr>
<tr><td>Oligo fwd</td><td>CACCGTGACAGAGTTTCTGTGATG</td><td></td><td></td><td></td><td></td><td></td><td></td><td></td><td></td><td></td><td></td></tr>
<tr><td>Oligo rev</td><td>AAACCATCACAGAAACTCTGTCAC</td><td></td><td></td><td></td><td></td><td></td><td></td><td></td><td></td><td></td><td></td></tr>
<tr><td>Chromosome</td><td>start</td><td>end</td><td>strand</td><td>MM</td><td>target_seq</td><td>PAM</td><td>alignment</td><td>distance</td><td>position</td><td>gene name</td><td>gene id</td></tr>
<tr><td>GU305914.1</td><td>194093</td><td>194115</td><td>+</td><td>0</td><td>GTGACAGAGTTTCTGTGATG</td><td>CGG</td><td>||||||||[||||||||||||]PAM</td><td>NA</td><td>Intergenic</td><td>NA</td><td>NA</td></tr>
<tr><td>T14743</td><td>AGAGTTTCTGTGATGCGGCACGG</td><td>1000</td><td></td><td></td><td></td><td></td><td></td><td></td><td></td><td></td><td></td></tr>
<tr><td>Oligo fwd</td><td>CACCAGAGTTTCTGTGATGCGGCA</td><td></td><td></td><td></td><td></td><td></td><td></td><td></td><td></td><td></td><td></td></tr>
<tr><td>Oligo rev</td><td>AAACTGCCGCATCACAGAAACTCT</td><td></td><td></td><td></td><td></td><td></td><td></td><td></td><td></td><td></td><td></td></tr>
<tr><td>Chromosome</td><td>start</td><td>end</td><td>strand</td><td>MM</td><td>target_seq</td><td>PAM</td><td>alignment</td><td>distance</td><td>position</td><td>gene name</td><td>gene id</td></tr>
</table>

<table>
<tr><td>GU305914.1</td><td>194098</td><td>194120</td><td>+</td><td>0</td><td>AGAGTTTCTGTGATGCGGCA</td><td>CGG</td><td>||||||||[||||||||||||]PAM</td><td>NA</td><td>Intergenic</td><td>NA</td><td>NA</td></tr>
<tr><td></td><td></td><td></td><td></td><td></td><td></td><td></td><td></td><td></td><td></td><td></td><td></td></tr>
<tr><td>T14744</td><td>GCACGGCTTTTATAGACTGATGG</td><td>1000</td><td></td><td></td><td></td><td></td><td></td><td></td><td></td><td></td><td></td></tr>
<tr><td>Oligo fwd</td><td>CACCGCACGGCTTTTATAGACTGA</td><td></td><td></td><td></td><td></td><td></td><td></td><td></td><td></td><td></td><td></td></tr>
<tr><td>Oligo rev</td><td>AAACTCAGTCTATAAAAGCCGTGC</td><td></td><td></td><td></td><td></td><td></td><td></td><td></td><td></td><td></td><td></td></tr>
<tr><td>Chromosome</td><td>start</td><td>end</td><td>strand</td><td>MM</td><td>target_seq</td><td>PAM</td><td>alignment</td><td>distance</td><td>position</td><td>gene name</td><td>gene id</td></tr>
<tr><td>GU305914.1</td><td>194115</td><td>194137</td><td>+</td><td>0</td><td>GCACGGCTTTTATAGACTGA</td><td>TGG</td><td>||||||||[||||||||||||]PAM</td><td>NA</td><td>Intergenic</td><td>NA</td><td>NA</td></tr>
<tr><td></td><td></td><td></td><td></td><td></td><td></td><td></td><td></td><td></td><td></td><td></td><td></td></tr>
<tr><td>T14745</td><td>GCTGCGCGCCCCCCCGCGAGCGG</td><td>1000</td><td></td><td></td><td></td><td></td><td></td><td></td><td></td><td></td><td></td></tr>
<tr><td>Oligo fwd</td><td>CACCGCTGCGCGCCCCCCCGCGAG</td><td></td><td></td><td></td><td></td><td></td><td></td><td></td><td></td><td></td><td></td></tr>
<tr><td>Oligo rev</td><td>AAACCTCGCGGGGGGGCGCGCAGC</td><td></td><td></td><td></td><td></td><td></td><td></td><td></td><td></td><td></td><td></td></tr>
<tr><td>Chromosome</td><td>start</td><td>end</td><td>strand</td><td>MM</td><td>target_seq</td><td>PAM</td><td>alignment</td><td>distance</td><td>position</td><td>gene name</td><td>gene id</td></tr>
<tr><td>GU305914.1</td><td>194140</td><td>194162</td><td>+</td><td>0</td><td>GCTGCGCGCCCCCCCGCGAG</td><td>CGG</td><td>||||||||[||||||||||||]PAM</td><td>NA</td><td>Intergenic</td><td>NA</td><td>NA</td></tr>
<tr><td></td><td></td><td></td><td></td><td></td><td></td><td></td><td></td><td></td><td></td><td></td><td></td></tr>
<tr><td>T14746</td><td>AGCGGCTGACACAGCGCCGCTGG</td><td>1000</td><td></td><td></td><td></td><td></td><td></td><td></td><td></td><td></td><td></td></tr>
<tr><td>Oligo fwd</td><td>CACCAGCGGCTGACACAGCGCCGC</td><td></td><td></td><td></td><td></td><td></td><td></td><td></td><td></td><td></td><td></td></tr>
<tr><td>Oligo rev</td><td>AAACGCGGCGCTGTGTCAGCCGCT</td><td></td><td></td><td></td><td></td><td></td><td></td><td></td><td></td><td></td><td></td></tr>
<tr><td>Chromosome</td><td>start</td><td>end</td><td>strand</td><td>MM</td><td>target_seq</td><td>PAM</td><td>alignment</td><td>distance</td><td>position</td><td>gene name</td><td>gene id</td></tr>
<tr><td>GU305914.1</td><td>194158</td><td>194180</td><td>+</td><td>0</td><td>AGCGGCTGACACAGCGCCGC</td><td>TGG</td><td>||||||||[||||||||||||]PAM</td><td>NA</td><td>Intergenic</td><td>NA</td><td>NA</td></tr>
<tr><td></td><td></td><td></td><td></td><td></td><td></td><td></td><td></td><td></td><td></td><td></td><td></td></tr>
<tr><td>T14747</td><td>AGCGCCGCTGGACTAGCGATCGG</td><td>1000</td><td></td><td></td><td></td><td></td><td></td><td></td><td></td><td></td><td></td></tr>
<tr><td>Oligo fwd</td><td>CACCAGCGCCGCTGGACTAGCGAT</td><td></td><td></td><td></td><td></td><td></td><td></td><td></td><td></td><td></td><td></td></tr>
<tr><td>Oligo rev</td><td>AAACATCGCTAGTCCAGCGGCGCT</td><td></td><td></td><td></td><td></td><td></td><td></td><td></td><td></td><td></td><td></td></tr>
<tr><td>Chromosome</td><td>start</td><td>end</td><td>strand</td><td>MM</td><td>target_seq</td><td>PAM</td><td>alignment</td><td>distance</td><td>position</td><td>gene name</td><td>gene id</td></tr>
<tr><td>GU305914.1</td><td>194170</td><td>194192</td><td>+</td><td>0</td><td>AGCGCCGCTGGACTAGCGAT</td><td>CGG</td><td>||||||||[||||||||||||]PAM</td><td>NA</td><td>Intergenic</td><td>NA</td><td>NA</td></tr>
<tr><td></td><td></td><td></td><td></td><td></td><td></td><td></td><td></td><td></td><td></td><td></td><td></td></tr>
<tr><td>T14748</td><td>GCTGGACTAGCGATCGGAGCCGG</td><td>1000</td><td></td><td></td><td></td><td></td><td></td><td></td><td></td><td></td><td></td></tr>
<tr><td>Oligo fwd</td><td>CACCGCTGGACTAGCGATCGGAGC</td><td></td><td></td><td></td><td></td><td></td><td></td><td></td><td></td><td></td><td></td></tr>
<tr><td>Oligo rev</td><td>AAACGCTCCGATCGCTAGTCCAGC</td><td></td><td></td><td></td><td></td><td></td><td></td><td></td><td></td><td></td><td></td></tr>
<tr><td>Chromosome</td><td>start</td><td>end</td><td>strand</td><td>MM</td><td>target_seq</td><td>PAM</td><td>alignment</td><td>distance</td><td>position</td><td>gene name</td><td>gene id</td></tr>
<tr><td>GU305914.1</td><td>194176</td><td>194198</td><td>+</td><td>0</td><td>GCTGGACTAGCGATCGGAGC</td><td>CGG</td><td>||||||||[||||||||||||]PAM</td><td>NA</td><td>Intergenic</td><td>NA</td><td>NA</td></tr>
<tr><td></td><td></td><td></td><td></td><td></td><td></td><td></td><td></td><td></td><td></td><td></td><td></td></tr>
<tr><td>T14749</td><td>TCGATCCCGCCTGAGTCGCGCGG</td><td>1000</td><td></td><td></td><td></td><td></td><td></td><td></td><td></td><td></td><td></td></tr>
<tr><td>Oligo fwd</td><td>CACCTCGATCCCGCCTGAGTCGCG</td><td></td><td></td><td></td><td></td><td></td><td></td><td></td><td></td><td></td><td></td></tr>
<tr><td>Oligo rev</td><td>AAACCGCGACTCAGGCGGGATCGA</td><td></td><td></td><td></td><td></td><td></td><td></td><td></td><td></td><td></td><td></td></tr>
<tr><td>Chromosome</td><td>start</td><td>end</td><td>strand</td><td>MM</td><td>target_seq</td><td>PAM</td><td>alignment</td><td>distance</td><td>position</td><td>gene name</td><td>gene id</td></tr>
<tr><td>GU305914.1</td><td>194217</td><td>194239</td><td>+</td><td>0</td><td>TCGATCCCGCCTGAGTCGCG</td><td>CGG</td><td>||||||||[||||||||||||]PAM</td><td>NA</td><td>Intergenic</td><td>NA</td><td>NA</td></tr>
<tr><td></td><td></td><td></td><td></td><td></td><td></td><td></td><td></td><td></td><td></td><td></td><td></td></tr>
<tr><td>T14750</td><td>GCCTGAGTCGCGCGGACCAGAGG</td><td>1000</td><td></td><td></td><td></td><td></td><td></td><td></td><td></td><td></td><td></td></tr>
<tr><td>Oligo fwd</td><td>CACCGCCTGAGTCGCGCGGACCAG</td><td></td><td></td><td></td><td></td><td></td><td></td><td></td><td></td><td></td><td></td></tr>
<tr><td>Oligo rev</td><td>AAACCTGGTCCGCGCGACTCAGGC</td><td></td><td></td><td></td><td></td><td></td><td></td><td></td><td></td><td></td><td></td></tr>
<tr><td>Chromosome</td><td>start</td><td>end</td><td>strand</td><td>MM</td><td>target_seq</td><td>PAM</td><td>alignment</td><td>distance</td><td>position</td><td>gene name</td><td>gene id</td></tr>
<tr><td>GU305914.1</td><td>194225</td><td>194247</td><td>+</td><td>0</td><td>GCCTGAGTCGCGCGGACCAG</td><td>AGG</td><td>||||||||[||||||||||||]PAM</td><td>NA</td><td>Intergenic</td><td>NA</td><td>NA</td></tr>
<tr><td></td><td></td><td></td><td></td><td></td><td></td><td></td><td></td><td></td><td></td><td></td><td></td></tr>
<tr><td>T14751</td><td>TGAGTCGCGCGGACCAGAGGTGG</td><td>1000</td><td></td><td></td><td></td><td></td><td></td><td></td><td></td><td></td><td></td></tr>
<tr><td>Oligo fwd</td><td>CACCTGAGTCGCGCGGACCAGAGG</td><td></td><td></td><td></td><td></td><td></td><td></td><td></td><td></td><td></td><td></td></tr>
<tr><td>Oligo rev</td><td>AAACCCTCTGGTCCGCGCGACTCA</td><td></td><td></td><td></td><td></td><td></td><td></td><td></td><td></td><td></td><td></td></tr>
<tr><td>Chromosome</td><td>start</td><td>end</td><td>strand</td><td>MM</td><td>target_seq</td><td>PAM</td><td>alignment</td><td>distance</td><td>position</td><td>gene name</td><td>gene id</td></tr>
<tr><td>GU305914.1</td><td>194228</td><td>194250</td><td>+</td><td>0</td><td>TGAGTCGCGCGGACCAGAGG</td><td>TGG</td><td>||||||||[||||||||||||]PAM</td><td>NA</td><td>Intergenic</td><td>NA</td><td>NA</td></tr>
<tr><td></td><td></td><td></td><td></td><td></td><td></td><td></td><td></td><td></td><td></td><td></td><td></td></tr>
<tr><td>T14752</td><td>ACCAGAGGTGGCAGTTTCAATGG</td><td>1000</td><td></td><td></td><td></td><td></td><td></td><td></td><td></td><td></td><td></td></tr>
<tr><td>Oligo fwd</td><td>CACCACCAGAGGTGGCAGTTTCAA</td><td></td><td></td><td></td><td></td><td></td><td></td><td></td><td></td><td></td><td></td></tr>
<tr><td>Oligo rev</td><td>AAACTTGAAACTGCCACCTCTGGT</td><td></td><td></td><td></td><td></td><td></td><td></td><td></td><td></td><td></td><td></td></tr>
<tr><td>Chromosome</td><td>start</td><td>end</td><td>strand</td><td>MM</td><td>target_seq</td><td>PAM</td><td>alignment</td><td>distance</td><td>position</td><td>gene name</td><td>gene id</td></tr>
<tr><td>GU305914.1</td><td>194240</td><td>194262</td><td>+</td><td>0</td><td>ACCAGAGGTGGCAGTTTCAA</td><td>TGG</td><td>||||||||[||||||||||||]PAM</td><td>NA</td><td>Intergenic</td><td>NA</td><td>NA</td></tr>
<tr><td></td><td></td><td></td><td></td><td></td><td></td><td></td><td></td><td></td><td></td><td></td><td></td></tr>
<tr><td>T14753</td><td>CCAGAGGTGGCAGTTTCAATGGG</td><td>1000</td><td></td><td></td><td></td><td></td><td></td><td></td><td></td><td></td><td></td></tr>
<tr><td>Oligo fwd</td><td>CACCCCAGAGGTGGCAGTTTCAAT</td><td></td><td></td><td></td><td></td><td></td><td></td><td></td><td></td><td></td><td></td></tr>
<tr><td>Oligo rev</td><td>AAACATTGAAACTGCCACCTCTGG</td><td></td><td></td><td></td><td></td><td></td><td></td><td></td><td></td><td></td><td></td></tr>
<tr><td>Chromosome</td><td>start</td><td>end</td><td>strand</td><td>MM</td><td>target_seq</td><td>PAM</td><td>alignment</td><td>distance</td><td>position</td><td>gene name</td><td>gene id</td></tr>
<tr><td>GU305914.1</td><td>194241</td><td>194263</td><td>+</td><td>0</td><td>CCAGAGGTGGCAGTTTCAAT</td><td>GGG</td><td>||||||||[||||||||||||]PAM</td><td>NA</td><td>Intergenic</td><td>NA</td><td>NA</td></tr>
<tr><td></td><td></td><td></td><td></td><td></td><td></td><td></td><td></td><td></td><td></td><td></td><td></td></tr>
<tr><td>T14754</td><td>CAGAGGTGGCAGTTTCAATGGGG</td><td>1000</td><td></td><td></td><td></td><td></td><td></td><td></td><td></td><td></td><td></td></tr>
<tr><td>Oligo fwd</td><td>CACCCAGAGGTGGCAGTTTCAATG</td><td></td><td></td><td></td><td></td><td></td><td></td><td></td><td></td><td></td><td></td></tr>
<tr><td>Oligo rev</td><td>AAACCATTGAAACTGCCACCTCTG</td><td></td><td></td><td></td><td></td><td></td><td></td><td></td><td></td><td></td><td></td></tr>
<tr><td>Chromosome</td><td>start</td><td>end</td><td>strand</td><td>MM</td><td>target_seq</td><td>PAM</td><td>alignment</td><td>distance</td><td>position</td><td>gene name</td><td>gene id</td></tr>
<tr><td>GU305914.1</td><td>194242</td><td>194264</td><td>+</td><td>0</td><td>CAGAGGTGGCAGTTTCAATG</td><td>GGG</td><td>||||||||[||||||||||||]PAM</td><td>NA</td><td>Intergenic</td><td>NA</td><td>NA</td></tr>
<tr><td></td><td></td><td></td><td></td><td></td><td></td><td></td><td></td><td></td><td></td><td></td><td></td></tr>
<tr><td>T14755</td><td>TCAATGGGGTTCCCGTCTTCTGG</td><td>1000</td><td></td><td></td><td></td><td></td><td></td><td></td><td></td><td></td><td></td></tr>
<tr><td>Oligo fwd</td><td>CACCTCAATGGGGTTCCCGTCTTC</td><td></td><td></td><td></td><td></td><td></td><td></td><td></td><td></td><td></td><td></td></tr>
<tr><td>Oligo rev</td><td>AAACGAAGACGGGAACCCCATTGA</td><td></td><td></td><td></td><td></td><td></td><td></td><td></td><td></td><td></td><td></td></tr>
<tr><td>Chromosome</td><td>start</td><td>end</td><td>strand</td><td>MM</td><td>target_seq</td><td>PAM</td><td>alignment</td><td>distance</td><td>position</td><td>gene name</td><td>gene id</td></tr>
<tr><td>GU305914.1</td><td>194256</td><td>194278</td><td>+</td><td>0</td><td>TCAATGGGGTTCCCGTCTTC</td><td>TGG</td><td>||||||||[||||||||||||]PAM</td><td>NA</td><td>Intergenic</td><td>NA</td><td>NA</td></tr>
<tr><td></td><td></td><td></td><td></td><td></td><td></td><td></td><td></td><td></td><td></td><td></td><td></td></tr>
<tr><td>T14756</td><td>CTGGAGCGAGCCGACCCTCTCGG</td><td>1000</td><td></td><td></td><td></td><td></td><td></td><td></td><td></td><td></td><td></td></tr>
<tr><td>Oligo fwd</td><td>CACCCTGGAGCGAGCCGACCCTCT</td><td></td><td></td><td></td><td></td><td></td><td></td><td></td><td></td><td></td><td></td></tr>
<tr><td>Oligo rev</td><td>AAACAGAGGGTCGGCTCGCTCCAG</td><td></td><td></td><td></td><td></td><td></td><td></td><td></td><td></td><td></td><td></td></tr>
<tr><td>Chromosome</td><td>start</td><td>end</td><td>strand</td><td>MM</td><td>target_seq</td><td>PAM</td><td>alignment</td><td>distance</td><td>position</td><td>gene name</td><td>gene id</td></tr>
<tr><td>GU305914.1</td><td>194275</td><td>194297</td><td>+</td><td>0</td><td>CTGGAGCGAGCCGACCCTCT</td><td>CGG</td><td>||||||||[||||||||||||]PAM</td><td>NA</td><td>Intergenic</td><td>NA</td><td>NA</td></tr>
<tr><td></td><td></td><td></td><td></td><td></td><td></td><td></td><td></td><td></td><td></td><td></td><td></td></tr>
<tr><td>T14757</td><td>GCCGCCGTCGAACATGTCCATGG</td><td>1000</td><td></td><td></td><td></td><td></td><td></td><td></td><td></td><td></td><td></td></tr>
<tr><td>Oligo fwd</td><td>CACCGCCGCCGTCGAACATGTCCA</td><td></td><td></td><td></td><td></td><td></td><td></td><td></td><td></td><td></td><td></td></tr>
<tr><td>Oligo rev</td><td>AAACTGGACATGTTCGACGGCGGC</td><td></td><td></td><td></td><td></td><td></td><td></td><td></td><td></td><td></td><td></td></tr>
<tr><td>Chromosome</td><td>start</td><td>end</td><td>strand</td><td>MM</td><td>target_seq</td><td>PAM</td><td>alignment</td><td>distance</td><td>position</td><td>gene name</td><td>gene id</td></tr>
<tr><td>GU305914.1</td><td>194335</td><td>194357</td><td>+</td><td>0</td><td>GCCGCCGTCGAACATGTCCA</td><td>TGG</td><td>||||||||[||||||||||||]PAM</td><td>NA</td><td>Intergenic</td><td>NA</td><td>NA</td></tr>
<tr><td></td><td></td><td></td><td></td><td></td><td></td><td></td><td></td><td></td><td></td><td></td><td></td></tr>
<tr><td>T14758</td><td>CGTCGAACATGTCCATGGCGCGG</td><td>1000</td><td></td><td></td><td></td><td></td><td></td><td></td><td></td><td></td><td></td></tr>
<tr><td>Oligo fwd</td><td>CACCCGTCGAACATGTCCATGGCG</td><td></td><td></td><td></td><td></td><td></td><td></td><td></td><td></td><td></td><td></td></tr>
<tr><td>Oligo rev</td><td>AAACCGCCATGGACATGTTCGACG</td><td></td><td></td><td></td><td></td><td></td><td></td><td></td><td></td><td></td><td></td></tr>
</table>

<table>
<tr><td>Chromosome</td><td>start</td><td>end</td><td>strand</td><td>MM</td><td>target_seq</td><td>PAM</td><td>alignment</td><td>distance</td><td>position</td><td>gene name</td><td>gene id</td></tr>
<tr><td>GU305914.1</td><td>194340</td><td>194362</td><td>+</td><td>0</td><td>CGTCGAACATGTCCATGGCG</td><td>CGG</td><td>||||||||[||||||||||||]PAM</td><td>NA</td><td>Intergenic</td><td>NA</td><td>NA</td></tr>
<tr><td>T14759</td><td>CGTAACGACGATAGAAGTCGCGG</td><td>1000</td><td></td><td></td><td></td><td></td><td></td><td></td><td></td><td></td><td></td></tr>
<tr><td>Oligo fwd</td><td>CACCCGTAACGACGATAGAAGTCG</td><td></td><td></td><td></td><td></td><td></td><td></td><td></td><td></td><td></td><td></td></tr>
<tr><td>Oligo rev</td><td>AAACCGACTTCTATCGTCGTTACG</td><td></td><td></td><td></td><td></td><td></td><td></td><td></td><td></td><td></td><td></td></tr>
<tr><td>Chromosome</td><td>start</td><td>end</td><td>strand</td><td>MM</td><td>target_seq</td><td>PAM</td><td>alignment</td><td>distance</td><td>position</td><td>gene name</td><td>gene id</td></tr>
<tr><td>GU305914.1</td><td>194364</td><td>194386</td><td>+</td><td>0</td><td>CGTAACGACGATAGAAGTCG</td><td>CGG</td><td>||||||||[||||||||||||]PAM</td><td>NA</td><td>Intergenic</td><td>NA</td><td>NA</td></tr>
<tr><td>T14760</td><td>ACGACGATAGAAGTCGCGGTCGG</td><td>1000</td><td></td><td></td><td></td><td></td><td></td><td></td><td></td><td></td><td></td></tr>
<tr><td>Oligo fwd</td><td>CACCACGACGATAGAAGTCGCGGT</td><td></td><td></td><td></td><td></td><td></td><td></td><td></td><td></td><td></td><td></td></tr>
<tr><td>Oligo rev</td><td>AAACACCGCGACTTCTATCGTCGT</td><td></td><td></td><td></td><td></td><td></td><td></td><td></td><td></td><td></td><td></td></tr>
<tr><td>Chromosome</td><td>start</td><td>end</td><td>strand</td><td>MM</td><td>target_seq</td><td>PAM</td><td>alignment</td><td>distance</td><td>position</td><td>gene name</td><td>gene id</td></tr>
<tr><td>GU305914.1</td><td>194368</td><td>194390</td><td>+</td><td>0</td><td>ACGACGATAGAAGTCGCGGT</td><td>CGG</td><td>||||||||[||||||||||||]PAM</td><td>NA</td><td>Intergenic</td><td>NA</td><td>NA</td></tr>
<tr><td>T14761</td><td>AGTCGCGGTCGGAGCACGACAGG</td><td>1000</td><td></td><td></td><td></td><td></td><td></td><td></td><td></td><td></td><td></td></tr>
<tr><td>Oligo fwd</td><td>CACCAGTCGCGGTCGGAGCACGAC</td><td></td><td></td><td></td><td></td><td></td><td></td><td></td><td></td><td></td><td></td></tr>
<tr><td>Oligo rev</td><td>AAACGTCGTGCTCCGACCGCGACT</td><td></td><td></td><td></td><td></td><td></td><td></td><td></td><td></td><td></td><td></td></tr>
<tr><td>Chromosome</td><td>start</td><td>end</td><td>strand</td><td>MM</td><td>target_seq</td><td>PAM</td><td>alignment</td><td>distance</td><td>position</td><td>gene name</td><td>gene id</td></tr>
<tr><td>GU305914.1</td><td>194379</td><td>194401</td><td>+</td><td>0</td><td>AGTCGCGGTCGGAGCACGAC</td><td>AGG</td><td>||||||||[||||||||||||]PAM</td><td>NA</td><td>Intergenic</td><td>NA</td><td>NA</td></tr>
<tr><td>T14762</td><td>GTCGCGGTCGGAGCACGACAGGG</td><td>1000</td><td></td><td></td><td></td><td></td><td></td><td></td><td></td><td></td><td></td></tr>
<tr><td>Oligo fwd</td><td>CACCGTCGCGGTCGGAGCACGACA</td><td></td><td></td><td></td><td></td><td></td><td></td><td></td><td></td><td></td><td></td></tr>
<tr><td>Oligo rev</td><td>AAACTGTCGTGCTCCGACCGCGAC</td><td></td><td></td><td></td><td></td><td></td><td></td><td></td><td></td><td></td><td></td></tr>
<tr><td>Chromosome</td><td>start</td><td>end</td><td>strand</td><td>MM</td><td>target_seq</td><td>PAM</td><td>alignment</td><td>distance</td><td>position</td><td>gene name</td><td>gene id</td></tr>
<tr><td>GU305914.1</td><td>194380</td><td>194402</td><td>+</td><td>0</td><td>GTCGCGGTCGGAGCACGACA</td><td>GGG</td><td>||||||||[||||||||||||]PAM</td><td>NA</td><td>Intergenic</td><td>NA</td><td>NA</td></tr>
<tr><td>T14763</td><td>GCGGTCGGAGCACGACAGGGAGG</td><td>1000</td><td></td><td></td><td></td><td></td><td></td><td></td><td></td><td></td><td></td></tr>
<tr><td>Oligo fwd</td><td>CACCGCGGTCGGAGCACGACAGGG</td><td></td><td></td><td></td><td></td><td></td><td></td><td></td><td></td><td></td><td></td></tr>
<tr><td>Oligo rev</td><td>AAACCCCTGTCGTGCTCCGACCGC</td><td></td><td></td><td></td><td></td><td></td><td></td><td></td><td></td><td></td><td></td></tr>
<tr><td>Chromosome</td><td>start</td><td>end</td><td>strand</td><td>MM</td><td>target_seq</td><td>PAM</td><td>alignment</td><td>distance</td><td>position</td><td>gene name</td><td>gene id</td></tr>
<tr><td>GU305914.1</td><td>194383</td><td>194405</td><td>+</td><td>0</td><td>GCGGTCGGAGCACGACAGGG</td><td>AGG</td><td>||||||||[||||||||||||]PAM</td><td>NA</td><td>Intergenic</td><td>NA</td><td>NA</td></tr>
<tr><td>T14764</td><td>AGGGAGGCGTCGCAGACCCGCGG</td><td>1000</td><td></td><td></td><td></td><td></td><td></td><td></td><td></td><td></td><td></td></tr>
<tr><td>Oligo fwd</td><td>CACCAGGGAGGCGTCGCAGACCCG</td><td></td><td></td><td></td><td></td><td></td><td></td><td></td><td></td><td></td><td></td></tr>
<tr><td>Oligo rev</td><td>AAACCGGGTCTGCGACGCCTCCCT</td><td></td><td></td><td></td><td></td><td></td><td></td><td></td><td></td><td></td><td></td></tr>
<tr><td>Chromosome</td><td>start</td><td>end</td><td>strand</td><td>MM</td><td>target_seq</td><td>PAM</td><td>alignment</td><td>distance</td><td>position</td><td>gene name</td><td>gene id</td></tr>
<tr><td>GU305914.1</td><td>194399</td><td>194421</td><td>+</td><td>0</td><td>AGGGAGGCGTCGCAGACCCG</td><td>CGG</td><td>||||||||[||||||||||||]PAM</td><td>NA</td><td>Intergenic</td><td>NA</td><td>NA</td></tr>
<tr><td>T14765</td><td>GGGAGGCGTCGCAGACCCGCGGG</td><td>1000</td><td></td><td></td><td></td><td></td><td></td><td></td><td></td><td></td><td></td></tr>
<tr><td>Oligo fwd</td><td>CACCGGGAGGCGTCGCAGACCCGC</td><td></td><td></td><td></td><td></td><td></td><td></td><td></td><td></td><td></td><td></td></tr>
<tr><td>Oligo rev</td><td>AAACGCGGGTCTGCGACGCCTCCC</td><td></td><td></td><td></td><td></td><td></td><td></td><td></td><td></td><td></td><td></td></tr>
<tr><td>Chromosome</td><td>start</td><td>end</td><td>strand</td><td>MM</td><td>target_seq</td><td>PAM</td><td>alignment</td><td>distance</td><td>position</td><td>gene name</td><td>gene id</td></tr>
<tr><td>GU305914.1</td><td>194400</td><td>194422</td><td>+</td><td>0</td><td>GGGAGGCGTCGCAGACCCGC</td><td>GGG</td><td>||||||||[||||||||||||]PAM</td><td>NA</td><td>Intergenic</td><td>NA</td><td>NA</td></tr>
<tr><td>T14766</td><td>CGTCGCAGACCCGCGGGTATCGG</td><td>1000</td><td></td><td></td><td></td><td></td><td></td><td></td><td></td><td></td><td></td></tr>
<tr><td>Oligo fwd</td><td>CACCCGTCGCAGACCCGCGGGTAT</td><td></td><td></td><td></td><td></td><td></td><td></td><td></td><td></td><td></td><td></td></tr>
<tr><td>Oligo rev</td><td>AAACATACCCGCGGGTCTGCGACG</td><td></td><td></td><td></td><td></td><td></td><td></td><td></td><td></td><td></td><td></td></tr>
<tr><td>Chromosome</td><td>start</td><td>end</td><td>strand</td><td>MM</td><td>target_seq</td><td>PAM</td><td>alignment</td><td>distance</td><td>position</td><td>gene name</td><td>gene id</td></tr>
<tr><td>GU305914.1</td><td>194406</td><td>194428</td><td>+</td><td>0</td><td>CGTCGCAGACCCGCGGGTAT</td><td>CGG</td><td>||||||||[||||||||||||]PAM</td><td>NA</td><td>Intergenic</td><td>NA</td><td>NA</td></tr>
<tr><td>T14767</td><td>CAGACCCGCGGGTATCGGTACGG</td><td>1000</td><td></td><td></td><td></td><td></td><td></td><td></td><td></td><td></td><td></td></tr>
<tr><td>Oligo fwd</td><td>CACCCAGACCCGCGGGTATCGGTA</td><td></td><td></td><td></td><td></td><td></td><td></td><td></td><td></td><td></td><td></td></tr>
<tr><td>Oligo rev</td><td>AAACTACCGATACCCGCGGGTCTG</td><td></td><td></td><td></td><td></td><td></td><td></td><td></td><td></td><td></td><td></td></tr>
<tr><td>Chromosome</td><td>start</td><td>end</td><td>strand</td><td>MM</td><td>target_seq</td><td>PAM</td><td>alignment</td><td>distance</td><td>position</td><td>gene name</td><td>gene id</td></tr>
<tr><td>GU305914.1</td><td>194411</td><td>194433</td><td>+</td><td>0</td><td>CAGACCCGCGGGTATCGGTA</td><td>CGG</td><td>||||||||[||||||||||||]PAM</td><td>NA</td><td>Intergenic</td><td>NA</td><td>NA</td></tr>
<tr><td>T14768</td><td>ACGGCCTCGTCTGTCCCGACAGG</td><td>1000</td><td></td><td></td><td></td><td></td><td></td><td></td><td></td><td></td><td></td></tr>
<tr><td>Oligo fwd</td><td>CACCACGGCCTCGTCTGTCCCGAC</td><td></td><td></td><td></td><td></td><td></td><td></td><td></td><td></td><td></td><td></td></tr>
<tr><td>Oligo rev</td><td>AAACGTCGGGACAGACGAGGCCGT</td><td></td><td></td><td></td><td></td><td></td><td></td><td></td><td></td><td></td><td></td></tr>
<tr><td>Chromosome</td><td>start</td><td>end</td><td>strand</td><td>MM</td><td>target_seq</td><td>PAM</td><td>alignment</td><td>distance</td><td>position</td><td>gene name</td><td>gene id</td></tr>
<tr><td>GU305914.1</td><td>194430</td><td>194452</td><td>+</td><td>0</td><td>ACGGCCTCGTCTGTCCCGAC</td><td>AGG</td><td>||||||||[||||||||||||]PAM</td><td>NA</td><td>Intergenic</td><td>NA</td><td>NA</td></tr>
<tr><td>T14769</td><td>CGGCCTCGTCTGTCCCGACAGGG</td><td>1000</td><td></td><td></td><td></td><td></td><td></td><td></td><td></td><td></td><td></td></tr>
<tr><td>Oligo fwd</td><td>CACCCGGCCTCGTCTGTCCCGACA</td><td></td><td></td><td></td><td></td><td></td><td></td><td></td><td></td><td></td><td></td></tr>
<tr><td>Oligo rev</td><td>AAACTGTCGGGACAGACGAGGCCG</td><td></td><td></td><td></td><td></td><td></td><td></td><td></td><td></td><td></td><td></td></tr>
<tr><td>Chromosome</td><td>start</td><td>end</td><td>strand</td><td>MM</td><td>target_seq</td><td>PAM</td><td>alignment</td><td>distance</td><td>position</td><td>gene name</td><td>gene id</td></tr>
<tr><td>GU305914.1</td><td>194431</td><td>194453</td><td>+</td><td>0</td><td>CGGCCTCGTCTGTCCCGACA</td><td>GGG</td><td>||||||||[||||||||||||]PAM</td><td>NA</td><td>Intergenic</td><td>NA</td><td>NA</td></tr>
<tr><td>T14770</td><td>CTGTCCCGACAGGGATTCGAAGG</td><td>1000</td><td></td><td></td><td></td><td></td><td></td><td></td><td></td><td></td><td></td></tr>
<tr><td>Oligo fwd</td><td>CACCCTGTCCCGACAGGGATTCGA</td><td></td><td></td><td></td><td></td><td></td><td></td><td></td><td></td><td></td><td></td></tr>
<tr><td>Oligo rev</td><td>AAACTCGAATCCCTGTCGGGACAG</td><td></td><td></td><td></td><td></td><td></td><td></td><td></td><td></td><td></td><td></td></tr>
<tr><td>Chromosome</td><td>start</td><td>end</td><td>strand</td><td>MM</td><td>target_seq</td><td>PAM</td><td>alignment</td><td>distance</td><td>position</td><td>gene name</td><td>gene id</td></tr>
<tr><td>GU305914.1</td><td>194440</td><td>194462</td><td>+</td><td>0</td><td>CTGTCCCGACAGGGATTCGA</td><td>AGG</td><td>||||||||[||||||||||||]PAM</td><td>NA</td><td>Intergenic</td><td>NA</td><td>NA</td></tr>
<tr><td>T14771</td><td>CAGGGATTCGAAGGCTTCCTCGG</td><td>1000</td><td></td><td></td><td></td><td></td><td></td><td></td><td></td><td></td><td></td></tr>
<tr><td>Oligo fwd</td><td>CACCCAGGGATTCGAAGGCTTCCT</td><td></td><td></td><td></td><td></td><td></td><td></td><td></td><td></td><td></td><td></td></tr>
<tr><td>Oligo rev</td><td>AAACAGGAAGCCTTCGAATCCCTG</td><td></td><td></td><td></td><td></td><td></td><td></td><td></td><td></td><td></td><td></td></tr>
<tr><td>Chromosome</td><td>start</td><td>end</td><td>strand</td><td>MM</td><td>target_seq</td><td>PAM</td><td>alignment</td><td>distance</td><td>position</td><td>gene name</td><td>gene id</td></tr>
<tr><td>GU305914.1</td><td>194449</td><td>194471</td><td>+</td><td>0</td><td>CAGGGATTCGAAGGCTTCCT</td><td>CGG</td><td>||||||||[||||||||||||]PAM</td><td>NA</td><td>Intergenic</td><td>NA</td><td>NA</td></tr>
<tr><td>T14772</td><td>AGGGATTCGAAGGCTTCCTCGGG</td><td>1000</td><td></td><td></td><td></td><td></td><td></td><td></td><td></td><td></td><td></td></tr>
<tr><td>Oligo fwd</td><td>CACCAGGGATTCGAAGGCTTCCTC</td><td></td><td></td><td></td><td></td><td></td><td></td><td></td><td></td><td></td><td></td></tr>
<tr><td>Oligo rev</td><td>AAACGAGGAAGCCTTCGAATCCCT</td><td></td><td></td><td></td><td></td><td></td><td></td><td></td><td></td><td></td><td></td></tr>
<tr><td>Chromosome</td><td>start</td><td>end</td><td>strand</td><td>MM</td><td>target_seq</td><td>PAM</td><td>alignment</td><td>distance</td><td>position</td><td>gene name</td><td>gene id</td></tr>
<tr><td>GU305914.1</td><td>194450</td><td>194472</td><td>+</td><td>0</td><td>AGGGATTCGAAGGCTTCCTC</td><td>GGG</td><td>||||||||[||||||||||||]PAM</td><td>NA</td><td>Intergenic</td><td>NA</td><td>NA</td></tr>
<tr><td>T14773</td><td>AGGCTTCCTCGGGATCCCCCCGG</td><td>1000</td><td></td><td></td><td></td><td></td><td></td><td></td><td></td><td></td><td></td></tr>
<tr><td>Oligo fwd</td><td>CACCAGGCTTCCTCGGGATCCCCC</td><td></td><td></td><td></td><td></td><td></td><td></td><td></td><td></td><td></td><td></td></tr>
</table>

<table>
<tr><td>Oligo rev</td><td>AAACGGGGGATCCCGAGGAAGCCT</td><td></td><td></td><td></td><td></td><td></td><td></td><td></td><td></td><td></td><td></td></tr>
<tr><td>Chromosome</td><td>start</td><td>end</td><td>strand</td><td>MM</td><td>target_seq</td><td>PAM</td><td>alignment</td><td>distance</td><td>position</td><td>gene name</td><td>gene id</td></tr>
<tr><td>GU305914.1</td><td>194460</td><td>194482</td><td>+</td><td>0</td><td>AGGCTTCCTCGGGATCCCCC</td><td>CGG</td><td>||||||||[||||||||||||]PAM</td><td>NA</td><td>Intergenic</td><td>NA</td><td>NA</td></tr>
<tr><td>T14774</td><td>GGCTTCCTCGGGATCCCCCGGG</td><td>1000</td><td></td><td></td><td></td><td></td><td></td><td></td><td></td><td></td><td></td></tr>
<tr><td>Oligo fwd</td><td>CACCGGCTTCCTCGGGATCCCCC</td><td></td><td></td><td></td><td></td><td></td><td></td><td></td><td></td><td></td><td></td></tr>
<tr><td>Oligo rev</td><td>AAACGGGGGGATCCCGAGGAAGCC</td><td></td><td></td><td></td><td></td><td></td><td></td><td></td><td></td><td></td><td></td></tr>
<tr><td>Chromosome</td><td>start</td><td>end</td><td>strand</td><td>MM</td><td>target_seq</td><td>PAM</td><td>alignment</td><td>distance</td><td>position</td><td>gene name</td><td>gene id</td></tr>
<tr><td>GU305914.1</td><td>194461</td><td>194483</td><td>+</td><td>0</td><td>GGCTTCCTCGGGATCCCCC</td><td>GGG</td><td>||||||||[||||||||||||]PAM</td><td>NA</td><td>Intergenic</td><td>NA</td><td>NA</td></tr>
<tr><td>T14775</td><td>GCTTCCTCGGGATCCCCCGGGG</td><td>1000</td><td></td><td></td><td></td><td></td><td></td><td></td><td></td><td></td><td></td></tr>
<tr><td>Oligo fwd</td><td>CACCGCTTCCTCGGGATCCCCCG</td><td></td><td></td><td></td><td></td><td></td><td></td><td></td><td></td><td></td><td></td></tr>
<tr><td>Oligo rev</td><td>AAACCGGGGGGATCCCGAGGAAGC</td><td></td><td></td><td></td><td></td><td></td><td></td><td></td><td></td><td></td><td></td></tr>
<tr><td>Chromosome</td><td>start</td><td>end</td><td>strand</td><td>MM</td><td>target_seq</td><td>PAM</td><td>alignment</td><td>distance</td><td>position</td><td>gene name</td><td>gene id</td></tr>
<tr><td>GU305914.1</td><td>194462</td><td>194484</td><td>+</td><td>0</td><td>GCTTCCTCGGGATCCCCCG</td><td>GGG</td><td>||||||||[||||||||||||]PAM</td><td>NA</td><td>Intergenic</td><td>NA</td><td>NA</td></tr>
<tr><td>T14776</td><td>CTTCCTCGGGATCCCCCGGGGG</td><td>1000</td><td></td><td></td><td></td><td></td><td></td><td></td><td></td><td></td><td></td></tr>
<tr><td>Oligo fwd</td><td>CACCCTTCCTCGGGATCCCCCGG</td><td></td><td></td><td></td><td></td><td></td><td></td><td></td><td></td><td></td><td></td></tr>
<tr><td>Oligo rev</td><td>AAACCCGGGGGGATCCCGAGGAAG</td><td></td><td></td><td></td><td></td><td></td><td></td><td></td><td></td><td></td><td></td></tr>
<tr><td>Chromosome</td><td>start</td><td>end</td><td>strand</td><td>MM</td><td>target_seq</td><td>PAM</td><td>alignment</td><td>distance</td><td>position</td><td>gene name</td><td>gene id</td></tr>
<tr><td>GU305914.1</td><td>194463</td><td>194485</td><td>+</td><td>0</td><td>CTTCCTCGGGATCCCCCGG</td><td>GGG</td><td>||||||||[||||||||||||]PAM</td><td>NA</td><td>Intergenic</td><td>NA</td><td>NA</td></tr>
<tr><td>T14777</td><td>CCTCGGGATCCCCCGGGGGTGG</td><td>1000</td><td></td><td></td><td></td><td></td><td></td><td></td><td></td><td></td><td></td></tr>
<tr><td>Oligo fwd</td><td>CACCCCTCGGGATCCCCCGGGGG</td><td></td><td></td><td></td><td></td><td></td><td></td><td></td><td></td><td></td><td></td></tr>
<tr><td>Oligo rev</td><td>AAACCCCCCGGGGGATCCCGAGG</td><td></td><td></td><td></td><td></td><td></td><td></td><td></td><td></td><td></td><td></td></tr>
<tr><td>Chromosome</td><td>start</td><td>end</td><td>strand</td><td>MM</td><td>target_seq</td><td>PAM</td><td>alignment</td><td>distance</td><td>position</td><td>gene name</td><td>gene id</td></tr>
<tr><td>GU305914.1</td><td>194466</td><td>194488</td><td>+</td><td>0</td><td>CCTCGGGATCCCCCGGGGG</td><td>TGG</td><td>||||||||[||||||||||||]PAM</td><td>NA</td><td>Intergenic</td><td>NA</td><td>NA</td></tr>
<tr><td>T14778</td><td>GTAAGACTCCGCCGTCACGATGG</td><td>1000</td><td></td><td></td><td></td><td></td><td></td><td></td><td></td><td></td><td></td></tr>
<tr><td>Oligo fwd</td><td>CACCGTAAGACTCCGCCGTCACGA</td><td></td><td></td><td></td><td></td><td></td><td></td><td></td><td></td><td></td><td></td></tr>
<tr><td>Oligo rev</td><td>AAACTCGTGACGGCGGAGTCTTAC</td><td></td><td></td><td></td><td></td><td></td><td></td><td></td><td></td><td></td><td></td></tr>
<tr><td>Chromosome</td><td>start</td><td>end</td><td>strand</td><td>MM</td><td>target_seq</td><td>PAM</td><td>alignment</td><td>distance</td><td>position</td><td>gene name</td><td>gene id</td></tr>
<tr><td>GU305914.1</td><td>194488</td><td>194510</td><td>+</td><td>0</td><td>GTAAGACTCCGCCGTCACGA</td><td>TGG</td><td>||||||||[||||||||||||]PAM</td><td>NA</td><td>Intergenic</td><td>NA</td><td>NA</td></tr>
<tr><td>T14779</td><td>CTCCGCCGTCACGATGGAGACGG</td><td>1000</td><td></td><td></td><td></td><td></td><td></td><td></td><td></td><td></td><td></td></tr>
<tr><td>Oligo fwd</td><td>CACCCTCCGCCGTCACGATGGAGA</td><td></td><td></td><td></td><td></td><td></td><td></td><td></td><td></td><td></td><td></td></tr>
<tr><td>Oligo rev</td><td>AAACTCTCCATCGTGACGGCGGAG</td><td></td><td></td><td></td><td></td><td></td><td></td><td></td><td></td><td></td><td></td></tr>
<tr><td>Chromosome</td><td>start</td><td>end</td><td>strand</td><td>MM</td><td>target_seq</td><td>PAM</td><td>alignment</td><td>distance</td><td>position</td><td>gene name</td><td>gene id</td></tr>
<tr><td>GU305914.1</td><td>194494</td><td>194516</td><td>+</td><td>0</td><td>CTCCGCCGTCACGATGGAGA</td><td>CGG</td><td>||||||||[||||||||||||]PAM</td><td>NA</td><td>Intergenic</td><td>NA</td><td>NA</td></tr>
<tr><td>T14780</td><td>TCCGCCGTCACGATGGAGACGGG</td><td>1000</td><td></td><td></td><td></td><td></td><td></td><td></td><td></td><td></td><td></td></tr>
<tr><td>Oligo fwd</td><td>CACCTCCGCCGTCACGATGGAGAC</td><td></td><td></td><td></td><td></td><td></td><td></td><td></td><td></td><td></td><td></td></tr>
<tr><td>Oligo rev</td><td>AAACGTCTCCATCGTGACGGCGGA</td><td></td><td></td><td></td><td></td><td></td><td></td><td></td><td></td><td></td><td></td></tr>
<tr><td>Chromosome</td><td>start</td><td>end</td><td>strand</td><td>MM</td><td>target_seq</td><td>PAM</td><td>alignment</td><td>distance</td><td>position</td><td>gene name</td><td>gene id</td></tr>
<tr><td>GU305914.1</td><td>194495</td><td>194517</td><td>+</td><td>0</td><td>TCCGCCGTCACGATGGAGAC</td><td>GGG</td><td>||||||||[||||||||||||]PAM</td><td>NA</td><td>Intergenic</td><td>NA</td><td>NA</td></tr>
<tr><td>T14781</td><td>CCGCCGTCACGATGGAGACGGGG</td><td>1000</td><td></td><td></td><td></td><td></td><td></td><td></td><td></td><td></td><td></td></tr>
<tr><td>Oligo fwd</td><td>CACCCCGCCGTCACGATGGAGACG</td><td></td><td></td><td></td><td></td><td></td><td></td><td></td><td></td><td></td><td></td></tr>
<tr><td>Oligo rev</td><td>AAACCGTCTCCATCGTGACGGCGG</td><td></td><td></td><td></td><td></td><td></td><td></td><td></td><td></td><td></td><td></td></tr>
<tr><td>Chromosome</td><td>start</td><td>end</td><td>strand</td><td>MM</td><td>target_seq</td><td>PAM</td><td>alignment</td><td>distance</td><td>position</td><td>gene name</td><td>gene id</td></tr>
<tr><td>GU305914.1</td><td>194496</td><td>194518</td><td>+</td><td>0</td><td>CCGCCGTCACGATGGAGACG</td><td>GGG</td><td>||||||||[||||||||||||]PAM</td><td>NA</td><td>Intergenic</td><td>NA</td><td>NA</td></tr>
<tr><td>T14782</td><td>GATGGAGACGGGGATCCCTTTGG</td><td>1000</td><td></td><td></td><td></td><td></td><td></td><td></td><td></td><td></td><td></td></tr>
<tr><td>Oligo fwd</td><td>CACCGATGGAGACGGGGATCCCTT</td><td></td><td></td><td></td><td></td><td></td><td></td><td></td><td></td><td></td><td></td></tr>
<tr><td>Oligo rev</td><td>AAACAAGGGATCCCCGTCTCCATC</td><td></td><td></td><td></td><td></td><td></td><td></td><td></td><td></td><td></td><td></td></tr>
<tr><td>Chromosome</td><td>start</td><td>end</td><td>strand</td><td>MM</td><td>target_seq</td><td>PAM</td><td>alignment</td><td>distance</td><td>position</td><td>gene name</td><td>gene id</td></tr>
<tr><td>GU305914.1</td><td>194506</td><td>194528</td><td>+</td><td>0</td><td>GATGGAGACGGGGATCCCTT</td><td>TGG</td><td>||||||||[||||||||||||]PAM</td><td>NA</td><td>Intergenic</td><td>NA</td><td>NA</td></tr>
<tr><td>GU305914.1</td><td>114172</td><td>114194</td><td>+</td><td>3</td><td>GACGGAGACGCCGATCCCTT</td><td>CGG</td><td>||-|||||[||--||||||||]PAM</td><td>NA</td><td>Intergenic</td><td>NA</td><td>NA</td></tr>
<tr><td>T14783</td><td>GGAGACGGGGATCCCTTTGGCGG</td><td>1000</td><td></td><td></td><td></td><td></td><td></td><td></td><td></td><td></td><td></td></tr>
<tr><td>Oligo fwd</td><td>CACCGGAGACGGGGATCCCTTTGG</td><td></td><td></td><td></td><td></td><td></td><td></td><td></td><td></td><td></td><td></td></tr>
<tr><td>Oligo rev</td><td>AAACCCAAAGGGATCCCCGTCTCC</td><td></td><td></td><td></td><td></td><td></td><td></td><td></td><td></td><td></td><td></td></tr>
<tr><td>Chromosome</td><td>start</td><td>end</td><td>strand</td><td>MM</td><td>target_seq</td><td>PAM</td><td>alignment</td><td>distance</td><td>position</td><td>gene name</td><td>gene id</td></tr>
<tr><td>GU305914.1</td><td>194509</td><td>194531</td><td>+</td><td>0</td><td>GGAGACGGGGATCCCTTTGG</td><td>CGG</td><td>||||||||[||||||||||||]PAM</td><td>NA</td><td>Intergenic</td><td>NA</td><td>NA</td></tr>
<tr><td>T14784</td><td>GAGACGGGGATCCCTTTGGCGGG</td><td>1000</td><td></td><td></td><td></td><td></td><td></td><td></td><td></td><td></td><td></td></tr>
<tr><td>Oligo fwd</td><td>CACCGAGACGGGGATCCCTTTGGC</td><td></td><td></td><td></td><td></td><td></td><td></td><td></td><td></td><td></td><td></td></tr>
<tr><td>Oligo rev</td><td>AAACGCCAAAGGGATCCCCGTCTC</td><td></td><td></td><td></td><td></td><td></td><td></td><td></td><td></td><td></td><td></td></tr>
<tr><td>Chromosome</td><td>start</td><td>end</td><td>strand</td><td>MM</td><td>target_seq</td><td>PAM</td><td>alignment</td><td>distance</td><td>position</td><td>gene name</td><td>gene id</td></tr>
<tr><td>GU305914.1</td><td>194510</td><td>194532</td><td>+</td><td>0</td><td>GAGACGGGGATCCCTTTGGC</td><td>GGG</td><td>||||||||[||||||||||||]PAM</td><td>NA</td><td>Intergenic</td><td>NA</td><td>NA</td></tr>
<tr><td>T14785</td><td>CGGGGATCCCTTTGGCGGGATGG</td><td>1000</td><td></td><td></td><td></td><td></td><td></td><td></td><td></td><td></td><td></td></tr>
<tr><td>Oligo fwd</td><td>CACCCGGGGATCCCTTTGGCGGGA</td><td></td><td></td><td></td><td></td><td></td><td></td><td></td><td></td><td></td><td></td></tr>
<tr><td>Oligo rev</td><td>AAACTCCCGCCAAAGGGATCCCCG</td><td></td><td></td><td></td><td></td><td></td><td></td><td></td><td></td><td></td><td></td></tr>
<tr><td>Chromosome</td><td>start</td><td>end</td><td>strand</td><td>MM</td><td>target_seq</td><td>PAM</td><td>alignment</td><td>distance</td><td>position</td><td>gene name</td><td>gene id</td></tr>
<tr><td>GU305914.1</td><td>194514</td><td>194536</td><td>+</td><td>0</td><td>CGGGGATCCCTTTGGCGGGA</td><td>TGG</td><td>||||||||[||||||||||||]PAM</td><td>NA</td><td>Intergenic</td><td>NA</td><td>NA</td></tr>
<tr><td>T14786</td><td>GGGGATCCCTTTGGCGGGATGGG</td><td>1000</td><td></td><td></td><td></td><td></td><td></td><td></td><td></td><td></td><td></td></tr>
<tr><td>Oligo fwd</td><td>CACCGGGGATCCCTTTGGCGGGAT</td><td></td><td></td><td></td><td></td><td></td><td></td><td></td><td></td><td></td><td></td></tr>
<tr><td>Oligo rev</td><td>AAACATCCCGCCAAAGGGATCCCC</td><td></td><td></td><td></td><td></td><td></td><td></td><td></td><td></td><td></td><td></td></tr>
<tr><td>Chromosome</td><td>start</td><td>end</td><td>strand</td><td>MM</td><td>target_seq</td><td>PAM</td><td>alignment</td><td>distance</td><td>position</td><td>gene name</td><td>gene id</td></tr>
<tr><td>GU305914.1</td><td>194515</td><td>194537</td><td>+</td><td>0</td><td>GGGGATCCCTTTGGCGGGAT</td><td>GGG</td><td>||||||||[||||||||||||]PAM</td><td>NA</td><td>Intergenic</td><td>NA</td><td>NA</td></tr>
<tr><td>T14787</td><td>TCGCCTCGCTCACGTTCGACAGG</td><td>1000</td><td></td><td></td><td></td><td></td><td></td><td></td><td></td><td></td><td></td></tr>
<tr><td>Oligo fwd</td><td>CACCTCGCCTCGCTCACGTTCGAC</td><td></td><td></td><td></td><td></td><td></td><td></td><td></td><td></td><td></td><td></td></tr>
<tr><td>Oligo rev</td><td>AAACGTCGAACGTGAGCGAGGCGA</td><td></td><td></td><td></td><td></td><td></td><td></td><td></td><td></td><td></td><td></td></tr>
<tr><td>Chromosome</td><td>start</td><td>end</td><td>strand</td><td>MM</td><td>target_seq</td><td>PAM</td><td>alignment</td><td>distance</td><td>position</td><td>gene name</td><td>gene id</td></tr>
<tr><td>GU305914.1</td><td>194550</td><td>194572</td><td>+</td><td>0</td><td>TCGCCTCGCTCACGTTCGAC</td><td>AGG</td><td>||||||||[||||||||||||]PAM</td><td>NA</td><td>Intergenic</td><td>NA</td><td>NA</td></tr>
</table>

T14788 CGCCTCGCTCACGTTCGACAGGG 1000
Oligo fwd CACCCGCCTCGCTCACGTTCGACA
Oligo rev AAACTGTCGAACGTGAGCGAGGCG
Chromosome start end strand MM target_seq PAM alignment distance position gene name gene id
GU305914.1 194551 194573 + 0 CGCCTCGCTCACGTTCGACA GGG ||||||||[||||||||||||]PAM NA Intergenic NA NA

T14789 GCCGTCCTGCAACCGCGATCTGG 1000
Oligo fwd CACCGCCGTCCTGCAACCGCGATC
Oligo rev AAACGATCGCGGTTGCAGGACGGC
Chromosome start end strand MM target_seq PAM alignment distance position gene name gene id
GU305914.1 194578 194600 + 0 GCCGTCCTGCAACCGCGATC TGG ||||||||[||||||||||||]PAM NA Intergenic NA NA

T14790 ATCTGGCGCCGAGTCGCCAGCGG 1000
Oligo fwd CACCATCTGGCGCCGAGTCGCCAG
Oligo rev AAACCTGGCGACTCGGCGCCAGAT
Chromosome start end strand MM target_seq PAM alignment distance position gene name gene id
GU305914.1 194595 194617 + 0 ATCTGGCGCCGAGTCGCCAG CGG ||||||||[||||||||||||]PAM NA Intergenic NA NA

T14791 GCCAGCGGTACCAGTCTTCCAGG 1000
Oligo fwd CACCGCCAGCGGTACCAGTCTTCC
Oligo rev AAACGGAAGACTGGTACCGCTGGC
Chromosome start end strand MM target_seq PAM alignment distance position gene name gene id
GU305914.1 194610 194632 + 0 GCCAGCGGTACCAGTCTTCC AGG ||||||||[||||||||||||]PAM NA Intergenic NA NA

T14792 GGTCGCGACGCGCGATCTCCTGG 1000
Oligo fwd CACCGGTCGCGACGCGCGATCTCC
Oligo rev AAACGGAGATCGCGCGTCGCGACC
Chromosome start end strand MM target_seq PAM alignment distance position gene name gene id
GU305914.1 194631 194653 + 0 GGTCGCGACGCGCGATCTCC TGG ||||||||[||||||||||||]PAM NA Intergenic NA NA

T14793 CTCCTGGCTCGCGAGCGACGTGG 1000
Oligo fwd CACCCTCCTGGCTCGCGAGCGACG
Oligo rev AAACCGTCGCTCGCGAGCCAGGAG
Chromosome start end strand MM target_seq PAM alignment distance position gene name gene id
GU305914.1 194647 194669 + 0 CTCCTGGCTCGCGAGCGACG TGG ||||||||[||||||||||||]PAM NA Intergenic NA NA

T14794 CTCGCGAGCGACGTGGCGATCGG 1000
Oligo fwd CACCCTCGCGAGCGACGTGGCGAT
Oligo rev AAACATCGCCACGTCGCTCGCGAG
Chromosome start end strand MM target_seq PAM alignment distance position gene name gene id
GU305914.1 194654 194676 + 0 CTCGCGAGCGACGTGGCGAT CGG ||||||||[||||||||||||]PAM NA Intergenic NA NA

T14795 GCGAGCGACGTGGCGATCGGCGG 1000
Oligo fwd CACCGCGAGCGACGTGGCGATCGG
Oligo rev AAACCCGATCGCCACGTCGCTCGC
Chromosome start end strand MM target_seq PAM alignment distance position gene name gene id
GU305914.1 194657 194679 + 0 GCGAGCGACGTGGCGATCGG CGG ||||||||[||||||||||||]PAM NA Intergenic NA NA

T14796 GCGACGTGGCGATCGGCGGCCGG 1000
Oligo fwd CACCGCGACGTGGCGATCGGCGGC
Oligo rev AAACGCCGCCGATCGCCACGTCGC
Chromosome start end strand MM target_seq PAM alignment distance position gene name gene id
GU305914.1 194661 194683 + 0 GCGACGTGGCGATCGGCGGC CGG ||||||||[||||||||||||]PAM NA Intergenic NA NA
GU305914.1 114149 114171 + 4 ACGACGCGGCGCTCGGGGGC TGG -|||||-|[|||-||||-|||]PAM NA Intergenic NA NA

T14797 GCGGCCGGTCTTCGTCCCTGTGG 1000
Oligo fwd CACCGCGGCCGGTCTTCGTCCCTG
Oligo rev AAACCAGGGACGAAGACCGGCCGC
Chromosome start end strand MM target_seq PAM alignment distance position gene name gene id
GU305914.1 194676 194698 + 0 GCGGCCGGTCTTCGTCCCTG TGG ||||||||[||||||||||||]PAM NA Intergenic NA NA

T14798 CGGCCGGTCTTCGTCCCTGTGGG 1000
Oligo fwd CACCCGGCCGGTCTTCGTCCCTGT
Oligo rev AAACACAGGGACGAAGACCGGCCG
Chromosome start end strand MM target_seq PAM alignment distance position gene name gene id
GU305914.1 194677 194699 + 0 CGGCCGGTCTTCGTCCCTGT GGG ||||||||[||||||||||||]PAM NA Intergenic NA NA

T14799 GGCCGGTCTTCGTCCCTGTGGGG 1000
Oligo fwd CACCGGCCGGTCTTCGTCCCTGTG
Oligo rev AAACCACAGGGACGAAGACCGGCC
Chromosome start end strand MM target_seq PAM alignment distance position gene name gene id
GU305914.1 194678 194700 + 0 GGCCGGTCTTCGTCCCTGTG GGG ||||||||[||||||||||||]PAM NA Intergenic NA NA

T14800 TCGTCCCTGTGGGGACAGAGCGG 1000
Oligo fwd CACCTCGTCCCTGTGGGGACAGAG
Oligo rev AAACCTCTGTCCCCACAGGGACGA
Chromosome start end strand MM target_seq PAM alignment distance position gene name gene id
GU305914.1 194687 194709 + 0 TCGTCCCTGTGGGGACAGAG CGG ||||||||[||||||||||||]PAM NA Intergenic NA NA

T14801 TCCCTGTGGGGACAGAGCGGCGG 1000
Oligo fwd CACCTCCCTGTGGGGACAGAGCGG
Oligo rev AAACCCGCTCTGTCCCCACAGGGA
Chromosome start end strand MM target_seq PAM alignment distance position gene name gene id
GU305914.1 194690 194712 + 0 TCCCTGTGGGGACAGAGCGG CGG ||||||||[||||||||||||]PAM NA Intergenic NA NA

T14802 CCCTGTGGGGACAGAGCGGCGGG 1000
Oligo fwd CACCCCCTGTGGGGACAGAGCGGC
Oligo rev AAACGCCGCTCTGTCCCCACAGGG
Chromosome start end strand MM target_seq PAM alignment distance position gene name gene id

GU305914.1 194691 194713 + 0 CCCTGTGGGGACAGAGCGGC GGG ||||||||[||||||||||||]PAM NA Intergenic NA NA

T14803 AGAGCGGCGGGCGTTCCAGACGG 1000
Oligo fwd CACCAGAGCGGCGGGCGTTCCAGA
Oligo rev AAACTCTGGAACGCCCGCCGCTCT
Chromosome start end strand MM target_seq PAM alignment distance position gene name gene id
GU305914.1 194703 194725 + 0 AGAGCGGCGGGCGTTCCAGA CGG ||||||||[||||||||||||]PAM NA Intergenic NA NA

T14804 GTTCCAGACGGAGTTCTCCCCGG 1000
Oligo fwd CACCGTTCCAGACGGAGTTCTCCC
Oligo rev AAACGGGAGAACTCCGTCTGGAAC
Chromosome start end strand MM target_seq PAM alignment distance position gene name gene id
GU305914.1 194715 194737 + 0 GTTCCAGACGGAGTTCTCCC CGG ||||||||[||||||||||||]PAM NA Intergenic NA NA

T14805 TTCCAGACGGAGTTCTCCCCGGG 1000
Oligo fwd CACCTTCCAGACGGAGTTCTCCCC
Oligo rev AAACGGGGAGAACTCCGTCTGGAA
Chromosome start end strand MM target_seq PAM alignment distance position gene name gene id
GU305914.1 194716 194738 + 0 TTCCAGACGGAGTTCTCCCC GGG ||||||||[||||||||||||]PAM NA Intergenic NA NA

T14806 GGAGTTCTCCCCGGGCGCCTCGG 1000
Oligo fwd CACCGGAGTTCTCCCCGGGCGCCT
Oligo rev AAACAGGCGCCCGGGGAGAACTCC
Chromosome start end strand MM target_seq PAM alignment distance position gene name gene id
GU305914.1 194724 194746 + 0 GGAGTTCTCCCCGGGCGCCT CGG ||||||||[||||||||||||]PAM NA Intergenic NA NA

T14807 CCCGGGCGCCTCGGTCGAAGCGG 1000
Oligo fwd CACCCCCGGGCGCCTCGGTCGAAG
Oligo rev AAACCTTCGACCGAGGCGCCCGGG
Chromosome start end strand MM target_seq PAM alignment distance position gene name gene id
GU305914.1 194733 194755 + 0 CCCGGGCGCCTCGGTCGAAG CGG ||||||||[||||||||||||]PAM NA Intergenic NA NA

T14808 GGCGCCTCGGTCGAAGCGGTTGG 1000
Oligo fwd CACCGGCGCCTCGGTCGAAGCGGT
Oligo rev AAACACCGCTTCGACCGAGGCGCC
Chromosome start end strand MM target_seq PAM alignment distance position gene name gene id
GU305914.1 194737 194759 + 0 GGCGCCTCGGTCGAAGCGGT TGG ||||||||[||||||||||||]PAM NA Intergenic NA NA

T14809 GCGCCTCGGTCGAAGCGGTTGGG 1000
Oligo fwd CACCGCGCCTCGGTCGAAGCGGTT
Oligo rev AAACAACCGCTTCGACCGAGGCGC
Chromosome start end strand MM target_seq PAM alignment distance position gene name gene id
GU305914.1 194738 194760 + 0 GCGCCTCGGTCGAAGCGGTT GGG ||||||||[||||||||||||]PAM NA Intergenic NA NA

T14810 TCGGTCGAAGCGGTTGGGCACGG 1000
Oligo fwd CACCTCGGTCGAAGCGGTTGGGCA
Oligo rev AAACTGCCCAACCGCTTCGACCGA
Chromosome start end strand MM target_seq PAM alignment distance position gene name gene id
GU305914.1 194743 194765 + 0 TCGGTCGAAGCGGTTGGGCA CGG ||||||||[||||||||||||]PAM NA Intergenic NA NA

T14811 GTCGAAGCGGTTGGGCACGGCGG 1000
Oligo fwd CACCGTCGAAGCGGTTGGGCACGG
Oligo rev AAACCCGTGCCCAACCGCTTCGAC
Chromosome start end strand MM target_seq PAM alignment distance position gene name gene id
GU305914.1 194746 194768 + 0 GTCGAAGCGGTTGGGCACGG CGG ||||||||[||||||||||||]PAM NA Intergenic NA NA

T14812 GAAGAAGACCGAGATGTCCTCGG 1000
Oligo fwd CACCGAAGAAGACCGAGATGTCCT
Oligo rev AAACAGGACATCTCGGTCTTCTTC
Chromosome start end strand MM target_seq PAM alignment distance position gene name gene id
GU305914.1 194794 194816 + 0 GAAGAAGACCGAGATGTCCT CGG ||||||||[||||||||||||]PAM NA Intergenic NA NA

T14813 TGAAGACGTCGTTACTCAGCAGG 1000
Oligo fwd CACCTGAAGACGTCGTTACTCAGC
Oligo rev AAACGCTGAGTAACGACGTCTTCA
Chromosome start end strand MM target_seq PAM alignment distance position gene name gene id
GU305914.1 194823 194845 + 0 TGAAGACGTCGTTACTCAGC AGG ||||||||[||||||||||||]PAM NA Intergenic NA NA

T14814 AGAAACCGTACACCGTTCCGCGG 1000
Oligo fwd CACCAGAAACCGTACACCGTTCCG
Oligo rev AAACCGGAACGGTGTACGGTTTCT
Chromosome start end strand MM target_seq PAM alignment distance position gene name gene id
GU305914.1 194847 194869 + 0 AGAAACCGTACACCGTTCCG CGG ||||||||[||||||||||||]PAM NA Intergenic NA NA

T14815 GAAACCGTACACCGTTCCGCGGG 1000
Oligo fwd CACCGAAACCGTACACCGTTCCGC
Oligo rev AAACGCGGAACGGTGTACGGTTTC
Chromosome start end strand MM target_seq PAM alignment distance position gene name gene id
GU305914.1 194848 194870 + 0 GAAACCGTACACCGTTCCGC GGG ||||||||[||||||||||||]PAM NA Intergenic NA NA

T14816 GCGGGCTCCGACCAGCACGACGG 1000
Oligo fwd CACCGCGGGCTCCGACCAGCACGA
Oligo rev AAACTCGTGCTGGTCGGAGCCCGC
Chromosome start end strand MM target_seq PAM alignment distance position gene name gene id
GU305914.1 194866 194888 + 0 GCGGGCTCCGACCAGCACGA CGG ||||||||[||||||||||||]PAM NA Intergenic NA NA

T14817 GGCTCCGACCAGCACGACGGTGG 1000
Oligo fwd CACCGGCTCCGACCAGCACGACGG
Oligo rev AAACCCGTCGTGCTGGTCGGAGCC

| | | | | | | | | | | | |
|---|---|---|---|---|---|---|---|---|---|---|---|
| Chromosome | start | end | strand | MM | target_seq | PAM | alignment | distance | position | gene name | gene id |
| GU305914.1 | 194869 | 194891 | + | 0 | GGCTCCGACCAGCACGACGG | TGG | \|\|\|\|\|\|\|\|[\|\|\|\|\|\|\|\|\|\|\|\|]PAM | NA | Intergenic | NA | NA |
| T14818 | CGAAGACGAAGCCGTCGTCCCGG | 1000 | | | | | | | | | |
| Oligo fwd | CACCCGAAGACGAAGCCGTCGTCC | | | | | | | | | | |
| Oligo rev | AAACGGACGACGGCTTCGTCTTCG | | | | | | | | | | |
| Chromosome | start | end | strand | MM | target_seq | PAM | alignment | distance | position | gene name | gene id |
| GU305914.1 | 194898 | 194920 | + | 0 | CGAAGACGAAGCCGTCGTCC | CGG | \|\|\|\|\|\|\|\|[\|\|\|\|\|\|\|\|\|\|\|\|]PAM | NA | Intergenic | NA | NA |
| T14819 | GTCGTCCCGGTACTCTCCCACGG | 1000 | | | | | | | | | |
| Oligo fwd | CACCGTCGTCCCGGTACTCTCCCA | | | | | | | | | | |
| Oligo rev | AAACTGGGAGAGTACCGGGACGAC | | | | | | | | | | |
| Chromosome | start | end | strand | MM | target_seq | PAM | alignment | distance | position | gene name | gene id |
| GU305914.1 | 194911 | 194933 | + | 0 | GTCGTCCCGGTACTCTCCCA | CGG | \|\|\|\|\|\|\|\|[\|\|\|\|\|\|\|\|\|\|\|\|]PAM | NA | Intergenic | NA | NA |
| T14820 | CTCTCCCACGGCGCCGATGACGG | 1000 | | | | | | | | | |
| Oligo fwd | CACCCTCTCCCACGGCGCCGATGA | | | | | | | | | | |
| Oligo rev | AAACTCATCGGCGCCGTGGGAGAG | | | | | | | | | | |
| Chromosome | start | end | strand | MM | target_seq | PAM | alignment | distance | position | gene name | gene id |
| GU305914.1 | 194923 | 194945 | + | 0 | CTCTCCCACGGCGCCGATGA | CGG | \|\|\|\|\|\|\|\|[\|\|\|\|\|\|\|\|\|\|\|\|]PAM | NA | Intergenic | NA | NA |
| T14821 | GCCGATGACGGTCCACGCCTCGG | 1000 | | | | | | | | | |
| Oligo fwd | CACCGCCGATGACGGTCCACGCCT | | | | | | | | | | |
| Oligo rev | AAACAGGCGTGGACCGTCATCGGC | | | | | | | | | | |
| Chromosome | start | end | strand | MM | target_seq | PAM | alignment | distance | position | gene name | gene id |
| GU305914.1 | 194935 | 194957 | + | 0 | GCCGATGACGGTCCACGCCT | CGG | \|\|\|\|\|\|\|\|[\|\|\|\|\|\|\|\|\|\|\|\|]PAM | NA | Intergenic | NA | NA |
| T14822 | CCACGCCTCGGCCAGCCGAAAGG | 1000 | | | | | | | | | |
| Oligo fwd | CACCCCACGCCTCGGCCAGCCGAA | | | | | | | | | | |
| Oligo rev | AAACTTCGGCTGGCCGAGGCGTGG | | | | | | | | | | |
| Chromosome | start | end | strand | MM | target_seq | PAM | alignment | distance | position | gene name | gene id |
| GU305914.1 | 194947 | 194969 | + | 0 | CCACGCCTCGGCCAGCCGAA | AGG | \|\|\|\|\|\|\|\|[\|\|\|\|\|\|\|\|\|\|\|\|]PAM | NA | Intergenic | NA | NA |
| T14823 | CGTGTCGTCCATCCGCGAGAAGG | 1000 | | | | | | | | | |
| Oligo fwd | CACCCGTGTCGTCCATCCGCGAGA | | | | | | | | | | |
| Oligo rev | AAACTCTCGCGGATGGACGACACG | | | | | | | | | | |
| Chromosome | start | end | strand | MM | target_seq | PAM | alignment | distance | position | gene name | gene id |
| GU305914.1 | 194998 | 195020 | + | 0 | CGTGTCGTCCATCCGCGAGA | AGG | \|\|\|\|\|\|\|\|[\|\|\|\|\|\|\|\|\|\|\|\|]PAM | NA | Intergenic | NA | NA |
| GU305914.1 | 199254 | 199276 | + | 4 | CGTGGCCACCAGCCGCGAGA | AGG | \|\|\|\|-\|--[\|\|\|-\|\|\|\|\|\|\|\|]PAM | NA | Intergenic | NA | NA |
| T14824 | GTGTCGTCCATCCGCGAGAAGGG | 1000 | | | | | | | | | |
| Oligo fwd | CACCGTGTCGTCCATCCGCGAGAA | | | | | | | | | | |
| Oligo rev | AAACTTCTCGCGGATGGACGACAC | | | | | | | | | | |
| Chromosome | start | end | strand | MM | target_seq | PAM | alignment | distance | position | gene name | gene id |
| GU305914.1 | 194999 | 195021 | + | 0 | GTGTCGTCCATCCGCGAGAA | GGG | \|\|\|\|\|\|\|\|[\|\|\|\|\|\|\|\|\|\|\|\|]PAM | NA | Intergenic | NA | NA |
| T14825 | TCGTCCATCCGCGAGAAGGGCGG | 1000 | | | | | | | | | |
| Oligo fwd | CACCTCGTCCATCCGCGAGAAGGG | | | | | | | | | | |
| Oligo rev | AAACCCCTTCTCGCGGATGGACGA | | | | | | | | | | |
| Chromosome | start | end | strand | MM | target_seq | PAM | alignment | distance | position | gene name | gene id |
| GU305914.1 | 195002 | 195024 | + | 0 | TCGTCCATCCGCGAGAAGGG | CGG | \|\|\|\|\|\|\|\|[\|\|\|\|\|\|\|\|\|\|\|\|]PAM | NA | Intergenic | NA | NA |
| T14826 | AGAAGGGCGGCTTCCTCTTCAGG | 1000 | | | | | | | | | |
| Oligo fwd | CACCAGAAGGGCGGCTTCCTCTTC | | | | | | | | | | |
| Oligo rev | AAACGAAGAGGAAGCCGCCCTTCT | | | | | | | | | | |
| Chromosome | start | end | strand | MM | target_seq | PAM | alignment | distance | position | gene name | gene id |
| GU305914.1 | 195015 | 195037 | + | 0 | AGAAGGGCGGCTTCCTCTTC | AGG | \|\|\|\|\|\|\|\|[\|\|\|\|\|\|\|\|\|\|\|\|]PAM | NA | Intergenic | NA | NA |
| GU305914.1 | 97356 | 97378 | + | 4 | GGAGGGGCGGGTTCCACTTC | GGG | -\|\|-\|\|\|\|[\|\|-\|\|\|\|-\|\|\|\|]PAM | NA | Intergenic | NA | NA |
| T14827 | GTCTCGCGCCGTCCCGAGACCGG | 1000 | | | | | | | | | |
| Oligo fwd | CACCGTCTCGCGCCGTCCCGAGAC | | | | | | | | | | |
| Oligo rev | AAACGTCTCGGGACGGCGCGAGAC | | | | | | | | | | |
| Chromosome | start | end | strand | MM | target_seq | PAM | alignment | distance | position | gene name | gene id |
| GU305914.1 | 195062 | 195084 | + | 0 | GTCTCGCGCCGTCCCGAGAC | CGG | \|\|\|\|\|\|\|\|[\|\|\|\|\|\|\|\|\|\|\|\|]PAM | NA | Intergenic | NA | NA |
| T14828 | TCTCGCGCCGTCCCGAGACCGGG | 1000 | | | | | | | | | |
| Oligo fwd | CACCTCTCGCGCCGTCCCGAGACC | | | | | | | | | | |
| Oligo rev | AAACGGTCTCGGGACGGCGCGAGA | | | | | | | | | | |
| Chromosome | start | end | strand | MM | target_seq | PAM | alignment | distance | position | gene name | gene id |
| GU305914.1 | 195063 | 195085 | + | 0 | TCTCGCGCCGTCCCGAGACC | GGG | \|\|\|\|\|\|\|\|[\|\|\|\|\|\|\|\|\|\|\|\|]PAM | NA | Intergenic | NA | NA |
| T14829 | GTCCCGAGACCGGGTCGCGCAGG | 1000 | | | | | | | | | |
| Oligo fwd | CACCGTCCCGAGACCGGGTCGCGC | | | | | | | | | | |
| Oligo rev | AAACGCGCGACCCGGTCTCGGGAC | | | | | | | | | | |
| Chromosome | start | end | strand | MM | target_seq | PAM | alignment | distance | position | gene name | gene id |
| GU305914.1 | 195072 | 195094 | + | 0 | GTCCCGAGACCGGGTCGCGC | AGG | \|\|\|\|\|\|\|\|[\|\|\|\|\|\|\|\|\|\|\|\|]PAM | NA | Intergenic | NA | NA |
| T14830 | TCCCGAGACCGGGTCGCGCAGGG | 1000 | | | | | | | | | |
| Oligo fwd | CACCTCCCGAGACCGGGTCGCGCA | | | | | | | | | | |
| Oligo rev | AAACTGCGCGACCCGGTCTCGGGA | | | | | | | | | | |
| Chromosome | start | end | strand | MM | target_seq | PAM | alignment | distance | position | gene name | gene id |
| GU305914.1 | 195073 | 195095 | + | 0 | TCCCGAGACCGGGTCGCGCA | GGG | \|\|\|\|\|\|\|\|[\|\|\|\|\|\|\|\|\|\|\|\|]PAM | NA | Intergenic | NA | NA |
| T14831 | CGTCGAGCCCCTGAACGCCGTGG | 1000 | | | | | | | | | |
| Oligo fwd | CACCCGTCGAGCCCCTGAACGCCG | | | | | | | | | | |
| Oligo rev | AAACCGGCGTTCAGGGGCTCGACG | | | | | | | | | | |
| Chromosome | start | end | strand | MM | target_seq | PAM | alignment | distance | position | gene name | gene id |
| GU305914.1 | 195100 | 195122 | + | 0 | CGTCGAGCCCCTGAACGCCG | TGG | \|\|\|\|\|\|\|\|[\|\|\|\|\|\|\|\|\|\|\|\|]PAM | NA | Intergenic | NA | NA |

| | | | | | | | | | | | |
|---|---|---|---|---|---|---|---|---|---|---|---|
| T14832 | CGAGCCCCTGAACGCCGTGGCGG | 1000 | | | | | | | | | |
| Oligo fwd | CACCCGAGCCCCTGAACGCCGTGG | | | | | | | | | | |
| Oligo rev | AAACCCACGGCGTTCAGGGGCTCG | | | | | | | | | | |
| Chromosome | start | end | strand | MM | target_seq | PAM | alignment | distance | position | gene name | gene id |
| GU305914.1 | 195103 | 195125 | + | 0 | CGAGCCCCTGAACGCCGTGG | CGG | \|\|\|\|\|\|\|\|[\|\|\|\|\|\|\|\|\|\|\|\|]PAM | NA | Intergenic | NA | NA |
| T14833 | GCCCCTGAACGCCGTGGCGGTGG | 1000 | | | | | | | | | |
| Oligo fwd | CACCGCCCCTGAACGCCGTGGCGG | | | | | | | | | | |
| Oligo rev | AAACCCGCCACGGCGTTCAGGGGC | | | | | | | | | | |
| Chromosome | start | end | strand | MM | target_seq | PAM | alignment | distance | position | gene name | gene id |
| GU305914.1 | 195106 | 195128 | + | 0 | GCCCCTGAACGCCGTGGCGG | TGG | \|\|\|\|\|\|\|\|[\|\|\|\|\|\|\|\|\|\|\|\|]PAM | NA | Intergenic | NA | NA |
| T14834 | CGGTGGCCGCCACGTTCTCCAGG | 1000 | | | | | | | | | |
| Oligo fwd | CACCCGGTGGCCGCCACGTTCTCC | | | | | | | | | | |
| Oligo rev | AAACGGAGAACGTGGCGGCCACCG | | | | | | | | | | |
| Chromosome | start | end | strand | MM | target_seq | PAM | alignment | distance | position | gene name | gene id |
| GU305914.1 | 195123 | 195145 | + | 0 | CGGTGGCCGCCACGTTCTCC | AGG | \|\|\|\|\|\|\|\|[\|\|\|\|\|\|\|\|\|\|\|\|]PAM | NA | Intergenic | NA | NA |
| T14835 | CACGTTCTCCAGGCCGACCACGG | 1000 | | | | | | | | | |
| Oligo fwd | CACCCACGTTCTCCAGGCCGACCA | | | | | | | | | | |
| Oligo rev | AAACTGGTCGGCCTGGAGAACGTG | | | | | | | | | | |
| Chromosome | start | end | strand | MM | target_seq | PAM | alignment | distance | position | gene name | gene id |
| GU305914.1 | 195133 | 195155 | + | 0 | CACGTTCTCCAGGCCGACCA | CGG | \|\|\|\|\|\|\|\|[\|\|\|\|\|\|\|\|\|\|\|\|]PAM | NA | Intergenic | NA | NA |
| T14836 | GTTCTCCAGGCCGACCACGGCGG | 1000 | | | | | | | | | |
| Oligo fwd | CACCGTTCTCCAGGCCGACCACGG | | | | | | | | | | |
| Oligo rev | AAACCCGTGGTCGGCCTGGAGAAC | | | | | | | | | | |
| Chromosome | start | end | strand | MM | target_seq | PAM | alignment | distance | position | gene name | gene id |
| GU305914.1 | 195136 | 195158 | + | 0 | GTTCTCCAGGCCGACCACGG | CGG | \|\|\|\|\|\|\|\|[\|\|\|\|\|\|\|\|\|\|\|\|]PAM | NA | Intergenic | NA | NA |
| T14837 | CTCCAGGCCGACCACGGCGGCGG | 1000 | | | | | | | | | |
| Oligo fwd | CACCCTCCAGGCCGACCACGGCGG | | | | | | | | | | |
| Oligo rev | AAACCCGCCGTGGTCGGCCTGGAG | | | | | | | | | | |
| Chromosome | start | end | strand | MM | target_seq | PAM | alignment | distance | position | gene name | gene id |
| GU305914.1 | 195139 | 195161 | + | 0 | CTCCAGGCCGACCACGGCGG | CGG | \|\|\|\|\|\|\|\|[\|\|\|\|\|\|\|\|\|\|\|\|]PAM | NA | Intergenic | NA | NA |
| T14838 | CGGACACGATCTTCCTGACCAGG | 1000 | | | | | | | | | |
| Oligo fwd | CACCCGGACACGATCTTCCTGACC | | | | | | | | | | |
| Oligo rev | AAACGGTCAGGAAGATCGTGTCCG | | | | | | | | | | |
| Chromosome | start | end | strand | MM | target_seq | PAM | alignment | distance | position | gene name | gene id |
| GU305914.1 | 195159 | 195181 | + | 0 | CGGACACGATCTTCCTGACC | AGG | \|\|\|\|\|\|\|\|[\|\|\|\|\|\|\|\|\|\|\|\|]PAM | NA | Intergenic | NA | NA |
| T14839 | GGACACGATCTTCCTGACCAGGG | 1000 | | | | | | | | | |
| Oligo fwd | CACCGGACACGATCTTCCTGACCA | | | | | | | | | | |
| Oligo rev | AAACTGGTCAGGAAGATCGTGTCC | | | | | | | | | | |
| Chromosome | start | end | strand | MM | target_seq | PAM | alignment | distance | position | gene name | gene id |
| GU305914.1 | 195160 | 195182 | + | 0 | GGACACGATCTTCCTGACCA | GGG | \|\|\|\|\|\|\|\|[\|\|\|\|\|\|\|\|\|\|\|\|]PAM | NA | Intergenic | NA | NA |
| T14840 | GATCTTCCTGACCAGGGTCTTGG | 1000 | | | | | | | | | |
| Oligo fwd | CACCGATCTTCCTGACCAGGGTCT | | | | | | | | | | |
| Oligo rev | AAACAGACCCTGGTCAGGAAGATC | | | | | | | | | | |
| Chromosome | start | end | strand | MM | target_seq | PAM | alignment | distance | position | gene name | gene id |
| GU305914.1 | 195166 | 195188 | + | 0 | GATCTTCCTGACCAGGGTCT | TGG | \|\|\|\|\|\|\|\|[\|\|\|\|\|\|\|\|\|\|\|\|]PAM | NA | Intergenic | NA | NA |
| T14841 | ATCTTCCTGACCAGGGTCTTGGG | 1000 | | | | | | | | | |
| Oligo fwd | CACCATCTTCCTGACCAGGGTCTT | | | | | | | | | | |
| Oligo rev | AAACAAGACCCTGGTCAGGAAGAT | | | | | | | | | | |
| Chromosome | start | end | strand | MM | target_seq | PAM | alignment | distance | position | gene name | gene id |
| GU305914.1 | 195167 | 195189 | + | 0 | ATCTTCCTGACCAGGGTCTT | GGG | \|\|\|\|\|\|\|\|[\|\|\|\|\|\|\|\|\|\|\|\|]PAM | NA | Intergenic | NA | NA |
| T14842 | CTGACCAGGGTCTTGGGCTTCGG | 1000 | | | | | | | | | |
| Oligo fwd | CACCCTGACCAGGGTCTTGGGCTT | | | | | | | | | | |
| Oligo rev | AAACAAGCCCAAGACCCTGGTCAG | | | | | | | | | | |
| Chromosome | start | end | strand | MM | target_seq | PAM | alignment | distance | position | gene name | gene id |
| GU305914.1 | 195173 | 195195 | + | 0 | CTGACCAGGGTCTTGGGCTT | CGG | \|\|\|\|\|\|\|\|[\|\|\|\|\|\|\|\|\|\|\|\|]PAM | NA | Intergenic | NA | NA |
| T14843 | GGGCTTCGGCATCGAGACCACGG | 1000 | | | | | | | | | |
| Oligo fwd | CACCGGGCTTCGGCATCGAGACCA | | | | | | | | | | |
| Oligo rev | AAACTGGTCTCGATGCCGAAGCCC | | | | | | | | | | |
| Chromosome | start | end | strand | MM | target_seq | PAM | alignment | distance | position | gene name | gene id |
| GU305914.1 | 195187 | 195209 | + | 0 | GGGCTTCGGCATCGAGACCA | CGG | \|\|\|\|\|\|\|\|[\|\|\|\|\|\|\|\|\|\|\|\|]PAM | NA | Intergenic | NA | NA |
| GU305914.1 | 136810 | 136832 | + | 4 | GGGTTCCGGCATCGTGACCG | TGG | \|\|\|-\|-\|\|[\|\|\|\|\|\|-\|\|\|\|-]PAM | NA | Intergenic | NA | NA |
| T14844 | GCGCCGCCGTCTCGATACACCGG | 1000 | | | | | | | | | |
| Oligo fwd | CACCGCGCCGCCGTCTCGATACAC | | | | | | | | | | |
| Oligo rev | AAACGTGTATCGAGACGGCGGCGC | | | | | | | | | | |
| Chromosome | start | end | strand | MM | target_seq | PAM | alignment | distance | position | gene name | gene id |
| GU305914.1 | 195209 | 195231 | + | 0 | GCGCCGCCGTCTCGATACAC | CGG | \|\|\|\|\|\|\|\|[\|\|\|\|\|\|\|\|\|\|\|\|]PAM | NA | Intergenic | NA | NA |
| T14845 | ACAGCCCCCGTCTCGCCAGCTGG | 1000 | | | | | | | | | |
| Oligo fwd | CACCACAGCCCCCGTCTCGCCAGC | | | | | | | | | | |
| Oligo rev | AAACGCTGGCGAGACGGGGGCTGT | | | | | | | | | | |
| Chromosome | start | end | strand | MM | target_seq | PAM | alignment | distance | position | gene name | gene id |
| GU305914.1 | 195243 | 195265 | + | 0 | ACAGCCCCCGTCTCGCCAGC | TGG | \|\|\|\|\|\|\|\|[\|\|\|\|\|\|\|\|\|\|\|\|]PAM | NA | Intergenic | NA | NA |
| T14846 | CGCCAGCTGGTCGATGTTCCTGG | 1000 | | | | | | | | | |
| Oligo fwd | CACCCGCCAGCTGGTCGATGTTCC | | | | | | | | | | |
| Oligo rev | AAACGGAACATCGACCAGCTGGCG | | | | | | | | | | |
| Chromosome | start | end | strand | MM | target_seq | PAM | alignment | distance | position | gene name | gene id |

GU305914.1 195256 195278 + 0 CGCCAGCTGGTCGATGTTCC TGG ||||||||[||||||||||||]PAM NA Intergenic NA NA

T14847 GGCCGCGTAGTACAGCCCGTCGG 1000
Oligo fwd CACCGGCCGCGTAGTACAGCCCGT
Oligo rev AAACACGGGCTGTACTACGCGGCC
Chromosome start end strand MM target_seq PAM alignment distance position gene name gene id
GU305914.1 195277 195299 + 0 GGCCGCGTAGTACAGCCCGT CGG ||||||||[||||||||||||]PAM NA Intergenic NA NA

T14848 GTAGTACAGCCCGTCGGACTCGG 1000
Oligo fwd CACCGTAGTACAGCCCGTCGGACT
Oligo rev AAACAGTCCGACGGGCTGTACTAC
Chromosome start end strand MM target_seq PAM alignment distance position gene name gene id
GU305914.1 195283 195305 + 0 GTAGTACAGCCCGTCGGACT CGG ||||||||[||||||||||||]PAM NA Intergenic NA NA

T14849 CGTCGGACTCGGCGTCGTACAGG 1000
Oligo fwd CACCCGTCGGACTCGGCGTCGTAC
Oligo rev AAACGTACGACGCCGAGTCCGACG
Chromosome start end strand MM target_seq PAM alignment distance position gene name gene id
GU305914.1 195294 195316 + 0 CGTCGGACTCGGCGTCGTAC AGG ||||||||[||||||||||||]PAM NA Intergenic NA NA

T14850 CGGCGTCGTACAGGAAGAAGCGG 1000
Oligo fwd CACCCGGCGTCGTACAGGAAGAAG
Oligo rev AAACCTTCTTCCTGTACGACGCCG
Chromosome start end strand MM target_seq PAM alignment distance position gene name gene id
GU305914.1 195303 195325 + 0 CGGCGTCGTACAGGAAGAAG CGG ||||||||[||||||||||||]PAM NA Intergenic NA NA

T14851 GGCCGCGCTCGTCCATGACCAGG 1000
Oligo fwd CACCGGCCGCGCTCGTCCATGACC
Oligo rev AAACGGTCATGGACGAGCGCGGCC
Chromosome start end strand MM target_seq PAM alignment distance position gene name gene id
GU305914.1 195324 195346 + 0 GGCCGCGCTCGTCCATGACC AGG ||||||||[||||||||||||]PAM NA Intergenic NA NA

T14852 GCCGCGCTCGTCCATGACCAGGG 1000
Oligo fwd CACCGCCGCGCTCGTCCATGACCA
Oligo rev AAACTGGTCATGGACGAGCGCGGC
Chromosome start end strand MM target_seq PAM alignment distance position gene name gene id
GU305914.1 195325 195347 + 0 GCCGCGCTCGTCCATGACCA GGG ||||||||[||||||||||||]PAM NA Intergenic NA NA

T14853 TCGTCCATGACCAGGGCGCACGG 1000
Oligo fwd CACCTCGTCCATGACCAGGGCGCA
Oligo rev AAACTGCGCCCTGGTCATGGACGA
Chromosome start end strand MM target_seq PAM alignment distance position gene name gene id
GU305914.1 195332 195354 + 0 TCGTCCATGACCAGGGCGCA CGG ||||||||[||||||||||||]PAM NA Intergenic NA NA

T14854 ACGGCGTCTCTCTGAAGTGCAGG 1000
Oligo fwd CACCACGGCGTCTCTCTGAAGTGC
Oligo rev AAACGCACTTCAGAGAGACGCCGT
Chromosome start end strand MM target_seq PAM alignment distance position gene name gene id
GU305914.1 195351 195373 + 0 ACGGCGTCTCTCTGAAGTGC AGG ||||||||[||||||||||||]PAM NA Intergenic NA NA

T14855 CGTCTCTCTGAAGTGCAGGTCGG 1000
Oligo fwd CACCCGTCTCTCTGAAGTGCAGGT
Oligo rev AAACACCTGCACTTCAGAGAGACG
Chromosome start end strand MM target_seq PAM alignment distance position gene name gene id
GU305914.1 195355 195377 + 0 CGTCTCTCTGAAGTGCAGGT CGG ||||||||[||||||||||||]PAM NA Intergenic NA NA

T14856 GTCTCTCTGAAGTGCAGGTCGGG 1000
Oligo fwd CACCGTCTCTCTGAAGTGCAGGTC
Oligo rev AAACGACCTGCACTTCAGAGAGAC
Chromosome start end strand MM target_seq PAM alignment distance position gene name gene id
GU305914.1 195356 195378 + 0 GTCTCTCTGAAGTGCAGGTC GGG ||||||||[||||||||||||]PAM NA Intergenic NA NA

T14857 GTGCAGGTCGGGCGAGCGCGCGG 1000
Oligo fwd CACCGTGCAGGTCGGGCGAGCGCG
Oligo rev AAACCGCGCTCGCCCGACCTGCAC
Chromosome start end strand MM target_seq PAM alignment distance position gene name gene id
GU305914.1 195367 195389 + 0 GTGCAGGTCGGGCGAGCGCG CGG ||||||||[||||||||||||]PAM NA Intergenic NA NA

T14858 AGCGCGCGGCGCAGACCCCCAGG 1000
Oligo fwd CACCAGCGCGCGGCGCAGACCCCC
Oligo rev AAACGGGGGTCTGCGCCGCGCGCT
Chromosome start end strand MM target_seq PAM alignment distance position gene name gene id
GU305914.1 195381 195403 + 0 AGCGCGCGGCGCAGACCCCC AGG ||||||||[||||||||||||]PAM NA Intergenic NA NA

T14859 GCGCGCGGCGCAGACCCCCAGGG 1000
Oligo fwd CACCGCGCGCGGCGCAGACCCCCA
Oligo rev AAACTGGGGGTCTGCGCCGCGCGC
Chromosome start end strand MM target_seq PAM alignment distance position gene name gene id
GU305914.1 195382 195404 + 0 GCGCGCGGCGCAGACCCCCA GGG ||||||||[||||||||||||]PAM NA Intergenic NA NA

T14860 CGCGCGGCGCAGACCCCCAGGGG 1000
Oligo fwd CACCCGCGCGGCGCAGACCCCCAG
Oligo rev AAACCTGGGGGTCTGCGCCGCGCG
Chromosome start end strand MM target_seq PAM alignment distance position gene name gene id
GU305914.1 195383 195405 + 0 CGCGCGGCGCAGACCCCCAG GGG ||||||||[||||||||||||]PAM NA Intergenic NA NA

T14861 CCCCAGGGGCGTCAGCTCGCCGG 1000
Oligo fwd CACCCCCCAGGGGCGTCAGCTCGC
Oligo rev AAACGCGAGCTGACGCCCCTGGGG

<table>
<tr><td>Chromosome</td><td>start</td><td>end</td><td>strand</td><td>MM</td><td>target_seq</td><td>PAM</td><td>alignment</td><td>distance</td><td>position</td><td>gene name</td><td>gene id</td></tr>
<tr><td>GU305914.1</td><td>195397</td><td>195419</td><td>+</td><td>0</td><td>CCCCAGGGGCGTCAGCTCGC</td><td>CGG</td><td>||||||||[||||||||||||]PAM</td><td>NA</td><td>Intergenic</td><td>NA</td><td>NA</td></tr>
<tr><td></td><td></td><td></td><td></td><td></td><td></td><td></td><td></td><td></td><td></td><td></td><td></td></tr>
<tr><td>T14862</td><td>CAGGGGCGTCAGCTCGCCGGCGG</td><td>1000</td><td></td><td></td><td></td><td></td><td></td><td></td><td></td><td></td><td></td></tr>
<tr><td>Oligo fwd</td><td>CACCCAGGGGCGTCAGCTCGCCGG</td><td></td><td></td><td></td><td></td><td></td><td></td><td></td><td></td><td></td><td></td></tr>
<tr><td>Oligo rev</td><td>AAACCCGGCGAGCTGACGCCCCTG</td><td></td><td></td><td></td><td></td><td></td><td></td><td></td><td></td><td></td><td></td></tr>
<tr><td>Chromosome</td><td>start</td><td>end</td><td>strand</td><td>MM</td><td>target_seq</td><td>PAM</td><td>alignment</td><td>distance</td><td>position</td><td>gene name</td><td>gene id</td></tr>
<tr><td>GU305914.1</td><td>195400</td><td>195422</td><td>+</td><td>0</td><td>CAGGGGCGTCAGCTCGCCGG</td><td>CGG</td><td>||||||||[||||||||||||]PAM</td><td>NA</td><td>Intergenic</td><td>NA</td><td>NA</td></tr>
<tr><td></td><td></td><td></td><td></td><td></td><td></td><td></td><td></td><td></td><td></td><td></td><td></td></tr>
<tr><td>T14863</td><td>GTCGCCGCAGCACAGATACTTGG</td><td>1000</td><td></td><td></td><td></td><td></td><td></td><td></td><td></td><td></td><td></td></tr>
<tr><td>Oligo fwd</td><td>CACCGTCGCCGCAGCACAGATACT</td><td></td><td></td><td></td><td></td><td></td><td></td><td></td><td></td><td></td><td></td></tr>
<tr><td>Oligo rev</td><td>AAACAGTATCTGTGCTGCGGCGAC</td><td></td><td></td><td></td><td></td><td></td><td></td><td></td><td></td><td></td><td></td></tr>
<tr><td>Chromosome</td><td>start</td><td>end</td><td>strand</td><td>MM</td><td>target_seq</td><td>PAM</td><td>alignment</td><td>distance</td><td>position</td><td>gene name</td><td>gene id</td></tr>
<tr><td>GU305914.1</td><td>195424</td><td>195446</td><td>+</td><td>0</td><td>GTCGCCGCAGCACAGATACT</td><td>TGG</td><td>||||||||[||||||||||||]PAM</td><td>NA</td><td>Intergenic</td><td>NA</td><td>NA</td></tr>
<tr><td></td><td></td><td></td><td></td><td></td><td></td><td></td><td></td><td></td><td></td><td></td><td></td></tr>
<tr><td>T14864</td><td>CGCAGCACAGATACTTGGACAGG</td><td>1000</td><td></td><td></td><td></td><td></td><td></td><td></td><td></td><td></td><td></td></tr>
<tr><td>Oligo fwd</td><td>CACCCGCAGCACAGATACTTGGAC</td><td></td><td></td><td></td><td></td><td></td><td></td><td></td><td></td><td></td><td></td></tr>
<tr><td>Oligo rev</td><td>AAACGTCCAAGTATCTGTGCTGCG</td><td></td><td></td><td></td><td></td><td></td><td></td><td></td><td></td><td></td><td></td></tr>
<tr><td>Chromosome</td><td>start</td><td>end</td><td>strand</td><td>MM</td><td>target_seq</td><td>PAM</td><td>alignment</td><td>distance</td><td>position</td><td>gene name</td><td>gene id</td></tr>
<tr><td>GU305914.1</td><td>195429</td><td>195451</td><td>+</td><td>0</td><td>CGCAGCACAGATACTTGGAC</td><td>AGG</td><td>||||||||[||||||||||||]PAM</td><td>NA</td><td>Intergenic</td><td>NA</td><td>NA</td></tr>
<tr><td></td><td></td><td></td><td></td><td></td><td></td><td></td><td></td><td></td><td></td><td></td><td></td></tr>
<tr><td>T14865</td><td>GGTCCACGCCGCGCAGCACGCGG</td><td>1000</td><td></td><td></td><td></td><td></td><td></td><td></td><td></td><td></td><td></td></tr>
<tr><td>Oligo fwd</td><td>CACCGGTCCACGCCGCGCAGCACG</td><td></td><td></td><td></td><td></td><td></td><td></td><td></td><td></td><td></td><td></td></tr>
<tr><td>Oligo rev</td><td>AAACCGTGCTGCGCGGCGTGGACC</td><td></td><td></td><td></td><td></td><td></td><td></td><td></td><td></td><td></td><td></td></tr>
<tr><td>Chromosome</td><td>start</td><td>end</td><td>strand</td><td>MM</td><td>target_seq</td><td>PAM</td><td>alignment</td><td>distance</td><td>position</td><td>gene name</td><td>gene id</td></tr>
<tr><td>GU305914.1</td><td>195450</td><td>195472</td><td>+</td><td>0</td><td>GGTCCACGCCGCGCAGCACG</td><td>CGG</td><td>||||||||[||||||||||||]PAM</td><td>NA</td><td>Intergenic</td><td>NA</td><td>NA</td></tr>
<tr><td></td><td></td><td></td><td></td><td></td><td></td><td></td><td></td><td></td><td></td><td></td><td></td></tr>
<tr><td>T14866</td><td>GCCGCGCAGCACGCGGATCTCGG</td><td>1000</td><td></td><td></td><td></td><td></td><td></td><td></td><td></td><td></td><td></td></tr>
<tr><td>Oligo fwd</td><td>CACCGCCGCGCAGCACGCGGATCT</td><td></td><td></td><td></td><td></td><td></td><td></td><td></td><td></td><td></td><td></td></tr>
<tr><td>Oligo rev</td><td>AAACAGATCCGCGTGCTGCGCGGC</td><td></td><td></td><td></td><td></td><td></td><td></td><td></td><td></td><td></td><td></td></tr>
<tr><td>Chromosome</td><td>start</td><td>end</td><td>strand</td><td>MM</td><td>target_seq</td><td>PAM</td><td>alignment</td><td>distance</td><td>position</td><td>gene name</td><td>gene id</td></tr>
<tr><td>GU305914.1</td><td>195457</td><td>195479</td><td>+</td><td>0</td><td>GCCGCGCAGCACGCGGATCT</td><td>CGG</td><td>||||||||[||||||||||||]PAM</td><td>NA</td><td>Intergenic</td><td>NA</td><td>NA</td></tr>
<tr><td></td><td></td><td></td><td></td><td></td><td></td><td></td><td></td><td></td><td></td><td></td><td></td></tr>
<tr><td>T14867</td><td>CGGATCTCGGAGCGTCTCTGCGG</td><td>1000</td><td></td><td></td><td></td><td></td><td></td><td></td><td></td><td></td><td></td></tr>
<tr><td>Oligo fwd</td><td>CACCCGGATCTCGGAGCGTCTCTG</td><td></td><td></td><td></td><td></td><td></td><td></td><td></td><td></td><td></td><td></td></tr>
<tr><td>Oligo rev</td><td>AAACCAGAGACGCTCCGAGATCCG</td><td></td><td></td><td></td><td></td><td></td><td></td><td></td><td></td><td></td><td></td></tr>
<tr><td>Chromosome</td><td>start</td><td>end</td><td>strand</td><td>MM</td><td>target_seq</td><td>PAM</td><td>alignment</td><td>distance</td><td>position</td><td>gene name</td><td>gene id</td></tr>
<tr><td>GU305914.1</td><td>195470</td><td>195492</td><td>+</td><td>0</td><td>CGGATCTCGGAGCGTCTCTG</td><td>CGG</td><td>||||||||[||||||||||||]PAM</td><td>NA</td><td>Intergenic</td><td>NA</td><td>NA</td></tr>
<tr><td></td><td></td><td></td><td></td><td></td><td></td><td></td><td></td><td></td><td></td><td></td><td></td></tr>
<tr><td>T14868</td><td>GGATCTCGGAGCGTCTCTGCGGG</td><td>1000</td><td></td><td></td><td></td><td></td><td></td><td></td><td></td><td></td><td></td></tr>
<tr><td>Oligo fwd</td><td>CACCGGATCTCGGAGCGTCTCTGC</td><td></td><td></td><td></td><td></td><td></td><td></td><td></td><td></td><td></td><td></td></tr>
<tr><td>Oligo rev</td><td>AAACGCAGAGACGCTCCGAGATCC</td><td></td><td></td><td></td><td></td><td></td><td></td><td></td><td></td><td></td><td></td></tr>
<tr><td>Chromosome</td><td>start</td><td>end</td><td>strand</td><td>MM</td><td>target_seq</td><td>PAM</td><td>alignment</td><td>distance</td><td>position</td><td>gene name</td><td>gene id</td></tr>
<tr><td>GU305914.1</td><td>195471</td><td>195493</td><td>+</td><td>0</td><td>GGATCTCGGAGCGTCTCTGC</td><td>GGG</td><td>||||||||[||||||||||||]PAM</td><td>NA</td><td>Intergenic</td><td>NA</td><td>NA</td></tr>
<tr><td></td><td></td><td></td><td></td><td></td><td></td><td></td><td></td><td></td><td></td><td></td><td></td></tr>
<tr><td>T14869</td><td>CGGAGCGTCTCTGCGGGATCAGG</td><td>1000</td><td></td><td></td><td></td><td></td><td></td><td></td><td></td><td></td><td></td></tr>
<tr><td>Oligo fwd</td><td>CACCCGGAGCGTCTCTGCGGGATC</td><td></td><td></td><td></td><td></td><td></td><td></td><td></td><td></td><td></td><td></td></tr>
<tr><td>Oligo rev</td><td>AAACGATCCCGCAGAGACGCTCCG</td><td></td><td></td><td></td><td></td><td></td><td></td><td></td><td></td><td></td><td></td></tr>
<tr><td>Chromosome</td><td>start</td><td>end</td><td>strand</td><td>MM</td><td>target_seq</td><td>PAM</td><td>alignment</td><td>distance</td><td>position</td><td>gene name</td><td>gene id</td></tr>
<tr><td>GU305914.1</td><td>195477</td><td>195499</td><td>+</td><td>0</td><td>CGGAGCGTCTCTGCGGGATC</td><td>AGG</td><td>||||||||[||||||||||||]PAM</td><td>NA</td><td>Intergenic</td><td>NA</td><td>NA</td></tr>
<tr><td></td><td></td><td></td><td></td><td></td><td></td><td></td><td></td><td></td><td></td><td></td><td></td></tr>
<tr><td>T14870</td><td>GGGATCAGGTACCAGTTCTCCGG</td><td>1000</td><td></td><td></td><td></td><td></td><td></td><td></td><td></td><td></td><td></td></tr>
<tr><td>Oligo fwd</td><td>CACCGGGATCAGGTACCAGTTCTC</td><td></td><td></td><td></td><td></td><td></td><td></td><td></td><td></td><td></td><td></td></tr>
<tr><td>Oligo rev</td><td>AAACGAGAACTGGTACCTGATCCC</td><td></td><td></td><td></td><td></td><td></td><td></td><td></td><td></td><td></td><td></td></tr>
<tr><td>Chromosome</td><td>start</td><td>end</td><td>strand</td><td>MM</td><td>target_seq</td><td>PAM</td><td>alignment</td><td>distance</td><td>position</td><td>gene name</td><td>gene id</td></tr>
<tr><td>GU305914.1</td><td>195491</td><td>195513</td><td>+</td><td>0</td><td>GGGATCAGGTACCAGTTCTC</td><td>CGG</td><td>||||||||[||||||||||||]PAM</td><td>NA</td><td>Intergenic</td><td>NA</td><td>NA</td></tr>
<tr><td></td><td></td><td></td><td></td><td></td><td></td><td></td><td></td><td></td><td></td><td></td><td></td></tr>
<tr><td>T14871</td><td>GGATCAGGTACCAGTTCTCCGGG</td><td>1000</td><td></td><td></td><td></td><td></td><td></td><td></td><td></td><td></td><td></td></tr>
<tr><td>Oligo fwd</td><td>CACCGGATCAGGTACCAGTTCTCC</td><td></td><td></td><td></td><td></td><td></td><td></td><td></td><td></td><td></td><td></td></tr>
<tr><td>Oligo rev</td><td>AAACGGAGAACTGGTACCTGATCC</td><td></td><td></td><td></td><td></td><td></td><td></td><td></td><td></td><td></td><td></td></tr>
<tr><td>Chromosome</td><td>start</td><td>end</td><td>strand</td><td>MM</td><td>target_seq</td><td>PAM</td><td>alignment</td><td>distance</td><td>position</td><td>gene name</td><td>gene id</td></tr>
<tr><td>GU305914.1</td><td>195492</td><td>195514</td><td>+</td><td>0</td><td>GGATCAGGTACCAGTTCTCC</td><td>GGG</td><td>||||||||[||||||||||||]PAM</td><td>NA</td><td>Intergenic</td><td>NA</td><td>NA</td></tr>
<tr><td></td><td></td><td></td><td></td><td></td><td></td><td></td><td></td><td></td><td></td><td></td><td></td></tr>
<tr><td>T14872</td><td>GGTACCAGTTCTCCGGGTAGCGG</td><td>1000</td><td></td><td></td><td></td><td></td><td></td><td></td><td></td><td></td><td></td></tr>
<tr><td>Oligo fwd</td><td>CACCGGTACCAGTTCTCCGGGTAG</td><td></td><td></td><td></td><td></td><td></td><td></td><td></td><td></td><td></td><td></td></tr>
<tr><td>Oligo rev</td><td>AAACCTACCCGGAGAACTGGTACC</td><td></td><td></td><td></td><td></td><td></td><td></td><td></td><td></td><td></td><td></td></tr>
<tr><td>Chromosome</td><td>start</td><td>end</td><td>strand</td><td>MM</td><td>target_seq</td><td>PAM</td><td>alignment</td><td>distance</td><td>position</td><td>gene name</td><td>gene id</td></tr>
<tr><td>GU305914.1</td><td>195498</td><td>195520</td><td>+</td><td>0</td><td>GGTACCAGTTCTCCGGGTAG</td><td>CGG</td><td>||||||||[||||||||||||]PAM</td><td>NA</td><td>Intergenic</td><td>NA</td><td>NA</td></tr>
<tr><td></td><td></td><td></td><td></td><td></td><td></td><td></td><td></td><td></td><td></td><td></td><td></td></tr>
<tr><td>T14873</td><td>TCTCCGGGTAGCGGAGCACGAGG</td><td>1000</td><td></td><td></td><td></td><td></td><td></td><td></td><td></td><td></td><td></td></tr>
<tr><td>Oligo fwd</td><td>CACCTCTCCGGGTAGCGGAGCACG</td><td></td><td></td><td></td><td></td><td></td><td></td><td></td><td></td><td></td><td></td></tr>
<tr><td>Oligo rev</td><td>AAACCGTGCTCCGCTACCCGGAGA</td><td></td><td></td><td></td><td></td><td></td><td></td><td></td><td></td><td></td><td></td></tr>
<tr><td>Chromosome</td><td>start</td><td>end</td><td>strand</td><td>MM</td><td>target_seq</td><td>PAM</td><td>alignment</td><td>distance</td><td>position</td><td>gene name</td><td>gene id</td></tr>
<tr><td>GU305914.1</td><td>195507</td><td>195529</td><td>+</td><td>0</td><td>TCTCCGGGTAGCGGAGCACG</td><td>AGG</td><td>||||||||[||||||||||||]PAM</td><td>NA</td><td>Intergenic</td><td>NA</td><td>NA</td></tr>
<tr><td></td><td></td><td></td><td></td><td></td><td></td><td></td><td></td><td></td><td></td><td></td><td></td></tr>
<tr><td>T14874</td><td>CTCCGGGTAGCGGAGCACGAGGG</td><td>1000</td><td></td><td></td><td></td><td></td><td></td><td></td><td></td><td></td><td></td></tr>
<tr><td>Oligo fwd</td><td>CACCCTCCGGGTAGCGGAGCACGA</td><td></td><td></td><td></td><td></td><td></td><td></td><td></td><td></td><td></td><td></td></tr>
<tr><td>Oligo rev</td><td>AAACTCGTGCTCCGCTACCCGGAG</td><td></td><td></td><td></td><td></td><td></td><td></td><td></td><td></td><td></td><td></td></tr>
<tr><td>Chromosome</td><td>start</td><td>end</td><td>strand</td><td>MM</td><td>target_seq</td><td>PAM</td><td>alignment</td><td>distance</td><td>position</td><td>gene name</td><td>gene id</td></tr>
<tr><td>GU305914.1</td><td>195508</td><td>195530</td><td>+</td><td>0</td><td>CTCCGGGTAGCGGAGCACGA</td><td>GGG</td><td>||||||||[||||||||||||]PAM</td><td>NA</td><td>Intergenic</td><td>NA</td><td>NA</td></tr>
<tr><td></td><td></td><td></td><td></td><td></td><td></td><td></td><td></td><td></td><td></td><td></td><td></td></tr>
<tr><td>T14875</td><td>TCCGGGTAGCGGAGCACGAGGGG</td><td>1000</td><td></td><td></td><td></td><td></td><td></td><td></td><td></td><td></td><td></td></tr>
<tr><td>Oligo fwd</td><td>CACCTCCGGGTAGCGGAGCACGAG</td><td></td><td></td><td></td><td></td><td></td><td></td><td></td><td></td><td></td><td></td></tr>
<tr><td>Oligo rev</td><td>AAACCTCGTGCTCCGCTACCCGGA</td><td></td><td></td><td></td><td></td><td></td><td></td><td></td><td></td><td></td><td></td></tr>
<tr><td>Chromosome</td><td>start</td><td>end</td><td>strand</td><td>MM</td><td>target_seq</td><td>PAM</td><td>alignment</td><td>distance</td><td>position</td><td>gene name</td><td>gene id</td></tr>
<tr><td>GU305914.1</td><td>195509</td><td>195531</td><td>+</td><td>0</td><td>TCCGGGTAGCGGAGCACGAG</td><td>GGG</td><td>||||||||[||||||||||||]PAM</td><td>NA</td><td>Intergenic</td><td>NA</td><td>NA</td></tr>
<tr><td></td><td></td><td></td><td></td><td></td><td></td><td></td><td></td><td></td><td></td><td></td><td></td></tr>
<tr><td>T14876</td><td>GGAGCACGAGGGGCTTGCCGAGG</td><td>1000</td><td></td><td></td><td></td><td></td><td></td><td></td><td></td><td></td><td></td></tr>
<tr><td>Oligo fwd</td><td>CACCGGAGCACGAGGGGCTTGCCG</td><td></td><td></td><td></td><td></td><td></td><td></td><td></td><td></td><td></td><td></td></tr>
</table>

| | | | | | | | | | | | |
|---|---|---|---|---|---|---|---|---|---|---|---|
| Oligo rev | AAACCGGCAAGCCCCTCGTGCTCC | | | | | | | | | | |
| Chromosome | start | end | strand | MM | target_seq | PAM | alignment | distance | position | gene name | gene id |
| GU305914.1 | 195519 | 195541 | + | 0 | GGAGCACGAGGGGCTTGCCG | AGG | \|\|\|\|\|\|\|\|[\|\|\|\|\|\|\|\|\|\|\|\|]PAM | NA | Intergenic | NA | NA |
| T14877 | GCCGAGGTAGTGCACCACCGCGG | 1000 | | | | | | | | | |
| Oligo fwd | CACCGCCGAGGTAGTGCACCACCG | | | | | | | | | | |
| Oligo rev | AAACCGGTGGTGCACTACCTCGGC | | | | | | | | | | |
| Chromosome | start | end | strand | MM | target_seq | PAM | alignment | distance | position | gene name | gene id |
| GU305914.1 | 195535 | 195557 | + | 0 | GCCGAGGTAGTGCACCACCG | CGG | \|\|\|\|\|\|\|\|[\|\|\|\|\|\|\|\|\|\|\|\|]PAM | NA | Intergenic | NA | NA |
| T14878 | GCACCACCGCGGCCGCGATCTGG | 1000 | | | | | | | | | |
| Oligo fwd | CACCGCACCACCGCGGCCGCGATC | | | | | | | | | | |
| Oligo rev | AAACGATCGCGGCCGCGGTGGTGC | | | | | | | | | | |
| Chromosome | start | end | strand | MM | target_seq | PAM | alignment | distance | position | gene name | gene id |
| GU305914.1 | 195546 | 195568 | + | 0 | GCACCACCGCGGCCGCGATC | TGG | \|\|\|\|\|\|\|\|[\|\|\|\|\|\|\|\|\|\|\|\|]PAM | NA | Intergenic | NA | NA |
| T14879 | ACCGCGGCCGCGATCTGGCCCGG | 1000 | | | | | | | | | |
| Oligo fwd | CACCACCGCGGCCGCGATCTGGCC | | | | | | | | | | |
| Oligo rev | AAACGGCCAGATCGCGGCCGCGGT | | | | | | | | | | |
| Chromosome | start | end | strand | MM | target_seq | PAM | alignment | distance | position | gene name | gene id |
| GU305914.1 | 195551 | 195573 | + | 0 | ACCGCGGCCGCGATCTGGCC | CGG | \|\|\|\|\|\|\|\|[\|\|\|\|\|\|\|\|\|\|\|\|]PAM | NA | Intergenic | NA | NA |
| T14880 | GCGGCCGCGATCTGGCCCGGCGG | 1000 | | | | | | | | | |
| Oligo fwd | CACCGCGGCCGCGATCTGGCCCGG | | | | | | | | | | |
| Oligo rev | AAACCCGGGCCAGATCGCGGCCGC | | | | | | | | | | |
| Chromosome | start | end | strand | MM | target_seq | PAM | alignment | distance | position | gene name | gene id |
| GU305914.1 | 195554 | 195576 | + | 0 | GCGGCCGCGATCTGGCCCGG | CGG | \|\|\|\|\|\|\|\|[\|\|\|\|\|\|\|\|\|\|\|\|]PAM | NA | Intergenic | NA | NA |
| GU305914.1 | 162792 | 162814 | - | 4 | CCGTCCGAGACCTGGCCCGG | CGG | -\|\|-\|\|\|-[\|\|-\|\|\|\|\|\|\|\|\|]PAM | NA | Intergenic | NA | NA |
| T14881 | CGGCCGCGATCTGGCCCGGCGGG | 1000 | | | | | | | | | |
| Oligo fwd | CACCCGGCCGCGATCTGGCCCGGC | | | | | | | | | | |
| Oligo rev | AAACGCCGGGCCAGATCGCGGCCG | | | | | | | | | | |
| Chromosome | start | end | strand | MM | target_seq | PAM | alignment | distance | position | gene name | gene id |
| GU305914.1 | 195555 | 195577 | + | 0 | CGGCCGCGATCTGGCCCGGC | GGG | \|\|\|\|\|\|\|\|[\|\|\|\|\|\|\|\|\|\|\|\|]PAM | NA | Intergenic | NA | NA |
| T14882 | TCTGGCCCGGCGGGAAACTGAGG | 1000 | | | | | | | | | |
| Oligo fwd | CACCTCTGGCCCGGCGGGAAACTG | | | | | | | | | | |
| Oligo rev | AAACCAGTTTCCCGCCGGGCCAGA | | | | | | | | | | |
| Chromosome | start | end | strand | MM | target_seq | PAM | alignment | distance | position | gene name | gene id |
| GU305914.1 | 195564 | 195586 | + | 0 | TCTGGCCCGGCGGGAAACTG | AGG | \|\|\|\|\|\|\|\|[\|\|\|\|\|\|\|\|\|\|\|\|]PAM | NA | Intergenic | NA | NA |
| T14883 | CTGGCCCGGCGGGAAACTGAGGG | 1000 | | | | | | | | | |
| Oligo fwd | CACCCTGGCCCGGCGGGAAACTGA | | | | | | | | | | |
| Oligo rev | AAACTCAGTTTCCCGCCGGGCCAG | | | | | | | | | | |
| Chromosome | start | end | strand | MM | target_seq | PAM | alignment | distance | position | gene name | gene id |
| GU305914.1 | 195565 | 195587 | + | 0 | CTGGCCCGGCGGGAAACTGA | GGG | \|\|\|\|\|\|\|\|[\|\|\|\|\|\|\|\|\|\|\|\|]PAM | NA | Intergenic | NA | NA |
| T14884 | CCGGCGGGAAACTGAGGGCCAGG | 1000 | | | | | | | | | |
| Oligo fwd | CACCCCGGCGGGAAACTGAGGGCC | | | | | | | | | | |
| Oligo rev | AAACGGCCCTCAGTTTCCCGCCGG | | | | | | | | | | |
| Chromosome | start | end | strand | MM | target_seq | PAM | alignment | distance | position | gene name | gene id |
| GU305914.1 | 195570 | 195592 | + | 0 | CCGGCGGGAAACTGAGGGCC | AGG | \|\|\|\|\|\|\|\|[\|\|\|\|\|\|\|\|\|\|\|\|]PAM | NA | Intergenic | NA | NA |
| T14885 | GCGGGAAACTGAGGGCCAGGTGG | 1000 | | | | | | | | | |
| Oligo fwd | CACCGCGGGAAACTGAGGGCCAGG | | | | | | | | | | |
| Oligo rev | AAACCCTGGCCCTCAGTTTCCCGC | | | | | | | | | | |
| Chromosome | start | end | strand | MM | target_seq | PAM | alignment | distance | position | gene name | gene id |
| GU305914.1 | 195573 | 195595 | + | 0 | GCGGGAAACTGAGGGCCAGG | TGG | \|\|\|\|\|\|\|\|[\|\|\|\|\|\|\|\|\|\|\|\|]PAM | NA | Intergenic | NA | NA |
| T14886 | CTTCTTCTCGTATTTCTCGACGG | 1000 | | | | | | | | | |
| Oligo fwd | CACCCTTCTTCTCGTATTTCTCGA | | | | | | | | | | |
| Oligo rev | AAACTCGAGAAATACGAGAAGAAG | | | | | | | | | | |
| Chromosome | start | end | strand | MM | target_seq | PAM | alignment | distance | position | gene name | gene id |
| GU305914.1 | 195646 | 195668 | + | 0 | CTTCTTCTCGTATTTCTCGA | CGG | \|\|\|\|\|\|\|\|[\|\|\|\|\|\|\|\|\|\|\|\|]PAM | NA | Intergenic | NA | NA |
| T14887 | GTATTTCTCGACGGCCTTCCTGG | 1000 | | | | | | | | | |
| Oligo fwd | CACCGTATTTCTCGACGGCCTTCC | | | | | | | | | | |
| Oligo rev | AAACGGAAGGCCGTCGAGAAATAC | | | | | | | | | | |
| Chromosome | start | end | strand | MM | target_seq | PAM | alignment | distance | position | gene name | gene id |
| GU305914.1 | 195655 | 195677 | + | 0 | GTATTTCTCGACGGCCTTCC | TGG | \|\|\|\|\|\|\|\|[\|\|\|\|\|\|\|\|\|\|\|\|]PAM | NA | Intergenic | NA | NA |
| T14888 | GGCCTTCCTGGACACCGTGACGG | 1000 | | | | | | | | | |
| Oligo fwd | CACCGGCCTTCCTGGACACCGTGA | | | | | | | | | | |
| Oligo rev | AAACTCACGGTGTCCAGGAAGGCC | | | | | | | | | | |
| Chromosome | start | end | strand | MM | target_seq | PAM | alignment | distance | position | gene name | gene id |
| GU305914.1 | 195667 | 195689 | + | 0 | GGCCTTCCTGGACACCGTGA | CGG | \|\|\|\|\|\|\|\|[\|\|\|\|\|\|\|\|\|\|\|\|]PAM | NA | Intergenic | NA | NA |
| GU305914.1 | 70558 | 70580 | + | 3 | GGCGTTCAGGGACACCGTGA | AGG | \|\|\|-\|\|\|-[-\|\|\|\|\|\|\|\|\|\|\|]PAM | NA | Intergenic | NA | NA |
| T14889 | CCTGGACACCGTGACGGCCGAGG | 1000 | | | | | | | | | |
| Oligo fwd | CACCCCTGGACACCGTGACGGCCG | | | | | | | | | | |
| Oligo rev | AAACCGGCCGTCACGGTGTCCAGG | | | | | | | | | | |
| Chromosome | start | end | strand | MM | target_seq | PAM | alignment | distance | position | gene name | gene id |
| GU305914.1 | 195673 | 195695 | + | 0 | CCTGGACACCGTGACGGCCG | AGG | \|\|\|\|\|\|\|\|[\|\|\|\|\|\|\|\|\|\|\|\|]PAM | NA | Intergenic | NA | NA |
| T14890 | GTGACGGCCGAGGTCTTCCACGG | 1000 | | | | | | | | | |
| Oligo fwd | CACCGTGACGGCCGAGGTCTTCCA | | | | | | | | | | |
| Oligo rev | AAACTGGAAGACCTCGGCCGTCAC | | | | | | | | | | |
| Chromosome | start | end | strand | MM | target_seq | PAM | alignment | distance | position | gene name | gene id |
| GU305914.1 | 195683 | 195705 | + | 0 | GTGACGGCCGAGGTCTTCCA | CGG | \|\|\|\|\|\|\|\|[\|\|\|\|\|\|\|\|\|\|\|\|]PAM | NA | Intergenic | NA | NA |

<table>
<tr><td>T14891</td><td>TCCCGCCGCGTCGCCGTCGTCGG</td><td>1000</td><td></td><td></td><td></td><td></td><td></td><td></td><td></td><td></td><td></td></tr>
<tr><td>Oligo fwd</td><td>CACCTCCCGCCGCGTCGCCGTCGT</td><td></td><td></td><td></td><td></td><td></td><td></td><td></td><td></td><td></td><td></td></tr>
<tr><td>Oligo rev</td><td>AAACACGACGGCGACGCGGCGGGA</td><td></td><td></td><td></td><td></td><td></td><td></td><td></td><td></td><td></td><td></td></tr>
<tr><td>Chromosome</td><td>start</td><td>end</td><td>strand</td><td>MM</td><td>target_seq</td><td>PAM</td><td>alignment</td><td>distance</td><td>position</td><td>gene name</td><td>gene id</td></tr>
<tr><td>GU305914.1</td><td>195724</td><td>195746</td><td>+</td><td>0</td><td>TCCCGCCGCGTCGCCGTCGT</td><td>CGG</td><td>||||||||[||||||||||||]PAM</td><td>NA</td><td>Intergenic</td><td>NA</td><td>NA</td></tr>
<tr><td>T14892</td><td>CCGCGTCGCCGTCGTCGGACAGG</td><td>1000</td><td></td><td></td><td></td><td></td><td></td><td></td><td></td><td></td><td></td></tr>
<tr><td>Oligo fwd</td><td>CACCCCGCGTCGCCGTCGTCGGAC</td><td></td><td></td><td></td><td></td><td></td><td></td><td></td><td></td><td></td><td></td></tr>
<tr><td>Oligo rev</td><td>AAACGTCCGACGACGGCGACGCGG</td><td></td><td></td><td></td><td></td><td></td><td></td><td></td><td></td><td></td><td></td></tr>
<tr><td>Chromosome</td><td>start</td><td>end</td><td>strand</td><td>MM</td><td>target_seq</td><td>PAM</td><td>alignment</td><td>distance</td><td>position</td><td>gene name</td><td>gene id</td></tr>
<tr><td>GU305914.1</td><td>195729</td><td>195751</td><td>+</td><td>0</td><td>CCGCGTCGCCGTCGTCGGAC</td><td>AGG</td><td>||||||||[||||||||||||]PAM</td><td>NA</td><td>Intergenic</td><td>NA</td><td>NA</td></tr>
<tr><td>T14893</td><td>CTGAGTCGCTCTCGCTGTCGCGG</td><td>1000</td><td></td><td></td><td></td><td></td><td></td><td></td><td></td><td></td><td></td></tr>
<tr><td>Oligo fwd</td><td>CACCCTGAGTCGCTCTCGCTGTCG</td><td></td><td></td><td></td><td></td><td></td><td></td><td></td><td></td><td></td><td></td></tr>
<tr><td>Oligo rev</td><td>AAACCGACAGCGAGAGCGACTCAG</td><td></td><td></td><td></td><td></td><td></td><td></td><td></td><td></td><td></td><td></td></tr>
<tr><td>Chromosome</td><td>start</td><td>end</td><td>strand</td><td>MM</td><td>target_seq</td><td>PAM</td><td>alignment</td><td>distance</td><td>position</td><td>gene name</td><td>gene id</td></tr>
<tr><td>GU305914.1</td><td>195777</td><td>195799</td><td>+</td><td>0</td><td>CTGAGTCGCTCTCGCTGTCG</td><td>CGG</td><td>||||||||[||||||||||||]PAM</td><td>NA</td><td>Intergenic</td><td>NA</td><td>NA</td></tr>
<tr><td>T14894</td><td>TGAGTCGCTCTCGCTGTCGCGGG</td><td>1000</td><td></td><td></td><td></td><td></td><td></td><td></td><td></td><td></td><td></td></tr>
<tr><td>Oligo fwd</td><td>CACCTGAGTCGCTCTCGCTGTCGC</td><td></td><td></td><td></td><td></td><td></td><td></td><td></td><td></td><td></td><td></td></tr>
<tr><td>Oligo rev</td><td>AAACGCGACAGCGAGAGCGACTCA</td><td></td><td></td><td></td><td></td><td></td><td></td><td></td><td></td><td></td><td></td></tr>
<tr><td>Chromosome</td><td>start</td><td>end</td><td>strand</td><td>MM</td><td>target_seq</td><td>PAM</td><td>alignment</td><td>distance</td><td>position</td><td>gene name</td><td>gene id</td></tr>
<tr><td>GU305914.1</td><td>195778</td><td>195800</td><td>+</td><td>0</td><td>TGAGTCGCTCTCGCTGTCGC</td><td>GGG</td><td>||||||||[||||||||||||]PAM</td><td>NA</td><td>Intergenic</td><td>NA</td><td>NA</td></tr>
<tr><td>T14895</td><td>GGAGCCCGCGTCAGAGCGCCCGG</td><td>1000</td><td></td><td></td><td></td><td></td><td></td><td></td><td></td><td></td><td></td></tr>
<tr><td>Oligo fwd</td><td>CACCGGAGCCCGCGTCAGAGCGCC</td><td></td><td></td><td></td><td></td><td></td><td></td><td></td><td></td><td></td><td></td></tr>
<tr><td>Oligo rev</td><td>AAACGGCGCTCTGACGCGGGCTCC</td><td></td><td></td><td></td><td></td><td></td><td></td><td></td><td></td><td></td><td></td></tr>
<tr><td>Chromosome</td><td>start</td><td>end</td><td>strand</td><td>MM</td><td>target_seq</td><td>PAM</td><td>alignment</td><td>distance</td><td>position</td><td>gene name</td><td>gene id</td></tr>
<tr><td>GU305914.1</td><td>195799</td><td>195821</td><td>+</td><td>0</td><td>GGAGCCCGCGTCAGAGCGCC</td><td>CGG</td><td>||||||||[||||||||||||]PAM</td><td>NA</td><td>Intergenic</td><td>NA</td><td>NA</td></tr>
<tr><td>T14896</td><td>GCGCCCGGTCCCGTCCAGCGCGG</td><td>1000</td><td></td><td></td><td></td><td></td><td></td><td></td><td></td><td></td><td></td></tr>
<tr><td>Oligo fwd</td><td>CACCGCGCCCGGTCCCGTCCAGCG</td><td></td><td></td><td></td><td></td><td></td><td></td><td></td><td></td><td></td><td></td></tr>
<tr><td>Oligo rev</td><td>AAACCGCTGGACGGGACCGGGCGC</td><td></td><td></td><td></td><td></td><td></td><td></td><td></td><td></td><td></td><td></td></tr>
<tr><td>Chromosome</td><td>start</td><td>end</td><td>strand</td><td>MM</td><td>target_seq</td><td>PAM</td><td>alignment</td><td>distance</td><td>position</td><td>gene name</td><td>gene id</td></tr>
<tr><td>GU305914.1</td><td>195814</td><td>195836</td><td>+</td><td>0</td><td>GCGCCCGGTCCCGTCCAGCG</td><td>CGG</td><td>||||||||[||||||||||||]PAM</td><td>NA</td><td>Intergenic</td><td>NA</td><td>NA</td></tr>
<tr><td>T14897</td><td>GCGCGGCGCGATCTATGAACAGG</td><td>1000</td><td></td><td></td><td></td><td></td><td></td><td></td><td></td><td></td><td></td></tr>
<tr><td>Oligo fwd</td><td>CACCGCGCGGCGCGATCTATGAAC</td><td></td><td></td><td></td><td></td><td></td><td></td><td></td><td></td><td></td><td></td></tr>
<tr><td>Oligo rev</td><td>AAACGTTCATAGATCGCGCCGCGC</td><td></td><td></td><td></td><td></td><td></td><td></td><td></td><td></td><td></td><td></td></tr>
<tr><td>Chromosome</td><td>start</td><td>end</td><td>strand</td><td>MM</td><td>target_seq</td><td>PAM</td><td>alignment</td><td>distance</td><td>position</td><td>gene name</td><td>gene id</td></tr>
<tr><td>GU305914.1</td><td>195831</td><td>195853</td><td>+</td><td>0</td><td>GCGCGGCGCGATCTATGAAC</td><td>AGG</td><td>||||||||[||||||||||||]PAM</td><td>NA</td><td>Intergenic</td><td>NA</td><td>NA</td></tr>
<tr><td>T14898</td><td>AGACTCCGCCATCGCGCCGCAGG</td><td>1000</td><td></td><td></td><td></td><td></td><td></td><td></td><td></td><td></td><td></td></tr>
<tr><td>Oligo fwd</td><td>CACCAGACTCCGCCATCGCGCCGC</td><td></td><td></td><td></td><td></td><td></td><td></td><td></td><td></td><td></td><td></td></tr>
<tr><td>Oligo rev</td><td>AAACGCGGCGCGATGGCGGAGTCT</td><td></td><td></td><td></td><td></td><td></td><td></td><td></td><td></td><td></td><td></td></tr>
<tr><td>Chromosome</td><td>start</td><td>end</td><td>strand</td><td>MM</td><td>target_seq</td><td>PAM</td><td>alignment</td><td>distance</td><td>position</td><td>gene name</td><td>gene id</td></tr>
<tr><td>GU305914.1</td><td>195856</td><td>195878</td><td>+</td><td>0</td><td>AGACTCCGCCATCGCGCCGC</td><td>AGG</td><td>||||||||[||||||||||||]PAM</td><td>NA</td><td>Intergenic</td><td>NA</td><td>NA</td></tr>
<tr><td>T14899</td><td>GCCGCAGGATTCGCGTCCCCCGG</td><td>1000</td><td></td><td></td><td></td><td></td><td></td><td></td><td></td><td></td><td></td></tr>
<tr><td>Oligo fwd</td><td>CACCGCCGCAGGATTCGCGTCCCC</td><td></td><td></td><td></td><td></td><td></td><td></td><td></td><td></td><td></td><td></td></tr>
<tr><td>Oligo rev</td><td>AAACGGGGACGCGAATCCTGCGGC</td><td></td><td></td><td></td><td></td><td></td><td></td><td></td><td></td><td></td><td></td></tr>
<tr><td>Chromosome</td><td>start</td><td>end</td><td>strand</td><td>MM</td><td>target_seq</td><td>PAM</td><td>alignment</td><td>distance</td><td>position</td><td>gene name</td><td>gene id</td></tr>
<tr><td>GU305914.1</td><td>195871</td><td>195893</td><td>+</td><td>0</td><td>GCCGCAGGATTCGCGTCCCC</td><td>CGG</td><td>||||||||[||||||||||||]PAM</td><td>NA</td><td>Intergenic</td><td>NA</td><td>NA</td></tr>
<tr><td>T14900</td><td>TCCCCCGGCCGTGCGATCCGTGG</td><td>1000</td><td></td><td></td><td></td><td></td><td></td><td></td><td></td><td></td><td></td></tr>
<tr><td>Oligo fwd</td><td>CACCTCCCCCGGCCGTGCGATCCG</td><td></td><td></td><td></td><td></td><td></td><td></td><td></td><td></td><td></td><td></td></tr>
<tr><td>Oligo rev</td><td>AAACCGGATCGCACGGCCGGGGGA</td><td></td><td></td><td></td><td></td><td></td><td></td><td></td><td></td><td></td><td></td></tr>
<tr><td>Chromosome</td><td>start</td><td>end</td><td>strand</td><td>MM</td><td>target_seq</td><td>PAM</td><td>alignment</td><td>distance</td><td>position</td><td>gene name</td><td>gene id</td></tr>
<tr><td>GU305914.1</td><td>195886</td><td>195908</td><td>+</td><td>0</td><td>TCCCCCGGCCGTGCGATCCG</td><td>TGG</td><td>||||||||[||||||||||||]PAM</td><td>NA</td><td>Intergenic</td><td>NA</td><td>NA</td></tr>
<tr><td>T14901</td><td>GTGCGATCCGTGGAGTCCAATGG</td><td>1000</td><td></td><td></td><td></td><td></td><td></td><td></td><td></td><td></td><td></td></tr>
<tr><td>Oligo fwd</td><td>CACCGTGCGATCCGTGGAGTCCAA</td><td></td><td></td><td></td><td></td><td></td><td></td><td></td><td></td><td></td><td></td></tr>
<tr><td>Oligo rev</td><td>AAACTTGGACTCCACGGATCGCAC</td><td></td><td></td><td></td><td></td><td></td><td></td><td></td><td></td><td></td><td></td></tr>
<tr><td>Chromosome</td><td>start</td><td>end</td><td>strand</td><td>MM</td><td>target_seq</td><td>PAM</td><td>alignment</td><td>distance</td><td>position</td><td>gene name</td><td>gene id</td></tr>
<tr><td>GU305914.1</td><td>195896</td><td>195918</td><td>+</td><td>0</td><td>GTGCGATCCGTGGAGTCCAA</td><td>TGG</td><td>||||||||[||||||||||||]PAM</td><td>NA</td><td>Intergenic</td><td>NA</td><td>NA</td></tr>
<tr><td>T14902</td><td>GATCCGTGGAGTCCAATGGCAGG</td><td>1000</td><td></td><td></td><td></td><td></td><td></td><td></td><td></td><td></td><td></td></tr>
<tr><td>Oligo fwd</td><td>CACCGATCCGTGGAGTCCAATGGC</td><td></td><td></td><td></td><td></td><td></td><td></td><td></td><td></td><td></td><td></td></tr>
<tr><td>Oligo rev</td><td>AAACGCCATTGGACTCCACGGATC</td><td></td><td></td><td></td><td></td><td></td><td></td><td></td><td></td><td></td><td></td></tr>
<tr><td>Chromosome</td><td>start</td><td>end</td><td>strand</td><td>MM</td><td>target_seq</td><td>PAM</td><td>alignment</td><td>distance</td><td>position</td><td>gene name</td><td>gene id</td></tr>
<tr><td>GU305914.1</td><td>195900</td><td>195922</td><td>+</td><td>0</td><td>GATCCGTGGAGTCCAATGGC</td><td>AGG</td><td>||||||||[||||||||||||]PAM</td><td>NA</td><td>Intergenic</td><td>NA</td><td>NA</td></tr>
<tr><td>T14903</td><td>GGAGTCCAATGGCAGGCCGAAGG</td><td>1000</td><td></td><td></td><td></td><td></td><td></td><td></td><td></td><td></td><td></td></tr>
<tr><td>Oligo fwd</td><td>CACCGGAGTCCAATGGCAGGCCGA</td><td></td><td></td><td></td><td></td><td></td><td></td><td></td><td></td><td></td><td></td></tr>
<tr><td>Oligo rev</td><td>AAACTCGGCCTGCCATTGGACTCC</td><td></td><td></td><td></td><td></td><td></td><td></td><td></td><td></td><td></td><td></td></tr>
<tr><td>Chromosome</td><td>start</td><td>end</td><td>strand</td><td>MM</td><td>target_seq</td><td>PAM</td><td>alignment</td><td>distance</td><td>position</td><td>gene name</td><td>gene id</td></tr>
<tr><td>GU305914.1</td><td>195907</td><td>195929</td><td>+</td><td>0</td><td>GGAGTCCAATGGCAGGCCGA</td><td>AGG</td><td>||||||||[||||||||||||]PAM</td><td>NA</td><td>Intergenic</td><td>NA</td><td>NA</td></tr>
<tr><td>T14904</td><td>ATGGCAGGCCGAAGGAAAACTGG</td><td>1000</td><td></td><td></td><td></td><td></td><td></td><td></td><td></td><td></td><td></td></tr>
<tr><td>Oligo fwd</td><td>CACCATGGCAGGCCGAAGGAAAAC</td><td></td><td></td><td></td><td></td><td></td><td></td><td></td><td></td><td></td><td></td></tr>
<tr><td>Oligo rev</td><td>AAACGTTTTCCTTCGGCCTGCCAT</td><td></td><td></td><td></td><td></td><td></td><td></td><td></td><td></td><td></td><td></td></tr>
<tr><td>Chromosome</td><td>start</td><td>end</td><td>strand</td><td>MM</td><td>target_seq</td><td>PAM</td><td>alignment</td><td>distance</td><td>position</td><td>gene name</td><td>gene id</td></tr>
<tr><td>GU305914.1</td><td>195915</td><td>195937</td><td>+</td><td>0</td><td>ATGGCAGGCCGAAGGAAAAC</td><td>TGG</td><td>||||||||[||||||||||||]PAM</td><td>NA</td><td>Intergenic</td><td>NA</td><td>NA</td></tr>
<tr><td>T14905</td><td>TGGTCATTTATAAAGCTTCTTGG</td><td>1000</td><td></td><td></td><td></td><td></td><td></td><td></td><td></td><td></td><td></td></tr>
<tr><td>Oligo fwd</td><td>CACCTGGTCATTTATAAAGCTTCT</td><td></td><td></td><td></td><td></td><td></td><td></td><td></td><td></td><td></td><td></td></tr>
<tr><td>Oligo rev</td><td>AAACAGAAGCTTTATAAATGACCA</td><td></td><td></td><td></td><td></td><td></td><td></td><td></td><td></td><td></td><td></td></tr>
<tr><td>Chromosome</td><td>start</td><td>end</td><td>strand</td><td>MM</td><td>target_seq</td><td>PAM</td><td>alignment</td><td>distance</td><td>position</td><td>gene name</td><td>gene id</td></tr>
</table>

<table>
<tr><td>GU305914.1</td><td>195935</td><td>195957</td><td>+</td><td>0</td><td>TGGTCATTTATAAAGCTTCT</td><td>TGG</td><td>||||||||[||||||||||||]PAM</td><td>NA</td><td>Intergenic</td><td>NA</td><td>NA</td></tr>
<tr><td>T14906</td><td>GGTCATTTATAAAGCTTCTTGGG</td><td>1000</td><td></td><td></td><td></td><td></td><td></td><td></td><td></td><td></td><td></td></tr>
<tr><td>Oligo fwd</td><td>CACCGGTCATTTATAAAGCTTCTT</td><td></td><td></td><td></td><td></td><td></td><td></td><td></td><td></td><td></td><td></td></tr>
<tr><td>Oligo rev</td><td>AAACAAGAAGCTTTATAAATGACC</td><td></td><td></td><td></td><td></td><td></td><td></td><td></td><td></td><td></td><td></td></tr>
<tr><td>Chromosome</td><td>start</td><td>end</td><td>strand</td><td>MM</td><td>target_seq</td><td>PAM</td><td>alignment</td><td>distance</td><td>position</td><td>gene name</td><td>gene id</td></tr>
<tr><td>GU305914.1</td><td>195936</td><td>195958</td><td>+</td><td>0</td><td>GGTCATTTATAAAGCTTCTT</td><td>GGG</td><td>||||||||[||||||||||||]PAM</td><td>NA</td><td>Intergenic</td><td>NA</td><td>NA</td></tr>
<tr><td>T14907</td><td>GTCATTTATAAAGCTTCTTGGGG</td><td>1000</td><td></td><td></td><td></td><td></td><td></td><td></td><td></td><td></td><td></td></tr>
<tr><td>Oligo fwd</td><td>CACCGTCATTTATAAAGCTTCTTG</td><td></td><td></td><td></td><td></td><td></td><td></td><td></td><td></td><td></td><td></td></tr>
<tr><td>Oligo rev</td><td>AAACCAAGAAGCTTTATAAATGAC</td><td></td><td></td><td></td><td></td><td></td><td></td><td></td><td></td><td></td><td></td></tr>
<tr><td>Chromosome</td><td>start</td><td>end</td><td>strand</td><td>MM</td><td>target_seq</td><td>PAM</td><td>alignment</td><td>distance</td><td>position</td><td>gene name</td><td>gene id</td></tr>
<tr><td>GU305914.1</td><td>195937</td><td>195959</td><td>+</td><td>0</td><td>GTCATTTATAAAGCTTCTTG</td><td>GGG</td><td>||||||||[||||||||||||]PAM</td><td>NA</td><td>Intergenic</td><td>NA</td><td>NA</td></tr>
<tr><td>T14908</td><td>TCATTTATAAAGCTTCTTGGGGG</td><td>1000</td><td></td><td></td><td></td><td></td><td></td><td></td><td></td><td></td><td></td></tr>
<tr><td>Oligo fwd</td><td>CACCTCATTTATAAAGCTTCTTGG</td><td></td><td></td><td></td><td></td><td></td><td></td><td></td><td></td><td></td><td></td></tr>
<tr><td>Oligo rev</td><td>AAACCCAAGAAGCTTTATAAATGA</td><td></td><td></td><td></td><td></td><td></td><td></td><td></td><td></td><td></td><td></td></tr>
<tr><td>Chromosome</td><td>start</td><td>end</td><td>strand</td><td>MM</td><td>target_seq</td><td>PAM</td><td>alignment</td><td>distance</td><td>position</td><td>gene name</td><td>gene id</td></tr>
<tr><td>GU305914.1</td><td>195938</td><td>195960</td><td>+</td><td>0</td><td>TCATTTATAAAGCTTCTTGG</td><td>GGG</td><td>||||||||[||||||||||||]PAM</td><td>NA</td><td>Intergenic</td><td>NA</td><td>NA</td></tr>
<tr><td>T14909</td><td>CATTTATAAAGCTTCTTGGGGGG</td><td>1000</td><td></td><td></td><td></td><td></td><td></td><td></td><td></td><td></td><td></td></tr>
<tr><td>Oligo fwd</td><td>CACCCATTTATAAAGCTTCTTGGG</td><td></td><td></td><td></td><td></td><td></td><td></td><td></td><td></td><td></td><td></td></tr>
<tr><td>Oligo rev</td><td>AAACCCCAAGAAGCTTTATAAATG</td><td></td><td></td><td></td><td></td><td></td><td></td><td></td><td></td><td></td><td></td></tr>
<tr><td>Chromosome</td><td>start</td><td>end</td><td>strand</td><td>MM</td><td>target_seq</td><td>PAM</td><td>alignment</td><td>distance</td><td>position</td><td>gene name</td><td>gene id</td></tr>
<tr><td>GU305914.1</td><td>195939</td><td>195961</td><td>+</td><td>0</td><td>CATTTATAAAGCTTCTTGGG</td><td>GGG</td><td>||||||||[||||||||||||]PAM</td><td>NA</td><td>Intergenic</td><td>NA</td><td>NA</td></tr>
<tr><td>T14910</td><td>AAGCTTCTTGGGGGGTAAACTGG</td><td>1000</td><td></td><td></td><td></td><td></td><td></td><td></td><td></td><td></td><td></td></tr>
<tr><td>Oligo fwd</td><td>CACCAAGCTTCTTGGGGGGTAAAC</td><td></td><td></td><td></td><td></td><td></td><td></td><td></td><td></td><td></td><td></td></tr>
<tr><td>Oligo rev</td><td>AAACGTTTACCCCCCAAGAAGCTT</td><td></td><td></td><td></td><td></td><td></td><td></td><td></td><td></td><td></td><td></td></tr>
<tr><td>Chromosome</td><td>start</td><td>end</td><td>strand</td><td>MM</td><td>target_seq</td><td>PAM</td><td>alignment</td><td>distance</td><td>position</td><td>gene name</td><td>gene id</td></tr>
<tr><td>GU305914.1</td><td>195947</td><td>195969</td><td>+</td><td>0</td><td>AAGCTTCTTGGGGGGTAAAC</td><td>TGG</td><td>||||||||[||||||||||||]PAM</td><td>NA</td><td>Intergenic</td><td>NA</td><td>NA</td></tr>
<tr><td>T14911</td><td>GGGTAAACTGGTGACCTGATCGG</td><td>1000</td><td></td><td></td><td></td><td></td><td></td><td></td><td></td><td></td><td></td></tr>
<tr><td>Oligo fwd</td><td>CACCGGGTAAACTGGTGACCTGAT</td><td></td><td></td><td></td><td></td><td></td><td></td><td></td><td></td><td></td><td></td></tr>
<tr><td>Oligo rev</td><td>AAACATCAGGTCACCAGTTTACCC</td><td></td><td></td><td></td><td></td><td></td><td></td><td></td><td></td><td></td><td></td></tr>
<tr><td>Chromosome</td><td>start</td><td>end</td><td>strand</td><td>MM</td><td>target_seq</td><td>PAM</td><td>alignment</td><td>distance</td><td>position</td><td>gene name</td><td>gene id</td></tr>
<tr><td>GU305914.1</td><td>195959</td><td>195981</td><td>+</td><td>0</td><td>GGGTAAACTGGTGACCTGAT</td><td>CGG</td><td>||||||||[||||||||||||]PAM</td><td>NA</td><td>Intergenic</td><td>NA</td><td>NA</td></tr>
<tr><td>T14912</td><td>GGTAAACTGGTGACCTGATCGGG</td><td>1000</td><td></td><td></td><td></td><td></td><td></td><td></td><td></td><td></td><td></td></tr>
<tr><td>Oligo fwd</td><td>CACCGGTAAACTGGTGACCTGATC</td><td></td><td></td><td></td><td></td><td></td><td></td><td></td><td></td><td></td><td></td></tr>
<tr><td>Oligo rev</td><td>AAACGATCAGGTCACCAGTTTACC</td><td></td><td></td><td></td><td></td><td></td><td></td><td></td><td></td><td></td><td></td></tr>
<tr><td>Chromosome</td><td>start</td><td>end</td><td>strand</td><td>MM</td><td>target_seq</td><td>PAM</td><td>alignment</td><td>distance</td><td>position</td><td>gene name</td><td>gene id</td></tr>
<tr><td>GU305914.1</td><td>195960</td><td>195982</td><td>+</td><td>0</td><td>GGTAAACTGGTGACCTGATC</td><td>GGG</td><td>||||||||[||||||||||||]PAM</td><td>NA</td><td>Intergenic</td><td>NA</td><td>NA</td></tr>
<tr><td>T14913</td><td>GTAAACTGGTGACCTGATCGGGG</td><td>1000</td><td></td><td></td><td></td><td></td><td></td><td></td><td></td><td></td><td></td></tr>
<tr><td>Oligo fwd</td><td>CACCGTAAACTGGTGACCTGATCG</td><td></td><td></td><td></td><td></td><td></td><td></td><td></td><td></td><td></td><td></td></tr>
<tr><td>Oligo rev</td><td>AAACCGATCAGGTCACCAGTTTAC</td><td></td><td></td><td></td><td></td><td></td><td></td><td></td><td></td><td></td><td></td></tr>
<tr><td>Chromosome</td><td>start</td><td>end</td><td>strand</td><td>MM</td><td>target_seq</td><td>PAM</td><td>alignment</td><td>distance</td><td>position</td><td>gene name</td><td>gene id</td></tr>
<tr><td>GU305914.1</td><td>195961</td><td>195983</td><td>+</td><td>0</td><td>GTAAACTGGTGACCTGATCG</td><td>GGG</td><td>||||||||[||||||||||||]PAM</td><td>NA</td><td>Intergenic</td><td>NA</td><td>NA</td></tr>
<tr><td>T14914</td><td>TAAACTGGTGACCTGATCGGGGG</td><td>1000</td><td></td><td></td><td></td><td></td><td></td><td></td><td></td><td></td><td></td></tr>
<tr><td>Oligo fwd</td><td>CACCTAAACTGGTGACCTGATCGG</td><td></td><td></td><td></td><td></td><td></td><td></td><td></td><td></td><td></td><td></td></tr>
<tr><td>Oligo rev</td><td>AAACCCGATCAGGTCACCAGTTTA</td><td></td><td></td><td></td><td></td><td></td><td></td><td></td><td></td><td></td><td></td></tr>
<tr><td>Chromosome</td><td>start</td><td>end</td><td>strand</td><td>MM</td><td>target_seq</td><td>PAM</td><td>alignment</td><td>distance</td><td>position</td><td>gene name</td><td>gene id</td></tr>
<tr><td>GU305914.1</td><td>195962</td><td>195984</td><td>+</td><td>0</td><td>TAAACTGGTGACCTGATCGG</td><td>GGG</td><td>||||||||[||||||||||||]PAM</td><td>NA</td><td>Intergenic</td><td>NA</td><td>NA</td></tr>
<tr><td>T14915</td><td>CTGGTGACCTGATCGGGGGAAGG</td><td>1000</td><td></td><td></td><td></td><td></td><td></td><td></td><td></td><td></td><td></td></tr>
<tr><td>Oligo fwd</td><td>CACCCTGGTGACCTGATCGGGGGA</td><td></td><td></td><td></td><td></td><td></td><td></td><td></td><td></td><td></td><td></td></tr>
<tr><td>Oligo rev</td><td>AAACTCCCCCGATCAGGTCACCAG</td><td></td><td></td><td></td><td></td><td></td><td></td><td></td><td></td><td></td><td></td></tr>
<tr><td>Chromosome</td><td>start</td><td>end</td><td>strand</td><td>MM</td><td>target_seq</td><td>PAM</td><td>alignment</td><td>distance</td><td>position</td><td>gene name</td><td>gene id</td></tr>
<tr><td>GU305914.1</td><td>195966</td><td>195988</td><td>+</td><td>0</td><td>CTGGTGACCTGATCGGGGGA</td><td>AGG</td><td>||||||||[||||||||||||]PAM</td><td>NA</td><td>Intergenic</td><td>NA</td><td>NA</td></tr>
<tr><td>T14916</td><td>CGCCGTATTCTCCGAACTTCTGG</td><td>1000</td><td></td><td></td><td></td><td></td><td></td><td></td><td></td><td></td><td></td></tr>
<tr><td>Oligo fwd</td><td>CACCCGCCGTATTCTCCGAACTTC</td><td></td><td></td><td></td><td></td><td></td><td></td><td></td><td></td><td></td><td></td></tr>
<tr><td>Oligo rev</td><td>AAACGAAGTTCGGAGAATACGGCG</td><td></td><td></td><td></td><td></td><td></td><td></td><td></td><td></td><td></td><td></td></tr>
<tr><td>Chromosome</td><td>start</td><td>end</td><td>strand</td><td>MM</td><td>target_seq</td><td>PAM</td><td>alignment</td><td>distance</td><td>position</td><td>gene name</td><td>gene id</td></tr>
<tr><td>GU305914.1</td><td>196005</td><td>196027</td><td>+</td><td>0</td><td>CGCCGTATTCTCCGAACTTC</td><td>TGG</td><td>||||||||[||||||||||||]PAM</td><td>NA</td><td>Intergenic</td><td>NA</td><td>NA</td></tr>
<tr><td>T14917</td><td>CGTCCTGACCATCGAAACGCAGG</td><td>1000</td><td></td><td></td><td></td><td></td><td></td><td></td><td></td><td></td><td></td></tr>
<tr><td>Oligo fwd</td><td>CACCCGTCCTGACCATCGAAACGC</td><td></td><td></td><td></td><td></td><td></td><td></td><td></td><td></td><td></td><td></td></tr>
<tr><td>Oligo rev</td><td>AAACGCGTTTCGATGGTCAGGACG</td><td></td><td></td><td></td><td></td><td></td><td></td><td></td><td></td><td></td><td></td></tr>
<tr><td>Chromosome</td><td>start</td><td>end</td><td>strand</td><td>MM</td><td>target_seq</td><td>PAM</td><td>alignment</td><td>distance</td><td>position</td><td>gene name</td><td>gene id</td></tr>
<tr><td>GU305914.1</td><td>196062</td><td>196084</td><td>+</td><td>0</td><td>CGTCCTGACCATCGAAACGC</td><td>AGG</td><td>||||||||[||||||||||||]PAM</td><td>NA</td><td>Intergenic</td><td>NA</td><td>NA</td></tr>
<tr><td>T14918</td><td>CGAAACGCAGGTCGCGTCTCTGG</td><td>1000</td><td></td><td></td><td></td><td></td><td></td><td></td><td></td><td></td><td></td></tr>
<tr><td>Oligo fwd</td><td>CACCCGAAACGCAGGTCGCGTCTC</td><td></td><td></td><td></td><td></td><td></td><td></td><td></td><td></td><td></td><td></td></tr>
<tr><td>Oligo rev</td><td>AAACGAGACGCGACCTGCGTTTCG</td><td></td><td></td><td></td><td></td><td></td><td></td><td></td><td></td><td></td><td></td></tr>
<tr><td>Chromosome</td><td>start</td><td>end</td><td>strand</td><td>MM</td><td>target_seq</td><td>PAM</td><td>alignment</td><td>distance</td><td>position</td><td>gene name</td><td>gene id</td></tr>
<tr><td>GU305914.1</td><td>196074</td><td>196096</td><td>+</td><td>0</td><td>CGAAACGCAGGTCGCGTCTC</td><td>TGG</td><td>||||||||[||||||||||||]PAM</td><td>NA</td><td>Intergenic</td><td>NA</td><td>NA</td></tr>
<tr><td>T14919</td><td>GAAACGCAGGTCGCGTCTCTGGG</td><td>1000</td><td></td><td></td><td></td><td></td><td></td><td></td><td></td><td></td><td></td></tr>
<tr><td>Oligo fwd</td><td>CACCGAAACGCAGGTCGCGTCTCT</td><td></td><td></td><td></td><td></td><td></td><td></td><td></td><td></td><td></td><td></td></tr>
<tr><td>Oligo rev</td><td>AAACAGAGACGCGACCTGCGTTTC</td><td></td><td></td><td></td><td></td><td></td><td></td><td></td><td></td><td></td><td></td></tr>
<tr><td>Chromosome</td><td>start</td><td>end</td><td>strand</td><td>MM</td><td>target_seq</td><td>PAM</td><td>alignment</td><td>distance</td><td>position</td><td>gene name</td><td>gene id</td></tr>
<tr><td>GU305914.1</td><td>196075</td><td>196097</td><td>+</td><td>0</td><td>GAAACGCAGGTCGCGTCTCT</td><td>GGG</td><td>||||||||[||||||||||||]PAM</td><td>NA</td><td>Intergenic</td><td>NA</td><td>NA</td></tr>
<tr><td>T14920</td><td>GGTCGCGTCTCTGGGCGCCGCGG</td><td>1000</td><td></td><td></td><td></td><td></td><td></td><td></td><td></td><td></td><td></td></tr>
<tr><td>Oligo fwd</td><td>CACCGGTCGCGTCTCTGGGCGCCG</td><td></td><td></td><td></td><td></td><td></td><td></td><td></td><td></td><td></td><td></td></tr>
<tr><td>Oligo rev</td><td>AAACCGGCGCCCAGAGACGCGACC</td><td></td><td></td><td></td><td></td><td></td><td></td><td></td><td></td><td></td><td></td></tr>
</table>

Chromosome start end strand MM target_seq PAM alignment distance position gene name gene id
GU305914.1 196083 196105 + 0 GGTCGCGTCTCTGGGCGCCG CGG ||||||||[||||||||||||]PAM NA Intergenic NA NA

T14921 CTCTCTCTGTCTGCTCCGTCTGG 1000
Oligo fwd CACCCTCTCTCTGTCTGCTCCGTC
Oligo rev AAACGACGGAGCAGACAGAGAGAG
Chromosome start end strand MM target_seq PAM alignment distance position gene name gene id
GU305914.1 196123 196145 + 0 CTCTCTCTGTCTGCTCCGTC TGG ||||||||[||||||||||||]PAM NA Intergenic NA NA

T14922 CTGTCTGCTCCGTCTGGCGCCGG 1000
Oligo fwd CACCCTGTCTGCTCCGTCTGGCGC
Oligo rev AAACGCGCCAGACGGAGCAGACAG
Chromosome start end strand MM target_seq PAM alignment distance position gene name gene id
GU305914.1 196129 196151 + 0 CTGTCTGCTCCGTCTGGCGC CGG ||||||||[||||||||||||]PAM NA Intergenic NA NA

T14923 GCGCCGGCGCCGTCAACTCACGG 1000
Oligo fwd CACCGCGCCGGCGCCGTCAACTCA
Oligo rev AAACTGAGTTGACGGCGCCGGCGC
Chromosome start end strand MM target_seq PAM alignment distance position gene name gene id
GU305914.1 196145 196167 + 0 GCGCCGGCGCCGTCAACTCA CGG ||||||||[||||||||||||]PAM NA Intergenic NA NA

T14924 CACGGTCTGCAGAGTCTGAGCGG 1000
Oligo fwd CACCCACGGTCTGCAGAGTCTGAG
Oligo rev AAACCTCAGACTCTGCAGACCGTG
Chromosome start end strand MM target_seq PAM alignment distance position gene name gene id
GU305914.1 196163 196185 + 0 CACGGTCTGCAGAGTCTGAG CGG ||||||||[||||||||||||]PAM NA Intergenic NA NA

T14925 TTGCCTCTCCAAGAGCCTTCTGG 1000
Oligo fwd CACCTTGCCTCTCCAAGAGCCTTC
Oligo rev AAACGAAGGCTCTTGGAGAGGCAA
Chromosome start end strand MM target_seq PAM alignment distance position gene name gene id
GU305914.1 196210 196232 + 0 TTGCCTCTCCAAGAGCCTTC TGG ||||||||[||||||||||||]PAM NA Intergenic NA NA

T14926 TCTCTGACATGTAACGCAACAGG 1000
Oligo fwd CACCTCTCTGACATGTAACGCAAC
Oligo rev AAACGTTGCGTTACATGTCAGAGA
Chromosome start end strand MM target_seq PAM alignment distance position gene name gene id
GU305914.1 196260 196282 + 0 TCTCTGACATGTAACGCAAC AGG ||||||||[||||||||||||]PAM NA Intergenic NA NA

T14927 CTCTGACATGTAACGCAACAGGG 1000
Oligo fwd CACCCTCTGACATGTAACGCAACA
Oligo rev AAACTGTTGCGTTACATGTCAGAG
Chromosome start end strand MM target_seq PAM alignment distance position gene name gene id
GU305914.1 196261 196283 + 0 CTCTGACATGTAACGCAACA GGG ||||||||[||||||||||||]PAM NA Intergenic NA NA

T14928 ACATGTAACGCAACAGGGTTCGG 1000
Oligo fwd CACCACATGTAACGCAACAGGGTT
Oligo rev AAACAACCCTGTTGCGTTACATGT
Chromosome start end strand MM target_seq PAM alignment distance position gene name gene id
GU305914.1 196266 196288 + 0 ACATGTAACGCAACAGGGTT CGG ||||||||[||||||||||||]PAM NA Intergenic NA NA

T14929 CATGTAACGCAACAGGGTTCGGG 1000
Oligo fwd CACCCATGTAACGCAACAGGGTTC
Oligo rev AAACGAACCCTGTTGCGTTACATG
Chromosome start end strand MM target_seq PAM alignment distance position gene name gene id
GU305914.1 196267 196289 + 0 CATGTAACGCAACAGGGTTC GGG ||||||||[||||||||||||]PAM NA Intergenic NA NA

T14930 GCAACAGGGTTCGGGTGTCTCGG 1000
Oligo fwd CACCGCAACAGGGTTCGGGTGTCT
Oligo rev AAACAGACACCCGAACCCTGTTGC
Chromosome start end strand MM target_seq PAM alignment distance position gene name gene id
GU305914.1 196275 196297 + 0 GCAACAGGGTTCGGGTGTCT CGG ||||||||[||||||||||||]PAM NA Intergenic NA NA

T14931 CGGGTGTCTCGGTTGCGATCCGG 1000
Oligo fwd CACCCGGGTGTCTCGGTTGCGATC
Oligo rev AAACGATCGCAACCGAGACACCCG
Chromosome start end strand MM target_seq PAM alignment distance position gene name gene id
GU305914.1 196286 196308 + 0 CGGGTGTCTCGGTTGCGATC CGG ||||||||[||||||||||||]PAM NA Intergenic NA NA

T14932 GGGTGTCTCGGTTGCGATCCGGG 1000
Oligo fwd CACCGGGTGTCTCGGTTGCGATCC
Oligo rev AAACGGATCGCAACCGAGACACCC
Chromosome start end strand MM target_seq PAM alignment distance position gene name gene id
GU305914.1 196287 196309 + 0 GGGTGTCTCGGTTGCGATCC GGG ||||||||[||||||||||||]PAM NA Intergenic NA NA

T14933 GCGTGTCGTTAGCACGCTCTCGG 1000
Oligo fwd CACCGCGTGTCGTTAGCACGCTCT
Oligo rev AAACAGAGCGTGCTAACGACACGC
Chromosome start end strand MM target_seq PAM alignment distance position gene name gene id
GU305914.1 196309 196331 + 0 GCGTGTCGTTAGCACGCTCT CGG ||||||||[||||||||||||]PAM NA Intergenic NA NA

T14934 CGTGTCGTTAGCACGCTCTCGGG 1000
Oligo fwd CACCCGTGTCGTTAGCACGCTCTC
Oligo rev AAACGAGAGCGTGCTAACGACACG
Chromosome start end strand MM target_seq PAM alignment distance position gene name gene id
GU305914.1 196310 196332 + 0 CGTGTCGTTAGCACGCTCTC GGG ||||||||[||||||||||||]PAM NA Intergenic NA NA

T14935 GCATGAGCGCGAGCCATCGTCGG 1000
Oligo fwd CACCGCATGAGCGCGAGCCATCGT

<table>
<tr><td>Oligo rev</td><td>AAACACGATGGCTCGCGCTCATGC</td><td></td><td></td><td></td><td></td><td></td><td></td><td></td><td></td><td></td><td></td></tr>
<tr><td>Chromosome</td><td>start</td><td>end</td><td>strand</td><td>MM</td><td>target_seq</td><td>PAM</td><td>alignment</td><td>distance</td><td>position</td><td>gene name</td><td>gene id</td></tr>
<tr><td>GU305914.1</td><td>196342</td><td>196364</td><td>+</td><td>0</td><td>GCATGAGCGCGAGCCATCGT</td><td>CGG</td><td>||||||||[||||||||||||]PAM</td><td>NA</td><td>Intergenic</td><td>NA</td><td>NA</td></tr>
<tr><td></td><td></td><td></td><td></td><td></td><td></td><td></td><td></td><td></td><td></td><td></td><td></td></tr>
<tr><td>T14936</td><td>CATGAGCGCGAGCCATCGTCGGG</td><td>1000</td><td></td><td></td><td></td><td></td><td></td><td></td><td></td><td></td><td></td></tr>
<tr><td>Oligo fwd</td><td>CACCCATGAGCGCGAGCCATCGTC</td><td></td><td></td><td></td><td></td><td></td><td></td><td></td><td></td><td></td><td></td></tr>
<tr><td>Oligo rev</td><td>AAACGACGATGGCTCGCGCTCATG</td><td></td><td></td><td></td><td></td><td></td><td></td><td></td><td></td><td></td><td></td></tr>
<tr><td>Chromosome</td><td>start</td><td>end</td><td>strand</td><td>MM</td><td>target_seq</td><td>PAM</td><td>alignment</td><td>distance</td><td>position</td><td>gene name</td><td>gene id</td></tr>
<tr><td>GU305914.1</td><td>196343</td><td>196365</td><td>+</td><td>0</td><td>CATGAGCGCGAGCCATCGTC</td><td>GGG</td><td>||||||||[||||||||||||]PAM</td><td>NA</td><td>Intergenic</td><td>NA</td><td>NA</td></tr>
<tr><td></td><td></td><td></td><td></td><td></td><td></td><td></td><td></td><td></td><td></td><td></td><td></td></tr>
<tr><td>T14937</td><td>ATGAGCGCGAGCCATCGTCGGGG</td><td>1000</td><td></td><td></td><td></td><td></td><td></td><td></td><td></td><td></td><td></td></tr>
<tr><td>Oligo fwd</td><td>CACCATGAGCGCGAGCCATCGTCG</td><td></td><td></td><td></td><td></td><td></td><td></td><td></td><td></td><td></td><td></td></tr>
<tr><td>Oligo rev</td><td>AAACCGACGATGGCTCGCGCTCAT</td><td></td><td></td><td></td><td></td><td></td><td></td><td></td><td></td><td></td><td></td></tr>
<tr><td>Chromosome</td><td>start</td><td>end</td><td>strand</td><td>MM</td><td>target_seq</td><td>PAM</td><td>alignment</td><td>distance</td><td>position</td><td>gene name</td><td>gene id</td></tr>
<tr><td>GU305914.1</td><td>196344</td><td>196366</td><td>+</td><td>0</td><td>ATGAGCGCGAGCCATCGTCG</td><td>GGG</td><td>||||||||[||||||||||||]PAM</td><td>NA</td><td>Intergenic</td><td>NA</td><td>NA</td></tr>
<tr><td></td><td></td><td></td><td></td><td></td><td></td><td></td><td></td><td></td><td></td><td></td><td></td></tr>
<tr><td>T14938</td><td>GTCCTCCTGTTCCCACTCGTTGG</td><td>1000</td><td></td><td></td><td></td><td></td><td></td><td></td><td></td><td></td><td></td></tr>
<tr><td>Oligo fwd</td><td>CACCGTCCTCCTGTTCCCACTCGT</td><td></td><td></td><td></td><td></td><td></td><td></td><td></td><td></td><td></td><td></td></tr>
<tr><td>Oligo rev</td><td>AAACACGAGTGGGAACAGGAGGAC</td><td></td><td></td><td></td><td></td><td></td><td></td><td></td><td></td><td></td><td></td></tr>
<tr><td>Chromosome</td><td>start</td><td>end</td><td>strand</td><td>MM</td><td>target_seq</td><td>PAM</td><td>alignment</td><td>distance</td><td>position</td><td>gene name</td><td>gene id</td></tr>
<tr><td>GU305914.1</td><td>196411</td><td>196433</td><td>+</td><td>0</td><td>GTCCTCCTGTTCCCACTCGT</td><td>TGG</td><td>||||||||[||||||||||||]PAM</td><td>NA</td><td>Intergenic</td><td>NA</td><td>NA</td></tr>
<tr><td></td><td></td><td></td><td></td><td></td><td></td><td></td><td></td><td></td><td></td><td></td><td></td></tr>
<tr><td>T14939</td><td>CTCGTTCGTGTACATCTCCCCGG</td><td>1000</td><td></td><td></td><td></td><td></td><td></td><td></td><td></td><td></td><td></td></tr>
<tr><td>Oligo fwd</td><td>CACCCTCGTTCGTGTACATCTCCC</td><td></td><td></td><td></td><td></td><td></td><td></td><td></td><td></td><td></td><td></td></tr>
<tr><td>Oligo rev</td><td>AAACGGGAGATGTACACGAACGAG</td><td></td><td></td><td></td><td></td><td></td><td></td><td></td><td></td><td></td><td></td></tr>
<tr><td>Chromosome</td><td>start</td><td>end</td><td>strand</td><td>MM</td><td>target_seq</td><td>PAM</td><td>alignment</td><td>distance</td><td>position</td><td>gene name</td><td>gene id</td></tr>
<tr><td>GU305914.1</td><td>196453</td><td>196475</td><td>+</td><td>0</td><td>CTCGTTCGTGTACATCTCCC</td><td>CGG</td><td>||||||||[||||||||||||]PAM</td><td>NA</td><td>Intergenic</td><td>NA</td><td>NA</td></tr>
<tr><td></td><td></td><td></td><td></td><td></td><td></td><td></td><td></td><td></td><td></td><td></td><td></td></tr>
<tr><td>T14940</td><td>CTCCCCGGCCTTTCTCTTCATGG</td><td>1000</td><td></td><td></td><td></td><td></td><td></td><td></td><td></td><td></td><td></td></tr>
<tr><td>Oligo fwd</td><td>CACCCTCCCCGGCCTTTCTCTTCA</td><td></td><td></td><td></td><td></td><td></td><td></td><td></td><td></td><td></td><td></td></tr>
<tr><td>Oligo rev</td><td>AAACTGAAGAGAAAGGCCGGGGAG</td><td></td><td></td><td></td><td></td><td></td><td></td><td></td><td></td><td></td><td></td></tr>
<tr><td>Chromosome</td><td>start</td><td>end</td><td>strand</td><td>MM</td><td>target_seq</td><td>PAM</td><td>alignment</td><td>distance</td><td>position</td><td>gene name</td><td>gene id</td></tr>
<tr><td>GU305914.1</td><td>196468</td><td>196490</td><td>+</td><td>0</td><td>CTCCCCGGCCTTTCTCTTCA</td><td>TGG</td><td>||||||||[||||||||||||]PAM</td><td>NA</td><td>Intergenic</td><td>NA</td><td>NA</td></tr>
<tr><td></td><td></td><td></td><td></td><td></td><td></td><td></td><td></td><td></td><td></td><td></td><td></td></tr>
<tr><td>T14941</td><td>CTTTCTCTTCATGGCGCGCACGG</td><td>1000</td><td></td><td></td><td></td><td></td><td></td><td></td><td></td><td></td><td></td></tr>
<tr><td>Oligo fwd</td><td>CACCCTTTCTCTTCATGGCGCGCA</td><td></td><td></td><td></td><td></td><td></td><td></td><td></td><td></td><td></td><td></td></tr>
<tr><td>Oligo rev</td><td>AAACTGCGCGCCATGAAGAGAAAG</td><td></td><td></td><td></td><td></td><td></td><td></td><td></td><td></td><td></td><td></td></tr>
<tr><td>Chromosome</td><td>start</td><td>end</td><td>strand</td><td>MM</td><td>target_seq</td><td>PAM</td><td>alignment</td><td>distance</td><td>position</td><td>gene name</td><td>gene id</td></tr>
<tr><td>GU305914.1</td><td>196477</td><td>196499</td><td>+</td><td>0</td><td>CTTTCTCTTCATGGCGCGCA</td><td>CGG</td><td>||||||||[||||||||||||]PAM</td><td>NA</td><td>Intergenic</td><td>NA</td><td>NA</td></tr>
<tr><td></td><td></td><td></td><td></td><td></td><td></td><td></td><td></td><td></td><td></td><td></td><td></td></tr>
<tr><td>T14942</td><td>TCTTCATGGCGCGCACGGCGTGG</td><td>1000</td><td></td><td></td><td></td><td></td><td></td><td></td><td></td><td></td><td></td></tr>
<tr><td>Oligo fwd</td><td>CACCTCTTCATGGCGCGCACGGCG</td><td></td><td></td><td></td><td></td><td></td><td></td><td></td><td></td><td></td><td></td></tr>
<tr><td>Oligo rev</td><td>AAACCGCCGTGCGCGCCATGAAGA</td><td></td><td></td><td></td><td></td><td></td><td></td><td></td><td></td><td></td><td></td></tr>
<tr><td>Chromosome</td><td>start</td><td>end</td><td>strand</td><td>MM</td><td>target_seq</td><td>PAM</td><td>alignment</td><td>distance</td><td>position</td><td>gene name</td><td>gene id</td></tr>
<tr><td>GU305914.1</td><td>196482</td><td>196504</td><td>+</td><td>0</td><td>TCTTCATGGCGCGCACGGCG</td><td>TGG</td><td>||||||||[||||||||||||]PAM</td><td>NA</td><td>Intergenic</td><td>NA</td><td>NA</td></tr>
<tr><td></td><td></td><td></td><td></td><td></td><td></td><td></td><td></td><td></td><td></td><td></td><td></td></tr>
<tr><td>T14943</td><td>GGCGCGCACGGCGTGGTTGATGG</td><td>1000</td><td></td><td></td><td></td><td></td><td></td><td></td><td></td><td></td><td></td></tr>
<tr><td>Oligo fwd</td><td>CACCGGCGCGCACGGCGTGGTTGA</td><td></td><td></td><td></td><td></td><td></td><td></td><td></td><td></td><td></td><td></td></tr>
<tr><td>Oligo rev</td><td>AAACTCAACCACGCCGTGCGCGCC</td><td></td><td></td><td></td><td></td><td></td><td></td><td></td><td></td><td></td><td></td></tr>
<tr><td>Chromosome</td><td>start</td><td>end</td><td>strand</td><td>MM</td><td>target_seq</td><td>PAM</td><td>alignment</td><td>distance</td><td>position</td><td>gene name</td><td>gene id</td></tr>
<tr><td>GU305914.1</td><td>196489</td><td>196511</td><td>+</td><td>0</td><td>GGCGCGCACGGCGTGGTTGA</td><td>TGG</td><td>||||||||[||||||||||||]PAM</td><td>NA</td><td>Intergenic</td><td>NA</td><td>NA</td></tr>
<tr><td></td><td></td><td></td><td></td><td></td><td></td><td></td><td></td><td></td><td></td><td></td><td></td></tr>
<tr><td>T14944</td><td>GTCGTCCCACGTCCGCATGTCGG</td><td>1000</td><td></td><td></td><td></td><td></td><td></td><td></td><td></td><td></td><td></td></tr>
<tr><td>Oligo fwd</td><td>CACCGTCGTCCCACGTCCGCATGT</td><td></td><td></td><td></td><td></td><td></td><td></td><td></td><td></td><td></td><td></td></tr>
<tr><td>Oligo rev</td><td>AAACACATGCGGACGTGGGACGAC</td><td></td><td></td><td></td><td></td><td></td><td></td><td></td><td></td><td></td><td></td></tr>
<tr><td>Chromosome</td><td>start</td><td>end</td><td>strand</td><td>MM</td><td>target_seq</td><td>PAM</td><td>alignment</td><td>distance</td><td>position</td><td>gene name</td><td>gene id</td></tr>
<tr><td>GU305914.1</td><td>196525</td><td>196547</td><td>+</td><td>0</td><td>GTCGTCCCACGTCCGCATGT</td><td>CGG</td><td>||||||||[||||||||||||]PAM</td><td>NA</td><td>Intergenic</td><td>NA</td><td>NA</td></tr>
<tr><td></td><td></td><td></td><td></td><td></td><td></td><td></td><td></td><td></td><td></td><td></td><td></td></tr>
<tr><td>T14945</td><td>TCGTCCCACGTCCGCATGTCGGG</td><td>1000</td><td></td><td></td><td></td><td></td><td></td><td></td><td></td><td></td><td></td></tr>
<tr><td>Oligo fwd</td><td>CACCTCGTCCCACGTCCGCATGTC</td><td></td><td></td><td></td><td></td><td></td><td></td><td></td><td></td><td></td><td></td></tr>
<tr><td>Oligo rev</td><td>AAACGACATGCGGACGTGGGACGA</td><td></td><td></td><td></td><td></td><td></td><td></td><td></td><td></td><td></td><td></td></tr>
<tr><td>Chromosome</td><td>start</td><td>end</td><td>strand</td><td>MM</td><td>target_seq</td><td>PAM</td><td>alignment</td><td>distance</td><td>position</td><td>gene name</td><td>gene id</td></tr>
<tr><td>GU305914.1</td><td>196526</td><td>196548</td><td>+</td><td>0</td><td>TCGTCCCACGTCCGCATGTC</td><td>GGG</td><td>||||||||[||||||||||||]PAM</td><td>NA</td><td>Intergenic</td><td>NA</td><td>NA</td></tr>
<tr><td></td><td></td><td></td><td></td><td></td><td></td><td></td><td></td><td></td><td></td><td></td><td></td></tr>
<tr><td>T14946</td><td>TCCGCATGTCGGGCCGAAAGCGG</td><td>1000</td><td></td><td></td><td></td><td></td><td></td><td></td><td></td><td></td><td></td></tr>
<tr><td>Oligo fwd</td><td>CACCTCCGCATGTCGGGCCGAAAG</td><td></td><td></td><td></td><td></td><td></td><td></td><td></td><td></td><td></td><td></td></tr>
<tr><td>Oligo rev</td><td>AAACCTTTCGGCCCGACATGCGGA</td><td></td><td></td><td></td><td></td><td></td><td></td><td></td><td></td><td></td><td></td></tr>
<tr><td>Chromosome</td><td>start</td><td>end</td><td>strand</td><td>MM</td><td>target_seq</td><td>PAM</td><td>alignment</td><td>distance</td><td>position</td><td>gene name</td><td>gene id</td></tr>
<tr><td>GU305914.1</td><td>196536</td><td>196558</td><td>+</td><td>0</td><td>TCCGCATGTCGGGCCGAAAG</td><td>CGG</td><td>||||||||[||||||||||||]PAM</td><td>NA</td><td>Intergenic</td><td>NA</td><td>NA</td></tr>
<tr><td></td><td></td><td></td><td></td><td></td><td></td><td></td><td></td><td></td><td></td><td></td><td></td></tr>
<tr><td>T14947</td><td>TGTCGGGCCGAAAGCGGTCCCGG</td><td>1000</td><td></td><td></td><td></td><td></td><td></td><td></td><td></td><td></td><td></td></tr>
<tr><td>Oligo fwd</td><td>CACCTGTCGGGCCGAAAGCGGTCC</td><td></td><td></td><td></td><td></td><td></td><td></td><td></td><td></td><td></td><td></td></tr>
<tr><td>Oligo rev</td><td>AAACGGACCGCTTTCGGCCCGACA</td><td></td><td></td><td></td><td></td><td></td><td></td><td></td><td></td><td></td><td></td></tr>
<tr><td>Chromosome</td><td>start</td><td>end</td><td>strand</td><td>MM</td><td>target_seq</td><td>PAM</td><td>alignment</td><td>distance</td><td>position</td><td>gene name</td><td>gene id</td></tr>
<tr><td>GU305914.1</td><td>196542</td><td>196564</td><td>+</td><td>0</td><td>TGTCGGGCCGAAAGCGGTCC</td><td>CGG</td><td>||||||||[||||||||||||]PAM</td><td>NA</td><td>Intergenic</td><td>NA</td><td>NA</td></tr>
<tr><td></td><td></td><td></td><td></td><td></td><td></td><td></td><td></td><td></td><td></td><td></td><td></td></tr>
<tr><td>T14948</td><td>ACACAGCCACTCGTGTTCGTTGG</td><td>1000</td><td></td><td></td><td></td><td></td><td></td><td></td><td></td><td></td><td></td></tr>
<tr><td>Oligo fwd</td><td>CACCACACAGCCACTCGTGTTCGT</td><td></td><td></td><td></td><td></td><td></td><td></td><td></td><td></td><td></td><td></td></tr>
<tr><td>Oligo rev</td><td>AAACACGAACACGAGTGGCTGTGT</td><td></td><td></td><td></td><td></td><td></td><td></td><td></td><td></td><td></td><td></td></tr>
<tr><td>Chromosome</td><td>start</td><td>end</td><td>strand</td><td>MM</td><td>target_seq</td><td>PAM</td><td>alignment</td><td>distance</td><td>position</td><td>gene name</td><td>gene id</td></tr>
<tr><td>GU305914.1</td><td>196567</td><td>196589</td><td>+</td><td>0</td><td>ACACAGCCACTCGTGTTCGT</td><td>TGG</td><td>||||||||[||||||||||||]PAM</td><td>NA</td><td>Intergenic</td><td>NA</td><td>NA</td></tr>
<tr><td>GU305914.1</td><td>160967</td><td>160989</td><td>-</td><td>4</td><td>ACACAGAGCATCGTGTTCGT</td><td>CGG</td><td>||||||--[--||||||||||]PAM</td><td>NA</td><td>Intergenic</td><td>NA</td><td>NA</td></tr>
<tr><td></td><td></td><td></td><td></td><td></td><td></td><td></td><td></td><td></td><td></td><td></td><td></td></tr>
<tr><td>T14949</td><td>CCTCGCTGCGCTCCGACTTGCGG</td><td>1000</td><td></td><td></td><td></td><td></td><td></td><td></td><td></td><td></td><td></td></tr>
<tr><td>Oligo fwd</td><td>CACCCCTCGCTGCGCTCCGACTTG</td><td></td><td></td><td></td><td></td><td></td><td></td><td></td><td></td><td></td><td></td></tr>
<tr><td>Oligo rev</td><td>AAACCAAGTCGGAGCGCAGCGAGG</td><td></td><td></td><td></td><td></td><td></td><td></td><td></td><td></td><td></td><td></td></tr>
<tr><td>Chromosome</td><td>start</td><td>end</td><td>strand</td><td>MM</td><td>target_seq</td><td>PAM</td><td>alignment</td><td>distance</td><td>position</td><td>gene name</td><td>gene id</td></tr>
<tr><td>GU305914.1</td><td>196599</td><td>196621</td><td>+</td><td>0</td><td>CCTCGCTGCGCTCCGACTTG</td><td>CGG</td><td>||||||||[||||||||||||]PAM</td><td>NA</td><td>Intergenic</td><td>NA</td><td>NA</td></tr>
</table>

| | | | | | | | | | | | |
|---|---|---|---|---|---|---|---|---|---|---|---|
| T14950 | GCGCTCCGACTTGCGGTGATCGG | 1000 | | | | | | | | | |
| Oligo fwd | CACCGCGCTCCGACTTGCGGTGAT | | | | | | | | | | |
| Oligo rev | AAACATCACCGCAAGTCGGAGCGC | | | | | | | | | | |
| Chromosome | start | end | strand | MM | target_seq | PAM | alignment | distance | position | gene name | gene id |
| GU305914.1 | 196606 | 196628 | + | 0 | GCGCTCCGACTTGCGGTGAT | CGG | \|\|\|\|\|\|\|\|[\|\|\|\|\|\|\|\|\|\|\|\|]PAM | NA | Intergenic | NA | NA |
| T14951 | CGCTCCGACTTGCGGTGATCGGG | 1000 | | | | | | | | | |
| Oligo fwd | CACCCGCTCCGACTTGCGGTGATC | | | | | | | | | | |
| Oligo rev | AAACGATCACCGCAAGTCGGAGCG | | | | | | | | | | |
| Chromosome | start | end | strand | MM | target_seq | PAM | alignment | distance | position | gene name | gene id |
| GU305914.1 | 196607 | 196629 | + | 0 | CGCTCCGACTTGCGGTGATC | GGG | \|\|\|\|\|\|\|\|[\|\|\|\|\|\|\|\|\|\|\|\|]PAM | NA | Intergenic | NA | NA |
| T14952 | CCGACTTGCGGTGATCGGGCAGG | 1000 | | | | | | | | | |
| Oligo fwd | CACCCCGACTTGCGGTGATCGGGC | | | | | | | | | | |
| Oligo rev | AAACGCCCGATCACCGCAAGTCGG | | | | | | | | | | |
| Chromosome | start | end | strand | MM | target_seq | PAM | alignment | distance | position | gene name | gene id |
| GU305914.1 | 196611 | 196633 | + | 0 | CCGACTTGCGGTGATCGGGC | AGG | \|\|\|\|\|\|\|\|[\|\|\|\|\|\|\|\|\|\|\|\|]PAM | NA | Intergenic | NA | NA |
| T14953 | ACTTGCGGTGATCGGGCAGGTGG | 1000 | | | | | | | | | |
| Oligo fwd | CACCACTTGCGGTGATCGGGCAGG | | | | | | | | | | |
| Oligo rev | AAACCCTGCCCGATCACCGCAAGT | | | | | | | | | | |
| Chromosome | start | end | strand | MM | target_seq | PAM | alignment | distance | position | gene name | gene id |
| GU305914.1 | 196614 | 196636 | + | 0 | ACTTGCGGTGATCGGGCAGG | TGG | \|\|\|\|\|\|\|\|[\|\|\|\|\|\|\|\|\|\|\|\|]PAM | NA | Intergenic | NA | NA |
| T14954 | CTTGCGGTGATCGGGCAGGTGGG | 1000 | | | | | | | | | |
| Oligo fwd | CACCCTTGCGGTGATCGGGCAGGT | | | | | | | | | | |
| Oligo rev | AAACACCTGCCCGATCACCGCAAG | | | | | | | | | | |
| Chromosome | start | end | strand | MM | target_seq | PAM | alignment | distance | position | gene name | gene id |
| GU305914.1 | 196615 | 196637 | + | 0 | CTTGCGGTGATCGGGCAGGT | GGG | \|\|\|\|\|\|\|\|[\|\|\|\|\|\|\|\|\|\|\|\|]PAM | NA | Intergenic | NA | NA |
| T14955 | TTGCGGTGATCGGGCAGGTGGGG | 1000 | | | | | | | | | |
| Oligo fwd | CACCTTGCGGTGATCGGGCAGGTG | | | | | | | | | | |
| Oligo rev | AAACCACCTGCCCGATCACCGCAA | | | | | | | | | | |
| Chromosome | start | end | strand | MM | target_seq | PAM | alignment | distance | position | gene name | gene id |
| GU305914.1 | 196616 | 196638 | + | 0 | TTGCGGTGATCGGGCAGGTG | GGG | \|\|\|\|\|\|\|\|[\|\|\|\|\|\|\|\|\|\|\|\|]PAM | NA | Intergenic | NA | NA |
| T14956 | AGATGACTCGCATCTCGCAGCGG | 1000 | | | | | | | | | |
| Oligo fwd | CACCAGATGACTCGCATCTCGCAG | | | | | | | | | | |
| Oligo rev | AAACCTGCGAGATGCGAGTCATCT | | | | | | | | | | |
| Chromosome | start | end | strand | MM | target_seq | PAM | alignment | distance | position | gene name | gene id |
| GU305914.1 | 196641 | 196663 | + | 0 | AGATGACTCGCATCTCGCAG | CGG | \|\|\|\|\|\|\|\|[\|\|\|\|\|\|\|\|\|\|\|\|]PAM | NA | Intergenic | NA | NA |
| T14957 | CAGCGGCCCTCCCCGCGCGCCGG | 1000 | | | | | | | | | |
| Oligo fwd | CACCCAGCGGCCCTCCCCGCGCGC | | | | | | | | | | |
| Oligo rev | AAACGCGCGCGGGGAGGGCCGCTG | | | | | | | | | | |
| Chromosome | start | end | strand | MM | target_seq | PAM | alignment | distance | position | gene name | gene id |
| GU305914.1 | 196658 | 196680 | + | 0 | CAGCGGCCCTCCCCGCGCGC | CGG | \|\|\|\|\|\|\|\|[\|\|\|\|\|\|\|\|\|\|\|\|]PAM | NA | Intergenic | NA | NA |
| T14958 | GGCCCTCCCCGCGCGCCGGACGG | 1000 | | | | | | | | | |
| Oligo fwd | CACCGGCCCTCCCCGCGCGCCGGA | | | | | | | | | | |
| Oligo rev | AAACTCCGGCGCGCGGGGAGGGCC | | | | | | | | | | |
| Chromosome | start | end | strand | MM | target_seq | PAM | alignment | distance | position | gene name | gene id |
| GU305914.1 | 196662 | 196684 | + | 0 | GGCCCTCCCCGCGCGCCGGA | CGG | \|\|\|\|\|\|\|\|[\|\|\|\|\|\|\|\|\|\|\|\|]PAM | NA | Intergenic | NA | NA |
| T14959 | GAAGCGCGCCCCGAACGCGATGG | 1000 | | | | | | | | | |
| Oligo fwd | CACCGAAGCGCGCCCCGAACGCGA | | | | | | | | | | |
| Oligo rev | AAACTCGCGTTCGGGGCGCGCTTC | | | | | | | | | | |
| Chromosome | start | end | strand | MM | target_seq | PAM | alignment | distance | position | gene name | gene id |
| GU305914.1 | 196687 | 196709 | + | 0 | GAAGCGCGCCCCGAACGCGA | TGG | \|\|\|\|\|\|\|\|[\|\|\|\|\|\|\|\|\|\|\|\|]PAM | NA | Intergenic | NA | NA |
| T14960 | CGCCCCGAACGCGATGGCCTTGG | 1000 | | | | | | | | | |
| Oligo fwd | CACCCGCCCCGAACGCGATGGCCT | | | | | | | | | | |
| Oligo rev | AAACAGGCCATCGCGTTCGGGGCG | | | | | | | | | | |
| Chromosome | start | end | strand | MM | target_seq | PAM | alignment | distance | position | gene name | gene id |
| GU305914.1 | 196693 | 196715 | + | 0 | CGCCCCGAACGCGATGGCCT | TGG | \|\|\|\|\|\|\|\|[\|\|\|\|\|\|\|\|\|\|\|\|]PAM | NA | Intergenic | NA | NA |
| GU305914.1 | 170366 | 170388 | + | 4 | CTCCGGGAACGCGATGGGCT | CGG | \|-\|\|--\|\|[\|\|\|\|\|\|\|\|\|-\|\|]PAM | NA | Intergenic | NA | NA |
| T14961 | CGCGATGGCCTTGGCCATGCCGG | 1000 | | | | | | | | | |
| Oligo fwd | CACCCGCGATGGCCTTGGCCATGC | | | | | | | | | | |
| Oligo rev | AAACGCATGGCCAAGGCCATCGCG | | | | | | | | | | |
| Chromosome | start | end | strand | MM | target_seq | PAM | alignment | distance | position | gene name | gene id |
| GU305914.1 | 196702 | 196724 | + | 0 | CGCGATGGCCTTGGCCATGC | CGG | \|\|\|\|\|\|\|\|[\|\|\|\|\|\|\|\|\|\|\|\|]PAM | NA | Intergenic | NA | NA |
| T14962 | TCGTGATGCCGTCCGCCACGCGG | 1000 | | | | | | | | | |
| Oligo fwd | CACCTCGTGATGCCGTCCGCCACG | | | | | | | | | | |
| Oligo rev | AAACCGTGGCGGACGGCATCACGA | | | | | | | | | | |
| Chromosome | start | end | strand | MM | target_seq | PAM | alignment | distance | position | gene name | gene id |
| GU305914.1 | 196737 | 196759 | + | 0 | TCGTGATGCCGTCCGCCACG | CGG | \|\|\|\|\|\|\|\|[\|\|\|\|\|\|\|\|\|\|\|\|]PAM | NA | Intergenic | NA | NA |
| T14963 | CGAACGTGTCGATGACGAGCAGG | 1000 | | | | | | | | | |
| Oligo fwd | CACCCGAACGTGTCGATGACGAGC | | | | | | | | | | |
| Oligo rev | AAACGCTCGTCATCGACACGTTCG | | | | | | | | | | |
| Chromosome | start | end | strand | MM | target_seq | PAM | alignment | distance | position | gene name | gene id |
| GU305914.1 | 196800 | 196822 | + | 0 | CGAACGTGTCGATGACGAGC | AGG | \|\|\|\|\|\|\|\|[\|\|\|\|\|\|\|\|\|\|\|\|]PAM | NA | Intergenic | NA | NA |
| T14964 | ACGTGTCGATGACGAGCAGGTGG | 1000 | | | | | | | | | |
| Oligo fwd | CACCACGTGTCGATGACGAGCAGG | | | | | | | | | | |
| Oligo rev | AAACCCTGCTCGTCATCGACACGT | | | | | | | | | | |
| Chromosome | start | end | strand | MM | target_seq | PAM | alignment | distance | position | gene name | gene id |

| | | | | | | | | | | | |
|---|---|---|---|---|---|---|---|---|---|---|---|
| GU305914.1 | 196803 | 196825 | + | 0 | ACGTGTCGATGACGAGCAGG | TGG | \|\|\|\|\|\|\|\|[\|\|\|\|\|\|\|\|\|\|\|\|]PAM | NA | Intergenic | NA | NA |

| | | | | | | | | | | | |
|---|---|---|---|---|---|---|---|---|---|---|---|
| T14965 | TGTCGATGACGAGCAGGTGGTGG | 1000 | | | | | | | | | |
| Oligo fwd | CACCTGTCGATGACGAGCAGGTGG | | | | | | | | | | |
| Oligo rev | AAACCCACCTGCTCGTCATCGACA | | | | | | | | | | |
| Chromosome | start | end | strand | MM | target_seq | PAM | alignment | distance | position | gene name | gene id |
| GU305914.1 | 196806 | 196828 | + | 0 | TGTCGATGACGAGCAGGTGG | TGG | \|\|\|\|\|\|\|\|[\|\|\|\|\|\|\|\|\|\|\|\|]PAM | NA | Intergenic | NA | NA |
| GU305914.1 | 54556 | 54578 | + | 4 | CGTCGAGGACGAGGTGGTGG | AGG | -\|\|\|\|\|-\|[\|\|\|\|\|--\|\|\|\|\|]PAM | NA | Intergenic | NA | NA |

| | | | | | | | | | | | |
|---|---|---|---|---|---|---|---|---|---|---|---|
| T14966 | TGGATGCTGAAGACGCCTGCCGG | 1000 | | | | | | | | | |
| Oligo fwd | CACCTGGATGCTGAAGACGCCTGC | | | | | | | | | | |
| Oligo rev | AAACGCAGGCGTCTTCAGCATCCA | | | | | | | | | | |
| Chromosome | start | end | strand | MM | target_seq | PAM | alignment | distance | position | gene name | gene id |
| GU305914.1 | 196826 | 196848 | + | 0 | TGGATGCTGAAGACGCCTGC | CGG | \|\|\|\|\|\|\|\|[\|\|\|\|\|\|\|\|\|\|\|\|]PAM | NA | Intergenic | NA | NA |

| | | | | | | | | | | | |
|---|---|---|---|---|---|---|---|---|---|---|---|
| T14967 | GAAGACGCCTGCCGGCGCGTAGG | 1000 | | | | | | | | | |
| Oligo fwd | CACCGAAGACGCCTGCCGGCGCGT | | | | | | | | | | |
| Oligo rev | AAACACGCGCCGGCAGGCGTCTTC | | | | | | | | | | |
| Chromosome | start | end | strand | MM | target_seq | PAM | alignment | distance | position | gene name | gene id |
| GU305914.1 | 196834 | 196856 | + | 0 | GAAGACGCCTGCCGGCGCGT | AGG | \|\|\|\|\|\|\|\|[\|\|\|\|\|\|\|\|\|\|\|\|]PAM | NA | Intergenic | NA | NA |

| | | | | | | | | | | | |
|---|---|---|---|---|---|---|---|---|---|---|---|
| T14968 | TGCCGGCGCGTAGGCGCCCACGG | 1000 | | | | | | | | | |
| Oligo fwd | CACCTGCCGGCGCGTAGGCGCCCA | | | | | | | | | | |
| Oligo rev | AAACTGGGCGCCTACGCGCCGGCA | | | | | | | | | | |
| Chromosome | start | end | strand | MM | target_seq | PAM | alignment | distance | position | gene name | gene id |
| GU305914.1 | 196843 | 196865 | + | 0 | TGCCGGCGCGTAGGCGCCCA | CGG | \|\|\|\|\|\|\|\|[\|\|\|\|\|\|\|\|\|\|\|\|]PAM | NA | Intergenic | NA | NA |

| | | | | | | | | | | | |
|---|---|---|---|---|---|---|---|---|---|---|---|
| T14969 | GTCGCCCCGTGATCTCGTCCAGG | 1000 | | | | | | | | | |
| Oligo fwd | CACCGTCGCCCCGTGATCTCGTCC | | | | | | | | | | |
| Oligo rev | AAACGGACGAGATCACGGGGCGAC | | | | | | | | | | |
| Chromosome | start | end | strand | MM | target_seq | PAM | alignment | distance | position | gene name | gene id |
| GU305914.1 | 196893 | 196915 | + | 0 | GTCGCCCCGTGATCTCGTCC | AGG | \|\|\|\|\|\|\|\|[\|\|\|\|\|\|\|\|\|\|\|\|]PAM | NA | Intergenic | NA | NA |

| | | | | | | | | | | | |
|---|---|---|---|---|---|---|---|---|---|---|---|
| T14970 | CCCCGTGATCTCGTCCAGGCAGG | 1000 | | | | | | | | | |
| Oligo fwd | CACCCCCCGTGATCTCGTCCAGGC | | | | | | | | | | |
| Oligo rev | AAACGCCTGGACGAGATCACGGGG | | | | | | | | | | |
| Chromosome | start | end | strand | MM | target_seq | PAM | alignment | distance | position | gene name | gene id |
| GU305914.1 | 196897 | 196919 | + | 0 | CCCCGTGATCTCGTCCAGGC | AGG | \|\|\|\|\|\|\|\|[\|\|\|\|\|\|\|\|\|\|\|\|]PAM | NA | Intergenic | NA | NA |

| | | | | | | | | | | | |
|---|---|---|---|---|---|---|---|---|---|---|---|
| T14971 | TCTCGTCCAGGCAGGCCTTCAGG | 1000 | | | | | | | | | |
| Oligo fwd | CACCTCTCGTCCAGGCAGGCCTTC | | | | | | | | | | |
| Oligo rev | AAACGAAGGCCTGCCTGGACGAGA | | | | | | | | | | |
| Chromosome | start | end | strand | MM | target_seq | PAM | alignment | distance | position | gene name | gene id |
| GU305914.1 | 196905 | 196927 | + | 0 | TCTCGTCCAGGCAGGCCTTC | AGG | \|\|\|\|\|\|\|\|[\|\|\|\|\|\|\|\|\|\|\|\|]PAM | NA | Intergenic | NA | NA |

| | | | | | | | | | | | |
|---|---|---|---|---|---|---|---|---|---|---|---|
| T14972 | AGGCAGGCCTTCAGGTTCGACGG | 1000 | | | | | | | | | |
| Oligo fwd | CACCAGGCAGGCCTTCAGGTTCGA | | | | | | | | | | |
| Oligo rev | AAACTCGAACCTGAAGGCCTGCCT | | | | | | | | | | |
| Chromosome | start | end | strand | MM | target_seq | PAM | alignment | distance | position | gene name | gene id |
| GU305914.1 | 196913 | 196935 | + | 0 | AGGCAGGCCTTCAGGTTCGA | CGG | \|\|\|\|\|\|\|\|[\|\|\|\|\|\|\|\|\|\|\|\|]PAM | NA | Intergenic | NA | NA |
| GU305914.1 | 90649 | 90671 | - | 4 | AAGAAGACCATCAGGTTCGA | GGG | \|-\|-\|\|-\|[\|-\|\|\|\|\|\|\|\|\|\|]PAM | NA | Intergenic | NA | NA |

| | | | | | | | | | | | |
|---|---|---|---|---|---|---|---|---|---|---|---|
| T14973 | GGCAGGCCTTCAGGTTCGACGGG | 1000 | | | | | | | | | |
| Oligo fwd | CACCGGCAGGCCTTCAGGTTCGAC | | | | | | | | | | |
| Oligo rev | AAACGTCGAACCTGAAGGCCTGCC | | | | | | | | | | |
| Chromosome | start | end | strand | MM | target_seq | PAM | alignment | distance | position | gene name | gene id |
| GU305914.1 | 196914 | 196936 | + | 0 | GGCAGGCCTTCAGGTTCGAC | GGG | \|\|\|\|\|\|\|\|[\|\|\|\|\|\|\|\|\|\|\|\|]PAM | NA | Intergenic | NA | NA |

| | | | | | | | | | | | |
|---|---|---|---|---|---|---|---|---|---|---|---|
| T14974 | TCGACGGGTTCATGATATCCAGG | 1000 | | | | | | | | | |
| Oligo fwd | CACCTCGACGGGTTCATGATATCC | | | | | | | | | | |
| Oligo rev | AAACGGATATCATGAACCCGTCGA | | | | | | | | | | |
| Chromosome | start | end | strand | MM | target_seq | PAM | alignment | distance | position | gene name | gene id |
| GU305914.1 | 196929 | 196951 | + | 0 | TCGACGGGTTCATGATATCC | AGG | \|\|\|\|\|\|\|\|[\|\|\|\|\|\|\|\|\|\|\|\|]PAM | NA | Intergenic | NA | NA |

| | | | | | | | | | | | |
|---|---|---|---|---|---|---|---|---|---|---|---|
| T14975 | CGACGGGTTCATGATATCCAGGG | 1000 | | | | | | | | | |
| Oligo fwd | CACCCGACGGGTTCATGATATCCA | | | | | | | | | | |
| Oligo rev | AAACTGGATATCATGAACCCGTCG | | | | | | | | | | |
| Chromosome | start | end | strand | MM | target_seq | PAM | alignment | distance | position | gene name | gene id |
| GU305914.1 | 196930 | 196952 | + | 0 | CGACGGGTTCATGATATCCA | GGG | \|\|\|\|\|\|\|\|[\|\|\|\|\|\|\|\|\|\|\|\|]PAM | NA | Intergenic | NA | NA |

| | | | | | | | | | | | |
|---|---|---|---|---|---|---|---|---|---|---|---|
| T14976 | GACGGGTTCATGATATCCAGGGG | 1000 | | | | | | | | | |
| Oligo fwd | CACCGACGGGTTCATGATATCCAG | | | | | | | | | | |
| Oligo rev | AAACCTGGATATCATGAACCCGTC | | | | | | | | | | |
| Chromosome | start | end | strand | MM | target_seq | PAM | alignment | distance | position | gene name | gene id |
| GU305914.1 | 196931 | 196953 | + | 0 | GACGGGTTCATGATATCCAG | GGG | \|\|\|\|\|\|\|\|[\|\|\|\|\|\|\|\|\|\|\|\|]PAM | NA | Intergenic | NA | NA |

| | | | | | | | | | | | |
|---|---|---|---|---|---|---|---|---|---|---|---|
| T14977 | ATCCAGGGGCCAGTAGCGCGCGG | 1000 | | | | | | | | | |
| Oligo fwd | CACCATCCAGGGGCCAGTAGCGCG | | | | | | | | | | |
| Oligo rev | AAACCGCGCTACTGGCCCCTGGAT | | | | | | | | | | |
| Chromosome | start | end | strand | MM | target_seq | PAM | alignment | distance | position | gene name | gene id |
| GU305914.1 | 196945 | 196967 | + | 0 | ATCCAGGGGCCAGTAGCGCG | CGG | \|\|\|\|\|\|\|\|[\|\|\|\|\|\|\|\|\|\|\|\|]PAM | NA | Intergenic | NA | NA |

| | | | | | | | | | | | |
|---|---|---|---|---|---|---|---|---|---|---|---|
| T14978 | ACGCCACCCAGCAGTTTCATCGG | 1000 | | | | | | | | | |
| Oligo fwd | CACCACGCCACCCAGCAGTTTCAT | | | | | | | | | | |
| Oligo rev | AAACATGAAACTGCTGGGTGGCGT | | | | | | | | | | |
| Chromosome | start | end | strand | MM | target_seq | PAM | alignment | distance | position | gene name | gene id |
| GU305914.1 | 196976 | 196998 | + | 0 | ACGCCACCCAGCAGTTTCAT | CGG | \|\|\|\|\|\|\|\|[\|\|\|\|\|\|\|\|\|\|\|\|]PAM | NA | Intergenic | NA | NA |

| | | |
|---|---|---|
| T14979 | CACCCAGCAGTTTCATCGGCTGG | 1000 |

<table>
<tr><td>Oligo fwd</td><td>CACCCACCCAGCAGTTTCATCGGC</td><td></td><td></td><td></td><td></td><td></td><td></td><td></td><td></td><td></td><td></td></tr>
<tr><td>Oligo rev</td><td>AAACGCCGATGAAACTGCTGGGTG</td><td></td><td></td><td></td><td></td><td></td><td></td><td></td><td></td><td></td><td></td></tr>
<tr><td>Chromosome</td><td>start</td><td>end</td><td>strand</td><td>MM</td><td>target_seq</td><td>PAM</td><td>alignment</td><td>distance</td><td>position</td><td>gene name</td><td>gene id</td></tr>
<tr><td>GU305914.1</td><td>196980</td><td>197002</td><td>+</td><td>0</td><td>CACCCAGCAGTTTCATCGGC</td><td>TGG</td><td>||||||||[||||||||||||]PAM</td><td>NA</td><td>Intergenic</td><td>NA</td><td>NA</td></tr>
<tr><td></td><td></td><td></td><td></td><td></td><td></td><td></td><td></td><td></td><td></td><td></td><td></td></tr>
<tr><td>T14980</td><td>CATCGGCTGGTACTTGAATCCGG</td><td>1000</td><td></td><td></td><td></td><td></td><td></td><td></td><td></td><td></td><td></td></tr>
<tr><td>Oligo fwd</td><td>CACCCATCGGCTGGTACTTGAATC</td><td></td><td></td><td></td><td></td><td></td><td></td><td></td><td></td><td></td><td></td></tr>
<tr><td>Oligo rev</td><td>AAACGATTCAAGTACCAGCCGATG</td><td></td><td></td><td></td><td></td><td></td><td></td><td></td><td></td><td></td><td></td></tr>
<tr><td>Chromosome</td><td>start</td><td>end</td><td>strand</td><td>MM</td><td>target_seq</td><td>PAM</td><td>alignment</td><td>distance</td><td>position</td><td>gene name</td><td>gene id</td></tr>
<tr><td>GU305914.1</td><td>196993</td><td>197015</td><td>+</td><td>0</td><td>CATCGGCTGGTACTTGAATC</td><td>CGG</td><td>||||||||[||||||||||||]PAM</td><td>NA</td><td>Intergenic</td><td>NA</td><td>NA</td></tr>
<tr><td></td><td></td><td></td><td></td><td></td><td></td><td></td><td></td><td></td><td></td><td></td><td></td></tr>
<tr><td>T14981</td><td>ATCGGCTGGTACTTGAATCCGGG</td><td>1000</td><td></td><td></td><td></td><td></td><td></td><td></td><td></td><td></td><td></td></tr>
<tr><td>Oligo fwd</td><td>CACCATCGGCTGGTACTTGAATCC</td><td></td><td></td><td></td><td></td><td></td><td></td><td></td><td></td><td></td><td></td></tr>
<tr><td>Oligo rev</td><td>AAACGGATTCAAGTACCAGCCGAT</td><td></td><td></td><td></td><td></td><td></td><td></td><td></td><td></td><td></td><td></td></tr>
<tr><td>Chromosome</td><td>start</td><td>end</td><td>strand</td><td>MM</td><td>target_seq</td><td>PAM</td><td>alignment</td><td>distance</td><td>position</td><td>gene name</td><td>gene id</td></tr>
<tr><td>GU305914.1</td><td>196994</td><td>197016</td><td>+</td><td>0</td><td>ATCGGCTGGTACTTGAATCC</td><td>GGG</td><td>||||||||[||||||||||||]PAM</td><td>NA</td><td>Intergenic</td><td>NA</td><td>NA</td></tr>
<tr><td></td><td></td><td></td><td></td><td></td><td></td><td></td><td></td><td></td><td></td><td></td><td></td></tr>
<tr><td>T14982</td><td>TCGGCTGGTACTTGAATCCGGGG</td><td>1000</td><td></td><td></td><td></td><td></td><td></td><td></td><td></td><td></td><td></td></tr>
<tr><td>Oligo fwd</td><td>CACCTCGGCTGGTACTTGAATCCG</td><td></td><td></td><td></td><td></td><td></td><td></td><td></td><td></td><td></td><td></td></tr>
<tr><td>Oligo rev</td><td>AAACCGGATTCAAGTACCAGCCGA</td><td></td><td></td><td></td><td></td><td></td><td></td><td></td><td></td><td></td><td></td></tr>
<tr><td>Chromosome</td><td>start</td><td>end</td><td>strand</td><td>MM</td><td>target_seq</td><td>PAM</td><td>alignment</td><td>distance</td><td>position</td><td>gene name</td><td>gene id</td></tr>
<tr><td>GU305914.1</td><td>196995</td><td>197017</td><td>+</td><td>0</td><td>TCGGCTGGTACTTGAATCCG</td><td>GGG</td><td>||||||||[||||||||||||]PAM</td><td>NA</td><td>Intergenic</td><td>NA</td><td>NA</td></tr>
<tr><td></td><td></td><td></td><td></td><td></td><td></td><td></td><td></td><td></td><td></td><td></td><td></td></tr>
<tr><td>T14983</td><td>CGGCTGGTACTTGAATCCGGGGG</td><td>1000</td><td></td><td></td><td></td><td></td><td></td><td></td><td></td><td></td><td></td></tr>
<tr><td>Oligo fwd</td><td>CACCCGGCTGGTACTTGAATCCGG</td><td></td><td></td><td></td><td></td><td></td><td></td><td></td><td></td><td></td><td></td></tr>
<tr><td>Oligo rev</td><td>AAACCCGGATTCAAGTACCAGCCG</td><td></td><td></td><td></td><td></td><td></td><td></td><td></td><td></td><td></td><td></td></tr>
<tr><td>Chromosome</td><td>start</td><td>end</td><td>strand</td><td>MM</td><td>target_seq</td><td>PAM</td><td>alignment</td><td>distance</td><td>position</td><td>gene name</td><td>gene id</td></tr>
<tr><td>GU305914.1</td><td>196996</td><td>197018</td><td>+</td><td>0</td><td>CGGCTGGTACTTGAATCCGG</td><td>GGG</td><td>||||||||[||||||||||||]PAM</td><td>NA</td><td>Intergenic</td><td>NA</td><td>NA</td></tr>
<tr><td></td><td></td><td></td><td></td><td></td><td></td><td></td><td></td><td></td><td></td><td></td><td></td></tr>
<tr><td>T14984</td><td>CGTCGCGCCCGCTGTTCTCCTGG</td><td>1000</td><td></td><td></td><td></td><td></td><td></td><td></td><td></td><td></td><td></td></tr>
<tr><td>Oligo fwd</td><td>CACCCGTCGCGCCCGCTGTTCTCC</td><td></td><td></td><td></td><td></td><td></td><td></td><td></td><td></td><td></td><td></td></tr>
<tr><td>Oligo rev</td><td>AAACGGAGAACAGCGGGCGCGACG</td><td></td><td></td><td></td><td></td><td></td><td></td><td></td><td></td><td></td><td></td></tr>
<tr><td>Chromosome</td><td>start</td><td>end</td><td>strand</td><td>MM</td><td>target_seq</td><td>PAM</td><td>alignment</td><td>distance</td><td>position</td><td>gene name</td><td>gene id</td></tr>
<tr><td>GU305914.1</td><td>197031</td><td>197053</td><td>+</td><td>0</td><td>CGTCGCGCCCGCTGTTCTCC</td><td>TGG</td><td>||||||||[||||||||||||]PAM</td><td>NA</td><td>Intergenic</td><td>NA</td><td>NA</td></tr>
<tr><td>GU305914.1</td><td>60651</td><td>60673</td><td>+</td><td>4</td><td>CGGGGCGCCCGATGTTCTCG</td><td>GGG</td><td>||--||||[|||-|||||||-]PAM</td><td>NA</td><td>Intergenic</td><td>NA</td><td>NA</td></tr>
<tr><td></td><td></td><td></td><td></td><td></td><td></td><td></td><td></td><td></td><td></td><td></td><td></td></tr>
<tr><td>T14985</td><td>GCTGTTCTCCTGGCAGTAATCGG</td><td>1000</td><td></td><td></td><td></td><td></td><td></td><td></td><td></td><td></td><td></td></tr>
<tr><td>Oligo fwd</td><td>CACCGCTGTTCTCCTGGCAGTAAT</td><td></td><td></td><td></td><td></td><td></td><td></td><td></td><td></td><td></td><td></td></tr>
<tr><td>Oligo rev</td><td>AAACATTACTGCCAGGAGAACAGC</td><td></td><td></td><td></td><td></td><td></td><td></td><td></td><td></td><td></td><td></td></tr>
<tr><td>Chromosome</td><td>start</td><td>end</td><td>strand</td><td>MM</td><td>target_seq</td><td>PAM</td><td>alignment</td><td>distance</td><td>position</td><td>gene name</td><td>gene id</td></tr>
<tr><td>GU305914.1</td><td>197041</td><td>197063</td><td>+</td><td>0</td><td>GCTGTTCTCCTGGCAGTAAT</td><td>CGG</td><td>||||||||[||||||||||||]PAM</td><td>NA</td><td>Intergenic</td><td>NA</td><td>NA</td></tr>
<tr><td></td><td></td><td></td><td></td><td></td><td></td><td></td><td></td><td></td><td></td><td></td><td></td></tr>
<tr><td>T14986</td><td>CCTGGCAGTAATCGGACACCCGG</td><td>1000</td><td></td><td></td><td></td><td></td><td></td><td></td><td></td><td></td><td></td></tr>
<tr><td>Oligo fwd</td><td>CACCCCTGGCAGTAATCGGACACC</td><td></td><td></td><td></td><td></td><td></td><td></td><td></td><td></td><td></td><td></td></tr>
<tr><td>Oligo rev</td><td>AAACGGTGTCCGATTACTGCCAGG</td><td></td><td></td><td></td><td></td><td></td><td></td><td></td><td></td><td></td><td></td></tr>
<tr><td>Chromosome</td><td>start</td><td>end</td><td>strand</td><td>MM</td><td>target_seq</td><td>PAM</td><td>alignment</td><td>distance</td><td>position</td><td>gene name</td><td>gene id</td></tr>
<tr><td>GU305914.1</td><td>197049</td><td>197071</td><td>+</td><td>0</td><td>CCTGGCAGTAATCGGACACC</td><td>CGG</td><td>||||||||[||||||||||||]PAM</td><td>NA</td><td>Intergenic</td><td>NA</td><td>NA</td></tr>
<tr><td></td><td></td><td></td><td></td><td></td><td></td><td></td><td></td><td></td><td></td><td></td><td></td></tr>
<tr><td>T14987</td><td>ACACCCGGTCGAGATCGTCCTGG</td><td>1000</td><td></td><td></td><td></td><td></td><td></td><td></td><td></td><td></td><td></td></tr>
<tr><td>Oligo fwd</td><td>CACCACACCCGGTCGAGATCGTCC</td><td></td><td></td><td></td><td></td><td></td><td></td><td></td><td></td><td></td><td></td></tr>
<tr><td>Oligo rev</td><td>AAACGGACGATCTCGACCGGGTGT</td><td></td><td></td><td></td><td></td><td></td><td></td><td></td><td></td><td></td><td></td></tr>
<tr><td>Chromosome</td><td>start</td><td>end</td><td>strand</td><td>MM</td><td>target_seq</td><td>PAM</td><td>alignment</td><td>distance</td><td>position</td><td>gene name</td><td>gene id</td></tr>
<tr><td>GU305914.1</td><td>197064</td><td>197086</td><td>+</td><td>0</td><td>ACACCCGGTCGAGATCGTCC</td><td>TGG</td><td>||||||||[||||||||||||]PAM</td><td>NA</td><td>Intergenic</td><td>NA</td><td>NA</td></tr>
<tr><td></td><td></td><td></td><td></td><td></td><td></td><td></td><td></td><td></td><td></td><td></td><td></td></tr>
<tr><td>T14988</td><td>ACATGATCAGCCCCACGACGAGG</td><td>1000</td><td></td><td></td><td></td><td></td><td></td><td></td><td></td><td></td><td></td></tr>
<tr><td>Oligo fwd</td><td>CACCACATGATCAGCCCCACGACG</td><td></td><td></td><td></td><td></td><td></td><td></td><td></td><td></td><td></td><td></td></tr>
<tr><td>Oligo rev</td><td>AAACCGTCGTGGGGCTGATCATGT</td><td></td><td></td><td></td><td></td><td></td><td></td><td></td><td></td><td></td><td></td></tr>
<tr><td>Chromosome</td><td>start</td><td>end</td><td>strand</td><td>MM</td><td>target_seq</td><td>PAM</td><td>alignment</td><td>distance</td><td>position</td><td>gene name</td><td>gene id</td></tr>
<tr><td>GU305914.1</td><td>197088</td><td>197110</td><td>+</td><td>0</td><td>ACATGATCAGCCCCACGACG</td><td>AGG</td><td>||||||||[||||||||||||]PAM</td><td>NA</td><td>Intergenic</td><td>NA</td><td>NA</td></tr>
<tr><td></td><td></td><td></td><td></td><td></td><td></td><td></td><td></td><td></td><td></td><td></td><td></td></tr>
<tr><td>T14989</td><td>TGATCAGCCCCACGACGAGGCGG</td><td>1000</td><td></td><td></td><td></td><td></td><td></td><td></td><td></td><td></td><td></td></tr>
<tr><td>Oligo fwd</td><td>CACCTGATCAGCCCCACGACGAGG</td><td></td><td></td><td></td><td></td><td></td><td></td><td></td><td></td><td></td><td></td></tr>
<tr><td>Oligo rev</td><td>AAACCCTCGTCGTGGGGCTGATCA</td><td></td><td></td><td></td><td></td><td></td><td></td><td></td><td></td><td></td><td></td></tr>
<tr><td>Chromosome</td><td>start</td><td>end</td><td>strand</td><td>MM</td><td>target_seq</td><td>PAM</td><td>alignment</td><td>distance</td><td>position</td><td>gene name</td><td>gene id</td></tr>
<tr><td>GU305914.1</td><td>197091</td><td>197113</td><td>+</td><td>0</td><td>TGATCAGCCCCACGACGAGG</td><td>CGG</td><td>||||||||[||||||||||||]PAM</td><td>NA</td><td>Intergenic</td><td>NA</td><td>NA</td></tr>
<tr><td></td><td></td><td></td><td></td><td></td><td></td><td></td><td></td><td></td><td></td><td></td><td></td></tr>
<tr><td>T14990</td><td>TCAGCCCCACGACGAGGCGGTGG</td><td>1000</td><td></td><td></td><td></td><td></td><td></td><td></td><td></td><td></td><td></td></tr>
<tr><td>Oligo fwd</td><td>CACCTCAGCCCCACGACGAGGCGG</td><td></td><td></td><td></td><td></td><td></td><td></td><td></td><td></td><td></td><td></td></tr>
<tr><td>Oligo rev</td><td>AAACCCGCCTCGTCGTGGGGCTGA</td><td></td><td></td><td></td><td></td><td></td><td></td><td></td><td></td><td></td><td></td></tr>
<tr><td>Chromosome</td><td>start</td><td>end</td><td>strand</td><td>MM</td><td>target_seq</td><td>PAM</td><td>alignment</td><td>distance</td><td>position</td><td>gene name</td><td>gene id</td></tr>
<tr><td>GU305914.1</td><td>197094</td><td>197116</td><td>+</td><td>0</td><td>TCAGCCCCACGACGAGGCGG</td><td>TGG</td><td>||||||||[||||||||||||]PAM</td><td>NA</td><td>Intergenic</td><td>NA</td><td>NA</td></tr>
<tr><td></td><td></td><td></td><td></td><td></td><td></td><td></td><td></td><td></td><td></td><td></td><td></td></tr>
<tr><td>T14991</td><td>CAGCCCCACGACGAGGCGGTGGG</td><td>1000</td><td></td><td></td><td></td><td></td><td></td><td></td><td></td><td></td><td></td></tr>
<tr><td>Oligo fwd</td><td>CACCCAGCCCCACGACGAGGCGGT</td><td></td><td></td><td></td><td></td><td></td><td></td><td></td><td></td><td></td><td></td></tr>
<tr><td>Oligo rev</td><td>AAACACCGCCTCGTCGTGGGGCTG</td><td></td><td></td><td></td><td></td><td></td><td></td><td></td><td></td><td></td><td></td></tr>
<tr><td>Chromosome</td><td>start</td><td>end</td><td>strand</td><td>MM</td><td>target_seq</td><td>PAM</td><td>alignment</td><td>distance</td><td>position</td><td>gene name</td><td>gene id</td></tr>
<tr><td>GU305914.1</td><td>197095</td><td>197117</td><td>+</td><td>0</td><td>CAGCCCCACGACGAGGCGGT</td><td>GGG</td><td>||||||||[||||||||||||]PAM</td><td>NA</td><td>Intergenic</td><td>NA</td><td>NA</td></tr>
<tr><td></td><td></td><td></td><td></td><td></td><td></td><td></td><td></td><td></td><td></td><td></td><td></td></tr>
<tr><td>T14992</td><td>AGCCCCACGACGAGGCGGTGGGG</td><td>1000</td><td></td><td></td><td></td><td></td><td></td><td></td><td></td><td></td><td></td></tr>
<tr><td>Oligo fwd</td><td>CACCAGCCCCACGACGAGGCGGTG</td><td></td><td></td><td></td><td></td><td></td><td></td><td></td><td></td><td></td><td></td></tr>
<tr><td>Oligo rev</td><td>AAACCACCGCCTCGTCGTGGGGCT</td><td></td><td></td><td></td><td></td><td></td><td></td><td></td><td></td><td></td><td></td></tr>
<tr><td>Chromosome</td><td>start</td><td>end</td><td>strand</td><td>MM</td><td>target_seq</td><td>PAM</td><td>alignment</td><td>distance</td><td>position</td><td>gene name</td><td>gene id</td></tr>
<tr><td>GU305914.1</td><td>197096</td><td>197118</td><td>+</td><td>0</td><td>AGCCCCACGACGAGGCGGTG</td><td>GGG</td><td>||||||||[||||||||||||]PAM</td><td>NA</td><td>Intergenic</td><td>NA</td><td>NA</td></tr>
<tr><td></td><td></td><td></td><td></td><td></td><td></td><td></td><td></td><td></td><td></td><td></td><td></td></tr>
<tr><td>T14993</td><td>GCCCCACGACGAGGCGGTGGGGG</td><td>1000</td><td></td><td></td><td></td><td></td><td></td><td></td><td></td><td></td><td></td></tr>
<tr><td>Oligo fwd</td><td>CACCGCCCCACGACGAGGCGGTGG</td><td></td><td></td><td></td><td></td><td></td><td></td><td></td><td></td><td></td><td></td></tr>
<tr><td>Oligo rev</td><td>AAACCCACCGCCTCGTCGTGGGGC</td><td></td><td></td><td></td><td></td><td></td><td></td><td></td><td></td><td></td><td></td></tr>
<tr><td>Chromosome</td><td>start</td><td>end</td><td>strand</td><td>MM</td><td>target_seq</td><td>PAM</td><td>alignment</td><td>distance</td><td>position</td><td>gene name</td><td>gene id</td></tr>
<tr><td>GU305914.1</td><td>197097</td><td>197119</td><td>+</td><td>0</td><td>GCCCCACGACGAGGCGGTGG</td><td>GGG</td><td>||||||||[||||||||||||]PAM</td><td>NA</td><td>Intergenic</td><td>NA</td><td>NA</td></tr>
</table>

T14994 CCCCACGACGAGGCGGTGGGGGG 1000
Oligo fwd CACCCCCCACGACGAGGCGGTGGG
Oligo rev AAACCCCACCGCCTCGTCGTGGGG
Chromosome start end strand MM target_seq PAM alignment distance position gene name gene id
GU305914.1 197098 197120 + 0 CCCCACGACGAGGCGGTGGG GGG ||||||||[||||||||||||]PAM NA Intergenic NA NA

T14995 GACGAGGCGGTGGGGGGTCGTGG 1000
Oligo fwd CACCGACGAGGCGGTGGGGGGTCG
Oligo rev AAACCGACCCCCCACCGCCTCGTC
Chromosome start end strand MM target_seq PAM alignment distance position gene name gene id
GU305914.1 197104 197126 + 0 GACGAGGCGGTGGGGGGTCG TGG ||||||||[||||||||||||]PAM NA Intergenic NA NA

T14996 TGGGGGGTCGTGGTCACCTGCGG 1000
Oligo fwd CACCTGGGGGGTCGTGGTCACCTG
Oligo rev AAACCAGGTGACCACGACCCCCCA
Chromosome start end strand MM target_seq PAM alignment distance position gene name gene id
GU305914.1 197114 197136 + 0 TGGGGGGTCGTGGTCACCTG CGG ||||||||[||||||||||||]PAM NA Intergenic NA NA

T14997 GGGTCGTGGTCACCTGCGGCAGG 1000
Oligo fwd CACCGGGTCGTGGTCACCTGCGGC
Oligo rev AAACGCCGCAGGTGACCACGACCC
Chromosome start end strand MM target_seq PAM alignment distance position gene name gene id
GU305914.1 197118 197140 + 0 GGGTCGTGGTCACCTGCGGC AGG ||||||||[||||||||||||]PAM NA Intergenic NA NA

T14998 GTGGTCACCTGCGGCAGGTGCGG 1000
Oligo fwd CACCGTGGTCACCTGCGGCAGGTG
Oligo rev AAACCACCTGCCGCAGGTGACCAC
Chromosome start end strand MM target_seq PAM alignment distance position gene name gene id
GU305914.1 197123 197145 + 0 GTGGTCACCTGCGGCAGGTG CGG ||||||||[||||||||||||]PAM NA Intergenic NA NA

T14999 TGGTCACCTGCGGCAGGTGCGGG 1000
Oligo fwd CACCTGGTCACCTGCGGCAGGTGC
Oligo rev AAACGCACCTGCCGCAGGTGACCA
Chromosome start end strand MM target_seq PAM alignment distance position gene name gene id
GU305914.1 197124 197146 + 0 TGGTCACCTGCGGCAGGTGC GGG ||||||||[||||||||||||]PAM NA Intergenic NA NA

T15000 CCTGCGGCAGGTGCGGGCGACGG 1000
Oligo fwd CACCCCTGCGGCAGGTGCGGGCGA
Oligo rev AAACTCGCCCGCACCTGCCGCAGG
Chromosome start end strand MM target_seq PAM alignment distance position gene name gene id
GU305914.1 197130 197152 + 0 CCTGCGGCAGGTGCGGGCGA CGG ||||||||[||||||||||||]PAM NA Intergenic NA NA

T15001 CGAAACGCGCCAGTTTATCCAGG 1000
Oligo fwd CACCCGAAACGCGCCAGTTTATCC
Oligo rev AAACGGATAAACTGGCGCGTTTCG
Chromosome start end strand MM target_seq PAM alignment distance position gene name gene id
GU305914.1 197178 197200 + 0 CGAAACGCGCCAGTTTATCC AGG ||||||||[||||||||||||]PAM NA Intergenic NA NA

T15002 CGCGCCAGTTTATCCAGGTCCGG 1000
Oligo fwd CACCCGCGCCAGTTTATCCAGGTC
Oligo rev AAACGACCTGGATAAACTGGCGCG
Chromosome start end strand MM target_seq PAM alignment distance position gene name gene id
GU305914.1 197183 197205 + 0 CGCGCCAGTTTATCCAGGTC CGG ||||||||[||||||||||||]PAM NA Intergenic NA NA

T15003 GCGCCAGTTTATCCAGGTCCGGG 1000
Oligo fwd CACCGCGCCAGTTTATCCAGGTCC
Oligo rev AAACGGACCTGGATAAACTGGCGC
Chromosome start end strand MM target_seq PAM alignment distance position gene name gene id
GU305914.1 197184 197206 + 0 GCGCCAGTTTATCCAGGTCC GGG ||||||||[||||||||||||]PAM NA Intergenic NA NA

T15004 CGCCAGTTTATCCAGGTCCGGGG 1000
Oligo fwd CACCCGCCAGTTTATCCAGGTCCG
Oligo rev AAACCGGACCTGGATAAACTGGCG
Chromosome start end strand MM target_seq PAM alignment distance position gene name gene id
GU305914.1 197185 197207 + 0 CGCCAGTTTATCCAGGTCCG GGG ||||||||[||||||||||||]PAM NA Intergenic NA NA

T15005 TATCCAGGTCCGGGGCCACCAGG 1000
Oligo fwd CACCTATCCAGGTCCGGGGCCACC
Oligo rev AAACGGTGGCCCCGGACCTGGATA
Chromosome start end strand MM target_seq PAM alignment distance position gene name gene id
GU305914.1 197193 197215 + 0 TATCCAGGTCCGGGGCCACC AGG ||||||||[||||||||||||]PAM NA Intergenic NA NA

T15006 GTCCGGGGCCACCAGGATCATGG 1000
Oligo fwd CACCGTCCGGGGCCACCAGGATCA
Oligo rev AAACTGATCCTGGTGGCCCCGGAC
Chromosome start end strand MM target_seq PAM alignment distance position gene name gene id
GU305914.1 197200 197222 + 0 GTCCGGGGCCACCAGGATCA TGG ||||||||[||||||||||||]PAM NA Intergenic NA NA

T15007 GAATCTCGTAGACGAAGACCCGG 1000
Oligo fwd CACCGAATCTCGTAGACGAAGACC
Oligo rev AAACGGTCTTCGTCTACGAGATTC
Chromosome start end strand MM target_seq PAM alignment distance position gene name gene id
GU305914.1 197232 197254 + 0 GAATCTCGTAGACGAAGACC CGG ||||||||[||||||||||||]PAM NA Intergenic NA NA
GU305914.1 31660 31682 + 4 TGATCTCGTAGGTGAAGACC TGG --||||||[|||--|||||||]PAM NA Intergenic NA NA

T15008 AATCTCGTAGACGAAGACCCGGG 1000
Oligo fwd CACCAATCTCGTAGACGAAGACCC
Oligo rev AAACGGGTCTTCGTCTACGAGATT

| | | | | | | | | | | | |
|---|---|---|---|---|---|---|---|---|---|---|---|
| Chromosome | start | end | strand | MM | target_seq | PAM | alignment | distance | position | gene name | gene id |
| GU305914.1 | 197233 | 197255 | + | 0 | AATCTCGTAGACGAAGACCC | GGG | \|\|\|\|\|\|\|\|[\|\|\|\|\|\|\|\|\|\|\|\|]PAM | NA | Intergenic | NA | NA |
| T15009 | GGTCCACGCTCCCAGACACAAGG | 1000 | | | | | | | | | |
| Oligo fwd | CACCGGTCCACGCTCCCAGACACA | | | | | | | | | | |
| Oligo rev | AAACTGTGTCTGGGAGCGTGGACC | | | | | | | | | | |
| Chromosome | start | end | strand | MM | target_seq | PAM | alignment | distance | position | gene name | gene id |
| GU305914.1 | 197254 | 197276 | + | 0 | GGTCCACGCTCCCAGACACA | AGG | \|\|\|\|\|\|\|\|[\|\|\|\|\|\|\|\|\|\|\|\|]PAM | NA | Intergenic | NA | NA |
| T15010 | CCACGCTCCCAGACACAAGGCGG | 1000 | | | | | | | | | |
| Oligo fwd | CACCCCACGCTCCCAGACACAAGG | | | | | | | | | | |
| Oligo rev | AAACCCTTGTGTCTGGGAGCGTGG | | | | | | | | | | |
| Chromosome | start | end | strand | MM | target_seq | PAM | alignment | distance | position | gene name | gene id |
| GU305914.1 | 197257 | 197279 | + | 0 | CCACGCTCCCAGACACAAGG | CGG | \|\|\|\|\|\|\|\|[\|\|\|\|\|\|\|\|\|\|\|\|]PAM | NA | Intergenic | NA | NA |
| T15011 | CCTCGTATCCGTCGTTGTTGCGG | 1000 | | | | | | | | | |
| Oligo fwd | CACCCCTCGTATCCGTCGTTGTTG | | | | | | | | | | |
| Oligo rev | AAACCAACAACGACGGATACGAGG | | | | | | | | | | |
| Chromosome | start | end | strand | MM | target_seq | PAM | alignment | distance | position | gene name | gene id |
| GU305914.1 | 197289 | 197311 | + | 0 | CCTCGTATCCGTCGTTGTTG | CGG | \|\|\|\|\|\|\|\|[\|\|\|\|\|\|\|\|\|\|\|\|]PAM | NA | Intergenic | NA | NA |
| T15012 | CGATCCGACCGACGACCGTCAGG | 1000 | | | | | | | | | |
| Oligo fwd | CACCCGATCCGACCGACGACCGTC | | | | | | | | | | |
| Oligo rev | AAACGACGGTCGTCGGTCGGATCG | | | | | | | | | | |
| Chromosome | start | end | strand | MM | target_seq | PAM | alignment | distance | position | gene name | gene id |
| GU305914.1 | 197316 | 197338 | + | 0 | CGATCCGACCGACGACCGTC | AGG | \|\|\|\|\|\|\|\|[\|\|\|\|\|\|\|\|\|\|\|\|]PAM | NA | Intergenic | NA | NA |
| T15013 | GATCCGACCGACGACCGTCAGGG | 1000 | | | | | | | | | |
| Oligo fwd | CACCGATCCGACCGACGACCGTCA | | | | | | | | | | |
| Oligo rev | AAACTGACGGTCGTCGGTCGGATC | | | | | | | | | | |
| Chromosome | start | end | strand | MM | target_seq | PAM | alignment | distance | position | gene name | gene id |
| GU305914.1 | 197317 | 197339 | + | 0 | GATCCGACCGACGACCGTCA | GGG | \|\|\|\|\|\|\|\|[\|\|\|\|\|\|\|\|\|\|\|\|]PAM | NA | Intergenic | NA | NA |
| T15014 | ATCCGACCGACGACCGTCAGGGG | 1000 | | | | | | | | | |
| Oligo fwd | CACCATCCGACCGACGACCGTCAG | | | | | | | | | | |
| Oligo rev | AAACCTGACGGTCGTCGGTCGGAT | | | | | | | | | | |
| Chromosome | start | end | strand | MM | target_seq | PAM | alignment | distance | position | gene name | gene id |
| GU305914.1 | 197318 | 197340 | + | 0 | ATCCGACCGACGACCGTCAG | GGG | \|\|\|\|\|\|\|\|[\|\|\|\|\|\|\|\|\|\|\|\|]PAM | NA | Intergenic | NA | NA |
| T15015 | CGCCTTTCAGTTCCTCGTAGCGG | 1000 | | | | | | | | | |
| Oligo fwd | CACCCGCCTTTCAGTTCCTCGTAG | | | | | | | | | | |
| Oligo rev | AAACCTACGAGGAACTGAAAGGCG | | | | | | | | | | |
| Chromosome | start | end | strand | MM | target_seq | PAM | alignment | distance | position | gene name | gene id |
| GU305914.1 | 197373 | 197395 | + | 0 | CGCCTTTCAGTTCCTCGTAG | CGG | \|\|\|\|\|\|\|\|[\|\|\|\|\|\|\|\|\|\|\|\|]PAM | NA | Intergenic | NA | NA |
| T15016 | CAGTTCCTCGTAGCGGCTCGCGG | 1000 | | | | | | | | | |
| Oligo fwd | CACCCAGTTCCTCGTAGCGGCTCG | | | | | | | | | | |
| Oligo rev | AAACCGAGCCGCTACGAGGAACTG | | | | | | | | | | |
| Chromosome | start | end | strand | MM | target_seq | PAM | alignment | distance | position | gene name | gene id |
| GU305914.1 | 197380 | 197402 | + | 0 | CAGTTCCTCGTAGCGGCTCG | CGG | \|\|\|\|\|\|\|\|[\|\|\|\|\|\|\|\|\|\|\|\|]PAM | NA | Intergenic | NA | NA |
| T15017 | AGTTCCTCGTAGCGGCTCGCGGG | 1000 | | | | | | | | | |
| Oligo fwd | CACCAGTTCCTCGTAGCGGCTCGC | | | | | | | | | | |
| Oligo rev | AAACGCGAGCCGCTACGAGGAACT | | | | | | | | | | |
| Chromosome | start | end | strand | MM | target_seq | PAM | alignment | distance | position | gene name | gene id |
| GU305914.1 | 197381 | 197403 | + | 0 | AGTTCCTCGTAGCGGCTCGC | GGG | \|\|\|\|\|\|\|\|[\|\|\|\|\|\|\|\|\|\|\|\|]PAM | NA | Intergenic | NA | NA |
| T15018 | GTTCCTCGTAGCGGCTCGCGGGG | 1000 | | | | | | | | | |
| Oligo fwd | CACCGTTCCTCGTAGCGGCTCGCG | | | | | | | | | | |
| Oligo rev | AAACCGCGAGCCGCTACGAGGAAC | | | | | | | | | | |
| Chromosome | start | end | strand | MM | target_seq | PAM | alignment | distance | position | gene name | gene id |
| GU305914.1 | 197382 | 197404 | + | 0 | GTTCCTCGTAGCGGCTCGCG | GGG | \|\|\|\|\|\|\|\|[\|\|\|\|\|\|\|\|\|\|\|\|]PAM | NA | Intergenic | NA | NA |
| GU305914.1 | 151991 | 152013 | - | 4 | GGTGGTCGTAGCGGCTCTCG | AGG | \|-\|--\|\|\|[\|\|\|\|\|\|\|\|\|-\|\|]PAM | NA | Intergenic | NA | NA |
| T15019 | TTCCTCGTAGCGGCTCGCGGGGG | 1000 | | | | | | | | | |
| Oligo fwd | CACCTTCCTCGTAGCGGCTCGCGG | | | | | | | | | | |
| Oligo rev | AAACCCGCGAGCCGCTACGAGGAA | | | | | | | | | | |
| Chromosome | start | end | strand | MM | target_seq | PAM | alignment | distance | position | gene name | gene id |
| GU305914.1 | 197383 | 197405 | + | 0 | TTCCTCGTAGCGGCTCGCGG | GGG | \|\|\|\|\|\|\|\|[\|\|\|\|\|\|\|\|\|\|\|\|]PAM | NA | Intergenic | NA | NA |
| T15020 | TCAGCTCCACGAACCAGTTGTGG | 1000 | | | | | | | | | |
| Oligo fwd | CACCTCAGCTCCACGAACCAGTTG | | | | | | | | | | |
| Oligo rev | AAACCAACTGGTTCGTGGAGCTGA | | | | | | | | | | |
| Chromosome | start | end | strand | MM | target_seq | PAM | alignment | distance | position | gene name | gene id |
| GU305914.1 | 197406 | 197428 | + | 0 | TCAGCTCCACGAACCAGTTG | TGG | \|\|\|\|\|\|\|\|[\|\|\|\|\|\|\|\|\|\|\|\|]PAM | NA | Intergenic | NA | NA |
| T15021 | CAGCTCCACGAACCAGTTGTGGG | 1000 | | | | | | | | | |
| Oligo fwd | CACCCAGCTCCACGAACCAGTTGT | | | | | | | | | | |
| Oligo rev | AAACACAACTGGTTCGTGGAGCTG | | | | | | | | | | |
| Chromosome | start | end | strand | MM | target_seq | PAM | alignment | distance | position | gene name | gene id |
| GU305914.1 | 197407 | 197429 | + | 0 | CAGCTCCACGAACCAGTTGT | GGG | \|\|\|\|\|\|\|\|[\|\|\|\|\|\|\|\|\|\|\|\|]PAM | NA | Intergenic | NA | NA |
| T15022 | AGCTCCACGAACCAGTTGTGGGG | 1000 | | | | | | | | | |
| Oligo fwd | CACCAGCTCCACGAACCAGTTGTG | | | | | | | | | | |
| Oligo rev | AAACCACAACTGGTTCGTGGAGCT | | | | | | | | | | |
| Chromosome | start | end | strand | MM | target_seq | PAM | alignment | distance | position | gene name | gene id |
| GU305914.1 | 197408 | 197430 | + | 0 | AGCTCCACGAACCAGTTGTG | GGG | \|\|\|\|\|\|\|\|[\|\|\|\|\|\|\|\|\|\|\|\|]PAM | NA | Intergenic | NA | NA |
| T15023 | GCTCCACGAACCAGTTGTGGGGG | 1000 | | | | | | | | | |

| | | | | | | | | | | | |
|---|---|---|---|---|---|---|---|---|---|---|---|
| Oligo fwd | CACCGCTCCACGAACCAGTTGTGG | | | | | | | | | | |
| Oligo rev | AAACCCACAACTGGTTCGTGGAGC | | | | | | | | | | |
| Chromosome | start | end | strand | MM | target_seq | PAM | alignment | distance | position | gene name | gene id |
| GU305914.1 | 197409 | 197431 | + | 0 | GCTCCACGAACCAGTTGTGG | GGG | \|\|\|\|\|\|\|\|[\|\|\|\|\|\|\|\|\|\|\|\|]PAM | NA | Intergenic | NA | NA |
| T15024 | CAGTTGTGGGGGATGCCCAGCGG | 1000 | | | | | | | | | |
| Oligo fwd | CACCCAGTTGTGGGGGATGCCCAG | | | | | | | | | | |
| Oligo rev | AAACCTGGGCATCCCCCACAACTG | | | | | | | | | | |
| Chromosome | start | end | strand | MM | target_seq | PAM | alignment | distance | position | gene name | gene id |
| GU305914.1 | 197420 | 197442 | + | 0 | CAGTTGTGGGGGATGCCCAG | CGG | \|\|\|\|\|\|\|\|[\|\|\|\|\|\|\|\|\|\|\|\|]PAM | NA | Intergenic | NA | NA |
| T15025 | GCCCAGCGGCAGACGCTCGCCGG | 1000 | | | | | | | | | |
| Oligo fwd | CACCGCCCAGCGGCAGACGCTCGC | | | | | | | | | | |
| Oligo rev | AAACGCGAGCGTCTGCCGCTGGGC | | | | | | | | | | |
| Chromosome | start | end | strand | MM | target_seq | PAM | alignment | distance | position | gene name | gene id |
| GU305914.1 | 197434 | 197456 | + | 0 | GCCCAGCGGCAGACGCTCGC | CGG | \|\|\|\|\|\|\|\|[\|\|\|\|\|\|\|\|\|\|\|\|]PAM | NA | Intergenic | NA | NA |
| T15026 | CGCCGGCGCGCCTCTCGACCAGG | 1000 | | | | | | | | | |
| Oligo fwd | CACCCGCCGGCGCGCCTCTCGACC | | | | | | | | | | |
| Oligo rev | AAACGGTCGAGAGGCGCGCCGGCG | | | | | | | | | | |
| Chromosome | start | end | strand | MM | target_seq | PAM | alignment | distance | position | gene name | gene id |
| GU305914.1 | 197451 | 197473 | + | 0 | CGCCGGCGCGCCTCTCGACC | AGG | \|\|\|\|\|\|\|\|[\|\|\|\|\|\|\|\|\|\|\|\|]PAM | NA | Intergenic | NA | NA |
| T15027 | GCCGGCGCGCCTCTCGACCAGGG | 1000 | | | | | | | | | |
| Oligo fwd | CACCGCCGGCGCGCCTCTCGACCA | | | | | | | | | | |
| Oligo rev | AAACTGGTCGAGAGGCGCGCCGGC | | | | | | | | | | |
| Chromosome | start | end | strand | MM | target_seq | PAM | alignment | distance | position | gene name | gene id |
| GU305914.1 | 197452 | 197474 | + | 0 | GCCGGCGCGCCTCTCGACCA | GGG | \|\|\|\|\|\|\|\|[\|\|\|\|\|\|\|\|\|\|\|\|]PAM | NA | Intergenic | NA | NA |
| T15028 | CCAGGGACGCCAGCTCCCCGCGG | 1000 | | | | | | | | | |
| Oligo fwd | CACCCCAGGGACGCCAGCTCCCCG | | | | | | | | | | |
| Oligo rev | AAACCGGGGAGCTGGCGTCCCTGG | | | | | | | | | | |
| Chromosome | start | end | strand | MM | target_seq | PAM | alignment | distance | position | gene name | gene id |
| GU305914.1 | 197469 | 197491 | + | 0 | CCAGGGACGCCAGCTCCCCG | CGG | \|\|\|\|\|\|\|\|[\|\|\|\|\|\|\|\|\|\|\|\|]PAM | NA | Intergenic | NA | NA |
| T15029 | CCAGCTCCCCGCGGTTCGACTGG | 1000 | | | | | | | | | |
| Oligo fwd | CACCCCAGCTCCCCGCGGTTCGAC | | | | | | | | | | |
| Oligo rev | AAACGTCGAACCGCGGGGAGCTGG | | | | | | | | | | |
| Chromosome | start | end | strand | MM | target_seq | PAM | alignment | distance | position | gene name | gene id |
| GU305914.1 | 197478 | 197500 | + | 0 | CCAGCTCCCCGCGGTTCGAC | TGG | \|\|\|\|\|\|\|\|[\|\|\|\|\|\|\|\|\|\|\|\|]PAM | NA | Intergenic | NA | NA |
| T15030 | GCTCCCCGCGGTTCGACTGGTGG | 1000 | | | | | | | | | |
| Oligo fwd | CACCGCTCCCCGCGGTTCGACTGG | | | | | | | | | | |
| Oligo rev | AAACCCAGTCGAACCGCGGGGAGC | | | | | | | | | | |
| Chromosome | start | end | strand | MM | target_seq | PAM | alignment | distance | position | gene name | gene id |
| GU305914.1 | 197481 | 197503 | + | 0 | GCTCCCCGCGGTTCGACTGG | TGG | \|\|\|\|\|\|\|\|[\|\|\|\|\|\|\|\|\|\|\|\|]PAM | NA | Intergenic | NA | NA |
| T15031 | CCCCGCGGTTCGACTGGTGGCGG | 1000 | | | | | | | | | |
| Oligo fwd | CACCCCCCGCGGTTCGACTGGTGG | | | | | | | | | | |
| Oligo rev | AAACCCACCAGTCGAACCGCGGGG | | | | | | | | | | |
| Chromosome | start | end | strand | MM | target_seq | PAM | alignment | distance | position | gene name | gene id |
| GU305914.1 | 197484 | 197506 | + | 0 | CCCCGCGGTTCGACTGGTGG | CGG | \|\|\|\|\|\|\|\|[\|\|\|\|\|\|\|\|\|\|\|\|]PAM | NA | Intergenic | NA | NA |
| T15032 | ACTGGTGGCGGTAGAAGTCGCGG | 1000 | | | | | | | | | |
| Oligo fwd | CACCACTGGTGGCGGTAGAAGTCG | | | | | | | | | | |
| Oligo rev | AAACCGACTTCTACCGCCACCAGT | | | | | | | | | | |
| Chromosome | start | end | strand | MM | target_seq | PAM | alignment | distance | position | gene name | gene id |
| GU305914.1 | 197496 | 197518 | + | 0 | ACTGGTGGCGGTAGAAGTCG | CGG | \|\|\|\|\|\|\|\|[\|\|\|\|\|\|\|\|\|\|\|\|]PAM | NA | Intergenic | NA | NA |
| T15033 | CGCGGAACGCGAGCATCACGCGG | 1000 | | | | | | | | | |
| Oligo fwd | CACCCGCGGAACGCGAGCATCACG | | | | | | | | | | |
| Oligo rev | AAACCGTGATGCTCGCGTTCCGCG | | | | | | | | | | |
| Chromosome | start | end | strand | MM | target_seq | PAM | alignment | distance | position | gene name | gene id |
| GU305914.1 | 197514 | 197536 | + | 0 | CGCGGAACGCGAGCATCACG | CGG | \|\|\|\|\|\|\|\|[\|\|\|\|\|\|\|\|\|\|\|\|]PAM | NA | Intergenic | NA | NA |
| T15034 | TCACGCGGAGAAACGATGAGCGG | 1000 | | | | | | | | | |
| Oligo fwd | CACCTCACGCGGAGAAACGATGAG | | | | | | | | | | |
| Oligo rev | AAACCTCATCGTTTCTCCGCGTGA | | | | | | | | | | |
| Chromosome | start | end | strand | MM | target_seq | PAM | alignment | distance | position | gene name | gene id |
| GU305914.1 | 197529 | 197551 | + | 0 | TCACGCGGAGAAACGATGAG | CGG | \|\|\|\|\|\|\|\|[\|\|\|\|\|\|\|\|\|\|\|\|]PAM | NA | Intergenic | NA | NA |
| T15035 | CACGCGGAGAAACGATGAGCGGG | 1000 | | | | | | | | | |
| Oligo fwd | CACCCACGCGGAGAAACGATGAGC | | | | | | | | | | |
| Oligo rev | AAACGCTCATCGTTTCTCCGCGTG | | | | | | | | | | |
| Chromosome | start | end | strand | MM | target_seq | PAM | alignment | distance | position | gene name | gene id |
| GU305914.1 | 197530 | 197552 | + | 0 | CACGCGGAGAAACGATGAGC | GGG | \|\|\|\|\|\|\|\|[\|\|\|\|\|\|\|\|\|\|\|\|]PAM | NA | Intergenic | NA | NA |
| T15036 | GCGGAGAAACGATGAGCGGGTGG | 1000 | | | | | | | | | |
| Oligo fwd | CACCGCGGAGAAACGATGAGCGGG | | | | | | | | | | |
| Oligo rev | AAACCCCGCTCATCGTTTCTCCGC | | | | | | | | | | |
| Chromosome | start | end | strand | MM | target_seq | PAM | alignment | distance | position | gene name | gene id |
| GU305914.1 | 197533 | 197555 | + | 0 | GCGGAGAAACGATGAGCGGG | TGG | \|\|\|\|\|\|\|\|[\|\|\|\|\|\|\|\|\|\|\|\|]PAM | NA | Intergenic | NA | NA |
| T15037 | TGAGCGGGTGGCGTGCCGATAGG | 1000 | | | | | | | | | |
| Oligo fwd | CACCTGAGCGGGTGGCGTGCCGAT | | | | | | | | | | |
| Oligo rev | AAACATCGGCACGCCACCCGCTCA | | | | | | | | | | |
| Chromosome | start | end | strand | MM | target_seq | PAM | alignment | distance | position | gene name | gene id |
| GU305914.1 | 197545 | 197567 | + | 0 | TGAGCGGGTGGCGTGCCGAT | AGG | \|\|\|\|\|\|\|\|[\|\|\|\|\|\|\|\|\|\|\|\|]PAM | NA | Intergenic | NA | NA |

<table>
<tr><td>T15038</td><td>GAGCGGGTGGCGTGCCGATAGGG</td><td>1000</td><td></td><td></td><td></td><td></td><td></td><td></td><td></td><td></td><td></td></tr>
<tr><td>Oligo fwd</td><td>CACCGAGCGGGTGGCGTGCCGATA</td><td></td><td></td><td></td><td></td><td></td><td></td><td></td><td></td><td></td><td></td></tr>
<tr><td>Oligo rev</td><td>AAACTATCGGCACGCCACCCGCTC</td><td></td><td></td><td></td><td></td><td></td><td></td><td></td><td></td><td></td><td></td></tr>
<tr><td>Chromosome</td><td>start</td><td>end</td><td>strand</td><td>MM</td><td>target_seq</td><td>PAM</td><td>alignment</td><td>distance</td><td>position</td><td>gene name</td><td>gene id</td></tr>
<tr><td>GU305914.1</td><td>197546</td><td>197568</td><td>+</td><td>0</td><td>GAGCGGGTGGCGTGCCGATA</td><td>GGG</td><td>||||||||[||||||||||||]PAM</td><td>NA</td><td>Intergenic</td><td>NA</td><td>NA</td></tr>
<tr><td></td><td></td><td></td><td></td><td></td><td></td><td></td><td></td><td></td><td></td><td></td><td></td></tr>
<tr><td>T15039</td><td>GTGGCGTGCCGATAGGGTGCCGG</td><td>1000</td><td></td><td></td><td></td><td></td><td></td><td></td><td></td><td></td><td></td></tr>
<tr><td>Oligo fwd</td><td>CACCGTGGCGTGCCGATAGGGTGC</td><td></td><td></td><td></td><td></td><td></td><td></td><td></td><td></td><td></td><td></td></tr>
<tr><td>Oligo rev</td><td>AAACGCACCCTATCGGCACGCCAC</td><td></td><td></td><td></td><td></td><td></td><td></td><td></td><td></td><td></td><td></td></tr>
<tr><td>Chromosome</td><td>start</td><td>end</td><td>strand</td><td>MM</td><td>target_seq</td><td>PAM</td><td>alignment</td><td>distance</td><td>position</td><td>gene name</td><td>gene id</td></tr>
<tr><td>GU305914.1</td><td>197552</td><td>197574</td><td>+</td><td>0</td><td>GTGGCGTGCCGATAGGGTGC</td><td>CGG</td><td>||||||||[||||||||||||]PAM</td><td>NA</td><td>Intergenic</td><td>NA</td><td>NA</td></tr>
<tr><td>GU305914.1</td><td>69202</td><td>69224</td><td>+</td><td>4</td><td>GCGGCGCGCCGACAAGGTGC</td><td>TGG</td><td>|-||||-|[||||-|-|||||]PAM</td><td>NA</td><td>Intergenic</td><td>NA</td><td>NA</td></tr>
<tr><td></td><td></td><td></td><td></td><td></td><td></td><td></td><td></td><td></td><td></td><td></td><td></td></tr>
<tr><td>T15040</td><td>CCGATAGGGTGCCGGATCATCGG</td><td>1000</td><td></td><td></td><td></td><td></td><td></td><td></td><td></td><td></td><td></td></tr>
<tr><td>Oligo fwd</td><td>CACCCCGATAGGGTGCCGGATCAT</td><td></td><td></td><td></td><td></td><td></td><td></td><td></td><td></td><td></td><td></td></tr>
<tr><td>Oligo rev</td><td>AAACATGATCCGGCACCCTATCGG</td><td></td><td></td><td></td><td></td><td></td><td></td><td></td><td></td><td></td><td></td></tr>
<tr><td>Chromosome</td><td>start</td><td>end</td><td>strand</td><td>MM</td><td>target_seq</td><td>PAM</td><td>alignment</td><td>distance</td><td>position</td><td>gene name</td><td>gene id</td></tr>
<tr><td>GU305914.1</td><td>197560</td><td>197582</td><td>+</td><td>0</td><td>CCGATAGGGTGCCGGATCAT</td><td>CGG</td><td>||||||||[||||||||||||]PAM</td><td>NA</td><td>Intergenic</td><td>NA</td><td>NA</td></tr>
<tr><td></td><td></td><td></td><td></td><td></td><td></td><td></td><td></td><td></td><td></td><td></td><td></td></tr>
<tr><td>T15041</td><td>GGGTGCCGGATCATCGGTCACGG</td><td>1000</td><td></td><td></td><td></td><td></td><td></td><td></td><td></td><td></td><td></td></tr>
<tr><td>Oligo fwd</td><td>CACCGGGTGCCGGATCATCGGTCA</td><td></td><td></td><td></td><td></td><td></td><td></td><td></td><td></td><td></td><td></td></tr>
<tr><td>Oligo rev</td><td>AAACTGACCGATGATCCGGCACCC</td><td></td><td></td><td></td><td></td><td></td><td></td><td></td><td></td><td></td><td></td></tr>
<tr><td>Chromosome</td><td>start</td><td>end</td><td>strand</td><td>MM</td><td>target_seq</td><td>PAM</td><td>alignment</td><td>distance</td><td>position</td><td>gene name</td><td>gene id</td></tr>
<tr><td>GU305914.1</td><td>197566</td><td>197588</td><td>+</td><td>0</td><td>GGGTGCCGGATCATCGGTCA</td><td>CGG</td><td>||||||||[||||||||||||]PAM</td><td>NA</td><td>Intergenic</td><td>NA</td><td>NA</td></tr>
<tr><td></td><td></td><td></td><td></td><td></td><td></td><td></td><td></td><td></td><td></td><td></td><td></td></tr>
<tr><td>T15042</td><td>GGTGCCGGATCATCGGTCACGGG</td><td>1000</td><td></td><td></td><td></td><td></td><td></td><td></td><td></td><td></td><td></td></tr>
<tr><td>Oligo fwd</td><td>CACCGGTGCCGGATCATCGGTCAC</td><td></td><td></td><td></td><td></td><td></td><td></td><td></td><td></td><td></td><td></td></tr>
<tr><td>Oligo rev</td><td>AAACGTGACCGATGATCCGGCACC</td><td></td><td></td><td></td><td></td><td></td><td></td><td></td><td></td><td></td><td></td></tr>
<tr><td>Chromosome</td><td>start</td><td>end</td><td>strand</td><td>MM</td><td>target_seq</td><td>PAM</td><td>alignment</td><td>distance</td><td>position</td><td>gene name</td><td>gene id</td></tr>
<tr><td>GU305914.1</td><td>197567</td><td>197589</td><td>+</td><td>0</td><td>GGTGCCGGATCATCGGTCAC</td><td>GGG</td><td>||||||||[||||||||||||]PAM</td><td>NA</td><td>Intergenic</td><td>NA</td><td>NA</td></tr>
<tr><td></td><td></td><td></td><td></td><td></td><td></td><td></td><td></td><td></td><td></td><td></td><td></td></tr>
<tr><td>T15043</td><td>GTGCCGGATCATCGGTCACGGGG</td><td>1000</td><td></td><td></td><td></td><td></td><td></td><td></td><td></td><td></td><td></td></tr>
<tr><td>Oligo fwd</td><td>CACCGTGCCGGATCATCGGTCACG</td><td></td><td></td><td></td><td></td><td></td><td></td><td></td><td></td><td></td><td></td></tr>
<tr><td>Oligo rev</td><td>AAACCGTGACCGATGATCCGGCAC</td><td></td><td></td><td></td><td></td><td></td><td></td><td></td><td></td><td></td><td></td></tr>
<tr><td>Chromosome</td><td>start</td><td>end</td><td>strand</td><td>MM</td><td>target_seq</td><td>PAM</td><td>alignment</td><td>distance</td><td>position</td><td>gene name</td><td>gene id</td></tr>
<tr><td>GU305914.1</td><td>197568</td><td>197590</td><td>+</td><td>0</td><td>GTGCCGGATCATCGGTCACG</td><td>GGG</td><td>||||||||[||||||||||||]PAM</td><td>NA</td><td>Intergenic</td><td>NA</td><td>NA</td></tr>
<tr><td></td><td></td><td></td><td></td><td></td><td></td><td></td><td></td><td></td><td></td><td></td><td></td></tr>
<tr><td>T15044</td><td>CAGTGTGCTGTCCATCTCGCCGG</td><td>1000</td><td></td><td></td><td></td><td></td><td></td><td></td><td></td><td></td><td></td></tr>
<tr><td>Oligo fwd</td><td>CACCCAGTGTGCTGTCCATCTCGC</td><td></td><td></td><td></td><td></td><td></td><td></td><td></td><td></td><td></td><td></td></tr>
<tr><td>Oligo rev</td><td>AAACGCGAGATGGACAGCACACTG</td><td></td><td></td><td></td><td></td><td></td><td></td><td></td><td></td><td></td><td></td></tr>
<tr><td>Chromosome</td><td>start</td><td>end</td><td>strand</td><td>MM</td><td>target_seq</td><td>PAM</td><td>alignment</td><td>distance</td><td>position</td><td>gene name</td><td>gene id</td></tr>
<tr><td>GU305914.1</td><td>197602</td><td>197624</td><td>+</td><td>0</td><td>CAGTGTGCTGTCCATCTCGC</td><td>CGG</td><td>||||||||[||||||||||||]PAM</td><td>NA</td><td>Intergenic</td><td>NA</td><td>NA</td></tr>
<tr><td>GU305914.1</td><td>144773</td><td>144795</td><td>-</td><td>4</td><td>CATGGTGCTGGCCATGTCGC</td><td>TGG</td><td>||--||||[||-||||-||||]PAM</td><td>NA</td><td>Intergenic</td><td>NA</td><td>NA</td></tr>
<tr><td></td><td></td><td></td><td></td><td></td><td></td><td></td><td></td><td></td><td></td><td></td><td></td></tr>
<tr><td>T15045</td><td>CATCTCGCCGGATGAGATGATGG</td><td>1000</td><td></td><td></td><td></td><td></td><td></td><td></td><td></td><td></td><td></td></tr>
<tr><td>Oligo fwd</td><td>CACCCATCTCGCCGGATGAGATGA</td><td></td><td></td><td></td><td></td><td></td><td></td><td></td><td></td><td></td><td></td></tr>
<tr><td>Oligo rev</td><td>AAACTCATCTCATCCGGCGAGATG</td><td></td><td></td><td></td><td></td><td></td><td></td><td></td><td></td><td></td><td></td></tr>
<tr><td>Chromosome</td><td>start</td><td>end</td><td>strand</td><td>MM</td><td>target_seq</td><td>PAM</td><td>alignment</td><td>distance</td><td>position</td><td>gene name</td><td>gene id</td></tr>
<tr><td>GU305914.1</td><td>197614</td><td>197636</td><td>+</td><td>0</td><td>CATCTCGCCGGATGAGATGA</td><td>TGG</td><td>||||||||[||||||||||||]PAM</td><td>NA</td><td>Intergenic</td><td>NA</td><td>NA</td></tr>
<tr><td></td><td></td><td></td><td></td><td></td><td></td><td></td><td></td><td></td><td></td><td></td><td></td></tr>
<tr><td>T15046</td><td>GGATGAGATGATGGTGATAATGG</td><td>1000</td><td></td><td></td><td></td><td></td><td></td><td></td><td></td><td></td><td></td></tr>
<tr><td>Oligo fwd</td><td>CACCGGATGAGATGATGGTGATAA</td><td></td><td></td><td></td><td></td><td></td><td></td><td></td><td></td><td></td><td></td></tr>
<tr><td>Oligo rev</td><td>AAACTTATCACCATCATCTCATCC</td><td></td><td></td><td></td><td></td><td></td><td></td><td></td><td></td><td></td><td></td></tr>
<tr><td>Chromosome</td><td>start</td><td>end</td><td>strand</td><td>MM</td><td>target_seq</td><td>PAM</td><td>alignment</td><td>distance</td><td>position</td><td>gene name</td><td>gene id</td></tr>
<tr><td>GU305914.1</td><td>197623</td><td>197645</td><td>+</td><td>0</td><td>GGATGAGATGATGGTGATAA</td><td>TGG</td><td>||||||||[||||||||||||]PAM</td><td>NA</td><td>Intergenic</td><td>NA</td><td>NA</td></tr>
<tr><td></td><td></td><td></td><td></td><td></td><td></td><td></td><td></td><td></td><td></td><td></td><td></td></tr>
<tr><td>T15047</td><td>GATAATGGTCTAGAGAAAAACGG</td><td>1000</td><td></td><td></td><td></td><td></td><td></td><td></td><td></td><td></td><td></td></tr>
<tr><td>Oligo fwd</td><td>CACCGATAATGGTCTAGAGAAAAA</td><td></td><td></td><td></td><td></td><td></td><td></td><td></td><td></td><td></td><td></td></tr>
<tr><td>Oligo rev</td><td>AAACTTTTTCTCTAGACCATTATC</td><td></td><td></td><td></td><td></td><td></td><td></td><td></td><td></td><td></td><td></td></tr>
<tr><td>Chromosome</td><td>start</td><td>end</td><td>strand</td><td>MM</td><td>target_seq</td><td>PAM</td><td>alignment</td><td>distance</td><td>position</td><td>gene name</td><td>gene id</td></tr>
<tr><td>GU305914.1</td><td>197638</td><td>197660</td><td>+</td><td>0</td><td>GATAATGGTCTAGAGAAAAA</td><td>CGG</td><td>||||||||[||||||||||||]PAM</td><td>NA</td><td>Intergenic</td><td>NA</td><td>NA</td></tr>
<tr><td></td><td></td><td></td><td></td><td></td><td></td><td></td><td></td><td></td><td></td><td></td><td></td></tr>
<tr><td>T15048</td><td>ATAATGGTCTAGAGAAAAACGGG</td><td>1000</td><td></td><td></td><td></td><td></td><td></td><td></td><td></td><td></td><td></td></tr>
<tr><td>Oligo fwd</td><td>CACCATAATGGTCTAGAGAAAAAC</td><td></td><td></td><td></td><td></td><td></td><td></td><td></td><td></td><td></td><td></td></tr>
<tr><td>Oligo rev</td><td>AAACGTTTTTCTCTAGACCATTAT</td><td></td><td></td><td></td><td></td><td></td><td></td><td></td><td></td><td></td><td></td></tr>
<tr><td>Chromosome</td><td>start</td><td>end</td><td>strand</td><td>MM</td><td>target_seq</td><td>PAM</td><td>alignment</td><td>distance</td><td>position</td><td>gene name</td><td>gene id</td></tr>
<tr><td>GU305914.1</td><td>197639</td><td>197661</td><td>+</td><td>0</td><td>ATAATGGTCTAGAGAAAAAC</td><td>GGG</td><td>||||||||[||||||||||||]PAM</td><td>NA</td><td>Intergenic</td><td>NA</td><td>NA</td></tr>
<tr><td></td><td></td><td></td><td></td><td></td><td></td><td></td><td></td><td></td><td></td><td></td><td></td></tr>
<tr><td>T15049</td><td>AACGGGCCGATTCTCCGCCGCGG</td><td>1000</td><td></td><td></td><td></td><td></td><td></td><td></td><td></td><td></td><td></td></tr>
<tr><td>Oligo fwd</td><td>CACCAACGGGCCGATTCTCCGCCG</td><td></td><td></td><td></td><td></td><td></td><td></td><td></td><td></td><td></td><td></td></tr>
<tr><td>Oligo rev</td><td>AAACCGGCGGAGAATCGGCCCGTT</td><td></td><td></td><td></td><td></td><td></td><td></td><td></td><td></td><td></td><td></td></tr>
<tr><td>Chromosome</td><td>start</td><td>end</td><td>strand</td><td>MM</td><td>target_seq</td><td>PAM</td><td>alignment</td><td>distance</td><td>position</td><td>gene name</td><td>gene id</td></tr>
<tr><td>GU305914.1</td><td>197656</td><td>197678</td><td>+</td><td>0</td><td>AACGGGCCGATTCTCCGCCG</td><td>CGG</td><td>||||||||[||||||||||||]PAM</td><td>NA</td><td>Intergenic</td><td>NA</td><td>NA</td></tr>
<tr><td></td><td></td><td></td><td></td><td></td><td></td><td></td><td></td><td></td><td></td><td></td><td></td></tr>
<tr><td>T15050</td><td>ACGGGCCGATTCTCCGCCGCGGG</td><td>1000</td><td></td><td></td><td></td><td></td><td></td><td></td><td></td><td></td><td></td></tr>
<tr><td>Oligo fwd</td><td>CACCACGGGCCGATTCTCCGCCGC</td><td></td><td></td><td></td><td></td><td></td><td></td><td></td><td></td><td></td><td></td></tr>
<tr><td>Oligo rev</td><td>AAACGCGGCGGAGAATCGGCCCGT</td><td></td><td></td><td></td><td></td><td></td><td></td><td></td><td></td><td></td><td></td></tr>
<tr><td>Chromosome</td><td>start</td><td>end</td><td>strand</td><td>MM</td><td>target_seq</td><td>PAM</td><td>alignment</td><td>distance</td><td>position</td><td>gene name</td><td>gene id</td></tr>
<tr><td>GU305914.1</td><td>197657</td><td>197679</td><td>+</td><td>0</td><td>ACGGGCCGATTCTCCGCCGC</td><td>GGG</td><td>||||||||[||||||||||||]PAM</td><td>NA</td><td>Intergenic</td><td>NA</td><td>NA</td></tr>
<tr><td></td><td></td><td></td><td></td><td></td><td></td><td></td><td></td><td></td><td></td><td></td><td></td></tr>
<tr><td>T15051</td><td>CGATTCTCCGCCGCGGGCCGTGG</td><td>1000</td><td></td><td></td><td></td><td></td><td></td><td></td><td></td><td></td><td></td></tr>
<tr><td>Oligo fwd</td><td>CACCCGATTCTCCGCCGCGGGCCG</td><td></td><td></td><td></td><td></td><td></td><td></td><td></td><td></td><td></td><td></td></tr>
<tr><td>Oligo rev</td><td>AAACCGGCCCGCGGCGGAGAATCG</td><td></td><td></td><td></td><td></td><td></td><td></td><td></td><td></td><td></td><td></td></tr>
<tr><td>Chromosome</td><td>start</td><td>end</td><td>strand</td><td>MM</td><td>target_seq</td><td>PAM</td><td>alignment</td><td>distance</td><td>position</td><td>gene name</td><td>gene id</td></tr>
<tr><td>GU305914.1</td><td>197663</td><td>197685</td><td>+</td><td>0</td><td>CGATTCTCCGCCGCGGGCCG</td><td>TGG</td><td>||||||||[||||||||||||]PAM</td><td>NA</td><td>Intergenic</td><td>NA</td><td>NA</td></tr>
<tr><td>GU305914.1</td><td>64951</td><td>64973</td><td>-</td><td>4</td><td>CTCTTCTCCGACGCGCGCCG</td><td>CGG</td><td>|--|||||[||-||||-||||]PAM</td><td>NA</td><td>Intergenic</td><td>NA</td><td>NA</td></tr>
<tr><td></td><td></td><td></td><td></td><td></td><td></td><td></td><td></td><td></td><td></td><td></td><td></td></tr>
<tr><td>T15052</td><td>CGGGCCGTGGCGATGTTGACAGG</td><td>1000</td><td></td><td></td><td></td><td></td><td></td><td></td><td></td><td></td><td></td></tr>
<tr><td>Oligo fwd</td><td>CACCCGGGCCGTGGCGATGTTGAC</td><td></td><td></td><td></td><td></td><td></td><td></td><td></td><td></td><td></td><td></td></tr>
</table>

<table>
<tr><td>Oligo rev</td><td>AAACGTCAACATCGCCACGGCCCG</td><td></td><td></td><td></td><td></td><td></td><td></td><td></td><td></td><td></td><td></td></tr>
<tr><td>Chromosome</td><td>start</td><td>end</td><td>strand</td><td>MM</td><td>target_seq</td><td>PAM</td><td>alignment</td><td>distance</td><td>position</td><td>gene name</td><td>gene id</td></tr>
<tr><td>GU305914.1</td><td>197676</td><td>197698</td><td>+</td><td>0</td><td>CGGGCCGTGGCGATGTTGAC</td><td>AGG</td><td>||||||||[||||||||||||]PAM</td><td>NA</td><td>Intergenic</td><td>NA</td><td>NA</td></tr>
<tr><td>T15053</td><td>GCGATGTTGACAGGCAGTCCCGG</td><td>1000</td><td></td><td></td><td></td><td></td><td></td><td></td><td></td><td></td><td></td></tr>
<tr><td>Oligo fwd</td><td>CACCGCGATGTTGACAGGCAGTCC</td><td></td><td></td><td></td><td></td><td></td><td></td><td></td><td></td><td></td><td></td></tr>
<tr><td>Oligo rev</td><td>AAACGGACTGCCTGTCAACATCGC</td><td></td><td></td><td></td><td></td><td></td><td></td><td></td><td></td><td></td><td></td></tr>
<tr><td>Chromosome</td><td>start</td><td>end</td><td>strand</td><td>MM</td><td>target_seq</td><td>PAM</td><td>alignment</td><td>distance</td><td>position</td><td>gene name</td><td>gene id</td></tr>
<tr><td>GU305914.1</td><td>197685</td><td>197707</td><td>+</td><td>0</td><td>GCGATGTTGACAGGCAGTCC</td><td>CGG</td><td>||||||||[||||||||||||]PAM</td><td>NA</td><td>Intergenic</td><td>NA</td><td>NA</td></tr>
<tr><td>T15054</td><td>TTGACAGGCAGTCCCGGTGACGG</td><td>1000</td><td></td><td></td><td></td><td></td><td></td><td></td><td></td><td></td><td></td></tr>
<tr><td>Oligo fwd</td><td>CACCTTGACAGGCAGTCCCGGTGA</td><td></td><td></td><td></td><td></td><td></td><td></td><td></td><td></td><td></td><td></td></tr>
<tr><td>Oligo rev</td><td>AAACTCACCGGGACTGCCTGTCAA</td><td></td><td></td><td></td><td></td><td></td><td></td><td></td><td></td><td></td><td></td></tr>
<tr><td>Chromosome</td><td>start</td><td>end</td><td>strand</td><td>MM</td><td>target_seq</td><td>PAM</td><td>alignment</td><td>distance</td><td>position</td><td>gene name</td><td>gene id</td></tr>
<tr><td>GU305914.1</td><td>197691</td><td>197713</td><td>+</td><td>0</td><td>TTGACAGGCAGTCCCGGTGA</td><td>CGG</td><td>||||||||[||||||||||||]PAM</td><td>NA</td><td>Intergenic</td><td>NA</td><td>NA</td></tr>
<tr><td>T15055</td><td>GGCAGTCCCGGTGACGGCCGAGG</td><td>1000</td><td></td><td></td><td></td><td></td><td></td><td></td><td></td><td></td><td></td></tr>
<tr><td>Oligo fwd</td><td>CACCGGCAGTCCCGGTGACGGCCG</td><td></td><td></td><td></td><td></td><td></td><td></td><td></td><td></td><td></td><td></td></tr>
<tr><td>Oligo rev</td><td>AAACCGGCCGTCACCGGGACTGCC</td><td></td><td></td><td></td><td></td><td></td><td></td><td></td><td></td><td></td><td></td></tr>
<tr><td>Chromosome</td><td>start</td><td>end</td><td>strand</td><td>MM</td><td>target_seq</td><td>PAM</td><td>alignment</td><td>distance</td><td>position</td><td>gene name</td><td>gene id</td></tr>
<tr><td>GU305914.1</td><td>197697</td><td>197719</td><td>+</td><td>0</td><td>GGCAGTCCCGGTGACGGCCG</td><td>AGG</td><td>||||||||[||||||||||||]PAM</td><td>NA</td><td>Intergenic</td><td>NA</td><td>NA</td></tr>
<tr><td>T15056</td><td>GCAGTCCCGGTGACGGCCGAGGG</td><td>1000</td><td></td><td></td><td></td><td></td><td></td><td></td><td></td><td></td><td></td></tr>
<tr><td>Oligo fwd</td><td>CACCGCAGTCCCGGTGACGGCCGA</td><td></td><td></td><td></td><td></td><td></td><td></td><td></td><td></td><td></td><td></td></tr>
<tr><td>Oligo rev</td><td>AAACTCGGCCGTCACCGGGACTGC</td><td></td><td></td><td></td><td></td><td></td><td></td><td></td><td></td><td></td><td></td></tr>
<tr><td>Chromosome</td><td>start</td><td>end</td><td>strand</td><td>MM</td><td>target_seq</td><td>PAM</td><td>alignment</td><td>distance</td><td>position</td><td>gene name</td><td>gene id</td></tr>
<tr><td>GU305914.1</td><td>197698</td><td>197720</td><td>+</td><td>0</td><td>GCAGTCCCGGTGACGGCCGA</td><td>GGG</td><td>||||||||[||||||||||||]PAM</td><td>NA</td><td>Intergenic</td><td>NA</td><td>NA</td></tr>
<tr><td>T15057</td><td>TCTGATGCTGTTAGTGAGCTAGG</td><td>1000</td><td></td><td></td><td></td><td></td><td></td><td></td><td></td><td></td><td></td></tr>
<tr><td>Oligo fwd</td><td>CACCTCTGATGCTGTTAGTGAGCT</td><td></td><td></td><td></td><td></td><td></td><td></td><td></td><td></td><td></td><td></td></tr>
<tr><td>Oligo rev</td><td>AAACAGCTCACTAACAGCATCAGA</td><td></td><td></td><td></td><td></td><td></td><td></td><td></td><td></td><td></td><td></td></tr>
<tr><td>Chromosome</td><td>start</td><td>end</td><td>strand</td><td>MM</td><td>target_seq</td><td>PAM</td><td>alignment</td><td>distance</td><td>position</td><td>gene name</td><td>gene id</td></tr>
<tr><td>GU305914.1</td><td>197729</td><td>197751</td><td>+</td><td>0</td><td>TCTGATGCTGTTAGTGAGCT</td><td>AGG</td><td>||||||||[||||||||||||]PAM</td><td>NA</td><td>Intergenic</td><td>NA</td><td>NA</td></tr>
<tr><td>T15058</td><td>GATGCTGTTAGTGAGCTAGGTGG</td><td>1000</td><td></td><td></td><td></td><td></td><td></td><td></td><td></td><td></td><td></td></tr>
<tr><td>Oligo fwd</td><td>CACCGATGCTGTTAGTGAGCTAGG</td><td></td><td></td><td></td><td></td><td></td><td></td><td></td><td></td><td></td><td></td></tr>
<tr><td>Oligo rev</td><td>AAACCCTAGCTCACTAACAGCATC</td><td></td><td></td><td></td><td></td><td></td><td></td><td></td><td></td><td></td><td></td></tr>
<tr><td>Chromosome</td><td>start</td><td>end</td><td>strand</td><td>MM</td><td>target_seq</td><td>PAM</td><td>alignment</td><td>distance</td><td>position</td><td>gene name</td><td>gene id</td></tr>
<tr><td>GU305914.1</td><td>197732</td><td>197754</td><td>+</td><td>0</td><td>GATGCTGTTAGTGAGCTAGG</td><td>TGG</td><td>||||||||[||||||||||||]PAM</td><td>NA</td><td>Intergenic</td><td>NA</td><td>NA</td></tr>
<tr><td>T15059</td><td>GGTGGAGATGATAATGCCGCTGG</td><td>1000</td><td></td><td></td><td></td><td></td><td></td><td></td><td></td><td></td><td></td></tr>
<tr><td>Oligo fwd</td><td>CACCGGTGGAGATGATAATGCCGC</td><td></td><td></td><td></td><td></td><td></td><td></td><td></td><td></td><td></td><td></td></tr>
<tr><td>Oligo rev</td><td>AAACGCGGCATTATCATCTCCACC</td><td></td><td></td><td></td><td></td><td></td><td></td><td></td><td></td><td></td><td></td></tr>
<tr><td>Chromosome</td><td>start</td><td>end</td><td>strand</td><td>MM</td><td>target_seq</td><td>PAM</td><td>alignment</td><td>distance</td><td>position</td><td>gene name</td><td>gene id</td></tr>
<tr><td>GU305914.1</td><td>197750</td><td>197772</td><td>+</td><td>0</td><td>GGTGGAGATGATAATGCCGC</td><td>TGG</td><td>||||||||[||||||||||||]PAM</td><td>NA</td><td>Intergenic</td><td>NA</td><td>NA</td></tr>
<tr><td>T15060</td><td>GGAGATGATAATGCCGCTGGTGG</td><td>1000</td><td></td><td></td><td></td><td></td><td></td><td></td><td></td><td></td><td></td></tr>
<tr><td>Oligo fwd</td><td>CACCGGAGATGATAATGCCGCTGG</td><td></td><td></td><td></td><td></td><td></td><td></td><td></td><td></td><td></td><td></td></tr>
<tr><td>Oligo rev</td><td>AAACCCAGCGGCATTATCATCTCC</td><td></td><td></td><td></td><td></td><td></td><td></td><td></td><td></td><td></td><td></td></tr>
<tr><td>Chromosome</td><td>start</td><td>end</td><td>strand</td><td>MM</td><td>target_seq</td><td>PAM</td><td>alignment</td><td>distance</td><td>position</td><td>gene name</td><td>gene id</td></tr>
<tr><td>GU305914.1</td><td>197753</td><td>197775</td><td>+</td><td>0</td><td>GGAGATGATAATGCCGCTGG</td><td>TGG</td><td>||||||||[||||||||||||]PAM</td><td>NA</td><td>Intergenic</td><td>NA</td><td>NA</td></tr>
<tr><td>T15061</td><td>GAGATGATAATGCCGCTGGTGGG</td><td>1000</td><td></td><td></td><td></td><td></td><td></td><td></td><td></td><td></td><td></td></tr>
<tr><td>Oligo fwd</td><td>CACCGAGATGATAATGCCGCTGGT</td><td></td><td></td><td></td><td></td><td></td><td></td><td></td><td></td><td></td><td></td></tr>
<tr><td>Oligo rev</td><td>AAACACCAGCGGCATTATCATCTC</td><td></td><td></td><td></td><td></td><td></td><td></td><td></td><td></td><td></td><td></td></tr>
<tr><td>Chromosome</td><td>start</td><td>end</td><td>strand</td><td>MM</td><td>target_seq</td><td>PAM</td><td>alignment</td><td>distance</td><td>position</td><td>gene name</td><td>gene id</td></tr>
<tr><td>GU305914.1</td><td>197754</td><td>197776</td><td>+</td><td>0</td><td>GAGATGATAATGCCGCTGGT</td><td>GGG</td><td>||||||||[||||||||||||]PAM</td><td>NA</td><td>Intergenic</td><td>NA</td><td>NA</td></tr>
<tr><td>T15062</td><td>AGATGATAATGCCGCTGGTGGGG</td><td>1000</td><td></td><td></td><td></td><td></td><td></td><td></td><td></td><td></td><td></td></tr>
<tr><td>Oligo fwd</td><td>CACCAGATGATAATGCCGCTGGTG</td><td></td><td></td><td></td><td></td><td></td><td></td><td></td><td></td><td></td><td></td></tr>
<tr><td>Oligo rev</td><td>AAACCACCAGCGGCATTATCATCT</td><td></td><td></td><td></td><td></td><td></td><td></td><td></td><td></td><td></td><td></td></tr>
<tr><td>Chromosome</td><td>start</td><td>end</td><td>strand</td><td>MM</td><td>target_seq</td><td>PAM</td><td>alignment</td><td>distance</td><td>position</td><td>gene name</td><td>gene id</td></tr>
<tr><td>GU305914.1</td><td>197755</td><td>197777</td><td>+</td><td>0</td><td>AGATGATAATGCCGCTGGTG</td><td>GGG</td><td>||||||||[||||||||||||]PAM</td><td>NA</td><td>Intergenic</td><td>NA</td><td>NA</td></tr>
<tr><td>T15063</td><td>TGATAATGCCGCTGGTGGGGTGG</td><td>1000</td><td></td><td></td><td></td><td></td><td></td><td></td><td></td><td></td><td></td></tr>
<tr><td>Oligo fwd</td><td>CACCTGATAATGCCGCTGGTGGGG</td><td></td><td></td><td></td><td></td><td></td><td></td><td></td><td></td><td></td><td></td></tr>
<tr><td>Oligo rev</td><td>AAACCCCCACCAGCGGCATTATCA</td><td></td><td></td><td></td><td></td><td></td><td></td><td></td><td></td><td></td><td></td></tr>
<tr><td>Chromosome</td><td>start</td><td>end</td><td>strand</td><td>MM</td><td>target_seq</td><td>PAM</td><td>alignment</td><td>distance</td><td>position</td><td>gene name</td><td>gene id</td></tr>
<tr><td>GU305914.1</td><td>197758</td><td>197780</td><td>+</td><td>0</td><td>TGATAATGCCGCTGGTGGGG</td><td>TGG</td><td>||||||||[||||||||||||]PAM</td><td>NA</td><td>Intergenic</td><td>NA</td><td>NA</td></tr>
<tr><td>T15064</td><td>GATAATGCCGCTGGTGGGGTGGG</td><td>1000</td><td></td><td></td><td></td><td></td><td></td><td></td><td></td><td></td><td></td></tr>
<tr><td>Oligo fwd</td><td>CACCGATAATGCCGCTGGTGGGGT</td><td></td><td></td><td></td><td></td><td></td><td></td><td></td><td></td><td></td><td></td></tr>
<tr><td>Oligo rev</td><td>AAACACCCCACCAGCGGCATTATC</td><td></td><td></td><td></td><td></td><td></td><td></td><td></td><td></td><td></td><td></td></tr>
<tr><td>Chromosome</td><td>start</td><td>end</td><td>strand</td><td>MM</td><td>target_seq</td><td>PAM</td><td>alignment</td><td>distance</td><td>position</td><td>gene name</td><td>gene id</td></tr>
<tr><td>GU305914.1</td><td>197759</td><td>197781</td><td>+</td><td>0</td><td>GATAATGCCGCTGGTGGGGT</td><td>GGG</td><td>||||||||[||||||||||||]PAM</td><td>NA</td><td>Intergenic</td><td>NA</td><td>NA</td></tr>
<tr><td>T15065</td><td>ATAATGCCGCTGGTGGGGTGGGG</td><td>1000</td><td></td><td></td><td></td><td></td><td></td><td></td><td></td><td></td><td></td></tr>
<tr><td>Oligo fwd</td><td>CACCATAATGCCGCTGGTGGGGTG</td><td></td><td></td><td></td><td></td><td></td><td></td><td></td><td></td><td></td><td></td></tr>
<tr><td>Oligo rev</td><td>AAACCACCCCACCAGCGGCATTAT</td><td></td><td></td><td></td><td></td><td></td><td></td><td></td><td></td><td></td><td></td></tr>
<tr><td>Chromosome</td><td>start</td><td>end</td><td>strand</td><td>MM</td><td>target_seq</td><td>PAM</td><td>alignment</td><td>distance</td><td>position</td><td>gene name</td><td>gene id</td></tr>
<tr><td>GU305914.1</td><td>197760</td><td>197782</td><td>+</td><td>0</td><td>ATAATGCCGCTGGTGGGGTG</td><td>GGG</td><td>||||||||[||||||||||||]PAM</td><td>NA</td><td>Intergenic</td><td>NA</td><td>NA</td></tr>
<tr><td>T15066</td><td>TAATGCCGCTGGTGGGGTGGGGG</td><td>1000</td><td></td><td></td><td></td><td></td><td></td><td></td><td></td><td></td><td></td></tr>
<tr><td>Oligo fwd</td><td>CACCTAATGCCGCTGGTGGGGTGG</td><td></td><td></td><td></td><td></td><td></td><td></td><td></td><td></td><td></td><td></td></tr>
<tr><td>Oligo rev</td><td>AAACCCACCCCACCAGCGGCATTA</td><td></td><td></td><td></td><td></td><td></td><td></td><td></td><td></td><td></td><td></td></tr>
<tr><td>Chromosome</td><td>start</td><td>end</td><td>strand</td><td>MM</td><td>target_seq</td><td>PAM</td><td>alignment</td><td>distance</td><td>position</td><td>gene name</td><td>gene id</td></tr>
<tr><td>GU305914.1</td><td>197761</td><td>197783</td><td>+</td><td>0</td><td>TAATGCCGCTGGTGGGGTGG</td><td>GGG</td><td>||||||||[||||||||||||]PAM</td><td>NA</td><td>Intergenic</td><td>NA</td><td>NA</td></tr>
<tr><td>T15067</td><td>AATGCCGCTGGTGGGGTGGGGGG</td><td>1000</td><td></td><td></td><td></td><td></td><td></td><td></td><td></td><td></td><td></td></tr>
</table>

<table>
<tr><td>Oligo fwd</td><td>CACCAATGCCGCTGGTGGGGTGGG</td><td></td><td></td><td></td><td></td><td></td><td></td><td></td><td></td><td></td><td></td></tr>
<tr><td>Oligo rev</td><td>AAACCCCACCCCACCAGCGGCATT</td><td></td><td></td><td></td><td></td><td></td><td></td><td></td><td></td><td></td><td></td></tr>
<tr><td>Chromosome</td><td>start</td><td>end</td><td>strand</td><td>MM</td><td>target_seq</td><td>PAM</td><td>alignment</td><td>distance</td><td>position</td><td>gene name</td><td>gene id</td></tr>
<tr><td>GU305914.1</td><td>197762</td><td>197784</td><td>+</td><td>0</td><td>AATGCCGCTGGTGGGGTGGG</td><td>GGG</td><td>||||||||[||||||||||||]PAM</td><td>NA</td><td>Intergenic</td><td>NA</td><td>NA</td></tr>
<tr><td></td><td></td><td></td><td></td><td></td><td></td><td></td><td></td><td></td><td></td><td></td><td></td></tr>
<tr><td>T15068</td><td>ATGCCGCTGGTGGGGTGGGGGGG</td><td>1000</td><td></td><td></td><td></td><td></td><td></td><td></td><td></td><td></td><td></td></tr>
<tr><td>Oligo fwd</td><td>CACCATGCCGCTGGTGGGGTGGGG</td><td></td><td></td><td></td><td></td><td></td><td></td><td></td><td></td><td></td><td></td></tr>
<tr><td>Oligo rev</td><td>AAACCCCCACCCCACCAGCGGCAT</td><td></td><td></td><td></td><td></td><td></td><td></td><td></td><td></td><td></td><td></td></tr>
<tr><td>Chromosome</td><td>start</td><td>end</td><td>strand</td><td>MM</td><td>target_seq</td><td>PAM</td><td>alignment</td><td>distance</td><td>position</td><td>gene name</td><td>gene id</td></tr>
<tr><td>GU305914.1</td><td>197763</td><td>197785</td><td>+</td><td>0</td><td>ATGCCGCTGGTGGGGTGGGG</td><td>GGG</td><td>||||||||[||||||||||||]PAM</td><td>NA</td><td>Intergenic</td><td>NA</td><td>NA</td></tr>
<tr><td></td><td></td><td></td><td></td><td></td><td></td><td></td><td></td><td></td><td></td><td></td><td></td></tr>
<tr><td>T15069</td><td>TGCCGCTGGTGGGGTGGGGGGGG</td><td>1000</td><td></td><td></td><td></td><td></td><td></td><td></td><td></td><td></td><td></td></tr>
<tr><td>Oligo fwd</td><td>CACCTGCCGCTGGTGGGGTGGGGG</td><td></td><td></td><td></td><td></td><td></td><td></td><td></td><td></td><td></td><td></td></tr>
<tr><td>Oligo rev</td><td>AAACCCCCCACCCCACCAGCGGCA</td><td></td><td></td><td></td><td></td><td></td><td></td><td></td><td></td><td></td><td></td></tr>
<tr><td>Chromosome</td><td>start</td><td>end</td><td>strand</td><td>MM</td><td>target_seq</td><td>PAM</td><td>alignment</td><td>distance</td><td>position</td><td>gene name</td><td>gene id</td></tr>
<tr><td>GU305914.1</td><td>197764</td><td>197786</td><td>+</td><td>0</td><td>TGCCGCTGGTGGGGTGGGGG</td><td>GGG</td><td>||||||||[||||||||||||]PAM</td><td>NA</td><td>Intergenic</td><td>NA</td><td>NA</td></tr>
<tr><td></td><td></td><td></td><td></td><td></td><td></td><td></td><td></td><td></td><td></td><td></td><td></td></tr>
<tr><td>T15070</td><td>GCCGCTGGTGGGGTGGGGGGGGG</td><td>1000</td><td></td><td></td><td></td><td></td><td></td><td></td><td></td><td></td><td></td></tr>
<tr><td>Oligo fwd</td><td>CACCGCCGCTGGTGGGGTGGGGGG</td><td></td><td></td><td></td><td></td><td></td><td></td><td></td><td></td><td></td><td></td></tr>
<tr><td>Oligo rev</td><td>AAACCCCCCCACCCCACCAGCGGC</td><td></td><td></td><td></td><td></td><td></td><td></td><td></td><td></td><td></td><td></td></tr>
<tr><td>Chromosome</td><td>start</td><td>end</td><td>strand</td><td>MM</td><td>target_seq</td><td>PAM</td><td>alignment</td><td>distance</td><td>position</td><td>gene name</td><td>gene id</td></tr>
<tr><td>GU305914.1</td><td>197765</td><td>197787</td><td>+</td><td>0</td><td>GCCGCTGGTGGGGTGGGGGG</td><td>GGG</td><td>||||||||[||||||||||||]PAM</td><td>NA</td><td>Intergenic</td><td>NA</td><td>NA</td></tr>
<tr><td></td><td></td><td></td><td></td><td></td><td></td><td></td><td></td><td></td><td></td><td></td><td></td></tr>
<tr><td>T15071</td><td>GGTGGGGTGGGGGGGGGTGATGG</td><td>1000</td><td></td><td></td><td></td><td></td><td></td><td></td><td></td><td></td><td></td></tr>
<tr><td>Oligo fwd</td><td>CACCGGTGGGGTGGGGGGGGGTGA</td><td></td><td></td><td></td><td></td><td></td><td></td><td></td><td></td><td></td><td></td></tr>
<tr><td>Oligo rev</td><td>AAACTCACCCCCCCCCACCCCACC</td><td></td><td></td><td></td><td></td><td></td><td></td><td></td><td></td><td></td><td></td></tr>
<tr><td>Chromosome</td><td>start</td><td>end</td><td>strand</td><td>MM</td><td>target_seq</td><td>PAM</td><td>alignment</td><td>distance</td><td>position</td><td>gene name</td><td>gene id</td></tr>
<tr><td>GU305914.1</td><td>197771</td><td>197793</td><td>+</td><td>0</td><td>GGTGGGGTGGGGGGGGGTGA</td><td>TGG</td><td>||||||||[||||||||||||]PAM</td><td>NA</td><td>Intergenic</td><td>NA</td><td>NA</td></tr>
<tr><td></td><td></td><td></td><td></td><td></td><td></td><td></td><td></td><td></td><td></td><td></td><td></td></tr>
<tr><td>T15072</td><td>GGGGTGGGGGGGGGTGATGGTGG</td><td>1000</td><td></td><td></td><td></td><td></td><td></td><td></td><td></td><td></td><td></td></tr>
<tr><td>Oligo fwd</td><td>CACCGGGGTGGGGGGGGGTGATGG</td><td></td><td></td><td></td><td></td><td></td><td></td><td></td><td></td><td></td><td></td></tr>
<tr><td>Oligo rev</td><td>AAACCCATCACCCCCCCCCACCCC</td><td></td><td></td><td></td><td></td><td></td><td></td><td></td><td></td><td></td><td></td></tr>
<tr><td>Chromosome</td><td>start</td><td>end</td><td>strand</td><td>MM</td><td>target_seq</td><td>PAM</td><td>alignment</td><td>distance</td><td>position</td><td>gene name</td><td>gene id</td></tr>
<tr><td>GU305914.1</td><td>197774</td><td>197796</td><td>+</td><td>0</td><td>GGGGTGGGGGGGGGTGATGG</td><td>TGG</td><td>||||||||[||||||||||||]PAM</td><td>NA</td><td>Intergenic</td><td>NA</td><td>NA</td></tr>
<tr><td></td><td></td><td></td><td></td><td></td><td></td><td></td><td></td><td></td><td></td><td></td><td></td></tr>
<tr><td>T15073</td><td>GGGTGGGGGGGGGTGATGGTGGG</td><td>1000</td><td></td><td></td><td></td><td></td><td></td><td></td><td></td><td></td><td></td></tr>
<tr><td>Oligo fwd</td><td>CACCGGGTGGGGGGGGGTGATGGT</td><td></td><td></td><td></td><td></td><td></td><td></td><td></td><td></td><td></td><td></td></tr>
<tr><td>Oligo rev</td><td>AAACACCATCACCCCCCCCCACCC</td><td></td><td></td><td></td><td></td><td></td><td></td><td></td><td></td><td></td><td></td></tr>
<tr><td>Chromosome</td><td>start</td><td>end</td><td>strand</td><td>MM</td><td>target_seq</td><td>PAM</td><td>alignment</td><td>distance</td><td>position</td><td>gene name</td><td>gene id</td></tr>
<tr><td>GU305914.1</td><td>197775</td><td>197797</td><td>+</td><td>0</td><td>GGGTGGGGGGGGGTGATGGT</td><td>GGG</td><td>||||||||[||||||||||||]PAM</td><td>NA</td><td>Intergenic</td><td>NA</td><td>NA</td></tr>
<tr><td></td><td></td><td></td><td></td><td></td><td></td><td></td><td></td><td></td><td></td><td></td><td></td></tr>
<tr><td>T15074</td><td>GGTGGGGGGGGGTGATGGTGGGG</td><td>1000</td><td></td><td></td><td></td><td></td><td></td><td></td><td></td><td></td><td></td></tr>
<tr><td>Oligo fwd</td><td>CACCGGTGGGGGGGGGTGATGGTG</td><td></td><td></td><td></td><td></td><td></td><td></td><td></td><td></td><td></td><td></td></tr>
<tr><td>Oligo rev</td><td>AAACCACCATCACCCCCCCCCACC</td><td></td><td></td><td></td><td></td><td></td><td></td><td></td><td></td><td></td><td></td></tr>
<tr><td>Chromosome</td><td>start</td><td>end</td><td>strand</td><td>MM</td><td>target_seq</td><td>PAM</td><td>alignment</td><td>distance</td><td>position</td><td>gene name</td><td>gene id</td></tr>
<tr><td>GU305914.1</td><td>197776</td><td>197798</td><td>+</td><td>0</td><td>GGTGGGGGGGGGTGATGGTG</td><td>GGG</td><td>||||||||[||||||||||||]PAM</td><td>NA</td><td>Intergenic</td><td>NA</td><td>NA</td></tr>
<tr><td></td><td></td><td></td><td></td><td></td><td></td><td></td><td></td><td></td><td></td><td></td><td></td></tr>
<tr><td>T15075</td><td>GTGGGGGGGGGTGATGGTGGGGG</td><td>1000</td><td></td><td></td><td></td><td></td><td></td><td></td><td></td><td></td><td></td></tr>
<tr><td>Oligo fwd</td><td>CACCGTGGGGGGGGGTGATGGTGG</td><td></td><td></td><td></td><td></td><td></td><td></td><td></td><td></td><td></td><td></td></tr>
<tr><td>Oligo rev</td><td>AAACCCACCATCACCCCCCCCCAC</td><td></td><td></td><td></td><td></td><td></td><td></td><td></td><td></td><td></td><td></td></tr>
<tr><td>Chromosome</td><td>start</td><td>end</td><td>strand</td><td>MM</td><td>target_seq</td><td>PAM</td><td>alignment</td><td>distance</td><td>position</td><td>gene name</td><td>gene id</td></tr>
<tr><td>GU305914.1</td><td>197777</td><td>197799</td><td>+</td><td>0</td><td>GTGGGGGGGGGTGATGGTGG</td><td>GGG</td><td>||||||||[||||||||||||]PAM</td><td>NA</td><td>Intergenic</td><td>NA</td><td>NA</td></tr>
<tr><td></td><td></td><td></td><td></td><td></td><td></td><td></td><td></td><td></td><td></td><td></td><td></td></tr>
<tr><td>T15076</td><td>GGGGGGTGATGGTGGGGGTGTGG</td><td>1000</td><td></td><td></td><td></td><td></td><td></td><td></td><td></td><td></td><td></td></tr>
<tr><td>Oligo fwd</td><td>CACCGGGGGGTGATGGTGGGGGTG</td><td></td><td></td><td></td><td></td><td></td><td></td><td></td><td></td><td></td><td></td></tr>
<tr><td>Oligo rev</td><td>AAACCACCCCCACCATCACCCCCC</td><td></td><td></td><td></td><td></td><td></td><td></td><td></td><td></td><td></td><td></td></tr>
<tr><td>Chromosome</td><td>start</td><td>end</td><td>strand</td><td>MM</td><td>target_seq</td><td>PAM</td><td>alignment</td><td>distance</td><td>position</td><td>gene name</td><td>gene id</td></tr>
<tr><td>GU305914.1</td><td>197782</td><td>197804</td><td>+</td><td>0</td><td>GGGGGGTGATGGTGGGGGTG</td><td>TGG</td><td>||||||||[||||||||||||]PAM</td><td>NA</td><td>Intergenic</td><td>NA</td><td>NA</td></tr>
<tr><td></td><td></td><td></td><td></td><td></td><td></td><td></td><td></td><td></td><td></td><td></td><td></td></tr>
<tr><td>T15077</td><td>GTGATGGTGGGGGTGTGGCTAGG</td><td>1000</td><td></td><td></td><td></td><td></td><td></td><td></td><td></td><td></td><td></td></tr>
<tr><td>Oligo fwd</td><td>CACCGTGATGGTGGGGGTGTGGCT</td><td></td><td></td><td></td><td></td><td></td><td></td><td></td><td></td><td></td><td></td></tr>
<tr><td>Oligo rev</td><td>AAACAGCCACACCCCCACCATCAC</td><td></td><td></td><td></td><td></td><td></td><td></td><td></td><td></td><td></td><td></td></tr>
<tr><td>Chromosome</td><td>start</td><td>end</td><td>strand</td><td>MM</td><td>target_seq</td><td>PAM</td><td>alignment</td><td>distance</td><td>position</td><td>gene name</td><td>gene id</td></tr>
<tr><td>GU305914.1</td><td>197787</td><td>197809</td><td>+</td><td>0</td><td>GTGATGGTGGGGGTGTGGCT</td><td>AGG</td><td>||||||||[||||||||||||]PAM</td><td>NA</td><td>Intergenic</td><td>NA</td><td>NA</td></tr>
<tr><td></td><td></td><td></td><td></td><td></td><td></td><td></td><td></td><td></td><td></td><td></td><td></td></tr>
<tr><td>T15078</td><td>GGGGTGTGGCTAGGTCTCCGAGG</td><td>1000</td><td></td><td></td><td></td><td></td><td></td><td></td><td></td><td></td><td></td></tr>
<tr><td>Oligo fwd</td><td>CACCGGGGTGTGGCTAGGTCTCCG</td><td></td><td></td><td></td><td></td><td></td><td></td><td></td><td></td><td></td><td></td></tr>
<tr><td>Oligo rev</td><td>AAACCGGAGACCTAGCCACACCCC</td><td></td><td></td><td></td><td></td><td></td><td></td><td></td><td></td><td></td><td></td></tr>
<tr><td>Chromosome</td><td>start</td><td>end</td><td>strand</td><td>MM</td><td>target_seq</td><td>PAM</td><td>alignment</td><td>distance</td><td>position</td><td>gene name</td><td>gene id</td></tr>
<tr><td>GU305914.1</td><td>197796</td><td>197818</td><td>+</td><td>0</td><td>GGGGTGTGGCTAGGTCTCCG</td><td>AGG</td><td>||||||||[||||||||||||]PAM</td><td>NA</td><td>Intergenic</td><td>NA</td><td>NA</td></tr>
<tr><td></td><td></td><td></td><td></td><td></td><td></td><td></td><td></td><td></td><td></td><td></td><td></td></tr>
<tr><td>T15079</td><td>GGGTGTGGCTAGGTCTCCGAGGG</td><td>1000</td><td></td><td></td><td></td><td></td><td></td><td></td><td></td><td></td><td></td></tr>
<tr><td>Oligo fwd</td><td>CACCGGGTGTGGCTAGGTCTCCGA</td><td></td><td></td><td></td><td></td><td></td><td></td><td></td><td></td><td></td><td></td></tr>
<tr><td>Oligo rev</td><td>AAACTCGGAGACCTAGCCACACCC</td><td></td><td></td><td></td><td></td><td></td><td></td><td></td><td></td><td></td><td></td></tr>
<tr><td>Chromosome</td><td>start</td><td>end</td><td>strand</td><td>MM</td><td>target_seq</td><td>PAM</td><td>alignment</td><td>distance</td><td>position</td><td>gene name</td><td>gene id</td></tr>
<tr><td>GU305914.1</td><td>197797</td><td>197819</td><td>+</td><td>0</td><td>GGGTGTGGCTAGGTCTCCGA</td><td>GGG</td><td>||||||||[||||||||||||]PAM</td><td>NA</td><td>Intergenic</td><td>NA</td><td>NA</td></tr>
<tr><td></td><td></td><td></td><td></td><td></td><td></td><td></td><td></td><td></td><td></td><td></td><td></td></tr>
<tr><td>T15080</td><td>GGCTAGGTCTCCGAGGGACCCGG</td><td>1000</td><td></td><td></td><td></td><td></td><td></td><td></td><td></td><td></td><td></td></tr>
<tr><td>Oligo fwd</td><td>CACCGGCTAGGTCTCCGAGGGACC</td><td></td><td></td><td></td><td></td><td></td><td></td><td></td><td></td><td></td><td></td></tr>
<tr><td>Oligo rev</td><td>AAACGGTCCCTCGGAGACCTAGCC</td><td></td><td></td><td></td><td></td><td></td><td></td><td></td><td></td><td></td><td></td></tr>
<tr><td>Chromosome</td><td>start</td><td>end</td><td>strand</td><td>MM</td><td>target_seq</td><td>PAM</td><td>alignment</td><td>distance</td><td>position</td><td>gene name</td><td>gene id</td></tr>
<tr><td>GU305914.1</td><td>197803</td><td>197825</td><td>+</td><td>0</td><td>GGCTAGGTCTCCGAGGGACC</td><td>CGG</td><td>||||||||[||||||||||||]PAM</td><td>NA</td><td>Intergenic</td><td>NA</td><td>NA</td></tr>
<tr><td></td><td></td><td></td><td></td><td></td><td></td><td></td><td></td><td></td><td></td><td></td><td></td></tr>
<tr><td>T15081</td><td>CCGAGGGACCCGGCCGCGCGCGG</td><td>1000</td><td></td><td></td><td></td><td></td><td></td><td></td><td></td><td></td><td></td></tr>
<tr><td>Oligo fwd</td><td>CACCCCGAGGGACCCGGCCGCGCG</td><td></td><td></td><td></td><td></td><td></td><td></td><td></td><td></td><td></td><td></td></tr>
<tr><td>Oligo rev</td><td>AAACCGCGCGGCCGGGTCCCTCGG</td><td></td><td></td><td></td><td></td><td></td><td></td><td></td><td></td><td></td><td></td></tr>
<tr><td>Chromosome</td><td>start</td><td>end</td><td>strand</td><td>MM</td><td>target_seq</td><td>PAM</td><td>alignment</td><td>distance</td><td>position</td><td>gene name</td><td>gene id</td></tr>
<tr><td>GU305914.1</td><td>197813</td><td>197835</td><td>+</td><td>0</td><td>CCGAGGGACCCGGCCGCGCG</td><td>CGG</td><td>||||||||[||||||||||||]PAM</td><td>NA</td><td>Intergenic</td><td>NA</td><td>NA</td></tr>
<tr><td>GU305914.1</td><td>207959</td><td>207981</td><td>-</td><td>4</td><td>GAGAGGGAGCCGGCCGCGTG</td><td>CGG</td><td>--||||||[-|||||||||-|]PAM</td><td>NA</td><td>Intergenic</td><td>NA</td><td>NA</td></tr>
</table>

<table>
<tr><td>T15082</td><td>ACCCGGCCGCGCGCGGTAGAAGG</td><td>1000</td><td></td><td></td><td></td><td></td><td></td><td></td><td></td><td></td><td></td></tr>
<tr><td>Oligo fwd</td><td>CACCACCCGGCCGCGCGCGGTAGA</td><td></td><td></td><td></td><td></td><td></td><td></td><td></td><td></td><td></td><td></td></tr>
<tr><td>Oligo rev</td><td>AAACTCTACCGCGCGCGGCCGGGT</td><td></td><td></td><td></td><td></td><td></td><td></td><td></td><td></td><td></td><td></td></tr>
<tr><td>Chromosome</td><td>start</td><td>end</td><td>strand</td><td>MM</td><td>target_seq</td><td>PAM</td><td>alignment</td><td>distance</td><td>position</td><td>gene name</td><td>gene id</td></tr>
<tr><td>GU305914.1</td><td>197820</td><td>197842</td><td>+</td><td>0</td><td>ACCCGGCCGCGCGCGGTAGA</td><td>AGG</td><td>||||||||[||||||||||||]PAM</td><td>NA</td><td>Intergenic</td><td>NA</td><td>NA</td></tr>
<tr><td>T15083</td><td>CCCGGCCGCGCGCGGTAGAAGGG</td><td>1000</td><td></td><td></td><td></td><td></td><td></td><td></td><td></td><td></td><td></td></tr>
<tr><td>Oligo fwd</td><td>CACCCCCGGCCGCGCGCGGTAGAA</td><td></td><td></td><td></td><td></td><td></td><td></td><td></td><td></td><td></td><td></td></tr>
<tr><td>Oligo rev</td><td>AAACTTCTACCGCGCGCGGCCGGG</td><td></td><td></td><td></td><td></td><td></td><td></td><td></td><td></td><td></td><td></td></tr>
<tr><td>Chromosome</td><td>start</td><td>end</td><td>strand</td><td>MM</td><td>target_seq</td><td>PAM</td><td>alignment</td><td>distance</td><td>position</td><td>gene name</td><td>gene id</td></tr>
<tr><td>GU305914.1</td><td>197821</td><td>197843</td><td>+</td><td>0</td><td>CCCGGCCGCGCGCGGTAGAA</td><td>GGG</td><td>||||||||[||||||||||||]PAM</td><td>NA</td><td>Intergenic</td><td>NA</td><td>NA</td></tr>
<tr><td>T15084</td><td>CCGGCCGCGCGCGGTAGAAGGGG</td><td>1000</td><td></td><td></td><td></td><td></td><td></td><td></td><td></td><td></td><td></td></tr>
<tr><td>Oligo fwd</td><td>CACCCCGGCCGCGCGCGGTAGAAG</td><td></td><td></td><td></td><td></td><td></td><td></td><td></td><td></td><td></td><td></td></tr>
<tr><td>Oligo rev</td><td>AAACCTTCTACCGCGCGCGGCCGG</td><td></td><td></td><td></td><td></td><td></td><td></td><td></td><td></td><td></td><td></td></tr>
<tr><td>Chromosome</td><td>start</td><td>end</td><td>strand</td><td>MM</td><td>target_seq</td><td>PAM</td><td>alignment</td><td>distance</td><td>position</td><td>gene name</td><td>gene id</td></tr>
<tr><td>GU305914.1</td><td>197822</td><td>197844</td><td>+</td><td>0</td><td>CCGGCCGCGCGCGGTAGAAG</td><td>GGG</td><td>||||||||[||||||||||||]PAM</td><td>NA</td><td>Intergenic</td><td>NA</td><td>NA</td></tr>
<tr><td>T15085</td><td>TATCGCAGAAAAACTGCCGTCGG</td><td>1000</td><td></td><td></td><td></td><td></td><td></td><td></td><td></td><td></td><td></td></tr>
<tr><td>Oligo fwd</td><td>CACCTATCGCAGAAAAACTGCCGT</td><td></td><td></td><td></td><td></td><td></td><td></td><td></td><td></td><td></td><td></td></tr>
<tr><td>Oligo rev</td><td>AAACACGGCAGTTTTTCTGCGATA</td><td></td><td></td><td></td><td></td><td></td><td></td><td></td><td></td><td></td><td></td></tr>
<tr><td>Chromosome</td><td>start</td><td>end</td><td>strand</td><td>MM</td><td>target_seq</td><td>PAM</td><td>alignment</td><td>distance</td><td>position</td><td>gene name</td><td>gene id</td></tr>
<tr><td>GU305914.1</td><td>197852</td><td>197874</td><td>+</td><td>0</td><td>TATCGCAGAAAAACTGCCGT</td><td>CGG</td><td>||||||||[||||||||||||]PAM</td><td>NA</td><td>Intergenic</td><td>NA</td><td>NA</td></tr>
<tr><td>T15086</td><td>CGCAGAAAAACTGCCGTCGGCGG</td><td>1000</td><td></td><td></td><td></td><td></td><td></td><td></td><td></td><td></td><td></td></tr>
<tr><td>Oligo fwd</td><td>CACCCGCAGAAAAACTGCCGTCGG</td><td></td><td></td><td></td><td></td><td></td><td></td><td></td><td></td><td></td><td></td></tr>
<tr><td>Oligo rev</td><td>AAACCCGACGGCAGTTTTTCTGCG</td><td></td><td></td><td></td><td></td><td></td><td></td><td></td><td></td><td></td><td></td></tr>
<tr><td>Chromosome</td><td>start</td><td>end</td><td>strand</td><td>MM</td><td>target_seq</td><td>PAM</td><td>alignment</td><td>distance</td><td>position</td><td>gene name</td><td>gene id</td></tr>
<tr><td>GU305914.1</td><td>197855</td><td>197877</td><td>+</td><td>0</td><td>CGCAGAAAAACTGCCGTCGG</td><td>CGG</td><td>||||||||[||||||||||||]PAM</td><td>NA</td><td>Intergenic</td><td>NA</td><td>NA</td></tr>
<tr><td>T15087</td><td>TCGTCATAGACTCCGTCTCGTGG</td><td>1000</td><td></td><td></td><td></td><td></td><td></td><td></td><td></td><td></td><td></td></tr>
<tr><td>Oligo fwd</td><td>CACCTCGTCATAGACTCCGTCTCG</td><td></td><td></td><td></td><td></td><td></td><td></td><td></td><td></td><td></td><td></td></tr>
<tr><td>Oligo rev</td><td>AAACCGAGACGGAGTCTATGACGA</td><td></td><td></td><td></td><td></td><td></td><td></td><td></td><td></td><td></td><td></td></tr>
<tr><td>Chromosome</td><td>start</td><td>end</td><td>strand</td><td>MM</td><td>target_seq</td><td>PAM</td><td>alignment</td><td>distance</td><td>position</td><td>gene name</td><td>gene id</td></tr>
<tr><td>GU305914.1</td><td>197897</td><td>197919</td><td>+</td><td>0</td><td>TCGTCATAGACTCCGTCTCG</td><td>TGG</td><td>||||||||[||||||||||||]PAM</td><td>NA</td><td>Intergenic</td><td>NA</td><td>NA</td></tr>
<tr><td>GU305914.1</td><td>96496</td><td>96518</td><td>+</td><td>4</td><td>TCCACACGGACTCCGTCTCG</td><td>GGG</td><td>||--||--[||||||||||||]PAM</td><td>NA</td><td>Intergenic</td><td>NA</td><td>NA</td></tr>
<tr><td>T15088</td><td>CATAGACTCCGTCTCGTGGCAGG</td><td>1000</td><td></td><td></td><td></td><td></td><td></td><td></td><td></td><td></td><td></td></tr>
<tr><td>Oligo fwd</td><td>CACCCATAGACTCCGTCTCGTGGC</td><td></td><td></td><td></td><td></td><td></td><td></td><td></td><td></td><td></td><td></td></tr>
<tr><td>Oligo rev</td><td>AAACGCCACGAGACGGAGTCTATG</td><td></td><td></td><td></td><td></td><td></td><td></td><td></td><td></td><td></td><td></td></tr>
<tr><td>Chromosome</td><td>start</td><td>end</td><td>strand</td><td>MM</td><td>target_seq</td><td>PAM</td><td>alignment</td><td>distance</td><td>position</td><td>gene name</td><td>gene id</td></tr>
<tr><td>GU305914.1</td><td>197901</td><td>197923</td><td>+</td><td>0</td><td>CATAGACTCCGTCTCGTGGC</td><td>AGG</td><td>||||||||[||||||||||||]PAM</td><td>NA</td><td>Intergenic</td><td>NA</td><td>NA</td></tr>
<tr><td>GU305914.1</td><td>199262</td><td>199284</td><td>-</td><td>4</td><td>GATAGACGCCTTCTCGCGGC</td><td>TGG</td><td>-||||||-[||-|||||-|||]PAM</td><td>NA</td><td>Intergenic</td><td>NA</td><td>NA</td></tr>
<tr><td>T15089</td><td>CAGGTCAACGCGAAGTACGTCGG</td><td>1000</td><td></td><td></td><td></td><td></td><td></td><td></td><td></td><td></td><td></td></tr>
<tr><td>Oligo fwd</td><td>CACCCAGGTCAACGCGAAGTACGT</td><td></td><td></td><td></td><td></td><td></td><td></td><td></td><td></td><td></td><td></td></tr>
<tr><td>Oligo rev</td><td>AAACACGTACTTCGCGTTGACCTG</td><td></td><td></td><td></td><td></td><td></td><td></td><td></td><td></td><td></td><td></td></tr>
<tr><td>Chromosome</td><td>start</td><td>end</td><td>strand</td><td>MM</td><td>target_seq</td><td>PAM</td><td>alignment</td><td>distance</td><td>position</td><td>gene name</td><td>gene id</td></tr>
<tr><td>GU305914.1</td><td>197920</td><td>197942</td><td>+</td><td>0</td><td>CAGGTCAACGCGAAGTACGT</td><td>CGG</td><td>||||||||[||||||||||||]PAM</td><td>NA</td><td>Intergenic</td><td>NA</td><td>NA</td></tr>
<tr><td>T15090</td><td>GTCAACGCGAAGTACGTCGGCGG</td><td>1000</td><td></td><td></td><td></td><td></td><td></td><td></td><td></td><td></td><td></td></tr>
<tr><td>Oligo fwd</td><td>CACCGTCAACGCGAAGTACGTCGG</td><td></td><td></td><td></td><td></td><td></td><td></td><td></td><td></td><td></td><td></td></tr>
<tr><td>Oligo rev</td><td>AAACCCGACGTACTTCGCGTTGAC</td><td></td><td></td><td></td><td></td><td></td><td></td><td></td><td></td><td></td><td></td></tr>
<tr><td>Chromosome</td><td>start</td><td>end</td><td>strand</td><td>MM</td><td>target_seq</td><td>PAM</td><td>alignment</td><td>distance</td><td>position</td><td>gene name</td><td>gene id</td></tr>
<tr><td>GU305914.1</td><td>197923</td><td>197945</td><td>+</td><td>0</td><td>GTCAACGCGAAGTACGTCGG</td><td>CGG</td><td>||||||||[||||||||||||]PAM</td><td>NA</td><td>Intergenic</td><td>NA</td><td>NA</td></tr>
<tr><td>T15091</td><td>GCGAAGTACGTCGGCGGCACCGG</td><td>1000</td><td></td><td></td><td></td><td></td><td></td><td></td><td></td><td></td><td></td></tr>
<tr><td>Oligo fwd</td><td>CACCGCGAAGTACGTCGGCGGCAC</td><td></td><td></td><td></td><td></td><td></td><td></td><td></td><td></td><td></td><td></td></tr>
<tr><td>Oligo rev</td><td>AAACGTGCCGCCGACGTACTTCGC</td><td></td><td></td><td></td><td></td><td></td><td></td><td></td><td></td><td></td><td></td></tr>
<tr><td>Chromosome</td><td>start</td><td>end</td><td>strand</td><td>MM</td><td>target_seq</td><td>PAM</td><td>alignment</td><td>distance</td><td>position</td><td>gene name</td><td>gene id</td></tr>
<tr><td>GU305914.1</td><td>197929</td><td>197951</td><td>+</td><td>0</td><td>GCGAAGTACGTCGGCGGCAC</td><td>CGG</td><td>||||||||[||||||||||||]PAM</td><td>NA</td><td>Intergenic</td><td>NA</td><td>NA</td></tr>
<tr><td>T15092</td><td>CGAAGTACGTCGGCGGCACCGGG</td><td>1000</td><td></td><td></td><td></td><td></td><td></td><td></td><td></td><td></td><td></td></tr>
<tr><td>Oligo fwd</td><td>CACCCGAAGTACGTCGGCGGCACC</td><td></td><td></td><td></td><td></td><td></td><td></td><td></td><td></td><td></td><td></td></tr>
<tr><td>Oligo rev</td><td>AAACGGTGCCGCCGACGTACTTCG</td><td></td><td></td><td></td><td></td><td></td><td></td><td></td><td></td><td></td><td></td></tr>
<tr><td>Chromosome</td><td>start</td><td>end</td><td>strand</td><td>MM</td><td>target_seq</td><td>PAM</td><td>alignment</td><td>distance</td><td>position</td><td>gene name</td><td>gene id</td></tr>
<tr><td>GU305914.1</td><td>197930</td><td>197952</td><td>+</td><td>0</td><td>CGAAGTACGTCGGCGGCACC</td><td>GGG</td><td>||||||||[||||||||||||]PAM</td><td>NA</td><td>Intergenic</td><td>NA</td><td>NA</td></tr>
<tr><td>T15093</td><td>CGGGCTCAGCTTGTACACGTAGG</td><td>1000</td><td></td><td></td><td></td><td></td><td></td><td></td><td></td><td></td><td></td></tr>
<tr><td>Oligo fwd</td><td>CACCCGGGCTCAGCTTGTACACGT</td><td></td><td></td><td></td><td></td><td></td><td></td><td></td><td></td><td></td><td></td></tr>
<tr><td>Oligo rev</td><td>AAACACGTGTACAAGCTGAGCCCG</td><td></td><td></td><td></td><td></td><td></td><td></td><td></td><td></td><td></td><td></td></tr>
<tr><td>Chromosome</td><td>start</td><td>end</td><td>strand</td><td>MM</td><td>target_seq</td><td>PAM</td><td>alignment</td><td>distance</td><td>position</td><td>gene name</td><td>gene id</td></tr>
<tr><td>GU305914.1</td><td>197949</td><td>197971</td><td>+</td><td>0</td><td>CGGGCTCAGCTTGTACACGT</td><td>AGG</td><td>||||||||[||||||||||||]PAM</td><td>NA</td><td>Intergenic</td><td>NA</td><td>NA</td></tr>
<tr><td>T15094</td><td>GGGCTCAGCTTGTACACGTAGGG</td><td>1000</td><td></td><td></td><td></td><td></td><td></td><td></td><td></td><td></td><td></td></tr>
<tr><td>Oligo fwd</td><td>CACCGGGCTCAGCTTGTACACGTA</td><td></td><td></td><td></td><td></td><td></td><td></td><td></td><td></td><td></td><td></td></tr>
<tr><td>Oligo rev</td><td>AAACTACGTGTACAAGCTGAGCCC</td><td></td><td></td><td></td><td></td><td></td><td></td><td></td><td></td><td></td><td></td></tr>
<tr><td>Chromosome</td><td>start</td><td>end</td><td>strand</td><td>MM</td><td>target_seq</td><td>PAM</td><td>alignment</td><td>distance</td><td>position</td><td>gene name</td><td>gene id</td></tr>
<tr><td>GU305914.1</td><td>197950</td><td>197972</td><td>+</td><td>0</td><td>GGGCTCAGCTTGTACACGTA</td><td>GGG</td><td>||||||||[||||||||||||]PAM</td><td>NA</td><td>Intergenic</td><td>NA</td><td>NA</td></tr>
<tr><td>T15095</td><td>CTCGACGTCCGACGCCGTCGCGG</td><td>1000</td><td></td><td></td><td></td><td></td><td></td><td></td><td></td><td></td><td></td></tr>
<tr><td>Oligo fwd</td><td>CACCCTCGACGTCCGACGCCGTCG</td><td></td><td></td><td></td><td></td><td></td><td></td><td></td><td></td><td></td><td></td></tr>
<tr><td>Oligo rev</td><td>AAACCGACGGCGTCGGACGTCGAG</td><td></td><td></td><td></td><td></td><td></td><td></td><td></td><td></td><td></td><td></td></tr>
<tr><td>Chromosome</td><td>start</td><td>end</td><td>strand</td><td>MM</td><td>target_seq</td><td>PAM</td><td>alignment</td><td>distance</td><td>position</td><td>gene name</td><td>gene id</td></tr>
<tr><td>GU305914.1</td><td>198000</td><td>198022</td><td>+</td><td>0</td><td>CTCGACGTCCGACGCCGTCG</td><td>CGG</td><td>||||||||[||||||||||||]PAM</td><td>NA</td><td>Intergenic</td><td>NA</td><td>NA</td></tr>
<tr><td>GU305914.1</td><td>118869</td><td>118891</td><td>-</td><td>4</td><td>GTCGTCGTCCGACGCCGAGG</td><td>AGG</td><td>-|||-|||[|||||||||--|]PAM</td><td>NA</td><td>Intergenic</td><td>NA</td><td>NA</td></tr>
<tr><td>T15096</td><td>GGCCGCCGTGAGCTTCACCTTGG</td><td>1000</td><td></td><td></td><td></td><td></td><td></td><td></td><td></td><td></td><td></td></tr>
</table>

<table>
<tr><td>Oligo fwd</td><td>CACCGGCCGCCGTGAGCTTCACCT</td><td></td><td></td><td></td><td></td><td></td><td></td><td></td><td></td><td></td><td></td><td></td></tr>
<tr><td>Oligo rev</td><td>AAACAGGTGAAGCTCACGGCGGCC</td><td></td><td></td><td></td><td></td><td></td><td></td><td></td><td></td><td></td><td></td><td></td></tr>
<tr><td>Chromosome</td><td>start</td><td></td><td>end</td><td>strand</td><td>MM</td><td>target_seq</td><td>PAM</td><td>alignment</td><td>distance</td><td>position</td><td>gene name</td><td>gene id</td></tr>
<tr><td>GU305914.1</td><td></td><td>198021</td><td>198043</td><td>+</td><td>0</td><td>GGCCGCCGTGAGCTTCACCT</td><td>TGG</td><td>||||||||[||||||||||||]PAM</td><td>NA</td><td>Intergenic</td><td>NA</td><td>NA</td></tr>
<tr><td></td><td></td><td></td><td></td><td></td><td></td><td></td><td></td><td></td><td></td><td></td><td></td><td></td></tr>
<tr><td>T15097</td><td>CCTTGGCGATCGCCGACTCCAGG</td><td>1000</td><td></td><td></td><td></td><td></td><td></td><td></td><td></td><td></td><td></td><td></td></tr>
<tr><td>Oligo fwd</td><td>CACCCCTTGGCGATCGCCGACTCC</td><td></td><td></td><td></td><td></td><td></td><td></td><td></td><td></td><td></td><td></td><td></td></tr>
<tr><td>Oligo rev</td><td>AAACGGAGTCGGCGATCGCCAAGG</td><td></td><td></td><td></td><td></td><td></td><td></td><td></td><td></td><td></td><td></td><td></td></tr>
<tr><td>Chromosome</td><td>start</td><td></td><td>end</td><td>strand</td><td>MM</td><td>target_seq</td><td>PAM</td><td>alignment</td><td>distance</td><td>position</td><td>gene name</td><td>gene id</td></tr>
<tr><td>GU305914.1</td><td></td><td>198038</td><td>198060</td><td>+</td><td>0</td><td>CCTTGGCGATCGCCGACTCC</td><td>AGG</td><td>||||||||[||||||||||||]PAM</td><td>NA</td><td>Intergenic</td><td>NA</td><td>NA</td></tr>
<tr><td></td><td></td><td></td><td></td><td></td><td></td><td></td><td></td><td></td><td></td><td></td><td></td><td></td></tr>
<tr><td>T15098</td><td>TGGCGATCGCCGACTCCAGGCGG</td><td>1000</td><td></td><td></td><td></td><td></td><td></td><td></td><td></td><td></td><td></td><td></td></tr>
<tr><td>Oligo fwd</td><td>CACCTGGCGATCGCCGACTCCAGG</td><td></td><td></td><td></td><td></td><td></td><td></td><td></td><td></td><td></td><td></td><td></td></tr>
<tr><td>Oligo rev</td><td>AAACCCTGGAGTCGGCGATCGCCA</td><td></td><td></td><td></td><td></td><td></td><td></td><td></td><td></td><td></td><td></td><td></td></tr>
<tr><td>Chromosome</td><td>start</td><td></td><td>end</td><td>strand</td><td>MM</td><td>target_seq</td><td>PAM</td><td>alignment</td><td>distance</td><td>position</td><td>gene name</td><td>gene id</td></tr>
<tr><td>GU305914.1</td><td></td><td>198041</td><td>198063</td><td>+</td><td>0</td><td>TGGCGATCGCCGACTCCAGG</td><td>CGG</td><td>||||||||[||||||||||||]PAM</td><td>NA</td><td>Intergenic</td><td>NA</td><td>NA</td></tr>
<tr><td></td><td></td><td></td><td></td><td></td><td></td><td></td><td></td><td></td><td></td><td></td><td></td><td></td></tr>
<tr><td>T15099</td><td>CGACTCCAGGCGGTCCCAGTAGG</td><td>1000</td><td></td><td></td><td></td><td></td><td></td><td></td><td></td><td></td><td></td><td></td></tr>
<tr><td>Oligo fwd</td><td>CACCCGACTCCAGGCGGTCCCAGT</td><td></td><td></td><td></td><td></td><td></td><td></td><td></td><td></td><td></td><td></td><td></td></tr>
<tr><td>Oligo rev</td><td>AAACACTGGGACCGCCTGGAGTCG</td><td></td><td></td><td></td><td></td><td></td><td></td><td></td><td></td><td></td><td></td><td></td></tr>
<tr><td>Chromosome</td><td>start</td><td></td><td>end</td><td>strand</td><td>MM</td><td>target_seq</td><td>PAM</td><td>alignment</td><td>distance</td><td>position</td><td>gene name</td><td>gene id</td></tr>
<tr><td>GU305914.1</td><td></td><td>198051</td><td>198073</td><td>+</td><td>0</td><td>CGACTCCAGGCGGTCCCAGT</td><td>AGG</td><td>||||||||[||||||||||||]PAM</td><td>NA</td><td>Intergenic</td><td>NA</td><td>NA</td></tr>
<tr><td></td><td></td><td></td><td></td><td></td><td></td><td></td><td></td><td></td><td></td><td></td><td></td><td></td></tr>
<tr><td>T15100</td><td>CGGTCCCAGTAGGTCTTCATCGG</td><td>1000</td><td></td><td></td><td></td><td></td><td></td><td></td><td></td><td></td><td></td><td></td></tr>
<tr><td>Oligo fwd</td><td>CACCCGGTCCCAGTAGGTCTTCAT</td><td></td><td></td><td></td><td></td><td></td><td></td><td></td><td></td><td></td><td></td><td></td></tr>
<tr><td>Oligo rev</td><td>AAACATGAAGACCTACTGGGACCG</td><td></td><td></td><td></td><td></td><td></td><td></td><td></td><td></td><td></td><td></td><td></td></tr>
<tr><td>Chromosome</td><td>start</td><td></td><td>end</td><td>strand</td><td>MM</td><td>target_seq</td><td>PAM</td><td>alignment</td><td>distance</td><td>position</td><td>gene name</td><td>gene id</td></tr>
<tr><td>GU305914.1</td><td></td><td>198061</td><td>198083</td><td>+</td><td>0</td><td>CGGTCCCAGTAGGTCTTCAT</td><td>CGG</td><td>||||||||[||||||||||||]PAM</td><td>NA</td><td>Intergenic</td><td>NA</td><td>NA</td></tr>
<tr><td></td><td></td><td></td><td></td><td></td><td></td><td></td><td></td><td></td><td></td><td></td><td></td><td></td></tr>
<tr><td>T15101</td><td>GTAGGTCTTCATCGGATCGCCGG</td><td>1000</td><td></td><td></td><td></td><td></td><td></td><td></td><td></td><td></td><td></td><td></td></tr>
<tr><td>Oligo fwd</td><td>CACCGTAGGTCTTCATCGGATCGC</td><td></td><td></td><td></td><td></td><td></td><td></td><td></td><td></td><td></td><td></td><td></td></tr>
<tr><td>Oligo rev</td><td>AAACGCGATCCGATGAAGACCTAC</td><td></td><td></td><td></td><td></td><td></td><td></td><td></td><td></td><td></td><td></td><td></td></tr>
<tr><td>Chromosome</td><td>start</td><td></td><td>end</td><td>strand</td><td>MM</td><td>target_seq</td><td>PAM</td><td>alignment</td><td>distance</td><td>position</td><td>gene name</td><td>gene id</td></tr>
<tr><td>GU305914.1</td><td></td><td>198069</td><td>198091</td><td>+</td><td>0</td><td>GTAGGTCTTCATCGGATCGC</td><td>CGG</td><td>||||||||[||||||||||||]PAM</td><td>NA</td><td>Intergenic</td><td>NA</td><td>NA</td></tr>
<tr><td></td><td></td><td></td><td></td><td></td><td></td><td></td><td></td><td></td><td></td><td></td><td></td><td></td></tr>
<tr><td>T15102</td><td>CGGATCGCCGGTGTCCGCGTTGG</td><td>1000</td><td></td><td></td><td></td><td></td><td></td><td></td><td></td><td></td><td></td><td></td></tr>
<tr><td>Oligo fwd</td><td>CACCCGGATCGCCGGTGTCCGCGT</td><td></td><td></td><td></td><td></td><td></td><td></td><td></td><td></td><td></td><td></td><td></td></tr>
<tr><td>Oligo rev</td><td>AAACACGCGGACACCGGCGATCCG</td><td></td><td></td><td></td><td></td><td></td><td></td><td></td><td></td><td></td><td></td><td></td></tr>
<tr><td>Chromosome</td><td>start</td><td></td><td>end</td><td>strand</td><td>MM</td><td>target_seq</td><td>PAM</td><td>alignment</td><td>distance</td><td>position</td><td>gene name</td><td>gene id</td></tr>
<tr><td>GU305914.1</td><td></td><td>198081</td><td>198103</td><td>+</td><td>0</td><td>CGGATCGCCGGTGTCCGCGT</td><td>TGG</td><td>||||||||[||||||||||||]PAM</td><td>NA</td><td>Intergenic</td><td>NA</td><td>NA</td></tr>
<tr><td></td><td></td><td></td><td></td><td></td><td></td><td></td><td></td><td></td><td></td><td></td><td></td><td></td></tr>
<tr><td>T15103</td><td>GCGTTGGTCACCAGCGCCCGCGG</td><td>1000</td><td></td><td></td><td></td><td></td><td></td><td></td><td></td><td></td><td></td><td></td></tr>
<tr><td>Oligo fwd</td><td>CACCGCGTTGGTCACCAGCGCCCG</td><td></td><td></td><td></td><td></td><td></td><td></td><td></td><td></td><td></td><td></td><td></td></tr>
<tr><td>Oligo rev</td><td>AAACCGGGCGCTGGTGACCAACGC</td><td></td><td></td><td></td><td></td><td></td><td></td><td></td><td></td><td></td><td></td><td></td></tr>
<tr><td>Chromosome</td><td>start</td><td></td><td>end</td><td>strand</td><td>MM</td><td>target_seq</td><td>PAM</td><td>alignment</td><td>distance</td><td>position</td><td>gene name</td><td>gene id</td></tr>
<tr><td>GU305914.1</td><td></td><td>198097</td><td>198119</td><td>+</td><td>0</td><td>GCGTTGGTCACCAGCGCCCG</td><td>CGG</td><td>||||||||[||||||||||||]PAM</td><td>NA</td><td>Intergenic</td><td>NA</td><td>NA</td></tr>
<tr><td></td><td></td><td></td><td></td><td></td><td></td><td></td><td></td><td></td><td></td><td></td><td></td><td></td></tr>
<tr><td>T15104</td><td>CAGCGCCCGCGGCGCCACGTTGG</td><td>1000</td><td></td><td></td><td></td><td></td><td></td><td></td><td></td><td></td><td></td><td></td></tr>
<tr><td>Oligo fwd</td><td>CACCCAGCGCCCGCGGCGCCACGT</td><td></td><td></td><td></td><td></td><td></td><td></td><td></td><td></td><td></td><td></td><td></td></tr>
<tr><td>Oligo rev</td><td>AAACACGTGGCGCCGCGGGCGCTG</td><td></td><td></td><td></td><td></td><td></td><td></td><td></td><td></td><td></td><td></td><td></td></tr>
<tr><td>Chromosome</td><td>start</td><td></td><td>end</td><td>strand</td><td>MM</td><td>target_seq</td><td>PAM</td><td>alignment</td><td>distance</td><td>position</td><td>gene name</td><td>gene id</td></tr>
<tr><td>GU305914.1</td><td></td><td>198108</td><td>198130</td><td>+</td><td>0</td><td>CAGCGCCCGCGGCGCCACGT</td><td>TGG</td><td>||||||||[||||||||||||]PAM</td><td>NA</td><td>Intergenic</td><td>NA</td><td>NA</td></tr>
<tr><td></td><td></td><td></td><td></td><td></td><td></td><td></td><td></td><td></td><td></td><td></td><td></td><td></td></tr>
<tr><td>T15105</td><td>AGCGCCCGCGGCGCCACGTTGGG</td><td>1000</td><td></td><td></td><td></td><td></td><td></td><td></td><td></td><td></td><td></td><td></td></tr>
<tr><td>Oligo fwd</td><td>CACCAGCGCCCGCGGCGCCACGTT</td><td></td><td></td><td></td><td></td><td></td><td></td><td></td><td></td><td></td><td></td><td></td></tr>
<tr><td>Oligo rev</td><td>AAACAACGTGGCGCCGCGGGCGCT</td><td></td><td></td><td></td><td></td><td></td><td></td><td></td><td></td><td></td><td></td><td></td></tr>
<tr><td>Chromosome</td><td>start</td><td></td><td>end</td><td>strand</td><td>MM</td><td>target_seq</td><td>PAM</td><td>alignment</td><td>distance</td><td>position</td><td>gene name</td><td>gene id</td></tr>
<tr><td>GU305914.1</td><td></td><td>198109</td><td>198131</td><td>+</td><td>0</td><td>AGCGCCCGCGGCGCCACGTT</td><td>GGG</td><td>||||||||[||||||||||||]PAM</td><td>NA</td><td>Intergenic</td><td>NA</td><td>NA</td></tr>
<tr><td></td><td></td><td></td><td></td><td></td><td></td><td></td><td></td><td></td><td></td><td></td><td></td><td></td></tr>
<tr><td>T15106</td><td>GCGCCCGCGGCGCCACGTTGGGG</td><td>1000</td><td></td><td></td><td></td><td></td><td></td><td></td><td></td><td></td><td></td><td></td></tr>
<tr><td>Oligo fwd</td><td>CACCGCGCCCGCGGCGCCACGTTG</td><td></td><td></td><td></td><td></td><td></td><td></td><td></td><td></td><td></td><td></td><td></td></tr>
<tr><td>Oligo rev</td><td>AAACCAACGTGGCGCCGCGGGCGC</td><td></td><td></td><td></td><td></td><td></td><td></td><td></td><td></td><td></td><td></td><td></td></tr>
<tr><td>Chromosome</td><td>start</td><td></td><td>end</td><td>strand</td><td>MM</td><td>target_seq</td><td>PAM</td><td>alignment</td><td>distance</td><td>position</td><td>gene name</td><td>gene id</td></tr>
<tr><td>GU305914.1</td><td></td><td>198110</td><td>198132</td><td>+</td><td>0</td><td>GCGCCCGCGGCGCCACGTTG</td><td>GGG</td><td>||||||||[||||||||||||]PAM</td><td>NA</td><td>Intergenic</td><td>NA</td><td>NA</td></tr>
<tr><td></td><td></td><td></td><td></td><td></td><td></td><td></td><td></td><td></td><td></td><td></td><td></td><td></td></tr>
<tr><td>T15107</td><td>GCGCCACGTTGGGGTCGTCCAGG</td><td>1000</td><td></td><td></td><td></td><td></td><td></td><td></td><td></td><td></td><td></td><td></td></tr>
<tr><td>Oligo fwd</td><td>CACCGCGCCACGTTGGGGTCGTCC</td><td></td><td></td><td></td><td></td><td></td><td></td><td></td><td></td><td></td><td></td><td></td></tr>
<tr><td>Oligo rev</td><td>AAACGGACGACCCCAACGTGGCGC</td><td></td><td></td><td></td><td></td><td></td><td></td><td></td><td></td><td></td><td></td><td></td></tr>
<tr><td>Chromosome</td><td>start</td><td></td><td>end</td><td>strand</td><td>MM</td><td>target_seq</td><td>PAM</td><td>alignment</td><td>distance</td><td>position</td><td>gene name</td><td>gene id</td></tr>
<tr><td>GU305914.1</td><td></td><td>198119</td><td>198141</td><td>+</td><td>0</td><td>GCGCCACGTTGGGGTCGTCC</td><td>AGG</td><td>||||||||[||||||||||||]PAM</td><td>NA</td><td>Intergenic</td><td>NA</td><td>NA</td></tr>
<tr><td></td><td></td><td></td><td></td><td></td><td></td><td></td><td></td><td></td><td></td><td></td><td></td><td></td></tr>
<tr><td>T15108</td><td>GTTGGGGTCGTCCAGGTCCCAGG</td><td>1000</td><td></td><td></td><td></td><td></td><td></td><td></td><td></td><td></td><td></td><td></td></tr>
<tr><td>Oligo fwd</td><td>CACCGTTGGGGTCGTCCAGGTCCC</td><td></td><td></td><td></td><td></td><td></td><td></td><td></td><td></td><td></td><td></td><td></td></tr>
<tr><td>Oligo rev</td><td>AAACGGGACCTGGACGACCCCAAC</td><td></td><td></td><td></td><td></td><td></td><td></td><td></td><td></td><td></td><td></td><td></td></tr>
<tr><td>Chromosome</td><td>start</td><td></td><td>end</td><td>strand</td><td>MM</td><td>target_seq</td><td>PAM</td><td>alignment</td><td>distance</td><td>position</td><td>gene name</td><td>gene id</td></tr>
<tr><td>GU305914.1</td><td></td><td>198126</td><td>198148</td><td>+</td><td>0</td><td>GTTGGGGTCGTCCAGGTCCC</td><td>AGG</td><td>||||||||[||||||||||||]PAM</td><td>NA</td><td>Intergenic</td><td>NA</td><td>NA</td></tr>
<tr><td></td><td></td><td></td><td></td><td></td><td></td><td></td><td></td><td></td><td></td><td></td><td></td><td></td></tr>
<tr><td>T15109</td><td>TTGGGGTCGTCCAGGTCCCAGGG</td><td>1000</td><td></td><td></td><td></td><td></td><td></td><td></td><td></td><td></td><td></td><td></td></tr>
<tr><td>Oligo fwd</td><td>CACCTTGGGGTCGTCCAGGTCCCA</td><td></td><td></td><td></td><td></td><td></td><td></td><td></td><td></td><td></td><td></td><td></td></tr>
<tr><td>Oligo rev</td><td>AAACTGGGACCTGGACGACCCCAA</td><td></td><td></td><td></td><td></td><td></td><td></td><td></td><td></td><td></td><td></td><td></td></tr>
<tr><td>Chromosome</td><td>start</td><td></td><td>end</td><td>strand</td><td>MM</td><td>target_seq</td><td>PAM</td><td>alignment</td><td>distance</td><td>position</td><td>gene name</td><td>gene id</td></tr>
<tr><td>GU305914.1</td><td></td><td>198127</td><td>198149</td><td>+</td><td>0</td><td>TTGGGGTCGTCCAGGTCCCA</td><td>GGG</td><td>||||||||[||||||||||||]PAM</td><td>NA</td><td>Intergenic</td><td>NA</td><td>NA</td></tr>
<tr><td></td><td></td><td></td><td></td><td></td><td></td><td></td><td></td><td></td><td></td><td></td><td></td><td></td></tr>
<tr><td>T15110</td><td>TGGGGTCGTCCAGGTCCCAGGGG</td><td>1000</td><td></td><td></td><td></td><td></td><td></td><td></td><td></td><td></td><td></td><td></td></tr>
<tr><td>Oligo fwd</td><td>CACCTGGGGTCGTCCAGGTCCCAG</td><td></td><td></td><td></td><td></td><td></td><td></td><td></td><td></td><td></td><td></td><td></td></tr>
<tr><td>Oligo rev</td><td>AAACCTGGGACCTGGACGACCCCA</td><td></td><td></td><td></td><td></td><td></td><td></td><td></td><td></td><td></td><td></td><td></td></tr>
<tr><td>Chromosome</td><td>start</td><td></td><td>end</td><td>strand</td><td>MM</td><td>target_seq</td><td>PAM</td><td>alignment</td><td>distance</td><td>position</td><td>gene name</td><td>gene id</td></tr>
<tr><td>GU305914.1</td><td></td><td>198128</td><td>198150</td><td>+</td><td>0</td><td>TGGGGTCGTCCAGGTCCCAG</td><td>GGG</td><td>||||||||[||||||||||||]PAM</td><td>NA</td><td>Intergenic</td><td>NA</td><td>NA</td></tr>
</table>

<table>
<tr><td>T15111</td><td>GGTCGTCCAGGTCCCAGGGGCGG</td><td>1000</td><td></td><td></td><td></td><td></td><td></td><td></td><td></td><td></td><td></td></tr>
<tr><td>Oligo fwd</td><td>CACCGGTCGTCCAGGTCCCAGGGG</td><td></td><td></td><td></td><td></td><td></td><td></td><td></td><td></td><td></td><td></td></tr>
<tr><td>Oligo rev</td><td>AAACCCCCTGGGACCTGGACGACC</td><td></td><td></td><td></td><td></td><td></td><td></td><td></td><td></td><td></td><td></td></tr>
<tr><td>Chromosome</td><td>start</td><td>end</td><td>strand</td><td>MM</td><td>target_seq</td><td>PAM</td><td>alignment</td><td>distance</td><td>position</td><td>gene name</td><td>gene id</td></tr>
<tr><td>GU305914.1</td><td>198131</td><td>198153</td><td>+</td><td>0</td><td>GGTCGTCCAGGTCCCAGGGG</td><td>CGG</td><td>||||||||[||||||||||||]PAM</td><td>NA</td><td>Intergenic</td><td>NA</td><td>NA</td></tr>
<tr><td>T15112</td><td>GTCCAGGTCCCAGGGGCGGCAGG</td><td>1000</td><td></td><td></td><td></td><td></td><td></td><td></td><td></td><td></td><td></td></tr>
<tr><td>Oligo fwd</td><td>CACCGTCCAGGTCCCAGGGGCGGC</td><td></td><td></td><td></td><td></td><td></td><td></td><td></td><td></td><td></td><td></td></tr>
<tr><td>Oligo rev</td><td>AAACGCCGCCCCTGGGACCTGGAC</td><td></td><td></td><td></td><td></td><td></td><td></td><td></td><td></td><td></td><td></td></tr>
<tr><td>Chromosome</td><td>start</td><td>end</td><td>strand</td><td>MM</td><td>target_seq</td><td>PAM</td><td>alignment</td><td>distance</td><td>position</td><td>gene name</td><td>gene id</td></tr>
<tr><td>GU305914.1</td><td>198135</td><td>198157</td><td>+</td><td>0</td><td>GTCCAGGTCCCAGGGGCGGC</td><td>AGG</td><td>||||||||[||||||||||||]PAM</td><td>NA</td><td>Intergenic</td><td>NA</td><td>NA</td></tr>
<tr><td>T15113</td><td>GCGGCAGGAGTGCACCTTGACGG</td><td>1000</td><td></td><td></td><td></td><td></td><td></td><td></td><td></td><td></td><td></td></tr>
<tr><td>Oligo fwd</td><td>CACCGCGGCAGGAGTGCACCTTGA</td><td></td><td></td><td></td><td></td><td></td><td></td><td></td><td></td><td></td><td></td></tr>
<tr><td>Oligo rev</td><td>AAACTCAAGGTGCACTCCTGCCGC</td><td></td><td></td><td></td><td></td><td></td><td></td><td></td><td></td><td></td><td></td></tr>
<tr><td>Chromosome</td><td>start</td><td>end</td><td>strand</td><td>MM</td><td>target_seq</td><td>PAM</td><td>alignment</td><td>distance</td><td>position</td><td>gene name</td><td>gene id</td></tr>
<tr><td>GU305914.1</td><td>198150</td><td>198172</td><td>+</td><td>0</td><td>GCGGCAGGAGTGCACCTTGA</td><td>CGG</td><td>||||||||[||||||||||||]PAM</td><td>NA</td><td>Intergenic</td><td>NA</td><td>NA</td></tr>
<tr><td>T15114</td><td>GCAGGAGTGCACCTTGACGGTGG</td><td>1000</td><td></td><td></td><td></td><td></td><td></td><td></td><td></td><td></td><td></td></tr>
<tr><td>Oligo fwd</td><td>CACCGCAGGAGTGCACCTTGACGG</td><td></td><td></td><td></td><td></td><td></td><td></td><td></td><td></td><td></td><td></td></tr>
<tr><td>Oligo rev</td><td>AAACCCGTCAAGGTGCACTCCTGC</td><td></td><td></td><td></td><td></td><td></td><td></td><td></td><td></td><td></td><td></td></tr>
<tr><td>Chromosome</td><td>start</td><td>end</td><td>strand</td><td>MM</td><td>target_seq</td><td>PAM</td><td>alignment</td><td>distance</td><td>position</td><td>gene name</td><td>gene id</td></tr>
<tr><td>GU305914.1</td><td>198153</td><td>198175</td><td>+</td><td>0</td><td>GCAGGAGTGCACCTTGACGG</td><td>TGG</td><td>||||||||[||||||||||||]PAM</td><td>NA</td><td>Intergenic</td><td>NA</td><td>NA</td></tr>
<tr><td>T15115</td><td>GACGGTGGACTGCACCAGCCAGG</td><td>1000</td><td></td><td></td><td></td><td></td><td></td><td></td><td></td><td></td><td></td></tr>
<tr><td>Oligo fwd</td><td>CACCGACGGTGGACTGCACCAGCC</td><td></td><td></td><td></td><td></td><td></td><td></td><td></td><td></td><td></td><td></td></tr>
<tr><td>Oligo rev</td><td>AAACGGCTGGTGCAGTCCACCGTC</td><td></td><td></td><td></td><td></td><td></td><td></td><td></td><td></td><td></td><td></td></tr>
<tr><td>Chromosome</td><td>start</td><td>end</td><td>strand</td><td>MM</td><td>target_seq</td><td>PAM</td><td>alignment</td><td>distance</td><td>position</td><td>gene name</td><td>gene id</td></tr>
<tr><td>GU305914.1</td><td>198168</td><td>198190</td><td>+</td><td>0</td><td>GACGGTGGACTGCACCAGCC</td><td>AGG</td><td>||||||||[||||||||||||]PAM</td><td>NA</td><td>Intergenic</td><td>NA</td><td>NA</td></tr>
<tr><td>T15116</td><td>CACCAGCCAGGCGTAGTGCTCGG</td><td>1000</td><td></td><td></td><td></td><td></td><td></td><td></td><td></td><td></td><td></td></tr>
<tr><td>Oligo fwd</td><td>CACCCACCAGCCAGGCGTAGTGCT</td><td></td><td></td><td></td><td></td><td></td><td></td><td></td><td></td><td></td><td></td></tr>
<tr><td>Oligo rev</td><td>AAACAGCACTACGCCTGGCTGGTG</td><td></td><td></td><td></td><td></td><td></td><td></td><td></td><td></td><td></td><td></td></tr>
<tr><td>Chromosome</td><td>start</td><td>end</td><td>strand</td><td>MM</td><td>target_seq</td><td>PAM</td><td>alignment</td><td>distance</td><td>position</td><td>gene name</td><td>gene id</td></tr>
<tr><td>GU305914.1</td><td>198180</td><td>198202</td><td>+</td><td>0</td><td>CACCAGCCAGGCGTAGTGCT</td><td>CGG</td><td>||||||||[||||||||||||]PAM</td><td>NA</td><td>Intergenic</td><td>NA</td><td>NA</td></tr>
<tr><td>T15117</td><td>GTAGTGCTCGGCCATGCAATCGG</td><td>1000</td><td></td><td></td><td></td><td></td><td></td><td></td><td></td><td></td><td></td></tr>
<tr><td>Oligo fwd</td><td>CACCGTAGTGCTCGGCCATGCAAT</td><td></td><td></td><td></td><td></td><td></td><td></td><td></td><td></td><td></td><td></td></tr>
<tr><td>Oligo rev</td><td>AAACATTGCATGGCCGAGCACTAC</td><td></td><td></td><td></td><td></td><td></td><td></td><td></td><td></td><td></td><td></td></tr>
<tr><td>Chromosome</td><td>start</td><td>end</td><td>strand</td><td>MM</td><td>target_seq</td><td>PAM</td><td>alignment</td><td>distance</td><td>position</td><td>gene name</td><td>gene id</td></tr>
<tr><td>GU305914.1</td><td>198192</td><td>198214</td><td>+</td><td>0</td><td>GTAGTGCTCGGCCATGCAAT</td><td>CGG</td><td>||||||||[||||||||||||]PAM</td><td>NA</td><td>Intergenic</td><td>NA</td><td>NA</td></tr>
<tr><td>T15118</td><td>GTGCTCGGCCATGCAATCGGCGG</td><td>1000</td><td></td><td></td><td></td><td></td><td></td><td></td><td></td><td></td><td></td></tr>
<tr><td>Oligo fwd</td><td>CACCGTGCTCGGCCATGCAATCGG</td><td></td><td></td><td></td><td></td><td></td><td></td><td></td><td></td><td></td><td></td></tr>
<tr><td>Oligo rev</td><td>AAACCCGATTGCATGGCCGAGCAC</td><td></td><td></td><td></td><td></td><td></td><td></td><td></td><td></td><td></td><td></td></tr>
<tr><td>Chromosome</td><td>start</td><td>end</td><td>strand</td><td>MM</td><td>target_seq</td><td>PAM</td><td>alignment</td><td>distance</td><td>position</td><td>gene name</td><td>gene id</td></tr>
<tr><td>GU305914.1</td><td>198195</td><td>198217</td><td>+</td><td>0</td><td>GTGCTCGGCCATGCAATCGG</td><td>CGG</td><td>||||||||[||||||||||||]PAM</td><td>NA</td><td>Intergenic</td><td>NA</td><td>NA</td></tr>
<tr><td>T15119</td><td>TGCTCGGCCATGCAATCGGCGGG</td><td>1000</td><td></td><td></td><td></td><td></td><td></td><td></td><td></td><td></td><td></td></tr>
<tr><td>Oligo fwd</td><td>CACCTGCTCGGCCATGCAATCGGC</td><td></td><td></td><td></td><td></td><td></td><td></td><td></td><td></td><td></td><td></td></tr>
<tr><td>Oligo rev</td><td>AAACGCCGATTGCATGGCCGAGCA</td><td></td><td></td><td></td><td></td><td></td><td></td><td></td><td></td><td></td><td></td></tr>
<tr><td>Chromosome</td><td>start</td><td>end</td><td>strand</td><td>MM</td><td>target_seq</td><td>PAM</td><td>alignment</td><td>distance</td><td>position</td><td>gene name</td><td>gene id</td></tr>
<tr><td>GU305914.1</td><td>198196</td><td>198218</td><td>+</td><td>0</td><td>TGCTCGGCCATGCAATCGGC</td><td>GGG</td><td>||||||||[||||||||||||]PAM</td><td>NA</td><td>Intergenic</td><td>NA</td><td>NA</td></tr>
<tr><td>T15120</td><td>GGGCACGTAGTCCTCATCGTCGG</td><td>1000</td><td></td><td></td><td></td><td></td><td></td><td></td><td></td><td></td><td></td></tr>
<tr><td>Oligo fwd</td><td>CACCGGGCACGTAGTCCTCATCGT</td><td></td><td></td><td></td><td></td><td></td><td></td><td></td><td></td><td></td><td></td></tr>
<tr><td>Oligo rev</td><td>AAACACGATGAGGACTACGTGCCC</td><td></td><td></td><td></td><td></td><td></td><td></td><td></td><td></td><td></td><td></td></tr>
<tr><td>Chromosome</td><td>start</td><td>end</td><td>strand</td><td>MM</td><td>target_seq</td><td>PAM</td><td>alignment</td><td>distance</td><td>position</td><td>gene name</td><td>gene id</td></tr>
<tr><td>GU305914.1</td><td>198216</td><td>198238</td><td>+</td><td>0</td><td>GGGCACGTAGTCCTCATCGT</td><td>CGG</td><td>||||||||[||||||||||||]PAM</td><td>NA</td><td>Intergenic</td><td>NA</td><td>NA</td></tr>
<tr><td>T15121</td><td>ACGTAGTCCTCATCGTCGGTCGG</td><td>1000</td><td></td><td></td><td></td><td></td><td></td><td></td><td></td><td></td><td></td></tr>
<tr><td>Oligo fwd</td><td>CACCACGTAGTCCTCATCGTCGGT</td><td></td><td></td><td></td><td></td><td></td><td></td><td></td><td></td><td></td><td></td></tr>
<tr><td>Oligo rev</td><td>AAACACCGACGATGAGGACTACGT</td><td></td><td></td><td></td><td></td><td></td><td></td><td></td><td></td><td></td><td></td></tr>
<tr><td>Chromosome</td><td>start</td><td>end</td><td>strand</td><td>MM</td><td>target_seq</td><td>PAM</td><td>alignment</td><td>distance</td><td>position</td><td>gene name</td><td>gene id</td></tr>
<tr><td>GU305914.1</td><td>198220</td><td>198242</td><td>+</td><td>0</td><td>ACGTAGTCCTCATCGTCGGT</td><td>CGG</td><td>||||||||[||||||||||||]PAM</td><td>NA</td><td>Intergenic</td><td>NA</td><td>NA</td></tr>
<tr><td>T15122</td><td>CGTCGGTCGGCGAGTCGAAGCGG</td><td>1000</td><td></td><td></td><td></td><td></td><td></td><td></td><td></td><td></td><td></td></tr>
<tr><td>Oligo fwd</td><td>CACCCGTCGGTCGGCGAGTCGAAG</td><td></td><td></td><td></td><td></td><td></td><td></td><td></td><td></td><td></td><td></td></tr>
<tr><td>Oligo rev</td><td>AAACCTTCGACTCGCCGACCGACG</td><td></td><td></td><td></td><td></td><td></td><td></td><td></td><td></td><td></td><td></td></tr>
<tr><td>Chromosome</td><td>start</td><td>end</td><td>strand</td><td>MM</td><td>target_seq</td><td>PAM</td><td>alignment</td><td>distance</td><td>position</td><td>gene name</td><td>gene id</td></tr>
<tr><td>GU305914.1</td><td>198233</td><td>198255</td><td>+</td><td>0</td><td>CGTCGGTCGGCGAGTCGAAG</td><td>CGG</td><td>||||||||[||||||||||||]PAM</td><td>NA</td><td>Intergenic</td><td>NA</td><td>NA</td></tr>
<tr><td>T15123</td><td>TCGGCGAGTCGAAGCGGTCGCGG</td><td>1000</td><td></td><td></td><td></td><td></td><td></td><td></td><td></td><td></td><td></td></tr>
<tr><td>Oligo fwd</td><td>CACCTCGGCGAGTCGAAGCGGTCG</td><td></td><td></td><td></td><td></td><td></td><td></td><td></td><td></td><td></td><td></td></tr>
<tr><td>Oligo rev</td><td>AAACCGACCGCTTCGACTCGCCGA</td><td></td><td></td><td></td><td></td><td></td><td></td><td></td><td></td><td></td><td></td></tr>
<tr><td>Chromosome</td><td>start</td><td>end</td><td>strand</td><td>MM</td><td>target_seq</td><td>PAM</td><td>alignment</td><td>distance</td><td>position</td><td>gene name</td><td>gene id</td></tr>
<tr><td>GU305914.1</td><td>198239</td><td>198261</td><td>+</td><td>0</td><td>TCGGCGAGTCGAAGCGGTCG</td><td>CGG</td><td>||||||||[||||||||||||]PAM</td><td>NA</td><td>Intergenic</td><td>NA</td><td>NA</td></tr>
<tr><td>T15124</td><td>GATGTACTTGAACTTCTGCTCGG</td><td>1000</td><td></td><td></td><td></td><td></td><td></td><td></td><td></td><td></td><td></td></tr>
<tr><td>Oligo fwd</td><td>CACCGATGTACTTGAACTTCTGCT</td><td></td><td></td><td></td><td></td><td></td><td></td><td></td><td></td><td></td><td></td></tr>
<tr><td>Oligo rev</td><td>AAACAGCAGAAGTTCAAGTACATC</td><td></td><td></td><td></td><td></td><td></td><td></td><td></td><td></td><td></td><td></td></tr>
<tr><td>Chromosome</td><td>start</td><td>end</td><td>strand</td><td>MM</td><td>target_seq</td><td>PAM</td><td>alignment</td><td>distance</td><td>position</td><td>gene name</td><td>gene id</td></tr>
<tr><td>GU305914.1</td><td>198273</td><td>198295</td><td>+</td><td>0</td><td>GATGTACTTGAACTTCTGCT</td><td>CGG</td><td>||||||||[||||||||||||]PAM</td><td>NA</td><td>Intergenic</td><td>NA</td><td>NA</td></tr>
<tr><td>T15125</td><td>ATGTACTTGAACTTCTGCTCGGG</td><td>1000</td><td></td><td></td><td></td><td></td><td></td><td></td><td></td><td></td><td></td></tr>
<tr><td>Oligo fwd</td><td>CACCATGTACTTGAACTTCTGCTC</td><td></td><td></td><td></td><td></td><td></td><td></td><td></td><td></td><td></td><td></td></tr>
<tr><td>Oligo rev</td><td>AAACGAGCAGAAGTTCAAGTACAT</td><td></td><td></td><td></td><td></td><td></td><td></td><td></td><td></td><td></td><td></td></tr>
<tr><td>Chromosome</td><td>start</td><td>end</td><td>strand</td><td>MM</td><td>target_seq</td><td>PAM</td><td>alignment</td><td>distance</td><td>position</td><td>gene name</td><td>gene id</td></tr>
<tr><td>GU305914.1</td><td>198274</td><td>198296</td><td>+</td><td>0</td><td>ATGTACTTGAACTTCTGCTC</td><td>GGG</td><td>||||||||[||||||||||||]PAM</td><td>NA</td><td>Intergenic</td><td>NA</td><td>NA</td></tr>
</table>

T15126 AACTTCTGCTCGGGCCAGTGCGG 1000
Oligo fwd CACCAACTTCTGCTCGGGCCAGTG
Oligo rev AAACCACTGGCCCGAGCAGAAGTT
Chromosome start end strand MM target_seq PAM alignment distance position gene name gene id
GU305914.1 198283 198305 + 0 AACTTCTGCTCGGGCCAGTG CGG ||||||||[||||||||||||]PAM NA Intergenic NA NA

T15127 ACTTCTGCTCGGGCCAGTGCGGG 1000
Oligo fwd CACCACTTCTGCTCGGGCCAGTGC
Oligo rev AAACGCACTGGCCCGAGCAGAAGT
Chromosome start end strand MM target_seq PAM alignment distance position gene name gene id
GU305914.1 198284 198306 + 0 ACTTCTGCTCGGGCCAGTGC GGG ||||||||[||||||||||||]PAM NA Intergenic NA NA

T15128 CTCGGGCCAGTGCGGGCACCTGG 1000
Oligo fwd CACCCTCGGGCCAGTGCGGGCACC
Oligo rev AAACGGTGCCCGCACTGGCCCGAG
Chromosome start end strand MM target_seq PAM alignment distance position gene name gene id
GU305914.1 198291 198313 + 0 CTCGGGCCAGTGCGGGCACC TGG ||||||||[||||||||||||]PAM NA Intergenic NA NA

T15129 TCGGGCCAGTGCGGGCACCTGGG 1000
Oligo fwd CACCTCGGGCCAGTGCGGGCACCT
Oligo rev AAACAGGTGCCCGCACTGGCCCGA
Chromosome start end strand MM target_seq PAM alignment distance position gene name gene id
GU305914.1 198292 198314 + 0 TCGGGCCAGTGCGGGCACCT GGG ||||||||[||||||||||||]PAM NA Intergenic NA NA

T15130 TGCTCTCCAGACGCTCCTTGCGG 1000
Oligo fwd CACCTGCTCTCCAGACGCTCCTTG
Oligo rev AAACCAAGGAGCGTCTGGAGAGCA
Chromosome start end strand MM target_seq PAM alignment distance position gene name gene id
GU305914.1 198317 198339 + 0 TGCTCTCCAGACGCTCCTTG CGG ||||||||[||||||||||||]PAM NA Intergenic NA NA

T15131 CCCTCTTCAGCATGCCCATCTGG 1000
Oligo fwd CACCCCCTCTTCAGCATGCCCATC
Oligo rev AAACGATGGGCATGCTGAAGAGGG
Chromosome start end strand MM target_seq PAM alignment distance position gene name gene id
GU305914.1 198377 198399 + 0 CCCTCTTCAGCATGCCCATC TGG ||||||||[||||||||||||]PAM NA Intergenic NA NA

T15132 CATGCCCATCTGGAAGAATTCGG 1000
Oligo fwd CACCCATGCCCATCTGGAAGAATT
Oligo rev AAACAATTCTTCCAGATGGGCATG
Chromosome start end strand MM target_seq PAM alignment distance position gene name gene id
GU305914.1 198387 198409 + 0 CATGCCCATCTGGAAGAATT CGG ||||||||[||||||||||||]PAM NA Intergenic NA NA

T15133 GAAGAATTCGGCCACGCTGTCGG 1000
Oligo fwd CACCGAAGAATTCGGCCACGCTGT
Oligo rev AAACACAGCGTGGCCGAATTCTTC
Chromosome start end strand MM target_seq PAM alignment distance position gene name gene id
GU305914.1 198399 198421 + 0 GAAGAATTCGGCCACGCTGT CGG ||||||||[||||||||||||]PAM NA Intergenic NA NA

T15134 TGTCGGCGACGCGCCGCGTCAGG 1000
Oligo fwd CACCTGTCGGCGACGCGCCGCGTC
Oligo rev AAACGACGCGGCGCGTCGCCGACA
Chromosome start end strand MM target_seq PAM alignment distance position gene name gene id
GU305914.1 198416 198438 + 0 TGTCGGCGACGCGCCGCGTC AGG ||||||||[||||||||||||]PAM NA Intergenic NA NA
GU305914.1 43392 43414 - 3 TGTCGGGGACGCGGCGCGTG GGG ||||||-|[|||||-|||||-]PAM NA Intergenic NA NA

T15135 CCGCTCTCGAGTACACCAGCAGG 1000
Oligo fwd CACCCCGCTCTCGAGTACACCAGC
Oligo rev AAACGCTGGTGTACTCGAGAGCGG
Chromosome start end strand MM target_seq PAM alignment distance position gene name gene id
GU305914.1 198440 198462 + 0 CCGCTCTCGAGTACACCAGC AGG ||||||||[||||||||||||]PAM NA Intergenic NA NA

T15136 GTACACCAGCAGGCTGTACACGG 1000
Oligo fwd CACCGTACACCAGCAGGCTGTACA
Oligo rev AAACTGTACAGCCTGCTGGTGTAC
Chromosome start end strand MM target_seq PAM alignment distance position gene name gene id
GU305914.1 198450 198472 + 0 GTACACCAGCAGGCTGTACA CGG ||||||||[||||||||||||]PAM NA Intergenic NA NA

T15137 GTACACGGCCCCGAACTTGTCGG 1000
Oligo fwd CACCGTACACGGCCCCGAACTTGT
Oligo rev AAACACAAGTTCGGGGCCGTGTAC
Chromosome start end strand MM target_seq PAM alignment distance position gene name gene id
GU305914.1 198465 198487 + 0 GTACACGGCCCCGAACTTGT CGG ||||||||[||||||||||||]PAM NA Intergenic NA NA

T15138 ACTTGTCGGCCACGATCACGTGG 1000
Oligo fwd CACCACTTGTCGGCCACGATCACG
Oligo rev AAACCGTGATCGTGGCCGACAAGT
Chromosome start end strand MM target_seq PAM alignment distance position gene name gene id
GU305914.1 198479 198501 + 0 ACTTGTCGGCCACGATCACG TGG ||||||||[||||||||||||]PAM NA Intergenic NA NA

T15139 GTCGGCCACGATCACGTGGTCGG 1000
Oligo fwd CACCGTCGGCCACGATCACGTGGT
Oligo rev AAACACCACGTGATCGTGGCCGAC
Chromosome start end strand MM target_seq PAM alignment distance position gene name gene id
GU305914.1 198483 198505 + 0 GTCGGCCACGATCACGTGGT CGG ||||||||[||||||||||||]PAM NA Intergenic NA NA

T15140 TCGGCCCTGAACTCCGCGTCCGG 1000
Oligo fwd CACCTCGGCCCTGAACTCCGCGTC
Oligo rev AAACGACGCGGAGTTCAGGGCCGA

| | | | | | | | | | | | |
|---|---|---|---|---|---|---|---|---|---|---|---|
| Chromosome | start | end | strand | MM | target_seq | PAM | alignment | distance | position | gene name | gene id |
| GU305914.1 | 198502 | 198524 | + | 0 | TCGGCCCTGAACTCCGCGTC | CGG | \|\|\|\|\|\|\|\|[\|\|\|\|\|\|\|\|\|\|\|\|]PAM | NA | Intergenic | NA | NA |
| T15141 | CCCTGAACTCCGCGTCCGGCAGG | 1000 | | | | | | | | | |
| Oligo fwd | CACCCCCTGAACTCCGCGTCCGGC | | | | | | | | | | |
| Oligo rev | AAACGCCGGACGCGGAGTTCAGGG | | | | | | | | | | |
| Chromosome | start | end | strand | MM | target_seq | PAM | alignment | distance | position | gene name | gene id |
| GU305914.1 | 198506 | 198528 | + | 0 | CCCTGAACTCCGCGTCCGGC | AGG | \|\|\|\|\|\|\|\|[\|\|\|\|\|\|\|\|\|\|\|\|]PAM | NA | Intergenic | NA | NA |
| T15142 | TCCGCGTCCGGCAGGCCCCCCGG | 1000 | | | | | | | | | |
| Oligo fwd | CACCTCCGCGTCCGGCAGGCCCCC | | | | | | | | | | |
| Oligo rev | AAACGGGGGCCTGCCGGACGCGGA | | | | | | | | | | |
| Chromosome | start | end | strand | MM | target_seq | PAM | alignment | distance | position | gene name | gene id |
| GU305914.1 | 198514 | 198536 | + | 0 | TCCGCGTCCGGCAGGCCCCC | CGG | \|\|\|\|\|\|\|\|[\|\|\|\|\|\|\|\|\|\|\|\|]PAM | NA | Intergenic | NA | NA |
| T15143 | GTTCCAGCACCGCTTGAACTCGG | 1000 | | | | | | | | | |
| Oligo fwd | CACCGTTCCAGCACCGCTTGAACT | | | | | | | | | | |
| Oligo rev | AAACAGTTCAAGCGGTGCTGGAAC | | | | | | | | | | |
| Chromosome | start | end | strand | MM | target_seq | PAM | alignment | distance | position | gene name | gene id |
| GU305914.1 | 198594 | 198616 | + | 0 | GTTCCAGCACCGCTTGAACT | CGG | \|\|\|\|\|\|\|\|[\|\|\|\|\|\|\|\|\|\|\|\|]PAM | NA | Intergenic | NA | NA |
| T15144 | TTCCAGCACCGCTTGAACTCGGG | 1000 | | | | | | | | | |
| Oligo fwd | CACCTTCCAGCACCGCTTGAACTC | | | | | | | | | | |
| Oligo rev | AAACGAGTTCAAGCGGTGCTGGAA | | | | | | | | | | |
| Chromosome | start | end | strand | MM | target_seq | PAM | alignment | distance | position | gene name | gene id |
| GU305914.1 | 198595 | 198617 | + | 0 | TTCCAGCACCGCTTGAACTC | GGG | \|\|\|\|\|\|\|\|[\|\|\|\|\|\|\|\|\|\|\|\|]PAM | NA | Intergenic | NA | NA |
| T15145 | TCCAGCACCGCTTGAACTCGGGG | 1000 | | | | | | | | | |
| Oligo fwd | CACCTCCAGCACCGCTTGAACTCG | | | | | | | | | | |
| Oligo rev | AAACCGAGTTCAAGCGGTGCTGGA | | | | | | | | | | |
| Chromosome | start | end | strand | MM | target_seq | PAM | alignment | distance | position | gene name | gene id |
| GU305914.1 | 198596 | 198618 | + | 0 | TCCAGCACCGCTTGAACTCG | GGG | \|\|\|\|\|\|\|\|[\|\|\|\|\|\|\|\|\|\|\|\|]PAM | NA | Intergenic | NA | NA |
| T15146 | ACTCGGGGTCGCACATGCCGCGG | 1000 | | | | | | | | | |
| Oligo fwd | CACCACTCGGGGTCGCACATGCCG | | | | | | | | | | |
| Oligo rev | AAACCGGCATGTGCGACCCCGAGT | | | | | | | | | | |
| Chromosome | start | end | strand | MM | target_seq | PAM | alignment | distance | position | gene name | gene id |
| GU305914.1 | 198611 | 198633 | + | 0 | ACTCGGGGTCGCACATGCCG | CGG | \|\|\|\|\|\|\|\|[\|\|\|\|\|\|\|\|\|\|\|\|]PAM | NA | Intergenic | NA | NA |
| T15147 | GGGTCGCACATGCCGCGGAACGG | 1000 | | | | | | | | | |
| Oligo fwd | CACCGGGTCGCACATGCCGCGGAA | | | | | | | | | | |
| Oligo rev | AAACTTCCGCGGCATGTGCGACCC | | | | | | | | | | |
| Chromosome | start | end | strand | MM | target_seq | PAM | alignment | distance | position | gene name | gene id |
| GU305914.1 | 198616 | 198638 | + | 0 | GGGTCGCACATGCCGCGGAA | CGG | \|\|\|\|\|\|\|\|[\|\|\|\|\|\|\|\|\|\|\|\|]PAM | NA | Intergenic | NA | NA |
| T15148 | CCAGATCCCCCTCAGCTCGCCGG | 1000 | | | | | | | | | |
| Oligo fwd | CACCCCAGATCCCCCTCAGCTCGC | | | | | | | | | | |
| Oligo rev | AAACGCGAGCTGAGGGGGATCTGG | | | | | | | | | | |
| Chromosome | start | end | strand | MM | target_seq | PAM | alignment | distance | position | gene name | gene id |
| GU305914.1 | 198639 | 198661 | + | 0 | CCAGATCCCCCTCAGCTCGC | CGG | \|\|\|\|\|\|\|\|[\|\|\|\|\|\|\|\|\|\|\|\|]PAM | NA | Intergenic | NA | NA |
| T15149 | CAGATCCCCCTCAGCTCGCCGGG | 1000 | | | | | | | | | |
| Oligo fwd | CACCCAGATCCCCCTCAGCTCGCC | | | | | | | | | | |
| Oligo rev | AAACGGCGAGCTGAGGGGGATCTG | | | | | | | | | | |
| Chromosome | start | end | strand | MM | target_seq | PAM | alignment | distance | position | gene name | gene id |
| GU305914.1 | 198640 | 198662 | + | 0 | CAGATCCCCCTCAGCTCGCC | GGG | \|\|\|\|\|\|\|\|[\|\|\|\|\|\|\|\|\|\|\|\|]PAM | NA | Intergenic | NA | NA |
| T15150 | AGATCCCCCTCAGCTCGCCGGGG | 1000 | | | | | | | | | |
| Oligo fwd | CACCAGATCCCCCTCAGCTCGCCG | | | | | | | | | | |
| Oligo rev | AAACCGGCGAGCTGAGGGGGATCT | | | | | | | | | | |
| Chromosome | start | end | strand | MM | target_seq | PAM | alignment | distance | position | gene name | gene id |
| GU305914.1 | 198641 | 198663 | + | 0 | AGATCCCCCTCAGCTCGCCG | GGG | \|\|\|\|\|\|\|\|[\|\|\|\|\|\|\|\|\|\|\|\|]PAM | NA | Intergenic | NA | NA |
| T15151 | GCTCGCCGGGGTCCTTCGCCAGG | 1000 | | | | | | | | | |
| Oligo fwd | CACCGCTCGCCGGGGTCCTTCGCC | | | | | | | | | | |
| Oligo rev | AAACGGCGAAGGACCCCGGCGAGC | | | | | | | | | | |
| Chromosome | start | end | strand | MM | target_seq | PAM | alignment | distance | position | gene name | gene id |
| GU305914.1 | 198653 | 198675 | + | 0 | GCTCGCCGGGGTCCTTCGCC | AGG | \|\|\|\|\|\|\|\|[\|\|\|\|\|\|\|\|\|\|\|\|]PAM | NA | Intergenic | NA | NA |
| T15152 | AGCGCCGACGTCCCGTAGCCCGG | 1000 | | | | | | | | | |
| Oligo fwd | CACCAGCGCCGACGTCCCGTAGCC | | | | | | | | | | |
| Oligo rev | AAACGGCTACGGGACGTCGGCGCT | | | | | | | | | | |
| Chromosome | start | end | strand | MM | target_seq | PAM | alignment | distance | position | gene name | gene id |
| GU305914.1 | 198679 | 198701 | + | 0 | AGCGCCGACGTCCCGTAGCC | CGG | \|\|\|\|\|\|\|\|[\|\|\|\|\|\|\|\|\|\|\|\|]PAM | NA | Intergenic | NA | NA |
| T15153 | GACGTCGCCGCGTCCGCGTTCGG | 1000 | | | | | | | | | |
| Oligo fwd | CACCGACGTCGCCGCGTCCGCGTT | | | | | | | | | | |
| Oligo rev | AAACAACGCGGACGCGGCGACGTC | | | | | | | | | | |
| Chromosome | start | end | strand | MM | target_seq | PAM | alignment | distance | position | gene name | gene id |
| GU305914.1 | 198751 | 198773 | + | 0 | GACGTCGCCGCGTCCGCGTT | CGG | \|\|\|\|\|\|\|\|[\|\|\|\|\|\|\|\|\|\|\|\|]PAM | NA | Intergenic | NA | NA |
| GU305914.1 | 110088 | 110110 | + | 3 | GGCGTCGCCGCGGCCGCGTC | GGG | \|-\|\|\|\|\|\|[\|\|\|\|-\|\|\|\|\|\|-]PAM | NA | Intergenic | NA | NA |
| T15154 | ACGTCGCCGCGTCCGCGTTCGGG | 1000 | | | | | | | | | |
| Oligo fwd | CACCACGTCGCCGCGTCCGCGTTC | | | | | | | | | | |
| Oligo rev | AAACGAACGCGGACGCGGCGACGT | | | | | | | | | | |
| Chromosome | start | end | strand | MM | target_seq | PAM | alignment | distance | position | gene name | gene id |
| GU305914.1 | 198752 | 198774 | + | 0 | ACGTCGCCGCGTCCGCGTTC | GGG | \|\|\|\|\|\|\|\|[\|\|\|\|\|\|\|\|\|\|\|\|]PAM | NA | Intergenic | NA | NA |
| T15155 | CGCCGCGTCCGCGTTCGGGTCGG | 1000 | | | | | | | | | |

Oligo fwd CACCCGCCGCGTCCGCGTTCGGGT
Oligo rev AAACACCCGAACGCGGACGCGGCG
Chromosome start end strand MM target_seq PAM alignment distance position gene name gene id
GU305914.1 198756 198778 + 0 CGCCGCGTCCGCGTTCGGGT CGG ||||||||[|||||||||||||]PAM NA Intergenic NA NA

T15156 CACCACCAGCAGCTCCATGCCGG 1000
Oligo fwd CACCCACCACCAGCAGCTCCATGC
Oligo rev AAACGCATGGAGCTGCTGGTGGTG
Chromosome start end strand MM target_seq PAM alignment distance position gene name gene id
GU305914.1 198783 198805 + 0 CACCACCAGCAGCTCCATGC CGG ||||||||[|||||||||||||]PAM NA Intergenic NA NA

T15157 ACCACCAGCAGCTCCATGCCGGG 1000
Oligo fwd CACCACCACCAGCAGCTCCATGCC
Oligo rev AAACGGCATGGAGCTGCTGGTGGT
Chromosome start end strand MM target_seq PAM alignment distance position gene name gene id
GU305914.1 198784 198806 + 0 ACCACCAGCAGCTCCATGCC GGG ||||||||[|||||||||||||]PAM NA Intergenic NA NA

T15158 CCACCAGCAGCTCCATGCCGGGG 1000
Oligo fwd CACCCCACCAGCAGCTCCATGCCG
Oligo rev AAACCGGCATGGAGCTGCTGGTGG
Chromosome start end strand MM target_seq PAM alignment distance position gene name gene id
GU305914.1 198785 198807 + 0 CCACCAGCAGCTCCATGCCG GGG ||||||||[|||||||||||||]PAM NA Intergenic NA NA

T15159 TCCATGCCGGGGTAGCCCCGCGG 1000
Oligo fwd CACCTCCATGCCGGGGTAGCCCCG
Oligo rev AAACCGGGGCTACCCCGGCATGGA
Chromosome start end strand MM target_seq PAM alignment distance position gene name gene id
GU305914.1 198796 198818 + 0 TCCATGCCGGGGTAGCCCCG CGG ||||||||[|||||||||||||]PAM NA Intergenic NA NA

T15160 GCCCCGCGGCACGTCGCACGAGG 1000
Oligo fwd CACCGCCCCGCGGCACGTCGCACG
Oligo rev AAACCGTGCGACGTGCCGCGGGGC
Chromosome start end strand MM target_seq PAM alignment distance position gene name gene id
GU305914.1 198810 198832 + 0 GCCCCGCGGCACGTCGCACG AGG ||||||||[|||||||||||||]PAM NA Intergenic NA NA

T15161 CGATGTACCTCAGCCGCACCAGG 1000
Oligo fwd CACCCGATGTACCTCAGCCGCACC
Oligo rev AAACGGTGCGGCTGAGGTACATCG
Chromosome start end strand MM target_seq PAM alignment distance position gene name gene id
GU305914.1 198833 198855 + 0 CGATGTACCTCAGCCGCACC AGG ||||||||[|||||||||||||]PAM NA Intergenic NA NA

T15162 AGCCGCACCAGGTCCGCGCGCGG 1000
Oligo fwd CACCAGCCGCACCAGGTCCGCGCG
Oligo rev AAACCGCGCGGACCTGGTGCGGCT
Chromosome start end strand MM target_seq PAM alignment distance position gene name gene id
GU305914.1 198844 198866 + 0 AGCCGCACCAGGTCCGCGCG CGG ||||||||[|||||||||||||]PAM NA Intergenic NA NA

T15163 CACCAGGTCCGCGCGCGGCACGG 1000
Oligo fwd CACCCACCAGGTCCGCGCGCGGCA
Oligo rev AAACTGCCGCGCGCGGACCTGGTG
Chromosome start end strand MM target_seq PAM alignment distance position gene name gene id
GU305914.1 198849 198871 + 0 CACCAGGTCCGCGCGCGGCA CGG ||||||||[|||||||||||||]PAM NA Intergenic NA NA

T15164 CCGCTGCATGTCTTCTTTGTAGG 1000
Oligo fwd CACCCCGCTGCATGTCTTCTTTGT
Oligo rev AAACACAAAGAAGACATGCAGCGG
Chromosome start end strand MM target_seq PAM alignment distance position gene name gene id
GU305914.1 198873 198895 + 0 CCGCTGCATGTCTTCTTTGT AGG ||||||||[|||||||||||||]PAM NA Intergenic NA NA

T15165 CTTTGTAGGCGAACGATCGTCGG 1000
Oligo fwd CACCCTTTGTAGGCGAACGATCGT
Oligo rev AAACACGATCGTTCGCCTACAAAG
Chromosome start end strand MM target_seq PAM alignment distance position gene name gene id
GU305914.1 198887 198909 + 0 CTTTGTAGGCGAACGATCGT CGG ||||||||[|||||||||||||]PAM NA Intergenic NA NA

T15166 TTTGTAGGCGAACGATCGTCGGG 1000
Oligo fwd CACCTTTGTAGGCGAACGATCGTC
Oligo rev AAACGACGATCGTTCGCCTACAAA
Chromosome start end strand MM target_seq PAM alignment distance position gene name gene id
GU305914.1 198888 198910 + 0 TTTGTAGGCGAACGATCGTC GGG ||||||||[|||||||||||||]PAM NA Intergenic NA NA

T15167 ATCGTCGGGAGATCCCGAAACGG 1000
Oligo fwd CACCATCGTCGGGAGATCCCGAAA
Oligo rev AAACTTTCGGGATCTCCCGACGAT
Chromosome start end strand MM target_seq PAM alignment distance position gene name gene id
GU305914.1 198902 198924 + 0 ATCGTCGGGAGATCCCGAAA CGG ||||||||[|||||||||||||]PAM NA Intergenic NA NA

T15168 CGAAACGGCCGAGCTCCTCCAGG 1000
Oligo fwd CACCCGAAACGGCCGAGCTCCTCC
Oligo rev AAACGGAGGAGCTCGGCCGTTTCG
Chromosome start end strand MM target_seq PAM alignment distance position gene name gene id
GU305914.1 198917 198939 + 0 CGAAACGGCCGAGCTCCTCC AGG ||||||||[|||||||||||||]PAM NA Intergenic NA NA

T15169 CCTCCAGGTTGCGCGCCACCAGG 1000
Oligo fwd CACCCCTCCAGGTTGCGCGCCACC
Oligo rev AAACGGTGGCGCGCAACCTGGAGG
Chromosome start end strand MM target_seq PAM alignment distance position gene name gene id
GU305914.1 198932 198954 + 0 CCTCCAGGTTGCGCGCCACC AGG ||||||||[|||||||||||||]PAM NA Intergenic NA NA

T15170 CGTACTCGCCCGACAGCACGAGG 1000
Oligo fwd CACCCGTACTCGCCCGACAGCACG
Oligo rev AAACCGTGCTGTCGGGCGAGTACG
Chromosome start end strand MM target_seq PAM alignment distance position gene name gene id
GU305914.1 198995 199017 + 0 CGTACTCGCCCGACAGCACG AGG ||||||||[||||||||||||]PAM NA Intergenic NA NA

T15171 GTACTCGCCCGACAGCACGAGGG 1000
Oligo fwd CACCGTACTCGCCCGACAGCACGA
Oligo rev AAACTCGTGCTGTCGGGCGAGTAC
Chromosome start end strand MM target_seq PAM alignment distance position gene name gene id
GU305914.1 198996 199018 + 0 GTACTCGCCCGACAGCACGA GGG ||||||||[||||||||||||]PAM NA Intergenic NA NA

T15172 CGCAACCCTCGCGTGTCGTACGG 1000
Oligo fwd CACCCGCAACCCTCGCGTGTCGTA
Oligo rev AAACTACGACACGCGAGGGTTGCG
Chromosome start end strand MM target_seq PAM alignment distance position gene name gene id
GU305914.1 199033 199055 + 0 CGCAACCCTCGCGTGTCGTA CGG ||||||||[||||||||||||]PAM NA Intergenic NA NA

T15173 GCAACCCTCGCGTGTCGTACGGG 1000
Oligo fwd CACCGCAACCCTCGCGTGTCGTAC
Oligo rev AAACGTACGACACGCGAGGGTTGC
Chromosome start end strand MM target_seq PAM alignment distance position gene name gene id
GU305914.1 199034 199056 + 0 GCAACCCTCGCGTGTCGTAC GGG ||||||||[||||||||||||]PAM NA Intergenic NA NA

T15174 CCCTCGCGTGTCGTACGGGTCGG 1000
Oligo fwd CACCCCCTCGCGTGTCGTACGGGT
Oligo rev AAACACCCGTACGACACGCGAGGG
Chromosome start end strand MM target_seq PAM alignment distance position gene name gene id
GU305914.1 199038 199060 + 0 CCCTCGCGTGTCGTACGGGT CGG ||||||||[||||||||||||]PAM NA Intergenic NA NA

T15175 TCGCGTGTCGTACGGGTCGGAGG 1000
Oligo fwd CACCTCGCGTGTCGTACGGGTCGG
Oligo rev AAACCCGACCCGTACGACACGCGA
Chromosome start end strand MM target_seq PAM alignment distance position gene name gene id
GU305914.1 199041 199063 + 0 TCGCGTGTCGTACGGGTCGG AGG ||||||||[||||||||||||]PAM NA Intergenic NA NA

T15176 GTCCGTCCGCACAGACCCAACGG 1000
Oligo fwd CACCGTCCGTCCGCACAGACCCAA
Oligo rev AAACTTGGGTCTGTGCGGACGGAC
Chromosome start end strand MM target_seq PAM alignment distance position gene name gene id
GU305914.1 199063 199085 + 0 GTCCGTCCGCACAGACCCAA CGG ||||||||[||||||||||||]PAM NA Intergenic NA NA

T15177 TCCGTCCGCACAGACCCAACGGG 1000
Oligo fwd CACCTCCGTCCGCACAGACCCAAC
Oligo rev AAACGTTGGGTCTGTGCGGACGGA
Chromosome start end strand MM target_seq PAM alignment distance position gene name gene id
GU305914.1 199064 199086 + 0 TCCGTCCGCACAGACCCAAC GGG ||||||||[||||||||||||]PAM NA Intergenic NA NA

T15178 CGCACAGACCCAACGGGATCAGG 1000
Oligo fwd CACCCGCACAGACCCAACGGGATC
Oligo rev AAACGATCCCGTTGGGTCTGTGCG
Chromosome start end strand MM target_seq PAM alignment distance position gene name gene id
GU305914.1 199070 199092 + 0 CGCACAGACCCAACGGGATC AGG ||||||||[||||||||||||]PAM NA Intergenic NA NA

T15179 GCACAGACCCAACGGGATCAGGG 1000
Oligo fwd CACCGCACAGACCCAACGGGATCA
Oligo rev AAACTGATCCCGTTGGGTCTGTGC
Chromosome start end strand MM target_seq PAM alignment distance position gene name gene id
GU305914.1 199071 199093 + 0 GCACAGACCCAACGGGATCA GGG ||||||||[||||||||||||]PAM NA Intergenic NA NA

T15180 TCAGGGCCTCTTCGCAACACAGG 1000
Oligo fwd CACCTCAGGGCCTCTTCGCAACAC
Oligo rev AAACGTGTTGCGAAGAGGCCCTGA
Chromosome start end strand MM target_seq PAM alignment distance position gene name gene id
GU305914.1 199088 199110 + 0 TCAGGGCCTCTTCGCAACAC AGG ||||||||[||||||||||||]PAM NA Intergenic NA NA

T15181 GTTCGCCCTGTCCGTCAGAACGG 1000
Oligo fwd CACCGTTCGCCCTGTCCGTCAGAA
Oligo rev AAACTTCTGACGGACAGGGCGAAC
Chromosome start end strand MM target_seq PAM alignment distance position gene name gene id
GU305914.1 199116 199138 + 0 GTTCGCCCTGTCCGTCAGAA CGG ||||||||[||||||||||||]PAM NA Intergenic NA NA

T15182 CACACCACGTACCAGTCCTGCGG 1000
Oligo fwd CACCCACACCACGTACCAGTCCTG
Oligo rev AAACCAGGACTGGTACGTGGTGTG
Chromosome start end strand MM target_seq PAM alignment distance position gene name gene id
GU305914.1 199162 199184 + 0 CACACCACGTACCAGTCCTG CGG ||||||||[||||||||||||]PAM NA Intergenic NA NA

T15183 ACCAGTCCTGCGGAAACCCCAGG 1000
Oligo fwd CACCACCAGTCCTGCGGAAACCCC
Oligo rev AAACGGGGTTTCCGCAGGACTGGT
Chromosome start end strand MM target_seq PAM alignment distance position gene name gene id
GU305914.1 199172 199194 + 0 ACCAGTCCTGCGGAAACCCC AGG ||||||||[||||||||||||]PAM NA Intergenic NA NA

T15184 CCAGTCCTGCGGAAACCCCAGGG 1000
Oligo fwd CACCCCAGTCCTGCGGAAACCCCA
Oligo rev AAACTGGGGTTTCCGCAGGACTGG
Chromosome start end strand MM target_seq PAM alignment distance position gene name gene id
GU305914.1 199173 199195 + 0 CCAGTCCTGCGGAAACCCCA GGG ||||||||[||||||||||||]PAM NA Intergenic NA NA

| | | | | | | | | | | | |
|---|---|---|---|---|---|---|---|---|---|---|---|
| T15185 | CAGTCCTGCGGAAACCCCAGGGG | 1000 | | | | | | | | | |
| Oligo fwd | CACCCAGTCCTGCGGAAACCCCAG | | | | | | | | | | |
| Oligo rev | AAACCTGGGGTTTCCGCAGGACTG | | | | | | | | | | |
| Chromosome | start | end | strand | MM | target_seq | PAM | alignment | distance | position | gene name | gene id |
| GU305914.1 | 199174 | 199196 | + | 0 | CAGTCCTGCGGAAACCCCAG | GGG | \|\|\|\|\|\|\|\|[\|\|\|\|\|\|\|\|\|\|\|\|]PAM | NA | Intergenic | NA | NA |

| | | | | | | | | | | | |
|---|---|---|---|---|---|---|---|---|---|---|---|
| T15186 | AGTCCTGCGGAAACCCCAGGGGG | 1000 | | | | | | | | | |
| Oligo fwd | CACCAGTCCTGCGGAAACCCCAGG | | | | | | | | | | |
| Oligo rev | AAACCCTGGGGTTTCCGCAGGACT | | | | | | | | | | |
| Chromosome | start | end | strand | MM | target_seq | PAM | alignment | distance | position | gene name | gene id |
| GU305914.1 | 199175 | 199197 | + | 0 | AGTCCTGCGGAAACCCCAGG | GGG | \|\|\|\|\|\|\|\|[\|\|\|\|\|\|\|\|\|\|\|\|]PAM | NA | Intergenic | NA | NA |

| | | | | | | | | | | | |
|---|---|---|---|---|---|---|---|---|---|---|---|
| T15187 | GCCTCTCGATCGCGCGCCGCCGG | 1000 | | | | | | | | | |
| Oligo fwd | CACCGCCTCTCGATCGCGCGCCGC | | | | | | | | | | |
| Oligo rev | AAACGCGGCGCGCGATCGAGAGGC | | | | | | | | | | |
| Chromosome | start | end | strand | MM | target_seq | PAM | alignment | distance | position | gene name | gene id |
| GU305914.1 | 199214 | 199236 | + | 0 | GCCTCTCGATCGCGCGCCGC | CGG | \|\|\|\|\|\|\|\|[\|\|\|\|\|\|\|\|\|\|\|\|]PAM | NA | Intergenic | NA | NA |

| | | | | | | | | | | | |
|---|---|---|---|---|---|---|---|---|---|---|---|
| T15188 | CCTCTCGATCGCGCGCCGCCGGG | 1000 | | | | | | | | | |
| Oligo fwd | CACCCCTCTCGATCGCGCGCCGCC | | | | | | | | | | |
| Oligo rev | AAACGGCGGCGCGCGATCGAGAGG | | | | | | | | | | |
| Chromosome | start | end | strand | MM | target_seq | PAM | alignment | distance | position | gene name | gene id |
| GU305914.1 | 199215 | 199237 | + | 0 | CCTCTCGATCGCGCGCCGCC | GGG | \|\|\|\|\|\|\|\|[\|\|\|\|\|\|\|\|\|\|\|\|]PAM | NA | Intergenic | NA | NA |

| | | | | | | | | | | | |
|---|---|---|---|---|---|---|---|---|---|---|---|
| T15189 | GGCGTTCTGATCGCCGCCCGTGG | 1000 | | | | | | | | | |
| Oligo fwd | CACCGGCGTTCTGATCGCCGCCCG | | | | | | | | | | |
| Oligo rev | AAACCGGGCGGCGATCAGAACGCC | | | | | | | | | | |
| Chromosome | start | end | strand | MM | target_seq | PAM | alignment | distance | position | gene name | gene id |
| GU305914.1 | 199236 | 199258 | + | 0 | GGCGTTCTGATCGCCGCCCG | TGG | \|\|\|\|\|\|\|\|[\|\|\|\|\|\|\|\|\|\|\|\|]PAM | NA | Intergenic | NA | NA |

| | | | | | | | | | | | |
|---|---|---|---|---|---|---|---|---|---|---|---|
| T15190 | CGTGGCCACCAGCCGCGAGAAGG | 1000 | | | | | | | | | |
| Oligo fwd | CACCCGTGGCCACCAGCCGCGAGA | | | | | | | | | | |
| Oligo rev | AAACTCTCGCGGCTGGTGGCCACG | | | | | | | | | | |
| Chromosome | start | end | strand | MM | target_seq | PAM | alignment | distance | position | gene name | gene id |
| GU305914.1 | 199254 | 199276 | + | 0 | CGTGGCCACCAGCCGCGAGA | AGG | \|\|\|\|\|\|\|\|[\|\|\|\|\|\|\|\|\|\|\|\|]PAM | NA | Intergenic | NA | NA |
| GU305914.1 | 194998 | 195020 | + | 4 | CGTGTCGTCCATCCGCGAGA | AGG | \|\|\|\|-\|--[\|\|\|-\|\|\|\|\|\|\|\|]PAM | NA | Intergenic | NA | NA |

| | | | | | | | | | | | |
|---|---|---|---|---|---|---|---|---|---|---|---|
| T15191 | GTCTATCACCCGCTTGTACGCGG | 1000 | | | | | | | | | |
| Oligo fwd | CACCGTCTATCACCCGCTTGTACG | | | | | | | | | | |
| Oligo rev | AAACCGTACAAGCGGGTGATAGAC | | | | | | | | | | |
| Chromosome | start | end | strand | MM | target_seq | PAM | alignment | distance | position | gene name | gene id |
| GU305914.1 | 199278 | 199300 | + | 0 | GTCTATCACCCGCTTGTACG | CGG | \|\|\|\|\|\|\|\|[\|\|\|\|\|\|\|\|\|\|\|\|]PAM | NA | Intergenic | NA | NA |

| | | | | | | | | | | | |
|---|---|---|---|---|---|---|---|---|---|---|---|
| T15192 | CACCCGCTTGTACGCGGAGATGG | 1000 | | | | | | | | | |
| Oligo fwd | CACCCACCCGCTTGTACGCGGAGA | | | | | | | | | | |
| Oligo rev | AAACTCTCCGCGTACAAGCGGGTG | | | | | | | | | | |
| Chromosome | start | end | strand | MM | target_seq | PAM | alignment | distance | position | gene name | gene id |
| GU305914.1 | 199284 | 199306 | + | 0 | CACCCGCTTGTACGCGGAGA | TGG | \|\|\|\|\|\|\|\|[\|\|\|\|\|\|\|\|\|\|\|\|]PAM | NA | Intergenic | NA | NA |

| | | | | | | | | | | | |
|---|---|---|---|---|---|---|---|---|---|---|---|
| T15193 | GAGATGGCGTTCGCCGTCACCGG | 1000 | | | | | | | | | |
| Oligo fwd | CACCGAGATGGCGTTCGCCGTCAC | | | | | | | | | | |
| Oligo rev | AAACGTGACGGCGAACGCCATCTC | | | | | | | | | | |
| Chromosome | start | end | strand | MM | target_seq | PAM | alignment | distance | position | gene name | gene id |
| GU305914.1 | 199300 | 199322 | + | 0 | GAGATGGCGTTCGCCGTCAC | CGG | \|\|\|\|\|\|\|\|[\|\|\|\|\|\|\|\|\|\|\|\|]PAM | NA | Intergenic | NA | NA |

| | | | | | | | | | | | |
|---|---|---|---|---|---|---|---|---|---|---|---|
| T15194 | GCCTCTCCATCTCGCCACCCCGG | 1000 | | | | | | | | | |
| Oligo fwd | CACCGCCTCTCCATCTCGCCACCC | | | | | | | | | | |
| Oligo rev | AAACGGGTGGCGAGATGGAGAGGC | | | | | | | | | | |
| Chromosome | start | end | strand | MM | target_seq | PAM | alignment | distance | position | gene name | gene id |
| GU305914.1 | 199322 | 199344 | + | 0 | GCCTCTCCATCTCGCCACCC | CGG | \|\|\|\|\|\|\|\|[\|\|\|\|\|\|\|\|\|\|\|\|]PAM | NA | Intergenic | NA | NA |

| | | | | | | | | | | | |
|---|---|---|---|---|---|---|---|---|---|---|---|
| T15195 | ACCCTTCCTCTCCTCTGCCTAGG | 1000 | | | | | | | | | |
| Oligo fwd | CACCACCCTTCCTCTCCTCTGCCT | | | | | | | | | | |
| Oligo rev | AAACAGGCAGAGGAGAGGAAGGGT | | | | | | | | | | |
| Chromosome | start | end | strand | MM | target_seq | PAM | alignment | distance | position | gene name | gene id |
| GU305914.1 | 199363 | 199385 | + | 0 | ACCCTTCCTCTCCTCTGCCT | AGG | \|\|\|\|\|\|\|\|[\|\|\|\|\|\|\|\|\|\|\|\|]PAM | NA | Intergenic | NA | NA |

| | | | | | | | | | | | |
|---|---|---|---|---|---|---|---|---|---|---|---|
| T15196 | CTCTGCCTAGGACACCCGAGAGG | 1000 | | | | | | | | | |
| Oligo fwd | CACCCTCTGCCTAGGACACCCGAG | | | | | | | | | | |
| Oligo rev | AAACCTCGGGTGTCCTAGGCAGAG | | | | | | | | | | |
| Chromosome | start | end | strand | MM | target_seq | PAM | alignment | distance | position | gene name | gene id |
| GU305914.1 | 199375 | 199397 | + | 0 | CTCTGCCTAGGACACCCGAG | AGG | \|\|\|\|\|\|\|\|[\|\|\|\|\|\|\|\|\|\|\|\|]PAM | NA | Intergenic | NA | NA |

| | | | | | | | | | | | |
|---|---|---|---|---|---|---|---|---|---|---|---|
| T15197 | CCAGAAGTGATCTGAGAAGATGG | 1000 | | | | | | | | | |
| Oligo fwd | CACCCCAGAAGTGATCTGAGAAGA | | | | | | | | | | |
| Oligo rev | AAACTCTTCTCAGATCACTTCTGG | | | | | | | | | | |
| Chromosome | start | end | strand | MM | target_seq | PAM | alignment | distance | position | gene name | gene id |
| GU305914.1 | 199437 | 199459 | + | 0 | CCAGAAGTGATCTGAGAAGA | TGG | \|\|\|\|\|\|\|\|[\|\|\|\|\|\|\|\|\|\|\|\|]PAM | NA | Intergenic | NA | NA |

| | | | | | | | | | | | |
|---|---|---|---|---|---|---|---|---|---|---|---|
| T15198 | TATTGTTTATTTATCAATACTGG | 1000 | | | | | | | | | |
| Oligo fwd | CACCTATTGTTTATTTATCAATAC | | | | | | | | | | |
| Oligo rev | AAACGTATTGATAAATAAACAATA | | | | | | | | | | |
| Chromosome | start | end | strand | MM | target_seq | PAM | alignment | distance | position | gene name | gene id |
| GU305914.1 | 199466 | 199488 | + | 0 | TATTGTTTATTTATCAATAC | TGG | \|\|\|\|\|\|\|\|[\|\|\|\|\|\|\|\|\|\|\|\|]PAM | NA | Intergenic | NA | NA |

| | | |
|---|---|---|
| T15199 | TTTATTTATCAATACTGGAGCGG | 1000 |
| Oligo fwd | CACCTTTATTTATCAATACTGGAG | |
| Oligo rev | AAACCTCCAGTATTGATAAATAAA | |

<table>
<tr><td>Chromosome</td><td>start</td><td>end</td><td>strand</td><td>MM</td><td>target_seq</td><td>PAM</td><td>alignment</td><td>distance</td><td>position</td><td>gene name</td><td>gene id</td></tr>
<tr><td>GU305914.1</td><td>199471</td><td>199493</td><td>+</td><td>0</td><td>TTTATTTATCAATACTGGAG</td><td>CGG</td><td>||||||||[||||||||||||]PAM</td><td>NA</td><td>Intergenic</td><td>NA</td><td>NA</td></tr>
<tr><td>T15200</td><td>TTATTTATCAATACTGGAGCGGG</td><td>1000</td><td></td><td></td><td></td><td></td><td></td><td></td><td></td><td></td><td></td></tr>
<tr><td>Oligo fwd</td><td>CACCTTATTTATCAATACTGGAGC</td><td></td><td></td><td></td><td></td><td></td><td></td><td></td><td></td><td></td><td></td></tr>
<tr><td>Oligo rev</td><td>AAACGCTCCAGTATTGATAAATAA</td><td></td><td></td><td></td><td></td><td></td><td></td><td></td><td></td><td></td><td></td></tr>
<tr><td>Chromosome</td><td>start</td><td>end</td><td>strand</td><td>MM</td><td>target_seq</td><td>PAM</td><td>alignment</td><td>distance</td><td>position</td><td>gene name</td><td>gene id</td></tr>
<tr><td>GU305914.1</td><td>199472</td><td>199494</td><td>+</td><td>0</td><td>TTATTTATCAATACTGGAGC</td><td>GGG</td><td>||||||||[||||||||||||]PAM</td><td>NA</td><td>Intergenic</td><td>NA</td><td>NA</td></tr>
<tr><td>T15201</td><td>TATTTATCAATACTGGAGCGGGG</td><td>1000</td><td></td><td></td><td></td><td></td><td></td><td></td><td></td><td></td><td></td></tr>
<tr><td>Oligo fwd</td><td>CACCTATTTATCAATACTGGAGCG</td><td></td><td></td><td></td><td></td><td></td><td></td><td></td><td></td><td></td><td></td></tr>
<tr><td>Oligo rev</td><td>AAACCGCTCCAGTATTGATAAATA</td><td></td><td></td><td></td><td></td><td></td><td></td><td></td><td></td><td></td><td></td></tr>
<tr><td>Chromosome</td><td>start</td><td>end</td><td>strand</td><td>MM</td><td>target_seq</td><td>PAM</td><td>alignment</td><td>distance</td><td>position</td><td>gene name</td><td>gene id</td></tr>
<tr><td>GU305914.1</td><td>199473</td><td>199495</td><td>+</td><td>0</td><td>TATTTATCAATACTGGAGCG</td><td>GGG</td><td>||||||||[||||||||||||]PAM</td><td>NA</td><td>Intergenic</td><td>NA</td><td>NA</td></tr>
<tr><td>T15202</td><td>TTATCAATACTGGAGCGGGGAGG</td><td>1000</td><td></td><td></td><td></td><td></td><td></td><td></td><td></td><td></td><td></td></tr>
<tr><td>Oligo fwd</td><td>CACCTTATCAATACTGGAGCGGGG</td><td></td><td></td><td></td><td></td><td></td><td></td><td></td><td></td><td></td><td></td></tr>
<tr><td>Oligo rev</td><td>AAACCCCCGCTCCAGTATTGATAA</td><td></td><td></td><td></td><td></td><td></td><td></td><td></td><td></td><td></td><td></td></tr>
<tr><td>Chromosome</td><td>start</td><td>end</td><td>strand</td><td>MM</td><td>target_seq</td><td>PAM</td><td>alignment</td><td>distance</td><td>position</td><td>gene name</td><td>gene id</td></tr>
<tr><td>GU305914.1</td><td>199476</td><td>199498</td><td>+</td><td>0</td><td>TTATCAATACTGGAGCGGGG</td><td>AGG</td><td>||||||||[||||||||||||]PAM</td><td>NA</td><td>Intergenic</td><td>NA</td><td>NA</td></tr>
<tr><td>T15203</td><td>TATCAATACTGGAGCGGGGAGGG</td><td>1000</td><td></td><td></td><td></td><td></td><td></td><td></td><td></td><td></td><td></td></tr>
<tr><td>Oligo fwd</td><td>CACCTATCAATACTGGAGCGGGGA</td><td></td><td></td><td></td><td></td><td></td><td></td><td></td><td></td><td></td><td></td></tr>
<tr><td>Oligo rev</td><td>AAACTCCCCGCTCCAGTATTGATA</td><td></td><td></td><td></td><td></td><td></td><td></td><td></td><td></td><td></td><td></td></tr>
<tr><td>Chromosome</td><td>start</td><td>end</td><td>strand</td><td>MM</td><td>target_seq</td><td>PAM</td><td>alignment</td><td>distance</td><td>position</td><td>gene name</td><td>gene id</td></tr>
<tr><td>GU305914.1</td><td>199477</td><td>199499</td><td>+</td><td>0</td><td>TATCAATACTGGAGCGGGGA</td><td>GGG</td><td>||||||||[||||||||||||]PAM</td><td>NA</td><td>Intergenic</td><td>NA</td><td>NA</td></tr>
<tr><td>T15204</td><td>ATCAATACTGGAGCGGGGAGGGG</td><td>1000</td><td></td><td></td><td></td><td></td><td></td><td></td><td></td><td></td><td></td></tr>
<tr><td>Oligo fwd</td><td>CACCATCAATACTGGAGCGGGGAG</td><td></td><td></td><td></td><td></td><td></td><td></td><td></td><td></td><td></td><td></td></tr>
<tr><td>Oligo rev</td><td>AAACCTCCCCGCTCCAGTATTGAT</td><td></td><td></td><td></td><td></td><td></td><td></td><td></td><td></td><td></td><td></td></tr>
<tr><td>Chromosome</td><td>start</td><td>end</td><td>strand</td><td>MM</td><td>target_seq</td><td>PAM</td><td>alignment</td><td>distance</td><td>position</td><td>gene name</td><td>gene id</td></tr>
<tr><td>GU305914.1</td><td>199478</td><td>199500</td><td>+</td><td>0</td><td>ATCAATACTGGAGCGGGGAG</td><td>GGG</td><td>||||||||[||||||||||||]PAM</td><td>NA</td><td>Intergenic</td><td>NA</td><td>NA</td></tr>
<tr><td>T15205</td><td>AATACTGGAGCGGGGAGGGGAGG</td><td>1000</td><td></td><td></td><td></td><td></td><td></td><td></td><td></td><td></td><td></td></tr>
<tr><td>Oligo fwd</td><td>CACCAATACTGGAGCGGGGAGGGG</td><td></td><td></td><td></td><td></td><td></td><td></td><td></td><td></td><td></td><td></td></tr>
<tr><td>Oligo rev</td><td>AAACCCCCTCCCCGCTCCAGTATT</td><td></td><td></td><td></td><td></td><td></td><td></td><td></td><td></td><td></td><td></td></tr>
<tr><td>Chromosome</td><td>start</td><td>end</td><td>strand</td><td>MM</td><td>target_seq</td><td>PAM</td><td>alignment</td><td>distance</td><td>position</td><td>gene name</td><td>gene id</td></tr>
<tr><td>GU305914.1</td><td>199481</td><td>199503</td><td>+</td><td>0</td><td>AATACTGGAGCGGGGAGGGG</td><td>AGG</td><td>||||||||[||||||||||||]PAM</td><td>NA</td><td>Intergenic</td><td>NA</td><td>NA</td></tr>
<tr><td>T15206</td><td>ATACTGGAGCGGGGAGGGGAGGG</td><td>1000</td><td></td><td></td><td></td><td></td><td></td><td></td><td></td><td></td><td></td></tr>
<tr><td>Oligo fwd</td><td>CACCATACTGGAGCGGGGAGGGGA</td><td></td><td></td><td></td><td></td><td></td><td></td><td></td><td></td><td></td><td></td></tr>
<tr><td>Oligo rev</td><td>AAACTCCCCTCCCCGCTCCAGTAT</td><td></td><td></td><td></td><td></td><td></td><td></td><td></td><td></td><td></td><td></td></tr>
<tr><td>Chromosome</td><td>start</td><td>end</td><td>strand</td><td>MM</td><td>target_seq</td><td>PAM</td><td>alignment</td><td>distance</td><td>position</td><td>gene name</td><td>gene id</td></tr>
<tr><td>GU305914.1</td><td>199482</td><td>199504</td><td>+</td><td>0</td><td>ATACTGGAGCGGGGAGGGGA</td><td>GGG</td><td>||||||||[||||||||||||]PAM</td><td>NA</td><td>Intergenic</td><td>NA</td><td>NA</td></tr>
<tr><td>T15207</td><td>TACTGGAGCGGGGAGGGGAGGGG</td><td>1000</td><td></td><td></td><td></td><td></td><td></td><td></td><td></td><td></td><td></td></tr>
<tr><td>Oligo fwd</td><td>CACCTACTGGAGCGGGGAGGGGAG</td><td></td><td></td><td></td><td></td><td></td><td></td><td></td><td></td><td></td><td></td></tr>
<tr><td>Oligo rev</td><td>AAACCTCCCCTCCCCGCTCCAGTA</td><td></td><td></td><td></td><td></td><td></td><td></td><td></td><td></td><td></td><td></td></tr>
<tr><td>Chromosome</td><td>start</td><td>end</td><td>strand</td><td>MM</td><td>target_seq</td><td>PAM</td><td>alignment</td><td>distance</td><td>position</td><td>gene name</td><td>gene id</td></tr>
<tr><td>GU305914.1</td><td>199483</td><td>199505</td><td>+</td><td>0</td><td>TACTGGAGCGGGGAGGGGAG</td><td>GGG</td><td>||||||||[||||||||||||]PAM</td><td>NA</td><td>Intergenic</td><td>NA</td><td>NA</td></tr>
<tr><td>T15208</td><td>GAGGGGAGGGGTTTATGTGATGG</td><td>1000</td><td></td><td></td><td></td><td></td><td></td><td></td><td></td><td></td><td></td></tr>
<tr><td>Oligo fwd</td><td>CACCGAGGGGAGGGGTTTATGTGA</td><td></td><td></td><td></td><td></td><td></td><td></td><td></td><td></td><td></td><td></td></tr>
<tr><td>Oligo rev</td><td>AAACTCACATAAACCCCTCCCCTC</td><td></td><td></td><td></td><td></td><td></td><td></td><td></td><td></td><td></td><td></td></tr>
<tr><td>Chromosome</td><td>start</td><td>end</td><td>strand</td><td>MM</td><td>target_seq</td><td>PAM</td><td>alignment</td><td>distance</td><td>position</td><td>gene name</td><td>gene id</td></tr>
<tr><td>GU305914.1</td><td>199495</td><td>199517</td><td>+</td><td>0</td><td>GAGGGGAGGGGTTTATGTGA</td><td>TGG</td><td>||||||||[||||||||||||]PAM</td><td>NA</td><td>Intergenic</td><td>NA</td><td>NA</td></tr>
<tr><td>T15209</td><td>GAGGGGTTTATGTGATGGCGAGG</td><td>1000</td><td></td><td></td><td></td><td></td><td></td><td></td><td></td><td></td><td></td></tr>
<tr><td>Oligo fwd</td><td>CACCGAGGGGTTTATGTGATGGCG</td><td></td><td></td><td></td><td></td><td></td><td></td><td></td><td></td><td></td><td></td></tr>
<tr><td>Oligo rev</td><td>AAACCGCCATCACATAAACCCCTC</td><td></td><td></td><td></td><td></td><td></td><td></td><td></td><td></td><td></td><td></td></tr>
<tr><td>Chromosome</td><td>start</td><td>end</td><td>strand</td><td>MM</td><td>target_seq</td><td>PAM</td><td>alignment</td><td>distance</td><td>position</td><td>gene name</td><td>gene id</td></tr>
<tr><td>GU305914.1</td><td>199500</td><td>199522</td><td>+</td><td>0</td><td>GAGGGGTTTATGTGATGGCG</td><td>AGG</td><td>||||||||[||||||||||||]PAM</td><td>NA</td><td>Intergenic</td><td>NA</td><td>NA</td></tr>
<tr><td>T15210</td><td>CGTCCGTCAGTCGTCATCGTCGG</td><td>1000</td><td></td><td></td><td></td><td></td><td></td><td></td><td></td><td></td><td></td></tr>
<tr><td>Oligo fwd</td><td>CACCCGTCCGTCAGTCGTCATCGT</td><td></td><td></td><td></td><td></td><td></td><td></td><td></td><td></td><td></td><td></td></tr>
<tr><td>Oligo rev</td><td>AAACACGATGACGACTGACGGACG</td><td></td><td></td><td></td><td></td><td></td><td></td><td></td><td></td><td></td><td></td></tr>
<tr><td>Chromosome</td><td>start</td><td>end</td><td>strand</td><td>MM</td><td>target_seq</td><td>PAM</td><td>alignment</td><td>distance</td><td>position</td><td>gene name</td><td>gene id</td></tr>
<tr><td>GU305914.1</td><td>199535</td><td>199557</td><td>+</td><td>0</td><td>CGTCCGTCAGTCGTCATCGT</td><td>CGG</td><td>||||||||[||||||||||||]PAM</td><td>NA</td><td>Intergenic</td><td>NA</td><td>NA</td></tr>
<tr><td>T15211</td><td>GCGTCCGCGTCCGCCGACCTCGG</td><td>1000</td><td></td><td></td><td></td><td></td><td></td><td></td><td></td><td></td><td></td></tr>
<tr><td>Oligo fwd</td><td>CACCGCGTCCGCGTCCGCCGACCT</td><td></td><td></td><td></td><td></td><td></td><td></td><td></td><td></td><td></td><td></td></tr>
<tr><td>Oligo rev</td><td>AAACAGGTCGGCGGACGCGGACGC</td><td></td><td></td><td></td><td></td><td></td><td></td><td></td><td></td><td></td><td></td></tr>
<tr><td>Chromosome</td><td>start</td><td>end</td><td>strand</td><td>MM</td><td>target_seq</td><td>PAM</td><td>alignment</td><td>distance</td><td>position</td><td>gene name</td><td>gene id</td></tr>
<tr><td>GU305914.1</td><td>199557</td><td>199579</td><td>+</td><td>0</td><td>GCGTCCGCGTCCGCCGACCT</td><td>CGG</td><td>||||||||[||||||||||||]PAM</td><td>NA</td><td>Intergenic</td><td>NA</td><td>NA</td></tr>
<tr><td>T15212</td><td>TCCGCCGACCTCGGATTCCGAGG</td><td>1000</td><td></td><td></td><td></td><td></td><td></td><td></td><td></td><td></td><td></td></tr>
<tr><td>Oligo fwd</td><td>CACCTCCGCCGACCTCGGATTCCG</td><td></td><td></td><td></td><td></td><td></td><td></td><td></td><td></td><td></td><td></td></tr>
<tr><td>Oligo rev</td><td>AAACCGGAATCCGAGGTCGGCGGA</td><td></td><td></td><td></td><td></td><td></td><td></td><td></td><td></td><td></td><td></td></tr>
<tr><td>Chromosome</td><td>start</td><td>end</td><td>strand</td><td>MM</td><td>target_seq</td><td>PAM</td><td>alignment</td><td>distance</td><td>position</td><td>gene name</td><td>gene id</td></tr>
<tr><td>GU305914.1</td><td>199566</td><td>199588</td><td>+</td><td>0</td><td>TCCGCCGACCTCGGATTCCG</td><td>AGG</td><td>||||||||[||||||||||||]PAM</td><td>NA</td><td>Intergenic</td><td>NA</td><td>NA</td></tr>
<tr><td>T15213</td><td>CCGCCGACCTCGGATTCCGAGGG</td><td>1000</td><td></td><td></td><td></td><td></td><td></td><td></td><td></td><td></td><td></td></tr>
<tr><td>Oligo fwd</td><td>CACCCCGCCGACCTCGGATTCCGA</td><td></td><td></td><td></td><td></td><td></td><td></td><td></td><td></td><td></td><td></td></tr>
<tr><td>Oligo rev</td><td>AAACTCGGAATCCGAGGTCGGCGG</td><td></td><td></td><td></td><td></td><td></td><td></td><td></td><td></td><td></td><td></td></tr>
<tr><td>Chromosome</td><td>start</td><td>end</td><td>strand</td><td>MM</td><td>target_seq</td><td>PAM</td><td>alignment</td><td>distance</td><td>position</td><td>gene name</td><td>gene id</td></tr>
<tr><td>GU305914.1</td><td>199567</td><td>199589</td><td>+</td><td>0</td><td>CCGCCGACCTCGGATTCCGA</td><td>GGG</td><td>||||||||[||||||||||||]PAM</td><td>NA</td><td>Intergenic</td><td>NA</td><td>NA</td></tr>
<tr><td>T15214</td><td>ATTCCGAGGGCAGCCTCTGAAGG</td><td>1000</td><td></td><td></td><td></td><td></td><td></td><td></td><td></td><td></td><td></td></tr>
<tr><td>Oligo fwd</td><td>CACCATTCCGAGGGCAGCCTCTGA</td><td></td><td></td><td></td><td></td><td></td><td></td><td></td><td></td><td></td><td></td></tr>
</table>

<table>
<tr><td>Oligo rev</td><td>AAACTCAGAGGCTGCCCTCGGAAT</td><td></td><td></td><td></td><td></td><td></td><td></td><td></td><td></td><td></td><td></td></tr>
<tr><td>Chromosome</td><td>start</td><td>end</td><td>strand</td><td>MM</td><td>target_seq</td><td>PAM</td><td>alignment</td><td>distance</td><td>position</td><td>gene name</td><td>gene id</td></tr>
<tr><td>GU305914.1</td><td>199580</td><td>199602</td><td>+</td><td>0</td><td>ATTCCGAGGGCAGCCTCTGA</td><td>AGG</td><td>||||||||[||||||||||||]PAM</td><td>NA</td><td>Intergenic</td><td>NA</td><td>NA</td></tr>
<tr><td>T15215</td><td>CCGAGGGCAGCCTCTGAAGGTGG</td><td>1000</td><td></td><td></td><td></td><td></td><td></td><td></td><td></td><td></td><td></td></tr>
<tr><td>Oligo fwd</td><td>CACCCCGAGGGCAGCCTCTGAAGG</td><td></td><td></td><td></td><td></td><td></td><td></td><td></td><td></td><td></td><td></td></tr>
<tr><td>Oligo rev</td><td>AAACCCTTCAGAGGCTGCCCTCGG</td><td></td><td></td><td></td><td></td><td></td><td></td><td></td><td></td><td></td><td></td></tr>
<tr><td>Chromosome</td><td>start</td><td>end</td><td>strand</td><td>MM</td><td>target_seq</td><td>PAM</td><td>alignment</td><td>distance</td><td>position</td><td>gene name</td><td>gene id</td></tr>
<tr><td>GU305914.1</td><td>199583</td><td>199605</td><td>+</td><td>0</td><td>CCGAGGGCAGCCTCTGAAGG</td><td>TGG</td><td>||||||||[||||||||||||]PAM</td><td>NA</td><td>Intergenic</td><td>NA</td><td>NA</td></tr>
<tr><td>T15216</td><td>CCGCCACCTCGACGTCGCGACGG</td><td>1000</td><td></td><td></td><td></td><td></td><td></td><td></td><td></td><td></td><td></td></tr>
<tr><td>Oligo fwd</td><td>CACCCCGCCACCTCGACGTCGCGA</td><td></td><td></td><td></td><td></td><td></td><td></td><td></td><td></td><td></td><td></td></tr>
<tr><td>Oligo rev</td><td>AAACTCGCGACGTCGAGGTGGCGG</td><td></td><td></td><td></td><td></td><td></td><td></td><td></td><td></td><td></td><td></td></tr>
<tr><td>Chromosome</td><td>start</td><td>end</td><td>strand</td><td>MM</td><td>target_seq</td><td>PAM</td><td>alignment</td><td>distance</td><td>position</td><td>gene name</td><td>gene id</td></tr>
<tr><td>GU305914.1</td><td>199627</td><td>199649</td><td>+</td><td>0</td><td>CCGCCACCTCGACGTCGCGA</td><td>CGG</td><td>||||||||[||||||||||||]PAM</td><td>NA</td><td>Intergenic</td><td>NA</td><td>NA</td></tr>
<tr><td>T15217</td><td>CGCGTCGAACGTCTTCCTCATGG</td><td>1000</td><td></td><td></td><td></td><td></td><td></td><td></td><td></td><td></td><td></td></tr>
<tr><td>Oligo fwd</td><td>CACCCGCGTCGAACGTCTTCCTCA</td><td></td><td></td><td></td><td></td><td></td><td></td><td></td><td></td><td></td><td></td></tr>
<tr><td>Oligo rev</td><td>AAACTGAGGAAGACGTTCGACGCG</td><td></td><td></td><td></td><td></td><td></td><td></td><td></td><td></td><td></td><td></td></tr>
<tr><td>Chromosome</td><td>start</td><td>end</td><td>strand</td><td>MM</td><td>target_seq</td><td>PAM</td><td>alignment</td><td>distance</td><td>position</td><td>gene name</td><td>gene id</td></tr>
<tr><td>GU305914.1</td><td>199667</td><td>199689</td><td>+</td><td>0</td><td>CGCGTCGAACGTCTTCCTCA</td><td>TGG</td><td>||||||||[||||||||||||]PAM</td><td>NA</td><td>Intergenic</td><td>NA</td><td>NA</td></tr>
<tr><td>T15218</td><td>GGCGTCGTCCATCTCGTCCCAGG</td><td>1000</td><td></td><td></td><td></td><td></td><td></td><td></td><td></td><td></td><td></td></tr>
<tr><td>Oligo fwd</td><td>CACCGGCGTCGTCCATCTCGTCCC</td><td></td><td></td><td></td><td></td><td></td><td></td><td></td><td></td><td></td><td></td></tr>
<tr><td>Oligo rev</td><td>AAACGGGACGAGATGGACGACGCC</td><td></td><td></td><td></td><td></td><td></td><td></td><td></td><td></td><td></td><td></td></tr>
<tr><td>Chromosome</td><td>start</td><td>end</td><td>strand</td><td>MM</td><td>target_seq</td><td>PAM</td><td>alignment</td><td>distance</td><td>position</td><td>gene name</td><td>gene id</td></tr>
<tr><td>GU305914.1</td><td>199688</td><td>199710</td><td>+</td><td>0</td><td>GGCGTCGTCCATCTCGTCCC</td><td>AGG</td><td>||||||||[||||||||||||]PAM</td><td>NA</td><td>Intergenic</td><td>NA</td><td>NA</td></tr>
<tr><td>T15219</td><td>CTCTCATCAGCCACCCCAGCTGG</td><td>1000</td><td></td><td></td><td></td><td></td><td></td><td></td><td></td><td></td><td></td></tr>
<tr><td>Oligo fwd</td><td>CACCCTCTCATCAGCCACCCCAGC</td><td></td><td></td><td></td><td></td><td></td><td></td><td></td><td></td><td></td><td></td></tr>
<tr><td>Oligo rev</td><td>AAACGCTGGGGTGGCTGATGAGAG</td><td></td><td></td><td></td><td></td><td></td><td></td><td></td><td></td><td></td><td></td></tr>
<tr><td>Chromosome</td><td>start</td><td>end</td><td>strand</td><td>MM</td><td>target_seq</td><td>PAM</td><td>alignment</td><td>distance</td><td>position</td><td>gene name</td><td>gene id</td></tr>
<tr><td>GU305914.1</td><td>199738</td><td>199760</td><td>+</td><td>0</td><td>CTCTCATCAGCCACCCCAGC</td><td>TGG</td><td>||||||||[||||||||||||]PAM</td><td>NA</td><td>Intergenic</td><td>NA</td><td>NA</td></tr>
<tr><td>T15220</td><td>TCTCATCAGCCACCCCAGCTGGG</td><td>1000</td><td></td><td></td><td></td><td></td><td></td><td></td><td></td><td></td><td></td></tr>
<tr><td>Oligo fwd</td><td>CACCTCTCATCAGCCACCCCAGCT</td><td></td><td></td><td></td><td></td><td></td><td></td><td></td><td></td><td></td><td></td></tr>
<tr><td>Oligo rev</td><td>AAACAGCTGGGGTGGCTGATGAGA</td><td></td><td></td><td></td><td></td><td></td><td></td><td></td><td></td><td></td><td></td></tr>
<tr><td>Chromosome</td><td>start</td><td>end</td><td>strand</td><td>MM</td><td>target_seq</td><td>PAM</td><td>alignment</td><td>distance</td><td>position</td><td>gene name</td><td>gene id</td></tr>
<tr><td>GU305914.1</td><td>199739</td><td>199761</td><td>+</td><td>0</td><td>TCTCATCAGCCACCCCAGCT</td><td>GGG</td><td>||||||||[||||||||||||]PAM</td><td>NA</td><td>Intergenic</td><td>NA</td><td>NA</td></tr>
<tr><td>T15221</td><td>CCAGCTGGGACGCGAAGTCGCGG</td><td>1000</td><td></td><td></td><td></td><td></td><td></td><td></td><td></td><td></td><td></td></tr>
<tr><td>Oligo fwd</td><td>CACCCCAGCTGGGACGCGAAGTCG</td><td></td><td></td><td></td><td></td><td></td><td></td><td></td><td></td><td></td><td></td></tr>
<tr><td>Oligo rev</td><td>AAACCGACTTCGCGTCCCAGCTGG</td><td></td><td></td><td></td><td></td><td></td><td></td><td></td><td></td><td></td><td></td></tr>
<tr><td>Chromosome</td><td>start</td><td>end</td><td>strand</td><td>MM</td><td>target_seq</td><td>PAM</td><td>alignment</td><td>distance</td><td>position</td><td>gene name</td><td>gene id</td></tr>
<tr><td>GU305914.1</td><td>199753</td><td>199775</td><td>+</td><td>0</td><td>CCAGCTGGGACGCGAAGTCG</td><td>CGG</td><td>||||||||[||||||||||||]PAM</td><td>NA</td><td>Intergenic</td><td>NA</td><td>NA</td></tr>
<tr><td>T15222</td><td>ACGCGAAGTCGCGGACGATGCGG</td><td>1000</td><td></td><td></td><td></td><td></td><td></td><td></td><td></td><td></td><td></td></tr>
<tr><td>Oligo fwd</td><td>CACCACGCGAAGTCGCGGACGATG</td><td></td><td></td><td></td><td></td><td></td><td></td><td></td><td></td><td></td><td></td></tr>
<tr><td>Oligo rev</td><td>AAACCATCGTCCGCGACTTCGCGT</td><td></td><td></td><td></td><td></td><td></td><td></td><td></td><td></td><td></td><td></td></tr>
<tr><td>Chromosome</td><td>start</td><td>end</td><td>strand</td><td>MM</td><td>target_seq</td><td>PAM</td><td>alignment</td><td>distance</td><td>position</td><td>gene name</td><td>gene id</td></tr>
<tr><td>GU305914.1</td><td>199762</td><td>199784</td><td>+</td><td>0</td><td>ACGCGAAGTCGCGGACGATG</td><td>CGG</td><td>||||||||[||||||||||||]PAM</td><td>NA</td><td>Intergenic</td><td>NA</td><td>NA</td></tr>
<tr><td>T15223</td><td>CACGTACGTCTCTTCCCACGCGG</td><td>1000</td><td></td><td></td><td></td><td></td><td></td><td></td><td></td><td></td><td></td></tr>
<tr><td>Oligo fwd</td><td>CACCCACGTACGTCTCTTCCCACG</td><td></td><td></td><td></td><td></td><td></td><td></td><td></td><td></td><td></td><td></td></tr>
<tr><td>Oligo rev</td><td>AAACCGTGGGAAGAGACGTACGTG</td><td></td><td></td><td></td><td></td><td></td><td></td><td></td><td></td><td></td><td></td></tr>
<tr><td>Chromosome</td><td>start</td><td>end</td><td>strand</td><td>MM</td><td>target_seq</td><td>PAM</td><td>alignment</td><td>distance</td><td>position</td><td>gene name</td><td>gene id</td></tr>
<tr><td>GU305914.1</td><td>199796</td><td>199818</td><td>+</td><td>0</td><td>CACGTACGTCTCTTCCCACG</td><td>CGG</td><td>||||||||[||||||||||||]PAM</td><td>NA</td><td>Intergenic</td><td>NA</td><td>NA</td></tr>
<tr><td>T15224</td><td>GTACGTCTCTTCCCACGCGGCGG</td><td>1000</td><td></td><td></td><td></td><td></td><td></td><td></td><td></td><td></td><td></td></tr>
<tr><td>Oligo fwd</td><td>CACCGTACGTCTCTTCCCACGCGG</td><td></td><td></td><td></td><td></td><td></td><td></td><td></td><td></td><td></td><td></td></tr>
<tr><td>Oligo rev</td><td>AAACCCGCGTGGGAAGAGACGTAC</td><td></td><td></td><td></td><td></td><td></td><td></td><td></td><td></td><td></td><td></td></tr>
<tr><td>Chromosome</td><td>start</td><td>end</td><td>strand</td><td>MM</td><td>target_seq</td><td>PAM</td><td>alignment</td><td>distance</td><td>position</td><td>gene name</td><td>gene id</td></tr>
<tr><td>GU305914.1</td><td>199799</td><td>199821</td><td>+</td><td>0</td><td>GTACGTCTCTTCCCACGCGG</td><td>CGG</td><td>||||||||[||||||||||||]PAM</td><td>NA</td><td>Intergenic</td><td>NA</td><td>NA</td></tr>
<tr><td>T15225</td><td>GGCCGCCACGTGCGCGCAGCCGG</td><td>1000</td><td></td><td></td><td></td><td></td><td></td><td></td><td></td><td></td><td></td></tr>
<tr><td>Oligo fwd</td><td>CACCGGCCGCCACGTGCGCGCAGC</td><td></td><td></td><td></td><td></td><td></td><td></td><td></td><td></td><td></td><td></td></tr>
<tr><td>Oligo rev</td><td>AAACGCTGCGCGCACGTGGCGGCC</td><td></td><td></td><td></td><td></td><td></td><td></td><td></td><td></td><td></td><td></td></tr>
<tr><td>Chromosome</td><td>start</td><td>end</td><td>strand</td><td>MM</td><td>target_seq</td><td>PAM</td><td>alignment</td><td>distance</td><td>position</td><td>gene name</td><td>gene id</td></tr>
<tr><td>GU305914.1</td><td>199820</td><td>199842</td><td>+</td><td>0</td><td>GGCCGCCACGTGCGCGCAGC</td><td>CGG</td><td>||||||||[||||||||||||]PAM</td><td>NA</td><td>Intergenic</td><td>NA</td><td>NA</td></tr>
<tr><td>T15226</td><td>GCCGCCACGTGCGCGCAGCCGGG</td><td>1000</td><td></td><td></td><td></td><td></td><td></td><td></td><td></td><td></td><td></td></tr>
<tr><td>Oligo fwd</td><td>CACCGCCGCCACGTGCGCGCAGCC</td><td></td><td></td><td></td><td></td><td></td><td></td><td></td><td></td><td></td><td></td></tr>
<tr><td>Oligo rev</td><td>AAACGGCTGCGCGCACGTGGCGGC</td><td></td><td></td><td></td><td></td><td></td><td></td><td></td><td></td><td></td><td></td></tr>
<tr><td>Chromosome</td><td>start</td><td>end</td><td>strand</td><td>MM</td><td>target_seq</td><td>PAM</td><td>alignment</td><td>distance</td><td>position</td><td>gene name</td><td>gene id</td></tr>
<tr><td>GU305914.1</td><td>199821</td><td>199843</td><td>+</td><td>0</td><td>GCCGCCACGTGCGCGCAGCC</td><td>GGG</td><td>||||||||[||||||||||||]PAM</td><td>NA</td><td>Intergenic</td><td>NA</td><td>NA</td></tr>
<tr><td>T15227</td><td>CGCAGCCGGGCGCCTCCAGCCGG</td><td>1000</td><td></td><td></td><td></td><td></td><td></td><td></td><td></td><td></td><td></td></tr>
<tr><td>Oligo fwd</td><td>CACCCGCAGCCGGGCGCCTCCAGC</td><td></td><td></td><td></td><td></td><td></td><td></td><td></td><td></td><td></td><td></td></tr>
<tr><td>Oligo rev</td><td>AAACGCTGGAGGCGCCCGGCTGCG</td><td></td><td></td><td></td><td></td><td></td><td></td><td></td><td></td><td></td><td></td></tr>
<tr><td>Chromosome</td><td>start</td><td>end</td><td>strand</td><td>MM</td><td>target_seq</td><td>PAM</td><td>alignment</td><td>distance</td><td>position</td><td>gene name</td><td>gene id</td></tr>
<tr><td>GU305914.1</td><td>199834</td><td>199856</td><td>+</td><td>0</td><td>CGCAGCCGGGCGCCTCCAGC</td><td>CGG</td><td>||||||||[||||||||||||]PAM</td><td>NA</td><td>Intergenic</td><td>NA</td><td>NA</td></tr>
<tr><td>T15228</td><td>GCAGCCGGGCGCCTCCAGCCGGG</td><td>1000</td><td></td><td></td><td></td><td></td><td></td><td></td><td></td><td></td><td></td></tr>
<tr><td>Oligo fwd</td><td>CACCGCAGCCGGGCGCCTCCAGCC</td><td></td><td></td><td></td><td></td><td></td><td></td><td></td><td></td><td></td><td></td></tr>
<tr><td>Oligo rev</td><td>AAACGGCTGGAGGCGCCCGGCTGC</td><td></td><td></td><td></td><td></td><td></td><td></td><td></td><td></td><td></td><td></td></tr>
<tr><td>Chromosome</td><td>start</td><td>end</td><td>strand</td><td>MM</td><td>target_seq</td><td>PAM</td><td>alignment</td><td>distance</td><td>position</td><td>gene name</td><td>gene id</td></tr>
<tr><td>GU305914.1</td><td>199835</td><td>199857</td><td>+</td><td>0</td><td>GCAGCCGGGCGCCTCCAGCC</td><td>GGG</td><td>||||||||[||||||||||||]PAM</td><td>NA</td><td>Intergenic</td><td>NA</td><td>NA</td></tr>
<tr><td>T15229</td><td>GGGCGCGCCCCATCGTGCCGCGG</td><td>1000</td><td></td><td></td><td></td><td></td><td></td><td></td><td></td><td></td><td></td></tr>
</table>

<table>
<tr><td>Oligo fwd</td><td>CACCGGGCGCGCCCCATCGTGCCG</td><td></td><td></td><td></td><td></td><td></td><td></td><td></td><td></td><td></td><td></td></tr>
<tr><td>Oligo rev</td><td>AAACCGGCACGATGGGGCGCGCCC</td><td></td><td></td><td></td><td></td><td></td><td></td><td></td><td></td><td></td><td></td></tr>
<tr><td>Chromosome</td><td>start</td><td>end</td><td>strand</td><td>MM</td><td>target_seq</td><td>PAM</td><td>alignment</td><td>distance</td><td>position</td><td>gene name</td><td>gene id</td></tr>
<tr><td>GU305914.1</td><td>199855</td><td>199877</td><td>+</td><td>0</td><td>GGGCGCGCCCCATCGTGCCG</td><td>CGG</td><td>||||||||[||||||||||||]PAM</td><td>NA</td><td>Intergenic</td><td>NA</td><td>NA</td></tr>
<tr><td>T15230</td><td>CCGCGGTCCAGACGCCGCTTCGG</td><td>1000</td><td></td><td></td><td></td><td></td><td></td><td></td><td></td><td></td><td></td></tr>
<tr><td>Oligo fwd</td><td>CACCCCGCGGTCCAGACGCCGCTT</td><td></td><td></td><td></td><td></td><td></td><td></td><td></td><td></td><td></td><td></td></tr>
<tr><td>Oligo rev</td><td>AAACAAGCGGCGTCTGGACCGCGG</td><td></td><td></td><td></td><td></td><td></td><td></td><td></td><td></td><td></td><td></td></tr>
<tr><td>Chromosome</td><td>start</td><td>end</td><td>strand</td><td>MM</td><td>target_seq</td><td>PAM</td><td>alignment</td><td>distance</td><td>position</td><td>gene name</td><td>gene id</td></tr>
<tr><td>GU305914.1</td><td>199872</td><td>199894</td><td>+</td><td>0</td><td>CCGCGGTCCAGACGCCGCTT</td><td>CGG</td><td>||||||||[||||||||||||]PAM</td><td>NA</td><td>Intergenic</td><td>NA</td><td>NA</td></tr>
<tr><td>T15231</td><td>GGTCCAGACGCCGCTTCGGCAGG</td><td>1000</td><td></td><td></td><td></td><td></td><td></td><td></td><td></td><td></td><td></td></tr>
<tr><td>Oligo fwd</td><td>CACCGGTCCAGACGCCGCTTCGGC</td><td></td><td></td><td></td><td></td><td></td><td></td><td></td><td></td><td></td><td></td></tr>
<tr><td>Oligo rev</td><td>AAACGCCGAAGCGGCGTCTGGACC</td><td></td><td></td><td></td><td></td><td></td><td></td><td></td><td></td><td></td><td></td></tr>
<tr><td>Chromosome</td><td>start</td><td>end</td><td>strand</td><td>MM</td><td>target_seq</td><td>PAM</td><td>alignment</td><td>distance</td><td>position</td><td>gene name</td><td>gene id</td></tr>
<tr><td>GU305914.1</td><td>199876</td><td>199898</td><td>+</td><td>0</td><td>GGTCCAGACGCCGCTTCGGC</td><td>AGG</td><td>||||||||[||||||||||||]PAM</td><td>NA</td><td>Intergenic</td><td>NA</td><td>NA</td></tr>
<tr><td>T15232</td><td>CCGCTTCGGCAGGTACAGCTTGG</td><td>1000</td><td></td><td></td><td></td><td></td><td></td><td></td><td></td><td></td><td></td></tr>
<tr><td>Oligo fwd</td><td>CACCCCGCTTCGGCAGGTACAGCT</td><td></td><td></td><td></td><td></td><td></td><td></td><td></td><td></td><td></td><td></td></tr>
<tr><td>Oligo rev</td><td>AAACAGCTGTACCTGCCGAAGCGG</td><td></td><td></td><td></td><td></td><td></td><td></td><td></td><td></td><td></td><td></td></tr>
<tr><td>Chromosome</td><td>start</td><td>end</td><td>strand</td><td>MM</td><td>target_seq</td><td>PAM</td><td>alignment</td><td>distance</td><td>position</td><td>gene name</td><td>gene id</td></tr>
<tr><td>GU305914.1</td><td>199886</td><td>199908</td><td>+</td><td>0</td><td>CCGCTTCGGCAGGTACAGCT</td><td>TGG</td><td>||||||||[||||||||||||]PAM</td><td>NA</td><td>Intergenic</td><td>NA</td><td>NA</td></tr>
<tr><td>T15233</td><td>CACGTTGTCCGCGATCCGCCAGG</td><td>1000</td><td></td><td></td><td></td><td></td><td></td><td></td><td></td><td></td><td></td></tr>
<tr><td>Oligo fwd</td><td>CACCCACGTTGTCCGCGATCCGCC</td><td></td><td></td><td></td><td></td><td></td><td></td><td></td><td></td><td></td><td></td></tr>
<tr><td>Oligo rev</td><td>AAACGGCGGATCGCGGACAACGTG</td><td></td><td></td><td></td><td></td><td></td><td></td><td></td><td></td><td></td><td></td></tr>
<tr><td>Chromosome</td><td>start</td><td>end</td><td>strand</td><td>MM</td><td>target_seq</td><td>PAM</td><td>alignment</td><td>distance</td><td>position</td><td>gene name</td><td>gene id</td></tr>
<tr><td>GU305914.1</td><td>199934</td><td>199956</td><td>+</td><td>0</td><td>CACGTTGTCCGCGATCCGCC</td><td>AGG</td><td>||||||||[||||||||||||]PAM</td><td>NA</td><td>Intergenic</td><td>NA</td><td>NA</td></tr>
<tr><td>T15234</td><td>CGCCAGGTCACGTCCATGTCCGG</td><td>1000</td><td></td><td></td><td></td><td></td><td></td><td></td><td></td><td></td><td></td></tr>
<tr><td>Oligo fwd</td><td>CACCCGCCAGGTCACGTCCATGTC</td><td></td><td></td><td></td><td></td><td></td><td></td><td></td><td></td><td></td><td></td></tr>
<tr><td>Oligo rev</td><td>AAACGACATGGACGTGACCTGGCG</td><td></td><td></td><td></td><td></td><td></td><td></td><td></td><td></td><td></td><td></td></tr>
<tr><td>Chromosome</td><td>start</td><td>end</td><td>strand</td><td>MM</td><td>target_seq</td><td>PAM</td><td>alignment</td><td>distance</td><td>position</td><td>gene name</td><td>gene id</td></tr>
<tr><td>GU305914.1</td><td>199950</td><td>199972</td><td>+</td><td>0</td><td>CGCCAGGTCACGTCCATGTC</td><td>CGG</td><td>||||||||[||||||||||||]PAM</td><td>NA</td><td>Intergenic</td><td>NA</td><td>NA</td></tr>
<tr><td>T15235</td><td>GCCAGGTCACGTCCATGTCCGGG</td><td>1000</td><td></td><td></td><td></td><td></td><td></td><td></td><td></td><td></td><td></td></tr>
<tr><td>Oligo fwd</td><td>CACCGCCAGGTCACGTCCATGTCC</td><td></td><td></td><td></td><td></td><td></td><td></td><td></td><td></td><td></td><td></td></tr>
<tr><td>Oligo rev</td><td>AAACGGACATGGACGTGACCTGGC</td><td></td><td></td><td></td><td></td><td></td><td></td><td></td><td></td><td></td><td></td></tr>
<tr><td>Chromosome</td><td>start</td><td>end</td><td>strand</td><td>MM</td><td>target_seq</td><td>PAM</td><td>alignment</td><td>distance</td><td>position</td><td>gene name</td><td>gene id</td></tr>
<tr><td>GU305914.1</td><td>199951</td><td>199973</td><td>+</td><td>0</td><td>GCCAGGTCACGTCCATGTCC</td><td>GGG</td><td>||||||||[||||||||||||]PAM</td><td>NA</td><td>Intergenic</td><td>NA</td><td>NA</td></tr>
<tr><td>T15236</td><td>CGTCCATGTCCGGGTCCGTGTGG</td><td>1000</td><td></td><td></td><td></td><td></td><td></td><td></td><td></td><td></td><td></td></tr>
<tr><td>Oligo fwd</td><td>CACCCGTCCATGTCCGGGTCCGTG</td><td></td><td></td><td></td><td></td><td></td><td></td><td></td><td></td><td></td><td></td></tr>
<tr><td>Oligo rev</td><td>AAACCACGGACCCGGACATGGACG</td><td></td><td></td><td></td><td></td><td></td><td></td><td></td><td></td><td></td><td></td></tr>
<tr><td>Chromosome</td><td>start</td><td>end</td><td>strand</td><td>MM</td><td>target_seq</td><td>PAM</td><td>alignment</td><td>distance</td><td>position</td><td>gene name</td><td>gene id</td></tr>
<tr><td>GU305914.1</td><td>199960</td><td>199982</td><td>+</td><td>0</td><td>CGTCCATGTCCGGGTCCGTG</td><td>TGG</td><td>||||||||[||||||||||||]PAM</td><td>NA</td><td>Intergenic</td><td>NA</td><td>NA</td></tr>
<tr><td>T15237</td><td>AGCACCGACTCGACGCGAAAAGG</td><td>1000</td><td></td><td></td><td></td><td></td><td></td><td></td><td></td><td></td><td></td></tr>
<tr><td>Oligo fwd</td><td>CACCAGCACCGACTCGACGCGAAA</td><td></td><td></td><td></td><td></td><td></td><td></td><td></td><td></td><td></td><td></td></tr>
<tr><td>Oligo rev</td><td>AAACTTTCGCGTCGAGTCGGTGCT</td><td></td><td></td><td></td><td></td><td></td><td></td><td></td><td></td><td></td><td></td></tr>
<tr><td>Chromosome</td><td>start</td><td>end</td><td>strand</td><td>MM</td><td>target_seq</td><td>PAM</td><td>alignment</td><td>distance</td><td>position</td><td>gene name</td><td>gene id</td></tr>
<tr><td>GU305914.1</td><td>200010</td><td>200032</td><td>+</td><td>0</td><td>AGCACCGACTCGACGCGAAA</td><td>AGG</td><td>||||||||[||||||||||||]PAM</td><td>NA</td><td>Intergenic</td><td>NA</td><td>NA</td></tr>
<tr><td>T15238</td><td>AAGGATCCTGCCCGACCCGCAGG</td><td>1000</td><td></td><td></td><td></td><td></td><td></td><td></td><td></td><td></td><td></td></tr>
<tr><td>Oligo fwd</td><td>CACCAAGGATCCTGCCCGACCCGC</td><td></td><td></td><td></td><td></td><td></td><td></td><td></td><td></td><td></td><td></td></tr>
<tr><td>Oligo rev</td><td>AAACGCGGGTCGGGCAGGATCCTT</td><td></td><td></td><td></td><td></td><td></td><td></td><td></td><td></td><td></td><td></td></tr>
<tr><td>Chromosome</td><td>start</td><td>end</td><td>strand</td><td>MM</td><td>target_seq</td><td>PAM</td><td>alignment</td><td>distance</td><td>position</td><td>gene name</td><td>gene id</td></tr>
<tr><td>GU305914.1</td><td>200029</td><td>200051</td><td>+</td><td>0</td><td>AAGGATCCTGCCCGACCCGC</td><td>AGG</td><td>||||||||[||||||||||||]PAM</td><td>NA</td><td>Intergenic</td><td>NA</td><td>NA</td></tr>
<tr><td>T15239</td><td>CGACCCCGAAGTGCTTCCAGTGG</td><td>1000</td><td></td><td></td><td></td><td></td><td></td><td></td><td></td><td></td><td></td></tr>
<tr><td>Oligo fwd</td><td>CACCCGACCCCGAAGTGCTTCCAG</td><td></td><td></td><td></td><td></td><td></td><td></td><td></td><td></td><td></td><td></td></tr>
<tr><td>Oligo rev</td><td>AAACCTGGAAGCACTTCGGGGTCG</td><td></td><td></td><td></td><td></td><td></td><td></td><td></td><td></td><td></td><td></td></tr>
<tr><td>Chromosome</td><td>start</td><td>end</td><td>strand</td><td>MM</td><td>target_seq</td><td>PAM</td><td>alignment</td><td>distance</td><td>position</td><td>gene name</td><td>gene id</td></tr>
<tr><td>GU305914.1</td><td>200056</td><td>200078</td><td>+</td><td>0</td><td>CGACCCCGAAGTGCTTCCAG</td><td>TGG</td><td>||||||||[||||||||||||]PAM</td><td>NA</td><td>Intergenic</td><td>NA</td><td>NA</td></tr>
<tr><td>T15240</td><td>GCTTCCAGTGGCTCGTCTTGAGG</td><td>1000</td><td></td><td></td><td></td><td></td><td></td><td></td><td></td><td></td><td></td></tr>
<tr><td>Oligo fwd</td><td>CACCGCTTCCAGTGGCTCGTCTTG</td><td></td><td></td><td></td><td></td><td></td><td></td><td></td><td></td><td></td><td></td></tr>
<tr><td>Oligo rev</td><td>AAACCAAGACGAGCCACTGGAAGC</td><td></td><td></td><td></td><td></td><td></td><td></td><td></td><td></td><td></td><td></td></tr>
<tr><td>Chromosome</td><td>start</td><td>end</td><td>strand</td><td>MM</td><td>target_seq</td><td>PAM</td><td>alignment</td><td>distance</td><td>position</td><td>gene name</td><td>gene id</td></tr>
<tr><td>GU305914.1</td><td>200068</td><td>200090</td><td>+</td><td>0</td><td>GCTTCCAGTGGCTCGTCTTG</td><td>AGG</td><td>||||||||[||||||||||||]PAM</td><td>NA</td><td>Intergenic</td><td>NA</td><td>NA</td></tr>
<tr><td>T15241</td><td>AGTGGCTCGTCTTGAGGTTTCGG</td><td>1000</td><td></td><td></td><td></td><td></td><td></td><td></td><td></td><td></td><td></td></tr>
<tr><td>Oligo fwd</td><td>CACCAGTGGCTCGTCTTGAGGTTT</td><td></td><td></td><td></td><td></td><td></td><td></td><td></td><td></td><td></td><td></td></tr>
<tr><td>Oligo rev</td><td>AAACAAACCTCAAGACGAGCCACT</td><td></td><td></td><td></td><td></td><td></td><td></td><td></td><td></td><td></td><td></td></tr>
<tr><td>Chromosome</td><td>start</td><td>end</td><td>strand</td><td>MM</td><td>target_seq</td><td>PAM</td><td>alignment</td><td>distance</td><td>position</td><td>gene name</td><td>gene id</td></tr>
<tr><td>GU305914.1</td><td>200074</td><td>200096</td><td>+</td><td>0</td><td>AGTGGCTCGTCTTGAGGTTT</td><td>CGG</td><td>||||||||[||||||||||||]PAM</td><td>NA</td><td>Intergenic</td><td>NA</td><td>NA</td></tr>
<tr><td>T15242</td><td>GTGGCTCGTCTTGAGGTTTCGGG</td><td>1000</td><td></td><td></td><td></td><td></td><td></td><td></td><td></td><td></td><td></td></tr>
<tr><td>Oligo fwd</td><td>CACCGTGGCTCGTCTTGAGGTTTC</td><td></td><td></td><td></td><td></td><td></td><td></td><td></td><td></td><td></td><td></td></tr>
<tr><td>Oligo rev</td><td>AAACGAAACCTCAAGACGAGCCAC</td><td></td><td></td><td></td><td></td><td></td><td></td><td></td><td></td><td></td><td></td></tr>
<tr><td>Chromosome</td><td>start</td><td>end</td><td>strand</td><td>MM</td><td>target_seq</td><td>PAM</td><td>alignment</td><td>distance</td><td>position</td><td>gene name</td><td>gene id</td></tr>
<tr><td>GU305914.1</td><td>200075</td><td>200097</td><td>+</td><td>0</td><td>GTGGCTCGTCTTGAGGTTTC</td><td>GGG</td><td>||||||||[||||||||||||]PAM</td><td>NA</td><td>Intergenic</td><td>NA</td><td>NA</td></tr>
<tr><td>T15243</td><td>TCTTGAGGTTTCGGGTGACGCGG</td><td>1000</td><td></td><td></td><td></td><td></td><td></td><td></td><td></td><td></td><td></td></tr>
<tr><td>Oligo fwd</td><td>CACCTCTTGAGGTTTCGGGTGACG</td><td></td><td></td><td></td><td></td><td></td><td></td><td></td><td></td><td></td><td></td></tr>
<tr><td>Oligo rev</td><td>AAACCGTCACCCGAAACCTCAAGA</td><td></td><td></td><td></td><td></td><td></td><td></td><td></td><td></td><td></td><td></td></tr>
<tr><td>Chromosome</td><td>start</td><td>end</td><td>strand</td><td>MM</td><td>target_seq</td><td>PAM</td><td>alignment</td><td>distance</td><td>position</td><td>gene name</td><td>gene id</td></tr>
<tr><td>GU305914.1</td><td>200083</td><td>200105</td><td>+</td><td>0</td><td>TCTTGAGGTTTCGGGTGACG</td><td>CGG</td><td>||||||||[||||||||||||]PAM</td><td>NA</td><td>Intergenic</td><td>NA</td><td>NA</td></tr>
</table>

| | | | | | | | | | | | |
|---|---|---|---|---|---|---|---|---|---|---|---|
| T15244 | GAGGTTTCGGGTGACGCGGTCGG | 1000 | | | | | | | | | |
| Oligo fwd | CACCGAGGTTTCGGGTGACGCGGT | | | | | | | | | | |
| Oligo rev | AAACACCGCGTCACCCGAAACCTC | | | | | | | | | | |
| Chromosome | start | end | strand | MM | target_seq | PAM | alignment | distance | position | gene name | gene id |
| GU305914.1 | 200087 | 200109 | + | 0 | GAGGTTTCGGGTGACGCGGT | CGG | \|\|\|\|\|\|\|\|[\|\|\|\|\|\|\|\|\|\|\|\|]PAM | NA | Intergenic | NA | NA |
| T15245 | GGTGACGCGGTCGGTGAACTTGG | 1000 | | | | | | | | | |
| Oligo fwd | CACCGGTGACGCGGTCGGTGAACT | | | | | | | | | | |
| Oligo rev | AAACAGTTCACCGACCGCGTCACC | | | | | | | | | | |
| Chromosome | start | end | strand | MM | target_seq | PAM | alignment | distance | position | gene name | gene id |
| GU305914.1 | 200096 | 200118 | + | 0 | GGTGACGCGGTCGGTGAACT | TGG | \|\|\|\|\|\|\|\|[\|\|\|\|\|\|\|\|\|\|\|\|]PAM | NA | Intergenic | NA | NA |
| T15246 | GAACTTGGCGAACTCTCCGTCGG | 1000 | | | | | | | | | |
| Oligo fwd | CACCGAACTTGGCGAACTCTCCGT | | | | | | | | | | |
| Oligo rev | AAACACGGAGAGTTCGCCAAGTTC | | | | | | | | | | |
| Chromosome | start | end | strand | MM | target_seq | PAM | alignment | distance | position | gene name | gene id |
| GU305914.1 | 200111 | 200133 | + | 0 | GAACTTGGCGAACTCTCCGT | CGG | \|\|\|\|\|\|\|\|[\|\|\|\|\|\|\|\|\|\|\|\|]PAM | NA | Intergenic | NA | NA |
| T15247 | CCGTCGGACATCCTGCAGAACGG | 1000 | | | | | | | | | |
| Oligo fwd | CACCCCGTCGGACATCCTGCAGAA | | | | | | | | | | |
| Oligo rev | AAACTTCTGCAGGATGTCCGACGG | | | | | | | | | | |
| Chromosome | start | end | strand | MM | target_seq | PAM | alignment | distance | position | gene name | gene id |
| GU305914.1 | 200127 | 200149 | + | 0 | CCGTCGGACATCCTGCAGAA | CGG | \|\|\|\|\|\|\|\|[\|\|\|\|\|\|\|\|\|\|\|\|]PAM | NA | Intergenic | NA | NA |
| T15248 | AGAACGGCCACCACCGCCGCAGG | 1000 | | | | | | | | | |
| Oligo fwd | CACCAGAACGGCCACCACCGCCGC | | | | | | | | | | |
| Oligo rev | AAACGCGGCGGTGGTGGCCGTTCT | | | | | | | | | | |
| Chromosome | start | end | strand | MM | target_seq | PAM | alignment | distance | position | gene name | gene id |
| GU305914.1 | 200143 | 200165 | + | 0 | AGAACGGCCACCACCGCCGC | AGG | \|\|\|\|\|\|\|\|[\|\|\|\|\|\|\|\|\|\|\|\|]PAM | NA | Intergenic | NA | NA |
| GU305914.1 | 36165 | 36187 | - | 4 | ACAACCACCACAACCGCCGC | CGG | \|-\|\|\|--\|[\|\|\|-\|\|\|\|\|\|\|\|]PAM | NA | Intergenic | NA | NA |
| T15249 | CCACCGCCGCAGGCACTCGTCGG | 1000 | | | | | | | | | |
| Oligo fwd | CACCCCACCGCCGCAGGCACTCGT | | | | | | | | | | |
| Oligo rev | AAACACGAGTGCCTGCGGCGGTGG | | | | | | | | | | |
| Chromosome | start | end | strand | MM | target_seq | PAM | alignment | distance | position | gene name | gene id |
| GU305914.1 | 200153 | 200175 | + | 0 | CCACCGCCGCAGGCACTCGT | CGG | \|\|\|\|\|\|\|\|[\|\|\|\|\|\|\|\|\|\|\|\|]PAM | NA | Intergenic | NA | NA |
| T15250 | TCGGTGCCGCACAGCACGAGCGG | 1000 | | | | | | | | | |
| Oligo fwd | CACCTCGGTGCCGCACAGCACGAG | | | | | | | | | | |
| Oligo rev | AAACCTCGTGCTGTGCGGCACCGA | | | | | | | | | | |
| Chromosome | start | end | strand | MM | target_seq | PAM | alignment | distance | position | gene name | gene id |
| GU305914.1 | 200172 | 200194 | + | 0 | TCGGTGCCGCACAGCACGAG | CGG | \|\|\|\|\|\|\|\|[\|\|\|\|\|\|\|\|\|\|\|\|]PAM | NA | Intergenic | NA | NA |
| T15251 | TGCCGCACAGCACGAGCGGCCGG | 1000 | | | | | | | | | |
| Oligo fwd | CACCTGCCGCACAGCACGAGCGGC | | | | | | | | | | |
| Oligo rev | AAACGCCGCTCGTGCTGTGCGGCA | | | | | | | | | | |
| Chromosome | start | end | strand | MM | target_seq | PAM | alignment | distance | position | gene name | gene id |
| GU305914.1 | 200176 | 200198 | + | 0 | TGCCGCACAGCACGAGCGGC | CGG | \|\|\|\|\|\|\|\|[\|\|\|\|\|\|\|\|\|\|\|\|]PAM | NA | Intergenic | NA | NA |
| T15252 | GAGCGGCCGGTCGCCCTCCCCGG | 1000 | | | | | | | | | |
| Oligo fwd | CACCGAGCGGCCGGTCGCCCTCCC | | | | | | | | | | |
| Oligo rev | AAACGGGAGGGCGACCGGCCGCTC | | | | | | | | | | |
| Chromosome | start | end | strand | MM | target_seq | PAM | alignment | distance | position | gene name | gene id |
| GU305914.1 | 200189 | 200211 | + | 0 | GAGCGGCCGGTCGCCCTCCC | CGG | \|\|\|\|\|\|\|\|[\|\|\|\|\|\|\|\|\|\|\|\|]PAM | NA | Intergenic | NA | NA |
| T15253 | AGCGGCCGGTCGCCCTCCCCGGG | 1000 | | | | | | | | | |
| Oligo fwd | CACCAGCGGCCGGTCGCCCTCCCC | | | | | | | | | | |
| Oligo rev | AAACGGGGAGGGCGACCGGCCGCT | | | | | | | | | | |
| Chromosome | start | end | strand | MM | target_seq | PAM | alignment | distance | position | gene name | gene id |
| GU305914.1 | 200190 | 200212 | + | 0 | AGCGGCCGGTCGCCCTCCCC | GGG | \|\|\|\|\|\|\|\|[\|\|\|\|\|\|\|\|\|\|\|\|]PAM | NA | Intergenic | NA | NA |
| T15254 | GCGGCCGGTCGCCCTCCCCGGGG | 1000 | | | | | | | | | |
| Oligo fwd | CACCGCGGCCGGTCGCCCTCCCCG | | | | | | | | | | |
| Oligo rev | AAACCGGGGAGGGCGACCGGCCGC | | | | | | | | | | |
| Chromosome | start | end | strand | MM | target_seq | PAM | alignment | distance | position | gene name | gene id |
| GU305914.1 | 200191 | 200213 | + | 0 | GCGGCCGGTCGCCCTCCCCG | GGG | \|\|\|\|\|\|\|\|[\|\|\|\|\|\|\|\|\|\|\|\|]PAM | NA | Intergenic | NA | NA |
| GU305914.1 | 200845 | 200867 | - | 4 | GCCGCCCGTCGCCCTCGCCA | TGG | \|\|-\|\|\|-\|[\|\|\|\|\|\|\|\|-\|\|-]PAM | NA | Intergenic | NA | NA |
| T15255 | CGGCCGGTCGCCCTCCCCGGGGG | 1000 | | | | | | | | | |
| Oligo fwd | CACCCGGCCGGTCGCCCTCCCCGG | | | | | | | | | | |
| Oligo rev | AAACCCGGGGAGGGCGACCGGCCG | | | | | | | | | | |
| Chromosome | start | end | strand | MM | target_seq | PAM | alignment | distance | position | gene name | gene id |
| GU305914.1 | 200192 | 200214 | + | 0 | CGGCCGGTCGCCCTCCCCGG | GGG | \|\|\|\|\|\|\|\|[\|\|\|\|\|\|\|\|\|\|\|\|]PAM | NA | Intergenic | NA | NA |
| T15256 | CCTCCCCGGGGGTGTGTAGCAGG | 1000 | | | | | | | | | |
| Oligo fwd | CACCCCTCCCCGGGGGTGTGTAGC | | | | | | | | | | |
| Oligo rev | AAACGCTACACACCCCCGGGGAGG | | | | | | | | | | |
| Chromosome | start | end | strand | MM | target_seq | PAM | alignment | distance | position | gene name | gene id |
| GU305914.1 | 200203 | 200225 | + | 0 | CCTCCCCGGGGGTGTGTAGC | AGG | \|\|\|\|\|\|\|\|[\|\|\|\|\|\|\|\|\|\|\|\|]PAM | NA | Intergenic | NA | NA |
| T15257 | CATCAGCGCGCGCTCCGCCACGG | 1000 | | | | | | | | | |
| Oligo fwd | CACCCATCAGCGCGCGCTCCGCCA | | | | | | | | | | |
| Oligo rev | AAACTGGCGGAGCGCGCGCTGATG | | | | | | | | | | |
| Chromosome | start | end | strand | MM | target_seq | PAM | alignment | distance | position | gene name | gene id |
| GU305914.1 | 200243 | 200265 | + | 0 | CATCAGCGCGCGCTCCGCCA | CGG | \|\|\|\|\|\|\|\|[\|\|\|\|\|\|\|\|\|\|\|\|]PAM | NA | Intergenic | NA | NA |
| T15258 | CAGCGCGCGCTCCGCCACGGCGG | 1000 | | | | | | | | | |
| Oligo fwd | CACCCAGCGCGCGCTCCGCCACGG | | | | | | | | | | |
| Oligo rev | AAACCCGTGGCGGAGCGCGCGCTG | | | | | | | | | | |

Chromosome start end strand MM target_seq PAM alignment distance position gene name gene id
GU305914.1 200246 200268 + 0 CAGCGCGCGCTCCGCCACGG CGG ||||||||[||||||||||||]PAM NA Intergenic NA NA

T15259 CGCCACGGCGGTCCCGTCTCTGG 1000
Oligo fwd CACCCGCCACGGCGGTCCCGTCTC
Oligo rev AAACGAGACGGGACCGCCGTGGCG
Chromosome start end strand MM target_seq PAM alignment distance position gene name gene id
GU305914.1 200258 200280 + 0 CGCCACGGCGGTCCCGTCTC TGG ||||||||[||||||||||||]PAM NA Intergenic NA NA

T15260 GCCACGGCGGTCCCGTCTCTGGG 1000
Oligo fwd CACCGCCACGGCGGTCCCGTCTCT
Oligo rev AAACAGAGACGGGACCGCCGTGGC
Chromosome start end strand MM target_seq PAM alignment distance position gene name gene id
GU305914.1 200259 200281 + 0 GCCACGGCGGTCCCGTCTCT GGG ||||||||[||||||||||||]PAM NA Intergenic NA NA

T15261 GAACTCCGCCACCAGCTTGACGG 1000
Oligo fwd CACCGAACTCCGCCACCAGCTTGA
Oligo rev AAACTCAAGCTGGTGGCGGAGTTC
Chromosome start end strand MM target_seq PAM alignment distance position gene name gene id
GU305914.1 200294 200316 + 0 GAACTCCGCCACCAGCTTGA CGG ||||||||[||||||||||||]PAM NA Intergenic NA NA

T15262 AACTCCGCCACCAGCTTGACGGG 1000
Oligo fwd CACCAACTCCGCCACCAGCTTGAC
Oligo rev AAACGTCAAGCTGGTGGCGGAGTT
Chromosome start end strand MM target_seq PAM alignment distance position gene name gene id
GU305914.1 200295 200317 + 0 AACTCCGCCACCAGCTTGAC GGG ||||||||[||||||||||||]PAM NA Intergenic NA NA

T15263 CACCAGCTTGACGGGCACCGCGG 1000
Oligo fwd CACCCACCAGCTTGACGGGCACCG
Oligo rev AAACCGGTGCCCGTCAAGCTGGTG
Chromosome start end strand MM target_seq PAM alignment distance position gene name gene id
GU305914.1 200303 200325 + 0 CACCAGCTTGACGGGCACCG CGG ||||||||[||||||||||||]PAM NA Intergenic NA NA

T15264 TGACGGGCACCGCGGTCGTCAGG 1000
Oligo fwd CACCTGACGGGCACCGCGGTCGTC
Oligo rev AAACGACGACCGCGGTGCCCGTCA
Chromosome start end strand MM target_seq PAM alignment distance position gene name gene id
GU305914.1 200311 200333 + 0 TGACGGGCACCGCGGTCGTC AGG ||||||||[||||||||||||]PAM NA Intergenic NA NA

T15265 GACGGGCACCGCGGTCGTCAGGG 1000
Oligo fwd CACCGACGGGCACCGCGGTCGTCA
Oligo rev AAACTGACGACCGCGGTGCCCGTC
Chromosome start end strand MM target_seq PAM alignment distance position gene name gene id
GU305914.1 200312 200334 + 0 GACGGGCACCGCGGTCGTCA GGG ||||||||[||||||||||||]PAM NA Intergenic NA NA

T15266 ACGGGCACCGCGGTCGTCAGGGG 1000
Oligo fwd CACCACGGGCACCGCGGTCGTCAG
Oligo rev AAACCTGACGACCGCGGTGCCCGT
Chromosome start end strand MM target_seq PAM alignment distance position gene name gene id
GU305914.1 200313 200335 + 0 ACGGGCACCGCGGTCGTCAG GGG ||||||||[||||||||||||]PAM NA Intergenic NA NA

T15267 GCACCGCGGTCGTCAGGGGCAGG 1000
Oligo fwd CACCGCACCGCGGTCGTCAGGGGC
Oligo rev AAACGCCCCTGACGACCGCGGTGC
Chromosome start end strand MM target_seq PAM alignment distance position gene name gene id
GU305914.1 200317 200339 + 0 GCACCGCGGTCGTCAGGGGC AGG ||||||||[||||||||||||]PAM NA Intergenic NA NA

T15268 GCAGGTGCCGTTCCGCGTACAGG 1000
Oligo fwd CACCGCAGGTGCCGTTCCGCGTAC
Oligo rev AAACGTACGCGGAACGGCACCTGC
Chromosome start end strand MM target_seq PAM alignment distance position gene name gene id
GU305914.1 200335 200357 + 0 GCAGGTGCCGTTCCGCGTAC AGG ||||||||[||||||||||||]PAM NA Intergenic NA NA

T15269 CAGGTGCCGTTCCGCGTACAGGG 1000
Oligo fwd CACCCAGGTGCCGTTCCGCGTACA
Oligo rev AAACTGTACGCGGAACGGCACCTG
Chromosome start end strand MM target_seq PAM alignment distance position gene name gene id
GU305914.1 200336 200358 + 0 CAGGTGCCGTTCCGCGTACA GGG ||||||||[||||||||||||]PAM NA Intergenic NA NA

T15270 AGGTGCCGTTCCGCGTACAGGGG 1000
Oligo fwd CACCAGGTGCCGTTCCGCGTACAG
Oligo rev AAACCTGTACGCGGAACGGCACCT
Chromosome start end strand MM target_seq PAM alignment distance position gene name gene id
GU305914.1 200337 200359 + 0 AGGTGCCGTTCCGCGTACAG GGG ||||||||[||||||||||||]PAM NA Intergenic NA NA

T15271 ACAGGGGTTCGAAGTTCACCAGG 1000
Oligo fwd CACCACAGGGGTTCGAAGTTCACC
Oligo rev AAACGGTGAACTTCGAACCCCTGT
Chromosome start end strand MM target_seq PAM alignment distance position gene name gene id
GU305914.1 200353 200375 + 0 ACAGGGGTTCGAAGTTCACC AGG ||||||||[||||||||||||]PAM NA Intergenic NA NA

T15272 GCTCAGACTCATGACGAAGACGG 1000
Oligo fwd CACCGCTCAGACTCATGACGAAGA
Oligo rev AAACTCTTCGTCATGAGTCTGAGC
Chromosome start end strand MM target_seq PAM alignment distance position gene name gene id
GU305914.1 200399 200421 + 0 GCTCAGACTCATGACGAAGA CGG ||||||||[||||||||||||]PAM NA Intergenic NA NA

T15273 ACTCATGACGAAGACGGCGTCGG 1000
Oligo fwd CACCACTCATGACGAAGACGGCGT

<table>
<tr><td>Oligo rev</td><td>AAACACGCCGTCTTCGTCATGAGT</td><td></td><td></td><td></td><td></td><td></td><td></td><td></td><td></td><td></td><td></td><td></td></tr>
<tr><td>Chromosome</td><td>start</td><td></td><td>end</td><td>strand</td><td>MM</td><td>target_seq</td><td>PAM</td><td>alignment</td><td>distance</td><td>position</td><td>gene name</td><td>gene id</td></tr>
<tr><td>GU305914.1</td><td></td><td>200405</td><td>200427</td><td>+</td><td>0</td><td>ACTCATGACGAAGACGGCGT</td><td>CGG</td><td>||||||||[||||||||||||]PAM</td><td>NA</td><td>Intergenic</td><td>NA</td><td>NA</td></tr>
<tr><td>T15274</td><td>AGACGGCGTCGGTCCGCTCCTGG</td><td></td><td>1000</td><td></td><td></td><td></td><td></td><td></td><td></td><td></td><td></td><td></td></tr>
<tr><td>Oligo fwd</td><td>CACCAGACGGCGTCGGTCCGCTCC</td><td></td><td></td><td></td><td></td><td></td><td></td><td></td><td></td><td></td><td></td><td></td></tr>
<tr><td>Oligo rev</td><td>AAACGGAGCGGACCGACGCCGTCT</td><td></td><td></td><td></td><td></td><td></td><td></td><td></td><td></td><td></td><td></td><td></td></tr>
<tr><td>Chromosome</td><td>start</td><td></td><td>end</td><td>strand</td><td>MM</td><td>target_seq</td><td>PAM</td><td>alignment</td><td>distance</td><td>position</td><td>gene name</td><td>gene id</td></tr>
<tr><td>GU305914.1</td><td></td><td>200416</td><td>200438</td><td>+</td><td>0</td><td>AGACGGCGTCGGTCCGCTCC</td><td>TGG</td><td>||||||||[||||||||||||]PAM</td><td>NA</td><td>Intergenic</td><td>NA</td><td>NA</td></tr>
<tr><td>T15275</td><td>GCTCCTGGTGACAGAGCAGCTGG</td><td></td><td>1000</td><td></td><td></td><td></td><td></td><td></td><td></td><td></td><td></td><td></td></tr>
<tr><td>Oligo fwd</td><td>CACCGCTCCTGGTGACAGAGCAGC</td><td></td><td></td><td></td><td></td><td></td><td></td><td></td><td></td><td></td><td></td><td></td></tr>
<tr><td>Oligo rev</td><td>AAACGCTGCTCTGTCACCAGGAGC</td><td></td><td></td><td></td><td></td><td></td><td></td><td></td><td></td><td></td><td></td><td></td></tr>
<tr><td>Chromosome</td><td>start</td><td></td><td>end</td><td>strand</td><td>MM</td><td>target_seq</td><td>PAM</td><td>alignment</td><td>distance</td><td>position</td><td>gene name</td><td>gene id</td></tr>
<tr><td>GU305914.1</td><td></td><td>200431</td><td>200453</td><td>+</td><td>0</td><td>GCTCCTGGTGACAGAGCAGC</td><td>TGG</td><td>||||||||[||||||||||||]PAM</td><td>NA</td><td>Intergenic</td><td>NA</td><td>NA</td></tr>
<tr><td>T15276</td><td>GGTGACAGAGCAGCTGGCCCTGG</td><td></td><td>1000</td><td></td><td></td><td></td><td></td><td></td><td></td><td></td><td></td><td></td></tr>
<tr><td>Oligo fwd</td><td>CACCGGTGACAGAGCAGCTGGCCC</td><td></td><td></td><td></td><td></td><td></td><td></td><td></td><td></td><td></td><td></td><td></td></tr>
<tr><td>Oligo rev</td><td>AAACGGGCCAGCTGCTCTGTCACC</td><td></td><td></td><td></td><td></td><td></td><td></td><td></td><td></td><td></td><td></td><td></td></tr>
<tr><td>Chromosome</td><td>start</td><td></td><td>end</td><td>strand</td><td>MM</td><td>target_seq</td><td>PAM</td><td>alignment</td><td>distance</td><td>position</td><td>gene name</td><td>gene id</td></tr>
<tr><td>GU305914.1</td><td></td><td>200437</td><td>200459</td><td>+</td><td>0</td><td>GGTGACAGAGCAGCTGGCCC</td><td>TGG</td><td>||||||||[||||||||||||]PAM</td><td>NA</td><td>Intergenic</td><td>NA</td><td>NA</td></tr>
<tr><td>T15277</td><td>GTGACAGAGCAGCTGGCCCTGGG</td><td></td><td>1000</td><td></td><td></td><td></td><td></td><td></td><td></td><td></td><td></td><td></td></tr>
<tr><td>Oligo fwd</td><td>CACCGTGACAGAGCAGCTGGCCCT</td><td></td><td></td><td></td><td></td><td></td><td></td><td></td><td></td><td></td><td></td><td></td></tr>
<tr><td>Oligo rev</td><td>AAACAGGGCCAGCTGCTCTGTCAC</td><td></td><td></td><td></td><td></td><td></td><td></td><td></td><td></td><td></td><td></td><td></td></tr>
<tr><td>Chromosome</td><td>start</td><td></td><td>end</td><td>strand</td><td>MM</td><td>target_seq</td><td>PAM</td><td>alignment</td><td>distance</td><td>position</td><td>gene name</td><td>gene id</td></tr>
<tr><td>GU305914.1</td><td></td><td>200438</td><td>200460</td><td>+</td><td>0</td><td>GTGACAGAGCAGCTGGCCCT</td><td>GGG</td><td>||||||||[||||||||||||]PAM</td><td>NA</td><td>Intergenic</td><td>NA</td><td>NA</td></tr>
<tr><td>T15278</td><td>TGACAGAGCAGCTGGCCCTGGGG</td><td></td><td>1000</td><td></td><td></td><td></td><td></td><td></td><td></td><td></td><td></td><td></td></tr>
<tr><td>Oligo fwd</td><td>CACCTGACAGAGCAGCTGGCCCTG</td><td></td><td></td><td></td><td></td><td></td><td></td><td></td><td></td><td></td><td></td><td></td></tr>
<tr><td>Oligo rev</td><td>AAACCAGGGCCAGCTGCTCTGTCA</td><td></td><td></td><td></td><td></td><td></td><td></td><td></td><td></td><td></td><td></td><td></td></tr>
<tr><td>Chromosome</td><td>start</td><td></td><td>end</td><td>strand</td><td>MM</td><td>target_seq</td><td>PAM</td><td>alignment</td><td>distance</td><td>position</td><td>gene name</td><td>gene id</td></tr>
<tr><td>GU305914.1</td><td></td><td>200439</td><td>200461</td><td>+</td><td>0</td><td>TGACAGAGCAGCTGGCCCTG</td><td>GGG</td><td>||||||||[||||||||||||]PAM</td><td>NA</td><td>Intergenic</td><td>NA</td><td>NA</td></tr>
<tr><td>T15279</td><td>GCAGCTGGCCCTGGGGACCCAGG</td><td></td><td>1000</td><td></td><td></td><td></td><td></td><td></td><td></td><td></td><td></td><td></td></tr>
<tr><td>Oligo fwd</td><td>CACCGCAGCTGGCCCTGGGGACCC</td><td></td><td></td><td></td><td></td><td></td><td></td><td></td><td></td><td></td><td></td><td></td></tr>
<tr><td>Oligo rev</td><td>AAACGGGTCCCCAGGGCCAGCTGC</td><td></td><td></td><td></td><td></td><td></td><td></td><td></td><td></td><td></td><td></td><td></td></tr>
<tr><td>Chromosome</td><td>start</td><td></td><td>end</td><td>strand</td><td>MM</td><td>target_seq</td><td>PAM</td><td>alignment</td><td>distance</td><td>position</td><td>gene name</td><td>gene id</td></tr>
<tr><td>GU305914.1</td><td></td><td>200446</td><td>200468</td><td>+</td><td>0</td><td>GCAGCTGGCCCTGGGGACCC</td><td>AGG</td><td>||||||||[||||||||||||]PAM</td><td>NA</td><td>Intergenic</td><td>NA</td><td>NA</td></tr>
<tr><td>T15280</td><td>CACGAACCATCTCAGCTCCTCGG</td><td></td><td>1000</td><td></td><td></td><td></td><td></td><td></td><td></td><td></td><td></td><td></td></tr>
<tr><td>Oligo fwd</td><td>CACCCACGAACCATCTCAGCTCCT</td><td></td><td></td><td></td><td></td><td></td><td></td><td></td><td></td><td></td><td></td><td></td></tr>
<tr><td>Oligo rev</td><td>AAACAGGAGCTGAGATGGTTCGTG</td><td></td><td></td><td></td><td></td><td></td><td></td><td></td><td></td><td></td><td></td><td></td></tr>
<tr><td>Chromosome</td><td>start</td><td></td><td>end</td><td>strand</td><td>MM</td><td>target_seq</td><td>PAM</td><td>alignment</td><td>distance</td><td>position</td><td>gene name</td><td>gene id</td></tr>
<tr><td>GU305914.1</td><td></td><td>200471</td><td>200493</td><td>+</td><td>0</td><td>CACGAACCATCTCAGCTCCT</td><td>CGG</td><td>||||||||[||||||||||||]PAM</td><td>NA</td><td>Intergenic</td><td>NA</td><td>NA</td></tr>
<tr><td>T15281</td><td>AACCATCTCAGCTCCTCGGTCGG</td><td></td><td>1000</td><td></td><td></td><td></td><td></td><td></td><td></td><td></td><td></td><td></td></tr>
<tr><td>Oligo fwd</td><td>CACCAACCATCTCAGCTCCTCGGT</td><td></td><td></td><td></td><td></td><td></td><td></td><td></td><td></td><td></td><td></td><td></td></tr>
<tr><td>Oligo rev</td><td>AAACACCGAGGAGCTGAGATGGTT</td><td></td><td></td><td></td><td></td><td></td><td></td><td></td><td></td><td></td><td></td><td></td></tr>
<tr><td>Chromosome</td><td>start</td><td></td><td>end</td><td>strand</td><td>MM</td><td>target_seq</td><td>PAM</td><td>alignment</td><td>distance</td><td>position</td><td>gene name</td><td>gene id</td></tr>
<tr><td>GU305914.1</td><td></td><td>200475</td><td>200497</td><td>+</td><td>0</td><td>AACCATCTCAGCTCCTCGGT</td><td>CGG</td><td>||||||||[||||||||||||]PAM</td><td>NA</td><td>Intergenic</td><td>NA</td><td>NA</td></tr>
<tr><td>T15282</td><td>CCTCGGTCGGAACGACGCCTCGG</td><td></td><td>1000</td><td></td><td></td><td></td><td></td><td></td><td></td><td></td><td></td><td></td></tr>
<tr><td>Oligo fwd</td><td>CACCCCTCGGTCGGAACGACGCCT</td><td></td><td></td><td></td><td></td><td></td><td></td><td></td><td></td><td></td><td></td><td></td></tr>
<tr><td>Oligo rev</td><td>AAACAGGCGTCGTTCCGACCGAGG</td><td></td><td></td><td></td><td></td><td></td><td></td><td></td><td></td><td></td><td></td><td></td></tr>
<tr><td>Chromosome</td><td>start</td><td></td><td>end</td><td>strand</td><td>MM</td><td>target_seq</td><td>PAM</td><td>alignment</td><td>distance</td><td>position</td><td>gene name</td><td>gene id</td></tr>
<tr><td>GU305914.1</td><td></td><td>200488</td><td>200510</td><td>+</td><td>0</td><td>CCTCGGTCGGAACGACGCCT</td><td>CGG</td><td>||||||||[||||||||||||]PAM</td><td>NA</td><td>Intergenic</td><td>NA</td><td>NA</td></tr>
<tr><td>T15283</td><td>GAACGACGCCTCGGAATCTCCGG</td><td></td><td>1000</td><td></td><td></td><td></td><td></td><td></td><td></td><td></td><td></td><td></td></tr>
<tr><td>Oligo fwd</td><td>CACCGAACGACGCCTCGGAATCTC</td><td></td><td></td><td></td><td></td><td></td><td></td><td></td><td></td><td></td><td></td><td></td></tr>
<tr><td>Oligo rev</td><td>AAACGAGATTCCGAGGCGTCGTTC</td><td></td><td></td><td></td><td></td><td></td><td></td><td></td><td></td><td></td><td></td><td></td></tr>
<tr><td>Chromosome</td><td>start</td><td></td><td>end</td><td>strand</td><td>MM</td><td>target_seq</td><td>PAM</td><td>alignment</td><td>distance</td><td>position</td><td>gene name</td><td>gene id</td></tr>
<tr><td>GU305914.1</td><td></td><td>200497</td><td>200519</td><td>+</td><td>0</td><td>GAACGACGCCTCGGAATCTC</td><td>CGG</td><td>||||||||[||||||||||||]PAM</td><td>NA</td><td>Intergenic</td><td>NA</td><td>NA</td></tr>
<tr><td>T15284</td><td>CTCGGAATCTCCGGACGACCTGG</td><td></td><td>1000</td><td></td><td></td><td></td><td></td><td></td><td></td><td></td><td></td><td></td></tr>
<tr><td>Oligo fwd</td><td>CACCCTCGGAATCTCCGGACGACC</td><td></td><td></td><td></td><td></td><td></td><td></td><td></td><td></td><td></td><td></td><td></td></tr>
<tr><td>Oligo rev</td><td>AAACGGTCGTCCGGAGATTCCGAG</td><td></td><td></td><td></td><td></td><td></td><td></td><td></td><td></td><td></td><td></td><td></td></tr>
<tr><td>Chromosome</td><td>start</td><td></td><td>end</td><td>strand</td><td>MM</td><td>target_seq</td><td>PAM</td><td>alignment</td><td>distance</td><td>position</td><td>gene name</td><td>gene id</td></tr>
<tr><td>GU305914.1</td><td></td><td>200506</td><td>200528</td><td>+</td><td>0</td><td>CTCGGAATCTCCGGACGACC</td><td>TGG</td><td>||||||||[||||||||||||]PAM</td><td>NA</td><td>Intergenic</td><td>NA</td><td>NA</td></tr>
<tr><td>T15285</td><td>CCTGGCCGAGCAGCGTGTAACGG</td><td></td><td>1000</td><td></td><td></td><td></td><td></td><td></td><td></td><td></td><td></td><td></td></tr>
<tr><td>Oligo fwd</td><td>CACCCCTGGCCGAGCAGCGTGTAA</td><td></td><td></td><td></td><td></td><td></td><td></td><td></td><td></td><td></td><td></td><td></td></tr>
<tr><td>Oligo rev</td><td>AAACTTACACGCTGCTCGGCCAGG</td><td></td><td></td><td></td><td></td><td></td><td></td><td></td><td></td><td></td><td></td><td></td></tr>
<tr><td>Chromosome</td><td>start</td><td></td><td>end</td><td>strand</td><td>MM</td><td>target_seq</td><td>PAM</td><td>alignment</td><td>distance</td><td>position</td><td>gene name</td><td>gene id</td></tr>
<tr><td>GU305914.1</td><td></td><td>200524</td><td>200546</td><td>+</td><td>0</td><td>CCTGGCCGAGCAGCGTGTAA</td><td>CGG</td><td>||||||||[||||||||||||]PAM</td><td>NA</td><td>Intergenic</td><td>NA</td><td>NA</td></tr>
<tr><td>T15286</td><td>AGCAGCGTGTAACGGCCCGCCGG</td><td></td><td>1000</td><td></td><td></td><td></td><td></td><td></td><td></td><td></td><td></td><td></td></tr>
<tr><td>Oligo fwd</td><td>CACCAGCAGCGTGTAACGGCCCGC</td><td></td><td></td><td></td><td></td><td></td><td></td><td></td><td></td><td></td><td></td><td></td></tr>
<tr><td>Oligo rev</td><td>AAACGCGGGCCGTTACACGCTGCT</td><td></td><td></td><td></td><td></td><td></td><td></td><td></td><td></td><td></td><td></td><td></td></tr>
<tr><td>Chromosome</td><td>start</td><td></td><td>end</td><td>strand</td><td>MM</td><td>target_seq</td><td>PAM</td><td>alignment</td><td>distance</td><td>position</td><td>gene name</td><td>gene id</td></tr>
<tr><td>GU305914.1</td><td></td><td>200532</td><td>200554</td><td>+</td><td>0</td><td>AGCAGCGTGTAACGGCCCGC</td><td>CGG</td><td>||||||||[||||||||||||]PAM</td><td>NA</td><td>Intergenic</td><td>NA</td><td>NA</td></tr>
<tr><td>T15287</td><td>GCCCCCGTACGTCTCCCAGACGG</td><td></td><td>1000</td><td></td><td></td><td></td><td></td><td></td><td></td><td></td><td></td><td></td></tr>
<tr><td>Oligo fwd</td><td>CACCGCCCCCGTACGTCTCCCAGA</td><td></td><td></td><td></td><td></td><td></td><td></td><td></td><td></td><td></td><td></td><td></td></tr>
<tr><td>Oligo rev</td><td>AAACTCTGGGAGACGTACGGGGGC</td><td></td><td></td><td></td><td></td><td></td><td></td><td></td><td></td><td></td><td></td><td></td></tr>
<tr><td>Chromosome</td><td>start</td><td></td><td>end</td><td>strand</td><td>MM</td><td>target_seq</td><td>PAM</td><td>alignment</td><td>distance</td><td>position</td><td>gene name</td><td>gene id</td></tr>
<tr><td>GU305914.1</td><td></td><td>200558</td><td>200580</td><td>+</td><td>0</td><td>GCCCCCGTACGTCTCCCAGA</td><td>CGG</td><td>||||||||[||||||||||||]PAM</td><td>NA</td><td>Intergenic</td><td>NA</td><td>NA</td></tr>
<tr><td>T15288</td><td>CCCCCGTACGTCTCCCAGACGGG</td><td></td><td>1000</td><td></td><td></td><td></td><td></td><td></td><td></td><td></td><td></td><td></td></tr>
</table>

<table>
<tr><td>Oligo fwd</td><td>CACCCCCCCGTACGTCTCCCAGAC</td><td></td><td></td><td></td><td></td><td></td><td></td><td></td><td></td><td></td><td></td></tr>
<tr><td>Oligo rev</td><td>AAACGTCTGGGAGACGTACGGGGG</td><td></td><td></td><td></td><td></td><td></td><td></td><td></td><td></td><td></td><td></td></tr>
<tr><td>Chromosome</td><td>start</td><td>end</td><td>strand</td><td>MM</td><td>target_seq</td><td>PAM</td><td>alignment</td><td>distance</td><td>position</td><td>gene name</td><td>gene id</td></tr>
<tr><td>GU305914.1</td><td>200559</td><td>200581</td><td>+</td><td>0</td><td>CCCCCGTACGTCTCCCAGAC</td><td>GGG</td><td>||||||||[||||||||||||]PAM</td><td>NA</td><td>Intergenic</td><td>NA</td><td>NA</td></tr>
<tr><td></td><td></td><td></td><td></td><td></td><td></td><td></td><td></td><td></td><td></td><td></td><td></td></tr>
<tr><td>T15289</td><td>CGTCTCCCAGACGGGATAGCCGG</td><td>1000</td><td></td><td></td><td></td><td></td><td></td><td></td><td></td><td></td><td></td></tr>
<tr><td>Oligo fwd</td><td>CACCCGTCTCCCAGACGGGATAGC</td><td></td><td></td><td></td><td></td><td></td><td></td><td></td><td></td><td></td><td></td></tr>
<tr><td>Oligo rev</td><td>AAACGCTATCCCGTCTGGGAGACG</td><td></td><td></td><td></td><td></td><td></td><td></td><td></td><td></td><td></td><td></td></tr>
<tr><td>Chromosome</td><td>start</td><td>end</td><td>strand</td><td>MM</td><td>target_seq</td><td>PAM</td><td>alignment</td><td>distance</td><td>position</td><td>gene name</td><td>gene id</td></tr>
<tr><td>GU305914.1</td><td>200567</td><td>200589</td><td>+</td><td>0</td><td>CGTCTCCCAGACGGGATAGC</td><td>CGG</td><td>||||||||[||||||||||||]PAM</td><td>NA</td><td>Intergenic</td><td>NA</td><td>NA</td></tr>
<tr><td></td><td></td><td></td><td></td><td></td><td></td><td></td><td></td><td></td><td></td><td></td><td></td></tr>
<tr><td>T15290</td><td>GTCTCCCAGACGGGATAGCCGGG</td><td>1000</td><td></td><td></td><td></td><td></td><td></td><td></td><td></td><td></td><td></td></tr>
<tr><td>Oligo fwd</td><td>CACCGTCTCCCAGACGGGATAGCC</td><td></td><td></td><td></td><td></td><td></td><td></td><td></td><td></td><td></td><td></td></tr>
<tr><td>Oligo rev</td><td>AAACGGCTATCCCGTCTGGGAGAC</td><td></td><td></td><td></td><td></td><td></td><td></td><td></td><td></td><td></td><td></td></tr>
<tr><td>Chromosome</td><td>start</td><td>end</td><td>strand</td><td>MM</td><td>target_seq</td><td>PAM</td><td>alignment</td><td>distance</td><td>position</td><td>gene name</td><td>gene id</td></tr>
<tr><td>GU305914.1</td><td>200568</td><td>200590</td><td>+</td><td>0</td><td>GTCTCCCAGACGGGATAGCC</td><td>GGG</td><td>||||||||[||||||||||||]PAM</td><td>NA</td><td>Intergenic</td><td>NA</td><td>NA</td></tr>
<tr><td></td><td></td><td></td><td></td><td></td><td></td><td></td><td></td><td></td><td></td><td></td><td></td></tr>
<tr><td>T15291</td><td>TCTCCCAGACGGGATAGCCGGGG</td><td>1000</td><td></td><td></td><td></td><td></td><td></td><td></td><td></td><td></td><td></td></tr>
<tr><td>Oligo fwd</td><td>CACCTCTCCCAGACGGGATAGCCG</td><td></td><td></td><td></td><td></td><td></td><td></td><td></td><td></td><td></td><td></td></tr>
<tr><td>Oligo rev</td><td>AAACCGGCTATCCCGTCTGGGAGA</td><td></td><td></td><td></td><td></td><td></td><td></td><td></td><td></td><td></td><td></td></tr>
<tr><td>Chromosome</td><td>start</td><td>end</td><td>strand</td><td>MM</td><td>target_seq</td><td>PAM</td><td>alignment</td><td>distance</td><td>position</td><td>gene name</td><td>gene id</td></tr>
<tr><td>GU305914.1</td><td>200569</td><td>200591</td><td>+</td><td>0</td><td>TCTCCCAGACGGGATAGCCG</td><td>GGG</td><td>||||||||[||||||||||||]PAM</td><td>NA</td><td>Intergenic</td><td>NA</td><td>NA</td></tr>
<tr><td></td><td></td><td></td><td></td><td></td><td></td><td></td><td></td><td></td><td></td><td></td><td></td></tr>
<tr><td>T15292</td><td>GGGGAAGCTCTCGACCTGATCGG</td><td>1000</td><td></td><td></td><td></td><td></td><td></td><td></td><td></td><td></td><td></td></tr>
<tr><td>Oligo fwd</td><td>CACCGGGGAAGCTCTCGACCTGAT</td><td></td><td></td><td></td><td></td><td></td><td></td><td></td><td></td><td></td><td></td></tr>
<tr><td>Oligo rev</td><td>AAACATCAGGTCGAGAGCTTCCCC</td><td></td><td></td><td></td><td></td><td></td><td></td><td></td><td></td><td></td><td></td></tr>
<tr><td>Chromosome</td><td>start</td><td>end</td><td>strand</td><td>MM</td><td>target_seq</td><td>PAM</td><td>alignment</td><td>distance</td><td>position</td><td>gene name</td><td>gene id</td></tr>
<tr><td>GU305914.1</td><td>200588</td><td>200610</td><td>+</td><td>0</td><td>GGGGAAGCTCTCGACCTGAT</td><td>CGG</td><td>||||||||[||||||||||||]PAM</td><td>NA</td><td>Intergenic</td><td>NA</td><td>NA</td></tr>
<tr><td></td><td></td><td></td><td></td><td></td><td></td><td></td><td></td><td></td><td></td><td></td><td></td></tr>
<tr><td>T15293</td><td>GAAGCTCTCGACCTGATCGGCGG</td><td>1000</td><td></td><td></td><td></td><td></td><td></td><td></td><td></td><td></td><td></td></tr>
<tr><td>Oligo fwd</td><td>CACCGAAGCTCTCGACCTGATCGG</td><td></td><td></td><td></td><td></td><td></td><td></td><td></td><td></td><td></td><td></td></tr>
<tr><td>Oligo rev</td><td>AAACCCGATCAGGTCGAGAGCTTC</td><td></td><td></td><td></td><td></td><td></td><td></td><td></td><td></td><td></td><td></td></tr>
<tr><td>Chromosome</td><td>start</td><td>end</td><td>strand</td><td>MM</td><td>target_seq</td><td>PAM</td><td>alignment</td><td>distance</td><td>position</td><td>gene name</td><td>gene id</td></tr>
<tr><td>GU305914.1</td><td>200591</td><td>200613</td><td>+</td><td>0</td><td>GAAGCTCTCGACCTGATCGG</td><td>CGG</td><td>||||||||[||||||||||||]PAM</td><td>NA</td><td>Intergenic</td><td>NA</td><td>NA</td></tr>
<tr><td></td><td></td><td></td><td></td><td></td><td></td><td></td><td></td><td></td><td></td><td></td><td></td></tr>
<tr><td>T15294</td><td>CGACCTGATCGGCGGAACGCAGG</td><td>1000</td><td></td><td></td><td></td><td></td><td></td><td></td><td></td><td></td><td></td></tr>
<tr><td>Oligo fwd</td><td>CACCCGACCTGATCGGCGGAACGC</td><td></td><td></td><td></td><td></td><td></td><td></td><td></td><td></td><td></td><td></td></tr>
<tr><td>Oligo rev</td><td>AAACGCGTTCCGCCGATCAGGTCG</td><td></td><td></td><td></td><td></td><td></td><td></td><td></td><td></td><td></td><td></td></tr>
<tr><td>Chromosome</td><td>start</td><td>end</td><td>strand</td><td>MM</td><td>target_seq</td><td>PAM</td><td>alignment</td><td>distance</td><td>position</td><td>gene name</td><td>gene id</td></tr>
<tr><td>GU305914.1</td><td>200599</td><td>200621</td><td>+</td><td>0</td><td>CGACCTGATCGGCGGAACGC</td><td>AGG</td><td>||||||||[||||||||||||]PAM</td><td>NA</td><td>Intergenic</td><td>NA</td><td>NA</td></tr>
<tr><td></td><td></td><td></td><td></td><td></td><td></td><td></td><td></td><td></td><td></td><td></td><td></td></tr>
<tr><td>T15295</td><td>GATCGGCGGAACGCAGGCAGAGG</td><td>1000</td><td></td><td></td><td></td><td></td><td></td><td></td><td></td><td></td><td></td></tr>
<tr><td>Oligo fwd</td><td>CACCGATCGGCGGAACGCAGGCAG</td><td></td><td></td><td></td><td></td><td></td><td></td><td></td><td></td><td></td><td></td></tr>
<tr><td>Oligo rev</td><td>AAACCTGCCTGCGTTCCGCCGATC</td><td></td><td></td><td></td><td></td><td></td><td></td><td></td><td></td><td></td><td></td></tr>
<tr><td>Chromosome</td><td>start</td><td>end</td><td>strand</td><td>MM</td><td>target_seq</td><td>PAM</td><td>alignment</td><td>distance</td><td>position</td><td>gene name</td><td>gene id</td></tr>
<tr><td>GU305914.1</td><td>200605</td><td>200627</td><td>+</td><td>0</td><td>GATCGGCGGAACGCAGGCAG</td><td>AGG</td><td>||||||||[||||||||||||]PAM</td><td>NA</td><td>Intergenic</td><td>NA</td><td>NA</td></tr>
<tr><td></td><td></td><td></td><td></td><td></td><td></td><td></td><td></td><td></td><td></td><td></td><td></td></tr>
<tr><td>T15296</td><td>AGGCAGAGGTACCAGCCCGTCGG</td><td>1000</td><td></td><td></td><td></td><td></td><td></td><td></td><td></td><td></td><td></td></tr>
<tr><td>Oligo fwd</td><td>CACCAGGCAGAGGTACCAGCCCGT</td><td></td><td></td><td></td><td></td><td></td><td></td><td></td><td></td><td></td><td></td></tr>
<tr><td>Oligo rev</td><td>AAACACGGGCTGGTACCTCTGCCT</td><td></td><td></td><td></td><td></td><td></td><td></td><td></td><td></td><td></td><td></td></tr>
<tr><td>Chromosome</td><td>start</td><td>end</td><td>strand</td><td>MM</td><td>target_seq</td><td>PAM</td><td>alignment</td><td>distance</td><td>position</td><td>gene name</td><td>gene id</td></tr>
<tr><td>GU305914.1</td><td>200619</td><td>200641</td><td>+</td><td>0</td><td>AGGCAGAGGTACCAGCCCGT</td><td>CGG</td><td>||||||||[||||||||||||]PAM</td><td>NA</td><td>Intergenic</td><td>NA</td><td>NA</td></tr>
<tr><td></td><td></td><td></td><td></td><td></td><td></td><td></td><td></td><td></td><td></td><td></td><td></td></tr>
<tr><td>T15297</td><td>GGCAGAGGTACCAGCCCGTCGGG</td><td>1000</td><td></td><td></td><td></td><td></td><td></td><td></td><td></td><td></td><td></td></tr>
<tr><td>Oligo fwd</td><td>CACCGGCAGAGGTACCAGCCCGTC</td><td></td><td></td><td></td><td></td><td></td><td></td><td></td><td></td><td></td><td></td></tr>
<tr><td>Oligo rev</td><td>AAACGACGGGCTGGTACCTCTGCC</td><td></td><td></td><td></td><td></td><td></td><td></td><td></td><td></td><td></td><td></td></tr>
<tr><td>Chromosome</td><td>start</td><td>end</td><td>strand</td><td>MM</td><td>target_seq</td><td>PAM</td><td>alignment</td><td>distance</td><td>position</td><td>gene name</td><td>gene id</td></tr>
<tr><td>GU305914.1</td><td>200620</td><td>200642</td><td>+</td><td>0</td><td>GGCAGAGGTACCAGCCCGTC</td><td>GGG</td><td>||||||||[||||||||||||]PAM</td><td>NA</td><td>Intergenic</td><td>NA</td><td>NA</td></tr>
<tr><td></td><td></td><td></td><td></td><td></td><td></td><td></td><td></td><td></td><td></td><td></td><td></td></tr>
<tr><td>T15298</td><td>GCAGAGGTACCAGCCCGTCGGGG</td><td>1000</td><td></td><td></td><td></td><td></td><td></td><td></td><td></td><td></td><td></td></tr>
<tr><td>Oligo fwd</td><td>CACCGCAGAGGTACCAGCCCGTCG</td><td></td><td></td><td></td><td></td><td></td><td></td><td></td><td></td><td></td><td></td></tr>
<tr><td>Oligo rev</td><td>AAACCGACGGGCTGGTACCTCTGC</td><td></td><td></td><td></td><td></td><td></td><td></td><td></td><td></td><td></td><td></td></tr>
<tr><td>Chromosome</td><td>start</td><td>end</td><td>strand</td><td>MM</td><td>target_seq</td><td>PAM</td><td>alignment</td><td>distance</td><td>position</td><td>gene name</td><td>gene id</td></tr>
<tr><td>GU305914.1</td><td>200621</td><td>200643</td><td>+</td><td>0</td><td>GCAGAGGTACCAGCCCGTCG</td><td>GGG</td><td>||||||||[||||||||||||]PAM</td><td>NA</td><td>Intergenic</td><td>NA</td><td>NA</td></tr>
<tr><td></td><td></td><td></td><td></td><td></td><td></td><td></td><td></td><td></td><td></td><td></td><td></td></tr>
<tr><td>T15299</td><td>ACCAGCCCGTCGGGGCGCCGAGG</td><td>1000</td><td></td><td></td><td></td><td></td><td></td><td></td><td></td><td></td><td></td></tr>
<tr><td>Oligo fwd</td><td>CACCACCAGCCCGTCGGGGCGCCG</td><td></td><td></td><td></td><td></td><td></td><td></td><td></td><td></td><td></td><td></td></tr>
<tr><td>Oligo rev</td><td>AAACCGGCGCCCCGACGGGCTGGT</td><td></td><td></td><td></td><td></td><td></td><td></td><td></td><td></td><td></td><td></td></tr>
<tr><td>Chromosome</td><td>start</td><td>end</td><td>strand</td><td>MM</td><td>target_seq</td><td>PAM</td><td>alignment</td><td>distance</td><td>position</td><td>gene name</td><td>gene id</td></tr>
<tr><td>GU305914.1</td><td>200629</td><td>200651</td><td>+</td><td>0</td><td>ACCAGCCCGTCGGGGCGCCG</td><td>AGG</td><td>||||||||[||||||||||||]PAM</td><td>NA</td><td>Intergenic</td><td>NA</td><td>NA</td></tr>
<tr><td></td><td></td><td></td><td></td><td></td><td></td><td></td><td></td><td></td><td></td><td></td><td></td></tr>
<tr><td>T15300</td><td>GCCGAGGCTCAGCAACCTGCCGG</td><td>1000</td><td></td><td></td><td></td><td></td><td></td><td></td><td></td><td></td><td></td></tr>
<tr><td>Oligo fwd</td><td>CACCGCCGAGGCTCAGCAACCTGC</td><td></td><td></td><td></td><td></td><td></td><td></td><td></td><td></td><td></td><td></td></tr>
<tr><td>Oligo rev</td><td>AAACGCAGGTTGCTGAGCCTCGGC</td><td></td><td></td><td></td><td></td><td></td><td></td><td></td><td></td><td></td><td></td></tr>
<tr><td>Chromosome</td><td>start</td><td>end</td><td>strand</td><td>MM</td><td>target_seq</td><td>PAM</td><td>alignment</td><td>distance</td><td>position</td><td>gene name</td><td>gene id</td></tr>
<tr><td>GU305914.1</td><td>200645</td><td>200667</td><td>+</td><td>0</td><td>GCCGAGGCTCAGCAACCTGC</td><td>CGG</td><td>||||||||[||||||||||||]PAM</td><td>NA</td><td>Intergenic</td><td>NA</td><td>NA</td></tr>
<tr><td></td><td></td><td></td><td></td><td></td><td></td><td></td><td></td><td></td><td></td><td></td><td></td></tr>
<tr><td>T15301</td><td>TTTCCTGACCACGTTCTCGACGG</td><td>1000</td><td></td><td></td><td></td><td></td><td></td><td></td><td></td><td></td><td></td></tr>
<tr><td>Oligo fwd</td><td>CACCTTTCCTGACCACGTTCTCGA</td><td></td><td></td><td></td><td></td><td></td><td></td><td></td><td></td><td></td><td></td></tr>
<tr><td>Oligo rev</td><td>AAACTCGAGAACGTGGTCAGGAAA</td><td></td><td></td><td></td><td></td><td></td><td></td><td></td><td></td><td></td><td></td></tr>
<tr><td>Chromosome</td><td>start</td><td>end</td><td>strand</td><td>MM</td><td>target_seq</td><td>PAM</td><td>alignment</td><td>distance</td><td>position</td><td>gene name</td><td>gene id</td></tr>
<tr><td>GU305914.1</td><td>200672</td><td>200694</td><td>+</td><td>0</td><td>TTTCCTGACCACGTTCTCGA</td><td>CGG</td><td>||||||||[||||||||||||]PAM</td><td>NA</td><td>Intergenic</td><td>NA</td><td>NA</td></tr>
<tr><td></td><td></td><td></td><td></td><td></td><td></td><td></td><td></td><td></td><td></td><td></td><td></td></tr>
<tr><td>T15302</td><td>CCACGTTCTCGACGGACGTGTGG</td><td>1000</td><td></td><td></td><td></td><td></td><td></td><td></td><td></td><td></td><td></td></tr>
<tr><td>Oligo fwd</td><td>CACCCCACGTTCTCGACGGACGTG</td><td></td><td></td><td></td><td></td><td></td><td></td><td></td><td></td><td></td><td></td></tr>
<tr><td>Oligo rev</td><td>AAACCACGTCCGTCGAGAACGTGG</td><td></td><td></td><td></td><td></td><td></td><td></td><td></td><td></td><td></td><td></td></tr>
<tr><td>Chromosome</td><td>start</td><td>end</td><td>strand</td><td>MM</td><td>target_seq</td><td>PAM</td><td>alignment</td><td>distance</td><td>position</td><td>gene name</td><td>gene id</td></tr>
<tr><td>GU305914.1</td><td>200680</td><td>200702</td><td>+</td><td>0</td><td>CCACGTTCTCGACGGACGTG</td><td>TGG</td><td>||||||||[||||||||||||]PAM</td><td>NA</td><td>Intergenic</td><td>NA</td><td>NA</td></tr>
</table>

| | | | | | | | | | | | |
|---|---|---|---|---|---|---|---|---|---|---|---|
| T15303 | ACGTGTGGTCGTTCTGCGCCAGG | 1000 | | | | | | | | | |
| Oligo fwd | CACCACGTGTGGTCGTTCTGCGCC | | | | | | | | | | |
| Oligo rev | AAACGGCGCAGAACGACCACACGT | | | | | | | | | | |
| Chromosome | start | end | strand | MM | target_seq | PAM | alignment | distance | position | gene name | gene id |
| GU305914.1 | 200695 | 200717 | + | 0 | ACGTGTGGTCGTTCTGCGCC | AGG | \|\|\|\|\|\|\|\|[\|\|\|\|\|\|\|\|\|\|\|\|]PAM | NA | Intergenic | NA | NA |
| T15304 | AGATGTCCGCGAAGTCCTGCAGG | 1000 | | | | | | | | | |
| Oligo fwd | CACCAGATGTCCGCGAAGTCCTGC | | | | | | | | | | |
| Oligo rev | AAACGCAGGACTTCGCGGACATCT | | | | | | | | | | |
| Chromosome | start | end | strand | MM | target_seq | PAM | alignment | distance | position | gene name | gene id |
| GU305914.1 | 200719 | 200741 | + | 0 | AGATGTCCGCGAAGTCCTGC | AGG | \|\|\|\|\|\|\|\|[\|\|\|\|\|\|\|\|\|\|\|\|]PAM | NA | Intergenic | NA | NA |
| T15305 | CGCGAAGTCCTGCAGGACCGCGG | 1000 | | | | | | | | | |
| Oligo fwd | CACCCGCGAAGTCCTGCAGGACCG | | | | | | | | | | |
| Oligo rev | AAACCGGTCCTGCAGGACTTCGCG | | | | | | | | | | |
| Chromosome | start | end | strand | MM | target_seq | PAM | alignment | distance | position | gene name | gene id |
| GU305914.1 | 200726 | 200748 | + | 0 | CGCGAAGTCCTGCAGGACCG | CGG | \|\|\|\|\|\|\|\|[\|\|\|\|\|\|\|\|\|\|\|\|]PAM | NA | Intergenic | NA | NA |
| T15306 | CTGCAGGACCGCGGTCACCGCGG | 1000 | | | | | | | | | |
| Oligo fwd | CACCCTGCAGGACCGCGGTCACCG | | | | | | | | | | |
| Oligo rev | AAACCGGTGACCGCGGTCCTGCAG | | | | | | | | | | |
| Chromosome | start | end | strand | MM | target_seq | PAM | alignment | distance | position | gene name | gene id |
| GU305914.1 | 200735 | 200757 | + | 0 | CTGCAGGACCGCGGTCACCG | CGG | \|\|\|\|\|\|\|\|[\|\|\|\|\|\|\|\|\|\|\|\|]PAM | NA | Intergenic | NA | NA |
| GU305914.1 | 163696 | 163718 | - | 4 | CTGCACGGCGGCGGCCACCG | CGG | \|\|\|\|\|-\|-[\|-\|\|\|\|-\|\|\|\|\|]PAM | NA | Intergenic | NA | NA |
| T15307 | CCGCGGCCGCCGTCAGATAGCGG | 1000 | | | | | | | | | |
| Oligo fwd | CACCCCGCGGCCGCCGTCAGATAG | | | | | | | | | | |
| Oligo rev | AAACCTATCTGACGGCGGCCGCGG | | | | | | | | | | |
| Chromosome | start | end | strand | MM | target_seq | PAM | alignment | distance | position | gene name | gene id |
| GU305914.1 | 200752 | 200774 | + | 0 | CCGCGGCCGCCGTCAGATAG | CGG | \|\|\|\|\|\|\|\|[\|\|\|\|\|\|\|\|\|\|\|\|]PAM | NA | Intergenic | NA | NA |
| T15308 | CGCGGCCGCCGTCAGATAGCGGG | 1000 | | | | | | | | | |
| Oligo fwd | CACCCGCGGCCGCCGTCAGATAGC | | | | | | | | | | |
| Oligo rev | AAACGCTATCTGACGGCGGCCGCG | | | | | | | | | | |
| Chromosome | start | end | strand | MM | target_seq | PAM | alignment | distance | position | gene name | gene id |
| GU305914.1 | 200753 | 200775 | + | 0 | CGCGGCCGCCGTCAGATAGC | GGG | \|\|\|\|\|\|\|\|[\|\|\|\|\|\|\|\|\|\|\|\|]PAM | NA | Intergenic | NA | NA |
| T15309 | GCCGTCAGATAGCGGGAGCGCGG | 1000 | | | | | | | | | |
| Oligo fwd | CACCGCCGTCAGATAGCGGGAGCG | | | | | | | | | | |
| Oligo rev | AAACCGCTCCCGCTATCTGACGGC | | | | | | | | | | |
| Chromosome | start | end | strand | MM | target_seq | PAM | alignment | distance | position | gene name | gene id |
| GU305914.1 | 200760 | 200782 | + | 0 | GCCGTCAGATAGCGGGAGCG | CGG | \|\|\|\|\|\|\|\|[\|\|\|\|\|\|\|\|\|\|\|\|]PAM | NA | Intergenic | NA | NA |
| T15310 | GAGCGCGGCCTGATGCCGACAGG | 1000 | | | | | | | | | |
| Oligo fwd | CACCGAGCGCGGCCTGATGCCGAC | | | | | | | | | | |
| Oligo rev | AAACGTCGGCATCAGGCCGCGCTC | | | | | | | | | | |
| Chromosome | start | end | strand | MM | target_seq | PAM | alignment | distance | position | gene name | gene id |
| GU305914.1 | 200775 | 200797 | + | 0 | GAGCGCGGCCTGATGCCGAC | AGG | \|\|\|\|\|\|\|\|[\|\|\|\|\|\|\|\|\|\|\|\|]PAM | NA | Intergenic | NA | NA |
| T15311 | GCCTGATGCCGACAGGCCTCTGG | 1000 | | | | | | | | | |
| Oligo fwd | CACCGCCTGATGCCGACAGGCCTC | | | | | | | | | | |
| Oligo rev | AAACGAGGCCTGTCGGCATCAGGC | | | | | | | | | | |
| Chromosome | start | end | strand | MM | target_seq | PAM | alignment | distance | position | gene name | gene id |
| GU305914.1 | 200782 | 200804 | + | 0 | GCCTGATGCCGACAGGCCTC | TGG | \|\|\|\|\|\|\|\|[\|\|\|\|\|\|\|\|\|\|\|\|]PAM | NA | Intergenic | NA | NA |
| T15312 | GGTCTCTGAAGTGATCCGATCGG | 1000 | | | | | | | | | |
| Oligo fwd | CACCGGTCTCTGAAGTGATCCGAT | | | | | | | | | | |
| Oligo rev | AAACATCGGATCACTTCAGAGACC | | | | | | | | | | |
| Chromosome | start | end | strand | MM | target_seq | PAM | alignment | distance | position | gene name | gene id |
| GU305914.1 | 200803 | 200825 | + | 0 | GGTCTCTGAAGTGATCCGAT | CGG | \|\|\|\|\|\|\|\|[\|\|\|\|\|\|\|\|\|\|\|\|]PAM | NA | Intergenic | NA | NA |
| T15313 | CCGATCGGATCGCCGCGCACAGG | 1000 | | | | | | | | | |
| Oligo fwd | CACCCCGATCGGATCGCCGCGCAC | | | | | | | | | | |
| Oligo rev | AAACGTGCGCGGCGATCCGATCGG | | | | | | | | | | |
| Chromosome | start | end | strand | MM | target_seq | PAM | alignment | distance | position | gene name | gene id |
| GU305914.1 | 200818 | 200840 | + | 0 | CCGATCGGATCGCCGCGCAC | AGG | \|\|\|\|\|\|\|\|[\|\|\|\|\|\|\|\|\|\|\|\|]PAM | NA | Intergenic | NA | NA |
| T15314 | CGATCGGATCGCCGCGCACAGGG | 1000 | | | | | | | | | |
| Oligo fwd | CACCCGATCGGATCGCCGCGCACA | | | | | | | | | | |
| Oligo rev | AAACTGTGCGCGGCGATCCGATCG | | | | | | | | | | |
| Chromosome | start | end | strand | MM | target_seq | PAM | alignment | distance | position | gene name | gene id |
| GU305914.1 | 200819 | 200841 | + | 0 | CGATCGGATCGCCGCGCACA | GGG | \|\|\|\|\|\|\|\|[\|\|\|\|\|\|\|\|\|\|\|\|]PAM | NA | Intergenic | NA | NA |
| T15315 | CGCCGCGCACAGGGCGTCCATGG | 1000 | | | | | | | | | |
| Oligo fwd | CACCCGCCGCGCACAGGGCGTCCA | | | | | | | | | | |
| Oligo rev | AAACTGGACGCCCTGTGCGCGGCG | | | | | | | | | | |
| Chromosome | start | end | strand | MM | target_seq | PAM | alignment | distance | position | gene name | gene id |
| GU305914.1 | 200828 | 200850 | + | 0 | CGCCGCGCACAGGGCGTCCA | TGG | \|\|\|\|\|\|\|\|[\|\|\|\|\|\|\|\|\|\|\|\|]PAM | NA | Intergenic | NA | NA |
| T15316 | CGCACAGGGCGTCCATGGCGAGG | 1000 | | | | | | | | | |
| Oligo fwd | CACCCGCACAGGGCGTCCATGGCG | | | | | | | | | | |
| Oligo rev | AAACCGCCATGGACGCCCTGTGCG | | | | | | | | | | |
| Chromosome | start | end | strand | MM | target_seq | PAM | alignment | distance | position | gene name | gene id |
| GU305914.1 | 200833 | 200855 | + | 0 | CGCACAGGGCGTCCATGGCG | AGG | \|\|\|\|\|\|\|\|[\|\|\|\|\|\|\|\|\|\|\|\|]PAM | NA | Intergenic | NA | NA |
| T15317 | GCACAGGGCGTCCATGGCGAGGG | 1000 | | | | | | | | | |
| Oligo fwd | CACCGCACAGGGCGTCCATGGCGA | | | | | | | | | | |
| Oligo rev | AAACTCGCCATGGACGCCCTGTGC | | | | | | | | | | |
| Chromosome | start | end | strand | MM | target_seq | PAM | alignment | distance | position | gene name | gene id |

| | | | | | | | | | | | |
|---|---|---|---|---|---|---|---|---|---|---|---|
| GU305914.1 | 200834 | 200856 | + | 0 | GCACAGGGCGTCCATGGCGA | GGG | \|\|\|\|\|\|\|\|[\|\|\|\|\|\|\|\|\|\|\|\|]PAM | NA | Intergenic | NA | NA |
| T15318 | GGCGTCCATGGCGAGGGCGACGG | 1000 | | | | | | | | | |
| Oligo fwd | CACCGGCGTCCATGGCGAGGGCGA | | | | | | | | | | |
| Oligo rev | AAACTCGCCCTCGCCATGGACGCC | | | | | | | | | | |
| Chromosome | start | end | strand | MM | target_seq | PAM | alignment | distance | position | gene name | gene id |
| GU305914.1 | 200840 | 200862 | + | 0 | GGCGTCCATGGCGAGGGCGA | CGG | \|\|\|\|\|\|\|\|[\|\|\|\|\|\|\|\|\|\|\|\|]PAM | NA | Intergenic | NA | NA |
| GU305914.1 | 129134 | 129156 | + | 4 | GGCGTGCGACGCGAGGGCGA | GGG | \|\|\|\|\|-\|-[--\|\|\|\|\|\|\|\|\|\|]PAM | NA | Intergenic | NA | NA |
| T15319 | GCGTCCATGGCGAGGGCGACGGG | 1000 | | | | | | | | | |
| Oligo fwd | CACCGCGTCCATGGCGAGGGCGAC | | | | | | | | | | |
| Oligo rev | AAACGTCGCCCTCGCCATGGACGC | | | | | | | | | | |
| Chromosome | start | end | strand | MM | target_seq | PAM | alignment | distance | position | gene name | gene id |
| GU305914.1 | 200841 | 200863 | + | 0 | GCGTCCATGGCGAGGGCGAC | GGG | \|\|\|\|\|\|\|\|[\|\|\|\|\|\|\|\|\|\|\|\|]PAM | NA | Intergenic | NA | NA |
| T15320 | TCCATGGCGAGGGCGACGGGCGG | 1000 | | | | | | | | | |
| Oligo fwd | CACCTCCATGGCGAGGGCGACGGG | | | | | | | | | | |
| Oligo rev | AAACCCCGTCGCCCTCGCCATGGA | | | | | | | | | | |
| Chromosome | start | end | strand | MM | target_seq | PAM | alignment | distance | position | gene name | gene id |
| GU305914.1 | 200844 | 200866 | + | 0 | TCCATGGCGAGGGCGACGGG | CGG | \|\|\|\|\|\|\|\|[\|\|\|\|\|\|\|\|\|\|\|\|]PAM | NA | Intergenic | NA | NA |
| T15321 | GCGAGGGCGACGGGCGGCAGCGG | 1000 | | | | | | | | | |
| Oligo fwd | CACCGCGAGGGCGACGGGCGGCAG | | | | | | | | | | |
| Oligo rev | AAACCTGCCGCCCGTCGCCCTCGC | | | | | | | | | | |
| Chromosome | start | end | strand | MM | target_seq | PAM | alignment | distance | position | gene name | gene id |
| GU305914.1 | 200850 | 200872 | + | 0 | GCGAGGGCGACGGGCGGCAG | CGG | \|\|\|\|\|\|\|\|[\|\|\|\|\|\|\|\|\|\|\|\|]PAM | NA | Intergenic | NA | NA |
| T15322 | CGAGGGCGACGGGCGGCAGCGGG | 1000 | | | | | | | | | |
| Oligo fwd | CACCCGAGGGCGACGGGCGGCAGC | | | | | | | | | | |
| Oligo rev | AAACGCTGCCGCCCGTCGCCCTCG | | | | | | | | | | |
| Chromosome | start | end | strand | MM | target_seq | PAM | alignment | distance | position | gene name | gene id |
| GU305914.1 | 200851 | 200873 | + | 0 | CGAGGGCGACGGGCGGCAGC | GGG | \|\|\|\|\|\|\|\|[\|\|\|\|\|\|\|\|\|\|\|\|]PAM | NA | Intergenic | NA | NA |
| T15323 | GACGGGCGGCAGCGGGAACACGG | 1000 | | | | | | | | | |
| Oligo fwd | CACCGACGGGCGGCAGCGGGAACA | | | | | | | | | | |
| Oligo rev | AAACTGTTCCCGCTGCCGCCCGTC | | | | | | | | | | |
| Chromosome | start | end | strand | MM | target_seq | PAM | alignment | distance | position | gene name | gene id |
| GU305914.1 | 200858 | 200880 | + | 0 | GACGGGCGGCAGCGGGAACA | CGG | \|\|\|\|\|\|\|\|[\|\|\|\|\|\|\|\|\|\|\|\|]PAM | NA | Intergenic | NA | NA |
| T15324 | ACGGGCGGCAGCGGGAACACGGG | 1000 | | | | | | | | | |
| Oligo fwd | CACCACGGGCGGCAGCGGGAACAC | | | | | | | | | | |
| Oligo rev | AAACGTGTTCCCGCTGCCGCCCGT | | | | | | | | | | |
| Chromosome | start | end | strand | MM | target_seq | PAM | alignment | distance | position | gene name | gene id |
| GU305914.1 | 200859 | 200881 | + | 0 | ACGGGCGGCAGCGGGAACAC | GGG | \|\|\|\|\|\|\|\|[\|\|\|\|\|\|\|\|\|\|\|\|]PAM | NA | Intergenic | NA | NA |
| T15325 | GGCGGCAGCGGGAACACGGGTGG | 1000 | | | | | | | | | |
| Oligo fwd | CACCGGCGGCAGCGGGAACACGGG | | | | | | | | | | |
| Oligo rev | AAACCCCGTGTTCCCGCTGCCGCC | | | | | | | | | | |
| Chromosome | start | end | strand | MM | target_seq | PAM | alignment | distance | position | gene name | gene id |
| GU305914.1 | 200862 | 200884 | + | 0 | GGCGGCAGCGGGAACACGGG | TGG | \|\|\|\|\|\|\|\|[\|\|\|\|\|\|\|\|\|\|\|\|]PAM | NA | Intergenic | NA | NA |
| T15326 | AGCGGGAACACGGGTGGAGAAGG | 1000 | | | | | | | | | |
| Oligo fwd | CACCAGCGGGAACACGGGTGGAGA | | | | | | | | | | |
| Oligo rev | AAACTCTCCACCCGTGTTCCCGCT | | | | | | | | | | |
| Chromosome | start | end | strand | MM | target_seq | PAM | alignment | distance | position | gene name | gene id |
| GU305914.1 | 200868 | 200890 | + | 0 | AGCGGGAACACGGGTGGAGA | AGG | \|\|\|\|\|\|\|\|[\|\|\|\|\|\|\|\|\|\|\|\|]PAM | NA | Intergenic | NA | NA |
| T15327 | GCGGGAACACGGGTGGAGAAGGG | 1000 | | | | | | | | | |
| Oligo fwd | CACCGCGGGAACACGGGTGGAGAA | | | | | | | | | | |
| Oligo rev | AAACTTCTCCACCCGTGTTCCCGC | | | | | | | | | | |
| Chromosome | start | end | strand | MM | target_seq | PAM | alignment | distance | position | gene name | gene id |
| GU305914.1 | 200869 | 200891 | + | 0 | GCGGGAACACGGGTGGAGAA | GGG | \|\|\|\|\|\|\|\|[\|\|\|\|\|\|\|\|\|\|\|\|]PAM | NA | Intergenic | NA | NA |
| T15328 | GGAACACGGGTGGAGAAGGGTGG | 1000 | | | | | | | | | |
| Oligo fwd | CACCGGAACACGGGTGGAGAAGGG | | | | | | | | | | |
| Oligo rev | AAACCCCTTCTCCACCCGTGTTCC | | | | | | | | | | |
| Chromosome | start | end | strand | MM | target_seq | PAM | alignment | distance | position | gene name | gene id |
| GU305914.1 | 200872 | 200894 | + | 0 | GGAACACGGGTGGAGAAGGG | TGG | \|\|\|\|\|\|\|\|[\|\|\|\|\|\|\|\|\|\|\|\|]PAM | NA | Intergenic | NA | NA |
| T15329 | CGGGTGGAGAAGGGTGGTCGCGG | 1000 | | | | | | | | | |
| Oligo fwd | CACCCGGGTGGAGAAGGGTGGTCG | | | | | | | | | | |
| Oligo rev | AAACCGACCACCCTTCTCCACCCG | | | | | | | | | | |
| Chromosome | start | end | strand | MM | target_seq | PAM | alignment | distance | position | gene name | gene id |
| GU305914.1 | 200878 | 200900 | + | 0 | CGGGTGGAGAAGGGTGGTCG | CGG | \|\|\|\|\|\|\|\|[\|\|\|\|\|\|\|\|\|\|\|\|]PAM | NA | Intergenic | NA | NA |
| T15330 | GGGTGGAGAAGGGTGGTCGCGGG | 1000 | | | | | | | | | |
| Oligo fwd | CACCGGGTGGAGAAGGGTGGTCGC | | | | | | | | | | |
| Oligo rev | AAACGCGACCACCCTTCTCCACCC | | | | | | | | | | |
| Chromosome | start | end | strand | MM | target_seq | PAM | alignment | distance | position | gene name | gene id |
| GU305914.1 | 200879 | 200901 | + | 0 | GGGTGGAGAAGGGTGGTCGC | GGG | \|\|\|\|\|\|\|\|[\|\|\|\|\|\|\|\|\|\|\|\|]PAM | NA | Intergenic | NA | NA |
| T15331 | TCACACACTCCGAATACCGATGG | 1000 | | | | | | | | | |
| Oligo fwd | CACCTCACACACTCCGAATACCGA | | | | | | | | | | |
| Oligo rev | AAACTCGGTATTCGGAGTGTGTGA | | | | | | | | | | |
| Chromosome | start | end | strand | MM | target_seq | PAM | alignment | distance | position | gene name | gene id |
| GU305914.1 | 200919 | 200941 | + | 0 | TCACACACTCCGAATACCGA | TGG | \|\|\|\|\|\|\|\|[\|\|\|\|\|\|\|\|\|\|\|\|]PAM | NA | Intergenic | NA | NA |
| T15332 | TACCGATGGCGTCGCGCTCGAGG | 1000 | | | | | | | | | |
| Oligo fwd | CACCTACCGATGGCGTCGCGCTCG | | | | | | | | | | |

<table>
<tr><td>Oligo rev</td><td>AAACCGAGCGCGACGCCATCGGTA</td><td></td><td></td><td></td><td></td><td></td><td></td><td></td><td></td><td></td><td></td></tr>
<tr><td>Chromosome</td><td>start</td><td>end</td><td>strand</td><td>MM</td><td>target_seq</td><td>PAM</td><td>alignment</td><td>distance</td><td>position</td><td>gene name</td><td>gene id</td></tr>
<tr><td>GU305914.1</td><td>200933</td><td>200955</td><td>+</td><td>0</td><td>TACCGATGGCGTCGCGCTCG</td><td>AGG</td><td>||||||||[||||||||||||]PAM</td><td>NA</td><td>Intergenic</td><td>NA</td><td>NA</td></tr>
<tr><td>GU305914.1</td><td>63577</td><td>63599</td><td>+</td><td>4</td><td>TGCGGATGGCCTCGCCCTCG</td><td>CGG</td><td>|-|-||||[||-||||-||||]PAM</td><td>NA</td><td>Intergenic</td><td>NA</td><td>NA</td></tr>
<tr><td></td><td></td><td></td><td></td><td></td><td></td><td></td><td></td><td></td><td></td><td></td><td></td></tr>
<tr><td>T15333</td><td>CGTCGCGCTCGAGGCGAGATCGG</td><td>1000</td><td></td><td></td><td></td><td></td><td></td><td></td><td></td><td></td><td></td></tr>
<tr><td>Oligo fwd</td><td>CACCCGTCGCGCTCGAGGCGAGAT</td><td></td><td></td><td></td><td></td><td></td><td></td><td></td><td></td><td></td><td></td></tr>
<tr><td>Oligo rev</td><td>AAACATCTCGCCTCGAGCGCGACG</td><td></td><td></td><td></td><td></td><td></td><td></td><td></td><td></td><td></td><td></td></tr>
<tr><td>Chromosome</td><td>start</td><td>end</td><td>strand</td><td>MM</td><td>target_seq</td><td>PAM</td><td>alignment</td><td>distance</td><td>position</td><td>gene name</td><td>gene id</td></tr>
<tr><td>GU305914.1</td><td>200942</td><td>200964</td><td>+</td><td>0</td><td>CGTCGCGCTCGAGGCGAGAT</td><td>CGG</td><td>||||||||[||||||||||||]PAM</td><td>NA</td><td>Intergenic</td><td>NA</td><td>NA</td></tr>
<tr><td></td><td></td><td></td><td></td><td></td><td></td><td></td><td></td><td></td><td></td><td></td><td></td></tr>
<tr><td>T15334</td><td>TTATCGCCCAAGTGTGAGAGAGG</td><td>1000</td><td></td><td></td><td></td><td></td><td></td><td></td><td></td><td></td><td></td></tr>
<tr><td>Oligo fwd</td><td>CACCTTATCGCCCAAGTGTGAGAG</td><td></td><td></td><td></td><td></td><td></td><td></td><td></td><td></td><td></td><td></td></tr>
<tr><td>Oligo rev</td><td>AAACCTCTCACACTTGGGCGATAA</td><td></td><td></td><td></td><td></td><td></td><td></td><td></td><td></td><td></td><td></td></tr>
<tr><td>Chromosome</td><td>start</td><td>end</td><td>strand</td><td>MM</td><td>target_seq</td><td>PAM</td><td>alignment</td><td>distance</td><td>position</td><td>gene name</td><td>gene id</td></tr>
<tr><td>GU305914.1</td><td>200969</td><td>200991</td><td>+</td><td>0</td><td>TTATCGCCCAAGTGTGAGAG</td><td>AGG</td><td>||||||||[||||||||||||]PAM</td><td>NA</td><td>Intergenic</td><td>NA</td><td>NA</td></tr>
<tr><td></td><td></td><td></td><td></td><td></td><td></td><td></td><td></td><td></td><td></td><td></td><td></td></tr>
<tr><td>T15335</td><td>TATCGCCCAAGTGTGAGAGAGGG</td><td>1000</td><td></td><td></td><td></td><td></td><td></td><td></td><td></td><td></td><td></td></tr>
<tr><td>Oligo fwd</td><td>CACCTATCGCCCAAGTGTGAGAGA</td><td></td><td></td><td></td><td></td><td></td><td></td><td></td><td></td><td></td><td></td></tr>
<tr><td>Oligo rev</td><td>AAACTCTCTCACACTTGGGCGATA</td><td></td><td></td><td></td><td></td><td></td><td></td><td></td><td></td><td></td><td></td></tr>
<tr><td>Chromosome</td><td>start</td><td>end</td><td>strand</td><td>MM</td><td>target_seq</td><td>PAM</td><td>alignment</td><td>distance</td><td>position</td><td>gene name</td><td>gene id</td></tr>
<tr><td>GU305914.1</td><td>200970</td><td>200992</td><td>+</td><td>0</td><td>TATCGCCCAAGTGTGAGAGA</td><td>GGG</td><td>||||||||[||||||||||||]PAM</td><td>NA</td><td>Intergenic</td><td>NA</td><td>NA</td></tr>
<tr><td></td><td></td><td></td><td></td><td></td><td></td><td></td><td></td><td></td><td></td><td></td><td></td></tr>
<tr><td>T15336</td><td>ATCGCCCAAGTGTGAGAGAGGGG</td><td>1000</td><td></td><td></td><td></td><td></td><td></td><td></td><td></td><td></td><td></td></tr>
<tr><td>Oligo fwd</td><td>CACCATCGCCCAAGTGTGAGAGAG</td><td></td><td></td><td></td><td></td><td></td><td></td><td></td><td></td><td></td><td></td></tr>
<tr><td>Oligo rev</td><td>AAACCTCTCTCACACTTGGGCGAT</td><td></td><td></td><td></td><td></td><td></td><td></td><td></td><td></td><td></td><td></td></tr>
<tr><td>Chromosome</td><td>start</td><td>end</td><td>strand</td><td>MM</td><td>target_seq</td><td>PAM</td><td>alignment</td><td>distance</td><td>position</td><td>gene name</td><td>gene id</td></tr>
<tr><td>GU305914.1</td><td>200971</td><td>200993</td><td>+</td><td>0</td><td>ATCGCCCAAGTGTGAGAGAG</td><td>GGG</td><td>||||||||[||||||||||||]PAM</td><td>NA</td><td>Intergenic</td><td>NA</td><td>NA</td></tr>
<tr><td></td><td></td><td></td><td></td><td></td><td></td><td></td><td></td><td></td><td></td><td></td><td></td></tr>
<tr><td>T15337</td><td>GCCCAAGTGTGAGAGAGGGGCGG</td><td>1000</td><td></td><td></td><td></td><td></td><td></td><td></td><td></td><td></td><td></td></tr>
<tr><td>Oligo fwd</td><td>CACCGCCCAAGTGTGAGAGAGGGG</td><td></td><td></td><td></td><td></td><td></td><td></td><td></td><td></td><td></td><td></td></tr>
<tr><td>Oligo rev</td><td>AAACCCCCTCTCTCACACTTGGGC</td><td></td><td></td><td></td><td></td><td></td><td></td><td></td><td></td><td></td><td></td></tr>
<tr><td>Chromosome</td><td>start</td><td>end</td><td>strand</td><td>MM</td><td>target_seq</td><td>PAM</td><td>alignment</td><td>distance</td><td>position</td><td>gene name</td><td>gene id</td></tr>
<tr><td>GU305914.1</td><td>200974</td><td>200996</td><td>+</td><td>0</td><td>GCCCAAGTGTGAGAGAGGGG</td><td>CGG</td><td>||||||||[||||||||||||]PAM</td><td>NA</td><td>Intergenic</td><td>NA</td><td>NA</td></tr>
<tr><td></td><td></td><td></td><td></td><td></td><td></td><td></td><td></td><td></td><td></td><td></td><td></td></tr>
<tr><td>T15338</td><td>CCCAAGTGTGAGAGAGGGGCGGG</td><td>1000</td><td></td><td></td><td></td><td></td><td></td><td></td><td></td><td></td><td></td></tr>
<tr><td>Oligo fwd</td><td>CACCCCCAAGTGTGAGAGAGGGGC</td><td></td><td></td><td></td><td></td><td></td><td></td><td></td><td></td><td></td><td></td></tr>
<tr><td>Oligo rev</td><td>AAACGCCCCTCTCTCACACTTGGG</td><td></td><td></td><td></td><td></td><td></td><td></td><td></td><td></td><td></td><td></td></tr>
<tr><td>Chromosome</td><td>start</td><td>end</td><td>strand</td><td>MM</td><td>target_seq</td><td>PAM</td><td>alignment</td><td>distance</td><td>position</td><td>gene name</td><td>gene id</td></tr>
<tr><td>GU305914.1</td><td>200975</td><td>200997</td><td>+</td><td>0</td><td>CCCAAGTGTGAGAGAGGGGC</td><td>GGG</td><td>||||||||[||||||||||||]PAM</td><td>NA</td><td>Intergenic</td><td>NA</td><td>NA</td></tr>
<tr><td></td><td></td><td></td><td></td><td></td><td></td><td></td><td></td><td></td><td></td><td></td><td></td></tr>
<tr><td>T15339</td><td>GAGAGATGACATGTCGTCACAGG</td><td>1000</td><td></td><td></td><td></td><td></td><td></td><td></td><td></td><td></td><td></td></tr>
<tr><td>Oligo fwd</td><td>CACCGAGAGATGACATGTCGTCAC</td><td></td><td></td><td></td><td></td><td></td><td></td><td></td><td></td><td></td><td></td></tr>
<tr><td>Oligo rev</td><td>AAACGTGACGACATGTCATCTCTC</td><td></td><td></td><td></td><td></td><td></td><td></td><td></td><td></td><td></td><td></td></tr>
<tr><td>Chromosome</td><td>start</td><td>end</td><td>strand</td><td>MM</td><td>target_seq</td><td>PAM</td><td>alignment</td><td>distance</td><td>position</td><td>gene name</td><td>gene id</td></tr>
<tr><td>GU305914.1</td><td>201004</td><td>201026</td><td>+</td><td>0</td><td>GAGAGATGACATGTCGTCAC</td><td>AGG</td><td>||||||||[||||||||||||]PAM</td><td>NA</td><td>Intergenic</td><td>NA</td><td>NA</td></tr>
<tr><td></td><td></td><td></td><td></td><td></td><td></td><td></td><td></td><td></td><td></td><td></td><td></td></tr>
<tr><td>T15340</td><td>AGAGATGACATGTCGTCACAGGG</td><td>1000</td><td></td><td></td><td></td><td></td><td></td><td></td><td></td><td></td><td></td></tr>
<tr><td>Oligo fwd</td><td>CACCAGAGATGACATGTCGTCACA</td><td></td><td></td><td></td><td></td><td></td><td></td><td></td><td></td><td></td><td></td></tr>
<tr><td>Oligo rev</td><td>AAACTGTGACGACATGTCATCTCT</td><td></td><td></td><td></td><td></td><td></td><td></td><td></td><td></td><td></td><td></td></tr>
<tr><td>Chromosome</td><td>start</td><td>end</td><td>strand</td><td>MM</td><td>target_seq</td><td>PAM</td><td>alignment</td><td>distance</td><td>position</td><td>gene name</td><td>gene id</td></tr>
<tr><td>GU305914.1</td><td>201005</td><td>201027</td><td>+</td><td>0</td><td>AGAGATGACATGTCGTCACA</td><td>GGG</td><td>||||||||[||||||||||||]PAM</td><td>NA</td><td>Intergenic</td><td>NA</td><td>NA</td></tr>
<tr><td></td><td></td><td></td><td></td><td></td><td></td><td></td><td></td><td></td><td></td><td></td><td></td></tr>
<tr><td>T15341</td><td>GAGATGACATGTCGTCACAGGGG</td><td>1000</td><td></td><td></td><td></td><td></td><td></td><td></td><td></td><td></td><td></td></tr>
<tr><td>Oligo fwd</td><td>CACCGAGATGACATGTCGTCACAG</td><td></td><td></td><td></td><td></td><td></td><td></td><td></td><td></td><td></td><td></td></tr>
<tr><td>Oligo rev</td><td>AAACCTGTGACGACATGTCATCTC</td><td></td><td></td><td></td><td></td><td></td><td></td><td></td><td></td><td></td><td></td></tr>
<tr><td>Chromosome</td><td>start</td><td>end</td><td>strand</td><td>MM</td><td>target_seq</td><td>PAM</td><td>alignment</td><td>distance</td><td>position</td><td>gene name</td><td>gene id</td></tr>
<tr><td>GU305914.1</td><td>201006</td><td>201028</td><td>+</td><td>0</td><td>GAGATGACATGTCGTCACAG</td><td>GGG</td><td>||||||||[||||||||||||]PAM</td><td>NA</td><td>Intergenic</td><td>NA</td><td>NA</td></tr>
<tr><td></td><td></td><td></td><td></td><td></td><td></td><td></td><td></td><td></td><td></td><td></td><td></td></tr>
<tr><td>T15342</td><td>AGGGGAAAACCGCCCCGTCGTGG</td><td>1000</td><td></td><td></td><td></td><td></td><td></td><td></td><td></td><td></td><td></td></tr>
<tr><td>Oligo fwd</td><td>CACCAGGGGAAAACCGCCCCGTCG</td><td></td><td></td><td></td><td></td><td></td><td></td><td></td><td></td><td></td><td></td></tr>
<tr><td>Oligo rev</td><td>AAACCGACGGGGCGGTTTTCCCCT</td><td></td><td></td><td></td><td></td><td></td><td></td><td></td><td></td><td></td><td></td></tr>
<tr><td>Chromosome</td><td>start</td><td>end</td><td>strand</td><td>MM</td><td>target_seq</td><td>PAM</td><td>alignment</td><td>distance</td><td>position</td><td>gene name</td><td>gene id</td></tr>
<tr><td>GU305914.1</td><td>201024</td><td>201046</td><td>+</td><td>0</td><td>AGGGGAAAACCGCCCCGTCG</td><td>TGG</td><td>||||||||[||||||||||||]PAM</td><td>NA</td><td>Intergenic</td><td>NA</td><td>NA</td></tr>
<tr><td></td><td></td><td></td><td></td><td></td><td></td><td></td><td></td><td></td><td></td><td></td><td></td></tr>
<tr><td>T15343</td><td>CCCCGTCGTGGACCTCGACGAGG</td><td>1000</td><td></td><td></td><td></td><td></td><td></td><td></td><td></td><td></td><td></td></tr>
<tr><td>Oligo fwd</td><td>CACCCCCCGTCGTGGACCTCGACG</td><td></td><td></td><td></td><td></td><td></td><td></td><td></td><td></td><td></td><td></td></tr>
<tr><td>Oligo rev</td><td>AAACCGTCGAGGTCCACGACGGGG</td><td></td><td></td><td></td><td></td><td></td><td></td><td></td><td></td><td></td><td></td></tr>
<tr><td>Chromosome</td><td>start</td><td>end</td><td>strand</td><td>MM</td><td>target_seq</td><td>PAM</td><td>alignment</td><td>distance</td><td>position</td><td>gene name</td><td>gene id</td></tr>
<tr><td>GU305914.1</td><td>201036</td><td>201058</td><td>+</td><td>0</td><td>CCCCGTCGTGGACCTCGACG</td><td>AGG</td><td>||||||||[||||||||||||]PAM</td><td>NA</td><td>Intergenic</td><td>NA</td><td>NA</td></tr>
<tr><td></td><td></td><td></td><td></td><td></td><td></td><td></td><td></td><td></td><td></td><td></td><td></td></tr>
<tr><td>T15344</td><td>CGTCGTGGACCTCGACGAGGCGG</td><td>1000</td><td></td><td></td><td></td><td></td><td></td><td></td><td></td><td></td><td></td></tr>
<tr><td>Oligo fwd</td><td>CACCCGTCGTGGACCTCGACGAGG</td><td></td><td></td><td></td><td></td><td></td><td></td><td></td><td></td><td></td><td></td></tr>
<tr><td>Oligo rev</td><td>AAACCCTCGTCGAGGTCCACGACG</td><td></td><td></td><td></td><td></td><td></td><td></td><td></td><td></td><td></td><td></td></tr>
<tr><td>Chromosome</td><td>start</td><td>end</td><td>strand</td><td>MM</td><td>target_seq</td><td>PAM</td><td>alignment</td><td>distance</td><td>position</td><td>gene name</td><td>gene id</td></tr>
<tr><td>GU305914.1</td><td>201039</td><td>201061</td><td>+</td><td>0</td><td>CGTCGTGGACCTCGACGAGG</td><td>CGG</td><td>||||||||[||||||||||||]PAM</td><td>NA</td><td>Intergenic</td><td>NA</td><td>NA</td></tr>
<tr><td></td><td></td><td></td><td></td><td></td><td></td><td></td><td></td><td></td><td></td><td></td><td></td></tr>
<tr><td>T15345</td><td>CGACGAGGCGGCTCACGCGTCGG</td><td>1000</td><td></td><td></td><td></td><td></td><td></td><td></td><td></td><td></td><td></td></tr>
<tr><td>Oligo fwd</td><td>CACCCGACGAGGCGGCTCACGCGT</td><td></td><td></td><td></td><td></td><td></td><td></td><td></td><td></td><td></td><td></td></tr>
<tr><td>Oligo rev</td><td>AAACACGCGTGAGCCGCCTCGTCG</td><td></td><td></td><td></td><td></td><td></td><td></td><td></td><td></td><td></td><td></td></tr>
<tr><td>Chromosome</td><td>start</td><td>end</td><td>strand</td><td>MM</td><td>target_seq</td><td>PAM</td><td>alignment</td><td>distance</td><td>position</td><td>gene name</td><td>gene id</td></tr>
<tr><td>GU305914.1</td><td>201051</td><td>201073</td><td>+</td><td>0</td><td>CGACGAGGCGGCTCACGCGT</td><td>CGG</td><td>||||||||[||||||||||||]PAM</td><td>NA</td><td>Intergenic</td><td>NA</td><td>NA</td></tr>
<tr><td></td><td></td><td></td><td></td><td></td><td></td><td></td><td></td><td></td><td></td><td></td><td></td></tr>
<tr><td>T15346</td><td>TCGGTCGCTCTCTCGTCGCGCGG</td><td>1000</td><td></td><td></td><td></td><td></td><td></td><td></td><td></td><td></td><td></td></tr>
<tr><td>Oligo fwd</td><td>CACCTCGGTCGCTCTCTCGTCGCG</td><td></td><td></td><td></td><td></td><td></td><td></td><td></td><td></td><td></td><td></td></tr>
<tr><td>Oligo rev</td><td>AAACCGCGACGAGAGAGCGACCGA</td><td></td><td></td><td></td><td></td><td></td><td></td><td></td><td></td><td></td><td></td></tr>
<tr><td>Chromosome</td><td>start</td><td>end</td><td>strand</td><td>MM</td><td>target_seq</td><td>PAM</td><td>alignment</td><td>distance</td><td>position</td><td>gene name</td><td>gene id</td></tr>
<tr><td>GU305914.1</td><td>201070</td><td>201092</td><td>+</td><td>0</td><td>TCGGTCGCTCTCTCGTCGCG</td><td>CGG</td><td>||||||||[||||||||||||]PAM</td><td>NA</td><td>Intergenic</td><td>NA</td><td>NA</td></tr>
</table>

<table>
<tr><td>T15347</td><td>GTCGCTCTCTCGTCGCGCGGCGG</td><td>1000</td><td></td><td></td><td></td><td></td><td></td><td></td><td></td><td></td><td></td></tr>
<tr><td>Oligo fwd</td><td>CACCGTCGCTCTCTCGTCGCGCGG</td><td></td><td></td><td></td><td></td><td></td><td></td><td></td><td></td><td></td><td></td></tr>
<tr><td>Oligo rev</td><td>AAACCCGCGCGACGAGAGAGCGAC</td><td></td><td></td><td></td><td></td><td></td><td></td><td></td><td></td><td></td><td></td></tr>
<tr><td>Chromosome</td><td>start</td><td>end</td><td>strand</td><td>MM</td><td>target_seq</td><td>PAM</td><td>alignment</td><td>distance</td><td>position</td><td>gene name</td><td>gene id</td></tr>
<tr><td>GU305914.1</td><td>201073</td><td>201095</td><td>+</td><td>0</td><td>GTCGCTCTCTCGTCGCGCGG</td><td>CGG</td><td>||||||||[||||||||||||]PAM</td><td>NA</td><td>Intergenic</td><td>NA</td><td>NA</td></tr>
<tr><td>T15348</td><td>GCTCTCTCGTCGCGCGGCGGTGG</td><td>1000</td><td></td><td></td><td></td><td></td><td></td><td></td><td></td><td></td><td></td></tr>
<tr><td>Oligo fwd</td><td>CACCGCTCTCTCGTCGCGCGGCGG</td><td></td><td></td><td></td><td></td><td></td><td></td><td></td><td></td><td></td><td></td></tr>
<tr><td>Oligo rev</td><td>AAACCCGCCGCGCGACGAGAGAGC</td><td></td><td></td><td></td><td></td><td></td><td></td><td></td><td></td><td></td><td></td></tr>
<tr><td>Chromosome</td><td>start</td><td>end</td><td>strand</td><td>MM</td><td>target_seq</td><td>PAM</td><td>alignment</td><td>distance</td><td>position</td><td>gene name</td><td>gene id</td></tr>
<tr><td>GU305914.1</td><td>201076</td><td>201098</td><td>+</td><td>0</td><td>GCTCTCTCGTCGCGCGGCGG</td><td>TGG</td><td>||||||||[||||||||||||]PAM</td><td>NA</td><td>Intergenic</td><td>NA</td><td>NA</td></tr>
<tr><td>GU305914.1</td><td>99735</td><td>99757</td><td>+</td><td>4</td><td>GCGCTCCCGTCTCGCCGCGG</td><td>CGG</td><td>||-|||-|[|||-|||-||||]PAM</td><td>NA</td><td>Intergenic</td><td>NA</td><td>NA</td></tr>
<tr><td>T15349</td><td>CTCTCGTCGCGCGGCGGTGGCGG</td><td>1000</td><td></td><td></td><td></td><td></td><td></td><td></td><td></td><td></td><td></td></tr>
<tr><td>Oligo fwd</td><td>CACCCTCTCGTCGCGCGGCGGTGG</td><td></td><td></td><td></td><td></td><td></td><td></td><td></td><td></td><td></td><td></td></tr>
<tr><td>Oligo rev</td><td>AAACCCACCGCCGCGCGACGAGAG</td><td></td><td></td><td></td><td></td><td></td><td></td><td></td><td></td><td></td><td></td></tr>
<tr><td>Chromosome</td><td>start</td><td>end</td><td>strand</td><td>MM</td><td>target_seq</td><td>PAM</td><td>alignment</td><td>distance</td><td>position</td><td>gene name</td><td>gene id</td></tr>
<tr><td>GU305914.1</td><td>201079</td><td>201101</td><td>+</td><td>0</td><td>CTCTCGTCGCGCGGCGGTGG</td><td>CGG</td><td>||||||||[||||||||||||]PAM</td><td>NA</td><td>Intergenic</td><td>NA</td><td>NA</td></tr>
<tr><td>GU305914.1</td><td>146006</td><td>146028</td><td>+</td><td>4</td><td>ATCGCGTCGCGCGCCGGTCG</td><td>CGG</td><td>-||-||||[|||||-||||-|]PAM</td><td>NA</td><td>Intergenic</td><td>NA</td><td>NA</td></tr>
<tr><td>T15350</td><td>TCTCGTCGCGCGGCGGTGGCGGG</td><td>1000</td><td></td><td></td><td></td><td></td><td></td><td></td><td></td><td></td><td></td></tr>
<tr><td>Oligo fwd</td><td>CACCTCTCGTCGCGCGGCGGTGGC</td><td></td><td></td><td></td><td></td><td></td><td></td><td></td><td></td><td></td><td></td></tr>
<tr><td>Oligo rev</td><td>AAACGCCACCGCCGCGCGACGAGA</td><td></td><td></td><td></td><td></td><td></td><td></td><td></td><td></td><td></td><td></td></tr>
<tr><td>Chromosome</td><td>start</td><td>end</td><td>strand</td><td>MM</td><td>target_seq</td><td>PAM</td><td>alignment</td><td>distance</td><td>position</td><td>gene name</td><td>gene id</td></tr>
<tr><td>GU305914.1</td><td>201080</td><td>201102</td><td>+</td><td>0</td><td>TCTCGTCGCGCGGCGGTGGC</td><td>GGG</td><td>||||||||[||||||||||||]PAM</td><td>NA</td><td>Intergenic</td><td>NA</td><td>NA</td></tr>
<tr><td>GU305914.1</td><td>146007</td><td>146029</td><td>+</td><td>3</td><td>TCGCGTCGCGCGCCGGTCGC</td><td>GGG</td><td>||-|||||[||||-||||-||]PAM</td><td>NA</td><td>Intergenic</td><td>NA</td><td>NA</td></tr>
<tr><td>T15351</td><td>CGCGCGGCGGTGGCGGGCGACGG</td><td>1000</td><td></td><td></td><td></td><td></td><td></td><td></td><td></td><td></td><td></td></tr>
<tr><td>Oligo fwd</td><td>CACCCGCGCGGCGGTGGCGGGCGA</td><td></td><td></td><td></td><td></td><td></td><td></td><td></td><td></td><td></td><td></td></tr>
<tr><td>Oligo rev</td><td>AAACTCGCCCGCCACCGCCGCGCG</td><td></td><td></td><td></td><td></td><td></td><td></td><td></td><td></td><td></td><td></td></tr>
<tr><td>Chromosome</td><td>start</td><td>end</td><td>strand</td><td>MM</td><td>target_seq</td><td>PAM</td><td>alignment</td><td>distance</td><td>position</td><td>gene name</td><td>gene id</td></tr>
<tr><td>GU305914.1</td><td>201086</td><td>201108</td><td>+</td><td>0</td><td>CGCGCGGCGGTGGCGGGCGA</td><td>CGG</td><td>||||||||[||||||||||||]PAM</td><td>NA</td><td>Intergenic</td><td>NA</td><td>NA</td></tr>
<tr><td>T15352</td><td>GGTGGCGGGCGACGGCGTCTCGG</td><td>1000</td><td></td><td></td><td></td><td></td><td></td><td></td><td></td><td></td><td></td></tr>
<tr><td>Oligo fwd</td><td>CACCGGTGGCGGGCGACGGCGTCT</td><td></td><td></td><td></td><td></td><td></td><td></td><td></td><td></td><td></td><td></td></tr>
<tr><td>Oligo rev</td><td>AAACAGACGCCGTCGCCCGCCACC</td><td></td><td></td><td></td><td></td><td></td><td></td><td></td><td></td><td></td><td></td></tr>
<tr><td>Chromosome</td><td>start</td><td>end</td><td>strand</td><td>MM</td><td>target_seq</td><td>PAM</td><td>alignment</td><td>distance</td><td>position</td><td>gene name</td><td>gene id</td></tr>
<tr><td>GU305914.1</td><td>201094</td><td>201116</td><td>+</td><td>0</td><td>GGTGGCGGGCGACGGCGTCT</td><td>CGG</td><td>||||||||[||||||||||||]PAM</td><td>NA</td><td>Intergenic</td><td>NA</td><td>NA</td></tr>
<tr><td>GU305914.1</td><td>131531</td><td>131553</td><td>+</td><td>4</td><td>GTTGGTGAGCGACGGCGTCA</td><td>AGG</td><td>|-|||-|-[|||||||||||-]PAM</td><td>NA</td><td>Intergenic</td><td>NA</td><td>NA</td></tr>
<tr><td>T15353</td><td>CGGGCGACGGCGTCTCGGCGCGG</td><td>1000</td><td></td><td></td><td></td><td></td><td></td><td></td><td></td><td></td><td></td></tr>
<tr><td>Oligo fwd</td><td>CACCCGGGCGACGGCGTCTCGGCG</td><td></td><td></td><td></td><td></td><td></td><td></td><td></td><td></td><td></td><td></td></tr>
<tr><td>Oligo rev</td><td>AAACCGCCGAGACGCCGTCGCCCG</td><td></td><td></td><td></td><td></td><td></td><td></td><td></td><td></td><td></td><td></td></tr>
<tr><td>Chromosome</td><td>start</td><td>end</td><td>strand</td><td>MM</td><td>target_seq</td><td>PAM</td><td>alignment</td><td>distance</td><td>position</td><td>gene name</td><td>gene id</td></tr>
<tr><td>GU305914.1</td><td>201099</td><td>201121</td><td>+</td><td>0</td><td>CGGGCGACGGCGTCTCGGCG</td><td>CGG</td><td>||||||||[||||||||||||]PAM</td><td>NA</td><td>Intergenic</td><td>NA</td><td>NA</td></tr>
<tr><td>T15354</td><td>GGGCGACGGCGTCTCGGCGCGGG</td><td>1000</td><td></td><td></td><td></td><td></td><td></td><td></td><td></td><td></td><td></td></tr>
<tr><td>Oligo fwd</td><td>CACCGGGCGACGGCGTCTCGGCGC</td><td></td><td></td><td></td><td></td><td></td><td></td><td></td><td></td><td></td><td></td></tr>
<tr><td>Oligo rev</td><td>AAACGCGCCGAGACGCCGTCGCCC</td><td></td><td></td><td></td><td></td><td></td><td></td><td></td><td></td><td></td><td></td></tr>
<tr><td>Chromosome</td><td>start</td><td>end</td><td>strand</td><td>MM</td><td>target_seq</td><td>PAM</td><td>alignment</td><td>distance</td><td>position</td><td>gene name</td><td>gene id</td></tr>
<tr><td>GU305914.1</td><td>201100</td><td>201122</td><td>+</td><td>0</td><td>GGGCGACGGCGTCTCGGCGC</td><td>GGG</td><td>||||||||[||||||||||||]PAM</td><td>NA</td><td>Intergenic</td><td>NA</td><td>NA</td></tr>
<tr><td>GU305914.1</td><td>150925</td><td>150947</td><td>-</td><td>4</td><td>GGACGAGGGCGGCGCGGCGC</td><td>TGG</td><td>||-|||-|[|||-|-||||||]PAM</td><td>NA</td><td>Intergenic</td><td>NA</td><td>NA</td></tr>
<tr><td>T15355</td><td>GGCGACGGCGTCTCGGCGCGGGG</td><td>1000</td><td></td><td></td><td></td><td></td><td></td><td></td><td></td><td></td><td></td></tr>
<tr><td>Oligo fwd</td><td>CACCGGCGACGGCGTCTCGGCGCG</td><td></td><td></td><td></td><td></td><td></td><td></td><td></td><td></td><td></td><td></td></tr>
<tr><td>Oligo rev</td><td>AAACCGCGCCGAGACGCCGTCGCC</td><td></td><td></td><td></td><td></td><td></td><td></td><td></td><td></td><td></td><td></td></tr>
<tr><td>Chromosome</td><td>start</td><td>end</td><td>strand</td><td>MM</td><td>target_seq</td><td>PAM</td><td>alignment</td><td>distance</td><td>position</td><td>gene name</td><td>gene id</td></tr>
<tr><td>GU305914.1</td><td>201101</td><td>201123</td><td>+</td><td>0</td><td>GGCGACGGCGTCTCGGCGCG</td><td>GGG</td><td>||||||||[||||||||||||]PAM</td><td>NA</td><td>Intergenic</td><td>NA</td><td>NA</td></tr>
<tr><td>T15356</td><td>CGGCGTCTCGGCGCGGGGATCGG</td><td>1000</td><td></td><td></td><td></td><td></td><td></td><td></td><td></td><td></td><td></td></tr>
<tr><td>Oligo fwd</td><td>CACCCGGCGTCTCGGCGCGGGGAT</td><td></td><td></td><td></td><td></td><td></td><td></td><td></td><td></td><td></td><td></td></tr>
<tr><td>Oligo rev</td><td>AAACATCCCCGCGCCGAGACGCCG</td><td></td><td></td><td></td><td></td><td></td><td></td><td></td><td></td><td></td><td></td></tr>
<tr><td>Chromosome</td><td>start</td><td>end</td><td>strand</td><td>MM</td><td>target_seq</td><td>PAM</td><td>alignment</td><td>distance</td><td>position</td><td>gene name</td><td>gene id</td></tr>
<tr><td>GU305914.1</td><td>201106</td><td>201128</td><td>+</td><td>0</td><td>CGGCGTCTCGGCGCGGGGAT</td><td>CGG</td><td>||||||||[||||||||||||]PAM</td><td>NA</td><td>Intergenic</td><td>NA</td><td>NA</td></tr>
<tr><td>T15357</td><td>GTCTCGGCGCGGGGATCGGACGG</td><td>1000</td><td></td><td></td><td></td><td></td><td></td><td></td><td></td><td></td><td></td></tr>
<tr><td>Oligo fwd</td><td>CACCGTCTCGGCGCGGGGATCGGA</td><td></td><td></td><td></td><td></td><td></td><td></td><td></td><td></td><td></td><td></td></tr>
<tr><td>Oligo rev</td><td>AAACTCCGATCCCCGCGCCGAGAC</td><td></td><td></td><td></td><td></td><td></td><td></td><td></td><td></td><td></td><td></td></tr>
<tr><td>Chromosome</td><td>start</td><td>end</td><td>strand</td><td>MM</td><td>target_seq</td><td>PAM</td><td>alignment</td><td>distance</td><td>position</td><td>gene name</td><td>gene id</td></tr>
<tr><td>GU305914.1</td><td>201110</td><td>201132</td><td>+</td><td>0</td><td>GTCTCGGCGCGGGGATCGGA</td><td>CGG</td><td>||||||||[||||||||||||]PAM</td><td>NA</td><td>Intergenic</td><td>NA</td><td>NA</td></tr>
<tr><td>T15358</td><td>TCTCGGCGCGGGGATCGGACGGG</td><td>1000</td><td></td><td></td><td></td><td></td><td></td><td></td><td></td><td></td><td></td></tr>
<tr><td>Oligo fwd</td><td>CACCTCTCGGCGCGGGGATCGGAC</td><td></td><td></td><td></td><td></td><td></td><td></td><td></td><td></td><td></td><td></td></tr>
<tr><td>Oligo rev</td><td>AAACGTCCGATCCCCGCGCCGAGA</td><td></td><td></td><td></td><td></td><td></td><td></td><td></td><td></td><td></td><td></td></tr>
<tr><td>Chromosome</td><td>start</td><td>end</td><td>strand</td><td>MM</td><td>target_seq</td><td>PAM</td><td>alignment</td><td>distance</td><td>position</td><td>gene name</td><td>gene id</td></tr>
<tr><td>GU305914.1</td><td>201111</td><td>201133</td><td>+</td><td>0</td><td>TCTCGGCGCGGGGATCGGAC</td><td>GGG</td><td>||||||||[||||||||||||]PAM</td><td>NA</td><td>Intergenic</td><td>NA</td><td>NA</td></tr>
<tr><td>T15359</td><td>CTCGGCGCGGGGATCGGACGGGG</td><td>1000</td><td></td><td></td><td></td><td></td><td></td><td></td><td></td><td></td><td></td></tr>
<tr><td>Oligo fwd</td><td>CACCCTCGGCGCGGGGATCGGACG</td><td></td><td></td><td></td><td></td><td></td><td></td><td></td><td></td><td></td><td></td></tr>
<tr><td>Oligo rev</td><td>AAACCGTCCGATCCCCGCGCCGAG</td><td></td><td></td><td></td><td></td><td></td><td></td><td></td><td></td><td></td><td></td></tr>
<tr><td>Chromosome</td><td>start</td><td>end</td><td>strand</td><td>MM</td><td>target_seq</td><td>PAM</td><td>alignment</td><td>distance</td><td>position</td><td>gene name</td><td>gene id</td></tr>
<tr><td>GU305914.1</td><td>201112</td><td>201134</td><td>+</td><td>0</td><td>CTCGGCGCGGGGATCGGACG</td><td>GGG</td><td>||||||||[||||||||||||]PAM</td><td>NA</td><td>Intergenic</td><td>NA</td><td>NA</td></tr>
<tr><td>T15360</td><td>GCGGGGATCGGACGGGGCGACGG</td><td>1000</td><td></td><td></td><td></td><td></td><td></td><td></td><td></td><td></td><td></td></tr>
<tr><td>Oligo fwd</td><td>CACCGCGGGGATCGGACGGGGCGA</td><td></td><td></td><td></td><td></td><td></td><td></td><td></td><td></td><td></td><td></td></tr>
<tr><td>Oligo rev</td><td>AAACTCGCCCCGTCCGATCCCCGC</td><td></td><td></td><td></td><td></td><td></td><td></td><td></td><td></td><td></td><td></td></tr>
<tr><td>Chromosome</td><td>start</td><td>end</td><td>strand</td><td>MM</td><td>target_seq</td><td>PAM</td><td>alignment</td><td>distance</td><td>position</td><td>gene name</td><td>gene id</td></tr>
<tr><td>GU305914.1</td><td>201118</td><td>201140</td><td>+</td><td>0</td><td>GCGGGGATCGGACGGGGCGA</td><td>CGG</td><td>||||||||[||||||||||||]PAM</td><td>NA</td><td>Intergenic</td><td>NA</td><td>NA</td></tr>
</table>

T15361 CGGGGATCGGACGGGGCGACGGG 1000
Oligo fwd CACCCGGGGATCGGACGGGGCGAC
Oligo rev AAACGTCGCCCCGTCCGATCCCCG
Chromosome start end strand MM target_seq PAM alignment distance position gene name gene id
GU305914.1 201119 201141 + 0 CGGGGATCGGACGGGGCGAC GGG ||||||||[||||||||||||]PAM NA Intergenic NA NA

T15362 GGGGATCGGACGGGGCGACGGGG 1000
Oligo fwd CACCGGGGATCGGACGGGGCGACG
Oligo rev AAACCGTCGCCCCGTCCGATCCCC
Chromosome start end strand MM target_seq PAM alignment distance position gene name gene id
GU305914.1 201120 201142 + 0 GGGGATCGGACGGGGCGACG GGG ||||||||[||||||||||||]PAM NA Intergenic NA NA

T15363 ACGGGGCGACGGGGACGACCCGG 1000
Oligo fwd CACCACGGGGCGACGGGGACGACC
Oligo rev AAACGGTCGTCCCCGTCGCCCCGT
Chromosome start end strand MM target_seq PAM alignment distance position gene name gene id
GU305914.1 201129 201151 + 0 ACGGGGCGACGGGGACGACC CGG ||||||||[||||||||||||]PAM NA Intergenic NA NA

T15364 GGGCGACGGGGACGACCCGGTGG 1000
Oligo fwd CACCGGGCGACGGGGACGACCCGG
Oligo rev AAACCCGGGTCGTCCCCGTCGCCC
Chromosome start end strand MM target_seq PAM alignment distance position gene name gene id
GU305914.1 201132 201154 + 0 GGGCGACGGGGACGACCCGG TGG ||||||||[||||||||||||]PAM NA Intergenic NA NA
GU305914.1 122463 122485 - 4 CGGCGACAGGAACGACCCGC CGG -||||||-[||-||||||||-]PAM NA Intergenic NA NA

T15365 GGGGACGACCCGGTGGAGTTCGG 1000
Oligo fwd CACCGGGGACGACCCGGTGGAGTT
Oligo rev AAACAACTCCACCGGGTCGTCCCC
Chromosome start end strand MM target_seq PAM alignment distance position gene name gene id
GU305914.1 201139 201161 + 0 GGGGACGACCCGGTGGAGTT CGG ||||||||[||||||||||||]PAM NA Intergenic NA NA

T15366 CCGGTGGAGTTCGGCGTCGACGG 1000
Oligo fwd CACCCCGGTGGAGTTCGGCGTCGA
Oligo rev AAACTCGACGCCGAACTCCACCGG
Chromosome start end strand MM target_seq PAM alignment distance position gene name gene id
GU305914.1 201148 201170 + 0 CCGGTGGAGTTCGGCGTCGA CGG ||||||||[||||||||||||]PAM NA Intergenic NA NA

T15367 CGGTGGAGTTCGGCGTCGACGGG 1000
Oligo fwd CACCCGGTGGAGTTCGGCGTCGAC
Oligo rev AAACGTCGACGCCGAACTCCACCG
Chromosome start end strand MM target_seq PAM alignment distance position gene name gene id
GU305914.1 201149 201171 + 0 CGGTGGAGTTCGGCGTCGAC GGG ||||||||[||||||||||||]PAM NA Intergenic NA NA

T15368 GGTGGAGTTCGGCGTCGACGGGG 1000
Oligo fwd CACCGGTGGAGTTCGGCGTCGACG
Oligo rev AAACCGTCGACGCCGAACTCCACC
Chromosome start end strand MM target_seq PAM alignment distance position gene name gene id
GU305914.1 201150 201172 + 0 GGTGGAGTTCGGCGTCGACG GGG ||||||||[||||||||||||]PAM NA Intergenic NA NA

T15369 GTGGAGTTCGGCGTCGACGGGGG 1000
Oligo fwd CACCGTGGAGTTCGGCGTCGACGG
Oligo rev AAACCCGTCGACGCCGAACTCCAC
Chromosome start end strand MM target_seq PAM alignment distance position gene name gene id
GU305914.1 201151 201173 + 0 GTGGAGTTCGGCGTCGACGG GGG ||||||||[||||||||||||]PAM NA Intergenic NA NA
GU305914.1 61376 61398 - 4 GTCGAGTGCGAGGTCGACGG GGG ||-||||-[||--||||||||]PAM NA Intergenic NA NA

T15370 GTTCGGCGTCGACGGGGGCGCGG 1000
Oligo fwd CACCGTTCGGCGTCGACGGGGGCG
Oligo rev AAACCGCCCCCGTCGACGCCGAAC
Chromosome start end strand MM target_seq PAM alignment distance position gene name gene id
GU305914.1 201156 201178 + 0 GTTCGGCGTCGACGGGGGCG CGG ||||||||[||||||||||||]PAM NA Intergenic NA NA
GU305914.1 159954 159976 - 4 TATCGGCGGCGACGGCGGCG AGG --||||||[-||||||-||||]PAM NA Intergenic NA NA

T15371 TTCGGCGTCGACGGGGGCGCGGG 1000
Oligo fwd CACCTTCGGCGTCGACGGGGGCGC
Oligo rev AAACGCGCCCCCGTCGACGCCGAA
Chromosome start end strand MM target_seq PAM alignment distance position gene name gene id
GU305914.1 201157 201179 + 0 TTCGGCGTCGACGGGGGCGC GGG ||||||||[||||||||||||]PAM NA Intergenic NA NA

T15372 GGCGTCGACGGGGGCGCGGGCGG 1000
Oligo fwd CACCGGCGTCGACGGGGGCGCGGG
Oligo rev AAACCCCGCGCCCCCGTCGACGCC
Chromosome start end strand MM target_seq PAM alignment distance position gene name gene id
GU305914.1 201160 201182 + 0 GGCGTCGACGGGGGCGCGGG CGG ||||||||[||||||||||||]PAM NA Intergenic NA NA

T15373 GTCGACGGGGGCGCGGGCGGTGG 1000
Oligo fwd CACCGTCGACGGGGGCGCGGGCGG
Oligo rev AAACCCGCCCGCGCCCCCGTCGAC
Chromosome start end strand MM target_seq PAM alignment distance position gene name gene id
GU305914.1 201163 201185 + 0 GTCGACGGGGGCGCGGGCGG TGG ||||||||[||||||||||||]PAM NA Intergenic NA NA

T15374 CGCGGGCGGTGGCGAATCGTCGG 1000
Oligo fwd CACCCGCGGGCGGTGGCGAATCGT
Oligo rev AAACACGATTCGCCACCGCCCGCG
Chromosome start end strand MM target_seq PAM alignment distance position gene name gene id
GU305914.1 201174 201196 + 0 CGCGGGCGGTGGCGAATCGT CGG ||||||||[||||||||||||]PAM NA Intergenic NA NA

T15375 GCGAATCGTCGGTCGTGTAGCGG 1000
Oligo fwd CACCGCGAATCGTCGGTCGTGTAG

| | | | | | | | | | | | |
|---|---|---|---|---|---|---|---|---|---|---|---|
| Oligo rev | AAACCTACACGACCGACGATTCGC | | | | | | | | | | |
| Chromosome | start | end | strand | MM | target_seq | PAM | alignment | distance | position | gene name | gene id |
| GU305914.1 | 201185 | 201207 | + | 0 | GCGAATCGTCGGTCGTGTAG | CGG | \|\|\|\|\|\|\|\|[\|\|\|\|\|\|\|\|\|\|\|\|]PAM | NA | Intergenic | NA | NA |
| T15376 | TGTAGCGGTACTGCCGCTCTCGG | 1000 | | | | | | | | | |
| Oligo fwd | CACCTGTAGCGGTACTGCCGCTCT | | | | | | | | | | |
| Oligo rev | AAACAGAGCGGCAGTACCGCTACA | | | | | | | | | | |
| Chromosome | start | end | strand | MM | target_seq | PAM | alignment | distance | position | gene name | gene id |
| GU305914.1 | 201200 | 201222 | + | 0 | TGTAGCGGTACTGCCGCTCT | CGG | \|\|\|\|\|\|\|\|[\|\|\|\|\|\|\|\|\|\|\|\|]PAM | NA | Intergenic | NA | NA |
| T15377 | AGCGGTACTGCCGCTCTCGGAGG | 1000 | | | | | | | | | |
| Oligo fwd | CACCAGCGGTACTGCCGCTCTCGG | | | | | | | | | | |
| Oligo rev | AAACCCGAGAGCGGCAGTACCGCT | | | | | | | | | | |
| Chromosome | start | end | strand | MM | target_seq | PAM | alignment | distance | position | gene name | gene id |
| GU305914.1 | 201203 | 201225 | + | 0 | AGCGGTACTGCCGCTCTCGG | AGG | \|\|\|\|\|\|\|\|[\|\|\|\|\|\|\|\|\|\|\|\|]PAM | NA | Intergenic | NA | NA |
| T15378 | GTGACAATCTCCCTCCGCCTCGG | 1000 | | | | | | | | | |
| Oligo fwd | CACCGTGACAATCTCCCTCCGCCT | | | | | | | | | | |
| Oligo rev | AAACAGGCGGAGGGAGATTGTCAC | | | | | | | | | | |
| Chromosome | start | end | strand | MM | target_seq | PAM | alignment | distance | position | gene name | gene id |
| GU305914.1 | 201231 | 201253 | + | 0 | GTGACAATCTCCCTCCGCCT | CGG | \|\|\|\|\|\|\|\|[\|\|\|\|\|\|\|\|\|\|\|\|]PAM | NA | Intergenic | NA | NA |
| T15379 | CTCCCTCCGCCTCGGAGCGCAGG | 1000 | | | | | | | | | |
| Oligo fwd | CACCCTCCCTCCGCCTCGGAGCGC | | | | | | | | | | |
| Oligo rev | AAACGCGCTCCGAGGCGGAGGGAG | | | | | | | | | | |
| Chromosome | start | end | strand | MM | target_seq | PAM | alignment | distance | position | gene name | gene id |
| GU305914.1 | 201239 | 201261 | + | 0 | CTCCCTCCGCCTCGGAGCGC | AGG | \|\|\|\|\|\|\|\|[\|\|\|\|\|\|\|\|\|\|\|\|]PAM | NA | Intergenic | NA | NA |
| T15380 | TCCCTCCGCCTCGGAGCGCAGGG | 1000 | | | | | | | | | |
| Oligo fwd | CACCTCCCTCCGCCTCGGAGCGCA | | | | | | | | | | |
| Oligo rev | AAACTGCGCTCCGAGGCGGAGGGA | | | | | | | | | | |
| Chromosome | start | end | strand | MM | target_seq | PAM | alignment | distance | position | gene name | gene id |
| GU305914.1 | 201240 | 201262 | + | 0 | TCCCTCCGCCTCGGAGCGCA | GGG | \|\|\|\|\|\|\|\|[\|\|\|\|\|\|\|\|\|\|\|\|]PAM | NA | Intergenic | NA | NA |
| T15381 | GCCTCGGAGCGCAGGGAGCGCGG | 1000 | | | | | | | | | |
| Oligo fwd | CACCGCCTCGGAGCGCAGGGAGCG | | | | | | | | | | |
| Oligo rev | AAACCGCTCCCTGCGCTCCGAGGC | | | | | | | | | | |
| Chromosome | start | end | strand | MM | target_seq | PAM | alignment | distance | position | gene name | gene id |
| GU305914.1 | 201247 | 201269 | + | 0 | GCCTCGGAGCGCAGGGAGCG | CGG | \|\|\|\|\|\|\|\|[\|\|\|\|\|\|\|\|\|\|\|\|]PAM | NA | Intergenic | NA | NA |
| T15382 | TCGGAGCGCAGGGAGCGCGGCGG | 1000 | | | | | | | | | |
| Oligo fwd | CACCTCGGAGCGCAGGGAGCGCGG | | | | | | | | | | |
| Oligo rev | AAACCCGCGCTCCCTGCGCTCCGA | | | | | | | | | | |
| Chromosome | start | end | strand | MM | target_seq | PAM | alignment | distance | position | gene name | gene id |
| GU305914.1 | 201250 | 201272 | + | 0 | TCGGAGCGCAGGGAGCGCGG | CGG | \|\|\|\|\|\|\|\|[\|\|\|\|\|\|\|\|\|\|\|\|]PAM | NA | Intergenic | NA | NA |
| T15383 | GCAGGGAGCGCGGCGGCTCCAGG | 1000 | | | | | | | | | |
| Oligo fwd | CACCGCAGGGAGCGCGGCGGCTCC | | | | | | | | | | |
| Oligo rev | AAACGGAGCCGCCGCGCTCCCTGC | | | | | | | | | | |
| Chromosome | start | end | strand | MM | target_seq | PAM | alignment | distance | position | gene name | gene id |
| GU305914.1 | 201257 | 201279 | + | 0 | GCAGGGAGCGCGGCGGCTCC | AGG | \|\|\|\|\|\|\|\|[\|\|\|\|\|\|\|\|\|\|\|\|]PAM | NA | Intergenic | NA | NA |
| T15384 | GAGCGCGGCGGCTCCAGGATCGG | 1000 | | | | | | | | | |
| Oligo fwd | CACCGAGCGCGGCGGCTCCAGGAT | | | | | | | | | | |
| Oligo rev | AAACATCCTGGAGCCGCCGCGCTC | | | | | | | | | | |
| Chromosome | start | end | strand | MM | target_seq | PAM | alignment | distance | position | gene name | gene id |
| GU305914.1 | 201262 | 201284 | + | 0 | GAGCGCGGCGGCTCCAGGAT | CGG | \|\|\|\|\|\|\|\|[\|\|\|\|\|\|\|\|\|\|\|\|]PAM | NA | Intergenic | NA | NA |
| T15385 | GCGGCTCCAGGATCGGTGCGTGG | 1000 | | | | | | | | | |
| Oligo fwd | CACCGCGGCTCCAGGATCGGTGCG | | | | | | | | | | |
| Oligo rev | AAACCGCACCGATCCTGGAGCCGC | | | | | | | | | | |
| Chromosome | start | end | strand | MM | target_seq | PAM | alignment | distance | position | gene name | gene id |
| GU305914.1 | 201269 | 201291 | + | 0 | GCGGCTCCAGGATCGGTGCG | TGG | \|\|\|\|\|\|\|\|[\|\|\|\|\|\|\|\|\|\|\|\|]PAM | NA | Intergenic | NA | NA |
| T15386 | CCAGGATCGGTGCGTGGTGACGG | 1000 | | | | | | | | | |
| Oligo fwd | CACCCCAGGATCGGTGCGTGGTGA | | | | | | | | | | |
| Oligo rev | AAACTCACCACGCACCGATCCTGG | | | | | | | | | | |
| Chromosome | start | end | strand | MM | target_seq | PAM | alignment | distance | position | gene name | gene id |
| GU305914.1 | 201275 | 201297 | + | 0 | CCAGGATCGGTGCGTGGTGA | CGG | \|\|\|\|\|\|\|\|[\|\|\|\|\|\|\|\|\|\|\|\|]PAM | NA | Intergenic | NA | NA |
| T15387 | CAGGATCGGTGCGTGGTGACGGG | 1000 | | | | | | | | | |
| Oligo fwd | CACCCAGGATCGGTGCGTGGTGAC | | | | | | | | | | |
| Oligo rev | AAACGTCACCACGCACCGATCCTG | | | | | | | | | | |
| Chromosome | start | end | strand | MM | target_seq | PAM | alignment | distance | position | gene name | gene id |
| GU305914.1 | 201276 | 201298 | + | 0 | CAGGATCGGTGCGTGGTGAC | GGG | \|\|\|\|\|\|\|\|[\|\|\|\|\|\|\|\|\|\|\|\|]PAM | NA | Intergenic | NA | NA |
| T15388 | GTGGTGACGGGTCGCGACGCCGG | 1000 | | | | | | | | | |
| Oligo fwd | CACCGTGGTGACGGGTCGCGACGC | | | | | | | | | | |
| Oligo rev | AAACGCGTCGCGACCCGTCACCAC | | | | | | | | | | |
| Chromosome | start | end | strand | MM | target_seq | PAM | alignment | distance | position | gene name | gene id |
| GU305914.1 | 201288 | 201310 | + | 0 | GTGGTGACGGGTCGCGACGC | CGG | \|\|\|\|\|\|\|\|[\|\|\|\|\|\|\|\|\|\|\|\|]PAM | NA | Intergenic | NA | NA |
| T15389 | CGCGACGCCGGTCTGACTCGAGG | 1000 | | | | | | | | | |
| Oligo fwd | CACCCGCGACGCCGGTCTGACTCG | | | | | | | | | | |
| Oligo rev | AAACCGAGTCAGACCGGCGTCGCG | | | | | | | | | | |
| Chromosome | start | end | strand | MM | target_seq | PAM | alignment | distance | position | gene name | gene id |
| GU305914.1 | 201300 | 201322 | + | 0 | CGCGACGCCGGTCTGACTCG | AGG | \|\|\|\|\|\|\|\|[\|\|\|\|\|\|\|\|\|\|\|\|]PAM | NA | Intergenic | NA | NA |
| T15390 | GACTCGAGGCCGCGTGCGCGCGG | 1000 | | | | | | | | | |

| | | | | | | | | | | | |
|---|---|---|---|---|---|---|---|---|---|---|---|
| Oligo fwd | CACCGACTCGAGGCCGCGTGCGCG | | | | | | | | | | |
| Oligo rev | AAACCGCGCACGCGGCCTCGAGTC | | | | | | | | | | |
| Chromosome | start | end | strand | MM | target_seq | PAM | alignment | distance | position | gene name | gene id |
| GU305914.1 | 201314 | 201336 | + | 0 | GACTCGAGGCCGCGTGCGCG | CGG | \|\|\|\|\|\|\|\|[\|\|\|\|\|\|\|\|\|\|\|\|]PAM | NA | Intergenic | NA | NA |
| T15391 | CGGCGCCTCGCCACCGCCGCCGG | 1000 | | | | | | | | | |
| Oligo fwd | CACCCGGCGCCTCGCCACCGCCGC | | | | | | | | | | |
| Oligo rev | AAACGCGGCGGTGGCGAGGCGCCG | | | | | | | | | | |
| Chromosome | start | end | strand | MM | target_seq | PAM | alignment | distance | position | gene name | gene id |
| GU305914.1 | 201334 | 201356 | + | 0 | CGGCGCCTCGCCACCGCCGC | CGG | \|\|\|\|\|\|\|\|[\|\|\|\|\|\|\|\|\|\|\|\|]PAM | NA | Intergenic | NA | NA |
| T15392 | GTGTCGTAGCGCCCCGCCTCCGG | 1000 | | | | | | | | | |
| Oligo fwd | CACCGTGTCGTAGCGCCCCGCCTC | | | | | | | | | | |
| Oligo rev | AAACGAGGCGGGGCGCTACGACAC | | | | | | | | | | |
| Chromosome | start | end | strand | MM | target_seq | PAM | alignment | distance | position | gene name | gene id |
| GU305914.1 | 201367 | 201389 | + | 0 | GTGTCGTAGCGCCCCGCCTC | CGG | \|\|\|\|\|\|\|\|[\|\|\|\|\|\|\|\|\|\|\|\|]PAM | NA | Intergenic | NA | NA |
| T15393 | TAGCGCCCCGCCTCCGGCGACGG | 1000 | | | | | | | | | |
| Oligo fwd | CACCTAGCGCCCCGCCTCCGGCGA | | | | | | | | | | |
| Oligo rev | AAACTCGCCGGAGGCGGGGCGCTA | | | | | | | | | | |
| Chromosome | start | end | strand | MM | target_seq | PAM | alignment | distance | position | gene name | gene id |
| GU305914.1 | 201373 | 201395 | + | 0 | TAGCGCCCCGCCTCCGGCGA | CGG | \|\|\|\|\|\|\|\|[\|\|\|\|\|\|\|\|\|\|\|\|]PAM | NA | Intergenic | NA | NA |
| T15394 | CGCCCCGCCTCCGGCGACGGAGG | 1000 | | | | | | | | | |
| Oligo fwd | CACCCGCCCCGCCTCCGGCGACGG | | | | | | | | | | |
| Oligo rev | AAACCCGTCGCCGGAGGCGGGGCG | | | | | | | | | | |
| Chromosome | start | end | strand | MM | target_seq | PAM | alignment | distance | position | gene name | gene id |
| GU305914.1 | 201376 | 201398 | + | 0 | CGCCCCGCCTCCGGCGACGG | AGG | \|\|\|\|\|\|\|\|[\|\|\|\|\|\|\|\|\|\|\|\|]PAM | NA | Intergenic | NA | NA |
| T15395 | CCCGCCTCCGGCGACGGAGGCGG | 1000 | | | | | | | | | |
| Oligo fwd | CACCCCCGCCTCCGGCGACGGAGG | | | | | | | | | | |
| Oligo rev | AAACCCTCCGTCGCCGGAGGCGGG | | | | | | | | | | |
| Chromosome | start | end | strand | MM | target_seq | PAM | alignment | distance | position | gene name | gene id |
| GU305914.1 | 201379 | 201401 | + | 0 | CCCGCCTCCGGCGACGGAGG | CGG | \|\|\|\|\|\|\|\|[\|\|\|\|\|\|\|\|\|\|\|\|]PAM | NA | Intergenic | NA | NA |
| GU305914.1 | 201380 | 201402 | - | 3 | GCCGCCTCCGTCGCCGGAGG | CGG | -\|\|\|\|\|\|\|[\|\|-\|\|-\|\|\|\|\|\|]PAM | NA | Intergenic | NA | NA |
| T15396 | GCCTCCGGCGACGGAGGCGGCGG | 1000 | | | | | | | | | |
| Oligo fwd | CACCGCCTCCGGCGACGGAGGCGG | | | | | | | | | | |
| Oligo rev | AAACCCGCCTCCGTCGCCGGAGGC | | | | | | | | | | |
| Chromosome | start | end | strand | MM | target_seq | PAM | alignment | distance | position | gene name | gene id |
| GU305914.1 | 201382 | 201404 | + | 0 | GCCTCCGGCGACGGAGGCGG | CGG | \|\|\|\|\|\|\|\|[\|\|\|\|\|\|\|\|\|\|\|\|]PAM | NA | Intergenic | NA | NA |
| T15397 | GGCGGCTTCTCCTTGAACCGAGG | 1000 | | | | | | | | | |
| Oligo fwd | CACCGGCGGCTTCTCCTTGAACCG | | | | | | | | | | |
| Oligo rev | AAACCGGTTCAAGGAGAAGCCGCC | | | | | | | | | | |
| Chromosome | start | end | strand | MM | target_seq | PAM | alignment | distance | position | gene name | gene id |
| GU305914.1 | 201400 | 201422 | + | 0 | GGCGGCTTCTCCTTGAACCG | AGG | \|\|\|\|\|\|\|\|[\|\|\|\|\|\|\|\|\|\|\|\|]PAM | NA | Intergenic | NA | NA |
| T15398 | GTCCGCCTTGTCCGCCGACTCGG | 1000 | | | | | | | | | |
| Oligo fwd | CACCGTCCGCCTTGTCCGCCGACT | | | | | | | | | | |
| Oligo rev | AAACAGTCGGCGGACAAGGCGGAC | | | | | | | | | | |
| Chromosome | start | end | strand | MM | target_seq | PAM | alignment | distance | position | gene name | gene id |
| GU305914.1 | 201429 | 201451 | + | 0 | GTCCGCCTTGTCCGCCGACT | CGG | \|\|\|\|\|\|\|\|[\|\|\|\|\|\|\|\|\|\|\|\|]PAM | NA | Intergenic | NA | NA |
| T15399 | TCCGCCTTGTCCGCCGACTCGGG | 1000 | | | | | | | | | |
| Oligo fwd | CACCTCCGCCTTGTCCGCCGACTC | | | | | | | | | | |
| Oligo rev | AAACGAGTCGGCGGACAAGGCGGA | | | | | | | | | | |
| Chromosome | start | end | strand | MM | target_seq | PAM | alignment | distance | position | gene name | gene id |
| GU305914.1 | 201430 | 201452 | + | 0 | TCCGCCTTGTCCGCCGACTC | GGG | \|\|\|\|\|\|\|\|[\|\|\|\|\|\|\|\|\|\|\|\|]PAM | NA | Intergenic | NA | NA |
| T15400 | CCGCCTTGTCCGCCGACTCGGGG | 1000 | | | | | | | | | |
| Oligo fwd | CACCCCGCCTTGTCCGCCGACTCG | | | | | | | | | | |
| Oligo rev | AAACCGAGTCGGCGGACAAGGCGG | | | | | | | | | | |
| Chromosome | start | end | strand | MM | target_seq | PAM | alignment | distance | position | gene name | gene id |
| GU305914.1 | 201431 | 201453 | + | 0 | CCGCCTTGTCCGCCGACTCG | GGG | \|\|\|\|\|\|\|\|[\|\|\|\|\|\|\|\|\|\|\|\|]PAM | NA | Intergenic | NA | NA |
| T15401 | GCAGCAGCGCCGACACCAAGCGG | 1000 | | | | | | | | | |
| Oligo fwd | CACCGCAGCAGCGCCGACACCAAG | | | | | | | | | | |
| Oligo rev | AAACCTTGGTGTCGGCGCTGCTGC | | | | | | | | | | |
| Chromosome | start | end | strand | MM | target_seq | PAM | alignment | distance | position | gene name | gene id |
| GU305914.1 | 201473 | 201495 | + | 0 | GCAGCAGCGCCGACACCAAG | CGG | \|\|\|\|\|\|\|\|[\|\|\|\|\|\|\|\|\|\|\|\|]PAM | NA | Intergenic | NA | NA |
| T15402 | GACACCAAGCGGCGCGTCCGCGG | 1000 | | | | | | | | | |
| Oligo fwd | CACCGACACCAAGCGGCGCGTCCG | | | | | | | | | | |
| Oligo rev | AAACCGGACGCGCCGCTTGGTGTC | | | | | | | | | | |
| Chromosome | start | end | strand | MM | target_seq | PAM | alignment | distance | position | gene name | gene id |
| GU305914.1 | 201484 | 201506 | + | 0 | GACACCAAGCGGCGCGTCCG | CGG | \|\|\|\|\|\|\|\|[\|\|\|\|\|\|\|\|\|\|\|\|]PAM | NA | Intergenic | NA | NA |
| T15403 | CAAGCGGCGCGTCCGCGGCTCGG | 1000 | | | | | | | | | |
| Oligo fwd | CACCCAAGCGGCGCGTCCGCGGCT | | | | | | | | | | |
| Oligo rev | AAACAGCCGCGGACGCGCCGCTTG | | | | | | | | | | |
| Chromosome | start | end | strand | MM | target_seq | PAM | alignment | distance | position | gene name | gene id |
| GU305914.1 | 201489 | 201511 | + | 0 | CAAGCGGCGCGTCCGCGGCT | CGG | \|\|\|\|\|\|\|\|[\|\|\|\|\|\|\|\|\|\|\|\|]PAM | NA | Intergenic | NA | NA |
| T15404 | AAGCGGCGCGTCCGCGGCTCGGG | 1000 | | | | | | | | | |
| Oligo fwd | CACCAAGCGGCGCGTCCGCGGCTC | | | | | | | | | | |
| Oligo rev | AAACGAGCCGCGGACGCGCCGCTT | | | | | | | | | | |
| Chromosome | start | end | strand | MM | target_seq | PAM | alignment | distance | position | gene name | gene id |
| GU305914.1 | 201490 | 201512 | + | 0 | AAGCGGCGCGTCCGCGGCTC | GGG | \|\|\|\|\|\|\|\|[\|\|\|\|\|\|\|\|\|\|\|\|]PAM | NA | Intergenic | NA | NA |

<table>
<tr><td>T15405</td><td>CGGCGCGTCCGCGGCTCGGGCGG</td><td>1000</td><td></td><td></td><td></td><td></td><td></td><td></td><td></td><td></td><td></td></tr>
<tr><td>Oligo fwd</td><td>CACCCGGCGCGTCCGCGGCTCGGG</td><td></td><td></td><td></td><td></td><td></td><td></td><td></td><td></td><td></td><td></td></tr>
<tr><td>Oligo rev</td><td>AAACCCCGAGCCGCGGACGCGCCG</td><td></td><td></td><td></td><td></td><td></td><td></td><td></td><td></td><td></td><td></td></tr>
<tr><td>Chromosome</td><td>start</td><td>end</td><td>strand</td><td>MM</td><td>target_seq</td><td>PAM</td><td>alignment</td><td>distance</td><td>position</td><td>gene name</td><td>gene id</td></tr>
<tr><td>GU305914.1</td><td>201493</td><td>201515</td><td>+</td><td>0</td><td>CGGCGCGTCCGCGGCTCGGG</td><td>CGG</td><td>||||||||[||||||||||||]PAM</td><td>NA</td><td>Intergenic</td><td>NA</td><td>NA</td></tr>
<tr><td>T15406</td><td>CGTCCGCGGCTCGGGCGGCGCGG</td><td>1000</td><td></td><td></td><td></td><td></td><td></td><td></td><td></td><td></td><td></td></tr>
<tr><td>Oligo fwd</td><td>CACCCGTCCGCGGCTCGGGCGGCG</td><td></td><td></td><td></td><td></td><td></td><td></td><td></td><td></td><td></td><td></td></tr>
<tr><td>Oligo rev</td><td>AAACCGCCGCCCGAGCCGCGGACG</td><td></td><td></td><td></td><td></td><td></td><td></td><td></td><td></td><td></td><td></td></tr>
<tr><td>Chromosome</td><td>start</td><td>end</td><td>strand</td><td>MM</td><td>target_seq</td><td>PAM</td><td>alignment</td><td>distance</td><td>position</td><td>gene name</td><td>gene id</td></tr>
<tr><td>GU305914.1</td><td>201498</td><td>201520</td><td>+</td><td>0</td><td>CGTCCGCGGCTCGGGCGGCG</td><td>CGG</td><td>||||||||[||||||||||||]PAM</td><td>NA</td><td>Intergenic</td><td>NA</td><td>NA</td></tr>
<tr><td>T15407</td><td>GCGCGGCGACGTGCGTGCAGAGG</td><td>1000</td><td></td><td></td><td></td><td></td><td></td><td></td><td></td><td></td><td></td></tr>
<tr><td>Oligo fwd</td><td>CACCGCGCGGCGACGTGCGTGCAG</td><td></td><td></td><td></td><td></td><td></td><td></td><td></td><td></td><td></td><td></td></tr>
<tr><td>Oligo rev</td><td>AAACCTGCACGCACGTCGCCGCGC</td><td></td><td></td><td></td><td></td><td></td><td></td><td></td><td></td><td></td><td></td></tr>
<tr><td>Chromosome</td><td>start</td><td>end</td><td>strand</td><td>MM</td><td>target_seq</td><td>PAM</td><td>alignment</td><td>distance</td><td>position</td><td>gene name</td><td>gene id</td></tr>
<tr><td>GU305914.1</td><td>201515</td><td>201537</td><td>+</td><td>0</td><td>GCGCGGCGACGTGCGTGCAG</td><td>AGG</td><td>||||||||[||||||||||||]PAM</td><td>NA</td><td>Intergenic</td><td>NA</td><td>NA</td></tr>
<tr><td>T15408</td><td>CCGTCACCTTGAGCCTCGACAGG</td><td>1000</td><td></td><td></td><td></td><td></td><td></td><td></td><td></td><td></td><td></td></tr>
<tr><td>Oligo fwd</td><td>CACCCCGTCACCTTGAGCCTCGAC</td><td></td><td></td><td></td><td></td><td></td><td></td><td></td><td></td><td></td><td></td></tr>
<tr><td>Oligo rev</td><td>AAACGTCGAGGCTCAAGGTGACGG</td><td></td><td></td><td></td><td></td><td></td><td></td><td></td><td></td><td></td><td></td></tr>
<tr><td>Chromosome</td><td>start</td><td>end</td><td>strand</td><td>MM</td><td>target_seq</td><td>PAM</td><td>alignment</td><td>distance</td><td>position</td><td>gene name</td><td>gene id</td></tr>
<tr><td>GU305914.1</td><td>201563</td><td>201585</td><td>+</td><td>0</td><td>CCGTCACCTTGAGCCTCGAC</td><td>AGG</td><td>||||||||[||||||||||||]PAM</td><td>NA</td><td>Intergenic</td><td>NA</td><td>NA</td></tr>
<tr><td>T15409</td><td>CCTTGAGCCTCGACAGGCGCAGG</td><td>1000</td><td></td><td></td><td></td><td></td><td></td><td></td><td></td><td></td><td></td></tr>
<tr><td>Oligo fwd</td><td>CACCCCTTGAGCCTCGACAGGCGC</td><td></td><td></td><td></td><td></td><td></td><td></td><td></td><td></td><td></td><td></td></tr>
<tr><td>Oligo rev</td><td>AAACGCGCCTGTCGAGGCTCAAGG</td><td></td><td></td><td></td><td></td><td></td><td></td><td></td><td></td><td></td><td></td></tr>
<tr><td>Chromosome</td><td>start</td><td>end</td><td>strand</td><td>MM</td><td>target_seq</td><td>PAM</td><td>alignment</td><td>distance</td><td>position</td><td>gene name</td><td>gene id</td></tr>
<tr><td>GU305914.1</td><td>201569</td><td>201591</td><td>+</td><td>0</td><td>CCTTGAGCCTCGACAGGCGC</td><td>AGG</td><td>||||||||[||||||||||||]PAM</td><td>NA</td><td>Intergenic</td><td>NA</td><td>NA</td></tr>
<tr><td>T15410</td><td>GAGCCTCGACAGGCGCAGGAAGG</td><td>1000</td><td></td><td></td><td></td><td></td><td></td><td></td><td></td><td></td><td></td></tr>
<tr><td>Oligo fwd</td><td>CACCGAGCCTCGACAGGCGCAGGA</td><td></td><td></td><td></td><td></td><td></td><td></td><td></td><td></td><td></td><td></td></tr>
<tr><td>Oligo rev</td><td>AAACTCCTGCGCCTGTCGAGGCTC</td><td></td><td></td><td></td><td></td><td></td><td></td><td></td><td></td><td></td><td></td></tr>
<tr><td>Chromosome</td><td>start</td><td>end</td><td>strand</td><td>MM</td><td>target_seq</td><td>PAM</td><td>alignment</td><td>distance</td><td>position</td><td>gene name</td><td>gene id</td></tr>
<tr><td>GU305914.1</td><td>201573</td><td>201595</td><td>+</td><td>0</td><td>GAGCCTCGACAGGCGCAGGA</td><td>AGG</td><td>||||||||[||||||||||||]PAM</td><td>NA</td><td>Intergenic</td><td>NA</td><td>NA</td></tr>
<tr><td>T15411</td><td>CAGGAAGGCGTCCGCCACCGCGG</td><td>1000</td><td></td><td></td><td></td><td></td><td></td><td></td><td></td><td></td><td></td></tr>
<tr><td>Oligo fwd</td><td>CACCCAGGAAGGCGTCCGCCACCG</td><td></td><td></td><td></td><td></td><td></td><td></td><td></td><td></td><td></td><td></td></tr>
<tr><td>Oligo rev</td><td>AAACCGGTGGCGGACGCCTTCCTG</td><td></td><td></td><td></td><td></td><td></td><td></td><td></td><td></td><td></td><td></td></tr>
<tr><td>Chromosome</td><td>start</td><td>end</td><td>strand</td><td>MM</td><td>target_seq</td><td>PAM</td><td>alignment</td><td>distance</td><td>position</td><td>gene name</td><td>gene id</td></tr>
<tr><td>GU305914.1</td><td>201588</td><td>201610</td><td>+</td><td>0</td><td>CAGGAAGGCGTCCGCCACCG</td><td>CGG</td><td>||||||||[||||||||||||]PAM</td><td>NA</td><td>Intergenic</td><td>NA</td><td>NA</td></tr>
<tr><td>T15412</td><td>CGCCACCGCGGTCTCCTCCTCGG</td><td>1000</td><td></td><td></td><td></td><td></td><td></td><td></td><td></td><td></td><td></td></tr>
<tr><td>Oligo fwd</td><td>CACCCGCCACCGCGGTCTCCTCCT</td><td></td><td></td><td></td><td></td><td></td><td></td><td></td><td></td><td></td><td></td></tr>
<tr><td>Oligo rev</td><td>AAACAGGAGGAGACCGCGGTGGCG</td><td></td><td></td><td></td><td></td><td></td><td></td><td></td><td></td><td></td><td></td></tr>
<tr><td>Chromosome</td><td>start</td><td>end</td><td>strand</td><td>MM</td><td>target_seq</td><td>PAM</td><td>alignment</td><td>distance</td><td>position</td><td>gene name</td><td>gene id</td></tr>
<tr><td>GU305914.1</td><td>201600</td><td>201622</td><td>+</td><td>0</td><td>CGCCACCGCGGTCTCCTCCT</td><td>CGG</td><td>||||||||[||||||||||||]PAM</td><td>NA</td><td>Intergenic</td><td>NA</td><td>NA</td></tr>
<tr><td>GU305914.1</td><td>165100</td><td>165122</td><td>-</td><td>3</td><td>GGCCATCGCGGTTTCCTCCT</td><td>CGG</td><td>-||||-||[||||-|||||||]PAM</td><td>NA</td><td>Intergenic</td><td>NA</td><td>NA</td></tr>
<tr><td>GU305914.1</td><td>32322</td><td>32344</td><td>+</td><td>4</td><td>CGCGTCCTCGGTCTCCTCGT</td><td>CGG</td><td>|||--||-[||||||||||-|]PAM</td><td>NA</td><td>Intergenic</td><td>NA</td><td>NA</td></tr>
<tr><td>T15413</td><td>CCGCGGTCTCCTCCTCGGACAGG</td><td>1000</td><td></td><td></td><td></td><td></td><td></td><td></td><td></td><td></td><td></td></tr>
<tr><td>Oligo fwd</td><td>CACCCCGCGGTCTCCTCCTCGGAC</td><td></td><td></td><td></td><td></td><td></td><td></td><td></td><td></td><td></td><td></td></tr>
<tr><td>Oligo rev</td><td>AAACGTCCGAGGAGGAGACCGCGG</td><td></td><td></td><td></td><td></td><td></td><td></td><td></td><td></td><td></td><td></td></tr>
<tr><td>Chromosome</td><td>start</td><td>end</td><td>strand</td><td>MM</td><td>target_seq</td><td>PAM</td><td>alignment</td><td>distance</td><td>position</td><td>gene name</td><td>gene id</td></tr>
<tr><td>GU305914.1</td><td>201605</td><td>201627</td><td>+</td><td>0</td><td>CCGCGGTCTCCTCCTCGGAC</td><td>AGG</td><td>||||||||[||||||||||||]PAM</td><td>NA</td><td>Intergenic</td><td>NA</td><td>NA</td></tr>
<tr><td>T15414</td><td>GGACAGGATCTCCGCCCTGCCGG</td><td>1000</td><td></td><td></td><td></td><td></td><td></td><td></td><td></td><td></td><td></td></tr>
<tr><td>Oligo fwd</td><td>CACCGGACAGGATCTCCGCCCTGC</td><td></td><td></td><td></td><td></td><td></td><td></td><td></td><td></td><td></td><td></td></tr>
<tr><td>Oligo rev</td><td>AAACGCAGGGCGGAGATCCTGTCC</td><td></td><td></td><td></td><td></td><td></td><td></td><td></td><td></td><td></td><td></td></tr>
<tr><td>Chromosome</td><td>start</td><td>end</td><td>strand</td><td>MM</td><td>target_seq</td><td>PAM</td><td>alignment</td><td>distance</td><td>position</td><td>gene name</td><td>gene id</td></tr>
<tr><td>GU305914.1</td><td>201621</td><td>201643</td><td>+</td><td>0</td><td>GGACAGGATCTCCGCCCTGC</td><td>CGG</td><td>||||||||[||||||||||||]PAM</td><td>NA</td><td>Intergenic</td><td>NA</td><td>NA</td></tr>
<tr><td>T15415</td><td>GACAGGATCTCCGCCCTGCCGGG</td><td>1000</td><td></td><td></td><td></td><td></td><td></td><td></td><td></td><td></td><td></td></tr>
<tr><td>Oligo fwd</td><td>CACCGACAGGATCTCCGCCCTGCC</td><td></td><td></td><td></td><td></td><td></td><td></td><td></td><td></td><td></td><td></td></tr>
<tr><td>Oligo rev</td><td>AAACGGCAGGGCGGAGATCCTGTC</td><td></td><td></td><td></td><td></td><td></td><td></td><td></td><td></td><td></td><td></td></tr>
<tr><td>Chromosome</td><td>start</td><td>end</td><td>strand</td><td>MM</td><td>target_seq</td><td>PAM</td><td>alignment</td><td>distance</td><td>position</td><td>gene name</td><td>gene id</td></tr>
<tr><td>GU305914.1</td><td>201622</td><td>201644</td><td>+</td><td>0</td><td>GACAGGATCTCCGCCCTGCC</td><td>GGG</td><td>||||||||[||||||||||||]PAM</td><td>NA</td><td>Intergenic</td><td>NA</td><td>NA</td></tr>
<tr><td>T15416</td><td>CTGCGCGAGCTGCGCCTCGCCGG</td><td>1000</td><td></td><td></td><td></td><td></td><td></td><td></td><td></td><td></td><td></td></tr>
<tr><td>Oligo fwd</td><td>CACCCTGCGCGAGCTGCGCCTCGC</td><td></td><td></td><td></td><td></td><td></td><td></td><td></td><td></td><td></td><td></td></tr>
<tr><td>Oligo rev</td><td>AAACGCGAGGCGCAGCTCGCGCAG</td><td></td><td></td><td></td><td></td><td></td><td></td><td></td><td></td><td></td><td></td></tr>
<tr><td>Chromosome</td><td>start</td><td>end</td><td>strand</td><td>MM</td><td>target_seq</td><td>PAM</td><td>alignment</td><td>distance</td><td>position</td><td>gene name</td><td>gene id</td></tr>
<tr><td>GU305914.1</td><td>201666</td><td>201688</td><td>+</td><td>0</td><td>CTGCGCGAGCTGCGCCTCGC</td><td>CGG</td><td>||||||||[||||||||||||]PAM</td><td>NA</td><td>Intergenic</td><td>NA</td><td>NA</td></tr>
<tr><td>T15417</td><td>TGCGCGAGCTGCGCCTCGCCGGG</td><td>1000</td><td></td><td></td><td></td><td></td><td></td><td></td><td></td><td></td><td></td></tr>
<tr><td>Oligo fwd</td><td>CACCTGCGCGAGCTGCGCCTCGCC</td><td></td><td></td><td></td><td></td><td></td><td></td><td></td><td></td><td></td><td></td></tr>
<tr><td>Oligo rev</td><td>AAACGGCGAGGCGCAGCTCGCGCA</td><td></td><td></td><td></td><td></td><td></td><td></td><td></td><td></td><td></td><td></td></tr>
<tr><td>Chromosome</td><td>start</td><td>end</td><td>strand</td><td>MM</td><td>target_seq</td><td>PAM</td><td>alignment</td><td>distance</td><td>position</td><td>gene name</td><td>gene id</td></tr>
<tr><td>GU305914.1</td><td>201667</td><td>201689</td><td>+</td><td>0</td><td>TGCGCGAGCTGCGCCTCGCC</td><td>GGG</td><td>||||||||[||||||||||||]PAM</td><td>NA</td><td>Intergenic</td><td>NA</td><td>NA</td></tr>
<tr><td>T15418</td><td>GCGCGAGCTGCGCCTCGCCGGGG</td><td>1000</td><td></td><td></td><td></td><td></td><td></td><td></td><td></td><td></td><td></td></tr>
<tr><td>Oligo fwd</td><td>CACCGCGCGAGCTGCGCCTCGCCG</td><td></td><td></td><td></td><td></td><td></td><td></td><td></td><td></td><td></td><td></td></tr>
<tr><td>Oligo rev</td><td>AAACCGGCGAGGCGCAGCTCGCGC</td><td></td><td></td><td></td><td></td><td></td><td></td><td></td><td></td><td></td><td></td></tr>
<tr><td>Chromosome</td><td>start</td><td>end</td><td>strand</td><td>MM</td><td>target_seq</td><td>PAM</td><td>alignment</td><td>distance</td><td>position</td><td>gene name</td><td>gene id</td></tr>
<tr><td>GU305914.1</td><td>201668</td><td>201690</td><td>+</td><td>0</td><td>GCGCGAGCTGCGCCTCGCCG</td><td>GGG</td><td>||||||||[||||||||||||]PAM</td><td>NA</td><td>Intergenic</td><td>NA</td><td>NA</td></tr>
<tr><td>T15419</td><td>CGCGAGCTGCGCCTCGCCGGGGG</td><td>1000</td><td></td><td></td><td></td><td></td><td></td><td></td><td></td><td></td><td></td></tr>
<tr><td>Oligo fwd</td><td>CACCCGCGAGCTGCGCCTCGCCGG</td><td></td><td></td><td></td><td></td><td></td><td></td><td></td><td></td><td></td><td></td></tr>
</table>

Oligo rev AAACCCGGCGAGGCGCAGCTCGCG
Chromosome start end strand MM target_seq PAM alignment distance position gene name gene id
GU305914.1 201669 201691 + 0 CGCGAGCTGCGCCTCGCCGG GGG ||||||||[||||||||||||]PAM NA Intergenic NA NA

T15420 GCGAGCTGCGCCTCGCCGGGGGG 1000
Oligo fwd CACCGCGAGCTGCGCCTCGCCGGG
Oligo rev AAACCCCGGCGAGGCGCAGCTCGC
Chromosome start end strand MM target_seq PAM alignment distance position gene name gene id
GU305914.1 201670 201692 + 0 GCGAGCTGCGCCTCGCCGGG GGG ||||||||[||||||||||||]PAM NA Intergenic NA NA

T15421 GTCGACGCACAGATGCGTCGCGG 1000
Oligo fwd CACCGTCGACGCACAGATGCGTCG
Oligo rev AAACCGACGCATCTGTGCGTCGAC
Chromosome start end strand MM target_seq PAM alignment distance position gene name gene id
GU305914.1 201699 201721 + 0 GTCGACGCACAGATGCGTCG CGG ||||||||[||||||||||||]PAM NA Intergenic NA NA

T15422 GACGCACAGATGCGTCGCGGCGG 1000
Oligo fwd CACCGACGCACAGATGCGTCGCGG
Oligo rev AAACCCGCGACGCATCTGTGCGTC
Chromosome start end strand MM target_seq PAM alignment distance position gene name gene id
GU305914.1 201702 201724 + 0 GACGCACAGATGCGTCGCGG CGG ||||||||[||||||||||||]PAM NA Intergenic NA NA

T15423 ACGCACAGATGCGTCGCGGCGGG 1000
Oligo fwd CACCACGCACAGATGCGTCGCGGC
Oligo rev AAACGCCGCGACGCATCTGTGCGT
Chromosome start end strand MM target_seq PAM alignment distance position gene name gene id
GU305914.1 201703 201725 + 0 ACGCACAGATGCGTCGCGGC GGG ||||||||[||||||||||||]PAM NA Intergenic NA NA

T15424 ACAGATGCGTCGCGGCGGGTCGG 1000
Oligo fwd CACCACAGATGCGTCGCGGCGGGT
Oligo rev AAACACCCGCCGCGACGCATCTGT
Chromosome start end strand MM target_seq PAM alignment distance position gene name gene id
GU305914.1 201707 201729 + 0 ACAGATGCGTCGCGGCGGGT CGG ||||||||[||||||||||||]PAM NA Intergenic NA NA

T15425 CAGATGCGTCGCGGCGGGTCGGG 1000
Oligo fwd CACCCAGATGCGTCGCGGCGGGTC
Oligo rev AAACGACCCGCCGCGACGCATCTG
Chromosome start end strand MM target_seq PAM alignment distance position gene name gene id
GU305914.1 201708 201730 + 0 CAGATGCGTCGCGGCGGGTC GGG ||||||||[||||||||||||]PAM NA Intergenic NA NA

T15426 GCGTCGCGGCGGGTCGGGAGTGG 1000
Oligo fwd CACCGCGTCGCGGCGGGTCGGGAG
Oligo rev AAACCTCCCGACCCGCCGCGACGC
Chromosome start end strand MM target_seq PAM alignment distance position gene name gene id
GU305914.1 201713 201735 + 0 GCGTCGCGGCGGGTCGGGAG TGG ||||||||[||||||||||||]PAM NA Intergenic NA NA

T15427 GGGAGTGGAACCACAGCTCCTGG 1000
Oligo fwd CACCGGGAGTGGAACCACAGCTCC
Oligo rev AAACGGAGCTGTGGTTCCACTCCC
Chromosome start end strand MM target_seq PAM alignment distance position gene name gene id
GU305914.1 201728 201750 + 0 GGGAGTGGAACCACAGCTCC TGG ||||||||[||||||||||||]PAM NA Intergenic NA NA

T15428 GTGGAACCACAGCTCCTGGTCGG 1000
Oligo fwd CACCGTGGAACCACAGCTCCTGGT
Oligo rev AAACACCAGGAGCTGTGGTTCCAC
Chromosome start end strand MM target_seq PAM alignment distance position gene name gene id
GU305914.1 201732 201754 + 0 GTGGAACCACAGCTCCTGGT CGG ||||||||[||||||||||||]PAM NA Intergenic NA NA

T15429 TGGAACCACAGCTCCTGGTCGGG 1000
Oligo fwd CACCTGGAACCACAGCTCCTGGTC
Oligo rev AAACGACCAGGAGCTGTGGTTCCA
Chromosome start end strand MM target_seq PAM alignment distance position gene name gene id
GU305914.1 201733 201755 + 0 TGGAACCACAGCTCCTGGTC GGG ||||||||[||||||||||||]PAM NA Intergenic NA NA

T15430 AACCACAGCTCCTGGTCGGGTGG 1000
Oligo fwd CACCAACCACAGCTCCTGGTCGGG
Oligo rev AAACCCCGACCAGGAGCTGTGGTT
Chromosome start end strand MM target_seq PAM alignment distance position gene name gene id
GU305914.1 201736 201758 + 0 AACCACAGCTCCTGGTCGGG TGG ||||||||[||||||||||||]PAM NA Intergenic NA NA

T15431 ACCACAGCTCCTGGTCGGGTGGG 1000
Oligo fwd CACCACCACAGCTCCTGGTCGGGT
Oligo rev AAACACCCGACCAGGAGCTGTGGT
Chromosome start end strand MM target_seq PAM alignment distance position gene name gene id
GU305914.1 201737 201759 + 0 ACCACAGCTCCTGGTCGGGT GGG ||||||||[||||||||||||]PAM NA Intergenic NA NA

T15432 CCACAGCTCCTGGTCGGGTGGGG 1000
Oligo fwd CACCCCACAGCTCCTGGTCGGGTG
Oligo rev AAACCACCCGACCAGGAGCTGTGG
Chromosome start end strand MM target_seq PAM alignment distance position gene name gene id
GU305914.1 201738 201760 + 0 CCACAGCTCCTGGTCGGGTG GGG ||||||||[||||||||||||]PAM NA Intergenic NA NA

T15433 CACAGCTCCTGGTCGGGTGGGGG 1000
Oligo fwd CACCCACAGCTCCTGGTCGGGTGG
Oligo rev AAACCCACCCGACCAGGAGCTGTG
Chromosome start end strand MM target_seq PAM alignment distance position gene name gene id
GU305914.1 201739 201761 + 0 CACAGCTCCTGGTCGGGTGG GGG ||||||||[||||||||||||]PAM NA Intergenic NA NA

T15434 GGTCGGGTGGGGGCTCGAGCCGG 1000

<table>
<tr><td>Oligo fwd</td><td>CACCGGTCGGGTGGGGGCTCGAGC</td><td></td><td></td><td></td><td></td><td></td><td></td><td></td><td></td><td></td><td></td></tr>
<tr><td>Oligo rev</td><td>AAACGCTCGAGCCCCCACCCGACC</td><td></td><td></td><td></td><td></td><td></td><td></td><td></td><td></td><td></td><td></td></tr>
<tr><td>Chromosome</td><td>start</td><td>end</td><td>strand</td><td>MM</td><td>target_seq</td><td>PAM</td><td>alignment</td><td>distance</td><td>position</td><td>gene name</td><td>gene id</td></tr>
<tr><td>GU305914.1</td><td>201749</td><td>201771</td><td>+</td><td>0</td><td>GGTCGGGTGGGGGCTCGAGC</td><td>CGG</td><td>||||||||[||||||||||||]PAM</td><td>NA</td><td>Intergenic</td><td>NA</td><td>NA</td></tr>
<tr><td>T15435</td><td>GGACGCTCCCCGCGAGATCCCGG</td><td>1000</td><td></td><td></td><td></td><td></td><td></td><td></td><td></td><td></td><td></td></tr>
<tr><td>Oligo fwd</td><td>CACCGGACGCTCCCCGCGAGATCC</td><td></td><td></td><td></td><td></td><td></td><td></td><td></td><td></td><td></td><td></td></tr>
<tr><td>Oligo rev</td><td>AAACGGATCTCGCGGGGAGCGTCC</td><td></td><td></td><td></td><td></td><td></td><td></td><td></td><td></td><td></td><td></td></tr>
<tr><td>Chromosome</td><td>start</td><td>end</td><td>strand</td><td>MM</td><td>target_seq</td><td>PAM</td><td>alignment</td><td>distance</td><td>position</td><td>gene name</td><td>gene id</td></tr>
<tr><td>GU305914.1</td><td>201770</td><td>201792</td><td>+</td><td>0</td><td>GGACGCTCCCCGCGAGATCC</td><td>CGG</td><td>||||||||[||||||||||||]PAM</td><td>NA</td><td>Intergenic</td><td>NA</td><td>NA</td></tr>
<tr><td>T15436</td><td>CGAGATCCCGGTCGACCCGACGG</td><td>1000</td><td></td><td></td><td></td><td></td><td></td><td></td><td></td><td></td><td></td></tr>
<tr><td>Oligo fwd</td><td>CACCCGAGATCCCGGTCGACCCGA</td><td></td><td></td><td></td><td></td><td></td><td></td><td></td><td></td><td></td><td></td></tr>
<tr><td>Oligo rev</td><td>AAACTCGGGTCGACCGGGATCTCG</td><td></td><td></td><td></td><td></td><td></td><td></td><td></td><td></td><td></td><td></td></tr>
<tr><td>Chromosome</td><td>start</td><td>end</td><td>strand</td><td>MM</td><td>target_seq</td><td>PAM</td><td>alignment</td><td>distance</td><td>position</td><td>gene name</td><td>gene id</td></tr>
<tr><td>GU305914.1</td><td>201782</td><td>201804</td><td>+</td><td>0</td><td>CGAGATCCCGGTCGACCCGA</td><td>CGG</td><td>||||||||[||||||||||||]PAM</td><td>NA</td><td>Intergenic</td><td>NA</td><td>NA</td></tr>
<tr><td>T15437</td><td>CCGGTCGACCCGACGGCCCGCGG</td><td>1000</td><td></td><td></td><td></td><td></td><td></td><td></td><td></td><td></td><td></td></tr>
<tr><td>Oligo fwd</td><td>CACCCCGGTCGACCCGACGGCCCG</td><td></td><td></td><td></td><td></td><td></td><td></td><td></td><td></td><td></td><td></td></tr>
<tr><td>Oligo rev</td><td>AAACCGGGCCGTCGGGTCGACCGG</td><td></td><td></td><td></td><td></td><td></td><td></td><td></td><td></td><td></td><td></td></tr>
<tr><td>Chromosome</td><td>start</td><td>end</td><td>strand</td><td>MM</td><td>target_seq</td><td>PAM</td><td>alignment</td><td>distance</td><td>position</td><td>gene name</td><td>gene id</td></tr>
<tr><td>GU305914.1</td><td>201789</td><td>201811</td><td>+</td><td>0</td><td>CCGGTCGACCCGACGGCCCG</td><td>CGG</td><td>||||||||[||||||||||||]PAM</td><td>NA</td><td>Intergenic</td><td>NA</td><td>NA</td></tr>
<tr><td>T15438</td><td>TCTTCAGCAGTCCCATGCGACGG</td><td>1000</td><td></td><td></td><td></td><td></td><td></td><td></td><td></td><td></td><td></td></tr>
<tr><td>Oligo fwd</td><td>CACCTCTTCAGCAGTCCCATGCGA</td><td></td><td></td><td></td><td></td><td></td><td></td><td></td><td></td><td></td><td></td></tr>
<tr><td>Oligo rev</td><td>AAACTCGCATGGGACTGCTGAAGA</td><td></td><td></td><td></td><td></td><td></td><td></td><td></td><td></td><td></td><td></td></tr>
<tr><td>Chromosome</td><td>start</td><td>end</td><td>strand</td><td>MM</td><td>target_seq</td><td>PAM</td><td>alignment</td><td>distance</td><td>position</td><td>gene name</td><td>gene id</td></tr>
<tr><td>GU305914.1</td><td>201818</td><td>201840</td><td>+</td><td>0</td><td>TCTTCAGCAGTCCCATGCGA</td><td>CGG</td><td>||||||||[||||||||||||]PAM</td><td>NA</td><td>Intergenic</td><td>NA</td><td>NA</td></tr>
<tr><td>T15439</td><td>TCCCATGCGACGGAGCTCCGCGG</td><td>1000</td><td></td><td></td><td></td><td></td><td></td><td></td><td></td><td></td><td></td></tr>
<tr><td>Oligo fwd</td><td>CACCTCCCATGCGACGGAGCTCCG</td><td></td><td></td><td></td><td></td><td></td><td></td><td></td><td></td><td></td><td></td></tr>
<tr><td>Oligo rev</td><td>AAACCGGAGCTCCGTCGCATGGGA</td><td></td><td></td><td></td><td></td><td></td><td></td><td></td><td></td><td></td><td></td></tr>
<tr><td>Chromosome</td><td>start</td><td>end</td><td>strand</td><td>MM</td><td>target_seq</td><td>PAM</td><td>alignment</td><td>distance</td><td>position</td><td>gene name</td><td>gene id</td></tr>
<tr><td>GU305914.1</td><td>201828</td><td>201850</td><td>+</td><td>0</td><td>TCCCATGCGACGGAGCTCCG</td><td>CGG</td><td>||||||||[||||||||||||]PAM</td><td>NA</td><td>Intergenic</td><td>NA</td><td>NA</td></tr>
<tr><td>T15440</td><td>ACGGAGCTCCGCGGCGCTGTCGG</td><td>1000</td><td></td><td></td><td></td><td></td><td></td><td></td><td></td><td></td><td></td></tr>
<tr><td>Oligo fwd</td><td>CACCACGGAGCTCCGCGGCGCTGT</td><td></td><td></td><td></td><td></td><td></td><td></td><td></td><td></td><td></td><td></td></tr>
<tr><td>Oligo rev</td><td>AAACACAGCGCCGCGGAGCTCCGT</td><td></td><td></td><td></td><td></td><td></td><td></td><td></td><td></td><td></td><td></td></tr>
<tr><td>Chromosome</td><td>start</td><td>end</td><td>strand</td><td>MM</td><td>target_seq</td><td>PAM</td><td>alignment</td><td>distance</td><td>position</td><td>gene name</td><td>gene id</td></tr>
<tr><td>GU305914.1</td><td>201837</td><td>201859</td><td>+</td><td>0</td><td>ACGGAGCTCCGCGGCGCTGT</td><td>CGG</td><td>||||||||[||||||||||||]PAM</td><td>NA</td><td>Intergenic</td><td>NA</td><td>NA</td></tr>
<tr><td>T15441</td><td>TGCCCGAGTCGTCGAAGACCAGG</td><td>1000</td><td></td><td></td><td></td><td></td><td></td><td></td><td></td><td></td><td></td></tr>
<tr><td>Oligo fwd</td><td>CACCTGCCCGAGTCGTCGAAGACC</td><td></td><td></td><td></td><td></td><td></td><td></td><td></td><td></td><td></td><td></td></tr>
<tr><td>Oligo rev</td><td>AAACGGTCTTCGACGACTCGGGCA</td><td></td><td></td><td></td><td></td><td></td><td></td><td></td><td></td><td></td><td></td></tr>
<tr><td>Chromosome</td><td>start</td><td>end</td><td>strand</td><td>MM</td><td>target_seq</td><td>PAM</td><td>alignment</td><td>distance</td><td>position</td><td>gene name</td><td>gene id</td></tr>
<tr><td>GU305914.1</td><td>201905</td><td>201927</td><td>+</td><td>0</td><td>TGCCCGAGTCGTCGAAGACC</td><td>AGG</td><td>||||||||[||||||||||||]PAM</td><td>NA</td><td>Intergenic</td><td>NA</td><td>NA</td></tr>
<tr><td>GU305914.1</td><td>97642</td><td>97664</td><td>+</td><td>4</td><td>CGCCCGTGTCGTCGAAGTCG</td><td>CGG</td><td>-|||||-|[|||||||||-|-]PAM</td><td>NA</td><td>Intergenic</td><td>NA</td><td>NA</td></tr>
<tr><td>T15442</td><td>AGACCAGGAGCTGCGCCCCGTGG</td><td>1000</td><td></td><td></td><td></td><td></td><td></td><td></td><td></td><td></td><td></td></tr>
<tr><td>Oligo fwd</td><td>CACCAGACCAGGAGCTGCGCCCCG</td><td></td><td></td><td></td><td></td><td></td><td></td><td></td><td></td><td></td><td></td></tr>
<tr><td>Oligo rev</td><td>AAACCGGGGCGCAGCTCCTGGTCT</td><td></td><td></td><td></td><td></td><td></td><td></td><td></td><td></td><td></td><td></td></tr>
<tr><td>Chromosome</td><td>start</td><td>end</td><td>strand</td><td>MM</td><td>target_seq</td><td>PAM</td><td>alignment</td><td>distance</td><td>position</td><td>gene name</td><td>gene id</td></tr>
<tr><td>GU305914.1</td><td>201920</td><td>201942</td><td>+</td><td>0</td><td>AGACCAGGAGCTGCGCCCCG</td><td>TGG</td><td>||||||||[||||||||||||]PAM</td><td>NA</td><td>Intergenic</td><td>NA</td><td>NA</td></tr>
<tr><td>T15443</td><td>CTGCGCCCCGTGGAACACCCCGG</td><td>1000</td><td></td><td></td><td></td><td></td><td></td><td></td><td></td><td></td><td></td></tr>
<tr><td>Oligo fwd</td><td>CACCCTGCGCCCCGTGGAACACCC</td><td></td><td></td><td></td><td></td><td></td><td></td><td></td><td></td><td></td><td></td></tr>
<tr><td>Oligo rev</td><td>AAACGGGTGTTCCACGGGGCGCAG</td><td></td><td></td><td></td><td></td><td></td><td></td><td></td><td></td><td></td><td></td></tr>
<tr><td>Chromosome</td><td>start</td><td>end</td><td>strand</td><td>MM</td><td>target_seq</td><td>PAM</td><td>alignment</td><td>distance</td><td>position</td><td>gene name</td><td>gene id</td></tr>
<tr><td>GU305914.1</td><td>201930</td><td>201952</td><td>+</td><td>0</td><td>CTGCGCCCCGTGGAACACCC</td><td>CGG</td><td>||||||||[||||||||||||]PAM</td><td>NA</td><td>Intergenic</td><td>NA</td><td>NA</td></tr>
<tr><td>T15444</td><td>CGTGGAACACCCCGGTGTCCCGG</td><td>1000</td><td></td><td></td><td></td><td></td><td></td><td></td><td></td><td></td><td></td></tr>
<tr><td>Oligo fwd</td><td>CACCCGTGGAACACCCCGGTGTCC</td><td></td><td></td><td></td><td></td><td></td><td></td><td></td><td></td><td></td><td></td></tr>
<tr><td>Oligo rev</td><td>AAACGGACACCGGGGTGTTCCACG</td><td></td><td></td><td></td><td></td><td></td><td></td><td></td><td></td><td></td><td></td></tr>
<tr><td>Chromosome</td><td>start</td><td>end</td><td>strand</td><td>MM</td><td>target_seq</td><td>PAM</td><td>alignment</td><td>distance</td><td>position</td><td>gene name</td><td>gene id</td></tr>
<tr><td>GU305914.1</td><td>201938</td><td>201960</td><td>+</td><td>0</td><td>CGTGGAACACCCCGGTGTCC</td><td>CGG</td><td>||||||||[||||||||||||]PAM</td><td>NA</td><td>Intergenic</td><td>NA</td><td>NA</td></tr>
<tr><td>T15445</td><td>TCCGAGCGTGTACCAGCGCGAGG</td><td>1000</td><td></td><td></td><td></td><td></td><td></td><td></td><td></td><td></td><td></td></tr>
<tr><td>Oligo fwd</td><td>CACCTCCGAGCGTGTACCAGCGCG</td><td></td><td></td><td></td><td></td><td></td><td></td><td></td><td></td><td></td><td></td></tr>
<tr><td>Oligo rev</td><td>AAACCGCGCTGGTACACGCTCGGA</td><td></td><td></td><td></td><td></td><td></td><td></td><td></td><td></td><td></td><td></td></tr>
<tr><td>Chromosome</td><td>start</td><td>end</td><td>strand</td><td>MM</td><td>target_seq</td><td>PAM</td><td>alignment</td><td>distance</td><td>position</td><td>gene name</td><td>gene id</td></tr>
<tr><td>GU305914.1</td><td>201975</td><td>201997</td><td>+</td><td>0</td><td>TCCGAGCGTGTACCAGCGCG</td><td>AGG</td><td>||||||||[||||||||||||]PAM</td><td>NA</td><td>Intergenic</td><td>NA</td><td>NA</td></tr>
<tr><td>T15446</td><td>ACATCTTTCGAGACGCGTATCGG</td><td>1000</td><td></td><td></td><td></td><td></td><td></td><td></td><td></td><td></td><td></td></tr>
<tr><td>Oligo fwd</td><td>CACCACATCTTTCGAGACGCGTAT</td><td></td><td></td><td></td><td></td><td></td><td></td><td></td><td></td><td></td><td></td></tr>
<tr><td>Oligo rev</td><td>AAACATACGCGTCTCGAAAGATGT</td><td></td><td></td><td></td><td></td><td></td><td></td><td></td><td></td><td></td><td></td></tr>
<tr><td>Chromosome</td><td>start</td><td>end</td><td>strand</td><td>MM</td><td>target_seq</td><td>PAM</td><td>alignment</td><td>distance</td><td>position</td><td>gene name</td><td>gene id</td></tr>
<tr><td>GU305914.1</td><td>201998</td><td>202020</td><td>+</td><td>0</td><td>ACATCTTTCGAGACGCGTAT</td><td>CGG</td><td>||||||||[||||||||||||]PAM</td><td>NA</td><td>Intergenic</td><td>NA</td><td>NA</td></tr>
<tr><td>T15447</td><td>TTCGAGACGCGTATCGGATGAGG</td><td>1000</td><td></td><td></td><td></td><td></td><td></td><td></td><td></td><td></td><td></td></tr>
<tr><td>Oligo fwd</td><td>CACCTTCGAGACGCGTATCGGATG</td><td></td><td></td><td></td><td></td><td></td><td></td><td></td><td></td><td></td><td></td></tr>
<tr><td>Oligo rev</td><td>AAACCATCCGATACGCGTCTCGAA</td><td></td><td></td><td></td><td></td><td></td><td></td><td></td><td></td><td></td><td></td></tr>
<tr><td>Chromosome</td><td>start</td><td>end</td><td>strand</td><td>MM</td><td>target_seq</td><td>PAM</td><td>alignment</td><td>distance</td><td>position</td><td>gene name</td><td>gene id</td></tr>
<tr><td>GU305914.1</td><td>202004</td><td>202026</td><td>+</td><td>0</td><td>TTCGAGACGCGTATCGGATG</td><td>AGG</td><td>||||||||[||||||||||||]PAM</td><td>NA</td><td>Intergenic</td><td>NA</td><td>NA</td></tr>
<tr><td>T15448</td><td>GATTTGTCCATCGCCACGAGCGG</td><td>1000</td><td></td><td></td><td></td><td></td><td></td><td></td><td></td><td></td><td></td></tr>
<tr><td>Oligo fwd</td><td>CACCGATTTGTCCATCGCCACGAG</td><td></td><td></td><td></td><td></td><td></td><td></td><td></td><td></td><td></td><td></td></tr>
<tr><td>Oligo rev</td><td>AAACCTCGTGGCGATGGACAAATC</td><td></td><td></td><td></td><td></td><td></td><td></td><td></td><td></td><td></td><td></td></tr>
<tr><td>Chromosome</td><td>start</td><td>end</td><td>strand</td><td>MM</td><td>target_seq</td><td>PAM</td><td>alignment</td><td>distance</td><td>position</td><td>gene name</td><td>gene id</td></tr>
<tr><td>GU305914.1</td><td>202039</td><td>202061</td><td>+</td><td>0</td><td>GATTTGTCCATCGCCACGAG</td><td>CGG</td><td>||||||||[||||||||||||]PAM</td><td>NA</td><td>Intergenic</td><td>NA</td><td>NA</td></tr>
</table>

| | | | | | | | | | | | |
|---|---|---|---|---|---|---|---|---|---|---|---|
| T15449 | ACTTGCAGCAGAGTTCGAGCCGG | 1000 | | | | | | | | | |
| Oligo fwd | CACCACTTGCAGCAGAGTTCGAGC | | | | | | | | | | |
| Oligo rev | AAACGCTCGAACTCTGCTGCAAGT | | | | | | | | | | |
| Chromosome | start | end | strand | MM | target_seq | PAM | alignment | distance | position | gene name | gene id |
| GU305914.1 | 202067 | 202089 | + | 0 | ACTTGCAGCAGAGTTCGAGC | CGG | ||||||||[||||||||||||]PAM | NA | Intergenic | NA | NA |
| T15450 | AGAGTTCGAGCCGGTCCAGTAGG | 1000 | | | | | | | | | |
| Oligo fwd | CACCAGAGTTCGAGCCGGTCCAGT | | | | | | | | | | |
| Oligo rev | AAACACTGGACCGGCTCGAACTCT | | | | | | | | | | |
| Chromosome | start | end | strand | MM | target_seq | PAM | alignment | distance | position | gene name | gene id |
| GU305914.1 | 202076 | 202098 | + | 0 | AGAGTTCGAGCCGGTCCAGT | AGG | ||||||||[||||||||||||]PAM | NA | Intergenic | NA | NA |
| T15451 | CGAGCCGGTCCAGTAGGATCAGG | 1000 | | | | | | | | | |
| Oligo fwd | CACCCGAGCCGGTCCAGTAGGATC | | | | | | | | | | |
| Oligo rev | AAACGATCCTACTGGACCGGCTCG | | | | | | | | | | |
| Chromosome | start | end | strand | MM | target_seq | PAM | alignment | distance | position | gene name | gene id |
| GU305914.1 | 202082 | 202104 | + | 0 | CGAGCCGGTCCAGTAGGATC | AGG | ||||||||[||||||||||||]PAM | NA | Intergenic | NA | NA |
| T15452 | GAGCCGGTCCAGTAGGATCAGGG | 1000 | | | | | | | | | |
| Oligo fwd | CACCGAGCCGGTCCAGTAGGATCA | | | | | | | | | | |
| Oligo rev | AAACTGATCCTACTGGACCGGCTC | | | | | | | | | | |
| Chromosome | start | end | strand | MM | target_seq | PAM | alignment | distance | position | gene name | gene id |
| GU305914.1 | 202083 | 202105 | + | 0 | GAGCCGGTCCAGTAGGATCA | GGG | ||||||||[||||||||||||]PAM | NA | Intergenic | NA | NA |
| T15453 | AGCCGGTCCAGTAGGATCAGGGG | 1000 | | | | | | | | | |
| Oligo fwd | CACCAGCCGGTCCAGTAGGATCAG | | | | | | | | | | |
| Oligo rev | AAACCTGATCCTACTGGACCGGCT | | | | | | | | | | |
| Chromosome | start | end | strand | MM | target_seq | PAM | alignment | distance | position | gene name | gene id |
| GU305914.1 | 202084 | 202106 | + | 0 | AGCCGGTCCAGTAGGATCAG | GGG | ||||||||[||||||||||||]PAM | NA | Intergenic | NA | NA |
| T15454 | GCCGGTCCAGTAGGATCAGGGGG | 1000 | | | | | | | | | |
| Oligo fwd | CACCGCCGGTCCAGTAGGATCAGG | | | | | | | | | | |
| Oligo rev | AAACCCTGATCCTACTGGACCGGC | | | | | | | | | | |
| Chromosome | start | end | strand | MM | target_seq | PAM | alignment | distance | position | gene name | gene id |
| GU305914.1 | 202085 | 202107 | + | 0 | GCCGGTCCAGTAGGATCAGG | GGG | ||||||||[||||||||||||]PAM | NA | Intergenic | NA | NA |
| T15455 | GTCCAGTAGGATCAGGGGGTTGG | 1000 | | | | | | | | | |
| Oligo fwd | CACCGTCCAGTAGGATCAGGGGGT | | | | | | | | | | |
| Oligo rev | AAACACCCCCTGATCCTACTGGAC | | | | | | | | | | |
| Chromosome | start | end | strand | MM | target_seq | PAM | alignment | distance | position | gene name | gene id |
| GU305914.1 | 202089 | 202111 | + | 0 | GTCCAGTAGGATCAGGGGGT | TGG | ||||||||[||||||||||||]PAM | NA | Intergenic | NA | NA |
| T15456 | GGGGTTGGTCTCCTGCTTGCCGG | 1000 | | | | | | | | | |
| Oligo fwd | CACCGGGGTTGGTCTCCTGCTTGC | | | | | | | | | | |
| Oligo rev | AAACGCAAGCAGGAGACCAACCCC | | | | | | | | | | |
| Chromosome | start | end | strand | MM | target_seq | PAM | alignment | distance | position | gene name | gene id |
| GU305914.1 | 202104 | 202126 | + | 0 | GGGGTTGGTCTCCTGCTTGC | CGG | ||||||||[||||||||||||]PAM | NA | Intergenic | NA | NA |
| T15457 | GGGTTGGTCTCCTGCTTGCCGGG | 1000 | | | | | | | | | |
| Oligo fwd | CACCGGGTTGGTCTCCTGCTTGCC | | | | | | | | | | |
| Oligo rev | AAACGGCAAGCAGGAGACCAACCC | | | | | | | | | | |
| Chromosome | start | end | strand | MM | target_seq | PAM | alignment | distance | position | gene name | gene id |
| GU305914.1 | 202105 | 202127 | + | 0 | GGGTTGGTCTCCTGCTTGCC | GGG | ||||||||[||||||||||||]PAM | NA | Intergenic | NA | NA |
| T15458 | GCAGCTCGATCACCGAGTTGCGG | 1000 | | | | | | | | | |
| Oligo fwd | CACCGCAGCTCGATCACCGAGTTG | | | | | | | | | | |
| Oligo rev | AAACCAACTCGGTGATCGAGCTGC | | | | | | | | | | |
| Chromosome | start | end | strand | MM | target_seq | PAM | alignment | distance | position | gene name | gene id |
| GU305914.1 | 202133 | 202155 | + | 0 | GCAGCTCGATCACCGAGTTG | CGG | ||||||||[||||||||||||]PAM | NA | Intergenic | NA | NA |
| T15459 | GATCACCGAGTTGCGGTACGTGG | 1000 | | | | | | | | | |
| Oligo fwd | CACCGATCACCGAGTTGCGGTACG | | | | | | | | | | |
| Oligo rev | AAACCGTACCGCAACTCGGTGATC | | | | | | | | | | |
| Chromosome | start | end | strand | MM | target_seq | PAM | alignment | distance | position | gene name | gene id |
| GU305914.1 | 202140 | 202162 | + | 0 | GATCACCGAGTTGCGGTACG | TGG | ||||||||[||||||||||||]PAM | NA | Intergenic | NA | NA |
| T15460 | AGTTGCGGTACGTGGACAGATGG | 1000 | | | | | | | | | |
| Oligo fwd | CACCAGTTGCGGTACGTGGACAGA | | | | | | | | | | |
| Oligo rev | AAACTCTGTCCACGTACCGCAACT | | | | | | | | | | |
| Chromosome | start | end | strand | MM | target_seq | PAM | alignment | distance | position | gene name | gene id |
| GU305914.1 | 202148 | 202170 | + | 0 | AGTTGCGGTACGTGGACAGA | TGG | ||||||||[||||||||||||]PAM | NA | Intergenic | NA | NA |
| T15461 | TGCGGTACGTGGACAGATGGCGG | 1000 | | | | | | | | | |
| Oligo fwd | CACCTGCGGTACGTGGACAGATGG | | | | | | | | | | |
| Oligo rev | AAACCCATCTGTCCACGTACCGCA | | | | | | | | | | |
| Chromosome | start | end | strand | MM | target_seq | PAM | alignment | distance | position | gene name | gene id |
| GU305914.1 | 202151 | 202173 | + | 0 | TGCGGTACGTGGACAGATGG | CGG | ||||||||[||||||||||||]PAM | NA | Intergenic | NA | NA |
| T15462 | ACGTGGACAGATGGCGGTTCAGG | 1000 | | | | | | | | | |
| Oligo fwd | CACCACGTGGACAGATGGCGGTTC | | | | | | | | | | |
| Oligo rev | AAACGAACCGCCATCTGTCCACGT | | | | | | | | | | |
| Chromosome | start | end | strand | MM | target_seq | PAM | alignment | distance | position | gene name | gene id |
| GU305914.1 | 202157 | 202179 | + | 0 | ACGTGGACAGATGGCGGTTC | AGG | ||||||||[||||||||||||]PAM | NA | Intergenic | NA | NA |
| T15463 | ATGGCGGTTCAGGTCGTAGCCGG | 1000 | | | | | | | | | |
| Oligo fwd | CACCATGGCGGTTCAGGTCGTAGC | | | | | | | | | | |
| Oligo rev | AAACGCTACGACCTGAACCGCCAT | | | | | | | | | | |
| Chromosome | start | end | strand | MM | target_seq | PAM | alignment | distance | position | gene name | gene id |

| | | | | | | | | | | | |
|---|---|---|---|---|---|---|---|---|---|---|---|
| GU305914.1 | 202167 | 202189 | + | 0 | ATGGCGGTTCAGGTCGTAGC | CGG | \|\|\|\|\|\|\|\|[\|\|\|\|\|\|\|\|\|\|\|\|]PAM | NA | Intergenic | NA | NA |
| T15464 | GGTCGTAGCCGGTCACGCAGAGG | 1000 | | | | | | | | | |
| Oligo fwd | CACCGGTCGTAGCCGGTCACGCAG | | | | | | | | | | |
| Oligo rev | AAACCTGCGTGACCGGCTACGACC | | | | | | | | | | |
| Chromosome | start | end | strand | MM | target_seq | PAM | alignment | distance | position | gene name | gene id |
| GU305914.1 | 202178 | 202200 | + | 0 | GGTCGTAGCCGGTCACGCAG | AGG | \|\|\|\|\|\|\|\|[\|\|\|\|\|\|\|\|\|\|\|\|]PAM | NA | Intergenic | NA | NA |
| T15465 | GCCGGTCACGCAGAGGCCGATGG | 1000 | | | | | | | | | |
| Oligo fwd | CACCGCCGGTCACGCAGAGGCCGA | | | | | | | | | | |
| Oligo rev | AAACTCGGCCTCTGCGTGACCGGC | | | | | | | | | | |
| Chromosome | start | end | strand | MM | target_seq | PAM | alignment | distance | position | gene name | gene id |
| GU305914.1 | 202185 | 202207 | + | 0 | GCCGGTCACGCAGAGGCCGA | TGG | \|\|\|\|\|\|\|\|[\|\|\|\|\|\|\|\|\|\|\|\|]PAM | NA | Intergenic | NA | NA |
| T15466 | AGGCCGATGGCGTCCCAGACCGG | 1000 | | | | | | | | | |
| Oligo fwd | CACCAGGCCGATGGCGTCCCAGAC | | | | | | | | | | |
| Oligo rev | AAACGTCTGGGACGCCATCGGCCT | | | | | | | | | | |
| Chromosome | start | end | strand | MM | target_seq | PAM | alignment | distance | position | gene name | gene id |
| GU305914.1 | 202198 | 202220 | + | 0 | AGGCCGATGGCGTCCCAGAC | CGG | \|\|\|\|\|\|\|\|[\|\|\|\|\|\|\|\|\|\|\|\|]PAM | NA | Intergenic | NA | NA |
| T15467 | GGCCGATGGCGTCCCAGACCGGG | 1000 | | | | | | | | | |
| Oligo fwd | CACCGGCCGATGGCGTCCCAGACC | | | | | | | | | | |
| Oligo rev | AAACGGTCTGGGACGCCATCGGCC | | | | | | | | | | |
| Chromosome | start | end | strand | MM | target_seq | PAM | alignment | distance | position | gene name | gene id |
| GU305914.1 | 202199 | 202221 | + | 0 | GGCCGATGGCGTCCCAGACC | GGG | \|\|\|\|\|\|\|\|[\|\|\|\|\|\|\|\|\|\|\|\|]PAM | NA | Intergenic | NA | NA |
| T15468 | CAGACCGGGTCTTTCTTCAACGG | 1000 | | | | | | | | | |
| Oligo fwd | CACCCAGACCGGGTCTTTCTTCAA | | | | | | | | | | |
| Oligo rev | AAACTTGAAGAAAGACCCGGTCTG | | | | | | | | | | |
| Chromosome | start | end | strand | MM | target_seq | PAM | alignment | distance | position | gene name | gene id |
| GU305914.1 | 202213 | 202235 | + | 0 | CAGACCGGGTCTTTCTTCAA | CGG | \|\|\|\|\|\|\|\|[\|\|\|\|\|\|\|\|\|\|\|\|]PAM | NA | Intergenic | NA | NA |
| T15469 | TTCTTCAACGGTATCGACAGCGG | 1000 | | | | | | | | | |
| Oligo fwd | CACCTTCTTCAACGGTATCGACAG | | | | | | | | | | |
| Oligo rev | AAACCTGTCGATACCGTTGAAGAA | | | | | | | | | | |
| Chromosome | start | end | strand | MM | target_seq | PAM | alignment | distance | position | gene name | gene id |
| GU305914.1 | 202225 | 202247 | + | 0 | TTCTTCAACGGTATCGACAG | CGG | \|\|\|\|\|\|\|\|[\|\|\|\|\|\|\|\|\|\|\|\|]PAM | NA | Intergenic | NA | NA |
| GU305914.1 | 228991 | 229013 | + | 4 | TTCTTCGGCGTTAGCGACAG | AGG | \|\|\|\|\|\|--[\|\|-\|\|-\|\|\|\|\|\|]PAM | NA | Intergenic | NA | NA |
| T15470 | CTATCCGCGAGAACTCGTCGAGG | 1000 | | | | | | | | | |
| Oligo fwd | CACCCTATCCGCGAGAACTCGTCG | | | | | | | | | | |
| Oligo rev | AAACCGACGAGTTCTCGCGGATAG | | | | | | | | | | |
| Chromosome | start | end | strand | MM | target_seq | PAM | alignment | distance | position | gene name | gene id |
| GU305914.1 | 202280 | 202302 | + | 0 | CTATCCGCGAGAACTCGTCG | AGG | \|\|\|\|\|\|\|\|[\|\|\|\|\|\|\|\|\|\|\|\|]PAM | NA | Intergenic | NA | NA |
| T15471 | CGAGAACTCGTCGAGGTCGTTGG | 1000 | | | | | | | | | |
| Oligo fwd | CACCCGAGAACTCGTCGAGGTCGT | | | | | | | | | | |
| Oligo rev | AAACACGACCTCGACGAGTTCTCG | | | | | | | | | | |
| Chromosome | start | end | strand | MM | target_seq | PAM | alignment | distance | position | gene name | gene id |
| GU305914.1 | 202287 | 202309 | + | 0 | CGAGAACTCGTCGAGGTCGT | TGG | \|\|\|\|\|\|\|\|[\|\|\|\|\|\|\|\|\|\|\|\|]PAM | NA | Intergenic | NA | NA |
| T15472 | GTCGTTGGCCGCCAACGCGATGG | 1000 | | | | | | | | | |
| Oligo fwd | CACCGTCGTTGGCCGCCAACGCGA | | | | | | | | | | |
| Oligo rev | AAACTCGCGTTGGCGGCCAACGAC | | | | | | | | | | |
| Chromosome | start | end | strand | MM | target_seq | PAM | alignment | distance | position | gene name | gene id |
| GU305914.1 | 202302 | 202324 | + | 0 | GTCGTTGGCCGCCAACGCGA | TGG | \|\|\|\|\|\|\|\|[\|\|\|\|\|\|\|\|\|\|\|\|]PAM | NA | Intergenic | NA | NA |
| T15473 | CAACGCGATGGCGTCCTCCTCGG | 1000 | | | | | | | | | |
| Oligo fwd | CACCCAACGCGATGGCGTCCTCCT | | | | | | | | | | |
| Oligo rev | AAACAGGAGGACGCCATCGCGTTG | | | | | | | | | | |
| Chromosome | start | end | strand | MM | target_seq | PAM | alignment | distance | position | gene name | gene id |
| GU305914.1 | 202314 | 202336 | + | 0 | CAACGCGATGGCGTCCTCCT | CGG | \|\|\|\|\|\|\|\|[\|\|\|\|\|\|\|\|\|\|\|\|]PAM | NA | Intergenic | NA | NA |
| T15474 | CATCCTCGACTCCTTGCCCATGG | 1000 | | | | | | | | | |
| Oligo fwd | CACCCATCCTCGACTCCTTGCCCA | | | | | | | | | | |
| Oligo rev | AAACTGGGCAAGGAGTCGAGGATG | | | | | | | | | | |
| Chromosome | start | end | strand | MM | target_seq | PAM | alignment | distance | position | gene name | gene id |
| GU305914.1 | 202350 | 202372 | + | 0 | CATCCTCGACTCCTTGCCCA | TGG | \|\|\|\|\|\|\|\|[\|\|\|\|\|\|\|\|\|\|\|\|]PAM | NA | Intergenic | NA | NA |
| T15475 | TCGACTCCTTGCCCATGGCGAGG | 1000 | | | | | | | | | |
| Oligo fwd | CACCTCGACTCCTTGCCCATGGCG | | | | | | | | | | |
| Oligo rev | AAACCGCCATGGGCAAGGAGTCGA | | | | | | | | | | |
| Chromosome | start | end | strand | MM | target_seq | PAM | alignment | distance | position | gene name | gene id |
| GU305914.1 | 202355 | 202377 | + | 0 | TCGACTCCTTGCCCATGGCG | AGG | \|\|\|\|\|\|\|\|[\|\|\|\|\|\|\|\|\|\|\|\|]PAM | NA | Intergenic | NA | NA |
| T15476 | CCTTGCCCATGGCGAGGTACAGG | 1000 | | | | | | | | | |
| Oligo fwd | CACCCCTTGCCCATGGCGAGGTAC | | | | | | | | | | |
| Oligo rev | AAACGTACCTCGCCATGGGCAAGG | | | | | | | | | | |
| Chromosome | start | end | strand | MM | target_seq | PAM | alignment | distance | position | gene name | gene id |
| GU305914.1 | 202361 | 202383 | + | 0 | CCTTGCCCATGGCGAGGTAC | AGG | \|\|\|\|\|\|\|\|[\|\|\|\|\|\|\|\|\|\|\|\|]PAM | NA | Intergenic | NA | NA |
| T15477 | CTTGCCCATGGCGAGGTACAGGG | 1000 | | | | | | | | | |
| Oligo fwd | CACCCTTGCCCATGGCGAGGTACA | | | | | | | | | | |
| Oligo rev | AAACTGTACCTCGCCATGGGCAAG | | | | | | | | | | |
| Chromosome | start | end | strand | MM | target_seq | PAM | alignment | distance | position | gene name | gene id |
| GU305914.1 | 202362 | 202384 | + | 0 | CTTGCCCATGGCGAGGTACA | GGG | \|\|\|\|\|\|\|\|[\|\|\|\|\|\|\|\|\|\|\|\|]PAM | NA | Intergenic | NA | NA |
| T15478 | AGGGACGTGTGCTGCACCATCGG | 1000 | | | | | | | | | |
| Oligo fwd | CACCAGGGACGTGTGCTGCACCAT | | | | | | | | | | |

Oligo rev AAACATGGTGCAGCACACGTCCCT
Chromosome start end strand MM target_seq PAM alignment distance position gene name gene id
GU305914.1 202381 202403 + 0 AGGGACGTGTGCTGCACCAT CGG ||||||||[||||||||||||]PAM NA Intergenic NA NA

T15479 ACGACCACCCACTCGCCCGTCGG 1000
Oligo fwd CACCACGACCACCCACTCGCCCGT
Oligo rev AAACACGGGCGAGTGGGTGGTCGT
Chromosome start end strand MM target_seq PAM alignment distance position gene name gene id
GU305914.1 202423 202445 + 0 ACGACCACCCACTCGCCCGT CGG ||||||||[||||||||||||]PAM NA Intergenic NA NA

T15480 CCACCCACTCGCCCGTCGGCAGG 1000
Oligo fwd CACCCCACCCACTCGCCCGTCGGC
Oligo rev AAACGCCGACGGGCGAGTGGGTGG
Chromosome start end strand MM target_seq PAM alignment distance position gene name gene id
GU305914.1 202427 202449 + 0 CCACCCACTCGCCCGTCGGC AGG ||||||||[||||||||||||]PAM NA Intergenic NA NA

T15481 CACTCGCCCGTCGGCAGGATCGG 1000
Oligo fwd CACCCACTCGCCCGTCGGCAGGAT
Oligo rev AAACATCCTGCCGACGGGCGAGTG
Chromosome start end strand MM target_seq PAM alignment distance position gene name gene id
GU305914.1 202432 202454 + 0 CACTCGCCCGTCGGCAGGAT CGG ||||||||[||||||||||||]PAM NA Intergenic NA NA

T15482 CGCCCGTCGGCAGGATCGGACGG 1000
Oligo fwd CACCCGCCCGTCGGCAGGATCGGA
Oligo rev AAACTCCGATCCTGCCGACGGGCG
Chromosome start end strand MM target_seq PAM alignment distance position gene name gene id
GU305914.1 202436 202458 + 0 CGCCCGTCGGCAGGATCGGA CGG ||||||||[||||||||||||]PAM NA Intergenic NA NA

T15483 CCGTCGGCAGGATCGGACGGAGG 1000
Oligo fwd CACCCCGTCGGCAGGATCGGACGG
Oligo rev AAACCCGTCCGATCCTGCCGACGG
Chromosome start end strand MM target_seq PAM alignment distance position gene name gene id
GU305914.1 202439 202461 + 0 CCGTCGGCAGGATCGGACGG AGG ||||||||[||||||||||||]PAM NA Intergenic NA NA

T15484 GCAGGATCGGACGGAGGTCCCGG 1000
Oligo fwd CACCGCAGGATCGGACGGAGGTCC
Oligo rev AAACGGACCTCCGTCCGATCCTGC
Chromosome start end strand MM target_seq PAM alignment distance position gene name gene id
GU305914.1 202445 202467 + 0 GCAGGATCGGACGGAGGTCC CGG ||||||||[||||||||||||]PAM NA Intergenic NA NA

T15485 ATCTCGACCAGTTCGCCCCGCGG 1000
Oligo fwd CACCATCTCGACCAGTTCGCCCCG
Oligo rev AAACCGGGGCGAACTGGTCGAGAT
Chromosome start end strand MM target_seq PAM alignment distance position gene name gene id
GU305914.1 202474 202496 + 0 ATCTCGACCAGTTCGCCCCG CGG ||||||||[||||||||||||]PAM NA Intergenic NA NA

T15486 CAGTTCGCCCCGCGGCACCACGG 1000
Oligo fwd CACCCAGTTCGCCCCGCGGCACCA
Oligo rev AAACTGGTGCCGCGGGGCGAACTG
Chromosome start end strand MM target_seq PAM alignment distance position gene name gene id
GU305914.1 202482 202504 + 0 CAGTTCGCCCCGCGGCACCA CGG ||||||||[||||||||||||]PAM NA Intergenic NA NA

T15487 ACCACGGCCAGCCACCACCCCGG 1000
Oligo fwd CACCACCACGGCCAGCCACCACCC
Oligo rev AAACGGGTGGTGGCTGGCCGTGGT
Chromosome start end strand MM target_seq PAM alignment distance position gene name gene id
GU305914.1 202498 202520 + 0 ACCACGGCCAGCCACCACCC CGG ||||||||[||||||||||||]PAM NA Intergenic NA NA

T15488 CCACGGCCAGCCACCACCCCGGG 1000
Oligo fwd CACCCCACGGCCAGCCACCACCCC
Oligo rev AAACGGGGTGGTGGCTGGCCGTGG
Chromosome start end strand MM target_seq PAM alignment distance position gene name gene id
GU305914.1 202499 202521 + 0 CCACGGCCAGCCACCACCCC GGG ||||||||[||||||||||||]PAM NA Intergenic NA NA

T15489 CACGGCCAGCCACCACCCCGGGG 1000
Oligo fwd CACCCACGGCCAGCCACCACCCCG
Oligo rev AAACCGGGGTGGTGGCTGGCCGTG
Chromosome start end strand MM target_seq PAM alignment distance position gene name gene id
GU305914.1 202500 202522 + 0 CACGGCCAGCCACCACCCCG GGG ||||||||[||||||||||||]PAM NA Intergenic NA NA

T15490 ACGGCCAGCCACCACCCCGGGGG 1000
Oligo fwd CACCACGGCCAGCCACCACCCCGG
Oligo rev AAACCCGGGGTGGTGGCTGGCCGT
Chromosome start end strand MM target_seq PAM alignment distance position gene name gene id
GU305914.1 202501 202523 + 0 ACGGCCAGCCACCACCCCGG GGG ||||||||[||||||||||||]PAM NA Intergenic NA NA

T15491 CGGCCAGCCACCACCCCGGGGGG 1000
Oligo fwd CACCCGGCCAGCCACCACCCCGGG
Oligo rev AAACCCCGGGGTGGTGGCTGGCCG
Chromosome start end strand MM target_seq PAM alignment distance position gene name gene id
GU305914.1 202502 202524 + 0 CGGCCAGCCACCACCCCGGG GGG ||||||||[||||||||||||]PAM NA Intergenic NA NA

T15492 GGGGTACCCGACACAGAGCTTGG 1000
Oligo fwd CACCGGGGTACCCGACACAGAGCT
Oligo rev AAACAGCTCTGTGTCGGGTACCCC
Chromosome start end strand MM target_seq PAM alignment distance position gene name gene id
GU305914.1 202521 202543 + 0 GGGGTACCCGACACAGAGCT TGG ||||||||[||||||||||||]PAM NA Intergenic NA NA

T15493 AACGTTCCGACGTCTGCGTGAGG 1000

<table>
<tr><td>Oligo fwd</td><td>CACCAACGTTCCGACGTCTGCGTG</td><td></td><td></td><td></td><td></td><td></td><td></td><td></td><td></td><td></td><td></td></tr>
<tr><td>Oligo rev</td><td>AAACCACGCAGACGTCGGAACGTT</td><td></td><td></td><td></td><td></td><td></td><td></td><td></td><td></td><td></td><td></td></tr>
<tr><td>Chromosome</td><td>start</td><td>end</td><td>strand</td><td>MM</td><td>target_seq</td><td>PAM</td><td>alignment</td><td>distance</td><td>position</td><td>gene name</td><td>gene id</td></tr>
<tr><td>GU305914.1</td><td>202577</td><td>202599</td><td>+</td><td>0</td><td>AACGTTCCGACGTCTGCGTG</td><td>AGG</td><td>||||||||[||||||||||||]PAM</td><td>NA</td><td>Intergenic</td><td>NA</td><td>NA</td></tr>
<tr><td>T15494</td><td>AAAAACTCTTGAAGTCTCTCAGG</td><td>1000</td><td></td><td></td><td></td><td></td><td></td><td></td><td></td><td></td><td></td></tr>
<tr><td>Oligo fwd</td><td>CACCAAAAACTCTTGAAGTCTCTC</td><td></td><td></td><td></td><td></td><td></td><td></td><td></td><td></td><td></td><td></td></tr>
<tr><td>Oligo rev</td><td>AAACGAGAGACTTCAAGAGTTTTT</td><td></td><td></td><td></td><td></td><td></td><td></td><td></td><td></td><td></td><td></td></tr>
<tr><td>Chromosome</td><td>start</td><td>end</td><td>strand</td><td>MM</td><td>target_seq</td><td>PAM</td><td>alignment</td><td>distance</td><td>position</td><td>gene name</td><td>gene id</td></tr>
<tr><td>GU305914.1</td><td>202601</td><td>202623</td><td>+</td><td>0</td><td>AAAAACTCTTGAAGTCTCTC</td><td>AGG</td><td>||||||||[||||||||||||]PAM</td><td>NA</td><td>Intergenic</td><td>NA</td><td>NA</td></tr>
<tr><td>T15495</td><td>AGTCTCTCAGGATCTGCTCTCGG</td><td>1000</td><td></td><td></td><td></td><td></td><td></td><td></td><td></td><td></td><td></td></tr>
<tr><td>Oligo fwd</td><td>CACCAGTCTCTCAGGATCTGCTCT</td><td></td><td></td><td></td><td></td><td></td><td></td><td></td><td></td><td></td><td></td></tr>
<tr><td>Oligo rev</td><td>AAACAGAGCAGATCCTGAGAGACT</td><td></td><td></td><td></td><td></td><td></td><td></td><td></td><td></td><td></td><td></td></tr>
<tr><td>Chromosome</td><td>start</td><td>end</td><td>strand</td><td>MM</td><td>target_seq</td><td>PAM</td><td>alignment</td><td>distance</td><td>position</td><td>gene name</td><td>gene id</td></tr>
<tr><td>GU305914.1</td><td>202613</td><td>202635</td><td>+</td><td>0</td><td>AGTCTCTCAGGATCTGCTCT</td><td>CGG</td><td>||||||||[||||||||||||]PAM</td><td>NA</td><td>Intergenic</td><td>NA</td><td>NA</td></tr>
<tr><td>T15496</td><td>AGGATCTGCTCTCGGTCCGCCGG</td><td>1000</td><td></td><td></td><td></td><td></td><td></td><td></td><td></td><td></td><td></td></tr>
<tr><td>Oligo fwd</td><td>CACCAGGATCTGCTCTCGGTCCGC</td><td></td><td></td><td></td><td></td><td></td><td></td><td></td><td></td><td></td><td></td></tr>
<tr><td>Oligo rev</td><td>AAACGCGGACCGAGAGCAGATCCT</td><td></td><td></td><td></td><td></td><td></td><td></td><td></td><td></td><td></td><td></td></tr>
<tr><td>Chromosome</td><td>start</td><td>end</td><td>strand</td><td>MM</td><td>target_seq</td><td>PAM</td><td>alignment</td><td>distance</td><td>position</td><td>gene name</td><td>gene id</td></tr>
<tr><td>GU305914.1</td><td>202621</td><td>202643</td><td>+</td><td>0</td><td>AGGATCTGCTCTCGGTCCGC</td><td>CGG</td><td>||||||||[||||||||||||]PAM</td><td>NA</td><td>Intergenic</td><td>NA</td><td>NA</td></tr>
<tr><td>T15497</td><td>GGATCTGCTCTCGGTCCGCCGGG</td><td>1000</td><td></td><td></td><td></td><td></td><td></td><td></td><td></td><td></td><td></td></tr>
<tr><td>Oligo fwd</td><td>CACCGGATCTGCTCTCGGTCCGCC</td><td></td><td></td><td></td><td></td><td></td><td></td><td></td><td></td><td></td><td></td></tr>
<tr><td>Oligo rev</td><td>AAACGGCGGACCGAGAGCAGATCC</td><td></td><td></td><td></td><td></td><td></td><td></td><td></td><td></td><td></td><td></td></tr>
<tr><td>Chromosome</td><td>start</td><td>end</td><td>strand</td><td>MM</td><td>target_seq</td><td>PAM</td><td>alignment</td><td>distance</td><td>position</td><td>gene name</td><td>gene id</td></tr>
<tr><td>GU305914.1</td><td>202622</td><td>202644</td><td>+</td><td>0</td><td>GGATCTGCTCTCGGTCCGCC</td><td>GGG</td><td>||||||||[||||||||||||]PAM</td><td>NA</td><td>Intergenic</td><td>NA</td><td>NA</td></tr>
<tr><td>T15498</td><td>TCTGCTCTCGGTCCGCCGGGCGG</td><td>1000</td><td></td><td></td><td></td><td></td><td></td><td></td><td></td><td></td><td></td></tr>
<tr><td>Oligo fwd</td><td>CACCTCTGCTCTCGGTCCGCCGGG</td><td></td><td></td><td></td><td></td><td></td><td></td><td></td><td></td><td></td><td></td></tr>
<tr><td>Oligo rev</td><td>AAACCCCGGCGGACCGAGAGCAGA</td><td></td><td></td><td></td><td></td><td></td><td></td><td></td><td></td><td></td><td></td></tr>
<tr><td>Chromosome</td><td>start</td><td>end</td><td>strand</td><td>MM</td><td>target_seq</td><td>PAM</td><td>alignment</td><td>distance</td><td>position</td><td>gene name</td><td>gene id</td></tr>
<tr><td>GU305914.1</td><td>202625</td><td>202647</td><td>+</td><td>0</td><td>TCTGCTCTCGGTCCGCCGGG</td><td>CGG</td><td>||||||||[||||||||||||]PAM</td><td>NA</td><td>Intergenic</td><td>NA</td><td>NA</td></tr>
<tr><td>T15499</td><td>GCCGGGCGGATCAGAGCCTCCGG</td><td>1000</td><td></td><td></td><td></td><td></td><td></td><td></td><td></td><td></td><td></td></tr>
<tr><td>Oligo fwd</td><td>CACCGCCGGGCGGATCAGAGCCTC</td><td></td><td></td><td></td><td></td><td></td><td></td><td></td><td></td><td></td><td></td></tr>
<tr><td>Oligo rev</td><td>AAACGAGGCTCTGATCCGCCCGGC</td><td></td><td></td><td></td><td></td><td></td><td></td><td></td><td></td><td></td><td></td></tr>
<tr><td>Chromosome</td><td>start</td><td>end</td><td>strand</td><td>MM</td><td>target_seq</td><td>PAM</td><td>alignment</td><td>distance</td><td>position</td><td>gene name</td><td>gene id</td></tr>
<tr><td>GU305914.1</td><td>202639</td><td>202661</td><td>+</td><td>0</td><td>GCCGGGCGGATCAGAGCCTC</td><td>CGG</td><td>||||||||[||||||||||||]PAM</td><td>NA</td><td>Intergenic</td><td>NA</td><td>NA</td></tr>
<tr><td>T15500</td><td>GATCAGAGCCTCCGGCCTCCAGG</td><td>1000</td><td></td><td></td><td></td><td></td><td></td><td></td><td></td><td></td><td></td></tr>
<tr><td>Oligo fwd</td><td>CACCGATCAGAGCCTCCGGCCTCC</td><td></td><td></td><td></td><td></td><td></td><td></td><td></td><td></td><td></td><td></td></tr>
<tr><td>Oligo rev</td><td>AAACGGAGGCCGGAGGCTCTGATC</td><td></td><td></td><td></td><td></td><td></td><td></td><td></td><td></td><td></td><td></td></tr>
<tr><td>Chromosome</td><td>start</td><td>end</td><td>strand</td><td>MM</td><td>target_seq</td><td>PAM</td><td>alignment</td><td>distance</td><td>position</td><td>gene name</td><td>gene id</td></tr>
<tr><td>GU305914.1</td><td>202647</td><td>202669</td><td>+</td><td>0</td><td>GATCAGAGCCTCCGGCCTCC</td><td>AGG</td><td>||||||||[||||||||||||]PAM</td><td>NA</td><td>Intergenic</td><td>NA</td><td>NA</td></tr>
<tr><td>T15501</td><td>AGAGCCTCCGGCCTCCAGGCCGG</td><td>1000</td><td></td><td></td><td></td><td></td><td></td><td></td><td></td><td></td><td></td></tr>
<tr><td>Oligo fwd</td><td>CACCAGAGCCTCCGGCCTCCAGGC</td><td></td><td></td><td></td><td></td><td></td><td></td><td></td><td></td><td></td><td></td></tr>
<tr><td>Oligo rev</td><td>AAACGCCTGGAGGCCGGAGGCTCT</td><td></td><td></td><td></td><td></td><td></td><td></td><td></td><td></td><td></td><td></td></tr>
<tr><td>Chromosome</td><td>start</td><td>end</td><td>strand</td><td>MM</td><td>target_seq</td><td>PAM</td><td>alignment</td><td>distance</td><td>position</td><td>gene name</td><td>gene id</td></tr>
<tr><td>GU305914.1</td><td>202651</td><td>202673</td><td>+</td><td>0</td><td>AGAGCCTCCGGCCTCCAGGC</td><td>CGG</td><td>||||||||[||||||||||||]PAM</td><td>NA</td><td>Intergenic</td><td>NA</td><td>NA</td></tr>
<tr><td>T15502</td><td>CGGCCTCCAGGCCGGATCTCCGG</td><td>1000</td><td></td><td></td><td></td><td></td><td></td><td></td><td></td><td></td><td></td></tr>
<tr><td>Oligo fwd</td><td>CACCCGGCCTCCAGGCCGGATCTC</td><td></td><td></td><td></td><td></td><td></td><td></td><td></td><td></td><td></td><td></td></tr>
<tr><td>Oligo rev</td><td>AAACGAGATCCGGCCTGGAGGCCG</td><td></td><td></td><td></td><td></td><td></td><td></td><td></td><td></td><td></td><td></td></tr>
<tr><td>Chromosome</td><td>start</td><td>end</td><td>strand</td><td>MM</td><td>target_seq</td><td>PAM</td><td>alignment</td><td>distance</td><td>position</td><td>gene name</td><td>gene id</td></tr>
<tr><td>GU305914.1</td><td>202659</td><td>202681</td><td>+</td><td>0</td><td>CGGCCTCCAGGCCGGATCTC</td><td>CGG</td><td>||||||||[||||||||||||]PAM</td><td>NA</td><td>Intergenic</td><td>NA</td><td>NA</td></tr>
<tr><td>T15503</td><td>CCAAGACATCGCGCACGAGACGG</td><td>1000</td><td></td><td></td><td></td><td></td><td></td><td></td><td></td><td></td><td></td></tr>
<tr><td>Oligo fwd</td><td>CACCCCAAGACATCGCGCACGAGA</td><td></td><td></td><td></td><td></td><td></td><td></td><td></td><td></td><td></td><td></td></tr>
<tr><td>Oligo rev</td><td>AAACTCTCGTGCGCGATGTCTTGG</td><td></td><td></td><td></td><td></td><td></td><td></td><td></td><td></td><td></td><td></td></tr>
<tr><td>Chromosome</td><td>start</td><td>end</td><td>strand</td><td>MM</td><td>target_seq</td><td>PAM</td><td>alignment</td><td>distance</td><td>position</td><td>gene name</td><td>gene id</td></tr>
<tr><td>GU305914.1</td><td>202686</td><td>202708</td><td>+</td><td>0</td><td>CCAAGACATCGCGCACGAGA</td><td>CGG</td><td>||||||||[||||||||||||]PAM</td><td>NA</td><td>Intergenic</td><td>NA</td><td>NA</td></tr>
<tr><td>T15504</td><td>CAAGACATCGCGCACGAGACGGG</td><td>1000</td><td></td><td></td><td></td><td></td><td></td><td></td><td></td><td></td><td></td></tr>
<tr><td>Oligo fwd</td><td>CACCCAAGACATCGCGCACGAGAC</td><td></td><td></td><td></td><td></td><td></td><td></td><td></td><td></td><td></td><td></td></tr>
<tr><td>Oligo rev</td><td>AAACGTCTCGTGCGCGATGTCTTG</td><td></td><td></td><td></td><td></td><td></td><td></td><td></td><td></td><td></td><td></td></tr>
<tr><td>Chromosome</td><td>start</td><td>end</td><td>strand</td><td>MM</td><td>target_seq</td><td>PAM</td><td>alignment</td><td>distance</td><td>position</td><td>gene name</td><td>gene id</td></tr>
<tr><td>GU305914.1</td><td>202687</td><td>202709</td><td>+</td><td>0</td><td>CAAGACATCGCGCACGAGAC</td><td>GGG</td><td>||||||||[||||||||||||]PAM</td><td>NA</td><td>Intergenic</td><td>NA</td><td>NA</td></tr>
<tr><td>T15505</td><td>GACATCGCGCACGAGACGGGCGG</td><td>1000</td><td></td><td></td><td></td><td></td><td></td><td></td><td></td><td></td><td></td></tr>
<tr><td>Oligo fwd</td><td>CACCGACATCGCGCACGAGACGGG</td><td></td><td></td><td></td><td></td><td></td><td></td><td></td><td></td><td></td><td></td></tr>
<tr><td>Oligo rev</td><td>AAACCCCGTCTCGTGCGCGATGTC</td><td></td><td></td><td></td><td></td><td></td><td></td><td></td><td></td><td></td><td></td></tr>
<tr><td>Chromosome</td><td>start</td><td>end</td><td>strand</td><td>MM</td><td>target_seq</td><td>PAM</td><td>alignment</td><td>distance</td><td>position</td><td>gene name</td><td>gene id</td></tr>
<tr><td>GU305914.1</td><td>202690</td><td>202712</td><td>+</td><td>0</td><td>GACATCGCGCACGAGACGGG</td><td>CGG</td><td>||||||||[||||||||||||]PAM</td><td>NA</td><td>Intergenic</td><td>NA</td><td>NA</td></tr>
<tr><td>T15506</td><td>CACGAGACGGGCGGACGAAGCGG</td><td>1000</td><td></td><td></td><td></td><td></td><td></td><td></td><td></td><td></td><td></td></tr>
<tr><td>Oligo fwd</td><td>CACCCACGAGACGGGCGGACGAAG</td><td></td><td></td><td></td><td></td><td></td><td></td><td></td><td></td><td></td><td></td></tr>
<tr><td>Oligo rev</td><td>AAACCTTCGTCCGCCCGTCTCGTG</td><td></td><td></td><td></td><td></td><td></td><td></td><td></td><td></td><td></td><td></td></tr>
<tr><td>Chromosome</td><td>start</td><td>end</td><td>strand</td><td>MM</td><td>target_seq</td><td>PAM</td><td>alignment</td><td>distance</td><td>position</td><td>gene name</td><td>gene id</td></tr>
<tr><td>GU305914.1</td><td>202699</td><td>202721</td><td>+</td><td>0</td><td>CACGAGACGGGCGGACGAAG</td><td>CGG</td><td>||||||||[||||||||||||]PAM</td><td>NA</td><td>Intergenic</td><td>NA</td><td>NA</td></tr>
<tr><td>T15507</td><td>GACGGGCGGACGAAGCGGAGCGG</td><td>1000</td><td></td><td></td><td></td><td></td><td></td><td></td><td></td><td></td><td></td></tr>
<tr><td>Oligo fwd</td><td>CACCGACGGGCGGACGAAGCGGAG</td><td></td><td></td><td></td><td></td><td></td><td></td><td></td><td></td><td></td><td></td></tr>
<tr><td>Oligo rev</td><td>AAACCTCCGCTTCGTCCGCCCGTC</td><td></td><td></td><td></td><td></td><td></td><td></td><td></td><td></td><td></td><td></td></tr>
<tr><td>Chromosome</td><td>start</td><td>end</td><td>strand</td><td>MM</td><td>target_seq</td><td>PAM</td><td>alignment</td><td>distance</td><td>position</td><td>gene name</td><td>gene id</td></tr>
<tr><td>GU305914.1</td><td>202704</td><td>202726</td><td>+</td><td>0</td><td>GACGGGCGGACGAAGCGGAG</td><td>CGG</td><td>||||||||[||||||||||||]PAM</td><td>NA</td><td>Intergenic</td><td>NA</td><td>NA</td></tr>
</table>

<table>
<tr><td>T15508</td><td>GACGAAGCGGAGCGGAGCCGAGG</td><td>1000</td><td colspan="9"></td></tr>
<tr><td>Oligo fwd</td><td>CACCGACGAAGCGGAGCGGAGCCG</td><td colspan="10"></td></tr>
<tr><td>Oligo rev</td><td>AAACCGGCTCCGCTCCGCTTCGTC</td><td colspan="10"></td></tr>
<tr><td>Chromosome</td><td>start</td><td>end</td><td>strand</td><td>MM</td><td>target_seq</td><td>PAM</td><td>alignment</td><td>distance</td><td>position</td><td>gene name</td><td>gene id</td></tr>
<tr><td>GU305914.1</td><td>202712</td><td>202734</td><td>+</td><td>0</td><td>GACGAAGCGGAGCGGAGCCG</td><td>AGG</td><td>||||||||[||||||||||||]PAM</td><td>NA</td><td>Intergenic</td><td>NA</td><td>NA</td></tr>
<tr><td>T15509</td><td>AGGCAACCACCTCTCTGCTGTGG</td><td>1000</td><td colspan="9"></td></tr>
<tr><td>Oligo fwd</td><td>CACCAGGCAACCACCTCTCTGCTG</td><td colspan="10"></td></tr>
<tr><td>Oligo rev</td><td>AAACCAGCAGAGAGGTGGTTGCCT</td><td colspan="10"></td></tr>
<tr><td>Chromosome</td><td>start</td><td>end</td><td>strand</td><td>MM</td><td>target_seq</td><td>PAM</td><td>alignment</td><td>distance</td><td>position</td><td>gene name</td><td>gene id</td></tr>
<tr><td>GU305914.1</td><td>202732</td><td>202754</td><td>+</td><td>0</td><td>AGGCAACCACCTCTCTGCTG</td><td>TGG</td><td>||||||||[||||||||||||]PAM</td><td>NA</td><td>Intergenic</td><td>NA</td><td>NA</td></tr>
<tr><td>T15510</td><td>AGAGATCTGTATACCGTTTCTGG</td><td>1000</td><td colspan="9"></td></tr>
<tr><td>Oligo fwd</td><td>CACCAGAGATCTGTATACCGTTTC</td><td colspan="10"></td></tr>
<tr><td>Oligo rev</td><td>AAACGAAACGGTATACAGATCTCT</td><td colspan="10"></td></tr>
<tr><td>Chromosome</td><td>start</td><td>end</td><td>strand</td><td>MM</td><td>target_seq</td><td>PAM</td><td>alignment</td><td>distance</td><td>position</td><td>gene name</td><td>gene id</td></tr>
<tr><td>GU305914.1</td><td>202759</td><td>202781</td><td>+</td><td>0</td><td>AGAGATCTGTATACCGTTTC</td><td>TGG</td><td>||||||||[||||||||||||]PAM</td><td>NA</td><td>Intergenic</td><td>NA</td><td>NA</td></tr>
<tr><td>T15511</td><td>ATACCGTTTCTGGAGTTACGCGG</td><td>1000</td><td colspan="9"></td></tr>
<tr><td>Oligo fwd</td><td>CACCATACCGTTTCTGGAGTTACG</td><td colspan="10"></td></tr>
<tr><td>Oligo rev</td><td>AAACCGTAACTCCAGAAACGGTAT</td><td colspan="10"></td></tr>
<tr><td>Chromosome</td><td>start</td><td>end</td><td>strand</td><td>MM</td><td>target_seq</td><td>PAM</td><td>alignment</td><td>distance</td><td>position</td><td>gene name</td><td>gene id</td></tr>
<tr><td>GU305914.1</td><td>202769</td><td>202791</td><td>+</td><td>0</td><td>ATACCGTTTCTGGAGTTACG</td><td>CGG</td><td>||||||||[||||||||||||]PAM</td><td>NA</td><td>Intergenic</td><td>NA</td><td>NA</td></tr>
<tr><td>T15512</td><td>CGGCCCTCTTTTACACCATCCGG</td><td>1000</td><td colspan="9"></td></tr>
<tr><td>Oligo fwd</td><td>CACCCGGCCCTCTTTTACACCATC</td><td colspan="10"></td></tr>
<tr><td>Oligo rev</td><td>AAACGATGGTGTAAAAGAGGGCCG</td><td colspan="10"></td></tr>
<tr><td>Chromosome</td><td>start</td><td>end</td><td>strand</td><td>MM</td><td>target_seq</td><td>PAM</td><td>alignment</td><td>distance</td><td>position</td><td>gene name</td><td>gene id</td></tr>
<tr><td>GU305914.1</td><td>202789</td><td>202811</td><td>+</td><td>0</td><td>CGGCCCTCTTTTACACCATC</td><td>CGG</td><td>||||||||[||||||||||||]PAM</td><td>NA</td><td>Intergenic</td><td>NA</td><td>NA</td></tr>
<tr><td>T15513</td><td>GGCCCTCTTTTACACCATCCGGG</td><td>1000</td><td colspan="9"></td></tr>
<tr><td>Oligo fwd</td><td>CACCGGCCCTCTTTTACACCATCC</td><td colspan="10"></td></tr>
<tr><td>Oligo rev</td><td>AAACGGATGGTGTAAAAGAGGGCC</td><td colspan="10"></td></tr>
<tr><td>Chromosome</td><td>start</td><td>end</td><td>strand</td><td>MM</td><td>target_seq</td><td>PAM</td><td>alignment</td><td>distance</td><td>position</td><td>gene name</td><td>gene id</td></tr>
<tr><td>GU305914.1</td><td>202790</td><td>202812</td><td>+</td><td>0</td><td>GGCCCTCTTTTACACCATCC</td><td>GGG</td><td>||||||||[||||||||||||]PAM</td><td>NA</td><td>Intergenic</td><td>NA</td><td>NA</td></tr>
<tr><td>T15514</td><td>ACCACACCCAGACAGATCAGAGG</td><td>1000</td><td colspan="9"></td></tr>
<tr><td>Oligo fwd</td><td>CACCACCACACCCAGACAGATCAG</td><td colspan="10"></td></tr>
<tr><td>Oligo rev</td><td>AAACCTGATCTGTCTGGGTGTGGT</td><td colspan="10"></td></tr>
<tr><td>Chromosome</td><td>start</td><td>end</td><td>strand</td><td>MM</td><td>target_seq</td><td>PAM</td><td>alignment</td><td>distance</td><td>position</td><td>gene name</td><td>gene id</td></tr>
<tr><td>GU305914.1</td><td>202815</td><td>202837</td><td>+</td><td>0</td><td>ACCACACCCAGACAGATCAG</td><td>AGG</td><td>||||||||[||||||||||||]PAM</td><td>NA</td><td>Intergenic</td><td>NA</td><td>NA</td></tr>
<tr><td>T15515</td><td>TCAGAGGCGCGCTCAAATGCTGG</td><td>1000</td><td colspan="9"></td></tr>
<tr><td>Oligo fwd</td><td>CACCTCAGAGGCGCGCTCAAATGC</td><td colspan="10"></td></tr>
<tr><td>Oligo rev</td><td>AAACGCATTTGAGCGCGCCTCTGA</td><td colspan="10"></td></tr>
<tr><td>Chromosome</td><td>start</td><td>end</td><td>strand</td><td>MM</td><td>target_seq</td><td>PAM</td><td>alignment</td><td>distance</td><td>position</td><td>gene name</td><td>gene id</td></tr>
<tr><td>GU305914.1</td><td>202831</td><td>202853</td><td>+</td><td>0</td><td>TCAGAGGCGCGCTCAAATGC</td><td>TGG</td><td>||||||||[||||||||||||]PAM</td><td>NA</td><td>Intergenic</td><td>NA</td><td>NA</td></tr>
<tr><td>T15516</td><td>CAGAGGCGCGCTCAAATGCTGGG</td><td>1000</td><td colspan="9"></td></tr>
<tr><td>Oligo fwd</td><td>CACCCAGAGGCGCGCTCAAATGCT</td><td colspan="10"></td></tr>
<tr><td>Oligo rev</td><td>AAACAGCATTTGAGCGCGCCTCTG</td><td colspan="10"></td></tr>
<tr><td>Chromosome</td><td>start</td><td>end</td><td>strand</td><td>MM</td><td>target_seq</td><td>PAM</td><td>alignment</td><td>distance</td><td>position</td><td>gene name</td><td>gene id</td></tr>
<tr><td>GU305914.1</td><td>202832</td><td>202854</td><td>+</td><td>0</td><td>CAGAGGCGCGCTCAAATGCT</td><td>GGG</td><td>||||||||[||||||||||||]PAM</td><td>NA</td><td>Intergenic</td><td>NA</td><td>NA</td></tr>
<tr><td>T15517</td><td>CGCGCTCAAATGCTGGGATCCGG</td><td>1000</td><td colspan="9"></td></tr>
<tr><td>Oligo fwd</td><td>CACCCGCGCTCAAATGCTGGGATC</td><td colspan="10"></td></tr>
<tr><td>Oligo rev</td><td>AAACGATCCCAGCATTTGAGCGCG</td><td colspan="10"></td></tr>
<tr><td>Chromosome</td><td>start</td><td>end</td><td>strand</td><td>MM</td><td>target_seq</td><td>PAM</td><td>alignment</td><td>distance</td><td>position</td><td>gene name</td><td>gene id</td></tr>
<tr><td>GU305914.1</td><td>202838</td><td>202860</td><td>+</td><td>0</td><td>CGCGCTCAAATGCTGGGATC</td><td>CGG</td><td>||||||||[||||||||||||]PAM</td><td>NA</td><td>Intergenic</td><td>NA</td><td>NA</td></tr>
<tr><td>T15518</td><td>GCGCTCAAATGCTGGGATCCGGG</td><td>1000</td><td colspan="9"></td></tr>
<tr><td>Oligo fwd</td><td>CACCGCGCTCAAATGCTGGGATCC</td><td colspan="10"></td></tr>
<tr><td>Oligo rev</td><td>AAACGGATCCCAGCATTTGAGCGC</td><td colspan="10"></td></tr>
<tr><td>Chromosome</td><td>start</td><td>end</td><td>strand</td><td>MM</td><td>target_seq</td><td>PAM</td><td>alignment</td><td>distance</td><td>position</td><td>gene name</td><td>gene id</td></tr>
<tr><td>GU305914.1</td><td>202839</td><td>202861</td><td>+</td><td>0</td><td>GCGCTCAAATGCTGGGATCC</td><td>GGG</td><td>||||||||[||||||||||||]PAM</td><td>NA</td><td>Intergenic</td><td>NA</td><td>NA</td></tr>
<tr><td>T15519</td><td>GGATCCGGGACCGTGACGATCGG</td><td>1000</td><td colspan="9"></td></tr>
<tr><td>Oligo fwd</td><td>CACCGGATCCGGGACCGTGACGAT</td><td colspan="10"></td></tr>
<tr><td>Oligo rev</td><td>AAACATCGTCACGGTCCCGGATCC</td><td colspan="10"></td></tr>
<tr><td>Chromosome</td><td>start</td><td>end</td><td>strand</td><td>MM</td><td>target_seq</td><td>PAM</td><td>alignment</td><td>distance</td><td>position</td><td>gene name</td><td>gene id</td></tr>
<tr><td>GU305914.1</td><td>202853</td><td>202875</td><td>+</td><td>0</td><td>GGATCCGGGACCGTGACGAT</td><td>CGG</td><td>||||||||[||||||||||||]PAM</td><td>NA</td><td>Intergenic</td><td>NA</td><td>NA</td></tr>
<tr><td>T15520</td><td>CGGACCGTCCCCGTTCGACCAGG</td><td>1000</td><td colspan="9"></td></tr>
<tr><td>Oligo fwd</td><td>CACCCGGACCGTCCCCGTTCGACC</td><td colspan="10"></td></tr>
<tr><td>Oligo rev</td><td>AAACGGTCGAACGGGGACGGTCCG</td><td colspan="10"></td></tr>
<tr><td>Chromosome</td><td>start</td><td>end</td><td>strand</td><td>MM</td><td>target_seq</td><td>PAM</td><td>alignment</td><td>distance</td><td>position</td><td>gene name</td><td>gene id</td></tr>
<tr><td>GU305914.1</td><td>202873</td><td>202895</td><td>+</td><td>0</td><td>CGGACCGTCCCCGTTCGACC</td><td>AGG</td><td>||||||||[||||||||||||]PAM</td><td>NA</td><td>Intergenic</td><td>NA</td><td>NA</td></tr>
<tr><td>T15521</td><td>ACCAGGCGTCCGACAACTCGCGG</td><td>1000</td><td colspan="9"></td></tr>
<tr><td>Oligo fwd</td><td>CACCACCAGGCGTCCGACAACTCG</td><td colspan="10"></td></tr>
<tr><td>Oligo rev</td><td>AAACCGAGTTGTCGGACGCCTGGT</td><td colspan="10"></td></tr>
<tr><td>Chromosome</td><td>start</td><td>end</td><td>strand</td><td>MM</td><td>target_seq</td><td>PAM</td><td>alignment</td><td>distance</td><td>position</td><td>gene name</td><td>gene id</td></tr>
<tr><td>GU305914.1</td><td>202890</td><td>202912</td><td>+</td><td>0</td><td>ACCAGGCGTCCGACAACTCG</td><td>CGG</td><td>||||||||[||||||||||||]PAM</td><td>NA</td><td>Intergenic</td><td>NA</td><td>NA</td></tr>
<tr><td>T15522</td><td>CCAGGCGTCCGACAACTCGCGGG</td><td>1000</td><td colspan="9"></td></tr>
<tr><td>Oligo fwd</td><td>CACCCCAGGCGTCCGACAACTCGC</td><td colspan="10"></td></tr>
<tr><td>Oligo rev</td><td>AAACGCGAGTTGTCGGACGCCTGG</td><td colspan="10"></td></tr>
<tr><td>Chromosome</td><td>start</td><td>end</td><td>strand</td><td>MM</td><td>target_seq</td><td>PAM</td><td>alignment</td><td>distance</td><td>position</td><td>gene name</td><td>gene id</td></tr>
<tr><td>GU305914.1</td><td>202891</td><td>202913</td><td>+</td><td>0</td><td>CCAGGCGTCCGACAACTCGC</td><td>GGG</td><td>||||||||[||||||||||||]PAM</td><td>NA</td><td>Intergenic</td><td>NA</td><td>NA</td></tr>
</table>

<table>
<tr><td>T15523</td><td>GGCGTCCGACAACTCGCGGGCGG</td><td>1000</td><td></td><td></td><td></td><td></td><td></td><td></td><td></td><td></td><td></td></tr>
<tr><td>Oligo fwd</td><td>CACCGGCGTCCGACAACTCGCGGG</td><td></td><td></td><td></td><td></td><td></td><td></td><td></td><td></td><td></td><td></td></tr>
<tr><td>Oligo rev</td><td>AAACCCCGCGAGTTGTCGGACGCC</td><td></td><td></td><td></td><td></td><td></td><td></td><td></td><td></td><td></td><td></td></tr>
<tr><td>Chromosome</td><td>start</td><td>end</td><td>strand</td><td>MM</td><td>target_seq</td><td>PAM</td><td>alignment</td><td>distance</td><td>position</td><td>gene name</td><td>gene id</td></tr>
<tr><td>GU305914.1</td><td>202894</td><td>202916</td><td>+</td><td>0</td><td>GGCGTCCGACAACTCGCGGG</td><td>CGG</td><td>||||||||[||||||||||||]PAM</td><td>NA</td><td>Intergenic</td><td>NA</td><td>NA</td></tr>
</table>

<table>
<tr><td>T15524</td><td>CTCGCGGGCGGACATCGTGACGG</td><td>1000</td><td></td><td></td><td></td><td></td><td></td><td></td><td></td><td></td><td></td></tr>
<tr><td>Oligo fwd</td><td>CACCCTCGCGGGCGGACATCGTGA</td><td></td><td></td><td></td><td></td><td></td><td></td><td></td><td></td><td></td><td></td></tr>
<tr><td>Oligo rev</td><td>AAACTCACGATGTCCGCCCGCGAG</td><td></td><td></td><td></td><td></td><td></td><td></td><td></td><td></td><td></td><td></td></tr>
<tr><td>Chromosome</td><td>start</td><td>end</td><td>strand</td><td>MM</td><td>target_seq</td><td>PAM</td><td>alignment</td><td>distance</td><td>position</td><td>gene name</td><td>gene id</td></tr>
<tr><td>GU305914.1</td><td>202906</td><td>202928</td><td>+</td><td>0</td><td>CTCGCGGGCGGACATCGTGA</td><td>CGG</td><td>||||||||[||||||||||||]PAM</td><td>NA</td><td>Intergenic</td><td>NA</td><td>NA</td></tr>
</table>

<table>
<tr><td>T15525</td><td>GACATCGTGACGGCATACGCCGG</td><td>1000</td><td></td><td></td><td></td><td></td><td></td><td></td><td></td><td></td><td></td></tr>
<tr><td>Oligo fwd</td><td>CACCGACATCGTGACGGCATACGC</td><td></td><td></td><td></td><td></td><td></td><td></td><td></td><td></td><td></td><td></td></tr>
<tr><td>Oligo rev</td><td>AAACGCGTATGCCGTCACGATGTC</td><td></td><td></td><td></td><td></td><td></td><td></td><td></td><td></td><td></td><td></td></tr>
<tr><td>Chromosome</td><td>start</td><td>end</td><td>strand</td><td>MM</td><td>target_seq</td><td>PAM</td><td>alignment</td><td>distance</td><td>position</td><td>gene name</td><td>gene id</td></tr>
<tr><td>GU305914.1</td><td>202916</td><td>202938</td><td>+</td><td>0</td><td>GACATCGTGACGGCATACGC</td><td>CGG</td><td>||||||||[||||||||||||]PAM</td><td>NA</td><td>Intergenic</td><td>NA</td><td>NA</td></tr>
</table>

<table>
<tr><td>T15526</td><td>CGTGACGGCATACGCCGGACAGG</td><td>1000</td><td></td><td></td><td></td><td></td><td></td><td></td><td></td><td></td><td></td></tr>
<tr><td>Oligo fwd</td><td>CACCCGTGACGGCATACGCCGGAC</td><td></td><td></td><td></td><td></td><td></td><td></td><td></td><td></td><td></td><td></td></tr>
<tr><td>Oligo rev</td><td>AAACGTCCGGCGTATGCCGTCACG</td><td></td><td></td><td></td><td></td><td></td><td></td><td></td><td></td><td></td><td></td></tr>
<tr><td>Chromosome</td><td>start</td><td>end</td><td>strand</td><td>MM</td><td>target_seq</td><td>PAM</td><td>alignment</td><td>distance</td><td>position</td><td>gene name</td><td>gene id</td></tr>
<tr><td>GU305914.1</td><td>202921</td><td>202943</td><td>+</td><td>0</td><td>CGTGACGGCATACGCCGGAC</td><td>AGG</td><td>||||||||[||||||||||||]PAM</td><td>NA</td><td>Intergenic</td><td>NA</td><td>NA</td></tr>
</table>

<table>
<tr><td>T15527</td><td>TGCTCATCACGTTGTCGCCGCGG</td><td>1000</td><td></td><td></td><td></td><td></td><td></td><td></td><td></td><td></td><td></td></tr>
<tr><td>Oligo fwd</td><td>CACCTGCTCATCACGTTGTCGCCG</td><td></td><td></td><td></td><td></td><td></td><td></td><td></td><td></td><td></td><td></td></tr>
<tr><td>Oligo rev</td><td>AAACCGGCGACAACGTGATGAGCA</td><td></td><td></td><td></td><td></td><td></td><td></td><td></td><td></td><td></td><td></td></tr>
<tr><td>Chromosome</td><td>start</td><td>end</td><td>strand</td><td>MM</td><td>target_seq</td><td>PAM</td><td>alignment</td><td>distance</td><td>position</td><td>gene name</td><td>gene id</td></tr>
<tr><td>GU305914.1</td><td>202956</td><td>202978</td><td>+</td><td>0</td><td>TGCTCATCACGTTGTCGCCG</td><td>CGG</td><td>||||||||[||||||||||||]PAM</td><td>NA</td><td>Intergenic</td><td>NA</td><td>NA</td></tr>
</table>

<table>
<tr><td>T15528</td><td>GTTGTCGCCGCGGAACCAGATGG</td><td>1000</td><td></td><td></td><td></td><td></td><td></td><td></td><td></td><td></td><td></td></tr>
<tr><td>Oligo fwd</td><td>CACCGTTGTCGCCGCGGAACCAGA</td><td></td><td></td><td></td><td></td><td></td><td></td><td></td><td></td><td></td><td></td></tr>
<tr><td>Oligo rev</td><td>AAACTCTGGTTCCGCGGCGACAAC</td><td></td><td></td><td></td><td></td><td></td><td></td><td></td><td></td><td></td><td></td></tr>
<tr><td>Chromosome</td><td>start</td><td>end</td><td>strand</td><td>MM</td><td>target_seq</td><td>PAM</td><td>alignment</td><td>distance</td><td>position</td><td>gene name</td><td>gene id</td></tr>
<tr><td>GU305914.1</td><td>202966</td><td>202988</td><td>+</td><td>0</td><td>GTTGTCGCCGCGGAACCAGA</td><td>TGG</td><td>||||||||[||||||||||||]PAM</td><td>NA</td><td>Intergenic</td><td>NA</td><td>NA</td></tr>
</table>

<table>
<tr><td>T15529</td><td>CCGCGGAACCAGATGGTCATCGG</td><td>1000</td><td></td><td></td><td></td><td></td><td></td><td></td><td></td><td></td><td></td></tr>
<tr><td>Oligo fwd</td><td>CACCCCGCGGAACCAGATGGTCAT</td><td></td><td></td><td></td><td></td><td></td><td></td><td></td><td></td><td></td><td></td></tr>
<tr><td>Oligo rev</td><td>AAACATGACCATCTGGTTCCGCGG</td><td></td><td></td><td></td><td></td><td></td><td></td><td></td><td></td><td></td><td></td></tr>
<tr><td>Chromosome</td><td>start</td><td>end</td><td>strand</td><td>MM</td><td>target_seq</td><td>PAM</td><td>alignment</td><td>distance</td><td>position</td><td>gene name</td><td>gene id</td></tr>
<tr><td>GU305914.1</td><td>202973</td><td>202995</td><td>+</td><td>0</td><td>CCGCGGAACCAGATGGTCAT</td><td>CGG</td><td>||||||||[||||||||||||]PAM</td><td>NA</td><td>Intergenic</td><td>NA</td><td>NA</td></tr>
</table>

<table>
<tr><td>T15530</td><td>GGAACCAGATGGTCATCGGTAGG</td><td>1000</td><td></td><td></td><td></td><td></td><td></td><td></td><td></td><td></td><td></td></tr>
<tr><td>Oligo fwd</td><td>CACCGGAACCAGATGGTCATCGGT</td><td></td><td></td><td></td><td></td><td></td><td></td><td></td><td></td><td></td><td></td></tr>
<tr><td>Oligo rev</td><td>AAACACCGATGACCATCTGGTTCC</td><td></td><td></td><td></td><td></td><td></td><td></td><td></td><td></td><td></td><td></td></tr>
<tr><td>Chromosome</td><td>start</td><td>end</td><td>strand</td><td>MM</td><td>target_seq</td><td>PAM</td><td>alignment</td><td>distance</td><td>position</td><td>gene name</td><td>gene id</td></tr>
<tr><td>GU305914.1</td><td>202977</td><td>202999</td><td>+</td><td>0</td><td>GGAACCAGATGGTCATCGGT</td><td>AGG</td><td>||||||||[||||||||||||]PAM</td><td>NA</td><td>Intergenic</td><td>NA</td><td>NA</td></tr>
<tr><td>GU305914.1</td><td>193700</td><td>193722</td><td>+</td><td>4</td><td>TGCCCCAGATCGTCATCGGT</td><td>GGG</td><td>-|--||||[||-|||||||||]PAM</td><td>NA</td><td>Intergenic</td><td>NA</td><td>NA</td></tr>
</table>

<table>
<tr><td>T15531</td><td>ATGGTCATCGGTAGGTCGCATGG</td><td>1000</td><td></td><td></td><td></td><td></td><td></td><td></td><td></td><td></td><td></td></tr>
<tr><td>Oligo fwd</td><td>CACCATGGTCATCGGTAGGTCGCA</td><td></td><td></td><td></td><td></td><td></td><td></td><td></td><td></td><td></td><td></td></tr>
<tr><td>Oligo rev</td><td>AAACTGCGACCTACCGATGACCAT</td><td></td><td></td><td></td><td></td><td></td><td></td><td></td><td></td><td></td><td></td></tr>
<tr><td>Chromosome</td><td>start</td><td>end</td><td>strand</td><td>MM</td><td>target_seq</td><td>PAM</td><td>alignment</td><td>distance</td><td>position</td><td>gene name</td><td>gene id</td></tr>
<tr><td>GU305914.1</td><td>202985</td><td>203007</td><td>+</td><td>0</td><td>ATGGTCATCGGTAGGTCGCA</td><td>TGG</td><td>||||||||[||||||||||||]PAM</td><td>NA</td><td>Intergenic</td><td>NA</td><td>NA</td></tr>
</table>

<table>
<tr><td>T15532</td><td>GTCATCGGTAGGTCGCATGGCGG</td><td>1000</td><td></td><td></td><td></td><td></td><td></td><td></td><td></td><td></td><td></td></tr>
<tr><td>Oligo fwd</td><td>CACCGTCATCGGTAGGTCGCATGG</td><td></td><td></td><td></td><td></td><td></td><td></td><td></td><td></td><td></td><td></td></tr>
<tr><td>Oligo rev</td><td>AAACCCATGCGACCTACCGATGAC</td><td></td><td></td><td></td><td></td><td></td><td></td><td></td><td></td><td></td><td></td></tr>
<tr><td>Chromosome</td><td>start</td><td>end</td><td>strand</td><td>MM</td><td>target_seq</td><td>PAM</td><td>alignment</td><td>distance</td><td>position</td><td>gene name</td><td>gene id</td></tr>
<tr><td>GU305914.1</td><td>202988</td><td>203010</td><td>+</td><td>0</td><td>GTCATCGGTAGGTCGCATGG</td><td>CGG</td><td>||||||||[||||||||||||]PAM</td><td>NA</td><td>Intergenic</td><td>NA</td><td>NA</td></tr>
</table>

<table>
<tr><td>T15533</td><td>GTTCCGCGCGTCTCTTCACCTGG</td><td>1000</td><td></td><td></td><td></td><td></td><td></td><td></td><td></td><td></td><td></td></tr>
<tr><td>Oligo fwd</td><td>CACCGTTCCGCGCGTCTCTTCACC</td><td></td><td></td><td></td><td></td><td></td><td></td><td></td><td></td><td></td><td></td></tr>
<tr><td>Oligo rev</td><td>AAACGGTGAAGAGACGCGCGGAAC</td><td></td><td></td><td></td><td></td><td></td><td></td><td></td><td></td><td></td><td></td></tr>
<tr><td>Chromosome</td><td>start</td><td>end</td><td>strand</td><td>MM</td><td>target_seq</td><td>PAM</td><td>alignment</td><td>distance</td><td>position</td><td>gene name</td><td>gene id</td></tr>
<tr><td>GU305914.1</td><td>203010</td><td>203032</td><td>+</td><td>0</td><td>GTTCCGCGCGTCTCTTCACC</td><td>TGG</td><td>||||||||[||||||||||||]PAM</td><td>NA</td><td>Intergenic</td><td>NA</td><td>NA</td></tr>
</table>

<table>
<tr><td>T15534</td><td>CGCGCGTCTCTTCACCTGGTAGG</td><td>1000</td><td></td><td></td><td></td><td></td><td></td><td></td><td></td><td></td><td></td></tr>
<tr><td>Oligo fwd</td><td>CACCCGCGCGTCTCTTCACCTGGT</td><td></td><td></td><td></td><td></td><td></td><td></td><td></td><td></td><td></td><td></td></tr>
<tr><td>Oligo rev</td><td>AAACACCAGGTGAAGAGACGCGCG</td><td></td><td></td><td></td><td></td><td></td><td></td><td></td><td></td><td></td><td></td></tr>
<tr><td>Chromosome</td><td>start</td><td>end</td><td>strand</td><td>MM</td><td>target_seq</td><td>PAM</td><td>alignment</td><td>distance</td><td>position</td><td>gene name</td><td>gene id</td></tr>
<tr><td>GU305914.1</td><td>203014</td><td>203036</td><td>+</td><td>0</td><td>CGCGCGTCTCTTCACCTGGT</td><td>AGG</td><td>||||||||[||||||||||||]PAM</td><td>NA</td><td>Intergenic</td><td>NA</td><td>NA</td></tr>
</table>

<table>
<tr><td>T15535</td><td>CGTCTCTTCACCTGGTAGGACGG</td><td>1000</td><td></td><td></td><td></td><td></td><td></td><td></td><td></td><td></td><td></td></tr>
<tr><td>Oligo fwd</td><td>CACCCGTCTCTTCACCTGGTAGGA</td><td></td><td></td><td></td><td></td><td></td><td></td><td></td><td></td><td></td><td></td></tr>
<tr><td>Oligo rev</td><td>AAACTCCTACCAGGTGAAGAGACG</td><td></td><td></td><td></td><td></td><td></td><td></td><td></td><td></td><td></td><td></td></tr>
<tr><td>Chromosome</td><td>start</td><td>end</td><td>strand</td><td>MM</td><td>target_seq</td><td>PAM</td><td>alignment</td><td>distance</td><td>position</td><td>gene name</td><td>gene id</td></tr>
<tr><td>GU305914.1</td><td>203018</td><td>203040</td><td>+</td><td>0</td><td>CGTCTCTTCACCTGGTAGGA</td><td>CGG</td><td>||||||||[||||||||||||]PAM</td><td>NA</td><td>Intergenic</td><td>NA</td><td>NA</td></tr>
</table>

<table>
<tr><td>T15536</td><td>ACCTGGTAGGACGGCCGTTTCGG</td><td>1000</td><td></td><td></td><td></td><td></td><td></td><td></td><td></td><td></td><td></td></tr>
<tr><td>Oligo fwd</td><td>CACCACCTGGTAGGACGGCCGTTT</td><td></td><td></td><td></td><td></td><td></td><td></td><td></td><td></td><td></td><td></td></tr>
<tr><td>Oligo rev</td><td>AAACAAACGGCCGTCCTACCAGGT</td><td></td><td></td><td></td><td></td><td></td><td></td><td></td><td></td><td></td><td></td></tr>
<tr><td>Chromosome</td><td>start</td><td>end</td><td>strand</td><td>MM</td><td>target_seq</td><td>PAM</td><td>alignment</td><td>distance</td><td>position</td><td>gene name</td><td>gene id</td></tr>
<tr><td>GU305914.1</td><td>203027</td><td>203049</td><td>+</td><td>0</td><td>ACCTGGTAGGACGGCCGTTT</td><td>CGG</td><td>||||||||[||||||||||||]PAM</td><td>NA</td><td>Intergenic</td><td>NA</td><td>NA</td></tr>
</table>

<table>
<tr><td>T15537</td><td>GACCGCCGCCATGACGCTCGCGG</td><td>1000</td></tr>
<tr><td>Oligo fwd</td><td>CACCGACCGCCGCCATGACGCTCG</td><td></td></tr>
<tr><td>Oligo rev</td><td>AAACCGAGCGTCATGGCGGCGGTC</td><td></td></tr>
</table>

<table>
<tr><td>Chromosome</td><td>start</td><td>end</td><td>strand</td><td>MM</td><td>target_seq</td><td>PAM</td><td>alignment</td><td>distance</td><td>position</td><td>gene name</td><td>gene id</td></tr>
<tr><td>GU305914.1</td><td>203134</td><td>203156</td><td>+</td><td>0</td><td>GACCGCCGCCATGACGCTCG</td><td>CGG</td><td>||||||||[||||||||||||]PAM</td><td>NA</td><td>Intergenic</td><td>NA</td><td>NA</td></tr>
<tr><td></td><td></td><td></td><td></td><td></td><td></td><td></td><td></td><td></td><td></td><td></td><td></td></tr>
<tr><td>T15538</td><td>ACCGCCGCCATGACGCTCGCGGG</td><td>1000</td><td></td><td></td><td></td><td></td><td></td><td></td><td></td><td></td><td></td></tr>
<tr><td>Oligo fwd</td><td>CACCACCGCCGCCATGACGCTCGC</td><td></td><td></td><td></td><td></td><td></td><td></td><td></td><td></td><td></td><td></td></tr>
<tr><td>Oligo rev</td><td>AAACGCGAGCGTCATGGCGGCGGT</td><td></td><td></td><td></td><td></td><td></td><td></td><td></td><td></td><td></td><td></td></tr>
<tr><td>Chromosome</td><td>start</td><td>end</td><td>strand</td><td>MM</td><td>target_seq</td><td>PAM</td><td>alignment</td><td>distance</td><td>position</td><td>gene name</td><td>gene id</td></tr>
<tr><td>GU305914.1</td><td>203135</td><td>203157</td><td>+</td><td>0</td><td>ACCGCCGCCATGACGCTCGC</td><td>GGG</td><td>||||||||[||||||||||||]PAM</td><td>NA</td><td>Intergenic</td><td>NA</td><td>NA</td></tr>
<tr><td></td><td></td><td></td><td></td><td></td><td></td><td></td><td></td><td></td><td></td><td></td><td></td></tr>
<tr><td>T15539</td><td>TGACGCTCGCGGGTATGAACAGG</td><td>1000</td><td></td><td></td><td></td><td></td><td></td><td></td><td></td><td></td><td></td></tr>
<tr><td>Oligo fwd</td><td>CACCTGACGCTCGCGGGTATGAAC</td><td></td><td></td><td></td><td></td><td></td><td></td><td></td><td></td><td></td><td></td></tr>
<tr><td>Oligo rev</td><td>AAACGTTCATACCCGCGAGCGTCA</td><td></td><td></td><td></td><td></td><td></td><td></td><td></td><td></td><td></td><td></td></tr>
<tr><td>Chromosome</td><td>start</td><td>end</td><td>strand</td><td>MM</td><td>target_seq</td><td>PAM</td><td>alignment</td><td>distance</td><td>position</td><td>gene name</td><td>gene id</td></tr>
<tr><td>GU305914.1</td><td>203145</td><td>203167</td><td>+</td><td>0</td><td>TGACGCTCGCGGGTATGAAC</td><td>AGG</td><td>||||||||[||||||||||||]PAM</td><td>NA</td><td>Intergenic</td><td>NA</td><td>NA</td></tr>
<tr><td></td><td></td><td></td><td></td><td></td><td></td><td></td><td></td><td></td><td></td><td></td><td></td></tr>
<tr><td>T15540</td><td>GACGCTCGCGGGTATGAACAGGG</td><td>1000</td><td></td><td></td><td></td><td></td><td></td><td></td><td></td><td></td><td></td></tr>
<tr><td>Oligo fwd</td><td>CACCGACGCTCGCGGGTATGAACA</td><td></td><td></td><td></td><td></td><td></td><td></td><td></td><td></td><td></td><td></td></tr>
<tr><td>Oligo rev</td><td>AAACTGTTCATACCCGCGAGCGTC</td><td></td><td></td><td></td><td></td><td></td><td></td><td></td><td></td><td></td><td></td></tr>
<tr><td>Chromosome</td><td>start</td><td>end</td><td>strand</td><td>MM</td><td>target_seq</td><td>PAM</td><td>alignment</td><td>distance</td><td>position</td><td>gene name</td><td>gene id</td></tr>
<tr><td>GU305914.1</td><td>203146</td><td>203168</td><td>+</td><td>0</td><td>GACGCTCGCGGGTATGAACA</td><td>GGG</td><td>||||||||[||||||||||||]PAM</td><td>NA</td><td>Intergenic</td><td>NA</td><td>NA</td></tr>
<tr><td></td><td></td><td></td><td></td><td></td><td></td><td></td><td></td><td></td><td></td><td></td><td></td></tr>
<tr><td>T15541</td><td>CGGGTATGAACAGGGCCAACAGG</td><td>1000</td><td></td><td></td><td></td><td></td><td></td><td></td><td></td><td></td><td></td></tr>
<tr><td>Oligo fwd</td><td>CACCCGGGTATGAACAGGGCCAAC</td><td></td><td></td><td></td><td></td><td></td><td></td><td></td><td></td><td></td><td></td></tr>
<tr><td>Oligo rev</td><td>AAACGTTGGCCCTGTTCATACCCG</td><td></td><td></td><td></td><td></td><td></td><td></td><td></td><td></td><td></td><td></td></tr>
<tr><td>Chromosome</td><td>start</td><td>end</td><td>strand</td><td>MM</td><td>target_seq</td><td>PAM</td><td>alignment</td><td>distance</td><td>position</td><td>gene name</td><td>gene id</td></tr>
<tr><td>GU305914.1</td><td>203154</td><td>203176</td><td>+</td><td>0</td><td>CGGGTATGAACAGGGCCAAC</td><td>AGG</td><td>||||||||[||||||||||||]PAM</td><td>NA</td><td>Intergenic</td><td>NA</td><td>NA</td></tr>
<tr><td></td><td></td><td></td><td></td><td></td><td></td><td></td><td></td><td></td><td></td><td></td><td></td></tr>
<tr><td>T15542</td><td>CACGATCTGCAGTTCGAGCCCGG</td><td>1000</td><td></td><td></td><td></td><td></td><td></td><td></td><td></td><td></td><td></td></tr>
<tr><td>Oligo fwd</td><td>CACCCACGATCTGCAGTTCGAGCC</td><td></td><td></td><td></td><td></td><td></td><td></td><td></td><td></td><td></td><td></td></tr>
<tr><td>Oligo rev</td><td>AAACGGCTCGAACTGCAGATCGTG</td><td></td><td></td><td></td><td></td><td></td><td></td><td></td><td></td><td></td><td></td></tr>
<tr><td>Chromosome</td><td>start</td><td>end</td><td>strand</td><td>MM</td><td>target_seq</td><td>PAM</td><td>alignment</td><td>distance</td><td>position</td><td>gene name</td><td>gene id</td></tr>
<tr><td>GU305914.1</td><td>203218</td><td>203240</td><td>+</td><td>0</td><td>CACGATCTGCAGTTCGAGCC</td><td>CGG</td><td>||||||||[||||||||||||]PAM</td><td>NA</td><td>Intergenic</td><td>NA</td><td>NA</td></tr>
<tr><td></td><td></td><td></td><td></td><td></td><td></td><td></td><td></td><td></td><td></td><td></td><td></td></tr>
<tr><td>T15543</td><td>CATAGACGTCGATATGTTTTTGG</td><td>1000</td><td></td><td></td><td></td><td></td><td></td><td></td><td></td><td></td><td></td></tr>
<tr><td>Oligo fwd</td><td>CACCCATAGACGTCGATATGTTTT</td><td></td><td></td><td></td><td></td><td></td><td></td><td></td><td></td><td></td><td></td></tr>
<tr><td>Oligo rev</td><td>AAACAAAACATATCGACGTCTATG</td><td></td><td></td><td></td><td></td><td></td><td></td><td></td><td></td><td></td><td></td></tr>
<tr><td>Chromosome</td><td>start</td><td>end</td><td>strand</td><td>MM</td><td>target_seq</td><td>PAM</td><td>alignment</td><td>distance</td><td>position</td><td>gene name</td><td>gene id</td></tr>
<tr><td>GU305914.1</td><td>203292</td><td>203314</td><td>+</td><td>0</td><td>CATAGACGTCGATATGTTTT</td><td>TGG</td><td>||||||||[||||||||||||]PAM</td><td>NA</td><td>Intergenic</td><td>NA</td><td>NA</td></tr>
<tr><td></td><td></td><td></td><td></td><td></td><td></td><td></td><td></td><td></td><td></td><td></td><td></td></tr>
<tr><td>T15544</td><td>TTTTTGGTAGAGCCCGTTACCGG</td><td>1000</td><td></td><td></td><td></td><td></td><td></td><td></td><td></td><td></td><td></td></tr>
<tr><td>Oligo fwd</td><td>CACCTTTTTGGTAGAGCCCGTTAC</td><td></td><td></td><td></td><td></td><td></td><td></td><td></td><td></td><td></td><td></td></tr>
<tr><td>Oligo rev</td><td>AAACGTAACGGGCTCTACCAAAAA</td><td></td><td></td><td></td><td></td><td></td><td></td><td></td><td></td><td></td><td></td></tr>
<tr><td>Chromosome</td><td>start</td><td>end</td><td>strand</td><td>MM</td><td>target_seq</td><td>PAM</td><td>alignment</td><td>distance</td><td>position</td><td>gene name</td><td>gene id</td></tr>
<tr><td>GU305914.1</td><td>203308</td><td>203330</td><td>+</td><td>0</td><td>TTTTTGGTAGAGCCCGTTAC</td><td>CGG</td><td>||||||||[||||||||||||]PAM</td><td>NA</td><td>Intergenic</td><td>NA</td><td>NA</td></tr>
<tr><td></td><td></td><td></td><td></td><td></td><td></td><td></td><td></td><td></td><td></td><td></td><td></td></tr>
<tr><td>T15545</td><td>TGGTAGAGCCCGTTACCGGACGG</td><td>1000</td><td></td><td></td><td></td><td></td><td></td><td></td><td></td><td></td><td></td></tr>
<tr><td>Oligo fwd</td><td>CACCTGGTAGAGCCCGTTACCGGA</td><td></td><td></td><td></td><td></td><td></td><td></td><td></td><td></td><td></td><td></td></tr>
<tr><td>Oligo rev</td><td>AAACTCCGGTAACGGGCTCTACCA</td><td></td><td></td><td></td><td></td><td></td><td></td><td></td><td></td><td></td><td></td></tr>
<tr><td>Chromosome</td><td>start</td><td>end</td><td>strand</td><td>MM</td><td>target_seq</td><td>PAM</td><td>alignment</td><td>distance</td><td>position</td><td>gene name</td><td>gene id</td></tr>
<tr><td>GU305914.1</td><td>203312</td><td>203334</td><td>+</td><td>0</td><td>TGGTAGAGCCCGTTACCGGA</td><td>CGG</td><td>||||||||[||||||||||||]PAM</td><td>NA</td><td>Intergenic</td><td>NA</td><td>NA</td></tr>
<tr><td></td><td></td><td></td><td></td><td></td><td></td><td></td><td></td><td></td><td></td><td></td><td></td></tr>
<tr><td>T15546</td><td>AGCCCGTTACCGGACGGTAGCGG</td><td>1000</td><td></td><td></td><td></td><td></td><td></td><td></td><td></td><td></td><td></td></tr>
<tr><td>Oligo fwd</td><td>CACCAGCCCGTTACCGGACGGTAG</td><td></td><td></td><td></td><td></td><td></td><td></td><td></td><td></td><td></td><td></td></tr>
<tr><td>Oligo rev</td><td>AAACCTACCGTCCGGTAACGGGCT</td><td></td><td></td><td></td><td></td><td></td><td></td><td></td><td></td><td></td><td></td></tr>
<tr><td>Chromosome</td><td>start</td><td>end</td><td>strand</td><td>MM</td><td>target_seq</td><td>PAM</td><td>alignment</td><td>distance</td><td>position</td><td>gene name</td><td>gene id</td></tr>
<tr><td>GU305914.1</td><td>203318</td><td>203340</td><td>+</td><td>0</td><td>AGCCCGTTACCGGACGGTAG</td><td>CGG</td><td>||||||||[||||||||||||]PAM</td><td>NA</td><td>Intergenic</td><td>NA</td><td>NA</td></tr>
<tr><td></td><td></td><td></td><td></td><td></td><td></td><td></td><td></td><td></td><td></td><td></td><td></td></tr>
<tr><td>T15547</td><td>GCCCGTTACCGGACGGTAGCGGG</td><td>1000</td><td></td><td></td><td></td><td></td><td></td><td></td><td></td><td></td><td></td></tr>
<tr><td>Oligo fwd</td><td>CACCGCCCGTTACCGGACGGTAGC</td><td></td><td></td><td></td><td></td><td></td><td></td><td></td><td></td><td></td><td></td></tr>
<tr><td>Oligo rev</td><td>AAACGCTACCGTCCGGTAACGGGC</td><td></td><td></td><td></td><td></td><td></td><td></td><td></td><td></td><td></td><td></td></tr>
<tr><td>Chromosome</td><td>start</td><td>end</td><td>strand</td><td>MM</td><td>target_seq</td><td>PAM</td><td>alignment</td><td>distance</td><td>position</td><td>gene name</td><td>gene id</td></tr>
<tr><td>GU305914.1</td><td>203319</td><td>203341</td><td>+</td><td>0</td><td>GCCCGTTACCGGACGGTAGC</td><td>GGG</td><td>||||||||[||||||||||||]PAM</td><td>NA</td><td>Intergenic</td><td>NA</td><td>NA</td></tr>
<tr><td></td><td></td><td></td><td></td><td></td><td></td><td></td><td></td><td></td><td></td><td></td><td></td></tr>
<tr><td>T15548</td><td>TACCGGACGGTAGCGGGTCGTGG</td><td>1000</td><td></td><td></td><td></td><td></td><td></td><td></td><td></td><td></td><td></td></tr>
<tr><td>Oligo fwd</td><td>CACCTACCGGACGGTAGCGGGTCG</td><td></td><td></td><td></td><td></td><td></td><td></td><td></td><td></td><td></td><td></td></tr>
<tr><td>Oligo rev</td><td>AAACCGACCCGCTACCGTCCGGTA</td><td></td><td></td><td></td><td></td><td></td><td></td><td></td><td></td><td></td><td></td></tr>
<tr><td>Chromosome</td><td>start</td><td>end</td><td>strand</td><td>MM</td><td>target_seq</td><td>PAM</td><td>alignment</td><td>distance</td><td>position</td><td>gene name</td><td>gene id</td></tr>
<tr><td>GU305914.1</td><td>203325</td><td>203347</td><td>+</td><td>0</td><td>TACCGGACGGTAGCGGGTCG</td><td>TGG</td><td>||||||||[||||||||||||]PAM</td><td>NA</td><td>Intergenic</td><td>NA</td><td>NA</td></tr>
<tr><td></td><td></td><td></td><td></td><td></td><td></td><td></td><td></td><td></td><td></td><td></td><td></td></tr>
<tr><td>T15549</td><td>TGGTGTTGTTTTGACGTTTGAGG</td><td>1000</td><td></td><td></td><td></td><td></td><td></td><td></td><td></td><td></td><td></td></tr>
<tr><td>Oligo fwd</td><td>CACCTGGTGTTGTTTTGACGTTTG</td><td></td><td></td><td></td><td></td><td></td><td></td><td></td><td></td><td></td><td></td></tr>
<tr><td>Oligo rev</td><td>AAACCAAACGTCAAAACAACACCA</td><td></td><td></td><td></td><td></td><td></td><td></td><td></td><td></td><td></td><td></td></tr>
<tr><td>Chromosome</td><td>start</td><td>end</td><td>strand</td><td>MM</td><td>target_seq</td><td>PAM</td><td>alignment</td><td>distance</td><td>position</td><td>gene name</td><td>gene id</td></tr>
<tr><td>GU305914.1</td><td>203345</td><td>203367</td><td>+</td><td>0</td><td>TGGTGTTGTTTTGACGTTTG</td><td>AGG</td><td>||||||||[||||||||||||]PAM</td><td>NA</td><td>Intergenic</td><td>NA</td><td>NA</td></tr>
<tr><td></td><td></td><td></td><td></td><td></td><td></td><td></td><td></td><td></td><td></td><td></td><td></td></tr>
<tr><td>T15550</td><td>CCACCACCGAATCTCTAAATCGG</td><td>1000</td><td></td><td></td><td></td><td></td><td></td><td></td><td></td><td></td><td></td></tr>
<tr><td>Oligo fwd</td><td>CACCCCACCACCGAATCTCTAAAT</td><td></td><td></td><td></td><td></td><td></td><td></td><td></td><td></td><td></td><td></td></tr>
<tr><td>Oligo rev</td><td>AAACATTTAGAGATTCGGTGGTGG</td><td></td><td></td><td></td><td></td><td></td><td></td><td></td><td></td><td></td><td></td></tr>
<tr><td>Chromosome</td><td>start</td><td>end</td><td>strand</td><td>MM</td><td>target_seq</td><td>PAM</td><td>alignment</td><td>distance</td><td>position</td><td>gene name</td><td>gene id</td></tr>
<tr><td>GU305914.1</td><td>203395</td><td>203417</td><td>+</td><td>0</td><td>CCACCACCGAATCTCTAAAT</td><td>CGG</td><td>||||||||[||||||||||||]PAM</td><td>NA</td><td>Intergenic</td><td>NA</td><td>NA</td></tr>
<tr><td></td><td></td><td></td><td></td><td></td><td></td><td></td><td></td><td></td><td></td><td></td><td></td></tr>
<tr><td>T15551</td><td>CCACCGAATCTCTAAATCGGCGG</td><td>1000</td><td></td><td></td><td></td><td></td><td></td><td></td><td></td><td></td><td></td></tr>
<tr><td>Oligo fwd</td><td>CACCCCACCGAATCTCTAAATCGG</td><td></td><td></td><td></td><td></td><td></td><td></td><td></td><td></td><td></td><td></td></tr>
<tr><td>Oligo rev</td><td>AAACCCGATTTAGAGATTCGGTGG</td><td></td><td></td><td></td><td></td><td></td><td></td><td></td><td></td><td></td><td></td></tr>
<tr><td>Chromosome</td><td>start</td><td>end</td><td>strand</td><td>MM</td><td>target_seq</td><td>PAM</td><td>alignment</td><td>distance</td><td>position</td><td>gene name</td><td>gene id</td></tr>
<tr><td>GU305914.1</td><td>203398</td><td>203420</td><td>+</td><td>0</td><td>CCACCGAATCTCTAAATCGG</td><td>CGG</td><td>||||||||[||||||||||||]PAM</td><td>NA</td><td>Intergenic</td><td>NA</td><td>NA</td></tr>
<tr><td></td><td></td><td></td><td></td><td></td><td></td><td></td><td></td><td></td><td></td><td></td><td></td></tr>
<tr><td>T15552</td><td>CACCGAATCTCTAAATCGGCGGG</td><td>1000</td><td></td><td></td><td></td><td></td><td></td><td></td><td></td><td></td><td></td></tr>
<tr><td>Oligo fwd</td><td>CACCCACCGAATCTCTAAATCGGC</td><td></td><td></td><td></td><td></td><td></td><td></td><td></td><td></td><td></td><td></td></tr>
</table>

<table>
<tr><td>Oligo rev</td><td>AAACGCCGATTTAGAGATTCGGTG</td><td></td><td></td><td></td><td></td><td></td><td></td><td></td><td></td><td></td><td></td></tr>
<tr><td>Chromosome</td><td>start</td><td>end</td><td>strand</td><td>MM</td><td>target_seq</td><td>PAM</td><td>alignment</td><td>distance</td><td>position</td><td>gene name</td><td>gene id</td></tr>
<tr><td>GU305914.1</td><td>203399</td><td>203421</td><td>+</td><td>0</td><td>CACCGAATCTCTAAATCGGC</td><td>GGG</td><td>||||||||[||||||||||||]PAM</td><td>NA</td><td>Intergenic</td><td>NA</td><td>NA</td></tr>
<tr><td></td><td></td><td></td><td></td><td></td><td></td><td></td><td></td><td></td><td></td><td></td><td></td></tr>
<tr><td>T15553</td><td>CATGCCGCCCTCGCGACCCGAGG</td><td>1000</td><td></td><td></td><td></td><td></td><td></td><td></td><td></td><td></td><td></td></tr>
<tr><td>Oligo fwd</td><td>CACCCATGCCGCCCTCGCGACCCG</td><td></td><td></td><td></td><td></td><td></td><td></td><td></td><td></td><td></td><td></td></tr>
<tr><td>Oligo rev</td><td>AAACCGGGTCGCGAGGGCGGCATG</td><td></td><td></td><td></td><td></td><td></td><td></td><td></td><td></td><td></td><td></td></tr>
<tr><td>Chromosome</td><td>start</td><td>end</td><td>strand</td><td>MM</td><td>target_seq</td><td>PAM</td><td>alignment</td><td>distance</td><td>position</td><td>gene name</td><td>gene id</td></tr>
<tr><td>GU305914.1</td><td>203452</td><td>203474</td><td>+</td><td>0</td><td>CATGCCGCCCTCGCGACCCG</td><td>AGG</td><td>||||||||[||||||||||||]PAM</td><td>NA</td><td>Intergenic</td><td>NA</td><td>NA</td></tr>
<tr><td></td><td></td><td></td><td></td><td></td><td></td><td></td><td></td><td></td><td></td><td></td><td></td></tr>
<tr><td>T15554</td><td>CGTCTCTCGACTTCCAGCTGTGG</td><td>1000</td><td></td><td></td><td></td><td></td><td></td><td></td><td></td><td></td><td></td></tr>
<tr><td>Oligo fwd</td><td>CACCCGTCTCTCGACTTCCAGCTG</td><td></td><td></td><td></td><td></td><td></td><td></td><td></td><td></td><td></td><td></td></tr>
<tr><td>Oligo rev</td><td>AAACCAGCTGGAAGTCGAGAGACG</td><td></td><td></td><td></td><td></td><td></td><td></td><td></td><td></td><td></td><td></td></tr>
<tr><td>Chromosome</td><td>start</td><td>end</td><td>strand</td><td>MM</td><td>target_seq</td><td>PAM</td><td>alignment</td><td>distance</td><td>position</td><td>gene name</td><td>gene id</td></tr>
<tr><td>GU305914.1</td><td>203477</td><td>203499</td><td>+</td><td>0</td><td>CGTCTCTCGACTTCCAGCTG</td><td>TGG</td><td>||||||||[||||||||||||]PAM</td><td>NA</td><td>Intergenic</td><td>NA</td><td>NA</td></tr>
<tr><td></td><td></td><td></td><td></td><td></td><td></td><td></td><td></td><td></td><td></td><td></td><td></td></tr>
<tr><td>T15555</td><td>GACTTCCAGCTGTGGTGATACGG</td><td>1000</td><td></td><td></td><td></td><td></td><td></td><td></td><td></td><td></td><td></td></tr>
<tr><td>Oligo fwd</td><td>CACCGACTTCCAGCTGTGGTGATA</td><td></td><td></td><td></td><td></td><td></td><td></td><td></td><td></td><td></td><td></td></tr>
<tr><td>Oligo rev</td><td>AAACTATCACCACAGCTGGAAGTC</td><td></td><td></td><td></td><td></td><td></td><td></td><td></td><td></td><td></td><td></td></tr>
<tr><td>Chromosome</td><td>start</td><td>end</td><td>strand</td><td>MM</td><td>target_seq</td><td>PAM</td><td>alignment</td><td>distance</td><td>position</td><td>gene name</td><td>gene id</td></tr>
<tr><td>GU305914.1</td><td>203485</td><td>203507</td><td>+</td><td>0</td><td>GACTTCCAGCTGTGGTGATA</td><td>CGG</td><td>||||||||[||||||||||||]PAM</td><td>NA</td><td>Intergenic</td><td>NA</td><td>NA</td></tr>
<tr><td></td><td></td><td></td><td></td><td></td><td></td><td></td><td></td><td></td><td></td><td></td><td></td></tr>
<tr><td>T15556</td><td>TTCCAGCTGTGGTGATACGGCGG</td><td>1000</td><td></td><td></td><td></td><td></td><td></td><td></td><td></td><td></td><td></td></tr>
<tr><td>Oligo fwd</td><td>CACCTTCCAGCTGTGGTGATACGG</td><td></td><td></td><td></td><td></td><td></td><td></td><td></td><td></td><td></td><td></td></tr>
<tr><td>Oligo rev</td><td>AAACCCGTATCACCACAGCTGGAA</td><td></td><td></td><td></td><td></td><td></td><td></td><td></td><td></td><td></td><td></td></tr>
<tr><td>Chromosome</td><td>start</td><td>end</td><td>strand</td><td>MM</td><td>target_seq</td><td>PAM</td><td>alignment</td><td>distance</td><td>position</td><td>gene name</td><td>gene id</td></tr>
<tr><td>GU305914.1</td><td>203488</td><td>203510</td><td>+</td><td>0</td><td>TTCCAGCTGTGGTGATACGG</td><td>CGG</td><td>||||||||[||||||||||||]PAM</td><td>NA</td><td>Intergenic</td><td>NA</td><td>NA</td></tr>
<tr><td></td><td></td><td></td><td></td><td></td><td></td><td></td><td></td><td></td><td></td><td></td><td></td></tr>
<tr><td>T15557</td><td>TTAGTTTTTTCTGACAGTCACGG</td><td>1000</td><td></td><td></td><td></td><td></td><td></td><td></td><td></td><td></td><td></td></tr>
<tr><td>Oligo fwd</td><td>CACCTTAGTTTTTTCTGACAGTCA</td><td></td><td></td><td></td><td></td><td></td><td></td><td></td><td></td><td></td><td></td></tr>
<tr><td>Oligo rev</td><td>AAACTGACTGTCAGAAAAAACTAA</td><td></td><td></td><td></td><td></td><td></td><td></td><td></td><td></td><td></td><td></td></tr>
<tr><td>Chromosome</td><td>start</td><td>end</td><td>strand</td><td>MM</td><td>target_seq</td><td>PAM</td><td>alignment</td><td>distance</td><td>position</td><td>gene name</td><td>gene id</td></tr>
<tr><td>GU305914.1</td><td>203538</td><td>203560</td><td>+</td><td>0</td><td>TTAGTTTTTTCTGACAGTCA</td><td>CGG</td><td>||||||||[||||||||||||]PAM</td><td>NA</td><td>Intergenic</td><td>NA</td><td>NA</td></tr>
<tr><td></td><td></td><td></td><td></td><td></td><td></td><td></td><td></td><td></td><td></td><td></td><td></td></tr>
<tr><td>T15558</td><td>ACAGTCACGGTCTCGTTCACTGG</td><td>1000</td><td></td><td></td><td></td><td></td><td></td><td></td><td></td><td></td><td></td></tr>
<tr><td>Oligo fwd</td><td>CACCACAGTCACGGTCTCGTTCAC</td><td></td><td></td><td></td><td></td><td></td><td></td><td></td><td></td><td></td><td></td></tr>
<tr><td>Oligo rev</td><td>AAACGTGAACGAGACCGTGACTGT</td><td></td><td></td><td></td><td></td><td></td><td></td><td></td><td></td><td></td><td></td></tr>
<tr><td>Chromosome</td><td>start</td><td>end</td><td>strand</td><td>MM</td><td>target_seq</td><td>PAM</td><td>alignment</td><td>distance</td><td>position</td><td>gene name</td><td>gene id</td></tr>
<tr><td>GU305914.1</td><td>203551</td><td>203573</td><td>+</td><td>0</td><td>ACAGTCACGGTCTCGTTCAC</td><td>TGG</td><td>||||||||[||||||||||||]PAM</td><td>NA</td><td>Intergenic</td><td>NA</td><td>NA</td></tr>
<tr><td></td><td></td><td></td><td></td><td></td><td></td><td></td><td></td><td></td><td></td><td></td><td></td></tr>
<tr><td>T15559</td><td>GTCACGGTCTCGTTCACTGGCGG</td><td>1000</td><td></td><td></td><td></td><td></td><td></td><td></td><td></td><td></td><td></td></tr>
<tr><td>Oligo fwd</td><td>CACCGTCACGGTCTCGTTCACTGG</td><td></td><td></td><td></td><td></td><td></td><td></td><td></td><td></td><td></td><td></td></tr>
<tr><td>Oligo rev</td><td>AAACCCAGTGAACGAGACCGTGAC</td><td></td><td></td><td></td><td></td><td></td><td></td><td></td><td></td><td></td><td></td></tr>
<tr><td>Chromosome</td><td>start</td><td>end</td><td>strand</td><td>MM</td><td>target_seq</td><td>PAM</td><td>alignment</td><td>distance</td><td>position</td><td>gene name</td><td>gene id</td></tr>
<tr><td>GU305914.1</td><td>203554</td><td>203576</td><td>+</td><td>0</td><td>GTCACGGTCTCGTTCACTGG</td><td>CGG</td><td>||||||||[||||||||||||]PAM</td><td>NA</td><td>Intergenic</td><td>NA</td><td>NA</td></tr>
<tr><td></td><td></td><td></td><td></td><td></td><td></td><td></td><td></td><td></td><td></td><td></td><td></td></tr>
<tr><td>T15560</td><td>GGACGTAGCCGTATCATCCTCGG</td><td>1000</td><td></td><td></td><td></td><td></td><td></td><td></td><td></td><td></td><td></td></tr>
<tr><td>Oligo fwd</td><td>CACCGGACGTAGCCGTATCATCCT</td><td></td><td></td><td></td><td></td><td></td><td></td><td></td><td></td><td></td><td></td></tr>
<tr><td>Oligo rev</td><td>AAACAGGATGATACGGCTACGTCC</td><td></td><td></td><td></td><td></td><td></td><td></td><td></td><td></td><td></td><td></td></tr>
<tr><td>Chromosome</td><td>start</td><td>end</td><td>strand</td><td>MM</td><td>target_seq</td><td>PAM</td><td>alignment</td><td>distance</td><td>position</td><td>gene name</td><td>gene id</td></tr>
<tr><td>GU305914.1</td><td>203575</td><td>203597</td><td>+</td><td>0</td><td>GGACGTAGCCGTATCATCCT</td><td>CGG</td><td>||||||||[||||||||||||]PAM</td><td>NA</td><td>Intergenic</td><td>NA</td><td>NA</td></tr>
<tr><td></td><td></td><td></td><td></td><td></td><td></td><td></td><td></td><td></td><td></td><td></td><td></td></tr>
<tr><td>T15561</td><td>ACAACCGACGTCTAGTTGTACGG</td><td>1000</td><td></td><td></td><td></td><td></td><td></td><td></td><td></td><td></td><td></td></tr>
<tr><td>Oligo fwd</td><td>CACCACAACCGACGTCTAGTTGTA</td><td></td><td></td><td></td><td></td><td></td><td></td><td></td><td></td><td></td><td></td></tr>
<tr><td>Oligo rev</td><td>AAACTACAACTAGACGTCGGTTGT</td><td></td><td></td><td></td><td></td><td></td><td></td><td></td><td></td><td></td><td></td></tr>
<tr><td>Chromosome</td><td>start</td><td>end</td><td>strand</td><td>MM</td><td>target_seq</td><td>PAM</td><td>alignment</td><td>distance</td><td>position</td><td>gene name</td><td>gene id</td></tr>
<tr><td>GU305914.1</td><td>203641</td><td>203663</td><td>+</td><td>0</td><td>ACAACCGACGTCTAGTTGTA</td><td>CGG</td><td>||||||||[||||||||||||]PAM</td><td>NA</td><td>Intergenic</td><td>NA</td><td>NA</td></tr>
<tr><td></td><td></td><td></td><td></td><td></td><td></td><td></td><td></td><td></td><td></td><td></td><td></td></tr>
<tr><td>T15562</td><td>GTTTTGCATTTTATCTAGCTCGG</td><td>1000</td><td></td><td></td><td></td><td></td><td></td><td></td><td></td><td></td><td></td></tr>
<tr><td>Oligo fwd</td><td>CACCGTTTTGCATTTTATCTAGCT</td><td></td><td></td><td></td><td></td><td></td><td></td><td></td><td></td><td></td><td></td></tr>
<tr><td>Oligo rev</td><td>AAACAGCTAGATAAAATGCAAAAC</td><td></td><td></td><td></td><td></td><td></td><td></td><td></td><td></td><td></td><td></td></tr>
<tr><td>Chromosome</td><td>start</td><td>end</td><td>strand</td><td>MM</td><td>target_seq</td><td>PAM</td><td>alignment</td><td>distance</td><td>position</td><td>gene name</td><td>gene id</td></tr>
<tr><td>GU305914.1</td><td>203689</td><td>203711</td><td>+</td><td>0</td><td>GTTTTGCATTTTATCTAGCT</td><td>CGG</td><td>||||||||[||||||||||||]PAM</td><td>NA</td><td>Intergenic</td><td>NA</td><td>NA</td></tr>
<tr><td></td><td></td><td></td><td></td><td></td><td></td><td></td><td></td><td></td><td></td><td></td><td></td></tr>
<tr><td>T15563</td><td>GATCCCGTGTGTCTTGTTATTGG</td><td>1000</td><td></td><td></td><td></td><td></td><td></td><td></td><td></td><td></td><td></td></tr>
<tr><td>Oligo fwd</td><td>CACCGATCCCGTGTGTCTTGTTAT</td><td></td><td></td><td></td><td></td><td></td><td></td><td></td><td></td><td></td><td></td></tr>
<tr><td>Oligo rev</td><td>AAACATAACAAGACACACGGGATC</td><td></td><td></td><td></td><td></td><td></td><td></td><td></td><td></td><td></td><td></td></tr>
<tr><td>Chromosome</td><td>start</td><td>end</td><td>strand</td><td>MM</td><td>target_seq</td><td>PAM</td><td>alignment</td><td>distance</td><td>position</td><td>gene name</td><td>gene id</td></tr>
<tr><td>GU305914.1</td><td>203785</td><td>203807</td><td>+</td><td>0</td><td>GATCCCGTGTGTCTTGTTAT</td><td>TGG</td><td>||||||||[||||||||||||]PAM</td><td>NA</td><td>Intergenic</td><td>NA</td><td>NA</td></tr>
<tr><td></td><td></td><td></td><td></td><td></td><td></td><td></td><td></td><td></td><td></td><td></td><td></td></tr>
<tr><td>T15564</td><td>GTTATTGGAGTATCCCACAAAGG</td><td>1000</td><td></td><td></td><td></td><td></td><td></td><td></td><td></td><td></td><td></td></tr>
<tr><td>Oligo fwd</td><td>CACCGTTATTGGAGTATCCCACAA</td><td></td><td></td><td></td><td></td><td></td><td></td><td></td><td></td><td></td><td></td></tr>
<tr><td>Oligo rev</td><td>AAACTTGTGGGATACTCCAATAAC</td><td></td><td></td><td></td><td></td><td></td><td></td><td></td><td></td><td></td><td></td></tr>
<tr><td>Chromosome</td><td>start</td><td>end</td><td>strand</td><td>MM</td><td>target_seq</td><td>PAM</td><td>alignment</td><td>distance</td><td>position</td><td>gene name</td><td>gene id</td></tr>
<tr><td>GU305914.1</td><td>203800</td><td>203822</td><td>+</td><td>0</td><td>GTTATTGGAGTATCCCACAA</td><td>AGG</td><td>||||||||[||||||||||||]PAM</td><td>NA</td><td>Intergenic</td><td>NA</td><td>NA</td></tr>
<tr><td></td><td></td><td></td><td></td><td></td><td></td><td></td><td></td><td></td><td></td><td></td><td></td></tr>
<tr><td>T15565</td><td>GGCTTCCCCGTCGACGTGTCCGG</td><td>1000</td><td></td><td></td><td></td><td></td><td></td><td></td><td></td><td></td><td></td></tr>
<tr><td>Oligo fwd</td><td>CACCGGCTTCCCCGTCGACGTGTC</td><td></td><td></td><td></td><td></td><td></td><td></td><td></td><td></td><td></td><td></td></tr>
<tr><td>Oligo rev</td><td>AAACGACACGTCGACGGGGAAGCC</td><td></td><td></td><td></td><td></td><td></td><td></td><td></td><td></td><td></td><td></td></tr>
<tr><td>Chromosome</td><td>start</td><td>end</td><td>strand</td><td>MM</td><td>target_seq</td><td>PAM</td><td>alignment</td><td>distance</td><td>position</td><td>gene name</td><td>gene id</td></tr>
<tr><td>GU305914.1</td><td>203821</td><td>203843</td><td>+</td><td>0</td><td>GGCTTCCCCGTCGACGTGTC</td><td>CGG</td><td>||||||||[||||||||||||]PAM</td><td>NA</td><td>Intergenic</td><td>NA</td><td>NA</td></tr>
<tr><td>GU305914.1</td><td>73003</td><td>73025</td><td>+</td><td>4</td><td>GGCGCCCCCGGCGACGAGTC</td><td>GGG</td><td>|||--|||[||-|||||-|||]PAM</td><td>NA</td><td>Intergenic</td><td>NA</td><td>NA</td></tr>
<tr><td></td><td></td><td></td><td></td><td></td><td></td><td></td><td></td><td></td><td></td><td></td><td></td></tr>
<tr><td>T15566</td><td>CGTGTCCGGTCCCAAAACAACGG</td><td>1000</td><td></td><td></td><td></td><td></td><td></td><td></td><td></td><td></td><td></td></tr>
<tr><td>Oligo fwd</td><td>CACCCGTGTCCGGTCCCAAAACAA</td><td></td><td></td><td></td><td></td><td></td><td></td><td></td><td></td><td></td><td></td></tr>
<tr><td>Oligo rev</td><td>AAACTTGTTTTGGGACCGGACACG</td><td></td><td></td><td></td><td></td><td></td><td></td><td></td><td></td><td></td><td></td></tr>
<tr><td>Chromosome</td><td>start</td><td>end</td><td>strand</td><td>MM</td><td>target_seq</td><td>PAM</td><td>alignment</td><td>distance</td><td>position</td><td>gene name</td><td>gene id</td></tr>
<tr><td>GU305914.1</td><td>203835</td><td>203857</td><td>+</td><td>0</td><td>CGTGTCCGGTCCCAAAACAA</td><td>CGG</td><td>||||||||[||||||||||||]PAM</td><td>NA</td><td>Intergenic</td><td>NA</td><td>NA</td></tr>
</table>

<table>
<tr><td>T15567</td><td>AAAACAACGGCGCGTATTTGCGG</td><td>1000</td><td></td><td></td><td></td><td></td><td></td><td></td><td></td><td></td><td></td></tr>
<tr><td>Oligo fwd</td><td>CACCAAAACAACGGCGCGTATTTG</td><td></td><td></td><td></td><td></td><td></td><td></td><td></td><td></td><td></td><td></td></tr>
<tr><td>Oligo rev</td><td>AAACCAAATACGCGCCGTTGTTTT</td><td></td><td></td><td></td><td></td><td></td><td></td><td></td><td></td><td></td><td></td></tr>
<tr><td>Chromosome</td><td>start</td><td>end</td><td>strand</td><td>MM</td><td>target_seq</td><td>PAM</td><td>alignment</td><td>distance</td><td>position</td><td>gene name</td><td>gene id</td></tr>
<tr><td>GU305914.1</td><td>203848</td><td>203870</td><td>+</td><td>0</td><td>AAAACAACGGCGCGTATTTG</td><td>CGG</td><td>||||||||[||||||||||||]PAM</td><td>NA</td><td>Intergenic</td><td>NA</td><td>NA</td></tr>
<tr><td></td><td></td><td></td><td></td><td></td><td></td><td></td><td></td><td></td><td></td><td></td><td></td></tr>
<tr><td>T15568</td><td>CGTGTATGCGTACCGCAGTCCGG</td><td>1000</td><td></td><td></td><td></td><td></td><td></td><td></td><td></td><td></td><td></td></tr>
<tr><td>Oligo fwd</td><td>CACCCGTGTATGCGTACCGCAGTC</td><td></td><td></td><td></td><td></td><td></td><td></td><td></td><td></td><td></td><td></td></tr>
<tr><td>Oligo rev</td><td>AAACGACTGCGGTACGCATACACG</td><td></td><td></td><td></td><td></td><td></td><td></td><td></td><td></td><td></td><td></td></tr>
<tr><td>Chromosome</td><td>start</td><td>end</td><td>strand</td><td>MM</td><td>target_seq</td><td>PAM</td><td>alignment</td><td>distance</td><td>position</td><td>gene name</td><td>gene id</td></tr>
<tr><td>GU305914.1</td><td>203887</td><td>203909</td><td>+</td><td>0</td><td>CGTGTATGCGTACCGCAGTC</td><td>CGG</td><td>||||||||[||||||||||||]PAM</td><td>NA</td><td>Intergenic</td><td>NA</td><td>NA</td></tr>
<tr><td></td><td></td><td></td><td></td><td></td><td></td><td></td><td></td><td></td><td></td><td></td><td></td></tr>
<tr><td>T15569</td><td>TGCGTACCGCAGTCCGGACTCGG</td><td>1000</td><td></td><td></td><td></td><td></td><td></td><td></td><td></td><td></td><td></td></tr>
<tr><td>Oligo fwd</td><td>CACCTGCGTACCGCAGTCCGGACT</td><td></td><td></td><td></td><td></td><td></td><td></td><td></td><td></td><td></td><td></td></tr>
<tr><td>Oligo rev</td><td>AAACAGTCCGGACTGCGGTACGCA</td><td></td><td></td><td></td><td></td><td></td><td></td><td></td><td></td><td></td><td></td></tr>
<tr><td>Chromosome</td><td>start</td><td>end</td><td>strand</td><td>MM</td><td>target_seq</td><td>PAM</td><td>alignment</td><td>distance</td><td>position</td><td>gene name</td><td>gene id</td></tr>
<tr><td>GU305914.1</td><td>203893</td><td>203915</td><td>+</td><td>0</td><td>TGCGTACCGCAGTCCGGACT</td><td>CGG</td><td>||||||||[||||||||||||]PAM</td><td>NA</td><td>Intergenic</td><td>NA</td><td>NA</td></tr>
<tr><td></td><td></td><td></td><td></td><td></td><td></td><td></td><td></td><td></td><td></td><td></td><td></td></tr>
<tr><td>T15570</td><td>CGGACGAGTCCTCTGTCCCGTGG</td><td>1000</td><td></td><td></td><td></td><td></td><td></td><td></td><td></td><td></td><td></td></tr>
<tr><td>Oligo fwd</td><td>CACCCGGACGAGTCCTCTGTCCCG</td><td></td><td></td><td></td><td></td><td></td><td></td><td></td><td></td><td></td><td></td></tr>
<tr><td>Oligo rev</td><td>AAACCGGGACAGAGGACTCGTCCG</td><td></td><td></td><td></td><td></td><td></td><td></td><td></td><td></td><td></td><td></td></tr>
<tr><td>Chromosome</td><td>start</td><td>end</td><td>strand</td><td>MM</td><td>target_seq</td><td>PAM</td><td>alignment</td><td>distance</td><td>position</td><td>gene name</td><td>gene id</td></tr>
<tr><td>GU305914.1</td><td>203913</td><td>203935</td><td>+</td><td>0</td><td>CGGACGAGTCCTCTGTCCCG</td><td>TGG</td><td>||||||||[||||||||||||]PAM</td><td>NA</td><td>Intergenic</td><td>NA</td><td>NA</td></tr>
<tr><td>GU305914.1</td><td>78082</td><td>78104</td><td>-</td><td>4</td><td>CGAACGGCTCCTCCGTCCCG</td><td>GGG</td><td>||-|||--[|||||-||||||]PAM</td><td>NA</td><td>Intergenic</td><td>NA</td><td>NA</td></tr>
<tr><td></td><td></td><td></td><td></td><td></td><td></td><td></td><td></td><td></td><td></td><td></td><td></td></tr>
<tr><td>T15571</td><td>GCGACATCTCACATTCATGTCGG</td><td>1000</td><td></td><td></td><td></td><td></td><td></td><td></td><td></td><td></td><td></td></tr>
<tr><td>Oligo fwd</td><td>CACCGCGACATCTCACATTCATGT</td><td></td><td></td><td></td><td></td><td></td><td></td><td></td><td></td><td></td><td></td></tr>
<tr><td>Oligo rev</td><td>AAACACATGAATGTGAGATGTCGC</td><td></td><td></td><td></td><td></td><td></td><td></td><td></td><td></td><td></td><td></td></tr>
<tr><td>Chromosome</td><td>start</td><td>end</td><td>strand</td><td>MM</td><td>target_seq</td><td>PAM</td><td>alignment</td><td>distance</td><td>position</td><td>gene name</td><td>gene id</td></tr>
<tr><td>GU305914.1</td><td>204023</td><td>204045</td><td>+</td><td>0</td><td>GCGACATCTCACATTCATGT</td><td>CGG</td><td>||||||||[||||||||||||]PAM</td><td>NA</td><td>Intergenic</td><td>NA</td><td>NA</td></tr>
<tr><td></td><td></td><td></td><td></td><td></td><td></td><td></td><td></td><td></td><td></td><td></td><td></td></tr>
<tr><td>T15572</td><td>GAAGAAATATTACGATGACACGG</td><td>1000</td><td></td><td></td><td></td><td></td><td></td><td></td><td></td><td></td><td></td></tr>
<tr><td>Oligo fwd</td><td>CACCGAAGAAATATTACGATGACA</td><td></td><td></td><td></td><td></td><td></td><td></td><td></td><td></td><td></td><td></td></tr>
<tr><td>Oligo rev</td><td>AAACTGTCATCGTAATATTTCTTC</td><td></td><td></td><td></td><td></td><td></td><td></td><td></td><td></td><td></td><td></td></tr>
<tr><td>Chromosome</td><td>start</td><td>end</td><td>strand</td><td>MM</td><td>target_seq</td><td>PAM</td><td>alignment</td><td>distance</td><td>position</td><td>gene name</td><td>gene id</td></tr>
<tr><td>GU305914.1</td><td>204049</td><td>204071</td><td>+</td><td>0</td><td>GAAGAAATATTACGATGACA</td><td>CGG</td><td>||||||||[||||||||||||]PAM</td><td>NA</td><td>Intergenic</td><td>NA</td><td>NA</td></tr>
<tr><td></td><td></td><td></td><td></td><td></td><td></td><td></td><td></td><td></td><td></td><td></td><td></td></tr>
<tr><td>T15573</td><td>AAGAAATATTACGATGACACGGG</td><td>1000</td><td></td><td></td><td></td><td></td><td></td><td></td><td></td><td></td><td></td></tr>
<tr><td>Oligo fwd</td><td>CACCAAGAAATATTACGATGACAC</td><td></td><td></td><td></td><td></td><td></td><td></td><td></td><td></td><td></td><td></td></tr>
<tr><td>Oligo rev</td><td>AAACGTGTCATCGTAATATTTCTT</td><td></td><td></td><td></td><td></td><td></td><td></td><td></td><td></td><td></td><td></td></tr>
<tr><td>Chromosome</td><td>start</td><td>end</td><td>strand</td><td>MM</td><td>target_seq</td><td>PAM</td><td>alignment</td><td>distance</td><td>position</td><td>gene name</td><td>gene id</td></tr>
<tr><td>GU305914.1</td><td>204050</td><td>204072</td><td>+</td><td>0</td><td>AAGAAATATTACGATGACAC</td><td>GGG</td><td>||||||||[||||||||||||]PAM</td><td>NA</td><td>Intergenic</td><td>NA</td><td>NA</td></tr>
<tr><td></td><td></td><td></td><td></td><td></td><td></td><td></td><td></td><td></td><td></td><td></td><td></td></tr>
<tr><td>T15574</td><td>AGAAATATTACGATGACACGGGG</td><td>1000</td><td></td><td></td><td></td><td></td><td></td><td></td><td></td><td></td><td></td></tr>
<tr><td>Oligo fwd</td><td>CACCAGAAATATTACGATGACACG</td><td></td><td></td><td></td><td></td><td></td><td></td><td></td><td></td><td></td><td></td></tr>
<tr><td>Oligo rev</td><td>AAACCGTGTCATCGTAATATTTCT</td><td></td><td></td><td></td><td></td><td></td><td></td><td></td><td></td><td></td><td></td></tr>
<tr><td>Chromosome</td><td>start</td><td>end</td><td>strand</td><td>MM</td><td>target_seq</td><td>PAM</td><td>alignment</td><td>distance</td><td>position</td><td>gene name</td><td>gene id</td></tr>
<tr><td>GU305914.1</td><td>204051</td><td>204073</td><td>+</td><td>0</td><td>AGAAATATTACGATGACACG</td><td>GGG</td><td>||||||||[||||||||||||]PAM</td><td>NA</td><td>Intergenic</td><td>NA</td><td>NA</td></tr>
<tr><td></td><td></td><td></td><td></td><td></td><td></td><td></td><td></td><td></td><td></td><td></td><td></td></tr>
<tr><td>T15575</td><td>TATTACGATGACACGGGGTGAGG</td><td>1000</td><td></td><td></td><td></td><td></td><td></td><td></td><td></td><td></td><td></td></tr>
<tr><td>Oligo fwd</td><td>CACCTATTACGATGACACGGGGTG</td><td></td><td></td><td></td><td></td><td></td><td></td><td></td><td></td><td></td><td></td></tr>
<tr><td>Oligo rev</td><td>AAACCACCCCGTGTCATCGTAATA</td><td></td><td></td><td></td><td></td><td></td><td></td><td></td><td></td><td></td><td></td></tr>
<tr><td>Chromosome</td><td>start</td><td>end</td><td>strand</td><td>MM</td><td>target_seq</td><td>PAM</td><td>alignment</td><td>distance</td><td>position</td><td>gene name</td><td>gene id</td></tr>
<tr><td>GU305914.1</td><td>204056</td><td>204078</td><td>+</td><td>0</td><td>TATTACGATGACACGGGGTG</td><td>AGG</td><td>||||||||[||||||||||||]PAM</td><td>NA</td><td>Intergenic</td><td>NA</td><td>NA</td></tr>
<tr><td></td><td></td><td></td><td></td><td></td><td></td><td></td><td></td><td></td><td></td><td></td><td></td></tr>
<tr><td>T15576</td><td>ATTACGATGACACGGGGTGAGGG</td><td>1000</td><td></td><td></td><td></td><td></td><td></td><td></td><td></td><td></td><td></td></tr>
<tr><td>Oligo fwd</td><td>CACCATTACGATGACACGGGGTGA</td><td></td><td></td><td></td><td></td><td></td><td></td><td></td><td></td><td></td><td></td></tr>
<tr><td>Oligo rev</td><td>AAACTCACCCCGTGTCATCGTAAT</td><td></td><td></td><td></td><td></td><td></td><td></td><td></td><td></td><td></td><td></td></tr>
<tr><td>Chromosome</td><td>start</td><td>end</td><td>strand</td><td>MM</td><td>target_seq</td><td>PAM</td><td>alignment</td><td>distance</td><td>position</td><td>gene name</td><td>gene id</td></tr>
<tr><td>GU305914.1</td><td>204057</td><td>204079</td><td>+</td><td>0</td><td>ATTACGATGACACGGGGTGA</td><td>GGG</td><td>||||||||[||||||||||||]PAM</td><td>NA</td><td>Intergenic</td><td>NA</td><td>NA</td></tr>
<tr><td></td><td></td><td></td><td></td><td></td><td></td><td></td><td></td><td></td><td></td><td></td><td></td></tr>
<tr><td>T15577</td><td>TTACGATGACACGGGGTGAGGGG</td><td>1000</td><td></td><td></td><td></td><td></td><td></td><td></td><td></td><td></td><td></td></tr>
<tr><td>Oligo fwd</td><td>CACCTTACGATGACACGGGGTGAG</td><td></td><td></td><td></td><td></td><td></td><td></td><td></td><td></td><td></td><td></td></tr>
<tr><td>Oligo rev</td><td>AAACCTCACCCCGTGTCATCGTAA</td><td></td><td></td><td></td><td></td><td></td><td></td><td></td><td></td><td></td><td></td></tr>
<tr><td>Chromosome</td><td>start</td><td>end</td><td>strand</td><td>MM</td><td>target_seq</td><td>PAM</td><td>alignment</td><td>distance</td><td>position</td><td>gene name</td><td>gene id</td></tr>
<tr><td>GU305914.1</td><td>204058</td><td>204080</td><td>+</td><td>0</td><td>TTACGATGACACGGGGTGAG</td><td>GGG</td><td>||||||||[||||||||||||]PAM</td><td>NA</td><td>Intergenic</td><td>NA</td><td>NA</td></tr>
<tr><td></td><td></td><td></td><td></td><td></td><td></td><td></td><td></td><td></td><td></td><td></td><td></td></tr>
<tr><td>T15578</td><td>ACGACAGACATACAGAGATTCGG</td><td>1000</td><td></td><td></td><td></td><td></td><td></td><td></td><td></td><td></td><td></td></tr>
<tr><td>Oligo fwd</td><td>CACCACGACAGACATACAGAGATT</td><td></td><td></td><td></td><td></td><td></td><td></td><td></td><td></td><td></td><td></td></tr>
<tr><td>Oligo rev</td><td>AAACAATCTCTGTATGTCTGTCGT</td><td></td><td></td><td></td><td></td><td></td><td></td><td></td><td></td><td></td><td></td></tr>
<tr><td>Chromosome</td><td>start</td><td>end</td><td>strand</td><td>MM</td><td>target_seq</td><td>PAM</td><td>alignment</td><td>distance</td><td>position</td><td>gene name</td><td>gene id</td></tr>
<tr><td>GU305914.1</td><td>204103</td><td>204125</td><td>+</td><td>0</td><td>ACGACAGACATACAGAGATT</td><td>CGG</td><td>||||||||[||||||||||||]PAM</td><td>NA</td><td>Intergenic</td><td>NA</td><td>NA</td></tr>
<tr><td></td><td></td><td></td><td></td><td></td><td></td><td></td><td></td><td></td><td></td><td></td><td></td></tr>
<tr><td>T15579</td><td>CTCCCGAACGTAGTTTTGTGCGG</td><td>1000</td><td></td><td></td><td></td><td></td><td></td><td></td><td></td><td></td><td></td></tr>
<tr><td>Oligo fwd</td><td>CACCCTCCCGAACGTAGTTTTGTG</td><td></td><td></td><td></td><td></td><td></td><td></td><td></td><td></td><td></td><td></td></tr>
<tr><td>Oligo rev</td><td>AAACCACAAAACTACGTTCGGGAG</td><td></td><td></td><td></td><td></td><td></td><td></td><td></td><td></td><td></td><td></td></tr>
<tr><td>Chromosome</td><td>start</td><td>end</td><td>strand</td><td>MM</td><td>target_seq</td><td>PAM</td><td>alignment</td><td>distance</td><td>position</td><td>gene name</td><td>gene id</td></tr>
<tr><td>GU305914.1</td><td>204207</td><td>204229</td><td>+</td><td>0</td><td>CTCCCGAACGTAGTTTTGTG</td><td>CGG</td><td>||||||||[||||||||||||]PAM</td><td>NA</td><td>Intergenic</td><td>NA</td><td>NA</td></tr>
<tr><td></td><td></td><td></td><td></td><td></td><td></td><td></td><td></td><td></td><td></td><td></td><td></td></tr>
<tr><td>T15580</td><td>ATACCATTGCACAGATGCACAGG</td><td>1000</td><td></td><td></td><td></td><td></td><td></td><td></td><td></td><td></td><td></td></tr>
<tr><td>Oligo fwd</td><td>CACCATACCATTGCACAGATGCAC</td><td></td><td></td><td></td><td></td><td></td><td></td><td></td><td></td><td></td><td></td></tr>
<tr><td>Oligo rev</td><td>AAACGTGCATCTGTGCAATGGTAT</td><td></td><td></td><td></td><td></td><td></td><td></td><td></td><td></td><td></td><td></td></tr>
<tr><td>Chromosome</td><td>start</td><td>end</td><td>strand</td><td>MM</td><td>target_seq</td><td>PAM</td><td>alignment</td><td>distance</td><td>position</td><td>gene name</td><td>gene id</td></tr>
<tr><td>GU305914.1</td><td>204255</td><td>204277</td><td>+</td><td>0</td><td>ATACCATTGCACAGATGCAC</td><td>AGG</td><td>||||||||[||||||||||||]PAM</td><td>NA</td><td>Intergenic</td><td>NA</td><td>NA</td></tr>
<tr><td></td><td></td><td></td><td></td><td></td><td></td><td></td><td></td><td></td><td></td><td></td><td></td></tr>
<tr><td>T15581</td><td>TACCATTGCACAGATGCACAGGG</td><td>1000</td><td></td><td></td><td></td><td></td><td></td><td></td><td></td><td></td><td></td></tr>
<tr><td>Oligo fwd</td><td>CACCTACCATTGCACAGATGCACA</td><td></td><td></td><td></td><td></td><td></td><td></td><td></td><td></td><td></td><td></td></tr>
<tr><td>Oligo rev</td><td>AAACTGTGCATCTGTGCAATGGTA</td><td></td><td></td><td></td><td></td><td></td><td></td><td></td><td></td><td></td><td></td></tr>
<tr><td>Chromosome</td><td>start</td><td>end</td><td>strand</td><td>MM</td><td>target_seq</td><td>PAM</td><td>alignment</td><td>distance</td><td>position</td><td>gene name</td><td>gene id</td></tr>
</table>

GU305914.1 204256 204278 + 0 TACCATTGCACAGATGCACA GGG ||||||||[||||||||||||]PAM NA Intergenic NA NA

T15582 GGCCGTTACCGCGACGAACATGG 1000
Oligo fwd CACCGGCCGTTACCGCGACGAACA
Oligo rev AAACTGTTCGTCGCGGTAACGGCC
Chromosome start end strand MM target_seq PAM alignment distance position gene name gene id
GU305914.1 204277 204299 + 0 GGCCGTTACCGCGACGAACA TGG ||||||||[||||||||||||]PAM NA Intergenic NA NA

T15583 ACGAACATGGCGACTAAAGTCGG 1000
Oligo fwd CACCACGAACATGGCGACTAAAGT
Oligo rev AAACACTTTAGTCGCCATGTTCGT
Chromosome start end strand MM target_seq PAM alignment distance position gene name gene id
GU305914.1 204290 204312 + 0 ACGAACATGGCGACTAAAGT CGG ||||||||[||||||||||||]PAM NA Intergenic NA NA

T15584 CATTAAACTGATGTCGTCCGAGG 1000
Oligo fwd CACCCATTAAACTGATGTCGTCCG
Oligo rev AAACCGGACGACATCAGTTTAATG
Chromosome start end strand MM target_seq PAM alignment distance position gene name gene id
GU305914.1 204382 204404 + 0 CATTAAACTGATGTCGTCCG AGG ||||||||[||||||||||||]PAM NA Intergenic NA NA

T15585 TAAACTGATGTCGTCCGAGGAGG 1000
Oligo fwd CACCTAAACTGATGTCGTCCGAGG
Oligo rev AAACCCTCGGACGACATCAGTTTA
Chromosome start end strand MM target_seq PAM alignment distance position gene name gene id
GU305914.1 204385 204407 + 0 TAAACTGATGTCGTCCGAGG AGG ||||||||[||||||||||||]PAM NA Intergenic NA NA

T15586 GATGTCGTCCGAGGAGGACGTGG 1000
Oligo fwd CACCGATGTCGTCCGAGGAGGACG
Oligo rev AAACCGTCCTCCTCGGACGACATC
Chromosome start end strand MM target_seq PAM alignment distance position gene name gene id
GU305914.1 204391 204413 + 0 GATGTCGTCCGAGGAGGACG TGG ||||||||[||||||||||||]PAM NA Intergenic NA NA

T15587 GATGTTCGTAGAGACCTCAACGG 1000
Oligo fwd CACCGATGTTCGTAGAGACCTCAA
Oligo rev AAACTTGAGGTCTCTACGAACATC
Chromosome start end strand MM target_seq PAM alignment distance position gene name gene id
GU305914.1 204427 204449 + 0 GATGTTCGTAGAGACCTCAA CGG ||||||||[||||||||||||]PAM NA Intergenic NA NA

T15588 CGTAGAGACCTCAACGGTCGCGG 1000
Oligo fwd CACCCGTAGAGACCTCAACGGTCG
Oligo rev AAACCGACCGTTGAGGTCTCTACG
Chromosome start end strand MM target_seq PAM alignment distance position gene name gene id
GU305914.1 204433 204455 + 0 CGTAGAGACCTCAACGGTCG CGG ||||||||[||||||||||||]PAM NA Intergenic NA NA

T15589 GACCTCAACGGTCGCGGTCTTGG 1000
Oligo fwd CACCGACCTCAACGGTCGCGGTCT
Oligo rev AAACAGACCGCGACCGTTGAGGTC
Chromosome start end strand MM target_seq PAM alignment distance position gene name gene id
GU305914.1 204439 204461 + 0 GACCTCAACGGTCGCGGTCT TGG ||||||||[||||||||||||]PAM NA Intergenic NA NA

T15590 AACGGTCGCGGTCTTGGTAGCGG 1000
Oligo fwd CACCAACGGTCGCGGTCTTGGTAG
Oligo rev AAACCTACCAAGACCGCGACCGTT
Chromosome start end strand MM target_seq PAM alignment distance position gene name gene id
GU305914.1 204445 204467 + 0 AACGGTCGCGGTCTTGGTAG CGG ||||||||[||||||||||||]PAM NA Intergenic NA NA

T15591 ACTCGATATCATGTCAAGCGCGG 1000
Oligo fwd CACCACTCGATATCATGTCAAGCG
Oligo rev AAACCGCTTGACATGATATCGAGT
Chromosome start end strand MM target_seq PAM alignment distance position gene name gene id
GU305914.1 204478 204500 + 0 ACTCGATATCATGTCAAGCG CGG ||||||||[||||||||||||]PAM NA Intergenic NA NA

T15592 AGCGCGGTACTCGAGACCGTCGG 1000
Oligo fwd CACCAGCGCGGTACTCGAGACCGT
Oligo rev AAACACGGTCTCGAGTACCGCGCT
Chromosome start end strand MM target_seq PAM alignment distance position gene name gene id
GU305914.1 204494 204516 + 0 AGCGCGGTACTCGAGACCGT CGG ||||||||[||||||||||||]PAM NA Intergenic NA NA

T15593 GCGCGGTACTCGAGACCGTCGGG 1000
Oligo fwd CACCGCGCGGTACTCGAGACCGTC
Oligo rev AAACGACGGTCTCGAGTACCGCGC
Chromosome start end strand MM target_seq PAM alignment distance position gene name gene id
GU305914.1 204495 204517 + 0 GCGCGGTACTCGAGACCGTC GGG ||||||||[||||||||||||]PAM NA Intergenic NA NA

T15594 GAGACCGTCGGGATGATAGTCGG 1000
Oligo fwd CACCGAGACCGTCGGGATGATAGT
Oligo rev AAACACTATCATCCCGACGGTCTC
Chromosome start end strand MM target_seq PAM alignment distance position gene name gene id
GU305914.1 204506 204528 + 0 GAGACCGTCGGGATGATAGT CGG ||||||||[||||||||||||]PAM NA Intergenic NA NA

T15595 GGGATGATAGTCGGCCTTGTCGG 1000
Oligo fwd CACCGGGATGATAGTCGGCCTTGT
Oligo rev AAACACAAGGCCGACTATCATCCC
Chromosome start end strand MM target_seq PAM alignment distance position gene name gene id
GU305914.1 204515 204537 + 0 GGGATGATAGTCGGCCTTGT CGG ||||||||[||||||||||||]PAM NA Intergenic NA NA

T15596 TGTCGGTCCTAGAGTACTGACGG 1000
Oligo fwd CACCTGTCGGTCCTAGAGTACTGA
Oligo rev AAACTCAGTACTCTAGGACCGACA

| | | | | | | | | | | | |
|---|---|---|---|---|---|---|---|---|---|---|---|
| Chromosome | start | end | strand | MM | target_seq | PAM | alignment | distance | position | gene name | gene id |
| GU305914.1 | 204532 | 204554 | + | 0 | TGTCGGTCCTAGAGTACTGA | CGG | \|\|\|\|\|\|\|\|[\|\|\|\|\|\|\|\|\|\|\|\|]PAM | NA | Intergenic | NA | NA |
| T15597 | GTCGTGTTCATACTCGAGAACGG | 1000 | | | | | | | | | |
| Oligo fwd | CACCGTCGTGTTCATACTCGAGAA | | | | | | | | | | |
| Oligo rev | AAACTTCTCGAGTATGAACACGAC | | | | | | | | | | |
| Chromosome | start | end | strand | MM | target_seq | PAM | alignment | distance | position | gene name | gene id |
| GU305914.1 | 204563 | 204585 | + | 0 | GTCGTGTTCATACTCGAGAA | CGG | \|\|\|\|\|\|\|\|[\|\|\|\|\|\|\|\|\|\|\|\|]PAM | NA | Intergenic | NA | NA |
| T15598 | ATACTCGAGAACGGTGTTTCTGG | 1000 | | | | | | | | | |
| Oligo fwd | CACCATACTCGAGAACGGTGTTTC | | | | | | | | | | |
| Oligo rev | AAACGAAACACCGTTCTCGAGTAT | | | | | | | | | | |
| Chromosome | start | end | strand | MM | target_seq | PAM | alignment | distance | position | gene name | gene id |
| GU305914.1 | 204572 | 204594 | + | 0 | ATACTCGAGAACGGTGTTTC | TGG | \|\|\|\|\|\|\|\|[\|\|\|\|\|\|\|\|\|\|\|\|]PAM | NA | Intergenic | NA | NA |
| T15599 | GTTTCTGGCGTAGTATGAAACGG | 1000 | | | | | | | | | |
| Oligo fwd | CACCGTTTCTGGCGTAGTATGAAA | | | | | | | | | | |
| Oligo rev | AAACTTTCATACTACGCCAGAAAC | | | | | | | | | | |
| Chromosome | start | end | strand | MM | target_seq | PAM | alignment | distance | position | gene name | gene id |
| GU305914.1 | 204587 | 204609 | + | 0 | GTTTCTGGCGTAGTATGAAA | CGG | \|\|\|\|\|\|\|\|[\|\|\|\|\|\|\|\|\|\|\|\|]PAM | NA | Intergenic | NA | NA |
| T15600 | CGTAGTAGTCGAGAGCGCCGTGG | 1000 | | | | | | | | | |
| Oligo fwd | CACCCGTAGTAGTCGAGAGCGCCG | | | | | | | | | | |
| Oligo rev | AAACCGGCGCTCTCGACTACTACG | | | | | | | | | | |
| Chromosome | start | end | strand | MM | target_seq | PAM | alignment | distance | position | gene name | gene id |
| GU305914.1 | 204628 | 204650 | + | 0 | CGTAGTAGTCGAGAGCGCCG | TGG | \|\|\|\|\|\|\|\|[\|\|\|\|\|\|\|\|\|\|\|\|]PAM | NA | Intergenic | NA | NA |
| T15601 | GTCGAGAGCGCCGTGGTAGTCGG | 1000 | | | | | | | | | |
| Oligo fwd | CACCGTCGAGAGCGCCGTGGTAGT | | | | | | | | | | |
| Oligo rev | AAACACTACCACGGCGCTCTCGAC | | | | | | | | | | |
| Chromosome | start | end | strand | MM | target_seq | PAM | alignment | distance | position | gene name | gene id |
| GU305914.1 | 204635 | 204657 | + | 0 | GTCGAGAGCGCCGTGGTAGT | CGG | \|\|\|\|\|\|\|\|[\|\|\|\|\|\|\|\|\|\|\|\|]PAM | NA | Intergenic | NA | NA |
| T15602 | TCGAGAGCGCCGTGGTAGTCGGG | 1000 | | | | | | | | | |
| Oligo fwd | CACCTCGAGAGCGCCGTGGTAGTC | | | | | | | | | | |
| Oligo rev | AAACGACTACCACGGCGCTCTCGA | | | | | | | | | | |
| Chromosome | start | end | strand | MM | target_seq | PAM | alignment | distance | position | gene name | gene id |
| GU305914.1 | 204636 | 204658 | + | 0 | TCGAGAGCGCCGTGGTAGTC | GGG | \|\|\|\|\|\|\|\|[\|\|\|\|\|\|\|\|\|\|\|\|]PAM | NA | Intergenic | NA | NA |
| T15603 | CGAGAGCGCCGTGGTAGTCGGGG | 1000 | | | | | | | | | |
| Oligo fwd | CACCCGAGAGCGCCGTGGTAGTCG | | | | | | | | | | |
| Oligo rev | AAACCGACTACCACGGCGCTCTCG | | | | | | | | | | |
| Chromosome | start | end | strand | MM | target_seq | PAM | alignment | distance | position | gene name | gene id |
| GU305914.1 | 204637 | 204659 | + | 0 | CGAGAGCGCCGTGGTAGTCG | GGG | \|\|\|\|\|\|\|\|[\|\|\|\|\|\|\|\|\|\|\|\|]PAM | NA | Intergenic | NA | NA |
| T15604 | GGTAGTCGGGGACGTCGTGATGG | 1000 | | | | | | | | | |
| Oligo fwd | CACCGGTAGTCGGGGACGTCGTGA | | | | | | | | | | |
| Oligo rev | AAACTCACGACGTCCCCGACTACC | | | | | | | | | | |
| Chromosome | start | end | strand | MM | target_seq | PAM | alignment | distance | position | gene name | gene id |
| GU305914.1 | 204649 | 204671 | + | 0 | GGTAGTCGGGGACGTCGTGA | TGG | \|\|\|\|\|\|\|\|[\|\|\|\|\|\|\|\|\|\|\|\|]PAM | NA | Intergenic | NA | NA |
| T15605 | GTAGTCGGGGACGTCGTGATGGG | 1000 | | | | | | | | | |
| Oligo fwd | CACCGTAGTCGGGGACGTCGTGAT | | | | | | | | | | |
| Oligo rev | AAACATCACGACGTCCCCGACTAC | | | | | | | | | | |
| Chromosome | start | end | strand | MM | target_seq | PAM | alignment | distance | position | gene name | gene id |
| GU305914.1 | 204650 | 204672 | + | 0 | GTAGTCGGGGACGTCGTGAT | GGG | \|\|\|\|\|\|\|\|[\|\|\|\|\|\|\|\|\|\|\|\|]PAM | NA | Intergenic | NA | NA |
| T15606 | TAGTCGGGGACGTCGTGATGGGG | 1000 | | | | | | | | | |
| Oligo fwd | CACCTAGTCGGGGACGTCGTGATG | | | | | | | | | | |
| Oligo rev | AAACCATCACGACGTCCCCGACTA | | | | | | | | | | |
| Chromosome | start | end | strand | MM | target_seq | PAM | alignment | distance | position | gene name | gene id |
| GU305914.1 | 204651 | 204673 | + | 0 | TAGTCGGGGACGTCGTGATG | GGG | \|\|\|\|\|\|\|\|[\|\|\|\|\|\|\|\|\|\|\|\|]PAM | NA | Intergenic | NA | NA |
| T15607 | AGTCGGGGACGTCGTGATGGGGG | 1000 | | | | | | | | | |
| Oligo fwd | CACCAGTCGGGGACGTCGTGATGG | | | | | | | | | | |
| Oligo rev | AAACCCATCACGACGTCCCCGACT | | | | | | | | | | |
| Chromosome | start | end | strand | MM | target_seq | PAM | alignment | distance | position | gene name | gene id |
| GU305914.1 | 204652 | 204674 | + | 0 | AGTCGGGGACGTCGTGATGG | GGG | \|\|\|\|\|\|\|\|[\|\|\|\|\|\|\|\|\|\|\|\|]PAM | NA | Intergenic | NA | NA |
| T15608 | GTCGGGGACGTCGTGATGGGGGG | 1000 | | | | | | | | | |
| Oligo fwd | CACCGTCGGGGACGTCGTGATGGG | | | | | | | | | | |
| Oligo rev | AAACCCCATCACGACGTCCCCGAC | | | | | | | | | | |
| Chromosome | start | end | strand | MM | target_seq | PAM | alignment | distance | position | gene name | gene id |
| GU305914.1 | 204653 | 204675 | + | 0 | GTCGGGGACGTCGTGATGGG | GGG | \|\|\|\|\|\|\|\|[\|\|\|\|\|\|\|\|\|\|\|\|]PAM | NA | Intergenic | NA | NA |
| T15609 | GGGGACGTCGTGATGGGGGGAGG | 1000 | | | | | | | | | |
| Oligo fwd | CACCGGGGACGTCGTGATGGGGGG | | | | | | | | | | |
| Oligo rev | AAACCCCCCCATCACGACGTCCCC | | | | | | | | | | |
| Chromosome | start | end | strand | MM | target_seq | PAM | alignment | distance | position | gene name | gene id |
| GU305914.1 | 204656 | 204678 | + | 0 | GGGGACGTCGTGATGGGGGG | AGG | \|\|\|\|\|\|\|\|[\|\|\|\|\|\|\|\|\|\|\|\|]PAM | NA | Intergenic | NA | NA |
| T15610 | ATGGGGGGAGGCACGTAGCGAGG | 1000 | | | | | | | | | |
| Oligo fwd | CACCATGGGGGGAGGCACGTAGCG | | | | | | | | | | |
| Oligo rev | AAACCGCTACGTGCCTCCCCCCAT | | | | | | | | | | |
| Chromosome | start | end | strand | MM | target_seq | PAM | alignment | distance | position | gene name | gene id |
| GU305914.1 | 204668 | 204690 | + | 0 | ATGGGGGGAGGCACGTAGCG | AGG | \|\|\|\|\|\|\|\|[\|\|\|\|\|\|\|\|\|\|\|\|]PAM | NA | Intergenic | NA | NA |
| T15611 | AAGCTCCGCGAACCGAAGACCGG | 1000 | | | | | | | | | |
| Oligo fwd | CACCAAGCTCCGCGAACCGAAGAC | | | | | | | | | | |

Oligo rev AAACGTCTTCGGTTCGCGGAGCTT
Chromosome start end strand MM target_seq PAM alignment distance position gene name gene id
GU305914.1 204708 204730 + 0 AAGCTCCGCGAACCGAAGAC CGG ||||||||[||||||||||||]PAM NA Intergenic NA NA

T15612 ACAGATGCTCTTAGACAGATCGG 1000
Oligo fwd CACCACAGATGCTCTTAGACAGAT
Oligo rev AAACATCTGTCTAAGAGCATCTGT
Chromosome start end strand MM target_seq PAM alignment distance position gene name gene id
GU305914.1 204736 204758 + 0 ACAGATGCTCTTAGACAGAT CGG ||||||||[||||||||||||]PAM NA Intergenic NA NA

T15613 ACAGATCGGAGTCTGATGTTTGG 1000
Oligo fwd CACCACAGATCGGAGTCTGATGTT
Oligo rev AAACAACATCAGACTCCGATCTGT
Chromosome start end strand MM target_seq PAM alignment distance position gene name gene id
GU305914.1 204750 204772 + 0 ACAGATCGGAGTCTGATGTT TGG ||||||||[||||||||||||]PAM NA Intergenic NA NA

T15614 ATATATAGTGAACCAGCTCTTGG 1000
Oligo fwd CACCATATATAGTGAACCAGCTCT
Oligo rev AAACAGAGCTGGTTCACTATATAT
Chromosome start end strand MM target_seq PAM alignment distance position gene name gene id
GU305914.1 204787 204809 + 0 ATATATAGTGAACCAGCTCT TGG ||||||||[||||||||||||]PAM NA Intergenic NA NA

T15615 CTTGGATACTTCGCCTTTTACGG 1000
Oligo fwd CACCCTTGGATACTTCGCCTTTTA
Oligo rev AAACTAAAAGGCGAAGTATCCAAG
Chromosome start end strand MM target_seq PAM alignment distance position gene name gene id
GU305914.1 204805 204827 + 0 CTTGGATACTTCGCCTTTTA CGG ||||||||[||||||||||||]PAM NA Intergenic NA NA

T15616 TTGGATACTTCGCCTTTTACGGG 1000
Oligo fwd CACCTTGGATACTTCGCCTTTTAC
Oligo rev AAACGTAAAAGGCGAAGTATCCAA
Chromosome start end strand MM target_seq PAM alignment distance position gene name gene id
GU305914.1 204806 204828 + 0 TTGGATACTTCGCCTTTTAC GGG ||||||||[||||||||||||]PAM NA Intergenic NA NA

T15617 GATACTTCGCCTTTTACGGGAGG 1000
Oligo fwd CACCGATACTTCGCCTTTTACGGG
Oligo rev AAACCCCGTAAAAGGCGAAGTATC
Chromosome start end strand MM target_seq PAM alignment distance position gene name gene id
GU305914.1 204809 204831 + 0 GATACTTCGCCTTTTACGGG AGG ||||||||[||||||||||||]PAM NA Intergenic NA NA

T15618 ATACTTCGCCTTTTACGGGAGGG 1000
Oligo fwd CACCATACTTCGCCTTTTACGGGA
Oligo rev AAACTCCCGTAAAAGGCGAAGTAT
Chromosome start end strand MM target_seq PAM alignment distance position gene name gene id
GU305914.1 204810 204832 + 0 ATACTTCGCCTTTTACGGGA GGG ||||||||[||||||||||||]PAM NA Intergenic NA NA

T15619 GCCTTTTACGGGAGGGAGTCCGG 1000
Oligo fwd CACCGCCTTTTACGGGAGGGAGTC
Oligo rev AAACGACTCCCTCCCGTAAAAGGC
Chromosome start end strand MM target_seq PAM alignment distance position gene name gene id
GU305914.1 204817 204839 + 0 GCCTTTTACGGGAGGGAGTC CGG ||||||||[||||||||||||]PAM NA Intergenic NA NA

T15620 ATCTCTATAGAAAAATAGAGCGG 1000
Oligo fwd CACCATCTCTATAGAAAAATAGAG
Oligo rev AAACCTCTATTTTTCTATAGAGAT
Chromosome start end strand MM target_seq PAM alignment distance position gene name gene id
GU305914.1 204848 204870 + 0 ATCTCTATAGAAAAATAGAG CGG ||||||||[||||||||||||]PAM NA Intergenic NA NA

T15621 TCGTTGTGTTGTTCGACGTGCGG 1000
Oligo fwd CACCTCGTTGTGTTGTTCGACGTG
Oligo rev AAACCACGTCGAACAACACAACGA
Chromosome start end strand MM target_seq PAM alignment distance position gene name gene id
GU305914.1 204894 204916 + 0 TCGTTGTGTTGTTCGACGTG CGG ||||||||[||||||||||||]PAM NA Intergenic NA NA

T15622 TGTTGTTCGACGTGCGGAATCGG 1000
Oligo fwd CACCTGTTGTTCGACGTGCGGAAT
Oligo rev AAACATTCCGCACGTCGAACAACA
Chromosome start end strand MM target_seq PAM alignment distance position gene name gene id
GU305914.1 204900 204922 + 0 TGTTGTTCGACGTGCGGAAT CGG ||||||||[||||||||||||]PAM NA Intergenic NA NA

T15623 GTTGTTCGACGTGCGGAATCGGG 1000
Oligo fwd CACCGTTGTTCGACGTGCGGAATC
Oligo rev AAACGATTCCGCACGTCGAACAAC
Chromosome start end strand MM target_seq PAM alignment distance position gene name gene id
GU305914.1 204901 204923 + 0 GTTGTTCGACGTGCGGAATC GGG ||||||||[||||||||||||]PAM NA Intergenic NA NA

T15624 ACGTGCGGAATCGGGTTTTCAGG 1000
Oligo fwd CACCACGTGCGGAATCGGGTTTTC
Oligo rev AAACGAAAACCCGATTCCGCACGT
Chromosome start end strand MM target_seq PAM alignment distance position gene name gene id
GU305914.1 204909 204931 + 0 ACGTGCGGAATCGGGTTTTC AGG ||||||||[||||||||||||]PAM NA Intergenic NA NA

T15625 AATCGGGTTTTCAGGCTGTTCGG 1000
Oligo fwd CACCAATCGGGTTTTCAGGCTGTT
Oligo rev AAACAACAGCCTGAAAACCCGATT
Chromosome start end strand MM target_seq PAM alignment distance position gene name gene id
GU305914.1 204917 204939 + 0 AATCGGGTTTTCAGGCTGTT CGG ||||||||[||||||||||||]PAM NA Intergenic NA NA

T15626 GCTGTTCGGACCGCCGAGTTTGG 1000

<table>
<tr><td>Oligo fwd</td><td>CACCGCTGTTCGGACCGCCGAGTT</td><td></td><td></td><td></td><td></td><td></td><td></td><td></td><td></td><td></td><td></td></tr>
<tr><td>Oligo rev</td><td>AAACAACTCGGCGGTCCGAACAGC</td><td></td><td></td><td></td><td></td><td></td><td></td><td></td><td></td><td></td><td></td></tr>
<tr><td>Chromosome</td><td>start</td><td>end</td><td>strand</td><td>MM</td><td>target_seq</td><td>PAM</td><td>alignment</td><td>distance</td><td>position</td><td>gene name</td><td>gene id</td></tr>
<tr><td>GU305914.1</td><td>204931</td><td>204953</td><td>+</td><td>0</td><td>GCTGTTCGGACCGCCGAGTT</td><td>TGG</td><td>||||||||[||||||||||||]PAM</td><td>NA</td><td>Intergenic</td><td>NA</td><td>NA</td></tr>
<tr><td>T15627</td><td>CGGACCGCCGAGTTTGGCAGCGG</td><td>1000</td><td></td><td></td><td></td><td></td><td></td><td></td><td></td><td></td><td></td></tr>
<tr><td>Oligo fwd</td><td>CACCCGGACCGCCGAGTTTGGCAG</td><td></td><td></td><td></td><td></td><td></td><td></td><td></td><td></td><td></td><td></td></tr>
<tr><td>Oligo rev</td><td>AAACCTGCCAAACTCGGCGGTCCG</td><td></td><td></td><td></td><td></td><td></td><td></td><td></td><td></td><td></td><td></td></tr>
<tr><td>Chromosome</td><td>start</td><td>end</td><td>strand</td><td>MM</td><td>target_seq</td><td>PAM</td><td>alignment</td><td>distance</td><td>position</td><td>gene name</td><td>gene id</td></tr>
<tr><td>GU305914.1</td><td>204937</td><td>204959</td><td>+</td><td>0</td><td>CGGACCGCCGAGTTTGGCAG</td><td>CGG</td><td>||||||||[||||||||||||]PAM</td><td>NA</td><td>Intergenic</td><td>NA</td><td>NA</td></tr>
<tr><td>T15628</td><td>TTTGTAGACTTTCAACCACCTGG</td><td>1000</td><td></td><td></td><td></td><td></td><td></td><td></td><td></td><td></td><td></td></tr>
<tr><td>Oligo fwd</td><td>CACCTTTGTAGACTTTCAACCACC</td><td></td><td></td><td></td><td></td><td></td><td></td><td></td><td></td><td></td><td></td></tr>
<tr><td>Oligo rev</td><td>AAACGGTGGTTGAAAGTCTACAAA</td><td></td><td></td><td></td><td></td><td></td><td></td><td></td><td></td><td></td><td></td></tr>
<tr><td>Chromosome</td><td>start</td><td>end</td><td>strand</td><td>MM</td><td>target_seq</td><td>PAM</td><td>alignment</td><td>distance</td><td>position</td><td>gene name</td><td>gene id</td></tr>
<tr><td>GU305914.1</td><td>204961</td><td>204983</td><td>+</td><td>0</td><td>TTTGTAGACTTTCAACCACC</td><td>TGG</td><td>||||||||[||||||||||||]PAM</td><td>NA</td><td>Intergenic</td><td>NA</td><td>NA</td></tr>
<tr><td>T15629</td><td>TTGTAGACTTTCAACCACCTGGG</td><td>1000</td><td></td><td></td><td></td><td></td><td></td><td></td><td></td><td></td><td></td></tr>
<tr><td>Oligo fwd</td><td>CACCTTGTAGACTTTCAACCACCT</td><td></td><td></td><td></td><td></td><td></td><td></td><td></td><td></td><td></td><td></td></tr>
<tr><td>Oligo rev</td><td>AAACAGGTGGTTGAAAGTCTACAA</td><td></td><td></td><td></td><td></td><td></td><td></td><td></td><td></td><td></td><td></td></tr>
<tr><td>Chromosome</td><td>start</td><td>end</td><td>strand</td><td>MM</td><td>target_seq</td><td>PAM</td><td>alignment</td><td>distance</td><td>position</td><td>gene name</td><td>gene id</td></tr>
<tr><td>GU305914.1</td><td>204962</td><td>204984</td><td>+</td><td>0</td><td>TTGTAGACTTTCAACCACCT</td><td>GGG</td><td>||||||||[||||||||||||]PAM</td><td>NA</td><td>Intergenic</td><td>NA</td><td>NA</td></tr>
<tr><td>T15630</td><td>TGTAGACTTTCAACCACCTGGGG</td><td>1000</td><td></td><td></td><td></td><td></td><td></td><td></td><td></td><td></td><td></td></tr>
<tr><td>Oligo fwd</td><td>CACCTGTAGACTTTCAACCACCTG</td><td></td><td></td><td></td><td></td><td></td><td></td><td></td><td></td><td></td><td></td></tr>
<tr><td>Oligo rev</td><td>AAACCAGGTGGTTGAAAGTCTACA</td><td></td><td></td><td></td><td></td><td></td><td></td><td></td><td></td><td></td><td></td></tr>
<tr><td>Chromosome</td><td>start</td><td>end</td><td>strand</td><td>MM</td><td>target_seq</td><td>PAM</td><td>alignment</td><td>distance</td><td>position</td><td>gene name</td><td>gene id</td></tr>
<tr><td>GU305914.1</td><td>204963</td><td>204985</td><td>+</td><td>0</td><td>TGTAGACTTTCAACCACCTG</td><td>GGG</td><td>||||||||[||||||||||||]PAM</td><td>NA</td><td>Intergenic</td><td>NA</td><td>NA</td></tr>
<tr><td>T15631</td><td>GACCTTGTTTTTCAGTAGATCGG</td><td>1000</td><td></td><td></td><td></td><td></td><td></td><td></td><td></td><td></td><td></td></tr>
<tr><td>Oligo fwd</td><td>CACCGACCTTGTTTTTCAGTAGAT</td><td></td><td></td><td></td><td></td><td></td><td></td><td></td><td></td><td></td><td></td></tr>
<tr><td>Oligo rev</td><td>AAACATCTACTGAAAAACAAGGTC</td><td></td><td></td><td></td><td></td><td></td><td></td><td></td><td></td><td></td><td></td></tr>
<tr><td>Chromosome</td><td>start</td><td>end</td><td>strand</td><td>MM</td><td>target_seq</td><td>PAM</td><td>alignment</td><td>distance</td><td>position</td><td>gene name</td><td>gene id</td></tr>
<tr><td>GU305914.1</td><td>204997</td><td>205019</td><td>+</td><td>0</td><td>GACCTTGTTTTTCAGTAGAT</td><td>CGG</td><td>||||||||[||||||||||||]PAM</td><td>NA</td><td>Intergenic</td><td>NA</td><td>NA</td></tr>
<tr><td>T15632</td><td>TCTGTCCCATTCACAAAAGCCGG</td><td>1000</td><td></td><td></td><td></td><td></td><td></td><td></td><td></td><td></td><td></td></tr>
<tr><td>Oligo fwd</td><td>CACCTCTGTCCCATTCACAAAAGC</td><td></td><td></td><td></td><td></td><td></td><td></td><td></td><td></td><td></td><td></td></tr>
<tr><td>Oligo rev</td><td>AAACGCTTTTGTGAATGGGACAGA</td><td></td><td></td><td></td><td></td><td></td><td></td><td></td><td></td><td></td><td></td></tr>
<tr><td>Chromosome</td><td>start</td><td>end</td><td>strand</td><td>MM</td><td>target_seq</td><td>PAM</td><td>alignment</td><td>distance</td><td>position</td><td>gene name</td><td>gene id</td></tr>
<tr><td>GU305914.1</td><td>205082</td><td>205104</td><td>+</td><td>0</td><td>TCTGTCCCATTCACAAAAGC</td><td>CGG</td><td>||||||||[||||||||||||]PAM</td><td>NA</td><td>Intergenic</td><td>NA</td><td>NA</td></tr>
<tr><td>GU305914.1</td><td>193576</td><td>193598</td><td>-</td><td>4</td><td>TCTGTCAACTTCACAAAACC</td><td>CGG</td><td>||||||--[-|||||||||-|]PAM</td><td>NA</td><td>Intergenic</td><td>NA</td><td>NA</td></tr>
<tr><td>T15633</td><td>CGGATTATATTTCAACAAAACGG</td><td>1000</td><td></td><td></td><td></td><td></td><td></td><td></td><td></td><td></td><td></td></tr>
<tr><td>Oligo fwd</td><td>CACCCGGATTATATTTCAACAAAA</td><td></td><td></td><td></td><td></td><td></td><td></td><td></td><td></td><td></td><td></td></tr>
<tr><td>Oligo rev</td><td>AAACTTTTGTTGAAATATAATCCG</td><td></td><td></td><td></td><td></td><td></td><td></td><td></td><td></td><td></td><td></td></tr>
<tr><td>Chromosome</td><td>start</td><td>end</td><td>strand</td><td>MM</td><td>target_seq</td><td>PAM</td><td>alignment</td><td>distance</td><td>position</td><td>gene name</td><td>gene id</td></tr>
<tr><td>GU305914.1</td><td>205102</td><td>205124</td><td>+</td><td>0</td><td>CGGATTATATTTCAACAAAA</td><td>CGG</td><td>||||||||[||||||||||||]PAM</td><td>NA</td><td>Intergenic</td><td>NA</td><td>NA</td></tr>
<tr><td>T15634</td><td>AACGGTTTCGTTCAGTACGATGG</td><td>1000</td><td></td><td></td><td></td><td></td><td></td><td></td><td></td><td></td><td></td></tr>
<tr><td>Oligo fwd</td><td>CACCAACGGTTTCGTTCAGTACGA</td><td></td><td></td><td></td><td></td><td></td><td></td><td></td><td></td><td></td><td></td></tr>
<tr><td>Oligo rev</td><td>AAACTCGTACTGAACGAAACCGTT</td><td></td><td></td><td></td><td></td><td></td><td></td><td></td><td></td><td></td><td></td></tr>
<tr><td>Chromosome</td><td>start</td><td>end</td><td>strand</td><td>MM</td><td>target_seq</td><td>PAM</td><td>alignment</td><td>distance</td><td>position</td><td>gene name</td><td>gene id</td></tr>
<tr><td>GU305914.1</td><td>205120</td><td>205142</td><td>+</td><td>0</td><td>AACGGTTTCGTTCAGTACGA</td><td>TGG</td><td>||||||||[||||||||||||]PAM</td><td>NA</td><td>Intergenic</td><td>NA</td><td>NA</td></tr>
<tr><td>T15635</td><td>GTACCAGAATGTAGCGTTGAAGG</td><td>1000</td><td></td><td></td><td></td><td></td><td></td><td></td><td></td><td></td><td></td></tr>
<tr><td>Oligo fwd</td><td>CACCGTACCAGAATGTAGCGTTGA</td><td></td><td></td><td></td><td></td><td></td><td></td><td></td><td></td><td></td><td></td></tr>
<tr><td>Oligo rev</td><td>AAACTCAACGCTACATTCTGGTAC</td><td></td><td></td><td></td><td></td><td></td><td></td><td></td><td></td><td></td><td></td></tr>
<tr><td>Chromosome</td><td>start</td><td>end</td><td>strand</td><td>MM</td><td>target_seq</td><td>PAM</td><td>alignment</td><td>distance</td><td>position</td><td>gene name</td><td>gene id</td></tr>
<tr><td>GU305914.1</td><td>205156</td><td>205178</td><td>+</td><td>0</td><td>GTACCAGAATGTAGCGTTGA</td><td>AGG</td><td>||||||||[||||||||||||]PAM</td><td>NA</td><td>Intergenic</td><td>NA</td><td>NA</td></tr>
<tr><td>T15636</td><td>CCGTTTTGAGAAATTCTCGCTGG</td><td>1000</td><td></td><td></td><td></td><td></td><td></td><td></td><td></td><td></td><td></td></tr>
<tr><td>Oligo fwd</td><td>CACCCCGTTTTGAGAAATTCTCGC</td><td></td><td></td><td></td><td></td><td></td><td></td><td></td><td></td><td></td><td></td></tr>
<tr><td>Oligo rev</td><td>AAACGCGAGAATTTCTCAAAACGG</td><td></td><td></td><td></td><td></td><td></td><td></td><td></td><td></td><td></td><td></td></tr>
<tr><td>Chromosome</td><td>start</td><td>end</td><td>strand</td><td>MM</td><td>target_seq</td><td>PAM</td><td>alignment</td><td>distance</td><td>position</td><td>gene name</td><td>gene id</td></tr>
<tr><td>GU305914.1</td><td>205233</td><td>205255</td><td>+</td><td>0</td><td>CCGTTTTGAGAAATTCTCGC</td><td>TGG</td><td>||||||||[||||||||||||]PAM</td><td>NA</td><td>Intergenic</td><td>NA</td><td>NA</td></tr>
<tr><td>T15637</td><td>TTAAAAAAATCATATTCTGTTGG</td><td>1000</td><td></td><td></td><td></td><td></td><td></td><td></td><td></td><td></td><td></td></tr>
<tr><td>Oligo fwd</td><td>CACCTTAAAAAAATCATATTCTGT</td><td></td><td></td><td></td><td></td><td></td><td></td><td></td><td></td><td></td><td></td></tr>
<tr><td>Oligo rev</td><td>AAACACAGAATATGATTTTTTTAA</td><td></td><td></td><td></td><td></td><td></td><td></td><td></td><td></td><td></td><td></td></tr>
<tr><td>Chromosome</td><td>start</td><td>end</td><td>strand</td><td>MM</td><td>target_seq</td><td>PAM</td><td>alignment</td><td>distance</td><td>position</td><td>gene name</td><td>gene id</td></tr>
<tr><td>GU305914.1</td><td>205259</td><td>205281</td><td>+</td><td>0</td><td>TTAAAAAAATCATATTCTGT</td><td>TGG</td><td>||||||||[||||||||||||]PAM</td><td>NA</td><td>Intergenic</td><td>NA</td><td>NA</td></tr>
<tr><td>T15638</td><td>CCATCCGTAACTGTCAGTGACGG</td><td>1000</td><td></td><td></td><td></td><td></td><td></td><td></td><td></td><td></td><td></td></tr>
<tr><td>Oligo fwd</td><td>CACCCCATCCGTAACTGTCAGTGA</td><td></td><td></td><td></td><td></td><td></td><td></td><td></td><td></td><td></td><td></td></tr>
<tr><td>Oligo rev</td><td>AAACTCACTGACAGTTACGGATGG</td><td></td><td></td><td></td><td></td><td></td><td></td><td></td><td></td><td></td><td></td></tr>
<tr><td>Chromosome</td><td>start</td><td>end</td><td>strand</td><td>MM</td><td>target_seq</td><td>PAM</td><td>alignment</td><td>distance</td><td>position</td><td>gene name</td><td>gene id</td></tr>
<tr><td>GU305914.1</td><td>205300</td><td>205322</td><td>+</td><td>0</td><td>CCATCCGTAACTGTCAGTGA</td><td>CGG</td><td>||||||||[||||||||||||]PAM</td><td>NA</td><td>Intergenic</td><td>NA</td><td>NA</td></tr>
<tr><td>T15639</td><td>TCATCAGTATGATCCCATCTCGG</td><td>1000</td><td></td><td></td><td></td><td></td><td></td><td></td><td></td><td></td><td></td></tr>
<tr><td>Oligo fwd</td><td>CACCTCATCAGTATGATCCCATCT</td><td></td><td></td><td></td><td></td><td></td><td></td><td></td><td></td><td></td><td></td></tr>
<tr><td>Oligo rev</td><td>AAACAGATGGGATCATACTGATGA</td><td></td><td></td><td></td><td></td><td></td><td></td><td></td><td></td><td></td><td></td></tr>
<tr><td>Chromosome</td><td>start</td><td>end</td><td>strand</td><td>MM</td><td>target_seq</td><td>PAM</td><td>alignment</td><td>distance</td><td>position</td><td>gene name</td><td>gene id</td></tr>
<tr><td>GU305914.1</td><td>205328</td><td>205350</td><td>+</td><td>0</td><td>TCATCAGTATGATCCCATCT</td><td>CGG</td><td>||||||||[||||||||||||]PAM</td><td>NA</td><td>Intergenic</td><td>NA</td><td>NA</td></tr>
<tr><td>T15640</td><td>CCATTTGACCGTCAGAAAACTGG</td><td>1000</td><td></td><td></td><td></td><td></td><td></td><td></td><td></td><td></td><td></td></tr>
<tr><td>Oligo fwd</td><td>CACCCCATTTGACCGTCAGAAAAC</td><td></td><td></td><td></td><td></td><td></td><td></td><td></td><td></td><td></td><td></td></tr>
<tr><td>Oligo rev</td><td>AAACGTTTTCTGACGGTCAAATGG</td><td></td><td></td><td></td><td></td><td></td><td></td><td></td><td></td><td></td><td></td></tr>
<tr><td>Chromosome</td><td>start</td><td>end</td><td>strand</td><td>MM</td><td>target_seq</td><td>PAM</td><td>alignment</td><td>distance</td><td>position</td><td>gene name</td><td>gene id</td></tr>
<tr><td>GU305914.1</td><td>205360</td><td>205382</td><td>+</td><td>0</td><td>CCATTTGACCGTCAGAAAAC</td><td>TGG</td><td>||||||||[||||||||||||]PAM</td><td>NA</td><td>Intergenic</td><td>NA</td><td>NA</td></tr>
</table>

<table>
<tr><td>T15641</td><td>CATCCAACGCGATCGTAGCTAGG</td><td>1000</td><td></td><td></td><td></td><td></td><td></td><td></td><td></td><td></td><td></td></tr>
<tr><td>Oligo fwd</td><td>CACCCATCCAACGCGATCGTAGCT</td><td></td><td></td><td></td><td></td><td></td><td></td><td></td><td></td><td></td><td></td></tr>
<tr><td>Oligo rev</td><td>AAACAGCTACGATCGCGTTGGATG</td><td></td><td></td><td></td><td></td><td></td><td></td><td></td><td></td><td></td><td></td></tr>
<tr><td>Chromosome</td><td>start</td><td>end</td><td>strand</td><td>MM</td><td>target_seq</td><td>PAM</td><td>alignment</td><td>distance</td><td>position</td><td>gene name</td><td>gene id</td></tr>
<tr><td>GU305914.1</td><td>205389</td><td>205411</td><td>+</td><td>0</td><td>CATCCAACGCGATCGTAGCT</td><td>AGG</td><td>||||||||[||||||||||||]PAM</td><td>NA</td><td>Intergenic</td><td>NA</td><td>NA</td></tr>
<tr><td>T15642</td><td>GCGATCGTAGCTAGGAAGTGAGG</td><td>1000</td><td></td><td></td><td></td><td></td><td></td><td></td><td></td><td></td><td></td></tr>
<tr><td>Oligo fwd</td><td>CACCGCGATCGTAGCTAGGAAGTG</td><td></td><td></td><td></td><td></td><td></td><td></td><td></td><td></td><td></td><td></td></tr>
<tr><td>Oligo rev</td><td>AAACCACTTCCTAGCTACGATCGC</td><td></td><td></td><td></td><td></td><td></td><td></td><td></td><td></td><td></td><td></td></tr>
<tr><td>Chromosome</td><td>start</td><td>end</td><td>strand</td><td>MM</td><td>target_seq</td><td>PAM</td><td>alignment</td><td>distance</td><td>position</td><td>gene name</td><td>gene id</td></tr>
<tr><td>GU305914.1</td><td>205397</td><td>205419</td><td>+</td><td>0</td><td>GCGATCGTAGCTAGGAAGTG</td><td>AGG</td><td>||||||||[||||||||||||]PAM</td><td>NA</td><td>Intergenic</td><td>NA</td><td>NA</td></tr>
<tr><td>T15643</td><td>GCATCGATAAAACCAGTGTCAGG</td><td>1000</td><td></td><td></td><td></td><td></td><td></td><td></td><td></td><td></td><td></td></tr>
<tr><td>Oligo fwd</td><td>CACCGCATCGATAAAACCAGTGTC</td><td></td><td></td><td></td><td></td><td></td><td></td><td></td><td></td><td></td><td></td></tr>
<tr><td>Oligo rev</td><td>AAACGACACTGGTTTTATCGATGC</td><td></td><td></td><td></td><td></td><td></td><td></td><td></td><td></td><td></td><td></td></tr>
<tr><td>Chromosome</td><td>start</td><td>end</td><td>strand</td><td>MM</td><td>target_seq</td><td>PAM</td><td>alignment</td><td>distance</td><td>position</td><td>gene name</td><td>gene id</td></tr>
<tr><td>GU305914.1</td><td>205437</td><td>205459</td><td>+</td><td>0</td><td>GCATCGATAAAACCAGTGTC</td><td>AGG</td><td>||||||||[||||||||||||]PAM</td><td>NA</td><td>Intergenic</td><td>NA</td><td>NA</td></tr>
<tr><td>T15644</td><td>AAACCAGTGTCAGGTTCACGCGG</td><td>1000</td><td></td><td></td><td></td><td></td><td></td><td></td><td></td><td></td><td></td></tr>
<tr><td>Oligo fwd</td><td>CACCAAACCAGTGTCAGGTTCACG</td><td></td><td></td><td></td><td></td><td></td><td></td><td></td><td></td><td></td><td></td></tr>
<tr><td>Oligo rev</td><td>AAACCGTGAACCTGACACTGGTTT</td><td></td><td></td><td></td><td></td><td></td><td></td><td></td><td></td><td></td><td></td></tr>
<tr><td>Chromosome</td><td>start</td><td>end</td><td>strand</td><td>MM</td><td>target_seq</td><td>PAM</td><td>alignment</td><td>distance</td><td>position</td><td>gene name</td><td>gene id</td></tr>
<tr><td>GU305914.1</td><td>205446</td><td>205468</td><td>+</td><td>0</td><td>AAACCAGTGTCAGGTTCACG</td><td>CGG</td><td>||||||||[||||||||||||]PAM</td><td>NA</td><td>Intergenic</td><td>NA</td><td>NA</td></tr>
<tr><td>T15645</td><td>TTCGAACCCGCGAGCACATTCGG</td><td>1000</td><td></td><td></td><td></td><td></td><td></td><td></td><td></td><td></td><td></td></tr>
<tr><td>Oligo fwd</td><td>CACCTTCGAACCCGCGAGCACATT</td><td></td><td></td><td></td><td></td><td></td><td></td><td></td><td></td><td></td><td></td></tr>
<tr><td>Oligo rev</td><td>AAACAATGTGCTCGCGGGTTCGAA</td><td></td><td></td><td></td><td></td><td></td><td></td><td></td><td></td><td></td><td></td></tr>
<tr><td>Chromosome</td><td>start</td><td>end</td><td>strand</td><td>MM</td><td>target_seq</td><td>PAM</td><td>alignment</td><td>distance</td><td>position</td><td>gene name</td><td>gene id</td></tr>
<tr><td>GU305914.1</td><td>205483</td><td>205505</td><td>+</td><td>0</td><td>TTCGAACCCGCGAGCACATT</td><td>CGG</td><td>||||||||[||||||||||||]PAM</td><td>NA</td><td>Intergenic</td><td>NA</td><td>NA</td></tr>
<tr><td>T15646</td><td>TCGAACCCGCGAGCACATTCGGG</td><td>1000</td><td></td><td></td><td></td><td></td><td></td><td></td><td></td><td></td><td></td></tr>
<tr><td>Oligo fwd</td><td>CACCTCGAACCCGCGAGCACATTC</td><td></td><td></td><td></td><td></td><td></td><td></td><td></td><td></td><td></td><td></td></tr>
<tr><td>Oligo rev</td><td>AAACGAATGTGCTCGCGGGTTCGA</td><td></td><td></td><td></td><td></td><td></td><td></td><td></td><td></td><td></td><td></td></tr>
<tr><td>Chromosome</td><td>start</td><td>end</td><td>strand</td><td>MM</td><td>target_seq</td><td>PAM</td><td>alignment</td><td>distance</td><td>position</td><td>gene name</td><td>gene id</td></tr>
<tr><td>GU305914.1</td><td>205484</td><td>205506</td><td>+</td><td>0</td><td>TCGAACCCGCGAGCACATTC</td><td>GGG</td><td>||||||||[||||||||||||]PAM</td><td>NA</td><td>Intergenic</td><td>NA</td><td>NA</td></tr>
<tr><td>T15647</td><td>ACATTCGGGATCTACGTCGCCGG</td><td>1000</td><td></td><td></td><td></td><td></td><td></td><td></td><td></td><td></td><td></td></tr>
<tr><td>Oligo fwd</td><td>CACCACATTCGGGATCTACGTCGC</td><td></td><td></td><td></td><td></td><td></td><td></td><td></td><td></td><td></td><td></td></tr>
<tr><td>Oligo rev</td><td>AAACGCGACGTAGATCCCGAATGT</td><td></td><td></td><td></td><td></td><td></td><td></td><td></td><td></td><td></td><td></td></tr>
<tr><td>Chromosome</td><td>start</td><td>end</td><td>strand</td><td>MM</td><td>target_seq</td><td>PAM</td><td>alignment</td><td>distance</td><td>position</td><td>gene name</td><td>gene id</td></tr>
<tr><td>GU305914.1</td><td>205498</td><td>205520</td><td>+</td><td>0</td><td>ACATTCGGGATCTACGTCGC</td><td>CGG</td><td>||||||||[||||||||||||]PAM</td><td>NA</td><td>Intergenic</td><td>NA</td><td>NA</td></tr>
<tr><td>T15648</td><td>CATTCGGGATCTACGTCGCCGGG</td><td>1000</td><td></td><td></td><td></td><td></td><td></td><td></td><td></td><td></td><td></td></tr>
<tr><td>Oligo fwd</td><td>CACCCATTCGGGATCTACGTCGCC</td><td></td><td></td><td></td><td></td><td></td><td></td><td></td><td></td><td></td><td></td></tr>
<tr><td>Oligo rev</td><td>AAACGGCGACGTAGATCCCGAATG</td><td></td><td></td><td></td><td></td><td></td><td></td><td></td><td></td><td></td><td></td></tr>
<tr><td>Chromosome</td><td>start</td><td>end</td><td>strand</td><td>MM</td><td>target_seq</td><td>PAM</td><td>alignment</td><td>distance</td><td>position</td><td>gene name</td><td>gene id</td></tr>
<tr><td>GU305914.1</td><td>205499</td><td>205521</td><td>+</td><td>0</td><td>CATTCGGGATCTACGTCGCC</td><td>GGG</td><td>||||||||[||||||||||||]PAM</td><td>NA</td><td>Intergenic</td><td>NA</td><td>NA</td></tr>
<tr><td>T15649</td><td>ATTCGGGATCTACGTCGCCGGGG</td><td>1000</td><td></td><td></td><td></td><td></td><td></td><td></td><td></td><td></td><td></td></tr>
<tr><td>Oligo fwd</td><td>CACCATTCGGGATCTACGTCGCCG</td><td></td><td></td><td></td><td></td><td></td><td></td><td></td><td></td><td></td><td></td></tr>
<tr><td>Oligo rev</td><td>AAACCGGCGACGTAGATCCCGAAT</td><td></td><td></td><td></td><td></td><td></td><td></td><td></td><td></td><td></td><td></td></tr>
<tr><td>Chromosome</td><td>start</td><td>end</td><td>strand</td><td>MM</td><td>target_seq</td><td>PAM</td><td>alignment</td><td>distance</td><td>position</td><td>gene name</td><td>gene id</td></tr>
<tr><td>GU305914.1</td><td>205500</td><td>205522</td><td>+</td><td>0</td><td>ATTCGGGATCTACGTCGCCG</td><td>GGG</td><td>||||||||[||||||||||||]PAM</td><td>NA</td><td>Intergenic</td><td>NA</td><td>NA</td></tr>
<tr><td>T15650</td><td>TTCGGGATCTACGTCGCCGGGGG</td><td>1000</td><td></td><td></td><td></td><td></td><td></td><td></td><td></td><td></td><td></td></tr>
<tr><td>Oligo fwd</td><td>CACCTTCGGGATCTACGTCGCCGG</td><td></td><td></td><td></td><td></td><td></td><td></td><td></td><td></td><td></td><td></td></tr>
<tr><td>Oligo rev</td><td>AAACCCGGCGACGTAGATCCCGAA</td><td></td><td></td><td></td><td></td><td></td><td></td><td></td><td></td><td></td><td></td></tr>
<tr><td>Chromosome</td><td>start</td><td>end</td><td>strand</td><td>MM</td><td>target_seq</td><td>PAM</td><td>alignment</td><td>distance</td><td>position</td><td>gene name</td><td>gene id</td></tr>
<tr><td>GU305914.1</td><td>205501</td><td>205523</td><td>+</td><td>0</td><td>TTCGGGATCTACGTCGCCGG</td><td>GGG</td><td>||||||||[||||||||||||]PAM</td><td>NA</td><td>Intergenic</td><td>NA</td><td>NA</td></tr>
<tr><td>T15651</td><td>GCTAGCGAACGAAACCGCCAAGG</td><td>1000</td><td></td><td></td><td></td><td></td><td></td><td></td><td></td><td></td><td></td></tr>
<tr><td>Oligo fwd</td><td>CACCGCTAGCGAACGAAACCGCCA</td><td></td><td></td><td></td><td></td><td></td><td></td><td></td><td></td><td></td><td></td></tr>
<tr><td>Oligo rev</td><td>AAACTGGCGGTTTCGTTCGCTAGC</td><td></td><td></td><td></td><td></td><td></td><td></td><td></td><td></td><td></td><td></td></tr>
<tr><td>Chromosome</td><td>start</td><td>end</td><td>strand</td><td>MM</td><td>target_seq</td><td>PAM</td><td>alignment</td><td>distance</td><td>position</td><td>gene name</td><td>gene id</td></tr>
<tr><td>GU305914.1</td><td>205537</td><td>205559</td><td>+</td><td>0</td><td>GCTAGCGAACGAAACCGCCA</td><td>AGG</td><td>||||||||[||||||||||||]PAM</td><td>NA</td><td>Intergenic</td><td>NA</td><td>NA</td></tr>
<tr><td>T15652</td><td>AAACCGCCAAGGCGCACATTAGG</td><td>1000</td><td></td><td></td><td></td><td></td><td></td><td></td><td></td><td></td><td></td></tr>
<tr><td>Oligo fwd</td><td>CACCAAACCGCCAAGGCGCACATT</td><td></td><td></td><td></td><td></td><td></td><td></td><td></td><td></td><td></td><td></td></tr>
<tr><td>Oligo rev</td><td>AAACAATGTGCGCCTTGGCGGTTT</td><td></td><td></td><td></td><td></td><td></td><td></td><td></td><td></td><td></td><td></td></tr>
<tr><td>Chromosome</td><td>start</td><td>end</td><td>strand</td><td>MM</td><td>target_seq</td><td>PAM</td><td>alignment</td><td>distance</td><td>position</td><td>gene name</td><td>gene id</td></tr>
<tr><td>GU305914.1</td><td>205548</td><td>205570</td><td>+</td><td>0</td><td>AAACCGCCAAGGCGCACATT</td><td>AGG</td><td>||||||||[||||||||||||]PAM</td><td>NA</td><td>Intergenic</td><td>NA</td><td>NA</td></tr>
<tr><td>T15653</td><td>TTAGGTATGAGACCACCCGACGG</td><td>1000</td><td></td><td></td><td></td><td></td><td></td><td></td><td></td><td></td><td></td></tr>
<tr><td>Oligo fwd</td><td>CACCTTAGGTATGAGACCACCCGA</td><td></td><td></td><td></td><td></td><td></td><td></td><td></td><td></td><td></td><td></td></tr>
<tr><td>Oligo rev</td><td>AAACTCGGGTGGTCTCATACCTAA</td><td></td><td></td><td></td><td></td><td></td><td></td><td></td><td></td><td></td><td></td></tr>
<tr><td>Chromosome</td><td>start</td><td>end</td><td>strand</td><td>MM</td><td>target_seq</td><td>PAM</td><td>alignment</td><td>distance</td><td>position</td><td>gene name</td><td>gene id</td></tr>
<tr><td>GU305914.1</td><td>205566</td><td>205588</td><td>+</td><td>0</td><td>TTAGGTATGAGACCACCCGA</td><td>CGG</td><td>||||||||[||||||||||||]PAM</td><td>NA</td><td>Intergenic</td><td>NA</td><td>NA</td></tr>
<tr><td>T15654</td><td>GACCACCCGACGGTCCATCGCGG</td><td>1000</td><td></td><td></td><td></td><td></td><td></td><td></td><td></td><td></td><td></td></tr>
<tr><td>Oligo fwd</td><td>CACCGACCACCCGACGGTCCATCG</td><td></td><td></td><td></td><td></td><td></td><td></td><td></td><td></td><td></td><td></td></tr>
<tr><td>Oligo rev</td><td>AAACCGATGGACCGTCGGGTGGTC</td><td></td><td></td><td></td><td></td><td></td><td></td><td></td><td></td><td></td><td></td></tr>
<tr><td>Chromosome</td><td>start</td><td>end</td><td>strand</td><td>MM</td><td>target_seq</td><td>PAM</td><td>alignment</td><td>distance</td><td>position</td><td>gene name</td><td>gene id</td></tr>
<tr><td>GU305914.1</td><td>205576</td><td>205598</td><td>+</td><td>0</td><td>GACCACCCGACGGTCCATCG</td><td>CGG</td><td>||||||||[||||||||||||]PAM</td><td>NA</td><td>Intergenic</td><td>NA</td><td>NA</td></tr>
<tr><td>T15655</td><td>CCATCGCGGAGTTGAAACACCGG</td><td>1000</td><td></td><td></td><td></td><td></td><td></td><td></td><td></td><td></td><td></td></tr>
<tr><td>Oligo fwd</td><td>CACCCCATCGCGGAGTTGAAACAC</td><td></td><td></td><td></td><td></td><td></td><td></td><td></td><td></td><td></td><td></td></tr>
<tr><td>Oligo rev</td><td>AAACGTGTTTCAACTCCGCGATGG</td><td></td><td></td><td></td><td></td><td></td><td></td><td></td><td></td><td></td><td></td></tr>
<tr><td>Chromosome</td><td>start</td><td>end</td><td>strand</td><td>MM</td><td>target_seq</td><td>PAM</td><td>alignment</td><td>distance</td><td>position</td><td>gene name</td><td>gene id</td></tr>
</table>

<table>
<tr><td>GU305914.1</td><td></td><td>205590</td><td>205612</td><td>+</td><td>0</td><td>CCATCGCGGAGTTGAAACAC</td><td>CGG</td><td>||||||||[||||||||||||]PAM</td><td>NA</td><td>Intergenic</td><td>NA</td><td>NA</td></tr>
<tr><td>T15656</td><td>TCTGCCCCTACATCTGTGTGCGG</td><td></td><td>1000</td><td></td><td></td><td></td><td></td><td></td><td></td><td></td><td></td><td></td></tr>
<tr><td>Oligo fwd</td><td>CACCTCTGCCCCTACATCTGTGTG</td><td></td><td></td><td></td><td></td><td></td><td></td><td></td><td></td><td></td><td></td><td></td></tr>
<tr><td>Oligo rev</td><td>AAACCACACAGATGTAGGGGCAGA</td><td></td><td></td><td></td><td></td><td></td><td></td><td></td><td></td><td></td><td></td><td></td></tr>
<tr><td>Chromosome</td><td>start</td><td></td><td>end</td><td>strand</td><td>MM</td><td>target_seq</td><td>PAM</td><td>alignment</td><td>distance</td><td>position</td><td>gene name</td><td>gene id</td></tr>
<tr><td>GU305914.1</td><td></td><td>205622</td><td>205644</td><td>+</td><td>0</td><td>TCTGCCCCTACATCTGTGTG</td><td>CGG</td><td>||||||||[||||||||||||]PAM</td><td>NA</td><td>Intergenic</td><td>NA</td><td>NA</td></tr>
<tr><td>T15657</td><td>GCAGTTCATCAAAGAAAACACGG</td><td></td><td>1000</td><td></td><td></td><td></td><td></td><td></td><td></td><td></td><td></td><td></td></tr>
<tr><td>Oligo fwd</td><td>CACCGCAGTTCATCAAAGAAAACA</td><td></td><td></td><td></td><td></td><td></td><td></td><td></td><td></td><td></td><td></td><td></td></tr>
<tr><td>Oligo rev</td><td>AAACTGTTTTCTTTGATGAACTGC</td><td></td><td></td><td></td><td></td><td></td><td></td><td></td><td></td><td></td><td></td><td></td></tr>
<tr><td>Chromosome</td><td>start</td><td></td><td>end</td><td>strand</td><td>MM</td><td>target_seq</td><td>PAM</td><td>alignment</td><td>distance</td><td>position</td><td>gene name</td><td>gene id</td></tr>
<tr><td>GU305914.1</td><td></td><td>205688</td><td>205710</td><td>+</td><td>0</td><td>GCAGTTCATCAAAGAAAACA</td><td>CGG</td><td>||||||||[||||||||||||]PAM</td><td>NA</td><td>Intergenic</td><td>NA</td><td>NA</td></tr>
<tr><td>T15658</td><td>CAGTTCATCAAAGAAAACACGGG</td><td></td><td>1000</td><td></td><td></td><td></td><td></td><td></td><td></td><td></td><td></td><td></td></tr>
<tr><td>Oligo fwd</td><td>CACCCAGTTCATCAAAGAAAACAC</td><td></td><td></td><td></td><td></td><td></td><td></td><td></td><td></td><td></td><td></td><td></td></tr>
<tr><td>Oligo rev</td><td>AAACGTGTTTTCTTTGATGAACTG</td><td></td><td></td><td></td><td></td><td></td><td></td><td></td><td></td><td></td><td></td><td></td></tr>
<tr><td>Chromosome</td><td>start</td><td></td><td>end</td><td>strand</td><td>MM</td><td>target_seq</td><td>PAM</td><td>alignment</td><td>distance</td><td>position</td><td>gene name</td><td>gene id</td></tr>
<tr><td>GU305914.1</td><td></td><td>205689</td><td>205711</td><td>+</td><td>0</td><td>CAGTTCATCAAAGAAAACAC</td><td>GGG</td><td>||||||||[||||||||||||]PAM</td><td>NA</td><td>Intergenic</td><td>NA</td><td>NA</td></tr>
<tr><td>T15659</td><td>AGTTCATCAAAGAAAACACGGGG</td><td></td><td>1000</td><td></td><td></td><td></td><td></td><td></td><td></td><td></td><td></td><td></td></tr>
<tr><td>Oligo fwd</td><td>CACCAGTTCATCAAAGAAAACACG</td><td></td><td></td><td></td><td></td><td></td><td></td><td></td><td></td><td></td><td></td><td></td></tr>
<tr><td>Oligo rev</td><td>AAACCGTGTTTTCTTTGATGAACT</td><td></td><td></td><td></td><td></td><td></td><td></td><td></td><td></td><td></td><td></td><td></td></tr>
<tr><td>Chromosome</td><td>start</td><td></td><td>end</td><td>strand</td><td>MM</td><td>target_seq</td><td>PAM</td><td>alignment</td><td>distance</td><td>position</td><td>gene name</td><td>gene id</td></tr>
<tr><td>GU305914.1</td><td></td><td>205690</td><td>205712</td><td>+</td><td>0</td><td>AGTTCATCAAAGAAAACACG</td><td>GGG</td><td>||||||||[||||||||||||]PAM</td><td>NA</td><td>Intergenic</td><td>NA</td><td>NA</td></tr>
<tr><td>T15660</td><td>GGGGTTTATTTTTATTGCATTGG</td><td></td><td>1000</td><td></td><td></td><td></td><td></td><td></td><td></td><td></td><td></td><td></td></tr>
<tr><td>Oligo fwd</td><td>CACCGGGGTTTATTTTTATTGCAT</td><td></td><td></td><td></td><td></td><td></td><td></td><td></td><td></td><td></td><td></td><td></td></tr>
<tr><td>Oligo rev</td><td>AAACATGCAATAAAAATAAACCCC</td><td></td><td></td><td></td><td></td><td></td><td></td><td></td><td></td><td></td><td></td><td></td></tr>
<tr><td>Chromosome</td><td>start</td><td></td><td>end</td><td>strand</td><td>MM</td><td>target_seq</td><td>PAM</td><td>alignment</td><td>distance</td><td>position</td><td>gene name</td><td>gene id</td></tr>
<tr><td>GU305914.1</td><td></td><td>205709</td><td>205731</td><td>+</td><td>0</td><td>GGGGTTTATTTTTATTGCAT</td><td>TGG</td><td>||||||||[||||||||||||]PAM</td><td>NA</td><td>Intergenic</td><td>NA</td><td>NA</td></tr>
<tr><td>T15661</td><td>GACTCTTACACGCACACGCAAGG</td><td></td><td>1000</td><td></td><td></td><td></td><td></td><td></td><td></td><td></td><td></td><td></td></tr>
<tr><td>Oligo fwd</td><td>CACCGACTCTTACACGCACACGCA</td><td></td><td></td><td></td><td></td><td></td><td></td><td></td><td></td><td></td><td></td><td></td></tr>
<tr><td>Oligo rev</td><td>AAACTGCGTGTGCGTGTAAGAGTC</td><td></td><td></td><td></td><td></td><td></td><td></td><td></td><td></td><td></td><td></td><td></td></tr>
<tr><td>Chromosome</td><td>start</td><td></td><td>end</td><td>strand</td><td>MM</td><td>target_seq</td><td>PAM</td><td>alignment</td><td>distance</td><td>position</td><td>gene name</td><td>gene id</td></tr>
<tr><td>GU305914.1</td><td></td><td>205738</td><td>205760</td><td>+</td><td>0</td><td>GACTCTTACACGCACACGCA</td><td>AGG</td><td>||||||||[||||||||||||]PAM</td><td>NA</td><td>Intergenic</td><td>NA</td><td>NA</td></tr>
<tr><td>T15662</td><td>TACACGCACACGCAAGGCATTGG</td><td></td><td>1000</td><td></td><td></td><td></td><td></td><td></td><td></td><td></td><td></td><td></td></tr>
<tr><td>Oligo fwd</td><td>CACCTACACGCACACGCAAGGCAT</td><td></td><td></td><td></td><td></td><td></td><td></td><td></td><td></td><td></td><td></td><td></td></tr>
<tr><td>Oligo rev</td><td>AAACATGCCTTGCGTGTGCGTGTA</td><td></td><td></td><td></td><td></td><td></td><td></td><td></td><td></td><td></td><td></td><td></td></tr>
<tr><td>Chromosome</td><td>start</td><td></td><td>end</td><td>strand</td><td>MM</td><td>target_seq</td><td>PAM</td><td>alignment</td><td>distance</td><td>position</td><td>gene name</td><td>gene id</td></tr>
<tr><td>GU305914.1</td><td></td><td>205744</td><td>205766</td><td>+</td><td>0</td><td>TACACGCACACGCAAGGCAT</td><td>TGG</td><td>||||||||[||||||||||||]PAM</td><td>NA</td><td>Intergenic</td><td>NA</td><td>NA</td></tr>
<tr><td>T15663</td><td>ATGCCCACGATGACCGCCCTCGG</td><td></td><td>1000</td><td></td><td></td><td></td><td></td><td></td><td></td><td></td><td></td><td></td></tr>
<tr><td>Oligo fwd</td><td>CACCATGCCCACGATGACCGCCCT</td><td></td><td></td><td></td><td></td><td></td><td></td><td></td><td></td><td></td><td></td><td></td></tr>
<tr><td>Oligo rev</td><td>AAACAGGGCGGTCATCGTGGGCAT</td><td></td><td></td><td></td><td></td><td></td><td></td><td></td><td></td><td></td><td></td><td></td></tr>
<tr><td>Chromosome</td><td>start</td><td></td><td>end</td><td>strand</td><td>MM</td><td>target_seq</td><td>PAM</td><td>alignment</td><td>distance</td><td>position</td><td>gene name</td><td>gene id</td></tr>
<tr><td>GU305914.1</td><td></td><td>205828</td><td>205850</td><td>+</td><td>0</td><td>ATGCCCACGATGACCGCCCT</td><td>CGG</td><td>||||||||[||||||||||||]PAM</td><td>NA</td><td>Intergenic</td><td>NA</td><td>NA</td></tr>
<tr><td>T15664</td><td>ATGACCGCCCTCGGATCGAAAGG</td><td></td><td>1000</td><td></td><td></td><td></td><td></td><td></td><td></td><td></td><td></td><td></td></tr>
<tr><td>Oligo fwd</td><td>CACCATGACCGCCCTCGGATCGAA</td><td></td><td></td><td></td><td></td><td></td><td></td><td></td><td></td><td></td><td></td><td></td></tr>
<tr><td>Oligo rev</td><td>AAACTTCGATCCGAGGGCGGTCAT</td><td></td><td></td><td></td><td></td><td></td><td></td><td></td><td></td><td></td><td></td><td></td></tr>
<tr><td>Chromosome</td><td>start</td><td></td><td>end</td><td>strand</td><td>MM</td><td>target_seq</td><td>PAM</td><td>alignment</td><td>distance</td><td>position</td><td>gene name</td><td>gene id</td></tr>
<tr><td>GU305914.1</td><td></td><td>205837</td><td>205859</td><td>+</td><td>0</td><td>ATGACCGCCCTCGGATCGAA</td><td>AGG</td><td>||||||||[||||||||||||]PAM</td><td>NA</td><td>Intergenic</td><td>NA</td><td>NA</td></tr>
<tr><td>T15665</td><td>GTTTCGATCTTCTCGATCTGTGG</td><td></td><td>1000</td><td></td><td></td><td></td><td></td><td></td><td></td><td></td><td></td><td></td></tr>
<tr><td>Oligo fwd</td><td>CACCGTTTCGATCTTCTCGATCTG</td><td></td><td></td><td></td><td></td><td></td><td></td><td></td><td></td><td></td><td></td><td></td></tr>
<tr><td>Oligo rev</td><td>AAACCAGATCGAGAAGATCGAAAC</td><td></td><td></td><td></td><td></td><td></td><td></td><td></td><td></td><td></td><td></td><td></td></tr>
<tr><td>Chromosome</td><td>start</td><td></td><td>end</td><td>strand</td><td>MM</td><td>target_seq</td><td>PAM</td><td>alignment</td><td>distance</td><td>position</td><td>gene name</td><td>gene id</td></tr>
<tr><td>GU305914.1</td><td></td><td>205859</td><td>205881</td><td>+</td><td>0</td><td>GTTTCGATCTTCTCGATCTG</td><td>TGG</td><td>||||||||[||||||||||||]PAM</td><td>NA</td><td>Intergenic</td><td>NA</td><td>NA</td></tr>
<tr><td>T15666</td><td>TCTGTGGTAATACGACCGCGTGG</td><td></td><td>1000</td><td></td><td></td><td></td><td></td><td></td><td></td><td></td><td></td><td></td></tr>
<tr><td>Oligo fwd</td><td>CACCTCTGTGGTAATACGACCGCG</td><td></td><td></td><td></td><td></td><td></td><td></td><td></td><td></td><td></td><td></td><td></td></tr>
<tr><td>Oligo rev</td><td>AAACCGCGGTCGTATTACCACAGA</td><td></td><td></td><td></td><td></td><td></td><td></td><td></td><td></td><td></td><td></td><td></td></tr>
<tr><td>Chromosome</td><td>start</td><td></td><td>end</td><td>strand</td><td>MM</td><td>target_seq</td><td>PAM</td><td>alignment</td><td>distance</td><td>position</td><td>gene name</td><td>gene id</td></tr>
<tr><td>GU305914.1</td><td></td><td>205875</td><td>205897</td><td>+</td><td>0</td><td>TCTGTGGTAATACGACCGCG</td><td>TGG</td><td>||||||||[||||||||||||]PAM</td><td>NA</td><td>Intergenic</td><td>NA</td><td>NA</td></tr>
<tr><td>T15667</td><td>TGTGACACGCGCAGCAATGTTGG</td><td></td><td>1000</td><td></td><td></td><td></td><td></td><td></td><td></td><td></td><td></td><td></td></tr>
<tr><td>Oligo fwd</td><td>CACCTGTGACACGCGCAGCAATGT</td><td></td><td></td><td></td><td></td><td></td><td></td><td></td><td></td><td></td><td></td><td></td></tr>
<tr><td>Oligo rev</td><td>AAACACATTGCTGCGCGTGTCACA</td><td></td><td></td><td></td><td></td><td></td><td></td><td></td><td></td><td></td><td></td><td></td></tr>
<tr><td>Chromosome</td><td>start</td><td></td><td>end</td><td>strand</td><td>MM</td><td>target_seq</td><td>PAM</td><td>alignment</td><td>distance</td><td>position</td><td>gene name</td><td>gene id</td></tr>
<tr><td>GU305914.1</td><td></td><td>205903</td><td>205925</td><td>+</td><td>0</td><td>TGTGACACGCGCAGCAATGT</td><td>TGG</td><td>||||||||[||||||||||||]PAM</td><td>NA</td><td>Intergenic</td><td>NA</td><td>NA</td></tr>
<tr><td>T15668</td><td>AACATCACAACTTAGCCATACGG</td><td></td><td>1000</td><td></td><td></td><td></td><td></td><td></td><td></td><td></td><td></td><td></td></tr>
<tr><td>Oligo fwd</td><td>CACCAACATCACAACTTAGCCATA</td><td></td><td></td><td></td><td></td><td></td><td></td><td></td><td></td><td></td><td></td><td></td></tr>
<tr><td>Oligo rev</td><td>AAACTATGGCTAAGTTGTGATGTT</td><td></td><td></td><td></td><td></td><td></td><td></td><td></td><td></td><td></td><td></td><td></td></tr>
<tr><td>Chromosome</td><td>start</td><td></td><td>end</td><td>strand</td><td>MM</td><td>target_seq</td><td>PAM</td><td>alignment</td><td>distance</td><td>position</td><td>gene name</td><td>gene id</td></tr>
<tr><td>GU305914.1</td><td></td><td>206016</td><td>206038</td><td>+</td><td>0</td><td>AACATCACAACTTAGCCATA</td><td>CGG</td><td>||||||||[||||||||||||]PAM</td><td>NA</td><td>Intergenic</td><td>NA</td><td>NA</td></tr>
<tr><td>T15669</td><td>ACATCACAACTTAGCCATACGGG</td><td></td><td>1000</td><td></td><td></td><td></td><td></td><td></td><td></td><td></td><td></td><td></td></tr>
<tr><td>Oligo fwd</td><td>CACCACATCACAACTTAGCCATAC</td><td></td><td></td><td></td><td></td><td></td><td></td><td></td><td></td><td></td><td></td><td></td></tr>
<tr><td>Oligo rev</td><td>AAACGTATGGCTAAGTTGTGATGT</td><td></td><td></td><td></td><td></td><td></td><td></td><td></td><td></td><td></td><td></td><td></td></tr>
<tr><td>Chromosome</td><td>start</td><td></td><td>end</td><td>strand</td><td>MM</td><td>target_seq</td><td>PAM</td><td>alignment</td><td>distance</td><td>position</td><td>gene name</td><td>gene id</td></tr>
<tr><td>GU305914.1</td><td></td><td>206017</td><td>206039</td><td>+</td><td>0</td><td>ACATCACAACTTAGCCATAC</td><td>GGG</td><td>||||||||[||||||||||||]PAM</td><td>NA</td><td>Intergenic</td><td>NA</td><td>NA</td></tr>
<tr><td>T15670</td><td>ACGGGATTATGACCAAACTTAGG</td><td></td><td>1000</td><td></td><td></td><td></td><td></td><td></td><td></td><td></td><td></td><td></td></tr>
<tr><td>Oligo fwd</td><td>CACCACGGGATTATGACCAAACTT</td><td></td><td></td><td></td><td></td><td></td><td></td><td></td><td></td><td></td><td></td><td></td></tr>
<tr><td>Oligo rev</td><td>AAACAAGTTTGGTCATAATCCCGT</td><td></td><td></td><td></td><td></td><td></td><td></td><td></td><td></td><td></td><td></td><td></td></tr>
</table>

Chromosome start end strand MM target_seq PAM alignment distance position gene name gene id
GU305914.1 206035 206057 + 0 ACGGGATTATGACCAAACTT AGG ||||||||[||||||||||||]PAM NA Intergenic NA NA

T15671 CGATTGTACAACGTCCTCCCCGG 1000
Oligo fwd CACCCGATTGTACAACGTCCTCCC
Oligo rev AAACGGGAGGACGTTGTACAATCG
Chromosome start end strand MM target_seq PAM alignment distance position gene name gene id
GU305914.1 206100 206122 + 0 CGATTGTACAACGTCCTCCC CGG ||||||||[||||||||||||]PAM NA Intergenic NA NA

T15672 ACGTCCTCCCCGGATGATGTCGG 1000
Oligo fwd CACCACGTCCTCCCCGGATGATGT
Oligo rev AAACACATCATCCGGGGAGGACGT
Chromosome start end strand MM target_seq PAM alignment distance position gene name gene id
GU305914.1 206110 206132 + 0 ACGTCCTCCCCGGATGATGT CGG ||||||||[||||||||||||]PAM NA Intergenic NA NA

T15673 AAACATACATTTAAAGTTAAAGG 1000
Oligo fwd CACCAAACATACATTTAAAGTTAA
Oligo rev AAACTTAACTTTAAATGTATGTTT
Chromosome start end strand MM target_seq PAM alignment distance position gene name gene id
GU305914.1 206179 206201 + 0 AAACATACATTTAAAGTTAA AGG ||||||||[||||||||||||]PAM NA Intergenic NA NA

T15674 CACAATTATTGAGAATGTTTCGG 1000
Oligo fwd CACCCACAATTATTGAGAATGTTT
Oligo rev AAACAAACATTCTCAATAATTGTG
Chromosome start end strand MM target_seq PAM alignment distance position gene name gene id
GU305914.1 206285 206307 + 0 CACAATTATTGAGAATGTTT CGG ||||||||[||||||||||||]PAM NA Intergenic NA NA

T15675 TTCGGTGTATGCTCGATAGCCGG 1000
Oligo fwd CACCTTCGGTGTATGCTCGATAGC
Oligo rev AAACGCTATCGAGCATACACCGAA
Chromosome start end strand MM target_seq PAM alignment distance position gene name gene id
GU305914.1 206303 206325 + 0 TTCGGTGTATGCTCGATAGC CGG ||||||||[||||||||||||]PAM NA Intergenic NA NA

T15676 GCTGTAAACTGTAGAATAATCGG 1000
Oligo fwd CACCGCTGTAAACTGTAGAATAAT
Oligo rev AAACATTATTCTACAGTTTACAGC
Chromosome start end strand MM target_seq PAM alignment distance position gene name gene id
GU305914.1 206325 206347 + 0 GCTGTAAACTGTAGAATAAT CGG ||||||||[||||||||||||]PAM NA Intergenic NA NA

T15677 CTCTGTCCGTCTCCGTTTTCAGG 1000
Oligo fwd CACCCTCTGTCCGTCTCCGTTTTC
Oligo rev AAACGAAAACGGAGACGGACAGAG
Chromosome start end strand MM target_seq PAM alignment distance position gene name gene id
GU305914.1 206392 206414 + 0 CTCTGTCCGTCTCCGTTTTC AGG ||||||||[||||||||||||]PAM NA Intergenic NA NA

T15678 TCTGTCCGTCTCCGTTTTCAGGG 1000
Oligo fwd CACCTCTGTCCGTCTCCGTTTTCA
Oligo rev AAACTGAAAACGGAGACGGACAGA
Chromosome start end strand MM target_seq PAM alignment distance position gene name gene id
GU305914.1 206393 206415 + 0 TCTGTCCGTCTCCGTTTTCA GGG ||||||||[||||||||||||]PAM NA Intergenic NA NA

T15679 GAGTCGACCATCTGTAACGCCGG 1000
Oligo fwd CACCGAGTCGACCATCTGTAACGC
Oligo rev AAACGCGTTACAGATGGTCGACTC
Chromosome start end strand MM target_seq PAM alignment distance position gene name gene id
GU305914.1 206439 206461 + 0 GAGTCGACCATCTGTAACGC CGG ||||||||[||||||||||||]PAM NA Intergenic NA NA

T15680 GATACGCACTCGAGCGTATCAGG 1000
Oligo fwd CACCGATACGCACTCGAGCGTATC
Oligo rev AAACGATACGCTCGAGTGCGTATC
Chromosome start end strand MM target_seq PAM alignment distance position gene name gene id
GU305914.1 206509 206531 + 0 GATACGCACTCGAGCGTATC AGG ||||||||[||||||||||||]PAM NA Intergenic NA NA

T15681 TGAGATCTTCCCTATGTAAACGG 1000
Oligo fwd CACCTGAGATCTTCCCTATGTAAA
Oligo rev AAACTTTACATAGGGAAGATCTCA
Chromosome start end strand MM target_seq PAM alignment distance position gene name gene id
GU305914.1 206597 206619 + 0 TGAGATCTTCCCTATGTAAA CGG ||||||||[||||||||||||]PAM NA Intergenic NA NA

T15682 TCCCTATGTAAACGGAGAACCGG 1000
Oligo fwd CACCTCCCTATGTAAACGGAGAAC
Oligo rev AAACGTTCTCCGTTTACATAGGGA
Chromosome start end strand MM target_seq PAM alignment distance position gene name gene id
GU305914.1 206605 206627 + 0 TCCCTATGTAAACGGAGAAC CGG ||||||||[||||||||||||]PAM NA Intergenic NA NA

T15683 CGCGAGTTCAGAGACGCTATAGG 1000
Oligo fwd CACCCGCGAGTTCAGAGACGCTAT
Oligo rev AAACATAGCGTCTCTGAACTCGCG
Chromosome start end strand MM target_seq PAM alignment distance position gene name gene id
GU305914.1 206686 206708 + 0 CGCGAGTTCAGAGACGCTAT AGG ||||||||[||||||||||||]PAM NA Intergenic NA NA

T15684 GACGCTATAGGAAAAGAATAAGG 1000
Oligo fwd CACCGACGCTATAGGAAAAGAATA
Oligo rev AAACTATTCTTTTCCTATAGCGTC
Chromosome start end strand MM target_seq PAM alignment distance position gene name gene id
GU305914.1 206698 206720 + 0 GACGCTATAGGAAAAGAATA AGG ||||||||[||||||||||||]PAM NA Intergenic NA NA

T15685 AGAATAAGGAAAAGAAATATCGG 1000
Oligo fwd CACCAGAATAAGGAAAAGAAATAT

<table>
<tr><td>Oligo rev</td><td>AAACATATTTCTTTTCCTTATTCT</td><td></td><td></td><td></td><td></td><td></td><td></td><td></td><td></td><td></td><td></td></tr>
<tr><td>Chromosome</td><td>start</td><td>end</td><td>strand</td><td>MM</td><td>target_seq</td><td>PAM</td><td>alignment</td><td>distance</td><td>position</td><td>gene name</td><td>gene id</td></tr>
<tr><td>GU305914.1</td><td>206712</td><td>206734</td><td>+</td><td>0</td><td>AGAATAAGGAAAAGAAATAT</td><td>CGG</td><td>||||||||[||||||||||||]PAM</td><td>NA</td><td>Intergenic</td><td>NA</td><td>NA</td></tr>
<tr><td>T15686</td><td>GTACGACGAAAGAGACATCCTGG</td><td>1000</td><td></td><td></td><td></td><td></td><td></td><td></td><td></td><td></td><td></td></tr>
<tr><td>Oligo fwd</td><td>CACCGTACGACGAAAGAGACATCC</td><td></td><td></td><td></td><td></td><td></td><td></td><td></td><td></td><td></td><td></td></tr>
<tr><td>Oligo rev</td><td>AAACGGATGTCTCTTTCGTCGTAC</td><td></td><td></td><td></td><td></td><td></td><td></td><td></td><td></td><td></td><td></td></tr>
<tr><td>Chromosome</td><td>start</td><td>end</td><td>strand</td><td>MM</td><td>target_seq</td><td>PAM</td><td>alignment</td><td>distance</td><td>position</td><td>gene name</td><td>gene id</td></tr>
<tr><td>GU305914.1</td><td>206771</td><td>206793</td><td>+</td><td>0</td><td>GTACGACGAAAGAGACATCC</td><td>TGG</td><td>||||||||[||||||||||||]PAM</td><td>NA</td><td>Intergenic</td><td>NA</td><td>NA</td></tr>
<tr><td>T15687</td><td>TACGACGAAAGAGACATCCTGGG</td><td>1000</td><td></td><td></td><td></td><td></td><td></td><td></td><td></td><td></td><td></td></tr>
<tr><td>Oligo fwd</td><td>CACCTACGACGAAAGAGACATCCT</td><td></td><td></td><td></td><td></td><td></td><td></td><td></td><td></td><td></td><td></td></tr>
<tr><td>Oligo rev</td><td>AAACAGGATGTCTCTTTCGTCGTA</td><td></td><td></td><td></td><td></td><td></td><td></td><td></td><td></td><td></td><td></td></tr>
<tr><td>Chromosome</td><td>start</td><td>end</td><td>strand</td><td>MM</td><td>target_seq</td><td>PAM</td><td>alignment</td><td>distance</td><td>position</td><td>gene name</td><td>gene id</td></tr>
<tr><td>GU305914.1</td><td>206772</td><td>206794</td><td>+</td><td>0</td><td>TACGACGAAAGAGACATCCT</td><td>GGG</td><td>||||||||[||||||||||||]PAM</td><td>NA</td><td>Intergenic</td><td>NA</td><td>NA</td></tr>
<tr><td>T15688</td><td>AAGCGAGACCACGCACCAGACGG</td><td>1000</td><td></td><td></td><td></td><td></td><td></td><td></td><td></td><td></td><td></td></tr>
<tr><td>Oligo fwd</td><td>CACCAAGCGAGACCACGCACCAGA</td><td></td><td></td><td></td><td></td><td></td><td></td><td></td><td></td><td></td><td></td></tr>
<tr><td>Oligo rev</td><td>AAACTCTGGTGCGTGGTCTCGCTT</td><td></td><td></td><td></td><td></td><td></td><td></td><td></td><td></td><td></td><td></td></tr>
<tr><td>Chromosome</td><td>start</td><td>end</td><td>strand</td><td>MM</td><td>target_seq</td><td>PAM</td><td>alignment</td><td>distance</td><td>position</td><td>gene name</td><td>gene id</td></tr>
<tr><td>GU305914.1</td><td>206817</td><td>206839</td><td>+</td><td>0</td><td>AAGCGAGACCACGCACCAGA</td><td>CGG</td><td>||||||||[||||||||||||]PAM</td><td>NA</td><td>Intergenic</td><td>NA</td><td>NA</td></tr>
<tr><td>T15689</td><td>CGCACCAGACGGCGATAGCGCGG</td><td>1000</td><td></td><td></td><td></td><td></td><td></td><td></td><td></td><td></td><td></td></tr>
<tr><td>Oligo fwd</td><td>CACCCGCACCAGACGGCGATAGCG</td><td></td><td></td><td></td><td></td><td></td><td></td><td></td><td></td><td></td><td></td></tr>
<tr><td>Oligo rev</td><td>AAACCGCTATCGCCGTCTGGTGCG</td><td></td><td></td><td></td><td></td><td></td><td></td><td></td><td></td><td></td><td></td></tr>
<tr><td>Chromosome</td><td>start</td><td>end</td><td>strand</td><td>MM</td><td>target_seq</td><td>PAM</td><td>alignment</td><td>distance</td><td>position</td><td>gene name</td><td>gene id</td></tr>
<tr><td>GU305914.1</td><td>206828</td><td>206850</td><td>+</td><td>0</td><td>CGCACCAGACGGCGATAGCG</td><td>CGG</td><td>||||||||[||||||||||||]PAM</td><td>NA</td><td>Intergenic</td><td>NA</td><td>NA</td></tr>
<tr><td>T15690</td><td>GCGATAGCGCGGACGCGAATCGG</td><td>1000</td><td></td><td></td><td></td><td></td><td></td><td></td><td></td><td></td><td></td></tr>
<tr><td>Oligo fwd</td><td>CACCGCGATAGCGCGGACGCGAAT</td><td></td><td></td><td></td><td></td><td></td><td></td><td></td><td></td><td></td><td></td></tr>
<tr><td>Oligo rev</td><td>AAACATTCGCGTCCGCGCTATCGC</td><td></td><td></td><td></td><td></td><td></td><td></td><td></td><td></td><td></td><td></td></tr>
<tr><td>Chromosome</td><td>start</td><td>end</td><td>strand</td><td>MM</td><td>target_seq</td><td>PAM</td><td>alignment</td><td>distance</td><td>position</td><td>gene name</td><td>gene id</td></tr>
<tr><td>GU305914.1</td><td>206839</td><td>206861</td><td>+</td><td>0</td><td>GCGATAGCGCGGACGCGAAT</td><td>CGG</td><td>||||||||[||||||||||||]PAM</td><td>NA</td><td>Intergenic</td><td>NA</td><td>NA</td></tr>
<tr><td>T15691</td><td>GCGCGGACGCGAATCGGACTCGG</td><td>1000</td><td></td><td></td><td></td><td></td><td></td><td></td><td></td><td></td><td></td></tr>
<tr><td>Oligo fwd</td><td>CACCGCGCGGACGCGAATCGGACT</td><td></td><td></td><td></td><td></td><td></td><td></td><td></td><td></td><td></td><td></td></tr>
<tr><td>Oligo rev</td><td>AAACAGTCCGATTCGCGTCCGCGC</td><td></td><td></td><td></td><td></td><td></td><td></td><td></td><td></td><td></td><td></td></tr>
<tr><td>Chromosome</td><td>start</td><td>end</td><td>strand</td><td>MM</td><td>target_seq</td><td>PAM</td><td>alignment</td><td>distance</td><td>position</td><td>gene name</td><td>gene id</td></tr>
<tr><td>GU305914.1</td><td>206845</td><td>206867</td><td>+</td><td>0</td><td>GCGCGGACGCGAATCGGACT</td><td>CGG</td><td>||||||||[||||||||||||]PAM</td><td>NA</td><td>Intergenic</td><td>NA</td><td>NA</td></tr>
<tr><td>T15692</td><td>GTCGCGAGCCGCGCGCGATACGG</td><td>1000</td><td></td><td></td><td></td><td></td><td></td><td></td><td></td><td></td><td></td></tr>
<tr><td>Oligo fwd</td><td>CACCGTCGCGAGCCGCGCGCGATA</td><td></td><td></td><td></td><td></td><td></td><td></td><td></td><td></td><td></td><td></td></tr>
<tr><td>Oligo rev</td><td>AAACTATCGCGCGCGGCTCGCGAC</td><td></td><td></td><td></td><td></td><td></td><td></td><td></td><td></td><td></td><td></td></tr>
<tr><td>Chromosome</td><td>start</td><td>end</td><td>strand</td><td>MM</td><td>target_seq</td><td>PAM</td><td>alignment</td><td>distance</td><td>position</td><td>gene name</td><td>gene id</td></tr>
<tr><td>GU305914.1</td><td>206878</td><td>206900</td><td>+</td><td>0</td><td>GTCGCGAGCCGCGCGCGATA</td><td>CGG</td><td>||||||||[||||||||||||]PAM</td><td>NA</td><td>Intergenic</td><td>NA</td><td>NA</td></tr>
<tr><td>T15693</td><td>ATCACACAAGTTTATTGAGATGG</td><td>1000</td><td></td><td></td><td></td><td></td><td></td><td></td><td></td><td></td><td></td></tr>
<tr><td>Oligo fwd</td><td>CACCATCACACAAGTTTATTGAGA</td><td></td><td></td><td></td><td></td><td></td><td></td><td></td><td></td><td></td><td></td></tr>
<tr><td>Oligo rev</td><td>AAACTCTCAATAAACTTGTGTGAT</td><td></td><td></td><td></td><td></td><td></td><td></td><td></td><td></td><td></td><td></td></tr>
<tr><td>Chromosome</td><td>start</td><td>end</td><td>strand</td><td>MM</td><td>target_seq</td><td>PAM</td><td>alignment</td><td>distance</td><td>position</td><td>gene name</td><td>gene id</td></tr>
<tr><td>GU305914.1</td><td>206919</td><td>206941</td><td>+</td><td>0</td><td>ATCACACAAGTTTATTGAGA</td><td>TGG</td><td>||||||||[||||||||||||]PAM</td><td>NA</td><td>Intergenic</td><td>NA</td><td>NA</td></tr>
<tr><td>T15694</td><td>TCACACAAGTTTATTGAGATGGG</td><td>1000</td><td></td><td></td><td></td><td></td><td></td><td></td><td></td><td></td><td></td></tr>
<tr><td>Oligo fwd</td><td>CACCTCACACAAGTTTATTGAGAT</td><td></td><td></td><td></td><td></td><td></td><td></td><td></td><td></td><td></td><td></td></tr>
<tr><td>Oligo rev</td><td>AAACATCTCAATAAACTTGTGTGA</td><td></td><td></td><td></td><td></td><td></td><td></td><td></td><td></td><td></td><td></td></tr>
<tr><td>Chromosome</td><td>start</td><td>end</td><td>strand</td><td>MM</td><td>target_seq</td><td>PAM</td><td>alignment</td><td>distance</td><td>position</td><td>gene name</td><td>gene id</td></tr>
<tr><td>GU305914.1</td><td>206920</td><td>206942</td><td>+</td><td>0</td><td>TCACACAAGTTTATTGAGAT</td><td>GGG</td><td>||||||||[||||||||||||]PAM</td><td>NA</td><td>Intergenic</td><td>NA</td><td>NA</td></tr>
<tr><td>T15695</td><td>CAAGTTTATTGAGATGGGACCGG</td><td>1000</td><td></td><td></td><td></td><td></td><td></td><td></td><td></td><td></td><td></td></tr>
<tr><td>Oligo fwd</td><td>CACCCAAGTTTATTGAGATGGGAC</td><td></td><td></td><td></td><td></td><td></td><td></td><td></td><td></td><td></td><td></td></tr>
<tr><td>Oligo rev</td><td>AAACGTCCCATCTCAATAAACTTG</td><td></td><td></td><td></td><td></td><td></td><td></td><td></td><td></td><td></td><td></td></tr>
<tr><td>Chromosome</td><td>start</td><td>end</td><td>strand</td><td>MM</td><td>target_seq</td><td>PAM</td><td>alignment</td><td>distance</td><td>position</td><td>gene name</td><td>gene id</td></tr>
<tr><td>GU305914.1</td><td>206925</td><td>206947</td><td>+</td><td>0</td><td>CAAGTTTATTGAGATGGGAC</td><td>CGG</td><td>||||||||[||||||||||||]PAM</td><td>NA</td><td>Intergenic</td><td>NA</td><td>NA</td></tr>
<tr><td>T15696</td><td>TTATTGAGATGGGACCGGAAAGG</td><td>1000</td><td></td><td></td><td></td><td></td><td></td><td></td><td></td><td></td><td></td></tr>
<tr><td>Oligo fwd</td><td>CACCTTATTGAGATGGGACCGGAA</td><td></td><td></td><td></td><td></td><td></td><td></td><td></td><td></td><td></td><td></td></tr>
<tr><td>Oligo rev</td><td>AAACTTCCGGTCCCATCTCAATAA</td><td></td><td></td><td></td><td></td><td></td><td></td><td></td><td></td><td></td><td></td></tr>
<tr><td>Chromosome</td><td>start</td><td>end</td><td>strand</td><td>MM</td><td>target_seq</td><td>PAM</td><td>alignment</td><td>distance</td><td>position</td><td>gene name</td><td>gene id</td></tr>
<tr><td>GU305914.1</td><td>206930</td><td>206952</td><td>+</td><td>0</td><td>TTATTGAGATGGGACCGGAA</td><td>AGG</td><td>||||||||[||||||||||||]PAM</td><td>NA</td><td>Intergenic</td><td>NA</td><td>NA</td></tr>
<tr><td>T15697</td><td>TGAATTTTTTTATTCATCGTCGG</td><td>1000</td><td></td><td></td><td></td><td></td><td></td><td></td><td></td><td></td><td></td></tr>
<tr><td>Oligo fwd</td><td>CACCTGAATTTTTTTATTCATCGT</td><td></td><td></td><td></td><td></td><td></td><td></td><td></td><td></td><td></td><td></td></tr>
<tr><td>Oligo rev</td><td>AAACACGATGAATAAAAAAATTCA</td><td></td><td></td><td></td><td></td><td></td><td></td><td></td><td></td><td></td><td></td></tr>
<tr><td>Chromosome</td><td>start</td><td>end</td><td>strand</td><td>MM</td><td>target_seq</td><td>PAM</td><td>alignment</td><td>distance</td><td>position</td><td>gene name</td><td>gene id</td></tr>
<tr><td>GU305914.1</td><td>206954</td><td>206976</td><td>+</td><td>0</td><td>TGAATTTTTTTATTCATCGT</td><td>CGG</td><td>||||||||[||||||||||||]PAM</td><td>NA</td><td>Intergenic</td><td>NA</td><td>NA</td></tr>
<tr><td>T15698</td><td>GAATTTTTTTATTCATCGTCGGG</td><td>1000</td><td></td><td></td><td></td><td></td><td></td><td></td><td></td><td></td><td></td></tr>
<tr><td>Oligo fwd</td><td>CACCGAATTTTTTTATTCATCGTC</td><td></td><td></td><td></td><td></td><td></td><td></td><td></td><td></td><td></td><td></td></tr>
<tr><td>Oligo rev</td><td>AAACGACGATGAATAAAAAAATTC</td><td></td><td></td><td></td><td></td><td></td><td></td><td></td><td></td><td></td><td></td></tr>
<tr><td>Chromosome</td><td>start</td><td>end</td><td>strand</td><td>MM</td><td>target_seq</td><td>PAM</td><td>alignment</td><td>distance</td><td>position</td><td>gene name</td><td>gene id</td></tr>
<tr><td>GU305914.1</td><td>206955</td><td>206977</td><td>+</td><td>0</td><td>GAATTTTTTTATTCATCGTC</td><td>GGG</td><td>||||||||[||||||||||||]PAM</td><td>NA</td><td>Intergenic</td><td>NA</td><td>NA</td></tr>
<tr><td>GU305914.1</td><td>214493</td><td>214515</td><td>+</td><td>4</td><td>TAATTGTTTTATTAATCGAC</td><td>GGG</td><td>-||||-||[|||||-||||-|]PAM</td><td>NA</td><td>Intergenic</td><td>NA</td><td>NA</td></tr>
<tr><td>T15699</td><td>TTATTCATCGTCGGGACCGCAGG</td><td>1000</td><td></td><td></td><td></td><td></td><td></td><td></td><td></td><td></td><td></td></tr>
<tr><td>Oligo fwd</td><td>CACCTTATTCATCGTCGGGACCGC</td><td></td><td></td><td></td><td></td><td></td><td></td><td></td><td></td><td></td><td></td></tr>
<tr><td>Oligo rev</td><td>AAACGCGGTCCCGACGATGAATAA</td><td></td><td></td><td></td><td></td><td></td><td></td><td></td><td></td><td></td><td></td></tr>
<tr><td>Chromosome</td><td>start</td><td>end</td><td>strand</td><td>MM</td><td>target_seq</td><td>PAM</td><td>alignment</td><td>distance</td><td>position</td><td>gene name</td><td>gene id</td></tr>
<tr><td>GU305914.1</td><td>206963</td><td>206985</td><td>+</td><td>0</td><td>TTATTCATCGTCGGGACCGC</td><td>AGG</td><td>||||||||[||||||||||||]PAM</td><td>NA</td><td>Intergenic</td><td>NA</td><td>NA</td></tr>
</table>

<table>
<tr><td>T15700</td><td>TATTCATCGTCGGGACCGCAGGG</td><td></td><td>1000</td><td></td><td></td><td></td><td></td><td></td><td></td><td></td><td></td><td></td></tr>
<tr><td>Oligo fwd</td><td>CACCTATTCATCGTCGGGACCGCA</td><td></td><td></td><td></td><td></td><td></td><td></td><td></td><td></td><td></td><td></td><td></td></tr>
<tr><td>Oligo rev</td><td>AAACTGCGGTCCCGACGATGAATA</td><td></td><td></td><td></td><td></td><td></td><td></td><td></td><td></td><td></td><td></td><td></td></tr>
<tr><td>Chromosome</td><td>start</td><td></td><td>end</td><td>strand</td><td>MM</td><td>target_seq</td><td>PAM</td><td>alignment</td><td>distance</td><td>position</td><td>gene name</td><td>gene id</td></tr>
<tr><td>GU305914.1</td><td></td><td>206964</td><td>206986</td><td>+</td><td>0</td><td>TATTCATCGTCGGGACCGCA</td><td>GGG</td><td>||||||||[||||||||||||]PAM</td><td>NA</td><td>Intergenic</td><td>NA</td><td>NA</td></tr>
<tr><td>T15701</td><td>CATATCTGTTAGCTATCCCACGG</td><td></td><td>1000</td><td></td><td></td><td></td><td></td><td></td><td></td><td></td><td></td><td></td></tr>
<tr><td>Oligo fwd</td><td>CACCCATATCTGTTAGCTATCCCA</td><td></td><td></td><td></td><td></td><td></td><td></td><td></td><td></td><td></td><td></td><td></td></tr>
<tr><td>Oligo rev</td><td>AAACTGGGATAGCTAACAGATATG</td><td></td><td></td><td></td><td></td><td></td><td></td><td></td><td></td><td></td><td></td><td></td></tr>
<tr><td>Chromosome</td><td>start</td><td></td><td>end</td><td>strand</td><td>MM</td><td>target_seq</td><td>PAM</td><td>alignment</td><td>distance</td><td>position</td><td>gene name</td><td>gene id</td></tr>
<tr><td>GU305914.1</td><td></td><td>206995</td><td>207017</td><td>+</td><td>0</td><td>CATATCTGTTAGCTATCCCA</td><td>CGG</td><td>||||||||[||||||||||||]PAM</td><td>NA</td><td>Intergenic</td><td>NA</td><td>NA</td></tr>
<tr><td>T15702</td><td>ATCTGTTAGCTATCCCACGGAGG</td><td></td><td>1000</td><td></td><td></td><td></td><td></td><td></td><td></td><td></td><td></td><td></td></tr>
<tr><td>Oligo fwd</td><td>CACCATCTGTTAGCTATCCCACGG</td><td></td><td></td><td></td><td></td><td></td><td></td><td></td><td></td><td></td><td></td><td></td></tr>
<tr><td>Oligo rev</td><td>AAACCCGTGGGATAGCTAACAGAT</td><td></td><td></td><td></td><td></td><td></td><td></td><td></td><td></td><td></td><td></td><td></td></tr>
<tr><td>Chromosome</td><td>start</td><td></td><td>end</td><td>strand</td><td>MM</td><td>target_seq</td><td>PAM</td><td>alignment</td><td>distance</td><td>position</td><td>gene name</td><td>gene id</td></tr>
<tr><td>GU305914.1</td><td></td><td>206998</td><td>207020</td><td>+</td><td>0</td><td>ATCTGTTAGCTATCCCACGG</td><td>AGG</td><td>||||||||[||||||||||||]PAM</td><td>NA</td><td>Intergenic</td><td>NA</td><td>NA</td></tr>
<tr><td>T15703</td><td>TCTGTTAGCTATCCCACGGAGGG</td><td></td><td>1000</td><td></td><td></td><td></td><td></td><td></td><td></td><td></td><td></td><td></td></tr>
<tr><td>Oligo fwd</td><td>CACCTCTGTTAGCTATCCCACGGA</td><td></td><td></td><td></td><td></td><td></td><td></td><td></td><td></td><td></td><td></td><td></td></tr>
<tr><td>Oligo rev</td><td>AAACTCCGTGGGATAGCTAACAGA</td><td></td><td></td><td></td><td></td><td></td><td></td><td></td><td></td><td></td><td></td><td></td></tr>
<tr><td>Chromosome</td><td>start</td><td></td><td>end</td><td>strand</td><td>MM</td><td>target_seq</td><td>PAM</td><td>alignment</td><td>distance</td><td>position</td><td>gene name</td><td>gene id</td></tr>
<tr><td>GU305914.1</td><td></td><td>206999</td><td>207021</td><td>+</td><td>0</td><td>TCTGTTAGCTATCCCACGGA</td><td>GGG</td><td>||||||||[||||||||||||]PAM</td><td>NA</td><td>Intergenic</td><td>NA</td><td>NA</td></tr>
<tr><td>T15704</td><td>ACGGAGGGAATATGCATGCACGG</td><td></td><td>1000</td><td></td><td></td><td></td><td></td><td></td><td></td><td></td><td></td><td></td></tr>
<tr><td>Oligo fwd</td><td>CACCACGGAGGGAATATGCATGCA</td><td></td><td></td><td></td><td></td><td></td><td></td><td></td><td></td><td></td><td></td><td></td></tr>
<tr><td>Oligo rev</td><td>AAACTGCATGCATATTCCCTCCGT</td><td></td><td></td><td></td><td></td><td></td><td></td><td></td><td></td><td></td><td></td><td></td></tr>
<tr><td>Chromosome</td><td>start</td><td></td><td>end</td><td>strand</td><td>MM</td><td>target_seq</td><td>PAM</td><td>alignment</td><td>distance</td><td>position</td><td>gene name</td><td>gene id</td></tr>
<tr><td>GU305914.1</td><td></td><td>207014</td><td>207036</td><td>+</td><td>0</td><td>ACGGAGGGAATATGCATGCA</td><td>CGG</td><td>||||||||[||||||||||||]PAM</td><td>NA</td><td>Intergenic</td><td>NA</td><td>NA</td></tr>
<tr><td>T15705</td><td>GGTCAACACGCCGATAACGAAGG</td><td></td><td>1000</td><td></td><td></td><td></td><td></td><td></td><td></td><td></td><td></td><td></td></tr>
<tr><td>Oligo fwd</td><td>CACCGGTCAACACGCCGATAACGA</td><td></td><td></td><td></td><td></td><td></td><td></td><td></td><td></td><td></td><td></td><td></td></tr>
<tr><td>Oligo rev</td><td>AAACTCGTTATCGGCGTGTTGACC</td><td></td><td></td><td></td><td></td><td></td><td></td><td></td><td></td><td></td><td></td><td></td></tr>
<tr><td>Chromosome</td><td>start</td><td></td><td>end</td><td>strand</td><td>MM</td><td>target_seq</td><td>PAM</td><td>alignment</td><td>distance</td><td>position</td><td>gene name</td><td>gene id</td></tr>
<tr><td>GU305914.1</td><td></td><td>207035</td><td>207057</td><td>+</td><td>0</td><td>GGTCAACACGCCGATAACGA</td><td>AGG</td><td>||||||||[||||||||||||]PAM</td><td>NA</td><td>Intergenic</td><td>NA</td><td>NA</td></tr>
<tr><td>T15706</td><td>CACGCCGATAACGAAGGCCGCGG</td><td></td><td>1000</td><td></td><td></td><td></td><td></td><td></td><td></td><td></td><td></td><td></td></tr>
<tr><td>Oligo fwd</td><td>CACCCACGCCGATAACGAAGGCCG</td><td></td><td></td><td></td><td></td><td></td><td></td><td></td><td></td><td></td><td></td><td></td></tr>
<tr><td>Oligo rev</td><td>AAACCGGCCTTCGTTATCGGCGTG</td><td></td><td></td><td></td><td></td><td></td><td></td><td></td><td></td><td></td><td></td><td></td></tr>
<tr><td>Chromosome</td><td>start</td><td></td><td>end</td><td>strand</td><td>MM</td><td>target_seq</td><td>PAM</td><td>alignment</td><td>distance</td><td>position</td><td>gene name</td><td>gene id</td></tr>
<tr><td>GU305914.1</td><td></td><td>207041</td><td>207063</td><td>+</td><td>0</td><td>CACGCCGATAACGAAGGCCG</td><td>CGG</td><td>||||||||[||||||||||||]PAM</td><td>NA</td><td>Intergenic</td><td>NA</td><td>NA</td></tr>
<tr><td>T15707</td><td>GATAACGAAGGCCGCGGCTACGG</td><td></td><td>1000</td><td></td><td></td><td></td><td></td><td></td><td></td><td></td><td></td><td></td></tr>
<tr><td>Oligo fwd</td><td>CACCGATAACGAAGGCCGCGGCTA</td><td></td><td></td><td></td><td></td><td></td><td></td><td></td><td></td><td></td><td></td><td></td></tr>
<tr><td>Oligo rev</td><td>AAACTAGCCGCGGCCTTCGTTATC</td><td></td><td></td><td></td><td></td><td></td><td></td><td></td><td></td><td></td><td></td><td></td></tr>
<tr><td>Chromosome</td><td>start</td><td></td><td>end</td><td>strand</td><td>MM</td><td>target_seq</td><td>PAM</td><td>alignment</td><td>distance</td><td>position</td><td>gene name</td><td>gene id</td></tr>
<tr><td>GU305914.1</td><td></td><td>207047</td><td>207069</td><td>+</td><td>0</td><td>GATAACGAAGGCCGCGGCTA</td><td>CGG</td><td>||||||||[||||||||||||]PAM</td><td>NA</td><td>Intergenic</td><td>NA</td><td>NA</td></tr>
<tr><td>T15708</td><td>ATAACGAAGGCCGCGGCTACGGG</td><td></td><td>1000</td><td></td><td></td><td></td><td></td><td></td><td></td><td></td><td></td><td></td></tr>
<tr><td>Oligo fwd</td><td>CACCATAACGAAGGCCGCGGCTAC</td><td></td><td></td><td></td><td></td><td></td><td></td><td></td><td></td><td></td><td></td><td></td></tr>
<tr><td>Oligo rev</td><td>AAACGTAGCCGCGGCCTTCGTTAT</td><td></td><td></td><td></td><td></td><td></td><td></td><td></td><td></td><td></td><td></td><td></td></tr>
<tr><td>Chromosome</td><td>start</td><td></td><td>end</td><td>strand</td><td>MM</td><td>target_seq</td><td>PAM</td><td>alignment</td><td>distance</td><td>position</td><td>gene name</td><td>gene id</td></tr>
<tr><td>GU305914.1</td><td></td><td>207048</td><td>207070</td><td>+</td><td>0</td><td>ATAACGAAGGCCGCGGCTAC</td><td>GGG</td><td>||||||||[||||||||||||]PAM</td><td>NA</td><td>Intergenic</td><td>NA</td><td>NA</td></tr>
<tr><td>T15709</td><td>GGGCGCCAGTCGTACCGCGTCGG</td><td></td><td>1000</td><td></td><td></td><td></td><td></td><td></td><td></td><td></td><td></td><td></td></tr>
<tr><td>Oligo fwd</td><td>CACCGGGCGCCAGTCGTACCGCGT</td><td></td><td></td><td></td><td></td><td></td><td></td><td></td><td></td><td></td><td></td><td></td></tr>
<tr><td>Oligo rev</td><td>AAACACGCGGTACGACTGGCGCCC</td><td></td><td></td><td></td><td></td><td></td><td></td><td></td><td></td><td></td><td></td><td></td></tr>
<tr><td>Chromosome</td><td>start</td><td></td><td>end</td><td>strand</td><td>MM</td><td>target_seq</td><td>PAM</td><td>alignment</td><td>distance</td><td>position</td><td>gene name</td><td>gene id</td></tr>
<tr><td>GU305914.1</td><td></td><td>207068</td><td>207090</td><td>+</td><td>0</td><td>GGGCGCCAGTCGTACCGCGT</td><td>CGG</td><td>||||||||[||||||||||||]PAM</td><td>NA</td><td>Intergenic</td><td>NA</td><td>NA</td></tr>
<tr><td>T15710</td><td>CCAGAGTCCGTGCGCCTGCACGG</td><td></td><td>1000</td><td></td><td></td><td></td><td></td><td></td><td></td><td></td><td></td><td></td></tr>
<tr><td>Oligo fwd</td><td>CACCCCAGAGTCCGTGCGCCTGCA</td><td></td><td></td><td></td><td></td><td></td><td></td><td></td><td></td><td></td><td></td><td></td></tr>
<tr><td>Oligo rev</td><td>AAACTGCAGGCGCACGGACTCTGG</td><td></td><td></td><td></td><td></td><td></td><td></td><td></td><td></td><td></td><td></td><td></td></tr>
<tr><td>Chromosome</td><td>start</td><td></td><td>end</td><td>strand</td><td>MM</td><td>target_seq</td><td>PAM</td><td>alignment</td><td>distance</td><td>position</td><td>gene name</td><td>gene id</td></tr>
<tr><td>GU305914.1</td><td></td><td>207119</td><td>207141</td><td>+</td><td>0</td><td>CCAGAGTCCGTGCGCCTGCA</td><td>CGG</td><td>||||||||[||||||||||||]PAM</td><td>NA</td><td>Intergenic</td><td>NA</td><td>NA</td></tr>
<tr><td>T15711</td><td>GGTCAGATACACCAGCCGAAAGG</td><td></td><td>1000</td><td></td><td></td><td></td><td></td><td></td><td></td><td></td><td></td><td></td></tr>
<tr><td>Oligo fwd</td><td>CACCGGTCAGATACACCAGCCGAA</td><td></td><td></td><td></td><td></td><td></td><td></td><td></td><td></td><td></td><td></td><td></td></tr>
<tr><td>Oligo rev</td><td>AAACTTCGGCTGGTGTATCTGACC</td><td></td><td></td><td></td><td></td><td></td><td></td><td></td><td></td><td></td><td></td><td></td></tr>
<tr><td>Chromosome</td><td>start</td><td></td><td>end</td><td>strand</td><td>MM</td><td>target_seq</td><td>PAM</td><td>alignment</td><td>distance</td><td>position</td><td>gene name</td><td>gene id</td></tr>
<tr><td>GU305914.1</td><td></td><td>207140</td><td>207162</td><td>+</td><td>0</td><td>GGTCAGATACACCAGCCGAA</td><td>AGG</td><td>||||||||[||||||||||||]PAM</td><td>NA</td><td>Intergenic</td><td>NA</td><td>NA</td></tr>
<tr><td>T15712</td><td>AGGCGATCACAAAGTAGATGAGG</td><td></td><td>1000</td><td></td><td></td><td></td><td></td><td></td><td></td><td></td><td></td><td></td></tr>
<tr><td>Oligo fwd</td><td>CACCAGGCGATCACAAAGTAGATG</td><td></td><td></td><td></td><td></td><td></td><td></td><td></td><td></td><td></td><td></td><td></td></tr>
<tr><td>Oligo rev</td><td>AAACCATCTACTTTGTGATCGCCT</td><td></td><td></td><td></td><td></td><td></td><td></td><td></td><td></td><td></td><td></td><td></td></tr>
<tr><td>Chromosome</td><td>start</td><td></td><td>end</td><td>strand</td><td>MM</td><td>target_seq</td><td>PAM</td><td>alignment</td><td>distance</td><td>position</td><td>gene name</td><td>gene id</td></tr>
<tr><td>GU305914.1</td><td></td><td>207160</td><td>207182</td><td>+</td><td>0</td><td>AGGCGATCACAAAGTAGATG</td><td>AGG</td><td>||||||||[||||||||||||]PAM</td><td>NA</td><td>Intergenic</td><td>NA</td><td>NA</td></tr>
<tr><td>T15713</td><td>AAAGTAGATGAGGATGCAGATGG</td><td></td><td>1000</td><td></td><td></td><td></td><td></td><td></td><td></td><td></td><td></td><td></td></tr>
<tr><td>Oligo fwd</td><td>CACCAAAGTAGATGAGGATGCAGA</td><td></td><td></td><td></td><td></td><td></td><td></td><td></td><td></td><td></td><td></td><td></td></tr>
<tr><td>Oligo rev</td><td>AAACTCTGCATCCTCATCTACTTT</td><td></td><td></td><td></td><td></td><td></td><td></td><td></td><td></td><td></td><td></td><td></td></tr>
<tr><td>Chromosome</td><td>start</td><td></td><td>end</td><td>strand</td><td>MM</td><td>target_seq</td><td>PAM</td><td>alignment</td><td>distance</td><td>position</td><td>gene name</td><td>gene id</td></tr>
<tr><td>GU305914.1</td><td></td><td>207170</td><td>207192</td><td>+</td><td>0</td><td>AAAGTAGATGAGGATGCAGA</td><td>TGG</td><td>||||||||[||||||||||||]PAM</td><td>NA</td><td>Intergenic</td><td>NA</td><td>NA</td></tr>
<tr><td>T15714</td><td>AAGTAGATGAGGATGCAGATGGG</td><td></td><td>1000</td><td></td><td></td><td></td><td></td><td></td><td></td><td></td><td></td><td></td></tr>
<tr><td>Oligo fwd</td><td>CACCAAGTAGATGAGGATGCAGAT</td><td></td><td></td><td></td><td></td><td></td><td></td><td></td><td></td><td></td><td></td><td></td></tr>
<tr><td>Oligo rev</td><td>AAACATCTGCATCCTCATCTACTT</td><td></td><td></td><td></td><td></td><td></td><td></td><td></td><td></td><td></td><td></td><td></td></tr>
<tr><td>Chromosome</td><td>start</td><td></td><td>end</td><td>strand</td><td>MM</td><td>target_seq</td><td>PAM</td><td>alignment</td><td>distance</td><td>position</td><td>gene name</td><td>gene id</td></tr>
<tr><td>GU305914.1</td><td></td><td>207171</td><td>207193</td><td>+</td><td>0</td><td>AAGTAGATGAGGATGCAGAT</td><td>GGG</td><td>||||||||[||||||||||||]PAM</td><td>NA</td><td>Intergenic</td><td>NA</td><td>NA</td></tr>
</table>

<table>
<tr><td>T15715</td><td>TAGATGAGGATGCAGATGGGCGG</td><td>1000</td><td></td><td></td><td></td><td></td><td></td><td></td><td></td><td></td><td></td></tr>
<tr><td>Oligo fwd</td><td>CACCTAGATGAGGATGCAGATGGG</td><td></td><td></td><td></td><td></td><td></td><td></td><td></td><td></td><td></td><td></td></tr>
<tr><td>Oligo rev</td><td>AAACCCCATCTGCATCCTCATCTA</td><td></td><td></td><td></td><td></td><td></td><td></td><td></td><td></td><td></td><td></td></tr>
<tr><td>Chromosome</td><td>start</td><td>end</td><td>strand</td><td>MM</td><td>target_seq</td><td>PAM</td><td>alignment</td><td>distance</td><td>position</td><td>gene name</td><td>gene id</td></tr>
<tr><td>GU305914.1</td><td>207174</td><td>207196</td><td>+</td><td>0</td><td>TAGATGAGGATGCAGATGGG</td><td>CGG</td><td>||||||||[||||||||||||]PAM</td><td>NA</td><td>Intergenic</td><td>NA</td><td>NA</td></tr>
<tr><td>T15716</td><td>AATCCATTTCAAGACTATCGCGG</td><td>1000</td><td></td><td></td><td></td><td></td><td></td><td></td><td></td><td></td><td></td></tr>
<tr><td>Oligo fwd</td><td>CACCAATCCATTTCAAGACTATCG</td><td></td><td></td><td></td><td></td><td></td><td></td><td></td><td></td><td></td><td></td></tr>
<tr><td>Oligo rev</td><td>AAACCGATAGTCTTGAAATGGATT</td><td></td><td></td><td></td><td></td><td></td><td></td><td></td><td></td><td></td><td></td></tr>
<tr><td>Chromosome</td><td>start</td><td>end</td><td>strand</td><td>MM</td><td>target_seq</td><td>PAM</td><td>alignment</td><td>distance</td><td>position</td><td>gene name</td><td>gene id</td></tr>
<tr><td>GU305914.1</td><td>207197</td><td>207219</td><td>+</td><td>0</td><td>AATCCATTTCAAGACTATCG</td><td>CGG</td><td>||||||||[||||||||||||]PAM</td><td>NA</td><td>Intergenic</td><td>NA</td><td>NA</td></tr>
<tr><td>T15717</td><td>CCATTTCAAGACTATCGCGGTGG</td><td>1000</td><td></td><td></td><td></td><td></td><td></td><td></td><td></td><td></td><td></td></tr>
<tr><td>Oligo fwd</td><td>CACCCCATTTCAAGACTATCGCGG</td><td></td><td></td><td></td><td></td><td></td><td></td><td></td><td></td><td></td><td></td></tr>
<tr><td>Oligo rev</td><td>AAACCCGCGATAGTCTTGAAATGG</td><td></td><td></td><td></td><td></td><td></td><td></td><td></td><td></td><td></td><td></td></tr>
<tr><td>Chromosome</td><td>start</td><td>end</td><td>strand</td><td>MM</td><td>target_seq</td><td>PAM</td><td>alignment</td><td>distance</td><td>position</td><td>gene name</td><td>gene id</td></tr>
<tr><td>GU305914.1</td><td>207200</td><td>207222</td><td>+</td><td>0</td><td>CCATTTCAAGACTATCGCGG</td><td>TGG</td><td>||||||||[||||||||||||]PAM</td><td>NA</td><td>Intergenic</td><td>NA</td><td>NA</td></tr>
<tr><td>T15718</td><td>ACTATCGCGGTGGACCAGTCCGG</td><td>1000</td><td></td><td></td><td></td><td></td><td></td><td></td><td></td><td></td><td></td></tr>
<tr><td>Oligo fwd</td><td>CACCACTATCGCGGTGGACCAGTC</td><td></td><td></td><td></td><td></td><td></td><td></td><td></td><td></td><td></td><td></td></tr>
<tr><td>Oligo rev</td><td>AAACGACTGGTCCACCGCGATAGT</td><td></td><td></td><td></td><td></td><td></td><td></td><td></td><td></td><td></td><td></td></tr>
<tr><td>Chromosome</td><td>start</td><td>end</td><td>strand</td><td>MM</td><td>target_seq</td><td>PAM</td><td>alignment</td><td>distance</td><td>position</td><td>gene name</td><td>gene id</td></tr>
<tr><td>GU305914.1</td><td>207210</td><td>207232</td><td>+</td><td>0</td><td>ACTATCGCGGTGGACCAGTC</td><td>CGG</td><td>||||||||[||||||||||||]PAM</td><td>NA</td><td>Intergenic</td><td>NA</td><td>NA</td></tr>
<tr><td>T15719</td><td>GTGGACCAGTCCGGCCTTTTCGG</td><td>1000</td><td></td><td></td><td></td><td></td><td></td><td></td><td></td><td></td><td></td></tr>
<tr><td>Oligo fwd</td><td>CACCGTGGACCAGTCCGGCCTTTT</td><td></td><td></td><td></td><td></td><td></td><td></td><td></td><td></td><td></td><td></td></tr>
<tr><td>Oligo rev</td><td>AAACAAAAGGCCGGACTGGTCCAC</td><td></td><td></td><td></td><td></td><td></td><td></td><td></td><td></td><td></td><td></td></tr>
<tr><td>Chromosome</td><td>start</td><td>end</td><td>strand</td><td>MM</td><td>target_seq</td><td>PAM</td><td>alignment</td><td>distance</td><td>position</td><td>gene name</td><td>gene id</td></tr>
<tr><td>GU305914.1</td><td>207219</td><td>207241</td><td>+</td><td>0</td><td>GTGGACCAGTCCGGCCTTTT</td><td>CGG</td><td>||||||||[||||||||||||]PAM</td><td>NA</td><td>Intergenic</td><td>NA</td><td>NA</td></tr>
<tr><td>T15720</td><td>TCGGTACCTCGTCCTCCTCTCGG</td><td>1000</td><td></td><td></td><td></td><td></td><td></td><td></td><td></td><td></td><td></td></tr>
<tr><td>Oligo fwd</td><td>CACCTCGGTACCTCGTCCTCCTCT</td><td></td><td></td><td></td><td></td><td></td><td></td><td></td><td></td><td></td><td></td></tr>
<tr><td>Oligo rev</td><td>AAACAGAGGAGGACGAGGTACCGA</td><td></td><td></td><td></td><td></td><td></td><td></td><td></td><td></td><td></td><td></td></tr>
<tr><td>Chromosome</td><td>start</td><td>end</td><td>strand</td><td>MM</td><td>target_seq</td><td>PAM</td><td>alignment</td><td>distance</td><td>position</td><td>gene name</td><td>gene id</td></tr>
<tr><td>GU305914.1</td><td>207238</td><td>207260</td><td>+</td><td>0</td><td>TCGGTACCTCGTCCTCCTCT</td><td>CGG</td><td>||||||||[||||||||||||]PAM</td><td>NA</td><td>Intergenic</td><td>NA</td><td>NA</td></tr>
<tr><td>T15721</td><td>GATCCTTCACTCTGAACGTCAGG</td><td>1000</td><td></td><td></td><td></td><td></td><td></td><td></td><td></td><td></td><td></td></tr>
<tr><td>Oligo fwd</td><td>CACCGATCCTTCACTCTGAACGTC</td><td></td><td></td><td></td><td></td><td></td><td></td><td></td><td></td><td></td><td></td></tr>
<tr><td>Oligo rev</td><td>AAACGACGTTCAGAGTGAAGGATC</td><td></td><td></td><td></td><td></td><td></td><td></td><td></td><td></td><td></td><td></td></tr>
<tr><td>Chromosome</td><td>start</td><td>end</td><td>strand</td><td>MM</td><td>target_seq</td><td>PAM</td><td>alignment</td><td>distance</td><td>position</td><td>gene name</td><td>gene id</td></tr>
<tr><td>GU305914.1</td><td>207274</td><td>207296</td><td>+</td><td>0</td><td>GATCCTTCACTCTGAACGTC</td><td>AGG</td><td>||||||||[||||||||||||]PAM</td><td>NA</td><td>Intergenic</td><td>NA</td><td>NA</td></tr>
<tr><td>T15722</td><td>TAACTTTTTGTTTCCGATCCCGG</td><td>1000</td><td></td><td></td><td></td><td></td><td></td><td></td><td></td><td></td><td></td></tr>
<tr><td>Oligo fwd</td><td>CACCTAACTTTTTGTTTCCGATCC</td><td></td><td></td><td></td><td></td><td></td><td></td><td></td><td></td><td></td><td></td></tr>
<tr><td>Oligo rev</td><td>AAACGGATCGGAAACAAAAAGTTA</td><td></td><td></td><td></td><td></td><td></td><td></td><td></td><td></td><td></td><td></td></tr>
<tr><td>Chromosome</td><td>start</td><td>end</td><td>strand</td><td>MM</td><td>target_seq</td><td>PAM</td><td>alignment</td><td>distance</td><td>position</td><td>gene name</td><td>gene id</td></tr>
<tr><td>GU305914.1</td><td>207312</td><td>207334</td><td>+</td><td>0</td><td>TAACTTTTTGTTTCCGATCC</td><td>CGG</td><td>||||||||[||||||||||||]PAM</td><td>NA</td><td>Intergenic</td><td>NA</td><td>NA</td></tr>
<tr><td>T15723</td><td>GTTCTAGATATCTACCCAGCAGG</td><td>1000</td><td></td><td></td><td></td><td></td><td></td><td></td><td></td><td></td><td></td></tr>
<tr><td>Oligo fwd</td><td>CACCGTTCTAGATATCTACCCAGC</td><td></td><td></td><td></td><td></td><td></td><td></td><td></td><td></td><td></td><td></td></tr>
<tr><td>Oligo rev</td><td>AAACGCTGGGTAGATATCTAGAAC</td><td></td><td></td><td></td><td></td><td></td><td></td><td></td><td></td><td></td><td></td></tr>
<tr><td>Chromosome</td><td>start</td><td>end</td><td>strand</td><td>MM</td><td>target_seq</td><td>PAM</td><td>alignment</td><td>distance</td><td>position</td><td>gene name</td><td>gene id</td></tr>
<tr><td>GU305914.1</td><td>207404</td><td>207426</td><td>+</td><td>0</td><td>GTTCTAGATATCTACCCAGC</td><td>AGG</td><td>||||||||[||||||||||||]PAM</td><td>NA</td><td>Intergenic</td><td>NA</td><td>NA</td></tr>
<tr><td>T15724</td><td>ACCGTCCCAGACTCCGTCGTCGG</td><td>1000</td><td></td><td></td><td></td><td></td><td></td><td></td><td></td><td></td><td></td></tr>
<tr><td>Oligo fwd</td><td>CACCACCGTCCCAGACTCCGTCGT</td><td></td><td></td><td></td><td></td><td></td><td></td><td></td><td></td><td></td><td></td></tr>
<tr><td>Oligo rev</td><td>AAACACGACGGAGTCTGGGACGGT</td><td></td><td></td><td></td><td></td><td></td><td></td><td></td><td></td><td></td><td></td></tr>
<tr><td>Chromosome</td><td>start</td><td>end</td><td>strand</td><td>MM</td><td>target_seq</td><td>PAM</td><td>alignment</td><td>distance</td><td>position</td><td>gene name</td><td>gene id</td></tr>
<tr><td>GU305914.1</td><td>207490</td><td>207512</td><td>+</td><td>0</td><td>ACCGTCCCAGACTCCGTCGT</td><td>CGG</td><td>||||||||[||||||||||||]PAM</td><td>NA</td><td>Intergenic</td><td>NA</td><td>NA</td></tr>
<tr><td>T15725</td><td>TCGTCGGTGACGTACACGAGCGG</td><td>1000</td><td></td><td></td><td></td><td></td><td></td><td></td><td></td><td></td><td></td></tr>
<tr><td>Oligo fwd</td><td>CACCTCGTCGGTGACGTACACGAG</td><td></td><td></td><td></td><td></td><td></td><td></td><td></td><td></td><td></td><td></td></tr>
<tr><td>Oligo rev</td><td>AAACCTCGTGTACGTCACCGACGA</td><td></td><td></td><td></td><td></td><td></td><td></td><td></td><td></td><td></td><td></td></tr>
<tr><td>Chromosome</td><td>start</td><td>end</td><td>strand</td><td>MM</td><td>target_seq</td><td>PAM</td><td>alignment</td><td>distance</td><td>position</td><td>gene name</td><td>gene id</td></tr>
<tr><td>GU305914.1</td><td>207506</td><td>207528</td><td>+</td><td>0</td><td>TCGTCGGTGACGTACACGAG</td><td>CGG</td><td>||||||||[||||||||||||]PAM</td><td>NA</td><td>Intergenic</td><td>NA</td><td>NA</td></tr>
<tr><td>T15726</td><td>ACCACCCCCACAGAGCCGCACGG</td><td>1000</td><td></td><td></td><td></td><td></td><td></td><td></td><td></td><td></td><td></td></tr>
<tr><td>Oligo fwd</td><td>CACCACCACCCCCACAGAGCCGCA</td><td></td><td></td><td></td><td></td><td></td><td></td><td></td><td></td><td></td><td></td></tr>
<tr><td>Oligo rev</td><td>AAACTGCGGCTCTGTGGGGGTGGT</td><td></td><td></td><td></td><td></td><td></td><td></td><td></td><td></td><td></td><td></td></tr>
<tr><td>Chromosome</td><td>start</td><td>end</td><td>strand</td><td>MM</td><td>target_seq</td><td>PAM</td><td>alignment</td><td>distance</td><td>position</td><td>gene name</td><td>gene id</td></tr>
<tr><td>GU305914.1</td><td>207538</td><td>207560</td><td>+</td><td>0</td><td>ACCACCCCCACAGAGCCGCA</td><td>CGG</td><td>||||||||[||||||||||||]PAM</td><td>NA</td><td>Intergenic</td><td>NA</td><td>NA</td></tr>
<tr><td>T15727</td><td>CATCAAGAAGACGCATCCTCCGG</td><td>1000</td><td></td><td></td><td></td><td></td><td></td><td></td><td></td><td></td><td></td></tr>
<tr><td>Oligo fwd</td><td>CACCCATCAAGAAGACGCATCCTC</td><td></td><td></td><td></td><td></td><td></td><td></td><td></td><td></td><td></td><td></td></tr>
<tr><td>Oligo rev</td><td>AAACGAGGATGCGTCTTCTTGATG</td><td></td><td></td><td></td><td></td><td></td><td></td><td></td><td></td><td></td><td></td></tr>
<tr><td>Chromosome</td><td>start</td><td>end</td><td>strand</td><td>MM</td><td>target_seq</td><td>PAM</td><td>alignment</td><td>distance</td><td>position</td><td>gene name</td><td>gene id</td></tr>
<tr><td>GU305914.1</td><td>207568</td><td>207590</td><td>+</td><td>0</td><td>CATCAAGAAGACGCATCCTC</td><td>CGG</td><td>||||||||[||||||||||||]PAM</td><td>NA</td><td>Intergenic</td><td>NA</td><td>NA</td></tr>
<tr><td>T15728</td><td>ATCAAGAAGACGCATCCTCCGGG</td><td>1000</td><td></td><td></td><td></td><td></td><td></td><td></td><td></td><td></td><td></td></tr>
<tr><td>Oligo fwd</td><td>CACCATCAAGAAGACGCATCCTCC</td><td></td><td></td><td></td><td></td><td></td><td></td><td></td><td></td><td></td><td></td></tr>
<tr><td>Oligo rev</td><td>AAACGGAGGATGCGTCTTCTTGAT</td><td></td><td></td><td></td><td></td><td></td><td></td><td></td><td></td><td></td><td></td></tr>
<tr><td>Chromosome</td><td>start</td><td>end</td><td>strand</td><td>MM</td><td>target_seq</td><td>PAM</td><td>alignment</td><td>distance</td><td>position</td><td>gene name</td><td>gene id</td></tr>
<tr><td>GU305914.1</td><td>207569</td><td>207591</td><td>+</td><td>0</td><td>ATCAAGAAGACGCATCCTCC</td><td>GGG</td><td>||||||||[||||||||||||]PAM</td><td>NA</td><td>Intergenic</td><td>NA</td><td>NA</td></tr>
<tr><td>T15729</td><td>TCAAGAAGACGCATCCTCCGGGG</td><td>1000</td><td></td><td></td><td></td><td></td><td></td><td></td><td></td><td></td><td></td></tr>
<tr><td>Oligo fwd</td><td>CACCTCAAGAAGACGCATCCTCCG</td><td></td><td></td><td></td><td></td><td></td><td></td><td></td><td></td><td></td><td></td></tr>
<tr><td>Oligo rev</td><td>AAACCGGAGGATGCGTCTTCTTGA</td><td></td><td></td><td></td><td></td><td></td><td></td><td></td><td></td><td></td><td></td></tr>
<tr><td>Chromosome</td><td>start</td><td>end</td><td>strand</td><td>MM</td><td>target_seq</td><td>PAM</td><td>alignment</td><td>distance</td><td>position</td><td>gene name</td><td>gene id</td></tr>
</table>

| | | | | | | | | | | | |
|---|---|---|---|---|---|---|---|---|---|---|---|
| GU305914.1 | 207570 | 207592 | + | 0 | TCAAGAAGACGCATCCTCCG | GGG | \|\|\|\|\|\|\|\|[\|\|\|\|\|\|\|\|\|\|\|\|]PAM | NA | Intergenic | NA | NA |
| T15730 | TCACTCATGAGCCATCTGAAAGG | 1000 | | | | | | | | | |
| Oligo fwd | CACCTCACTCATGAGCCATCTGAA | | | | | | | | | | |
| Oligo rev | AAACTTCAGATGGCTCATGAGTGA | | | | | | | | | | |
| Chromosome | start | end | strand | MM | target_seq | PAM | alignment | distance | position | gene name | gene id |
| GU305914.1 | 207596 | 207618 | + | 0 | TCACTCATGAGCCATCTGAA | AGG | \|\|\|\|\|\|\|\|[\|\|\|\|\|\|\|\|\|\|\|\|]PAM | NA | Intergenic | NA | NA |
| T15731 | TGAGCCATCTGAAAGGTCTGAGG | 1000 | | | | | | | | | |
| Oligo fwd | CACCTGAGCCATCTGAAAGGTCTG | | | | | | | | | | |
| Oligo rev | AAACCAGACCTTTCAGATGGCTCA | | | | | | | | | | |
| Chromosome | start | end | strand | MM | target_seq | PAM | alignment | distance | position | gene name | gene id |
| GU305914.1 | 207603 | 207625 | + | 0 | TGAGCCATCTGAAAGGTCTG | AGG | \|\|\|\|\|\|\|\|[\|\|\|\|\|\|\|\|\|\|\|\|]PAM | NA | Intergenic | NA | NA |
| T15732 | GCCATCTGAAAGGTCTGAGGTGG | 1000 | | | | | | | | | |
| Oligo fwd | CACCGCCATCTGAAAGGTCTGAGG | | | | | | | | | | |
| Oligo rev | AAACCCTCAGACCTTTCAGATGGC | | | | | | | | | | |
| Chromosome | start | end | strand | MM | target_seq | PAM | alignment | distance | position | gene name | gene id |
| GU305914.1 | 207606 | 207628 | + | 0 | GCCATCTGAAAGGTCTGAGG | TGG | \|\|\|\|\|\|\|\|[\|\|\|\|\|\|\|\|\|\|\|\|]PAM | NA | Intergenic | NA | NA |
| T15733 | CTTTATATACAAAGTCTAAGCGG | 1000 | | | | | | | | | |
| Oligo fwd | CACCCTTTATATACAAAGTCTAAG | | | | | | | | | | |
| Oligo rev | AAACCTTAGACTTTGTATATAAAG | | | | | | | | | | |
| Chromosome | start | end | strand | MM | target_seq | PAM | alignment | distance | position | gene name | gene id |
| GU305914.1 | 207647 | 207669 | + | 0 | CTTTATATACAAAGTCTAAG | CGG | \|\|\|\|\|\|\|\|[\|\|\|\|\|\|\|\|\|\|\|\|]PAM | NA | Intergenic | NA | NA |
| T15734 | TATACAAAGTCTAAGCGGAGCGG | 1000 | | | | | | | | | |
| Oligo fwd | CACCTATACAAAGTCTAAGCGGAG | | | | | | | | | | |
| Oligo rev | AAACCTCCGCTTAGACTTTGTATA | | | | | | | | | | |
| Chromosome | start | end | strand | MM | target_seq | PAM | alignment | distance | position | gene name | gene id |
| GU305914.1 | 207652 | 207674 | + | 0 | TATACAAAGTCTAAGCGGAG | CGG | \|\|\|\|\|\|\|\|[\|\|\|\|\|\|\|\|\|\|\|\|]PAM | NA | Intergenic | NA | NA |
| T15735 | ATACAAAGTCTAAGCGGAGCGGG | 1000 | | | | | | | | | |
| Oligo fwd | CACCATACAAAGTCTAAGCGGAGC | | | | | | | | | | |
| Oligo rev | AAACGCTCCGCTTAGACTTTGTAT | | | | | | | | | | |
| Chromosome | start | end | strand | MM | target_seq | PAM | alignment | distance | position | gene name | gene id |
| GU305914.1 | 207653 | 207675 | + | 0 | ATACAAAGTCTAAGCGGAGC | GGG | \|\|\|\|\|\|\|\|[\|\|\|\|\|\|\|\|\|\|\|\|]PAM | NA | Intergenic | NA | NA |
| T15736 | TAAGCGGAGCGGGTGTCACGCGG | 1000 | | | | | | | | | |
| Oligo fwd | CACCTAAGCGGAGCGGGTGTCACG | | | | | | | | | | |
| Oligo rev | AAACCGTGACACCCGCTCCGCTTA | | | | | | | | | | |
| Chromosome | start | end | strand | MM | target_seq | PAM | alignment | distance | position | gene name | gene id |
| GU305914.1 | 207663 | 207685 | + | 0 | TAAGCGGAGCGGGTGTCACG | CGG | \|\|\|\|\|\|\|\|[\|\|\|\|\|\|\|\|\|\|\|\|]PAM | NA | Intergenic | NA | NA |
| T15737 | AGCGGGTGTCACGCGGCATCCGG | 1000 | | | | | | | | | |
| Oligo fwd | CACCAGCGGGTGTCACGCGGCATC | | | | | | | | | | |
| Oligo rev | AAACGATGCCGCGTGACACCCGCT | | | | | | | | | | |
| Chromosome | start | end | strand | MM | target_seq | PAM | alignment | distance | position | gene name | gene id |
| GU305914.1 | 207670 | 207692 | + | 0 | AGCGGGTGTCACGCGGCATC | CGG | \|\|\|\|\|\|\|\|[\|\|\|\|\|\|\|\|\|\|\|\|]PAM | NA | Intergenic | NA | NA |
| T15738 | GCTTCCTCGTGTCAATCGACCGG | 1000 | | | | | | | | | |
| Oligo fwd | CACCGCTTCCTCGTGTCAATCGAC | | | | | | | | | | |
| Oligo rev | AAACGTCGATTGACACGAGGAAGC | | | | | | | | | | |
| Chromosome | start | end | strand | MM | target_seq | PAM | alignment | distance | position | gene name | gene id |
| GU305914.1 | 207697 | 207719 | + | 0 | GCTTCCTCGTGTCAATCGAC | CGG | \|\|\|\|\|\|\|\|[\|\|\|\|\|\|\|\|\|\|\|\|]PAM | NA | Intergenic | NA | NA |
| T15739 | ACGTCCGTGCTCGACTCGCCAGG | 1000 | | | | | | | | | |
| Oligo fwd | CACCACGTCCGTGCTCGACTCGCC | | | | | | | | | | |
| Oligo rev | AAACGGCGAGTCGAGCACGGACGT | | | | | | | | | | |
| Chromosome | start | end | strand | MM | target_seq | PAM | alignment | distance | position | gene name | gene id |
| GU305914.1 | 207728 | 207750 | + | 0 | ACGTCCGTGCTCGACTCGCC | AGG | \|\|\|\|\|\|\|\|[\|\|\|\|\|\|\|\|\|\|\|\|]PAM | NA | Intergenic | NA | NA |
| T15740 | TCCGTGCTCGACTCGCCAGGCGG | 1000 | | | | | | | | | |
| Oligo fwd | CACCTCCGTGCTCGACTCGCCAGG | | | | | | | | | | |
| Oligo rev | AAACCCTGGCGAGTCGAGCACGGA | | | | | | | | | | |
| Chromosome | start | end | strand | MM | target_seq | PAM | alignment | distance | position | gene name | gene id |
| GU305914.1 | 207731 | 207753 | + | 0 | TCCGTGCTCGACTCGCCAGG | CGG | \|\|\|\|\|\|\|\|[\|\|\|\|\|\|\|\|\|\|\|\|]PAM | NA | Intergenic | NA | NA |
| T15741 | CTGATCTGTCTGTGCCGCGTCGG | 1000 | | | | | | | | | |
| Oligo fwd | CACCCTGATCTGTCTGTGCCGCGT | | | | | | | | | | |
| Oligo rev | AAACACGCGGCACAGACAGATCAG | | | | | | | | | | |
| Chromosome | start | end | strand | MM | target_seq | PAM | alignment | distance | position | gene name | gene id |
| GU305914.1 | 207776 | 207798 | + | 0 | CTGATCTGTCTGTGCCGCGT | CGG | \|\|\|\|\|\|\|\|[\|\|\|\|\|\|\|\|\|\|\|\|]PAM | NA | Intergenic | NA | NA |
| T15742 | AGTCAGCTCGTCCAGACTCGCGG | 1000 | | | | | | | | | |
| Oligo fwd | CACCAGTCAGCTCGTCCAGACTCG | | | | | | | | | | |
| Oligo rev | AAACCGAGTCTGGACGAGCTGACT | | | | | | | | | | |
| Chromosome | start | end | strand | MM | target_seq | PAM | alignment | distance | position | gene name | gene id |
| GU305914.1 | 207799 | 207821 | + | 0 | AGTCAGCTCGTCCAGACTCG | CGG | \|\|\|\|\|\|\|\|[\|\|\|\|\|\|\|\|\|\|\|\|]PAM | NA | Intergenic | NA | NA |
| T15743 | CCAGACTCGCGGCGCGATAGCGG | 1000 | | | | | | | | | |
| Oligo fwd | CACCCCAGACTCGCGGCGCGATAG | | | | | | | | | | |
| Oligo rev | AAACCTATCGCGCCGCGAGTCTGG | | | | | | | | | | |
| Chromosome | start | end | strand | MM | target_seq | PAM | alignment | distance | position | gene name | gene id |
| GU305914.1 | 207810 | 207832 | + | 0 | CCAGACTCGCGGCGCGATAG | CGG | \|\|\|\|\|\|\|\|[\|\|\|\|\|\|\|\|\|\|\|\|]PAM | NA | Intergenic | NA | NA |
| T15744 | CGCGGCGCGATAGCGGTCGTCGG | 1000 | | | | | | | | | |
| Oligo fwd | CACCCGCGGCGCGATAGCGGTCGT | | | | | | | | | | |
| Oligo rev | AAACACGACCGCTATCGCGCCGCG | | | | | | | | | | |

<table>
<tr><td>Chromosome</td><td>start</td><td>end</td><td>strand</td><td>MM</td><td>target_seq</td><td>PAM</td><td>alignment</td><td>distance</td><td>position</td><td>gene name</td><td>gene id</td></tr>
<tr><td>GU305914.1</td><td>207817</td><td>207839</td><td>+</td><td>0</td><td>CGCGGCGCGATAGCGGTCGT</td><td>CGG</td><td>||||||||[||||||||||||]PAM</td><td>NA</td><td>Intergenic</td><td>NA</td><td>NA</td></tr>
<tr><td>T15745</td><td>GTCGCGCCTGAGAAACACATAGG</td><td>1000</td><td></td><td></td><td></td><td></td><td></td><td></td><td></td><td></td><td></td></tr>
<tr><td>Oligo fwd</td><td>CACCGTCGCGCCTGAGAAACACAT</td><td></td><td></td><td></td><td></td><td></td><td></td><td></td><td></td><td></td><td></td></tr>
<tr><td>Oligo rev</td><td>AAACATGTGTTTCTCAGGCGCGAC</td><td></td><td></td><td></td><td></td><td></td><td></td><td></td><td></td><td></td><td></td></tr>
<tr><td>Chromosome</td><td>start</td><td>end</td><td>strand</td><td>MM</td><td>target_seq</td><td>PAM</td><td>alignment</td><td>distance</td><td>position</td><td>gene name</td><td>gene id</td></tr>
<tr><td>GU305914.1</td><td>207865</td><td>207887</td><td>+</td><td>0</td><td>GTCGCGCCTGAGAAACACAT</td><td>AGG</td><td>||||||||[||||||||||||]PAM</td><td>NA</td><td>Intergenic</td><td>NA</td><td>NA</td></tr>
<tr><td>T15746</td><td>CGTCGATACGATCGCCACGACGG</td><td>1000</td><td></td><td></td><td></td><td></td><td></td><td></td><td></td><td></td><td></td></tr>
<tr><td>Oligo fwd</td><td>CACCCGTCGATACGATCGCCACGA</td><td></td><td></td><td></td><td></td><td></td><td></td><td></td><td></td><td></td><td></td></tr>
<tr><td>Oligo rev</td><td>AAACTCGTGGCGATCGTATCGACG</td><td></td><td></td><td></td><td></td><td></td><td></td><td></td><td></td><td></td><td></td></tr>
<tr><td>Chromosome</td><td>start</td><td>end</td><td>strand</td><td>MM</td><td>target_seq</td><td>PAM</td><td>alignment</td><td>distance</td><td>position</td><td>gene name</td><td>gene id</td></tr>
<tr><td>GU305914.1</td><td>207916</td><td>207938</td><td>+</td><td>0</td><td>CGTCGATACGATCGCCACGA</td><td>CGG</td><td>||||||||[||||||||||||]PAM</td><td>NA</td><td>Intergenic</td><td>NA</td><td>NA</td></tr>
<tr><td>T15747</td><td>CGATCGCCACGACGGTGACCAGG</td><td>1000</td><td></td><td></td><td></td><td></td><td></td><td></td><td></td><td></td><td></td></tr>
<tr><td>Oligo fwd</td><td>CACCCGATCGCCACGACGGTGACC</td><td></td><td></td><td></td><td></td><td></td><td></td><td></td><td></td><td></td><td></td></tr>
<tr><td>Oligo rev</td><td>AAACGGTCACCGTCGTGGCGATCG</td><td></td><td></td><td></td><td></td><td></td><td></td><td></td><td></td><td></td><td></td></tr>
<tr><td>Chromosome</td><td>start</td><td>end</td><td>strand</td><td>MM</td><td>target_seq</td><td>PAM</td><td>alignment</td><td>distance</td><td>position</td><td>gene name</td><td>gene id</td></tr>
<tr><td>GU305914.1</td><td>207924</td><td>207946</td><td>+</td><td>0</td><td>CGATCGCCACGACGGTGACC</td><td>AGG</td><td>||||||||[||||||||||||]PAM</td><td>NA</td><td>Intergenic</td><td>NA</td><td>NA</td></tr>
<tr><td>T15748</td><td>GATCGCCACGACGGTGACCAGGG</td><td>1000</td><td></td><td></td><td></td><td></td><td></td><td></td><td></td><td></td><td></td></tr>
<tr><td>Oligo fwd</td><td>CACCGATCGCCACGACGGTGACCA</td><td></td><td></td><td></td><td></td><td></td><td></td><td></td><td></td><td></td><td></td></tr>
<tr><td>Oligo rev</td><td>AAACTGGTCACCGTCGTGGCGATC</td><td></td><td></td><td></td><td></td><td></td><td></td><td></td><td></td><td></td><td></td></tr>
<tr><td>Chromosome</td><td>start</td><td>end</td><td>strand</td><td>MM</td><td>target_seq</td><td>PAM</td><td>alignment</td><td>distance</td><td>position</td><td>gene name</td><td>gene id</td></tr>
<tr><td>GU305914.1</td><td>207925</td><td>207947</td><td>+</td><td>0</td><td>GATCGCCACGACGGTGACCA</td><td>GGG</td><td>||||||||[||||||||||||]PAM</td><td>NA</td><td>Intergenic</td><td>NA</td><td>NA</td></tr>
<tr><td>T15749</td><td>GGACAGCTCCATGCCGCACGCGG</td><td>1000</td><td></td><td></td><td></td><td></td><td></td><td></td><td></td><td></td><td></td></tr>
<tr><td>Oligo fwd</td><td>CACCGGACAGCTCCATGCCGCACG</td><td></td><td></td><td></td><td></td><td></td><td></td><td></td><td></td><td></td><td></td></tr>
<tr><td>Oligo rev</td><td>AAACCGTGCGGCATGGAGCTGTCC</td><td></td><td></td><td></td><td></td><td></td><td></td><td></td><td></td><td></td><td></td></tr>
<tr><td>Chromosome</td><td>start</td><td>end</td><td>strand</td><td>MM</td><td>target_seq</td><td>PAM</td><td>alignment</td><td>distance</td><td>position</td><td>gene name</td><td>gene id</td></tr>
<tr><td>GU305914.1</td><td>207946</td><td>207968</td><td>+</td><td>0</td><td>GGACAGCTCCATGCCGCACG</td><td>CGG</td><td>||||||||[||||||||||||]PAM</td><td>NA</td><td>Intergenic</td><td>NA</td><td>NA</td></tr>
<tr><td>T15750</td><td>AGCTCCATGCCGCACGCGGCCGG</td><td>1000</td><td></td><td></td><td></td><td></td><td></td><td></td><td></td><td></td><td></td></tr>
<tr><td>Oligo fwd</td><td>CACCAGCTCCATGCCGCACGCGGC</td><td></td><td></td><td></td><td></td><td></td><td></td><td></td><td></td><td></td><td></td></tr>
<tr><td>Oligo rev</td><td>AAACGCCGCGTGCGGCATGGAGCT</td><td></td><td></td><td></td><td></td><td></td><td></td><td></td><td></td><td></td><td></td></tr>
<tr><td>Chromosome</td><td>start</td><td>end</td><td>strand</td><td>MM</td><td>target_seq</td><td>PAM</td><td>alignment</td><td>distance</td><td>position</td><td>gene name</td><td>gene id</td></tr>
<tr><td>GU305914.1</td><td>207950</td><td>207972</td><td>+</td><td>0</td><td>AGCTCCATGCCGCACGCGGC</td><td>CGG</td><td>||||||||[||||||||||||]PAM</td><td>NA</td><td>Intergenic</td><td>NA</td><td>NA</td></tr>
<tr><td>T15751</td><td>CGTTGCCGTTCTCAAGATCTTGG</td><td>1000</td><td></td><td></td><td></td><td></td><td></td><td></td><td></td><td></td><td></td></tr>
<tr><td>Oligo fwd</td><td>CACCCGTTGCCGTTCTCAAGATCT</td><td></td><td></td><td></td><td></td><td></td><td></td><td></td><td></td><td></td><td></td></tr>
<tr><td>Oligo rev</td><td>AAACAGATCTTGAGAACGGCAACG</td><td></td><td></td><td></td><td></td><td></td><td></td><td></td><td></td><td></td><td></td></tr>
<tr><td>Chromosome</td><td>start</td><td>end</td><td>strand</td><td>MM</td><td>target_seq</td><td>PAM</td><td>alignment</td><td>distance</td><td>position</td><td>gene name</td><td>gene id</td></tr>
<tr><td>GU305914.1</td><td>207988</td><td>208010</td><td>+</td><td>0</td><td>CGTTGCCGTTCTCAAGATCT</td><td>TGG</td><td>||||||||[||||||||||||]PAM</td><td>NA</td><td>Intergenic</td><td>NA</td><td>NA</td></tr>
<tr><td>T15752</td><td>ATCTTGGCCTCGTCGTGCGACGG</td><td>1000</td><td></td><td></td><td></td><td></td><td></td><td></td><td></td><td></td><td></td></tr>
<tr><td>Oligo fwd</td><td>CACCATCTTGGCCTCGTCGTGCGA</td><td></td><td></td><td></td><td></td><td></td><td></td><td></td><td></td><td></td><td></td></tr>
<tr><td>Oligo rev</td><td>AAACTCGCACGACGAGGCCAAGAT</td><td></td><td></td><td></td><td></td><td></td><td></td><td></td><td></td><td></td><td></td></tr>
<tr><td>Chromosome</td><td>start</td><td>end</td><td>strand</td><td>MM</td><td>target_seq</td><td>PAM</td><td>alignment</td><td>distance</td><td>position</td><td>gene name</td><td>gene id</td></tr>
<tr><td>GU305914.1</td><td>208004</td><td>208026</td><td>+</td><td>0</td><td>ATCTTGGCCTCGTCGTGCGA</td><td>CGG</td><td>||||||||[||||||||||||]PAM</td><td>NA</td><td>Intergenic</td><td>NA</td><td>NA</td></tr>
<tr><td>T15753</td><td>TCTTGGCCTCGTCGTGCGACGGG</td><td>1000</td><td></td><td></td><td></td><td></td><td></td><td></td><td></td><td></td><td></td></tr>
<tr><td>Oligo fwd</td><td>CACCTCTTGGCCTCGTCGTGCGAC</td><td></td><td></td><td></td><td></td><td></td><td></td><td></td><td></td><td></td><td></td></tr>
<tr><td>Oligo rev</td><td>AAACGTCGCACGACGAGGCCAAGA</td><td></td><td></td><td></td><td></td><td></td><td></td><td></td><td></td><td></td><td></td></tr>
<tr><td>Chromosome</td><td>start</td><td>end</td><td>strand</td><td>MM</td><td>target_seq</td><td>PAM</td><td>alignment</td><td>distance</td><td>position</td><td>gene name</td><td>gene id</td></tr>
<tr><td>GU305914.1</td><td>208005</td><td>208027</td><td>+</td><td>0</td><td>TCTTGGCCTCGTCGTGCGAC</td><td>GGG</td><td>||||||||[||||||||||||]PAM</td><td>NA</td><td>Intergenic</td><td>NA</td><td>NA</td></tr>
<tr><td>T15754</td><td>CTTGGCCTCGTCGTGCGACGGGG</td><td>1000</td><td></td><td></td><td></td><td></td><td></td><td></td><td></td><td></td><td></td></tr>
<tr><td>Oligo fwd</td><td>CACCCTTGGCCTCGTCGTGCGACG</td><td></td><td></td><td></td><td></td><td></td><td></td><td></td><td></td><td></td><td></td></tr>
<tr><td>Oligo rev</td><td>AAACCGTCGCACGACGAGGCCAAG</td><td></td><td></td><td></td><td></td><td></td><td></td><td></td><td></td><td></td><td></td></tr>
<tr><td>Chromosome</td><td>start</td><td>end</td><td>strand</td><td>MM</td><td>target_seq</td><td>PAM</td><td>alignment</td><td>distance</td><td>position</td><td>gene name</td><td>gene id</td></tr>
<tr><td>GU305914.1</td><td>208006</td><td>208028</td><td>+</td><td>0</td><td>CTTGGCCTCGTCGTGCGACG</td><td>GGG</td><td>||||||||[||||||||||||]PAM</td><td>NA</td><td>Intergenic</td><td>NA</td><td>NA</td></tr>
<tr><td>T15755</td><td>AATGACACAGACCAGATCCGAGG</td><td>1000</td><td></td><td></td><td></td><td></td><td></td><td></td><td></td><td></td><td></td></tr>
<tr><td>Oligo fwd</td><td>CACCAATGACACAGACCAGATCCG</td><td></td><td></td><td></td><td></td><td></td><td></td><td></td><td></td><td></td><td></td></tr>
<tr><td>Oligo rev</td><td>AAACCGGATCTGGTCTGTGTCATT</td><td></td><td></td><td></td><td></td><td></td><td></td><td></td><td></td><td></td><td></td></tr>
<tr><td>Chromosome</td><td>start</td><td>end</td><td>strand</td><td>MM</td><td>target_seq</td><td>PAM</td><td>alignment</td><td>distance</td><td>position</td><td>gene name</td><td>gene id</td></tr>
<tr><td>GU305914.1</td><td>208063</td><td>208085</td><td>+</td><td>0</td><td>AATGACACAGACCAGATCCG</td><td>AGG</td><td>||||||||[||||||||||||]PAM</td><td>NA</td><td>Intergenic</td><td>NA</td><td>NA</td></tr>
<tr><td>T15756</td><td>ATGACACAGACCAGATCCGAGGG</td><td>1000</td><td></td><td></td><td></td><td></td><td></td><td></td><td></td><td></td><td></td></tr>
<tr><td>Oligo fwd</td><td>CACCATGACACAGACCAGATCCGA</td><td></td><td></td><td></td><td></td><td></td><td></td><td></td><td></td><td></td><td></td></tr>
<tr><td>Oligo rev</td><td>AAACTCGGATCTGGTCTGTGTCAT</td><td></td><td></td><td></td><td></td><td></td><td></td><td></td><td></td><td></td><td></td></tr>
<tr><td>Chromosome</td><td>start</td><td>end</td><td>strand</td><td>MM</td><td>target_seq</td><td>PAM</td><td>alignment</td><td>distance</td><td>position</td><td>gene name</td><td>gene id</td></tr>
<tr><td>GU305914.1</td><td>208064</td><td>208086</td><td>+</td><td>0</td><td>ATGACACAGACCAGATCCGA</td><td>GGG</td><td>||||||||[||||||||||||]PAM</td><td>NA</td><td>Intergenic</td><td>NA</td><td>NA</td></tr>
<tr><td>T15757</td><td>ATCCGAGGGCTCGAAGCTCGAGG</td><td>1000</td><td></td><td></td><td></td><td></td><td></td><td></td><td></td><td></td><td></td></tr>
<tr><td>Oligo fwd</td><td>CACCATCCGAGGGCTCGAAGCTCG</td><td></td><td></td><td></td><td></td><td></td><td></td><td></td><td></td><td></td><td></td></tr>
<tr><td>Oligo rev</td><td>AAACCGAGCTTCGAGCCCTCGGAT</td><td></td><td></td><td></td><td></td><td></td><td></td><td></td><td></td><td></td><td></td></tr>
<tr><td>Chromosome</td><td>start</td><td>end</td><td>strand</td><td>MM</td><td>target_seq</td><td>PAM</td><td>alignment</td><td>distance</td><td>position</td><td>gene name</td><td>gene id</td></tr>
<tr><td>GU305914.1</td><td>208078</td><td>208100</td><td>+</td><td>0</td><td>ATCCGAGGGCTCGAAGCTCG</td><td>AGG</td><td>||||||||[||||||||||||]PAM</td><td>NA</td><td>Intergenic</td><td>NA</td><td>NA</td></tr>
<tr><td>T15758</td><td>CCACCGTATCCGAAAAGAACCGG</td><td>1000</td><td></td><td></td><td></td><td></td><td></td><td></td><td></td><td></td><td></td></tr>
<tr><td>Oligo fwd</td><td>CACCCCACCGTATCCGAAAAGAAC</td><td></td><td></td><td></td><td></td><td></td><td></td><td></td><td></td><td></td><td></td></tr>
<tr><td>Oligo rev</td><td>AAACGTTCTTTTCGGATACGGTGG</td><td></td><td></td><td></td><td></td><td></td><td></td><td></td><td></td><td></td><td></td></tr>
<tr><td>Chromosome</td><td>start</td><td>end</td><td>strand</td><td>MM</td><td>target_seq</td><td>PAM</td><td>alignment</td><td>distance</td><td>position</td><td>gene name</td><td>gene id</td></tr>
<tr><td>GU305914.1</td><td>208122</td><td>208144</td><td>+</td><td>0</td><td>CCACCGTATCCGAAAAGAAC</td><td>CGG</td><td>||||||||[||||||||||||]PAM</td><td>NA</td><td>Intergenic</td><td>NA</td><td>NA</td></tr>
<tr><td>T15759</td><td>CCGTATCCGAAAAGAACCGGCGG</td><td>1000</td><td></td><td></td><td></td><td></td><td></td><td></td><td></td><td></td><td></td></tr>
<tr><td>Oligo fwd</td><td>CACCCCGTATCCGAAAAGAACCGG</td><td></td><td></td><td></td><td></td><td></td><td></td><td></td><td></td><td></td><td></td></tr>
</table>

<table>
<tr><td>Oligo rev</td><td>AAACCCGGTTCTTTTCGGATACGG</td><td></td><td></td><td></td><td></td><td></td><td></td><td></td><td></td><td></td><td></td></tr>
<tr><td>Chromosome</td><td>start</td><td>end</td><td>strand</td><td>MM</td><td>target_seq</td><td>PAM</td><td>alignment</td><td>distance</td><td>position</td><td>gene name</td><td>gene id</td></tr>
<tr><td>GU305914.1</td><td>208125</td><td>208147</td><td>+</td><td>0</td><td>CCGTATCCGAAAAGAACCGG</td><td>CGG</td><td>||||||||[||||||||||||]PAM</td><td>NA</td><td>Intergenic</td><td>NA</td><td>NA</td></tr>
<tr><td>GU305914.1</td><td>124574</td><td>124596</td><td>+</td><td>4</td><td>CCGGGTCCGAGTAGAACCGG</td><td>TGG</td><td>|||--|||[||--||||||||]PAM</td><td>NA</td><td>Intergenic</td><td>NA</td><td>NA</td></tr>
<tr><td></td><td></td><td></td><td></td><td></td><td></td><td></td><td></td><td></td><td></td><td></td><td></td></tr>
<tr><td>T15760</td><td>CGTATCCGAAAAGAACCGGCGGG</td><td>1000</td><td></td><td></td><td></td><td></td><td></td><td></td><td></td><td></td><td></td></tr>
<tr><td>Oligo fwd</td><td>CACCCGTATCCGAAAAGAACCGGC</td><td></td><td></td><td></td><td></td><td></td><td></td><td></td><td></td><td></td><td></td></tr>
<tr><td>Oligo rev</td><td>AAACGCCGGTTCTTTTCGGATACG</td><td></td><td></td><td></td><td></td><td></td><td></td><td></td><td></td><td></td><td></td></tr>
<tr><td>Chromosome</td><td>start</td><td>end</td><td>strand</td><td>MM</td><td>target_seq</td><td>PAM</td><td>alignment</td><td>distance</td><td>position</td><td>gene name</td><td>gene id</td></tr>
<tr><td>GU305914.1</td><td>208126</td><td>208148</td><td>+</td><td>0</td><td>CGTATCCGAAAAGAACCGGC</td><td>GGG</td><td>||||||||[||||||||||||]PAM</td><td>NA</td><td>Intergenic</td><td>NA</td><td>NA</td></tr>
<tr><td></td><td></td><td></td><td></td><td></td><td></td><td></td><td></td><td></td><td></td><td></td><td></td></tr>
<tr><td>T15761</td><td>GCCCCTCACCGTCACGTCGAAGG</td><td>1000</td><td></td><td></td><td></td><td></td><td></td><td></td><td></td><td></td><td></td></tr>
<tr><td>Oligo fwd</td><td>CACCGCCCCTCACCGTCACGTCGA</td><td></td><td></td><td></td><td></td><td></td><td></td><td></td><td></td><td></td><td></td></tr>
<tr><td>Oligo rev</td><td>AAACTCGACGTGACGGTGAGGGGC</td><td></td><td></td><td></td><td></td><td></td><td></td><td></td><td></td><td></td><td></td></tr>
<tr><td>Chromosome</td><td>start</td><td>end</td><td>strand</td><td>MM</td><td>target_seq</td><td>PAM</td><td>alignment</td><td>distance</td><td>position</td><td>gene name</td><td>gene id</td></tr>
<tr><td>GU305914.1</td><td>208180</td><td>208202</td><td>+</td><td>0</td><td>GCCCCTCACCGTCACGTCGA</td><td>AGG</td><td>||||||||[||||||||||||]PAM</td><td>NA</td><td>Intergenic</td><td>NA</td><td>NA</td></tr>
<tr><td></td><td></td><td></td><td></td><td></td><td></td><td></td><td></td><td></td><td></td><td></td><td></td></tr>
<tr><td>T15762</td><td>CCCCTCACCGTCACGTCGAAGGG</td><td>1000</td><td></td><td></td><td></td><td></td><td></td><td></td><td></td><td></td><td></td></tr>
<tr><td>Oligo fwd</td><td>CACCCCCCTCACCGTCACGTCGAA</td><td></td><td></td><td></td><td></td><td></td><td></td><td></td><td></td><td></td><td></td></tr>
<tr><td>Oligo rev</td><td>AAACTTCGACGTGACGGTGAGGGG</td><td></td><td></td><td></td><td></td><td></td><td></td><td></td><td></td><td></td><td></td></tr>
<tr><td>Chromosome</td><td>start</td><td>end</td><td>strand</td><td>MM</td><td>target_seq</td><td>PAM</td><td>alignment</td><td>distance</td><td>position</td><td>gene name</td><td>gene id</td></tr>
<tr><td>GU305914.1</td><td>208181</td><td>208203</td><td>+</td><td>0</td><td>CCCCTCACCGTCACGTCGAA</td><td>GGG</td><td>||||||||[||||||||||||]PAM</td><td>NA</td><td>Intergenic</td><td>NA</td><td>NA</td></tr>
<tr><td></td><td></td><td></td><td></td><td></td><td></td><td></td><td></td><td></td><td></td><td></td><td></td></tr>
<tr><td>T15763</td><td>ACCGTCACGTCGAAGGGTAGCGG</td><td>1000</td><td></td><td></td><td></td><td></td><td></td><td></td><td></td><td></td><td></td></tr>
<tr><td>Oligo fwd</td><td>CACCACCGTCACGTCGAAGGGTAG</td><td></td><td></td><td></td><td></td><td></td><td></td><td></td><td></td><td></td><td></td></tr>
<tr><td>Oligo rev</td><td>AAACCTACCCTTCGACGTGACGGT</td><td></td><td></td><td></td><td></td><td></td><td></td><td></td><td></td><td></td><td></td></tr>
<tr><td>Chromosome</td><td>start</td><td>end</td><td>strand</td><td>MM</td><td>target_seq</td><td>PAM</td><td>alignment</td><td>distance</td><td>position</td><td>gene name</td><td>gene id</td></tr>
<tr><td>GU305914.1</td><td>208187</td><td>208209</td><td>+</td><td>0</td><td>ACCGTCACGTCGAAGGGTAG</td><td>CGG</td><td>||||||||[||||||||||||]PAM</td><td>NA</td><td>Intergenic</td><td>NA</td><td>NA</td></tr>
<tr><td></td><td></td><td></td><td></td><td></td><td></td><td></td><td></td><td></td><td></td><td></td><td></td></tr>
<tr><td>T15764</td><td>CCGTCACGTCGAAGGGTAGCGGG</td><td>1000</td><td></td><td></td><td></td><td></td><td></td><td></td><td></td><td></td><td></td></tr>
<tr><td>Oligo fwd</td><td>CACCCCGTCACGTCGAAGGGTAGC</td><td></td><td></td><td></td><td></td><td></td><td></td><td></td><td></td><td></td><td></td></tr>
<tr><td>Oligo rev</td><td>AAACGCTACCCTTCGACGTGACGG</td><td></td><td></td><td></td><td></td><td></td><td></td><td></td><td></td><td></td><td></td></tr>
<tr><td>Chromosome</td><td>start</td><td>end</td><td>strand</td><td>MM</td><td>target_seq</td><td>PAM</td><td>alignment</td><td>distance</td><td>position</td><td>gene name</td><td>gene id</td></tr>
<tr><td>GU305914.1</td><td>208188</td><td>208210</td><td>+</td><td>0</td><td>CCGTCACGTCGAAGGGTAGC</td><td>GGG</td><td>||||||||[||||||||||||]PAM</td><td>NA</td><td>Intergenic</td><td>NA</td><td>NA</td></tr>
<tr><td></td><td></td><td></td><td></td><td></td><td></td><td></td><td></td><td></td><td></td><td></td><td></td></tr>
<tr><td>T15765</td><td>CGTCACGTCGAAGGGTAGCGGGG</td><td>1000</td><td></td><td></td><td></td><td></td><td></td><td></td><td></td><td></td><td></td></tr>
<tr><td>Oligo fwd</td><td>CACCCGTCACGTCGAAGGGTAGCG</td><td></td><td></td><td></td><td></td><td></td><td></td><td></td><td></td><td></td><td></td></tr>
<tr><td>Oligo rev</td><td>AAACCGCTACCCTTCGACGTGACG</td><td></td><td></td><td></td><td></td><td></td><td></td><td></td><td></td><td></td><td></td></tr>
<tr><td>Chromosome</td><td>start</td><td>end</td><td>strand</td><td>MM</td><td>target_seq</td><td>PAM</td><td>alignment</td><td>distance</td><td>position</td><td>gene name</td><td>gene id</td></tr>
<tr><td>GU305914.1</td><td>208189</td><td>208211</td><td>+</td><td>0</td><td>CGTCACGTCGAAGGGTAGCG</td><td>GGG</td><td>||||||||[||||||||||||]PAM</td><td>NA</td><td>Intergenic</td><td>NA</td><td>NA</td></tr>
<tr><td></td><td></td><td></td><td></td><td></td><td></td><td></td><td></td><td></td><td></td><td></td><td></td></tr>
<tr><td>T15766</td><td>ACGTCGAAGGGTAGCGGGGCCGG</td><td>1000</td><td></td><td></td><td></td><td></td><td></td><td></td><td></td><td></td><td></td></tr>
<tr><td>Oligo fwd</td><td>CACCACGTCGAAGGGTAGCGGGGC</td><td></td><td></td><td></td><td></td><td></td><td></td><td></td><td></td><td></td><td></td></tr>
<tr><td>Oligo rev</td><td>AAACGCCCCGCTACCCTTCGACGT</td><td></td><td></td><td></td><td></td><td></td><td></td><td></td><td></td><td></td><td></td></tr>
<tr><td>Chromosome</td><td>start</td><td>end</td><td>strand</td><td>MM</td><td>target_seq</td><td>PAM</td><td>alignment</td><td>distance</td><td>position</td><td>gene name</td><td>gene id</td></tr>
<tr><td>GU305914.1</td><td>208193</td><td>208215</td><td>+</td><td>0</td><td>ACGTCGAAGGGTAGCGGGGC</td><td>CGG</td><td>||||||||[||||||||||||]PAM</td><td>NA</td><td>Intergenic</td><td>NA</td><td>NA</td></tr>
<tr><td></td><td></td><td></td><td></td><td></td><td></td><td></td><td></td><td></td><td></td><td></td><td></td></tr>
<tr><td>T15767</td><td>CGTCGAAGGGTAGCGGGGCCGGG</td><td>1000</td><td></td><td></td><td></td><td></td><td></td><td></td><td></td><td></td><td></td></tr>
<tr><td>Oligo fwd</td><td>CACCCGTCGAAGGGTAGCGGGGCC</td><td></td><td></td><td></td><td></td><td></td><td></td><td></td><td></td><td></td><td></td></tr>
<tr><td>Oligo rev</td><td>AAACGGCCCCGCTACCCTTCGACG</td><td></td><td></td><td></td><td></td><td></td><td></td><td></td><td></td><td></td><td></td></tr>
<tr><td>Chromosome</td><td>start</td><td>end</td><td>strand</td><td>MM</td><td>target_seq</td><td>PAM</td><td>alignment</td><td>distance</td><td>position</td><td>gene name</td><td>gene id</td></tr>
<tr><td>GU305914.1</td><td>208194</td><td>208216</td><td>+</td><td>0</td><td>CGTCGAAGGGTAGCGGGGCC</td><td>GGG</td><td>||||||||[||||||||||||]PAM</td><td>NA</td><td>Intergenic</td><td>NA</td><td>NA</td></tr>
<tr><td></td><td></td><td></td><td></td><td></td><td></td><td></td><td></td><td></td><td></td><td></td><td></td></tr>
<tr><td>T15768</td><td>GAAGAATCCGTGTCTTCCCACGG</td><td>1000</td><td></td><td></td><td></td><td></td><td></td><td></td><td></td><td></td><td></td></tr>
<tr><td>Oligo fwd</td><td>CACCGAAGAATCCGTGTCTTCCCA</td><td></td><td></td><td></td><td></td><td></td><td></td><td></td><td></td><td></td><td></td></tr>
<tr><td>Oligo rev</td><td>AAACTGGGAAGACACGGATTCTTC</td><td></td><td></td><td></td><td></td><td></td><td></td><td></td><td></td><td></td><td></td></tr>
<tr><td>Chromosome</td><td>start</td><td>end</td><td>strand</td><td>MM</td><td>target_seq</td><td>PAM</td><td>alignment</td><td>distance</td><td>position</td><td>gene name</td><td>gene id</td></tr>
<tr><td>GU305914.1</td><td>208234</td><td>208256</td><td>+</td><td>0</td><td>GAAGAATCCGTGTCTTCCCA</td><td>CGG</td><td>||||||||[||||||||||||]PAM</td><td>NA</td><td>Intergenic</td><td>NA</td><td>NA</td></tr>
<tr><td></td><td></td><td></td><td></td><td></td><td></td><td></td><td></td><td></td><td></td><td></td><td></td></tr>
<tr><td>T15769</td><td>AAAACTAACCTCCAAGTGCTCGG</td><td>1000</td><td></td><td></td><td></td><td></td><td></td><td></td><td></td><td></td><td></td></tr>
<tr><td>Oligo fwd</td><td>CACCAAAACTAACCTCCAAGTGCT</td><td></td><td></td><td></td><td></td><td></td><td></td><td></td><td></td><td></td><td></td></tr>
<tr><td>Oligo rev</td><td>AAACAGCACTTGGAGGTTAGTTTT</td><td></td><td></td><td></td><td></td><td></td><td></td><td></td><td></td><td></td><td></td></tr>
<tr><td>Chromosome</td><td>start</td><td>end</td><td>strand</td><td>MM</td><td>target_seq</td><td>PAM</td><td>alignment</td><td>distance</td><td>position</td><td>gene name</td><td>gene id</td></tr>
<tr><td>GU305914.1</td><td>208264</td><td>208286</td><td>+</td><td>0</td><td>AAAACTAACCTCCAAGTGCT</td><td>CGG</td><td>||||||||[||||||||||||]PAM</td><td>NA</td><td>Intergenic</td><td>NA</td><td>NA</td></tr>
<tr><td></td><td></td><td></td><td></td><td></td><td></td><td></td><td></td><td></td><td></td><td></td><td></td></tr>
<tr><td>T15770</td><td>AAACTAACCTCCAAGTGCTCGGG</td><td>1000</td><td></td><td></td><td></td><td></td><td></td><td></td><td></td><td></td><td></td></tr>
<tr><td>Oligo fwd</td><td>CACCAAACTAACCTCCAAGTGCTC</td><td></td><td></td><td></td><td></td><td></td><td></td><td></td><td></td><td></td><td></td></tr>
<tr><td>Oligo rev</td><td>AAACGAGCACTTGGAGGTTAGTTT</td><td></td><td></td><td></td><td></td><td></td><td></td><td></td><td></td><td></td><td></td></tr>
<tr><td>Chromosome</td><td>start</td><td>end</td><td>strand</td><td>MM</td><td>target_seq</td><td>PAM</td><td>alignment</td><td>distance</td><td>position</td><td>gene name</td><td>gene id</td></tr>
<tr><td>GU305914.1</td><td>208265</td><td>208287</td><td>+</td><td>0</td><td>AAACTAACCTCCAAGTGCTC</td><td>GGG</td><td>||||||||[||||||||||||]PAM</td><td>NA</td><td>Intergenic</td><td>NA</td><td>NA</td></tr>
<tr><td></td><td></td><td></td><td></td><td></td><td></td><td></td><td></td><td></td><td></td><td></td><td></td></tr>
<tr><td>T15771</td><td>GACGCAGCACGCTCAGATCGAGG</td><td>1000</td><td></td><td></td><td></td><td></td><td></td><td></td><td></td><td></td><td></td></tr>
<tr><td>Oligo fwd</td><td>CACCGACGCAGCACGCTCAGATCG</td><td></td><td></td><td></td><td></td><td></td><td></td><td></td><td></td><td></td><td></td></tr>
<tr><td>Oligo rev</td><td>AAACCGATCTGAGCGTGCTGCGTC</td><td></td><td></td><td></td><td></td><td></td><td></td><td></td><td></td><td></td><td></td></tr>
<tr><td>Chromosome</td><td>start</td><td>end</td><td>strand</td><td>MM</td><td>target_seq</td><td>PAM</td><td>alignment</td><td>distance</td><td>position</td><td>gene name</td><td>gene id</td></tr>
<tr><td>GU305914.1</td><td>208353</td><td>208375</td><td>+</td><td>0</td><td>GACGCAGCACGCTCAGATCG</td><td>AGG</td><td>||||||||[||||||||||||]PAM</td><td>NA</td><td>Intergenic</td><td>NA</td><td>NA</td></tr>
<tr><td></td><td></td><td></td><td></td><td></td><td></td><td></td><td></td><td></td><td></td><td></td><td></td></tr>
<tr><td>T15772</td><td>CAGCACGCTCAGATCGAGGTCGG</td><td>1000</td><td></td><td></td><td></td><td></td><td></td><td></td><td></td><td></td><td></td></tr>
<tr><td>Oligo fwd</td><td>CACCCAGCACGCTCAGATCGAGGT</td><td></td><td></td><td></td><td></td><td></td><td></td><td></td><td></td><td></td><td></td></tr>
<tr><td>Oligo rev</td><td>AAACACCTCGATCTGAGCGTGCTG</td><td></td><td></td><td></td><td></td><td></td><td></td><td></td><td></td><td></td><td></td></tr>
<tr><td>Chromosome</td><td>start</td><td>end</td><td>strand</td><td>MM</td><td>target_seq</td><td>PAM</td><td>alignment</td><td>distance</td><td>position</td><td>gene name</td><td>gene id</td></tr>
<tr><td>GU305914.1</td><td>208357</td><td>208379</td><td>+</td><td>0</td><td>CAGCACGCTCAGATCGAGGT</td><td>CGG</td><td>||||||||[||||||||||||]PAM</td><td>NA</td><td>Intergenic</td><td>NA</td><td>NA</td></tr>
<tr><td></td><td></td><td></td><td></td><td></td><td></td><td></td><td></td><td></td><td></td><td></td><td></td></tr>
<tr><td>T15773</td><td>CACGCTCAGATCGAGGTCGGTGG</td><td>1000</td><td></td><td></td><td></td><td></td><td></td><td></td><td></td><td></td><td></td></tr>
<tr><td>Oligo fwd</td><td>CACCCACGCTCAGATCGAGGTCGG</td><td></td><td></td><td></td><td></td><td></td><td></td><td></td><td></td><td></td><td></td></tr>
<tr><td>Oligo rev</td><td>AAACCCGACCTCGATCTGAGCGTG</td><td></td><td></td><td></td><td></td><td></td><td></td><td></td><td></td><td></td><td></td></tr>
<tr><td>Chromosome</td><td>start</td><td>end</td><td>strand</td><td>MM</td><td>target_seq</td><td>PAM</td><td>alignment</td><td>distance</td><td>position</td><td>gene name</td><td>gene id</td></tr>
<tr><td>GU305914.1</td><td>208360</td><td>208382</td><td>+</td><td>0</td><td>CACGCTCAGATCGAGGTCGG</td><td>TGG</td><td>||||||||[||||||||||||]PAM</td><td>NA</td><td>Intergenic</td><td>NA</td><td>NA</td></tr>
</table>

<table>
<tr><td>T15774</td><td>GGTCGGTGGTGAACTTTGTCAGG</td><td>1000</td><td></td><td></td><td></td><td></td><td></td><td></td><td></td><td></td><td></td></tr>
<tr><td>Oligo fwd</td><td>CACCGGTCGGTGGTGAACTTTGTC</td><td></td><td></td><td></td><td></td><td></td><td></td><td></td><td></td><td></td><td></td></tr>
<tr><td>Oligo rev</td><td>AAACGACAAAGTTCACCACCGACC</td><td></td><td></td><td></td><td></td><td></td><td></td><td></td><td></td><td></td><td></td></tr>
<tr><td>Chromosome</td><td>start</td><td>end</td><td>strand</td><td>MM</td><td>target_seq</td><td>PAM</td><td>alignment</td><td>distance</td><td>position</td><td>gene name</td><td>gene id</td></tr>
<tr><td>GU305914.1</td><td>208374</td><td>208396</td><td>+</td><td>0</td><td>GGTCGGTGGTGAACTTTGTC</td><td>AGG</td><td>||||||||[||||||||||||]PAM</td><td>NA</td><td>Intergenic</td><td>NA</td><td>NA</td></tr>
<tr><td>T15775</td><td>TCTAACGTTCGTACCGTCCGCGG</td><td>1000</td><td></td><td></td><td></td><td></td><td></td><td></td><td></td><td></td><td></td></tr>
<tr><td>Oligo fwd</td><td>CACCTCTAACGTTCGTACCGTCCG</td><td></td><td></td><td></td><td></td><td></td><td></td><td></td><td></td><td></td><td></td></tr>
<tr><td>Oligo rev</td><td>AAACCGGACGGTACGAACGTTAGA</td><td></td><td></td><td></td><td></td><td></td><td></td><td></td><td></td><td></td><td></td></tr>
<tr><td>Chromosome</td><td>start</td><td>end</td><td>strand</td><td>MM</td><td>target_seq</td><td>PAM</td><td>alignment</td><td>distance</td><td>position</td><td>gene name</td><td>gene id</td></tr>
<tr><td>GU305914.1</td><td>208459</td><td>208481</td><td>+</td><td>0</td><td>TCTAACGTTCGTACCGTCCG</td><td>CGG</td><td>||||||||[||||||||||||]PAM</td><td>NA</td><td>Intergenic</td><td>NA</td><td>NA</td></tr>
<tr><td>T15776</td><td>CGTCCGCGGTGTGTACAACGAGG</td><td>1000</td><td></td><td></td><td></td><td></td><td></td><td></td><td></td><td></td><td></td></tr>
<tr><td>Oligo fwd</td><td>CACCCGTCCGCGGTGTGTACAACG</td><td></td><td></td><td></td><td></td><td></td><td></td><td></td><td></td><td></td><td></td></tr>
<tr><td>Oligo rev</td><td>AAACCGTTGTACACACCGCGGACG</td><td></td><td></td><td></td><td></td><td></td><td></td><td></td><td></td><td></td><td></td></tr>
<tr><td>Chromosome</td><td>start</td><td>end</td><td>strand</td><td>MM</td><td>target_seq</td><td>PAM</td><td>alignment</td><td>distance</td><td>position</td><td>gene name</td><td>gene id</td></tr>
<tr><td>GU305914.1</td><td>208473</td><td>208495</td><td>+</td><td>0</td><td>CGTCCGCGGTGTGTACAACG</td><td>AGG</td><td>||||||||[||||||||||||]PAM</td><td>NA</td><td>Intergenic</td><td>NA</td><td>NA</td></tr>
<tr><td>T15777</td><td>ACGAGGCATCCGACCAACTGCGG</td><td>1000</td><td></td><td></td><td></td><td></td><td></td><td></td><td></td><td></td><td></td></tr>
<tr><td>Oligo fwd</td><td>CACCACGAGGCATCCGACCAACTG</td><td></td><td></td><td></td><td></td><td></td><td></td><td></td><td></td><td></td><td></td></tr>
<tr><td>Oligo rev</td><td>AAACCAGTTGGTCGGATGCCTCGT</td><td></td><td></td><td></td><td></td><td></td><td></td><td></td><td></td><td></td><td></td></tr>
<tr><td>Chromosome</td><td>start</td><td>end</td><td>strand</td><td>MM</td><td>target_seq</td><td>PAM</td><td>alignment</td><td>distance</td><td>position</td><td>gene name</td><td>gene id</td></tr>
<tr><td>GU305914.1</td><td>208490</td><td>208512</td><td>+</td><td>0</td><td>ACGAGGCATCCGACCAACTG</td><td>CGG</td><td>||||||||[||||||||||||]PAM</td><td>NA</td><td>Intergenic</td><td>NA</td><td>NA</td></tr>
<tr><td>T15778</td><td>GAACAGCCGCGCTTTGACCGAGG</td><td>1000</td><td></td><td></td><td></td><td></td><td></td><td></td><td></td><td></td><td></td></tr>
<tr><td>Oligo fwd</td><td>CACCGAACAGCCGCGCTTTGACCG</td><td></td><td></td><td></td><td></td><td></td><td></td><td></td><td></td><td></td><td></td></tr>
<tr><td>Oligo rev</td><td>AAACCGGTCAAAGCGCGGCTGTTC</td><td></td><td></td><td></td><td></td><td></td><td></td><td></td><td></td><td></td><td></td></tr>
<tr><td>Chromosome</td><td>start</td><td>end</td><td>strand</td><td>MM</td><td>target_seq</td><td>PAM</td><td>alignment</td><td>distance</td><td>position</td><td>gene name</td><td>gene id</td></tr>
<tr><td>GU305914.1</td><td>208542</td><td>208564</td><td>+</td><td>0</td><td>GAACAGCCGCGCTTTGACCG</td><td>AGG</td><td>||||||||[||||||||||||]PAM</td><td>NA</td><td>Intergenic</td><td>NA</td><td>NA</td></tr>
<tr><td>T15779</td><td>CGCTTTGACCGAGGAGCCGACGG</td><td>1000</td><td></td><td></td><td></td><td></td><td></td><td></td><td></td><td></td><td></td></tr>
<tr><td>Oligo fwd</td><td>CACCCGCTTTGACCGAGGAGCCGA</td><td></td><td></td><td></td><td></td><td></td><td></td><td></td><td></td><td></td><td></td></tr>
<tr><td>Oligo rev</td><td>AAACTCGGCTCCTCGGTCAAAGCG</td><td></td><td></td><td></td><td></td><td></td><td></td><td></td><td></td><td></td><td></td></tr>
<tr><td>Chromosome</td><td>start</td><td>end</td><td>strand</td><td>MM</td><td>target_seq</td><td>PAM</td><td>alignment</td><td>distance</td><td>position</td><td>gene name</td><td>gene id</td></tr>
<tr><td>GU305914.1</td><td>208551</td><td>208573</td><td>+</td><td>0</td><td>CGCTTTGACCGAGGAGCCGA</td><td>CGG</td><td>||||||||[||||||||||||]PAM</td><td>NA</td><td>Intergenic</td><td>NA</td><td>NA</td></tr>
<tr><td>T15780</td><td>ACCGAGGAGCCGACGGAACGCGG</td><td>1000</td><td></td><td></td><td></td><td></td><td></td><td></td><td></td><td></td><td></td></tr>
<tr><td>Oligo fwd</td><td>CACCACCGAGGAGCCGACGGAACG</td><td></td><td></td><td></td><td></td><td></td><td></td><td></td><td></td><td></td><td></td></tr>
<tr><td>Oligo rev</td><td>AAACCGTTCCGTCGGCTCCTCGGT</td><td></td><td></td><td></td><td></td><td></td><td></td><td></td><td></td><td></td><td></td></tr>
<tr><td>Chromosome</td><td>start</td><td>end</td><td>strand</td><td>MM</td><td>target_seq</td><td>PAM</td><td>alignment</td><td>distance</td><td>position</td><td>gene name</td><td>gene id</td></tr>
<tr><td>GU305914.1</td><td>208558</td><td>208580</td><td>+</td><td>0</td><td>ACCGAGGAGCCGACGGAACG</td><td>CGG</td><td>||||||||[||||||||||||]PAM</td><td>NA</td><td>Intergenic</td><td>NA</td><td>NA</td></tr>
<tr><td>T15781</td><td>GGAGCCGACGGAACGCGGTCAGG</td><td>1000</td><td></td><td></td><td></td><td></td><td></td><td></td><td></td><td></td><td></td></tr>
<tr><td>Oligo fwd</td><td>CACCGGAGCCGACGGAACGCGGTC</td><td></td><td></td><td></td><td></td><td></td><td></td><td></td><td></td><td></td><td></td></tr>
<tr><td>Oligo rev</td><td>AAACGACCGCGTTCCGTCGGCTCC</td><td></td><td></td><td></td><td></td><td></td><td></td><td></td><td></td><td></td><td></td></tr>
<tr><td>Chromosome</td><td>start</td><td>end</td><td>strand</td><td>MM</td><td>target_seq</td><td>PAM</td><td>alignment</td><td>distance</td><td>position</td><td>gene name</td><td>gene id</td></tr>
<tr><td>GU305914.1</td><td>208563</td><td>208585</td><td>+</td><td>0</td><td>GGAGCCGACGGAACGCGGTC</td><td>AGG</td><td>||||||||[||||||||||||]PAM</td><td>NA</td><td>Intergenic</td><td>NA</td><td>NA</td></tr>
<tr><td>T15782</td><td>GCCGACGGAACGCGGTCAGGTGG</td><td>1000</td><td></td><td></td><td></td><td></td><td></td><td></td><td></td><td></td><td></td></tr>
<tr><td>Oligo fwd</td><td>CACCGCCGACGGAACGCGGTCAGG</td><td></td><td></td><td></td><td></td><td></td><td></td><td></td><td></td><td></td><td></td></tr>
<tr><td>Oligo rev</td><td>AAACCCTGACCGCGTTCCGTCGGC</td><td></td><td></td><td></td><td></td><td></td><td></td><td></td><td></td><td></td><td></td></tr>
<tr><td>Chromosome</td><td>start</td><td>end</td><td>strand</td><td>MM</td><td>target_seq</td><td>PAM</td><td>alignment</td><td>distance</td><td>position</td><td>gene name</td><td>gene id</td></tr>
<tr><td>GU305914.1</td><td>208566</td><td>208588</td><td>+</td><td>0</td><td>GCCGACGGAACGCGGTCAGG</td><td>TGG</td><td>||||||||[||||||||||||]PAM</td><td>NA</td><td>Intergenic</td><td>NA</td><td>NA</td></tr>
<tr><td>T15783</td><td>ATCCCACAGAAAATGTACTTCGG</td><td>1000</td><td></td><td></td><td></td><td></td><td></td><td></td><td></td><td></td><td></td></tr>
<tr><td>Oligo fwd</td><td>CACCATCCCACAGAAAATGTACTT</td><td></td><td></td><td></td><td></td><td></td><td></td><td></td><td></td><td></td><td></td></tr>
<tr><td>Oligo rev</td><td>AAACAAGTACATTTTCTGTGGGAT</td><td></td><td></td><td></td><td></td><td></td><td></td><td></td><td></td><td></td><td></td></tr>
<tr><td>Chromosome</td><td>start</td><td>end</td><td>strand</td><td>MM</td><td>target_seq</td><td>PAM</td><td>alignment</td><td>distance</td><td>position</td><td>gene name</td><td>gene id</td></tr>
<tr><td>GU305914.1</td><td>208597</td><td>208619</td><td>+</td><td>0</td><td>ATCCCACAGAAAATGTACTT</td><td>CGG</td><td>||||||||[||||||||||||]PAM</td><td>NA</td><td>Intergenic</td><td>NA</td><td>NA</td></tr>
<tr><td>T15784</td><td>TCCCACAGAAAATGTACTTCGGG</td><td>1000</td><td></td><td></td><td></td><td></td><td></td><td></td><td></td><td></td><td></td></tr>
<tr><td>Oligo fwd</td><td>CACCTCCCACAGAAAATGTACTTC</td><td></td><td></td><td></td><td></td><td></td><td></td><td></td><td></td><td></td><td></td></tr>
<tr><td>Oligo rev</td><td>AAACGAAGTACATTTTCTGTGGGA</td><td></td><td></td><td></td><td></td><td></td><td></td><td></td><td></td><td></td><td></td></tr>
<tr><td>Chromosome</td><td>start</td><td>end</td><td>strand</td><td>MM</td><td>target_seq</td><td>PAM</td><td>alignment</td><td>distance</td><td>position</td><td>gene name</td><td>gene id</td></tr>
<tr><td>GU305914.1</td><td>208598</td><td>208620</td><td>+</td><td>0</td><td>TCCCACAGAAAATGTACTTC</td><td>GGG</td><td>||||||||[||||||||||||]PAM</td><td>NA</td><td>Intergenic</td><td>NA</td><td>NA</td></tr>
<tr><td>T15785</td><td>CTTCGGGATCATCGACCTTCAGG</td><td>1000</td><td></td><td></td><td></td><td></td><td></td><td></td><td></td><td></td><td></td></tr>
<tr><td>Oligo fwd</td><td>CACCCTTCGGGATCATCGACCTTC</td><td></td><td></td><td></td><td></td><td></td><td></td><td></td><td></td><td></td><td></td></tr>
<tr><td>Oligo rev</td><td>AAACGAAGGTCGATGATCCCGAAG</td><td></td><td></td><td></td><td></td><td></td><td></td><td></td><td></td><td></td><td></td></tr>
<tr><td>Chromosome</td><td>start</td><td>end</td><td>strand</td><td>MM</td><td>target_seq</td><td>PAM</td><td>alignment</td><td>distance</td><td>position</td><td>gene name</td><td>gene id</td></tr>
<tr><td>GU305914.1</td><td>208614</td><td>208636</td><td>+</td><td>0</td><td>CTTCGGGATCATCGACCTTC</td><td>AGG</td><td>||||||||[||||||||||||]PAM</td><td>NA</td><td>Intergenic</td><td>NA</td><td>NA</td></tr>
<tr><td>T15786</td><td>GGGATCATCGACCTTCAGGTCGG</td><td>1000</td><td></td><td></td><td></td><td></td><td></td><td></td><td></td><td></td><td></td></tr>
<tr><td>Oligo fwd</td><td>CACCGGGATCATCGACCTTCAGGT</td><td></td><td></td><td></td><td></td><td></td><td></td><td></td><td></td><td></td><td></td></tr>
<tr><td>Oligo rev</td><td>AAACACCTGAAGGTCGATGATCCC</td><td></td><td></td><td></td><td></td><td></td><td></td><td></td><td></td><td></td><td></td></tr>
<tr><td>Chromosome</td><td>start</td><td>end</td><td>strand</td><td>MM</td><td>target_seq</td><td>PAM</td><td>alignment</td><td>distance</td><td>position</td><td>gene name</td><td>gene id</td></tr>
<tr><td>GU305914.1</td><td>208618</td><td>208640</td><td>+</td><td>0</td><td>GGGATCATCGACCTTCAGGT</td><td>CGG</td><td>||||||||[||||||||||||]PAM</td><td>NA</td><td>Intergenic</td><td>NA</td><td>NA</td></tr>
<tr><td>T15787</td><td>GACCTTCAGGTCGGATACAGAGG</td><td>1000</td><td></td><td></td><td></td><td></td><td></td><td></td><td></td><td></td><td></td></tr>
<tr><td>Oligo fwd</td><td>CACCGACCTTCAGGTCGGATACAG</td><td></td><td></td><td></td><td></td><td></td><td></td><td></td><td></td><td></td><td></td></tr>
<tr><td>Oligo rev</td><td>AAACCTGTATCCGACCTGAAGGTC</td><td></td><td></td><td></td><td></td><td></td><td></td><td></td><td></td><td></td><td></td></tr>
<tr><td>Chromosome</td><td>start</td><td>end</td><td>strand</td><td>MM</td><td>target_seq</td><td>PAM</td><td>alignment</td><td>distance</td><td>position</td><td>gene name</td><td>gene id</td></tr>
<tr><td>GU305914.1</td><td>208627</td><td>208649</td><td>+</td><td>0</td><td>GACCTTCAGGTCGGATACAG</td><td>AGG</td><td>||||||||[||||||||||||]PAM</td><td>NA</td><td>Intergenic</td><td>NA</td><td>NA</td></tr>
<tr><td>T15788</td><td>ACCTTCAGGTCGGATACAGAGGG</td><td>1000</td><td></td><td></td><td></td><td></td><td></td><td></td><td></td><td></td><td></td></tr>
<tr><td>Oligo fwd</td><td>CACCACCTTCAGGTCGGATACAGA</td><td></td><td></td><td></td><td></td><td></td><td></td><td></td><td></td><td></td><td></td></tr>
<tr><td>Oligo rev</td><td>AAACTCTGTATCCGACCTGAAGGT</td><td></td><td></td><td></td><td></td><td></td><td></td><td></td><td></td><td></td><td></td></tr>
<tr><td>Chromosome</td><td>start</td><td>end</td><td>strand</td><td>MM</td><td>target_seq</td><td>PAM</td><td>alignment</td><td>distance</td><td>position</td><td>gene name</td><td>gene id</td></tr>
<tr><td>GU305914.1</td><td>208628</td><td>208650</td><td>+</td><td>0</td><td>ACCTTCAGGTCGGATACAGA</td><td>GGG</td><td>||||||||[||||||||||||]PAM</td><td>NA</td><td>Intergenic</td><td>NA</td><td>NA</td></tr>
</table>

T15789 GGTCGGATACAGAGGGATCGCGG 1000
Oligo fwd CACCGGTCGGATACAGAGGGATCG
Oligo rev AAACCGATCCCTCTGTATCCGACC
Chromosome start end strand MM target_seq PAM alignment distance position gene name gene id
GU305914.1 208635 208657 + 0 GGTCGGATACAGAGGGATCG CGG ||||||||[||||||||||||]PAM NA Intergenic NA NA

T15790 CGCGACCGTCTATTCTTACGAGG 1000
Oligo fwd CACCCGCGACCGTCTATTCTTACG
Oligo rev AAACCGTAAGAATAGACGGTCGCG
Chromosome start end strand MM target_seq PAM alignment distance position gene name gene id
GU305914.1 208671 208693 + 0 CGCGACCGTCTATTCTTACG AGG ||||||||[||||||||||||]PAM NA Intergenic NA NA

T15791 GCGACCGTCTATTCTTACGAGGG 1000
Oligo fwd CACCGCGACCGTCTATTCTTACGA
Oligo rev AAACTCGTAAGAATAGACGGTCGC
Chromosome start end strand MM target_seq PAM alignment distance position gene name gene id
GU305914.1 208672 208694 + 0 GCGACCGTCTATTCTTACGA GGG ||||||||[||||||||||||]PAM NA Intergenic NA NA

T15792 CGACCGTCTATTCTTACGAGGGG 1000
Oligo fwd CACCCGACCGTCTATTCTTACGAG
Oligo rev AAACCTCGTAAGAATAGACGGTCG
Chromosome start end strand MM target_seq PAM alignment distance position gene name gene id
GU305914.1 208673 208695 + 0 CGACCGTCTATTCTTACGAG GGG ||||||||[||||||||||||]PAM NA Intergenic NA NA

T15793 CGAGGGGAAAGCTATCGTTCAGG 1000
Oligo fwd CACCCGAGGGGAAAGCTATCGTTC
Oligo rev AAACGAACGATAGCTTTCCCCTCG
Chromosome start end strand MM target_seq PAM alignment distance position gene name gene id
GU305914.1 208689 208711 + 0 CGAGGGGAAAGCTATCGTTC AGG ||||||||[||||||||||||]PAM NA Intergenic NA NA

T15794 GAGGGGAAAGCTATCGTTCAGGG 1000
Oligo fwd CACCGAGGGGAAAGCTATCGTTCA
Oligo rev AAACTGAACGATAGCTTTCCCCTC
Chromosome start end strand MM target_seq PAM alignment distance position gene name gene id
GU305914.1 208690 208712 + 0 GAGGGGAAAGCTATCGTTCA GGG ||||||||[||||||||||||]PAM NA Intergenic NA NA

T15795 GCACGATGAAATAACACACCAGG 1000
Oligo fwd CACCGCACGATGAAATAACACACC
Oligo rev AAACGGTGTGTTATTTCATCGTGC
Chromosome start end strand MM target_seq PAM alignment distance position gene name gene id
GU305914.1 208785 208807 + 0 GCACGATGAAATAACACACC AGG ||||||||[||||||||||||]PAM NA Intergenic NA NA

T15796 TAACCCCCCACGTTCGCACACGG 1000
Oligo fwd CACCTAACCCCCCACGTTCGCACA
Oligo rev AAACTGTGCGAACGTGGGGGGTTA
Chromosome start end strand MM target_seq PAM alignment distance position gene name gene id
GU305914.1 208809 208831 + 0 TAACCCCCCACGTTCGCACA CGG ||||||||[||||||||||||]PAM NA Intergenic NA NA

T15797 CACCGCCATCACGACAGACGTGG 1000
Oligo fwd CACCCACCGCCATCACGACAGACG
Oligo rev AAACCGTCTGTCGTGATGGCGGTG
Chromosome start end strand MM target_seq PAM alignment distance position gene name gene id
GU305914.1 208834 208856 + 0 CACCGCCATCACGACAGACG TGG ||||||||[||||||||||||]PAM NA Intergenic NA NA

T15798 ATCGAAAGCTGAGCTCGCTGTGG 1000
Oligo fwd CACCATCGAAAGCTGAGCTCGCTG
Oligo rev AAACCAGCGAGCTCAGCTTTCGAT
Chromosome start end strand MM target_seq PAM alignment distance position gene name gene id
GU305914.1 208917 208939 + 0 ATCGAAAGCTGAGCTCGCTG TGG ||||||||[||||||||||||]PAM NA Intergenic NA NA

T15799 TCGAAAGCTGAGCTCGCTGTGGG 1000
Oligo fwd CACCTCGAAAGCTGAGCTCGCTGT
Oligo rev AAACACAGCGAGCTCAGCTTTCGA
Chromosome start end strand MM target_seq PAM alignment distance position gene name gene id
GU305914.1 208918 208940 + 0 TCGAAAGCTGAGCTCGCTGT GGG ||||||||[||||||||||||]PAM NA Intergenic NA NA

T15800 ACGTCGATTGCGTTCTCTTGCGG 1000
Oligo fwd CACCACGTCGATTGCGTTCTCTTG
Oligo rev AAACCAAGAGAACGCAATCGACGT
Chromosome start end strand MM target_seq PAM alignment distance position gene name gene id
GU305914.1 208941 208963 + 0 ACGTCGATTGCGTTCTCTTG CGG ||||||||[||||||||||||]PAM NA Intergenic NA NA

T15801 TTGCGTTCTCTTGCGGAACCTGG 1000
Oligo fwd CACCTTGCGTTCTCTTGCGGAACC
Oligo rev AAACGGTTCCGCAAGAGAACGCAA
Chromosome start end strand MM target_seq PAM alignment distance position gene name gene id
GU305914.1 208948 208970 + 0 TTGCGTTCTCTTGCGGAACC TGG ||||||||[||||||||||||]PAM NA Intergenic NA NA

T15802 GCAGCAGTCGTATCTTGTGTCGG 1000
Oligo fwd CACCGCAGCAGTCGTATCTTGTGT
Oligo rev AAACACACAAGATACGACTGCTGC
Chromosome start end strand MM target_seq PAM alignment distance position gene name gene id
GU305914.1 208989 209011 + 0 GCAGCAGTCGTATCTTGTGT CGG ||||||||[||||||||||||]PAM NA Intergenic NA NA

T15803 CTTGTGTCGGAAGATGACCACGG 1000
Oligo fwd CACCCTTGTGTCGGAAGATGACCA
Oligo rev AAACTGGTCATCTTCCGACACAAG
Chromosome start end strand MM target_seq PAM alignment distance position gene name gene id

GU305914.1 209002 209024 + 0 CTTGTGTCGGAAGATGACCA CGG ||||||||[||||||||||||]PAM NA Intergenic NA NA

T15804 GTGTCGGAAGATGACCACGGAGG 1000
Oligo fwd CACCGTGTCGGAAGATGACCACGG
Oligo rev AAACCCGTGGTCATCTTCCGACAC
Chromosome start end strand MM target_seq PAM alignment distance position gene name gene id
GU305914.1 209005 209027 + 0 GTGTCGGAAGATGACCACGG AGG ||||||||[||||||||||||]PAM NA Intergenic NA NA

T15805 TAGCACGACGAGTAGAACCGCGG 1000
Oligo fwd CACCTAGCACGACGAGTAGAACCG
Oligo rev AAACCGGTTCTACTCGTCGTGCTA
Chromosome start end strand MM target_seq PAM alignment distance position gene name gene id
GU305914.1 209029 209051 + 0 TAGCACGACGAGTAGAACCG CGG ||||||||[||||||||||||]PAM NA Intergenic NA NA

T15806 AGCACGACGAGTAGAACCGCGGG 1000
Oligo fwd CACCAGCACGACGAGTAGAACCGC
Oligo rev AAACGCGGTTCTACTCGTCGTGCT
Chromosome start end strand MM target_seq PAM alignment distance position gene name gene id
GU305914.1 209030 209052 + 0 AGCACGACGAGTAGAACCGC GGG ||||||||[||||||||||||]PAM NA Intergenic NA NA

T15807 GACGAGTAGAACCGCGGGAACGG 1000
Oligo fwd CACCGACGAGTAGAACCGCGGGAA
Oligo rev AAACTTCCCGCGGTTCTACTCGTC
Chromosome start end strand MM target_seq PAM alignment distance position gene name gene id
GU305914.1 209035 209057 + 0 GACGAGTAGAACCGCGGGAA CGG ||||||||[||||||||||||]PAM NA Intergenic NA NA
GU305914.1 124579 124601 + 4 TCCGAGTAGAACCGGTGGAA GGG --||||||[||||||--||||]PAM NA Intergenic NA NA

T15808 AACGGCGAGCAGATAGAGCAAGG 1000
Oligo fwd CACCAACGGCGAGCAGATAGAGCA
Oligo rev AAACTGCTCTATCTGCTCGCCGTT
Chromosome start end strand MM target_seq PAM alignment distance position gene name gene id
GU305914.1 209053 209075 + 0 AACGGCGAGCAGATAGAGCA AGG ||||||||[||||||||||||]PAM NA Intergenic NA NA

T15809 GACATGCGATGTCGCCTCTTCGG 1000
Oligo fwd CACCGACATGCGATGTCGCCTCTT
Oligo rev AAACAAGAGGCGACATCGCATGTC
Chromosome start end strand MM target_seq PAM alignment distance position gene name gene id
GU305914.1 209095 209117 + 0 GACATGCGATGTCGCCTCTT CGG ||||||||[||||||||||||]PAM NA Intergenic NA NA

T15810 GCTCACGACGATCGACCGATTGG 1000
Oligo fwd CACCGCTCACGACGATCGACCGAT
Oligo rev AAACATCGGTCGATCGTCGTGAGC
Chromosome start end strand MM target_seq PAM alignment distance position gene name gene id
GU305914.1 209122 209144 + 0 GCTCACGACGATCGACCGAT TGG ||||||||[||||||||||||]PAM NA Intergenic NA NA

T15811 GATCGACCGATTGGTGTATATGG 1000
Oligo fwd CACCGATCGACCGATTGGTGTATA
Oligo rev AAACTATACACCAATCGGTCGATC
Chromosome start end strand MM target_seq PAM alignment distance position gene name gene id
GU305914.1 209131 209153 + 0 GATCGACCGATTGGTGTATA TGG ||||||||[||||||||||||]PAM NA Intergenic NA NA

T15812 ATCGACCGATTGGTGTATATGGG 1000
Oligo fwd CACCATCGACCGATTGGTGTATAT
Oligo rev AAACATATACACCAATCGGTCGAT
Chromosome start end strand MM target_seq PAM alignment distance position gene name gene id
GU305914.1 209132 209154 + 0 ATCGACCGATTGGTGTATAT GGG ||||||||[||||||||||||]PAM NA Intergenic NA NA

T15813 CAGTTCTGAGCCGTTCGAGACGG 1000
Oligo fwd CACCCAGTTCTGAGCCGTTCGAGA
Oligo rev AAACTCTCGAACGGCTCAGAACTG
Chromosome start end strand MM target_seq PAM alignment distance position gene name gene id
GU305914.1 209164 209186 + 0 CAGTTCTGAGCCGTTCGAGA CGG ||||||||[||||||||||||]PAM NA Intergenic NA NA

T15814 CTGAGCCGTTCGAGACGGAGAGG 1000
Oligo fwd CACCCTGAGCCGTTCGAGACGGAG
Oligo rev AAACCTCCGTCTCGAACGGCTCAG
Chromosome start end strand MM target_seq PAM alignment distance position gene name gene id
GU305914.1 209169 209191 + 0 CTGAGCCGTTCGAGACGGAG AGG ||||||||[||||||||||||]PAM NA Intergenic NA NA

T15815 GCCGTTCGAGACGGAGAGGACGG 1000
Oligo fwd CACCGCCGTTCGAGACGGAGAGGA
Oligo rev AAACTCCTCTCCGTCTCGAACGGC
Chromosome start end strand MM target_seq PAM alignment distance position gene name gene id
GU305914.1 209173 209195 + 0 GCCGTTCGAGACGGAGAGGA CGG ||||||||[||||||||||||]PAM NA Intergenic NA NA

T15816 GAGACGGAGAGGACGGTGCGCGG 1000
Oligo fwd CACCGAGACGGAGAGGACGGTGCG
Oligo rev AAACCGCACCGTCCTCTCCGTCTC
Chromosome start end strand MM target_seq PAM alignment distance position gene name gene id
GU305914.1 209180 209202 + 0 GAGACGGAGAGGACGGTGCG CGG ||||||||[||||||||||||]PAM NA Intergenic NA NA

T15817 GAGAGGACGGTGCGCGGTGACGG 1000
Oligo fwd CACCGAGAGGACGGTGCGCGGTGA
Oligo rev AAACTCACCGCGCACCGTCCTCTC
Chromosome start end strand MM target_seq PAM alignment distance position gene name gene id
GU305914.1 209186 209208 + 0 GAGAGGACGGTGCGCGGTGA CGG ||||||||[||||||||||||]PAM NA Intergenic NA NA

T15818 CGGTGACGGACTGAGTTCTCCGG 1000
Oligo fwd CACCCGGTGACGGACTGAGTTCTC

<table>
<tr><td>Oligo rev</td><td>AAACGAGAACTCAGTCCGTCACCG</td><td></td><td></td><td></td><td></td><td></td><td></td><td></td><td></td><td></td><td></td></tr>
<tr><td>Chromosome</td><td>start</td><td>end</td><td>strand</td><td>MM</td><td>target_seq</td><td>PAM</td><td>alignment</td><td>distance</td><td>position</td><td>gene name</td><td>gene id</td></tr>
<tr><td>GU305914.1</td><td>209200</td><td>209222</td><td>+</td><td>0</td><td>CGGTGACGGACTGAGTTCTC</td><td>CGG</td><td>||||||||[||||||||||||]PAM</td><td>NA</td><td>Intergenic</td><td>NA</td><td>NA</td></tr>
<tr><td>T15819</td><td>GGTGACGGACTGAGTTCTCCGGG</td><td>1000</td><td></td><td></td><td></td><td></td><td></td><td></td><td></td><td></td><td></td></tr>
<tr><td>Oligo fwd</td><td>CACCGGTGACGGACTGAGTTCTCC</td><td></td><td></td><td></td><td></td><td></td><td></td><td></td><td></td><td></td><td></td></tr>
<tr><td>Oligo rev</td><td>AAACGGAGAACTCAGTCCGTCACC</td><td></td><td></td><td></td><td></td><td></td><td></td><td></td><td></td><td></td><td></td></tr>
<tr><td>Chromosome</td><td>start</td><td>end</td><td>strand</td><td>MM</td><td>target_seq</td><td>PAM</td><td>alignment</td><td>distance</td><td>position</td><td>gene name</td><td>gene id</td></tr>
<tr><td>GU305914.1</td><td>209201</td><td>209223</td><td>+</td><td>0</td><td>GGTGACGGACTGAGTTCTCC</td><td>GGG</td><td>||||||||[||||||||||||]PAM</td><td>NA</td><td>Intergenic</td><td>NA</td><td>NA</td></tr>
<tr><td>T15820</td><td>TGCCACACGTCTCTCCGTCGTGG</td><td>1000</td><td></td><td></td><td></td><td></td><td></td><td></td><td></td><td></td><td></td></tr>
<tr><td>Oligo fwd</td><td>CACCTGCCACACGTCTCTCCGTCG</td><td></td><td></td><td></td><td></td><td></td><td></td><td></td><td></td><td></td><td></td></tr>
<tr><td>Oligo rev</td><td>AAACCGACGGAGAGACGTGTGGCA</td><td></td><td></td><td></td><td></td><td></td><td></td><td></td><td></td><td></td><td></td></tr>
<tr><td>Chromosome</td><td>start</td><td>end</td><td>strand</td><td>MM</td><td>target_seq</td><td>PAM</td><td>alignment</td><td>distance</td><td>position</td><td>gene name</td><td>gene id</td></tr>
<tr><td>GU305914.1</td><td>209236</td><td>209258</td><td>+</td><td>0</td><td>TGCCACACGTCTCTCCGTCG</td><td>TGG</td><td>||||||||[||||||||||||]PAM</td><td>NA</td><td>Intergenic</td><td>NA</td><td>NA</td></tr>
<tr><td>GU305914.1</td><td>102791</td><td>102813</td><td>-</td><td>4</td><td>TCCCAGACGTCGCTTCGTCG</td><td>CGG</td><td>|-|||-||[|||-||-|||||]PAM</td><td>NA</td><td>Intergenic</td><td>NA</td><td>NA</td></tr>
<tr><td>T15821</td><td>CACACGTCTCTCCGTCGTGGCGG</td><td>1000</td><td></td><td></td><td></td><td></td><td></td><td></td><td></td><td></td><td></td></tr>
<tr><td>Oligo fwd</td><td>CACCCACACGTCTCTCCGTCGTGG</td><td></td><td></td><td></td><td></td><td></td><td></td><td></td><td></td><td></td><td></td></tr>
<tr><td>Oligo rev</td><td>AAACCCACGACGGAGAGACGTGTG</td><td></td><td></td><td></td><td></td><td></td><td></td><td></td><td></td><td></td><td></td></tr>
<tr><td>Chromosome</td><td>start</td><td>end</td><td>strand</td><td>MM</td><td>target_seq</td><td>PAM</td><td>alignment</td><td>distance</td><td>position</td><td>gene name</td><td>gene id</td></tr>
<tr><td>GU305914.1</td><td>209239</td><td>209261</td><td>+</td><td>0</td><td>CACACGTCTCTCCGTCGTGG</td><td>CGG</td><td>||||||||[||||||||||||]PAM</td><td>NA</td><td>Intergenic</td><td>NA</td><td>NA</td></tr>
<tr><td>T15822</td><td>GTCGTGGCGGCCGAACTGTTCGG</td><td>1000</td><td></td><td></td><td></td><td></td><td></td><td></td><td></td><td></td><td></td></tr>
<tr><td>Oligo fwd</td><td>CACCGTCGTGGCGGCCGAACTGTT</td><td></td><td></td><td></td><td></td><td></td><td></td><td></td><td></td><td></td><td></td></tr>
<tr><td>Oligo rev</td><td>AAACAACAGTTCGGCCGCCACGAC</td><td></td><td></td><td></td><td></td><td></td><td></td><td></td><td></td><td></td><td></td></tr>
<tr><td>Chromosome</td><td>start</td><td>end</td><td>strand</td><td>MM</td><td>target_seq</td><td>PAM</td><td>alignment</td><td>distance</td><td>position</td><td>gene name</td><td>gene id</td></tr>
<tr><td>GU305914.1</td><td>209252</td><td>209274</td><td>+</td><td>0</td><td>GTCGTGGCGGCCGAACTGTT</td><td>CGG</td><td>||||||||[||||||||||||]PAM</td><td>NA</td><td>Intergenic</td><td>NA</td><td>NA</td></tr>
<tr><td>T15823</td><td>CGGCCGAACTGTTCGGCTGCCGG</td><td>1000</td><td></td><td></td><td></td><td></td><td></td><td></td><td></td><td></td><td></td></tr>
<tr><td>Oligo fwd</td><td>CACCCGGCCGAACTGTTCGGCTGC</td><td></td><td></td><td></td><td></td><td></td><td></td><td></td><td></td><td></td><td></td></tr>
<tr><td>Oligo rev</td><td>AAACGCAGCCGAACAGTTCGGCCG</td><td></td><td></td><td></td><td></td><td></td><td></td><td></td><td></td><td></td><td></td></tr>
<tr><td>Chromosome</td><td>start</td><td>end</td><td>strand</td><td>MM</td><td>target_seq</td><td>PAM</td><td>alignment</td><td>distance</td><td>position</td><td>gene name</td><td>gene id</td></tr>
<tr><td>GU305914.1</td><td>209259</td><td>209281</td><td>+</td><td>0</td><td>CGGCCGAACTGTTCGGCTGC</td><td>CGG</td><td>||||||||[||||||||||||]PAM</td><td>NA</td><td>Intergenic</td><td>NA</td><td>NA</td></tr>
<tr><td>T15824</td><td>CGAACTGTTCGGCTGCCGGATGG</td><td>1000</td><td></td><td></td><td></td><td></td><td></td><td></td><td></td><td></td><td></td></tr>
<tr><td>Oligo fwd</td><td>CACCCGAACTGTTCGGCTGCCGGA</td><td></td><td></td><td></td><td></td><td></td><td></td><td></td><td></td><td></td><td></td></tr>
<tr><td>Oligo rev</td><td>AAACTCCGGCAGCCGAACAGTTCG</td><td></td><td></td><td></td><td></td><td></td><td></td><td></td><td></td><td></td><td></td></tr>
<tr><td>Chromosome</td><td>start</td><td>end</td><td>strand</td><td>MM</td><td>target_seq</td><td>PAM</td><td>alignment</td><td>distance</td><td>position</td><td>gene name</td><td>gene id</td></tr>
<tr><td>GU305914.1</td><td>209263</td><td>209285</td><td>+</td><td>0</td><td>CGAACTGTTCGGCTGCCGGA</td><td>TGG</td><td>||||||||[||||||||||||]PAM</td><td>NA</td><td>Intergenic</td><td>NA</td><td>NA</td></tr>
<tr><td>T15825</td><td>GATGGTGACGTTCCAGCCCGCGG</td><td>1000</td><td></td><td></td><td></td><td></td><td></td><td></td><td></td><td></td><td></td></tr>
<tr><td>Oligo fwd</td><td>CACCGATGGTGACGTTCCAGCCCG</td><td></td><td></td><td></td><td></td><td></td><td></td><td></td><td></td><td></td><td></td></tr>
<tr><td>Oligo rev</td><td>AAACCGGGCTGGAACGTCACCATC</td><td></td><td></td><td></td><td></td><td></td><td></td><td></td><td></td><td></td><td></td></tr>
<tr><td>Chromosome</td><td>start</td><td>end</td><td>strand</td><td>MM</td><td>target_seq</td><td>PAM</td><td>alignment</td><td>distance</td><td>position</td><td>gene name</td><td>gene id</td></tr>
<tr><td>GU305914.1</td><td>209281</td><td>209303</td><td>+</td><td>0</td><td>GATGGTGACGTTCCAGCCCG</td><td>CGG</td><td>||||||||[||||||||||||]PAM</td><td>NA</td><td>Intergenic</td><td>NA</td><td>NA</td></tr>
<tr><td>T15826</td><td>GAGCATATTATCTCGCACCGCGG</td><td>1000</td><td></td><td></td><td></td><td></td><td></td><td></td><td></td><td></td><td></td></tr>
<tr><td>Oligo fwd</td><td>CACCGAGCATATTATCTCGCACCG</td><td></td><td></td><td></td><td></td><td></td><td></td><td></td><td></td><td></td><td></td></tr>
<tr><td>Oligo rev</td><td>AAACCGGTGCGAGATAATATGCTC</td><td></td><td></td><td></td><td></td><td></td><td></td><td></td><td></td><td></td><td></td></tr>
<tr><td>Chromosome</td><td>start</td><td>end</td><td>strand</td><td>MM</td><td>target_seq</td><td>PAM</td><td>alignment</td><td>distance</td><td>position</td><td>gene name</td><td>gene id</td></tr>
<tr><td>GU305914.1</td><td>209303</td><td>209325</td><td>+</td><td>0</td><td>GAGCATATTATCTCGCACCG</td><td>CGG</td><td>||||||||[||||||||||||]PAM</td><td>NA</td><td>Intergenic</td><td>NA</td><td>NA</td></tr>
<tr><td>T15827</td><td>TCGCACCGCGGTGAGTTGTTCGG</td><td>1000</td><td></td><td></td><td></td><td></td><td></td><td></td><td></td><td></td><td></td></tr>
<tr><td>Oligo fwd</td><td>CACCTCGCACCGCGGTGAGTTGTT</td><td></td><td></td><td></td><td></td><td></td><td></td><td></td><td></td><td></td><td></td></tr>
<tr><td>Oligo rev</td><td>AAACAACAACTCACCGCGGTGCGA</td><td></td><td></td><td></td><td></td><td></td><td></td><td></td><td></td><td></td><td></td></tr>
<tr><td>Chromosome</td><td>start</td><td>end</td><td>strand</td><td>MM</td><td>target_seq</td><td>PAM</td><td>alignment</td><td>distance</td><td>position</td><td>gene name</td><td>gene id</td></tr>
<tr><td>GU305914.1</td><td>209315</td><td>209337</td><td>+</td><td>0</td><td>TCGCACCGCGGTGAGTTGTT</td><td>CGG</td><td>||||||||[||||||||||||]PAM</td><td>NA</td><td>Intergenic</td><td>NA</td><td>NA</td></tr>
<tr><td>T15828</td><td>CGCACCGCGGTGAGTTGTTCGGG</td><td>1000</td><td></td><td></td><td></td><td></td><td></td><td></td><td></td><td></td><td></td></tr>
<tr><td>Oligo fwd</td><td>CACCCGCACCGCGGTGAGTTGTTC</td><td></td><td></td><td></td><td></td><td></td><td></td><td></td><td></td><td></td><td></td></tr>
<tr><td>Oligo rev</td><td>AAACGAACAACTCACCGCGGTGCG</td><td></td><td></td><td></td><td></td><td></td><td></td><td></td><td></td><td></td><td></td></tr>
<tr><td>Chromosome</td><td>start</td><td>end</td><td>strand</td><td>MM</td><td>target_seq</td><td>PAM</td><td>alignment</td><td>distance</td><td>position</td><td>gene name</td><td>gene id</td></tr>
<tr><td>GU305914.1</td><td>209316</td><td>209338</td><td>+</td><td>0</td><td>CGCACCGCGGTGAGTTGTTC</td><td>GGG</td><td>||||||||[||||||||||||]PAM</td><td>NA</td><td>Intergenic</td><td>NA</td><td>NA</td></tr>
<tr><td>T15829</td><td>GCACCGCGGTGAGTTGTTCGGGG</td><td>1000</td><td></td><td></td><td></td><td></td><td></td><td></td><td></td><td></td><td></td></tr>
<tr><td>Oligo fwd</td><td>CACCGCACCGCGGTGAGTTGTTCG</td><td></td><td></td><td></td><td></td><td></td><td></td><td></td><td></td><td></td><td></td></tr>
<tr><td>Oligo rev</td><td>AAACCGAACAACTCACCGCGGTGC</td><td></td><td></td><td></td><td></td><td></td><td></td><td></td><td></td><td></td><td></td></tr>
<tr><td>Chromosome</td><td>start</td><td>end</td><td>strand</td><td>MM</td><td>target_seq</td><td>PAM</td><td>alignment</td><td>distance</td><td>position</td><td>gene name</td><td>gene id</td></tr>
<tr><td>GU305914.1</td><td>209317</td><td>209339</td><td>+</td><td>0</td><td>GCACCGCGGTGAGTTGTTCG</td><td>GGG</td><td>||||||||[||||||||||||]PAM</td><td>NA</td><td>Intergenic</td><td>NA</td><td>NA</td></tr>
<tr><td>T15830</td><td>TGATGCATTCAGCTGAACATCGG</td><td>1000</td><td></td><td></td><td></td><td></td><td></td><td></td><td></td><td></td><td></td></tr>
<tr><td>Oligo fwd</td><td>CACCTGATGCATTCAGCTGAACAT</td><td></td><td></td><td></td><td></td><td></td><td></td><td></td><td></td><td></td><td></td></tr>
<tr><td>Oligo rev</td><td>AAACATGTTCAGCTGAATGCATCA</td><td></td><td></td><td></td><td></td><td></td><td></td><td></td><td></td><td></td><td></td></tr>
<tr><td>Chromosome</td><td>start</td><td>end</td><td>strand</td><td>MM</td><td>target_seq</td><td>PAM</td><td>alignment</td><td>distance</td><td>position</td><td>gene name</td><td>gene id</td></tr>
<tr><td>GU305914.1</td><td>209395</td><td>209417</td><td>+</td><td>0</td><td>TGATGCATTCAGCTGAACAT</td><td>CGG</td><td>||||||||[||||||||||||]PAM</td><td>NA</td><td>Intergenic</td><td>NA</td><td>NA</td></tr>
<tr><td>T15831</td><td>ATAGTCATCCACAGTATCACCGG</td><td>1000</td><td></td><td></td><td></td><td></td><td></td><td></td><td></td><td></td><td></td></tr>
<tr><td>Oligo fwd</td><td>CACCATAGTCATCCACAGTATCAC</td><td></td><td></td><td></td><td></td><td></td><td></td><td></td><td></td><td></td><td></td></tr>
<tr><td>Oligo rev</td><td>AAACGTGATACTGTGGATGACTAT</td><td></td><td></td><td></td><td></td><td></td><td></td><td></td><td></td><td></td><td></td></tr>
<tr><td>Chromosome</td><td>start</td><td>end</td><td>strand</td><td>MM</td><td>target_seq</td><td>PAM</td><td>alignment</td><td>distance</td><td>position</td><td>gene name</td><td>gene id</td></tr>
<tr><td>GU305914.1</td><td>209426</td><td>209448</td><td>+</td><td>0</td><td>ATAGTCATCCACAGTATCAC</td><td>CGG</td><td>||||||||[||||||||||||]PAM</td><td>NA</td><td>Intergenic</td><td>NA</td><td>NA</td></tr>
<tr><td>T15832</td><td>GTTCGTCATACTAGAGAAACTGG</td><td>1000</td><td></td><td></td><td></td><td></td><td></td><td></td><td></td><td></td><td></td></tr>
<tr><td>Oligo fwd</td><td>CACCGTTCGTCATACTAGAGAAAC</td><td></td><td></td><td></td><td></td><td></td><td></td><td></td><td></td><td></td><td></td></tr>
<tr><td>Oligo rev</td><td>AAACGTTTCTCTAGTATGACGAAC</td><td></td><td></td><td></td><td></td><td></td><td></td><td></td><td></td><td></td><td></td></tr>
<tr><td>Chromosome</td><td>start</td><td>end</td><td>strand</td><td>MM</td><td>target_seq</td><td>PAM</td><td>alignment</td><td>distance</td><td>position</td><td>gene name</td><td>gene id</td></tr>
<tr><td>GU305914.1</td><td>209479</td><td>209501</td><td>+</td><td>0</td><td>GTTCGTCATACTAGAGAAAC</td><td>TGG</td><td>||||||||[||||||||||||]PAM</td><td>NA</td><td>Intergenic</td><td>NA</td><td>NA</td></tr>
</table>

| | | | | | | | | | | | |
|---|---|---|---|---|---|---|---|---|---|---|---|
| T15833 | ACTGGACACCGACCGTTCTTTGG | 1000 | | | | | | | | | |
| Oligo fwd | CACCACTGGACACCGACCGTTCTT | | | | | | | | | | |
| Oligo rev | AAACAAGAACGGTCGGTGTCCAGT | | | | | | | | | | |
| Chromosome | start | end | strand | MM | target_seq | PAM | alignment | distance | position | gene name | gene id |
| GU305914.1 | 209497 | 209519 | + | 0 | ACTGGACACCGACCGTTCTT | TGG | \|\|\|\|\|\|\|\|[\|\|\|\|\|\|\|\|\|\|\|\|]PAM | NA | Intergenic | NA | NA |
| T15834 | TCGAATATCACCACTGTGTGTGG | 1000 | | | | | | | | | |
| Oligo fwd | CACCTCGAATATCACCACTGTGTG | | | | | | | | | | |
| Oligo rev | AAACCACACAGTGGTGATATTCGA | | | | | | | | | | |
| Chromosome | start | end | strand | MM | target_seq | PAM | alignment | distance | position | gene name | gene id |
| GU305914.1 | 209563 | 209585 | + | 0 | TCGAATATCACCACTGTGTG | TGG | \|\|\|\|\|\|\|\|[\|\|\|\|\|\|\|\|\|\|\|\|]PAM | NA | Intergenic | NA | NA |
| T15835 | TATCACCACTGTGTGTGGTTAGG | 1000 | | | | | | | | | |
| Oligo fwd | CACCTATCACCACTGTGTGTGGTT | | | | | | | | | | |
| Oligo rev | AAACAACCACACACAGTGGTGATA | | | | | | | | | | |
| Chromosome | start | end | strand | MM | target_seq | PAM | alignment | distance | position | gene name | gene id |
| GU305914.1 | 209568 | 209590 | + | 0 | TATCACCACTGTGTGTGGTT | AGG | \|\|\|\|\|\|\|\|[\|\|\|\|\|\|\|\|\|\|\|\|]PAM | NA | Intergenic | NA | NA |
| T15836 | TACCCCATACCCTGAAGCCATGG | 1000 | | | | | | | | | |
| Oligo fwd | CACCTACCCCATACCCTGAAGCCA | | | | | | | | | | |
| Oligo rev | AAACTGGCTTCAGGGTATGGGGTA | | | | | | | | | | |
| Chromosome | start | end | strand | MM | target_seq | PAM | alignment | distance | position | gene name | gene id |
| GU305914.1 | 209596 | 209618 | + | 0 | TACCCCATACCCTGAAGCCA | TGG | \|\|\|\|\|\|\|\|[\|\|\|\|\|\|\|\|\|\|\|\|]PAM | NA | Intergenic | NA | NA |
| T15837 | TCACCATCATCACTGTCGAAAGG | 1000 | | | | | | | | | |
| Oligo fwd | CACCTCACCATCATCACTGTCGAA | | | | | | | | | | |
| Oligo rev | AAACTTCGACAGTGATGATGGTGA | | | | | | | | | | |
| Chromosome | start | end | strand | MM | target_seq | PAM | alignment | distance | position | gene name | gene id |
| GU305914.1 | 209651 | 209673 | + | 0 | TCACCATCATCACTGTCGAA | AGG | \|\|\|\|\|\|\|\|[\|\|\|\|\|\|\|\|\|\|\|\|]PAM | NA | Intergenic | NA | NA |
| T15838 | ATCATCACTGTCGAAAGGCGAGG | 1000 | | | | | | | | | |
| Oligo fwd | CACCATCATCACTGTCGAAAGGCG | | | | | | | | | | |
| Oligo rev | AAACCGCCTTTCGACAGTGATGAT | | | | | | | | | | |
| Chromosome | start | end | strand | MM | target_seq | PAM | alignment | distance | position | gene name | gene id |
| GU305914.1 | 209656 | 209678 | + | 0 | ATCATCACTGTCGAAAGGCG | AGG | \|\|\|\|\|\|\|\|[\|\|\|\|\|\|\|\|\|\|\|\|]PAM | NA | Intergenic | NA | NA |
| T15839 | GAAAGGCGAGGATATTAAGTTGG | 1000 | | | | | | | | | |
| Oligo fwd | CACCGAAAGGCGAGGATATTAAGT | | | | | | | | | | |
| Oligo rev | AAACACTTAATATCCTCGCCTTTC | | | | | | | | | | |
| Chromosome | start | end | strand | MM | target_seq | PAM | alignment | distance | position | gene name | gene id |
| GU305914.1 | 209668 | 209690 | + | 0 | GAAAGGCGAGGATATTAAGT | TGG | \|\|\|\|\|\|\|\|[\|\|\|\|\|\|\|\|\|\|\|\|]PAM | NA | Intergenic | NA | NA |
| T15840 | ATTATAATATAACACATCCGTGG | 1000 | | | | | | | | | |
| Oligo fwd | CACCATTATAATATAACACATCCG | | | | | | | | | | |
| Oligo rev | AAACCGGATGTGTTATATTATAAT | | | | | | | | | | |
| Chromosome | start | end | strand | MM | target_seq | PAM | alignment | distance | position | gene name | gene id |
| GU305914.1 | 209701 | 209723 | + | 0 | ATTATAATATAACACATCCG | TGG | \|\|\|\|\|\|\|\|[\|\|\|\|\|\|\|\|\|\|\|\|]PAM | NA | Intergenic | NA | NA |
| T15841 | AGTAGAAATTCACCGATAGATGG | 1000 | | | | | | | | | |
| Oligo fwd | CACCAGTAGAAATTCACCGATAGA | | | | | | | | | | |
| Oligo rev | AAACTCTATCGGTGAATTTCTACT | | | | | | | | | | |
| Chromosome | start | end | strand | MM | target_seq | PAM | alignment | distance | position | gene name | gene id |
| GU305914.1 | 209748 | 209770 | + | 0 | AGTAGAAATTCACCGATAGA | TGG | \|\|\|\|\|\|\|\|[\|\|\|\|\|\|\|\|\|\|\|\|]PAM | NA | Intergenic | NA | NA |
| T15842 | CACACTTATCTCACCAGCCTCGG | 1000 | | | | | | | | | |
| Oligo fwd | CACCCACACTTATCTCACCAGCCT | | | | | | | | | | |
| Oligo rev | AAACAGGCTGGTGAGATAAGTGTG | | | | | | | | | | |
| Chromosome | start | end | strand | MM | target_seq | PAM | alignment | distance | position | gene name | gene id |
| GU305914.1 | 209782 | 209804 | + | 0 | CACACTTATCTCACCAGCCT | CGG | \|\|\|\|\|\|\|\|[\|\|\|\|\|\|\|\|\|\|\|\|]PAM | NA | Intergenic | NA | NA |
| T15843 | CAGCCTCGGAAAGCAGTCCTCGG | 1000 | | | | | | | | | |
| Oligo fwd | CACCCAGCCTCGGAAAGCAGTCCT | | | | | | | | | | |
| Oligo rev | AAACAGGACTGCTTTCCGAGGCTG | | | | | | | | | | |
| Chromosome | start | end | strand | MM | target_seq | PAM | alignment | distance | position | gene name | gene id |
| GU305914.1 | 209796 | 209818 | + | 0 | CAGCCTCGGAAAGCAGTCCT | CGG | \|\|\|\|\|\|\|\|[\|\|\|\|\|\|\|\|\|\|\|\|]PAM | NA | Intergenic | NA | NA |
| T15844 | CGGAAAGCAGTCCTCGGACTTGG | 1000 | | | | | | | | | |
| Oligo fwd | CACCCGGAAAGCAGTCCTCGGACT | | | | | | | | | | |
| Oligo rev | AAACAGTCCGAGGACTGCTTTCCG | | | | | | | | | | |
| Chromosome | start | end | strand | MM | target_seq | PAM | alignment | distance | position | gene name | gene id |
| GU305914.1 | 209802 | 209824 | + | 0 | CGGAAAGCAGTCCTCGGACT | TGG | \|\|\|\|\|\|\|\|[\|\|\|\|\|\|\|\|\|\|\|\|]PAM | NA | Intergenic | NA | NA |
| T15845 | GGAAAGCAGTCCTCGGACTTGGG | 1000 | | | | | | | | | |
| Oligo fwd | CACCGGAAAGCAGTCCTCGGACTT | | | | | | | | | | |
| Oligo rev | AAACAAGTCCGAGGACTGCTTTCC | | | | | | | | | | |
| Chromosome | start | end | strand | MM | target_seq | PAM | alignment | distance | position | gene name | gene id |
| GU305914.1 | 209803 | 209825 | + | 0 | GGAAAGCAGTCCTCGGACTT | GGG | \|\|\|\|\|\|\|\|[\|\|\|\|\|\|\|\|\|\|\|\|]PAM | NA | Intergenic | NA | NA |
| T15846 | CCTTGCATGAAAATGAATTCAGG | 1000 | | | | | | | | | |
| Oligo fwd | CACCCCTTGCATGAAAATGAATTC | | | | | | | | | | |
| Oligo rev | AAACGAATTCATTTTCATGCAAGG | | | | | | | | | | |
| Chromosome | start | end | strand | MM | target_seq | PAM | alignment | distance | position | gene name | gene id |
| GU305914.1 | 209843 | 209865 | + | 0 | CCTTGCATGAAAATGAATTC | AGG | \|\|\|\|\|\|\|\|[\|\|\|\|\|\|\|\|\|\|\|\|]PAM | NA | Intergenic | NA | NA |
| T15847 | GACCCACCCTGTACCACAGACGG | 1000 | | | | | | | | | |
| Oligo fwd | CACCGACCCACCCTGTACCACAGA | | | | | | | | | | |
| Oligo rev | AAACTCTGTGGTACAGGGTGGGTC | | | | | | | | | | |
| Chromosome | start | end | strand | MM | target_seq | PAM | alignment | distance | position | gene name | gene id |
| GU305914.1 | 209884 | 209906 | + | 0 | GACCCACCCTGTACCACAGA | CGG | \|\|\|\|\|\|\|\|[\|\|\|\|\|\|\|\|\|\|\|\|]PAM | NA | Intergenic | NA | NA |

<table>
<tr><td>T15848</td><td>TTATCGTACACCGTCAACAGCGG</td><td>1000</td><td></td><td></td><td></td><td></td><td></td><td></td><td></td><td></td><td></td></tr>
<tr><td>Oligo fwd</td><td>CACCTTATCGTACACCGTCAACAG</td><td></td><td></td><td></td><td></td><td></td><td></td><td></td><td></td><td></td><td></td></tr>
<tr><td>Oligo rev</td><td>AAACCTGTTGACGGTGTACGATAA</td><td></td><td></td><td></td><td></td><td></td><td></td><td></td><td></td><td></td><td></td></tr>
<tr><td>Chromosome</td><td>start</td><td>end</td><td>strand</td><td>MM</td><td>target_seq</td><td>PAM</td><td>alignment</td><td>distance</td><td>position</td><td>gene name</td><td>gene id</td></tr>
<tr><td>GU305914.1</td><td>209912</td><td>209934</td><td>+</td><td>0</td><td>TTATCGTACACCGTCAACAG</td><td>CGG</td><td>||||||||[||||||||||||]PAM</td><td>NA</td><td>Intergenic</td><td>NA</td><td>NA</td></tr>
<tr><td>T15849</td><td>CACTGTCTGCTTTAGACCGACGG</td><td>1000</td><td></td><td></td><td></td><td></td><td></td><td></td><td></td><td></td><td></td></tr>
<tr><td>Oligo fwd</td><td>CACCCACTGTCTGCTTTAGACCGA</td><td></td><td></td><td></td><td></td><td></td><td></td><td></td><td></td><td></td><td></td></tr>
<tr><td>Oligo rev</td><td>AAACTCGGTCTAAAGCAGACAGTG</td><td></td><td></td><td></td><td></td><td></td><td></td><td></td><td></td><td></td><td></td></tr>
<tr><td>Chromosome</td><td>start</td><td>end</td><td>strand</td><td>MM</td><td>target_seq</td><td>PAM</td><td>alignment</td><td>distance</td><td>position</td><td>gene name</td><td>gene id</td></tr>
<tr><td>GU305914.1</td><td>209935</td><td>209957</td><td>+</td><td>0</td><td>CACTGTCTGCTTTAGACCGA</td><td>CGG</td><td>||||||||[||||||||||||]PAM</td><td>NA</td><td>Intergenic</td><td>NA</td><td>NA</td></tr>
<tr><td>T15850</td><td>TGTGACCGCATCATGAAGCCCGG</td><td>1000</td><td></td><td></td><td></td><td></td><td></td><td></td><td></td><td></td><td></td></tr>
<tr><td>Oligo fwd</td><td>CACCTGTGACCGCATCATGAAGCC</td><td></td><td></td><td></td><td></td><td></td><td></td><td></td><td></td><td></td><td></td></tr>
<tr><td>Oligo rev</td><td>AAACGGCTTCATGATGCGGTCACA</td><td></td><td></td><td></td><td></td><td></td><td></td><td></td><td></td><td></td><td></td></tr>
<tr><td>Chromosome</td><td>start</td><td>end</td><td>strand</td><td>MM</td><td>target_seq</td><td>PAM</td><td>alignment</td><td>distance</td><td>position</td><td>gene name</td><td>gene id</td></tr>
<tr><td>GU305914.1</td><td>209962</td><td>209984</td><td>+</td><td>0</td><td>TGTGACCGCATCATGAAGCC</td><td>CGG</td><td>||||||||[||||||||||||]PAM</td><td>NA</td><td>Intergenic</td><td>NA</td><td>NA</td></tr>
<tr><td>T15851</td><td>CCGTCGACGCCGTAAGAACGAGG</td><td>1000</td><td></td><td></td><td></td><td></td><td></td><td></td><td></td><td></td><td></td></tr>
<tr><td>Oligo fwd</td><td>CACCCCGTCGACGCCGTAAGAACG</td><td></td><td></td><td></td><td></td><td></td><td></td><td></td><td></td><td></td><td></td></tr>
<tr><td>Oligo rev</td><td>AAACCGTTCTTACGGCGTCGACGG</td><td></td><td></td><td></td><td></td><td></td><td></td><td></td><td></td><td></td><td></td></tr>
<tr><td>Chromosome</td><td>start</td><td>end</td><td>strand</td><td>MM</td><td>target_seq</td><td>PAM</td><td>alignment</td><td>distance</td><td>position</td><td>gene name</td><td>gene id</td></tr>
<tr><td>GU305914.1</td><td>209985</td><td>210007</td><td>+</td><td>0</td><td>CCGTCGACGCCGTAAGAACG</td><td>AGG</td><td>||||||||[||||||||||||]PAM</td><td>NA</td><td>Intergenic</td><td>NA</td><td>NA</td></tr>
<tr><td>T15852</td><td>GACGCCGTAAGAACGAGGACCGG</td><td>1000</td><td></td><td></td><td></td><td></td><td></td><td></td><td></td><td></td><td></td></tr>
<tr><td>Oligo fwd</td><td>CACCGACGCCGTAAGAACGAGGAC</td><td></td><td></td><td></td><td></td><td></td><td></td><td></td><td></td><td></td><td></td></tr>
<tr><td>Oligo rev</td><td>AAACGTCCTCGTTCTTACGGCGTC</td><td></td><td></td><td></td><td></td><td></td><td></td><td></td><td></td><td></td><td></td></tr>
<tr><td>Chromosome</td><td>start</td><td>end</td><td>strand</td><td>MM</td><td>target_seq</td><td>PAM</td><td>alignment</td><td>distance</td><td>position</td><td>gene name</td><td>gene id</td></tr>
<tr><td>GU305914.1</td><td>209990</td><td>210012</td><td>+</td><td>0</td><td>GACGCCGTAAGAACGAGGAC</td><td>CGG</td><td>||||||||[||||||||||||]PAM</td><td>NA</td><td>Intergenic</td><td>NA</td><td>NA</td></tr>
<tr><td>T15853</td><td>GACCGGCGACTCACACACAGAGG</td><td>1000</td><td></td><td></td><td></td><td></td><td></td><td></td><td></td><td></td><td></td></tr>
<tr><td>Oligo fwd</td><td>CACCGACCGGCGACTCACACACAG</td><td></td><td></td><td></td><td></td><td></td><td></td><td></td><td></td><td></td><td></td></tr>
<tr><td>Oligo rev</td><td>AAACCTGTGTGTGAGTCGCCGGTC</td><td></td><td></td><td></td><td></td><td></td><td></td><td></td><td></td><td></td><td></td></tr>
<tr><td>Chromosome</td><td>start</td><td>end</td><td>strand</td><td>MM</td><td>target_seq</td><td>PAM</td><td>alignment</td><td>distance</td><td>position</td><td>gene name</td><td>gene id</td></tr>
<tr><td>GU305914.1</td><td>210007</td><td>210029</td><td>+</td><td>0</td><td>GACCGGCGACTCACACACAG</td><td>AGG</td><td>||||||||[||||||||||||]PAM</td><td>NA</td><td>Intergenic</td><td>NA</td><td>NA</td></tr>
<tr><td>T15854</td><td>CGACTCACACACAGAGGCTGCGG</td><td>1000</td><td></td><td></td><td></td><td></td><td></td><td></td><td></td><td></td><td></td></tr>
<tr><td>Oligo fwd</td><td>CACCCGACTCACACACAGAGGCTG</td><td></td><td></td><td></td><td></td><td></td><td></td><td></td><td></td><td></td><td></td></tr>
<tr><td>Oligo rev</td><td>AAACCAGCCTCTGTGTGTGAGTCG</td><td></td><td></td><td></td><td></td><td></td><td></td><td></td><td></td><td></td><td></td></tr>
<tr><td>Chromosome</td><td>start</td><td>end</td><td>strand</td><td>MM</td><td>target_seq</td><td>PAM</td><td>alignment</td><td>distance</td><td>position</td><td>gene name</td><td>gene id</td></tr>
<tr><td>GU305914.1</td><td>210013</td><td>210035</td><td>+</td><td>0</td><td>CGACTCACACACAGAGGCTG</td><td>CGG</td><td>||||||||[||||||||||||]PAM</td><td>NA</td><td>Intergenic</td><td>NA</td><td>NA</td></tr>
<tr><td>T15855</td><td>CGTTCAATGAGTCACCACAGCGG</td><td>1000</td><td></td><td></td><td></td><td></td><td></td><td></td><td></td><td></td><td></td></tr>
<tr><td>Oligo fwd</td><td>CACCCGTTCAATGAGTCACCACAG</td><td></td><td></td><td></td><td></td><td></td><td></td><td></td><td></td><td></td><td></td></tr>
<tr><td>Oligo rev</td><td>AAACCTGTGGTGACTCATTGAACG</td><td></td><td></td><td></td><td></td><td></td><td></td><td></td><td></td><td></td><td></td></tr>
<tr><td>Chromosome</td><td>start</td><td>end</td><td>strand</td><td>MM</td><td>target_seq</td><td>PAM</td><td>alignment</td><td>distance</td><td>position</td><td>gene name</td><td>gene id</td></tr>
<tr><td>GU305914.1</td><td>210124</td><td>210146</td><td>+</td><td>0</td><td>CGTTCAATGAGTCACCACAG</td><td>CGG</td><td>||||||||[||||||||||||]PAM</td><td>NA</td><td>Intergenic</td><td>NA</td><td>NA</td></tr>
<tr><td>T15856</td><td>TTATCCCTGAGCCCATTCCCAGG</td><td>1000</td><td></td><td></td><td></td><td></td><td></td><td></td><td></td><td></td><td></td></tr>
<tr><td>Oligo fwd</td><td>CACCTTATCCCTGAGCCCATTCCC</td><td></td><td></td><td></td><td></td><td></td><td></td><td></td><td></td><td></td><td></td></tr>
<tr><td>Oligo rev</td><td>AAACGGGAATGGGCTCAGGGATAA</td><td></td><td></td><td></td><td></td><td></td><td></td><td></td><td></td><td></td><td></td></tr>
<tr><td>Chromosome</td><td>start</td><td>end</td><td>strand</td><td>MM</td><td>target_seq</td><td>PAM</td><td>alignment</td><td>distance</td><td>position</td><td>gene name</td><td>gene id</td></tr>
<tr><td>GU305914.1</td><td>210195</td><td>210217</td><td>+</td><td>0</td><td>TTATCCCTGAGCCCATTCCC</td><td>AGG</td><td>||||||||[||||||||||||]PAM</td><td>NA</td><td>Intergenic</td><td>NA</td><td>NA</td></tr>
<tr><td>T15857</td><td>CCGCCCCAGATTCCCCACCGAGG</td><td>1000</td><td></td><td></td><td></td><td></td><td></td><td></td><td></td><td></td><td></td></tr>
<tr><td>Oligo fwd</td><td>CACCCCGCCCCAGATTCCCCACCG</td><td></td><td></td><td></td><td></td><td></td><td></td><td></td><td></td><td></td><td></td></tr>
<tr><td>Oligo rev</td><td>AAACCGGTGGGGAATCTGGGGCGG</td><td></td><td></td><td></td><td></td><td></td><td></td><td></td><td></td><td></td><td></td></tr>
<tr><td>Chromosome</td><td>start</td><td>end</td><td>strand</td><td>MM</td><td>target_seq</td><td>PAM</td><td>alignment</td><td>distance</td><td>position</td><td>gene name</td><td>gene id</td></tr>
<tr><td>GU305914.1</td><td>210223</td><td>210245</td><td>+</td><td>0</td><td>CCGCCCCAGATTCCCCACCG</td><td>AGG</td><td>||||||||[||||||||||||]PAM</td><td>NA</td><td>Intergenic</td><td>NA</td><td>NA</td></tr>
<tr><td>T15858</td><td>CATTGTCCAGTTAGTGTATGTGG</td><td>1000</td><td></td><td></td><td></td><td></td><td></td><td></td><td></td><td></td><td></td></tr>
<tr><td>Oligo fwd</td><td>CACCCATTGTCCAGTTAGTGTATG</td><td></td><td></td><td></td><td></td><td></td><td></td><td></td><td></td><td></td><td></td></tr>
<tr><td>Oligo rev</td><td>AAACCATACACTAACTGGACAATG</td><td></td><td></td><td></td><td></td><td></td><td></td><td></td><td></td><td></td><td></td></tr>
<tr><td>Chromosome</td><td>start</td><td>end</td><td>strand</td><td>MM</td><td>target_seq</td><td>PAM</td><td>alignment</td><td>distance</td><td>position</td><td>gene name</td><td>gene id</td></tr>
<tr><td>GU305914.1</td><td>210251</td><td>210273</td><td>+</td><td>0</td><td>CATTGTCCAGTTAGTGTATG</td><td>TGG</td><td>||||||||[||||||||||||]PAM</td><td>NA</td><td>Intergenic</td><td>NA</td><td>NA</td></tr>
<tr><td>T15859</td><td>AGTTAGTGTATGTGGCCCGACGG</td><td>1000</td><td></td><td></td><td></td><td></td><td></td><td></td><td></td><td></td><td></td></tr>
<tr><td>Oligo fwd</td><td>CACCAGTTAGTGTATGTGGCCCGA</td><td></td><td></td><td></td><td></td><td></td><td></td><td></td><td></td><td></td><td></td></tr>
<tr><td>Oligo rev</td><td>AAACTCGGGCCACATACACTAACT</td><td></td><td></td><td></td><td></td><td></td><td></td><td></td><td></td><td></td><td></td></tr>
<tr><td>Chromosome</td><td>start</td><td>end</td><td>strand</td><td>MM</td><td>target_seq</td><td>PAM</td><td>alignment</td><td>distance</td><td>position</td><td>gene name</td><td>gene id</td></tr>
<tr><td>GU305914.1</td><td>210259</td><td>210281</td><td>+</td><td>0</td><td>AGTTAGTGTATGTGGCCCGA</td><td>CGG</td><td>||||||||[||||||||||||]PAM</td><td>NA</td><td>Intergenic</td><td>NA</td><td>NA</td></tr>
<tr><td>T15860</td><td>GTTAGTGTATGTGGCCCGACGGG</td><td>1000</td><td></td><td></td><td></td><td></td><td></td><td></td><td></td><td></td><td></td></tr>
<tr><td>Oligo fwd</td><td>CACCGTTAGTGTATGTGGCCCGAC</td><td></td><td></td><td></td><td></td><td></td><td></td><td></td><td></td><td></td><td></td></tr>
<tr><td>Oligo rev</td><td>AAACGTCGGGCCACATACACTAAC</td><td></td><td></td><td></td><td></td><td></td><td></td><td></td><td></td><td></td><td></td></tr>
<tr><td>Chromosome</td><td>start</td><td>end</td><td>strand</td><td>MM</td><td>target_seq</td><td>PAM</td><td>alignment</td><td>distance</td><td>position</td><td>gene name</td><td>gene id</td></tr>
<tr><td>GU305914.1</td><td>210260</td><td>210282</td><td>+</td><td>0</td><td>GTTAGTGTATGTGGCCCGAC</td><td>GGG</td><td>||||||||[||||||||||||]PAM</td><td>NA</td><td>Intergenic</td><td>NA</td><td>NA</td></tr>
<tr><td>T15861</td><td>AGTGTATGTGGCCCGACGGGCGG</td><td>1000</td><td></td><td></td><td></td><td></td><td></td><td></td><td></td><td></td><td></td></tr>
<tr><td>Oligo fwd</td><td>CACCAGTGTATGTGGCCCGACGGG</td><td></td><td></td><td></td><td></td><td></td><td></td><td></td><td></td><td></td><td></td></tr>
<tr><td>Oligo rev</td><td>AAACCCCGTCGGGCCACATACACT</td><td></td><td></td><td></td><td></td><td></td><td></td><td></td><td></td><td></td><td></td></tr>
<tr><td>Chromosome</td><td>start</td><td>end</td><td>strand</td><td>MM</td><td>target_seq</td><td>PAM</td><td>alignment</td><td>distance</td><td>position</td><td>gene name</td><td>gene id</td></tr>
<tr><td>GU305914.1</td><td>210263</td><td>210285</td><td>+</td><td>0</td><td>AGTGTATGTGGCCCGACGGG</td><td>CGG</td><td>||||||||[||||||||||||]PAM</td><td>NA</td><td>Intergenic</td><td>NA</td><td>NA</td></tr>
<tr><td>T15862</td><td>TCTGACATTGTCCAGTGTGCCGG</td><td>1000</td><td></td><td></td><td></td><td></td><td></td><td></td><td></td><td></td><td></td></tr>
<tr><td>Oligo fwd</td><td>CACCTCTGACATTGTCCAGTGTGC</td><td></td><td></td><td></td><td></td><td></td><td></td><td></td><td></td><td></td><td></td></tr>
<tr><td>Oligo rev</td><td>AAACGCACACTGGACAATGTCAGA</td><td></td><td></td><td></td><td></td><td></td><td></td><td></td><td></td><td></td><td></td></tr>
<tr><td>Chromosome</td><td>start</td><td>end</td><td>strand</td><td>MM</td><td>target_seq</td><td>PAM</td><td>alignment</td><td>distance</td><td>position</td><td>gene name</td><td>gene id</td></tr>
</table>

GU305914.1 210308 210330 + 0 TCTGACATTGTCCAGTGTGC CGG ||||||||[||||||||||||]PAM NA Intergenic NA NA

T15863 CGCACGAACATCACCACACGCGG 1000
Oligo fwd CACCCGCACGAACATCACCACACG
Oligo rev AAACCGTGTGGTGATGTTCGTGCG
Chromosome start end strand MM target_seq PAM alignment distance position gene name gene id
GU305914.1 210332 210354 + 0 CGCACGAACATCACCACACG CGG ||||||||[||||||||||||]PAM NA Intergenic NA NA

T15864 CGCGGCAGTTGATGTAGACCAGG 1000
Oligo fwd CACCCGCGGCAGTTGATGTAGACC
Oligo rev AAACGGTCTACATCAACTGCCGCG
Chromosome start end strand MM target_seq PAM alignment distance position gene name gene id
GU305914.1 210350 210372 + 0 CGCGGCAGTTGATGTAGACC AGG ||||||||[||||||||||||]PAM NA Intergenic NA NA

T15865 ACCAGGCGATACCGCATACGAGG 1000
Oligo fwd CACCACCAGGCGATACCGCATACG
Oligo rev AAACCGTATGCGGTATCGCCTGGT
Chromosome start end strand MM target_seq PAM alignment distance position gene name gene id
GU305914.1 210367 210389 + 0 ACCAGGCGATACCGCATACG AGG ||||||||[||||||||||||]PAM NA Intergenic NA NA

T15866 CACGACTCCGATCGCGATCAAGG 1000
Oligo fwd CACCCACGACTCCGATCGCGATCA
Oligo rev AAACTGATCGCGATCGGAGTCGTG
Chromosome start end strand MM target_seq PAM alignment distance position gene name gene id
GU305914.1 210450 210472 + 0 CACGACTCCGATCGCGATCA AGG ||||||||[||||||||||||]PAM NA Intergenic NA NA

T15867 CCACACGTCCATGAAGAAGATGG 1000
Oligo fwd CACCCCACACGTCCATGAAGAAGA
Oligo rev AAACTCTTCTTCATGGACGTGTGG
Chromosome start end strand MM target_seq PAM alignment distance position gene name gene id
GU305914.1 210474 210496 + 0 CCACACGTCCATGAAGAAGA TGG ||||||||[||||||||||||]PAM NA Intergenic NA NA

T15868 GAAGAAGATGGAGTACTCCGCGG 1000
Oligo fwd CACCGAAGAAGATGGAGTACTCCG
Oligo rev AAACCGGAGTACTCCATCTTCTTC
Chromosome start end strand MM target_seq PAM alignment distance position gene name gene id
GU305914.1 210486 210508 + 0 GAAGAAGATGGAGTACTCCG CGG ||||||||[||||||||||||]PAM NA Intergenic NA NA

T15869 GATGGAGTACTCCGCGGAGACGG 1000
Oligo fwd CACCGATGGAGTACTCCGCGGAGA
Oligo rev AAACTCTCCGCGGAGTACTCCATC
Chromosome start end strand MM target_seq PAM alignment distance position gene name gene id
GU305914.1 210492 210514 + 0 GATGGAGTACTCCGCGGAGA CGG ||||||||[||||||||||||]PAM NA Intergenic NA NA

T15870 ATGGAGTACTCCGCGGAGACGGG 1000
Oligo fwd CACCATGGAGTACTCCGCGGAGAC
Oligo rev AAACGTCTCCGCGGAGTACTCCAT
Chromosome start end strand MM target_seq PAM alignment distance position gene name gene id
GU305914.1 210493 210515 + 0 ATGGAGTACTCCGCGGAGAC GGG ||||||||[||||||||||||]PAM NA Intergenic NA NA

T15871 GAGTACTCCGCGGAGACGGGAGG 1000
Oligo fwd CACCGAGTACTCCGCGGAGACGGG
Oligo rev AAACCCCGTCTCCGCGGAGTACTC
Chromosome start end strand MM target_seq PAM alignment distance position gene name gene id
GU305914.1 210496 210518 + 0 GAGTACTCCGCGGAGACGGG AGG ||||||||[||||||||||||]PAM NA Intergenic NA NA

T15872 CCTGTGCCACGAGACGACCGCGG 1000
Oligo fwd CACCCCTGTGCCACGAGACGACCG
Oligo rev AAACCGGTCGTCTCGTGGCACAGG
Chromosome start end strand MM target_seq PAM alignment distance position gene name gene id
GU305914.1 210540 210562 + 0 CCTGTGCCACGAGACGACCG CGG ||||||||[||||||||||||]PAM NA Intergenic NA NA

T15873 GATCGACGACAACAGCATGCCGG 1000
Oligo fwd CACCGATCGACGACAACAGCATGC
Oligo rev AAACGCATGCTGTTGTCGTCGATC
Chromosome start end strand MM target_seq PAM alignment distance position gene name gene id
GU305914.1 210582 210604 + 0 GATCGACGACAACAGCATGC CGG ||||||||[||||||||||||]PAM NA Intergenic NA NA

T15874 ATCGACGACAACAGCATGCCGGG 1000
Oligo fwd CACCATCGACGACAACAGCATGCC
Oligo rev AAACGGCATGCTGTTGTCGTCGAT
Chromosome start end strand MM target_seq PAM alignment distance position gene name gene id
GU305914.1 210583 210605 + 0 ATCGACGACAACAGCATGCC GGG ||||||||[||||||||||||]PAM NA Intergenic NA NA

T15875 AGCATGCCGGGACTCTGTTGAGG 1000
Oligo fwd CACCAGCATGCCGGGACTCTGTTG
Oligo rev AAACCAACAGAGTCCCGGCATGCT
Chromosome start end strand MM target_seq PAM alignment distance position gene name gene id
GU305914.1 210595 210617 + 0 AGCATGCCGGGACTCTGTTG AGG ||||||||[||||||||||||]PAM NA Intergenic NA NA

T15876 GCATGCCGGGACTCTGTTGAGGG 1000
Oligo fwd CACCGCATGCCGGGACTCTGTTGA
Oligo rev AAACTCAACAGAGTCCCGGCATGC
Chromosome start end strand MM target_seq PAM alignment distance position gene name gene id
GU305914.1 210596 210618 + 0 GCATGCCGGGACTCTGTTGA GGG ||||||||[||||||||||||]PAM NA Intergenic NA NA

T15877 ACCGTAGCGTACCATCCCGTCGG 1000
Oligo fwd CACCACCGTAGCGTACCATCCCGT
Oligo rev AAACACGGGATGGTACGCTACGGT

| Chromosome | start | end | strand | MM | target_seq | PAM | alignment | distance | position | gene name | gene id |
|---|---|---|---|---|---|---|---|---|---|---|---|
| GU305914.1 | 210634 | 210656 | + | 0 | ACCGTAGCGTACCATCCCGT | CGG | \|\|\|\|\|\|\|\|[\|\|\|\|\|\|\|\|\|\|\|\|]PAM | NA | Intergenic | NA | NA |
| T15878 | CCGTAGCGTACCATCCCGTCGGG | 1000 | | | | | | | | | |
| Oligo fwd | CACCCCGTAGCGTACCATCCCGTC | | | | | | | | | | |
| Oligo rev | AAACGACGGGATGGTACGCTACGG | | | | | | | | | | |
| Chromosome | start | end | strand | MM | target_seq | PAM | alignment | distance | position | gene name | gene id |
| GU305914.1 | 210635 | 210657 | + | 0 | CCGTAGCGTACCATCCCGTC | GGG | \|\|\|\|\|\|\|\|[\|\|\|\|\|\|\|\|\|\|\|\|]PAM | NA | Intergenic | NA | NA |
| T15879 | CGTAGCGTACCATCCCGTCGGGG | 1000 | | | | | | | | | |
| Oligo fwd | CACCCGTAGCGTACCATCCCGTCG | | | | | | | | | | |
| Oligo rev | AAACCGACGGGATGGTACGCTACG | | | | | | | | | | |
| Chromosome | start | end | strand | MM | target_seq | PAM | alignment | distance | position | gene name | gene id |
| GU305914.1 | 210636 | 210658 | + | 0 | CGTAGCGTACCATCCCGTCG | GGG | \|\|\|\|\|\|\|\|[\|\|\|\|\|\|\|\|\|\|\|\|]PAM | NA | Intergenic | NA | NA |
| T15880 | AGAAATCACGAACGCGAAGTCGG | 1000 | | | | | | | | | |
| Oligo fwd | CACCAGAAATCACGAACGCGAAGT | | | | | | | | | | |
| Oligo rev | AAACACTTCGCGTTCGTGATTTCT | | | | | | | | | | |
| Chromosome | start | end | strand | MM | target_seq | PAM | alignment | distance | position | gene name | gene id |
| GU305914.1 | 210660 | 210682 | + | 0 | AGAAATCACGAACGCGAAGT | CGG | \|\|\|\|\|\|\|\|[\|\|\|\|\|\|\|\|\|\|\|\|]PAM | NA | Intergenic | NA | NA |
| T15881 | CGAAGTCGGTACTACCGTTCAGG | 1000 | | | | | | | | | |
| Oligo fwd | CACCCGAAGTCGGTACTACCGTTC | | | | | | | | | | |
| Oligo rev | AAACGAACGGTAGTACCGACTTCG | | | | | | | | | | |
| Chromosome | start | end | strand | MM | target_seq | PAM | alignment | distance | position | gene name | gene id |
| GU305914.1 | 210674 | 210696 | + | 0 | CGAAGTCGGTACTACCGTTC | AGG | \|\|\|\|\|\|\|\|[\|\|\|\|\|\|\|\|\|\|\|\|]PAM | NA | Intergenic | NA | NA |
| T15882 | AGACTGATGTTGTACGCCAACGG | 1000 | | | | | | | | | |
| Oligo fwd | CACCAGACTGATGTTGTACGCCAA | | | | | | | | | | |
| Oligo rev | AAACTTGGCGTACAACATCAGTCT | | | | | | | | | | |
| Chromosome | start | end | strand | MM | target_seq | PAM | alignment | distance | position | gene name | gene id |
| GU305914.1 | 210781 | 210803 | + | 0 | AGACTGATGTTGTACGCCAA | CGG | \|\|\|\|\|\|\|\|[\|\|\|\|\|\|\|\|\|\|\|\|]PAM | NA | Intergenic | NA | NA |
| T15883 | AACCTGAGTGCGCATCGTACAGG | 1000 | | | | | | | | | |
| Oligo fwd | CACCAACCTGAGTGCGCATCGTAC | | | | | | | | | | |
| Oligo rev | AAACGTACGATGCGCACTCAGGTT | | | | | | | | | | |
| Chromosome | start | end | strand | MM | target_seq | PAM | alignment | distance | position | gene name | gene id |
| GU305914.1 | 210804 | 210826 | + | 0 | AACCTGAGTGCGCATCGTAC | AGG | \|\|\|\|\|\|\|\|[\|\|\|\|\|\|\|\|\|\|\|\|]PAM | NA | Intergenic | NA | NA |
| T15884 | CAGCGAGTATTCGTTGTGTCTGG | 1000 | | | | | | | | | |
| Oligo fwd | CACCCAGCGAGTATTCGTTGTGTC | | | | | | | | | | |
| Oligo rev | AAACGACACAACGAATACTCGCTG | | | | | | | | | | |
| Chromosome | start | end | strand | MM | target_seq | PAM | alignment | distance | position | gene name | gene id |
| GU305914.1 | 210855 | 210877 | + | 0 | CAGCGAGTATTCGTTGTGTC | TGG | \|\|\|\|\|\|\|\|[\|\|\|\|\|\|\|\|\|\|\|\|]PAM | NA | Intergenic | NA | NA |
| T15885 | AGCGAGTATTCGTTGTGTCTGGG | 1000 | | | | | | | | | |
| Oligo fwd | CACCAGCGAGTATTCGTTGTGTCT | | | | | | | | | | |
| Oligo rev | AAACAGACACAACGAATACTCGCT | | | | | | | | | | |
| Chromosome | start | end | strand | MM | target_seq | PAM | alignment | distance | position | gene name | gene id |
| GU305914.1 | 210856 | 210878 | + | 0 | AGCGAGTATTCGTTGTGTCT | GGG | \|\|\|\|\|\|\|\|[\|\|\|\|\|\|\|\|\|\|\|\|]PAM | NA | Intergenic | NA | NA |
| T15886 | GCGAGTATTCGTTGTGTCTGGGG | 1000 | | | | | | | | | |
| Oligo fwd | CACCGCGAGTATTCGTTGTGTCTG | | | | | | | | | | |
| Oligo rev | AAACCAGACACAACGAATACTCGC | | | | | | | | | | |
| Chromosome | start | end | strand | MM | target_seq | PAM | alignment | distance | position | gene name | gene id |
| GU305914.1 | 210857 | 210879 | + | 0 | GCGAGTATTCGTTGTGTCTG | GGG | \|\|\|\|\|\|\|\|[\|\|\|\|\|\|\|\|\|\|\|\|]PAM | NA | Intergenic | NA | NA |
| T15887 | GTTGTGTCTGGGGTTGTCCATGG | 1000 | | | | | | | | | |
| Oligo fwd | CACCGTTGTGTCTGGGGTTGTCCA | | | | | | | | | | |
| Oligo rev | AAACTGGACAACCCCAGACACAAC | | | | | | | | | | |
| Chromosome | start | end | strand | MM | target_seq | PAM | alignment | distance | position | gene name | gene id |
| GU305914.1 | 210867 | 210889 | + | 0 | GTTGTGTCTGGGGTTGTCCA | TGG | \|\|\|\|\|\|\|\|[\|\|\|\|\|\|\|\|\|\|\|\|]PAM | NA | Intergenic | NA | NA |
| T15888 | GGCGACGATCTTCCGACAGAAGG | 1000 | | | | | | | | | |
| Oligo fwd | CACCGGCGACGATCTTCCGACAGA | | | | | | | | | | |
| Oligo rev | AAACTCTGTCGGAAGATCGTCGCC | | | | | | | | | | |
| Chromosome | start | end | strand | MM | target_seq | PAM | alignment | distance | position | gene name | gene id |
| GU305914.1 | 210888 | 210910 | + | 0 | GGCGACGATCTTCCGACAGA | AGG | \|\|\|\|\|\|\|\|[\|\|\|\|\|\|\|\|\|\|\|\|]PAM | NA | Intergenic | NA | NA |
| T15889 | AGAAGGACGTCCAGCGTTTCTGG | 1000 | | | | | | | | | |
| Oligo fwd | CACCAGAAGGACGTCCAGCGTTTC | | | | | | | | | | |
| Oligo rev | AAACGAAACGCTGGACGTCCTTCT | | | | | | | | | | |
| Chromosome | start | end | strand | MM | target_seq | PAM | alignment | distance | position | gene name | gene id |
| GU305914.1 | 210905 | 210927 | + | 0 | AGAAGGACGTCCAGCGTTTC | TGG | \|\|\|\|\|\|\|\|[\|\|\|\|\|\|\|\|\|\|\|\|]PAM | NA | Intergenic | NA | NA |
| T15890 | GTCCCTCAAGATAGAGATTCCGG | 1000 | | | | | | | | | |
| Oligo fwd | CACCGTCCCTCAAGATAGAGATTC | | | | | | | | | | |
| Oligo rev | AAACGAATCTCTATCTTGAGGGAC | | | | | | | | | | |
| Chromosome | start | end | strand | MM | target_seq | PAM | alignment | distance | position | gene name | gene id |
| GU305914.1 | 210942 | 210964 | + | 0 | GTCCCTCAAGATAGAGATTC | CGG | \|\|\|\|\|\|\|\|[\|\|\|\|\|\|\|\|\|\|\|\|]PAM | NA | Intergenic | NA | NA |
| T15891 | GTAGCGCGCGATGTCATCCTCGG | 1000 | | | | | | | | | |
| Oligo fwd | CACCGTAGCGCGCGATGTCATCCT | | | | | | | | | | |
| Oligo rev | AAACAGGATGACATCGCGCGCTAC | | | | | | | | | | |
| Chromosome | start | end | strand | MM | target_seq | PAM | alignment | distance | position | gene name | gene id |
| GU305914.1 | 210975 | 210997 | + | 0 | GTAGCGCGCGATGTCATCCT | CGG | \|\|\|\|\|\|\|\|[\|\|\|\|\|\|\|\|\|\|\|\|]PAM | NA | Intergenic | NA | NA |
| T15892 | GATGTCATCCTCGGATACGTCGG | 1000 | | | | | | | | | |
| Oligo fwd | CACCGATGTCATCCTCGGATACGT | | | | | | | | | | |

<table>
<tr><td>Oligo rev</td><td>AAACACGTATCCGAGGATGACATC</td><td></td><td></td><td></td><td></td><td></td><td></td><td></td><td></td><td></td><td></td></tr>
<tr><td>Chromosome</td><td>start</td><td>end</td><td>strand</td><td>MM</td><td>target_seq</td><td>PAM</td><td>alignment</td><td>distance</td><td>position</td><td>gene name</td><td>gene id</td></tr>
<tr><td>GU305914.1</td><td>210984</td><td>211006</td><td>+</td><td>0</td><td>GATGTCATCCTCGGATACGT</td><td>CGG</td><td>||||||||[||||||||||||]PAM</td><td>NA</td><td>Intergenic</td><td>NA</td><td>NA</td></tr>
<tr><td>GU305914.1</td><td>132646</td><td>132668</td><td>+</td><td>4</td><td>GATGTGCTCCTCGCACACGT</td><td>AGG</td><td>|||||--|[|||||-|-||||]PAM</td><td>NA</td><td>Intergenic</td><td>NA</td><td>NA</td></tr>
<tr><td></td><td></td><td></td><td></td><td></td><td></td><td></td><td></td><td></td><td></td><td></td><td></td></tr>
<tr><td>T15893</td><td>ATGTCATCCTCGGATACGTCGGG</td><td>1000</td><td></td><td></td><td></td><td></td><td></td><td></td><td></td><td></td><td></td></tr>
<tr><td>Oligo fwd</td><td>CACCATGTCATCCTCGGATACGTC</td><td></td><td></td><td></td><td></td><td></td><td></td><td></td><td></td><td></td><td></td></tr>
<tr><td>Oligo rev</td><td>AAACGACGTATCCGAGGATGACAT</td><td></td><td></td><td></td><td></td><td></td><td></td><td></td><td></td><td></td><td></td></tr>
<tr><td>Chromosome</td><td>start</td><td>end</td><td>strand</td><td>MM</td><td>target_seq</td><td>PAM</td><td>alignment</td><td>distance</td><td>position</td><td>gene name</td><td>gene id</td></tr>
<tr><td>GU305914.1</td><td>210985</td><td>211007</td><td>+</td><td>0</td><td>ATGTCATCCTCGGATACGTC</td><td>GGG</td><td>||||||||[||||||||||||]PAM</td><td>NA</td><td>Intergenic</td><td>NA</td><td>NA</td></tr>
<tr><td></td><td></td><td></td><td></td><td></td><td></td><td></td><td></td><td></td><td></td><td></td><td></td></tr>
<tr><td>T15894</td><td>TCGGATACGTCGGGAGAGTACGG</td><td>1000</td><td></td><td></td><td></td><td></td><td></td><td></td><td></td><td></td><td></td></tr>
<tr><td>Oligo fwd</td><td>CACCTCGGATACGTCGGGAGAGTA</td><td></td><td></td><td></td><td></td><td></td><td></td><td></td><td></td><td></td><td></td></tr>
<tr><td>Oligo rev</td><td>AAACTACTCTCCCGACGTATCCGA</td><td></td><td></td><td></td><td></td><td></td><td></td><td></td><td></td><td></td><td></td></tr>
<tr><td>Chromosome</td><td>start</td><td>end</td><td>strand</td><td>MM</td><td>target_seq</td><td>PAM</td><td>alignment</td><td>distance</td><td>position</td><td>gene name</td><td>gene id</td></tr>
<tr><td>GU305914.1</td><td>210994</td><td>211016</td><td>+</td><td>0</td><td>TCGGATACGTCGGGAGAGTA</td><td>CGG</td><td>||||||||[||||||||||||]PAM</td><td>NA</td><td>Intergenic</td><td>NA</td><td>NA</td></tr>
<tr><td></td><td></td><td></td><td></td><td></td><td></td><td></td><td></td><td></td><td></td><td></td><td></td></tr>
<tr><td>T15895</td><td>GATACGTCGGGAGAGTACGGCGG</td><td>1000</td><td></td><td></td><td></td><td></td><td></td><td></td><td></td><td></td><td></td></tr>
<tr><td>Oligo fwd</td><td>CACCGATACGTCGGGAGAGTACGG</td><td></td><td></td><td></td><td></td><td></td><td></td><td></td><td></td><td></td><td></td></tr>
<tr><td>Oligo rev</td><td>AAACCCGTACTCTCCCGACGTATC</td><td></td><td></td><td></td><td></td><td></td><td></td><td></td><td></td><td></td><td></td></tr>
<tr><td>Chromosome</td><td>start</td><td>end</td><td>strand</td><td>MM</td><td>target_seq</td><td>PAM</td><td>alignment</td><td>distance</td><td>position</td><td>gene name</td><td>gene id</td></tr>
<tr><td>GU305914.1</td><td>210997</td><td>211019</td><td>+</td><td>0</td><td>GATACGTCGGGAGAGTACGG</td><td>CGG</td><td>||||||||[||||||||||||]PAM</td><td>NA</td><td>Intergenic</td><td>NA</td><td>NA</td></tr>
<tr><td></td><td></td><td></td><td></td><td></td><td></td><td></td><td></td><td></td><td></td><td></td><td></td></tr>
<tr><td>T15896</td><td>CGTTTCGACGCCTTTGTCTACGG</td><td>1000</td><td></td><td></td><td></td><td></td><td></td><td></td><td></td><td></td><td></td></tr>
<tr><td>Oligo fwd</td><td>CACCCGTTTCGACGCCTTTGTCTA</td><td></td><td></td><td></td><td></td><td></td><td></td><td></td><td></td><td></td><td></td></tr>
<tr><td>Oligo rev</td><td>AAACTAGACAAAGGCGTCGAAACG</td><td></td><td></td><td></td><td></td><td></td><td></td><td></td><td></td><td></td><td></td></tr>
<tr><td>Chromosome</td><td>start</td><td>end</td><td>strand</td><td>MM</td><td>target_seq</td><td>PAM</td><td>alignment</td><td>distance</td><td>position</td><td>gene name</td><td>gene id</td></tr>
<tr><td>GU305914.1</td><td>211029</td><td>211051</td><td>+</td><td>0</td><td>CGTTTCGACGCCTTTGTCTA</td><td>CGG</td><td>||||||||[||||||||||||]PAM</td><td>NA</td><td>Intergenic</td><td>NA</td><td>NA</td></tr>
<tr><td></td><td></td><td></td><td></td><td></td><td></td><td></td><td></td><td></td><td></td><td></td><td></td></tr>
<tr><td>T15897</td><td>GCCGTCCATCATCTGAAGTACGG</td><td>1000</td><td></td><td></td><td></td><td></td><td></td><td></td><td></td><td></td><td></td></tr>
<tr><td>Oligo fwd</td><td>CACCGCCGTCCATCATCTGAAGTA</td><td></td><td></td><td></td><td></td><td></td><td></td><td></td><td></td><td></td><td></td></tr>
<tr><td>Oligo rev</td><td>AAACTACTTCAGATGATGGACGGC</td><td></td><td></td><td></td><td></td><td></td><td></td><td></td><td></td><td></td><td></td></tr>
<tr><td>Chromosome</td><td>start</td><td>end</td><td>strand</td><td>MM</td><td>target_seq</td><td>PAM</td><td>alignment</td><td>distance</td><td>position</td><td>gene name</td><td>gene id</td></tr>
<tr><td>GU305914.1</td><td>211071</td><td>211093</td><td>+</td><td>0</td><td>GCCGTCCATCATCTGAAGTA</td><td>CGG</td><td>||||||||[||||||||||||]PAM</td><td>NA</td><td>Intergenic</td><td>NA</td><td>NA</td></tr>
<tr><td></td><td></td><td></td><td></td><td></td><td></td><td></td><td></td><td></td><td></td><td></td><td></td></tr>
<tr><td>T15898</td><td>TGAAGTACGGTACATTCGAATGG</td><td>1000</td><td></td><td></td><td></td><td></td><td></td><td></td><td></td><td></td><td></td></tr>
<tr><td>Oligo fwd</td><td>CACCTGAAGTACGGTACATTCGAA</td><td></td><td></td><td></td><td></td><td></td><td></td><td></td><td></td><td></td><td></td></tr>
<tr><td>Oligo rev</td><td>AAACTTCGAATGTACCGTACTTCA</td><td></td><td></td><td></td><td></td><td></td><td></td><td></td><td></td><td></td><td></td></tr>
<tr><td>Chromosome</td><td>start</td><td>end</td><td>strand</td><td>MM</td><td>target_seq</td><td>PAM</td><td>alignment</td><td>distance</td><td>position</td><td>gene name</td><td>gene id</td></tr>
<tr><td>GU305914.1</td><td>211084</td><td>211106</td><td>+</td><td>0</td><td>TGAAGTACGGTACATTCGAA</td><td>TGG</td><td>||||||||[||||||||||||]PAM</td><td>NA</td><td>Intergenic</td><td>NA</td><td>NA</td></tr>
<tr><td></td><td></td><td></td><td></td><td></td><td></td><td></td><td></td><td></td><td></td><td></td><td></td></tr>
<tr><td>T15899</td><td>CATCTATATTCGAAGACTACCGG</td><td>1000</td><td></td><td></td><td></td><td></td><td></td><td></td><td></td><td></td><td></td></tr>
<tr><td>Oligo fwd</td><td>CACCCATCTATATTCGAAGACTAC</td><td></td><td></td><td></td><td></td><td></td><td></td><td></td><td></td><td></td><td></td></tr>
<tr><td>Oligo rev</td><td>AAACGTAGTCTTCGAATATAGATG</td><td></td><td></td><td></td><td></td><td></td><td></td><td></td><td></td><td></td><td></td></tr>
<tr><td>Chromosome</td><td>start</td><td>end</td><td>strand</td><td>MM</td><td>target_seq</td><td>PAM</td><td>alignment</td><td>distance</td><td>position</td><td>gene name</td><td>gene id</td></tr>
<tr><td>GU305914.1</td><td>211117</td><td>211139</td><td>+</td><td>0</td><td>CATCTATATTCGAAGACTAC</td><td>CGG</td><td>||||||||[||||||||||||]PAM</td><td>NA</td><td>Intergenic</td><td>NA</td><td>NA</td></tr>
<tr><td></td><td></td><td></td><td></td><td></td><td></td><td></td><td></td><td></td><td></td><td></td><td></td></tr>
<tr><td>T15900</td><td>TTCGAAGACTACCGGCCGTCCGG</td><td>1000</td><td></td><td></td><td></td><td></td><td></td><td></td><td></td><td></td><td></td></tr>
<tr><td>Oligo fwd</td><td>CACCTTCGAAGACTACCGGCCGTC</td><td></td><td></td><td></td><td></td><td></td><td></td><td></td><td></td><td></td><td></td></tr>
<tr><td>Oligo rev</td><td>AAACGACGGCCGGTAGTCTTCGAA</td><td></td><td></td><td></td><td></td><td></td><td></td><td></td><td></td><td></td><td></td></tr>
<tr><td>Chromosome</td><td>start</td><td>end</td><td>strand</td><td>MM</td><td>target_seq</td><td>PAM</td><td>alignment</td><td>distance</td><td>position</td><td>gene name</td><td>gene id</td></tr>
<tr><td>GU305914.1</td><td>211125</td><td>211147</td><td>+</td><td>0</td><td>TTCGAAGACTACCGGCCGTC</td><td>CGG</td><td>||||||||[||||||||||||]PAM</td><td>NA</td><td>Intergenic</td><td>NA</td><td>NA</td></tr>
<tr><td></td><td></td><td></td><td></td><td></td><td></td><td></td><td></td><td></td><td></td><td></td><td></td></tr>
<tr><td>T15901</td><td>AGCTCGCTCACGTTCATCCAAGG</td><td>1000</td><td></td><td></td><td></td><td></td><td></td><td></td><td></td><td></td><td></td></tr>
<tr><td>Oligo fwd</td><td>CACCAGCTCGCTCACGTTCATCCA</td><td></td><td></td><td></td><td></td><td></td><td></td><td></td><td></td><td></td><td></td></tr>
<tr><td>Oligo rev</td><td>AAACTGGATGAACGTGAGCGAGCT</td><td></td><td></td><td></td><td></td><td></td><td></td><td></td><td></td><td></td><td></td></tr>
<tr><td>Chromosome</td><td>start</td><td>end</td><td>strand</td><td>MM</td><td>target_seq</td><td>PAM</td><td>alignment</td><td>distance</td><td>position</td><td>gene name</td><td>gene id</td></tr>
<tr><td>GU305914.1</td><td>211219</td><td>211241</td><td>+</td><td>0</td><td>AGCTCGCTCACGTTCATCCA</td><td>AGG</td><td>||||||||[||||||||||||]PAM</td><td>NA</td><td>Intergenic</td><td>NA</td><td>NA</td></tr>
<tr><td></td><td></td><td></td><td></td><td></td><td></td><td></td><td></td><td></td><td></td><td></td><td></td></tr>
<tr><td>T15902</td><td>AGGAAAATGTTTCACCTCTCCGG</td><td>1000</td><td></td><td></td><td></td><td></td><td></td><td></td><td></td><td></td><td></td></tr>
<tr><td>Oligo fwd</td><td>CACCAGGAAAATGTTTCACCTCTC</td><td></td><td></td><td></td><td></td><td></td><td></td><td></td><td></td><td></td><td></td></tr>
<tr><td>Oligo rev</td><td>AAACGAGAGGTGAAACATTTTCCT</td><td></td><td></td><td></td><td></td><td></td><td></td><td></td><td></td><td></td><td></td></tr>
<tr><td>Chromosome</td><td>start</td><td>end</td><td>strand</td><td>MM</td><td>target_seq</td><td>PAM</td><td>alignment</td><td>distance</td><td>position</td><td>gene name</td><td>gene id</td></tr>
<tr><td>GU305914.1</td><td>211239</td><td>211261</td><td>+</td><td>0</td><td>AGGAAAATGTTTCACCTCTC</td><td>CGG</td><td>||||||||[||||||||||||]PAM</td><td>NA</td><td>Intergenic</td><td>NA</td><td>NA</td></tr>
<tr><td></td><td></td><td></td><td></td><td></td><td></td><td></td><td></td><td></td><td></td><td></td><td></td></tr>
<tr><td>T15903</td><td>AAAATGTTTCACCTCTCCGGAGG</td><td>1000</td><td></td><td></td><td></td><td></td><td></td><td></td><td></td><td></td><td></td></tr>
<tr><td>Oligo fwd</td><td>CACCAAAATGTTTCACCTCTCCGG</td><td></td><td></td><td></td><td></td><td></td><td></td><td></td><td></td><td></td><td></td></tr>
<tr><td>Oligo rev</td><td>AAACCCGGAGAGGTGAAACATTTT</td><td></td><td></td><td></td><td></td><td></td><td></td><td></td><td></td><td></td><td></td></tr>
<tr><td>Chromosome</td><td>start</td><td>end</td><td>strand</td><td>MM</td><td>target_seq</td><td>PAM</td><td>alignment</td><td>distance</td><td>position</td><td>gene name</td><td>gene id</td></tr>
<tr><td>GU305914.1</td><td>211242</td><td>211264</td><td>+</td><td>0</td><td>AAAATGTTTCACCTCTCCGG</td><td>AGG</td><td>||||||||[||||||||||||]PAM</td><td>NA</td><td>Intergenic</td><td>NA</td><td>NA</td></tr>
<tr><td></td><td></td><td></td><td></td><td></td><td></td><td></td><td></td><td></td><td></td><td></td><td></td></tr>
<tr><td>T15904</td><td>ACTGCTCTAGATTGATACTGTGG</td><td>1000</td><td></td><td></td><td></td><td></td><td></td><td></td><td></td><td></td><td></td></tr>
<tr><td>Oligo fwd</td><td>CACCACTGCTCTAGATTGATACTG</td><td></td><td></td><td></td><td></td><td></td><td></td><td></td><td></td><td></td><td></td></tr>
<tr><td>Oligo rev</td><td>AAACCAGTATCAATCTAGAGCAGT</td><td></td><td></td><td></td><td></td><td></td><td></td><td></td><td></td><td></td><td></td></tr>
<tr><td>Chromosome</td><td>start</td><td>end</td><td>strand</td><td>MM</td><td>target_seq</td><td>PAM</td><td>alignment</td><td>distance</td><td>position</td><td>gene name</td><td>gene id</td></tr>
<tr><td>GU305914.1</td><td>211289</td><td>211311</td><td>+</td><td>0</td><td>ACTGCTCTAGATTGATACTG</td><td>TGG</td><td>||||||||[||||||||||||]PAM</td><td>NA</td><td>Intergenic</td><td>NA</td><td>NA</td></tr>
<tr><td></td><td></td><td></td><td></td><td></td><td></td><td></td><td></td><td></td><td></td><td></td><td></td></tr>
<tr><td>T15905</td><td>GCTCTAGATTGATACTGTGGCGG</td><td>1000</td><td></td><td></td><td></td><td></td><td></td><td></td><td></td><td></td><td></td></tr>
<tr><td>Oligo fwd</td><td>CACCGCTCTAGATTGATACTGTGG</td><td></td><td></td><td></td><td></td><td></td><td></td><td></td><td></td><td></td><td></td></tr>
<tr><td>Oligo rev</td><td>AAACCCACAGTATCAATCTAGAGC</td><td></td><td></td><td></td><td></td><td></td><td></td><td></td><td></td><td></td><td></td></tr>
<tr><td>Chromosome</td><td>start</td><td>end</td><td>strand</td><td>MM</td><td>target_seq</td><td>PAM</td><td>alignment</td><td>distance</td><td>position</td><td>gene name</td><td>gene id</td></tr>
<tr><td>GU305914.1</td><td>211292</td><td>211314</td><td>+</td><td>0</td><td>GCTCTAGATTGATACTGTGG</td><td>CGG</td><td>||||||||[||||||||||||]PAM</td><td>NA</td><td>Intergenic</td><td>NA</td><td>NA</td></tr>
<tr><td></td><td></td><td></td><td></td><td></td><td></td><td></td><td></td><td></td><td></td><td></td><td></td></tr>
<tr><td>T15906</td><td>ATACTGTGGCGGAACATCTCCGG</td><td>1000</td><td></td><td></td><td></td><td></td><td></td><td></td><td></td><td></td><td></td></tr>
<tr><td>Oligo fwd</td><td>CACCATACTGTGGCGGAACATCTC</td><td></td><td></td><td></td><td></td><td></td><td></td><td></td><td></td><td></td><td></td></tr>
<tr><td>Oligo rev</td><td>AAACGAGATGTTCCGCCACAGTAT</td><td></td><td></td><td></td><td></td><td></td><td></td><td></td><td></td><td></td><td></td></tr>
<tr><td>Chromosome</td><td>start</td><td>end</td><td>strand</td><td>MM</td><td>target_seq</td><td>PAM</td><td>alignment</td><td>distance</td><td>position</td><td>gene name</td><td>gene id</td></tr>
<tr><td>GU305914.1</td><td>211303</td><td>211325</td><td>+</td><td>0</td><td>ATACTGTGGCGGAACATCTC</td><td>CGG</td><td>||||||||[||||||||||||]PAM</td><td>NA</td><td>Intergenic</td><td>NA</td><td>NA</td></tr>
</table>

| | | | | | | | | | | | |
|---|---|---|---|---|---|---|---|---|---|---|---|
| T15907 | ACGAGATGAACGTCGCCGTCAGG | 1000 | | | | | | | | | |
| Oligo fwd | CACCACGAGATGAACGTCGCCGTC | | | | | | | | | | |
| Oligo rev | AAACGACGGCGACGTTCATCTCGT | | | | | | | | | | |
| Chromosome | start | end | strand | MM | target_seq | PAM | alignment | distance | position | gene name | gene id |
| GU305914.1 | 211337 | 211359 | + | 0 | ACGAGATGAACGTCGCCGTC | AGG | ||||||||[||||||||||||]PAM | NA | Intergenic | NA | NA |
| T15908 | TGCTGCCCGCCGTCTCGCCTCGG | 1000 | | | | | | | | | |
| Oligo fwd | CACCTGCTGCCCGCCGTCTCGCCT | | | | | | | | | | |
| Oligo rev | AAACAGGCGAGACGGCGGGCAGCA | | | | | | | | | | |
| Chromosome | start | end | strand | MM | target_seq | PAM | alignment | distance | position | gene name | gene id |
| GU305914.1 | 211412 | 211434 | + | 0 | TGCTGCCCGCCGTCTCGCCT | CGG | ||||||||[||||||||||||]PAM | NA | Intergenic | NA | NA |
| T15909 | CTCGGTTTATGAGAACCGAGAGG | 1000 | | | | | | | | | |
| Oligo fwd | CACCCTCGGTTTATGAGAACCGAG | | | | | | | | | | |
| Oligo rev | AAACCTCGGTTCTCATAAACCGAG | | | | | | | | | | |
| Chromosome | start | end | strand | MM | target_seq | PAM | alignment | distance | position | gene name | gene id |
| GU305914.1 | 211430 | 211452 | + | 0 | CTCGGTTTATGAGAACCGAG | AGG | ||||||||[||||||||||||]PAM | NA | Intergenic | NA | NA |
| T15910 | GAGAACCGAGAGGAGCAAGTAGG | 1000 | | | | | | | | | |
| Oligo fwd | CACCGAGAACCGAGAGGAGCAAGT | | | | | | | | | | |
| Oligo rev | AAACACTTGCTCCTCTCGGTTCTC | | | | | | | | | | |
| Chromosome | start | end | strand | MM | target_seq | PAM | alignment | distance | position | gene name | gene id |
| GU305914.1 | 211440 | 211462 | + | 0 | GAGAACCGAGAGGAGCAAGT | AGG | ||||||||[||||||||||||]PAM | NA | Intergenic | NA | NA |
| T15911 | GATAGCGCCCAGCATTGTACAGG | 1000 | | | | | | | | | |
| Oligo fwd | CACCGATAGCGCCCAGCATTGTAC | | | | | | | | | | |
| Oligo rev | AAACGTACAATGCTGGGCGCTATC | | | | | | | | | | |
| Chromosome | start | end | strand | MM | target_seq | PAM | alignment | distance | position | gene name | gene id |
| GU305914.1 | 211464 | 211486 | + | 0 | GATAGCGCCCAGCATTGTAC | AGG | ||||||||[||||||||||||]PAM | NA | Intergenic | NA | NA |
| T15912 | GCATTGTACAGGCTAGCGATCGG | 1000 | | | | | | | | | |
| Oligo fwd | CACCGCATTGTACAGGCTAGCGAT | | | | | | | | | | |
| Oligo rev | AAACATCGCTAGCCTGTACAATGC | | | | | | | | | | |
| Chromosome | start | end | strand | MM | target_seq | PAM | alignment | distance | position | gene name | gene id |
| GU305914.1 | 211475 | 211497 | + | 0 | GCATTGTACAGGCTAGCGAT | CGG | ||||||||[||||||||||||]PAM | NA | Intergenic | NA | NA |
| T15913 | CATTGTACAGGCTAGCGATCGGG | 1000 | | | | | | | | | |
| Oligo fwd | CACCCATTGTACAGGCTAGCGATC | | | | | | | | | | |
| Oligo rev | AAACGATCGCTAGCCTGTACAATG | | | | | | | | | | |
| Chromosome | start | end | strand | MM | target_seq | PAM | alignment | distance | position | gene name | gene id |
| GU305914.1 | 211476 | 211498 | + | 0 | CATTGTACAGGCTAGCGATC | GGG | ||||||||[||||||||||||]PAM | NA | Intergenic | NA | NA |
| T15914 | GCTAGCGATCGGGAGACCGACGG | 1000 | | | | | | | | | |
| Oligo fwd | CACCGCTAGCGATCGGGAGACCGA | | | | | | | | | | |
| Oligo rev | AAACTCGGTCTCCCGATCGCTAGC | | | | | | | | | | |
| Chromosome | start | end | strand | MM | target_seq | PAM | alignment | distance | position | gene name | gene id |
| GU305914.1 | 211486 | 211508 | + | 0 | GCTAGCGATCGGGAGACCGA | CGG | ||||||||[||||||||||||]PAM | NA | Intergenic | NA | NA |
| T15915 | GGGAGACCGACGGTGAAACGTGG | 1000 | | | | | | | | | |
| Oligo fwd | CACCGGGAGACCGACGGTGAAACG | | | | | | | | | | |
| Oligo rev | AAACCGTTTCACCGTCGGTCTCCC | | | | | | | | | | |
| Chromosome | start | end | strand | MM | target_seq | PAM | alignment | distance | position | gene name | gene id |
| GU305914.1 | 211496 | 211518 | + | 0 | GGGAGACCGACGGTGAAACG | TGG | ||||||||[||||||||||||]PAM | NA | Intergenic | NA | NA |
| T15916 | ACCGACGGTGAAACGTGGAGCGG | 1000 | | | | | | | | | |
| Oligo fwd | CACCACCGACGGTGAAACGTGGAG | | | | | | | | | | |
| Oligo rev | AAACCTCCACGTTTCACCGTCGGT | | | | | | | | | | |
| Chromosome | start | end | strand | MM | target_seq | PAM | alignment | distance | position | gene name | gene id |
| GU305914.1 | 211501 | 211523 | + | 0 | ACCGACGGTGAAACGTGGAG | CGG | ||||||||[||||||||||||]PAM | NA | Intergenic | NA | NA |
| T15917 | TGAAACGTGGAGCGGTGACACGG | 1000 | | | | | | | | | |
| Oligo fwd | CACCTGAAACGTGGAGCGGTGACA | | | | | | | | | | |
| Oligo rev | AAACTGTCACCGCTCCACGTTTCA | | | | | | | | | | |
| Chromosome | start | end | strand | MM | target_seq | PAM | alignment | distance | position | gene name | gene id |
| GU305914.1 | 211509 | 211531 | + | 0 | TGAAACGTGGAGCGGTGACA | CGG | ||||||||[||||||||||||]PAM | NA | Intergenic | NA | NA |
| T15918 | GGCTTGTGATGTTACTCTTTCGG | 1000 | | | | | | | | | |
| Oligo fwd | CACCGGCTTGTGATGTTACTCTTT | | | | | | | | | | |
| Oligo rev | AAACAAAGAGTAACATCACAAGCC | | | | | | | | | | |
| Chromosome | start | end | strand | MM | target_seq | PAM | alignment | distance | position | gene name | gene id |
| GU305914.1 | 211530 | 211552 | + | 0 | GGCTTGTGATGTTACTCTTT | CGG | ||||||||[||||||||||||]PAM | NA | Intergenic | NA | NA |
| T15919 | GCTTGTGATGTTACTCTTTCGGG | 1000 | | | | | | | | | |
| Oligo fwd | CACCGCTTGTGATGTTACTCTTTC | | | | | | | | | | |
| Oligo rev | AAACGAAAGAGTAACATCACAAGC | | | | | | | | | | |
| Chromosome | start | end | strand | MM | target_seq | PAM | alignment | distance | position | gene name | gene id |
| GU305914.1 | 211531 | 211553 | + | 0 | GCTTGTGATGTTACTCTTTC | GGG | ||||||||[||||||||||||]PAM | NA | Intergenic | NA | NA |
| T15920 | TAACATCGAAAGCGAAAGTCCGG | 1000 | | | | | | | | | |
| Oligo fwd | CACCTAACATCGAAAGCGAAAGTC | | | | | | | | | | |
| Oligo rev | AAACGACTTTCGCTTTCGATGTTA | | | | | | | | | | |
| Chromosome | start | end | strand | MM | target_seq | PAM | alignment | distance | position | gene name | gene id |
| GU305914.1 | 211620 | 211642 | + | 0 | TAACATCGAAAGCGAAAGTC | CGG | ||||||||[||||||||||||]PAM | NA | Intergenic | NA | NA |
| T15921 | GAAAGTCCGGACTTTCCCAAAGG | 1000 | | | | | | | | | |
| Oligo fwd | CACCGAAAGTCCGGACTTTCCCAA | | | | | | | | | | |
| Oligo rev | AAACTTGGGAAAGTCCGGACTTTC | | | | | | | | | | |
| Chromosome | start | end | strand | MM | target_seq | PAM | alignment | distance | position | gene name | gene id |
| GU305914.1 | 211633 | 211655 | + | 0 | GAAAGTCCGGACTTTCCCAA | AGG | ||||||||[||||||||||||]PAM | NA | Intergenic | NA | NA |

| | | | | | | | | | | | |
|---|---|---|---|---|---|---|---|---|---|---|---|
| T15922 | CCAAAGGCCTGAGCCGACCGAGG | 1000 | | | | | | | | | |
| Oligo fwd | CACCCCAAAGGCCTGAGCCGACCG | | | | | | | | | | |
| Oligo rev | AAACCGGTCGGCTCAGGCCTTTGG | | | | | | | | | | |
| Chromosome | start | end | strand | MM | target_seq | PAM | alignment | distance | position | gene name | gene id |
| GU305914.1 | 211649 | 211671 | + | 0 | CCAAAGGCCTGAGCCGACCG | AGG | ||||||||[||||||||||||]PAM | NA | Intergenic | NA | NA |

| | | | | | | | | | | | |
|---|---|---|---|---|---|---|---|---|---|---|---|
| T15923 | AAGGCCTGAGCCGACCGAGGCGG | 1000 | | | | | | | | | |
| Oligo fwd | CACCAAGGCCTGAGCCGACCGAGG | | | | | | | | | | |
| Oligo rev | AAACCCTCGGTCGGCTCAGGCCTT | | | | | | | | | | |
| Chromosome | start | end | strand | MM | target_seq | PAM | alignment | distance | position | gene name | gene id |
| GU305914.1 | 211652 | 211674 | + | 0 | AAGGCCTGAGCCGACCGAGG | CGG | ||||||||[||||||||||||]PAM | NA | Intergenic | NA | NA |

| | | | | | | | | | | | |
|---|---|---|---|---|---|---|---|---|---|---|---|
| T15924 | TTTCATCCTCGCGCGTGTAACGG | 1000 | | | | | | | | | |
| Oligo fwd | CACCTTTCATCCTCGCGCGTGTAA | | | | | | | | | | |
| Oligo rev | AAACTTACACGCGCGAGGATGAAA | | | | | | | | | | |
| Chromosome | start | end | strand | MM | target_seq | PAM | alignment | distance | position | gene name | gene id |
| GU305914.1 | 211750 | 211772 | + | 0 | TTTCATCCTCGCGCGTGTAA | CGG | ||||||||[||||||||||||]PAM | NA | Intergenic | NA | NA |

| | | | | | | | | | | | |
|---|---|---|---|---|---|---|---|---|---|---|---|
| T15925 | GATCGAATATCGCAGCCGTTCGG | 1000 | | | | | | | | | |
| Oligo fwd | CACCGATCGAATATCGCAGCCGTT | | | | | | | | | | |
| Oligo rev | AAACAACGGCTGCGATATTCGATC | | | | | | | | | | |
| Chromosome | start | end | strand | MM | target_seq | PAM | alignment | distance | position | gene name | gene id |
| GU305914.1 | 211772 | 211794 | + | 0 | GATCGAATATCGCAGCCGTT | CGG | ||||||||[||||||||||||]PAM | NA | Intergenic | NA | NA |

| | | | | | | | | | | | |
|---|---|---|---|---|---|---|---|---|---|---|---|
| T15926 | GCCGTTCGGCCATGCGAGCCAGG | 1000 | | | | | | | | | |
| Oligo fwd | CACCGCCGTTCGGCCATGCGAGCC | | | | | | | | | | |
| Oligo rev | AAACGGCTCGCATGGCCGAACGGC | | | | | | | | | | |
| Chromosome | start | end | strand | MM | target_seq | PAM | alignment | distance | position | gene name | gene id |
| GU305914.1 | 211786 | 211808 | + | 0 | GCCGTTCGGCCATGCGAGCC | AGG | ||||||||[||||||||||||]PAM | NA | Intergenic | NA | NA |

| | | | | | | | | | | | |
|---|---|---|---|---|---|---|---|---|---|---|---|
| T15927 | GGCCATGCGAGCCAGGCGCCCGG | 1000 | | | | | | | | | |
| Oligo fwd | CACCGGCCATGCGAGCCAGGCGCC | | | | | | | | | | |
| Oligo rev | AAACGGCGCCTGGCTCGCATGGCC | | | | | | | | | | |
| Chromosome | start | end | strand | MM | target_seq | PAM | alignment | distance | position | gene name | gene id |
| GU305914.1 | 211793 | 211815 | + | 0 | GGCCATGCGAGCCAGGCGCC | CGG | ||||||||[||||||||||||]PAM | NA | Intergenic | NA | NA |

| | | | | | | | | | | | |
|---|---|---|---|---|---|---|---|---|---|---|---|
| T15928 | CGGTGACGATCCCGTGCACCAGG | 1000 | | | | | | | | | |
| Oligo fwd | CACCCGGTGACGATCCCGTGCACC | | | | | | | | | | |
| Oligo rev | AAACGGTGCACGGGATCGTCACCG | | | | | | | | | | |
| Chromosome | start | end | strand | MM | target_seq | PAM | alignment | distance | position | gene name | gene id |
| GU305914.1 | 211813 | 211835 | + | 0 | CGGTGACGATCCCGTGCACC | AGG | ||||||||[||||||||||||]PAM | NA | Intergenic | NA | NA |

| | | | | | | | | | | | |
|---|---|---|---|---|---|---|---|---|---|---|---|
| T15929 | CACGAATAACGCGACGAATGCGG | 1000 | | | | | | | | | |
| Oligo fwd | CACCCACGAATAACGCGACGAATG | | | | | | | | | | |
| Oligo rev | AAACCATTCGTCGCGTTATTCGTG | | | | | | | | | | |
| Chromosome | start | end | strand | MM | target_seq | PAM | alignment | distance | position | gene name | gene id |
| GU305914.1 | 211847 | 211869 | + | 0 | CACGAATAACGCGACGAATG | CGG | ||||||||[||||||||||||]PAM | NA | Intergenic | NA | NA |

| | | | | | | | | | | | |
|---|---|---|---|---|---|---|---|---|---|---|---|
| T15930 | TAACGCGACGAATGCGGCGACGG | 1000 | | | | | | | | | |
| Oligo fwd | CACCTAACGCGACGAATGCGGCGA | | | | | | | | | | |
| Oligo rev | AAACTCGCCGCATTCGTCGCGTTA | | | | | | | | | | |
| Chromosome | start | end | strand | MM | target_seq | PAM | alignment | distance | position | gene name | gene id |
| GU305914.1 | 211853 | 211875 | + | 0 | TAACGCGACGAATGCGGCGA | CGG | ||||||||[||||||||||||]PAM | NA | Intergenic | NA | NA |

| | | | | | | | | | | | |
|---|---|---|---|---|---|---|---|---|---|---|---|
| T15931 | TGCGGCGACGGTGATCGCGATGG | 1000 | | | | | | | | | |
| Oligo fwd | CACCTGCGGCGACGGTGATCGCGA | | | | | | | | | | |
| Oligo rev | AAACTCGCGATCACCGTCGCCGCA | | | | | | | | | | |
| Chromosome | start | end | strand | MM | target_seq | PAM | alignment | distance | position | gene name | gene id |
| GU305914.1 | 211865 | 211887 | + | 0 | TGCGGCGACGGTGATCGCGA | TGG | ||||||||[||||||||||||]PAM | NA | Intergenic | NA | NA |

| | | | | | | | | | | | |
|---|---|---|---|---|---|---|---|---|---|---|---|
| T15932 | GGTGATCGCGATGGTCACCGTGG | 1000 | | | | | | | | | |
| Oligo fwd | CACCGGTGATCGCGATGGTCACCG | | | | | | | | | | |
| Oligo rev | AAACCGGTGACCATCGCGATCACC | | | | | | | | | | |
| Chromosome | start | end | strand | MM | target_seq | PAM | alignment | distance | position | gene name | gene id |
| GU305914.1 | 211874 | 211896 | + | 0 | GGTGATCGCGATGGTCACCG | TGG | ||||||||[||||||||||||]PAM | NA | Intergenic | NA | NA |

| | | | | | | | | | | | |
|---|---|---|---|---|---|---|---|---|---|---|---|
| T15933 | TCGAATCGTTGATCGTCCTCTGG | 1000 | | | | | | | | | |
| Oligo fwd | CACCTCGAATCGTTGATCGTCCTC | | | | | | | | | | |
| Oligo rev | AAACGAGGACGATCAACGATTCGA | | | | | | | | | | |
| Chromosome | start | end | strand | MM | target_seq | PAM | alignment | distance | position | gene name | gene id |
| GU305914.1 | 211903 | 211925 | + | 0 | TCGAATCGTTGATCGTCCTC | TGG | ||||||||[||||||||||||]PAM | NA | Intergenic | NA | NA |

| | | | | | | | | | | | |
|---|---|---|---|---|---|---|---|---|---|---|---|
| T15934 | GTTGATCGTCCTCTGGACGAAGG | 1000 | | | | | | | | | |
| Oligo fwd | CACCGTTGATCGTCCTCTGGACGA | | | | | | | | | | |
| Oligo rev | AAACTCGTCCAGAGGACGATCAAC | | | | | | | | | | |
| Chromosome | start | end | strand | MM | target_seq | PAM | alignment | distance | position | gene name | gene id |
| GU305914.1 | 211910 | 211932 | + | 0 | GTTGATCGTCCTCTGGACGA | AGG | ||||||||[||||||||||||]PAM | NA | Intergenic | NA | NA |

| | | | | | | | | | | | |
|---|---|---|---|---|---|---|---|---|---|---|---|
| T15935 | ATCGTCCTCTGGACGAAGGCCGG | 1000 | | | | | | | | | |
| Oligo fwd | CACCATCGTCCTCTGGACGAAGGC | | | | | | | | | | |
| Oligo rev | AAACGCCTTCGTCCAGAGGACGAT | | | | | | | | | | |
| Chromosome | start | end | strand | MM | target_seq | PAM | alignment | distance | position | gene name | gene id |
| GU305914.1 | 211914 | 211936 | + | 0 | ATCGTCCTCTGGACGAAGGC | CGG | ||||||||[||||||||||||]PAM | NA | Intergenic | NA | NA |

| | | | | | | | | | | | |
|---|---|---|---|---|---|---|---|---|---|---|---|
| T15936 | GAAGGCCGGACCGTGTTCATCGG | 1000 | | | | | | | | | |
| Oligo fwd | CACCGAAGGCCGGACCGTGTTCAT | | | | | | | | | | |
| Oligo rev | AAACATGAACACGGTCCGGCCTTC | | | | | | | | | | |
| Chromosome | start | end | strand | MM | target_seq | PAM | alignment | distance | position | gene name | gene id |

<table>
<tr><td>GU305914.1</td><td>211928</td><td>211950</td><td>+</td><td>0</td><td>GAAGGCCGGACCGTGTTCAT</td><td>CGG</td><td>||||||||[||||||||||||]PAM</td><td>NA</td><td>Intergenic</td><td>NA</td><td>NA</td></tr>
<tr><td colspan="12"></td></tr>
<tr><td>T15937</td><td>TCATCGGACATGAATCCGTGCGG</td><td>1000</td><td colspan="9"></td></tr>
<tr><td>Oligo fwd</td><td>CACCTCATCGGACATGAATCCGTG</td><td colspan="10"></td></tr>
<tr><td>Oligo rev</td><td>AAACCACGGATTCATGTCCGATGA</td><td colspan="10"></td></tr>
<tr><td>Chromosome</td><td>start</td><td>end</td><td>strand</td><td>MM</td><td>target_seq</td><td>PAM</td><td>alignment</td><td>distance</td><td>position</td><td>gene name</td><td>gene id</td></tr>
<tr><td>GU305914.1</td><td>211944</td><td>211966</td><td>+</td><td>0</td><td>TCATCGGACATGAATCCGTG</td><td>CGG</td><td>||||||||[||||||||||||]PAM</td><td>NA</td><td>Intergenic</td><td>NA</td><td>NA</td></tr>
<tr><td colspan="12"></td></tr>
<tr><td>T15938</td><td>CATGAATCCGTGCGGTTCTACGG</td><td>1000</td><td colspan="9"></td></tr>
<tr><td>Oligo fwd</td><td>CACCCATGAATCCGTGCGGTTCTA</td><td colspan="10"></td></tr>
<tr><td>Oligo rev</td><td>AAACTAGAACCGCACGGATTCATG</td><td colspan="10"></td></tr>
<tr><td>Chromosome</td><td>start</td><td>end</td><td>strand</td><td>MM</td><td>target_seq</td><td>PAM</td><td>alignment</td><td>distance</td><td>position</td><td>gene name</td><td>gene id</td></tr>
<tr><td>GU305914.1</td><td>211952</td><td>211974</td><td>+</td><td>0</td><td>CATGAATCCGTGCGGTTCTA</td><td>CGG</td><td>||||||||[||||||||||||]PAM</td><td>NA</td><td>Intergenic</td><td>NA</td><td>NA</td></tr>
<tr><td colspan="12"></td></tr>
<tr><td>T15939</td><td>ATGAATCCGTGCGGTTCTACGGG</td><td>1000</td><td colspan="9"></td></tr>
<tr><td>Oligo fwd</td><td>CACCATGAATCCGTGCGGTTCTAC</td><td colspan="10"></td></tr>
<tr><td>Oligo rev</td><td>AAACGTAGAACCGCACGGATTCAT</td><td colspan="10"></td></tr>
<tr><td>Chromosome</td><td>start</td><td>end</td><td>strand</td><td>MM</td><td>target_seq</td><td>PAM</td><td>alignment</td><td>distance</td><td>position</td><td>gene name</td><td>gene id</td></tr>
<tr><td>GU305914.1</td><td>211953</td><td>211975</td><td>+</td><td>0</td><td>ATGAATCCGTGCGGTTCTAC</td><td>GGG</td><td>||||||||[||||||||||||]PAM</td><td>NA</td><td>Intergenic</td><td>NA</td><td>NA</td></tr>
<tr><td colspan="12"></td></tr>
<tr><td>T15940</td><td>TGAATCCGTGCGGTTCTACGGGG</td><td>1000</td><td colspan="9"></td></tr>
<tr><td>Oligo fwd</td><td>CACCTGAATCCGTGCGGTTCTACG</td><td colspan="10"></td></tr>
<tr><td>Oligo rev</td><td>AAACCGTAGAACCGCACGGATTCA</td><td colspan="10"></td></tr>
<tr><td>Chromosome</td><td>start</td><td>end</td><td>strand</td><td>MM</td><td>target_seq</td><td>PAM</td><td>alignment</td><td>distance</td><td>position</td><td>gene name</td><td>gene id</td></tr>
<tr><td>GU305914.1</td><td>211954</td><td>211976</td><td>+</td><td>0</td><td>TGAATCCGTGCGGTTCTACG</td><td>GGG</td><td>||||||||[||||||||||||]PAM</td><td>NA</td><td>Intergenic</td><td>NA</td><td>NA</td></tr>
<tr><td colspan="12"></td></tr>
<tr><td>T15941</td><td>TGCGGTTCTACGGGGAGCAGCGG</td><td>1000</td><td colspan="9"></td></tr>
<tr><td>Oligo fwd</td><td>CACCTGCGGTTCTACGGGGAGCAG</td><td colspan="10"></td></tr>
<tr><td>Oligo rev</td><td>AAACCTGCTCCCCGTAGAACCGCA</td><td colspan="10"></td></tr>
<tr><td>Chromosome</td><td>start</td><td>end</td><td>strand</td><td>MM</td><td>target_seq</td><td>PAM</td><td>alignment</td><td>distance</td><td>position</td><td>gene name</td><td>gene id</td></tr>
<tr><td>GU305914.1</td><td>211962</td><td>211984</td><td>+</td><td>0</td><td>TGCGGTTCTACGGGGAGCAG</td><td>CGG</td><td>||||||||[||||||||||||]PAM</td><td>NA</td><td>Intergenic</td><td>NA</td><td>NA</td></tr>
<tr><td colspan="12"></td></tr>
<tr><td>T15942</td><td>TTGAGCCTGCCTCGCTTTCGAGG</td><td>1000</td><td colspan="9"></td></tr>
<tr><td>Oligo fwd</td><td>CACCTTGAGCCTGCCTCGCTTTCG</td><td colspan="10"></td></tr>
<tr><td>Oligo rev</td><td>AAACCGAAAGCGAGGCAGGCTCAA</td><td colspan="10"></td></tr>
<tr><td>Chromosome</td><td>start</td><td>end</td><td>strand</td><td>MM</td><td>target_seq</td><td>PAM</td><td>alignment</td><td>distance</td><td>position</td><td>gene name</td><td>gene id</td></tr>
<tr><td>GU305914.1</td><td>211991</td><td>212013</td><td>+</td><td>0</td><td>TTGAGCCTGCCTCGCTTTCG</td><td>AGG</td><td>||||||||[||||||||||||]PAM</td><td>NA</td><td>Intergenic</td><td>NA</td><td>NA</td></tr>
<tr><td colspan="12"></td></tr>
<tr><td>T15943</td><td>CGAGGCCGCTACCATCTCCTCGG</td><td>1000</td><td colspan="9"></td></tr>
<tr><td>Oligo fwd</td><td>CACCCGAGGCCGCTACCATCTCCT</td><td colspan="10"></td></tr>
<tr><td>Oligo rev</td><td>AAACAGGAGATGGTAGCGGCCTCG</td><td colspan="10"></td></tr>
<tr><td>Chromosome</td><td>start</td><td>end</td><td>strand</td><td>MM</td><td>target_seq</td><td>PAM</td><td>alignment</td><td>distance</td><td>position</td><td>gene name</td><td>gene id</td></tr>
<tr><td>GU305914.1</td><td>212009</td><td>212031</td><td>+</td><td>0</td><td>CGAGGCCGCTACCATCTCCT</td><td>CGG</td><td>||||||||[||||||||||||]PAM</td><td>NA</td><td>Intergenic</td><td>NA</td><td>NA</td></tr>
<tr><td colspan="12"></td></tr>
<tr><td>T15944</td><td>TCTCCTCGGAGAACCGAAGACGG</td><td>1000</td><td colspan="9"></td></tr>
<tr><td>Oligo fwd</td><td>CACCTCTCCTCGGAGAACCGAAGA</td><td colspan="10"></td></tr>
<tr><td>Oligo rev</td><td>AAACTCTTCGGTTCTCCGAGGAGA</td><td colspan="10"></td></tr>
<tr><td>Chromosome</td><td>start</td><td>end</td><td>strand</td><td>MM</td><td>target_seq</td><td>PAM</td><td>alignment</td><td>distance</td><td>position</td><td>gene name</td><td>gene id</td></tr>
<tr><td>GU305914.1</td><td>212023</td><td>212045</td><td>+</td><td>0</td><td>TCTCCTCGGAGAACCGAAGA</td><td>CGG</td><td>||||||||[||||||||||||]PAM</td><td>NA</td><td>Intergenic</td><td>NA</td><td>NA</td></tr>
<tr><td colspan="12"></td></tr>
<tr><td>T15945</td><td>GACCGTCCAGCCGCTCGACGAGG</td><td>1000</td><td colspan="9"></td></tr>
<tr><td>Oligo fwd</td><td>CACCGACCGTCCAGCCGCTCGACG</td><td colspan="10"></td></tr>
<tr><td>Oligo rev</td><td>AAACCGTCGAGCGGCTGGACGGTC</td><td colspan="10"></td></tr>
<tr><td>Chromosome</td><td>start</td><td>end</td><td>strand</td><td>MM</td><td>target_seq</td><td>PAM</td><td>alignment</td><td>distance</td><td>position</td><td>gene name</td><td>gene id</td></tr>
<tr><td>GU305914.1</td><td>212045</td><td>212067</td><td>+</td><td>0</td><td>GACCGTCCAGCCGCTCGACG</td><td>AGG</td><td>||||||||[||||||||||||]PAM</td><td>NA</td><td>Intergenic</td><td>NA</td><td>NA</td></tr>
<tr><td>GU305914.1</td><td>147735</td><td>147757</td><td>+</td><td>3</td><td>GGCCGTCCAGCCCCTGGACG</td><td>GGG</td><td>|-||||||[||||-||-||||]PAM</td><td>NA</td><td>Intergenic</td><td>NA</td><td>NA</td></tr>
<tr><td colspan="12"></td></tr>
<tr><td>T15946</td><td>GCTCGACGAGGTGATCTCGCAGG</td><td>1000</td><td colspan="9"></td></tr>
<tr><td>Oligo fwd</td><td>CACCGCTCGACGAGGTGATCTCGC</td><td colspan="10"></td></tr>
<tr><td>Oligo rev</td><td>AAACGCGAGATCACCTCGTCGAGC</td><td colspan="10"></td></tr>
<tr><td>Chromosome</td><td>start</td><td>end</td><td>strand</td><td>MM</td><td>target_seq</td><td>PAM</td><td>alignment</td><td>distance</td><td>position</td><td>gene name</td><td>gene id</td></tr>
<tr><td>GU305914.1</td><td>212057</td><td>212079</td><td>+</td><td>0</td><td>GCTCGACGAGGTGATCTCGC</td><td>AGG</td><td>||||||||[||||||||||||]PAM</td><td>NA</td><td>Intergenic</td><td>NA</td><td>NA</td></tr>
<tr><td colspan="12"></td></tr>
<tr><td>T15947</td><td>AAGATCTGACTTCTGTTGACCGG</td><td>1000</td><td colspan="9"></td></tr>
<tr><td>Oligo fwd</td><td>CACCAAGATCTGACTTCTGTTGAC</td><td colspan="10"></td></tr>
<tr><td>Oligo rev</td><td>AAACGTCAACAGAAGTCAGATCTT</td><td colspan="10"></td></tr>
<tr><td>Chromosome</td><td>start</td><td>end</td><td>strand</td><td>MM</td><td>target_seq</td><td>PAM</td><td>alignment</td><td>distance</td><td>position</td><td>gene name</td><td>gene id</td></tr>
<tr><td>GU305914.1</td><td>212088</td><td>212110</td><td>+</td><td>0</td><td>AAGATCTGACTTCTGTTGAC</td><td>CGG</td><td>||||||||[||||||||||||]PAM</td><td>NA</td><td>Intergenic</td><td>NA</td><td>NA</td></tr>
<tr><td colspan="12"></td></tr>
<tr><td>T15948</td><td>ACTTCTGTTGACCGGCAGATAGG</td><td>1000</td><td colspan="9"></td></tr>
<tr><td>Oligo fwd</td><td>CACCACTTCTGTTGACCGGCAGAT</td><td colspan="10"></td></tr>
<tr><td>Oligo rev</td><td>AAACATCTGCCGGTCAACAGAAGT</td><td colspan="10"></td></tr>
<tr><td>Chromosome</td><td>start</td><td>end</td><td>strand</td><td>MM</td><td>target_seq</td><td>PAM</td><td>alignment</td><td>distance</td><td>position</td><td>gene name</td><td>gene id</td></tr>
<tr><td>GU305914.1</td><td>212096</td><td>212118</td><td>+</td><td>0</td><td>ACTTCTGTTGACCGGCAGAT</td><td>AGG</td><td>||||||||[||||||||||||]PAM</td><td>NA</td><td>Intergenic</td><td>NA</td><td>NA</td></tr>
<tr><td colspan="12"></td></tr>
<tr><td>T15949</td><td>GATGTCCGTGTAGTAGTCCGCGG</td><td>1000</td><td colspan="9"></td></tr>
<tr><td>Oligo fwd</td><td>CACCGATGTCCGTGTAGTAGTCCG</td><td colspan="10"></td></tr>
<tr><td>Oligo rev</td><td>AAACCGGACTACTACACGGACATC</td><td colspan="10"></td></tr>
<tr><td>Chromosome</td><td>start</td><td>end</td><td>strand</td><td>MM</td><td>target_seq</td><td>PAM</td><td>alignment</td><td>distance</td><td>position</td><td>gene name</td><td>gene id</td></tr>
<tr><td>GU305914.1</td><td>212129</td><td>212151</td><td>+</td><td>0</td><td>GATGTCCGTGTAGTAGTCCG</td><td>CGG</td><td>||||||||[||||||||||||]PAM</td><td>NA</td><td>Intergenic</td><td>NA</td><td>NA</td></tr>
<tr><td colspan="12"></td></tr>
<tr><td>T15950</td><td>CGTGTAGTAGTCCGCGGACAAGG</td><td>1000</td><td colspan="9"></td></tr>
<tr><td>Oligo fwd</td><td>CACCCGTGTAGTAGTCCGCGGACA</td><td colspan="10"></td></tr>
<tr><td>Oligo rev</td><td>AAACTGTCCGCGGACTACTACACG</td><td colspan="10"></td></tr>
<tr><td>Chromosome</td><td>start</td><td>end</td><td>strand</td><td>MM</td><td>target_seq</td><td>PAM</td><td>alignment</td><td>distance</td><td>position</td><td>gene name</td><td>gene id</td></tr>
<tr><td>GU305914.1</td><td>212135</td><td>212157</td><td>+</td><td>0</td><td>CGTGTAGTAGTCCGCGGACA</td><td>AGG</td><td>||||||||[||||||||||||]PAM</td><td>NA</td><td>Intergenic</td><td>NA</td><td>NA</td></tr>
<tr><td colspan="12"></td></tr>
<tr><td>T15951</td><td>CCGCGGACAAGGCTATCGCACGG</td><td>1000</td><td colspan="9"></td></tr>
<tr><td>Oligo fwd</td><td>CACCCCGCGGACAAGGCTATCGCA</td><td colspan="10"></td></tr>
</table>

Oligo rev AAACTGCGATAGCCTTGTCCGCGG
Chromosome start end strand MM target_seq PAM alignment distance position gene name gene id
GU305914.1 212146 212168 + 0 CCGCGGACAAGGCTATCGCA CGG ||||||||[||||||||||||]PAM NA Intergenic NA NA

T15952 CGCGGACAAGGCTATCGCACGGG 1000
Oligo fwd CACCCGCGGACAAGGCTATCGCAC
Oligo rev AAACGTGCGATAGCCTTGTCCGCG
Chromosome start end strand MM target_seq PAM alignment distance position gene name gene id
GU305914.1 212147 212169 + 0 CGCGGACAAGGCTATCGCAC GGG ||||||||[||||||||||||]PAM NA Intergenic NA NA

T15953 AGGCTATCGCACGGGAGTCCCGG 1000
Oligo fwd CACCAGGCTATCGCACGGGAGTCC
Oligo rev AAACGGACTCCCGTGCGATAGCCT
Chromosome start end strand MM target_seq PAM alignment distance position gene name gene id
GU305914.1 212155 212177 + 0 AGGCTATCGCACGGGAGTCC CGG ||||||||[||||||||||||]PAM NA Intergenic NA NA

T15954 TATCGCACGGGAGTCCCGGCCGG 1000
Oligo fwd CACCTATCGCACGGGAGTCCCGGC
Oligo rev AAACGCCGGGACTCCCGTGCGATA
Chromosome start end strand MM target_seq PAM alignment distance position gene name gene id
GU305914.1 212159 212181 + 0 TATCGCACGGGAGTCCCGGC CGG ||||||||[||||||||||||]PAM NA Intergenic NA NA

T15955 GTCCCGGCCGGCGCTCCACACGG 1000
Oligo fwd CACCGTCCCGGCCGGCGCTCCACA
Oligo rev AAACTGTGGAGCGCCGGCCGGGAC
Chromosome start end strand MM target_seq PAM alignment distance position gene name gene id
GU305914.1 212171 212193 + 0 GTCCCGGCCGGCGCTCCACA CGG ||||||||[||||||||||||]PAM NA Intergenic NA NA

T15956 CGGCGCTCCACACGGTCATATGG 1000
Oligo fwd CACCCGGCGCTCCACACGGTCATA
Oligo rev AAACTATGACCGTGTGGAGCGCCG
Chromosome start end strand MM target_seq PAM alignment distance position gene name gene id
GU305914.1 212179 212201 + 0 CGGCGCTCCACACGGTCATA TGG ||||||||[||||||||||||]PAM NA Intergenic NA NA

T15957 ACGGTCATATGGTATTTCAAAGG 1000
Oligo fwd CACCACGGTCATATGGTATTTCAA
Oligo rev AAACTTGAAATACCATATGACCGT
Chromosome start end strand MM target_seq PAM alignment distance position gene name gene id
GU305914.1 212190 212212 + 0 ACGGTCATATGGTATTTCAA AGG ||||||||[||||||||||||]PAM NA Intergenic NA NA

T15958 CGGTCATATGGTATTTCAAAGGG 1000
Oligo fwd CACCCGGTCATATGGTATTTCAAA
Oligo rev AAACTTTGAAATACCATATGACCG
Chromosome start end strand MM target_seq PAM alignment distance position gene name gene id
GU305914.1 212191 212213 + 0 CGGTCATATGGTATTTCAAA GGG ||||||||[||||||||||||]PAM NA Intergenic NA NA

T15959 TATGGTATTTCAAAGGGACCTGG 1000
Oligo fwd CACCTATGGTATTTCAAAGGGACC
Oligo rev AAACGGTCCCTTTGAAATACCATA
Chromosome start end strand MM target_seq PAM alignment distance position gene name gene id
GU305914.1 212197 212219 + 0 TATGGTATTTCAAAGGGACC TGG ||||||||[||||||||||||]PAM NA Intergenic NA NA

T15960 ATGGTATTTCAAAGGGACCTGGG 1000
Oligo fwd CACCATGGTATTTCAAAGGGACCT
Oligo rev AAACAGGTCCCTTTGAAATACCAT
Chromosome start end strand MM target_seq PAM alignment distance position gene name gene id
GU305914.1 212198 212220 + 0 ATGGTATTTCAAAGGGACCT GGG ||||||||[||||||||||||]PAM NA Intergenic NA NA

T15961 TGTCATCTTCTAGAGTATACTGG 1000
Oligo fwd CACCTGTCATCTTCTAGAGTATAC
Oligo rev AAACGTATACTCTAGAAGATGACA
Chromosome start end strand MM target_seq PAM alignment distance position gene name gene id
GU305914.1 212269 212291 + 0 TGTCATCTTCTAGAGTATAC TGG ||||||||[||||||||||||]PAM NA Intergenic NA NA

T15962 AGAGTATACTGGTTGTCGTCCGG 1000
Oligo fwd CACCAGAGTATACTGGTTGTCGTC
Oligo rev AAACGACGACAACCAGTATACTCT
Chromosome start end strand MM target_seq PAM alignment distance position gene name gene id
GU305914.1 212280 212302 + 0 AGAGTATACTGGTTGTCGTC CGG ||||||||[||||||||||||]PAM NA Intergenic NA NA

T15963 TATACTGGTTGTCGTCCGGACGG 1000
Oligo fwd CACCTATACTGGTTGTCGTCCGGA
Oligo rev AAACTCCGGACGACAACCAGTATA
Chromosome start end strand MM target_seq PAM alignment distance position gene name gene id
GU305914.1 212284 212306 + 0 TATACTGGTTGTCGTCCGGA CGG ||||||||[||||||||||||]PAM NA Intergenic NA NA

T15964 CTGGTTGTCGTCCGGACGGTCGG 1000
Oligo fwd CACCCTGGTTGTCGTCCGGACGGT
Oligo rev AAACACCGTCCGGACGACAACCAG
Chromosome start end strand MM target_seq PAM alignment distance position gene name gene id
GU305914.1 212288 212310 + 0 CTGGTTGTCGTCCGGACGGT CGG ||||||||[||||||||||||]PAM NA Intergenic NA NA

T15965 TCTTACATATCTCAGTCCACCGG 1000
Oligo fwd CACCTCTTACATATCTCAGTCCAC
Oligo rev AAACGTGGACTGAGATATGTAAGA
Chromosome start end strand MM target_seq PAM alignment distance position gene name gene id
GU305914.1 212323 212345 + 0 TCTTACATATCTCAGTCCAC CGG ||||||||[||||||||||||]PAM NA Intergenic NA NA

T15966 ACCTCTGATATACTTCACCAAGG 1000

<table>
<tr><td>Oligo fwd</td><td>CACCACCTCTGATATACTTCACCA</td><td></td><td></td><td></td><td></td><td></td><td></td><td></td><td></td><td></td><td></td></tr>
<tr><td>Oligo rev</td><td>AAACTGGTGAAGTATATCAGAGGT</td><td></td><td></td><td></td><td></td><td></td><td></td><td></td><td></td><td></td><td></td></tr>
<tr><td>Chromosome</td><td>start</td><td>end</td><td>strand</td><td>MM</td><td>target_seq</td><td>PAM</td><td>alignment</td><td>distance</td><td>position</td><td>gene name</td><td>gene id</td></tr>
<tr><td>GU305914.1</td><td>212390</td><td>212412</td><td>+</td><td>0</td><td>ACCTCTGATATACTTCACCA</td><td>AGG</td><td>||||||||[||||||||||||]PAM</td><td>NA</td><td>Intergenic</td><td>NA</td><td>NA</td></tr>
<tr><td>T15967</td><td>GTAATCGTACAGTTGAACTGAGG</td><td>1000</td><td></td><td></td><td></td><td></td><td></td><td></td><td></td><td></td><td></td></tr>
<tr><td>Oligo fwd</td><td>CACCGTAATCGTACAGTTGAACTG</td><td></td><td></td><td></td><td></td><td></td><td></td><td></td><td></td><td></td><td></td></tr>
<tr><td>Oligo rev</td><td>AAACCAGTTCAACTGTACGATTAC</td><td></td><td></td><td></td><td></td><td></td><td></td><td></td><td></td><td></td><td></td></tr>
<tr><td>Chromosome</td><td>start</td><td>end</td><td>strand</td><td>MM</td><td>target_seq</td><td>PAM</td><td>alignment</td><td>distance</td><td>position</td><td>gene name</td><td>gene id</td></tr>
<tr><td>GU305914.1</td><td>212508</td><td>212530</td><td>+</td><td>0</td><td>GTAATCGTACAGTTGAACTG</td><td>AGG</td><td>||||||||[||||||||||||]PAM</td><td>NA</td><td>Intergenic</td><td>NA</td><td>NA</td></tr>
<tr><td>T15968</td><td>TAATCGTACAGTTGAACTGAGGG</td><td>1000</td><td></td><td></td><td></td><td></td><td></td><td></td><td></td><td></td><td></td></tr>
<tr><td>Oligo fwd</td><td>CACCTAATCGTACAGTTGAACTGA</td><td></td><td></td><td></td><td></td><td></td><td></td><td></td><td></td><td></td><td></td></tr>
<tr><td>Oligo rev</td><td>AAACTCAGTTCAACTGTACGATTA</td><td></td><td></td><td></td><td></td><td></td><td></td><td></td><td></td><td></td><td></td></tr>
<tr><td>Chromosome</td><td>start</td><td>end</td><td>strand</td><td>MM</td><td>target_seq</td><td>PAM</td><td>alignment</td><td>distance</td><td>position</td><td>gene name</td><td>gene id</td></tr>
<tr><td>GU305914.1</td><td>212509</td><td>212531</td><td>+</td><td>0</td><td>TAATCGTACAGTTGAACTGA</td><td>GGG</td><td>||||||||[||||||||||||]PAM</td><td>NA</td><td>Intergenic</td><td>NA</td><td>NA</td></tr>
<tr><td>T15969</td><td>GTACAGTTGAACTGAGGGACCGG</td><td>1000</td><td></td><td></td><td></td><td></td><td></td><td></td><td></td><td></td><td></td></tr>
<tr><td>Oligo fwd</td><td>CACCGTACAGTTGAACTGAGGGAC</td><td></td><td></td><td></td><td></td><td></td><td></td><td></td><td></td><td></td><td></td></tr>
<tr><td>Oligo rev</td><td>AAACGTCCCTCAGTTCAACTGTAC</td><td></td><td></td><td></td><td></td><td></td><td></td><td></td><td></td><td></td><td></td></tr>
<tr><td>Chromosome</td><td>start</td><td>end</td><td>strand</td><td>MM</td><td>target_seq</td><td>PAM</td><td>alignment</td><td>distance</td><td>position</td><td>gene name</td><td>gene id</td></tr>
<tr><td>GU305914.1</td><td>212514</td><td>212536</td><td>+</td><td>0</td><td>GTACAGTTGAACTGAGGGAC</td><td>CGG</td><td>||||||||[||||||||||||]PAM</td><td>NA</td><td>Intergenic</td><td>NA</td><td>NA</td></tr>
<tr><td>T15970</td><td>ATACTCTACGACCAGCTCCCCGG</td><td>1000</td><td></td><td></td><td></td><td></td><td></td><td></td><td></td><td></td><td></td></tr>
<tr><td>Oligo fwd</td><td>CACCATACTCTACGACCAGCTCCC</td><td></td><td></td><td></td><td></td><td></td><td></td><td></td><td></td><td></td><td></td></tr>
<tr><td>Oligo rev</td><td>AAACGGGAGCTGGTCGTAGAGTAT</td><td></td><td></td><td></td><td></td><td></td><td></td><td></td><td></td><td></td><td></td></tr>
<tr><td>Chromosome</td><td>start</td><td>end</td><td>strand</td><td>MM</td><td>target_seq</td><td>PAM</td><td>alignment</td><td>distance</td><td>position</td><td>gene name</td><td>gene id</td></tr>
<tr><td>GU305914.1</td><td>212543</td><td>212565</td><td>+</td><td>0</td><td>ATACTCTACGACCAGCTCCC</td><td>CGG</td><td>||||||||[||||||||||||]PAM</td><td>NA</td><td>Intergenic</td><td>NA</td><td>NA</td></tr>
<tr><td>T15971</td><td>TACTCTACGACCAGCTCCCCGGG</td><td>1000</td><td></td><td></td><td></td><td></td><td></td><td></td><td></td><td></td><td></td></tr>
<tr><td>Oligo fwd</td><td>CACCTACTCTACGACCAGCTCCCC</td><td></td><td></td><td></td><td></td><td></td><td></td><td></td><td></td><td></td><td></td></tr>
<tr><td>Oligo rev</td><td>AAACGGGGAGCTGGTCGTAGAGTA</td><td></td><td></td><td></td><td></td><td></td><td></td><td></td><td></td><td></td><td></td></tr>
<tr><td>Chromosome</td><td>start</td><td>end</td><td>strand</td><td>MM</td><td>target_seq</td><td>PAM</td><td>alignment</td><td>distance</td><td>position</td><td>gene name</td><td>gene id</td></tr>
<tr><td>GU305914.1</td><td>212544</td><td>212566</td><td>+</td><td>0</td><td>TACTCTACGACCAGCTCCCC</td><td>GGG</td><td>||||||||[||||||||||||]PAM</td><td>NA</td><td>Intergenic</td><td>NA</td><td>NA</td></tr>
<tr><td>T15972</td><td>ACTCTACGACCAGCTCCCCGGGG</td><td>1000</td><td></td><td></td><td></td><td></td><td></td><td></td><td></td><td></td><td></td></tr>
<tr><td>Oligo fwd</td><td>CACCACTCTACGACCAGCTCCCCG</td><td></td><td></td><td></td><td></td><td></td><td></td><td></td><td></td><td></td><td></td></tr>
<tr><td>Oligo rev</td><td>AAACCGGGGAGCTGGTCGTAGAGT</td><td></td><td></td><td></td><td></td><td></td><td></td><td></td><td></td><td></td><td></td></tr>
<tr><td>Chromosome</td><td>start</td><td>end</td><td>strand</td><td>MM</td><td>target_seq</td><td>PAM</td><td>alignment</td><td>distance</td><td>position</td><td>gene name</td><td>gene id</td></tr>
<tr><td>GU305914.1</td><td>212545</td><td>212567</td><td>+</td><td>0</td><td>ACTCTACGACCAGCTCCCCG</td><td>GGG</td><td>||||||||[||||||||||||]PAM</td><td>NA</td><td>Intergenic</td><td>NA</td><td>NA</td></tr>
<tr><td>T15973</td><td>CTCTACGACCAGCTCCCCGGGGG</td><td>1000</td><td></td><td></td><td></td><td></td><td></td><td></td><td></td><td></td><td></td></tr>
<tr><td>Oligo fwd</td><td>CACCCTCTACGACCAGCTCCCCGG</td><td></td><td></td><td></td><td></td><td></td><td></td><td></td><td></td><td></td><td></td></tr>
<tr><td>Oligo rev</td><td>AAACCCGGGGAGCTGGTCGTAGAG</td><td></td><td></td><td></td><td></td><td></td><td></td><td></td><td></td><td></td><td></td></tr>
<tr><td>Chromosome</td><td>start</td><td>end</td><td>strand</td><td>MM</td><td>target_seq</td><td>PAM</td><td>alignment</td><td>distance</td><td>position</td><td>gene name</td><td>gene id</td></tr>
<tr><td>GU305914.1</td><td>212546</td><td>212568</td><td>+</td><td>0</td><td>CTCTACGACCAGCTCCCCGG</td><td>GGG</td><td>||||||||[||||||||||||]PAM</td><td>NA</td><td>Intergenic</td><td>NA</td><td>NA</td></tr>
<tr><td>T15974</td><td>CGACCAGCTCCCCGGGGGTCTGG</td><td>1000</td><td></td><td></td><td></td><td></td><td></td><td></td><td></td><td></td><td></td></tr>
<tr><td>Oligo fwd</td><td>CACCCGACCAGCTCCCCGGGGGTC</td><td></td><td></td><td></td><td></td><td></td><td></td><td></td><td></td><td></td><td></td></tr>
<tr><td>Oligo rev</td><td>AAACGACCCCCGGGGAGCTGGTCG</td><td></td><td></td><td></td><td></td><td></td><td></td><td></td><td></td><td></td><td></td></tr>
<tr><td>Chromosome</td><td>start</td><td>end</td><td>strand</td><td>MM</td><td>target_seq</td><td>PAM</td><td>alignment</td><td>distance</td><td>position</td><td>gene name</td><td>gene id</td></tr>
<tr><td>GU305914.1</td><td>212551</td><td>212573</td><td>+</td><td>0</td><td>CGACCAGCTCCCCGGGGGTC</td><td>TGG</td><td>||||||||[||||||||||||]PAM</td><td>NA</td><td>Intergenic</td><td>NA</td><td>NA</td></tr>
<tr><td>T15975</td><td>AGTACTTCGAGACGAACTCGCGG</td><td>1000</td><td></td><td></td><td></td><td></td><td></td><td></td><td></td><td></td><td></td></tr>
<tr><td>Oligo fwd</td><td>CACCAGTACTTCGAGACGAACTCG</td><td></td><td></td><td></td><td></td><td></td><td></td><td></td><td></td><td></td><td></td></tr>
<tr><td>Oligo rev</td><td>AAACCGAGTTCGTCTCGAAGTACT</td><td></td><td></td><td></td><td></td><td></td><td></td><td></td><td></td><td></td><td></td></tr>
<tr><td>Chromosome</td><td>start</td><td>end</td><td>strand</td><td>MM</td><td>target_seq</td><td>PAM</td><td>alignment</td><td>distance</td><td>position</td><td>gene name</td><td>gene id</td></tr>
<tr><td>GU305914.1</td><td>212587</td><td>212609</td><td>+</td><td>0</td><td>AGTACTTCGAGACGAACTCG</td><td>CGG</td><td>||||||||[||||||||||||]PAM</td><td>NA</td><td>Intergenic</td><td>NA</td><td>NA</td></tr>
<tr><td>T15976</td><td>CTGTAGTAGAAACTGTATCTCGG</td><td>1000</td><td></td><td></td><td></td><td></td><td></td><td></td><td></td><td></td><td></td></tr>
<tr><td>Oligo fwd</td><td>CACCCTGTAGTAGAAACTGTATCT</td><td></td><td></td><td></td><td></td><td></td><td></td><td></td><td></td><td></td><td></td></tr>
<tr><td>Oligo rev</td><td>AAACAGATACAGTTTCTACTACAG</td><td></td><td></td><td></td><td></td><td></td><td></td><td></td><td></td><td></td><td></td></tr>
<tr><td>Chromosome</td><td>start</td><td>end</td><td>strand</td><td>MM</td><td>target_seq</td><td>PAM</td><td>alignment</td><td>distance</td><td>position</td><td>gene name</td><td>gene id</td></tr>
<tr><td>GU305914.1</td><td>212612</td><td>212634</td><td>+</td><td>0</td><td>CTGTAGTAGAAACTGTATCT</td><td>CGG</td><td>||||||||[||||||||||||]PAM</td><td>NA</td><td>Intergenic</td><td>NA</td><td>NA</td></tr>
<tr><td>T15977</td><td>TGTAGTAGAAACTGTATCTCGGG</td><td>1000</td><td></td><td></td><td></td><td></td><td></td><td></td><td></td><td></td><td></td></tr>
<tr><td>Oligo fwd</td><td>CACCTGTAGTAGAAACTGTATCTC</td><td></td><td></td><td></td><td></td><td></td><td></td><td></td><td></td><td></td><td></td></tr>
<tr><td>Oligo rev</td><td>AAACGAGATACAGTTTCTACTACA</td><td></td><td></td><td></td><td></td><td></td><td></td><td></td><td></td><td></td><td></td></tr>
<tr><td>Chromosome</td><td>start</td><td>end</td><td>strand</td><td>MM</td><td>target_seq</td><td>PAM</td><td>alignment</td><td>distance</td><td>position</td><td>gene name</td><td>gene id</td></tr>
<tr><td>GU305914.1</td><td>212613</td><td>212635</td><td>+</td><td>0</td><td>TGTAGTAGAAACTGTATCTC</td><td>GGG</td><td>||||||||[||||||||||||]PAM</td><td>NA</td><td>Intergenic</td><td>NA</td><td>NA</td></tr>
<tr><td>T15978</td><td>CTCGGGCCACGACTCGTTCGCGG</td><td>1000</td><td></td><td></td><td></td><td></td><td></td><td></td><td></td><td></td><td></td></tr>
<tr><td>Oligo fwd</td><td>CACCCTCGGGCCACGACTCGTTCG</td><td></td><td></td><td></td><td></td><td></td><td></td><td></td><td></td><td></td><td></td></tr>
<tr><td>Oligo rev</td><td>AAACCGAACGAGTCGTGGCCCGAG</td><td></td><td></td><td></td><td></td><td></td><td></td><td></td><td></td><td></td><td></td></tr>
<tr><td>Chromosome</td><td>start</td><td>end</td><td>strand</td><td>MM</td><td>target_seq</td><td>PAM</td><td>alignment</td><td>distance</td><td>position</td><td>gene name</td><td>gene id</td></tr>
<tr><td>GU305914.1</td><td>212630</td><td>212652</td><td>+</td><td>0</td><td>CTCGGGCCACGACTCGTTCG</td><td>CGG</td><td>||||||||[||||||||||||]PAM</td><td>NA</td><td>Intergenic</td><td>NA</td><td>NA</td></tr>
<tr><td>T15979</td><td>TCGGGCCACGACTCGTTCGCGGG</td><td>1000</td><td></td><td></td><td></td><td></td><td></td><td></td><td></td><td></td><td></td></tr>
<tr><td>Oligo fwd</td><td>CACCTCGGGCCACGACTCGTTCGC</td><td></td><td></td><td></td><td></td><td></td><td></td><td></td><td></td><td></td><td></td></tr>
<tr><td>Oligo rev</td><td>AAACGCGAACGAGTCGTGGCCCGA</td><td></td><td></td><td></td><td></td><td></td><td></td><td></td><td></td><td></td><td></td></tr>
<tr><td>Chromosome</td><td>start</td><td>end</td><td>strand</td><td>MM</td><td>target_seq</td><td>PAM</td><td>alignment</td><td>distance</td><td>position</td><td>gene name</td><td>gene id</td></tr>
<tr><td>GU305914.1</td><td>212631</td><td>212653</td><td>+</td><td>0</td><td>TCGGGCCACGACTCGTTCGC</td><td>GGG</td><td>||||||||[||||||||||||]PAM</td><td>NA</td><td>Intergenic</td><td>NA</td><td>NA</td></tr>
<tr><td>T15980</td><td>AACGACCACTCATACGTGCCAGG</td><td>1000</td><td></td><td></td><td></td><td></td><td></td><td></td><td></td><td></td><td></td></tr>
<tr><td>Oligo fwd</td><td>CACCAACGACCACTCATACGTGCC</td><td></td><td></td><td></td><td></td><td></td><td></td><td></td><td></td><td></td><td></td></tr>
<tr><td>Oligo rev</td><td>AAACGGCACGTATGAGTGGTCGTT</td><td></td><td></td><td></td><td></td><td></td><td></td><td></td><td></td><td></td><td></td></tr>
<tr><td>Chromosome</td><td>start</td><td>end</td><td>strand</td><td>MM</td><td>target_seq</td><td>PAM</td><td>alignment</td><td>distance</td><td>position</td><td>gene name</td><td>gene id</td></tr>
<tr><td>GU305914.1</td><td>212655</td><td>212677</td><td>+</td><td>0</td><td>AACGACCACTCATACGTGCC</td><td>AGG</td><td>||||||||[||||||||||||]PAM</td><td>NA</td><td>Intergenic</td><td>NA</td><td>NA</td></tr>
</table>

<table>
<tr><td>T15981</td><td>TCTCCATGCTGACAATCGAGAGG</td><td>1000</td></tr>
<tr><td>Oligo fwd</td><td>CACCTCTCCATGCTGACAATCGAG</td></tr>
<tr><td>Oligo rev</td><td>AAACCTCGATTGTCAGCATGGAGA</td></tr>
<tr><td>Chromosome</td><td>start</td><td>end</td><td>strand</td><td>MM</td><td>target_seq</td><td>PAM</td><td>alignment</td><td>distance</td><td>position</td><td>gene name</td><td>gene id</td></tr>
<tr><td>GU305914.1</td><td>212689</td><td>212711</td><td>+</td><td>0</td><td>TCTCCATGCTGACAATCGAG</td><td>AGG</td><td>||||||||[||||||||||||]PAM</td><td>NA</td><td>Intergenic</td><td>NA</td><td>NA</td></tr>
<tr><td>T15982</td><td>CTCCATGCTGACAATCGAGAGGG</td><td>1000</td></tr>
<tr><td>Oligo fwd</td><td>CACCCTCCATGCTGACAATCGAGA</td></tr>
<tr><td>Oligo rev</td><td>AAACTCTCGATTGTCAGCATGGAG</td></tr>
<tr><td>Chromosome</td><td>start</td><td>end</td><td>strand</td><td>MM</td><td>target_seq</td><td>PAM</td><td>alignment</td><td>distance</td><td>position</td><td>gene name</td><td>gene id</td></tr>
<tr><td>GU305914.1</td><td>212690</td><td>212712</td><td>+</td><td>0</td><td>CTCCATGCTGACAATCGAGA</td><td>GGG</td><td>||||||||[||||||||||||]PAM</td><td>NA</td><td>Intergenic</td><td>NA</td><td>NA</td></tr>
<tr><td>T15983</td><td>CGTCGCCCCGCTCGATGTTCTGG</td><td>1000</td></tr>
<tr><td>Oligo fwd</td><td>CACCCGTCGCCCCGCTCGATGTTC</td></tr>
<tr><td>Oligo rev</td><td>AAACGAACATCGAGCGGGGCGACG</td></tr>
<tr><td>Chromosome</td><td>start</td><td>end</td><td>strand</td><td>MM</td><td>target_seq</td><td>PAM</td><td>alignment</td><td>distance</td><td>position</td><td>gene name</td><td>gene id</td></tr>
<tr><td>GU305914.1</td><td>212738</td><td>212760</td><td>+</td><td>0</td><td>CGTCGCCCCGCTCGATGTTC</td><td>TGG</td><td>||||||||[||||||||||||]PAM</td><td>NA</td><td>Intergenic</td><td>NA</td><td>NA</td></tr>
<tr><td>T15984</td><td>CCCGCTCGATGTTCTGGAGACGG</td><td>1000</td></tr>
<tr><td>Oligo fwd</td><td>CACCCCCGCTCGATGTTCTGGAGA</td></tr>
<tr><td>Oligo rev</td><td>AAACTCTCCAGAACATCGAGCGGG</td></tr>
<tr><td>Chromosome</td><td>start</td><td>end</td><td>strand</td><td>MM</td><td>target_seq</td><td>PAM</td><td>alignment</td><td>distance</td><td>position</td><td>gene name</td><td>gene id</td></tr>
<tr><td>GU305914.1</td><td>212744</td><td>212766</td><td>+</td><td>0</td><td>CCCGCTCGATGTTCTGGAGA</td><td>CGG</td><td>||||||||[||||||||||||]PAM</td><td>NA</td><td>Intergenic</td><td>NA</td><td>NA</td></tr>
<tr><td>T15985</td><td>ACGGAAAAGATGAGCTCCATAGG</td><td>1000</td></tr>
<tr><td>Oligo fwd</td><td>CACCACGGAAAAGATGAGCTCCAT</td></tr>
<tr><td>Oligo rev</td><td>AAACATGGAGCTCATCTTTTCCGT</td></tr>
<tr><td>Chromosome</td><td>start</td><td>end</td><td>strand</td><td>MM</td><td>target_seq</td><td>PAM</td><td>alignment</td><td>distance</td><td>position</td><td>gene name</td><td>gene id</td></tr>
<tr><td>GU305914.1</td><td>212763</td><td>212785</td><td>+</td><td>0</td><td>ACGGAAAAGATGAGCTCCAT</td><td>AGG</td><td>||||||||[||||||||||||]PAM</td><td>NA</td><td>Intergenic</td><td>NA</td><td>NA</td></tr>
<tr><td>T15986</td><td>AGATGAGCTCCATAGGTCCCTGG</td><td>1000</td></tr>
<tr><td>Oligo fwd</td><td>CACCAGATGAGCTCCATAGGTCCC</td></tr>
<tr><td>Oligo rev</td><td>AAACGGGACCTATGGAGCTCATCT</td></tr>
<tr><td>Chromosome</td><td>start</td><td>end</td><td>strand</td><td>MM</td><td>target_seq</td><td>PAM</td><td>alignment</td><td>distance</td><td>position</td><td>gene name</td><td>gene id</td></tr>
<tr><td>GU305914.1</td><td>212770</td><td>212792</td><td>+</td><td>0</td><td>AGATGAGCTCCATAGGTCCC</td><td>TGG</td><td>||||||||[||||||||||||]PAM</td><td>NA</td><td>Intergenic</td><td>NA</td><td>NA</td></tr>
<tr><td>T15987</td><td>CGTTCCCGTCCTGTTAACTTCGG</td><td>1000</td></tr>
<tr><td>Oligo fwd</td><td>CACCCGTTCCCGTCCTGTTAACTT</td></tr>
<tr><td>Oligo rev</td><td>AAACAAGTTAACAGGACGGGAACG</td></tr>
<tr><td>Chromosome</td><td>start</td><td>end</td><td>strand</td><td>MM</td><td>target_seq</td><td>PAM</td><td>alignment</td><td>distance</td><td>position</td><td>gene name</td><td>gene id</td></tr>
<tr><td>GU305914.1</td><td>212804</td><td>212826</td><td>+</td><td>0</td><td>CGTTCCCGTCCTGTTAACTT</td><td>CGG</td><td>||||||||[||||||||||||]PAM</td><td>NA</td><td>Intergenic</td><td>NA</td><td>NA</td></tr>
<tr><td>T15988</td><td>GTTCCCGTCCTGTTAACTTCGGG</td><td>1000</td></tr>
<tr><td>Oligo fwd</td><td>CACCGTTCCCGTCCTGTTAACTTC</td></tr>
<tr><td>Oligo rev</td><td>AAACGAAGTTAACAGGACGGGAAC</td></tr>
<tr><td>Chromosome</td><td>start</td><td>end</td><td>strand</td><td>MM</td><td>target_seq</td><td>PAM</td><td>alignment</td><td>distance</td><td>position</td><td>gene name</td><td>gene id</td></tr>
<tr><td>GU305914.1</td><td>212805</td><td>212827</td><td>+</td><td>0</td><td>GTTCCCGTCCTGTTAACTTC</td><td>GGG</td><td>||||||||[||||||||||||]PAM</td><td>NA</td><td>Intergenic</td><td>NA</td><td>NA</td></tr>
<tr><td>T15989</td><td>CCTCCTGAGATGAGAACCAACGG</td><td>1000</td></tr>
<tr><td>Oligo fwd</td><td>CACCCCTCCTGAGATGAGAACCAA</td></tr>
<tr><td>Oligo rev</td><td>AAACTTGGTTCTCATCTCAGGAGG</td></tr>
<tr><td>Chromosome</td><td>start</td><td>end</td><td>strand</td><td>MM</td><td>target_seq</td><td>PAM</td><td>alignment</td><td>distance</td><td>position</td><td>gene name</td><td>gene id</td></tr>
<tr><td>GU305914.1</td><td>212862</td><td>212884</td><td>+</td><td>0</td><td>CCTCCTGAGATGAGAACCAA</td><td>CGG</td><td>||||||||[||||||||||||]PAM</td><td>NA</td><td>Intergenic</td><td>NA</td><td>NA</td></tr>
<tr><td>T15990</td><td>AGATGAGAACCAACGGCAGAAGG</td><td>1000</td></tr>
<tr><td>Oligo fwd</td><td>CACCAGATGAGAACCAACGGCAGA</td></tr>
<tr><td>Oligo rev</td><td>AAACTCTGCCGTTGGTTCTCATCT</td></tr>
<tr><td>Chromosome</td><td>start</td><td>end</td><td>strand</td><td>MM</td><td>target_seq</td><td>PAM</td><td>alignment</td><td>distance</td><td>position</td><td>gene name</td><td>gene id</td></tr>
<tr><td>GU305914.1</td><td>212869</td><td>212891</td><td>+</td><td>0</td><td>AGATGAGAACCAACGGCAGA</td><td>AGG</td><td>||||||||[||||||||||||]PAM</td><td>NA</td><td>Intergenic</td><td>NA</td><td>NA</td></tr>
<tr><td>T15991</td><td>GAACCAACGGCAGAAGGAGAAGG</td><td>1000</td></tr>
<tr><td>Oligo fwd</td><td>CACCGAACCAACGGCAGAAGGAGA</td></tr>
<tr><td>Oligo rev</td><td>AAACTCTCCTTCTGCCGTTGGTTC</td></tr>
<tr><td>Chromosome</td><td>start</td><td>end</td><td>strand</td><td>MM</td><td>target_seq</td><td>PAM</td><td>alignment</td><td>distance</td><td>position</td><td>gene name</td><td>gene id</td></tr>
<tr><td>GU305914.1</td><td>212875</td><td>212897</td><td>+</td><td>0</td><td>GAACCAACGGCAGAAGGAGA</td><td>AGG</td><td>||||||||[||||||||||||]PAM</td><td>NA</td><td>Intergenic</td><td>NA</td><td>NA</td></tr>
<tr><td>T15992</td><td>AACCAACGGCAGAAGGAGAAGGG</td><td>1000</td></tr>
<tr><td>Oligo fwd</td><td>CACCAACCAACGGCAGAAGGAGAA</td></tr>
<tr><td>Oligo rev</td><td>AAACTTCTCCTTCTGCCGTTGGTT</td></tr>
<tr><td>Chromosome</td><td>start</td><td>end</td><td>strand</td><td>MM</td><td>target_seq</td><td>PAM</td><td>alignment</td><td>distance</td><td>position</td><td>gene name</td><td>gene id</td></tr>
<tr><td>GU305914.1</td><td>212876</td><td>212898</td><td>+</td><td>0</td><td>AACCAACGGCAGAAGGAGAA</td><td>GGG</td><td>||||||||[||||||||||||]PAM</td><td>NA</td><td>Intergenic</td><td>NA</td><td>NA</td></tr>
<tr><td>T15993</td><td>ACCAACGGCAGAAGGAGAAGGGG</td><td>1000</td></tr>
<tr><td>Oligo fwd</td><td>CACCACCAACGGCAGAAGGAGAAG</td></tr>
<tr><td>Oligo rev</td><td>AAACCTTCTCCTTCTGCCGTTGGT</td></tr>
<tr><td>Chromosome</td><td>start</td><td>end</td><td>strand</td><td>MM</td><td>target_seq</td><td>PAM</td><td>alignment</td><td>distance</td><td>position</td><td>gene name</td><td>gene id</td></tr>
<tr><td>GU305914.1</td><td>212877</td><td>212899</td><td>+</td><td>0</td><td>ACCAACGGCAGAAGGAGAAG</td><td>GGG</td><td>||||||||[||||||||||||]PAM</td><td>NA</td><td>Intergenic</td><td>NA</td><td>NA</td></tr>
<tr><td>T15994</td><td>AAGGGGAATCATCGTTCTCCCGG</td><td>1000</td></tr>
<tr><td>Oligo fwd</td><td>CACCAAGGGGAATCATCGTTCTCC</td></tr>
<tr><td>Oligo rev</td><td>AAACGGAGAACGATGATTCCCCTT</td></tr>
<tr><td>Chromosome</td><td>start</td><td>end</td><td>strand</td><td>MM</td><td>target_seq</td><td>PAM</td><td>alignment</td><td>distance</td><td>position</td><td>gene name</td><td>gene id</td></tr>
<tr><td>GU305914.1</td><td>212894</td><td>212916</td><td>+</td><td>0</td><td>AAGGGGAATCATCGTTCTCC</td><td>CGG</td><td>||||||||[||||||||||||]PAM</td><td>NA</td><td>Intergenic</td><td>NA</td><td>NA</td></tr>
<tr><td>T15995</td><td>CCGGTACCCAGCTTTGTGTGTGG</td><td>1000</td></tr>
<tr><td>Oligo fwd</td><td>CACCCCGGTACCCAGCTTTGTGTG</td></tr>
<tr><td>Oligo rev</td><td>AAACCACACAAAGCTGGGTACCGG</td></tr>
<tr><td>Chromosome</td><td>start</td><td>end</td><td>strand</td><td>MM</td><td>target_seq</td><td>PAM</td><td>alignment</td><td>distance</td><td>position</td><td>gene name</td><td>gene id</td></tr>
<tr><td>GU305914.1</td><td>212913</td><td>212935</td><td>+</td><td>0</td><td>CCGGTACCCAGCTTTGTGTG</td><td>TGG</td><td>||||||||[||||||||||||]PAM</td><td>NA</td><td>Intergenic</td><td>NA</td><td>NA</td></tr>
</table>

<table>
<tr><td>T15996</td><td>CGGTACCCAGCTTTGTGTGTGGG</td><td>1000</td><td></td><td></td><td></td><td></td><td></td><td></td><td></td><td></td><td></td></tr>
<tr><td>Oligo fwd</td><td>CACCCGGTACCCAGCTTTGTGTGT</td><td></td><td></td><td></td><td></td><td></td><td></td><td></td><td></td><td></td><td></td></tr>
<tr><td>Oligo rev</td><td>AAACACACACAAAGCTGGGTACCG</td><td></td><td></td><td></td><td></td><td></td><td></td><td></td><td></td><td></td><td></td></tr>
<tr><td>Chromosome</td><td>start</td><td>end</td><td>strand</td><td>MM</td><td>target_seq</td><td>PAM</td><td>alignment</td><td>distance</td><td>position</td><td>gene name</td><td>gene id</td></tr>
<tr><td>GU305914.1</td><td>212914</td><td>212936</td><td>+</td><td>0</td><td>CGGTACCCAGCTTTGTGTGT</td><td>GGG</td><td>||||||||[||||||||||||]PAM</td><td>NA</td><td>Intergenic</td><td>NA</td><td>NA</td></tr>
<tr><td>T15997</td><td>GGTACCCAGCTTTGTGTGTGGGG</td><td>1000</td><td></td><td></td><td></td><td></td><td></td><td></td><td></td><td></td><td></td></tr>
<tr><td>Oligo fwd</td><td>CACCGGTACCCAGCTTTGTGTGTG</td><td></td><td></td><td></td><td></td><td></td><td></td><td></td><td></td><td></td><td></td></tr>
<tr><td>Oligo rev</td><td>AAACCACACACAAAGCTGGGTACC</td><td></td><td></td><td></td><td></td><td></td><td></td><td></td><td></td><td></td><td></td></tr>
<tr><td>Chromosome</td><td>start</td><td>end</td><td>strand</td><td>MM</td><td>target_seq</td><td>PAM</td><td>alignment</td><td>distance</td><td>position</td><td>gene name</td><td>gene id</td></tr>
<tr><td>GU305914.1</td><td>212915</td><td>212937</td><td>+</td><td>0</td><td>GGTACCCAGCTTTGTGTGTG</td><td>GGG</td><td>||||||||[||||||||||||]PAM</td><td>NA</td><td>Intergenic</td><td>NA</td><td>NA</td></tr>
<tr><td>T15998</td><td>GCCGATAAATCACGCCCTTATGG</td><td>1000</td><td></td><td></td><td></td><td></td><td></td><td></td><td></td><td></td><td></td></tr>
<tr><td>Oligo fwd</td><td>CACCGCCGATAAATCACGCCCTTA</td><td></td><td></td><td></td><td></td><td></td><td></td><td></td><td></td><td></td><td></td></tr>
<tr><td>Oligo rev</td><td>AAACTAAGGGCGTGATTTATCGGC</td><td></td><td></td><td></td><td></td><td></td><td></td><td></td><td></td><td></td><td></td></tr>
<tr><td>Chromosome</td><td>start</td><td>end</td><td>strand</td><td>MM</td><td>target_seq</td><td>PAM</td><td>alignment</td><td>distance</td><td>position</td><td>gene name</td><td>gene id</td></tr>
<tr><td>GU305914.1</td><td>212955</td><td>212977</td><td>+</td><td>0</td><td>GCCGATAAATCACGCCCTTA</td><td>TGG</td><td>||||||||[||||||||||||]PAM</td><td>NA</td><td>Intergenic</td><td>NA</td><td>NA</td></tr>
<tr><td>T15999</td><td>CACCCTTATCGTCTTCCCAACGG</td><td>1000</td><td></td><td></td><td></td><td></td><td></td><td></td><td></td><td></td><td></td></tr>
<tr><td>Oligo fwd</td><td>CACCCACCCTTATCGTCTTCCCAA</td><td></td><td></td><td></td><td></td><td></td><td></td><td></td><td></td><td></td><td></td></tr>
<tr><td>Oligo rev</td><td>AAACTTGGGAAGACGATAAGGGTG</td><td></td><td></td><td></td><td></td><td></td><td></td><td></td><td></td><td></td><td></td></tr>
<tr><td>Chromosome</td><td>start</td><td>end</td><td>strand</td><td>MM</td><td>target_seq</td><td>PAM</td><td>alignment</td><td>distance</td><td>position</td><td>gene name</td><td>gene id</td></tr>
<tr><td>GU305914.1</td><td>213087</td><td>213109</td><td>+</td><td>0</td><td>CACCCTTATCGTCTTCCCAA</td><td>CGG</td><td>||||||||[||||||||||||]PAM</td><td>NA</td><td>Intergenic</td><td>NA</td><td>NA</td></tr>
<tr><td>T16000</td><td>CTTCCCAACGGTGTGTGATCAGG</td><td>1000</td><td></td><td></td><td></td><td></td><td></td><td></td><td></td><td></td><td></td></tr>
<tr><td>Oligo fwd</td><td>CACCCTTCCCAACGGTGTGTGATC</td><td></td><td></td><td></td><td></td><td></td><td></td><td></td><td></td><td></td><td></td></tr>
<tr><td>Oligo rev</td><td>AAACGATCACACACCGTTGGGAAG</td><td></td><td></td><td></td><td></td><td></td><td></td><td></td><td></td><td></td><td></td></tr>
<tr><td>Chromosome</td><td>start</td><td>end</td><td>strand</td><td>MM</td><td>target_seq</td><td>PAM</td><td>alignment</td><td>distance</td><td>position</td><td>gene name</td><td>gene id</td></tr>
<tr><td>GU305914.1</td><td>213099</td><td>213121</td><td>+</td><td>0</td><td>CTTCCCAACGGTGTGTGATC</td><td>AGG</td><td>||||||||[||||||||||||]PAM</td><td>NA</td><td>Intergenic</td><td>NA</td><td>NA</td></tr>
<tr><td>T16001</td><td>TGTGATCAGGAGTACTCCCGCGG</td><td>1000</td><td></td><td></td><td></td><td></td><td></td><td></td><td></td><td></td><td></td></tr>
<tr><td>Oligo fwd</td><td>CACCTGTGATCAGGAGTACTCCCG</td><td></td><td></td><td></td><td></td><td></td><td></td><td></td><td></td><td></td><td></td></tr>
<tr><td>Oligo rev</td><td>AAACCGGGAGTACTCCTGATCACA</td><td></td><td></td><td></td><td></td><td></td><td></td><td></td><td></td><td></td><td></td></tr>
<tr><td>Chromosome</td><td>start</td><td>end</td><td>strand</td><td>MM</td><td>target_seq</td><td>PAM</td><td>alignment</td><td>distance</td><td>position</td><td>gene name</td><td>gene id</td></tr>
<tr><td>GU305914.1</td><td>213112</td><td>213134</td><td>+</td><td>0</td><td>TGTGATCAGGAGTACTCCCG</td><td>CGG</td><td>||||||||[||||||||||||]PAM</td><td>NA</td><td>Intergenic</td><td>NA</td><td>NA</td></tr>
<tr><td>T16002</td><td>GATCCCAGATAAAATCACAACGG</td><td>1000</td><td></td><td></td><td></td><td></td><td></td><td></td><td></td><td></td><td></td></tr>
<tr><td>Oligo fwd</td><td>CACCGATCCCAGATAAAATCACAA</td><td></td><td></td><td></td><td></td><td></td><td></td><td></td><td></td><td></td><td></td></tr>
<tr><td>Oligo rev</td><td>AAACTTGTGATTTTATCTGGGATC</td><td></td><td></td><td></td><td></td><td></td><td></td><td></td><td></td><td></td><td></td></tr>
<tr><td>Chromosome</td><td>start</td><td>end</td><td>strand</td><td>MM</td><td>target_seq</td><td>PAM</td><td>alignment</td><td>distance</td><td>position</td><td>gene name</td><td>gene id</td></tr>
<tr><td>GU305914.1</td><td>213136</td><td>213158</td><td>+</td><td>0</td><td>GATCCCAGATAAAATCACAA</td><td>CGG</td><td>||||||||[||||||||||||]PAM</td><td>NA</td><td>Intergenic</td><td>NA</td><td>NA</td></tr>
<tr><td>T16003</td><td>ATAAAATCACAACGGTGATCAGG</td><td>1000</td><td></td><td></td><td></td><td></td><td></td><td></td><td></td><td></td><td></td></tr>
<tr><td>Oligo fwd</td><td>CACCATAAAATCACAACGGTGATC</td><td></td><td></td><td></td><td></td><td></td><td></td><td></td><td></td><td></td><td></td></tr>
<tr><td>Oligo rev</td><td>AAACGATCACCGTTGTGATTTTAT</td><td></td><td></td><td></td><td></td><td></td><td></td><td></td><td></td><td></td><td></td></tr>
<tr><td>Chromosome</td><td>start</td><td>end</td><td>strand</td><td>MM</td><td>target_seq</td><td>PAM</td><td>alignment</td><td>distance</td><td>position</td><td>gene name</td><td>gene id</td></tr>
<tr><td>GU305914.1</td><td>213144</td><td>213166</td><td>+</td><td>0</td><td>ATAAAATCACAACGGTGATC</td><td>AGG</td><td>||||||||[||||||||||||]PAM</td><td>NA</td><td>Intergenic</td><td>NA</td><td>NA</td></tr>
<tr><td>T16004</td><td>AGTATCGTGTCGAGCCGATTCGG</td><td>1000</td><td></td><td></td><td></td><td></td><td></td><td></td><td></td><td></td><td></td></tr>
<tr><td>Oligo fwd</td><td>CACCAGTATCGTGTCGAGCCGATT</td><td></td><td></td><td></td><td></td><td></td><td></td><td></td><td></td><td></td><td></td></tr>
<tr><td>Oligo rev</td><td>AAACAATCGGCTCGACACGATACT</td><td></td><td></td><td></td><td></td><td></td><td></td><td></td><td></td><td></td><td></td></tr>
<tr><td>Chromosome</td><td>start</td><td>end</td><td>strand</td><td>MM</td><td>target_seq</td><td>PAM</td><td>alignment</td><td>distance</td><td>position</td><td>gene name</td><td>gene id</td></tr>
<tr><td>GU305914.1</td><td>213173</td><td>213195</td><td>+</td><td>0</td><td>AGTATCGTGTCGAGCCGATT</td><td>CGG</td><td>||||||||[||||||||||||]PAM</td><td>NA</td><td>Intergenic</td><td>NA</td><td>NA</td></tr>
<tr><td>T16005</td><td>GTATCGTGTCGAGCCGATTCGGG</td><td>1000</td><td></td><td></td><td></td><td></td><td></td><td></td><td></td><td></td><td></td></tr>
<tr><td>Oligo fwd</td><td>CACCGTATCGTGTCGAGCCGATTC</td><td></td><td></td><td></td><td></td><td></td><td></td><td></td><td></td><td></td><td></td></tr>
<tr><td>Oligo rev</td><td>AAACGAATCGGCTCGACACGATAC</td><td></td><td></td><td></td><td></td><td></td><td></td><td></td><td></td><td></td><td></td></tr>
<tr><td>Chromosome</td><td>start</td><td>end</td><td>strand</td><td>MM</td><td>target_seq</td><td>PAM</td><td>alignment</td><td>distance</td><td>position</td><td>gene name</td><td>gene id</td></tr>
<tr><td>GU305914.1</td><td>213174</td><td>213196</td><td>+</td><td>0</td><td>GTATCGTGTCGAGCCGATTC</td><td>GGG</td><td>||||||||[||||||||||||]PAM</td><td>NA</td><td>Intergenic</td><td>NA</td><td>NA</td></tr>
<tr><td>T16006</td><td>CGAGCCGATTCGGGCCTATCTGG</td><td>1000</td><td></td><td></td><td></td><td></td><td></td><td></td><td></td><td></td><td></td></tr>
<tr><td>Oligo fwd</td><td>CACCCGAGCCGATTCGGGCCTATC</td><td></td><td></td><td></td><td></td><td></td><td></td><td></td><td></td><td></td><td></td></tr>
<tr><td>Oligo rev</td><td>AAACGATAGGCCCGAATCGGCTCG</td><td></td><td></td><td></td><td></td><td></td><td></td><td></td><td></td><td></td><td></td></tr>
<tr><td>Chromosome</td><td>start</td><td>end</td><td>strand</td><td>MM</td><td>target_seq</td><td>PAM</td><td>alignment</td><td>distance</td><td>position</td><td>gene name</td><td>gene id</td></tr>
<tr><td>GU305914.1</td><td>213183</td><td>213205</td><td>+</td><td>0</td><td>CGAGCCGATTCGGGCCTATC</td><td>TGG</td><td>||||||||[||||||||||||]PAM</td><td>NA</td><td>Intergenic</td><td>NA</td><td>NA</td></tr>
<tr><td>T16007</td><td>GGTTTTCCGTATCCTCATCCCGG</td><td>1000</td><td></td><td></td><td></td><td></td><td></td><td></td><td></td><td></td><td></td></tr>
<tr><td>Oligo fwd</td><td>CACCGGTTTTCCGTATCCTCATCC</td><td></td><td></td><td></td><td></td><td></td><td></td><td></td><td></td><td></td><td></td></tr>
<tr><td>Oligo rev</td><td>AAACGGATGAGGATACGGAAAACC</td><td></td><td></td><td></td><td></td><td></td><td></td><td></td><td></td><td></td><td></td></tr>
<tr><td>Chromosome</td><td>start</td><td>end</td><td>strand</td><td>MM</td><td>target_seq</td><td>PAM</td><td>alignment</td><td>distance</td><td>position</td><td>gene name</td><td>gene id</td></tr>
<tr><td>GU305914.1</td><td>213204</td><td>213226</td><td>+</td><td>0</td><td>GGTTTTCCGTATCCTCATCC</td><td>CGG</td><td>||||||||[||||||||||||]PAM</td><td>NA</td><td>Intergenic</td><td>NA</td><td>NA</td></tr>
<tr><td>T16008</td><td>CCCGGTCGATCAGCGTCGTGAGG</td><td>1000</td><td></td><td></td><td></td><td></td><td></td><td></td><td></td><td></td><td></td></tr>
<tr><td>Oligo fwd</td><td>CACCCCCGGTCGATCAGCGTCGTG</td><td></td><td></td><td></td><td></td><td></td><td></td><td></td><td></td><td></td><td></td></tr>
<tr><td>Oligo rev</td><td>AAACCACGACGCTGATCGACCGGG</td><td></td><td></td><td></td><td></td><td></td><td></td><td></td><td></td><td></td><td></td></tr>
<tr><td>Chromosome</td><td>start</td><td>end</td><td>strand</td><td>MM</td><td>target_seq</td><td>PAM</td><td>alignment</td><td>distance</td><td>position</td><td>gene name</td><td>gene id</td></tr>
<tr><td>GU305914.1</td><td>213222</td><td>213244</td><td>+</td><td>0</td><td>CCCGGTCGATCAGCGTCGTG</td><td>AGG</td><td>||||||||[||||||||||||]PAM</td><td>NA</td><td>Intergenic</td><td>NA</td><td>NA</td></tr>
<tr><td>T16009</td><td>CCGGTCGATCAGCGTCGTGAGGG</td><td>1000</td><td></td><td></td><td></td><td></td><td></td><td></td><td></td><td></td><td></td></tr>
<tr><td>Oligo fwd</td><td>CACCCCGGTCGATCAGCGTCGTGA</td><td></td><td></td><td></td><td></td><td></td><td></td><td></td><td></td><td></td><td></td></tr>
<tr><td>Oligo rev</td><td>AAACTCACGACGCTGATCGACCGG</td><td></td><td></td><td></td><td></td><td></td><td></td><td></td><td></td><td></td><td></td></tr>
<tr><td>Chromosome</td><td>start</td><td>end</td><td>strand</td><td>MM</td><td>target_seq</td><td>PAM</td><td>alignment</td><td>distance</td><td>position</td><td>gene name</td><td>gene id</td></tr>
<tr><td>GU305914.1</td><td>213223</td><td>213245</td><td>+</td><td>0</td><td>CCGGTCGATCAGCGTCGTGA</td><td>GGG</td><td>||||||||[||||||||||||]PAM</td><td>NA</td><td>Intergenic</td><td>NA</td><td>NA</td></tr>
<tr><td>T16010</td><td>GTCGATCAGCGTCGTGAGGGTGG</td><td>1000</td><td></td><td></td><td></td><td></td><td></td><td></td><td></td><td></td><td></td></tr>
<tr><td>Oligo fwd</td><td>CACCGTCGATCAGCGTCGTGAGGG</td><td></td><td></td><td></td><td></td><td></td><td></td><td></td><td></td><td></td><td></td></tr>
<tr><td>Oligo rev</td><td>AAACCCCTCACGACGCTGATCGAC</td><td></td><td></td><td></td><td></td><td></td><td></td><td></td><td></td><td></td><td></td></tr>
<tr><td>Chromosome</td><td>start</td><td>end</td><td>strand</td><td>MM</td><td>target_seq</td><td>PAM</td><td>alignment</td><td>distance</td><td>position</td><td>gene name</td><td>gene id</td></tr>
</table>

| | | | | | | | | | | | |
|---|---|---|---|---|---|---|---|---|---|---|---|
| GU305914.1 | 213226 | 213248 | + | 0 | GTCGATCAGCGTCGTGAGGG | TGG | ||||||||[||||||||||||]PAM | NA | Intergenic | NA | NA |
| T16011 | GGTGATCGTGTCACCCTCGTCGG | 1000 | | | | | | | | | |
| Oligo fwd | CACCGGTGATCGTGTCACCCTCGT | | | | | | | | | | |
| Oligo rev | AAACACGAGGGTGACACGATCACC | | | | | | | | | | |
| Chromosome | start | end | strand | MM | target_seq | PAM | alignment | distance | position | gene name | gene id |
| GU305914.1 | 213247 | 213269 | + | 0 | GGTGATCGTGTCACCCTCGT | CGG | ||||||||[||||||||||||]PAM | NA | Intergenic | NA | NA |
| T16012 | GATCGTGTCACCCTCGTCGGAGG | 1000 | | | | | | | | | |
| Oligo fwd | CACCGATCGTGTCACCCTCGTCGG | | | | | | | | | | |
| Oligo rev | AAACCCGACGAGGGTGACACGATC | | | | | | | | | | |
| Chromosome | start | end | strand | MM | target_seq | PAM | alignment | distance | position | gene name | gene id |
| GU305914.1 | 213250 | 213272 | + | 0 | GATCGTGTCACCCTCGTCGG | AGG | ||||||||[||||||||||||]PAM | NA | Intergenic | NA | NA |
| T16013 | CGTGTCACCCTCGTCGGAGGTGG | 1000 | | | | | | | | | |
| Oligo fwd | CACCCGTGTCACCCTCGTCGGAGG | | | | | | | | | | |
| Oligo rev | AAACCCTCCGACGAGGGTGACACG | | | | | | | | | | |
| Chromosome | start | end | strand | MM | target_seq | PAM | alignment | distance | position | gene name | gene id |
| GU305914.1 | 213253 | 213275 | + | 0 | CGTGTCACCCTCGTCGGAGG | TGG | ||||||||[||||||||||||]PAM | NA | Intergenic | NA | NA |
| T16014 | GTCACCCTCGTCGGAGGTGGCGG | 1000 | | | | | | | | | |
| Oligo fwd | CACCGTCACCCTCGTCGGAGGTGG | | | | | | | | | | |
| Oligo rev | AAACCCACCTCCGACGAGGGTGAC | | | | | | | | | | |
| Chromosome | start | end | strand | MM | target_seq | PAM | alignment | distance | position | gene name | gene id |
| GU305914.1 | 213256 | 213278 | + | 0 | GTCACCCTCGTCGGAGGTGG | CGG | ||||||||[||||||||||||]PAM | NA | Intergenic | NA | NA |
| T16015 | CCCTCGTCGGAGGTGGCGGTCGG | 1000 | | | | | | | | | |
| Oligo fwd | CACCCCCTCGTCGGAGGTGGCGGT | | | | | | | | | | |
| Oligo rev | AAACACCGCCACCTCCGACGAGGG | | | | | | | | | | |
| Chromosome | start | end | strand | MM | target_seq | PAM | alignment | distance | position | gene name | gene id |
| GU305914.1 | 213260 | 213282 | + | 0 | CCCTCGTCGGAGGTGGCGGT | CGG | ||||||||[||||||||||||]PAM | NA | Intergenic | NA | NA |
| T16016 | GTCGGAGGTGGCGGTCGGAGAGG | 1000 | | | | | | | | | |
| Oligo fwd | CACCGTCGGAGGTGGCGGTCGGAG | | | | | | | | | | |
| Oligo rev | AAACCTCCGACCGCCACCTCCGAC | | | | | | | | | | |
| Chromosome | start | end | strand | MM | target_seq | PAM | alignment | distance | position | gene name | gene id |
| GU305914.1 | 213265 | 213287 | + | 0 | GTCGGAGGTGGCGGTCGGAG | AGG | ||||||||[||||||||||||]PAM | NA | Intergenic | NA | NA |
| GU305914.1 | 75857 | 75879 | + | 3 | GGCGGAGGCGGCGGTCGGCG | AGG | |-||||||[-|||||||||-|]PAM | NA | Intergenic | NA | NA |
| T16017 | GGAGGTGGCGGTCGGAGAGGCGG | 1000 | | | | | | | | | |
| Oligo fwd | CACCGGAGGTGGCGGTCGGAGAGG | | | | | | | | | | |
| Oligo rev | AAACCCTCTCCGACCGCCACCTCC | | | | | | | | | | |
| Chromosome | start | end | strand | MM | target_seq | PAM | alignment | distance | position | gene name | gene id |
| GU305914.1 | 213268 | 213290 | + | 0 | GGAGGTGGCGGTCGGAGAGG | CGG | ||||||||[||||||||||||]PAM | NA | Intergenic | NA | NA |
| GU305914.1 | 75860 | 75882 | + | 2 | GGAGGCGGCGGTCGGCGAGG | GGG | |||||-||[|||||||-||||]PAM | NA | Intergenic | NA | NA |
| GU305914.1 | 75736 | 75758 | + | 3 | GGAAGTGGAGGTCGGCGAGG | CGG | |||-||||[-||||||-||||]PAM | NA | Intergenic | NA | NA |
| GU305914.1 | 173582 | 173604 | + | 4 | CGAGGTGTCGGTCGGCGACG | AGG | -||||||-[|||||||-||-|]PAM | NA | Intergenic | NA | NA |
| T16018 | GGTAGTCCCTGTTCTCAGCCTGG | 1000 | | | | | | | | | |
| Oligo fwd | CACCGGTAGTCCCTGTTCTCAGCC | | | | | | | | | | |
| Oligo rev | AAACGGCTGAGAACAGGGACTACC | | | | | | | | | | |
| Chromosome | start | end | strand | MM | target_seq | PAM | alignment | distance | position | gene name | gene id |
| GU305914.1 | 213289 | 213311 | + | 0 | GGTAGTCCCTGTTCTCAGCC | TGG | ||||||||[||||||||||||]PAM | NA | Intergenic | NA | NA |
| T16019 | TCTCAGCCTGGTCGTACTCAAGG | 1000 | | | | | | | | | |
| Oligo fwd | CACCTCTCAGCCTGGTCGTACTCA | | | | | | | | | | |
| Oligo rev | AAACTGAGTACGACCAGGCTGAGA | | | | | | | | | | |
| Chromosome | start | end | strand | MM | target_seq | PAM | alignment | distance | position | gene name | gene id |
| GU305914.1 | 213301 | 213323 | + | 0 | TCTCAGCCTGGTCGTACTCA | AGG | ||||||||[||||||||||||]PAM | NA | Intergenic | NA | NA |
| T16020 | CCTGGTCGTACTCAAGGTCGTGG | 1000 | | | | | | | | | |
| Oligo fwd | CACCCCTGGTCGTACTCAAGGTCG | | | | | | | | | | |
| Oligo rev | AAACCGACCTTGAGTACGACCAGG | | | | | | | | | | |
| Chromosome | start | end | strand | MM | target_seq | PAM | alignment | distance | position | gene name | gene id |
| GU305914.1 | 213307 | 213329 | + | 0 | CCTGGTCGTACTCAAGGTCG | TGG | ||||||||[||||||||||||]PAM | NA | Intergenic | NA | NA |
| T16021 | GTCGTACTCAAGGTCGTGGTCGG | 1000 | | | | | | | | | |
| Oligo fwd | CACCGTCGTACTCAAGGTCGTGGT | | | | | | | | | | |
| Oligo rev | AAACACCACGACCTTGAGTACGAC | | | | | | | | | | |
| Chromosome | start | end | strand | MM | target_seq | PAM | alignment | distance | position | gene name | gene id |
| GU305914.1 | 213311 | 213333 | + | 0 | GTCGTACTCAAGGTCGTGGT | CGG | ||||||||[||||||||||||]PAM | NA | Intergenic | NA | NA |
| T16022 | TCGTACTCAAGGTCGTGGTCGGG | 1000 | | | | | | | | | |
| Oligo fwd | CACCTCGTACTCAAGGTCGTGGTC | | | | | | | | | | |
| Oligo rev | AAACGACCACGACCTTGAGTACGA | | | | | | | | | | |
| Chromosome | start | end | strand | MM | target_seq | PAM | alignment | distance | position | gene name | gene id |
| GU305914.1 | 213312 | 213334 | + | 0 | TCGTACTCAAGGTCGTGGTC | GGG | ||||||||[||||||||||||]PAM | NA | Intergenic | NA | NA |
| T16023 | ACTCAAGGTCGTGGTCGGGATGG | 1000 | | | | | | | | | |
| Oligo fwd | CACCACTCAAGGTCGTGGTCGGGA | | | | | | | | | | |
| Oligo rev | AAACTCCCGACCACGACCTTGAGT | | | | | | | | | | |
| Chromosome | start | end | strand | MM | target_seq | PAM | alignment | distance | position | gene name | gene id |
| GU305914.1 | 213316 | 213338 | + | 0 | ACTCAAGGTCGTGGTCGGGA | TGG | ||||||||[||||||||||||]PAM | NA | Intergenic | NA | NA |
| T16024 | GTCGGGATGGATGTCTTAGACGG | 1000 | | | | | | | | | |
| Oligo fwd | CACCGTCGGGATGGATGTCTTAGA | | | | | | | | | | |
| Oligo rev | AAACTCTAAGACATCCATCCCGAC | | | | | | | | | | |
| Chromosome | start | end | strand | MM | target_seq | PAM | alignment | distance | position | gene name | gene id |
| GU305914.1 | 213329 | 213351 | + | 0 | GTCGGGATGGATGTCTTAGA | CGG | ||||||||[||||||||||||]PAM | NA | Intergenic | NA | NA |

<table>
<tr><td>T16025</td><td>TGTCTTAGACGGTGTGCTCCCGG</td><td></td><td>1000</td><td></td><td></td><td></td><td></td><td></td><td></td><td></td><td></td></tr>
<tr><td>Oligo fwd</td><td>CACCTGTCTTAGACGGTGTGCTCC</td><td></td><td></td><td></td><td></td><td></td><td></td><td></td><td></td><td></td><td></td></tr>
<tr><td>Oligo rev</td><td>AAACGGAGCACACCGTCTAAGACA</td><td></td><td></td><td></td><td></td><td></td><td></td><td></td><td></td><td></td><td></td></tr>
<tr><td>Chromosome</td><td>start</td><td>end</td><td>strand</td><td>MM</td><td>target_seq</td><td>PAM</td><td>alignment</td><td>distance</td><td>position</td><td>gene name</td><td>gene id</td></tr>
<tr><td>GU305914.1</td><td>213340</td><td>213362</td><td>+</td><td>0</td><td>TGTCTTAGACGGTGTGCTCC</td><td>CGG</td><td>||||||||[||||||||||||]PAM</td><td>NA</td><td>Intergenic</td><td>NA</td><td>NA</td></tr>
<tr><td>T16026</td><td>CCCGGTAGACCTCCTCGAAACGG</td><td></td><td>1000</td><td></td><td></td><td></td><td></td><td></td><td></td><td></td><td></td></tr>
<tr><td>Oligo fwd</td><td>CACCCCCGGTAGACCTCCTCGAAA</td><td></td><td></td><td></td><td></td><td></td><td></td><td></td><td></td><td></td><td></td></tr>
<tr><td>Oligo rev</td><td>AAACTTTCGAGGAGGTCTACCGGG</td><td></td><td></td><td></td><td></td><td></td><td></td><td></td><td></td><td></td><td></td></tr>
<tr><td>Chromosome</td><td>start</td><td>end</td><td>strand</td><td>MM</td><td>target_seq</td><td>PAM</td><td>alignment</td><td>distance</td><td>position</td><td>gene name</td><td>gene id</td></tr>
<tr><td>GU305914.1</td><td>213358</td><td>213380</td><td>+</td><td>0</td><td>CCCGGTAGACCTCCTCGAAA</td><td>CGG</td><td>||||||||[||||||||||||]PAM</td><td>NA</td><td>Intergenic</td><td>NA</td><td>NA</td></tr>
<tr><td>GU305914.1</td><td>146589</td><td>146611</td><td>-</td><td>4</td><td>CGCCGTAGACCTCCTTGAAG</td><td>AGG</td><td>|-|-||||[|||||||-|||-]PAM</td><td>NA</td><td>Intergenic</td><td>NA</td><td>NA</td></tr>
<tr><td>T16027</td><td>ACGGCCGTCGTGCTATTCATCGG</td><td></td><td>1000</td><td></td><td></td><td></td><td></td><td></td><td></td><td></td><td></td></tr>
<tr><td>Oligo fwd</td><td>CACCACGGCCGTCGTGCTATTCAT</td><td></td><td></td><td></td><td></td><td></td><td></td><td></td><td></td><td></td><td></td></tr>
<tr><td>Oligo rev</td><td>AAACATGAATAGCACGACGGCCGT</td><td></td><td></td><td></td><td></td><td></td><td></td><td></td><td></td><td></td><td></td></tr>
<tr><td>Chromosome</td><td>start</td><td>end</td><td>strand</td><td>MM</td><td>target_seq</td><td>PAM</td><td>alignment</td><td>distance</td><td>position</td><td>gene name</td><td>gene id</td></tr>
<tr><td>GU305914.1</td><td>213377</td><td>213399</td><td>+</td><td>0</td><td>ACGGCCGTCGTGCTATTCAT</td><td>CGG</td><td>||||||||[||||||||||||]PAM</td><td>NA</td><td>Intergenic</td><td>NA</td><td>NA</td></tr>
<tr><td>T16028</td><td>CGGCATCGTAGTCGCGATACTGG</td><td></td><td>1000</td><td></td><td></td><td></td><td></td><td></td><td></td><td></td><td></td></tr>
<tr><td>Oligo fwd</td><td>CACCCGGCATCGTAGTCGCGATAC</td><td></td><td></td><td></td><td></td><td></td><td></td><td></td><td></td><td></td><td></td></tr>
<tr><td>Oligo rev</td><td>AAACGTATCGCGACTACGATGCCG</td><td></td><td></td><td></td><td></td><td></td><td></td><td></td><td></td><td></td><td></td></tr>
<tr><td>Chromosome</td><td>start</td><td>end</td><td>strand</td><td>MM</td><td>target_seq</td><td>PAM</td><td>alignment</td><td>distance</td><td>position</td><td>gene name</td><td>gene id</td></tr>
<tr><td>GU305914.1</td><td>213397</td><td>213419</td><td>+</td><td>0</td><td>CGGCATCGTAGTCGCGATAC</td><td>TGG</td><td>||||||||[||||||||||||]PAM</td><td>NA</td><td>Intergenic</td><td>NA</td><td>NA</td></tr>
<tr><td>T16029</td><td>GCTAGTCGTGTATCCCTGCGTGG</td><td></td><td>1000</td><td></td><td></td><td></td><td></td><td></td><td></td><td></td><td></td></tr>
<tr><td>Oligo fwd</td><td>CACCGCTAGTCGTGTATCCCTGCG</td><td></td><td></td><td></td><td></td><td></td><td></td><td></td><td></td><td></td><td></td></tr>
<tr><td>Oligo rev</td><td>AAACCGCAGGGATACACGACTAGC</td><td></td><td></td><td></td><td></td><td></td><td></td><td></td><td></td><td></td><td></td></tr>
<tr><td>Chromosome</td><td>start</td><td>end</td><td>strand</td><td>MM</td><td>target_seq</td><td>PAM</td><td>alignment</td><td>distance</td><td>position</td><td>gene name</td><td>gene id</td></tr>
<tr><td>GU305914.1</td><td>213433</td><td>213455</td><td>+</td><td>0</td><td>GCTAGTCGTGTATCCCTGCG</td><td>TGG</td><td>||||||||[||||||||||||]PAM</td><td>NA</td><td>Intergenic</td><td>NA</td><td>NA</td></tr>
<tr><td>T16030</td><td>TCCCTGCGTGGTCACCGTAGTGG</td><td></td><td>1000</td><td></td><td></td><td></td><td></td><td></td><td></td><td></td><td></td></tr>
<tr><td>Oligo fwd</td><td>CACCTCCCTGCGTGGTCACCGTAG</td><td></td><td></td><td></td><td></td><td></td><td></td><td></td><td></td><td></td><td></td></tr>
<tr><td>Oligo rev</td><td>AAACCTACGGTGACCACGCAGGGA</td><td></td><td></td><td></td><td></td><td></td><td></td><td></td><td></td><td></td><td></td></tr>
<tr><td>Chromosome</td><td>start</td><td>end</td><td>strand</td><td>MM</td><td>target_seq</td><td>PAM</td><td>alignment</td><td>distance</td><td>position</td><td>gene name</td><td>gene id</td></tr>
<tr><td>GU305914.1</td><td>213445</td><td>213467</td><td>+</td><td>0</td><td>TCCCTGCGTGGTCACCGTAG</td><td>TGG</td><td>||||||||[||||||||||||]PAM</td><td>NA</td><td>Intergenic</td><td>NA</td><td>NA</td></tr>
<tr><td>T16031</td><td>CGTGGTCACCGTAGTGGCGTTGG</td><td></td><td>1000</td><td></td><td></td><td></td><td></td><td></td><td></td><td></td><td></td></tr>
<tr><td>Oligo fwd</td><td>CACCCGTGGTCACCGTAGTGGCGT</td><td></td><td></td><td></td><td></td><td></td><td></td><td></td><td></td><td></td><td></td></tr>
<tr><td>Oligo rev</td><td>AAACACGCCACTACGGTGACCACG</td><td></td><td></td><td></td><td></td><td></td><td></td><td></td><td></td><td></td><td></td></tr>
<tr><td>Chromosome</td><td>start</td><td>end</td><td>strand</td><td>MM</td><td>target_seq</td><td>PAM</td><td>alignment</td><td>distance</td><td>position</td><td>gene name</td><td>gene id</td></tr>
<tr><td>GU305914.1</td><td>213451</td><td>213473</td><td>+</td><td>0</td><td>CGTGGTCACCGTAGTGGCGT</td><td>TGG</td><td>||||||||[||||||||||||]PAM</td><td>NA</td><td>Intergenic</td><td>NA</td><td>NA</td></tr>
<tr><td>T16032</td><td>GTGGTCACCGTAGTGGCGTTGGG</td><td></td><td>1000</td><td></td><td></td><td></td><td></td><td></td><td></td><td></td><td></td></tr>
<tr><td>Oligo fwd</td><td>CACCGTGGTCACCGTAGTGGCGTT</td><td></td><td></td><td></td><td></td><td></td><td></td><td></td><td></td><td></td><td></td></tr>
<tr><td>Oligo rev</td><td>AAACAACGCCACTACGGTGACCAC</td><td></td><td></td><td></td><td></td><td></td><td></td><td></td><td></td><td></td><td></td></tr>
<tr><td>Chromosome</td><td>start</td><td>end</td><td>strand</td><td>MM</td><td>target_seq</td><td>PAM</td><td>alignment</td><td>distance</td><td>position</td><td>gene name</td><td>gene id</td></tr>
<tr><td>GU305914.1</td><td>213452</td><td>213474</td><td>+</td><td>0</td><td>GTGGTCACCGTAGTGGCGTT</td><td>GGG</td><td>||||||||[||||||||||||]PAM</td><td>NA</td><td>Intergenic</td><td>NA</td><td>NA</td></tr>
<tr><td>T16033</td><td>TGGTCACCGTAGTGGCGTTGGGG</td><td></td><td>1000</td><td></td><td></td><td></td><td></td><td></td><td></td><td></td><td></td></tr>
<tr><td>Oligo fwd</td><td>CACCTGGTCACCGTAGTGGCGTTG</td><td></td><td></td><td></td><td></td><td></td><td></td><td></td><td></td><td></td><td></td></tr>
<tr><td>Oligo rev</td><td>AAACCAACGCCACTACGGTGACCA</td><td></td><td></td><td></td><td></td><td></td><td></td><td></td><td></td><td></td><td></td></tr>
<tr><td>Chromosome</td><td>start</td><td>end</td><td>strand</td><td>MM</td><td>target_seq</td><td>PAM</td><td>alignment</td><td>distance</td><td>position</td><td>gene name</td><td>gene id</td></tr>
<tr><td>GU305914.1</td><td>213453</td><td>213475</td><td>+</td><td>0</td><td>TGGTCACCGTAGTGGCGTTG</td><td>GGG</td><td>||||||||[||||||||||||]PAM</td><td>NA</td><td>Intergenic</td><td>NA</td><td>NA</td></tr>
<tr><td>T16034</td><td>CACCGTAGTGGCGTTGGGGCTGG</td><td></td><td>1000</td><td></td><td></td><td></td><td></td><td></td><td></td><td></td><td></td></tr>
<tr><td>Oligo fwd</td><td>CACCCACCGTAGTGGCGTTGGGGC</td><td></td><td></td><td></td><td></td><td></td><td></td><td></td><td></td><td></td><td></td></tr>
<tr><td>Oligo rev</td><td>AAACGCCCCAACGCCACTACGGTG</td><td></td><td></td><td></td><td></td><td></td><td></td><td></td><td></td><td></td><td></td></tr>
<tr><td>Chromosome</td><td>start</td><td>end</td><td>strand</td><td>MM</td><td>target_seq</td><td>PAM</td><td>alignment</td><td>distance</td><td>position</td><td>gene name</td><td>gene id</td></tr>
<tr><td>GU305914.1</td><td>213457</td><td>213479</td><td>+</td><td>0</td><td>CACCGTAGTGGCGTTGGGGC</td><td>TGG</td><td>||||||||[||||||||||||]PAM</td><td>NA</td><td>Intergenic</td><td>NA</td><td>NA</td></tr>
<tr><td>T16035</td><td>CTCATACTCGAAGTGATCATCGG</td><td></td><td>1000</td><td></td><td></td><td></td><td></td><td></td><td></td><td></td><td></td></tr>
<tr><td>Oligo fwd</td><td>CACCCTCATACTCGAAGTGATCAT</td><td></td><td></td><td></td><td></td><td></td><td></td><td></td><td></td><td></td><td></td></tr>
<tr><td>Oligo rev</td><td>AAACATGATCACTTCGAGTATGAG</td><td></td><td></td><td></td><td></td><td></td><td></td><td></td><td></td><td></td><td></td></tr>
<tr><td>Chromosome</td><td>start</td><td>end</td><td>strand</td><td>MM</td><td>target_seq</td><td>PAM</td><td>alignment</td><td>distance</td><td>position</td><td>gene name</td><td>gene id</td></tr>
<tr><td>GU305914.1</td><td>213482</td><td>213504</td><td>+</td><td>0</td><td>CTCATACTCGAAGTGATCAT</td><td>CGG</td><td>||||||||[||||||||||||]PAM</td><td>NA</td><td>Intergenic</td><td>NA</td><td>NA</td></tr>
<tr><td>T16036</td><td>GTGATCATCGGTGTCGTGCTAGG</td><td></td><td>1000</td><td></td><td></td><td></td><td></td><td></td><td></td><td></td><td></td></tr>
<tr><td>Oligo fwd</td><td>CACCGTGATCATCGGTGTCGTGCT</td><td></td><td></td><td></td><td></td><td></td><td></td><td></td><td></td><td></td><td></td></tr>
<tr><td>Oligo rev</td><td>AAACAGCACGACACCGATGATCAC</td><td></td><td></td><td></td><td></td><td></td><td></td><td></td><td></td><td></td><td></td></tr>
<tr><td>Chromosome</td><td>start</td><td>end</td><td>strand</td><td>MM</td><td>target_seq</td><td>PAM</td><td>alignment</td><td>distance</td><td>position</td><td>gene name</td><td>gene id</td></tr>
<tr><td>GU305914.1</td><td>213494</td><td>213516</td><td>+</td><td>0</td><td>GTGATCATCGGTGTCGTGCT</td><td>AGG</td><td>||||||||[||||||||||||]PAM</td><td>NA</td><td>Intergenic</td><td>NA</td><td>NA</td></tr>
<tr><td>T16037</td><td>TTTCGAAGATGTATGAACGACGG</td><td></td><td>1000</td><td></td><td></td><td></td><td></td><td></td><td></td><td></td><td></td></tr>
<tr><td>Oligo fwd</td><td>CACCTTTCGAAGATGTATGAACGA</td><td></td><td></td><td></td><td></td><td></td><td></td><td></td><td></td><td></td><td></td></tr>
<tr><td>Oligo rev</td><td>AAACTCGTTCATACATCTTCGAAA</td><td></td><td></td><td></td><td></td><td></td><td></td><td></td><td></td><td></td><td></td></tr>
<tr><td>Chromosome</td><td>start</td><td>end</td><td>strand</td><td>MM</td><td>target_seq</td><td>PAM</td><td>alignment</td><td>distance</td><td>position</td><td>gene name</td><td>gene id</td></tr>
<tr><td>GU305914.1</td><td>213517</td><td>213539</td><td>+</td><td>0</td><td>TTTCGAAGATGTATGAACGA</td><td>CGG</td><td>||||||||[||||||||||||]PAM</td><td>NA</td><td>Intergenic</td><td>NA</td><td>NA</td></tr>
<tr><td>T16038</td><td>CGAAGATGTATGAACGACGGCGG</td><td></td><td>1000</td><td></td><td></td><td></td><td></td><td></td><td></td><td></td><td></td></tr>
<tr><td>Oligo fwd</td><td>CACCCGAAGATGTATGAACGACGG</td><td></td><td></td><td></td><td></td><td></td><td></td><td></td><td></td><td></td><td></td></tr>
<tr><td>Oligo rev</td><td>AAACCCGTCGTTCATACATCTTCG</td><td></td><td></td><td></td><td></td><td></td><td></td><td></td><td></td><td></td><td></td></tr>
<tr><td>Chromosome</td><td>start</td><td>end</td><td>strand</td><td>MM</td><td>target_seq</td><td>PAM</td><td>alignment</td><td>distance</td><td>position</td><td>gene name</td><td>gene id</td></tr>
<tr><td>GU305914.1</td><td>213520</td><td>213542</td><td>+</td><td>0</td><td>CGAAGATGTATGAACGACGG</td><td>CGG</td><td>||||||||[||||||||||||]PAM</td><td>NA</td><td>Intergenic</td><td>NA</td><td>NA</td></tr>
<tr><td>T16039</td><td>TGTATGAACGACGGCGGCCGTGG</td><td></td><td>1000</td><td></td><td></td><td></td><td></td><td></td><td></td><td></td><td></td></tr>
<tr><td>Oligo fwd</td><td>CACCTGTATGAACGACGGCGGCCG</td><td></td><td></td><td></td><td></td><td></td><td></td><td></td><td></td><td></td><td></td></tr>
<tr><td>Oligo rev</td><td>AAACCGGCCGCCGTCGTTCATACA</td><td></td><td></td><td></td><td></td><td></td><td></td><td></td><td></td><td></td><td></td></tr>
</table>

Chromosome start end strand MM target_seq PAM alignment distance position gene name gene id
GU305914.1 213526 213548 + 0 TGTATGAACGACGGCGGCCG TGG ||||||||[||||||||||||]PAM NA Intergenic NA NA

T16040 ACGACGGCGGCCGTGGACGATGG 1000
Oligo fwd CACCACGACGGCGGCCGTGGACGA
Oligo rev AAACTCGTCCACGGCCGCCGTCGT
Chromosome start end strand MM target_seq PAM alignment distance position gene name gene id
GU305914.1 213533 213555 + 0 ACGACGGCGGCCGTGGACGA TGG ||||||||[||||||||||||]PAM NA Intergenic NA NA

T16041 GGCGGCCGTGGACGATGGTTCGG 1000
Oligo fwd CACCGGCGGCCGTGGACGATGGTT
Oligo rev AAACAACCATCGTCCACGGCCGCC
Chromosome start end strand MM target_seq PAM alignment distance position gene name gene id
GU305914.1 213538 213560 + 0 GGCGGCCGTGGACGATGGTT CGG ||||||||[||||||||||||]PAM NA Intergenic NA NA
GU305914.1 213607 213629 + 3 GGTGACCGTGGACGACGGTT CGG ||-|-|||[|||||||-||||]PAM NA Intergenic NA NA

T16042 GTGGACGATGGTTCGGTCGTCGG 1000
Oligo fwd CACCGTGGACGATGGTTCGGTCGT
Oligo rev AAACACGACCGAACCATCGTCCAC
Chromosome start end strand MM target_seq PAM alignment distance position gene name gene id
GU305914.1 213545 213567 + 0 GTGGACGATGGTTCGGTCGT CGG ||||||||[||||||||||||]PAM NA Intergenic NA NA
GU305914.1 213614 213636 + 1 GTGGACGACGGTTCGGTCGT CGG ||||||||[-|||||||||||]PAM NA Intergenic NA NA
GU305914.1 24982 25004 + 4 CGAGACGATGGGTCGGTCGT GGG ---|||||[|||-||||||||]PAM NA Intergenic NA NA

T16043 CGTCGGTAGCACAGTAGAAGTGG 1000
Oligo fwd CACCCGTCGGTAGCACAGTAGAAG
Oligo rev AAACCTTCTACTGTGCTACCGACG
Chromosome start end strand MM target_seq PAM alignment distance position gene name gene id
GU305914.1 213562 213584 + 0 CGTCGGTAGCACAGTAGAAG TGG ||||||||[||||||||||||]PAM NA Intergenic NA NA
GU305914.1 213631 213653 + 2 CGTCGGTAGCGCAGTAGTAG TGG ||||||||[||-||||||-||]PAM NA Intergenic NA NA

T16044 CGGTAGCACAGTAGAAGTGGTGG 1000
Oligo fwd CACCCGGTAGCACAGTAGAAGTGG
Oligo rev AAACCCACTTCTACTGTGCTACCG
Chromosome start end strand MM target_seq PAM alignment distance position gene name gene id
GU305914.1 213565 213587 + 0 CGGTAGCACAGTAGAAGTGG TGG ||||||||[||||||||||||]PAM NA Intergenic NA NA
GU305914.1 213634 213656 + 2 CGGTAGCGCAGTAGTAGTGG CGG |||||||-[||||||-|||||]PAM NA Intergenic NA NA

T16045 TAGCACAGTAGAAGTGGTGGTGG 1000
Oligo fwd CACCTAGCACAGTAGAAGTGGTGG
Oligo rev AAACCCACCACTTCTACTGTGCTA
Chromosome start end strand MM target_seq PAM alignment distance position gene name gene id
GU305914.1 213568 213590 + 0 TAGCACAGTAGAAGTGGTGG TGG ||||||||[||||||||||||]PAM NA Intergenic NA NA
GU305914.1 213637 213659 + 3 TAGCGCAGTAGTAGTGGCGG TGG ||||-|||[|||-|||||-||]PAM NA Intergenic NA NA

T16046 CACAGTAGAAGTGGTGGTGGTGG 1000
Oligo fwd CACCCACAGTAGAAGTGGTGGTGG
Oligo rev AAACCCACCACCACTTCTACTGTG
Chromosome start end strand MM target_seq PAM alignment distance position gene name gene id
GU305914.1 213571 213593 + 0 CACAGTAGAAGTGGTGGTGG TGG ||||||||[||||||||||||]PAM NA Intergenic NA NA
GU305914.1 213640 213662 + 3 CGCAGTAGTAGTGGCGGTGG TGG |-||||||[-|||||-|||||]PAM NA Intergenic NA NA

T16047 AGTAGAAGTGGTGGTGGTGGTGG 1000
Oligo fwd CACCAGTAGAAGTGGTGGTGGTGG
Oligo rev AAACCCACCACCACCACTTCTACT
Chromosome start end strand MM target_seq PAM alignment distance position gene name gene id
GU305914.1 213574 213596 + 0 AGTAGAAGTGGTGGTGGTGG TGG ||||||||[||||||||||||]PAM NA Intergenic NA NA
GU305914.1 213643 213665 + 2 AGTAGTAGTGGCGGTGGTGG TGG |||||-||[|||-||||||||]PAM NA Intergenic NA NA
GU305914.1 213580 213602 + 4 AGTGGTGGTGGTGGTGGTAG TGG |||-|--|[||||||||||-|]PAM NA Intergenic NA NA

T16048 AGTGGTGGTGGTGGTGGTAGTGG 1000
Oligo fwd CACCAGTGGTGGTGGTGGTGGTAG
Oligo rev AAACCTACCACCACCACCACCACT
Chromosome start end strand MM target_seq PAM alignment distance position gene name gene id
GU305914.1 213580 213602 + 0 AGTGGTGGTGGTGGTGGTAG TGG ||||||||[||||||||||||]PAM NA Intergenic NA NA
GU305914.1 213586 213608 + 2 GGTGGTGGTGGTAGTGGTAG TGG -|||||||[||||-|||||||]PAM NA Intergenic NA NA
GU305914.1 213574 213596 + 4 AGTAGAAGTGGTGGTGGTGG TGG |||-|--|[||||||||||-|]PAM NA Intergenic NA NA
GU305914.1 213643 213665 + 4 AGTAGTAGTGGCGGTGGTGG TGG |||-||-|[|||-||||||-|]PAM NA Intergenic NA NA

T16049 GGTGGTGGTGGTAGTGGTAGTGG 1000
Oligo fwd CACCGGTGGTGGTGGTAGTGGTAG
Oligo rev AAACCTACCACTACCACCACCACC
Chromosome start end strand MM target_seq PAM alignment distance position gene name gene id
GU305914.1 213586 213608 + 0 GGTGGTGGTGGTAGTGGTAG TGG ||||||||[||||||||||||]PAM NA Intergenic NA NA
GU305914.1 213580 213602 + 2 AGTGGTGGTGGTGGTGGTAG TGG -|||||||[||||-|||||||]PAM NA Intergenic NA NA
GU305914.1 192597 192619 + 4 TGTGGTCGTCGTAGTAGTAG TGG -|||||-|[|-|||||-||||]PAM NA Intergenic NA NA
GU305914.1 174642 174664 + 4 GGAGATGGTGGTAGTGGTGA CGG ||-|-|||[||||||||||--]PAM NA Intergenic NA NA

T16050 GGTAGTGGTAGTGGTGACCGTGG 1000
Oligo fwd CACCGGTAGTGGTAGTGGTGACCG
Oligo rev AAACCGGTCACCACTACCACTACC
Chromosome start end strand MM target_seq PAM alignment distance position gene name gene id
GU305914.1 213595 213617 + 0 GGTAGTGGTAGTGGTGACCG TGG ||||||||[||||||||||||]PAM NA Intergenic NA NA

T16051 GTAGTGGTGACCGTGGACGACGG 1000
Oligo fwd CACCGTAGTGGTGACCGTGGACGA
Oligo rev AAACTCGTCCACGGTCACCACTAC
Chromosome start end strand MM target_seq PAM alignment distance position gene name gene id
GU305914.1 213602 213624 + 0 GTAGTGGTGACCGTGGACGA CGG ||||||||[||||||||||||]PAM NA Intergenic NA NA

T16052 GGTGACCGTGGACGACGGTTCGG 1000
Oligo fwd CACCGGTGACCGTGGACGACGGTT
Oligo rev AAACAACCGTCGTCCACGGTCACC

| Chromosome | start | end | strand | MM | target_seq | PAM | alignment | distance | position | gene name | gene id |
|---|---|---|---|---|---|---|---|---|---|---|---|
| GU305914.1 | 213607 | 213629 | + | 0 | GGTGACCGTGGACGACGGTT | CGG | \|\|\|\|\|\|\|\|[\|\|\|\|\|\|\|\|\|\|\|\|]PAM | NA | Intergenic | NA | NA |
| GU305914.1 | 213538 | 213560 | + | 3 | GGCGGCCGTGGACGATGGTT | CGG | \|\|-\|-\|\|\|[\|\|\|\|\|\|\|-\|\|\|\|]PAM | NA | Intergenic | NA | NA |

T16053 GTGGACGACGGTTCGGTCGTCGG 1000
Oligo fwd CACCGTGGACGACGGTTCGGTCGT
Oligo rev AAACACGACCGAACCGTCGTCCAC

| Chromosome | start | end | strand | MM | target_seq | PAM | alignment | distance | position | gene name | gene id |
|---|---|---|---|---|---|---|---|---|---|---|---|
| GU305914.1 | 213614 | 213636 | + | 0 | GTGGACGACGGTTCGGTCGT | CGG | \|\|\|\|\|\|\|\|[\|\|\|\|\|\|\|\|\|\|\|\|]PAM | NA | Intergenic | NA | NA |
| GU305914.1 | 213545 | 213567 | + | 1 | GTGGACGATGGTTCGGTCGT | CGG | \|\|\|\|\|\|\|\|[-\|\|\|\|\|\|\|\|\|\|\|]PAM | NA | Intergenic | NA | NA |

T16054 CGTCGGTAGCGCAGTAGTAGTGG 1000
Oligo fwd CACCCGTCGGTAGCGCAGTAGTAG
Oligo rev AAACCTACTACTGCGCTACCGACG

| Chromosome | start | end | strand | MM | target_seq | PAM | alignment | distance | position | gene name | gene id |
|---|---|---|---|---|---|---|---|---|---|---|---|
| GU305914.1 | 213631 | 213653 | + | 0 | CGTCGGTAGCGCAGTAGTAG | TGG | \|\|\|\|\|\|\|\|[\|\|\|\|\|\|\|\|\|\|\|\|]PAM | NA | Intergenic | NA | NA |
| GU305914.1 | 213562 | 213584 | + | 2 | CGTCGGTAGCACAGTAGAAG | TGG | \|\|\|\|\|\|\|\|[\|\|-\|\|\|\|\|\|-\|\|]PAM | NA | Intergenic | NA | NA |

T16055 CGGTAGCGCAGTAGTAGTGGCGG 1000
Oligo fwd CACCCGGTAGCGCAGTAGTAGTGG
Oligo rev AAACCCACTACTACTGCGCTACCG

| Chromosome | start | end | strand | MM | target_seq | PAM | alignment | distance | position | gene name | gene id |
|---|---|---|---|---|---|---|---|---|---|---|---|
| GU305914.1 | 213634 | 213656 | + | 0 | CGGTAGCGCAGTAGTAGTGG | CGG | \|\|\|\|\|\|\|\|[\|\|\|\|\|\|\|\|\|\|\|\|]PAM | NA | Intergenic | NA | NA |
| GU305914.1 | 213565 | 213587 | + | 2 | CGGTAGCACAGTAGAAGTGG | TGG | \|\|\|\|\|\|\|-[\|\|\|\|\|\|-\|\|\|\|\|]PAM | NA | Intergenic | NA | NA |

T16056 TAGCGCAGTAGTAGTGGCGGTGG 1000
Oligo fwd CACCTAGCGCAGTAGTAGTGGCGG
Oligo rev AAACCCGCCACTACTACTGCGCTA

| Chromosome | start | end | strand | MM | target_seq | PAM | alignment | distance | position | gene name | gene id |
|---|---|---|---|---|---|---|---|---|---|---|---|
| GU305914.1 | 213637 | 213659 | + | 0 | TAGCGCAGTAGTAGTGGCGG | TGG | \|\|\|\|\|\|\|\|[\|\|\|\|\|\|\|\|\|\|\|\|]PAM | NA | Intergenic | NA | NA |
| GU305914.1 | 213568 | 213590 | + | 3 | TAGCACAGTAGAAGTGGTGG | TGG | \|\|\|\|-\|\|\|[\|\|\|-\|\|\|\|\|-\|\|]PAM | NA | Intergenic | NA | NA |

T16057 CGCAGTAGTAGTGGCGGTGGTGG 1000
Oligo fwd CACCCGCAGTAGTAGTGGCGGTGG
Oligo rev AAACCCACCGCCACTACTACTGCG

| Chromosome | start | end | strand | MM | target_seq | PAM | alignment | distance | position | gene name | gene id |
|---|---|---|---|---|---|---|---|---|---|---|---|
| GU305914.1 | 213640 | 213662 | + | 0 | CGCAGTAGTAGTGGCGGTGG | TGG | \|\|\|\|\|\|\|\|[\|\|\|\|\|\|\|\|\|\|\|\|]PAM | NA | Intergenic | NA | NA |
| GU305914.1 | 213571 | 213593 | + | 3 | CACAGTAGAAGTGGTGGTGG | TGG | \|-\|\|\|\|\|\|[-\|\|\|\|\|-\|\|\|\|\|]PAM | NA | Intergenic | NA | NA |
| GU305914.1 | 192603 | 192625 | + | 4 | CGTCGTAGTAGTAGTGGTGG | TGG | \|\|--\|\|\|\|[\|\|\|\|-\|-\|\|\|\|\|]PAM | NA | Intergenic | NA | NA |

T16058 AGTAGTAGTGGCGGTGGTGGTGG 1000
Oligo fwd CACCAGTAGTAGTGGCGGTGGTGG
Oligo rev AAACCCACCACCGCCACTACTACT

| Chromosome | start | end | strand | MM | target_seq | PAM | alignment | distance | position | gene name | gene id |
|---|---|---|---|---|---|---|---|---|---|---|---|
| GU305914.1 | 213643 | 213665 | + | 0 | AGTAGTAGTGGCGGTGGTGG | TGG | \|\|\|\|\|\|\|\|[\|\|\|\|\|\|\|\|\|\|\|\|]PAM | NA | Intergenic | NA | NA |
| GU305914.1 | 213574 | 213596 | + | 2 | AGTAGAAGTGGTGGTGGTGG | TGG | \|\|\|\|\|-\|\|[\|\|\|-\|\|\|\|\|\|\|\|]PAM | NA | Intergenic | NA | NA |
| GU305914.1 | 213580 | 213602 | + | 4 | AGTGGTGGTGGTGGTGGTAG | TGG | \|\|\|-\|\|-\|[\|\|\|-\|\|\|\|\|\|-\|]PAM | NA | Intergenic | NA | NA |

T16059 GGTGGTATTTTGTCCCTTCGTGG 1000
Oligo fwd CACCGGTGGTATTTTGTCCCTTCG
Oligo rev AAACCGAAGGGACAAAATACCACC

| Chromosome | start | end | strand | MM | target_seq | PAM | alignment | distance | position | gene name | gene id |
|---|---|---|---|---|---|---|---|---|---|---|---|
| GU305914.1 | 213661 | 213683 | + | 0 | GGTGGTATTTTGTCCCTTCG | TGG | \|\|\|\|\|\|\|\|[\|\|\|\|\|\|\|\|\|\|\|\|]PAM | NA | Intergenic | NA | NA |

T16060 TTTTGTCCCTTCGTGGTAACAGG 1000
Oligo fwd CACCTTTTGTCCCTTCGTGGTAAC
Oligo rev AAACGTTACCACGAAGGGACAAAA

| Chromosome | start | end | strand | MM | target_seq | PAM | alignment | distance | position | gene name | gene id |
|---|---|---|---|---|---|---|---|---|---|---|---|
| GU305914.1 | 213668 | 213690 | + | 0 | TTTTGTCCCTTCGTGGTAAC | AGG | \|\|\|\|\|\|\|\|[\|\|\|\|\|\|\|\|\|\|\|\|]PAM | NA | Intergenic | NA | NA |

T16061 TTTGTCCCTTCGTGGTAACAGGG 1000
Oligo fwd CACCTTTGTCCCTTCGTGGTAACA
Oligo rev AAACTGTTACCACGAAGGGACAAA

| Chromosome | start | end | strand | MM | target_seq | PAM | alignment | distance | position | gene name | gene id |
|---|---|---|---|---|---|---|---|---|---|---|---|
| GU305914.1 | 213669 | 213691 | + | 0 | TTTGTCCCTTCGTGGTAACA | GGG | \|\|\|\|\|\|\|\|[\|\|\|\|\|\|\|\|\|\|\|\|]PAM | NA | Intergenic | NA | NA |

T16062 CCCTTCGTGGTAACAGGGCTTGG 1000
Oligo fwd CACCCCCTTCGTGGTAACAGGGCT
Oligo rev AAACAGCCCTGTTACCACGAAGGG

| Chromosome | start | end | strand | MM | target_seq | PAM | alignment | distance | position | gene name | gene id |
|---|---|---|---|---|---|---|---|---|---|---|---|
| GU305914.1 | 213674 | 213696 | + | 0 | CCCTTCGTGGTAACAGGGCT | TGG | \|\|\|\|\|\|\|\|[\|\|\|\|\|\|\|\|\|\|\|\|]PAM | NA | Intergenic | NA | NA |

T16063 AGGGCTTGGAGTAGTAACAACGG 1000
Oligo fwd CACCAGGGCTTGGAGTAGTAACAA
Oligo rev AAACTTGTTACTACTCCAAGCCCT

| Chromosome | start | end | strand | MM | target_seq | PAM | alignment | distance | position | gene name | gene id |
|---|---|---|---|---|---|---|---|---|---|---|---|
| GU305914.1 | 213688 | 213710 | + | 0 | AGGGCTTGGAGTAGTAACAA | CGG | \|\|\|\|\|\|\|\|[\|\|\|\|\|\|\|\|\|\|\|\|]PAM | NA | Intergenic | NA | NA |

T16064 GTAACAACGGCACCTGCACTAGG 1000
Oligo fwd CACCGTAACAACGGCACCTGCACT
Oligo rev AAACAGTGCAGGTGCCGTTGTTAC

| Chromosome | start | end | strand | MM | target_seq | PAM | alignment | distance | position | gene name | gene id |
|---|---|---|---|---|---|---|---|---|---|---|---|
| GU305914.1 | 213701 | 213723 | + | 0 | GTAACAACGGCACCTGCACT | AGG | \|\|\|\|\|\|\|\|[\|\|\|\|\|\|\|\|\|\|\|\|]PAM | NA | Intergenic | NA | NA |

T16065 TAACAACGGCACCTGCACTAGGG 1000

| | | | | | | | | | | | |
|---|---|---|---|---|---|---|---|---|---|---|---|
| Oligo fwd | CACCTAACAACGGCACCTGCACTA | | | | | | | | | | |
| Oligo rev | AAACTAGTGCAGGTGCCGTTGTTA | | | | | | | | | | |
| Chromosome | start | end | strand | MM | target_seq | PAM | alignment | distance | position | gene name | gene id |
| GU305914.1 | 213702 | 213724 | + | 0 | TAACAACGGCACCTGCACTA | GGG | \|\|\|\|\|\|\|\|[\|\|\|\|\|\|\|\|\|\|\|\|]PAM | NA | Intergenic | NA | NA |
| T16066 | AACAACGGCACCTGCACTAGGGG | 1000 | | | | | | | | | |
| Oligo fwd | CACCAACAACGGCACCTGCACTAG | | | | | | | | | | |
| Oligo rev | AAACCTAGTGCAGGTGCCGTTGTT | | | | | | | | | | |
| Chromosome | start | end | strand | MM | target_seq | PAM | alignment | distance | position | gene name | gene id |
| GU305914.1 | 213703 | 213725 | + | 0 | AACAACGGCACCTGCACTAG | GGG | \|\|\|\|\|\|\|\|[\|\|\|\|\|\|\|\|\|\|\|\|]PAM | NA | Intergenic | NA | NA |
| T16067 | CGAAGAGATCGCGTCCGCAGCGG | 1000 | | | | | | | | | |
| Oligo fwd | CACCCGAAGAGATCGCGTCCGCAG | | | | | | | | | | |
| Oligo rev | AAACCTGCGGACGCGATCTCTTCG | | | | | | | | | | |
| Chromosome | start | end | strand | MM | target_seq | PAM | alignment | distance | position | gene name | gene id |
| GU305914.1 | 213733 | 213755 | + | 0 | CGAAGAGATCGCGTCCGCAG | CGG | \|\|\|\|\|\|\|\|[\|\|\|\|\|\|\|\|\|\|\|\|]PAM | NA | Intergenic | NA | NA |
| T16068 | GATCGCGTCCGCAGCGGTAGAGG | 1000 | | | | | | | | | |
| Oligo fwd | CACCGATCGCGTCCGCAGCGGTAG | | | | | | | | | | |
| Oligo rev | AAACCTACCGCTGCGGACGCGATC | | | | | | | | | | |
| Chromosome | start | end | strand | MM | target_seq | PAM | alignment | distance | position | gene name | gene id |
| GU305914.1 | 213739 | 213761 | + | 0 | GATCGCGTCCGCAGCGGTAG | AGG | \|\|\|\|\|\|\|\|[\|\|\|\|\|\|\|\|\|\|\|\|]PAM | NA | Intergenic | NA | NA |
| T16069 | GTCCGCAGCGGTAGAGGTATCGG | 1000 | | | | | | | | | |
| Oligo fwd | CACCGTCCGCAGCGGTAGAGGTAT | | | | | | | | | | |
| Oligo rev | AAACATACCTCTACCGCTGCGGAC | | | | | | | | | | |
| Chromosome | start | end | strand | MM | target_seq | PAM | alignment | distance | position | gene name | gene id |
| GU305914.1 | 213745 | 213767 | + | 0 | GTCCGCAGCGGTAGAGGTAT | CGG | \|\|\|\|\|\|\|\|[\|\|\|\|\|\|\|\|\|\|\|\|]PAM | NA | Intergenic | NA | NA |
| T16070 | TCCGCAGCGGTAGAGGTATCGGG | 1000 | | | | | | | | | |
| Oligo fwd | CACCTCCGCAGCGGTAGAGGTATC | | | | | | | | | | |
| Oligo rev | AAACGATACCTCTACCGCTGCGGA | | | | | | | | | | |
| Chromosome | start | end | strand | MM | target_seq | PAM | alignment | distance | position | gene name | gene id |
| GU305914.1 | 213746 | 213768 | + | 0 | TCCGCAGCGGTAGAGGTATC | GGG | \|\|\|\|\|\|\|\|[\|\|\|\|\|\|\|\|\|\|\|\|]PAM | NA | Intergenic | NA | NA |
| T16071 | GGTATCGGGCTCCGTAGTCTCGG | 1000 | | | | | | | | | |
| Oligo fwd | CACCGGTATCGGGCTCCGTAGTCT | | | | | | | | | | |
| Oligo rev | AAACAGACTACGGAGCCCGATACC | | | | | | | | | | |
| Chromosome | start | end | strand | MM | target_seq | PAM | alignment | distance | position | gene name | gene id |
| GU305914.1 | 213760 | 213782 | + | 0 | GGTATCGGGCTCCGTAGTCT | CGG | \|\|\|\|\|\|\|\|[\|\|\|\|\|\|\|\|\|\|\|\|]PAM | NA | Intergenic | NA | NA |
| T16072 | GTATCGGGCTCCGTAGTCTCGGG | 1000 | | | | | | | | | |
| Oligo fwd | CACCGTATCGGGCTCCGTAGTCTC | | | | | | | | | | |
| Oligo rev | AAACGAGACTACGGAGCCCGATAC | | | | | | | | | | |
| Chromosome | start | end | strand | MM | target_seq | PAM | alignment | distance | position | gene name | gene id |
| GU305914.1 | 213761 | 213783 | + | 0 | GTATCGGGCTCCGTAGTCTC | GGG | \|\|\|\|\|\|\|\|[\|\|\|\|\|\|\|\|\|\|\|\|]PAM | NA | Intergenic | NA | NA |
| T16073 | GGCTCCGTAGTCTCGGGCTGAGG | 1000 | | | | | | | | | |
| Oligo fwd | CACCGGCTCCGTAGTCTCGGGCTG | | | | | | | | | | |
| Oligo rev | AAACCAGCCCGAGACTACGGAGCC | | | | | | | | | | |
| Chromosome | start | end | strand | MM | target_seq | PAM | alignment | distance | position | gene name | gene id |
| GU305914.1 | 213767 | 213789 | + | 0 | GGCTCCGTAGTCTCGGGCTG | AGG | \|\|\|\|\|\|\|\|[\|\|\|\|\|\|\|\|\|\|\|\|]PAM | NA | Intergenic | NA | NA |
| GU305914.1 | 66685 | 66707 | + | 4 | GGATCCGGAGCCGCGGGCTG | CGG | \|\|-\|\|\|\|-[\|\|-\|-\|\|\|\|\|\|\|]PAM | NA | Intergenic | NA | NA |
| T16074 | GTCTCGGGCTGAGGTGTAGTCGG | 1000 | | | | | | | | | |
| Oligo fwd | CACCGTCTCGGGCTGAGGTGTAGT | | | | | | | | | | |
| Oligo rev | AAACACTACACCTCAGCCCGAGAC | | | | | | | | | | |
| Chromosome | start | end | strand | MM | target_seq | PAM | alignment | distance | position | gene name | gene id |
| GU305914.1 | 213776 | 213798 | + | 0 | GTCTCGGGCTGAGGTGTAGT | CGG | \|\|\|\|\|\|\|\|[\|\|\|\|\|\|\|\|\|\|\|\|]PAM | NA | Intergenic | NA | NA |
| T16075 | GGGCTGAGGTGTAGTCGGAGCGG | 1000 | | | | | | | | | |
| Oligo fwd | CACCGGGCTGAGGTGTAGTCGGAG | | | | | | | | | | |
| Oligo rev | AAACCTCCGACTACACCTCAGCCC | | | | | | | | | | |
| Chromosome | start | end | strand | MM | target_seq | PAM | alignment | distance | position | gene name | gene id |
| GU305914.1 | 213781 | 213803 | + | 0 | GGGCTGAGGTGTAGTCGGAG | CGG | \|\|\|\|\|\|\|\|[\|\|\|\|\|\|\|\|\|\|\|\|]PAM | NA | Intergenic | NA | NA |
| T16076 | GACAAGCGTAGTCAGCAGTTCGG | 1000 | | | | | | | | | |
| Oligo fwd | CACCGACAAGCGTAGTCAGCAGTT | | | | | | | | | | |
| Oligo rev | AAACAACTGCTGACTACGCTTGTC | | | | | | | | | | |
| Chromosome | start | end | strand | MM | target_seq | PAM | alignment | distance | position | gene name | gene id |
| GU305914.1 | 213862 | 213884 | + | 0 | GACAAGCGTAGTCAGCAGTT | CGG | \|\|\|\|\|\|\|\|[\|\|\|\|\|\|\|\|\|\|\|\|]PAM | NA | Intergenic | NA | NA |
| T16077 | CAGTTCGGTCGTAGCGAACGTGG | 1000 | | | | | | | | | |
| Oligo fwd | CACCCAGTTCGGTCGTAGCGAACG | | | | | | | | | | |
| Oligo rev | AAACCGTTCGCTACGACCGAACTG | | | | | | | | | | |
| Chromosome | start | end | strand | MM | target_seq | PAM | alignment | distance | position | gene name | gene id |
| GU305914.1 | 213877 | 213899 | + | 0 | CAGTTCGGTCGTAGCGAACG | TGG | \|\|\|\|\|\|\|\|[\|\|\|\|\|\|\|\|\|\|\|\|]PAM | NA | Intergenic | NA | NA |
| T16078 | GGTCGTAGCGAACGTGGTACCGG | 1000 | | | | | | | | | |
| Oligo fwd | CACCGGTCGTAGCGAACGTGGTAC | | | | | | | | | | |
| Oligo rev | AAACGTACCACGTTCGCTACGACC | | | | | | | | | | |
| Chromosome | start | end | strand | MM | target_seq | PAM | alignment | distance | position | gene name | gene id |
| GU305914.1 | 213883 | 213905 | + | 0 | GGTCGTAGCGAACGTGGTAC | CGG | \|\|\|\|\|\|\|\|[\|\|\|\|\|\|\|\|\|\|\|\|]PAM | NA | Intergenic | NA | NA |
| GU305914.1 | 191313 | 191335 | - | 4 | GATCGTCGCGACCGTGCTAC | TGG | \|-\|\|\|\|-\|[\|\|\|-\|\|\|\|-\|\|\|]PAM | NA | Intergenic | NA | NA |
| T16079 | GTCGTAGCGAACGTGGTACCGGG | 1000 | | | | | | | | | |
| Oligo fwd | CACCGTCGTAGCGAACGTGGTACC | | | | | | | | | | |
| Oligo rev | AAACGGTACCACGTTCGCTACGAC | | | | | | | | | | |
| Chromosome | start | end | strand | MM | target_seq | PAM | alignment | distance | position | gene name | gene id |

GU305914.1 213884 213906 + 0 GTCGTAGCGAACGTGGTACC GGG ||||||||[||||||||||||]PAM NA Intergenic NA NA

T16080 CGTGGTACCGGGCTCGCTAGTGG 1000
Oligo fwd CACCCGTGGTACCGGGCTCGCTAG
Oligo rev AAACCTAGCGAGCCCGGTACCACG
Chromosome start end strand MM target_seq PAM alignment distance position gene name gene id
GU305914.1 213895 213917 + 0 CGTGGTACCGGGCTCGCTAG TGG ||||||||[||||||||||||]PAM NA Intergenic NA NA

T16081 GTGGTACCGGGCTCGCTAGTGGG 1000
Oligo fwd CACCGTGGTACCGGGCTCGCTAGT
Oligo rev AAACACTAGCGAGCCCGGTACCAC
Chromosome start end strand MM target_seq PAM alignment distance position gene name gene id
GU305914.1 213896 213918 + 0 GTGGTACCGGGCTCGCTAGT GGG ||||||||[||||||||||||]PAM NA Intergenic NA NA

T16082 ACCGGGCTCGCTAGTGGGCTCGG 1000
Oligo fwd CACCACCGGGCTCGCTAGTGGGCT
Oligo rev AAACAGCCCACTAGCGAGCCCGGT
Chromosome start end strand MM target_seq PAM alignment distance position gene name gene id
GU305914.1 213901 213923 + 0 ACCGGGCTCGCTAGTGGGCT CGG ||||||||[||||||||||||]PAM NA Intergenic NA NA

T16083 CCGGGCTCGCTAGTGGGCTCGGG 1000
Oligo fwd CACCCCGGGCTCGCTAGTGGGCTC
Oligo rev AAACGAGCCCACTAGCGAGCCCGG
Chromosome start end strand MM target_seq PAM alignment distance position gene name gene id
GU305914.1 213902 213924 + 0 CCGGGCTCGCTAGTGGGCTC GGG ||||||||[||||||||||||]PAM NA Intergenic NA NA

T16084 CTAGTGGGCTCGGGCGTCGTAGG 1000
Oligo fwd CACCCTAGTGGGCTCGGGCGTCGT
Oligo rev AAACACGACGCCCGAGCCCACTAG
Chromosome start end strand MM target_seq PAM alignment distance position gene name gene id
GU305914.1 213911 213933 + 0 CTAGTGGGCTCGGGCGTCGT AGG ||||||||[||||||||||||]PAM NA Intergenic NA NA

T16085 CTCGGGCGTCGTAGGATCTTCGG 1000
Oligo fwd CACCCTCGGGCGTCGTAGGATCTT
Oligo rev AAACAAGATCCTACGACGCCCGAG
Chromosome start end strand MM target_seq PAM alignment distance position gene name gene id
GU305914.1 213919 213941 + 0 CTCGGGCGTCGTAGGATCTT CGG ||||||||[||||||||||||]PAM NA Intergenic NA NA

T16086 TTCGGATGTCATAAGATCCCCGG 1000
Oligo fwd CACCTTCGGATGTCATAAGATCCC
Oligo rev AAACGGGATCTTATGACATCCGAA
Chromosome start end strand MM target_seq PAM alignment distance position gene name gene id
GU305914.1 213937 213959 + 0 TTCGGATGTCATAAGATCCC CGG ||||||||[||||||||||||]PAM NA Intergenic NA NA

T16087 CCCCGGACCCCGTCATCTCTAGG 1000
Oligo fwd CACCCCCCGGACCCCGTCATCTCT
Oligo rev AAACAGAGATGACGGGGTCCGGGG
Chromosome start end strand MM target_seq PAM alignment distance position gene name gene id
GU305914.1 213954 213976 + 0 CCCCGGACCCCGTCATCTCT AGG ||||||||[||||||||||||]PAM NA Intergenic NA NA

T16088 CCCGGACCCCGTCATCTCTAGGG 1000
Oligo fwd CACCCCCGGACCCCGTCATCTCTA
Oligo rev AAACTAGAGATGACGGGGTCCGGG
Chromosome start end strand MM target_seq PAM alignment distance position gene name gene id
GU305914.1 213955 213977 + 0 CCCGGACCCCGTCATCTCTA GGG ||||||||[||||||||||||]PAM NA Intergenic NA NA

T16089 AGTAGCGTCGAGAGATACCGTGG 1000
Oligo fwd CACCAGTAGCGTCGAGAGATACCG
Oligo rev AAACCGGTATCTCTCGACGCTACT
Chromosome start end strand MM target_seq PAM alignment distance position gene name gene id
GU305914.1 213988 214010 + 0 AGTAGCGTCGAGAGATACCG TGG ||||||||[||||||||||||]PAM NA Intergenic NA NA

T16090 GAGAGATACCGTGGTAGAAGTGG 1000
Oligo fwd CACCGAGAGATACCGTGGTAGAAG
Oligo rev AAACCTTCTACCACGGTATCTCTC
Chromosome start end strand MM target_seq PAM alignment distance position gene name gene id
GU305914.1 213997 214019 + 0 GAGAGATACCGTGGTAGAAG TGG ||||||||[||||||||||||]PAM NA Intergenic NA NA

T16091 AGAAAGTTCCATTCCTACCGAGG 1000
Oligo fwd CACCAGAAAGTTCCATTCCTACCG
Oligo rev AAACCGGTAGGAATGGAACTTTCT
Chromosome start end strand MM target_seq PAM alignment distance position gene name gene id
GU305914.1 214027 214049 + 0 AGAAAGTTCCATTCCTACCG AGG ||||||||[||||||||||||]PAM NA Intergenic NA NA

T16092 TAAACGACCCAATGCGACGACGG 1000
Oligo fwd CACCTAAACGACCCAATGCGACGA
Oligo rev AAACTCGTCGCATTGGGTCGTTTA
Chromosome start end strand MM target_seq PAM alignment distance position gene name gene id
GU305914.1 214069 214091 + 0 TAAACGACCCAATGCGACGA CGG ||||||||[||||||||||||]PAM NA Intergenic NA NA

T16093 AAACGACCCAATGCGACGACGGG 1000
Oligo fwd CACCAAACGACCCAATGCGACGAC
Oligo rev AAACGTCGTCGCATTGGGTCGTTT
Chromosome start end strand MM target_seq PAM alignment distance position gene name gene id
GU305914.1 214070 214092 + 0 AAACGACCCAATGCGACGAC GGG ||||||||[||||||||||||]PAM NA Intergenic NA NA

T16094 GACCCAATGCGACGACGGGATGG 1000
Oligo fwd CACCGACCCAATGCGACGACGGGA
Oligo rev AAACTCCCGTCGTCGCATTGGGTC

Chromosome start end strand MM target_seq PAM alignment distance position gene name gene id
GU305914.1 214074 214096 + 0 GACCCAATGCGACGACGGGA TGG ||||||||[||||||||||||]PAM NA Intergenic NA NA

T16095 CCGTAACATCGCCCGCATCGTGG 1000
Oligo fwd CACCCCGTAACATCGCCCGCATCG
Oligo rev AAACCGATGCGGGCGATGTTACGG
Chromosome start end strand MM target_seq PAM alignment distance position gene name gene id
GU305914.1 214132 214154 + 0 CCGTAACATCGCCCGCATCG TGG ||||||||[||||||||||||]PAM NA Intergenic NA NA

T16096 TCGTGGTCGAGACCGTATCTCGG 1000
Oligo fwd CACCTCGTGGTCGAGACCGTATCT
Oligo rev AAACAGATACGGTCTCGACCACGA
Chromosome start end strand MM target_seq PAM alignment distance position gene name gene id
GU305914.1 214149 214171 + 0 TCGTGGTCGAGACCGTATCT CGG ||||||||[||||||||||||]PAM NA Intergenic NA NA

T16097 CACCCCCTCGTGAATGTGATTGG 1000
Oligo fwd CACCCACCCCCTCGTGAATGTGAT
Oligo rev AAACATCACATTCACGAGGGGGTG
Chromosome start end strand MM target_seq PAM alignment distance position gene name gene id
GU305914.1 214174 214196 + 0 CACCCCCTCGTGAATGTGAT TGG ||||||||[||||||||||||]PAM NA Intergenic NA NA

T16098 CCCCTCGTGAATGTGATTGGCGG 1000
Oligo fwd CACCCCCCTCGTGAATGTGATTGG
Oligo rev AAACCCAATCACATTCACGAGGGG
Chromosome start end strand MM target_seq PAM alignment distance position gene name gene id
GU305914.1 214177 214199 + 0 CCCCTCGTGAATGTGATTGG CGG ||||||||[||||||||||||]PAM NA Intergenic NA NA

T16099 CCCTCGTGAATGTGATTGGCGGG 1000
Oligo fwd CACCCCCTCGTGAATGTGATTGGC
Oligo rev AAACGCCAATCACATTCACGAGGG
Chromosome start end strand MM target_seq PAM alignment distance position gene name gene id
GU305914.1 214178 214200 + 0 CCCTCGTGAATGTGATTGGC GGG ||||||||[||||||||||||]PAM NA Intergenic NA NA

T16100 ATGTGATTGGCGGGCAGACGCGG 1000
Oligo fwd CACCATGTGATTGGCGGGCAGACG
Oligo rev AAACCGTCTGCCCGCCAATCACAT
Chromosome start end strand MM target_seq PAM alignment distance position gene name gene id
GU305914.1 214187 214209 + 0 ATGTGATTGGCGGGCAGACG CGG ||||||||[||||||||||||]PAM NA Intergenic NA NA

T16101 TGTGATTGGCGGGCAGACGCGGG 1000
Oligo fwd CACCTGTGATTGGCGGGCAGACGC
Oligo rev AAACGCGTCTGCCCGCCAATCACA
Chromosome start end strand MM target_seq PAM alignment distance position gene name gene id
GU305914.1 214188 214210 + 0 TGTGATTGGCGGGCAGACGC GGG ||||||||[||||||||||||]PAM NA Intergenic NA NA

T16102 GTGATTGGCGGGCAGACGCGGGG 1000
Oligo fwd CACCGTGATTGGCGGGCAGACGCG
Oligo rev AAACCGCGTCTGCCCGCCAATCAC
Chromosome start end strand MM target_seq PAM alignment distance position gene name gene id
GU305914.1 214189 214211 + 0 GTGATTGGCGGGCAGACGCG GGG ||||||||[||||||||||||]PAM NA Intergenic NA NA

T16103 GGCGGGCAGACGCGGGGTTCTGG 1000
Oligo fwd CACCGGCGGGCAGACGCGGGGTTC
Oligo rev AAACGAACCCCGCGTCTGCCCGCC
Chromosome start end strand MM target_seq PAM alignment distance position gene name gene id
GU305914.1 214195 214217 + 0 GGCGGGCAGACGCGGGGTTC TGG ||||||||[||||||||||||]PAM NA Intergenic NA NA

T16104 GCAGACGCGGGGTTCTGGTCTGG 1000
Oligo fwd CACCGCAGACGCGGGGTTCTGGTC
Oligo rev AAACGACCAGAACCCCGCGTCTGC
Chromosome start end strand MM target_seq PAM alignment distance position gene name gene id
GU305914.1 214200 214222 + 0 GCAGACGCGGGGTTCTGGTC TGG ||||||||[||||||||||||]PAM NA Intergenic NA NA

T16105 CAGACGCGGGGTTCTGGTCTGGG 1000
Oligo fwd CACCCAGACGCGGGGTTCTGGTCT
Oligo rev AAACAGACCAGAACCCCGCGTCTG
Chromosome start end strand MM target_seq PAM alignment distance position gene name gene id
GU305914.1 214201 214223 + 0 CAGACGCGGGGTTCTGGTCT GGG ||||||||[||||||||||||]PAM NA Intergenic NA NA

T16106 TATTGCCTTTCATGTTTGAGTGG 1000
Oligo fwd CACCTATTGCCTTTCATGTTTGAG
Oligo rev AAACCTCAAACATGAAAGGCAATA
Chromosome start end strand MM target_seq PAM alignment distance position gene name gene id
GU305914.1 214242 214264 + 0 TATTGCCTTTCATGTTTGAG TGG ||||||||[||||||||||||]PAM NA Intergenic NA NA

T16107 TGGCCACACCCAGATATCTACGG 1000
Oligo fwd CACCTGGCCACACCCAGATATCTA
Oligo rev AAACTAGATATCTGGGTGTGGCCA
Chromosome start end strand MM target_seq PAM alignment distance position gene name gene id
GU305914.1 214262 214284 + 0 TGGCCACACCCAGATATCTA CGG ||||||||[||||||||||||]PAM NA Intergenic NA NA

T16108 CTACACCACGATAATCTGTCAGG 1000
Oligo fwd CACCCTACACCACGATAATCTGTC
Oligo rev AAACGACAGATTATCGTGGTGTAG
Chromosome start end strand MM target_seq PAM alignment distance position gene name gene id
GU305914.1 214332 214354 + 0 CTACACCACGATAATCTGTC AGG ||||||||[||||||||||||]PAM NA Intergenic NA NA

T16109 GTTTAACTCGATGTTTTTTTAGG 1000
Oligo fwd CACCGTTTAACTCGATGTTTTTTT

<table>
<tr><td>Oligo rev</td><td>AAACAAAAAAACATCGAGTTAAAC</td><td></td><td></td><td></td><td></td><td></td><td></td><td></td><td></td><td></td><td></td></tr>
<tr><td>Chromosome</td><td>start</td><td>end</td><td>strand</td><td>MM</td><td>target_seq</td><td>PAM</td><td>alignment</td><td>distance</td><td>position</td><td>gene name</td><td>gene id</td></tr>
<tr><td>GU305914.1</td><td>214364</td><td>214386</td><td>+</td><td>0</td><td>GTTTAACTCGATGTTTTTTT</td><td>AGG</td><td>||||||||[||||||||||||]PAM</td><td>NA</td><td>Intergenic</td><td>NA</td><td>NA</td></tr>
<tr><td></td><td></td><td></td><td></td><td></td><td></td><td></td><td></td><td></td><td></td><td></td><td></td></tr>
<tr><td>T16110</td><td>CGAGAGAAATCTGACTCGTGCGG</td><td>1000</td><td></td><td></td><td></td><td></td><td></td><td></td><td></td><td></td><td></td></tr>
<tr><td>Oligo fwd</td><td>CACCCGAGAGAAATCTGACTCGTG</td><td></td><td></td><td></td><td></td><td></td><td></td><td></td><td></td><td></td><td></td></tr>
<tr><td>Oligo rev</td><td>AAACCACGAGTCAGATTTCTCTCG</td><td></td><td></td><td></td><td></td><td></td><td></td><td></td><td></td><td></td><td></td></tr>
<tr><td>Chromosome</td><td>start</td><td>end</td><td>strand</td><td>MM</td><td>target_seq</td><td>PAM</td><td>alignment</td><td>distance</td><td>position</td><td>gene name</td><td>gene id</td></tr>
<tr><td>GU305914.1</td><td>214412</td><td>214434</td><td>+</td><td>0</td><td>CGAGAGAAATCTGACTCGTG</td><td>CGG</td><td>||||||||[||||||||||||]PAM</td><td>NA</td><td>Intergenic</td><td>NA</td><td>NA</td></tr>
<tr><td></td><td></td><td></td><td></td><td></td><td></td><td></td><td></td><td></td><td></td><td></td><td></td></tr>
<tr><td>T16111</td><td>AGAAATCTGACTCGTGCGGCCGG</td><td>1000</td><td></td><td></td><td></td><td></td><td></td><td></td><td></td><td></td><td></td></tr>
<tr><td>Oligo fwd</td><td>CACCAGAAATCTGACTCGTGCGGC</td><td></td><td></td><td></td><td></td><td></td><td></td><td></td><td></td><td></td><td></td></tr>
<tr><td>Oligo rev</td><td>AAACGCCGCACGAGTCAGATTTCT</td><td></td><td></td><td></td><td></td><td></td><td></td><td></td><td></td><td></td><td></td></tr>
<tr><td>Chromosome</td><td>start</td><td>end</td><td>strand</td><td>MM</td><td>target_seq</td><td>PAM</td><td>alignment</td><td>distance</td><td>position</td><td>gene name</td><td>gene id</td></tr>
<tr><td>GU305914.1</td><td>214416</td><td>214438</td><td>+</td><td>0</td><td>AGAAATCTGACTCGTGCGGC</td><td>CGG</td><td>||||||||[||||||||||||]PAM</td><td>NA</td><td>Intergenic</td><td>NA</td><td>NA</td></tr>
<tr><td></td><td></td><td></td><td></td><td></td><td></td><td></td><td></td><td></td><td></td><td></td><td></td></tr>
<tr><td>T16112</td><td>TCGCACGCGCCGACAATAAAAGG</td><td>1000</td><td></td><td></td><td></td><td></td><td></td><td></td><td></td><td></td><td></td></tr>
<tr><td>Oligo fwd</td><td>CACCTCGCACGCGCCGACAATAAA</td><td></td><td></td><td></td><td></td><td></td><td></td><td></td><td></td><td></td><td></td></tr>
<tr><td>Oligo rev</td><td>AAACTTTATTGTCGGCGCGTGCGA</td><td></td><td></td><td></td><td></td><td></td><td></td><td></td><td></td><td></td><td></td></tr>
<tr><td>Chromosome</td><td>start</td><td>end</td><td>strand</td><td>MM</td><td>target_seq</td><td>PAM</td><td>alignment</td><td>distance</td><td>position</td><td>gene name</td><td>gene id</td></tr>
<tr><td>GU305914.1</td><td>214439</td><td>214461</td><td>+</td><td>0</td><td>TCGCACGCGCCGACAATAAA</td><td>AGG</td><td>||||||||[||||||||||||]PAM</td><td>NA</td><td>Intergenic</td><td>NA</td><td>NA</td></tr>
<tr><td></td><td></td><td></td><td></td><td></td><td></td><td></td><td></td><td></td><td></td><td></td><td></td></tr>
<tr><td>T16113</td><td>AAAAGGACGCGATACACGTTTGG</td><td>1000</td><td></td><td></td><td></td><td></td><td></td><td></td><td></td><td></td><td></td></tr>
<tr><td>Oligo fwd</td><td>CACCAAAAGGACGCGATACACGTT</td><td></td><td></td><td></td><td></td><td></td><td></td><td></td><td></td><td></td><td></td></tr>
<tr><td>Oligo rev</td><td>AAACAACGTGTATCGCGTCCTTTT</td><td></td><td></td><td></td><td></td><td></td><td></td><td></td><td></td><td></td><td></td></tr>
<tr><td>Chromosome</td><td>start</td><td>end</td><td>strand</td><td>MM</td><td>target_seq</td><td>PAM</td><td>alignment</td><td>distance</td><td>position</td><td>gene name</td><td>gene id</td></tr>
<tr><td>GU305914.1</td><td>214456</td><td>214478</td><td>+</td><td>0</td><td>AAAAGGACGCGATACACGTT</td><td>TGG</td><td>||||||||[||||||||||||]PAM</td><td>NA</td><td>Intergenic</td><td>NA</td><td>NA</td></tr>
<tr><td></td><td></td><td></td><td></td><td></td><td></td><td></td><td></td><td></td><td></td><td></td><td></td></tr>
<tr><td>T16114</td><td>AAAGGACGCGATACACGTTTGGG</td><td>1000</td><td></td><td></td><td></td><td></td><td></td><td></td><td></td><td></td><td></td></tr>
<tr><td>Oligo fwd</td><td>CACCAAAGGACGCGATACACGTTT</td><td></td><td></td><td></td><td></td><td></td><td></td><td></td><td></td><td></td><td></td></tr>
<tr><td>Oligo rev</td><td>AAACAAACGTGTATCGCGTCCTTT</td><td></td><td></td><td></td><td></td><td></td><td></td><td></td><td></td><td></td><td></td></tr>
<tr><td>Chromosome</td><td>start</td><td>end</td><td>strand</td><td>MM</td><td>target_seq</td><td>PAM</td><td>alignment</td><td>distance</td><td>position</td><td>gene name</td><td>gene id</td></tr>
<tr><td>GU305914.1</td><td>214457</td><td>214479</td><td>+</td><td>0</td><td>AAAGGACGCGATACACGTTT</td><td>GGG</td><td>||||||||[||||||||||||]PAM</td><td>NA</td><td>Intergenic</td><td>NA</td><td>NA</td></tr>
<tr><td></td><td></td><td></td><td></td><td></td><td></td><td></td><td></td><td></td><td></td><td></td><td></td></tr>
<tr><td>T16115</td><td>ATAATTGTTTTATTAATCGACGG</td><td>1000</td><td></td><td></td><td></td><td></td><td></td><td></td><td></td><td></td><td></td></tr>
<tr><td>Oligo fwd</td><td>CACCATAATTGTTTTATTAATCGA</td><td></td><td></td><td></td><td></td><td></td><td></td><td></td><td></td><td></td><td></td></tr>
<tr><td>Oligo rev</td><td>AAACTCGATTAATAAAACAATTAT</td><td></td><td></td><td></td><td></td><td></td><td></td><td></td><td></td><td></td><td></td></tr>
<tr><td>Chromosome</td><td>start</td><td>end</td><td>strand</td><td>MM</td><td>target_seq</td><td>PAM</td><td>alignment</td><td>distance</td><td>position</td><td>gene name</td><td>gene id</td></tr>
<tr><td>GU305914.1</td><td>214492</td><td>214514</td><td>+</td><td>0</td><td>ATAATTGTTTTATTAATCGA</td><td>CGG</td><td>||||||||[||||||||||||]PAM</td><td>NA</td><td>Intergenic</td><td>NA</td><td>NA</td></tr>
<tr><td></td><td></td><td></td><td></td><td></td><td></td><td></td><td></td><td></td><td></td><td></td><td></td></tr>
<tr><td>T16116</td><td>TAATTGTTTTATTAATCGACGGG</td><td>1000</td><td></td><td></td><td></td><td></td><td></td><td></td><td></td><td></td><td></td></tr>
<tr><td>Oligo fwd</td><td>CACCTAATTGTTTTATTAATCGAC</td><td></td><td></td><td></td><td></td><td></td><td></td><td></td><td></td><td></td><td></td></tr>
<tr><td>Oligo rev</td><td>AAACGTCGATTAATAAAACAATTA</td><td></td><td></td><td></td><td></td><td></td><td></td><td></td><td></td><td></td><td></td></tr>
<tr><td>Chromosome</td><td>start</td><td>end</td><td>strand</td><td>MM</td><td>target_seq</td><td>PAM</td><td>alignment</td><td>distance</td><td>position</td><td>gene name</td><td>gene id</td></tr>
<tr><td>GU305914.1</td><td>214493</td><td>214515</td><td>+</td><td>0</td><td>TAATTGTTTTATTAATCGAC</td><td>GGG</td><td>||||||||[||||||||||||]PAM</td><td>NA</td><td>Intergenic</td><td>NA</td><td>NA</td></tr>
<tr><td>GU305914.1</td><td>206955</td><td>206977</td><td>+</td><td>4</td><td>GAATTTTTTTATTCATCGTC</td><td>GGG</td><td>-||||-||[|||||-||||-|]PAM</td><td>NA</td><td>Intergenic</td><td>NA</td><td>NA</td></tr>
<tr><td></td><td></td><td></td><td></td><td></td><td></td><td></td><td></td><td></td><td></td><td></td><td></td></tr>
<tr><td>T16117</td><td>GTTTTATTAATCGACGGGAGCGG</td><td>1000</td><td></td><td></td><td></td><td></td><td></td><td></td><td></td><td></td><td></td></tr>
<tr><td>Oligo fwd</td><td>CACCGTTTTATTAATCGACGGGAG</td><td></td><td></td><td></td><td></td><td></td><td></td><td></td><td></td><td></td><td></td></tr>
<tr><td>Oligo rev</td><td>AAACCTCCCGTCGATTAATAAAAC</td><td></td><td></td><td></td><td></td><td></td><td></td><td></td><td></td><td></td><td></td></tr>
<tr><td>Chromosome</td><td>start</td><td>end</td><td>strand</td><td>MM</td><td>target_seq</td><td>PAM</td><td>alignment</td><td>distance</td><td>position</td><td>gene name</td><td>gene id</td></tr>
<tr><td>GU305914.1</td><td>214498</td><td>214520</td><td>+</td><td>0</td><td>GTTTTATTAATCGACGGGAG</td><td>CGG</td><td>||||||||[||||||||||||]PAM</td><td>NA</td><td>Intergenic</td><td>NA</td><td>NA</td></tr>
<tr><td></td><td></td><td></td><td></td><td></td><td></td><td></td><td></td><td></td><td></td><td></td><td></td></tr>
<tr><td>T16118</td><td>TTTTATTAATCGACGGGAGCGGG</td><td>1000</td><td></td><td></td><td></td><td></td><td></td><td></td><td></td><td></td><td></td></tr>
<tr><td>Oligo fwd</td><td>CACCTTTTATTAATCGACGGGAGC</td><td></td><td></td><td></td><td></td><td></td><td></td><td></td><td></td><td></td><td></td></tr>
<tr><td>Oligo rev</td><td>AAACGCTCCCGTCGATTAATAAAA</td><td></td><td></td><td></td><td></td><td></td><td></td><td></td><td></td><td></td><td></td></tr>
<tr><td>Chromosome</td><td>start</td><td>end</td><td>strand</td><td>MM</td><td>target_seq</td><td>PAM</td><td>alignment</td><td>distance</td><td>position</td><td>gene name</td><td>gene id</td></tr>
<tr><td>GU305914.1</td><td>214499</td><td>214521</td><td>+</td><td>0</td><td>TTTTATTAATCGACGGGAGC</td><td>GGG</td><td>||||||||[||||||||||||]PAM</td><td>NA</td><td>Intergenic</td><td>NA</td><td>NA</td></tr>
<tr><td></td><td></td><td></td><td></td><td></td><td></td><td></td><td></td><td></td><td></td><td></td><td></td></tr>
<tr><td>T16119</td><td>TTTATTAATCGACGGGAGCGGGG</td><td>1000</td><td></td><td></td><td></td><td></td><td></td><td></td><td></td><td></td><td></td></tr>
<tr><td>Oligo fwd</td><td>CACCTTTATTAATCGACGGGAGCG</td><td></td><td></td><td></td><td></td><td></td><td></td><td></td><td></td><td></td><td></td></tr>
<tr><td>Oligo rev</td><td>AAACCGCTCCCGTCGATTAATAAA</td><td></td><td></td><td></td><td></td><td></td><td></td><td></td><td></td><td></td><td></td></tr>
<tr><td>Chromosome</td><td>start</td><td>end</td><td>strand</td><td>MM</td><td>target_seq</td><td>PAM</td><td>alignment</td><td>distance</td><td>position</td><td>gene name</td><td>gene id</td></tr>
<tr><td>GU305914.1</td><td>214500</td><td>214522</td><td>+</td><td>0</td><td>TTTATTAATCGACGGGAGCG</td><td>GGG</td><td>||||||||[||||||||||||]PAM</td><td>NA</td><td>Intergenic</td><td>NA</td><td>NA</td></tr>
<tr><td></td><td></td><td></td><td></td><td></td><td></td><td></td><td></td><td></td><td></td><td></td><td></td></tr>
<tr><td>T16120</td><td>TTATTAATCGACGGGAGCGGGGG</td><td>1000</td><td></td><td></td><td></td><td></td><td></td><td></td><td></td><td></td><td></td></tr>
<tr><td>Oligo fwd</td><td>CACCTTATTAATCGACGGGAGCGG</td><td></td><td></td><td></td><td></td><td></td><td></td><td></td><td></td><td></td><td></td></tr>
<tr><td>Oligo rev</td><td>AAACCCGCTCCCGTCGATTAATAA</td><td></td><td></td><td></td><td></td><td></td><td></td><td></td><td></td><td></td><td></td></tr>
<tr><td>Chromosome</td><td>start</td><td>end</td><td>strand</td><td>MM</td><td>target_seq</td><td>PAM</td><td>alignment</td><td>distance</td><td>position</td><td>gene name</td><td>gene id</td></tr>
<tr><td>GU305914.1</td><td>214501</td><td>214523</td><td>+</td><td>0</td><td>TTATTAATCGACGGGAGCGG</td><td>GGG</td><td>||||||||[||||||||||||]PAM</td><td>NA</td><td>Intergenic</td><td>NA</td><td>NA</td></tr>
<tr><td></td><td></td><td></td><td></td><td></td><td></td><td></td><td></td><td></td><td></td><td></td><td></td></tr>
<tr><td>T16121</td><td>TATTAATCGACGGGAGCGGGGGG</td><td>1000</td><td></td><td></td><td></td><td></td><td></td><td></td><td></td><td></td><td></td></tr>
<tr><td>Oligo fwd</td><td>CACCTATTAATCGACGGGAGCGGG</td><td></td><td></td><td></td><td></td><td></td><td></td><td></td><td></td><td></td><td></td></tr>
<tr><td>Oligo rev</td><td>AAACCCCGCTCCCGTCGATTAATA</td><td></td><td></td><td></td><td></td><td></td><td></td><td></td><td></td><td></td><td></td></tr>
<tr><td>Chromosome</td><td>start</td><td>end</td><td>strand</td><td>MM</td><td>target_seq</td><td>PAM</td><td>alignment</td><td>distance</td><td>position</td><td>gene name</td><td>gene id</td></tr>
<tr><td>GU305914.1</td><td>214502</td><td>214524</td><td>+</td><td>0</td><td>TATTAATCGACGGGAGCGGG</td><td>GGG</td><td>||||||||[||||||||||||]PAM</td><td>NA</td><td>Intergenic</td><td>NA</td><td>NA</td></tr>
<tr><td></td><td></td><td></td><td></td><td></td><td></td><td></td><td></td><td></td><td></td><td></td><td></td></tr>
<tr><td>T16122</td><td>ATCGACGGGAGCGGGGGGACCGG</td><td>1000</td><td></td><td></td><td></td><td></td><td></td><td></td><td></td><td></td><td></td></tr>
<tr><td>Oligo fwd</td><td>CACCATCGACGGGAGCGGGGGGAC</td><td></td><td></td><td></td><td></td><td></td><td></td><td></td><td></td><td></td><td></td></tr>
<tr><td>Oligo rev</td><td>AAACGTCCCCCCGCTCCCGTCGAT</td><td></td><td></td><td></td><td></td><td></td><td></td><td></td><td></td><td></td><td></td></tr>
<tr><td>Chromosome</td><td>start</td><td>end</td><td>strand</td><td>MM</td><td>target_seq</td><td>PAM</td><td>alignment</td><td>distance</td><td>position</td><td>gene name</td><td>gene id</td></tr>
<tr><td>GU305914.1</td><td>214507</td><td>214529</td><td>+</td><td>0</td><td>ATCGACGGGAGCGGGGGGAC</td><td>CGG</td><td>||||||||[||||||||||||]PAM</td><td>NA</td><td>Intergenic</td><td>NA</td><td>NA</td></tr>
<tr><td></td><td></td><td></td><td></td><td></td><td></td><td></td><td></td><td></td><td></td><td></td><td></td></tr>
<tr><td>T16123</td><td>TCGACGGGAGCGGGGGGACCGGG</td><td>1000</td><td></td><td></td><td></td><td></td><td></td><td></td><td></td><td></td><td></td></tr>
<tr><td>Oligo fwd</td><td>CACCTCGACGGGAGCGGGGGGACC</td><td></td><td></td><td></td><td></td><td></td><td></td><td></td><td></td><td></td><td></td></tr>
<tr><td>Oligo rev</td><td>AAACGGTCCCCCCGCTCCCGTCGA</td><td></td><td></td><td></td><td></td><td></td><td></td><td></td><td></td><td></td><td></td></tr>
<tr><td>Chromosome</td><td>start</td><td>end</td><td>strand</td><td>MM</td><td>target_seq</td><td>PAM</td><td>alignment</td><td>distance</td><td>position</td><td>gene name</td><td>gene id</td></tr>
<tr><td>GU305914.1</td><td>214508</td><td>214530</td><td>+</td><td>0</td><td>TCGACGGGAGCGGGGGGACC</td><td>GGG</td><td>||||||||[||||||||||||]PAM</td><td>NA</td><td>Intergenic</td><td>NA</td><td>NA</td></tr>
</table>

| | | | | | | | | | | | |
|---|---|---|---|---|---|---|---|---|---|---|---|
| T16124 | CGACGGGAGCGGGGGGACCGGGG | 1000 | | | | | | | | | |
| Oligo fwd | CACCCGACGGGAGCGGGGGGACCG | | | | | | | | | | |
| Oligo rev | AAACCGGTCCCCCCGCTCCCGTCG | | | | | | | | | | |
| Chromosome | start | end | strand | MM | target_seq | PAM | alignment | distance | position | gene name | gene id |
| GU305914.1 | 214509 | 214531 | + | 0 | CGACGGGAGCGGGGGGACCG | GGG | \|\|\|\|\|\|\|\|[\|\|\|\|\|\|\|\|\|\|\|\|]PAM | NA | Intergenic | NA | NA |

| | | | | | | | | | | | |
|---|---|---|---|---|---|---|---|---|---|---|---|
| T16125 | CCGGGGTGATCATTTGTAGACGG | 1000 | | | | | | | | | |
| Oligo fwd | CACCCCGGGGTGATCATTTGTAGA | | | | | | | | | | |
| Oligo rev | AAACTCTACAAATGATCACCCCGG | | | | | | | | | | |
| Chromosome | start | end | strand | MM | target_seq | PAM | alignment | distance | position | gene name | gene id |
| GU305914.1 | 214526 | 214548 | + | 0 | CCGGGGTGATCATTTGTAGA | CGG | \|\|\|\|\|\|\|\|[\|\|\|\|\|\|\|\|\|\|\|\|]PAM | NA | Intergenic | NA | NA |

| | | | | | | | | | | | |
|---|---|---|---|---|---|---|---|---|---|---|---|
| T16126 | CGGGGTGATCATTTGTAGACGGG | 1000 | | | | | | | | | |
| Oligo fwd | CACCCGGGGTGATCATTTGTAGAC | | | | | | | | | | |
| Oligo rev | AAACGTCTACAAATGATCACCCCG | | | | | | | | | | |
| Chromosome | start | end | strand | MM | target_seq | PAM | alignment | distance | position | gene name | gene id |
| GU305914.1 | 214527 | 214549 | + | 0 | CGGGGTGATCATTTGTAGAC | GGG | \|\|\|\|\|\|\|\|[\|\|\|\|\|\|\|\|\|\|\|\|]PAM | NA | Intergenic | NA | NA |

| | | | | | | | | | | | |
|---|---|---|---|---|---|---|---|---|---|---|---|
| T16127 | GGTGATCATTTGTAGACGGGCGG | 1000 | | | | | | | | | |
| Oligo fwd | CACCGGTGATCATTTGTAGACGGG | | | | | | | | | | |
| Oligo rev | AAACCCCGTCTACAAATGATCACC | | | | | | | | | | |
| Chromosome | start | end | strand | MM | target_seq | PAM | alignment | distance | position | gene name | gene id |
| GU305914.1 | 214530 | 214552 | + | 0 | GGTGATCATTTGTAGACGGG | CGG | \|\|\|\|\|\|\|\|[\|\|\|\|\|\|\|\|\|\|\|\|]PAM | NA | Intergenic | NA | NA |

| | | | | | | | | | | | |
|---|---|---|---|---|---|---|---|---|---|---|---|
| T16128 | GTGATCATTTGTAGACGGGCGGG | 1000 | | | | | | | | | |
| Oligo fwd | CACCGTGATCATTTGTAGACGGGC | | | | | | | | | | |
| Oligo rev | AAACGCCCGTCTACAAATGATCAC | | | | | | | | | | |
| Chromosome | start | end | strand | MM | target_seq | PAM | alignment | distance | position | gene name | gene id |
| GU305914.1 | 214531 | 214553 | + | 0 | GTGATCATTTGTAGACGGGC | GGG | \|\|\|\|\|\|\|\|[\|\|\|\|\|\|\|\|\|\|\|\|]PAM | NA | Intergenic | NA | NA |

| | | | | | | | | | | | |
|---|---|---|---|---|---|---|---|---|---|---|---|
| T16129 | GCTCTTCTTCTCCCCTTCACAGG | 1000 | | | | | | | | | |
| Oligo fwd | CACCGCTCTTCTTCTCCCCTTCAC | | | | | | | | | | |
| Oligo rev | AAACGTGAAGGGGAGAAGAAGAGC | | | | | | | | | | |
| Chromosome | start | end | strand | MM | target_seq | PAM | alignment | distance | position | gene name | gene id |
| GU305914.1 | 214556 | 214578 | + | 0 | GCTCTTCTTCTCCCCTTCAC | AGG | \|\|\|\|\|\|\|\|[\|\|\|\|\|\|\|\|\|\|\|\|]PAM | NA | Intergenic | NA | NA |

| | | | | | | | | | | | |
|---|---|---|---|---|---|---|---|---|---|---|---|
| T16130 | CTTCTTCTCCCCTTCACAGGAGG | 1000 | | | | | | | | | |
| Oligo fwd | CACCCTTCTTCTCCCCTTCACAGG | | | | | | | | | | |
| Oligo rev | AAACCCTGTGAAGGGGAGAAGAAG | | | | | | | | | | |
| Chromosome | start | end | strand | MM | target_seq | PAM | alignment | distance | position | gene name | gene id |
| GU305914.1 | 214559 | 214581 | + | 0 | CTTCTTCTCCCCTTCACAGG | AGG | \|\|\|\|\|\|\|\|[\|\|\|\|\|\|\|\|\|\|\|\|]PAM | NA | Intergenic | NA | NA |

| | | | | | | | | | | | |
|---|---|---|---|---|---|---|---|---|---|---|---|
| T16131 | CTTCTCCCCTTCACAGGAGGCGG | 1000 | | | | | | | | | |
| Oligo fwd | CACCCTTCTCCCCTTCACAGGAGG | | | | | | | | | | |
| Oligo rev | AAACCCTCCTGTGAAGGGGAGAAG | | | | | | | | | | |
| Chromosome | start | end | strand | MM | target_seq | PAM | alignment | distance | position | gene name | gene id |
| GU305914.1 | 214562 | 214584 | + | 0 | CTTCTCCCCTTCACAGGAGG | CGG | \|\|\|\|\|\|\|\|[\|\|\|\|\|\|\|\|\|\|\|\|]PAM | NA | Intergenic | NA | NA |

| | | | | | | | | | | | |
|---|---|---|---|---|---|---|---|---|---|---|---|
| T16132 | AGGAGGCGGCGAAGCACGAACGG | 1000 | | | | | | | | | |
| Oligo fwd | CACCAGGAGGCGGCGAAGCACGAA | | | | | | | | | | |
| Oligo rev | AAACTTCGTGCTTCGCCGCCTCCT | | | | | | | | | | |
| Chromosome | start | end | strand | MM | target_seq | PAM | alignment | distance | position | gene name | gene id |
| GU305914.1 | 214576 | 214598 | + | 0 | AGGAGGCGGCGAAGCACGAA | CGG | \|\|\|\|\|\|\|\|[\|\|\|\|\|\|\|\|\|\|\|\|]PAM | NA | Intergenic | NA | NA |

| | | | | | | | | | | | |
|---|---|---|---|---|---|---|---|---|---|---|---|
| T16133 | GCATCCAGCACATCACGACGAGG | 1000 | | | | | | | | | |
| Oligo fwd | CACCGCATCCAGCACATCACGACG | | | | | | | | | | |
| Oligo rev | AAACCGTCGTGATGTGCTGGATGC | | | | | | | | | | |
| Chromosome | start | end | strand | MM | target_seq | PAM | alignment | distance | position | gene name | gene id |
| GU305914.1 | 214612 | 214634 | + | 0 | GCATCCAGCACATCACGACG | AGG | \|\|\|\|\|\|\|\|[\|\|\|\|\|\|\|\|\|\|\|\|]PAM | NA | Intergenic | NA | NA |

| | | | | | | | | | | | |
|---|---|---|---|---|---|---|---|---|---|---|---|
| T16134 | GTACGCAGAGCACGACCAGTAGG | 1000 | | | | | | | | | |
| Oligo fwd | CACCGTACGCAGAGCACGACCAGT | | | | | | | | | | |
| Oligo rev | AAACACTGGTCGTGCTCTGCGTAC | | | | | | | | | | |
| Chromosome | start | end | strand | MM | target_seq | PAM | alignment | distance | position | gene name | gene id |
| GU305914.1 | 214636 | 214658 | + | 0 | GTACGCAGAGCACGACCAGT | AGG | \|\|\|\|\|\|\|\|[\|\|\|\|\|\|\|\|\|\|\|\|]PAM | NA | Intergenic | NA | NA |

| | | | | | | | | | | | |
|---|---|---|---|---|---|---|---|---|---|---|---|
| T16135 | TACGCAGAGCACGACCAGTAGGG | 1000 | | | | | | | | | |
| Oligo fwd | CACCTACGCAGAGCACGACCAGTA | | | | | | | | | | |
| Oligo rev | AAACTACTGGTCGTGCTCTGCGTA | | | | | | | | | | |
| Chromosome | start | end | strand | MM | target_seq | PAM | alignment | distance | position | gene name | gene id |
| GU305914.1 | 214637 | 214659 | + | 0 | TACGCAGAGCACGACCAGTA | GGG | \|\|\|\|\|\|\|\|[\|\|\|\|\|\|\|\|\|\|\|\|]PAM | NA | Intergenic | NA | NA |

| | | | | | | | | | | | |
|---|---|---|---|---|---|---|---|---|---|---|---|
| T16136 | CGTGACGTTGATGATGAGTTCGG | 1000 | | | | | | | | | |
| Oligo fwd | CACCCGTGACGTTGATGATGAGTT | | | | | | | | | | |
| Oligo rev | AAACAACTCATCATCAACGTCACG | | | | | | | | | | |
| Chromosome | start | end | strand | MM | target_seq | PAM | alignment | distance | position | gene name | gene id |
| GU305914.1 | 214664 | 214686 | + | 0 | CGTGACGTTGATGATGAGTT | CGG | \|\|\|\|\|\|\|\|[\|\|\|\|\|\|\|\|\|\|\|\|]PAM | NA | Intergenic | NA | NA |

| | | | | | | | | | | | |
|---|---|---|---|---|---|---|---|---|---|---|---|
| T16137 | TGATGAGTTCGGCGTCGTTTAGG | 1000 | | | | | | | | | |
| Oligo fwd | CACCTGATGAGTTCGGCGTCGTTT | | | | | | | | | | |
| Oligo rev | AAACAAACGACGCCGAACTCATCA | | | | | | | | | | |
| Chromosome | start | end | strand | MM | target_seq | PAM | alignment | distance | position | gene name | gene id |
| GU305914.1 | 214675 | 214697 | + | 0 | TGATGAGTTCGGCGTCGTTT | AGG | \|\|\|\|\|\|\|\|[\|\|\|\|\|\|\|\|\|\|\|\|]PAM | NA | Intergenic | NA | NA |

| | | | | | | | | | | | |
|---|---|---|---|---|---|---|---|---|---|---|---|
| T16138 | AGTTCGGCGTCGTTTAGGTACGG | 1000 | | | | | | | | | |
| Oligo fwd | CACCAGTTCGGCGTCGTTTAGGTA | | | | | | | | | | |
| Oligo rev | AAACTACCTAAACGACGCCGAACT | | | | | | | | | | |
| Chromosome | start | end | strand | MM | target_seq | PAM | alignment | distance | position | gene name | gene id |
| GU305914.1 | 214680 | 214702 | + | 0 | AGTTCGGCGTCGTTTAGGTA | CGG | \|\|\|\|\|\|\|\|[\|\|\|\|\|\|\|\|\|\|\|\|]PAM | NA | Intergenic | NA | NA |

| | | | | | | | | | | | | |
|---|---|---|---|---|---|---|---|---|---|---|---|---|
| T16139 | GTTTAGGTACGGAACCACTTTGG | | 1000 | | | | | | | | | |
| Oligo fwd | CACCGTTTAGGTACGGAACCACTT | | | | | | | | | | | |
| Oligo rev | AAACAAGTGGTTCCGTACCTAAAC | | | | | | | | | | | |
| Chromosome | start | | end | strand | MM | target_seq | PAM | alignment | distance | position | gene name | gene id |
| GU305914.1 | | 214691 | 214713 | + | 0 | GTTTAGGTACGGAACCACTT | TGG | \|\|\|\|\|\|\|\|[\|\|\|\|\|\|\|\|\|\|\|\|]PAM | NA | Intergenic | NA | NA |
| T16140 | ACCACTTTGGTATGATTCTTCGG | | 1000 | | | | | | | | | |
| Oligo fwd | CACCACCACTTTGGTATGATTCTT | | | | | | | | | | | |
| Oligo rev | AAACAAGAATCATACCAAAGTGGT | | | | | | | | | | | |
| Chromosome | start | | end | strand | MM | target_seq | PAM | alignment | distance | position | gene name | gene id |
| GU305914.1 | | 214704 | 214726 | + | 0 | ACCACTTTGGTATGATTCTT | CGG | \|\|\|\|\|\|\|\|[\|\|\|\|\|\|\|\|\|\|\|\|]PAM | NA | Intergenic | NA | NA |
| T16141 | GGTATGATTCTTCGGCGTCGAGG | | 1000 | | | | | | | | | |
| Oligo fwd | CACCGGTATGATTCTTCGGCGTCG | | | | | | | | | | | |
| Oligo rev | AAACCGACGCCGAAGAATCATACC | | | | | | | | | | | |
| Chromosome | start | | end | strand | MM | target_seq | PAM | alignment | distance | position | gene name | gene id |
| GU305914.1 | | 214712 | 214734 | + | 0 | GGTATGATTCTTCGGCGTCG | AGG | \|\|\|\|\|\|\|\|[\|\|\|\|\|\|\|\|\|\|\|\|]PAM | NA | Intergenic | NA | NA |
| T16142 | CGGCGTCGAGGTGACCTTGACGG | | 1000 | | | | | | | | | |
| Oligo fwd | CACCCGGCGTCGAGGTGACCTTGA | | | | | | | | | | | |
| Oligo rev | AAACTCAAGGTCACCTCGACGCCG | | | | | | | | | | | |
| Chromosome | start | | end | strand | MM | target_seq | PAM | alignment | distance | position | gene name | gene id |
| GU305914.1 | | 214724 | 214746 | + | 0 | CGGCGTCGAGGTGACCTTGA | CGG | \|\|\|\|\|\|\|\|[\|\|\|\|\|\|\|\|\|\|\|\|]PAM | NA | Intergenic | NA | NA |
| T16143 | GAGGTGACCTTGACGGATGATGG | | 1000 | | | | | | | | | |
| Oligo fwd | CACCGAGGTGACCTTGACGGATGA | | | | | | | | | | | |
| Oligo rev | AAACTCATCCGTCAAGGTCACCTC | | | | | | | | | | | |
| Chromosome | start | | end | strand | MM | target_seq | PAM | alignment | distance | position | gene name | gene id |
| GU305914.1 | | 214731 | 214753 | + | 0 | GAGGTGACCTTGACGGATGA | TGG | \|\|\|\|\|\|\|\|[\|\|\|\|\|\|\|\|\|\|\|\|]PAM | NA | Intergenic | NA | NA |
| T16144 | GAGCGAACCCTCCTGTTGAGCGG | | 1000 | | | | | | | | | |
| Oligo fwd | CACCGAGCGAACCCTCCTGTTGAG | | | | | | | | | | | |
| Oligo rev | AAACCTCAACAGGAGGGTTCGCTC | | | | | | | | | | | |
| Chromosome | start | | end | strand | MM | target_seq | PAM | alignment | distance | position | gene name | gene id |
| GU305914.1 | | 214758 | 214780 | + | 0 | GAGCGAACCCTCCTGTTGAG | CGG | \|\|\|\|\|\|\|\|[\|\|\|\|\|\|\|\|\|\|\|\|]PAM | NA | Intergenic | NA | NA |
| T16145 | AGCGAACCCTCCTGTTGAGCGGG | | 1000 | | | | | | | | | |
| Oligo fwd | CACCAGCGAACCCTCCTGTTGAGC | | | | | | | | | | | |
| Oligo rev | AAACGCTCAACAGGAGGGTTCGCT | | | | | | | | | | | |
| Chromosome | start | | end | strand | MM | target_seq | PAM | alignment | distance | position | gene name | gene id |
| GU305914.1 | | 214759 | 214781 | + | 0 | AGCGAACCCTCCTGTTGAGC | GGG | \|\|\|\|\|\|\|\|[\|\|\|\|\|\|\|\|\|\|\|\|]PAM | NA | Intergenic | NA | NA |
| T16146 | CCTCCTGTTGAGCGGGCATCTGG | | 1000 | | | | | | | | | |
| Oligo fwd | CACCCCTCCTGTTGAGCGGGCATC | | | | | | | | | | | |
| Oligo rev | AAACGATGCCCGCTCAACAGGAGG | | | | | | | | | | | |
| Chromosome | start | | end | strand | MM | target_seq | PAM | alignment | distance | position | gene name | gene id |
| GU305914.1 | | 214766 | 214788 | + | 0 | CCTCCTGTTGAGCGGGCATC | TGG | \|\|\|\|\|\|\|\|[\|\|\|\|\|\|\|\|\|\|\|\|]PAM | NA | Intergenic | NA | NA |
| T16147 | CTCCTGTTGAGCGGGCATCTGGG | | 1000 | | | | | | | | | |
| Oligo fwd | CACCCTCCTGTTGAGCGGGCATCT | | | | | | | | | | | |
| Oligo rev | AAACAGATGCCCGCTCAACAGGAG | | | | | | | | | | | |
| Chromosome | start | | end | strand | MM | target_seq | PAM | alignment | distance | position | gene name | gene id |
| GU305914.1 | | 214767 | 214789 | + | 0 | CTCCTGTTGAGCGGGCATCT | GGG | \|\|\|\|\|\|\|\|[\|\|\|\|\|\|\|\|\|\|\|\|]PAM | NA | Intergenic | NA | NA |
| T16148 | ACGACGCACGCGTCCCGAAGTGG | | 1000 | | | | | | | | | |
| Oligo fwd | CACCACGACGCACGCGTCCCGAAG | | | | | | | | | | | |
| Oligo rev | AAACCTTCGGGACGCGTGCGTCGT | | | | | | | | | | | |
| Chromosome | start | | end | strand | MM | target_seq | PAM | alignment | distance | position | gene name | gene id |
| GU305914.1 | | 214808 | 214830 | + | 0 | ACGACGCACGCGTCCCGAAG | TGG | \|\|\|\|\|\|\|\|[\|\|\|\|\|\|\|\|\|\|\|\|]PAM | NA | Intergenic | NA | NA |
| T16149 | ATTCCTCACCGAGCCCAGCCAGG | | 1000 | | | | | | | | | |
| Oligo fwd | CACCATTCCTCACCGAGCCCAGCC | | | | | | | | | | | |
| Oligo rev | AAACGGCTGGGCTCGGTGAGGAAT | | | | | | | | | | | |
| Chromosome | start | | end | strand | MM | target_seq | PAM | alignment | distance | position | gene name | gene id |
| GU305914.1 | | 214832 | 214854 | + | 0 | ATTCCTCACCGAGCCCAGCC | AGG | \|\|\|\|\|\|\|\|[\|\|\|\|\|\|\|\|\|\|\|\|]PAM | NA | Intergenic | NA | NA |
| T16150 | TCGCACCGTGCATACGAGACCGG | | 1000 | | | | | | | | | |
| Oligo fwd | CACCTCGCACCGTGCATACGAGAC | | | | | | | | | | | |
| Oligo rev | AAACGTCTCGTATGCACGGTGCGA | | | | | | | | | | | |
| Chromosome | start | | end | strand | MM | target_seq | PAM | alignment | distance | position | gene name | gene id |
| GU305914.1 | | 214859 | 214881 | + | 0 | TCGCACCGTGCATACGAGAC | CGG | \|\|\|\|\|\|\|\|[\|\|\|\|\|\|\|\|\|\|\|\|]PAM | NA | Intergenic | NA | NA |
| T16151 | CGCACCGTGCATACGAGACCGGG | | 1000 | | | | | | | | | |
| Oligo fwd | CACCCGCACCGTGCATACGAGACC | | | | | | | | | | | |
| Oligo rev | AAACGGTCTCGTATGCACGGTGCG | | | | | | | | | | | |
| Chromosome | start | | end | strand | MM | target_seq | PAM | alignment | distance | position | gene name | gene id |
| GU305914.1 | | 214860 | 214882 | + | 0 | CGCACCGTGCATACGAGACC | GGG | \|\|\|\|\|\|\|\|[\|\|\|\|\|\|\|\|\|\|\|\|]PAM | NA | Intergenic | NA | NA |
| T16152 | ACCGGGTGCACTCTCCGTCACGG | | 1000 | | | | | | | | | |
| Oligo fwd | CACCACCGGGTGCACTCTCCGTCA | | | | | | | | | | | |
| Oligo rev | AAACTGACGGAGAGTGCACCCGGT | | | | | | | | | | | |
| Chromosome | start | | end | strand | MM | target_seq | PAM | alignment | distance | position | gene name | gene id |
| GU305914.1 | | 214877 | 214899 | + | 0 | ACCGGGTGCACTCTCCGTCA | CGG | \|\|\|\|\|\|\|\|[\|\|\|\|\|\|\|\|\|\|\|\|]PAM | NA | Intergenic | NA | NA |
| T16153 | CGTGTAATTGAGCCGACCGCCGG | | 1000 | | | | | | | | | |
| Oligo fwd | CACCCGTGTAATTGAGCCGACCGC | | | | | | | | | | | |
| Oligo rev | AAACGCGGTCGGCTCAATTACACG | | | | | | | | | | | |
| Chromosome | start | | end | strand | MM | target_seq | PAM | alignment | distance | position | gene name | gene id |

<table>
<tr><td>GU305914.1</td><td></td><td>214922</td><td>214944</td><td>+</td><td>0</td><td>CGTGTAATTGAGCCGACCGC</td><td>CGG</td><td>||||||||[||||||||||||]PAM</td><td>NA</td><td>Intergenic</td><td>NA</td><td>NA</td></tr>
<tr><td></td></tr>
<tr><td>T16154</td><td>GTTACCGCATTCGTCCGTGATGG</td><td></td><td>1000</td></tr>
<tr><td>Oligo fwd</td><td>CACCGTTACCGCATTCGTCCGTGA</td></tr>
<tr><td>Oligo rev</td><td>AAACTCACGGACGAATGCGGTAAC</td></tr>
<tr><td>Chromosome</td><td>start</td><td></td><td>end</td><td>strand</td><td>MM</td><td>target_seq</td><td>PAM</td><td>alignment</td><td>distance</td><td>position</td><td>gene name</td><td>gene id</td></tr>
<tr><td>GU305914.1</td><td></td><td>214955</td><td>214977</td><td>+</td><td>0</td><td>GTTACCGCATTCGTCCGTGA</td><td>TGG</td><td>||||||||[||||||||||||]PAM</td><td>NA</td><td>Intergenic</td><td>NA</td><td>NA</td></tr>
<tr><td></td></tr>
<tr><td>T16155</td><td>TAGCCGACCTGACGTTCCACAGG</td><td></td><td>1000</td></tr>
<tr><td>Oligo fwd</td><td>CACCTAGCCGACCTGACGTTCCAC</td></tr>
<tr><td>Oligo rev</td><td>AAACGTGGAACGTCAGGTCGGCTA</td></tr>
<tr><td>Chromosome</td><td>start</td><td></td><td>end</td><td>strand</td><td>MM</td><td>target_seq</td><td>PAM</td><td>alignment</td><td>distance</td><td>position</td><td>gene name</td><td>gene id</td></tr>
<tr><td>GU305914.1</td><td></td><td>215080</td><td>215102</td><td>+</td><td>0</td><td>TAGCCGACCTGACGTTCCAC</td><td>AGG</td><td>||||||||[||||||||||||]PAM</td><td>NA</td><td>Intergenic</td><td>NA</td><td>NA</td></tr>
<tr><td></td></tr>
<tr><td>T16156</td><td>TCGTTCAACGAACAGTTATTAGG</td><td></td><td>1000</td></tr>
<tr><td>Oligo fwd</td><td>CACCTCGTTCAACGAACAGTTATT</td></tr>
<tr><td>Oligo rev</td><td>AAACAATAACTGTTCGTTGAACGA</td></tr>
<tr><td>Chromosome</td><td>start</td><td></td><td>end</td><td>strand</td><td>MM</td><td>target_seq</td><td>PAM</td><td>alignment</td><td>distance</td><td>position</td><td>gene name</td><td>gene id</td></tr>
<tr><td>GU305914.1</td><td></td><td>215190</td><td>215212</td><td>+</td><td>0</td><td>TCGTTCAACGAACAGTTATT</td><td>AGG</td><td>||||||||[||||||||||||]PAM</td><td>NA</td><td>Intergenic</td><td>NA</td><td>NA</td></tr>
<tr><td></td></tr>
<tr><td>T16157</td><td>AACGAACAGTTATTAGGCTTTGG</td><td></td><td>1000</td></tr>
<tr><td>Oligo fwd</td><td>CACCAACGAACAGTTATTAGGCTT</td></tr>
<tr><td>Oligo rev</td><td>AAACAAGCCTAATAACTGTTCGTT</td></tr>
<tr><td>Chromosome</td><td>start</td><td></td><td>end</td><td>strand</td><td>MM</td><td>target_seq</td><td>PAM</td><td>alignment</td><td>distance</td><td>position</td><td>gene name</td><td>gene id</td></tr>
<tr><td>GU305914.1</td><td></td><td>215196</td><td>215218</td><td>+</td><td>0</td><td>AACGAACAGTTATTAGGCTT</td><td>TGG</td><td>||||||||[||||||||||||]PAM</td><td>NA</td><td>Intergenic</td><td>NA</td><td>NA</td></tr>
<tr><td></td></tr>
<tr><td>T16158</td><td>ATTAGGCTTTGGTTCTCCGTAGG</td><td></td><td>1000</td></tr>
<tr><td>Oligo fwd</td><td>CACCATTAGGCTTTGGTTCTCCGT</td></tr>
<tr><td>Oligo rev</td><td>AAACACGGAGAACCAAAGCCTAAT</td></tr>
<tr><td>Chromosome</td><td>start</td><td></td><td>end</td><td>strand</td><td>MM</td><td>target_seq</td><td>PAM</td><td>alignment</td><td>distance</td><td>position</td><td>gene name</td><td>gene id</td></tr>
<tr><td>GU305914.1</td><td></td><td>215207</td><td>215229</td><td>+</td><td>0</td><td>ATTAGGCTTTGGTTCTCCGT</td><td>AGG</td><td>||||||||[||||||||||||]PAM</td><td>NA</td><td>Intergenic</td><td>NA</td><td>NA</td></tr>
<tr><td></td></tr>
<tr><td>T16159</td><td>CTTTGGTTCTCCGTAGGTTATGG</td><td></td><td>1000</td></tr>
<tr><td>Oligo fwd</td><td>CACCCTTTGGTTCTCCGTAGGTTA</td></tr>
<tr><td>Oligo rev</td><td>AAACTAACCTACGGAGAACCAAAG</td></tr>
<tr><td>Chromosome</td><td>start</td><td></td><td>end</td><td>strand</td><td>MM</td><td>target_seq</td><td>PAM</td><td>alignment</td><td>distance</td><td>position</td><td>gene name</td><td>gene id</td></tr>
<tr><td>GU305914.1</td><td></td><td>215213</td><td>215235</td><td>+</td><td>0</td><td>CTTTGGTTCTCCGTAGGTTA</td><td>TGG</td><td>||||||||[||||||||||||]PAM</td><td>NA</td><td>Intergenic</td><td>NA</td><td>NA</td></tr>
<tr><td></td></tr>
<tr><td>T16160</td><td>GGTTATGGTACGATTCTTGATGG</td><td></td><td>1000</td></tr>
<tr><td>Oligo fwd</td><td>CACCGGTTATGGTACGATTCTTGA</td></tr>
<tr><td>Oligo rev</td><td>AAACTCAAGAATCGTACCATAACC</td></tr>
<tr><td>Chromosome</td><td>start</td><td></td><td>end</td><td>strand</td><td>MM</td><td>target_seq</td><td>PAM</td><td>alignment</td><td>distance</td><td>position</td><td>gene name</td><td>gene id</td></tr>
<tr><td>GU305914.1</td><td></td><td>215228</td><td>215250</td><td>+</td><td>0</td><td>GGTTATGGTACGATTCTTGA</td><td>TGG</td><td>||||||||[||||||||||||]PAM</td><td>NA</td><td>Intergenic</td><td>NA</td><td>NA</td></tr>
<tr><td></td></tr>
<tr><td>T16161</td><td>GGAGTTATAGTACAGACGACAGG</td><td></td><td>1000</td></tr>
<tr><td>Oligo fwd</td><td>CACCGGAGTTATAGTACAGACGAC</td></tr>
<tr><td>Oligo rev</td><td>AAACGTCGTCTGTACTATAACTCC</td></tr>
<tr><td>Chromosome</td><td>start</td><td></td><td>end</td><td>strand</td><td>MM</td><td>target_seq</td><td>PAM</td><td>alignment</td><td>distance</td><td>position</td><td>gene name</td><td>gene id</td></tr>
<tr><td>GU305914.1</td><td></td><td>215249</td><td>215271</td><td>+</td><td>0</td><td>GGAGTTATAGTACAGACGAC</td><td>AGG</td><td>||||||||[||||||||||||]PAM</td><td>NA</td><td>Intergenic</td><td>NA</td><td>NA</td></tr>
<tr><td></td></tr>
<tr><td>T16162</td><td>GTAGTAATCCGACATGTTGTTGG</td><td></td><td>1000</td></tr>
<tr><td>Oligo fwd</td><td>CACCGTAGTAATCCGACATGTTGT</td></tr>
<tr><td>Oligo rev</td><td>AAACACAACATGTCGGATTACTAC</td></tr>
<tr><td>Chromosome</td><td>start</td><td></td><td>end</td><td>strand</td><td>MM</td><td>target_seq</td><td>PAM</td><td>alignment</td><td>distance</td><td>position</td><td>gene name</td><td>gene id</td></tr>
<tr><td>GU305914.1</td><td></td><td>215291</td><td>215313</td><td>+</td><td>0</td><td>GTAGTAATCCGACATGTTGT</td><td>TGG</td><td>||||||||[||||||||||||]PAM</td><td>NA</td><td>Intergenic</td><td>NA</td><td>NA</td></tr>
<tr><td></td></tr>
<tr><td>T16163</td><td>TAGTAATCCGACATGTTGTTGGG</td><td></td><td>1000</td></tr>
<tr><td>Oligo fwd</td><td>CACCTAGTAATCCGACATGTTGTT</td></tr>
<tr><td>Oligo rev</td><td>AAACAACAACATGTCGGATTACTA</td></tr>
<tr><td>Chromosome</td><td>start</td><td></td><td>end</td><td>strand</td><td>MM</td><td>target_seq</td><td>PAM</td><td>alignment</td><td>distance</td><td>position</td><td>gene name</td><td>gene id</td></tr>
<tr><td>GU305914.1</td><td></td><td>215292</td><td>215314</td><td>+</td><td>0</td><td>TAGTAATCCGACATGTTGTT</td><td>GGG</td><td>||||||||[||||||||||||]PAM</td><td>NA</td><td>Intergenic</td><td>NA</td><td>NA</td></tr>
<tr><td></td></tr>
<tr><td>T16164</td><td>CCGACATGTTGTTGGGTCTCAGG</td><td></td><td>1000</td></tr>
<tr><td>Oligo fwd</td><td>CACCCCGACATGTTGTTGGGTCTC</td></tr>
<tr><td>Oligo rev</td><td>AAACGAGACCCAACAACATGTCGG</td></tr>
<tr><td>Chromosome</td><td>start</td><td></td><td>end</td><td>strand</td><td>MM</td><td>target_seq</td><td>PAM</td><td>alignment</td><td>distance</td><td>position</td><td>gene name</td><td>gene id</td></tr>
<tr><td>GU305914.1</td><td></td><td>215299</td><td>215321</td><td>+</td><td>0</td><td>CCGACATGTTGTTGGGTCTC</td><td>AGG</td><td>||||||||[||||||||||||]PAM</td><td>NA</td><td>Intergenic</td><td>NA</td><td>NA</td></tr>
<tr><td></td></tr>
<tr><td>T16165</td><td>AACGTATATTCCAATTCCAGAGG</td><td></td><td>1000</td></tr>
<tr><td>Oligo fwd</td><td>CACCAACGTATATTCCAATTCCAG</td></tr>
<tr><td>Oligo rev</td><td>AAACCTGGAATTGGAATATACGTT</td></tr>
<tr><td>Chromosome</td><td>start</td><td></td><td>end</td><td>strand</td><td>MM</td><td>target_seq</td><td>PAM</td><td>alignment</td><td>distance</td><td>position</td><td>gene name</td><td>gene id</td></tr>
<tr><td>GU305914.1</td><td></td><td>215325</td><td>215347</td><td>+</td><td>0</td><td>AACGTATATTCCAATTCCAG</td><td>AGG</td><td>||||||||[||||||||||||]PAM</td><td>NA</td><td>Intergenic</td><td>NA</td><td>NA</td></tr>
<tr><td></td></tr>
<tr><td>T16166</td><td>ACGTATATTCCAATTCCAGAGGG</td><td></td><td>1000</td></tr>
<tr><td>Oligo fwd</td><td>CACCACGTATATTCCAATTCCAGA</td></tr>
<tr><td>Oligo rev</td><td>AAACTCTGGAATTGGAATATACGT</td></tr>
<tr><td>Chromosome</td><td>start</td><td></td><td>end</td><td>strand</td><td>MM</td><td>target_seq</td><td>PAM</td><td>alignment</td><td>distance</td><td>position</td><td>gene name</td><td>gene id</td></tr>
<tr><td>GU305914.1</td><td></td><td>215326</td><td>215348</td><td>+</td><td>0</td><td>ACGTATATTCCAATTCCAGA</td><td>GGG</td><td>||||||||[||||||||||||]PAM</td><td>NA</td><td>Intergenic</td><td>NA</td><td>NA</td></tr>
<tr><td></td></tr>
<tr><td>T16167</td><td>GTCACCCACGAGCTCCCTCACGG</td><td></td><td>1000</td></tr>
<tr><td>Oligo fwd</td><td>CACCGTCACCCACGAGCTCCCTCA</td></tr>
<tr><td>Oligo rev</td><td>AAACTGAGGGAGCTCGTGGGTGAC</td></tr>
<tr><td>Chromosome</td><td>start</td><td></td><td>end</td><td>strand</td><td>MM</td><td>target_seq</td><td>PAM</td><td>alignment</td><td>distance</td><td>position</td><td>gene name</td><td>gene id</td></tr>
<tr><td>GU305914.1</td><td></td><td>215351</td><td>215373</td><td>+</td><td>0</td><td>GTCACCCACGAGCTCCCTCA</td><td>CGG</td><td>||||||||[||||||||||||]PAM</td><td>NA</td><td>Intergenic</td><td>NA</td><td>NA</td></tr>
<tr><td></td></tr>
<tr><td>T16168</td><td>CGAGCTCCCTCACGGTCTGCAGG</td><td></td><td>1000</td></tr>
<tr><td>Oligo fwd</td><td>CACCCGAGCTCCCTCACGGTCTGC</td></tr>
<tr><td>Oligo rev</td><td>AAACGCAGACCGTGAGGGAGCTCG</td></tr>
</table>

<table>
<tr><td>Chromosome</td><td>start</td><td>end</td><td>strand</td><td>MM</td><td>target_seq</td><td>PAM</td><td>alignment</td><td>distance</td><td>position</td><td>gene name</td><td>gene id</td></tr>
<tr><td>GU305914.1</td><td>215359</td><td>215381</td><td>+</td><td>0</td><td>CGAGCTCCCTCACGGTCTGC</td><td>AGG</td><td>||||||||[||||||||||||]PAM</td><td>NA</td><td>Intergenic</td><td>NA</td><td>NA</td></tr>
<tr><td></td><td></td><td></td><td></td><td></td><td></td><td></td><td></td><td></td><td></td><td></td><td></td></tr>
<tr><td>T16169</td><td>ATTCGCGATCAGAGCCAGTTCGG</td><td>1000</td><td></td><td></td><td></td><td></td><td></td><td></td><td></td><td></td><td></td></tr>
<tr><td>Oligo fwd</td><td>CACCATTCGCGATCAGAGCCAGTT</td><td></td><td></td><td></td><td></td><td></td><td></td><td></td><td></td><td></td><td></td></tr>
<tr><td>Oligo rev</td><td>AAACAACTGGCTCTGATCGCGAAT</td><td></td><td></td><td></td><td></td><td></td><td></td><td></td><td></td><td></td><td></td></tr>
<tr><td>Chromosome</td><td>start</td><td>end</td><td>strand</td><td>MM</td><td>target_seq</td><td>PAM</td><td>alignment</td><td>distance</td><td>position</td><td>gene name</td><td>gene id</td></tr>
<tr><td>GU305914.1</td><td>215393</td><td>215415</td><td>+</td><td>0</td><td>ATTCGCGATCAGAGCCAGTT</td><td>CGG</td><td>||||||||[||||||||||||]PAM</td><td>NA</td><td>Intergenic</td><td>NA</td><td>NA</td></tr>
<tr><td></td><td></td><td></td><td></td><td></td><td></td><td></td><td></td><td></td><td></td><td></td><td></td></tr>
<tr><td>T16170</td><td>GACCCTAGATCCGTAGCGTTAGG</td><td>1000</td><td></td><td></td><td></td><td></td><td></td><td></td><td></td><td></td><td></td></tr>
<tr><td>Oligo fwd</td><td>CACCGACCCTAGATCCGTAGCGTT</td><td></td><td></td><td></td><td></td><td></td><td></td><td></td><td></td><td></td><td></td></tr>
<tr><td>Oligo rev</td><td>AAACAACGCTACGGATCTAGGGTC</td><td></td><td></td><td></td><td></td><td></td><td></td><td></td><td></td><td></td><td></td></tr>
<tr><td>Chromosome</td><td>start</td><td>end</td><td>strand</td><td>MM</td><td>target_seq</td><td>PAM</td><td>alignment</td><td>distance</td><td>position</td><td>gene name</td><td>gene id</td></tr>
<tr><td>GU305914.1</td><td>215415</td><td>215437</td><td>+</td><td>0</td><td>GACCCTAGATCCGTAGCGTT</td><td>AGG</td><td>||||||||[||||||||||||]PAM</td><td>NA</td><td>Intergenic</td><td>NA</td><td>NA</td></tr>
<tr><td></td><td></td><td></td><td></td><td></td><td></td><td></td><td></td><td></td><td></td><td></td><td></td></tr>
<tr><td>T16171</td><td>ACCCTAGATCCGTAGCGTTAGGG</td><td>1000</td><td></td><td></td><td></td><td></td><td></td><td></td><td></td><td></td><td></td></tr>
<tr><td>Oligo fwd</td><td>CACCACCCTAGATCCGTAGCGTTA</td><td></td><td></td><td></td><td></td><td></td><td></td><td></td><td></td><td></td><td></td></tr>
<tr><td>Oligo rev</td><td>AAACTAACGCTACGGATCTAGGGT</td><td></td><td></td><td></td><td></td><td></td><td></td><td></td><td></td><td></td><td></td></tr>
<tr><td>Chromosome</td><td>start</td><td>end</td><td>strand</td><td>MM</td><td>target_seq</td><td>PAM</td><td>alignment</td><td>distance</td><td>position</td><td>gene name</td><td>gene id</td></tr>
<tr><td>GU305914.1</td><td>215416</td><td>215438</td><td>+</td><td>0</td><td>ACCCTAGATCCGTAGCGTTA</td><td>GGG</td><td>||||||||[||||||||||||]PAM</td><td>NA</td><td>Intergenic</td><td>NA</td><td>NA</td></tr>
<tr><td></td><td></td><td></td><td></td><td></td><td></td><td></td><td></td><td></td><td></td><td></td><td></td></tr>
<tr><td>T16172</td><td>ACATCGTGATCTGTCCGTCTTGG</td><td>1000</td><td></td><td></td><td></td><td></td><td></td><td></td><td></td><td></td><td></td></tr>
<tr><td>Oligo fwd</td><td>CACCACATCGTGATCTGTCCGTCT</td><td></td><td></td><td></td><td></td><td></td><td></td><td></td><td></td><td></td><td></td></tr>
<tr><td>Oligo rev</td><td>AAACAGACGGACAGATCACGATGT</td><td></td><td></td><td></td><td></td><td></td><td></td><td></td><td></td><td></td><td></td></tr>
<tr><td>Chromosome</td><td>start</td><td>end</td><td>strand</td><td>MM</td><td>target_seq</td><td>PAM</td><td>alignment</td><td>distance</td><td>position</td><td>gene name</td><td>gene id</td></tr>
<tr><td>GU305914.1</td><td>215452</td><td>215474</td><td>+</td><td>0</td><td>ACATCGTGATCTGTCCGTCT</td><td>TGG</td><td>||||||||[||||||||||||]PAM</td><td>NA</td><td>Intergenic</td><td>NA</td><td>NA</td></tr>
<tr><td></td><td></td><td></td><td></td><td></td><td></td><td></td><td></td><td></td><td></td><td></td><td></td></tr>
<tr><td>T16173</td><td>CGTGATCTGTCCGTCTTGGATGG</td><td>1000</td><td></td><td></td><td></td><td></td><td></td><td></td><td></td><td></td><td></td></tr>
<tr><td>Oligo fwd</td><td>CACCCGTGATCTGTCCGTCTTGGA</td><td></td><td></td><td></td><td></td><td></td><td></td><td></td><td></td><td></td><td></td></tr>
<tr><td>Oligo rev</td><td>AAACTCCAAGACGGACAGATCACG</td><td></td><td></td><td></td><td></td><td></td><td></td><td></td><td></td><td></td><td></td></tr>
<tr><td>Chromosome</td><td>start</td><td>end</td><td>strand</td><td>MM</td><td>target_seq</td><td>PAM</td><td>alignment</td><td>distance</td><td>position</td><td>gene name</td><td>gene id</td></tr>
<tr><td>GU305914.1</td><td>215456</td><td>215478</td><td>+</td><td>0</td><td>CGTGATCTGTCCGTCTTGGA</td><td>TGG</td><td>||||||||[||||||||||||]PAM</td><td>NA</td><td>Intergenic</td><td>NA</td><td>NA</td></tr>
<tr><td></td><td></td><td></td><td></td><td></td><td></td><td></td><td></td><td></td><td></td><td></td><td></td></tr>
<tr><td>T16174</td><td>TTGGATGGTCATGAGATCGAAGG</td><td>1000</td><td></td><td></td><td></td><td></td><td></td><td></td><td></td><td></td><td></td></tr>
<tr><td>Oligo fwd</td><td>CACCTTGGATGGTCATGAGATCGA</td><td></td><td></td><td></td><td></td><td></td><td></td><td></td><td></td><td></td><td></td></tr>
<tr><td>Oligo rev</td><td>AAACTCGATCTCATGACCATCCAA</td><td></td><td></td><td></td><td></td><td></td><td></td><td></td><td></td><td></td><td></td></tr>
<tr><td>Chromosome</td><td>start</td><td>end</td><td>strand</td><td>MM</td><td>target_seq</td><td>PAM</td><td>alignment</td><td>distance</td><td>position</td><td>gene name</td><td>gene id</td></tr>
<tr><td>GU305914.1</td><td>215471</td><td>215493</td><td>+</td><td>0</td><td>TTGGATGGTCATGAGATCGA</td><td>AGG</td><td>||||||||[||||||||||||]PAM</td><td>NA</td><td>Intergenic</td><td>NA</td><td>NA</td></tr>
<tr><td></td><td></td><td></td><td></td><td></td><td></td><td></td><td></td><td></td><td></td><td></td><td></td></tr>
<tr><td>T16175</td><td>TCTGATACGAGCGACGTAATCGG</td><td>1000</td><td></td><td></td><td></td><td></td><td></td><td></td><td></td><td></td><td></td></tr>
<tr><td>Oligo fwd</td><td>CACCTCTGATACGAGCGACGTAAT</td><td></td><td></td><td></td><td></td><td></td><td></td><td></td><td></td><td></td><td></td></tr>
<tr><td>Oligo rev</td><td>AAACATTACGTCGCTCGTATCAGA</td><td></td><td></td><td></td><td></td><td></td><td></td><td></td><td></td><td></td><td></td></tr>
<tr><td>Chromosome</td><td>start</td><td>end</td><td>strand</td><td>MM</td><td>target_seq</td><td>PAM</td><td>alignment</td><td>distance</td><td>position</td><td>gene name</td><td>gene id</td></tr>
<tr><td>GU305914.1</td><td>215504</td><td>215526</td><td>+</td><td>0</td><td>TCTGATACGAGCGACGTAAT</td><td>CGG</td><td>||||||||[||||||||||||]PAM</td><td>NA</td><td>Intergenic</td><td>NA</td><td>NA</td></tr>
<tr><td></td><td></td><td></td><td></td><td></td><td></td><td></td><td></td><td></td><td></td><td></td><td></td></tr>
<tr><td>T16176</td><td>CGGTCGAATTCTCTGAATTCAGG</td><td>1000</td><td></td><td></td><td></td><td></td><td></td><td></td><td></td><td></td><td></td></tr>
<tr><td>Oligo fwd</td><td>CACCCGGTCGAATTCTCTGAATTC</td><td></td><td></td><td></td><td></td><td></td><td></td><td></td><td></td><td></td><td></td></tr>
<tr><td>Oligo rev</td><td>AAACGAATTCAGAGAATTCGACCG</td><td></td><td></td><td></td><td></td><td></td><td></td><td></td><td></td><td></td><td></td></tr>
<tr><td>Chromosome</td><td>start</td><td>end</td><td>strand</td><td>MM</td><td>target_seq</td><td>PAM</td><td>alignment</td><td>distance</td><td>position</td><td>gene name</td><td>gene id</td></tr>
<tr><td>GU305914.1</td><td>215524</td><td>215546</td><td>+</td><td>0</td><td>CGGTCGAATTCTCTGAATTC</td><td>AGG</td><td>||||||||[||||||||||||]PAM</td><td>NA</td><td>Intergenic</td><td>NA</td><td>NA</td></tr>
<tr><td></td><td></td><td></td><td></td><td></td><td></td><td></td><td></td><td></td><td></td><td></td><td></td></tr>
<tr><td>T16177</td><td>CTGAATTCAGGTAGATTGTCAGG</td><td>1000</td><td></td><td></td><td></td><td></td><td></td><td></td><td></td><td></td><td></td></tr>
<tr><td>Oligo fwd</td><td>CACCCTGAATTCAGGTAGATTGTC</td><td></td><td></td><td></td><td></td><td></td><td></td><td></td><td></td><td></td><td></td></tr>
<tr><td>Oligo rev</td><td>AAACGACAATCTACCTGAATTCAG</td><td></td><td></td><td></td><td></td><td></td><td></td><td></td><td></td><td></td><td></td></tr>
<tr><td>Chromosome</td><td>start</td><td>end</td><td>strand</td><td>MM</td><td>target_seq</td><td>PAM</td><td>alignment</td><td>distance</td><td>position</td><td>gene name</td><td>gene id</td></tr>
<tr><td>GU305914.1</td><td>215536</td><td>215558</td><td>+</td><td>0</td><td>CTGAATTCAGGTAGATTGTC</td><td>AGG</td><td>||||||||[||||||||||||]PAM</td><td>NA</td><td>Intergenic</td><td>NA</td><td>NA</td></tr>
<tr><td></td><td></td><td></td><td></td><td></td><td></td><td></td><td></td><td></td><td></td><td></td><td></td></tr>
<tr><td>T16178</td><td>ATTCAGGTAGATTGTCAGGACGG</td><td>1000</td><td></td><td></td><td></td><td></td><td></td><td></td><td></td><td></td><td></td></tr>
<tr><td>Oligo fwd</td><td>CACCATTCAGGTAGATTGTCAGGA</td><td></td><td></td><td></td><td></td><td></td><td></td><td></td><td></td><td></td><td></td></tr>
<tr><td>Oligo rev</td><td>AAACTCCTGACAATCTACCTGAAT</td><td></td><td></td><td></td><td></td><td></td><td></td><td></td><td></td><td></td><td></td></tr>
<tr><td>Chromosome</td><td>start</td><td>end</td><td>strand</td><td>MM</td><td>target_seq</td><td>PAM</td><td>alignment</td><td>distance</td><td>position</td><td>gene name</td><td>gene id</td></tr>
<tr><td>GU305914.1</td><td>215540</td><td>215562</td><td>+</td><td>0</td><td>ATTCAGGTAGATTGTCAGGA</td><td>CGG</td><td>||||||||[||||||||||||]PAM</td><td>NA</td><td>Intergenic</td><td>NA</td><td>NA</td></tr>
<tr><td></td><td></td><td></td><td></td><td></td><td></td><td></td><td></td><td></td><td></td><td></td><td></td></tr>
<tr><td>T16179</td><td>ATAGTCTTGTAAATCGATGTAGG</td><td>1000</td><td></td><td></td><td></td><td></td><td></td><td></td><td></td><td></td><td></td></tr>
<tr><td>Oligo fwd</td><td>CACCATAGTCTTGTAAATCGATGT</td><td></td><td></td><td></td><td></td><td></td><td></td><td></td><td></td><td></td><td></td></tr>
<tr><td>Oligo rev</td><td>AAACACATCGATTTACAAGACTAT</td><td></td><td></td><td></td><td></td><td></td><td></td><td></td><td></td><td></td><td></td></tr>
<tr><td>Chromosome</td><td>start</td><td>end</td><td>strand</td><td>MM</td><td>target_seq</td><td>PAM</td><td>alignment</td><td>distance</td><td>position</td><td>gene name</td><td>gene id</td></tr>
<tr><td>GU305914.1</td><td>215597</td><td>215619</td><td>+</td><td>0</td><td>ATAGTCTTGTAAATCGATGT</td><td>AGG</td><td>||||||||[||||||||||||]PAM</td><td>NA</td><td>Intergenic</td><td>NA</td><td>NA</td></tr>
<tr><td></td><td></td><td></td><td></td><td></td><td></td><td></td><td></td><td></td><td></td><td></td><td></td></tr>
<tr><td>T16180</td><td>TTGTAAATCGATGTAGGCGTAGG</td><td>1000</td><td></td><td></td><td></td><td></td><td></td><td></td><td></td><td></td><td></td></tr>
<tr><td>Oligo fwd</td><td>CACCTTGTAAATCGATGTAGGCGT</td><td></td><td></td><td></td><td></td><td></td><td></td><td></td><td></td><td></td><td></td></tr>
<tr><td>Oligo rev</td><td>AAACACGCCTACATCGATTTACAA</td><td></td><td></td><td></td><td></td><td></td><td></td><td></td><td></td><td></td><td></td></tr>
<tr><td>Chromosome</td><td>start</td><td>end</td><td>strand</td><td>MM</td><td>target_seq</td><td>PAM</td><td>alignment</td><td>distance</td><td>position</td><td>gene name</td><td>gene id</td></tr>
<tr><td>GU305914.1</td><td>215603</td><td>215625</td><td>+</td><td>0</td><td>TTGTAAATCGATGTAGGCGT</td><td>AGG</td><td>||||||||[||||||||||||]PAM</td><td>NA</td><td>Intergenic</td><td>NA</td><td>NA</td></tr>
<tr><td></td><td></td><td></td><td></td><td></td><td></td><td></td><td></td><td></td><td></td><td></td><td></td></tr>
<tr><td>T16181</td><td>AGGCACAAGCGATCAAGAGCAGG</td><td>1000</td><td></td><td></td><td></td><td></td><td></td><td></td><td></td><td></td><td></td></tr>
<tr><td>Oligo fwd</td><td>CACCAGGCACAAGCGATCAAGAGC</td><td></td><td></td><td></td><td></td><td></td><td></td><td></td><td></td><td></td><td></td></tr>
<tr><td>Oligo rev</td><td>AAACGCTCTTGATCGCTTGTGCCT</td><td></td><td></td><td></td><td></td><td></td><td></td><td></td><td></td><td></td><td></td></tr>
<tr><td>Chromosome</td><td>start</td><td>end</td><td>strand</td><td>MM</td><td>target_seq</td><td>PAM</td><td>alignment</td><td>distance</td><td>position</td><td>gene name</td><td>gene id</td></tr>
<tr><td>GU305914.1</td><td>215623</td><td>215645</td><td>+</td><td>0</td><td>AGGCACAAGCGATCAAGAGC</td><td>AGG</td><td>||||||||[||||||||||||]PAM</td><td>NA</td><td>Intergenic</td><td>NA</td><td>NA</td></tr>
<tr><td></td><td></td><td></td><td></td><td></td><td></td><td></td><td></td><td></td><td></td><td></td><td></td></tr>
<tr><td>T16182</td><td>AGCTGTCGATGATACGTAGTCGG</td><td>1000</td><td></td><td></td><td></td><td></td><td></td><td></td><td></td><td></td><td></td></tr>
<tr><td>Oligo fwd</td><td>CACCAGCTGTCGATGATACGTAGT</td><td></td><td></td><td></td><td></td><td></td><td></td><td></td><td></td><td></td><td></td></tr>
<tr><td>Oligo rev</td><td>AAACACTACGTATCATCGACAGCT</td><td></td><td></td><td></td><td></td><td></td><td></td><td></td><td></td><td></td><td></td></tr>
<tr><td>Chromosome</td><td>start</td><td>end</td><td>strand</td><td>MM</td><td>target_seq</td><td>PAM</td><td>alignment</td><td>distance</td><td>position</td><td>gene name</td><td>gene id</td></tr>
<tr><td>GU305914.1</td><td>215720</td><td>215742</td><td>+</td><td>0</td><td>AGCTGTCGATGATACGTAGT</td><td>CGG</td><td>||||||||[||||||||||||]PAM</td><td>NA</td><td>Intergenic</td><td>NA</td><td>NA</td></tr>
<tr><td></td><td></td><td></td><td></td><td></td><td></td><td></td><td></td><td></td><td></td><td></td><td></td></tr>
<tr><td>T16183</td><td>TGATACGTAGTCGGACAATGAGG</td><td>1000</td><td></td><td></td><td></td><td></td><td></td><td></td><td></td><td></td><td></td></tr>
<tr><td>Oligo fwd</td><td>CACCTGATACGTAGTCGGACAATG</td><td></td><td></td><td></td><td></td><td></td><td></td><td></td><td></td><td></td><td></td></tr>
</table>

<table>
<tr><td>Oligo rev</td><td>AAACCATTGTCCGACTACGTATCA</td><td></td><td></td><td></td><td></td><td></td><td></td><td></td><td></td><td></td><td></td></tr>
<tr><td>Chromosome</td><td>start</td><td>end</td><td>strand</td><td>MM</td><td>target_seq</td><td>PAM</td><td>alignment</td><td>distance</td><td>position</td><td>gene name</td><td>gene id</td></tr>
<tr><td>GU305914.1</td><td>215729</td><td>215751</td><td>+</td><td>0</td><td>TGATACGTAGTCGGACAATG</td><td>AGG</td><td>||||||||[||||||||||||]PAM</td><td>NA</td><td>Intergenic</td><td>NA</td><td>NA</td></tr>
<tr><td></td><td></td><td></td><td></td><td></td><td></td><td></td><td></td><td></td><td></td><td></td><td></td></tr>
<tr><td>T16184</td><td>GATACGTAGTCGGACAATGAGGG</td><td>1000</td><td></td><td></td><td></td><td></td><td></td><td></td><td></td><td></td><td></td></tr>
<tr><td>Oligo fwd</td><td>CACCGATACGTAGTCGGACAATGA</td><td></td><td></td><td></td><td></td><td></td><td></td><td></td><td></td><td></td><td></td></tr>
<tr><td>Oligo rev</td><td>AAACTCATTGTCCGACTACGTATC</td><td></td><td></td><td></td><td></td><td></td><td></td><td></td><td></td><td></td><td></td></tr>
<tr><td>Chromosome</td><td>start</td><td>end</td><td>strand</td><td>MM</td><td>target_seq</td><td>PAM</td><td>alignment</td><td>distance</td><td>position</td><td>gene name</td><td>gene id</td></tr>
<tr><td>GU305914.1</td><td>215730</td><td>215752</td><td>+</td><td>0</td><td>GATACGTAGTCGGACAATGA</td><td>GGG</td><td>||||||||[||||||||||||]PAM</td><td>NA</td><td>Intergenic</td><td>NA</td><td>NA</td></tr>
<tr><td></td><td></td><td></td><td></td><td></td><td></td><td></td><td></td><td></td><td></td><td></td><td></td></tr>
<tr><td>T16185</td><td>ATACGTAGTCGGACAATGAGGGG</td><td>1000</td><td></td><td></td><td></td><td></td><td></td><td></td><td></td><td></td><td></td></tr>
<tr><td>Oligo fwd</td><td>CACCATACGTAGTCGGACAATGAG</td><td></td><td></td><td></td><td></td><td></td><td></td><td></td><td></td><td></td><td></td></tr>
<tr><td>Oligo rev</td><td>AAACCTCATTGTCCGACTACGTAT</td><td></td><td></td><td></td><td></td><td></td><td></td><td></td><td></td><td></td><td></td></tr>
<tr><td>Chromosome</td><td>start</td><td>end</td><td>strand</td><td>MM</td><td>target_seq</td><td>PAM</td><td>alignment</td><td>distance</td><td>position</td><td>gene name</td><td>gene id</td></tr>
<tr><td>GU305914.1</td><td>215731</td><td>215753</td><td>+</td><td>0</td><td>ATACGTAGTCGGACAATGAG</td><td>GGG</td><td>||||||||[||||||||||||]PAM</td><td>NA</td><td>Intergenic</td><td>NA</td><td>NA</td></tr>
<tr><td>GU305914.1</td><td>101773</td><td>101795</td><td>+</td><td>4</td><td>ACACGTATTCGGACAGGGAG</td><td>AGG</td><td>|-|||||-[|||||||--|||]PAM</td><td>NA</td><td>Intergenic</td><td>NA</td><td>NA</td></tr>
<tr><td></td><td></td><td></td><td></td><td></td><td></td><td></td><td></td><td></td><td></td><td></td><td></td></tr>
<tr><td>T16186</td><td>TACGTAGTCGGACAATGAGGGGG</td><td>1000</td><td></td><td></td><td></td><td></td><td></td><td></td><td></td><td></td><td></td></tr>
<tr><td>Oligo fwd</td><td>CACCTACGTAGTCGGACAATGAGG</td><td></td><td></td><td></td><td></td><td></td><td></td><td></td><td></td><td></td><td></td></tr>
<tr><td>Oligo rev</td><td>AAACCCTCATTGTCCGACTACGTA</td><td></td><td></td><td></td><td></td><td></td><td></td><td></td><td></td><td></td><td></td></tr>
<tr><td>Chromosome</td><td>start</td><td>end</td><td>strand</td><td>MM</td><td>target_seq</td><td>PAM</td><td>alignment</td><td>distance</td><td>position</td><td>gene name</td><td>gene id</td></tr>
<tr><td>GU305914.1</td><td>215732</td><td>215754</td><td>+</td><td>0</td><td>TACGTAGTCGGACAATGAGG</td><td>GGG</td><td>||||||||[||||||||||||]PAM</td><td>NA</td><td>Intergenic</td><td>NA</td><td>NA</td></tr>
<tr><td></td><td></td><td></td><td></td><td></td><td></td><td></td><td></td><td></td><td></td><td></td><td></td></tr>
<tr><td>T16187</td><td>CTGTGTGCCAAAGATCTTTCTGG</td><td>1000</td><td></td><td></td><td></td><td></td><td></td><td></td><td></td><td></td><td></td></tr>
<tr><td>Oligo fwd</td><td>CACCCTGTGTGCCAAAGATCTTTC</td><td></td><td></td><td></td><td></td><td></td><td></td><td></td><td></td><td></td><td></td></tr>
<tr><td>Oligo rev</td><td>AAACGAAAGATCTTTGGCACACAG</td><td></td><td></td><td></td><td></td><td></td><td></td><td></td><td></td><td></td><td></td></tr>
<tr><td>Chromosome</td><td>start</td><td>end</td><td>strand</td><td>MM</td><td>target_seq</td><td>PAM</td><td>alignment</td><td>distance</td><td>position</td><td>gene name</td><td>gene id</td></tr>
<tr><td>GU305914.1</td><td>215755</td><td>215777</td><td>+</td><td>0</td><td>CTGTGTGCCAAAGATCTTTC</td><td>TGG</td><td>||||||||[||||||||||||]PAM</td><td>NA</td><td>Intergenic</td><td>NA</td><td>NA</td></tr>
<tr><td></td><td></td><td></td><td></td><td></td><td></td><td></td><td></td><td></td><td></td><td></td><td></td></tr>
<tr><td>T16188</td><td>AAGATCTTTCTGGCACTGAATGG</td><td>1000</td><td></td><td></td><td></td><td></td><td></td><td></td><td></td><td></td><td></td></tr>
<tr><td>Oligo fwd</td><td>CACCAAGATCTTTCTGGCACTGAA</td><td></td><td></td><td></td><td></td><td></td><td></td><td></td><td></td><td></td><td></td></tr>
<tr><td>Oligo rev</td><td>AAACTTCAGTGCCAGAAAGATCTT</td><td></td><td></td><td></td><td></td><td></td><td></td><td></td><td></td><td></td><td></td></tr>
<tr><td>Chromosome</td><td>start</td><td>end</td><td>strand</td><td>MM</td><td>target_seq</td><td>PAM</td><td>alignment</td><td>distance</td><td>position</td><td>gene name</td><td>gene id</td></tr>
<tr><td>GU305914.1</td><td>215765</td><td>215787</td><td>+</td><td>0</td><td>AAGATCTTTCTGGCACTGAA</td><td>TGG</td><td>||||||||[||||||||||||]PAM</td><td>NA</td><td>Intergenic</td><td>NA</td><td>NA</td></tr>
<tr><td></td><td></td><td></td><td></td><td></td><td></td><td></td><td></td><td></td><td></td><td></td><td></td></tr>
<tr><td>T16189</td><td>CTGGCACTGAATGGACCAGACGG</td><td>1000</td><td></td><td></td><td></td><td></td><td></td><td></td><td></td><td></td><td></td></tr>
<tr><td>Oligo fwd</td><td>CACCCTGGCACTGAATGGACCAGA</td><td></td><td></td><td></td><td></td><td></td><td></td><td></td><td></td><td></td><td></td></tr>
<tr><td>Oligo rev</td><td>AAACTCTGGTCCATTCAGTGCCAG</td><td></td><td></td><td></td><td></td><td></td><td></td><td></td><td></td><td></td><td></td></tr>
<tr><td>Chromosome</td><td>start</td><td>end</td><td>strand</td><td>MM</td><td>target_seq</td><td>PAM</td><td>alignment</td><td>distance</td><td>position</td><td>gene name</td><td>gene id</td></tr>
<tr><td>GU305914.1</td><td>215774</td><td>215796</td><td>+</td><td>0</td><td>CTGGCACTGAATGGACCAGA</td><td>CGG</td><td>||||||||[||||||||||||]PAM</td><td>NA</td><td>Intergenic</td><td>NA</td><td>NA</td></tr>
<tr><td></td><td></td><td></td><td></td><td></td><td></td><td></td><td></td><td></td><td></td><td></td><td></td></tr>
<tr><td>T16190</td><td>TGGCACTGAATGGACCAGACGGG</td><td>1000</td><td></td><td></td><td></td><td></td><td></td><td></td><td></td><td></td><td></td></tr>
<tr><td>Oligo fwd</td><td>CACCTGGCACTGAATGGACCAGAC</td><td></td><td></td><td></td><td></td><td></td><td></td><td></td><td></td><td></td><td></td></tr>
<tr><td>Oligo rev</td><td>AAACGTCTGGTCCATTCAGTGCCA</td><td></td><td></td><td></td><td></td><td></td><td></td><td></td><td></td><td></td><td></td></tr>
<tr><td>Chromosome</td><td>start</td><td>end</td><td>strand</td><td>MM</td><td>target_seq</td><td>PAM</td><td>alignment</td><td>distance</td><td>position</td><td>gene name</td><td>gene id</td></tr>
<tr><td>GU305914.1</td><td>215775</td><td>215797</td><td>+</td><td>0</td><td>TGGCACTGAATGGACCAGAC</td><td>GGG</td><td>||||||||[||||||||||||]PAM</td><td>NA</td><td>Intergenic</td><td>NA</td><td>NA</td></tr>
<tr><td></td><td></td><td></td><td></td><td></td><td></td><td></td><td></td><td></td><td></td><td></td><td></td></tr>
<tr><td>T16191</td><td>TGTCACAATACACGTCATACCGG</td><td>1000</td><td></td><td></td><td></td><td></td><td></td><td></td><td></td><td></td><td></td></tr>
<tr><td>Oligo fwd</td><td>CACCTGTCACAATACACGTCATAC</td><td></td><td></td><td></td><td></td><td></td><td></td><td></td><td></td><td></td><td></td></tr>
<tr><td>Oligo rev</td><td>AAACGTATGACGTGTATTGTGACA</td><td></td><td></td><td></td><td></td><td></td><td></td><td></td><td></td><td></td><td></td></tr>
<tr><td>Chromosome</td><td>start</td><td>end</td><td>strand</td><td>MM</td><td>target_seq</td><td>PAM</td><td>alignment</td><td>distance</td><td>position</td><td>gene name</td><td>gene id</td></tr>
<tr><td>GU305914.1</td><td>215821</td><td>215843</td><td>+</td><td>0</td><td>TGTCACAATACACGTCATAC</td><td>CGG</td><td>||||||||[||||||||||||]PAM</td><td>NA</td><td>Intergenic</td><td>NA</td><td>NA</td></tr>
<tr><td></td><td></td><td></td><td></td><td></td><td></td><td></td><td></td><td></td><td></td><td></td><td></td></tr>
<tr><td>T16192</td><td>GTCACAATACACGTCATACCGGG</td><td>1000</td><td></td><td></td><td></td><td></td><td></td><td></td><td></td><td></td><td></td></tr>
<tr><td>Oligo fwd</td><td>CACCGTCACAATACACGTCATACC</td><td></td><td></td><td></td><td></td><td></td><td></td><td></td><td></td><td></td><td></td></tr>
<tr><td>Oligo rev</td><td>AAACGGTATGACGTGTATTGTGAC</td><td></td><td></td><td></td><td></td><td></td><td></td><td></td><td></td><td></td><td></td></tr>
<tr><td>Chromosome</td><td>start</td><td>end</td><td>strand</td><td>MM</td><td>target_seq</td><td>PAM</td><td>alignment</td><td>distance</td><td>position</td><td>gene name</td><td>gene id</td></tr>
<tr><td>GU305914.1</td><td>215822</td><td>215844</td><td>+</td><td>0</td><td>GTCACAATACACGTCATACC</td><td>GGG</td><td>||||||||[||||||||||||]PAM</td><td>NA</td><td>Intergenic</td><td>NA</td><td>NA</td></tr>
<tr><td></td><td></td><td></td><td></td><td></td><td></td><td></td><td></td><td></td><td></td><td></td><td></td></tr>
<tr><td>T16193</td><td>TCACAATACACGTCATACCGGGG</td><td>1000</td><td></td><td></td><td></td><td></td><td></td><td></td><td></td><td></td><td></td></tr>
<tr><td>Oligo fwd</td><td>CACCTCACAATACACGTCATACCG</td><td></td><td></td><td></td><td></td><td></td><td></td><td></td><td></td><td></td><td></td></tr>
<tr><td>Oligo rev</td><td>AAACCGGTATGACGTGTATTGTGA</td><td></td><td></td><td></td><td></td><td></td><td></td><td></td><td></td><td></td><td></td></tr>
<tr><td>Chromosome</td><td>start</td><td>end</td><td>strand</td><td>MM</td><td>target_seq</td><td>PAM</td><td>alignment</td><td>distance</td><td>position</td><td>gene name</td><td>gene id</td></tr>
<tr><td>GU305914.1</td><td>215823</td><td>215845</td><td>+</td><td>0</td><td>TCACAATACACGTCATACCG</td><td>GGG</td><td>||||||||[||||||||||||]PAM</td><td>NA</td><td>Intergenic</td><td>NA</td><td>NA</td></tr>
<tr><td></td><td></td><td></td><td></td><td></td><td></td><td></td><td></td><td></td><td></td><td></td><td></td></tr>
<tr><td>T16194</td><td>CGTCATACCGGGGACTACATCGG</td><td>1000</td><td></td><td></td><td></td><td></td><td></td><td></td><td></td><td></td><td></td></tr>
<tr><td>Oligo fwd</td><td>CACCCGTCATACCGGGGACTACAT</td><td></td><td></td><td></td><td></td><td></td><td></td><td></td><td></td><td></td><td></td></tr>
<tr><td>Oligo rev</td><td>AAACATGTAGTCCCCGGTATGACG</td><td></td><td></td><td></td><td></td><td></td><td></td><td></td><td></td><td></td><td></td></tr>
<tr><td>Chromosome</td><td>start</td><td>end</td><td>strand</td><td>MM</td><td>target_seq</td><td>PAM</td><td>alignment</td><td>distance</td><td>position</td><td>gene name</td><td>gene id</td></tr>
<tr><td>GU305914.1</td><td>215833</td><td>215855</td><td>+</td><td>0</td><td>CGTCATACCGGGGACTACAT</td><td>CGG</td><td>||||||||[||||||||||||]PAM</td><td>NA</td><td>Intergenic</td><td>NA</td><td>NA</td></tr>
<tr><td>GU305914.1</td><td>30766</td><td>30788</td><td>-</td><td>4</td><td>CGCCCTACGGGGGACTATAT</td><td>AGG</td><td>||-|-|||[-||||||||-||]PAM</td><td>NA</td><td>Intergenic</td><td>NA</td><td>NA</td></tr>
<tr><td></td><td></td><td></td><td></td><td></td><td></td><td></td><td></td><td></td><td></td><td></td><td></td></tr>
<tr><td>T16195</td><td>ATACCGGGGACTACATCGGTCGG</td><td>1000</td><td></td><td></td><td></td><td></td><td></td><td></td><td></td><td></td><td></td></tr>
<tr><td>Oligo fwd</td><td>CACCATACCGGGGACTACATCGGT</td><td></td><td></td><td></td><td></td><td></td><td></td><td></td><td></td><td></td><td></td></tr>
<tr><td>Oligo rev</td><td>AAACACCGATGTAGTCCCCGGTAT</td><td></td><td></td><td></td><td></td><td></td><td></td><td></td><td></td><td></td><td></td></tr>
<tr><td>Chromosome</td><td>start</td><td>end</td><td>strand</td><td>MM</td><td>target_seq</td><td>PAM</td><td>alignment</td><td>distance</td><td>position</td><td>gene name</td><td>gene id</td></tr>
<tr><td>GU305914.1</td><td>215837</td><td>215859</td><td>+</td><td>0</td><td>ATACCGGGGACTACATCGGT</td><td>CGG</td><td>||||||||[||||||||||||]PAM</td><td>NA</td><td>Intergenic</td><td>NA</td><td>NA</td></tr>
<tr><td></td><td></td><td></td><td></td><td></td><td></td><td></td><td></td><td></td><td></td><td></td><td></td></tr>
<tr><td>T16196</td><td>ACATCGGTCGGTATTGAATTAGG</td><td>1000</td><td></td><td></td><td></td><td></td><td></td><td></td><td></td><td></td><td></td></tr>
<tr><td>Oligo fwd</td><td>CACCACATCGGTCGGTATTGAATT</td><td></td><td></td><td></td><td></td><td></td><td></td><td></td><td></td><td></td><td></td></tr>
<tr><td>Oligo rev</td><td>AAACAATTCAATACCGACCGATGT</td><td></td><td></td><td></td><td></td><td></td><td></td><td></td><td></td><td></td><td></td></tr>
<tr><td>Chromosome</td><td>start</td><td>end</td><td>strand</td><td>MM</td><td>target_seq</td><td>PAM</td><td>alignment</td><td>distance</td><td>position</td><td>gene name</td><td>gene id</td></tr>
<tr><td>GU305914.1</td><td>215849</td><td>215871</td><td>+</td><td>0</td><td>ACATCGGTCGGTATTGAATT</td><td>AGG</td><td>||||||||[||||||||||||]PAM</td><td>NA</td><td>Intergenic</td><td>NA</td><td>NA</td></tr>
<tr><td></td><td></td><td></td><td></td><td></td><td></td><td></td><td></td><td></td><td></td><td></td><td></td></tr>
<tr><td>T16197</td><td>CATCGGTCGGTATTGAATTAGGG</td><td>1000</td><td></td><td></td><td></td><td></td><td></td><td></td><td></td><td></td><td></td></tr>
<tr><td>Oligo fwd</td><td>CACCCATCGGTCGGTATTGAATTA</td><td></td><td></td><td></td><td></td><td></td><td></td><td></td><td></td><td></td><td></td></tr>
<tr><td>Oligo rev</td><td>AAACTAATTCAATACCGACCGATG</td><td></td><td></td><td></td><td></td><td></td><td></td><td></td><td></td><td></td><td></td></tr>
<tr><td>Chromosome</td><td>start</td><td>end</td><td>strand</td><td>MM</td><td>target_seq</td><td>PAM</td><td>alignment</td><td>distance</td><td>position</td><td>gene name</td><td>gene id</td></tr>
<tr><td>GU305914.1</td><td>215850</td><td>215872</td><td>+</td><td>0</td><td>CATCGGTCGGTATTGAATTA</td><td>GGG</td><td>||||||||[||||||||||||]PAM</td><td>NA</td><td>Intergenic</td><td>NA</td><td>NA</td></tr>
</table>

<table>
<tr><td>T16198</td><td>AGCTCATCTCACGTAGAACATGG</td><td>1000</td><td></td><td></td><td></td><td></td><td></td><td></td><td></td><td></td><td></td></tr>
<tr><td>Oligo fwd</td><td>CACCAGCTCATCTCACGTAGAACA</td><td></td><td></td><td></td><td></td><td></td><td></td><td></td><td></td><td></td><td></td></tr>
<tr><td>Oligo rev</td><td>AAACTGTTCTACGTGAGATGAGCT</td><td></td><td></td><td></td><td></td><td></td><td></td><td></td><td></td><td></td><td></td></tr>
<tr><td>Chromosome</td><td>start</td><td>end</td><td>strand</td><td>MM</td><td>target_seq</td><td>PAM</td><td>alignment</td><td>distance</td><td>position</td><td>gene name</td><td>gene id</td></tr>
<tr><td>GU305914.1</td><td>215874</td><td>215896</td><td>+</td><td>0</td><td>AGCTCATCTCACGTAGAACA</td><td>TGG</td><td>||||||||[||||||||||||]PAM</td><td>NA</td><td>Intergenic</td><td>NA</td><td>NA</td></tr>
<tr><td>T16199</td><td>GACGCATAAATACCACACATAGG</td><td>1000</td><td></td><td></td><td></td><td></td><td></td><td></td><td></td><td></td><td></td></tr>
<tr><td>Oligo fwd</td><td>CACCGACGCATAAATACCACACAT</td><td></td><td></td><td></td><td></td><td></td><td></td><td></td><td></td><td></td><td></td></tr>
<tr><td>Oligo rev</td><td>AAACATGTGTGGTATTTATGCGTC</td><td></td><td></td><td></td><td></td><td></td><td></td><td></td><td></td><td></td><td></td></tr>
<tr><td>Chromosome</td><td>start</td><td>end</td><td>strand</td><td>MM</td><td>target_seq</td><td>PAM</td><td>alignment</td><td>distance</td><td>position</td><td>gene name</td><td>gene id</td></tr>
<tr><td>GU305914.1</td><td>216007</td><td>216029</td><td>+</td><td>0</td><td>GACGCATAAATACCACACAT</td><td>AGG</td><td>||||||||[||||||||||||]PAM</td><td>NA</td><td>Intergenic</td><td>NA</td><td>NA</td></tr>
<tr><td>T16200</td><td>ACCACACATAGGATGAGTATCGG</td><td>1000</td><td></td><td></td><td></td><td></td><td></td><td></td><td></td><td></td><td></td></tr>
<tr><td>Oligo fwd</td><td>CACCACCACACATAGGATGAGTAT</td><td></td><td></td><td></td><td></td><td></td><td></td><td></td><td></td><td></td><td></td></tr>
<tr><td>Oligo rev</td><td>AAACATACTCATCCTATGTGTGGT</td><td></td><td></td><td></td><td></td><td></td><td></td><td></td><td></td><td></td><td></td></tr>
<tr><td>Chromosome</td><td>start</td><td>end</td><td>strand</td><td>MM</td><td>target_seq</td><td>PAM</td><td>alignment</td><td>distance</td><td>position</td><td>gene name</td><td>gene id</td></tr>
<tr><td>GU305914.1</td><td>216018</td><td>216040</td><td>+</td><td>0</td><td>ACCACACATAGGATGAGTAT</td><td>CGG</td><td>||||||||[||||||||||||]PAM</td><td>NA</td><td>Intergenic</td><td>NA</td><td>NA</td></tr>
<tr><td>T16201</td><td>AGGATGAGTATCGGTAAGTTCGG</td><td>1000</td><td></td><td></td><td></td><td></td><td></td><td></td><td></td><td></td><td></td></tr>
<tr><td>Oligo fwd</td><td>CACCAGGATGAGTATCGGTAAGTT</td><td></td><td></td><td></td><td></td><td></td><td></td><td></td><td></td><td></td><td></td></tr>
<tr><td>Oligo rev</td><td>AAACAACTTACCGATACTCATCCT</td><td></td><td></td><td></td><td></td><td></td><td></td><td></td><td></td><td></td><td></td></tr>
<tr><td>Chromosome</td><td>start</td><td>end</td><td>strand</td><td>MM</td><td>target_seq</td><td>PAM</td><td>alignment</td><td>distance</td><td>position</td><td>gene name</td><td>gene id</td></tr>
<tr><td>GU305914.1</td><td>216027</td><td>216049</td><td>+</td><td>0</td><td>AGGATGAGTATCGGTAAGTT</td><td>CGG</td><td>||||||||[||||||||||||]PAM</td><td>NA</td><td>Intergenic</td><td>NA</td><td>NA</td></tr>
<tr><td>T16202</td><td>TAAGTTCGGTAAATGAGAAGCGG</td><td>1000</td><td></td><td></td><td></td><td></td><td></td><td></td><td></td><td></td><td></td></tr>
<tr><td>Oligo fwd</td><td>CACCTAAGTTCGGTAAATGAGAAG</td><td></td><td></td><td></td><td></td><td></td><td></td><td></td><td></td><td></td><td></td></tr>
<tr><td>Oligo rev</td><td>AAACCTTCTCATTTACCGAACTTA</td><td></td><td></td><td></td><td></td><td></td><td></td><td></td><td></td><td></td><td></td></tr>
<tr><td>Chromosome</td><td>start</td><td>end</td><td>strand</td><td>MM</td><td>target_seq</td><td>PAM</td><td>alignment</td><td>distance</td><td>position</td><td>gene name</td><td>gene id</td></tr>
<tr><td>GU305914.1</td><td>216041</td><td>216063</td><td>+</td><td>0</td><td>TAAGTTCGGTAAATGAGAAG</td><td>CGG</td><td>||||||||[||||||||||||]PAM</td><td>NA</td><td>Intergenic</td><td>NA</td><td>NA</td></tr>
<tr><td>T16203</td><td>GTAAATGAGAAGCGGTAGAACGG</td><td>1000</td><td></td><td></td><td></td><td></td><td></td><td></td><td></td><td></td><td></td></tr>
<tr><td>Oligo fwd</td><td>CACCGTAAATGAGAAGCGGTAGAA</td><td></td><td></td><td></td><td></td><td></td><td></td><td></td><td></td><td></td><td></td></tr>
<tr><td>Oligo rev</td><td>AAACTTCTACCGCTTCTCATTTAC</td><td></td><td></td><td></td><td></td><td></td><td></td><td></td><td></td><td></td><td></td></tr>
<tr><td>Chromosome</td><td>start</td><td>end</td><td>strand</td><td>MM</td><td>target_seq</td><td>PAM</td><td>alignment</td><td>distance</td><td>position</td><td>gene name</td><td>gene id</td></tr>
<tr><td>GU305914.1</td><td>216049</td><td>216071</td><td>+</td><td>0</td><td>GTAAATGAGAAGCGGTAGAA</td><td>CGG</td><td>||||||||[||||||||||||]PAM</td><td>NA</td><td>Intergenic</td><td>NA</td><td>NA</td></tr>
<tr><td>T16204</td><td>TTCGAAGAGCTGATATTGACAGG</td><td>1000</td><td></td><td></td><td></td><td></td><td></td><td></td><td></td><td></td><td></td></tr>
<tr><td>Oligo fwd</td><td>CACCTTCGAAGAGCTGATATTGAC</td><td></td><td></td><td></td><td></td><td></td><td></td><td></td><td></td><td></td><td></td></tr>
<tr><td>Oligo rev</td><td>AAACGTCAATATCAGCTCTTCGAA</td><td></td><td></td><td></td><td></td><td></td><td></td><td></td><td></td><td></td><td></td></tr>
<tr><td>Chromosome</td><td>start</td><td>end</td><td>strand</td><td>MM</td><td>target_seq</td><td>PAM</td><td>alignment</td><td>distance</td><td>position</td><td>gene name</td><td>gene id</td></tr>
<tr><td>GU305914.1</td><td>216108</td><td>216130</td><td>+</td><td>0</td><td>TTCGAAGAGCTGATATTGAC</td><td>AGG</td><td>||||||||[||||||||||||]PAM</td><td>NA</td><td>Intergenic</td><td>NA</td><td>NA</td></tr>
<tr><td>T16205</td><td>CGTTTCGTGCGATATTGTGACGG</td><td>1000</td><td></td><td></td><td></td><td></td><td></td><td></td><td></td><td></td><td></td></tr>
<tr><td>Oligo fwd</td><td>CACCCGTTTCGTGCGATATTGTGA</td><td></td><td></td><td></td><td></td><td></td><td></td><td></td><td></td><td></td><td></td></tr>
<tr><td>Oligo rev</td><td>AAACTCACAATATCGCACGAAACG</td><td></td><td></td><td></td><td></td><td></td><td></td><td></td><td></td><td></td><td></td></tr>
<tr><td>Chromosome</td><td>start</td><td>end</td><td>strand</td><td>MM</td><td>target_seq</td><td>PAM</td><td>alignment</td><td>distance</td><td>position</td><td>gene name</td><td>gene id</td></tr>
<tr><td>GU305914.1</td><td>216135</td><td>216157</td><td>+</td><td>0</td><td>CGTTTCGTGCGATATTGTGA</td><td>CGG</td><td>||||||||[||||||||||||]PAM</td><td>NA</td><td>Intergenic</td><td>NA</td><td>NA</td></tr>
<tr><td>T16206</td><td>TTCGTGCGATATTGTGACGGAGG</td><td>1000</td><td></td><td></td><td></td><td></td><td></td><td></td><td></td><td></td><td></td></tr>
<tr><td>Oligo fwd</td><td>CACCTTCGTGCGATATTGTGACGG</td><td></td><td></td><td></td><td></td><td></td><td></td><td></td><td></td><td></td><td></td></tr>
<tr><td>Oligo rev</td><td>AAACCCGTCACAATATCGCACGAA</td><td></td><td></td><td></td><td></td><td></td><td></td><td></td><td></td><td></td><td></td></tr>
<tr><td>Chromosome</td><td>start</td><td>end</td><td>strand</td><td>MM</td><td>target_seq</td><td>PAM</td><td>alignment</td><td>distance</td><td>position</td><td>gene name</td><td>gene id</td></tr>
<tr><td>GU305914.1</td><td>216138</td><td>216160</td><td>+</td><td>0</td><td>TTCGTGCGATATTGTGACGG</td><td>AGG</td><td>||||||||[||||||||||||]PAM</td><td>NA</td><td>Intergenic</td><td>NA</td><td>NA</td></tr>
<tr><td>T16207</td><td>CACGACGCCCTTCCATCCCAAGG</td><td>1000</td><td></td><td></td><td></td><td></td><td></td><td></td><td></td><td></td><td></td></tr>
<tr><td>Oligo fwd</td><td>CACCCACGACGCCCTTCCATCCCA</td><td></td><td></td><td></td><td></td><td></td><td></td><td></td><td></td><td></td><td></td></tr>
<tr><td>Oligo rev</td><td>AAACTGGGATGGAAGGGCGTCGTG</td><td></td><td></td><td></td><td></td><td></td><td></td><td></td><td></td><td></td><td></td></tr>
<tr><td>Chromosome</td><td>start</td><td>end</td><td>strand</td><td>MM</td><td>target_seq</td><td>PAM</td><td>alignment</td><td>distance</td><td>position</td><td>gene name</td><td>gene id</td></tr>
<tr><td>GU305914.1</td><td>216164</td><td>216186</td><td>+</td><td>0</td><td>CACGACGCCCTTCCATCCCA</td><td>AGG</td><td>||||||||[||||||||||||]PAM</td><td>NA</td><td>Intergenic</td><td>NA</td><td>NA</td></tr>
<tr><td>T16208</td><td>CATCTTTGAGCGTGACTACCCGG</td><td>1000</td><td></td><td></td><td></td><td></td><td></td><td></td><td></td><td></td><td></td></tr>
<tr><td>Oligo fwd</td><td>CACCCATCTTTGAGCGTGACTACC</td><td></td><td></td><td></td><td></td><td></td><td></td><td></td><td></td><td></td><td></td></tr>
<tr><td>Oligo rev</td><td>AAACGGTAGTCACGCTCAAAGATG</td><td></td><td></td><td></td><td></td><td></td><td></td><td></td><td></td><td></td><td></td></tr>
<tr><td>Chromosome</td><td>start</td><td>end</td><td>strand</td><td>MM</td><td>target_seq</td><td>PAM</td><td>alignment</td><td>distance</td><td>position</td><td>gene name</td><td>gene id</td></tr>
<tr><td>GU305914.1</td><td>216201</td><td>216223</td><td>+</td><td>0</td><td>CATCTTTGAGCGTGACTACC</td><td>CGG</td><td>||||||||[||||||||||||]PAM</td><td>NA</td><td>Intergenic</td><td>NA</td><td>NA</td></tr>
<tr><td>T16209</td><td>ATCTTTGAGCGTGACTACCCGGG</td><td>1000</td><td></td><td></td><td></td><td></td><td></td><td></td><td></td><td></td><td></td></tr>
<tr><td>Oligo fwd</td><td>CACCATCTTTGAGCGTGACTACCC</td><td></td><td></td><td></td><td></td><td></td><td></td><td></td><td></td><td></td><td></td></tr>
<tr><td>Oligo rev</td><td>AAACGGGTAGTCACGCTCAAAGAT</td><td></td><td></td><td></td><td></td><td></td><td></td><td></td><td></td><td></td><td></td></tr>
<tr><td>Chromosome</td><td>start</td><td>end</td><td>strand</td><td>MM</td><td>target_seq</td><td>PAM</td><td>alignment</td><td>distance</td><td>position</td><td>gene name</td><td>gene id</td></tr>
<tr><td>GU305914.1</td><td>216202</td><td>216224</td><td>+</td><td>0</td><td>ATCTTTGAGCGTGACTACCC</td><td>GGG</td><td>||||||||[||||||||||||]PAM</td><td>NA</td><td>Intergenic</td><td>NA</td><td>NA</td></tr>
<tr><td>T16210</td><td>TCTTTGAGCGTGACTACCCGGGG</td><td>1000</td><td></td><td></td><td></td><td></td><td></td><td></td><td></td><td></td><td></td></tr>
<tr><td>Oligo fwd</td><td>CACCTCTTTGAGCGTGACTACCCG</td><td></td><td></td><td></td><td></td><td></td><td></td><td></td><td></td><td></td><td></td></tr>
<tr><td>Oligo rev</td><td>AAACCGGGTAGTCACGCTCAAAGA</td><td></td><td></td><td></td><td></td><td></td><td></td><td></td><td></td><td></td><td></td></tr>
<tr><td>Chromosome</td><td>start</td><td>end</td><td>strand</td><td>MM</td><td>target_seq</td><td>PAM</td><td>alignment</td><td>distance</td><td>position</td><td>gene name</td><td>gene id</td></tr>
<tr><td>GU305914.1</td><td>216203</td><td>216225</td><td>+</td><td>0</td><td>TCTTTGAGCGTGACTACCCG</td><td>GGG</td><td>||||||||[||||||||||||]PAM</td><td>NA</td><td>Intergenic</td><td>NA</td><td>NA</td></tr>
<tr><td>T16211</td><td>GTGACTACCCGGGGCCTTCACGG</td><td>1000</td><td></td><td></td><td></td><td></td><td></td><td></td><td></td><td></td><td></td></tr>
<tr><td>Oligo fwd</td><td>CACCGTGACTACCCGGGGCCTTCA</td><td></td><td></td><td></td><td></td><td></td><td></td><td></td><td></td><td></td><td></td></tr>
<tr><td>Oligo rev</td><td>AAACTGAAGGCCCCGGGTAGTCAC</td><td></td><td></td><td></td><td></td><td></td><td></td><td></td><td></td><td></td><td></td></tr>
<tr><td>Chromosome</td><td>start</td><td>end</td><td>strand</td><td>MM</td><td>target_seq</td><td>PAM</td><td>alignment</td><td>distance</td><td>position</td><td>gene name</td><td>gene id</td></tr>
<tr><td>GU305914.1</td><td>216212</td><td>216234</td><td>+</td><td>0</td><td>GTGACTACCCGGGGCCTTCA</td><td>CGG</td><td>||||||||[||||||||||||]PAM</td><td>NA</td><td>Intergenic</td><td>NA</td><td>NA</td></tr>
<tr><td>T16212</td><td>TACCCGGGGCCTTCACGGTAAGG</td><td>1000</td><td></td><td></td><td></td><td></td><td></td><td></td><td></td><td></td><td></td></tr>
<tr><td>Oligo fwd</td><td>CACCTACCCGGGGCCTTCACGGTA</td><td></td><td></td><td></td><td></td><td></td><td></td><td></td><td></td><td></td><td></td></tr>
<tr><td>Oligo rev</td><td>AAACTACCGTGAAGGCCCCGGGTA</td><td></td><td></td><td></td><td></td><td></td><td></td><td></td><td></td><td></td><td></td></tr>
<tr><td>Chromosome</td><td>start</td><td>end</td><td>strand</td><td>MM</td><td>target_seq</td><td>PAM</td><td>alignment</td><td>distance</td><td>position</td><td>gene name</td><td>gene id</td></tr>
</table>

<table>
<tr><td>GU305914.1</td><td>216217</td><td>216239</td><td>+</td><td>0</td><td>TACCCGGGGCCTTCACGGTA</td><td>AGG</td><td>||||||||[||||||||||||]PAM</td><td>NA</td><td>Intergenic</td><td>NA</td><td>NA</td></tr>
<tr><td>T16213</td><td>CGCAGAGCTGTTAGTGTTCAAGG</td><td>1000</td><td></td><td></td><td></td><td></td><td></td><td></td><td></td><td></td><td></td></tr>
<tr><td>Oligo fwd</td><td>CACCCGCAGAGCTGTTAGTGTTCA</td><td></td><td></td><td></td><td></td><td></td><td></td><td></td><td></td><td></td><td></td></tr>
<tr><td>Oligo rev</td><td>AAACTGAACACTAACAGCTCTGCG</td><td></td><td></td><td></td><td></td><td></td><td></td><td></td><td></td><td></td><td></td></tr>
<tr><td>Chromosome</td><td>start</td><td>end</td><td>strand</td><td>MM</td><td>target_seq</td><td>PAM</td><td>alignment</td><td>distance</td><td>position</td><td>gene name</td><td>gene id</td></tr>
<tr><td>GU305914.1</td><td>216275</td><td>216297</td><td>+</td><td>0</td><td>CGCAGAGCTGTTAGTGTTCA</td><td>AGG</td><td>||||||||[||||||||||||]PAM</td><td>NA</td><td>Intergenic</td><td>NA</td><td>NA</td></tr>
<tr><td>T16214</td><td>AGTGTTCAAGGATAATTCTACGG</td><td>1000</td><td></td><td></td><td></td><td></td><td></td><td></td><td></td><td></td><td></td></tr>
<tr><td>Oligo fwd</td><td>CACCAGTGTTCAAGGATAATTCTA</td><td></td><td></td><td></td><td></td><td></td><td></td><td></td><td></td><td></td><td></td></tr>
<tr><td>Oligo rev</td><td>AAACTAGAATTATCCTTGAACACT</td><td></td><td></td><td></td><td></td><td></td><td></td><td></td><td></td><td></td><td></td></tr>
<tr><td>Chromosome</td><td>start</td><td>end</td><td>strand</td><td>MM</td><td>target_seq</td><td>PAM</td><td>alignment</td><td>distance</td><td>position</td><td>gene name</td><td>gene id</td></tr>
<tr><td>GU305914.1</td><td>216287</td><td>216309</td><td>+</td><td>0</td><td>AGTGTTCAAGGATAATTCTA</td><td>CGG</td><td>||||||||[||||||||||||]PAM</td><td>NA</td><td>Intergenic</td><td>NA</td><td>NA</td></tr>
<tr><td>T16215</td><td>AAGGATAATTCTACGGTGAACGG</td><td>1000</td><td></td><td></td><td></td><td></td><td></td><td></td><td></td><td></td><td></td></tr>
<tr><td>Oligo fwd</td><td>CACCAAGGATAATTCTACGGTGAA</td><td></td><td></td><td></td><td></td><td></td><td></td><td></td><td></td><td></td><td></td></tr>
<tr><td>Oligo rev</td><td>AAACTTCACCGTAGAATTATCCTT</td><td></td><td></td><td></td><td></td><td></td><td></td><td></td><td></td><td></td><td></td></tr>
<tr><td>Chromosome</td><td>start</td><td>end</td><td>strand</td><td>MM</td><td>target_seq</td><td>PAM</td><td>alignment</td><td>distance</td><td>position</td><td>gene name</td><td>gene id</td></tr>
<tr><td>GU305914.1</td><td>216294</td><td>216316</td><td>+</td><td>0</td><td>AAGGATAATTCTACGGTGAA</td><td>CGG</td><td>||||||||[||||||||||||]PAM</td><td>NA</td><td>Intergenic</td><td>NA</td><td>NA</td></tr>
<tr><td>T16216</td><td>CCTCGATGTGACCACACACGTGG</td><td>1000</td><td></td><td></td><td></td><td></td><td></td><td></td><td></td><td></td><td></td></tr>
<tr><td>Oligo fwd</td><td>CACCCCTCGATGTGACCACACACG</td><td></td><td></td><td></td><td></td><td></td><td></td><td></td><td></td><td></td><td></td></tr>
<tr><td>Oligo rev</td><td>AAACCGTGTGTGGTCACATCGAGG</td><td></td><td></td><td></td><td></td><td></td><td></td><td></td><td></td><td></td><td></td></tr>
<tr><td>Chromosome</td><td>start</td><td>end</td><td>strand</td><td>MM</td><td>target_seq</td><td>PAM</td><td>alignment</td><td>distance</td><td>position</td><td>gene name</td><td>gene id</td></tr>
<tr><td>GU305914.1</td><td>216326</td><td>216348</td><td>+</td><td>0</td><td>CCTCGATGTGACCACACACG</td><td>TGG</td><td>||||||||[||||||||||||]PAM</td><td>NA</td><td>Intergenic</td><td>NA</td><td>NA</td></tr>
<tr><td>T16217</td><td>GACCACACACGTGGTCAAGCCGG</td><td>1000</td><td></td><td></td><td></td><td></td><td></td><td></td><td></td><td></td><td></td></tr>
<tr><td>Oligo fwd</td><td>CACCGACCACACACGTGGTCAAGC</td><td></td><td></td><td></td><td></td><td></td><td></td><td></td><td></td><td></td><td></td></tr>
<tr><td>Oligo rev</td><td>AAACGCTTGACCACGTGTGTGGTC</td><td></td><td></td><td></td><td></td><td></td><td></td><td></td><td></td><td></td><td></td></tr>
<tr><td>Chromosome</td><td>start</td><td>end</td><td>strand</td><td>MM</td><td>target_seq</td><td>PAM</td><td>alignment</td><td>distance</td><td>position</td><td>gene name</td><td>gene id</td></tr>
<tr><td>GU305914.1</td><td>216335</td><td>216357</td><td>+</td><td>0</td><td>GACCACACACGTGGTCAAGC</td><td>CGG</td><td>||||||||[||||||||||||]PAM</td><td>NA</td><td>Intergenic</td><td>NA</td><td>NA</td></tr>
<tr><td>T16218</td><td>GTTGTACCAGAACTCGACTTCGG</td><td>1000</td><td></td><td></td><td></td><td></td><td></td><td></td><td></td><td></td><td></td></tr>
<tr><td>Oligo fwd</td><td>CACCGTTGTACCAGAACTCGACTT</td><td></td><td></td><td></td><td></td><td></td><td></td><td></td><td></td><td></td><td></td></tr>
<tr><td>Oligo rev</td><td>AAACAAGTCGAGTTCTGGTACAAC</td><td></td><td></td><td></td><td></td><td></td><td></td><td></td><td></td><td></td><td></td></tr>
<tr><td>Chromosome</td><td>start</td><td>end</td><td>strand</td><td>MM</td><td>target_seq</td><td>PAM</td><td>alignment</td><td>distance</td><td>position</td><td>gene name</td><td>gene id</td></tr>
<tr><td>GU305914.1</td><td>216362</td><td>216384</td><td>+</td><td>0</td><td>GTTGTACCAGAACTCGACTT</td><td>CGG</td><td>||||||||[||||||||||||]PAM</td><td>NA</td><td>Intergenic</td><td>NA</td><td>NA</td></tr>
<tr><td>T16219</td><td>ATGCGAGTTGCTTTAACTTTTGG</td><td>1000</td><td></td><td></td><td></td><td></td><td></td><td></td><td></td><td></td><td></td></tr>
<tr><td>Oligo fwd</td><td>CACCATGCGAGTTGCTTTAACTTT</td><td></td><td></td><td></td><td></td><td></td><td></td><td></td><td></td><td></td><td></td></tr>
<tr><td>Oligo rev</td><td>AAACAAAGTTAAAGCAACTCGCAT</td><td></td><td></td><td></td><td></td><td></td><td></td><td></td><td></td><td></td><td></td></tr>
<tr><td>Chromosome</td><td>start</td><td>end</td><td>strand</td><td>MM</td><td>target_seq</td><td>PAM</td><td>alignment</td><td>distance</td><td>position</td><td>gene name</td><td>gene id</td></tr>
<tr><td>GU305914.1</td><td>216397</td><td>216419</td><td>+</td><td>0</td><td>ATGCGAGTTGCTTTAACTTT</td><td>TGG</td><td>||||||||[||||||||||||]PAM</td><td>NA</td><td>Intergenic</td><td>NA</td><td>NA</td></tr>
<tr><td>T16220</td><td>AACGTTGCTTTTTATCATATAGG</td><td>1000</td><td></td><td></td><td></td><td></td><td></td><td></td><td></td><td></td><td></td></tr>
<tr><td>Oligo fwd</td><td>CACCAACGTTGCTTTTTATCATAT</td><td></td><td></td><td></td><td></td><td></td><td></td><td></td><td></td><td></td><td></td></tr>
<tr><td>Oligo rev</td><td>AAACATATGATAAAAAGCAACGTT</td><td></td><td></td><td></td><td></td><td></td><td></td><td></td><td></td><td></td><td></td></tr>
<tr><td>Chromosome</td><td>start</td><td>end</td><td>strand</td><td>MM</td><td>target_seq</td><td>PAM</td><td>alignment</td><td>distance</td><td>position</td><td>gene name</td><td>gene id</td></tr>
<tr><td>GU305914.1</td><td>216452</td><td>216474</td><td>+</td><td>0</td><td>AACGTTGCTTTTTATCATAT</td><td>AGG</td><td>||||||||[||||||||||||]PAM</td><td>NA</td><td>Intergenic</td><td>NA</td><td>NA</td></tr>
<tr><td>T16221</td><td>GGTACCAATTTCTTTTGCGTCGG</td><td>1000</td><td></td><td></td><td></td><td></td><td></td><td></td><td></td><td></td><td></td></tr>
<tr><td>Oligo fwd</td><td>CACCGGTACCAATTTCTTTTGCGT</td><td></td><td></td><td></td><td></td><td></td><td></td><td></td><td></td><td></td><td></td></tr>
<tr><td>Oligo rev</td><td>AAACACGCAAAAGAAATTGGTACC</td><td></td><td></td><td></td><td></td><td></td><td></td><td></td><td></td><td></td><td></td></tr>
<tr><td>Chromosome</td><td>start</td><td>end</td><td>strand</td><td>MM</td><td>target_seq</td><td>PAM</td><td>alignment</td><td>distance</td><td>position</td><td>gene name</td><td>gene id</td></tr>
<tr><td>GU305914.1</td><td>216473</td><td>216495</td><td>+</td><td>0</td><td>GGTACCAATTTCTTTTGCGT</td><td>CGG</td><td>||||||||[||||||||||||]PAM</td><td>NA</td><td>Intergenic</td><td>NA</td><td>NA</td></tr>
<tr><td>T16222</td><td>ATACGCTACCCTGACCGTATCGG</td><td>1000</td><td></td><td></td><td></td><td></td><td></td><td></td><td></td><td></td><td></td></tr>
<tr><td>Oligo fwd</td><td>CACCATACGCTACCCTGACCGTAT</td><td></td><td></td><td></td><td></td><td></td><td></td><td></td><td></td><td></td><td></td></tr>
<tr><td>Oligo rev</td><td>AAACATACGGTCAGGGTAGCGTAT</td><td></td><td></td><td></td><td></td><td></td><td></td><td></td><td></td><td></td><td></td></tr>
<tr><td>Chromosome</td><td>start</td><td>end</td><td>strand</td><td>MM</td><td>target_seq</td><td>PAM</td><td>alignment</td><td>distance</td><td>position</td><td>gene name</td><td>gene id</td></tr>
<tr><td>GU305914.1</td><td>216548</td><td>216570</td><td>+</td><td>0</td><td>ATACGCTACCCTGACCGTAT</td><td>CGG</td><td>||||||||[||||||||||||]PAM</td><td>NA</td><td>Intergenic</td><td>NA</td><td>NA</td></tr>
<tr><td>T16223</td><td>CCCAGAGACTTTGTGTCAATAGG</td><td>1000</td><td></td><td></td><td></td><td></td><td></td><td></td><td></td><td></td><td></td></tr>
<tr><td>Oligo fwd</td><td>CACCCCCAGAGACTTTGTGTCAAT</td><td></td><td></td><td></td><td></td><td></td><td></td><td></td><td></td><td></td><td></td></tr>
<tr><td>Oligo rev</td><td>AAACATTGACACAAAGTCTCTGGG</td><td></td><td></td><td></td><td></td><td></td><td></td><td></td><td></td><td></td><td></td></tr>
<tr><td>Chromosome</td><td>start</td><td>end</td><td>strand</td><td>MM</td><td>target_seq</td><td>PAM</td><td>alignment</td><td>distance</td><td>position</td><td>gene name</td><td>gene id</td></tr>
<tr><td>GU305914.1</td><td>216646</td><td>216668</td><td>+</td><td>0</td><td>CCCAGAGACTTTGTGTCAAT</td><td>AGG</td><td>||||||||[||||||||||||]PAM</td><td>NA</td><td>Intergenic</td><td>NA</td><td>NA</td></tr>
<tr><td>T16224</td><td>ATATCAGTCGAACTGACATCCGG</td><td>1000</td><td></td><td></td><td></td><td></td><td></td><td></td><td></td><td></td><td></td></tr>
<tr><td>Oligo fwd</td><td>CACCATATCAGTCGAACTGACATC</td><td></td><td></td><td></td><td></td><td></td><td></td><td></td><td></td><td></td><td></td></tr>
<tr><td>Oligo rev</td><td>AAACGATGTCAGTTCGACTGATAT</td><td></td><td></td><td></td><td></td><td></td><td></td><td></td><td></td><td></td><td></td></tr>
<tr><td>Chromosome</td><td>start</td><td>end</td><td>strand</td><td>MM</td><td>target_seq</td><td>PAM</td><td>alignment</td><td>distance</td><td>position</td><td>gene name</td><td>gene id</td></tr>
<tr><td>GU305914.1</td><td>216711</td><td>216733</td><td>+</td><td>0</td><td>ATATCAGTCGAACTGACATC</td><td>CGG</td><td>||||||||[||||||||||||]PAM</td><td>NA</td><td>Intergenic</td><td>NA</td><td>NA</td></tr>
<tr><td>T16225</td><td>CGGACAGATTTTACATATCTTGG</td><td>1000</td><td></td><td></td><td></td><td></td><td></td><td></td><td></td><td></td><td></td></tr>
<tr><td>Oligo fwd</td><td>CACCCGGACAGATTTTACATATCT</td><td></td><td></td><td></td><td></td><td></td><td></td><td></td><td></td><td></td><td></td></tr>
<tr><td>Oligo rev</td><td>AAACAGATATGTAAAATCTGTCCG</td><td></td><td></td><td></td><td></td><td></td><td></td><td></td><td></td><td></td><td></td></tr>
<tr><td>Chromosome</td><td>start</td><td>end</td><td>strand</td><td>MM</td><td>target_seq</td><td>PAM</td><td>alignment</td><td>distance</td><td>position</td><td>gene name</td><td>gene id</td></tr>
<tr><td>GU305914.1</td><td>216731</td><td>216753</td><td>+</td><td>0</td><td>CGGACAGATTTTACATATCT</td><td>TGG</td><td>||||||||[||||||||||||]PAM</td><td>NA</td><td>Intergenic</td><td>NA</td><td>NA</td></tr>
<tr><td>T16226</td><td>GGACAGATTTTACATATCTTGGG</td><td>1000</td><td></td><td></td><td></td><td></td><td></td><td></td><td></td><td></td><td></td></tr>
<tr><td>Oligo fwd</td><td>CACCGGACAGATTTTACATATCTT</td><td></td><td></td><td></td><td></td><td></td><td></td><td></td><td></td><td></td><td></td></tr>
<tr><td>Oligo rev</td><td>AAACAAGATATGTAAAATCTGTCC</td><td></td><td></td><td></td><td></td><td></td><td></td><td></td><td></td><td></td><td></td></tr>
<tr><td>Chromosome</td><td>start</td><td>end</td><td>strand</td><td>MM</td><td>target_seq</td><td>PAM</td><td>alignment</td><td>distance</td><td>position</td><td>gene name</td><td>gene id</td></tr>
<tr><td>GU305914.1</td><td>216732</td><td>216754</td><td>+</td><td>0</td><td>GGACAGATTTTACATATCTT</td><td>GGG</td><td>||||||||[||||||||||||]PAM</td><td>NA</td><td>Intergenic</td><td>NA</td><td>NA</td></tr>
<tr><td>T16227</td><td>GACAGATTTTACATATCTTGGGG</td><td>1000</td><td></td><td></td><td></td><td></td><td></td><td></td><td></td><td></td><td></td></tr>
<tr><td>Oligo fwd</td><td>CACCGACAGATTTTACATATCTTG</td><td></td><td></td><td></td><td></td><td></td><td></td><td></td><td></td><td></td><td></td></tr>
<tr><td>Oligo rev</td><td>AAACCAAGATATGTAAAATCTGTC</td><td></td><td></td><td></td><td></td><td></td><td></td><td></td><td></td><td></td><td></td></tr>
</table>

<table>
<tr><td>Chromosome</td><td>start</td><td>end</td><td>strand</td><td>MM</td><td>target_seq</td><td>PAM</td><td>alignment</td><td>distance</td><td>position</td><td>gene name</td><td>gene id</td></tr>
<tr><td>GU305914.1</td><td>216733</td><td>216755</td><td>+</td><td>0</td><td>GACAGATTTTACATATCTTG</td><td>GGG</td><td>||||||||[||||||||||||]PAM</td><td>NA</td><td>Intergenic</td><td>NA</td><td>NA</td></tr>
<tr><td>T16228</td><td>CCAACGTCCCACACGACGACTGG</td><td>1000</td><td></td><td></td><td></td><td></td><td></td><td></td><td></td><td></td><td></td></tr>
<tr><td>Oligo fwd</td><td>CACCCCAACGTCCCACACGACGAC</td><td></td><td></td><td></td><td></td><td></td><td></td><td></td><td></td><td></td><td></td></tr>
<tr><td>Oligo rev</td><td>AAACGTCGTCGTGTGGGACGTTGG</td><td></td><td></td><td></td><td></td><td></td><td></td><td></td><td></td><td></td><td></td></tr>
<tr><td>Chromosome</td><td>start</td><td>end</td><td>strand</td><td>MM</td><td>target_seq</td><td>PAM</td><td>alignment</td><td>distance</td><td>position</td><td>gene name</td><td>gene id</td></tr>
<tr><td>GU305914.1</td><td>216765</td><td>216787</td><td>+</td><td>0</td><td>CCAACGTCCCACACGACGAC</td><td>TGG</td><td>||||||||[||||||||||||]PAM</td><td>NA</td><td>Intergenic</td><td>NA</td><td>NA</td></tr>
<tr><td>T16229</td><td>CTACAGAAATCGTACGCTTATGG</td><td>1000</td><td></td><td></td><td></td><td></td><td></td><td></td><td></td><td></td><td></td></tr>
<tr><td>Oligo fwd</td><td>CACCCTACAGAAATCGTACGCTTA</td><td></td><td></td><td></td><td></td><td></td><td></td><td></td><td></td><td></td><td></td></tr>
<tr><td>Oligo rev</td><td>AAACTAAGCGTACGATTTCTGTAG</td><td></td><td></td><td></td><td></td><td></td><td></td><td></td><td></td><td></td><td></td></tr>
<tr><td>Chromosome</td><td>start</td><td>end</td><td>strand</td><td>MM</td><td>target_seq</td><td>PAM</td><td>alignment</td><td>distance</td><td>position</td><td>gene name</td><td>gene id</td></tr>
<tr><td>GU305914.1</td><td>216799</td><td>216821</td><td>+</td><td>0</td><td>CTACAGAAATCGTACGCTTA</td><td>TGG</td><td>||||||||[||||||||||||]PAM</td><td>NA</td><td>Intergenic</td><td>NA</td><td>NA</td></tr>
<tr><td>T16230</td><td>ATGGACGCCACCTCTTGTAACGG</td><td>1000</td><td></td><td></td><td></td><td></td><td></td><td></td><td></td><td></td><td></td></tr>
<tr><td>Oligo fwd</td><td>CACCATGGACGCCACCTCTTGTAA</td><td></td><td></td><td></td><td></td><td></td><td></td><td></td><td></td><td></td><td></td></tr>
<tr><td>Oligo rev</td><td>AAACTTACAAGAGGTGGCGTCCAT</td><td></td><td></td><td></td><td></td><td></td><td></td><td></td><td></td><td></td><td></td></tr>
<tr><td>Chromosome</td><td>start</td><td>end</td><td>strand</td><td>MM</td><td>target_seq</td><td>PAM</td><td>alignment</td><td>distance</td><td>position</td><td>gene name</td><td>gene id</td></tr>
<tr><td>GU305914.1</td><td>216818</td><td>216840</td><td>+</td><td>0</td><td>ATGGACGCCACCTCTTGTAA</td><td>CGG</td><td>||||||||[||||||||||||]PAM</td><td>NA</td><td>Intergenic</td><td>NA</td><td>NA</td></tr>
<tr><td>T16231</td><td>TTCACAATATATGTGTCATCAGG</td><td>1000</td><td></td><td></td><td></td><td></td><td></td><td></td><td></td><td></td><td></td></tr>
<tr><td>Oligo fwd</td><td>CACCTTCACAATATATGTGTCATC</td><td></td><td></td><td></td><td></td><td></td><td></td><td></td><td></td><td></td><td></td></tr>
<tr><td>Oligo rev</td><td>AAACGATGACACATATATTGTGAA</td><td></td><td></td><td></td><td></td><td></td><td></td><td></td><td></td><td></td><td></td></tr>
<tr><td>Chromosome</td><td>start</td><td>end</td><td>strand</td><td>MM</td><td>target_seq</td><td>PAM</td><td>alignment</td><td>distance</td><td>position</td><td>gene name</td><td>gene id</td></tr>
<tr><td>GU305914.1</td><td>216885</td><td>216907</td><td>+</td><td>0</td><td>TTCACAATATATGTGTCATC</td><td>AGG</td><td>||||||||[||||||||||||]PAM</td><td>NA</td><td>Intergenic</td><td>NA</td><td>NA</td></tr>
<tr><td>T16232</td><td>ATATATGTGTCATCAGGATCAGG</td><td>1000</td><td></td><td></td><td></td><td></td><td></td><td></td><td></td><td></td><td></td></tr>
<tr><td>Oligo fwd</td><td>CACCATATATGTGTCATCAGGATC</td><td></td><td></td><td></td><td></td><td></td><td></td><td></td><td></td><td></td><td></td></tr>
<tr><td>Oligo rev</td><td>AAACGATCCTGATGACACATATAT</td><td></td><td></td><td></td><td></td><td></td><td></td><td></td><td></td><td></td><td></td></tr>
<tr><td>Chromosome</td><td>start</td><td>end</td><td>strand</td><td>MM</td><td>target_seq</td><td>PAM</td><td>alignment</td><td>distance</td><td>position</td><td>gene name</td><td>gene id</td></tr>
<tr><td>GU305914.1</td><td>216891</td><td>216913</td><td>+</td><td>0</td><td>ATATATGTGTCATCAGGATC</td><td>AGG</td><td>||||||||[||||||||||||]PAM</td><td>NA</td><td>Intergenic</td><td>NA</td><td>NA</td></tr>
<tr><td>T16233</td><td>CAATCAAGAGTAATTTTACTAGG</td><td>1000</td><td></td><td></td><td></td><td></td><td></td><td></td><td></td><td></td><td></td></tr>
<tr><td>Oligo fwd</td><td>CACCCAATCAAGAGTAATTTTACT</td><td></td><td></td><td></td><td></td><td></td><td></td><td></td><td></td><td></td><td></td></tr>
<tr><td>Oligo rev</td><td>AAACAGTAAAATTACTCTTGATTG</td><td></td><td></td><td></td><td></td><td></td><td></td><td></td><td></td><td></td><td></td></tr>
<tr><td>Chromosome</td><td>start</td><td>end</td><td>strand</td><td>MM</td><td>target_seq</td><td>PAM</td><td>alignment</td><td>distance</td><td>position</td><td>gene name</td><td>gene id</td></tr>
<tr><td>GU305914.1</td><td>216949</td><td>216971</td><td>+</td><td>0</td><td>CAATCAAGAGTAATTTTACT</td><td>AGG</td><td>||||||||[||||||||||||]PAM</td><td>NA</td><td>Intergenic</td><td>NA</td><td>NA</td></tr>
<tr><td>T16234</td><td>AATCAAGAGTAATTTTACTAGGG</td><td>1000</td><td></td><td></td><td></td><td></td><td></td><td></td><td></td><td></td><td></td></tr>
<tr><td>Oligo fwd</td><td>CACCAATCAAGAGTAATTTTACTA</td><td></td><td></td><td></td><td></td><td></td><td></td><td></td><td></td><td></td><td></td></tr>
<tr><td>Oligo rev</td><td>AAACTAGTAAAATTACTCTTGATT</td><td></td><td></td><td></td><td></td><td></td><td></td><td></td><td></td><td></td><td></td></tr>
<tr><td>Chromosome</td><td>start</td><td>end</td><td>strand</td><td>MM</td><td>target_seq</td><td>PAM</td><td>alignment</td><td>distance</td><td>position</td><td>gene name</td><td>gene id</td></tr>
<tr><td>GU305914.1</td><td>216950</td><td>216972</td><td>+</td><td>0</td><td>AATCAAGAGTAATTTTACTA</td><td>GGG</td><td>||||||||[||||||||||||]PAM</td><td>NA</td><td>Intergenic</td><td>NA</td><td>NA</td></tr>
<tr><td>T16235</td><td>ATCAAGAGTAATTTTACTAGGGG</td><td>1000</td><td></td><td></td><td></td><td></td><td></td><td></td><td></td><td></td><td></td></tr>
<tr><td>Oligo fwd</td><td>CACCATCAAGAGTAATTTTACTAG</td><td></td><td></td><td></td><td></td><td></td><td></td><td></td><td></td><td></td><td></td></tr>
<tr><td>Oligo rev</td><td>AAACCTAGTAAAATTACTCTTGAT</td><td></td><td></td><td></td><td></td><td></td><td></td><td></td><td></td><td></td><td></td></tr>
<tr><td>Chromosome</td><td>start</td><td>end</td><td>strand</td><td>MM</td><td>target_seq</td><td>PAM</td><td>alignment</td><td>distance</td><td>position</td><td>gene name</td><td>gene id</td></tr>
<tr><td>GU305914.1</td><td>216951</td><td>216973</td><td>+</td><td>0</td><td>ATCAAGAGTAATTTTACTAG</td><td>GGG</td><td>||||||||[||||||||||||]PAM</td><td>NA</td><td>Intergenic</td><td>NA</td><td>NA</td></tr>
<tr><td>T16236</td><td>TCAAGAGTAATTTTACTAGGGGG</td><td>1000</td><td></td><td></td><td></td><td></td><td></td><td></td><td></td><td></td><td></td></tr>
<tr><td>Oligo fwd</td><td>CACCTCAAGAGTAATTTTACTAGG</td><td></td><td></td><td></td><td></td><td></td><td></td><td></td><td></td><td></td><td></td></tr>
<tr><td>Oligo rev</td><td>AAACCCTAGTAAAATTACTCTTGA</td><td></td><td></td><td></td><td></td><td></td><td></td><td></td><td></td><td></td><td></td></tr>
<tr><td>Chromosome</td><td>start</td><td>end</td><td>strand</td><td>MM</td><td>target_seq</td><td>PAM</td><td>alignment</td><td>distance</td><td>position</td><td>gene name</td><td>gene id</td></tr>
<tr><td>GU305914.1</td><td>216952</td><td>216974</td><td>+</td><td>0</td><td>TCAAGAGTAATTTTACTAGG</td><td>GGG</td><td>||||||||[||||||||||||]PAM</td><td>NA</td><td>Intergenic</td><td>NA</td><td>NA</td></tr>
<tr><td>T16237</td><td>GGGACGATGACCATGTCAAGAGG</td><td>1000</td><td></td><td></td><td></td><td></td><td></td><td></td><td></td><td></td><td></td></tr>
<tr><td>Oligo fwd</td><td>CACCGGGACGATGACCATGTCAAG</td><td></td><td></td><td></td><td></td><td></td><td></td><td></td><td></td><td></td><td></td></tr>
<tr><td>Oligo rev</td><td>AAACCTTGACATGGTCATCGTCCC</td><td></td><td></td><td></td><td></td><td></td><td></td><td></td><td></td><td></td><td></td></tr>
<tr><td>Chromosome</td><td>start</td><td>end</td><td>strand</td><td>MM</td><td>target_seq</td><td>PAM</td><td>alignment</td><td>distance</td><td>position</td><td>gene name</td><td>gene id</td></tr>
<tr><td>GU305914.1</td><td>216972</td><td>216994</td><td>+</td><td>0</td><td>GGGACGATGACCATGTCAAG</td><td>AGG</td><td>||||||||[||||||||||||]PAM</td><td>NA</td><td>Intergenic</td><td>NA</td><td>NA</td></tr>
<tr><td>T16238</td><td>TGTCAAGAGGTACTGAATATCGG</td><td>1000</td><td></td><td></td><td></td><td></td><td></td><td></td><td></td><td></td><td></td></tr>
<tr><td>Oligo fwd</td><td>CACCTGTCAAGAGGTACTGAATAT</td><td></td><td></td><td></td><td></td><td></td><td></td><td></td><td></td><td></td><td></td></tr>
<tr><td>Oligo rev</td><td>AAACATATTCAGTACCTCTTGACA</td><td></td><td></td><td></td><td></td><td></td><td></td><td></td><td></td><td></td><td></td></tr>
<tr><td>Chromosome</td><td>start</td><td>end</td><td>strand</td><td>MM</td><td>target_seq</td><td>PAM</td><td>alignment</td><td>distance</td><td>position</td><td>gene name</td><td>gene id</td></tr>
<tr><td>GU305914.1</td><td>216985</td><td>217007</td><td>+</td><td>0</td><td>TGTCAAGAGGTACTGAATAT</td><td>CGG</td><td>||||||||[||||||||||||]PAM</td><td>NA</td><td>Intergenic</td><td>NA</td><td>NA</td></tr>
<tr><td>T16239</td><td>GTCAAGAGGTACTGAATATCGGG</td><td>1000</td><td></td><td></td><td></td><td></td><td></td><td></td><td></td><td></td><td></td></tr>
<tr><td>Oligo fwd</td><td>CACCGTCAAGAGGTACTGAATATC</td><td></td><td></td><td></td><td></td><td></td><td></td><td></td><td></td><td></td><td></td></tr>
<tr><td>Oligo rev</td><td>AAACGATATTCAGTACCTCTTGAC</td><td></td><td></td><td></td><td></td><td></td><td></td><td></td><td></td><td></td><td></td></tr>
<tr><td>Chromosome</td><td>start</td><td>end</td><td>strand</td><td>MM</td><td>target_seq</td><td>PAM</td><td>alignment</td><td>distance</td><td>position</td><td>gene name</td><td>gene id</td></tr>
<tr><td>GU305914.1</td><td>216986</td><td>217008</td><td>+</td><td>0</td><td>GTCAAGAGGTACTGAATATC</td><td>GGG</td><td>||||||||[||||||||||||]PAM</td><td>NA</td><td>Intergenic</td><td>NA</td><td>NA</td></tr>
<tr><td>T16240</td><td>TCAAGAGGTACTGAATATCGGGG</td><td>1000</td><td></td><td></td><td></td><td></td><td></td><td></td><td></td><td></td><td></td></tr>
<tr><td>Oligo fwd</td><td>CACCTCAAGAGGTACTGAATATCG</td><td></td><td></td><td></td><td></td><td></td><td></td><td></td><td></td><td></td><td></td></tr>
<tr><td>Oligo rev</td><td>AAACCGATATTCAGTACCTCTTGA</td><td></td><td></td><td></td><td></td><td></td><td></td><td></td><td></td><td></td><td></td></tr>
<tr><td>Chromosome</td><td>start</td><td>end</td><td>strand</td><td>MM</td><td>target_seq</td><td>PAM</td><td>alignment</td><td>distance</td><td>position</td><td>gene name</td><td>gene id</td></tr>
<tr><td>GU305914.1</td><td>216987</td><td>217009</td><td>+</td><td>0</td><td>TCAAGAGGTACTGAATATCG</td><td>GGG</td><td>||||||||[||||||||||||]PAM</td><td>NA</td><td>Intergenic</td><td>NA</td><td>NA</td></tr>
<tr><td>T16241</td><td>GGGTACACTTTCTCAAATATCGG</td><td>1000</td><td></td><td></td><td></td><td></td><td></td><td></td><td></td><td></td><td></td></tr>
<tr><td>Oligo fwd</td><td>CACCGGGTACACTTTCTCAAATAT</td><td></td><td></td><td></td><td></td><td></td><td></td><td></td><td></td><td></td><td></td></tr>
<tr><td>Oligo rev</td><td>AAACATATTTGAGAAAGTGTACCC</td><td></td><td></td><td></td><td></td><td></td><td></td><td></td><td></td><td></td><td></td></tr>
<tr><td>Chromosome</td><td>start</td><td>end</td><td>strand</td><td>MM</td><td>target_seq</td><td>PAM</td><td>alignment</td><td>distance</td><td>position</td><td>gene name</td><td>gene id</td></tr>
<tr><td>GU305914.1</td><td>217007</td><td>217029</td><td>+</td><td>0</td><td>GGGTACACTTTCTCAAATAT</td><td>CGG</td><td>||||||||[||||||||||||]PAM</td><td>NA</td><td>Intergenic</td><td>NA</td><td>NA</td></tr>
<tr><td>T16242</td><td>GGTACACTTTCTCAAATATCGGG</td><td>1000</td><td></td><td></td><td></td><td></td><td></td><td></td><td></td><td></td><td></td></tr>
<tr><td>Oligo fwd</td><td>CACCGGTACACTTTCTCAAATATC</td><td></td><td></td><td></td><td></td><td></td><td></td><td></td><td></td><td></td><td></td></tr>
</table>

Oligo rev AAACGATATTTGAGAAAGTGTACC
Chromosome start end strand MM target_seq PAM alignment distance position gene name gene id
GU305914.1 217008 217030 + 0 GGTACACTTTCTCAAATATC GGG ||||||||[||||||||||||]PAM NA Intergenic NA NA

T16243 GTACACTTTCTCAAATATCGGGG 1000
Oligo fwd CACCGTACACTTTCTCAAATATCG
Oligo rev AAACCGATATTTGAGAAAGTGTAC
Chromosome start end strand MM target_seq PAM alignment distance position gene name gene id
GU305914.1 217009 217031 + 0 GTACACTTTCTCAAATATCG GGG ||||||||[||||||||||||]PAM NA Intergenic NA NA

T16244 AAATAAGCACCCACAGACACCGG 1000
Oligo fwd CACCAAATAAGCACCCACAGACAC
Oligo rev AAACGTGTCTGTGGGTGCTTATTT
Chromosome start end strand MM target_seq PAM alignment distance position gene name gene id
GU305914.1 217070 217092 + 0 AAATAAGCACCCACAGACAC CGG ||||||||[||||||||||||]PAM NA Intergenic NA NA

T16245 ACCACTAACACGTCTAACCACGG 1000
Oligo fwd CACCACCACTAACACGTCTAACCA
Oligo rev AAACTGGTTAGACGTGTTAGTGGT
Chromosome start end strand MM target_seq PAM alignment distance position gene name gene id
GU305914.1 217109 217131 + 0 ACCACTAACACGTCTAACCA CGG ||||||||[||||||||||||]PAM NA Intergenic NA NA

T16246 CGTCTAACCACGGAGTCCGCAGG 1000
Oligo fwd CACCCGTCTAACCACGGAGTCCGC
Oligo rev AAACGCGGACTCCGTGGTTAGACG
Chromosome start end strand MM target_seq PAM alignment distance position gene name gene id
GU305914.1 217119 217141 + 0 CGTCTAACCACGGAGTCCGC AGG ||||||||[||||||||||||]PAM NA Intergenic NA NA

T16247 GTCTAACCACGGAGTCCGCAGGG 1000
Oligo fwd CACCGTCTAACCACGGAGTCCGCA
Oligo rev AAACTGCGGACTCCGTGGTTAGAC
Chromosome start end strand MM target_seq PAM alignment distance position gene name gene id
GU305914.1 217120 217142 + 0 GTCTAACCACGGAGTCCGCA GGG ||||||||[||||||||||||]PAM NA Intergenic NA NA

T16248 ATCATTATCCTCTACAGAGACGG 1000
Oligo fwd CACCATCATTATCCTCTACAGAGA
Oligo rev AAACTCTCTGTAGAGGATAATGAT
Chromosome start end strand MM target_seq PAM alignment distance position gene name gene id
GU305914.1 217147 217169 + 0 ATCATTATCCTCTACAGAGA CGG ||||||||[||||||||||||]PAM NA Intergenic NA NA

T16249 CAGAGACGGACGCCGCTGTACGG 1000
Oligo fwd CACCCAGAGACGGACGCCGCTGTA
Oligo rev AAACTACAGCGGCGTCCGTCTCTG
Chromosome start end strand MM target_seq PAM alignment distance position gene name gene id
GU305914.1 217161 217183 + 0 CAGAGACGGACGCCGCTGTA CGG ||||||||[||||||||||||]PAM NA Intergenic NA NA

T16250 GCCGCTGTACGGTTTCTCACCGG 1000
Oligo fwd CACCGCCGCTGTACGGTTTCTCAC
Oligo rev AAACGTGAGAAACCGTACAGCGGC
Chromosome start end strand MM target_seq PAM alignment distance position gene name gene id
GU305914.1 217172 217194 + 0 GCCGCTGTACGGTTTCTCAC CGG ||||||||[||||||||||||]PAM NA Intergenic NA NA

T16251 ACGGTTTCTCACCGGTTTGACGG 1000
Oligo fwd CACCACGGTTTCTCACCGGTTTGA
Oligo rev AAACTCAAACCGGTGAGAAACCGT
Chromosome start end strand MM target_seq PAM alignment distance position gene name gene id
GU305914.1 217180 217202 + 0 ACGGTTTCTCACCGGTTTGA CGG ||||||||[||||||||||||]PAM NA Intergenic NA NA

T16252 ACCGGTTTGACGGTCGTTCGCGG 1000
Oligo fwd CACCACCGGTTTGACGGTCGTTCG
Oligo rev AAACCGAACGACCGTCAAACCGGT
Chromosome start end strand MM target_seq PAM alignment distance position gene name gene id
GU305914.1 217190 217212 + 0 ACCGGTTTGACGGTCGTTCG CGG ||||||||[||||||||||||]PAM NA Intergenic NA NA

T16253 GGTTTGACGGTCGTTCGCGGCGG 1000
Oligo fwd CACCGGTTTGACGGTCGTTCGCGG
Oligo rev AAACCCGCGAACGACCGTCAAACC
Chromosome start end strand MM target_seq PAM alignment distance position gene name gene id
GU305914.1 217193 217215 + 0 GGTTTGACGGTCGTTCGCGG CGG ||||||||[||||||||||||]PAM NA Intergenic NA NA

T16254 CGGTCGTTCGCGGCGGCGCGAGG 1000
Oligo fwd CACCCGGTCGTTCGCGGCGGCGCG
Oligo rev AAACCGCGCCGCCGCGAACGACCG
Chromosome start end strand MM target_seq PAM alignment distance position gene name gene id
GU305914.1 217200 217222 + 0 CGGTCGTTCGCGGCGGCGCG AGG ||||||||[||||||||||||]PAM NA Intergenic NA NA

T16255 ATGCTCCAAGACTTCGTCCTCGG 1000
Oligo fwd CACCATGCTCCAAGACTTCGTCCT
Oligo rev AAACAGGACGAAGTCTTGGAGCAT
Chromosome start end strand MM target_seq PAM alignment distance position gene name gene id
GU305914.1 217240 217262 + 0 ATGCTCCAAGACTTCGTCCT CGG ||||||||[||||||||||||]PAM NA Intergenic NA NA
GU305914.1 216181 216203 - 4 ATGTTACGTGACTTCGTCCT TGG |||-|-|-[-|||||||||||]PAM NA Intergenic NA NA

T16256 TCGTCCTCGGACGCGACACCCGG 1000
Oligo fwd CACCTCGTCCTCGGACGCGACACC
Oligo rev AAACGGTGTCGCGTCCGAGGACGA
Chromosome start end strand MM target_seq PAM alignment distance position gene name gene id
GU305914.1 217253 217275 + 0 TCGTCCTCGGACGCGACACC CGG ||||||||[||||||||||||]PAM NA Intergenic NA NA

T16257 CGTCCTCGGACGCGACACCCGGG 1000
Oligo fwd CACCCGTCCTCGGACGCGACACCC
Oligo rev AAACGGGTGTCGCGTCCGAGGACG
Chromosome start end strand MM target_seq PAM alignment distance position gene name gene id
GU305914.1 217254 217276 + 0 CGTCCTCGGACGCGACACCC GGG ||||||||[||||||||||||]PAM NA Intergenic NA NA

T16258 ACGACGTCCTCGTAAGTCAGCGG 1000
Oligo fwd CACCACGACGTCCTCGTAAGTCAG
Oligo rev AAACCTGACTTACGAGGACGTCGT
Chromosome start end strand MM target_seq PAM alignment distance position gene name gene id
GU305914.1 217277 217299 + 0 ACGACGTCCTCGTAAGTCAG CGG ||||||||[||||||||||||]PAM NA Intergenic NA NA

T16259 CCTCGTAAGTCAGCGGTCTACGG 1000
Oligo fwd CACCCCTCGTAAGTCAGCGGTCTA
Oligo rev AAACTAGACCGCTGACTTACGAGG
Chromosome start end strand MM target_seq PAM alignment distance position gene name gene id
GU305914.1 217284 217306 + 0 CCTCGTAAGTCAGCGGTCTA CGG ||||||||[||||||||||||]PAM NA Intergenic NA NA

T16260 GTCTACGGATACAACCGTGTTGG 1000
Oligo fwd CACCGTCTACGGATACAACCGTGT
Oligo rev AAACACACGGTTGTATCCGTAGAC
Chromosome start end strand MM target_seq PAM alignment distance position gene name gene id
GU305914.1 217299 217321 + 0 GTCTACGGATACAACCGTGT TGG ||||||||[||||||||||||]PAM NA Intergenic NA NA

T16261 CTTCACTGACGCCAACCAGCCGG 1000
Oligo fwd CACCCTTCACTGACGCCAACCAGC
Oligo rev AAACGCTGGTTGGCGTCAGTGAAG
Chromosome start end strand MM target_seq PAM alignment distance position gene name gene id
GU305914.1 217336 217358 + 0 CTTCACTGACGCCAACCAGC CGG ||||||||[||||||||||||]PAM NA Intergenic NA NA

T16262 AGCAGAGTAGATCTTACAGACGG 1000
Oligo fwd CACCAGCAGAGTAGATCTTACAGA
Oligo rev AAACTCTGTAAGATCTACTCTGCT
Chromosome start end strand MM target_seq PAM alignment distance position gene name gene id
GU305914.1 217360 217382 + 0 AGCAGAGTAGATCTTACAGA CGG ||||||||[||||||||||||]PAM NA Intergenic NA NA

T16263 GCAGAGTAGATCTTACAGACGGG 1000
Oligo fwd CACCGCAGAGTAGATCTTACAGAC
Oligo rev AAACGTCTGTAAGATCTACTCTGC
Chromosome start end strand MM target_seq PAM alignment distance position gene name gene id
GU305914.1 217361 217383 + 0 GCAGAGTAGATCTTACAGAC GGG ||||||||[||||||||||||]PAM NA Intergenic NA NA

T16264 GTTTCCAGCCGCCGTCACGCTGG 1000
Oligo fwd CACCGTTTCCAGCCGCCGTCACGC
Oligo rev AAACGCGTGACGGCGGCTGGAAAC
Chromosome start end strand MM target_seq PAM alignment distance position gene name gene id
GU305914.1 217396 217418 + 0 GTTTCCAGCCGCCGTCACGC TGG ||||||||[||||||||||||]PAM NA Intergenic NA NA

T16265 GGCCTTCGCGTAATGTAACACGG 1000
Oligo fwd CACCGGCCTTCGCGTAATGTAACA
Oligo rev AAACTGTTACATTACGCGAAGGCC
Chromosome start end strand MM target_seq PAM alignment distance position gene name gene id
GU305914.1 217417 217439 + 0 GGCCTTCGCGTAATGTAACA CGG ||||||||[||||||||||||]PAM NA Intergenic NA NA

T16266 GCCTTCGCGTAATGTAACACGGG 1000
Oligo fwd CACCGCCTTCGCGTAATGTAACAC
Oligo rev AAACGTGTTACATTACGCGAAGGC
Chromosome start end strand MM target_seq PAM alignment distance position gene name gene id
GU305914.1 217418 217440 + 0 GCCTTCGCGTAATGTAACAC GGG ||||||||[||||||||||||]PAM NA Intergenic NA NA

T16267 TATCAACGAGAAAGTTTGTGCGG 1000
Oligo fwd CACCTATCAACGAGAAAGTTTGTG
Oligo rev AAACCACAAACTTTCTCGTTGATA
Chromosome start end strand MM target_seq PAM alignment distance position gene name gene id
GU305914.1 217446 217468 + 0 TATCAACGAGAAAGTTTGTG CGG ||||||||[||||||||||||]PAM NA Intergenic NA NA

T16268 CGCAGTGCAGCGAAAGCCGCCGG 1000
Oligo fwd CACCCGCAGTGCAGCGAAAGCCGC
Oligo rev AAACGCGGCTTTCGCTGCACTGCG
Chromosome start end strand MM target_seq PAM alignment distance position gene name gene id
GU305914.1 217513 217535 + 0 CGCAGTGCAGCGAAAGCCGC CGG ||||||||[||||||||||||]PAM NA Intergenic NA NA

T16269 CACATTACAGAATTCTACCTCGG 1000
Oligo fwd CACCCACATTACAGAATTCTACCT
Oligo rev AAACAGGTAGAATTCTGTAATGTG
Chromosome start end strand MM target_seq PAM alignment distance position gene name gene id
GU305914.1 217537 217559 + 0 CACATTACAGAATTCTACCT CGG ||||||||[||||||||||||]PAM NA Intergenic NA NA

T16270 CAGAATTCTACCTCGGTATGCGG 1000
Oligo fwd CACCCAGAATTCTACCTCGGTATG
Oligo rev AAACCATACCGAGGTAGAATTCTG
Chromosome start end strand MM target_seq PAM alignment distance position gene name gene id
GU305914.1 217544 217566 + 0 CAGAATTCTACCTCGGTATG CGG ||||||||[||||||||||||]PAM NA Intergenic NA NA

T16271 AATTCTACCTCGGTATGCGGCGG 1000
Oligo fwd CACCAATTCTACCTCGGTATGCGG
Oligo rev AAACCCGCATACCGAGGTAGAATT
Chromosome start end strand MM target_seq PAM alignment distance position gene name gene id
GU305914.1 217547 217569 + 0 AATTCTACCTCGGTATGCGG CGG ||||||||[||||||||||||]PAM NA Intergenic NA NA

<table>
<tr><td>T16272</td><td>TGTATTAATTTATCAAGCACAGG</td><td>1000</td><td></td><td></td><td></td><td></td><td></td><td></td><td></td><td></td><td></td></tr>
<tr><td>Oligo fwd</td><td>CACCTGTATTAATTTATCAAGCAC</td><td></td><td></td><td></td><td></td><td></td><td></td><td></td><td></td><td></td><td></td></tr>
<tr><td>Oligo rev</td><td>AAACGTGCTTGATAAATTAATACA</td><td></td><td></td><td></td><td></td><td></td><td></td><td></td><td></td><td></td><td></td></tr>
<tr><td>Chromosome</td><td>start</td><td>end</td><td>strand</td><td>MM</td><td>target_seq</td><td>PAM</td><td>alignment</td><td>distance</td><td>position</td><td>gene name</td><td>gene id</td></tr>
<tr><td>GU305914.1</td><td>217580</td><td>217602</td><td>+</td><td>0</td><td>TGTATTAATTTATCAAGCAC</td><td>AGG</td><td>||||||||[||||||||||||]PAM</td><td>NA</td><td>Intergenic</td><td>NA</td><td>NA</td></tr>
<tr><td>T16273</td><td>GTATTAATTTATCAAGCACAGGG</td><td>1000</td><td></td><td></td><td></td><td></td><td></td><td></td><td></td><td></td><td></td></tr>
<tr><td>Oligo fwd</td><td>CACCGTATTAATTTATCAAGCACA</td><td></td><td></td><td></td><td></td><td></td><td></td><td></td><td></td><td></td><td></td></tr>
<tr><td>Oligo rev</td><td>AAACTGTGCTTGATAAATTAATAC</td><td></td><td></td><td></td><td></td><td></td><td></td><td></td><td></td><td></td><td></td></tr>
<tr><td>Chromosome</td><td>start</td><td>end</td><td>strand</td><td>MM</td><td>target_seq</td><td>PAM</td><td>alignment</td><td>distance</td><td>position</td><td>gene name</td><td>gene id</td></tr>
<tr><td>GU305914.1</td><td>217581</td><td>217603</td><td>+</td><td>0</td><td>GTATTAATTTATCAAGCACA</td><td>GGG</td><td>||||||||[||||||||||||]PAM</td><td>NA</td><td>Intergenic</td><td>NA</td><td>NA</td></tr>
<tr><td>T16274</td><td>TATTAATTTATCAAGCACAGGGG</td><td>1000</td><td></td><td></td><td></td><td></td><td></td><td></td><td></td><td></td><td></td></tr>
<tr><td>Oligo fwd</td><td>CACCTATTAATTTATCAAGCACAG</td><td></td><td></td><td></td><td></td><td></td><td></td><td></td><td></td><td></td><td></td></tr>
<tr><td>Oligo rev</td><td>AAACCTGTGCTTGATAAATTAATA</td><td></td><td></td><td></td><td></td><td></td><td></td><td></td><td></td><td></td><td></td></tr>
<tr><td>Chromosome</td><td>start</td><td>end</td><td>strand</td><td>MM</td><td>target_seq</td><td>PAM</td><td>alignment</td><td>distance</td><td>position</td><td>gene name</td><td>gene id</td></tr>
<tr><td>GU305914.1</td><td>217582</td><td>217604</td><td>+</td><td>0</td><td>TATTAATTTATCAAGCACAG</td><td>GGG</td><td>||||||||[||||||||||||]PAM</td><td>NA</td><td>Intergenic</td><td>NA</td><td>NA</td></tr>
<tr><td>T16275</td><td>CCAGACTCTTTTGATACGCTCGG</td><td>1000</td><td></td><td></td><td></td><td></td><td></td><td></td><td></td><td></td><td></td></tr>
<tr><td>Oligo fwd</td><td>CACCCCAGACTCTTTTGATACGCT</td><td></td><td></td><td></td><td></td><td></td><td></td><td></td><td></td><td></td><td></td></tr>
<tr><td>Oligo rev</td><td>AAACAGCGTATCAAAAGAGTCTGG</td><td></td><td></td><td></td><td></td><td></td><td></td><td></td><td></td><td></td><td></td></tr>
<tr><td>Chromosome</td><td>start</td><td>end</td><td>strand</td><td>MM</td><td>target_seq</td><td>PAM</td><td>alignment</td><td>distance</td><td>position</td><td>gene name</td><td>gene id</td></tr>
<tr><td>GU305914.1</td><td>217606</td><td>217628</td><td>+</td><td>0</td><td>CCAGACTCTTTTGATACGCT</td><td>CGG</td><td>||||||||[||||||||||||]PAM</td><td>NA</td><td>Intergenic</td><td>NA</td><td>NA</td></tr>
<tr><td>T16276</td><td>TTTTGATACGCTCGGAATTTTGG</td><td>1000</td><td></td><td></td><td></td><td></td><td></td><td></td><td></td><td></td><td></td></tr>
<tr><td>Oligo fwd</td><td>CACCTTTTGATACGCTCGGAATTT</td><td></td><td></td><td></td><td></td><td></td><td></td><td></td><td></td><td></td><td></td></tr>
<tr><td>Oligo rev</td><td>AAACAAATTCCGAGCGTATCAAAA</td><td></td><td></td><td></td><td></td><td></td><td></td><td></td><td></td><td></td><td></td></tr>
<tr><td>Chromosome</td><td>start</td><td>end</td><td>strand</td><td>MM</td><td>target_seq</td><td>PAM</td><td>alignment</td><td>distance</td><td>position</td><td>gene name</td><td>gene id</td></tr>
<tr><td>GU305914.1</td><td>217614</td><td>217636</td><td>+</td><td>0</td><td>TTTTGATACGCTCGGAATTT</td><td>TGG</td><td>||||||||[||||||||||||]PAM</td><td>NA</td><td>Intergenic</td><td>NA</td><td>NA</td></tr>
<tr><td>T16277</td><td>TGGAGTAGTACATCATATCCAGG</td><td>1000</td><td></td><td></td><td></td><td></td><td></td><td></td><td></td><td></td><td></td></tr>
<tr><td>Oligo fwd</td><td>CACCTGGAGTAGTACATCATATCC</td><td></td><td></td><td></td><td></td><td></td><td></td><td></td><td></td><td></td><td></td></tr>
<tr><td>Oligo rev</td><td>AAACGGATATGATGTACTACTCCA</td><td></td><td></td><td></td><td></td><td></td><td></td><td></td><td></td><td></td><td></td></tr>
<tr><td>Chromosome</td><td>start</td><td>end</td><td>strand</td><td>MM</td><td>target_seq</td><td>PAM</td><td>alignment</td><td>distance</td><td>position</td><td>gene name</td><td>gene id</td></tr>
<tr><td>GU305914.1</td><td>217634</td><td>217656</td><td>+</td><td>0</td><td>TGGAGTAGTACATCATATCC</td><td>AGG</td><td>||||||||[||||||||||||]PAM</td><td>NA</td><td>Intergenic</td><td>NA</td><td>NA</td></tr>
<tr><td>T16278</td><td>GTATGATCCAATCTGATGTCCGG</td><td>1000</td><td></td><td></td><td></td><td></td><td></td><td></td><td></td><td></td><td></td></tr>
<tr><td>Oligo fwd</td><td>CACCGTATGATCCAATCTGATGTC</td><td></td><td></td><td></td><td></td><td></td><td></td><td></td><td></td><td></td><td></td></tr>
<tr><td>Oligo rev</td><td>AAACGACATCAGATTGGATCATAC</td><td></td><td></td><td></td><td></td><td></td><td></td><td></td><td></td><td></td><td></td></tr>
<tr><td>Chromosome</td><td>start</td><td>end</td><td>strand</td><td>MM</td><td>target_seq</td><td>PAM</td><td>alignment</td><td>distance</td><td>position</td><td>gene name</td><td>gene id</td></tr>
<tr><td>GU305914.1</td><td>217661</td><td>217683</td><td>+</td><td>0</td><td>GTATGATCCAATCTGATGTC</td><td>CGG</td><td>||||||||[||||||||||||]PAM</td><td>NA</td><td>Intergenic</td><td>NA</td><td>NA</td></tr>
<tr><td>T16279</td><td>CCGGATCAGAAGCATCAGCGAGG</td><td>1000</td><td></td><td></td><td></td><td></td><td></td><td></td><td></td><td></td><td></td></tr>
<tr><td>Oligo fwd</td><td>CACCCCGGATCAGAAGCATCAGCG</td><td></td><td></td><td></td><td></td><td></td><td></td><td></td><td></td><td></td><td></td></tr>
<tr><td>Oligo rev</td><td>AAACCGCTGATGCTTCTGATCCGG</td><td></td><td></td><td></td><td></td><td></td><td></td><td></td><td></td><td></td><td></td></tr>
<tr><td>Chromosome</td><td>start</td><td>end</td><td>strand</td><td>MM</td><td>target_seq</td><td>PAM</td><td>alignment</td><td>distance</td><td>position</td><td>gene name</td><td>gene id</td></tr>
<tr><td>GU305914.1</td><td>217680</td><td>217702</td><td>+</td><td>0</td><td>CCGGATCAGAAGCATCAGCG</td><td>AGG</td><td>||||||||[||||||||||||]PAM</td><td>NA</td><td>Intergenic</td><td>NA</td><td>NA</td></tr>
<tr><td>T16280</td><td>TCAGAAGCATCAGCGAGGAGCGG</td><td>1000</td><td></td><td></td><td></td><td></td><td></td><td></td><td></td><td></td><td></td></tr>
<tr><td>Oligo fwd</td><td>CACCTCAGAAGCATCAGCGAGGAG</td><td></td><td></td><td></td><td></td><td></td><td></td><td></td><td></td><td></td><td></td></tr>
<tr><td>Oligo rev</td><td>AAACCTCCTCGCTGATGCTTCTGA</td><td></td><td></td><td></td><td></td><td></td><td></td><td></td><td></td><td></td><td></td></tr>
<tr><td>Chromosome</td><td>start</td><td>end</td><td>strand</td><td>MM</td><td>target_seq</td><td>PAM</td><td>alignment</td><td>distance</td><td>position</td><td>gene name</td><td>gene id</td></tr>
<tr><td>GU305914.1</td><td>217685</td><td>217707</td><td>+</td><td>0</td><td>TCAGAAGCATCAGCGAGGAG</td><td>CGG</td><td>||||||||[||||||||||||]PAM</td><td>NA</td><td>Intergenic</td><td>NA</td><td>NA</td></tr>
<tr><td>T16281</td><td>CAGAAGCATCAGCGAGGAGCGGG</td><td>1000</td><td></td><td></td><td></td><td></td><td></td><td></td><td></td><td></td><td></td></tr>
<tr><td>Oligo fwd</td><td>CACCCAGAAGCATCAGCGAGGAGC</td><td></td><td></td><td></td><td></td><td></td><td></td><td></td><td></td><td></td><td></td></tr>
<tr><td>Oligo rev</td><td>AAACGCTCCTCGCTGATGCTTCTG</td><td></td><td></td><td></td><td></td><td></td><td></td><td></td><td></td><td></td><td></td></tr>
<tr><td>Chromosome</td><td>start</td><td>end</td><td>strand</td><td>MM</td><td>target_seq</td><td>PAM</td><td>alignment</td><td>distance</td><td>position</td><td>gene name</td><td>gene id</td></tr>
<tr><td>GU305914.1</td><td>217686</td><td>217708</td><td>+</td><td>0</td><td>CAGAAGCATCAGCGAGGAGC</td><td>GGG</td><td>||||||||[||||||||||||]PAM</td><td>NA</td><td>Intergenic</td><td>NA</td><td>NA</td></tr>
<tr><td>T16282</td><td>AGAAGCATCAGCGAGGAGCGGGG</td><td>1000</td><td></td><td></td><td></td><td></td><td></td><td></td><td></td><td></td><td></td></tr>
<tr><td>Oligo fwd</td><td>CACCAGAAGCATCAGCGAGGAGCG</td><td></td><td></td><td></td><td></td><td></td><td></td><td></td><td></td><td></td><td></td></tr>
<tr><td>Oligo rev</td><td>AAACCGCTCCTCGCTGATGCTTCT</td><td></td><td></td><td></td><td></td><td></td><td></td><td></td><td></td><td></td><td></td></tr>
<tr><td>Chromosome</td><td>start</td><td>end</td><td>strand</td><td>MM</td><td>target_seq</td><td>PAM</td><td>alignment</td><td>distance</td><td>position</td><td>gene name</td><td>gene id</td></tr>
<tr><td>GU305914.1</td><td>217687</td><td>217709</td><td>+</td><td>0</td><td>AGAAGCATCAGCGAGGAGCG</td><td>GGG</td><td>||||||||[||||||||||||]PAM</td><td>NA</td><td>Intergenic</td><td>NA</td><td>NA</td></tr>
<tr><td>T16283</td><td>CATCAGCGAGGAGCGGGGATAGG</td><td>1000</td><td></td><td></td><td></td><td></td><td></td><td></td><td></td><td></td><td></td></tr>
<tr><td>Oligo fwd</td><td>CACCCATCAGCGAGGAGCGGGGAT</td><td></td><td></td><td></td><td></td><td></td><td></td><td></td><td></td><td></td><td></td></tr>
<tr><td>Oligo rev</td><td>AAACATCCCCGCTCCTCGCTGATG</td><td></td><td></td><td></td><td></td><td></td><td></td><td></td><td></td><td></td><td></td></tr>
<tr><td>Chromosome</td><td>start</td><td>end</td><td>strand</td><td>MM</td><td>target_seq</td><td>PAM</td><td>alignment</td><td>distance</td><td>position</td><td>gene name</td><td>gene id</td></tr>
<tr><td>GU305914.1</td><td>217692</td><td>217714</td><td>+</td><td>0</td><td>CATCAGCGAGGAGCGGGGAT</td><td>AGG</td><td>||||||||[||||||||||||]PAM</td><td>NA</td><td>Intergenic</td><td>NA</td><td>NA</td></tr>
<tr><td>T16284</td><td>GCGAGGAGCGGGGATAGGAGCGG</td><td>1000</td><td></td><td></td><td></td><td></td><td></td><td></td><td></td><td></td><td></td></tr>
<tr><td>Oligo fwd</td><td>CACCGCGAGGAGCGGGGATAGGAG</td><td></td><td></td><td></td><td></td><td></td><td></td><td></td><td></td><td></td><td></td></tr>
<tr><td>Oligo rev</td><td>AAACCTCCTATCCCCGCTCCTCGC</td><td></td><td></td><td></td><td></td><td></td><td></td><td></td><td></td><td></td><td></td></tr>
<tr><td>Chromosome</td><td>start</td><td>end</td><td>strand</td><td>MM</td><td>target_seq</td><td>PAM</td><td>alignment</td><td>distance</td><td>position</td><td>gene name</td><td>gene id</td></tr>
<tr><td>GU305914.1</td><td>217697</td><td>217719</td><td>+</td><td>0</td><td>GCGAGGAGCGGGGATAGGAG</td><td>CGG</td><td>||||||||[||||||||||||]PAM</td><td>NA</td><td>Intergenic</td><td>NA</td><td>NA</td></tr>
<tr><td>T16285</td><td>CACCACGTTCAAGATCACGCAGG</td><td>1000</td><td></td><td></td><td></td><td></td><td></td><td></td><td></td><td></td><td></td></tr>
<tr><td>Oligo fwd</td><td>CACCCACCACGTTCAAGATCACGC</td><td></td><td></td><td></td><td></td><td></td><td></td><td></td><td></td><td></td><td></td></tr>
<tr><td>Oligo rev</td><td>AAACGCGTGATCTTGAACGTGGTG</td><td></td><td></td><td></td><td></td><td></td><td></td><td></td><td></td><td></td><td></td></tr>
<tr><td>Chromosome</td><td>start</td><td>end</td><td>strand</td><td>MM</td><td>target_seq</td><td>PAM</td><td>alignment</td><td>distance</td><td>position</td><td>gene name</td><td>gene id</td></tr>
<tr><td>GU305914.1</td><td>217800</td><td>217822</td><td>+</td><td>0</td><td>CACCACGTTCAAGATCACGC</td><td>AGG</td><td>||||||||[||||||||||||]PAM</td><td>NA</td><td>Intergenic</td><td>NA</td><td>NA</td></tr>
<tr><td>T16286</td><td>GTTCGCTAGATAAAGTCCTTCGG</td><td>1000</td><td></td><td></td><td></td><td></td><td></td><td></td><td></td><td></td><td></td></tr>
<tr><td>Oligo fwd</td><td>CACCGTTCGCTAGATAAAGTCCTT</td><td></td><td></td><td></td><td></td><td></td><td></td><td></td><td></td><td></td><td></td></tr>
<tr><td>Oligo rev</td><td>AAACAAGGACTTTATCTAGCGAAC</td><td></td><td></td><td></td><td></td><td></td><td></td><td></td><td></td><td></td><td></td></tr>
<tr><td>Chromosome</td><td>start</td><td>end</td><td>strand</td><td>MM</td><td>target_seq</td><td>PAM</td><td>alignment</td><td>distance</td><td>position</td><td>gene name</td><td>gene id</td></tr>
</table>

GU305914.1 217822 217844 + 0 GTTCGCTAGATAAAGTCCTT CGG ||||||||[||||||||||||]PAM NA Intergenic NA NA

T16287 GGTCTCGAAAAAGAGTATCTCGG 1000
Oligo fwd CACCGGTCTCGAAAAAGAGTATCT
Oligo rev AAACAGATACTCTTTTTCGAGACC
Chromosome start end strand MM target_seq PAM alignment distance position gene name gene id
GU305914.1 217843 217865 + 0 GGTCTCGAAAAAGAGTATCT CGG ||||||||[||||||||||||]PAM NA Intergenic NA NA

T16288 TATCTCGGTCGACAACTCGTCGG 1000
Oligo fwd CACCTATCTCGGTCGACAACTCGT
Oligo rev AAACACGAGTTGTCGACCGAGATA
Chromosome start end strand MM target_seq PAM alignment distance position gene name gene id
GU305914.1 217858 217880 + 0 TATCTCGGTCGACAACTCGT CGG ||||||||[||||||||||||]PAM NA Intergenic NA NA

T16289 CGACAACTCGTCGGTATCGTTGG 1000
Oligo fwd CACCCGACAACTCGTCGGTATCGT
Oligo rev AAACACGATACCGACGAGTTGTCG
Chromosome start end strand MM target_seq PAM alignment distance position gene name gene id
GU305914.1 217867 217889 + 0 CGACAACTCGTCGGTATCGT TGG ||||||||[||||||||||||]PAM NA Intergenic NA NA

T16290 GACAACTCGTCGGTATCGTTGGG 1000
Oligo fwd CACCGACAACTCGTCGGTATCGTT
Oligo rev AAACAACGATACCGACGAGTTGTC
Chromosome start end strand MM target_seq PAM alignment distance position gene name gene id
GU305914.1 217868 217890 + 0 GACAACTCGTCGGTATCGTT GGG ||||||||[||||||||||||]PAM NA Intergenic NA NA

T16291 GGTATCGTTGGGTACGACCCAGG 1000
Oligo fwd CACCGGTATCGTTGGGTACGACCC
Oligo rev AAACGGGTCGTACCCAACGATACC
Chromosome start end strand MM target_seq PAM alignment distance position gene name gene id
GU305914.1 217879 217901 + 0 GGTATCGTTGGGTACGACCC AGG ||||||||[||||||||||||]PAM NA Intergenic NA NA

T16292 TTGGGTACGACCCAGGTTAGAGG 1000
Oligo fwd CACCTTGGGTACGACCCAGGTTAG
Oligo rev AAACCTAACCTGGGTCGTACCCAA
Chromosome start end strand MM target_seq PAM alignment distance position gene name gene id
GU305914.1 217886 217908 + 0 TTGGGTACGACCCAGGTTAG AGG ||||||||[||||||||||||]PAM NA Intergenic NA NA

T16293 GATCGCCTCTCTGCCATTCGAGG 1000
Oligo fwd CACCGATCGCCTCTCTGCCATTCG
Oligo rev AAACCGAATGGCAGAGAGGCGATC
Chromosome start end strand MM target_seq PAM alignment distance position gene name gene id
GU305914.1 217914 217936 + 0 GATCGCCTCTCTGCCATTCG AGG ||||||||[||||||||||||]PAM NA Intergenic NA NA

T16294 CTCTCTGCCATTCGAGGAGACGG 1000
Oligo fwd CACCCTCTCTGCCATTCGAGGAGA
Oligo rev AAACTCTCCTCGAATGGCAGAGAG
Chromosome start end strand MM target_seq PAM alignment distance position gene name gene id
GU305914.1 217920 217942 + 0 CTCTCTGCCATTCGAGGAGA CGG ||||||||[||||||||||||]PAM NA Intergenic NA NA

T16295 CGAACTGAACGTTCCTGTCGAGG 1000
Oligo fwd CACCCGAACTGAACGTTCCTGTCG
Oligo rev AAACCGACAGGAACGTTCAGTTCG
Chromosome start end strand MM target_seq PAM alignment distance position gene name gene id
GU305914.1 217971 217993 + 0 CGAACTGAACGTTCCTGTCG AGG ||||||||[||||||||||||]PAM NA Intergenic NA NA

T16296 TGTCGAGGTATCCGATGATCTGG 1000
Oligo fwd CACCTGTCGAGGTATCCGATGATC
Oligo rev AAACGATCATCGGATACCTCGACA
Chromosome start end strand MM target_seq PAM alignment distance position gene name gene id
GU305914.1 217986 218008 + 0 TGTCGAGGTATCCGATGATC TGG ||||||||[||||||||||||]PAM NA Intergenic NA NA

T16297 CCGATGATCTGGATACTCAGCGG 1000
Oligo fwd CACCCCGATGATCTGGATACTCAG
Oligo rev AAACCTGAGTATCCAGATCATCGG
Chromosome start end strand MM target_seq PAM alignment distance position gene name gene id
GU305914.1 217997 218019 + 0 CCGATGATCTGGATACTCAG CGG ||||||||[||||||||||||]PAM NA Intergenic NA NA

T16298 TTGAGCAGAGAAGAGAGCTGCGG 1000
Oligo fwd CACCTTGAGCAGAGAAGAGAGCTG
Oligo rev AAACCAGCTCTCTTCTCTGCTCAA
Chromosome start end strand MM target_seq PAM alignment distance position gene name gene id
GU305914.1 218041 218063 + 0 TTGAGCAGAGAAGAGAGCTG CGG ||||||||[||||||||||||]PAM NA Intergenic NA NA

T16299 AGAGAGCTGCGGAGACCAGAAGG 1000
Oligo fwd CACCAGAGAGCTGCGGAGACCAGA
Oligo rev AAACTCTGGTCTCCGCAGCTCTCT
Chromosome start end strand MM target_seq PAM alignment distance position gene name gene id
GU305914.1 218052 218074 + 0 AGAGAGCTGCGGAGACCAGA AGG ||||||||[||||||||||||]PAM NA Intergenic NA NA

T16300 CGAACGCGCGATAGAAATAGCGG 1000
Oligo fwd CACCCGAACGCGCGATAGAAATAG
Oligo rev AAACCTATTTCTATCGCGCGTTCG
Chromosome start end strand MM target_seq PAM alignment distance position gene name gene id
GU305914.1 218124 218146 + 0 CGAACGCGCGATAGAAATAG CGG ||||||||[||||||||||||]PAM NA Intergenic NA NA

T16301 CATTGTTAATACTGACAAACTGG 1000
Oligo fwd CACCCATTGTTAATACTGACAAAC
Oligo rev AAACGTTTGTCAGTATTAACAATG

| Chromosome | start | end | strand | MM | target_seq | PAM | alignment | distance | position | gene name | gene id |
|---|---|---|---|---|---|---|---|---|---|---|---|
| GU305914.1 | 218207 | 218229 | + | 0 | CATTGTTAATACTGACAAAC | TGG | \|\|\|\|\|\|\|\|[\|\|\|\|\|\|\|\|\|\|\|\|]PAM | NA | Intergenic | NA | NA |
| T16302 | TATAGTTAGCACCGTTAGAGAGG | 1000 | | | | | | | | | |
| Oligo fwd | CACCTATAGTTAGCACCGTTAGAG | | | | | | | | | | |
| Oligo rev | AAACCTCTAACGGTGCTAACTATA | | | | | | | | | | |
| Chromosome | start | end | strand | MM | target_seq | PAM | alignment | distance | position | gene name | gene id |
| GU305914.1 | 218234 | 218256 | + | 0 | TATAGTTAGCACCGTTAGAG | AGG | \|\|\|\|\|\|\|\|[\|\|\|\|\|\|\|\|\|\|\|\|]PAM | NA | Intergenic | NA | NA |
| T16303 | ATAGTTAGCACCGTTAGAGAGGG | 1000 | | | | | | | | | |
| Oligo fwd | CACCATAGTTAGCACCGTTAGAGA | | | | | | | | | | |
| Oligo rev | AAACTCTCTAACGGTGCTAACTAT | | | | | | | | | | |
| Chromosome | start | end | strand | MM | target_seq | PAM | alignment | distance | position | gene name | gene id |
| GU305914.1 | 218235 | 218257 | + | 0 | ATAGTTAGCACCGTTAGAGA | GGG | \|\|\|\|\|\|\|\|[\|\|\|\|\|\|\|\|\|\|\|\|]PAM | NA | Intergenic | NA | NA |
| T16304 | CACTAAGACCGCGACGATCGCGG | 1000 | | | | | | | | | |
| Oligo fwd | CACCCACTAAGACCGCGACGATCG | | | | | | | | | | |
| Oligo rev | AAACCGATCGTCGCGGTCTTAGTG | | | | | | | | | | |
| Chromosome | start | end | strand | MM | target_seq | PAM | alignment | distance | position | gene name | gene id |
| GU305914.1 | 218352 | 218374 | + | 0 | CACTAAGACCGCGACGATCG | CGG | \|\|\|\|\|\|\|\|[\|\|\|\|\|\|\|\|\|\|\|\|]PAM | NA | Intergenic | NA | NA |
| GU305914.1 | 121271 | 121293 | - | 4 | GACCAAGTACGCGACGATCG | CGG | -\|\|-\|\|\|-[-\|\|\|\|\|\|\|\|\|\|\|]PAM | NA | Intergenic | NA | NA |
| GU305914.1 | 62760 | 62782 | - | 4 | CCCCAAGACCGCGACCAACG | CGG | \|-\|-\|\|\|\|[\|\|\|\|\|\|\|-\|-\|\|]PAM | NA | Intergenic | NA | NA |
| T16305 | ATCGCGGCCACCAGCATCACCGG | 1000 | | | | | | | | | |
| Oligo fwd | CACCATCGCGGCCACCAGCATCAC | | | | | | | | | | |
| Oligo rev | AAACGTGATGCTGGTGGCCGCGAT | | | | | | | | | | |
| Chromosome | start | end | strand | MM | target_seq | PAM | alignment | distance | position | gene name | gene id |
| GU305914.1 | 218368 | 218390 | + | 0 | ATCGCGGCCACCAGCATCAC | CGG | \|\|\|\|\|\|\|\|[\|\|\|\|\|\|\|\|\|\|\|\|]PAM | NA | Intergenic | NA | NA |
| T16306 | CACCGGAACCTCGCTGCCCATGG | 1000 | | | | | | | | | |
| Oligo fwd | CACCCACCGGAACCTCGCTGCCCA | | | | | | | | | | |
| Oligo rev | AAACTGGGCAGCGAGGTTCCGGTG | | | | | | | | | | |
| Chromosome | start | end | strand | MM | target_seq | PAM | alignment | distance | position | gene name | gene id |
| GU305914.1 | 218385 | 218407 | + | 0 | CACCGGAACCTCGCTGCCCA | TGG | \|\|\|\|\|\|\|\|[\|\|\|\|\|\|\|\|\|\|\|\|]PAM | NA | Intergenic | NA | NA |
| T16307 | ACCGGAACCTCGCTGCCCATGGG | 1000 | | | | | | | | | |
| Oligo fwd | CACCACCGGAACCTCGCTGCCCAT | | | | | | | | | | |
| Oligo rev | AAACATGGGCAGCGAGGTTCCGGT | | | | | | | | | | |
| Chromosome | start | end | strand | MM | target_seq | PAM | alignment | distance | position | gene name | gene id |
| GU305914.1 | 218386 | 218408 | + | 0 | ACCGGAACCTCGCTGCCCAT | GGG | \|\|\|\|\|\|\|\|[\|\|\|\|\|\|\|\|\|\|\|\|]PAM | NA | Intergenic | NA | NA |
| T16308 | ACCTCGCTGCCCATGGGTAGCGG | 1000 | | | | | | | | | |
| Oligo fwd | CACCACCTCGCTGCCCATGGGTAG | | | | | | | | | | |
| Oligo rev | AAACCTACCCATGGGCAGCGAGGT | | | | | | | | | | |
| Chromosome | start | end | strand | MM | target_seq | PAM | alignment | distance | position | gene name | gene id |
| GU305914.1 | 218392 | 218414 | + | 0 | ACCTCGCTGCCCATGGGTAG | CGG | \|\|\|\|\|\|\|\|[\|\|\|\|\|\|\|\|\|\|\|\|]PAM | NA | Intergenic | NA | NA |
| T16309 | CCCATGGGTAGCGGTCCGTAAGG | 1000 | | | | | | | | | |
| Oligo fwd | CACCCCCATGGGTAGCGGTCCGTA | | | | | | | | | | |
| Oligo rev | AAACTACGGACCGCTACCCATGGG | | | | | | | | | | |
| Chromosome | start | end | strand | MM | target_seq | PAM | alignment | distance | position | gene name | gene id |
| GU305914.1 | 218401 | 218423 | + | 0 | CCCATGGGTAGCGGTCCGTA | AGG | \|\|\|\|\|\|\|\|[\|\|\|\|\|\|\|\|\|\|\|\|]PAM | NA | Intergenic | NA | NA |
| T16310 | GGGTAGCGGTCCGTAAGGATAGG | 1000 | | | | | | | | | |
| Oligo fwd | CACCGGGTAGCGGTCCGTAAGGAT | | | | | | | | | | |
| Oligo rev | AAACATCCTTACGGACCGCTACCC | | | | | | | | | | |
| Chromosome | start | end | strand | MM | target_seq | PAM | alignment | distance | position | gene name | gene id |
| GU305914.1 | 218406 | 218428 | + | 0 | GGGTAGCGGTCCGTAAGGAT | AGG | \|\|\|\|\|\|\|\|[\|\|\|\|\|\|\|\|\|\|\|\|]PAM | NA | Intergenic | NA | NA |
| T16311 | GGTAGCGGTCCGTAAGGATAGGG | 1000 | | | | | | | | | |
| Oligo fwd | CACCGGTAGCGGTCCGTAAGGATA | | | | | | | | | | |
| Oligo rev | AAACTATCCTTACGGACCGCTACC | | | | | | | | | | |
| Chromosome | start | end | strand | MM | target_seq | PAM | alignment | distance | position | gene name | gene id |
| GU305914.1 | 218407 | 218429 | + | 0 | GGTAGCGGTCCGTAAGGATA | GGG | \|\|\|\|\|\|\|\|[\|\|\|\|\|\|\|\|\|\|\|\|]PAM | NA | Intergenic | NA | NA |
| T16312 | CATGAAAGACGCCCACTTTCCGG | 1000 | | | | | | | | | |
| Oligo fwd | CACCCATGAAAGACGCCCACTTTC | | | | | | | | | | |
| Oligo rev | AAACGAAAGTGGGCGTCTTTCATG | | | | | | | | | | |
| Chromosome | start | end | strand | MM | target_seq | PAM | alignment | distance | position | gene name | gene id |
| GU305914.1 | 218436 | 218458 | + | 0 | CATGAAAGACGCCCACTTTC | CGG | \|\|\|\|\|\|\|\|[\|\|\|\|\|\|\|\|\|\|\|\|]PAM | NA | Intergenic | NA | NA |
| T16313 | ATGAAAGACGCCCACTTTCCGGG | 1000 | | | | | | | | | |
| Oligo fwd | CACCATGAAAGACGCCCACTTTCC | | | | | | | | | | |
| Oligo rev | AAACGGAAAGTGGGCGTCTTTCAT | | | | | | | | | | |
| Chromosome | start | end | strand | MM | target_seq | PAM | alignment | distance | position | gene name | gene id |
| GU305914.1 | 218437 | 218459 | + | 0 | ATGAAAGACGCCCACTTTCC | GGG | \|\|\|\|\|\|\|\|[\|\|\|\|\|\|\|\|\|\|\|\|]PAM | NA | Intergenic | NA | NA |
| T16314 | AAGACACTAGACAGCTCAGACGG | 1000 | | | | | | | | | |
| Oligo fwd | CACCAAGACACTAGACAGCTCAGA | | | | | | | | | | |
| Oligo rev | AAACTCTGAGCTGTCTAGTGTCTT | | | | | | | | | | |
| Chromosome | start | end | strand | MM | target_seq | PAM | alignment | distance | position | gene name | gene id |
| GU305914.1 | 218462 | 218484 | + | 0 | AAGACACTAGACAGCTCAGA | CGG | \|\|\|\|\|\|\|\|[\|\|\|\|\|\|\|\|\|\|\|\|]PAM | NA | Intergenic | NA | NA |
| T16315 | CAGACGGTGAACATCATCCGTGG | 1000 | | | | | | | | | |
| Oligo fwd | CACCCAGACGGTGAACATCATCCG | | | | | | | | | | |
| Oligo rev | AAACCGGATGATGTTCACCGTCTG | | | | | | | | | | |
| Chromosome | start | end | strand | MM | target_seq | PAM | alignment | distance | position | gene name | gene id |
| GU305914.1 | 218478 | 218500 | + | 0 | CAGACGGTGAACATCATCCG | TGG | \|\|\|\|\|\|\|\|[\|\|\|\|\|\|\|\|\|\|\|\|]PAM | NA | Intergenic | NA | NA |

<table>
<tr><td>T16316</td><td>AGACGGTGAACATCATCCGTGGG</td><td>1000</td><td></td><td></td><td></td><td></td><td></td><td></td><td></td><td></td><td></td></tr>
<tr><td>Oligo fwd</td><td>CACCAGACGGTGAACATCATCCGT</td><td></td><td></td><td></td><td></td><td></td><td></td><td></td><td></td><td></td><td></td></tr>
<tr><td>Oligo rev</td><td>AAACACGGATGATGTTCACCGTCT</td><td></td><td></td><td></td><td></td><td></td><td></td><td></td><td></td><td></td><td></td></tr>
<tr><td>Chromosome</td><td>start</td><td>end</td><td>strand</td><td>MM</td><td>target_seq</td><td>PAM</td><td>alignment</td><td>distance</td><td>position</td><td>gene name</td><td>gene id</td></tr>
<tr><td>GU305914.1</td><td>218479</td><td>218501</td><td>+</td><td>0</td><td>AGACGGTGAACATCATCCGT</td><td>GGG</td><td>||||||||[||||||||||||]PAM</td><td>NA</td><td>Intergenic</td><td>NA</td><td>NA</td></tr>
<tr><td>T16317</td><td>GGTGAACATCATCCGTGGGAAGG</td><td>1000</td><td></td><td></td><td></td><td></td><td></td><td></td><td></td><td></td><td></td></tr>
<tr><td>Oligo fwd</td><td>CACCGGTGAACATCATCCGTGGGA</td><td></td><td></td><td></td><td></td><td></td><td></td><td></td><td></td><td></td><td></td></tr>
<tr><td>Oligo rev</td><td>AAACTCCCACGGATGATGTTCACC</td><td></td><td></td><td></td><td></td><td></td><td></td><td></td><td></td><td></td><td></td></tr>
<tr><td>Chromosome</td><td>start</td><td>end</td><td>strand</td><td>MM</td><td>target_seq</td><td>PAM</td><td>alignment</td><td>distance</td><td>position</td><td>gene name</td><td>gene id</td></tr>
<tr><td>GU305914.1</td><td>218483</td><td>218505</td><td>+</td><td>0</td><td>GGTGAACATCATCCGTGGGA</td><td>AGG</td><td>||||||||[||||||||||||]PAM</td><td>NA</td><td>Intergenic</td><td>NA</td><td>NA</td></tr>
<tr><td>T16318</td><td>GTGAACATCATCCGTGGGAAGGG</td><td>1000</td><td></td><td></td><td></td><td></td><td></td><td></td><td></td><td></td><td></td></tr>
<tr><td>Oligo fwd</td><td>CACCGTGAACATCATCCGTGGGAA</td><td></td><td></td><td></td><td></td><td></td><td></td><td></td><td></td><td></td><td></td></tr>
<tr><td>Oligo rev</td><td>AAACTTCCCACGGATGATGTTCAC</td><td></td><td></td><td></td><td></td><td></td><td></td><td></td><td></td><td></td><td></td></tr>
<tr><td>Chromosome</td><td>start</td><td>end</td><td>strand</td><td>MM</td><td>target_seq</td><td>PAM</td><td>alignment</td><td>distance</td><td>position</td><td>gene name</td><td>gene id</td></tr>
<tr><td>GU305914.1</td><td>218484</td><td>218506</td><td>+</td><td>0</td><td>GTGAACATCATCCGTGGGAA</td><td>GGG</td><td>||||||||[||||||||||||]PAM</td><td>NA</td><td>Intergenic</td><td>NA</td><td>NA</td></tr>
<tr><td>T16319</td><td>ACCCCGCCACCGACTCCGTCCGG</td><td>1000</td><td></td><td></td><td></td><td></td><td></td><td></td><td></td><td></td><td></td></tr>
<tr><td>Oligo fwd</td><td>CACCACCCCGCCACCGACTCCGTC</td><td></td><td></td><td></td><td></td><td></td><td></td><td></td><td></td><td></td><td></td></tr>
<tr><td>Oligo rev</td><td>AAACGACGGAGTCGGTGGCGGGGT</td><td></td><td></td><td></td><td></td><td></td><td></td><td></td><td></td><td></td><td></td></tr>
<tr><td>Chromosome</td><td>start</td><td>end</td><td>strand</td><td>MM</td><td>target_seq</td><td>PAM</td><td>alignment</td><td>distance</td><td>position</td><td>gene name</td><td>gene id</td></tr>
<tr><td>GU305914.1</td><td>218524</td><td>218546</td><td>+</td><td>0</td><td>ACCCCGCCACCGACTCCGTC</td><td>CGG</td><td>||||||||[||||||||||||]PAM</td><td>NA</td><td>Intergenic</td><td>NA</td><td>NA</td></tr>
<tr><td>T16320</td><td>GGCATCCAATAAAAACCCCTAGG</td><td>1000</td><td></td><td></td><td></td><td></td><td></td><td></td><td></td><td></td><td></td></tr>
<tr><td>Oligo fwd</td><td>CACCGGCATCCAATAAAAACCCCT</td><td></td><td></td><td></td><td></td><td></td><td></td><td></td><td></td><td></td><td></td></tr>
<tr><td>Oligo rev</td><td>AAACAGGGGTTTTTATTGGATGCC</td><td></td><td></td><td></td><td></td><td></td><td></td><td></td><td></td><td></td><td></td></tr>
<tr><td>Chromosome</td><td>start</td><td>end</td><td>strand</td><td>MM</td><td>target_seq</td><td>PAM</td><td>alignment</td><td>distance</td><td>position</td><td>gene name</td><td>gene id</td></tr>
<tr><td>GU305914.1</td><td>218545</td><td>218567</td><td>+</td><td>0</td><td>GGCATCCAATAAAAACCCCT</td><td>AGG</td><td>||||||||[||||||||||||]PAM</td><td>NA</td><td>Intergenic</td><td>NA</td><td>NA</td></tr>
<tr><td>T16321</td><td>CATCCAGTAACCATGACATCTGG</td><td>1000</td><td></td><td></td><td></td><td></td><td></td><td></td><td></td><td></td><td></td></tr>
<tr><td>Oligo fwd</td><td>CACCCATCCAGTAACCATGACATC</td><td></td><td></td><td></td><td></td><td></td><td></td><td></td><td></td><td></td><td></td></tr>
<tr><td>Oligo rev</td><td>AAACGATGTCATGGTTACTGGATG</td><td></td><td></td><td></td><td></td><td></td><td></td><td></td><td></td><td></td><td></td></tr>
<tr><td>Chromosome</td><td>start</td><td>end</td><td>strand</td><td>MM</td><td>target_seq</td><td>PAM</td><td>alignment</td><td>distance</td><td>position</td><td>gene name</td><td>gene id</td></tr>
<tr><td>GU305914.1</td><td>218582</td><td>218604</td><td>+</td><td>0</td><td>CATCCAGTAACCATGACATC</td><td>TGG</td><td>||||||||[||||||||||||]PAM</td><td>NA</td><td>Intergenic</td><td>NA</td><td>NA</td></tr>
<tr><td>T16322</td><td>CATGACATCTGGATGTCGAAAGG</td><td>1000</td><td></td><td></td><td></td><td></td><td></td><td></td><td></td><td></td><td></td></tr>
<tr><td>Oligo fwd</td><td>CACCCATGACATCTGGATGTCGAA</td><td></td><td></td><td></td><td></td><td></td><td></td><td></td><td></td><td></td><td></td></tr>
<tr><td>Oligo rev</td><td>AAACTTCGACATCCAGATGTCATG</td><td></td><td></td><td></td><td></td><td></td><td></td><td></td><td></td><td></td><td></td></tr>
<tr><td>Chromosome</td><td>start</td><td>end</td><td>strand</td><td>MM</td><td>target_seq</td><td>PAM</td><td>alignment</td><td>distance</td><td>position</td><td>gene name</td><td>gene id</td></tr>
<tr><td>GU305914.1</td><td>218593</td><td>218615</td><td>+</td><td>0</td><td>CATGACATCTGGATGTCGAA</td><td>AGG</td><td>||||||||[||||||||||||]PAM</td><td>NA</td><td>Intergenic</td><td>NA</td><td>NA</td></tr>
<tr><td>T16323</td><td>ATGACATCTGGATGTCGAAAGGG</td><td>1000</td><td></td><td></td><td></td><td></td><td></td><td></td><td></td><td></td><td></td></tr>
<tr><td>Oligo fwd</td><td>CACCATGACATCTGGATGTCGAAA</td><td></td><td></td><td></td><td></td><td></td><td></td><td></td><td></td><td></td><td></td></tr>
<tr><td>Oligo rev</td><td>AAACTTTCGACATCCAGATGTCAT</td><td></td><td></td><td></td><td></td><td></td><td></td><td></td><td></td><td></td><td></td></tr>
<tr><td>Chromosome</td><td>start</td><td>end</td><td>strand</td><td>MM</td><td>target_seq</td><td>PAM</td><td>alignment</td><td>distance</td><td>position</td><td>gene name</td><td>gene id</td></tr>
<tr><td>GU305914.1</td><td>218594</td><td>218616</td><td>+</td><td>0</td><td>ATGACATCTGGATGTCGAAA</td><td>GGG</td><td>||||||||[||||||||||||]PAM</td><td>NA</td><td>Intergenic</td><td>NA</td><td>NA</td></tr>
<tr><td>T16324</td><td>CATCTGGATGTCGAAAGGGCTGG</td><td>1000</td><td></td><td></td><td></td><td></td><td></td><td></td><td></td><td></td><td></td></tr>
<tr><td>Oligo fwd</td><td>CACCCATCTGGATGTCGAAAGGGC</td><td></td><td></td><td></td><td></td><td></td><td></td><td></td><td></td><td></td><td></td></tr>
<tr><td>Oligo rev</td><td>AAACGCCCTTTCGACATCCAGATG</td><td></td><td></td><td></td><td></td><td></td><td></td><td></td><td></td><td></td><td></td></tr>
<tr><td>Chromosome</td><td>start</td><td>end</td><td>strand</td><td>MM</td><td>target_seq</td><td>PAM</td><td>alignment</td><td>distance</td><td>position</td><td>gene name</td><td>gene id</td></tr>
<tr><td>GU305914.1</td><td>218598</td><td>218620</td><td>+</td><td>0</td><td>CATCTGGATGTCGAAAGGGC</td><td>TGG</td><td>||||||||[||||||||||||]PAM</td><td>NA</td><td>Intergenic</td><td>NA</td><td>NA</td></tr>
<tr><td>T16325</td><td>CTTGACCGTAGAATTTCTTTCGG</td><td>1000</td><td></td><td></td><td></td><td></td><td></td><td></td><td></td><td></td><td></td></tr>
<tr><td>Oligo fwd</td><td>CACCCTTGACCGTAGAATTTCTTT</td><td></td><td></td><td></td><td></td><td></td><td></td><td></td><td></td><td></td><td></td></tr>
<tr><td>Oligo rev</td><td>AAACAAAGAAATTCTACGGTCAAG</td><td></td><td></td><td></td><td></td><td></td><td></td><td></td><td></td><td></td><td></td></tr>
<tr><td>Chromosome</td><td>start</td><td>end</td><td>strand</td><td>MM</td><td>target_seq</td><td>PAM</td><td>alignment</td><td>distance</td><td>position</td><td>gene name</td><td>gene id</td></tr>
<tr><td>GU305914.1</td><td>218637</td><td>218659</td><td>+</td><td>0</td><td>CTTGACCGTAGAATTTCTTT</td><td>CGG</td><td>||||||||[||||||||||||]PAM</td><td>NA</td><td>Intergenic</td><td>NA</td><td>NA</td></tr>
<tr><td>T16326</td><td>GACCGAGAGTATTGAGTCTTCGG</td><td>1000</td><td></td><td></td><td></td><td></td><td></td><td></td><td></td><td></td><td></td></tr>
<tr><td>Oligo fwd</td><td>CACCGACCGAGAGTATTGAGTCTT</td><td></td><td></td><td></td><td></td><td></td><td></td><td></td><td></td><td></td><td></td></tr>
<tr><td>Oligo rev</td><td>AAACAAGACTCAATACTCTCGGTC</td><td></td><td></td><td></td><td></td><td></td><td></td><td></td><td></td><td></td><td></td></tr>
<tr><td>Chromosome</td><td>start</td><td>end</td><td>strand</td><td>MM</td><td>target_seq</td><td>PAM</td><td>alignment</td><td>distance</td><td>position</td><td>gene name</td><td>gene id</td></tr>
<tr><td>GU305914.1</td><td>218693</td><td>218715</td><td>+</td><td>0</td><td>GACCGAGAGTATTGAGTCTT</td><td>CGG</td><td>||||||||[||||||||||||]PAM</td><td>NA</td><td>Intergenic</td><td>NA</td><td>NA</td></tr>
<tr><td>T16327</td><td>GCTACTCCAACGCCCCCCGATGG</td><td>1000</td><td></td><td></td><td></td><td></td><td></td><td></td><td></td><td></td><td></td></tr>
<tr><td>Oligo fwd</td><td>CACCGCTACTCCAACGCCCCCCGA</td><td></td><td></td><td></td><td></td><td></td><td></td><td></td><td></td><td></td><td></td></tr>
<tr><td>Oligo rev</td><td>AAACTCGGGGGGCGTTGGAGTAGC</td><td></td><td></td><td></td><td></td><td></td><td></td><td></td><td></td><td></td><td></td></tr>
<tr><td>Chromosome</td><td>start</td><td>end</td><td>strand</td><td>MM</td><td>target_seq</td><td>PAM</td><td>alignment</td><td>distance</td><td>position</td><td>gene name</td><td>gene id</td></tr>
<tr><td>GU305914.1</td><td>218730</td><td>218752</td><td>+</td><td>0</td><td>GCTACTCCAACGCCCCCCGA</td><td>TGG</td><td>||||||||[||||||||||||]PAM</td><td>NA</td><td>Intergenic</td><td>NA</td><td>NA</td></tr>
<tr><td>T16328</td><td>CCCCCGATGGTATTCATGTGCGG</td><td>1000</td><td></td><td></td><td></td><td></td><td></td><td></td><td></td><td></td><td></td></tr>
<tr><td>Oligo fwd</td><td>CACCCCCCCGATGGTATTCATGTG</td><td></td><td></td><td></td><td></td><td></td><td></td><td></td><td></td><td></td><td></td></tr>
<tr><td>Oligo rev</td><td>AAACCACATGAATACCATCGGGGG</td><td></td><td></td><td></td><td></td><td></td><td></td><td></td><td></td><td></td><td></td></tr>
<tr><td>Chromosome</td><td>start</td><td>end</td><td>strand</td><td>MM</td><td>target_seq</td><td>PAM</td><td>alignment</td><td>distance</td><td>position</td><td>gene name</td><td>gene id</td></tr>
<tr><td>GU305914.1</td><td>218743</td><td>218765</td><td>+</td><td>0</td><td>CCCCCGATGGTATTCATGTG</td><td>CGG</td><td>||||||||[||||||||||||]PAM</td><td>NA</td><td>Intergenic</td><td>NA</td><td>NA</td></tr>
<tr><td>T16329</td><td>GTGCGGCCATAAACTAGTCATGG</td><td>1000</td><td></td><td></td><td></td><td></td><td></td><td></td><td></td><td></td><td></td></tr>
<tr><td>Oligo fwd</td><td>CACCGTGCGGCCATAAACTAGTCA</td><td></td><td></td><td></td><td></td><td></td><td></td><td></td><td></td><td></td><td></td></tr>
<tr><td>Oligo rev</td><td>AAACTGACTAGTTTATGGCCGCAC</td><td></td><td></td><td></td><td></td><td></td><td></td><td></td><td></td><td></td><td></td></tr>
<tr><td>Chromosome</td><td>start</td><td>end</td><td>strand</td><td>MM</td><td>target_seq</td><td>PAM</td><td>alignment</td><td>distance</td><td>position</td><td>gene name</td><td>gene id</td></tr>
<tr><td>GU305914.1</td><td>218760</td><td>218782</td><td>+</td><td>0</td><td>GTGCGGCCATAAACTAGTCA</td><td>TGG</td><td>||||||||[||||||||||||]PAM</td><td>NA</td><td>Intergenic</td><td>NA</td><td>NA</td></tr>
<tr><td>T16330</td><td>GGCCGTGTTGTTGATCCACACGG</td><td>1000</td><td></td><td></td><td></td><td></td><td></td><td></td><td></td><td></td><td></td></tr>
<tr><td>Oligo fwd</td><td>CACCGGCCGTGTTGTTGATCCACA</td><td></td><td></td><td></td><td></td><td></td><td></td><td></td><td></td><td></td><td></td></tr>
<tr><td>Oligo rev</td><td>AAACTGTGGATCAACAACACGGCC</td><td></td><td></td><td></td><td></td><td></td><td></td><td></td><td></td><td></td><td></td></tr>
<tr><td>Chromosome</td><td>start</td><td>end</td><td>strand</td><td>MM</td><td>target_seq</td><td>PAM</td><td>alignment</td><td>distance</td><td>position</td><td>gene name</td><td>gene id</td></tr>
<tr><td>GU305914.1</td><td>218781</td><td>218803</td><td>+</td><td>0</td><td>GGCCGTGTTGTTGATCCACA</td><td>CGG</td><td>||||||||[||||||||||||]PAM</td><td>NA</td><td>Intergenic</td><td>NA</td><td>NA</td></tr>
</table>

<table>
<tr><td>T16331</td><td>GGAAAAAGTACCGTTATCATCGG</td><td>1000</td><td></td><td></td><td></td><td></td><td></td><td></td><td></td><td></td><td></td></tr>
<tr><td>Oligo fwd</td><td>CACCGGAAAAAGTACCGTTATCAT</td><td></td><td></td><td></td><td></td><td></td><td></td><td></td><td></td><td></td><td></td></tr>
<tr><td>Oligo rev</td><td>AAACATGATAACGGTACTTTTTCC</td><td></td><td></td><td></td><td></td><td></td><td></td><td></td><td></td><td></td><td></td></tr>
<tr><td>Chromosome</td><td>start</td><td>end</td><td>strand</td><td>MM</td><td>target_seq</td><td>PAM</td><td>alignment</td><td>distance</td><td>position</td><td>gene name</td><td>gene id</td></tr>
<tr><td>GU305914.1</td><td>218802</td><td>218824</td><td>+</td><td>0</td><td>GGAAAAAGTACCGTTATCAT</td><td>CGG</td><td>||||||||[||||||||||||]PAM</td><td>NA</td><td>Intergenic</td><td>NA</td><td>NA</td></tr>
<tr><td>T16332</td><td>GTTGTTAAACTTAAAATCATCGG</td><td>1000</td><td></td><td></td><td></td><td></td><td></td><td></td><td></td><td></td><td></td></tr>
<tr><td>Oligo fwd</td><td>CACCGTTGTTAAACTTAAAATCAT</td><td></td><td></td><td></td><td></td><td></td><td></td><td></td><td></td><td></td><td></td></tr>
<tr><td>Oligo rev</td><td>AAACATGATTTTAAGTTTAACAAC</td><td></td><td></td><td></td><td></td><td></td><td></td><td></td><td></td><td></td><td></td></tr>
<tr><td>Chromosome</td><td>start</td><td>end</td><td>strand</td><td>MM</td><td>target_seq</td><td>PAM</td><td>alignment</td><td>distance</td><td>position</td><td>gene name</td><td>gene id</td></tr>
<tr><td>GU305914.1</td><td>218841</td><td>218863</td><td>+</td><td>0</td><td>GTTGTTAAACTTAAAATCAT</td><td>CGG</td><td>||||||||[||||||||||||]PAM</td><td>NA</td><td>Intergenic</td><td>NA</td><td>NA</td></tr>
<tr><td>T16333</td><td>CCTCTGTCGAACCCAGCAGACGG</td><td>1000</td><td></td><td></td><td></td><td></td><td></td><td></td><td></td><td></td><td></td></tr>
<tr><td>Oligo fwd</td><td>CACCCCTCTGTCGAACCCAGCAGA</td><td></td><td></td><td></td><td></td><td></td><td></td><td></td><td></td><td></td><td></td></tr>
<tr><td>Oligo rev</td><td>AAACTCTGCTGGGTTCGACAGAGG</td><td></td><td></td><td></td><td></td><td></td><td></td><td></td><td></td><td></td><td></td></tr>
<tr><td>Chromosome</td><td>start</td><td>end</td><td>strand</td><td>MM</td><td>target_seq</td><td>PAM</td><td>alignment</td><td>distance</td><td>position</td><td>gene name</td><td>gene id</td></tr>
<tr><td>GU305914.1</td><td>218880</td><td>218902</td><td>+</td><td>0</td><td>CCTCTGTCGAACCCAGCAGA</td><td>CGG</td><td>||||||||[||||||||||||]PAM</td><td>NA</td><td>Intergenic</td><td>NA</td><td>NA</td></tr>
<tr><td>T16334</td><td>ACAGTCGTACGTGATCTGAACGG</td><td>1000</td><td></td><td></td><td></td><td></td><td></td><td></td><td></td><td></td><td></td></tr>
<tr><td>Oligo fwd</td><td>CACCACAGTCGTACGTGATCTGAA</td><td></td><td></td><td></td><td></td><td></td><td></td><td></td><td></td><td></td><td></td></tr>
<tr><td>Oligo rev</td><td>AAACTTCAGATCACGTACGACTGT</td><td></td><td></td><td></td><td></td><td></td><td></td><td></td><td></td><td></td><td></td></tr>
<tr><td>Chromosome</td><td>start</td><td>end</td><td>strand</td><td>MM</td><td>target_seq</td><td>PAM</td><td>alignment</td><td>distance</td><td>position</td><td>gene name</td><td>gene id</td></tr>
<tr><td>GU305914.1</td><td>218925</td><td>218947</td><td>+</td><td>0</td><td>ACAGTCGTACGTGATCTGAA</td><td>CGG</td><td>||||||||[||||||||||||]PAM</td><td>NA</td><td>Intergenic</td><td>NA</td><td>NA</td></tr>
<tr><td>T16335</td><td>AGATCATGCCCGTCTCGTCGAGG</td><td>1000</td><td></td><td></td><td></td><td></td><td></td><td></td><td></td><td></td><td></td></tr>
<tr><td>Oligo fwd</td><td>CACCAGATCATGCCCGTCTCGTCG</td><td></td><td></td><td></td><td></td><td></td><td></td><td></td><td></td><td></td><td></td></tr>
<tr><td>Oligo rev</td><td>AAACCGACGAGACGGGCATGATCT</td><td></td><td></td><td></td><td></td><td></td><td></td><td></td><td></td><td></td><td></td></tr>
<tr><td>Chromosome</td><td>start</td><td>end</td><td>strand</td><td>MM</td><td>target_seq</td><td>PAM</td><td>alignment</td><td>distance</td><td>position</td><td>gene name</td><td>gene id</td></tr>
<tr><td>GU305914.1</td><td>218948</td><td>218970</td><td>+</td><td>0</td><td>AGATCATGCCCGTCTCGTCG</td><td>AGG</td><td>||||||||[||||||||||||]PAM</td><td>NA</td><td>Intergenic</td><td>NA</td><td>NA</td></tr>
<tr><td>T16336</td><td>GTTCGAAGACCTGTTCTGAGCGG</td><td>1000</td><td></td><td></td><td></td><td></td><td></td><td></td><td></td><td></td><td></td></tr>
<tr><td>Oligo fwd</td><td>CACCGTTCGAAGACCTGTTCTGAG</td><td></td><td></td><td></td><td></td><td></td><td></td><td></td><td></td><td></td><td></td></tr>
<tr><td>Oligo rev</td><td>AAACCTCAGAACAGGTCTTCGAAC</td><td></td><td></td><td></td><td></td><td></td><td></td><td></td><td></td><td></td><td></td></tr>
<tr><td>Chromosome</td><td>start</td><td>end</td><td>strand</td><td>MM</td><td>target_seq</td><td>PAM</td><td>alignment</td><td>distance</td><td>position</td><td>gene name</td><td>gene id</td></tr>
<tr><td>GU305914.1</td><td>218973</td><td>218995</td><td>+</td><td>0</td><td>GTTCGAAGACCTGTTCTGAG</td><td>CGG</td><td>||||||||[||||||||||||]PAM</td><td>NA</td><td>Intergenic</td><td>NA</td><td>NA</td></tr>
<tr><td>T16337</td><td>GCATAAAGTCTCGATGTTGTTGG</td><td>1000</td><td></td><td></td><td></td><td></td><td></td><td></td><td></td><td></td><td></td></tr>
<tr><td>Oligo fwd</td><td>CACCGCATAAAGTCTCGATGTTGT</td><td></td><td></td><td></td><td></td><td></td><td></td><td></td><td></td><td></td><td></td></tr>
<tr><td>Oligo rev</td><td>AAACACAACATCGAGACTTTATGC</td><td></td><td></td><td></td><td></td><td></td><td></td><td></td><td></td><td></td><td></td></tr>
<tr><td>Chromosome</td><td>start</td><td>end</td><td>strand</td><td>MM</td><td>target_seq</td><td>PAM</td><td>alignment</td><td>distance</td><td>position</td><td>gene name</td><td>gene id</td></tr>
<tr><td>GU305914.1</td><td>219002</td><td>219024</td><td>+</td><td>0</td><td>GCATAAAGTCTCGATGTTGT</td><td>TGG</td><td>||||||||[||||||||||||]PAM</td><td>NA</td><td>Intergenic</td><td>NA</td><td>NA</td></tr>
<tr><td>T16338</td><td>GTCTCGATGTTGTTGGAGAAAGG</td><td>1000</td><td></td><td></td><td></td><td></td><td></td><td></td><td></td><td></td><td></td></tr>
<tr><td>Oligo fwd</td><td>CACCGTCTCGATGTTGTTGGAGAA</td><td></td><td></td><td></td><td></td><td></td><td></td><td></td><td></td><td></td><td></td></tr>
<tr><td>Oligo rev</td><td>AAACTTCTCCAACAACATCGAGAC</td><td></td><td></td><td></td><td></td><td></td><td></td><td></td><td></td><td></td><td></td></tr>
<tr><td>Chromosome</td><td>start</td><td>end</td><td>strand</td><td>MM</td><td>target_seq</td><td>PAM</td><td>alignment</td><td>distance</td><td>position</td><td>gene name</td><td>gene id</td></tr>
<tr><td>GU305914.1</td><td>219009</td><td>219031</td><td>+</td><td>0</td><td>GTCTCGATGTTGTTGGAGAA</td><td>AGG</td><td>||||||||[||||||||||||]PAM</td><td>NA</td><td>Intergenic</td><td>NA</td><td>NA</td></tr>
<tr><td>T16339</td><td>GTTGGAGAAAGGTCACCTCTTGG</td><td>1000</td><td></td><td></td><td></td><td></td><td></td><td></td><td></td><td></td><td></td></tr>
<tr><td>Oligo fwd</td><td>CACCGTTGGAGAAAGGTCACCTCT</td><td></td><td></td><td></td><td></td><td></td><td></td><td></td><td></td><td></td><td></td></tr>
<tr><td>Oligo rev</td><td>AAACAGAGGTGACCTTTCTCCAAC</td><td></td><td></td><td></td><td></td><td></td><td></td><td></td><td></td><td></td><td></td></tr>
<tr><td>Chromosome</td><td>start</td><td>end</td><td>strand</td><td>MM</td><td>target_seq</td><td>PAM</td><td>alignment</td><td>distance</td><td>position</td><td>gene name</td><td>gene id</td></tr>
<tr><td>GU305914.1</td><td>219020</td><td>219042</td><td>+</td><td>0</td><td>GTTGGAGAAAGGTCACCTCT</td><td>TGG</td><td>||||||||[||||||||||||]PAM</td><td>NA</td><td>Intergenic</td><td>NA</td><td>NA</td></tr>
<tr><td>T16340</td><td>TTGGAGAAAGGTCACCTCTTGGG</td><td>1000</td><td></td><td></td><td></td><td></td><td></td><td></td><td></td><td></td><td></td></tr>
<tr><td>Oligo fwd</td><td>CACCTTGGAGAAAGGTCACCTCTT</td><td></td><td></td><td></td><td></td><td></td><td></td><td></td><td></td><td></td><td></td></tr>
<tr><td>Oligo rev</td><td>AAACAAGAGGTGACCTTTCTCCAA</td><td></td><td></td><td></td><td></td><td></td><td></td><td></td><td></td><td></td><td></td></tr>
<tr><td>Chromosome</td><td>start</td><td>end</td><td>strand</td><td>MM</td><td>target_seq</td><td>PAM</td><td>alignment</td><td>distance</td><td>position</td><td>gene name</td><td>gene id</td></tr>
<tr><td>GU305914.1</td><td>219021</td><td>219043</td><td>+</td><td>0</td><td>TTGGAGAAAGGTCACCTCTT</td><td>GGG</td><td>||||||||[||||||||||||]PAM</td><td>NA</td><td>Intergenic</td><td>NA</td><td>NA</td></tr>
<tr><td>T16341</td><td>GTAGCCGTACGTCCACGTCGTGG</td><td>1000</td><td></td><td></td><td></td><td></td><td></td><td></td><td></td><td></td><td></td></tr>
<tr><td>Oligo fwd</td><td>CACCGTAGCCGTACGTCCACGTCG</td><td></td><td></td><td></td><td></td><td></td><td></td><td></td><td></td><td></td><td></td></tr>
<tr><td>Oligo rev</td><td>AAACCGACGTGGACGTACGGCTAC</td><td></td><td></td><td></td><td></td><td></td><td></td><td></td><td></td><td></td><td></td></tr>
<tr><td>Chromosome</td><td>start</td><td>end</td><td>strand</td><td>MM</td><td>target_seq</td><td>PAM</td><td>alignment</td><td>distance</td><td>position</td><td>gene name</td><td>gene id</td></tr>
<tr><td>GU305914.1</td><td>219060</td><td>219082</td><td>+</td><td>0</td><td>GTAGCCGTACGTCCACGTCG</td><td>TGG</td><td>||||||||[||||||||||||]PAM</td><td>NA</td><td>Intergenic</td><td>NA</td><td>NA</td></tr>
<tr><td>T16342</td><td>TATCGTTCACGCTGAGTAAGAGG</td><td>1000</td><td></td><td></td><td></td><td></td><td></td><td></td><td></td><td></td><td></td></tr>
<tr><td>Oligo fwd</td><td>CACCTATCGTTCACGCTGAGTAAG</td><td></td><td></td><td></td><td></td><td></td><td></td><td></td><td></td><td></td><td></td></tr>
<tr><td>Oligo rev</td><td>AAACCTTACTCAGCGTGAACGATA</td><td></td><td></td><td></td><td></td><td></td><td></td><td></td><td></td><td></td><td></td></tr>
<tr><td>Chromosome</td><td>start</td><td>end</td><td>strand</td><td>MM</td><td>target_seq</td><td>PAM</td><td>alignment</td><td>distance</td><td>position</td><td>gene name</td><td>gene id</td></tr>
<tr><td>GU305914.1</td><td>219089</td><td>219111</td><td>+</td><td>0</td><td>TATCGTTCACGCTGAGTAAG</td><td>AGG</td><td>||||||||[||||||||||||]PAM</td><td>NA</td><td>Intergenic</td><td>NA</td><td>NA</td></tr>
<tr><td>T16343</td><td>ATCGTTCACGCTGAGTAAGAGGG</td><td>1000</td><td></td><td></td><td></td><td></td><td></td><td></td><td></td><td></td><td></td></tr>
<tr><td>Oligo fwd</td><td>CACCATCGTTCACGCTGAGTAAGA</td><td></td><td></td><td></td><td></td><td></td><td></td><td></td><td></td><td></td><td></td></tr>
<tr><td>Oligo rev</td><td>AAACTCTTACTCAGCGTGAACGAT</td><td></td><td></td><td></td><td></td><td></td><td></td><td></td><td></td><td></td><td></td></tr>
<tr><td>Chromosome</td><td>start</td><td>end</td><td>strand</td><td>MM</td><td>target_seq</td><td>PAM</td><td>alignment</td><td>distance</td><td>position</td><td>gene name</td><td>gene id</td></tr>
<tr><td>GU305914.1</td><td>219090</td><td>219112</td><td>+</td><td>0</td><td>ATCGTTCACGCTGAGTAAGA</td><td>GGG</td><td>||||||||[||||||||||||]PAM</td><td>NA</td><td>Intergenic</td><td>NA</td><td>NA</td></tr>
<tr><td>T16344</td><td>GAGGGCATCCCTCTCATTGATGG</td><td>1000</td><td></td><td></td><td></td><td></td><td></td><td></td><td></td><td></td><td></td></tr>
<tr><td>Oligo fwd</td><td>CACCGAGGGCATCCCTCTCATTGA</td><td></td><td></td><td></td><td></td><td></td><td></td><td></td><td></td><td></td><td></td></tr>
<tr><td>Oligo rev</td><td>AAACTCAATGAGAGGGATGCCCTC</td><td></td><td></td><td></td><td></td><td></td><td></td><td></td><td></td><td></td><td></td></tr>
<tr><td>Chromosome</td><td>start</td><td>end</td><td>strand</td><td>MM</td><td>target_seq</td><td>PAM</td><td>alignment</td><td>distance</td><td>position</td><td>gene name</td><td>gene id</td></tr>
<tr><td>GU305914.1</td><td>219108</td><td>219130</td><td>+</td><td>0</td><td>GAGGGCATCCCTCTCATTGA</td><td>TGG</td><td>||||||||[||||||||||||]PAM</td><td>NA</td><td>Intergenic</td><td>NA</td><td>NA</td></tr>
<tr><td>T16345</td><td>ACCCCAACTGAAATTATAAAAGG</td><td>1000</td><td></td><td></td><td></td><td></td><td></td><td></td><td></td><td></td><td></td></tr>
<tr><td>Oligo fwd</td><td>CACCACCCCAACTGAAATTATAAA</td><td></td><td></td><td></td><td></td><td></td><td></td><td></td><td></td><td></td><td></td></tr>
<tr><td>Oligo rev</td><td>AAACTTTATAATTTCAGTTGGGGT</td><td></td><td></td><td></td><td></td><td></td><td></td><td></td><td></td><td></td><td></td></tr>
<tr><td>Chromosome</td><td>start</td><td>end</td><td>strand</td><td>MM</td><td>target_seq</td><td>PAM</td><td>alignment</td><td>distance</td><td>position</td><td>gene name</td><td>gene id</td></tr>
</table>

<table>
<tr><td>GU305914.1</td><td></td><td>219135</td><td>219157</td><td>+</td><td>0</td><td>ACCCCAACTGAAATTATAAA</td><td>AGG</td><td>||||||||[||||||||||||]PAM</td><td>NA</td><td>Intergenic</td><td>NA</td><td>NA</td></tr>
<tr><td>T16346</td><td>TTATAAAAGGTTCGACACGAAGG</td><td></td><td>1000</td><td></td><td></td><td></td><td></td><td></td><td></td><td></td><td></td><td></td></tr>
<tr><td>Oligo fwd</td><td>CACCTTATAAAAGGTTCGACACGA</td><td></td><td></td><td></td><td></td><td></td><td></td><td></td><td></td><td></td><td></td><td></td></tr>
<tr><td>Oligo rev</td><td>AAACTCGTGTCGAACCTTTTATAA</td><td></td><td></td><td></td><td></td><td></td><td></td><td></td><td></td><td></td><td></td><td></td></tr>
<tr><td>Chromosome</td><td>start</td><td></td><td>end</td><td>strand</td><td>MM</td><td>target_seq</td><td>PAM</td><td>alignment</td><td>distance</td><td>position</td><td>gene name</td><td>gene id</td></tr>
<tr><td>GU305914.1</td><td></td><td>219148</td><td>219170</td><td>+</td><td>0</td><td>TTATAAAAGGTTCGACACGA</td><td>AGG</td><td>||||||||[||||||||||||]PAM</td><td>NA</td><td>Intergenic</td><td>NA</td><td>NA</td></tr>
<tr><td>T16347</td><td>AAAGGTTCGACACGAAGGAACGG</td><td></td><td>1000</td><td></td><td></td><td></td><td></td><td></td><td></td><td></td><td></td><td></td></tr>
<tr><td>Oligo fwd</td><td>CACCAAAGGTTCGACACGAAGGAA</td><td></td><td></td><td></td><td></td><td></td><td></td><td></td><td></td><td></td><td></td><td></td></tr>
<tr><td>Oligo rev</td><td>AAACTTCCTTCGTGTCGAACCTTT</td><td></td><td></td><td></td><td></td><td></td><td></td><td></td><td></td><td></td><td></td><td></td></tr>
<tr><td>Chromosome</td><td>start</td><td></td><td>end</td><td>strand</td><td>MM</td><td>target_seq</td><td>PAM</td><td>alignment</td><td>distance</td><td>position</td><td>gene name</td><td>gene id</td></tr>
<tr><td>GU305914.1</td><td></td><td>219153</td><td>219175</td><td>+</td><td>0</td><td>AAAGGTTCGACACGAAGGAA</td><td>CGG</td><td>||||||||[||||||||||||]PAM</td><td>NA</td><td>Intergenic</td><td>NA</td><td>NA</td></tr>
<tr><td>T16348</td><td>GGTTCGACACGAAGGAACGGCGG</td><td></td><td>1000</td><td></td><td></td><td></td><td></td><td></td><td></td><td></td><td></td><td></td></tr>
<tr><td>Oligo fwd</td><td>CACCGGTTCGACACGAAGGAACGG</td><td></td><td></td><td></td><td></td><td></td><td></td><td></td><td></td><td></td><td></td><td></td></tr>
<tr><td>Oligo rev</td><td>AAACCCGTTCCTTCGTGTCGAACC</td><td></td><td></td><td></td><td></td><td></td><td></td><td></td><td></td><td></td><td></td><td></td></tr>
<tr><td>Chromosome</td><td>start</td><td></td><td>end</td><td>strand</td><td>MM</td><td>target_seq</td><td>PAM</td><td>alignment</td><td>distance</td><td>position</td><td>gene name</td><td>gene id</td></tr>
<tr><td>GU305914.1</td><td></td><td>219156</td><td>219178</td><td>+</td><td>0</td><td>GGTTCGACACGAAGGAACGG</td><td>CGG</td><td>||||||||[||||||||||||]PAM</td><td>NA</td><td>Intergenic</td><td>NA</td><td>NA</td></tr>
<tr><td>T16349</td><td>GGAACGGCGGTTATGAGCTCTGG</td><td></td><td>1000</td><td></td><td></td><td></td><td></td><td></td><td></td><td></td><td></td><td></td></tr>
<tr><td>Oligo fwd</td><td>CACCGGAACGGCGGTTATGAGCTC</td><td></td><td></td><td></td><td></td><td></td><td></td><td></td><td></td><td></td><td></td><td></td></tr>
<tr><td>Oligo rev</td><td>AAACGAGCTCATAACCGCCGTTCC</td><td></td><td></td><td></td><td></td><td></td><td></td><td></td><td></td><td></td><td></td><td></td></tr>
<tr><td>Chromosome</td><td>start</td><td></td><td>end</td><td>strand</td><td>MM</td><td>target_seq</td><td>PAM</td><td>alignment</td><td>distance</td><td>position</td><td>gene name</td><td>gene id</td></tr>
<tr><td>GU305914.1</td><td></td><td>219169</td><td>219191</td><td>+</td><td>0</td><td>GGAACGGCGGTTATGAGCTC</td><td>TGG</td><td>||||||||[||||||||||||]PAM</td><td>NA</td><td>Intergenic</td><td>NA</td><td>NA</td></tr>
<tr><td>T16350</td><td>ATGAGCTCTGGTGTCGTAGACGG</td><td></td><td>1000</td><td></td><td></td><td></td><td></td><td></td><td></td><td></td><td></td><td></td></tr>
<tr><td>Oligo fwd</td><td>CACCATGAGCTCTGGTGTCGTAGA</td><td></td><td></td><td></td><td></td><td></td><td></td><td></td><td></td><td></td><td></td><td></td></tr>
<tr><td>Oligo rev</td><td>AAACTCTACGACACCAGAGCTCAT</td><td></td><td></td><td></td><td></td><td></td><td></td><td></td><td></td><td></td><td></td><td></td></tr>
<tr><td>Chromosome</td><td>start</td><td></td><td>end</td><td>strand</td><td>MM</td><td>target_seq</td><td>PAM</td><td>alignment</td><td>distance</td><td>position</td><td>gene name</td><td>gene id</td></tr>
<tr><td>GU305914.1</td><td></td><td>219181</td><td>219203</td><td>+</td><td>0</td><td>ATGAGCTCTGGTGTCGTAGA</td><td>CGG</td><td>||||||||[||||||||||||]PAM</td><td>NA</td><td>Intergenic</td><td>NA</td><td>NA</td></tr>
<tr><td>T16351</td><td>TGGTGTCGTAGACGGCAGAGTGG</td><td></td><td>1000</td><td></td><td></td><td></td><td></td><td></td><td></td><td></td><td></td><td></td></tr>
<tr><td>Oligo fwd</td><td>CACCTGGTGTCGTAGACGGCAGAG</td><td></td><td></td><td></td><td></td><td></td><td></td><td></td><td></td><td></td><td></td><td></td></tr>
<tr><td>Oligo rev</td><td>AAACCTCTGCCGTCTACGACACCA</td><td></td><td></td><td></td><td></td><td></td><td></td><td></td><td></td><td></td><td></td><td></td></tr>
<tr><td>Chromosome</td><td>start</td><td></td><td>end</td><td>strand</td><td>MM</td><td>target_seq</td><td>PAM</td><td>alignment</td><td>distance</td><td>position</td><td>gene name</td><td>gene id</td></tr>
<tr><td>GU305914.1</td><td></td><td>219189</td><td>219211</td><td>+</td><td>0</td><td>TGGTGTCGTAGACGGCAGAG</td><td>TGG</td><td>||||||||[||||||||||||]PAM</td><td>NA</td><td>Intergenic</td><td>NA</td><td>NA</td></tr>
<tr><td>T16352</td><td>CAGAGTGGTTAGATTCTGCAAGG</td><td></td><td>1000</td><td></td><td></td><td></td><td></td><td></td><td></td><td></td><td></td><td></td></tr>
<tr><td>Oligo fwd</td><td>CACCCAGAGTGGTTAGATTCTGCA</td><td></td><td></td><td></td><td></td><td></td><td></td><td></td><td></td><td></td><td></td><td></td></tr>
<tr><td>Oligo rev</td><td>AAACTGCAGAATCTAACCACTCTG</td><td></td><td></td><td></td><td></td><td></td><td></td><td></td><td></td><td></td><td></td><td></td></tr>
<tr><td>Chromosome</td><td>start</td><td></td><td>end</td><td>strand</td><td>MM</td><td>target_seq</td><td>PAM</td><td>alignment</td><td>distance</td><td>position</td><td>gene name</td><td>gene id</td></tr>
<tr><td>GU305914.1</td><td></td><td>219204</td><td>219226</td><td>+</td><td>0</td><td>CAGAGTGGTTAGATTCTGCA</td><td>AGG</td><td>||||||||[||||||||||||]PAM</td><td>NA</td><td>Intergenic</td><td>NA</td><td>NA</td></tr>
<tr><td>T16353</td><td>GGTACTATTGTGTATCGACGCGG</td><td></td><td>1000</td><td></td><td></td><td></td><td></td><td></td><td></td><td></td><td></td><td></td></tr>
<tr><td>Oligo fwd</td><td>CACCGGTACTATTGTGTATCGACG</td><td></td><td></td><td></td><td></td><td></td><td></td><td></td><td></td><td></td><td></td><td></td></tr>
<tr><td>Oligo rev</td><td>AAACCGTCGATACACAATAGTACC</td><td></td><td></td><td></td><td></td><td></td><td></td><td></td><td></td><td></td><td></td><td></td></tr>
<tr><td>Chromosome</td><td>start</td><td></td><td>end</td><td>strand</td><td>MM</td><td>target_seq</td><td>PAM</td><td>alignment</td><td>distance</td><td>position</td><td>gene name</td><td>gene id</td></tr>
<tr><td>GU305914.1</td><td></td><td>219225</td><td>219247</td><td>+</td><td>0</td><td>GGTACTATTGTGTATCGACG</td><td>CGG</td><td>||||||||[||||||||||||]PAM</td><td>NA</td><td>Intergenic</td><td>NA</td><td>NA</td></tr>
<tr><td>T16354</td><td>GTTGAAACGTTCGCAGAACCAGG</td><td></td><td>1000</td><td></td><td></td><td></td><td></td><td></td><td></td><td></td><td></td><td></td></tr>
<tr><td>Oligo fwd</td><td>CACCGTTGAAACGTTCGCAGAACC</td><td></td><td></td><td></td><td></td><td></td><td></td><td></td><td></td><td></td><td></td><td></td></tr>
<tr><td>Oligo rev</td><td>AAACGGTTCTGCGAACGTTTCAAC</td><td></td><td></td><td></td><td></td><td></td><td></td><td></td><td></td><td></td><td></td><td></td></tr>
<tr><td>Chromosome</td><td>start</td><td></td><td>end</td><td>strand</td><td>MM</td><td>target_seq</td><td>PAM</td><td>alignment</td><td>distance</td><td>position</td><td>gene name</td><td>gene id</td></tr>
<tr><td>GU305914.1</td><td></td><td>219250</td><td>219272</td><td>+</td><td>0</td><td>GTTGAAACGTTCGCAGAACC</td><td>AGG</td><td>||||||||[||||||||||||]PAM</td><td>NA</td><td>Intergenic</td><td>NA</td><td>NA</td></tr>
<tr><td>T16355</td><td>GAACCAGGTGTCGTGACGAGCGG</td><td></td><td>1000</td><td></td><td></td><td></td><td></td><td></td><td></td><td></td><td></td><td></td></tr>
<tr><td>Oligo fwd</td><td>CACCGAACCAGGTGTCGTGACGAG</td><td></td><td></td><td></td><td></td><td></td><td></td><td></td><td></td><td></td><td></td><td></td></tr>
<tr><td>Oligo rev</td><td>AAACCTCGTCACGACACCTGGTTC</td><td></td><td></td><td></td><td></td><td></td><td></td><td></td><td></td><td></td><td></td><td></td></tr>
<tr><td>Chromosome</td><td>start</td><td></td><td>end</td><td>strand</td><td>MM</td><td>target_seq</td><td>PAM</td><td>alignment</td><td>distance</td><td>position</td><td>gene name</td><td>gene id</td></tr>
<tr><td>GU305914.1</td><td></td><td>219265</td><td>219287</td><td>+</td><td>0</td><td>GAACCAGGTGTCGTGACGAG</td><td>CGG</td><td>||||||||[||||||||||||]PAM</td><td>NA</td><td>Intergenic</td><td>NA</td><td>NA</td></tr>
<tr><td>T16356</td><td>GACATCACCGATGCTACCGTAGG</td><td></td><td>1000</td><td></td><td></td><td></td><td></td><td></td><td></td><td></td><td></td><td></td></tr>
<tr><td>Oligo fwd</td><td>CACCGACATCACCGATGCTACCGT</td><td></td><td></td><td></td><td></td><td></td><td></td><td></td><td></td><td></td><td></td><td></td></tr>
<tr><td>Oligo rev</td><td>AAACACGGTAGCATCGGTGATGTC</td><td></td><td></td><td></td><td></td><td></td><td></td><td></td><td></td><td></td><td></td><td></td></tr>
<tr><td>Chromosome</td><td>start</td><td></td><td>end</td><td>strand</td><td>MM</td><td>target_seq</td><td>PAM</td><td>alignment</td><td>distance</td><td>position</td><td>gene name</td><td>gene id</td></tr>
<tr><td>GU305914.1</td><td></td><td>219307</td><td>219329</td><td>+</td><td>0</td><td>GACATCACCGATGCTACCGT</td><td>AGG</td><td>||||||||[||||||||||||]PAM</td><td>NA</td><td>Intergenic</td><td>NA</td><td>NA</td></tr>
<tr><td>T16357</td><td>CGATGCTACCGTAGGAGCCGCGG</td><td></td><td>1000</td><td></td><td></td><td></td><td></td><td></td><td></td><td></td><td></td><td></td></tr>
<tr><td>Oligo fwd</td><td>CACCCGATGCTACCGTAGGAGCCG</td><td></td><td></td><td></td><td></td><td></td><td></td><td></td><td></td><td></td><td></td><td></td></tr>
<tr><td>Oligo rev</td><td>AAACCGGCTCCTACGGTAGCATCG</td><td></td><td></td><td></td><td></td><td></td><td></td><td></td><td></td><td></td><td></td><td></td></tr>
<tr><td>Chromosome</td><td>start</td><td></td><td>end</td><td>strand</td><td>MM</td><td>target_seq</td><td>PAM</td><td>alignment</td><td>distance</td><td>position</td><td>gene name</td><td>gene id</td></tr>
<tr><td>GU305914.1</td><td></td><td>219315</td><td>219337</td><td>+</td><td>0</td><td>CGATGCTACCGTAGGAGCCG</td><td>CGG</td><td>||||||||[||||||||||||]PAM</td><td>NA</td><td>Intergenic</td><td>NA</td><td>NA</td></tr>
<tr><td>T16358</td><td>ATTATATATAGATTTATCTGAGG</td><td></td><td>1000</td><td></td><td></td><td></td><td></td><td></td><td></td><td></td><td></td><td></td></tr>
<tr><td>Oligo fwd</td><td>CACCATTATATATAGATTTATCTG</td><td></td><td></td><td></td><td></td><td></td><td></td><td></td><td></td><td></td><td></td><td></td></tr>
<tr><td>Oligo rev</td><td>AAACCAGATAAATCTATATATAAT</td><td></td><td></td><td></td><td></td><td></td><td></td><td></td><td></td><td></td><td></td><td></td></tr>
<tr><td>Chromosome</td><td>start</td><td></td><td>end</td><td>strand</td><td>MM</td><td>target_seq</td><td>PAM</td><td>alignment</td><td>distance</td><td>position</td><td>gene name</td><td>gene id</td></tr>
<tr><td>GU305914.1</td><td></td><td>219367</td><td>219389</td><td>+</td><td>0</td><td>ATTATATATAGATTTATCTG</td><td>AGG</td><td>||||||||[||||||||||||]PAM</td><td>NA</td><td>Intergenic</td><td>NA</td><td>NA</td></tr>
<tr><td>T16359</td><td>AAAATACGACGTCGTCCACATGG</td><td></td><td>1000</td><td></td><td></td><td></td><td></td><td></td><td></td><td></td><td></td><td></td></tr>
<tr><td>Oligo fwd</td><td>CACCAAAATACGACGTCGTCCACA</td><td></td><td></td><td></td><td></td><td></td><td></td><td></td><td></td><td></td><td></td><td></td></tr>
<tr><td>Oligo rev</td><td>AAACTGTGGACGACGTCGTATTTT</td><td></td><td></td><td></td><td></td><td></td><td></td><td></td><td></td><td></td><td></td><td></td></tr>
<tr><td>Chromosome</td><td>start</td><td></td><td>end</td><td>strand</td><td>MM</td><td>target_seq</td><td>PAM</td><td>alignment</td><td>distance</td><td>position</td><td>gene name</td><td>gene id</td></tr>
<tr><td>GU305914.1</td><td></td><td>219447</td><td>219469</td><td>+</td><td>0</td><td>AAAATACGACGTCGTCCACA</td><td>TGG</td><td>||||||||[||||||||||||]PAM</td><td>NA</td><td>Intergenic</td><td>NA</td><td>NA</td></tr>
<tr><td>T16360</td><td>CGTCCACATGGTCGCTCCTGCGG</td><td></td><td>1000</td><td></td><td></td><td></td><td></td><td></td><td></td><td></td><td></td><td></td></tr>
<tr><td>Oligo fwd</td><td>CACCCGTCCACATGGTCGCTCCTG</td><td></td><td></td><td></td><td></td><td></td><td></td><td></td><td></td><td></td><td></td><td></td></tr>
<tr><td>Oligo rev</td><td>AAACCAGGAGCGACCATGTGGACG</td><td></td><td></td><td></td><td></td><td></td><td></td><td></td><td></td><td></td><td></td><td></td></tr>
</table>

<table>
<tr><td>Chromosome</td><td>start</td><td>end</td><td>strand</td><td>MM</td><td>target_seq</td><td>PAM</td><td>alignment</td><td>distance</td><td>position</td><td>gene name</td><td>gene id</td></tr>
<tr><td>GU305914.1</td><td>219459</td><td>219481</td><td>+</td><td>0</td><td>CGTCCACATGGTCGCTCCTG</td><td>CGG</td><td>||||||||[||||||||||||]PAM</td><td>NA</td><td>Intergenic</td><td>NA</td><td>NA</td></tr>
<tr><td>T16361</td><td>CAAAAGTTTATTACATATAACGG</td><td>1000</td><td></td><td></td><td></td><td></td><td></td><td></td><td></td><td></td><td></td></tr>
<tr><td>Oligo fwd</td><td>CACCCAAAAGTTTATTACATATAA</td><td></td><td></td><td></td><td></td><td></td><td></td><td></td><td></td><td></td><td></td></tr>
<tr><td>Oligo rev</td><td>AAACTTATATGTAATAAACTTTTG</td><td></td><td></td><td></td><td></td><td></td><td></td><td></td><td></td><td></td><td></td></tr>
<tr><td>Chromosome</td><td>start</td><td>end</td><td>strand</td><td>MM</td><td>target_seq</td><td>PAM</td><td>alignment</td><td>distance</td><td>position</td><td>gene name</td><td>gene id</td></tr>
<tr><td>GU305914.1</td><td>219570</td><td>219592</td><td>+</td><td>0</td><td>CAAAAGTTTATTACATATAA</td><td>CGG</td><td>||||||||[||||||||||||]PAM</td><td>NA</td><td>Intergenic</td><td>NA</td><td>NA</td></tr>
<tr><td>T16362</td><td>AATGTTTCGTATAGAAAGCATGG</td><td>1000</td><td></td><td></td><td></td><td></td><td></td><td></td><td></td><td></td><td></td></tr>
<tr><td>Oligo fwd</td><td>CACCAATGTTTCGTATAGAAAGCA</td><td></td><td></td><td></td><td></td><td></td><td></td><td></td><td></td><td></td><td></td></tr>
<tr><td>Oligo rev</td><td>AAACTGCTTTCTATACGAAACATT</td><td></td><td></td><td></td><td></td><td></td><td></td><td></td><td></td><td></td><td></td></tr>
<tr><td>Chromosome</td><td>start</td><td>end</td><td>strand</td><td>MM</td><td>target_seq</td><td>PAM</td><td>alignment</td><td>distance</td><td>position</td><td>gene name</td><td>gene id</td></tr>
<tr><td>GU305914.1</td><td>219595</td><td>219617</td><td>+</td><td>0</td><td>AATGTTTCGTATAGAAAGCA</td><td>TGG</td><td>||||||||[||||||||||||]PAM</td><td>NA</td><td>Intergenic</td><td>NA</td><td>NA</td></tr>
<tr><td>T16363</td><td>ATGTTTCGTATAGAAAGCATGGG</td><td>1000</td><td></td><td></td><td></td><td></td><td></td><td></td><td></td><td></td><td></td></tr>
<tr><td>Oligo fwd</td><td>CACCATGTTTCGTATAGAAAGCAT</td><td></td><td></td><td></td><td></td><td></td><td></td><td></td><td></td><td></td><td></td></tr>
<tr><td>Oligo rev</td><td>AAACATGCTTTCTATACGAAACAT</td><td></td><td></td><td></td><td></td><td></td><td></td><td></td><td></td><td></td><td></td></tr>
<tr><td>Chromosome</td><td>start</td><td>end</td><td>strand</td><td>MM</td><td>target_seq</td><td>PAM</td><td>alignment</td><td>distance</td><td>position</td><td>gene name</td><td>gene id</td></tr>
<tr><td>GU305914.1</td><td>219596</td><td>219618</td><td>+</td><td>0</td><td>ATGTTTCGTATAGAAAGCAT</td><td>GGG</td><td>||||||||[||||||||||||]PAM</td><td>NA</td><td>Intergenic</td><td>NA</td><td>NA</td></tr>
<tr><td>T16364</td><td>TGTTTCGTATAGAAAGCATGGGG</td><td>1000</td><td></td><td></td><td></td><td></td><td></td><td></td><td></td><td></td><td></td></tr>
<tr><td>Oligo fwd</td><td>CACCTGTTTCGTATAGAAAGCATG</td><td></td><td></td><td></td><td></td><td></td><td></td><td></td><td></td><td></td><td></td></tr>
<tr><td>Oligo rev</td><td>AAACCATGCTTTCTATACGAAACA</td><td></td><td></td><td></td><td></td><td></td><td></td><td></td><td></td><td></td><td></td></tr>
<tr><td>Chromosome</td><td>start</td><td>end</td><td>strand</td><td>MM</td><td>target_seq</td><td>PAM</td><td>alignment</td><td>distance</td><td>position</td><td>gene name</td><td>gene id</td></tr>
<tr><td>GU305914.1</td><td>219597</td><td>219619</td><td>+</td><td>0</td><td>TGTTTCGTATAGAAAGCATG</td><td>GGG</td><td>||||||||[||||||||||||]PAM</td><td>NA</td><td>Intergenic</td><td>NA</td><td>NA</td></tr>
<tr><td>T16365</td><td>GTTTCGTATAGAAAGCATGGGGG</td><td>1000</td><td></td><td></td><td></td><td></td><td></td><td></td><td></td><td></td><td></td></tr>
<tr><td>Oligo fwd</td><td>CACCGTTTCGTATAGAAAGCATGG</td><td></td><td></td><td></td><td></td><td></td><td></td><td></td><td></td><td></td><td></td></tr>
<tr><td>Oligo rev</td><td>AAACCCATGCTTTCTATACGAAAC</td><td></td><td></td><td></td><td></td><td></td><td></td><td></td><td></td><td></td><td></td></tr>
<tr><td>Chromosome</td><td>start</td><td>end</td><td>strand</td><td>MM</td><td>target_seq</td><td>PAM</td><td>alignment</td><td>distance</td><td>position</td><td>gene name</td><td>gene id</td></tr>
<tr><td>GU305914.1</td><td>219598</td><td>219620</td><td>+</td><td>0</td><td>GTTTCGTATAGAAAGCATGG</td><td>GGG</td><td>||||||||[||||||||||||]PAM</td><td>NA</td><td>Intergenic</td><td>NA</td><td>NA</td></tr>
<tr><td>T16366</td><td>TAAGCTCGCAGATCGATTGAAGG</td><td>1000</td><td></td><td></td><td></td><td></td><td></td><td></td><td></td><td></td><td></td></tr>
<tr><td>Oligo fwd</td><td>CACCTAAGCTCGCAGATCGATTGA</td><td></td><td></td><td></td><td></td><td></td><td></td><td></td><td></td><td></td><td></td></tr>
<tr><td>Oligo rev</td><td>AAACTCAATCGATCTGCGAGCTTA</td><td></td><td></td><td></td><td></td><td></td><td></td><td></td><td></td><td></td><td></td></tr>
<tr><td>Chromosome</td><td>start</td><td>end</td><td>strand</td><td>MM</td><td>target_seq</td><td>PAM</td><td>alignment</td><td>distance</td><td>position</td><td>gene name</td><td>gene id</td></tr>
<tr><td>GU305914.1</td><td>219629</td><td>219651</td><td>+</td><td>0</td><td>TAAGCTCGCAGATCGATTGA</td><td>AGG</td><td>||||||||[||||||||||||]PAM</td><td>NA</td><td>Intergenic</td><td>NA</td><td>NA</td></tr>
<tr><td>T16367</td><td>GATTGAAGGTTCGATCTGACAGG</td><td>1000</td><td></td><td></td><td></td><td></td><td></td><td></td><td></td><td></td><td></td></tr>
<tr><td>Oligo fwd</td><td>CACCGATTGAAGGTTCGATCTGAC</td><td></td><td></td><td></td><td></td><td></td><td></td><td></td><td></td><td></td><td></td></tr>
<tr><td>Oligo rev</td><td>AAACGTCAGATCGAACCTTCAATC</td><td></td><td></td><td></td><td></td><td></td><td></td><td></td><td></td><td></td><td></td></tr>
<tr><td>Chromosome</td><td>start</td><td>end</td><td>strand</td><td>MM</td><td>target_seq</td><td>PAM</td><td>alignment</td><td>distance</td><td>position</td><td>gene name</td><td>gene id</td></tr>
<tr><td>GU305914.1</td><td>219643</td><td>219665</td><td>+</td><td>0</td><td>GATTGAAGGTTCGATCTGAC</td><td>AGG</td><td>||||||||[||||||||||||]PAM</td><td>NA</td><td>Intergenic</td><td>NA</td><td>NA</td></tr>
<tr><td>T16368</td><td>AGGTTCGATCTGACAGGAACAGG</td><td>1000</td><td></td><td></td><td></td><td></td><td></td><td></td><td></td><td></td><td></td></tr>
<tr><td>Oligo fwd</td><td>CACCAGGTTCGATCTGACAGGAAC</td><td></td><td></td><td></td><td></td><td></td><td></td><td></td><td></td><td></td><td></td></tr>
<tr><td>Oligo rev</td><td>AAACGTTCCTGTCAGATCGAACCT</td><td></td><td></td><td></td><td></td><td></td><td></td><td></td><td></td><td></td><td></td></tr>
<tr><td>Chromosome</td><td>start</td><td>end</td><td>strand</td><td>MM</td><td>target_seq</td><td>PAM</td><td>alignment</td><td>distance</td><td>position</td><td>gene name</td><td>gene id</td></tr>
<tr><td>GU305914.1</td><td>219649</td><td>219671</td><td>+</td><td>0</td><td>AGGTTCGATCTGACAGGAAC</td><td>AGG</td><td>||||||||[||||||||||||]PAM</td><td>NA</td><td>Intergenic</td><td>NA</td><td>NA</td></tr>
<tr><td>T16369</td><td>ATTTGAGTTAGCGTGTTATTAGG</td><td>1000</td><td></td><td></td><td></td><td></td><td></td><td></td><td></td><td></td><td></td></tr>
<tr><td>Oligo fwd</td><td>CACCATTTGAGTTAGCGTGTTATT</td><td></td><td></td><td></td><td></td><td></td><td></td><td></td><td></td><td></td><td></td></tr>
<tr><td>Oligo rev</td><td>AAACAATAACACGCTAACTCAAAT</td><td></td><td></td><td></td><td></td><td></td><td></td><td></td><td></td><td></td><td></td></tr>
<tr><td>Chromosome</td><td>start</td><td>end</td><td>strand</td><td>MM</td><td>target_seq</td><td>PAM</td><td>alignment</td><td>distance</td><td>position</td><td>gene name</td><td>gene id</td></tr>
<tr><td>GU305914.1</td><td>219692</td><td>219714</td><td>+</td><td>0</td><td>ATTTGAGTTAGCGTGTTATT</td><td>AGG</td><td>||||||||[||||||||||||]PAM</td><td>NA</td><td>Intergenic</td><td>NA</td><td>NA</td></tr>
<tr><td>T16370</td><td>GTGTTATTAGGAAGACTCCCAGG</td><td>1000</td><td></td><td></td><td></td><td></td><td></td><td></td><td></td><td></td><td></td></tr>
<tr><td>Oligo fwd</td><td>CACCGTGTTATTAGGAAGACTCCC</td><td></td><td></td><td></td><td></td><td></td><td></td><td></td><td></td><td></td><td></td></tr>
<tr><td>Oligo rev</td><td>AAACGGGAGTCTTCCTAATAACAC</td><td></td><td></td><td></td><td></td><td></td><td></td><td></td><td></td><td></td><td></td></tr>
<tr><td>Chromosome</td><td>start</td><td>end</td><td>strand</td><td>MM</td><td>target_seq</td><td>PAM</td><td>alignment</td><td>distance</td><td>position</td><td>gene name</td><td>gene id</td></tr>
<tr><td>GU305914.1</td><td>219704</td><td>219726</td><td>+</td><td>0</td><td>GTGTTATTAGGAAGACTCCC</td><td>AGG</td><td>||||||||[||||||||||||]PAM</td><td>NA</td><td>Intergenic</td><td>NA</td><td>NA</td></tr>
<tr><td>T16371</td><td>AGCATCTGTTGCCACGATGATGG</td><td>1000</td><td></td><td></td><td></td><td></td><td></td><td></td><td></td><td></td><td></td></tr>
<tr><td>Oligo fwd</td><td>CACCAGCATCTGTTGCCACGATGA</td><td></td><td></td><td></td><td></td><td></td><td></td><td></td><td></td><td></td><td></td></tr>
<tr><td>Oligo rev</td><td>AAACTCATCGTGGCAACAGATGCT</td><td></td><td></td><td></td><td></td><td></td><td></td><td></td><td></td><td></td><td></td></tr>
<tr><td>Chromosome</td><td>start</td><td>end</td><td>strand</td><td>MM</td><td>target_seq</td><td>PAM</td><td>alignment</td><td>distance</td><td>position</td><td>gene name</td><td>gene id</td></tr>
<tr><td>GU305914.1</td><td>219736</td><td>219758</td><td>+</td><td>0</td><td>AGCATCTGTTGCCACGATGA</td><td>TGG</td><td>||||||||[||||||||||||]PAM</td><td>NA</td><td>Intergenic</td><td>NA</td><td>NA</td></tr>
<tr><td>T16372</td><td>TCTGTTGCCACGATGATGGCTGG</td><td>1000</td><td></td><td></td><td></td><td></td><td></td><td></td><td></td><td></td><td></td></tr>
<tr><td>Oligo fwd</td><td>CACCTCTGTTGCCACGATGATGGC</td><td></td><td></td><td></td><td></td><td></td><td></td><td></td><td></td><td></td><td></td></tr>
<tr><td>Oligo rev</td><td>AAACGCCATCATCGTGGCAACAGA</td><td></td><td></td><td></td><td></td><td></td><td></td><td></td><td></td><td></td><td></td></tr>
<tr><td>Chromosome</td><td>start</td><td>end</td><td>strand</td><td>MM</td><td>target_seq</td><td>PAM</td><td>alignment</td><td>distance</td><td>position</td><td>gene name</td><td>gene id</td></tr>
<tr><td>GU305914.1</td><td>219740</td><td>219762</td><td>+</td><td>0</td><td>TCTGTTGCCACGATGATGGC</td><td>TGG</td><td>||||||||[||||||||||||]PAM</td><td>NA</td><td>Intergenic</td><td>NA</td><td>NA</td></tr>
<tr><td>T16373</td><td>ACGATGATGGCTGGATGCGATGG</td><td>1000</td><td></td><td></td><td></td><td></td><td></td><td></td><td></td><td></td><td></td></tr>
<tr><td>Oligo fwd</td><td>CACCACGATGATGGCTGGATGCGA</td><td></td><td></td><td></td><td></td><td></td><td></td><td></td><td></td><td></td><td></td></tr>
<tr><td>Oligo rev</td><td>AAACTCGCATCCAGCCATCATCGT</td><td></td><td></td><td></td><td></td><td></td><td></td><td></td><td></td><td></td><td></td></tr>
<tr><td>Chromosome</td><td>start</td><td>end</td><td>strand</td><td>MM</td><td>target_seq</td><td>PAM</td><td>alignment</td><td>distance</td><td>position</td><td>gene name</td><td>gene id</td></tr>
<tr><td>GU305914.1</td><td>219749</td><td>219771</td><td>+</td><td>0</td><td>ACGATGATGGCTGGATGCGA</td><td>TGG</td><td>||||||||[||||||||||||]PAM</td><td>NA</td><td>Intergenic</td><td>NA</td><td>NA</td></tr>
<tr><td>T16374</td><td>ATGGCCTGATGAGTAATAAGCGG</td><td>1000</td><td></td><td></td><td></td><td></td><td></td><td></td><td></td><td></td><td></td></tr>
<tr><td>Oligo fwd</td><td>CACCATGGCCTGATGAGTAATAAG</td><td></td><td></td><td></td><td></td><td></td><td></td><td></td><td></td><td></td><td></td></tr>
<tr><td>Oligo rev</td><td>AAACCTTATTACTCATCAGGCCAT</td><td></td><td></td><td></td><td></td><td></td><td></td><td></td><td></td><td></td><td></td></tr>
<tr><td>Chromosome</td><td>start</td><td>end</td><td>strand</td><td>MM</td><td>target_seq</td><td>PAM</td><td>alignment</td><td>distance</td><td>position</td><td>gene name</td><td>gene id</td></tr>
<tr><td>GU305914.1</td><td>219768</td><td>219790</td><td>+</td><td>0</td><td>ATGGCCTGATGAGTAATAAG</td><td>CGG</td><td>||||||||[||||||||||||]PAM</td><td>NA</td><td>Intergenic</td><td>NA</td><td>NA</td></tr>
<tr><td>T16375</td><td>TCCACATGTTTCTACTTTACAGG</td><td>1000</td><td></td><td></td><td></td><td></td><td></td><td></td><td></td><td></td><td></td></tr>
<tr><td>Oligo fwd</td><td>CACCTCCACATGTTTCTACTTTAC</td><td></td><td></td><td></td><td></td><td></td><td></td><td></td><td></td><td></td><td></td></tr>
</table>

| | | | | | | | | | | | |
|---|---|---|---|---|---|---|---|---|---|---|---|
| Oligo rev | AAACGTAAAGTAGAAACATGTGGA | | | | | | | | | | |
| Chromosome | start | end | strand | MM | target_seq | PAM | alignment | distance | position | gene name | gene id |
| GU305914.1 | 219828 | 219850 | + | 0 | TCCACATGTTTCTACTTTAC | AGG | ||||||||[||||||||||||]PAM | NA | Intergenic | NA | NA |
| T16376 | GTACGCGTACTCAAATTCGTCGG | 1000 | | | | | | | | | |
| Oligo fwd | CACCGTACGCGTACTCAAATTCGT | | | | | | | | | | |
| Oligo rev | AAACACGAATTTGAGTACGCGTAC | | | | | | | | | | |
| Chromosome | start | end | strand | MM | target_seq | PAM | alignment | distance | position | gene name | gene id |
| GU305914.1 | 219858 | 219880 | + | 0 | GTACGCGTACTCAAATTCGT | CGG | ||||||||[||||||||||||]PAM | NA | Intergenic | NA | NA |
| T16377 | TCCACAGCGATCTGTTAACTTGG | 1000 | | | | | | | | | |
| Oligo fwd | CACCTCCACAGCGATCTGTTAACT | | | | | | | | | | |
| Oligo rev | AAACAGTTAACAGATCGCTGTGGA | | | | | | | | | | |
| Chromosome | start | end | strand | MM | target_seq | PAM | alignment | distance | position | gene name | gene id |
| GU305914.1 | 219881 | 219903 | + | 0 | TCCACAGCGATCTGTTAACT | TGG | ||||||||[||||||||||||]PAM | NA | Intergenic | NA | NA |
| T16378 | TAACTTGGACAGATCCGTCATGG | 1000 | | | | | | | | | |
| Oligo fwd | CACCTAACTTGGACAGATCCGTCA | | | | | | | | | | |
| Oligo rev | AAACTGACGGATCTGTCCAAGTTA | | | | | | | | | | |
| Chromosome | start | end | strand | MM | target_seq | PAM | alignment | distance | position | gene name | gene id |
| GU305914.1 | 219896 | 219918 | + | 0 | TAACTTGGACAGATCCGTCA | TGG | ||||||||[||||||||||||]PAM | NA | Intergenic | NA | NA |
| T16379 | CGTATTAGATGAATGATACCAGG | 1000 | | | | | | | | | |
| Oligo fwd | CACCCGTATTAGATGAATGATACC | | | | | | | | | | |
| Oligo rev | AAACGGTATCATTCATCTAATACG | | | | | | | | | | |
| Chromosome | start | end | strand | MM | target_seq | PAM | alignment | distance | position | gene name | gene id |
| GU305914.1 | 220008 | 220030 | + | 0 | CGTATTAGATGAATGATACC | AGG | ||||||||[||||||||||||]PAM | NA | Intergenic | NA | NA |
| T16380 | TACGTGCCACACATCCTCTGTGG | 1000 | | | | | | | | | |
| Oligo fwd | CACCTACGTGCCACACATCCTCTG | | | | | | | | | | |
| Oligo rev | AAACCAGAGGATGTGTGGCACGTA | | | | | | | | | | |
| Chromosome | start | end | strand | MM | target_seq | PAM | alignment | distance | position | gene name | gene id |
| GU305914.1 | 220039 | 220061 | + | 0 | TACGTGCCACACATCCTCTG | TGG | ||||||||[||||||||||||]PAM | NA | Intergenic | NA | NA |
| T16381 | CTGTACTCCGAGCGTTTGTCCGG | 1000 | | | | | | | | | |
| Oligo fwd | CACCCTGTACTCCGAGCGTTTGTC | | | | | | | | | | |
| Oligo rev | AAACGACAAACGCTCGGAGTACAG | | | | | | | | | | |
| Chromosome | start | end | strand | MM | target_seq | PAM | alignment | distance | position | gene name | gene id |
| GU305914.1 | 220074 | 220096 | + | 0 | CTGTACTCCGAGCGTTTGTC | CGG | ||||||||[||||||||||||]PAM | NA | Intergenic | NA | NA |
| T16382 | GCGTTTGTCCGGTCCGCGTTCGG | 1000 | | | | | | | | | |
| Oligo fwd | CACCGCGTTTGTCCGGTCCGCGTT | | | | | | | | | | |
| Oligo rev | AAACAACGCGGACCGGACAAACGC | | | | | | | | | | |
| Chromosome | start | end | strand | MM | target_seq | PAM | alignment | distance | position | gene name | gene id |
| GU305914.1 | 220085 | 220107 | + | 0 | GCGTTTGTCCGGTCCGCGTT | CGG | ||||||||[||||||||||||]PAM | NA | Intergenic | NA | NA |
| T16383 | CGTGTATGTGTTGTGATCAAAGG | 1000 | | | | | | | | | |
| Oligo fwd | CACCCGTGTATGTGTTGTGATCAA | | | | | | | | | | |
| Oligo rev | AAACTTGATCACAACACATACACG | | | | | | | | | | |
| Chromosome | start | end | strand | MM | target_seq | PAM | alignment | distance | position | gene name | gene id |
| GU305914.1 | 220111 | 220133 | + | 0 | CGTGTATGTGTTGTGATCAA | AGG | ||||||||[||||||||||||]PAM | NA | Intergenic | NA | NA |
| T16384 | GTGATCAAAGGTCCCAACACCGG | 1000 | | | | | | | | | |
| Oligo fwd | CACCGTGATCAAAGGTCCCAACAC | | | | | | | | | | |
| Oligo rev | AAACGTGTTGGGACCTTTGATCAC | | | | | | | | | | |
| Chromosome | start | end | strand | MM | target_seq | PAM | alignment | distance | position | gene name | gene id |
| GU305914.1 | 220123 | 220145 | + | 0 | GTGATCAAAGGTCCCAACAC | CGG | ||||||||[||||||||||||]PAM | NA | Intergenic | NA | NA |
| T16385 | AGGTCCCAACACCGGAAGCGCGG | 1000 | | | | | | | | | |
| Oligo fwd | CACCAGGTCCCAACACCGGAAGCG | | | | | | | | | | |
| Oligo rev | AAACCGCTTCCGGTGTTGGGACCT | | | | | | | | | | |
| Chromosome | start | end | strand | MM | target_seq | PAM | alignment | distance | position | gene name | gene id |
| GU305914.1 | 220131 | 220153 | + | 0 | AGGTCCCAACACCGGAAGCG | CGG | ||||||||[||||||||||||]PAM | NA | Intergenic | NA | NA |
| T16386 | CGGAAGCGCGGAAGAACGACCGG | 1000 | | | | | | | | | |
| Oligo fwd | CACCCGGAAGCGCGGAAGAACGAC | | | | | | | | | | |
| Oligo rev | AAACGTCGTTCTTCCGCGCTTCCG | | | | | | | | | | |
| Chromosome | start | end | strand | MM | target_seq | PAM | alignment | distance | position | gene name | gene id |
| GU305914.1 | 220143 | 220165 | + | 0 | CGGAAGCGCGGAAGAACGAC | CGG | ||||||||[||||||||||||]PAM | NA | Intergenic | NA | NA |
| T16387 | GGAAGAACGACCGGTAACATCGG | 1000 | | | | | | | | | |
| Oligo fwd | CACCGGAAGAACGACCGGTAACAT | | | | | | | | | | |
| Oligo rev | AAACATGTTACCGGTCGTTCTTCC | | | | | | | | | | |
| Chromosome | start | end | strand | MM | target_seq | PAM | alignment | distance | position | gene name | gene id |
| GU305914.1 | 220152 | 220174 | + | 0 | GGAAGAACGACCGGTAACAT | CGG | ||||||||[||||||||||||]PAM | NA | Intergenic | NA | NA |
| T16388 | GAGATAAGTCCGTCACACAAAGG | 1000 | | | | | | | | | |
| Oligo fwd | CACCGAGATAAGTCCGTCACACAA | | | | | | | | | | |
| Oligo rev | AAACTTGTGTGACGGACTTATCTC | | | | | | | | | | |
| Chromosome | start | end | strand | MM | target_seq | PAM | alignment | distance | position | gene name | gene id |
| GU305914.1 | 220198 | 220220 | + | 0 | GAGATAAGTCCGTCACACAA | AGG | ||||||||[||||||||||||]PAM | NA | Intergenic | NA | NA |
| T16389 | TTCATTTTAAGTGTTGAAGATGG | 1000 | | | | | | | | | |
| Oligo fwd | CACCTTCATTTTAAGTGTTGAAGA | | | | | | | | | | |
| Oligo rev | AAACTCTTCAACACTTAAAATGAA | | | | | | | | | | |
| Chromosome | start | end | strand | MM | target_seq | PAM | alignment | distance | position | gene name | gene id |
| GU305914.1 | 220222 | 220244 | + | 0 | TTCATTTTAAGTGTTGAAGA | TGG | ||||||||[||||||||||||]PAM | NA | Intergenic | NA | NA |
| T16390 | TCATTTTAAGTGTTGAAGATGGG | 1000 | | | | | | | | | |

<table>
<tr><td>Oligo fwd</td><td>CACCTCATTTTAAGTGTTGAAGAT</td><td></td><td></td><td></td><td></td><td></td><td></td><td></td><td></td><td></td><td></td></tr>
<tr><td>Oligo rev</td><td>AAACATCTTCAACACTTAAAATGA</td><td></td><td></td><td></td><td></td><td></td><td></td><td></td><td></td><td></td><td></td></tr>
<tr><td>Chromosome</td><td>start</td><td>end</td><td>strand</td><td>MM</td><td>target_seq</td><td>PAM</td><td>alignment</td><td>distance</td><td>position</td><td>gene name</td><td>gene id</td></tr>
<tr><td>GU305914.1</td><td>220223</td><td>220245</td><td>+</td><td>0</td><td>TCATTTTAAGTGTTGAAGAT</td><td>GGG</td><td>||||||||[||||||||||||]PAM</td><td>NA</td><td>Intergenic</td><td>NA</td><td>NA</td></tr>
<tr><td>T16391</td><td>AGATGGGCCTGATTTTGAAGTGG</td><td>1000</td><td></td><td></td><td></td><td></td><td></td><td></td><td></td><td></td><td></td></tr>
<tr><td>Oligo fwd</td><td>CACCAGATGGGCCTGATTTTGAAG</td><td></td><td></td><td></td><td></td><td></td><td></td><td></td><td></td><td></td><td></td></tr>
<tr><td>Oligo rev</td><td>AAACCTTCAAAATCAGGCCCATCT</td><td></td><td></td><td></td><td></td><td></td><td></td><td></td><td></td><td></td><td></td></tr>
<tr><td>Chromosome</td><td>start</td><td>end</td><td>strand</td><td>MM</td><td>target_seq</td><td>PAM</td><td>alignment</td><td>distance</td><td>position</td><td>gene name</td><td>gene id</td></tr>
<tr><td>GU305914.1</td><td>220239</td><td>220261</td><td>+</td><td>0</td><td>AGATGGGCCTGATTTTGAAG</td><td>TGG</td><td>||||||||[||||||||||||]PAM</td><td>NA</td><td>Intergenic</td><td>NA</td><td>NA</td></tr>
<tr><td>T16392</td><td>CCTCTGACGATTCTGAATCGCGG</td><td>1000</td><td></td><td></td><td></td><td></td><td></td><td></td><td></td><td></td><td></td></tr>
<tr><td>Oligo fwd</td><td>CACCCCTCTGACGATTCTGAATCG</td><td></td><td></td><td></td><td></td><td></td><td></td><td></td><td></td><td></td><td></td></tr>
<tr><td>Oligo rev</td><td>AAACCGATTCAGAATCGTCAGAGG</td><td></td><td></td><td></td><td></td><td></td><td></td><td></td><td></td><td></td><td></td></tr>
<tr><td>Chromosome</td><td>start</td><td>end</td><td>strand</td><td>MM</td><td>target_seq</td><td>PAM</td><td>alignment</td><td>distance</td><td>position</td><td>gene name</td><td>gene id</td></tr>
<tr><td>GU305914.1</td><td>220292</td><td>220314</td><td>+</td><td>0</td><td>CCTCTGACGATTCTGAATCG</td><td>CGG</td><td>||||||||[||||||||||||]PAM</td><td>NA</td><td>Intergenic</td><td>NA</td><td>NA</td></tr>
<tr><td>T16393</td><td>TCGCGGTTAGCGCTACGTGAAGG</td><td>1000</td><td></td><td></td><td></td><td></td><td></td><td></td><td></td><td></td><td></td></tr>
<tr><td>Oligo fwd</td><td>CACCTCGCGGTTAGCGCTACGTGA</td><td></td><td></td><td></td><td></td><td></td><td></td><td></td><td></td><td></td><td></td></tr>
<tr><td>Oligo rev</td><td>AAACTCACGTAGCGCTAACCGCGA</td><td></td><td></td><td></td><td></td><td></td><td></td><td></td><td></td><td></td><td></td></tr>
<tr><td>Chromosome</td><td>start</td><td>end</td><td>strand</td><td>MM</td><td>target_seq</td><td>PAM</td><td>alignment</td><td>distance</td><td>position</td><td>gene name</td><td>gene id</td></tr>
<tr><td>GU305914.1</td><td>220309</td><td>220331</td><td>+</td><td>0</td><td>TCGCGGTTAGCGCTACGTGA</td><td>AGG</td><td>||||||||[||||||||||||]PAM</td><td>NA</td><td>Intergenic</td><td>NA</td><td>NA</td></tr>
<tr><td>T16394</td><td>GATGATCCACTAAGAACATATGG</td><td>1000</td><td></td><td></td><td></td><td></td><td></td><td></td><td></td><td></td><td></td></tr>
<tr><td>Oligo fwd</td><td>CACCGATGATCCACTAAGAACATA</td><td></td><td></td><td></td><td></td><td></td><td></td><td></td><td></td><td></td><td></td></tr>
<tr><td>Oligo rev</td><td>AAACTATGTTCTTAGTGGATCATC</td><td></td><td></td><td></td><td></td><td></td><td></td><td></td><td></td><td></td><td></td></tr>
<tr><td>Chromosome</td><td>start</td><td>end</td><td>strand</td><td>MM</td><td>target_seq</td><td>PAM</td><td>alignment</td><td>distance</td><td>position</td><td>gene name</td><td>gene id</td></tr>
<tr><td>GU305914.1</td><td>220357</td><td>220379</td><td>+</td><td>0</td><td>GATGATCCACTAAGAACATA</td><td>TGG</td><td>||||||||[||||||||||||]PAM</td><td>NA</td><td>Intergenic</td><td>NA</td><td>NA</td></tr>
<tr><td>T16395</td><td>AAAAGTAGCTAGAATAGTCTCGG</td><td>1000</td><td></td><td></td><td></td><td></td><td></td><td></td><td></td><td></td><td></td></tr>
<tr><td>Oligo fwd</td><td>CACCAAAAGTAGCTAGAATAGTCT</td><td></td><td></td><td></td><td></td><td></td><td></td><td></td><td></td><td></td><td></td></tr>
<tr><td>Oligo rev</td><td>AAACAGACTATTCTAGCTACTTTT</td><td></td><td></td><td></td><td></td><td></td><td></td><td></td><td></td><td></td><td></td></tr>
<tr><td>Chromosome</td><td>start</td><td>end</td><td>strand</td><td>MM</td><td>target_seq</td><td>PAM</td><td>alignment</td><td>distance</td><td>position</td><td>gene name</td><td>gene id</td></tr>
<tr><td>GU305914.1</td><td>220413</td><td>220435</td><td>+</td><td>0</td><td>AAAAGTAGCTAGAATAGTCT</td><td>CGG</td><td>||||||||[||||||||||||]PAM</td><td>NA</td><td>Intergenic</td><td>NA</td><td>NA</td></tr>
<tr><td>T16396</td><td>AATAGTCTCGGTCTTATTACCGG</td><td>1000</td><td></td><td></td><td></td><td></td><td></td><td></td><td></td><td></td><td></td></tr>
<tr><td>Oligo fwd</td><td>CACCAATAGTCTCGGTCTTATTAC</td><td></td><td></td><td></td><td></td><td></td><td></td><td></td><td></td><td></td><td></td></tr>
<tr><td>Oligo rev</td><td>AAACGTAATAAGACCGAGACTATT</td><td></td><td></td><td></td><td></td><td></td><td></td><td></td><td></td><td></td><td></td></tr>
<tr><td>Chromosome</td><td>start</td><td>end</td><td>strand</td><td>MM</td><td>target_seq</td><td>PAM</td><td>alignment</td><td>distance</td><td>position</td><td>gene name</td><td>gene id</td></tr>
<tr><td>GU305914.1</td><td>220425</td><td>220447</td><td>+</td><td>0</td><td>AATAGTCTCGGTCTTATTAC</td><td>CGG</td><td>||||||||[||||||||||||]PAM</td><td>NA</td><td>Intergenic</td><td>NA</td><td>NA</td></tr>
<tr><td>T16397</td><td>ATCAGAACCAGATGACGTACAGG</td><td>1000</td><td></td><td></td><td></td><td></td><td></td><td></td><td></td><td></td><td></td></tr>
<tr><td>Oligo fwd</td><td>CACCATCAGAACCAGATGACGTAC</td><td></td><td></td><td></td><td></td><td></td><td></td><td></td><td></td><td></td><td></td></tr>
<tr><td>Oligo rev</td><td>AAACGTACGTCATCTGGTTCTGAT</td><td></td><td></td><td></td><td></td><td></td><td></td><td></td><td></td><td></td><td></td></tr>
<tr><td>Chromosome</td><td>start</td><td>end</td><td>strand</td><td>MM</td><td>target_seq</td><td>PAM</td><td>alignment</td><td>distance</td><td>position</td><td>gene name</td><td>gene id</td></tr>
<tr><td>GU305914.1</td><td>220488</td><td>220510</td><td>+</td><td>0</td><td>ATCAGAACCAGATGACGTAC</td><td>AGG</td><td>||||||||[||||||||||||]PAM</td><td>NA</td><td>Intergenic</td><td>NA</td><td>NA</td></tr>
<tr><td>T16398</td><td>ATATTCGAGTGAATTGACATCGG</td><td>1000</td><td></td><td></td><td></td><td></td><td></td><td></td><td></td><td></td><td></td></tr>
<tr><td>Oligo fwd</td><td>CACCATATTCGAGTGAATTGACAT</td><td></td><td></td><td></td><td></td><td></td><td></td><td></td><td></td><td></td><td></td></tr>
<tr><td>Oligo rev</td><td>AAACATGTCAATTCACTCGAATAT</td><td></td><td></td><td></td><td></td><td></td><td></td><td></td><td></td><td></td><td></td></tr>
<tr><td>Chromosome</td><td>start</td><td>end</td><td>strand</td><td>MM</td><td>target_seq</td><td>PAM</td><td>alignment</td><td>distance</td><td>position</td><td>gene name</td><td>gene id</td></tr>
<tr><td>GU305914.1</td><td>220512</td><td>220534</td><td>+</td><td>0</td><td>ATATTCGAGTGAATTGACAT</td><td>CGG</td><td>||||||||[||||||||||||]PAM</td><td>NA</td><td>Intergenic</td><td>NA</td><td>NA</td></tr>
<tr><td>T16399</td><td>TATTCGAGTGAATTGACATCGGG</td><td>1000</td><td></td><td></td><td></td><td></td><td></td><td></td><td></td><td></td><td></td></tr>
<tr><td>Oligo fwd</td><td>CACCTATTCGAGTGAATTGACATC</td><td></td><td></td><td></td><td></td><td></td><td></td><td></td><td></td><td></td><td></td></tr>
<tr><td>Oligo rev</td><td>AAACGATGTCAATTCACTCGAATA</td><td></td><td></td><td></td><td></td><td></td><td></td><td></td><td></td><td></td><td></td></tr>
<tr><td>Chromosome</td><td>start</td><td>end</td><td>strand</td><td>MM</td><td>target_seq</td><td>PAM</td><td>alignment</td><td>distance</td><td>position</td><td>gene name</td><td>gene id</td></tr>
<tr><td>GU305914.1</td><td>220513</td><td>220535</td><td>+</td><td>0</td><td>TATTCGAGTGAATTGACATC</td><td>GGG</td><td>||||||||[||||||||||||]PAM</td><td>NA</td><td>Intergenic</td><td>NA</td><td>NA</td></tr>
<tr><td>T16400</td><td>ATTCGAGTGAATTGACATCGGGG</td><td>1000</td><td></td><td></td><td></td><td></td><td></td><td></td><td></td><td></td><td></td></tr>
<tr><td>Oligo fwd</td><td>CACCATTCGAGTGAATTGACATCG</td><td></td><td></td><td></td><td></td><td></td><td></td><td></td><td></td><td></td><td></td></tr>
<tr><td>Oligo rev</td><td>AAACCGATGTCAATTCACTCGAAT</td><td></td><td></td><td></td><td></td><td></td><td></td><td></td><td></td><td></td><td></td></tr>
<tr><td>Chromosome</td><td>start</td><td>end</td><td>strand</td><td>MM</td><td>target_seq</td><td>PAM</td><td>alignment</td><td>distance</td><td>position</td><td>gene name</td><td>gene id</td></tr>
<tr><td>GU305914.1</td><td>220514</td><td>220536</td><td>+</td><td>0</td><td>ATTCGAGTGAATTGACATCG</td><td>GGG</td><td>||||||||[||||||||||||]PAM</td><td>NA</td><td>Intergenic</td><td>NA</td><td>NA</td></tr>
<tr><td>T16401</td><td>CTTTAGCTGTGAGATGAACTTGG</td><td>1000</td><td></td><td></td><td></td><td></td><td></td><td></td><td></td><td></td><td></td></tr>
<tr><td>Oligo fwd</td><td>CACCCTTTAGCTGTGAGATGAACT</td><td></td><td></td><td></td><td></td><td></td><td></td><td></td><td></td><td></td><td></td></tr>
<tr><td>Oligo rev</td><td>AAACAGTTCATCTCACAGCTAAAG</td><td></td><td></td><td></td><td></td><td></td><td></td><td></td><td></td><td></td><td></td></tr>
<tr><td>Chromosome</td><td>start</td><td>end</td><td>strand</td><td>MM</td><td>target_seq</td><td>PAM</td><td>alignment</td><td>distance</td><td>position</td><td>gene name</td><td>gene id</td></tr>
<tr><td>GU305914.1</td><td>220538</td><td>220560</td><td>+</td><td>0</td><td>CTTTAGCTGTGAGATGAACT</td><td>TGG</td><td>||||||||[||||||||||||]PAM</td><td>NA</td><td>Intergenic</td><td>NA</td><td>NA</td></tr>
<tr><td>T16402</td><td>CCGCGTTGTTAATTTGTATCAGG</td><td>1000</td><td></td><td></td><td></td><td></td><td></td><td></td><td></td><td></td><td></td></tr>
<tr><td>Oligo fwd</td><td>CACCCCGCGTTGTTAATTTGTATC</td><td></td><td></td><td></td><td></td><td></td><td></td><td></td><td></td><td></td><td></td></tr>
<tr><td>Oligo rev</td><td>AAACGATACAAATTAACAACGCGG</td><td></td><td></td><td></td><td></td><td></td><td></td><td></td><td></td><td></td><td></td></tr>
<tr><td>Chromosome</td><td>start</td><td>end</td><td>strand</td><td>MM</td><td>target_seq</td><td>PAM</td><td>alignment</td><td>distance</td><td>position</td><td>gene name</td><td>gene id</td></tr>
<tr><td>GU305914.1</td><td>220565</td><td>220587</td><td>+</td><td>0</td><td>CCGCGTTGTTAATTTGTATC</td><td>AGG</td><td>||||||||[||||||||||||]PAM</td><td>NA</td><td>Intergenic</td><td>NA</td><td>NA</td></tr>
<tr><td>T16403</td><td>GAGCTTCGTTATGAGACGCATGG</td><td>1000</td><td></td><td></td><td></td><td></td><td></td><td></td><td></td><td></td><td></td></tr>
<tr><td>Oligo fwd</td><td>CACCGAGCTTCGTTATGAGACGCA</td><td></td><td></td><td></td><td></td><td></td><td></td><td></td><td></td><td></td><td></td></tr>
<tr><td>Oligo rev</td><td>AAACTGCGTCTCATAACGAAGCTC</td><td></td><td></td><td></td><td></td><td></td><td></td><td></td><td></td><td></td><td></td></tr>
<tr><td>Chromosome</td><td>start</td><td>end</td><td>strand</td><td>MM</td><td>target_seq</td><td>PAM</td><td>alignment</td><td>distance</td><td>position</td><td>gene name</td><td>gene id</td></tr>
<tr><td>GU305914.1</td><td>220605</td><td>220627</td><td>+</td><td>0</td><td>GAGCTTCGTTATGAGACGCA</td><td>TGG</td><td>||||||||[||||||||||||]PAM</td><td>NA</td><td>Intergenic</td><td>NA</td><td>NA</td></tr>
<tr><td>T16404</td><td>CGATCGTCGTTTGATGTTATCGG</td><td>1000</td><td></td><td></td><td></td><td></td><td></td><td></td><td></td><td></td><td></td></tr>
<tr><td>Oligo fwd</td><td>CACCCGATCGTCGTTTGATGTTAT</td><td></td><td></td><td></td><td></td><td></td><td></td><td></td><td></td><td></td><td></td></tr>
<tr><td>Oligo rev</td><td>AAACATAACATCAAACGACGATCG</td><td></td><td></td><td></td><td></td><td></td><td></td><td></td><td></td><td></td><td></td></tr>
<tr><td>Chromosome</td><td>start</td><td>end</td><td>strand</td><td>MM</td><td>target_seq</td><td>PAM</td><td>alignment</td><td>distance</td><td>position</td><td>gene name</td><td>gene id</td></tr>
<tr><td>GU305914.1</td><td>220634</td><td>220656</td><td>+</td><td>0</td><td>CGATCGTCGTTTGATGTTAT</td><td>CGG</td><td>||||||||[||||||||||||]PAM</td><td>NA</td><td>Intergenic</td><td>NA</td><td>NA</td></tr>
</table>

<table>
<tr><td>T16405</td><td>GATCGTCGTTTGATGTTATCGGG</td><td>1000</td><td colspan="9"></td></tr>
<tr><td>Oligo fwd</td><td>CACCGATCGTCGTTTGATGTTATC</td><td colspan="10"></td></tr>
<tr><td>Oligo rev</td><td>AAACGATAACATCAAACGACGATC</td><td colspan="10"></td></tr>
<tr><td>Chromosome</td><td>start</td><td>end</td><td>strand</td><td>MM</td><td>target_seq</td><td>PAM</td><td>alignment</td><td>distance</td><td>position</td><td>gene name</td><td>gene id</td></tr>
<tr><td>GU305914.1</td><td>220635</td><td>220657</td><td>+</td><td>0</td><td>GATCGTCGTTTGATGTTATC</td><td>GGG</td><td>||||||||[||||||||||||]PAM</td><td>NA</td><td>Intergenic</td><td>NA</td><td>NA</td></tr>
<tr><td>T16406</td><td>ATCGTCGTTTGATGTTATCGGGG</td><td>1000</td><td colspan="9"></td></tr>
<tr><td>Oligo fwd</td><td>CACCATCGTCGTTTGATGTTATCG</td><td colspan="10"></td></tr>
<tr><td>Oligo rev</td><td>AAACCGATAACATCAAACGACGAT</td><td colspan="10"></td></tr>
<tr><td>Chromosome</td><td>start</td><td>end</td><td>strand</td><td>MM</td><td>target_seq</td><td>PAM</td><td>alignment</td><td>distance</td><td>position</td><td>gene name</td><td>gene id</td></tr>
<tr><td>GU305914.1</td><td>220636</td><td>220658</td><td>+</td><td>0</td><td>ATCGTCGTTTGATGTTATCG</td><td>GGG</td><td>||||||||[||||||||||||]PAM</td><td>NA</td><td>Intergenic</td><td>NA</td><td>NA</td></tr>
<tr><td>T16407</td><td>GCTGATGACAAAAAGAACCACGG</td><td>1000</td><td colspan="9"></td></tr>
<tr><td>Oligo fwd</td><td>CACCGCTGATGACAAAAAGAACCA</td><td colspan="10"></td></tr>
<tr><td>Oligo rev</td><td>AAACTGGTTCTTTTTGTCATCAGC</td><td colspan="10"></td></tr>
<tr><td>Chromosome</td><td>start</td><td>end</td><td>strand</td><td>MM</td><td>target_seq</td><td>PAM</td><td>alignment</td><td>distance</td><td>position</td><td>gene name</td><td>gene id</td></tr>
<tr><td>GU305914.1</td><td>220705</td><td>220727</td><td>+</td><td>0</td><td>GCTGATGACAAAAAGAACCA</td><td>CGG</td><td>||||||||[||||||||||||]PAM</td><td>NA</td><td>Intergenic</td><td>NA</td><td>NA</td></tr>
<tr><td>T16408</td><td>CGACGAGATTTTCCAATCTCTGG</td><td>1000</td><td colspan="9"></td></tr>
<tr><td>Oligo fwd</td><td>CACCCGACGAGATTTTCCAATCTC</td><td colspan="10"></td></tr>
<tr><td>Oligo rev</td><td>AAACGAGATTGGAAAATCTCGTCG</td><td colspan="10"></td></tr>
<tr><td>Chromosome</td><td>start</td><td>end</td><td>strand</td><td>MM</td><td>target_seq</td><td>PAM</td><td>alignment</td><td>distance</td><td>position</td><td>gene name</td><td>gene id</td></tr>
<tr><td>GU305914.1</td><td>220729</td><td>220751</td><td>+</td><td>0</td><td>CGACGAGATTTTCCAATCTC</td><td>TGG</td><td>||||||||[||||||||||||]PAM</td><td>NA</td><td>Intergenic</td><td>NA</td><td>NA</td></tr>
<tr><td>T16409</td><td>GACGAGATTTTCCAATCTCTGGG</td><td>1000</td><td colspan="9"></td></tr>
<tr><td>Oligo fwd</td><td>CACCGACGAGATTTTCCAATCTCT</td><td colspan="10"></td></tr>
<tr><td>Oligo rev</td><td>AAACAGAGATTGGAAAATCTCGTC</td><td colspan="10"></td></tr>
<tr><td>Chromosome</td><td>start</td><td>end</td><td>strand</td><td>MM</td><td>target_seq</td><td>PAM</td><td>alignment</td><td>distance</td><td>position</td><td>gene name</td><td>gene id</td></tr>
<tr><td>GU305914.1</td><td>220730</td><td>220752</td><td>+</td><td>0</td><td>GACGAGATTTTCCAATCTCT</td><td>GGG</td><td>||||||||[||||||||||||]PAM</td><td>NA</td><td>Intergenic</td><td>NA</td><td>NA</td></tr>
<tr><td>T16410</td><td>TTCCAATCTCTGGGATCAGCTGG</td><td>1000</td><td colspan="9"></td></tr>
<tr><td>Oligo fwd</td><td>CACCTTCCAATCTCTGGGATCAGC</td><td colspan="10"></td></tr>
<tr><td>Oligo rev</td><td>AAACGCTGATCCCAGAGATTGGAA</td><td colspan="10"></td></tr>
<tr><td>Chromosome</td><td>start</td><td>end</td><td>strand</td><td>MM</td><td>target_seq</td><td>PAM</td><td>alignment</td><td>distance</td><td>position</td><td>gene name</td><td>gene id</td></tr>
<tr><td>GU305914.1</td><td>220739</td><td>220761</td><td>+</td><td>0</td><td>TTCCAATCTCTGGGATCAGC</td><td>TGG</td><td>||||||||[||||||||||||]PAM</td><td>NA</td><td>Intergenic</td><td>NA</td><td>NA</td></tr>
<tr><td>T16411</td><td>TGGCACGTAAACGTCAAAGACGG</td><td>1000</td><td colspan="9"></td></tr>
<tr><td>Oligo fwd</td><td>CACCTGGCACGTAAACGTCAAAGA</td><td colspan="10"></td></tr>
<tr><td>Oligo rev</td><td>AAACTCTTTGACGTTTACGTGCCA</td><td colspan="10"></td></tr>
<tr><td>Chromosome</td><td>start</td><td>end</td><td>strand</td><td>MM</td><td>target_seq</td><td>PAM</td><td>alignment</td><td>distance</td><td>position</td><td>gene name</td><td>gene id</td></tr>
<tr><td>GU305914.1</td><td>220759</td><td>220781</td><td>+</td><td>0</td><td>TGGCACGTAAACGTCAAAGA</td><td>CGG</td><td>||||||||[||||||||||||]PAM</td><td>NA</td><td>Intergenic</td><td>NA</td><td>NA</td></tr>
<tr><td>T16412</td><td>GGCACGTAAACGTCAAAGACGGG</td><td>1000</td><td colspan="9"></td></tr>
<tr><td>Oligo fwd</td><td>CACCGGCACGTAAACGTCAAAGAC</td><td colspan="10"></td></tr>
<tr><td>Oligo rev</td><td>AAACGTCTTTGACGTTTACGTGCC</td><td colspan="10"></td></tr>
<tr><td>Chromosome</td><td>start</td><td>end</td><td>strand</td><td>MM</td><td>target_seq</td><td>PAM</td><td>alignment</td><td>distance</td><td>position</td><td>gene name</td><td>gene id</td></tr>
<tr><td>GU305914.1</td><td>220760</td><td>220782</td><td>+</td><td>0</td><td>GGCACGTAAACGTCAAAGAC</td><td>GGG</td><td>||||||||[||||||||||||]PAM</td><td>NA</td><td>Intergenic</td><td>NA</td><td>NA</td></tr>
<tr><td>T16413</td><td>CAAAGACGGGATCGTCGAAACGG</td><td>1000</td><td colspan="9"></td></tr>
<tr><td>Oligo fwd</td><td>CACCCAAAGACGGGATCGTCGAAA</td><td colspan="10"></td></tr>
<tr><td>Oligo rev</td><td>AAACTTTCGACGATCCCGTCTTTG</td><td colspan="10"></td></tr>
<tr><td>Chromosome</td><td>start</td><td>end</td><td>strand</td><td>MM</td><td>target_seq</td><td>PAM</td><td>alignment</td><td>distance</td><td>position</td><td>gene name</td><td>gene id</td></tr>
<tr><td>GU305914.1</td><td>220773</td><td>220795</td><td>+</td><td>0</td><td>CAAAGACGGGATCGTCGAAA</td><td>CGG</td><td>||||||||[||||||||||||]PAM</td><td>NA</td><td>Intergenic</td><td>NA</td><td>NA</td></tr>
<tr><td>T16414</td><td>GGGATCGTCGAAACGGATCTCGG</td><td>1000</td><td colspan="9"></td></tr>
<tr><td>Oligo fwd</td><td>CACCGGGATCGTCGAAACGGATCT</td><td colspan="10"></td></tr>
<tr><td>Oligo rev</td><td>AAACAGATCCGTTTCGACGATCCC</td><td colspan="10"></td></tr>
<tr><td>Chromosome</td><td>start</td><td>end</td><td>strand</td><td>MM</td><td>target_seq</td><td>PAM</td><td>alignment</td><td>distance</td><td>position</td><td>gene name</td><td>gene id</td></tr>
<tr><td>GU305914.1</td><td>220780</td><td>220802</td><td>+</td><td>0</td><td>GGGATCGTCGAAACGGATCT</td><td>CGG</td><td>||||||||[||||||||||||]PAM</td><td>NA</td><td>Intergenic</td><td>NA</td><td>NA</td></tr>
<tr><td>T16415</td><td>GGATCGTCGAAACGGATCTCGGG</td><td>1000</td><td colspan="9"></td></tr>
<tr><td>Oligo fwd</td><td>CACCGGATCGTCGAAACGGATCTC</td><td colspan="10"></td></tr>
<tr><td>Oligo rev</td><td>AAACGAGATCCGTTTCGACGATCC</td><td colspan="10"></td></tr>
<tr><td>Chromosome</td><td>start</td><td>end</td><td>strand</td><td>MM</td><td>target_seq</td><td>PAM</td><td>alignment</td><td>distance</td><td>position</td><td>gene name</td><td>gene id</td></tr>
<tr><td>GU305914.1</td><td>220781</td><td>220803</td><td>+</td><td>0</td><td>GGATCGTCGAAACGGATCTC</td><td>GGG</td><td>||||||||[||||||||||||]PAM</td><td>NA</td><td>Intergenic</td><td>NA</td><td>NA</td></tr>
<tr><td>T16416</td><td>GAAACGGATCTCGGGAGAGACGG</td><td>1000</td><td colspan="9"></td></tr>
<tr><td>Oligo fwd</td><td>CACCGAAACGGATCTCGGGAGAGA</td><td colspan="10"></td></tr>
<tr><td>Oligo rev</td><td>AAACTCTCTCCCGAGATCCGTTTC</td><td colspan="10"></td></tr>
<tr><td>Chromosome</td><td>start</td><td>end</td><td>strand</td><td>MM</td><td>target_seq</td><td>PAM</td><td>alignment</td><td>distance</td><td>position</td><td>gene name</td><td>gene id</td></tr>
<tr><td>GU305914.1</td><td>220789</td><td>220811</td><td>+</td><td>0</td><td>GAAACGGATCTCGGGAGAGA</td><td>CGG</td><td>||||||||[||||||||||||]PAM</td><td>NA</td><td>Intergenic</td><td>NA</td><td>NA</td></tr>
<tr><td>T16417</td><td>ATCTACACGTACAGTGCGTGCGG</td><td>1000</td><td colspan="9"></td></tr>
<tr><td>Oligo fwd</td><td>CACCATCTACACGTACAGTGCGTG</td><td colspan="10"></td></tr>
<tr><td>Oligo rev</td><td>AAACCACGCACTGTACGTGTAGAT</td><td colspan="10"></td></tr>
<tr><td>Chromosome</td><td>start</td><td>end</td><td>strand</td><td>MM</td><td>target_seq</td><td>PAM</td><td>alignment</td><td>distance</td><td>position</td><td>gene name</td><td>gene id</td></tr>
<tr><td>GU305914.1</td><td>220827</td><td>220849</td><td>+</td><td>0</td><td>ATCTACACGTACAGTGCGTG</td><td>CGG</td><td>||||||||[||||||||||||]PAM</td><td>NA</td><td>Intergenic</td><td>NA</td><td>NA</td></tr>
<tr><td>T16418</td><td>AACCGCGAGAAAAGTAGAGCCGG</td><td>1000</td><td colspan="9"></td></tr>
<tr><td>Oligo fwd</td><td>CACCAACCGCGAGAAAAGTAGAGC</td><td colspan="10"></td></tr>
<tr><td>Oligo rev</td><td>AAACGCTCTACTTTTCTCGCGGTT</td><td colspan="10"></td></tr>
<tr><td>Chromosome</td><td>start</td><td>end</td><td>strand</td><td>MM</td><td>target_seq</td><td>PAM</td><td>alignment</td><td>distance</td><td>position</td><td>gene name</td><td>gene id</td></tr>
<tr><td>GU305914.1</td><td>220866</td><td>220888</td><td>+</td><td>0</td><td>AACCGCGAGAAAAGTAGAGC</td><td>CGG</td><td>||||||||[||||||||||||]PAM</td><td>NA</td><td>Intergenic</td><td>NA</td><td>NA</td></tr>
<tr><td>T16419</td><td>GTATCTTGCACTATGCCGTCCGG</td><td>1000</td><td colspan="9"></td></tr>
<tr><td>Oligo fwd</td><td>CACCGTATCTTGCACTATGCCGTC</td><td colspan="10"></td></tr>
<tr><td>Oligo rev</td><td>AAACGACGGCATAGTGCAAGATAC</td><td colspan="10"></td></tr>
<tr><td>Chromosome</td><td>start</td><td>end</td><td>strand</td><td>MM</td><td>target_seq</td><td>PAM</td><td>alignment</td><td>distance</td><td>position</td><td>gene name</td><td>gene id</td></tr>
<tr><td>GU305914.1</td><td>220937</td><td>220959</td><td>+</td><td>0</td><td>GTATCTTGCACTATGCCGTC</td><td>CGG</td><td>||||||||[||||||||||||]PAM</td><td>NA</td><td>Intergenic</td><td>NA</td><td>NA</td></tr>
</table>

<table>
<tr><td>T16420</td><td>AGAGTCCACATACCAAGAGATGG</td><td>1000</td><td></td><td></td><td></td><td></td><td></td><td></td><td></td><td></td><td></td></tr>
<tr><td>Oligo fwd</td><td>CACCAGAGTCCACATACCAAGAGA</td><td></td><td></td><td></td><td></td><td></td><td></td><td></td><td></td><td></td><td></td></tr>
<tr><td>Oligo rev</td><td>AAACTCTCTTGGTATGTGGACTCT</td><td></td><td></td><td></td><td></td><td></td><td></td><td></td><td></td><td></td><td></td></tr>
<tr><td>Chromosome</td><td>start</td><td>end</td><td>strand</td><td>MM</td><td>target_seq</td><td>PAM</td><td>alignment</td><td>distance</td><td>position</td><td>gene name</td><td>gene id</td></tr>
<tr><td>GU305914.1</td><td>220987</td><td>221009</td><td>+</td><td>0</td><td>AGAGTCCACATACCAAGAGA</td><td>TGG</td><td>||||||||[||||||||||||]PAM</td><td>NA</td><td>Intergenic</td><td>NA</td><td>NA</td></tr>
<tr><td>T16421</td><td>ACCAAGAGATGGTATGATCTAGG</td><td>1000</td><td></td><td></td><td></td><td></td><td></td><td></td><td></td><td></td><td></td></tr>
<tr><td>Oligo fwd</td><td>CACCACCAAGAGATGGTATGATCT</td><td></td><td></td><td></td><td></td><td></td><td></td><td></td><td></td><td></td><td></td></tr>
<tr><td>Oligo rev</td><td>AAACAGATCATACCATCTCTTGGT</td><td></td><td></td><td></td><td></td><td></td><td></td><td></td><td></td><td></td><td></td></tr>
<tr><td>Chromosome</td><td>start</td><td>end</td><td>strand</td><td>MM</td><td>target_seq</td><td>PAM</td><td>alignment</td><td>distance</td><td>position</td><td>gene name</td><td>gene id</td></tr>
<tr><td>GU305914.1</td><td>220998</td><td>221020</td><td>+</td><td>0</td><td>ACCAAGAGATGGTATGATCT</td><td>AGG</td><td>||||||||[||||||||||||]PAM</td><td>NA</td><td>Intergenic</td><td>NA</td><td>NA</td></tr>
<tr><td>T16422</td><td>TGATCTAGGCGATTGCCAAAAGG</td><td>1000</td><td></td><td></td><td></td><td></td><td></td><td></td><td></td><td></td><td></td></tr>
<tr><td>Oligo fwd</td><td>CACCTGATCTAGGCGATTGCCAAA</td><td></td><td></td><td></td><td></td><td></td><td></td><td></td><td></td><td></td><td></td></tr>
<tr><td>Oligo rev</td><td>AAACTTTGGCAATCGCCTAGATCA</td><td></td><td></td><td></td><td></td><td></td><td></td><td></td><td></td><td></td><td></td></tr>
<tr><td>Chromosome</td><td>start</td><td>end</td><td>strand</td><td>MM</td><td>target_seq</td><td>PAM</td><td>alignment</td><td>distance</td><td>position</td><td>gene name</td><td>gene id</td></tr>
<tr><td>GU305914.1</td><td>221012</td><td>221034</td><td>+</td><td>0</td><td>TGATCTAGGCGATTGCCAAA</td><td>AGG</td><td>||||||||[||||||||||||]PAM</td><td>NA</td><td>Intergenic</td><td>NA</td><td>NA</td></tr>
<tr><td>T16423</td><td>GGCGATTGCCAAAAGGACAATGG</td><td>1000</td><td></td><td></td><td></td><td></td><td></td><td></td><td></td><td></td><td></td></tr>
<tr><td>Oligo fwd</td><td>CACCGGCGATTGCCAAAAGGACAA</td><td></td><td></td><td></td><td></td><td></td><td></td><td></td><td></td><td></td><td></td></tr>
<tr><td>Oligo rev</td><td>AAACTTGTCCTTTTGGCAATCGCC</td><td></td><td></td><td></td><td></td><td></td><td></td><td></td><td></td><td></td><td></td></tr>
<tr><td>Chromosome</td><td>start</td><td>end</td><td>strand</td><td>MM</td><td>target_seq</td><td>PAM</td><td>alignment</td><td>distance</td><td>position</td><td>gene name</td><td>gene id</td></tr>
<tr><td>GU305914.1</td><td>221019</td><td>221041</td><td>+</td><td>0</td><td>GGCGATTGCCAAAAGGACAA</td><td>TGG</td><td>||||||||[||||||||||||]PAM</td><td>NA</td><td>Intergenic</td><td>NA</td><td>NA</td></tr>
<tr><td>T16424</td><td>ATTGCCAAAAGGACAATGGCAGG</td><td>1000</td><td></td><td></td><td></td><td></td><td></td><td></td><td></td><td></td><td></td></tr>
<tr><td>Oligo fwd</td><td>CACCATTGCCAAAAGGACAATGGC</td><td></td><td></td><td></td><td></td><td></td><td></td><td></td><td></td><td></td><td></td></tr>
<tr><td>Oligo rev</td><td>AAACGCCATTGTCCTTTTGGCAAT</td><td></td><td></td><td></td><td></td><td></td><td></td><td></td><td></td><td></td><td></td></tr>
<tr><td>Chromosome</td><td>start</td><td>end</td><td>strand</td><td>MM</td><td>target_seq</td><td>PAM</td><td>alignment</td><td>distance</td><td>position</td><td>gene name</td><td>gene id</td></tr>
<tr><td>GU305914.1</td><td>221023</td><td>221045</td><td>+</td><td>0</td><td>ATTGCCAAAAGGACAATGGC</td><td>AGG</td><td>||||||||[||||||||||||]PAM</td><td>NA</td><td>Intergenic</td><td>NA</td><td>NA</td></tr>
<tr><td>T16425</td><td>ACTAGCGATGATGACGCGTGCGG</td><td>1000</td><td></td><td></td><td></td><td></td><td></td><td></td><td></td><td></td><td></td></tr>
<tr><td>Oligo fwd</td><td>CACCACTAGCGATGATGACGCGTG</td><td></td><td></td><td></td><td></td><td></td><td></td><td></td><td></td><td></td><td></td></tr>
<tr><td>Oligo rev</td><td>AAACCACGCGTCATCATCGCTAGT</td><td></td><td></td><td></td><td></td><td></td><td></td><td></td><td></td><td></td><td></td></tr>
<tr><td>Chromosome</td><td>start</td><td>end</td><td>strand</td><td>MM</td><td>target_seq</td><td>PAM</td><td>alignment</td><td>distance</td><td>position</td><td>gene name</td><td>gene id</td></tr>
<tr><td>GU305914.1</td><td>221074</td><td>221096</td><td>+</td><td>0</td><td>ACTAGCGATGATGACGCGTG</td><td>CGG</td><td>||||||||[||||||||||||]PAM</td><td>NA</td><td>Intergenic</td><td>NA</td><td>NA</td></tr>
<tr><td>T16426</td><td>GATGATGACGCGTGCGGTTAAGG</td><td>1000</td><td></td><td></td><td></td><td></td><td></td><td></td><td></td><td></td><td></td></tr>
<tr><td>Oligo fwd</td><td>CACCGATGATGACGCGTGCGGTTA</td><td></td><td></td><td></td><td></td><td></td><td></td><td></td><td></td><td></td><td></td></tr>
<tr><td>Oligo rev</td><td>AAACTAACCGCACGCGTCATCATC</td><td></td><td></td><td></td><td></td><td></td><td></td><td></td><td></td><td></td><td></td></tr>
<tr><td>Chromosome</td><td>start</td><td>end</td><td>strand</td><td>MM</td><td>target_seq</td><td>PAM</td><td>alignment</td><td>distance</td><td>position</td><td>gene name</td><td>gene id</td></tr>
<tr><td>GU305914.1</td><td>221080</td><td>221102</td><td>+</td><td>0</td><td>GATGATGACGCGTGCGGTTA</td><td>AGG</td><td>||||||||[||||||||||||]PAM</td><td>NA</td><td>Intergenic</td><td>NA</td><td>NA</td></tr>
<tr><td>T16427</td><td>CGTGCGGTTAAGGAAAACCCAGG</td><td>1000</td><td></td><td></td><td></td><td></td><td></td><td></td><td></td><td></td><td></td></tr>
<tr><td>Oligo fwd</td><td>CACCCGTGCGGTTAAGGAAAACCC</td><td></td><td></td><td></td><td></td><td></td><td></td><td></td><td></td><td></td><td></td></tr>
<tr><td>Oligo rev</td><td>AAACGGGTTTTCCTTAACCGCACG</td><td></td><td></td><td></td><td></td><td></td><td></td><td></td><td></td><td></td><td></td></tr>
<tr><td>Chromosome</td><td>start</td><td>end</td><td>strand</td><td>MM</td><td>target_seq</td><td>PAM</td><td>alignment</td><td>distance</td><td>position</td><td>gene name</td><td>gene id</td></tr>
<tr><td>GU305914.1</td><td>221090</td><td>221112</td><td>+</td><td>0</td><td>CGTGCGGTTAAGGAAAACCC</td><td>AGG</td><td>||||||||[||||||||||||]PAM</td><td>NA</td><td>Intergenic</td><td>NA</td><td>NA</td></tr>
<tr><td>T16428</td><td>AACCCAGGCTCCGACCCCGACGG</td><td>1000</td><td></td><td></td><td></td><td></td><td></td><td></td><td></td><td></td><td></td></tr>
<tr><td>Oligo fwd</td><td>CACCAACCCAGGCTCCGACCCCGA</td><td></td><td></td><td></td><td></td><td></td><td></td><td></td><td></td><td></td><td></td></tr>
<tr><td>Oligo rev</td><td>AAACTCGGGGTCGGAGCCTGGGTT</td><td></td><td></td><td></td><td></td><td></td><td></td><td></td><td></td><td></td><td></td></tr>
<tr><td>Chromosome</td><td>start</td><td>end</td><td>strand</td><td>MM</td><td>target_seq</td><td>PAM</td><td>alignment</td><td>distance</td><td>position</td><td>gene name</td><td>gene id</td></tr>
<tr><td>GU305914.1</td><td>221105</td><td>221127</td><td>+</td><td>0</td><td>AACCCAGGCTCCGACCCCGA</td><td>CGG</td><td>||||||||[||||||||||||]PAM</td><td>NA</td><td>Intergenic</td><td>NA</td><td>NA</td></tr>
<tr><td>T16429</td><td>AGACTGTTCTATACATCTTAAGG</td><td>1000</td><td></td><td></td><td></td><td></td><td></td><td></td><td></td><td></td><td></td></tr>
<tr><td>Oligo fwd</td><td>CACCAGACTGTTCTATACATCTTA</td><td></td><td></td><td></td><td></td><td></td><td></td><td></td><td></td><td></td><td></td></tr>
<tr><td>Oligo rev</td><td>AAACTAAGATGTATAGAACAGTCT</td><td></td><td></td><td></td><td></td><td></td><td></td><td></td><td></td><td></td><td></td></tr>
<tr><td>Chromosome</td><td>start</td><td>end</td><td>strand</td><td>MM</td><td>target_seq</td><td>PAM</td><td>alignment</td><td>distance</td><td>position</td><td>gene name</td><td>gene id</td></tr>
<tr><td>GU305914.1</td><td>221128</td><td>221150</td><td>+</td><td>0</td><td>AGACTGTTCTATACATCTTA</td><td>AGG</td><td>||||||||[||||||||||||]PAM</td><td>NA</td><td>Intergenic</td><td>NA</td><td>NA</td></tr>
<tr><td>T16430</td><td>TTAAGGACATGTCGTCGCTCCGG</td><td>1000</td><td></td><td></td><td></td><td></td><td></td><td></td><td></td><td></td><td></td></tr>
<tr><td>Oligo fwd</td><td>CACCTTAAGGACATGTCGTCGCTC</td><td></td><td></td><td></td><td></td><td></td><td></td><td></td><td></td><td></td><td></td></tr>
<tr><td>Oligo rev</td><td>AAACGAGCGACGACATGTCCTTAA</td><td></td><td></td><td></td><td></td><td></td><td></td><td></td><td></td><td></td><td></td></tr>
<tr><td>Chromosome</td><td>start</td><td>end</td><td>strand</td><td>MM</td><td>target_seq</td><td>PAM</td><td>alignment</td><td>distance</td><td>position</td><td>gene name</td><td>gene id</td></tr>
<tr><td>GU305914.1</td><td>221145</td><td>221167</td><td>+</td><td>0</td><td>TTAAGGACATGTCGTCGCTC</td><td>CGG</td><td>||||||||[||||||||||||]PAM</td><td>NA</td><td>Intergenic</td><td>NA</td><td>NA</td></tr>
<tr><td>T16431</td><td>TAAGGACATGTCGTCGCTCCGGG</td><td>1000</td><td></td><td></td><td></td><td></td><td></td><td></td><td></td><td></td><td></td></tr>
<tr><td>Oligo fwd</td><td>CACCTAAGGACATGTCGTCGCTCC</td><td></td><td></td><td></td><td></td><td></td><td></td><td></td><td></td><td></td><td></td></tr>
<tr><td>Oligo rev</td><td>AAACGGAGCGACGACATGTCCTTA</td><td></td><td></td><td></td><td></td><td></td><td></td><td></td><td></td><td></td><td></td></tr>
<tr><td>Chromosome</td><td>start</td><td>end</td><td>strand</td><td>MM</td><td>target_seq</td><td>PAM</td><td>alignment</td><td>distance</td><td>position</td><td>gene name</td><td>gene id</td></tr>
<tr><td>GU305914.1</td><td>221146</td><td>221168</td><td>+</td><td>0</td><td>TAAGGACATGTCGTCGCTCC</td><td>GGG</td><td>||||||||[||||||||||||]PAM</td><td>NA</td><td>Intergenic</td><td>NA</td><td>NA</td></tr>
<tr><td>T16432</td><td>TGTATGCATGAAAAGCATAAAGG</td><td>1000</td><td></td><td></td><td></td><td></td><td></td><td></td><td></td><td></td><td></td></tr>
<tr><td>Oligo fwd</td><td>CACCTGTATGCATGAAAAGCATAA</td><td></td><td></td><td></td><td></td><td></td><td></td><td></td><td></td><td></td><td></td></tr>
<tr><td>Oligo rev</td><td>AAACTTATGCTTTTCATGCATACA</td><td></td><td></td><td></td><td></td><td></td><td></td><td></td><td></td><td></td><td></td></tr>
<tr><td>Chromosome</td><td>start</td><td>end</td><td>strand</td><td>MM</td><td>target_seq</td><td>PAM</td><td>alignment</td><td>distance</td><td>position</td><td>gene name</td><td>gene id</td></tr>
<tr><td>GU305914.1</td><td>221188</td><td>221210</td><td>+</td><td>0</td><td>TGTATGCATGAAAAGCATAA</td><td>AGG</td><td>||||||||[||||||||||||]PAM</td><td>NA</td><td>Intergenic</td><td>NA</td><td>NA</td></tr>
<tr><td>T16433</td><td>CGATGATGTGTATCATCGTGAGG</td><td>1000</td><td></td><td></td><td></td><td></td><td></td><td></td><td></td><td></td><td></td></tr>
<tr><td>Oligo fwd</td><td>CACCCGATGATGTGTATCATCGTG</td><td></td><td></td><td></td><td></td><td></td><td></td><td></td><td></td><td></td><td></td></tr>
<tr><td>Oligo rev</td><td>AAACCACGATGATACACATCATCG</td><td></td><td></td><td></td><td></td><td></td><td></td><td></td><td></td><td></td><td></td></tr>
<tr><td>Chromosome</td><td>start</td><td>end</td><td>strand</td><td>MM</td><td>target_seq</td><td>PAM</td><td>alignment</td><td>distance</td><td>position</td><td>gene name</td><td>gene id</td></tr>
<tr><td>GU305914.1</td><td>221211</td><td>221233</td><td>+</td><td>0</td><td>CGATGATGTGTATCATCGTG</td><td>AGG</td><td>||||||||[||||||||||||]PAM</td><td>NA</td><td>Intergenic</td><td>NA</td><td>NA</td></tr>
<tr><td>T16434</td><td>GTGTATCATCGTGAGGTGATAGG</td><td>1000</td><td></td><td></td><td></td><td></td><td></td><td></td><td></td><td></td><td></td></tr>
<tr><td>Oligo fwd</td><td>CACCGTGTATCATCGTGAGGTGAT</td><td></td><td></td><td></td><td></td><td></td><td></td><td></td><td></td><td></td><td></td></tr>
<tr><td>Oligo rev</td><td>AAACATCACCTCACGATGATACAC</td><td></td><td></td><td></td><td></td><td></td><td></td><td></td><td></td><td></td><td></td></tr>
<tr><td>Chromosome</td><td>start</td><td>end</td><td>strand</td><td>MM</td><td>target_seq</td><td>PAM</td><td>alignment</td><td>distance</td><td>position</td><td>gene name</td><td>gene id</td></tr>
</table>

<table>
<tr><td>GU305914.1</td><td>221218</td><td>221240</td><td>+</td><td>0</td><td>GTGTATCATCGTGAGGTGAT</td><td>AGG</td><td>||||||||[||||||||||||]PAM</td><td>NA</td><td>Intergenic</td><td>NA</td><td>NA</td></tr>
<tr><td>T16435</td><td>TAGGAAAGCCCCATGTCTATCGG</td><td>1000</td><td></td><td></td><td></td><td></td><td></td><td></td><td></td><td></td><td></td></tr>
<tr><td>Oligo fwd</td><td>CACCTAGGAAAGCCCCATGTCTAT</td><td></td><td></td><td></td><td></td><td></td><td></td><td></td><td></td><td></td><td></td></tr>
<tr><td>Oligo rev</td><td>AAACATAGACATGGGGCTTTCCTA</td><td></td><td></td><td></td><td></td><td></td><td></td><td></td><td></td><td></td><td></td></tr>
<tr><td>Chromosome</td><td>start</td><td>end</td><td>strand</td><td>MM</td><td>target_seq</td><td>PAM</td><td>alignment</td><td>distance</td><td>position</td><td>gene name</td><td>gene id</td></tr>
<tr><td>GU305914.1</td><td>221237</td><td>221259</td><td>+</td><td>0</td><td>TAGGAAAGCCCCATGTCTAT</td><td>CGG</td><td>||||||||[||||||||||||]PAM</td><td>NA</td><td>Intergenic</td><td>NA</td><td>NA</td></tr>
<tr><td>T16436</td><td>TGTCTATCGGAAACACACTACGG</td><td>1000</td><td></td><td></td><td></td><td></td><td></td><td></td><td></td><td></td><td></td></tr>
<tr><td>Oligo fwd</td><td>CACCTGTCTATCGGAAACACACTA</td><td></td><td></td><td></td><td></td><td></td><td></td><td></td><td></td><td></td><td></td></tr>
<tr><td>Oligo rev</td><td>AAACTAGTGTGTTTCCGATAGACA</td><td></td><td></td><td></td><td></td><td></td><td></td><td></td><td></td><td></td><td></td></tr>
<tr><td>Chromosome</td><td>start</td><td>end</td><td>strand</td><td>MM</td><td>target_seq</td><td>PAM</td><td>alignment</td><td>distance</td><td>position</td><td>gene name</td><td>gene id</td></tr>
<tr><td>GU305914.1</td><td>221250</td><td>221272</td><td>+</td><td>0</td><td>TGTCTATCGGAAACACACTA</td><td>CGG</td><td>||||||||[||||||||||||]PAM</td><td>NA</td><td>Intergenic</td><td>NA</td><td>NA</td></tr>
<tr><td>T16437</td><td>AACACACTACGGTTCAAGAGAGG</td><td>1000</td><td></td><td></td><td></td><td></td><td></td><td></td><td></td><td></td><td></td></tr>
<tr><td>Oligo fwd</td><td>CACCAACACACTACGGTTCAAGAG</td><td></td><td></td><td></td><td></td><td></td><td></td><td></td><td></td><td></td><td></td></tr>
<tr><td>Oligo rev</td><td>AAACCTCTTGAACCGTAGTGTGTT</td><td></td><td></td><td></td><td></td><td></td><td></td><td></td><td></td><td></td><td></td></tr>
<tr><td>Chromosome</td><td>start</td><td>end</td><td>strand</td><td>MM</td><td>target_seq</td><td>PAM</td><td>alignment</td><td>distance</td><td>position</td><td>gene name</td><td>gene id</td></tr>
<tr><td>GU305914.1</td><td>221261</td><td>221283</td><td>+</td><td>0</td><td>AACACACTACGGTTCAAGAG</td><td>AGG</td><td>||||||||[||||||||||||]PAM</td><td>NA</td><td>Intergenic</td><td>NA</td><td>NA</td></tr>
<tr><td>T16438</td><td>ACACACTACGGTTCAAGAGAGGG</td><td>1000</td><td></td><td></td><td></td><td></td><td></td><td></td><td></td><td></td><td></td></tr>
<tr><td>Oligo fwd</td><td>CACCACACACTACGGTTCAAGAGA</td><td></td><td></td><td></td><td></td><td></td><td></td><td></td><td></td><td></td><td></td></tr>
<tr><td>Oligo rev</td><td>AAACTCTCTTGAACCGTAGTGTGT</td><td></td><td></td><td></td><td></td><td></td><td></td><td></td><td></td><td></td><td></td></tr>
<tr><td>Chromosome</td><td>start</td><td>end</td><td>strand</td><td>MM</td><td>target_seq</td><td>PAM</td><td>alignment</td><td>distance</td><td>position</td><td>gene name</td><td>gene id</td></tr>
<tr><td>GU305914.1</td><td>221262</td><td>221284</td><td>+</td><td>0</td><td>ACACACTACGGTTCAAGAGA</td><td>GGG</td><td>||||||||[||||||||||||]PAM</td><td>NA</td><td>Intergenic</td><td>NA</td><td>NA</td></tr>
<tr><td>T16439</td><td>AGATTCCACCATCTCAGATTCGG</td><td>1000</td><td></td><td></td><td></td><td></td><td></td><td></td><td></td><td></td><td></td></tr>
<tr><td>Oligo fwd</td><td>CACCAGATTCCACCATCTCAGATT</td><td></td><td></td><td></td><td></td><td></td><td></td><td></td><td></td><td></td><td></td></tr>
<tr><td>Oligo rev</td><td>AAACAATCTGAGATGGTGGAATCT</td><td></td><td></td><td></td><td></td><td></td><td></td><td></td><td></td><td></td><td></td></tr>
<tr><td>Chromosome</td><td>start</td><td>end</td><td>strand</td><td>MM</td><td>target_seq</td><td>PAM</td><td>alignment</td><td>distance</td><td>position</td><td>gene name</td><td>gene id</td></tr>
<tr><td>GU305914.1</td><td>221296</td><td>221318</td><td>+</td><td>0</td><td>AGATTCCACCATCTCAGATT</td><td>CGG</td><td>||||||||[||||||||||||]PAM</td><td>NA</td><td>Intergenic</td><td>NA</td><td>NA</td></tr>
<tr><td>T16440</td><td>GATTCCACCATCTCAGATTCGGG</td><td>1000</td><td></td><td></td><td></td><td></td><td></td><td></td><td></td><td></td><td></td></tr>
<tr><td>Oligo fwd</td><td>CACCGATTCCACCATCTCAGATTC</td><td></td><td></td><td></td><td></td><td></td><td></td><td></td><td></td><td></td><td></td></tr>
<tr><td>Oligo rev</td><td>AAACGAATCTGAGATGGTGGAATC</td><td></td><td></td><td></td><td></td><td></td><td></td><td></td><td></td><td></td><td></td></tr>
<tr><td>Chromosome</td><td>start</td><td>end</td><td>strand</td><td>MM</td><td>target_seq</td><td>PAM</td><td>alignment</td><td>distance</td><td>position</td><td>gene name</td><td>gene id</td></tr>
<tr><td>GU305914.1</td><td>221297</td><td>221319</td><td>+</td><td>0</td><td>GATTCCACCATCTCAGATTC</td><td>GGG</td><td>||||||||[||||||||||||]PAM</td><td>NA</td><td>Intergenic</td><td>NA</td><td>NA</td></tr>
<tr><td>T16441</td><td>CACCATCTCAGATTCGGGAATGG</td><td>1000</td><td></td><td></td><td></td><td></td><td></td><td></td><td></td><td></td><td></td></tr>
<tr><td>Oligo fwd</td><td>CACCCACCATCTCAGATTCGGGAA</td><td></td><td></td><td></td><td></td><td></td><td></td><td></td><td></td><td></td><td></td></tr>
<tr><td>Oligo rev</td><td>AAACTTCCCGAATCTGAGATGGTG</td><td></td><td></td><td></td><td></td><td></td><td></td><td></td><td></td><td></td><td></td></tr>
<tr><td>Chromosome</td><td>start</td><td>end</td><td>strand</td><td>MM</td><td>target_seq</td><td>PAM</td><td>alignment</td><td>distance</td><td>position</td><td>gene name</td><td>gene id</td></tr>
<tr><td>GU305914.1</td><td>221302</td><td>221324</td><td>+</td><td>0</td><td>CACCATCTCAGATTCGGGAA</td><td>TGG</td><td>||||||||[||||||||||||]PAM</td><td>NA</td><td>Intergenic</td><td>NA</td><td>NA</td></tr>
<tr><td>T16442</td><td>CAGATTCGGGAATGGTGCGAAGG</td><td>1000</td><td></td><td></td><td></td><td></td><td></td><td></td><td></td><td></td><td></td></tr>
<tr><td>Oligo fwd</td><td>CACCCAGATTCGGGAATGGTGCGA</td><td></td><td></td><td></td><td></td><td></td><td></td><td></td><td></td><td></td><td></td></tr>
<tr><td>Oligo rev</td><td>AAACTCGCACCATTCCCGAATCTG</td><td></td><td></td><td></td><td></td><td></td><td></td><td></td><td></td><td></td><td></td></tr>
<tr><td>Chromosome</td><td>start</td><td>end</td><td>strand</td><td>MM</td><td>target_seq</td><td>PAM</td><td>alignment</td><td>distance</td><td>position</td><td>gene name</td><td>gene id</td></tr>
<tr><td>GU305914.1</td><td>221310</td><td>221332</td><td>+</td><td>0</td><td>CAGATTCGGGAATGGTGCGA</td><td>AGG</td><td>||||||||[||||||||||||]PAM</td><td>NA</td><td>Intergenic</td><td>NA</td><td>NA</td></tr>
<tr><td>T16443</td><td>AAGGACCGCCCGCAGTTGCAAGG</td><td>1000</td><td></td><td></td><td></td><td></td><td></td><td></td><td></td><td></td><td></td></tr>
<tr><td>Oligo fwd</td><td>CACCAAGGACCGCCCGCAGTTGCA</td><td></td><td></td><td></td><td></td><td></td><td></td><td></td><td></td><td></td><td></td></tr>
<tr><td>Oligo rev</td><td>AAACTGCAACTGCGGGCGGTCCTT</td><td></td><td></td><td></td><td></td><td></td><td></td><td></td><td></td><td></td><td></td></tr>
<tr><td>Chromosome</td><td>start</td><td>end</td><td>strand</td><td>MM</td><td>target_seq</td><td>PAM</td><td>alignment</td><td>distance</td><td>position</td><td>gene name</td><td>gene id</td></tr>
<tr><td>GU305914.1</td><td>221329</td><td>221351</td><td>+</td><td>0</td><td>AAGGACCGCCCGCAGTTGCA</td><td>AGG</td><td>||||||||[||||||||||||]PAM</td><td>NA</td><td>Intergenic</td><td>NA</td><td>NA</td></tr>
<tr><td>T16444</td><td>AGGACCGCCCGCAGTTGCAAGGG</td><td>1000</td><td></td><td></td><td></td><td></td><td></td><td></td><td></td><td></td><td></td></tr>
<tr><td>Oligo fwd</td><td>CACCAGGACCGCCCGCAGTTGCAA</td><td></td><td></td><td></td><td></td><td></td><td></td><td></td><td></td><td></td><td></td></tr>
<tr><td>Oligo rev</td><td>AAACTTGCAACTGCGGGCGGTCCT</td><td></td><td></td><td></td><td></td><td></td><td></td><td></td><td></td><td></td><td></td></tr>
<tr><td>Chromosome</td><td>start</td><td>end</td><td>strand</td><td>MM</td><td>target_seq</td><td>PAM</td><td>alignment</td><td>distance</td><td>position</td><td>gene name</td><td>gene id</td></tr>
<tr><td>GU305914.1</td><td>221330</td><td>221352</td><td>+</td><td>0</td><td>AGGACCGCCCGCAGTTGCAA</td><td>GGG</td><td>||||||||[||||||||||||]PAM</td><td>NA</td><td>Intergenic</td><td>NA</td><td>NA</td></tr>
<tr><td>T16445</td><td>GGACCGCCCGCAGTTGCAAGGGG</td><td>1000</td><td></td><td></td><td></td><td></td><td></td><td></td><td></td><td></td><td></td></tr>
<tr><td>Oligo fwd</td><td>CACCGGACCGCCCGCAGTTGCAAG</td><td></td><td></td><td></td><td></td><td></td><td></td><td></td><td></td><td></td><td></td></tr>
<tr><td>Oligo rev</td><td>AAACCTTGCAACTGCGGGCGGTCC</td><td></td><td></td><td></td><td></td><td></td><td></td><td></td><td></td><td></td><td></td></tr>
<tr><td>Chromosome</td><td>start</td><td>end</td><td>strand</td><td>MM</td><td>target_seq</td><td>PAM</td><td>alignment</td><td>distance</td><td>position</td><td>gene name</td><td>gene id</td></tr>
<tr><td>GU305914.1</td><td>221331</td><td>221353</td><td>+</td><td>0</td><td>GGACCGCCCGCAGTTGCAAG</td><td>GGG</td><td>||||||||[||||||||||||]PAM</td><td>NA</td><td>Intergenic</td><td>NA</td><td>NA</td></tr>
<tr><td>T16446</td><td>GACCGCCCGCAGTTGCAAGGGGG</td><td>1000</td><td></td><td></td><td></td><td></td><td></td><td></td><td></td><td></td><td></td></tr>
<tr><td>Oligo fwd</td><td>CACCGACCGCCCGCAGTTGCAAGG</td><td></td><td></td><td></td><td></td><td></td><td></td><td></td><td></td><td></td><td></td></tr>
<tr><td>Oligo rev</td><td>AAACCCTTGCAACTGCGGGCGGTC</td><td></td><td></td><td></td><td></td><td></td><td></td><td></td><td></td><td></td><td></td></tr>
<tr><td>Chromosome</td><td>start</td><td>end</td><td>strand</td><td>MM</td><td>target_seq</td><td>PAM</td><td>alignment</td><td>distance</td><td>position</td><td>gene name</td><td>gene id</td></tr>
<tr><td>GU305914.1</td><td>221332</td><td>221354</td><td>+</td><td>0</td><td>GACCGCCCGCAGTTGCAAGG</td><td>GGG</td><td>||||||||[||||||||||||]PAM</td><td>NA</td><td>Intergenic</td><td>NA</td><td>NA</td></tr>
<tr><td>GU305914.1</td><td>64060</td><td>64082</td><td>-</td><td>4</td><td>CACCGTCTGCAGTTGCACGG</td><td>CGG</td><td>-||||-|-[|||||||||-||]PAM</td><td>NA</td><td>Intergenic</td><td>NA</td><td>NA</td></tr>
<tr><td>T16447</td><td>CGCAGTTGCAAGGGGGTACCCGG</td><td>1000</td><td></td><td></td><td></td><td></td><td></td><td></td><td></td><td></td><td></td></tr>
<tr><td>Oligo fwd</td><td>CACCCGCAGTTGCAAGGGGGTACC</td><td></td><td></td><td></td><td></td><td></td><td></td><td></td><td></td><td></td><td></td></tr>
<tr><td>Oligo rev</td><td>AAACGGTACCCCCTTGCAACTGCG</td><td></td><td></td><td></td><td></td><td></td><td></td><td></td><td></td><td></td><td></td></tr>
<tr><td>Chromosome</td><td>start</td><td>end</td><td>strand</td><td>MM</td><td>target_seq</td><td>PAM</td><td>alignment</td><td>distance</td><td>position</td><td>gene name</td><td>gene id</td></tr>
<tr><td>GU305914.1</td><td>221339</td><td>221361</td><td>+</td><td>0</td><td>CGCAGTTGCAAGGGGGTACC</td><td>CGG</td><td>||||||||[||||||||||||]PAM</td><td>NA</td><td>Intergenic</td><td>NA</td><td>NA</td></tr>
<tr><td>T16448</td><td>GCAGTTGCAAGGGGGTACCCGGG</td><td>1000</td><td></td><td></td><td></td><td></td><td></td><td></td><td></td><td></td><td></td></tr>
<tr><td>Oligo fwd</td><td>CACCGCAGTTGCAAGGGGGTACCC</td><td></td><td></td><td></td><td></td><td></td><td></td><td></td><td></td><td></td><td></td></tr>
<tr><td>Oligo rev</td><td>AAACGGGTACCCCCTTGCAACTGC</td><td></td><td></td><td></td><td></td><td></td><td></td><td></td><td></td><td></td><td></td></tr>
<tr><td>Chromosome</td><td>start</td><td>end</td><td>strand</td><td>MM</td><td>target_seq</td><td>PAM</td><td>alignment</td><td>distance</td><td>position</td><td>gene name</td><td>gene id</td></tr>
<tr><td>GU305914.1</td><td>221340</td><td>221362</td><td>+</td><td>0</td><td>GCAGTTGCAAGGGGGTACCC</td><td>GGG</td><td>||||||||[||||||||||||]PAM</td><td>NA</td><td>Intergenic</td><td>NA</td><td>NA</td></tr>
<tr><td>T16449</td><td>TCTCAGACTGACTTCGTCTGAGG</td><td>1000</td><td></td><td></td><td></td><td></td><td></td><td></td><td></td><td></td><td></td></tr>
<tr><td>Oligo fwd</td><td>CACCTCTCAGACTGACTTCGTCTG</td><td></td><td></td><td></td><td></td><td></td><td></td><td></td><td></td><td></td><td></td></tr>
</table>

<table>
<tr><td>Oligo rev</td><td>AAACCAGACGAAGTCAGTCTGAGA</td><td></td><td></td><td></td><td></td><td></td><td></td><td></td><td></td><td></td><td></td></tr>
<tr><td>Chromosome</td><td>start</td><td>end</td><td>strand</td><td>MM</td><td>target_seq</td><td>PAM</td><td>alignment</td><td>distance</td><td>position</td><td>gene name</td><td>gene id</td></tr>
<tr><td>GU305914.1</td><td>221391</td><td>221413</td><td>+</td><td>0</td><td>TCTCAGACTGACTTCGTCTG</td><td>AGG</td><td>||||||||[||||||||||||]PAM</td><td>NA</td><td>Intergenic</td><td>NA</td><td>NA</td></tr>
<tr><td>T16450</td><td>CGTAGATAAGCGCGAACAGTAGG</td><td>1000</td><td></td><td></td><td></td><td></td><td></td><td></td><td></td><td></td><td></td></tr>
<tr><td>Oligo fwd</td><td>CACCCGTAGATAAGCGCGAACAGT</td><td></td><td></td><td></td><td></td><td></td><td></td><td></td><td></td><td></td><td></td></tr>
<tr><td>Oligo rev</td><td>AAACACTGTTCGCGCTTATCTACG</td><td></td><td></td><td></td><td></td><td></td><td></td><td></td><td></td><td></td><td></td></tr>
<tr><td>Chromosome</td><td>start</td><td>end</td><td>strand</td><td>MM</td><td>target_seq</td><td>PAM</td><td>alignment</td><td>distance</td><td>position</td><td>gene name</td><td>gene id</td></tr>
<tr><td>GU305914.1</td><td>221430</td><td>221452</td><td>+</td><td>0</td><td>CGTAGATAAGCGCGAACAGT</td><td>AGG</td><td>||||||||[||||||||||||]PAM</td><td>NA</td><td>Intergenic</td><td>NA</td><td>NA</td></tr>
<tr><td>GU305914.1</td><td>65573</td><td>65595</td><td>-</td><td>4</td><td>CGTAGCGCAGCGCGAACAGC</td><td>CGG</td><td>|||||---[|||||||||||-]PAM</td><td>NA</td><td>Intergenic</td><td>NA</td><td>NA</td></tr>
<tr><td>T16451</td><td>GATAAGCGCGAACAGTAGGCTGG</td><td>1000</td><td></td><td></td><td></td><td></td><td></td><td></td><td></td><td></td><td></td></tr>
<tr><td>Oligo fwd</td><td>CACCGATAAGCGCGAACAGTAGGC</td><td></td><td></td><td></td><td></td><td></td><td></td><td></td><td></td><td></td><td></td></tr>
<tr><td>Oligo rev</td><td>AAACGCCTACTGTTCGCGCTTATC</td><td></td><td></td><td></td><td></td><td></td><td></td><td></td><td></td><td></td><td></td></tr>
<tr><td>Chromosome</td><td>start</td><td>end</td><td>strand</td><td>MM</td><td>target_seq</td><td>PAM</td><td>alignment</td><td>distance</td><td>position</td><td>gene name</td><td>gene id</td></tr>
<tr><td>GU305914.1</td><td>221434</td><td>221456</td><td>+</td><td>0</td><td>GATAAGCGCGAACAGTAGGC</td><td>TGG</td><td>||||||||[||||||||||||]PAM</td><td>NA</td><td>Intergenic</td><td>NA</td><td>NA</td></tr>
<tr><td>T16452</td><td>CGAACAGTAGGCTGGCTGCCAGG</td><td>1000</td><td></td><td></td><td></td><td></td><td></td><td></td><td></td><td></td><td></td></tr>
<tr><td>Oligo fwd</td><td>CACCCGAACAGTAGGCTGGCTGCC</td><td></td><td></td><td></td><td></td><td></td><td></td><td></td><td></td><td></td><td></td></tr>
<tr><td>Oligo rev</td><td>AAACGGCAGCCAGCCTACTGTTCG</td><td></td><td></td><td></td><td></td><td></td><td></td><td></td><td></td><td></td><td></td></tr>
<tr><td>Chromosome</td><td>start</td><td>end</td><td>strand</td><td>MM</td><td>target_seq</td><td>PAM</td><td>alignment</td><td>distance</td><td>position</td><td>gene name</td><td>gene id</td></tr>
<tr><td>GU305914.1</td><td>221442</td><td>221464</td><td>+</td><td>0</td><td>CGAACAGTAGGCTGGCTGCC</td><td>AGG</td><td>||||||||[||||||||||||]PAM</td><td>NA</td><td>Intergenic</td><td>NA</td><td>NA</td></tr>
<tr><td>T16453</td><td>GTAGGCTGGCTGCCAGGCAGAGG</td><td>1000</td><td></td><td></td><td></td><td></td><td></td><td></td><td></td><td></td><td></td></tr>
<tr><td>Oligo fwd</td><td>CACCGTAGGCTGGCTGCCAGGCAG</td><td></td><td></td><td></td><td></td><td></td><td></td><td></td><td></td><td></td><td></td></tr>
<tr><td>Oligo rev</td><td>AAACCTGCCTGGCAGCCAGCCTAC</td><td></td><td></td><td></td><td></td><td></td><td></td><td></td><td></td><td></td><td></td></tr>
<tr><td>Chromosome</td><td>start</td><td>end</td><td>strand</td><td>MM</td><td>target_seq</td><td>PAM</td><td>alignment</td><td>distance</td><td>position</td><td>gene name</td><td>gene id</td></tr>
<tr><td>GU305914.1</td><td>221448</td><td>221470</td><td>+</td><td>0</td><td>GTAGGCTGGCTGCCAGGCAG</td><td>AGG</td><td>||||||||[||||||||||||]PAM</td><td>NA</td><td>Intergenic</td><td>NA</td><td>NA</td></tr>
<tr><td>T16454</td><td>GGCAGAGGATCACAAGACTCAGG</td><td>1000</td><td></td><td></td><td></td><td></td><td></td><td></td><td></td><td></td><td></td></tr>
<tr><td>Oligo fwd</td><td>CACCGGCAGAGGATCACAAGACTC</td><td></td><td></td><td></td><td></td><td></td><td></td><td></td><td></td><td></td><td></td></tr>
<tr><td>Oligo rev</td><td>AAACGAGTCTTGTGATCCTCTGCC</td><td></td><td></td><td></td><td></td><td></td><td></td><td></td><td></td><td></td><td></td></tr>
<tr><td>Chromosome</td><td>start</td><td>end</td><td>strand</td><td>MM</td><td>target_seq</td><td>PAM</td><td>alignment</td><td>distance</td><td>position</td><td>gene name</td><td>gene id</td></tr>
<tr><td>GU305914.1</td><td>221463</td><td>221485</td><td>+</td><td>0</td><td>GGCAGAGGATCACAAGACTC</td><td>AGG</td><td>||||||||[||||||||||||]PAM</td><td>NA</td><td>Intergenic</td><td>NA</td><td>NA</td></tr>
<tr><td>T16455</td><td>GCAGAGGATCACAAGACTCAGGG</td><td>1000</td><td></td><td></td><td></td><td></td><td></td><td></td><td></td><td></td><td></td></tr>
<tr><td>Oligo fwd</td><td>CACCGCAGAGGATCACAAGACTCA</td><td></td><td></td><td></td><td></td><td></td><td></td><td></td><td></td><td></td><td></td></tr>
<tr><td>Oligo rev</td><td>AAACTGAGTCTTGTGATCCTCTGC</td><td></td><td></td><td></td><td></td><td></td><td></td><td></td><td></td><td></td><td></td></tr>
<tr><td>Chromosome</td><td>start</td><td>end</td><td>strand</td><td>MM</td><td>target_seq</td><td>PAM</td><td>alignment</td><td>distance</td><td>position</td><td>gene name</td><td>gene id</td></tr>
<tr><td>GU305914.1</td><td>221464</td><td>221486</td><td>+</td><td>0</td><td>GCAGAGGATCACAAGACTCA</td><td>GGG</td><td>||||||||[||||||||||||]PAM</td><td>NA</td><td>Intergenic</td><td>NA</td><td>NA</td></tr>
<tr><td>T16456</td><td>GATCACAAGACTCAGGGCCACGG</td><td>1000</td><td></td><td></td><td></td><td></td><td></td><td></td><td></td><td></td><td></td></tr>
<tr><td>Oligo fwd</td><td>CACCGATCACAAGACTCAGGGCCA</td><td></td><td></td><td></td><td></td><td></td><td></td><td></td><td></td><td></td><td></td></tr>
<tr><td>Oligo rev</td><td>AAACTGGCCCTGAGTCTTGTGATC</td><td></td><td></td><td></td><td></td><td></td><td></td><td></td><td></td><td></td><td></td></tr>
<tr><td>Chromosome</td><td>start</td><td>end</td><td>strand</td><td>MM</td><td>target_seq</td><td>PAM</td><td>alignment</td><td>distance</td><td>position</td><td>gene name</td><td>gene id</td></tr>
<tr><td>GU305914.1</td><td>221470</td><td>221492</td><td>+</td><td>0</td><td>GATCACAAGACTCAGGGCCA</td><td>CGG</td><td>||||||||[||||||||||||]PAM</td><td>NA</td><td>Intergenic</td><td>NA</td><td>NA</td></tr>
<tr><td>T16457</td><td>AATCACCGAGCCGAGACCGTTGG</td><td>1000</td><td></td><td></td><td></td><td></td><td></td><td></td><td></td><td></td><td></td></tr>
<tr><td>Oligo fwd</td><td>CACCAATCACCGAGCCGAGACCGT</td><td></td><td></td><td></td><td></td><td></td><td></td><td></td><td></td><td></td><td></td></tr>
<tr><td>Oligo rev</td><td>AAACACGGTCTCGGCTCGGTGATT</td><td></td><td></td><td></td><td></td><td></td><td></td><td></td><td></td><td></td><td></td></tr>
<tr><td>Chromosome</td><td>start</td><td>end</td><td>strand</td><td>MM</td><td>target_seq</td><td>PAM</td><td>alignment</td><td>distance</td><td>position</td><td>gene name</td><td>gene id</td></tr>
<tr><td>GU305914.1</td><td>221521</td><td>221543</td><td>+</td><td>0</td><td>AATCACCGAGCCGAGACCGT</td><td>TGG</td><td>||||||||[||||||||||||]PAM</td><td>NA</td><td>Intergenic</td><td>NA</td><td>NA</td></tr>
<tr><td>T16458</td><td>CGAGCCGAGACCGTTGGTTGTGG</td><td>1000</td><td></td><td></td><td></td><td></td><td></td><td></td><td></td><td></td><td></td></tr>
<tr><td>Oligo fwd</td><td>CACCCGAGCCGAGACCGTTGGTTG</td><td></td><td></td><td></td><td></td><td></td><td></td><td></td><td></td><td></td><td></td></tr>
<tr><td>Oligo rev</td><td>AAACCAACCAACGGTCTCGGCTCG</td><td></td><td></td><td></td><td></td><td></td><td></td><td></td><td></td><td></td><td></td></tr>
<tr><td>Chromosome</td><td>start</td><td>end</td><td>strand</td><td>MM</td><td>target_seq</td><td>PAM</td><td>alignment</td><td>distance</td><td>position</td><td>gene name</td><td>gene id</td></tr>
<tr><td>GU305914.1</td><td>221527</td><td>221549</td><td>+</td><td>0</td><td>CGAGCCGAGACCGTTGGTTG</td><td>TGG</td><td>||||||||[||||||||||||]PAM</td><td>NA</td><td>Intergenic</td><td>NA</td><td>NA</td></tr>
<tr><td>T16459</td><td>AGACCGTTGGTTGTGGTGATCGG</td><td>1000</td><td></td><td></td><td></td><td></td><td></td><td></td><td></td><td></td><td></td></tr>
<tr><td>Oligo fwd</td><td>CACCAGACCGTTGGTTGTGGTGAT</td><td></td><td></td><td></td><td></td><td></td><td></td><td></td><td></td><td></td><td></td></tr>
<tr><td>Oligo rev</td><td>AAACATCACCACAACCAACGGTCT</td><td></td><td></td><td></td><td></td><td></td><td></td><td></td><td></td><td></td><td></td></tr>
<tr><td>Chromosome</td><td>start</td><td>end</td><td>strand</td><td>MM</td><td>target_seq</td><td>PAM</td><td>alignment</td><td>distance</td><td>position</td><td>gene name</td><td>gene id</td></tr>
<tr><td>GU305914.1</td><td>221534</td><td>221556</td><td>+</td><td>0</td><td>AGACCGTTGGTTGTGGTGAT</td><td>CGG</td><td>||||||||[||||||||||||]PAM</td><td>NA</td><td>Intergenic</td><td>NA</td><td>NA</td></tr>
<tr><td>T16460</td><td>CCGTTGGTTGTGGTGATCGGCGG</td><td>1000</td><td></td><td></td><td></td><td></td><td></td><td></td><td></td><td></td><td></td></tr>
<tr><td>Oligo fwd</td><td>CACCCCGTTGGTTGTGGTGATCGG</td><td></td><td></td><td></td><td></td><td></td><td></td><td></td><td></td><td></td><td></td></tr>
<tr><td>Oligo rev</td><td>AAACCCGATCACCACAACCAACGG</td><td></td><td></td><td></td><td></td><td></td><td></td><td></td><td></td><td></td><td></td></tr>
<tr><td>Chromosome</td><td>start</td><td>end</td><td>strand</td><td>MM</td><td>target_seq</td><td>PAM</td><td>alignment</td><td>distance</td><td>position</td><td>gene name</td><td>gene id</td></tr>
<tr><td>GU305914.1</td><td>221537</td><td>221559</td><td>+</td><td>0</td><td>CCGTTGGTTGTGGTGATCGG</td><td>CGG</td><td>||||||||[||||||||||||]PAM</td><td>NA</td><td>Intergenic</td><td>NA</td><td>NA</td></tr>
<tr><td>T16461</td><td>GGACCTCCCAAACCGTTCGAAGG</td><td>1000</td><td></td><td></td><td></td><td></td><td></td><td></td><td></td><td></td><td></td></tr>
<tr><td>Oligo fwd</td><td>CACCGGACCTCCCAAACCGTTCGA</td><td></td><td></td><td></td><td></td><td></td><td></td><td></td><td></td><td></td><td></td></tr>
<tr><td>Oligo rev</td><td>AAACTCGAACGGTTTGGGAGGTCC</td><td></td><td></td><td></td><td></td><td></td><td></td><td></td><td></td><td></td><td></td></tr>
<tr><td>Chromosome</td><td>start</td><td>end</td><td>strand</td><td>MM</td><td>target_seq</td><td>PAM</td><td>alignment</td><td>distance</td><td>position</td><td>gene name</td><td>gene id</td></tr>
<tr><td>GU305914.1</td><td>221558</td><td>221580</td><td>+</td><td>0</td><td>GGACCTCCCAAACCGTTCGA</td><td>AGG</td><td>||||||||[||||||||||||]PAM</td><td>NA</td><td>Intergenic</td><td>NA</td><td>NA</td></tr>
<tr><td>T16462</td><td>AACCGTTCGAAGGATATGCCAGG</td><td>1000</td><td></td><td></td><td></td><td></td><td></td><td></td><td></td><td></td><td></td></tr>
<tr><td>Oligo fwd</td><td>CACCAACCGTTCGAAGGATATGCC</td><td></td><td></td><td></td><td></td><td></td><td></td><td></td><td></td><td></td><td></td></tr>
<tr><td>Oligo rev</td><td>AAACGGCATATCCTTCGAACGGTT</td><td></td><td></td><td></td><td></td><td></td><td></td><td></td><td></td><td></td><td></td></tr>
<tr><td>Chromosome</td><td>start</td><td>end</td><td>strand</td><td>MM</td><td>target_seq</td><td>PAM</td><td>alignment</td><td>distance</td><td>position</td><td>gene name</td><td>gene id</td></tr>
<tr><td>GU305914.1</td><td>221568</td><td>221590</td><td>+</td><td>0</td><td>AACCGTTCGAAGGATATGCC</td><td>AGG</td><td>||||||||[||||||||||||]PAM</td><td>NA</td><td>Intergenic</td><td>NA</td><td>NA</td></tr>
<tr><td>T16463</td><td>GAACCCTGCCGCCGCCACCACGG</td><td>1000</td><td></td><td></td><td></td><td></td><td></td><td></td><td></td><td></td><td></td></tr>
<tr><td>Oligo fwd</td><td>CACCGAACCCTGCCGCCGCCACCA</td><td></td><td></td><td></td><td></td><td></td><td></td><td></td><td></td><td></td><td></td></tr>
<tr><td>Oligo rev</td><td>AAACTGGTGGCGGCGGCAGGGTTC</td><td></td><td></td><td></td><td></td><td></td><td></td><td></td><td></td><td></td><td></td></tr>
<tr><td>Chromosome</td><td>start</td><td>end</td><td>strand</td><td>MM</td><td>target_seq</td><td>PAM</td><td>alignment</td><td>distance</td><td>position</td><td>gene name</td><td>gene id</td></tr>
<tr><td>GU305914.1</td><td>221595</td><td>221617</td><td>+</td><td>0</td><td>GAACCCTGCCGCCGCCACCA</td><td>CGG</td><td>||||||||[||||||||||||]PAM</td><td>NA</td><td>Intergenic</td><td>NA</td><td>NA</td></tr>
</table>

<table>
<tr><td>T16464</td><td>GCCGCCGCCACCACGGCCTCCGG</td><td>1000</td><td></td><td></td><td></td><td></td><td></td><td></td><td></td><td></td><td></td></tr>
<tr><td>Oligo fwd</td><td>CACCGCCGCCGCCACCACGGCCTC</td><td></td><td></td><td></td><td></td><td></td><td></td><td></td><td></td><td></td><td></td></tr>
<tr><td>Oligo rev</td><td>AAACGAGGCCGTGGTGGCGGCGGC</td><td></td><td></td><td></td><td></td><td></td><td></td><td></td><td></td><td></td><td></td></tr>
<tr><td>Chromosome</td><td>start</td><td>end</td><td>strand</td><td>MM</td><td>target_seq</td><td>PAM</td><td>alignment</td><td>distance</td><td>position</td><td>gene name</td><td>gene id</td></tr>
<tr><td>GU305914.1</td><td>221602</td><td>221624</td><td>+</td><td>0</td><td>GCCGCCGCCACCACGGCCTC</td><td>CGG</td><td>||||||||[||||||||||||]PAM</td><td>NA</td><td>Intergenic</td><td>NA</td><td>NA</td></tr>
<tr><td>GU305914.1</td><td>161759</td><td>161781</td><td>+</td><td>2</td><td>GCCGCCGCCTCCTCGGCCTC</td><td>CGG</td><td>||||||||[|-||-|||||||]PAM</td><td>NA</td><td>Intergenic</td><td>NA</td><td>NA</td></tr>
<tr><td></td><td></td><td></td><td></td><td></td><td></td><td></td><td></td><td></td><td></td><td></td><td></td></tr>
<tr><td>T16465</td><td>GTCGTCACGTTTTCGATAACAGG</td><td>1000</td><td></td><td></td><td></td><td></td><td></td><td></td><td></td><td></td><td></td></tr>
<tr><td>Oligo fwd</td><td>CACCGTCGTCACGTTTTCGATAAC</td><td></td><td></td><td></td><td></td><td></td><td></td><td></td><td></td><td></td><td></td></tr>
<tr><td>Oligo rev</td><td>AAACGTTATCGAAAACGTGACGAC</td><td></td><td></td><td></td><td></td><td></td><td></td><td></td><td></td><td></td><td></td></tr>
<tr><td>Chromosome</td><td>start</td><td>end</td><td>strand</td><td>MM</td><td>target_seq</td><td>PAM</td><td>alignment</td><td>distance</td><td>position</td><td>gene name</td><td>gene id</td></tr>
<tr><td>GU305914.1</td><td>221624</td><td>221646</td><td>+</td><td>0</td><td>GTCGTCACGTTTTCGATAAC</td><td>AGG</td><td>||||||||[||||||||||||]PAM</td><td>NA</td><td>Intergenic</td><td>NA</td><td>NA</td></tr>
<tr><td></td><td></td><td></td><td></td><td></td><td></td><td></td><td></td><td></td><td></td><td></td><td></td></tr>
<tr><td>T16466</td><td>AGGCCCCCCTAACGCGCTACTGG</td><td>1000</td><td></td><td></td><td></td><td></td><td></td><td></td><td></td><td></td><td></td></tr>
<tr><td>Oligo fwd</td><td>CACCAGGCCCCCCTAACGCGCTAC</td><td></td><td></td><td></td><td></td><td></td><td></td><td></td><td></td><td></td><td></td></tr>
<tr><td>Oligo rev</td><td>AAACGTAGCGCGTTAGGGGGGCCT</td><td></td><td></td><td></td><td></td><td></td><td></td><td></td><td></td><td></td><td></td></tr>
<tr><td>Chromosome</td><td>start</td><td>end</td><td>strand</td><td>MM</td><td>target_seq</td><td>PAM</td><td>alignment</td><td>distance</td><td>position</td><td>gene name</td><td>gene id</td></tr>
<tr><td>GU305914.1</td><td>221644</td><td>221666</td><td>+</td><td>0</td><td>AGGCCCCCCTAACGCGCTAC</td><td>TGG</td><td>||||||||[||||||||||||]PAM</td><td>NA</td><td>Intergenic</td><td>NA</td><td>NA</td></tr>
<tr><td></td><td></td><td></td><td></td><td></td><td></td><td></td><td></td><td></td><td></td><td></td><td></td></tr>
<tr><td>T16467</td><td>CGACGTTAAAATCTGCGACGTGG</td><td>1000</td><td></td><td></td><td></td><td></td><td></td><td></td><td></td><td></td><td></td></tr>
<tr><td>Oligo fwd</td><td>CACCCGACGTTAAAATCTGCGACG</td><td></td><td></td><td></td><td></td><td></td><td></td><td></td><td></td><td></td><td></td></tr>
<tr><td>Oligo rev</td><td>AAACCGTCGCAGATTTTAACGTCG</td><td></td><td></td><td></td><td></td><td></td><td></td><td></td><td></td><td></td><td></td></tr>
<tr><td>Chromosome</td><td>start</td><td>end</td><td>strand</td><td>MM</td><td>target_seq</td><td>PAM</td><td>alignment</td><td>distance</td><td>position</td><td>gene name</td><td>gene id</td></tr>
<tr><td>GU305914.1</td><td>221677</td><td>221699</td><td>+</td><td>0</td><td>CGACGTTAAAATCTGCGACG</td><td>TGG</td><td>||||||||[||||||||||||]PAM</td><td>NA</td><td>Intergenic</td><td>NA</td><td>NA</td></tr>
<tr><td></td><td></td><td></td><td></td><td></td><td></td><td></td><td></td><td></td><td></td><td></td><td></td></tr>
<tr><td>T16468</td><td>CTGCGACGTGGTAGCCAAGACGG</td><td>1000</td><td></td><td></td><td></td><td></td><td></td><td></td><td></td><td></td><td></td></tr>
<tr><td>Oligo fwd</td><td>CACCCTGCGACGTGGTAGCCAAGA</td><td></td><td></td><td></td><td></td><td></td><td></td><td></td><td></td><td></td><td></td></tr>
<tr><td>Oligo rev</td><td>AAACTCTTGGCTACCACGTCGCAG</td><td></td><td></td><td></td><td></td><td></td><td></td><td></td><td></td><td></td><td></td></tr>
<tr><td>Chromosome</td><td>start</td><td>end</td><td>strand</td><td>MM</td><td>target_seq</td><td>PAM</td><td>alignment</td><td>distance</td><td>position</td><td>gene name</td><td>gene id</td></tr>
<tr><td>GU305914.1</td><td>221689</td><td>221711</td><td>+</td><td>0</td><td>CTGCGACGTGGTAGCCAAGA</td><td>CGG</td><td>||||||||[||||||||||||]PAM</td><td>NA</td><td>Intergenic</td><td>NA</td><td>NA</td></tr>
<tr><td></td><td></td><td></td><td></td><td></td><td></td><td></td><td></td><td></td><td></td><td></td><td></td></tr>
<tr><td>T16469</td><td>GGTAGCCAAGACGGTGTGATCGG</td><td>1000</td><td></td><td></td><td></td><td></td><td></td><td></td><td></td><td></td><td></td></tr>
<tr><td>Oligo fwd</td><td>CACCGGTAGCCAAGACGGTGTGAT</td><td></td><td></td><td></td><td></td><td></td><td></td><td></td><td></td><td></td><td></td></tr>
<tr><td>Oligo rev</td><td>AAACATCACACCGTCTTGGCTACC</td><td></td><td></td><td></td><td></td><td></td><td></td><td></td><td></td><td></td><td></td></tr>
<tr><td>Chromosome</td><td>start</td><td>end</td><td>strand</td><td>MM</td><td>target_seq</td><td>PAM</td><td>alignment</td><td>distance</td><td>position</td><td>gene name</td><td>gene id</td></tr>
<tr><td>GU305914.1</td><td>221698</td><td>221720</td><td>+</td><td>0</td><td>GGTAGCCAAGACGGTGTGAT</td><td>CGG</td><td>||||||||[||||||||||||]PAM</td><td>NA</td><td>Intergenic</td><td>NA</td><td>NA</td></tr>
<tr><td></td><td></td><td></td><td></td><td></td><td></td><td></td><td></td><td></td><td></td><td></td><td></td></tr>
<tr><td>T16470</td><td>GGTGTGATCGGAATAATTGTCGG</td><td>1000</td><td></td><td></td><td></td><td></td><td></td><td></td><td></td><td></td><td></td></tr>
<tr><td>Oligo fwd</td><td>CACCGGTGTGATCGGAATAATTGT</td><td></td><td></td><td></td><td></td><td></td><td></td><td></td><td></td><td></td><td></td></tr>
<tr><td>Oligo rev</td><td>AAACACAATTATTCCGATCACACC</td><td></td><td></td><td></td><td></td><td></td><td></td><td></td><td></td><td></td><td></td></tr>
<tr><td>Chromosome</td><td>start</td><td>end</td><td>strand</td><td>MM</td><td>target_seq</td><td>PAM</td><td>alignment</td><td>distance</td><td>position</td><td>gene name</td><td>gene id</td></tr>
<tr><td>GU305914.1</td><td>221710</td><td>221732</td><td>+</td><td>0</td><td>GGTGTGATCGGAATAATTGT</td><td>CGG</td><td>||||||||[||||||||||||]PAM</td><td>NA</td><td>Intergenic</td><td>NA</td><td>NA</td></tr>
<tr><td></td><td></td><td></td><td></td><td></td><td></td><td></td><td></td><td></td><td></td><td></td><td></td></tr>
<tr><td>T16471</td><td>TGATCGGAATAATTGTCGGCAGG</td><td>1000</td><td></td><td></td><td></td><td></td><td></td><td></td><td></td><td></td><td></td></tr>
<tr><td>Oligo fwd</td><td>CACCTGATCGGAATAATTGTCGGC</td><td></td><td></td><td></td><td></td><td></td><td></td><td></td><td></td><td></td><td></td></tr>
<tr><td>Oligo rev</td><td>AAACGCCGACAATTATTCCGATCA</td><td></td><td></td><td></td><td></td><td></td><td></td><td></td><td></td><td></td><td></td></tr>
<tr><td>Chromosome</td><td>start</td><td>end</td><td>strand</td><td>MM</td><td>target_seq</td><td>PAM</td><td>alignment</td><td>distance</td><td>position</td><td>gene name</td><td>gene id</td></tr>
<tr><td>GU305914.1</td><td>221714</td><td>221736</td><td>+</td><td>0</td><td>TGATCGGAATAATTGTCGGC</td><td>AGG</td><td>||||||||[||||||||||||]PAM</td><td>NA</td><td>Intergenic</td><td>NA</td><td>NA</td></tr>
<tr><td></td><td></td><td></td><td></td><td></td><td></td><td></td><td></td><td></td><td></td><td></td><td></td></tr>
<tr><td>T16472</td><td>GATCGGAATAATTGTCGGCAGGG</td><td>1000</td><td></td><td></td><td></td><td></td><td></td><td></td><td></td><td></td><td></td></tr>
<tr><td>Oligo fwd</td><td>CACCGATCGGAATAATTGTCGGCA</td><td></td><td></td><td></td><td></td><td></td><td></td><td></td><td></td><td></td><td></td></tr>
<tr><td>Oligo rev</td><td>AAACTGCCGACAATTATTCCGATC</td><td></td><td></td><td></td><td></td><td></td><td></td><td></td><td></td><td></td><td></td></tr>
<tr><td>Chromosome</td><td>start</td><td>end</td><td>strand</td><td>MM</td><td>target_seq</td><td>PAM</td><td>alignment</td><td>distance</td><td>position</td><td>gene name</td><td>gene id</td></tr>
<tr><td>GU305914.1</td><td>221715</td><td>221737</td><td>+</td><td>0</td><td>GATCGGAATAATTGTCGGCA</td><td>GGG</td><td>||||||||[||||||||||||]PAM</td><td>NA</td><td>Intergenic</td><td>NA</td><td>NA</td></tr>
<tr><td></td><td></td><td></td><td></td><td></td><td></td><td></td><td></td><td></td><td></td><td></td><td></td></tr>
<tr><td>T16473</td><td>ATAATTGTCGGCAGGGACGTAGG</td><td>1000</td><td></td><td></td><td></td><td></td><td></td><td></td><td></td><td></td><td></td></tr>
<tr><td>Oligo fwd</td><td>CACCATAATTGTCGGCAGGGACGT</td><td></td><td></td><td></td><td></td><td></td><td></td><td></td><td></td><td></td><td></td></tr>
<tr><td>Oligo rev</td><td>AAACACGTCCCTGCCGACAATTAT</td><td></td><td></td><td></td><td></td><td></td><td></td><td></td><td></td><td></td><td></td></tr>
<tr><td>Chromosome</td><td>start</td><td>end</td><td>strand</td><td>MM</td><td>target_seq</td><td>PAM</td><td>alignment</td><td>distance</td><td>position</td><td>gene name</td><td>gene id</td></tr>
<tr><td>GU305914.1</td><td>221722</td><td>221744</td><td>+</td><td>0</td><td>ATAATTGTCGGCAGGGACGT</td><td>AGG</td><td>||||||||[||||||||||||]PAM</td><td>NA</td><td>Intergenic</td><td>NA</td><td>NA</td></tr>
<tr><td></td><td></td><td></td><td></td><td></td><td></td><td></td><td></td><td></td><td></td><td></td><td></td></tr>
<tr><td>T16474</td><td>ATTGTCGGCAGGGACGTAGGTGG</td><td>1000</td><td></td><td></td><td></td><td></td><td></td><td></td><td></td><td></td><td></td></tr>
<tr><td>Oligo fwd</td><td>CACCATTGTCGGCAGGGACGTAGG</td><td></td><td></td><td></td><td></td><td></td><td></td><td></td><td></td><td></td><td></td></tr>
<tr><td>Oligo rev</td><td>AAACCCTACGTCCCTGCCGACAAT</td><td></td><td></td><td></td><td></td><td></td><td></td><td></td><td></td><td></td><td></td></tr>
<tr><td>Chromosome</td><td>start</td><td>end</td><td>strand</td><td>MM</td><td>target_seq</td><td>PAM</td><td>alignment</td><td>distance</td><td>position</td><td>gene name</td><td>gene id</td></tr>
<tr><td>GU305914.1</td><td>221725</td><td>221747</td><td>+</td><td>0</td><td>ATTGTCGGCAGGGACGTAGG</td><td>TGG</td><td>||||||||[||||||||||||]PAM</td><td>NA</td><td>Intergenic</td><td>NA</td><td>NA</td></tr>
<tr><td></td><td></td><td></td><td></td><td></td><td></td><td></td><td></td><td></td><td></td><td></td><td></td></tr>
<tr><td>T16475</td><td>TAGAGATACATACCGCGATTCGG</td><td>1000</td><td></td><td></td><td></td><td></td><td></td><td></td><td></td><td></td><td></td></tr>
<tr><td>Oligo fwd</td><td>CACCTAGAGATACATACCGCGATT</td><td></td><td></td><td></td><td></td><td></td><td></td><td></td><td></td><td></td><td></td></tr>
<tr><td>Oligo rev</td><td>AAACAATCGCGGTATGTATCTCTA</td><td></td><td></td><td></td><td></td><td></td><td></td><td></td><td></td><td></td><td></td></tr>
<tr><td>Chromosome</td><td>start</td><td>end</td><td>strand</td><td>MM</td><td>target_seq</td><td>PAM</td><td>alignment</td><td>distance</td><td>position</td><td>gene name</td><td>gene id</td></tr>
<tr><td>GU305914.1</td><td>221756</td><td>221778</td><td>+</td><td>0</td><td>TAGAGATACATACCGCGATT</td><td>CGG</td><td>||||||||[||||||||||||]PAM</td><td>NA</td><td>Intergenic</td><td>NA</td><td>NA</td></tr>
<tr><td></td><td></td><td></td><td></td><td></td><td></td><td></td><td></td><td></td><td></td><td></td><td></td></tr>
<tr><td>T16476</td><td>GATACATACCGCGATTCGGTAGG</td><td>1000</td><td></td><td></td><td></td><td></td><td></td><td></td><td></td><td></td><td></td></tr>
<tr><td>Oligo fwd</td><td>CACCGATACATACCGCGATTCGGT</td><td></td><td></td><td></td><td></td><td></td><td></td><td></td><td></td><td></td><td></td></tr>
<tr><td>Oligo rev</td><td>AAACACCGAATCGCGGTATGTATC</td><td></td><td></td><td></td><td></td><td></td><td></td><td></td><td></td><td></td><td></td></tr>
<tr><td>Chromosome</td><td>start</td><td>end</td><td>strand</td><td>MM</td><td>target_seq</td><td>PAM</td><td>alignment</td><td>distance</td><td>position</td><td>gene name</td><td>gene id</td></tr>
<tr><td>GU305914.1</td><td>221760</td><td>221782</td><td>+</td><td>0</td><td>GATACATACCGCGATTCGGT</td><td>AGG</td><td>||||||||[||||||||||||]PAM</td><td>NA</td><td>Intergenic</td><td>NA</td><td>NA</td></tr>
<tr><td></td><td></td><td></td><td></td><td></td><td></td><td></td><td></td><td></td><td></td><td></td><td></td></tr>
<tr><td>T16477</td><td>CCGCGATCCACCGTTCGAGCCGG</td><td>1000</td><td></td><td></td><td></td><td></td><td></td><td></td><td></td><td></td><td></td></tr>
<tr><td>Oligo fwd</td><td>CACCCCGCGATCCACCGTTCGAGC</td><td></td><td></td><td></td><td></td><td></td><td></td><td></td><td></td><td></td><td></td></tr>
<tr><td>Oligo rev</td><td>AAACGCTCGAACGGTGGATCGCGG</td><td></td><td></td><td></td><td></td><td></td><td></td><td></td><td></td><td></td><td></td></tr>
<tr><td>Chromosome</td><td>start</td><td>end</td><td>strand</td><td>MM</td><td>target_seq</td><td>PAM</td><td>alignment</td><td>distance</td><td>position</td><td>gene name</td><td>gene id</td></tr>
<tr><td>GU305914.1</td><td>221814</td><td>221836</td><td>+</td><td>0</td><td>CCGCGATCCACCGTTCGAGC</td><td>CGG</td><td>||||||||[||||||||||||]PAM</td><td>NA</td><td>Intergenic</td><td>NA</td><td>NA</td></tr>
<tr><td></td><td></td><td></td><td></td><td></td><td></td><td></td><td></td><td></td><td></td><td></td><td></td></tr>
<tr><td>T16478</td><td>GAAAGCCACCCGACAGTGAACGG</td><td>1000</td><td></td><td></td><td></td><td></td><td></td><td></td><td></td><td></td><td></td></tr>
<tr><td>Oligo fwd</td><td>CACCGAAAGCCACCCGACAGTGAA</td><td></td><td></td><td></td><td></td><td></td><td></td><td></td><td></td><td></td><td></td></tr>
<tr><td>Oligo rev</td><td>AAACTTCACTGTCGGGTGGCTTTC</td><td></td><td></td><td></td><td></td><td></td><td></td><td></td><td></td><td></td><td></td></tr>
<tr><td>Chromosome</td><td>start</td><td>end</td><td>strand</td><td>MM</td><td>target_seq</td><td>PAM</td><td>alignment</td><td>distance</td><td>position</td><td>gene name</td><td>gene id</td></tr>
</table>

<table>
<tr><td>GU305914.1</td><td>221839</td><td>221861</td><td>+</td><td>0</td><td>GAAAGCCACCCGACAGTGAA</td><td>CGG</td><td>||||||||[||||||||||||]PAM</td><td>NA</td><td>Intergenic</td><td>NA</td><td>NA</td></tr>
<tr><td>T16479</td><td>GCCACCCGACAGTGAACGGCCGG</td><td>1000</td><td></td><td></td><td></td><td></td><td></td><td></td><td></td><td></td><td></td></tr>
<tr><td>Oligo fwd</td><td>CACCGCCACCCGACAGTGAACGGC</td><td></td><td></td><td></td><td></td><td></td><td></td><td></td><td></td><td></td><td></td></tr>
<tr><td>Oligo rev</td><td>AAACGCCGTTCACTGTCGGGTGGC</td><td></td><td></td><td></td><td></td><td></td><td></td><td></td><td></td><td></td><td></td></tr>
<tr><td>Chromosome</td><td>start</td><td>end</td><td>strand</td><td>MM</td><td>target_seq</td><td>PAM</td><td>alignment</td><td>distance</td><td>position</td><td>gene name</td><td>gene id</td></tr>
<tr><td>GU305914.1</td><td>221843</td><td>221865</td><td>+</td><td>0</td><td>GCCACCCGACAGTGAACGGC</td><td>CGG</td><td>||||||||[||||||||||||]PAM</td><td>NA</td><td>Intergenic</td><td>NA</td><td>NA</td></tr>
<tr><td>T16480</td><td>CCACCCGACAGTGAACGGCCGGG</td><td>1000</td><td></td><td></td><td></td><td></td><td></td><td></td><td></td><td></td><td></td></tr>
<tr><td>Oligo fwd</td><td>CACCCCACCCGACAGTGAACGGCC</td><td></td><td></td><td></td><td></td><td></td><td></td><td></td><td></td><td></td><td></td></tr>
<tr><td>Oligo rev</td><td>AAACGGCCGTTCACTGTCGGGTGG</td><td></td><td></td><td></td><td></td><td></td><td></td><td></td><td></td><td></td><td></td></tr>
<tr><td>Chromosome</td><td>start</td><td>end</td><td>strand</td><td>MM</td><td>target_seq</td><td>PAM</td><td>alignment</td><td>distance</td><td>position</td><td>gene name</td><td>gene id</td></tr>
<tr><td>GU305914.1</td><td>221844</td><td>221866</td><td>+</td><td>0</td><td>CCACCCGACAGTGAACGGCC</td><td>GGG</td><td>||||||||[||||||||||||]PAM</td><td>NA</td><td>Intergenic</td><td>NA</td><td>NA</td></tr>
<tr><td>T16481</td><td>TGCCTAACGAGTGACACTTAAGG</td><td>1000</td><td></td><td></td><td></td><td></td><td></td><td></td><td></td><td></td><td></td></tr>
<tr><td>Oligo fwd</td><td>CACCTGCCTAACGAGTGACACTTA</td><td></td><td></td><td></td><td></td><td></td><td></td><td></td><td></td><td></td><td></td></tr>
<tr><td>Oligo rev</td><td>AAACTAAGTGTCACTCGTTAGGCA</td><td></td><td></td><td></td><td></td><td></td><td></td><td></td><td></td><td></td><td></td></tr>
<tr><td>Chromosome</td><td>start</td><td>end</td><td>strand</td><td>MM</td><td>target_seq</td><td>PAM</td><td>alignment</td><td>distance</td><td>position</td><td>gene name</td><td>gene id</td></tr>
<tr><td>GU305914.1</td><td>221867</td><td>221889</td><td>+</td><td>0</td><td>TGCCTAACGAGTGACACTTA</td><td>AGG</td><td>||||||||[||||||||||||]PAM</td><td>NA</td><td>Intergenic</td><td>NA</td><td>NA</td></tr>
<tr><td>T16482</td><td>GTGACACTTAAGGTCGCGAGCGG</td><td>1000</td><td></td><td></td><td></td><td></td><td></td><td></td><td></td><td></td><td></td></tr>
<tr><td>Oligo fwd</td><td>CACCGTGACACTTAAGGTCGCGAG</td><td></td><td></td><td></td><td></td><td></td><td></td><td></td><td></td><td></td><td></td></tr>
<tr><td>Oligo rev</td><td>AAACCTCGCGACCTTAAGTGTCAC</td><td></td><td></td><td></td><td></td><td></td><td></td><td></td><td></td><td></td><td></td></tr>
<tr><td>Chromosome</td><td>start</td><td>end</td><td>strand</td><td>MM</td><td>target_seq</td><td>PAM</td><td>alignment</td><td>distance</td><td>position</td><td>gene name</td><td>gene id</td></tr>
<tr><td>GU305914.1</td><td>221877</td><td>221899</td><td>+</td><td>0</td><td>GTGACACTTAAGGTCGCGAG</td><td>CGG</td><td>||||||||[||||||||||||]PAM</td><td>NA</td><td>Intergenic</td><td>NA</td><td>NA</td></tr>
<tr><td>T16483</td><td>TGACACTTAAGGTCGCGAGCGGG</td><td>1000</td><td></td><td></td><td></td><td></td><td></td><td></td><td></td><td></td><td></td></tr>
<tr><td>Oligo fwd</td><td>CACCTGACACTTAAGGTCGCGAGC</td><td></td><td></td><td></td><td></td><td></td><td></td><td></td><td></td><td></td><td></td></tr>
<tr><td>Oligo rev</td><td>AAACGCTCGCGACCTTAAGTGTCA</td><td></td><td></td><td></td><td></td><td></td><td></td><td></td><td></td><td></td><td></td></tr>
<tr><td>Chromosome</td><td>start</td><td>end</td><td>strand</td><td>MM</td><td>target_seq</td><td>PAM</td><td>alignment</td><td>distance</td><td>position</td><td>gene name</td><td>gene id</td></tr>
<tr><td>GU305914.1</td><td>221878</td><td>221900</td><td>+</td><td>0</td><td>TGACACTTAAGGTCGCGAGC</td><td>GGG</td><td>||||||||[||||||||||||]PAM</td><td>NA</td><td>Intergenic</td><td>NA</td><td>NA</td></tr>
<tr><td>T16484</td><td>GACACTTAAGGTCGCGAGCGGGG</td><td>1000</td><td></td><td></td><td></td><td></td><td></td><td></td><td></td><td></td><td></td></tr>
<tr><td>Oligo fwd</td><td>CACCGACACTTAAGGTCGCGAGCG</td><td></td><td></td><td></td><td></td><td></td><td></td><td></td><td></td><td></td><td></td></tr>
<tr><td>Oligo rev</td><td>AAACCGCTCGCGACCTTAAGTGTC</td><td></td><td></td><td></td><td></td><td></td><td></td><td></td><td></td><td></td><td></td></tr>
<tr><td>Chromosome</td><td>start</td><td>end</td><td>strand</td><td>MM</td><td>target_seq</td><td>PAM</td><td>alignment</td><td>distance</td><td>position</td><td>gene name</td><td>gene id</td></tr>
<tr><td>GU305914.1</td><td>221879</td><td>221901</td><td>+</td><td>0</td><td>GACACTTAAGGTCGCGAGCG</td><td>GGG</td><td>||||||||[||||||||||||]PAM</td><td>NA</td><td>Intergenic</td><td>NA</td><td>NA</td></tr>
<tr><td>T16485</td><td>ACACTTAAGGTCGCGAGCGGGGG</td><td>1000</td><td></td><td></td><td></td><td></td><td></td><td></td><td></td><td></td><td></td></tr>
<tr><td>Oligo fwd</td><td>CACCACACTTAAGGTCGCGAGCGG</td><td></td><td></td><td></td><td></td><td></td><td></td><td></td><td></td><td></td><td></td></tr>
<tr><td>Oligo rev</td><td>AAACCCGCTCGCGACCTTAAGTGT</td><td></td><td></td><td></td><td></td><td></td><td></td><td></td><td></td><td></td><td></td></tr>
<tr><td>Chromosome</td><td>start</td><td>end</td><td>strand</td><td>MM</td><td>target_seq</td><td>PAM</td><td>alignment</td><td>distance</td><td>position</td><td>gene name</td><td>gene id</td></tr>
<tr><td>GU305914.1</td><td>221880</td><td>221902</td><td>+</td><td>0</td><td>ACACTTAAGGTCGCGAGCGG</td><td>GGG</td><td>||||||||[||||||||||||]PAM</td><td>NA</td><td>Intergenic</td><td>NA</td><td>NA</td></tr>
<tr><td>T16486</td><td>GGTCGCGAGCGGGGGAAGAAAGG</td><td>1000</td><td></td><td></td><td></td><td></td><td></td><td></td><td></td><td></td><td></td></tr>
<tr><td>Oligo fwd</td><td>CACCGGTCGCGAGCGGGGGAAGAA</td><td></td><td></td><td></td><td></td><td></td><td></td><td></td><td></td><td></td><td></td></tr>
<tr><td>Oligo rev</td><td>AAACTTCTTCCCCCGCTCGCGACC</td><td></td><td></td><td></td><td></td><td></td><td></td><td></td><td></td><td></td><td></td></tr>
<tr><td>Chromosome</td><td>start</td><td>end</td><td>strand</td><td>MM</td><td>target_seq</td><td>PAM</td><td>alignment</td><td>distance</td><td>position</td><td>gene name</td><td>gene id</td></tr>
<tr><td>GU305914.1</td><td>221888</td><td>221910</td><td>+</td><td>0</td><td>GGTCGCGAGCGGGGGAAGAA</td><td>AGG</td><td>||||||||[||||||||||||]PAM</td><td>NA</td><td>Intergenic</td><td>NA</td><td>NA</td></tr>
<tr><td>T16487</td><td>CGGGGGAAGAAAGGAGAACACGG</td><td>1000</td><td></td><td></td><td></td><td></td><td></td><td></td><td></td><td></td><td></td></tr>
<tr><td>Oligo fwd</td><td>CACCCGGGGGAAGAAAGGAGAACA</td><td></td><td></td><td></td><td></td><td></td><td></td><td></td><td></td><td></td><td></td></tr>
<tr><td>Oligo rev</td><td>AAACTGTTCTCCTTTCTTCCCCCG</td><td></td><td></td><td></td><td></td><td></td><td></td><td></td><td></td><td></td><td></td></tr>
<tr><td>Chromosome</td><td>start</td><td>end</td><td>strand</td><td>MM</td><td>target_seq</td><td>PAM</td><td>alignment</td><td>distance</td><td>position</td><td>gene name</td><td>gene id</td></tr>
<tr><td>GU305914.1</td><td>221897</td><td>221919</td><td>+</td><td>0</td><td>CGGGGGAAGAAAGGAGAACA</td><td>CGG</td><td>||||||||[||||||||||||]PAM</td><td>NA</td><td>Intergenic</td><td>NA</td><td>NA</td></tr>
<tr><td>T16488</td><td>ACGTTAATAAAAATTATACCCGG</td><td>1000</td><td></td><td></td><td></td><td></td><td></td><td></td><td></td><td></td><td></td></tr>
<tr><td>Oligo fwd</td><td>CACCACGTTAATAAAAATTATACC</td><td></td><td></td><td></td><td></td><td></td><td></td><td></td><td></td><td></td><td></td></tr>
<tr><td>Oligo rev</td><td>AAACGGTATAATTTTTATTAACGT</td><td></td><td></td><td></td><td></td><td></td><td></td><td></td><td></td><td></td><td></td></tr>
<tr><td>Chromosome</td><td>start</td><td>end</td><td>strand</td><td>MM</td><td>target_seq</td><td>PAM</td><td>alignment</td><td>distance</td><td>position</td><td>gene name</td><td>gene id</td></tr>
<tr><td>GU305914.1</td><td>221963</td><td>221985</td><td>+</td><td>0</td><td>ACGTTAATAAAAATTATACC</td><td>CGG</td><td>||||||||[||||||||||||]PAM</td><td>NA</td><td>Intergenic</td><td>NA</td><td>NA</td></tr>
<tr><td>T16489</td><td>TAAAAATTATACCCGGTTATAGG</td><td>1000</td><td></td><td></td><td></td><td></td><td></td><td></td><td></td><td></td><td></td></tr>
<tr><td>Oligo fwd</td><td>CACCTAAAAATTATACCCGGTTAT</td><td></td><td></td><td></td><td></td><td></td><td></td><td></td><td></td><td></td><td></td></tr>
<tr><td>Oligo rev</td><td>AAACATAACCGGGTATAATTTTTA</td><td></td><td></td><td></td><td></td><td></td><td></td><td></td><td></td><td></td><td></td></tr>
<tr><td>Chromosome</td><td>start</td><td>end</td><td>strand</td><td>MM</td><td>target_seq</td><td>PAM</td><td>alignment</td><td>distance</td><td>position</td><td>gene name</td><td>gene id</td></tr>
<tr><td>GU305914.1</td><td>221970</td><td>221992</td><td>+</td><td>0</td><td>TAAAAATTATACCCGGTTAT</td><td>AGG</td><td>||||||||[||||||||||||]PAM</td><td>NA</td><td>Intergenic</td><td>NA</td><td>NA</td></tr>
<tr><td>T16490</td><td>AAAAATTATACCCGGTTATAGGG</td><td>1000</td><td></td><td></td><td></td><td></td><td></td><td></td><td></td><td></td><td></td></tr>
<tr><td>Oligo fwd</td><td>CACCAAAAATTATACCCGGTTATA</td><td></td><td></td><td></td><td></td><td></td><td></td><td></td><td></td><td></td><td></td></tr>
<tr><td>Oligo rev</td><td>AAACTATAACCGGGTATAATTTTT</td><td></td><td></td><td></td><td></td><td></td><td></td><td></td><td></td><td></td><td></td></tr>
<tr><td>Chromosome</td><td>start</td><td>end</td><td>strand</td><td>MM</td><td>target_seq</td><td>PAM</td><td>alignment</td><td>distance</td><td>position</td><td>gene name</td><td>gene id</td></tr>
<tr><td>GU305914.1</td><td>221971</td><td>221993</td><td>+</td><td>0</td><td>AAAAATTATACCCGGTTATA</td><td>GGG</td><td>||||||||[||||||||||||]PAM</td><td>NA</td><td>Intergenic</td><td>NA</td><td>NA</td></tr>
<tr><td>T16491</td><td>TTATACCCGGTTATAGGGAACGG</td><td>1000</td><td></td><td></td><td></td><td></td><td></td><td></td><td></td><td></td><td></td></tr>
<tr><td>Oligo fwd</td><td>CACCTTATACCCGGTTATAGGGAA</td><td></td><td></td><td></td><td></td><td></td><td></td><td></td><td></td><td></td><td></td></tr>
<tr><td>Oligo rev</td><td>AAACTTCCCTATAACCGGGTATAA</td><td></td><td></td><td></td><td></td><td></td><td></td><td></td><td></td><td></td><td></td></tr>
<tr><td>Chromosome</td><td>start</td><td>end</td><td>strand</td><td>MM</td><td>target_seq</td><td>PAM</td><td>alignment</td><td>distance</td><td>position</td><td>gene name</td><td>gene id</td></tr>
<tr><td>GU305914.1</td><td>221976</td><td>221998</td><td>+</td><td>0</td><td>TTATACCCGGTTATAGGGAA</td><td>CGG</td><td>||||||||[||||||||||||]PAM</td><td>NA</td><td>Intergenic</td><td>NA</td><td>NA</td></tr>
<tr><td>T16492</td><td>TATACCCGGTTATAGGGAACGGG</td><td>1000</td><td></td><td></td><td></td><td></td><td></td><td></td><td></td><td></td><td></td></tr>
<tr><td>Oligo fwd</td><td>CACCTATACCCGGTTATAGGGAAC</td><td></td><td></td><td></td><td></td><td></td><td></td><td></td><td></td><td></td><td></td></tr>
<tr><td>Oligo rev</td><td>AAACGTTCCCTATAACCGGGTATA</td><td></td><td></td><td></td><td></td><td></td><td></td><td></td><td></td><td></td><td></td></tr>
<tr><td>Chromosome</td><td>start</td><td>end</td><td>strand</td><td>MM</td><td>target_seq</td><td>PAM</td><td>alignment</td><td>distance</td><td>position</td><td>gene name</td><td>gene id</td></tr>
<tr><td>GU305914.1</td><td>221977</td><td>221999</td><td>+</td><td>0</td><td>TATACCCGGTTATAGGGAAC</td><td>GGG</td><td>||||||||[||||||||||||]PAM</td><td>NA</td><td>Intergenic</td><td>NA</td><td>NA</td></tr>
<tr><td>T16493</td><td>CAGTCTCTCTTTGTCACCGCCGG</td><td>1000</td><td></td><td></td><td></td><td></td><td></td><td></td><td></td><td></td><td></td></tr>
<tr><td>Oligo fwd</td><td>CACCCAGTCTCTCTTTGTCACCGC</td><td></td><td></td><td></td><td></td><td></td><td></td><td></td><td></td><td></td><td></td></tr>
<tr><td>Oligo rev</td><td>AAACGCGGTGACAAAGAGAGACTG</td><td></td><td></td><td></td><td></td><td></td><td></td><td></td><td></td><td></td><td></td></tr>
</table>

<table>
<tr><td>Chromosome</td><td>start</td><td></td><td>end</td><td>strand</td><td>MM</td><td>target_seq</td><td>PAM</td><td>alignment</td><td>distance</td><td>position</td><td>gene name</td><td>gene id</td></tr>
<tr><td>GU305914.1</td><td></td><td>222003</td><td>222025</td><td>+</td><td>0</td><td>CAGTCTCTCTTTGTCACCGC</td><td>CGG</td><td>||||||||[||||||||||||]PAM</td><td>NA</td><td>Intergenic</td><td>NA</td><td>NA</td></tr>
<tr><td></td><td></td><td></td><td></td><td></td><td></td><td></td><td></td><td></td><td></td><td></td><td></td><td></td></tr>
<tr><td>T16494</td><td>CCGCCGGATCGCGATCCGTTCGG</td><td></td><td>1000</td><td></td><td></td><td></td><td></td><td></td><td></td><td></td><td></td><td></td></tr>
<tr><td>Oligo fwd</td><td>CACCCCGCCGGATCGCGATCCGTT</td><td></td><td></td><td></td><td></td><td></td><td></td><td></td><td></td><td></td><td></td><td></td></tr>
<tr><td>Oligo rev</td><td>AAACAACGGATCGCGATCCGGCGG</td><td></td><td></td><td></td><td></td><td></td><td></td><td></td><td></td><td></td><td></td><td></td></tr>
<tr><td>Chromosome</td><td>start</td><td></td><td>end</td><td>strand</td><td>MM</td><td>target_seq</td><td>PAM</td><td>alignment</td><td>distance</td><td>position</td><td>gene name</td><td>gene id</td></tr>
<tr><td>GU305914.1</td><td></td><td>222019</td><td>222041</td><td>+</td><td>0</td><td>CCGCCGGATCGCGATCCGTT</td><td>CGG</td><td>||||||||[||||||||||||]PAM</td><td>NA</td><td>Intergenic</td><td>NA</td><td>NA</td></tr>
<tr><td></td><td></td><td></td><td></td><td></td><td></td><td></td><td></td><td></td><td></td><td></td><td></td><td></td></tr>
<tr><td>T16495</td><td>GCGATCCGTTCGGCCGTCTGAGG</td><td></td><td>1000</td><td></td><td></td><td></td><td></td><td></td><td></td><td></td><td></td><td></td></tr>
<tr><td>Oligo fwd</td><td>CACCGCGATCCGTTCGGCCGTCTG</td><td></td><td></td><td></td><td></td><td></td><td></td><td></td><td></td><td></td><td></td><td></td></tr>
<tr><td>Oligo rev</td><td>AAACCAGACGGCCGAACGGATCGC</td><td></td><td></td><td></td><td></td><td></td><td></td><td></td><td></td><td></td><td></td><td></td></tr>
<tr><td>Chromosome</td><td>start</td><td></td><td>end</td><td>strand</td><td>MM</td><td>target_seq</td><td>PAM</td><td>alignment</td><td>distance</td><td>position</td><td>gene name</td><td>gene id</td></tr>
<tr><td>GU305914.1</td><td></td><td>222029</td><td>222051</td><td>+</td><td>0</td><td>GCGATCCGTTCGGCCGTCTG</td><td>AGG</td><td>||||||||[||||||||||||]PAM</td><td>NA</td><td>Intergenic</td><td>NA</td><td>NA</td></tr>
<tr><td></td><td></td><td></td><td></td><td></td><td></td><td></td><td></td><td></td><td></td><td></td><td></td><td></td></tr>
<tr><td>T16496</td><td>GTACACGACCGCTAACACCGCGG</td><td></td><td>1000</td><td></td><td></td><td></td><td></td><td></td><td></td><td></td><td></td><td></td></tr>
<tr><td>Oligo fwd</td><td>CACCGTACACGACCGCTAACACCG</td><td></td><td></td><td></td><td></td><td></td><td></td><td></td><td></td><td></td><td></td><td></td></tr>
<tr><td>Oligo rev</td><td>AAACCGGTGTTAGCGGTCGTGTAC</td><td></td><td></td><td></td><td></td><td></td><td></td><td></td><td></td><td></td><td></td><td></td></tr>
<tr><td>Chromosome</td><td>start</td><td></td><td>end</td><td>strand</td><td>MM</td><td>target_seq</td><td>PAM</td><td>alignment</td><td>distance</td><td>position</td><td>gene name</td><td>gene id</td></tr>
<tr><td>GU305914.1</td><td></td><td>222051</td><td>222073</td><td>+</td><td>0</td><td>GTACACGACCGCTAACACCG</td><td>CGG</td><td>||||||||[||||||||||||]PAM</td><td>NA</td><td>Intergenic</td><td>NA</td><td>NA</td></tr>
<tr><td></td><td></td><td></td><td></td><td></td><td></td><td></td><td></td><td></td><td></td><td></td><td></td><td></td></tr>
<tr><td>T16497</td><td>GATCAGTCCGATGAGCAGAAAGG</td><td></td><td>1000</td><td></td><td></td><td></td><td></td><td></td><td></td><td></td><td></td><td></td></tr>
<tr><td>Oligo fwd</td><td>CACCGATCAGTCCGATGAGCAGAA</td><td></td><td></td><td></td><td></td><td></td><td></td><td></td><td></td><td></td><td></td><td></td></tr>
<tr><td>Oligo rev</td><td>AAACTTCTGCTCATCGGACTGATC</td><td></td><td></td><td></td><td></td><td></td><td></td><td></td><td></td><td></td><td></td><td></td></tr>
<tr><td>Chromosome</td><td>start</td><td></td><td>end</td><td>strand</td><td>MM</td><td>target_seq</td><td>PAM</td><td>alignment</td><td>distance</td><td>position</td><td>gene name</td><td>gene id</td></tr>
<tr><td>GU305914.1</td><td></td><td>222078</td><td>222100</td><td>+</td><td>0</td><td>GATCAGTCCGATGAGCAGAA</td><td>AGG</td><td>||||||||[||||||||||||]PAM</td><td>NA</td><td>Intergenic</td><td>NA</td><td>NA</td></tr>
<tr><td></td><td></td><td></td><td></td><td></td><td></td><td></td><td></td><td></td><td></td><td></td><td></td><td></td></tr>
<tr><td>T16498</td><td>TCCGATGAGCAGAAAGGTCACGG</td><td></td><td>1000</td><td></td><td></td><td></td><td></td><td></td><td></td><td></td><td></td><td></td></tr>
<tr><td>Oligo fwd</td><td>CACCTCCGATGAGCAGAAAGGTCA</td><td></td><td></td><td></td><td></td><td></td><td></td><td></td><td></td><td></td><td></td><td></td></tr>
<tr><td>Oligo rev</td><td>AAACTGACCTTTCTGCTCATCGGA</td><td></td><td></td><td></td><td></td><td></td><td></td><td></td><td></td><td></td><td></td><td></td></tr>
<tr><td>Chromosome</td><td>start</td><td></td><td>end</td><td>strand</td><td>MM</td><td>target_seq</td><td>PAM</td><td>alignment</td><td>distance</td><td>position</td><td>gene name</td><td>gene id</td></tr>
<tr><td>GU305914.1</td><td></td><td>222084</td><td>222106</td><td>+</td><td>0</td><td>TCCGATGAGCAGAAAGGTCA</td><td>CGG</td><td>||||||||[||||||||||||]PAM</td><td>NA</td><td>Intergenic</td><td>NA</td><td>NA</td></tr>
<tr><td></td><td></td><td></td><td></td><td></td><td></td><td></td><td></td><td></td><td></td><td></td><td></td><td></td></tr>
<tr><td>T16499</td><td>AAGGTCACGGCCGCGACGAAAGG</td><td></td><td>1000</td><td></td><td></td><td></td><td></td><td></td><td></td><td></td><td></td><td></td></tr>
<tr><td>Oligo fwd</td><td>CACCAAGGTCACGGCCGCGACGAA</td><td></td><td></td><td></td><td></td><td></td><td></td><td></td><td></td><td></td><td></td><td></td></tr>
<tr><td>Oligo rev</td><td>AAACTTCGTCGCGGCCGTGACCTT</td><td></td><td></td><td></td><td></td><td></td><td></td><td></td><td></td><td></td><td></td><td></td></tr>
<tr><td>Chromosome</td><td>start</td><td></td><td>end</td><td>strand</td><td>MM</td><td>target_seq</td><td>PAM</td><td>alignment</td><td>distance</td><td>position</td><td>gene name</td><td>gene id</td></tr>
<tr><td>GU305914.1</td><td></td><td>222097</td><td>222119</td><td>+</td><td>0</td><td>AAGGTCACGGCCGCGACGAA</td><td>AGG</td><td>||||||||[||||||||||||]PAM</td><td>NA</td><td>Intergenic</td><td>NA</td><td>NA</td></tr>
<tr><td></td><td></td><td></td><td></td><td></td><td></td><td></td><td></td><td></td><td></td><td></td><td></td><td></td></tr>
<tr><td>T16500</td><td>ATCGTCGTTTATCAGAAGCGAGG</td><td></td><td>1000</td><td></td><td></td><td></td><td></td><td></td><td></td><td></td><td></td><td></td></tr>
<tr><td>Oligo fwd</td><td>CACCATCGTCGTTTATCAGAAGCG</td><td></td><td></td><td></td><td></td><td></td><td></td><td></td><td></td><td></td><td></td><td></td></tr>
<tr><td>Oligo rev</td><td>AAACCGCTTCTGATAAACGACGAT</td><td></td><td></td><td></td><td></td><td></td><td></td><td></td><td></td><td></td><td></td><td></td></tr>
<tr><td>Chromosome</td><td>start</td><td></td><td>end</td><td>strand</td><td>MM</td><td>target_seq</td><td>PAM</td><td>alignment</td><td>distance</td><td>position</td><td>gene name</td><td>gene id</td></tr>
<tr><td>GU305914.1</td><td></td><td>222144</td><td>222166</td><td>+</td><td>0</td><td>ATCGTCGTTTATCAGAAGCG</td><td>AGG</td><td>||||||||[||||||||||||]PAM</td><td>NA</td><td>Intergenic</td><td>NA</td><td>NA</td></tr>
<tr><td></td><td></td><td></td><td></td><td></td><td></td><td></td><td></td><td></td><td></td><td></td><td></td><td></td></tr>
<tr><td>T16501</td><td>TCGTTTATCAGAAGCGAGGTTGG</td><td></td><td>1000</td><td></td><td></td><td></td><td></td><td></td><td></td><td></td><td></td><td></td></tr>
<tr><td>Oligo fwd</td><td>CACCTCGTTTATCAGAAGCGAGGT</td><td></td><td></td><td></td><td></td><td></td><td></td><td></td><td></td><td></td><td></td><td></td></tr>
<tr><td>Oligo rev</td><td>AAACACCTCGCTTCTGATAAACGA</td><td></td><td></td><td></td><td></td><td></td><td></td><td></td><td></td><td></td><td></td><td></td></tr>
<tr><td>Chromosome</td><td>start</td><td></td><td>end</td><td>strand</td><td>MM</td><td>target_seq</td><td>PAM</td><td>alignment</td><td>distance</td><td>position</td><td>gene name</td><td>gene id</td></tr>
<tr><td>GU305914.1</td><td></td><td>222148</td><td>222170</td><td>+</td><td>0</td><td>TCGTTTATCAGAAGCGAGGT</td><td>TGG</td><td>||||||||[||||||||||||]PAM</td><td>NA</td><td>Intergenic</td><td>NA</td><td>NA</td></tr>
<tr><td></td><td></td><td></td><td></td><td></td><td></td><td></td><td></td><td></td><td></td><td></td><td></td><td></td></tr>
<tr><td>T16502</td><td>CGTTTATCAGAAGCGAGGTTGGG</td><td></td><td>1000</td><td></td><td></td><td></td><td></td><td></td><td></td><td></td><td></td><td></td></tr>
<tr><td>Oligo fwd</td><td>CACCCGTTTATCAGAAGCGAGGTT</td><td></td><td></td><td></td><td></td><td></td><td></td><td></td><td></td><td></td><td></td><td></td></tr>
<tr><td>Oligo rev</td><td>AAACAACCTCGCTTCTGATAAACG</td><td></td><td></td><td></td><td></td><td></td><td></td><td></td><td></td><td></td><td></td><td></td></tr>
<tr><td>Chromosome</td><td>start</td><td></td><td>end</td><td>strand</td><td>MM</td><td>target_seq</td><td>PAM</td><td>alignment</td><td>distance</td><td>position</td><td>gene name</td><td>gene id</td></tr>
<tr><td>GU305914.1</td><td></td><td>222149</td><td>222171</td><td>+</td><td>0</td><td>CGTTTATCAGAAGCGAGGTT</td><td>GGG</td><td>||||||||[||||||||||||]PAM</td><td>NA</td><td>Intergenic</td><td>NA</td><td>NA</td></tr>
<tr><td></td><td></td><td></td><td></td><td></td><td></td><td></td><td></td><td></td><td></td><td></td><td></td><td></td></tr>
<tr><td>T16503</td><td>GTTTATCAGAAGCGAGGTTGGGG</td><td></td><td>1000</td><td></td><td></td><td></td><td></td><td></td><td></td><td></td><td></td><td></td></tr>
<tr><td>Oligo fwd</td><td>CACCGTTTATCAGAAGCGAGGTTG</td><td></td><td></td><td></td><td></td><td></td><td></td><td></td><td></td><td></td><td></td><td></td></tr>
<tr><td>Oligo rev</td><td>AAACCAACCTCGCTTCTGATAAAC</td><td></td><td></td><td></td><td></td><td></td><td></td><td></td><td></td><td></td><td></td><td></td></tr>
<tr><td>Chromosome</td><td>start</td><td></td><td>end</td><td>strand</td><td>MM</td><td>target_seq</td><td>PAM</td><td>alignment</td><td>distance</td><td>position</td><td>gene name</td><td>gene id</td></tr>
<tr><td>GU305914.1</td><td></td><td>222150</td><td>222172</td><td>+</td><td>0</td><td>GTTTATCAGAAGCGAGGTTG</td><td>GGG</td><td>||||||||[||||||||||||]PAM</td><td>NA</td><td>Intergenic</td><td>NA</td><td>NA</td></tr>
<tr><td></td><td></td><td></td><td></td><td></td><td></td><td></td><td></td><td></td><td></td><td></td><td></td><td></td></tr>
<tr><td>T16504</td><td>CAGAAGCGAGGTTGGGGCGTCGG</td><td></td><td>1000</td><td></td><td></td><td></td><td></td><td></td><td></td><td></td><td></td><td></td></tr>
<tr><td>Oligo fwd</td><td>CACCCAGAAGCGAGGTTGGGGCGT</td><td></td><td></td><td></td><td></td><td></td><td></td><td></td><td></td><td></td><td></td><td></td></tr>
<tr><td>Oligo rev</td><td>AAACACGCCCCAACCTCGCTTCTG</td><td></td><td></td><td></td><td></td><td></td><td></td><td></td><td></td><td></td><td></td><td></td></tr>
<tr><td>Chromosome</td><td>start</td><td></td><td>end</td><td>strand</td><td>MM</td><td>target_seq</td><td>PAM</td><td>alignment</td><td>distance</td><td>position</td><td>gene name</td><td>gene id</td></tr>
<tr><td>GU305914.1</td><td></td><td>222156</td><td>222178</td><td>+</td><td>0</td><td>CAGAAGCGAGGTTGGGGCGT</td><td>CGG</td><td>||||||||[||||||||||||]PAM</td><td>NA</td><td>Intergenic</td><td>NA</td><td>NA</td></tr>
<tr><td></td><td></td><td></td><td></td><td></td><td></td><td></td><td></td><td></td><td></td><td></td><td></td><td></td></tr>
<tr><td>T16505</td><td>AGAAGCGAGGTTGGGGCGTCGGG</td><td></td><td>1000</td><td></td><td></td><td></td><td></td><td></td><td></td><td></td><td></td><td></td></tr>
<tr><td>Oligo fwd</td><td>CACCAGAAGCGAGGTTGGGGCGTC</td><td></td><td></td><td></td><td></td><td></td><td></td><td></td><td></td><td></td><td></td><td></td></tr>
<tr><td>Oligo rev</td><td>AAACGACGCCCCAACCTCGCTTCT</td><td></td><td></td><td></td><td></td><td></td><td></td><td></td><td></td><td></td><td></td><td></td></tr>
<tr><td>Chromosome</td><td>start</td><td></td><td>end</td><td>strand</td><td>MM</td><td>target_seq</td><td>PAM</td><td>alignment</td><td>distance</td><td>position</td><td>gene name</td><td>gene id</td></tr>
<tr><td>GU305914.1</td><td></td><td>222157</td><td>222179</td><td>+</td><td>0</td><td>AGAAGCGAGGTTGGGGCGTC</td><td>GGG</td><td>||||||||[||||||||||||]PAM</td><td>NA</td><td>Intergenic</td><td>NA</td><td>NA</td></tr>
<tr><td></td><td></td><td></td><td></td><td></td><td></td><td></td><td></td><td></td><td></td><td></td><td></td><td></td></tr>
<tr><td>T16506</td><td>AGCGAGGTTGGGGCGTCGGGTGG</td><td></td><td>1000</td><td></td><td></td><td></td><td></td><td></td><td></td><td></td><td></td><td></td></tr>
<tr><td>Oligo fwd</td><td>CACCAGCGAGGTTGGGGCGTCGGG</td><td></td><td></td><td></td><td></td><td></td><td></td><td></td><td></td><td></td><td></td><td></td></tr>
<tr><td>Oligo rev</td><td>AAACCCCGACGCCCCAACCTCGCT</td><td></td><td></td><td></td><td></td><td></td><td></td><td></td><td></td><td></td><td></td><td></td></tr>
<tr><td>Chromosome</td><td>start</td><td></td><td>end</td><td>strand</td><td>MM</td><td>target_seq</td><td>PAM</td><td>alignment</td><td>distance</td><td>position</td><td>gene name</td><td>gene id</td></tr>
<tr><td>GU305914.1</td><td></td><td>222160</td><td>222182</td><td>+</td><td>0</td><td>AGCGAGGTTGGGGCGTCGGG</td><td>TGG</td><td>||||||||[||||||||||||]PAM</td><td>NA</td><td>Intergenic</td><td>NA</td><td>NA</td></tr>
<tr><td></td><td></td><td></td><td></td><td></td><td></td><td></td><td></td><td></td><td></td><td></td><td></td><td></td></tr>
<tr><td>T16507</td><td>GCGAGGTTGGGGCGTCGGGTGGG</td><td></td><td>1000</td><td></td><td></td><td></td><td></td><td></td><td></td><td></td><td></td><td></td></tr>
<tr><td>Oligo fwd</td><td>CACCGCGAGGTTGGGGCGTCGGGT</td><td></td><td></td><td></td><td></td><td></td><td></td><td></td><td></td><td></td><td></td><td></td></tr>
<tr><td>Oligo rev</td><td>AAACACCCGACGCCCCAACCTCGC</td><td></td><td></td><td></td><td></td><td></td><td></td><td></td><td></td><td></td><td></td><td></td></tr>
<tr><td>Chromosome</td><td>start</td><td></td><td>end</td><td>strand</td><td>MM</td><td>target_seq</td><td>PAM</td><td>alignment</td><td>distance</td><td>position</td><td>gene name</td><td>gene id</td></tr>
<tr><td>GU305914.1</td><td></td><td>222161</td><td>222183</td><td>+</td><td>0</td><td>GCGAGGTTGGGGCGTCGGGT</td><td>GGG</td><td>||||||||[||||||||||||]PAM</td><td>NA</td><td>Intergenic</td><td>NA</td><td>NA</td></tr>
<tr><td></td><td></td><td></td><td></td><td></td><td></td><td></td><td></td><td></td><td></td><td></td><td></td><td></td></tr>
<tr><td>T16508</td><td>AGGTTGGGGCGTCGGGTGGGTGG</td><td></td><td>1000</td><td></td><td></td><td></td><td></td><td></td><td></td><td></td><td></td><td></td></tr>
<tr><td>Oligo fwd</td><td>CACCAGGTTGGGGCGTCGGGTGGG</td><td></td><td></td><td></td><td></td><td></td><td></td><td></td><td></td><td></td><td></td><td></td></tr>
</table>

Oligo rev AAACCCCACCCGACGCCCCAACCT
Chromosome start end strand MM target_seq PAM alignment distance position gene name gene id
GU305914.1 222164 222186 + 0 AGGTTGGGGCGTCGGGTGGG TGG ||||||||[||||||||||||]PAM NA Intergenic NA NA

T16509 GGGCGTCGGGTGGGTGGTGCTGG 1000
Oligo fwd CACCGGGCGTCGGGTGGGTGGTGC
Oligo rev AAACGCACCACCCACCCGACGCCC
Chromosome start end strand MM target_seq PAM alignment distance position gene name gene id
GU305914.1 222170 222192 + 0 GGGCGTCGGGTGGGTGGTGC TGG ||||||||[||||||||||||]PAM NA Intergenic NA NA

T16510 CGTCGGGTGGGTGGTGCTGGTGG 1000
Oligo fwd CACCCGTCGGGTGGGTGGTGCTGG
Oligo rev AAACCCAGCACCACCCACCCGACG
Chromosome start end strand MM target_seq PAM alignment distance position gene name gene id
GU305914.1 222173 222195 + 0 CGTCGGGTGGGTGGTGCTGG TGG ||||||||[||||||||||||]PAM NA Intergenic NA NA

T16511 GTCGGGTGGGTGGTGCTGGTGGG 1000
Oligo fwd CACCGTCGGGTGGGTGGTGCTGGT
Oligo rev AAACACCAGCACCACCCACCCGAC
Chromosome start end strand MM target_seq PAM alignment distance position gene name gene id
GU305914.1 222174 222196 + 0 GTCGGGTGGGTGGTGCTGGT GGG ||||||||[||||||||||||]PAM NA Intergenic NA NA

T16512 GGTGGGAATCGTTCTTCAGATGG 1000
Oligo fwd CACCGGTGGGAATCGTTCTTCAGA
Oligo rev AAACTCTGAAGAACGATTCCCACC
Chromosome start end strand MM target_seq PAM alignment distance position gene name gene id
GU305914.1 222191 222213 + 0 GGTGGGAATCGTTCTTCAGA TGG ||||||||[||||||||||||]PAM NA Intergenic NA NA

T16513 GCCTTCTGCGTCGACCTCGACGG 1000
Oligo fwd CACCGCCTTCTGCGTCGACCTCGA
Oligo rev AAACTCGAGGTCGACGCAGAAGGC
Chromosome start end strand MM target_seq PAM alignment distance position gene name gene id
GU305914.1 222213 222235 + 0 GCCTTCTGCGTCGACCTCGA CGG ||||||||[||||||||||||]PAM NA Intergenic NA NA

T16514 GCGTCGACCTCGACGGCGCCCGG 1000
Oligo fwd CACCGCGTCGACCTCGACGGCGCC
Oligo rev AAACGGCGCCGTCGAGGTCGACGC
Chromosome start end strand MM target_seq PAM alignment distance position gene name gene id
GU305914.1 222220 222242 + 0 GCGTCGACCTCGACGGCGCC CGG ||||||||[||||||||||||]PAM NA Intergenic NA NA

T16515 ACCTCGACGGCGCCCGGCTCTGG 1000
Oligo fwd CACCACCTCGACGGCGCCCGGCTC
Oligo rev AAACGAGCCGGGCGCCGTCGAGGT
Chromosome start end strand MM target_seq PAM alignment distance position gene name gene id
GU305914.1 222226 222248 + 0 ACCTCGACGGCGCCCGGCTC TGG ||||||||[||||||||||||]PAM NA Intergenic NA NA

T16516 CCTCGACGGCGCCCGGCTCTGGG 1000
Oligo fwd CACCCCTCGACGGCGCCCGGCTCT
Oligo rev AAACAGAGCCGGGCGCCGTCGAGG
Chromosome start end strand MM target_seq PAM alignment distance position gene name gene id
GU305914.1 222227 222249 + 0 CCTCGACGGCGCCCGGCTCT GGG ||||||||[||||||||||||]PAM NA Intergenic NA NA

T16517 CTCGACGGCGCCCGGCTCTGGGG 1000
Oligo fwd CACCCTCGACGGCGCCCGGCTCTG
Oligo rev AAACCAGAGCCGGGCGCCGTCGAG
Chromosome start end strand MM target_seq PAM alignment distance position gene name gene id
GU305914.1 222228 222250 + 0 CTCGACGGCGCCCGGCTCTG GGG ||||||||[||||||||||||]PAM NA Intergenic NA NA

T16518 CCGGCTCTGGGGCGACTTGTTGG 1000
Oligo fwd CACCCCGGCTCTGGGGCGACTTGT
Oligo rev AAACACAAGTCGCCCCAGAGCCGG
Chromosome start end strand MM target_seq PAM alignment distance position gene name gene id
GU305914.1 222239 222261 + 0 CCGGCTCTGGGGCGACTTGT TGG ||||||||[||||||||||||]PAM NA Intergenic NA NA

T16519 CGGCTCTGGGGCGACTTGTTGGG 1000
Oligo fwd CACCCGGCTCTGGGGCGACTTGTT
Oligo rev AAACAACAAGTCGCCCCAGAGCCG
Chromosome start end strand MM target_seq PAM alignment distance position gene name gene id
GU305914.1 222240 222262 + 0 CGGCTCTGGGGCGACTTGTT GGG ||||||||[||||||||||||]PAM NA Intergenic NA NA

T16520 GTATATCTATCATCTCCGTGAGG 1000
Oligo fwd CACCGTATATCTATCATCTCCGTG
Oligo rev AAACCACGGAGATGATAGATATAC
Chromosome start end strand MM target_seq PAM alignment distance position gene name gene id
GU305914.1 222281 222303 + 0 GTATATCTATCATCTCCGTG AGG ||||||||[||||||||||||]PAM NA Intergenic NA NA

T16521 ATCTATCATCTCCGTGAGGTAGG 1000
Oligo fwd CACCATCTATCATCTCCGTGAGGT
Oligo rev AAACACCTCACGGAGATGATAGAT
Chromosome start end strand MM target_seq PAM alignment distance position gene name gene id
GU305914.1 222285 222307 + 0 ATCTATCATCTCCGTGAGGT AGG ||||||||[||||||||||||]PAM NA Intergenic NA NA

T16522 CTCCGTGAGGTAGGCGACGTCGG 1000
Oligo fwd CACCCTCCGTGAGGTAGGCGACGT
Oligo rev AAACACGTCGCCTACCTCACGGAG
Chromosome start end strand MM target_seq PAM alignment distance position gene name gene id
GU305914.1 222294 222316 + 0 CTCCGTGAGGTAGGCGACGT CGG ||||||||[||||||||||||]PAM NA Intergenic NA NA

T16523 TCCGTGAGGTAGGCGACGTCGGG 1000

<table>
<tr><td>Oligo fwd</td><td>CACCTCCGTGAGGTAGGCGACGTC</td><td colspan="10"></td></tr>
<tr><td>Oligo rev</td><td>AAACGACGTCGCCTACCTCACGGA</td><td colspan="10"></td></tr>
<tr><td>Chromosome</td><td>start</td><td>end</td><td>strand</td><td>MM</td><td>target_seq</td><td>PAM</td><td>alignment</td><td>distance</td><td>position</td><td>gene name</td><td>gene id</td></tr>
<tr><td>GU305914.1</td><td>222295</td><td>222317</td><td>+</td><td>0</td><td>TCCGTGAGGTAGGCGACGTC</td><td>GGG</td><td>||||||||[||||||||||||]PAM</td><td>NA</td><td>Intergenic</td><td>NA</td><td>NA</td></tr>
<tr><td>T16524</td><td>GACCCGAAGACGATACCGCGTGG</td><td>1000</td><td colspan="9"></td></tr>
<tr><td>Oligo fwd</td><td>CACCGACCCGAAGACGATACCGCG</td><td colspan="10"></td></tr>
<tr><td>Oligo rev</td><td>AAACCGCGGTATCGTCTTCGGGTC</td><td colspan="10"></td></tr>
<tr><td>Chromosome</td><td>start</td><td>end</td><td>strand</td><td>MM</td><td>target_seq</td><td>PAM</td><td>alignment</td><td>distance</td><td>position</td><td>gene name</td><td>gene id</td></tr>
<tr><td>GU305914.1</td><td>222317</td><td>222339</td><td>+</td><td>0</td><td>GACCCGAAGACGATACCGCG</td><td>TGG</td><td>||||||||[||||||||||||]PAM</td><td>NA</td><td>Intergenic</td><td>NA</td><td>NA</td></tr>
<tr><td>T16525</td><td>ACCCGAAGACGATACCGCGTGGG</td><td>1000</td><td colspan="9"></td></tr>
<tr><td>Oligo fwd</td><td>CACCACCCGAAGACGATACCGCGT</td><td colspan="10"></td></tr>
<tr><td>Oligo rev</td><td>AAACACGCGGTATCGTCTTCGGGT</td><td colspan="10"></td></tr>
<tr><td>Chromosome</td><td>start</td><td>end</td><td>strand</td><td>MM</td><td>target_seq</td><td>PAM</td><td>alignment</td><td>distance</td><td>position</td><td>gene name</td><td>gene id</td></tr>
<tr><td>GU305914.1</td><td>222318</td><td>222340</td><td>+</td><td>0</td><td>ACCCGAAGACGATACCGCGT</td><td>GGG</td><td>||||||||[||||||||||||]PAM</td><td>NA</td><td>Intergenic</td><td>NA</td><td>NA</td></tr>
<tr><td>T16526</td><td>CTGCAGCGCCTCTTGAACGTCGG</td><td>1000</td><td colspan="9"></td></tr>
<tr><td>Oligo fwd</td><td>CACCCTGCAGCGCCTCTTGAACGT</td><td colspan="10"></td></tr>
<tr><td>Oligo rev</td><td>AAACACGTTCAAGAGGCGCTGCAG</td><td colspan="10"></td></tr>
<tr><td>Chromosome</td><td>start</td><td>end</td><td>strand</td><td>MM</td><td>target_seq</td><td>PAM</td><td>alignment</td><td>distance</td><td>position</td><td>gene name</td><td>gene id</td></tr>
<tr><td>GU305914.1</td><td>222342</td><td>222364</td><td>+</td><td>0</td><td>CTGCAGCGCCTCTTGAACGT</td><td>CGG</td><td>||||||||[||||||||||||]PAM</td><td>NA</td><td>Intergenic</td><td>NA</td><td>NA</td></tr>
<tr><td>T16527</td><td>TGCAGCGCCTCTTGAACGTCGGG</td><td>1000</td><td colspan="9"></td></tr>
<tr><td>Oligo fwd</td><td>CACCTGCAGCGCCTCTTGAACGTC</td><td colspan="10"></td></tr>
<tr><td>Oligo rev</td><td>AAACGACGTTCAAGAGGCGCTGCA</td><td colspan="10"></td></tr>
<tr><td>Chromosome</td><td>start</td><td>end</td><td>strand</td><td>MM</td><td>target_seq</td><td>PAM</td><td>alignment</td><td>distance</td><td>position</td><td>gene name</td><td>gene id</td></tr>
<tr><td>GU305914.1</td><td>222343</td><td>222365</td><td>+</td><td>0</td><td>TGCAGCGCCTCTTGAACGTC</td><td>GGG</td><td>||||||||[||||||||||||]PAM</td><td>NA</td><td>Intergenic</td><td>NA</td><td>NA</td></tr>
<tr><td>T16528</td><td>AACGTCGGGTTGTTCGCTTATGG</td><td>1000</td><td colspan="9"></td></tr>
<tr><td>Oligo fwd</td><td>CACCAACGTCGGGTTGTTCGCTTA</td><td colspan="10"></td></tr>
<tr><td>Oligo rev</td><td>AAACTAAGCGAACAACCCGACGTT</td><td colspan="10"></td></tr>
<tr><td>Chromosome</td><td>start</td><td>end</td><td>strand</td><td>MM</td><td>target_seq</td><td>PAM</td><td>alignment</td><td>distance</td><td>position</td><td>gene name</td><td>gene id</td></tr>
<tr><td>GU305914.1</td><td>222357</td><td>222379</td><td>+</td><td>0</td><td>AACGTCGGGTTGTTCGCTTA</td><td>TGG</td><td>||||||||[||||||||||||]PAM</td><td>NA</td><td>Intergenic</td><td>NA</td><td>NA</td></tr>
<tr><td>T16529</td><td>TCGGGTTGTTCGCTTATGGTCGG</td><td>1000</td><td colspan="9"></td></tr>
<tr><td>Oligo fwd</td><td>CACCTCGGGTTGTTCGCTTATGGT</td><td colspan="10"></td></tr>
<tr><td>Oligo rev</td><td>AAACACCATAAGCGAACAACCCGA</td><td colspan="10"></td></tr>
<tr><td>Chromosome</td><td>start</td><td>end</td><td>strand</td><td>MM</td><td>target_seq</td><td>PAM</td><td>alignment</td><td>distance</td><td>position</td><td>gene name</td><td>gene id</td></tr>
<tr><td>GU305914.1</td><td>222361</td><td>222383</td><td>+</td><td>0</td><td>TCGGGTTGTTCGCTTATGGT</td><td>CGG</td><td>||||||||[||||||||||||]PAM</td><td>NA</td><td>Intergenic</td><td>NA</td><td>NA</td></tr>
<tr><td>T16530</td><td>GCTTATGGTCGGTCCGTCACCGG</td><td>1000</td><td colspan="9"></td></tr>
<tr><td>Oligo fwd</td><td>CACCGCTTATGGTCGGTCCGTCAC</td><td colspan="10"></td></tr>
<tr><td>Oligo rev</td><td>AAACGTGACGGACCGACCATAAGC</td><td colspan="10"></td></tr>
<tr><td>Chromosome</td><td>start</td><td>end</td><td>strand</td><td>MM</td><td>target_seq</td><td>PAM</td><td>alignment</td><td>distance</td><td>position</td><td>gene name</td><td>gene id</td></tr>
<tr><td>GU305914.1</td><td>222372</td><td>222394</td><td>+</td><td>0</td><td>GCTTATGGTCGGTCCGTCAC</td><td>CGG</td><td>||||||||[||||||||||||]PAM</td><td>NA</td><td>Intergenic</td><td>NA</td><td>NA</td></tr>
<tr><td>T16531</td><td>CCGTCACCGGCACCCTCCGTCGG</td><td>1000</td><td colspan="9"></td></tr>
<tr><td>Oligo fwd</td><td>CACCCCGTCACCGGCACCCTCCGT</td><td colspan="10"></td></tr>
<tr><td>Oligo rev</td><td>AAACACGGAGGGTGCCGGTGACGG</td><td colspan="10"></td></tr>
<tr><td>Chromosome</td><td>start</td><td>end</td><td>strand</td><td>MM</td><td>target_seq</td><td>PAM</td><td>alignment</td><td>distance</td><td>position</td><td>gene name</td><td>gene id</td></tr>
<tr><td>GU305914.1</td><td>222385</td><td>222407</td><td>+</td><td>0</td><td>CCGTCACCGGCACCCTCCGT</td><td>CGG</td><td>||||||||[||||||||||||]PAM</td><td>NA</td><td>Intergenic</td><td>NA</td><td>NA</td></tr>
<tr><td>T16532</td><td>CGTCACCGGCACCCTCCGTCGGG</td><td>1000</td><td colspan="9"></td></tr>
<tr><td>Oligo fwd</td><td>CACCCGTCACCGGCACCCTCCGTC</td><td colspan="10"></td></tr>
<tr><td>Oligo rev</td><td>AAACGACGGAGGGTGCCGGTGACG</td><td colspan="10"></td></tr>
<tr><td>Chromosome</td><td>start</td><td>end</td><td>strand</td><td>MM</td><td>target_seq</td><td>PAM</td><td>alignment</td><td>distance</td><td>position</td><td>gene name</td><td>gene id</td></tr>
<tr><td>GU305914.1</td><td>222386</td><td>222408</td><td>+</td><td>0</td><td>CGTCACCGGCACCCTCCGTC</td><td>GGG</td><td>||||||||[||||||||||||]PAM</td><td>NA</td><td>Intergenic</td><td>NA</td><td>NA</td></tr>
<tr><td>T16533</td><td>GCACCCTCCGTCGGGTACGAAGG</td><td>1000</td><td colspan="9"></td></tr>
<tr><td>Oligo fwd</td><td>CACCGCACCCTCCGTCGGGTACGA</td><td colspan="10"></td></tr>
<tr><td>Oligo rev</td><td>AAACTCGTACCCGACGGAGGGTGC</td><td colspan="10"></td></tr>
<tr><td>Chromosome</td><td>start</td><td>end</td><td>strand</td><td>MM</td><td>target_seq</td><td>PAM</td><td>alignment</td><td>distance</td><td>position</td><td>gene name</td><td>gene id</td></tr>
<tr><td>GU305914.1</td><td>222394</td><td>222416</td><td>+</td><td>0</td><td>GCACCCTCCGTCGGGTACGA</td><td>AGG</td><td>||||||||[||||||||||||]PAM</td><td>NA</td><td>Intergenic</td><td>NA</td><td>NA</td></tr>
<tr><td>T16534</td><td>TGCTTGTCGCGCGATCGACGAGG</td><td>1000</td><td colspan="9"></td></tr>
<tr><td>Oligo fwd</td><td>CACCTGCTTGTCGCGCGATCGACG</td><td colspan="10"></td></tr>
<tr><td>Oligo rev</td><td>AAACCGTCGATCGCGCGACAAGCA</td><td colspan="10"></td></tr>
<tr><td>Chromosome</td><td>start</td><td>end</td><td>strand</td><td>MM</td><td>target_seq</td><td>PAM</td><td>alignment</td><td>distance</td><td>position</td><td>gene name</td><td>gene id</td></tr>
<tr><td>GU305914.1</td><td>222431</td><td>222453</td><td>+</td><td>0</td><td>TGCTTGTCGCGCGATCGACG</td><td>AGG</td><td>||||||||[||||||||||||]PAM</td><td>NA</td><td>Intergenic</td><td>NA</td><td>NA</td></tr>
<tr><td>T16535</td><td>GCGCGACGAAGCGTTCGTTGCGG</td><td>1000</td><td colspan="9"></td></tr>
<tr><td>Oligo fwd</td><td>CACCGCGCGACGAAGCGTTCGTTG</td><td colspan="10"></td></tr>
<tr><td>Oligo rev</td><td>AAACCAACGAACGCTTCGTCGCGC</td><td colspan="10"></td></tr>
<tr><td>Chromosome</td><td>start</td><td>end</td><td>strand</td><td>MM</td><td>target_seq</td><td>PAM</td><td>alignment</td><td>distance</td><td>position</td><td>gene name</td><td>gene id</td></tr>
<tr><td>GU305914.1</td><td>222485</td><td>222507</td><td>+</td><td>0</td><td>GCGCGACGAAGCGTTCGTTG</td><td>CGG</td><td>||||||||[||||||||||||]PAM</td><td>NA</td><td>Intergenic</td><td>NA</td><td>NA</td></tr>
<tr><td>T16536</td><td>GCGTTCGTTGCGGCCCTCCATGG</td><td>1000</td><td colspan="9"></td></tr>
<tr><td>Oligo fwd</td><td>CACCGCGTTCGTTGCGGCCCTCCA</td><td colspan="10"></td></tr>
<tr><td>Oligo rev</td><td>AAACTGGAGGGCCGCAACGAACGC</td><td colspan="10"></td></tr>
<tr><td>Chromosome</td><td>start</td><td>end</td><td>strand</td><td>MM</td><td>target_seq</td><td>PAM</td><td>alignment</td><td>distance</td><td>position</td><td>gene name</td><td>gene id</td></tr>
<tr><td>GU305914.1</td><td>222495</td><td>222517</td><td>+</td><td>0</td><td>GCGTTCGTTGCGGCCCTCCA</td><td>TGG</td><td>||||||||[||||||||||||]PAM</td><td>NA</td><td>Intergenic</td><td>NA</td><td>NA</td></tr>
<tr><td>T16537</td><td>TTGCGGCCCTCCATGGCTGATGG</td><td>1000</td><td colspan="9"></td></tr>
<tr><td>Oligo fwd</td><td>CACCTTGCGGCCCTCCATGGCTGA</td><td colspan="10"></td></tr>
<tr><td>Oligo rev</td><td>AAACTCAGCCATGGAGGGCCGCAA</td><td colspan="10"></td></tr>
<tr><td>Chromosome</td><td>start</td><td>end</td><td>strand</td><td>MM</td><td>target_seq</td><td>PAM</td><td>alignment</td><td>distance</td><td>position</td><td>gene name</td><td>gene id</td></tr>
<tr><td>GU305914.1</td><td>222502</td><td>222524</td><td>+</td><td>0</td><td>TTGCGGCCCTCCATGGCTGA</td><td>TGG</td><td>||||||||[||||||||||||]PAM</td><td>NA</td><td>Intergenic</td><td>NA</td><td>NA</td></tr>
</table>

<table>
<tr><td>T16538</td><td>CGCGACACGACCCCATTCACCGG</td><td>1000</td><td></td><td></td><td></td><td></td><td></td><td></td><td></td><td></td><td></td></tr>
<tr><td>Oligo fwd</td><td>CACCCGCGACACGACCCCATTCAC</td><td></td><td></td><td></td><td></td><td></td><td></td><td></td><td></td><td></td><td></td></tr>
<tr><td>Oligo rev</td><td>AAACGTGAATGGGGTCGTGTCGCG</td><td></td><td></td><td></td><td></td><td></td><td></td><td></td><td></td><td></td><td></td></tr>
<tr><td>Chromosome</td><td>start</td><td>end</td><td>strand</td><td>MM</td><td>target_seq</td><td>PAM</td><td>alignment</td><td>distance</td><td>position</td><td>gene name</td><td>gene id</td></tr>
<tr><td>GU305914.1</td><td>222528</td><td>222550</td><td>+</td><td>0</td><td>CGCGACACGACCCCATTCAC</td><td>CGG</td><td>||||||||[||||||||||||]PAM</td><td>NA</td><td>Intergenic</td><td>NA</td><td>NA</td></tr>
<tr><td></td><td></td><td></td><td></td><td></td><td></td><td></td><td></td><td></td><td></td><td></td><td></td></tr>
<tr><td>T16539</td><td>GCACGCCACCAGTAAAATACCGG</td><td>1000</td><td></td><td></td><td></td><td></td><td></td><td></td><td></td><td></td><td></td></tr>
<tr><td>Oligo fwd</td><td>CACCGCACGCCACCAGTAAAATAC</td><td></td><td></td><td></td><td></td><td></td><td></td><td></td><td></td><td></td><td></td></tr>
<tr><td>Oligo rev</td><td>AAACGTATTTTACTGGTGGCGTGC</td><td></td><td></td><td></td><td></td><td></td><td></td><td></td><td></td><td></td><td></td></tr>
<tr><td>Chromosome</td><td>start</td><td>end</td><td>strand</td><td>MM</td><td>target_seq</td><td>PAM</td><td>alignment</td><td>distance</td><td>position</td><td>gene name</td><td>gene id</td></tr>
<tr><td>GU305914.1</td><td>222550</td><td>222572</td><td>+</td><td>0</td><td>GCACGCCACCAGTAAAATAC</td><td>CGG</td><td>||||||||[||||||||||||]PAM</td><td>NA</td><td>Intergenic</td><td>NA</td><td>NA</td></tr>
<tr><td></td><td></td><td></td><td></td><td></td><td></td><td></td><td></td><td></td><td></td><td></td><td></td></tr>
<tr><td>T16540</td><td>AAATACCGGATTTCGAACGTCGG</td><td>1000</td><td></td><td></td><td></td><td></td><td></td><td></td><td></td><td></td><td></td></tr>
<tr><td>Oligo fwd</td><td>CACCAAATACCGGATTTCGAACGT</td><td></td><td></td><td></td><td></td><td></td><td></td><td></td><td></td><td></td><td></td></tr>
<tr><td>Oligo rev</td><td>AAACACGTTCGAAATCCGGTATTT</td><td></td><td></td><td></td><td></td><td></td><td></td><td></td><td></td><td></td><td></td></tr>
<tr><td>Chromosome</td><td>start</td><td>end</td><td>strand</td><td>MM</td><td>target_seq</td><td>PAM</td><td>alignment</td><td>distance</td><td>position</td><td>gene name</td><td>gene id</td></tr>
<tr><td>GU305914.1</td><td>222564</td><td>222586</td><td>+</td><td>0</td><td>AAATACCGGATTTCGAACGT</td><td>CGG</td><td>||||||||[||||||||||||]PAM</td><td>NA</td><td>Intergenic</td><td>NA</td><td>NA</td></tr>
<tr><td></td><td></td><td></td><td></td><td></td><td></td><td></td><td></td><td></td><td></td><td></td><td></td></tr>
<tr><td>T16541</td><td>AATACCGGATTTCGAACGTCGGG</td><td>1000</td><td></td><td></td><td></td><td></td><td></td><td></td><td></td><td></td><td></td></tr>
<tr><td>Oligo fwd</td><td>CACCAATACCGGATTTCGAACGTC</td><td></td><td></td><td></td><td></td><td></td><td></td><td></td><td></td><td></td><td></td></tr>
<tr><td>Oligo rev</td><td>AAACGACGTTCGAAATCCGGTATT</td><td></td><td></td><td></td><td></td><td></td><td></td><td></td><td></td><td></td><td></td></tr>
<tr><td>Chromosome</td><td>start</td><td>end</td><td>strand</td><td>MM</td><td>target_seq</td><td>PAM</td><td>alignment</td><td>distance</td><td>position</td><td>gene name</td><td>gene id</td></tr>
<tr><td>GU305914.1</td><td>222565</td><td>222587</td><td>+</td><td>0</td><td>AATACCGGATTTCGAACGTC</td><td>GGG</td><td>||||||||[||||||||||||]PAM</td><td>NA</td><td>Intergenic</td><td>NA</td><td>NA</td></tr>
<tr><td></td><td></td><td></td><td></td><td></td><td></td><td></td><td></td><td></td><td></td><td></td><td></td></tr>
<tr><td>T16542</td><td>TCGAACGTCGGGAACAGCGACGG</td><td>1000</td><td></td><td></td><td></td><td></td><td></td><td></td><td></td><td></td><td></td></tr>
<tr><td>Oligo fwd</td><td>CACCTCGAACGTCGGGAACAGCGA</td><td></td><td></td><td></td><td></td><td></td><td></td><td></td><td></td><td></td><td></td></tr>
<tr><td>Oligo rev</td><td>AAACTCGCTGTTCCCGACGTTCGA</td><td></td><td></td><td></td><td></td><td></td><td></td><td></td><td></td><td></td><td></td></tr>
<tr><td>Chromosome</td><td>start</td><td>end</td><td>strand</td><td>MM</td><td>target_seq</td><td>PAM</td><td>alignment</td><td>distance</td><td>position</td><td>gene name</td><td>gene id</td></tr>
<tr><td>GU305914.1</td><td>222576</td><td>222598</td><td>+</td><td>0</td><td>TCGAACGTCGGGAACAGCGA</td><td>CGG</td><td>||||||||[||||||||||||]PAM</td><td>NA</td><td>Intergenic</td><td>NA</td><td>NA</td></tr>
<tr><td></td><td></td><td></td><td></td><td></td><td></td><td></td><td></td><td></td><td></td><td></td><td></td></tr>
<tr><td>T16543</td><td>ACGGCCTGACGTCCGAGACCAGG</td><td>1000</td><td></td><td></td><td></td><td></td><td></td><td></td><td></td><td></td><td></td></tr>
<tr><td>Oligo fwd</td><td>CACCACGGCCTGACGTCCGAGACC</td><td></td><td></td><td></td><td></td><td></td><td></td><td></td><td></td><td></td><td></td></tr>
<tr><td>Oligo rev</td><td>AAACGGTCTCGGACGTCAGGCCGT</td><td></td><td></td><td></td><td></td><td></td><td></td><td></td><td></td><td></td><td></td></tr>
<tr><td>Chromosome</td><td>start</td><td>end</td><td>strand</td><td>MM</td><td>target_seq</td><td>PAM</td><td>alignment</td><td>distance</td><td>position</td><td>gene name</td><td>gene id</td></tr>
<tr><td>GU305914.1</td><td>222595</td><td>222617</td><td>+</td><td>0</td><td>ACGGCCTGACGTCCGAGACC</td><td>AGG</td><td>||||||||[||||||||||||]PAM</td><td>NA</td><td>Intergenic</td><td>NA</td><td>NA</td></tr>
<tr><td></td><td></td><td></td><td></td><td></td><td></td><td></td><td></td><td></td><td></td><td></td><td></td></tr>
<tr><td>T16544</td><td>GTACACGACGATCGTGACGATGG</td><td>1000</td><td></td><td></td><td></td><td></td><td></td><td></td><td></td><td></td><td></td></tr>
<tr><td>Oligo fwd</td><td>CACCGTACACGACGATCGTGACGA</td><td></td><td></td><td></td><td></td><td></td><td></td><td></td><td></td><td></td><td></td></tr>
<tr><td>Oligo rev</td><td>AAACTCGTCACGATCGTCGTGTAC</td><td></td><td></td><td></td><td></td><td></td><td></td><td></td><td></td><td></td><td></td></tr>
<tr><td>Chromosome</td><td>start</td><td>end</td><td>strand</td><td>MM</td><td>target_seq</td><td>PAM</td><td>alignment</td><td>distance</td><td>position</td><td>gene name</td><td>gene id</td></tr>
<tr><td>GU305914.1</td><td>222623</td><td>222645</td><td>+</td><td>0</td><td>GTACACGACGATCGTGACGA</td><td>TGG</td><td>||||||||[||||||||||||]PAM</td><td>NA</td><td>Intergenic</td><td>NA</td><td>NA</td></tr>
<tr><td>GU305914.1</td><td>124214</td><td>124236</td><td>-</td><td>4</td><td>CAACACGCCGATCGTGACCA</td><td>AGG</td><td>--|||||-[||||||||||-|]PAM</td><td>NA</td><td>Intergenic</td><td>NA</td><td>NA</td></tr>
<tr><td></td><td></td><td></td><td></td><td></td><td></td><td></td><td></td><td></td><td></td><td></td><td></td></tr>
<tr><td>T16545</td><td>AATACACAGCATCACCACGAAGG</td><td>1000</td><td></td><td></td><td></td><td></td><td></td><td></td><td></td><td></td><td></td></tr>
<tr><td>Oligo fwd</td><td>CACCAATACACAGCATCACCACGA</td><td></td><td></td><td></td><td></td><td></td><td></td><td></td><td></td><td></td><td></td></tr>
<tr><td>Oligo rev</td><td>AAACTCGTGGTGATGCTGTGTATT</td><td></td><td></td><td></td><td></td><td></td><td></td><td></td><td></td><td></td><td></td></tr>
<tr><td>Chromosome</td><td>start</td><td>end</td><td>strand</td><td>MM</td><td>target_seq</td><td>PAM</td><td>alignment</td><td>distance</td><td>position</td><td>gene name</td><td>gene id</td></tr>
<tr><td>GU305914.1</td><td>222647</td><td>222669</td><td>+</td><td>0</td><td>AATACACAGCATCACCACGA</td><td>AGG</td><td>||||||||[||||||||||||]PAM</td><td>NA</td><td>Intergenic</td><td>NA</td><td>NA</td></tr>
<tr><td></td><td></td><td></td><td></td><td></td><td></td><td></td><td></td><td></td><td></td><td></td><td></td></tr>
<tr><td>T16546</td><td>ACACAGCATCACCACGAAGGCGG</td><td>1000</td><td></td><td></td><td></td><td></td><td></td><td></td><td></td><td></td><td></td></tr>
<tr><td>Oligo fwd</td><td>CACCACACAGCATCACCACGAAGG</td><td></td><td></td><td></td><td></td><td></td><td></td><td></td><td></td><td></td><td></td></tr>
<tr><td>Oligo rev</td><td>AAACCCTTCGTGGTGATGCTGTGT</td><td></td><td></td><td></td><td></td><td></td><td></td><td></td><td></td><td></td><td></td></tr>
<tr><td>Chromosome</td><td>start</td><td>end</td><td>strand</td><td>MM</td><td>target_seq</td><td>PAM</td><td>alignment</td><td>distance</td><td>position</td><td>gene name</td><td>gene id</td></tr>
<tr><td>GU305914.1</td><td>222650</td><td>222672</td><td>+</td><td>0</td><td>ACACAGCATCACCACGAAGG</td><td>CGG</td><td>||||||||[||||||||||||]PAM</td><td>NA</td><td>Intergenic</td><td>NA</td><td>NA</td></tr>
<tr><td></td><td></td><td></td><td></td><td></td><td></td><td></td><td></td><td></td><td></td><td></td><td></td></tr>
<tr><td>T16547</td><td>GCATCACCACGAAGGCGGCGAGG</td><td>1000</td><td></td><td></td><td></td><td></td><td></td><td></td><td></td><td></td><td></td></tr>
<tr><td>Oligo fwd</td><td>CACCGCATCACCACGAAGGCGGCG</td><td></td><td></td><td></td><td></td><td></td><td></td><td></td><td></td><td></td><td></td></tr>
<tr><td>Oligo rev</td><td>AAACCGCCGCCTTCGTGGTGATGC</td><td></td><td></td><td></td><td></td><td></td><td></td><td></td><td></td><td></td><td></td></tr>
<tr><td>Chromosome</td><td>start</td><td>end</td><td>strand</td><td>MM</td><td>target_seq</td><td>PAM</td><td>alignment</td><td>distance</td><td>position</td><td>gene name</td><td>gene id</td></tr>
<tr><td>GU305914.1</td><td>222655</td><td>222677</td><td>+</td><td>0</td><td>GCATCACCACGAAGGCGGCG</td><td>AGG</td><td>||||||||[||||||||||||]PAM</td><td>NA</td><td>Intergenic</td><td>NA</td><td>NA</td></tr>
<tr><td></td><td></td><td></td><td></td><td></td><td></td><td></td><td></td><td></td><td></td><td></td><td></td></tr>
<tr><td>T16548</td><td>ACAGTCGCAGCTCTTCGATCCGG</td><td>1000</td><td></td><td></td><td></td><td></td><td></td><td></td><td></td><td></td><td></td></tr>
<tr><td>Oligo fwd</td><td>CACCACAGTCGCAGCTCTTCGATC</td><td></td><td></td><td></td><td></td><td></td><td></td><td></td><td></td><td></td><td></td></tr>
<tr><td>Oligo rev</td><td>AAACGATCGAAGAGCTGCGACTGT</td><td></td><td></td><td></td><td></td><td></td><td></td><td></td><td></td><td></td><td></td></tr>
<tr><td>Chromosome</td><td>start</td><td>end</td><td>strand</td><td>MM</td><td>target_seq</td><td>PAM</td><td>alignment</td><td>distance</td><td>position</td><td>gene name</td><td>gene id</td></tr>
<tr><td>GU305914.1</td><td>222692</td><td>222714</td><td>+</td><td>0</td><td>ACAGTCGCAGCTCTTCGATC</td><td>CGG</td><td>||||||||[||||||||||||]PAM</td><td>NA</td><td>Intergenic</td><td>NA</td><td>NA</td></tr>
<tr><td></td><td></td><td></td><td></td><td></td><td></td><td></td><td></td><td></td><td></td><td></td><td></td></tr>
<tr><td>T16549</td><td>CGGTTGAACTGCGCGCCACGTGG</td><td>1000</td><td></td><td></td><td></td><td></td><td></td><td></td><td></td><td></td><td></td></tr>
<tr><td>Oligo fwd</td><td>CACCCGGTTGAACTGCGCGCCACG</td><td></td><td></td><td></td><td></td><td></td><td></td><td></td><td></td><td></td><td></td></tr>
<tr><td>Oligo rev</td><td>AAACCGTGGCGCGCAGTTCAACCG</td><td></td><td></td><td></td><td></td><td></td><td></td><td></td><td></td><td></td><td></td></tr>
<tr><td>Chromosome</td><td>start</td><td>end</td><td>strand</td><td>MM</td><td>target_seq</td><td>PAM</td><td>alignment</td><td>distance</td><td>position</td><td>gene name</td><td>gene id</td></tr>
<tr><td>GU305914.1</td><td>222712</td><td>222734</td><td>+</td><td>0</td><td>CGGTTGAACTGCGCGCCACG</td><td>TGG</td><td>||||||||[||||||||||||]PAM</td><td>NA</td><td>Intergenic</td><td>NA</td><td>NA</td></tr>
<tr><td></td><td></td><td></td><td></td><td></td><td></td><td></td><td></td><td></td><td></td><td></td><td></td></tr>
<tr><td>T16550</td><td>TGAACTGCGCGCCACGTGGACGG</td><td>1000</td><td></td><td></td><td></td><td></td><td></td><td></td><td></td><td></td><td></td></tr>
<tr><td>Oligo fwd</td><td>CACCTGAACTGCGCGCCACGTGGA</td><td></td><td></td><td></td><td></td><td></td><td></td><td></td><td></td><td></td><td></td></tr>
<tr><td>Oligo rev</td><td>AAACTCCACGTGGCGCGCAGTTCA</td><td></td><td></td><td></td><td></td><td></td><td></td><td></td><td></td><td></td><td></td></tr>
<tr><td>Chromosome</td><td>start</td><td>end</td><td>strand</td><td>MM</td><td>target_seq</td><td>PAM</td><td>alignment</td><td>distance</td><td>position</td><td>gene name</td><td>gene id</td></tr>
<tr><td>GU305914.1</td><td>222716</td><td>222738</td><td>+</td><td>0</td><td>TGAACTGCGCGCCACGTGGA</td><td>CGG</td><td>||||||||[||||||||||||]PAM</td><td>NA</td><td>Intergenic</td><td>NA</td><td>NA</td></tr>
<tr><td></td><td></td><td></td><td></td><td></td><td></td><td></td><td></td><td></td><td></td><td></td><td></td></tr>
<tr><td>T16551</td><td>GAACTGCGCGCCACGTGGACGGG</td><td>1000</td><td></td><td></td><td></td><td></td><td></td><td></td><td></td><td></td><td></td></tr>
<tr><td>Oligo fwd</td><td>CACCGAACTGCGCGCCACGTGGAC</td><td></td><td></td><td></td><td></td><td></td><td></td><td></td><td></td><td></td><td></td></tr>
<tr><td>Oligo rev</td><td>AAACGTCCACGTGGCGCGCAGTTC</td><td></td><td></td><td></td><td></td><td></td><td></td><td></td><td></td><td></td><td></td></tr>
<tr><td>Chromosome</td><td>start</td><td>end</td><td>strand</td><td>MM</td><td>target_seq</td><td>PAM</td><td>alignment</td><td>distance</td><td>position</td><td>gene name</td><td>gene id</td></tr>
<tr><td>GU305914.1</td><td>222717</td><td>222739</td><td>+</td><td>0</td><td>GAACTGCGCGCCACGTGGAC</td><td>GGG</td><td>||||||||[||||||||||||]PAM</td><td>NA</td><td>Intergenic</td><td>NA</td><td>NA</td></tr>
<tr><td></td><td></td><td></td><td></td><td></td><td></td><td></td><td></td><td></td><td></td><td></td><td></td></tr>
<tr><td>T16552</td><td>ACGGGTCGACATTCGATCGTCGG</td><td>1000</td><td></td><td></td><td></td><td></td><td></td><td></td><td></td><td></td><td></td></tr>
<tr><td>Oligo fwd</td><td>CACCACGGGTCGACATTCGATCGT</td><td></td><td></td><td></td><td></td><td></td><td></td><td></td><td></td><td></td><td></td></tr>
<tr><td>Oligo rev</td><td>AAACACGATCGAATGTCGACCCGT</td><td></td><td></td><td></td><td></td><td></td><td></td><td></td><td></td><td></td><td></td></tr>
<tr><td>Chromosome</td><td>start</td><td>end</td><td>strand</td><td>MM</td><td>target_seq</td><td>PAM</td><td>alignment</td><td>distance</td><td>position</td><td>gene name</td><td>gene id</td></tr>
</table>

<table>
<tr><td>GU305914.1</td><td>222735</td><td>222757</td><td>+</td><td>0</td><td>ACGGGTCGACATTCGATCGT</td><td>CGG</td><td>||||||||[||||||||||||]PAM</td><td>NA</td><td>Intergenic</td><td>NA</td><td>NA</td></tr>
<tr><td>T16553</td><td>TCGACATTCGATCGTCGGCAAGG</td><td>1000</td><td></td><td></td><td></td><td></td><td></td><td></td><td></td><td></td><td></td></tr>
<tr><td>Oligo fwd</td><td>CACCTCGACATTCGATCGTCGGCA</td><td></td><td></td><td></td><td></td><td></td><td></td><td></td><td></td><td></td><td></td></tr>
<tr><td>Oligo rev</td><td>AAACTGCCGACGATCGAATGTCGA</td><td></td><td></td><td></td><td></td><td></td><td></td><td></td><td></td><td></td><td></td></tr>
<tr><td>Chromosome</td><td>start</td><td>end</td><td>strand</td><td>MM</td><td>target_seq</td><td>PAM</td><td>alignment</td><td>distance</td><td>position</td><td>gene name</td><td>gene id</td></tr>
<tr><td>GU305914.1</td><td>222740</td><td>222762</td><td>+</td><td>0</td><td>TCGACATTCGATCGTCGGCA</td><td>AGG</td><td>||||||||[||||||||||||]PAM</td><td>NA</td><td>Intergenic</td><td>NA</td><td>NA</td></tr>
<tr><td>T16554</td><td>CGATCGTCGGCAAGGATAGCAGG</td><td>1000</td><td></td><td></td><td></td><td></td><td></td><td></td><td></td><td></td><td></td></tr>
<tr><td>Oligo fwd</td><td>CACCCGATCGTCGGCAAGGATAGC</td><td></td><td></td><td></td><td></td><td></td><td></td><td></td><td></td><td></td><td></td></tr>
<tr><td>Oligo rev</td><td>AAACGCTATCCTTGCCGACGATCG</td><td></td><td></td><td></td><td></td><td></td><td></td><td></td><td></td><td></td><td></td></tr>
<tr><td>Chromosome</td><td>start</td><td>end</td><td>strand</td><td>MM</td><td>target_seq</td><td>PAM</td><td>alignment</td><td>distance</td><td>position</td><td>gene name</td><td>gene id</td></tr>
<tr><td>GU305914.1</td><td>222748</td><td>222770</td><td>+</td><td>0</td><td>CGATCGTCGGCAAGGATAGC</td><td>AGG</td><td>||||||||[||||||||||||]PAM</td><td>NA</td><td>Intergenic</td><td>NA</td><td>NA</td></tr>
<tr><td>T16555</td><td>CAAGGATAGCAGGTTACCGCCGG</td><td>1000</td><td></td><td></td><td></td><td></td><td></td><td></td><td></td><td></td><td></td></tr>
<tr><td>Oligo fwd</td><td>CACCCAAGGATAGCAGGTTACCGC</td><td></td><td></td><td></td><td></td><td></td><td></td><td></td><td></td><td></td><td></td></tr>
<tr><td>Oligo rev</td><td>AAACGCGGTAACCTGCTATCCTTG</td><td></td><td></td><td></td><td></td><td></td><td></td><td></td><td></td><td></td><td></td></tr>
<tr><td>Chromosome</td><td>start</td><td>end</td><td>strand</td><td>MM</td><td>target_seq</td><td>PAM</td><td>alignment</td><td>distance</td><td>position</td><td>gene name</td><td>gene id</td></tr>
<tr><td>GU305914.1</td><td>222758</td><td>222780</td><td>+</td><td>0</td><td>CAAGGATAGCAGGTTACCGC</td><td>CGG</td><td>||||||||[||||||||||||]PAM</td><td>NA</td><td>Intergenic</td><td>NA</td><td>NA</td></tr>
<tr><td>T16556</td><td>GACGACGCACGATACGCAGACGG</td><td>1000</td><td></td><td></td><td></td><td></td><td></td><td></td><td></td><td></td><td></td></tr>
<tr><td>Oligo fwd</td><td>CACCGACGACGCACGATACGCAGA</td><td></td><td></td><td></td><td></td><td></td><td></td><td></td><td></td><td></td><td></td></tr>
<tr><td>Oligo rev</td><td>AAACTCTGCGTATCGTGCGTCGTC</td><td></td><td></td><td></td><td></td><td></td><td></td><td></td><td></td><td></td><td></td></tr>
<tr><td>Chromosome</td><td>start</td><td>end</td><td>strand</td><td>MM</td><td>target_seq</td><td>PAM</td><td>alignment</td><td>distance</td><td>position</td><td>gene name</td><td>gene id</td></tr>
<tr><td>GU305914.1</td><td>222785</td><td>222807</td><td>+</td><td>0</td><td>GACGACGCACGATACGCAGA</td><td>CGG</td><td>||||||||[||||||||||||]PAM</td><td>NA</td><td>Intergenic</td><td>NA</td><td>NA</td></tr>
<tr><td>T16557</td><td>ACGACGCACGATACGCAGACGGG</td><td>1000</td><td></td><td></td><td></td><td></td><td></td><td></td><td></td><td></td><td></td></tr>
<tr><td>Oligo fwd</td><td>CACCACGACGCACGATACGCAGAC</td><td></td><td></td><td></td><td></td><td></td><td></td><td></td><td></td><td></td><td></td></tr>
<tr><td>Oligo rev</td><td>AAACGTCTGCGTATCGTGCGTCGT</td><td></td><td></td><td></td><td></td><td></td><td></td><td></td><td></td><td></td><td></td></tr>
<tr><td>Chromosome</td><td>start</td><td>end</td><td>strand</td><td>MM</td><td>target_seq</td><td>PAM</td><td>alignment</td><td>distance</td><td>position</td><td>gene name</td><td>gene id</td></tr>
<tr><td>GU305914.1</td><td>222786</td><td>222808</td><td>+</td><td>0</td><td>ACGACGCACGATACGCAGAC</td><td>GGG</td><td>||||||||[||||||||||||]PAM</td><td>NA</td><td>Intergenic</td><td>NA</td><td>NA</td></tr>
<tr><td>T16558</td><td>CGACGCACGATACGCAGACGGGG</td><td>1000</td><td></td><td></td><td></td><td></td><td></td><td></td><td></td><td></td><td></td></tr>
<tr><td>Oligo fwd</td><td>CACCCGACGCACGATACGCAGACG</td><td></td><td></td><td></td><td></td><td></td><td></td><td></td><td></td><td></td><td></td></tr>
<tr><td>Oligo rev</td><td>AAACCGTCTGCGTATCGTGCGTCG</td><td></td><td></td><td></td><td></td><td></td><td></td><td></td><td></td><td></td><td></td></tr>
<tr><td>Chromosome</td><td>start</td><td>end</td><td>strand</td><td>MM</td><td>target_seq</td><td>PAM</td><td>alignment</td><td>distance</td><td>position</td><td>gene name</td><td>gene id</td></tr>
<tr><td>GU305914.1</td><td>222787</td><td>222809</td><td>+</td><td>0</td><td>CGACGCACGATACGCAGACG</td><td>GGG</td><td>||||||||[||||||||||||]PAM</td><td>NA</td><td>Intergenic</td><td>NA</td><td>NA</td></tr>
<tr><td>T16559</td><td>CGCACGATACGCAGACGGGGAGG</td><td>1000</td><td></td><td></td><td></td><td></td><td></td><td></td><td></td><td></td><td></td></tr>
<tr><td>Oligo fwd</td><td>CACCCGCACGATACGCAGACGGGG</td><td></td><td></td><td></td><td></td><td></td><td></td><td></td><td></td><td></td><td></td></tr>
<tr><td>Oligo rev</td><td>AAACCCCCGTCTGCGTATCGTGCG</td><td></td><td></td><td></td><td></td><td></td><td></td><td></td><td></td><td></td><td></td></tr>
<tr><td>Chromosome</td><td>start</td><td>end</td><td>strand</td><td>MM</td><td>target_seq</td><td>PAM</td><td>alignment</td><td>distance</td><td>position</td><td>gene name</td><td>gene id</td></tr>
<tr><td>GU305914.1</td><td>222790</td><td>222812</td><td>+</td><td>0</td><td>CGCACGATACGCAGACGGGG</td><td>AGG</td><td>||||||||[||||||||||||]PAM</td><td>NA</td><td>Intergenic</td><td>NA</td><td>NA</td></tr>
<tr><td>T16560</td><td>CGATACGCAGACGGGGAGGTCGG</td><td>1000</td><td></td><td></td><td></td><td></td><td></td><td></td><td></td><td></td><td></td></tr>
<tr><td>Oligo fwd</td><td>CACCCGATACGCAGACGGGGAGGT</td><td></td><td></td><td></td><td></td><td></td><td></td><td></td><td></td><td></td><td></td></tr>
<tr><td>Oligo rev</td><td>AAACACCTCCCCGTCTGCGTATCG</td><td></td><td></td><td></td><td></td><td></td><td></td><td></td><td></td><td></td><td></td></tr>
<tr><td>Chromosome</td><td>start</td><td>end</td><td>strand</td><td>MM</td><td>target_seq</td><td>PAM</td><td>alignment</td><td>distance</td><td>position</td><td>gene name</td><td>gene id</td></tr>
<tr><td>GU305914.1</td><td>222794</td><td>222816</td><td>+</td><td>0</td><td>CGATACGCAGACGGGGAGGT</td><td>CGG</td><td>||||||||[||||||||||||]PAM</td><td>NA</td><td>Intergenic</td><td>NA</td><td>NA</td></tr>
<tr><td>T16561</td><td>CGCAGACGGGGAGGTCGGAGCGG</td><td>1000</td><td></td><td></td><td></td><td></td><td></td><td></td><td></td><td></td><td></td></tr>
<tr><td>Oligo fwd</td><td>CACCCGCAGACGGGGAGGTCGGAG</td><td></td><td></td><td></td><td></td><td></td><td></td><td></td><td></td><td></td><td></td></tr>
<tr><td>Oligo rev</td><td>AAACCTCCGACCTCCCCGTCTGCG</td><td></td><td></td><td></td><td></td><td></td><td></td><td></td><td></td><td></td><td></td></tr>
<tr><td>Chromosome</td><td>start</td><td>end</td><td>strand</td><td>MM</td><td>target_seq</td><td>PAM</td><td>alignment</td><td>distance</td><td>position</td><td>gene name</td><td>gene id</td></tr>
<tr><td>GU305914.1</td><td>222799</td><td>222821</td><td>+</td><td>0</td><td>CGCAGACGGGGAGGTCGGAG</td><td>CGG</td><td>||||||||[||||||||||||]PAM</td><td>NA</td><td>Intergenic</td><td>NA</td><td>NA</td></tr>
<tr><td>T16562</td><td>ACGGGGAGGTCGGAGCGGATCGG</td><td>1000</td><td></td><td></td><td></td><td></td><td></td><td></td><td></td><td></td><td></td></tr>
<tr><td>Oligo fwd</td><td>CACCACGGGGAGGTCGGAGCGGAT</td><td></td><td></td><td></td><td></td><td></td><td></td><td></td><td></td><td></td><td></td></tr>
<tr><td>Oligo rev</td><td>AAACATCCGCTCCGACCTCCCCGT</td><td></td><td></td><td></td><td></td><td></td><td></td><td></td><td></td><td></td><td></td></tr>
<tr><td>Chromosome</td><td>start</td><td>end</td><td>strand</td><td>MM</td><td>target_seq</td><td>PAM</td><td>alignment</td><td>distance</td><td>position</td><td>gene name</td><td>gene id</td></tr>
<tr><td>GU305914.1</td><td>222804</td><td>222826</td><td>+</td><td>0</td><td>ACGGGGAGGTCGGAGCGGAT</td><td>CGG</td><td>||||||||[||||||||||||]PAM</td><td>NA</td><td>Intergenic</td><td>NA</td><td>NA</td></tr>
<tr><td>T16563</td><td>CGGAGCGGATCGGAAAACTGAGG</td><td>1000</td><td></td><td></td><td></td><td></td><td></td><td></td><td></td><td></td><td></td></tr>
<tr><td>Oligo fwd</td><td>CACCCGGAGCGGATCGGAAAACTG</td><td></td><td></td><td></td><td></td><td></td><td></td><td></td><td></td><td></td><td></td></tr>
<tr><td>Oligo rev</td><td>AAACCAGTTTTCCGATCCGCTCCG</td><td></td><td></td><td></td><td></td><td></td><td></td><td></td><td></td><td></td><td></td></tr>
<tr><td>Chromosome</td><td>start</td><td>end</td><td>strand</td><td>MM</td><td>target_seq</td><td>PAM</td><td>alignment</td><td>distance</td><td>position</td><td>gene name</td><td>gene id</td></tr>
<tr><td>GU305914.1</td><td>222814</td><td>222836</td><td>+</td><td>0</td><td>CGGAGCGGATCGGAAAACTG</td><td>AGG</td><td>||||||||[||||||||||||]PAM</td><td>NA</td><td>Intergenic</td><td>NA</td><td>NA</td></tr>
<tr><td>T16564</td><td>GGAGCGGATCGGAAAACTGAGGG</td><td>1000</td><td></td><td></td><td></td><td></td><td></td><td></td><td></td><td></td><td></td></tr>
<tr><td>Oligo fwd</td><td>CACCGGAGCGGATCGGAAAACTGA</td><td></td><td></td><td></td><td></td><td></td><td></td><td></td><td></td><td></td><td></td></tr>
<tr><td>Oligo rev</td><td>AAACTCAGTTTTCCGATCCGCTCC</td><td></td><td></td><td></td><td></td><td></td><td></td><td></td><td></td><td></td><td></td></tr>
<tr><td>Chromosome</td><td>start</td><td>end</td><td>strand</td><td>MM</td><td>target_seq</td><td>PAM</td><td>alignment</td><td>distance</td><td>position</td><td>gene name</td><td>gene id</td></tr>
<tr><td>GU305914.1</td><td>222815</td><td>222837</td><td>+</td><td>0</td><td>GGAGCGGATCGGAAAACTGA</td><td>GGG</td><td>||||||||[||||||||||||]PAM</td><td>NA</td><td>Intergenic</td><td>NA</td><td>NA</td></tr>
<tr><td>T16565</td><td>GGATCGGAAAACTGAGGGCCAGG</td><td>1000</td><td></td><td></td><td></td><td></td><td></td><td></td><td></td><td></td><td></td></tr>
<tr><td>Oligo fwd</td><td>CACCGGATCGGAAAACTGAGGGCC</td><td></td><td></td><td></td><td></td><td></td><td></td><td></td><td></td><td></td><td></td></tr>
<tr><td>Oligo rev</td><td>AAACGGCCCTCAGTTTTCCGATCC</td><td></td><td></td><td></td><td></td><td></td><td></td><td></td><td></td><td></td><td></td></tr>
<tr><td>Chromosome</td><td>start</td><td>end</td><td>strand</td><td>MM</td><td>target_seq</td><td>PAM</td><td>alignment</td><td>distance</td><td>position</td><td>gene name</td><td>gene id</td></tr>
<tr><td>GU305914.1</td><td>222820</td><td>222842</td><td>+</td><td>0</td><td>GGATCGGAAAACTGAGGGCC</td><td>AGG</td><td>||||||||[||||||||||||]PAM</td><td>NA</td><td>Intergenic</td><td>NA</td><td>NA</td></tr>
<tr><td>T16566</td><td>CGGAAAACTGAGGGCCAGGTCGG</td><td>1000</td><td></td><td></td><td></td><td></td><td></td><td></td><td></td><td></td><td></td></tr>
<tr><td>Oligo fwd</td><td>CACCCGGAAAACTGAGGGCCAGGT</td><td></td><td></td><td></td><td></td><td></td><td></td><td></td><td></td><td></td><td></td></tr>
<tr><td>Oligo rev</td><td>AAACACCTGGCCCTCAGTTTTCCG</td><td></td><td></td><td></td><td></td><td></td><td></td><td></td><td></td><td></td><td></td></tr>
<tr><td>Chromosome</td><td>start</td><td>end</td><td>strand</td><td>MM</td><td>target_seq</td><td>PAM</td><td>alignment</td><td>distance</td><td>position</td><td>gene name</td><td>gene id</td></tr>
<tr><td>GU305914.1</td><td>222824</td><td>222846</td><td>+</td><td>0</td><td>CGGAAAACTGAGGGCCAGGT</td><td>CGG</td><td>||||||||[||||||||||||]PAM</td><td>NA</td><td>Intergenic</td><td>NA</td><td>NA</td></tr>
<tr><td>T16567</td><td>GGCCAGGTCGGATGCGTTGACGG</td><td>1000</td><td></td><td></td><td></td><td></td><td></td><td></td><td></td><td></td><td></td></tr>
<tr><td>Oligo fwd</td><td>CACCGGCCAGGTCGGATGCGTTGA</td><td></td><td></td><td></td><td></td><td></td><td></td><td></td><td></td><td></td><td></td></tr>
<tr><td>Oligo rev</td><td>AAACTCAACGCATCCGACCTGGCC</td><td></td><td></td><td></td><td></td><td></td><td></td><td></td><td></td><td></td><td></td></tr>
</table>

<table>
<tr><td>Chromosome</td><td>start</td><td>end</td><td>strand</td><td>MM</td><td>target_seq</td><td>PAM</td><td>alignment</td><td>distance</td><td>position</td><td>gene name</td><td>gene id</td></tr>
<tr><td>GU305914.1</td><td>222836</td><td>222858</td><td>+</td><td>0</td><td>GGCCAGGTCGGATGCGTTGA</td><td>CGG</td><td>||||||||[||||||||||||]PAM</td><td>NA</td><td>Intergenic</td><td>NA</td><td>NA</td></tr>
<tr><td>T16568</td><td>TGACGGTAAAACCCTCGAGCCGG</td><td>1000</td><td></td><td></td><td></td><td></td><td></td><td></td><td></td><td></td><td></td></tr>
<tr><td>Oligo fwd</td><td>CACCTGACGGTAAAACCCTCGAGC</td><td></td><td></td><td></td><td></td><td></td><td></td><td></td><td></td><td></td><td></td></tr>
<tr><td>Oligo rev</td><td>AAACGCTCGAGGGTTTTACCGTCA</td><td></td><td></td><td></td><td></td><td></td><td></td><td></td><td></td><td></td><td></td></tr>
<tr><td>Chromosome</td><td>start</td><td>end</td><td>strand</td><td>MM</td><td>target_seq</td><td>PAM</td><td>alignment</td><td>distance</td><td>position</td><td>gene name</td><td>gene id</td></tr>
<tr><td>GU305914.1</td><td>222853</td><td>222875</td><td>+</td><td>0</td><td>TGACGGTAAAACCCTCGAGC</td><td>CGG</td><td>||||||||[||||||||||||]PAM</td><td>NA</td><td>Intergenic</td><td>NA</td><td>NA</td></tr>
<tr><td>T16569</td><td>GTTCGACATCCACAAGAAGTCGG</td><td>1000</td><td></td><td></td><td></td><td></td><td></td><td></td><td></td><td></td><td></td></tr>
<tr><td>Oligo fwd</td><td>CACCGTTCGACATCCACAAGAAGT</td><td></td><td></td><td></td><td></td><td></td><td></td><td></td><td></td><td></td><td></td></tr>
<tr><td>Oligo rev</td><td>AAACACTTCTTGTGGATGTCGAAC</td><td></td><td></td><td></td><td></td><td></td><td></td><td></td><td></td><td></td><td></td></tr>
<tr><td>Chromosome</td><td>start</td><td>end</td><td>strand</td><td>MM</td><td>target_seq</td><td>PAM</td><td>alignment</td><td>distance</td><td>position</td><td>gene name</td><td>gene id</td></tr>
<tr><td>GU305914.1</td><td>222875</td><td>222897</td><td>+</td><td>0</td><td>GTTCGACATCCACAAGAAGT</td><td>CGG</td><td>||||||||[||||||||||||]PAM</td><td>NA</td><td>Intergenic</td><td>NA</td><td>NA</td></tr>
<tr><td>T16570</td><td>CGACATCCACAAGAAGTCGGTGG</td><td>1000</td><td></td><td></td><td></td><td></td><td></td><td></td><td></td><td></td><td></td></tr>
<tr><td>Oligo fwd</td><td>CACCCGACATCCACAAGAAGTCGG</td><td></td><td></td><td></td><td></td><td></td><td></td><td></td><td></td><td></td><td></td></tr>
<tr><td>Oligo rev</td><td>AAACCCGACTTCTTGTGGATGTCG</td><td></td><td></td><td></td><td></td><td></td><td></td><td></td><td></td><td></td><td></td></tr>
<tr><td>Chromosome</td><td>start</td><td>end</td><td>strand</td><td>MM</td><td>target_seq</td><td>PAM</td><td>alignment</td><td>distance</td><td>position</td><td>gene name</td><td>gene id</td></tr>
<tr><td>GU305914.1</td><td>222878</td><td>222900</td><td>+</td><td>0</td><td>CGACATCCACAAGAAGTCGG</td><td>TGG</td><td>||||||||[||||||||||||]PAM</td><td>NA</td><td>Intergenic</td><td>NA</td><td>NA</td></tr>
<tr><td>T16571</td><td>CAAGAAGTCGGTGGTGCCCAAGG</td><td>1000</td><td></td><td></td><td></td><td></td><td></td><td></td><td></td><td></td><td></td></tr>
<tr><td>Oligo fwd</td><td>CACCCAAGAAGTCGGTGGTGCCCA</td><td></td><td></td><td></td><td></td><td></td><td></td><td></td><td></td><td></td><td></td></tr>
<tr><td>Oligo rev</td><td>AAACTGGGCACCACCGACTTCTTG</td><td></td><td></td><td></td><td></td><td></td><td></td><td></td><td></td><td></td><td></td></tr>
<tr><td>Chromosome</td><td>start</td><td>end</td><td>strand</td><td>MM</td><td>target_seq</td><td>PAM</td><td>alignment</td><td>distance</td><td>position</td><td>gene name</td><td>gene id</td></tr>
<tr><td>GU305914.1</td><td>222887</td><td>222909</td><td>+</td><td>0</td><td>CAAGAAGTCGGTGGTGCCCA</td><td>AGG</td><td>||||||||[||||||||||||]PAM</td><td>NA</td><td>Intergenic</td><td>NA</td><td>NA</td></tr>
<tr><td>T16572</td><td>CTCCTCGCCCGCTGCCACGATGG</td><td>1000</td><td></td><td></td><td></td><td></td><td></td><td></td><td></td><td></td><td></td></tr>
<tr><td>Oligo fwd</td><td>CACCCTCCTCGCCCGCTGCCACGA</td><td></td><td></td><td></td><td></td><td></td><td></td><td></td><td></td><td></td><td></td></tr>
<tr><td>Oligo rev</td><td>AAACTCGTGGCAGCGGGCGAGGAG</td><td></td><td></td><td></td><td></td><td></td><td></td><td></td><td></td><td></td><td></td></tr>
<tr><td>Chromosome</td><td>start</td><td>end</td><td>strand</td><td>MM</td><td>target_seq</td><td>PAM</td><td>alignment</td><td>distance</td><td>position</td><td>gene name</td><td>gene id</td></tr>
<tr><td>GU305914.1</td><td>222911</td><td>222933</td><td>+</td><td>0</td><td>CTCCTCGCCCGCTGCCACGA</td><td>TGG</td><td>||||||||[||||||||||||]PAM</td><td>NA</td><td>Intergenic</td><td>NA</td><td>NA</td></tr>
<tr><td>T16573</td><td>CGCCCGCTGCCACGATGGCCTGG</td><td>1000</td><td></td><td></td><td></td><td></td><td></td><td></td><td></td><td></td><td></td></tr>
<tr><td>Oligo fwd</td><td>CACCCGCCCGCTGCCACGATGGCC</td><td></td><td></td><td></td><td></td><td></td><td></td><td></td><td></td><td></td><td></td></tr>
<tr><td>Oligo rev</td><td>AAACGGCCATCGTGGCAGCGGGCG</td><td></td><td></td><td></td><td></td><td></td><td></td><td></td><td></td><td></td><td></td></tr>
<tr><td>Chromosome</td><td>start</td><td>end</td><td>strand</td><td>MM</td><td>target_seq</td><td>PAM</td><td>alignment</td><td>distance</td><td>position</td><td>gene name</td><td>gene id</td></tr>
<tr><td>GU305914.1</td><td>222916</td><td>222938</td><td>+</td><td>0</td><td>CGCCCGCTGCCACGATGGCC</td><td>TGG</td><td>||||||||[||||||||||||]PAM</td><td>NA</td><td>Intergenic</td><td>NA</td><td>NA</td></tr>
<tr><td>T16574</td><td>CACGATGGCCTGGTTGTTGACGG</td><td>1000</td><td></td><td></td><td></td><td></td><td></td><td></td><td></td><td></td><td></td></tr>
<tr><td>Oligo fwd</td><td>CACCCACGATGGCCTGGTTGTTGA</td><td></td><td></td><td></td><td></td><td></td><td></td><td></td><td></td><td></td><td></td></tr>
<tr><td>Oligo rev</td><td>AAACTCAACAACCAGGCCATCGTG</td><td></td><td></td><td></td><td></td><td></td><td></td><td></td><td></td><td></td><td></td></tr>
<tr><td>Chromosome</td><td>start</td><td>end</td><td>strand</td><td>MM</td><td>target_seq</td><td>PAM</td><td>alignment</td><td>distance</td><td>position</td><td>gene name</td><td>gene id</td></tr>
<tr><td>GU305914.1</td><td>222926</td><td>222948</td><td>+</td><td>0</td><td>CACGATGGCCTGGTTGTTGA</td><td>CGG</td><td>||||||||[||||||||||||]PAM</td><td>NA</td><td>Intergenic</td><td>NA</td><td>NA</td></tr>
<tr><td>T16575</td><td>CAGAAGATGCGCGAGTACCGAGG</td><td>1000</td><td></td><td></td><td></td><td></td><td></td><td></td><td></td><td></td><td></td></tr>
<tr><td>Oligo fwd</td><td>CACCCAGAAGATGCGCGAGTACCG</td><td></td><td></td><td></td><td></td><td></td><td></td><td></td><td></td><td></td><td></td></tr>
<tr><td>Oligo rev</td><td>AAACCGGTACTCGCGCATCTTCTG</td><td></td><td></td><td></td><td></td><td></td><td></td><td></td><td></td><td></td><td></td></tr>
<tr><td>Chromosome</td><td>start</td><td>end</td><td>strand</td><td>MM</td><td>target_seq</td><td>PAM</td><td>alignment</td><td>distance</td><td>position</td><td>gene name</td><td>gene id</td></tr>
<tr><td>GU305914.1</td><td>222951</td><td>222973</td><td>+</td><td>0</td><td>CAGAAGATGCGCGAGTACCG</td><td>AGG</td><td>||||||||[||||||||||||]PAM</td><td>NA</td><td>Intergenic</td><td>NA</td><td>NA</td></tr>
<tr><td>T16576</td><td>AAGATGCGCGAGTACCGAGGAGG</td><td>1000</td><td></td><td></td><td></td><td></td><td></td><td></td><td></td><td></td><td></td></tr>
<tr><td>Oligo fwd</td><td>CACCAAGATGCGCGAGTACCGAGG</td><td></td><td></td><td></td><td></td><td></td><td></td><td></td><td></td><td></td><td></td></tr>
<tr><td>Oligo rev</td><td>AAACCCTCGGTACTCGCGCATCTT</td><td></td><td></td><td></td><td></td><td></td><td></td><td></td><td></td><td></td><td></td></tr>
<tr><td>Chromosome</td><td>start</td><td>end</td><td>strand</td><td>MM</td><td>target_seq</td><td>PAM</td><td>alignment</td><td>distance</td><td>position</td><td>gene name</td><td>gene id</td></tr>
<tr><td>GU305914.1</td><td>222954</td><td>222976</td><td>+</td><td>0</td><td>AAGATGCGCGAGTACCGAGG</td><td>AGG</td><td>||||||||[||||||||||||]PAM</td><td>NA</td><td>Intergenic</td><td>NA</td><td>NA</td></tr>
<tr><td>T16577</td><td>AGATGCGCGAGTACCGAGGAGGG</td><td>1000</td><td></td><td></td><td></td><td></td><td></td><td></td><td></td><td></td><td></td></tr>
<tr><td>Oligo fwd</td><td>CACCAGATGCGCGAGTACCGAGGA</td><td></td><td></td><td></td><td></td><td></td><td></td><td></td><td></td><td></td><td></td></tr>
<tr><td>Oligo rev</td><td>AAACTCCTCGGTACTCGCGCATCT</td><td></td><td></td><td></td><td></td><td></td><td></td><td></td><td></td><td></td><td></td></tr>
<tr><td>Chromosome</td><td>start</td><td>end</td><td>strand</td><td>MM</td><td>target_seq</td><td>PAM</td><td>alignment</td><td>distance</td><td>position</td><td>gene name</td><td>gene id</td></tr>
<tr><td>GU305914.1</td><td>222955</td><td>222977</td><td>+</td><td>0</td><td>AGATGCGCGAGTACCGAGGA</td><td>GGG</td><td>||||||||[||||||||||||]PAM</td><td>NA</td><td>Intergenic</td><td>NA</td><td>NA</td></tr>
<tr><td>T16578</td><td>CGACAGTAGTCCTTTCCGAACGG</td><td>1000</td><td></td><td></td><td></td><td></td><td></td><td></td><td></td><td></td><td></td></tr>
<tr><td>Oligo fwd</td><td>CACCCGACAGTAGTCCTTTCCGAA</td><td></td><td></td><td></td><td></td><td></td><td></td><td></td><td></td><td></td><td></td></tr>
<tr><td>Oligo rev</td><td>AAACTTCGGAAAGGACTACTGTCG</td><td></td><td></td><td></td><td></td><td></td><td></td><td></td><td></td><td></td><td></td></tr>
<tr><td>Chromosome</td><td>start</td><td>end</td><td>strand</td><td>MM</td><td>target_seq</td><td>PAM</td><td>alignment</td><td>distance</td><td>position</td><td>gene name</td><td>gene id</td></tr>
<tr><td>GU305914.1</td><td>222999</td><td>223021</td><td>+</td><td>0</td><td>CGACAGTAGTCCTTTCCGAA</td><td>CGG</td><td>||||||||[||||||||||||]PAM</td><td>NA</td><td>Intergenic</td><td>NA</td><td>NA</td></tr>
<tr><td>T16579</td><td>CCGAACGGCTCCGCCGCGTCCGG</td><td>1000</td><td></td><td></td><td></td><td></td><td></td><td></td><td></td><td></td><td></td></tr>
<tr><td>Oligo fwd</td><td>CACCCCGAACGGCTCCGCCGCGTC</td><td></td><td></td><td></td><td></td><td></td><td></td><td></td><td></td><td></td><td></td></tr>
<tr><td>Oligo rev</td><td>AAACGACGCGGCGGAGCCGTTCGG</td><td></td><td></td><td></td><td></td><td></td><td></td><td></td><td></td><td></td><td></td></tr>
<tr><td>Chromosome</td><td>start</td><td>end</td><td>strand</td><td>MM</td><td>target_seq</td><td>PAM</td><td>alignment</td><td>distance</td><td>position</td><td>gene name</td><td>gene id</td></tr>
<tr><td>GU305914.1</td><td>223014</td><td>223036</td><td>+</td><td>0</td><td>CCGAACGGCTCCGCCGCGTC</td><td>CGG</td><td>||||||||[||||||||||||]PAM</td><td>NA</td><td>Intergenic</td><td>NA</td><td>NA</td></tr>
<tr><td>T16580</td><td>TCCGCCGCGTCCGGTACCACCGG</td><td>1000</td><td></td><td></td><td></td><td></td><td></td><td></td><td></td><td></td><td></td></tr>
<tr><td>Oligo fwd</td><td>CACCTCCGCCGCGTCCGGTACCAC</td><td></td><td></td><td></td><td></td><td></td><td></td><td></td><td></td><td></td><td></td></tr>
<tr><td>Oligo rev</td><td>AAACGTGGTACCGGACGCGGCGGA</td><td></td><td></td><td></td><td></td><td></td><td></td><td></td><td></td><td></td><td></td></tr>
<tr><td>Chromosome</td><td>start</td><td>end</td><td>strand</td><td>MM</td><td>target_seq</td><td>PAM</td><td>alignment</td><td>distance</td><td>position</td><td>gene name</td><td>gene id</td></tr>
<tr><td>GU305914.1</td><td>223023</td><td>223045</td><td>+</td><td>0</td><td>TCCGCCGCGTCCGGTACCAC</td><td>CGG</td><td>||||||||[||||||||||||]PAM</td><td>NA</td><td>Intergenic</td><td>NA</td><td>NA</td></tr>
<tr><td>T16581</td><td>GTACCACCGGCTCGATCGTCAGG</td><td>1000</td><td></td><td></td><td></td><td></td><td></td><td></td><td></td><td></td><td></td></tr>
<tr><td>Oligo fwd</td><td>CACCGTACCACCGGCTCGATCGTC</td><td></td><td></td><td></td><td></td><td></td><td></td><td></td><td></td><td></td><td></td></tr>
<tr><td>Oligo rev</td><td>AAACGACGATCGAGCCGGTGGTAC</td><td></td><td></td><td></td><td></td><td></td><td></td><td></td><td></td><td></td><td></td></tr>
<tr><td>Chromosome</td><td>start</td><td>end</td><td>strand</td><td>MM</td><td>target_seq</td><td>PAM</td><td>alignment</td><td>distance</td><td>position</td><td>gene name</td><td>gene id</td></tr>
<tr><td>GU305914.1</td><td>223036</td><td>223058</td><td>+</td><td>0</td><td>GTACCACCGGCTCGATCGTC</td><td>AGG</td><td>||||||||[||||||||||||]PAM</td><td>NA</td><td>Intergenic</td><td>NA</td><td>NA</td></tr>
<tr><td>T16582</td><td>ATCGTCAGGAGCTCCTCGCCCGG</td><td>1000</td><td></td><td></td><td></td><td></td><td></td><td></td><td></td><td></td><td></td></tr>
<tr><td>Oligo fwd</td><td>CACCATCGTCAGGAGCTCCTCGCC</td><td></td><td></td><td></td><td></td><td></td><td></td><td></td><td></td><td></td><td></td></tr>
</table>

<table>
<tr><td>Oligo rev</td><td>AAACGGCGAGGAGCTCCTGACGAT</td><td></td><td></td><td></td><td></td><td></td><td></td><td></td><td></td><td></td><td></td><td></td></tr>
<tr><td>Chromosome</td><td>start</td><td></td><td>end</td><td>strand</td><td>MM</td><td>target_seq</td><td>PAM</td><td>alignment</td><td>distance</td><td>position</td><td>gene name</td><td>gene id</td></tr>
<tr><td>GU305914.1</td><td></td><td>223050</td><td>223072</td><td>+</td><td>0</td><td>ATCGTCAGGAGCTCCTCGCC</td><td>CGG</td><td>||||||||[||||||||||||]PAM</td><td>NA</td><td>Intergenic</td><td>NA</td><td>NA</td></tr>
<tr><td>GU305914.1</td><td></td><td>122521</td><td>122543</td><td>-</td><td>4</td><td>AATGTCAGGAGATCCTCGGC</td><td>CGG</td><td>|--|||||[|||-||||||-|]PAM</td><td>NA</td><td>Intergenic</td><td>NA</td><td>NA</td></tr>
<tr><td></td><td></td><td></td><td></td><td></td><td></td><td></td><td></td><td></td><td></td><td></td><td></td><td></td></tr>
<tr><td>T16583</td><td>TCGTCAGGAGCTCCTCGCCCGGG</td><td></td><td>1000</td><td></td><td></td><td></td><td></td><td></td><td></td><td></td><td></td><td></td></tr>
<tr><td>Oligo fwd</td><td>CACCTCGTCAGGAGCTCCTCGCCC</td><td></td><td></td><td></td><td></td><td></td><td></td><td></td><td></td><td></td><td></td><td></td></tr>
<tr><td>Oligo rev</td><td>AAACGGGCGAGGAGCTCCTGACGA</td><td></td><td></td><td></td><td></td><td></td><td></td><td></td><td></td><td></td><td></td><td></td></tr>
<tr><td>Chromosome</td><td>start</td><td></td><td>end</td><td>strand</td><td>MM</td><td>target_seq</td><td>PAM</td><td>alignment</td><td>distance</td><td>position</td><td>gene name</td><td>gene id</td></tr>
<tr><td>GU305914.1</td><td></td><td>223051</td><td>223073</td><td>+</td><td>0</td><td>TCGTCAGGAGCTCCTCGCCC</td><td>GGG</td><td>||||||||[||||||||||||]PAM</td><td>NA</td><td>Intergenic</td><td>NA</td><td>NA</td></tr>
<tr><td></td><td></td><td></td><td></td><td></td><td></td><td></td><td></td><td></td><td></td><td></td><td></td><td></td></tr>
<tr><td>T16584</td><td>CCTCGCCCGGGACCCAACGTAGG</td><td></td><td>1000</td><td></td><td></td><td></td><td></td><td></td><td></td><td></td><td></td><td></td></tr>
<tr><td>Oligo fwd</td><td>CACCCCTCGCCCGGGACCCAACGT</td><td></td><td></td><td></td><td></td><td></td><td></td><td></td><td></td><td></td><td></td><td></td></tr>
<tr><td>Oligo rev</td><td>AAACACGTTGGGTCCCGGGCGAGG</td><td></td><td></td><td></td><td></td><td></td><td></td><td></td><td></td><td></td><td></td><td></td></tr>
<tr><td>Chromosome</td><td>start</td><td></td><td>end</td><td>strand</td><td>MM</td><td>target_seq</td><td>PAM</td><td>alignment</td><td>distance</td><td>position</td><td>gene name</td><td>gene id</td></tr>
<tr><td>GU305914.1</td><td></td><td>223063</td><td>223085</td><td>+</td><td>0</td><td>CCTCGCCCGGGACCCAACGT</td><td>AGG</td><td>||||||||[||||||||||||]PAM</td><td>NA</td><td>Intergenic</td><td>NA</td><td>NA</td></tr>
<tr><td></td><td></td><td></td><td></td><td></td><td></td><td></td><td></td><td></td><td></td><td></td><td></td><td></td></tr>
<tr><td>T16585</td><td>CGCCCGGGACCCAACGTAGGAGG</td><td></td><td>1000</td><td></td><td></td><td></td><td></td><td></td><td></td><td></td><td></td><td></td></tr>
<tr><td>Oligo fwd</td><td>CACCCGCCCGGGACCCAACGTAGG</td><td></td><td></td><td></td><td></td><td></td><td></td><td></td><td></td><td></td><td></td><td></td></tr>
<tr><td>Oligo rev</td><td>AAACCCTACGTTGGGTCCCGGGCG</td><td></td><td></td><td></td><td></td><td></td><td></td><td></td><td></td><td></td><td></td><td></td></tr>
<tr><td>Chromosome</td><td>start</td><td></td><td>end</td><td>strand</td><td>MM</td><td>target_seq</td><td>PAM</td><td>alignment</td><td>distance</td><td>position</td><td>gene name</td><td>gene id</td></tr>
<tr><td>GU305914.1</td><td></td><td>223066</td><td>223088</td><td>+</td><td>0</td><td>CGCCCGGGACCCAACGTAGG</td><td>AGG</td><td>||||||||[||||||||||||]PAM</td><td>NA</td><td>Intergenic</td><td>NA</td><td>NA</td></tr>
<tr><td></td><td></td><td></td><td></td><td></td><td></td><td></td><td></td><td></td><td></td><td></td><td></td><td></td></tr>
<tr><td>T16586</td><td>GCCCGGGACCCAACGTAGGAGGG</td><td></td><td>1000</td><td></td><td></td><td></td><td></td><td></td><td></td><td></td><td></td><td></td></tr>
<tr><td>Oligo fwd</td><td>CACCGCCCGGGACCCAACGTAGGA</td><td></td><td></td><td></td><td></td><td></td><td></td><td></td><td></td><td></td><td></td><td></td></tr>
<tr><td>Oligo rev</td><td>AAACTCCTACGTTGGGTCCCGGGC</td><td></td><td></td><td></td><td></td><td></td><td></td><td></td><td></td><td></td><td></td><td></td></tr>
<tr><td>Chromosome</td><td>start</td><td></td><td>end</td><td>strand</td><td>MM</td><td>target_seq</td><td>PAM</td><td>alignment</td><td>distance</td><td>position</td><td>gene name</td><td>gene id</td></tr>
<tr><td>GU305914.1</td><td></td><td>223067</td><td>223089</td><td>+</td><td>0</td><td>GCCCGGGACCCAACGTAGGA</td><td>GGG</td><td>||||||||[||||||||||||]PAM</td><td>NA</td><td>Intergenic</td><td>NA</td><td>NA</td></tr>
<tr><td></td><td></td><td></td><td></td><td></td><td></td><td></td><td></td><td></td><td></td><td></td><td></td><td></td></tr>
<tr><td>T16587</td><td>CCCGGGACCCAACGTAGGAGGGG</td><td></td><td>1000</td><td></td><td></td><td></td><td></td><td></td><td></td><td></td><td></td><td></td></tr>
<tr><td>Oligo fwd</td><td>CACCCCCGGGACCCAACGTAGGAG</td><td></td><td></td><td></td><td></td><td></td><td></td><td></td><td></td><td></td><td></td><td></td></tr>
<tr><td>Oligo rev</td><td>AAACCTCCTACGTTGGGTCCCGGG</td><td></td><td></td><td></td><td></td><td></td><td></td><td></td><td></td><td></td><td></td><td></td></tr>
<tr><td>Chromosome</td><td>start</td><td></td><td>end</td><td>strand</td><td>MM</td><td>target_seq</td><td>PAM</td><td>alignment</td><td>distance</td><td>position</td><td>gene name</td><td>gene id</td></tr>
<tr><td>GU305914.1</td><td></td><td>223068</td><td>223090</td><td>+</td><td>0</td><td>CCCGGGACCCAACGTAGGAG</td><td>GGG</td><td>||||||||[||||||||||||]PAM</td><td>NA</td><td>Intergenic</td><td>NA</td><td>NA</td></tr>
<tr><td></td><td></td><td></td><td></td><td></td><td></td><td></td><td></td><td></td><td></td><td></td><td></td><td></td></tr>
<tr><td>T16588</td><td>GGGACCCAACGTAGGAGGGGAGG</td><td></td><td>1000</td><td></td><td></td><td></td><td></td><td></td><td></td><td></td><td></td><td></td></tr>
<tr><td>Oligo fwd</td><td>CACCGGGACCCAACGTAGGAGGGG</td><td></td><td></td><td></td><td></td><td></td><td></td><td></td><td></td><td></td><td></td><td></td></tr>
<tr><td>Oligo rev</td><td>AAACCCCCTCCTACGTTGGGTCCC</td><td></td><td></td><td></td><td></td><td></td><td></td><td></td><td></td><td></td><td></td><td></td></tr>
<tr><td>Chromosome</td><td>start</td><td></td><td>end</td><td>strand</td><td>MM</td><td>target_seq</td><td>PAM</td><td>alignment</td><td>distance</td><td>position</td><td>gene name</td><td>gene id</td></tr>
<tr><td>GU305914.1</td><td></td><td>223071</td><td>223093</td><td>+</td><td>0</td><td>GGGACCCAACGTAGGAGGGG</td><td>AGG</td><td>||||||||[||||||||||||]PAM</td><td>NA</td><td>Intergenic</td><td>NA</td><td>NA</td></tr>
<tr><td></td><td></td><td></td><td></td><td></td><td></td><td></td><td></td><td></td><td></td><td></td><td></td><td></td></tr>
<tr><td>T16589</td><td>ACCCAACGTAGGAGGGGAGGCGG</td><td></td><td>1000</td><td></td><td></td><td></td><td></td><td></td><td></td><td></td><td></td><td></td></tr>
<tr><td>Oligo fwd</td><td>CACCACCCAACGTAGGAGGGGAGG</td><td></td><td></td><td></td><td></td><td></td><td></td><td></td><td></td><td></td><td></td><td></td></tr>
<tr><td>Oligo rev</td><td>AAACCCTCCCCTCCTACGTTGGGT</td><td></td><td></td><td></td><td></td><td></td><td></td><td></td><td></td><td></td><td></td><td></td></tr>
<tr><td>Chromosome</td><td>start</td><td></td><td>end</td><td>strand</td><td>MM</td><td>target_seq</td><td>PAM</td><td>alignment</td><td>distance</td><td>position</td><td>gene name</td><td>gene id</td></tr>
<tr><td>GU305914.1</td><td></td><td>223074</td><td>223096</td><td>+</td><td>0</td><td>ACCCAACGTAGGAGGGGAGG</td><td>CGG</td><td>||||||||[||||||||||||]PAM</td><td>NA</td><td>Intergenic</td><td>NA</td><td>NA</td></tr>
<tr><td></td><td></td><td></td><td></td><td></td><td></td><td></td><td></td><td></td><td></td><td></td><td></td><td></td></tr>
<tr><td>T16590</td><td>GCGGAATCAGACACATAGCCTGG</td><td></td><td>1000</td><td></td><td></td><td></td><td></td><td></td><td></td><td></td><td></td><td></td></tr>
<tr><td>Oligo fwd</td><td>CACCGCGGAATCAGACACATAGCC</td><td></td><td></td><td></td><td></td><td></td><td></td><td></td><td></td><td></td><td></td><td></td></tr>
<tr><td>Oligo rev</td><td>AAACGGCTATGTGTCTGATTCCGC</td><td></td><td></td><td></td><td></td><td></td><td></td><td></td><td></td><td></td><td></td><td></td></tr>
<tr><td>Chromosome</td><td>start</td><td></td><td>end</td><td>strand</td><td>MM</td><td>target_seq</td><td>PAM</td><td>alignment</td><td>distance</td><td>position</td><td>gene name</td><td>gene id</td></tr>
<tr><td>GU305914.1</td><td></td><td>223093</td><td>223115</td><td>+</td><td>0</td><td>GCGGAATCAGACACATAGCC</td><td>TGG</td><td>||||||||[||||||||||||]PAM</td><td>NA</td><td>Intergenic</td><td>NA</td><td>NA</td></tr>
<tr><td></td><td></td><td></td><td></td><td></td><td></td><td></td><td></td><td></td><td></td><td></td><td></td><td></td></tr>
<tr><td>T16591</td><td>GACACATAGCCTGGCTCACATGG</td><td></td><td>1000</td><td></td><td></td><td></td><td></td><td></td><td></td><td></td><td></td><td></td></tr>
<tr><td>Oligo fwd</td><td>CACCGACACATAGCCTGGCTCACA</td><td></td><td></td><td></td><td></td><td></td><td></td><td></td><td></td><td></td><td></td><td></td></tr>
<tr><td>Oligo rev</td><td>AAACTGTGAGCCAGGCTATGTGTC</td><td></td><td></td><td></td><td></td><td></td><td></td><td></td><td></td><td></td><td></td><td></td></tr>
<tr><td>Chromosome</td><td>start</td><td></td><td>end</td><td>strand</td><td>MM</td><td>target_seq</td><td>PAM</td><td>alignment</td><td>distance</td><td>position</td><td>gene name</td><td>gene id</td></tr>
<tr><td>GU305914.1</td><td></td><td>223102</td><td>223124</td><td>+</td><td>0</td><td>GACACATAGCCTGGCTCACA</td><td>TGG</td><td>||||||||[||||||||||||]PAM</td><td>NA</td><td>Intergenic</td><td>NA</td><td>NA</td></tr>
<tr><td></td><td></td><td></td><td></td><td></td><td></td><td></td><td></td><td></td><td></td><td></td><td></td><td></td></tr>
<tr><td>T16592</td><td>GGTCGATGTCGCACACGACGAGG</td><td></td><td>1000</td><td></td><td></td><td></td><td></td><td></td><td></td><td></td><td></td><td></td></tr>
<tr><td>Oligo fwd</td><td>CACCGGTCGATGTCGCACACGACG</td><td></td><td></td><td></td><td></td><td></td><td></td><td></td><td></td><td></td><td></td><td></td></tr>
<tr><td>Oligo rev</td><td>AAACCGTCGTGTGCGACATCGACC</td><td></td><td></td><td></td><td></td><td></td><td></td><td></td><td></td><td></td><td></td><td></td></tr>
<tr><td>Chromosome</td><td>start</td><td></td><td>end</td><td>strand</td><td>MM</td><td>target_seq</td><td>PAM</td><td>alignment</td><td>distance</td><td>position</td><td>gene name</td><td>gene id</td></tr>
<tr><td>GU305914.1</td><td></td><td>223123</td><td>223145</td><td>+</td><td>0</td><td>GGTCGATGTCGCACACGACG</td><td>AGG</td><td>||||||||[||||||||||||]PAM</td><td>NA</td><td>Intergenic</td><td>NA</td><td>NA</td></tr>
<tr><td>GU305914.1</td><td></td><td>130982</td><td>131004</td><td>-</td><td>4</td><td>GCTCGCGGTCGCCCACGACG</td><td>AGG</td><td>|-|||--|[||||-|||||||]PAM</td><td>NA</td><td>Intergenic</td><td>NA</td><td>NA</td></tr>
<tr><td></td><td></td><td></td><td></td><td></td><td></td><td></td><td></td><td></td><td></td><td></td><td></td><td></td></tr>
<tr><td>T16593</td><td>ACGAGGCCCGAGAAGCCGCTAGG</td><td></td><td>1000</td><td></td><td></td><td></td><td></td><td></td><td></td><td></td><td></td><td></td></tr>
<tr><td>Oligo fwd</td><td>CACCACGAGGCCCGAGAAGCCGCT</td><td></td><td></td><td></td><td></td><td></td><td></td><td></td><td></td><td></td><td></td><td></td></tr>
<tr><td>Oligo rev</td><td>AAACAGCGGCTTCTCGGGCCTCGT</td><td></td><td></td><td></td><td></td><td></td><td></td><td></td><td></td><td></td><td></td><td></td></tr>
<tr><td>Chromosome</td><td>start</td><td></td><td>end</td><td>strand</td><td>MM</td><td>target_seq</td><td>PAM</td><td>alignment</td><td>distance</td><td>position</td><td>gene name</td><td>gene id</td></tr>
<tr><td>GU305914.1</td><td></td><td>223140</td><td>223162</td><td>+</td><td>0</td><td>ACGAGGCCCGAGAAGCCGCT</td><td>AGG</td><td>||||||||[||||||||||||]PAM</td><td>NA</td><td>Intergenic</td><td>NA</td><td>NA</td></tr>
<tr><td></td><td></td><td></td><td></td><td></td><td></td><td></td><td></td><td></td><td></td><td></td><td></td><td></td></tr>
<tr><td>T16594</td><td>AGGCCCGAGAAGCCGCTAGGCGG</td><td></td><td>1000</td><td></td><td></td><td></td><td></td><td></td><td></td><td></td><td></td><td></td></tr>
<tr><td>Oligo fwd</td><td>CACCAGGCCCGAGAAGCCGCTAGG</td><td></td><td></td><td></td><td></td><td></td><td></td><td></td><td></td><td></td><td></td><td></td></tr>
<tr><td>Oligo rev</td><td>AAACCCTAGCGGCTTCTCGGGCCT</td><td></td><td></td><td></td><td></td><td></td><td></td><td></td><td></td><td></td><td></td><td></td></tr>
<tr><td>Chromosome</td><td>start</td><td></td><td>end</td><td>strand</td><td>MM</td><td>target_seq</td><td>PAM</td><td>alignment</td><td>distance</td><td>position</td><td>gene name</td><td>gene id</td></tr>
<tr><td>GU305914.1</td><td></td><td>223143</td><td>223165</td><td>+</td><td>0</td><td>AGGCCCGAGAAGCCGCTAGG</td><td>CGG</td><td>||||||||[||||||||||||]PAM</td><td>NA</td><td>Intergenic</td><td>NA</td><td>NA</td></tr>
<tr><td></td><td></td><td></td><td></td><td></td><td></td><td></td><td></td><td></td><td></td><td></td><td></td><td></td></tr>
<tr><td>T16595</td><td>GCCGCTAGGCGGCCACAGCTCGG</td><td></td><td>1000</td><td></td><td></td><td></td><td></td><td></td><td></td><td></td><td></td><td></td></tr>
<tr><td>Oligo fwd</td><td>CACCGCCGCTAGGCGGCCACAGCT</td><td></td><td></td><td></td><td></td><td></td><td></td><td></td><td></td><td></td><td></td><td></td></tr>
<tr><td>Oligo rev</td><td>AAACAGCTGTGGCCGCCTAGCGGC</td><td></td><td></td><td></td><td></td><td></td><td></td><td></td><td></td><td></td><td></td><td></td></tr>
<tr><td>Chromosome</td><td>start</td><td></td><td>end</td><td>strand</td><td>MM</td><td>target_seq</td><td>PAM</td><td>alignment</td><td>distance</td><td>position</td><td>gene name</td><td>gene id</td></tr>
<tr><td>GU305914.1</td><td></td><td>223154</td><td>223176</td><td>+</td><td>0</td><td>GCCGCTAGGCGGCCACAGCT</td><td>CGG</td><td>||||||||[||||||||||||]PAM</td><td>NA</td><td>Intergenic</td><td>NA</td><td>NA</td></tr>
<tr><td></td><td></td><td></td><td></td><td></td><td></td><td></td><td></td><td></td><td></td><td></td><td></td><td></td></tr>
<tr><td>T16596</td><td>GTCCGCCATCCCCGTGAAACCGG</td><td></td><td>1000</td><td></td><td></td><td></td><td></td><td></td><td></td><td></td><td></td><td></td></tr>
<tr><td>Oligo fwd</td><td>CACCGTCCGCCATCCCCGTGAAAC</td><td></td><td></td><td></td><td></td><td></td><td></td><td></td><td></td><td></td><td></td><td></td></tr>
<tr><td>Oligo rev</td><td>AAACGTTTCACGGGGATGGCGGAC</td><td></td><td></td><td></td><td></td><td></td><td></td><td></td><td></td><td></td><td></td><td></td></tr>
<tr><td>Chromosome</td><td>start</td><td></td><td>end</td><td>strand</td><td>MM</td><td>target_seq</td><td>PAM</td><td>alignment</td><td>distance</td><td>position</td><td>gene name</td><td>gene id</td></tr>
<tr><td>GU305914.1</td><td></td><td>223193</td><td>223215</td><td>+</td><td>0</td><td>GTCCGCCATCCCCGTGAAAC</td><td>CGG</td><td>||||||||[||||||||||||]PAM</td><td>NA</td><td>Intergenic</td><td>NA</td><td>NA</td></tr>
</table>

T16597 TCCCCGTGAAACCGGCGACGTGG 1000
Oligo fwd CACCTCCCCGTGAAACCGGCGACG
Oligo rev AAACCGTCGCCGGTTTCACGGGGA
Chromosome start end strand MM target_seq PAM alignment distance position gene name gene id
GU305914.1 223201 223223 + 0 TCCCCGTGAAACCGGCGACG TGG ||||||||[||||||||||||]PAM NA Intergenic NA NA

T16598 GACGTGGATGTGAAAGCTCTCGG 1000
Oligo fwd CACCGACGTGGATGTGAAAGCTCT
Oligo rev AAACAGAGCTTTCACATCCACGTC
Chromosome start end strand MM target_seq PAM alignment distance position gene name gene id
GU305914.1 223217 223239 + 0 GACGTGGATGTGAAAGCTCT CGG ||||||||[||||||||||||]PAM NA Intergenic NA NA

T16599 AAAGCTCTCGGCATCGAACGTGG 1000
Oligo fwd CACCAAAGCTCTCGGCATCGAACG
Oligo rev AAACCGTTCGATGCCGAGAGCTTT
Chromosome start end strand MM target_seq PAM alignment distance position gene name gene id
GU305914.1 223229 223251 + 0 AAAGCTCTCGGCATCGAACG TGG ||||||||[||||||||||||]PAM NA Intergenic NA NA

T16600 ACGTGGCGATCGTCGTCGTCTGG 1000
Oligo fwd CACCACGTGGCGATCGTCGTCGTC
Oligo rev AAACGACGACGACGATCGCCACGT
Chromosome start end strand MM target_seq PAM alignment distance position gene name gene id
GU305914.1 223246 223268 + 0 ACGTGGCGATCGTCGTCGTC TGG ||||||||[||||||||||||]PAM NA Intergenic NA NA

T16601 TCGTCGTCGTCTGGAAATCACGG 1000
Oligo fwd CACCTCGTCGTCGTCTGGAAATCA
Oligo rev AAACTGATTTCCAGACGACGACGA
Chromosome start end strand MM target_seq PAM alignment distance position gene name gene id
GU305914.1 223255 223277 + 0 TCGTCGTCGTCTGGAAATCA CGG ||||||||[||||||||||||]PAM NA Intergenic NA NA

T16602 CGTCTGGAAATCACGGCCTTCGG 1000
Oligo fwd CACCCGTCTGGAAATCACGGCCTT
Oligo rev AAACAAGGCCGTGATTTCCAGACG
Chromosome start end strand MM target_seq PAM alignment distance position gene name gene id
GU305914.1 223262 223284 + 0 CGTCTGGAAATCACGGCCTT CGG ||||||||[||||||||||||]PAM NA Intergenic NA NA

T16603 GTCTGGAAATCACGGCCTTCGGG 1000
Oligo fwd CACCGTCTGGAAATCACGGCCTTC
Oligo rev AAACGAAGGCCGTGATTTCCAGAC
Chromosome start end strand MM target_seq PAM alignment distance position gene name gene id
GU305914.1 223263 223285 + 0 GTCTGGAAATCACGGCCTTC GGG ||||||||[||||||||||||]PAM NA Intergenic NA NA

T16604 GACAAACTTCTTCACCGAACCGG 1000
Oligo fwd CACCGACAAACTTCTTCACCGAAC
Oligo rev AAACGTTCGGTGAAGAAGTTTGTC
Chromosome start end strand MM target_seq PAM alignment distance position gene name gene id
GU305914.1 223307 223329 + 0 GACAAACTTCTTCACCGAAC CGG ||||||||[||||||||||||]PAM NA Intergenic NA NA

T16605 ACAAACTTCTTCACCGAACCGGG 1000
Oligo fwd CACCACAAACTTCTTCACCGAACC
Oligo rev AAACGGTTCGGTGAAGAAGTTTGT
Chromosome start end strand MM target_seq PAM alignment distance position gene name gene id
GU305914.1 223308 223330 + 0 ACAAACTTCTTCACCGAACC GGG ||||||||[||||||||||||]PAM NA Intergenic NA NA

T16606 TCACCGAACCGGGTTTCACGTGG 1000
Oligo fwd CACCTCACCGAACCGGGTTTCACG
Oligo rev AAACCGTGAAACCCGGTTCGGTGA
Chromosome start end strand MM target_seq PAM alignment distance position gene name gene id
GU305914.1 223318 223340 + 0 TCACCGAACCGGGTTTCACG TGG ||||||||[||||||||||||]PAM NA Intergenic NA NA

T16607 GGGTTTCACGTGGCAGATCACGG 1000
Oligo fwd CACCGGGTTTCACGTGGCAGATCA
Oligo rev AAACTGATCTGCCACGTGAAACCC
Chromosome start end strand MM target_seq PAM alignment distance position gene name gene id
GU305914.1 223328 223350 + 0 GGGTTTCACGTGGCAGATCA CGG ||||||||[||||||||||||]PAM NA Intergenic NA NA

T16608 TTTCACGTGGCAGATCACGGCGG 1000
Oligo fwd CACCTTTCACGTGGCAGATCACGG
Oligo rev AAACCCGTGATCTGCCACGTGAAA
Chromosome start end strand MM target_seq PAM alignment distance position gene name gene id
GU305914.1 223331 223353 + 0 TTTCACGTGGCAGATCACGG CGG ||||||||[||||||||||||]PAM NA Intergenic NA NA

T16609 AGATCACGGCGGCTTTGTCCTGG 1000
Oligo fwd CACCAGATCACGGCGGCTTTGTCC
Oligo rev AAACGGACAAAGCCGCCGTGATCT
Chromosome start end strand MM target_seq PAM alignment distance position gene name gene id
GU305914.1 223342 223364 + 0 AGATCACGGCGGCTTTGTCC TGG ||||||||[||||||||||||]PAM NA Intergenic NA NA

T16610 CACGGCGGCTTTGTCCTGGTCGG 1000
Oligo fwd CACCCACGGCGGCTTTGTCCTGGT
Oligo rev AAACACCAGGACAAAGCCGCCGTG
Chromosome start end strand MM target_seq PAM alignment distance position gene name gene id
GU305914.1 223346 223368 + 0 CACGGCGGCTTTGTCCTGGT CGG ||||||||[||||||||||||]PAM NA Intergenic NA NA

T16611 ACGGCGGCTTTGTCCTGGTCGGG 1000
Oligo fwd CACCACGGCGGCTTTGTCCTGGTC
Oligo rev AAACGACCAGGACAAAGCCGCCGT
Chromosome start end strand MM target_seq PAM alignment distance position gene name gene id

<table>
<tr><td>GU305914.1</td><td></td><td>223347</td><td>223369</td><td>+</td><td>0</td><td>ACGGCGGCTTTGTCCTGGTC</td><td>GGG</td><td>||||||||[||||||||||||]PAM</td><td>NA</td><td>Intergenic</td><td>NA</td><td>NA</td></tr>
<tr><td>T16612</td><td>TGTTCATGCAGTTATCTGAGAGG</td><td></td><td>1000</td><td></td><td></td><td></td><td></td><td></td><td></td><td></td><td></td><td></td></tr>
<tr><td>Oligo fwd</td><td>CACCTGTTCATGCAGTTATCTGAG</td><td></td><td></td><td></td><td></td><td></td><td></td><td></td><td></td><td></td><td></td><td></td></tr>
<tr><td>Oligo rev</td><td>AAACCTCAGATAACTGCATGAACA</td><td></td><td></td><td></td><td></td><td></td><td></td><td></td><td></td><td></td><td></td><td></td></tr>
<tr><td>Chromosome</td><td>start</td><td></td><td>end</td><td>strand</td><td>MM</td><td>target_seq</td><td>PAM</td><td>alignment</td><td>distance</td><td>position</td><td>gene name</td><td>gene id</td></tr>
<tr><td>GU305914.1</td><td></td><td>223378</td><td>223400</td><td>+</td><td>0</td><td>TGTTCATGCAGTTATCTGAG</td><td>AGG</td><td>||||||||[||||||||||||]PAM</td><td>NA</td><td>Intergenic</td><td>NA</td><td>NA</td></tr>
<tr><td>T16613</td><td>TCCGTCGACGCAGCTTCCGACGG</td><td></td><td>1000</td><td></td><td></td><td></td><td></td><td></td><td></td><td></td><td></td><td></td></tr>
<tr><td>Oligo fwd</td><td>CACCTCCGTCGACGCAGCTTCCGA</td><td></td><td></td><td></td><td></td><td></td><td></td><td></td><td></td><td></td><td></td><td></td></tr>
<tr><td>Oligo rev</td><td>AAACTCGGAAGCTGCGTCGACGGA</td><td></td><td></td><td></td><td></td><td></td><td></td><td></td><td></td><td></td><td></td><td></td></tr>
<tr><td>Chromosome</td><td>start</td><td></td><td>end</td><td>strand</td><td>MM</td><td>target_seq</td><td>PAM</td><td>alignment</td><td>distance</td><td>position</td><td>gene name</td><td>gene id</td></tr>
<tr><td>GU305914.1</td><td></td><td>223403</td><td>223425</td><td>+</td><td>0</td><td>TCCGTCGACGCAGCTTCCGA</td><td>CGG</td><td>||||||||[||||||||||||]PAM</td><td>NA</td><td>Intergenic</td><td>NA</td><td>NA</td></tr>
<tr><td>T16614</td><td>GTCGACGCAGCTTCCGACGGCGG</td><td></td><td>1000</td><td></td><td></td><td></td><td></td><td></td><td></td><td></td><td></td><td></td></tr>
<tr><td>Oligo fwd</td><td>CACCGTCGACGCAGCTTCCGACGG</td><td></td><td></td><td></td><td></td><td></td><td></td><td></td><td></td><td></td><td></td><td></td></tr>
<tr><td>Oligo rev</td><td>AAACCCGTCGGAAGCTGCGTCGAC</td><td></td><td></td><td></td><td></td><td></td><td></td><td></td><td></td><td></td><td></td><td></td></tr>
<tr><td>Chromosome</td><td>start</td><td></td><td>end</td><td>strand</td><td>MM</td><td>target_seq</td><td>PAM</td><td>alignment</td><td>distance</td><td>position</td><td>gene name</td><td>gene id</td></tr>
<tr><td>GU305914.1</td><td></td><td>223406</td><td>223428</td><td>+</td><td>0</td><td>GTCGACGCAGCTTCCGACGG</td><td>CGG</td><td>||||||||[||||||||||||]PAM</td><td>NA</td><td>Intergenic</td><td>NA</td><td>NA</td></tr>
<tr><td>T16615</td><td>GCTTCCGACGGCGGACGCCGAGG</td><td></td><td>1000</td><td></td><td></td><td></td><td></td><td></td><td></td><td></td><td></td><td></td></tr>
<tr><td>Oligo fwd</td><td>CACCGCTTCCGACGGCGGACGCCG</td><td></td><td></td><td></td><td></td><td></td><td></td><td></td><td></td><td></td><td></td><td></td></tr>
<tr><td>Oligo rev</td><td>AAACCGGCGTCCGCCGTCGGAAGC</td><td></td><td></td><td></td><td></td><td></td><td></td><td></td><td></td><td></td><td></td><td></td></tr>
<tr><td>Chromosome</td><td>start</td><td></td><td>end</td><td>strand</td><td>MM</td><td>target_seq</td><td>PAM</td><td>alignment</td><td>distance</td><td>position</td><td>gene name</td><td>gene id</td></tr>
<tr><td>GU305914.1</td><td></td><td>223415</td><td>223437</td><td>+</td><td>0</td><td>GCTTCCGACGGCGGACGCCG</td><td>AGG</td><td>||||||||[||||||||||||]PAM</td><td>NA</td><td>Intergenic</td><td>NA</td><td>NA</td></tr>
<tr><td>T16616</td><td>CTGAAGCCGCCGCGTAGATGAGG</td><td></td><td>1000</td><td></td><td></td><td></td><td></td><td></td><td></td><td></td><td></td><td></td></tr>
<tr><td>Oligo fwd</td><td>CACCCTGAAGCCGCCGCGTAGATG</td><td></td><td></td><td></td><td></td><td></td><td></td><td></td><td></td><td></td><td></td><td></td></tr>
<tr><td>Oligo rev</td><td>AAACCATCTACGCGGCGGCTTCAG</td><td></td><td></td><td></td><td></td><td></td><td></td><td></td><td></td><td></td><td></td><td></td></tr>
<tr><td>Chromosome</td><td>start</td><td></td><td>end</td><td>strand</td><td>MM</td><td>target_seq</td><td>PAM</td><td>alignment</td><td>distance</td><td>position</td><td>gene name</td><td>gene id</td></tr>
<tr><td>GU305914.1</td><td></td><td>223450</td><td>223472</td><td>+</td><td>0</td><td>CTGAAGCCGCCGCGTAGATG</td><td>AGG</td><td>||||||||[||||||||||||]PAM</td><td>NA</td><td>Intergenic</td><td>NA</td><td>NA</td></tr>
<tr><td>T16617</td><td>TGAAGCCGCCGCGTAGATGAGGG</td><td></td><td>1000</td><td></td><td></td><td></td><td></td><td></td><td></td><td></td><td></td><td></td></tr>
<tr><td>Oligo fwd</td><td>CACCTGAAGCCGCCGCGTAGATGA</td><td></td><td></td><td></td><td></td><td></td><td></td><td></td><td></td><td></td><td></td><td></td></tr>
<tr><td>Oligo rev</td><td>AAACTCATCTACGCGGCGGCTTCA</td><td></td><td></td><td></td><td></td><td></td><td></td><td></td><td></td><td></td><td></td><td></td></tr>
<tr><td>Chromosome</td><td>start</td><td></td><td>end</td><td>strand</td><td>MM</td><td>target_seq</td><td>PAM</td><td>alignment</td><td>distance</td><td>position</td><td>gene name</td><td>gene id</td></tr>
<tr><td>GU305914.1</td><td></td><td>223451</td><td>223473</td><td>+</td><td>0</td><td>TGAAGCCGCCGCGTAGATGA</td><td>GGG</td><td>||||||||[||||||||||||]PAM</td><td>NA</td><td>Intergenic</td><td>NA</td><td>NA</td></tr>
<tr><td>T16618</td><td>GTAGATGAGGGCGAGCATCCCGG</td><td></td><td>1000</td><td></td><td></td><td></td><td></td><td></td><td></td><td></td><td></td><td></td></tr>
<tr><td>Oligo fwd</td><td>CACCGTAGATGAGGGCGAGCATCC</td><td></td><td></td><td></td><td></td><td></td><td></td><td></td><td></td><td></td><td></td><td></td></tr>
<tr><td>Oligo rev</td><td>AAACGGATGCTCGCCCTCATCTAC</td><td></td><td></td><td></td><td></td><td></td><td></td><td></td><td></td><td></td><td></td><td></td></tr>
<tr><td>Chromosome</td><td>start</td><td></td><td>end</td><td>strand</td><td>MM</td><td>target_seq</td><td>PAM</td><td>alignment</td><td>distance</td><td>position</td><td>gene name</td><td>gene id</td></tr>
<tr><td>GU305914.1</td><td></td><td>223463</td><td>223485</td><td>+</td><td>0</td><td>GTAGATGAGGGCGAGCATCC</td><td>CGG</td><td>||||||||[||||||||||||]PAM</td><td>NA</td><td>Intergenic</td><td>NA</td><td>NA</td></tr>
<tr><td>T16619</td><td>TCCCGGCCTTGAAGAGTCGTCGG</td><td></td><td>1000</td><td></td><td></td><td></td><td></td><td></td><td></td><td></td><td></td><td></td></tr>
<tr><td>Oligo fwd</td><td>CACCTCCCGGCCTTGAAGAGTCGT</td><td></td><td></td><td></td><td></td><td></td><td></td><td></td><td></td><td></td><td></td><td></td></tr>
<tr><td>Oligo rev</td><td>AAACACGACTCTTCAAGGCCGGGA</td><td></td><td></td><td></td><td></td><td></td><td></td><td></td><td></td><td></td><td></td><td></td></tr>
<tr><td>Chromosome</td><td>start</td><td></td><td>end</td><td>strand</td><td>MM</td><td>target_seq</td><td>PAM</td><td>alignment</td><td>distance</td><td>position</td><td>gene name</td><td>gene id</td></tr>
<tr><td>GU305914.1</td><td></td><td>223480</td><td>223502</td><td>+</td><td>0</td><td>TCCCGGCCTTGAAGAGTCGT</td><td>CGG</td><td>||||||||[||||||||||||]PAM</td><td>NA</td><td>Intergenic</td><td>NA</td><td>NA</td></tr>
<tr><td>T16620</td><td>CCCGGCCTTGAAGAGTCGTCGGG</td><td></td><td>1000</td><td></td><td></td><td></td><td></td><td></td><td></td><td></td><td></td><td></td></tr>
<tr><td>Oligo fwd</td><td>CACCCCCGGCCTTGAAGAGTCGTC</td><td></td><td></td><td></td><td></td><td></td><td></td><td></td><td></td><td></td><td></td><td></td></tr>
<tr><td>Oligo rev</td><td>AAACGACGACTCTTCAAGGCCGGG</td><td></td><td></td><td></td><td></td><td></td><td></td><td></td><td></td><td></td><td></td><td></td></tr>
<tr><td>Chromosome</td><td>start</td><td></td><td>end</td><td>strand</td><td>MM</td><td>target_seq</td><td>PAM</td><td>alignment</td><td>distance</td><td>position</td><td>gene name</td><td>gene id</td></tr>
<tr><td>GU305914.1</td><td></td><td>223481</td><td>223503</td><td>+</td><td>0</td><td>CCCGGCCTTGAAGAGTCGTC</td><td>GGG</td><td>||||||||[||||||||||||]PAM</td><td>NA</td><td>Intergenic</td><td>NA</td><td>NA</td></tr>
<tr><td>T16621</td><td>TGAAGAGTCGTCGGGATCGACGG</td><td></td><td>1000</td><td></td><td></td><td></td><td></td><td></td><td></td><td></td><td></td><td></td></tr>
<tr><td>Oligo fwd</td><td>CACCTGAAGAGTCGTCGGGATCGA</td><td></td><td></td><td></td><td></td><td></td><td></td><td></td><td></td><td></td><td></td><td></td></tr>
<tr><td>Oligo rev</td><td>AAACTCGATCCCGACGACTCTTCA</td><td></td><td></td><td></td><td></td><td></td><td></td><td></td><td></td><td></td><td></td><td></td></tr>
<tr><td>Chromosome</td><td>start</td><td></td><td>end</td><td>strand</td><td>MM</td><td>target_seq</td><td>PAM</td><td>alignment</td><td>distance</td><td>position</td><td>gene name</td><td>gene id</td></tr>
<tr><td>GU305914.1</td><td></td><td>223489</td><td>223511</td><td>+</td><td>0</td><td>TGAAGAGTCGTCGGGATCGA</td><td>CGG</td><td>||||||||[||||||||||||]PAM</td><td>NA</td><td>Intergenic</td><td>NA</td><td>NA</td></tr>
<tr><td>T16622</td><td>TCGGGATCGACGGTCCTCCTCGG</td><td></td><td>1000</td><td></td><td></td><td></td><td></td><td></td><td></td><td></td><td></td><td></td></tr>
<tr><td>Oligo fwd</td><td>CACCTCGGGATCGACGGTCCTCCT</td><td></td><td></td><td></td><td></td><td></td><td></td><td></td><td></td><td></td><td></td><td></td></tr>
<tr><td>Oligo rev</td><td>AAACAGGAGGACCGTCGATCCCGA</td><td></td><td></td><td></td><td></td><td></td><td></td><td></td><td></td><td></td><td></td><td></td></tr>
<tr><td>Chromosome</td><td>start</td><td></td><td>end</td><td>strand</td><td>MM</td><td>target_seq</td><td>PAM</td><td>alignment</td><td>distance</td><td>position</td><td>gene name</td><td>gene id</td></tr>
<tr><td>GU305914.1</td><td></td><td>223499</td><td>223521</td><td>+</td><td>0</td><td>TCGGGATCGACGGTCCTCCT</td><td>CGG</td><td>||||||||[||||||||||||]PAM</td><td>NA</td><td>Intergenic</td><td>NA</td><td>NA</td></tr>
<tr><td>T16623</td><td>TCGACGGTCCTCCTCGGTCCAGG</td><td></td><td>1000</td><td></td><td></td><td></td><td></td><td></td><td></td><td></td><td></td><td></td></tr>
<tr><td>Oligo fwd</td><td>CACCTCGACGGTCCTCCTCGGTCC</td><td></td><td></td><td></td><td></td><td></td><td></td><td></td><td></td><td></td><td></td><td></td></tr>
<tr><td>Oligo rev</td><td>AAACGGACCGAGGAGGACCGTCGA</td><td></td><td></td><td></td><td></td><td></td><td></td><td></td><td></td><td></td><td></td><td></td></tr>
<tr><td>Chromosome</td><td>start</td><td></td><td>end</td><td>strand</td><td>MM</td><td>target_seq</td><td>PAM</td><td>alignment</td><td>distance</td><td>position</td><td>gene name</td><td>gene id</td></tr>
<tr><td>GU305914.1</td><td></td><td>223505</td><td>223527</td><td>+</td><td>0</td><td>TCGACGGTCCTCCTCGGTCC</td><td>AGG</td><td>||||||||[||||||||||||]PAM</td><td>NA</td><td>Intergenic</td><td>NA</td><td>NA</td></tr>
<tr><td>T16624</td><td>CGACGGTCCTCCTCGGTCCAGGG</td><td></td><td>1000</td><td></td><td></td><td></td><td></td><td></td><td></td><td></td><td></td><td></td></tr>
<tr><td>Oligo fwd</td><td>CACCCGACGGTCCTCCTCGGTCCA</td><td></td><td></td><td></td><td></td><td></td><td></td><td></td><td></td><td></td><td></td><td></td></tr>
<tr><td>Oligo rev</td><td>AAACTGGACCGAGGAGGACCGTCG</td><td></td><td></td><td></td><td></td><td></td><td></td><td></td><td></td><td></td><td></td><td></td></tr>
<tr><td>Chromosome</td><td>start</td><td></td><td>end</td><td>strand</td><td>MM</td><td>target_seq</td><td>PAM</td><td>alignment</td><td>distance</td><td>position</td><td>gene name</td><td>gene id</td></tr>
<tr><td>GU305914.1</td><td></td><td>223506</td><td>223528</td><td>+</td><td>0</td><td>CGACGGTCCTCCTCGGTCCA</td><td>GGG</td><td>||||||||[||||||||||||]PAM</td><td>NA</td><td>Intergenic</td><td>NA</td><td>NA</td></tr>
<tr><td>T16625</td><td>GACGGTCCTCCTCGGTCCAGGGG</td><td></td><td>1000</td><td></td><td></td><td></td><td></td><td></td><td></td><td></td><td></td><td></td></tr>
<tr><td>Oligo fwd</td><td>CACCGACGGTCCTCCTCGGTCCAG</td><td></td><td></td><td></td><td></td><td></td><td></td><td></td><td></td><td></td><td></td><td></td></tr>
<tr><td>Oligo rev</td><td>AAACCTGGACCGAGGAGGACCGTC</td><td></td><td></td><td></td><td></td><td></td><td></td><td></td><td></td><td></td><td></td><td></td></tr>
<tr><td>Chromosome</td><td>start</td><td></td><td>end</td><td>strand</td><td>MM</td><td>target_seq</td><td>PAM</td><td>alignment</td><td>distance</td><td>position</td><td>gene name</td><td>gene id</td></tr>
<tr><td>GU305914.1</td><td></td><td>223507</td><td>223529</td><td>+</td><td>0</td><td>GACGGTCCTCCTCGGTCCAG</td><td>GGG</td><td>||||||||[||||||||||||]PAM</td><td>NA</td><td>Intergenic</td><td>NA</td><td>NA</td></tr>
<tr><td>T16626</td><td>TCGGTCCAGGGGATACAGAACGG</td><td></td><td>1000</td><td></td><td></td><td></td><td></td><td></td><td></td><td></td><td></td><td></td></tr>
<tr><td>Oligo fwd</td><td>CACCTCGGTCCAGGGGATACAGAA</td><td></td><td></td><td></td><td></td><td></td><td></td><td></td><td></td><td></td><td></td><td></td></tr>
<tr><td>Oligo rev</td><td>AAACTTCTGTATCCCCTGGACCGA</td><td></td><td></td><td></td><td></td><td></td><td></td><td></td><td></td><td></td><td></td><td></td></tr>
</table>

Chromosome start end strand MM target_seq PAM alignment distance position gene name gene id
GU305914.1 223518 223540 + 0 TCGGTCCAGGGGATACAGAA CGG ||||||||[||||||||||||]PAM NA Intergenic NA NA

T16627 CGGTCCAGGGGATACAGAACGGG 1000
Oligo fwd CACCCGGTCCAGGGGATACAGAAC
Oligo rev AAACGTTCTGTATCCCCTGGACCG
Chromosome start end strand MM target_seq PAM alignment distance position gene name gene id
GU305914.1 223519 223541 + 0 CGGTCCAGGGGATACAGAAC GGG ||||||||[||||||||||||]PAM NA Intergenic NA NA

T16628 GGATACAGAACGGGAGTCGTCGG 1000
Oligo fwd CACCGGATACAGAACGGGAGTCGT
Oligo rev AAACACGACTCCCGTTCTGTATCC
Chromosome start end strand MM target_seq PAM alignment distance position gene name gene id
GU305914.1 223528 223550 + 0 GGATACAGAACGGGAGTCGT CGG ||||||||[||||||||||||]PAM NA Intergenic NA NA

T16629 TACAGAACGGGAGTCGTCGGCGG 1000
Oligo fwd CACCTACAGAACGGGAGTCGTCGG
Oligo rev AAACCCGACGACTCCCGTTCTGTA
Chromosome start end strand MM target_seq PAM alignment distance position gene name gene id
GU305914.1 223531 223553 + 0 TACAGAACGGGAGTCGTCGG CGG ||||||||[||||||||||||]PAM NA Intergenic NA NA

T16630 ACGGGAGTCGTCGGCGGTAGCGG 1000
Oligo fwd CACCACGGGAGTCGTCGGCGGTAG
Oligo rev AAACCTACCGCCGACGACTCCCGT
Chromosome start end strand MM target_seq PAM alignment distance position gene name gene id
GU305914.1 223537 223559 + 0 ACGGGAGTCGTCGGCGGTAG CGG ||||||||[||||||||||||]PAM NA Intergenic NA NA
GU305914.1 109437 109459 + 2 ACGGGAGACGGCGGCGGTAG CGG |||||||-[||-|||||||||]PAM NA Intergenic NA NA

T16631 CGTCGGCGGTAGCGGCGACGCGG 1000
Oligo fwd CACCCGTCGGCGGTAGCGGCGACG
Oligo rev AAACCGTCGCCGCTACCGCCGACG
Chromosome start end strand MM target_seq PAM alignment distance position gene name gene id
GU305914.1 223545 223567 + 0 CGTCGGCGGTAGCGGCGACG CGG ||||||||[||||||||||||]PAM NA Intergenic NA NA

T16632 GTAGCGGCGACGCGGTCTTCTGG 1000
Oligo fwd CACCGTAGCGGCGACGCGGTCTTC
Oligo rev AAACGAAGACCGCGTCGCCGCTAC
Chromosome start end strand MM target_seq PAM alignment distance position gene name gene id
GU305914.1 223553 223575 + 0 GTAGCGGCGACGCGGTCTTC TGG ||||||||[||||||||||||]PAM NA Intergenic NA NA

T16633 GCGACGCGGTCTTCTGGAAGAGG 1000
Oligo fwd CACCGCGACGCGGTCTTCTGGAAG
Oligo rev AAACCTTCCAGAAGACCGCGTCGC
Chromosome start end strand MM target_seq PAM alignment distance position gene name gene id
GU305914.1 223559 223581 + 0 GCGACGCGGTCTTCTGGAAG AGG ||||||||[||||||||||||]PAM NA Intergenic NA NA

T16634 CGCGGTCTTCTGGAAGAGGTCGG 1000
Oligo fwd CACCCGCGGTCTTCTGGAAGAGGT
Oligo rev AAACACCTCTTCCAGAAGACCGCG
Chromosome start end strand MM target_seq PAM alignment distance position gene name gene id
GU305914.1 223563 223585 + 0 CGCGGTCTTCTGGAAGAGGT CGG ||||||||[||||||||||||]PAM NA Intergenic NA NA

T16635 GAAGAGGTCGGTACACAGCGAGG 1000
Oligo fwd CACCGAAGAGGTCGGTACACAGCG
Oligo rev AAACCGCTGTGTACCGACCTCTTC
Chromosome start end strand MM target_seq PAM alignment distance position gene name gene id
GU305914.1 223575 223597 + 0 GAAGAGGTCGGTACACAGCG AGG ||||||||[||||||||||||]PAM NA Intergenic NA NA

T16636 GGTACACAGCGAGGACCACGAGG 1000
Oligo fwd CACCGGTACACAGCGAGGACCACG
Oligo rev AAACCGTGGTCCTCGCTGTGTACC
Chromosome start end strand MM target_seq PAM alignment distance position gene name gene id
GU305914.1 223584 223606 + 0 GGTACACAGCGAGGACCACG AGG ||||||||[||||||||||||]PAM NA Intergenic NA NA

T16637 GACCGACCCGAGTGCCGCCGCGG 1000
Oligo fwd CACCGACCGACCCGAGTGCCGCCG
Oligo rev AAACCGGCGGCACTCGGGTCGGTC
Chromosome start end strand MM target_seq PAM alignment distance position gene name gene id
GU305914.1 223606 223628 + 0 GACCGACCCGAGTGCCGCCG CGG ||||||||[||||||||||||]PAM NA Intergenic NA NA

T16638 CGCGGCTACCCCGATCGAAATGG 1000
Oligo fwd CACCCGCGGCTACCCCGATCGAAA
Oligo rev AAACTTTCGATCGGGGTAGCCGCG
Chromosome start end strand MM target_seq PAM alignment distance position gene name gene id
GU305914.1 223624 223646 + 0 CGCGGCTACCCCGATCGAAA TGG ||||||||[||||||||||||]PAM NA Intergenic NA NA

T16639 GGCTACACGCAACGCGTCGTCGG 1000
Oligo fwd CACCGGCTACACGCAACGCGTCGT
Oligo rev AAACACGACGCGTTGCGTGTAGCC
Chromosome start end strand MM target_seq PAM alignment distance position gene name gene id
GU305914.1 223645 223667 + 0 GGCTACACGCAACGCGTCGT CGG ||||||||[||||||||||||]PAM NA Intergenic NA NA

T16640 GCAACGCGTCGTCGGCACATCGG 1000
Oligo fwd CACCGCAACGCGTCGTCGGCACAT
Oligo rev AAACATGTGCCGACGACGCGTTGC
Chromosome start end strand MM target_seq PAM alignment distance position gene name gene id
GU305914.1 223653 223675 + 0 GCAACGCGTCGTCGGCACAT CGG ||||||||[||||||||||||]PAM NA Intergenic NA NA

T16641 TCGTCGGCACATCGGCAACTCGG 1000

| | | | | | | | | | | | |
|---|---|---|---|---|---|---|---|---|---|---|---|
| Oligo fwd | CACCTCGTCGGCACATCGGCAACT | | | | | | | | | | |
| Oligo rev | AAACAGTTGCCGATGTGCCGACGA | | | | | | | | | | |
| Chromosome | start | end | strand | MM | target_seq | PAM | alignment | distance | position | gene name | gene id |
| GU305914.1 | 223661 | 223683 | + | 0 | TCGTCGGCACATCGGCAACT | CGG | ||||||||[||||||||||||]PAM | NA | Intergenic | NA | NA |
| T16642 | CGTCGGCACATCGGCAACTCGGG | 1000 | | | | | | | | | |
| Oligo fwd | CACCCGTCGGCACATCGGCAACTC | | | | | | | | | | |
| Oligo rev | AAACGAGTTGCCGATGTGCCGACG | | | | | | | | | | |
| Chromosome | start | end | strand | MM | target_seq | PAM | alignment | distance | position | gene name | gene id |
| GU305914.1 | 223662 | 223684 | + | 0 | CGTCGGCACATCGGCAACTC | GGG | ||||||||[||||||||||||]PAM | NA | Intergenic | NA | NA |
| T16643 | CGGGTTCCCCGCATCCTGCCAGG | 1000 | | | | | | | | | |
| Oligo fwd | CACCCGGGTTCCCCGCATCCTGCC | | | | | | | | | | |
| Oligo rev | AAACGGCAGGATGCGGGGAACCCG | | | | | | | | | | |
| Chromosome | start | end | strand | MM | target_seq | PAM | alignment | distance | position | gene name | gene id |
| GU305914.1 | 223681 | 223703 | + | 0 | CGGGTTCCCCGCATCCTGCC | AGG | ||||||||[||||||||||||]PAM | NA | Intergenic | NA | NA |
| T16644 | GGGTTCCCCGCATCCTGCCAGGG | 1000 | | | | | | | | | |
| Oligo fwd | CACCGGGTTCCCCGCATCCTGCCA | | | | | | | | | | |
| Oligo rev | AAACTGGCAGGATGCGGGGAACCC | | | | | | | | | | |
| Chromosome | start | end | strand | MM | target_seq | PAM | alignment | distance | position | gene name | gene id |
| GU305914.1 | 223682 | 223704 | + | 0 | GGGTTCCCCGCATCCTGCCA | GGG | ||||||||[||||||||||||]PAM | NA | Intergenic | NA | NA |
| T16645 | CCCGCATCCTGCCAGGGAGACGG | 1000 | | | | | | | | | |
| Oligo fwd | CACCCCCGCATCCTGCCAGGGAGA | | | | | | | | | | |
| Oligo rev | AAACTCTCCCTGGCAGGATGCGGG | | | | | | | | | | |
| Chromosome | start | end | strand | MM | target_seq | PAM | alignment | distance | position | gene name | gene id |
| GU305914.1 | 223688 | 223710 | + | 0 | CCCGCATCCTGCCAGGGAGA | CGG | ||||||||[||||||||||||]PAM | NA | Intergenic | NA | NA |
| T16646 | GTCTCCCCCTCCGACAGCCGCGG | 1000 | | | | | | | | | |
| Oligo fwd | CACCGTCTCCCCCTCCGACAGCCG | | | | | | | | | | |
| Oligo rev | AAACCGGCTGTCGGAGGGGGAGAC | | | | | | | | | | |
| Chromosome | start | end | strand | MM | target_seq | PAM | alignment | distance | position | gene name | gene id |
| GU305914.1 | 223720 | 223742 | + | 0 | GTCTCCCCCTCCGACAGCCG | CGG | ||||||||[||||||||||||]PAM | NA | Intergenic | NA | NA |
| T16647 | ACATCGCCTTCCCACGTGTGTGG | 1000 | | | | | | | | | |
| Oligo fwd | CACCACATCGCCTTCCCACGTGTG | | | | | | | | | | |
| Oligo rev | AAACCACACGTGGGAAGGCGATGT | | | | | | | | | | |
| Chromosome | start | end | strand | MM | target_seq | PAM | alignment | distance | position | gene name | gene id |
| GU305914.1 | 223747 | 223769 | + | 0 | ACATCGCCTTCCCACGTGTG | TGG | ||||||||[||||||||||||]PAM | NA | Intergenic | NA | NA |
| T16648 | CCCACGTGTGTGGCGACCCGAGG | 1000 | | | | | | | | | |
| Oligo fwd | CACCCCCACGTGTGTGGCGACCCG | | | | | | | | | | |
| Oligo rev | AAACCGGGTCGCCACACACGTGGG | | | | | | | | | | |
| Chromosome | start | end | strand | MM | target_seq | PAM | alignment | distance | position | gene name | gene id |
| GU305914.1 | 223757 | 223779 | + | 0 | CCCACGTGTGTGGCGACCCG | AGG | ||||||||[||||||||||||]PAM | NA | Intergenic | NA | NA |
| T16649 | CGACCCGAGGCCCCAAGATCTGG | 1000 | | | | | | | | | |
| Oligo fwd | CACCCGACCCGAGGCCCCAAGATC | | | | | | | | | | |
| Oligo rev | AAACGATCTTGGGGCCTCGGGTCG | | | | | | | | | | |
| Chromosome | start | end | strand | MM | target_seq | PAM | alignment | distance | position | gene name | gene id |
| GU305914.1 | 223770 | 223792 | + | 0 | CGACCCGAGGCCCCAAGATC | TGG | ||||||||[||||||||||||]PAM | NA | Intergenic | NA | NA |
| T16650 | CCGAGGCCCCAAGATCTGGCAGG | 1000 | | | | | | | | | |
| Oligo fwd | CACCCCGAGGCCCCAAGATCTGGC | | | | | | | | | | |
| Oligo rev | AAACGCCAGATCTTGGGGCCTCGG | | | | | | | | | | |
| Chromosome | start | end | strand | MM | target_seq | PAM | alignment | distance | position | gene name | gene id |
| GU305914.1 | 223774 | 223796 | + | 0 | CCGAGGCCCCAAGATCTGGC | AGG | ||||||||[||||||||||||]PAM | NA | Intergenic | NA | NA |
| T16651 | AGATCTGGCAGGAGAAGCACCGG | 1000 | | | | | | | | | |
| Oligo fwd | CACCAGATCTGGCAGGAGAAGCAC | | | | | | | | | | |
| Oligo rev | AAACGTGCTTCTCCTGCCAGATCT | | | | | | | | | | |
| Chromosome | start | end | strand | MM | target_seq | PAM | alignment | distance | position | gene name | gene id |
| GU305914.1 | 223785 | 223807 | + | 0 | AGATCTGGCAGGAGAAGCAC | CGG | ||||||||[||||||||||||]PAM | NA | Intergenic | NA | NA |
| T16652 | CACCGGTCCCGAGACCGTCTCGG | 1000 | | | | | | | | | |
| Oligo fwd | CACCCACCGGTCCCGAGACCGTCT | | | | | | | | | | |
| Oligo rev | AAACAGACGGTCTCGGGACCGGTG | | | | | | | | | | |
| Chromosome | start | end | strand | MM | target_seq | PAM | alignment | distance | position | gene name | gene id |
| GU305914.1 | 223802 | 223824 | + | 0 | CACCGGTCCCGAGACCGTCT | CGG | ||||||||[||||||||||||]PAM | NA | Intergenic | NA | NA |
| GU305914.1 | 223810 | 223832 | - | 4 | TGCTGGTCCCGAGACGGTCT | CGG | --|-||||[|||||||-||||]PAM | NA | Intergenic | NA | NA |
| T16653 | ACCGGTCCCGAGACCGTCTCGGG | 1000 | | | | | | | | | |
| Oligo fwd | CACCACCGGTCCCGAGACCGTCTC | | | | | | | | | | |
| Oligo rev | AAACGAGACGGTCTCGGGACCGGT | | | | | | | | | | |
| Chromosome | start | end | strand | MM | target_seq | PAM | alignment | distance | position | gene name | gene id |
| GU305914.1 | 223803 | 223825 | + | 0 | ACCGGTCCCGAGACCGTCTC | GGG | ||||||||[||||||||||||]PAM | NA | Intergenic | NA | NA |
| GU305914.1 | 223809 | 223831 | - | 3 | GCTGGTCCCGAGACGGTCTC | GGG | -|-|||||[||||||-|||||]PAM | NA | Intergenic | NA | NA |
| T16654 | TCTCGGGACCAGCAGATCGCCGG | 1000 | | | | | | | | | |
| Oligo fwd | CACCTCTCGGGACCAGCAGATCGC | | | | | | | | | | |
| Oligo rev | AAACGCGATCTGCTGGTCCCGAGA | | | | | | | | | | |
| Chromosome | start | end | strand | MM | target_seq | PAM | alignment | distance | position | gene name | gene id |
| GU305914.1 | 223819 | 223841 | + | 0 | TCTCGGGACCAGCAGATCGC | CGG | ||||||||[||||||||||||]PAM | NA | Intergenic | NA | NA |
| T16655 | GGACCAGCAGATCGCCGGTCAGG | 1000 | | | | | | | | | |
| Oligo fwd | CACCGGACCAGCAGATCGCCGGTC | | | | | | | | | | |
| Oligo rev | AAACGACCGGCGATCTGCTGGTCC | | | | | | | | | | |
| Chromosome | start | end | strand | MM | target_seq | PAM | alignment | distance | position | gene name | gene id |

GU305914.1 223824 223846 + 0 GGACCAGCAGATCGCCGGTC AGG ||||||||[||||||||||||]PAM NA Intergenic NA NA

T16656 GACCAGCAGATCGCCGGTCAGGG 1000
Oligo fwd CACCGACCAGCAGATCGCCGGTCA
Oligo rev AAACTGACCGGCGATCTGCTGGTC
Chromosome start end strand MM target_seq PAM alignment distance position gene name gene id
GU305914.1 223825 223847 + 0 GACCAGCAGATCGCCGGTCA GGG ||||||||[||||||||||||]PAM NA Intergenic NA NA

T16657 ACCAGCAGATCGCCGGTCAGGGG 1000
Oligo fwd CACCACCAGCAGATCGCCGGTCAG
Oligo rev AAACCTGACCGGCGATCTGCTGGT
Chromosome start end strand MM target_seq PAM alignment distance position gene name gene id
GU305914.1 223826 223848 + 0 ACCAGCAGATCGCCGGTCAG GGG ||||||||[||||||||||||]PAM NA Intergenic NA NA

T16658 CCGGTCAGGGGATCTAGTGCCGG 1000
Oligo fwd CACCCCGGTCAGGGGATCTAGTGC
Oligo rev AAACGCACTAGATCCCCTGACCGG
Chromosome start end strand MM target_seq PAM alignment distance position gene name gene id
GU305914.1 223838 223860 + 0 CCGGTCAGGGGATCTAGTGC CGG ||||||||[||||||||||||]PAM NA Intergenic NA NA

T16659 CCGGCCGCGACCCGACCGTGCGG 1000
Oligo fwd CACCCCGGCCGCGACCCGACCGTG
Oligo rev AAACCACGGTCGGGTCGCGGCCGG
Chromosome start end strand MM target_seq PAM alignment distance position gene name gene id
GU305914.1 223857 223879 + 0 CCGGCCGCGACCCGACCGTG CGG ||||||||[||||||||||||]PAM NA Intergenic NA NA

T16660 ATTCGCTAGCCGTCTCAGCCTGG 1000
Oligo fwd CACCATTCGCTAGCCGTCTCAGCC
Oligo rev AAACGGCTGAGACGGCTAGCGAAT
Chromosome start end strand MM target_seq PAM alignment distance position gene name gene id
GU305914.1 224041 224063 + 0 ATTCGCTAGCCGTCTCAGCC TGG ||||||||[||||||||||||]PAM NA Intergenic NA NA

T16661 TTCGCTAGCCGTCTCAGCCTGGG 1000
Oligo fwd CACCTTCGCTAGCCGTCTCAGCCT
Oligo rev AAACAGGCTGAGACGGCTAGCGAA
Chromosome start end strand MM target_seq PAM alignment distance position gene name gene id
GU305914.1 224042 224064 + 0 TTCGCTAGCCGTCTCAGCCT GGG ||||||||[||||||||||||]PAM NA Intergenic NA NA

T16662 TAGCCGTCTCAGCCTGGGACCGG 1000
Oligo fwd CACCTAGCCGTCTCAGCCTGGGAC
Oligo rev AAACGTCCCAGGCTGAGACGGCTA
Chromosome start end strand MM target_seq PAM alignment distance position gene name gene id
GU305914.1 224047 224069 + 0 TAGCCGTCTCAGCCTGGGAC CGG ||||||||[||||||||||||]PAM NA Intergenic NA NA

T16663 GCGACGACCGAGTACTTCGTCGG 1000
Oligo fwd CACCGCGACGACCGAGTACTTCGT
Oligo rev AAACACGAAGTACTCGGTCGTCGC
Chromosome start end strand MM target_seq PAM alignment distance position gene name gene id
GU305914.1 224069 224091 + 0 GCGACGACCGAGTACTTCGT CGG ||||||||[||||||||||||]PAM NA Intergenic NA NA

T16664 TCTCGTCCCCGCCGCACGACAGG 1000
Oligo fwd CACCTCTCGTCCCCGCCGCACGAC
Oligo rev AAACGTCGTGCGGCGGGGACGAGA
Chromosome start end strand MM target_seq PAM alignment distance position gene name gene id
GU305914.1 224097 224119 + 0 TCTCGTCCCCGCCGCACGAC AGG ||||||||[||||||||||||]PAM NA Intergenic NA NA

T16665 ACGACAGGACTACCTCCAGCCGG 1000
Oligo fwd CACCACGACAGGACTACCTCCAGC
Oligo rev AAACGCTGGAGGTAGTCCTGTCGT
Chromosome start end strand MM target_seq PAM alignment distance position gene name gene id
GU305914.1 224112 224134 + 0 ACGACAGGACTACCTCCAGC CGG ||||||||[||||||||||||]PAM NA Intergenic NA NA

T16666 AGGACTACCTCCAGCCGGTGCGG 1000
Oligo fwd CACCAGGACTACCTCCAGCCGGTG
Oligo rev AAACCACCGGCTGGAGGTAGTCCT
Chromosome start end strand MM target_seq PAM alignment distance position gene name gene id
GU305914.1 224117 224139 + 0 AGGACTACCTCCAGCCGGTG CGG ||||||||[||||||||||||]PAM NA Intergenic NA NA

T16667 CAGATAGAACACGACCCTGTCGG 1000
Oligo fwd CACCCAGATAGAACACGACCCTGT
Oligo rev AAACACAGGGTCGTGTTCTATCTG
Chromosome start end strand MM target_seq PAM alignment distance position gene name gene id
GU305914.1 224140 224162 + 0 CAGATAGAACACGACCCTGT CGG ||||||||[||||||||||||]PAM NA Intergenic NA NA

T16668 ATAGAACACGACCCTGTCGGAGG 1000
Oligo fwd CACCATAGAACACGACCCTGTCGG
Oligo rev AAACCCGACAGGGTCGTGTTCTAT
Chromosome start end strand MM target_seq PAM alignment distance position gene name gene id
GU305914.1 224143 224165 + 0 ATAGAACACGACCCTGTCGG AGG ||||||||[||||||||||||]PAM NA Intergenic NA NA

T16669 CGCGAGCGACAACTCGTACTCGG 1000
Oligo fwd CACCCGCGAGCGACAACTCGTACT
Oligo rev AAACAGTACGAGTTGTCGCTCGCG
Chromosome start end strand MM target_seq PAM alignment distance position gene name gene id
GU305914.1 224167 224189 + 0 CGCGAGCGACAACTCGTACT CGG ||||||||[||||||||||||]PAM NA Intergenic NA NA

T16670 ACAACTCGTACTCGGCCGCCCGG 1000
Oligo fwd CACCACAACTCGTACTCGGCCGCC
Oligo rev AAACGGCGGCCGAGTACGAGTTGT

| Chromosome | start | end | strand | MM | target_seq | PAM | alignment | distance | position | gene name | gene id |
|---|---|---|---|---|---|---|---|---|---|---|---|
| GU305914.1 | 224175 | 224197 | + | 0 | ACAACTCGTACTCGGCCGCC | CGG | ||||||||[||||||||||||]PAM | NA | Intergenic | NA | NA |
| T16671 | TCGTACTCGGCCGCCCGGCCCGG | 1000 | | | | | | | | | |
| Oligo fwd | CACCTCGTACTCGGCCGCCCGGCC | | | | | | | | | | |
| Oligo rev | AAACGGCCGGGCGGCCGAGTACGA | | | | | | | | | | |
| Chromosome | start | end | strand | MM | target_seq | PAM | alignment | distance | position | gene name | gene id |
| GU305914.1 | 224180 | 224202 | + | 0 | TCGTACTCGGCCGCCCGGCC | CGG | ||||||||[||||||||||||]PAM | NA | Intergenic | NA | NA |
| T16672 | GCCGCCCGGCCCGGCCATCTCGG | 1000 | | | | | | | | | |
| Oligo fwd | CACCGCCGCCCGGCCCGGCCATCT | | | | | | | | | | |
| Oligo rev | AAACAGATGGCCGGGCCGGGCGGC | | | | | | | | | | |
| Chromosome | start | end | strand | MM | target_seq | PAM | alignment | distance | position | gene name | gene id |
| GU305914.1 | 224189 | 224211 | + | 0 | GCCGCCCGGCCCGGCCATCT | CGG | ||||||||[||||||||||||]PAM | NA | Intergenic | NA | NA |
| T16673 | CCCGGCCCGGCCATCTCGGCCGG | 1000 | | | | | | | | | |
| Oligo fwd | CACCCCCGGCCCGGCCATCTCGGC | | | | | | | | | | |
| Oligo rev | AAACGCCGAGATGGCCGGGCCGGG | | | | | | | | | | |
| Chromosome | start | end | strand | MM | target_seq | PAM | alignment | distance | position | gene name | gene id |
| GU305914.1 | 224193 | 224215 | + | 0 | CCCGGCCCGGCCATCTCGGC | CGG | ||||||||[||||||||||||]PAM | NA | Intergenic | NA | NA |
| T16674 | GAGATGTAGTCCGCTCGCGAAGG | 1000 | | | | | | | | | |
| Oligo fwd | CACCGAGATGTAGTCCGCTCGCGA | | | | | | | | | | |
| Oligo rev | AAACTCGCGAGCGGACTACATCTC | | | | | | | | | | |
| Chromosome | start | end | strand | MM | target_seq | PAM | alignment | distance | position | gene name | gene id |
| GU305914.1 | 224219 | 224241 | + | 0 | GAGATGTAGTCCGCTCGCGA | AGG | ||||||||[||||||||||||]PAM | NA | Intergenic | NA | NA |
| T16675 | CTCGCGAAGGCCGAGACCAGCGG | 1000 | | | | | | | | | |
| Oligo fwd | CACCCTCGCGAAGGCCGAGACCAG | | | | | | | | | | |
| Oligo rev | AAACCTGGTCTCGGCCTTCGCGAG | | | | | | | | | | |
| Chromosome | start | end | strand | MM | target_seq | PAM | alignment | distance | position | gene name | gene id |
| GU305914.1 | 224232 | 224254 | + | 0 | CTCGCGAAGGCCGAGACCAG | CGG | ||||||||[||||||||||||]PAM | NA | Intergenic | NA | NA |
| T16676 | GCCGAGACCAGCGGCAGCCCAGG | 1000 | | | | | | | | | |
| Oligo fwd | CACCGCCGAGACCAGCGGCAGCCC | | | | | | | | | | |
| Oligo rev | AAACGGGCTGCCGCTGGTCTCGGC | | | | | | | | | | |
| Chromosome | start | end | strand | MM | target_seq | PAM | alignment | distance | position | gene name | gene id |
| GU305914.1 | 224241 | 224263 | + | 0 | GCCGAGACCAGCGGCAGCCC | AGG | ||||||||[||||||||||||]PAM | NA | Intergenic | NA | NA |
| T16677 | GAGACCAGCGGCAGCCCAGGAGG | 1000 | | | | | | | | | |
| Oligo fwd | CACCGAGACCAGCGGCAGCCCAGG | | | | | | | | | | |
| Oligo rev | AAACCCTGGGCTGCCGCTGGTCTC | | | | | | | | | | |
| Chromosome | start | end | strand | MM | target_seq | PAM | alignment | distance | position | gene name | gene id |
| GU305914.1 | 224244 | 224266 | + | 0 | GAGACCAGCGGCAGCCCAGG | AGG | ||||||||[||||||||||||]PAM | NA | Intergenic | NA | NA |
| T16678 | GCGGCAGCCCAGGAGGAGACCGG | 1000 | | | | | | | | | |
| Oligo fwd | CACCGCGGCAGCCCAGGAGGAGAC | | | | | | | | | | |
| Oligo rev | AAACGTCTCCTCCTGGGCTGCCGC | | | | | | | | | | |
| Chromosome | start | end | strand | MM | target_seq | PAM | alignment | distance | position | gene name | gene id |
| GU305914.1 | 224251 | 224273 | + | 0 | GCGGCAGCCCAGGAGGAGAC | CGG | ||||||||[||||||||||||]PAM | NA | Intergenic | NA | NA |
| T16679 | CGGCAGCCCAGGAGGAGACCGGG | 1000 | | | | | | | | | |
| Oligo fwd | CACCCGGCAGCCCAGGAGGAGACC | | | | | | | | | | |
| Oligo rev | AAACGGTCTCCTCCTGGGCTGCCG | | | | | | | | | | |
| Chromosome | start | end | strand | MM | target_seq | PAM | alignment | distance | position | gene name | gene id |
| GU305914.1 | 224252 | 224274 | + | 0 | CGGCAGCCCAGGAGGAGACC | GGG | ||||||||[||||||||||||]PAM | NA | Intergenic | NA | NA |
| T16680 | AGGAGACCGGGTCTCGTCAGCGG | 1000 | | | | | | | | | |
| Oligo fwd | CACCAGGAGACCGGGTCTCGTCAG | | | | | | | | | | |
| Oligo rev | AAACCTGACGAGACCCGGTCTCCT | | | | | | | | | | |
| Chromosome | start | end | strand | MM | target_seq | PAM | alignment | distance | position | gene name | gene id |
| GU305914.1 | 224264 | 224286 | + | 0 | AGGAGACCGGGTCTCGTCAG | CGG | ||||||||[||||||||||||]PAM | NA | Intergenic | NA | NA |
| T16681 | GACCGGGTCTCGTCAGCGGCAGG | 1000 | | | | | | | | | |
| Oligo fwd | CACCGACCGGGTCTCGTCAGCGGC | | | | | | | | | | |
| Oligo rev | AAACGCCGCTGACGAGACCCGGTC | | | | | | | | | | |
| Chromosome | start | end | strand | MM | target_seq | PAM | alignment | distance | position | gene name | gene id |
| GU305914.1 | 224268 | 224290 | + | 0 | GACCGGGTCTCGTCAGCGGC | AGG | ||||||||[||||||||||||]PAM | NA | Intergenic | NA | NA |
| T16682 | ACCGGGTCTCGTCAGCGGCAGGG | 1000 | | | | | | | | | |
| Oligo fwd | CACCACCGGGTCTCGTCAGCGGCA | | | | | | | | | | |
| Oligo rev | AAACTGCCGCTGACGAGACCCGGT | | | | | | | | | | |
| Chromosome | start | end | strand | MM | target_seq | PAM | alignment | distance | position | gene name | gene id |
| GU305914.1 | 224269 | 224291 | + | 0 | ACCGGGTCTCGTCAGCGGCA | GGG | ||||||||[||||||||||||]PAM | NA | Intergenic | NA | NA |
| T16683 | TCGTCAGCGGCAGGGTCACCAGG | 1000 | | | | | | | | | |
| Oligo fwd | CACCTCGTCAGCGGCAGGGTCACC | | | | | | | | | | |
| Oligo rev | AAACGGTGACCCTGCCGCTGACGA | | | | | | | | | | |
| Chromosome | start | end | strand | MM | target_seq | PAM | alignment | distance | position | gene name | gene id |
| GU305914.1 | 224277 | 224299 | + | 0 | TCGTCAGCGGCAGGGTCACC | AGG | ||||||||[||||||||||||]PAM | NA | Intergenic | NA | NA |
| T16684 | CAGCGGCAGGGTCACCAGGACGG | 1000 | | | | | | | | | |
| Oligo fwd | CACCCAGCGGCAGGGTCACCAGGA | | | | | | | | | | |
| Oligo rev | AAACTCCTGGTGACCCTGCCGCTG | | | | | | | | | | |
| Chromosome | start | end | strand | MM | target_seq | PAM | alignment | distance | position | gene name | gene id |
| GU305914.1 | 224281 | 224303 | + | 0 | CAGCGGCAGGGTCACCAGGA | CGG | ||||||||[||||||||||||]PAM | NA | Intergenic | NA | NA |
| T16685 | GGCAGGGTCACCAGGACGGATGG | 1000 | | | | | | | | | |
| Oligo fwd | CACCGGCAGGGTCACCAGGACGGA | | | | | | | | | | |

| | | | | | | | | | | | |
|---|---|---|---|---|---|---|---|---|---|---|---|
| Oligo rev | AAACTCCGTCCTGGTGACCCTGCC | | | | | | | | | | |
| Chromosome | start | end | strand | MM | target_seq | PAM | alignment | distance | position | gene name | gene id |
| GU305914.1 | 224285 | 224307 | + | 0 | GGCAGGGTCACCAGGACGGA | TGG | \|\|\|\|\|\|\|\|[\|\|\|\|\|\|\|\|\|\|\|\|]PAM | NA | Intergenic | NA | NA |
| T16686 | GATGGCCGTCCCGCTCGTGTAGG | 1000 | | | | | | | | | |
| Oligo fwd | CACCGATGGCCGTCCCGCTCGTGT | | | | | | | | | | |
| Oligo rev | AAACACACGAGCGGGACGGCCATC | | | | | | | | | | |
| Chromosome | start | end | strand | MM | target_seq | PAM | alignment | distance | position | gene name | gene id |
| GU305914.1 | 224303 | 224325 | + | 0 | GATGGCCGTCCCGCTCGTGT | AGG | \|\|\|\|\|\|\|\|[\|\|\|\|\|\|\|\|\|\|\|\|]PAM | NA | Intergenic | NA | NA |
| GU305914.1 | 73095 | 73117 | - | 4 | GAAGGGCGTCTCGCTCGTGG | CGG | \|\|-\|\|-\|\|[\|\|-\|\|\|\|\|\|\|\|-]PAM | NA | Intergenic | NA | NA |
| T16687 | ATGGCCGTCCCGCTCGTGTAGGG | 1000 | | | | | | | | | |
| Oligo fwd | CACCATGGCCGTCCCGCTCGTGTA | | | | | | | | | | |
| Oligo rev | AAACTACACGAGCGGGACGGCCAT | | | | | | | | | | |
| Chromosome | start | end | strand | MM | target_seq | PAM | alignment | distance | position | gene name | gene id |
| GU305914.1 | 224304 | 224326 | + | 0 | ATGGCCGTCCCGCTCGTGTA | GGG | \|\|\|\|\|\|\|\|[\|\|\|\|\|\|\|\|\|\|\|\|]PAM | NA | Intergenic | NA | NA |
| T16688 | TGGCCGTCCCGCTCGTGTAGGGG | 1000 | | | | | | | | | |
| Oligo fwd | CACCTGGCCGTCCCGCTCGTGTAG | | | | | | | | | | |
| Oligo rev | AAACCTACACGAGCGGGACGGCCA | | | | | | | | | | |
| Chromosome | start | end | strand | MM | target_seq | PAM | alignment | distance | position | gene name | gene id |
| GU305914.1 | 224305 | 224327 | + | 0 | TGGCCGTCCCGCTCGTGTAG | GGG | \|\|\|\|\|\|\|\|[\|\|\|\|\|\|\|\|\|\|\|\|]PAM | NA | Intergenic | NA | NA |
| T16689 | GGGACATCAACAGCCACGTCAGG | 1000 | | | | | | | | | |
| Oligo fwd | CACCGGGACATCAACAGCCACGTC | | | | | | | | | | |
| Oligo rev | AAACGACGTGGCTGTTGATGTCCC | | | | | | | | | | |
| Chromosome | start | end | strand | MM | target_seq | PAM | alignment | distance | position | gene name | gene id |
| GU305914.1 | 224325 | 224347 | + | 0 | GGGACATCAACAGCCACGTC | AGG | \|\|\|\|\|\|\|\|[\|\|\|\|\|\|\|\|\|\|\|\|]PAM | NA | Intergenic | NA | NA |
| T16690 | TCAACAGCCACGTCAGGAGCAGG | 1000 | | | | | | | | | |
| Oligo fwd | CACCTCAACAGCCACGTCAGGAGC | | | | | | | | | | |
| Oligo rev | AAACGCTCCTGACGTGGCTGTTGA | | | | | | | | | | |
| Chromosome | start | end | strand | MM | target_seq | PAM | alignment | distance | position | gene name | gene id |
| GU305914.1 | 224331 | 224353 | + | 0 | TCAACAGCCACGTCAGGAGC | AGG | \|\|\|\|\|\|\|\|[\|\|\|\|\|\|\|\|\|\|\|\|]PAM | NA | Intergenic | NA | NA |
| T16691 | CAACAGCCACGTCAGGAGCAGGG | 1000 | | | | | | | | | |
| Oligo fwd | CACCCAACAGCCACGTCAGGAGCA | | | | | | | | | | |
| Oligo rev | AAACTGCTCCTGACGTGGCTGTTG | | | | | | | | | | |
| Chromosome | start | end | strand | MM | target_seq | PAM | alignment | distance | position | gene name | gene id |
| GU305914.1 | 224332 | 224354 | + | 0 | CAACAGCCACGTCAGGAGCA | GGG | \|\|\|\|\|\|\|\|[\|\|\|\|\|\|\|\|\|\|\|\|]PAM | NA | Intergenic | NA | NA |
| T16692 | AGGAGCAGGGTCAGCATCGTAGG | 1000 | | | | | | | | | |
| Oligo fwd | CACCAGGAGCAGGGTCAGCATCGT | | | | | | | | | | |
| Oligo rev | AAACACGATGCTGACCCTGCTCCT | | | | | | | | | | |
| Chromosome | start | end | strand | MM | target_seq | PAM | alignment | distance | position | gene name | gene id |
| GU305914.1 | 224345 | 224367 | + | 0 | AGGAGCAGGGTCAGCATCGT | AGG | \|\|\|\|\|\|\|\|[\|\|\|\|\|\|\|\|\|\|\|\|]PAM | NA | Intergenic | NA | NA |
| T16693 | GGGTCAGCATCGTAGGCCCGAGG | 1000 | | | | | | | | | |
| Oligo fwd | CACCGGGTCAGCATCGTAGGCCCG | | | | | | | | | | |
| Oligo rev | AAACCGGGCCTACGATGCTGACCC | | | | | | | | | | |
| Chromosome | start | end | strand | MM | target_seq | PAM | alignment | distance | position | gene name | gene id |
| GU305914.1 | 224352 | 224374 | + | 0 | GGGTCAGCATCGTAGGCCCG | AGG | \|\|\|\|\|\|\|\|[\|\|\|\|\|\|\|\|\|\|\|\|]PAM | NA | Intergenic | NA | NA |
| T16694 | GGTCAGCATCGTAGGCCCGAGGG | 1000 | | | | | | | | | |
| Oligo fwd | CACCGGTCAGCATCGTAGGCCCGA | | | | | | | | | | |
| Oligo rev | AAACTCGGGCCTACGATGCTGACC | | | | | | | | | | |
| Chromosome | start | end | strand | MM | target_seq | PAM | alignment | distance | position | gene name | gene id |
| GU305914.1 | 224353 | 224375 | + | 0 | GGTCAGCATCGTAGGCCCGA | GGG | \|\|\|\|\|\|\|\|[\|\|\|\|\|\|\|\|\|\|\|\|]PAM | NA | Intergenic | NA | NA |
| T16695 | CGTAGGCCCGAGGGCCATGACGG | 1000 | | | | | | | | | |
| Oligo fwd | CACCCGTAGGCCCGAGGGCCATGA | | | | | | | | | | |
| Oligo rev | AAACTCATGGCCCTCGGGCCTACG | | | | | | | | | | |
| Chromosome | start | end | strand | MM | target_seq | PAM | alignment | distance | position | gene name | gene id |
| GU305914.1 | 224362 | 224384 | + | 0 | CGTAGGCCCGAGGGCCATGA | CGG | \|\|\|\|\|\|\|\|[\|\|\|\|\|\|\|\|\|\|\|\|]PAM | NA | Intergenic | NA | NA |
| T16696 | GGCCCGAGGGCCATGACGGCCGG | 1000 | | | | | | | | | |
| Oligo fwd | CACCGGCCCGAGGGCCATGACGGC | | | | | | | | | | |
| Oligo rev | AAACGCCGTCATGGCCCTCGGGCC | | | | | | | | | | |
| Chromosome | start | end | strand | MM | target_seq | PAM | alignment | distance | position | gene name | gene id |
| GU305914.1 | 224366 | 224388 | + | 0 | GGCCCGAGGGCCATGACGGC | CGG | \|\|\|\|\|\|\|\|[\|\|\|\|\|\|\|\|\|\|\|\|]PAM | NA | Intergenic | NA | NA |
| T16697 | CCGAGGGCCATGACGGCCGGTGG | 1000 | | | | | | | | | |
| Oligo fwd | CACCCCGAGGGCCATGACGGCCGG | | | | | | | | | | |
| Oligo rev | AAACCCGGCCGTCATGGCCCTCGG | | | | | | | | | | |
| Chromosome | start | end | strand | MM | target_seq | PAM | alignment | distance | position | gene name | gene id |
| GU305914.1 | 224369 | 224391 | + | 0 | CCGAGGGCCATGACGGCCGG | TGG | \|\|\|\|\|\|\|\|[\|\|\|\|\|\|\|\|\|\|\|\|]PAM | NA | Intergenic | NA | NA |
| T16698 | GGCCGGTGGCCGCGAGAAAAAGG | 1000 | | | | | | | | | |
| Oligo fwd | CACCGGCCGGTGGCCGCGAGAAAA | | | | | | | | | | |
| Oligo rev | AAACTTTTCTCGCGGCCACCGGCC | | | | | | | | | | |
| Chromosome | start | end | strand | MM | target_seq | PAM | alignment | distance | position | gene name | gene id |
| GU305914.1 | 224383 | 224405 | + | 0 | GGCCGGTGGCCGCGAGAAAA | AGG | \|\|\|\|\|\|\|\|[\|\|\|\|\|\|\|\|\|\|\|\|]PAM | NA | Intergenic | NA | NA |
| T16699 | AAGCAGAAAGAGAACTTTATTGG | 1000 | | | | | | | | | |
| Oligo fwd | CACCAAGCAGAAAGAGAACTTTAT | | | | | | | | | | |
| Oligo rev | AAACATAAAGTTCTCTTTCTGCTT | | | | | | | | | | |
| Chromosome | start | end | strand | MM | target_seq | PAM | alignment | distance | position | gene name | gene id |
| GU305914.1 | 224434 | 224456 | + | 0 | AAGCAGAAAGAGAACTTTAT | TGG | \|\|\|\|\|\|\|\|[\|\|\|\|\|\|\|\|\|\|\|\|]PAM | NA | Intergenic | NA | NA |

<table>
<tr><td>T16700</td><td>AGCAGAAAGAGAACTTTATTGGG</td><td>1000</td><td></td><td></td><td></td><td></td><td></td><td></td><td></td><td></td><td></td></tr>
<tr><td>Oligo fwd</td><td>CACCAGCAGAAAGAGAACTTTATT</td><td></td><td></td><td></td><td></td><td></td><td></td><td></td><td></td><td></td><td></td></tr>
<tr><td>Oligo rev</td><td>AAACAATAAAGTTCTCTTTCTGCT</td><td></td><td></td><td></td><td></td><td></td><td></td><td></td><td></td><td></td><td></td></tr>
<tr><td>Chromosome</td><td>start</td><td>end</td><td>strand</td><td>MM</td><td>target_seq</td><td>PAM</td><td>alignment</td><td>distance</td><td>position</td><td>gene name</td><td>gene id</td></tr>
<tr><td>GU305914.1</td><td>224435</td><td>224457</td><td>+</td><td>0</td><td>AGCAGAAAGAGAACTTTATT</td><td>GGG</td><td>|||||||[|||||||||||||]PAM</td><td>NA</td><td>Intergenic</td><td>NA</td><td>NA</td></tr>
<tr><td>T16701</td><td>GCAGAAAGAGAACTTTATTGGGG</td><td>1000</td><td></td><td></td><td></td><td></td><td></td><td></td><td></td><td></td><td></td></tr>
<tr><td>Oligo fwd</td><td>CACCGCAGAAAGAGAACTTTATTG</td><td></td><td></td><td></td><td></td><td></td><td></td><td></td><td></td><td></td><td></td></tr>
<tr><td>Oligo rev</td><td>AAACCAATAAAGTTCTCTTTCTGC</td><td></td><td></td><td></td><td></td><td></td><td></td><td></td><td></td><td></td><td></td></tr>
<tr><td>Chromosome</td><td>start</td><td>end</td><td>strand</td><td>MM</td><td>target_seq</td><td>PAM</td><td>alignment</td><td>distance</td><td>position</td><td>gene name</td><td>gene id</td></tr>
<tr><td>GU305914.1</td><td>224436</td><td>224458</td><td>+</td><td>0</td><td>GCAGAAAGAGAACTTTATTG</td><td>GGG</td><td>|||||||[|||||||||||||]PAM</td><td>NA</td><td>Intergenic</td><td>NA</td><td>NA</td></tr>
<tr><td>T16702</td><td>CAGAAAGAGAACTTTATTGGGGG</td><td>1000</td><td></td><td></td><td></td><td></td><td></td><td></td><td></td><td></td><td></td></tr>
<tr><td>Oligo fwd</td><td>CACCCAGAAAGAGAACTTTATTGG</td><td></td><td></td><td></td><td></td><td></td><td></td><td></td><td></td><td></td><td></td></tr>
<tr><td>Oligo rev</td><td>AAACCCAATAAAGTTCTCTTTCTG</td><td></td><td></td><td></td><td></td><td></td><td></td><td></td><td></td><td></td><td></td></tr>
<tr><td>Chromosome</td><td>start</td><td>end</td><td>strand</td><td>MM</td><td>target_seq</td><td>PAM</td><td>alignment</td><td>distance</td><td>position</td><td>gene name</td><td>gene id</td></tr>
<tr><td>GU305914.1</td><td>224437</td><td>224459</td><td>+</td><td>0</td><td>CAGAAAGAGAACTTTATTGG</td><td>GGG</td><td>|||||||[|||||||||||||]PAM</td><td>NA</td><td>Intergenic</td><td>NA</td><td>NA</td></tr>
<tr><td>T16703</td><td>AAAGAGAACTTTATTGGGGGAGG</td><td>1000</td><td></td><td></td><td></td><td></td><td></td><td></td><td></td><td></td><td></td></tr>
<tr><td>Oligo fwd</td><td>CACCAAAGAGAACTTTATTGGGGG</td><td></td><td></td><td></td><td></td><td></td><td></td><td></td><td></td><td></td><td></td></tr>
<tr><td>Oligo rev</td><td>AAACCCCCCAATAAAGTTCTCTTT</td><td></td><td></td><td></td><td></td><td></td><td></td><td></td><td></td><td></td><td></td></tr>
<tr><td>Chromosome</td><td>start</td><td>end</td><td>strand</td><td>MM</td><td>target_seq</td><td>PAM</td><td>alignment</td><td>distance</td><td>position</td><td>gene name</td><td>gene id</td></tr>
<tr><td>GU305914.1</td><td>224440</td><td>224462</td><td>+</td><td>0</td><td>AAAGAGAACTTTATTGGGGG</td><td>AGG</td><td>|||||||[|||||||||||||]PAM</td><td>NA</td><td>Intergenic</td><td>NA</td><td>NA</td></tr>
<tr><td>T16704</td><td>GGGAGGTTGTAACATATCAACGG</td><td>1000</td><td></td><td></td><td></td><td></td><td></td><td></td><td></td><td></td><td></td></tr>
<tr><td>Oligo fwd</td><td>CACCGGGAGGTTGTAACATATCAA</td><td></td><td></td><td></td><td></td><td></td><td></td><td></td><td></td><td></td><td></td></tr>
<tr><td>Oligo rev</td><td>AAACTTGATATGTTACAACCTCCC</td><td></td><td></td><td></td><td></td><td></td><td></td><td></td><td></td><td></td><td></td></tr>
<tr><td>Chromosome</td><td>start</td><td>end</td><td>strand</td><td>MM</td><td>target_seq</td><td>PAM</td><td>alignment</td><td>distance</td><td>position</td><td>gene name</td><td>gene id</td></tr>
<tr><td>GU305914.1</td><td>224457</td><td>224479</td><td>+</td><td>0</td><td>GGGAGGTTGTAACATATCAA</td><td>CGG</td><td>|||||||[|||||||||||||]PAM</td><td>NA</td><td>Intergenic</td><td>NA</td><td>NA</td></tr>
<tr><td>T16705</td><td>CAACGGTTTAATAATAGAAAAGG</td><td>1000</td><td></td><td></td><td></td><td></td><td></td><td></td><td></td><td></td><td></td></tr>
<tr><td>Oligo fwd</td><td>CACCCAACGGTTTAATAATAGAAA</td><td></td><td></td><td></td><td></td><td></td><td></td><td></td><td></td><td></td><td></td></tr>
<tr><td>Oligo rev</td><td>AAACTTTCTATTATTAAACCGTTG</td><td></td><td></td><td></td><td></td><td></td><td></td><td></td><td></td><td></td><td></td></tr>
<tr><td>Chromosome</td><td>start</td><td>end</td><td>strand</td><td>MM</td><td>target_seq</td><td>PAM</td><td>alignment</td><td>distance</td><td>position</td><td>gene name</td><td>gene id</td></tr>
<tr><td>GU305914.1</td><td>224474</td><td>224496</td><td>+</td><td>0</td><td>CAACGGTTTAATAATAGAAA</td><td>AGG</td><td>|||||||[|||||||||||||]PAM</td><td>NA</td><td>Intergenic</td><td>NA</td><td>NA</td></tr>
<tr><td>T16706</td><td>TTAATAATAGAAAAGGAAATCGG</td><td>1000</td><td></td><td></td><td></td><td></td><td></td><td></td><td></td><td></td><td></td></tr>
<tr><td>Oligo fwd</td><td>CACCTTAATAATAGAAAAGGAAAT</td><td></td><td></td><td></td><td></td><td></td><td></td><td></td><td></td><td></td><td></td></tr>
<tr><td>Oligo rev</td><td>AAACATTTCCTTTTCTATTATTAA</td><td></td><td></td><td></td><td></td><td></td><td></td><td></td><td></td><td></td><td></td></tr>
<tr><td>Chromosome</td><td>start</td><td>end</td><td>strand</td><td>MM</td><td>target_seq</td><td>PAM</td><td>alignment</td><td>distance</td><td>position</td><td>gene name</td><td>gene id</td></tr>
<tr><td>GU305914.1</td><td>224481</td><td>224503</td><td>+</td><td>0</td><td>TTAATAATAGAAAAGGAAAT</td><td>CGG</td><td>|||||||[|||||||||||||]PAM</td><td>NA</td><td>Intergenic</td><td>NA</td><td>NA</td></tr>
<tr><td>T16707</td><td>TAATAATAGAAAAGGAAATCGGG</td><td>1000</td><td></td><td></td><td></td><td></td><td></td><td></td><td></td><td></td><td></td></tr>
<tr><td>Oligo fwd</td><td>CACCTAATAATAGAAAAGGAAATC</td><td></td><td></td><td></td><td></td><td></td><td></td><td></td><td></td><td></td><td></td></tr>
<tr><td>Oligo rev</td><td>AAACGATTTCCTTTTCTATTATTA</td><td></td><td></td><td></td><td></td><td></td><td></td><td></td><td></td><td></td><td></td></tr>
<tr><td>Chromosome</td><td>start</td><td>end</td><td>strand</td><td>MM</td><td>target_seq</td><td>PAM</td><td>alignment</td><td>distance</td><td>position</td><td>gene name</td><td>gene id</td></tr>
<tr><td>GU305914.1</td><td>224482</td><td>224504</td><td>+</td><td>0</td><td>TAATAATAGAAAAGGAAATC</td><td>GGG</td><td>|||||||[|||||||||||||]PAM</td><td>NA</td><td>Intergenic</td><td>NA</td><td>NA</td></tr>
<tr><td>T16708</td><td>CTCGTTCACGAGTATCTCATAGG</td><td>1000</td><td></td><td></td><td></td><td></td><td></td><td></td><td></td><td></td><td></td></tr>
<tr><td>Oligo fwd</td><td>CACCCTCGTTCACGAGTATCTCAT</td><td></td><td></td><td></td><td></td><td></td><td></td><td></td><td></td><td></td><td></td></tr>
<tr><td>Oligo rev</td><td>AAACATGAGATACTCGTGAACGAG</td><td></td><td></td><td></td><td></td><td></td><td></td><td></td><td></td><td></td><td></td></tr>
<tr><td>Chromosome</td><td>start</td><td>end</td><td>strand</td><td>MM</td><td>target_seq</td><td>PAM</td><td>alignment</td><td>distance</td><td>position</td><td>gene name</td><td>gene id</td></tr>
<tr><td>GU305914.1</td><td>224526</td><td>224548</td><td>+</td><td>0</td><td>CTCGTTCACGAGTATCTCAT</td><td>AGG</td><td>|||||||[|||||||||||||]PAM</td><td>NA</td><td>Intergenic</td><td>NA</td><td>NA</td></tr>
<tr><td>T16709</td><td>TTCACGAGTATCTCATAGGCCGG</td><td>1000</td><td></td><td></td><td></td><td></td><td></td><td></td><td></td><td></td><td></td></tr>
<tr><td>Oligo fwd</td><td>CACCTTCACGAGTATCTCATAGGC</td><td></td><td></td><td></td><td></td><td></td><td></td><td></td><td></td><td></td><td></td></tr>
<tr><td>Oligo rev</td><td>AAACGCCTATGAGATACTCGTGAA</td><td></td><td></td><td></td><td></td><td></td><td></td><td></td><td></td><td></td><td></td></tr>
<tr><td>Chromosome</td><td>start</td><td>end</td><td>strand</td><td>MM</td><td>target_seq</td><td>PAM</td><td>alignment</td><td>distance</td><td>position</td><td>gene name</td><td>gene id</td></tr>
<tr><td>GU305914.1</td><td>224530</td><td>224552</td><td>+</td><td>0</td><td>TTCACGAGTATCTCATAGGC</td><td>CGG</td><td>|||||||[|||||||||||||]PAM</td><td>NA</td><td>Intergenic</td><td>NA</td><td>NA</td></tr>
<tr><td>T16710</td><td>CCTTTTTTTCCTCCTGCGCCCGG</td><td>1000</td><td></td><td></td><td></td><td></td><td></td><td></td><td></td><td></td><td></td></tr>
<tr><td>Oligo fwd</td><td>CACCCCTTTTTTTCCTCCTGCGCC</td><td></td><td></td><td></td><td></td><td></td><td></td><td></td><td></td><td></td><td></td></tr>
<tr><td>Oligo rev</td><td>AAACGGCGCAGGAGGAAAAAAAGG</td><td></td><td></td><td></td><td></td><td></td><td></td><td></td><td></td><td></td><td></td></tr>
<tr><td>Chromosome</td><td>start</td><td>end</td><td>strand</td><td>MM</td><td>target_seq</td><td>PAM</td><td>alignment</td><td>distance</td><td>position</td><td>gene name</td><td>gene id</td></tr>
<tr><td>GU305914.1</td><td>224557</td><td>224579</td><td>+</td><td>0</td><td>CCTTTTTTTCCTCCTGCGCC</td><td>CGG</td><td>|||||||[|||||||||||||]PAM</td><td>NA</td><td>Intergenic</td><td>NA</td><td>NA</td></tr>
<tr><td>T16711</td><td>TTCCTCCTGCGCCCGGCGAGCGG</td><td>1000</td><td></td><td></td><td></td><td></td><td></td><td></td><td></td><td></td><td></td></tr>
<tr><td>Oligo fwd</td><td>CACCTTCCTCCTGCGCCCGGCGAG</td><td></td><td></td><td></td><td></td><td></td><td></td><td></td><td></td><td></td><td></td></tr>
<tr><td>Oligo rev</td><td>AAACCTCGCCGGGCGCAGGAGGAA</td><td></td><td></td><td></td><td></td><td></td><td></td><td></td><td></td><td></td><td></td></tr>
<tr><td>Chromosome</td><td>start</td><td>end</td><td>strand</td><td>MM</td><td>target_seq</td><td>PAM</td><td>alignment</td><td>distance</td><td>position</td><td>gene name</td><td>gene id</td></tr>
<tr><td>GU305914.1</td><td>224564</td><td>224586</td><td>+</td><td>0</td><td>TTCCTCCTGCGCCCGGCGAG</td><td>CGG</td><td>|||||||[|||||||||||||]PAM</td><td>NA</td><td>Intergenic</td><td>NA</td><td>NA</td></tr>
<tr><td>T16712</td><td>TCCTCCTGCGCCCGGCGAGCGGG</td><td>1000</td><td></td><td></td><td></td><td></td><td></td><td></td><td></td><td></td><td></td></tr>
<tr><td>Oligo fwd</td><td>CACCTCCTCCTGCGCCCGGCGAGC</td><td></td><td></td><td></td><td></td><td></td><td></td><td></td><td></td><td></td><td></td></tr>
<tr><td>Oligo rev</td><td>AAACGCTCGCCGGGCGCAGGAGGA</td><td></td><td></td><td></td><td></td><td></td><td></td><td></td><td></td><td></td><td></td></tr>
<tr><td>Chromosome</td><td>start</td><td>end</td><td>strand</td><td>MM</td><td>target_seq</td><td>PAM</td><td>alignment</td><td>distance</td><td>position</td><td>gene name</td><td>gene id</td></tr>
<tr><td>GU305914.1</td><td>224565</td><td>224587</td><td>+</td><td>0</td><td>TCCTCCTGCGCCCGGCGAGC</td><td>GGG</td><td>|||||||[|||||||||||||]PAM</td><td>NA</td><td>Intergenic</td><td>NA</td><td>NA</td></tr>
<tr><td>T16713</td><td>TCCTGCGCCCGGCGAGCGGGAGG</td><td>1000</td><td></td><td></td><td></td><td></td><td></td><td></td><td></td><td></td><td></td></tr>
<tr><td>Oligo fwd</td><td>CACCTCCTGCGCCCGGCGAGCGGG</td><td></td><td></td><td></td><td></td><td></td><td></td><td></td><td></td><td></td><td></td></tr>
<tr><td>Oligo rev</td><td>AAACCCCGCTCGCCGGGCGCAGGA</td><td></td><td></td><td></td><td></td><td></td><td></td><td></td><td></td><td></td><td></td></tr>
<tr><td>Chromosome</td><td>start</td><td>end</td><td>strand</td><td>MM</td><td>target_seq</td><td>PAM</td><td>alignment</td><td>distance</td><td>position</td><td>gene name</td><td>gene id</td></tr>
<tr><td>GU305914.1</td><td>224568</td><td>224590</td><td>+</td><td>0</td><td>TCCTGCGCCCGGCGAGCGGG</td><td>AGG</td><td>|||||||[|||||||||||||]PAM</td><td>NA</td><td>Intergenic</td><td>NA</td><td>NA</td></tr>
<tr><td>GU305914.1</td><td>113525</td><td>113547</td><td>-</td><td>4</td><td>TCCCGCGTCCGGCGCGCGAG</td><td>CGG</td><td>|||-|||-[||||||-|||-|]PAM</td><td>NA</td><td>Intergenic</td><td>NA</td><td>NA</td></tr>
<tr><td>T16714</td><td>CCTGCGCCCGGCGAGCGGGAGGG</td><td>1000</td><td></td><td></td><td></td><td></td><td></td><td></td><td></td><td></td><td></td></tr>
<tr><td>Oligo fwd</td><td>CACCCCTGCGCCCGGCGAGCGGGA</td><td></td><td></td><td></td><td></td><td></td><td></td><td></td><td></td><td></td><td></td></tr>
<tr><td>Oligo rev</td><td>AAACTCCCGCTCGCCGGGCGCAGG</td><td></td><td></td><td></td><td></td><td></td><td></td><td></td><td></td><td></td><td></td></tr>
<tr><td>Chromosome</td><td>start</td><td>end</td><td>strand</td><td>MM</td><td>target_seq</td><td>PAM</td><td>alignment</td><td>distance</td><td>position</td><td>gene name</td><td>gene id</td></tr>
</table>

<table>
<tr><td>GU305914.1</td><td>224569</td><td>224591</td><td>+</td><td>0</td><td>CCTGCGCCCGGCGAGCGGGA</td><td>GGG</td><td>||||||||[||||||||||||]PAM</td><td>NA</td><td>Intergenic</td><td>NA</td><td>NA</td></tr>
<tr><td>T16715</td><td>CTGCGCCCGGCGAGCGGGAGGGG</td><td>1000</td><td></td><td></td><td></td><td></td><td></td><td></td><td></td><td></td><td></td></tr>
<tr><td>Oligo fwd</td><td>CACCCTGCGCCCGGCGAGCGGGAG</td><td></td><td></td><td></td><td></td><td></td><td></td><td></td><td></td><td></td><td></td></tr>
<tr><td>Oligo rev</td><td>AAACCTCCCGCTCGCCGGGCGCAG</td><td></td><td></td><td></td><td></td><td></td><td></td><td></td><td></td><td></td><td></td></tr>
<tr><td>Chromosome</td><td>start</td><td>end</td><td>strand</td><td>MM</td><td>target_seq</td><td>PAM</td><td>alignment</td><td>distance</td><td>position</td><td>gene name</td><td>gene id</td></tr>
<tr><td>GU305914.1</td><td>224570</td><td>224592</td><td>+</td><td>0</td><td>CTGCGCCCGGCGAGCGGGAG</td><td>GGG</td><td>||||||||[||||||||||||]PAM</td><td>NA</td><td>Intergenic</td><td>NA</td><td>NA</td></tr>
<tr><td>GU305914.1</td><td>75865</td><td>75887</td><td>+</td><td>4</td><td>CGGCGGTCGGCGAGGGGGAG</td><td>CGG</td><td>|-|||--|[||||||-|||||]PAM</td><td>NA</td><td>Intergenic</td><td>NA</td><td>NA</td></tr>
<tr><td>T16716</td><td>GGGAGGGGAAGCAACAGCTGCGG</td><td>1000</td><td></td><td></td><td></td><td></td><td></td><td></td><td></td><td></td><td></td></tr>
<tr><td>Oligo fwd</td><td>CACCGGGAGGGGAAGCAACAGCTG</td><td></td><td></td><td></td><td></td><td></td><td></td><td></td><td></td><td></td><td></td></tr>
<tr><td>Oligo rev</td><td>AAACCAGCTGTTGCTTCCCCTCCC</td><td></td><td></td><td></td><td></td><td></td><td></td><td></td><td></td><td></td><td></td></tr>
<tr><td>Chromosome</td><td>start</td><td>end</td><td>strand</td><td>MM</td><td>target_seq</td><td>PAM</td><td>alignment</td><td>distance</td><td>position</td><td>gene name</td><td>gene id</td></tr>
<tr><td>GU305914.1</td><td>224585</td><td>224607</td><td>+</td><td>0</td><td>GGGAGGGGAAGCAACAGCTG</td><td>CGG</td><td>||||||||[||||||||||||]PAM</td><td>NA</td><td>Intergenic</td><td>NA</td><td>NA</td></tr>
<tr><td>T16717</td><td>TGCCGATCAGCACGATCCCCAGG</td><td>1000</td><td></td><td></td><td></td><td></td><td></td><td></td><td></td><td></td><td></td></tr>
<tr><td>Oligo fwd</td><td>CACCTGCCGATCAGCACGATCCCC</td><td></td><td></td><td></td><td></td><td></td><td></td><td></td><td></td><td></td><td></td></tr>
<tr><td>Oligo rev</td><td>AAACGGGGATCGTGCTGATCGGCA</td><td></td><td></td><td></td><td></td><td></td><td></td><td></td><td></td><td></td><td></td></tr>
<tr><td>Chromosome</td><td>start</td><td>end</td><td>strand</td><td>MM</td><td>target_seq</td><td>PAM</td><td>alignment</td><td>distance</td><td>position</td><td>gene name</td><td>gene id</td></tr>
<tr><td>GU305914.1</td><td>224624</td><td>224646</td><td>+</td><td>0</td><td>TGCCGATCAGCACGATCCCC</td><td>AGG</td><td>||||||||[||||||||||||]PAM</td><td>NA</td><td>Intergenic</td><td>NA</td><td>NA</td></tr>
<tr><td>T16718</td><td>GCCGATCAGCACGATCCCCAGGG</td><td>1000</td><td></td><td></td><td></td><td></td><td></td><td></td><td></td><td></td><td></td></tr>
<tr><td>Oligo fwd</td><td>CACCGCCGATCAGCACGATCCCCA</td><td></td><td></td><td></td><td></td><td></td><td></td><td></td><td></td><td></td><td></td></tr>
<tr><td>Oligo rev</td><td>AAACTGGGGATCGTGCTGATCGGC</td><td></td><td></td><td></td><td></td><td></td><td></td><td></td><td></td><td></td><td></td></tr>
<tr><td>Chromosome</td><td>start</td><td>end</td><td>strand</td><td>MM</td><td>target_seq</td><td>PAM</td><td>alignment</td><td>distance</td><td>position</td><td>gene name</td><td>gene id</td></tr>
<tr><td>GU305914.1</td><td>224625</td><td>224647</td><td>+</td><td>0</td><td>GCCGATCAGCACGATCCCCA</td><td>GGG</td><td>||||||||[||||||||||||]PAM</td><td>NA</td><td>Intergenic</td><td>NA</td><td>NA</td></tr>
<tr><td>T16719</td><td>TCAGCACGATCCCCAGGGCCAGG</td><td>1000</td><td></td><td></td><td></td><td></td><td></td><td></td><td></td><td></td><td></td></tr>
<tr><td>Oligo fwd</td><td>CACCTCAGCACGATCCCCAGGGCC</td><td></td><td></td><td></td><td></td><td></td><td></td><td></td><td></td><td></td><td></td></tr>
<tr><td>Oligo rev</td><td>AAACGGCCCTGGGGATCGTGCTGA</td><td></td><td></td><td></td><td></td><td></td><td></td><td></td><td></td><td></td><td></td></tr>
<tr><td>Chromosome</td><td>start</td><td>end</td><td>strand</td><td>MM</td><td>target_seq</td><td>PAM</td><td>alignment</td><td>distance</td><td>position</td><td>gene name</td><td>gene id</td></tr>
<tr><td>GU305914.1</td><td>224630</td><td>224652</td><td>+</td><td>0</td><td>TCAGCACGATCCCCAGGGCC</td><td>AGG</td><td>||||||||[||||||||||||]PAM</td><td>NA</td><td>Intergenic</td><td>NA</td><td>NA</td></tr>
<tr><td>T16720</td><td>CCACGTGATGACGATCAGCACGG</td><td>1000</td><td></td><td></td><td></td><td></td><td></td><td></td><td></td><td></td><td></td></tr>
<tr><td>Oligo fwd</td><td>CACCCCACGTGATGACGATCAGCA</td><td></td><td></td><td></td><td></td><td></td><td></td><td></td><td></td><td></td><td></td></tr>
<tr><td>Oligo rev</td><td>AAACTGCTGATCGTCATCACGTGG</td><td></td><td></td><td></td><td></td><td></td><td></td><td></td><td></td><td></td><td></td></tr>
<tr><td>Chromosome</td><td>start</td><td>end</td><td>strand</td><td>MM</td><td>target_seq</td><td>PAM</td><td>alignment</td><td>distance</td><td>position</td><td>gene name</td><td>gene id</td></tr>
<tr><td>GU305914.1</td><td>224673</td><td>224695</td><td>+</td><td>0</td><td>CCACGTGATGACGATCAGCA</td><td>CGG</td><td>||||||||[||||||||||||]PAM</td><td>NA</td><td>Intergenic</td><td>NA</td><td>NA</td></tr>
<tr><td>T16721</td><td>TGACGATCAGCACGGTGTGCAGG</td><td>1000</td><td></td><td></td><td></td><td></td><td></td><td></td><td></td><td></td><td></td></tr>
<tr><td>Oligo fwd</td><td>CACCTGACGATCAGCACGGTGTGC</td><td></td><td></td><td></td><td></td><td></td><td></td><td></td><td></td><td></td><td></td></tr>
<tr><td>Oligo rev</td><td>AAACGCACACCGTGCTGATCGTCA</td><td></td><td></td><td></td><td></td><td></td><td></td><td></td><td></td><td></td><td></td></tr>
<tr><td>Chromosome</td><td>start</td><td>end</td><td>strand</td><td>MM</td><td>target_seq</td><td>PAM</td><td>alignment</td><td>distance</td><td>position</td><td>gene name</td><td>gene id</td></tr>
<tr><td>GU305914.1</td><td>224681</td><td>224703</td><td>+</td><td>0</td><td>TGACGATCAGCACGGTGTGC</td><td>AGG</td><td>||||||||[||||||||||||]PAM</td><td>NA</td><td>Intergenic</td><td>NA</td><td>NA</td></tr>
<tr><td>T16722</td><td>CGGTGTGCAGGATGTCCATGTGG</td><td>1000</td><td></td><td></td><td></td><td></td><td></td><td></td><td></td><td></td><td></td></tr>
<tr><td>Oligo fwd</td><td>CACCCGGTGTGCAGGATGTCCATG</td><td></td><td></td><td></td><td></td><td></td><td></td><td></td><td></td><td></td><td></td></tr>
<tr><td>Oligo rev</td><td>AAACCATGGACATCCTGCACACCG</td><td></td><td></td><td></td><td></td><td></td><td></td><td></td><td></td><td></td><td></td></tr>
<tr><td>Chromosome</td><td>start</td><td>end</td><td>strand</td><td>MM</td><td>target_seq</td><td>PAM</td><td>alignment</td><td>distance</td><td>position</td><td>gene name</td><td>gene id</td></tr>
<tr><td>GU305914.1</td><td>224693</td><td>224715</td><td>+</td><td>0</td><td>CGGTGTGCAGGATGTCCATG</td><td>TGG</td><td>||||||||[||||||||||||]PAM</td><td>NA</td><td>Intergenic</td><td>NA</td><td>NA</td></tr>
<tr><td>T16723</td><td>TGTGCAGGATGTCCATGTGGCGG</td><td>1000</td><td></td><td></td><td></td><td></td><td></td><td></td><td></td><td></td><td></td></tr>
<tr><td>Oligo fwd</td><td>CACCTGTGCAGGATGTCCATGTGG</td><td></td><td></td><td></td><td></td><td></td><td></td><td></td><td></td><td></td><td></td></tr>
<tr><td>Oligo rev</td><td>AAACCCACATGGACATCCTGCACA</td><td></td><td></td><td></td><td></td><td></td><td></td><td></td><td></td><td></td><td></td></tr>
<tr><td>Chromosome</td><td>start</td><td>end</td><td>strand</td><td>MM</td><td>target_seq</td><td>PAM</td><td>alignment</td><td>distance</td><td>position</td><td>gene name</td><td>gene id</td></tr>
<tr><td>GU305914.1</td><td>224696</td><td>224718</td><td>+</td><td>0</td><td>TGTGCAGGATGTCCATGTGG</td><td>CGG</td><td>||||||||[||||||||||||]PAM</td><td>NA</td><td>Intergenic</td><td>NA</td><td>NA</td></tr>
<tr><td>T16724</td><td>TCCATGTGGCGGCGTTCCGTCGG</td><td>1000</td><td></td><td></td><td></td><td></td><td></td><td></td><td></td><td></td><td></td></tr>
<tr><td>Oligo fwd</td><td>CACCTCCATGTGGCGGCGTTCCGT</td><td></td><td></td><td></td><td></td><td></td><td></td><td></td><td></td><td></td><td></td></tr>
<tr><td>Oligo rev</td><td>AAACACGGAACGCCGCCACATGGA</td><td></td><td></td><td></td><td></td><td></td><td></td><td></td><td></td><td></td><td></td></tr>
<tr><td>Chromosome</td><td>start</td><td>end</td><td>strand</td><td>MM</td><td>target_seq</td><td>PAM</td><td>alignment</td><td>distance</td><td>position</td><td>gene name</td><td>gene id</td></tr>
<tr><td>GU305914.1</td><td>224707</td><td>224729</td><td>+</td><td>0</td><td>TCCATGTGGCGGCGTTCCGT</td><td>CGG</td><td>||||||||[||||||||||||]PAM</td><td>NA</td><td>Intergenic</td><td>NA</td><td>NA</td></tr>
<tr><td>GU305914.1</td><td>208567</td><td>208589</td><td>-</td><td>4</td><td>TCCACCTGACCGCGTTCCGT</td><td>CGG</td><td>||||--||[-|-|||||||||]PAM</td><td>NA</td><td>Intergenic</td><td>NA</td><td>NA</td></tr>
<tr><td>T16725</td><td>GCGTTCCGTCGGCGTCACCGTGG</td><td>1000</td><td></td><td></td><td></td><td></td><td></td><td></td><td></td><td></td><td></td></tr>
<tr><td>Oligo fwd</td><td>CACCGCGTTCCGTCGGCGTCACCG</td><td></td><td></td><td></td><td></td><td></td><td></td><td></td><td></td><td></td><td></td></tr>
<tr><td>Oligo rev</td><td>AAACCGGTGACGCCGACGGAACGC</td><td></td><td></td><td></td><td></td><td></td><td></td><td></td><td></td><td></td><td></td></tr>
<tr><td>Chromosome</td><td>start</td><td>end</td><td>strand</td><td>MM</td><td>target_seq</td><td>PAM</td><td>alignment</td><td>distance</td><td>position</td><td>gene name</td><td>gene id</td></tr>
<tr><td>GU305914.1</td><td>224718</td><td>224740</td><td>+</td><td>0</td><td>GCGTTCCGTCGGCGTCACCG</td><td>TGG</td><td>||||||||[||||||||||||]PAM</td><td>NA</td><td>Intergenic</td><td>NA</td><td>NA</td></tr>
<tr><td>T16726</td><td>GTATCCTGCCCGAAGTCGAACGG</td><td>1000</td><td></td><td></td><td></td><td></td><td></td><td></td><td></td><td></td><td></td></tr>
<tr><td>Oligo fwd</td><td>CACCGTATCCTGCCCGAAGTCGAA</td><td></td><td></td><td></td><td></td><td></td><td></td><td></td><td></td><td></td><td></td></tr>
<tr><td>Oligo rev</td><td>AAACTTCGACTTCGGGCAGGATAC</td><td></td><td></td><td></td><td></td><td></td><td></td><td></td><td></td><td></td><td></td></tr>
<tr><td>Chromosome</td><td>start</td><td>end</td><td>strand</td><td>MM</td><td>target_seq</td><td>PAM</td><td>alignment</td><td>distance</td><td>position</td><td>gene name</td><td>gene id</td></tr>
<tr><td>GU305914.1</td><td>224740</td><td>224762</td><td>+</td><td>0</td><td>GTATCCTGCCCGAAGTCGAA</td><td>CGG</td><td>||||||||[||||||||||||]PAM</td><td>NA</td><td>Intergenic</td><td>NA</td><td>NA</td></tr>
<tr><td>T16727</td><td>GCAGCGCTATAGTTTTCGCCCGG</td><td>1000</td><td></td><td></td><td></td><td></td><td></td><td></td><td></td><td></td><td></td></tr>
<tr><td>Oligo fwd</td><td>CACCGCAGCGCTATAGTTTTCGCC</td><td></td><td></td><td></td><td></td><td></td><td></td><td></td><td></td><td></td><td></td></tr>
<tr><td>Oligo rev</td><td>AAACGGCGAAAACTATAGCGCTGC</td><td></td><td></td><td></td><td></td><td></td><td></td><td></td><td></td><td></td><td></td></tr>
<tr><td>Chromosome</td><td>start</td><td>end</td><td>strand</td><td>MM</td><td>target_seq</td><td>PAM</td><td>alignment</td><td>distance</td><td>position</td><td>gene name</td><td>gene id</td></tr>
<tr><td>GU305914.1</td><td>224762</td><td>224784</td><td>+</td><td>0</td><td>GCAGCGCTATAGTTTTCGCC</td><td>CGG</td><td>||||||||[||||||||||||]PAM</td><td>NA</td><td>Intergenic</td><td>NA</td><td>NA</td></tr>
<tr><td>T16728</td><td>CAGCGCTATAGTTTTCGCCCGGG</td><td>1000</td><td></td><td></td><td></td><td></td><td></td><td></td><td></td><td></td><td></td></tr>
<tr><td>Oligo fwd</td><td>CACCCAGCGCTATAGTTTTCGCCC</td><td></td><td></td><td></td><td></td><td></td><td></td><td></td><td></td><td></td><td></td></tr>
<tr><td>Oligo rev</td><td>AAACGGGCGAAAACTATAGCGCTG</td><td></td><td></td><td></td><td></td><td></td><td></td><td></td><td></td><td></td><td></td></tr>
<tr><td>Chromosome</td><td>start</td><td>end</td><td>strand</td><td>MM</td><td>target_seq</td><td>PAM</td><td>alignment</td><td>distance</td><td>position</td><td>gene name</td><td>gene id</td></tr>
<tr><td>GU305914.1</td><td>224763</td><td>224785</td><td>+</td><td>0</td><td>CAGCGCTATAGTTTTCGCCC</td><td>GGG</td><td>||||||||[||||||||||||]PAM</td><td>NA</td><td>Intergenic</td><td>NA</td><td>NA</td></tr>
<tr><td>T16729</td><td>GCTATAGTTTTCGCCCGGGCCGG</td><td>1000</td><td></td><td></td><td></td><td></td><td></td><td></td><td></td><td></td><td></td></tr>
</table>

<table>
<tr><td>Oligo fwd</td><td>CACCGCTATAGTTTTCGCCCGGGC</td><td></td><td></td><td></td><td></td><td></td><td></td><td></td><td></td><td></td><td></td></tr>
<tr><td>Oligo rev</td><td>AAACGCCCGGGCGAAAACTATAGC</td><td></td><td></td><td></td><td></td><td></td><td></td><td></td><td></td><td></td><td></td></tr>
<tr><td>Chromosome</td><td>start</td><td>end</td><td>strand</td><td>MM</td><td>target_seq</td><td>PAM</td><td>alignment</td><td>distance</td><td>position</td><td>gene name</td><td>gene id</td></tr>
<tr><td>GU305914.1</td><td>224767</td><td>224789</td><td>+</td><td>0</td><td>GCTATAGTTTTCGCCCGGGC</td><td>CGG</td><td>||||||||[||||||||||||]PAM</td><td>NA</td><td>Intergenic</td><td>NA</td><td>NA</td></tr>
<tr><td></td><td></td><td></td><td></td><td></td><td></td><td></td><td></td><td></td><td></td><td></td><td></td></tr>
<tr><td>T16730</td><td>CTATAGTTTTCGCCCGGGCCGGG</td><td>1000</td><td></td><td></td><td></td><td></td><td></td><td></td><td></td><td></td><td></td></tr>
<tr><td>Oligo fwd</td><td>CACCCTATAGTTTTCGCCCGGGCC</td><td></td><td></td><td></td><td></td><td></td><td></td><td></td><td></td><td></td><td></td></tr>
<tr><td>Oligo rev</td><td>AAACGGCCCGGGCGAAAACTATAG</td><td></td><td></td><td></td><td></td><td></td><td></td><td></td><td></td><td></td><td></td></tr>
<tr><td>Chromosome</td><td>start</td><td>end</td><td>strand</td><td>MM</td><td>target_seq</td><td>PAM</td><td>alignment</td><td>distance</td><td>position</td><td>gene name</td><td>gene id</td></tr>
<tr><td>GU305914.1</td><td>224768</td><td>224790</td><td>+</td><td>0</td><td>CTATAGTTTTCGCCCGGGCC</td><td>GGG</td><td>||||||||[||||||||||||]PAM</td><td>NA</td><td>Intergenic</td><td>NA</td><td>NA</td></tr>
<tr><td></td><td></td><td></td><td></td><td></td><td></td><td></td><td></td><td></td><td></td><td></td><td></td></tr>
<tr><td>T16731</td><td>CCCGGGCCGGGCAGCTCAACCGG</td><td>1000</td><td></td><td></td><td></td><td></td><td></td><td></td><td></td><td></td><td></td></tr>
<tr><td>Oligo fwd</td><td>CACCCCCGGGCCGGGCAGCTCAAC</td><td></td><td></td><td></td><td></td><td></td><td></td><td></td><td></td><td></td><td></td></tr>
<tr><td>Oligo rev</td><td>AAACGTTGAGCTGCCCGGCCCGGG</td><td></td><td></td><td></td><td></td><td></td><td></td><td></td><td></td><td></td><td></td></tr>
<tr><td>Chromosome</td><td>start</td><td>end</td><td>strand</td><td>MM</td><td>target_seq</td><td>PAM</td><td>alignment</td><td>distance</td><td>position</td><td>gene name</td><td>gene id</td></tr>
<tr><td>GU305914.1</td><td>224780</td><td>224802</td><td>+</td><td>0</td><td>CCCGGGCCGGGCAGCTCAAC</td><td>CGG</td><td>||||||||[||||||||||||]PAM</td><td>NA</td><td>Intergenic</td><td>NA</td><td>NA</td></tr>
<tr><td></td><td></td><td></td><td></td><td></td><td></td><td></td><td></td><td></td><td></td><td></td><td></td></tr>
<tr><td>T16732</td><td>CCGGGCCGGGCAGCTCAACCGGG</td><td>1000</td><td></td><td></td><td></td><td></td><td></td><td></td><td></td><td></td><td></td></tr>
<tr><td>Oligo fwd</td><td>CACCCCGGGCCGGGCAGCTCAACC</td><td></td><td></td><td></td><td></td><td></td><td></td><td></td><td></td><td></td><td></td></tr>
<tr><td>Oligo rev</td><td>AAACGGTTGAGCTGCCCGGCCCGG</td><td></td><td></td><td></td><td></td><td></td><td></td><td></td><td></td><td></td><td></td></tr>
<tr><td>Chromosome</td><td>start</td><td>end</td><td>strand</td><td>MM</td><td>target_seq</td><td>PAM</td><td>alignment</td><td>distance</td><td>position</td><td>gene name</td><td>gene id</td></tr>
<tr><td>GU305914.1</td><td>224781</td><td>224803</td><td>+</td><td>0</td><td>CCGGGCCGGGCAGCTCAACC</td><td>GGG</td><td>||||||||[||||||||||||]PAM</td><td>NA</td><td>Intergenic</td><td>NA</td><td>NA</td></tr>
<tr><td></td><td></td><td></td><td></td><td></td><td></td><td></td><td></td><td></td><td></td><td></td><td></td></tr>
<tr><td>T16733</td><td>CGGGCAGCTCAACCGGGAGCTGG</td><td>1000</td><td></td><td></td><td></td><td></td><td></td><td></td><td></td><td></td><td></td></tr>
<tr><td>Oligo fwd</td><td>CACCCGGGCAGCTCAACCGGGAGC</td><td></td><td></td><td></td><td></td><td></td><td></td><td></td><td></td><td></td><td></td></tr>
<tr><td>Oligo rev</td><td>AAACGCTCCCGGTTGAGCTGCCCG</td><td></td><td></td><td></td><td></td><td></td><td></td><td></td><td></td><td></td><td></td></tr>
<tr><td>Chromosome</td><td>start</td><td>end</td><td>strand</td><td>MM</td><td>target_seq</td><td>PAM</td><td>alignment</td><td>distance</td><td>position</td><td>gene name</td><td>gene id</td></tr>
<tr><td>GU305914.1</td><td>224787</td><td>224809</td><td>+</td><td>0</td><td>CGGGCAGCTCAACCGGGAGC</td><td>TGG</td><td>||||||||[||||||||||||]PAM</td><td>NA</td><td>Intergenic</td><td>NA</td><td>NA</td></tr>
<tr><td></td><td></td><td></td><td></td><td></td><td></td><td></td><td></td><td></td><td></td><td></td><td></td></tr>
<tr><td>T16734</td><td>TCGTGCACTGCAGACATGCCAGG</td><td>1000</td><td></td><td></td><td></td><td></td><td></td><td></td><td></td><td></td><td></td></tr>
<tr><td>Oligo fwd</td><td>CACCTCGTGCACTGCAGACATGCC</td><td></td><td></td><td></td><td></td><td></td><td></td><td></td><td></td><td></td><td></td></tr>
<tr><td>Oligo rev</td><td>AAACGGCATGTCTGCAGTGCACGA</td><td></td><td></td><td></td><td></td><td></td><td></td><td></td><td></td><td></td><td></td></tr>
<tr><td>Chromosome</td><td>start</td><td>end</td><td>strand</td><td>MM</td><td>target_seq</td><td>PAM</td><td>alignment</td><td>distance</td><td>position</td><td>gene name</td><td>gene id</td></tr>
<tr><td>GU305914.1</td><td>224831</td><td>224853</td><td>+</td><td>0</td><td>TCGTGCACTGCAGACATGCC</td><td>AGG</td><td>||||||||[||||||||||||]PAM</td><td>NA</td><td>Intergenic</td><td>NA</td><td>NA</td></tr>
<tr><td></td><td></td><td></td><td></td><td></td><td></td><td></td><td></td><td></td><td></td><td></td><td></td></tr>
<tr><td>T16735</td><td>ACTGCAGACATGCCAGGCGCTGG</td><td>1000</td><td></td><td></td><td></td><td></td><td></td><td></td><td></td><td></td><td></td></tr>
<tr><td>Oligo fwd</td><td>CACCACTGCAGACATGCCAGGCGC</td><td></td><td></td><td></td><td></td><td></td><td></td><td></td><td></td><td></td><td></td></tr>
<tr><td>Oligo rev</td><td>AAACGCGCCTGGCATGTCTGCAGT</td><td></td><td></td><td></td><td></td><td></td><td></td><td></td><td></td><td></td><td></td></tr>
<tr><td>Chromosome</td><td>start</td><td>end</td><td>strand</td><td>MM</td><td>target_seq</td><td>PAM</td><td>alignment</td><td>distance</td><td>position</td><td>gene name</td><td>gene id</td></tr>
<tr><td>GU305914.1</td><td>224837</td><td>224859</td><td>+</td><td>0</td><td>ACTGCAGACATGCCAGGCGC</td><td>TGG</td><td>||||||||[||||||||||||]PAM</td><td>NA</td><td>Intergenic</td><td>NA</td><td>NA</td></tr>
<tr><td></td><td></td><td></td><td></td><td></td><td></td><td></td><td></td><td></td><td></td><td></td><td></td></tr>
<tr><td>T16736</td><td>CCAGGCGCTGGATCGTGCTGCGG</td><td>1000</td><td></td><td></td><td></td><td></td><td></td><td></td><td></td><td></td><td></td></tr>
<tr><td>Oligo fwd</td><td>CACCCCAGGCGCTGGATCGTGCTG</td><td></td><td></td><td></td><td></td><td></td><td></td><td></td><td></td><td></td><td></td></tr>
<tr><td>Oligo rev</td><td>AAACCAGCACGATCCAGCGCCTGG</td><td></td><td></td><td></td><td></td><td></td><td></td><td></td><td></td><td></td><td></td></tr>
<tr><td>Chromosome</td><td>start</td><td>end</td><td>strand</td><td>MM</td><td>target_seq</td><td>PAM</td><td>alignment</td><td>distance</td><td>position</td><td>gene name</td><td>gene id</td></tr>
<tr><td>GU305914.1</td><td>224849</td><td>224871</td><td>+</td><td>0</td><td>CCAGGCGCTGGATCGTGCTG</td><td>CGG</td><td>||||||||[||||||||||||]PAM</td><td>NA</td><td>Intergenic</td><td>NA</td><td>NA</td></tr>
<tr><td></td><td></td><td></td><td></td><td></td><td></td><td></td><td></td><td></td><td></td><td></td><td></td></tr>
<tr><td>T16737</td><td>CAGGCGCTGGATCGTGCTGCGGG</td><td>1000</td><td></td><td></td><td></td><td></td><td></td><td></td><td></td><td></td><td></td></tr>
<tr><td>Oligo fwd</td><td>CACCCAGGCGCTGGATCGTGCTGC</td><td></td><td></td><td></td><td></td><td></td><td></td><td></td><td></td><td></td><td></td></tr>
<tr><td>Oligo rev</td><td>AAACGCAGCACGATCCAGCGCCTG</td><td></td><td></td><td></td><td></td><td></td><td></td><td></td><td></td><td></td><td></td></tr>
<tr><td>Chromosome</td><td>start</td><td>end</td><td>strand</td><td>MM</td><td>target_seq</td><td>PAM</td><td>alignment</td><td>distance</td><td>position</td><td>gene name</td><td>gene id</td></tr>
<tr><td>GU305914.1</td><td>224850</td><td>224872</td><td>+</td><td>0</td><td>CAGGCGCTGGATCGTGCTGC</td><td>GGG</td><td>||||||||[||||||||||||]PAM</td><td>NA</td><td>Intergenic</td><td>NA</td><td>NA</td></tr>
<tr><td></td><td></td><td></td><td></td><td></td><td></td><td></td><td></td><td></td><td></td><td></td><td></td></tr>
<tr><td>T16738</td><td>TCGTGCTGCGGGCGAAGACGTGG</td><td>1000</td><td></td><td></td><td></td><td></td><td></td><td></td><td></td><td></td><td></td></tr>
<tr><td>Oligo fwd</td><td>CACCTCGTGCTGCGGGCGAAGACG</td><td></td><td></td><td></td><td></td><td></td><td></td><td></td><td></td><td></td><td></td></tr>
<tr><td>Oligo rev</td><td>AAACCGTCTTCGCCCGCAGCACGA</td><td></td><td></td><td></td><td></td><td></td><td></td><td></td><td></td><td></td><td></td></tr>
<tr><td>Chromosome</td><td>start</td><td>end</td><td>strand</td><td>MM</td><td>target_seq</td><td>PAM</td><td>alignment</td><td>distance</td><td>position</td><td>gene name</td><td>gene id</td></tr>
<tr><td>GU305914.1</td><td>224861</td><td>224883</td><td>+</td><td>0</td><td>TCGTGCTGCGGGCGAAGACG</td><td>TGG</td><td>||||||||[||||||||||||]PAM</td><td>NA</td><td>Intergenic</td><td>NA</td><td>NA</td></tr>
<tr><td></td><td></td><td></td><td></td><td></td><td></td><td></td><td></td><td></td><td></td><td></td><td></td></tr>
<tr><td>T16739</td><td>GACGTGGATGTCGCCCGTGTAGG</td><td>1000</td><td></td><td></td><td></td><td></td><td></td><td></td><td></td><td></td><td></td></tr>
<tr><td>Oligo fwd</td><td>CACCGACGTGGATGTCGCCCGTGT</td><td></td><td></td><td></td><td></td><td></td><td></td><td></td><td></td><td></td><td></td></tr>
<tr><td>Oligo rev</td><td>AAACACACGGGCGACATCCACGTC</td><td></td><td></td><td></td><td></td><td></td><td></td><td></td><td></td><td></td><td></td></tr>
<tr><td>Chromosome</td><td>start</td><td>end</td><td>strand</td><td>MM</td><td>target_seq</td><td>PAM</td><td>alignment</td><td>distance</td><td>position</td><td>gene name</td><td>gene id</td></tr>
<tr><td>GU305914.1</td><td>224877</td><td>224899</td><td>+</td><td>0</td><td>GACGTGGATGTCGCCCGTGT</td><td>AGG</td><td>||||||||[||||||||||||]PAM</td><td>NA</td><td>Intergenic</td><td>NA</td><td>NA</td></tr>
<tr><td></td><td></td><td></td><td></td><td></td><td></td><td></td><td></td><td></td><td></td><td></td><td></td></tr>
<tr><td>T16740</td><td>GTAGTACGTCCGATCGCCGTCGG</td><td>1000</td><td></td><td></td><td></td><td></td><td></td><td></td><td></td><td></td><td></td></tr>
<tr><td>Oligo fwd</td><td>CACCGTAGTACGTCCGATCGCCGT</td><td></td><td></td><td></td><td></td><td></td><td></td><td></td><td></td><td></td><td></td></tr>
<tr><td>Oligo rev</td><td>AAACACGGCGATCGGACGTACTAC</td><td></td><td></td><td></td><td></td><td></td><td></td><td></td><td></td><td></td><td></td></tr>
<tr><td>Chromosome</td><td>start</td><td>end</td><td>strand</td><td>MM</td><td>target_seq</td><td>PAM</td><td>alignment</td><td>distance</td><td>position</td><td>gene name</td><td>gene id</td></tr>
<tr><td>GU305914.1</td><td>224907</td><td>224929</td><td>+</td><td>0</td><td>GTAGTACGTCCGATCGCCGT</td><td>CGG</td><td>||||||||[||||||||||||]PAM</td><td>NA</td><td>Intergenic</td><td>NA</td><td>NA</td></tr>
<tr><td></td><td></td><td></td><td></td><td></td><td></td><td></td><td></td><td></td><td></td><td></td><td></td></tr>
<tr><td>T16741</td><td>TAGTACGTCCGATCGCCGTCGGG</td><td>1000</td><td></td><td></td><td></td><td></td><td></td><td></td><td></td><td></td><td></td></tr>
<tr><td>Oligo fwd</td><td>CACCTAGTACGTCCGATCGCCGTC</td><td></td><td></td><td></td><td></td><td></td><td></td><td></td><td></td><td></td><td></td></tr>
<tr><td>Oligo rev</td><td>AAACGACGGCGATCGGACGTACTA</td><td></td><td></td><td></td><td></td><td></td><td></td><td></td><td></td><td></td><td></td></tr>
<tr><td>Chromosome</td><td>start</td><td>end</td><td>strand</td><td>MM</td><td>target_seq</td><td>PAM</td><td>alignment</td><td>distance</td><td>position</td><td>gene name</td><td>gene id</td></tr>
<tr><td>GU305914.1</td><td>224908</td><td>224930</td><td>+</td><td>0</td><td>TAGTACGTCCGATCGCCGTC</td><td>GGG</td><td>||||||||[||||||||||||]PAM</td><td>NA</td><td>Intergenic</td><td>NA</td><td>NA</td></tr>
<tr><td></td><td></td><td></td><td></td><td></td><td></td><td></td><td></td><td></td><td></td><td></td><td></td></tr>
<tr><td>T16742</td><td>AGTACGTCCGATCGCCGTCGGGG</td><td>1000</td><td></td><td></td><td></td><td></td><td></td><td></td><td></td><td></td><td></td></tr>
<tr><td>Oligo fwd</td><td>CACCAGTACGTCCGATCGCCGTCG</td><td></td><td></td><td></td><td></td><td></td><td></td><td></td><td></td><td></td><td></td></tr>
<tr><td>Oligo rev</td><td>AAACCGACGGCGATCGGACGTACT</td><td></td><td></td><td></td><td></td><td></td><td></td><td></td><td></td><td></td><td></td></tr>
<tr><td>Chromosome</td><td>start</td><td>end</td><td>strand</td><td>MM</td><td>target_seq</td><td>PAM</td><td>alignment</td><td>distance</td><td>position</td><td>gene name</td><td>gene id</td></tr>
<tr><td>GU305914.1</td><td>224909</td><td>224931</td><td>+</td><td>0</td><td>AGTACGTCCGATCGCCGTCG</td><td>GGG</td><td>||||||||[||||||||||||]PAM</td><td>NA</td><td>Intergenic</td><td>NA</td><td>NA</td></tr>
<tr><td></td><td></td><td></td><td></td><td></td><td></td><td></td><td></td><td></td><td></td><td></td><td></td></tr>
<tr><td>T16743</td><td>GGAGTTGTTCCCAACGCACGCGG</td><td>1000</td><td></td><td></td><td></td><td></td><td></td><td></td><td></td><td></td><td></td></tr>
<tr><td>Oligo fwd</td><td>CACCGGAGTTGTTCCCAACGCACG</td><td></td><td></td><td></td><td></td><td></td><td></td><td></td><td></td><td></td><td></td></tr>
<tr><td>Oligo rev</td><td>AAACCGTGCGTTGGGAACAACTCC</td><td></td><td></td><td></td><td></td><td></td><td></td><td></td><td></td><td></td><td></td></tr>
<tr><td>Chromosome</td><td>start</td><td>end</td><td>strand</td><td>MM</td><td>target_seq</td><td>PAM</td><td>alignment</td><td>distance</td><td>position</td><td>gene name</td><td>gene id</td></tr>
<tr><td>GU305914.1</td><td>224930</td><td>224952</td><td>+</td><td>0</td><td>GGAGTTGTTCCCAACGCACG</td><td>CGG</td><td>||||||||[||||||||||||]PAM</td><td>NA</td><td>Intergenic</td><td>NA</td><td>NA</td></tr>
</table>

<table>
<tr><td>T16744</td><td>CCAACGCACGCGGCCCTCCACGG</td><td>1000</td><td></td><td></td><td></td><td></td><td></td><td></td><td></td><td></td><td></td></tr>
<tr><td>Oligo fwd</td><td>CACCCCAACGCACGCGGCCCTCCA</td><td></td><td></td><td></td><td></td><td></td><td></td><td></td><td></td><td></td><td></td></tr>
<tr><td>Oligo rev</td><td>AAACTGGAGGGCCGCGTGCGTTGG</td><td></td><td></td><td></td><td></td><td></td><td></td><td></td><td></td><td></td><td></td></tr>
<tr><td>Chromosome</td><td>start</td><td>end</td><td>strand</td><td>MM</td><td>target_seq</td><td>PAM</td><td>alignment</td><td>distance</td><td>position</td><td>gene name</td><td>gene id</td></tr>
<tr><td>GU305914.1</td><td>224940</td><td>224962</td><td>+</td><td>0</td><td>CCAACGCACGCGGCCCTCCA</td><td>CGG</td><td>||||||||[||||||||||||]PAM</td><td>NA</td><td>Intergenic</td><td>NA</td><td>NA</td></tr>
<tr><td>T16745</td><td>CAACGCACGCGGCCCTCCACGGG</td><td>1000</td><td></td><td></td><td></td><td></td><td></td><td></td><td></td><td></td><td></td></tr>
<tr><td>Oligo fwd</td><td>CACCCAACGCACGCGGCCCTCCAC</td><td></td><td></td><td></td><td></td><td></td><td></td><td></td><td></td><td></td><td></td></tr>
<tr><td>Oligo rev</td><td>AAACGTGGAGGGCCGCGTGCGTTG</td><td></td><td></td><td></td><td></td><td></td><td></td><td></td><td></td><td></td><td></td></tr>
<tr><td>Chromosome</td><td>start</td><td>end</td><td>strand</td><td>MM</td><td>target_seq</td><td>PAM</td><td>alignment</td><td>distance</td><td>position</td><td>gene name</td><td>gene id</td></tr>
<tr><td>GU305914.1</td><td>224941</td><td>224963</td><td>+</td><td>0</td><td>CAACGCACGCGGCCCTCCAC</td><td>GGG</td><td>||||||||[||||||||||||]PAM</td><td>NA</td><td>Intergenic</td><td>NA</td><td>NA</td></tr>
<tr><td>T16746</td><td>AACGCACGCGGCCCTCCACGGGG</td><td>1000</td><td></td><td></td><td></td><td></td><td></td><td></td><td></td><td></td><td></td></tr>
<tr><td>Oligo fwd</td><td>CACCAACGCACGCGGCCCTCCACG</td><td></td><td></td><td></td><td></td><td></td><td></td><td></td><td></td><td></td><td></td></tr>
<tr><td>Oligo rev</td><td>AAACCGTGGAGGGCCGCGTGCGTT</td><td></td><td></td><td></td><td></td><td></td><td></td><td></td><td></td><td></td><td></td></tr>
<tr><td>Chromosome</td><td>start</td><td>end</td><td>strand</td><td>MM</td><td>target_seq</td><td>PAM</td><td>alignment</td><td>distance</td><td>position</td><td>gene name</td><td>gene id</td></tr>
<tr><td>GU305914.1</td><td>224942</td><td>224964</td><td>+</td><td>0</td><td>AACGCACGCGGCCCTCCACG</td><td>GGG</td><td>||||||||[||||||||||||]PAM</td><td>NA</td><td>Intergenic</td><td>NA</td><td>NA</td></tr>
<tr><td>T16747</td><td>CGCTCCACTCGATGCTCACCAGG</td><td>1000</td><td></td><td></td><td></td><td></td><td></td><td></td><td></td><td></td><td></td></tr>
<tr><td>Oligo fwd</td><td>CACCCGCTCCACTCGATGCTCACC</td><td></td><td></td><td></td><td></td><td></td><td></td><td></td><td></td><td></td><td></td></tr>
<tr><td>Oligo rev</td><td>AAACGGTGAGCATCGAGTGGAGCG</td><td></td><td></td><td></td><td></td><td></td><td></td><td></td><td></td><td></td><td></td></tr>
<tr><td>Chromosome</td><td>start</td><td>end</td><td>strand</td><td>MM</td><td>target_seq</td><td>PAM</td><td>alignment</td><td>distance</td><td>position</td><td>gene name</td><td>gene id</td></tr>
<tr><td>GU305914.1</td><td>224972</td><td>224994</td><td>+</td><td>0</td><td>CGCTCCACTCGATGCTCACC</td><td>AGG</td><td>||||||||[||||||||||||]PAM</td><td>NA</td><td>Intergenic</td><td>NA</td><td>NA</td></tr>
<tr><td>T16748</td><td>GCTCCACTCGATGCTCACCAGGG</td><td>1000</td><td></td><td></td><td></td><td></td><td></td><td></td><td></td><td></td><td></td></tr>
<tr><td>Oligo fwd</td><td>CACCGCTCCACTCGATGCTCACCA</td><td></td><td></td><td></td><td></td><td></td><td></td><td></td><td></td><td></td><td></td></tr>
<tr><td>Oligo rev</td><td>AAACTGGTGAGCATCGAGTGGAGC</td><td></td><td></td><td></td><td></td><td></td><td></td><td></td><td></td><td></td><td></td></tr>
<tr><td>Chromosome</td><td>start</td><td>end</td><td>strand</td><td>MM</td><td>target_seq</td><td>PAM</td><td>alignment</td><td>distance</td><td>position</td><td>gene name</td><td>gene id</td></tr>
<tr><td>GU305914.1</td><td>224973</td><td>224995</td><td>+</td><td>0</td><td>GCTCCACTCGATGCTCACCA</td><td>GGG</td><td>||||||||[||||||||||||]PAM</td><td>NA</td><td>Intergenic</td><td>NA</td><td>NA</td></tr>
<tr><td>T16749</td><td>CTCCACTCGATGCTCACCAGGGG</td><td>1000</td><td></td><td></td><td></td><td></td><td></td><td></td><td></td><td></td><td></td></tr>
<tr><td>Oligo fwd</td><td>CACCCTCCACTCGATGCTCACCAG</td><td></td><td></td><td></td><td></td><td></td><td></td><td></td><td></td><td></td><td></td></tr>
<tr><td>Oligo rev</td><td>AAACCTGGTGAGCATCGAGTGGAG</td><td></td><td></td><td></td><td></td><td></td><td></td><td></td><td></td><td></td><td></td></tr>
<tr><td>Chromosome</td><td>start</td><td>end</td><td>strand</td><td>MM</td><td>target_seq</td><td>PAM</td><td>alignment</td><td>distance</td><td>position</td><td>gene name</td><td>gene id</td></tr>
<tr><td>GU305914.1</td><td>224974</td><td>224996</td><td>+</td><td>0</td><td>CTCCACTCGATGCTCACCAG</td><td>GGG</td><td>||||||||[||||||||||||]PAM</td><td>NA</td><td>Intergenic</td><td>NA</td><td>NA</td></tr>
<tr><td>T16750</td><td>CTCGATGCTCACCAGGGGATAGG</td><td>1000</td><td></td><td></td><td></td><td></td><td></td><td></td><td></td><td></td><td></td></tr>
<tr><td>Oligo fwd</td><td>CACCCTCGATGCTCACCAGGGGAT</td><td></td><td></td><td></td><td></td><td></td><td></td><td></td><td></td><td></td><td></td></tr>
<tr><td>Oligo rev</td><td>AAACATCCCCTGGTGAGCATCGAG</td><td></td><td></td><td></td><td></td><td></td><td></td><td></td><td></td><td></td><td></td></tr>
<tr><td>Chromosome</td><td>start</td><td>end</td><td>strand</td><td>MM</td><td>target_seq</td><td>PAM</td><td>alignment</td><td>distance</td><td>position</td><td>gene name</td><td>gene id</td></tr>
<tr><td>GU305914.1</td><td>224979</td><td>225001</td><td>+</td><td>0</td><td>CTCGATGCTCACCAGGGGAT</td><td>AGG</td><td>||||||||[||||||||||||]PAM</td><td>NA</td><td>Intergenic</td><td>NA</td><td>NA</td></tr>
<tr><td>T16751</td><td>TCGATGCTCACCAGGGGATAGGG</td><td>1000</td><td></td><td></td><td></td><td></td><td></td><td></td><td></td><td></td><td></td></tr>
<tr><td>Oligo fwd</td><td>CACCTCGATGCTCACCAGGGGATA</td><td></td><td></td><td></td><td></td><td></td><td></td><td></td><td></td><td></td><td></td></tr>
<tr><td>Oligo rev</td><td>AAACTATCCCCTGGTGAGCATCGA</td><td></td><td></td><td></td><td></td><td></td><td></td><td></td><td></td><td></td><td></td></tr>
<tr><td>Chromosome</td><td>start</td><td>end</td><td>strand</td><td>MM</td><td>target_seq</td><td>PAM</td><td>alignment</td><td>distance</td><td>position</td><td>gene name</td><td>gene id</td></tr>
<tr><td>GU305914.1</td><td>224980</td><td>225002</td><td>+</td><td>0</td><td>TCGATGCTCACCAGGGGATA</td><td>GGG</td><td>||||||||[||||||||||||]PAM</td><td>NA</td><td>Intergenic</td><td>NA</td><td>NA</td></tr>
<tr><td>T16752</td><td>GACCGTGCCACCAGCGCATCAGG</td><td>1000</td><td></td><td></td><td></td><td></td><td></td><td></td><td></td><td></td><td></td></tr>
<tr><td>Oligo fwd</td><td>CACCGACCGTGCCACCAGCGCATC</td><td></td><td></td><td></td><td></td><td></td><td></td><td></td><td></td><td></td><td></td></tr>
<tr><td>Oligo rev</td><td>AAACGATGCGCTGGTGGCACGGTC</td><td></td><td></td><td></td><td></td><td></td><td></td><td></td><td></td><td></td><td></td></tr>
<tr><td>Chromosome</td><td>start</td><td>end</td><td>strand</td><td>MM</td><td>target_seq</td><td>PAM</td><td>alignment</td><td>distance</td><td>position</td><td>gene name</td><td>gene id</td></tr>
<tr><td>GU305914.1</td><td>225002</td><td>225024</td><td>+</td><td>0</td><td>GACCGTGCCACCAGCGCATC</td><td>AGG</td><td>||||||||[||||||||||||]PAM</td><td>NA</td><td>Intergenic</td><td>NA</td><td>NA</td></tr>
<tr><td>T16753</td><td>ACCGTGCCACCAGCGCATCAGGG</td><td>1000</td><td></td><td></td><td></td><td></td><td></td><td></td><td></td><td></td><td></td></tr>
<tr><td>Oligo fwd</td><td>CACCACCGTGCCACCAGCGCATCA</td><td></td><td></td><td></td><td></td><td></td><td></td><td></td><td></td><td></td><td></td></tr>
<tr><td>Oligo rev</td><td>AAACTGATGCGCTGGTGGCACGGT</td><td></td><td></td><td></td><td></td><td></td><td></td><td></td><td></td><td></td><td></td></tr>
<tr><td>Chromosome</td><td>start</td><td>end</td><td>strand</td><td>MM</td><td>target_seq</td><td>PAM</td><td>alignment</td><td>distance</td><td>position</td><td>gene name</td><td>gene id</td></tr>
<tr><td>GU305914.1</td><td>225003</td><td>225025</td><td>+</td><td>0</td><td>ACCGTGCCACCAGCGCATCA</td><td>GGG</td><td>||||||||[||||||||||||]PAM</td><td>NA</td><td>Intergenic</td><td>NA</td><td>NA</td></tr>
<tr><td>T16754</td><td>TGCCACCAGCGCATCAGGGTAGG</td><td>1000</td><td></td><td></td><td></td><td></td><td></td><td></td><td></td><td></td><td></td></tr>
<tr><td>Oligo fwd</td><td>CACCTGCCACCAGCGCATCAGGGT</td><td></td><td></td><td></td><td></td><td></td><td></td><td></td><td></td><td></td><td></td></tr>
<tr><td>Oligo rev</td><td>AAACACCCTGATGCGCTGGTGGCA</td><td></td><td></td><td></td><td></td><td></td><td></td><td></td><td></td><td></td><td></td></tr>
<tr><td>Chromosome</td><td>start</td><td>end</td><td>strand</td><td>MM</td><td>target_seq</td><td>PAM</td><td>alignment</td><td>distance</td><td>position</td><td>gene name</td><td>gene id</td></tr>
<tr><td>GU305914.1</td><td>225007</td><td>225029</td><td>+</td><td>0</td><td>TGCCACCAGCGCATCAGGGT</td><td>AGG</td><td>||||||||[||||||||||||]PAM</td><td>NA</td><td>Intergenic</td><td>NA</td><td>NA</td></tr>
<tr><td>T16755</td><td>AGCGCATCAGGGTAGGCTGCAGG</td><td>1000</td><td></td><td></td><td></td><td></td><td></td><td></td><td></td><td></td><td></td></tr>
<tr><td>Oligo fwd</td><td>CACCAGCGCATCAGGGTAGGCTGC</td><td></td><td></td><td></td><td></td><td></td><td></td><td></td><td></td><td></td><td></td></tr>
<tr><td>Oligo rev</td><td>AAACGCAGCCTACCCTGATGCGCT</td><td></td><td></td><td></td><td></td><td></td><td></td><td></td><td></td><td></td><td></td></tr>
<tr><td>Chromosome</td><td>start</td><td>end</td><td>strand</td><td>MM</td><td>target_seq</td><td>PAM</td><td>alignment</td><td>distance</td><td>position</td><td>gene name</td><td>gene id</td></tr>
<tr><td>GU305914.1</td><td>225014</td><td>225036</td><td>+</td><td>0</td><td>AGCGCATCAGGGTAGGCTGC</td><td>AGG</td><td>||||||||[||||||||||||]PAM</td><td>NA</td><td>Intergenic</td><td>NA</td><td>NA</td></tr>
<tr><td>T16756</td><td>GCGCATCAGGGTAGGCTGCAGGG</td><td>1000</td><td></td><td></td><td></td><td></td><td></td><td></td><td></td><td></td><td></td></tr>
<tr><td>Oligo fwd</td><td>CACCGCGCATCAGGGTAGGCTGCA</td><td></td><td></td><td></td><td></td><td></td><td></td><td></td><td></td><td></td><td></td></tr>
<tr><td>Oligo rev</td><td>AAACTGCAGCCTACCCTGATGCGC</td><td></td><td></td><td></td><td></td><td></td><td></td><td></td><td></td><td></td><td></td></tr>
<tr><td>Chromosome</td><td>start</td><td>end</td><td>strand</td><td>MM</td><td>target_seq</td><td>PAM</td><td>alignment</td><td>distance</td><td>position</td><td>gene name</td><td>gene id</td></tr>
<tr><td>GU305914.1</td><td>225015</td><td>225037</td><td>+</td><td>0</td><td>GCGCATCAGGGTAGGCTGCA</td><td>GGG</td><td>||||||||[||||||||||||]PAM</td><td>NA</td><td>Intergenic</td><td>NA</td><td>NA</td></tr>
<tr><td>T16757</td><td>CGCATCAGGGTAGGCTGCAGGGG</td><td>1000</td><td></td><td></td><td></td><td></td><td></td><td></td><td></td><td></td><td></td></tr>
<tr><td>Oligo fwd</td><td>CACCCGCATCAGGGTAGGCTGCAG</td><td></td><td></td><td></td><td></td><td></td><td></td><td></td><td></td><td></td><td></td></tr>
<tr><td>Oligo rev</td><td>AAACCTGCAGCCTACCCTGATGCG</td><td></td><td></td><td></td><td></td><td></td><td></td><td></td><td></td><td></td><td></td></tr>
<tr><td>Chromosome</td><td>start</td><td>end</td><td>strand</td><td>MM</td><td>target_seq</td><td>PAM</td><td>alignment</td><td>distance</td><td>position</td><td>gene name</td><td>gene id</td></tr>
<tr><td>GU305914.1</td><td>225016</td><td>225038</td><td>+</td><td>0</td><td>CGCATCAGGGTAGGCTGCAG</td><td>GGG</td><td>||||||||[||||||||||||]PAM</td><td>NA</td><td>Intergenic</td><td>NA</td><td>NA</td></tr>
<tr><td>T16758</td><td>GCTGCAGGGGTGTCGACCCACGG</td><td>1000</td><td></td><td></td><td></td><td></td><td></td><td></td><td></td><td></td><td></td></tr>
<tr><td>Oligo fwd</td><td>CACCGCTGCAGGGGTGTCGACCCA</td><td></td><td></td><td></td><td></td><td></td><td></td><td></td><td></td><td></td><td></td></tr>
<tr><td>Oligo rev</td><td>AAACTGGGTCGACACCCCTGCAGC</td><td></td><td></td><td></td><td></td><td></td><td></td><td></td><td></td><td></td><td></td></tr>
<tr><td>Chromosome</td><td>start</td><td>end</td><td>strand</td><td>MM</td><td>target_seq</td><td>PAM</td><td>alignment</td><td>distance</td><td>position</td><td>gene name</td><td>gene id</td></tr>
<tr><td>GU305914.1</td><td>225029</td><td>225051</td><td>+</td><td>0</td><td>GCTGCAGGGGTGTCGACCCA</td><td>CGG</td><td>||||||||[||||||||||||]PAM</td><td>NA</td><td>Intergenic</td><td>NA</td><td>NA</td></tr>
</table>

T16759 GGGTGTCGACCCACGGATGTTGG 1000
Oligo fwd CACCGGGTGTCGACCCACGGATGT
Oligo rev AAACACATCCGTGGGTCGACACCC
Chromosome start end strand MM target_seq PAM alignment distance position gene name gene id
GU305914.1 225036 225058 + 0 GGGTGTCGACCCACGGATGT TGG ||||||||[||||||||||||]PAM NA Intergenic NA NA

T16760 TCGACCCACGGATGTTGGCCAGG 1000
Oligo fwd CACCTCGACCCACGGATGTTGGCC
Oligo rev AAACGGCCAACATCCGTGGGTCGA
Chromosome start end strand MM target_seq PAM alignment distance position gene name gene id
GU305914.1 225041 225063 + 0 TCGACCCACGGATGTTGGCC AGG ||||||||[||||||||||||]PAM NA Intergenic NA NA

T16761 TGTTGGCCAGGCAACGAAACAGG 1000
Oligo fwd CACCTGTTGGCCAGGCAACGAAAC
Oligo rev AAACGTTTCGTTGCCTGGCCAACA
Chromosome start end strand MM target_seq PAM alignment distance position gene name gene id
GU305914.1 225053 225075 + 0 TGTTGGCCAGGCAACGAAAC AGG ||||||||[||||||||||||]PAM NA Intergenic NA NA

T16762 GCCAGGCAACGAAACAGGAACGG 1000
Oligo fwd CACCGCCAGGCAACGAAACAGGAA
Oligo rev AAACTTCCTGTTTCGTTGCCTGGC
Chromosome start end strand MM target_seq PAM alignment distance position gene name gene id
GU305914.1 225058 225080 + 0 GCCAGGCAACGAAACAGGAA CGG ||||||||[||||||||||||]PAM NA Intergenic NA NA

T16763 CGGAGCGAGCGTGTCTCTGTCGG 1000
Oligo fwd CACCCGGAGCGAGCGTGTCTCTGT
Oligo rev AAACACAGAGACACGCTCGCTCCG
Chromosome start end strand MM target_seq PAM alignment distance position gene name gene id
GU305914.1 225078 225100 + 0 CGGAGCGAGCGTGTCTCTGT CGG ||||||||[||||||||||||]PAM NA Intergenic NA NA

T16764 CTGCAACAGCGCCACCTGAGCGG 1000
Oligo fwd CACCCTGCAACAGCGCCACCTGAG
Oligo rev AAACCTCAGGTGGCGCTGTTGCAG
Chromosome start end strand MM target_seq PAM alignment distance position gene name gene id
GU305914.1 225105 225127 + 0 CTGCAACAGCGCCACCTGAG CGG ||||||||[||||||||||||]PAM NA Intergenic NA NA

T16765 TGCAACAGCGCCACCTGAGCGGG 1000
Oligo fwd CACCTGCAACAGCGCCACCTGAGC
Oligo rev AAACGCTCAGGTGGCGCTGTTGCA
Chromosome start end strand MM target_seq PAM alignment distance position gene name gene id
GU305914.1 225106 225128 + 0 TGCAACAGCGCCACCTGAGC GGG ||||||||[||||||||||||]PAM NA Intergenic NA NA

T16766 AACAGCGCCACCTGAGCGGGCGG 1000
Oligo fwd CACCAACAGCGCCACCTGAGCGGG
Oligo rev AAACCCCGCTCAGGTGGCGCTGTT
Chromosome start end strand MM target_seq PAM alignment distance position gene name gene id
GU305914.1 225109 225131 + 0 AACAGCGCCACCTGAGCGGG CGG ||||||||[||||||||||||]PAM NA Intergenic NA NA

T16767 CACCTGAGCGGGCGGCAGCACGG 1000
Oligo fwd CACCCACCTGAGCGGGCGGCAGCA
Oligo rev AAACTGCTGCCGCCCGCTCAGGTG
Chromosome start end strand MM target_seq PAM alignment distance position gene name gene id
GU305914.1 225117 225139 + 0 CACCTGAGCGGGCGGCAGCA CGG ||||||||[||||||||||||]PAM NA Intergenic NA NA

T16768 CTTGATCACGATTTCCCCGACGG 1000
Oligo fwd CACCCTTGATCACGATTTCCCCGA
Oligo rev AAACTCGGGGAAATCGTGATCAAG
Chromosome start end strand MM target_seq PAM alignment distance position gene name gene id
GU305914.1 225165 225187 + 0 CTTGATCACGATTTCCCCGA CGG ||||||||[||||||||||||]PAM NA Intergenic NA NA

T16769 ACGATTTCCCCGACGGTCTGCGG 1000
Oligo fwd CACCACGATTTCCCCGACGGTCTG
Oligo rev AAACCAGACCGTCGGGGAAATCGT
Chromosome start end strand MM target_seq PAM alignment distance position gene name gene id
GU305914.1 225172 225194 + 0 ACGATTTCCCCGACGGTCTG CGG ||||||||[||||||||||||]PAM NA Intergenic NA NA

T16770 ATTTCCCCGACGGTCTGCGGTGG 1000
Oligo fwd CACCATTTCCCCGACGGTCTGCGG
Oligo rev AAACCCGCAGACCGTCGGGGAAAT
Chromosome start end strand MM target_seq PAM alignment distance position gene name gene id
GU305914.1 225175 225197 + 0 ATTTCCCCGACGGTCTGCGG TGG ||||||||[||||||||||||]PAM NA Intergenic NA NA

T16771 CGGTGGCAGTTTGAGCCATCCGG 1000
Oligo fwd CACCCGGTGGCAGTTTGAGCCATC
Oligo rev AAACGATGGCTCAAACTGCCACCG
Chromosome start end strand MM target_seq PAM alignment distance position gene name gene id
GU305914.1 225192 225214 + 0 CGGTGGCAGTTTGAGCCATC CGG ||||||||[||||||||||||]PAM NA Intergenic NA NA

T16772 TCCGGACGCGTACACGTCCACGG 1000
Oligo fwd CACCTCCGGACGCGTACACGTCCA
Oligo rev AAACTGGACGTGTACGCGTCCGGA
Chromosome start end strand MM target_seq PAM alignment distance position gene name gene id
GU305914.1 225210 225232 + 0 TCCGGACGCGTACACGTCCA CGG ||||||||[||||||||||||]PAM NA Intergenic NA NA

T16773 GTACACGTCCACGGTGTCGTCGG 1000
Oligo fwd CACCGTACACGTCCACGGTGTCGT
Oligo rev AAACACGACACCGTGGACGTGTAC
Chromosome start end strand MM target_seq PAM alignment distance position gene name gene id

<table>
<tr><td>GU305914.1</td><td>225219</td><td>225241</td><td>+</td><td>0</td><td>GTACACGTCCACGGTGTCGT</td><td>CGG</td><td>||||||||[||||||||||||]PAM</td><td>NA</td><td>Intergenic</td><td>NA</td><td>NA</td></tr>
<tr><td></td><td></td><td></td><td></td><td></td><td></td><td></td><td></td><td></td><td></td><td></td><td></td></tr>
<tr><td>T16774</td><td>TCCACGGTGTCGTCGGTGAGCGG</td><td>1000</td><td></td><td></td><td></td><td></td><td></td><td></td><td></td><td></td><td></td></tr>
<tr><td>Oligo fwd</td><td>CACCTCCACGGTGTCGTCGGTGAG</td><td></td><td></td><td></td><td></td><td></td><td></td><td></td><td></td><td></td><td></td></tr>
<tr><td>Oligo rev</td><td>AAACCTCACCGACGACACCGTGGA</td><td></td><td></td><td></td><td></td><td></td><td></td><td></td><td></td><td></td><td></td></tr>
<tr><td>Chromosome</td><td>start</td><td>end</td><td>strand</td><td>MM</td><td>target_seq</td><td>PAM</td><td>alignment</td><td>distance</td><td>position</td><td>gene name</td><td>gene id</td></tr>
<tr><td>GU305914.1</td><td>225226</td><td>225248</td><td>+</td><td>0</td><td>TCCACGGTGTCGTCGGTGAG</td><td>CGG</td><td>||||||||[||||||||||||]PAM</td><td>NA</td><td>Intergenic</td><td>NA</td><td>NA</td></tr>
<tr><td>GU305914.1</td><td>128762</td><td>128784</td><td>+</td><td>4</td><td>TCCGCGGCGTCGACGGAGAG</td><td>TGG</td><td>|||-|||-[||||-|||-|||]PAM</td><td>NA</td><td>Intergenic</td><td>NA</td><td>NA</td></tr>
<tr><td></td><td></td><td></td><td></td><td></td><td></td><td></td><td></td><td></td><td></td><td></td><td></td></tr>
<tr><td>T16775</td><td>ACGGTGTCGTCGGTGAGCGGAGG</td><td>1000</td><td></td><td></td><td></td><td></td><td></td><td></td><td></td><td></td><td></td></tr>
<tr><td>Oligo fwd</td><td>CACCACGGTGTCGTCGGTGAGCGG</td><td></td><td></td><td></td><td></td><td></td><td></td><td></td><td></td><td></td><td></td></tr>
<tr><td>Oligo rev</td><td>AAACCCGCTCACCGACGACACCGT</td><td></td><td></td><td></td><td></td><td></td><td></td><td></td><td></td><td></td><td></td></tr>
<tr><td>Chromosome</td><td>start</td><td>end</td><td>strand</td><td>MM</td><td>target_seq</td><td>PAM</td><td>alignment</td><td>distance</td><td>position</td><td>gene name</td><td>gene id</td></tr>
<tr><td>GU305914.1</td><td>225229</td><td>225251</td><td>+</td><td>0</td><td>ACGGTGTCGTCGGTGAGCGG</td><td>AGG</td><td>||||||||[||||||||||||]PAM</td><td>NA</td><td>Intergenic</td><td>NA</td><td>NA</td></tr>
<tr><td>GU305914.1</td><td>67730</td><td>67752</td><td>+</td><td>4</td><td>TCGGTGATGTCGGGGAGCGG</td><td>CGG</td><td>-|||||--[|||||-||||||]PAM</td><td>NA</td><td>Intergenic</td><td>NA</td><td>NA</td></tr>
<tr><td></td><td></td><td></td><td></td><td></td><td></td><td></td><td></td><td></td><td></td><td></td><td></td></tr>
<tr><td>T16776</td><td>CGGAGGCCCGTGAAAGATCAAGG</td><td>1000</td><td></td><td></td><td></td><td></td><td></td><td></td><td></td><td></td><td></td></tr>
<tr><td>Oligo fwd</td><td>CACCCGGAGGCCCGTGAAAGATCA</td><td></td><td></td><td></td><td></td><td></td><td></td><td></td><td></td><td></td><td></td></tr>
<tr><td>Oligo rev</td><td>AAACTGATCTTTCACGGGCCTCCG</td><td></td><td></td><td></td><td></td><td></td><td></td><td></td><td></td><td></td><td></td></tr>
<tr><td>Chromosome</td><td>start</td><td>end</td><td>strand</td><td>MM</td><td>target_seq</td><td>PAM</td><td>alignment</td><td>distance</td><td>position</td><td>gene name</td><td>gene id</td></tr>
<tr><td>GU305914.1</td><td>225246</td><td>225268</td><td>+</td><td>0</td><td>CGGAGGCCCGTGAAAGATCA</td><td>AGG</td><td>||||||||[||||||||||||]PAM</td><td>NA</td><td>Intergenic</td><td>NA</td><td>NA</td></tr>
<tr><td></td><td></td><td></td><td></td><td></td><td></td><td></td><td></td><td></td><td></td><td></td><td></td></tr>
<tr><td>T16777</td><td>TCAAGGCGTCTCCGACGTAGAGG</td><td>1000</td><td></td><td></td><td></td><td></td><td></td><td></td><td></td><td></td><td></td></tr>
<tr><td>Oligo fwd</td><td>CACCTCAAGGCGTCTCCGACGTAG</td><td></td><td></td><td></td><td></td><td></td><td></td><td></td><td></td><td></td><td></td></tr>
<tr><td>Oligo rev</td><td>AAACCTACGTCGGAGACGCCTTGA</td><td></td><td></td><td></td><td></td><td></td><td></td><td></td><td></td><td></td><td></td></tr>
<tr><td>Chromosome</td><td>start</td><td>end</td><td>strand</td><td>MM</td><td>target_seq</td><td>PAM</td><td>alignment</td><td>distance</td><td>position</td><td>gene name</td><td>gene id</td></tr>
<tr><td>GU305914.1</td><td>225263</td><td>225285</td><td>+</td><td>0</td><td>TCAAGGCGTCTCCGACGTAG</td><td>AGG</td><td>||||||||[||||||||||||]PAM</td><td>NA</td><td>Intergenic</td><td>NA</td><td>NA</td></tr>
<tr><td></td><td></td><td></td><td></td><td></td><td></td><td></td><td></td><td></td><td></td><td></td><td></td></tr>
<tr><td>T16778</td><td>CGTAGAGGTCGCTGTTATCTCGG</td><td>1000</td><td></td><td></td><td></td><td></td><td></td><td></td><td></td><td></td><td></td></tr>
<tr><td>Oligo fwd</td><td>CACCCGTAGAGGTCGCTGTTATCT</td><td></td><td></td><td></td><td></td><td></td><td></td><td></td><td></td><td></td><td></td></tr>
<tr><td>Oligo rev</td><td>AAACAGATAACAGCGACCTCTACG</td><td></td><td></td><td></td><td></td><td></td><td></td><td></td><td></td><td></td><td></td></tr>
<tr><td>Chromosome</td><td>start</td><td>end</td><td>strand</td><td>MM</td><td>target_seq</td><td>PAM</td><td>alignment</td><td>distance</td><td>position</td><td>gene name</td><td>gene id</td></tr>
<tr><td>GU305914.1</td><td>225278</td><td>225300</td><td>+</td><td>0</td><td>CGTAGAGGTCGCTGTTATCT</td><td>CGG</td><td>||||||||[||||||||||||]PAM</td><td>NA</td><td>Intergenic</td><td>NA</td><td>NA</td></tr>
<tr><td></td><td></td><td></td><td></td><td></td><td></td><td></td><td></td><td></td><td></td><td></td><td></td></tr>
<tr><td>T16779</td><td>GCTGACCCAAGTACACCGTCTGG</td><td>1000</td><td></td><td></td><td></td><td></td><td></td><td></td><td></td><td></td><td></td></tr>
<tr><td>Oligo fwd</td><td>CACCGCTGACCCAAGTACACCGTC</td><td></td><td></td><td></td><td></td><td></td><td></td><td></td><td></td><td></td><td></td></tr>
<tr><td>Oligo rev</td><td>AAACGACGGTGTACTTGGGTCAGC</td><td></td><td></td><td></td><td></td><td></td><td></td><td></td><td></td><td></td><td></td></tr>
<tr><td>Chromosome</td><td>start</td><td>end</td><td>strand</td><td>MM</td><td>target_seq</td><td>PAM</td><td>alignment</td><td>distance</td><td>position</td><td>gene name</td><td>gene id</td></tr>
<tr><td>GU305914.1</td><td>225326</td><td>225348</td><td>+</td><td>0</td><td>GCTGACCCAAGTACACCGTC</td><td>TGG</td><td>||||||||[||||||||||||]PAM</td><td>NA</td><td>Intergenic</td><td>NA</td><td>NA</td></tr>
<tr><td></td><td></td><td></td><td></td><td></td><td></td><td></td><td></td><td></td><td></td><td></td><td></td></tr>
<tr><td>T16780</td><td>AGTACACCGTCTGGTCGTCGTGG</td><td>1000</td><td></td><td></td><td></td><td></td><td></td><td></td><td></td><td></td><td></td></tr>
<tr><td>Oligo fwd</td><td>CACCAGTACACCGTCTGGTCGTCG</td><td></td><td></td><td></td><td></td><td></td><td></td><td></td><td></td><td></td><td></td></tr>
<tr><td>Oligo rev</td><td>AAACCGACGACCAGACGGTGTACT</td><td></td><td></td><td></td><td></td><td></td><td></td><td></td><td></td><td></td><td></td></tr>
<tr><td>Chromosome</td><td>start</td><td>end</td><td>strand</td><td>MM</td><td>target_seq</td><td>PAM</td><td>alignment</td><td>distance</td><td>position</td><td>gene name</td><td>gene id</td></tr>
<tr><td>GU305914.1</td><td>225335</td><td>225357</td><td>+</td><td>0</td><td>AGTACACCGTCTGGTCGTCG</td><td>TGG</td><td>||||||||[||||||||||||]PAM</td><td>NA</td><td>Intergenic</td><td>NA</td><td>NA</td></tr>
<tr><td></td><td></td><td></td><td></td><td></td><td></td><td></td><td></td><td></td><td></td><td></td><td></td></tr>
<tr><td>T16781</td><td>CTGGTCGTCGTGGTTGCGATAGG</td><td>1000</td><td></td><td></td><td></td><td></td><td></td><td></td><td></td><td></td><td></td></tr>
<tr><td>Oligo fwd</td><td>CACCCTGGTCGTCGTGGTTGCGAT</td><td></td><td></td><td></td><td></td><td></td><td></td><td></td><td></td><td></td><td></td></tr>
<tr><td>Oligo rev</td><td>AAACATCGCAACCACGACGACCAG</td><td></td><td></td><td></td><td></td><td></td><td></td><td></td><td></td><td></td><td></td></tr>
<tr><td>Chromosome</td><td>start</td><td>end</td><td>strand</td><td>MM</td><td>target_seq</td><td>PAM</td><td>alignment</td><td>distance</td><td>position</td><td>gene name</td><td>gene id</td></tr>
<tr><td>GU305914.1</td><td>225345</td><td>225367</td><td>+</td><td>0</td><td>CTGGTCGTCGTGGTTGCGAT</td><td>AGG</td><td>||||||||[||||||||||||]PAM</td><td>NA</td><td>Intergenic</td><td>NA</td><td>NA</td></tr>
<tr><td></td><td></td><td></td><td></td><td></td><td></td><td></td><td></td><td></td><td></td><td></td><td></td></tr>
<tr><td>T16782</td><td>AGCCCGAGCGCACGTCGTGATGG</td><td>1000</td><td></td><td></td><td></td><td></td><td></td><td></td><td></td><td></td><td></td></tr>
<tr><td>Oligo fwd</td><td>CACCAGCCCGAGCGCACGTCGTGA</td><td></td><td></td><td></td><td></td><td></td><td></td><td></td><td></td><td></td><td></td></tr>
<tr><td>Oligo rev</td><td>AAACTCACGACGTGCGCTCGGGCT</td><td></td><td></td><td></td><td></td><td></td><td></td><td></td><td></td><td></td><td></td></tr>
<tr><td>Chromosome</td><td>start</td><td>end</td><td>strand</td><td>MM</td><td>target_seq</td><td>PAM</td><td>alignment</td><td>distance</td><td>position</td><td>gene name</td><td>gene id</td></tr>
<tr><td>GU305914.1</td><td>225377</td><td>225399</td><td>+</td><td>0</td><td>AGCCCGAGCGCACGTCGTGA</td><td>TGG</td><td>||||||||[||||||||||||]PAM</td><td>NA</td><td>Intergenic</td><td>NA</td><td>NA</td></tr>
<tr><td></td><td></td><td></td><td></td><td></td><td></td><td></td><td></td><td></td><td></td><td></td><td></td></tr>
<tr><td>T16783</td><td>CCGAGCGCACGTCGTGATGGCGG</td><td>1000</td><td></td><td></td><td></td><td></td><td></td><td></td><td></td><td></td><td></td></tr>
<tr><td>Oligo fwd</td><td>CACCCCGAGCGCACGTCGTGATGG</td><td></td><td></td><td></td><td></td><td></td><td></td><td></td><td></td><td></td><td></td></tr>
<tr><td>Oligo rev</td><td>AAACCCATCACGACGTGCGCTCGG</td><td></td><td></td><td></td><td></td><td></td><td></td><td></td><td></td><td></td><td></td></tr>
<tr><td>Chromosome</td><td>start</td><td>end</td><td>strand</td><td>MM</td><td>target_seq</td><td>PAM</td><td>alignment</td><td>distance</td><td>position</td><td>gene name</td><td>gene id</td></tr>
<tr><td>GU305914.1</td><td>225380</td><td>225402</td><td>+</td><td>0</td><td>CCGAGCGCACGTCGTGATGG</td><td>CGG</td><td>||||||||[||||||||||||]PAM</td><td>NA</td><td>Intergenic</td><td>NA</td><td>NA</td></tr>
<tr><td></td><td></td><td></td><td></td><td></td><td></td><td></td><td></td><td></td><td></td><td></td><td></td></tr>
<tr><td>T16784</td><td>GTCGTGATGGCGGCCGCTCCAGG</td><td>1000</td><td></td><td></td><td></td><td></td><td></td><td></td><td></td><td></td><td></td></tr>
<tr><td>Oligo fwd</td><td>CACCGTCGTGATGGCGGCCGCTCC</td><td></td><td></td><td></td><td></td><td></td><td></td><td></td><td></td><td></td><td></td></tr>
<tr><td>Oligo rev</td><td>AAACGGAGCGGCCGCCATCACGAC</td><td></td><td></td><td></td><td></td><td></td><td></td><td></td><td></td><td></td><td></td></tr>
<tr><td>Chromosome</td><td>start</td><td>end</td><td>strand</td><td>MM</td><td>target_seq</td><td>PAM</td><td>alignment</td><td>distance</td><td>position</td><td>gene name</td><td>gene id</td></tr>
<tr><td>GU305914.1</td><td>225390</td><td>225412</td><td>+</td><td>0</td><td>GTCGTGATGGCGGCCGCTCC</td><td>AGG</td><td>||||||||[||||||||||||]PAM</td><td>NA</td><td>Intergenic</td><td>NA</td><td>NA</td></tr>
<tr><td></td><td></td><td></td><td></td><td></td><td></td><td></td><td></td><td></td><td></td><td></td><td></td></tr>
<tr><td>T16785</td><td>GGCGGCCGCTCCAGGAGAACTGG</td><td>1000</td><td></td><td></td><td></td><td></td><td></td><td></td><td></td><td></td><td></td></tr>
<tr><td>Oligo fwd</td><td>CACCGGCGGCCGCTCCAGGAGAAC</td><td></td><td></td><td></td><td></td><td></td><td></td><td></td><td></td><td></td><td></td></tr>
<tr><td>Oligo rev</td><td>AAACGTTCTCCTGGAGCGGCCGCC</td><td></td><td></td><td></td><td></td><td></td><td></td><td></td><td></td><td></td><td></td></tr>
<tr><td>Chromosome</td><td>start</td><td>end</td><td>strand</td><td>MM</td><td>target_seq</td><td>PAM</td><td>alignment</td><td>distance</td><td>position</td><td>gene name</td><td>gene id</td></tr>
<tr><td>GU305914.1</td><td>225398</td><td>225420</td><td>+</td><td>0</td><td>GGCGGCCGCTCCAGGAGAAC</td><td>TGG</td><td>||||||||[||||||||||||]PAM</td><td>NA</td><td>Intergenic</td><td>NA</td><td>NA</td></tr>
<tr><td></td><td></td><td></td><td></td><td></td><td></td><td></td><td></td><td></td><td></td><td></td><td></td></tr>
<tr><td>T16786</td><td>AGGAGAACTGGATCACGTTCTGG</td><td>1000</td><td></td><td></td><td></td><td></td><td></td><td></td><td></td><td></td><td></td></tr>
<tr><td>Oligo fwd</td><td>CACCAGGAGAACTGGATCACGTTC</td><td></td><td></td><td></td><td></td><td></td><td></td><td></td><td></td><td></td><td></td></tr>
<tr><td>Oligo rev</td><td>AAACGAACGTGATCCAGTTCTCCT</td><td></td><td></td><td></td><td></td><td></td><td></td><td></td><td></td><td></td><td></td></tr>
<tr><td>Chromosome</td><td>start</td><td>end</td><td>strand</td><td>MM</td><td>target_seq</td><td>PAM</td><td>alignment</td><td>distance</td><td>position</td><td>gene name</td><td>gene id</td></tr>
<tr><td>GU305914.1</td><td>225410</td><td>225432</td><td>+</td><td>0</td><td>AGGAGAACTGGATCACGTTC</td><td>TGG</td><td>||||||||[||||||||||||]PAM</td><td>NA</td><td>Intergenic</td><td>NA</td><td>NA</td></tr>
<tr><td></td><td></td><td></td><td></td><td></td><td></td><td></td><td></td><td></td><td></td><td></td><td></td></tr>
<tr><td>T16787</td><td>ACTGGATCACGTTCTGGTCTTGG</td><td>1000</td><td></td><td></td><td></td><td></td><td></td><td></td><td></td><td></td><td></td></tr>
<tr><td>Oligo fwd</td><td>CACCACTGGATCACGTTCTGGTCT</td><td></td><td></td><td></td><td></td><td></td><td></td><td></td><td></td><td></td><td></td></tr>
<tr><td>Oligo rev</td><td>AAACAGACCAGAACGTGATCCAGT</td><td></td><td></td><td></td><td></td><td></td><td></td><td></td><td></td><td></td><td></td></tr>
<tr><td>Chromosome</td><td>start</td><td>end</td><td>strand</td><td>MM</td><td>target_seq</td><td>PAM</td><td>alignment</td><td>distance</td><td>position</td><td>gene name</td><td>gene id</td></tr>
<tr><td>GU305914.1</td><td>225416</td><td>225438</td><td>+</td><td>0</td><td>ACTGGATCACGTTCTGGTCT</td><td>TGG</td><td>||||||||[||||||||||||]PAM</td><td>NA</td><td>Intergenic</td><td>NA</td><td>NA</td></tr>
<tr><td>GU305914.1</td><td>132042</td><td>132064</td><td>+</td><td>4</td><td>GTTGGATCTCGTTCTGGTCC</td><td>GGG</td><td>--||||||[-||||||||||-]PAM</td><td>NA</td><td>Intergenic</td><td>NA</td><td>NA</td></tr>
</table>

| | | | | | | | | | | | |
|---|---|---|---|---|---|---|---|---|---|---|---|
| T16788 | CTGGATCACGTTCTGGTCTTGGG | 1000 | | | | | | | | | |
| Oligo fwd | CACCCTGGATCACGTTCTGGTCTT | | | | | | | | | | |
| Oligo rev | AAACAAGACCAGAACGTGATCCAG | | | | | | | | | | |
| Chromosome | start | end | strand | MM | target_seq | PAM | alignment | distance | position | gene name | gene id |
| GU305914.1 | 225417 | 225439 | + | 0 | CTGGATCACGTTCTGGTCTT | GGG | \|\|\|\|\|\|\|\|[\|\|\|\|\|\|\|\|\|\|\|\|]PAM | NA | Intergenic | NA | NA |
| T16789 | TCTGGTCTTGGGTATCGTACAGG | 1000 | | | | | | | | | |
| Oligo fwd | CACCTCTGGTCTTGGGTATCGTAC | | | | | | | | | | |
| Oligo rev | AAACGTACGATACCCAAGACCAGA | | | | | | | | | | |
| Chromosome | start | end | strand | MM | target_seq | PAM | alignment | distance | position | gene name | gene id |
| GU305914.1 | 225428 | 225450 | + | 0 | TCTGGTCTTGGGTATCGTAC | AGG | \|\|\|\|\|\|\|\|[\|\|\|\|\|\|\|\|\|\|\|\|]PAM | NA | Intergenic | NA | NA |
| T16790 | ATCGTACAGGAAGATCTTGATGG | 1000 | | | | | | | | | |
| Oligo fwd | CACCATCGTACAGGAAGATCTTGA | | | | | | | | | | |
| Oligo rev | AAACTCAAGATCTTCCTGTACGAT | | | | | | | | | | |
| Chromosome | start | end | strand | MM | target_seq | PAM | alignment | distance | position | gene name | gene id |
| GU305914.1 | 225441 | 225463 | + | 0 | ATCGTACAGGAAGATCTTGA | TGG | \|\|\|\|\|\|\|\|[\|\|\|\|\|\|\|\|\|\|\|\|]PAM | NA | Intergenic | NA | NA |
| T16791 | TCGTACAGGAAGATCTTGATGGG | 1000 | | | | | | | | | |
| Oligo fwd | CACCTCGTACAGGAAGATCTTGAT | | | | | | | | | | |
| Oligo rev | AAACATCAAGATCTTCCTGTACGA | | | | | | | | | | |
| Chromosome | start | end | strand | MM | target_seq | PAM | alignment | distance | position | gene name | gene id |
| GU305914.1 | 225442 | 225464 | + | 0 | TCGTACAGGAAGATCTTGAT | GGG | \|\|\|\|\|\|\|\|[\|\|\|\|\|\|\|\|\|\|\|\|]PAM | NA | Intergenic | NA | NA |
| T16792 | CAGGAAGATCTTGATGGGCTCGG | 1000 | | | | | | | | | |
| Oligo fwd | CACCCAGGAAGATCTTGATGGGCT | | | | | | | | | | |
| Oligo rev | AAACAGCCCATCAAGATCTTCCTG | | | | | | | | | | |
| Chromosome | start | end | strand | MM | target_seq | PAM | alignment | distance | position | gene name | gene id |
| GU305914.1 | 225447 | 225469 | + | 0 | CAGGAAGATCTTGATGGGCT | CGG | \|\|\|\|\|\|\|\|[\|\|\|\|\|\|\|\|\|\|\|\|]PAM | NA | Intergenic | NA | NA |
| T16793 | AGGAAGATCTTGATGGGCTCGGG | 1000 | | | | | | | | | |
| Oligo fwd | CACCAGGAAGATCTTGATGGGCTC | | | | | | | | | | |
| Oligo rev | AAACGAGCCCATCAAGATCTTCCT | | | | | | | | | | |
| Chromosome | start | end | strand | MM | target_seq | PAM | alignment | distance | position | gene name | gene id |
| GU305914.1 | 225448 | 225470 | + | 0 | AGGAAGATCTTGATGGGCTC | GGG | \|\|\|\|\|\|\|\|[\|\|\|\|\|\|\|\|\|\|\|\|]PAM | NA | Intergenic | NA | NA |
| T16794 | ATCTTGATGGGCTCGGGACGCGG | 1000 | | | | | | | | | |
| Oligo fwd | CACCATCTTGATGGGCTCGGGACG | | | | | | | | | | |
| Oligo rev | AAACCGTCCCGAGCCCATCAAGAT | | | | | | | | | | |
| Chromosome | start | end | strand | MM | target_seq | PAM | alignment | distance | position | gene name | gene id |
| GU305914.1 | 225454 | 225476 | + | 0 | ATCTTGATGGGCTCGGGACG | CGG | \|\|\|\|\|\|\|\|[\|\|\|\|\|\|\|\|\|\|\|\|]PAM | NA | Intergenic | NA | NA |
| T16795 | CGGGACGCGGACGCTCCTCCTGG | 1000 | | | | | | | | | |
| Oligo fwd | CACCCGGGACGCGGACGCTCCTCC | | | | | | | | | | |
| Oligo rev | AAACGGAGGAGCGTCCGCGTCCCG | | | | | | | | | | |
| Chromosome | start | end | strand | MM | target_seq | PAM | alignment | distance | position | gene name | gene id |
| GU305914.1 | 225467 | 225489 | + | 0 | CGGGACGCGGACGCTCCTCC | TGG | \|\|\|\|\|\|\|\|[\|\|\|\|\|\|\|\|\|\|\|\|]PAM | NA | Intergenic | NA | NA |
| T16796 | GGGACGCGGACGCTCCTCCTGGG | 1000 | | | | | | | | | |
| Oligo fwd | CACCGGGACGCGGACGCTCCTCCT | | | | | | | | | | |
| Oligo rev | AAACAGGAGGAGCGTCCGCGTCCC | | | | | | | | | | |
| Chromosome | start | end | strand | MM | target_seq | PAM | alignment | distance | position | gene name | gene id |
| GU305914.1 | 225468 | 225490 | + | 0 | GGGACGCGGACGCTCCTCCT | GGG | \|\|\|\|\|\|\|\|[\|\|\|\|\|\|\|\|\|\|\|\|]PAM | NA | Intergenic | NA | NA |
| T16797 | CCTCCTGGGTGACCGAGCCGAGG | 1000 | | | | | | | | | |
| Oligo fwd | CACCCCTCCTGGGTGACCGAGCCG | | | | | | | | | | |
| Oligo rev | AAACCGGCTCGGTCACCCAGGAGG | | | | | | | | | | |
| Chromosome | start | end | strand | MM | target_seq | PAM | alignment | distance | position | gene name | gene id |
| GU305914.1 | 225482 | 225504 | + | 0 | CCTCCTGGGTGACCGAGCCG | AGG | \|\|\|\|\|\|\|\|[\|\|\|\|\|\|\|\|\|\|\|\|]PAM | NA | Intergenic | NA | NA |
| T16798 | CTCCTGGGTGACCGAGCCGAGGG | 1000 | | | | | | | | | |
| Oligo fwd | CACCCTCCTGGGTGACCGAGCCGA | | | | | | | | | | |
| Oligo rev | AAACTCGGCTCGGTCACCCAGGAG | | | | | | | | | | |
| Chromosome | start | end | strand | MM | target_seq | PAM | alignment | distance | position | gene name | gene id |
| GU305914.1 | 225483 | 225505 | + | 0 | CTCCTGGGTGACCGAGCCGA | GGG | \|\|\|\|\|\|\|\|[\|\|\|\|\|\|\|\|\|\|\|\|]PAM | NA | Intergenic | NA | NA |
| T16799 | CTGGGTGACCGAGCCGAGGGCGG | 1000 | | | | | | | | | |
| Oligo fwd | CACCCTGGGTGACCGAGCCGAGGG | | | | | | | | | | |
| Oligo rev | AAACCCCTCGGCTCGGTCACCCAG | | | | | | | | | | |
| Chromosome | start | end | strand | MM | target_seq | PAM | alignment | distance | position | gene name | gene id |
| GU305914.1 | 225486 | 225508 | + | 0 | CTGGGTGACCGAGCCGAGGG | CGG | \|\|\|\|\|\|\|\|[\|\|\|\|\|\|\|\|\|\|\|\|]PAM | NA | Intergenic | NA | NA |
| T16800 | GGTGACCGAGCCGAGGGCGGCGG | 1000 | | | | | | | | | |
| Oligo fwd | CACCGGTGACCGAGCCGAGGGCGG | | | | | | | | | | |
| Oligo rev | AAACCCGCCCTCGGCTCGGTCACC | | | | | | | | | | |
| Chromosome | start | end | strand | MM | target_seq | PAM | alignment | distance | position | gene name | gene id |
| GU305914.1 | 225489 | 225511 | + | 0 | GGTGACCGAGCCGAGGGCGG | CGG | \|\|\|\|\|\|\|\|[\|\|\|\|\|\|\|\|\|\|\|\|]PAM | NA | Intergenic | NA | NA |
| T16801 | GACCGAGCCGAGGGCGGCGGTGG | 1000 | | | | | | | | | |
| Oligo fwd | CACCGACCGAGCCGAGGGCGGCGG | | | | | | | | | | |
| Oligo rev | AAACCCGCCGCCCTCGGCTCGGTC | | | | | | | | | | |
| Chromosome | start | end | strand | MM | target_seq | PAM | alignment | distance | position | gene name | gene id |
| GU305914.1 | 225492 | 225514 | + | 0 | GACCGAGCCGAGGGCGGCGG | TGG | \|\|\|\|\|\|\|\|[\|\|\|\|\|\|\|\|\|\|\|\|]PAM | NA | Intergenic | NA | NA |
| GU305914.1 | 135539 | 135561 | - | 4 | GAGCGGGCCGAAGTCGGCGG | CGG | \|\|-\|\|-\|\|[\|\|\|-\|-\|\|\|\|\|\|]PAM | NA | Intergenic | NA | NA |
| T16802 | CGAGCCGAGGGCGGCGGTGGTGG | 1000 | | | | | | | | | |
| Oligo fwd | CACCCGAGCCGAGGGCGGCGGTGG | | | | | | | | | | |
| Oligo rev | AAACCCACCGCCGCCCTCGGCTCG | | | | | | | | | | |
| Chromosome | start | end | strand | MM | target_seq | PAM | alignment | distance | position | gene name | gene id |

| | | | | | | | | | | | |
|---|---|---|---|---|---|---|---|---|---|---|---|
| GU305914.1 | 225495 | 225517 | + | 0 | CGAGCCGAGGGCGGCGGTGG | TGG | \|\|\|\|\|\|\|\|[\|\|\|\|\|\|\|\|\|\|\|\|]PAM | NA | Intergenic | NA | NA |
| T16803 | GCCGAGGGCGGCGGTGGTGGTGG | 1000 | | | | | | | | | |
| Oligo fwd | CACCGCCGAGGGCGGCGGTGGTGG | | | | | | | | | | |
| Oligo rev | AAACCCACCACCGCCGCCCTCGGC | | | | | | | | | | |
| Chromosome | start | end | strand | MM | target_seq | PAM | alignment | distance | position | gene name | gene id |
| GU305914.1 | 225498 | 225520 | + | 0 | GCCGAGGGCGGCGGTGGTGG | TGG | \|\|\|\|\|\|\|\|[\|\|\|\|\|\|\|\|\|\|\|\|]PAM | NA | Intergenic | NA | NA |
| GU305914.1 | 62437 | 62459 | + | 4 | GGCGGTGGCGGCGGGGGTGG | TGG | \|-\|\|--\|\|[\|\|\|\|\|\|-\|\|\|\|\|]PAM | NA | Intergenic | NA | NA |
| GU305914.1 | 36161 | 36183 | + | 4 | GCTGCCGGCGGCGGTTGTGG | TGG | \|\|-\|--\|\|[\|\|\|\|\|\|\|-\|\|\|\|]PAM | NA | Intergenic | NA | NA |
| GU305914.1 | 52719 | 52741 | + | 4 | GCCGCTGGCGGTGGCGGTGG | CGG | \|\|\|\|--\|\|[\|\|\|-\|\|-\|\|\|\|\|]PAM | NA | Intergenic | NA | NA |
| GU305914.1 | 111175 | 111197 | - | 4 | GCCGACGACGGCGAAGGTGG | AGG | \|\|\|\|\|-\|-[\|\|\|\|\|--\|\|\|\|\|]PAM | NA | Intergenic | NA | NA |
| GU305914.1 | 149482 | 149504 | - | 4 | GGCGACGGCGGCGGCGGCGG | CGG | \|-\|\|\|-\|\|[\|\|\|\|\|\|-\|\|-\|\|]PAM | NA | Intergenic | NA | NA |
| T16804 | GCGGTGGTGGTGGTTTTCTTCGG | 1000 | | | | | | | | | |
| Oligo fwd | CACCGCGGTGGTGGTGGTTTTCTT | | | | | | | | | | |
| Oligo rev | AAACAAGAAAACCACCACCACCGC | | | | | | | | | | |
| Chromosome | start | end | strand | MM | target_seq | PAM | alignment | distance | position | gene name | gene id |
| GU305914.1 | 225508 | 225530 | + | 0 | GCGGTGGTGGTGGTTTTCTT | CGG | \|\|\|\|\|\|\|\|[\|\|\|\|\|\|\|\|\|\|\|\|]PAM | NA | Intergenic | NA | NA |
| T16805 | TTTGCTACAGTCTACGAGCGCGG | 1000 | | | | | | | | | |
| Oligo fwd | CACCTTTGCTACAGTCTACGAGCG | | | | | | | | | | |
| Oligo rev | AAACCGCTCGTAGACTGTAGCAAA | | | | | | | | | | |
| Chromosome | start | end | strand | MM | target_seq | PAM | alignment | distance | position | gene name | gene id |
| GU305914.1 | 225531 | 225553 | + | 0 | TTTGCTACAGTCTACGAGCG | CGG | \|\|\|\|\|\|\|\|[\|\|\|\|\|\|\|\|\|\|\|\|]PAM | NA | Intergenic | NA | NA |
| T16806 | TTGCTACAGTCTACGAGCGCGGG | 1000 | | | | | | | | | |
| Oligo fwd | CACCTTGCTACAGTCTACGAGCGC | | | | | | | | | | |
| Oligo rev | AAACGCGCTCGTAGACTGTAGCAA | | | | | | | | | | |
| Chromosome | start | end | strand | MM | target_seq | PAM | alignment | distance | position | gene name | gene id |
| GU305914.1 | 225532 | 225554 | + | 0 | TTGCTACAGTCTACGAGCGC | GGG | \|\|\|\|\|\|\|\|[\|\|\|\|\|\|\|\|\|\|\|\|]PAM | NA | Intergenic | NA | NA |
| T16807 | TGCTACAGTCTACGAGCGCGGGG | 1000 | | | | | | | | | |
| Oligo fwd | CACCTGCTACAGTCTACGAGCGCG | | | | | | | | | | |
| Oligo rev | AAACCGCGCTCGTAGACTGTAGCA | | | | | | | | | | |
| Chromosome | start | end | strand | MM | target_seq | PAM | alignment | distance | position | gene name | gene id |
| GU305914.1 | 225533 | 225555 | + | 0 | TGCTACAGTCTACGAGCGCG | GGG | \|\|\|\|\|\|\|\|[\|\|\|\|\|\|\|\|\|\|\|\|]PAM | NA | Intergenic | NA | NA |
| T16808 | TACAGTCTACGAGCGCGGGGCGG | 1000 | | | | | | | | | |
| Oligo fwd | CACCTACAGTCTACGAGCGCGGGG | | | | | | | | | | |
| Oligo rev | AAACCCCCGCGCTCGTAGACTGTA | | | | | | | | | | |
| Chromosome | start | end | strand | MM | target_seq | PAM | alignment | distance | position | gene name | gene id |
| GU305914.1 | 225536 | 225558 | + | 0 | TACAGTCTACGAGCGCGGGG | CGG | \|\|\|\|\|\|\|\|[\|\|\|\|\|\|\|\|\|\|\|\|]PAM | NA | Intergenic | NA | NA |
| T16809 | CGAGCCACGCGAGACACCAGAGG | 1000 | | | | | | | | | |
| Oligo fwd | CACCCGAGCCACGCGAGACACCAG | | | | | | | | | | |
| Oligo rev | AAACCTGGTGTCTCGCGTGGCTCG | | | | | | | | | | |
| Chromosome | start | end | strand | MM | target_seq | PAM | alignment | distance | position | gene name | gene id |
| GU305914.1 | 225563 | 225585 | + | 0 | CGAGCCACGCGAGACACCAG | AGG | \|\|\|\|\|\|\|\|[\|\|\|\|\|\|\|\|\|\|\|\|]PAM | NA | Intergenic | NA | NA |
| T16810 | ACGCGAGACACCAGAGGCAGAGG | 1000 | | | | | | | | | |
| Oligo fwd | CACCACGCGAGACACCAGAGGCAG | | | | | | | | | | |
| Oligo rev | AAACCTGCCTCTGGTGTCTCGCGT | | | | | | | | | | |
| Chromosome | start | end | strand | MM | target_seq | PAM | alignment | distance | position | gene name | gene id |
| GU305914.1 | 225569 | 225591 | + | 0 | ACGCGAGACACCAGAGGCAG | AGG | \|\|\|\|\|\|\|\|[\|\|\|\|\|\|\|\|\|\|\|\|]PAM | NA | Intergenic | NA | NA |
| T16811 | CGAGACACCAGAGGCAGAGGAGG | 1000 | | | | | | | | | |
| Oligo fwd | CACCCGAGACACCAGAGGCAGAGG | | | | | | | | | | |
| Oligo rev | AAACCCTCTGCCTCTGGTGTCTCG | | | | | | | | | | |
| Chromosome | start | end | strand | MM | target_seq | PAM | alignment | distance | position | gene name | gene id |
| GU305914.1 | 225572 | 225594 | + | 0 | CGAGACACCAGAGGCAGAGG | AGG | \|\|\|\|\|\|\|\|[\|\|\|\|\|\|\|\|\|\|\|\|]PAM | NA | Intergenic | NA | NA |
| T16812 | GCAGAGGAGGACAGAGAAGCTGG | 1000 | | | | | | | | | |
| Oligo fwd | CACCGCAGAGGAGGACAGAGAAGC | | | | | | | | | | |
| Oligo rev | AAACGCTTCTCTGTCCTCCTCTGC | | | | | | | | | | |
| Chromosome | start | end | strand | MM | target_seq | PAM | alignment | distance | position | gene name | gene id |
| GU305914.1 | 225585 | 225607 | + | 0 | GCAGAGGAGGACAGAGAAGC | TGG | \|\|\|\|\|\|\|\|[\|\|\|\|\|\|\|\|\|\|\|\|]PAM | NA | Intergenic | NA | NA |
| T16813 | GAAGCTGGCCGCACGTAGCCTGG | 1000 | | | | | | | | | |
| Oligo fwd | CACCGAAGCTGGCCGCACGTAGCC | | | | | | | | | | |
| Oligo rev | AAACGGCTACGTGCGGCCAGCTTC | | | | | | | | | | |
| Chromosome | start | end | strand | MM | target_seq | PAM | alignment | distance | position | gene name | gene id |
| GU305914.1 | 225600 | 225622 | + | 0 | GAAGCTGGCCGCACGTAGCC | TGG | \|\|\|\|\|\|\|\|[\|\|\|\|\|\|\|\|\|\|\|\|]PAM | NA | Intergenic | NA | NA |
| T16814 | GCCGCACGTAGCCTGGCGACTGG | 1000 | | | | | | | | | |
| Oligo fwd | CACCGCCGCACGTAGCCTGGCGAC | | | | | | | | | | |
| Oligo rev | AAACGTCGCCAGGCTACGTGCGGC | | | | | | | | | | |
| Chromosome | start | end | strand | MM | target_seq | PAM | alignment | distance | position | gene name | gene id |
| GU305914.1 | 225607 | 225629 | + | 0 | GCCGCACGTAGCCTGGCGAC | TGG | \|\|\|\|\|\|\|\|[\|\|\|\|\|\|\|\|\|\|\|\|]PAM | NA | Intergenic | NA | NA |
| T16815 | GCGACTGGCGATAGCTGTTGAGG | 1000 | | | | | | | | | |
| Oligo fwd | CACCGCGACTGGCGATAGCTGTTG | | | | | | | | | | |
| Oligo rev | AAACCAACAGCTATCGCCAGTCGC | | | | | | | | | | |
| Chromosome | start | end | strand | MM | target_seq | PAM | alignment | distance | position | gene name | gene id |
| GU305914.1 | 225622 | 225644 | + | 0 | GCGACTGGCGATAGCTGTTG | AGG | \|\|\|\|\|\|\|\|[\|\|\|\|\|\|\|\|\|\|\|\|]PAM | NA | Intergenic | NA | NA |
| T16816 | GATAGCTGTTGAGGAATTGCTGG | 1000 | | | | | | | | | |
| Oligo fwd | CACCGATAGCTGTTGAGGAATTGC | | | | | | | | | | |
| Oligo rev | AAACGCAATTCCTCAACAGCTATC | | | | | | | | | | |
| Chromosome | start | end | strand | MM | target_seq | PAM | alignment | distance | position | gene name | gene id |

GU305914.1 225631 225653 + 0 GATAGCTGTTGAGGAATTGC TGG ||||||||[||||||||||||]PAM NA Intergenic NA NA

T16817 AATTGCTGGAAGCGCCATCCCGG 1000
Oligo fwd CACCAATTGCTGGAAGCGCCATCC
Oligo rev AAACGGATGGCGCTTCCAGCAATT
Chromosome start end strand MM target_seq PAM alignment distance position gene name gene id
GU305914.1 225645 225667 + 0 AATTGCTGGAAGCGCCATCC CGG ||||||||[||||||||||||]PAM NA Intergenic NA NA

T16818 GGATCGTTAGTGTCCGCCGCCGG 1000
Oligo fwd CACCGGATCGTTAGTGTCCGCCGC
Oligo rev AAACGCGGCGGACACTAACGATCC
Chromosome start end strand MM target_seq PAM alignment distance position gene name gene id
GU305914.1 225666 225688 + 0 GGATCGTTAGTGTCCGCCGC CGG ||||||||[||||||||||||]PAM NA Intergenic NA NA

T16819 CCGCCGGAGTCCTCTGTCTCTGG 1000
Oligo fwd CACCCCGCCGGAGTCCTCTGTCTC
Oligo rev AAACGAGACAGAGGACTCCGGCGG
Chromosome start end strand MM target_seq PAM alignment distance position gene name gene id
GU305914.1 225682 225704 + 0 CCGCCGGAGTCCTCTGTCTC TGG ||||||||[||||||||||||]PAM NA Intergenic NA NA

T16820 CCTCTGTCTCTGGACGCCCGCGG 1000
Oligo fwd CACCCCTCTGTCTCTGGACGCCCG
Oligo rev AAACCGGGCGTCCAGAGACAGAGG
Chromosome start end strand MM target_seq PAM alignment distance position gene name gene id
GU305914.1 225692 225714 + 0 CCTCTGTCTCTGGACGCCCG CGG ||||||||[||||||||||||]PAM NA Intergenic NA NA

T16821 CTGTCTCTGGACGCCCGCGGCGG 1000
Oligo fwd CACCCTGTCTCTGGACGCCCGCGG
Oligo rev AAACCCGCGGGCGTCCAGAGACAG
Chromosome start end strand MM target_seq PAM alignment distance position gene name gene id
GU305914.1 225695 225717 + 0 CTGTCTCTGGACGCCCGCGG CGG ||||||||[||||||||||||]PAM NA Intergenic NA NA

T16822 TGTCTCTGGACGCCCGCGGCGGG 1000
Oligo fwd CACCTGTCTCTGGACGCCCGCGGC
Oligo rev AAACGCCGCGGGCGTCCAGAGACA
Chromosome start end strand MM target_seq PAM alignment distance position gene name gene id
GU305914.1 225696 225718 + 0 TGTCTCTGGACGCCCGCGGC GGG ||||||||[||||||||||||]PAM NA Intergenic NA NA

T16823 GTCTCTGGACGCCCGCGGCGGGG 1000
Oligo fwd CACCGTCTCTGGACGCCCGCGGCG
Oligo rev AAACCGCCGCGGGCGTCCAGAGAC
Chromosome start end strand MM target_seq PAM alignment distance position gene name gene id
GU305914.1 225697 225719 + 0 GTCTCTGGACGCCCGCGGCG GGG ||||||||[||||||||||||]PAM NA Intergenic NA NA

T16824 TCTCTGGACGCCCGCGGCGGGGG 1000
Oligo fwd CACCTCTCTGGACGCCCGCGGCGG
Oligo rev AAACCCGCCGCGGGCGTCCAGAGA
Chromosome start end strand MM target_seq PAM alignment distance position gene name gene id
GU305914.1 225698 225720 + 0 TCTCTGGACGCCCGCGGCGG GGG ||||||||[||||||||||||]PAM NA Intergenic NA NA

T16825 GACGCCCGCGGCGGGGGCGCTGG 1000
Oligo fwd CACCGACGCCCGCGGCGGGGGCGC
Oligo rev AAACGCGCCCCCGCCGCGGGCGTC
Chromosome start end strand MM target_seq PAM alignment distance position gene name gene id
GU305914.1 225704 225726 + 0 GACGCCCGCGGCGGGGGCGC TGG ||||||||[||||||||||||]PAM NA Intergenic NA NA
GU305914.1 58680 58702 + 3 GACGCGCGCGGCGGAGTCGC CGG |||||-||[||||||-|-|||]PAM NA Intergenic NA NA

T16826 GGCGGGGGCGCTGGCGCTGATGG 1000
Oligo fwd CACCGGCGGGGGCGCTGGCGCTGA
Oligo rev AAACTCAGCGCCAGCGCCCCCGCC
Chromosome start end strand MM target_seq PAM alignment distance position gene name gene id
GU305914.1 225713 225735 + 0 GGCGGGGGCGCTGGCGCTGA TGG ||||||||[||||||||||||]PAM NA Intergenic NA NA
GU305914.1 66716 66738 + 3 GGCGGACGCGCTGACGCTGA TGG |||||--|[|||||-|||||||]PAM NA Intergenic NA NA

T16827 ATGGTGCCGCTCCCTCTCGAAGG 1000
Oligo fwd CACCATGGTGCCGCTCCCTCTCGA
Oligo rev AAACTCGAGAGGGAGCGGCACCAT
Chromosome start end strand MM target_seq PAM alignment distance position gene name gene id
GU305914.1 225732 225754 + 0 ATGGTGCCGCTCCCTCTCGA AGG ||||||||[||||||||||||]PAM NA Intergenic NA NA

T16828 TGGTGCCGCTCCCTCTCGAAGGG 1000
Oligo fwd CACCTGGTGCCGCTCCCTCTCGAA
Oligo rev AAACTTCGAGAGGGAGCGGCACCA
Chromosome start end strand MM target_seq PAM alignment distance position gene name gene id
GU305914.1 225733 225755 + 0 TGGTGCCGCTCCCTCTCGAA GGG ||||||||[||||||||||||]PAM NA Intergenic NA NA

T16829 CGCTCCCTCTCGAAGGGTCGTGG 1000
Oligo fwd CACCCGCTCCCTCTCGAAGGGTCG
Oligo rev AAACCGACCCTTCGAGAGGGAGCG
Chromosome start end strand MM target_seq PAM alignment distance position gene name gene id
GU305914.1 225739 225761 + 0 CGCTCCCTCTCGAAGGGTCG TGG ||||||||[||||||||||||]PAM NA Intergenic NA NA

T16830 CGAAGGGTCGTGGCGAATGCTGG 1000
Oligo fwd CACCCGAAGGGTCGTGGCGAATGC
Oligo rev AAACGCATTCGCCACGACCCTTCG
Chromosome start end strand MM target_seq PAM alignment distance position gene name gene id
GU305914.1 225749 225771 + 0 CGAAGGGTCGTGGCGAATGC TGG ||||||||[||||||||||||]PAM NA Intergenic NA NA

T16831 GCTCTTTTCTAGATCTCCTGAGG 1000

Oligo fwd CACCGCTCTTTTCTAGATCTCCTG
Oligo rev AAACCAGGAGATCTAGAAAAGAGC
Chromosome start end strand MM target_seq PAM alignment distance position gene name gene id
GU305914.1 225791 225813 + 0 GCTCTTTTCTAGATCTCCTG AGG ||||||||[||||||||||||]PAM NA Intergenic NA NA

T16832 CTCTTTTCTAGATCTCCTGAGGG 1000
Oligo fwd CACCCTCTTTTCTAGATCTCCTGA
Oligo rev AAACTCAGGAGATCTAGAAAAGAG
Chromosome start end strand MM target_seq PAM alignment distance position gene name gene id
GU305914.1 225792 225814 + 0 CTCTTTTCTAGATCTCCTGA GGG ||||||||[||||||||||||]PAM NA Intergenic NA NA

T16833 TCTTTTCTAGATCTCCTGAGGGG 1000
Oligo fwd CACCTCTTTTCTAGATCTCCTGAG
Oligo rev AAACCTCAGGAGATCTAGAAAAGA
Chromosome start end strand MM target_seq PAM alignment distance position gene name gene id
GU305914.1 225793 225815 + 0 TCTTTTCTAGATCTCCTGAG GGG ||||||||[||||||||||||]PAM NA Intergenic NA NA

T16834 CTTTTCTAGATCTCCTGAGGGGG 1000
Oligo fwd CACCCTTTTCTAGATCTCCTGAGG
Oligo rev AAACCCTCAGGAGATCTAGAAAAG
Chromosome start end strand MM target_seq PAM alignment distance position gene name gene id
GU305914.1 225794 225816 + 0 CTTTTCTAGATCTCCTGAGG GGG ||||||||[||||||||||||]PAM NA Intergenic NA NA

T16835 TTCTAGATCTCCTGAGGGGGAGG 1000
Oligo fwd CACCTTCTAGATCTCCTGAGGGGG
Oligo rev AAACCCCCCTCAGGAGATCTAGAA
Chromosome start end strand MM target_seq PAM alignment distance position gene name gene id
GU305914.1 225797 225819 + 0 TTCTAGATCTCCTGAGGGGG AGG ||||||||[||||||||||||]PAM NA Intergenic NA NA

T16836 TCTTCCGCACGCGCAGCTGCTGG 1000
Oligo fwd CACCTCTTCCGCACGCGCAGCTGC
Oligo rev AAACGCAGCTGCGCGTGCGGAAGA
Chromosome start end strand MM target_seq PAM alignment distance position gene name gene id
GU305914.1 225854 225876 + 0 TCTTCCGCACGCGCAGCTGC TGG ||||||||[||||||||||||]PAM NA Intergenic NA NA

T16837 GCGCAGCTGCTGGCCCTCAATGG 1000
Oligo fwd CACCGCGCAGCTGCTGGCCCTCAA
Oligo rev AAACTTGAGGGCCAGCAGCTGCGC
Chromosome start end strand MM target_seq PAM alignment distance position gene name gene id
GU305914.1 225864 225886 + 0 GCGCAGCTGCTGGCCCTCAA TGG ||||||||[||||||||||||]PAM NA Intergenic NA NA

T16838 GCCCTCAATGGCGACTGTTGCGG 1000
Oligo fwd CACCGCCCTCAATGGCGACTGTTG
Oligo rev AAACCAACAGTCGCCATTGAGGGC
Chromosome start end strand MM target_seq PAM alignment distance position gene name gene id
GU305914.1 225876 225898 + 0 GCCCTCAATGGCGACTGTTG CGG ||||||||[||||||||||||]PAM NA Intergenic NA NA

T16839 ACCGACCCGCGACGAGCTCGAGG 1000
Oligo fwd CACCACCGACCCGCGACGAGCTCG
Oligo rev AAACCGAGCTCGTCGCGGGTCGGT
Chromosome start end strand MM target_seq PAM alignment distance position gene name gene id
GU305914.1 225905 225927 + 0 ACCGACCCGCGACGAGCTCG AGG ||||||||[||||||||||||]PAM NA Intergenic NA NA

T16840 CGCGACGAGCTCGAGGCCCGAGG 1000
Oligo fwd CACCCGCGACGAGCTCGAGGCCCG
Oligo rev AAACCGGGCCTCGAGCTCGTCGCG
Chromosome start end strand MM target_seq PAM alignment distance position gene name gene id
GU305914.1 225912 225934 + 0 CGCGACGAGCTCGAGGCCCG AGG ||||||||[||||||||||||]PAM NA Intergenic NA NA

T16841 GCGACGAGCTCGAGGCCCGAGGG 1000
Oligo fwd CACCGCGACGAGCTCGAGGCCCGA
Oligo rev AAACTCGGGCCTCGAGCTCGTCGC
Chromosome start end strand MM target_seq PAM alignment distance position gene name gene id
GU305914.1 225913 225935 + 0 GCGACGAGCTCGAGGCCCGA GGG ||||||||[||||||||||||]PAM NA Intergenic NA NA

T16842 GAGGCCCGAGGGTCAGCGTCAGG 1000
Oligo fwd CACCGAGGCCCGAGGGTCAGCGTC
Oligo rev AAACGACGCTGACCCTCGGGCCTC
Chromosome start end strand MM target_seq PAM alignment distance position gene name gene id
GU305914.1 225924 225946 + 0 GAGGCCCGAGGGTCAGCGTC AGG ||||||||[||||||||||||]PAM NA Intergenic NA NA

T16843 GGTCAGCGTCAGGCCCGCCGTGG 1000
Oligo fwd CACCGGTCAGCGTCAGGCCCGCCG
Oligo rev AAACCGGCGGGCCTGACGCTGACC
Chromosome start end strand MM target_seq PAM alignment distance position gene name gene id
GU305914.1 225934 225956 + 0 GGTCAGCGTCAGGCCCGCCG TGG ||||||||[||||||||||||]PAM NA Intergenic NA NA

T16844 CACCGCTCCGAACAGCACTCCGG 1000
Oligo fwd CACCCACCGCTCCGAACAGCACTC
Oligo rev AAACGAGTGCTGTTCGGAGCGGTG
Chromosome start end strand MM target_seq PAM alignment distance position gene name gene id
GU305914.1 225958 225980 + 0 CACCGCTCCGAACAGCACTC CGG ||||||||[||||||||||||]PAM NA Intergenic NA NA

T16845 GCGCTCCACGCGCAGCCGTTTGG 1000
Oligo fwd CACCGCGCTCCACGCGCAGCCGTT
Oligo rev AAACAACGGCTGCGCGTGGAGCGC
Chromosome start end strand MM target_seq PAM alignment distance position gene name gene id
GU305914.1 225998 226020 + 0 GCGCTCCACGCGCAGCCGTT TGG ||||||||[||||||||||||]PAM NA Intergenic NA NA

T16846 CGCTCCACGCGCAGCCGTTTGGG 1000
Oligo fwd CACCCGCTCCACGCGCAGCCGTTT
Oligo rev AAACAAACGGCTGCGCGTGGAGCG
Chromosome start end strand MM target_seq PAM alignment distance position gene name gene id
GU305914.1 225999 226021 + 0 CGCTCCACGCGCAGCCGTTT GGG ||||||||[||||||||||||]PAM NA Intergenic NA NA

T16847 GCTCCACGCGCAGCCGTTTGGGG 1000
Oligo fwd CACCGCTCCACGCGCAGCCGTTTG
Oligo rev AAACCAAACGGCTGCGCGTGGAGC
Chromosome start end strand MM target_seq PAM alignment distance position gene name gene id
GU305914.1 226000 226022 + 0 GCTCCACGCGCAGCCGTTTG GGG ||||||||[||||||||||||]PAM NA Intergenic NA NA

T16848 GCCGTTTGGGGTCGACCGTTTGG 1000
Oligo fwd CACCGCCGTTTGGGGTCGACCGTT
Oligo rev AAACAACGGTCGACCCCAAACGGC
Chromosome start end strand MM target_seq PAM alignment distance position gene name gene id
GU305914.1 226012 226034 + 0 GCCGTTTGGGGTCGACCGTT TGG ||||||||[||||||||||||]PAM NA Intergenic NA NA

T16849 GGTCGACCGTTTGGAAACGACGG 1000
Oligo fwd CACCGGTCGACCGTTTGGAAACGA
Oligo rev AAACTCGTTTCCAAACGGTCGACC
Chromosome start end strand MM target_seq PAM alignment distance position gene name gene id
GU305914.1 226021 226043 + 0 GGTCGACCGTTTGGAAACGA CGG ||||||||[||||||||||||]PAM NA Intergenic NA NA

T16850 GTCGACCGTTTGGAAACGACGGG 1000
Oligo fwd CACCGTCGACCGTTTGGAAACGAC
Oligo rev AAACGTCGTTTCCAAACGGTCGAC
Chromosome start end strand MM target_seq PAM alignment distance position gene name gene id
GU305914.1 226022 226044 + 0 GTCGACCGTTTGGAAACGAC GGG ||||||||[||||||||||||]PAM NA Intergenic NA NA

T16851 TCGACCGTTTGGAAACGACGGGG 1000
Oligo fwd CACCTCGACCGTTTGGAAACGACG
Oligo rev AAACCGTCGTTTCCAAACGGTCGA
Chromosome start end strand MM target_seq PAM alignment distance position gene name gene id
GU305914.1 226023 226045 + 0 TCGACCGTTTGGAAACGACG GGG ||||||||[||||||||||||]PAM NA Intergenic NA NA

T16852 GGAAACGACGGGGAGTCTCCTGG 1000
Oligo fwd CACCGGAAACGACGGGGAGTCTCC
Oligo rev AAACGGAGACTCCCCGTCGTTTCC
Chromosome start end strand MM target_seq PAM alignment distance position gene name gene id
GU305914.1 226033 226055 + 0 GGAAACGACGGGGAGTCTCC TGG ||||||||[||||||||||||]PAM NA Intergenic NA NA

T16853 ACGACGGGGAGTCTCCTGGAAGG 1000
Oligo fwd CACCACGACGGGGAGTCTCCTGGA
Oligo rev AAACTCCAGGAGACTCCCCGTCGT
Chromosome start end strand MM target_seq PAM alignment distance position gene name gene id
GU305914.1 226037 226059 + 0 ACGACGGGGAGTCTCCTGGA AGG ||||||||[||||||||||||]PAM NA Intergenic NA NA

T16854 ACGGGGAGTCTCCTGGAAGGAGG 1000
Oligo fwd CACCACGGGGAGTCTCCTGGAAGG
Oligo rev AAACCCTTCCAGGAGACTCCCCGT
Chromosome start end strand MM target_seq PAM alignment distance position gene name gene id
GU305914.1 226040 226062 + 0 ACGGGGAGTCTCCTGGAAGG AGG ||||||||[||||||||||||]PAM NA Intergenic NA NA

T16855 ATGAGAACGACACGCCGTCGTGG 1000
Oligo fwd CACCATGAGAACGACACGCCGTCG
Oligo rev AAACCGACGGCGTGTCGTTCTCAT
Chromosome start end strand MM target_seq PAM alignment distance position gene name gene id
GU305914.1 226063 226085 + 0 ATGAGAACGACACGCCGTCG TGG ||||||||[||||||||||||]PAM NA Intergenic NA NA

T16856 GTGGCTACCCCCAGCAGTCTCGG 1000
Oligo fwd CACCGTGGCTACCCCCAGCAGTCT
Oligo rev AAACAGACTGCTGGGGGTAGCCAC
Chromosome start end strand MM target_seq PAM alignment distance position gene name gene id
GU305914.1 226082 226104 + 0 GTGGCTACCCCCAGCAGTCT CGG ||||||||[||||||||||||]PAM NA Intergenic NA NA

T16857 CTACCCCCAGCAGTCTCGGTTGG 1000
Oligo fwd CACCCTACCCCCAGCAGTCTCGGT
Oligo rev AAACACCGAGACTGCTGGGGGTAG
Chromosome start end strand MM target_seq PAM alignment distance position gene name gene id
GU305914.1 226086 226108 + 0 CTACCCCCAGCAGTCTCGGT TGG ||||||||[||||||||||||]PAM NA Intergenic NA NA

T16858 TGACCGTGCATGAAACGCAGAGG 1000
Oligo fwd CACCTGACCGTGCATGAAACGCAG
Oligo rev AAACCTGCGTTTCATGCACGGTCA
Chromosome start end strand MM target_seq PAM alignment distance position gene name gene id
GU305914.1 226110 226132 + 0 TGACCGTGCATGAAACGCAG AGG ||||||||[||||||||||||]PAM NA Intergenic NA NA

T16859 GACCGTGCATGAAACGCAGAGGG 1000
Oligo fwd CACCGACCGTGCATGAAACGCAGA
Oligo rev AAACTCTGCGTTTCATGCACGGTC
Chromosome start end strand MM target_seq PAM alignment distance position gene name gene id
GU305914.1 226111 226133 + 0 GACCGTGCATGAAACGCAGA GGG ||||||||[||||||||||||]PAM NA Intergenic NA NA

T16860 TGAAACGCAGAGGGTCCCCGCGG 1000
Oligo fwd CACCTGAAACGCAGAGGGTCCCCG
Oligo rev AAACCGGGGACCCTCTGCGTTTCA
Chromosome start end strand MM target_seq PAM alignment distance position gene name gene id
GU305914.1 226120 226142 + 0 TGAAACGCAGAGGGTCCCCG CGG ||||||||[||||||||||||]PAM NA Intergenic NA NA

<table>
<tr><td>T16861</td><td>GCAGAGGGTCCCCGCGGCCCAGG</td><td>1000</td><td></td><td></td><td></td><td></td><td></td><td></td><td></td><td></td><td></td></tr>
<tr><td>Oligo fwd</td><td>CACCGCAGAGGGTCCCCGCGGCCC</td><td></td><td></td><td></td><td></td><td></td><td></td><td></td><td></td><td></td><td></td></tr>
<tr><td>Oligo rev</td><td>AAACGGGCCGCGGGGACCCTCTGC</td><td></td><td></td><td></td><td></td><td></td><td></td><td></td><td></td><td></td><td></td></tr>
<tr><td>Chromosome</td><td>start</td><td>end</td><td>strand</td><td>MM</td><td>target_seq</td><td>PAM</td><td>alignment</td><td>distance</td><td>position</td><td>gene name</td><td>gene id</td></tr>
<tr><td>GU305914.1</td><td>226126</td><td>226148</td><td>+</td><td>0</td><td>GCAGAGGGTCCCCGCGGCCC</td><td>AGG</td><td>||||||||[||||||||||||]PAM</td><td>NA</td><td>Intergenic</td><td>NA</td><td>NA</td></tr>
<tr><td>T16862</td><td>GGGTCCCCGCGGCCCAGGCCTGG</td><td>1000</td><td></td><td></td><td></td><td></td><td></td><td></td><td></td><td></td><td></td></tr>
<tr><td>Oligo fwd</td><td>CACCGGGTCCCCGCGGCCCAGGCC</td><td></td><td></td><td></td><td></td><td></td><td></td><td></td><td></td><td></td><td></td></tr>
<tr><td>Oligo rev</td><td>AAACGGCCTGGGCCGCGGGGACCC</td><td></td><td></td><td></td><td></td><td></td><td></td><td></td><td></td><td></td><td></td></tr>
<tr><td>Chromosome</td><td>start</td><td>end</td><td>strand</td><td>MM</td><td>target_seq</td><td>PAM</td><td>alignment</td><td>distance</td><td>position</td><td>gene name</td><td>gene id</td></tr>
<tr><td>GU305914.1</td><td>226131</td><td>226153</td><td>+</td><td>0</td><td>GGGTCCCCGCGGCCCAGGCC</td><td>TGG</td><td>||||||||[||||||||||||]PAM</td><td>NA</td><td>Intergenic</td><td>NA</td><td>NA</td></tr>
<tr><td>T16863</td><td>ACCGTTTCGATCTCGAACCACGG</td><td>1000</td><td></td><td></td><td></td><td></td><td></td><td></td><td></td><td></td><td></td></tr>
<tr><td>Oligo fwd</td><td>CACCACCGTTTCGATCTCGAACCA</td><td></td><td></td><td></td><td></td><td></td><td></td><td></td><td></td><td></td><td></td></tr>
<tr><td>Oligo rev</td><td>AAACTGGTTCGAGATCGAAACGGT</td><td></td><td></td><td></td><td></td><td></td><td></td><td></td><td></td><td></td><td></td></tr>
<tr><td>Chromosome</td><td>start</td><td>end</td><td>strand</td><td>MM</td><td>target_seq</td><td>PAM</td><td>alignment</td><td>distance</td><td>position</td><td>gene name</td><td>gene id</td></tr>
<tr><td>GU305914.1</td><td>226178</td><td>226200</td><td>+</td><td>0</td><td>ACCGTTTCGATCTCGAACCA</td><td>CGG</td><td>||||||||[||||||||||||]PAM</td><td>NA</td><td>Intergenic</td><td>NA</td><td>NA</td></tr>
<tr><td>T16864</td><td>CCGTTTCGATCTCGAACCACGGG</td><td>1000</td><td></td><td></td><td></td><td></td><td></td><td></td><td></td><td></td><td></td></tr>
<tr><td>Oligo fwd</td><td>CACCCCGTTTCGATCTCGAACCAC</td><td></td><td></td><td></td><td></td><td></td><td></td><td></td><td></td><td></td><td></td></tr>
<tr><td>Oligo rev</td><td>AAACGTGGTTCGAGATCGAAACGG</td><td></td><td></td><td></td><td></td><td></td><td></td><td></td><td></td><td></td><td></td></tr>
<tr><td>Chromosome</td><td>start</td><td>end</td><td>strand</td><td>MM</td><td>target_seq</td><td>PAM</td><td>alignment</td><td>distance</td><td>position</td><td>gene name</td><td>gene id</td></tr>
<tr><td>GU305914.1</td><td>226179</td><td>226201</td><td>+</td><td>0</td><td>CCGTTTCGATCTCGAACCAC</td><td>GGG</td><td>||||||||[||||||||||||]PAM</td><td>NA</td><td>Intergenic</td><td>NA</td><td>NA</td></tr>
<tr><td>T16865</td><td>TTTCGATCTCGAACCACGGGCGG</td><td>1000</td><td></td><td></td><td></td><td></td><td></td><td></td><td></td><td></td><td></td></tr>
<tr><td>Oligo fwd</td><td>CACCTTTCGATCTCGAACCACGGG</td><td></td><td></td><td></td><td></td><td></td><td></td><td></td><td></td><td></td><td></td></tr>
<tr><td>Oligo rev</td><td>AAACCCCGTGGTTCGAGATCGAAA</td><td></td><td></td><td></td><td></td><td></td><td></td><td></td><td></td><td></td><td></td></tr>
<tr><td>Chromosome</td><td>start</td><td>end</td><td>strand</td><td>MM</td><td>target_seq</td><td>PAM</td><td>alignment</td><td>distance</td><td>position</td><td>gene name</td><td>gene id</td></tr>
<tr><td>GU305914.1</td><td>226182</td><td>226204</td><td>+</td><td>0</td><td>TTTCGATCTCGAACCACGGG</td><td>CGG</td><td>||||||||[||||||||||||]PAM</td><td>NA</td><td>Intergenic</td><td>NA</td><td>NA</td></tr>
<tr><td>T16866</td><td>CGATCTCGAACCACGGGCGGAGG</td><td>1000</td><td></td><td></td><td></td><td></td><td></td><td></td><td></td><td></td><td></td></tr>
<tr><td>Oligo fwd</td><td>CACCCGATCTCGAACCACGGGCGG</td><td></td><td></td><td></td><td></td><td></td><td></td><td></td><td></td><td></td><td></td></tr>
<tr><td>Oligo rev</td><td>AAACCCGCCCGTGGTTCGAGATCG</td><td></td><td></td><td></td><td></td><td></td><td></td><td></td><td></td><td></td><td></td></tr>
<tr><td>Chromosome</td><td>start</td><td>end</td><td>strand</td><td>MM</td><td>target_seq</td><td>PAM</td><td>alignment</td><td>distance</td><td>position</td><td>gene name</td><td>gene id</td></tr>
<tr><td>GU305914.1</td><td>226185</td><td>226207</td><td>+</td><td>0</td><td>CGATCTCGAACCACGGGCGG</td><td>AGG</td><td>||||||||[||||||||||||]PAM</td><td>NA</td><td>Intergenic</td><td>NA</td><td>NA</td></tr>
<tr><td>T16867</td><td>TCGAACCACGGGCGGAGGTCCGG</td><td>1000</td><td></td><td></td><td></td><td></td><td></td><td></td><td></td><td></td><td></td></tr>
<tr><td>Oligo fwd</td><td>CACCTCGAACCACGGGCGGAGGTC</td><td></td><td></td><td></td><td></td><td></td><td></td><td></td><td></td><td></td><td></td></tr>
<tr><td>Oligo rev</td><td>AAACGACCTCCGCCCGTGGTTCGA</td><td></td><td></td><td></td><td></td><td></td><td></td><td></td><td></td><td></td><td></td></tr>
<tr><td>Chromosome</td><td>start</td><td>end</td><td>strand</td><td>MM</td><td>target_seq</td><td>PAM</td><td>alignment</td><td>distance</td><td>position</td><td>gene name</td><td>gene id</td></tr>
<tr><td>GU305914.1</td><td>226190</td><td>226212</td><td>+</td><td>0</td><td>TCGAACCACGGGCGGAGGTC</td><td>CGG</td><td>||||||||[||||||||||||]PAM</td><td>NA</td><td>Intergenic</td><td>NA</td><td>NA</td></tr>
<tr><td>T16868</td><td>CACGGGCGGAGGTCCGGACTTGG</td><td>1000</td><td></td><td></td><td></td><td></td><td></td><td></td><td></td><td></td><td></td></tr>
<tr><td>Oligo fwd</td><td>CACCCACGGGCGGAGGTCCGGACT</td><td></td><td></td><td></td><td></td><td></td><td></td><td></td><td></td><td></td><td></td></tr>
<tr><td>Oligo rev</td><td>AAACAGTCCGGACCTCCGCCCGTG</td><td></td><td></td><td></td><td></td><td></td><td></td><td></td><td></td><td></td><td></td></tr>
<tr><td>Chromosome</td><td>start</td><td>end</td><td>strand</td><td>MM</td><td>target_seq</td><td>PAM</td><td>alignment</td><td>distance</td><td>position</td><td>gene name</td><td>gene id</td></tr>
<tr><td>GU305914.1</td><td>226196</td><td>226218</td><td>+</td><td>0</td><td>CACGGGCGGAGGTCCGGACT</td><td>TGG</td><td>||||||||[||||||||||||]PAM</td><td>NA</td><td>Intergenic</td><td>NA</td><td>NA</td></tr>
<tr><td>GU305914.1</td><td>89992</td><td>90014</td><td>-</td><td>4</td><td>GACGTGCGGAGGTCCGGGCA</td><td>CGG</td><td>-|||-|||[|||||||||-|-]PAM</td><td>NA</td><td>Intergenic</td><td>NA</td><td>NA</td></tr>
<tr><td>T16869</td><td>GGAGGTCCGGACTTGGTCTTCGG</td><td>1000</td><td></td><td></td><td></td><td></td><td></td><td></td><td></td><td></td><td></td></tr>
<tr><td>Oligo fwd</td><td>CACCGGAGGTCCGGACTTGGTCTT</td><td></td><td></td><td></td><td></td><td></td><td></td><td></td><td></td><td></td><td></td></tr>
<tr><td>Oligo rev</td><td>AAACAAGACCAAGTCCGGACCTCC</td><td></td><td></td><td></td><td></td><td></td><td></td><td></td><td></td><td></td><td></td></tr>
<tr><td>Chromosome</td><td>start</td><td>end</td><td>strand</td><td>MM</td><td>target_seq</td><td>PAM</td><td>alignment</td><td>distance</td><td>position</td><td>gene name</td><td>gene id</td></tr>
<tr><td>GU305914.1</td><td>226203</td><td>226225</td><td>+</td><td>0</td><td>GGAGGTCCGGACTTGGTCTT</td><td>CGG</td><td>||||||||[||||||||||||]PAM</td><td>NA</td><td>Intergenic</td><td>NA</td><td>NA</td></tr>
<tr><td>T16870</td><td>GAGGTCCGGACTTGGTCTTCGGG</td><td>1000</td><td></td><td></td><td></td><td></td><td></td><td></td><td></td><td></td><td></td></tr>
<tr><td>Oligo fwd</td><td>CACCGAGGTCCGGACTTGGTCTTC</td><td></td><td></td><td></td><td></td><td></td><td></td><td></td><td></td><td></td><td></td></tr>
<tr><td>Oligo rev</td><td>AAACGAAGACCAAGTCCGGACCTC</td><td></td><td></td><td></td><td></td><td></td><td></td><td></td><td></td><td></td><td></td></tr>
<tr><td>Chromosome</td><td>start</td><td>end</td><td>strand</td><td>MM</td><td>target_seq</td><td>PAM</td><td>alignment</td><td>distance</td><td>position</td><td>gene name</td><td>gene id</td></tr>
<tr><td>GU305914.1</td><td>226204</td><td>226226</td><td>+</td><td>0</td><td>GAGGTCCGGACTTGGTCTTC</td><td>GGG</td><td>||||||||[||||||||||||]PAM</td><td>NA</td><td>Intergenic</td><td>NA</td><td>NA</td></tr>
<tr><td>T16871</td><td>GGACTTGGTCTTCGGGTCTGAGG</td><td>1000</td><td></td><td></td><td></td><td></td><td></td><td></td><td></td><td></td><td></td></tr>
<tr><td>Oligo fwd</td><td>CACCGGACTTGGTCTTCGGGTCTG</td><td></td><td></td><td></td><td></td><td></td><td></td><td></td><td></td><td></td><td></td></tr>
<tr><td>Oligo rev</td><td>AAACCAGACCCGAAGACCAAGTCC</td><td></td><td></td><td></td><td></td><td></td><td></td><td></td><td></td><td></td><td></td></tr>
<tr><td>Chromosome</td><td>start</td><td>end</td><td>strand</td><td>MM</td><td>target_seq</td><td>PAM</td><td>alignment</td><td>distance</td><td>position</td><td>gene name</td><td>gene id</td></tr>
<tr><td>GU305914.1</td><td>226211</td><td>226233</td><td>+</td><td>0</td><td>GGACTTGGTCTTCGGGTCTG</td><td>AGG</td><td>||||||||[||||||||||||]PAM</td><td>NA</td><td>Intergenic</td><td>NA</td><td>NA</td></tr>
<tr><td>T16872</td><td>GACTTGGTCTTCGGGTCTGAGGG</td><td>1000</td><td></td><td></td><td></td><td></td><td></td><td></td><td></td><td></td><td></td></tr>
<tr><td>Oligo fwd</td><td>CACCGACTTGGTCTTCGGGTCTGA</td><td></td><td></td><td></td><td></td><td></td><td></td><td></td><td></td><td></td><td></td></tr>
<tr><td>Oligo rev</td><td>AAACTCAGACCCGAAGACCAAGTC</td><td></td><td></td><td></td><td></td><td></td><td></td><td></td><td></td><td></td><td></td></tr>
<tr><td>Chromosome</td><td>start</td><td>end</td><td>strand</td><td>MM</td><td>target_seq</td><td>PAM</td><td>alignment</td><td>distance</td><td>position</td><td>gene name</td><td>gene id</td></tr>
<tr><td>GU305914.1</td><td>226212</td><td>226234</td><td>+</td><td>0</td><td>GACTTGGTCTTCGGGTCTGA</td><td>GGG</td><td>||||||||[||||||||||||]PAM</td><td>NA</td><td>Intergenic</td><td>NA</td><td>NA</td></tr>
<tr><td>T16873</td><td>TGGTCTTCGGGTCTGAGGGACGG</td><td>1000</td><td></td><td></td><td></td><td></td><td></td><td></td><td></td><td></td><td></td></tr>
<tr><td>Oligo fwd</td><td>CACCTGGTCTTCGGGTCTGAGGGA</td><td></td><td></td><td></td><td></td><td></td><td></td><td></td><td></td><td></td><td></td></tr>
<tr><td>Oligo rev</td><td>AAACTCCCTCAGACCCGAAGACCA</td><td></td><td></td><td></td><td></td><td></td><td></td><td></td><td></td><td></td><td></td></tr>
<tr><td>Chromosome</td><td>start</td><td>end</td><td>strand</td><td>MM</td><td>target_seq</td><td>PAM</td><td>alignment</td><td>distance</td><td>position</td><td>gene name</td><td>gene id</td></tr>
<tr><td>GU305914.1</td><td>226216</td><td>226238</td><td>+</td><td>0</td><td>TGGTCTTCGGGTCTGAGGGA</td><td>CGG</td><td>||||||||[||||||||||||]PAM</td><td>NA</td><td>Intergenic</td><td>NA</td><td>NA</td></tr>
<tr><td>T16874</td><td>TCTTCGGGTCTGAGGGACGGTGG</td><td>1000</td><td></td><td></td><td></td><td></td><td></td><td></td><td></td><td></td><td></td></tr>
<tr><td>Oligo fwd</td><td>CACCTCTTCGGGTCTGAGGGACGG</td><td></td><td></td><td></td><td></td><td></td><td></td><td></td><td></td><td></td><td></td></tr>
<tr><td>Oligo rev</td><td>AAACCCGTCCCTCAGACCCGAAGA</td><td></td><td></td><td></td><td></td><td></td><td></td><td></td><td></td><td></td><td></td></tr>
<tr><td>Chromosome</td><td>start</td><td>end</td><td>strand</td><td>MM</td><td>target_seq</td><td>PAM</td><td>alignment</td><td>distance</td><td>position</td><td>gene name</td><td>gene id</td></tr>
<tr><td>GU305914.1</td><td>226219</td><td>226241</td><td>+</td><td>0</td><td>TCTTCGGGTCTGAGGGACGG</td><td>TGG</td><td>||||||||[||||||||||||]PAM</td><td>NA</td><td>Intergenic</td><td>NA</td><td>NA</td></tr>
<tr><td>T16875</td><td>CCGCTTCCCGTCGTCTCCCGCGG</td><td>1000</td><td></td><td></td><td></td><td></td><td></td><td></td><td></td><td></td><td></td></tr>
<tr><td>Oligo fwd</td><td>CACCCCGCTTCCCGTCGTCTCCCG</td><td></td><td></td><td></td><td></td><td></td><td></td><td></td><td></td><td></td><td></td></tr>
<tr><td>Oligo rev</td><td>AAACCGGGAGACGACGGGAAGCGG</td><td></td><td></td><td></td><td></td><td></td><td></td><td></td><td></td><td></td><td></td></tr>
</table>

| | | | | | | | | | | | |
|---|---|---|---|---|---|---|---|---|---|---|---|
| Chromosome | start | end | strand | MM | target_seq | PAM | alignment | distance | position | gene name | gene id |
| GU305914.1 | 226256 | 226278 | + | 0 | CCGCTTCCCGTCGTCTCCCG | CGG | \|\|\|\|\|\|\|\|[\|\|\|\|\|\|\|\|\|\|\|\|]PAM | NA | Intergenic | NA | NA |
| T16876 | CTTCCCGTCGTCTCCCGCGGCGG | 1000 | | | | | | | | | |
| Oligo fwd | CACCCTTCCCGTCGTCTCCCGCGG | | | | | | | | | | |
| Oligo rev | AAACCCGCGGGAGACGACGGGAAG | | | | | | | | | | |
| Chromosome | start | end | strand | MM | target_seq | PAM | alignment | distance | position | gene name | gene id |
| GU305914.1 | 226259 | 226281 | + | 0 | CTTCCCGTCGTCTCCCGCGG | CGG | \|\|\|\|\|\|\|\|[\|\|\|\|\|\|\|\|\|\|\|\|]PAM | NA | Intergenic | NA | NA |
| T16877 | TTCCCGTCGTCTCCCGCGGCGGG | 1000 | | | | | | | | | |
| Oligo fwd | CACCTTCCCGTCGTCTCCCGCGGC | | | | | | | | | | |
| Oligo rev | AAACGCCGCGGGAGACGACGGGAA | | | | | | | | | | |
| Chromosome | start | end | strand | MM | target_seq | PAM | alignment | distance | position | gene name | gene id |
| GU305914.1 | 226260 | 226282 | + | 0 | TTCCCGTCGTCTCCCGCGGC | GGG | \|\|\|\|\|\|\|\|[\|\|\|\|\|\|\|\|\|\|\|\|]PAM | NA | Intergenic | NA | NA |
| T16878 | TACGAGAAGCTGCTTGAACCAGG | 1000 | | | | | | | | | |
| Oligo fwd | CACCTACGAGAAGCTGCTTGAACC | | | | | | | | | | |
| Oligo rev | AAACGGTTCAAGCAGCTTCTCGTA | | | | | | | | | | |
| Chromosome | start | end | strand | MM | target_seq | PAM | alignment | distance | position | gene name | gene id |
| GU305914.1 | 226292 | 226314 | + | 0 | TACGAGAAGCTGCTTGAACC | AGG | \|\|\|\|\|\|\|\|[\|\|\|\|\|\|\|\|\|\|\|\|]PAM | NA | Intergenic | NA | NA |
| T16879 | GCTGCTTGAACCAGGTCCCGAGG | 1000 | | | | | | | | | |
| Oligo fwd | CACCGCTGCTTGAACCAGGTCCCG | | | | | | | | | | |
| Oligo rev | AAACCGGGACCTGGTTCAAGCAGC | | | | | | | | | | |
| Chromosome | start | end | strand | MM | target_seq | PAM | alignment | distance | position | gene name | gene id |
| GU305914.1 | 226300 | 226322 | + | 0 | GCTGCTTGAACCAGGTCCCG | AGG | \|\|\|\|\|\|\|\|[\|\|\|\|\|\|\|\|\|\|\|\|]PAM | NA | Intergenic | NA | NA |
| T16880 | TCGCCGTTCTGTAGATCGACAGG | 1000 | | | | | | | | | |
| Oligo fwd | CACCTCGCCGTTCTGTAGATCGAC | | | | | | | | | | |
| Oligo rev | AAACGTCGATCTACAGAACGGCGA | | | | | | | | | | |
| Chromosome | start | end | strand | MM | target_seq | PAM | alignment | distance | position | gene name | gene id |
| GU305914.1 | 226329 | 226351 | + | 0 | TCGCCGTTCTGTAGATCGAC | AGG | \|\|\|\|\|\|\|\|[\|\|\|\|\|\|\|\|\|\|\|\|]PAM | NA | Intergenic | NA | NA |
| T16881 | CGCCGTTCTGTAGATCGACAGGG | 1000 | | | | | | | | | |
| Oligo fwd | CACCCGCCGTTCTGTAGATCGACA | | | | | | | | | | |
| Oligo rev | AAACTGTCGATCTACAGAACGGCG | | | | | | | | | | |
| Chromosome | start | end | strand | MM | target_seq | PAM | alignment | distance | position | gene name | gene id |
| GU305914.1 | 226330 | 226352 | + | 0 | CGCCGTTCTGTAGATCGACA | GGG | \|\|\|\|\|\|\|\|[\|\|\|\|\|\|\|\|\|\|\|\|]PAM | NA | Intergenic | NA | NA |
| T16882 | CTGTAGATCGACAGGGTCTGAGG | 1000 | | | | | | | | | |
| Oligo fwd | CACCCTGTAGATCGACAGGGTCTG | | | | | | | | | | |
| Oligo rev | AAACCAGACCCTGTCGATCTACAG | | | | | | | | | | |
| Chromosome | start | end | strand | MM | target_seq | PAM | alignment | distance | position | gene name | gene id |
| GU305914.1 | 226337 | 226359 | + | 0 | CTGTAGATCGACAGGGTCTG | AGG | \|\|\|\|\|\|\|\|[\|\|\|\|\|\|\|\|\|\|\|\|]PAM | NA | Intergenic | NA | NA |
| T16883 | CGACAGGGTCTGAGGTCCCGAGG | 1000 | | | | | | | | | |
| Oligo fwd | CACCCGACAGGGTCTGAGGTCCCG | | | | | | | | | | |
| Oligo rev | AAACCGGGACCTCAGACCCTGTCG | | | | | | | | | | |
| Chromosome | start | end | strand | MM | target_seq | PAM | alignment | distance | position | gene name | gene id |
| GU305914.1 | 226345 | 226367 | + | 0 | CGACAGGGTCTGAGGTCCCG | AGG | \|\|\|\|\|\|\|\|[\|\|\|\|\|\|\|\|\|\|\|\|]PAM | NA | Intergenic | NA | NA |
| T16884 | GGTCTGAGGTCCCGAGGACTCGG | 1000 | | | | | | | | | |
| Oligo fwd | CACCGGTCTGAGGTCCCGAGGACT | | | | | | | | | | |
| Oligo rev | AAACAGTCCTCGGGACCTCAGACC | | | | | | | | | | |
| Chromosome | start | end | strand | MM | target_seq | PAM | alignment | distance | position | gene name | gene id |
| GU305914.1 | 226351 | 226373 | + | 0 | GGTCTGAGGTCCCGAGGACT | CGG | \|\|\|\|\|\|\|\|[\|\|\|\|\|\|\|\|\|\|\|\|]PAM | NA | Intergenic | NA | NA |
| T16885 | GTCTGAGGTCCCGAGGACTCGGG | 1000 | | | | | | | | | |
| Oligo fwd | CACCGTCTGAGGTCCCGAGGACTC | | | | | | | | | | |
| Oligo rev | AAACGAGTCCTCGGGACCTCAGAC | | | | | | | | | | |
| Chromosome | start | end | strand | MM | target_seq | PAM | alignment | distance | position | gene name | gene id |
| GU305914.1 | 226352 | 226374 | + | 0 | GTCTGAGGTCCCGAGGACTC | GGG | \|\|\|\|\|\|\|\|[\|\|\|\|\|\|\|\|\|\|\|\|]PAM | NA | Intergenic | NA | NA |
| T16886 | TGAGGTCCCGAGGACTCGGGCGG | 1000 | | | | | | | | | |
| Oligo fwd | CACCTGAGGTCCCGAGGACTCGGG | | | | | | | | | | |
| Oligo rev | AAACCCCGAGTCCTCGGGACCTCA | | | | | | | | | | |
| Chromosome | start | end | strand | MM | target_seq | PAM | alignment | distance | position | gene name | gene id |
| GU305914.1 | 226355 | 226377 | + | 0 | TGAGGTCCCGAGGACTCGGG | CGG | \|\|\|\|\|\|\|\|[\|\|\|\|\|\|\|\|\|\|\|\|]PAM | NA | Intergenic | NA | NA |
| T16887 | CACTGACCCGTTTAACCACCAGG | 1000 | | | | | | | | | |
| Oligo fwd | CACCCACTGACCCGTTTAACCACC | | | | | | | | | | |
| Oligo rev | AAACGGTGGTTAAACGGGTCAGTG | | | | | | | | | | |
| Chromosome | start | end | strand | MM | target_seq | PAM | alignment | distance | position | gene name | gene id |
| GU305914.1 | 226396 | 226418 | + | 0 | CACTGACCCGTTTAACCACC | AGG | \|\|\|\|\|\|\|\|[\|\|\|\|\|\|\|\|\|\|\|\|]PAM | NA | Intergenic | NA | NA |
| T16888 | AACCACCAGGCCAAGCCGTACGG | 1000 | | | | | | | | | |
| Oligo fwd | CACCAACCACCAGGCCAAGCCGTA | | | | | | | | | | |
| Oligo rev | AAACTACGGCTTGGCCTGGTGGTT | | | | | | | | | | |
| Chromosome | start | end | strand | MM | target_seq | PAM | alignment | distance | position | gene name | gene id |
| GU305914.1 | 226409 | 226431 | + | 0 | AACCACCAGGCCAAGCCGTA | CGG | \|\|\|\|\|\|\|\|[\|\|\|\|\|\|\|\|\|\|\|\|]PAM | NA | Intergenic | NA | NA |
| T16889 | GAGCCGAACGATTCGAAAGCAGG | 1000 | | | | | | | | | |
| Oligo fwd | CACCGAGCCGAACGATTCGAAAGC | | | | | | | | | | |
| Oligo rev | AAACGCTTTCGAATCGTTCGGCTC | | | | | | | | | | |
| Chromosome | start | end | strand | MM | target_seq | PAM | alignment | distance | position | gene name | gene id |
| GU305914.1 | 226431 | 226453 | + | 0 | GAGCCGAACGATTCGAAAGC | AGG | \|\|\|\|\|\|\|\|[\|\|\|\|\|\|\|\|\|\|\|\|]PAM | NA | Intergenic | NA | NA |
| T16890 | CCGAACGATTCGAAAGCAGGCGG | 1000 | | | | | | | | | |
| Oligo fwd | CACCCCGAACGATTCGAAAGCAGG | | | | | | | | | | |

| | | | | | | | | | | | | |
|---|---|---|---|---|---|---|---|---|---|---|---|---|
| Oligo rev | AAACCCTGCTTTCGAATCGTTCGG | | | | | | | | | | | |
| Chromosome | start | | end | strand | MM | target_seq | PAM | alignment | distance | position | gene name | gene id |
| GU305914.1 | | 226434 | 226456 | + | 0 | CCGAACGATTCGAAAGCAGG | CGG | \|\|\|\|\|\|\|\|[\|\|\|\|\|\|\|\|\|\|\|\|]PAM | NA | Intergenic | NA | NA |
| T16891 | CGAACGATTCGAAAGCAGGCGGG | | 1000 | | | | | | | | | |
| Oligo fwd | CACCCGAACGATTCGAAAGCAGGC | | | | | | | | | | | |
| Oligo rev | AAACGCCTGCTTTCGAATCGTTCG | | | | | | | | | | | |
| Chromosome | start | | end | strand | MM | target_seq | PAM | alignment | distance | position | gene name | gene id |
| GU305914.1 | | 226435 | 226457 | + | 0 | CGAACGATTCGAAAGCAGGC | GGG | \|\|\|\|\|\|\|\|[\|\|\|\|\|\|\|\|\|\|\|\|]PAM | NA | Intergenic | NA | NA |
| T16892 | GAACGATTCGAAAGCAGGCGGGG | | 1000 | | | | | | | | | |
| Oligo fwd | CACCGAACGATTCGAAAGCAGGCG | | | | | | | | | | | |
| Oligo rev | AAACCGCCTGCTTTCGAATCGTTC | | | | | | | | | | | |
| Chromosome | start | | end | strand | MM | target_seq | PAM | alignment | distance | position | gene name | gene id |
| GU305914.1 | | 226436 | 226458 | + | 0 | GAACGATTCGAAAGCAGGCG | GGG | \|\|\|\|\|\|\|\|[\|\|\|\|\|\|\|\|\|\|\|\|]PAM | NA | Intergenic | NA | NA |
| T16893 | AACGATTCGAAAGCAGGCGGGGG | | 1000 | | | | | | | | | |
| Oligo fwd | CACCAACGATTCGAAAGCAGGCGG | | | | | | | | | | | |
| Oligo rev | AAACCCGCCTGCTTTCGAATCGTT | | | | | | | | | | | |
| Chromosome | start | | end | strand | MM | target_seq | PAM | alignment | distance | position | gene name | gene id |
| GU305914.1 | | 226437 | 226459 | + | 0 | AACGATTCGAAAGCAGGCGG | GGG | \|\|\|\|\|\|\|\|[\|\|\|\|\|\|\|\|\|\|\|\|]PAM | NA | Intergenic | NA | NA |
| T16894 | AGCAGGCGGGGGTCGCCGCGTGG | | 1000 | | | | | | | | | |
| Oligo fwd | CACCAGCAGGCGGGGGTCGCCGCG | | | | | | | | | | | |
| Oligo rev | AAACCGCGGCGACCCCCGCCTGCT | | | | | | | | | | | |
| Chromosome | start | | end | strand | MM | target_seq | PAM | alignment | distance | position | gene name | gene id |
| GU305914.1 | | 226448 | 226470 | + | 0 | AGCAGGCGGGGGTCGCCGCG | TGG | \|\|\|\|\|\|\|\|[\|\|\|\|\|\|\|\|\|\|\|\|]PAM | NA | Intergenic | NA | NA |
| T16895 | AGGCGGGGGTCGCCGCGTGGAGG | | 1000 | | | | | | | | | |
| Oligo fwd | CACCAGGCGGGGGTCGCCGCGTGG | | | | | | | | | | | |
| Oligo rev | AAACCCACGCGGCGACCCCCGCCT | | | | | | | | | | | |
| Chromosome | start | | end | strand | MM | target_seq | PAM | alignment | distance | position | gene name | gene id |
| GU305914.1 | | 226451 | 226473 | + | 0 | AGGCGGGGGTCGCCGCGTGG | AGG | \|\|\|\|\|\|\|\|[\|\|\|\|\|\|\|\|\|\|\|\|]PAM | NA | Intergenic | NA | NA |
| T16896 | GGGGGTCGCCGCGTGGAGGCTGG | | 1000 | | | | | | | | | |
| Oligo fwd | CACCGGGGGTCGCCGCGTGGAGGC | | | | | | | | | | | |
| Oligo rev | AAACGCCTCCACGCGGCGACCCCC | | | | | | | | | | | |
| Chromosome | start | | end | strand | MM | target_seq | PAM | alignment | distance | position | gene name | gene id |
| GU305914.1 | | 226455 | 226477 | + | 0 | GGGGGTCGCCGCGTGGAGGC | TGG | \|\|\|\|\|\|\|\|[\|\|\|\|\|\|\|\|\|\|\|\|]PAM | NA | Intergenic | NA | NA |
| T16897 | CATCGCACAGCCCAGCTTTCTGG | | 1000 | | | | | | | | | |
| Oligo fwd | CACCCATCGCACAGCCCAGCTTTC | | | | | | | | | | | |
| Oligo rev | AAACGAAAGCTGGGCTGTGCGATG | | | | | | | | | | | |
| Chromosome | start | | end | strand | MM | target_seq | PAM | alignment | distance | position | gene name | gene id |
| GU305914.1 | | 226478 | 226500 | + | 0 | CATCGCACAGCCCAGCTTTC | TGG | \|\|\|\|\|\|\|\|[\|\|\|\|\|\|\|\|\|\|\|\|]PAM | NA | Intergenic | NA | NA |
| T16898 | CGCTTCTTCCCAGCTTAACGAGG | | 1000 | | | | | | | | | |
| Oligo fwd | CACCCGCTTCTTCCCAGCTTAACG | | | | | | | | | | | |
| Oligo rev | AAACCGTTAAGCTGGGAAGAAGCG | | | | | | | | | | | |
| Chromosome | start | | end | strand | MM | target_seq | PAM | alignment | distance | position | gene name | gene id |
| GU305914.1 | | 226504 | 226526 | + | 0 | CGCTTCTTCCCAGCTTAACG | AGG | \|\|\|\|\|\|\|\|[\|\|\|\|\|\|\|\|\|\|\|\|]PAM | NA | Intergenic | NA | NA |
| T16899 | GCTTCTTCCCAGCTTAACGAGGG | | 1000 | | | | | | | | | |
| Oligo fwd | CACCGCTTCTTCCCAGCTTAACGA | | | | | | | | | | | |
| Oligo rev | AAACTCGTTAAGCTGGGAAGAAGC | | | | | | | | | | | |
| Chromosome | start | | end | strand | MM | target_seq | PAM | alignment | distance | position | gene name | gene id |
| GU305914.1 | | 226505 | 226527 | + | 0 | GCTTCTTCCCAGCTTAACGA | GGG | \|\|\|\|\|\|\|\|[\|\|\|\|\|\|\|\|\|\|\|\|]PAM | NA | Intergenic | NA | NA |
| T16900 | TTCCCAGCTTAACGAGGGCGCGG | | 1000 | | | | | | | | | |
| Oligo fwd | CACCTTCCCAGCTTAACGAGGGCG | | | | | | | | | | | |
| Oligo rev | AAACCGCCCTCGTTAAGCTGGGAA | | | | | | | | | | | |
| Chromosome | start | | end | strand | MM | target_seq | PAM | alignment | distance | position | gene name | gene id |
| GU305914.1 | | 226510 | 226532 | + | 0 | TTCCCAGCTTAACGAGGGCG | CGG | \|\|\|\|\|\|\|\|[\|\|\|\|\|\|\|\|\|\|\|\|]PAM | NA | Intergenic | NA | NA |
| T16901 | TCCCAGCTTAACGAGGGCGCGGG | | 1000 | | | | | | | | | |
| Oligo fwd | CACCTCCCAGCTTAACGAGGGCGC | | | | | | | | | | | |
| Oligo rev | AAACGCGCCCTCGTTAAGCTGGGA | | | | | | | | | | | |
| Chromosome | start | | end | strand | MM | target_seq | PAM | alignment | distance | position | gene name | gene id |
| GU305914.1 | | 226511 | 226533 | + | 0 | TCCCAGCTTAACGAGGGCGC | GGG | \|\|\|\|\|\|\|\|[\|\|\|\|\|\|\|\|\|\|\|\|]PAM | NA | Intergenic | NA | NA |
| T16902 | TAACGAGGGCGCGGGCTCCACGG | | 1000 | | | | | | | | | |
| Oligo fwd | CACCTAACGAGGGCGCGGGCTCCA | | | | | | | | | | | |
| Oligo rev | AAACTGGAGCCCGCGCCCTCGTTA | | | | | | | | | | | |
| Chromosome | start | | end | strand | MM | target_seq | PAM | alignment | distance | position | gene name | gene id |
| GU305914.1 | | 226519 | 226541 | + | 0 | TAACGAGGGCGCGGGCTCCA | CGG | \|\|\|\|\|\|\|\|[\|\|\|\|\|\|\|\|\|\|\|\|]PAM | NA | Intergenic | NA | NA |
| T16903 | CGCGGGCTCCACGGAGACCTCGG | | 1000 | | | | | | | | | |
| Oligo fwd | CACCCGCGGGCTCCACGGAGACCT | | | | | | | | | | | |
| Oligo rev | AAACAGGTCTCCGTGGAGCCCGCG | | | | | | | | | | | |
| Chromosome | start | | end | strand | MM | target_seq | PAM | alignment | distance | position | gene name | gene id |
| GU305914.1 | | 226528 | 226550 | + | 0 | CGCGGGCTCCACGGAGACCT | CGG | \|\|\|\|\|\|\|\|[\|\|\|\|\|\|\|\|\|\|\|\|]PAM | NA | Intergenic | NA | NA |
| T16904 | CTCGGAAGACGAGTTGAACACGG | | 1000 | | | | | | | | | |
| Oligo fwd | CACCCTCGGAAGACGAGTTGAACA | | | | | | | | | | | |
| Oligo rev | AAACTGTTCAACTCGTCTTCCGAG | | | | | | | | | | | |
| Chromosome | start | | end | strand | MM | target_seq | PAM | alignment | distance | position | gene name | gene id |
| GU305914.1 | | 226546 | 226568 | + | 0 | CTCGGAAGACGAGTTGAACA | CGG | \|\|\|\|\|\|\|\|[\|\|\|\|\|\|\|\|\|\|\|\|]PAM | NA | Intergenic | NA | NA |
| T16905 | AGTTGAACACGGCACACGCCAGG | | 1000 | | | | | | | | | |

Oligo fwd CACCAGTTGAACACGGCACACGCC
Oligo rev AAACGGCGTGTGCCGTGTTCAACT
Chromosome start end strand MM target_seq PAM alignment distance position gene name gene id
GU305914.1 226557 226579 + 0 AGTTGAACACGGCACACGCC AGG ||||||||[||||||||||||]PAM NA Intergenic NA NA

T16906 CAGTACTCCTTCGTCTTCCTCGG 1000
Oligo fwd CACCCAGTACTCCTTCGTCTTCCT
Oligo rev AAACAGGAAGACGAAGGAGTACTG
Chromosome start end strand MM target_seq PAM alignment distance position gene name gene id
GU305914.1 226622 226644 + 0 CAGTACTCCTTCGTCTTCCT CGG ||||||||[||||||||||||]PAM NA Intergenic NA NA

T16907 TACTCCTTCGTCTTCCTCGGAGG 1000
Oligo fwd CACCTACTCCTTCGTCTTCCTCGG
Oligo rev AAACCCGAGGAAGACGAAGGAGTA
Chromosome start end strand MM target_seq PAM alignment distance position gene name gene id
GU305914.1 226625 226647 + 0 TACTCCTTCGTCTTCCTCGG AGG ||||||||[||||||||||||]PAM NA Intergenic NA NA

T16908 TTCGTCTTCCTCGGAGGCTTCGG 1000
Oligo fwd CACCTTCGTCTTCCTCGGAGGCTT
Oligo rev AAACAAGCCTCCGAGGAAGACGAA
Chromosome start end strand MM target_seq PAM alignment distance position gene name gene id
GU305914.1 226631 226653 + 0 TTCGTCTTCCTCGGAGGCTT CGG ||||||||[||||||||||||]PAM NA Intergenic NA NA

T16909 GGAGTCTGAGTCTGTTTTTGAGG 1000
Oligo fwd CACCGGAGTCTGAGTCTGTTTTTG
Oligo rev AAACCAAAAACAGACTCAGACTCC
Chromosome start end strand MM target_seq PAM alignment distance position gene name gene id
GU305914.1 226652 226674 + 0 GGAGTCTGAGTCTGTTTTTG AGG ||||||||[||||||||||||]PAM NA Intergenic NA NA

T16910 GAGTCTGAGTCTGTTTTTGAGGG 1000
Oligo fwd CACCGAGTCTGAGTCTGTTTTTGA
Oligo rev AAACTCAAAAACAGACTCAGACTC
Chromosome start end strand MM target_seq PAM alignment distance position gene name gene id
GU305914.1 226653 226675 + 0 GAGTCTGAGTCTGTTTTTGA GGG ||||||||[||||||||||||]PAM NA Intergenic NA NA

T16911 ACACCTCGCCTCGAGCCACTCGG 1000
Oligo fwd CACCACACCTCGCCTCGAGCCACT
Oligo rev AAACAGTGGCTCGAGGCGAGGTGT
Chromosome start end strand MM target_seq PAM alignment distance position gene name gene id
GU305914.1 226703 226725 + 0 ACACCTCGCCTCGAGCCACT CGG ||||||||[||||||||||||]PAM NA Intergenic NA NA

T16912 CTCGAGCCACTCGGAGAGCGCGG 1000
Oligo fwd CACCCTCGAGCCACTCGGAGAGCG
Oligo rev AAACCGCTCTCCGAGTGGCTCGAG
Chromosome start end strand MM target_seq PAM alignment distance position gene name gene id
GU305914.1 226712 226734 + 0 CTCGAGCCACTCGGAGAGCG CGG ||||||||[||||||||||||]PAM NA Intergenic NA NA

T16913 GAGCCACTCGGAGAGCGCGGAGG 1000
Oligo fwd CACCGAGCCACTCGGAGAGCGCGG
Oligo rev AAACCCGCGCTCTCCGAGTGGCTC
Chromosome start end strand MM target_seq PAM alignment distance position gene name gene id
GU305914.1 226715 226737 + 0 GAGCCACTCGGAGAGCGCGG AGG ||||||||[||||||||||||]PAM NA Intergenic NA NA

T16914 CACTCGGAGAGCGCGGAGGTCGG 1000
Oligo fwd CACCCACTCGGAGAGCGCGGAGGT
Oligo rev AAACACCTCCGCGCTCTCCGAGTG
Chromosome start end strand MM target_seq PAM alignment distance position gene name gene id
GU305914.1 226719 226741 + 0 CACTCGGAGAGCGCGGAGGT CGG ||||||||[||||||||||||]PAM NA Intergenic NA NA

T16915 ACTCGGAGAGCGCGGAGGTCGGG 1000
Oligo fwd CACCACTCGGAGAGCGCGGAGGTC
Oligo rev AAACGACCTCCGCGCTCTCCGAGT
Chromosome start end strand MM target_seq PAM alignment distance position gene name gene id
GU305914.1 226720 226742 + 0 ACTCGGAGAGCGCGGAGGTC GGG ||||||||[||||||||||||]PAM NA Intergenic NA NA

T16916 GGAGGTCGGGACGAAGAATCTGG 1000
Oligo fwd CACCGGAGGTCGGGACGAAGAATC
Oligo rev AAACGATTCTTCGTCCCGACCTCC
Chromosome start end strand MM target_seq PAM alignment distance position gene name gene id
GU305914.1 226733 226755 + 0 GGAGGTCGGGACGAAGAATC TGG ||||||||[||||||||||||]PAM NA Intergenic NA NA

T16917 GAGGTCGGGACGAAGAATCTGGG 1000
Oligo fwd CACCGAGGTCGGGACGAAGAATCT
Oligo rev AAACAGATTCTTCGTCCCGACCTC
Chromosome start end strand MM target_seq PAM alignment distance position gene name gene id
GU305914.1 226734 226756 + 0 GAGGTCGGGACGAAGAATCT GGG ||||||||[||||||||||||]PAM NA Intergenic NA NA

T16918 GTCGGGACGAAGAATCTGGGCGG 1000
Oligo fwd CACCGTCGGGACGAAGAATCTGGG
Oligo rev AAACCCCAGATTCTTCGTCCCGAC
Chromosome start end strand MM target_seq PAM alignment distance position gene name gene id
GU305914.1 226737 226759 + 0 GTCGGGACGAAGAATCTGGG CGG ||||||||[||||||||||||]PAM NA Intergenic NA NA

T16919 GAATCTGGGCGGCCGTCTCCAGG 1000
Oligo fwd CACCGAATCTGGGCGGCCGTCTCC
Oligo rev AAACGGAGACGGCCGCCCAGATTC
Chromosome start end strand MM target_seq PAM alignment distance position gene name gene id
GU305914.1 226748 226770 + 0 GAATCTGGGCGGCCGTCTCC AGG ||||||||[||||||||||||]PAM NA Intergenic NA NA

| | | | | | | | | | | | | |
|---|---|---|---|---|---|---|---|---|---|---|---|---|
| T16920 | GAGTCCTCGCCGATCGAGACCGG | | 1000 | | | | | | | | | |
| Oligo fwd | CACCGAGTCCTCGCCGATCGAGAC | | | | | | | | | | | |
| Oligo rev | AAACGTCTCGATCGGCGAGGACTC | | | | | | | | | | | |
| Chromosome | start | | end | strand | MM | target_seq | PAM | alignment | distance | position | gene name | gene id |
| GU305914.1 | | 226772 | 226794 | + | 0 | GAGTCCTCGCCGATCGAGAC | CGG | \|\|\|\|\|\|\|\|[\|\|\|\|\|\|\|\|\|\|\|\|]PAM | NA | Intergenic | NA | NA |
| T16921 | AGTCCTCGCCGATCGAGACCGGG | | 1000 | | | | | | | | | |
| Oligo fwd | CACCAGTCCTCGCCGATCGAGACC | | | | | | | | | | | |
| Oligo rev | AAACGGTCTCGATCGGCGAGGACT | | | | | | | | | | | |
| Chromosome | start | | end | strand | MM | target_seq | PAM | alignment | distance | position | gene name | gene id |
| GU305914.1 | | 226773 | 226795 | + | 0 | AGTCCTCGCCGATCGAGACC | GGG | \|\|\|\|\|\|\|\|[\|\|\|\|\|\|\|\|\|\|\|\|]PAM | NA | Intergenic | NA | NA |
| T16922 | GCGCCGCTGCGATTCTTTCGAGG | | 1000 | | | | | | | | | |
| Oligo fwd | CACCGCGCCGCTGCGATTCTTTCG | | | | | | | | | | | |
| Oligo rev | AAACCGAAAGAATCGCAGCGGCGC | | | | | | | | | | | |
| Chromosome | start | | end | strand | MM | target_seq | PAM | alignment | distance | position | gene name | gene id |
| GU305914.1 | | 226810 | 226832 | + | 0 | GCGCCGCTGCGATTCTTTCG | AGG | \|\|\|\|\|\|\|\|[\|\|\|\|\|\|\|\|\|\|\|\|]PAM | NA | Intergenic | NA | NA |
| T16923 | TTTCGAGGTCTCGCTCTCCCAGG | | 1000 | | | | | | | | | |
| Oligo fwd | CACCTTTCGAGGTCTCGCTCTCCC | | | | | | | | | | | |
| Oligo rev | AAACGGGAGAGCGAGACCTCGAAA | | | | | | | | | | | |
| Chromosome | start | | end | strand | MM | target_seq | PAM | alignment | distance | position | gene name | gene id |
| GU305914.1 | | 226825 | 226847 | + | 0 | TTTCGAGGTCTCGCTCTCCC | AGG | \|\|\|\|\|\|\|\|[\|\|\|\|\|\|\|\|\|\|\|\|]PAM | NA | Intergenic | NA | NA |
| GU305914.1 | | 81208 | 81230 | + | 4 | GTTCGAGTTCTCGCCCTCCG | AGG | -\|\|\|\|\|\|-[\|\|\|\|\|\|-\|\|\|\|-]PAM | NA | Intergenic | NA | NA |
| T16924 | GGTCTCGCTCTCCCAGGCGAAGG | | 1000 | | | | | | | | | |
| Oligo fwd | CACCGGTCTCGCTCTCCCAGGCGA | | | | | | | | | | | |
| Oligo rev | AAACTCGCCTGGGAGAGCGAGACC | | | | | | | | | | | |
| Chromosome | start | | end | strand | MM | target_seq | PAM | alignment | distance | position | gene name | gene id |
| GU305914.1 | | 226831 | 226853 | + | 0 | GGTCTCGCTCTCCCAGGCGA | AGG | \|\|\|\|\|\|\|\|[\|\|\|\|\|\|\|\|\|\|\|\|]PAM | NA | Intergenic | NA | NA |
| T16925 | AGGCCCCTCCGTCTCTGAGCAGG | | 1000 | | | | | | | | | |
| Oligo fwd | CACCAGGCCCCTCCGTCTCTGAGC | | | | | | | | | | | |
| Oligo rev | AAACGCTCAGAGACGGAGGGGCCT | | | | | | | | | | | |
| Chromosome | start | | end | strand | MM | target_seq | PAM | alignment | distance | position | gene name | gene id |
| GU305914.1 | | 226851 | 226873 | + | 0 | AGGCCCCTCCGTCTCTGAGC | AGG | \|\|\|\|\|\|\|\|[\|\|\|\|\|\|\|\|\|\|\|\|]PAM | NA | Intergenic | NA | NA |
| GU305914.1 | | 64998 | 65020 | - | 4 | AGGCGCCGTCGTCTCTGAGG | CGG | \|\|\|\|-\|\|-[-\|\|\|\|\|\|\|\|\|\|-]PAM | NA | Intergenic | NA | NA |
| T16926 | GGCCCCTCCGTCTCTGAGCAGGG | | 1000 | | | | | | | | | |
| Oligo fwd | CACCGGCCCCTCCGTCTCTGAGCA | | | | | | | | | | | |
| Oligo rev | AAACTGCTCAGAGACGGAGGGGCC | | | | | | | | | | | |
| Chromosome | start | | end | strand | MM | target_seq | PAM | alignment | distance | position | gene name | gene id |
| GU305914.1 | | 226852 | 226874 | + | 0 | GGCCCCTCCGTCTCTGAGCA | GGG | \|\|\|\|\|\|\|\|[\|\|\|\|\|\|\|\|\|\|\|\|]PAM | NA | Intergenic | NA | NA |
| T16927 | CTCTGAGCAGGGTGATCCATTGG | | 1000 | | | | | | | | | |
| Oligo fwd | CACCCTCTGAGCAGGGTGATCCAT | | | | | | | | | | | |
| Oligo rev | AAACATGGATCACCCTGCTCAGAG | | | | | | | | | | | |
| Chromosome | start | | end | strand | MM | target_seq | PAM | alignment | distance | position | gene name | gene id |
| GU305914.1 | | 226863 | 226885 | + | 0 | CTCTGAGCAGGGTGATCCAT | TGG | \|\|\|\|\|\|\|\|[\|\|\|\|\|\|\|\|\|\|\|\|]PAM | NA | Intergenic | NA | NA |
| T16928 | CGAACCTTCGATCTGCCACCAGG | | 1000 | | | | | | | | | |
| Oligo fwd | CACCCGAACCTTCGATCTGCCACC | | | | | | | | | | | |
| Oligo rev | AAACGGTGGCAGATCGAAGGTTCG | | | | | | | | | | | |
| Chromosome | start | | end | strand | MM | target_seq | PAM | alignment | distance | position | gene name | gene id |
| GU305914.1 | | 226924 | 226946 | + | 0 | CGAACCTTCGATCTGCCACC | AGG | \|\|\|\|\|\|\|\|[\|\|\|\|\|\|\|\|\|\|\|\|]PAM | NA | Intergenic | NA | NA |
| T16929 | CACCAGGTCCCGTTGATCCGAGG | | 1000 | | | | | | | | | |
| Oligo fwd | CACCCACCAGGTCCCGTTGATCCG | | | | | | | | | | | |
| Oligo rev | AAACCGGATCAACGGGACCTGGTG | | | | | | | | | | | |
| Chromosome | start | | end | strand | MM | target_seq | PAM | alignment | distance | position | gene name | gene id |
| GU305914.1 | | 226940 | 226962 | + | 0 | CACCAGGTCCCGTTGATCCG | AGG | \|\|\|\|\|\|\|\|[\|\|\|\|\|\|\|\|\|\|\|\|]PAM | NA | Intergenic | NA | NA |
| T16930 | CGAAACGACGTTGTTATATACGG | | 1000 | | | | | | | | | |
| Oligo fwd | CACCCGAAACGACGTTGTTATATA | | | | | | | | | | | |
| Oligo rev | AAACTATATAACAACGTCGTTTCG | | | | | | | | | | | |
| Chromosome | start | | end | strand | MM | target_seq | PAM | alignment | distance | position | gene name | gene id |
| GU305914.1 | | 227006 | 227028 | + | 0 | CGAAACGACGTTGTTATATA | CGG | \|\|\|\|\|\|\|\|[\|\|\|\|\|\|\|\|\|\|\|\|]PAM | NA | Intergenic | NA | NA |
| T16931 | GGAACCTTTGTTTCCACCCACGG | | 1000 | | | | | | | | | |
| Oligo fwd | CACCGGAACCTTTGTTTCCACCCA | | | | | | | | | | | |
| Oligo rev | AAACTGGGTGGAAACAAAGGTTCC | | | | | | | | | | | |
| Chromosome | start | | end | strand | MM | target_seq | PAM | alignment | distance | position | gene name | gene id |
| GU305914.1 | | 227027 | 227049 | + | 0 | GGAACCTTTGTTTCCACCCA | CGG | \|\|\|\|\|\|\|\|[\|\|\|\|\|\|\|\|\|\|\|\|]PAM | NA | Intergenic | NA | NA |
| T16932 | ACCTTTGTTTCCACCCACGGTGG | | 1000 | | | | | | | | | |
| Oligo fwd | CACCACCTTTGTTTCCACCCACGG | | | | | | | | | | | |
| Oligo rev | AAACCCGTGGGTGGAAACAAAGGT | | | | | | | | | | | |
| Chromosome | start | | end | strand | MM | target_seq | PAM | alignment | distance | position | gene name | gene id |
| GU305914.1 | | 227030 | 227052 | + | 0 | ACCTTTGTTTCCACCCACGG | TGG | \|\|\|\|\|\|\|\|[\|\|\|\|\|\|\|\|\|\|\|\|]PAM | NA | Intergenic | NA | NA |
| T16933 | GTCACAACACCACCCCAGAATGG | | 1000 | | | | | | | | | |
| Oligo fwd | CACCGTCACAACACCACCCCAGAA | | | | | | | | | | | |
| Oligo rev | AAACTTCTGGGGTGGTGTTGTGAC | | | | | | | | | | | |
| Chromosome | start | | end | strand | MM | target_seq | PAM | alignment | distance | position | gene name | gene id |
| GU305914.1 | | 227086 | 227108 | + | 0 | GTCACAACACCACCCCAGAA | TGG | \|\|\|\|\|\|\|\|[\|\|\|\|\|\|\|\|\|\|\|\|]PAM | NA | Intergenic | NA | NA |
| T16934 | ACCCCAGAATGGTTACACGCAGG | | 1000 | | | | | | | | | |
| Oligo fwd | CACCACCCCAGAATGGTTACACGC | | | | | | | | | | | |
| Oligo rev | AAACGCGTGTAACCATTCTGGGGT | | | | | | | | | | | |

Chromosome start end strand MM target_seq PAM alignment distance position gene name gene id
GU305914.1 227097 227119 + 0 ACCCCAGAATGGTTACACGC AGG ||||||||[||||||||||||]PAM NA Intergenic NA NA

T16935 CCAACCACCACAAACCACGTCGG 1000
Oligo fwd CACCCCAACCACCACAAACCACGT
Oligo rev AAACACGTGGTTTGTGGTGGTTGG
Chromosome start end strand MM target_seq PAM alignment distance position gene name gene id
GU305914.1 227121 227143 + 0 CCAACCACCACAAACCACGT CGG ||||||||[||||||||||||]PAM NA Intergenic NA NA

T16936 CAACCACCACAAACCACGTCGGG 1000
Oligo fwd CACCCAACCACCACAAACCACGTC
Oligo rev AAACGACGTGGTTTGTGGTGGTTG
Chromosome start end strand MM target_seq PAM alignment distance position gene name gene id
GU305914.1 227122 227144 + 0 CAACCACCACAAACCACGTC GGG ||||||||[||||||||||||]PAM NA Intergenic NA NA

T16937 ATGATGTCTCATGCCACTTGAGG 1000
Oligo fwd CACCATGATGTCTCATGCCACTTG
Oligo rev AAACCAAGTGGCATGAGACATCAT
Chromosome start end strand MM target_seq PAM alignment distance position gene name gene id
GU305914.1 227189 227211 + 0 ATGATGTCTCATGCCACTTG AGG ||||||||[||||||||||||]PAM NA Intergenic NA NA

T16938 TGCCACTTGAGGTAGAAACTTGG 1000
Oligo fwd CACCTGCCACTTGAGGTAGAAACT
Oligo rev AAACAGTTTCTACCTCAAGTGGCA
Chromosome start end strand MM target_seq PAM alignment distance position gene name gene id
GU305914.1 227200 227222 + 0 TGCCACTTGAGGTAGAAACT TGG ||||||||[||||||||||||]PAM NA Intergenic NA NA

T16939 CTTGAGGTAGAAACTTGGCCCGG 1000
Oligo fwd CACCCTTGAGGTAGAAACTTGGCC
Oligo rev AAACGGCCAAGTTTCTACCTCAAG
Chromosome start end strand MM target_seq PAM alignment distance position gene name gene id
GU305914.1 227205 227227 + 0 CTTGAGGTAGAAACTTGGCC CGG ||||||||[||||||||||||]PAM NA Intergenic NA NA

T16940 TTGAGGTAGAAACTTGGCCCGGG 1000
Oligo fwd CACCTTGAGGTAGAAACTTGGCCC
Oligo rev AAACGGGCCAAGTTTCTACCTCAA
Chromosome start end strand MM target_seq PAM alignment distance position gene name gene id
GU305914.1 227206 227228 + 0 TTGAGGTAGAAACTTGGCCC GGG ||||||||[||||||||||||]PAM NA Intergenic NA NA

T16941 AGAAACTTGGCCCGGGACCGCGG 1000
Oligo fwd CACCAGAAACTTGGCCCGGGACCG
Oligo rev AAACCGGTCCCGGGCCAAGTTTCT
Chromosome start end strand MM target_seq PAM alignment distance position gene name gene id
GU305914.1 227213 227235 + 0 AGAAACTTGGCCCGGGACCG CGG ||||||||[||||||||||||]PAM NA Intergenic NA NA

T16942 CCCAAACTTTCCACACTCAAAGG 1000
Oligo fwd CACCCCCAAACTTTCCACACTCAA
Oligo rev AAACTTGAGTGTGGAAAGTTTGGG
Chromosome start end strand MM target_seq PAM alignment distance position gene name gene id
GU305914.1 227242 227264 + 0 CCCAAACTTTCCACACTCAA AGG ||||||||[||||||||||||]PAM NA Intergenic NA NA

T16943 CCAAAACTAATACCACTTTCCGG 1000
Oligo fwd CACCCCAAAACTAATACCACTTTC
Oligo rev AAACGAAAGTGGTATTAGTTTTGG
Chromosome start end strand MM target_seq PAM alignment distance position gene name gene id
GU305914.1 227285 227307 + 0 CCAAAACTAATACCACTTTC CGG ||||||||[||||||||||||]PAM NA Intergenic NA NA

T16944 TACCACTTTCCGGAACGAGACGG 1000
Oligo fwd CACCTACCACTTTCCGGAACGAGA
Oligo rev AAACTCTCGTTCCGGAAAGTGGTA
Chromosome start end strand MM target_seq PAM alignment distance position gene name gene id
GU305914.1 227295 227317 + 0 TACCACTTTCCGGAACGAGA CGG ||||||||[||||||||||||]PAM NA Intergenic NA NA

T16945 ACCACTTTCCGGAACGAGACGGG 1000
Oligo fwd CACCACCACTTTCCGGAACGAGAC
Oligo rev AAACGTCTCGTTCCGGAAAGTGGT
Chromosome start end strand MM target_seq PAM alignment distance position gene name gene id
GU305914.1 227296 227318 + 0 ACCACTTTCCGGAACGAGAC GGG ||||||||[||||||||||||]PAM NA Intergenic NA NA

T16946 CCAAGTTATTGTCCTCCCGATGG 1000
Oligo fwd CACCCCAAGTTATTGTCCTCCCGA
Oligo rev AAACTCGGGAGGACAATAACTTGG
Chromosome start end strand MM target_seq PAM alignment distance position gene name gene id
GU305914.1 227348 227370 + 0 CCAAGTTATTGTCCTCCCGA TGG ||||||||[||||||||||||]PAM NA Intergenic NA NA

T16947 TTGTCCTCCCGATGGTTCGTCGG 1000
Oligo fwd CACCTTGTCCTCCCGATGGTTCGT
Oligo rev AAACACGAACCATCGGGAGGACAA
Chromosome start end strand MM target_seq PAM alignment distance position gene name gene id
GU305914.1 227356 227378 + 0 TTGTCCTCCCGATGGTTCGT CGG ||||||||[||||||||||||]PAM NA Intergenic NA NA

T16948 ATGGTTCGTCGGCGCCAGAGAGG 1000
Oligo fwd CACCATGGTTCGTCGGCGCCAGAG
Oligo rev AAACCTCTGGCGCCGACGAACCAT
Chromosome start end strand MM target_seq PAM alignment distance position gene name gene id
GU305914.1 227367 227389 + 0 ATGGTTCGTCGGCGCCAGAG AGG ||||||||[||||||||||||]PAM NA Intergenic NA NA

T16949 GCCAGAGAGGTCGCTTTCCCAGG 1000
Oligo fwd CACCGCCAGAGAGGTCGCTTTCCC

<table>
<tr><td>Oligo rev</td><td>AAACGGGAAAGCGACCTCTCTGGC</td><td></td><td></td><td></td><td></td><td></td><td></td><td></td><td></td><td></td><td></td></tr>
<tr><td>Chromosome</td><td>start</td><td>end</td><td>strand</td><td>MM</td><td>target_seq</td><td>PAM</td><td>alignment</td><td>distance</td><td>position</td><td>gene name</td><td>gene id</td></tr>
<tr><td>GU305914.1</td><td>227380</td><td>227402</td><td>+</td><td>0</td><td>GCCAGAGAGGTCGCTTTCCC</td><td>AGG</td><td>||||||||[||||||||||||]PAM</td><td>NA</td><td>Intergenic</td><td>NA</td><td>NA</td></tr>
<tr><td></td><td></td><td></td><td></td><td></td><td></td><td></td><td></td><td></td><td></td><td></td><td></td></tr>
<tr><td>T16950</td><td>AGAGAGGTCGCTTTCCCAGGAGG</td><td>1000</td><td></td><td></td><td></td><td></td><td></td><td></td><td></td><td></td><td></td></tr>
<tr><td>Oligo fwd</td><td>CACCAGAGAGGTCGCTTTCCCAGG</td><td></td><td></td><td></td><td></td><td></td><td></td><td></td><td></td><td></td><td></td></tr>
<tr><td>Oligo rev</td><td>AAACCCTGGGAAAGCGACCTCTCT</td><td></td><td></td><td></td><td></td><td></td><td></td><td></td><td></td><td></td><td></td></tr>
<tr><td>Chromosome</td><td>start</td><td>end</td><td>strand</td><td>MM</td><td>target_seq</td><td>PAM</td><td>alignment</td><td>distance</td><td>position</td><td>gene name</td><td>gene id</td></tr>
<tr><td>GU305914.1</td><td>227383</td><td>227405</td><td>+</td><td>0</td><td>AGAGAGGTCGCTTTCCCAGG</td><td>AGG</td><td>||||||||[||||||||||||]PAM</td><td>NA</td><td>Intergenic</td><td>NA</td><td>NA</td></tr>
<tr><td></td><td></td><td></td><td></td><td></td><td></td><td></td><td></td><td></td><td></td><td></td><td></td></tr>
<tr><td>T16951</td><td>GTCGCTTTCCCAGGAGGTCTTGG</td><td>1000</td><td></td><td></td><td></td><td></td><td></td><td></td><td></td><td></td><td></td></tr>
<tr><td>Oligo fwd</td><td>CACCGTCGCTTTCCCAGGAGGTCT</td><td></td><td></td><td></td><td></td><td></td><td></td><td></td><td></td><td></td><td></td></tr>
<tr><td>Oligo rev</td><td>AAACAGACCTCCTGGGAAAGCGAC</td><td></td><td></td><td></td><td></td><td></td><td></td><td></td><td></td><td></td><td></td></tr>
<tr><td>Chromosome</td><td>start</td><td>end</td><td>strand</td><td>MM</td><td>target_seq</td><td>PAM</td><td>alignment</td><td>distance</td><td>position</td><td>gene name</td><td>gene id</td></tr>
<tr><td>GU305914.1</td><td>227389</td><td>227411</td><td>+</td><td>0</td><td>GTCGCTTTCCCAGGAGGTCT</td><td>TGG</td><td>||||||||[||||||||||||]PAM</td><td>NA</td><td>Intergenic</td><td>NA</td><td>NA</td></tr>
<tr><td></td><td></td><td></td><td></td><td></td><td></td><td></td><td></td><td></td><td></td><td></td><td></td></tr>
<tr><td>T16952</td><td>TGGCTCTCGAGAGCCAAGCCCGG</td><td>1000</td><td></td><td></td><td></td><td></td><td></td><td></td><td></td><td></td><td></td></tr>
<tr><td>Oligo fwd</td><td>CACCTGGCTCTCGAGAGCCAAGCC</td><td></td><td></td><td></td><td></td><td></td><td></td><td></td><td></td><td></td><td></td></tr>
<tr><td>Oligo rev</td><td>AAACGGCTTGGCTCTCGAGAGCCA</td><td></td><td></td><td></td><td></td><td></td><td></td><td></td><td></td><td></td><td></td></tr>
<tr><td>Chromosome</td><td>start</td><td>end</td><td>strand</td><td>MM</td><td>target_seq</td><td>PAM</td><td>alignment</td><td>distance</td><td>position</td><td>gene name</td><td>gene id</td></tr>
<tr><td>GU305914.1</td><td>227409</td><td>227431</td><td>+</td><td>0</td><td>TGGCTCTCGAGAGCCAAGCC</td><td>CGG</td><td>||||||||[||||||||||||]PAM</td><td>NA</td><td>Intergenic</td><td>NA</td><td>NA</td></tr>
<tr><td></td><td></td><td></td><td></td><td></td><td></td><td></td><td></td><td></td><td></td><td></td><td></td></tr>
<tr><td>T16953</td><td>CGAGAGCCAAGCCCGGTGCCAGG</td><td>1000</td><td></td><td></td><td></td><td></td><td></td><td></td><td></td><td></td><td></td></tr>
<tr><td>Oligo fwd</td><td>CACCCGAGAGCCAAGCCCGGTGCC</td><td></td><td></td><td></td><td></td><td></td><td></td><td></td><td></td><td></td><td></td></tr>
<tr><td>Oligo rev</td><td>AAACGGCACCGGGCTTGGCTCTCG</td><td></td><td></td><td></td><td></td><td></td><td></td><td></td><td></td><td></td><td></td></tr>
<tr><td>Chromosome</td><td>start</td><td>end</td><td>strand</td><td>MM</td><td>target_seq</td><td>PAM</td><td>alignment</td><td>distance</td><td>position</td><td>gene name</td><td>gene id</td></tr>
<tr><td>GU305914.1</td><td>227416</td><td>227438</td><td>+</td><td>0</td><td>CGAGAGCCAAGCCCGGTGCC</td><td>AGG</td><td>||||||||[||||||||||||]PAM</td><td>NA</td><td>Intergenic</td><td>NA</td><td>NA</td></tr>
<tr><td></td><td></td><td></td><td></td><td></td><td></td><td></td><td></td><td></td><td></td><td></td><td></td></tr>
<tr><td>T16954</td><td>GCCAGGAACCTTGTACCCACAGG</td><td>1000</td><td></td><td></td><td></td><td></td><td></td><td></td><td></td><td></td><td></td></tr>
<tr><td>Oligo fwd</td><td>CACCGCCAGGAACCTTGTACCCAC</td><td></td><td></td><td></td><td></td><td></td><td></td><td></td><td></td><td></td><td></td></tr>
<tr><td>Oligo rev</td><td>AAACGTGGGTACAAGGTTCCTGGC</td><td></td><td></td><td></td><td></td><td></td><td></td><td></td><td></td><td></td><td></td></tr>
<tr><td>Chromosome</td><td>start</td><td>end</td><td>strand</td><td>MM</td><td>target_seq</td><td>PAM</td><td>alignment</td><td>distance</td><td>position</td><td>gene name</td><td>gene id</td></tr>
<tr><td>GU305914.1</td><td>227433</td><td>227455</td><td>+</td><td>0</td><td>GCCAGGAACCTTGTACCCAC</td><td>AGG</td><td>||||||||[||||||||||||]PAM</td><td>NA</td><td>Intergenic</td><td>NA</td><td>NA</td></tr>
<tr><td></td><td></td><td></td><td></td><td></td><td></td><td></td><td></td><td></td><td></td><td></td><td></td></tr>
<tr><td>T16955</td><td>GGAACCTTGTACCCACAGGATGG</td><td>1000</td><td></td><td></td><td></td><td></td><td></td><td></td><td></td><td></td><td></td></tr>
<tr><td>Oligo fwd</td><td>CACCGGAACCTTGTACCCACAGGA</td><td></td><td></td><td></td><td></td><td></td><td></td><td></td><td></td><td></td><td></td></tr>
<tr><td>Oligo rev</td><td>AAACTCCTGTGGGTACAAGGTTCC</td><td></td><td></td><td></td><td></td><td></td><td></td><td></td><td></td><td></td><td></td></tr>
<tr><td>Chromosome</td><td>start</td><td>end</td><td>strand</td><td>MM</td><td>target_seq</td><td>PAM</td><td>alignment</td><td>distance</td><td>position</td><td>gene name</td><td>gene id</td></tr>
<tr><td>GU305914.1</td><td>227437</td><td>227459</td><td>+</td><td>0</td><td>GGAACCTTGTACCCACAGGA</td><td>TGG</td><td>||||||||[||||||||||||]PAM</td><td>NA</td><td>Intergenic</td><td>NA</td><td>NA</td></tr>
<tr><td></td><td></td><td></td><td></td><td></td><td></td><td></td><td></td><td></td><td></td><td></td><td></td></tr>
<tr><td>T16956</td><td>ACCCACCCCTCCATGTGACGCGG</td><td>1000</td><td></td><td></td><td></td><td></td><td></td><td></td><td></td><td></td><td></td></tr>
<tr><td>Oligo fwd</td><td>CACCACCCACCCCTCCATGTGACG</td><td></td><td></td><td></td><td></td><td></td><td></td><td></td><td></td><td></td><td></td></tr>
<tr><td>Oligo rev</td><td>AAACCGTCACATGGAGGGGTGGGT</td><td></td><td></td><td></td><td></td><td></td><td></td><td></td><td></td><td></td><td></td></tr>
<tr><td>Chromosome</td><td>start</td><td>end</td><td>strand</td><td>MM</td><td>target_seq</td><td>PAM</td><td>alignment</td><td>distance</td><td>position</td><td>gene name</td><td>gene id</td></tr>
<tr><td>GU305914.1</td><td>227468</td><td>227490</td><td>+</td><td>0</td><td>ACCCACCCCTCCATGTGACG</td><td>CGG</td><td>||||||||[||||||||||||]PAM</td><td>NA</td><td>Intergenic</td><td>NA</td><td>NA</td></tr>
<tr><td></td><td></td><td></td><td></td><td></td><td></td><td></td><td></td><td></td><td></td><td></td><td></td></tr>
<tr><td>T16957</td><td>CCCACCCCTCCATGTGACGCGGG</td><td>1000</td><td></td><td></td><td></td><td></td><td></td><td></td><td></td><td></td><td></td></tr>
<tr><td>Oligo fwd</td><td>CACCCCCACCCCTCCATGTGACGC</td><td></td><td></td><td></td><td></td><td></td><td></td><td></td><td></td><td></td><td></td></tr>
<tr><td>Oligo rev</td><td>AAACGCGTCACATGGAGGGGTGGG</td><td></td><td></td><td></td><td></td><td></td><td></td><td></td><td></td><td></td><td></td></tr>
<tr><td>Chromosome</td><td>start</td><td>end</td><td>strand</td><td>MM</td><td>target_seq</td><td>PAM</td><td>alignment</td><td>distance</td><td>position</td><td>gene name</td><td>gene id</td></tr>
<tr><td>GU305914.1</td><td>227469</td><td>227491</td><td>+</td><td>0</td><td>CCCACCCCTCCATGTGACGC</td><td>GGG</td><td>||||||||[||||||||||||]PAM</td><td>NA</td><td>Intergenic</td><td>NA</td><td>NA</td></tr>
<tr><td></td><td></td><td></td><td></td><td></td><td></td><td></td><td></td><td></td><td></td><td></td><td></td></tr>
<tr><td>T16958</td><td>TAGCTTTTATTGATTCGAACTGG</td><td>1000</td><td></td><td></td><td></td><td></td><td></td><td></td><td></td><td></td><td></td></tr>
<tr><td>Oligo fwd</td><td>CACCTAGCTTTTATTGATTCGAAC</td><td></td><td></td><td></td><td></td><td></td><td></td><td></td><td></td><td></td><td></td></tr>
<tr><td>Oligo rev</td><td>AAACGTTCGAATCAATAAAAGCTA</td><td></td><td></td><td></td><td></td><td></td><td></td><td></td><td></td><td></td><td></td></tr>
<tr><td>Chromosome</td><td>start</td><td>end</td><td>strand</td><td>MM</td><td>target_seq</td><td>PAM</td><td>alignment</td><td>distance</td><td>position</td><td>gene name</td><td>gene id</td></tr>
<tr><td>GU305914.1</td><td>227542</td><td>227564</td><td>+</td><td>0</td><td>TAGCTTTTATTGATTCGAAC</td><td>TGG</td><td>||||||||[||||||||||||]PAM</td><td>NA</td><td>Intergenic</td><td>NA</td><td>NA</td></tr>
<tr><td></td><td></td><td></td><td></td><td></td><td></td><td></td><td></td><td></td><td></td><td></td><td></td></tr>
<tr><td>T16959</td><td>CGAACTGGCGAAAGCTCAAGCGG</td><td>1000</td><td></td><td></td><td></td><td></td><td></td><td></td><td></td><td></td><td></td></tr>
<tr><td>Oligo fwd</td><td>CACCCGAACTGGCGAAAGCTCAAG</td><td></td><td></td><td></td><td></td><td></td><td></td><td></td><td></td><td></td><td></td></tr>
<tr><td>Oligo rev</td><td>AAACCTTGAGCTTTCGCCAGTTCG</td><td></td><td></td><td></td><td></td><td></td><td></td><td></td><td></td><td></td><td></td></tr>
<tr><td>Chromosome</td><td>start</td><td>end</td><td>strand</td><td>MM</td><td>target_seq</td><td>PAM</td><td>alignment</td><td>distance</td><td>position</td><td>gene name</td><td>gene id</td></tr>
<tr><td>GU305914.1</td><td>227557</td><td>227579</td><td>+</td><td>0</td><td>CGAACTGGCGAAAGCTCAAG</td><td>CGG</td><td>||||||||[||||||||||||]PAM</td><td>NA</td><td>Intergenic</td><td>NA</td><td>NA</td></tr>
<tr><td></td><td></td><td></td><td></td><td></td><td></td><td></td><td></td><td></td><td></td><td></td><td></td></tr>
<tr><td>T16960</td><td>GAACTGGCGAAAGCTCAAGCGGG</td><td>1000</td><td></td><td></td><td></td><td></td><td></td><td></td><td></td><td></td><td></td></tr>
<tr><td>Oligo fwd</td><td>CACCGAACTGGCGAAAGCTCAAGC</td><td></td><td></td><td></td><td></td><td></td><td></td><td></td><td></td><td></td><td></td></tr>
<tr><td>Oligo rev</td><td>AAACGCTTGAGCTTTCGCCAGTTC</td><td></td><td></td><td></td><td></td><td></td><td></td><td></td><td></td><td></td><td></td></tr>
<tr><td>Chromosome</td><td>start</td><td>end</td><td>strand</td><td>MM</td><td>target_seq</td><td>PAM</td><td>alignment</td><td>distance</td><td>position</td><td>gene name</td><td>gene id</td></tr>
<tr><td>GU305914.1</td><td>227558</td><td>227580</td><td>+</td><td>0</td><td>GAACTGGCGAAAGCTCAAGC</td><td>GGG</td><td>||||||||[||||||||||||]PAM</td><td>NA</td><td>Intergenic</td><td>NA</td><td>NA</td></tr>
<tr><td></td><td></td><td></td><td></td><td></td><td></td><td></td><td></td><td></td><td></td><td></td><td></td></tr>
<tr><td>T16961</td><td>TGGCGAAAGCTCAAGCGGGAAGG</td><td>1000</td><td></td><td></td><td></td><td></td><td></td><td></td><td></td><td></td><td></td></tr>
<tr><td>Oligo fwd</td><td>CACCTGGCGAAAGCTCAAGCGGGA</td><td></td><td></td><td></td><td></td><td></td><td></td><td></td><td></td><td></td><td></td></tr>
<tr><td>Oligo rev</td><td>AAACTCCCGCTTGAGCTTTCGCCA</td><td></td><td></td><td></td><td></td><td></td><td></td><td></td><td></td><td></td><td></td></tr>
<tr><td>Chromosome</td><td>start</td><td>end</td><td>strand</td><td>MM</td><td>target_seq</td><td>PAM</td><td>alignment</td><td>distance</td><td>position</td><td>gene name</td><td>gene id</td></tr>
<tr><td>GU305914.1</td><td>227562</td><td>227584</td><td>+</td><td>0</td><td>TGGCGAAAGCTCAAGCGGGA</td><td>AGG</td><td>||||||||[||||||||||||]PAM</td><td>NA</td><td>Intergenic</td><td>NA</td><td>NA</td></tr>
<tr><td></td><td></td><td></td><td></td><td></td><td></td><td></td><td></td><td></td><td></td><td></td><td></td></tr>
<tr><td>T16962</td><td>TCAAGCGGGAAGGAGATGAGAGG</td><td>1000</td><td></td><td></td><td></td><td></td><td></td><td></td><td></td><td></td><td></td></tr>
<tr><td>Oligo fwd</td><td>CACCTCAAGCGGGAAGGAGATGAG</td><td></td><td></td><td></td><td></td><td></td><td></td><td></td><td></td><td></td><td></td></tr>
<tr><td>Oligo rev</td><td>AAACCTCATCTCCTTCCCGCTTGA</td><td></td><td></td><td></td><td></td><td></td><td></td><td></td><td></td><td></td><td></td></tr>
<tr><td>Chromosome</td><td>start</td><td>end</td><td>strand</td><td>MM</td><td>target_seq</td><td>PAM</td><td>alignment</td><td>distance</td><td>position</td><td>gene name</td><td>gene id</td></tr>
<tr><td>GU305914.1</td><td>227572</td><td>227594</td><td>+</td><td>0</td><td>TCAAGCGGGAAGGAGATGAG</td><td>AGG</td><td>||||||||[||||||||||||]PAM</td><td>NA</td><td>Intergenic</td><td>NA</td><td>NA</td></tr>
<tr><td></td><td></td><td></td><td></td><td></td><td></td><td></td><td></td><td></td><td></td><td></td><td></td></tr>
<tr><td>T16963</td><td>AAGGAGATGAGAGGTCGTCCCGG</td><td>1000</td><td></td><td></td><td></td><td></td><td></td><td></td><td></td><td></td><td></td></tr>
<tr><td>Oligo fwd</td><td>CACCAAGGAGATGAGAGGTCGTCC</td><td></td><td></td><td></td><td></td><td></td><td></td><td></td><td></td><td></td><td></td></tr>
<tr><td>Oligo rev</td><td>AAACGGACGACCTCTCATCTCCTT</td><td></td><td></td><td></td><td></td><td></td><td></td><td></td><td></td><td></td><td></td></tr>
<tr><td>Chromosome</td><td>start</td><td>end</td><td>strand</td><td>MM</td><td>target_seq</td><td>PAM</td><td>alignment</td><td>distance</td><td>position</td><td>gene name</td><td>gene id</td></tr>
<tr><td>GU305914.1</td><td>227581</td><td>227603</td><td>+</td><td>0</td><td>AAGGAGATGAGAGGTCGTCC</td><td>CGG</td><td>||||||||[||||||||||||]PAM</td><td>NA</td><td>Intergenic</td><td>NA</td><td>NA</td></tr>
<tr><td></td><td></td><td></td><td></td><td></td><td></td><td></td><td></td><td></td><td></td><td></td><td></td></tr>
<tr><td>T16964</td><td>CCGACCCCCCGACGTCCGCTCGG</td><td>1000</td><td></td><td></td><td></td><td></td><td></td><td></td><td></td><td></td><td></td></tr>
</table>

Oligo fwd CACCCCGACCCCCCGACGTCCGCT
Oligo rev AAACAGCGGACGTCGGGGGGTCGG
Chromosome start end strand MM target_seq PAM alignment distance position gene name gene id
GU305914.1 227610 227632 + 0 CCGACCCCCCGACGTCCGCT CGG ||||||||[||||||||||||]PAM NA Intergenic NA NA

T16965 TCGGAACAGAAAAACAACCCCGG 1000
Oligo fwd CACCTCGGAACAGAAAAACAACCC
Oligo rev AAACGGGTTGTTTTTCTGTTCCGA
Chromosome start end strand MM target_seq PAM alignment distance position gene name gene id
GU305914.1 227629 227651 + 0 TCGGAACAGAAAAACAACCC CGG ||||||||[||||||||||||]PAM NA Intergenic NA NA

T16966 CGGAACAGAAAAACAACCCCGGG 1000
Oligo fwd CACCCGGAACAGAAAAACAACCCC
Oligo rev AAACGGGGTTGTTTTTCTGTTCCG
Chromosome start end strand MM target_seq PAM alignment distance position gene name gene id
GU305914.1 227630 227652 + 0 CGGAACAGAAAAACAACCCC GGG ||||||||[||||||||||||]PAM NA Intergenic NA NA

T16967 CTGACCGTTCGATGTGACGAAGG 1000
Oligo fwd CACCCTGACCGTTCGATGTGACGA
Oligo rev AAACTCGTCACATCGAACGGTCAG
Chromosome start end strand MM target_seq PAM alignment distance position gene name gene id
GU305914.1 227659 227681 + 0 CTGACCGTTCGATGTGACGA AGG ||||||||[||||||||||||]PAM NA Intergenic NA NA

T16968 TGACCGTTCGATGTGACGAAGGG 1000
Oligo fwd CACCTGACCGTTCGATGTGACGAA
Oligo rev AAACTTCGTCACATCGAACGGTCA
Chromosome start end strand MM target_seq PAM alignment distance position gene name gene id
GU305914.1 227660 227682 + 0 TGACCGTTCGATGTGACGAA GGG ||||||||[||||||||||||]PAM NA Intergenic NA NA

T16969 GTTCGATGTGACGAAGGGCGAGG 1000
Oligo fwd CACCGTTCGATGTGACGAAGGGCG
Oligo rev AAACCGCCCTTCGTCACATCGAAC
Chromosome start end strand MM target_seq PAM alignment distance position gene name gene id
GU305914.1 227665 227687 + 0 GTTCGATGTGACGAAGGGCG AGG ||||||||[||||||||||||]PAM NA Intergenic NA NA

T16970 TTCGATGTGACGAAGGGCGAGGG 1000
Oligo fwd CACCTTCGATGTGACGAAGGGCGA
Oligo rev AAACTCGCCCTTCGTCACATCGAA
Chromosome start end strand MM target_seq PAM alignment distance position gene name gene id
GU305914.1 227666 227688 + 0 TTCGATGTGACGAAGGGCGA GGG ||||||||[||||||||||||]PAM NA Intergenic NA NA

T16971 TCGATGTGACGAAGGGCGAGGGG 1000
Oligo fwd CACCTCGATGTGACGAAGGGCGAG
Oligo rev AAACCTCGCCCTTCGTCACATCGA
Chromosome start end strand MM target_seq PAM alignment distance position gene name gene id
GU305914.1 227667 227689 + 0 TCGATGTGACGAAGGGCGAG GGG ||||||||[||||||||||||]PAM NA Intergenic NA NA

T16972 AAGGGCGAGGGGAAAAACATCGG 1000
Oligo fwd CACCAAGGGCGAGGGGAAAAACAT
Oligo rev AAACATGTTTTTCCCCTCGCCCTT
Chromosome start end strand MM target_seq PAM alignment distance position gene name gene id
GU305914.1 227678 227700 + 0 AAGGGCGAGGGGAAAAACAT CGG ||||||||[||||||||||||]PAM NA Intergenic NA NA

T16973 AGGGCGAGGGGAAAAACATCGGG 1000
Oligo fwd CACCAGGGCGAGGGGAAAAACATC
Oligo rev AAACGATGTTTTTCCCCTCGCCCT
Chromosome start end strand MM target_seq PAM alignment distance position gene name gene id
GU305914.1 227679 227701 + 0 AGGGCGAGGGGAAAAACATC GGG ||||||||[||||||||||||]PAM NA Intergenic NA NA

T16974 GGGCGAGGGGAAAAACATCGGGG 1000
Oligo fwd CACCGGGCGAGGGGAAAAACATCG
Oligo rev AAACCGATGTTTTTCCCCTCGCCC
Chromosome start end strand MM target_seq PAM alignment distance position gene name gene id
GU305914.1 227680 227702 + 0 GGGCGAGGGGAAAAACATCG GGG ||||||||[||||||||||||]PAM NA Intergenic NA NA

T16975 GAGGGGAAAAACATCGGGGAAGG 1000
Oligo fwd CACCGAGGGGAAAAACATCGGGGA
Oligo rev AAACTCCCCGATGTTTTTCCCCTC
Chromosome start end strand MM target_seq PAM alignment distance position gene name gene id
GU305914.1 227684 227706 + 0 GAGGGGAAAAACATCGGGGA AGG ||||||||[||||||||||||]PAM NA Intergenic NA NA

T16976 AAACATCGGGGAAGGACAAAAGG 1000
Oligo fwd CACCAAACATCGGGGAAGGACAAA
Oligo rev AAACTTTGTCCTTCCCCGATGTTT
Chromosome start end strand MM target_seq PAM alignment distance position gene name gene id
GU305914.1 227692 227714 + 0 AAACATCGGGGAAGGACAAA AGG ||||||||[||||||||||||]PAM NA Intergenic NA NA

T16977 GGGAAGGACAAAAGGTCATTCGG 1000
Oligo fwd CACCGGGAAGGACAAAAGGTCATT
Oligo rev AAACAATGACCTTTTGTCCTTCCC
Chromosome start end strand MM target_seq PAM alignment distance position gene name gene id
GU305914.1 227700 227722 + 0 GGGAAGGACAAAAGGTCATT CGG ||||||||[||||||||||||]PAM NA Intergenic NA NA

T16978 GGAAGGACAAAAGGTCATTCGGG 1000
Oligo fwd CACCGGAAGGACAAAAGGTCATTC
Oligo rev AAACGAATGACCTTTTGTCCTTCC
Chromosome start end strand MM target_seq PAM alignment distance position gene name gene id
GU305914.1 227701 227723 + 0 GGAAGGACAAAAGGTCATTC GGG ||||||||[||||||||||||]PAM NA Intergenic NA NA

<table>
<tr><td>T16979</td><td>ACAAAAGGTCATTCGGGAGCTGG</td><td>1000</td><td></td><td></td><td></td><td></td><td></td><td></td><td></td><td></td><td></td></tr>
<tr><td>Oligo fwd</td><td>CACCACAAAAGGTCATTCGGGAGC</td><td></td><td></td><td></td><td></td><td></td><td></td><td></td><td></td><td></td><td></td></tr>
<tr><td>Oligo rev</td><td>AAACGCTCCCGAATGACCTTTTGT</td><td></td><td></td><td></td><td></td><td></td><td></td><td></td><td></td><td></td><td></td></tr>
<tr><td>Chromosome</td><td>start</td><td>end</td><td>strand</td><td>MM</td><td>target_seq</td><td>PAM</td><td>alignment</td><td>distance</td><td>position</td><td>gene name</td><td>gene id</td></tr>
<tr><td>GU305914.1</td><td>227707</td><td>227729</td><td>+</td><td>0</td><td>ACAAAAGGTCATTCGGGAGC</td><td>TGG</td><td>||||||||[||||||||||||]PAM</td><td>NA</td><td>Intergenic</td><td>NA</td><td>NA</td></tr>
<tr><td>T16980</td><td>CAAAAGGTCATTCGGGAGCTGGG</td><td>1000</td><td></td><td></td><td></td><td></td><td></td><td></td><td></td><td></td><td></td></tr>
<tr><td>Oligo fwd</td><td>CACCCAAAAGGTCATTCGGGAGCT</td><td></td><td></td><td></td><td></td><td></td><td></td><td></td><td></td><td></td><td></td></tr>
<tr><td>Oligo rev</td><td>AAACAGCTCCCGAATGACCTTTTG</td><td></td><td></td><td></td><td></td><td></td><td></td><td></td><td></td><td></td><td></td></tr>
<tr><td>Chromosome</td><td>start</td><td>end</td><td>strand</td><td>MM</td><td>target_seq</td><td>PAM</td><td>alignment</td><td>distance</td><td>position</td><td>gene name</td><td>gene id</td></tr>
<tr><td>GU305914.1</td><td>227708</td><td>227730</td><td>+</td><td>0</td><td>CAAAAGGTCATTCGGGAGCT</td><td>GGG</td><td>||||||||[||||||||||||]PAM</td><td>NA</td><td>Intergenic</td><td>NA</td><td>NA</td></tr>
<tr><td>T16981</td><td>AAAAGGTCATTCGGGAGCTGGGG</td><td>1000</td><td></td><td></td><td></td><td></td><td></td><td></td><td></td><td></td><td></td></tr>
<tr><td>Oligo fwd</td><td>CACCAAAAGGTCATTCGGGAGCTG</td><td></td><td></td><td></td><td></td><td></td><td></td><td></td><td></td><td></td><td></td></tr>
<tr><td>Oligo rev</td><td>AAACCAGCTCCCGAATGACCTTTT</td><td></td><td></td><td></td><td></td><td></td><td></td><td></td><td></td><td></td><td></td></tr>
<tr><td>Chromosome</td><td>start</td><td>end</td><td>strand</td><td>MM</td><td>target_seq</td><td>PAM</td><td>alignment</td><td>distance</td><td>position</td><td>gene name</td><td>gene id</td></tr>
<tr><td>GU305914.1</td><td>227709</td><td>227731</td><td>+</td><td>0</td><td>AAAAGGTCATTCGGGAGCTG</td><td>GGG</td><td>||||||||[||||||||||||]PAM</td><td>NA</td><td>Intergenic</td><td>NA</td><td>NA</td></tr>
<tr><td>T16982</td><td>TCATTCGGGAGCTGGGGTCATGG</td><td>1000</td><td></td><td></td><td></td><td></td><td></td><td></td><td></td><td></td><td></td></tr>
<tr><td>Oligo fwd</td><td>CACCTCATTCGGGAGCTGGGGTCA</td><td></td><td></td><td></td><td></td><td></td><td></td><td></td><td></td><td></td><td></td></tr>
<tr><td>Oligo rev</td><td>AAACTGACCCCAGCTCCCGAATGA</td><td></td><td></td><td></td><td></td><td></td><td></td><td></td><td></td><td></td><td></td></tr>
<tr><td>Chromosome</td><td>start</td><td>end</td><td>strand</td><td>MM</td><td>target_seq</td><td>PAM</td><td>alignment</td><td>distance</td><td>position</td><td>gene name</td><td>gene id</td></tr>
<tr><td>GU305914.1</td><td>227715</td><td>227737</td><td>+</td><td>0</td><td>TCATTCGGGAGCTGGGGTCA</td><td>TGG</td><td>||||||||[||||||||||||]PAM</td><td>NA</td><td>Intergenic</td><td>NA</td><td>NA</td></tr>
<tr><td>T16983</td><td>CGGGAGCTGGGGTCATGGAACGG</td><td>1000</td><td></td><td></td><td></td><td></td><td></td><td></td><td></td><td></td><td></td></tr>
<tr><td>Oligo fwd</td><td>CACCCGGGAGCTGGGGTCATGGAA</td><td></td><td></td><td></td><td></td><td></td><td></td><td></td><td></td><td></td><td></td></tr>
<tr><td>Oligo rev</td><td>AAACTTCCATGACCCCAGCTCCCG</td><td></td><td></td><td></td><td></td><td></td><td></td><td></td><td></td><td></td><td></td></tr>
<tr><td>Chromosome</td><td>start</td><td>end</td><td>strand</td><td>MM</td><td>target_seq</td><td>PAM</td><td>alignment</td><td>distance</td><td>position</td><td>gene name</td><td>gene id</td></tr>
<tr><td>GU305914.1</td><td>227720</td><td>227742</td><td>+</td><td>0</td><td>CGGGAGCTGGGGTCATGGAA</td><td>CGG</td><td>||||||||[||||||||||||]PAM</td><td>NA</td><td>Intergenic</td><td>NA</td><td>NA</td></tr>
<tr><td>T16984</td><td>CTGGGGTCATGGAACGGTCAAGG</td><td>1000</td><td></td><td></td><td></td><td></td><td></td><td></td><td></td><td></td><td></td></tr>
<tr><td>Oligo fwd</td><td>CACCCTGGGGTCATGGAACGGTCA</td><td></td><td></td><td></td><td></td><td></td><td></td><td></td><td></td><td></td><td></td></tr>
<tr><td>Oligo rev</td><td>AAACTGACCGTTCCATGACCCCAG</td><td></td><td></td><td></td><td></td><td></td><td></td><td></td><td></td><td></td><td></td></tr>
<tr><td>Chromosome</td><td>start</td><td>end</td><td>strand</td><td>MM</td><td>target_seq</td><td>PAM</td><td>alignment</td><td>distance</td><td>position</td><td>gene name</td><td>gene id</td></tr>
<tr><td>GU305914.1</td><td>227726</td><td>227748</td><td>+</td><td>0</td><td>CTGGGGTCATGGAACGGTCA</td><td>AGG</td><td>||||||||[||||||||||||]PAM</td><td>NA</td><td>Intergenic</td><td>NA</td><td>NA</td></tr>
<tr><td>T16985</td><td>ATGGAACGGTCAAGGTTACGAGG</td><td>1000</td><td></td><td></td><td></td><td></td><td></td><td></td><td></td><td></td><td></td></tr>
<tr><td>Oligo fwd</td><td>CACCATGGAACGGTCAAGGTTACG</td><td></td><td></td><td></td><td></td><td></td><td></td><td></td><td></td><td></td><td></td></tr>
<tr><td>Oligo rev</td><td>AAACCGTAACCTTGACCGTTCCAT</td><td></td><td></td><td></td><td></td><td></td><td></td><td></td><td></td><td></td><td></td></tr>
<tr><td>Chromosome</td><td>start</td><td>end</td><td>strand</td><td>MM</td><td>target_seq</td><td>PAM</td><td>alignment</td><td>distance</td><td>position</td><td>gene name</td><td>gene id</td></tr>
<tr><td>GU305914.1</td><td>227734</td><td>227756</td><td>+</td><td>0</td><td>ATGGAACGGTCAAGGTTACG</td><td>AGG</td><td>||||||||[||||||||||||]PAM</td><td>NA</td><td>Intergenic</td><td>NA</td><td>NA</td></tr>
<tr><td>T16986</td><td>GGTCAAGGTTACGAGGAAAAAGG</td><td>1000</td><td></td><td></td><td></td><td></td><td></td><td></td><td></td><td></td><td></td></tr>
<tr><td>Oligo fwd</td><td>CACCGGTCAAGGTTACGAGGAAAA</td><td></td><td></td><td></td><td></td><td></td><td></td><td></td><td></td><td></td><td></td></tr>
<tr><td>Oligo rev</td><td>AAACTTTTCCTCGTAACCTTGACC</td><td></td><td></td><td></td><td></td><td></td><td></td><td></td><td></td><td></td><td></td></tr>
<tr><td>Chromosome</td><td>start</td><td>end</td><td>strand</td><td>MM</td><td>target_seq</td><td>PAM</td><td>alignment</td><td>distance</td><td>position</td><td>gene name</td><td>gene id</td></tr>
<tr><td>GU305914.1</td><td>227741</td><td>227763</td><td>+</td><td>0</td><td>GGTCAAGGTTACGAGGAAAA</td><td>AGG</td><td>||||||||[||||||||||||]PAM</td><td>NA</td><td>Intergenic</td><td>NA</td><td>NA</td></tr>
<tr><td>T16987</td><td>AAGGTTACGAGGAAAAAGGTCGG</td><td>1000</td><td></td><td></td><td></td><td></td><td></td><td></td><td></td><td></td><td></td></tr>
<tr><td>Oligo fwd</td><td>CACCAAGGTTACGAGGAAAAAGGT</td><td></td><td></td><td></td><td></td><td></td><td></td><td></td><td></td><td></td><td></td></tr>
<tr><td>Oligo rev</td><td>AAACACCTTTTTCCTCGTAACCTT</td><td></td><td></td><td></td><td></td><td></td><td></td><td></td><td></td><td></td><td></td></tr>
<tr><td>Chromosome</td><td>start</td><td>end</td><td>strand</td><td>MM</td><td>target_seq</td><td>PAM</td><td>alignment</td><td>distance</td><td>position</td><td>gene name</td><td>gene id</td></tr>
<tr><td>GU305914.1</td><td>227745</td><td>227767</td><td>+</td><td>0</td><td>AAGGTTACGAGGAAAAAGGT</td><td>CGG</td><td>||||||||[||||||||||||]PAM</td><td>NA</td><td>Intergenic</td><td>NA</td><td>NA</td></tr>
<tr><td>T16988</td><td>AGGTTACGAGGAAAAAGGTCGGG</td><td>1000</td><td></td><td></td><td></td><td></td><td></td><td></td><td></td><td></td><td></td></tr>
<tr><td>Oligo fwd</td><td>CACCAGGTTACGAGGAAAAAGGTC</td><td></td><td></td><td></td><td></td><td></td><td></td><td></td><td></td><td></td><td></td></tr>
<tr><td>Oligo rev</td><td>AAACGACCTTTTTCCTCGTAACCT</td><td></td><td></td><td></td><td></td><td></td><td></td><td></td><td></td><td></td><td></td></tr>
<tr><td>Chromosome</td><td>start</td><td>end</td><td>strand</td><td>MM</td><td>target_seq</td><td>PAM</td><td>alignment</td><td>distance</td><td>position</td><td>gene name</td><td>gene id</td></tr>
<tr><td>GU305914.1</td><td>227746</td><td>227768</td><td>+</td><td>0</td><td>AGGTTACGAGGAAAAAGGTC</td><td>GGG</td><td>||||||||[||||||||||||]PAM</td><td>NA</td><td>Intergenic</td><td>NA</td><td>NA</td></tr>
<tr><td>T16989</td><td>ACGAGGAAAAAGGTCGGGACAGG</td><td>1000</td><td></td><td></td><td></td><td></td><td></td><td></td><td></td><td></td><td></td></tr>
<tr><td>Oligo fwd</td><td>CACCACGAGGAAAAAGGTCGGGAC</td><td></td><td></td><td></td><td></td><td></td><td></td><td></td><td></td><td></td><td></td></tr>
<tr><td>Oligo rev</td><td>AAACGTCCCGACCTTTTTCCTCGT</td><td></td><td></td><td></td><td></td><td></td><td></td><td></td><td></td><td></td><td></td></tr>
<tr><td>Chromosome</td><td>start</td><td>end</td><td>strand</td><td>MM</td><td>target_seq</td><td>PAM</td><td>alignment</td><td>distance</td><td>position</td><td>gene name</td><td>gene id</td></tr>
<tr><td>GU305914.1</td><td>227751</td><td>227773</td><td>+</td><td>0</td><td>ACGAGGAAAAAGGTCGGGAC</td><td>AGG</td><td>||||||||[||||||||||||]PAM</td><td>NA</td><td>Intergenic</td><td>NA</td><td>NA</td></tr>
<tr><td>T16990</td><td>GAAAAAGGTCGGGACAGGTCAGG</td><td>1000</td><td></td><td></td><td></td><td></td><td></td><td></td><td></td><td></td><td></td></tr>
<tr><td>Oligo fwd</td><td>CACCGAAAAAGGTCGGGACAGGTC</td><td></td><td></td><td></td><td></td><td></td><td></td><td></td><td></td><td></td><td></td></tr>
<tr><td>Oligo rev</td><td>AAACGACCTGTCCCGACCTTTTTC</td><td></td><td></td><td></td><td></td><td></td><td></td><td></td><td></td><td></td><td></td></tr>
<tr><td>Chromosome</td><td>start</td><td>end</td><td>strand</td><td>MM</td><td>target_seq</td><td>PAM</td><td>alignment</td><td>distance</td><td>position</td><td>gene name</td><td>gene id</td></tr>
<tr><td>GU305914.1</td><td>227756</td><td>227778</td><td>+</td><td>0</td><td>GAAAAAGGTCGGGACAGGTC</td><td>AGG</td><td>||||||||[||||||||||||]PAM</td><td>NA</td><td>Intergenic</td><td>NA</td><td>NA</td></tr>
<tr><td>T16991</td><td>AAAAAGGTCGGGACAGGTCAGGG</td><td>1000</td><td></td><td></td><td></td><td></td><td></td><td></td><td></td><td></td><td></td></tr>
<tr><td>Oligo fwd</td><td>CACCAAAAAGGTCGGGACAGGTCA</td><td></td><td></td><td></td><td></td><td></td><td></td><td></td><td></td><td></td><td></td></tr>
<tr><td>Oligo rev</td><td>AAACTGACCTGTCCCGACCTTTTT</td><td></td><td></td><td></td><td></td><td></td><td></td><td></td><td></td><td></td><td></td></tr>
<tr><td>Chromosome</td><td>start</td><td>end</td><td>strand</td><td>MM</td><td>target_seq</td><td>PAM</td><td>alignment</td><td>distance</td><td>position</td><td>gene name</td><td>gene id</td></tr>
<tr><td>GU305914.1</td><td>227757</td><td>227779</td><td>+</td><td>0</td><td>AAAAAGGTCGGGACAGGTCA</td><td>GGG</td><td>||||||||[||||||||||||]PAM</td><td>NA</td><td>Intergenic</td><td>NA</td><td>NA</td></tr>
<tr><td>T16992</td><td>TCAGGGATCTTAATGCTAAGTGG</td><td>1000</td><td></td><td></td><td></td><td></td><td></td><td></td><td></td><td></td><td></td></tr>
<tr><td>Oligo fwd</td><td>CACCTCAGGGATCTTAATGCTAAG</td><td></td><td></td><td></td><td></td><td></td><td></td><td></td><td></td><td></td><td></td></tr>
<tr><td>Oligo rev</td><td>AAACCTTAGCATTAAGATCCCTGA</td><td></td><td></td><td></td><td></td><td></td><td></td><td></td><td></td><td></td><td></td></tr>
<tr><td>Chromosome</td><td>start</td><td>end</td><td>strand</td><td>MM</td><td>target_seq</td><td>PAM</td><td>alignment</td><td>distance</td><td>position</td><td>gene name</td><td>gene id</td></tr>
<tr><td>GU305914.1</td><td>227774</td><td>227796</td><td>+</td><td>0</td><td>TCAGGGATCTTAATGCTAAG</td><td>TGG</td><td>||||||||[||||||||||||]PAM</td><td>NA</td><td>Intergenic</td><td>NA</td><td>NA</td></tr>
<tr><td>T16993</td><td>TTAATGCTAAGTGGATAACACGG</td><td>1000</td><td></td><td></td><td></td><td></td><td></td><td></td><td></td><td></td><td></td></tr>
<tr><td>Oligo fwd</td><td>CACCTTAATGCTAAGTGGATAACA</td><td></td><td></td><td></td><td></td><td></td><td></td><td></td><td></td><td></td><td></td></tr>
<tr><td>Oligo rev</td><td>AAACTGTTATCCACTTAGCATTAA</td><td></td><td></td><td></td><td></td><td></td><td></td><td></td><td></td><td></td><td></td></tr>
<tr><td>Chromosome</td><td>start</td><td>end</td><td>strand</td><td>MM</td><td>target_seq</td><td>PAM</td><td>alignment</td><td>distance</td><td>position</td><td>gene name</td><td>gene id</td></tr>
<tr><td>GU305914.1</td><td>227783</td><td>227805</td><td>+</td><td>0</td><td>TTAATGCTAAGTGGATAACA</td><td>CGG</td><td>||||||||[||||||||||||]PAM</td><td>NA</td><td>Intergenic</td><td>NA</td><td>NA</td></tr>
</table>

<table>
<tr><td>T16994</td><td>TAAGTGGATAACACGGAAGACGG</td><td>1000</td><td></td><td></td><td></td><td></td><td></td><td></td><td></td><td></td><td></td></tr>
<tr><td>Oligo fwd</td><td>CACCTAAGTGGATAACACGGAAGA</td><td></td><td></td><td></td><td></td><td></td><td></td><td></td><td></td><td></td><td></td></tr>
<tr><td>Oligo rev</td><td>AAACTCTTCCGTGTTATCCACTTA</td><td></td><td></td><td></td><td></td><td></td><td></td><td></td><td></td><td></td><td></td></tr>
<tr><td>Chromosome</td><td>start</td><td>end</td><td>strand</td><td>MM</td><td>target_seq</td><td>PAM</td><td>alignment</td><td>distance</td><td>position</td><td>gene name</td><td>gene id</td></tr>
<tr><td>GU305914.1</td><td>227790</td><td>227812</td><td>+</td><td>0</td><td>TAAGTGGATAACACGGAAGA</td><td>CGG</td><td>||||||||[||||||||||||]PAM</td><td>NA</td><td>Intergenic</td><td>NA</td><td>NA</td></tr>
<tr><td>T16995</td><td>TGGATAACACGGAAGACGGACGG</td><td>1000</td><td></td><td></td><td></td><td></td><td></td><td></td><td></td><td></td><td></td></tr>
<tr><td>Oligo fwd</td><td>CACCTGGATAACACGGAAGACGGA</td><td></td><td></td><td></td><td></td><td></td><td></td><td></td><td></td><td></td><td></td></tr>
<tr><td>Oligo rev</td><td>AAACTCCGTCTTCCGTGTTATCCA</td><td></td><td></td><td></td><td></td><td></td><td></td><td></td><td></td><td></td><td></td></tr>
<tr><td>Chromosome</td><td>start</td><td>end</td><td>strand</td><td>MM</td><td>target_seq</td><td>PAM</td><td>alignment</td><td>distance</td><td>position</td><td>gene name</td><td>gene id</td></tr>
<tr><td>GU305914.1</td><td>227794</td><td>227816</td><td>+</td><td>0</td><td>TGGATAACACGGAAGACGGA</td><td>CGG</td><td>||||||||[||||||||||||]PAM</td><td>NA</td><td>Intergenic</td><td>NA</td><td>NA</td></tr>
<tr><td>T16996</td><td>ACGGACGGCCGTCGACCGATCGG</td><td>1000</td><td></td><td></td><td></td><td></td><td></td><td></td><td></td><td></td><td></td></tr>
<tr><td>Oligo fwd</td><td>CACCACGGACGGCCGTCGACCGAT</td><td></td><td></td><td></td><td></td><td></td><td></td><td></td><td></td><td></td><td></td></tr>
<tr><td>Oligo rev</td><td>AAACATCGGTCGACGGCCGTCCGT</td><td></td><td></td><td></td><td></td><td></td><td></td><td></td><td></td><td></td><td></td></tr>
<tr><td>Chromosome</td><td>start</td><td>end</td><td>strand</td><td>MM</td><td>target_seq</td><td>PAM</td><td>alignment</td><td>distance</td><td>position</td><td>gene name</td><td>gene id</td></tr>
<tr><td>GU305914.1</td><td>227809</td><td>227831</td><td>+</td><td>0</td><td>ACGGACGGCCGTCGACCGAT</td><td>CGG</td><td>||||||||[||||||||||||]PAM</td><td>NA</td><td>Intergenic</td><td>NA</td><td>NA</td></tr>
<tr><td>T16997</td><td>CGACCGATCGGACCAGCCGACGG</td><td>1000</td><td></td><td></td><td></td><td></td><td></td><td></td><td></td><td></td><td></td></tr>
<tr><td>Oligo fwd</td><td>CACCCGACCGATCGGACCAGCCGA</td><td></td><td></td><td></td><td></td><td></td><td></td><td></td><td></td><td></td><td></td></tr>
<tr><td>Oligo rev</td><td>AAACTCGGCTGGTCCGATCGGTCG</td><td></td><td></td><td></td><td></td><td></td><td></td><td></td><td></td><td></td><td></td></tr>
<tr><td>Chromosome</td><td>start</td><td>end</td><td>strand</td><td>MM</td><td>target_seq</td><td>PAM</td><td>alignment</td><td>distance</td><td>position</td><td>gene name</td><td>gene id</td></tr>
<tr><td>GU305914.1</td><td>227821</td><td>227843</td><td>+</td><td>0</td><td>CGACCGATCGGACCAGCCGA</td><td>CGG</td><td>||||||||[||||||||||||]PAM</td><td>NA</td><td>Intergenic</td><td>NA</td><td>NA</td></tr>
<tr><td>T16998</td><td>TCGGACCAGCCGACGGTTGACGG</td><td>1000</td><td></td><td></td><td></td><td></td><td></td><td></td><td></td><td></td><td></td></tr>
<tr><td>Oligo fwd</td><td>CACCTCGGACCAGCCGACGGTTGA</td><td></td><td></td><td></td><td></td><td></td><td></td><td></td><td></td><td></td><td></td></tr>
<tr><td>Oligo rev</td><td>AAACTCAACCGTCGGCTGGTCCGA</td><td></td><td></td><td></td><td></td><td></td><td></td><td></td><td></td><td></td><td></td></tr>
<tr><td>Chromosome</td><td>start</td><td>end</td><td>strand</td><td>MM</td><td>target_seq</td><td>PAM</td><td>alignment</td><td>distance</td><td>position</td><td>gene name</td><td>gene id</td></tr>
<tr><td>GU305914.1</td><td>227828</td><td>227850</td><td>+</td><td>0</td><td>TCGGACCAGCCGACGGTTGA</td><td>CGG</td><td>||||||||[||||||||||||]PAM</td><td>NA</td><td>Intergenic</td><td>NA</td><td>NA</td></tr>
<tr><td>T16999</td><td>CCAGCCGACGGTTGACGGTGAGG</td><td>1000</td><td></td><td></td><td></td><td></td><td></td><td></td><td></td><td></td><td></td></tr>
<tr><td>Oligo fwd</td><td>CACCCCAGCCGACGGTTGACGGTG</td><td></td><td></td><td></td><td></td><td></td><td></td><td></td><td></td><td></td><td></td></tr>
<tr><td>Oligo rev</td><td>AAACCACCGTCAACCGTCGGCTGG</td><td></td><td></td><td></td><td></td><td></td><td></td><td></td><td></td><td></td><td></td></tr>
<tr><td>Chromosome</td><td>start</td><td>end</td><td>strand</td><td>MM</td><td>target_seq</td><td>PAM</td><td>alignment</td><td>distance</td><td>position</td><td>gene name</td><td>gene id</td></tr>
<tr><td>GU305914.1</td><td>227833</td><td>227855</td><td>+</td><td>0</td><td>CCAGCCGACGGTTGACGGTG</td><td>AGG</td><td>||||||||[||||||||||||]PAM</td><td>NA</td><td>Intergenic</td><td>NA</td><td>NA</td></tr>
<tr><td>T17000</td><td>GGTTGACGGTGAGGATTTAGTGG</td><td>1000</td><td></td><td></td><td></td><td></td><td></td><td></td><td></td><td></td><td></td></tr>
<tr><td>Oligo fwd</td><td>CACCGGTTGACGGTGAGGATTTAG</td><td></td><td></td><td></td><td></td><td></td><td></td><td></td><td></td><td></td><td></td></tr>
<tr><td>Oligo rev</td><td>AAACCTAAATCCTCACCGTCAACC</td><td></td><td></td><td></td><td></td><td></td><td></td><td></td><td></td><td></td><td></td></tr>
<tr><td>Chromosome</td><td>start</td><td>end</td><td>strand</td><td>MM</td><td>target_seq</td><td>PAM</td><td>alignment</td><td>distance</td><td>position</td><td>gene name</td><td>gene id</td></tr>
<tr><td>GU305914.1</td><td>227842</td><td>227864</td><td>+</td><td>0</td><td>GGTTGACGGTGAGGATTTAG</td><td>TGG</td><td>||||||||[||||||||||||]PAM</td><td>NA</td><td>Intergenic</td><td>NA</td><td>NA</td></tr>
<tr><td>T17001</td><td>GGCATGTTAGCACCGCGACCCGG</td><td>1000</td><td></td><td></td><td></td><td></td><td></td><td></td><td></td><td></td><td></td></tr>
<tr><td>Oligo fwd</td><td>CACCGGCATGTTAGCACCGCGACC</td><td></td><td></td><td></td><td></td><td></td><td></td><td></td><td></td><td></td><td></td></tr>
<tr><td>Oligo rev</td><td>AAACGGTCGCGGTGCTAACATGCC</td><td></td><td></td><td></td><td></td><td></td><td></td><td></td><td></td><td></td><td></td></tr>
<tr><td>Chromosome</td><td>start</td><td>end</td><td>strand</td><td>MM</td><td>target_seq</td><td>PAM</td><td>alignment</td><td>distance</td><td>position</td><td>gene name</td><td>gene id</td></tr>
<tr><td>GU305914.1</td><td>227863</td><td>227885</td><td>+</td><td>0</td><td>GGCATGTTAGCACCGCGACC</td><td>CGG</td><td>||||||||[||||||||||||]PAM</td><td>NA</td><td>Intergenic</td><td>NA</td><td>NA</td></tr>
<tr><td>T17002</td><td>ATGTTAGCACCGCGACCCGGAGG</td><td>1000</td><td></td><td></td><td></td><td></td><td></td><td></td><td></td><td></td><td></td></tr>
<tr><td>Oligo fwd</td><td>CACCATGTTAGCACCGCGACCCGG</td><td></td><td></td><td></td><td></td><td></td><td></td><td></td><td></td><td></td><td></td></tr>
<tr><td>Oligo rev</td><td>AAACCCGGGTCGCGGTGCTAACAT</td><td></td><td></td><td></td><td></td><td></td><td></td><td></td><td></td><td></td><td></td></tr>
<tr><td>Chromosome</td><td>start</td><td>end</td><td>strand</td><td>MM</td><td>target_seq</td><td>PAM</td><td>alignment</td><td>distance</td><td>position</td><td>gene name</td><td>gene id</td></tr>
<tr><td>GU305914.1</td><td>227866</td><td>227888</td><td>+</td><td>0</td><td>ATGTTAGCACCGCGACCCGG</td><td>AGG</td><td>||||||||[||||||||||||]PAM</td><td>NA</td><td>Intergenic</td><td>NA</td><td>NA</td></tr>
<tr><td>T17003</td><td>CCGGAGGTCTTAGAACATATAGG</td><td>1000</td><td></td><td></td><td></td><td></td><td></td><td></td><td></td><td></td><td></td></tr>
<tr><td>Oligo fwd</td><td>CACCCCGGAGGTCTTAGAACATAT</td><td></td><td></td><td></td><td></td><td></td><td></td><td></td><td></td><td></td><td></td></tr>
<tr><td>Oligo rev</td><td>AAACATATGTTCTAAGACCTCCGG</td><td></td><td></td><td></td><td></td><td></td><td></td><td></td><td></td><td></td><td></td></tr>
<tr><td>Chromosome</td><td>start</td><td>end</td><td>strand</td><td>MM</td><td>target_seq</td><td>PAM</td><td>alignment</td><td>distance</td><td>position</td><td>gene name</td><td>gene id</td></tr>
<tr><td>GU305914.1</td><td>227882</td><td>227904</td><td>+</td><td>0</td><td>CCGGAGGTCTTAGAACATAT</td><td>AGG</td><td>||||||||[||||||||||||]PAM</td><td>NA</td><td>Intergenic</td><td>NA</td><td>NA</td></tr>
<tr><td>T17004</td><td>CGGAGGTCTTAGAACATATAGGG</td><td>1000</td><td></td><td></td><td></td><td></td><td></td><td></td><td></td><td></td><td></td></tr>
<tr><td>Oligo fwd</td><td>CACCCGGAGGTCTTAGAACATATA</td><td></td><td></td><td></td><td></td><td></td><td></td><td></td><td></td><td></td><td></td></tr>
<tr><td>Oligo rev</td><td>AAACTATATGTTCTAAGACCTCCG</td><td></td><td></td><td></td><td></td><td></td><td></td><td></td><td></td><td></td><td></td></tr>
<tr><td>Chromosome</td><td>start</td><td>end</td><td>strand</td><td>MM</td><td>target_seq</td><td>PAM</td><td>alignment</td><td>distance</td><td>position</td><td>gene name</td><td>gene id</td></tr>
<tr><td>GU305914.1</td><td>227883</td><td>227905</td><td>+</td><td>0</td><td>CGGAGGTCTTAGAACATATA</td><td>GGG</td><td>||||||||[||||||||||||]PAM</td><td>NA</td><td>Intergenic</td><td>NA</td><td>NA</td></tr>
<tr><td>T17005</td><td>ACATATAGGGAACCACGATGAGG</td><td>1000</td><td></td><td></td><td></td><td></td><td></td><td></td><td></td><td></td><td></td></tr>
<tr><td>Oligo fwd</td><td>CACCACATATAGGGAACCACGATG</td><td></td><td></td><td></td><td></td><td></td><td></td><td></td><td></td><td></td><td></td></tr>
<tr><td>Oligo rev</td><td>AAACCATCGTGGTTCCCTATATGT</td><td></td><td></td><td></td><td></td><td></td><td></td><td></td><td></td><td></td><td></td></tr>
<tr><td>Chromosome</td><td>start</td><td>end</td><td>strand</td><td>MM</td><td>target_seq</td><td>PAM</td><td>alignment</td><td>distance</td><td>position</td><td>gene name</td><td>gene id</td></tr>
<tr><td>GU305914.1</td><td>227896</td><td>227918</td><td>+</td><td>0</td><td>ACATATAGGGAACCACGATG</td><td>AGG</td><td>||||||||[||||||||||||]PAM</td><td>NA</td><td>Intergenic</td><td>NA</td><td>NA</td></tr>
<tr><td>T17006</td><td>CATATAGGGAACCACGATGAGGG</td><td>1000</td><td></td><td></td><td></td><td></td><td></td><td></td><td></td><td></td><td></td></tr>
<tr><td>Oligo fwd</td><td>CACCCATATAGGGAACCACGATGA</td><td></td><td></td><td></td><td></td><td></td><td></td><td></td><td></td><td></td><td></td></tr>
<tr><td>Oligo rev</td><td>AAACTCATCGTGGTTCCCTATATG</td><td></td><td></td><td></td><td></td><td></td><td></td><td></td><td></td><td></td><td></td></tr>
<tr><td>Chromosome</td><td>start</td><td>end</td><td>strand</td><td>MM</td><td>target_seq</td><td>PAM</td><td>alignment</td><td>distance</td><td>position</td><td>gene name</td><td>gene id</td></tr>
<tr><td>GU305914.1</td><td>227897</td><td>227919</td><td>+</td><td>0</td><td>CATATAGGGAACCACGATGA</td><td>GGG</td><td>||||||||[||||||||||||]PAM</td><td>NA</td><td>Intergenic</td><td>NA</td><td>NA</td></tr>
<tr><td>T17007</td><td>GGGAACCACGATGAGGGTCTCGG</td><td>1000</td><td></td><td></td><td></td><td></td><td></td><td></td><td></td><td></td><td></td></tr>
<tr><td>Oligo fwd</td><td>CACCGGGAACCACGATGAGGGTCT</td><td></td><td></td><td></td><td></td><td></td><td></td><td></td><td></td><td></td><td></td></tr>
<tr><td>Oligo rev</td><td>AAACAGACCCTCATCGTGGTTCCC</td><td></td><td></td><td></td><td></td><td></td><td></td><td></td><td></td><td></td><td></td></tr>
<tr><td>Chromosome</td><td>start</td><td>end</td><td>strand</td><td>MM</td><td>target_seq</td><td>PAM</td><td>alignment</td><td>distance</td><td>position</td><td>gene name</td><td>gene id</td></tr>
<tr><td>GU305914.1</td><td>227903</td><td>227925</td><td>+</td><td>0</td><td>GGGAACCACGATGAGGGTCT</td><td>CGG</td><td>||||||||[||||||||||||]PAM</td><td>NA</td><td>Intergenic</td><td>NA</td><td>NA</td></tr>
<tr><td>T17008</td><td>ACCACGATGAGGGTCTCGGACGG</td><td>1000</td><td></td><td></td><td></td><td></td><td></td><td></td><td></td><td></td><td></td></tr>
<tr><td>Oligo fwd</td><td>CACCACCACGATGAGGGTCTCGGA</td><td></td><td></td><td></td><td></td><td></td><td></td><td></td><td></td><td></td><td></td></tr>
<tr><td>Oligo rev</td><td>AAACTCCGAGACCCTCATCGTGGT</td><td></td><td></td><td></td><td></td><td></td><td></td><td></td><td></td><td></td><td></td></tr>
<tr><td>Chromosome</td><td>start</td><td>end</td><td>strand</td><td>MM</td><td>target_seq</td><td>PAM</td><td>alignment</td><td>distance</td><td>position</td><td>gene name</td><td>gene id</td></tr>
</table>

| | | | | | | | | | | | |
|---|---|---|---|---|---|---|---|---|---|---|---|
| GU305914.1 | 227907 | 227929 | + | 0 | ACCACGATGAGGGTCTCGGA | CGG | \|\|\|\|\|\|\|\|[\|\|\|\|\|\|\|\|\|\|\|\|]PAM | NA | Intergenic | NA | NA |
| T17009 | GGTCTCGGACGGAGCTATAGCGG | 1000 | | | | | | | | | |
| Oligo fwd | CACCGGTCTCGGACGGAGCTATAG | | | | | | | | | | |
| Oligo rev | AAACCTATAGCTCCGTCCGAGACC | | | | | | | | | | |
| Chromosome | start | end | strand | MM | target_seq | PAM | alignment | distance | position | gene name | gene id |
| GU305914.1 | 227918 | 227940 | + | 0 | GGTCTCGGACGGAGCTATAG | CGG | \|\|\|\|\|\|\|\|[\|\|\|\|\|\|\|\|\|\|\|\|]PAM | NA | Intergenic | NA | NA |
| T17010 | GTCTCGGACGGAGCTATAGCGGG | 1000 | | | | | | | | | |
| Oligo fwd | CACCGTCTCGGACGGAGCTATAGC | | | | | | | | | | |
| Oligo rev | AAACGCTATAGCTCCGTCCGAGAC | | | | | | | | | | |
| Chromosome | start | end | strand | MM | target_seq | PAM | alignment | distance | position | gene name | gene id |
| GU305914.1 | 227919 | 227941 | + | 0 | GTCTCGGACGGAGCTATAGC | GGG | \|\|\|\|\|\|\|\|[\|\|\|\|\|\|\|\|\|\|\|\|]PAM | NA | Intergenic | NA | NA |
| T17011 | TCTCGGACGGAGCTATAGCGGGG | 1000 | | | | | | | | | |
| Oligo fwd | CACCTCTCGGACGGAGCTATAGCG | | | | | | | | | | |
| Oligo rev | AAACCGCTATAGCTCCGTCCGAGA | | | | | | | | | | |
| Chromosome | start | end | strand | MM | target_seq | PAM | alignment | distance | position | gene name | gene id |
| GU305914.1 | 227920 | 227942 | + | 0 | TCTCGGACGGAGCTATAGCG | GGG | \|\|\|\|\|\|\|\|[\|\|\|\|\|\|\|\|\|\|\|\|]PAM | NA | Intergenic | NA | NA |
| T17012 | TTACACAGAAACGAAACCTAAGG | 1000 | | | | | | | | | |
| Oligo fwd | CACCTTACACAGAAACGAAACCTA | | | | | | | | | | |
| Oligo rev | AAACTAGGTTTCGTTTCTGTGTAA | | | | | | | | | | |
| Chromosome | start | end | strand | MM | target_seq | PAM | alignment | distance | position | gene name | gene id |
| GU305914.1 | 227973 | 227995 | + | 0 | TTACACAGAAACGAAACCTA | AGG | \|\|\|\|\|\|\|\|[\|\|\|\|\|\|\|\|\|\|\|\|]PAM | NA | Intergenic | NA | NA |
| T17013 | ACAGAAACGAAACCTAAGGACGG | 1000 | | | | | | | | | |
| Oligo fwd | CACCACAGAAACGAAACCTAAGGA | | | | | | | | | | |
| Oligo rev | AAACTCCTTAGGTTTCGTTTCTGT | | | | | | | | | | |
| Chromosome | start | end | strand | MM | target_seq | PAM | alignment | distance | position | gene name | gene id |
| GU305914.1 | 227977 | 227999 | + | 0 | ACAGAAACGAAACCTAAGGA | CGG | \|\|\|\|\|\|\|\|[\|\|\|\|\|\|\|\|\|\|\|\|]PAM | NA | Intergenic | NA | NA |
| T17014 | CAGAAACGAAACCTAAGGACGGG | 1000 | | | | | | | | | |
| Oligo fwd | CACCCAGAAACGAAACCTAAGGAC | | | | | | | | | | |
| Oligo rev | AAACGTCCTTAGGTTTCGTTTCTG | | | | | | | | | | |
| Chromosome | start | end | strand | MM | target_seq | PAM | alignment | distance | position | gene name | gene id |
| GU305914.1 | 227978 | 228000 | + | 0 | CAGAAACGAAACCTAAGGAC | GGG | \|\|\|\|\|\|\|\|[\|\|\|\|\|\|\|\|\|\|\|\|]PAM | NA | Intergenic | NA | NA |
| T17015 | GGACGGGTCATCTACGAACTAGG | 1000 | | | | | | | | | |
| Oligo fwd | CACCGGACGGGTCATCTACGAACT | | | | | | | | | | |
| Oligo rev | AAACAGTTCGTAGATGACCCGTCC | | | | | | | | | | |
| Chromosome | start | end | strand | MM | target_seq | PAM | alignment | distance | position | gene name | gene id |
| GU305914.1 | 227994 | 228016 | + | 0 | GGACGGGTCATCTACGAACT | AGG | \|\|\|\|\|\|\|\|[\|\|\|\|\|\|\|\|\|\|\|\|]PAM | NA | Intergenic | NA | NA |
| T17016 | GACGGGTCATCTACGAACTAGGG | 1000 | | | | | | | | | |
| Oligo fwd | CACCGACGGGTCATCTACGAACTA | | | | | | | | | | |
| Oligo rev | AAACTAGTTCGTAGATGACCCGTC | | | | | | | | | | |
| Chromosome | start | end | strand | MM | target_seq | PAM | alignment | distance | position | gene name | gene id |
| GU305914.1 | 227995 | 228017 | + | 0 | GACGGGTCATCTACGAACTA | GGG | \|\|\|\|\|\|\|\|[\|\|\|\|\|\|\|\|\|\|\|\|]PAM | NA | Intergenic | NA | NA |
| T17017 | TACGAACTAGGGAGCATGCGAGG | 1000 | | | | | | | | | |
| Oligo fwd | CACCTACGAACTAGGGAGCATGCG | | | | | | | | | | |
| Oligo rev | AAACCGCATGCTCCCTAGTTCGTA | | | | | | | | | | |
| Chromosome | start | end | strand | MM | target_seq | PAM | alignment | distance | position | gene name | gene id |
| GU305914.1 | 228006 | 228028 | + | 0 | TACGAACTAGGGAGCATGCG | AGG | \|\|\|\|\|\|\|\|[\|\|\|\|\|\|\|\|\|\|\|\|]PAM | NA | Intergenic | NA | NA |
| T17018 | AACTAGGGAGCATGCGAGGACGG | 1000 | | | | | | | | | |
| Oligo fwd | CACCAACTAGGGAGCATGCGAGGA | | | | | | | | | | |
| Oligo rev | AAACTCCTCGCATGCTCCCTAGTT | | | | | | | | | | |
| Chromosome | start | end | strand | MM | target_seq | PAM | alignment | distance | position | gene name | gene id |
| GU305914.1 | 228010 | 228032 | + | 0 | AACTAGGGAGCATGCGAGGA | CGG | \|\|\|\|\|\|\|\|[\|\|\|\|\|\|\|\|\|\|\|\|]PAM | NA | Intergenic | NA | NA |
| T17019 | CATGCGAGGACGGACACACGAGG | 1000 | | | | | | | | | |
| Oligo fwd | CACCCATGCGAGGACGGACACACG | | | | | | | | | | |
| Oligo rev | AAACCGTGTGTCCGTCCTCGCATG | | | | | | | | | | |
| Chromosome | start | end | strand | MM | target_seq | PAM | alignment | distance | position | gene name | gene id |
| GU305914.1 | 228020 | 228042 | + | 0 | CATGCGAGGACGGACACACG | AGG | \|\|\|\|\|\|\|\|[\|\|\|\|\|\|\|\|\|\|\|\|]PAM | NA | Intergenic | NA | NA |
| T17020 | ATGCGAGGACGGACACACGAGGG | 1000 | | | | | | | | | |
| Oligo fwd | CACCATGCGAGGACGGACACACGA | | | | | | | | | | |
| Oligo rev | AAACTCGTGTGTCCGTCCTCGCAT | | | | | | | | | | |
| Chromosome | start | end | strand | MM | target_seq | PAM | alignment | distance | position | gene name | gene id |
| GU305914.1 | 228021 | 228043 | + | 0 | ATGCGAGGACGGACACACGA | GGG | \|\|\|\|\|\|\|\|[\|\|\|\|\|\|\|\|\|\|\|\|]PAM | NA | Intergenic | NA | NA |
| T17021 | GAGGACGGACACACGAGGGAAGG | 1000 | | | | | | | | | |
| Oligo fwd | CACCGAGGACGGACACACGAGGGA | | | | | | | | | | |
| Oligo rev | AAACTCCCTCGTGTGTCCGTCCTC | | | | | | | | | | |
| Chromosome | start | end | strand | MM | target_seq | PAM | alignment | distance | position | gene name | gene id |
| GU305914.1 | 228025 | 228047 | + | 0 | GAGGACGGACACACGAGGGA | AGG | \|\|\|\|\|\|\|\|[\|\|\|\|\|\|\|\|\|\|\|\|]PAM | NA | Intergenic | NA | NA |
| T17022 | TCAAGTAGCATCTCCCGATGAGG | 1000 | | | | | | | | | |
| Oligo fwd | CACCTCAAGTAGCATCTCCCGATG | | | | | | | | | | |
| Oligo rev | AAACCATCGGGAGATGCTACTTGA | | | | | | | | | | |
| Chromosome | start | end | strand | MM | target_seq | PAM | alignment | distance | position | gene name | gene id |
| GU305914.1 | 228101 | 228123 | + | 0 | TCAAGTAGCATCTCCCGATG | AGG | \|\|\|\|\|\|\|\|[\|\|\|\|\|\|\|\|\|\|\|\|]PAM | NA | Intergenic | NA | NA |
| T17023 | CCGATGAGGAGACCGTTCTCAGG | 1000 | | | | | | | | | |
| Oligo fwd | CACCCCGATGAGGAGACCGTTCTC | | | | | | | | | | |
| Oligo rev | AAACGAGAACGGTCTCCTCATCGG | | | | | | | | | | |

<table>
<tr><td>Chromosome</td><td>start</td><td>end</td><td>strand</td><td>MM</td><td>target_seq</td><td>PAM</td><td>alignment</td><td>distance</td><td>position</td><td>gene name</td><td>gene id</td></tr>
<tr><td>GU305914.1</td><td>228115</td><td>228137</td><td>+</td><td>0</td><td>CCGATGAGGAGACCGTTCTC</td><td>AGG</td><td>||||||||[||||||||||||]PAM</td><td>NA</td><td>Intergenic</td><td>NA</td><td>NA</td></tr>
<tr><td></td><td></td><td></td><td></td><td></td><td></td><td></td><td></td><td></td><td></td><td></td><td></td></tr>
<tr><td>T17024</td><td>GTTCTCAGGCAGACATCCGAAGG</td><td>1000</td><td></td><td></td><td></td><td></td><td></td><td></td><td></td><td></td><td></td></tr>
<tr><td>Oligo fwd</td><td>CACCGTTCTCAGGCAGACATCCGA</td><td></td><td></td><td></td><td></td><td></td><td></td><td></td><td></td><td></td><td></td></tr>
<tr><td>Oligo rev</td><td>AAACTCGGATGTCTGCCTGAGAAC</td><td></td><td></td><td></td><td></td><td></td><td></td><td></td><td></td><td></td><td></td></tr>
<tr><td>Chromosome</td><td>start</td><td>end</td><td>strand</td><td>MM</td><td>target_seq</td><td>PAM</td><td>alignment</td><td>distance</td><td>position</td><td>gene name</td><td>gene id</td></tr>
<tr><td>GU305914.1</td><td>228129</td><td>228151</td><td>+</td><td>0</td><td>GTTCTCAGGCAGACATCCGA</td><td>AGG</td><td>||||||||[||||||||||||]PAM</td><td>NA</td><td>Intergenic</td><td>NA</td><td>NA</td></tr>
<tr><td></td><td></td><td></td><td></td><td></td><td></td><td></td><td></td><td></td><td></td><td></td><td></td></tr>
<tr><td>T17025</td><td>TTCTCAGGCAGACATCCGAAGGG</td><td>1000</td><td></td><td></td><td></td><td></td><td></td><td></td><td></td><td></td><td></td></tr>
<tr><td>Oligo fwd</td><td>CACCTTCTCAGGCAGACATCCGAA</td><td></td><td></td><td></td><td></td><td></td><td></td><td></td><td></td><td></td><td></td></tr>
<tr><td>Oligo rev</td><td>AAACTTCGGATGTCTGCCTGAGAA</td><td></td><td></td><td></td><td></td><td></td><td></td><td></td><td></td><td></td><td></td></tr>
<tr><td>Chromosome</td><td>start</td><td>end</td><td>strand</td><td>MM</td><td>target_seq</td><td>PAM</td><td>alignment</td><td>distance</td><td>position</td><td>gene name</td><td>gene id</td></tr>
<tr><td>GU305914.1</td><td>228130</td><td>228152</td><td>+</td><td>0</td><td>TTCTCAGGCAGACATCCGAA</td><td>GGG</td><td>||||||||[||||||||||||]PAM</td><td>NA</td><td>Intergenic</td><td>NA</td><td>NA</td></tr>
<tr><td></td><td></td><td></td><td></td><td></td><td></td><td></td><td></td><td></td><td></td><td></td><td></td></tr>
<tr><td>T17026</td><td>TTATTCCGCAGATCGTCGTCTGG</td><td>1000</td><td></td><td></td><td></td><td></td><td></td><td></td><td></td><td></td><td></td></tr>
<tr><td>Oligo fwd</td><td>CACCTTATTCCGCAGATCGTCGTC</td><td></td><td></td><td></td><td></td><td></td><td></td><td></td><td></td><td></td><td></td></tr>
<tr><td>Oligo rev</td><td>AAACGACGACGATCTGCGGAATAA</td><td></td><td></td><td></td><td></td><td></td><td></td><td></td><td></td><td></td><td></td></tr>
<tr><td>Chromosome</td><td>start</td><td>end</td><td>strand</td><td>MM</td><td>target_seq</td><td>PAM</td><td>alignment</td><td>distance</td><td>position</td><td>gene name</td><td>gene id</td></tr>
<tr><td>GU305914.1</td><td>228171</td><td>228193</td><td>+</td><td>0</td><td>TTATTCCGCAGATCGTCGTC</td><td>TGG</td><td>||||||||[||||||||||||]PAM</td><td>NA</td><td>Intergenic</td><td>NA</td><td>NA</td></tr>
<tr><td></td><td></td><td></td><td></td><td></td><td></td><td></td><td></td><td></td><td></td><td></td><td></td></tr>
<tr><td>T17027</td><td>TATTCCGCAGATCGTCGTCTGGG</td><td>1000</td><td></td><td></td><td></td><td></td><td></td><td></td><td></td><td></td><td></td></tr>
<tr><td>Oligo fwd</td><td>CACCTATTCCGCAGATCGTCGTCT</td><td></td><td></td><td></td><td></td><td></td><td></td><td></td><td></td><td></td><td></td></tr>
<tr><td>Oligo rev</td><td>AAACAGACGACGATCTGCGGAATA</td><td></td><td></td><td></td><td></td><td></td><td></td><td></td><td></td><td></td><td></td></tr>
<tr><td>Chromosome</td><td>start</td><td>end</td><td>strand</td><td>MM</td><td>target_seq</td><td>PAM</td><td>alignment</td><td>distance</td><td>position</td><td>gene name</td><td>gene id</td></tr>
<tr><td>GU305914.1</td><td>228172</td><td>228194</td><td>+</td><td>0</td><td>TATTCCGCAGATCGTCGTCT</td><td>GGG</td><td>||||||||[||||||||||||]PAM</td><td>NA</td><td>Intergenic</td><td>NA</td><td>NA</td></tr>
<tr><td></td><td></td><td></td><td></td><td></td><td></td><td></td><td></td><td></td><td></td><td></td><td></td></tr>
<tr><td>T17028</td><td>ATTCCGCAGATCGTCGTCTGGGG</td><td>1000</td><td></td><td></td><td></td><td></td><td></td><td></td><td></td><td></td><td></td></tr>
<tr><td>Oligo fwd</td><td>CACCATTCCGCAGATCGTCGTCTG</td><td></td><td></td><td></td><td></td><td></td><td></td><td></td><td></td><td></td><td></td></tr>
<tr><td>Oligo rev</td><td>AAACCAGACGACGATCTGCGGAAT</td><td></td><td></td><td></td><td></td><td></td><td></td><td></td><td></td><td></td><td></td></tr>
<tr><td>Chromosome</td><td>start</td><td>end</td><td>strand</td><td>MM</td><td>target_seq</td><td>PAM</td><td>alignment</td><td>distance</td><td>position</td><td>gene name</td><td>gene id</td></tr>
<tr><td>GU305914.1</td><td>228173</td><td>228195</td><td>+</td><td>0</td><td>ATTCCGCAGATCGTCGTCTG</td><td>GGG</td><td>||||||||[||||||||||||]PAM</td><td>NA</td><td>Intergenic</td><td>NA</td><td>NA</td></tr>
<tr><td>GU305914.1</td><td>193699</td><td>193721</td><td>+</td><td>4</td><td>ATGCCCCAGATCGTCATCGG</td><td>TGG</td><td>||-||-||[|||||||-||-|]PAM</td><td>NA</td><td>Intergenic</td><td>NA</td><td>NA</td></tr>
<tr><td></td><td></td><td></td><td></td><td></td><td></td><td></td><td></td><td></td><td></td><td></td><td></td></tr>
<tr><td>T17029</td><td>TTCCGCAGATCGTCGTCTGGGGG</td><td>1000</td><td></td><td></td><td></td><td></td><td></td><td></td><td></td><td></td><td></td></tr>
<tr><td>Oligo fwd</td><td>CACCTTCCGCAGATCGTCGTCTGG</td><td></td><td></td><td></td><td></td><td></td><td></td><td></td><td></td><td></td><td></td></tr>
<tr><td>Oligo rev</td><td>AAACCCAGACGACGATCTGCGGAA</td><td></td><td></td><td></td><td></td><td></td><td></td><td></td><td></td><td></td><td></td></tr>
<tr><td>Chromosome</td><td>start</td><td>end</td><td>strand</td><td>MM</td><td>target_seq</td><td>PAM</td><td>alignment</td><td>distance</td><td>position</td><td>gene name</td><td>gene id</td></tr>
<tr><td>GU305914.1</td><td>228174</td><td>228196</td><td>+</td><td>0</td><td>TTCCGCAGATCGTCGTCTGG</td><td>GGG</td><td>||||||||[||||||||||||]PAM</td><td>NA</td><td>Intergenic</td><td>NA</td><td>NA</td></tr>
<tr><td></td><td></td><td></td><td></td><td></td><td></td><td></td><td></td><td></td><td></td><td></td><td></td></tr>
<tr><td>T17030</td><td>TCGTCGTCTGGGGGAGATCACGG</td><td>1000</td><td></td><td></td><td></td><td></td><td></td><td></td><td></td><td></td><td></td></tr>
<tr><td>Oligo fwd</td><td>CACCTCGTCGTCTGGGGGAGATCA</td><td></td><td></td><td></td><td></td><td></td><td></td><td></td><td></td><td></td><td></td></tr>
<tr><td>Oligo rev</td><td>AAACTGATCTCCCCCAGACGACGA</td><td></td><td></td><td></td><td></td><td></td><td></td><td></td><td></td><td></td><td></td></tr>
<tr><td>Chromosome</td><td>start</td><td>end</td><td>strand</td><td>MM</td><td>target_seq</td><td>PAM</td><td>alignment</td><td>distance</td><td>position</td><td>gene name</td><td>gene id</td></tr>
<tr><td>GU305914.1</td><td>228183</td><td>228205</td><td>+</td><td>0</td><td>TCGTCGTCTGGGGGAGATCA</td><td>CGG</td><td>||||||||[||||||||||||]PAM</td><td>NA</td><td>Intergenic</td><td>NA</td><td>NA</td></tr>
<tr><td></td><td></td><td></td><td></td><td></td><td></td><td></td><td></td><td></td><td></td><td></td><td></td></tr>
<tr><td>T17031</td><td>CGTCTGGGGGAGATCACGGACGG</td><td>1000</td><td></td><td></td><td></td><td></td><td></td><td></td><td></td><td></td><td></td></tr>
<tr><td>Oligo fwd</td><td>CACCCGTCTGGGGGAGATCACGGA</td><td></td><td></td><td></td><td></td><td></td><td></td><td></td><td></td><td></td><td></td></tr>
<tr><td>Oligo rev</td><td>AAACTCCGTGATCTCCCCCAGACG</td><td></td><td></td><td></td><td></td><td></td><td></td><td></td><td></td><td></td><td></td></tr>
<tr><td>Chromosome</td><td>start</td><td>end</td><td>strand</td><td>MM</td><td>target_seq</td><td>PAM</td><td>alignment</td><td>distance</td><td>position</td><td>gene name</td><td>gene id</td></tr>
<tr><td>GU305914.1</td><td>228187</td><td>228209</td><td>+</td><td>0</td><td>CGTCTGGGGGAGATCACGGA</td><td>CGG</td><td>||||||||[||||||||||||]PAM</td><td>NA</td><td>Intergenic</td><td>NA</td><td>NA</td></tr>
<tr><td></td><td></td><td></td><td></td><td></td><td></td><td></td><td></td><td></td><td></td><td></td><td></td></tr>
<tr><td>T17032</td><td>GTCTGGGGGAGATCACGGACGGG</td><td>1000</td><td></td><td></td><td></td><td></td><td></td><td></td><td></td><td></td><td></td></tr>
<tr><td>Oligo fwd</td><td>CACCGTCTGGGGGAGATCACGGAC</td><td></td><td></td><td></td><td></td><td></td><td></td><td></td><td></td><td></td><td></td></tr>
<tr><td>Oligo rev</td><td>AAACGTCCGTGATCTCCCCCAGAC</td><td></td><td></td><td></td><td></td><td></td><td></td><td></td><td></td><td></td><td></td></tr>
<tr><td>Chromosome</td><td>start</td><td>end</td><td>strand</td><td>MM</td><td>target_seq</td><td>PAM</td><td>alignment</td><td>distance</td><td>position</td><td>gene name</td><td>gene id</td></tr>
<tr><td>GU305914.1</td><td>228188</td><td>228210</td><td>+</td><td>0</td><td>GTCTGGGGGAGATCACGGAC</td><td>GGG</td><td>||||||||[||||||||||||]PAM</td><td>NA</td><td>Intergenic</td><td>NA</td><td>NA</td></tr>
<tr><td></td><td></td><td></td><td></td><td></td><td></td><td></td><td></td><td></td><td></td><td></td><td></td></tr>
<tr><td>T17033</td><td>TCTGGGGGAGATCACGGACGGGG</td><td>1000</td><td></td><td></td><td></td><td></td><td></td><td></td><td></td><td></td><td></td></tr>
<tr><td>Oligo fwd</td><td>CACCTCTGGGGGAGATCACGGACG</td><td></td><td></td><td></td><td></td><td></td><td></td><td></td><td></td><td></td><td></td></tr>
<tr><td>Oligo rev</td><td>AAACCGTCCGTGATCTCCCCCAGA</td><td></td><td></td><td></td><td></td><td></td><td></td><td></td><td></td><td></td><td></td></tr>
<tr><td>Chromosome</td><td>start</td><td>end</td><td>strand</td><td>MM</td><td>target_seq</td><td>PAM</td><td>alignment</td><td>distance</td><td>position</td><td>gene name</td><td>gene id</td></tr>
<tr><td>GU305914.1</td><td>228189</td><td>228211</td><td>+</td><td>0</td><td>TCTGGGGGAGATCACGGACG</td><td>GGG</td><td>||||||||[||||||||||||]PAM</td><td>NA</td><td>Intergenic</td><td>NA</td><td>NA</td></tr>
<tr><td></td><td></td><td></td><td></td><td></td><td></td><td></td><td></td><td></td><td></td><td></td><td></td></tr>
<tr><td>T17034</td><td>CTGGGGGAGATCACGGACGGGGG</td><td>1000</td><td></td><td></td><td></td><td></td><td></td><td></td><td></td><td></td><td></td></tr>
<tr><td>Oligo fwd</td><td>CACCCTGGGGGAGATCACGGACGG</td><td></td><td></td><td></td><td></td><td></td><td></td><td></td><td></td><td></td><td></td></tr>
<tr><td>Oligo rev</td><td>AAACCCGTCCGTGATCTCCCCCAG</td><td></td><td></td><td></td><td></td><td></td><td></td><td></td><td></td><td></td><td></td></tr>
<tr><td>Chromosome</td><td>start</td><td>end</td><td>strand</td><td>MM</td><td>target_seq</td><td>PAM</td><td>alignment</td><td>distance</td><td>position</td><td>gene name</td><td>gene id</td></tr>
<tr><td>GU305914.1</td><td>228190</td><td>228212</td><td>+</td><td>0</td><td>CTGGGGGAGATCACGGACGG</td><td>GGG</td><td>||||||||[||||||||||||]PAM</td><td>NA</td><td>Intergenic</td><td>NA</td><td>NA</td></tr>
<tr><td></td><td></td><td></td><td></td><td></td><td></td><td></td><td></td><td></td><td></td><td></td><td></td></tr>
<tr><td>T17035</td><td>TGGGGGAGATCACGGACGGGGGG</td><td>1000</td><td></td><td></td><td></td><td></td><td></td><td></td><td></td><td></td><td></td></tr>
<tr><td>Oligo fwd</td><td>CACCTGGGGGAGATCACGGACGGG</td><td></td><td></td><td></td><td></td><td></td><td></td><td></td><td></td><td></td><td></td></tr>
<tr><td>Oligo rev</td><td>AAACCCCGTCCGTGATCTCCCCCA</td><td></td><td></td><td></td><td></td><td></td><td></td><td></td><td></td><td></td><td></td></tr>
<tr><td>Chromosome</td><td>start</td><td>end</td><td>strand</td><td>MM</td><td>target_seq</td><td>PAM</td><td>alignment</td><td>distance</td><td>position</td><td>gene name</td><td>gene id</td></tr>
<tr><td>GU305914.1</td><td>228191</td><td>228213</td><td>+</td><td>0</td><td>TGGGGGAGATCACGGACGGG</td><td>GGG</td><td>||||||||[||||||||||||]PAM</td><td>NA</td><td>Intergenic</td><td>NA</td><td>NA</td></tr>
<tr><td></td><td></td><td></td><td></td><td></td><td></td><td></td><td></td><td></td><td></td><td></td><td></td></tr>
<tr><td>T17036</td><td>AGATCACGGACGGGGGGAGACGG</td><td>1000</td><td></td><td></td><td></td><td></td><td></td><td></td><td></td><td></td><td></td></tr>
<tr><td>Oligo fwd</td><td>CACCAGATCACGGACGGGGGGAGA</td><td></td><td></td><td></td><td></td><td></td><td></td><td></td><td></td><td></td><td></td></tr>
<tr><td>Oligo rev</td><td>AAACTCTCCCCCCGTCCGTGATCT</td><td></td><td></td><td></td><td></td><td></td><td></td><td></td><td></td><td></td><td></td></tr>
<tr><td>Chromosome</td><td>start</td><td>end</td><td>strand</td><td>MM</td><td>target_seq</td><td>PAM</td><td>alignment</td><td>distance</td><td>position</td><td>gene name</td><td>gene id</td></tr>
<tr><td>GU305914.1</td><td>228197</td><td>228219</td><td>+</td><td>0</td><td>AGATCACGGACGGGGGGAGA</td><td>CGG</td><td>||||||||[||||||||||||]PAM</td><td>NA</td><td>Intergenic</td><td>NA</td><td>NA</td></tr>
<tr><td></td><td></td><td></td><td></td><td></td><td></td><td></td><td></td><td></td><td></td><td></td><td></td></tr>
<tr><td>T17037</td><td>TCACGGACGGGGGGAGACGGCGG</td><td>1000</td><td></td><td></td><td></td><td></td><td></td><td></td><td></td><td></td><td></td></tr>
<tr><td>Oligo fwd</td><td>CACCTCACGGACGGGGGGAGACGG</td><td></td><td></td><td></td><td></td><td></td><td></td><td></td><td></td><td></td><td></td></tr>
<tr><td>Oligo rev</td><td>AAACCCGTCTCCCCCCGTCCGTGA</td><td></td><td></td><td></td><td></td><td></td><td></td><td></td><td></td><td></td><td></td></tr>
<tr><td>Chromosome</td><td>start</td><td>end</td><td>strand</td><td>MM</td><td>target_seq</td><td>PAM</td><td>alignment</td><td>distance</td><td>position</td><td>gene name</td><td>gene id</td></tr>
<tr><td>GU305914.1</td><td>228200</td><td>228222</td><td>+</td><td>0</td><td>TCACGGACGGGGGGAGACGG</td><td>CGG</td><td>||||||||[||||||||||||]PAM</td><td>NA</td><td>Intergenic</td><td>NA</td><td>NA</td></tr>
<tr><td></td><td></td><td></td><td></td><td></td><td></td><td></td><td></td><td></td><td></td><td></td><td></td></tr>
<tr><td>T17038</td><td>ACGGGGGGAGACGGCGGACGAGG</td><td>1000</td><td></td><td></td><td></td><td></td><td></td><td></td><td></td><td></td><td></td></tr>
</table>

<table>
<tr><td>Oligo fwd</td><td>CACCACGGGGGGAGACGGCGGACG</td><td></td><td></td><td></td><td></td><td></td><td></td><td></td><td></td><td></td><td></td></tr>
<tr><td>Oligo rev</td><td>AAACCGTCCGCCGTCTCCCCCCGT</td><td></td><td></td><td></td><td></td><td></td><td></td><td></td><td></td><td></td><td></td></tr>
<tr><td>Chromosome</td><td>start</td><td>end</td><td>strand</td><td>MM</td><td>target_seq</td><td>PAM</td><td>alignment</td><td>distance</td><td>position</td><td>gene name</td><td>gene id</td></tr>
<tr><td>GU305914.1</td><td>228206</td><td>228228</td><td>+</td><td>0</td><td>ACGGGGGGAGACGGCGGACG</td><td>AGG</td><td>||||||||[||||||||||||]PAM</td><td>NA</td><td>Intergenic</td><td>NA</td><td>NA</td></tr>
<tr><td></td><td></td><td></td><td></td><td></td><td></td><td></td><td></td><td></td><td></td><td></td><td></td></tr>
<tr><td>T17039</td><td>GGCGGACGAGGATCGCTCCGAGG</td><td>1000</td><td></td><td></td><td></td><td></td><td></td><td></td><td></td><td></td><td></td></tr>
<tr><td>Oligo fwd</td><td>CACCGGCGGACGAGGATCGCTCCG</td><td></td><td></td><td></td><td></td><td></td><td></td><td></td><td></td><td></td><td></td></tr>
<tr><td>Oligo rev</td><td>AAACCGGAGCGATCCTCGTCCGCC</td><td></td><td></td><td></td><td></td><td></td><td></td><td></td><td></td><td></td><td></td></tr>
<tr><td>Chromosome</td><td>start</td><td>end</td><td>strand</td><td>MM</td><td>target_seq</td><td>PAM</td><td>alignment</td><td>distance</td><td>position</td><td>gene name</td><td>gene id</td></tr>
<tr><td>GU305914.1</td><td>228218</td><td>228240</td><td>+</td><td>0</td><td>GGCGGACGAGGATCGCTCCG</td><td>AGG</td><td>||||||||[||||||||||||]PAM</td><td>NA</td><td>Intergenic</td><td>NA</td><td>NA</td></tr>
<tr><td></td><td></td><td></td><td></td><td></td><td></td><td></td><td></td><td></td><td></td><td></td><td></td></tr>
<tr><td>T17040</td><td>GGATAGCGTTTCTACTCCGAAGG</td><td>1000</td><td></td><td></td><td></td><td></td><td></td><td></td><td></td><td></td><td></td></tr>
<tr><td>Oligo fwd</td><td>CACCGGATAGCGTTTCTACTCCGA</td><td></td><td></td><td></td><td></td><td></td><td></td><td></td><td></td><td></td><td></td></tr>
<tr><td>Oligo rev</td><td>AAACTCGGAGTAGAAACGCTATCC</td><td></td><td></td><td></td><td></td><td></td><td></td><td></td><td></td><td></td><td></td></tr>
<tr><td>Chromosome</td><td>start</td><td>end</td><td>strand</td><td>MM</td><td>target_seq</td><td>PAM</td><td>alignment</td><td>distance</td><td>position</td><td>gene name</td><td>gene id</td></tr>
<tr><td>GU305914.1</td><td>228239</td><td>228261</td><td>+</td><td>0</td><td>GGATAGCGTTTCTACTCCGA</td><td>AGG</td><td>||||||||[||||||||||||]PAM</td><td>NA</td><td>Intergenic</td><td>NA</td><td>NA</td></tr>
<tr><td></td><td></td><td></td><td></td><td></td><td></td><td></td><td></td><td></td><td></td><td></td><td></td></tr>
<tr><td>T17041</td><td>CGTTTCTACTCCGAAGGAGACGG</td><td>1000</td><td></td><td></td><td></td><td></td><td></td><td></td><td></td><td></td><td></td></tr>
<tr><td>Oligo fwd</td><td>CACCCGTTTCTACTCCGAAGGAGA</td><td></td><td></td><td></td><td></td><td></td><td></td><td></td><td></td><td></td><td></td></tr>
<tr><td>Oligo rev</td><td>AAACTCTCCTTCGGAGTAGAAACG</td><td></td><td></td><td></td><td></td><td></td><td></td><td></td><td></td><td></td><td></td></tr>
<tr><td>Chromosome</td><td>start</td><td>end</td><td>strand</td><td>MM</td><td>target_seq</td><td>PAM</td><td>alignment</td><td>distance</td><td>position</td><td>gene name</td><td>gene id</td></tr>
<tr><td>GU305914.1</td><td>228245</td><td>228267</td><td>+</td><td>0</td><td>CGTTTCTACTCCGAAGGAGA</td><td>CGG</td><td>||||||||[||||||||||||]PAM</td><td>NA</td><td>Intergenic</td><td>NA</td><td>NA</td></tr>
<tr><td></td><td></td><td></td><td></td><td></td><td></td><td></td><td></td><td></td><td></td><td></td><td></td></tr>
<tr><td>T17042</td><td>GTTTCTACTCCGAAGGAGACGGG</td><td>1000</td><td></td><td></td><td></td><td></td><td></td><td></td><td></td><td></td><td></td></tr>
<tr><td>Oligo fwd</td><td>CACCGTTTCTACTCCGAAGGAGAC</td><td></td><td></td><td></td><td></td><td></td><td></td><td></td><td></td><td></td><td></td></tr>
<tr><td>Oligo rev</td><td>AAACGTCTCCTTCGGAGTAGAAAC</td><td></td><td></td><td></td><td></td><td></td><td></td><td></td><td></td><td></td><td></td></tr>
<tr><td>Chromosome</td><td>start</td><td>end</td><td>strand</td><td>MM</td><td>target_seq</td><td>PAM</td><td>alignment</td><td>distance</td><td>position</td><td>gene name</td><td>gene id</td></tr>
<tr><td>GU305914.1</td><td>228246</td><td>228268</td><td>+</td><td>0</td><td>GTTTCTACTCCGAAGGAGAC</td><td>GGG</td><td>||||||||[||||||||||||]PAM</td><td>NA</td><td>Intergenic</td><td>NA</td><td>NA</td></tr>
<tr><td></td><td></td><td></td><td></td><td></td><td></td><td></td><td></td><td></td><td></td><td></td><td></td></tr>
<tr><td>T17043</td><td>CTCCGAAGGAGACGGGCAGTCGG</td><td>1000</td><td></td><td></td><td></td><td></td><td></td><td></td><td></td><td></td><td></td></tr>
<tr><td>Oligo fwd</td><td>CACCCTCCGAAGGAGACGGGCAGT</td><td></td><td></td><td></td><td></td><td></td><td></td><td></td><td></td><td></td><td></td></tr>
<tr><td>Oligo rev</td><td>AAACACTGCCCGTCTCCTTCGGAG</td><td></td><td></td><td></td><td></td><td></td><td></td><td></td><td></td><td></td><td></td></tr>
<tr><td>Chromosome</td><td>start</td><td>end</td><td>strand</td><td>MM</td><td>target_seq</td><td>PAM</td><td>alignment</td><td>distance</td><td>position</td><td>gene name</td><td>gene id</td></tr>
<tr><td>GU305914.1</td><td>228253</td><td>228275</td><td>+</td><td>0</td><td>CTCCGAAGGAGACGGGCAGT</td><td>CGG</td><td>||||||||[||||||||||||]PAM</td><td>NA</td><td>Intergenic</td><td>NA</td><td>NA</td></tr>
<tr><td></td><td></td><td></td><td></td><td></td><td></td><td></td><td></td><td></td><td></td><td></td><td></td></tr>
<tr><td>T17044</td><td>GAAGGAGACGGGCAGTCGGTCGG</td><td>1000</td><td></td><td></td><td></td><td></td><td></td><td></td><td></td><td></td><td></td></tr>
<tr><td>Oligo fwd</td><td>CACCGAAGGAGACGGGCAGTCGGT</td><td></td><td></td><td></td><td></td><td></td><td></td><td></td><td></td><td></td><td></td></tr>
<tr><td>Oligo rev</td><td>AAACACCGACTGCCCGTCTCCTTC</td><td></td><td></td><td></td><td></td><td></td><td></td><td></td><td></td><td></td><td></td></tr>
<tr><td>Chromosome</td><td>start</td><td>end</td><td>strand</td><td>MM</td><td>target_seq</td><td>PAM</td><td>alignment</td><td>distance</td><td>position</td><td>gene name</td><td>gene id</td></tr>
<tr><td>GU305914.1</td><td>228257</td><td>228279</td><td>+</td><td>0</td><td>GAAGGAGACGGGCAGTCGGT</td><td>CGG</td><td>||||||||[||||||||||||]PAM</td><td>NA</td><td>Intergenic</td><td>NA</td><td>NA</td></tr>
<tr><td></td><td></td><td></td><td></td><td></td><td></td><td></td><td></td><td></td><td></td><td></td><td></td></tr>
<tr><td>T17045</td><td>GGCAGTCGGTCGGATGTTTCAGG</td><td>1000</td><td></td><td></td><td></td><td></td><td></td><td></td><td></td><td></td><td></td></tr>
<tr><td>Oligo fwd</td><td>CACCGGCAGTCGGTCGGATGTTTC</td><td></td><td></td><td></td><td></td><td></td><td></td><td></td><td></td><td></td><td></td></tr>
<tr><td>Oligo rev</td><td>AAACGAAACATCCGACCGACTGCC</td><td></td><td></td><td></td><td></td><td></td><td></td><td></td><td></td><td></td><td></td></tr>
<tr><td>Chromosome</td><td>start</td><td>end</td><td>strand</td><td>MM</td><td>target_seq</td><td>PAM</td><td>alignment</td><td>distance</td><td>position</td><td>gene name</td><td>gene id</td></tr>
<tr><td>GU305914.1</td><td>228267</td><td>228289</td><td>+</td><td>0</td><td>GGCAGTCGGTCGGATGTTTC</td><td>AGG</td><td>||||||||[||||||||||||]PAM</td><td>NA</td><td>Intergenic</td><td>NA</td><td>NA</td></tr>
<tr><td></td><td></td><td></td><td></td><td></td><td></td><td></td><td></td><td></td><td></td><td></td><td></td></tr>
<tr><td>T17046</td><td>GTCGGTCGGATGTTTCAGGACGG</td><td>1000</td><td></td><td></td><td></td><td></td><td></td><td></td><td></td><td></td><td></td></tr>
<tr><td>Oligo fwd</td><td>CACCGTCGGTCGGATGTTTCAGGA</td><td></td><td></td><td></td><td></td><td></td><td></td><td></td><td></td><td></td><td></td></tr>
<tr><td>Oligo rev</td><td>AAACTCCTGAAACATCCGACCGAC</td><td></td><td></td><td></td><td></td><td></td><td></td><td></td><td></td><td></td><td></td></tr>
<tr><td>Chromosome</td><td>start</td><td>end</td><td>strand</td><td>MM</td><td>target_seq</td><td>PAM</td><td>alignment</td><td>distance</td><td>position</td><td>gene name</td><td>gene id</td></tr>
<tr><td>GU305914.1</td><td>228271</td><td>228293</td><td>+</td><td>0</td><td>GTCGGTCGGATGTTTCAGGA</td><td>CGG</td><td>||||||||[||||||||||||]PAM</td><td>NA</td><td>Intergenic</td><td>NA</td><td>NA</td></tr>
<tr><td></td><td></td><td></td><td></td><td></td><td></td><td></td><td></td><td></td><td></td><td></td><td></td></tr>
<tr><td>T17047</td><td>GGTCGGATGTTTCAGGACGGAGG</td><td>1000</td><td></td><td></td><td></td><td></td><td></td><td></td><td></td><td></td><td></td></tr>
<tr><td>Oligo fwd</td><td>CACCGGTCGGATGTTTCAGGACGG</td><td></td><td></td><td></td><td></td><td></td><td></td><td></td><td></td><td></td><td></td></tr>
<tr><td>Oligo rev</td><td>AAACCCGTCCTGAAACATCCGACC</td><td></td><td></td><td></td><td></td><td></td><td></td><td></td><td></td><td></td><td></td></tr>
<tr><td>Chromosome</td><td>start</td><td>end</td><td>strand</td><td>MM</td><td>target_seq</td><td>PAM</td><td>alignment</td><td>distance</td><td>position</td><td>gene name</td><td>gene id</td></tr>
<tr><td>GU305914.1</td><td>228274</td><td>228296</td><td>+</td><td>0</td><td>GGTCGGATGTTTCAGGACGG</td><td>AGG</td><td>||||||||[||||||||||||]PAM</td><td>NA</td><td>Intergenic</td><td>NA</td><td>NA</td></tr>
<tr><td></td><td></td><td></td><td></td><td></td><td></td><td></td><td></td><td></td><td></td><td></td><td></td></tr>
<tr><td>T17048</td><td>GTCGGATGTTTCAGGACGGAGGG</td><td>1000</td><td></td><td></td><td></td><td></td><td></td><td></td><td></td><td></td><td></td></tr>
<tr><td>Oligo fwd</td><td>CACCGTCGGATGTTTCAGGACGGA</td><td></td><td></td><td></td><td></td><td></td><td></td><td></td><td></td><td></td><td></td></tr>
<tr><td>Oligo rev</td><td>AAACTCCGTCCTGAAACATCCGAC</td><td></td><td></td><td></td><td></td><td></td><td></td><td></td><td></td><td></td><td></td></tr>
<tr><td>Chromosome</td><td>start</td><td>end</td><td>strand</td><td>MM</td><td>target_seq</td><td>PAM</td><td>alignment</td><td>distance</td><td>position</td><td>gene name</td><td>gene id</td></tr>
<tr><td>GU305914.1</td><td>228275</td><td>228297</td><td>+</td><td>0</td><td>GTCGGATGTTTCAGGACGGA</td><td>GGG</td><td>||||||||[||||||||||||]PAM</td><td>NA</td><td>Intergenic</td><td>NA</td><td>NA</td></tr>
<tr><td></td><td></td><td></td><td></td><td></td><td></td><td></td><td></td><td></td><td></td><td></td><td></td></tr>
<tr><td>T17049</td><td>TCGGATGTTTCAGGACGGAGGGG</td><td>1000</td><td></td><td></td><td></td><td></td><td></td><td></td><td></td><td></td><td></td></tr>
<tr><td>Oligo fwd</td><td>CACCTCGGATGTTTCAGGACGGAG</td><td></td><td></td><td></td><td></td><td></td><td></td><td></td><td></td><td></td><td></td></tr>
<tr><td>Oligo rev</td><td>AAACCTCCGTCCTGAAACATCCGA</td><td></td><td></td><td></td><td></td><td></td><td></td><td></td><td></td><td></td><td></td></tr>
<tr><td>Chromosome</td><td>start</td><td>end</td><td>strand</td><td>MM</td><td>target_seq</td><td>PAM</td><td>alignment</td><td>distance</td><td>position</td><td>gene name</td><td>gene id</td></tr>
<tr><td>GU305914.1</td><td>228276</td><td>228298</td><td>+</td><td>0</td><td>TCGGATGTTTCAGGACGGAG</td><td>GGG</td><td>||||||||[||||||||||||]PAM</td><td>NA</td><td>Intergenic</td><td>NA</td><td>NA</td></tr>
<tr><td></td><td></td><td></td><td></td><td></td><td></td><td></td><td></td><td></td><td></td><td></td><td></td></tr>
<tr><td>T17050</td><td>ATGTTTCAGGACGGAGGGGAAGG</td><td>1000</td><td></td><td></td><td></td><td></td><td></td><td></td><td></td><td></td><td></td></tr>
<tr><td>Oligo fwd</td><td>CACCATGTTTCAGGACGGAGGGGA</td><td></td><td></td><td></td><td></td><td></td><td></td><td></td><td></td><td></td><td></td></tr>
<tr><td>Oligo rev</td><td>AAACTCCCCTCCGTCCTGAAACAT</td><td></td><td></td><td></td><td></td><td></td><td></td><td></td><td></td><td></td><td></td></tr>
<tr><td>Chromosome</td><td>start</td><td>end</td><td>strand</td><td>MM</td><td>target_seq</td><td>PAM</td><td>alignment</td><td>distance</td><td>position</td><td>gene name</td><td>gene id</td></tr>
<tr><td>GU305914.1</td><td>228280</td><td>228302</td><td>+</td><td>0</td><td>ATGTTTCAGGACGGAGGGGA</td><td>AGG</td><td>||||||||[||||||||||||]PAM</td><td>NA</td><td>Intergenic</td><td>NA</td><td>NA</td></tr>
<tr><td></td><td></td><td></td><td></td><td></td><td></td><td></td><td></td><td></td><td></td><td></td><td></td></tr>
<tr><td>T17051</td><td>TTTCAGGACGGAGGGGAAGGAGG</td><td>1000</td><td></td><td></td><td></td><td></td><td></td><td></td><td></td><td></td><td></td></tr>
<tr><td>Oligo fwd</td><td>CACCTTTCAGGACGGAGGGGAAGG</td><td></td><td></td><td></td><td></td><td></td><td></td><td></td><td></td><td></td><td></td></tr>
<tr><td>Oligo rev</td><td>AAACCCTTCCCCTCCGTCCTGAAA</td><td></td><td></td><td></td><td></td><td></td><td></td><td></td><td></td><td></td><td></td></tr>
<tr><td>Chromosome</td><td>start</td><td>end</td><td>strand</td><td>MM</td><td>target_seq</td><td>PAM</td><td>alignment</td><td>distance</td><td>position</td><td>gene name</td><td>gene id</td></tr>
<tr><td>GU305914.1</td><td>228283</td><td>228305</td><td>+</td><td>0</td><td>TTTCAGGACGGAGGGGAAGG</td><td>AGG</td><td>||||||||[||||||||||||]PAM</td><td>NA</td><td>Intergenic</td><td>NA</td><td>NA</td></tr>
<tr><td></td><td></td><td></td><td></td><td></td><td></td><td></td><td></td><td></td><td></td><td></td><td></td></tr>
<tr><td>T17052</td><td>GGGAAGGAGGATTCACGTCGAGG</td><td>1000</td><td></td><td></td><td></td><td></td><td></td><td></td><td></td><td></td><td></td></tr>
<tr><td>Oligo fwd</td><td>CACCGGGAAGGAGGATTCACGTCG</td><td></td><td></td><td></td><td></td><td></td><td></td><td></td><td></td><td></td><td></td></tr>
<tr><td>Oligo rev</td><td>AAACCGACGTGAATCCTCCTTCCC</td><td></td><td></td><td></td><td></td><td></td><td></td><td></td><td></td><td></td><td></td></tr>
<tr><td>Chromosome</td><td>start</td><td>end</td><td>strand</td><td>MM</td><td>target_seq</td><td>PAM</td><td>alignment</td><td>distance</td><td>position</td><td>gene name</td><td>gene id</td></tr>
<tr><td>GU305914.1</td><td>228296</td><td>228318</td><td>+</td><td>0</td><td>GGGAAGGAGGATTCACGTCG</td><td>AGG</td><td>||||||||[||||||||||||]PAM</td><td>NA</td><td>Intergenic</td><td>NA</td><td>NA</td></tr>
</table>

T17053 GTCGAGGACCGAGAGCCAGCCGG 1000

Oligo fwd CACCGTCGAGGACCGAGAGCCAGC

Oligo rev AAACGCTGGCTCTCGGTCCTCGAC

Chromosome start end strand MM target_seq PAM alignment distance position gene name gene id

GU305914.1 228312 228334 + 0 GTCGAGGACCGAGAGCCAGC CGG ||||||||[||||||||||||]PAM NA Intergenic NA NA

T17054 GGACCGAGAGCCAGCCGGAGAGG 1000

Oligo fwd CACCGGACCGAGAGCCAGCCGGAG

Oligo rev AAACCTCCGGCTGGCTCTCGGTCC

Chromosome start end strand MM target_seq PAM alignment distance position gene name gene id

GU305914.1 228317 228339 + 0 GGACCGAGAGCCAGCCGGAG AGG ||||||||[||||||||||||]PAM NA Intergenic NA NA

T17055 ATCCGAGCCAGATCTAGCGTCGG 1000

Oligo fwd CACCATCCGAGCCAGATCTAGCGT

Oligo rev AAACACGCTAGATCTGGCTCGGAT

Chromosome start end strand MM target_seq PAM alignment distance position gene name gene id

GU305914.1 228378 228400 + 0 ATCCGAGCCAGATCTAGCGT CGG ||||||||[||||||||||||]PAM NA Intergenic NA NA

T17056 TCCGAGCCAGATCTAGCGTCGGG 1000

Oligo fwd CACCTCCGAGCCAGATCTAGCGTC

Oligo rev AAACGACGCTAGATCTGGCTCGGA

Chromosome start end strand MM target_seq PAM alignment distance position gene name gene id

GU305914.1 228379 228401 + 0 TCCGAGCCAGATCTAGCGTC GGG ||||||||[||||||||||||]PAM NA Intergenic NA NA

T17057 GCGTCGGGCCTCCCTCATCACGG 1000

Oligo fwd CACCGCGTCGGGCCTCCCTCATCA

Oligo rev AAACTGATGAGGGAGGCCCGACGC

Chromosome start end strand MM target_seq PAM alignment distance position gene name gene id

GU305914.1 228394 228416 + 0 GCGTCGGGCCTCCCTCATCA CGG ||||||||[||||||||||||]PAM NA Intergenic NA NA

T17058 CGTCGGGCCTCCCTCATCACGGG 1000

Oligo fwd CACCCGTCGGGCCTCCCTCATCAC

Oligo rev AAACGTGATGAGGGAGGCCCGACG

Chromosome start end strand MM target_seq PAM alignment distance position gene name gene id

GU305914.1 228395 228417 + 0 CGTCGGGCCTCCCTCATCAC GGG ||||||||[||||||||||||]PAM NA Intergenic NA NA

T17059 GTCGGGCCTCCCTCATCACGGGG 1000

Oligo fwd CACCGTCGGGCCTCCCTCATCACG

Oligo rev AAACCGTGATGAGGGAGGCCCGAC

Chromosome start end strand MM target_seq PAM alignment distance position gene name gene id

GU305914.1 228396 228418 + 0 GTCGGGCCTCCCTCATCACG GGG ||||||||[||||||||||||]PAM NA Intergenic NA NA

T17060 TCGGGCCTCCCTCATCACGGGGG 1000

Oligo fwd CACCTCGGGCCTCCCTCATCACGG

Oligo rev AAACCCGTGATGAGGGAGGCCCGA

Chromosome start end strand MM target_seq PAM alignment distance position gene name gene id

GU305914.1 228397 228419 + 0 TCGGGCCTCCCTCATCACGG GGG ||||||||[||||||||||||]PAM NA Intergenic NA NA

T17061 CGGGCCTCCCTCATCACGGGGGG 1000

Oligo fwd CACCCGGGCCTCCCTCATCACGGG

Oligo rev AAACCCCGTGATGAGGGAGGCCCG

Chromosome start end strand MM target_seq PAM alignment distance position gene name gene id

GU305914.1 228398 228420 + 0 CGGGCCTCCCTCATCACGGG GGG ||||||||[||||||||||||]PAM NA Intergenic NA NA

T17062 CGGGGGGCACCTACCTCTTTCGG 1000

Oligo fwd CACCCGGGGGGCACCTACCTCTTT

Oligo rev AAACAAAGAGGTAGGTGCCCCCCG

Chromosome start end strand MM target_seq PAM alignment distance position gene name gene id

GU305914.1 228414 228436 + 0 CGGGGGGCACCTACCTCTTT CGG ||||||||[||||||||||||]PAM NA Intergenic NA NA

T17063 GGGGCACCTACCTCTTTCGGTGG 1000

Oligo fwd CACCGGGGCACCTACCTCTTTCGG

Oligo rev AAACCCGAAAGAGGTAGGTGCCCC

Chromosome start end strand MM target_seq PAM alignment distance position gene name gene id

GU305914.1 228417 228439 + 0 GGGGCACCTACCTCTTTCGG TGG ||||||||[||||||||||||]PAM NA Intergenic NA NA

T17064 CTCCTGCGTATCCAAGATCTTGG 1000

Oligo fwd CACCCTCCTGCGTATCCAAGATCT

Oligo rev AAACAGATCTTGGATACGCAGGAG

Chromosome start end strand MM target_seq PAM alignment distance position gene name gene id

GU305914.1 228447 228469 + 0 CTCCTGCGTATCCAAGATCT TGG ||||||||[||||||||||||]PAM NA Intergenic NA NA

T17065 AACGACGTCTCCAACCGCGCTGG 1000

Oligo fwd CACCAACGACGTCTCCAACCGCGC

Oligo rev AAACGCGCGGTTGGAGACGTCGTT

Chromosome start end strand MM target_seq PAM alignment distance position gene name gene id

GU305914.1 228482 228504 + 0 AACGACGTCTCCAACCGCGC TGG ||||||||[||||||||||||]PAM NA Intergenic NA NA

T17066 ACGACGTCTCCAACCGCGCTGGG 1000

Oligo fwd CACCACGACGTCTCCAACCGCGCT

Oligo rev AAACAGCGCGGTTGGAGACGTCGT

Chromosome start end strand MM target_seq PAM alignment distance position gene name gene id

GU305914.1 228483 228505 + 0 ACGACGTCTCCAACCGCGCT GGG ||||||||[||||||||||||]PAM NA Intergenic NA NA

T17067 CAACCGCGCTGGGCGAACAGAGG 1000

Oligo fwd CACCCAACCGCGCTGGGCGAACAG

Oligo rev AAACCTGTTCGCCCAGCGCGGTTG

Chromosome start end strand MM target_seq PAM alignment distance position gene name gene id

GU305914.1 228493 228515 + 0 CAACCGCGCTGGGCGAACAG AGG ||||||||[||||||||||||]PAM NA Intergenic NA NA

GU305914.1 229798 229820 + 4 CAGCCCCGTTGCGCGAACAG GGG ||-||-||[-||-||||||||]PAM NA Intergenic NA NA

T17068 AACAGAGGACGCTCGCTGATCGG 1000
Oligo fwd CACCAACAGAGGACGCTCGCTGAT
Oligo rev AAACATCAGCGAGCGTCCTCTGTT
Chromosome start end strand MM target_seq PAM alignment distance position gene name gene id
GU305914.1 228508 228530 + 0 AACAGAGGACGCTCGCTGAT CGG ||||||||[||||||||||||]PAM NA Intergenic NA NA

T17069 GAGGACGCTCGCTGATCGGACGG 1000
Oligo fwd CACCGAGGACGCTCGCTGATCGGA
Oligo rev AAACTCCGATCAGCGAGCGTCCTC
Chromosome start end strand MM target_seq PAM alignment distance position gene name gene id
GU305914.1 228512 228534 + 0 GAGGACGCTCGCTGATCGGA CGG ||||||||[||||||||||||]PAM NA Intergenic NA NA

T17070 ACGGACGTTCGTGTAGTACACGG 1000
Oligo fwd CACCACGGACGTTCGTGTAGTACA
Oligo rev AAACTGTACTACACGAACGTCCGT
Chromosome start end strand MM target_seq PAM alignment distance position gene name gene id
GU305914.1 228531 228553 + 0 ACGGACGTTCGTGTAGTACA CGG ||||||||[||||||||||||]PAM NA Intergenic NA NA

T17071 CGGCGATGCGACGTGAATCCAGG 1000
Oligo fwd CACCCGGCGATGCGACGTGAATCC
Oligo rev AAACGGATTCACGTCGCATCGCCG
Chromosome start end strand MM target_seq PAM alignment distance position gene name gene id
GU305914.1 228551 228573 + 0 CGGCGATGCGACGTGAATCC AGG ||||||||[||||||||||||]PAM NA Intergenic NA NA

T17072 ATGCGACGTGAATCCAGGTCAGG 1000
Oligo fwd CACCATGCGACGTGAATCCAGGTC
Oligo rev AAACGACCTGGATTCACGTCGCAT
Chromosome start end strand MM target_seq PAM alignment distance position gene name gene id
GU305914.1 228556 228578 + 0 ATGCGACGTGAATCCAGGTC AGG ||||||||[||||||||||||]PAM NA Intergenic NA NA

T17073 TGCGACGTGAATCCAGGTCAGGG 1000
Oligo fwd CACCTGCGACGTGAATCCAGGTCA
Oligo rev AAACTGACCTGGATTCACGTCGCA
Chromosome start end strand MM target_seq PAM alignment distance position gene name gene id
GU305914.1 228557 228579 + 0 TGCGACGTGAATCCAGGTCA GGG ||||||||[||||||||||||]PAM NA Intergenic NA NA

T17074 GACGTGAATCCAGGTCAGGGAGG 1000
Oligo fwd CACCGACGTGAATCCAGGTCAGGG
Oligo rev AAACCCCTGACCTGGATTCACGTC
Chromosome start end strand MM target_seq PAM alignment distance position gene name gene id
GU305914.1 228560 228582 + 0 GACGTGAATCCAGGTCAGGG AGG ||||||||[||||||||||||]PAM NA Intergenic NA NA

T17075 AGGTAGAATGCGACACCCAGCGG 1000
Oligo fwd CACCAGGTAGAATGCGACACCCAG
Oligo rev AAACCTGGGTGTCGCATTCTACCT
Chromosome start end strand MM target_seq PAM alignment distance position gene name gene id
GU305914.1 228580 228602 + 0 AGGTAGAATGCGACACCCAG CGG ||||||||[||||||||||||]PAM NA Intergenic NA NA

T17076 GGTAGAATGCGACACCCAGCGGG 1000
Oligo fwd CACCGGTAGAATGCGACACCCAGC
Oligo rev AAACGCTGGGTGTCGCATTCTACC
Chromosome start end strand MM target_seq PAM alignment distance position gene name gene id
GU305914.1 228581 228603 + 0 GGTAGAATGCGACACCCAGC GGG ||||||||[||||||||||||]PAM NA Intergenic NA NA

T17077 AATGCGACACCCAGCGGGTGAGG 1000
Oligo fwd CACCAATGCGACACCCAGCGGGTG
Oligo rev AAACCACCCGCTGGGTGTCGCATT
Chromosome start end strand MM target_seq PAM alignment distance position gene name gene id
GU305914.1 228586 228608 + 0 AATGCGACACCCAGCGGGTG AGG ||||||||[||||||||||||]PAM NA Intergenic NA NA

T17078 ACCCAGCGGGTGAGGCCCGTAGG 1000
Oligo fwd CACCACCCAGCGGGTGAGGCCCGT
Oligo rev AAACACGGGCCTCACCCGCTGGGT
Chromosome start end strand MM target_seq PAM alignment distance position gene name gene id
GU305914.1 228594 228616 + 0 ACCCAGCGGGTGAGGCCCGT AGG ||||||||[||||||||||||]PAM NA Intergenic NA NA

T17079 CCCAGCGGGTGAGGCCCGTAGGG 1000
Oligo fwd CACCCCCAGCGGGTGAGGCCCGTA
Oligo rev AAACTACGGGCCTCACCCGCTGGG
Chromosome start end strand MM target_seq PAM alignment distance position gene name gene id
GU305914.1 228595 228617 + 0 CCCAGCGGGTGAGGCCCGTA GGG ||||||||[||||||||||||]PAM NA Intergenic NA NA

T17080 GGTGAGGCCCGTAGGGTTCGCGG 1000
Oligo fwd CACCGGTGAGGCCCGTAGGGTTCG
Oligo rev AAACCGAACCCTACGGGCCTCACC
Chromosome start end strand MM target_seq PAM alignment distance position gene name gene id
GU305914.1 228602 228624 + 0 GGTGAGGCCCGTAGGGTTCG CGG ||||||||[||||||||||||]PAM NA Intergenic NA NA

T17081 GTGAGGCCCGTAGGGTTCGCGGG 1000
Oligo fwd CACCGTGAGGCCCGTAGGGTTCGC
Oligo rev AAACGCGAACCCTACGGGCCTCAC
Chromosome start end strand MM target_seq PAM alignment distance position gene name gene id
GU305914.1 228603 228625 + 0 GTGAGGCCCGTAGGGTTCGC GGG ||||||||[||||||||||||]PAM NA Intergenic NA NA

T17082 TGAGGCCCGTAGGGTTCGCGGGG 1000
Oligo fwd CACCTGAGGCCCGTAGGGTTCGCG
Oligo rev AAACCGCGAACCCTACGGGCCTCA

| | | | | | | | | | | | |
|---|---|---|---|---|---|---|---|---|---|---|---|
| Chromosome | start | end | strand | MM | target_seq | PAM | alignment | distance | position | gene name | gene id |
| GU305914.1 | 228604 | 228626 | + | 0 | TGAGGCCCGTAGGGTTCGCG | GGG | \|\|\|\|\|\|\|\|[\|\|\|\|\|\|\|\|\|\|\|\|]PAM | NA | Intergenic | NA | NA |
| T17083 | GTAGGGTTCGCGGGGATCGTAGG | 1000 | | | | | | | | | |
| Oligo fwd | CACCGTAGGGTTCGCGGGGATCGT | | | | | | | | | | |
| Oligo rev | AAACACGATCCCCGCGAACCCTAC | | | | | | | | | | |
| Chromosome | start | end | strand | MM | target_seq | PAM | alignment | distance | position | gene name | gene id |
| GU305914.1 | 228612 | 228634 | + | 0 | GTAGGGTTCGCGGGGATCGT | AGG | \|\|\|\|\|\|\|\|[\|\|\|\|\|\|\|\|\|\|\|\|]PAM | NA | Intergenic | NA | NA |
| T17084 | TAGGGTTCGCGGGGATCGTAGGG | 1000 | | | | | | | | | |
| Oligo fwd | CACCTAGGGTTCGCGGGGATCGTA | | | | | | | | | | |
| Oligo rev | AAACTACGATCCCCGCGAACCCTA | | | | | | | | | | |
| Chromosome | start | end | strand | MM | target_seq | PAM | alignment | distance | position | gene name | gene id |
| GU305914.1 | 228613 | 228635 | + | 0 | TAGGGTTCGCGGGGATCGTA | GGG | \|\|\|\|\|\|\|\|[\|\|\|\|\|\|\|\|\|\|\|\|]PAM | NA | Intergenic | NA | NA |
| T17085 | AGGGACAGCAACAGAAAAGACGG | 1000 | | | | | | | | | |
| Oligo fwd | CACCAGGGACAGCAACAGAAAAGA | | | | | | | | | | |
| Oligo rev | AAACTCTTTTCTGTTGCTGTCCCT | | | | | | | | | | |
| Chromosome | start | end | strand | MM | target_seq | PAM | alignment | distance | position | gene name | gene id |
| GU305914.1 | 228632 | 228654 | + | 0 | AGGGACAGCAACAGAAAAGA | CGG | \|\|\|\|\|\|\|\|[\|\|\|\|\|\|\|\|\|\|\|\|]PAM | NA | Intergenic | NA | NA |
| T17086 | AGCAACAGAAAAGACGGCAACGG | 1000 | | | | | | | | | |
| Oligo fwd | CACCAGCAACAGAAAAGACGGCAA | | | | | | | | | | |
| Oligo rev | AAACTTGCCGTCTTTTCTGTTGCT | | | | | | | | | | |
| Chromosome | start | end | strand | MM | target_seq | PAM | alignment | distance | position | gene name | gene id |
| GU305914.1 | 228638 | 228660 | + | 0 | AGCAACAGAAAAGACGGCAA | CGG | \|\|\|\|\|\|\|\|[\|\|\|\|\|\|\|\|\|\|\|\|]PAM | NA | Intergenic | NA | NA |
| T17087 | CTATCGTCACCGAACACTTTCGG | 1000 | | | | | | | | | |
| Oligo fwd | CACCCTATCGTCACCGAACACTTT | | | | | | | | | | |
| Oligo rev | AAACAAAGTGTTCGGTGACGATAG | | | | | | | | | | |
| Chromosome | start | end | strand | MM | target_seq | PAM | alignment | distance | position | gene name | gene id |
| GU305914.1 | 228661 | 228683 | + | 0 | CTATCGTCACCGAACACTTT | CGG | \|\|\|\|\|\|\|\|[\|\|\|\|\|\|\|\|\|\|\|\|]PAM | NA | Intergenic | NA | NA |
| T17088 | CGGTGACATGAACGATGAAGTGG | 1000 | | | | | | | | | |
| Oligo fwd | CACCCGGTGACATGAACGATGAAG | | | | | | | | | | |
| Oligo rev | AAACCTTCATCGTTCATGTCACCG | | | | | | | | | | |
| Chromosome | start | end | strand | MM | target_seq | PAM | alignment | distance | position | gene name | gene id |
| GU305914.1 | 228681 | 228703 | + | 0 | CGGTGACATGAACGATGAAG | TGG | \|\|\|\|\|\|\|\|[\|\|\|\|\|\|\|\|\|\|\|\|]PAM | NA | Intergenic | NA | NA |
| T17089 | TGACATGAACGATGAAGTGGCGG | 1000 | | | | | | | | | |
| Oligo fwd | CACCTGACATGAACGATGAAGTGG | | | | | | | | | | |
| Oligo rev | AAACCCACTTCATCGTTCATGTCA | | | | | | | | | | |
| Chromosome | start | end | strand | MM | target_seq | PAM | alignment | distance | position | gene name | gene id |
| GU305914.1 | 228684 | 228706 | + | 0 | TGACATGAACGATGAAGTGG | CGG | \|\|\|\|\|\|\|\|[\|\|\|\|\|\|\|\|\|\|\|\|]PAM | NA | Intergenic | NA | NA |
| T17090 | GGCGGAGAAAGTAGCAAGCTCGG | 1000 | | | | | | | | | |
| Oligo fwd | CACCGGCGGAGAAAGTAGCAAGCT | | | | | | | | | | |
| Oligo rev | AAACAGCTTGCTACTTTCTCCGCC | | | | | | | | | | |
| Chromosome | start | end | strand | MM | target_seq | PAM | alignment | distance | position | gene name | gene id |
| GU305914.1 | 228702 | 228724 | + | 0 | GGCGGAGAAAGTAGCAAGCT | CGG | \|\|\|\|\|\|\|\|[\|\|\|\|\|\|\|\|\|\|\|\|]PAM | NA | Intergenic | NA | NA |
| T17091 | GAAAGTAGCAAGCTCGGTGTCGG | 1000 | | | | | | | | | |
| Oligo fwd | CACCGAAAGTAGCAAGCTCGGTGT | | | | | | | | | | |
| Oligo rev | AAACACACCGAGCTTGCTACTTTC | | | | | | | | | | |
| Chromosome | start | end | strand | MM | target_seq | PAM | alignment | distance | position | gene name | gene id |
| GU305914.1 | 228708 | 228730 | + | 0 | GAAAGTAGCAAGCTCGGTGT | CGG | \|\|\|\|\|\|\|\|[\|\|\|\|\|\|\|\|\|\|\|\|]PAM | NA | Intergenic | NA | NA |
| T17092 | GTAGCAAGCTCGGTGTCGGTCGG | 1000 | | | | | | | | | |
| Oligo fwd | CACCGTAGCAAGCTCGGTGTCGGT | | | | | | | | | | |
| Oligo rev | AAACACCGACACCGAGCTTGCTAC | | | | | | | | | | |
| Chromosome | start | end | strand | MM | target_seq | PAM | alignment | distance | position | gene name | gene id |
| GU305914.1 | 228712 | 228734 | + | 0 | GTAGCAAGCTCGGTGTCGGT | CGG | \|\|\|\|\|\|\|\|[\|\|\|\|\|\|\|\|\|\|\|\|]PAM | NA | Intergenic | NA | NA |
| T17093 | ATTGCAGCTCCCAGCGACGAAGG | 1000 | | | | | | | | | |
| Oligo fwd | CACCATTGCAGCTCCCAGCGACGA | | | | | | | | | | |
| Oligo rev | AAACTCGTCGCTGGGAGCTGCAAT | | | | | | | | | | |
| Chromosome | start | end | strand | MM | target_seq | PAM | alignment | distance | position | gene name | gene id |
| GU305914.1 | 228740 | 228762 | + | 0 | ATTGCAGCTCCCAGCGACGA | AGG | \|\|\|\|\|\|\|\|[\|\|\|\|\|\|\|\|\|\|\|\|]PAM | NA | Intergenic | NA | NA |
| T17094 | GCTCCCAGCGACGAAGGTTGTGG | 1000 | | | | | | | | | |
| Oligo fwd | CACCGCTCCCAGCGACGAAGGTTG | | | | | | | | | | |
| Oligo rev | AAACCAACCTTCGTCGCTGGGAGC | | | | | | | | | | |
| Chromosome | start | end | strand | MM | target_seq | PAM | alignment | distance | position | gene name | gene id |
| GU305914.1 | 228746 | 228768 | + | 0 | GCTCCCAGCGACGAAGGTTG | TGG | \|\|\|\|\|\|\|\|[\|\|\|\|\|\|\|\|\|\|\|\|]PAM | NA | Intergenic | NA | NA |
| T17095 | GTTGTGGCCGATACCCGCGTAGG | 1000 | | | | | | | | | |
| Oligo fwd | CACCGTTGTGGCCGATACCCGCGT | | | | | | | | | | |
| Oligo rev | AAACACGCGGGTATCGGCCACAAC | | | | | | | | | | |
| Chromosome | start | end | strand | MM | target_seq | PAM | alignment | distance | position | gene name | gene id |
| GU305914.1 | 228762 | 228784 | + | 0 | GTTGTGGCCGATACCCGCGT | AGG | \|\|\|\|\|\|\|\|[\|\|\|\|\|\|\|\|\|\|\|\|]PAM | NA | Intergenic | NA | NA |
| T17096 | TTGTGGCCGATACCCGCGTAGGG | 1000 | | | | | | | | | |
| Oligo fwd | CACCTTGTGGCCGATACCCGCGTA | | | | | | | | | | |
| Oligo rev | AAACTACGCGGGTATCGGCCACAA | | | | | | | | | | |
| Chromosome | start | end | strand | MM | target_seq | PAM | alignment | distance | position | gene name | gene id |
| GU305914.1 | 228763 | 228785 | + | 0 | TTGTGGCCGATACCCGCGTA | GGG | \|\|\|\|\|\|\|\|[\|\|\|\|\|\|\|\|\|\|\|\|]PAM | NA | Intergenic | NA | NA |
| T17097 | TGTGGCCGATACCCGCGTAGGGG | 1000 | | | | | | | | | |
| Oligo fwd | CACCTGTGGCCGATACCCGCGTAG | | | | | | | | | | |

Oligo rev AAACCTACGCGGGTATCGGCCACA
Chromosome start end strand MM target_seq PAM alignment distance position gene name gene id
GU305914.1 228764 228786 + 0 TGTGGCCGATACCCGCGTAG GGG ||||||||[||||||||||||]PAM NA Intergenic NA NA

T17098 ATACCCGCGTAGGGGACGAACGG 1000
Oligo fwd CACCATACCCGCGTAGGGGACGAA
Oligo rev AAACTTCGTCCCCTACGCGGGTAT
Chromosome start end strand MM target_seq PAM alignment distance position gene name gene id
GU305914.1 228772 228794 + 0 ATACCCGCGTAGGGGACGAA CGG ||||||||[||||||||||||]PAM NA Intergenic NA NA

T17099 TACCCGCGTAGGGGACGAACGGG 1000
Oligo fwd CACCTACCCGCGTAGGGGACGAAC
Oligo rev AAACGTTCGTCCCCTACGCGGGTA
Chromosome start end strand MM target_seq PAM alignment distance position gene name gene id
GU305914.1 228773 228795 + 0 TACCCGCGTAGGGGACGAAC GGG ||||||||[||||||||||||]PAM NA Intergenic NA NA

T17100 CGCGTTGCTCATCGCGAGTCCGG 1000
Oligo fwd CACCCGCGTTGCTCATCGCGAGTC
Oligo rev AAACGACTCGCGATGAGCAACGCG
Chromosome start end strand MM target_seq PAM alignment distance position gene name gene id
GU305914.1 228798 228820 + 0 CGCGTTGCTCATCGCGAGTC CGG ||||||||[||||||||||||]PAM NA Intergenic NA NA

T17101 TCATCGCGAGTCCGGCGTCGCGG 1000
Oligo fwd CACCTCATCGCGAGTCCGGCGTCG
Oligo rev AAACCGACGCCGGACTCGCGATGA
Chromosome start end strand MM target_seq PAM alignment distance position gene name gene id
GU305914.1 228806 228828 + 0 TCATCGCGAGTCCGGCGTCG CGG ||||||||[||||||||||||]PAM NA Intergenic NA NA

T17102 CATCGCGAGTCCGGCGTCGCGGG 1000
Oligo fwd CACCCATCGCGAGTCCGGCGTCGC
Oligo rev AAACGCGACGCCGGACTCGCGATG
Chromosome start end strand MM target_seq PAM alignment distance position gene name gene id
GU305914.1 228807 228829 + 0 CATCGCGAGTCCGGCGTCGC GGG ||||||||[||||||||||||]PAM NA Intergenic NA NA

T17103 ATCGCGAGTCCGGCGTCGCGGGG 1000
Oligo fwd CACCATCGCGAGTCCGGCGTCGCG
Oligo rev AAACCGCGACGCCGGACTCGCGAT
Chromosome start end strand MM target_seq PAM alignment distance position gene name gene id
GU305914.1 228808 228830 + 0 ATCGCGAGTCCGGCGTCGCG GGG ||||||||[||||||||||||]PAM NA Intergenic NA NA

T17104 GGAGCGAAAAGCTGATTAGATGG 1000
Oligo fwd CACCGGAGCGAAAAGCTGATTAGA
Oligo rev AAACTCTAATCAGCTTTTCGCTCC
Chromosome start end strand MM target_seq PAM alignment distance position gene name gene id
GU305914.1 228829 228851 + 0 GGAGCGAAAAGCTGATTAGA TGG ||||||||[||||||||||||]PAM NA Intergenic NA NA

T17105 CGCGAAGCTGACAGACCAGCAGG 1000
Oligo fwd CACCCGCGAAGCTGACAGACCAGC
Oligo rev AAACGCTGGTCTGTCAGCTTCGCG
Chromosome start end strand MM target_seq PAM alignment distance position gene name gene id
GU305914.1 228886 228908 + 0 CGCGAAGCTGACAGACCAGC AGG ||||||||[||||||||||||]PAM NA Intergenic NA NA

T17106 GCTGACAGACCAGCAGGACTTGG 1000
Oligo fwd CACCGCTGACAGACCAGCAGGACT
Oligo rev AAACAGTCCTGCTGGTCTGTCAGC
Chromosome start end strand MM target_seq PAM alignment distance position gene name gene id
GU305914.1 228892 228914 + 0 GCTGACAGACCAGCAGGACT TGG ||||||||[||||||||||||]PAM NA Intergenic NA NA

T17107 AGACCAGCAGGACTTGGTTTCGG 1000
Oligo fwd CACCAGACCAGCAGGACTTGGTTT
Oligo rev AAACAAACCAAGTCCTGCTGGTCT
Chromosome start end strand MM target_seq PAM alignment distance position gene name gene id
GU305914.1 228898 228920 + 0 AGACCAGCAGGACTTGGTTT CGG ||||||||[||||||||||||]PAM NA Intergenic NA NA

T17108 CTTGGTTTCGGTCTTACGCATGG 1000
Oligo fwd CACCCTTGGTTTCGGTCTTACGCA
Oligo rev AAACTGCGTAAGACCGAAACCAAG
Chromosome start end strand MM target_seq PAM alignment distance position gene name gene id
GU305914.1 228910 228932 + 0 CTTGGTTTCGGTCTTACGCA TGG ||||||||[||||||||||||]PAM NA Intergenic NA NA

T17109 CATGGTCATTCCACTGCACACGG 1000
Oligo fwd CACCCATGGTCATTCCACTGCACA
Oligo rev AAACTGTGCAGTGGAATGACCATG
Chromosome start end strand MM target_seq PAM alignment distance position gene name gene id
GU305914.1 228928 228950 + 0 CATGGTCATTCCACTGCACA CGG ||||||||[||||||||||||]PAM NA Intergenic NA NA

T17110 TCATTCCACTGCACACGGTGTGG 1000
Oligo fwd CACCTCATTCCACTGCACACGGTG
Oligo rev AAACCACCGTGTGCAGTGGAATGA
Chromosome start end strand MM target_seq PAM alignment distance position gene name gene id
GU305914.1 228933 228955 + 0 TCATTCCACTGCACACGGTG TGG ||||||||[||||||||||||]PAM NA Intergenic NA NA

T17111 CATTCCACTGCACACGGTGTGGG 1000
Oligo fwd CACCCATTCCACTGCACACGGTGT
Oligo rev AAACACACCGTGTGCAGTGGAATG
Chromosome start end strand MM target_seq PAM alignment distance position gene name gene id
GU305914.1 228934 228956 + 0 CATTCCACTGCACACGGTGT GGG ||||||||[||||||||||||]PAM NA Intergenic NA NA

T17112 ATTTTCGCAGAAATCTTCTTCGG 1000

<table>
<tr><td>Oligo fwd</td><td>CACCATTTTCGCAGAAATCTTCTT</td><td></td><td></td><td></td><td></td><td></td><td></td><td></td><td></td><td></td></tr>
<tr><td>Oligo rev</td><td>AAACAAGAAGATTTCTGCGAAAAT</td><td></td><td></td><td></td><td></td><td></td><td></td><td></td><td></td><td></td></tr>
<tr><td>Chromosome</td><td>start</td><td>end</td><td>strand</td><td>MM</td><td>target_seq</td><td>PAM</td><td>alignment</td><td>distance</td><td>position</td><td>gene name</td><td>gene id</td></tr>
<tr><td>GU305914.1</td><td>228976</td><td>228998</td><td>+</td><td>0</td><td>ATTTTCGCAGAAATCTTCTT</td><td>CGG</td><td>||||||||[||||||||||||]PAM</td><td>NA</td><td>Intergenic</td><td>NA</td><td>NA</td></tr>
<tr><td></td></tr>
<tr><td>T17113</td><td>TTCTTCGGCGTTAGCGACAGAGG</td><td>1000</td></tr>
<tr><td>Oligo fwd</td><td>CACCTTCTTCGGCGTTAGCGACAG</td></tr>
<tr><td>Oligo rev</td><td>AAACCTGTCGCTAACGCCGAAGAA</td></tr>
<tr><td>Chromosome</td><td>start</td><td>end</td><td>strand</td><td>MM</td><td>target_seq</td><td>PAM</td><td>alignment</td><td>distance</td><td>position</td><td>gene name</td><td>gene id</td></tr>
<tr><td>GU305914.1</td><td>228991</td><td>229013</td><td>+</td><td>0</td><td>TTCTTCGGCGTTAGCGACAG</td><td>AGG</td><td>||||||||[||||||||||||]PAM</td><td>NA</td><td>Intergenic</td><td>NA</td><td>NA</td></tr>
<tr><td>GU305914.1</td><td>202225</td><td>202247</td><td>+</td><td>4</td><td>TTCTTCAACGGTATCGACAG</td><td>CGG</td><td>||||||--[||-||-||||||]PAM</td><td>NA</td><td>Intergenic</td><td>NA</td><td>NA</td></tr>
<tr><td></td></tr>
<tr><td>T17114</td><td>CGTTAGCGACAGAGGTATGCAGG</td><td>1000</td></tr>
<tr><td>Oligo fwd</td><td>CACCCGTTAGCGACAGAGGTATGC</td></tr>
<tr><td>Oligo rev</td><td>AAACGCATACCTCTGTCGCTAACG</td></tr>
<tr><td>Chromosome</td><td>start</td><td>end</td><td>strand</td><td>MM</td><td>target_seq</td><td>PAM</td><td>alignment</td><td>distance</td><td>position</td><td>gene name</td><td>gene id</td></tr>
<tr><td>GU305914.1</td><td>228999</td><td>229021</td><td>+</td><td>0</td><td>CGTTAGCGACAGAGGTATGC</td><td>AGG</td><td>||||||||[||||||||||||]PAM</td><td>NA</td><td>Intergenic</td><td>NA</td><td>NA</td></tr>
<tr><td></td></tr>
<tr><td>T17115</td><td>GGTATGCAGGCAGTGTTCACTGG</td><td>1000</td></tr>
<tr><td>Oligo fwd</td><td>CACCGGTATGCAGGCAGTGTTCAC</td></tr>
<tr><td>Oligo rev</td><td>AAACGTGAACACTGCCTGCATACC</td></tr>
<tr><td>Chromosome</td><td>start</td><td>end</td><td>strand</td><td>MM</td><td>target_seq</td><td>PAM</td><td>alignment</td><td>distance</td><td>position</td><td>gene name</td><td>gene id</td></tr>
<tr><td>GU305914.1</td><td>229012</td><td>229034</td><td>+</td><td>0</td><td>GGTATGCAGGCAGTGTTCAC</td><td>TGG</td><td>||||||||[||||||||||||]PAM</td><td>NA</td><td>Intergenic</td><td>NA</td><td>NA</td></tr>
<tr><td></td></tr>
<tr><td>T17116</td><td>GCAGGCAGTGTTCACTGGAACGG</td><td>1000</td></tr>
<tr><td>Oligo fwd</td><td>CACCGCAGGCAGTGTTCACTGGAA</td></tr>
<tr><td>Oligo rev</td><td>AAACTTCCAGTGAACACTGCCTGC</td></tr>
<tr><td>Chromosome</td><td>start</td><td>end</td><td>strand</td><td>MM</td><td>target_seq</td><td>PAM</td><td>alignment</td><td>distance</td><td>position</td><td>gene name</td><td>gene id</td></tr>
<tr><td>GU305914.1</td><td>229017</td><td>229039</td><td>+</td><td>0</td><td>GCAGGCAGTGTTCACTGGAA</td><td>CGG</td><td>||||||||[||||||||||||]PAM</td><td>NA</td><td>Intergenic</td><td>NA</td><td>NA</td></tr>
<tr><td></td></tr>
<tr><td>T17117</td><td>GGCAGTGTTCACTGGAACGGAGG</td><td>1000</td></tr>
<tr><td>Oligo fwd</td><td>CACCGGCAGTGTTCACTGGAACGG</td></tr>
<tr><td>Oligo rev</td><td>AAACCCGTTCCAGTGAACACTGCC</td></tr>
<tr><td>Chromosome</td><td>start</td><td>end</td><td>strand</td><td>MM</td><td>target_seq</td><td>PAM</td><td>alignment</td><td>distance</td><td>position</td><td>gene name</td><td>gene id</td></tr>
<tr><td>GU305914.1</td><td>229020</td><td>229042</td><td>+</td><td>0</td><td>GGCAGTGTTCACTGGAACGG</td><td>AGG</td><td>||||||||[||||||||||||]PAM</td><td>NA</td><td>Intergenic</td><td>NA</td><td>NA</td></tr>
<tr><td></td></tr>
<tr><td>T17118</td><td>GCAGTGTTCACTGGAACGGAGGG</td><td>1000</td></tr>
<tr><td>Oligo fwd</td><td>CACCGCAGTGTTCACTGGAACGGA</td></tr>
<tr><td>Oligo rev</td><td>AAACTCCGTTCCAGTGAACACTGC</td></tr>
<tr><td>Chromosome</td><td>start</td><td>end</td><td>strand</td><td>MM</td><td>target_seq</td><td>PAM</td><td>alignment</td><td>distance</td><td>position</td><td>gene name</td><td>gene id</td></tr>
<tr><td>GU305914.1</td><td>229021</td><td>229043</td><td>+</td><td>0</td><td>GCAGTGTTCACTGGAACGGA</td><td>GGG</td><td>||||||||[||||||||||||]PAM</td><td>NA</td><td>Intergenic</td><td>NA</td><td>NA</td></tr>
<tr><td></td></tr>
<tr><td>T17119</td><td>GATACATGCACCGTTATCGATGG</td><td>1000</td></tr>
<tr><td>Oligo fwd</td><td>CACCGATACATGCACCGTTATCGA</td></tr>
<tr><td>Oligo rev</td><td>AAACTCGATAACGGTGCATGTATC</td></tr>
<tr><td>Chromosome</td><td>start</td><td>end</td><td>strand</td><td>MM</td><td>target_seq</td><td>PAM</td><td>alignment</td><td>distance</td><td>position</td><td>gene name</td><td>gene id</td></tr>
<tr><td>GU305914.1</td><td>229051</td><td>229073</td><td>+</td><td>0</td><td>GATACATGCACCGTTATCGA</td><td>TGG</td><td>||||||||[||||||||||||]PAM</td><td>NA</td><td>Intergenic</td><td>NA</td><td>NA</td></tr>
<tr><td></td></tr>
<tr><td>T17120</td><td>ATTTAGCGTGTCACACGACCTGG</td><td>1000</td></tr>
<tr><td>Oligo fwd</td><td>CACCATTTAGCGTGTCACACGACC</td></tr>
<tr><td>Oligo rev</td><td>AAACGGTCGTGTGACACGCTAAAT</td></tr>
<tr><td>Chromosome</td><td>start</td><td>end</td><td>strand</td><td>MM</td><td>target_seq</td><td>PAM</td><td>alignment</td><td>distance</td><td>position</td><td>gene name</td><td>gene id</td></tr>
<tr><td>GU305914.1</td><td>229078</td><td>229100</td><td>+</td><td>0</td><td>ATTTAGCGTGTCACACGACC</td><td>TGG</td><td>||||||||[||||||||||||]PAM</td><td>NA</td><td>Intergenic</td><td>NA</td><td>NA</td></tr>
<tr><td></td></tr>
<tr><td>T17121</td><td>ACACGACCTGGACCTCATGACGG</td><td>1000</td></tr>
<tr><td>Oligo fwd</td><td>CACCACACGACCTGGACCTCATGA</td></tr>
<tr><td>Oligo rev</td><td>AAACTCATGAGGTCCAGGTCGTGT</td></tr>
<tr><td>Chromosome</td><td>start</td><td>end</td><td>strand</td><td>MM</td><td>target_seq</td><td>PAM</td><td>alignment</td><td>distance</td><td>position</td><td>gene name</td><td>gene id</td></tr>
<tr><td>GU305914.1</td><td>229090</td><td>229112</td><td>+</td><td>0</td><td>ACACGACCTGGACCTCATGA</td><td>CGG</td><td>||||||||[||||||||||||]PAM</td><td>NA</td><td>Intergenic</td><td>NA</td><td>NA</td></tr>
<tr><td></td></tr>
<tr><td>T17122</td><td>CCTGGACCTCATGACGGACGCGG</td><td>1000</td></tr>
<tr><td>Oligo fwd</td><td>CACCCCTGGACCTCATGACGGACG</td></tr>
<tr><td>Oligo rev</td><td>AAACCGTCCGTCATGAGGTCCAGG</td></tr>
<tr><td>Chromosome</td><td>start</td><td>end</td><td>strand</td><td>MM</td><td>target_seq</td><td>PAM</td><td>alignment</td><td>distance</td><td>position</td><td>gene name</td><td>gene id</td></tr>
<tr><td>GU305914.1</td><td>229096</td><td>229118</td><td>+</td><td>0</td><td>CCTGGACCTCATGACGGACG</td><td>CGG</td><td>||||||||[||||||||||||]PAM</td><td>NA</td><td>Intergenic</td><td>NA</td><td>NA</td></tr>
<tr><td></td></tr>
<tr><td>T17123</td><td>CTGGACCTCATGACGGACGCGGG</td><td>1000</td></tr>
<tr><td>Oligo fwd</td><td>CACCCTGGACCTCATGACGGACGC</td></tr>
<tr><td>Oligo rev</td><td>AAACGCGTCCGTCATGAGGTCCAG</td></tr>
<tr><td>Chromosome</td><td>start</td><td>end</td><td>strand</td><td>MM</td><td>target_seq</td><td>PAM</td><td>alignment</td><td>distance</td><td>position</td><td>gene name</td><td>gene id</td></tr>
<tr><td>GU305914.1</td><td>229097</td><td>229119</td><td>+</td><td>0</td><td>CTGGACCTCATGACGGACGC</td><td>GGG</td><td>||||||||[||||||||||||]PAM</td><td>NA</td><td>Intergenic</td><td>NA</td><td>NA</td></tr>
<tr><td></td></tr>
<tr><td>T17124</td><td>ATGACGGACGCGGGAACCGCCGG</td><td>1000</td></tr>
<tr><td>Oligo fwd</td><td>CACCATGACGGACGCGGGAACCGC</td></tr>
<tr><td>Oligo rev</td><td>AAACGCGGTTCCCGCGTCCGTCAT</td></tr>
<tr><td>Chromosome</td><td>start</td><td>end</td><td>strand</td><td>MM</td><td>target_seq</td><td>PAM</td><td>alignment</td><td>distance</td><td>position</td><td>gene name</td><td>gene id</td></tr>
<tr><td>GU305914.1</td><td>229106</td><td>229128</td><td>+</td><td>0</td><td>ATGACGGACGCGGGAACCGC</td><td>CGG</td><td>||||||||[||||||||||||]PAM</td><td>NA</td><td>Intergenic</td><td>NA</td><td>NA</td></tr>
<tr><td></td></tr>
<tr><td>T17125</td><td>GAGCACAGCTCACCACAGACAGG</td><td>1000</td></tr>
<tr><td>Oligo fwd</td><td>CACCGAGCACAGCTCACCACAGAC</td></tr>
<tr><td>Oligo rev</td><td>AAACGTCTGTGGTGAGCTGTGCTC</td></tr>
<tr><td>Chromosome</td><td>start</td><td>end</td><td>strand</td><td>MM</td><td>target_seq</td><td>PAM</td><td>alignment</td><td>distance</td><td>position</td><td>gene name</td><td>gene id</td></tr>
<tr><td>GU305914.1</td><td>229128</td><td>229150</td><td>+</td><td>0</td><td>GAGCACAGCTCACCACAGAC</td><td>AGG</td><td>||||||||[||||||||||||]PAM</td><td>NA</td><td>Intergenic</td><td>NA</td><td>NA</td></tr>
<tr><td></td></tr>
<tr><td>T17126</td><td>GAGAGACCACCGAAAGCGACAGG</td><td>1000</td></tr>
<tr><td>Oligo fwd</td><td>CACCGAGAGACCACCGAAAGCGAC</td></tr>
<tr><td>Oligo rev</td><td>AAACGTCGCTTTCGGTGGTCTCTC</td></tr>
<tr><td>Chromosome</td><td>start</td><td>end</td><td>strand</td><td>MM</td><td>target_seq</td><td>PAM</td><td>alignment</td><td>distance</td><td>position</td><td>gene name</td><td>gene id</td></tr>
<tr><td>GU305914.1</td><td>229155</td><td>229177</td><td>+</td><td>0</td><td>GAGAGACCACCGAAAGCGAC</td><td>AGG</td><td>||||||||[||||||||||||]PAM</td><td>NA</td><td>Intergenic</td><td>NA</td><td>NA</td></tr>
</table>

T17127 CACCGAAAGCGACAGGTCACCGG 1000
Oligo fwd CACCCACCGAAAGCGACAGGTCAC
Oligo rev AAACGTGACCTGTCGCTTTCGGTG
Chromosome start end strand MM target_seq PAM alignment distance position gene name gene id
GU305914.1 229162 229184 + 0 CACCGAAAGCGACAGGTCAC CGG ||||||||[||||||||||||]PAM NA Intergenic NA NA

T17128 ACCGAAAGCGACAGGTCACCGGG 1000
Oligo fwd CACCACCGAAAGCGACAGGTCACC
Oligo rev AAACGGTGACCTGTCGCTTTCGGT
Chromosome start end strand MM target_seq PAM alignment distance position gene name gene id
GU305914.1 229163 229185 + 0 ACCGAAAGCGACAGGTCACC GGG ||||||||[||||||||||||]PAM NA Intergenic NA NA

T17129 ACACTTCAAGCGCACCCGCGCGG 1000
Oligo fwd CACCACACTTCAAGCGCACCCGCG
Oligo rev AAACCGCGGGTGCGCTTGAAGTGT
Chromosome start end strand MM target_seq PAM alignment distance position gene name gene id
GU305914.1 229229 229251 + 0 ACACTTCAAGCGCACCCGCG CGG ||||||||[||||||||||||]PAM NA Intergenic NA NA

T17130 CACTTCAAGCGCACCCGCGCGGG 1000
Oligo fwd CACCCACTTCAAGCGCACCCGCGC
Oligo rev AAACGCGCGGGTGCGCTTGAAGTG
Chromosome start end strand MM target_seq PAM alignment distance position gene name gene id
GU305914.1 229230 229252 + 0 CACTTCAAGCGCACCCGCGC GGG ||||||||[||||||||||||]PAM NA Intergenic NA NA

T17131 ACTTCAAGCGCACCCGCGCGGGG 1000
Oligo fwd CACCACTTCAAGCGCACCCGCGCG
Oligo rev AAACCGCGCGGGTGCGCTTGAAGT
Chromosome start end strand MM target_seq PAM alignment distance position gene name gene id
GU305914.1 229231 229253 + 0 ACTTCAAGCGCACCCGCGCG GGG ||||||||[||||||||||||]PAM NA Intergenic NA NA

T17132 CTTCAAGCGCACCCGCGCGGGGG 1000
Oligo fwd CACCCTTCAAGCGCACCCGCGCGG
Oligo rev AAACCCGCGCGGGTGCGCTTGAAG
Chromosome start end strand MM target_seq PAM alignment distance position gene name gene id
GU305914.1 229232 229254 + 0 CTTCAAGCGCACCCGCGCGG GGG ||||||||[||||||||||||]PAM NA Intergenic NA NA

T17133 CAAGCGCACCCGCGCGGGGGAGG 1000
Oligo fwd CACCCAAGCGCACCCGCGCGGGGG
Oligo rev AAACCCCCCGCGCGGGTGCGCTTG
Chromosome start end strand MM target_seq PAM alignment distance position gene name gene id
GU305914.1 229235 229257 + 0 CAAGCGCACCCGCGCGGGGG AGG ||||||||[||||||||||||]PAM NA Intergenic NA NA

T17134 CGGGGGAGGAGACAAAAGAGCGG 1000
Oligo fwd CACCCGGGGGAGGAGACAAAAGAG
Oligo rev AAACCTCTTTTGTCTCCTCCCCCG
Chromosome start end strand MM target_seq PAM alignment distance position gene name gene id
GU305914.1 229249 229271 + 0 CGGGGGAGGAGACAAAAGAG CGG ||||||||[||||||||||||]PAM NA Intergenic NA NA

T17135 GGGGGAGGAGACAAAAGAGCGGG 1000
Oligo fwd CACCGGGGGAGGAGACAAAAGAGC
Oligo rev AAACGCTCTTTTGTCTCCTCCCCC
Chromosome start end strand MM target_seq PAM alignment distance position gene name gene id
GU305914.1 229250 229272 + 0 GGGGGAGGAGACAAAAGAGC GGG ||||||||[||||||||||||]PAM NA Intergenic NA NA

T17136 GGGGAGGAGACAAAAGAGCGGGG 1000
Oligo fwd CACCGGGGAGGAGACAAAAGAGCG
Oligo rev AAACCGCTCTTTTGTCTCCTCCCC
Chromosome start end strand MM target_seq PAM alignment distance position gene name gene id
GU305914.1 229251 229273 + 0 GGGGAGGAGACAAAAGAGCG GGG ||||||||[||||||||||||]PAM NA Intergenic NA NA

T17137 CGAGAAGTCAGACCGATGAGCGG 1000
Oligo fwd CACCCGAGAAGTCAGACCGATGAG
Oligo rev AAACCTCATCGGTCTGACTTCTCG
Chromosome start end strand MM target_seq PAM alignment distance position gene name gene id
GU305914.1 229279 229301 + 0 CGAGAAGTCAGACCGATGAG CGG ||||||||[||||||||||||]PAM NA Intergenic NA NA

T17138 GAGAAGTCAGACCGATGAGCGGG 1000
Oligo fwd CACCGAGAAGTCAGACCGATGAGC
Oligo rev AAACGCTCATCGGTCTGACTTCTC
Chromosome start end strand MM target_seq PAM alignment distance position gene name gene id
GU305914.1 229280 229302 + 0 GAGAAGTCAGACCGATGAGC GGG ||||||||[||||||||||||]PAM NA Intergenic NA NA

T17139 GTCAGACCGATGAGCGGGAACGG 1000
Oligo fwd CACCGTCAGACCGATGAGCGGGAA
Oligo rev AAACTTCCCGCTCATCGGTCTGAC
Chromosome start end strand MM target_seq PAM alignment distance position gene name gene id
GU305914.1 229285 229307 + 0 GTCAGACCGATGAGCGGGAA CGG ||||||||[||||||||||||]PAM NA Intergenic NA NA

T17140 ACAGCGATCCGCGACACGAACGG 1000
Oligo fwd CACCACAGCGATCCGCGACACGAA
Oligo rev AAACTTCGTGTCGCGGATCGCTGT
Chromosome start end strand MM target_seq PAM alignment distance position gene name gene id
GU305914.1 229308 229330 + 0 ACAGCGATCCGCGACACGAA CGG ||||||||[||||||||||||]PAM NA Intergenic NA NA

T17141 GATCCGCGACACGAACGGCGAGG 1000
Oligo fwd CACCGATCCGCGACACGAACGGCG
Oligo rev AAACCGCCGTTCGTGTCGCGGATC
Chromosome start end strand MM target_seq PAM alignment distance position gene name gene id

GU305914.1    229313    229335    +    0    GATCCGCGACACGAACGGCG    AGG    ||||||||[||||||||||||]PAM    NA    Intergenic    NA    NA

T17142    ATCCGCGACACGAACGGCGAGGG    1000
Oligo fwd    CACCATCCGCGACACGAACGGCGA
Oligo rev    AAACTCGCCGTTCGTGTCGCGGAT
Chromosome    start    end    strand    MM    target_seq    PAM    alignment    distance    position    gene name    gene id
GU305914.1    229314    229336    +    0    ATCCGCGACACGAACGGCGA    GGG    ||||||||[||||||||||||]PAM    NA    Intergenic    NA    NA

T17143    ACGAACGGCGAGGGAGCGCGG    1000
Oligo fwd    CACCACGAACGGCGAGGGAGCGCG
Oligo rev    AAACCGCGCTCCCTCGCCGTTCGT
Chromosome    start    end    strand    MM    target_seq    PAM    alignment    distance    position    gene name    gene id
GU305914.1    229323    229345    +    0    ACGAACGGCGAGGGAGCGCG    CGG    ||||||||[||||||||||||]PAM    NA    Intergenic    NA    NA

T17144    GCGGAGCGAGATCGCGCACTCGG    1000
Oligo fwd    CACCGCGGAGCGAGATCGCGCACT
Oligo rev    AAACAGTGCGCGATCTCGCTCCGC
Chromosome    start    end    strand    MM    target_seq    PAM    alignment    distance    position    gene name    gene id
GU305914.1    229342    229364    +    0    GCGGAGCGAGATCGCGCACT    CGG    ||||||||[||||||||||||]PAM    NA    Intergenic    NA    NA

T17145    CGGCCTCCCGCAGCGCTCACAGG    1000
Oligo fwd    CACCCGGCCTCCCGCAGCGCTCAC
Oligo rev    AAACGTGAGCGCTGCGGGAGGCCG
Chromosome    start    end    strand    MM    target_seq    PAM    alignment    distance    position    gene name    gene id
GU305914.1    229362    229384    +    0    CGGCCTCCCGCAGCGCTCAC    AGG    ||||||||[||||||||||||]PAM    NA    Intergenic    NA    NA

T17146    GGCCTCCCGCAGCGCTCACAGGG    1000
Oligo fwd    CACCGGCCTCCCGCAGCGCTCACA
Oligo rev    AAACTGTGAGCGCTGCGGGAGGCC
Chromosome    start    end    strand    MM    target_seq    PAM    alignment    distance    position    gene name    gene id
GU305914.1    229363    229385    +    0    GGCCTCCCGCAGCGCTCACA    GGG    ||||||||[||||||||||||]PAM    NA    Intergenic    NA    NA

T17147    AAGTTGACCGATCACGAACTCGG    1000
Oligo fwd    CACCAAGTTGACCGATCACGAACT
Oligo rev    AAACAGTTCGTGATCGGTCAACTT
Chromosome    start    end    strand    MM    target_seq    PAM    alignment    distance    position    gene name    gene id
GU305914.1    229387    229409    +    0    AAGTTGACCGATCACGAACT    CGG    ||||||||[||||||||||||]PAM    NA    Intergenic    NA    NA

T17148    TGACCGATCACGAACTCGGCTGG    1000
Oligo fwd    CACCTGACCGATCACGAACTCGGC
Oligo rev    AAACGCCGAGTTCGTGATCGGTCA
Chromosome    start    end    strand    MM    target_seq    PAM    alignment    distance    position    gene name    gene id
GU305914.1    229391    229413    +    0    TGACCGATCACGAACTCGGC    TGG    ||||||||[||||||||||||]PAM    NA    Intergenic    NA    NA

T17149    ACACGAACATCGTTATTACCTGG    1000
Oligo fwd    CACCACACGAACATCGTTATTACC
Oligo rev    AAACGGTAATAACGATGTTCGTGT
Chromosome    start    end    strand    MM    target_seq    PAM    alignment    distance    position    gene name    gene id
GU305914.1    229425    229447    +    0    ACACGAACATCGTTATTACC    TGG    ||||||||[||||||||||||]PAM    NA    Intergenic    NA    NA

T17150    GTTATTACCTGGTACCTGCGCGG    1000
Oligo fwd    CACCGTTATTACCTGGTACCTGCG
Oligo rev    AAACCGCAGGTACCAGGTAATAAC
Chromosome    start    end    strand    MM    target_seq    PAM    alignment    distance    position    gene name    gene id
GU305914.1    229436    229458    +    0    GTTATTACCTGGTACCTGCG    CGG    ||||||||[||||||||||||]PAM    NA    Intergenic    NA    NA

T17151    ATTACCTGGTACCTGCGCGGCGG    1000
Oligo fwd    CACCATTACCTGGTACCTGCGCGG
Oligo rev    AAACCCGCGCAGGTACCAGGTAAT
Chromosome    start    end    strand    MM    target_seq    PAM    alignment    distance    position    gene name    gene id
GU305914.1    229439    229461    +    0    ATTACCTGGTACCTGCGCGG    CGG    ||||||||[||||||||||||]PAM    NA    Intergenic    NA    NA

T17152    CCGTAACGCCTGTAGCATGCCGG    1000
Oligo fwd    CACCCCGTAACGCCTGTAGCATGC
Oligo rev    AAACGCATGCTACAGGCGTTACGG
Chromosome    start    end    strand    MM    target_seq    PAM    alignment    distance    position    gene name    gene id
GU305914.1    229467    229489    +    0    CCGTAACGCCTGTAGCATGC    CGG    ||||||||[||||||||||||]PAM    NA    Intergenic    NA    NA

T17153    TAACGCCTGTAGCATGCCGGCGG    1000
Oligo fwd    CACCTAACGCCTGTAGCATGCCGG
Oligo rev    AAACCCGGCATGCTACAGGCGTTA
Chromosome    start    end    strand    MM    target_seq    PAM    alignment    distance    position    gene name    gene id
GU305914.1    229470    229492    +    0    TAACGCCTGTAGCATGCCGG    CGG    ||||||||[||||||||||||]PAM    NA    Intergenic    NA    NA

T17154    AACGCCTGTAGCATGCCGGCGGG    1000
Oligo fwd    CACCAACGCCTGTAGCATGCCGGC
Oligo rev    AAACGCCGGCATGCTACAGGCGTT
Chromosome    start    end    strand    MM    target_seq    PAM    alignment    distance    position    gene name    gene id
GU305914.1    229471    229493    +    0    AACGCCTGTAGCATGCCGGC    GGG    ||||||||[||||||||||||]PAM    NA    Intergenic    NA    NA

T17155    ACGCCTGTAGCATGCCGGCGGGG    1000
Oligo fwd    CACCACGCCTGTAGCATGCCGGCG
Oligo rev    AAACCGCCGGCATGCTACAGGCGT
Chromosome    start    end    strand    MM    target_seq    PAM    alignment    distance    position    gene name    gene id
GU305914.1    229472    229494    +    0    ACGCCTGTAGCATGCCGGCG    GGG    ||||||||[||||||||||||]PAM    NA    Intergenic    NA    NA

T17156    GCCGGCGGGGCGCCCTCGCCCGG    1000
Oligo fwd    CACCGCCGGCGGGGCGCCCTCGCC
Oligo rev    AAACGGCGAGGGCGCCCCGCCGGC

<table>
<tr><td>Chromosome</td><td>start</td><td>end</td><td>strand</td><td>MM</td><td>target_seq</td><td>PAM</td><td>alignment</td><td>distance</td><td>position</td><td>gene name</td><td>gene id</td></tr>
<tr><td>GU305914.1</td><td>229485</td><td>229507</td><td>+</td><td>0</td><td>GCCGGCGGGGCGCCCTCGCC</td><td>CGG</td><td>||||||||[||||||||||||]PAM</td><td>NA</td><td>Intergenic</td><td>NA</td><td>NA</td></tr>
<tr><td></td><td></td><td></td><td></td><td></td><td></td><td></td><td></td><td></td><td></td><td></td><td></td></tr>
<tr><td>T17157</td><td>CCTCCGTGAACGACGTTCTCCGG</td><td>1000</td><td></td><td></td><td></td><td></td><td></td><td></td><td></td><td></td><td></td></tr>
<tr><td>Oligo fwd</td><td>CACCCCTCCGTGAACGACGTTCTC</td><td></td><td></td><td></td><td></td><td></td><td></td><td></td><td></td><td></td><td></td></tr>
<tr><td>Oligo rev</td><td>AAACGAGAACGTCGTTCACGGAGG</td><td></td><td></td><td></td><td></td><td></td><td></td><td></td><td></td><td></td><td></td></tr>
<tr><td>Chromosome</td><td>start</td><td>end</td><td>strand</td><td>MM</td><td>target_seq</td><td>PAM</td><td>alignment</td><td>distance</td><td>position</td><td>gene name</td><td>gene id</td></tr>
<tr><td>GU305914.1</td><td>229530</td><td>229552</td><td>+</td><td>0</td><td>CCTCCGTGAACGACGTTCTC</td><td>CGG</td><td>||||||||[||||||||||||]PAM</td><td>NA</td><td>Intergenic</td><td>NA</td><td>NA</td></tr>
<tr><td></td><td></td><td></td><td></td><td></td><td></td><td></td><td></td><td></td><td></td><td></td><td></td></tr>
<tr><td>T17158</td><td>GAACGACGTTCTCCGGAGCGCGG</td><td>1000</td><td></td><td></td><td></td><td></td><td></td><td></td><td></td><td></td><td></td></tr>
<tr><td>Oligo fwd</td><td>CACCGAACGACGTTCTCCGGAGCG</td><td></td><td></td><td></td><td></td><td></td><td></td><td></td><td></td><td></td><td></td></tr>
<tr><td>Oligo rev</td><td>AAACCGCTCCGGAGAACGTCGTTC</td><td></td><td></td><td></td><td></td><td></td><td></td><td></td><td></td><td></td><td></td></tr>
<tr><td>Chromosome</td><td>start</td><td>end</td><td>strand</td><td>MM</td><td>target_seq</td><td>PAM</td><td>alignment</td><td>distance</td><td>position</td><td>gene name</td><td>gene id</td></tr>
<tr><td>GU305914.1</td><td>229537</td><td>229559</td><td>+</td><td>0</td><td>GAACGACGTTCTCCGGAGCG</td><td>CGG</td><td>||||||||[||||||||||||]PAM</td><td>NA</td><td>Intergenic</td><td>NA</td><td>NA</td></tr>
<tr><td></td><td></td><td></td><td></td><td></td><td></td><td></td><td></td><td></td><td></td><td></td><td></td></tr>
<tr><td>T17159</td><td>GTTCTCCGGAGCGCGGTACGTGG</td><td>1000</td><td></td><td></td><td></td><td></td><td></td><td></td><td></td><td></td><td></td></tr>
<tr><td>Oligo fwd</td><td>CACCGTTCTCCGGAGCGCGGTACG</td><td></td><td></td><td></td><td></td><td></td><td></td><td></td><td></td><td></td><td></td></tr>
<tr><td>Oligo rev</td><td>AAACCGTACCGCGCTCCGGAGAAC</td><td></td><td></td><td></td><td></td><td></td><td></td><td></td><td></td><td></td><td></td></tr>
<tr><td>Chromosome</td><td>start</td><td>end</td><td>strand</td><td>MM</td><td>target_seq</td><td>PAM</td><td>alignment</td><td>distance</td><td>position</td><td>gene name</td><td>gene id</td></tr>
<tr><td>GU305914.1</td><td>229544</td><td>229566</td><td>+</td><td>0</td><td>GTTCTCCGGAGCGCGGTACG</td><td>TGG</td><td>||||||||[||||||||||||]PAM</td><td>NA</td><td>Intergenic</td><td>NA</td><td>NA</td></tr>
<tr><td></td><td></td><td></td><td></td><td></td><td></td><td></td><td></td><td></td><td></td><td></td><td></td></tr>
<tr><td>T17160</td><td>ACGTGGCAACACTCGAACAGCGG</td><td>1000</td><td></td><td></td><td></td><td></td><td></td><td></td><td></td><td></td><td></td></tr>
<tr><td>Oligo fwd</td><td>CACCACGTGGCAACACTCGAACAG</td><td></td><td></td><td></td><td></td><td></td><td></td><td></td><td></td><td></td><td></td></tr>
<tr><td>Oligo rev</td><td>AAACCTGTTCGAGTGTTGCCACGT</td><td></td><td></td><td></td><td></td><td></td><td></td><td></td><td></td><td></td><td></td></tr>
<tr><td>Chromosome</td><td>start</td><td>end</td><td>strand</td><td>MM</td><td>target_seq</td><td>PAM</td><td>alignment</td><td>distance</td><td>position</td><td>gene name</td><td>gene id</td></tr>
<tr><td>GU305914.1</td><td>229561</td><td>229583</td><td>+</td><td>0</td><td>ACGTGGCAACACTCGAACAG</td><td>CGG</td><td>||||||||[||||||||||||]PAM</td><td>NA</td><td>Intergenic</td><td>NA</td><td>NA</td></tr>
<tr><td></td><td></td><td></td><td></td><td></td><td></td><td></td><td></td><td></td><td></td><td></td><td></td></tr>
<tr><td>T17161</td><td>TGGCAACACTCGAACAGCGGAGG</td><td>1000</td><td></td><td></td><td></td><td></td><td></td><td></td><td></td><td></td><td></td></tr>
<tr><td>Oligo fwd</td><td>CACCTGGCAACACTCGAACAGCGG</td><td></td><td></td><td></td><td></td><td></td><td></td><td></td><td></td><td></td><td></td></tr>
<tr><td>Oligo rev</td><td>AAACCCGCTGTTCGAGTGTTGCCA</td><td></td><td></td><td></td><td></td><td></td><td></td><td></td><td></td><td></td><td></td></tr>
<tr><td>Chromosome</td><td>start</td><td>end</td><td>strand</td><td>MM</td><td>target_seq</td><td>PAM</td><td>alignment</td><td>distance</td><td>position</td><td>gene name</td><td>gene id</td></tr>
<tr><td>GU305914.1</td><td>229564</td><td>229586</td><td>+</td><td>0</td><td>TGGCAACACTCGAACAGCGG</td><td>AGG</td><td>||||||||[||||||||||||]PAM</td><td>NA</td><td>Intergenic</td><td>NA</td><td>NA</td></tr>
<tr><td></td><td></td><td></td><td></td><td></td><td></td><td></td><td></td><td></td><td></td><td></td><td></td></tr>
<tr><td>T17162</td><td>AACAGCGGAGGAACGCTCGAAGG</td><td>1000</td><td></td><td></td><td></td><td></td><td></td><td></td><td></td><td></td><td></td></tr>
<tr><td>Oligo fwd</td><td>CACCAACAGCGGAGGAACGCTCGA</td><td></td><td></td><td></td><td></td><td></td><td></td><td></td><td></td><td></td><td></td></tr>
<tr><td>Oligo rev</td><td>AAACTCGAGCGTTCCTCCGCTGTT</td><td></td><td></td><td></td><td></td><td></td><td></td><td></td><td></td><td></td><td></td></tr>
<tr><td>Chromosome</td><td>start</td><td>end</td><td>strand</td><td>MM</td><td>target_seq</td><td>PAM</td><td>alignment</td><td>distance</td><td>position</td><td>gene name</td><td>gene id</td></tr>
<tr><td>GU305914.1</td><td>229576</td><td>229598</td><td>+</td><td>0</td><td>AACAGCGGAGGAACGCTCGA</td><td>AGG</td><td>||||||||[||||||||||||]PAM</td><td>NA</td><td>Intergenic</td><td>NA</td><td>NA</td></tr>
<tr><td></td><td></td><td></td><td></td><td></td><td></td><td></td><td></td><td></td><td></td><td></td><td></td></tr>
<tr><td>T17163</td><td>TCGAGCGCGAACCCGCGACGCGG</td><td>1000</td><td></td><td></td><td></td><td></td><td></td><td></td><td></td><td></td><td></td></tr>
<tr><td>Oligo fwd</td><td>CACCTCGAGCGCGAACCCGCGACG</td><td></td><td></td><td></td><td></td><td></td><td></td><td></td><td></td><td></td><td></td></tr>
<tr><td>Oligo rev</td><td>AAACCGTCGCGGGTTCGCGCTCGA</td><td></td><td></td><td></td><td></td><td></td><td></td><td></td><td></td><td></td><td></td></tr>
<tr><td>Chromosome</td><td>start</td><td>end</td><td>strand</td><td>MM</td><td>target_seq</td><td>PAM</td><td>alignment</td><td>distance</td><td>position</td><td>gene name</td><td>gene id</td></tr>
<tr><td>GU305914.1</td><td>229599</td><td>229621</td><td>+</td><td>0</td><td>TCGAGCGCGAACCCGCGACG</td><td>CGG</td><td>||||||||[||||||||||||]PAM</td><td>NA</td><td>Intergenic</td><td>NA</td><td>NA</td></tr>
<tr><td></td><td></td><td></td><td></td><td></td><td></td><td></td><td></td><td></td><td></td><td></td><td></td></tr>
<tr><td>T17164</td><td>GAACCCGCGACGCGGCACGCTGG</td><td>1000</td><td></td><td></td><td></td><td></td><td></td><td></td><td></td><td></td><td></td></tr>
<tr><td>Oligo fwd</td><td>CACCGAACCCGCGACGCGGCACGC</td><td></td><td></td><td></td><td></td><td></td><td></td><td></td><td></td><td></td><td></td></tr>
<tr><td>Oligo rev</td><td>AAACGCGTGCCGCGTCGCGGGTTC</td><td></td><td></td><td></td><td></td><td></td><td></td><td></td><td></td><td></td><td></td></tr>
<tr><td>Chromosome</td><td>start</td><td>end</td><td>strand</td><td>MM</td><td>target_seq</td><td>PAM</td><td>alignment</td><td>distance</td><td>position</td><td>gene name</td><td>gene id</td></tr>
<tr><td>GU305914.1</td><td>229607</td><td>229629</td><td>+</td><td>0</td><td>GAACCCGCGACGCGGCACGC</td><td>TGG</td><td>||||||||[||||||||||||]PAM</td><td>NA</td><td>Intergenic</td><td>NA</td><td>NA</td></tr>
<tr><td></td><td></td><td></td><td></td><td></td><td></td><td></td><td></td><td></td><td></td><td></td><td></td></tr>
<tr><td>T17165</td><td>CGCTGGTCGCGTTGCGTCACCGG</td><td>1000</td><td></td><td></td><td></td><td></td><td></td><td></td><td></td><td></td><td></td></tr>
<tr><td>Oligo fwd</td><td>CACCCGCTGGTCGCGTTGCGTCAC</td><td></td><td></td><td></td><td></td><td></td><td></td><td></td><td></td><td></td><td></td></tr>
<tr><td>Oligo rev</td><td>AAACGTGACGCAACGCGACCAGCG</td><td></td><td></td><td></td><td></td><td></td><td></td><td></td><td></td><td></td><td></td></tr>
<tr><td>Chromosome</td><td>start</td><td>end</td><td>strand</td><td>MM</td><td>target_seq</td><td>PAM</td><td>alignment</td><td>distance</td><td>position</td><td>gene name</td><td>gene id</td></tr>
<tr><td>GU305914.1</td><td>229624</td><td>229646</td><td>+</td><td>0</td><td>CGCTGGTCGCGTTGCGTCAC</td><td>CGG</td><td>||||||||[||||||||||||]PAM</td><td>NA</td><td>Intergenic</td><td>NA</td><td>NA</td></tr>
<tr><td></td><td></td><td></td><td></td><td></td><td></td><td></td><td></td><td></td><td></td><td></td><td></td></tr>
<tr><td>T17166</td><td>TCACCGGCGTCGCGTTACGACGG</td><td>1000</td><td></td><td></td><td></td><td></td><td></td><td></td><td></td><td></td><td></td></tr>
<tr><td>Oligo fwd</td><td>CACCTCACCGGCGTCGCGTTACGA</td><td></td><td></td><td></td><td></td><td></td><td></td><td></td><td></td><td></td><td></td></tr>
<tr><td>Oligo rev</td><td>AAACTCGTAACGCGACGCCGGTGA</td><td></td><td></td><td></td><td></td><td></td><td></td><td></td><td></td><td></td><td></td></tr>
<tr><td>Chromosome</td><td>start</td><td>end</td><td>strand</td><td>MM</td><td>target_seq</td><td>PAM</td><td>alignment</td><td>distance</td><td>position</td><td>gene name</td><td>gene id</td></tr>
<tr><td>GU305914.1</td><td>229640</td><td>229662</td><td>+</td><td>0</td><td>TCACCGGCGTCGCGTTACGA</td><td>CGG</td><td>||||||||[||||||||||||]PAM</td><td>NA</td><td>Intergenic</td><td>NA</td><td>NA</td></tr>
<tr><td></td><td></td><td></td><td></td><td></td><td></td><td></td><td></td><td></td><td></td><td></td><td></td></tr>
<tr><td>T17167</td><td>CACCGGCGTCGCGTTACGACGGG</td><td>1000</td><td></td><td></td><td></td><td></td><td></td><td></td><td></td><td></td><td></td></tr>
<tr><td>Oligo fwd</td><td>CACCCACCGGCGTCGCGTTACGAC</td><td></td><td></td><td></td><td></td><td></td><td></td><td></td><td></td><td></td><td></td></tr>
<tr><td>Oligo rev</td><td>AAACGTCGTAACGCGACGCCGGTG</td><td></td><td></td><td></td><td></td><td></td><td></td><td></td><td></td><td></td><td></td></tr>
<tr><td>Chromosome</td><td>start</td><td>end</td><td>strand</td><td>MM</td><td>target_seq</td><td>PAM</td><td>alignment</td><td>distance</td><td>position</td><td>gene name</td><td>gene id</td></tr>
<tr><td>GU305914.1</td><td>229641</td><td>229663</td><td>+</td><td>0</td><td>CACCGGCGTCGCGTTACGAC</td><td>GGG</td><td>||||||||[||||||||||||]PAM</td><td>NA</td><td>Intergenic</td><td>NA</td><td>NA</td></tr>
<tr><td></td><td></td><td></td><td></td><td></td><td></td><td></td><td></td><td></td><td></td><td></td><td></td></tr>
<tr><td>T17168</td><td>ACGACGGGCATCTCGAAACCCGG</td><td>1000</td><td></td><td></td><td></td><td></td><td></td><td></td><td></td><td></td><td></td></tr>
<tr><td>Oligo fwd</td><td>CACCACGACGGGCATCTCGAAACC</td><td></td><td></td><td></td><td></td><td></td><td></td><td></td><td></td><td></td><td></td></tr>
<tr><td>Oligo rev</td><td>AAACGGTTTCGAGATGCCCGTCGT</td><td></td><td></td><td></td><td></td><td></td><td></td><td></td><td></td><td></td><td></td></tr>
<tr><td>Chromosome</td><td>start</td><td>end</td><td>strand</td><td>MM</td><td>target_seq</td><td>PAM</td><td>alignment</td><td>distance</td><td>position</td><td>gene name</td><td>gene id</td></tr>
<tr><td>GU305914.1</td><td>229656</td><td>229678</td><td>+</td><td>0</td><td>ACGACGGGCATCTCGAAACC</td><td>CGG</td><td>||||||||[||||||||||||]PAM</td><td>NA</td><td>Intergenic</td><td>NA</td><td>NA</td></tr>
<tr><td></td><td></td><td></td><td></td><td></td><td></td><td></td><td></td><td></td><td></td><td></td><td></td></tr>
<tr><td>T17169</td><td>CGACGGGCATCTCGAAACCCGGG</td><td>1000</td><td></td><td></td><td></td><td></td><td></td><td></td><td></td><td></td><td></td></tr>
<tr><td>Oligo fwd</td><td>CACCCGACGGGCATCTCGAAACCC</td><td></td><td></td><td></td><td></td><td></td><td></td><td></td><td></td><td></td><td></td></tr>
<tr><td>Oligo rev</td><td>AAACGGGTTTCGAGATGCCCGTCG</td><td></td><td></td><td></td><td></td><td></td><td></td><td></td><td></td><td></td><td></td></tr>
<tr><td>Chromosome</td><td>start</td><td>end</td><td>strand</td><td>MM</td><td>target_seq</td><td>PAM</td><td>alignment</td><td>distance</td><td>position</td><td>gene name</td><td>gene id</td></tr>
<tr><td>GU305914.1</td><td>229657</td><td>229679</td><td>+</td><td>0</td><td>CGACGGGCATCTCGAAACCC</td><td>GGG</td><td>||||||||[||||||||||||]PAM</td><td>NA</td><td>Intergenic</td><td>NA</td><td>NA</td></tr>
<tr><td></td><td></td><td></td><td></td><td></td><td></td><td></td><td></td><td></td><td></td><td></td><td></td></tr>
<tr><td>T17170</td><td>CGGGCATCTCGAAACCCGGGCGG</td><td>1000</td><td></td><td></td><td></td><td></td><td></td><td></td><td></td><td></td><td></td></tr>
<tr><td>Oligo fwd</td><td>CACCCGGGCATCTCGAAACCCGGG</td><td></td><td></td><td></td><td></td><td></td><td></td><td></td><td></td><td></td><td></td></tr>
<tr><td>Oligo rev</td><td>AAACCCCGGGTTTCGAGATGCCCG</td><td></td><td></td><td></td><td></td><td></td><td></td><td></td><td></td><td></td><td></td></tr>
<tr><td>Chromosome</td><td>start</td><td>end</td><td>strand</td><td>MM</td><td>target_seq</td><td>PAM</td><td>alignment</td><td>distance</td><td>position</td><td>gene name</td><td>gene id</td></tr>
<tr><td>GU305914.1</td><td>229660</td><td>229682</td><td>+</td><td>0</td><td>CGGGCATCTCGAAACCCGGG</td><td>CGG</td><td>||||||||[||||||||||||]PAM</td><td>NA</td><td>Intergenic</td><td>NA</td><td>NA</td></tr>
<tr><td></td><td></td><td></td><td></td><td></td><td></td><td></td><td></td><td></td><td></td><td></td><td></td></tr>
<tr><td>T17171</td><td>GCGGTCGCTCGACCAGCGAGCGG</td><td>1000</td><td></td><td></td><td></td><td></td><td></td><td></td><td></td><td></td><td></td></tr>
<tr><td>Oligo fwd</td><td>CACCGCGGTCGCTCGACCAGCGAG</td><td></td><td></td><td></td><td></td><td></td><td></td><td></td><td></td><td></td><td></td></tr>
</table>

Oligo rev AAACCTCGCTGGTCGAGCGACCGC
Chromosome start end strand MM target_seq PAM alignment distance position gene name gene id
GU305914.1 229679 229701 + 0 GCGGTCGCTCGACCAGCGAG CGG ||||||||[||||||||||||]PAM NA Intergenic NA NA

T17172 CGGTCGCTCGACCAGCGAGCGGG 1000
Oligo fwd CACCCGGTCGCTCGACCAGCGAGC
Oligo rev AAACGCTCGCTGGTCGAGCGACCG
Chromosome start end strand MM target_seq PAM alignment distance position gene name gene id
GU305914.1 229680 229702 + 0 CGGTCGCTCGACCAGCGAGC GGG ||||||||[||||||||||||]PAM NA Intergenic NA NA
GU305914.1 88246 88268 - 4 CCGCCGCTCGTCCGGCGAGC GGG |-|-||||[||-||-||||||]PAM NA Intergenic NA NA

T17173 GAGCGGGTGCTCGCTACGCGCGG 1000
Oligo fwd CACCGAGCGGGTGCTCGCTACGCG
Oligo rev AAACCGCGTAGCGAGCACCCGCTC
Chromosome start end strand MM target_seq PAM alignment distance position gene name gene id
GU305914.1 229696 229718 + 0 GAGCGGGTGCTCGCTACGCG CGG ||||||||[||||||||||||]PAM NA Intergenic NA NA
GU305914.1 162263 162285 - 4 GAGCGGTCGGTCGCTACGAG GGG ||||||--[|-||||||||-|]PAM NA Intergenic NA NA

T17174 GGGTGCTCGCTACGCGCGGCCGG 1000
Oligo fwd CACCGGGTGCTCGCTACGCGCGGC
Oligo rev AAACGCCGCGCGTAGCGAGCACCC
Chromosome start end strand MM target_seq PAM alignment distance position gene name gene id
GU305914.1 229700 229722 + 0 GGGTGCTCGCTACGCGCGGC CGG ||||||||[||||||||||||]PAM NA Intergenic NA NA

T17175 GACGAGCCTGAGCGCGTAACTGG 1000
Oligo fwd CACCGACGAGCCTGAGCGCGTAAC
Oligo rev AAACGTTACGCGCTCAGGCTCGTC
Chromosome start end strand MM target_seq PAM alignment distance position gene name gene id
GU305914.1 229748 229770 + 0 GACGAGCCTGAGCGCGTAAC TGG ||||||||[||||||||||||]PAM NA Intergenic NA NA

T17176 GCGTAACTGGTCTGCCACCACGG 1000
Oligo fwd CACCGCGTAACTGGTCTGCCACCA
Oligo rev AAACTGGTGGCAGACCAGTTACGC
Chromosome start end strand MM target_seq PAM alignment distance position gene name gene id
GU305914.1 229761 229783 + 0 GCGTAACTGGTCTGCCACCA CGG ||||||||[||||||||||||]PAM NA Intergenic NA NA

T17177 CGTAACTGGTCTGCCACCACGGG 1000
Oligo fwd CACCCGTAACTGGTCTGCCACCAC
Oligo rev AAACGTGGTGGCAGACCAGTTACG
Chromosome start end strand MM target_seq PAM alignment distance position gene name gene id
GU305914.1 229762 229784 + 0 CGTAACTGGTCTGCCACCAC GGG ||||||||[||||||||||||]PAM NA Intergenic NA NA

T17178 GTAACTGGTCTGCCACCACGGGG 1000
Oligo fwd CACCGTAACTGGTCTGCCACCACG
Oligo rev AAACCGTGGTGGCAGACCAGTTAC
Chromosome start end strand MM target_seq PAM alignment distance position gene name gene id
GU305914.1 229763 229785 + 0 GTAACTGGTCTGCCACCACG GGG ||||||||[||||||||||||]PAM NA Intergenic NA NA

T17179 ACTGGTCTGCCACCACGGGGTGG 1000
Oligo fwd CACCACTGGTCTGCCACCACGGGG
Oligo rev AAACCCCCGTGGTGGCAGACCAGT
Chromosome start end strand MM target_seq PAM alignment distance position gene name gene id
GU305914.1 229766 229788 + 0 ACTGGTCTGCCACCACGGGG TGG ||||||||[||||||||||||]PAM NA Intergenic NA NA

T17180 GCCAGCCCCGTTGCGCGAACAGG 1000
Oligo fwd CACCGCCAGCCCCGTTGCGCGAAC
Oligo rev AAACGTTCGCGCAACGGGGCTGGC
Chromosome start end strand MM target_seq PAM alignment distance position gene name gene id
GU305914.1 229796 229818 + 0 GCCAGCCCCGTTGCGCGAAC AGG ||||||||[||||||||||||]PAM NA Intergenic NA NA

T17181 CCAGCCCCGTTGCGCGAACAGGG 1000
Oligo fwd CACCCCAGCCCCGTTGCGCGAACA
Oligo rev AAACTGTTCGCGCAACGGGGCTGG
Chromosome start end strand MM target_seq PAM alignment distance position gene name gene id
GU305914.1 229797 229819 + 0 CCAGCCCCGTTGCGCGAACA GGG ||||||||[||||||||||||]PAM NA Intergenic NA NA

T17182 CAGCCCCGTTGCGCGAACAGGGG 1000
Oligo fwd CACCCAGCCCCGTTGCGCGAACAG
Oligo rev AAACCTGTTCGCGCAACGGGGCTG
Chromosome start end strand MM target_seq PAM alignment distance position gene name gene id
GU305914.1 229798 229820 + 0 CAGCCCCGTTGCGCGAACAG GGG ||||||||[||||||||||||]PAM NA Intergenic NA NA
GU305914.1 228493 228515 + 4 CAACCGCGCTGGGCGAACAG AGG ||-||-||[-||-||||||||]PAM NA Intergenic NA NA

T17183 AGCCCCGTTGCGCGAACAGGGGG 1000
Oligo fwd CACCAGCCCCGTTGCGCGAACAGG
Oligo rev AAACCCTGTTCGCGCAACGGGGCT
Chromosome start end strand MM target_seq PAM alignment distance position gene name gene id
GU305914.1 229799 229821 + 0 AGCCCCGTTGCGCGAACAGG GGG ||||||||[||||||||||||]PAM NA Intergenic NA NA

T17184 CGAACAGGGGGCGCTCGCACAGG 1000
Oligo fwd CACCCGAACAGGGGGCGCTCGCAC
Oligo rev AAACGTGCGAGCGCCCCCTGTTCG
Chromosome start end strand MM target_seq PAM alignment distance position gene name gene id
GU305914.1 229811 229833 + 0 CGAACAGGGGGCGCTCGCAC AGG ||||||||[||||||||||||]PAM NA Intergenic NA NA

T17185 GAACAGGGGGCGCTCGCACAGGG 1000
Oligo fwd CACCGAACAGGGGGCGCTCGCACA
Oligo rev AAACTGTGCGAGCGCCCCCTGTTC
Chromosome start end strand MM target_seq PAM alignment distance position gene name gene id

GU305914.1 229812 229834 + 0 GAACAGGGGGCGCTCGCACA GGG ||||||||[||||||||||||]PAM NA Intergenic NA NA

T17186 AACAGGGGGCGCTCGCACAGGGG 1000
Oligo fwd CACCAACAGGGGGCGCTCGCACAG
Oligo rev AAACCTGTGCGAGCGCCCCCTGTT
Chromosome start end strand MM target_seq PAM alignment distance position gene name gene id
GU305914.1 229813 229835 + 0 AACAGGGGGCGCTCGCACAG GGG ||||||||[||||||||||||]PAM NA Intergenic NA NA

T17187 ACAGGGGGCGCTCGCACAGGGGG 1000
Oligo fwd CACCACAGGGGGCGCTCGCACAGG
Oligo rev AAACCCTGTGCGAGCGCCCCCTGT
Chromosome start end strand MM target_seq PAM alignment distance position gene name gene id
GU305914.1 229814 229836 + 0 ACAGGGGGCGCTCGCACAGG GGG ||||||||[||||||||||||]PAM NA Intergenic NA NA

T17188 CGTAGTAGACTCCCAGAAAACGG 1000
Oligo fwd CACCCGTAGTAGACTCCCAGAAAA
Oligo rev AAACTTTTCTGGGAGTCTACTACG
Chromosome start end strand MM target_seq PAM alignment distance position gene name gene id
GU305914.1 229847 229869 + 0 CGTAGTAGACTCCCAGAAAA CGG ||||||||[||||||||||||]PAM NA Intergenic NA NA

T17189 TCGCGTATAAAATGACGCCCAGG 1000
Oligo fwd CACCTCGCGTATAAAATGACGCCC
Oligo rev AAACGGGCGTCATTTTATACGCGA
Chromosome start end strand MM target_seq PAM alignment distance position gene name gene id
GU305914.1 229881 229903 + 0 TCGCGTATAAAATGACGCCC AGG ||||||||[||||||||||||]PAM NA Intergenic NA NA

T17190 CGCGTATAAAATGACGCCCAGGG 1000
Oligo fwd CACCCGCGTATAAAATGACGCCCA
Oligo rev AAACTGGGCGTCATTTTATACGCG
Chromosome start end strand MM target_seq PAM alignment distance position gene name gene id
GU305914.1 229882 229904 + 0 CGCGTATAAAATGACGCCCA GGG ||||||||[||||||||||||]PAM NA Intergenic NA NA

T17191 AAAATGACGCCCAGGGTCCCCGG 1000
Oligo fwd CACCAAAATGACGCCCAGGGTCCC
Oligo rev AAACGGGACCCTGGGCGTCATTTT
Chromosome start end strand MM target_seq PAM alignment distance position gene name gene id
GU305914.1 229889 229911 + 0 AAAATGACGCCCAGGGTCCC CGG ||||||||[||||||||||||]PAM NA Intergenic NA NA

T17192 TCCCCGGACGTTACGCCGCGAGG 1000
Oligo fwd CACCTCCCCGGACGTTACGCCGCG
Oligo rev AAACCGCGGCGTAACGTCCGGGGA
Chromosome start end strand MM target_seq PAM alignment distance position gene name gene id
GU305914.1 229905 229927 + 0 TCCCCGGACGTTACGCCGCG AGG ||||||||[||||||||||||]PAM NA Intergenic NA NA

T17193 CCCCGGACGTTACGCCGCGAGGG 1000
Oligo fwd CACCCCCCGGACGTTACGCCGCGA
Oligo rev AAACTCGCGGCGTAACGTCCGGGG
Chromosome start end strand MM target_seq PAM alignment distance position gene name gene id
GU305914.1 229906 229928 + 0 CCCCGGACGTTACGCCGCGA GGG ||||||||[||||||||||||]PAM NA Intergenic NA NA

T17194 CGGACGTTACGCCGCGAGGGCGG 1000
Oligo fwd CACCCGGACGTTACGCCGCGAGGG
Oligo rev AAACCCCTCGCGGCGTAACGTCCG
Chromosome start end strand MM target_seq PAM alignment distance position gene name gene id
GU305914.1 229909 229931 + 0 CGGACGTTACGCCGCGAGGG CGG ||||||||[||||||||||||]PAM NA Intergenic NA NA
GU305914.1 188571 188593 - 4 GGGAAGTTACAACGCGAGGG TGG -|||-|||[||--||||||||]PAM NA Intergenic NA NA

T17195 TACGCCGCGAGGGCGGCTCCAGG 1000
Oligo fwd CACCTACGCCGCGAGGGCGGCTCC
Oligo rev AAACGGAGCCGCCCTCGCGGCGTA
Chromosome start end strand MM target_seq PAM alignment distance position gene name gene id
GU305914.1 229916 229938 + 0 TACGCCGCGAGGGCGGCTCC AGG ||||||||[||||||||||||]PAM NA Intergenic NA NA

T17196 ACGCCGCGAGGGCGGCTCCAGGG 1000
Oligo fwd CACCACGCCGCGAGGGCGGCTCCA
Oligo rev AAACTGGAGCCGCCCTCGCGGCGT
Chromosome start end strand MM target_seq PAM alignment distance position gene name gene id
GU305914.1 229917 229939 + 0 ACGCCGCGAGGGCGGCTCCA GGG ||||||||[||||||||||||]PAM NA Intergenic NA NA

T17197 CGCCGCGAGGGCGGCTCCAGGGG 1000
Oligo fwd CACCCGCCGCGAGGGCGGCTCCAG
Oligo rev AAACCTGGAGCCGCCCTCGCGGCG
Chromosome start end strand MM target_seq PAM alignment distance position gene name gene id
GU305914.1 229918 229940 + 0 CGCCGCGAGGGCGGCTCCAG GGG ||||||||[||||||||||||]PAM NA Intergenic NA NA

T17198 GCCGCGAGGGCGGCTCCAGGGGG 1000
Oligo fwd CACCGCCGCGAGGGCGGCTCCAGG
Oligo rev AAACCCTGGAGCCGCCCTCGCGGC
Chromosome start end strand MM target_seq PAM alignment distance position gene name gene id
GU305914.1 229919 229941 + 0 GCCGCGAGGGCGGCTCCAGG GGG ||||||||[||||||||||||]PAM NA Intergenic NA NA

T17199 GCTCCAGGGGGGTTCACCCCCCGG 1000
Oligo fwd CACCGCTCCAGGGGGGTTCACCCCCC
Oligo rev AAACGGGGGTGAACCCCCTGGAGC
Chromosome start end strand MM target_seq PAM alignment distance position gene name gene id
GU305914.1 229931 229953 + 0 GCTCCAGGGGGGTTCACCCCC CGG ||||||||[||||||||||||]PAM NA Intergenic NA NA

T17200 CTCCAGGGGGGTTCACCCCCCGGG 1000
Oligo fwd CACCCTCCAGGGGGGTTCACCCCCC

<table>
<tr><td>Oligo rev</td><td>AAACGGGGGGTGAACCCCCTGGAG</td><td></td><td></td><td></td><td></td><td></td><td></td><td></td><td></td><td></td><td></td></tr>
<tr><td>Chromosome</td><td>start</td><td>end</td><td>strand</td><td>MM</td><td>target_seq</td><td>PAM</td><td>alignment</td><td>distance</td><td>position</td><td>gene name</td><td>gene id</td></tr>
<tr><td>GU305914.1</td><td>229932</td><td>229954</td><td>+</td><td>0</td><td>CTCCAGGGGGTTCACCCCCC</td><td>GGG</td><td>||||||||[||||||||||||]PAM</td><td>NA</td><td>Intergenic</td><td>NA</td><td>NA</td></tr>
<tr><td>T17201</td><td>GGGGGTTCACCCCCCGGGCGCGG</td><td>1000</td><td></td><td></td><td></td><td></td><td></td><td></td><td></td><td></td><td></td></tr>
<tr><td>Oligo fwd</td><td>CACCGGGGGTTCACCCCCCGGGCG</td><td></td><td></td><td></td><td></td><td></td><td></td><td></td><td></td><td></td><td></td></tr>
<tr><td>Oligo rev</td><td>AAACCGCCCGGGGGGTGAACCCCC</td><td></td><td></td><td></td><td></td><td></td><td></td><td></td><td></td><td></td><td></td></tr>
<tr><td>Chromosome</td><td>start</td><td>end</td><td>strand</td><td>MM</td><td>target_seq</td><td>PAM</td><td>alignment</td><td>distance</td><td>position</td><td>gene name</td><td>gene id</td></tr>
<tr><td>GU305914.1</td><td>229937</td><td>229959</td><td>+</td><td>0</td><td>GGGGGTTCACCCCCCGGGCG</td><td>CGG</td><td>||||||||[||||||||||||]PAM</td><td>NA</td><td>Intergenic</td><td>NA</td><td>NA</td></tr>
<tr><td>T17202</td><td>CACCGTCTCGTGCGCCATCCCGG</td><td>1000</td><td></td><td></td><td></td><td></td><td></td><td></td><td></td><td></td><td></td></tr>
<tr><td>Oligo fwd</td><td>CACCCACCGTCTCGTGCGCCATCC</td><td></td><td></td><td></td><td></td><td></td><td></td><td></td><td></td><td></td><td></td></tr>
<tr><td>Oligo rev</td><td>AAACGGATGGCGCACGAGACGGTG</td><td></td><td></td><td></td><td></td><td></td><td></td><td></td><td></td><td></td><td></td></tr>
<tr><td>Chromosome</td><td>start</td><td>end</td><td>strand</td><td>MM</td><td>target_seq</td><td>PAM</td><td>alignment</td><td>distance</td><td>position</td><td>gene name</td><td>gene id</td></tr>
<tr><td>GU305914.1</td><td>229965</td><td>229987</td><td>+</td><td>0</td><td>CACCGTCTCGTGCGCCATCC</td><td>CGG</td><td>||||||||[||||||||||||]PAM</td><td>NA</td><td>Intergenic</td><td>NA</td><td>NA</td></tr>
<tr><td>T17203</td><td>GTCTCGTGCGCCATCCCGGCCGG</td><td>1000</td><td></td><td></td><td></td><td></td><td></td><td></td><td></td><td></td><td></td></tr>
<tr><td>Oligo fwd</td><td>CACCGTCTCGTGCGCCATCCCGGC</td><td></td><td></td><td></td><td></td><td></td><td></td><td></td><td></td><td></td><td></td></tr>
<tr><td>Oligo rev</td><td>AAACGCCGGGATGGCGCACGAGAC</td><td></td><td></td><td></td><td></td><td></td><td></td><td></td><td></td><td></td><td></td></tr>
<tr><td>Chromosome</td><td>start</td><td>end</td><td>strand</td><td>MM</td><td>target_seq</td><td>PAM</td><td>alignment</td><td>distance</td><td>position</td><td>gene name</td><td>gene id</td></tr>
<tr><td>GU305914.1</td><td>229969</td><td>229991</td><td>+</td><td>0</td><td>GTCTCGTGCGCCATCCCGGC</td><td>CGG</td><td>||||||||[||||||||||||]PAM</td><td>NA</td><td>Intergenic</td><td>NA</td><td>NA</td></tr>
<tr><td>T17204</td><td>TCGTGCGCCATCCCGGCCGGCGG</td><td>1000</td><td></td><td></td><td></td><td></td><td></td><td></td><td></td><td></td><td></td></tr>
<tr><td>Oligo fwd</td><td>CACCTCGTGCGCCATCCCGGCCGG</td><td></td><td></td><td></td><td></td><td></td><td></td><td></td><td></td><td></td><td></td></tr>
<tr><td>Oligo rev</td><td>AAACCCGGCCGGGATGGCGCACGA</td><td></td><td></td><td></td><td></td><td></td><td></td><td></td><td></td><td></td><td></td></tr>
<tr><td>Chromosome</td><td>start</td><td>end</td><td>strand</td><td>MM</td><td>target_seq</td><td>PAM</td><td>alignment</td><td>distance</td><td>position</td><td>gene name</td><td>gene id</td></tr>
<tr><td>GU305914.1</td><td>229972</td><td>229994</td><td>+</td><td>0</td><td>TCGTGCGCCATCCCGGCCGG</td><td>CGG</td><td>||||||||[||||||||||||]PAM</td><td>NA</td><td>Intergenic</td><td>NA</td><td>NA</td></tr>
<tr><td>T17205</td><td>GCCATCCCGGCCGGCGGCAGTGG</td><td>1000</td><td></td><td></td><td></td><td></td><td></td><td></td><td></td><td></td><td></td></tr>
<tr><td>Oligo fwd</td><td>CACCGCCATCCCGGCCGGCGGCAG</td><td></td><td></td><td></td><td></td><td></td><td></td><td></td><td></td><td></td><td></td></tr>
<tr><td>Oligo rev</td><td>AAACCTGCCGCCGGCCGGGATGGC</td><td></td><td></td><td></td><td></td><td></td><td></td><td></td><td></td><td></td><td></td></tr>
<tr><td>Chromosome</td><td>start</td><td>end</td><td>strand</td><td>MM</td><td>target_seq</td><td>PAM</td><td>alignment</td><td>distance</td><td>position</td><td>gene name</td><td>gene id</td></tr>
<tr><td>GU305914.1</td><td>229978</td><td>230000</td><td>+</td><td>0</td><td>GCCATCCCGGCCGGCGGCAG</td><td>TGG</td><td>||||||||[||||||||||||]PAM</td><td>NA</td><td>Intergenic</td><td>NA</td><td>NA</td></tr>
<tr><td>T17206</td><td>TCCCGGCCGGCGGCAGTGGCAGG</td><td>1000</td><td></td><td></td><td></td><td></td><td></td><td></td><td></td><td></td><td></td></tr>
<tr><td>Oligo fwd</td><td>CACCTCCCGGCCGGCGGCAGTGGC</td><td></td><td></td><td></td><td></td><td></td><td></td><td></td><td></td><td></td><td></td></tr>
<tr><td>Oligo rev</td><td>AAACGCCACTGCCGCCGGCCGGGA</td><td></td><td></td><td></td><td></td><td></td><td></td><td></td><td></td><td></td><td></td></tr>
<tr><td>Chromosome</td><td>start</td><td>end</td><td>strand</td><td>MM</td><td>target_seq</td><td>PAM</td><td>alignment</td><td>distance</td><td>position</td><td>gene name</td><td>gene id</td></tr>
<tr><td>GU305914.1</td><td>229982</td><td>230004</td><td>+</td><td>0</td><td>TCCCGGCCGGCGGCAGTGGC</td><td>AGG</td><td>||||||||[||||||||||||]PAM</td><td>NA</td><td>Intergenic</td><td>NA</td><td>NA</td></tr>
<tr><td>T17207</td><td>CGGCCGGCGGCAGTGGCAGGTGG</td><td>1000</td><td></td><td></td><td></td><td></td><td></td><td></td><td></td><td></td><td></td></tr>
<tr><td>Oligo fwd</td><td>CACCCGGCCGGCGGCAGTGGCAGG</td><td></td><td></td><td></td><td></td><td></td><td></td><td></td><td></td><td></td><td></td></tr>
<tr><td>Oligo rev</td><td>AAACCCTGCCACTGCCGCCGGCCG</td><td></td><td></td><td></td><td></td><td></td><td></td><td></td><td></td><td></td><td></td></tr>
<tr><td>Chromosome</td><td>start</td><td>end</td><td>strand</td><td>MM</td><td>target_seq</td><td>PAM</td><td>alignment</td><td>distance</td><td>position</td><td>gene name</td><td>gene id</td></tr>
<tr><td>GU305914.1</td><td>229985</td><td>230007</td><td>+</td><td>0</td><td>CGGCCGGCGGCAGTGGCAGG</td><td>TGG</td><td>||||||||[||||||||||||]PAM</td><td>NA</td><td>Intergenic</td><td>NA</td><td>NA</td></tr>
<tr><td>T17208</td><td>GGCCGGCGGCAGTGGCAGGTGGG</td><td>1000</td><td></td><td></td><td></td><td></td><td></td><td></td><td></td><td></td><td></td></tr>
<tr><td>Oligo fwd</td><td>CACCGGCCGGCGGCAGTGGCAGGT</td><td></td><td></td><td></td><td></td><td></td><td></td><td></td><td></td><td></td><td></td></tr>
<tr><td>Oligo rev</td><td>AAACACCTGCCACTGCCGCCGGCC</td><td></td><td></td><td></td><td></td><td></td><td></td><td></td><td></td><td></td><td></td></tr>
<tr><td>Chromosome</td><td>start</td><td>end</td><td>strand</td><td>MM</td><td>target_seq</td><td>PAM</td><td>alignment</td><td>distance</td><td>position</td><td>gene name</td><td>gene id</td></tr>
<tr><td>GU305914.1</td><td>229986</td><td>230008</td><td>+</td><td>0</td><td>GGCCGGCGGCAGTGGCAGGT</td><td>GGG</td><td>||||||||[||||||||||||]PAM</td><td>NA</td><td>Intergenic</td><td>NA</td><td>NA</td></tr>
<tr><td>T17209</td><td>CGGCAGTGGCAGGTGGGCGTCGG</td><td>1000</td><td></td><td></td><td></td><td></td><td></td><td></td><td></td><td></td><td></td></tr>
<tr><td>Oligo fwd</td><td>CACCCGGCAGTGGCAGGTGGGCGT</td><td></td><td></td><td></td><td></td><td></td><td></td><td></td><td></td><td></td><td></td></tr>
<tr><td>Oligo rev</td><td>AAACACGCCCACCTGCCACTGCCG</td><td></td><td></td><td></td><td></td><td></td><td></td><td></td><td></td><td></td><td></td></tr>
<tr><td>Chromosome</td><td>start</td><td>end</td><td>strand</td><td>MM</td><td>target_seq</td><td>PAM</td><td>alignment</td><td>distance</td><td>position</td><td>gene name</td><td>gene id</td></tr>
<tr><td>GU305914.1</td><td>229992</td><td>230014</td><td>+</td><td>0</td><td>CGGCAGTGGCAGGTGGGCGT</td><td>CGG</td><td>||||||||[||||||||||||]PAM</td><td>NA</td><td>Intergenic</td><td>NA</td><td>NA</td></tr>
<tr><td>T17210</td><td>GGCAGTGGCAGGTGGGCGTCGGG</td><td>1000</td><td></td><td></td><td></td><td></td><td></td><td></td><td></td><td></td><td></td></tr>
<tr><td>Oligo fwd</td><td>CACCGGCAGTGGCAGGTGGGCGTC</td><td></td><td></td><td></td><td></td><td></td><td></td><td></td><td></td><td></td><td></td></tr>
<tr><td>Oligo rev</td><td>AAACGACGCCCACCTGCCACTGCC</td><td></td><td></td><td></td><td></td><td></td><td></td><td></td><td></td><td></td><td></td></tr>
<tr><td>Chromosome</td><td>start</td><td>end</td><td>strand</td><td>MM</td><td>target_seq</td><td>PAM</td><td>alignment</td><td>distance</td><td>position</td><td>gene name</td><td>gene id</td></tr>
<tr><td>GU305914.1</td><td>229993</td><td>230015</td><td>+</td><td>0</td><td>GGCAGTGGCAGGTGGGCGTC</td><td>GGG</td><td>||||||||[||||||||||||]PAM</td><td>NA</td><td>Intergenic</td><td>NA</td><td>NA</td></tr>
<tr><td>T17211</td><td>GCAGTGGCAGGTGGGCGTCGGGG</td><td>1000</td><td></td><td></td><td></td><td></td><td></td><td></td><td></td><td></td><td></td></tr>
<tr><td>Oligo fwd</td><td>CACCGCAGTGGCAGGTGGGCGTCG</td><td></td><td></td><td></td><td></td><td></td><td></td><td></td><td></td><td></td><td></td></tr>
<tr><td>Oligo rev</td><td>AAACCGACGCCCACCTGCCACTGC</td><td></td><td></td><td></td><td></td><td></td><td></td><td></td><td></td><td></td><td></td></tr>
<tr><td>Chromosome</td><td>start</td><td>end</td><td>strand</td><td>MM</td><td>target_seq</td><td>PAM</td><td>alignment</td><td>distance</td><td>position</td><td>gene name</td><td>gene id</td></tr>
<tr><td>GU305914.1</td><td>229994</td><td>230016</td><td>+</td><td>0</td><td>GCAGTGGCAGGTGGGCGTCG</td><td>GGG</td><td>||||||||[||||||||||||]PAM</td><td>NA</td><td>Intergenic</td><td>NA</td><td>NA</td></tr>
<tr><td>T17212</td><td>CGTCGGGGCGCGCGTCCGCTCGG</td><td>1000</td><td></td><td></td><td></td><td></td><td></td><td></td><td></td><td></td><td></td></tr>
<tr><td>Oligo fwd</td><td>CACCCGTCGGGGCGCGCGTCCGCT</td><td></td><td></td><td></td><td></td><td></td><td></td><td></td><td></td><td></td><td></td></tr>
<tr><td>Oligo rev</td><td>AAACAGCGGACGCGCGCCCCGACG</td><td></td><td></td><td></td><td></td><td></td><td></td><td></td><td></td><td></td><td></td></tr>
<tr><td>Chromosome</td><td>start</td><td>end</td><td>strand</td><td>MM</td><td>target_seq</td><td>PAM</td><td>alignment</td><td>distance</td><td>position</td><td>gene name</td><td>gene id</td></tr>
<tr><td>GU305914.1</td><td>230009</td><td>230031</td><td>+</td><td>0</td><td>CGTCGGGGCGCGCGTCCGCT</td><td>CGG</td><td>||||||||[||||||||||||]PAM</td><td>NA</td><td>Intergenic</td><td>NA</td><td>NA</td></tr>
<tr><td>T17213</td><td>CCCACGCCGTACACGTCTCGCGG</td><td>1000</td><td></td><td></td><td></td><td></td><td></td><td></td><td></td><td></td><td></td></tr>
<tr><td>Oligo fwd</td><td>CACCCCCACGCCGTACACGTCTCG</td><td></td><td></td><td></td><td></td><td></td><td></td><td></td><td></td><td></td><td></td></tr>
<tr><td>Oligo rev</td><td>AAACCGAGACGTGTACGGCGTGGG</td><td></td><td></td><td></td><td></td><td></td><td></td><td></td><td></td><td></td><td></td></tr>
<tr><td>Chromosome</td><td>start</td><td>end</td><td>strand</td><td>MM</td><td>target_seq</td><td>PAM</td><td>alignment</td><td>distance</td><td>position</td><td>gene name</td><td>gene id</td></tr>
<tr><td>GU305914.1</td><td>230044</td><td>230066</td><td>+</td><td>0</td><td>CCCACGCCGTACACGTCTCG</td><td>CGG</td><td>||||||||[||||||||||||]PAM</td><td>NA</td><td>Intergenic</td><td>NA</td><td>NA</td></tr>
<tr><td>T17214</td><td>CCACGCCGTACACGTCTCGCGGG</td><td>1000</td><td></td><td></td><td></td><td></td><td></td><td></td><td></td><td></td><td></td></tr>
<tr><td>Oligo fwd</td><td>CACCCCACGCCGTACACGTCTCGC</td><td></td><td></td><td></td><td></td><td></td><td></td><td></td><td></td><td></td><td></td></tr>
<tr><td>Oligo rev</td><td>AAACGCGAGACGTGTACGGCGTGG</td><td></td><td></td><td></td><td></td><td></td><td></td><td></td><td></td><td></td><td></td></tr>
<tr><td>Chromosome</td><td>start</td><td>end</td><td>strand</td><td>MM</td><td>target_seq</td><td>PAM</td><td>alignment</td><td>distance</td><td>position</td><td>gene name</td><td>gene id</td></tr>
<tr><td>GU305914.1</td><td>230045</td><td>230067</td><td>+</td><td>0</td><td>CCACGCCGTACACGTCTCGC</td><td>GGG</td><td>||||||||[||||||||||||]PAM</td><td>NA</td><td>Intergenic</td><td>NA</td><td>NA</td></tr>
<tr><td>T17215</td><td>CAGTTCCCAGTACACTCCCGCGG</td><td>1000</td><td></td><td></td><td></td><td></td><td></td><td></td><td></td><td></td><td></td></tr>
</table>

| | | | | | | | | | | | |
|---|---|---|---|---|---|---|---|---|---|---|---|
| Oligo fwd | CACCCAGTTCCCAGTACACTCCCG | | | | | | | | | | |
| Oligo rev | AAACCGGGAGTGTACTGGGAACTG | | | | | | | | | | |
| Chromosome | start | end | strand | MM | target_seq | PAM | alignment | distance | position | gene name | gene id |
| GU305914.1 | 230076 | 230098 | + | 0 | CAGTTCCCAGTACACTCCCG | CGG | \|\|\|\|\|\|\|\|[\|\|\|\|\|\|\|\|\|\|\|\|]PAM | NA | Intergenic | NA | NA |
| T17216 | CACTCCCGCGGTAGTCGTGCCGG | 1000 | | | | | | | | | |
| Oligo fwd | CACCCACTCCCGCGGTAGTCGTGC | | | | | | | | | | |
| Oligo rev | AAACGCACGACTACCGCGGGAGTG | | | | | | | | | | |
| Chromosome | start | end | strand | MM | target_seq | PAM | alignment | distance | position | gene name | gene id |
| GU305914.1 | 230088 | 230110 | + | 0 | CACTCCCGCGGTAGTCGTGC | CGG | \|\|\|\|\|\|\|\|[\|\|\|\|\|\|\|\|\|\|\|\|]PAM | NA | Intergenic | NA | NA |
| T17217 | CGGTAGTCGTGCCGGACAGCCGG | 1000 | | | | | | | | | |
| Oligo fwd | CACCCGGTAGTCGTGCCGGACAGC | | | | | | | | | | |
| Oligo rev | AAACGCTGTCCGGCACGACTACCG | | | | | | | | | | |
| Chromosome | start | end | strand | MM | target_seq | PAM | alignment | distance | position | gene name | gene id |
| GU305914.1 | 230096 | 230118 | + | 0 | CGGTAGTCGTGCCGGACAGC | CGG | \|\|\|\|\|\|\|\|[\|\|\|\|\|\|\|\|\|\|\|\|]PAM | NA | Intergenic | NA | NA |
| T17218 | CGTGCCGGACAGCCGGACGAAGG | 1000 | | | | | | | | | |
| Oligo fwd | CACCCGTGCCGGACAGCCGGACGA | | | | | | | | | | |
| Oligo rev | AAACTCGTCCGGCTGTCCGGCACG | | | | | | | | | | |
| Chromosome | start | end | strand | MM | target_seq | PAM | alignment | distance | position | gene name | gene id |
| GU305914.1 | 230103 | 230125 | + | 0 | CGTGCCGGACAGCCGGACGA | AGG | \|\|\|\|\|\|\|\|[\|\|\|\|\|\|\|\|\|\|\|\|]PAM | NA | Intergenic | NA | NA |
| T17219 | GACTCCTCCACACACATATGAGG | 1000 | | | | | | | | | |
| Oligo fwd | CACCGACTCCTCCACACACATATG | | | | | | | | | | |
| Oligo rev | AAACCATATGTGTGTGGAGGAGTC | | | | | | | | | | |
| Chromosome | start | end | strand | MM | target_seq | PAM | alignment | distance | position | gene name | gene id |
| GU305914.1 | 230159 | 230181 | + | 0 | GACTCCTCCACACACATATG | AGG | \|\|\|\|\|\|\|\|[\|\|\|\|\|\|\|\|\|\|\|\|]PAM | NA | Intergenic | NA | NA |
| T17220 | CTCCACGCCATTCACTTGCATGG | 1000 | | | | | | | | | |
| Oligo fwd | CACCCTCCACGCCATTCACTTGCA | | | | | | | | | | |
| Oligo rev | AAACTGCAAGTGAATGGCGTGGAG | | | | | | | | | | |
| Chromosome | start | end | strand | MM | target_seq | PAM | alignment | distance | position | gene name | gene id |
| GU305914.1 | 230242 | 230264 | + | 0 | CTCCACGCCATTCACTTGCA | TGG | \|\|\|\|\|\|\|\|[\|\|\|\|\|\|\|\|\|\|\|\|]PAM | NA | Intergenic | NA | NA |
| T17221 | TCCACGCCATTCACTTGCATGGG | 1000 | | | | | | | | | |
| Oligo fwd | CACCTCCACGCCATTCACTTGCAT | | | | | | | | | | |
| Oligo rev | AAACATGCAAGTGAATGGCGTGGA | | | | | | | | | | |
| Chromosome | start | end | strand | MM | target_seq | PAM | alignment | distance | position | gene name | gene id |
| GU305914.1 | 230243 | 230265 | + | 0 | TCCACGCCATTCACTTGCAT | GGG | \|\|\|\|\|\|\|\|[\|\|\|\|\|\|\|\|\|\|\|\|]PAM | NA | Intergenic | NA | NA |
| T17222 | ACGCCATTCACTTGCATGGGCGG | 1000 | | | | | | | | | |
| Oligo fwd | CACCACGCCATTCACTTGCATGGG | | | | | | | | | | |
| Oligo rev | AAACCCCATGCAAGTGAATGGCGT | | | | | | | | | | |
| Chromosome | start | end | strand | MM | target_seq | PAM | alignment | distance | position | gene name | gene id |
| GU305914.1 | 230246 | 230268 | + | 0 | ACGCCATTCACTTGCATGGG | CGG | \|\|\|\|\|\|\|\|[\|\|\|\|\|\|\|\|\|\|\|\|]PAM | NA | Intergenic | NA | NA |
| T17223 | CGCCATTCACTTGCATGGGCGGG | 1000 | | | | | | | | | |
| Oligo fwd | CACCCGCCATTCACTTGCATGGGC | | | | | | | | | | |
| Oligo rev | AAACGCCCATGCAAGTGAATGGCG | | | | | | | | | | |
| Chromosome | start | end | strand | MM | target_seq | PAM | alignment | distance | position | gene name | gene id |
| GU305914.1 | 230247 | 230269 | + | 0 | CGCCATTCACTTGCATGGGC | GGG | \|\|\|\|\|\|\|\|[\|\|\|\|\|\|\|\|\|\|\|\|]PAM | NA | Intergenic | NA | NA |
| T17224 | GCTGCACTTTTTTGCCGCCGAGG | 1000 | | | | | | | | | |
| Oligo fwd | CACCGCTGCACTTTTTTGCCGCCG | | | | | | | | | | |
| Oligo rev | AAACCGGCGGCAAAAAAGTGCAGC | | | | | | | | | | |
| Chromosome | start | end | strand | MM | target_seq | PAM | alignment | distance | position | gene name | gene id |
| GU305914.1 | 230269 | 230291 | + | 0 | GCTGCACTTTTTTGCCGCCG | AGG | \|\|\|\|\|\|\|\|[\|\|\|\|\|\|\|\|\|\|\|\|]PAM | NA | Intergenic | NA | NA |
| T17225 | CTGCACTTTTTTGCCGCCGAGGG | 1000 | | | | | | | | | |
| Oligo fwd | CACCCTGCACTTTTTTGCCGCCGA | | | | | | | | | | |
| Oligo rev | AAACTCGGCGGCAAAAAAGTGCAG | | | | | | | | | | |
| Chromosome | start | end | strand | MM | target_seq | PAM | alignment | distance | position | gene name | gene id |
| GU305914.1 | 230270 | 230292 | + | 0 | CTGCACTTTTTTGCCGCCGA | GGG | \|\|\|\|\|\|\|\|[\|\|\|\|\|\|\|\|\|\|\|\|]PAM | NA | Intergenic | NA | NA |
| T17226 | TTTTGCCGCCGAGGGCGCCGCGG | 1000 | | | | | | | | | |
| Oligo fwd | CACCTTTTGCCGCCGAGGGCGCCG | | | | | | | | | | |
| Oligo rev | AAACCGGCGCCCTCGGCGGCAAAA | | | | | | | | | | |
| Chromosome | start | end | strand | MM | target_seq | PAM | alignment | distance | position | gene name | gene id |
| GU305914.1 | 230278 | 230300 | + | 0 | TTTTGCCGCCGAGGGCGCCG | CGG | \|\|\|\|\|\|\|\|[\|\|\|\|\|\|\|\|\|\|\|\|]PAM | NA | Intergenic | NA | NA |
| T17227 | CACATACTTCTTTTTTTCGTAGG | 1000 | | | | | | | | | |
| Oligo fwd | CACCCACATACTTCTTTTTTTCGT | | | | | | | | | | |
| Oligo rev | AAACACGAAAAAAAGAAGTATGTG | | | | | | | | | | |
| Chromosome | start | end | strand | MM | target_seq | PAM | alignment | distance | position | gene name | gene id |
| GU305914.1 | 230301 | 230323 | + | 0 | CACATACTTCTTTTTTTCGT | AGG | \|\|\|\|\|\|\|\|[\|\|\|\|\|\|\|\|\|\|\|\|]PAM | NA | Intergenic | NA | NA |
| T17228 | ACATACTTCTTTTTTTCGTAGGG | 1000 | | | | | | | | | |
| Oligo fwd | CACCACATACTTCTTTTTTTCGTA | | | | | | | | | | |
| Oligo rev | AAACTACGAAAAAAAGAAGTATGT | | | | | | | | | | |
| Chromosome | start | end | strand | MM | target_seq | PAM | alignment | distance | position | gene name | gene id |
| GU305914.1 | 230302 | 230324 | + | 0 | ACATACTTCTTTTTTTCGTA | GGG | \|\|\|\|\|\|\|\|[\|\|\|\|\|\|\|\|\|\|\|\|]PAM | NA | Intergenic | NA | NA |
| T17229 | GTACCACTTTTTATCCCCCCCGG | 1000 | | | | | | | | | |
| Oligo fwd | CACCGTACCACTTTTTATCCCCCC | | | | | | | | | | |
| Oligo rev | AAACGGGGGGATAAAAAGTGGTAC | | | | | | | | | | |
| Chromosome | start | end | strand | MM | target_seq | PAM | alignment | distance | position | gene name | gene id |
| GU305914.1 | 230335 | 230357 | + | 0 | GTACCACTTTTTATCCCCCC | CGG | \|\|\|\|\|\|\|\|[\|\|\|\|\|\|\|\|\|\|\|\|]PAM | NA | Intergenic | NA | NA |

<table>
<tr><td>T17230</td><td>CCCGGCCGTCTGAGTGCGCGCGG</td><td>1000</td><td></td><td></td><td></td><td></td><td></td><td></td><td></td><td></td><td></td></tr>
<tr><td>Oligo fwd</td><td>CACCCCCGGCCGTCTGAGTGCGCG</td><td></td><td></td><td></td><td></td><td></td><td></td><td></td><td></td><td></td><td></td></tr>
<tr><td>Oligo rev</td><td>AAACCGCGCACTCAGACGGCCGGG</td><td></td><td></td><td></td><td></td><td></td><td></td><td></td><td></td><td></td><td></td></tr>
<tr><td>Chromosome</td><td>start</td><td>end</td><td>strand</td><td>MM</td><td>target_seq</td><td>PAM</td><td>alignment</td><td>distance</td><td>position</td><td>gene name</td><td>gene id</td></tr>
<tr><td>GU305914.1</td><td>230353</td><td>230375</td><td>+</td><td>0</td><td>CCCGGCCGTCTGAGTGCGCG</td><td>CGG</td><td>||||||||[||||||||||||]PAM</td><td>NA</td><td>Intergenic</td><td>NA</td><td>NA</td></tr>
<tr><td>GU305914.1</td><td>38</td><td>60</td><td>+</td><td>0</td><td>CCCGGCCGTCTGAGTGCGCG</td><td>CGG</td><td>||||||||[||||||||||||]PAM</td><td>NA</td><td>Intergenic</td><td>NA</td><td>NA</td></tr>
<tr><td>GU305914.1</td><td>149055</td><td>149077</td><td>+</td><td>4</td><td>ACCGGCAGTCCGAGTCCGCG</td><td>CGG</td><td>-|||||-|[||-||||-||||]PAM</td><td>NA</td><td>Intergenic</td><td>NA</td><td>NA</td></tr>
</table>

<table>
<tr><td>T17231</td><td>CCGGCCGTCTGAGTGCGCGCGGG</td><td>1000</td><td></td><td></td><td></td><td></td><td></td><td></td><td></td><td></td><td></td></tr>
<tr><td>Oligo fwd</td><td>CACCCCGGCCGTCTGAGTGCGCGC</td><td></td><td></td><td></td><td></td><td></td><td></td><td></td><td></td><td></td><td></td></tr>
<tr><td>Oligo rev</td><td>AAACGCGCGCACTCAGACGGCCGG</td><td></td><td></td><td></td><td></td><td></td><td></td><td></td><td></td><td></td><td></td></tr>
<tr><td>Chromosome</td><td>start</td><td>end</td><td>strand</td><td>MM</td><td>target_seq</td><td>PAM</td><td>alignment</td><td>distance</td><td>position</td><td>gene name</td><td>gene id</td></tr>
<tr><td>GU305914.1</td><td>230354</td><td>230376</td><td>+</td><td>0</td><td>CCGGCCGTCTGAGTGCGCGC</td><td>GGG</td><td>||||||||[||||||||||||]PAM</td><td>NA</td><td>Intergenic</td><td>NA</td><td>NA</td></tr>
<tr><td>GU305914.1</td><td>39</td><td>61</td><td>+</td><td>0</td><td>CCGGCCGTCTGAGTGCGCGC</td><td>GGG</td><td>||||||||[||||||||||||]PAM</td><td>NA</td><td>Intergenic</td><td>NA</td><td>NA</td></tr>
</table>

<table>
<tr><td>T17232</td><td>GATGAATCAGCTTGATTCGTTGG</td><td>1000</td><td></td><td></td><td></td><td></td><td></td><td></td><td></td><td></td><td></td></tr>
<tr><td>Oligo fwd</td><td>CACCGATGAATCAGCTTGATTCGT</td><td></td><td></td><td></td><td></td><td></td><td></td><td></td><td></td><td></td><td></td></tr>
<tr><td>Oligo rev</td><td>AAACACGAATCAAGCTGATTCATC</td><td></td><td></td><td></td><td></td><td></td><td></td><td></td><td></td><td></td><td></td></tr>
<tr><td>Chromosome</td><td>start</td><td>end</td><td>strand</td><td>MM</td><td>target_seq</td><td>PAM</td><td>alignment</td><td>distance</td><td>position</td><td>gene name</td><td>gene id</td></tr>
<tr><td>GU305914.1</td><td>230378</td><td>230400</td><td>-</td><td>0</td><td>GATGAATCAGCTTGATTCGT</td><td>TGG</td><td>||||||||[||||||||||||]PAM</td><td>NA</td><td>Intergenic</td><td>NA</td><td>NA</td></tr>
</table>

<table>
<tr><td>T17233</td><td>ATGAATCAGCTTGATTCGTTGGG</td><td>1000</td><td></td><td></td><td></td><td></td><td></td><td></td><td></td><td></td><td></td></tr>
<tr><td>Oligo fwd</td><td>CACCATGAATCAGCTTGATTCGTT</td><td></td><td></td><td></td><td></td><td></td><td></td><td></td><td></td><td></td><td></td></tr>
<tr><td>Oligo rev</td><td>AAACAACGAATCAAGCTGATTCAT</td><td></td><td></td><td></td><td></td><td></td><td></td><td></td><td></td><td></td><td></td></tr>
<tr><td>Chromosome</td><td>start</td><td>end</td><td>strand</td><td>MM</td><td>target_seq</td><td>PAM</td><td>alignment</td><td>distance</td><td>position</td><td>gene name</td><td>gene id</td></tr>
<tr><td>GU305914.1</td><td>230377</td><td>230399</td><td>-</td><td>0</td><td>ATGAATCAGCTTGATTCGTT</td><td>GGG</td><td>||||||||[||||||||||||]PAM</td><td>NA</td><td>Intergenic</td><td>NA</td><td>NA</td></tr>
</table>

<table>
<tr><td>T17234</td><td>TGGGCCCGCGCGCACTCAGACGG</td><td>1000</td><td></td><td></td><td></td><td></td><td></td><td></td><td></td><td></td><td></td></tr>
<tr><td>Oligo fwd</td><td>CACCTGGGCCCGCGCGCACTCAGA</td><td></td><td></td><td></td><td></td><td></td><td></td><td></td><td></td><td></td><td></td></tr>
<tr><td>Oligo rev</td><td>AAACTCTGAGTGCGCGCGGGCCCA</td><td></td><td></td><td></td><td></td><td></td><td></td><td></td><td></td><td></td><td></td></tr>
<tr><td>Chromosome</td><td>start</td><td>end</td><td>strand</td><td>MM</td><td>target_seq</td><td>PAM</td><td>alignment</td><td>distance</td><td>position</td><td>gene name</td><td>gene id</td></tr>
<tr><td>GU305914.1</td><td>230358</td><td>230380</td><td>-</td><td>0</td><td>TGGGCCCGCGCGCACTCAGA</td><td>CGG</td><td>||||||||[||||||||||||]PAM</td><td>NA</td><td>Intergenic</td><td>NA</td><td>NA</td></tr>
<tr><td>GU305914.1</td><td>43</td><td>65</td><td>-</td><td>1</td><td>GGGGCCCGCGCGCACTCAGA</td><td>CGG</td><td>-|||||||[||||||||||||]PAM</td><td>NA</td><td>Intergenic</td><td>NA</td><td>NA</td></tr>
</table>

<table>
<tr><td>T17235</td><td>CCCGCGCGCACTCAGACGGCCGG</td><td>1000</td><td></td><td></td><td></td><td></td><td></td><td></td><td></td><td></td><td></td></tr>
<tr><td>Oligo fwd</td><td>CACCCCCGCGCGCACTCAGACGGC</td><td></td><td></td><td></td><td></td><td></td><td></td><td></td><td></td><td></td><td></td></tr>
<tr><td>Oligo rev</td><td>AAACGCCGTCTGAGTGCGCGCGGG</td><td></td><td></td><td></td><td></td><td></td><td></td><td></td><td></td><td></td><td></td></tr>
<tr><td>Chromosome</td><td>start</td><td>end</td><td>strand</td><td>MM</td><td>target_seq</td><td>PAM</td><td>alignment</td><td>distance</td><td>position</td><td>gene name</td><td>gene id</td></tr>
<tr><td>GU305914.1</td><td>39</td><td>61</td><td>-</td><td>0</td><td>CCCGCGCGCACTCAGACGGC</td><td>CGG</td><td>||||||||[||||||||||||]PAM</td><td>NA</td><td>Intergenic</td><td>NA</td><td>NA</td></tr>
<tr><td>GU305914.1</td><td>230354</td><td>230376</td><td>-</td><td>0</td><td>CCCGCGCGCACTCAGACGGC</td><td>CGG</td><td>||||||||[||||||||||||]PAM</td><td>NA</td><td>Intergenic</td><td>NA</td><td>NA</td></tr>
</table>

<table>
<tr><td>T17236</td><td>CCGCGCGCACTCAGACGGCCGGG</td><td>1000</td><td></td><td></td><td></td><td></td><td></td><td></td><td></td><td></td><td></td></tr>
<tr><td>Oligo fwd</td><td>CACCCCGCGCGCACTCAGACGGCC</td><td></td><td></td><td></td><td></td><td></td><td></td><td></td><td></td><td></td><td></td></tr>
<tr><td>Oligo rev</td><td>AAACGGCCGTCTGAGTGCGCGCGG</td><td></td><td></td><td></td><td></td><td></td><td></td><td></td><td></td><td></td><td></td></tr>
<tr><td>Chromosome</td><td>start</td><td>end</td><td>strand</td><td>MM</td><td>target_seq</td><td>PAM</td><td>alignment</td><td>distance</td><td>position</td><td>gene name</td><td>gene id</td></tr>
<tr><td>GU305914.1</td><td>38</td><td>60</td><td>-</td><td>0</td><td>CCGCGCGCACTCAGACGGCC</td><td>GGG</td><td>||||||||[||||||||||||]PAM</td><td>NA</td><td>Intergenic</td><td>NA</td><td>NA</td></tr>
<tr><td>GU305914.1</td><td>230353</td><td>230375</td><td>-</td><td>0</td><td>CCGCGCGCACTCAGACGGCC</td><td>GGG</td><td>||||||||[||||||||||||]PAM</td><td>NA</td><td>Intergenic</td><td>NA</td><td>NA</td></tr>
</table>

<table>
<tr><td>T17237</td><td>CGCGCGCACTCAGACGGCCGGGG</td><td>1000</td><td></td><td></td><td></td><td></td><td></td><td></td><td></td><td></td><td></td></tr>
<tr><td>Oligo fwd</td><td>CACCCGCGCGCACTCAGACGGCCG</td><td></td><td></td><td></td><td></td><td></td><td></td><td></td><td></td><td></td><td></td></tr>
<tr><td>Oligo rev</td><td>AAACCGGCCGTCTGAGTGCGCGCG</td><td></td><td></td><td></td><td></td><td></td><td></td><td></td><td></td><td></td><td></td></tr>
<tr><td>Chromosome</td><td>start</td><td>end</td><td>strand</td><td>MM</td><td>target_seq</td><td>PAM</td><td>alignment</td><td>distance</td><td>position</td><td>gene name</td><td>gene id</td></tr>
<tr><td>GU305914.1</td><td>37</td><td>59</td><td>-</td><td>0</td><td>CGCGCGCACTCAGACGGCCG</td><td>GGG</td><td>||||||||[||||||||||||]PAM</td><td>NA</td><td>Intergenic</td><td>NA</td><td>NA</td></tr>
<tr><td>GU305914.1</td><td>230352</td><td>230374</td><td>-</td><td>0</td><td>CGCGCGCACTCAGACGGCCG</td><td>GGG</td><td>||||||||[||||||||||||]PAM</td><td>NA</td><td>Intergenic</td><td>NA</td><td>NA</td></tr>
</table>

<table>
<tr><td>T17238</td><td>GCGCGCACTCAGACGGCCGGGGG</td><td>1000</td><td></td><td></td><td></td><td></td><td></td><td></td><td></td><td></td><td></td></tr>
<tr><td>Oligo fwd</td><td>CACCGCGCGCACTCAGACGGCCGG</td><td></td><td></td><td></td><td></td><td></td><td></td><td></td><td></td><td></td><td></td></tr>
<tr><td>Oligo rev</td><td>AAACCCGGCCGTCTGAGTGCGCGC</td><td></td><td></td><td></td><td></td><td></td><td></td><td></td><td></td><td></td><td></td></tr>
<tr><td>Chromosome</td><td>start</td><td>end</td><td>strand</td><td>MM</td><td>target_seq</td><td>PAM</td><td>alignment</td><td>distance</td><td>position</td><td>gene name</td><td>gene id</td></tr>
<tr><td>GU305914.1</td><td>230351</td><td>230373</td><td>-</td><td>0</td><td>GCGCGCACTCAGACGGCCGG</td><td>GGG</td><td>||||||||[||||||||||||]PAM</td><td>NA</td><td>Intergenic</td><td>NA</td><td>NA</td></tr>
<tr><td>GU305914.1</td><td>36</td><td>58</td><td>-</td><td>0</td><td>GCGCGCACTCAGACGGCCGG</td><td>GGG</td><td>||||||||[||||||||||||]PAM</td><td>NA</td><td>Intergenic</td><td>NA</td><td>NA</td></tr>
<tr><td>GU305914.1</td><td>92153</td><td>92175</td><td>+</td><td>4</td><td>ACGCGCGCTCGGAGGGCCGG</td><td>CGG</td><td>-|||||-|[||-||-||||||]PAM</td><td>NA</td><td>Intergenic</td><td>NA</td><td>NA</td></tr>
</table>

<table>
<tr><td>T17239</td><td>CGCGCACTCAGACGGCCGGGGGG</td><td>1000</td><td></td><td></td><td></td><td></td><td></td><td></td><td></td><td></td><td></td></tr>
<tr><td>Oligo fwd</td><td>CACCCGCGCACTCAGACGGCCGGG</td><td></td><td></td><td></td><td></td><td></td><td></td><td></td><td></td><td></td><td></td></tr>
<tr><td>Oligo rev</td><td>AAACCCCGGCCGTCTGAGTGCGCG</td><td></td><td></td><td></td><td></td><td></td><td></td><td></td><td></td><td></td><td></td></tr>
<tr><td>Chromosome</td><td>start</td><td>end</td><td>strand</td><td>MM</td><td>target_seq</td><td>PAM</td><td>alignment</td><td>distance</td><td>position</td><td>gene name</td><td>gene id</td></tr>
<tr><td>GU305914.1</td><td>35</td><td>57</td><td>-</td><td>0</td><td>CGCGCACTCAGACGGCCGGG</td><td>GGG</td><td>||||||||[||||||||||||]PAM</td><td>NA</td><td>Intergenic</td><td>NA</td><td>NA</td></tr>
<tr><td>GU305914.1</td><td>230350</td><td>230372</td><td>-</td><td>0</td><td>CGCGCACTCAGACGGCCGGG</td><td>GGG</td><td>||||||||[||||||||||||]PAM</td><td>NA</td><td>Intergenic</td><td>NA</td><td>NA</td></tr>
</table>

<table>
<tr><td>T17240</td><td>GCGCACTCAGACGGCCGGGGGGG</td><td>1000</td><td></td><td></td><td></td><td></td><td></td><td></td><td></td><td></td><td></td></tr>
<tr><td>Oligo fwd</td><td>CACCGCGCACTCAGACGGCCGGGG</td><td></td><td></td><td></td><td></td><td></td><td></td><td></td><td></td><td></td><td></td></tr>
<tr><td>Oligo rev</td><td>AAACCCCCGGCCGTCTGAGTGCGC</td><td></td><td></td><td></td><td></td><td></td><td></td><td></td><td></td><td></td><td></td></tr>
<tr><td>Chromosome</td><td>start</td><td>end</td><td>strand</td><td>MM</td><td>target_seq</td><td>PAM</td><td>alignment</td><td>distance</td><td>position</td><td>gene name</td><td>gene id</td></tr>
<tr><td>GU305914.1</td><td>34</td><td>56</td><td>-</td><td>0</td><td>GCGCACTCAGACGGCCGGGG</td><td>GGG</td><td>||||||||[||||||||||||]PAM</td><td>NA</td><td>Intergenic</td><td>NA</td><td>NA</td></tr>
<tr><td>GU305914.1</td><td>230349</td><td>230371</td><td>-</td><td>0</td><td>GCGCACTCAGACGGCCGGGG</td><td>GGG</td><td>||||||||[||||||||||||]PAM</td><td>NA</td><td>Intergenic</td><td>NA</td><td>NA</td></tr>
</table>

<table>
<tr><td>T17241</td><td>CGGCCGGGGGGGATAAAAAGTGG</td><td>1000</td><td></td><td></td><td></td><td></td><td></td><td></td><td></td><td></td><td></td></tr>
<tr><td>Oligo fwd</td><td>CACCCGGCCGGGGGGGATAAAAAG</td><td></td><td></td><td></td><td></td><td></td><td></td><td></td><td></td><td></td><td></td></tr>
<tr><td>Oligo rev</td><td>AAACCTTTTTATCCCCCCCGGCCG</td><td></td><td></td><td></td><td></td><td></td><td></td><td></td><td></td><td></td><td></td></tr>
<tr><td>Chromosome</td><td>start</td><td>end</td><td>strand</td><td>MM</td><td>target_seq</td><td>PAM</td><td>alignment</td><td>distance</td><td>position</td><td>gene name</td><td>gene id</td></tr>
<tr><td>GU305914.1</td><td>230338</td><td>230360</td><td>-</td><td>0</td><td>CGGCCGGGGGGGATAAAAAG</td><td>TGG</td><td>||||||||[||||||||||||]PAM</td><td>NA</td><td>Intergenic</td><td>NA</td><td>NA</td></tr>
</table>

<table>
<tr><td>T17242</td><td>ATAAAAAGTGGTACTGAGCTAGG</td><td>1000</td><td></td><td></td><td></td><td></td><td></td><td></td><td></td><td></td><td></td></tr>
<tr><td>Oligo fwd</td><td>CACCATAAAAAGTGGTACTGAGCT</td><td></td><td></td><td></td><td></td><td></td><td></td><td></td><td></td><td></td><td></td></tr>
<tr><td>Oligo rev</td><td>AAACAGCTCAGTACCACTTTTTAT</td><td></td><td></td><td></td><td></td><td></td><td></td><td></td><td></td><td></td><td></td></tr>
<tr><td>Chromosome</td><td>start</td><td>end</td><td>strand</td><td>MM</td><td>target_seq</td><td>PAM</td><td>alignment</td><td>distance</td><td>position</td><td>gene name</td><td>gene id</td></tr>
<tr><td>GU305914.1</td><td>230326</td><td>230348</td><td>-</td><td>0</td><td>ATAAAAAGTGGTACTGAGCT</td><td>AGG</td><td>||||||||[||||||||||||]PAM</td><td>NA</td><td>Intergenic</td><td>NA</td><td>NA</td></tr>
</table>

<table>
<tr><td>T17243</td><td>AAAAAAAGAAGTATGTGCCGCGG</td><td>1000</td><td></td><td></td><td></td><td></td><td></td><td></td><td></td><td></td><td></td></tr>
<tr><td>Oligo fwd</td><td>CACCAAAAAAAGAAGTATGTGCCG</td><td></td><td></td><td></td><td></td><td></td><td></td><td></td><td></td><td></td><td></td></tr>
<tr><td>Oligo rev</td><td>AAACCGGCACATACTTCTTTTTTT</td><td></td><td></td><td></td><td></td><td></td><td></td><td></td><td></td><td></td><td></td></tr>
<tr><td>Chromosome</td><td>start</td><td>end</td><td>strand</td><td>MM</td><td>target_seq</td><td>PAM</td><td>alignment</td><td>distance</td><td>position</td><td>gene name</td><td>gene id</td></tr>
<tr><td>GU305914.1</td><td>230295</td><td>230317</td><td>-</td><td>0</td><td>AAAAAAAGAAGTATGTGCCG</td><td>CGG</td><td>||||||||[||||||||||||]PAM</td><td>NA</td><td>Intergenic</td><td>NA</td><td>NA</td></tr>
<tr><td>T17244</td><td>AGTATGTGCCGCGGCGCCCTCGG</td><td>1000</td><td></td><td></td><td></td><td></td><td></td><td></td><td></td><td></td><td></td></tr>
<tr><td>Oligo fwd</td><td>CACCAGTATGTGCCGCGGCGCCCT</td><td></td><td></td><td></td><td></td><td></td><td></td><td></td><td></td><td></td><td></td></tr>
<tr><td>Oligo rev</td><td>AAACAGGGCGCCGCGGCACATACT</td><td></td><td></td><td></td><td></td><td></td><td></td><td></td><td></td><td></td><td></td></tr>
<tr><td>Chromosome</td><td>start</td><td>end</td><td>strand</td><td>MM</td><td>target_seq</td><td>PAM</td><td>alignment</td><td>distance</td><td>position</td><td>gene name</td><td>gene id</td></tr>
<tr><td>GU305914.1</td><td>230286</td><td>230308</td><td>-</td><td>0</td><td>AGTATGTGCCGCGGCGCCCT</td><td>CGG</td><td>||||||||[||||||||||||]PAM</td><td>NA</td><td>Intergenic</td><td>NA</td><td>NA</td></tr>
<tr><td>T17245</td><td>ATGTGCCGCGGCGCCCTCGGCGG</td><td>1000</td><td></td><td></td><td></td><td></td><td></td><td></td><td></td><td></td><td></td></tr>
<tr><td>Oligo fwd</td><td>CACCATGTGCCGCGGCGCCCTCGG</td><td></td><td></td><td></td><td></td><td></td><td></td><td></td><td></td><td></td><td></td></tr>
<tr><td>Oligo rev</td><td>AAACCCGAGGGCGCCGCGGCACAT</td><td></td><td></td><td></td><td></td><td></td><td></td><td></td><td></td><td></td><td></td></tr>
<tr><td>Chromosome</td><td>start</td><td>end</td><td>strand</td><td>MM</td><td>target_seq</td><td>PAM</td><td>alignment</td><td>distance</td><td>position</td><td>gene name</td><td>gene id</td></tr>
<tr><td>GU305914.1</td><td>230283</td><td>230305</td><td>-</td><td>0</td><td>ATGTGCCGCGGCGCCCTCGG</td><td>CGG</td><td>||||||||[||||||||||||]PAM</td><td>NA</td><td>Intergenic</td><td>NA</td><td>NA</td></tr>
<tr><td>T17246</td><td>AGCCCGCCCATGCAAGTGAATGG</td><td>1000</td><td></td><td></td><td></td><td></td><td></td><td></td><td></td><td></td><td></td></tr>
<tr><td>Oligo fwd</td><td>CACCAGCCCGCCCATGCAAGTGAA</td><td></td><td></td><td></td><td></td><td></td><td></td><td></td><td></td><td></td><td></td></tr>
<tr><td>Oligo rev</td><td>AAACTTCACTTGCATGGGCGGGCT</td><td></td><td></td><td></td><td></td><td></td><td></td><td></td><td></td><td></td><td></td></tr>
<tr><td>Chromosome</td><td>start</td><td>end</td><td>strand</td><td>MM</td><td>target_seq</td><td>PAM</td><td>alignment</td><td>distance</td><td>position</td><td>gene name</td><td>gene id</td></tr>
<tr><td>GU305914.1</td><td>230249</td><td>230271</td><td>-</td><td>0</td><td>AGCCCGCCCATGCAAGTGAA</td><td>TGG</td><td>||||||||[||||||||||||]PAM</td><td>NA</td><td>Intergenic</td><td>NA</td><td>NA</td></tr>
<tr><td>T17247</td><td>GCCCATGCAAGTGAATGGCGTGG</td><td>1000</td><td></td><td></td><td></td><td></td><td></td><td></td><td></td><td></td><td></td></tr>
<tr><td>Oligo fwd</td><td>CACCGCCCATGCAAGTGAATGGCG</td><td></td><td></td><td></td><td></td><td></td><td></td><td></td><td></td><td></td><td></td></tr>
<tr><td>Oligo rev</td><td>AAACCGCCATTCACTTGCATGGGC</td><td></td><td></td><td></td><td></td><td></td><td></td><td></td><td></td><td></td><td></td></tr>
<tr><td>Chromosome</td><td>start</td><td>end</td><td>strand</td><td>MM</td><td>target_seq</td><td>PAM</td><td>alignment</td><td>distance</td><td>position</td><td>gene name</td><td>gene id</td></tr>
<tr><td>GU305914.1</td><td>230244</td><td>230266</td><td>-</td><td>0</td><td>GCCCATGCAAGTGAATGGCG</td><td>TGG</td><td>||||||||[||||||||||||]PAM</td><td>NA</td><td>Intergenic</td><td>NA</td><td>NA</td></tr>
<tr><td>T17248</td><td>TGCAAGTGAATGGCGTGGAGTGG</td><td>1000</td><td></td><td></td><td></td><td></td><td></td><td></td><td></td><td></td><td></td></tr>
<tr><td>Oligo fwd</td><td>CACCTGCAAGTGAATGGCGTGGAG</td><td></td><td></td><td></td><td></td><td></td><td></td><td></td><td></td><td></td><td></td></tr>
<tr><td>Oligo rev</td><td>AAACCTCCACGCCATTCACTTGCA</td><td></td><td></td><td></td><td></td><td></td><td></td><td></td><td></td><td></td><td></td></tr>
<tr><td>Chromosome</td><td>start</td><td>end</td><td>strand</td><td>MM</td><td>target_seq</td><td>PAM</td><td>alignment</td><td>distance</td><td>position</td><td>gene name</td><td>gene id</td></tr>
<tr><td>GU305914.1</td><td>230239</td><td>230261</td><td>-</td><td>0</td><td>TGCAAGTGAATGGCGTGGAG</td><td>TGG</td><td>||||||||[||||||||||||]PAM</td><td>NA</td><td>Intergenic</td><td>NA</td><td>NA</td></tr>
<tr><td>T17249</td><td>GCAAGTGAATGGCGTGGAGTGGG</td><td>1000</td><td></td><td></td><td></td><td></td><td></td><td></td><td></td><td></td><td></td></tr>
<tr><td>Oligo fwd</td><td>CACCGCAAGTGAATGGCGTGGAGT</td><td></td><td></td><td></td><td></td><td></td><td></td><td></td><td></td><td></td><td></td></tr>
<tr><td>Oligo rev</td><td>AAACACTCCACGCCATTCACTTGC</td><td></td><td></td><td></td><td></td><td></td><td></td><td></td><td></td><td></td><td></td></tr>
<tr><td>Chromosome</td><td>start</td><td>end</td><td>strand</td><td>MM</td><td>target_seq</td><td>PAM</td><td>alignment</td><td>distance</td><td>position</td><td>gene name</td><td>gene id</td></tr>
<tr><td>GU305914.1</td><td>230238</td><td>230260</td><td>-</td><td>0</td><td>GCAAGTGAATGGCGTGGAGT</td><td>GGG</td><td>||||||||[||||||||||||]PAM</td><td>NA</td><td>Intergenic</td><td>NA</td><td>NA</td></tr>
<tr><td>T17250</td><td>GAATGGCGTGGAGTGGGTGTTGG</td><td>1000</td><td></td><td></td><td></td><td></td><td></td><td></td><td></td><td></td><td></td></tr>
<tr><td>Oligo fwd</td><td>CACCGAATGGCGTGGAGTGGGTGT</td><td></td><td></td><td></td><td></td><td></td><td></td><td></td><td></td><td></td><td></td></tr>
<tr><td>Oligo rev</td><td>AAACACACCCACTCCACGCCATTC</td><td></td><td></td><td></td><td></td><td></td><td></td><td></td><td></td><td></td><td></td></tr>
<tr><td>Chromosome</td><td>start</td><td>end</td><td>strand</td><td>MM</td><td>target_seq</td><td>PAM</td><td>alignment</td><td>distance</td><td>position</td><td>gene name</td><td>gene id</td></tr>
<tr><td>GU305914.1</td><td>230232</td><td>230254</td><td>-</td><td>0</td><td>GAATGGCGTGGAGTGGGTGT</td><td>TGG</td><td>||||||||[||||||||||||]PAM</td><td>NA</td><td>Intergenic</td><td>NA</td><td>NA</td></tr>
<tr><td>T17251</td><td>AATGGCGTGGAGTGGGTGTTGGG</td><td>1000</td><td></td><td></td><td></td><td></td><td></td><td></td><td></td><td></td><td></td></tr>
<tr><td>Oligo fwd</td><td>CACCAATGGCGTGGAGTGGGTGTT</td><td></td><td></td><td></td><td></td><td></td><td></td><td></td><td></td><td></td><td></td></tr>
<tr><td>Oligo rev</td><td>AAACAACACCCACTCCACGCCATT</td><td></td><td></td><td></td><td></td><td></td><td></td><td></td><td></td><td></td><td></td></tr>
<tr><td>Chromosome</td><td>start</td><td>end</td><td>strand</td><td>MM</td><td>target_seq</td><td>PAM</td><td>alignment</td><td>distance</td><td>position</td><td>gene name</td><td>gene id</td></tr>
<tr><td>GU305914.1</td><td>230231</td><td>230253</td><td>-</td><td>0</td><td>AATGGCGTGGAGTGGGTGTT</td><td>GGG</td><td>||||||||[||||||||||||]PAM</td><td>NA</td><td>Intergenic</td><td>NA</td><td>NA</td></tr>
<tr><td>T17252</td><td>TGGGCTGCTAGTTGTTGCTGAGG</td><td>1000</td><td></td><td></td><td></td><td></td><td></td><td></td><td></td><td></td><td></td></tr>
<tr><td>Oligo fwd</td><td>CACCTGGGCTGCTAGTTGTTGCTG</td><td></td><td></td><td></td><td></td><td></td><td></td><td></td><td></td><td></td><td></td></tr>
<tr><td>Oligo rev</td><td>AAACCAGCAACAACTAGCAGCCCA</td><td></td><td></td><td></td><td></td><td></td><td></td><td></td><td></td><td></td><td></td></tr>
<tr><td>Chromosome</td><td>start</td><td>end</td><td>strand</td><td>MM</td><td>target_seq</td><td>PAM</td><td>alignment</td><td>distance</td><td>position</td><td>gene name</td><td>gene id</td></tr>
<tr><td>GU305914.1</td><td>230212</td><td>230234</td><td>-</td><td>0</td><td>TGGGCTGCTAGTTGTTGCTG</td><td>AGG</td><td>||||||||[||||||||||||]PAM</td><td>NA</td><td>Intergenic</td><td>NA</td><td>NA</td></tr>
<tr><td>T17253</td><td>GCTAGTTGTTGCTGAGGATGAGG</td><td>1000</td><td></td><td></td><td></td><td></td><td></td><td></td><td></td><td></td><td></td></tr>
<tr><td>Oligo fwd</td><td>CACCGCTAGTTGTTGCTGAGGATG</td><td></td><td></td><td></td><td></td><td></td><td></td><td></td><td></td><td></td><td></td></tr>
<tr><td>Oligo rev</td><td>AAACCATCCTCAGCAACAACTAGC</td><td></td><td></td><td></td><td></td><td></td><td></td><td></td><td></td><td></td><td></td></tr>
<tr><td>Chromosome</td><td>start</td><td>end</td><td>strand</td><td>MM</td><td>target_seq</td><td>PAM</td><td>alignment</td><td>distance</td><td>position</td><td>gene name</td><td>gene id</td></tr>
<tr><td>GU305914.1</td><td>230206</td><td>230228</td><td>-</td><td>0</td><td>GCTAGTTGTTGCTGAGGATG</td><td>AGG</td><td>||||||||[||||||||||||]PAM</td><td>NA</td><td>Intergenic</td><td>NA</td><td>NA</td></tr>
<tr><td>T17254</td><td>GTTGCTGAGGATGAGGTGTGCGG</td><td>1000</td><td></td><td></td><td></td><td></td><td></td><td></td><td></td><td></td><td></td></tr>
<tr><td>Oligo fwd</td><td>CACCGTTGCTGAGGATGAGGTGTG</td><td></td><td></td><td></td><td></td><td></td><td></td><td></td><td></td><td></td><td></td></tr>
<tr><td>Oligo rev</td><td>AAACCACACCTCATCCTCAGCAAC</td><td></td><td></td><td></td><td></td><td></td><td></td><td></td><td></td><td></td><td></td></tr>
<tr><td>Chromosome</td><td>start</td><td>end</td><td>strand</td><td>MM</td><td>target_seq</td><td>PAM</td><td>alignment</td><td>distance</td><td>position</td><td>gene name</td><td>gene id</td></tr>
<tr><td>GU305914.1</td><td>230199</td><td>230221</td><td>-</td><td>0</td><td>GTTGCTGAGGATGAGGTGTG</td><td>CGG</td><td>||||||||[||||||||||||]PAM</td><td>NA</td><td>Intergenic</td><td>NA</td><td>NA</td></tr>
<tr><td>T17255</td><td>ATCTAAACCTCATATGTGTGTGG</td><td>1000</td><td></td><td></td><td></td><td></td><td></td><td></td><td></td><td></td><td></td></tr>
<tr><td>Oligo fwd</td><td>CACCATCTAAACCTCATATGTGTG</td><td></td><td></td><td></td><td></td><td></td><td></td><td></td><td></td><td></td><td></td></tr>
<tr><td>Oligo rev</td><td>AAACCACACATATGAGGTTTAGAT</td><td></td><td></td><td></td><td></td><td></td><td></td><td></td><td></td><td></td><td></td></tr>
<tr><td>Chromosome</td><td>start</td><td>end</td><td>strand</td><td>MM</td><td>target_seq</td><td>PAM</td><td>alignment</td><td>distance</td><td>position</td><td>gene name</td><td>gene id</td></tr>
<tr><td>GU305914.1</td><td>230166</td><td>230188</td><td>-</td><td>0</td><td>ATCTAAACCTCATATGTGTG</td><td>TGG</td><td>||||||||[||||||||||||]PAM</td><td>NA</td><td>Intergenic</td><td>NA</td><td>NA</td></tr>
<tr><td>T17256</td><td>TAAACCTCATATGTGTGTGGAGG</td><td>1000</td><td></td><td></td><td></td><td></td><td></td><td></td><td></td><td></td><td></td></tr>
<tr><td>Oligo fwd</td><td>CACCTAAACCTCATATGTGTGTGG</td><td></td><td></td><td></td><td></td><td></td><td></td><td></td><td></td><td></td><td></td></tr>
<tr><td>Oligo rev</td><td>AAACCCACACACATATGAGGTTTA</td><td></td><td></td><td></td><td></td><td></td><td></td><td></td><td></td><td></td><td></td></tr>
<tr><td>Chromosome</td><td>start</td><td>end</td><td>strand</td><td>MM</td><td>target_seq</td><td>PAM</td><td>alignment</td><td>distance</td><td>position</td><td>gene name</td><td>gene id</td></tr>
<tr><td>GU305914.1</td><td>230163</td><td>230185</td><td>-</td><td>0</td><td>TAAACCTCATATGTGTGTGG</td><td>AGG</td><td>||||||||[||||||||||||]PAM</td><td>NA</td><td>Intergenic</td><td>NA</td><td>NA</td></tr>
<tr><td>T17257</td><td>TCGTTAAAAAAACATGTGCTCGG</td><td>1000</td><td></td><td></td><td></td><td></td><td></td><td></td><td></td><td></td><td></td></tr>
<tr><td>Oligo fwd</td><td>CACCTCGTTAAAAAAACATGTGCT</td><td></td><td></td><td></td><td></td><td></td><td></td><td></td><td></td><td></td><td></td></tr>
<tr><td>Oligo rev</td><td>AAACAGCACATGTTTTTTTAACGA</td><td></td><td></td><td></td><td></td><td></td><td></td><td></td><td></td><td></td><td></td></tr>
<tr><td>Chromosome</td><td>start</td><td>end</td><td>strand</td><td>MM</td><td>target_seq</td><td>PAM</td><td>alignment</td><td>distance</td><td>position</td><td>gene name</td><td>gene id</td></tr>
<tr><td>GU305914.1</td><td>230138</td><td>230160</td><td>-</td><td>0</td><td>TCGTTAAAAAAACATGTGCT</td><td>CGG</td><td>||||||||[||||||||||||]PAM</td><td>NA</td><td>Intergenic</td><td>NA</td><td>NA</td></tr>
</table>

<table>
<tr><td>T17258</td><td>CATGTGCTCGGTTAACGAGTTGG</td><td>1000</td><td></td><td></td><td></td><td></td><td></td><td></td><td></td><td></td><td></td></tr>
<tr><td>Oligo fwd</td><td>CACCCATGTGCTCGGTTAACGAGT</td><td></td><td></td><td></td><td></td><td></td><td></td><td></td><td></td><td></td><td></td></tr>
<tr><td>Oligo rev</td><td>AAACACTCGTTAACCGAGCACATG</td><td></td><td></td><td></td><td></td><td></td><td></td><td></td><td></td><td></td><td></td></tr>
<tr><td>Chromosome</td><td>start</td><td>end</td><td>strand</td><td>MM</td><td>target_seq</td><td>PAM</td><td>alignment</td><td>distance</td><td>position</td><td>gene name</td><td>gene id</td></tr>
<tr><td>GU305914.1</td><td>230126</td><td>230148</td><td>-</td><td>0</td><td>CATGTGCTCGGTTAACGAGT</td><td>TGG</td><td>||||||||[||||||||||||]PAM</td><td>NA</td><td>Intergenic</td><td>NA</td><td>NA</td></tr>
<tr><td>T17259</td><td>TTAACGAGTTGGCCTTCGTCCGG</td><td>1000</td><td></td><td></td><td></td><td></td><td></td><td></td><td></td><td></td><td></td></tr>
<tr><td>Oligo fwd</td><td>CACCTTAACGAGTTGGCCTTCGTC</td><td></td><td></td><td></td><td></td><td></td><td></td><td></td><td></td><td></td><td></td></tr>
<tr><td>Oligo rev</td><td>AAACGACGAAGGCCAACTCGTTAA</td><td></td><td></td><td></td><td></td><td></td><td></td><td></td><td></td><td></td><td></td></tr>
<tr><td>Chromosome</td><td>start</td><td>end</td><td>strand</td><td>MM</td><td>target_seq</td><td>PAM</td><td>alignment</td><td>distance</td><td>position</td><td>gene name</td><td>gene id</td></tr>
<tr><td>GU305914.1</td><td>230115</td><td>230137</td><td>-</td><td>0</td><td>TTAACGAGTTGGCCTTCGTC</td><td>CGG</td><td>||||||||[||||||||||||]PAM</td><td>NA</td><td>Intergenic</td><td>NA</td><td>NA</td></tr>
<tr><td>T17260</td><td>TTGGCCTTCGTCCGGCTGTCCGG</td><td>1000</td><td></td><td></td><td></td><td></td><td></td><td></td><td></td><td></td><td></td></tr>
<tr><td>Oligo fwd</td><td>CACCTTGGCCTTCGTCCGGCTGTC</td><td></td><td></td><td></td><td></td><td></td><td></td><td></td><td></td><td></td><td></td></tr>
<tr><td>Oligo rev</td><td>AAACGACAGCCGGACGAAGGCCAA</td><td></td><td></td><td></td><td></td><td></td><td></td><td></td><td></td><td></td><td></td></tr>
<tr><td>Chromosome</td><td>start</td><td>end</td><td>strand</td><td>MM</td><td>target_seq</td><td>PAM</td><td>alignment</td><td>distance</td><td>position</td><td>gene name</td><td>gene id</td></tr>
<tr><td>GU305914.1</td><td>230107</td><td>230129</td><td>-</td><td>0</td><td>TTGGCCTTCGTCCGGCTGTC</td><td>CGG</td><td>||||||||[||||||||||||]PAM</td><td>NA</td><td>Intergenic</td><td>NA</td><td>NA</td></tr>
<tr><td>T17261</td><td>GCTGTCCGGCACGACTACCGCGG</td><td>1000</td><td></td><td></td><td></td><td></td><td></td><td></td><td></td><td></td><td></td></tr>
<tr><td>Oligo fwd</td><td>CACCGCTGTCCGGCACGACTACCG</td><td></td><td></td><td></td><td></td><td></td><td></td><td></td><td></td><td></td><td></td></tr>
<tr><td>Oligo rev</td><td>AAACCGGTAGTCGTGCCGGACAGC</td><td></td><td></td><td></td><td></td><td></td><td></td><td></td><td></td><td></td><td></td></tr>
<tr><td>Chromosome</td><td>start</td><td>end</td><td>strand</td><td>MM</td><td>target_seq</td><td>PAM</td><td>alignment</td><td>distance</td><td>position</td><td>gene name</td><td>gene id</td></tr>
<tr><td>GU305914.1</td><td>230093</td><td>230115</td><td>-</td><td>0</td><td>GCTGTCCGGCACGACTACCG</td><td>CGG</td><td>||||||||[||||||||||||]PAM</td><td>NA</td><td>Intergenic</td><td>NA</td><td>NA</td></tr>
<tr><td>T17262</td><td>CTGTCCGGCACGACTACCGCGGG</td><td>1000</td><td></td><td></td><td></td><td></td><td></td><td></td><td></td><td></td><td></td></tr>
<tr><td>Oligo fwd</td><td>CACCCTGTCCGGCACGACTACCGC</td><td></td><td></td><td></td><td></td><td></td><td></td><td></td><td></td><td></td><td></td></tr>
<tr><td>Oligo rev</td><td>AAACGCGGTAGTCGTGCCGGACAG</td><td></td><td></td><td></td><td></td><td></td><td></td><td></td><td></td><td></td><td></td></tr>
<tr><td>Chromosome</td><td>start</td><td>end</td><td>strand</td><td>MM</td><td>target_seq</td><td>PAM</td><td>alignment</td><td>distance</td><td>position</td><td>gene name</td><td>gene id</td></tr>
<tr><td>GU305914.1</td><td>230092</td><td>230114</td><td>-</td><td>0</td><td>CTGTCCGGCACGACTACCGC</td><td>GGG</td><td>||||||||[||||||||||||]PAM</td><td>NA</td><td>Intergenic</td><td>NA</td><td>NA</td></tr>
<tr><td>GU305914.1</td><td>53506</td><td>53528</td><td>-</td><td>4</td><td>CTCTCGGGCACGACCACCGT</td><td>CGG</td><td>||-||-||[||||||-||||-]PAM</td><td>NA</td><td>Intergenic</td><td>NA</td><td>NA</td></tr>
<tr><td>T17263</td><td>CGACTACCGCGGGAGTGTACTGG</td><td>1000</td><td></td><td></td><td></td><td></td><td></td><td></td><td></td><td></td><td></td></tr>
<tr><td>Oligo fwd</td><td>CACCCGACTACCGCGGGAGTGTAC</td><td></td><td></td><td></td><td></td><td></td><td></td><td></td><td></td><td></td><td></td></tr>
<tr><td>Oligo rev</td><td>AAACGTACACTCCCGCGGTAGTCG</td><td></td><td></td><td></td><td></td><td></td><td></td><td></td><td></td><td></td><td></td></tr>
<tr><td>Chromosome</td><td>start</td><td>end</td><td>strand</td><td>MM</td><td>target_seq</td><td>PAM</td><td>alignment</td><td>distance</td><td>position</td><td>gene name</td><td>gene id</td></tr>
<tr><td>GU305914.1</td><td>230082</td><td>230104</td><td>-</td><td>0</td><td>CGACTACCGCGGGAGTGTAC</td><td>TGG</td><td>||||||||[||||||||||||]PAM</td><td>NA</td><td>Intergenic</td><td>NA</td><td>NA</td></tr>
<tr><td>T17264</td><td>GACTACCGCGGGAGTGTACTGGG</td><td>1000</td><td></td><td></td><td></td><td></td><td></td><td></td><td></td><td></td><td></td></tr>
<tr><td>Oligo fwd</td><td>CACCGACTACCGCGGGAGTGTACT</td><td></td><td></td><td></td><td></td><td></td><td></td><td></td><td></td><td></td><td></td></tr>
<tr><td>Oligo rev</td><td>AAACAGTACACTCCCGCGGTAGTC</td><td></td><td></td><td></td><td></td><td></td><td></td><td></td><td></td><td></td><td></td></tr>
<tr><td>Chromosome</td><td>start</td><td>end</td><td>strand</td><td>MM</td><td>target_seq</td><td>PAM</td><td>alignment</td><td>distance</td><td>position</td><td>gene name</td><td>gene id</td></tr>
<tr><td>GU305914.1</td><td>230081</td><td>230103</td><td>-</td><td>0</td><td>GACTACCGCGGGAGTGTACT</td><td>GGG</td><td>||||||||[||||||||||||]PAM</td><td>NA</td><td>Intergenic</td><td>NA</td><td>NA</td></tr>
<tr><td>T17265</td><td>GCGCGCCCGCGAGACGTGTACGG</td><td>1000</td><td></td><td></td><td></td><td></td><td></td><td></td><td></td><td></td><td></td></tr>
<tr><td>Oligo fwd</td><td>CACCGCGCGCCCGCGAGACGTGTA</td><td></td><td></td><td></td><td></td><td></td><td></td><td></td><td></td><td></td><td></td></tr>
<tr><td>Oligo rev</td><td>AAACTACACGTCTCGCGGGCGCGC</td><td></td><td></td><td></td><td></td><td></td><td></td><td></td><td></td><td></td><td></td></tr>
<tr><td>Chromosome</td><td>start</td><td>end</td><td>strand</td><td>MM</td><td>target_seq</td><td>PAM</td><td>alignment</td><td>distance</td><td>position</td><td>gene name</td><td>gene id</td></tr>
<tr><td>GU305914.1</td><td>230050</td><td>230072</td><td>-</td><td>0</td><td>GCGCGCCCGCGAGACGTGTA</td><td>CGG</td><td>||||||||[||||||||||||]PAM</td><td>NA</td><td>Intergenic</td><td>NA</td><td>NA</td></tr>
<tr><td>T17266</td><td>CCCGCGAGACGTGTACGGCGTGG</td><td>1000</td><td></td><td></td><td></td><td></td><td></td><td></td><td></td><td></td><td></td></tr>
<tr><td>Oligo fwd</td><td>CACCCCCGCGAGACGTGTACGGCG</td><td></td><td></td><td></td><td></td><td></td><td></td><td></td><td></td><td></td><td></td></tr>
<tr><td>Oligo rev</td><td>AAACCGCCGTACACGTCTCGCGGG</td><td></td><td></td><td></td><td></td><td></td><td></td><td></td><td></td><td></td><td></td></tr>
<tr><td>Chromosome</td><td>start</td><td>end</td><td>strand</td><td>MM</td><td>target_seq</td><td>PAM</td><td>alignment</td><td>distance</td><td>position</td><td>gene name</td><td>gene id</td></tr>
<tr><td>GU305914.1</td><td>230045</td><td>230067</td><td>-</td><td>0</td><td>CCCGCGAGACGTGTACGGCG</td><td>TGG</td><td>||||||||[||||||||||||]PAM</td><td>NA</td><td>Intergenic</td><td>NA</td><td>NA</td></tr>
<tr><td>T17267</td><td>CCGCGAGACGTGTACGGCGTGGG</td><td>1000</td><td></td><td></td><td></td><td></td><td></td><td></td><td></td><td></td><td></td></tr>
<tr><td>Oligo fwd</td><td>CACCCCGCGAGACGTGTACGGCGT</td><td></td><td></td><td></td><td></td><td></td><td></td><td></td><td></td><td></td><td></td></tr>
<tr><td>Oligo rev</td><td>AAACACGCCGTACACGTCTCGCGG</td><td></td><td></td><td></td><td></td><td></td><td></td><td></td><td></td><td></td><td></td></tr>
<tr><td>Chromosome</td><td>start</td><td>end</td><td>strand</td><td>MM</td><td>target_seq</td><td>PAM</td><td>alignment</td><td>distance</td><td>position</td><td>gene name</td><td>gene id</td></tr>
<tr><td>GU305914.1</td><td>230044</td><td>230066</td><td>-</td><td>0</td><td>CCGCGAGACGTGTACGGCGT</td><td>GGG</td><td>||||||||[||||||||||||]PAM</td><td>NA</td><td>Intergenic</td><td>NA</td><td>NA</td></tr>
<tr><td>T17268</td><td>GGGTGACTTAGATCGCCGAGCGG</td><td>1000</td><td></td><td></td><td></td><td></td><td></td><td></td><td></td><td></td><td></td></tr>
<tr><td>Oligo fwd</td><td>CACCGGGTGACTTAGATCGCCGAG</td><td></td><td></td><td></td><td></td><td></td><td></td><td></td><td></td><td></td><td></td></tr>
<tr><td>Oligo rev</td><td>AAACCTCGGCGATCTAAGTCACCC</td><td></td><td></td><td></td><td></td><td></td><td></td><td></td><td></td><td></td><td></td></tr>
<tr><td>Chromosome</td><td>start</td><td>end</td><td>strand</td><td>MM</td><td>target_seq</td><td>PAM</td><td>alignment</td><td>distance</td><td>position</td><td>gene name</td><td>gene id</td></tr>
<tr><td>GU305914.1</td><td>230024</td><td>230046</td><td>-</td><td>0</td><td>GGGTGACTTAGATCGCCGAG</td><td>CGG</td><td>||||||||[||||||||||||]PAM</td><td>NA</td><td>Intergenic</td><td>NA</td><td>NA</td></tr>
<tr><td>T17269</td><td>CGCCCACCTGCCACTGCCGCCGG</td><td>1000</td><td></td><td></td><td></td><td></td><td></td><td></td><td></td><td></td><td></td></tr>
<tr><td>Oligo fwd</td><td>CACCCGCCCACCTGCCACTGCCGC</td><td></td><td></td><td></td><td></td><td></td><td></td><td></td><td></td><td></td><td></td></tr>
<tr><td>Oligo rev</td><td>AAACGCGGCAGTGGCAGGTGGGCG</td><td></td><td></td><td></td><td></td><td></td><td></td><td></td><td></td><td></td><td></td></tr>
<tr><td>Chromosome</td><td>start</td><td>end</td><td>strand</td><td>MM</td><td>target_seq</td><td>PAM</td><td>alignment</td><td>distance</td><td>position</td><td>gene name</td><td>gene id</td></tr>
<tr><td>GU305914.1</td><td>229988</td><td>230010</td><td>-</td><td>0</td><td>CGCCCACCTGCCACTGCCGC</td><td>CGG</td><td>||||||||[||||||||||||]PAM</td><td>NA</td><td>Intergenic</td><td>NA</td><td>NA</td></tr>
<tr><td>T17270</td><td>CACCTGCCACTGCCGCCGGCCGG</td><td>1000</td><td></td><td></td><td></td><td></td><td></td><td></td><td></td><td></td><td></td></tr>
<tr><td>Oligo fwd</td><td>CACCCACCTGCCACTGCCGCCGGC</td><td></td><td></td><td></td><td></td><td></td><td></td><td></td><td></td><td></td><td></td></tr>
<tr><td>Oligo rev</td><td>AAACGCCGGCGGCAGTGGCAGGTG</td><td></td><td></td><td></td><td></td><td></td><td></td><td></td><td></td><td></td><td></td></tr>
<tr><td>Chromosome</td><td>start</td><td>end</td><td>strand</td><td>MM</td><td>target_seq</td><td>PAM</td><td>alignment</td><td>distance</td><td>position</td><td>gene name</td><td>gene id</td></tr>
<tr><td>GU305914.1</td><td>229984</td><td>230006</td><td>-</td><td>0</td><td>CACCTGCCACTGCCGCCGGC</td><td>CGG</td><td>||||||||[||||||||||||]PAM</td><td>NA</td><td>Intergenic</td><td>NA</td><td>NA</td></tr>
<tr><td>T17271</td><td>ACCTGCCACTGCCGCCGGCCGGG</td><td>1000</td><td></td><td></td><td></td><td></td><td></td><td></td><td></td><td></td><td></td></tr>
<tr><td>Oligo fwd</td><td>CACCACCTGCCACTGCCGCCGGCC</td><td></td><td></td><td></td><td></td><td></td><td></td><td></td><td></td><td></td><td></td></tr>
<tr><td>Oligo rev</td><td>AAACGGCCGGCGGCAGTGGCAGGT</td><td></td><td></td><td></td><td></td><td></td><td></td><td></td><td></td><td></td><td></td></tr>
<tr><td>Chromosome</td><td>start</td><td>end</td><td>strand</td><td>MM</td><td>target_seq</td><td>PAM</td><td>alignment</td><td>distance</td><td>position</td><td>gene name</td><td>gene id</td></tr>
<tr><td>GU305914.1</td><td>229983</td><td>230005</td><td>-</td><td>0</td><td>ACCTGCCACTGCCGCCGGCC</td><td>GGG</td><td>||||||||[||||||||||||]PAM</td><td>NA</td><td>Intergenic</td><td>NA</td><td>NA</td></tr>
<tr><td>T17272</td><td>GCCACTGCCGCCGGCCGGGATGG</td><td>1000</td><td></td><td></td><td></td><td></td><td></td><td></td><td></td><td></td><td></td></tr>
<tr><td>Oligo fwd</td><td>CACCGCCACTGCCGCCGGCCGGGA</td><td></td><td></td><td></td><td></td><td></td><td></td><td></td><td></td><td></td><td></td></tr>
<tr><td>Oligo rev</td><td>AAACTCCCGGCCGGCGGCAGTGGC</td><td></td><td></td><td></td><td></td><td></td><td></td><td></td><td></td><td></td><td></td></tr>
</table>

| Chromosome | start | end | strand | MM | target_seq | PAM | alignment | distance | position | gene name | gene id |
|---|---|---|---|---|---|---|---|---|---|---|---|
| GU305914.1 | 229979 | 230001 | - | 0 | GCCACTGCCGCCGGCCGGGA | TGG | \|\|\|\|\|\|\|\|[\|\|\|\|\|\|\|\|\|\|\|\|]PAM | NA | Intergenic | NA | NA |
| GU305914.1 | 62675 | 62697 | - | 4 | GCTGCTGCCGCCGGCGGGGG | CGG | \|\|--\|\|\|\|[\|\|\|\|\|\|\|-\|\|\|-]PAM | NA | Intergenic | NA | NA |
| T17273 | GGCCGGGATGGCGCACGAGACGG | 1000 | | | | | | | | | |
| Oligo fwd | CACCGGCCGGGATGGCGCACGAGA | | | | | | | | | | |
| Oligo rev | AAACTCTCGTGCGCCATCCCGGCC | | | | | | | | | | |
| Chromosome | start | end | strand | MM | target_seq | PAM | alignment | distance | position | gene name | gene id |
| GU305914.1 | 229967 | 229989 | - | 0 | GGCCGGGATGGCGCACGAGA | CGG | \|\|\|\|\|\|\|\|[\|\|\|\|\|\|\|\|\|\|\|\|]PAM | NA | Intergenic | NA | NA |
| GU305914.1 | 67219 | 67241 | + | 4 | GGACGCGATCGCGGACGAGA | AGG | \|\|-\|\|-\|\|[\|-\|\|\|-\|\|\|\|\|\|]PAM | NA | Intergenic | NA | NA |
| T17274 | AGACGGTGTTTTTCCGCGCCCGG | 1000 | | | | | | | | | |
| Oligo fwd | CACCAGACGGTGTTTTTCCGCGCC | | | | | | | | | | |
| Oligo rev | AAACGGCGCGGAAAAACACCGTCT | | | | | | | | | | |
| Chromosome | start | end | strand | MM | target_seq | PAM | alignment | distance | position | gene name | gene id |
| GU305914.1 | 229950 | 229972 | - | 0 | AGACGGTGTTTTTCCGCGCC | CGG | \|\|\|\|\|\|\|\|[\|\|\|\|\|\|\|\|\|\|\|\|]PAM | NA | Intergenic | NA | NA |
| T17275 | GACGGTGTTTTTCCGCGCCCGGG | 1000 | | | | | | | | | |
| Oligo fwd | CACCGACGGTGTTTTTCCGCGCCC | | | | | | | | | | |
| Oligo rev | AAACGGGCGCGGAAAAACACCGTC | | | | | | | | | | |
| Chromosome | start | end | strand | MM | target_seq | PAM | alignment | distance | position | gene name | gene id |
| GU305914.1 | 229949 | 229971 | - | 0 | GACGGTGTTTTTCCGCGCCC | GGG | \|\|\|\|\|\|\|\|[\|\|\|\|\|\|\|\|\|\|\|\|]PAM | NA | Intergenic | NA | NA |
| T17276 | ACGGTGTTTTTCCGCGCCCGGGG | 1000 | | | | | | | | | |
| Oligo fwd | CACCACGGTGTTTTTCCGCGCCCG | | | | | | | | | | |
| Oligo rev | AAACCGGGCGCGGAAAAACACCGT | | | | | | | | | | |
| Chromosome | start | end | strand | MM | target_seq | PAM | alignment | distance | position | gene name | gene id |
| GU305914.1 | 229948 | 229970 | - | 0 | ACGGTGTTTTTCCGCGCCCG | GGG | \|\|\|\|\|\|\|\|[\|\|\|\|\|\|\|\|\|\|\|\|]PAM | NA | Intergenic | NA | NA |
| T17277 | CGGTGTTTTTCCGCGCCCGGGGG | 1000 | | | | | | | | | |
| Oligo fwd | CACCCGGTGTTTTTCCGCGCCCGG | | | | | | | | | | |
| Oligo rev | AAACCCGGGCGCGGAAAAACACCG | | | | | | | | | | |
| Chromosome | start | end | strand | MM | target_seq | PAM | alignment | distance | position | gene name | gene id |
| GU305914.1 | 229947 | 229969 | - | 0 | CGGTGTTTTTCCGCGCCCGG | GGG | \|\|\|\|\|\|\|\|[\|\|\|\|\|\|\|\|\|\|\|\|]PAM | NA | Intergenic | NA | NA |
| T17278 | GGTGTTTTTCCGCGCCCGGGGGG | 1000 | | | | | | | | | |
| Oligo fwd | CACCGGTGTTTTTCCGCGCCCGGG | | | | | | | | | | |
| Oligo rev | AAACCCCGGGCGCGGAAAAACACC | | | | | | | | | | |
| Chromosome | start | end | strand | MM | target_seq | PAM | alignment | distance | position | gene name | gene id |
| GU305914.1 | 229946 | 229968 | - | 0 | GGTGTTTTTCCGCGCCCGGG | GGG | \|\|\|\|\|\|\|\|[\|\|\|\|\|\|\|\|\|\|\|\|]PAM | NA | Intergenic | NA | NA |
| T17279 | CGCCCGGGGGGTGAACCCCCTGG | 1000 | | | | | | | | | |
| Oligo fwd | CACCCGCCCGGGGGGTGAACCCCC | | | | | | | | | | |
| Oligo rev | AAACGGGGGTTCACCCCCCGGGCG | | | | | | | | | | |
| Chromosome | start | end | strand | MM | target_seq | PAM | alignment | distance | position | gene name | gene id |
| GU305914.1 | 229934 | 229956 | - | 0 | CGCCCGGGGGGTGAACCCCC | TGG | \|\|\|\|\|\|\|\|[\|\|\|\|\|\|\|\|\|\|\|\|]PAM | NA | Intergenic | NA | NA |
| T17280 | ACCCCCTGGAGCCGCCCTCGCGG | 1000 | | | | | | | | | |
| Oligo fwd | CACCACCCCCTGGAGCCGCCCTCG | | | | | | | | | | |
| Oligo rev | AAACCGAGGGCGGCTCCAGGGGGT | | | | | | | | | | |
| Chromosome | start | end | strand | MM | target_seq | PAM | alignment | distance | position | gene name | gene id |
| GU305914.1 | 229920 | 229942 | - | 0 | ACCCCCTGGAGCCGCCCTCG | CGG | \|\|\|\|\|\|\|\|[\|\|\|\|\|\|\|\|\|\|\|\|]PAM | NA | Intergenic | NA | NA |
| T17281 | CGCCCTCGCGGCGTAACGTCCGG | 1000 | | | | | | | | | |
| Oligo fwd | CACCCGCCCTCGCGGCGTAACGTC | | | | | | | | | | |
| Oligo rev | AAACGACGTTACGCCGCGAGGGCG | | | | | | | | | | |
| Chromosome | start | end | strand | MM | target_seq | PAM | alignment | distance | position | gene name | gene id |
| GU305914.1 | 229908 | 229930 | - | 0 | CGCCCTCGCGGCGTAACGTC | CGG | \|\|\|\|\|\|\|\|[\|\|\|\|\|\|\|\|\|\|\|\|]PAM | NA | Intergenic | NA | NA |
| T17282 | GCCCTCGCGGCGTAACGTCCGGG | 1000 | | | | | | | | | |
| Oligo fwd | CACCGCCCTCGCGGCGTAACGTCC | | | | | | | | | | |
| Oligo rev | AAACGGACGTTACGCCGCGAGGGC | | | | | | | | | | |
| Chromosome | start | end | strand | MM | target_seq | PAM | alignment | distance | position | gene name | gene id |
| GU305914.1 | 229907 | 229929 | - | 0 | GCCCTCGCGGCGTAACGTCC | GGG | \|\|\|\|\|\|\|\|[\|\|\|\|\|\|\|\|\|\|\|\|]PAM | NA | Intergenic | NA | NA |
| T17283 | CCCTCGCGGCGTAACGTCCGGGG | 1000 | | | | | | | | | |
| Oligo fwd | CACCCCCTCGCGGCGTAACGTCCG | | | | | | | | | | |
| Oligo rev | AAACCGGACGTTACGCCGCGAGGG | | | | | | | | | | |
| Chromosome | start | end | strand | MM | target_seq | PAM | alignment | distance | position | gene name | gene id |
| GU305914.1 | 229906 | 229928 | - | 0 | CCCTCGCGGCGTAACGTCCG | GGG | \|\|\|\|\|\|\|\|[\|\|\|\|\|\|\|\|\|\|\|\|]PAM | NA | Intergenic | NA | NA |
| T17284 | GGCGTAACGTCCGGGGACCCTGG | 1000 | | | | | | | | | |
| Oligo fwd | CACCGGCGTAACGTCCGGGGACCC | | | | | | | | | | |
| Oligo rev | AAACGGGTCCCCGGACGTTACGCC | | | | | | | | | | |
| Chromosome | start | end | strand | MM | target_seq | PAM | alignment | distance | position | gene name | gene id |
| GU305914.1 | 229899 | 229921 | - | 0 | GGCGTAACGTCCGGGGACCC | TGG | \|\|\|\|\|\|\|\|[\|\|\|\|\|\|\|\|\|\|\|\|]PAM | NA | Intergenic | NA | NA |
| T17285 | GCGTAACGTCCGGGGACCCTGGG | 1000 | | | | | | | | | |
| Oligo fwd | CACCGCGTAACGTCCGGGGACCCT | | | | | | | | | | |
| Oligo rev | AAACAGGGTCCCCGGACGTTACGC | | | | | | | | | | |
| Chromosome | start | end | strand | MM | target_seq | PAM | alignment | distance | position | gene name | gene id |
| GU305914.1 | 229898 | 229920 | - | 0 | GCGTAACGTCCGGGGACCCT | GGG | \|\|\|\|\|\|\|\|[\|\|\|\|\|\|\|\|\|\|\|\|]PAM | NA | Intergenic | NA | NA |
| T17286 | GGCGTCATTTTATACGCGAACGG | 1000 | | | | | | | | | |
| Oligo fwd | CACCGGCGTCATTTTATACGCGAA | | | | | | | | | | |
| Oligo rev | AAACTTCGCGTATAAAATGACGCC | | | | | | | | | | |
| Chromosome | start | end | strand | MM | target_seq | PAM | alignment | distance | position | gene name | gene id |
| GU305914.1 | 229877 | 229899 | - | 0 | GGCGTCATTTTATACGCGAA | CGG | \|\|\|\|\|\|\|\|[\|\|\|\|\|\|\|\|\|\|\|\|]PAM | NA | Intergenic | NA | NA |

T17287 GTCATTTTATACGCGAACGGTGG 1000
Oligo fwd CACCGTCATTTTATACGCGAACGG
Oligo rev AAACCCGTTCGCGTATAAAATGAC
Chromosome start end strand MM target_seq PAM alignment distance position gene name gene id
GU305914.1 229874 229896 - 0 GTCATTTTATACGCGAACGG TGG ||||||||[||||||||||||]PAM NA Intergenic NA NA

T17288 AACGGTGGACAGCCGTTTTCTGG 1000
Oligo fwd CACCAACGGTGGACAGCCGTTTTC
Oligo rev AAACGAAAACGGCTGTCCACCGTT
Chromosome start end strand MM target_seq PAM alignment distance position gene name gene id
GU305914.1 229859 229881 - 0 AACGGTGGACAGCCGTTTTC TGG ||||||||[||||||||||||]PAM NA Intergenic NA NA

T17289 ACGGTGGACAGCCGTTTTCTGGG 1000
Oligo fwd CACCACGGTGGACAGCCGTTTTCT
Oligo rev AAACAGAAAACGGCTGTCCACCGT
Chromosome start end strand MM target_seq PAM alignment distance position gene name gene id
GU305914.1 229858 229880 - 0 ACGGTGGACAGCCGTTTTCT GGG ||||||||[||||||||||||]PAM NA Intergenic NA NA

T17290 TTTTCTGGGAGTCTACTACGCGG 1000
Oligo fwd CACCTTTTCTGGGAGTCTACTACG
Oligo rev AAACCGTAGTAGACTCCCAGAAAA
Chromosome start end strand MM target_seq PAM alignment distance position gene name gene id
GU305914.1 229844 229866 - 0 TTTTCTGGGAGTCTACTACG CGG ||||||||[||||||||||||]PAM NA Intergenic NA NA

T17291 TCTGGGAGTCTACTACGCGGTGG 1000
Oligo fwd CACCTCTGGGAGTCTACTACGCGG
Oligo rev AAACCCGCGTAGTAGACTCCCAGA
Chromosome start end strand MM target_seq PAM alignment distance position gene name gene id
GU305914.1 229841 229863 - 0 TCTGGGAGTCTACTACGCGG TGG ||||||||[||||||||||||]PAM NA Intergenic NA NA

T17292 AGCGCCCCCTGTTCGCGCAACGG 1000
Oligo fwd CACCAGCGCCCCCTGTTCGCGCAA
Oligo rev AAACTTGCGCGAACAGGGGGCGCT
Chromosome start end strand MM target_seq PAM alignment distance position gene name gene id
GU305914.1 229803 229825 - 0 AGCGCCCCCTGTTCGCGCAA CGG ||||||||[||||||||||||]PAM NA Intergenic NA NA

T17293 GCGCCCCCTGTTCGCGCAACGGG 1000
Oligo fwd CACCGCGCCCCCTGTTCGCGCAAC
Oligo rev AAACGTTGCGCGAACAGGGGGCGC
Chromosome start end strand MM target_seq PAM alignment distance position gene name gene id
GU305914.1 229802 229824 - 0 GCGCCCCCTGTTCGCGCAAC GGG ||||||||[||||||||||||]PAM NA Intergenic NA NA

T17294 CGCCCCCTGTTCGCGCAACGGGG 1000
Oligo fwd CACCCGCCCCCTGTTCGCGCAACG
Oligo rev AAACCGTTGCGCGAACAGGGGGCG
Chromosome start end strand MM target_seq PAM alignment distance position gene name gene id
GU305914.1 229801 229823 - 0 CGCCCCCTGTTCGCGCAACG GGG ||||||||[||||||||||||]PAM NA Intergenic NA NA
GU305914.1 228496 228518 - 4 CGTCCTCTGTTCGCCCAGCG CGG ||-||-||[|||||||-||-||]PAM NA Intergenic NA NA

T17295 CCCTGTTCGCGCAACGGGGCTGG 1000
Oligo fwd CACCCCCTGTTCGCGCAACGGGGC
Oligo rev AAACGCCCCGTTGCGCGAACAGGG
Chromosome start end strand MM target_seq PAM alignment distance position gene name gene id
GU305914.1 229797 229819 - 0 CCCTGTTCGCGCAACGGGGC TGG ||||||||[||||||||||||]PAM NA Intergenic NA NA

T17296 TGTTCGCGCAACGGGGCTGGCGG 1000
Oligo fwd CACCTGTTCGCGCAACGGGGCTGG
Oligo rev AAACCCAGCCCCGTTGCGCGAACA
Chromosome start end strand MM target_seq PAM alignment distance position gene name gene id
GU305914.1 229794 229816 - 0 TGTTCGCGCAACGGGGCTGG CGG ||||||||[||||||||||||]PAM NA Intergenic NA NA

T17297 CTGGCGGCAGCGCCACCCCGTGG 1000
Oligo fwd CACCCTGGCGGCAGCGCCACCCCG
Oligo rev AAACCGGGGTGGCGCTGCCGCCAG
Chromosome start end strand MM target_seq PAM alignment distance position gene name gene id
GU305914.1 229778 229800 - 0 CTGGCGGCAGCGCCACCCCG TGG ||||||||[||||||||||||]PAM NA Intergenic NA NA

T17298 GCGGCAGCGCCACCCCGTGGTGG 1000
Oligo fwd CACCGCGGCAGCGCCACCCCGTGG
Oligo rev AAACCCACGGGGTGGCGCTGCCGC
Chromosome start end strand MM target_seq PAM alignment distance position gene name gene id
GU305914.1 229775 229797 - 0 GCGGCAGCGCCACCCCGTGG TGG ||||||||[||||||||||||]PAM NA Intergenic NA NA

T17299 GGCAGACCAGTTACGCGCTCAGG 1000
Oligo fwd CACCGGCAGACCAGTTACGCGCTC
Oligo rev AAACGAGCGCGTAACTGGTCTGCC
Chromosome start end strand MM target_seq PAM alignment distance position gene name gene id
GU305914.1 229754 229776 - 0 GGCAGACCAGTTACGCGCTC AGG ||||||||[||||||||||||]PAM NA Intergenic NA NA

T17300 GTTACGCGCTCAGGCTCGTCAGG 1000
Oligo fwd CACCGTTACGCGCTCAGGCTCGTC
Oligo rev AAACGACGAGCCTGAGCGCGTAAC
Chromosome start end strand MM target_seq PAM alignment distance position gene name gene id
GU305914.1 229745 229767 - 0 GTTACGCGCTCAGGCTCGTC AGG ||||||||[||||||||||||]PAM NA Intergenic NA NA

T17301 ACGCGCTCAGGCTCGTCAGGCGG 1000
Oligo fwd CACCACGCGCTCAGGCTCGTCAGG
Oligo rev AAACCCTGACGAGCCTGAGCGCGT
Chromosome start end strand MM target_seq PAM alignment distance position gene name gene id

GU305914.1 229742 229764 - 0 ACGCGCTCAGGCTCGTCAGG CGG ||||||||[||||||||||||]PAM NA Intergenic NA NA

T17302 CGCTCAGGCTCGTCAGGCGGCGG 1000
Oligo fwd CACCCGCTCAGGCTCGTCAGGCGG
Oligo rev AAACCCGCCTGACGAGCCTGAGCG
Chromosome start end strand MM target_seq PAM alignment distance position gene name gene id
GU305914.1 229739 229761 - 0 CGCTCAGGCTCGTCAGGCGG CGG ||||||||[||||||||||||]PAM NA Intergenic NA NA

T17303 GGCTCGTCAGGCGGCGGATGTGG 1000
Oligo fwd CACCGGCTCGTCAGGCGGCGGATG
Oligo rev AAACCATCCGCCGCCTGACGAGCC
Chromosome start end strand MM target_seq PAM alignment distance position gene name gene id
GU305914.1 229733 229755 - 0 GGCTCGTCAGGCGGCGGATG TGG ||||||||[||||||||||||]PAM NA Intergenic NA NA
GU305914.1 150580 150602 + 4 TGCTCTTCAGCCGGTGGATG CGG -||||-||[||-|||-|||||]PAM NA Intergenic NA NA

T17304 GCTCGTCAGGCGGCGGATGTGGG 1000
Oligo fwd CACCGCTCGTCAGGCGGCGGATGT
Oligo rev AAACACATCCGCCGCCTGACGAGC
Chromosome start end strand MM target_seq PAM alignment distance position gene name gene id
GU305914.1 229732 229754 - 0 GCTCGTCAGGCGGCGGATGT GGG ||||||||[||||||||||||]PAM NA Intergenic NA NA

T17305 CGGATGTGGGTGAGTGCGTCCGG 1000
Oligo fwd CACCCGGATGTGGGTGAGTGCGTC
Oligo rev AAACGACGCACTCACCCACATCCG
Chromosome start end strand MM target_seq PAM alignment distance position gene name gene id
GU305914.1 229719 229741 - 0 CGGATGTGGGTGAGTGCGTC CGG ||||||||[||||||||||||]PAM NA Intergenic NA NA

T17306 CGTAGCGAGCACCCGCTCGCTGG 1000
Oligo fwd CACCCGTAGCGAGCACCCGCTCGC
Oligo rev AAACGCGAGCGGGTGCTCGCTACG
Chromosome start end strand MM target_seq PAM alignment distance position gene name gene id
GU305914.1 229691 229713 - 0 CGTAGCGAGCACCCGCTCGC TGG ||||||||[||||||||||||]PAM NA Intergenic NA NA

T17307 TCGCTGGTCGAGCGACCGCCCGG 1000
Oligo fwd CACCTCGCTGGTCGAGCGACCGCC
Oligo rev AAACGGCGGTCGCTCGACCAGCGA
Chromosome start end strand MM target_seq PAM alignment distance position gene name gene id
GU305914.1 229675 229697 - 0 TCGCTGGTCGAGCGACCGCC CGG ||||||||[||||||||||||]PAM NA Intergenic NA NA

T17308 CGCTGGTCGAGCGACCGCCCGGG 1000
Oligo fwd CACCCGCTGGTCGAGCGACCGCCC
Oligo rev AAACGGGCGGTCGCTCGACCAGCG
Chromosome start end strand MM target_seq PAM alignment distance position gene name gene id
GU305914.1 229674 229696 - 0 CGCTGGTCGAGCGACCGCCC GGG ||||||||[||||||||||||]PAM NA Intergenic NA NA

T17309 TGCCCGTCGTAACGCGACGCCGG 1000
Oligo fwd CACCTGCCCGTCGTAACGCGACGC
Oligo rev AAACGCGTCGCGTTACGACGGGCA
Chromosome start end strand MM target_seq PAM alignment distance position gene name gene id
GU305914.1 229643 229665 - 0 TGCCCGTCGTAACGCGACGC CGG ||||||||[||||||||||||]PAM NA Intergenic NA NA

T17310 GCGACCAGCGTGCCGCGTCGCGG 1000
Oligo fwd CACCGCGACCAGCGTGCCGCGTCG
Oligo rev AAACCGACGCGGCACGCTGGTCGC
Chromosome start end strand MM target_seq PAM alignment distance position gene name gene id
GU305914.1 229611 229633 - 0 GCGACCAGCGTGCCGCGTCG CGG ||||||||[||||||||||||]PAM NA Intergenic NA NA

T17311 CGACCAGCGTGCCGCGTCGCGGG 1000
Oligo fwd CACCCGACCAGCGTGCCGCGTCGC
Oligo rev AAACGCGACGCGGCACGCTGGTCG
Chromosome start end strand MM target_seq PAM alignment distance position gene name gene id
GU305914.1 229610 229632 - 0 CGACCAGCGTGCCGCGTCGC GGG ||||||||[||||||||||||]PAM NA Intergenic NA NA

T17312 TGTTGCCACGTACCGCGCTCCGG 1000
Oligo fwd CACCTGTTGCCACGTACCGCGCTC
Oligo rev AAACGAGCGCGGTACGTGGCAACA
Chromosome start end strand MM target_seq PAM alignment distance position gene name gene id
GU305914.1 229549 229571 - 0 TGTTGCCACGTACCGCGCTC CGG ||||||||[||||||||||||]PAM NA Intergenic NA NA

T17313 GCTCCGGAGAACGTCGTTCACGG 1000
Oligo fwd CACCGCTCCGGAGAACGTCGTTCA
Oligo rev AAACTGAACGACGTTCTCCGGAGC
Chromosome start end strand MM target_seq PAM alignment distance position gene name gene id
GU305914.1 229533 229555 - 0 GCTCCGGAGAACGTCGTTCA CGG ||||||||[||||||||||||]PAM NA Intergenic NA NA

T17314 CCGGAGAACGTCGTTCACGGAGG 1000
Oligo fwd CACCCCGGAGAACGTCGTTCACGG
Oligo rev AAACCCGTGAACGACGTTCTCCGG
Chromosome start end strand MM target_seq PAM alignment distance position gene name gene id
GU305914.1 229530 229552 - 0 CCGGAGAACGTCGTTCACGG AGG ||||||||[||||||||||||]PAM NA Intergenic NA NA

T17315 TCGTTCACGGAGGTGTTACGCGG 1000
Oligo fwd CACCTCGTTCACGGAGGTGTTACG
Oligo rev AAACCGTAACACCTCCGTGAACGA
Chromosome start end strand MM target_seq PAM alignment distance position gene name gene id
GU305914.1 229520 229542 - 0 TCGTTCACGGAGGTGTTACG CGG ||||||||[||||||||||||]PAM NA Intergenic NA NA

T17316 TACGCGGAGCGCATGACTGCCGG 1000
Oligo fwd CACCTACGCGGAGCGCATGACTGC

Oligo rev AAACGCAGTCATGCGCTCCGCGTA
Chromosome start end strand MM target_seq PAM alignment distance position gene name gene id
GU305914.1 229504 229526 - 0 TACGCGGAGCGCATGACTGC CGG ||||||||[||||||||||||]PAM NA Intergenic NA NA

T17317 ACGCGGAGCGCATGACTGCCGGG 1000
Oligo fwd CACCACGCGGAGCGCATGACTGCC
Oligo rev AAACGGCAGTCATGCGCTCCGCGT
Chromosome start end strand MM target_seq PAM alignment distance position gene name gene id
GU305914.1 229503 229525 - 0 ACGCGGAGCGCATGACTGCC GGG ||||||||[||||||||||||]PAM NA Intergenic NA NA

T17318 GAGCGCATGACTGCCGGGCGAGG 1000
Oligo fwd CACCGAGCGCATGACTGCCGGGCG
Oligo rev AAACCGCCCGGCAGTCATGCGCTC
Chromosome start end strand MM target_seq PAM alignment distance position gene name gene id
GU305914.1 229498 229520 - 0 GAGCGCATGACTGCCGGGCG AGG ||||||||[||||||||||||]PAM NA Intergenic NA NA

T17319 AGCGCATGACTGCCGGGCGAGGG 1000
Oligo fwd CACCAGCGCATGACTGCCGGGCGA
Oligo rev AAACTCGCCCGGCAGTCATGCGCT
Chromosome start end strand MM target_seq PAM alignment distance position gene name gene id
GU305914.1 229497 229519 - 0 AGCGCATGACTGCCGGGCGA GGG ||||||||[||||||||||||]PAM NA Intergenic NA NA

T17320 GCCGGGCGAGGGCGCCCCGCCGG 1000
Oligo fwd CACCGCCGGGCGAGGGCGCCCCGC
Oligo rev AAACGCGGGGCGCCCTCGCCCGGC
Chromosome start end strand MM target_seq PAM alignment distance position gene name gene id
GU305914.1 229486 229508 - 0 GCCGGGCGAGGGCGCCCCGC CGG ||||||||[||||||||||||]PAM NA Intergenic NA NA

T17321 GCGCCCCGCCGGCATGCTACAGG 1000
Oligo fwd CACCGCGCCCCGCCGGCATGCTAC
Oligo rev AAACGTAGCATGCCGGCGGGGCGC
Chromosome start end strand MM target_seq PAM alignment distance position gene name gene id
GU305914.1 229475 229497 - 0 GCGCCCCGCCGGCATGCTAC AGG ||||||||[||||||||||||]PAM NA Intergenic NA NA

T17322 CCGGCATGCTACAGGCGTTACGG 1000
Oligo fwd CACCCCGGCATGCTACAGGCGTTA
Oligo rev AAACTAACGCCTGTAGCATGCCGG
Chromosome start end strand MM target_seq PAM alignment distance position gene name gene id
GU305914.1 229467 229489 - 0 CCGGCATGCTACAGGCGTTA CGG ||||||||[||||||||||||]PAM NA Intergenic NA NA

T17323 CGGCATGCTACAGGCGTTACGGG 1000
Oligo fwd CACCCGGCATGCTACAGGCGTTAC
Oligo rev AAACGTAACGCCTGTAGCATGCCG
Chromosome start end strand MM target_seq PAM alignment distance position gene name gene id
GU305914.1 229466 229488 - 0 CGGCATGCTACAGGCGTTAC GGG ||||||||[||||||||||||]PAM NA Intergenic NA NA

T17324 TTACGGGTTGACCGCCGCGCAGG 1000
Oligo fwd CACCTTACGGGTTGACCGCCGCGC
Oligo rev AAACGCGCGGCGGTCAACCCGTAA
Chromosome start end strand MM target_seq PAM alignment distance position gene name gene id
GU305914.1 229450 229472 - 0 TTACGGGTTGACCGCCGCGC AGG ||||||||[||||||||||||]PAM NA Intergenic NA NA

T17325 TTGACCGCCGCGCAGGTACCAGG 1000
Oligo fwd CACCTTGACCGCCGCGCAGGTACC
Oligo rev AAACGGTACCTGCGCGGCGGTCAA
Chromosome start end strand MM target_seq PAM alignment distance position gene name gene id
GU305914.1 229443 229465 - 0 TTGACCGCCGCGCAGGTACC AGG ||||||||[||||||||||||]PAM NA Intergenic NA NA

T17326 TTTCCAGCCGAGTTCGTGATCGG 1000
Oligo fwd CACCTTTCCAGCCGAGTTCGTGAT
Oligo rev AAACATCACGAACTCGGCTGGAAA
Chromosome start end strand MM target_seq PAM alignment distance position gene name gene id
GU305914.1 229394 229416 - 0 TTTCCAGCCGAGTTCGTGAT CGG ||||||||[||||||||||||]PAM NA Intergenic NA NA

T17327 AACTTTCCCTGTGAGCGCTGCGG 1000
Oligo fwd CACCAACTTTCCCTGTGAGCGCTG
Oligo rev AAACCAGCGCTCACAGGGAAAGTT
Chromosome start end strand MM target_seq PAM alignment distance position gene name gene id
GU305914.1 229369 229391 - 0 AACTTTCCCTGTGAGCGCTG CGG ||||||||[||||||||||||]PAM NA Intergenic NA NA

T17328 ACTTTCCCTGTGAGCGCTGCGGG 1000
Oligo fwd CACCACTTTCCCTGTGAGCGCTGC
Oligo rev AAACGCAGCGCTCACAGGGAAAGT
Chromosome start end strand MM target_seq PAM alignment distance position gene name gene id
GU305914.1 229368 229390 - 0 ACTTTCCCTGTGAGCGCTGC GGG ||||||||[||||||||||||]PAM NA Intergenic NA NA

T17329 TTCCCTGTGAGCGCTGCGGGAGG 1000
Oligo fwd CACCTTCCCTGTGAGCGCTGCGGG
Oligo rev AAACCCCGCAGCGCTCACAGGGAA
Chromosome start end strand MM target_seq PAM alignment distance position gene name gene id
GU305914.1 229365 229387 - 0 TTCCCTGTGAGCGCTGCGGG AGG ||||||||[||||||||||||]PAM NA Intergenic NA NA

T17330 CTCCCTCGCCGTTCGTGTCGCGG 1000
Oligo fwd CACCCTCCCTCGCCGTTCGTGTCG
Oligo rev AAACCGACACGAACGGCGAGGGAG
Chromosome start end strand MM target_seq PAM alignment distance position gene name gene id
GU305914.1 229316 229338 - 0 CTCCCTCGCCGTTCGTGTCG CGG ||||||||[||||||||||||]PAM NA Intergenic NA NA

T17331 CGCTGTCCGTTCCCGCTCATCGG 1000

| | | | | | | | | | | | | |
|---|---|---|---|---|---|---|---|---|---|---|---|---|
| Oligo fwd | CACCCGCTGTCCGTTCCCGCTCAT | | | | | | | | | | | |
| Oligo rev | AAACATGAGCGGGAACGGACAGCG | | | | | | | | | | | |
| Chromosome | start | | end | strand | MM | target_seq | PAM | alignment | distance | position | gene name | gene id |
| GU305914.1 | | 229291 | 229313 | - | 0 | CGCTGTCCGTTCCCGCTCAT | CGG | \|\|\|\|\|\|\|\|[\|\|\|\|\|\|\|\|\|\|\|\|]PAM | NA | Intergenic | NA | NA |
| T17332 | CGCTCATCGGTCTGACTTCTCGG | | 1000 | | | | | | | | | |
| Oligo fwd | CACCCGCTCATCGGTCTGACTTCT | | | | | | | | | | | |
| Oligo rev | AAACAGAAGTCAGACCGATGAGCG | | | | | | | | | | | |
| Chromosome | start | | end | strand | MM | target_seq | PAM | alignment | distance | position | gene name | gene id |
| GU305914.1 | | 229278 | 229300 | - | 0 | CGCTCATCGGTCTGACTTCT | CGG | \|\|\|\|\|\|\|\|[\|\|\|\|\|\|\|\|\|\|\|\|]PAM | NA | Intergenic | NA | NA |
| T17333 | CTTTTGTCTCCTCCCCCGCGCGG | | 1000 | | | | | | | | | |
| Oligo fwd | CACCCTTTTGTCTCCTCCCCCGCG | | | | | | | | | | | |
| Oligo rev | AAACCGCGGGGGAGGAGACAAAAG | | | | | | | | | | | |
| Chromosome | start | | end | strand | MM | target_seq | PAM | alignment | distance | position | gene name | gene id |
| GU305914.1 | | 229244 | 229266 | - | 0 | CTTTTGTCTCCTCCCCCGCG | CGG | \|\|\|\|\|\|\|\|[\|\|\|\|\|\|\|\|\|\|\|\|]PAM | NA | Intergenic | NA | NA |
| T17334 | TTTTGTCTCCTCCCCCGCGCGGG | | 1000 | | | | | | | | | |
| Oligo fwd | CACCTTTTGTCTCCTCCCCCGCGC | | | | | | | | | | | |
| Oligo rev | AAACGCGCGGGGGAGGAGACAAAA | | | | | | | | | | | |
| Chromosome | start | | end | strand | MM | target_seq | PAM | alignment | distance | position | gene name | gene id |
| GU305914.1 | | 229243 | 229265 | - | 0 | TTTTGTCTCCTCCCCCGCGC | GGG | \|\|\|\|\|\|\|\|[\|\|\|\|\|\|\|\|\|\|\|\|]PAM | NA | Intergenic | NA | NA |
| T17335 | AAGTGTATAAAAGCTGAGTGTGG | | 1000 | | | | | | | | | |
| Oligo fwd | CACCAAGTGTATAAAAGCTGAGTG | | | | | | | | | | | |
| Oligo rev | AAACCACTCAGCTTTTATACACTT | | | | | | | | | | | |
| Chromosome | start | | end | strand | MM | target_seq | PAM | alignment | distance | position | gene name | gene id |
| GU305914.1 | | 229212 | 229234 | - | 0 | AAGTGTATAAAAGCTGAGTG | TGG | \|\|\|\|\|\|\|\|[\|\|\|\|\|\|\|\|\|\|\|\|]PAM | NA | Intergenic | NA | NA |
| T17336 | TATAAAAGCTGAGTGTGGAGCGG | | 1000 | | | | | | | | | |
| Oligo fwd | CACCTATAAAAGCTGAGTGTGGAG | | | | | | | | | | | |
| Oligo rev | AAACCTCCACACTCAGCTTTTATA | | | | | | | | | | | |
| Chromosome | start | | end | strand | MM | target_seq | PAM | alignment | distance | position | gene name | gene id |
| GU305914.1 | | 229207 | 229229 | - | 0 | TATAAAAGCTGAGTGTGGAG | CGG | \|\|\|\|\|\|\|\|[\|\|\|\|\|\|\|\|\|\|\|\|]PAM | NA | Intergenic | NA | NA |
| T17337 | CAGTCGACAGCTTCACATCCCGG | | 1000 | | | | | | | | | |
| Oligo fwd | CACCCAGTCGACAGCTTCACATCC | | | | | | | | | | | |
| Oligo rev | AAACGGATGTGAAGCTGTCGACTG | | | | | | | | | | | |
| Chromosome | start | | end | strand | MM | target_seq | PAM | alignment | distance | position | gene name | gene id |
| GU305914.1 | | 229181 | 229203 | - | 0 | CAGTCGACAGCTTCACATCC | CGG | \|\|\|\|\|\|\|\|[\|\|\|\|\|\|\|\|\|\|\|\|]PAM | NA | Intergenic | NA | NA |
| T17338 | TCCCGGTGACCTGTCGCTTTCGG | | 1000 | | | | | | | | | |
| Oligo fwd | CACCTCCCGGTGACCTGTCGCTTT | | | | | | | | | | | |
| Oligo rev | AAACAAAGCGACAGGTCACCGGGA | | | | | | | | | | | |
| Chromosome | start | | end | strand | MM | target_seq | PAM | alignment | distance | position | gene name | gene id |
| GU305914.1 | | 229164 | 229186 | - | 0 | TCCCGGTGACCTGTCGCTTT | CGG | \|\|\|\|\|\|\|\|[\|\|\|\|\|\|\|\|\|\|\|\|]PAM | NA | Intergenic | NA | NA |
| T17339 | CGGTGACCTGTCGCTTTCGGTGG | | 1000 | | | | | | | | | |
| Oligo fwd | CACCCGGTGACCTGTCGCTTTCGG | | | | | | | | | | | |
| Oligo rev | AAACCCGAAAGCGACAGGTCACCG | | | | | | | | | | | |
| Chromosome | start | | end | strand | MM | target_seq | PAM | alignment | distance | position | gene name | gene id |
| GU305914.1 | | 229161 | 229183 | - | 0 | CGGTGACCTGTCGCTTTCGG | TGG | \|\|\|\|\|\|\|\|[\|\|\|\|\|\|\|\|\|\|\|\|]PAM | NA | Intergenic | NA | NA |
| T17340 | GGTCTCTCTCTGCCTGTCTGTGG | | 1000 | | | | | | | | | |
| Oligo fwd | CACCGGTCTCTCTCTGCCTGTCTG | | | | | | | | | | | |
| Oligo rev | AAACCAGACAGGCAGAGAGAGACC | | | | | | | | | | | |
| Chromosome | start | | end | strand | MM | target_seq | PAM | alignment | distance | position | gene name | gene id |
| GU305914.1 | | 229140 | 229162 | - | 0 | GGTCTCTCTCTGCCTGTCTG | TGG | \|\|\|\|\|\|\|\|[\|\|\|\|\|\|\|\|\|\|\|\|]PAM | NA | Intergenic | NA | NA |
| T17341 | GTCTGTGGTGAGCTGTGCTCCGG | | 1000 | | | | | | | | | |
| Oligo fwd | CACCGTCTGTGGTGAGCTGTGCTC | | | | | | | | | | | |
| Oligo rev | AAACGAGCACAGCTCACCACAGAC | | | | | | | | | | | |
| Chromosome | start | | end | strand | MM | target_seq | PAM | alignment | distance | position | gene name | gene id |
| GU305914.1 | | 229125 | 229147 | - | 0 | GTCTGTGGTGAGCTGTGCTC | CGG | \|\|\|\|\|\|\|\|[\|\|\|\|\|\|\|\|\|\|\|\|]PAM | NA | Intergenic | NA | NA |
| T17342 | TGTGGTGAGCTGTGCTCCGGCGG | | 1000 | | | | | | | | | |
| Oligo fwd | CACCTGTGGTGAGCTGTGCTCCGG | | | | | | | | | | | |
| Oligo rev | AAACCCGGAGCACAGCTCACCACA | | | | | | | | | | | |
| Chromosome | start | | end | strand | MM | target_seq | PAM | alignment | distance | position | gene name | gene id |
| GU305914.1 | | 229122 | 229144 | - | 0 | TGTGGTGAGCTGTGCTCCGG | CGG | \|\|\|\|\|\|\|\|[\|\|\|\|\|\|\|\|\|\|\|\|]PAM | NA | Intergenic | NA | NA |
| T17343 | CGGTTCCCGCGTCCGTCATGAGG | | 1000 | | | | | | | | | |
| Oligo fwd | CACCCGGTTCCCGCGTCCGTCATG | | | | | | | | | | | |
| Oligo rev | AAACCATGACGGACGCGGGAACCG | | | | | | | | | | | |
| Chromosome | start | | end | strand | MM | target_seq | PAM | alignment | distance | position | gene name | gene id |
| GU305914.1 | | 229102 | 229124 | - | 0 | CGGTTCCCGCGTCCGTCATG | AGG | \|\|\|\|\|\|\|\|[\|\|\|\|\|\|\|\|\|\|\|\|]PAM | NA | Intergenic | NA | NA |
| T17344 | CCGCGTCCGTCATGAGGTCCAGG | | 1000 | | | | | | | | | |
| Oligo fwd | CACCCCGCGTCCGTCATGAGGTCC | | | | | | | | | | | |
| Oligo rev | AAACGGACCTCATGACGGACGCGG | | | | | | | | | | | |
| Chromosome | start | | end | strand | MM | target_seq | PAM | alignment | distance | position | gene name | gene id |
| GU305914.1 | | 229096 | 229118 | - | 0 | CCGCGTCCGTCATGAGGTCC | AGG | \|\|\|\|\|\|\|\|[\|\|\|\|\|\|\|\|\|\|\|\|]PAM | NA | Intergenic | NA | NA |
| GU305914.1 | | 67399 | 67421 | - | 4 | TCGCGTACGTCAGGAAGTCC | AGG | -\|\|\|\|\|-\|[\|\|\|\|-\|\|-\|\|\|\|]PAM | NA | Intergenic | NA | NA |
| T17345 | GTCGTGTGACACGCTAAATATGG | | 1000 | | | | | | | | | |
| Oligo fwd | CACCGTCGTGTGACACGCTAAATA | | | | | | | | | | | |
| Oligo rev | AAACTATTTAGCGTGTCACACGAC | | | | | | | | | | | |
| Chromosome | start | | end | strand | MM | target_seq | PAM | alignment | distance | position | gene name | gene id |
| GU305914.1 | | 229074 | 229096 | - | 0 | GTCGTGTGACACGCTAAATA | TGG | \|\|\|\|\|\|\|\|[\|\|\|\|\|\|\|\|\|\|\|\|]PAM | NA | Intergenic | NA | NA |

<table>
<tr><td>T17346</td><td>CTAAATATGGCCATCGATAACGG</td><td>1000</td><td></td><td></td><td></td><td></td><td></td><td></td><td></td><td></td><td></td></tr>
<tr><td>Oligo fwd</td><td>CACCCTAAATATGGCCATCGATAA</td><td></td><td></td><td></td><td></td><td></td><td></td><td></td><td></td><td></td><td></td></tr>
<tr><td>Oligo rev</td><td>AAACTTATCGATGGCCATATTTAG</td><td></td><td></td><td></td><td></td><td></td><td></td><td></td><td></td><td></td><td></td></tr>
<tr><td>Chromosome</td><td>start</td><td>end</td><td>strand</td><td>MM</td><td>target_seq</td><td>PAM</td><td>alignment</td><td>distance</td><td>position</td><td>gene name</td><td>gene id</td></tr>
<tr><td>GU305914.1</td><td>229061</td><td>229083</td><td>-</td><td>0</td><td>CTAAATATGGCCATCGATAA</td><td>CGG</td><td>||||||||[||||||||||||]PAM</td><td>NA</td><td>Intergenic</td><td>NA</td><td>NA</td></tr>
<tr><td>T17347</td><td>AAATCAGCTGCTGCGTCGTCTGG</td><td>1000</td><td></td><td></td><td></td><td></td><td></td><td></td><td></td><td></td><td></td></tr>
<tr><td>Oligo fwd</td><td>CACCAAATCAGCTGCTGCGTCGTC</td><td></td><td></td><td></td><td></td><td></td><td></td><td></td><td></td><td></td><td></td></tr>
<tr><td>Oligo rev</td><td>AAACGACGACGCAGCAGCTGATTT</td><td></td><td></td><td></td><td></td><td></td><td></td><td></td><td></td><td></td><td></td></tr>
<tr><td>Chromosome</td><td>start</td><td>end</td><td>strand</td><td>MM</td><td>target_seq</td><td>PAM</td><td>alignment</td><td>distance</td><td>position</td><td>gene name</td><td>gene id</td></tr>
<tr><td>GU305914.1</td><td>228957</td><td>228979</td><td>-</td><td>0</td><td>AAATCAGCTGCTGCGTCGTC</td><td>TGG</td><td>||||||||[||||||||||||]PAM</td><td>NA</td><td>Intergenic</td><td>NA</td><td>NA</td></tr>
<tr><td>T17348</td><td>CTGGCCCACACCGTGTGCAGTGG</td><td>1000</td><td></td><td></td><td></td><td></td><td></td><td></td><td></td><td></td><td></td></tr>
<tr><td>Oligo fwd</td><td>CACCCTGGCCCACACCGTGTGCAG</td><td></td><td></td><td></td><td></td><td></td><td></td><td></td><td></td><td></td><td></td></tr>
<tr><td>Oligo rev</td><td>AAACCTGCACACGGTGTGGGCCAG</td><td></td><td></td><td></td><td></td><td></td><td></td><td></td><td></td><td></td><td></td></tr>
<tr><td>Chromosome</td><td>start</td><td>end</td><td>strand</td><td>MM</td><td>target_seq</td><td>PAM</td><td>alignment</td><td>distance</td><td>position</td><td>gene name</td><td>gene id</td></tr>
<tr><td>GU305914.1</td><td>228938</td><td>228960</td><td>-</td><td>0</td><td>CTGGCCCACACCGTGTGCAG</td><td>TGG</td><td>||||||||[||||||||||||]PAM</td><td>NA</td><td>Intergenic</td><td>NA</td><td>NA</td></tr>
<tr><td>T17349</td><td>AGACCGAAACCAAGTCCTGCTGG</td><td>1000</td><td></td><td></td><td></td><td></td><td></td><td></td><td></td><td></td><td></td></tr>
<tr><td>Oligo fwd</td><td>CACCAGACCGAAACCAAGTCCTGC</td><td></td><td></td><td></td><td></td><td></td><td></td><td></td><td></td><td></td><td></td></tr>
<tr><td>Oligo rev</td><td>AAACGCAGGACTTGGTTTCGGTCT</td><td></td><td></td><td></td><td></td><td></td><td></td><td></td><td></td><td></td><td></td></tr>
<tr><td>Chromosome</td><td>start</td><td>end</td><td>strand</td><td>MM</td><td>target_seq</td><td>PAM</td><td>alignment</td><td>distance</td><td>position</td><td>gene name</td><td>gene id</td></tr>
<tr><td>GU305914.1</td><td>228901</td><td>228923</td><td>-</td><td>0</td><td>AGACCGAAACCAAGTCCTGC</td><td>TGG</td><td>||||||||[||||||||||||]PAM</td><td>NA</td><td>Intergenic</td><td>NA</td><td>NA</td></tr>
<tr><td>T17350</td><td>CGCGCGCGACGAAGTTGCGATGG</td><td>1000</td><td></td><td></td><td></td><td></td><td></td><td></td><td></td><td></td><td></td></tr>
<tr><td>Oligo fwd</td><td>CACCCGCGCGCGACGAAGTTGCGA</td><td></td><td></td><td></td><td></td><td></td><td></td><td></td><td></td><td></td><td></td></tr>
<tr><td>Oligo rev</td><td>AAACTCGCAACTTCGTCGCGCGCG</td><td></td><td></td><td></td><td></td><td></td><td></td><td></td><td></td><td></td><td></td></tr>
<tr><td>Chromosome</td><td>start</td><td>end</td><td>strand</td><td>MM</td><td>target_seq</td><td>PAM</td><td>alignment</td><td>distance</td><td>position</td><td>gene name</td><td>gene id</td></tr>
<tr><td>GU305914.1</td><td>228867</td><td>228889</td><td>-</td><td>0</td><td>CGCGCGCGACGAAGTTGCGA</td><td>TGG</td><td>||||||||[||||||||||||]PAM</td><td>NA</td><td>Intergenic</td><td>NA</td><td>NA</td></tr>
<tr><td>T17351</td><td>CTTTTCGCTCCCCGCGACGCCGG</td><td>1000</td><td></td><td></td><td></td><td></td><td></td><td></td><td></td><td></td><td></td></tr>
<tr><td>Oligo fwd</td><td>CACCCTTTTCGCTCCCCGCGACGC</td><td></td><td></td><td></td><td></td><td></td><td></td><td></td><td></td><td></td><td></td></tr>
<tr><td>Oligo rev</td><td>AAACGCGTCGCGGGGAGCGAAAAG</td><td></td><td></td><td></td><td></td><td></td><td></td><td></td><td></td><td></td><td></td></tr>
<tr><td>Chromosome</td><td>start</td><td>end</td><td>strand</td><td>MM</td><td>target_seq</td><td>PAM</td><td>alignment</td><td>distance</td><td>position</td><td>gene name</td><td>gene id</td></tr>
<tr><td>GU305914.1</td><td>228817</td><td>228839</td><td>-</td><td>0</td><td>CTTTTCGCTCCCCGCGACGC</td><td>CGG</td><td>||||||||[||||||||||||]PAM</td><td>NA</td><td>Intergenic</td><td>NA</td><td>NA</td></tr>
<tr><td>T17352</td><td>CGGACTCGCGATGAGCAACGCGG</td><td>1000</td><td></td><td></td><td></td><td></td><td></td><td></td><td></td><td></td><td></td></tr>
<tr><td>Oligo fwd</td><td>CACCCGGACTCGCGATGAGCAACG</td><td></td><td></td><td></td><td></td><td></td><td></td><td></td><td></td><td></td><td></td></tr>
<tr><td>Oligo rev</td><td>AAACCGTTGCTCATCGCGAGTCCG</td><td></td><td></td><td></td><td></td><td></td><td></td><td></td><td></td><td></td><td></td></tr>
<tr><td>Chromosome</td><td>start</td><td>end</td><td>strand</td><td>MM</td><td>target_seq</td><td>PAM</td><td>alignment</td><td>distance</td><td>position</td><td>gene name</td><td>gene id</td></tr>
<tr><td>GU305914.1</td><td>228797</td><td>228819</td><td>-</td><td>0</td><td>CGGACTCGCGATGAGCAACG</td><td>CGG</td><td>||||||||[||||||||||||]PAM</td><td>NA</td><td>Intergenic</td><td>NA</td><td>NA</td></tr>
<tr><td>T17353</td><td>GGTCCCGTTCGTCCCCTACGCGG</td><td>1000</td><td></td><td></td><td></td><td></td><td></td><td></td><td></td><td></td><td></td></tr>
<tr><td>Oligo fwd</td><td>CACCGGTCCCGTTCGTCCCCTACG</td><td></td><td></td><td></td><td></td><td></td><td></td><td></td><td></td><td></td><td></td></tr>
<tr><td>Oligo rev</td><td>AAACCGTAGGGGACGAACGGGACC</td><td></td><td></td><td></td><td></td><td></td><td></td><td></td><td></td><td></td><td></td></tr>
<tr><td>Chromosome</td><td>start</td><td>end</td><td>strand</td><td>MM</td><td>target_seq</td><td>PAM</td><td>alignment</td><td>distance</td><td>position</td><td>gene name</td><td>gene id</td></tr>
<tr><td>GU305914.1</td><td>228776</td><td>228798</td><td>-</td><td>0</td><td>GGTCCCGTTCGTCCCCTACG</td><td>CGG</td><td>||||||||[||||||||||||]PAM</td><td>NA</td><td>Intergenic</td><td>NA</td><td>NA</td></tr>
<tr><td>T17354</td><td>GTCCCGTTCGTCCCCTACGCGGG</td><td>1000</td><td></td><td></td><td></td><td></td><td></td><td></td><td></td><td></td><td></td></tr>
<tr><td>Oligo fwd</td><td>CACCGTCCCGTTCGTCCCCTACGC</td><td></td><td></td><td></td><td></td><td></td><td></td><td></td><td></td><td></td><td></td></tr>
<tr><td>Oligo rev</td><td>AAACGCGTAGGGGACGAACGGGAC</td><td></td><td></td><td></td><td></td><td></td><td></td><td></td><td></td><td></td><td></td></tr>
<tr><td>Chromosome</td><td>start</td><td>end</td><td>strand</td><td>MM</td><td>target_seq</td><td>PAM</td><td>alignment</td><td>distance</td><td>position</td><td>gene name</td><td>gene id</td></tr>
<tr><td>GU305914.1</td><td>228775</td><td>228797</td><td>-</td><td>0</td><td>GTCCCGTTCGTCCCCTACGC</td><td>GGG</td><td>||||||||[||||||||||||]PAM</td><td>NA</td><td>Intergenic</td><td>NA</td><td>NA</td></tr>
<tr><td>T17355</td><td>TTCGTCCCCTACGCGGGTATCGG</td><td>1000</td><td></td><td></td><td></td><td></td><td></td><td></td><td></td><td></td><td></td></tr>
<tr><td>Oligo fwd</td><td>CACCTTCGTCCCCTACGCGGGTAT</td><td></td><td></td><td></td><td></td><td></td><td></td><td></td><td></td><td></td><td></td></tr>
<tr><td>Oligo rev</td><td>AAACATACCCGCGTAGGGGACGAA</td><td></td><td></td><td></td><td></td><td></td><td></td><td></td><td></td><td></td><td></td></tr>
<tr><td>Chromosome</td><td>start</td><td>end</td><td>strand</td><td>MM</td><td>target_seq</td><td>PAM</td><td>alignment</td><td>distance</td><td>position</td><td>gene name</td><td>gene id</td></tr>
<tr><td>GU305914.1</td><td>228769</td><td>228791</td><td>-</td><td>0</td><td>TTCGTCCCCTACGCGGGTAT</td><td>CGG</td><td>||||||||[||||||||||||]PAM</td><td>NA</td><td>Intergenic</td><td>NA</td><td>NA</td></tr>
<tr><td>T17356</td><td>TCGGCCACAACCTTCGTCGCTGG</td><td>1000</td><td></td><td></td><td></td><td></td><td></td><td></td><td></td><td></td><td></td></tr>
<tr><td>Oligo fwd</td><td>CACCTCGGCCACAACCTTCGTCGC</td><td></td><td></td><td></td><td></td><td></td><td></td><td></td><td></td><td></td><td></td></tr>
<tr><td>Oligo rev</td><td>AAACGCGACGAAGGTTGTGGCCGA</td><td></td><td></td><td></td><td></td><td></td><td></td><td></td><td></td><td></td><td></td></tr>
<tr><td>Chromosome</td><td>start</td><td>end</td><td>strand</td><td>MM</td><td>target_seq</td><td>PAM</td><td>alignment</td><td>distance</td><td>position</td><td>gene name</td><td>gene id</td></tr>
<tr><td>GU305914.1</td><td>228750</td><td>228772</td><td>-</td><td>0</td><td>TCGGCCACAACCTTCGTCGC</td><td>TGG</td><td>||||||||[||||||||||||]PAM</td><td>NA</td><td>Intergenic</td><td>NA</td><td>NA</td></tr>
<tr><td>T17357</td><td>CGGCCACAACCTTCGTCGCTGGG</td><td>1000</td><td></td><td></td><td></td><td></td><td></td><td></td><td></td><td></td><td></td></tr>
<tr><td>Oligo fwd</td><td>CACCCGGCCACAACCTTCGTCGCT</td><td></td><td></td><td></td><td></td><td></td><td></td><td></td><td></td><td></td><td></td></tr>
<tr><td>Oligo rev</td><td>AAACAGCGACGAAGGTTGTGGCCG</td><td></td><td></td><td></td><td></td><td></td><td></td><td></td><td></td><td></td><td></td></tr>
<tr><td>Chromosome</td><td>start</td><td>end</td><td>strand</td><td>MM</td><td>target_seq</td><td>PAM</td><td>alignment</td><td>distance</td><td>position</td><td>gene name</td><td>gene id</td></tr>
<tr><td>GU305914.1</td><td>228749</td><td>228771</td><td>-</td><td>0</td><td>CGGCCACAACCTTCGTCGCT</td><td>GGG</td><td>||||||||[||||||||||||]PAM</td><td>NA</td><td>Intergenic</td><td>NA</td><td>NA</td></tr>
<tr><td>T17358</td><td>TTCATGTCACCGAAAGTGTTCGG</td><td>1000</td><td></td><td></td><td></td><td></td><td></td><td></td><td></td><td></td><td></td></tr>
<tr><td>Oligo fwd</td><td>CACCTTCATGTCACCGAAAGTGTT</td><td></td><td></td><td></td><td></td><td></td><td></td><td></td><td></td><td></td><td></td></tr>
<tr><td>Oligo rev</td><td>AAACAACACTTTCGGTGACATGAA</td><td></td><td></td><td></td><td></td><td></td><td></td><td></td><td></td><td></td><td></td></tr>
<tr><td>Chromosome</td><td>start</td><td>end</td><td>strand</td><td>MM</td><td>target_seq</td><td>PAM</td><td>alignment</td><td>distance</td><td>position</td><td>gene name</td><td>gene id</td></tr>
<tr><td>GU305914.1</td><td>228670</td><td>228692</td><td>-</td><td>0</td><td>TTCATGTCACCGAAAGTGTT</td><td>CGG</td><td>||||||||[||||||||||||]PAM</td><td>NA</td><td>Intergenic</td><td>NA</td><td>NA</td></tr>
<tr><td>T17359</td><td>TACGATCCCCGCGAACCCTACGG</td><td>1000</td><td></td><td></td><td></td><td></td><td></td><td></td><td></td><td></td><td></td></tr>
<tr><td>Oligo fwd</td><td>CACCTACGATCCCCGCGAACCCTA</td><td></td><td></td><td></td><td></td><td></td><td></td><td></td><td></td><td></td><td></td></tr>
<tr><td>Oligo rev</td><td>AAACTAGGGTTCGCGGGGATCGTA</td><td></td><td></td><td></td><td></td><td></td><td></td><td></td><td></td><td></td><td></td></tr>
<tr><td>Chromosome</td><td>start</td><td>end</td><td>strand</td><td>MM</td><td>target_seq</td><td>PAM</td><td>alignment</td><td>distance</td><td>position</td><td>gene name</td><td>gene id</td></tr>
<tr><td>GU305914.1</td><td>228610</td><td>228632</td><td>-</td><td>0</td><td>TACGATCCCCGCGAACCCTA</td><td>CGG</td><td>||||||||[||||||||||||]PAM</td><td>NA</td><td>Intergenic</td><td>NA</td><td>NA</td></tr>
<tr><td>T17360</td><td>ACGATCCCCGCGAACCCTACGGG</td><td>1000</td><td></td><td></td><td></td><td></td><td></td><td></td><td></td><td></td><td></td></tr>
<tr><td>Oligo fwd</td><td>CACCACGATCCCCGCGAACCCTAC</td><td></td><td></td><td></td><td></td><td></td><td></td><td></td><td></td><td></td><td></td></tr>
<tr><td>Oligo rev</td><td>AAACGTAGGGTTCGCGGGGATCGT</td><td></td><td></td><td></td><td></td><td></td><td></td><td></td><td></td><td></td><td></td></tr>
<tr><td>Chromosome</td><td>start</td><td>end</td><td>strand</td><td>MM</td><td>target_seq</td><td>PAM</td><td>alignment</td><td>distance</td><td>position</td><td>gene name</td><td>gene id</td></tr>
</table>

GU305914.1 228609 228631 - 0 ACGATCCCCGCGAACCCTAC GGG ||||||||[||||||||||||]PAM NA Intergenic NA NA

T17361 ACCCTACGGGCCTCACCCGCTGG 1000
Oligo fwd CACCACCCTACGGGCCTCACCCGC
Oligo rev AAACGCGGGTGAGGCCCGTAGGGT
Chromosome start end strand MM target_seq PAM alignment distance position gene name gene id
GU305914.1 228596 228618 - 0 ACCCTACGGGCCTCACCCGC TGG ||||||||[||||||||||||]PAM NA Intergenic NA NA

T17362 CCCTACGGGCCTCACCCGCTGGG 1000
Oligo fwd CACCCCCTACGGGCCTCACCCGCT
Oligo rev AAACAGCGGGTGAGGCCCGTAGGG
Chromosome start end strand MM target_seq PAM alignment distance position gene name gene id
GU305914.1 228595 228617 - 0 CCCTACGGGCCTCACCCGCT GGG ||||||||[||||||||||||]PAM NA Intergenic NA NA

T17363 CGCATTCTACCTCCCTGACCTGG 1000
Oligo fwd CACCCGCATTCTACCTCCCTGACC
Oligo rev AAACGGTCAGGGAGGTAGAATGCG
Chromosome start end strand MM target_seq PAM alignment distance position gene name gene id
GU305914.1 228569 228591 - 0 CGCATTCTACCTCCCTGACC TGG ||||||||[||||||||||||]PAM NA Intergenic NA NA

T17364 CGTCCTCTGTTCGCCCAGCGCGG 1000
Oligo fwd CACCCGTCCTCTGTTCGCCCAGCG
Oligo rev AAACCGCTGGGCGAACAGAGGACG
Chromosome start end strand MM target_seq PAM alignment distance position gene name gene id
GU305914.1 228496 228518 - 0 CGTCCTCTGTTCGCCCAGCG CGG ||||||||[||||||||||||]PAM NA Intergenic NA NA
GU305914.1 229801 229823 - 4 CGCCCCCTGTTCGCGCAACG GGG ||-||-||[||||||-||-||]PAM NA Intergenic NA NA

T17365 CTCTGTTCGCCCAGCGCGGTTGG 1000
Oligo fwd CACCCTCTGTTCGCCCAGCGCGGT
Oligo rev AAACACCGCGCTGGGCGAACAGAG
Chromosome start end strand MM target_seq PAM alignment distance position gene name gene id
GU305914.1 228492 228514 - 0 CTCTGTTCGCCCAGCGCGGT TGG ||||||||[||||||||||||]PAM NA Intergenic NA NA

T17366 TCAGACGTTCGCCAAGATCTTGG 1000
Oligo fwd CACCTCAGACGTTCGCCAAGATCT
Oligo rev AAACAGATCTTGGCGAACGTCTGA
Chromosome start end strand MM target_seq PAM alignment distance position gene name gene id
GU305914.1 228458 228480 - 0 TCAGACGTTCGCCAAGATCT TGG ||||||||[||||||||||||]PAM NA Intergenic NA NA

T17367 CGCCAAGATCTTGGATACGCAGG 1000
Oligo fwd CACCCGCCAAGATCTTGGATACGC
Oligo rev AAACGCGTATCCAAGATCTTGGCG
Chromosome start end strand MM target_seq PAM alignment distance position gene name gene id
GU305914.1 228449 228471 - 0 CGCCAAGATCTTGGATACGC AGG ||||||||[||||||||||||]PAM NA Intergenic NA NA

T17368 AGATCTTGGATACGCAGGAGCGG 1000
Oligo fwd CACCAGATCTTGGATACGCAGGAG
Oligo rev AAACCTCCTGCGTATCCAAGATCT
Chromosome start end strand MM target_seq PAM alignment distance position gene name gene id
GU305914.1 228444 228466 - 0 AGATCTTGGATACGCAGGAG CGG ||||||||[||||||||||||]PAM NA Intergenic NA NA

T17369 TCTTGGATACGCAGGAGCGGCGG 1000
Oligo fwd CACCTCTTGGATACGCAGGAGCGG
Oligo rev AAACCCGCTCCTGCGTATCCAAGA
Chromosome start end strand MM target_seq PAM alignment distance position gene name gene id
GU305914.1 228441 228463 - 0 TCTTGGATACGCAGGAGCGG CGG ||||||||[||||||||||||]PAM NA Intergenic NA NA

T17370 CTTGGATACGCAGGAGCGGCGGG 1000
Oligo fwd CACCCTTGGATACGCAGGAGCGGC
Oligo rev AAACGCCGCTCCTGCGTATCCAAG
Chromosome start end strand MM target_seq PAM alignment distance position gene name gene id
GU305914.1 228440 228462 - 0 CTTGGATACGCAGGAGCGGC GGG ||||||||[||||||||||||]PAM NA Intergenic NA NA

T17371 GAGCGGCGGGCCACCGAAAGAGG 1000
Oligo fwd CACCGAGCGGCGGGCCACCGAAAG
Oligo rev AAACCTTTCGGTGGCCCGCCGCTC
Chromosome start end strand MM target_seq PAM alignment distance position gene name gene id
GU305914.1 228427 228449 - 0 GAGCGGCGGGCCACCGAAAG AGG ||||||||[||||||||||||]PAM NA Intergenic NA NA

T17372 GGCGGGCCACCGAAAGAGGTAGG 1000
Oligo fwd CACCGGCGGGCCACCGAAAGAGGT
Oligo rev AAACACCTCTTTCGGTGGCCCGCC
Chromosome start end strand MM target_seq PAM alignment distance position gene name gene id
GU305914.1 228423 228445 - 0 GGCGGGCCACCGAAAGAGGT AGG ||||||||[||||||||||||]PAM NA Intergenic NA NA

T17373 GGTAGGTGCCCCCCGTGATGAGG 1000
Oligo fwd CACCGGTAGGTGCCCCCCGTGATG
Oligo rev AAACCATCACGGGGGGCACCTACC
Chromosome start end strand MM target_seq PAM alignment distance position gene name gene id
GU305914.1 228406 228428 - 0 GGTAGGTGCCCCCCGTGATG AGG ||||||||[||||||||||||]PAM NA Intergenic NA NA

T17374 GTAGGTGCCCCCCGTGATGAGGG 1000
Oligo fwd CACCGTAGGTGCCCCCCGTGATGA
Oligo rev AAACTCATCACGGGGGGCACCTAC
Chromosome start end strand MM target_seq PAM alignment distance position gene name gene id
GU305914.1 228405 228427 - 0 GTAGGTGCCCCCCGTGATGA GGG ||||||||[||||||||||||]PAM NA Intergenic NA NA

T17375 GGTGCCCCCCGTGATGAGGGAGG 1000
Oligo fwd CACCGGTGCCCCCCGTGATGAGGG

| | | | | | | | | | | | |
|---|---|---|---|---|---|---|---|---|---|---|---|
| Oligo rev | AAACCCCTCATCACGGGGGGCACC | | | | | | | | | | |
| Chromosome | start | end | strand | MM | target_seq | PAM | alignment | distance | position | gene name | gene id |
| GU305914.1 | 228402 | 228424 | - | 0 | GGTGCCCCCCGTGATGAGGG | AGG | \|\|\|\|\|\|\|\|[\|\|\|\|\|\|\|\|\|\|\|\|]PAM | NA | Intergenic | NA | NA |
| T17376 | GGGAGGCCCGACGCTAGATCTGG | 1000 | | | | | | | | | |
| Oligo fwd | CACCGGGAGGCCCGACGCTAGATC | | | | | | | | | | |
| Oligo rev | AAACGATCTAGCGTCGGGCCTCCC | | | | | | | | | | |
| Chromosome | start | end | strand | MM | target_seq | PAM | alignment | distance | position | gene name | gene id |
| GU305914.1 | 228385 | 228407 | - | 0 | GGGAGGCCCGACGCTAGATC | TGG | \|\|\|\|\|\|\|\|[\|\|\|\|\|\|\|\|\|\|\|\|]PAM | NA | Intergenic | NA | NA |
| T17377 | GCCCGACGCTAGATCTGGCTCGG | 1000 | | | | | | | | | |
| Oligo fwd | CACCGCCCGACGCTAGATCTGGCT | | | | | | | | | | |
| Oligo rev | AAACAGCCAGATCTAGCGTCGGGC | | | | | | | | | | |
| Chromosome | start | end | strand | MM | target_seq | PAM | alignment | distance | position | gene name | gene id |
| GU305914.1 | 228380 | 228402 | - | 0 | GCCCGACGCTAGATCTGGCT | CGG | \|\|\|\|\|\|\|\|[\|\|\|\|\|\|\|\|\|\|\|\|]PAM | NA | Intergenic | NA | NA |
| T17378 | TAACGTTGTTTTTGTCTGACAGG | 1000 | | | | | | | | | |
| Oligo fwd | CACCTAACGTTGTTTTTGTCTGAC | | | | | | | | | | |
| Oligo rev | AAACGTCAGACAAAAACAACGTTA | | | | | | | | | | |
| Chromosome | start | end | strand | MM | target_seq | PAM | alignment | distance | position | gene name | gene id |
| GU305914.1 | 228346 | 228368 | - | 0 | TAACGTTGTTTTTGTCTGAC | AGG | \|\|\|\|\|\|\|\|[\|\|\|\|\|\|\|\|\|\|\|\|]PAM | NA | Intergenic | NA | NA |
| T17379 | CTGACAGGTGACAACCTCTCCGG | 1000 | | | | | | | | | |
| Oligo fwd | CACCCTGACAGGTGACAACCTCTC | | | | | | | | | | |
| Oligo rev | AAACGAGAGGTTGTCACCTGTCAG | | | | | | | | | | |
| Chromosome | start | end | strand | MM | target_seq | PAM | alignment | distance | position | gene name | gene id |
| GU305914.1 | 228331 | 228353 | - | 0 | CTGACAGGTGACAACCTCTC | CGG | \|\|\|\|\|\|\|\|[\|\|\|\|\|\|\|\|\|\|\|\|]PAM | NA | Intergenic | NA | NA |
| T17380 | CAGGTGACAACCTCTCCGGCTGG | 1000 | | | | | | | | | |
| Oligo fwd | CACCCAGGTGACAACCTCTCCGGC | | | | | | | | | | |
| Oligo rev | AAACGCCGGAGAGGTTGTCACCTG | | | | | | | | | | |
| Chromosome | start | end | strand | MM | target_seq | PAM | alignment | distance | position | gene name | gene id |
| GU305914.1 | 228327 | 228349 | - | 0 | CAGGTGACAACCTCTCCGGC | TGG | \|\|\|\|\|\|\|\|[\|\|\|\|\|\|\|\|\|\|\|\|]PAM | NA | Intergenic | NA | NA |
| T17381 | CAACCTCTCCGGCTGGCTCTCGG | 1000 | | | | | | | | | |
| Oligo fwd | CACCCAACCTCTCCGGCTGGCTCT | | | | | | | | | | |
| Oligo rev | AAACAGAGCCAGCCGGAGAGGTTG | | | | | | | | | | |
| Chromosome | start | end | strand | MM | target_seq | PAM | alignment | distance | position | gene name | gene id |
| GU305914.1 | 228320 | 228342 | - | 0 | CAACCTCTCCGGCTGGCTCT | CGG | \|\|\|\|\|\|\|\|[\|\|\|\|\|\|\|\|\|\|\|\|]PAM | NA | Intergenic | NA | NA |
| T17382 | GACCGACTGCCCGTCTCCTTCGG | 1000 | | | | | | | | | |
| Oligo fwd | CACCGACCGACTGCCCGTCTCCTT | | | | | | | | | | |
| Oligo rev | AAACAAGGAGACGGGCAGTCGGTC | | | | | | | | | | |
| Chromosome | start | end | strand | MM | target_seq | PAM | alignment | distance | position | gene name | gene id |
| GU305914.1 | 228255 | 228277 | - | 0 | GACCGACTGCCCGTCTCCTT | CGG | \|\|\|\|\|\|\|\|[\|\|\|\|\|\|\|\|\|\|\|\|]PAM | NA | Intergenic | NA | NA |
| T17383 | CGGAGTAGAAACGCTATCCTCGG | 1000 | | | | | | | | | |
| Oligo fwd | CACCCGGAGTAGAAACGCTATCCT | | | | | | | | | | |
| Oligo rev | AAACAGGATAGCGTTTCTACTCCG | | | | | | | | | | |
| Chromosome | start | end | strand | MM | target_seq | PAM | alignment | distance | position | gene name | gene id |
| GU305914.1 | 228235 | 228257 | - | 0 | CGGAGTAGAAACGCTATCCT | CGG | \|\|\|\|\|\|\|\|[\|\|\|\|\|\|\|\|\|\|\|\|]PAM | NA | Intergenic | NA | NA |
| T17384 | CTCCCCCAGACGACGATCTGCGG | 1000 | | | | | | | | | |
| Oligo fwd | CACCCTCCCCCAGACGACGATCTG | | | | | | | | | | |
| Oligo rev | AAACCAGATCGTCGTCTGGGGGAG | | | | | | | | | | |
| Chromosome | start | end | strand | MM | target_seq | PAM | alignment | distance | position | gene name | gene id |
| GU305914.1 | 228176 | 228198 | - | 0 | CTCCCCCAGACGACGATCTG | CGG | \|\|\|\|\|\|\|\|[\|\|\|\|\|\|\|\|\|\|\|\|]PAM | NA | Intergenic | NA | NA |
| T17385 | GCTGTGAAAAGAAGTCCCTTCGG | 1000 | | | | | | | | | |
| Oligo fwd | CACCGCTGTGAAAAGAAGTCCCTT | | | | | | | | | | |
| Oligo rev | AAACAAGGGACTTCTTTTCACAGC | | | | | | | | | | |
| Chromosome | start | end | strand | MM | target_seq | PAM | alignment | distance | position | gene name | gene id |
| GU305914.1 | 228145 | 228167 | - | 0 | GCTGTGAAAAGAAGTCCCTT | CGG | \|\|\|\|\|\|\|\|[\|\|\|\|\|\|\|\|\|\|\|\|]PAM | NA | Intergenic | NA | NA |
| T17386 | TTCGGATGTCTGCCTGAGAACGG | 1000 | | | | | | | | | |
| Oligo fwd | CACCTTCGGATGTCTGCCTGAGAA | | | | | | | | | | |
| Oligo rev | AAACTTCTCAGGCAGACATCCGAA | | | | | | | | | | |
| Chromosome | start | end | strand | MM | target_seq | PAM | alignment | distance | position | gene name | gene id |
| GU305914.1 | 228127 | 228149 | - | 0 | TTCGGATGTCTGCCTGAGAA | CGG | \|\|\|\|\|\|\|\|[\|\|\|\|\|\|\|\|\|\|\|\|]PAM | NA | Intergenic | NA | NA |
| T17387 | CCTGAGAACGGTCTCCTCATCGG | 1000 | | | | | | | | | |
| Oligo fwd | CACCCCTGAGAACGGTCTCCTCAT | | | | | | | | | | |
| Oligo rev | AAACATGAGGAGACCGTTCTCAGG | | | | | | | | | | |
| Chromosome | start | end | strand | MM | target_seq | PAM | alignment | distance | position | gene name | gene id |
| GU305914.1 | 228115 | 228137 | - | 0 | CCTGAGAACGGTCTCCTCAT | CGG | \|\|\|\|\|\|\|\|[\|\|\|\|\|\|\|\|\|\|\|\|]PAM | NA | Intergenic | NA | NA |
| T17388 | CTGAGAACGGTCTCCTCATCGGG | 1000 | | | | | | | | | |
| Oligo fwd | CACCCTGAGAACGGTCTCCTCATC | | | | | | | | | | |
| Oligo rev | AAACGATGAGGAGACCGTTCTCAG | | | | | | | | | | |
| Chromosome | start | end | strand | MM | target_seq | PAM | alignment | distance | position | gene name | gene id |
| GU305914.1 | 228114 | 228136 | - | 0 | CTGAGAACGGTCTCCTCATC | GGG | \|\|\|\|\|\|\|\|[\|\|\|\|\|\|\|\|\|\|\|\|]PAM | NA | Intergenic | NA | NA |
| T17389 | TTCGTAGATGACCCGTCCTTAGG | 1000 | | | | | | | | | |
| Oligo fwd | CACCTTCGTAGATGACCCGTCCTT | | | | | | | | | | |
| Oligo rev | AAACAAGGACGGGTCATCTACGAA | | | | | | | | | | |
| Chromosome | start | end | strand | MM | target_seq | PAM | alignment | distance | position | gene name | gene id |
| GU305914.1 | 227989 | 228011 | - | 0 | TTCGTAGATGACCCGTCCTT | AGG | \|\|\|\|\|\|\|\|[\|\|\|\|\|\|\|\|\|\|\|\|]PAM | NA | Intergenic | NA | NA |
| T17390 | AGGTTTCGTTTCTGTGTAAACGG | 1000 | | | | | | | | | |

<table>
<tr><td>Oligo fwd</td><td>CACCAGGTTTCGTTTCTGTGTAAA</td></tr>
<tr><td>Oligo rev</td><td>AAACTTTACACAGAAACGAAACCT</td></tr>
<tr><td>Chromosome</td><td>start</td><td>end</td><td>strand</td><td>MM</td><td>target_seq</td><td>PAM</td><td>alignment</td><td>distance</td><td>position</td><td>gene name</td><td>gene id</td></tr>
<tr><td>GU305914.1</td><td>227969</td><td>227991</td><td>-</td><td>0</td><td>AGGTTTCGTTTCTGTGTAAA</td><td>CGG</td><td>||||||||[||||||||||||]PAM</td><td>NA</td><td>Intergenic</td><td>NA</td><td>NA</td></tr>
<tr><td>T17391</td><td>CTCGTGAAGTTTCGTCTTCTCGG</td><td>1000</td></tr>
<tr><td>Oligo fwd</td><td>CACCCTCGTGAAGTTTCGTCTTCT</td></tr>
<tr><td>Oligo rev</td><td>AAACAGAAGACGAAACTTCACGAG</td></tr>
<tr><td>Chromosome</td><td>start</td><td>end</td><td>strand</td><td>MM</td><td>target_seq</td><td>PAM</td><td>alignment</td><td>distance</td><td>position</td><td>gene name</td><td>gene id</td></tr>
<tr><td>GU305914.1</td><td>227946</td><td>227968</td><td>-</td><td>0</td><td>CTCGTGAAGTTTCGTCTTCT</td><td>CGG</td><td>||||||||[||||||||||||]PAM</td><td>NA</td><td>Intergenic</td><td>NA</td><td>NA</td></tr>
<tr><td>T17392</td><td>TCCGTCCGAGACCCTCATCGTGG</td><td>1000</td></tr>
<tr><td>Oligo fwd</td><td>CACCTCCGTCCGAGACCCTCATCG</td></tr>
<tr><td>Oligo rev</td><td>AAACCGATGAGGGTCTCGGACGGA</td></tr>
<tr><td>Chromosome</td><td>start</td><td>end</td><td>strand</td><td>MM</td><td>target_seq</td><td>PAM</td><td>alignment</td><td>distance</td><td>position</td><td>gene name</td><td>gene id</td></tr>
<tr><td>GU305914.1</td><td>227908</td><td>227930</td><td>-</td><td>0</td><td>TCCGTCCGAGACCCTCATCG</td><td>TGG</td><td>||||||||[||||||||||||]PAM</td><td>NA</td><td>Intergenic</td><td>NA</td><td>NA</td></tr>
<tr><td>T17393</td><td>CCTATATGTTCTAAGACCTCCGG</td><td>1000</td></tr>
<tr><td>Oligo fwd</td><td>CACCCCTATATGTTCTAAGACCTC</td></tr>
<tr><td>Oligo rev</td><td>AAACGAGGTCTTAGAACATATAGG</td></tr>
<tr><td>Chromosome</td><td>start</td><td>end</td><td>strand</td><td>MM</td><td>target_seq</td><td>PAM</td><td>alignment</td><td>distance</td><td>position</td><td>gene name</td><td>gene id</td></tr>
<tr><td>GU305914.1</td><td>227882</td><td>227904</td><td>-</td><td>0</td><td>CCTATATGTTCTAAGACCTC</td><td>CGG</td><td>||||||||[||||||||||||]PAM</td><td>NA</td><td>Intergenic</td><td>NA</td><td>NA</td></tr>
<tr><td>T17394</td><td>CTATATGTTCTAAGACCTCCGGG</td><td>1000</td></tr>
<tr><td>Oligo fwd</td><td>CACCCTATATGTTCTAAGACCTCC</td></tr>
<tr><td>Oligo rev</td><td>AAACGGAGGTCTTAGAACATATAG</td></tr>
<tr><td>Chromosome</td><td>start</td><td>end</td><td>strand</td><td>MM</td><td>target_seq</td><td>PAM</td><td>alignment</td><td>distance</td><td>position</td><td>gene name</td><td>gene id</td></tr>
<tr><td>GU305914.1</td><td>227881</td><td>227903</td><td>-</td><td>0</td><td>CTATATGTTCTAAGACCTCC</td><td>GGG</td><td>||||||||[||||||||||||]PAM</td><td>NA</td><td>Intergenic</td><td>NA</td><td>NA</td></tr>
<tr><td>T17395</td><td>GTTCTAAGACCTCCGGGTCGCGG</td><td>1000</td></tr>
<tr><td>Oligo fwd</td><td>CACCGTTCTAAGACCTCCGGGTCG</td></tr>
<tr><td>Oligo rev</td><td>AAACCGACCCGGAGGTCTTAGAAC</td></tr>
<tr><td>Chromosome</td><td>start</td><td>end</td><td>strand</td><td>MM</td><td>target_seq</td><td>PAM</td><td>alignment</td><td>distance</td><td>position</td><td>gene name</td><td>gene id</td></tr>
<tr><td>GU305914.1</td><td>227875</td><td>227897</td><td>-</td><td>0</td><td>GTTCTAAGACCTCCGGGTCG</td><td>CGG</td><td>||||||||[||||||||||||]PAM</td><td>NA</td><td>Intergenic</td><td>NA</td><td>NA</td></tr>
<tr><td>T17396</td><td>AAATCCTCACCGTCAACCGTCGG</td><td>1000</td></tr>
<tr><td>Oligo fwd</td><td>CACCAAATCCTCACCGTCAACCGT</td></tr>
<tr><td>Oligo rev</td><td>AAACACGGTTGACGGTGAGGATTT</td></tr>
<tr><td>Chromosome</td><td>start</td><td>end</td><td>strand</td><td>MM</td><td>target_seq</td><td>PAM</td><td>alignment</td><td>distance</td><td>position</td><td>gene name</td><td>gene id</td></tr>
<tr><td>GU305914.1</td><td>227837</td><td>227859</td><td>-</td><td>0</td><td>AAATCCTCACCGTCAACCGT</td><td>CGG</td><td>||||||||[||||||||||||]PAM</td><td>NA</td><td>Intergenic</td><td>NA</td><td>NA</td></tr>
<tr><td>T17397</td><td>CCTCACCGTCAACCGTCGGCTGG</td><td>1000</td></tr>
<tr><td>Oligo fwd</td><td>CACCCCTCACCGTCAACCGTCGGC</td></tr>
<tr><td>Oligo rev</td><td>AAACGCCGACGGTTGACGGTGAGG</td></tr>
<tr><td>Chromosome</td><td>start</td><td>end</td><td>strand</td><td>MM</td><td>target_seq</td><td>PAM</td><td>alignment</td><td>distance</td><td>position</td><td>gene name</td><td>gene id</td></tr>
<tr><td>GU305914.1</td><td>227833</td><td>227855</td><td>-</td><td>0</td><td>CCTCACCGTCAACCGTCGGC</td><td>TGG</td><td>||||||||[||||||||||||]PAM</td><td>NA</td><td>Intergenic</td><td>NA</td><td>NA</td></tr>
<tr><td>T17398</td><td>CAACCGTCGGCTGGTCCGATCGG</td><td>1000</td></tr>
<tr><td>Oligo fwd</td><td>CACCCAACCGTCGGCTGGTCCGAT</td></tr>
<tr><td>Oligo rev</td><td>AAACATCGGACCAGCCGACGGTTG</td></tr>
<tr><td>Chromosome</td><td>start</td><td>end</td><td>strand</td><td>MM</td><td>target_seq</td><td>PAM</td><td>alignment</td><td>distance</td><td>position</td><td>gene name</td><td>gene id</td></tr>
<tr><td>GU305914.1</td><td>227824</td><td>227846</td><td>-</td><td>0</td><td>CAACCGTCGGCTGGTCCGAT</td><td>CGG</td><td>||||||||[||||||||||||]PAM</td><td>NA</td><td>Intergenic</td><td>NA</td><td>NA</td></tr>
<tr><td>T17399</td><td>CGGCTGGTCCGATCGGTCGACGG</td><td>1000</td></tr>
<tr><td>Oligo fwd</td><td>CACCCGGCTGGTCCGATCGGTCGA</td></tr>
<tr><td>Oligo rev</td><td>AAACTCGACCGATCGGACCAGCCG</td></tr>
<tr><td>Chromosome</td><td>start</td><td>end</td><td>strand</td><td>MM</td><td>target_seq</td><td>PAM</td><td>alignment</td><td>distance</td><td>position</td><td>gene name</td><td>gene id</td></tr>
<tr><td>GU305914.1</td><td>227817</td><td>227839</td><td>-</td><td>0</td><td>CGGCTGGTCCGATCGGTCGA</td><td>CGG</td><td>||||||||[||||||||||||]PAM</td><td>NA</td><td>Intergenic</td><td>NA</td><td>NA</td></tr>
<tr><td>T17400</td><td>TCGCCCTTCGTCACATCGAACGG</td><td>1000</td></tr>
<tr><td>Oligo fwd</td><td>CACCTCGCCCTTCGTCACATCGAA</td></tr>
<tr><td>Oligo rev</td><td>AAACTTCGATGTGACGAAGGGCGA</td></tr>
<tr><td>Chromosome</td><td>start</td><td>end</td><td>strand</td><td>MM</td><td>target_seq</td><td>PAM</td><td>alignment</td><td>distance</td><td>position</td><td>gene name</td><td>gene id</td></tr>
<tr><td>GU305914.1</td><td>227663</td><td>227685</td><td>-</td><td>0</td><td>TCGCCCTTCGTCACATCGAA</td><td>CGG</td><td>||||||||[||||||||||||]PAM</td><td>NA</td><td>Intergenic</td><td>NA</td><td>NA</td></tr>
<tr><td>T17401</td><td>TCGAACGGTCAGTTCGTTCCCGG</td><td>1000</td></tr>
<tr><td>Oligo fwd</td><td>CACCTCGAACGGTCAGTTCGTTCC</td></tr>
<tr><td>Oligo rev</td><td>AAACGGAACGAACTGACCGTTCGA</td></tr>
<tr><td>Chromosome</td><td>start</td><td>end</td><td>strand</td><td>MM</td><td>target_seq</td><td>PAM</td><td>alignment</td><td>distance</td><td>position</td><td>gene name</td><td>gene id</td></tr>
<tr><td>GU305914.1</td><td>227648</td><td>227670</td><td>-</td><td>0</td><td>TCGAACGGTCAGTTCGTTCC</td><td>CGG</td><td>||||||||[||||||||||||]PAM</td><td>NA</td><td>Intergenic</td><td>NA</td><td>NA</td></tr>
<tr><td>T17402</td><td>CGAACGGTCAGTTCGTTCCCGGG</td><td>1000</td></tr>
<tr><td>Oligo fwd</td><td>CACCCGAACGGTCAGTTCGTTCCC</td></tr>
<tr><td>Oligo rev</td><td>AAACGGGAACGAACTGACCGTTCG</td></tr>
<tr><td>Chromosome</td><td>start</td><td>end</td><td>strand</td><td>MM</td><td>target_seq</td><td>PAM</td><td>alignment</td><td>distance</td><td>position</td><td>gene name</td><td>gene id</td></tr>
<tr><td>GU305914.1</td><td>227647</td><td>227669</td><td>-</td><td>0</td><td>CGAACGGTCAGTTCGTTCCC</td><td>GGG</td><td>||||||||[||||||||||||]PAM</td><td>NA</td><td>Intergenic</td><td>NA</td><td>NA</td></tr>
<tr><td>T17403</td><td>GAACGGTCAGTTCGTTCCCGGGG</td><td>1000</td></tr>
<tr><td>Oligo fwd</td><td>CACCGAACGGTCAGTTCGTTCCCG</td></tr>
<tr><td>Oligo rev</td><td>AAACCGGGAACGAACTGACCGTTC</td></tr>
<tr><td>Chromosome</td><td>start</td><td>end</td><td>strand</td><td>MM</td><td>target_seq</td><td>PAM</td><td>alignment</td><td>distance</td><td>position</td><td>gene name</td><td>gene id</td></tr>
<tr><td>GU305914.1</td><td>227646</td><td>227668</td><td>-</td><td>0</td><td>GAACGGTCAGTTCGTTCCCG</td><td>GGG</td><td>||||||||[||||||||||||]PAM</td><td>NA</td><td>Intergenic</td><td>NA</td><td>NA</td></tr>
<tr><td>T17404</td><td>GGTTGTTTTTCTGTTCCGAGCGG</td><td>1000</td></tr>
<tr><td>Oligo fwd</td><td>CACCGGTTGTTTTTCTGTTCCGAG</td></tr>
<tr><td>Oligo rev</td><td>AAACCTCGGAACAGAAAAACAACC</td></tr>
<tr><td>Chromosome</td><td>start</td><td>end</td><td>strand</td><td>MM</td><td>target_seq</td><td>PAM</td><td>alignment</td><td>distance</td><td>position</td><td>gene name</td><td>gene id</td></tr>
<tr><td>GU305914.1</td><td>227625</td><td>227647</td><td>-</td><td>0</td><td>GGTTGTTTTTCTGTTCCGAG</td><td>CGG</td><td>||||||||[||||||||||||]PAM</td><td>NA</td><td>Intergenic</td><td>NA</td><td>NA</td></tr>
</table>

T17405 TTTCTGTTCCGAGCGGACGTCGG 1000
Oligo fwd CACCTTTCTGTTCCGAGCGGACGT
Oligo rev AAACACGTCCGCTCGGAACAGAAA
Chromosome start end strand MM target_seq PAM alignment distance position gene name gene id
GU305914.1 227618 227640 - 0 TTTCTGTTCCGAGCGGACGT CGG ||||||||[||||||||||||]PAM NA Intergenic NA NA

T17406 TTCTGTTCCGAGCGGACGTCGGG 1000
Oligo fwd CACCTTCTGTTCCGAGCGGACGTC
Oligo rev AAACGACGTCCGCTCGGAACAGAA
Chromosome start end strand MM target_seq PAM alignment distance position gene name gene id
GU305914.1 227617 227639 - 0 TTCTGTTCCGAGCGGACGTC GGG ||||||||[||||||||||||]PAM NA Intergenic NA NA

T17407 TCTGTTCCGAGCGGACGTCGGGG 1000
Oligo fwd CACCTCTGTTCCGAGCGGACGTCG
Oligo rev AAACCGACGTCCGCTCGGAACAGA
Chromosome start end strand MM target_seq PAM alignment distance position gene name gene id
GU305914.1 227616 227638 - 0 TCTGTTCCGAGCGGACGTCG GGG ||||||||[||||||||||||]PAM NA Intergenic NA NA

T17408 CTGTTCCGAGCGGACGTCGGGGG 1000
Oligo fwd CACCCTGTTCCGAGCGGACGTCGG
Oligo rev AAACCCGACGTCCGCTCGGAACAG
Chromosome start end strand MM target_seq PAM alignment distance position gene name gene id
GU305914.1 227615 227637 - 0 CTGTTCCGAGCGGACGTCGG GGG ||||||||[||||||||||||]PAM NA Intergenic NA NA

T17409 TGTTCCGAGCGGACGTCGGGGGG 1000
Oligo fwd CACCTGTTCCGAGCGGACGTCGGG
Oligo rev AAACCCCGACGTCCGCTCGGAACA
Chromosome start end strand MM target_seq PAM alignment distance position gene name gene id
GU305914.1 227614 227636 - 0 TGTTCCGAGCGGACGTCGGG GGG ||||||||[||||||||||||]PAM NA Intergenic NA NA

T17410 CCGAGCGGACGTCGGGGGGTCGG 1000
Oligo fwd CACCCCGAGCGGACGTCGGGGGGT
Oligo rev AAACACCCCCCGACGTCCGCTCGG
Chromosome start end strand MM target_seq PAM alignment distance position gene name gene id
GU305914.1 227610 227632 - 0 CCGAGCGGACGTCGGGGGGT CGG ||||||||[||||||||||||]PAM NA Intergenic NA NA

T17411 GTCGGGGGGTCGGTCTCATCCGG 1000
Oligo fwd CACCGTCGGGGGGTCGGTCTCATC
Oligo rev AAACGATGAGACCGACCCCCCGAC
Chromosome start end strand MM target_seq PAM alignment distance position gene name gene id
GU305914.1 227600 227622 - 0 GTCGGGGGGTCGGTCTCATC CGG ||||||||[||||||||||||]PAM NA Intergenic NA NA

T17412 TCGGGGGGTCGGTCTCATCCGGG 1000
Oligo fwd CACCTCGGGGGGTCGGTCTCATCC
Oligo rev AAACGGATGAGACCGACCCCCCGA
Chromosome start end strand MM target_seq PAM alignment distance position gene name gene id
GU305914.1 227599 227621 - 0 TCGGGGGGTCGGTCTCATCC GGG ||||||||[||||||||||||]PAM NA Intergenic NA NA

T17413 CAATGATTTGACATCTCGTCTGG 1000
Oligo fwd CACCCAATGATTTGACATCTCGTC
Oligo rev AAACGACGAGATGTCAAATCATTG
Chromosome start end strand MM target_seq PAM alignment distance position gene name gene id
GU305914.1 227517 227539 - 0 CAATGATTTGACATCTCGTC TGG ||||||||[||||||||||||]PAM NA Intergenic NA NA

T17414 TCTGGTACATCGTGTTTTTCTGG 1000
Oligo fwd CACCTCTGGTACATCGTGTTTTTC
Oligo rev AAACGAAAAACACGATGTACCAGA
Chromosome start end strand MM target_seq PAM alignment distance position gene name gene id
GU305914.1 227499 227521 - 0 TCTGGTACATCGTGTTTTTC TGG ||||||||[||||||||||||]PAM NA Intergenic NA NA

T17415 CTGGTACATCGTGTTTTTCTGGG 1000
Oligo fwd CACCCTGGTACATCGTGTTTTTCT
Oligo rev AAACAGAAAAACACGATGTACCAG
Chromosome start end strand MM target_seq PAM alignment distance position gene name gene id
GU305914.1 227498 227520 - 0 CTGGTACATCGTGTTTTTCT GGG ||||||||[||||||||||||]PAM NA Intergenic NA NA

T17416 GGGTGCTTACCCGCGTCACATGG 1000
Oligo fwd CACCGGGTGCTTACCCGCGTCACA
Oligo rev AAACTGTGACGCGGGTAAGCACCC
Chromosome start end strand MM target_seq PAM alignment distance position gene name gene id
GU305914.1 227478 227500 - 0 GGGTGCTTACCCGCGTCACA TGG ||||||||[||||||||||||]PAM NA Intergenic NA NA

T17417 TGCTTACCCGCGTCACATGGAGG 1000
Oligo fwd CACCTGCTTACCCGCGTCACATGG
Oligo rev AAACCCATGTGACGCGGGTAAGCA
Chromosome start end strand MM target_seq PAM alignment distance position gene name gene id
GU305914.1 227475 227497 - 0 TGCTTACCCGCGTCACATGG AGG ||||||||[||||||||||||]PAM NA Intergenic NA NA

T17418 GCTTACCCGCGTCACATGGAGGG 1000
Oligo fwd CACCGCTTACCCGCGTCACATGGA
Oligo rev AAACTCCATGTGACGCGGGTAAGC
Chromosome start end strand MM target_seq PAM alignment distance position gene name gene id
GU305914.1 227474 227496 - 0 GCTTACCCGCGTCACATGGA GGG ||||||||[||||||||||||]PAM NA Intergenic NA NA

T17419 CTTACCCGCGTCACATGGAGGGG 1000
Oligo fwd CACCCTTACCCGCGTCACATGGAG
Oligo rev AAACCTCCATGTGACGCGGGTAAG
Chromosome start end strand MM target_seq PAM alignment distance position gene name gene id
GU305914.1 227473 227495 - 0 CTTACCCGCGTCACATGGAG GGG ||||||||[||||||||||||]PAM NA Intergenic NA NA

<table>
<tr><td>T17420</td><td>ACCCGCGTCACATGGAGGGGTGG</td><td>1000</td><td></td><td></td><td></td><td></td><td></td><td></td><td></td><td></td><td></td></tr>
<tr><td>Oligo fwd</td><td>CACCACCCGCGTCACATGGAGGGG</td><td></td><td></td><td></td><td></td><td></td><td></td><td></td><td></td><td></td><td></td></tr>
<tr><td>Oligo rev</td><td>AAACCCCCTCCATGTGACGCGGGT</td><td></td><td></td><td></td><td></td><td></td><td></td><td></td><td></td><td></td><td></td></tr>
<tr><td>Chromosome</td><td>start</td><td>end</td><td>strand</td><td>MM</td><td>target_seq</td><td>PAM</td><td>alignment</td><td>distance</td><td>position</td><td>gene name</td><td>gene id</td></tr>
<tr><td>GU305914.1</td><td>227470</td><td>227492</td><td>-</td><td>0</td><td>ACCCGCGTCACATGGAGGGG</td><td>TGG</td><td>||||||||[||||||||||||]PAM</td><td>NA</td><td>Intergenic</td><td>NA</td><td>NA</td></tr>
<tr><td>T17421</td><td>CCCGCGTCACATGGAGGGGTGGG</td><td>1000</td><td></td><td></td><td></td><td></td><td></td><td></td><td></td><td></td><td></td></tr>
<tr><td>Oligo fwd</td><td>CACCCCCGCGTCACATGGAGGGGT</td><td></td><td></td><td></td><td></td><td></td><td></td><td></td><td></td><td></td><td></td></tr>
<tr><td>Oligo rev</td><td>AAACACCCCTCCATGTGACGCGGG</td><td></td><td></td><td></td><td></td><td></td><td></td><td></td><td></td><td></td><td></td></tr>
<tr><td>Chromosome</td><td>start</td><td>end</td><td>strand</td><td>MM</td><td>target_seq</td><td>PAM</td><td>alignment</td><td>distance</td><td>position</td><td>gene name</td><td>gene id</td></tr>
<tr><td>GU305914.1</td><td>227469</td><td>227491</td><td>-</td><td>0</td><td>CCCGCGTCACATGGAGGGGT</td><td>GGG</td><td>||||||||[||||||||||||]PAM</td><td>NA</td><td>Intergenic</td><td>NA</td><td>NA</td></tr>
<tr><td>T17422</td><td>GTCACATGGAGGGGTGGGTTCGG</td><td>1000</td><td></td><td></td><td></td><td></td><td></td><td></td><td></td><td></td><td></td></tr>
<tr><td>Oligo fwd</td><td>CACCGTCACATGGAGGGGTGGGTT</td><td></td><td></td><td></td><td></td><td></td><td></td><td></td><td></td><td></td><td></td></tr>
<tr><td>Oligo rev</td><td>AAACAACCCACCCCTCCATGTGAC</td><td></td><td></td><td></td><td></td><td></td><td></td><td></td><td></td><td></td><td></td></tr>
<tr><td>Chromosome</td><td>start</td><td>end</td><td>strand</td><td>MM</td><td>target_seq</td><td>PAM</td><td>alignment</td><td>distance</td><td>position</td><td>gene name</td><td>gene id</td></tr>
<tr><td>GU305914.1</td><td>227464</td><td>227486</td><td>-</td><td>0</td><td>GTCACATGGAGGGGTGGGTT</td><td>CGG</td><td>||||||||[||||||||||||]PAM</td><td>NA</td><td>Intergenic</td><td>NA</td><td>NA</td></tr>
<tr><td>T17423</td><td>CATGGAGGGGTGGGTTCGGCAGG</td><td>1000</td><td></td><td></td><td></td><td></td><td></td><td></td><td></td><td></td><td></td></tr>
<tr><td>Oligo fwd</td><td>CACCCATGGAGGGGTGGGTTCGGC</td><td></td><td></td><td></td><td></td><td></td><td></td><td></td><td></td><td></td><td></td></tr>
<tr><td>Oligo rev</td><td>AAACGCCGAACCCACCCCTCCATG</td><td></td><td></td><td></td><td></td><td></td><td></td><td></td><td></td><td></td><td></td></tr>
<tr><td>Chromosome</td><td>start</td><td>end</td><td>strand</td><td>MM</td><td>target_seq</td><td>PAM</td><td>alignment</td><td>distance</td><td>position</td><td>gene name</td><td>gene id</td></tr>
<tr><td>GU305914.1</td><td>227460</td><td>227482</td><td>-</td><td>0</td><td>CATGGAGGGGTGGGTTCGGC</td><td>AGG</td><td>||||||||[||||||||||||]PAM</td><td>NA</td><td>Intergenic</td><td>NA</td><td>NA</td></tr>
<tr><td>T17424</td><td>GGGTTCGGCAGGCCATCCTGTGG</td><td>1000</td><td></td><td></td><td></td><td></td><td></td><td></td><td></td><td></td><td></td></tr>
<tr><td>Oligo fwd</td><td>CACCGGGTTCGGCAGGCCATCCTG</td><td></td><td></td><td></td><td></td><td></td><td></td><td></td><td></td><td></td><td></td></tr>
<tr><td>Oligo rev</td><td>AAACCAGGATGGCCTGCCGAACCC</td><td></td><td></td><td></td><td></td><td></td><td></td><td></td><td></td><td></td><td></td></tr>
<tr><td>Chromosome</td><td>start</td><td>end</td><td>strand</td><td>MM</td><td>target_seq</td><td>PAM</td><td>alignment</td><td>distance</td><td>position</td><td>gene name</td><td>gene id</td></tr>
<tr><td>GU305914.1</td><td>227449</td><td>227471</td><td>-</td><td>0</td><td>GGGTTCGGCAGGCCATCCTG</td><td>TGG</td><td>||||||||[||||||||||||]PAM</td><td>NA</td><td>Intergenic</td><td>NA</td><td>NA</td></tr>
<tr><td>T17425</td><td>GGTTCGGCAGGCCATCCTGTGGG</td><td>1000</td><td></td><td></td><td></td><td></td><td></td><td></td><td></td><td></td><td></td></tr>
<tr><td>Oligo fwd</td><td>CACCGGTTCGGCAGGCCATCCTGT</td><td></td><td></td><td></td><td></td><td></td><td></td><td></td><td></td><td></td><td></td></tr>
<tr><td>Oligo rev</td><td>AAACACAGGATGGCCTGCCGAACC</td><td></td><td></td><td></td><td></td><td></td><td></td><td></td><td></td><td></td><td></td></tr>
<tr><td>Chromosome</td><td>start</td><td>end</td><td>strand</td><td>MM</td><td>target_seq</td><td>PAM</td><td>alignment</td><td>distance</td><td>position</td><td>gene name</td><td>gene id</td></tr>
<tr><td>GU305914.1</td><td>227448</td><td>227470</td><td>-</td><td>0</td><td>GGTTCGGCAGGCCATCCTGT</td><td>GGG</td><td>||||||||[||||||||||||]PAM</td><td>NA</td><td>Intergenic</td><td>NA</td><td>NA</td></tr>
<tr><td>T17426</td><td>CAGGCCATCCTGTGGGTACAAGG</td><td>1000</td><td></td><td></td><td></td><td></td><td></td><td></td><td></td><td></td><td></td></tr>
<tr><td>Oligo fwd</td><td>CACCCAGGCCATCCTGTGGGTACA</td><td></td><td></td><td></td><td></td><td></td><td></td><td></td><td></td><td></td><td></td></tr>
<tr><td>Oligo rev</td><td>AAACTGTACCCACAGGATGGCCTG</td><td></td><td></td><td></td><td></td><td></td><td></td><td></td><td></td><td></td><td></td></tr>
<tr><td>Chromosome</td><td>start</td><td>end</td><td>strand</td><td>MM</td><td>target_seq</td><td>PAM</td><td>alignment</td><td>distance</td><td>position</td><td>gene name</td><td>gene id</td></tr>
<tr><td>GU305914.1</td><td>227441</td><td>227463</td><td>-</td><td>0</td><td>CAGGCCATCCTGTGGGTACA</td><td>AGG</td><td>||||||||[||||||||||||]PAM</td><td>NA</td><td>Intergenic</td><td>NA</td><td>NA</td></tr>
<tr><td>T17427</td><td>TCCTGTGGGTACAAGGTTCCTGG</td><td>1000</td><td></td><td></td><td></td><td></td><td></td><td></td><td></td><td></td><td></td></tr>
<tr><td>Oligo fwd</td><td>CACCTCCTGTGGGTACAAGGTTCC</td><td></td><td></td><td></td><td></td><td></td><td></td><td></td><td></td><td></td><td></td></tr>
<tr><td>Oligo rev</td><td>AAACGGAACCTTGTACCCACAGGA</td><td></td><td></td><td></td><td></td><td></td><td></td><td></td><td></td><td></td><td></td></tr>
<tr><td>Chromosome</td><td>start</td><td>end</td><td>strand</td><td>MM</td><td>target_seq</td><td>PAM</td><td>alignment</td><td>distance</td><td>position</td><td>gene name</td><td>gene id</td></tr>
<tr><td>GU305914.1</td><td>227434</td><td>227456</td><td>-</td><td>0</td><td>TCCTGTGGGTACAAGGTTCC</td><td>TGG</td><td>||||||||[||||||||||||]PAM</td><td>NA</td><td>Intergenic</td><td>NA</td><td>NA</td></tr>
<tr><td>T17428</td><td>GGGTACAAGGTTCCTGGCACCGG</td><td>1000</td><td></td><td></td><td></td><td></td><td></td><td></td><td></td><td></td><td></td></tr>
<tr><td>Oligo fwd</td><td>CACCGGGTACAAGGTTCCTGGCAC</td><td></td><td></td><td></td><td></td><td></td><td></td><td></td><td></td><td></td><td></td></tr>
<tr><td>Oligo rev</td><td>AAACGTGCCAGGAACCTTGTACCC</td><td></td><td></td><td></td><td></td><td></td><td></td><td></td><td></td><td></td><td></td></tr>
<tr><td>Chromosome</td><td>start</td><td>end</td><td>strand</td><td>MM</td><td>target_seq</td><td>PAM</td><td>alignment</td><td>distance</td><td>position</td><td>gene name</td><td>gene id</td></tr>
<tr><td>GU305914.1</td><td>227428</td><td>227450</td><td>-</td><td>0</td><td>GGGTACAAGGTTCCTGGCAC</td><td>CGG</td><td>||||||||[||||||||||||]PAM</td><td>NA</td><td>Intergenic</td><td>NA</td><td>NA</td></tr>
<tr><td>T17429</td><td>GGTACAAGGTTCCTGGCACCGGG</td><td>1000</td><td></td><td></td><td></td><td></td><td></td><td></td><td></td><td></td><td></td></tr>
<tr><td>Oligo fwd</td><td>CACCGGTACAAGGTTCCTGGCACC</td><td></td><td></td><td></td><td></td><td></td><td></td><td></td><td></td><td></td><td></td></tr>
<tr><td>Oligo rev</td><td>AAACGGTGCCAGGAACCTTGTACC</td><td></td><td></td><td></td><td></td><td></td><td></td><td></td><td></td><td></td><td></td></tr>
<tr><td>Chromosome</td><td>start</td><td>end</td><td>strand</td><td>MM</td><td>target_seq</td><td>PAM</td><td>alignment</td><td>distance</td><td>position</td><td>gene name</td><td>gene id</td></tr>
<tr><td>GU305914.1</td><td>227427</td><td>227449</td><td>-</td><td>0</td><td>GGTACAAGGTTCCTGGCACC</td><td>GGG</td><td>||||||||[||||||||||||]PAM</td><td>NA</td><td>Intergenic</td><td>NA</td><td>NA</td></tr>
<tr><td>T17430</td><td>AAGGTTCCTGGCACCGGGCTTGG</td><td>1000</td><td></td><td></td><td></td><td></td><td></td><td></td><td></td><td></td><td></td></tr>
<tr><td>Oligo fwd</td><td>CACCAAGGTTCCTGGCACCGGGCT</td><td></td><td></td><td></td><td></td><td></td><td></td><td></td><td></td><td></td><td></td></tr>
<tr><td>Oligo rev</td><td>AAACAGCCCGGTGCCAGGAACCTT</td><td></td><td></td><td></td><td></td><td></td><td></td><td></td><td></td><td></td><td></td></tr>
<tr><td>Chromosome</td><td>start</td><td>end</td><td>strand</td><td>MM</td><td>target_seq</td><td>PAM</td><td>alignment</td><td>distance</td><td>position</td><td>gene name</td><td>gene id</td></tr>
<tr><td>GU305914.1</td><td>227422</td><td>227444</td><td>-</td><td>0</td><td>AAGGTTCCTGGCACCGGGCT</td><td>TGG</td><td>||||||||[||||||||||||]PAM</td><td>NA</td><td>Intergenic</td><td>NA</td><td>NA</td></tr>
<tr><td>T17431</td><td>TCTCGAGAGCCAAGACCTCCTGG</td><td>1000</td><td></td><td></td><td></td><td></td><td></td><td></td><td></td><td></td><td></td></tr>
<tr><td>Oligo fwd</td><td>CACCTCTCGAGAGCCAAGACCTCC</td><td></td><td></td><td></td><td></td><td></td><td></td><td></td><td></td><td></td><td></td></tr>
<tr><td>Oligo rev</td><td>AAACGGAGGTCTTGGCTCTCGAGA</td><td></td><td></td><td></td><td></td><td></td><td></td><td></td><td></td><td></td><td></td></tr>
<tr><td>Chromosome</td><td>start</td><td>end</td><td>strand</td><td>MM</td><td>target_seq</td><td>PAM</td><td>alignment</td><td>distance</td><td>position</td><td>gene name</td><td>gene id</td></tr>
<tr><td>GU305914.1</td><td>227398</td><td>227420</td><td>-</td><td>0</td><td>TCTCGAGAGCCAAGACCTCC</td><td>TGG</td><td>||||||||[||||||||||||]PAM</td><td>NA</td><td>Intergenic</td><td>NA</td><td>NA</td></tr>
<tr><td>T17432</td><td>CTCGAGAGCCAAGACCTCCTGGG</td><td>1000</td><td></td><td></td><td></td><td></td><td></td><td></td><td></td><td></td><td></td></tr>
<tr><td>Oligo fwd</td><td>CACCCTCGAGAGCCAAGACCTCCT</td><td></td><td></td><td></td><td></td><td></td><td></td><td></td><td></td><td></td><td></td></tr>
<tr><td>Oligo rev</td><td>AAACAGGAGGTCTTGGCTCTCGAG</td><td></td><td></td><td></td><td></td><td></td><td></td><td></td><td></td><td></td><td></td></tr>
<tr><td>Chromosome</td><td>start</td><td>end</td><td>strand</td><td>MM</td><td>target_seq</td><td>PAM</td><td>alignment</td><td>distance</td><td>position</td><td>gene name</td><td>gene id</td></tr>
<tr><td>GU305914.1</td><td>227397</td><td>227419</td><td>-</td><td>0</td><td>CTCGAGAGCCAAGACCTCCT</td><td>GGG</td><td>||||||||[||||||||||||]PAM</td><td>NA</td><td>Intergenic</td><td>NA</td><td>NA</td></tr>
<tr><td>T17433</td><td>TCCTGGGAAAGCGACCTCTCTGG</td><td>1000</td><td></td><td></td><td></td><td></td><td></td><td></td><td></td><td></td><td></td></tr>
<tr><td>Oligo fwd</td><td>CACCTCCTGGGAAAGCGACCTCTC</td><td></td><td></td><td></td><td></td><td></td><td></td><td></td><td></td><td></td><td></td></tr>
<tr><td>Oligo rev</td><td>AAACGAGAGGTCGCTTTCCCAGGA</td><td></td><td></td><td></td><td></td><td></td><td></td><td></td><td></td><td></td><td></td></tr>
<tr><td>Chromosome</td><td>start</td><td>end</td><td>strand</td><td>MM</td><td>target_seq</td><td>PAM</td><td>alignment</td><td>distance</td><td>position</td><td>gene name</td><td>gene id</td></tr>
<tr><td>GU305914.1</td><td>227381</td><td>227403</td><td>-</td><td>0</td><td>TCCTGGGAAAGCGACCTCTC</td><td>TGG</td><td>||||||||[||||||||||||]PAM</td><td>NA</td><td>Intergenic</td><td>NA</td><td>NA</td></tr>
<tr><td>T17434</td><td>CTCTGGCGCCGACGAACCATCGG</td><td>1000</td><td></td><td></td><td></td><td></td><td></td><td></td><td></td><td></td><td></td></tr>
<tr><td>Oligo fwd</td><td>CACCCTCTGGCGCCGACGAACCAT</td><td></td><td></td><td></td><td></td><td></td><td></td><td></td><td></td><td></td><td></td></tr>
<tr><td>Oligo rev</td><td>AAACATGGTTCGTCGGCGCCAGAG</td><td></td><td></td><td></td><td></td><td></td><td></td><td></td><td></td><td></td><td></td></tr>
<tr><td>Chromosome</td><td>start</td><td>end</td><td>strand</td><td>MM</td><td>target_seq</td><td>PAM</td><td>alignment</td><td>distance</td><td>position</td><td>gene name</td><td>gene id</td></tr>
</table>

GU305914.1 227364 227386 - 0 CTCTGGCGCCGACGAACCAT CGG ||||||||[||||||||||||]PAM NA Intergenic NA NA

T17435 TCTGGCGCCGACGAACCATCGGG 1000
Oligo fwd CACCTCTGGCGCCGACGAACCATC
Oligo rev AAACGATGGTTCGTCGGCGCCAGA
Chromosome start end strand MM target_seq PAM alignment distance position gene name gene id
GU305914.1 227363 227385 - 0 TCTGGCGCCGACGAACCATC GGG ||||||||[||||||||||||]PAM NA Intergenic NA NA

T17436 GGCGCCGACGAACCATCGGGAGG 1000
Oligo fwd CACCGGCGCCGACGAACCATCGGG
Oligo rev AAACCCCGATGGTTCGTCGGCGCC
Chromosome start end strand MM target_seq PAM alignment distance position gene name gene id
GU305914.1 227360 227382 - 0 GGCGCCGACGAACCATCGGG AGG ||||||||[||||||||||||]PAM NA Intergenic NA NA

T17437 CCATCGGGAGGACAATAACTTGG 1000
Oligo fwd CACCCCATCGGGAGGACAATAACT
Oligo rev AAACAGTTATTGTCCTCCCGATGG
Chromosome start end strand MM target_seq PAM alignment distance position gene name gene id
GU305914.1 227348 227370 - 0 CCATCGGGAGGACAATAACT TGG ||||||||[||||||||||||]PAM NA Intergenic NA NA

T17438 CAATAACTTGGCGTGAGTAACGG 1000
Oligo fwd CACCCAATAACTTGGCGTGAGTAA
Oligo rev AAACTTACTCACGCCAAGTTATTG
Chromosome start end strand MM target_seq PAM alignment distance position gene name gene id
GU305914.1 227336 227358 - 0 CAATAACTTGGCGTGAGTAA CGG ||||||||[||||||||||||]PAM NA Intergenic NA NA

T17439 GAGTAACGGTCGAACGACGACGG 1000
Oligo fwd CACCGAGTAACGGTCGAACGACGA
Oligo rev AAACTCGTCGTTCGACCGTTACTC
Chromosome start end strand MM target_seq PAM alignment distance position gene name gene id
GU305914.1 227322 227344 - 0 GAGTAACGGTCGAACGACGA CGG ||||||||[||||||||||||]PAM NA Intergenic NA NA

T17440 GACGGTTTCCCGTCTCGTTCCGG 1000
Oligo fwd CACCGACGGTTTCCCGTCTCGTTC
Oligo rev AAACGAACGAGACGGGAAACCGTC
Chromosome start end strand MM target_seq PAM alignment distance position gene name gene id
GU305914.1 227304 227326 - 0 GACGGTTTCCCGTCTCGTTC CGG ||||||||[||||||||||||]PAM NA Intergenic NA NA

T17441 TCCCGTCTCGTTCCGGAAAGTGG 1000
Oligo fwd CACCTCCCGTCTCGTTCCGGAAAG
Oligo rev AAACCTTTCCGGAACGAGACGGGA
Chromosome start end strand MM target_seq PAM alignment distance position gene name gene id
GU305914.1 227297 227319 - 0 TCCCGTCTCGTTCCGGAAAG TGG ||||||||[||||||||||||]PAM NA Intergenic NA NA

T17442 CCGGAAAGTGGTATTAGTTTTGG 1000
Oligo fwd CACCCCGGAAAGTGGTATTAGTTT
Oligo rev AAACAAACTAATACCACTTTCCGG
Chromosome start end strand MM target_seq PAM alignment distance position gene name gene id
GU305914.1 227285 227307 - 0 CCGGAAAGTGGTATTAGTTT TGG ||||||||[||||||||||||]PAM NA Intergenic NA NA

T17443 TCGAAACTGACCTTTGAGTGTGG 1000
Oligo fwd CACCTCGAAACTGACCTTTGAGTG
Oligo rev AAACCACTCAAAGGTCAGTTTCGA
Chromosome start end strand MM target_seq PAM alignment distance position gene name gene id
GU305914.1 227252 227274 - 0 TCGAAACTGACCTTTGAGTG TGG ||||||||[||||||||||||]PAM NA Intergenic NA NA

T17444 ACCTTTGAGTGTGGAAAGTTTGG 1000
Oligo fwd CACCACCTTTGAGTGTGGAAAGTT
Oligo rev AAACAACTTTCCACACTCAAAGGT
Chromosome start end strand MM target_seq PAM alignment distance position gene name gene id
GU305914.1 227243 227265 - 0 ACCTTTGAGTGTGGAAAGTT TGG ||||||||[||||||||||||]PAM NA Intergenic NA NA

T17445 CCTTTGAGTGTGGAAAGTTTGGG 1000
Oligo fwd CACCCCTTTGAGTGTGGAAAGTTT
Oligo rev AAACAAACTTTCCACACTCAAAGG
Chromosome start end strand MM target_seq PAM alignment distance position gene name gene id
GU305914.1 227242 227264 - 0 CCTTTGAGTGTGGAAAGTTT GGG ||||||||[||||||||||||]PAM NA Intergenic NA NA

T17446 CTTTGAGTGTGGAAAGTTTGGGG 1000
Oligo fwd CACCCTTTGAGTGTGGAAAGTTTG
Oligo rev AAACCAAACTTTCCACACTCAAAG
Chromosome start end strand MM target_seq PAM alignment distance position gene name gene id
GU305914.1 227241 227263 - 0 CTTTGAGTGTGGAAAGTTTG GGG ||||||||[||||||||||||]PAM NA Intergenic NA NA

T17447 GAAAGTTTGGGGACTTTCCGCGG 1000
Oligo fwd CACCGAAAGTTTGGGGACTTTCCG
Oligo rev AAACCGGAAAGTCCCCAAACTTTC
Chromosome start end strand MM target_seq PAM alignment distance position gene name gene id
GU305914.1 227230 227252 - 0 GAAAGTTTGGGGACTTTCCG CGG ||||||||[||||||||||||]PAM NA Intergenic NA NA

T17448 TTGGGGACTTTCCGCGGTCCCGG 1000
Oligo fwd CACCTTGGGGACTTTCCGCGGTCC
Oligo rev AAACGGACCGCGGAAAGTCCCCAA
Chromosome start end strand MM target_seq PAM alignment distance position gene name gene id
GU305914.1 227224 227246 - 0 TTGGGGACTTTCCGCGGTCC CGG ||||||||[||||||||||||]PAM NA Intergenic NA NA

T17449 TGGGGACTTTCCGCGGTCCCGGG 1000
Oligo fwd CACCTGGGGACTTTCCGCGGTCCC
Oligo rev AAACGGGACCGCGGAAAGTCCCCA

| | | | | | | | | | | | |
|---|---|---|---|---|---|---|---|---|---|---|---|
| Chromosome | start | end | strand | MM | target_seq | PAM | alignment | distance | position | gene name | gene id |
| GU305914.1 | 227223 | 227245 | - | 0 | TGGGGACTTTCCGCGGTCCC | GGG | \|\|\|\|\|\|\|\|[\|\|\|\|\|\|\|\|\|\|\|\|]PAM | NA | Intergenic | NA | NA |
| T17450 | GGCCAAGTTTCTACCTCAAGTGG | 1000 | | | | | | | | | |
| Oligo fwd | CACCGGCCAAGTTTCTACCTCAAG | | | | | | | | | | |
| Oligo rev | AAACCTTGAGGTAGAAACTTGGCC | | | | | | | | | | |
| Chromosome | start | end | strand | MM | target_seq | PAM | alignment | distance | position | gene name | gene id |
| GU305914.1 | 227202 | 227224 | - | 0 | GGCCAAGTTTCTACCTCAAG | TGG | \|\|\|\|\|\|\|\|[\|\|\|\|\|\|\|\|\|\|\|\|]PAM | NA | Intergenic | NA | NA |
| T17451 | CATGAGACATCATGCTCACGTGG | 1000 | | | | | | | | | |
| Oligo fwd | CACCCATGAGACATCATGCTCACG | | | | | | | | | | |
| Oligo rev | AAACCGTGAGCATGATGTCTCATG | | | | | | | | | | |
| Chromosome | start | end | strand | MM | target_seq | PAM | alignment | distance | position | gene name | gene id |
| GU305914.1 | 227179 | 227201 | - | 0 | CATGAGACATCATGCTCACG | TGG | \|\|\|\|\|\|\|\|[\|\|\|\|\|\|\|\|\|\|\|\|]PAM | NA | Intergenic | NA | NA |
| T17452 | CATGCTCACGTGGCAGAGTGTGG | 1000 | | | | | | | | | |
| Oligo fwd | CACCCATGCTCACGTGGCAGAGTG | | | | | | | | | | |
| Oligo rev | AAACCACTCTGCCACGTGAGCATG | | | | | | | | | | |
| Chromosome | start | end | strand | MM | target_seq | PAM | alignment | distance | position | gene name | gene id |
| GU305914.1 | 227169 | 227191 | - | 0 | CATGCTCACGTGGCAGAGTG | TGG | \|\|\|\|\|\|\|\|[\|\|\|\|\|\|\|\|\|\|\|\|]PAM | NA | Intergenic | NA | NA |
| T17453 | GAGTGTGGAAACTTTGACAGCGG | 1000 | | | | | | | | | |
| Oligo fwd | CACCGAGTGTGGAAACTTTGACAG | | | | | | | | | | |
| Oligo rev | AAACCTGTCAAAGTTTCCACACTC | | | | | | | | | | |
| Chromosome | start | end | strand | MM | target_seq | PAM | alignment | distance | position | gene name | gene id |
| GU305914.1 | 227154 | 227176 | - | 0 | GAGTGTGGAAACTTTGACAG | CGG | \|\|\|\|\|\|\|\|[\|\|\|\|\|\|\|\|\|\|\|\|]PAM | NA | Intergenic | NA | NA |
| T17454 | GCGGATGTTGCGACCCGACGTGG | 1000 | | | | | | | | | |
| Oligo fwd | CACCGCGGATGTTGCGACCCGACG | | | | | | | | | | |
| Oligo rev | AAACCGTCGGGTCGCAACATCCGC | | | | | | | | | | |
| Chromosome | start | end | strand | MM | target_seq | PAM | alignment | distance | position | gene name | gene id |
| GU305914.1 | 227135 | 227157 | - | 0 | GCGGATGTTGCGACCCGACG | TGG | \|\|\|\|\|\|\|\|[\|\|\|\|\|\|\|\|\|\|\|\|]PAM | NA | Intergenic | NA | NA |
| T17455 | TTGCGACCCGACGTGGTTTGTGG | 1000 | | | | | | | | | |
| Oligo fwd | CACCTTGCGACCCGACGTGGTTTG | | | | | | | | | | |
| Oligo rev | AAACCAAACCACGTCGGGTCGCAA | | | | | | | | | | |
| Chromosome | start | end | strand | MM | target_seq | PAM | alignment | distance | position | gene name | gene id |
| GU305914.1 | 227128 | 227150 | - | 0 | TTGCGACCCGACGTGGTTTG | TGG | \|\|\|\|\|\|\|\|[\|\|\|\|\|\|\|\|\|\|\|\|]PAM | NA | Intergenic | NA | NA |
| T17456 | CGACCCGACGTGGTTTGTGGTGG | 1000 | | | | | | | | | |
| Oligo fwd | CACCCGACCCGACGTGGTTTGTGG | | | | | | | | | | |
| Oligo rev | AAACCCACAAACCACGTCGGGTCG | | | | | | | | | | |
| Chromosome | start | end | strand | MM | target_seq | PAM | alignment | distance | position | gene name | gene id |
| GU305914.1 | 227125 | 227147 | - | 0 | CGACCCGACGTGGTTTGTGG | TGG | \|\|\|\|\|\|\|\|[\|\|\|\|\|\|\|\|\|\|\|\|]PAM | NA | Intergenic | NA | NA |
| T17457 | CCGACGTGGTTTGTGGTGGTTGG | 1000 | | | | | | | | | |
| Oligo fwd | CACCCCGACGTGGTTTGTGGTGGT | | | | | | | | | | |
| Oligo rev | AAACACCACCACAAACCACGTCGG | | | | | | | | | | |
| Chromosome | start | end | strand | MM | target_seq | PAM | alignment | distance | position | gene name | gene id |
| GU305914.1 | 227121 | 227143 | - | 0 | CCGACGTGGTTTGTGGTGGT | TGG | \|\|\|\|\|\|\|\|[\|\|\|\|\|\|\|\|\|\|\|\|]PAM | NA | Intergenic | NA | NA |
| T17458 | GGTCCTGCGTGTAACCATTCTGG | 1000 | | | | | | | | | |
| Oligo fwd | CACCGGTCCTGCGTGTAACCATTC | | | | | | | | | | |
| Oligo rev | AAACGAATGGTTACACGCAGGACC | | | | | | | | | | |
| Chromosome | start | end | strand | MM | target_seq | PAM | alignment | distance | position | gene name | gene id |
| GU305914.1 | 227100 | 227122 | - | 0 | GGTCCTGCGTGTAACCATTC | TGG | \|\|\|\|\|\|\|\|[\|\|\|\|\|\|\|\|\|\|\|\|]PAM | NA | Intergenic | NA | NA |
| T17459 | GTCCTGCGTGTAACCATTCTGGG | 1000 | | | | | | | | | |
| Oligo fwd | CACCGTCCTGCGTGTAACCATTCT | | | | | | | | | | |
| Oligo rev | AAACAGAATGGTTACACGCAGGAC | | | | | | | | | | |
| Chromosome | start | end | strand | MM | target_seq | PAM | alignment | distance | position | gene name | gene id |
| GU305914.1 | 227099 | 227121 | - | 0 | GTCCTGCGTGTAACCATTCT | GGG | \|\|\|\|\|\|\|\|[\|\|\|\|\|\|\|\|\|\|\|\|]PAM | NA | Intergenic | NA | NA |
| T17460 | TCCTGCGTGTAACCATTCTGGGG | 1000 | | | | | | | | | |
| Oligo fwd | CACCTCCTGCGTGTAACCATTCTG | | | | | | | | | | |
| Oligo rev | AAACCAGAATGGTTACACGCAGGA | | | | | | | | | | |
| Chromosome | start | end | strand | MM | target_seq | PAM | alignment | distance | position | gene name | gene id |
| GU305914.1 | 227098 | 227120 | - | 0 | TCCTGCGTGTAACCATTCTG | GGG | \|\|\|\|\|\|\|\|[\|\|\|\|\|\|\|\|\|\|\|\|]PAM | NA | Intergenic | NA | NA |
| T17461 | TGCGTGTAACCATTCTGGGGTGG | 1000 | | | | | | | | | |
| Oligo fwd | CACCTGCGTGTAACCATTCTGGGG | | | | | | | | | | |
| Oligo rev | AAACCCCCAGAATGGTTACACGCA | | | | | | | | | | |
| Chromosome | start | end | strand | MM | target_seq | PAM | alignment | distance | position | gene name | gene id |
| GU305914.1 | 227095 | 227117 | - | 0 | TGCGTGTAACCATTCTGGGG | TGG | \|\|\|\|\|\|\|\|[\|\|\|\|\|\|\|\|\|\|\|\|]PAM | NA | Intergenic | NA | NA |
| T17462 | TGGTGTTGTGACTCGAGCAACGG | 1000 | | | | | | | | | |
| Oligo fwd | CACCTGGTGTTGTGACTCGAGCAA | | | | | | | | | | |
| Oligo rev | AAACTTGCTCGAGTCACAACACCA | | | | | | | | | | |
| Chromosome | start | end | strand | MM | target_seq | PAM | alignment | distance | position | gene name | gene id |
| GU305914.1 | 227075 | 227097 | - | 0 | TGGTGTTGTGACTCGAGCAA | CGG | \|\|\|\|\|\|\|\|[\|\|\|\|\|\|\|\|\|\|\|\|]PAM | NA | Intergenic | NA | NA |
| T17463 | GTGACTCGAGCAACGGTGATTGG | 1000 | | | | | | | | | |
| Oligo fwd | CACCGTGACTCGAGCAACGGTGAT | | | | | | | | | | |
| Oligo rev | AAACATCACCGTTGCTCGAGTCAC | | | | | | | | | | |
| Chromosome | start | end | strand | MM | target_seq | PAM | alignment | distance | position | gene name | gene id |
| GU305914.1 | 227068 | 227090 | - | 0 | GTGACTCGAGCAACGGTGAT | TGG | \|\|\|\|\|\|\|\|[\|\|\|\|\|\|\|\|\|\|\|\|]PAM | NA | Intergenic | NA | NA |
| T17464 | GCGTCGATAGCGTTCCACCGTGG | 1000 | | | | | | | | | |
| Oligo fwd | CACCGCGTCGATAGCGTTCCACCG | | | | | | | | | | |

| | | | | | | | | | | | | |
|---|---|---|---|---|---|---|---|---|---|---|---|---|
| Oligo rev | AAACCGGTGGAACGCTATCGACGC | | | | | | | | | | | |
| Chromosome | start | | end | strand | MM | target_seq | PAM | alignment | distance | position | gene name | gene id |
| GU305914.1 | | 227044 | 227066 | - | 0 | GCGTCGATAGCGTTCCACCG | TGG | \|\|\|\|\|\|\|\|[\|\|\|\|\|\|\|\|\|\|\|\|]PAM | NA | Intergenic | NA | NA |
| T17465 | CGTCGATAGCGTTCCACCGTGGG | | 1000 | | | | | | | | | |
| Oligo fwd | CACCCGTCGATAGCGTTCCACCGT | | | | | | | | | | | |
| Oligo rev | AAACACGGTGGAACGCTATCGACG | | | | | | | | | | | |
| Chromosome | start | | end | strand | MM | target_seq | PAM | alignment | distance | position | gene name | gene id |
| GU305914.1 | | 227043 | 227065 | - | 0 | CGTCGATAGCGTTCCACCGT | GGG | \|\|\|\|\|\|\|\|[\|\|\|\|\|\|\|\|\|\|\|\|]PAM | NA | Intergenic | NA | NA |
| T17466 | CGATAGCGTTCCACCGTGGGTGG | | 1000 | | | | | | | | | |
| Oligo fwd | CACCCGATAGCGTTCCACCGTGGG | | | | | | | | | | | |
| Oligo rev | AAACCCCACGGTGGAACGCTATCG | | | | | | | | | | | |
| Chromosome | start | | end | strand | MM | target_seq | PAM | alignment | distance | position | gene name | gene id |
| GU305914.1 | | 227040 | 227062 | - | 0 | CGATAGCGTTCCACCGTGGG | TGG | \|\|\|\|\|\|\|\|[\|\|\|\|\|\|\|\|\|\|\|\|]PAM | NA | Intergenic | NA | NA |
| T17467 | TCCACCGTGGGTGGAAACAAAGG | | 1000 | | | | | | | | | |
| Oligo fwd | CACCTCCACCGTGGGTGGAAACAA | | | | | | | | | | | |
| Oligo rev | AAACTTGTTTCCACCCACGGTGGA | | | | | | | | | | | |
| Chromosome | start | | end | strand | MM | target_seq | PAM | alignment | distance | position | gene name | gene id |
| GU305914.1 | | 227031 | 227053 | - | 0 | TCCACCGTGGGTGGAAACAA | AGG | \|\|\|\|\|\|\|\|[\|\|\|\|\|\|\|\|\|\|\|\|]PAM | NA | Intergenic | NA | NA |
| T17468 | CGTATATAACAACGTCGTTTCGG | | 1000 | | | | | | | | | |
| Oligo fwd | CACCCGTATATAACAACGTCGTTT | | | | | | | | | | | |
| Oligo rev | AAACAAACGACGTTGTTATATACG | | | | | | | | | | | |
| Chromosome | start | | end | strand | MM | target_seq | PAM | alignment | distance | position | gene name | gene id |
| GU305914.1 | | 227005 | 227027 | - | 0 | CGTATATAACAACGTCGTTT | CGG | \|\|\|\|\|\|\|\|[\|\|\|\|\|\|\|\|\|\|\|\|]PAM | NA | Intergenic | NA | NA |
| T17469 | GTATATAACAACGTCGTTTCGGG | | 1000 | | | | | | | | | |
| Oligo fwd | CACCGTATATAACAACGTCGTTTC | | | | | | | | | | | |
| Oligo rev | AAACGAAACGACGTTGTTATATAC | | | | | | | | | | | |
| Chromosome | start | | end | strand | MM | target_seq | PAM | alignment | distance | position | gene name | gene id |
| GU305914.1 | | 227004 | 227026 | - | 0 | GTATATAACAACGTCGTTTC | GGG | \|\|\|\|\|\|\|\|[\|\|\|\|\|\|\|\|\|\|\|\|]PAM | NA | Intergenic | NA | NA |
| T17470 | TATATAACAACGTCGTTTCGGGG | | 1000 | | | | | | | | | |
| Oligo fwd | CACCTATATAACAACGTCGTTTCG | | | | | | | | | | | |
| Oligo rev | AAACCGAAACGACGTTGTTATATA | | | | | | | | | | | |
| Chromosome | start | | end | strand | MM | target_seq | PAM | alignment | distance | position | gene name | gene id |
| GU305914.1 | | 227003 | 227025 | - | 0 | TATATAACAACGTCGTTTCG | GGG | \|\|\|\|\|\|\|\|[\|\|\|\|\|\|\|\|\|\|\|\|]PAM | NA | Intergenic | NA | NA |
| T17471 | ATAACAACGTCGTTTCGGGGCGG | | 1000 | | | | | | | | | |
| Oligo fwd | CACCATAACAACGTCGTTTCGGGG | | | | | | | | | | | |
| Oligo rev | AAACCCCCGAAACGACGTTGTTAT | | | | | | | | | | | |
| Chromosome | start | | end | strand | MM | target_seq | PAM | alignment | distance | position | gene name | gene id |
| GU305914.1 | | 227000 | 227022 | - | 0 | ATAACAACGTCGTTTCGGGG | CGG | \|\|\|\|\|\|\|\|[\|\|\|\|\|\|\|\|\|\|\|\|]PAM | NA | Intergenic | NA | NA |
| T17472 | GAACAGTTGCAGAGTGCGTGTGG | | 1000 | | | | | | | | | |
| Oligo fwd | CACCGAACAGTTGCAGAGTGCGTG | | | | | | | | | | | |
| Oligo rev | AAACCACGCACTCTGCAACTGTTC | | | | | | | | | | | |
| Chromosome | start | | end | strand | MM | target_seq | PAM | alignment | distance | position | gene name | gene id |
| GU305914.1 | | 226975 | 226997 | - | 0 | GAACAGTTGCAGAGTGCGTG | TGG | \|\|\|\|\|\|\|\|[\|\|\|\|\|\|\|\|\|\|\|\|]PAM | NA | Intergenic | NA | NA |
| T17473 | GTTGCAGAGTGCGTGTGGCACGG | | 1000 | | | | | | | | | |
| Oligo fwd | CACCGTTGCAGAGTGCGTGTGGCA | | | | | | | | | | | |
| Oligo rev | AAACTGCCACACGCACTCTGCAAC | | | | | | | | | | | |
| Chromosome | start | | end | strand | MM | target_seq | PAM | alignment | distance | position | gene name | gene id |
| GU305914.1 | | 226970 | 226992 | - | 0 | GTTGCAGAGTGCGTGTGGCA | CGG | \|\|\|\|\|\|\|\|[\|\|\|\|\|\|\|\|\|\|\|\|]PAM | NA | Intergenic | NA | NA |
| T17474 | AGAGTGCGTGTGGCACGGTTCGG | | 1000 | | | | | | | | | |
| Oligo fwd | CACCAGAGTGCGTGTGGCACGGTT | | | | | | | | | | | |
| Oligo rev | AAACAACCGTGCCACACGCACTCT | | | | | | | | | | | |
| Chromosome | start | | end | strand | MM | target_seq | PAM | alignment | distance | position | gene name | gene id |
| GU305914.1 | | 226965 | 226987 | - | 0 | AGAGTGCGTGTGGCACGGTT | CGG | \|\|\|\|\|\|\|\|[\|\|\|\|\|\|\|\|\|\|\|\|]PAM | NA | Intergenic | NA | NA |
| T17475 | TGTGGCACGGTTCGGCTCCTCGG | | 1000 | | | | | | | | | |
| Oligo fwd | CACCTGTGGCACGGTTCGGCTCCT | | | | | | | | | | | |
| Oligo rev | AAACAGGAGCCGAACCGTGCCACA | | | | | | | | | | | |
| Chromosome | start | | end | strand | MM | target_seq | PAM | alignment | distance | position | gene name | gene id |
| GU305914.1 | | 226957 | 226979 | - | 0 | TGTGGCACGGTTCGGCTCCT | CGG | \|\|\|\|\|\|\|\|[\|\|\|\|\|\|\|\|\|\|\|\|]PAM | NA | Intergenic | NA | NA |
| T17476 | GGTTCGGCTCCTCGGATCAACGG | | 1000 | | | | | | | | | |
| Oligo fwd | CACCGGTTCGGCTCCTCGGATCAA | | | | | | | | | | | |
| Oligo rev | AAACTTGATCCGAGGAGCCGAACC | | | | | | | | | | | |
| Chromosome | start | | end | strand | MM | target_seq | PAM | alignment | distance | position | gene name | gene id |
| GU305914.1 | | 226949 | 226971 | - | 0 | GGTTCGGCTCCTCGGATCAA | CGG | \|\|\|\|\|\|\|\|[\|\|\|\|\|\|\|\|\|\|\|\|]PAM | NA | Intergenic | NA | NA |
| T17477 | GTTCGGCTCCTCGGATCAACGGG | | 1000 | | | | | | | | | |
| Oligo fwd | CACCGTTCGGCTCCTCGGATCAAC | | | | | | | | | | | |
| Oligo rev | AAACGTTGATCCGAGGAGCCGAAC | | | | | | | | | | | |
| Chromosome | start | | end | strand | MM | target_seq | PAM | alignment | distance | position | gene name | gene id |
| GU305914.1 | | 226948 | 226970 | - | 0 | GTTCGGCTCCTCGGATCAAC | GGG | \|\|\|\|\|\|\|\|[\|\|\|\|\|\|\|\|\|\|\|\|]PAM | NA | Intergenic | NA | NA |
| T17478 | CTCCTCGGATCAACGGGACCTGG | | 1000 | | | | | | | | | |
| Oligo fwd | CACCCTCCTCGGATCAACGGGACC | | | | | | | | | | | |
| Oligo rev | AAACGGTCCCGTTGATCCGAGGAG | | | | | | | | | | | |
| Chromosome | start | | end | strand | MM | target_seq | PAM | alignment | distance | position | gene name | gene id |
| GU305914.1 | | 226942 | 226964 | - | 0 | CTCCTCGGATCAACGGGACC | TGG | \|\|\|\|\|\|\|\|[\|\|\|\|\|\|\|\|\|\|\|\|]PAM | NA | Intergenic | NA | NA |
| T17479 | CTCGGATCAACGGGACCTGGTGG | | 1000 | | | | | | | | | |

Oligo fwd CACCCTCGGATCAACGGGACCTGG
Oligo rev AAACCCAGGTCCCGTTGATCCGAG
Chromosome start end strand MM target_seq PAM alignment distance position gene name gene id
GU305914.1 226939 226961 - 0 CTCGGATCAACGGGACCTGG TGG ||||||||[||||||||||||]PAM NA Intergenic NA NA

T17480 GGGACCTGGTGGCAGATCGAAGG 1000
Oligo fwd CACCGGGACCTGGTGGCAGATCGA
Oligo rev AAACTCGATCTGCCACCAGGTCCC
Chromosome start end strand MM target_seq PAM alignment distance position gene name gene id
GU305914.1 226928 226950 - 0 GGGACCTGGTGGCAGATCGA AGG ||||||||[||||||||||||]PAM NA Intergenic NA NA

T17481 GGCAGATCGAAGGTTCGACTCGG 1000
Oligo fwd CACCGGCAGATCGAAGGTTCGACT
Oligo rev AAACAGTCGAACCTTCGATCTGCC
Chromosome start end strand MM target_seq PAM alignment distance position gene name gene id
GU305914.1 226918 226940 - 0 GGCAGATCGAAGGTTCGACT CGG ||||||||[||||||||||||]PAM NA Intergenic NA NA

T17482 GATCGAAGGTTCGACTCGGACGG 1000
Oligo fwd CACCGATCGAAGGTTCGACTCGGA
Oligo rev AAACTCCGAGTCGAACCTTCGATC
Chromosome start end strand MM target_seq PAM alignment distance position gene name gene id
GU305914.1 226914 226936 - 0 GATCGAAGGTTCGACTCGGA CGG ||||||||[||||||||||||]PAM NA Intergenic NA NA

T17483 ACGGAGATCGCGAAGTTCAACGG 1000
Oligo fwd CACCACGGAGATCGCGAAGTTCAA
Oligo rev AAACTTGAACTTCGCGATCTCCGT
Chromosome start end strand MM target_seq PAM alignment distance position gene name gene id
GU305914.1 226895 226917 - 0 ACGGAGATCGCGAAGTTCAA CGG ||||||||[||||||||||||]PAM NA Intergenic NA NA

T17484 TCGCGAAGTTCAACGGCTCGTGG 1000
Oligo fwd CACCTCGCGAAGTTCAACGGCTCG
Oligo rev AAACCGAGCCGTTGAACTTCGCGA
Chromosome start end strand MM target_seq PAM alignment distance position gene name gene id
GU305914.1 226888 226910 - 0 TCGCGAAGTTCAACGGCTCG TGG ||||||||[||||||||||||]PAM NA Intergenic NA NA

T17485 TCAACGGCTCGTGGTTCCAATGG 1000
Oligo fwd CACCTCAACGGCTCGTGGTTCCAA
Oligo rev AAACTTGGAACCACGAGCCGTTGA
Chromosome start end strand MM target_seq PAM alignment distance position gene name gene id
GU305914.1 226879 226901 - 0 TCAACGGCTCGTGGTTCCAA TGG ||||||||[||||||||||||]PAM NA Intergenic NA NA

T17486 TGGATCACCCTGCTCAGAGACGG 1000
Oligo fwd CACCTGGATCACCCTGCTCAGAGA
Oligo rev AAACTCTCTGAGCAGGGTGATCCA
Chromosome start end strand MM target_seq PAM alignment distance position gene name gene id
GU305914.1 226859 226881 - 0 TGGATCACCCTGCTCAGAGA CGG ||||||||[||||||||||||]PAM NA Intergenic NA NA

T17487 ATCACCCTGCTCAGAGACGGAGG 1000
Oligo fwd CACCATCACCCTGCTCAGAGACGG
Oligo rev AAACCCGTCTCTGAGCAGGGTGAT
Chromosome start end strand MM target_seq PAM alignment distance position gene name gene id
GU305914.1 226856 226878 - 0 ATCACCCTGCTCAGAGACGG AGG ||||||||[||||||||||||]PAM NA Intergenic NA NA

T17488 TCACCCTGCTCAGAGACGGAGGG 1000
Oligo fwd CACCTCACCCTGCTCAGAGACGGA
Oligo rev AAACTCCGTCTCTGAGCAGGGTGA
Chromosome start end strand MM target_seq PAM alignment distance position gene name gene id
GU305914.1 226855 226877 - 0 TCACCCTGCTCAGAGACGGA GGG ||||||||[||||||||||||]PAM NA Intergenic NA NA

T17489 CACCCTGCTCAGAGACGGAGGGG 1000
Oligo fwd CACCCACCCTGCTCAGAGACGGAG
Oligo rev AAACCTCCGTCTCTGAGCAGGGTG
Chromosome start end strand MM target_seq PAM alignment distance position gene name gene id
GU305914.1 226854 226876 - 0 CACCCTGCTCAGAGACGGAG GGG ||||||||[||||||||||||]PAM NA Intergenic NA NA

T17490 GAGACGGAGGGGCCTTCGCCTGG 1000
Oligo fwd CACCGAGACGGAGGGGCCTTCGCC
Oligo rev AAACGGCGAAGGCCCCTCCGTCTC
Chromosome start end strand MM target_seq PAM alignment distance position gene name gene id
GU305914.1 226843 226865 - 0 GAGACGGAGGGGCCTTCGCC TGG ||||||||[||||||||||||]PAM NA Intergenic NA NA

T17491 AGACGGAGGGGCCTTCGCCTGGG 1000
Oligo fwd CACCAGACGGAGGGGCCTTCGCCT
Oligo rev AAACAGGCGAAGGCCCCTCCGTCT
Chromosome start end strand MM target_seq PAM alignment distance position gene name gene id
GU305914.1 226842 226864 - 0 AGACGGAGGGGCCTTCGCCT GGG ||||||||[||||||||||||]PAM NA Intergenic NA NA

T17492 AGACCTCGAAAGAATCGCAGCGG 1000
Oligo fwd CACCAGACCTCGAAAGAATCGCAG
Oligo rev AAACCTGCGATTCTTTCGAGGTCT
Chromosome start end strand MM target_seq PAM alignment distance position gene name gene id
GU305914.1 226813 226835 - 0 AGACCTCGAAAGAATCGCAG CGG ||||||||[||||||||||||]PAM NA Intergenic NA NA

T17493 CGAAAGAATCGCAGCGGCGCTGG 1000
Oligo fwd CACCCGAAAGAATCGCAGCGGCGC
Oligo rev AAACGCGCCGCTGCGATTCTTTCG
Chromosome start end strand MM target_seq PAM alignment distance position gene name gene id
GU305914.1 226807 226829 - 0 CGAAAGAATCGCAGCGGCGC TGG ||||||||[||||||||||||]PAM NA Intergenic NA NA

<table>
<tr><td>T17494</td><td>ATCGCAGCGGCGCTGGCAGAAGG</td><td>1000</td><td></td><td></td><td></td><td></td><td></td><td></td><td></td><td></td><td></td></tr>
<tr><td>Oligo fwd</td><td>CACCATCGCAGCGGCGCTGGCAGA</td><td></td><td></td><td></td><td></td><td></td><td></td><td></td><td></td><td></td><td></td></tr>
<tr><td>Oligo rev</td><td>AAACTCTGCCAGCGCCGCTGCGAT</td><td></td><td></td><td></td><td></td><td></td><td></td><td></td><td></td><td></td><td></td></tr>
<tr><td>Chromosome</td><td>start</td><td>end</td><td>strand</td><td>MM</td><td>target_seq</td><td>PAM</td><td>alignment</td><td>distance</td><td>position</td><td>gene name</td><td>gene id</td></tr>
<tr><td>GU305914.1</td><td>226800</td><td>226822</td><td>-</td><td>0</td><td>ATCGCAGCGGCGCTGGCAGA</td><td>AGG</td><td>||||||||[||||||||||||]PAM</td><td>NA</td><td>Intergenic</td><td>NA</td><td>NA</td></tr>
<tr><td>T17495</td><td>GCGCTGGCAGAAGGCGAGCCCGG</td><td>1000</td><td></td><td></td><td></td><td></td><td></td><td></td><td></td><td></td><td></td></tr>
<tr><td>Oligo fwd</td><td>CACCGCGCTGGCAGAAGGCGAGCC</td><td></td><td></td><td></td><td></td><td></td><td></td><td></td><td></td><td></td><td></td></tr>
<tr><td>Oligo rev</td><td>AAACGGCTCGCCTTCTGCCAGCGC</td><td></td><td></td><td></td><td></td><td></td><td></td><td></td><td></td><td></td><td></td></tr>
<tr><td>Chromosome</td><td>start</td><td>end</td><td>strand</td><td>MM</td><td>target_seq</td><td>PAM</td><td>alignment</td><td>distance</td><td>position</td><td>gene name</td><td>gene id</td></tr>
<tr><td>GU305914.1</td><td>226791</td><td>226813</td><td>-</td><td>0</td><td>GCGCTGGCAGAAGGCGAGCC</td><td>CGG</td><td>||||||||[||||||||||||]PAM</td><td>NA</td><td>Intergenic</td><td>NA</td><td>NA</td></tr>
<tr><td>T17496</td><td>AAGGCGAGCCCGGTCTCGATCGG</td><td>1000</td><td></td><td></td><td></td><td></td><td></td><td></td><td></td><td></td><td></td></tr>
<tr><td>Oligo fwd</td><td>CACCAAGGCGAGCCCGGTCTCGAT</td><td></td><td></td><td></td><td></td><td></td><td></td><td></td><td></td><td></td><td></td></tr>
<tr><td>Oligo rev</td><td>AAACATCGAGACCGGGCTCGCCTT</td><td></td><td></td><td></td><td></td><td></td><td></td><td></td><td></td><td></td><td></td></tr>
<tr><td>Chromosome</td><td>start</td><td>end</td><td>strand</td><td>MM</td><td>target_seq</td><td>PAM</td><td>alignment</td><td>distance</td><td>position</td><td>gene name</td><td>gene id</td></tr>
<tr><td>GU305914.1</td><td>226781</td><td>226803</td><td>-</td><td>0</td><td>AAGGCGAGCCCGGTCTCGAT</td><td>CGG</td><td>||||||||[||||||||||||]PAM</td><td>NA</td><td>Intergenic</td><td>NA</td><td>NA</td></tr>
<tr><td>T17497</td><td>GAGCCCGGTCTCGATCGGCGAGG</td><td>1000</td><td></td><td></td><td></td><td></td><td></td><td></td><td></td><td></td><td></td></tr>
<tr><td>Oligo fwd</td><td>CACCGAGCCCGGTCTCGATCGGCG</td><td></td><td></td><td></td><td></td><td></td><td></td><td></td><td></td><td></td><td></td></tr>
<tr><td>Oligo rev</td><td>AAACCGCCGATCGAGACCGGGCTC</td><td></td><td></td><td></td><td></td><td></td><td></td><td></td><td></td><td></td><td></td></tr>
<tr><td>Chromosome</td><td>start</td><td>end</td><td>strand</td><td>MM</td><td>target_seq</td><td>PAM</td><td>alignment</td><td>distance</td><td>position</td><td>gene name</td><td>gene id</td></tr>
<tr><td>GU305914.1</td><td>226776</td><td>226798</td><td>-</td><td>0</td><td>GAGCCCGGTCTCGATCGGCG</td><td>AGG</td><td>||||||||[||||||||||||]PAM</td><td>NA</td><td>Intergenic</td><td>NA</td><td>NA</td></tr>
<tr><td>T17498</td><td>TCGATCGGCGAGGACTCTCCTGG</td><td>1000</td><td></td><td></td><td></td><td></td><td></td><td></td><td></td><td></td><td></td></tr>
<tr><td>Oligo fwd</td><td>CACCTCGATCGGCGAGGACTCTCC</td><td></td><td></td><td></td><td></td><td></td><td></td><td></td><td></td><td></td><td></td></tr>
<tr><td>Oligo rev</td><td>AAACGGAGAGTCCTCGCCGATCGA</td><td></td><td></td><td></td><td></td><td></td><td></td><td></td><td></td><td></td><td></td></tr>
<tr><td>Chromosome</td><td>start</td><td>end</td><td>strand</td><td>MM</td><td>target_seq</td><td>PAM</td><td>alignment</td><td>distance</td><td>position</td><td>gene name</td><td>gene id</td></tr>
<tr><td>GU305914.1</td><td>226766</td><td>226788</td><td>-</td><td>0</td><td>TCGATCGGCGAGGACTCTCC</td><td>TGG</td><td>||||||||[||||||||||||]PAM</td><td>NA</td><td>Intergenic</td><td>NA</td><td>NA</td></tr>
<tr><td>T17499</td><td>GGCGAGGACTCTCCTGGAGACGG</td><td>1000</td><td></td><td></td><td></td><td></td><td></td><td></td><td></td><td></td><td></td></tr>
<tr><td>Oligo fwd</td><td>CACCGGCGAGGACTCTCCTGGAGA</td><td></td><td></td><td></td><td></td><td></td><td></td><td></td><td></td><td></td><td></td></tr>
<tr><td>Oligo rev</td><td>AAACTCTCCAGGAGAGTCCTCGCC</td><td></td><td></td><td></td><td></td><td></td><td></td><td></td><td></td><td></td><td></td></tr>
<tr><td>Chromosome</td><td>start</td><td>end</td><td>strand</td><td>MM</td><td>target_seq</td><td>PAM</td><td>alignment</td><td>distance</td><td>position</td><td>gene name</td><td>gene id</td></tr>
<tr><td>GU305914.1</td><td>226760</td><td>226782</td><td>-</td><td>0</td><td>GGCGAGGACTCTCCTGGAGA</td><td>CGG</td><td>||||||||[||||||||||||]PAM</td><td>NA</td><td>Intergenic</td><td>NA</td><td>NA</td></tr>
<tr><td>T17500</td><td>CGACCTCCGCGCTCTCCGAGTGG</td><td>1000</td><td></td><td></td><td></td><td></td><td></td><td></td><td></td><td></td><td></td></tr>
<tr><td>Oligo fwd</td><td>CACCCGACCTCCGCGCTCTCCGAG</td><td></td><td></td><td></td><td></td><td></td><td></td><td></td><td></td><td></td><td></td></tr>
<tr><td>Oligo rev</td><td>AAACCTCGGAGAGCGCGGAGGTCG</td><td></td><td></td><td></td><td></td><td></td><td></td><td></td><td></td><td></td><td></td></tr>
<tr><td>Chromosome</td><td>start</td><td>end</td><td>strand</td><td>MM</td><td>target_seq</td><td>PAM</td><td>alignment</td><td>distance</td><td>position</td><td>gene name</td><td>gene id</td></tr>
<tr><td>GU305914.1</td><td>226718</td><td>226740</td><td>-</td><td>0</td><td>CGACCTCCGCGCTCTCCGAG</td><td>TGG</td><td>||||||||[||||||||||||]PAM</td><td>NA</td><td>Intergenic</td><td>NA</td><td>NA</td></tr>
<tr><td>GU305914.1</td><td>42901</td><td>42923</td><td>-</td><td>3</td><td>CGTCCCTCGCGCTCTCCGAG</td><td>GGG</td><td>||-||--|[||||||||||||]PAM</td><td>NA</td><td>Intergenic</td><td>NA</td><td>NA</td></tr>
<tr><td>T17501</td><td>CGCGCTCTCCGAGTGGCTCGAGG</td><td>1000</td><td></td><td></td><td></td><td></td><td></td><td></td><td></td><td></td><td></td></tr>
<tr><td>Oligo fwd</td><td>CACCCGCGCTCTCCGAGTGGCTCG</td><td></td><td></td><td></td><td></td><td></td><td></td><td></td><td></td><td></td><td></td></tr>
<tr><td>Oligo rev</td><td>AAACCGAGCCACTCGGAGAGCGCG</td><td></td><td></td><td></td><td></td><td></td><td></td><td></td><td></td><td></td><td></td></tr>
<tr><td>Chromosome</td><td>start</td><td>end</td><td>strand</td><td>MM</td><td>target_seq</td><td>PAM</td><td>alignment</td><td>distance</td><td>position</td><td>gene name</td><td>gene id</td></tr>
<tr><td>GU305914.1</td><td>226711</td><td>226733</td><td>-</td><td>0</td><td>CGCGCTCTCCGAGTGGCTCG</td><td>AGG</td><td>||||||||[||||||||||||]PAM</td><td>NA</td><td>Intergenic</td><td>NA</td><td>NA</td></tr>
<tr><td>T17502</td><td>TCTCCGAGTGGCTCGAGGCGAGG</td><td>1000</td><td></td><td></td><td></td><td></td><td></td><td></td><td></td><td></td><td></td></tr>
<tr><td>Oligo fwd</td><td>CACCTCTCCGAGTGGCTCGAGGCG</td><td></td><td></td><td></td><td></td><td></td><td></td><td></td><td></td><td></td><td></td></tr>
<tr><td>Oligo rev</td><td>AAACCGCCTCGAGCCACTCGGAGA</td><td></td><td></td><td></td><td></td><td></td><td></td><td></td><td></td><td></td><td></td></tr>
<tr><td>Chromosome</td><td>start</td><td>end</td><td>strand</td><td>MM</td><td>target_seq</td><td>PAM</td><td>alignment</td><td>distance</td><td>position</td><td>gene name</td><td>gene id</td></tr>
<tr><td>GU305914.1</td><td>226706</td><td>226728</td><td>-</td><td>0</td><td>TCTCCGAGTGGCTCGAGGCG</td><td>AGG</td><td>||||||||[||||||||||||]PAM</td><td>NA</td><td>Intergenic</td><td>NA</td><td>NA</td></tr>
<tr><td>T17503</td><td>GTCAGCAACAGCAGAAGAAGTGG</td><td>1000</td><td></td><td></td><td></td><td></td><td></td><td></td><td></td><td></td><td></td></tr>
<tr><td>Oligo fwd</td><td>CACCGTCAGCAACAGCAGAAGAAG</td><td></td><td></td><td></td><td></td><td></td><td></td><td></td><td></td><td></td><td></td></tr>
<tr><td>Oligo rev</td><td>AAACCTTCTTCTGCTGTTGCTGAC</td><td></td><td></td><td></td><td></td><td></td><td></td><td></td><td></td><td></td><td></td></tr>
<tr><td>Chromosome</td><td>start</td><td>end</td><td>strand</td><td>MM</td><td>target_seq</td><td>PAM</td><td>alignment</td><td>distance</td><td>position</td><td>gene name</td><td>gene id</td></tr>
<tr><td>GU305914.1</td><td>226682</td><td>226704</td><td>-</td><td>0</td><td>GTCAGCAACAGCAGAAGAAG</td><td>TGG</td><td>||||||||[||||||||||||]PAM</td><td>NA</td><td>Intergenic</td><td>NA</td><td>NA</td></tr>
<tr><td>GU305914.1</td><td>27408</td><td>27430</td><td>+</td><td>4</td><td>GGCAGCAGCAGCAGCAGCAG</td><td>CGG</td><td>|-|||||-[||||||-||-||]PAM</td><td>NA</td><td>Intergenic</td><td>NA</td><td>NA</td></tr>
<tr><td>T17504</td><td>AGCAACAGCAGAAGAAGTGGTGG</td><td>1000</td><td></td><td></td><td></td><td></td><td></td><td></td><td></td><td></td><td></td></tr>
<tr><td>Oligo fwd</td><td>CACCAGCAACAGCAGAAGAAGTGG</td><td></td><td></td><td></td><td></td><td></td><td></td><td></td><td></td><td></td><td></td></tr>
<tr><td>Oligo rev</td><td>AAACCCACTTCTTCTGCTGTTGCT</td><td></td><td></td><td></td><td></td><td></td><td></td><td></td><td></td><td></td><td></td></tr>
<tr><td>Chromosome</td><td>start</td><td>end</td><td>strand</td><td>MM</td><td>target_seq</td><td>PAM</td><td>alignment</td><td>distance</td><td>position</td><td>gene name</td><td>gene id</td></tr>
<tr><td>GU305914.1</td><td>226679</td><td>226701</td><td>-</td><td>0</td><td>AGCAACAGCAGAAGAAGTGG</td><td>TGG</td><td>||||||||[||||||||||||]PAM</td><td>NA</td><td>Intergenic</td><td>NA</td><td>NA</td></tr>
<tr><td>T17505</td><td>CTCAGACTCCGAAGCCTCCGAGG</td><td>1000</td><td></td><td></td><td></td><td></td><td></td><td></td><td></td><td></td><td></td></tr>
<tr><td>Oligo fwd</td><td>CACCCTCAGACTCCGAAGCCTCCG</td><td></td><td></td><td></td><td></td><td></td><td></td><td></td><td></td><td></td><td></td></tr>
<tr><td>Oligo rev</td><td>AAACCGGAGGCTTCGGAGTCTGAG</td><td></td><td></td><td></td><td></td><td></td><td></td><td></td><td></td><td></td><td></td></tr>
<tr><td>Chromosome</td><td>start</td><td>end</td><td>strand</td><td>MM</td><td>target_seq</td><td>PAM</td><td>alignment</td><td>distance</td><td>position</td><td>gene name</td><td>gene id</td></tr>
<tr><td>GU305914.1</td><td>226639</td><td>226661</td><td>-</td><td>0</td><td>CTCAGACTCCGAAGCCTCCG</td><td>AGG</td><td>||||||||[||||||||||||]PAM</td><td>NA</td><td>Intergenic</td><td>NA</td><td>NA</td></tr>
<tr><td>T17506</td><td>GAAGCCTCCGAGGAAGACGAAGG</td><td>1000</td><td></td><td></td><td></td><td></td><td></td><td></td><td></td><td></td><td></td></tr>
<tr><td>Oligo fwd</td><td>CACCGAAGCCTCCGAGGAAGACGA</td><td></td><td></td><td></td><td></td><td></td><td></td><td></td><td></td><td></td><td></td></tr>
<tr><td>Oligo rev</td><td>AAACTCGTCTTCCTCGGAGGCTTC</td><td></td><td></td><td></td><td></td><td></td><td></td><td></td><td></td><td></td><td></td></tr>
<tr><td>Chromosome</td><td>start</td><td>end</td><td>strand</td><td>MM</td><td>target_seq</td><td>PAM</td><td>alignment</td><td>distance</td><td>position</td><td>gene name</td><td>gene id</td></tr>
<tr><td>GU305914.1</td><td>226629</td><td>226651</td><td>-</td><td>0</td><td>GAAGCCTCCGAGGAAGACGA</td><td>AGG</td><td>||||||||[||||||||||||]PAM</td><td>NA</td><td>Intergenic</td><td>NA</td><td>NA</td></tr>
<tr><td>GU305914.1</td><td>164319</td><td>164341</td><td>+</td><td>4</td><td>GAATCCGAGGAGGAAGACGA</td><td>TGG</td><td>|||-||--[-|||||||||||]PAM</td><td>NA</td><td>Intergenic</td><td>NA</td><td>NA</td></tr>
<tr><td>T17507</td><td>CGAGGAAGACGAAGGAGTACTGG</td><td>1000</td><td></td><td></td><td></td><td></td><td></td><td></td><td></td><td></td><td></td></tr>
<tr><td>Oligo fwd</td><td>CACCCGAGGAAGACGAAGGAGTAC</td><td></td><td></td><td></td><td></td><td></td><td></td><td></td><td></td><td></td><td></td></tr>
<tr><td>Oligo rev</td><td>AAACGTACTCCTTCGTCTTCCTCG</td><td></td><td></td><td></td><td></td><td></td><td></td><td></td><td></td><td></td><td></td></tr>
<tr><td>Chromosome</td><td>start</td><td>end</td><td>strand</td><td>MM</td><td>target_seq</td><td>PAM</td><td>alignment</td><td>distance</td><td>position</td><td>gene name</td><td>gene id</td></tr>
<tr><td>GU305914.1</td><td>226621</td><td>226643</td><td>-</td><td>0</td><td>CGAGGAAGACGAAGGAGTAC</td><td>TGG</td><td>||||||||[||||||||||||]PAM</td><td>NA</td><td>Intergenic</td><td>NA</td><td>NA</td></tr>
<tr><td>T17508</td><td>GAGGAAGACGAAGGAGTACTGGG</td><td>1000</td><td></td><td></td><td></td><td></td><td></td><td></td><td></td><td></td><td></td></tr>
<tr><td>Oligo fwd</td><td>CACCGAGGAAGACGAAGGAGTACT</td><td></td><td></td><td></td><td></td><td></td><td></td><td></td><td></td><td></td><td></td></tr>
</table>

| | | | | | | | | | | | |
|---|---|---|---|---|---|---|---|---|---|---|---|
| Oligo rev | AAACAGTACTCCTTCGTCTTCCTC | | | | | | | | | | |
| Chromosome | start | end | strand | MM | target_seq | PAM | alignment | distance | position | gene name | gene id |
| GU305914.1 | 226620 | 226642 | - | 0 | GAGGAAGACGAAGGAGTACT | GGG | \|\|\|\|\|\|\|\|[\|\|\|\|\|\|\|\|\|\|\|\|]PAM | NA | Intergenic | NA | NA |
| GU305914.1 | 224076 | 224098 | - | 4 | GAGGGAGCCGACGAAGTACT | CGG | \|\|\|\|-\|\|-[\|\|\|-\|-\|\|\|\|\|\|]PAM | NA | Intergenic | NA | NA |
| T17509 | AGGAGTACTGGGTACATTCGTGG | 1000 | | | | | | | | | |
| Oligo fwd | CACCAGGAGTACTGGGTACATTCG | | | | | | | | | | |
| Oligo rev | AAACCGAATGTACCCAGTACTCCT | | | | | | | | | | |
| Chromosome | start | end | strand | MM | target_seq | PAM | alignment | distance | position | gene name | gene id |
| GU305914.1 | 226609 | 226631 | - | 0 | AGGAGTACTGGGTACATTCG | TGG | \|\|\|\|\|\|\|\|[\|\|\|\|\|\|\|\|\|\|\|\|]PAM | NA | Intergenic | NA | NA |
| T17510 | CGAGTTGACAGAAAATTTCGCGG | 1000 | | | | | | | | | |
| Oligo fwd | CACCCGAGTTGACAGAAAATTTCG | | | | | | | | | | |
| Oligo rev | AAACCGAAATTTTCTGTCAACTCG | | | | | | | | | | |
| Chromosome | start | end | strand | MM | target_seq | PAM | alignment | distance | position | gene name | gene id |
| GU305914.1 | 226584 | 226606 | - | 0 | CGAGTTGACAGAAAATTTCG | CGG | \|\|\|\|\|\|\|\|[\|\|\|\|\|\|\|\|\|\|\|\|]PAM | NA | Intergenic | NA | NA |
| T17511 | GAGTTGACAGAAAATTTCGCGGG | 1000 | | | | | | | | | |
| Oligo fwd | CACCGAGTTGACAGAAAATTTCGC | | | | | | | | | | |
| Oligo rev | AAACGCGAAATTTTCTGTCAACTC | | | | | | | | | | |
| Chromosome | start | end | strand | MM | target_seq | PAM | alignment | distance | position | gene name | gene id |
| GU305914.1 | 226583 | 226605 | - | 0 | GAGTTGACAGAAAATTTCGC | GGG | \|\|\|\|\|\|\|\|[\|\|\|\|\|\|\|\|\|\|\|\|]PAM | NA | Intergenic | NA | NA |
| T17512 | AGAAAATTTCGCGGGCAGCCTGG | 1000 | | | | | | | | | |
| Oligo fwd | CACCAGAAAATTTCGCGGGCAGCC | | | | | | | | | | |
| Oligo rev | AAACGGCTGCCCGCGAAATTTTCT | | | | | | | | | | |
| Chromosome | start | end | strand | MM | target_seq | PAM | alignment | distance | position | gene name | gene id |
| GU305914.1 | 226575 | 226597 | - | 0 | AGAAAATTTCGCGGGCAGCC | TGG | \|\|\|\|\|\|\|\|[\|\|\|\|\|\|\|\|\|\|\|\|]PAM | NA | Intergenic | NA | NA |
| T17513 | CGTGTTCAACTCGTCTTCCGAGG | 1000 | | | | | | | | | |
| Oligo fwd | CACCCGTGTTCAACTCGTCTTCCG | | | | | | | | | | |
| Oligo rev | AAACCGGAAGACGAGTTGAACACG | | | | | | | | | | |
| Chromosome | start | end | strand | MM | target_seq | PAM | alignment | distance | position | gene name | gene id |
| GU305914.1 | 226545 | 226567 | - | 0 | CGTGTTCAACTCGTCTTCCG | AGG | \|\|\|\|\|\|\|\|[\|\|\|\|\|\|\|\|\|\|\|\|]PAM | NA | Intergenic | NA | NA |
| T17514 | CTCGTCTTCCGAGGTCTCCGTGG | 1000 | | | | | | | | | |
| Oligo fwd | CACCCTCGTCTTCCGAGGTCTCCG | | | | | | | | | | |
| Oligo rev | AAACCGGAGACCTCGGAAGACGAG | | | | | | | | | | |
| Chromosome | start | end | strand | MM | target_seq | PAM | alignment | distance | position | gene name | gene id |
| GU305914.1 | 226536 | 226558 | - | 0 | CTCGTCTTCCGAGGTCTCCG | TGG | \|\|\|\|\|\|\|\|[\|\|\|\|\|\|\|\|\|\|\|\|]PAM | NA | Intergenic | NA | NA |
| T17515 | AGCCCGCGCCCTCGTTAAGCTGG | 1000 | | | | | | | | | |
| Oligo fwd | CACCAGCCCGCGCCCTCGTTAAGC | | | | | | | | | | |
| Oligo rev | AAACGCTTAACGAGGGCGCGGGCT | | | | | | | | | | |
| Chromosome | start | end | strand | MM | target_seq | PAM | alignment | distance | position | gene name | gene id |
| GU305914.1 | 226513 | 226535 | - | 0 | AGCCCGCGCCCTCGTTAAGC | TGG | \|\|\|\|\|\|\|\|[\|\|\|\|\|\|\|\|\|\|\|\|]PAM | NA | Intergenic | NA | NA |
| T17516 | GCCCGCGCCCTCGTTAAGCTGGG | 1000 | | | | | | | | | |
| Oligo fwd | CACCGCCCGCGCCCTCGTTAAGCT | | | | | | | | | | |
| Oligo rev | AAACAGCTTAACGAGGGCGCGGGC | | | | | | | | | | |
| Chromosome | start | end | strand | MM | target_seq | PAM | alignment | distance | position | gene name | gene id |
| GU305914.1 | 226512 | 226534 | - | 0 | GCCCGCGCCCTCGTTAAGCT | GGG | \|\|\|\|\|\|\|\|[\|\|\|\|\|\|\|\|\|\|\|\|]PAM | NA | Intergenic | NA | NA |
| T17517 | AAGAAGCGTCGCCAGAAAGCTGG | 1000 | | | | | | | | | |
| Oligo fwd | CACCAAGAAGCGTCGCCAGAAAGC | | | | | | | | | | |
| Oligo rev | AAACGCTTTCTGGCGACGCTTCTT | | | | | | | | | | |
| Chromosome | start | end | strand | MM | target_seq | PAM | alignment | distance | position | gene name | gene id |
| GU305914.1 | 226489 | 226511 | - | 0 | AAGAAGCGTCGCCAGAAAGC | TGG | \|\|\|\|\|\|\|\|[\|\|\|\|\|\|\|\|\|\|\|\|]PAM | NA | Intergenic | NA | NA |
| T17518 | AGAAGCGTCGCCAGAAAGCTGGG | 1000 | | | | | | | | | |
| Oligo fwd | CACCAGAAGCGTCGCCAGAAAGCT | | | | | | | | | | |
| Oligo rev | AAACAGCTTTCTGGCGACGCTTCT | | | | | | | | | | |
| Chromosome | start | end | strand | MM | target_seq | PAM | alignment | distance | position | gene name | gene id |
| GU305914.1 | 226488 | 226510 | - | 0 | AGAAGCGTCGCCAGAAAGCT | GGG | \|\|\|\|\|\|\|\|[\|\|\|\|\|\|\|\|\|\|\|\|]PAM | NA | Intergenic | NA | NA |
| T17519 | GTGCGATGCCAGCCTCCACGCGG | 1000 | | | | | | | | | |
| Oligo fwd | CACCGTGCGATGCCAGCCTCCACG | | | | | | | | | | |
| Oligo rev | AAACCGTGGAGGCTGGCATCGCAC | | | | | | | | | | |
| Chromosome | start | end | strand | MM | target_seq | PAM | alignment | distance | position | gene name | gene id |
| GU305914.1 | 226463 | 226485 | - | 0 | GTGCGATGCCAGCCTCCACG | CGG | \|\|\|\|\|\|\|\|[\|\|\|\|\|\|\|\|\|\|\|\|]PAM | NA | Intergenic | NA | NA |
| T17520 | CCGCCTGCTTTCGAATCGTTCGG | 1000 | | | | | | | | | |
| Oligo fwd | CACCCCGCCTGCTTTCGAATCGTT | | | | | | | | | | |
| Oligo rev | AAACAACGATTCGAAAGCAGGCGG | | | | | | | | | | |
| Chromosome | start | end | strand | MM | target_seq | PAM | alignment | distance | position | gene name | gene id |
| GU305914.1 | 226434 | 226456 | - | 0 | CCGCCTGCTTTCGAATCGTT | CGG | \|\|\|\|\|\|\|\|[\|\|\|\|\|\|\|\|\|\|\|\|]PAM | NA | Intergenic | NA | NA |
| T17521 | TCGAATCGTTCGGCTCCGTACGG | 1000 | | | | | | | | | |
| Oligo fwd | CACCTCGAATCGTTCGGCTCCGTA | | | | | | | | | | |
| Oligo rev | AAACTACGGAGCCGAACGATTCGA | | | | | | | | | | |
| Chromosome | start | end | strand | MM | target_seq | PAM | alignment | distance | position | gene name | gene id |
| GU305914.1 | 226424 | 226446 | - | 0 | TCGAATCGTTCGGCTCCGTA | CGG | \|\|\|\|\|\|\|\|[\|\|\|\|\|\|\|\|\|\|\|\|]PAM | NA | Intergenic | NA | NA |
| T17522 | TCGTTCGGCTCCGTACGGCTTGG | 1000 | | | | | | | | | |
| Oligo fwd | CACCTCGTTCGGCTCCGTACGGCT | | | | | | | | | | |
| Oligo rev | AAACAGCCGTACGGAGCCGAACGA | | | | | | | | | | |
| Chromosome | start | end | strand | MM | target_seq | PAM | alignment | distance | position | gene name | gene id |
| GU305914.1 | 226419 | 226441 | - | 0 | TCGTTCGGCTCCGTACGGCT | TGG | \|\|\|\|\|\|\|\|[\|\|\|\|\|\|\|\|\|\|\|\|]PAM | NA | Intergenic | NA | NA |

<table>
<tr><td>T17523</td><td>CGGCTCCGTACGGCTTGGCCTGG</td><td>1000</td><td></td><td></td><td></td><td></td><td></td><td></td><td></td><td></td><td></td></tr>
<tr><td>Oligo fwd</td><td>CACCCGGCTCCGTACGGCTTGGCC</td><td></td><td></td><td></td><td></td><td></td><td></td><td></td><td></td><td></td><td></td></tr>
<tr><td>Oligo rev</td><td>AAACGGCCAAGCCGTACGGAGCCG</td><td></td><td></td><td></td><td></td><td></td><td></td><td></td><td></td><td></td><td></td></tr>
<tr><td>Chromosome</td><td>start</td><td>end</td><td>strand</td><td>MM</td><td>target_seq</td><td>PAM</td><td>alignment</td><td>distance</td><td>position</td><td>gene name</td><td>gene id</td></tr>
<tr><td>GU305914.1</td><td>226414</td><td>226436</td><td>-</td><td>0</td><td>CGGCTCCGTACGGCTTGGCC</td><td>TGG</td><td>||||||||[||||||||||||]PAM</td><td>NA</td><td>Intergenic</td><td>NA</td><td>NA</td></tr>
<tr><td>T17524</td><td>CTCCGTACGGCTTGGCCTGGTGG</td><td>1000</td><td></td><td></td><td></td><td></td><td></td><td></td><td></td><td></td><td></td></tr>
<tr><td>Oligo fwd</td><td>CACCCTCCGTACGGCTTGGCCTGG</td><td></td><td></td><td></td><td></td><td></td><td></td><td></td><td></td><td></td><td></td></tr>
<tr><td>Oligo rev</td><td>AAACCCAGGCCAAGCCGTACGGAG</td><td></td><td></td><td></td><td></td><td></td><td></td><td></td><td></td><td></td><td></td></tr>
<tr><td>Chromosome</td><td>start</td><td>end</td><td>strand</td><td>MM</td><td>target_seq</td><td>PAM</td><td>alignment</td><td>distance</td><td>position</td><td>gene name</td><td>gene id</td></tr>
<tr><td>GU305914.1</td><td>226411</td><td>226433</td><td>-</td><td>0</td><td>CTCCGTACGGCTTGGCCTGG</td><td>TGG</td><td>||||||||[||||||||||||]PAM</td><td>NA</td><td>Intergenic</td><td>NA</td><td>NA</td></tr>
<tr><td>T17525</td><td>GGCTTGGCCTGGTGGTTAAACGG</td><td>1000</td><td></td><td></td><td></td><td></td><td></td><td></td><td></td><td></td><td></td></tr>
<tr><td>Oligo fwd</td><td>CACCGGCTTGGCCTGGTGGTTAAA</td><td></td><td></td><td></td><td></td><td></td><td></td><td></td><td></td><td></td><td></td></tr>
<tr><td>Oligo rev</td><td>AAACTTTAACCACCAGGCCAAGCC</td><td></td><td></td><td></td><td></td><td></td><td></td><td></td><td></td><td></td><td></td></tr>
<tr><td>Chromosome</td><td>start</td><td>end</td><td>strand</td><td>MM</td><td>target_seq</td><td>PAM</td><td>alignment</td><td>distance</td><td>position</td><td>gene name</td><td>gene id</td></tr>
<tr><td>GU305914.1</td><td>226403</td><td>226425</td><td>-</td><td>0</td><td>GGCTTGGCCTGGTGGTTAAA</td><td>CGG</td><td>||||||||[||||||||||||]PAM</td><td>NA</td><td>Intergenic</td><td>NA</td><td>NA</td></tr>
<tr><td>T17526</td><td>GCTTGGCCTGGTGGTTAAACGGG</td><td>1000</td><td></td><td></td><td></td><td></td><td></td><td></td><td></td><td></td><td></td></tr>
<tr><td>Oligo fwd</td><td>CACCGCTTGGCCTGGTGGTTAAAC</td><td></td><td></td><td></td><td></td><td></td><td></td><td></td><td></td><td></td><td></td></tr>
<tr><td>Oligo rev</td><td>AAACGTTTAACCACCAGGCCAAGC</td><td></td><td></td><td></td><td></td><td></td><td></td><td></td><td></td><td></td><td></td></tr>
<tr><td>Chromosome</td><td>start</td><td>end</td><td>strand</td><td>MM</td><td>target_seq</td><td>PAM</td><td>alignment</td><td>distance</td><td>position</td><td>gene name</td><td>gene id</td></tr>
<tr><td>GU305914.1</td><td>226402</td><td>226424</td><td>-</td><td>0</td><td>GCTTGGCCTGGTGGTTAAAC</td><td>GGG</td><td>||||||||[||||||||||||]PAM</td><td>NA</td><td>Intergenic</td><td>NA</td><td>NA</td></tr>
<tr><td>T17527</td><td>CTGGTGGTTAAACGGGTCAGTGG</td><td>1000</td><td></td><td></td><td></td><td></td><td></td><td></td><td></td><td></td><td></td></tr>
<tr><td>Oligo fwd</td><td>CACCCTGGTGGTTAAACGGGTCAG</td><td></td><td></td><td></td><td></td><td></td><td></td><td></td><td></td><td></td><td></td></tr>
<tr><td>Oligo rev</td><td>AAACCTGACCCGTTTAACCACCAG</td><td></td><td></td><td></td><td></td><td></td><td></td><td></td><td></td><td></td><td></td></tr>
<tr><td>Chromosome</td><td>start</td><td>end</td><td>strand</td><td>MM</td><td>target_seq</td><td>PAM</td><td>alignment</td><td>distance</td><td>position</td><td>gene name</td><td>gene id</td></tr>
<tr><td>GU305914.1</td><td>226395</td><td>226417</td><td>-</td><td>0</td><td>CTGGTGGTTAAACGGGTCAG</td><td>TGG</td><td>||||||||[||||||||||||]PAM</td><td>NA</td><td>Intergenic</td><td>NA</td><td>NA</td></tr>
<tr><td>T17528</td><td>GTGGTTAAACGGGTCAGTGGCGG</td><td>1000</td><td></td><td></td><td></td><td></td><td></td><td></td><td></td><td></td><td></td></tr>
<tr><td>Oligo fwd</td><td>CACCGTGGTTAAACGGGTCAGTGG</td><td></td><td></td><td></td><td></td><td></td><td></td><td></td><td></td><td></td><td></td></tr>
<tr><td>Oligo rev</td><td>AAACCCACTGACCCGTTTAACCAC</td><td></td><td></td><td></td><td></td><td></td><td></td><td></td><td></td><td></td><td></td></tr>
<tr><td>Chromosome</td><td>start</td><td>end</td><td>strand</td><td>MM</td><td>target_seq</td><td>PAM</td><td>alignment</td><td>distance</td><td>position</td><td>gene name</td><td>gene id</td></tr>
<tr><td>GU305914.1</td><td>226392</td><td>226414</td><td>-</td><td>0</td><td>GTGGTTAAACGGGTCAGTGG</td><td>CGG</td><td>||||||||[||||||||||||]PAM</td><td>NA</td><td>Intergenic</td><td>NA</td><td>NA</td></tr>
<tr><td>T17529</td><td>ATCGTATCCGCCCGAGTCCTCGG</td><td>1000</td><td></td><td></td><td></td><td></td><td></td><td></td><td></td><td></td><td></td></tr>
<tr><td>Oligo fwd</td><td>CACCATCGTATCCGCCCGAGTCCT</td><td></td><td></td><td></td><td></td><td></td><td></td><td></td><td></td><td></td><td></td></tr>
<tr><td>Oligo rev</td><td>AAACAGGACTCGGGCGGATACGAT</td><td></td><td></td><td></td><td></td><td></td><td></td><td></td><td></td><td></td><td></td></tr>
<tr><td>Chromosome</td><td>start</td><td>end</td><td>strand</td><td>MM</td><td>target_seq</td><td>PAM</td><td>alignment</td><td>distance</td><td>position</td><td>gene name</td><td>gene id</td></tr>
<tr><td>GU305914.1</td><td>226362</td><td>226384</td><td>-</td><td>0</td><td>ATCGTATCCGCCCGAGTCCT</td><td>CGG</td><td>||||||||[||||||||||||]PAM</td><td>NA</td><td>Intergenic</td><td>NA</td><td>NA</td></tr>
<tr><td>T17530</td><td>TCGTATCCGCCCGAGTCCTCGGG</td><td>1000</td><td></td><td></td><td></td><td></td><td></td><td></td><td></td><td></td><td></td></tr>
<tr><td>Oligo fwd</td><td>CACCTCGTATCCGCCCGAGTCCTC</td><td></td><td></td><td></td><td></td><td></td><td></td><td></td><td></td><td></td><td></td></tr>
<tr><td>Oligo rev</td><td>AAACGAGGACTCGGGCGGATACGA</td><td></td><td></td><td></td><td></td><td></td><td></td><td></td><td></td><td></td><td></td></tr>
<tr><td>Chromosome</td><td>start</td><td>end</td><td>strand</td><td>MM</td><td>target_seq</td><td>PAM</td><td>alignment</td><td>distance</td><td>position</td><td>gene name</td><td>gene id</td></tr>
<tr><td>GU305914.1</td><td>226361</td><td>226383</td><td>-</td><td>0</td><td>TCGTATCCGCCCGAGTCCTC</td><td>GGG</td><td>||||||||[||||||||||||]PAM</td><td>NA</td><td>Intergenic</td><td>NA</td><td>NA</td></tr>
<tr><td>T17531</td><td>GACCCTGTCGATCTACAGAACGG</td><td>1000</td><td></td><td></td><td></td><td></td><td></td><td></td><td></td><td></td><td></td></tr>
<tr><td>Oligo fwd</td><td>CACCGACCCTGTCGATCTACAGAA</td><td></td><td></td><td></td><td></td><td></td><td></td><td></td><td></td><td></td><td></td></tr>
<tr><td>Oligo rev</td><td>AAACTTCTGTAGATCGACAGGGTC</td><td></td><td></td><td></td><td></td><td></td><td></td><td></td><td></td><td></td><td></td></tr>
<tr><td>Chromosome</td><td>start</td><td>end</td><td>strand</td><td>MM</td><td>target_seq</td><td>PAM</td><td>alignment</td><td>distance</td><td>position</td><td>gene name</td><td>gene id</td></tr>
<tr><td>GU305914.1</td><td>226332</td><td>226354</td><td>-</td><td>0</td><td>GACCCTGTCGATCTACAGAA</td><td>CGG</td><td>||||||||[||||||||||||]PAM</td><td>NA</td><td>Intergenic</td><td>NA</td><td>NA</td></tr>
<tr><td>T17532</td><td>GTCGATCTACAGAACGGCGATGG</td><td>1000</td><td></td><td></td><td></td><td></td><td></td><td></td><td></td><td></td><td></td></tr>
<tr><td>Oligo fwd</td><td>CACCGTCGATCTACAGAACGGCGA</td><td></td><td></td><td></td><td></td><td></td><td></td><td></td><td></td><td></td><td></td></tr>
<tr><td>Oligo rev</td><td>AAACTCGCCGTTCTGTAGATCGAC</td><td></td><td></td><td></td><td></td><td></td><td></td><td></td><td></td><td></td><td></td></tr>
<tr><td>Chromosome</td><td>start</td><td>end</td><td>strand</td><td>MM</td><td>target_seq</td><td>PAM</td><td>alignment</td><td>distance</td><td>position</td><td>gene name</td><td>gene id</td></tr>
<tr><td>GU305914.1</td><td>226326</td><td>226348</td><td>-</td><td>0</td><td>GTCGATCTACAGAACGGCGA</td><td>TGG</td><td>||||||||[||||||||||||]PAM</td><td>NA</td><td>Intergenic</td><td>NA</td><td>NA</td></tr>
<tr><td>T17533</td><td>CAGAACGGCGATGGAGACCTCGG</td><td>1000</td><td></td><td></td><td></td><td></td><td></td><td></td><td></td><td></td><td></td></tr>
<tr><td>Oligo fwd</td><td>CACCCAGAACGGCGATGGAGACCT</td><td></td><td></td><td></td><td></td><td></td><td></td><td></td><td></td><td></td><td></td></tr>
<tr><td>Oligo rev</td><td>AAACAGGTCTCCATCGCCGTTCTG</td><td></td><td></td><td></td><td></td><td></td><td></td><td></td><td></td><td></td><td></td></tr>
<tr><td>Chromosome</td><td>start</td><td>end</td><td>strand</td><td>MM</td><td>target_seq</td><td>PAM</td><td>alignment</td><td>distance</td><td>position</td><td>gene name</td><td>gene id</td></tr>
<tr><td>GU305914.1</td><td>226317</td><td>226339</td><td>-</td><td>0</td><td>CAGAACGGCGATGGAGACCT</td><td>CGG</td><td>||||||||[||||||||||||]PAM</td><td>NA</td><td>Intergenic</td><td>NA</td><td>NA</td></tr>
<tr><td>T17534</td><td>AGAACGGCGATGGAGACCTCGGG</td><td>1000</td><td></td><td></td><td></td><td></td><td></td><td></td><td></td><td></td><td></td></tr>
<tr><td>Oligo fwd</td><td>CACCAGAACGGCGATGGAGACCTC</td><td></td><td></td><td></td><td></td><td></td><td></td><td></td><td></td><td></td><td></td></tr>
<tr><td>Oligo rev</td><td>AAACGAGGTCTCCATCGCCGTTCT</td><td></td><td></td><td></td><td></td><td></td><td></td><td></td><td></td><td></td><td></td></tr>
<tr><td>Chromosome</td><td>start</td><td>end</td><td>strand</td><td>MM</td><td>target_seq</td><td>PAM</td><td>alignment</td><td>distance</td><td>position</td><td>gene name</td><td>gene id</td></tr>
<tr><td>GU305914.1</td><td>226316</td><td>226338</td><td>-</td><td>0</td><td>AGAACGGCGATGGAGACCTC</td><td>GGG</td><td>||||||||[||||||||||||]PAM</td><td>NA</td><td>Intergenic</td><td>NA</td><td>NA</td></tr>
<tr><td>T17535</td><td>GCGATGGAGACCTCGGGACCTGG</td><td>1000</td><td></td><td></td><td></td><td></td><td></td><td></td><td></td><td></td><td></td></tr>
<tr><td>Oligo fwd</td><td>CACCGCGATGGAGACCTCGGGACC</td><td></td><td></td><td></td><td></td><td></td><td></td><td></td><td></td><td></td><td></td></tr>
<tr><td>Oligo rev</td><td>AAACGGTCCCGAGGTCTCCATCGC</td><td></td><td></td><td></td><td></td><td></td><td></td><td></td><td></td><td></td><td></td></tr>
<tr><td>Chromosome</td><td>start</td><td>end</td><td>strand</td><td>MM</td><td>target_seq</td><td>PAM</td><td>alignment</td><td>distance</td><td>position</td><td>gene name</td><td>gene id</td></tr>
<tr><td>GU305914.1</td><td>226310</td><td>226332</td><td>-</td><td>0</td><td>GCGATGGAGACCTCGGGACC</td><td>TGG</td><td>||||||||[||||||||||||]PAM</td><td>NA</td><td>Intergenic</td><td>NA</td><td>NA</td></tr>
<tr><td>T17536</td><td>CGTACGAGCAACACCCGCCGCGG</td><td>1000</td><td></td><td></td><td></td><td></td><td></td><td></td><td></td><td></td><td></td></tr>
<tr><td>Oligo fwd</td><td>CACCCGTACGAGCAACACCCGCCG</td><td></td><td></td><td></td><td></td><td></td><td></td><td></td><td></td><td></td><td></td></tr>
<tr><td>Oligo rev</td><td>AAACCGGCGGGTGTTGCTCGTACG</td><td></td><td></td><td></td><td></td><td></td><td></td><td></td><td></td><td></td><td></td></tr>
<tr><td>Chromosome</td><td>start</td><td>end</td><td>strand</td><td>MM</td><td>target_seq</td><td>PAM</td><td>alignment</td><td>distance</td><td>position</td><td>gene name</td><td>gene id</td></tr>
<tr><td>GU305914.1</td><td>226273</td><td>226295</td><td>-</td><td>0</td><td>CGTACGAGCAACACCCGCCG</td><td>CGG</td><td>||||||||[||||||||||||]PAM</td><td>NA</td><td>Intergenic</td><td>NA</td><td>NA</td></tr>
<tr><td>GU305914.1</td><td>27178</td><td>27200</td><td>+</td><td>3</td><td>GGTCCGAGCAACACACGCCG</td><td>CGG</td><td>-||-||||[||||||-|||||]PAM</td><td>NA</td><td>Intergenic</td><td>NA</td><td>NA</td></tr>
<tr><td>T17537</td><td>GTACGAGCAACACCCGCCGCGGG</td><td>1000</td><td></td><td></td><td></td><td></td><td></td><td></td><td></td><td></td><td></td></tr>
<tr><td>Oligo fwd</td><td>CACCGTACGAGCAACACCCGCCGC</td><td></td><td></td><td></td><td></td><td></td><td></td><td></td><td></td><td></td><td></td></tr>
<tr><td>Oligo rev</td><td>AAACGCGGCGGGTGTTGCTCGTAC</td><td></td><td></td><td></td><td></td><td></td><td></td><td></td><td></td><td></td><td></td></tr>
<tr><td>Chromosome</td><td>start</td><td>end</td><td>strand</td><td>MM</td><td>target_seq</td><td>PAM</td><td>alignment</td><td>distance</td><td>position</td><td>gene name</td><td>gene id</td></tr>
</table>

GU305914.1 226272 226294 - 0 GTACGAGCAACACCCGCCGC GGG ||||||||[||||||||||||]PAM NA Intergenic NA NA

T17538 ACACCCGCCGCGGGAGACGACGG 1000
Oligo fwd CACCACACCCGCCGCGGGAGACGA
Oligo rev AAACTCGTCTCCCGCGGCGGGTGT
Chromosome start end strand MM target_seq PAM alignment distance position gene name gene id
GU305914.1 226263 226285 - 0 ACACCCGCCGCGGGAGACGA CGG ||||||||[||||||||||||]PAM NA Intergenic NA NA

T17539 CACCCGCCGCGGGAGACGACGGG 1000
Oligo fwd CACCCACCCGCCGCGGGAGACGAC
Oligo rev AAACGTCGTCTCCCGCGGCGGGTG
Chromosome start end strand MM target_seq PAM alignment distance position gene name gene id
GU305914.1 226262 226284 - 0 CACCCGCCGCGGGAGACGAC GGG ||||||||[||||||||||||]PAM NA Intergenic NA NA

T17540 CCGCGGGAGACGACGGGAAGCGG 1000
Oligo fwd CACCCCGCGGGAGACGACGGGAAG
Oligo rev AAACCTTCCCGTCGTCTCCCGCGG
Chromosome start end strand MM target_seq PAM alignment distance position gene name gene id
GU305914.1 226256 226278 - 0 CCGCGGGAGACGACGGGAAG CGG ||||||||[||||||||||||]PAM NA Intergenic NA NA

T17541 TCAGACCCGAAGACCAAGTCCGG 1000
Oligo fwd CACCTCAGACCCGAAGACCAAGTC
Oligo rev AAACGACTTGGTCTTCGGGTCTGA
Chromosome start end strand MM target_seq PAM alignment distance position gene name gene id
GU305914.1 226209 226231 - 0 TCAGACCCGAAGACCAAGTC CGG ||||||||[||||||||||||]PAM NA Intergenic NA NA

T17542 CAAGTCCGGACCTCCGCCCGTGG 1000
Oligo fwd CACCCAAGTCCGGACCTCCGCCCG
Oligo rev AAACCGGGCGGAGGTCCGGACTTG
Chromosome start end strand MM target_seq PAM alignment distance position gene name gene id
GU305914.1 226195 226217 - 0 CAAGTCCGGACCTCCGCCCG TGG ||||||||[||||||||||||]PAM NA Intergenic NA NA

T17543 CCCGTGGTTCGAGATCGAAACGG 1000
Oligo fwd CACCCCCGTGGTTCGAGATCGAAA
Oligo rev AAACTTTCGATCTCGAACCACGGG
Chromosome start end strand MM target_seq PAM alignment distance position gene name gene id
GU305914.1 226179 226201 - 0 CCCGTGGTTCGAGATCGAAA CGG ||||||||[||||||||||||]PAM NA Intergenic NA NA

T17544 AGATCGAAACGGTGTCAGATCGG 1000
Oligo fwd CACCAGATCGAAACGGTGTCAGAT
Oligo rev AAACATCTGACACCGTTTCGATCT
Chromosome start end strand MM target_seq PAM alignment distance position gene name gene id
GU305914.1 226168 226190 - 0 AGATCGAAACGGTGTCAGAT CGG ||||||||[||||||||||||]PAM NA Intergenic NA NA

T17545 CGAAACGGTGTCAGATCGGATGG 1000
Oligo fwd CACCCGAAACGGTGTCAGATCGGA
Oligo rev AAACTCCGATCTGACACCGTTTCG
Chromosome start end strand MM target_seq PAM alignment distance position gene name gene id
GU305914.1 226164 226186 - 0 CGAAACGGTGTCAGATCGGA TGG ||||||||[||||||||||||]PAM NA Intergenic NA NA

T17546 GTCAGATCGGATGGTGCTTCAGG 1000
Oligo fwd CACCGTCAGATCGGATGGTGCTTC
Oligo rev AAACGAAGCACCATCCGATCTGAC
Chromosome start end strand MM target_seq PAM alignment distance position gene name gene id
GU305914.1 226155 226177 - 0 GTCAGATCGGATGGTGCTTC AGG ||||||||[||||||||||||]PAM NA Intergenic NA NA

T17547 TCGGATGGTGCTTCAGGACCAGG 1000
Oligo fwd CACCTCGGATGGTGCTTCAGGACC
Oligo rev AAACGGTCCTGAAGCACCATCCGA
Chromosome start end strand MM target_seq PAM alignment distance position gene name gene id
GU305914.1 226149 226171 - 0 TCGGATGGTGCTTCAGGACC AGG ||||||||[||||||||||||]PAM NA Intergenic NA NA

T17548 TGGTGCTTCAGGACCAGGCCTGG 1000
Oligo fwd CACCTGGTGCTTCAGGACCAGGCC
Oligo rev AAACGGCCTGGTCCTGAAGCACCA
Chromosome start end strand MM target_seq PAM alignment distance position gene name gene id
GU305914.1 226144 226166 - 0 TGGTGCTTCAGGACCAGGCC TGG ||||||||[||||||||||||]PAM NA Intergenic NA NA

T17549 GGTGCTTCAGGACCAGGCCTGGG 1000
Oligo fwd CACCGGTGCTTCAGGACCAGGCCT
Oligo rev AAACAGGCCTGGTCCTGAAGCACC
Chromosome start end strand MM target_seq PAM alignment distance position gene name gene id
GU305914.1 226143 226165 - 0 GGTGCTTCAGGACCAGGCCT GGG ||||||||[||||||||||||]PAM NA Intergenic NA NA

T17550 TCAGGACCAGGCCTGGGCCGCGG 1000
Oligo fwd CACCTCAGGACCAGGCCTGGGCCG
Oligo rev AAACCGGCCCAGGCCTGGTCCTGA
Chromosome start end strand MM target_seq PAM alignment distance position gene name gene id
GU305914.1 226137 226159 - 0 TCAGGACCAGGCCTGGGCCG CGG ||||||||[||||||||||||]PAM NA Intergenic NA NA

T17551 CAGGACCAGGCCTGGGCCGCGGG 1000
Oligo fwd CACCCAGGACCAGGCCTGGGCCGC
Oligo rev AAACGCGGCCCAGGCCTGGTCCTG
Chromosome start end strand MM target_seq PAM alignment distance position gene name gene id
GU305914.1 226136 226158 - 0 CAGGACCAGGCCTGGGCCGC GGG ||||||||[||||||||||||]PAM NA Intergenic NA NA

T17552 AGGACCAGGCCTGGGCCGCGGGG 1000
Oligo fwd CACCAGGACCAGGCCTGGGCCGCG
Oligo rev AAACCGCGGCCCAGGCCTGGTCCT

<table>
<tr><td>Chromosome</td><td>start</td><td>end</td><td>strand</td><td>MM</td><td>target_seq</td><td>PAM</td><td>alignment</td><td>distance</td><td>position</td><td>gene name</td><td>gene id</td></tr>
<tr><td>GU305914.1</td><td>226135</td><td>226157</td><td>-</td><td>0</td><td>AGGACCAGGCCTGGGCCGCG</td><td>GGG</td><td>||||||||[||||||||||||]PAM</td><td>NA</td><td>Intergenic</td><td>NA</td><td>NA</td></tr>
<tr><td>T17553</td><td>GACCCTCTGCGTTTCATGCACGG</td><td>1000</td><td></td><td></td><td></td><td></td><td></td><td></td><td></td><td></td><td></td></tr>
<tr><td>Oligo fwd</td><td>CACCGACCCTCTGCGTTTCATGCA</td><td></td><td></td><td></td><td></td><td></td><td></td><td></td><td></td><td></td><td></td></tr>
<tr><td>Oligo rev</td><td>AAACTGCATGAAACGCAGAGGGTC</td><td></td><td></td><td></td><td></td><td></td><td></td><td></td><td></td><td></td><td></td></tr>
<tr><td>Chromosome</td><td>start</td><td>end</td><td>strand</td><td>MM</td><td>target_seq</td><td>PAM</td><td>alignment</td><td>distance</td><td>position</td><td>gene name</td><td>gene id</td></tr>
<tr><td>GU305914.1</td><td>226113</td><td>226135</td><td>-</td><td>0</td><td>GACCCTCTGCGTTTCATGCA</td><td>CGG</td><td>||||||||[||||||||||||]PAM</td><td>NA</td><td>Intergenic</td><td>NA</td><td>NA</td></tr>
<tr><td>T17554</td><td>GGTCATCCAACCGAGACTGCTGG</td><td>1000</td><td></td><td></td><td></td><td></td><td></td><td></td><td></td><td></td><td></td></tr>
<tr><td>Oligo fwd</td><td>CACCGGTCATCCAACCGAGACTGC</td><td></td><td></td><td></td><td></td><td></td><td></td><td></td><td></td><td></td><td></td></tr>
<tr><td>Oligo rev</td><td>AAACGCAGTCTCGGTTGGATGACC</td><td></td><td></td><td></td><td></td><td></td><td></td><td></td><td></td><td></td><td></td></tr>
<tr><td>Chromosome</td><td>start</td><td>end</td><td>strand</td><td>MM</td><td>target_seq</td><td>PAM</td><td>alignment</td><td>distance</td><td>position</td><td>gene name</td><td>gene id</td></tr>
<tr><td>GU305914.1</td><td>226092</td><td>226114</td><td>-</td><td>0</td><td>GGTCATCCAACCGAGACTGC</td><td>TGG</td><td>||||||||[||||||||||||]PAM</td><td>NA</td><td>Intergenic</td><td>NA</td><td>NA</td></tr>
<tr><td>T17555</td><td>GTCATCCAACCGAGACTGCTGGG</td><td>1000</td><td></td><td></td><td></td><td></td><td></td><td></td><td></td><td></td><td></td></tr>
<tr><td>Oligo fwd</td><td>CACCGTCATCCAACCGAGACTGCT</td><td></td><td></td><td></td><td></td><td></td><td></td><td></td><td></td><td></td><td></td></tr>
<tr><td>Oligo rev</td><td>AAACAGCAGTCTCGGTTGGATGAC</td><td></td><td></td><td></td><td></td><td></td><td></td><td></td><td></td><td></td><td></td></tr>
<tr><td>Chromosome</td><td>start</td><td>end</td><td>strand</td><td>MM</td><td>target_seq</td><td>PAM</td><td>alignment</td><td>distance</td><td>position</td><td>gene name</td><td>gene id</td></tr>
<tr><td>GU305914.1</td><td>226091</td><td>226113</td><td>-</td><td>0</td><td>GTCATCCAACCGAGACTGCT</td><td>GGG</td><td>||||||||[||||||||||||]PAM</td><td>NA</td><td>Intergenic</td><td>NA</td><td>NA</td></tr>
<tr><td>T17556</td><td>TCATCCAACCGAGACTGCTGGGG</td><td>1000</td><td></td><td></td><td></td><td></td><td></td><td></td><td></td><td></td><td></td></tr>
<tr><td>Oligo fwd</td><td>CACCTCATCCAACCGAGACTGCTG</td><td></td><td></td><td></td><td></td><td></td><td></td><td></td><td></td><td></td><td></td></tr>
<tr><td>Oligo rev</td><td>AAACCAGCAGTCTCGGTTGGATGA</td><td></td><td></td><td></td><td></td><td></td><td></td><td></td><td></td><td></td><td></td></tr>
<tr><td>Chromosome</td><td>start</td><td>end</td><td>strand</td><td>MM</td><td>target_seq</td><td>PAM</td><td>alignment</td><td>distance</td><td>position</td><td>gene name</td><td>gene id</td></tr>
<tr><td>GU305914.1</td><td>226090</td><td>226112</td><td>-</td><td>0</td><td>TCATCCAACCGAGACTGCTG</td><td>GGG</td><td>||||||||[||||||||||||]PAM</td><td>NA</td><td>Intergenic</td><td>NA</td><td>NA</td></tr>
<tr><td>T17557</td><td>CATCCAACCGAGACTGCTGGGGG</td><td>1000</td><td></td><td></td><td></td><td></td><td></td><td></td><td></td><td></td><td></td></tr>
<tr><td>Oligo fwd</td><td>CACCCATCCAACCGAGACTGCTGG</td><td></td><td></td><td></td><td></td><td></td><td></td><td></td><td></td><td></td><td></td></tr>
<tr><td>Oligo rev</td><td>AAACCCAGCAGTCTCGGTTGGATG</td><td></td><td></td><td></td><td></td><td></td><td></td><td></td><td></td><td></td><td></td></tr>
<tr><td>Chromosome</td><td>start</td><td>end</td><td>strand</td><td>MM</td><td>target_seq</td><td>PAM</td><td>alignment</td><td>distance</td><td>position</td><td>gene name</td><td>gene id</td></tr>
<tr><td>GU305914.1</td><td>226089</td><td>226111</td><td>-</td><td>0</td><td>CATCCAACCGAGACTGCTGG</td><td>GGG</td><td>||||||||[||||||||||||]PAM</td><td>NA</td><td>Intergenic</td><td>NA</td><td>NA</td></tr>
<tr><td>T17558</td><td>ACTGCTGGGGGTAGCCACGACGG</td><td>1000</td><td></td><td></td><td></td><td></td><td></td><td></td><td></td><td></td><td></td></tr>
<tr><td>Oligo fwd</td><td>CACCACTGCTGGGGGTAGCCACGA</td><td></td><td></td><td></td><td></td><td></td><td></td><td></td><td></td><td></td><td></td></tr>
<tr><td>Oligo rev</td><td>AAACTCGTGGCTACCCCCAGCAGT</td><td></td><td></td><td></td><td></td><td></td><td></td><td></td><td></td><td></td><td></td></tr>
<tr><td>Chromosome</td><td>start</td><td>end</td><td>strand</td><td>MM</td><td>target_seq</td><td>PAM</td><td>alignment</td><td>distance</td><td>position</td><td>gene name</td><td>gene id</td></tr>
<tr><td>GU305914.1</td><td>226077</td><td>226099</td><td>-</td><td>0</td><td>ACTGCTGGGGGTAGCCACGA</td><td>CGG</td><td>||||||||[||||||||||||]PAM</td><td>NA</td><td>Intergenic</td><td>NA</td><td>NA</td></tr>
<tr><td>T17559</td><td>GTCGTTCTCATCCTCCTTCCAGG</td><td>1000</td><td></td><td></td><td></td><td></td><td></td><td></td><td></td><td></td><td></td></tr>
<tr><td>Oligo fwd</td><td>CACCGTCGTTCTCATCCTCCTTCC</td><td></td><td></td><td></td><td></td><td></td><td></td><td></td><td></td><td></td><td></td></tr>
<tr><td>Oligo rev</td><td>AAACGGAAGGAGGATGAGAACGAC</td><td></td><td></td><td></td><td></td><td></td><td></td><td></td><td></td><td></td><td></td></tr>
<tr><td>Chromosome</td><td>start</td><td>end</td><td>strand</td><td>MM</td><td>target_seq</td><td>PAM</td><td>alignment</td><td>distance</td><td>position</td><td>gene name</td><td>gene id</td></tr>
<tr><td>GU305914.1</td><td>226051</td><td>226073</td><td>-</td><td>0</td><td>GTCGTTCTCATCCTCCTTCC</td><td>AGG</td><td>||||||||[||||||||||||]PAM</td><td>NA</td><td>Intergenic</td><td>NA</td><td>NA</td></tr>
<tr><td>T17560</td><td>GACTCCCCGTCGTTTCCAAACGG</td><td>1000</td><td></td><td></td><td></td><td></td><td></td><td></td><td></td><td></td><td></td></tr>
<tr><td>Oligo fwd</td><td>CACCGACTCCCCGTCGTTTCCAAA</td><td></td><td></td><td></td><td></td><td></td><td></td><td></td><td></td><td></td><td></td></tr>
<tr><td>Oligo rev</td><td>AAACTTTGGAAACGACGGGGAGTC</td><td></td><td></td><td></td><td></td><td></td><td></td><td></td><td></td><td></td><td></td></tr>
<tr><td>Chromosome</td><td>start</td><td>end</td><td>strand</td><td>MM</td><td>target_seq</td><td>PAM</td><td>alignment</td><td>distance</td><td>position</td><td>gene name</td><td>gene id</td></tr>
<tr><td>GU305914.1</td><td>226027</td><td>226049</td><td>-</td><td>0</td><td>GACTCCCCGTCGTTTCCAAA</td><td>CGG</td><td>||||||||[||||||||||||]PAM</td><td>NA</td><td>Intergenic</td><td>NA</td><td>NA</td></tr>
<tr><td>T17561</td><td>TCCAAACGGTCGACCCCAAACGG</td><td>1000</td><td></td><td></td><td></td><td></td><td></td><td></td><td></td><td></td><td></td></tr>
<tr><td>Oligo fwd</td><td>CACCTCCAAACGGTCGACCCCAAA</td><td></td><td></td><td></td><td></td><td></td><td></td><td></td><td></td><td></td><td></td></tr>
<tr><td>Oligo rev</td><td>AAACTTTGGGGTCGACCGTTTGGA</td><td></td><td></td><td></td><td></td><td></td><td></td><td></td><td></td><td></td><td></td></tr>
<tr><td>Chromosome</td><td>start</td><td>end</td><td>strand</td><td>MM</td><td>target_seq</td><td>PAM</td><td>alignment</td><td>distance</td><td>position</td><td>gene name</td><td>gene id</td></tr>
<tr><td>GU305914.1</td><td>226013</td><td>226035</td><td>-</td><td>0</td><td>TCCAAACGGTCGACCCCAAA</td><td>CGG</td><td>||||||||[||||||||||||]PAM</td><td>NA</td><td>Intergenic</td><td>NA</td><td>NA</td></tr>
<tr><td>T17562</td><td>CGACCCCAAACGGCTGCGCGTGG</td><td>1000</td><td></td><td></td><td></td><td></td><td></td><td></td><td></td><td></td><td></td></tr>
<tr><td>Oligo fwd</td><td>CACCCGACCCCAAACGGCTGCGCG</td><td></td><td></td><td></td><td></td><td></td><td></td><td></td><td></td><td></td><td></td></tr>
<tr><td>Oligo rev</td><td>AAACCGCGCAGCCGTTTGGGGTCG</td><td></td><td></td><td></td><td></td><td></td><td></td><td></td><td></td><td></td><td></td></tr>
<tr><td>Chromosome</td><td>start</td><td>end</td><td>strand</td><td>MM</td><td>target_seq</td><td>PAM</td><td>alignment</td><td>distance</td><td>position</td><td>gene name</td><td>gene id</td></tr>
<tr><td>GU305914.1</td><td>226003</td><td>226025</td><td>-</td><td>0</td><td>CGACCCCAAACGGCTGCGCG</td><td>TGG</td><td>||||||||[||||||||||||]PAM</td><td>NA</td><td>Intergenic</td><td>NA</td><td>NA</td></tr>
<tr><td>T17563</td><td>CGGCTGCGCGTGGAGCGCGACGG</td><td>1000</td><td></td><td></td><td></td><td></td><td></td><td></td><td></td><td></td><td></td></tr>
<tr><td>Oligo fwd</td><td>CACCCGGCTGCGCGTGGAGCGCGA</td><td></td><td></td><td></td><td></td><td></td><td></td><td></td><td></td><td></td><td></td></tr>
<tr><td>Oligo rev</td><td>AAACTCGCGCTCCACGCGCAGCCG</td><td></td><td></td><td></td><td></td><td></td><td></td><td></td><td></td><td></td><td></td></tr>
<tr><td>Chromosome</td><td>start</td><td>end</td><td>strand</td><td>MM</td><td>target_seq</td><td>PAM</td><td>alignment</td><td>distance</td><td>position</td><td>gene name</td><td>gene id</td></tr>
<tr><td>GU305914.1</td><td>225993</td><td>226015</td><td>-</td><td>0</td><td>CGGCTGCGCGTGGAGCGCGA</td><td>CGG</td><td>||||||||[||||||||||||]PAM</td><td>NA</td><td>Intergenic</td><td>NA</td><td>NA</td></tr>
<tr><td>T17564</td><td>CTGCGCGTGGAGCGCGACGGCGG</td><td>1000</td><td></td><td></td><td></td><td></td><td></td><td></td><td></td><td></td><td></td></tr>
<tr><td>Oligo fwd</td><td>CACCCTGCGCGTGGAGCGCGACGG</td><td></td><td></td><td></td><td></td><td></td><td></td><td></td><td></td><td></td><td></td></tr>
<tr><td>Oligo rev</td><td>AAACCCGTCGCGCTCCACGCGCAG</td><td></td><td></td><td></td><td></td><td></td><td></td><td></td><td></td><td></td><td></td></tr>
<tr><td>Chromosome</td><td>start</td><td>end</td><td>strand</td><td>MM</td><td>target_seq</td><td>PAM</td><td>alignment</td><td>distance</td><td>position</td><td>gene name</td><td>gene id</td></tr>
<tr><td>GU305914.1</td><td>225990</td><td>226012</td><td>-</td><td>0</td><td>CTGCGCGTGGAGCGCGACGG</td><td>CGG</td><td>||||||||[||||||||||||]PAM</td><td>NA</td><td>Intergenic</td><td>NA</td><td>NA</td></tr>
<tr><td>T17565</td><td>CGCGTGGAGCGCGACGGCGGCGG</td><td>1000</td><td></td><td></td><td></td><td></td><td></td><td></td><td></td><td></td><td></td></tr>
<tr><td>Oligo fwd</td><td>CACCCGCGTGGAGCGCGACGGCGG</td><td></td><td></td><td></td><td></td><td></td><td></td><td></td><td></td><td></td><td></td></tr>
<tr><td>Oligo rev</td><td>AAACCCGCCGTCGCGCTCCACGCG</td><td></td><td></td><td></td><td></td><td></td><td></td><td></td><td></td><td></td><td></td></tr>
<tr><td>Chromosome</td><td>start</td><td>end</td><td>strand</td><td>MM</td><td>target_seq</td><td>PAM</td><td>alignment</td><td>distance</td><td>position</td><td>gene name</td><td>gene id</td></tr>
<tr><td>GU305914.1</td><td>225987</td><td>226009</td><td>-</td><td>0</td><td>CGCGTGGAGCGCGACGGCGG</td><td>CGG</td><td>||||||||[||||||||||||]PAM</td><td>NA</td><td>Intergenic</td><td>NA</td><td>NA</td></tr>
<tr><td>GU305914.1</td><td>96942</td><td>96964</td><td>+</td><td>4</td><td>GGCGAGCAGGGCGACGGCGG</td><td>AGG</td><td>-|||-|-|[|-||||||||||]PAM</td><td>NA</td><td>Intergenic</td><td>NA</td><td>NA</td></tr>
<tr><td>GU305914.1</td><td>90265</td><td>90287</td><td>-</td><td>4</td><td>CGGGAGGCGGGCGACGGCGG</td><td>AGG</td><td>||-|-||-[|-||||||||||]PAM</td><td>NA</td><td>Intergenic</td><td>NA</td><td>NA</td></tr>
<tr><td>T17566</td><td>GTGGAGCGCGACGGCGGCGGCGG</td><td>1000</td><td></td><td></td><td></td><td></td><td></td><td></td><td></td><td></td><td></td></tr>
<tr><td>Oligo fwd</td><td>CACCGTGGAGCGCGACGGCGGCGG</td><td></td><td></td><td></td><td></td><td></td><td></td><td></td><td></td><td></td><td></td></tr>
<tr><td>Oligo rev</td><td>AAACCCGCCGCCGTCGCGCTCCAC</td><td></td><td></td><td></td><td></td><td></td><td></td><td></td><td></td><td></td><td></td></tr>
<tr><td>Chromosome</td><td>start</td><td>end</td><td>strand</td><td>MM</td><td>target_seq</td><td>PAM</td><td>alignment</td><td>distance</td><td>position</td><td>gene name</td><td>gene id</td></tr>
<tr><td>GU305914.1</td><td>225984</td><td>226006</td><td>-</td><td>0</td><td>GTGGAGCGCGACGGCGGCGG</td><td>CGG</td><td>||||||||[||||||||||||]PAM</td><td>NA</td><td>Intergenic</td><td>NA</td><td>NA</td></tr>
<tr><td>GU305914.1</td><td>176454</td><td>176476</td><td>+</td><td>4</td><td>GACGATGGCGACGGCGGCGG</td><td>AGG</td><td>|--||--|[||||||||||||]PAM</td><td>NA</td><td>Intergenic</td><td>NA</td><td>NA</td></tr>
</table>

GU305914.1 134259 134281 + 4 GGGAAGTGCGATGGCGGCGG AGG |-|-||-|[|||-||||||||]PAM NA Intergenic NA NA
GU305914.1 110629 110651 - 4 GTGCAGGACGCCGGCGGCGG AGG |||-||--[||-|||||||||]PAM NA Intergenic NA NA
GU305914.1 96945 96967 + 4 GAGCAGGGCGACGGCGGAGG CGG |-|-||-|[|||||||||-||]PAM NA Intergenic NA NA
GU305914.1 90262 90284 - 4 GAGGCGGGCGACGGCGGAGG CGG |-||-|-|[|||||||||-||]PAM NA Intergenic NA NA

T17567 TGGAGCGCGACGGCGGCGGCGGG 1000
Oligo fwd CACCTGGAGCGCGACGGCGGCGGC
Oligo rev AAACGCCGCCGCCGTCGCGCTCCA
Chromosome start end strand MM target_seq PAM alignment distance position gene name gene id
GU305914.1 225983 226005 - 0 TGGAGCGCGACGGCGGCGGC GGG ||||||||[||||||||||||]PAM NA Intergenic NA NA

T17568 GCGACGGCGGCGGCGGGAGCCGG 1000
Oligo fwd CACCGCGACGGCGGCGGCGGGAGC
Oligo rev AAACGCTCCCGCCGCCGCCGTCGC
Chromosome start end strand MM target_seq PAM alignment distance position gene name gene id
GU305914.1 225977 225999 - 0 GCGACGGCGGCGGCGGGAGC CGG ||||||||[||||||||||||]PAM NA Intergenic NA NA
GU305914.1 149475 149497 - 2 GCGGCGGCGGCGGCGGCAGC GGG |||-||||[||||||||-|||]PAM NA Intergenic NA NA
GU305914.1 168830 168852 + 3 GCGGCGGCGGCGGCGACAGC GGG |||-||||[|||||||--|||]PAM NA Intergenic NA NA
GU305914.1 35654 35676 + 4 TCCACGGCGGCGGCGGCGGC GGG -|-|||||[||||||||--||]PAM NA Intergenic NA NA

T17569 GCGGGAGCCGGAGTGCTGTTCGG 1000
Oligo fwd CACCGCGGGAGCCGGAGTGCTGTT
Oligo rev AAACAACAGCACTCCGGCTCCCGC
Chromosome start end strand MM target_seq PAM alignment distance position gene name gene id
GU305914.1 225965 225987 - 0 GCGGGAGCCGGAGTGCTGTT CGG ||||||||[||||||||||||]PAM NA Intergenic NA NA

T17570 AGCCGGAGTGCTGTTCGGAGCGG 1000
Oligo fwd CACCAGCCGGAGTGCTGTTCGGAG
Oligo rev AAACCTCCGAACAGCACTCCGGCT
Chromosome start end strand MM target_seq PAM alignment distance position gene name gene id
GU305914.1 225960 225982 - 0 AGCCGGAGTGCTGTTCGGAG CGG ||||||||[||||||||||||]PAM NA Intergenic NA NA

T17571 CGGAGTGCTGTTCGGAGCGGTGG 1000
Oligo fwd CACCCGGAGTGCTGTTCGGAGCGG
Oligo rev AAACCCGCTCCGAACAGCACTCCG
Chromosome start end strand MM target_seq PAM alignment distance position gene name gene id
GU305914.1 225957 225979 - 0 CGGAGTGCTGTTCGGAGCGG TGG ||||||||[||||||||||||]PAM NA Intergenic NA NA

T17572 GCTGTTCGGAGCGGTGGCCACGG 1000
Oligo fwd CACCGCTGTTCGGAGCGGTGGCCA
Oligo rev AAACTGGCCACCGCTCCGAACAGC
Chromosome start end strand MM target_seq PAM alignment distance position gene name gene id
GU305914.1 225951 225973 - 0 GCTGTTCGGAGCGGTGGCCA CGG ||||||||[||||||||||||]PAM NA Intergenic NA NA

T17573 GTTCGGAGCGGTGGCCACGGCGG 1000
Oligo fwd CACCGTTCGGAGCGGTGGCCACGG
Oligo rev AAACCCGTGGCCACCGCTCCGAAC
Chromosome start end strand MM target_seq PAM alignment distance position gene name gene id
GU305914.1 225948 225970 - 0 GTTCGGAGCGGTGGCCACGG CGG ||||||||[||||||||||||]PAM NA Intergenic NA NA

T17574 TTCGGAGCGGTGGCCACGGCGGG 1000
Oligo fwd CACCTTCGGAGCGGTGGCCACGGC
Oligo rev AAACGCCGTGGCCACCGCTCCGAA
Chromosome start end strand MM target_seq PAM alignment distance position gene name gene id
GU305914.1 225947 225969 - 0 TTCGGAGCGGTGGCCACGGC GGG ||||||||[||||||||||||]PAM NA Intergenic NA NA

T17575 GCGGGCCTGACGCTGACCCTCGG 1000
Oligo fwd CACCGCGGGCCTGACGCTGACCCT
Oligo rev AAACAGGGTCAGCGTCAGGCCCGC
Chromosome start end strand MM target_seq PAM alignment distance position gene name gene id
GU305914.1 225929 225951 - 0 GCGGGCCTGACGCTGACCCT CGG ||||||||[||||||||||||]PAM NA Intergenic NA NA

T17576 CGGGCCTGACGCTGACCCTCGGG 1000
Oligo fwd CACCCGGGCCTGACGCTGACCCTC
Oligo rev AAACGAGGGTCAGCGTCAGGCCCG
Chromosome start end strand MM target_seq PAM alignment distance position gene name gene id
GU305914.1 225928 225950 - 0 CGGGCCTGACGCTGACCCTC GGG ||||||||[||||||||||||]PAM NA Intergenic NA NA

T17577 CTCGGGCCTCGAGCTCGTCGCGG 1000
Oligo fwd CACCCTCGGGCCTCGAGCTCGTCG
Oligo rev AAACCGACGAGCTCGAGGCCCGAG
Chromosome start end strand MM target_seq PAM alignment distance position gene name gene id
GU305914.1 225911 225933 - 0 CTCGGGCCTCGAGCTCGTCG CGG ||||||||[||||||||||||]PAM NA Intergenic NA NA

T17578 TCGGGCCTCGAGCTCGTCGCGGG 1000
Oligo fwd CACCTCGGGCCTCGAGCTCGTCGC
Oligo rev AAACGCGACGAGCTCGAGGCCCGA
Chromosome start end strand MM target_seq PAM alignment distance position gene name gene id
GU305914.1 225910 225932 - 0 TCGGGCCTCGAGCTCGTCGC GGG ||||||||[||||||||||||]PAM NA Intergenic NA NA

T17579 GCCTCGAGCTCGTCGCGGGTCGG 1000
Oligo fwd CACCGCCTCGAGCTCGTCGCGGGT
Oligo rev AAACACCCGCGACGAGCTCGAGGC
Chromosome start end strand MM target_seq PAM alignment distance position gene name gene id
GU305914.1 225906 225928 - 0 GCCTCGAGCTCGTCGCGGGT CGG ||||||||[||||||||||||]PAM NA Intergenic NA NA

T17580 GAGCTCGTCGCGGGTCGGTCAGG 1000
Oligo fwd CACCGAGCTCGTCGCGGGTCGGTC
Oligo rev AAACGACCGACCCGCGACGAGCTC

Chromosome start end strand MM target_seq PAM alignment distance position gene name gene id
GU305914.1 225901 225923 - 0 GAGCTCGTCGCGGGTCGGTC AGG ||||||||[||||||||||||]PAM NA Intergenic NA NA

T17581 TTCCGCAACAGTCGCCATTGAGG 1000
Oligo fwd CACCTTCCGCAACAGTCGCCATTG
Oligo rev AAACCAATGGCGACTGTTGCGGAA
Chromosome start end strand MM target_seq PAM alignment distance position gene name gene id
GU305914.1 225878 225900 - 0 TTCCGCAACAGTCGCCATTG AGG ||||||||[||||||||||||]PAM NA Intergenic NA NA

T17582 TCCGCAACAGTCGCCATTGAGGG 1000
Oligo fwd CACCTCCGCAACAGTCGCCATTGA
Oligo rev AAACTCAATGGCGACTGTTGCGGA
Chromosome start end strand MM target_seq PAM alignment distance position gene name gene id
GU305914.1 225877 225899 - 0 TCCGCAACAGTCGCCATTGA GGG ||||||||[||||||||||||]PAM NA Intergenic NA NA

T17583 AGGGCCAGCAGCTGCGCGTGCGG 1000
Oligo fwd CACCAGGGCCAGCAGCTGCGCGTG
Oligo rev AAACCACGCGCAGCTGCTGGCCCT
Chromosome start end strand MM target_seq PAM alignment distance position gene name gene id
GU305914.1 225858 225880 - 0 AGGGCCAGCAGCTGCGCGTG CGG ||||||||[||||||||||||]PAM NA Intergenic NA NA

T17584 AGCAGCTGCGCGTGCGGAAGAGG 1000
Oligo fwd CACCAGCAGCTGCGCGTGCGGAAG
Oligo rev AAACCTTCCGCACGCGCAGCTGCT
Chromosome start end strand MM target_seq PAM alignment distance position gene name gene id
GU305914.1 225852 225874 - 0 AGCAGCTGCGCGTGCGGAAG AGG ||||||||[||||||||||||]PAM NA Intergenic NA NA

T17585 GCAGCTGCGCGTGCGGAAGAGGG 1000
Oligo fwd CACCGCAGCTGCGCGTGCGGAAGA
Oligo rev AAACTCTTCCGCACGCGCAGCTGC
Chromosome start end strand MM target_seq PAM alignment distance position gene name gene id
GU305914.1 225851 225873 - 0 GCAGCTGCGCGTGCGGAAGA GGG ||||||||[||||||||||||]PAM NA Intergenic NA NA

T17586 CGCGTGCGGAAGAGGGTGTTTGG 1000
Oligo fwd CACCCGCGTGCGGAAGAGGGTGTT
Oligo rev AAACAACACCCTCTTCCGCACGCG
Chromosome start end strand MM target_seq PAM alignment distance position gene name gene id
GU305914.1 225844 225866 - 0 CGCGTGCGGAAGAGGGTGTT TGG ||||||||[||||||||||||]PAM NA Intergenic NA NA

T17587 GGATCGACGTTCGCTCTGTCAGG 1000
Oligo fwd CACCGGATCGACGTTCGCTCTGTC
Oligo rev AAACGACAGAGCGAACGTCGATCC
Chromosome start end strand MM target_seq PAM alignment distance position gene name gene id
GU305914.1 225823 225845 - 0 GGATCGACGTTCGCTCTGTC AGG ||||||||[||||||||||||]PAM NA Intergenic NA NA

T17588 GATCGACGTTCGCTCTGTCAGGG 1000
Oligo fwd CACCGATCGACGTTCGCTCTGTCA
Oligo rev AAACTGACAGAGCGAACGTCGATC
Chromosome start end strand MM target_seq PAM alignment distance position gene name gene id
GU305914.1 225822 225844 - 0 GATCGACGTTCGCTCTGTCA GGG ||||||||[||||||||||||]PAM NA Intergenic NA NA

T17589 TGTCAGGGCGCCTCCCCCTCAGG 1000
Oligo fwd CACCTGTCAGGGCGCCTCCCCCTC
Oligo rev AAACGAGGGGGAGGCGCCCTGACA
Chromosome start end strand MM target_seq PAM alignment distance position gene name gene id
GU305914.1 225807 225829 - 0 TGTCAGGGCGCCTCCCCCTC AGG ||||||||[||||||||||||]PAM NA Intergenic NA NA

T17590 AGGAGATCTAGAAAAGAGCGAGG 1000
Oligo fwd CACCAGGAGATCTAGAAAAGAGCG
Oligo rev AAACCGCTCTTTTCTAGATCTCCT
Chromosome start end strand MM target_seq PAM alignment distance position gene name gene id
GU305914.1 225787 225809 - 0 AGGAGATCTAGAAAAGAGCG AGG ||||||||[||||||||||||]PAM NA Intergenic NA NA

T17591 ATCTAGAAAAGAGCGAGGTCTGG 1000
Oligo fwd CACCATCTAGAAAAGAGCGAGGTC
Oligo rev AAACGACCTCGCTCTTTTCTAGAT
Chromosome start end strand MM target_seq PAM alignment distance position gene name gene id
GU305914.1 225782 225804 - 0 ATCTAGAAAAGAGCGAGGTC TGG ||||||||[||||||||||||]PAM NA Intergenic NA NA

T17592 ATTCGCCACGACCCTTCGAGAGG 1000
Oligo fwd CACCATTCGCCACGACCCTTCGAG
Oligo rev AAACCTCGAAGGGTCGTGGCGAAT
Chromosome start end strand MM target_seq PAM alignment distance position gene name gene id
GU305914.1 225744 225766 - 0 ATTCGCCACGACCCTTCGAG AGG ||||||||[||||||||||||]PAM NA Intergenic NA NA

T17593 TTCGCCACGACCCTTCGAGAGGG 1000
Oligo fwd CACCTTCGCCACGACCCTTCGAGA
Oligo rev AAACTCTCGAAGGGTCGTGGCGAA
Chromosome start end strand MM target_seq PAM alignment distance position gene name gene id
GU305914.1 225743 225765 - 0 TTCGCCACGACCCTTCGAGA GGG ||||||||[||||||||||||]PAM NA Intergenic NA NA

T17594 CACGACCCTTCGAGAGGGAGCGG 1000
Oligo fwd CACCCACGACCCTTCGAGAGGGAG
Oligo rev AAACCTCCCTCTCGAAGGGTCGTG
Chromosome start end strand MM target_seq PAM alignment distance position gene name gene id
GU305914.1 225738 225760 - 0 CACGACCCTTCGAGAGGGAG CGG ||||||||[||||||||||||]PAM NA Intergenic NA NA

T17595 CAGCGCCAGCGCCCCCGCCGCGG 1000
Oligo fwd CACCCAGCGCCAGCGCCCCCGCCG

<table>
<tr><td>Oligo rev</td><td>AAACCGGCGGGGGCGCTGGCGCTG</td><td></td><td></td><td></td><td></td><td></td><td></td><td></td><td></td><td></td><td></td></tr>
<tr><td>Chromosome</td><td>start</td><td>end</td><td>strand</td><td>MM</td><td>target_seq</td><td>PAM</td><td>alignment</td><td>distance</td><td>position</td><td>gene name</td><td>gene id</td></tr>
<tr><td>GU305914.1</td><td>225709</td><td>225731</td><td>-</td><td>0</td><td>CAGCGCCAGCGCCCCCGCCG</td><td>CGG</td><td>||||||||[||||||||||||]PAM</td><td>NA</td><td>Intergenic</td><td>NA</td><td>NA</td></tr>
<tr><td>T17596</td><td>AGCGCCAGCGCCCCCGCCGCGGG</td><td>1000</td><td></td><td></td><td></td><td></td><td></td><td></td><td></td><td></td><td></td></tr>
<tr><td>Oligo fwd</td><td>CACCAGCGCCAGCGCCCCCGCCGC</td><td></td><td></td><td></td><td></td><td></td><td></td><td></td><td></td><td></td><td></td></tr>
<tr><td>Oligo rev</td><td>AAACGCGGCGGGGGCGCTGGCGCT</td><td></td><td></td><td></td><td></td><td></td><td></td><td></td><td></td><td></td><td></td></tr>
<tr><td>Chromosome</td><td>start</td><td>end</td><td>strand</td><td>MM</td><td>target_seq</td><td>PAM</td><td>alignment</td><td>distance</td><td>position</td><td>gene name</td><td>gene id</td></tr>
<tr><td>GU305914.1</td><td>225708</td><td>225730</td><td>-</td><td>0</td><td>AGCGCCAGCGCCCCCGCCGC</td><td>GGG</td><td>||||||||[||||||||||||]PAM</td><td>NA</td><td>Intergenic</td><td>NA</td><td>NA</td></tr>
<tr><td>GU305914.1</td><td>62668</td><td>62690</td><td>+</td><td>3</td><td>AGCGCCGCCGCCCCCGCCGG</td><td>CGG</td><td>||||||--[|||||||||||-]PAM</td><td>NA</td><td>Intergenic</td><td>NA</td><td>NA</td></tr>
<tr><td>T17597</td><td>CCGCGGGCGTCCAGAGACAGAGG</td><td>1000</td><td></td><td></td><td></td><td></td><td></td><td></td><td></td><td></td><td></td></tr>
<tr><td>Oligo fwd</td><td>CACCCCGCGGGCGTCCAGAGACAG</td><td></td><td></td><td></td><td></td><td></td><td></td><td></td><td></td><td></td><td></td></tr>
<tr><td>Oligo rev</td><td>AAACCTGTCTCTGGACGCCCGCGG</td><td></td><td></td><td></td><td></td><td></td><td></td><td></td><td></td><td></td><td></td></tr>
<tr><td>Chromosome</td><td>start</td><td>end</td><td>strand</td><td>MM</td><td>target_seq</td><td>PAM</td><td>alignment</td><td>distance</td><td>position</td><td>gene name</td><td>gene id</td></tr>
<tr><td>GU305914.1</td><td>225692</td><td>225714</td><td>-</td><td>0</td><td>CCGCGGGCGTCCAGAGACAG</td><td>AGG</td><td>||||||||[||||||||||||]PAM</td><td>NA</td><td>Intergenic</td><td>NA</td><td>NA</td></tr>
<tr><td>T17598</td><td>CGTCCAGAGACAGAGGACTCCGG</td><td>1000</td><td></td><td></td><td></td><td></td><td></td><td></td><td></td><td></td><td></td></tr>
<tr><td>Oligo fwd</td><td>CACCCGTCCAGAGACAGAGGACTC</td><td></td><td></td><td></td><td></td><td></td><td></td><td></td><td></td><td></td><td></td></tr>
<tr><td>Oligo rev</td><td>AAACGAGTCCTCTGTCTCTGGACG</td><td></td><td></td><td></td><td></td><td></td><td></td><td></td><td></td><td></td><td></td></tr>
<tr><td>Chromosome</td><td>start</td><td>end</td><td>strand</td><td>MM</td><td>target_seq</td><td>PAM</td><td>alignment</td><td>distance</td><td>position</td><td>gene name</td><td>gene id</td></tr>
<tr><td>GU305914.1</td><td>225685</td><td>225707</td><td>-</td><td>0</td><td>CGTCCAGAGACAGAGGACTC</td><td>CGG</td><td>||||||||[||||||||||||]PAM</td><td>NA</td><td>Intergenic</td><td>NA</td><td>NA</td></tr>
<tr><td>T17599</td><td>CCAGAGACAGAGGACTCCGGCGG</td><td>1000</td><td></td><td></td><td></td><td></td><td></td><td></td><td></td><td></td><td></td></tr>
<tr><td>Oligo fwd</td><td>CACCCCAGAGACAGAGGACTCCGG</td><td></td><td></td><td></td><td></td><td></td><td></td><td></td><td></td><td></td><td></td></tr>
<tr><td>Oligo rev</td><td>AAACCCGGAGTCCTCTGTCTCTGG</td><td></td><td></td><td></td><td></td><td></td><td></td><td></td><td></td><td></td><td></td></tr>
<tr><td>Chromosome</td><td>start</td><td>end</td><td>strand</td><td>MM</td><td>target_seq</td><td>PAM</td><td>alignment</td><td>distance</td><td>position</td><td>gene name</td><td>gene id</td></tr>
<tr><td>GU305914.1</td><td>225682</td><td>225704</td><td>-</td><td>0</td><td>CCAGAGACAGAGGACTCCGG</td><td>CGG</td><td>||||||||[||||||||||||]PAM</td><td>NA</td><td>Intergenic</td><td>NA</td><td>NA</td></tr>
<tr><td>T17600</td><td>GAGACAGAGGACTCCGGCGGCGG</td><td>1000</td><td></td><td></td><td></td><td></td><td></td><td></td><td></td><td></td><td></td></tr>
<tr><td>Oligo fwd</td><td>CACCGAGACAGAGGACTCCGGCGG</td><td></td><td></td><td></td><td></td><td></td><td></td><td></td><td></td><td></td><td></td></tr>
<tr><td>Oligo rev</td><td>AAACCCGCCGGAGTCCTCTGTCTC</td><td></td><td></td><td></td><td></td><td></td><td></td><td></td><td></td><td></td><td></td></tr>
<tr><td>Chromosome</td><td>start</td><td>end</td><td>strand</td><td>MM</td><td>target_seq</td><td>PAM</td><td>alignment</td><td>distance</td><td>position</td><td>gene name</td><td>gene id</td></tr>
<tr><td>GU305914.1</td><td>225679</td><td>225701</td><td>-</td><td>0</td><td>GAGACAGAGGACTCCGGCGG</td><td>CGG</td><td>||||||||[||||||||||||]PAM</td><td>NA</td><td>Intergenic</td><td>NA</td><td>NA</td></tr>
<tr><td>T17601</td><td>GGCGGCGGACACTAACGATCCGG</td><td>1000</td><td></td><td></td><td></td><td></td><td></td><td></td><td></td><td></td><td></td></tr>
<tr><td>Oligo fwd</td><td>CACCGGCGGCGGACACTAACGATC</td><td></td><td></td><td></td><td></td><td></td><td></td><td></td><td></td><td></td><td></td></tr>
<tr><td>Oligo rev</td><td>AAACGATCGTTAGTGTCCGCCGCC</td><td></td><td></td><td></td><td></td><td></td><td></td><td></td><td></td><td></td><td></td></tr>
<tr><td>Chromosome</td><td>start</td><td>end</td><td>strand</td><td>MM</td><td>target_seq</td><td>PAM</td><td>alignment</td><td>distance</td><td>position</td><td>gene name</td><td>gene id</td></tr>
<tr><td>GU305914.1</td><td>225664</td><td>225686</td><td>-</td><td>0</td><td>GGCGGCGGACACTAACGATC</td><td>CGG</td><td>||||||||[||||||||||||]PAM</td><td>NA</td><td>Intergenic</td><td>NA</td><td>NA</td></tr>
<tr><td>T17602</td><td>GCGGCGGACACTAACGATCCGGG</td><td>1000</td><td></td><td></td><td></td><td></td><td></td><td></td><td></td><td></td><td></td></tr>
<tr><td>Oligo fwd</td><td>CACCGCGGCGGACACTAACGATCC</td><td></td><td></td><td></td><td></td><td></td><td></td><td></td><td></td><td></td><td></td></tr>
<tr><td>Oligo rev</td><td>AAACGGATCGTTAGTGTCCGCCGC</td><td></td><td></td><td></td><td></td><td></td><td></td><td></td><td></td><td></td><td></td></tr>
<tr><td>Chromosome</td><td>start</td><td>end</td><td>strand</td><td>MM</td><td>target_seq</td><td>PAM</td><td>alignment</td><td>distance</td><td>position</td><td>gene name</td><td>gene id</td></tr>
<tr><td>GU305914.1</td><td>225663</td><td>225685</td><td>-</td><td>0</td><td>GCGGCGGACACTAACGATCC</td><td>GGG</td><td>||||||||[||||||||||||]PAM</td><td>NA</td><td>Intergenic</td><td>NA</td><td>NA</td></tr>
<tr><td>T17603</td><td>CGGACACTAACGATCCGGGATGG</td><td>1000</td><td></td><td></td><td></td><td></td><td></td><td></td><td></td><td></td><td></td></tr>
<tr><td>Oligo fwd</td><td>CACCCGGACACTAACGATCCGGGA</td><td></td><td></td><td></td><td></td><td></td><td></td><td></td><td></td><td></td><td></td></tr>
<tr><td>Oligo rev</td><td>AAACTCCCGGATCGTTAGTGTCCG</td><td></td><td></td><td></td><td></td><td></td><td></td><td></td><td></td><td></td><td></td></tr>
<tr><td>Chromosome</td><td>start</td><td>end</td><td>strand</td><td>MM</td><td>target_seq</td><td>PAM</td><td>alignment</td><td>distance</td><td>position</td><td>gene name</td><td>gene id</td></tr>
<tr><td>GU305914.1</td><td>225659</td><td>225681</td><td>-</td><td>0</td><td>CGGACACTAACGATCCGGGA</td><td>TGG</td><td>||||||||[||||||||||||]PAM</td><td>NA</td><td>Intergenic</td><td>NA</td><td>NA</td></tr>
<tr><td>T17604</td><td>AACAGCTATCGCCAGTCGCCAGG</td><td>1000</td><td></td><td></td><td></td><td></td><td></td><td></td><td></td><td></td><td></td></tr>
<tr><td>Oligo fwd</td><td>CACCAACAGCTATCGCCAGTCGCC</td><td></td><td></td><td></td><td></td><td></td><td></td><td></td><td></td><td></td><td></td></tr>
<tr><td>Oligo rev</td><td>AAACGGCGACTGGCGATAGCTGTT</td><td></td><td></td><td></td><td></td><td></td><td></td><td></td><td></td><td></td><td></td></tr>
<tr><td>Chromosome</td><td>start</td><td>end</td><td>strand</td><td>MM</td><td>target_seq</td><td>PAM</td><td>alignment</td><td>distance</td><td>position</td><td>gene name</td><td>gene id</td></tr>
<tr><td>GU305914.1</td><td>225618</td><td>225640</td><td>-</td><td>0</td><td>AACAGCTATCGCCAGTCGCC</td><td>AGG</td><td>||||||||[||||||||||||]PAM</td><td>NA</td><td>Intergenic</td><td>NA</td><td>NA</td></tr>
<tr><td>T17605</td><td>GCCAGTCGCCAGGCTACGTGCGG</td><td>1000</td><td></td><td></td><td></td><td></td><td></td><td></td><td></td><td></td><td></td></tr>
<tr><td>Oligo fwd</td><td>CACCGCCAGTCGCCAGGCTACGTG</td><td></td><td></td><td></td><td></td><td></td><td></td><td></td><td></td><td></td><td></td></tr>
<tr><td>Oligo rev</td><td>AAACCACGTAGCCTGGCGACTGGC</td><td></td><td></td><td></td><td></td><td></td><td></td><td></td><td></td><td></td><td></td></tr>
<tr><td>Chromosome</td><td>start</td><td>end</td><td>strand</td><td>MM</td><td>target_seq</td><td>PAM</td><td>alignment</td><td>distance</td><td>position</td><td>gene name</td><td>gene id</td></tr>
<tr><td>GU305914.1</td><td>225608</td><td>225630</td><td>-</td><td>0</td><td>GCCAGTCGCCAGGCTACGTG</td><td>CGG</td><td>||||||||[||||||||||||]PAM</td><td>NA</td><td>Intergenic</td><td>NA</td><td>NA</td></tr>
<tr><td>T17606</td><td>TCTCTGTCCTCCTCTGCCTCTGG</td><td>1000</td><td></td><td></td><td></td><td></td><td></td><td></td><td></td><td></td><td></td></tr>
<tr><td>Oligo fwd</td><td>CACCTCTCTGTCCTCCTCTGCCTC</td><td></td><td></td><td></td><td></td><td></td><td></td><td></td><td></td><td></td><td></td></tr>
<tr><td>Oligo rev</td><td>AAACGAGGCAGAGGAGGACAGAGA</td><td></td><td></td><td></td><td></td><td></td><td></td><td></td><td></td><td></td><td></td></tr>
<tr><td>Chromosome</td><td>start</td><td>end</td><td>strand</td><td>MM</td><td>target_seq</td><td>PAM</td><td>alignment</td><td>distance</td><td>position</td><td>gene name</td><td>gene id</td></tr>
<tr><td>GU305914.1</td><td>225579</td><td>225601</td><td>-</td><td>0</td><td>TCTCTGTCCTCCTCTGCCTC</td><td>TGG</td><td>||||||||[||||||||||||]PAM</td><td>NA</td><td>Intergenic</td><td>NA</td><td>NA</td></tr>
<tr><td>T17607</td><td>TCTGCCTCTGGTGTCTCGCGTGG</td><td>1000</td><td></td><td></td><td></td><td></td><td></td><td></td><td></td><td></td><td></td></tr>
<tr><td>Oligo fwd</td><td>CACCTCTGCCTCTGGTGTCTCGCG</td><td></td><td></td><td></td><td></td><td></td><td></td><td></td><td></td><td></td><td></td></tr>
<tr><td>Oligo rev</td><td>AAACCGCGAGACACCAGAGGCAGA</td><td></td><td></td><td></td><td></td><td></td><td></td><td></td><td></td><td></td><td></td></tr>
<tr><td>Chromosome</td><td>start</td><td>end</td><td>strand</td><td>MM</td><td>target_seq</td><td>PAM</td><td>alignment</td><td>distance</td><td>position</td><td>gene name</td><td>gene id</td></tr>
<tr><td>GU305914.1</td><td>225567</td><td>225589</td><td>-</td><td>0</td><td>TCTGCCTCTGGTGTCTCGCG</td><td>TGG</td><td>||||||||[||||||||||||]PAM</td><td>NA</td><td>Intergenic</td><td>NA</td><td>NA</td></tr>
<tr><td>T17608</td><td>ACCACCACCACCGCCGCCCTCGG</td><td>1000</td><td></td><td></td><td></td><td></td><td></td><td></td><td></td><td></td><td></td></tr>
<tr><td>Oligo fwd</td><td>CACCACCACCACCACCGCCGCCCT</td><td></td><td></td><td></td><td></td><td></td><td></td><td></td><td></td><td></td><td></td></tr>
<tr><td>Oligo rev</td><td>AAACAGGGCGGCGGTGGTGGTGGT</td><td></td><td></td><td></td><td></td><td></td><td></td><td></td><td></td><td></td><td></td></tr>
<tr><td>Chromosome</td><td>start</td><td>end</td><td>strand</td><td>MM</td><td>target_seq</td><td>PAM</td><td>alignment</td><td>distance</td><td>position</td><td>gene name</td><td>gene id</td></tr>
<tr><td>GU305914.1</td><td>225499</td><td>225521</td><td>-</td><td>0</td><td>ACCACCACCACCGCCGCCCT</td><td>CGG</td><td>||||||||[||||||||||||]PAM</td><td>NA</td><td>Intergenic</td><td>NA</td><td>NA</td></tr>
<tr><td>T17609</td><td>CACCACCGCCGCCCTCGGCTCGG</td><td>1000</td><td></td><td></td><td></td><td></td><td></td><td></td><td></td><td></td><td></td></tr>
<tr><td>Oligo fwd</td><td>CACCCACCACCGCCGCCCTCGGCT</td><td></td><td></td><td></td><td></td><td></td><td></td><td></td><td></td><td></td><td></td></tr>
<tr><td>Oligo rev</td><td>AAACAGCCGAGGGCGGCGGTGGTG</td><td></td><td></td><td></td><td></td><td></td><td></td><td></td><td></td><td></td><td></td></tr>
<tr><td>Chromosome</td><td>start</td><td>end</td><td>strand</td><td>MM</td><td>target_seq</td><td>PAM</td><td>alignment</td><td>distance</td><td>position</td><td>gene name</td><td>gene id</td></tr>
<tr><td>GU305914.1</td><td>225494</td><td>225516</td><td>-</td><td>0</td><td>CACCACCGCCGCCCTCGGCT</td><td>CGG</td><td>||||||||[||||||||||||]PAM</td><td>NA</td><td>Intergenic</td><td>NA</td><td>NA</td></tr>
</table>

T17610 CGCCCTCGGCTCGGTCACCCAGG 1000
Oligo fwd CACCCGCCCTCGGCTCGGTCACCC
Oligo rev AAACGGGTGACCGAGCCGAGGGCG
Chromosome start end strand MM target_seq PAM alignment distance position gene name gene id
GU305914.1 225485 225507 - 0 CGCCCTCGGCTCGGTCACCC AGG ||||||||[||||||||||||]PAM NA Intergenic NA NA

T17611 CCTCGGCTCGGTCACCCAGGAGG 1000
Oligo fwd CACCCCTCGGCTCGGTCACCCAGG
Oligo rev AAACCCTGGGTGACCGAGCCGAGG
Chromosome start end strand MM target_seq PAM alignment distance position gene name gene id
GU305914.1 225482 225504 - 0 CCTCGGCTCGGTCACCCAGG AGG ||||||||[||||||||||||]PAM NA Intergenic NA NA

T17612 AGAACGTGATCCAGTTCTCCTGG 1000
Oligo fwd CACCAGAACGTGATCCAGTTCTCC
Oligo rev AAACGGAGAACTGGATCACGTTCT
Chromosome start end strand MM target_seq PAM alignment distance position gene name gene id
GU305914.1 225408 225430 - 0 AGAACGTGATCCAGTTCTCC TGG ||||||||[||||||||||||]PAM NA Intergenic NA NA

T17613 GTGATCCAGTTCTCCTGGAGCGG 1000
Oligo fwd CACCGTGATCCAGTTCTCCTGGAG
Oligo rev AAACCTCCAGGAGAACTGGATCAC
Chromosome start end strand MM target_seq PAM alignment distance position gene name gene id
GU305914.1 225403 225425 - 0 GTGATCCAGTTCTCCTGGAG CGG ||||||||[||||||||||||]PAM NA Intergenic NA NA

T17614 CCGCCATCACGACGTGCGCTCGG 1000
Oligo fwd CACCCCGCCATCACGACGTGCGCT
Oligo rev AAACAGCGCACGTCGTGATGGCGG
Chromosome start end strand MM target_seq PAM alignment distance position gene name gene id
GU305914.1 225380 225402 - 0 CCGCCATCACGACGTGCGCT CGG ||||||||[||||||||||||]PAM NA Intergenic NA NA

T17615 CGCCATCACGACGTGCGCTCGGG 1000
Oligo fwd CACCCGCCATCACGACGTGCGCTC
Oligo rev AAACGAGCGCACGTCGTGATGGCG
Chromosome start end strand MM target_seq PAM alignment distance position gene name gene id
GU305914.1 225379 225401 - 0 CGCCATCACGACGTGCGCTC GGG ||||||||[||||||||||||]PAM NA Intergenic NA NA

T17616 TCGCAACCACGACGACCAGACGG 1000
Oligo fwd CACCTCGCAACCACGACGACCAGA
Oligo rev AAACTCTGGTCGTCGTGGTTGCGA
Chromosome start end strand MM target_seq PAM alignment distance position gene name gene id
GU305914.1 225341 225363 - 0 TCGCAACCACGACGACCAGA CGG ||||||||[||||||||||||]PAM NA Intergenic NA NA

T17617 CGACGACCAGACGGTGTACTTGG 1000
Oligo fwd CACCCGACGACCAGACGGTGTACT
Oligo rev AAACAGTACACCGTCTGGTCGTCG
Chromosome start end strand MM target_seq PAM alignment distance position gene name gene id
GU305914.1 225332 225354 - 0 CGACGACCAGACGGTGTACT TGG ||||||||[||||||||||||]PAM NA Intergenic NA NA

T17618 GACGACCAGACGGTGTACTTGGG 1000
Oligo fwd CACCGACGACCAGACGGTGTACTT
Oligo rev AAACAAGTACACCGTCTGGTCGTC
Chromosome start end strand MM target_seq PAM alignment distance position gene name gene id
GU305914.1 225331 225353 - 0 GACGACCAGACGGTGTACTT GGG ||||||||[||||||||||||]PAM NA Intergenic NA NA

T17619 TGTACTTGGGTCAGCTCGACTGG 1000
Oligo fwd CACCTGTACTTGGGTCAGCTCGAC
Oligo rev AAACGTCGAGCTGACCCAAGTACA
Chromosome start end strand MM target_seq PAM alignment distance position gene name gene id
GU305914.1 225318 225340 - 0 TGTACTTGGGTCAGCTCGAC TGG ||||||||[||||||||||||]PAM NA Intergenic NA NA

T17620 TTGGGTCAGCTCGACTGGAGAGG 1000
Oligo fwd CACCTTGGGTCAGCTCGACTGGAG
Oligo rev AAACCTCCAGTCGAGCTGACCCAA
Chromosome start end strand MM target_seq PAM alignment distance position gene name gene id
GU305914.1 225313 225335 - 0 TTGGGTCAGCTCGACTGGAG AGG ||||||||[||||||||||||]PAM NA Intergenic NA NA

T17621 TCGACTGGAGAGGCTTCGTGAGG 1000
Oligo fwd CACCTCGACTGGAGAGGCTTCGTG
Oligo rev AAACCACGAAGCCTCTCCAGTCGA
Chromosome start end strand MM target_seq PAM alignment distance position gene name gene id
GU305914.1 225303 225325 - 0 TCGACTGGAGAGGCTTCGTG AGG ||||||||[||||||||||||]PAM NA Intergenic NA NA

T17622 GATAACAGCGACCTCTACGTCGG 1000
Oligo fwd CACCGATAACAGCGACCTCTACGT
Oligo rev AAACACGTAGAGGTCGCTGTTATC
Chromosome start end strand MM target_seq PAM alignment distance position gene name gene id
GU305914.1 225274 225296 - 0 GATAACAGCGACCTCTACGT CGG ||||||||[||||||||||||]PAM NA Intergenic NA NA

T17623 GGAGACGCCTTGATCTTTCACGG 1000
Oligo fwd CACCGGAGACGCCTTGATCTTTCA
Oligo rev AAACTGAAAGATCAAGGCGTCTCC
Chromosome start end strand MM target_seq PAM alignment distance position gene name gene id
GU305914.1 225253 225275 - 0 GGAGACGCCTTGATCTTTCA CGG ||||||||[||||||||||||]PAM NA Intergenic NA NA

T17624 GAGACGCCTTGATCTTTCACGGG 1000
Oligo fwd CACCGAGACGCCTTGATCTTTCAC
Oligo rev AAACGTGAAAGATCAAGGCGTCTC
Chromosome start end strand MM target_seq PAM alignment distance position gene name gene id
GU305914.1 225252 225274 - 0 GAGACGCCTTGATCTTTCAC GGG ||||||||[||||||||||||]PAM NA Intergenic NA NA

<table>
<tr><td>T17625</td><td>TCCGCTCACCGACGACACCGTGG</td><td>1000</td><td></td><td></td><td></td><td></td><td></td><td></td><td></td><td></td><td></td></tr>
<tr><td>Oligo fwd</td><td>CACCTCCGCTCACCGACGACACCG</td><td></td><td></td><td></td><td></td><td></td><td></td><td></td><td></td><td></td><td></td></tr>
<tr><td>Oligo rev</td><td>AAACCGGTGTCGTCGGTGAGCGGA</td><td></td><td></td><td></td><td></td><td></td><td></td><td></td><td></td><td></td><td></td></tr>
<tr><td>Chromosome</td><td>start</td><td>end</td><td>strand</td><td>MM</td><td>target_seq</td><td>PAM</td><td>alignment</td><td>distance</td><td>position</td><td>gene name</td><td>gene id</td></tr>
<tr><td>GU305914.1</td><td>225227</td><td>225249</td><td>-</td><td>0</td><td>TCCGCTCACCGACGACACCG</td><td>TGG</td><td>||||||||[||||||||||||]PAM</td><td>NA</td><td>Intergenic</td><td>NA</td><td>NA</td></tr>
<tr><td>T17626</td><td>ACCGTGGACGTGTACGCGTCCGG</td><td>1000</td><td></td><td></td><td></td><td></td><td></td><td></td><td></td><td></td><td></td></tr>
<tr><td>Oligo fwd</td><td>CACCACCGTGGACGTGTACGCGTC</td><td></td><td></td><td></td><td></td><td></td><td></td><td></td><td></td><td></td><td></td></tr>
<tr><td>Oligo rev</td><td>AAACGACGCGTACACGTCCACGGT</td><td></td><td></td><td></td><td></td><td></td><td></td><td></td><td></td><td></td><td></td></tr>
<tr><td>Chromosome</td><td>start</td><td>end</td><td>strand</td><td>MM</td><td>target_seq</td><td>PAM</td><td>alignment</td><td>distance</td><td>position</td><td>gene name</td><td>gene id</td></tr>
<tr><td>GU305914.1</td><td>225211</td><td>225233</td><td>-</td><td>0</td><td>ACCGTGGACGTGTACGCGTC</td><td>CGG</td><td>||||||||[||||||||||||]PAM</td><td>NA</td><td>Intergenic</td><td>NA</td><td>NA</td></tr>
<tr><td>T17627</td><td>TGGACGTGTACGCGTCCGGATGG</td><td>1000</td><td></td><td></td><td></td><td></td><td></td><td></td><td></td><td></td><td></td></tr>
<tr><td>Oligo fwd</td><td>CACCTGGACGTGTACGCGTCCGGA</td><td></td><td></td><td></td><td></td><td></td><td></td><td></td><td></td><td></td><td></td></tr>
<tr><td>Oligo rev</td><td>AAACTCCGGACGCGTACACGTCCA</td><td></td><td></td><td></td><td></td><td></td><td></td><td></td><td></td><td></td><td></td></tr>
<tr><td>Chromosome</td><td>start</td><td>end</td><td>strand</td><td>MM</td><td>target_seq</td><td>PAM</td><td>alignment</td><td>distance</td><td>position</td><td>gene name</td><td>gene id</td></tr>
<tr><td>GU305914.1</td><td>225207</td><td>225229</td><td>-</td><td>0</td><td>TGGACGTGTACGCGTCCGGA</td><td>TGG</td><td>||||||||[||||||||||||]PAM</td><td>NA</td><td>Intergenic</td><td>NA</td><td>NA</td></tr>
<tr><td>T17628</td><td>AAACTGCCACCGCAGACCGTCGG</td><td>1000</td><td></td><td></td><td></td><td></td><td></td><td></td><td></td><td></td><td></td></tr>
<tr><td>Oligo fwd</td><td>CACCAAACTGCCACCGCAGACCGT</td><td></td><td></td><td></td><td></td><td></td><td></td><td></td><td></td><td></td><td></td></tr>
<tr><td>Oligo rev</td><td>AAACACGGTCTGCGGTGGCAGTTT</td><td></td><td></td><td></td><td></td><td></td><td></td><td></td><td></td><td></td><td></td></tr>
<tr><td>Chromosome</td><td>start</td><td>end</td><td>strand</td><td>MM</td><td>target_seq</td><td>PAM</td><td>alignment</td><td>distance</td><td>position</td><td>gene name</td><td>gene id</td></tr>
<tr><td>GU305914.1</td><td>225181</td><td>225203</td><td>-</td><td>0</td><td>AAACTGCCACCGCAGACCGT</td><td>CGG</td><td>||||||||[||||||||||||]PAM</td><td>NA</td><td>Intergenic</td><td>NA</td><td>NA</td></tr>
<tr><td>T17629</td><td>AACTGCCACCGCAGACCGTCGGG</td><td>1000</td><td></td><td></td><td></td><td></td><td></td><td></td><td></td><td></td><td></td></tr>
<tr><td>Oligo fwd</td><td>CACCAACTGCCACCGCAGACCGTC</td><td></td><td></td><td></td><td></td><td></td><td></td><td></td><td></td><td></td><td></td></tr>
<tr><td>Oligo rev</td><td>AAACGACGGTCTGCGGTGGCAGTT</td><td></td><td></td><td></td><td></td><td></td><td></td><td></td><td></td><td></td><td></td></tr>
<tr><td>Chromosome</td><td>start</td><td>end</td><td>strand</td><td>MM</td><td>target_seq</td><td>PAM</td><td>alignment</td><td>distance</td><td>position</td><td>gene name</td><td>gene id</td></tr>
<tr><td>GU305914.1</td><td>225180</td><td>225202</td><td>-</td><td>0</td><td>AACTGCCACCGCAGACCGTC</td><td>GGG</td><td>||||||||[||||||||||||]PAM</td><td>NA</td><td>Intergenic</td><td>NA</td><td>NA</td></tr>
<tr><td>T17630</td><td>ACTGCCACCGCAGACCGTCGGGG</td><td>1000</td><td></td><td></td><td></td><td></td><td></td><td></td><td></td><td></td><td></td></tr>
<tr><td>Oligo fwd</td><td>CACCACTGCCACCGCAGACCGTCG</td><td></td><td></td><td></td><td></td><td></td><td></td><td></td><td></td><td></td><td></td></tr>
<tr><td>Oligo rev</td><td>AAACCGACGGTCTGCGGTGGCAGT</td><td></td><td></td><td></td><td></td><td></td><td></td><td></td><td></td><td></td><td></td></tr>
<tr><td>Chromosome</td><td>start</td><td>end</td><td>strand</td><td>MM</td><td>target_seq</td><td>PAM</td><td>alignment</td><td>distance</td><td>position</td><td>gene name</td><td>gene id</td></tr>
<tr><td>GU305914.1</td><td>225179</td><td>225201</td><td>-</td><td>0</td><td>ACTGCCACCGCAGACCGTCG</td><td>GGG</td><td>||||||||[||||||||||||]PAM</td><td>NA</td><td>Intergenic</td><td>NA</td><td>NA</td></tr>
<tr><td>T17631</td><td>CGTCGGGGAAATCGTGATCAAGG</td><td>1000</td><td></td><td></td><td></td><td></td><td></td><td></td><td></td><td></td><td></td></tr>
<tr><td>Oligo fwd</td><td>CACCCGTCGGGGAAATCGTGATCA</td><td></td><td></td><td></td><td></td><td></td><td></td><td></td><td></td><td></td><td></td></tr>
<tr><td>Oligo rev</td><td>AAACTGATCACGATTTCCCCGACG</td><td></td><td></td><td></td><td></td><td></td><td></td><td></td><td></td><td></td><td></td></tr>
<tr><td>Chromosome</td><td>start</td><td>end</td><td>strand</td><td>MM</td><td>target_seq</td><td>PAM</td><td>alignment</td><td>distance</td><td>position</td><td>gene name</td><td>gene id</td></tr>
<tr><td>GU305914.1</td><td>225164</td><td>225186</td><td>-</td><td>0</td><td>CGTCGGGGAAATCGTGATCA</td><td>AGG</td><td>||||||||[||||||||||||]PAM</td><td>NA</td><td>Intergenic</td><td>NA</td><td>NA</td></tr>
<tr><td>T17632</td><td>GTCGGGGAAATCGTGATCAAGGG</td><td>1000</td><td></td><td></td><td></td><td></td><td></td><td></td><td></td><td></td><td></td></tr>
<tr><td>Oligo fwd</td><td>CACCGTCGGGGAAATCGTGATCAA</td><td></td><td></td><td></td><td></td><td></td><td></td><td></td><td></td><td></td><td></td></tr>
<tr><td>Oligo rev</td><td>AAACTTGATCACGATTTCCCCGAC</td><td></td><td></td><td></td><td></td><td></td><td></td><td></td><td></td><td></td><td></td></tr>
<tr><td>Chromosome</td><td>start</td><td>end</td><td>strand</td><td>MM</td><td>target_seq</td><td>PAM</td><td>alignment</td><td>distance</td><td>position</td><td>gene name</td><td>gene id</td></tr>
<tr><td>GU305914.1</td><td>225163</td><td>225185</td><td>-</td><td>0</td><td>GTCGGGGAAATCGTGATCAA</td><td>GGG</td><td>||||||||[||||||||||||]PAM</td><td>NA</td><td>Intergenic</td><td>NA</td><td>NA</td></tr>
<tr><td>T17633</td><td>GAAATCGTGATCAAGGGCTACGG</td><td>1000</td><td></td><td></td><td></td><td></td><td></td><td></td><td></td><td></td><td></td></tr>
<tr><td>Oligo fwd</td><td>CACCGAAATCGTGATCAAGGGCTA</td><td></td><td></td><td></td><td></td><td></td><td></td><td></td><td></td><td></td><td></td></tr>
<tr><td>Oligo rev</td><td>AAACTAGCCCTTGATCACGATTTC</td><td></td><td></td><td></td><td></td><td></td><td></td><td></td><td></td><td></td><td></td></tr>
<tr><td>Chromosome</td><td>start</td><td>end</td><td>strand</td><td>MM</td><td>target_seq</td><td>PAM</td><td>alignment</td><td>distance</td><td>position</td><td>gene name</td><td>gene id</td></tr>
<tr><td>GU305914.1</td><td>225157</td><td>225179</td><td>-</td><td>0</td><td>GAAATCGTGATCAAGGGCTA</td><td>CGG</td><td>||||||||[||||||||||||]PAM</td><td>NA</td><td>Intergenic</td><td>NA</td><td>NA</td></tr>
<tr><td>T17634</td><td>GATCAAGGGCTACGGACAGACGG</td><td>1000</td><td></td><td></td><td></td><td></td><td></td><td></td><td></td><td></td><td></td></tr>
<tr><td>Oligo fwd</td><td>CACCGATCAAGGGCTACGGACAGA</td><td></td><td></td><td></td><td></td><td></td><td></td><td></td><td></td><td></td><td></td></tr>
<tr><td>Oligo rev</td><td>AAACTCTGTCCGTAGCCCTTGATC</td><td></td><td></td><td></td><td></td><td></td><td></td><td></td><td></td><td></td><td></td></tr>
<tr><td>Chromosome</td><td>start</td><td>end</td><td>strand</td><td>MM</td><td>target_seq</td><td>PAM</td><td>alignment</td><td>distance</td><td>position</td><td>gene name</td><td>gene id</td></tr>
<tr><td>GU305914.1</td><td>225149</td><td>225171</td><td>-</td><td>0</td><td>GATCAAGGGCTACGGACAGA</td><td>CGG</td><td>||||||||[||||||||||||]PAM</td><td>NA</td><td>Intergenic</td><td>NA</td><td>NA</td></tr>
<tr><td>T17635</td><td>CAAGGGCTACGGACAGACGGCGG</td><td>1000</td><td></td><td></td><td></td><td></td><td></td><td></td><td></td><td></td><td></td></tr>
<tr><td>Oligo fwd</td><td>CACCCAAGGGCTACGGACAGACGG</td><td></td><td></td><td></td><td></td><td></td><td></td><td></td><td></td><td></td><td></td></tr>
<tr><td>Oligo rev</td><td>AAACCCGTCTGTCCGTAGCCCTTG</td><td></td><td></td><td></td><td></td><td></td><td></td><td></td><td></td><td></td><td></td></tr>
<tr><td>Chromosome</td><td>start</td><td>end</td><td>strand</td><td>MM</td><td>target_seq</td><td>PAM</td><td>alignment</td><td>distance</td><td>position</td><td>gene name</td><td>gene id</td></tr>
<tr><td>GU305914.1</td><td>225146</td><td>225168</td><td>-</td><td>0</td><td>CAAGGGCTACGGACAGACGG</td><td>CGG</td><td>||||||||[||||||||||||]PAM</td><td>NA</td><td>Intergenic</td><td>NA</td><td>NA</td></tr>
<tr><td>T17636</td><td>GACCGTGCTGCCGCCCGCTCAGG</td><td>1000</td><td></td><td></td><td></td><td></td><td></td><td></td><td></td><td></td><td></td></tr>
<tr><td>Oligo fwd</td><td>CACCGACCGTGCTGCCGCCCGCTC</td><td></td><td></td><td></td><td></td><td></td><td></td><td></td><td></td><td></td><td></td></tr>
<tr><td>Oligo rev</td><td>AAACGAGCGGGCGGCAGCACGGTC</td><td></td><td></td><td></td><td></td><td></td><td></td><td></td><td></td><td></td><td></td></tr>
<tr><td>Chromosome</td><td>start</td><td>end</td><td>strand</td><td>MM</td><td>target_seq</td><td>PAM</td><td>alignment</td><td>distance</td><td>position</td><td>gene name</td><td>gene id</td></tr>
<tr><td>GU305914.1</td><td>225119</td><td>225141</td><td>-</td><td>0</td><td>GACCGTGCTGCCGCCCGCTC</td><td>AGG</td><td>||||||||[||||||||||||]PAM</td><td>NA</td><td>Intergenic</td><td>NA</td><td>NA</td></tr>
<tr><td>T17637</td><td>CGTGCTGCCGCCCGCTCAGGTGG</td><td>1000</td><td></td><td></td><td></td><td></td><td></td><td></td><td></td><td></td><td></td></tr>
<tr><td>Oligo fwd</td><td>CACCCGTGCTGCCGCCCGCTCAGG</td><td></td><td></td><td></td><td></td><td></td><td></td><td></td><td></td><td></td><td></td></tr>
<tr><td>Oligo rev</td><td>AAACCCTGAGCGGGCGGCAGCACG</td><td></td><td></td><td></td><td></td><td></td><td></td><td></td><td></td><td></td><td></td></tr>
<tr><td>Chromosome</td><td>start</td><td>end</td><td>strand</td><td>MM</td><td>target_seq</td><td>PAM</td><td>alignment</td><td>distance</td><td>position</td><td>gene name</td><td>gene id</td></tr>
<tr><td>GU305914.1</td><td>225116</td><td>225138</td><td>-</td><td>0</td><td>CGTGCTGCCGCCCGCTCAGG</td><td>TGG</td><td>||||||||[||||||||||||]PAM</td><td>NA</td><td>Intergenic</td><td>NA</td><td>NA</td></tr>
<tr><td>T17638</td><td>TCCGTTCCTGTTTCGTTGCCTGG</td><td>1000</td><td></td><td></td><td></td><td></td><td></td><td></td><td></td><td></td><td></td></tr>
<tr><td>Oligo fwd</td><td>CACCTCCGTTCCTGTTTCGTTGCC</td><td></td><td></td><td></td><td></td><td></td><td></td><td></td><td></td><td></td><td></td></tr>
<tr><td>Oligo rev</td><td>AAACGGCAACGAAACAGGAACGGA</td><td></td><td></td><td></td><td></td><td></td><td></td><td></td><td></td><td></td><td></td></tr>
<tr><td>Chromosome</td><td>start</td><td>end</td><td>strand</td><td>MM</td><td>target_seq</td><td>PAM</td><td>alignment</td><td>distance</td><td>position</td><td>gene name</td><td>gene id</td></tr>
<tr><td>GU305914.1</td><td>225059</td><td>225081</td><td>-</td><td>0</td><td>TCCGTTCCTGTTTCGTTGCC</td><td>TGG</td><td>||||||||[||||||||||||]PAM</td><td>NA</td><td>Intergenic</td><td>NA</td><td>NA</td></tr>
<tr><td>T17639</td><td>CGTTGCCTGGCCAACATCCGTGG</td><td>1000</td><td></td><td></td><td></td><td></td><td></td><td></td><td></td><td></td><td></td></tr>
<tr><td>Oligo fwd</td><td>CACCCGTTGCCTGGCCAACATCCG</td><td></td><td></td><td></td><td></td><td></td><td></td><td></td><td></td><td></td><td></td></tr>
<tr><td>Oligo rev</td><td>AAACCGGATGTTGGCCAGGCAACG</td><td></td><td></td><td></td><td></td><td></td><td></td><td></td><td></td><td></td><td></td></tr>
<tr><td>Chromosome</td><td>start</td><td>end</td><td>strand</td><td>MM</td><td>target_seq</td><td>PAM</td><td>alignment</td><td>distance</td><td>position</td><td>gene name</td><td>gene id</td></tr>
</table>

<table>
<tr><td>GU305914.1</td><td>225046</td><td>225068</td><td>-</td><td>0</td><td>CGTTGCCTGGCCAACATCCG</td><td>TGG</td><td>||||||||[||||||||||||]PAM</td><td>NA</td><td>Intergenic</td><td>NA</td><td>NA</td></tr>
<tr><td>T17640</td><td>GTTGCCTGGCCAACATCCGTGGG</td><td>1000</td><td></td><td></td><td></td><td></td><td></td><td></td><td></td><td></td><td></td></tr>
<tr><td>Oligo fwd</td><td>CACCGTTGCCTGGCCAACATCCGT</td><td></td><td></td><td></td><td></td><td></td><td></td><td></td><td></td><td></td><td></td></tr>
<tr><td>Oligo rev</td><td>AAACACGGATGTTGGCCAGGCAAC</td><td></td><td></td><td></td><td></td><td></td><td></td><td></td><td></td><td></td><td></td></tr>
<tr><td>Chromosome</td><td>start</td><td>end</td><td>strand</td><td>MM</td><td>target_seq</td><td>PAM</td><td>alignment</td><td>distance</td><td>position</td><td>gene name</td><td>gene id</td></tr>
<tr><td>GU305914.1</td><td>225045</td><td>225067</td><td>-</td><td>0</td><td>GTTGCCTGGCCAACATCCGT</td><td>GGG</td><td>||||||||[||||||||||||]PAM</td><td>NA</td><td>Intergenic</td><td>NA</td><td>NA</td></tr>
<tr><td>T17641</td><td>TGCAGCCTACCCTGATGCGCTGG</td><td>1000</td><td></td><td></td><td></td><td></td><td></td><td></td><td></td><td></td><td></td></tr>
<tr><td>Oligo fwd</td><td>CACCTGCAGCCTACCCTGATGCGC</td><td></td><td></td><td></td><td></td><td></td><td></td><td></td><td></td><td></td><td></td></tr>
<tr><td>Oligo rev</td><td>AAACGCGCATCAGGGTAGGCTGCA</td><td></td><td></td><td></td><td></td><td></td><td></td><td></td><td></td><td></td><td></td></tr>
<tr><td>Chromosome</td><td>start</td><td>end</td><td>strand</td><td>MM</td><td>target_seq</td><td>PAM</td><td>alignment</td><td>distance</td><td>position</td><td>gene name</td><td>gene id</td></tr>
<tr><td>GU305914.1</td><td>225012</td><td>225034</td><td>-</td><td>0</td><td>TGCAGCCTACCCTGATGCGC</td><td>TGG</td><td>||||||||[||||||||||||]PAM</td><td>NA</td><td>Intergenic</td><td>NA</td><td>NA</td></tr>
<tr><td>T17642</td><td>AGCCTACCCTGATGCGCTGGTGG</td><td>1000</td><td></td><td></td><td></td><td></td><td></td><td></td><td></td><td></td><td></td></tr>
<tr><td>Oligo fwd</td><td>CACCAGCCTACCCTGATGCGCTGG</td><td></td><td></td><td></td><td></td><td></td><td></td><td></td><td></td><td></td><td></td></tr>
<tr><td>Oligo rev</td><td>AAACCCAGCGCATCAGGGTAGGCT</td><td></td><td></td><td></td><td></td><td></td><td></td><td></td><td></td><td></td><td></td></tr>
<tr><td>Chromosome</td><td>start</td><td>end</td><td>strand</td><td>MM</td><td>target_seq</td><td>PAM</td><td>alignment</td><td>distance</td><td>position</td><td>gene name</td><td>gene id</td></tr>
<tr><td>GU305914.1</td><td>225009</td><td>225031</td><td>-</td><td>0</td><td>AGCCTACCCTGATGCGCTGG</td><td>TGG</td><td>||||||||[||||||||||||]PAM</td><td>NA</td><td>Intergenic</td><td>NA</td><td>NA</td></tr>
<tr><td>T17643</td><td>ACCCTGATGCGCTGGTGGCACGG</td><td>1000</td><td></td><td></td><td></td><td></td><td></td><td></td><td></td><td></td><td></td></tr>
<tr><td>Oligo fwd</td><td>CACCACCCTGATGCGCTGGTGGCA</td><td></td><td></td><td></td><td></td><td></td><td></td><td></td><td></td><td></td><td></td></tr>
<tr><td>Oligo rev</td><td>AAACTGCCACCAGCGCATCAGGGT</td><td></td><td></td><td></td><td></td><td></td><td></td><td></td><td></td><td></td><td></td></tr>
<tr><td>Chromosome</td><td>start</td><td>end</td><td>strand</td><td>MM</td><td>target_seq</td><td>PAM</td><td>alignment</td><td>distance</td><td>position</td><td>gene name</td><td>gene id</td></tr>
<tr><td>GU305914.1</td><td>225004</td><td>225026</td><td>-</td><td>0</td><td>ACCCTGATGCGCTGGTGGCA</td><td>CGG</td><td>||||||||[||||||||||||]PAM</td><td>NA</td><td>Intergenic</td><td>NA</td><td>NA</td></tr>
<tr><td>T17644</td><td>GTGGCACGGTCCCTATCCCCTGG</td><td>1000</td><td></td><td></td><td></td><td></td><td></td><td></td><td></td><td></td><td></td></tr>
<tr><td>Oligo fwd</td><td>CACCGTGGCACGGTCCCTATCCCC</td><td></td><td></td><td></td><td></td><td></td><td></td><td></td><td></td><td></td><td></td></tr>
<tr><td>Oligo rev</td><td>AAACGGGGATAGGGACCGTGCCAC</td><td></td><td></td><td></td><td></td><td></td><td></td><td></td><td></td><td></td><td></td></tr>
<tr><td>Chromosome</td><td>start</td><td>end</td><td>strand</td><td>MM</td><td>target_seq</td><td>PAM</td><td>alignment</td><td>distance</td><td>position</td><td>gene name</td><td>gene id</td></tr>
<tr><td>GU305914.1</td><td>224990</td><td>225012</td><td>-</td><td>0</td><td>GTGGCACGGTCCCTATCCCC</td><td>TGG</td><td>||||||||[||||||||||||]PAM</td><td>NA</td><td>Intergenic</td><td>NA</td><td>NA</td></tr>
<tr><td>T17645</td><td>ATCCCCTGGTGAGCATCGAGTGG</td><td>1000</td><td></td><td></td><td></td><td></td><td></td><td></td><td></td><td></td><td></td></tr>
<tr><td>Oligo fwd</td><td>CACCATCCCCTGGTGAGCATCGAG</td><td></td><td></td><td></td><td></td><td></td><td></td><td></td><td></td><td></td><td></td></tr>
<tr><td>Oligo rev</td><td>AAACCTCGATGCTCACCAGGGGAT</td><td></td><td></td><td></td><td></td><td></td><td></td><td></td><td></td><td></td><td></td></tr>
<tr><td>Chromosome</td><td>start</td><td>end</td><td>strand</td><td>MM</td><td>target_seq</td><td>PAM</td><td>alignment</td><td>distance</td><td>position</td><td>gene name</td><td>gene id</td></tr>
<tr><td>GU305914.1</td><td>224976</td><td>224998</td><td>-</td><td>0</td><td>ATCCCCTGGTGAGCATCGAG</td><td>TGG</td><td>||||||||[||||||||||||]PAM</td><td>NA</td><td>Intergenic</td><td>NA</td><td>NA</td></tr>
<tr><td>T17646</td><td>GTGGAGCGAGAACAACCCCGTGG</td><td>1000</td><td></td><td></td><td></td><td></td><td></td><td></td><td></td><td></td><td></td></tr>
<tr><td>Oligo fwd</td><td>CACCGTGGAGCGAGAACAACCCCG</td><td></td><td></td><td></td><td></td><td></td><td></td><td></td><td></td><td></td><td></td></tr>
<tr><td>Oligo rev</td><td>AAACCGGGGTTGTTCTCGCTCCAC</td><td></td><td></td><td></td><td></td><td></td><td></td><td></td><td></td><td></td><td></td></tr>
<tr><td>Chromosome</td><td>start</td><td>end</td><td>strand</td><td>MM</td><td>target_seq</td><td>PAM</td><td>alignment</td><td>distance</td><td>position</td><td>gene name</td><td>gene id</td></tr>
<tr><td>GU305914.1</td><td>224957</td><td>224979</td><td>-</td><td>0</td><td>GTGGAGCGAGAACAACCCCG</td><td>TGG</td><td>||||||||[||||||||||||]PAM</td><td>NA</td><td>Intergenic</td><td>NA</td><td>NA</td></tr>
<tr><td>T17647</td><td>GAGCGAGAACAACCCCGTGGAGG</td><td>1000</td><td></td><td></td><td></td><td></td><td></td><td></td><td></td><td></td><td></td></tr>
<tr><td>Oligo fwd</td><td>CACCGAGCGAGAACAACCCCGTGG</td><td></td><td></td><td></td><td></td><td></td><td></td><td></td><td></td><td></td><td></td></tr>
<tr><td>Oligo rev</td><td>AAACCCACGGGGTTGTTCTCGCTC</td><td></td><td></td><td></td><td></td><td></td><td></td><td></td><td></td><td></td><td></td></tr>
<tr><td>Chromosome</td><td>start</td><td>end</td><td>strand</td><td>MM</td><td>target_seq</td><td>PAM</td><td>alignment</td><td>distance</td><td>position</td><td>gene name</td><td>gene id</td></tr>
<tr><td>GU305914.1</td><td>224954</td><td>224976</td><td>-</td><td>0</td><td>GAGCGAGAACAACCCCGTGG</td><td>AGG</td><td>||||||||[||||||||||||]PAM</td><td>NA</td><td>Intergenic</td><td>NA</td><td>NA</td></tr>
<tr><td>T17648</td><td>AGCGAGAACAACCCCGTGGAGGG</td><td>1000</td><td></td><td></td><td></td><td></td><td></td><td></td><td></td><td></td><td></td></tr>
<tr><td>Oligo fwd</td><td>CACCAGCGAGAACAACCCCGTGGA</td><td></td><td></td><td></td><td></td><td></td><td></td><td></td><td></td><td></td><td></td></tr>
<tr><td>Oligo rev</td><td>AAACTCCACGGGGTTGTTCTCGCT</td><td></td><td></td><td></td><td></td><td></td><td></td><td></td><td></td><td></td><td></td></tr>
<tr><td>Chromosome</td><td>start</td><td>end</td><td>strand</td><td>MM</td><td>target_seq</td><td>PAM</td><td>alignment</td><td>distance</td><td>position</td><td>gene name</td><td>gene id</td></tr>
<tr><td>GU305914.1</td><td>224953</td><td>224975</td><td>-</td><td>0</td><td>AGCGAGAACAACCCCGTGGA</td><td>GGG</td><td>||||||||[||||||||||||]PAM</td><td>NA</td><td>Intergenic</td><td>NA</td><td>NA</td></tr>
<tr><td>T17649</td><td>CCGTGGAGGGCCGCGTGCGTTGG</td><td>1000</td><td></td><td></td><td></td><td></td><td></td><td></td><td></td><td></td><td></td></tr>
<tr><td>Oligo fwd</td><td>CACCCCGTGGAGGGCCGCGTGCGT</td><td></td><td></td><td></td><td></td><td></td><td></td><td></td><td></td><td></td><td></td></tr>
<tr><td>Oligo rev</td><td>AAACACGCACGCGGCCCTCCACGG</td><td></td><td></td><td></td><td></td><td></td><td></td><td></td><td></td><td></td><td></td></tr>
<tr><td>Chromosome</td><td>start</td><td>end</td><td>strand</td><td>MM</td><td>target_seq</td><td>PAM</td><td>alignment</td><td>distance</td><td>position</td><td>gene name</td><td>gene id</td></tr>
<tr><td>GU305914.1</td><td>224940</td><td>224962</td><td>-</td><td>0</td><td>CCGTGGAGGGCCGCGTGCGT</td><td>TGG</td><td>||||||||[||||||||||||]PAM</td><td>NA</td><td>Intergenic</td><td>NA</td><td>NA</td></tr>
<tr><td>T17650</td><td>CGTGGAGGGCCGCGTGCGTTGGG</td><td>1000</td><td></td><td></td><td></td><td></td><td></td><td></td><td></td><td></td><td></td></tr>
<tr><td>Oligo fwd</td><td>CACCCGTGGAGGGCCGCGTGCGTT</td><td></td><td></td><td></td><td></td><td></td><td></td><td></td><td></td><td></td><td></td></tr>
<tr><td>Oligo rev</td><td>AAACAACGCACGCGGCCCTCCACG</td><td></td><td></td><td></td><td></td><td></td><td></td><td></td><td></td><td></td><td></td></tr>
<tr><td>Chromosome</td><td>start</td><td>end</td><td>strand</td><td>MM</td><td>target_seq</td><td>PAM</td><td>alignment</td><td>distance</td><td>position</td><td>gene name</td><td>gene id</td></tr>
<tr><td>GU305914.1</td><td>224939</td><td>224961</td><td>-</td><td>0</td><td>CGTGGAGGGCCGCGTGCGTT</td><td>GGG</td><td>||||||||[||||||||||||]PAM</td><td>NA</td><td>Intergenic</td><td>NA</td><td>NA</td></tr>
<tr><td>T17651</td><td>CGTTGGGAACAACTCCCCGACGG</td><td>1000</td><td></td><td></td><td></td><td></td><td></td><td></td><td></td><td></td><td></td></tr>
<tr><td>Oligo fwd</td><td>CACCCGTTGGGAACAACTCCCCGA</td><td></td><td></td><td></td><td></td><td></td><td></td><td></td><td></td><td></td><td></td></tr>
<tr><td>Oligo rev</td><td>AAACTCGGGGAGTTGTTCCCAACG</td><td></td><td></td><td></td><td></td><td></td><td></td><td></td><td></td><td></td><td></td></tr>
<tr><td>Chromosome</td><td>start</td><td>end</td><td>strand</td><td>MM</td><td>target_seq</td><td>PAM</td><td>alignment</td><td>distance</td><td>position</td><td>gene name</td><td>gene id</td></tr>
<tr><td>GU305914.1</td><td>224923</td><td>224945</td><td>-</td><td>0</td><td>CGTTGGGAACAACTCCCCGA</td><td>CGG</td><td>||||||||[||||||||||||]PAM</td><td>NA</td><td>Intergenic</td><td>NA</td><td>NA</td></tr>
<tr><td>T17652</td><td>AACAACTCCCCGACGGCGATCGG</td><td>1000</td><td></td><td></td><td></td><td></td><td></td><td></td><td></td><td></td><td></td></tr>
<tr><td>Oligo fwd</td><td>CACCAACAACTCCCCGACGGCGAT</td><td></td><td></td><td></td><td></td><td></td><td></td><td></td><td></td><td></td><td></td></tr>
<tr><td>Oligo rev</td><td>AAACATCGCCGTCGGGGAGTTGTT</td><td></td><td></td><td></td><td></td><td></td><td></td><td></td><td></td><td></td><td></td></tr>
<tr><td>Chromosome</td><td>start</td><td>end</td><td>strand</td><td>MM</td><td>target_seq</td><td>PAM</td><td>alignment</td><td>distance</td><td>position</td><td>gene name</td><td>gene id</td></tr>
<tr><td>GU305914.1</td><td>224916</td><td>224938</td><td>-</td><td>0</td><td>AACAACTCCCCGACGGCGAT</td><td>CGG</td><td>||||||||[||||||||||||]PAM</td><td>NA</td><td>Intergenic</td><td>NA</td><td>NA</td></tr>
<tr><td>T17653</td><td>GTACTACGTGAACGCCTACACGG</td><td>1000</td><td></td><td></td><td></td><td></td><td></td><td></td><td></td><td></td><td></td></tr>
<tr><td>Oligo fwd</td><td>CACCGTACTACGTGAACGCCTACA</td><td></td><td></td><td></td><td></td><td></td><td></td><td></td><td></td><td></td><td></td></tr>
<tr><td>Oligo rev</td><td>AAACTGTAGGCGTTCACGTAGTAC</td><td></td><td></td><td></td><td></td><td></td><td></td><td></td><td></td><td></td><td></td></tr>
<tr><td>Chromosome</td><td>start</td><td>end</td><td>strand</td><td>MM</td><td>target_seq</td><td>PAM</td><td>alignment</td><td>distance</td><td>position</td><td>gene name</td><td>gene id</td></tr>
<tr><td>GU305914.1</td><td>224891</td><td>224913</td><td>-</td><td>0</td><td>GTACTACGTGAACGCCTACA</td><td>CGG</td><td>||||||||[||||||||||||]PAM</td><td>NA</td><td>Intergenic</td><td>NA</td><td>NA</td></tr>
<tr><td>T17654</td><td>TACTACGTGAACGCCTACACGGG</td><td>1000</td><td></td><td></td><td></td><td></td><td></td><td></td><td></td><td></td><td></td></tr>
<tr><td>Oligo fwd</td><td>CACCTACTACGTGAACGCCTACAC</td><td></td><td></td><td></td><td></td><td></td><td></td><td></td><td></td><td></td><td></td></tr>
<tr><td>Oligo rev</td><td>AAACGTGTAGGCGTTCACGTAGTA</td><td></td><td></td><td></td><td></td><td></td><td></td><td></td><td></td><td></td><td></td></tr>
</table>

Chromosome start end strand MM target_seq PAM alignment distance position gene name gene id
GU305914.1 224890 224912 - 0 TACTACGTGAACGCCTACAC GGG ||||||||[||||||||||||]PAM NA Intergenic NA NA

T17655 CCGCAGCACGATCCAGCGCCTGG 1000
Oligo fwd CACCCCGCAGCACGATCCAGCGCC
Oligo rev AAACGGCGCTGGATCGTGCTGCGG
Chromosome start end strand MM target_seq PAM alignment distance position gene name gene id
GU305914.1 224849 224871 - 0 CCGCAGCACGATCCAGCGCC TGG ||||||||[||||||||||||]PAM NA Intergenic NA NA

T17656 ATGTCTGCAGTGCACGATGCAGG 1000
Oligo fwd CACCATGTCTGCAGTGCACGATGC
Oligo rev AAACGCATCGTGCACTGCAGACAT
Chromosome start end strand MM target_seq PAM alignment distance position gene name gene id
GU305914.1 224825 224847 - 0 ATGTCTGCAGTGCACGATGC AGG ||||||||[||||||||||||]PAM NA Intergenic NA NA

T17657 TCTGCAGTGCACGATGCAGGCGG 1000
Oligo fwd CACCTCTGCAGTGCACGATGCAGG
Oligo rev AAACCCTGCATCGTGCACTGCAGA
Chromosome start end strand MM target_seq PAM alignment distance position gene name gene id
GU305914.1 224822 224844 - 0 TCTGCAGTGCACGATGCAGG CGG ||||||||[||||||||||||]PAM NA Intergenic NA NA

T17658 GTGCACGATGCAGGCGGACAAGG 1000
Oligo fwd CACCGTGCACGATGCAGGCGGACA
Oligo rev AAACTGTCCGCCTGCATCGTGCAC
Chromosome start end strand MM target_seq PAM alignment distance position gene name gene id
GU305914.1 224816 224838 - 0 GTGCACGATGCAGGCGGACA AGG ||||||||[||||||||||||]PAM NA Intergenic NA NA

T17659 ACAAGGTAGCTACCAGCTCCCGG 1000
Oligo fwd CACCACAAGGTAGCTACCAGCTCC
Oligo rev AAACGGAGCTGGTAGCTACCTTGT
Chromosome start end strand MM target_seq PAM alignment distance position gene name gene id
GU305914.1 224799 224821 - 0 ACAAGGTAGCTACCAGCTCC CGG ||||||||[||||||||||||]PAM NA Intergenic NA NA

T17660 CAGCTCCCGGTTGAGCTGCCCGG 1000
Oligo fwd CACCCAGCTCCCGGTTGAGCTGCC
Oligo rev AAACGGCAGCTCAACCGGGAGCTG
Chromosome start end strand MM target_seq PAM alignment distance position gene name gene id
GU305914.1 224786 224808 - 0 CAGCTCCCGGTTGAGCTGCC CGG ||||||||[||||||||||||]PAM NA Intergenic NA NA

T17661 CCCGGTTGAGCTGCCCGGCCCGG 1000
Oligo fwd CACCCCCGGTTGAGCTGCCCGGCC
Oligo rev AAACGGCCGGGCAGCTCAACCGGG
Chromosome start end strand MM target_seq PAM alignment distance position gene name gene id
GU305914.1 224781 224803 - 0 CCCGGTTGAGCTGCCCGGCC CGG ||||||||[||||||||||||]PAM NA Intergenic NA NA

T17662 CCGGTTGAGCTGCCCGGCCCGGG 1000
Oligo fwd CACCCCGGTTGAGCTGCCCGGCCC
Oligo rev AAACGGGCCGGGCAGCTCAACCGG
Chromosome start end strand MM target_seq PAM alignment distance position gene name gene id
GU305914.1 224780 224802 - 0 CCGGTTGAGCTGCCCGGCCC GGG ||||||||[||||||||||||]PAM NA Intergenic NA NA

T17663 ATAGCGCTGCCGTTCGACTTCGG 1000
Oligo fwd CACCATAGCGCTGCCGTTCGACTT
Oligo rev AAACAAGTCGAACGGCAGCGCTAT
Chromosome start end strand MM target_seq PAM alignment distance position gene name gene id
GU305914.1 224749 224771 - 0 ATAGCGCTGCCGTTCGACTT CGG ||||||||[||||||||||||]PAM NA Intergenic NA NA

T17664 TAGCGCTGCCGTTCGACTTCGGG 1000
Oligo fwd CACCTAGCGCTGCCGTTCGACTTC
Oligo rev AAACGAAGTCGAACGGCAGCGCTA
Chromosome start end strand MM target_seq PAM alignment distance position gene name gene id
GU305914.1 224748 224770 - 0 TAGCGCTGCCGTTCGACTTC GGG ||||||||[||||||||||||]PAM NA Intergenic NA NA

T17665 GCTGCCGTTCGACTTCGGGCAGG 1000
Oligo fwd CACCGCTGCCGTTCGACTTCGGGC
Oligo rev AAACGCCCGAAGTCGAACGGCAGC
Chromosome start end strand MM target_seq PAM alignment distance position gene name gene id
GU305914.1 224744 224766 - 0 GCTGCCGTTCGACTTCGGGC AGG ||||||||[||||||||||||]PAM NA Intergenic NA NA

T17666 CGACTTCGGGCAGGATACCACGG 1000
Oligo fwd CACCCGACTTCGGGCAGGATACCA
Oligo rev AAACTGGTATCCTGCCCGAAGTCG
Chromosome start end strand MM target_seq PAM alignment distance position gene name gene id
GU305914.1 224735 224757 - 0 CGACTTCGGGCAGGATACCA CGG ||||||||[||||||||||||]PAM NA Intergenic NA NA

T17667 GGATACCACGGTGACGCCGACGG 1000
Oligo fwd CACCGGATACCACGGTGACGCCGA
Oligo rev AAACTCGGCGTCACCGTGGTATCC
Chromosome start end strand MM target_seq PAM alignment distance position gene name gene id
GU305914.1 224723 224745 - 0 GGATACCACGGTGACGCCGA CGG ||||||||[||||||||||||]PAM NA Intergenic NA NA

T17668 GCCGACGGAACGCCGCCACATGG 1000
Oligo fwd CACCGCCGACGGAACGCCGCCACA
Oligo rev AAACTGTGGCGGCGTTCCGTCGGC
Chromosome start end strand MM target_seq PAM alignment distance position gene name gene id
GU305914.1 224708 224730 - 0 GCCGACGGAACGCCGCCACA TGG ||||||||[||||||||||||]PAM NA Intergenic NA NA

T17669 CCGTGCTGATCGTCATCACGTGG 1000
Oligo fwd CACCCCGTGCTGATCGTCATCACG

<table>
<tr><td>Oligo rev</td><td>AAACCGTGATGACGATCAGCACGG</td><td></td><td></td><td></td><td></td><td></td><td></td><td></td><td></td><td></td><td></td></tr>
<tr><td>Chromosome</td><td>start</td><td>end</td><td>strand</td><td>MM</td><td>target_seq</td><td>PAM</td><td>alignment</td><td>distance</td><td>position</td><td>gene name</td><td>gene id</td></tr>
<tr><td>GU305914.1</td><td>224673</td><td>224695</td><td>-</td><td>0</td><td>CCGTGCTGATCGTCATCACG</td><td>TGG</td><td>||||||||[||||||||||||]PAM</td><td>NA</td><td>Intergenic</td><td>NA</td><td>NA</td></tr>
<tr><td></td><td></td><td></td><td></td><td></td><td></td><td></td><td></td><td></td><td></td><td></td><td></td></tr>
<tr><td>T17670</td><td>GATCGTCATCACGTGGTTCGTGG</td><td>1000</td><td></td><td></td><td></td><td></td><td></td><td></td><td></td><td></td><td></td></tr>
<tr><td>Oligo fwd</td><td>CACCGATCGTCATCACGTGGTTCG</td><td></td><td></td><td></td><td></td><td></td><td></td><td></td><td></td><td></td><td></td></tr>
<tr><td>Oligo rev</td><td>AAACCGAACCACGTGATGACGATC</td><td></td><td></td><td></td><td></td><td></td><td></td><td></td><td></td><td></td><td></td></tr>
<tr><td>Chromosome</td><td>start</td><td>end</td><td>strand</td><td>MM</td><td>target_seq</td><td>PAM</td><td>alignment</td><td>distance</td><td>position</td><td>gene name</td><td>gene id</td></tr>
<tr><td>GU305914.1</td><td>224666</td><td>224688</td><td>-</td><td>0</td><td>GATCGTCATCACGTGGTTCG</td><td>TGG</td><td>||||||||[||||||||||||]PAM</td><td>NA</td><td>Intergenic</td><td>NA</td><td>NA</td></tr>
<tr><td></td><td></td><td></td><td></td><td></td><td></td><td></td><td></td><td></td><td></td><td></td><td></td></tr>
<tr><td>T17671</td><td>CGTCATCACGTGGTTCGTGGCGG</td><td>1000</td><td></td><td></td><td></td><td></td><td></td><td></td><td></td><td></td><td></td></tr>
<tr><td>Oligo fwd</td><td>CACCCGTCATCACGTGGTTCGTGG</td><td></td><td></td><td></td><td></td><td></td><td></td><td></td><td></td><td></td><td></td></tr>
<tr><td>Oligo rev</td><td>AAACCCACGAACCACGTGATGACG</td><td></td><td></td><td></td><td></td><td></td><td></td><td></td><td></td><td></td><td></td></tr>
<tr><td>Chromosome</td><td>start</td><td>end</td><td>strand</td><td>MM</td><td>target_seq</td><td>PAM</td><td>alignment</td><td>distance</td><td>position</td><td>gene name</td><td>gene id</td></tr>
<tr><td>GU305914.1</td><td>224663</td><td>224685</td><td>-</td><td>0</td><td>CGTCATCACGTGGTTCGTGG</td><td>CGG</td><td>||||||||[||||||||||||]PAM</td><td>NA</td><td>Intergenic</td><td>NA</td><td>NA</td></tr>
<tr><td>GU305914.1</td><td>96323</td><td>96345</td><td>+</td><td>4</td><td>CATGATCACGTCGTCCGTGG</td><td>GGG</td><td>|-|-||||[|||-||-|||||]PAM</td><td>NA</td><td>Intergenic</td><td>NA</td><td>NA</td></tr>
<tr><td></td><td></td><td></td><td></td><td></td><td></td><td></td><td></td><td></td><td></td><td></td><td></td></tr>
<tr><td>T17672</td><td>GTCATCACGTGGTTCGTGGCGGG</td><td>1000</td><td></td><td></td><td></td><td></td><td></td><td></td><td></td><td></td><td></td></tr>
<tr><td>Oligo fwd</td><td>CACCGTCATCACGTGGTTCGTGGC</td><td></td><td></td><td></td><td></td><td></td><td></td><td></td><td></td><td></td><td></td></tr>
<tr><td>Oligo rev</td><td>AAACGCCACGAACCACGTGATGAC</td><td></td><td></td><td></td><td></td><td></td><td></td><td></td><td></td><td></td><td></td></tr>
<tr><td>Chromosome</td><td>start</td><td>end</td><td>strand</td><td>MM</td><td>target_seq</td><td>PAM</td><td>alignment</td><td>distance</td><td>position</td><td>gene name</td><td>gene id</td></tr>
<tr><td>GU305914.1</td><td>224662</td><td>224684</td><td>-</td><td>0</td><td>GTCATCACGTGGTTCGTGGC</td><td>GGG</td><td>||||||||[||||||||||||]PAM</td><td>NA</td><td>Intergenic</td><td>NA</td><td>NA</td></tr>
<tr><td></td><td></td><td></td><td></td><td></td><td></td><td></td><td></td><td></td><td></td><td></td><td></td></tr>
<tr><td>T17673</td><td>ACGTGGTTCGTGGCGGGACTCGG</td><td>1000</td><td></td><td></td><td></td><td></td><td></td><td></td><td></td><td></td><td></td></tr>
<tr><td>Oligo fwd</td><td>CACCACGTGGTTCGTGGCGGGACT</td><td></td><td></td><td></td><td></td><td></td><td></td><td></td><td></td><td></td><td></td></tr>
<tr><td>Oligo rev</td><td>AAACAGTCCCGCCACGAACCACGT</td><td></td><td></td><td></td><td></td><td></td><td></td><td></td><td></td><td></td><td></td></tr>
<tr><td>Chromosome</td><td>start</td><td>end</td><td>strand</td><td>MM</td><td>target_seq</td><td>PAM</td><td>alignment</td><td>distance</td><td>position</td><td>gene name</td><td>gene id</td></tr>
<tr><td>GU305914.1</td><td>224656</td><td>224678</td><td>-</td><td>0</td><td>ACGTGGTTCGTGGCGGGACT</td><td>CGG</td><td>||||||||[||||||||||||]PAM</td><td>NA</td><td>Intergenic</td><td>NA</td><td>NA</td></tr>
<tr><td></td><td></td><td></td><td></td><td></td><td></td><td></td><td></td><td></td><td></td><td></td><td></td></tr>
<tr><td>T17674</td><td>CGTGGTTCGTGGCGGGACTCGGG</td><td>1000</td><td></td><td></td><td></td><td></td><td></td><td></td><td></td><td></td><td></td></tr>
<tr><td>Oligo fwd</td><td>CACCCGTGGTTCGTGGCGGGACTC</td><td></td><td></td><td></td><td></td><td></td><td></td><td></td><td></td><td></td><td></td></tr>
<tr><td>Oligo rev</td><td>AAACGAGTCCCGCCACGAACCACG</td><td></td><td></td><td></td><td></td><td></td><td></td><td></td><td></td><td></td><td></td></tr>
<tr><td>Chromosome</td><td>start</td><td>end</td><td>strand</td><td>MM</td><td>target_seq</td><td>PAM</td><td>alignment</td><td>distance</td><td>position</td><td>gene name</td><td>gene id</td></tr>
<tr><td>GU305914.1</td><td>224655</td><td>224677</td><td>-</td><td>0</td><td>CGTGGTTCGTGGCGGGACTC</td><td>GGG</td><td>||||||||[||||||||||||]PAM</td><td>NA</td><td>Intergenic</td><td>NA</td><td>NA</td></tr>
<tr><td></td><td></td><td></td><td></td><td></td><td></td><td></td><td></td><td></td><td></td><td></td><td></td></tr>
<tr><td>T17675</td><td>CGTGGCGGGACTCGGGATCCTGG</td><td>1000</td><td></td><td></td><td></td><td></td><td></td><td></td><td></td><td></td><td></td></tr>
<tr><td>Oligo fwd</td><td>CACCCGTGGCGGGACTCGGGATCC</td><td></td><td></td><td></td><td></td><td></td><td></td><td></td><td></td><td></td><td></td></tr>
<tr><td>Oligo rev</td><td>AAACGGATCCCGAGTCCCGCCACG</td><td></td><td></td><td></td><td></td><td></td><td></td><td></td><td></td><td></td><td></td></tr>
<tr><td>Chromosome</td><td>start</td><td>end</td><td>strand</td><td>MM</td><td>target_seq</td><td>PAM</td><td>alignment</td><td>distance</td><td>position</td><td>gene name</td><td>gene id</td></tr>
<tr><td>GU305914.1</td><td>224648</td><td>224670</td><td>-</td><td>0</td><td>CGTGGCGGGACTCGGGATCC</td><td>TGG</td><td>||||||||[||||||||||||]PAM</td><td>NA</td><td>Intergenic</td><td>NA</td><td>NA</td></tr>
<tr><td></td><td></td><td></td><td></td><td></td><td></td><td></td><td></td><td></td><td></td><td></td><td></td></tr>
<tr><td>T17676</td><td>GGGACTCGGGATCCTGGCCCTGG</td><td>1000</td><td></td><td></td><td></td><td></td><td></td><td></td><td></td><td></td><td></td></tr>
<tr><td>Oligo fwd</td><td>CACCGGGACTCGGGATCCTGGCCC</td><td></td><td></td><td></td><td></td><td></td><td></td><td></td><td></td><td></td><td></td></tr>
<tr><td>Oligo rev</td><td>AAACGGGCCAGGATCCCGAGTCCC</td><td></td><td></td><td></td><td></td><td></td><td></td><td></td><td></td><td></td><td></td></tr>
<tr><td>Chromosome</td><td>start</td><td>end</td><td>strand</td><td>MM</td><td>target_seq</td><td>PAM</td><td>alignment</td><td>distance</td><td>position</td><td>gene name</td><td>gene id</td></tr>
<tr><td>GU305914.1</td><td>224642</td><td>224664</td><td>-</td><td>0</td><td>GGGACTCGGGATCCTGGCCC</td><td>TGG</td><td>||||||||[||||||||||||]PAM</td><td>NA</td><td>Intergenic</td><td>NA</td><td>NA</td></tr>
<tr><td></td><td></td><td></td><td></td><td></td><td></td><td></td><td></td><td></td><td></td><td></td><td></td></tr>
<tr><td>T17677</td><td>GGACTCGGGATCCTGGCCCTGGG</td><td>1000</td><td></td><td></td><td></td><td></td><td></td><td></td><td></td><td></td><td></td></tr>
<tr><td>Oligo fwd</td><td>CACCGGACTCGGGATCCTGGCCCT</td><td></td><td></td><td></td><td></td><td></td><td></td><td></td><td></td><td></td><td></td></tr>
<tr><td>Oligo rev</td><td>AAACAGGGCCAGGATCCCGAGTCC</td><td></td><td></td><td></td><td></td><td></td><td></td><td></td><td></td><td></td><td></td></tr>
<tr><td>Chromosome</td><td>start</td><td>end</td><td>strand</td><td>MM</td><td>target_seq</td><td>PAM</td><td>alignment</td><td>distance</td><td>position</td><td>gene name</td><td>gene id</td></tr>
<tr><td>GU305914.1</td><td>224641</td><td>224663</td><td>-</td><td>0</td><td>GGACTCGGGATCCTGGCCCT</td><td>GGG</td><td>||||||||[||||||||||||]PAM</td><td>NA</td><td>Intergenic</td><td>NA</td><td>NA</td></tr>
<tr><td></td><td></td><td></td><td></td><td></td><td></td><td></td><td></td><td></td><td></td><td></td><td></td></tr>
<tr><td>T17678</td><td>GACTCGGGATCCTGGCCCTGGGG</td><td>1000</td><td></td><td></td><td></td><td></td><td></td><td></td><td></td><td></td><td></td></tr>
<tr><td>Oligo fwd</td><td>CACCGACTCGGGATCCTGGCCCTG</td><td></td><td></td><td></td><td></td><td></td><td></td><td></td><td></td><td></td><td></td></tr>
<tr><td>Oligo rev</td><td>AAACCAGGGCCAGGATCCCGAGTC</td><td></td><td></td><td></td><td></td><td></td><td></td><td></td><td></td><td></td><td></td></tr>
<tr><td>Chromosome</td><td>start</td><td>end</td><td>strand</td><td>MM</td><td>target_seq</td><td>PAM</td><td>alignment</td><td>distance</td><td>position</td><td>gene name</td><td>gene id</td></tr>
<tr><td>GU305914.1</td><td>224640</td><td>224662</td><td>-</td><td>0</td><td>GACTCGGGATCCTGGCCCTG</td><td>GGG</td><td>||||||||[||||||||||||]PAM</td><td>NA</td><td>Intergenic</td><td>NA</td><td>NA</td></tr>
<tr><td></td><td></td><td></td><td></td><td></td><td></td><td></td><td></td><td></td><td></td><td></td><td></td></tr>
<tr><td>T17679</td><td>GCCCTGGGGATCGTGCTGATCGG</td><td>1000</td><td></td><td></td><td></td><td></td><td></td><td></td><td></td><td></td><td></td></tr>
<tr><td>Oligo fwd</td><td>CACCGCCCTGGGGATCGTGCTGAT</td><td></td><td></td><td></td><td></td><td></td><td></td><td></td><td></td><td></td><td></td></tr>
<tr><td>Oligo rev</td><td>AAACATCAGCACGATCCCCAGGGC</td><td></td><td></td><td></td><td></td><td></td><td></td><td></td><td></td><td></td><td></td></tr>
<tr><td>Chromosome</td><td>start</td><td>end</td><td>strand</td><td>MM</td><td>target_seq</td><td>PAM</td><td>alignment</td><td>distance</td><td>position</td><td>gene name</td><td>gene id</td></tr>
<tr><td>GU305914.1</td><td>224626</td><td>224648</td><td>-</td><td>0</td><td>GCCCTGGGGATCGTGCTGAT</td><td>CGG</td><td>||||||||[||||||||||||]PAM</td><td>NA</td><td>Intergenic</td><td>NA</td><td>NA</td></tr>
<tr><td>GU305914.1</td><td>120529</td><td>120551</td><td>-</td><td>4</td><td>GCCCTAGAGATCGTGCCGAC</td><td>GGG</td><td>|||||-|-[||||||||-||-]PAM</td><td>NA</td><td>Intergenic</td><td>NA</td><td>NA</td></tr>
<tr><td></td><td></td><td></td><td></td><td></td><td></td><td></td><td></td><td></td><td></td><td></td><td></td></tr>
<tr><td>T17680</td><td>TTGCTTCCCCTCCCGCTCGCCGG</td><td>1000</td><td></td><td></td><td></td><td></td><td></td><td></td><td></td><td></td><td></td></tr>
<tr><td>Oligo fwd</td><td>CACCTTGCTTCCCCTCCCGCTCGC</td><td></td><td></td><td></td><td></td><td></td><td></td><td></td><td></td><td></td><td></td></tr>
<tr><td>Oligo rev</td><td>AAACGCGAGCGGGAGGGGAAGCAA</td><td></td><td></td><td></td><td></td><td></td><td></td><td></td><td></td><td></td><td></td></tr>
<tr><td>Chromosome</td><td>start</td><td>end</td><td>strand</td><td>MM</td><td>target_seq</td><td>PAM</td><td>alignment</td><td>distance</td><td>position</td><td>gene name</td><td>gene id</td></tr>
<tr><td>GU305914.1</td><td>224576</td><td>224598</td><td>-</td><td>0</td><td>TTGCTTCCCCTCCCGCTCGC</td><td>CGG</td><td>||||||||[||||||||||||]PAM</td><td>NA</td><td>Intergenic</td><td>NA</td><td>NA</td></tr>
<tr><td></td><td></td><td></td><td></td><td></td><td></td><td></td><td></td><td></td><td></td><td></td><td></td></tr>
<tr><td>T17681</td><td>TGCTTCCCCTCCCGCTCGCCGGG</td><td>1000</td><td></td><td></td><td></td><td></td><td></td><td></td><td></td><td></td><td></td></tr>
<tr><td>Oligo fwd</td><td>CACCTGCTTCCCCTCCCGCTCGCC</td><td></td><td></td><td></td><td></td><td></td><td></td><td></td><td></td><td></td><td></td></tr>
<tr><td>Oligo rev</td><td>AAACGGCGAGCGGGAGGGGAAGCA</td><td></td><td></td><td></td><td></td><td></td><td></td><td></td><td></td><td></td><td></td></tr>
<tr><td>Chromosome</td><td>start</td><td>end</td><td>strand</td><td>MM</td><td>target_seq</td><td>PAM</td><td>alignment</td><td>distance</td><td>position</td><td>gene name</td><td>gene id</td></tr>
<tr><td>GU305914.1</td><td>224575</td><td>224597</td><td>-</td><td>0</td><td>TGCTTCCCCTCCCGCTCGCC</td><td>GGG</td><td>||||||||[||||||||||||]PAM</td><td>NA</td><td>Intergenic</td><td>NA</td><td>NA</td></tr>
<tr><td></td><td></td><td></td><td></td><td></td><td></td><td></td><td></td><td></td><td></td><td></td><td></td></tr>
<tr><td>T17682</td><td>CCCTCCCGCTCGCCGGGCGCAGG</td><td>1000</td><td></td><td></td><td></td><td></td><td></td><td></td><td></td><td></td><td></td></tr>
<tr><td>Oligo fwd</td><td>CACCCCCTCCCGCTCGCCGGGCGC</td><td></td><td></td><td></td><td></td><td></td><td></td><td></td><td></td><td></td><td></td></tr>
<tr><td>Oligo rev</td><td>AAACGCGCCCGGCGAGCGGGAGGG</td><td></td><td></td><td></td><td></td><td></td><td></td><td></td><td></td><td></td><td></td></tr>
<tr><td>Chromosome</td><td>start</td><td>end</td><td>strand</td><td>MM</td><td>target_seq</td><td>PAM</td><td>alignment</td><td>distance</td><td>position</td><td>gene name</td><td>gene id</td></tr>
<tr><td>GU305914.1</td><td>224569</td><td>224591</td><td>-</td><td>0</td><td>CCCTCCCGCTCGCCGGGCGC</td><td>AGG</td><td>||||||||[||||||||||||]PAM</td><td>NA</td><td>Intergenic</td><td>NA</td><td>NA</td></tr>
<tr><td></td><td></td><td></td><td></td><td></td><td></td><td></td><td></td><td></td><td></td><td></td><td></td></tr>
<tr><td>T17683</td><td>TCCCGCTCGCCGGGCGCAGGAGG</td><td>1000</td><td></td><td></td><td></td><td></td><td></td><td></td><td></td><td></td><td></td></tr>
<tr><td>Oligo fwd</td><td>CACCTCCCGCTCGCCGGGCGCAGG</td><td></td><td></td><td></td><td></td><td></td><td></td><td></td><td></td><td></td><td></td></tr>
<tr><td>Oligo rev</td><td>AAACCCTGCGCCCGGCGAGCGGGA</td><td></td><td></td><td></td><td></td><td></td><td></td><td></td><td></td><td></td><td></td></tr>
<tr><td>Chromosome</td><td>start</td><td>end</td><td>strand</td><td>MM</td><td>target_seq</td><td>PAM</td><td>alignment</td><td>distance</td><td>position</td><td>gene name</td><td>gene id</td></tr>
<tr><td>GU305914.1</td><td>224566</td><td>224588</td><td>-</td><td>0</td><td>TCCCGCTCGCCGGGCGCAGG</td><td>AGG</td><td>||||||||[||||||||||||]PAM</td><td>NA</td><td>Intergenic</td><td>NA</td><td>NA</td></tr>
</table>

T17684 CCGGGCGCAGGAGGAAAAAAAGG 1000
Oligo fwd CACCCCGGGCGCAGGAGGAAAAAA
Oligo rev AAACTTTTTTCCTCCTGCGCCCGG
Chromosome start end strand MM target_seq PAM alignment distance position gene name gene id
GU305914.1 224557 224579 - 0 CCGGGCGCAGGAGGAAAAAA AGG ||||||||[||||||||||||]PAM NA Intergenic NA NA

T17685 GGCGCAGGAGGAAAAAAAGGCGG 1000
Oligo fwd CACCGGCGCAGGAGGAAAAAAAGG
Oligo rev AAACCCTTTTTTTCCTCCTGCGCC
Chromosome start end strand MM target_seq PAM alignment distance position gene name gene id
GU305914.1 224554 224576 - 0 GGCGCAGGAGGAAAAAAAGG CGG ||||||||[||||||||||||]PAM NA Intergenic NA NA

T17686 AGGAGGAAAAAAAGGCGGACCGG 1000
Oligo fwd CACCAGGAGGAAAAAAAGGCGGAC
Oligo rev AAACGTCCGCCTTTTTTTCCTCCT
Chromosome start end strand MM target_seq PAM alignment distance position gene name gene id
GU305914.1 224549 224571 - 0 AGGAGGAAAAAAAGGCGGAC CGG ||||||||[||||||||||||]PAM NA Intergenic NA NA

T17687 CTATGAGATACTCGTGAACGAGG 1000
Oligo fwd CACCCTATGAGATACTCGTGAACG
Oligo rev AAACCGTTCACGAGTATCTCATAG
Chromosome start end strand MM target_seq PAM alignment distance position gene name gene id
GU305914.1 224525 224547 - 0 CTATGAGATACTCGTGAACG AGG ||||||||[||||||||||||]PAM NA Intergenic NA NA

T17688 GATACTCGTGAACGAGGAGACGG 1000
Oligo fwd CACCGATACTCGTGAACGAGGAGA
Oligo rev AAACTCTCCTCGTTCACGAGTATC
Chromosome start end strand MM target_seq PAM alignment distance position gene name gene id
GU305914.1 224519 224541 - 0 GATACTCGTGAACGAGGAGA CGG ||||||||[||||||||||||]PAM NA Intergenic NA NA

T17689 GTGAACGAGGAGACGGCGTGAGG 1000
Oligo fwd CACCGTGAACGAGGAGACGGCGTG
Oligo rev AAACCACGCCGTCTCCTCGTTCAC
Chromosome start end strand MM target_seq PAM alignment distance position gene name gene id
GU305914.1 224512 224534 - 0 GTGAACGAGGAGACGGCGTG AGG ||||||||[||||||||||||]PAM NA Intergenic NA NA

T17690 CGTTCTCTTCCTTTTTCTCGCGG 1000
Oligo fwd CACCCGTTCTCTTCCTTTTTCTCG
Oligo rev AAACCGAGAAAAAGGAAGAGAACG
Chromosome start end strand MM target_seq PAM alignment distance position gene name gene id
GU305914.1 224392 224414 - 0 CGTTCTCTTCCTTTTTCTCG CGG ||||||||[||||||||||||]PAM NA Intergenic NA NA

T17691 TTCCTTTTTCTCGCGGCCACCGG 1000
Oligo fwd CACCTTCCTTTTTCTCGCGGCCAC
Oligo rev AAACGTGGCCGCGAGAAAAAGGAA
Chromosome start end strand MM target_seq PAM alignment distance position gene name gene id
GU305914.1 224385 224407 - 0 TTCCTTTTTCTCGCGGCCAC CGG ||||||||[||||||||||||]PAM NA Intergenic NA NA

T17692 CTCGCGGCCACCGGCCGTCATGG 1000
Oligo fwd CACCCTCGCGGCCACCGGCCGTCA
Oligo rev AAACTGACGGCCGGTGGCCGCGAG
Chromosome start end strand MM target_seq PAM alignment distance position gene name gene id
GU305914.1 224376 224398 - 0 CTCGCGGCCACCGGCCGTCA TGG ||||||||[||||||||||||]PAM NA Intergenic NA NA
GU305914.1 110498 110520 + 4 CTCGCGCGACCCGGCCGTCA CGG ||||||--[--||||||||||]PAM NA Intergenic NA NA

T17693 CCACCGGCCGTCATGGCCCTCGG 1000
Oligo fwd CACCCCACCGGCCGTCATGGCCCT
Oligo rev AAACAGGGCCATGACGGCCGGTGG
Chromosome start end strand MM target_seq PAM alignment distance position gene name gene id
GU305914.1 224369 224391 - 0 CCACCGGCCGTCATGGCCCT CGG ||||||||[||||||||||||]PAM NA Intergenic NA NA

T17694 CACCGGCCGTCATGGCCCTCGGG 1000
Oligo fwd CACCCACCGGCCGTCATGGCCCTC
Oligo rev AAACGAGGGCCATGACGGCCGGTG
Chromosome start end strand MM target_seq PAM alignment distance position gene name gene id
GU305914.1 224368 224390 - 0 CACCGGCCGTCATGGCCCTC GGG ||||||||[||||||||||||]PAM NA Intergenic NA NA

T17695 TGCTGACCCTGCTCCTGACGTGG 1000
Oligo fwd CACCTGCTGACCCTGCTCCTGACG
Oligo rev AAACCGTCAGGAGCAGGGTCAGCA
Chromosome start end strand MM target_seq PAM alignment distance position gene name gene id
GU305914.1 224338 224360 - 0 TGCTGACCCTGCTCCTGACG TGG ||||||||[||||||||||||]PAM NA Intergenic NA NA

T17696 GTTGATGTCCCCTACACGAGCGG 1000
Oligo fwd CACCGTTGATGTCCCCTACACGAG
Oligo rev AAACCTCGTGTAGGGGACATCAAC
Chromosome start end strand MM target_seq PAM alignment distance position gene name gene id
GU305914.1 224313 224335 - 0 GTTGATGTCCCCTACACGAG CGG ||||||||[||||||||||||]PAM NA Intergenic NA NA

T17697 TTGATGTCCCCTACACGAGCGGG 1000
Oligo fwd CACCTTGATGTCCCCTACACGAGC
Oligo rev AAACGCTCGTGTAGGGGACATCAA
Chromosome start end strand MM target_seq PAM alignment distance position gene name gene id
GU305914.1 224312 224334 - 0 TTGATGTCCCCTACACGAGC GGG ||||||||[||||||||||||]PAM NA Intergenic NA NA

T17698 TGTCCCCTACACGAGCGGGACGG 1000
Oligo fwd CACCTGTCCCCTACACGAGCGGGA
Oligo rev AAACTCCCGCTCGTGTAGGGGACA

<table>
<tr><td>Chromosome</td><td>start</td><td>end</td><td>strand</td><td>MM</td><td>target_seq</td><td>PAM</td><td>alignment</td><td>distance</td><td>position</td><td>gene name</td><td>gene id</td></tr>
<tr><td>GU305914.1</td><td>224308</td><td>224330</td><td>-</td><td>0</td><td>TGTCCCCTACACGAGCGGGA</td><td>CGG</td><td>||||||||[||||||||||||]PAM</td><td>NA</td><td>Intergenic</td><td>NA</td><td>NA</td></tr>
<tr><td>T17699</td><td>AGCGGGACGGCCATCCGTCCTGG</td><td>1000</td><td></td><td></td><td></td><td></td><td></td><td></td><td></td><td></td><td></td></tr>
<tr><td>Oligo fwd</td><td>CACCAGCGGGACGGCCATCCGTCC</td><td></td><td></td><td></td><td></td><td></td><td></td><td></td><td></td><td></td><td></td></tr>
<tr><td>Oligo rev</td><td>AAACGGACGGATGGCCGTCCCGCT</td><td></td><td></td><td></td><td></td><td></td><td></td><td></td><td></td><td></td><td></td></tr>
<tr><td>Chromosome</td><td>start</td><td>end</td><td>strand</td><td>MM</td><td>target_seq</td><td>PAM</td><td>alignment</td><td>distance</td><td>position</td><td>gene name</td><td>gene id</td></tr>
<tr><td>GU305914.1</td><td>224295</td><td>224317</td><td>-</td><td>0</td><td>AGCGGGACGGCCATCCGTCC</td><td>TGG</td><td>||||||||[||||||||||||]PAM</td><td>NA</td><td>Intergenic</td><td>NA</td><td>NA</td></tr>
<tr><td>T17700</td><td>ACCCTGCCGCTGACGAGACCCGG</td><td>1000</td><td></td><td></td><td></td><td></td><td></td><td></td><td></td><td></td><td></td></tr>
<tr><td>Oligo fwd</td><td>CACCACCCTGCCGCTGACGAGACC</td><td></td><td></td><td></td><td></td><td></td><td></td><td></td><td></td><td></td><td></td></tr>
<tr><td>Oligo rev</td><td>AAACGGTCTCGTCAGCGGCAGGGT</td><td></td><td></td><td></td><td></td><td></td><td></td><td></td><td></td><td></td><td></td></tr>
<tr><td>Chromosome</td><td>start</td><td>end</td><td>strand</td><td>MM</td><td>target_seq</td><td>PAM</td><td>alignment</td><td>distance</td><td>position</td><td>gene name</td><td>gene id</td></tr>
<tr><td>GU305914.1</td><td>224270</td><td>224292</td><td>-</td><td>0</td><td>ACCCTGCCGCTGACGAGACC</td><td>CGG</td><td>||||||||[||||||||||||]PAM</td><td>NA</td><td>Intergenic</td><td>NA</td><td>NA</td></tr>
<tr><td>T17701</td><td>GACGAGACCCGGTCTCCTCCTGG</td><td>1000</td><td></td><td></td><td></td><td></td><td></td><td></td><td></td><td></td><td></td></tr>
<tr><td>Oligo fwd</td><td>CACCGACGAGACCCGGTCTCCTCC</td><td></td><td></td><td></td><td></td><td></td><td></td><td></td><td></td><td></td><td></td></tr>
<tr><td>Oligo rev</td><td>AAACGGAGGAGACCGGGTCTCGTC</td><td></td><td></td><td></td><td></td><td></td><td></td><td></td><td></td><td></td><td></td></tr>
<tr><td>Chromosome</td><td>start</td><td>end</td><td>strand</td><td>MM</td><td>target_seq</td><td>PAM</td><td>alignment</td><td>distance</td><td>position</td><td>gene name</td><td>gene id</td></tr>
<tr><td>GU305914.1</td><td>224259</td><td>224281</td><td>-</td><td>0</td><td>GACGAGACCCGGTCTCCTCC</td><td>TGG</td><td>||||||||[||||||||||||]PAM</td><td>NA</td><td>Intergenic</td><td>NA</td><td>NA</td></tr>
<tr><td>T17702</td><td>ACGAGACCCGGTCTCCTCCTGGG</td><td>1000</td><td></td><td></td><td></td><td></td><td></td><td></td><td></td><td></td><td></td></tr>
<tr><td>Oligo fwd</td><td>CACCACGAGACCCGGTCTCCTCCT</td><td></td><td></td><td></td><td></td><td></td><td></td><td></td><td></td><td></td><td></td></tr>
<tr><td>Oligo rev</td><td>AAACAGGAGGAGACCGGGTCTCGT</td><td></td><td></td><td></td><td></td><td></td><td></td><td></td><td></td><td></td><td></td></tr>
<tr><td>Chromosome</td><td>start</td><td>end</td><td>strand</td><td>MM</td><td>target_seq</td><td>PAM</td><td>alignment</td><td>distance</td><td>position</td><td>gene name</td><td>gene id</td></tr>
<tr><td>GU305914.1</td><td>224258</td><td>224280</td><td>-</td><td>0</td><td>ACGAGACCCGGTCTCCTCCT</td><td>GGG</td><td>||||||||[||||||||||||]PAM</td><td>NA</td><td>Intergenic</td><td>NA</td><td>NA</td></tr>
<tr><td>T17703</td><td>GTCTCCTCCTGGGCTGCCGCTGG</td><td>1000</td><td></td><td></td><td></td><td></td><td></td><td></td><td></td><td></td><td></td></tr>
<tr><td>Oligo fwd</td><td>CACCGTCTCCTCCTGGGCTGCCGC</td><td></td><td></td><td></td><td></td><td></td><td></td><td></td><td></td><td></td><td></td></tr>
<tr><td>Oligo rev</td><td>AAACGCGGCAGCCCAGGAGGAGAC</td><td></td><td></td><td></td><td></td><td></td><td></td><td></td><td></td><td></td><td></td></tr>
<tr><td>Chromosome</td><td>start</td><td>end</td><td>strand</td><td>MM</td><td>target_seq</td><td>PAM</td><td>alignment</td><td>distance</td><td>position</td><td>gene name</td><td>gene id</td></tr>
<tr><td>GU305914.1</td><td>224248</td><td>224270</td><td>-</td><td>0</td><td>GTCTCCTCCTGGGCTGCCGC</td><td>TGG</td><td>||||||||[||||||||||||]PAM</td><td>NA</td><td>Intergenic</td><td>NA</td><td>NA</td></tr>
<tr><td>T17704</td><td>TCCTGGGCTGCCGCTGGTCTCGG</td><td>1000</td><td></td><td></td><td></td><td></td><td></td><td></td><td></td><td></td><td></td></tr>
<tr><td>Oligo fwd</td><td>CACCTCCTGGGCTGCCGCTGGTCT</td><td></td><td></td><td></td><td></td><td></td><td></td><td></td><td></td><td></td><td></td></tr>
<tr><td>Oligo rev</td><td>AAACAGACCAGCGGCAGCCCAGGA</td><td></td><td></td><td></td><td></td><td></td><td></td><td></td><td></td><td></td><td></td></tr>
<tr><td>Chromosome</td><td>start</td><td>end</td><td>strand</td><td>MM</td><td>target_seq</td><td>PAM</td><td>alignment</td><td>distance</td><td>position</td><td>gene name</td><td>gene id</td></tr>
<tr><td>GU305914.1</td><td>224242</td><td>224264</td><td>-</td><td>0</td><td>TCCTGGGCTGCCGCTGGTCT</td><td>CGG</td><td>||||||||[||||||||||||]PAM</td><td>NA</td><td>Intergenic</td><td>NA</td><td>NA</td></tr>
<tr><td>T17705</td><td>CTGGTCTCGGCCTTCGCGAGCGG</td><td>1000</td><td></td><td></td><td></td><td></td><td></td><td></td><td></td><td></td><td></td></tr>
<tr><td>Oligo fwd</td><td>CACCCTGGTCTCGGCCTTCGCGAG</td><td></td><td></td><td></td><td></td><td></td><td></td><td></td><td></td><td></td><td></td></tr>
<tr><td>Oligo rev</td><td>AAACCTCGCGAAGGCCGAGACCAG</td><td></td><td></td><td></td><td></td><td></td><td></td><td></td><td></td><td></td><td></td></tr>
<tr><td>Chromosome</td><td>start</td><td>end</td><td>strand</td><td>MM</td><td>target_seq</td><td>PAM</td><td>alignment</td><td>distance</td><td>position</td><td>gene name</td><td>gene id</td></tr>
<tr><td>GU305914.1</td><td>224229</td><td>224251</td><td>-</td><td>0</td><td>CTGGTCTCGGCCTTCGCGAG</td><td>CGG</td><td>||||||||[||||||||||||]PAM</td><td>NA</td><td>Intergenic</td><td>NA</td><td>NA</td></tr>
<tr><td>T17706</td><td>GAGCGGACTACATCTCGATCCGG</td><td>1000</td><td></td><td></td><td></td><td></td><td></td><td></td><td></td><td></td><td></td></tr>
<tr><td>Oligo fwd</td><td>CACCGAGCGGACTACATCTCGATC</td><td></td><td></td><td></td><td></td><td></td><td></td><td></td><td></td><td></td><td></td></tr>
<tr><td>Oligo rev</td><td>AAACGATCGAGATGTAGTCCGCTC</td><td></td><td></td><td></td><td></td><td></td><td></td><td></td><td></td><td></td><td></td></tr>
<tr><td>Chromosome</td><td>start</td><td>end</td><td>strand</td><td>MM</td><td>target_seq</td><td>PAM</td><td>alignment</td><td>distance</td><td>position</td><td>gene name</td><td>gene id</td></tr>
<tr><td>GU305914.1</td><td>224212</td><td>224234</td><td>-</td><td>0</td><td>GAGCGGACTACATCTCGATC</td><td>CGG</td><td>||||||||[||||||||||||]PAM</td><td>NA</td><td>Intergenic</td><td>NA</td><td>NA</td></tr>
<tr><td>T17707</td><td>ACATCTCGATCCGGCCGAGATGG</td><td>1000</td><td></td><td></td><td></td><td></td><td></td><td></td><td></td><td></td><td></td></tr>
<tr><td>Oligo fwd</td><td>CACCACATCTCGATCCGGCCGAGA</td><td></td><td></td><td></td><td></td><td></td><td></td><td></td><td></td><td></td><td></td></tr>
<tr><td>Oligo rev</td><td>AAACTCTCGGCCGGATCGAGATGT</td><td></td><td></td><td></td><td></td><td></td><td></td><td></td><td></td><td></td><td></td></tr>
<tr><td>Chromosome</td><td>start</td><td>end</td><td>strand</td><td>MM</td><td>target_seq</td><td>PAM</td><td>alignment</td><td>distance</td><td>position</td><td>gene name</td><td>gene id</td></tr>
<tr><td>GU305914.1</td><td>224203</td><td>224225</td><td>-</td><td>0</td><td>ACATCTCGATCCGGCCGAGA</td><td>TGG</td><td>||||||||[||||||||||||]PAM</td><td>NA</td><td>Intergenic</td><td>NA</td><td>NA</td></tr>
<tr><td>T17708</td><td>CTCGATCCGGCCGAGATGGCCGG</td><td>1000</td><td></td><td></td><td></td><td></td><td></td><td></td><td></td><td></td><td></td></tr>
<tr><td>Oligo fwd</td><td>CACCCTCGATCCGGCCGAGATGGC</td><td></td><td></td><td></td><td></td><td></td><td></td><td></td><td></td><td></td><td></td></tr>
<tr><td>Oligo rev</td><td>AAACGCCATCTCGGCCGGATCGAG</td><td></td><td></td><td></td><td></td><td></td><td></td><td></td><td></td><td></td><td></td></tr>
<tr><td>Chromosome</td><td>start</td><td>end</td><td>strand</td><td>MM</td><td>target_seq</td><td>PAM</td><td>alignment</td><td>distance</td><td>position</td><td>gene name</td><td>gene id</td></tr>
<tr><td>GU305914.1</td><td>224199</td><td>224221</td><td>-</td><td>0</td><td>CTCGATCCGGCCGAGATGGC</td><td>CGG</td><td>||||||||[||||||||||||]PAM</td><td>NA</td><td>Intergenic</td><td>NA</td><td>NA</td></tr>
<tr><td>T17709</td><td>TCGATCCGGCCGAGATGGCCGGG</td><td>1000</td><td></td><td></td><td></td><td></td><td></td><td></td><td></td><td></td><td></td></tr>
<tr><td>Oligo fwd</td><td>CACCTCGATCCGGCCGAGATGGCC</td><td></td><td></td><td></td><td></td><td></td><td></td><td></td><td></td><td></td><td></td></tr>
<tr><td>Oligo rev</td><td>AAACGGCCATCTCGGCCGGATCGA</td><td></td><td></td><td></td><td></td><td></td><td></td><td></td><td></td><td></td><td></td></tr>
<tr><td>Chromosome</td><td>start</td><td>end</td><td>strand</td><td>MM</td><td>target_seq</td><td>PAM</td><td>alignment</td><td>distance</td><td>position</td><td>gene name</td><td>gene id</td></tr>
<tr><td>GU305914.1</td><td>224198</td><td>224220</td><td>-</td><td>0</td><td>TCGATCCGGCCGAGATGGCC</td><td>GGG</td><td>||||||||[||||||||||||]PAM</td><td>NA</td><td>Intergenic</td><td>NA</td><td>NA</td></tr>
<tr><td>T17710</td><td>TCCGGCCGAGATGGCCGGGCCGG</td><td>1000</td><td></td><td></td><td></td><td></td><td></td><td></td><td></td><td></td><td></td></tr>
<tr><td>Oligo fwd</td><td>CACCTCCGGCCGAGATGGCCGGGC</td><td></td><td></td><td></td><td></td><td></td><td></td><td></td><td></td><td></td><td></td></tr>
<tr><td>Oligo rev</td><td>AAACGCCCGGCCATCTCGGCCGGA</td><td></td><td></td><td></td><td></td><td></td><td></td><td></td><td></td><td></td><td></td></tr>
<tr><td>Chromosome</td><td>start</td><td>end</td><td>strand</td><td>MM</td><td>target_seq</td><td>PAM</td><td>alignment</td><td>distance</td><td>position</td><td>gene name</td><td>gene id</td></tr>
<tr><td>GU305914.1</td><td>224194</td><td>224216</td><td>-</td><td>0</td><td>TCCGGCCGAGATGGCCGGGC</td><td>CGG</td><td>||||||||[||||||||||||]PAM</td><td>NA</td><td>Intergenic</td><td>NA</td><td>NA</td></tr>
<tr><td>T17711</td><td>CCGGCCGAGATGGCCGGGCCGGG</td><td>1000</td><td></td><td></td><td></td><td></td><td></td><td></td><td></td><td></td><td></td></tr>
<tr><td>Oligo fwd</td><td>CACCCCGGCCGAGATGGCCGGGCC</td><td></td><td></td><td></td><td></td><td></td><td></td><td></td><td></td><td></td><td></td></tr>
<tr><td>Oligo rev</td><td>AAACGGCCCGGCCATCTCGGCCGG</td><td></td><td></td><td></td><td></td><td></td><td></td><td></td><td></td><td></td><td></td></tr>
<tr><td>Chromosome</td><td>start</td><td>end</td><td>strand</td><td>MM</td><td>target_seq</td><td>PAM</td><td>alignment</td><td>distance</td><td>position</td><td>gene name</td><td>gene id</td></tr>
<tr><td>GU305914.1</td><td>224193</td><td>224215</td><td>-</td><td>0</td><td>CCGGCCGAGATGGCCGGGCC</td><td>GGG</td><td>||||||||[||||||||||||]PAM</td><td>NA</td><td>Intergenic</td><td>NA</td><td>NA</td></tr>
<tr><td>T17712</td><td>GCCGAGATGGCCGGGCCGGGCGG</td><td>1000</td><td></td><td></td><td></td><td></td><td></td><td></td><td></td><td></td><td></td></tr>
<tr><td>Oligo fwd</td><td>CACCGCCGAGATGGCCGGGCCGGG</td><td></td><td></td><td></td><td></td><td></td><td></td><td></td><td></td><td></td><td></td></tr>
<tr><td>Oligo rev</td><td>AAACCCCGGCCCGGCCATCTCGGC</td><td></td><td></td><td></td><td></td><td></td><td></td><td></td><td></td><td></td><td></td></tr>
<tr><td>Chromosome</td><td>start</td><td>end</td><td>strand</td><td>MM</td><td>target_seq</td><td>PAM</td><td>alignment</td><td>distance</td><td>position</td><td>gene name</td><td>gene id</td></tr>
<tr><td>GU305914.1</td><td>224190</td><td>224212</td><td>-</td><td>0</td><td>GCCGAGATGGCCGGGCCGGG</td><td>CGG</td><td>||||||||[||||||||||||]PAM</td><td>NA</td><td>Intergenic</td><td>NA</td><td>NA</td></tr>
<tr><td>T17713</td><td>TGTCGCTCGCGTCCTCCGACAGG</td><td>1000</td><td></td><td></td><td></td><td></td><td></td><td></td><td></td><td></td><td></td></tr>
<tr><td>Oligo fwd</td><td>CACCTGTCGCTCGCGTCCTCCGAC</td><td></td><td></td><td></td><td></td><td></td><td></td><td></td><td></td><td></td><td></td></tr>
</table>

<table>
<tr><td>Oligo rev</td><td>AAACGTCGGAGGACGCGAGCGACA</td><td></td><td></td><td></td><td></td><td></td><td></td><td></td><td></td><td></td><td></td></tr>
<tr><td>Chromosome</td><td>start</td><td>end</td><td>strand</td><td>MM</td><td>target_seq</td><td>PAM</td><td>alignment</td><td>distance</td><td>position</td><td>gene name</td><td>gene id</td></tr>
<tr><td>GU305914.1</td><td>224155</td><td>224177</td><td>-</td><td>0</td><td>TGTCGCTCGCGTCCTCCGAC</td><td>AGG</td><td>||||||||[||||||||||||]PAM</td><td>NA</td><td>Intergenic</td><td>NA</td><td>NA</td></tr>
<tr><td>T17714</td><td>GTCGCTCGCGTCCTCCGACAGGG</td><td>1000</td><td></td><td></td><td></td><td></td><td></td><td></td><td></td><td></td><td></td></tr>
<tr><td>Oligo fwd</td><td>CACCGTCGCTCGCGTCCTCCGACA</td><td></td><td></td><td></td><td></td><td></td><td></td><td></td><td></td><td></td><td></td></tr>
<tr><td>Oligo rev</td><td>AAACTGTCGGAGGACGCGAGCGAC</td><td></td><td></td><td></td><td></td><td></td><td></td><td></td><td></td><td></td><td></td></tr>
<tr><td>Chromosome</td><td>start</td><td>end</td><td>strand</td><td>MM</td><td>target_seq</td><td>PAM</td><td>alignment</td><td>distance</td><td>position</td><td>gene name</td><td>gene id</td></tr>
<tr><td>GU305914.1</td><td>224154</td><td>224176</td><td>-</td><td>0</td><td>GTCGCTCGCGTCCTCCGACA</td><td>GGG</td><td>||||||||[||||||||||||]PAM</td><td>NA</td><td>Intergenic</td><td>NA</td><td>NA</td></tr>
<tr><td>T17715</td><td>TCGTGTTCTATCTGCCGCACCGG</td><td>1000</td><td></td><td></td><td></td><td></td><td></td><td></td><td></td><td></td><td></td></tr>
<tr><td>Oligo fwd</td><td>CACCTCGTGTTCTATCTGCCGCAC</td><td></td><td></td><td></td><td></td><td></td><td></td><td></td><td></td><td></td><td></td></tr>
<tr><td>Oligo rev</td><td>AAACGTGCGGCAGATAGAACACGA</td><td></td><td></td><td></td><td></td><td></td><td></td><td></td><td></td><td></td><td></td></tr>
<tr><td>Chromosome</td><td>start</td><td>end</td><td>strand</td><td>MM</td><td>target_seq</td><td>PAM</td><td>alignment</td><td>distance</td><td>position</td><td>gene name</td><td>gene id</td></tr>
<tr><td>GU305914.1</td><td>224131</td><td>224153</td><td>-</td><td>0</td><td>TCGTGTTCTATCTGCCGCAC</td><td>CGG</td><td>||||||||[||||||||||||]PAM</td><td>NA</td><td>Intergenic</td><td>NA</td><td>NA</td></tr>
<tr><td>T17716</td><td>GTTCTATCTGCCGCACCGGCTGG</td><td>1000</td><td></td><td></td><td></td><td></td><td></td><td></td><td></td><td></td><td></td></tr>
<tr><td>Oligo fwd</td><td>CACCGTTCTATCTGCCGCACCGGC</td><td></td><td></td><td></td><td></td><td></td><td></td><td></td><td></td><td></td><td></td></tr>
<tr><td>Oligo rev</td><td>AAACGCCGGTGCGGCAGATAGAAC</td><td></td><td></td><td></td><td></td><td></td><td></td><td></td><td></td><td></td><td></td></tr>
<tr><td>Chromosome</td><td>start</td><td>end</td><td>strand</td><td>MM</td><td>target_seq</td><td>PAM</td><td>alignment</td><td>distance</td><td>position</td><td>gene name</td><td>gene id</td></tr>
<tr><td>GU305914.1</td><td>224127</td><td>224149</td><td>-</td><td>0</td><td>GTTCTATCTGCCGCACCGGC</td><td>TGG</td><td>||||||||[||||||||||||]PAM</td><td>NA</td><td>Intergenic</td><td>NA</td><td>NA</td></tr>
<tr><td>T17717</td><td>CTATCTGCCGCACCGGCTGGAGG</td><td>1000</td><td></td><td></td><td></td><td></td><td></td><td></td><td></td><td></td><td></td></tr>
<tr><td>Oligo fwd</td><td>CACCCTATCTGCCGCACCGGCTGG</td><td></td><td></td><td></td><td></td><td></td><td></td><td></td><td></td><td></td><td></td></tr>
<tr><td>Oligo rev</td><td>AAACCCAGCCGGTGCGGCAGATAG</td><td></td><td></td><td></td><td></td><td></td><td></td><td></td><td></td><td></td><td></td></tr>
<tr><td>Chromosome</td><td>start</td><td>end</td><td>strand</td><td>MM</td><td>target_seq</td><td>PAM</td><td>alignment</td><td>distance</td><td>position</td><td>gene name</td><td>gene id</td></tr>
<tr><td>GU305914.1</td><td>224124</td><td>224146</td><td>-</td><td>0</td><td>CTATCTGCCGCACCGGCTGG</td><td>AGG</td><td>||||||||[||||||||||||]PAM</td><td>NA</td><td>Intergenic</td><td>NA</td><td>NA</td></tr>
<tr><td>T17718</td><td>CTGGAGGTAGTCCTGTCGTGCGG</td><td>1000</td><td></td><td></td><td></td><td></td><td></td><td></td><td></td><td></td><td></td></tr>
<tr><td>Oligo fwd</td><td>CACCCTGGAGGTAGTCCTGTCGTG</td><td></td><td></td><td></td><td></td><td></td><td></td><td></td><td></td><td></td><td></td></tr>
<tr><td>Oligo rev</td><td>AAACCACGACAGGACTACCTCCAG</td><td></td><td></td><td></td><td></td><td></td><td></td><td></td><td></td><td></td><td></td></tr>
<tr><td>Chromosome</td><td>start</td><td>end</td><td>strand</td><td>MM</td><td>target_seq</td><td>PAM</td><td>alignment</td><td>distance</td><td>position</td><td>gene name</td><td>gene id</td></tr>
<tr><td>GU305914.1</td><td>224108</td><td>224130</td><td>-</td><td>0</td><td>CTGGAGGTAGTCCTGTCGTG</td><td>CGG</td><td>||||||||[||||||||||||]PAM</td><td>NA</td><td>Intergenic</td><td>NA</td><td>NA</td></tr>
<tr><td>T17719</td><td>GAGGTAGTCCTGTCGTGCGGCGG</td><td>1000</td><td></td><td></td><td></td><td></td><td></td><td></td><td></td><td></td><td></td></tr>
<tr><td>Oligo fwd</td><td>CACCGAGGTAGTCCTGTCGTGCGG</td><td></td><td></td><td></td><td></td><td></td><td></td><td></td><td></td><td></td><td></td></tr>
<tr><td>Oligo rev</td><td>AAACCCGCACGACAGGACTACCTC</td><td></td><td></td><td></td><td></td><td></td><td></td><td></td><td></td><td></td><td></td></tr>
<tr><td>Chromosome</td><td>start</td><td>end</td><td>strand</td><td>MM</td><td>target_seq</td><td>PAM</td><td>alignment</td><td>distance</td><td>position</td><td>gene name</td><td>gene id</td></tr>
<tr><td>GU305914.1</td><td>224105</td><td>224127</td><td>-</td><td>0</td><td>GAGGTAGTCCTGTCGTGCGG</td><td>CGG</td><td>||||||||[||||||||||||]PAM</td><td>NA</td><td>Intergenic</td><td>NA</td><td>NA</td></tr>
<tr><td>T17720</td><td>AGGTAGTCCTGTCGTGCGGCGGG</td><td>1000</td><td></td><td></td><td></td><td></td><td></td><td></td><td></td><td></td><td></td></tr>
<tr><td>Oligo fwd</td><td>CACCAGGTAGTCCTGTCGTGCGGC</td><td></td><td></td><td></td><td></td><td></td><td></td><td></td><td></td><td></td><td></td></tr>
<tr><td>Oligo rev</td><td>AAACGCCGCACGACAGGACTACCT</td><td></td><td></td><td></td><td></td><td></td><td></td><td></td><td></td><td></td><td></td></tr>
<tr><td>Chromosome</td><td>start</td><td>end</td><td>strand</td><td>MM</td><td>target_seq</td><td>PAM</td><td>alignment</td><td>distance</td><td>position</td><td>gene name</td><td>gene id</td></tr>
<tr><td>GU305914.1</td><td>224104</td><td>224126</td><td>-</td><td>0</td><td>AGGTAGTCCTGTCGTGCGGC</td><td>GGG</td><td>||||||||[||||||||||||]PAM</td><td>NA</td><td>Intergenic</td><td>NA</td><td>NA</td></tr>
<tr><td>T17721</td><td>GGTAGTCCTGTCGTGCGGCGGGG</td><td>1000</td><td></td><td></td><td></td><td></td><td></td><td></td><td></td><td></td><td></td></tr>
<tr><td>Oligo fwd</td><td>CACCGGTAGTCCTGTCGTGCGGCG</td><td></td><td></td><td></td><td></td><td></td><td></td><td></td><td></td><td></td><td></td></tr>
<tr><td>Oligo rev</td><td>AAACCGCCGCACGACAGGACTACC</td><td></td><td></td><td></td><td></td><td></td><td></td><td></td><td></td><td></td><td></td></tr>
<tr><td>Chromosome</td><td>start</td><td>end</td><td>strand</td><td>MM</td><td>target_seq</td><td>PAM</td><td>alignment</td><td>distance</td><td>position</td><td>gene name</td><td>gene id</td></tr>
<tr><td>GU305914.1</td><td>224103</td><td>224125</td><td>-</td><td>0</td><td>GGTAGTCCTGTCGTGCGGCG</td><td>GGG</td><td>||||||||[||||||||||||]PAM</td><td>NA</td><td>Intergenic</td><td>NA</td><td>NA</td></tr>
<tr><td>T17722</td><td>TGTCGTGCGGCGGGGACGAGAGG</td><td>1000</td><td></td><td></td><td></td><td></td><td></td><td></td><td></td><td></td><td></td></tr>
<tr><td>Oligo fwd</td><td>CACCTGTCGTGCGGCGGGGACGAG</td><td></td><td></td><td></td><td></td><td></td><td></td><td></td><td></td><td></td><td></td></tr>
<tr><td>Oligo rev</td><td>AAACCTCGTCCCCGCCGCACGACA</td><td></td><td></td><td></td><td></td><td></td><td></td><td></td><td></td><td></td><td></td></tr>
<tr><td>Chromosome</td><td>start</td><td>end</td><td>strand</td><td>MM</td><td>target_seq</td><td>PAM</td><td>alignment</td><td>distance</td><td>position</td><td>gene name</td><td>gene id</td></tr>
<tr><td>GU305914.1</td><td>224095</td><td>224117</td><td>-</td><td>0</td><td>TGTCGTGCGGCGGGGACGAG</td><td>AGG</td><td>||||||||[||||||||||||]PAM</td><td>NA</td><td>Intergenic</td><td>NA</td><td>NA</td></tr>
<tr><td>T17723</td><td>GTCGTGCGGCGGGGACGAGAGGG</td><td>1000</td><td></td><td></td><td></td><td></td><td></td><td></td><td></td><td></td><td></td></tr>
<tr><td>Oligo fwd</td><td>CACCGTCGTGCGGCGGGGACGAGA</td><td></td><td></td><td></td><td></td><td></td><td></td><td></td><td></td><td></td><td></td></tr>
<tr><td>Oligo rev</td><td>AAACTCTCGTCCCCGCCGCACGAC</td><td></td><td></td><td></td><td></td><td></td><td></td><td></td><td></td><td></td><td></td></tr>
<tr><td>Chromosome</td><td>start</td><td>end</td><td>strand</td><td>MM</td><td>target_seq</td><td>PAM</td><td>alignment</td><td>distance</td><td>position</td><td>gene name</td><td>gene id</td></tr>
<tr><td>GU305914.1</td><td>224094</td><td>224116</td><td>-</td><td>0</td><td>GTCGTGCGGCGGGGACGAGA</td><td>GGG</td><td>||||||||[||||||||||||]PAM</td><td>NA</td><td>Intergenic</td><td>NA</td><td>NA</td></tr>
<tr><td>T17724</td><td>GAGGGAGCCGACGAAGTACTCGG</td><td>1000</td><td></td><td></td><td></td><td></td><td></td><td></td><td></td><td></td><td></td></tr>
<tr><td>Oligo fwd</td><td>CACCGAGGGAGCCGACGAAGTACT</td><td></td><td></td><td></td><td></td><td></td><td></td><td></td><td></td><td></td><td></td></tr>
<tr><td>Oligo rev</td><td>AAACAGTACTTCGTCGGCTCCCTC</td><td></td><td></td><td></td><td></td><td></td><td></td><td></td><td></td><td></td><td></td></tr>
<tr><td>Chromosome</td><td>start</td><td>end</td><td>strand</td><td>MM</td><td>target_seq</td><td>PAM</td><td>alignment</td><td>distance</td><td>position</td><td>gene name</td><td>gene id</td></tr>
<tr><td>GU305914.1</td><td>224076</td><td>224098</td><td>-</td><td>0</td><td>GAGGGAGCCGACGAAGTACT</td><td>CGG</td><td>||||||||[||||||||||||]PAM</td><td>NA</td><td>Intergenic</td><td>NA</td><td>NA</td></tr>
<tr><td>GU305914.1</td><td>226620</td><td>226642</td><td>-</td><td>4</td><td>GAGGAAGACGAAGGAGTACT</td><td>GGG</td><td>||||-||-[|||-|-||||||]PAM</td><td>NA</td><td>Intergenic</td><td>NA</td><td>NA</td></tr>
<tr><td>T17725</td><td>ACGAAGTACTCGGTCGTCGCCGG</td><td>1000</td><td></td><td></td><td></td><td></td><td></td><td></td><td></td><td></td><td></td></tr>
<tr><td>Oligo fwd</td><td>CACCACGAAGTACTCGGTCGTCGC</td><td></td><td></td><td></td><td></td><td></td><td></td><td></td><td></td><td></td><td></td></tr>
<tr><td>Oligo rev</td><td>AAACGCGACGACCGAGTACTTCGT</td><td></td><td></td><td></td><td></td><td></td><td></td><td></td><td></td><td></td><td></td></tr>
<tr><td>Chromosome</td><td>start</td><td>end</td><td>strand</td><td>MM</td><td>target_seq</td><td>PAM</td><td>alignment</td><td>distance</td><td>position</td><td>gene name</td><td>gene id</td></tr>
<tr><td>GU305914.1</td><td>224066</td><td>224088</td><td>-</td><td>0</td><td>ACGAAGTACTCGGTCGTCGC</td><td>CGG</td><td>||||||||[||||||||||||]PAM</td><td>NA</td><td>Intergenic</td><td>NA</td><td>NA</td></tr>
<tr><td>T17726</td><td>ACTCGGTCGTCGCCGGTCCCAGG</td><td>1000</td><td></td><td></td><td></td><td></td><td></td><td></td><td></td><td></td><td></td></tr>
<tr><td>Oligo fwd</td><td>CACCACTCGGTCGTCGCCGGTCCC</td><td></td><td></td><td></td><td></td><td></td><td></td><td></td><td></td><td></td><td></td></tr>
<tr><td>Oligo rev</td><td>AAACGGGACCGGCGACGACCGAGT</td><td></td><td></td><td></td><td></td><td></td><td></td><td></td><td></td><td></td><td></td></tr>
<tr><td>Chromosome</td><td>start</td><td>end</td><td>strand</td><td>MM</td><td>target_seq</td><td>PAM</td><td>alignment</td><td>distance</td><td>position</td><td>gene name</td><td>gene id</td></tr>
<tr><td>GU305914.1</td><td>224059</td><td>224081</td><td>-</td><td>0</td><td>ACTCGGTCGTCGCCGGTCCC</td><td>AGG</td><td>||||||||[||||||||||||]PAM</td><td>NA</td><td>Intergenic</td><td>NA</td><td>NA</td></tr>
<tr><td>T17727</td><td>TCGCCGGTCCCAGGCTGAGACGG</td><td>1000</td><td></td><td></td><td></td><td></td><td></td><td></td><td></td><td></td><td></td></tr>
<tr><td>Oligo fwd</td><td>CACCTCGCCGGTCCCAGGCTGAGA</td><td></td><td></td><td></td><td></td><td></td><td></td><td></td><td></td><td></td><td></td></tr>
<tr><td>Oligo rev</td><td>AAACTCTCAGCCTGGGACCGGCGA</td><td></td><td></td><td></td><td></td><td></td><td></td><td></td><td></td><td></td><td></td></tr>
<tr><td>Chromosome</td><td>start</td><td>end</td><td>strand</td><td>MM</td><td>target_seq</td><td>PAM</td><td>alignment</td><td>distance</td><td>position</td><td>gene name</td><td>gene id</td></tr>
<tr><td>GU305914.1</td><td>224050</td><td>224072</td><td>-</td><td>0</td><td>TCGCCGGTCCCAGGCTGAGA</td><td>CGG</td><td>||||||||[||||||||||||]PAM</td><td>NA</td><td>Intergenic</td><td>NA</td><td>NA</td></tr>
</table>

<table>
<tr><td>T17728</td><td>CTGAGACGGCTAGCGAATCGCGG</td><td></td><td>1000</td></tr>
<tr><td>Oligo fwd</td><td>CACCCTGAGACGGCTAGCGAATCG</td></tr>
<tr><td>Oligo rev</td><td>AAACCGATTCGCTAGCCGTCTCAG</td></tr>
<tr><td>Chromosome</td><td>start</td><td></td><td>end</td><td>strand</td><td>MM</td><td>target_seq</td><td>PAM</td><td>alignment</td><td>distance</td><td>position</td><td>gene name</td><td>gene id</td></tr>
<tr><td>GU305914.1</td><td></td><td>224036</td><td>224058</td><td>-</td><td>0</td><td>CTGAGACGGCTAGCGAATCG</td><td>CGG</td><td>||||||||[||||||||||||]PAM</td><td>NA</td><td>Intergenic</td><td>NA</td><td>NA</td></tr>
<tr><td>T17729</td><td>GTTCTGACTTTTCTCGCGTGCGG</td><td></td><td>1000</td></tr>
<tr><td>Oligo fwd</td><td>CACCGTTCTGACTTTTCTCGCGTG</td></tr>
<tr><td>Oligo rev</td><td>AAACCACGCGAGAAAAGTCAGAAC</td></tr>
<tr><td>Chromosome</td><td>start</td><td></td><td>end</td><td>strand</td><td>MM</td><td>target_seq</td><td>PAM</td><td>alignment</td><td>distance</td><td>position</td><td>gene name</td><td>gene id</td></tr>
<tr><td>GU305914.1</td><td></td><td>224014</td><td>224036</td><td>-</td><td>0</td><td>GTTCTGACTTTTCTCGCGTG</td><td>CGG</td><td>||||||||[||||||||||||]PAM</td><td>NA</td><td>Intergenic</td><td>NA</td><td>NA</td></tr>
<tr><td>T17730</td><td>CGTGCGGTGTGAGCAGTATCTGG</td><td></td><td>1000</td></tr>
<tr><td>Oligo fwd</td><td>CACCCGTGCGGTGTGAGCAGTATC</td></tr>
<tr><td>Oligo rev</td><td>AAACGATACTGCTCACACCGCACG</td></tr>
<tr><td>Chromosome</td><td>start</td><td></td><td>end</td><td>strand</td><td>MM</td><td>target_seq</td><td>PAM</td><td>alignment</td><td>distance</td><td>position</td><td>gene name</td><td>gene id</td></tr>
<tr><td>GU305914.1</td><td></td><td>223998</td><td>224020</td><td>-</td><td>0</td><td>CGTGCGGTGTGAGCAGTATC</td><td>TGG</td><td>||||||||[||||||||||||]PAM</td><td>NA</td><td>Intergenic</td><td>NA</td><td>NA</td></tr>
<tr><td>T17731</td><td>GTGCGGTGTGAGCAGTATCTGGG</td><td></td><td>1000</td></tr>
<tr><td>Oligo fwd</td><td>CACCGTGCGGTGTGAGCAGTATCT</td></tr>
<tr><td>Oligo rev</td><td>AAACAGATACTGCTCACACCGCAC</td></tr>
<tr><td>Chromosome</td><td>start</td><td></td><td>end</td><td>strand</td><td>MM</td><td>target_seq</td><td>PAM</td><td>alignment</td><td>distance</td><td>position</td><td>gene name</td><td>gene id</td></tr>
<tr><td>GU305914.1</td><td></td><td>223997</td><td>224019</td><td>-</td><td>0</td><td>GTGCGGTGTGAGCAGTATCT</td><td>GGG</td><td>||||||||[||||||||||||]PAM</td><td>NA</td><td>Intergenic</td><td>NA</td><td>NA</td></tr>
<tr><td>T17732</td><td>TGTGAGCAGTATCTGGGTGACGG</td><td></td><td>1000</td></tr>
<tr><td>Oligo fwd</td><td>CACCTGTGAGCAGTATCTGGGTGA</td></tr>
<tr><td>Oligo rev</td><td>AAACTCACCCAGATACTGCTCACA</td></tr>
<tr><td>Chromosome</td><td>start</td><td></td><td>end</td><td>strand</td><td>MM</td><td>target_seq</td><td>PAM</td><td>alignment</td><td>distance</td><td>position</td><td>gene name</td><td>gene id</td></tr>
<tr><td>GU305914.1</td><td></td><td>223991</td><td>224013</td><td>-</td><td>0</td><td>TGTGAGCAGTATCTGGGTGA</td><td>CGG</td><td>||||||||[||||||||||||]PAM</td><td>NA</td><td>Intergenic</td><td>NA</td><td>NA</td></tr>
<tr><td>T17733</td><td>GTGAGCAGTATCTGGGTGACGGG</td><td></td><td>1000</td></tr>
<tr><td>Oligo fwd</td><td>CACCGTGAGCAGTATCTGGGTGAC</td></tr>
<tr><td>Oligo rev</td><td>AAACGTCACCCAGATACTGCTCAC</td></tr>
<tr><td>Chromosome</td><td>start</td><td></td><td>end</td><td>strand</td><td>MM</td><td>target_seq</td><td>PAM</td><td>alignment</td><td>distance</td><td>position</td><td>gene name</td><td>gene id</td></tr>
<tr><td>GU305914.1</td><td></td><td>223990</td><td>224012</td><td>-</td><td>0</td><td>GTGAGCAGTATCTGGGTGAC</td><td>GGG</td><td>||||||||[||||||||||||]PAM</td><td>NA</td><td>Intergenic</td><td>NA</td><td>NA</td></tr>
<tr><td>T17734</td><td>TGAGCAGTATCTGGGTGACGGGG</td><td></td><td>1000</td></tr>
<tr><td>Oligo fwd</td><td>CACCTGAGCAGTATCTGGGTGACG</td></tr>
<tr><td>Oligo rev</td><td>AAACCGTCACCCAGATACTGCTCA</td></tr>
<tr><td>Chromosome</td><td>start</td><td></td><td>end</td><td>strand</td><td>MM</td><td>target_seq</td><td>PAM</td><td>alignment</td><td>distance</td><td>position</td><td>gene name</td><td>gene id</td></tr>
<tr><td>GU305914.1</td><td></td><td>223989</td><td>224011</td><td>-</td><td>0</td><td>TGAGCAGTATCTGGGTGACG</td><td>GGG</td><td>||||||||[||||||||||||]PAM</td><td>NA</td><td>Intergenic</td><td>NA</td><td>NA</td></tr>
<tr><td>T17735</td><td>GAGCAGTATCTGGGTGACGGGGG</td><td></td><td>1000</td></tr>
<tr><td>Oligo fwd</td><td>CACCGAGCAGTATCTGGGTGACGG</td></tr>
<tr><td>Oligo rev</td><td>AAACCCGTCACCCAGATACTGCTC</td></tr>
<tr><td>Chromosome</td><td>start</td><td></td><td>end</td><td>strand</td><td>MM</td><td>target_seq</td><td>PAM</td><td>alignment</td><td>distance</td><td>position</td><td>gene name</td><td>gene id</td></tr>
<tr><td>GU305914.1</td><td></td><td>223988</td><td>224010</td><td>-</td><td>0</td><td>GAGCAGTATCTGGGTGACGG</td><td>GGG</td><td>||||||||[||||||||||||]PAM</td><td>NA</td><td>Intergenic</td><td>NA</td><td>NA</td></tr>
<tr><td>T17736</td><td>TCTGGGTGACGGGGGTAGTGTGG</td><td></td><td>1000</td></tr>
<tr><td>Oligo fwd</td><td>CACCTCTGGGTGACGGGGGTAGTG</td></tr>
<tr><td>Oligo rev</td><td>AAACCACTACCCCCGTCACCCAGA</td></tr>
<tr><td>Chromosome</td><td>start</td><td></td><td>end</td><td>strand</td><td>MM</td><td>target_seq</td><td>PAM</td><td>alignment</td><td>distance</td><td>position</td><td>gene name</td><td>gene id</td></tr>
<tr><td>GU305914.1</td><td></td><td>223980</td><td>224002</td><td>-</td><td>0</td><td>TCTGGGTGACGGGGGTAGTG</td><td>TGG</td><td>||||||||[||||||||||||]PAM</td><td>NA</td><td>Intergenic</td><td>NA</td><td>NA</td></tr>
<tr><td>T17737</td><td>CTGGGTGACGGGGGTAGTGTGGG</td><td></td><td>1000</td></tr>
<tr><td>Oligo fwd</td><td>CACCCTGGGTGACGGGGGTAGTGT</td></tr>
<tr><td>Oligo rev</td><td>AAACACACTACCCCCGTCACCCAG</td></tr>
<tr><td>Chromosome</td><td>start</td><td></td><td>end</td><td>strand</td><td>MM</td><td>target_seq</td><td>PAM</td><td>alignment</td><td>distance</td><td>position</td><td>gene name</td><td>gene id</td></tr>
<tr><td>GU305914.1</td><td></td><td>223979</td><td>224001</td><td>-</td><td>0</td><td>CTGGGTGACGGGGGTAGTGT</td><td>GGG</td><td>||||||||[||||||||||||]PAM</td><td>NA</td><td>Intergenic</td><td>NA</td><td>NA</td></tr>
<tr><td>T17738</td><td>GGTGACGGGGGTAGTGTGGGAGG</td><td></td><td>1000</td></tr>
<tr><td>Oligo fwd</td><td>CACCGGTGACGGGGGTAGTGTGGG</td></tr>
<tr><td>Oligo rev</td><td>AAACCCCACACTACCCCCGTCACC</td></tr>
<tr><td>Chromosome</td><td>start</td><td></td><td>end</td><td>strand</td><td>MM</td><td>target_seq</td><td>PAM</td><td>alignment</td><td>distance</td><td>position</td><td>gene name</td><td>gene id</td></tr>
<tr><td>GU305914.1</td><td></td><td>223976</td><td>223998</td><td>-</td><td>0</td><td>GGTGACGGGGGTAGTGTGGG</td><td>AGG</td><td>||||||||[||||||||||||]PAM</td><td>NA</td><td>Intergenic</td><td>NA</td><td>NA</td></tr>
<tr><td>T17739</td><td>GTGACGGGGGTAGTGTGGGAGGG</td><td></td><td>1000</td></tr>
<tr><td>Oligo fwd</td><td>CACCGTGACGGGGGTAGTGTGGGA</td></tr>
<tr><td>Oligo rev</td><td>AAACTCCCACACTACCCCCGTCAC</td></tr>
<tr><td>Chromosome</td><td>start</td><td></td><td>end</td><td>strand</td><td>MM</td><td>target_seq</td><td>PAM</td><td>alignment</td><td>distance</td><td>position</td><td>gene name</td><td>gene id</td></tr>
<tr><td>GU305914.1</td><td></td><td>223975</td><td>223997</td><td>-</td><td>0</td><td>GTGACGGGGGTAGTGTGGGA</td><td>GGG</td><td>||||||||[||||||||||||]PAM</td><td>NA</td><td>Intergenic</td><td>NA</td><td>NA</td></tr>
<tr><td>T17740</td><td>TGACGGGGGTAGTGTGGGAGGGG</td><td></td><td>1000</td></tr>
<tr><td>Oligo fwd</td><td>CACCTGACGGGGGTAGTGTGGGAG</td></tr>
<tr><td>Oligo rev</td><td>AAACCTCCCACACTACCCCCGTCA</td></tr>
<tr><td>Chromosome</td><td>start</td><td></td><td>end</td><td>strand</td><td>MM</td><td>target_seq</td><td>PAM</td><td>alignment</td><td>distance</td><td>position</td><td>gene name</td><td>gene id</td></tr>
<tr><td>GU305914.1</td><td></td><td>223974</td><td>223996</td><td>-</td><td>0</td><td>TGACGGGGGTAGTGTGGGAG</td><td>GGG</td><td>||||||||[||||||||||||]PAM</td><td>NA</td><td>Intergenic</td><td>NA</td><td>NA</td></tr>
<tr><td>T17741</td><td>CGGGGGTAGTGTGGGAGGGGTGG</td><td></td><td>1000</td></tr>
<tr><td>Oligo fwd</td><td>CACCCGGGGGTAGTGTGGGAGGGG</td></tr>
<tr><td>Oligo rev</td><td>AAACCCCCTCCCACACTACCCCCG</td></tr>
<tr><td>Chromosome</td><td>start</td><td></td><td>end</td><td>strand</td><td>MM</td><td>target_seq</td><td>PAM</td><td>alignment</td><td>distance</td><td>position</td><td>gene name</td><td>gene id</td></tr>
<tr><td>GU305914.1</td><td></td><td>223971</td><td>223993</td><td>-</td><td>0</td><td>CGGGGGTAGTGTGGGAGGGG</td><td>TGG</td><td>||||||||[||||||||||||]PAM</td><td>NA</td><td>Intergenic</td><td>NA</td><td>NA</td></tr>
<tr><td>T17742</td><td>GGTAGTGTGGGAGGGGTGGAAGG</td><td></td><td>1000</td></tr>
<tr><td>Oligo fwd</td><td>CACCGGTAGTGTGGGAGGGGTGGA</td></tr>
<tr><td>Oligo rev</td><td>AAACTCCACCCCTCCCACACTACC</td></tr>
<tr><td>Chromosome</td><td>start</td><td></td><td>end</td><td>strand</td><td>MM</td><td>target_seq</td><td>PAM</td><td>alignment</td><td>distance</td><td>position</td><td>gene name</td><td>gene id</td></tr>
<tr><td>GU305914.1</td><td></td><td>223967</td><td>223989</td><td>-</td><td>0</td><td>GGTAGTGTGGGAGGGGTGGA</td><td>AGG</td><td>||||||||[||||||||||||]PAM</td><td>NA</td><td>Intergenic</td><td>NA</td><td>NA</td></tr>
</table>

| | | | | | | | | | | | |
|---|---|---|---|---|---|---|---|---|---|---|---|
| T17743 | GTAGTGTGGGAGGGGTGGAAGGG | 1000 | | | | | | | | | |
| Oligo fwd | CACCGTAGTGTGGGAGGGGTGGAA | | | | | | | | | | |
| Oligo rev | AAACTTCCACCCCTCCCACACTAC | | | | | | | | | | |
| Chromosome | start | end | strand | MM | target_seq | PAM | alignment | distance | position | gene name | gene id |
| GU305914.1 | 223966 | 223988 | - | 0 | GTAGTGTGGGAGGGGTGGAA | GGG | \|\|\|\|\|\|\|\|[\|\|\|\|\|\|\|\|\|\|\|\|]PAM | NA | Intergenic | NA | NA |
| T17744 | TGGAAGGGATATATACAGTGAGG | 1000 | | | | | | | | | |
| Oligo fwd | CACCTGGAAGGGATATATACAGTG | | | | | | | | | | |
| Oligo rev | AAACCACTGTATATATCCCTTCCA | | | | | | | | | | |
| Chromosome | start | end | strand | MM | target_seq | PAM | alignment | distance | position | gene name | gene id |
| GU305914.1 | 223951 | 223973 | - | 0 | TGGAAGGGATATATACAGTG | AGG | \|\|\|\|\|\|\|\|[\|\|\|\|\|\|\|\|\|\|\|\|]PAM | NA | Intergenic | NA | NA |
| T17745 | AAGGGATATATACAGTGAGGTGG | 1000 | | | | | | | | | |
| Oligo fwd | CACCAAGGGATATATACAGTGAGG | | | | | | | | | | |
| Oligo rev | AAACCCTCACTGTATATATCCCTT | | | | | | | | | | |
| Chromosome | start | end | strand | MM | target_seq | PAM | alignment | distance | position | gene name | gene id |
| GU305914.1 | 223948 | 223970 | - | 0 | AAGGGATATATACAGTGAGG | TGG | \|\|\|\|\|\|\|\|[\|\|\|\|\|\|\|\|\|\|\|\|]PAM | NA | Intergenic | NA | NA |
| T17746 | GGATATATACAGTGAGGTGGTGG | 1000 | | | | | | | | | |
| Oligo fwd | CACCGGATATATACAGTGAGGTGG | | | | | | | | | | |
| Oligo rev | AAACCCACCTCACTGTATATATCC | | | | | | | | | | |
| Chromosome | start | end | strand | MM | target_seq | PAM | alignment | distance | position | gene name | gene id |
| GU305914.1 | 223945 | 223967 | - | 0 | GGATATATACAGTGAGGTGG | TGG | \|\|\|\|\|\|\|\|[\|\|\|\|\|\|\|\|\|\|\|\|]PAM | NA | Intergenic | NA | NA |
| T17747 | GTGAGGTGGTGGATAGATACAGG | 1000 | | | | | | | | | |
| Oligo fwd | CACCGTGAGGTGGTGGATAGATAC | | | | | | | | | | |
| Oligo rev | AAACGTATCTATCCACCACCTCAC | | | | | | | | | | |
| Chromosome | start | end | strand | MM | target_seq | PAM | alignment | distance | position | gene name | gene id |
| GU305914.1 | 223934 | 223956 | - | 0 | GTGAGGTGGTGGATAGATAC | AGG | \|\|\|\|\|\|\|\|[\|\|\|\|\|\|\|\|\|\|\|\|]PAM | NA | Intergenic | NA | NA |
| T17748 | TGGATAGATACAGGTTTAGTCGG | 1000 | | | | | | | | | |
| Oligo fwd | CACCTGGATAGATACAGGTTTAGT | | | | | | | | | | |
| Oligo rev | AAACACTAAACCTGTATCTATCCA | | | | | | | | | | |
| Chromosome | start | end | strand | MM | target_seq | PAM | alignment | distance | position | gene name | gene id |
| GU305914.1 | 223925 | 223947 | - | 0 | TGGATAGATACAGGTTTAGT | CGG | \|\|\|\|\|\|\|\|[\|\|\|\|\|\|\|\|\|\|\|\|]PAM | NA | Intergenic | NA | NA |
| T17749 | GGATAGATACAGGTTTAGTCGGG | 1000 | | | | | | | | | |
| Oligo fwd | CACCGGATAGATACAGGTTTAGTC | | | | | | | | | | |
| Oligo rev | AAACGACTAAACCTGTATCTATCC | | | | | | | | | | |
| Chromosome | start | end | strand | MM | target_seq | PAM | alignment | distance | position | gene name | gene id |
| GU305914.1 | 223924 | 223946 | - | 0 | GGATAGATACAGGTTTAGTC | GGG | \|\|\|\|\|\|\|\|[\|\|\|\|\|\|\|\|\|\|\|\|]PAM | NA | Intergenic | NA | NA |
| T17750 | GAGCGTGCAGCGTGTGCGAGAGG | 1000 | | | | | | | | | |
| Oligo fwd | CACCGAGCGTGCAGCGTGTGCGAG | | | | | | | | | | |
| Oligo rev | AAACCTCGCACACGCTGCACGCTC | | | | | | | | | | |
| Chromosome | start | end | strand | MM | target_seq | PAM | alignment | distance | position | gene name | gene id |
| GU305914.1 | 223902 | 223924 | - | 0 | GAGCGTGCAGCGTGTGCGAG | AGG | \|\|\|\|\|\|\|\|[\|\|\|\|\|\|\|\|\|\|\|\|]PAM | NA | Intergenic | NA | NA |
| T17751 | AGCGTGCAGCGTGTGCGAGAGGG | 1000 | | | | | | | | | |
| Oligo fwd | CACCAGCGTGCAGCGTGTGCGAGA | | | | | | | | | | |
| Oligo rev | AAACTCTCGCACACGCTGCACGCT | | | | | | | | | | |
| Chromosome | start | end | strand | MM | target_seq | PAM | alignment | distance | position | gene name | gene id |
| GU305914.1 | 223901 | 223923 | - | 0 | AGCGTGCAGCGTGTGCGAGA | GGG | \|\|\|\|\|\|\|\|[\|\|\|\|\|\|\|\|\|\|\|\|]PAM | NA | Intergenic | NA | NA |
| T17752 | TCGCGTTGTCGCGAACCGCACGG | 1000 | | | | | | | | | |
| Oligo fwd | CACCTCGCGTTGTCGCGAACCGCA | | | | | | | | | | |
| Oligo rev | AAACTGCGGTTCGCGACAACGCGA | | | | | | | | | | |
| Chromosome | start | end | strand | MM | target_seq | PAM | alignment | distance | position | gene name | gene id |
| GU305914.1 | 223872 | 223894 | - | 0 | TCGCGTTGTCGCGAACCGCA | CGG | \|\|\|\|\|\|\|\|[\|\|\|\|\|\|\|\|\|\|\|\|]PAM | NA | Intergenic | NA | NA |
| T17753 | GTTGTCGCGAACCGCACGGTCGG | 1000 | | | | | | | | | |
| Oligo fwd | CACCGTTGTCGCGAACCGCACGGT | | | | | | | | | | |
| Oligo rev | AAACACCGTGCGGTTCGCGACAAC | | | | | | | | | | |
| Chromosome | start | end | strand | MM | target_seq | PAM | alignment | distance | position | gene name | gene id |
| GU305914.1 | 223868 | 223890 | - | 0 | GTTGTCGCGAACCGCACGGT | CGG | \|\|\|\|\|\|\|\|[\|\|\|\|\|\|\|\|\|\|\|\|]PAM | NA | Intergenic | NA | NA |
| T17754 | TTGTCGCGAACCGCACGGTCGGG | 1000 | | | | | | | | | |
| Oligo fwd | CACCTTGTCGCGAACCGCACGGTC | | | | | | | | | | |
| Oligo rev | AAACGACCGTGCGGTTCGCGACAA | | | | | | | | | | |
| Chromosome | start | end | strand | MM | target_seq | PAM | alignment | distance | position | gene name | gene id |
| GU305914.1 | 223867 | 223889 | - | 0 | TTGTCGCGAACCGCACGGTC | GGG | \|\|\|\|\|\|\|\|[\|\|\|\|\|\|\|\|\|\|\|\|]PAM | NA | Intergenic | NA | NA |
| T17755 | CGAACCGCACGGTCGGGTCGCGG | 1000 | | | | | | | | | |
| Oligo fwd | CACCCGAACCGCACGGTCGGGTCG | | | | | | | | | | |
| Oligo rev | AAACCGACCCGACCGTGCGGTTCG | | | | | | | | | | |
| Chromosome | start | end | strand | MM | target_seq | PAM | alignment | distance | position | gene name | gene id |
| GU305914.1 | 223861 | 223883 | - | 0 | CGAACCGCACGGTCGGGTCG | CGG | \|\|\|\|\|\|\|\|[\|\|\|\|\|\|\|\|\|\|\|\|]PAM | NA | Intergenic | NA | NA |
| T17756 | CCGCACGGTCGGGTCGCGGCCGG | 1000 | | | | | | | | | |
| Oligo fwd | CACCCCGCACGGTCGGGTCGCGGC | | | | | | | | | | |
| Oligo rev | AAACGCCGCGACCCGACCGTGCGG | | | | | | | | | | |
| Chromosome | start | end | strand | MM | target_seq | PAM | alignment | distance | position | gene name | gene id |
| GU305914.1 | 223857 | 223879 | - | 0 | CCGCACGGTCGGGTCGCGGC | CGG | \|\|\|\|\|\|\|\|[\|\|\|\|\|\|\|\|\|\|\|\|]PAM | NA | Intergenic | NA | NA |
| T17757 | CCGGCACTAGATCCCCTGACCGG | 1000 | | | | | | | | | |
| Oligo fwd | CACCCCGGCACTAGATCCCCTGAC | | | | | | | | | | |
| Oligo rev | AAACGTCAGGGGATCTAGTGCCGG | | | | | | | | | | |
| Chromosome | start | end | strand | MM | target_seq | PAM | alignment | distance | position | gene name | gene id |

<table>
<tr><td>GU305914.1</td><td>223838</td><td>223860</td><td>-</td><td>0</td><td>CCGGCACTAGATCCCCTGAC</td><td>CGG</td><td>||||||||[||||||||||||]PAM</td><td>NA</td><td>Intergenic</td><td>NA</td><td>NA</td></tr>
<tr><td>T17758</td><td>TCCCCTGACCGGCGATCTGCTGG</td><td>1000</td><td></td><td></td><td></td><td></td><td></td><td></td><td></td><td></td><td></td></tr>
<tr><td>Oligo fwd</td><td>CACCTCCCCTGACCGGCGATCTGC</td><td></td><td></td><td></td><td></td><td></td><td></td><td></td><td></td><td></td><td></td></tr>
<tr><td>Oligo rev</td><td>AAACGCAGATCGCCGGTCAGGGGA</td><td></td><td></td><td></td><td></td><td></td><td></td><td></td><td></td><td></td><td></td></tr>
<tr><td>Chromosome</td><td>start</td><td>end</td><td>strand</td><td>MM</td><td>target_seq</td><td>PAM</td><td>alignment</td><td>distance</td><td>position</td><td>gene name</td><td>gene id</td></tr>
<tr><td>GU305914.1</td><td>223827</td><td>223849</td><td>-</td><td>0</td><td>TCCCCTGACCGGCGATCTGC</td><td>TGG</td><td>||||||||[||||||||||||]PAM</td><td>NA</td><td>Intergenic</td><td>NA</td><td>NA</td></tr>
<tr><td>T17759</td><td>GCGATCTGCTGGTCCCGAGACGG</td><td>1000</td><td></td><td></td><td></td><td></td><td></td><td></td><td></td><td></td><td></td></tr>
<tr><td>Oligo fwd</td><td>CACCGCGATCTGCTGGTCCCGAGA</td><td></td><td></td><td></td><td></td><td></td><td></td><td></td><td></td><td></td><td></td></tr>
<tr><td>Oligo rev</td><td>AAACTCTCGGGACCAGCAGATCGC</td><td></td><td></td><td></td><td></td><td></td><td></td><td></td><td></td><td></td><td></td></tr>
<tr><td>Chromosome</td><td>start</td><td>end</td><td>strand</td><td>MM</td><td>target_seq</td><td>PAM</td><td>alignment</td><td>distance</td><td>position</td><td>gene name</td><td>gene id</td></tr>
<tr><td>GU305914.1</td><td>223816</td><td>223838</td><td>-</td><td>0</td><td>GCGATCTGCTGGTCCCGAGA</td><td>CGG</td><td>||||||||[||||||||||||]PAM</td><td>NA</td><td>Intergenic</td><td>NA</td><td>NA</td></tr>
<tr><td>T17760</td><td>TGCTGGTCCCGAGACGGTCTCGG</td><td>1000</td><td></td><td></td><td></td><td></td><td></td><td></td><td></td><td></td><td></td></tr>
<tr><td>Oligo fwd</td><td>CACCTGCTGGTCCCGAGACGGTCT</td><td></td><td></td><td></td><td></td><td></td><td></td><td></td><td></td><td></td><td></td></tr>
<tr><td>Oligo rev</td><td>AAACAGACCGTCTCGGGACCAGCA</td><td></td><td></td><td></td><td></td><td></td><td></td><td></td><td></td><td></td><td></td></tr>
<tr><td>Chromosome</td><td>start</td><td>end</td><td>strand</td><td>MM</td><td>target_seq</td><td>PAM</td><td>alignment</td><td>distance</td><td>position</td><td>gene name</td><td>gene id</td></tr>
<tr><td>GU305914.1</td><td>223810</td><td>223832</td><td>-</td><td>0</td><td>TGCTGGTCCCGAGACGGTCT</td><td>CGG</td><td>||||||||[||||||||||||]PAM</td><td>NA</td><td>Intergenic</td><td>NA</td><td>NA</td></tr>
<tr><td>GU305914.1</td><td>223802</td><td>223824</td><td>+</td><td>4</td><td>CACCGGTCCCGAGACCGTCT</td><td>CGG</td><td>--|-||||[|||||||-||||]PAM</td><td>NA</td><td>Intergenic</td><td>NA</td><td>NA</td></tr>
<tr><td>T17761</td><td>GCTGGTCCCGAGACGGTCTCGGG</td><td>1000</td><td></td><td></td><td></td><td></td><td></td><td></td><td></td><td></td><td></td></tr>
<tr><td>Oligo fwd</td><td>CACCGCTGGTCCCGAGACGGTCTC</td><td></td><td></td><td></td><td></td><td></td><td></td><td></td><td></td><td></td><td></td></tr>
<tr><td>Oligo rev</td><td>AAACGAGACCGTCTCGGGACCAGC</td><td></td><td></td><td></td><td></td><td></td><td></td><td></td><td></td><td></td><td></td></tr>
<tr><td>Chromosome</td><td>start</td><td>end</td><td>strand</td><td>MM</td><td>target_seq</td><td>PAM</td><td>alignment</td><td>distance</td><td>position</td><td>gene name</td><td>gene id</td></tr>
<tr><td>GU305914.1</td><td>223809</td><td>223831</td><td>-</td><td>0</td><td>GCTGGTCCCGAGACGGTCTC</td><td>GGG</td><td>||||||||[||||||||||||]PAM</td><td>NA</td><td>Intergenic</td><td>NA</td><td>NA</td></tr>
<tr><td>GU305914.1</td><td>223803</td><td>223825</td><td>+</td><td>3</td><td>ACCGGTCCCGAGACCGTCTC</td><td>GGG</td><td>-|-|||||[||||||-|||||]PAM</td><td>NA</td><td>Intergenic</td><td>NA</td><td>NA</td></tr>
<tr><td>T17762</td><td>TCCCGAGACGGTCTCGGGACCGG</td><td>1000</td><td></td><td></td><td></td><td></td><td></td><td></td><td></td><td></td><td></td></tr>
<tr><td>Oligo fwd</td><td>CACCTCCCGAGACGGTCTCGGGAC</td><td></td><td></td><td></td><td></td><td></td><td></td><td></td><td></td><td></td><td></td></tr>
<tr><td>Oligo rev</td><td>AAACGTCCCGAGACCGTCTCGGGA</td><td></td><td></td><td></td><td></td><td></td><td></td><td></td><td></td><td></td><td></td></tr>
<tr><td>Chromosome</td><td>start</td><td>end</td><td>strand</td><td>MM</td><td>target_seq</td><td>PAM</td><td>alignment</td><td>distance</td><td>position</td><td>gene name</td><td>gene id</td></tr>
<tr><td>GU305914.1</td><td>223804</td><td>223826</td><td>-</td><td>0</td><td>TCCCGAGACGGTCTCGGGAC</td><td>CGG</td><td>||||||||[||||||||||||]PAM</td><td>NA</td><td>Intergenic</td><td>NA</td><td>NA</td></tr>
<tr><td>T17763</td><td>GTGCTTCTCCTGCCAGATCTTGG</td><td>1000</td><td></td><td></td><td></td><td></td><td></td><td></td><td></td><td></td><td></td></tr>
<tr><td>Oligo fwd</td><td>CACCGTGCTTCTCCTGCCAGATCT</td><td></td><td></td><td></td><td></td><td></td><td></td><td></td><td></td><td></td><td></td></tr>
<tr><td>Oligo rev</td><td>AAACAGATCTGGCAGGAGAAGCAC</td><td></td><td></td><td></td><td></td><td></td><td></td><td></td><td></td><td></td><td></td></tr>
<tr><td>Chromosome</td><td>start</td><td>end</td><td>strand</td><td>MM</td><td>target_seq</td><td>PAM</td><td>alignment</td><td>distance</td><td>position</td><td>gene name</td><td>gene id</td></tr>
<tr><td>GU305914.1</td><td>223782</td><td>223804</td><td>-</td><td>0</td><td>GTGCTTCTCCTGCCAGATCT</td><td>TGG</td><td>||||||||[||||||||||||]PAM</td><td>NA</td><td>Intergenic</td><td>NA</td><td>NA</td></tr>
<tr><td>T17764</td><td>TGCTTCTCCTGCCAGATCTTGGG</td><td>1000</td><td></td><td></td><td></td><td></td><td></td><td></td><td></td><td></td><td></td></tr>
<tr><td>Oligo fwd</td><td>CACCTGCTTCTCCTGCCAGATCTT</td><td></td><td></td><td></td><td></td><td></td><td></td><td></td><td></td><td></td><td></td></tr>
<tr><td>Oligo rev</td><td>AAACAAGATCTGGCAGGAGAAGCA</td><td></td><td></td><td></td><td></td><td></td><td></td><td></td><td></td><td></td><td></td></tr>
<tr><td>Chromosome</td><td>start</td><td>end</td><td>strand</td><td>MM</td><td>target_seq</td><td>PAM</td><td>alignment</td><td>distance</td><td>position</td><td>gene name</td><td>gene id</td></tr>
<tr><td>GU305914.1</td><td>223781</td><td>223803</td><td>-</td><td>0</td><td>TGCTTCTCCTGCCAGATCTT</td><td>GGG</td><td>||||||||[||||||||||||]PAM</td><td>NA</td><td>Intergenic</td><td>NA</td><td>NA</td></tr>
<tr><td>T17765</td><td>GCTTCTCCTGCCAGATCTTGGGG</td><td>1000</td><td></td><td></td><td></td><td></td><td></td><td></td><td></td><td></td><td></td></tr>
<tr><td>Oligo fwd</td><td>CACCGCTTCTCCTGCCAGATCTTG</td><td></td><td></td><td></td><td></td><td></td><td></td><td></td><td></td><td></td><td></td></tr>
<tr><td>Oligo rev</td><td>AAACCAAGATCTGGCAGGAGAAGC</td><td></td><td></td><td></td><td></td><td></td><td></td><td></td><td></td><td></td><td></td></tr>
<tr><td>Chromosome</td><td>start</td><td>end</td><td>strand</td><td>MM</td><td>target_seq</td><td>PAM</td><td>alignment</td><td>distance</td><td>position</td><td>gene name</td><td>gene id</td></tr>
<tr><td>GU305914.1</td><td>223780</td><td>223802</td><td>-</td><td>0</td><td>GCTTCTCCTGCCAGATCTTG</td><td>GGG</td><td>||||||||[||||||||||||]PAM</td><td>NA</td><td>Intergenic</td><td>NA</td><td>NA</td></tr>
<tr><td>T17766</td><td>CCTGCCAGATCTTGGGGCCTCGG</td><td>1000</td><td></td><td></td><td></td><td></td><td></td><td></td><td></td><td></td><td></td></tr>
<tr><td>Oligo fwd</td><td>CACCCCTGCCAGATCTTGGGGCCT</td><td></td><td></td><td></td><td></td><td></td><td></td><td></td><td></td><td></td><td></td></tr>
<tr><td>Oligo rev</td><td>AAACAGGCCCCAAGATCTGGCAGG</td><td></td><td></td><td></td><td></td><td></td><td></td><td></td><td></td><td></td><td></td></tr>
<tr><td>Chromosome</td><td>start</td><td>end</td><td>strand</td><td>MM</td><td>target_seq</td><td>PAM</td><td>alignment</td><td>distance</td><td>position</td><td>gene name</td><td>gene id</td></tr>
<tr><td>GU305914.1</td><td>223774</td><td>223796</td><td>-</td><td>0</td><td>CCTGCCAGATCTTGGGGCCT</td><td>CGG</td><td>||||||||[||||||||||||]PAM</td><td>NA</td><td>Intergenic</td><td>NA</td><td>NA</td></tr>
<tr><td>T17767</td><td>CTGCCAGATCTTGGGGCCTCGGG</td><td>1000</td><td></td><td></td><td></td><td></td><td></td><td></td><td></td><td></td><td></td></tr>
<tr><td>Oligo fwd</td><td>CACCCTGCCAGATCTTGGGGCCTC</td><td></td><td></td><td></td><td></td><td></td><td></td><td></td><td></td><td></td><td></td></tr>
<tr><td>Oligo rev</td><td>AAACGAGGCCCCAAGATCTGGCAG</td><td></td><td></td><td></td><td></td><td></td><td></td><td></td><td></td><td></td><td></td></tr>
<tr><td>Chromosome</td><td>start</td><td>end</td><td>strand</td><td>MM</td><td>target_seq</td><td>PAM</td><td>alignment</td><td>distance</td><td>position</td><td>gene name</td><td>gene id</td></tr>
<tr><td>GU305914.1</td><td>223773</td><td>223795</td><td>-</td><td>0</td><td>CTGCCAGATCTTGGGGCCTC</td><td>GGG</td><td>||||||||[||||||||||||]PAM</td><td>NA</td><td>Intergenic</td><td>NA</td><td>NA</td></tr>
<tr><td>T17768</td><td>GCCTCGGGTCGCCACACACGTGG</td><td>1000</td><td></td><td></td><td></td><td></td><td></td><td></td><td></td><td></td><td></td></tr>
<tr><td>Oligo fwd</td><td>CACCGCCTCGGGTCGCCACACACG</td><td></td><td></td><td></td><td></td><td></td><td></td><td></td><td></td><td></td><td></td></tr>
<tr><td>Oligo rev</td><td>AAACCGTGTGTGGCGACCCGAGGC</td><td></td><td></td><td></td><td></td><td></td><td></td><td></td><td></td><td></td><td></td></tr>
<tr><td>Chromosome</td><td>start</td><td>end</td><td>strand</td><td>MM</td><td>target_seq</td><td>PAM</td><td>alignment</td><td>distance</td><td>position</td><td>gene name</td><td>gene id</td></tr>
<tr><td>GU305914.1</td><td>223758</td><td>223780</td><td>-</td><td>0</td><td>GCCTCGGGTCGCCACACACG</td><td>TGG</td><td>||||||||[||||||||||||]PAM</td><td>NA</td><td>Intergenic</td><td>NA</td><td>NA</td></tr>
<tr><td>T17769</td><td>CCTCGGGTCGCCACACACGTGGG</td><td>1000</td><td></td><td></td><td></td><td></td><td></td><td></td><td></td><td></td><td></td></tr>
<tr><td>Oligo fwd</td><td>CACCCCTCGGGTCGCCACACACGT</td><td></td><td></td><td></td><td></td><td></td><td></td><td></td><td></td><td></td><td></td></tr>
<tr><td>Oligo rev</td><td>AAACACGTGTGTGGCGACCCGAGG</td><td></td><td></td><td></td><td></td><td></td><td></td><td></td><td></td><td></td><td></td></tr>
<tr><td>Chromosome</td><td>start</td><td>end</td><td>strand</td><td>MM</td><td>target_seq</td><td>PAM</td><td>alignment</td><td>distance</td><td>position</td><td>gene name</td><td>gene id</td></tr>
<tr><td>GU305914.1</td><td>223757</td><td>223779</td><td>-</td><td>0</td><td>CCTCGGGTCGCCACACACGT</td><td>GGG</td><td>||||||||[||||||||||||]PAM</td><td>NA</td><td>Intergenic</td><td>NA</td><td>NA</td></tr>
<tr><td>T17770</td><td>GGGTCGCCACACACGTGGGAAGG</td><td>1000</td><td></td><td></td><td></td><td></td><td></td><td></td><td></td><td></td><td></td></tr>
<tr><td>Oligo fwd</td><td>CACCGGGTCGCCACACACGTGGGA</td><td></td><td></td><td></td><td></td><td></td><td></td><td></td><td></td><td></td><td></td></tr>
<tr><td>Oligo rev</td><td>AAACTCCCACGTGTGTGGCGACCC</td><td></td><td></td><td></td><td></td><td></td><td></td><td></td><td></td><td></td><td></td></tr>
<tr><td>Chromosome</td><td>start</td><td>end</td><td>strand</td><td>MM</td><td>target_seq</td><td>PAM</td><td>alignment</td><td>distance</td><td>position</td><td>gene name</td><td>gene id</td></tr>
<tr><td>GU305914.1</td><td>223753</td><td>223775</td><td>-</td><td>0</td><td>GGGTCGCCACACACGTGGGA</td><td>AGG</td><td>||||||||[||||||||||||]PAM</td><td>NA</td><td>Intergenic</td><td>NA</td><td>NA</td></tr>
<tr><td>T17771</td><td>GGGAAGGCGATGTTTCTCCGCGG</td><td>1000</td><td></td><td></td><td></td><td></td><td></td><td></td><td></td><td></td><td></td></tr>
<tr><td>Oligo fwd</td><td>CACCGGGAAGGCGATGTTTCTCCG</td><td></td><td></td><td></td><td></td><td></td><td></td><td></td><td></td><td></td><td></td></tr>
<tr><td>Oligo rev</td><td>AAACCGGAGAAACATCGCCTTCCC</td><td></td><td></td><td></td><td></td><td></td><td></td><td></td><td></td><td></td><td></td></tr>
<tr><td>Chromosome</td><td>start</td><td>end</td><td>strand</td><td>MM</td><td>target_seq</td><td>PAM</td><td>alignment</td><td>distance</td><td>position</td><td>gene name</td><td>gene id</td></tr>
<tr><td>GU305914.1</td><td>223737</td><td>223759</td><td>-</td><td>0</td><td>GGGAAGGCGATGTTTCTCCG</td><td>CGG</td><td>||||||||[||||||||||||]PAM</td><td>NA</td><td>Intergenic</td><td>NA</td><td>NA</td></tr>
<tr><td>T17772</td><td>CGATGTTTCTCCGCGGCTGTCGG</td><td>1000</td><td></td><td></td><td></td><td></td><td></td><td></td><td></td><td></td><td></td></tr>
</table>

<table>
<tr><td>Oligo fwd</td><td>CACCCGATGTTTCTCCGCGGCTGT</td><td></td><td></td><td></td><td></td><td></td><td></td><td></td><td></td><td></td><td></td></tr>
<tr><td>Oligo rev</td><td>AAACACAGCCGCGGAGAAACATCG</td><td></td><td></td><td></td><td></td><td></td><td></td><td></td><td></td><td></td><td></td></tr>
<tr><td>Chromosome</td><td>start</td><td>end</td><td>strand</td><td>MM</td><td>target_seq</td><td>PAM</td><td>alignment</td><td>distance</td><td>position</td><td>gene name</td><td>gene id</td></tr>
<tr><td>GU305914.1</td><td>223730</td><td>223752</td><td>-</td><td>0</td><td>CGATGTTTCTCCGCGGCTGT</td><td>CGG</td><td>||||||||[||||||||||||]PAM</td><td>NA</td><td>Intergenic</td><td>NA</td><td>NA</td></tr>
<tr><td>T17773</td><td>TGTTTCTCCGCGGCTGTCGGAGG</td><td>1000</td><td></td><td></td><td></td><td></td><td></td><td></td><td></td><td></td><td></td></tr>
<tr><td>Oligo fwd</td><td>CACCTGTTTCTCCGCGGCTGTCGG</td><td></td><td></td><td></td><td></td><td></td><td></td><td></td><td></td><td></td><td></td></tr>
<tr><td>Oligo rev</td><td>AAACCCGACAGCCGCGGAGAAACA</td><td></td><td></td><td></td><td></td><td></td><td></td><td></td><td></td><td></td><td></td></tr>
<tr><td>Chromosome</td><td>start</td><td>end</td><td>strand</td><td>MM</td><td>target_seq</td><td>PAM</td><td>alignment</td><td>distance</td><td>position</td><td>gene name</td><td>gene id</td></tr>
<tr><td>GU305914.1</td><td>223727</td><td>223749</td><td>-</td><td>0</td><td>TGTTTCTCCGCGGCTGTCGG</td><td>AGG</td><td>||||||||[||||||||||||]PAM</td><td>NA</td><td>Intergenic</td><td>NA</td><td>NA</td></tr>
<tr><td>T17774</td><td>GTTTCTCCGCGGCTGTCGGAGGG</td><td>1000</td><td></td><td></td><td></td><td></td><td></td><td></td><td></td><td></td><td></td></tr>
<tr><td>Oligo fwd</td><td>CACCGTTTCTCCGCGGCTGTCGGA</td><td></td><td></td><td></td><td></td><td></td><td></td><td></td><td></td><td></td><td></td></tr>
<tr><td>Oligo rev</td><td>AAACTCCGACAGCCGCGGAGAAAC</td><td></td><td></td><td></td><td></td><td></td><td></td><td></td><td></td><td></td><td></td></tr>
<tr><td>Chromosome</td><td>start</td><td>end</td><td>strand</td><td>MM</td><td>target_seq</td><td>PAM</td><td>alignment</td><td>distance</td><td>position</td><td>gene name</td><td>gene id</td></tr>
<tr><td>GU305914.1</td><td>223726</td><td>223748</td><td>-</td><td>0</td><td>GTTTCTCCGCGGCTGTCGGA</td><td>GGG</td><td>||||||||[||||||||||||]PAM</td><td>NA</td><td>Intergenic</td><td>NA</td><td>NA</td></tr>
<tr><td>T17775</td><td>TTTCTCCGCGGCTGTCGGAGGGG</td><td>1000</td><td></td><td></td><td></td><td></td><td></td><td></td><td></td><td></td><td></td></tr>
<tr><td>Oligo fwd</td><td>CACCTTTCTCCGCGGCTGTCGGAG</td><td></td><td></td><td></td><td></td><td></td><td></td><td></td><td></td><td></td><td></td></tr>
<tr><td>Oligo rev</td><td>AAACCTCCGACAGCCGCGGAGAAA</td><td></td><td></td><td></td><td></td><td></td><td></td><td></td><td></td><td></td><td></td></tr>
<tr><td>Chromosome</td><td>start</td><td>end</td><td>strand</td><td>MM</td><td>target_seq</td><td>PAM</td><td>alignment</td><td>distance</td><td>position</td><td>gene name</td><td>gene id</td></tr>
<tr><td>GU305914.1</td><td>223725</td><td>223747</td><td>-</td><td>0</td><td>TTTCTCCGCGGCTGTCGGAG</td><td>GGG</td><td>||||||||[||||||||||||]PAM</td><td>NA</td><td>Intergenic</td><td>NA</td><td>NA</td></tr>
<tr><td>T17776</td><td>TTCTCCGCGGCTGTCGGAGGGGG</td><td>1000</td><td></td><td></td><td></td><td></td><td></td><td></td><td></td><td></td><td></td></tr>
<tr><td>Oligo fwd</td><td>CACCTTCTCCGCGGCTGTCGGAGG</td><td></td><td></td><td></td><td></td><td></td><td></td><td></td><td></td><td></td><td></td></tr>
<tr><td>Oligo rev</td><td>AAACCCTCCGACAGCCGCGGAGAA</td><td></td><td></td><td></td><td></td><td></td><td></td><td></td><td></td><td></td><td></td></tr>
<tr><td>Chromosome</td><td>start</td><td>end</td><td>strand</td><td>MM</td><td>target_seq</td><td>PAM</td><td>alignment</td><td>distance</td><td>position</td><td>gene name</td><td>gene id</td></tr>
<tr><td>GU305914.1</td><td>223724</td><td>223746</td><td>-</td><td>0</td><td>TTCTCCGCGGCTGTCGGAGG</td><td>GGG</td><td>||||||||[||||||||||||]PAM</td><td>NA</td><td>Intergenic</td><td>NA</td><td>NA</td></tr>
<tr><td>T17777</td><td>ACGCGCAGCTACCGTCTCCCTGG</td><td>1000</td><td></td><td></td><td></td><td></td><td></td><td></td><td></td><td></td><td></td></tr>
<tr><td>Oligo fwd</td><td>CACCACGCGCAGCTACCGTCTCCC</td><td></td><td></td><td></td><td></td><td></td><td></td><td></td><td></td><td></td><td></td></tr>
<tr><td>Oligo rev</td><td>AAACGGGAGACGGTAGCTGCGCGT</td><td></td><td></td><td></td><td></td><td></td><td></td><td></td><td></td><td></td><td></td></tr>
<tr><td>Chromosome</td><td>start</td><td>end</td><td>strand</td><td>MM</td><td>target_seq</td><td>PAM</td><td>alignment</td><td>distance</td><td>position</td><td>gene name</td><td>gene id</td></tr>
<tr><td>GU305914.1</td><td>223699</td><td>223721</td><td>-</td><td>0</td><td>ACGCGCAGCTACCGTCTCCC</td><td>TGG</td><td>||||||||[||||||||||||]PAM</td><td>NA</td><td>Intergenic</td><td>NA</td><td>NA</td></tr>
<tr><td>T17778</td><td>GCAGCTACCGTCTCCCTGGCAGG</td><td>1000</td><td></td><td></td><td></td><td></td><td></td><td></td><td></td><td></td><td></td></tr>
<tr><td>Oligo fwd</td><td>CACCGCAGCTACCGTCTCCCTGGC</td><td></td><td></td><td></td><td></td><td></td><td></td><td></td><td></td><td></td><td></td></tr>
<tr><td>Oligo rev</td><td>AAACGCCAGGGAGACGGTAGCTGC</td><td></td><td></td><td></td><td></td><td></td><td></td><td></td><td></td><td></td><td></td></tr>
<tr><td>Chromosome</td><td>start</td><td>end</td><td>strand</td><td>MM</td><td>target_seq</td><td>PAM</td><td>alignment</td><td>distance</td><td>position</td><td>gene name</td><td>gene id</td></tr>
<tr><td>GU305914.1</td><td>223695</td><td>223717</td><td>-</td><td>0</td><td>GCAGCTACCGTCTCCCTGGC</td><td>AGG</td><td>||||||||[||||||||||||]PAM</td><td>NA</td><td>Intergenic</td><td>NA</td><td>NA</td></tr>
<tr><td>T17779</td><td>ACCGTCTCCCTGGCAGGATGCGG</td><td>1000</td><td></td><td></td><td></td><td></td><td></td><td></td><td></td><td></td><td></td></tr>
<tr><td>Oligo fwd</td><td>CACCACCGTCTCCCTGGCAGGATG</td><td></td><td></td><td></td><td></td><td></td><td></td><td></td><td></td><td></td><td></td></tr>
<tr><td>Oligo rev</td><td>AAACCATCCTGCCAGGGAGACGGT</td><td></td><td></td><td></td><td></td><td></td><td></td><td></td><td></td><td></td><td></td></tr>
<tr><td>Chromosome</td><td>start</td><td>end</td><td>strand</td><td>MM</td><td>target_seq</td><td>PAM</td><td>alignment</td><td>distance</td><td>position</td><td>gene name</td><td>gene id</td></tr>
<tr><td>GU305914.1</td><td>223689</td><td>223711</td><td>-</td><td>0</td><td>ACCGTCTCCCTGGCAGGATG</td><td>CGG</td><td>||||||||[||||||||||||]PAM</td><td>NA</td><td>Intergenic</td><td>NA</td><td>NA</td></tr>
<tr><td>T17780</td><td>CCGTCTCCCTGGCAGGATGCGGG</td><td>1000</td><td></td><td></td><td></td><td></td><td></td><td></td><td></td><td></td><td></td></tr>
<tr><td>Oligo fwd</td><td>CACCCCGTCTCCCTGGCAGGATGC</td><td></td><td></td><td></td><td></td><td></td><td></td><td></td><td></td><td></td><td></td></tr>
<tr><td>Oligo rev</td><td>AAACGCATCCTGCCAGGGAGACGG</td><td></td><td></td><td></td><td></td><td></td><td></td><td></td><td></td><td></td><td></td></tr>
<tr><td>Chromosome</td><td>start</td><td>end</td><td>strand</td><td>MM</td><td>target_seq</td><td>PAM</td><td>alignment</td><td>distance</td><td>position</td><td>gene name</td><td>gene id</td></tr>
<tr><td>GU305914.1</td><td>223688</td><td>223710</td><td>-</td><td>0</td><td>CCGTCTCCCTGGCAGGATGC</td><td>GGG</td><td>||||||||[||||||||||||]PAM</td><td>NA</td><td>Intergenic</td><td>NA</td><td>NA</td></tr>
<tr><td>T17781</td><td>CGTCTCCCTGGCAGGATGCGGGG</td><td>1000</td><td></td><td></td><td></td><td></td><td></td><td></td><td></td><td></td><td></td></tr>
<tr><td>Oligo fwd</td><td>CACCCGTCTCCCTGGCAGGATGCG</td><td></td><td></td><td></td><td></td><td></td><td></td><td></td><td></td><td></td><td></td></tr>
<tr><td>Oligo rev</td><td>AAACCGCATCCTGCCAGGGAGACG</td><td></td><td></td><td></td><td></td><td></td><td></td><td></td><td></td><td></td><td></td></tr>
<tr><td>Chromosome</td><td>start</td><td>end</td><td>strand</td><td>MM</td><td>target_seq</td><td>PAM</td><td>alignment</td><td>distance</td><td>position</td><td>gene name</td><td>gene id</td></tr>
<tr><td>GU305914.1</td><td>223687</td><td>223709</td><td>-</td><td>0</td><td>CGTCTCCCTGGCAGGATGCG</td><td>GGG</td><td>||||||||[||||||||||||]PAM</td><td>NA</td><td>Intergenic</td><td>NA</td><td>NA</td></tr>
<tr><td>T17782</td><td>TTGCGTGTAGCCATTTCGATCGG</td><td>1000</td><td></td><td></td><td></td><td></td><td></td><td></td><td></td><td></td><td></td></tr>
<tr><td>Oligo fwd</td><td>CACCTTGCGTGTAGCCATTTCGAT</td><td></td><td></td><td></td><td></td><td></td><td></td><td></td><td></td><td></td><td></td></tr>
<tr><td>Oligo rev</td><td>AAACATCGAAATGGCTACACGCAA</td><td></td><td></td><td></td><td></td><td></td><td></td><td></td><td></td><td></td><td></td></tr>
<tr><td>Chromosome</td><td>start</td><td>end</td><td>strand</td><td>MM</td><td>target_seq</td><td>PAM</td><td>alignment</td><td>distance</td><td>position</td><td>gene name</td><td>gene id</td></tr>
<tr><td>GU305914.1</td><td>223634</td><td>223656</td><td>-</td><td>0</td><td>TTGCGTGTAGCCATTTCGAT</td><td>CGG</td><td>||||||||[||||||||||||]PAM</td><td>NA</td><td>Intergenic</td><td>NA</td><td>NA</td></tr>
<tr><td>T17783</td><td>TGCGTGTAGCCATTTCGATCGGG</td><td>1000</td><td></td><td></td><td></td><td></td><td></td><td></td><td></td><td></td><td></td></tr>
<tr><td>Oligo fwd</td><td>CACCTGCGTGTAGCCATTTCGATC</td><td></td><td></td><td></td><td></td><td></td><td></td><td></td><td></td><td></td><td></td></tr>
<tr><td>Oligo rev</td><td>AAACGATCGAAATGGCTACACGCA</td><td></td><td></td><td></td><td></td><td></td><td></td><td></td><td></td><td></td><td></td></tr>
<tr><td>Chromosome</td><td>start</td><td>end</td><td>strand</td><td>MM</td><td>target_seq</td><td>PAM</td><td>alignment</td><td>distance</td><td>position</td><td>gene name</td><td>gene id</td></tr>
<tr><td>GU305914.1</td><td>223633</td><td>223655</td><td>-</td><td>0</td><td>TGCGTGTAGCCATTTCGATC</td><td>GGG</td><td>||||||||[||||||||||||]PAM</td><td>NA</td><td>Intergenic</td><td>NA</td><td>NA</td></tr>
<tr><td>T17784</td><td>GCGTGTAGCCATTTCGATCGGGG</td><td>1000</td><td></td><td></td><td></td><td></td><td></td><td></td><td></td><td></td><td></td></tr>
<tr><td>Oligo fwd</td><td>CACCGCGTGTAGCCATTTCGATCG</td><td></td><td></td><td></td><td></td><td></td><td></td><td></td><td></td><td></td><td></td></tr>
<tr><td>Oligo rev</td><td>AAACCGATCGAAATGGCTACACGC</td><td></td><td></td><td></td><td></td><td></td><td></td><td></td><td></td><td></td><td></td></tr>
<tr><td>Chromosome</td><td>start</td><td>end</td><td>strand</td><td>MM</td><td>target_seq</td><td>PAM</td><td>alignment</td><td>distance</td><td>position</td><td>gene name</td><td>gene id</td></tr>
<tr><td>GU305914.1</td><td>223632</td><td>223654</td><td>-</td><td>0</td><td>GCGTGTAGCCATTTCGATCG</td><td>GGG</td><td>||||||||[||||||||||||]PAM</td><td>NA</td><td>Intergenic</td><td>NA</td><td>NA</td></tr>
<tr><td>T17785</td><td>CATTTCGATCGGGGTAGCCGCGG</td><td>1000</td><td></td><td></td><td></td><td></td><td></td><td></td><td></td><td></td><td></td></tr>
<tr><td>Oligo fwd</td><td>CACCCATTTCGATCGGGGTAGCCG</td><td></td><td></td><td></td><td></td><td></td><td></td><td></td><td></td><td></td><td></td></tr>
<tr><td>Oligo rev</td><td>AAACCGGCTACCCCGATCGAAATG</td><td></td><td></td><td></td><td></td><td></td><td></td><td></td><td></td><td></td><td></td></tr>
<tr><td>Chromosome</td><td>start</td><td>end</td><td>strand</td><td>MM</td><td>target_seq</td><td>PAM</td><td>alignment</td><td>distance</td><td>position</td><td>gene name</td><td>gene id</td></tr>
<tr><td>GU305914.1</td><td>223623</td><td>223645</td><td>-</td><td>0</td><td>CATTTCGATCGGGGTAGCCG</td><td>CGG</td><td>||||||||[||||||||||||]PAM</td><td>NA</td><td>Intergenic</td><td>NA</td><td>NA</td></tr>
<tr><td>T17786</td><td>TTCGATCGGGGTAGCCGCGGCGG</td><td>1000</td><td></td><td></td><td></td><td></td><td></td><td></td><td></td><td></td><td></td></tr>
<tr><td>Oligo fwd</td><td>CACCTTCGATCGGGGTAGCCGCGG</td><td></td><td></td><td></td><td></td><td></td><td></td><td></td><td></td><td></td><td></td></tr>
<tr><td>Oligo rev</td><td>AAACCCGCGGCTACCCCGATCGAA</td><td></td><td></td><td></td><td></td><td></td><td></td><td></td><td></td><td></td><td></td></tr>
<tr><td>Chromosome</td><td>start</td><td>end</td><td>strand</td><td>MM</td><td>target_seq</td><td>PAM</td><td>alignment</td><td>distance</td><td>position</td><td>gene name</td><td>gene id</td></tr>
<tr><td>GU305914.1</td><td>223620</td><td>223642</td><td>-</td><td>0</td><td>TTCGATCGGGGTAGCCGCGG</td><td>CGG</td><td>||||||||[||||||||||||]PAM</td><td>NA</td><td>Intergenic</td><td>NA</td><td>NA</td></tr>
</table>

| | | | | | | | | | | | |
|---|---|---|---|---|---|---|---|---|---|---|---|
| T17787 | GGGGTAGCCGCGGCGGCACTCGG | 1000 | | | | | | | | | |
| Oligo fwd | CACCGGGGTAGCCGCGGCGGCACT | | | | | | | | | | |
| Oligo rev | AAACAGTGCCGCCGCGGCTACCCC | | | | | | | | | | |
| Chromosome | start | end | strand | MM | target_seq | PAM | alignment | distance | position | gene name | gene id |
| GU305914.1 | 223613 | 223635 | - | 0 | GGGGTAGCCGCGGCGGCACT | CGG | \|\|\|\|\|\|\|\|[\|\|\|\|\|\|\|\|\|\|\|\|]PAM | NA | Intergenic | NA | NA |
| T17788 | GGGTAGCCGCGGCGGCACTCGGG | 1000 | | | | | | | | | |
| Oligo fwd | CACCGGGTAGCCGCGGCGGCACTC | | | | | | | | | | |
| Oligo rev | AAACGAGTGCCGCCGCGGCTACCC | | | | | | | | | | |
| Chromosome | start | end | strand | MM | target_seq | PAM | alignment | distance | position | gene name | gene id |
| GU305914.1 | 223612 | 223634 | - | 0 | GGGTAGCCGCGGCGGCACTC | GGG | \|\|\|\|\|\|\|\|[\|\|\|\|\|\|\|\|\|\|\|\|]PAM | NA | Intergenic | NA | NA |
| T17789 | AGCCGCGGCGGCACTCGGGTCGG | 1000 | | | | | | | | | |
| Oligo fwd | CACCAGCCGCGGCGGCACTCGGGT | | | | | | | | | | |
| Oligo rev | AAACACCCGAGTGCCGCCGCGGCT | | | | | | | | | | |
| Chromosome | start | end | strand | MM | target_seq | PAM | alignment | distance | position | gene name | gene id |
| GU305914.1 | 223608 | 223630 | - | 0 | AGCCGCGGCGGCACTCGGGT | CGG | \|\|\|\|\|\|\|\|[\|\|\|\|\|\|\|\|\|\|\|\|]PAM | NA | Intergenic | NA | NA |
| T17790 | GGCACTCGGGTCGGTCCTCGTGG | 1000 | | | | | | | | | |
| Oligo fwd | CACCGGCACTCGGGTCGGTCCTCG | | | | | | | | | | |
| Oligo rev | AAACCGAGGACCGACCCGAGTGCC | | | | | | | | | | |
| Chromosome | start | end | strand | MM | target_seq | PAM | alignment | distance | position | gene name | gene id |
| GU305914.1 | 223599 | 223621 | - | 0 | GGCACTCGGGTCGGTCCTCG | TGG | \|\|\|\|\|\|\|\|[\|\|\|\|\|\|\|\|\|\|\|\|]PAM | NA | Intergenic | NA | NA |
| T17791 | GACTCCCGTTCTGTATCCCCTGG | 1000 | | | | | | | | | |
| Oligo fwd | CACCGACTCCCGTTCTGTATCCCC | | | | | | | | | | |
| Oligo rev | AAACGGGGATACAGAACGGGAGTC | | | | | | | | | | |
| Chromosome | start | end | strand | MM | target_seq | PAM | alignment | distance | position | gene name | gene id |
| GU305914.1 | 223523 | 223545 | - | 0 | GACTCCCGTTCTGTATCCCC | TGG | \|\|\|\|\|\|\|\|[\|\|\|\|\|\|\|\|\|\|\|\|]PAM | NA | Intergenic | NA | NA |
| T17792 | GTTCTGTATCCCCTGGACCGAGG | 1000 | | | | | | | | | |
| Oligo fwd | CACCGTTCTGTATCCCCTGGACCG | | | | | | | | | | |
| Oligo rev | AAACCGGTCCAGGGGATACAGAAC | | | | | | | | | | |
| Chromosome | start | end | strand | MM | target_seq | PAM | alignment | distance | position | gene name | gene id |
| GU305914.1 | 223516 | 223538 | - | 0 | GTTCTGTATCCCCTGGACCG | AGG | \|\|\|\|\|\|\|\|[\|\|\|\|\|\|\|\|\|\|\|\|]PAM | NA | Intergenic | NA | NA |
| T17793 | CTGTATCCCCTGGACCGAGGAGG | 1000 | | | | | | | | | |
| Oligo fwd | CACCCTGTATCCCCTGGACCGAGG | | | | | | | | | | |
| Oligo rev | AAACCCTCGGTCCAGGGGATACAG | | | | | | | | | | |
| Chromosome | start | end | strand | MM | target_seq | PAM | alignment | distance | position | gene name | gene id |
| GU305914.1 | 223513 | 223535 | - | 0 | CTGTATCCCCTGGACCGAGG | AGG | \|\|\|\|\|\|\|\|[\|\|\|\|\|\|\|\|\|\|\|\|]PAM | NA | Intergenic | NA | NA |
| T17794 | TCGATCCCGACGACTCTTCAAGG | 1000 | | | | | | | | | |
| Oligo fwd | CACCTCGATCCCGACGACTCTTCA | | | | | | | | | | |
| Oligo rev | AAACTGAAGAGTCGTCGGGATCGA | | | | | | | | | | |
| Chromosome | start | end | strand | MM | target_seq | PAM | alignment | distance | position | gene name | gene id |
| GU305914.1 | 223486 | 223508 | - | 0 | TCGATCCCGACGACTCTTCA | AGG | \|\|\|\|\|\|\|\|[\|\|\|\|\|\|\|\|\|\|\|\|]PAM | NA | Intergenic | NA | NA |
| T17795 | TCCCGACGACTCTTCAAGGCCGG | 1000 | | | | | | | | | |
| Oligo fwd | CACCTCCCGACGACTCTTCAAGGC | | | | | | | | | | |
| Oligo rev | AAACGCCTTGAAGAGTCGTCGGGA | | | | | | | | | | |
| Chromosome | start | end | strand | MM | target_seq | PAM | alignment | distance | position | gene name | gene id |
| GU305914.1 | 223482 | 223504 | - | 0 | TCCCGACGACTCTTCAAGGC | CGG | \|\|\|\|\|\|\|\|[\|\|\|\|\|\|\|\|\|\|\|\|]PAM | NA | Intergenic | NA | NA |
| T17796 | CCCGACGACTCTTCAAGGCCGGG | 1000 | | | | | | | | | |
| Oligo fwd | CACCCCCGACGACTCTTCAAGGCC | | | | | | | | | | |
| Oligo rev | AAACGGCCTTGAAGAGTCGTCGGG | | | | | | | | | | |
| Chromosome | start | end | strand | MM | target_seq | PAM | alignment | distance | position | gene name | gene id |
| GU305914.1 | 223481 | 223503 | - | 0 | CCCGACGACTCTTCAAGGCC | GGG | \|\|\|\|\|\|\|\|[\|\|\|\|\|\|\|\|\|\|\|\|]PAM | NA | Intergenic | NA | NA |
| T17797 | GATGCTCGCCCTCATCTACGCGG | 1000 | | | | | | | | | |
| Oligo fwd | CACCGATGCTCGCCCTCATCTACG | | | | | | | | | | |
| Oligo rev | AAACCGTAGATGAGGGCGAGCATC | | | | | | | | | | |
| Chromosome | start | end | strand | MM | target_seq | PAM | alignment | distance | position | gene name | gene id |
| GU305914.1 | 223459 | 223481 | - | 0 | GATGCTCGCCCTCATCTACG | CGG | \|\|\|\|\|\|\|\|[\|\|\|\|\|\|\|\|\|\|\|\|]PAM | NA | Intergenic | NA | NA |
| T17798 | GCTCGCCCTCATCTACGCGGCGG | 1000 | | | | | | | | | |
| Oligo fwd | CACCGCTCGCCCTCATCTACGCGG | | | | | | | | | | |
| Oligo rev | AAACCCGCGTAGATGAGGGCGAGC | | | | | | | | | | |
| Chromosome | start | end | strand | MM | target_seq | PAM | alignment | distance | position | gene name | gene id |
| GU305914.1 | 223456 | 223478 | - | 0 | GCTCGCCCTCATCTACGCGG | CGG | \|\|\|\|\|\|\|\|[\|\|\|\|\|\|\|\|\|\|\|\|]PAM | NA | Intergenic | NA | NA |
| T17799 | CTACGCGGCGGCTTCAGCTGCGG | 1000 | | | | | | | | | |
| Oligo fwd | CACCCTACGCGGCGGCTTCAGCTG | | | | | | | | | | |
| Oligo rev | AAACCAGCTGAAGCCGCCGCGTAG | | | | | | | | | | |
| Chromosome | start | end | strand | MM | target_seq | PAM | alignment | distance | position | gene name | gene id |
| GU305914.1 | 223444 | 223466 | - | 0 | CTACGCGGCGGCTTCAGCTG | CGG | \|\|\|\|\|\|\|\|[\|\|\|\|\|\|\|\|\|\|\|\|]PAM | NA | Intergenic | NA | NA |
| T17800 | TACGCGGCGGCTTCAGCTGCGGG | 1000 | | | | | | | | | |
| Oligo fwd | CACCTACGCGGCGGCTTCAGCTGC | | | | | | | | | | |
| Oligo rev | AAACGCAGCTGAAGCCGCCGCGTA | | | | | | | | | | |
| Chromosome | start | end | strand | MM | target_seq | PAM | alignment | distance | position | gene name | gene id |
| GU305914.1 | 223443 | 223465 | - | 0 | TACGCGGCGGCTTCAGCTGC | GGG | \|\|\|\|\|\|\|\|[\|\|\|\|\|\|\|\|\|\|\|\|]PAM | NA | Intergenic | NA | NA |
| T17801 | GGCGGCTTCAGCTGCGGGCACGG | 1000 | | | | | | | | | |
| Oligo fwd | CACCGGCGGCTTCAGCTGCGGGCA | | | | | | | | | | |
| Oligo rev | AAACTGCCCGCAGCTGAAGCCGCC | | | | | | | | | | |
| Chromosome | start | end | strand | MM | target_seq | PAM | alignment | distance | position | gene name | gene id |
| GU305914.1 | 223438 | 223460 | - | 0 | GGCGGCTTCAGCTGCGGGCA | CGG | \|\|\|\|\|\|\|\|[\|\|\|\|\|\|\|\|\|\|\|\|]PAM | NA | Intergenic | NA | NA |

<table>
<tr><td>T17802</td><td>TTCAGCTGCGGGCACGGCCTCGG</td><td>1000</td></tr>
<tr><td>Oligo fwd</td><td>CACCTTCAGCTGCGGGCACGGCCT</td></tr>
<tr><td>Oligo rev</td><td>AAACAGGCCGTGCCCGCAGCTGAA</td></tr>
<tr><td>Chromosome</td><td>start</td><td>end</td><td>strand</td><td>MM</td><td>target_seq</td><td>PAM</td><td>alignment</td><td>distance</td><td>position</td><td>gene name</td><td>gene id</td></tr>
<tr><td>GU305914.1</td><td>223432</td><td>223454</td><td>-</td><td>0</td><td>TTCAGCTGCGGGCACGGCCT</td><td>CGG</td><td>||||||||[||||||||||||]PAM</td><td>NA</td><td>Intergenic</td><td>NA</td><td>NA</td></tr>
<tr><td>T17803</td><td>ACGGCCTCGGCGTCCGCCGTCGG</td><td>1000</td></tr>
<tr><td>Oligo fwd</td><td>CACCACGGCCTCGGCGTCCGCCGT</td></tr>
<tr><td>Oligo rev</td><td>AAACACGGCGGACGCCGAGGCCGT</td></tr>
<tr><td>Chromosome</td><td>start</td><td>end</td><td>strand</td><td>MM</td><td>target_seq</td><td>PAM</td><td>alignment</td><td>distance</td><td>position</td><td>gene name</td><td>gene id</td></tr>
<tr><td>GU305914.1</td><td>223419</td><td>223441</td><td>-</td><td>0</td><td>ACGGCCTCGGCGTCCGCCGT</td><td>CGG</td><td>||||||||[||||||||||||]PAM</td><td>NA</td><td>Intergenic</td><td>NA</td><td>NA</td></tr>
<tr><td>T17804</td><td>GCCGTCGGAAGCTGCGTCGACGG</td><td>1000</td></tr>
<tr><td>Oligo fwd</td><td>CACCGCCGTCGGAAGCTGCGTCGA</td></tr>
<tr><td>Oligo rev</td><td>AAACTCGACGCAGCTTCCGACGGC</td></tr>
<tr><td>Chromosome</td><td>start</td><td>end</td><td>strand</td><td>MM</td><td>target_seq</td><td>PAM</td><td>alignment</td><td>distance</td><td>position</td><td>gene name</td><td>gene id</td></tr>
<tr><td>GU305914.1</td><td>223404</td><td>223426</td><td>-</td><td>0</td><td>GCCGTCGGAAGCTGCGTCGA</td><td>CGG</td><td>||||||||[||||||||||||]PAM</td><td>NA</td><td>Intergenic</td><td>NA</td><td>NA</td></tr>
<tr><td>T17805</td><td>GAACAAGATCTATCCCGACCAGG</td><td>1000</td></tr>
<tr><td>Oligo fwd</td><td>CACCGAACAAGATCTATCCCGACC</td></tr>
<tr><td>Oligo rev</td><td>AAACGGTCGGGATAGATCTTGTTC</td></tr>
<tr><td>Chromosome</td><td>start</td><td>end</td><td>strand</td><td>MM</td><td>target_seq</td><td>PAM</td><td>alignment</td><td>distance</td><td>position</td><td>gene name</td><td>gene id</td></tr>
<tr><td>GU305914.1</td><td>223360</td><td>223382</td><td>-</td><td>0</td><td>GAACAAGATCTATCCCGACC</td><td>AGG</td><td>||||||||[||||||||||||]PAM</td><td>NA</td><td>Intergenic</td><td>NA</td><td>NA</td></tr>
<tr><td>T17806</td><td>GTGATCTGCCACGTGAAACCCGG</td><td>1000</td></tr>
<tr><td>Oligo fwd</td><td>CACCGTGATCTGCCACGTGAAACC</td></tr>
<tr><td>Oligo rev</td><td>AAACGGTTTCACGTGGCAGATCAC</td></tr>
<tr><td>Chromosome</td><td>start</td><td>end</td><td>strand</td><td>MM</td><td>target_seq</td><td>PAM</td><td>alignment</td><td>distance</td><td>position</td><td>gene name</td><td>gene id</td></tr>
<tr><td>GU305914.1</td><td>223326</td><td>223348</td><td>-</td><td>0</td><td>GTGATCTGCCACGTGAAACC</td><td>CGG</td><td>||||||||[||||||||||||]PAM</td><td>NA</td><td>Intergenic</td><td>NA</td><td>NA</td></tr>
<tr><td>T17807</td><td>CTGCCACGTGAAACCCGGTTCGG</td><td>1000</td></tr>
<tr><td>Oligo fwd</td><td>CACCCTGCCACGTGAAACCCGGTT</td></tr>
<tr><td>Oligo rev</td><td>AAACAACCGGGTTTCACGTGGCAG</td></tr>
<tr><td>Chromosome</td><td>start</td><td>end</td><td>strand</td><td>MM</td><td>target_seq</td><td>PAM</td><td>alignment</td><td>distance</td><td>position</td><td>gene name</td><td>gene id</td></tr>
<tr><td>GU305914.1</td><td>223321</td><td>223343</td><td>-</td><td>0</td><td>CTGCCACGTGAAACCCGGTT</td><td>CGG</td><td>||||||||[||||||||||||]PAM</td><td>NA</td><td>Intergenic</td><td>NA</td><td>NA</td></tr>
<tr><td>T17808</td><td>GGTTCGGTGAAGAAGTTTGTCGG</td><td>1000</td></tr>
<tr><td>Oligo fwd</td><td>CACCGGTTCGGTGAAGAAGTTTGT</td></tr>
<tr><td>Oligo rev</td><td>AAACACAAACTTCTTCACCGAACC</td></tr>
<tr><td>Chromosome</td><td>start</td><td>end</td><td>strand</td><td>MM</td><td>target_seq</td><td>PAM</td><td>alignment</td><td>distance</td><td>position</td><td>gene name</td><td>gene id</td></tr>
<tr><td>GU305914.1</td><td>223305</td><td>223327</td><td>-</td><td>0</td><td>GGTTCGGTGAAGAAGTTTGT</td><td>CGG</td><td>||||||||[||||||||||||]PAM</td><td>NA</td><td>Intergenic</td><td>NA</td><td>NA</td></tr>
<tr><td>T17809</td><td>GTTCGGTGAAGAAGTTTGTCGGG</td><td>1000</td></tr>
<tr><td>Oligo fwd</td><td>CACCGTTCGGTGAAGAAGTTTGTC</td></tr>
<tr><td>Oligo rev</td><td>AAACGACAAACTTCTTCACCGAAC</td></tr>
<tr><td>Chromosome</td><td>start</td><td>end</td><td>strand</td><td>MM</td><td>target_seq</td><td>PAM</td><td>alignment</td><td>distance</td><td>position</td><td>gene name</td><td>gene id</td></tr>
<tr><td>GU305914.1</td><td>223304</td><td>223326</td><td>-</td><td>0</td><td>GTTCGGTGAAGAAGTTTGTC</td><td>GGG</td><td>||||||||[||||||||||||]PAM</td><td>NA</td><td>Intergenic</td><td>NA</td><td>NA</td></tr>
<tr><td>T17810</td><td>TGAAGAAGTTTGTCGGGAGCTGG</td><td>1000</td></tr>
<tr><td>Oligo fwd</td><td>CACCTGAAGAAGTTTGTCGGGAGC</td></tr>
<tr><td>Oligo rev</td><td>AAACGCTCCCGACAAACTTCTTCA</td></tr>
<tr><td>Chromosome</td><td>start</td><td>end</td><td>strand</td><td>MM</td><td>target_seq</td><td>PAM</td><td>alignment</td><td>distance</td><td>position</td><td>gene name</td><td>gene id</td></tr>
<tr><td>GU305914.1</td><td>223298</td><td>223320</td><td>-</td><td>0</td><td>TGAAGAAGTTTGTCGGGAGC</td><td>TGG</td><td>||||||||[||||||||||||]PAM</td><td>NA</td><td>Intergenic</td><td>NA</td><td>NA</td></tr>
<tr><td>T17811</td><td>TGGTCGTTCGCGCTGCCCGAAGG</td><td>1000</td></tr>
<tr><td>Oligo fwd</td><td>CACCTGGTCGTTCGCGCTGCCCGA</td></tr>
<tr><td>Oligo rev</td><td>AAACTCGGGCAGCGCGAACGACCA</td></tr>
<tr><td>Chromosome</td><td>start</td><td>end</td><td>strand</td><td>MM</td><td>target_seq</td><td>PAM</td><td>alignment</td><td>distance</td><td>position</td><td>gene name</td><td>gene id</td></tr>
<tr><td>GU305914.1</td><td>223278</td><td>223300</td><td>-</td><td>0</td><td>TGGTCGTTCGCGCTGCCCGA</td><td>AGG</td><td>||||||||[||||||||||||]PAM</td><td>NA</td><td>Intergenic</td><td>NA</td><td>NA</td></tr>
<tr><td>T17812</td><td>AGCTTTCACATCCACGTCGCCGG</td><td>1000</td></tr>
<tr><td>Oligo fwd</td><td>CACCAGCTTTCACATCCACGTCGC</td></tr>
<tr><td>Oligo rev</td><td>AAACGCGACGTGGATGTGAAAGCT</td></tr>
<tr><td>Chromosome</td><td>start</td><td>end</td><td>strand</td><td>MM</td><td>target_seq</td><td>PAM</td><td>alignment</td><td>distance</td><td>position</td><td>gene name</td><td>gene id</td></tr>
<tr><td>GU305914.1</td><td>223212</td><td>223234</td><td>-</td><td>0</td><td>AGCTTTCACATCCACGTCGC</td><td>CGG</td><td>||||||||[||||||||||||]PAM</td><td>NA</td><td>Intergenic</td><td>NA</td><td>NA</td></tr>
<tr><td>T17813</td><td>CATCCACGTCGCCGGTTTCACGG</td><td>1000</td></tr>
<tr><td>Oligo fwd</td><td>CACCCATCCACGTCGCCGGTTTCA</td></tr>
<tr><td>Oligo rev</td><td>AAACTGAAACCGGCGACGTGGATG</td></tr>
<tr><td>Chromosome</td><td>start</td><td>end</td><td>strand</td><td>MM</td><td>target_seq</td><td>PAM</td><td>alignment</td><td>distance</td><td>position</td><td>gene name</td><td>gene id</td></tr>
<tr><td>GU305914.1</td><td>223204</td><td>223226</td><td>-</td><td>0</td><td>CATCCACGTCGCCGGTTTCA</td><td>CGG</td><td>||||||||[||||||||||||]PAM</td><td>NA</td><td>Intergenic</td><td>NA</td><td>NA</td></tr>
<tr><td>T17814</td><td>ATCCACGTCGCCGGTTTCACGGG</td><td>1000</td></tr>
<tr><td>Oligo fwd</td><td>CACCATCCACGTCGCCGGTTTCAC</td></tr>
<tr><td>Oligo rev</td><td>AAACGTGAAACCGGCGACGTGGAT</td></tr>
<tr><td>Chromosome</td><td>start</td><td>end</td><td>strand</td><td>MM</td><td>target_seq</td><td>PAM</td><td>alignment</td><td>distance</td><td>position</td><td>gene name</td><td>gene id</td></tr>
<tr><td>GU305914.1</td><td>223203</td><td>223225</td><td>-</td><td>0</td><td>ATCCACGTCGCCGGTTTCAC</td><td>GGG</td><td>||||||||[||||||||||||]PAM</td><td>NA</td><td>Intergenic</td><td>NA</td><td>NA</td></tr>
<tr><td>T17815</td><td>TCCACGTCGCCGGTTTCACGGGG</td><td>1000</td></tr>
<tr><td>Oligo fwd</td><td>CACCTCCACGTCGCCGGTTTCACG</td></tr>
<tr><td>Oligo rev</td><td>AAACCGTGAAACCGGCGACGTGGA</td></tr>
<tr><td>Chromosome</td><td>start</td><td>end</td><td>strand</td><td>MM</td><td>target_seq</td><td>PAM</td><td>alignment</td><td>distance</td><td>position</td><td>gene name</td><td>gene id</td></tr>
<tr><td>GU305914.1</td><td>223202</td><td>223224</td><td>-</td><td>0</td><td>TCCACGTCGCCGGTTTCACG</td><td>GGG</td><td>||||||||[||||||||||||]PAM</td><td>NA</td><td>Intergenic</td><td>NA</td><td>NA</td></tr>
<tr><td>T17816</td><td>CGTCGCCGGTTTCACGGGGATGG</td><td>1000</td></tr>
<tr><td>Oligo fwd</td><td>CACCCGTCGCCGGTTTCACGGGGA</td></tr>
<tr><td>Oligo rev</td><td>AAACTCCCCGTGAAACCGGCGACG</td></tr>
<tr><td>Chromosome</td><td>start</td><td>end</td><td>strand</td><td>MM</td><td>target_seq</td><td>PAM</td><td>alignment</td><td>distance</td><td>position</td><td>gene name</td><td>gene id</td></tr>
</table>

| | | | | | | | | | | | |
|---|---|---|---|---|---|---|---|---|---|---|---|
| GU305914.1 | 223198 | 223220 | - | 0 | CGTCGCCGGTTTCACGGGGA | TGG | ||||||||[||||||||||||]PAM | NA | Intergenic | NA | NA |
| GU305914.1 | 32020 | 32042 | - | 4 | CGCCGGCGGTTCCAAGGGGA | TGG | ||-||-||[|||-||-|||||]PAM | NA | Intergenic | NA | NA |
| T17817 | CGCCGGTTTCACGGGGATGGCGG | 1000 | | | | | | | | | |
| Oligo fwd | CACCCGCCGGTTTCACGGGGATGG | | | | | | | | | | |
| Oligo rev | AAACCCATCCCCGTGAAACCGGCG | | | | | | | | | | |
| Chromosome | start | end | strand | MM | target_seq | PAM | alignment | distance | position | gene name | gene id |
| GU305914.1 | 223195 | 223217 | - | 0 | CGCCGGTTTCACGGGGATGG | CGG | ||||||||[||||||||||||]PAM | NA | Intergenic | NA | NA |
| GU305914.1 | 80808 | 80830 | - | 4 | CTACGGTTTCACGGGGGTAG | CGG | |--|||||[||||||||-|-|]PAM | NA | Intergenic | NA | NA |
| T17818 | TGTCAATGACGACCGAGCTGTGG | 1000 | | | | | | | | | |
| Oligo fwd | CACCTGTCAATGACGACCGAGCTG | | | | | | | | | | |
| Oligo rev | AAACCAGCTCGGTCGTCATTGACA | | | | | | | | | | |
| Chromosome | start | end | strand | MM | target_seq | PAM | alignment | distance | position | gene name | gene id |
| GU305914.1 | 223166 | 223188 | - | 0 | TGTCAATGACGACCGAGCTG | TGG | ||||||||[||||||||||||]PAM | NA | Intergenic | NA | NA |
| T17819 | ACCGAGCTGTGGCCGCCTAGCGG | 1000 | | | | | | | | | |
| Oligo fwd | CACCACCGAGCTGTGGCCGCCTAG | | | | | | | | | | |
| Oligo rev | AAACCTAGGCGGCCACAGCTCGGT | | | | | | | | | | |
| Chromosome | start | end | strand | MM | target_seq | PAM | alignment | distance | position | gene name | gene id |
| GU305914.1 | 223155 | 223177 | - | 0 | ACCGAGCTGTGGCCGCCTAG | CGG | ||||||||[||||||||||||]PAM | NA | Intergenic | NA | NA |
| T17820 | GTGGCCGCCTAGCGGCTTCTCGG | 1000 | | | | | | | | | |
| Oligo fwd | CACCGTGGCCGCCTAGCGGCTTCT | | | | | | | | | | |
| Oligo rev | AAACAGAAGCCGCTAGGCGGCCAC | | | | | | | | | | |
| Chromosome | start | end | strand | MM | target_seq | PAM | alignment | distance | position | gene name | gene id |
| GU305914.1 | 223147 | 223169 | - | 0 | GTGGCCGCCTAGCGGCTTCT | CGG | ||||||||[||||||||||||]PAM | NA | Intergenic | NA | NA |
| T17821 | TGGCCGCCTAGCGGCTTCTCGGG | 1000 | | | | | | | | | |
| Oligo fwd | CACCTGGCCGCCTAGCGGCTTCTC | | | | | | | | | | |
| Oligo rev | AAACGAGAAGCCGCTAGGCGGCCA | | | | | | | | | | |
| Chromosome | start | end | strand | MM | target_seq | PAM | alignment | distance | position | gene name | gene id |
| GU305914.1 | 223146 | 223168 | - | 0 | TGGCCGCCTAGCGGCTTCTC | GGG | ||||||||[||||||||||||]PAM | NA | Intergenic | NA | NA |
| T17822 | CGACATCGACCATGTGAGCCAGG | 1000 | | | | | | | | | |
| Oligo fwd | CACCCGACATCGACCATGTGAGCC | | | | | | | | | | |
| Oligo rev | AAACGGCTCACATGGTCGATGTCG | | | | | | | | | | |
| Chromosome | start | end | strand | MM | target_seq | PAM | alignment | distance | position | gene name | gene id |
| GU305914.1 | 223111 | 223133 | - | 0 | CGACATCGACCATGTGAGCC | AGG | ||||||||[||||||||||||]PAM | NA | Intergenic | NA | NA |
| T17823 | TTCCGCCTCCCCTCCTACGTTGG | 1000 | | | | | | | | | |
| Oligo fwd | CACCTTCCGCCTCCCCTCCTACGT | | | | | | | | | | |
| Oligo rev | AAACACGTAGGAGGGGAGGCGGAA | | | | | | | | | | |
| Chromosome | start | end | strand | MM | target_seq | PAM | alignment | distance | position | gene name | gene id |
| GU305914.1 | 223076 | 223098 | - | 0 | TTCCGCCTCCCCTCCTACGT | TGG | ||||||||[||||||||||||]PAM | NA | Intergenic | NA | NA |
| T17824 | TCCGCCTCCCCTCCTACGTTGGG | 1000 | | | | | | | | | |
| Oligo fwd | CACCTCCGCCTCCCCTCCTACGTT | | | | | | | | | | |
| Oligo rev | AAACAACGTAGGAGGGGAGGCGGA | | | | | | | | | | |
| Chromosome | start | end | strand | MM | target_seq | PAM | alignment | distance | position | gene name | gene id |
| GU305914.1 | 223075 | 223097 | - | 0 | TCCGCCTCCCCTCCTACGTT | GGG | ||||||||[||||||||||||]PAM | NA | Intergenic | NA | NA |
| T17825 | TCCCCTCCTACGTTGGGTCCCGG | 1000 | | | | | | | | | |
| Oligo fwd | CACCTCCCCTCCTACGTTGGGTCC | | | | | | | | | | |
| Oligo rev | AAACGGACCCAACGTAGGAGGGGA | | | | | | | | | | |
| Chromosome | start | end | strand | MM | target_seq | PAM | alignment | distance | position | gene name | gene id |
| GU305914.1 | 223069 | 223091 | - | 0 | TCCCCTCCTACGTTGGGTCC | CGG | ||||||||[||||||||||||]PAM | NA | Intergenic | NA | NA |
| T17826 | CCCCTCCTACGTTGGGTCCCGGG | 1000 | | | | | | | | | |
| Oligo fwd | CACCCCCCTCCTACGTTGGGTCCC | | | | | | | | | | |
| Oligo rev | AAACGGGACCCAACGTAGGAGGGG | | | | | | | | | | |
| Chromosome | start | end | strand | MM | target_seq | PAM | alignment | distance | position | gene name | gene id |
| GU305914.1 | 223068 | 223090 | - | 0 | CCCCTCCTACGTTGGGTCCC | GGG | ||||||||[||||||||||||]PAM | NA | Intergenic | NA | NA |
| T17827 | CCTACGTTGGGTCCCGGGCGAGG | 1000 | | | | | | | | | |
| Oligo fwd | CACCCCTACGTTGGGTCCCGGGCG | | | | | | | | | | |
| Oligo rev | AAACCGCCCGGGACCCAACGTAGG | | | | | | | | | | |
| Chromosome | start | end | strand | MM | target_seq | PAM | alignment | distance | position | gene name | gene id |
| GU305914.1 | 223063 | 223085 | - | 0 | CCTACGTTGGGTCCCGGGCG | AGG | ||||||||[||||||||||||]PAM | NA | Intergenic | NA | NA |
| T17828 | GGAGCTCCTGACGATCGAGCCGG | 1000 | | | | | | | | | |
| Oligo fwd | CACCGGAGCTCCTGACGATCGAGC | | | | | | | | | | |
| Oligo rev | AAACGCTCGATCGTCAGGAGCTCC | | | | | | | | | | |
| Chromosome | start | end | strand | MM | target_seq | PAM | alignment | distance | position | gene name | gene id |
| GU305914.1 | 223042 | 223064 | - | 0 | GGAGCTCCTGACGATCGAGC | CGG | ||||||||[||||||||||||]PAM | NA | Intergenic | NA | NA |
| T17829 | GCTCCTGACGATCGAGCCGGTGG | 1000 | | | | | | | | | |
| Oligo fwd | CACCGCTCCTGACGATCGAGCCGG | | | | | | | | | | |
| Oligo rev | AAACCCGGCTCGATCGTCAGGAGC | | | | | | | | | | |
| Chromosome | start | end | strand | MM | target_seq | PAM | alignment | distance | position | gene name | gene id |
| GU305914.1 | 223039 | 223061 | - | 0 | GCTCCTGACGATCGAGCCGG | TGG | ||||||||[||||||||||||]PAM | NA | Intergenic | NA | NA |
| T17830 | GACGATCGAGCCGGTGGTACCGG | 1000 | | | | | | | | | |
| Oligo fwd | CACCGACGATCGAGCCGGTGGTAC | | | | | | | | | | |
| Oligo rev | AAACGTACCACCGGCTCGATCGTC | | | | | | | | | | |
| Chromosome | start | end | strand | MM | target_seq | PAM | alignment | distance | position | gene name | gene id |
| GU305914.1 | 223033 | 223055 | - | 0 | GACGATCGAGCCGGTGGTAC | CGG | ||||||||[||||||||||||]PAM | NA | Intergenic | NA | NA |
| T17831 | CGAGCCGGTGGTACCGGACGCGG | 1000 | | | | | | | | | |

| | | | | | | | | | | | | |
|---|---|---|---|---|---|---|---|---|---|---|---|---|
| Oligo fwd | CACCCGAGCCGGTGGTACCGGACG | | | | | | | | | | | |
| Oligo rev | AAACCGTCCGGTACCACCGGCTCG | | | | | | | | | | | |
| Chromosome | start | | end | strand | MM | target_seq | PAM | alignment | distance | position | gene name | gene id |
| GU305914.1 | | 223027 | 223049 | - | 0 | CGAGCCGGTGGTACCGGACG | CGG | \|\|\|\|\|\|\|\|[\|\|\|\|\|\|\|\|\|\|\|\|]PAM | NA | Intergenic | NA | NA |
| T17832 | GCCGGTGGTACCGGACGCGGCGG | 1000 | | | | | | | | | | |
| Oligo fwd | CACCGCCGGTGGTACCGGACGCGG | | | | | | | | | | | |
| Oligo rev | AAACCCGCGTCCGGTACCACCGGC | | | | | | | | | | | |
| Chromosome | start | | end | strand | MM | target_seq | PAM | alignment | distance | position | gene name | gene id |
| GU305914.1 | | 223024 | 223046 | - | 0 | GCCGGTGGTACCGGACGCGG | CGG | \|\|\|\|\|\|\|\|[\|\|\|\|\|\|\|\|\|\|\|\|]PAM | NA | Intergenic | NA | NA |
| T17833 | CCGGACGCGGCGGAGCCGTTCGG | 1000 | | | | | | | | | | |
| Oligo fwd | CACCCCGGACGCGGCGGAGCCGTT | | | | | | | | | | | |
| Oligo rev | AAACAACGGCTCCGCCGCGTCCGG | | | | | | | | | | | |
| Chromosome | start | | end | strand | MM | target_seq | PAM | alignment | distance | position | gene name | gene id |
| GU305914.1 | | 223014 | 223036 | - | 0 | CCGGACGCGGCGGAGCCGTT | CGG | \|\|\|\|\|\|\|\|[\|\|\|\|\|\|\|\|\|\|\|\|]PAM | NA | Intergenic | NA | NA |
| T17834 | CGCGGCGGAGCCGTTCGGAAAGG | 1000 | | | | | | | | | | |
| Oligo fwd | CACCCGCGGCGGAGCCGTTCGGAA | | | | | | | | | | | |
| Oligo rev | AAACTTCCGAACGGCTCCGCCGCG | | | | | | | | | | | |
| Chromosome | start | | end | strand | MM | target_seq | PAM | alignment | distance | position | gene name | gene id |
| GU305914.1 | | 223009 | 223031 | - | 0 | CGCGGCGGAGCCGTTCGGAA | AGG | \|\|\|\|\|\|\|\|[\|\|\|\|\|\|\|\|\|\|\|\|]PAM | NA | Intergenic | NA | NA |
| T17835 | CGTTCGGAAAGGACTACTGTCGG | 1000 | | | | | | | | | | |
| Oligo fwd | CACCCGTTCGGAAAGGACTACTGT | | | | | | | | | | | |
| Oligo rev | AAACACAGTAGTCCTTTCCGAACG | | | | | | | | | | | |
| Chromosome | start | | end | strand | MM | target_seq | PAM | alignment | distance | position | gene name | gene id |
| GU305914.1 | | 222998 | 223020 | - | 0 | CGTTCGGAAAGGACTACTGT | CGG | \|\|\|\|\|\|\|\|[\|\|\|\|\|\|\|\|\|\|\|\|]PAM | NA | Intergenic | NA | NA |
| T17836 | GAAAGGACTACTGTCGGACGTGG | 1000 | | | | | | | | | | |
| Oligo fwd | CACCGAAAGGACTACTGTCGGACG | | | | | | | | | | | |
| Oligo rev | AAACCGTCCGACAGTAGTCCTTTC | | | | | | | | | | | |
| Chromosome | start | | end | strand | MM | target_seq | PAM | alignment | distance | position | gene name | gene id |
| GU305914.1 | | 222992 | 223014 | - | 0 | GAAAGGACTACTGTCGGACG | TGG | \|\|\|\|\|\|\|\|[\|\|\|\|\|\|\|\|\|\|\|\|]PAM | NA | Intergenic | NA | NA |
| T17837 | AAAGGACTACTGTCGGACGTGGG | 1000 | | | | | | | | | | |
| Oligo fwd | CACCAAAGGACTACTGTCGGACGT | | | | | | | | | | | |
| Oligo rev | AAACACGTCCGACAGTAGTCCTTT | | | | | | | | | | | |
| Chromosome | start | | end | strand | MM | target_seq | PAM | alignment | distance | position | gene name | gene id |
| GU305914.1 | | 222991 | 223013 | - | 0 | AAAGGACTACTGTCGGACGT | GGG | \|\|\|\|\|\|\|\|[\|\|\|\|\|\|\|\|\|\|\|\|]PAM | NA | Intergenic | NA | NA |
| T17838 | AAGGACTACTGTCGGACGTGGGG | 1000 | | | | | | | | | | |
| Oligo fwd | CACCAAGGACTACTGTCGGACGTG | | | | | | | | | | | |
| Oligo rev | AAACCACGTCCGACAGTAGTCCTT | | | | | | | | | | | |
| Chromosome | start | | end | strand | MM | target_seq | PAM | alignment | distance | position | gene name | gene id |
| GU305914.1 | | 222990 | 223012 | - | 0 | AAGGACTACTGTCGGACGTG | GGG | \|\|\|\|\|\|\|\|[\|\|\|\|\|\|\|\|\|\|\|\|]PAM | NA | Intergenic | NA | NA |
| T17839 | TCGGACGTGGGGCGCTGACGCGG | 1000 | | | | | | | | | | |
| Oligo fwd | CACCTCGGACGTGGGGCGCTGACG | | | | | | | | | | | |
| Oligo rev | AAACCGTCAGCGCCCCACGTCCGA | | | | | | | | | | | |
| Chromosome | start | | end | strand | MM | target_seq | PAM | alignment | distance | position | gene name | gene id |
| GU305914.1 | | 222979 | 223001 | - | 0 | TCGGACGTGGGGCGCTGACG | CGG | \|\|\|\|\|\|\|\|[\|\|\|\|\|\|\|\|\|\|\|\|]PAM | NA | Intergenic | NA | NA |
| T17840 | CGGACGTGGGGCGCTGACGCGGG | 1000 | | | | | | | | | | |
| Oligo fwd | CACCCGGACGTGGGGCGCTGACGC | | | | | | | | | | | |
| Oligo rev | AAACGCGTCAGCGCCCCACGTCCG | | | | | | | | | | | |
| Chromosome | start | | end | strand | MM | target_seq | PAM | alignment | distance | position | gene name | gene id |
| GU305914.1 | | 222978 | 223000 | - | 0 | CGGACGTGGGGCGCTGACGC | GGG | \|\|\|\|\|\|\|\|[\|\|\|\|\|\|\|\|\|\|\|\|]PAM | NA | Intergenic | NA | NA |
| T17841 | GCGCTGACGCGGGCCCTCCTCGG | 1000 | | | | | | | | | | |
| Oligo fwd | CACCGCGCTGACGCGGGCCCTCCT | | | | | | | | | | | |
| Oligo rev | AAACAGGAGGGCCCGCGTCAGCGC | | | | | | | | | | | |
| Chromosome | start | | end | strand | MM | target_seq | PAM | alignment | distance | position | gene name | gene id |
| GU305914.1 | | 222968 | 222990 | - | 0 | GCGCTGACGCGGGCCCTCCT | CGG | \|\|\|\|\|\|\|\|[\|\|\|\|\|\|\|\|\|\|\|\|]PAM | NA | Intergenic | NA | NA |
| T17842 | CTCGGTACTCGCGCATCTTCTGG | 1000 | | | | | | | | | | |
| Oligo fwd | CACCCTCGGTACTCGCGCATCTTC | | | | | | | | | | | |
| Oligo rev | AAACGAAGATGCGCGAGTACCGAG | | | | | | | | | | | |
| Chromosome | start | | end | strand | MM | target_seq | PAM | alignment | distance | position | gene name | gene id |
| GU305914.1 | | 222950 | 222972 | - | 0 | CTCGGTACTCGCGCATCTTC | TGG | \|\|\|\|\|\|\|\|[\|\|\|\|\|\|\|\|\|\|\|\|]PAM | NA | Intergenic | NA | NA |
| T17843 | TCGGTACTCGCGCATCTTCTGGG | 1000 | | | | | | | | | | |
| Oligo fwd | CACCTCGGTACTCGCGCATCTTCT | | | | | | | | | | | |
| Oligo rev | AAACAGAAGATGCGCGAGTACCGA | | | | | | | | | | | |
| Chromosome | start | | end | strand | MM | target_seq | PAM | alignment | distance | position | gene name | gene id |
| GU305914.1 | | 222949 | 222971 | - | 0 | TCGGTACTCGCGCATCTTCT | GGG | \|\|\|\|\|\|\|\|[\|\|\|\|\|\|\|\|\|\|\|\|]PAM | NA | Intergenic | NA | NA |
| T17844 | CTTCTGGGCCGTCAACAACCAGG | 1000 | | | | | | | | | | |
| Oligo fwd | CACCCTTCTGGGCCGTCAACAACC | | | | | | | | | | | |
| Oligo rev | AAACGGTTGTTGACGGCCCAGAAG | | | | | | | | | | | |
| Chromosome | start | | end | strand | MM | target_seq | PAM | alignment | distance | position | gene name | gene id |
| GU305914.1 | | 222934 | 222956 | - | 0 | CTTCTGGGCCGTCAACAACC | AGG | \|\|\|\|\|\|\|\|[\|\|\|\|\|\|\|\|\|\|\|\|]PAM | NA | Intergenic | NA | NA |
| T17845 | CGTCAACAACCAGGCCATCGTGG | 1000 | | | | | | | | | | |
| Oligo fwd | CACCCGTCAACAACCAGGCCATCG | | | | | | | | | | | |
| Oligo rev | AAACCGATGGCCTGGTTGTTGACG | | | | | | | | | | | |
| Chromosome | start | | end | strand | MM | target_seq | PAM | alignment | distance | position | gene name | gene id |
| GU305914.1 | | 222925 | 222947 | - | 0 | CGTCAACAACCAGGCCATCG | TGG | \|\|\|\|\|\|\|\|[\|\|\|\|\|\|\|\|\|\|\|\|]PAM | NA | Intergenic | NA | NA |

T17846 CAACCAGGCCATCGTGGCAGCGG 1000
Oligo fwd CACCCAACCAGGCCATCGTGGCAG
Oligo rev AAACCTGCCACGATGGCCTGGTTG
Chromosome start end strand MM target_seq PAM alignment distance position gene name gene id
GU305914.1 222919 222941 - 0 CAACCAGGCCATCGTGGCAG CGG ||||||||[||||||||||||]PAM NA Intergenic NA NA

T17847 AACCAGGCCATCGTGGCAGCGGG 1000
Oligo fwd CACCAACCAGGCCATCGTGGCAGC
Oligo rev AAACGCTGCCACGATGGCCTGGTT
Chromosome start end strand MM target_seq PAM alignment distance position gene name gene id
GU305914.1 222918 222940 - 0 AACCAGGCCATCGTGGCAGC GGG ||||||||[||||||||||||]PAM NA Intergenic NA NA

T17848 GGCCATCGTGGCAGCGGGCGAGG 1000
Oligo fwd CACCGGCCATCGTGGCAGCGGGCG
Oligo rev AAACCGCCCGCTGCCACGATGGCC
Chromosome start end strand MM target_seq PAM alignment distance position gene name gene id
GU305914.1 222913 222935 - 0 GGCCATCGTGGCAGCGGGCG AGG ||||||||[||||||||||||]PAM NA Intergenic NA NA

T17849 GGCAGCGGGCGAGGAGACCTTGG 1000
Oligo fwd CACCGGCAGCGGGCGAGGAGACCT
Oligo rev AAACAGGTCTCCTCGCCCGCTGCC
Chromosome start end strand MM target_seq PAM alignment distance position gene name gene id
GU305914.1 222904 222926 - 0 GGCAGCGGGCGAGGAGACCT TGG ||||||||[||||||||||||]PAM NA Intergenic NA NA

T17850 GCAGCGGGCGAGGAGACCTTGGG 1000
Oligo fwd CACCGCAGCGGGCGAGGAGACCTT
Oligo rev AAACAAGGTCTCCTCGCCCGCTGC
Chromosome start end strand MM target_seq PAM alignment distance position gene name gene id
GU305914.1 222903 222925 - 0 GCAGCGGGCGAGGAGACCTT GGG ||||||||[||||||||||||]PAM NA Intergenic NA NA

T17851 TGGGCACCACCGACTTCTTGTGG 1000
Oligo fwd CACCTGGGCACCACCGACTTCTTG
Oligo rev AAACCAAGAAGTCGGTGGTGCCCA
Chromosome start end strand MM target_seq PAM alignment distance position gene name gene id
GU305914.1 222884 222906 - 0 TGGGCACCACCGACTTCTTG TGG ||||||||[||||||||||||]PAM NA Intergenic NA NA

T17852 ACTTCTTGTGGATGTCGAACCGG 1000
Oligo fwd CACCACTTCTTGTGGATGTCGAAC
Oligo rev AAACGTTCGACATCCACAAGAAGT
Chromosome start end strand MM target_seq PAM alignment distance position gene name gene id
GU305914.1 222872 222894 - 0 ACTTCTTGTGGATGTCGAAC CGG ||||||||[||||||||||||]PAM NA Intergenic NA NA

T17853 GTGGATGTCGAACCGGCTCGAGG 1000
Oligo fwd CACCGTGGATGTCGAACCGGCTCG
Oligo rev AAACCGAGCCGGTTCGACATCCAC
Chromosome start end strand MM target_seq PAM alignment distance position gene name gene id
GU305914.1 222865 222887 - 0 GTGGATGTCGAACCGGCTCG AGG ||||||||[||||||||||||]PAM NA Intergenic NA NA

T17854 TGGATGTCGAACCGGCTCGAGGG 1000
Oligo fwd CACCTGGATGTCGAACCGGCTCGA
Oligo rev AAACTCGAGCCGGTTCGACATCCA
Chromosome start end strand MM target_seq PAM alignment distance position gene name gene id
GU305914.1 222864 222886 - 0 TGGATGTCGAACCGGCTCGA GGG ||||||||[||||||||||||]PAM NA Intergenic NA NA

T17855 TACCGTCAACGCATCCGACCTGG 1000
Oligo fwd CACCTACCGTCAACGCATCCGACC
Oligo rev AAACGGTCGGATGCGTTGACGGTA
Chromosome start end strand MM target_seq PAM alignment distance position gene name gene id
GU305914.1 222838 222860 - 0 TACCGTCAACGCATCCGACC TGG ||||||||[||||||||||||]PAM NA Intergenic NA NA

T17856 GTCTGCGTATCGTGCGTCGTCGG 1000
Oligo fwd CACCGTCTGCGTATCGTGCGTCGT
Oligo rev AAACACGACGCACGATACGCAGAC
Chromosome start end strand MM target_seq PAM alignment distance position gene name gene id
GU305914.1 222783 222805 - 0 GTCTGCGTATCGTGCGTCGT CGG ||||||||[||||||||||||]PAM NA Intergenic NA NA

T17857 TCTGCGTATCGTGCGTCGTCGGG 1000
Oligo fwd CACCTCTGCGTATCGTGCGTCGTC
Oligo rev AAACGACGACGCACGATACGCAGA
Chromosome start end strand MM target_seq PAM alignment distance position gene name gene id
GU305914.1 222782 222804 - 0 TCTGCGTATCGTGCGTCGTC GGG ||||||||[||||||||||||]PAM NA Intergenic NA NA

T17858 GTATCGTGCGTCGTCGGGTCCGG 1000
Oligo fwd CACCGTATCGTGCGTCGTCGGGTC
Oligo rev AAACGACCCGACGACGCACGATAC
Chromosome start end strand MM target_seq PAM alignment distance position gene name gene id
GU305914.1 222777 222799 - 0 GTATCGTGCGTCGTCGGGTC CGG ||||||||[||||||||||||]PAM NA Intergenic NA NA

T17859 TCGTGCGTCGTCGGGTCCGGCGG 1000
Oligo fwd CACCTCGTGCGTCGTCGGGTCCGG
Oligo rev AAACCCGGACCCGACGACGCACGA
Chromosome start end strand MM target_seq PAM alignment distance position gene name gene id
GU305914.1 222774 222796 - 0 TCGTGCGTCGTCGGGTCCGG CGG ||||||||[||||||||||||]PAM NA Intergenic NA NA

T17860 CGAATGTCGACCCGTCCACGTGG 1000
Oligo fwd CACCCGAATGTCGACCCGTCCACG
Oligo rev AAACCGTGGACGGGTCGACATTCG
Chromosome start end strand MM target_seq PAM alignment distance position gene name gene id
GU305914.1 222727 222749 - 0 CGAATGTCGACCCGTCCACG TGG ||||||||[||||||||||||]PAM NA Intergenic NA NA

T17861 CACGTGGCGCGCAGTTCAACCGG 1000
Oligo fwd CACCCACGTGGCGCGCAGTTCAAC
Oligo rev AAACGTTGAACTGCGCGCCACGTG
Chromosome start end strand MM target_seq PAM alignment distance position gene name gene id
GU305914.1 222711 222733 - 0 CACGTGGCGCGCAGTTCAAC CGG ||||||||[||||||||||||]PAM NA Intergenic NA NA

T17862 GAAGAGCTGCGACTGTCAGTTGG 1000
Oligo fwd CACCGAAGAGCTGCGACTGTCAGT
Oligo rev AAACACTGACAGTCGCAGCTCTTC
Chromosome start end strand MM target_seq PAM alignment distance position gene name gene id
GU305914.1 222685 222707 - 0 GAAGAGCTGCGACTGTCAGT TGG ||||||||[||||||||||||]PAM NA Intergenic NA NA

T17863 GCGTTTCCTCGCCGCCTTCGTGG 1000
Oligo fwd CACCGCGTTTCCTCGCCGCCTTCG
Oligo rev AAACCGAAGGCGGCGAGGAAACGC
Chromosome start end strand MM target_seq PAM alignment distance position gene name gene id
GU305914.1 222661 222683 - 0 GCGTTTCCTCGCCGCCTTCG TGG ||||||||[||||||||||||]PAM NA Intergenic NA NA

T17864 GATCGTCGTGTACATCTACCTGG 1000
Oligo fwd CACCGATCGTCGTGTACATCTACC
Oligo rev AAACGGTAGATGTACACGACGATC
Chromosome start end strand MM target_seq PAM alignment distance position gene name gene id
GU305914.1 222613 222635 - 0 GATCGTCGTGTACATCTACC TGG ||||||||[||||||||||||]PAM NA Intergenic NA NA

T17865 CGTGTACATCTACCTGGTCTCGG 1000
Oligo fwd CACCCGTGTACATCTACCTGGTCT
Oligo rev AAACAGACCAGGTAGATGTACACG
Chromosome start end strand MM target_seq PAM alignment distance position gene name gene id
GU305914.1 222607 222629 - 0 CGTGTACATCTACCTGGTCT CGG ||||||||[||||||||||||]PAM NA Intergenic NA NA

T17866 TCTACCTGGTCTCGGACGTCAGG 1000
Oligo fwd CACCTCTACCTGGTCTCGGACGTC
Oligo rev AAACGACGTCCGAGACCAGGTAGA
Chromosome start end strand MM target_seq PAM alignment distance position gene name gene id
GU305914.1 222599 222621 - 0 TCTACCTGGTCTCGGACGTC AGG ||||||||[||||||||||||]PAM NA Intergenic NA NA

T17867 TGTTCCCGACGTTCGAAATCCGG 1000
Oligo fwd CACCTGTTCCCGACGTTCGAAATC
Oligo rev AAACGATTTCGAACGTCGGGAACA
Chromosome start end strand MM target_seq PAM alignment distance position gene name gene id
GU305914.1 222569 222591 - 0 TGTTCCCGACGTTCGAAATC CGG ||||||||[||||||||||||]PAM NA Intergenic NA NA

T17868 TTCGAAATCCGGTATTTTACTGG 1000
Oligo fwd CACCTTCGAAATCCGGTATTTTAC
Oligo rev AAACGTAAAATACCGGATTTCGAA
Chromosome start end strand MM target_seq PAM alignment distance position gene name gene id
GU305914.1 222558 222580 - 0 TTCGAAATCCGGTATTTTAC TGG ||||||||[||||||||||||]PAM NA Intergenic NA NA

T17869 GAAATCCGGTATTTTACTGGTGG 1000
Oligo fwd CACCGAAATCCGGTATTTTACTGG
Oligo rev AAACCCAGTAAAATACCGGATTTC
Chromosome start end strand MM target_seq PAM alignment distance position gene name gene id
GU305914.1 222555 222577 - 0 GAAATCCGGTATTTTACTGG TGG ||||||||[||||||||||||]PAM NA Intergenic NA NA

T17870 GTATTTTACTGGTGGCGTGCCGG 1000
Oligo fwd CACCGTATTTTACTGGTGGCGTGC
Oligo rev AAACGCACGCCACCAGTAAAATAC
Chromosome start end strand MM target_seq PAM alignment distance position gene name gene id
GU305914.1 222547 222569 - 0 GTATTTTACTGGTGGCGTGC CGG ||||||||[||||||||||||]PAM NA Intergenic NA NA

T17871 ACTGGTGGCGTGCCGGTGAATGG 1000
Oligo fwd CACCACTGGTGGCGTGCCGGTGAA
Oligo rev AAACTTCACCGGCACGCCACCAGT
Chromosome start end strand MM target_seq PAM alignment distance position gene name gene id
GU305914.1 222540 222562 - 0 ACTGGTGGCGTGCCGGTGAA TGG ||||||||[||||||||||||]PAM NA Intergenic NA NA

T17872 CTGGTGGCGTGCCGGTGAATGGG 1000
Oligo fwd CACCCTGGTGGCGTGCCGGTGAAT
Oligo rev AAACATTCACCGGCACGCCACCAG
Chromosome start end strand MM target_seq PAM alignment distance position gene name gene id
GU305914.1 222539 222561 - 0 CTGGTGGCGTGCCGGTGAAT GGG ||||||||[||||||||||||]PAM NA Intergenic NA NA

T17873 TGGTGGCGTGCCGGTGAATGGGG 1000
Oligo fwd CACCTGGTGGCGTGCCGGTGAATG
Oligo rev AAACCATTCACCGGCACGCCACCA
Chromosome start end strand MM target_seq PAM alignment distance position gene name gene id
GU305914.1 222538 222560 - 0 TGGTGGCGTGCCGGTGAATG GGG ||||||||[||||||||||||]PAM NA Intergenic NA NA

T17874 TGTCGCGCAACCATCAGCCATGG 1000
Oligo fwd CACCTGTCGCGCAACCATCAGCCA
Oligo rev AAACTGGCTGATGGTTGCGCGACA
Chromosome start end strand MM target_seq PAM alignment distance position gene name gene id
GU305914.1 222512 222534 - 0 TGTCGCGCAACCATCAGCCA TGG ||||||||[||||||||||||]PAM NA Intergenic NA NA

T17875 CGCGCAACCATCAGCCATGGAGG 1000
Oligo fwd CACCCGCGCAACCATCAGCCATGG
Oligo rev AAACCCATGGCTGATGGTTGCGCG
Chromosome start end strand MM target_seq PAM alignment distance position gene name gene id

| | | | | | | | | | | | |
|---|---|---|---|---|---|---|---|---|---|---|---|
| GU305914.1 | 222509 | 222531 | - | 0 | CGCGCAACCATCAGCCATGG | AGG | \|\|\|\|\|\|\|\|[\|\|\|\|\|\|\|\|\|\|\|\|]PAM | NA | Intergenic | NA | NA |
| T17876 | GCGCAACCATCAGCCATGGAGGG | 1000 | | | | | | | | | |
| Oligo fwd | CACCGCGCAACCATCAGCCATGGA | | | | | | | | | | |
| Oligo rev | AAACTCCATGGCTGATGGTTGCGC | | | | | | | | | | |
| Chromosome | start | end | strand | MM | target_seq | PAM | alignment | distance | position | gene name | gene id |
| GU305914.1 | 222508 | 222530 | - | 0 | GCGCAACCATCAGCCATGGA | GGG | \|\|\|\|\|\|\|\|[\|\|\|\|\|\|\|\|\|\|\|\|]PAM | NA | Intergenic | NA | NA |
| T17877 | ACGAACGCTTCGTCGCGCGTCGG | 1000 | | | | | | | | | |
| Oligo fwd | CACCACGAACGCTTCGTCGCGCGT | | | | | | | | | | |
| Oligo rev | AAACACGCGCGACGAAGCGTTCGT | | | | | | | | | | |
| Chromosome | start | end | strand | MM | target_seq | PAM | alignment | distance | position | gene name | gene id |
| GU305914.1 | 222480 | 222502 | - | 0 | ACGAACGCTTCGTCGCGCGT | CGG | \|\|\|\|\|\|\|\|[\|\|\|\|\|\|\|\|\|\|\|\|]PAM | NA | Intergenic | NA | NA |
| T17878 | TCGCGCGACAAGCAGTGTATCGG | 1000 | | | | | | | | | |
| Oligo fwd | CACCTCGCGCGACAAGCAGTGTAT | | | | | | | | | | |
| Oligo rev | AAACATACACTGCTTGTCGCGCGA | | | | | | | | | | |
| Chromosome | start | end | strand | MM | target_seq | PAM | alignment | distance | position | gene name | gene id |
| GU305914.1 | 222422 | 222444 | - | 0 | TCGCGCGACAAGCAGTGTAT | CGG | \|\|\|\|\|\|\|\|[\|\|\|\|\|\|\|\|\|\|\|\|]PAM | NA | Intergenic | NA | NA |
| T17879 | CGCGCGACAAGCAGTGTATCGGG | 1000 | | | | | | | | | |
| Oligo fwd | CACCCGCGCGACAAGCAGTGTATC | | | | | | | | | | |
| Oligo rev | AAACGATACACTGCTTGTCGCGCG | | | | | | | | | | |
| Chromosome | start | end | strand | MM | target_seq | PAM | alignment | distance | position | gene name | gene id |
| GU305914.1 | 222421 | 222443 | - | 0 | CGCGCGACAAGCAGTGTATC | GGG | \|\|\|\|\|\|\|\|[\|\|\|\|\|\|\|\|\|\|\|\|]PAM | NA | Intergenic | NA | NA |
| T17880 | GGGAACGCCTTCGTACCCGACGG | 1000 | | | | | | | | | |
| Oligo fwd | CACCGGGAACGCCTTCGTACCCGA | | | | | | | | | | |
| Oligo rev | AAACTCGGGTACGAAGGCGTTCCC | | | | | | | | | | |
| Chromosome | start | end | strand | MM | target_seq | PAM | alignment | distance | position | gene name | gene id |
| GU305914.1 | 222401 | 222423 | - | 0 | GGGAACGCCTTCGTACCCGA | CGG | \|\|\|\|\|\|\|\|[\|\|\|\|\|\|\|\|\|\|\|\|]PAM | NA | Intergenic | NA | NA |
| GU305914.1 | 17180 | 17202 | - | 4 | GGGGATGCCGTCGTACCTGA | TGG | \|\|\|-\|-\|\|[\|-\|\|\|\|\|\|\|-\|\|]PAM | NA | Intergenic | NA | NA |
| T17881 | AACGCCTTCGTACCCGACGGAGG | 1000 | | | | | | | | | |
| Oligo fwd | CACCAACGCCTTCGTACCCGACGG | | | | | | | | | | |
| Oligo rev | AAACCCGTCGGGTACGAAGGCGTT | | | | | | | | | | |
| Chromosome | start | end | strand | MM | target_seq | PAM | alignment | distance | position | gene name | gene id |
| GU305914.1 | 222398 | 222420 | - | 0 | AACGCCTTCGTACCCGACGG | AGG | \|\|\|\|\|\|\|\|[\|\|\|\|\|\|\|\|\|\|\|\|]PAM | NA | Intergenic | NA | NA |
| T17882 | ACGCCTTCGTACCCGACGGAGGG | 1000 | | | | | | | | | |
| Oligo fwd | CACCACGCCTTCGTACCCGACGGA | | | | | | | | | | |
| Oligo rev | AAACTCCGTCGGGTACGAAGGCGT | | | | | | | | | | |
| Chromosome | start | end | strand | MM | target_seq | PAM | alignment | distance | position | gene name | gene id |
| GU305914.1 | 222397 | 222419 | - | 0 | ACGCCTTCGTACCCGACGGA | GGG | \|\|\|\|\|\|\|\|[\|\|\|\|\|\|\|\|\|\|\|\|]PAM | NA | Intergenic | NA | NA |
| T17883 | TCGTACCCGACGGAGGGTGCCGG | 1000 | | | | | | | | | |
| Oligo fwd | CACCTCGTACCCGACGGAGGGTGC | | | | | | | | | | |
| Oligo rev | AAACGCACCCTCCGTCGGGTACGA | | | | | | | | | | |
| Chromosome | start | end | strand | MM | target_seq | PAM | alignment | distance | position | gene name | gene id |
| GU305914.1 | 222391 | 222413 | - | 0 | TCGTACCCGACGGAGGGTGC | CGG | \|\|\|\|\|\|\|\|[\|\|\|\|\|\|\|\|\|\|\|\|]PAM | NA | Intergenic | NA | NA |
| T17884 | CCGACGGAGGGTGCCGGTGACGG | 1000 | | | | | | | | | |
| Oligo fwd | CACCCCGACGGAGGGTGCCGGTGA | | | | | | | | | | |
| Oligo rev | AAACTCACCGGCACCCTCCGTCGG | | | | | | | | | | |
| Chromosome | start | end | strand | MM | target_seq | PAM | alignment | distance | position | gene name | gene id |
| GU305914.1 | 222385 | 222407 | - | 0 | CCGACGGAGGGTGCCGGTGA | CGG | \|\|\|\|\|\|\|\|[\|\|\|\|\|\|\|\|\|\|\|\|]PAM | NA | Intergenic | NA | NA |
| T17885 | CGAACAACCCGACGTTCAAGAGG | 1000 | | | | | | | | | |
| Oligo fwd | CACCCGAACAACCCGACGTTCAAG | | | | | | | | | | |
| Oligo rev | AAACCTTGAACGTCGGGTTGTTCG | | | | | | | | | | |
| Chromosome | start | end | strand | MM | target_seq | PAM | alignment | distance | position | gene name | gene id |
| GU305914.1 | 222350 | 222372 | - | 0 | CGAACAACCCGACGTTCAAG | AGG | \|\|\|\|\|\|\|\|[\|\|\|\|\|\|\|\|\|\|\|\|]PAM | NA | Intergenic | NA | NA |
| T17886 | AGAGGCGCTGCAGTCCCACGCGG | 1000 | | | | | | | | | |
| Oligo fwd | CACCAGAGGCGCTGCAGTCCCACG | | | | | | | | | | |
| Oligo rev | AAACCGTGGGACTGCAGCGCCTCT | | | | | | | | | | |
| Chromosome | start | end | strand | MM | target_seq | PAM | alignment | distance | position | gene name | gene id |
| GU305914.1 | 222332 | 222354 | - | 0 | AGAGGCGCTGCAGTCCCACG | CGG | \|\|\|\|\|\|\|\|[\|\|\|\|\|\|\|\|\|\|\|\|]PAM | NA | Intergenic | NA | NA |
| T17887 | GTCCCACGCGGTATCGTCTTCGG | 1000 | | | | | | | | | |
| Oligo fwd | CACCGTCCCACGCGGTATCGTCTT | | | | | | | | | | |
| Oligo rev | AAACAAGACGATACCGCGTGGGAC | | | | | | | | | | |
| Chromosome | start | end | strand | MM | target_seq | PAM | alignment | distance | position | gene name | gene id |
| GU305914.1 | 222320 | 222342 | - | 0 | GTCCCACGCGGTATCGTCTT | CGG | \|\|\|\|\|\|\|\|[\|\|\|\|\|\|\|\|\|\|\|\|]PAM | NA | Intergenic | NA | NA |
| T17888 | TCCCACGCGGTATCGTCTTCGGG | 1000 | | | | | | | | | |
| Oligo fwd | CACCTCCCACGCGGTATCGTCTTC | | | | | | | | | | |
| Oligo rev | AAACGAAGACGATACCGCGTGGGA | | | | | | | | | | |
| Chromosome | start | end | strand | MM | target_seq | PAM | alignment | distance | position | gene name | gene id |
| GU305914.1 | 222319 | 222341 | - | 0 | TCCCACGCGGTATCGTCTTC | GGG | \|\|\|\|\|\|\|\|[\|\|\|\|\|\|\|\|\|\|\|\|]PAM | NA | Intergenic | NA | NA |
| T17889 | TCCCGACGTCGCCTACCTCACGG | 1000 | | | | | | | | | |
| Oligo fwd | CACCTCCCGACGTCGCCTACCTCA | | | | | | | | | | |
| Oligo rev | AAACTGAGGTAGGCGACGTCGGGA | | | | | | | | | | |
| Chromosome | start | end | strand | MM | target_seq | PAM | alignment | distance | position | gene name | gene id |
| GU305914.1 | 222296 | 222318 | - | 0 | TCCCGACGTCGCCTACCTCA | CGG | \|\|\|\|\|\|\|\|[\|\|\|\|\|\|\|\|\|\|\|\|]PAM | NA | Intergenic | NA | NA |
| T17890 | ACGGAGATGATAGATATACTAGG | 1000 | | | | | | | | | |
| Oligo fwd | CACCACGGAGATGATAGATATACT | | | | | | | | | | |

| | | | | | | | | | | | |
|---|---|---|---|---|---|---|---|---|---|---|---|
| Oligo rev | AAACAGTATATCTATCATCTCCGT | | | | | | | | | | |
| Chromosome | start | end | strand | MM | target_seq | PAM | alignment | distance | position | gene name | gene id |
| GU305914.1 | 222277 | 222299 | - | 0 | ACGGAGATGATAGATATACT | AGG | \|\|\|\|\|\|\|\|[\|\|\|\|\|\|\|\|\|\|\|\|]PAM | NA | Intergenic | NA | NA |
| T17891 | GATAGATATACTAGGTCGCGAGG | 1000 | | | | | | | | | |
| Oligo fwd | CACCGATAGATATACTAGGTCGCG | | | | | | | | | | |
| Oligo rev | AAACCGCGACCTAGTATATCTATC | | | | | | | | | | |
| Chromosome | start | end | strand | MM | target_seq | PAM | alignment | distance | position | gene name | gene id |
| GU305914.1 | 222269 | 222291 | - | 0 | GATAGATATACTAGGTCGCG | AGG | \|\|\|\|\|\|\|\|[\|\|\|\|\|\|\|\|\|\|\|\|]PAM | NA | Intergenic | NA | NA |
| T17892 | CCAACAAGTCGCCCCAGAGCCGG | 1000 | | | | | | | | | |
| Oligo fwd | CACCCCAACAAGTCGCCCCAGAGC | | | | | | | | | | |
| Oligo rev | AAACGCTCTGGGGCGACTTGTTGG | | | | | | | | | | |
| Chromosome | start | end | strand | MM | target_seq | PAM | alignment | distance | position | gene name | gene id |
| GU305914.1 | 222239 | 222261 | - | 0 | CCAACAAGTCGCCCCAGAGC | CGG | \|\|\|\|\|\|\|\|[\|\|\|\|\|\|\|\|\|\|\|\|]PAM | NA | Intergenic | NA | NA |
| T17893 | CAACAAGTCGCCCCAGAGCCGGG | 1000 | | | | | | | | | |
| Oligo fwd | CACCCAACAAGTCGCCCCAGAGCC | | | | | | | | | | |
| Oligo rev | AAACGGCTCTGGGGCGACTTGTTG | | | | | | | | | | |
| Chromosome | start | end | strand | MM | target_seq | PAM | alignment | distance | position | gene name | gene id |
| GU305914.1 | 222238 | 222260 | - | 0 | CAACAAGTCGCCCCAGAGCC | GGG | \|\|\|\|\|\|\|\|[\|\|\|\|\|\|\|\|\|\|\|\|]PAM | NA | Intergenic | NA | NA |
| T17894 | CCCAGAGCCGGGCGCCGTCGAGG | 1000 | | | | | | | | | |
| Oligo fwd | CACCCCCAGAGCCGGGCGCCGTCG | | | | | | | | | | |
| Oligo rev | AAACCGACGGCGCCCGGCTCTGGG | | | | | | | | | | |
| Chromosome | start | end | strand | MM | target_seq | PAM | alignment | distance | position | gene name | gene id |
| GU305914.1 | 222227 | 222249 | - | 0 | CCCAGAGCCGGGCGCCGTCG | AGG | \|\|\|\|\|\|\|\|[\|\|\|\|\|\|\|\|\|\|\|\|]PAM | NA | Intergenic | NA | NA |
| T17895 | GCCGTCGAGGTCGACGCAGAAGG | 1000 | | | | | | | | | |
| Oligo fwd | CACCGCCGTCGAGGTCGACGCAGA | | | | | | | | | | |
| Oligo rev | AAACTCTGCGTCGACCTCGACGGC | | | | | | | | | | |
| Chromosome | start | end | strand | MM | target_seq | PAM | alignment | distance | position | gene name | gene id |
| GU305914.1 | 222214 | 222236 | - | 0 | GCCGTCGAGGTCGACGCAGA | AGG | \|\|\|\|\|\|\|\|[\|\|\|\|\|\|\|\|\|\|\|\|]PAM | NA | Intergenic | NA | NA |
| T17896 | GACGATCTCAACACGAGCATCGG | 1000 | | | | | | | | | |
| Oligo fwd | CACCGACGATCTCAACACGAGCAT | | | | | | | | | | |
| Oligo rev | AAACATGCTCGTGTTGAGATCGTC | | | | | | | | | | |
| Chromosome | start | end | strand | MM | target_seq | PAM | alignment | distance | position | gene name | gene id |
| GU305914.1 | 222127 | 222149 | - | 0 | GACGATCTCAACACGAGCAT | CGG | \|\|\|\|\|\|\|\|[\|\|\|\|\|\|\|\|\|\|\|\|]PAM | NA | Intergenic | NA | NA |
| T17897 | CTCAACACGAGCATCGGAAAAGG | 1000 | | | | | | | | | |
| Oligo fwd | CACCCTCAACACGAGCATCGGAAA | | | | | | | | | | |
| Oligo rev | AAACTTTCCGATGCTCGTGTTGAG | | | | | | | | | | |
| Chromosome | start | end | strand | MM | target_seq | PAM | alignment | distance | position | gene name | gene id |
| GU305914.1 | 222121 | 222143 | - | 0 | CTCAACACGAGCATCGGAAA | AGG | \|\|\|\|\|\|\|\|[\|\|\|\|\|\|\|\|\|\|\|\|]PAM | NA | Intergenic | NA | NA |
| T17898 | CGGAAAAGGTCCTTTCGTCGCGG | 1000 | | | | | | | | | |
| Oligo fwd | CACCCGGAAAAGGTCCTTTCGTCG | | | | | | | | | | |
| Oligo rev | AAACCGACGAAAGGACCTTTTCCG | | | | | | | | | | |
| Chromosome | start | end | strand | MM | target_seq | PAM | alignment | distance | position | gene name | gene id |
| GU305914.1 | 222107 | 222129 | - | 0 | CGGAAAAGGTCCTTTCGTCG | CGG | \|\|\|\|\|\|\|\|[\|\|\|\|\|\|\|\|\|\|\|\|]PAM | NA | Intergenic | NA | NA |
| T17899 | GCCGTGACCTTTCTGCTCATCGG | 1000 | | | | | | | | | |
| Oligo fwd | CACCGCCGTGACCTTTCTGCTCAT | | | | | | | | | | |
| Oligo rev | AAACATGAGCAGAAAGGTCACGGC | | | | | | | | | | |
| Chromosome | start | end | strand | MM | target_seq | PAM | alignment | distance | position | gene name | gene id |
| GU305914.1 | 222085 | 222107 | - | 0 | GCCGTGACCTTTCTGCTCAT | CGG | \|\|\|\|\|\|\|\|[\|\|\|\|\|\|\|\|\|\|\|\|]PAM | NA | Intergenic | NA | NA |
| T17900 | TTTCTGCTCATCGGACTGATCGG | 1000 | | | | | | | | | |
| Oligo fwd | CACCTTTCTGCTCATCGGACTGAT | | | | | | | | | | |
| Oligo rev | AAACATCAGTCCGATGAGCAGAAA | | | | | | | | | | |
| Chromosome | start | end | strand | MM | target_seq | PAM | alignment | distance | position | gene name | gene id |
| GU305914.1 | 222076 | 222098 | - | 0 | TTTCTGCTCATCGGACTGAT | CGG | \|\|\|\|\|\|\|\|[\|\|\|\|\|\|\|\|\|\|\|\|]PAM | NA | Intergenic | NA | NA |
| T17901 | TTCTGCTCATCGGACTGATCGGG | 1000 | | | | | | | | | |
| Oligo fwd | CACCTTCTGCTCATCGGACTGATC | | | | | | | | | | |
| Oligo rev | AAACGATCAGTCCGATGAGCAGAA | | | | | | | | | | |
| Chromosome | start | end | strand | MM | target_seq | PAM | alignment | distance | position | gene name | gene id |
| GU305914.1 | 222075 | 222097 | - | 0 | TTCTGCTCATCGGACTGATC | GGG | \|\|\|\|\|\|\|\|[\|\|\|\|\|\|\|\|\|\|\|\|]PAM | NA | Intergenic | NA | NA |
| T17902 | TCTGCTCATCGGACTGATCGGGG | 1000 | | | | | | | | | |
| Oligo fwd | CACCTCTGCTCATCGGACTGATCG | | | | | | | | | | |
| Oligo rev | AAACCGATCAGTCCGATGAGCAGA | | | | | | | | | | |
| Chromosome | start | end | strand | MM | target_seq | PAM | alignment | distance | position | gene name | gene id |
| GU305914.1 | 222074 | 222096 | - | 0 | TCTGCTCATCGGACTGATCG | GGG | \|\|\|\|\|\|\|\|[\|\|\|\|\|\|\|\|\|\|\|\|]PAM | NA | Intergenic | NA | NA |
| T17903 | CATCGGACTGATCGGGGCCGCGG | 1000 | | | | | | | | | |
| Oligo fwd | CACCCATCGGACTGATCGGGGCCG | | | | | | | | | | |
| Oligo rev | AAACCGGCCCCGATCAGTCCGATG | | | | | | | | | | |
| Chromosome | start | end | strand | MM | target_seq | PAM | alignment | distance | position | gene name | gene id |
| GU305914.1 | 222068 | 222090 | - | 0 | CATCGGACTGATCGGGGCCG | CGG | \|\|\|\|\|\|\|\|[\|\|\|\|\|\|\|\|\|\|\|\|]PAM | NA | Intergenic | NA | NA |
| T17904 | GATCGGGGCCGCGGTGTTAGCGG | 1000 | | | | | | | | | |
| Oligo fwd | CACCGATCGGGGCCGCGGTGTTAG | | | | | | | | | | |
| Oligo rev | AAACCTAACACCGCGGCCCCGATC | | | | | | | | | | |
| Chromosome | start | end | strand | MM | target_seq | PAM | alignment | distance | position | gene name | gene id |
| GU305914.1 | 222059 | 222081 | - | 0 | GATCGGGGCCGCGGTGTTAG | CGG | \|\|\|\|\|\|\|\|[\|\|\|\|\|\|\|\|\|\|\|\|]PAM | NA | Intergenic | NA | NA |
| T17905 | TAGCGGTCGTGTACCTCAGACGG | 1000 | | | | | | | | | |

| | | | | | | | | | | | |
|---|---|---|---|---|---|---|---|---|---|---|---|
| Oligo fwd | CACCTAGCGGTCGTGTACCTCAGA | | | | | | | | | | |
| Oligo rev | AAACTCTGAGGTACACGACCGCTA | | | | | | | | | | |
| Chromosome | start | end | strand | MM | target_seq | PAM | alignment | distance | position | gene name | gene id |
| GU305914.1 | 222042 | 222064 | - | 0 | TAGCGGTCGTGTACCTCAGA | CGG | \|\|\|\|\|\|\|\|[\|\|\|\|\|\|\|\|\|\|\|\|]PAM | NA | Intergenic | NA | NA |
| T17906 | GTGTACCTCAGACGGCCGAACGG | 1000 | | | | | | | | | |
| Oligo fwd | CACCGTGTACCTCAGACGGCCGAA | | | | | | | | | | |
| Oligo rev | AAACTTCGGCCGTCTGAGGTACAC | | | | | | | | | | |
| Chromosome | start | end | strand | MM | target_seq | PAM | alignment | distance | position | gene name | gene id |
| GU305914.1 | 222034 | 222056 | - | 0 | GTGTACCTCAGACGGCCGAA | CGG | \|\|\|\|\|\|\|\|[\|\|\|\|\|\|\|\|\|\|\|\|]PAM | NA | Intergenic | NA | NA |
| T17907 | CGGCCGAACGGATCGCGATCCGG | 1000 | | | | | | | | | |
| Oligo fwd | CACCCGGCCGAACGGATCGCGATC | | | | | | | | | | |
| Oligo rev | AAACGATCGCGATCCGTTCGGCCG | | | | | | | | | | |
| Chromosome | start | end | strand | MM | target_seq | PAM | alignment | distance | position | gene name | gene id |
| GU305914.1 | 222022 | 222044 | - | 0 | CGGCCGAACGGATCGCGATC | CGG | \|\|\|\|\|\|\|\|[\|\|\|\|\|\|\|\|\|\|\|\|]PAM | NA | Intergenic | NA | NA |
| GU305914.1 | 114787 | 114809 | - | 4 | CCGCCGCGCGGGTCGCGATC | AGG | \|-\|\|\|\|--[\|\|\|-\|\|\|\|\|\|\|\|]PAM | NA | Intergenic | NA | NA |
| T17908 | CCGAACGGATCGCGATCCGGCGG | 1000 | | | | | | | | | |
| Oligo fwd | CACCCCGAACGGATCGCGATCCGG | | | | | | | | | | |
| Oligo rev | AAACCCGGATCGCGATCCGTTCGG | | | | | | | | | | |
| Chromosome | start | end | strand | MM | target_seq | PAM | alignment | distance | position | gene name | gene id |
| GU305914.1 | 222019 | 222041 | - | 0 | CCGAACGGATCGCGATCCGG | CGG | \|\|\|\|\|\|\|\|[\|\|\|\|\|\|\|\|\|\|\|\|]PAM | NA | Intergenic | NA | NA |
| T17909 | CGGCGGTGACAAAGAGAGACTGG | 1000 | | | | | | | | | |
| Oligo fwd | CACCCGGCGGTGACAAAGAGAGAC | | | | | | | | | | |
| Oligo rev | AAACGTCTCTCTTTGTCACCGCCG | | | | | | | | | | |
| Chromosome | start | end | strand | MM | target_seq | PAM | alignment | distance | position | gene name | gene id |
| GU305914.1 | 222002 | 222024 | - | 0 | CGGCGGTGACAAAGAGAGAC | TGG | \|\|\|\|\|\|\|\|[\|\|\|\|\|\|\|\|\|\|\|\|]PAM | NA | Intergenic | NA | NA |
| T17910 | GGCGGTGACAAAGAGAGACTGGG | 1000 | | | | | | | | | |
| Oligo fwd | CACCGGCGGTGACAAAGAGAGACT | | | | | | | | | | |
| Oligo rev | AAACAGTCTCTCTTTGTCACCGCC | | | | | | | | | | |
| Chromosome | start | end | strand | MM | target_seq | PAM | alignment | distance | position | gene name | gene id |
| GU305914.1 | 222001 | 222023 | - | 0 | GGCGGTGACAAAGAGAGACT | GGG | \|\|\|\|\|\|\|\|[\|\|\|\|\|\|\|\|\|\|\|\|]PAM | NA | Intergenic | NA | NA |
| T17911 | TGGGTCCCGTTCCCTATAACCGG | 1000 | | | | | | | | | |
| Oligo fwd | CACCTGGGTCCCGTTCCCTATAAC | | | | | | | | | | |
| Oligo rev | AAACGTTATAGGGAACGGGACCCA | | | | | | | | | | |
| Chromosome | start | end | strand | MM | target_seq | PAM | alignment | distance | position | gene name | gene id |
| GU305914.1 | 221982 | 222004 | - | 0 | TGGGTCCCGTTCCCTATAAC | CGG | \|\|\|\|\|\|\|\|[\|\|\|\|\|\|\|\|\|\|\|\|]PAM | NA | Intergenic | NA | NA |
| T17912 | GGGTCCCGTTCCCTATAACCGGG | 1000 | | | | | | | | | |
| Oligo fwd | CACCGGGTCCCGTTCCCTATAACC | | | | | | | | | | |
| Oligo rev | AAACGGTTATAGGGAACGGGACCC | | | | | | | | | | |
| Chromosome | start | end | strand | MM | target_seq | PAM | alignment | distance | position | gene name | gene id |
| GU305914.1 | 221981 | 222003 | - | 0 | GGGTCCCGTTCCCTATAACC | GGG | \|\|\|\|\|\|\|\|[\|\|\|\|\|\|\|\|\|\|\|\|]PAM | NA | Intergenic | NA | NA |
| T17913 | ACGTGTTTCTAATAAAACACTGG | 1000 | | | | | | | | | |
| Oligo fwd | CACCACGTGTTTCTAATAAAACAC | | | | | | | | | | |
| Oligo rev | AAACGTGTTTTATTAGAAACACGT | | | | | | | | | | |
| Chromosome | start | end | strand | MM | target_seq | PAM | alignment | distance | position | gene name | gene id |
| GU305914.1 | 221944 | 221966 | - | 0 | ACGTGTTTCTAATAAAACAC | TGG | \|\|\|\|\|\|\|\|[\|\|\|\|\|\|\|\|\|\|\|\|]PAM | NA | Intergenic | NA | NA |
| T17914 | GACCTTAAGTGTCACTCGTTAGG | 1000 | | | | | | | | | |
| Oligo fwd | CACCGACCTTAAGTGTCACTCGTT | | | | | | | | | | |
| Oligo rev | AAACAACGAGTGACACTTAAGGTC | | | | | | | | | | |
| Chromosome | start | end | strand | MM | target_seq | PAM | alignment | distance | position | gene name | gene id |
| GU305914.1 | 221869 | 221891 | - | 0 | GACCTTAAGTGTCACTCGTT | AGG | \|\|\|\|\|\|\|\|[\|\|\|\|\|\|\|\|\|\|\|\|]PAM | NA | Intergenic | NA | NA |
| T17915 | AGTGTCACTCGTTAGGCACCCGG | 1000 | | | | | | | | | |
| Oligo fwd | CACCAGTGTCACTCGTTAGGCACC | | | | | | | | | | |
| Oligo rev | AAACGGTGCCTAACGAGTGACACT | | | | | | | | | | |
| Chromosome | start | end | strand | MM | target_seq | PAM | alignment | distance | position | gene name | gene id |
| GU305914.1 | 221862 | 221884 | - | 0 | AGTGTCACTCGTTAGGCACC | CGG | \|\|\|\|\|\|\|\|[\|\|\|\|\|\|\|\|\|\|\|\|]PAM | NA | Intergenic | NA | NA |
| T17916 | GGCACCCGGCCGTTCACTGTCGG | 1000 | | | | | | | | | |
| Oligo fwd | CACCGGCACCCGGCCGTTCACTGT | | | | | | | | | | |
| Oligo rev | AAACACAGTGAACGGCCGGGTGCC | | | | | | | | | | |
| Chromosome | start | end | strand | MM | target_seq | PAM | alignment | distance | position | gene name | gene id |
| GU305914.1 | 221848 | 221870 | - | 0 | GGCACCCGGCCGTTCACTGT | CGG | \|\|\|\|\|\|\|\|[\|\|\|\|\|\|\|\|\|\|\|\|]PAM | NA | Intergenic | NA | NA |
| T17917 | GCACCCGGCCGTTCACTGTCGGG | 1000 | | | | | | | | | |
| Oligo fwd | CACCGCACCCGGCCGTTCACTGTC | | | | | | | | | | |
| Oligo rev | AAACGACAGTGAACGGCCGGGTGC | | | | | | | | | | |
| Chromosome | start | end | strand | MM | target_seq | PAM | alignment | distance | position | gene name | gene id |
| GU305914.1 | 221847 | 221869 | - | 0 | GCACCCGGCCGTTCACTGTC | GGG | \|\|\|\|\|\|\|\|[\|\|\|\|\|\|\|\|\|\|\|\|]PAM | NA | Intergenic | NA | NA |
| T17918 | CCCGGCCGTTCACTGTCGGGTGG | 1000 | | | | | | | | | |
| Oligo fwd | CACCCCCGGCCGTTCACTGTCGGG | | | | | | | | | | |
| Oligo rev | AAACCCCGACAGTGAACGGCCGGG | | | | | | | | | | |
| Chromosome | start | end | strand | MM | target_seq | PAM | alignment | distance | position | gene name | gene id |
| GU305914.1 | 221844 | 221866 | - | 0 | CCCGGCCGTTCACTGTCGGG | TGG | \|\|\|\|\|\|\|\|[\|\|\|\|\|\|\|\|\|\|\|\|]PAM | NA | Intergenic | NA | NA |
| T17919 | ACTGTCGGGTGGCTTTCGACCGG | 1000 | | | | | | | | | |
| Oligo fwd | CACCACTGTCGGGTGGCTTTCGAC | | | | | | | | | | |
| Oligo rev | AAACGTCGAAAGCCACCCGACAGT | | | | | | | | | | |
| Chromosome | start | end | strand | MM | target_seq | PAM | alignment | distance | position | gene name | gene id |
| GU305914.1 | 221833 | 221855 | - | 0 | ACTGTCGGGTGGCTTTCGAC | CGG | \|\|\|\|\|\|\|\|[\|\|\|\|\|\|\|\|\|\|\|\|]PAM | NA | Intergenic | NA | NA |

<table>
<tr><td>T17920</td><td>TGGCTTTCGACCGGCTCGAACGG</td><td></td><td>1000</td><td></td><td></td><td></td><td></td><td></td><td></td><td></td><td></td><td></td></tr>
<tr><td>Oligo fwd</td><td>CACCTGGCTTTCGACCGGCTCGAA</td><td></td><td></td><td></td><td></td><td></td><td></td><td></td><td></td><td></td><td></td><td></td></tr>
<tr><td>Oligo rev</td><td>AAACTTCGAGCCGGTCGAAAGCCA</td><td></td><td></td><td></td><td></td><td></td><td></td><td></td><td></td><td></td><td></td><td></td></tr>
<tr><td>Chromosome</td><td>start</td><td></td><td>end</td><td>strand</td><td>MM</td><td>target_seq</td><td>PAM</td><td>alignment</td><td>distance</td><td>position</td><td>gene name</td><td>gene id</td></tr>
<tr><td>GU305914.1</td><td></td><td>221824</td><td>221846</td><td>-</td><td>0</td><td>TGGCTTTCGACCGGCTCGAA</td><td>CGG</td><td>||||||||[||||||||||||]PAM</td><td>NA</td><td>Intergenic</td><td>NA</td><td>NA</td></tr>
<tr><td>T17921</td><td>CTTTCGACCGGCTCGAACGGTGG</td><td></td><td>1000</td><td></td><td></td><td></td><td></td><td></td><td></td><td></td><td></td><td></td></tr>
<tr><td>Oligo fwd</td><td>CACCCTTTCGACCGGCTCGAACGG</td><td></td><td></td><td></td><td></td><td></td><td></td><td></td><td></td><td></td><td></td><td></td></tr>
<tr><td>Oligo rev</td><td>AAACCCGTTCGAGCCGGTCGAAAG</td><td></td><td></td><td></td><td></td><td></td><td></td><td></td><td></td><td></td><td></td><td></td></tr>
<tr><td>Chromosome</td><td>start</td><td></td><td>end</td><td>strand</td><td>MM</td><td>target_seq</td><td>PAM</td><td>alignment</td><td>distance</td><td>position</td><td>gene name</td><td>gene id</td></tr>
<tr><td>GU305914.1</td><td></td><td>221821</td><td>221843</td><td>-</td><td>0</td><td>CTTTCGACCGGCTCGAACGG</td><td>TGG</td><td>||||||||[||||||||||||]PAM</td><td>NA</td><td>Intergenic</td><td>NA</td><td>NA</td></tr>
<tr><td>T17922</td><td>CCGGCTCGAACGGTGGATCGCGG</td><td></td><td>1000</td><td></td><td></td><td></td><td></td><td></td><td></td><td></td><td></td><td></td></tr>
<tr><td>Oligo fwd</td><td>CACCCCGGCTCGAACGGTGGATCG</td><td></td><td></td><td></td><td></td><td></td><td></td><td></td><td></td><td></td><td></td><td></td></tr>
<tr><td>Oligo rev</td><td>AAACCGATCCACCGTTCGAGCCGG</td><td></td><td></td><td></td><td></td><td></td><td></td><td></td><td></td><td></td><td></td><td></td></tr>
<tr><td>Chromosome</td><td>start</td><td></td><td>end</td><td>strand</td><td>MM</td><td>target_seq</td><td>PAM</td><td>alignment</td><td>distance</td><td>position</td><td>gene name</td><td>gene id</td></tr>
<tr><td>GU305914.1</td><td></td><td>221814</td><td>221836</td><td>-</td><td>0</td><td>CCGGCTCGAACGGTGGATCG</td><td>CGG</td><td>||||||||[||||||||||||]PAM</td><td>NA</td><td>Intergenic</td><td>NA</td><td>NA</td></tr>
<tr><td>T17923</td><td>CTGTACGTCCTACCGAATCGCGG</td><td></td><td>1000</td><td></td><td></td><td></td><td></td><td></td><td></td><td></td><td></td><td></td></tr>
<tr><td>Oligo fwd</td><td>CACCCTGTACGTCCTACCGAATCG</td><td></td><td></td><td></td><td></td><td></td><td></td><td></td><td></td><td></td><td></td><td></td></tr>
<tr><td>Oligo rev</td><td>AAACCGATTCGGTAGGACGTACAG</td><td></td><td></td><td></td><td></td><td></td><td></td><td></td><td></td><td></td><td></td><td></td></tr>
<tr><td>Chromosome</td><td>start</td><td></td><td>end</td><td>strand</td><td>MM</td><td>target_seq</td><td>PAM</td><td>alignment</td><td>distance</td><td>position</td><td>gene name</td><td>gene id</td></tr>
<tr><td>GU305914.1</td><td></td><td>221768</td><td>221790</td><td>-</td><td>0</td><td>CTGTACGTCCTACCGAATCG</td><td>CGG</td><td>||||||||[||||||||||||]PAM</td><td>NA</td><td>Intergenic</td><td>NA</td><td>NA</td></tr>
<tr><td>T17924</td><td>TTATTCCGATCACACCGTCTTGG</td><td></td><td>1000</td><td></td><td></td><td></td><td></td><td></td><td></td><td></td><td></td><td></td></tr>
<tr><td>Oligo fwd</td><td>CACCTTATTCCGATCACACCGTCT</td><td></td><td></td><td></td><td></td><td></td><td></td><td></td><td></td><td></td><td></td><td></td></tr>
<tr><td>Oligo rev</td><td>AAACAGACGGTGTGATCGGAATAA</td><td></td><td></td><td></td><td></td><td></td><td></td><td></td><td></td><td></td><td></td><td></td></tr>
<tr><td>Chromosome</td><td>start</td><td></td><td>end</td><td>strand</td><td>MM</td><td>target_seq</td><td>PAM</td><td>alignment</td><td>distance</td><td>position</td><td>gene name</td><td>gene id</td></tr>
<tr><td>GU305914.1</td><td></td><td>221703</td><td>221725</td><td>-</td><td>0</td><td>TTATTCCGATCACACCGTCT</td><td>TGG</td><td>||||||||[||||||||||||]PAM</td><td>NA</td><td>Intergenic</td><td>NA</td><td>NA</td></tr>
<tr><td>T17925</td><td>GTCGCAGATTTTAACGTCGACGG</td><td></td><td>1000</td><td></td><td></td><td></td><td></td><td></td><td></td><td></td><td></td><td></td></tr>
<tr><td>Oligo fwd</td><td>CACCGTCGCAGATTTTAACGTCGA</td><td></td><td></td><td></td><td></td><td></td><td></td><td></td><td></td><td></td><td></td><td></td></tr>
<tr><td>Oligo rev</td><td>AAACTCGACGTTAAAATCTGCGAC</td><td></td><td></td><td></td><td></td><td></td><td></td><td></td><td></td><td></td><td></td><td></td></tr>
<tr><td>Chromosome</td><td>start</td><td></td><td>end</td><td>strand</td><td>MM</td><td>target_seq</td><td>PAM</td><td>alignment</td><td>distance</td><td>position</td><td>gene name</td><td>gene id</td></tr>
<tr><td>GU305914.1</td><td></td><td>221673</td><td>221695</td><td>-</td><td>0</td><td>GTCGCAGATTTTAACGTCGA</td><td>CGG</td><td>||||||||[||||||||||||]PAM</td><td>NA</td><td>Intergenic</td><td>NA</td><td>NA</td></tr>
<tr><td>T17926</td><td>GCAGATTTTAACGTCGACGGTGG</td><td></td><td>1000</td><td></td><td></td><td></td><td></td><td></td><td></td><td></td><td></td><td></td></tr>
<tr><td>Oligo fwd</td><td>CACCGCAGATTTTAACGTCGACGG</td><td></td><td></td><td></td><td></td><td></td><td></td><td></td><td></td><td></td><td></td><td></td></tr>
<tr><td>Oligo rev</td><td>AAACCCGTCGACGTTAAAATCTGC</td><td></td><td></td><td></td><td></td><td></td><td></td><td></td><td></td><td></td><td></td><td></td></tr>
<tr><td>Chromosome</td><td>start</td><td></td><td>end</td><td>strand</td><td>MM</td><td>target_seq</td><td>PAM</td><td>alignment</td><td>distance</td><td>position</td><td>gene name</td><td>gene id</td></tr>
<tr><td>GU305914.1</td><td></td><td>221670</td><td>221692</td><td>-</td><td>0</td><td>GCAGATTTTAACGTCGACGG</td><td>TGG</td><td>||||||||[||||||||||||]PAM</td><td>NA</td><td>Intergenic</td><td>NA</td><td>NA</td></tr>
<tr><td>T17927</td><td>GTGGTCGCCAGTAGCGCGTTAGG</td><td></td><td>1000</td><td></td><td></td><td></td><td></td><td></td><td></td><td></td><td></td><td></td></tr>
<tr><td>Oligo fwd</td><td>CACCGTGGTCGCCAGTAGCGCGTT</td><td></td><td></td><td></td><td></td><td></td><td></td><td></td><td></td><td></td><td></td><td></td></tr>
<tr><td>Oligo rev</td><td>AAACAACGCGCTACTGGCGACCAC</td><td></td><td></td><td></td><td></td><td></td><td></td><td></td><td></td><td></td><td></td><td></td></tr>
<tr><td>Chromosome</td><td>start</td><td></td><td>end</td><td>strand</td><td>MM</td><td>target_seq</td><td>PAM</td><td>alignment</td><td>distance</td><td>position</td><td>gene name</td><td>gene id</td></tr>
<tr><td>GU305914.1</td><td></td><td>221651</td><td>221673</td><td>-</td><td>0</td><td>GTGGTCGCCAGTAGCGCGTT</td><td>AGG</td><td>||||||||[||||||||||||]PAM</td><td>NA</td><td>Intergenic</td><td>NA</td><td>NA</td></tr>
<tr><td>T17928</td><td>TGGTCGCCAGTAGCGCGTTAGGG</td><td></td><td>1000</td><td></td><td></td><td></td><td></td><td></td><td></td><td></td><td></td><td></td></tr>
<tr><td>Oligo fwd</td><td>CACCTGGTCGCCAGTAGCGCGTTA</td><td></td><td></td><td></td><td></td><td></td><td></td><td></td><td></td><td></td><td></td><td></td></tr>
<tr><td>Oligo rev</td><td>AAACTAACGCGCTACTGGCGACCA</td><td></td><td></td><td></td><td></td><td></td><td></td><td></td><td></td><td></td><td></td><td></td></tr>
<tr><td>Chromosome</td><td>start</td><td></td><td>end</td><td>strand</td><td>MM</td><td>target_seq</td><td>PAM</td><td>alignment</td><td>distance</td><td>position</td><td>gene name</td><td>gene id</td></tr>
<tr><td>GU305914.1</td><td></td><td>221650</td><td>221672</td><td>-</td><td>0</td><td>TGGTCGCCAGTAGCGCGTTA</td><td>GGG</td><td>||||||||[||||||||||||]PAM</td><td>NA</td><td>Intergenic</td><td>NA</td><td>NA</td></tr>
<tr><td>T17929</td><td>GGTCGCCAGTAGCGCGTTAGGGG</td><td></td><td>1000</td><td></td><td></td><td></td><td></td><td></td><td></td><td></td><td></td><td></td></tr>
<tr><td>Oligo fwd</td><td>CACCGGTCGCCAGTAGCGCGTTAG</td><td></td><td></td><td></td><td></td><td></td><td></td><td></td><td></td><td></td><td></td><td></td></tr>
<tr><td>Oligo rev</td><td>AAACCTAACGCGCTACTGGCGACC</td><td></td><td></td><td></td><td></td><td></td><td></td><td></td><td></td><td></td><td></td><td></td></tr>
<tr><td>Chromosome</td><td>start</td><td></td><td>end</td><td>strand</td><td>MM</td><td>target_seq</td><td>PAM</td><td>alignment</td><td>distance</td><td>position</td><td>gene name</td><td>gene id</td></tr>
<tr><td>GU305914.1</td><td></td><td>221649</td><td>221671</td><td>-</td><td>0</td><td>GGTCGCCAGTAGCGCGTTAG</td><td>GGG</td><td>||||||||[||||||||||||]PAM</td><td>NA</td><td>Intergenic</td><td>NA</td><td>NA</td></tr>
<tr><td>T17930</td><td>GTCGCCAGTAGCGCGTTAGGGGG</td><td></td><td>1000</td><td></td><td></td><td></td><td></td><td></td><td></td><td></td><td></td><td></td></tr>
<tr><td>Oligo fwd</td><td>CACCGTCGCCAGTAGCGCGTTAGG</td><td></td><td></td><td></td><td></td><td></td><td></td><td></td><td></td><td></td><td></td><td></td></tr>
<tr><td>Oligo rev</td><td>AAACCCTAACGCGCTACTGGCGAC</td><td></td><td></td><td></td><td></td><td></td><td></td><td></td><td></td><td></td><td></td><td></td></tr>
<tr><td>Chromosome</td><td>start</td><td></td><td>end</td><td>strand</td><td>MM</td><td>target_seq</td><td>PAM</td><td>alignment</td><td>distance</td><td>position</td><td>gene name</td><td>gene id</td></tr>
<tr><td>GU305914.1</td><td></td><td>221648</td><td>221670</td><td>-</td><td>0</td><td>GTCGCCAGTAGCGCGTTAGG</td><td>GGG</td><td>||||||||[||||||||||||]PAM</td><td>NA</td><td>Intergenic</td><td>NA</td><td>NA</td></tr>
<tr><td>T17931</td><td>TCGCCAGTAGCGCGTTAGGGGGG</td><td></td><td>1000</td><td></td><td></td><td></td><td></td><td></td><td></td><td></td><td></td><td></td></tr>
<tr><td>Oligo fwd</td><td>CACCTCGCCAGTAGCGCGTTAGGG</td><td></td><td></td><td></td><td></td><td></td><td></td><td></td><td></td><td></td><td></td><td></td></tr>
<tr><td>Oligo rev</td><td>AAACCCCTAACGCGCTACTGGCGA</td><td></td><td></td><td></td><td></td><td></td><td></td><td></td><td></td><td></td><td></td><td></td></tr>
<tr><td>Chromosome</td><td>start</td><td></td><td>end</td><td>strand</td><td>MM</td><td>target_seq</td><td>PAM</td><td>alignment</td><td>distance</td><td>position</td><td>gene name</td><td>gene id</td></tr>
<tr><td>GU305914.1</td><td></td><td>221647</td><td>221669</td><td>-</td><td>0</td><td>TCGCCAGTAGCGCGTTAGGG</td><td>GGG</td><td>||||||||[||||||||||||]PAM</td><td>NA</td><td>Intergenic</td><td>NA</td><td>NA</td></tr>
<tr><td>T17932</td><td>GTTATCGAAAACGTGACGACCGG</td><td></td><td>1000</td><td></td><td></td><td></td><td></td><td></td><td></td><td></td><td></td><td></td></tr>
<tr><td>Oligo fwd</td><td>CACCGTTATCGAAAACGTGACGAC</td><td></td><td></td><td></td><td></td><td></td><td></td><td></td><td></td><td></td><td></td><td></td></tr>
<tr><td>Oligo rev</td><td>AAACGTCGTCACGTTTTCGATAAC</td><td></td><td></td><td></td><td></td><td></td><td></td><td></td><td></td><td></td><td></td><td></td></tr>
<tr><td>Chromosome</td><td>start</td><td></td><td>end</td><td>strand</td><td>MM</td><td>target_seq</td><td>PAM</td><td>alignment</td><td>distance</td><td>position</td><td>gene name</td><td>gene id</td></tr>
<tr><td>GU305914.1</td><td></td><td>221621</td><td>221643</td><td>-</td><td>0</td><td>GTTATCGAAAACGTGACGAC</td><td>CGG</td><td>||||||||[||||||||||||]PAM</td><td>NA</td><td>Intergenic</td><td>NA</td><td>NA</td></tr>
<tr><td>T17933</td><td>ATCGAAAACGTGACGACCGGAGG</td><td></td><td>1000</td><td></td><td></td><td></td><td></td><td></td><td></td><td></td><td></td><td></td></tr>
<tr><td>Oligo fwd</td><td>CACCATCGAAAACGTGACGACCGG</td><td></td><td></td><td></td><td></td><td></td><td></td><td></td><td></td><td></td><td></td><td></td></tr>
<tr><td>Oligo rev</td><td>AAACCCGGTCGTCACGTTTTCGAT</td><td></td><td></td><td></td><td></td><td></td><td></td><td></td><td></td><td></td><td></td><td></td></tr>
<tr><td>Chromosome</td><td>start</td><td></td><td>end</td><td>strand</td><td>MM</td><td>target_seq</td><td>PAM</td><td>alignment</td><td>distance</td><td>position</td><td>gene name</td><td>gene id</td></tr>
<tr><td>GU305914.1</td><td></td><td>221618</td><td>221640</td><td>-</td><td>0</td><td>ATCGAAAACGTGACGACCGG</td><td>AGG</td><td>||||||||[||||||||||||]PAM</td><td>NA</td><td>Intergenic</td><td>NA</td><td>NA</td></tr>
<tr><td>T17934</td><td>AACGTGACGACCGGAGGCCGTGG</td><td></td><td>1000</td><td></td><td></td><td></td><td></td><td></td><td></td><td></td><td></td><td></td></tr>
<tr><td>Oligo fwd</td><td>CACCAACGTGACGACCGGAGGCCG</td><td></td><td></td><td></td><td></td><td></td><td></td><td></td><td></td><td></td><td></td><td></td></tr>
<tr><td>Oligo rev</td><td>AAACCGGCCTCCGGTCGTCACGTT</td><td></td><td></td><td></td><td></td><td></td><td></td><td></td><td></td><td></td><td></td><td></td></tr>
<tr><td>Chromosome</td><td>start</td><td></td><td>end</td><td>strand</td><td>MM</td><td>target_seq</td><td>PAM</td><td>alignment</td><td>distance</td><td>position</td><td>gene name</td><td>gene id</td></tr>
</table>

| | | | | | | | | | | | |
|---|---|---|---|---|---|---|---|---|---|---|---|
| GU305914.1 | 221612 | 221634 | - | 0 | AACGTGACGACCGGAGGCCG | TGG | ||||||||[||||||||||||]PAM | NA | Intergenic | NA | NA |
| T17935 | GTGACGACCGGAGGCCGTGGTGG | 1000 | | | | | | | | | |
| Oligo fwd | CACCGTGACGACCGGAGGCCGTGG | | | | | | | | | | |
| Oligo rev | AAACCCACGGCCTCCGGTCGTCAC | | | | | | | | | | |
| Chromosome | start | end | strand | MM | target_seq | PAM | alignment | distance | position | gene name | gene id |
| GU305914.1 | 221609 | 221631 | - | 0 | GTGACGACCGGAGGCCGTGG | TGG | ||||||||[||||||||||||]PAM | NA | Intergenic | NA | NA |
| T17936 | ACGACCGGAGGCCGTGGTGGCGG | 1000 | | | | | | | | | |
| Oligo fwd | CACCACGACCGGAGGCCGTGGTGG | | | | | | | | | | |
| Oligo rev | AAACCCACCACGGCCTCCGGTCGT | | | | | | | | | | |
| Chromosome | start | end | strand | MM | target_seq | PAM | alignment | distance | position | gene name | gene id |
| GU305914.1 | 221606 | 221628 | - | 0 | ACGACCGGAGGCCGTGGTGG | CGG | ||||||||[||||||||||||]PAM | NA | Intergenic | NA | NA |
| T17937 | ACCGGAGGCCGTGGTGGCGGCGG | 1000 | | | | | | | | | |
| Oligo fwd | CACCACCGGAGGCCGTGGTGGCGG | | | | | | | | | | |
| Oligo rev | AAACCCGCCACCACGGCCTCCGGT | | | | | | | | | | |
| Chromosome | start | end | strand | MM | target_seq | PAM | alignment | distance | position | gene name | gene id |
| GU305914.1 | 221603 | 221625 | - | 0 | ACCGGAGGCCGTGGTGGCGG | CGG | ||||||||[||||||||||||]PAM | NA | Intergenic | NA | NA |
| GU305914.1 | 161760 | 161782 | - | 3 | CCCGGAGGCCGAGGAGGCGG | CGG | -|||||||[|||-||-|||||]PAM | NA | Intergenic | NA | NA |
| GU305914.1 | 164854 | 164876 | - | 4 | AGCGGCGGCCGTGGTGGCTC | TGG | |-|||-||[||||||||||--]PAM | NA | Intergenic | NA | NA |
| T17938 | GAGGCCGTGGTGGCGGCGGCAGG | 1000 | | | | | | | | | |
| Oligo fwd | CACCGAGGCCGTGGTGGCGGCGGC | | | | | | | | | | |
| Oligo rev | AAACGCCGCCGCCACCACGGCCTC | | | | | | | | | | |
| Chromosome | start | end | strand | MM | target_seq | PAM | alignment | distance | position | gene name | gene id |
| GU305914.1 | 221599 | 221621 | - | 0 | GAGGCCGTGGTGGCGGCGGC | AGG | ||||||||[||||||||||||]PAM | NA | Intergenic | NA | NA |
| GU305914.1 | 114657 | 114679 | - | 3 | GCGGTGGTGGTGGCGGCGGC | GGG | |-||--||[||||||||||||]PAM | NA | Intergenic | NA | NA |
| GU305914.1 | 48960 | 48982 | - | 4 | CATGACGGGGTGGCGGCGGC | GGG | -|-|-||-[||||||||||||]PAM | NA | Intergenic | NA | NA |
| GU305914.1 | 86780 | 86802 | - | 4 | GAGGGGGCGGGGGCGGCGGC | AGG | ||||--|-[||-|||||||||]PAM | NA | Intergenic | NA | NA |
| GU305914.1 | 62444 | 62466 | + | 4 | GCGGCGGGGGTGGTGGCGGC | GGG | |-|||-|-[|||||-||||||]PAM | NA | Intergenic | NA | NA |
| GU305914.1 | 90246 | 90268 | - | 4 | GAGGCGGCGGAGCCGGCGGC | AGG | |||||-|-[||-|-|||||||]PAM | NA | Intergenic | NA | NA |
| GU305914.1 | 48160 | 48182 | + | 4 | GCGGCGGTGGCGGCGGCGGT | GGG | |-|||-||[||-||||||||-]PAM | NA | Intergenic | NA | NA |
| T17939 | AGGCCGTGGTGGCGGCGGCAGGG | 1000 | | | | | | | | | |
| Oligo fwd | CACCAGGCCGTGGTGGCGGCGGCA | | | | | | | | | | |
| Oligo rev | AAACTGCCGCCGCCACCACGGCCT | | | | | | | | | | |
| Chromosome | start | end | strand | MM | target_seq | PAM | alignment | distance | position | gene name | gene id |
| GU305914.1 | 221598 | 221620 | - | 0 | AGGCCGTGGTGGCGGCGGCA | GGG | ||||||||[||||||||||||]PAM | NA | Intergenic | NA | NA |
| GU305914.1 | 86779 | 86801 | - | 4 | AGGGGGCGGGGGCGGCGGCA | GGG | |||--|-|[|-||||||||||]PAM | NA | Intergenic | NA | NA |
| GU305914.1 | 48959 | 48981 | - | 4 | ATGACGGGGTGGCGGCGGCG | GGG | |-|-||-|[|||||||||||-]PAM | NA | Intergenic | NA | NA |
| T17940 | CGGCGGCAGGGTTCATATCCTGG | 1000 | | | | | | | | | |
| Oligo fwd | CACCCGGCGGCAGGGTTCATATCC | | | | | | | | | | |
| Oligo rev | AAACGGATATGAACCCTGCCGCCG | | | | | | | | | | |
| Chromosome | start | end | strand | MM | target_seq | PAM | alignment | distance | position | gene name | gene id |
| GU305914.1 | 221586 | 221608 | - | 0 | CGGCGGCAGGGTTCATATCC | TGG | ||||||||[||||||||||||]PAM | NA | Intergenic | NA | NA |
| T17941 | ATCCTGGCATATCCTTCGAACGG | 1000 | | | | | | | | | |
| Oligo fwd | CACCATCCTGGCATATCCTTCGAA | | | | | | | | | | |
| Oligo rev | AAACTTCGAAGGATATGCCAGGAT | | | | | | | | | | |
| Chromosome | start | end | strand | MM | target_seq | PAM | alignment | distance | position | gene name | gene id |
| GU305914.1 | 221570 | 221592 | - | 0 | ATCCTGGCATATCCTTCGAA | CGG | ||||||||[||||||||||||]PAM | NA | Intergenic | NA | NA |
| T17942 | GGCATATCCTTCGAACGGTTTGG | 1000 | | | | | | | | | |
| Oligo fwd | CACCGGCATATCCTTCGAACGGTT | | | | | | | | | | |
| Oligo rev | AAACAACCGTTCGAAGGATATGCC | | | | | | | | | | |
| Chromosome | start | end | strand | MM | target_seq | PAM | alignment | distance | position | gene name | gene id |
| GU305914.1 | 221565 | 221587 | - | 0 | GGCATATCCTTCGAACGGTT | TGG | ||||||||[||||||||||||]PAM | NA | Intergenic | NA | NA |
| T17943 | GCATATCCTTCGAACGGTTTGGG | 1000 | | | | | | | | | |
| Oligo fwd | CACCGCATATCCTTCGAACGGTTT | | | | | | | | | | |
| Oligo rev | AAACAAACCGTTCGAAGGATATGC | | | | | | | | | | |
| Chromosome | start | end | strand | MM | target_seq | PAM | alignment | distance | position | gene name | gene id |
| GU305914.1 | 221564 | 221586 | - | 0 | GCATATCCTTCGAACGGTTT | GGG | ||||||||[||||||||||||]PAM | NA | Intergenic | NA | NA |
| T17944 | TATCCTTCGAACGGTTTGGGAGG | 1000 | | | | | | | | | |
| Oligo fwd | CACCTATCCTTCGAACGGTTTGGG | | | | | | | | | | |
| Oligo rev | AAACCCCAAACCGTTCGAAGGATA | | | | | | | | | | |
| Chromosome | start | end | strand | MM | target_seq | PAM | alignment | distance | position | gene name | gene id |
| GU305914.1 | 221561 | 221583 | - | 0 | TATCCTTCGAACGGTTTGGG | AGG | ||||||||[||||||||||||]PAM | NA | Intergenic | NA | NA |
| T17945 | CCGCCGATCACCACAACCAACGG | 1000 | | | | | | | | | |
| Oligo fwd | CACCCCGCCGATCACCACAACCAA | | | | | | | | | | |
| Oligo rev | AAACTTGGTTGTGGTGATCGGCGG | | | | | | | | | | |
| Chromosome | start | end | strand | MM | target_seq | PAM | alignment | distance | position | gene name | gene id |
| GU305914.1 | 221537 | 221559 | - | 0 | CCGCCGATCACCACAACCAA | CGG | ||||||||[||||||||||||]PAM | NA | Intergenic | NA | NA |
| T17946 | ATCACCACAACCAACGGTCTCGG | 1000 | | | | | | | | | |
| Oligo fwd | CACCATCACCACAACCAACGGTCT | | | | | | | | | | |
| Oligo rev | AAACAGACCGTTGGTTGTGGTGAT | | | | | | | | | | |
| Chromosome | start | end | strand | MM | target_seq | PAM | alignment | distance | position | gene name | gene id |
| GU305914.1 | 221531 | 221553 | - | 0 | ATCACCACAACCAACGGTCT | CGG | ||||||||[||||||||||||]PAM | NA | Intergenic | NA | NA |
| T17947 | CACAACCAACGGTCTCGGCTCGG | 1000 | | | | | | | | | |
| Oligo fwd | CACCCACAACCAACGGTCTCGGCT | | | | | | | | | | |
| Oligo rev | AAACAGCCGAGACCGTTGGTTGTG | | | | | | | | | | |
| Chromosome | start | end | strand | MM | target_seq | PAM | alignment | distance | position | gene name | gene id |
| GU305914.1 | 221526 | 221548 | - | 0 | CACAACCAACGGTCTCGGCT | CGG | ||||||||[||||||||||||]PAM | NA | Intergenic | NA | NA |

<table>
<tr><td>T17948</td><td>GGCTCGGTGATTTACACGAGCGG</td><td>1000</td><td></td><td></td><td></td><td></td><td></td><td></td><td></td><td></td><td></td></tr>
<tr><td>Oligo fwd</td><td>CACCGGCTCGGTGATTTACACGAG</td><td></td><td></td><td></td><td></td><td></td><td></td><td></td><td></td><td></td><td></td></tr>
<tr><td>Oligo rev</td><td>AAACCTCGTGTAAATCACCGAGCC</td><td></td><td></td><td></td><td></td><td></td><td></td><td></td><td></td><td></td><td></td></tr>
<tr><td>Chromosome</td><td>start</td><td>end</td><td>strand</td><td>MM</td><td>target_seq</td><td>PAM</td><td>alignment</td><td>distance</td><td>position</td><td>gene name</td><td>gene id</td></tr>
<tr><td>GU305914.1</td><td>221510</td><td>221532</td><td>-</td><td>0</td><td>GGCTCGGTGATTTACACGAG</td><td>CGG</td><td>||||||||[||||||||||||]PAM</td><td>NA</td><td>Intergenic</td><td>NA</td><td>NA</td></tr>
<tr><td>T17949</td><td>TTACACGAGCGGTAGTTACACGG</td><td>1000</td><td></td><td></td><td></td><td></td><td></td><td></td><td></td><td></td><td></td></tr>
<tr><td>Oligo fwd</td><td>CACCTTACACGAGCGGTAGTTACA</td><td></td><td></td><td></td><td></td><td></td><td></td><td></td><td></td><td></td><td></td></tr>
<tr><td>Oligo rev</td><td>AAACTGTAACTACCGCTCGTGTAA</td><td></td><td></td><td></td><td></td><td></td><td></td><td></td><td></td><td></td><td></td></tr>
<tr><td>Chromosome</td><td>start</td><td>end</td><td>strand</td><td>MM</td><td>target_seq</td><td>PAM</td><td>alignment</td><td>distance</td><td>position</td><td>gene name</td><td>gene id</td></tr>
<tr><td>GU305914.1</td><td>221499</td><td>221521</td><td>-</td><td>0</td><td>TTACACGAGCGGTAGTTACA</td><td>CGG</td><td>||||||||[||||||||||||]PAM</td><td>NA</td><td>Intergenic</td><td>NA</td><td>NA</td></tr>
<tr><td>T17950</td><td>GAGCGGTAGTTACACGGCACTGG</td><td>1000</td><td></td><td></td><td></td><td></td><td></td><td></td><td></td><td></td><td></td></tr>
<tr><td>Oligo fwd</td><td>CACCGAGCGGTAGTTACACGGCAC</td><td></td><td></td><td></td><td></td><td></td><td></td><td></td><td></td><td></td><td></td></tr>
<tr><td>Oligo rev</td><td>AAACGTGCCGTGTAACTACCGCTC</td><td></td><td></td><td></td><td></td><td></td><td></td><td></td><td></td><td></td><td></td></tr>
<tr><td>Chromosome</td><td>start</td><td>end</td><td>strand</td><td>MM</td><td>target_seq</td><td>PAM</td><td>alignment</td><td>distance</td><td>position</td><td>gene name</td><td>gene id</td></tr>
<tr><td>GU305914.1</td><td>221493</td><td>221515</td><td>-</td><td>0</td><td>GAGCGGTAGTTACACGGCAC</td><td>TGG</td><td>||||||||[||||||||||||]PAM</td><td>NA</td><td>Intergenic</td><td>NA</td><td>NA</td></tr>
<tr><td>T17951</td><td>TAGTTACACGGCACTGGCCGTGG</td><td>1000</td><td></td><td></td><td></td><td></td><td></td><td></td><td></td><td></td><td></td></tr>
<tr><td>Oligo fwd</td><td>CACCTAGTTACACGGCACTGGCCG</td><td></td><td></td><td></td><td></td><td></td><td></td><td></td><td></td><td></td><td></td></tr>
<tr><td>Oligo rev</td><td>AAACCGGCCAGTGCCGTGTAACTA</td><td></td><td></td><td></td><td></td><td></td><td></td><td></td><td></td><td></td><td></td></tr>
<tr><td>Chromosome</td><td>start</td><td>end</td><td>strand</td><td>MM</td><td>target_seq</td><td>PAM</td><td>alignment</td><td>distance</td><td>position</td><td>gene name</td><td>gene id</td></tr>
<tr><td>GU305914.1</td><td>221487</td><td>221509</td><td>-</td><td>0</td><td>TAGTTACACGGCACTGGCCG</td><td>TGG</td><td>||||||||[||||||||||||]PAM</td><td>NA</td><td>Intergenic</td><td>NA</td><td>NA</td></tr>
<tr><td>T17952</td><td>GAGTCTTGTGATCCTCTGCCTGG</td><td>1000</td><td></td><td></td><td></td><td></td><td></td><td></td><td></td><td></td><td></td></tr>
<tr><td>Oligo fwd</td><td>CACCGAGTCTTGTGATCCTCTGCC</td><td></td><td></td><td></td><td></td><td></td><td></td><td></td><td></td><td></td><td></td></tr>
<tr><td>Oligo rev</td><td>AAACGGCAGAGGATCACAAGACTC</td><td></td><td></td><td></td><td></td><td></td><td></td><td></td><td></td><td></td><td></td></tr>
<tr><td>Chromosome</td><td>start</td><td>end</td><td>strand</td><td>MM</td><td>target_seq</td><td>PAM</td><td>alignment</td><td>distance</td><td>position</td><td>gene name</td><td>gene id</td></tr>
<tr><td>GU305914.1</td><td>221460</td><td>221482</td><td>-</td><td>0</td><td>GAGTCTTGTGATCCTCTGCC</td><td>TGG</td><td>||||||||[||||||||||||]PAM</td><td>NA</td><td>Intergenic</td><td>NA</td><td>NA</td></tr>
<tr><td>T17953</td><td>TTCGCGCTTATCTACGCGCACGG</td><td>1000</td><td></td><td></td><td></td><td></td><td></td><td></td><td></td><td></td><td></td></tr>
<tr><td>Oligo fwd</td><td>CACCTTCGCGCTTATCTACGCGCA</td><td></td><td></td><td></td><td></td><td></td><td></td><td></td><td></td><td></td><td></td></tr>
<tr><td>Oligo rev</td><td>AAACTGCGCGTAGATAAGCGCGAA</td><td></td><td></td><td></td><td></td><td></td><td></td><td></td><td></td><td></td><td></td></tr>
<tr><td>Chromosome</td><td>start</td><td>end</td><td>strand</td><td>MM</td><td>target_seq</td><td>PAM</td><td>alignment</td><td>distance</td><td>position</td><td>gene name</td><td>gene id</td></tr>
<tr><td>GU305914.1</td><td>221423</td><td>221445</td><td>-</td><td>0</td><td>TTCGCGCTTATCTACGCGCA</td><td>CGG</td><td>||||||||[||||||||||||]PAM</td><td>NA</td><td>Intergenic</td><td>NA</td><td>NA</td></tr>
<tr><td>T17954</td><td>GTCAGTCTGAGACTCTGAACAGG</td><td>1000</td><td></td><td></td><td></td><td></td><td></td><td></td><td></td><td></td><td></td></tr>
<tr><td>Oligo fwd</td><td>CACCGTCAGTCTGAGACTCTGAAC</td><td></td><td></td><td></td><td></td><td></td><td></td><td></td><td></td><td></td><td></td></tr>
<tr><td>Oligo rev</td><td>AAACGTTCAGAGTCTCAGACTGAC</td><td></td><td></td><td></td><td></td><td></td><td></td><td></td><td></td><td></td><td></td></tr>
<tr><td>Chromosome</td><td>start</td><td>end</td><td>strand</td><td>MM</td><td>target_seq</td><td>PAM</td><td>alignment</td><td>distance</td><td>position</td><td>gene name</td><td>gene id</td></tr>
<tr><td>GU305914.1</td><td>221380</td><td>221402</td><td>-</td><td>0</td><td>GTCAGTCTGAGACTCTGAAC</td><td>AGG</td><td>||||||||[||||||||||||]PAM</td><td>NA</td><td>Intergenic</td><td>NA</td><td>NA</td></tr>
<tr><td>T17955</td><td>TCAGTCTGAGACTCTGAACAGGG</td><td>1000</td><td></td><td></td><td></td><td></td><td></td><td></td><td></td><td></td><td></td></tr>
<tr><td>Oligo fwd</td><td>CACCTCAGTCTGAGACTCTGAACA</td><td></td><td></td><td></td><td></td><td></td><td></td><td></td><td></td><td></td><td></td></tr>
<tr><td>Oligo rev</td><td>AAACTGTTCAGAGTCTCAGACTGA</td><td></td><td></td><td></td><td></td><td></td><td></td><td></td><td></td><td></td><td></td></tr>
<tr><td>Chromosome</td><td>start</td><td>end</td><td>strand</td><td>MM</td><td>target_seq</td><td>PAM</td><td>alignment</td><td>distance</td><td>position</td><td>gene name</td><td>gene id</td></tr>
<tr><td>GU305914.1</td><td>221379</td><td>221401</td><td>-</td><td>0</td><td>TCAGTCTGAGACTCTGAACA</td><td>GGG</td><td>||||||||[||||||||||||]PAM</td><td>NA</td><td>Intergenic</td><td>NA</td><td>NA</td></tr>
<tr><td>T17956</td><td>GAGACTCTGAACAGGGACAGTGG</td><td>1000</td><td></td><td></td><td></td><td></td><td></td><td></td><td></td><td></td><td></td></tr>
<tr><td>Oligo fwd</td><td>CACCGAGACTCTGAACAGGGACAG</td><td></td><td></td><td></td><td></td><td></td><td></td><td></td><td></td><td></td><td></td></tr>
<tr><td>Oligo rev</td><td>AAACCTGTCCCTGTTCAGAGTCTC</td><td></td><td></td><td></td><td></td><td></td><td></td><td></td><td></td><td></td><td></td></tr>
<tr><td>Chromosome</td><td>start</td><td>end</td><td>strand</td><td>MM</td><td>target_seq</td><td>PAM</td><td>alignment</td><td>distance</td><td>position</td><td>gene name</td><td>gene id</td></tr>
<tr><td>GU305914.1</td><td>221372</td><td>221394</td><td>-</td><td>0</td><td>GAGACTCTGAACAGGGACAG</td><td>TGG</td><td>||||||||[||||||||||||]PAM</td><td>NA</td><td>Intergenic</td><td>NA</td><td>NA</td></tr>
<tr><td>T17957</td><td>GGACAGTGGCGAGTTCGACCCGG</td><td>1000</td><td></td><td></td><td></td><td></td><td></td><td></td><td></td><td></td><td></td></tr>
<tr><td>Oligo fwd</td><td>CACCGGACAGTGGCGAGTTCGACC</td><td></td><td></td><td></td><td></td><td></td><td></td><td></td><td></td><td></td><td></td></tr>
<tr><td>Oligo rev</td><td>AAACGGTCGAACTCGCCACTGTCC</td><td></td><td></td><td></td><td></td><td></td><td></td><td></td><td></td><td></td><td></td></tr>
<tr><td>Chromosome</td><td>start</td><td>end</td><td>strand</td><td>MM</td><td>target_seq</td><td>PAM</td><td>alignment</td><td>distance</td><td>position</td><td>gene name</td><td>gene id</td></tr>
<tr><td>GU305914.1</td><td>221358</td><td>221380</td><td>-</td><td>0</td><td>GGACAGTGGCGAGTTCGACC</td><td>CGG</td><td>||||||||[||||||||||||]PAM</td><td>NA</td><td>Intergenic</td><td>NA</td><td>NA</td></tr>
<tr><td>T17958</td><td>GACAGTGGCGAGTTCGACCCGGG</td><td>1000</td><td></td><td></td><td></td><td></td><td></td><td></td><td></td><td></td><td></td></tr>
<tr><td>Oligo fwd</td><td>CACCGACAGTGGCGAGTTCGACCC</td><td></td><td></td><td></td><td></td><td></td><td></td><td></td><td></td><td></td><td></td></tr>
<tr><td>Oligo rev</td><td>AAACGGGTCGAACTCGCCACTGTC</td><td></td><td></td><td></td><td></td><td></td><td></td><td></td><td></td><td></td><td></td></tr>
<tr><td>Chromosome</td><td>start</td><td>end</td><td>strand</td><td>MM</td><td>target_seq</td><td>PAM</td><td>alignment</td><td>distance</td><td>position</td><td>gene name</td><td>gene id</td></tr>
<tr><td>GU305914.1</td><td>221357</td><td>221379</td><td>-</td><td>0</td><td>GACAGTGGCGAGTTCGACCC</td><td>GGG</td><td>||||||||[||||||||||||]PAM</td><td>NA</td><td>Intergenic</td><td>NA</td><td>NA</td></tr>
<tr><td>T17959</td><td>CGGGTACCCCCTTGCAACTGCGG</td><td>1000</td><td></td><td></td><td></td><td></td><td></td><td></td><td></td><td></td><td></td></tr>
<tr><td>Oligo fwd</td><td>CACCCGGGTACCCCCTTGCAACTG</td><td></td><td></td><td></td><td></td><td></td><td></td><td></td><td></td><td></td><td></td></tr>
<tr><td>Oligo rev</td><td>AAACCAGTTGCAAGGGGGTACCCG</td><td></td><td></td><td></td><td></td><td></td><td></td><td></td><td></td><td></td><td></td></tr>
<tr><td>Chromosome</td><td>start</td><td>end</td><td>strand</td><td>MM</td><td>target_seq</td><td>PAM</td><td>alignment</td><td>distance</td><td>position</td><td>gene name</td><td>gene id</td></tr>
<tr><td>GU305914.1</td><td>221338</td><td>221360</td><td>-</td><td>0</td><td>CGGGTACCCCCTTGCAACTG</td><td>CGG</td><td>||||||||[||||||||||||]PAM</td><td>NA</td><td>Intergenic</td><td>NA</td><td>NA</td></tr>
<tr><td>T17960</td><td>GGGTACCCCCTTGCAACTGCGGG</td><td>1000</td><td></td><td></td><td></td><td></td><td></td><td></td><td></td><td></td><td></td></tr>
<tr><td>Oligo fwd</td><td>CACCGGGTACCCCCTTGCAACTGC</td><td></td><td></td><td></td><td></td><td></td><td></td><td></td><td></td><td></td><td></td></tr>
<tr><td>Oligo rev</td><td>AAACGCAGTTGCAAGGGGGTACCC</td><td></td><td></td><td></td><td></td><td></td><td></td><td></td><td></td><td></td><td></td></tr>
<tr><td>Chromosome</td><td>start</td><td>end</td><td>strand</td><td>MM</td><td>target_seq</td><td>PAM</td><td>alignment</td><td>distance</td><td>position</td><td>gene name</td><td>gene id</td></tr>
<tr><td>GU305914.1</td><td>221337</td><td>221359</td><td>-</td><td>0</td><td>GGGTACCCCCTTGCAACTGC</td><td>GGG</td><td>||||||||[||||||||||||]PAM</td><td>NA</td><td>Intergenic</td><td>NA</td><td>NA</td></tr>
<tr><td>T17961</td><td>TACCCCCTTGCAACTGCGGGCGG</td><td>1000</td><td></td><td></td><td></td><td></td><td></td><td></td><td></td><td></td><td></td></tr>
<tr><td>Oligo fwd</td><td>CACCTACCCCCTTGCAACTGCGGG</td><td></td><td></td><td></td><td></td><td></td><td></td><td></td><td></td><td></td><td></td></tr>
<tr><td>Oligo rev</td><td>AAACCCCGCAGTTGCAAGGGGGTA</td><td></td><td></td><td></td><td></td><td></td><td></td><td></td><td></td><td></td><td></td></tr>
<tr><td>Chromosome</td><td>start</td><td>end</td><td>strand</td><td>MM</td><td>target_seq</td><td>PAM</td><td>alignment</td><td>distance</td><td>position</td><td>gene name</td><td>gene id</td></tr>
<tr><td>GU305914.1</td><td>221334</td><td>221356</td><td>-</td><td>0</td><td>TACCCCCTTGCAACTGCGGG</td><td>CGG</td><td>||||||||[||||||||||||]PAM</td><td>NA</td><td>Intergenic</td><td>NA</td><td>NA</td></tr>
<tr><td>T17962</td><td>CACCATTCCCGAATCTGAGATGG</td><td>1000</td><td></td><td></td><td></td><td></td><td></td><td></td><td></td><td></td><td></td></tr>
<tr><td>Oligo fwd</td><td>CACCCACCATTCCCGAATCTGAGA</td><td></td><td></td><td></td><td></td><td></td><td></td><td></td><td></td><td></td><td></td></tr>
<tr><td>Oligo rev</td><td>AAACTCTCAGATTCGGGAATGGTG</td><td></td><td></td><td></td><td></td><td></td><td></td><td></td><td></td><td></td><td></td></tr>
<tr><td>Chromosome</td><td>start</td><td>end</td><td>strand</td><td>MM</td><td>target_seq</td><td>PAM</td><td>alignment</td><td>distance</td><td>position</td><td>gene name</td><td>gene id</td></tr>
</table>

| | | | | | | | | | | | |
|---|---|---|---|---|---|---|---|---|---|---|---|
| GU305914.1 | | 221304 | 221326 | - | 0 | CACCATTCCCGAATCTGAGA | TGG | \|\|\|\|\|\|\|\|[\|\|\|\|\|\|\|\|\|\|\|\|]PAM | NA | Intergenic | NA | NA |
| T17963 | CATTCCCGAATCTGAGATGGTGG | | 1000 | | | | | | | | |
| Oligo fwd | CACCCATTCCCGAATCTGAGATGG | | | | | | | | | | |
| Oligo rev | AAACCCATCTCAGATTCGGGAATG | | | | | | | | | | |
| Chromosome | start | | end | strand | MM | target_seq | PAM | alignment | distance | position | gene name | gene id |
| GU305914.1 | | 221301 | 221323 | - | 0 | CATTCCCGAATCTGAGATGG | TGG | \|\|\|\|\|\|\|\|[\|\|\|\|\|\|\|\|\|\|\|\|]PAM | NA | Intergenic | NA | NA |
| T17964 | GATGGTGGAATCTAACGAGTAGG | | 1000 | | | | | | | | |
| Oligo fwd | CACCGATGGTGGAATCTAACGAGT | | | | | | | | | | |
| Oligo rev | AAACACTCGTTAGATTCCACCATC | | | | | | | | | | |
| Chromosome | start | | end | strand | MM | target_seq | PAM | alignment | distance | position | gene name | gene id |
| GU305914.1 | | 221286 | 221308 | - | 0 | GATGGTGGAATCTAACGAGT | AGG | \|\|\|\|\|\|\|\|[\|\|\|\|\|\|\|\|\|\|\|\|]PAM | NA | Intergenic | NA | NA |
| T17965 | ATGGTGGAATCTAACGAGTAGGG | | 1000 | | | | | | | | |
| Oligo fwd | CACCATGGTGGAATCTAACGAGTA | | | | | | | | | | |
| Oligo rev | AAACTACTCGTTAGATTCCACCAT | | | | | | | | | | |
| Chromosome | start | | end | strand | MM | target_seq | PAM | alignment | distance | position | gene name | gene id |
| GU305914.1 | | 221285 | 221307 | - | 0 | ATGGTGGAATCTAACGAGTA | GGG | \|\|\|\|\|\|\|\|[\|\|\|\|\|\|\|\|\|\|\|\|]PAM | NA | Intergenic | NA | NA |
| T17966 | TAGTGTGTTTCCGATAGACATGG | | 1000 | | | | | | | | |
| Oligo fwd | CACCTAGTGTGTTTCCGATAGACA | | | | | | | | | | |
| Oligo rev | AAACTGTCTATCGGAAACACACTA | | | | | | | | | | |
| Chromosome | start | | end | strand | MM | target_seq | PAM | alignment | distance | position | gene name | gene id |
| GU305914.1 | | 221247 | 221269 | - | 0 | TAGTGTGTTTCCGATAGACA | TGG | \|\|\|\|\|\|\|\|[\|\|\|\|\|\|\|\|\|\|\|\|]PAM | NA | Intergenic | NA | NA |
| T17967 | AGTGTGTTTCCGATAGACATGGG | | 1000 | | | | | | | | |
| Oligo fwd | CACCAGTGTGTTTCCGATAGACAT | | | | | | | | | | |
| Oligo rev | AAACATGTCTATCGGAAACACACT | | | | | | | | | | |
| Chromosome | start | | end | strand | MM | target_seq | PAM | alignment | distance | position | gene name | gene id |
| GU305914.1 | | 221246 | 221268 | - | 0 | AGTGTGTTTCCGATAGACAT | GGG | \|\|\|\|\|\|\|\|[\|\|\|\|\|\|\|\|\|\|\|\|]PAM | NA | Intergenic | NA | NA |
| T17968 | GTGTGTTTCCGATAGACATGGGG | | 1000 | | | | | | | | |
| Oligo fwd | CACCGTGTGTTTCCGATAGACATG | | | | | | | | | | |
| Oligo rev | AAACCATGTCTATCGGAAACACAC | | | | | | | | | | |
| Chromosome | start | | end | strand | MM | target_seq | PAM | alignment | distance | position | gene name | gene id |
| GU305914.1 | | 221245 | 221267 | - | 0 | GTGTGTTTCCGATAGACATG | GGG | \|\|\|\|\|\|\|\|[\|\|\|\|\|\|\|\|\|\|\|\|]PAM | NA | Intergenic | NA | NA |
| T17969 | AATCGAGACACGTCTGTTCCCGG | | 1000 | | | | | | | | |
| Oligo fwd | CACCAATCGAGACACGTCTGTTCC | | | | | | | | | | |
| Oligo rev | AAACGGAACAGACGTGTCTCGATT | | | | | | | | | | |
| Chromosome | start | | end | strand | MM | target_seq | PAM | alignment | distance | position | gene name | gene id |
| GU305914.1 | | 221164 | 221186 | - | 0 | AATCGAGACACGTCTGTTCC | CGG | \|\|\|\|\|\|\|\|[\|\|\|\|\|\|\|\|\|\|\|\|]PAM | NA | Intergenic | NA | NA |
| T17970 | ATGTATAGAACAGTCTCCGTCGG | | 1000 | | | | | | | | |
| Oligo fwd | CACCATGTATAGAACAGTCTCCGT | | | | | | | | | | |
| Oligo rev | AAACACGGAGACTGTTCTATACAT | | | | | | | | | | |
| Chromosome | start | | end | strand | MM | target_seq | PAM | alignment | distance | position | gene name | gene id |
| GU305914.1 | | 221121 | 221143 | - | 0 | ATGTATAGAACAGTCTCCGT | CGG | \|\|\|\|\|\|\|\|[\|\|\|\|\|\|\|\|\|\|\|\|]PAM | NA | Intergenic | NA | NA |
| T17971 | TGTATAGAACAGTCTCCGTCGGG | | 1000 | | | | | | | | |
| Oligo fwd | CACCTGTATAGAACAGTCTCCGTC | | | | | | | | | | |
| Oligo rev | AAACGACGGAGACTGTTCTATACA | | | | | | | | | | |
| Chromosome | start | | end | strand | MM | target_seq | PAM | alignment | distance | position | gene name | gene id |
| GU305914.1 | | 221120 | 221142 | - | 0 | TGTATAGAACAGTCTCCGTC | GGG | \|\|\|\|\|\|\|\|[\|\|\|\|\|\|\|\|\|\|\|\|]PAM | NA | Intergenic | NA | NA |
| T17972 | GTATAGAACAGTCTCCGTCGGGG | | 1000 | | | | | | | | |
| Oligo fwd | CACCGTATAGAACAGTCTCCGTCG | | | | | | | | | | |
| Oligo rev | AAACCGACGGAGACTGTTCTATAC | | | | | | | | | | |
| Chromosome | start | | end | strand | MM | target_seq | PAM | alignment | distance | position | gene name | gene id |
| GU305914.1 | | 221119 | 221141 | - | 0 | GTATAGAACAGTCTCCGTCG | GGG | \|\|\|\|\|\|\|\|[\|\|\|\|\|\|\|\|\|\|\|\|]PAM | NA | Intergenic | NA | NA |
| T17973 | AGAACAGTCTCCGTCGGGGTCGG | | 1000 | | | | | | | | |
| Oligo fwd | CACCAGAACAGTCTCCGTCGGGGT | | | | | | | | | | |
| Oligo rev | AAACACCCCGACGGAGACTGTTCT | | | | | | | | | | |
| Chromosome | start | | end | strand | MM | target_seq | PAM | alignment | distance | position | gene name | gene id |
| GU305914.1 | | 221115 | 221137 | - | 0 | AGAACAGTCTCCGTCGGGGT | CGG | \|\|\|\|\|\|\|\|[\|\|\|\|\|\|\|\|\|\|\|\|]PAM | NA | Intergenic | NA | NA |
| T17974 | TCTCCGTCGGGGTCGGAGCCTGG | | 1000 | | | | | | | | |
| Oligo fwd | CACCTCTCCGTCGGGGTCGGAGCC | | | | | | | | | | |
| Oligo rev | AAACGGCTCCGACCCCGACGGAGA | | | | | | | | | | |
| Chromosome | start | | end | strand | MM | target_seq | PAM | alignment | distance | position | gene name | gene id |
| GU305914.1 | | 221108 | 221130 | - | 0 | TCTCCGTCGGGGTCGGAGCC | TGG | \|\|\|\|\|\|\|\|[\|\|\|\|\|\|\|\|\|\|\|\|]PAM | NA | Intergenic | NA | NA |
| GU305914.1 | | 114835 | 114857 | - | 4 | TCCCCGGCGGGGTCGTAGCG | AGG | \|\|-\|\|\|-\|[\|\|\|\|\|\|\|-\|\|\|-]PAM | NA | Intergenic | NA | NA |
| T17975 | CTCCGTCGGGGTCGGAGCCTGGG | | 1000 | | | | | | | | |
| Oligo fwd | CACCCTCCGTCGGGGTCGGAGCCT | | | | | | | | | | |
| Oligo rev | AAACAGGCTCCGACCCCGACGGAG | | | | | | | | | | |
| Chromosome | start | | end | strand | MM | target_seq | PAM | alignment | distance | position | gene name | gene id |
| GU305914.1 | | 221107 | 221129 | - | 0 | CTCCGTCGGGGTCGGAGCCT | GGG | \|\|\|\|\|\|\|\|[\|\|\|\|\|\|\|\|\|\|\|\|]PAM | NA | Intergenic | NA | NA |
| T17976 | GTCGCCTGCCATTGTCCTTTTGG | | 1000 | | | | | | | | |
| Oligo fwd | CACCGTCGCCTGCCATTGTCCTTT | | | | | | | | | | |
| Oligo rev | AAACAAAGGACAATGGCAGGCGAC | | | | | | | | | | |
| Chromosome | start | | end | strand | MM | target_seq | PAM | alignment | distance | position | gene name | gene id |
| GU305914.1 | | 221027 | 221049 | - | 0 | GTCGCCTGCCATTGTCCTTT | TGG | \|\|\|\|\|\|\|\|[\|\|\|\|\|\|\|\|\|\|\|\|]PAM | NA | Intergenic | NA | NA |
| T17977 | GCCTAGATCATACCATCTCTTGG | | 1000 | | | | | | | | |
| Oligo fwd | CACCGCCTAGATCATACCATCTCT | | | | | | | | | | |

Oligo rev AAACAGAGATGGTATGATCTAGGC
Chromosome start end strand MM target_seq PAM alignment distance position gene name gene id
GU305914.1 220999 221021 - 0 GCCTAGATCATACCATCTCT TGG ||||||||[||||||||||||]PAM NA Intergenic NA NA

T17978 TCATACCATCTCTTGGTATGTGG 1000
Oligo fwd CACCTCATACCATCTCTTGGTATG
Oligo rev AAACCATACCAAGAGATGGTATGA
Chromosome start end strand MM target_seq PAM alignment distance position gene name gene id
GU305914.1 220992 221014 - 0 TCATACCATCTCTTGGTATG TGG ||||||||[||||||||||||]PAM NA Intergenic NA NA

T17979 GAACACGTAGCTACTGTCGACGG 1000
Oligo fwd CACCGAACACGTAGCTACTGTCGA
Oligo rev AAACTCGACAGTAGCTACGTGTTC
Chromosome start end strand MM target_seq PAM alignment distance position gene name gene id
GU305914.1 220964 220986 - 0 GAACACGTAGCTACTGTCGA CGG ||||||||[||||||||||||]PAM NA Intergenic NA NA

T17980 AGCTACTGTCGACGGTCGACCGG 1000
Oligo fwd CACCAGCTACTGTCGACGGTCGAC
Oligo rev AAACGTCGACCGTCGACAGTAGCT
Chromosome start end strand MM target_seq PAM alignment distance position gene name gene id
GU305914.1 220956 220978 - 0 AGCTACTGTCGACGGTCGAC CGG ||||||||[||||||||||||]PAM NA Intergenic NA NA

T17981 ACTGTCGACGGTCGACCGGACGG 1000
Oligo fwd CACCACTGTCGACGGTCGACCGGA
Oligo rev AAACTCCGGTCGACCGTCGACAGT
Chromosome start end strand MM target_seq PAM alignment distance position gene name gene id
GU305914.1 220952 220974 - 0 ACTGTCGACGGTCGACCGGA CGG ||||||||[||||||||||||]PAM NA Intergenic NA NA

T17982 GGACGGCATAGTGCAAGATACGG 1000
Oligo fwd CACCGGACGGCATAGTGCAAGATA
Oligo rev AAACTATCTTGCACTATGCCGTCC
Chromosome start end strand MM target_seq PAM alignment distance position gene name gene id
GU305914.1 220935 220957 - 0 GGACGGCATAGTGCAAGATA CGG ||||||||[||||||||||||]PAM NA Intergenic NA NA

T17983 CGGCATAGTGCAAGATACGGTGG 1000
Oligo fwd CACCCGGCATAGTGCAAGATACGG
Oligo rev AAACCCGTATCTTGCACTATGCCG
Chromosome start end strand MM target_seq PAM alignment distance position gene name gene id
GU305914.1 220932 220954 - 0 CGGCATAGTGCAAGATACGG TGG ||||||||[||||||||||||]PAM NA Intergenic NA NA

T17984 CAAGATACGGTGGAGATCAGCGG 1000
Oligo fwd CACCCAAGATACGGTGGAGATCAG
Oligo rev AAACCTGATCTCCACCGTATCTTG
Chromosome start end strand MM target_seq PAM alignment distance position gene name gene id
GU305914.1 220922 220944 - 0 CAAGATACGGTGGAGATCAG CGG ||||||||[||||||||||||]PAM NA Intergenic NA NA

T17985 CGGTGGAGATCAGCGGCACGTGG 1000
Oligo fwd CACCCGGTGGAGATCAGCGGCACG
Oligo rev AAACCGTGCCGCTGATCTCCACCG
Chromosome start end strand MM target_seq PAM alignment distance position gene name gene id
GU305914.1 220915 220937 - 0 CGGTGGAGATCAGCGGCACG TGG ||||||||[||||||||||||]PAM NA Intergenic NA NA

T17986 CGTGGCGAAACACGTCTCGTTGG 1000
Oligo fwd CACCCGTGGCGAAACACGTCTCGT
Oligo rev AAACACGAGACGTGTTTCGCCACG
Chromosome start end strand MM target_seq PAM alignment distance position gene name gene id
GU305914.1 220897 220919 - 0 CGTGGCGAAACACGTCTCGT TGG ||||||||[||||||||||||]PAM NA Intergenic NA NA

T17987 CGAAACACGTCTCGTTGGCGAGG 1000
Oligo fwd CACCCGAAACACGTCTCGTTGGCG
Oligo rev AAACCGCCAACGAGACGTGTTTCG
Chromosome start end strand MM target_seq PAM alignment distance position gene name gene id
GU305914.1 220892 220914 - 0 CGAAACACGTCTCGTTGGCG AGG ||||||||[||||||||||||]PAM NA Intergenic NA NA
GU305914.1 61818 61840 + 4 CGGAGCACTTCTCGTTGACG AGG ||-|-|||[-||||||||-||]PAM NA Intergenic NA NA

T17988 CGTCTCGTTGGCGAGGAGACCGG 1000
Oligo fwd CACCCGTCTCGTTGGCGAGGAGAC
Oligo rev AAACGTCTCCTCGCCAACGAGACG
Chromosome start end strand MM target_seq PAM alignment distance position gene name gene id
GU305914.1 220885 220907 - 0 CGTCTCGTTGGCGAGGAGAC CGG ||||||||[||||||||||||]PAM NA Intergenic NA NA

T17989 GACCGGCTCTACTTTTCTCGCGG 1000
Oligo fwd CACCGACCGGCTCTACTTTTCTCG
Oligo rev AAACCGAGAAAAGTAGAGCCGGTC
Chromosome start end strand MM target_seq PAM alignment distance position gene name gene id
GU305914.1 220868 220890 - 0 GACCGGCTCTACTTTTCTCG CGG ||||||||[||||||||||||]PAM NA Intergenic NA NA

T17990 TCTACTTTTCTCGCGGTTTACGG 1000
Oligo fwd CACCTCTACTTTTCTCGCGGTTTA
Oligo rev AAACTAAACCGCGAGAAAAGTAGA
Chromosome start end strand MM target_seq PAM alignment distance position gene name gene id
GU305914.1 220861 220883 - 0 TCTACTTTTCTCGCGGTTTA CGG ||||||||[||||||||||||]PAM NA Intergenic NA NA

T17991 TTCTCGCGGTTTACGGATATCGG 1000
Oligo fwd CACCTTCTCGCGGTTTACGGATAT
Oligo rev AAACATATCCGTAAACCGCGAGAA
Chromosome start end strand MM target_seq PAM alignment distance position gene name gene id
GU305914.1 220854 220876 - 0 TTCTCGCGGTTTACGGATAT CGG ||||||||[||||||||||||]PAM NA Intergenic NA NA

| | | | | | | | | | | | |
|---|---|---|---|---|---|---|---|---|---|---|---|
| T17992 | TCTCGCGGTTTACGGATATCGGG | 1000 | | | | | | | | | |
| Oligo fwd | CACCTCTCGCGGTTTACGGATATC | | | | | | | | | | |
| Oligo rev | AAACGATATCCGTAAACCGCGAGA | | | | | | | | | | |
| Chromosome | start | end | strand | MM | target_seq | PAM | alignment | distance | position | gene name | gene id |
| GU305914.1 | 220853 | 220875 | - | 0 | TCTCGCGGTTTACGGATATC | GGG | ||||||||[||||||||||||]PAM | NA | Intergenic | NA | NA |
| T17993 | TGCCAGCTGATCCCAGAGATTGG | 1000 | | | | | | | | | |
| Oligo fwd | CACCTGCCAGCTGATCCCAGAGAT | | | | | | | | | | |
| Oligo rev | AAACATCTCTGGGATCAGCTGGCA | | | | | | | | | | |
| Chromosome | start | end | strand | MM | target_seq | PAM | alignment | distance | position | gene name | gene id |
| GU305914.1 | 220741 | 220763 | - | 0 | TGCCAGCTGATCCCAGAGAT | TGG | ||||||||[||||||||||||]PAM | NA | Intergenic | NA | NA |
| T17994 | TTGGAAAATCTCGTCGACCGTGG | 1000 | | | | | | | | | |
| Oligo fwd | CACCTTGGAAAATCTCGTCGACCG | | | | | | | | | | |
| Oligo rev | AAACCGGTCGACGAGATTTTCCAA | | | | | | | | | | |
| Chromosome | start | end | strand | MM | target_seq | PAM | alignment | distance | position | gene name | gene id |
| GU305914.1 | 220722 | 220744 | - | 0 | TTGGAAAATCTCGTCGACCG | TGG | ||||||||[||||||||||||]PAM | NA | Intergenic | NA | NA |
| T17995 | TTTTCATATATCTATCTGCTTGG | 1000 | | | | | | | | | |
| Oligo fwd | CACCTTTTCATATATCTATCTGCT | | | | | | | | | | |
| Oligo rev | AAACAGCAGATAGATATATGAAAA | | | | | | | | | | |
| Chromosome | start | end | strand | MM | target_seq | PAM | alignment | distance | position | gene name | gene id |
| GU305914.1 | 220673 | 220695 | - | 0 | TTTTCATATATCTATCTGCT | TGG | ||||||||[||||||||||||]PAM | NA | Intergenic | NA | NA |
| T17996 | TCTGCTTGGAACTCTTGCTTCGG | 1000 | | | | | | | | | |
| Oligo fwd | CACCTCTGCTTGGAACTCTTGCTT | | | | | | | | | | |
| Oligo rev | AAACAAGCAAGAGTTCCAAGCAGA | | | | | | | | | | |
| Chromosome | start | end | strand | MM | target_seq | PAM | alignment | distance | position | gene name | gene id |
| GU305914.1 | 220659 | 220681 | - | 0 | TCTGCTTGGAACTCTTGCTT | CGG | ||||||||[||||||||||||]PAM | NA | Intergenic | NA | NA |
| T17997 | ACGAAGCTCTGTTTGCTTCTAGG | 1000 | | | | | | | | | |
| Oligo fwd | CACCACGAAGCTCTGTTTGCTTCT | | | | | | | | | | |
| Oligo rev | AAACAGAAGCAAACAGAGCTTCGT | | | | | | | | | | |
| Chromosome | start | end | strand | MM | target_seq | PAM | alignment | distance | position | gene name | gene id |
| GU305914.1 | 220591 | 220613 | - | 0 | ACGAAGCTCTGTTTGCTTCT | AGG | ||||||||[||||||||||||]PAM | NA | Intergenic | NA | NA |
| T17998 | CCTGATACAAATTAACAACGCGG | 1000 | | | | | | | | | |
| Oligo fwd | CACCCCTGATACAAATTAACAACG | | | | | | | | | | |
| Oligo rev | AAACCGTTGTTAATTTGTATCAGG | | | | | | | | | | |
| Chromosome | start | end | strand | MM | target_seq | PAM | alignment | distance | position | gene name | gene id |
| GU305914.1 | 220565 | 220587 | - | 0 | CCTGATACAAATTAACAACG | CGG | ||||||||[||||||||||||]PAM | NA | Intergenic | NA | NA |
| T17999 | GAATATACCTGTACGTCATCTGG | 1000 | | | | | | | | | |
| Oligo fwd | CACCGAATATACCTGTACGTCATC | | | | | | | | | | |
| Oligo rev | AAACGATGACGTACAGGTATATTC | | | | | | | | | | |
| Chromosome | start | end | strand | MM | target_seq | PAM | alignment | distance | position | gene name | gene id |
| GU305914.1 | 220495 | 220517 | - | 0 | GAATATACCTGTACGTCATC | TGG | ||||||||[||||||||||||]PAM | NA | Intergenic | NA | NA |
| T18000 | TGTACGTCATCTGGTTCTGATGG | 1000 | | | | | | | | | |
| Oligo fwd | CACCTGTACGTCATCTGGTTCTGA | | | | | | | | | | |
| Oligo rev | AAACTCAGAACCAGATGACGTACA | | | | | | | | | | |
| Chromosome | start | end | strand | MM | target_seq | PAM | alignment | distance | position | gene name | gene id |
| GU305914.1 | 220486 | 220508 | - | 0 | TGTACGTCATCTGGTTCTGA | TGG | ||||||||[||||||||||||]PAM | NA | Intergenic | NA | NA |
| T18001 | GATGGCTTTGAAAATGTTCAAGG | 1000 | | | | | | | | | |
| Oligo fwd | CACCGATGGCTTTGAAAATGTTCA | | | | | | | | | | |
| Oligo rev | AAACTGAACATTTTCAAAGCCATC | | | | | | | | | | |
| Chromosome | start | end | strand | MM | target_seq | PAM | alignment | distance | position | gene name | gene id |
| GU305914.1 | 220468 | 220490 | - | 0 | GATGGCTTTGAAAATGTTCA | AGG | ||||||||[||||||||||||]PAM | NA | Intergenic | NA | NA |
| T18002 | TTGAAAATGTTCAAGGATTTTGG | 1000 | | | | | | | | | |
| Oligo fwd | CACCTTGAAAATGTTCAAGGATTT | | | | | | | | | | |
| Oligo rev | AAACAAATCCTTGAACATTTTCAA | | | | | | | | | | |
| Chromosome | start | end | strand | MM | target_seq | PAM | alignment | distance | position | gene name | gene id |
| GU305914.1 | 220461 | 220483 | - | 0 | TTGAAAATGTTCAAGGATTT | TGG | ||||||||[||||||||||||]PAM | NA | Intergenic | NA | NA |
| T18003 | TTTTGGAAGATACTATGTACCGG | 1000 | | | | | | | | | |
| Oligo fwd | CACCTTTTGGAAGATACTATGTAC | | | | | | | | | | |
| Oligo rev | AAACGTACATAGTATCTTCCAAAA | | | | | | | | | | |
| Chromosome | start | end | strand | MM | target_seq | PAM | alignment | distance | position | gene name | gene id |
| GU305914.1 | 220444 | 220466 | - | 0 | TTTTGGAAGATACTATGTAC | CGG | ||||||||[||||||||||||]PAM | NA | Intergenic | NA | NA |
| T18004 | GAGACTATTCTAGCTACTTTTGG | 1000 | | | | | | | | | |
| Oligo fwd | CACCGAGACTATTCTAGCTACTTT | | | | | | | | | | |
| Oligo rev | AAACAAAGTAGCTAGAATAGTCTC | | | | | | | | | | |
| Chromosome | start | end | strand | MM | target_seq | PAM | alignment | distance | position | gene name | gene id |
| GU305914.1 | 220411 | 220433 | - | 0 | GAGACTATTCTAGCTACTTT | TGG | ||||||||[||||||||||||]PAM | NA | Intergenic | NA | NA |
| T18005 | ACTATTCTAGCTACTTTTGGTGG | 1000 | | | | | | | | | |
| Oligo fwd | CACCACTATTCTAGCTACTTTTGG | | | | | | | | | | |
| Oligo rev | AAACCCAAAAGTAGCTAGAATAGT | | | | | | | | | | |
| Chromosome | start | end | strand | MM | target_seq | PAM | alignment | distance | position | gene name | gene id |
| GU305914.1 | 220408 | 220430 | - | 0 | ACTATTCTAGCTACTTTTGG | TGG | ||||||||[||||||||||||]PAM | NA | Intergenic | NA | NA |
| T18006 | GCTACTTTTGGTGGCTATAGTGG | 1000 | | | | | | | | | |
| Oligo fwd | CACCGCTACTTTTGGTGGCTATAG | | | | | | | | | | |
| Oligo rev | AAACCTATAGCCACCAAAAGTAGC | | | | | | | | | | |
| Chromosome | start | end | strand | MM | target_seq | PAM | alignment | distance | position | gene name | gene id |
| GU305914.1 | 220399 | 220421 | - | 0 | GCTACTTTTGGTGGCTATAG | TGG | ||||||||[||||||||||||]PAM | NA | Intergenic | NA | NA |

| | | | | | | | | | | | |
|---|---|---|---|---|---|---|---|---|---|---|---|
| T18007 | ACTTTTGGTGGCTATAGTGGCGG | 1000 | | | | | | | | | |
| Oligo fwd | CACCACTTTTGGTGGCTATAGTGG | | | | | | | | | | |
| Oligo rev | AAACCCACTATAGCCACCAAAAGT | | | | | | | | | | |
| Chromosome | start | end | strand | MM | target_seq | PAM | alignment | distance | position | gene name | gene id |
| GU305914.1 | 220396 | 220418 | - | 0 | ACTTTTGGTGGCTATAGTGG | CGG | \|\|\|\|\|\|\|\|[\|\|\|\|\|\|\|\|\|\|\|\|]PAM | NA | Intergenic | NA | NA |
| T18008 | AAACATCCATATGTTCTTAGTGG | 1000 | | | | | | | | | |
| Oligo fwd | CACCAAACATCCATATGTTCTTAG | | | | | | | | | | |
| Oligo rev | AAACCTAAGAACATATGGATGTTT | | | | | | | | | | |
| Chromosome | start | end | strand | MM | target_seq | PAM | alignment | distance | position | gene name | gene id |
| GU305914.1 | 220363 | 220385 | - | 0 | AAACATCCATATGTTCTTAG | TGG | \|\|\|\|\|\|\|\|[\|\|\|\|\|\|\|\|\|\|\|\|]PAM | NA | Intergenic | NA | NA |
| T18009 | TGGATCATCGACTACAACTATGG | 1000 | | | | | | | | | |
| Oligo fwd | CACCTGGATCATCGACTACAACTA | | | | | | | | | | |
| Oligo rev | AAACTAGTTGTAGTCGATGATCCA | | | | | | | | | | |
| Chromosome | start | end | strand | MM | target_seq | PAM | alignment | distance | position | gene name | gene id |
| GU305914.1 | 220343 | 220365 | - | 0 | TGGATCATCGACTACAACTA | TGG | \|\|\|\|\|\|\|\|[\|\|\|\|\|\|\|\|\|\|\|\|]PAM | NA | Intergenic | NA | NA |
| GU305914.1 | 50240 | 50262 | - | 4 | TGTATCGTCGACTGCAAGTA | CGG | \|\|-\|\|\|-\|[\|\|\|\|\|-\|\|\|-\|\|]PAM | NA | Intergenic | NA | NA |
| T18010 | CCGCGATTCAGAATCGTCAGAGG | 1000 | | | | | | | | | |
| Oligo fwd | CACCCCGCGATTCAGAATCGTCAG | | | | | | | | | | |
| Oligo rev | AAACCTGACGATTCTGAATCGCGG | | | | | | | | | | |
| Chromosome | start | end | strand | MM | target_seq | PAM | alignment | distance | position | gene name | gene id |
| GU305914.1 | 220292 | 220314 | - | 0 | CCGCGATTCAGAATCGTCAG | AGG | \|\|\|\|\|\|\|\|[\|\|\|\|\|\|\|\|\|\|\|\|]PAM | NA | Intergenic | NA | NA |
| T18011 | ATCGTCAGAGGAAGATGAAGAGG | 1000 | | | | | | | | | |
| Oligo fwd | CACCATCGTCAGAGGAAGATGAAG | | | | | | | | | | |
| Oligo rev | AAACCTTCATCTTCCTCTGACGAT | | | | | | | | | | |
| Chromosome | start | end | strand | MM | target_seq | PAM | alignment | distance | position | gene name | gene id |
| GU305914.1 | 220280 | 220302 | - | 0 | ATCGTCAGAGGAAGATGAAG | AGG | \|\|\|\|\|\|\|\|[\|\|\|\|\|\|\|\|\|\|\|\|]PAM | NA | Intergenic | NA | NA |
| T18012 | AGAGGAAGATGAAGAGGAAGAGG | 1000 | | | | | | | | | |
| Oligo fwd | CACCAGAGGAAGATGAAGAGGAAG | | | | | | | | | | |
| Oligo rev | AAACCTTCCTCTTCATCTTCCTCT | | | | | | | | | | |
| Chromosome | start | end | strand | MM | target_seq | PAM | alignment | distance | position | gene name | gene id |
| GU305914.1 | 220274 | 220296 | - | 0 | AGAGGAAGATGAAGAGGAAG | AGG | \|\|\|\|\|\|\|\|[\|\|\|\|\|\|\|\|\|\|\|\|]PAM | NA | Intergenic | NA | NA |
| GU305914.1 | 45065 | 45087 | + | 4 | AGGAGAAGAGGAGGAGGAAG | AGG | \|\|--\|\|\|\|[\|-\|\|-\|\|\|\|\|\|\|]PAM | NA | Intergenic | NA | NA |
| GU305914.1 | 118851 | 118873 | - | 4 | GGAGGAGGAAGAAGACGAAG | AGG | -\|\|\|\|\|-\|[\|-\|\|\|\|\|-\|\|\|\|]PAM | NA | Intergenic | NA | NA |
| T18013 | GACGACACCACTTCAAAATCAGG | 1000 | | | | | | | | | |
| Oligo fwd | CACCGACGACACCACTTCAAAATC | | | | | | | | | | |
| Oligo rev | AAACGATTTTGAAGTGGTGTCGTC | | | | | | | | | | |
| Chromosome | start | end | strand | MM | target_seq | PAM | alignment | distance | position | gene name | gene id |
| GU305914.1 | 220246 | 220268 | - | 0 | GACGACACCACTTCAAAATC | AGG | \|\|\|\|\|\|\|\|[\|\|\|\|\|\|\|\|\|\|\|\|]PAM | NA | Intergenic | NA | NA |
| T18014 | AAAATGAAACCTTTGTGTGACGG | 1000 | | | | | | | | | |
| Oligo fwd | CACCAAAATGAAACCTTTGTGTGA | | | | | | | | | | |
| Oligo rev | AAACTCACACAAAGGTTTCATTTT | | | | | | | | | | |
| Chromosome | start | end | strand | MM | target_seq | PAM | alignment | distance | position | gene name | gene id |
| GU305914.1 | 220207 | 220229 | - | 0 | AAAATGAAACCTTTGTGTGA | CGG | \|\|\|\|\|\|\|\|[\|\|\|\|\|\|\|\|\|\|\|\|]PAM | NA | Intergenic | NA | NA |
| T18015 | GGACTTATCTCATGCAAACTTGG | 1000 | | | | | | | | | |
| Oligo fwd | CACCGGACTTATCTCATGCAAACT | | | | | | | | | | |
| Oligo rev | AAACAGTTTGCATGAGATAAGTCC | | | | | | | | | | |
| Chromosome | start | end | strand | MM | target_seq | PAM | alignment | distance | position | gene name | gene id |
| GU305914.1 | 220186 | 220208 | - | 0 | GGACTTATCTCATGCAAACT | TGG | \|\|\|\|\|\|\|\|[\|\|\|\|\|\|\|\|\|\|\|\|]PAM | NA | Intergenic | NA | NA |
| T18016 | TACAACAGTACCGATGTTACCGG | 1000 | | | | | | | | | |
| Oligo fwd | CACCTACAACAGTACCGATGTTAC | | | | | | | | | | |
| Oligo rev | AAACGTAACATCGGTACTGTTGTA | | | | | | | | | | |
| Chromosome | start | end | strand | MM | target_seq | PAM | alignment | distance | position | gene name | gene id |
| GU305914.1 | 220162 | 220184 | - | 0 | TACAACAGTACCGATGTTAC | CGG | \|\|\|\|\|\|\|\|[\|\|\|\|\|\|\|\|\|\|\|\|]PAM | NA | Intergenic | NA | NA |
| T18017 | CGGTCGTTCTTCCGCGCTTCCGG | 1000 | | | | | | | | | |
| Oligo fwd | CACCCGGTCGTTCTTCCGCGCTTC | | | | | | | | | | |
| Oligo rev | AAACGAAGCGCGGAAGAACGACCG | | | | | | | | | | |
| Chromosome | start | end | strand | MM | target_seq | PAM | alignment | distance | position | gene name | gene id |
| GU305914.1 | 220142 | 220164 | - | 0 | CGGTCGTTCTTCCGCGCTTC | CGG | \|\|\|\|\|\|\|\|[\|\|\|\|\|\|\|\|\|\|\|\|]PAM | NA | Intergenic | NA | NA |
| T18018 | TTCTTCCGCGCTTCCGGTGTTGG | 1000 | | | | | | | | | |
| Oligo fwd | CACCTTCTTCCGCGCTTCCGGTGT | | | | | | | | | | |
| Oligo rev | AAACACACCGGAAGCGCGGAAGAA | | | | | | | | | | |
| Chromosome | start | end | strand | MM | target_seq | PAM | alignment | distance | position | gene name | gene id |
| GU305914.1 | 220136 | 220158 | - | 0 | TTCTTCCGCGCTTCCGGTGT | TGG | \|\|\|\|\|\|\|\|[\|\|\|\|\|\|\|\|\|\|\|\|]PAM | NA | Intergenic | NA | NA |
| T18019 | TCTTCCGCGCTTCCGGTGTTGGG | 1000 | | | | | | | | | |
| Oligo fwd | CACCTCTTCCGCGCTTCCGGTGTT | | | | | | | | | | |
| Oligo rev | AAACAACACCGGAAGCGCGGAAGA | | | | | | | | | | |
| Chromosome | start | end | strand | MM | target_seq | PAM | alignment | distance | position | gene name | gene id |
| GU305914.1 | 220135 | 220157 | - | 0 | TCTTCCGCGCTTCCGGTGTT | GGG | \|\|\|\|\|\|\|\|[\|\|\|\|\|\|\|\|\|\|\|\|]PAM | NA | Intergenic | NA | NA |
| T18020 | CACATACACGCAACCGAACGCGG | 1000 | | | | | | | | | |
| Oligo fwd | CACCCACATACACGCAACCGAACG | | | | | | | | | | |
| Oligo rev | AAACCGTTCGGTTGCGTGTATGTG | | | | | | | | | | |
| Chromosome | start | end | strand | MM | target_seq | PAM | alignment | distance | position | gene name | gene id |
| GU305914.1 | 220098 | 220120 | - | 0 | CACATACACGCAACCGAACG | CGG | \|\|\|\|\|\|\|\|[\|\|\|\|\|\|\|\|\|\|\|\|]PAM | NA | Intergenic | NA | NA |
| T18021 | ACACGCAACCGAACGCGGACCGG | 1000 | | | | | | | | | |

<table>
<tr><td>Oligo fwd</td><td>CACCACACGCAACCGAACGCGGAC</td><td></td><td></td><td></td><td></td><td></td><td></td><td></td><td></td><td></td><td></td></tr>
<tr><td>Oligo rev</td><td>AAACGTCCGCGTTCGGTTGCGTGT</td><td></td><td></td><td></td><td></td><td></td><td></td><td></td><td></td><td></td><td></td></tr>
<tr><td>Chromosome</td><td>start</td><td>end</td><td>strand</td><td>MM</td><td>target_seq</td><td>PAM</td><td>alignment</td><td>distance</td><td>position</td><td>gene name</td><td>gene id</td></tr>
<tr><td>GU305914.1</td><td>220093</td><td>220115</td><td>-</td><td>0</td><td>ACACGCAACCGAACGCGGAC</td><td>CGG</td><td>||||||||[||||||||||||]PAM</td><td>NA</td><td>Intergenic</td><td>NA</td><td>NA</td></tr>
<tr><td>T18022</td><td>ACGCGGACCGGACAAACGCTCGG</td><td>1000</td><td></td><td></td><td></td><td></td><td></td><td></td><td></td><td></td><td></td></tr>
<tr><td>Oligo fwd</td><td>CACCACGCGGACCGGACAAACGCT</td><td></td><td></td><td></td><td></td><td></td><td></td><td></td><td></td><td></td><td></td></tr>
<tr><td>Oligo rev</td><td>AAACAGCGTTTGTCCGGTCCGCGT</td><td></td><td></td><td></td><td></td><td></td><td></td><td></td><td></td><td></td><td></td></tr>
<tr><td>Chromosome</td><td>start</td><td>end</td><td>strand</td><td>MM</td><td>target_seq</td><td>PAM</td><td>alignment</td><td>distance</td><td>position</td><td>gene name</td><td>gene id</td></tr>
<tr><td>GU305914.1</td><td>220081</td><td>220103</td><td>-</td><td>0</td><td>ACGCGGACCGGACAAACGCT</td><td>CGG</td><td>||||||||[||||||||||||]PAM</td><td>NA</td><td>Intergenic</td><td>NA</td><td>NA</td></tr>
<tr><td>T18023</td><td>AGATGTCATGCAGACCACAGAGG</td><td>1000</td><td></td><td></td><td></td><td></td><td></td><td></td><td></td><td></td><td></td></tr>
<tr><td>Oligo fwd</td><td>CACCAGATGTCATGCAGACCACAG</td><td></td><td></td><td></td><td></td><td></td><td></td><td></td><td></td><td></td><td></td></tr>
<tr><td>Oligo rev</td><td>AAACCTGTGGTCTGCATGACATCT</td><td></td><td></td><td></td><td></td><td></td><td></td><td></td><td></td><td></td><td></td></tr>
<tr><td>Chromosome</td><td>start</td><td>end</td><td>strand</td><td>MM</td><td>target_seq</td><td>PAM</td><td>alignment</td><td>distance</td><td>position</td><td>gene name</td><td>gene id</td></tr>
<tr><td>GU305914.1</td><td>220053</td><td>220075</td><td>-</td><td>0</td><td>AGATGTCATGCAGACCACAG</td><td>AGG</td><td>||||||||[||||||||||||]PAM</td><td>NA</td><td>Intergenic</td><td>NA</td><td>NA</td></tr>
<tr><td>T18024</td><td>TGCAGACCACAGAGGATGTGTGG</td><td>1000</td><td></td><td></td><td></td><td></td><td></td><td></td><td></td><td></td><td></td></tr>
<tr><td>Oligo fwd</td><td>CACCTGCAGACCACAGAGGATGTG</td><td></td><td></td><td></td><td></td><td></td><td></td><td></td><td></td><td></td><td></td></tr>
<tr><td>Oligo rev</td><td>AAACCACATCCTCTGTGGTCTGCA</td><td></td><td></td><td></td><td></td><td></td><td></td><td></td><td></td><td></td><td></td></tr>
<tr><td>Chromosome</td><td>start</td><td>end</td><td>strand</td><td>MM</td><td>target_seq</td><td>PAM</td><td>alignment</td><td>distance</td><td>position</td><td>gene name</td><td>gene id</td></tr>
<tr><td>GU305914.1</td><td>220045</td><td>220067</td><td>-</td><td>0</td><td>TGCAGACCACAGAGGATGTG</td><td>TGG</td><td>||||||||[||||||||||||]PAM</td><td>NA</td><td>Intergenic</td><td>NA</td><td>NA</td></tr>
<tr><td>T18025</td><td>GTGGCACGTACAATGTAACCTGG</td><td>1000</td><td></td><td></td><td></td><td></td><td></td><td></td><td></td><td></td><td></td></tr>
<tr><td>Oligo fwd</td><td>CACCGTGGCACGTACAATGTAACC</td><td></td><td></td><td></td><td></td><td></td><td></td><td></td><td></td><td></td><td></td></tr>
<tr><td>Oligo rev</td><td>AAACGGTTACATTGTACGTGCCAC</td><td></td><td></td><td></td><td></td><td></td><td></td><td></td><td></td><td></td><td></td></tr>
<tr><td>Chromosome</td><td>start</td><td>end</td><td>strand</td><td>MM</td><td>target_seq</td><td>PAM</td><td>alignment</td><td>distance</td><td>position</td><td>gene name</td><td>gene id</td></tr>
<tr><td>GU305914.1</td><td>220026</td><td>220048</td><td>-</td><td>0</td><td>GTGGCACGTACAATGTAACC</td><td>TGG</td><td>||||||||[||||||||||||]PAM</td><td>NA</td><td>Intergenic</td><td>NA</td><td>NA</td></tr>
<tr><td>T18026</td><td>TCTAATACGATTACAACGATAGG</td><td>1000</td><td></td><td></td><td></td><td></td><td></td><td></td><td></td><td></td><td></td></tr>
<tr><td>Oligo fwd</td><td>CACCTCTAATACGATTACAACGAT</td><td></td><td></td><td></td><td></td><td></td><td></td><td></td><td></td><td></td><td></td></tr>
<tr><td>Oligo rev</td><td>AAACATCGTTGTAATCGTATTAGA</td><td></td><td></td><td></td><td></td><td></td><td></td><td></td><td></td><td></td><td></td></tr>
<tr><td>Chromosome</td><td>start</td><td>end</td><td>strand</td><td>MM</td><td>target_seq</td><td>PAM</td><td>alignment</td><td>distance</td><td>position</td><td>gene name</td><td>gene id</td></tr>
<tr><td>GU305914.1</td><td>219994</td><td>220016</td><td>-</td><td>0</td><td>TCTAATACGATTACAACGAT</td><td>AGG</td><td>||||||||[||||||||||||]PAM</td><td>NA</td><td>Intergenic</td><td>NA</td><td>NA</td></tr>
<tr><td>T18027</td><td>ATACGATTACAACGATAGGTCGG</td><td>1000</td><td></td><td></td><td></td><td></td><td></td><td></td><td></td><td></td><td></td></tr>
<tr><td>Oligo fwd</td><td>CACCATACGATTACAACGATAGGT</td><td></td><td></td><td></td><td></td><td></td><td></td><td></td><td></td><td></td><td></td></tr>
<tr><td>Oligo rev</td><td>AAACACCTATCGTTGTAATCGTAT</td><td></td><td></td><td></td><td></td><td></td><td></td><td></td><td></td><td></td><td></td></tr>
<tr><td>Chromosome</td><td>start</td><td>end</td><td>strand</td><td>MM</td><td>target_seq</td><td>PAM</td><td>alignment</td><td>distance</td><td>position</td><td>gene name</td><td>gene id</td></tr>
<tr><td>GU305914.1</td><td>219990</td><td>220012</td><td>-</td><td>0</td><td>ATACGATTACAACGATAGGT</td><td>CGG</td><td>||||||||[||||||||||||]PAM</td><td>NA</td><td>Intergenic</td><td>NA</td><td>NA</td></tr>
<tr><td>T18028</td><td>TACGATTACAACGATAGGTCGGG</td><td>1000</td><td></td><td></td><td></td><td></td><td></td><td></td><td></td><td></td><td></td></tr>
<tr><td>Oligo fwd</td><td>CACCTACGATTACAACGATAGGTC</td><td></td><td></td><td></td><td></td><td></td><td></td><td></td><td></td><td></td><td></td></tr>
<tr><td>Oligo rev</td><td>AAACGACCTATCGTTGTAATCGTA</td><td></td><td></td><td></td><td></td><td></td><td></td><td></td><td></td><td></td><td></td></tr>
<tr><td>Chromosome</td><td>start</td><td>end</td><td>strand</td><td>MM</td><td>target_seq</td><td>PAM</td><td>alignment</td><td>distance</td><td>position</td><td>gene name</td><td>gene id</td></tr>
<tr><td>GU305914.1</td><td>219989</td><td>220011</td><td>-</td><td>0</td><td>TACGATTACAACGATAGGTC</td><td>GGG</td><td>||||||||[||||||||||||]PAM</td><td>NA</td><td>Intergenic</td><td>NA</td><td>NA</td></tr>
<tr><td>T18029</td><td>CGATAGGTCGGGCAACGTATTGG</td><td>1000</td><td></td><td></td><td></td><td></td><td></td><td></td><td></td><td></td><td></td></tr>
<tr><td>Oligo fwd</td><td>CACCCGATAGGTCGGGCAACGTAT</td><td></td><td></td><td></td><td></td><td></td><td></td><td></td><td></td><td></td><td></td></tr>
<tr><td>Oligo rev</td><td>AAACATACGTTGCCCGACCTATCG</td><td></td><td></td><td></td><td></td><td></td><td></td><td></td><td></td><td></td><td></td></tr>
<tr><td>Chromosome</td><td>start</td><td>end</td><td>strand</td><td>MM</td><td>target_seq</td><td>PAM</td><td>alignment</td><td>distance</td><td>position</td><td>gene name</td><td>gene id</td></tr>
<tr><td>GU305914.1</td><td>219978</td><td>220000</td><td>-</td><td>0</td><td>CGATAGGTCGGGCAACGTAT</td><td>TGG</td><td>||||||||[||||||||||||]PAM</td><td>NA</td><td>Intergenic</td><td>NA</td><td>NA</td></tr>
<tr><td>T18030</td><td>TGGAATGAATCTTATTCAGATGG</td><td>1000</td><td></td><td></td><td></td><td></td><td></td><td></td><td></td><td></td><td></td></tr>
<tr><td>Oligo fwd</td><td>CACCTGGAATGAATCTTATTCAGA</td><td></td><td></td><td></td><td></td><td></td><td></td><td></td><td></td><td></td><td></td></tr>
<tr><td>Oligo rev</td><td>AAACTCTGAATAAGATTCATTCCA</td><td></td><td></td><td></td><td></td><td></td><td></td><td></td><td></td><td></td><td></td></tr>
<tr><td>Chromosome</td><td>start</td><td>end</td><td>strand</td><td>MM</td><td>target_seq</td><td>PAM</td><td>alignment</td><td>distance</td><td>position</td><td>gene name</td><td>gene id</td></tr>
<tr><td>GU305914.1</td><td>219958</td><td>219980</td><td>-</td><td>0</td><td>TGGAATGAATCTTATTCAGA</td><td>TGG</td><td>||||||||[||||||||||||]PAM</td><td>NA</td><td>Intergenic</td><td>NA</td><td>NA</td></tr>
<tr><td>T18031</td><td>CAGATGGATGTAACACAACGTGG</td><td>1000</td><td></td><td></td><td></td><td></td><td></td><td></td><td></td><td></td><td></td></tr>
<tr><td>Oligo fwd</td><td>CACCCAGATGGATGTAACACAACG</td><td></td><td></td><td></td><td></td><td></td><td></td><td></td><td></td><td></td><td></td></tr>
<tr><td>Oligo rev</td><td>AAACCGTTGTGTTACATCCATCTG</td><td></td><td></td><td></td><td></td><td></td><td></td><td></td><td></td><td></td><td></td></tr>
<tr><td>Chromosome</td><td>start</td><td>end</td><td>strand</td><td>MM</td><td>target_seq</td><td>PAM</td><td>alignment</td><td>distance</td><td>position</td><td>gene name</td><td>gene id</td></tr>
<tr><td>GU305914.1</td><td>219942</td><td>219964</td><td>-</td><td>0</td><td>CAGATGGATGTAACACAACG</td><td>TGG</td><td>||||||||[||||||||||||]PAM</td><td>NA</td><td>Intergenic</td><td>NA</td><td>NA</td></tr>
<tr><td>T18032</td><td>AGATGGATGTAACACAACGTGGG</td><td>1000</td><td></td><td></td><td></td><td></td><td></td><td></td><td></td><td></td><td></td></tr>
<tr><td>Oligo fwd</td><td>CACCAGATGGATGTAACACAACGT</td><td></td><td></td><td></td><td></td><td></td><td></td><td></td><td></td><td></td><td></td></tr>
<tr><td>Oligo rev</td><td>AAACACGTTGTGTTACATCCATCT</td><td></td><td></td><td></td><td></td><td></td><td></td><td></td><td></td><td></td><td></td></tr>
<tr><td>Chromosome</td><td>start</td><td>end</td><td>strand</td><td>MM</td><td>target_seq</td><td>PAM</td><td>alignment</td><td>distance</td><td>position</td><td>gene name</td><td>gene id</td></tr>
<tr><td>GU305914.1</td><td>219941</td><td>219963</td><td>-</td><td>0</td><td>AGATGGATGTAACACAACGT</td><td>GGG</td><td>||||||||[||||||||||||]PAM</td><td>NA</td><td>Intergenic</td><td>NA</td><td>NA</td></tr>
<tr><td>T18033</td><td>GATGGATGTAACACAACGTGGGG</td><td>1000</td><td></td><td></td><td></td><td></td><td></td><td></td><td></td><td></td><td></td></tr>
<tr><td>Oligo fwd</td><td>CACCGATGGATGTAACACAACGTG</td><td></td><td></td><td></td><td></td><td></td><td></td><td></td><td></td><td></td><td></td></tr>
<tr><td>Oligo rev</td><td>AAACCACGTTGTGTTACATCCATC</td><td></td><td></td><td></td><td></td><td></td><td></td><td></td><td></td><td></td><td></td></tr>
<tr><td>Chromosome</td><td>start</td><td>end</td><td>strand</td><td>MM</td><td>target_seq</td><td>PAM</td><td>alignment</td><td>distance</td><td>position</td><td>gene name</td><td>gene id</td></tr>
<tr><td>GU305914.1</td><td>219940</td><td>219962</td><td>-</td><td>0</td><td>GATGGATGTAACACAACGTG</td><td>GGG</td><td>||||||||[||||||||||||]PAM</td><td>NA</td><td>Intergenic</td><td>NA</td><td>NA</td></tr>
<tr><td>T18034</td><td>GGGGCAGTGAGAAACTTAAATGG</td><td>1000</td><td></td><td></td><td></td><td></td><td></td><td></td><td></td><td></td><td></td></tr>
<tr><td>Oligo fwd</td><td>CACCGGGGCAGTGAGAAACTTAAA</td><td></td><td></td><td></td><td></td><td></td><td></td><td></td><td></td><td></td><td></td></tr>
<tr><td>Oligo rev</td><td>AAACTTTAAGTTTCTCACTGCCCC</td><td></td><td></td><td></td><td></td><td></td><td></td><td></td><td></td><td></td><td></td></tr>
<tr><td>Chromosome</td><td>start</td><td>end</td><td>strand</td><td>MM</td><td>target_seq</td><td>PAM</td><td>alignment</td><td>distance</td><td>position</td><td>gene name</td><td>gene id</td></tr>
<tr><td>GU305914.1</td><td>219921</td><td>219943</td><td>-</td><td>0</td><td>GGGGCAGTGAGAAACTTAAA</td><td>TGG</td><td>||||||||[||||||||||||]PAM</td><td>NA</td><td>Intergenic</td><td>NA</td><td>NA</td></tr>
<tr><td>T18035</td><td>GGGCAGTGAGAAACTTAAATGGG</td><td>1000</td><td></td><td></td><td></td><td></td><td></td><td></td><td></td><td></td><td></td></tr>
<tr><td>Oligo fwd</td><td>CACCGGGCAGTGAGAAACTTAAAT</td><td></td><td></td><td></td><td></td><td></td><td></td><td></td><td></td><td></td><td></td></tr>
<tr><td>Oligo rev</td><td>AAACATTTAAGTTTCTCACTGCCC</td><td></td><td></td><td></td><td></td><td></td><td></td><td></td><td></td><td></td><td></td></tr>
<tr><td>Chromosome</td><td>start</td><td>end</td><td>strand</td><td>MM</td><td>target_seq</td><td>PAM</td><td>alignment</td><td>distance</td><td>position</td><td>gene name</td><td>gene id</td></tr>
<tr><td>GU305914.1</td><td>219920</td><td>219942</td><td>-</td><td>0</td><td>GGGCAGTGAGAAACTTAAAT</td><td>GGG</td><td>||||||||[||||||||||||]PAM</td><td>NA</td><td>Intergenic</td><td>NA</td><td>NA</td></tr>
</table>

T18036 AAACTTAAATGGGACCATGACGG 1000
Oligo fwd CACCAAACTTAAATGGGACCATGA
Oligo rev AAACTCATGGTCCCATTTAAGTTT
Chromosome start end strand MM target_seq PAM alignment distance position gene name gene id
GU305914.1 219910 219932 - 0 AAACTTAAATGGGACCATGA CGG ||||||||[||||||||||||]PAM NA Intergenic NA NA

T18037 TCCAAGTTAACAGATCGCTGTGG 1000
Oligo fwd CACCTCCAAGTTAACAGATCGCTG
Oligo rev AAACCAGCGATCTGTTAACTTGGA
Chromosome start end strand MM target_seq PAM alignment distance position gene name gene id
GU305914.1 219882 219904 - 0 TCCAAGTTAACAGATCGCTG TGG ||||||||[||||||||||||]PAM NA Intergenic NA NA

T18038 TCCTGTAAAGTAGAAACATGTGG 1000
Oligo fwd CACCTCCTGTAAAGTAGAAACATG
Oligo rev AAACCATGTTTCTACTTTACAGGA
Chromosome start end strand MM target_seq PAM alignment distance position gene name gene id
GU305914.1 219829 219851 - 0 TCCTGTAAAGTAGAAACATG TGG ||||||||[||||||||||||]PAM NA Intergenic NA NA

T18039 AGTAGAAACATGTGGAGCTGCGG 1000
Oligo fwd CACCAGTAGAAACATGTGGAGCTG
Oligo rev AAACCAGCTCCACATGTTTCTACT
Chromosome start end strand MM target_seq PAM alignment distance position gene name gene id
GU305914.1 219821 219843 - 0 AGTAGAAACATGTGGAGCTG CGG ||||||||[||||||||||||]PAM NA Intergenic NA NA

T18040 AAACATGTGGAGCTGCGGTATGG 1000
Oligo fwd CACCAAACATGTGGAGCTGCGGTA
Oligo rev AAACTACCGCAGCTCCACATGTTT
Chromosome start end strand MM target_seq PAM alignment distance position gene name gene id
GU305914.1 219816 219838 - 0 AAACATGTGGAGCTGCGGTA TGG ||||||||[||||||||||||]PAM NA Intergenic NA NA

T18041 TGGAGCTGCGGTATGGAGTATGG 1000
Oligo fwd CACCTGGAGCTGCGGTATGGAGTA
Oligo rev AAACTACTCCATACCGCAGCTCCA
Chromosome start end strand MM target_seq PAM alignment distance position gene name gene id
GU305914.1 219809 219831 - 0 TGGAGCTGCGGTATGGAGTA TGG ||||||||[||||||||||||]PAM NA Intergenic NA NA

T18042 TGGAGTATGGTTTGTGATTCAGG 1000
Oligo fwd CACCTGGAGTATGGTTTGTGATTC
Oligo rev AAACGAATCACAAACCATACTCCA
Chromosome start end strand MM target_seq PAM alignment distance position gene name gene id
GU305914.1 219796 219818 - 0 TGGAGTATGGTTTGTGATTC AGG ||||||||[||||||||||||]PAM NA Intergenic NA NA

T18043 TACACCGCTTATTACTCATCAGG 1000
Oligo fwd CACCTACACCGCTTATTACTCATC
Oligo rev AAACGATGAGTAATAAGCGGTGTA
Chromosome start end strand MM target_seq PAM alignment distance position gene name gene id
GU305914.1 219772 219794 - 0 TACACCGCTTATTACTCATC AGG ||||||||[||||||||||||]PAM NA Intergenic NA NA

T18044 ATCGCATCCAGCCATCATCGTGG 1000
Oligo fwd CACCATCGCATCCAGCCATCATCG
Oligo rev AAACCGATGATGGCTGGATGCGAT
Chromosome start end strand MM target_seq PAM alignment distance position gene name gene id
GU305914.1 219747 219769 - 0 ATCGCATCCAGCCATCATCG TGG ||||||||[||||||||||||]PAM NA Intergenic NA NA

T18045 ACAGATGCTATTAGTATTCCTGG 1000
Oligo fwd CACCACAGATGCTATTAGTATTCC
Oligo rev AAACGGAATACTAATAGCATCTGT
Chromosome start end strand MM target_seq PAM alignment distance position gene name gene id
GU305914.1 219722 219744 - 0 ACAGATGCTATTAGTATTCC TGG ||||||||[||||||||||||]PAM NA Intergenic NA NA

T18046 CAGATGCTATTAGTATTCCTGGG 1000
Oligo fwd CACCCAGATGCTATTAGTATTCCT
Oligo rev AAACAGGAATACTAATAGCATCTG
Chromosome start end strand MM target_seq PAM alignment distance position gene name gene id
GU305914.1 219721 219743 - 0 CAGATGCTATTAGTATTCCT GGG ||||||||[||||||||||||]PAM NA Intergenic NA NA

T18047 TAATAACACGCTAACTCAAATGG 1000
Oligo fwd CACCTAATAACACGCTAACTCAAA
Oligo rev AAACTTTGAGTTAGCGTGTTATTA
Chromosome start end strand MM target_seq PAM alignment distance position gene name gene id
GU305914.1 219690 219712 - 0 TAATAACACGCTAACTCAAA TGG ||||||||[||||||||||||]PAM NA Intergenic NA NA

T18048 AACACGCTAACTCAAATGGCTGG 1000
Oligo fwd CACCAACACGCTAACTCAAATGGC
Oligo rev AAACGCCATTTGAGTTAGCGTGTT
Chromosome start end strand MM target_seq PAM alignment distance position gene name gene id
GU305914.1 219686 219708 - 0 AACACGCTAACTCAAATGGC TGG ||||||||[||||||||||||]PAM NA Intergenic NA NA

T18049 TACTGTATTTGTCATTGTTTTGG 1000
Oligo fwd CACCTACTGTATTTGTCATTGTTT
Oligo rev AAACAAACAATGACAAATACAGTA
Chromosome start end strand MM target_seq PAM alignment distance position gene name gene id
GU305914.1 219540 219562 - 0 TACTGTATTTGTCATTGTTT TGG ||||||||[||||||||||||]PAM NA Intergenic NA NA

T18050 GGCTCGCTTCTATACAGATAAGG 1000
Oligo fwd CACCGGCTCGCTTCTATACAGATA
Oligo rev AAACTATCTGTATAGAAGCGAGCC
Chromosome start end strand MM target_seq PAM alignment distance position gene name gene id
GU305914.1 219519 219541 - 0 GGCTCGCTTCTATACAGATA AGG ||||||||[||||||||||||]PAM NA Intergenic NA NA

T18051 GCTCGCTTCTATACAGATAAGGG 1000
Oligo fwd CACCGCTCGCTTCTATACAGATAA
Oligo rev AAACTTATCTGTATAGAAGCGAGC
Chromosome start end strand MM target_seq PAM alignment distance position gene name gene id
GU305914.1 219518 219540 - 0 GCTCGCTTCTATACAGATAA GGG ||||||||[||||||||||||]PAM NA Intergenic NA NA

T18052 CGCTTCTATACAGATAAGGGCGG 1000
Oligo fwd CACCCGCTTCTATACAGATAAGGG
Oligo rev AAACCCCTTATCTGTATAGAAGCG
Chromosome start end strand MM target_seq PAM alignment distance position gene name gene id
GU305914.1 219515 219537 - 0 CGCTTCTATACAGATAAGGG CGG ||||||||[||||||||||||]PAM NA Intergenic NA NA

T18053 GCTTCTATACAGATAAGGGCGGG 1000
Oligo fwd CACCGCTTCTATACAGATAAGGGC
Oligo rev AAACGCCCTTATCTGTATAGAAGC
Chromosome start end strand MM target_seq PAM alignment distance position gene name gene id
GU305914.1 219514 219536 - 0 GCTTCTATACAGATAAGGGC GGG ||||||||[||||||||||||]PAM NA Intergenic NA NA

T18054 ACAGATAAGGGCGGGTGTGTCGG 1000
Oligo fwd CACCACAGATAAGGGCGGGTGTGT
Oligo rev AAACACACACCCGCCCTTATCTGT
Chromosome start end strand MM target_seq PAM alignment distance position gene name gene id
GU305914.1 219506 219528 - 0 ACAGATAAGGGCGGGTGTGT CGG ||||||||[||||||||||||]PAM NA Intergenic NA NA

T18055 ATTATCTCGTGATTCTCCGCAGG 1000
Oligo fwd CACCATTATCTCGTGATTCTCCGC
Oligo rev AAACGCGGAGAATCACGAGATAAT
Chromosome start end strand MM target_seq PAM alignment distance position gene name gene id
GU305914.1 219475 219497 - 0 ATTATCTCGTGATTCTCCGC AGG ||||||||[||||||||||||]PAM NA Intergenic NA NA

T18056 TCTCCGCAGGAGCGACCATGTGG 1000
Oligo fwd CACCTCTCCGCAGGAGCGACCATG
Oligo rev AAACCATGGTCGCTCCTGCGGAGA
Chromosome start end strand MM target_seq PAM alignment distance position gene name gene id
GU305914.1 219462 219484 - 0 TCTCCGCAGGAGCGACCATG TGG ||||||||[||||||||||||]PAM NA Intergenic NA NA

T18057 ACGACGTCGTATTTTTTAATCGG 1000
Oligo fwd CACCACGACGTCGTATTTTTTAAT
Oligo rev AAACATTAAAAAATACGACGTCGT
Chromosome start end strand MM target_seq PAM alignment distance position gene name gene id
GU305914.1 219439 219461 - 0 ACGACGTCGTATTTTTTAAT CGG ||||||||[||||||||||||]PAM NA Intergenic NA NA

T18058 GTCGTATTTTTTAATCGGACTGG 1000
Oligo fwd CACCGTCGTATTTTTTAATCGGAC
Oligo rev AAACGTCCGATTAAAAAATACGAC
Chromosome start end strand MM target_seq PAM alignment distance position gene name gene id
GU305914.1 219434 219456 - 0 GTCGTATTTTTTAATCGGAC TGG ||||||||[||||||||||||]PAM NA Intergenic NA NA

T18059 TCGTATTTTTTAATCGGACTGGG 1000
Oligo fwd CACCTCGTATTTTTTAATCGGACT
Oligo rev AAACAGTCCGATTAAAAAATACGA
Chromosome start end strand MM target_seq PAM alignment distance position gene name gene id
GU305914.1 219433 219455 - 0 TCGTATTTTTTAATCGGACT GGG ||||||||[||||||||||||]PAM NA Intergenic NA NA

T18060 AGTTCTCTCAACGTTCAGTCAGG 1000
Oligo fwd CACCAGTTCTCTCAACGTTCAGTC
Oligo rev AAACGACTGAACGTTGAGAGAACT
Chromosome start end strand MM target_seq PAM alignment distance position gene name gene id
GU305914.1 219404 219426 - 0 AGTTCTCTCAACGTTCAGTC AGG ||||||||[||||||||||||]PAM NA Intergenic NA NA

T18061 CAGATAAATCTATATATAATTGG 1000
Oligo fwd CACCCAGATAAATCTATATATAAT
Oligo rev AAACATTATATATAGATTTATCTG
Chromosome start end strand MM target_seq PAM alignment distance position gene name gene id
GU305914.1 219364 219386 - 0 CAGATAAATCTATATATAAT TGG ||||||||[||||||||||||]PAM NA Intergenic NA NA

T18062 AAATTGTTCTTTTTCTACCGCGG 1000
Oligo fwd CACCAAATTGTTCTTTTTCTACCG
Oligo rev AAACCGGTAGAAAAAGAACAATTT
Chromosome start end strand MM target_seq PAM alignment distance position gene name gene id
GU305914.1 219332 219354 - 0 AAATTGTTCTTTTTCTACCG CGG ||||||||[||||||||||||]PAM NA Intergenic NA NA

T18063 TTTTTCTACCGCGGCTCCTACGG 1000
Oligo fwd CACCTTTTTCTACCGCGGCTCCTA
Oligo rev AAACTAGGAGCCGCGGTAGAAAAA
Chromosome start end strand MM target_seq PAM alignment distance position gene name gene id
GU305914.1 219323 219345 - 0 TTTTTCTACCGCGGCTCCTA CGG ||||||||[||||||||||||]PAM NA Intergenic NA NA

T18064 CGCGGCTCCTACGGTAGCATCGG 1000
Oligo fwd CACCCGCGGCTCCTACGGTAGCAT
Oligo rev AAACATGCTACCGTAGGAGCCGCG
Chromosome start end strand MM target_seq PAM alignment distance position gene name gene id
GU305914.1 219314 219336 - 0 CGCGGCTCCTACGGTAGCAT CGG ||||||||[||||||||||||]PAM NA Intergenic NA NA

T18065 AGCATCGGTGATGTCTACTGCGG 1000
Oligo fwd CACCAGCATCGGTGATGTCTACTG
Oligo rev AAACCAGTAGACATCACCGATGCT
Chromosome start end strand MM target_seq PAM alignment distance position gene name gene id

GU305914.1 219299 219321 - 0 AGCATCGGTGATGTCTACTG CGG ||||||||[||||||||||||]PAM NA Intergenic NA NA

T18066 ACACCGCTCGTCACGACACCTGG 1000
Oligo fwd CACCACACCGCTCGTCACGACACC
Oligo rev AAACGGTGTCGTGACGAGCGGTGT
Chromosome start end strand MM target_seq PAM alignment distance position gene name gene id
GU305914.1 219268 219290 - 0 ACACCGCTCGTCACGACACC TGG ||||||||[||||||||||||]PAM NA Intergenic NA NA

T18067 GAACCTTTTATAATTTCAGTTGG 1000
Oligo fwd CACCGAACCTTTTATAATTTCAGT
Oligo rev AAACACTGAAATTATAAAAGGTTC
Chromosome start end strand MM target_seq PAM alignment distance position gene name gene id
GU305914.1 219138 219160 - 0 GAACCTTTTATAATTTCAGT TGG ||||||||[||||||||||||]PAM NA Intergenic NA NA

T18068 AACCTTTTATAATTTCAGTTGGG 1000
Oligo fwd CACCAACCTTTTATAATTTCAGTT
Oligo rev AAACAACTGAAATTATAAAAGGTT
Chromosome start end strand MM target_seq PAM alignment distance position gene name gene id
GU305914.1 219137 219159 - 0 AACCTTTTATAATTTCAGTT GGG ||||||||[||||||||||||]PAM NA Intergenic NA NA

T18069 ACCTTTTATAATTTCAGTTGGGG 1000
Oligo fwd CACCACCTTTTATAATTTCAGTTG
Oligo rev AAACCAACTGAAATTATAAAAGGT
Chromosome start end strand MM target_seq PAM alignment distance position gene name gene id
GU305914.1 219136 219158 - 0 ACCTTTTATAATTTCAGTTG GGG ||||||||[||||||||||||]PAM NA Intergenic NA NA

T18070 GGGGTGCATCCATCAATGAGAGG 1000
Oligo fwd CACCGGGGTGCATCCATCAATGAG
Oligo rev AAACCTCATTGATGGATGCACCCC
Chromosome start end strand MM target_seq PAM alignment distance position gene name gene id
GU305914.1 219117 219139 - 0 GGGGTGCATCCATCAATGAG AGG ||||||||[||||||||||||]PAM NA Intergenic NA NA

T18071 GGGTGCATCCATCAATGAGAGGG 1000
Oligo fwd CACCGGGTGCATCCATCAATGAGA
Oligo rev AAACTCTCATTGATGGATGCACCC
Chromosome start end strand MM target_seq PAM alignment distance position gene name gene id
GU305914.1 219116 219138 - 0 GGGTGCATCCATCAATGAGA GGG ||||||||[||||||||||||]PAM NA Intergenic NA NA

T18072 ACGATAATTCGACCACGACGTGG 1000
Oligo fwd CACCACGATAATTCGACCACGACG
Oligo rev AAACCGTCGTGGTCGAATTATCGT
Chromosome start end strand MM target_seq PAM alignment distance position gene name gene id
GU305914.1 219072 219094 - 0 ACGATAATTCGACCACGACG TGG ||||||||[||||||||||||]PAM NA Intergenic NA NA

T18073 TCGACCACGACGTGGACGTACGG 1000
Oligo fwd CACCTCGACCACGACGTGGACGTA
Oligo rev AAACTACGTCCACGTCGTGGTCGA
Chromosome start end strand MM target_seq PAM alignment distance position gene name gene id
GU305914.1 219064 219086 - 0 TCGACCACGACGTGGACGTA CGG ||||||||[||||||||||||]PAM NA Intergenic NA NA

T18074 TGGACGTACGGCTACGATGTCGG 1000
Oligo fwd CACCTGGACGTACGGCTACGATGT
Oligo rev AAACACATCGTAGCCGTACGTCCA
Chromosome start end strand MM target_seq PAM alignment distance position gene name gene id
GU305914.1 219052 219074 - 0 TGGACGTACGGCTACGATGT CGG ||||||||[||||||||||||]PAM NA Intergenic NA NA

T18075 TGTCGGCAACACGTCCCAAGAGG 1000
Oligo fwd CACCTGTCGGCAACACGTCCCAAG
Oligo rev AAACCTTGGGACGTGTTGCCGACA
Chromosome start end strand MM target_seq PAM alignment distance position gene name gene id
GU305914.1 219035 219057 - 0 TGTCGGCAACACGTCCCAAG AGG ||||||||[||||||||||||]PAM NA Intergenic NA NA

T18076 TGCGAGATACCGCTCAGAACAGG 1000
Oligo fwd CACCTGCGAGATACCGCTCAGAAC
Oligo rev AAACGTTCTGAGCGGTATCTCGCA
Chromosome start end strand MM target_seq PAM alignment distance position gene name gene id
GU305914.1 218982 219004 - 0 TGCGAGATACCGCTCAGAAC AGG ||||||||[||||||||||||]PAM NA Intergenic NA NA

T18077 TTCGAACTACCTCGACGAGACGG 1000
Oligo fwd CACCTTCGAACTACCTCGACGAGA
Oligo rev AAACTCTCGTCGAGGTAGTTCGAA
Chromosome start end strand MM target_seq PAM alignment distance position gene name gene id
GU305914.1 218957 218979 - 0 TTCGAACTACCTCGACGAGA CGG ||||||||[||||||||||||]PAM NA Intergenic NA NA

T18078 TCGAACTACCTCGACGAGACGGG 1000
Oligo fwd CACCTCGAACTACCTCGACGAGAC
Oligo rev AAACGTCTCGTCGAGGTAGTTCGA
Chromosome start end strand MM target_seq PAM alignment distance position gene name gene id
GU305914.1 218956 218978 - 0 TCGAACTACCTCGACGAGAC GGG ||||||||[||||||||||||]PAM NA Intergenic NA NA

T18079 AGATCACGTACGACTGTTGTCGG 1000
Oligo fwd CACCAGATCACGTACGACTGTTGT
Oligo rev AAACACAACAGTCGTACGTGATCT
Chromosome start end strand MM target_seq PAM alignment distance position gene name gene id
GU305914.1 218919 218941 - 0 AGATCACGTACGACTGTTGT CGG ||||||||[||||||||||||]PAM NA Intergenic NA NA

T18080 GTACGACTGTTGTCGGATGCTGG 1000
Oligo fwd CACCGTACGACTGTTGTCGGATGC
Oligo rev AAACGCATCCGACAACAGTCGTAC

<table>
<tr><td>Chromosome</td><td>start</td><td>end</td><td>strand</td><td>MM</td><td>target_seq</td><td>PAM</td><td>alignment</td><td>distance</td><td>position</td><td>gene name</td><td>gene id</td></tr>
<tr><td>GU305914.1</td><td>218912</td><td>218934</td><td>-</td><td>0</td><td>GTACGACTGTTGTCGGATGC</td><td>TGG</td><td>||||||||[||||||||||||]PAM</td><td>NA</td><td>Intergenic</td><td>NA</td><td>NA</td></tr>
<tr><td>T18081</td><td>TACGACTGTTGTCGGATGCTGGG</td><td>1000</td><td></td><td></td><td></td><td></td><td></td><td></td><td></td><td></td><td></td></tr>
<tr><td>Oligo fwd</td><td>CACCTACGACTGTTGTCGGATGCT</td><td></td><td></td><td></td><td></td><td></td><td></td><td></td><td></td><td></td><td></td></tr>
<tr><td>Oligo rev</td><td>AAACAGCATCCGACAACAGTCGTA</td><td></td><td></td><td></td><td></td><td></td><td></td><td></td><td></td><td></td><td></td></tr>
<tr><td>Chromosome</td><td>start</td><td>end</td><td>strand</td><td>MM</td><td>target_seq</td><td>PAM</td><td>alignment</td><td>distance</td><td>position</td><td>gene name</td><td>gene id</td></tr>
<tr><td>GU305914.1</td><td>218911</td><td>218933</td><td>-</td><td>0</td><td>TACGACTGTTGTCGGATGCT</td><td>GGG</td><td>||||||||[||||||||||||]PAM</td><td>NA</td><td>Intergenic</td><td>NA</td><td>NA</td></tr>
<tr><td>T18082</td><td>TGTTGTCGGATGCTGGGCAATGG</td><td>1000</td><td></td><td></td><td></td><td></td><td></td><td></td><td></td><td></td><td></td></tr>
<tr><td>Oligo fwd</td><td>CACCTGTTGTCGGATGCTGGGCAA</td><td></td><td></td><td></td><td></td><td></td><td></td><td></td><td></td><td></td><td></td></tr>
<tr><td>Oligo rev</td><td>AAACTTGCCCAGCATCCGACAACA</td><td></td><td></td><td></td><td></td><td></td><td></td><td></td><td></td><td></td><td></td></tr>
<tr><td>Chromosome</td><td>start</td><td>end</td><td>strand</td><td>MM</td><td>target_seq</td><td>PAM</td><td>alignment</td><td>distance</td><td>position</td><td>gene name</td><td>gene id</td></tr>
<tr><td>GU305914.1</td><td>218905</td><td>218927</td><td>-</td><td>0</td><td>TGTTGTCGGATGCTGGGCAA</td><td>TGG</td><td>||||||||[||||||||||||]PAM</td><td>NA</td><td>Intergenic</td><td>NA</td><td>NA</td></tr>
<tr><td>T18083</td><td>GTTGTCGGATGCTGGGCAATGGG</td><td>1000</td><td></td><td></td><td></td><td></td><td></td><td></td><td></td><td></td><td></td></tr>
<tr><td>Oligo fwd</td><td>CACCGTTGTCGGATGCTGGGCAAT</td><td></td><td></td><td></td><td></td><td></td><td></td><td></td><td></td><td></td><td></td></tr>
<tr><td>Oligo rev</td><td>AAACATTGCCCAGCATCCGACAAC</td><td></td><td></td><td></td><td></td><td></td><td></td><td></td><td></td><td></td><td></td></tr>
<tr><td>Chromosome</td><td>start</td><td>end</td><td>strand</td><td>MM</td><td>target_seq</td><td>PAM</td><td>alignment</td><td>distance</td><td>position</td><td>gene name</td><td>gene id</td></tr>
<tr><td>GU305914.1</td><td>218904</td><td>218926</td><td>-</td><td>0</td><td>GTTGTCGGATGCTGGGCAAT</td><td>GGG</td><td>||||||||[||||||||||||]PAM</td><td>NA</td><td>Intergenic</td><td>NA</td><td>NA</td></tr>
<tr><td>T18084</td><td>TGGGCAATGGGACCGTCTGCTGG</td><td>1000</td><td></td><td></td><td></td><td></td><td></td><td></td><td></td><td></td><td></td></tr>
<tr><td>Oligo fwd</td><td>CACCTGGGCAATGGGACCGTCTGC</td><td></td><td></td><td></td><td></td><td></td><td></td><td></td><td></td><td></td><td></td></tr>
<tr><td>Oligo rev</td><td>AAACGCAGACGGTCCCATTGCCCA</td><td></td><td></td><td></td><td></td><td></td><td></td><td></td><td></td><td></td><td></td></tr>
<tr><td>Chromosome</td><td>start</td><td>end</td><td>strand</td><td>MM</td><td>target_seq</td><td>PAM</td><td>alignment</td><td>distance</td><td>position</td><td>gene name</td><td>gene id</td></tr>
<tr><td>GU305914.1</td><td>218892</td><td>218914</td><td>-</td><td>0</td><td>TGGGCAATGGGACCGTCTGC</td><td>TGG</td><td>||||||||[||||||||||||]PAM</td><td>NA</td><td>Intergenic</td><td>NA</td><td>NA</td></tr>
<tr><td>T18085</td><td>GGGCAATGGGACCGTCTGCTGGG</td><td>1000</td><td></td><td></td><td></td><td></td><td></td><td></td><td></td><td></td><td></td></tr>
<tr><td>Oligo fwd</td><td>CACCGGGCAATGGGACCGTCTGCT</td><td></td><td></td><td></td><td></td><td></td><td></td><td></td><td></td><td></td><td></td></tr>
<tr><td>Oligo rev</td><td>AAACAGCAGACGGTCCCATTGCCC</td><td></td><td></td><td></td><td></td><td></td><td></td><td></td><td></td><td></td><td></td></tr>
<tr><td>Chromosome</td><td>start</td><td>end</td><td>strand</td><td>MM</td><td>target_seq</td><td>PAM</td><td>alignment</td><td>distance</td><td>position</td><td>gene name</td><td>gene id</td></tr>
<tr><td>GU305914.1</td><td>218891</td><td>218913</td><td>-</td><td>0</td><td>GGGCAATGGGACCGTCTGCT</td><td>GGG</td><td>||||||||[||||||||||||]PAM</td><td>NA</td><td>Intergenic</td><td>NA</td><td>NA</td></tr>
<tr><td>T18086</td><td>CCGTCTGCTGGGTTCGACAGAGG</td><td>1000</td><td></td><td></td><td></td><td></td><td></td><td></td><td></td><td></td><td></td></tr>
<tr><td>Oligo fwd</td><td>CACCCCGTCTGCTGGGTTCGACAG</td><td></td><td></td><td></td><td></td><td></td><td></td><td></td><td></td><td></td><td></td></tr>
<tr><td>Oligo rev</td><td>AAACCTGTCGAACCCAGCAGACGG</td><td></td><td></td><td></td><td></td><td></td><td></td><td></td><td></td><td></td><td></td></tr>
<tr><td>Chromosome</td><td>start</td><td>end</td><td>strand</td><td>MM</td><td>target_seq</td><td>PAM</td><td>alignment</td><td>distance</td><td>position</td><td>gene name</td><td>gene id</td></tr>
<tr><td>GU305914.1</td><td>218880</td><td>218902</td><td>-</td><td>0</td><td>CCGTCTGCTGGGTTCGACAG</td><td>AGG</td><td>||||||||[||||||||||||]PAM</td><td>NA</td><td>Intergenic</td><td>NA</td><td>NA</td></tr>
<tr><td>T18087</td><td>TGGGTTCGACAGAGGATAAACGG</td><td>1000</td><td></td><td></td><td></td><td></td><td></td><td></td><td></td><td></td><td></td></tr>
<tr><td>Oligo fwd</td><td>CACCTGGGTTCGACAGAGGATAAA</td><td></td><td></td><td></td><td></td><td></td><td></td><td></td><td></td><td></td><td></td></tr>
<tr><td>Oligo rev</td><td>AAACTTTATCCTCTGTCGAACCCA</td><td></td><td></td><td></td><td></td><td></td><td></td><td></td><td></td><td></td><td></td></tr>
<tr><td>Chromosome</td><td>start</td><td>end</td><td>strand</td><td>MM</td><td>target_seq</td><td>PAM</td><td>alignment</td><td>distance</td><td>position</td><td>gene name</td><td>gene id</td></tr>
<tr><td>GU305914.1</td><td>218872</td><td>218894</td><td>-</td><td>0</td><td>TGGGTTCGACAGAGGATAAA</td><td>CGG</td><td>||||||||[||||||||||||]PAM</td><td>NA</td><td>Intergenic</td><td>NA</td><td>NA</td></tr>
<tr><td>T18088</td><td>GTCAGCGTCTCCGATGATAACGG</td><td>1000</td><td></td><td></td><td></td><td></td><td></td><td></td><td></td><td></td><td></td></tr>
<tr><td>Oligo fwd</td><td>CACCGTCAGCGTCTCCGATGATAA</td><td></td><td></td><td></td><td></td><td></td><td></td><td></td><td></td><td></td><td></td></tr>
<tr><td>Oligo rev</td><td>AAACTTATCATCGGAGACGCTGAC</td><td></td><td></td><td></td><td></td><td></td><td></td><td></td><td></td><td></td><td></td></tr>
<tr><td>Chromosome</td><td>start</td><td>end</td><td>strand</td><td>MM</td><td>target_seq</td><td>PAM</td><td>alignment</td><td>distance</td><td>position</td><td>gene name</td><td>gene id</td></tr>
<tr><td>GU305914.1</td><td>218812</td><td>218834</td><td>-</td><td>0</td><td>GTCAGCGTCTCCGATGATAA</td><td>CGG</td><td>||||||||[||||||||||||]PAM</td><td>NA</td><td>Intergenic</td><td>NA</td><td>NA</td></tr>
<tr><td>T18089</td><td>ATAACGGTACTTTTTCCGTGTGG</td><td>1000</td><td></td><td></td><td></td><td></td><td></td><td></td><td></td><td></td><td></td></tr>
<tr><td>Oligo fwd</td><td>CACCATAACGGTACTTTTTCCGTG</td><td></td><td></td><td></td><td></td><td></td><td></td><td></td><td></td><td></td><td></td></tr>
<tr><td>Oligo rev</td><td>AAACCACGGAAAAAGTACCGTTAT</td><td></td><td></td><td></td><td></td><td></td><td></td><td></td><td></td><td></td><td></td></tr>
<tr><td>Chromosome</td><td>start</td><td>end</td><td>strand</td><td>MM</td><td>target_seq</td><td>PAM</td><td>alignment</td><td>distance</td><td>position</td><td>gene name</td><td>gene id</td></tr>
<tr><td>GU305914.1</td><td>218796</td><td>218818</td><td>-</td><td>0</td><td>ATAACGGTACTTTTTCCGTG</td><td>TGG</td><td>||||||||[||||||||||||]PAM</td><td>NA</td><td>Intergenic</td><td>NA</td><td>NA</td></tr>
<tr><td>GU305914.1</td><td>157064</td><td>157086</td><td>+</td><td>3</td><td>ATGATGATACTTTTTCCGTG</td><td>TGG</td><td>||-|-|-|[||||||||||||]PAM</td><td>NA</td><td>Intergenic</td><td>NA</td><td>NA</td></tr>
<tr><td>T18090</td><td>TTCCGTGTGGATCAACAACACGG</td><td>1000</td><td></td><td></td><td></td><td></td><td></td><td></td><td></td><td></td><td></td></tr>
<tr><td>Oligo fwd</td><td>CACCTTCCGTGTGGATCAACAACA</td><td></td><td></td><td></td><td></td><td></td><td></td><td></td><td></td><td></td><td></td></tr>
<tr><td>Oligo rev</td><td>AAACTGTTGTTGATCCACACGGAA</td><td></td><td></td><td></td><td></td><td></td><td></td><td></td><td></td><td></td><td></td></tr>
<tr><td>Chromosome</td><td>start</td><td>end</td><td>strand</td><td>MM</td><td>target_seq</td><td>PAM</td><td>alignment</td><td>distance</td><td>position</td><td>gene name</td><td>gene id</td></tr>
<tr><td>GU305914.1</td><td>218783</td><td>218805</td><td>-</td><td>0</td><td>TTCCGTGTGGATCAACAACA</td><td>CGG</td><td>||||||||[||||||||||||]PAM</td><td>NA</td><td>Intergenic</td><td>NA</td><td>NA</td></tr>
<tr><td>T18091</td><td>ACACGGCCATGACTAGTTTATGG</td><td>1000</td><td></td><td></td><td></td><td></td><td></td><td></td><td></td><td></td><td></td></tr>
<tr><td>Oligo fwd</td><td>CACCACACGGCCATGACTAGTTTA</td><td></td><td></td><td></td><td></td><td></td><td></td><td></td><td></td><td></td><td></td></tr>
<tr><td>Oligo rev</td><td>AAACTAAACTAGTCATGGCCGTGT</td><td></td><td></td><td></td><td></td><td></td><td></td><td></td><td></td><td></td><td></td></tr>
<tr><td>Chromosome</td><td>start</td><td>end</td><td>strand</td><td>MM</td><td>target_seq</td><td>PAM</td><td>alignment</td><td>distance</td><td>position</td><td>gene name</td><td>gene id</td></tr>
<tr><td>GU305914.1</td><td>218766</td><td>218788</td><td>-</td><td>0</td><td>ACACGGCCATGACTAGTTTA</td><td>TGG</td><td>||||||||[||||||||||||]PAM</td><td>NA</td><td>Intergenic</td><td>NA</td><td>NA</td></tr>
<tr><td>T18092</td><td>TGGCCGCACATGAATACCATCGG</td><td>1000</td><td></td><td></td><td></td><td></td><td></td><td></td><td></td><td></td><td></td></tr>
<tr><td>Oligo fwd</td><td>CACCTGGCCGCACATGAATACCAT</td><td></td><td></td><td></td><td></td><td></td><td></td><td></td><td></td><td></td><td></td></tr>
<tr><td>Oligo rev</td><td>AAACATGGTATTCATGTGCGGCCA</td><td></td><td></td><td></td><td></td><td></td><td></td><td></td><td></td><td></td><td></td></tr>
<tr><td>Chromosome</td><td>start</td><td>end</td><td>strand</td><td>MM</td><td>target_seq</td><td>PAM</td><td>alignment</td><td>distance</td><td>position</td><td>gene name</td><td>gene id</td></tr>
<tr><td>GU305914.1</td><td>218746</td><td>218768</td><td>-</td><td>0</td><td>TGGCCGCACATGAATACCAT</td><td>CGG</td><td>||||||||[||||||||||||]PAM</td><td>NA</td><td>Intergenic</td><td>NA</td><td>NA</td></tr>
<tr><td>T18093</td><td>GGCCGCACATGAATACCATCGGG</td><td>1000</td><td></td><td></td><td></td><td></td><td></td><td></td><td></td><td></td><td></td></tr>
<tr><td>Oligo fwd</td><td>CACCGGCCGCACATGAATACCATC</td><td></td><td></td><td></td><td></td><td></td><td></td><td></td><td></td><td></td><td></td></tr>
<tr><td>Oligo rev</td><td>AAACGATGGTATTCATGTGCGGCC</td><td></td><td></td><td></td><td></td><td></td><td></td><td></td><td></td><td></td><td></td></tr>
<tr><td>Chromosome</td><td>start</td><td>end</td><td>strand</td><td>MM</td><td>target_seq</td><td>PAM</td><td>alignment</td><td>distance</td><td>position</td><td>gene name</td><td>gene id</td></tr>
<tr><td>GU305914.1</td><td>218745</td><td>218767</td><td>-</td><td>0</td><td>GGCCGCACATGAATACCATC</td><td>GGG</td><td>||||||||[||||||||||||]PAM</td><td>NA</td><td>Intergenic</td><td>NA</td><td>NA</td></tr>
<tr><td>T18094</td><td>GCCGCACATGAATACCATCGGGG</td><td>1000</td><td></td><td></td><td></td><td></td><td></td><td></td><td></td><td></td><td></td></tr>
<tr><td>Oligo fwd</td><td>CACCGCCGCACATGAATACCATCG</td><td></td><td></td><td></td><td></td><td></td><td></td><td></td><td></td><td></td><td></td></tr>
<tr><td>Oligo rev</td><td>AAACCGATGGTATTCATGTGCGGC</td><td></td><td></td><td></td><td></td><td></td><td></td><td></td><td></td><td></td><td></td></tr>
<tr><td>Chromosome</td><td>start</td><td>end</td><td>strand</td><td>MM</td><td>target_seq</td><td>PAM</td><td>alignment</td><td>distance</td><td>position</td><td>gene name</td><td>gene id</td></tr>
<tr><td>GU305914.1</td><td>218744</td><td>218766</td><td>-</td><td>0</td><td>GCCGCACATGAATACCATCG</td><td>GGG</td><td>||||||||[||||||||||||]PAM</td><td>NA</td><td>Intergenic</td><td>NA</td><td>NA</td></tr>
<tr><td>T18095</td><td>CCGCACATGAATACCATCGGGGG</td><td>1000</td><td></td><td></td><td></td><td></td><td></td><td></td><td></td><td></td><td></td></tr>
</table>

<table>
<tr><td>Oligo fwd</td><td>CACCCCGCACATGAATACCATCGG</td><td></td><td></td><td></td><td></td><td></td><td></td><td></td><td></td><td></td><td></td></tr>
<tr><td>Oligo rev</td><td>AAACCCGATGGTATTCATGTGCGG</td><td></td><td></td><td></td><td></td><td></td><td></td><td></td><td></td><td></td><td></td></tr>
<tr><td>Chromosome</td><td>start</td><td>end</td><td>strand</td><td>MM</td><td>target_seq</td><td>PAM</td><td>alignment</td><td>distance</td><td>position</td><td>gene name</td><td>gene id</td></tr>
<tr><td>GU305914.1</td><td>218743</td><td>218765</td><td>-</td><td>0</td><td>CCGCACATGAATACCATCGG</td><td>GGG</td><td>||||||||[||||||||||||]PAM</td><td>NA</td><td>Intergenic</td><td>NA</td><td>NA</td></tr>
<tr><td></td><td></td><td></td><td></td><td></td><td></td><td></td><td></td><td></td><td></td><td></td><td></td></tr>
<tr><td>T18096</td><td>CGCACATGAATACCATCGGGGGG</td><td>1000</td><td></td><td></td><td></td><td></td><td></td><td></td><td></td><td></td><td></td></tr>
<tr><td>Oligo fwd</td><td>CACCCGCACATGAATACCATCGGG</td><td></td><td></td><td></td><td></td><td></td><td></td><td></td><td></td><td></td><td></td></tr>
<tr><td>Oligo rev</td><td>AAACCCCGATGGTATTCATGTGCG</td><td></td><td></td><td></td><td></td><td></td><td></td><td></td><td></td><td></td><td></td></tr>
<tr><td>Chromosome</td><td>start</td><td>end</td><td>strand</td><td>MM</td><td>target_seq</td><td>PAM</td><td>alignment</td><td>distance</td><td>position</td><td>gene name</td><td>gene id</td></tr>
<tr><td>GU305914.1</td><td>218742</td><td>218764</td><td>-</td><td>0</td><td>CGCACATGAATACCATCGGG</td><td>GGG</td><td>||||||||[||||||||||||]PAM</td><td>NA</td><td>Intergenic</td><td>NA</td><td>NA</td></tr>
<tr><td></td><td></td><td></td><td></td><td></td><td></td><td></td><td></td><td></td><td></td><td></td><td></td></tr>
<tr><td>T18097</td><td>TGAATACCATCGGGGGGCGTTGG</td><td>1000</td><td></td><td></td><td></td><td></td><td></td><td></td><td></td><td></td><td></td></tr>
<tr><td>Oligo fwd</td><td>CACCTGAATACCATCGGGGGGCGT</td><td></td><td></td><td></td><td></td><td></td><td></td><td></td><td></td><td></td><td></td></tr>
<tr><td>Oligo rev</td><td>AAACACGCCCCCCGATGGTATTCA</td><td></td><td></td><td></td><td></td><td></td><td></td><td></td><td></td><td></td><td></td></tr>
<tr><td>Chromosome</td><td>start</td><td>end</td><td>strand</td><td>MM</td><td>target_seq</td><td>PAM</td><td>alignment</td><td>distance</td><td>position</td><td>gene name</td><td>gene id</td></tr>
<tr><td>GU305914.1</td><td>218736</td><td>218758</td><td>-</td><td>0</td><td>TGAATACCATCGGGGGGCGT</td><td>TGG</td><td>||||||||[||||||||||||]PAM</td><td>NA</td><td>Intergenic</td><td>NA</td><td>NA</td></tr>
<tr><td></td><td></td><td></td><td></td><td></td><td></td><td></td><td></td><td></td><td></td><td></td><td></td></tr>
<tr><td>T18098</td><td>ATCCGAAGACTCAATACTCTCGG</td><td>1000</td><td></td><td></td><td></td><td></td><td></td><td></td><td></td><td></td><td></td></tr>
<tr><td>Oligo fwd</td><td>CACCATCCGAAGACTCAATACTCT</td><td></td><td></td><td></td><td></td><td></td><td></td><td></td><td></td><td></td><td></td></tr>
<tr><td>Oligo rev</td><td>AAACAGAGTATTGAGTCTTCGGAT</td><td></td><td></td><td></td><td></td><td></td><td></td><td></td><td></td><td></td><td></td></tr>
<tr><td>Chromosome</td><td>start</td><td>end</td><td>strand</td><td>MM</td><td>target_seq</td><td>PAM</td><td>alignment</td><td>distance</td><td>position</td><td>gene name</td><td>gene id</td></tr>
<tr><td>GU305914.1</td><td>218695</td><td>218717</td><td>-</td><td>0</td><td>ATCCGAAGACTCAATACTCT</td><td>CGG</td><td>||||||||[||||||||||||]PAM</td><td>NA</td><td>Intergenic</td><td>NA</td><td>NA</td></tr>
<tr><td></td><td></td><td></td><td></td><td></td><td></td><td></td><td></td><td></td><td></td><td></td><td></td></tr>
<tr><td>T18099</td><td>AATACTCTCGGTCGCGCTGTCGG</td><td>1000</td><td></td><td></td><td></td><td></td><td></td><td></td><td></td><td></td><td></td></tr>
<tr><td>Oligo fwd</td><td>CACCAATACTCTCGGTCGCGCTGT</td><td></td><td></td><td></td><td></td><td></td><td></td><td></td><td></td><td></td><td></td></tr>
<tr><td>Oligo rev</td><td>AAACACAGCGCGACCGAGAGTATT</td><td></td><td></td><td></td><td></td><td></td><td></td><td></td><td></td><td></td><td></td></tr>
<tr><td>Chromosome</td><td>start</td><td>end</td><td>strand</td><td>MM</td><td>target_seq</td><td>PAM</td><td>alignment</td><td>distance</td><td>position</td><td>gene name</td><td>gene id</td></tr>
<tr><td>GU305914.1</td><td>218683</td><td>218705</td><td>-</td><td>0</td><td>AATACTCTCGGTCGCGCTGT</td><td>CGG</td><td>||||||||[||||||||||||]PAM</td><td>NA</td><td>Intergenic</td><td>NA</td><td>NA</td></tr>
<tr><td></td><td></td><td></td><td></td><td></td><td></td><td></td><td></td><td></td><td></td><td></td><td></td></tr>
<tr><td>T18100</td><td>TCGCGCTGTCGGCTACACGAAGG</td><td>1000</td><td></td><td></td><td></td><td></td><td></td><td></td><td></td><td></td><td></td></tr>
<tr><td>Oligo fwd</td><td>CACCTCGCGCTGTCGGCTACACGA</td><td></td><td></td><td></td><td></td><td></td><td></td><td></td><td></td><td></td><td></td></tr>
<tr><td>Oligo rev</td><td>AAACTCGTGTAGCCGACAGCGCGA</td><td></td><td></td><td></td><td></td><td></td><td></td><td></td><td></td><td></td><td></td></tr>
<tr><td>Chromosome</td><td>start</td><td>end</td><td>strand</td><td>MM</td><td>target_seq</td><td>PAM</td><td>alignment</td><td>distance</td><td>position</td><td>gene name</td><td>gene id</td></tr>
<tr><td>GU305914.1</td><td>218672</td><td>218694</td><td>-</td><td>0</td><td>TCGCGCTGTCGGCTACACGA</td><td>AGG</td><td>||||||||[||||||||||||]PAM</td><td>NA</td><td>Intergenic</td><td>NA</td><td>NA</td></tr>
<tr><td></td><td></td><td></td><td></td><td></td><td></td><td></td><td></td><td></td><td></td><td></td><td></td></tr>
<tr><td>T18101</td><td>CGCGACCGAAAGAAATTCTACGG</td><td>1000</td><td></td><td></td><td></td><td></td><td></td><td></td><td></td><td></td><td></td></tr>
<tr><td>Oligo fwd</td><td>CACCCGCGACCGAAAGAAATTCTA</td><td></td><td></td><td></td><td></td><td></td><td></td><td></td><td></td><td></td><td></td></tr>
<tr><td>Oligo rev</td><td>AAACTAGAATTTCTTTCGGTCGCG</td><td></td><td></td><td></td><td></td><td></td><td></td><td></td><td></td><td></td><td></td></tr>
<tr><td>Chromosome</td><td>start</td><td>end</td><td>strand</td><td>MM</td><td>target_seq</td><td>PAM</td><td>alignment</td><td>distance</td><td>position</td><td>gene name</td><td>gene id</td></tr>
<tr><td>GU305914.1</td><td>218642</td><td>218664</td><td>-</td><td>0</td><td>CGCGACCGAAAGAAATTCTA</td><td>CGG</td><td>||||||||[||||||||||||]PAM</td><td>NA</td><td>Intergenic</td><td>NA</td><td>NA</td></tr>
<tr><td></td><td></td><td></td><td></td><td></td><td></td><td></td><td></td><td></td><td></td><td></td><td></td></tr>
<tr><td>T18102</td><td>CTACGGTCAAGTGTTTTGTCAGG</td><td>1000</td><td></td><td></td><td></td><td></td><td></td><td></td><td></td><td></td><td></td></tr>
<tr><td>Oligo fwd</td><td>CACCCTACGGTCAAGTGTTTTGTC</td><td></td><td></td><td></td><td></td><td></td><td></td><td></td><td></td><td></td><td></td></tr>
<tr><td>Oligo rev</td><td>AAACGACAAAACACTTGACCGTAG</td><td></td><td></td><td></td><td></td><td></td><td></td><td></td><td></td><td></td><td></td></tr>
<tr><td>Chromosome</td><td>start</td><td>end</td><td>strand</td><td>MM</td><td>target_seq</td><td>PAM</td><td>alignment</td><td>distance</td><td>position</td><td>gene name</td><td>gene id</td></tr>
<tr><td>GU305914.1</td><td>218625</td><td>218647</td><td>-</td><td>0</td><td>CTACGGTCAAGTGTTTTGTC</td><td>AGG</td><td>||||||||[||||||||||||]PAM</td><td>NA</td><td>Intergenic</td><td>NA</td><td>NA</td></tr>
<tr><td></td><td></td><td></td><td></td><td></td><td></td><td></td><td></td><td></td><td></td><td></td><td></td></tr>
<tr><td>T18103</td><td>CTTTCGACATCCAGATGTCATGG</td><td>1000</td><td></td><td></td><td></td><td></td><td></td><td></td><td></td><td></td><td></td></tr>
<tr><td>Oligo fwd</td><td>CACCCTTTCGACATCCAGATGTCA</td><td></td><td></td><td></td><td></td><td></td><td></td><td></td><td></td><td></td><td></td></tr>
<tr><td>Oligo rev</td><td>AAACTGACATCTGGATGTCGAAAG</td><td></td><td></td><td></td><td></td><td></td><td></td><td></td><td></td><td></td><td></td></tr>
<tr><td>Chromosome</td><td>start</td><td>end</td><td>strand</td><td>MM</td><td>target_seq</td><td>PAM</td><td>alignment</td><td>distance</td><td>position</td><td>gene name</td><td>gene id</td></tr>
<tr><td>GU305914.1</td><td>218592</td><td>218614</td><td>-</td><td>0</td><td>CTTTCGACATCCAGATGTCA</td><td>TGG</td><td>||||||||[||||||||||||]PAM</td><td>NA</td><td>Intergenic</td><td>NA</td><td>NA</td></tr>
<tr><td></td><td></td><td></td><td></td><td></td><td></td><td></td><td></td><td></td><td></td><td></td><td></td></tr>
<tr><td>T18104</td><td>CATCCAGATGTCATGGTTACTGG</td><td>1000</td><td></td><td></td><td></td><td></td><td></td><td></td><td></td><td></td><td></td></tr>
<tr><td>Oligo fwd</td><td>CACCCATCCAGATGTCATGGTTAC</td><td></td><td></td><td></td><td></td><td></td><td></td><td></td><td></td><td></td><td></td></tr>
<tr><td>Oligo rev</td><td>AAACGTAACCATGACATCTGGATG</td><td></td><td></td><td></td><td></td><td></td><td></td><td></td><td></td><td></td><td></td></tr>
<tr><td>Chromosome</td><td>start</td><td>end</td><td>strand</td><td>MM</td><td>target_seq</td><td>PAM</td><td>alignment</td><td>distance</td><td>position</td><td>gene name</td><td>gene id</td></tr>
<tr><td>GU305914.1</td><td>218585</td><td>218607</td><td>-</td><td>0</td><td>CATCCAGATGTCATGGTTAC</td><td>TGG</td><td>||||||||[||||||||||||]PAM</td><td>NA</td><td>Intergenic</td><td>NA</td><td>NA</td></tr>
<tr><td></td><td></td><td></td><td></td><td></td><td></td><td></td><td></td><td></td><td></td><td></td><td></td></tr>
<tr><td>T18105</td><td>CAGATGTCATGGTTACTGGATGG</td><td>1000</td><td></td><td></td><td></td><td></td><td></td><td></td><td></td><td></td><td></td></tr>
<tr><td>Oligo fwd</td><td>CACCCAGATGTCATGGTTACTGGA</td><td></td><td></td><td></td><td></td><td></td><td></td><td></td><td></td><td></td><td></td></tr>
<tr><td>Oligo rev</td><td>AAACTCCAGTAACCATGACATCTG</td><td></td><td></td><td></td><td></td><td></td><td></td><td></td><td></td><td></td><td></td></tr>
<tr><td>Chromosome</td><td>start</td><td>end</td><td>strand</td><td>MM</td><td>target_seq</td><td>PAM</td><td>alignment</td><td>distance</td><td>position</td><td>gene name</td><td>gene id</td></tr>
<tr><td>GU305914.1</td><td>218581</td><td>218603</td><td>-</td><td>0</td><td>CAGATGTCATGGTTACTGGA</td><td>TGG</td><td>||||||||[||||||||||||]PAM</td><td>NA</td><td>Intergenic</td><td>NA</td><td>NA</td></tr>
<tr><td></td><td></td><td></td><td></td><td></td><td></td><td></td><td></td><td></td><td></td><td></td><td></td></tr>
<tr><td>T18106</td><td>AGATGTCATGGTTACTGGATGGG</td><td>1000</td><td></td><td></td><td></td><td></td><td></td><td></td><td></td><td></td><td></td></tr>
<tr><td>Oligo fwd</td><td>CACCAGATGTCATGGTTACTGGAT</td><td></td><td></td><td></td><td></td><td></td><td></td><td></td><td></td><td></td><td></td></tr>
<tr><td>Oligo rev</td><td>AAACATCCAGTAACCATGACATCT</td><td></td><td></td><td></td><td></td><td></td><td></td><td></td><td></td><td></td><td></td></tr>
<tr><td>Chromosome</td><td>start</td><td>end</td><td>strand</td><td>MM</td><td>target_seq</td><td>PAM</td><td>alignment</td><td>distance</td><td>position</td><td>gene name</td><td>gene id</td></tr>
<tr><td>GU305914.1</td><td>218580</td><td>218602</td><td>-</td><td>0</td><td>AGATGTCATGGTTACTGGAT</td><td>GGG</td><td>||||||||[||||||||||||]PAM</td><td>NA</td><td>Intergenic</td><td>NA</td><td>NA</td></tr>
<tr><td></td><td></td><td></td><td></td><td></td><td></td><td></td><td></td><td></td><td></td><td></td><td></td></tr>
<tr><td>T18107</td><td>ATGGGTACGAGATTAAACCTAGG</td><td>1000</td><td></td><td></td><td></td><td></td><td></td><td></td><td></td><td></td><td></td></tr>
<tr><td>Oligo fwd</td><td>CACCATGGGTACGAGATTAAACCT</td><td></td><td></td><td></td><td></td><td></td><td></td><td></td><td></td><td></td><td></td></tr>
<tr><td>Oligo rev</td><td>AAACAGGTTTAATCTCGTACCCAT</td><td></td><td></td><td></td><td></td><td></td><td></td><td></td><td></td><td></td><td></td></tr>
<tr><td>Chromosome</td><td>start</td><td>end</td><td>strand</td><td>MM</td><td>target_seq</td><td>PAM</td><td>alignment</td><td>distance</td><td>position</td><td>gene name</td><td>gene id</td></tr>
<tr><td>GU305914.1</td><td>218562</td><td>218584</td><td>-</td><td>0</td><td>ATGGGTACGAGATTAAACCT</td><td>AGG</td><td>||||||||[||||||||||||]PAM</td><td>NA</td><td>Intergenic</td><td>NA</td><td>NA</td></tr>
<tr><td></td><td></td><td></td><td></td><td></td><td></td><td></td><td></td><td></td><td></td><td></td><td></td></tr>
<tr><td>T18108</td><td>TGGGTACGAGATTAAACCTAGGG</td><td>1000</td><td></td><td></td><td></td><td></td><td></td><td></td><td></td><td></td><td></td></tr>
<tr><td>Oligo fwd</td><td>CACCTGGGTACGAGATTAAACCTA</td><td></td><td></td><td></td><td></td><td></td><td></td><td></td><td></td><td></td><td></td></tr>
<tr><td>Oligo rev</td><td>AAACTAGGTTTAATCTCGTACCCA</td><td></td><td></td><td></td><td></td><td></td><td></td><td></td><td></td><td></td><td></td></tr>
<tr><td>Chromosome</td><td>start</td><td>end</td><td>strand</td><td>MM</td><td>target_seq</td><td>PAM</td><td>alignment</td><td>distance</td><td>position</td><td>gene name</td><td>gene id</td></tr>
<tr><td>GU305914.1</td><td>218561</td><td>218583</td><td>-</td><td>0</td><td>TGGGTACGAGATTAAACCTA</td><td>GGG</td><td>||||||||[||||||||||||]PAM</td><td>NA</td><td>Intergenic</td><td>NA</td><td>NA</td></tr>
<tr><td></td><td></td><td></td><td></td><td></td><td></td><td></td><td></td><td></td><td></td><td></td><td></td></tr>
<tr><td>T18109</td><td>GGGTACGAGATTAAACCTAGGGG</td><td>1000</td><td></td><td></td><td></td><td></td><td></td><td></td><td></td><td></td><td></td></tr>
<tr><td>Oligo fwd</td><td>CACCGGGTACGAGATTAAACCTAG</td><td></td><td></td><td></td><td></td><td></td><td></td><td></td><td></td><td></td><td></td></tr>
<tr><td>Oligo rev</td><td>AAACCTAGGTTTAATCTCGTACCC</td><td></td><td></td><td></td><td></td><td></td><td></td><td></td><td></td><td></td><td></td></tr>
<tr><td>Chromosome</td><td>start</td><td>end</td><td>strand</td><td>MM</td><td>target_seq</td><td>PAM</td><td>alignment</td><td>distance</td><td>position</td><td>gene name</td><td>gene id</td></tr>
<tr><td>GU305914.1</td><td>218560</td><td>218582</td><td>-</td><td>0</td><td>GGGTACGAGATTAAACCTAG</td><td>GGG</td><td>||||||||[||||||||||||]PAM</td><td>NA</td><td>Intergenic</td><td>NA</td><td>NA</td></tr>
</table>

| | | | | | | | | | | | |
|---|---|---|---|---|---|---|---|---|---|---|---|
| T18110 | TTAAACCTAGGGGTTTTTATTGG | 1000 | | | | | | | | | |
| Oligo fwd | CACCTTAAACCTAGGGGTTTTTAT | | | | | | | | | | |
| Oligo rev | AAACATAAAAACCCCTAGGTTTAA | | | | | | | | | | |
| Chromosome | start | end | strand | MM | target_seq | PAM | alignment | distance | position | gene name | gene id |
| GU305914.1 | 218550 | 218572 | - | 0 | TTAAACCTAGGGGTTTTTAT | TGG | \|\|\|\|\|\|\|\|[\|\|\|\|\|\|\|\|\|\|\|\|]PAM | NA | Intergenic | NA | NA |
| T18111 | TAGGGGTTTTTATTGGATGCCGG | 1000 | | | | | | | | | |
| Oligo fwd | CACCTAGGGGTTTTTATTGGATGC | | | | | | | | | | |
| Oligo rev | AAACGCATCCAATAAAAACCCCTA | | | | | | | | | | |
| Chromosome | start | end | strand | MM | target_seq | PAM | alignment | distance | position | gene name | gene id |
| GU305914.1 | 218543 | 218565 | - | 0 | TAGGGGTTTTTATTGGATGC | CGG | \|\|\|\|\|\|\|\|[\|\|\|\|\|\|\|\|\|\|\|\|]PAM | NA | Intergenic | NA | NA |
| T18112 | GGTTTTTATTGGATGCCGGACGG | 1000 | | | | | | | | | |
| Oligo fwd | CACCGGTTTTTATTGGATGCCGGA | | | | | | | | | | |
| Oligo rev | AAACTCCGGCATCCAATAAAAACC | | | | | | | | | | |
| Chromosome | start | end | strand | MM | target_seq | PAM | alignment | distance | position | gene name | gene id |
| GU305914.1 | 218539 | 218561 | - | 0 | GGTTTTTATTGGATGCCGGA | CGG | \|\|\|\|\|\|\|\|[\|\|\|\|\|\|\|\|\|\|\|\|]PAM | NA | Intergenic | NA | NA |
| T18113 | TATTGGATGCCGGACGGAGTCGG | 1000 | | | | | | | | | |
| Oligo fwd | CACCTATTGGATGCCGGACGGAGT | | | | | | | | | | |
| Oligo rev | AAACACTCCGTCCGGCATCCAATA | | | | | | | | | | |
| Chromosome | start | end | strand | MM | target_seq | PAM | alignment | distance | position | gene name | gene id |
| GU305914.1 | 218533 | 218555 | - | 0 | TATTGGATGCCGGACGGAGT | CGG | \|\|\|\|\|\|\|\|[\|\|\|\|\|\|\|\|\|\|\|\|]PAM | NA | Intergenic | NA | NA |
| T18114 | TGGATGCCGGACGGAGTCGGTGG | 1000 | | | | | | | | | |
| Oligo fwd | CACCTGGATGCCGGACGGAGTCGG | | | | | | | | | | |
| Oligo rev | AAACCCGACTCCGTCCGGCATCCA | | | | | | | | | | |
| Chromosome | start | end | strand | MM | target_seq | PAM | alignment | distance | position | gene name | gene id |
| GU305914.1 | 218530 | 218552 | - | 0 | TGGATGCCGGACGGAGTCGG | TGG | \|\|\|\|\|\|\|\|[\|\|\|\|\|\|\|\|\|\|\|\|]PAM | NA | Intergenic | NA | NA |
| T18115 | ATGCCGGACGGAGTCGGTGGCGG | 1000 | | | | | | | | | |
| Oligo fwd | CACCATGCCGGACGGAGTCGGTGG | | | | | | | | | | |
| Oligo rev | AAACCCACCGACTCCGTCCGGCAT | | | | | | | | | | |
| Chromosome | start | end | strand | MM | target_seq | PAM | alignment | distance | position | gene name | gene id |
| GU305914.1 | 218527 | 218549 | - | 0 | ATGCCGGACGGAGTCGGTGG | CGG | \|\|\|\|\|\|\|\|[\|\|\|\|\|\|\|\|\|\|\|\|]PAM | NA | Intergenic | NA | NA |
| T18116 | TGCCGGACGGAGTCGGTGGCGGG | 1000 | | | | | | | | | |
| Oligo fwd | CACCTGCCGGACGGAGTCGGTGGC | | | | | | | | | | |
| Oligo rev | AAACGCCACCGACTCCGTCCGGCA | | | | | | | | | | |
| Chromosome | start | end | strand | MM | target_seq | PAM | alignment | distance | position | gene name | gene id |
| GU305914.1 | 218526 | 218548 | - | 0 | TGCCGGACGGAGTCGGTGGC | GGG | \|\|\|\|\|\|\|\|[\|\|\|\|\|\|\|\|\|\|\|\|]PAM | NA | Intergenic | NA | NA |
| T18117 | GCCGGACGGAGTCGGTGGCGGGG | 1000 | | | | | | | | | |
| Oligo fwd | CACCGCCGGACGGAGTCGGTGGCG | | | | | | | | | | |
| Oligo rev | AAACCGCCACCGACTCCGTCCGGC | | | | | | | | | | |
| Chromosome | start | end | strand | MM | target_seq | PAM | alignment | distance | position | gene name | gene id |
| GU305914.1 | 218525 | 218547 | - | 0 | GCCGGACGGAGTCGGTGGCG | GGG | \|\|\|\|\|\|\|\|[\|\|\|\|\|\|\|\|\|\|\|\|]PAM | NA | Intergenic | NA | NA |
| T18118 | GACGGAGTCGGTGGCGGGGTCGG | 1000 | | | | | | | | | |
| Oligo fwd | CACCGACGGAGTCGGTGGCGGGGT | | | | | | | | | | |
| Oligo rev | AAACACCCCGCCACCGACTCCGTC | | | | | | | | | | |
| Chromosome | start | end | strand | MM | target_seq | PAM | alignment | distance | position | gene name | gene id |
| GU305914.1 | 218521 | 218543 | - | 0 | GACGGAGTCGGTGGCGGGGT | CGG | \|\|\|\|\|\|\|\|[\|\|\|\|\|\|\|\|\|\|\|\|]PAM | NA | Intergenic | NA | NA |
| GU305914.1 | 109443 | 109465 | + | 4 | GACGGCGGCGGTAGCGGCGT | CGG | \|\|\|\|\|-\|-[\|\|\|\|-\|\|\|\|-\|\|]PAM | NA | Intergenic | NA | NA |
| T18119 | GAGTCGGTGGCGGGGTCGGACGG | 1000 | | | | | | | | | |
| Oligo fwd | CACCGAGTCGGTGGCGGGGTCGGA | | | | | | | | | | |
| Oligo rev | AAACTCCGACCCCGCCACCGACTC | | | | | | | | | | |
| Chromosome | start | end | strand | MM | target_seq | PAM | alignment | distance | position | gene name | gene id |
| GU305914.1 | 218517 | 218539 | - | 0 | GAGTCGGTGGCGGGGTCGGA | CGG | \|\|\|\|\|\|\|\|[\|\|\|\|\|\|\|\|\|\|\|\|]PAM | NA | Intergenic | NA | NA |
| T18120 | GATCAAGATCACCCTTCCCACGG | 1000 | | | | | | | | | |
| Oligo fwd | CACCGATCAAGATCACCCTTCCCA | | | | | | | | | | |
| Oligo rev | AAACTGGGAAGGGTGATCTTGATC | | | | | | | | | | |
| Chromosome | start | end | strand | MM | target_seq | PAM | alignment | distance | position | gene name | gene id |
| GU305914.1 | 218495 | 218517 | - | 0 | GATCAAGATCACCCTTCCCA | CGG | \|\|\|\|\|\|\|\|[\|\|\|\|\|\|\|\|\|\|\|\|]PAM | NA | Intergenic | NA | NA |
| T18121 | AGCTGTCTAGTGTCTTCGCCCGG | 1000 | | | | | | | | | |
| Oligo fwd | CACCAGCTGTCTAGTGTCTTCGCC | | | | | | | | | | |
| Oligo rev | AAACGGCGAAGACACTAGACAGCT | | | | | | | | | | |
| Chromosome | start | end | strand | MM | target_seq | PAM | alignment | distance | position | gene name | gene id |
| GU305914.1 | 218455 | 218477 | - | 0 | AGCTGTCTAGTGTCTTCGCC | CGG | \|\|\|\|\|\|\|\|[\|\|\|\|\|\|\|\|\|\|\|\|]PAM | NA | Intergenic | NA | NA |
| T18122 | TAGTGTCTTCGCCCGGAAAGTGG | 1000 | | | | | | | | | |
| Oligo fwd | CACCTAGTGTCTTCGCCCGGAAAG | | | | | | | | | | |
| Oligo rev | AAACCTTTCCGGGCGAAGACACTA | | | | | | | | | | |
| Chromosome | start | end | strand | MM | target_seq | PAM | alignment | distance | position | gene name | gene id |
| GU305914.1 | 218448 | 218470 | - | 0 | TAGTGTCTTCGCCCGGAAAG | TGG | \|\|\|\|\|\|\|\|[\|\|\|\|\|\|\|\|\|\|\|\|]PAM | NA | Intergenic | NA | NA |
| T18123 | AGTGTCTTCGCCCGGAAAGTGGG | 1000 | | | | | | | | | |
| Oligo fwd | CACCAGTGTCTTCGCCCGGAAAGT | | | | | | | | | | |
| Oligo rev | AAACACTTTCCGGGCGAAGACACT | | | | | | | | | | |
| Chromosome | start | end | strand | MM | target_seq | PAM | alignment | distance | position | gene name | gene id |
| GU305914.1 | 218447 | 218469 | - | 0 | AGTGTCTTCGCCCGGAAAGT | GGG | \|\|\|\|\|\|\|\|[\|\|\|\|\|\|\|\|\|\|\|\|]PAM | NA | Intergenic | NA | NA |
| T18124 | CGGAAAGTGGGCGTCTTTCATGG | 1000 | | | | | | | | | |
| Oligo fwd | CACCCGGAAAGTGGGCGTCTTTCA | | | | | | | | | | |
| Oligo rev | AAACTGAAAGACGCCCACTTTCCG | | | | | | | | | | |
| Chromosome | start | end | strand | MM | target_seq | PAM | alignment | distance | position | gene name | gene id |

<table>
<tr><td>GU305914.1</td><td>218435</td><td>218457</td><td>-</td><td>0</td><td>CGGAAAGTGGGCGTCTTTCA</td><td>TGG</td><td>||||||||[||||||||||||]PAM</td><td>NA</td><td>Intergenic</td><td>NA</td><td>NA</td></tr>
<tr><td>T18125</td><td>AAAGTGGGCGTCTTTCATGGTGG</td><td>1000</td><td></td><td></td><td></td><td></td><td></td><td></td><td></td><td></td><td></td></tr>
<tr><td>Oligo fwd</td><td>CACCAAAGTGGGCGTCTTTCATGG</td><td></td><td></td><td></td><td></td><td></td><td></td><td></td><td></td><td></td><td></td></tr>
<tr><td>Oligo rev</td><td>AAACCCATGAAAGACGCCCACTTT</td><td></td><td></td><td></td><td></td><td></td><td></td><td></td><td></td><td></td><td></td></tr>
<tr><td>Chromosome</td><td>start</td><td>end</td><td>strand</td><td>MM</td><td>target_seq</td><td>PAM</td><td>alignment</td><td>distance</td><td>position</td><td>gene name</td><td>gene id</td></tr>
<tr><td>GU305914.1</td><td>218432</td><td>218454</td><td>-</td><td>0</td><td>AAAGTGGGCGTCTTTCATGG</td><td>TGG</td><td>||||||||[||||||||||||]PAM</td><td>NA</td><td>Intergenic</td><td>NA</td><td>NA</td></tr>
<tr><td>T18126</td><td>ATGGTGGTGCCCTATCCTTACGG</td><td>1000</td><td></td><td></td><td></td><td></td><td></td><td></td><td></td><td></td><td></td></tr>
<tr><td>Oligo fwd</td><td>CACCATGGTGGTGCCCTATCCTTA</td><td></td><td></td><td></td><td></td><td></td><td></td><td></td><td></td><td></td><td></td></tr>
<tr><td>Oligo rev</td><td>AAACTAAGGATAGGGCACCACCAT</td><td></td><td></td><td></td><td></td><td></td><td></td><td></td><td></td><td></td><td></td></tr>
<tr><td>Chromosome</td><td>start</td><td>end</td><td>strand</td><td>MM</td><td>target_seq</td><td>PAM</td><td>alignment</td><td>distance</td><td>position</td><td>gene name</td><td>gene id</td></tr>
<tr><td>GU305914.1</td><td>218416</td><td>218438</td><td>-</td><td>0</td><td>ATGGTGGTGCCCTATCCTTA</td><td>CGG</td><td>||||||||[||||||||||||]PAM</td><td>NA</td><td>Intergenic</td><td>NA</td><td>NA</td></tr>
<tr><td>T18127</td><td>TCCTTACGGACCGCTACCCATGG</td><td>1000</td><td></td><td></td><td></td><td></td><td></td><td></td><td></td><td></td><td></td></tr>
<tr><td>Oligo fwd</td><td>CACCTCCTTACGGACCGCTACCCA</td><td></td><td></td><td></td><td></td><td></td><td></td><td></td><td></td><td></td><td></td></tr>
<tr><td>Oligo rev</td><td>AAACTGGGTAGCGGTCCGTAAGGA</td><td></td><td></td><td></td><td></td><td></td><td></td><td></td><td></td><td></td><td></td></tr>
<tr><td>Chromosome</td><td>start</td><td>end</td><td>strand</td><td>MM</td><td>target_seq</td><td>PAM</td><td>alignment</td><td>distance</td><td>position</td><td>gene name</td><td>gene id</td></tr>
<tr><td>GU305914.1</td><td>218402</td><td>218424</td><td>-</td><td>0</td><td>TCCTTACGGACCGCTACCCA</td><td>TGG</td><td>||||||||[||||||||||||]PAM</td><td>NA</td><td>Intergenic</td><td>NA</td><td>NA</td></tr>
<tr><td>T18128</td><td>CCTTACGGACCGCTACCCATGGG</td><td>1000</td><td></td><td></td><td></td><td></td><td></td><td></td><td></td><td></td><td></td></tr>
<tr><td>Oligo fwd</td><td>CACCCCTTACGGACCGCTACCCAT</td><td></td><td></td><td></td><td></td><td></td><td></td><td></td><td></td><td></td><td></td></tr>
<tr><td>Oligo rev</td><td>AAACATGGGTAGCGGTCCGTAAGG</td><td></td><td></td><td></td><td></td><td></td><td></td><td></td><td></td><td></td><td></td></tr>
<tr><td>Chromosome</td><td>start</td><td>end</td><td>strand</td><td>MM</td><td>target_seq</td><td>PAM</td><td>alignment</td><td>distance</td><td>position</td><td>gene name</td><td>gene id</td></tr>
<tr><td>GU305914.1</td><td>218401</td><td>218423</td><td>-</td><td>0</td><td>CCTTACGGACCGCTACCCAT</td><td>GGG</td><td>||||||||[||||||||||||]PAM</td><td>NA</td><td>Intergenic</td><td>NA</td><td>NA</td></tr>
<tr><td>T18129</td><td>ACCGCTACCCATGGGCAGCGAGG</td><td>1000</td><td></td><td></td><td></td><td></td><td></td><td></td><td></td><td></td><td></td></tr>
<tr><td>Oligo fwd</td><td>CACCACCGCTACCCATGGGCAGCG</td><td></td><td></td><td></td><td></td><td></td><td></td><td></td><td></td><td></td><td></td></tr>
<tr><td>Oligo rev</td><td>AAACCGCTGCCCATGGGTAGCGGT</td><td></td><td></td><td></td><td></td><td></td><td></td><td></td><td></td><td></td><td></td></tr>
<tr><td>Chromosome</td><td>start</td><td>end</td><td>strand</td><td>MM</td><td>target_seq</td><td>PAM</td><td>alignment</td><td>distance</td><td>position</td><td>gene name</td><td>gene id</td></tr>
<tr><td>GU305914.1</td><td>218393</td><td>218415</td><td>-</td><td>0</td><td>ACCGCTACCCATGGGCAGCG</td><td>AGG</td><td>||||||||[||||||||||||]PAM</td><td>NA</td><td>Intergenic</td><td>NA</td><td>NA</td></tr>
<tr><td>T18130</td><td>ACCCATGGGCAGCGAGGTTCCGG</td><td>1000</td><td></td><td></td><td></td><td></td><td></td><td></td><td></td><td></td><td></td></tr>
<tr><td>Oligo fwd</td><td>CACCACCCATGGGCAGCGAGGTTC</td><td></td><td></td><td></td><td></td><td></td><td></td><td></td><td></td><td></td><td></td></tr>
<tr><td>Oligo rev</td><td>AAACGAACCTCGCTGCCCATGGGT</td><td></td><td></td><td></td><td></td><td></td><td></td><td></td><td></td><td></td><td></td></tr>
<tr><td>Chromosome</td><td>start</td><td>end</td><td>strand</td><td>MM</td><td>target_seq</td><td>PAM</td><td>alignment</td><td>distance</td><td>position</td><td>gene name</td><td>gene id</td></tr>
<tr><td>GU305914.1</td><td>218387</td><td>218409</td><td>-</td><td>0</td><td>ACCCATGGGCAGCGAGGTTC</td><td>CGG</td><td>||||||||[||||||||||||]PAM</td><td>NA</td><td>Intergenic</td><td>NA</td><td>NA</td></tr>
<tr><td>T18131</td><td>CAGCGAGGTTCCGGTGATGCTGG</td><td>1000</td><td></td><td></td><td></td><td></td><td></td><td></td><td></td><td></td><td></td></tr>
<tr><td>Oligo fwd</td><td>CACCCAGCGAGGTTCCGGTGATGC</td><td></td><td></td><td></td><td></td><td></td><td></td><td></td><td></td><td></td><td></td></tr>
<tr><td>Oligo rev</td><td>AAACGCATCACCGGAACCTCGCTG</td><td></td><td></td><td></td><td></td><td></td><td></td><td></td><td></td><td></td><td></td></tr>
<tr><td>Chromosome</td><td>start</td><td>end</td><td>strand</td><td>MM</td><td>target_seq</td><td>PAM</td><td>alignment</td><td>distance</td><td>position</td><td>gene name</td><td>gene id</td></tr>
<tr><td>GU305914.1</td><td>218378</td><td>218400</td><td>-</td><td>0</td><td>CAGCGAGGTTCCGGTGATGC</td><td>TGG</td><td>||||||||[||||||||||||]PAM</td><td>NA</td><td>Intergenic</td><td>NA</td><td>NA</td></tr>
<tr><td>T18132</td><td>CGAGGTTCCGGTGATGCTGGTGG</td><td>1000</td><td></td><td></td><td></td><td></td><td></td><td></td><td></td><td></td><td></td></tr>
<tr><td>Oligo fwd</td><td>CACCCGAGGTTCCGGTGATGCTGG</td><td></td><td></td><td></td><td></td><td></td><td></td><td></td><td></td><td></td><td></td></tr>
<tr><td>Oligo rev</td><td>AAACCCAGCATCACCGGAACCTCG</td><td></td><td></td><td></td><td></td><td></td><td></td><td></td><td></td><td></td><td></td></tr>
<tr><td>Chromosome</td><td>start</td><td>end</td><td>strand</td><td>MM</td><td>target_seq</td><td>PAM</td><td>alignment</td><td>distance</td><td>position</td><td>gene name</td><td>gene id</td></tr>
<tr><td>GU305914.1</td><td>218375</td><td>218397</td><td>-</td><td>0</td><td>CGAGGTTCCGGTGATGCTGG</td><td>TGG</td><td>||||||||[||||||||||||]PAM</td><td>NA</td><td>Intergenic</td><td>NA</td><td>NA</td></tr>
<tr><td>T18133</td><td>GCTGGTGGCCGCGATCGTCGCGG</td><td>1000</td><td></td><td></td><td></td><td></td><td></td><td></td><td></td><td></td><td></td></tr>
<tr><td>Oligo fwd</td><td>CACCGCTGGTGGCCGCGATCGTCG</td><td></td><td></td><td></td><td></td><td></td><td></td><td></td><td></td><td></td><td></td></tr>
<tr><td>Oligo rev</td><td>AAACCGACGATCGCGGCCACCAGC</td><td></td><td></td><td></td><td></td><td></td><td></td><td></td><td></td><td></td><td></td></tr>
<tr><td>Chromosome</td><td>start</td><td>end</td><td>strand</td><td>MM</td><td>target_seq</td><td>PAM</td><td>alignment</td><td>distance</td><td>position</td><td>gene name</td><td>gene id</td></tr>
<tr><td>GU305914.1</td><td>218360</td><td>218382</td><td>-</td><td>0</td><td>GCTGGTGGCCGCGATCGTCG</td><td>CGG</td><td>||||||||[||||||||||||]PAM</td><td>NA</td><td>Intergenic</td><td>NA</td><td>NA</td></tr>
<tr><td>GU305914.1</td><td>138095</td><td>138117</td><td>-</td><td>4</td><td>GCTGTCGGCGGCGATCGACG</td><td>CGG</td><td>||||--||[|-|||||||-||]PAM</td><td>NA</td><td>Intergenic</td><td>NA</td><td>NA</td></tr>
<tr><td>T18134</td><td>CGCGATCGTCGCGGTCTTAGTGG</td><td>1000</td><td></td><td></td><td></td><td></td><td></td><td></td><td></td><td></td><td></td></tr>
<tr><td>Oligo fwd</td><td>CACCCGCGATCGTCGCGGTCTTAG</td><td></td><td></td><td></td><td></td><td></td><td></td><td></td><td></td><td></td><td></td></tr>
<tr><td>Oligo rev</td><td>AAACCTAAGACCGCGACGATCGCG</td><td></td><td></td><td></td><td></td><td></td><td></td><td></td><td></td><td></td><td></td></tr>
<tr><td>Chromosome</td><td>start</td><td>end</td><td>strand</td><td>MM</td><td>target_seq</td><td>PAM</td><td>alignment</td><td>distance</td><td>position</td><td>gene name</td><td>gene id</td></tr>
<tr><td>GU305914.1</td><td>218351</td><td>218373</td><td>-</td><td>0</td><td>CGCGATCGTCGCGGTCTTAG</td><td>TGG</td><td>||||||||[||||||||||||]PAM</td><td>NA</td><td>Intergenic</td><td>NA</td><td>NA</td></tr>
<tr><td>T18135</td><td>TTAGTGGTCTTCTTGATGATAGG</td><td>1000</td><td></td><td></td><td></td><td></td><td></td><td></td><td></td><td></td><td></td></tr>
<tr><td>Oligo fwd</td><td>CACCTTAGTGGTCTTCTTGATGAT</td><td></td><td></td><td></td><td></td><td></td><td></td><td></td><td></td><td></td><td></td></tr>
<tr><td>Oligo rev</td><td>AAACATCATCAAGAAGACCACTAA</td><td></td><td></td><td></td><td></td><td></td><td></td><td></td><td></td><td></td><td></td></tr>
<tr><td>Chromosome</td><td>start</td><td>end</td><td>strand</td><td>MM</td><td>target_seq</td><td>PAM</td><td>alignment</td><td>distance</td><td>position</td><td>gene name</td><td>gene id</td></tr>
<tr><td>GU305914.1</td><td>218335</td><td>218357</td><td>-</td><td>0</td><td>TTAGTGGTCTTCTTGATGAT</td><td>AGG</td><td>||||||||[||||||||||||]PAM</td><td>NA</td><td>Intergenic</td><td>NA</td><td>NA</td></tr>
<tr><td>T18136</td><td>TAGTGGTCTTCTTGATGATAGGG</td><td>1000</td><td></td><td></td><td></td><td></td><td></td><td></td><td></td><td></td><td></td></tr>
<tr><td>Oligo fwd</td><td>CACCTAGTGGTCTTCTTGATGATA</td><td></td><td></td><td></td><td></td><td></td><td></td><td></td><td></td><td></td><td></td></tr>
<tr><td>Oligo rev</td><td>AAACTATCATCAAGAAGACCACTA</td><td></td><td></td><td></td><td></td><td></td><td></td><td></td><td></td><td></td><td></td></tr>
<tr><td>Chromosome</td><td>start</td><td>end</td><td>strand</td><td>MM</td><td>target_seq</td><td>PAM</td><td>alignment</td><td>distance</td><td>position</td><td>gene name</td><td>gene id</td></tr>
<tr><td>GU305914.1</td><td>218334</td><td>218356</td><td>-</td><td>0</td><td>TAGTGGTCTTCTTGATGATA</td><td>GGG</td><td>||||||||[||||||||||||]PAM</td><td>NA</td><td>Intergenic</td><td>NA</td><td>NA</td></tr>
<tr><td>T18137</td><td>TTCTTGATGATAGGGTGCATCGG</td><td>1000</td><td></td><td></td><td></td><td></td><td></td><td></td><td></td><td></td><td></td></tr>
<tr><td>Oligo fwd</td><td>CACCTTCTTGATGATAGGGTGCAT</td><td></td><td></td><td></td><td></td><td></td><td></td><td></td><td></td><td></td><td></td></tr>
<tr><td>Oligo rev</td><td>AAACATGCACCCTATCATCAAGAA</td><td></td><td></td><td></td><td></td><td></td><td></td><td></td><td></td><td></td><td></td></tr>
<tr><td>Chromosome</td><td>start</td><td>end</td><td>strand</td><td>MM</td><td>target_seq</td><td>PAM</td><td>alignment</td><td>distance</td><td>position</td><td>gene name</td><td>gene id</td></tr>
<tr><td>GU305914.1</td><td>218326</td><td>218348</td><td>-</td><td>0</td><td>TTCTTGATGATAGGGTGCAT</td><td>CGG</td><td>||||||||[||||||||||||]PAM</td><td>NA</td><td>Intergenic</td><td>NA</td><td>NA</td></tr>
<tr><td>T18138</td><td>TTTGCTCGTTCGTCTGCGAGCGG</td><td>1000</td><td></td><td></td><td></td><td></td><td></td><td></td><td></td><td></td><td></td></tr>
<tr><td>Oligo fwd</td><td>CACCTTTGCTCGTTCGTCTGCGAG</td><td></td><td></td><td></td><td></td><td></td><td></td><td></td><td></td><td></td><td></td></tr>
<tr><td>Oligo rev</td><td>AAACCTCGCAGACGAACGAGCAAA</td><td></td><td></td><td></td><td></td><td></td><td></td><td></td><td></td><td></td><td></td></tr>
<tr><td>Chromosome</td><td>start</td><td>end</td><td>strand</td><td>MM</td><td>target_seq</td><td>PAM</td><td>alignment</td><td>distance</td><td>position</td><td>gene name</td><td>gene id</td></tr>
<tr><td>GU305914.1</td><td>218301</td><td>218323</td><td>-</td><td>0</td><td>TTTGCTCGTTCGTCTGCGAG</td><td>CGG</td><td>||||||||[||||||||||||]PAM</td><td>NA</td><td>Intergenic</td><td>NA</td><td>NA</td></tr>
<tr><td>T18139</td><td>AAATCTGTCGCCCTCTCTAACGG</td><td>1000</td><td></td><td></td><td></td><td></td><td></td><td></td><td></td><td></td><td></td></tr>
<tr><td>Oligo fwd</td><td>CACCAAATCTGTCGCCCTCTCTAA</td><td></td><td></td><td></td><td></td><td></td><td></td><td></td><td></td><td></td><td></td></tr>
</table>

<table>
<tr><td>Oligo rev</td><td>AAACTTAGAGAGGGCGACAGATTT</td><td></td><td></td><td></td><td></td><td></td><td></td><td></td><td></td><td></td><td></td></tr>
<tr><td>Chromosome</td><td>start</td><td>end</td><td>strand</td><td>MM</td><td>target_seq</td><td>PAM</td><td>alignment</td><td>distance</td><td>position</td><td>gene name</td><td>gene id</td></tr>
<tr><td>GU305914.1</td><td>218245</td><td>218267</td><td>-</td><td>0</td><td>AAATCTGTCGCCCTCTCTAA</td><td>CGG</td><td>||||||||[||||||||||||]PAM</td><td>NA</td><td>Intergenic</td><td>NA</td><td>NA</td></tr>
<tr><td>T18140</td><td>CAGTTTGTCAGTATTAACAATGG</td><td>1000</td><td></td><td></td><td></td><td></td><td></td><td></td><td></td><td></td><td></td></tr>
<tr><td>Oligo fwd</td><td>CACCCAGTTTGTCAGTATTAACAA</td><td></td><td></td><td></td><td></td><td></td><td></td><td></td><td></td><td></td><td></td></tr>
<tr><td>Oligo rev</td><td>AAACTTGTTAATACTGACAAACTG</td><td></td><td></td><td></td><td></td><td></td><td></td><td></td><td></td><td></td><td></td></tr>
<tr><td>Chromosome</td><td>start</td><td>end</td><td>strand</td><td>MM</td><td>target_seq</td><td>PAM</td><td>alignment</td><td>distance</td><td>position</td><td>gene name</td><td>gene id</td></tr>
<tr><td>GU305914.1</td><td>218206</td><td>218228</td><td>-</td><td>0</td><td>CAGTTTGTCAGTATTAACAA</td><td>TGG</td><td>||||||||[||||||||||||]PAM</td><td>NA</td><td>Intergenic</td><td>NA</td><td>NA</td></tr>
<tr><td>T18141</td><td>TTCTATCGAAATCTCTGTCGAGG</td><td>1000</td><td></td><td></td><td></td><td></td><td></td><td></td><td></td><td></td><td></td></tr>
<tr><td>Oligo fwd</td><td>CACCTTCTATCGAAATCTCTGTCG</td><td></td><td></td><td></td><td></td><td></td><td></td><td></td><td></td><td></td><td></td></tr>
<tr><td>Oligo rev</td><td>AAACCGACAGAGATTTCGATAGAA</td><td></td><td></td><td></td><td></td><td></td><td></td><td></td><td></td><td></td><td></td></tr>
<tr><td>Chromosome</td><td>start</td><td>end</td><td>strand</td><td>MM</td><td>target_seq</td><td>PAM</td><td>alignment</td><td>distance</td><td>position</td><td>gene name</td><td>gene id</td></tr>
<tr><td>GU305914.1</td><td>218180</td><td>218202</td><td>-</td><td>0</td><td>TTCTATCGAAATCTCTGTCG</td><td>AGG</td><td>||||||||[||||||||||||]PAM</td><td>NA</td><td>Intergenic</td><td>NA</td><td>NA</td></tr>
<tr><td>T18142</td><td>CGCTATTTCTATCGCGCGTTCGG</td><td>1000</td><td></td><td></td><td></td><td></td><td></td><td></td><td></td><td></td><td></td></tr>
<tr><td>Oligo fwd</td><td>CACCCGCTATTTCTATCGCGCGTT</td><td></td><td></td><td></td><td></td><td></td><td></td><td></td><td></td><td></td><td></td></tr>
<tr><td>Oligo rev</td><td>AAACAACGCGCGATAGAAATAGCG</td><td></td><td></td><td></td><td></td><td></td><td></td><td></td><td></td><td></td><td></td></tr>
<tr><td>Chromosome</td><td>start</td><td>end</td><td>strand</td><td>MM</td><td>target_seq</td><td>PAM</td><td>alignment</td><td>distance</td><td>position</td><td>gene name</td><td>gene id</td></tr>
<tr><td>GU305914.1</td><td>218123</td><td>218145</td><td>-</td><td>0</td><td>CGCTATTTCTATCGCGCGTT</td><td>CGG</td><td>||||||||[||||||||||||]PAM</td><td>NA</td><td>Intergenic</td><td>NA</td><td>NA</td></tr>
<tr><td>T18143</td><td>GCTATTTCTATCGCGCGTTCGGG</td><td>1000</td><td></td><td></td><td></td><td></td><td></td><td></td><td></td><td></td><td></td></tr>
<tr><td>Oligo fwd</td><td>CACCGCTATTTCTATCGCGCGTTC</td><td></td><td></td><td></td><td></td><td></td><td></td><td></td><td></td><td></td><td></td></tr>
<tr><td>Oligo rev</td><td>AAACGAACGCGCGATAGAAATAGC</td><td></td><td></td><td></td><td></td><td></td><td></td><td></td><td></td><td></td><td></td></tr>
<tr><td>Chromosome</td><td>start</td><td>end</td><td>strand</td><td>MM</td><td>target_seq</td><td>PAM</td><td>alignment</td><td>distance</td><td>position</td><td>gene name</td><td>gene id</td></tr>
<tr><td>GU305914.1</td><td>218122</td><td>218144</td><td>-</td><td>0</td><td>GCTATTTCTATCGCGCGTTC</td><td>GGG</td><td>||||||||[||||||||||||]PAM</td><td>NA</td><td>Intergenic</td><td>NA</td><td>NA</td></tr>
<tr><td>T18144</td><td>CTATTTCTATCGCGCGTTCGGGG</td><td>1000</td><td></td><td></td><td></td><td></td><td></td><td></td><td></td><td></td><td></td></tr>
<tr><td>Oligo fwd</td><td>CACCCTATTTCTATCGCGCGTTCG</td><td></td><td></td><td></td><td></td><td></td><td></td><td></td><td></td><td></td><td></td></tr>
<tr><td>Oligo rev</td><td>AAACCGAACGCGCGATAGAAATAG</td><td></td><td></td><td></td><td></td><td></td><td></td><td></td><td></td><td></td><td></td></tr>
<tr><td>Chromosome</td><td>start</td><td>end</td><td>strand</td><td>MM</td><td>target_seq</td><td>PAM</td><td>alignment</td><td>distance</td><td>position</td><td>gene name</td><td>gene id</td></tr>
<tr><td>GU305914.1</td><td>218121</td><td>218143</td><td>-</td><td>0</td><td>CTATTTCTATCGCGCGTTCG</td><td>GGG</td><td>||||||||[||||||||||||]PAM</td><td>NA</td><td>Intergenic</td><td>NA</td><td>NA</td></tr>
<tr><td>T18145</td><td>CGCGCGTTCGGGGAATAGTCAGG</td><td>1000</td><td></td><td></td><td></td><td></td><td></td><td></td><td></td><td></td><td></td></tr>
<tr><td>Oligo fwd</td><td>CACCCGCGCGTTCGGGGAATAGTC</td><td></td><td></td><td></td><td></td><td></td><td></td><td></td><td></td><td></td><td></td></tr>
<tr><td>Oligo rev</td><td>AAACGACTATTCCCCGAACGCGCG</td><td></td><td></td><td></td><td></td><td></td><td></td><td></td><td></td><td></td><td></td></tr>
<tr><td>Chromosome</td><td>start</td><td>end</td><td>strand</td><td>MM</td><td>target_seq</td><td>PAM</td><td>alignment</td><td>distance</td><td>position</td><td>gene name</td><td>gene id</td></tr>
<tr><td>GU305914.1</td><td>218111</td><td>218133</td><td>-</td><td>0</td><td>CGCGCGTTCGGGGAATAGTC</td><td>AGG</td><td>||||||||[||||||||||||]PAM</td><td>NA</td><td>Intergenic</td><td>NA</td><td>NA</td></tr>
<tr><td>T18146</td><td>CAGGTTGTGACATGTTGTCGAGG</td><td>1000</td><td></td><td></td><td></td><td></td><td></td><td></td><td></td><td></td><td></td></tr>
<tr><td>Oligo fwd</td><td>CACCCAGGTTGTGACATGTTGTCG</td><td></td><td></td><td></td><td></td><td></td><td></td><td></td><td></td><td></td><td></td></tr>
<tr><td>Oligo rev</td><td>AAACCGACAACATGTCACAACCTG</td><td></td><td></td><td></td><td></td><td></td><td></td><td></td><td></td><td></td><td></td></tr>
<tr><td>Chromosome</td><td>start</td><td>end</td><td>strand</td><td>MM</td><td>target_seq</td><td>PAM</td><td>alignment</td><td>distance</td><td>position</td><td>gene name</td><td>gene id</td></tr>
<tr><td>GU305914.1</td><td>218092</td><td>218114</td><td>-</td><td>0</td><td>CAGGTTGTGACATGTTGTCG</td><td>AGG</td><td>||||||||[||||||||||||]PAM</td><td>NA</td><td>Intergenic</td><td>NA</td><td>NA</td></tr>
<tr><td>T18147</td><td>GTTGTGACATGTTGTCGAGGTGG</td><td>1000</td><td></td><td></td><td></td><td></td><td></td><td></td><td></td><td></td><td></td></tr>
<tr><td>Oligo fwd</td><td>CACCGTTGTGACATGTTGTCGAGG</td><td></td><td></td><td></td><td></td><td></td><td></td><td></td><td></td><td></td><td></td></tr>
<tr><td>Oligo rev</td><td>AAACCCTCGACAACATGTCACAAC</td><td></td><td></td><td></td><td></td><td></td><td></td><td></td><td></td><td></td><td></td></tr>
<tr><td>Chromosome</td><td>start</td><td>end</td><td>strand</td><td>MM</td><td>target_seq</td><td>PAM</td><td>alignment</td><td>distance</td><td>position</td><td>gene name</td><td>gene id</td></tr>
<tr><td>GU305914.1</td><td>218089</td><td>218111</td><td>-</td><td>0</td><td>GTTGTGACATGTTGTCGAGG</td><td>TGG</td><td>||||||||[||||||||||||]PAM</td><td>NA</td><td>Intergenic</td><td>NA</td><td>NA</td></tr>
<tr><td>T18148</td><td>TTGTGACATGTTGTCGAGGTGGG</td><td>1000</td><td></td><td></td><td></td><td></td><td></td><td></td><td></td><td></td><td></td></tr>
<tr><td>Oligo fwd</td><td>CACCTTGTGACATGTTGTCGAGGT</td><td></td><td></td><td></td><td></td><td></td><td></td><td></td><td></td><td></td><td></td></tr>
<tr><td>Oligo rev</td><td>AAACACCTCGACAACATGTCACAA</td><td></td><td></td><td></td><td></td><td></td><td></td><td></td><td></td><td></td><td></td></tr>
<tr><td>Chromosome</td><td>start</td><td>end</td><td>strand</td><td>MM</td><td>target_seq</td><td>PAM</td><td>alignment</td><td>distance</td><td>position</td><td>gene name</td><td>gene id</td></tr>
<tr><td>GU305914.1</td><td>218088</td><td>218110</td><td>-</td><td>0</td><td>TTGTGACATGTTGTCGAGGT</td><td>GGG</td><td>||||||||[||||||||||||]PAM</td><td>NA</td><td>Intergenic</td><td>NA</td><td>NA</td></tr>
<tr><td>T18149</td><td>TGTGACATGTTGTCGAGGTGGGG</td><td>1000</td><td></td><td></td><td></td><td></td><td></td><td></td><td></td><td></td><td></td></tr>
<tr><td>Oligo fwd</td><td>CACCTGTGACATGTTGTCGAGGTG</td><td></td><td></td><td></td><td></td><td></td><td></td><td></td><td></td><td></td><td></td></tr>
<tr><td>Oligo rev</td><td>AAACCACCTCGACAACATGTCACA</td><td></td><td></td><td></td><td></td><td></td><td></td><td></td><td></td><td></td><td></td></tr>
<tr><td>Chromosome</td><td>start</td><td>end</td><td>strand</td><td>MM</td><td>target_seq</td><td>PAM</td><td>alignment</td><td>distance</td><td>position</td><td>gene name</td><td>gene id</td></tr>
<tr><td>GU305914.1</td><td>218087</td><td>218109</td><td>-</td><td>0</td><td>TGTGACATGTTGTCGAGGTG</td><td>GGG</td><td>||||||||[||||||||||||]PAM</td><td>NA</td><td>Intergenic</td><td>NA</td><td>NA</td></tr>
<tr><td>T18150</td><td>GTTGTCGAGGTGGGGAACGCTGG</td><td>1000</td><td></td><td></td><td></td><td></td><td></td><td></td><td></td><td></td><td></td></tr>
<tr><td>Oligo fwd</td><td>CACCGTTGTCGAGGTGGGGAACGC</td><td></td><td></td><td></td><td></td><td></td><td></td><td></td><td></td><td></td><td></td></tr>
<tr><td>Oligo rev</td><td>AAACGCGTTCCCCACCTCGACAAC</td><td></td><td></td><td></td><td></td><td></td><td></td><td></td><td></td><td></td><td></td></tr>
<tr><td>Chromosome</td><td>start</td><td>end</td><td>strand</td><td>MM</td><td>target_seq</td><td>PAM</td><td>alignment</td><td>distance</td><td>position</td><td>gene name</td><td>gene id</td></tr>
<tr><td>GU305914.1</td><td>218079</td><td>218101</td><td>-</td><td>0</td><td>GTTGTCGAGGTGGGGAACGC</td><td>TGG</td><td>||||||||[||||||||||||]PAM</td><td>NA</td><td>Intergenic</td><td>NA</td><td>NA</td></tr>
<tr><td>T18151</td><td>TTGTCGAGGTGGGGAACGCTGGG</td><td>1000</td><td></td><td></td><td></td><td></td><td></td><td></td><td></td><td></td><td></td></tr>
<tr><td>Oligo fwd</td><td>CACCTTGTCGAGGTGGGGAACGCT</td><td></td><td></td><td></td><td></td><td></td><td></td><td></td><td></td><td></td><td></td></tr>
<tr><td>Oligo rev</td><td>AAACAGCGTTCCCCACCTCGACAA</td><td></td><td></td><td></td><td></td><td></td><td></td><td></td><td></td><td></td><td></td></tr>
<tr><td>Chromosome</td><td>start</td><td>end</td><td>strand</td><td>MM</td><td>target_seq</td><td>PAM</td><td>alignment</td><td>distance</td><td>position</td><td>gene name</td><td>gene id</td></tr>
<tr><td>GU305914.1</td><td>218078</td><td>218100</td><td>-</td><td>0</td><td>TTGTCGAGGTGGGGAACGCT</td><td>GGG</td><td>||||||||[||||||||||||]PAM</td><td>NA</td><td>Intergenic</td><td>NA</td><td>NA</td></tr>
<tr><td>T18152</td><td>GGGAACGCTGGGCTGCCTTCTGG</td><td>1000</td><td></td><td></td><td></td><td></td><td></td><td></td><td></td><td></td><td></td></tr>
<tr><td>Oligo fwd</td><td>CACCGGGAACGCTGGGCTGCCTTC</td><td></td><td></td><td></td><td></td><td></td><td></td><td></td><td></td><td></td><td></td></tr>
<tr><td>Oligo rev</td><td>AAACGAAGGCAGCCCAGCGTTCCC</td><td></td><td></td><td></td><td></td><td></td><td></td><td></td><td></td><td></td><td></td></tr>
<tr><td>Chromosome</td><td>start</td><td>end</td><td>strand</td><td>MM</td><td>target_seq</td><td>PAM</td><td>alignment</td><td>distance</td><td>position</td><td>gene name</td><td>gene id</td></tr>
<tr><td>GU305914.1</td><td>218067</td><td>218089</td><td>-</td><td>0</td><td>GGGAACGCTGGGCTGCCTTC</td><td>TGG</td><td>||||||||[||||||||||||]PAM</td><td>NA</td><td>Intergenic</td><td>NA</td><td>NA</td></tr>
<tr><td>T18153</td><td>CCGCTGAGTATCCAGATCATCGG</td><td>1000</td><td></td><td></td><td></td><td></td><td></td><td></td><td></td><td></td><td></td></tr>
<tr><td>Oligo fwd</td><td>CACCCCGCTGAGTATCCAGATCAT</td><td></td><td></td><td></td><td></td><td></td><td></td><td></td><td></td><td></td><td></td></tr>
<tr><td>Oligo rev</td><td>AAACATGATCTGGATACTCAGCGG</td><td></td><td></td><td></td><td></td><td></td><td></td><td></td><td></td><td></td><td></td></tr>
<tr><td>Chromosome</td><td>start</td><td>end</td><td>strand</td><td>MM</td><td>target_seq</td><td>PAM</td><td>alignment</td><td>distance</td><td>position</td><td>gene name</td><td>gene id</td></tr>
<tr><td>GU305914.1</td><td>217997</td><td>218019</td><td>-</td><td>0</td><td>CCGCTGAGTATCCAGATCAT</td><td>CGG</td><td>||||||||[||||||||||||]PAM</td><td>NA</td><td>Intergenic</td><td>NA</td><td>NA</td></tr>
<tr><td>T18154</td><td>AGATCATCGGATACCTCGACAGG</td><td>1000</td><td></td><td></td><td></td><td></td><td></td><td></td><td></td><td></td><td></td></tr>
</table>

<table>
<tr><td>Oligo fwd</td><td>CACCAGATCATCGGATACCTCGAC</td></tr>
<tr><td>Oligo rev</td><td>AAACGTCGAGGTATCCGATGATCT</td></tr>
<tr><td>Chromosome</td><td>start</td><td>end</td><td>strand</td><td>MM</td><td>target_seq</td><td>PAM</td><td>alignment</td><td>distance</td><td>position</td><td>gene name</td><td>gene id</td></tr>
<tr><td>GU305914.1</td><td>217984</td><td>218006</td><td>-</td><td>0</td><td>AGATCATCGGATACCTCGAC</td><td>AGG</td><td>||||||||[||||||||||||]PAM</td><td>NA</td><td>Intergenic</td><td>NA</td><td>NA</td></tr>
<tr><td></td></tr>
<tr><td>T18155</td><td>CGACAGGAACGTTCAGTTCGCGG</td><td>1000</td></tr>
<tr><td>Oligo fwd</td><td>CACCCGACAGGAACGTTCAGTTCG</td></tr>
<tr><td>Oligo rev</td><td>AAACCGAACTGAACGTTCCTGTCG</td></tr>
<tr><td>Chromosome</td><td>start</td><td>end</td><td>strand</td><td>MM</td><td>target_seq</td><td>PAM</td><td>alignment</td><td>distance</td><td>position</td><td>gene name</td><td>gene id</td></tr>
<tr><td>GU305914.1</td><td>217968</td><td>217990</td><td>-</td><td>0</td><td>CGACAGGAACGTTCAGTTCG</td><td>CGG</td><td>||||||||[||||||||||||]PAM</td><td>NA</td><td>Intergenic</td><td>NA</td><td>NA</td></tr>
<tr><td></td></tr>
<tr><td>T18156</td><td>GCGGCGAGTCTCTCGATCAACGG</td><td>1000</td></tr>
<tr><td>Oligo fwd</td><td>CACCGCGGCGAGTCTCTCGATCAA</td></tr>
<tr><td>Oligo rev</td><td>AAACTTGATCGAGAGACTCGCCGC</td></tr>
<tr><td>Chromosome</td><td>start</td><td>end</td><td>strand</td><td>MM</td><td>target_seq</td><td>PAM</td><td>alignment</td><td>distance</td><td>position</td><td>gene name</td><td>gene id</td></tr>
<tr><td>GU305914.1</td><td>217949</td><td>217971</td><td>-</td><td>0</td><td>GCGGCGAGTCTCTCGATCAA</td><td>CGG</td><td>||||||||[||||||||||||]PAM</td><td>NA</td><td>Intergenic</td><td>NA</td><td>NA</td></tr>
<tr><td></td></tr>
<tr><td>T18157</td><td>GTAGCTACCGTCTCCTCGAATGG</td><td>1000</td></tr>
<tr><td>Oligo fwd</td><td>CACCGTAGCTACCGTCTCCTCGAA</td></tr>
<tr><td>Oligo rev</td><td>AAACTTCGAGGAGACGGTAGCTAC</td></tr>
<tr><td>Chromosome</td><td>start</td><td>end</td><td>strand</td><td>MM</td><td>target_seq</td><td>PAM</td><td>alignment</td><td>distance</td><td>position</td><td>gene name</td><td>gene id</td></tr>
<tr><td>GU305914.1</td><td>217927</td><td>217949</td><td>-</td><td>0</td><td>GTAGCTACCGTCTCCTCGAA</td><td>TGG</td><td>||||||||[||||||||||||]PAM</td><td>NA</td><td>Intergenic</td><td>NA</td><td>NA</td></tr>
<tr><td></td></tr>
<tr><td>T18158</td><td>CGTCTCCTCGAATGGCAGAGAGG</td><td>1000</td></tr>
<tr><td>Oligo fwd</td><td>CACCCGTCTCCTCGAATGGCAGAG</td></tr>
<tr><td>Oligo rev</td><td>AAACCTCTGCCATTCGAGGAGACG</td></tr>
<tr><td>Chromosome</td><td>start</td><td>end</td><td>strand</td><td>MM</td><td>target_seq</td><td>PAM</td><td>alignment</td><td>distance</td><td>position</td><td>gene name</td><td>gene id</td></tr>
<tr><td>GU305914.1</td><td>217919</td><td>217941</td><td>-</td><td>0</td><td>CGTCTCCTCGAATGGCAGAG</td><td>AGG</td><td>||||||||[||||||||||||]PAM</td><td>NA</td><td>Intergenic</td><td>NA</td><td>NA</td></tr>
<tr><td></td></tr>
<tr><td>T18159</td><td>GCGATCTCGTACCTCTAACCTGG</td><td>1000</td></tr>
<tr><td>Oligo fwd</td><td>CACCGCGATCTCGTACCTCTAACC</td></tr>
<tr><td>Oligo rev</td><td>AAACGGTTAGAGGTACGAGATCGC</td></tr>
<tr><td>Chromosome</td><td>start</td><td>end</td><td>strand</td><td>MM</td><td>target_seq</td><td>PAM</td><td>alignment</td><td>distance</td><td>position</td><td>gene name</td><td>gene id</td></tr>
<tr><td>GU305914.1</td><td>217897</td><td>217919</td><td>-</td><td>0</td><td>GCGATCTCGTACCTCTAACC</td><td>TGG</td><td>||||||||[||||||||||||]PAM</td><td>NA</td><td>Intergenic</td><td>NA</td><td>NA</td></tr>
<tr><td></td></tr>
<tr><td>T18160</td><td>CGATCTCGTACCTCTAACCTGGG</td><td>1000</td></tr>
<tr><td>Oligo fwd</td><td>CACCCGATCTCGTACCTCTAACCT</td></tr>
<tr><td>Oligo rev</td><td>AAACAGGTTAGAGGTACGAGATCG</td></tr>
<tr><td>Chromosome</td><td>start</td><td>end</td><td>strand</td><td>MM</td><td>target_seq</td><td>PAM</td><td>alignment</td><td>distance</td><td>position</td><td>gene name</td><td>gene id</td></tr>
<tr><td>GU305914.1</td><td>217896</td><td>217918</td><td>-</td><td>0</td><td>CGATCTCGTACCTCTAACCT</td><td>GGG</td><td>||||||||[||||||||||||]PAM</td><td>NA</td><td>Intergenic</td><td>NA</td><td>NA</td></tr>
<tr><td></td></tr>
<tr><td>T18161</td><td>ATACTCTTTTTCGAGACCGAAGG</td><td>1000</td></tr>
<tr><td>Oligo fwd</td><td>CACCATACTCTTTTTCGAGACCGA</td></tr>
<tr><td>Oligo rev</td><td>AAACTCGGTCTCGAAAAAGAGTAT</td></tr>
<tr><td>Chromosome</td><td>start</td><td>end</td><td>strand</td><td>MM</td><td>target_seq</td><td>PAM</td><td>alignment</td><td>distance</td><td>position</td><td>gene name</td><td>gene id</td></tr>
<tr><td>GU305914.1</td><td>217838</td><td>217860</td><td>-</td><td>0</td><td>ATACTCTTTTTCGAGACCGA</td><td>AGG</td><td>||||||||[||||||||||||]PAM</td><td>NA</td><td>Intergenic</td><td>NA</td><td>NA</td></tr>
<tr><td></td></tr>
<tr><td>T18162</td><td>AACCTGCGTGATCTTGAACGTGG</td><td>1000</td></tr>
<tr><td>Oligo fwd</td><td>CACCAACCTGCGTGATCTTGAACG</td></tr>
<tr><td>Oligo rev</td><td>AAACCGTTCAAGATCACGCAGGTT</td></tr>
<tr><td>Chromosome</td><td>start</td><td>end</td><td>strand</td><td>MM</td><td>target_seq</td><td>PAM</td><td>alignment</td><td>distance</td><td>position</td><td>gene name</td><td>gene id</td></tr>
<tr><td>GU305914.1</td><td>217802</td><td>217824</td><td>-</td><td>0</td><td>AACCTGCGTGATCTTGAACG</td><td>TGG</td><td>||||||||[||||||||||||]PAM</td><td>NA</td><td>Intergenic</td><td>NA</td><td>NA</td></tr>
<tr><td></td></tr>
<tr><td>T18163</td><td>GTCTCGTACACGTGTACGTTCGG</td><td>1000</td></tr>
<tr><td>Oligo fwd</td><td>CACCGTCTCGTACACGTGTACGTT</td></tr>
<tr><td>Oligo rev</td><td>AAACAACGTACACGTGTACGAGAC</td></tr>
<tr><td>Chromosome</td><td>start</td><td>end</td><td>strand</td><td>MM</td><td>target_seq</td><td>PAM</td><td>alignment</td><td>distance</td><td>position</td><td>gene name</td><td>gene id</td></tr>
<tr><td>GU305914.1</td><td>217763</td><td>217785</td><td>-</td><td>0</td><td>GTCTCGTACACGTGTACGTT</td><td>CGG</td><td>||||||||[||||||||||||]PAM</td><td>NA</td><td>Intergenic</td><td>NA</td><td>NA</td></tr>
<tr><td></td></tr>
<tr><td>T18164</td><td>GCGATGCGATGTTTCACGTACGG</td><td>1000</td></tr>
<tr><td>Oligo fwd</td><td>CACCGCGATGCGATGTTTCACGTA</td></tr>
<tr><td>Oligo rev</td><td>AAACTACGTGAAACATCGCATCGC</td></tr>
<tr><td>Chromosome</td><td>start</td><td>end</td><td>strand</td><td>MM</td><td>target_seq</td><td>PAM</td><td>alignment</td><td>distance</td><td>position</td><td>gene name</td><td>gene id</td></tr>
<tr><td>GU305914.1</td><td>217731</td><td>217753</td><td>-</td><td>0</td><td>GCGATGCGATGTTTCACGTA</td><td>CGG</td><td>||||||||[||||||||||||]PAM</td><td>NA</td><td>Intergenic</td><td>NA</td><td>NA</td></tr>
<tr><td></td></tr>
<tr><td>T18165</td><td>CGATGCGATGTTTCACGTACGGG</td><td>1000</td></tr>
<tr><td>Oligo fwd</td><td>CACCCGATGCGATGTTTCACGTAC</td></tr>
<tr><td>Oligo rev</td><td>AAACGTACGTGAAACATCGCATCG</td></tr>
<tr><td>Chromosome</td><td>start</td><td>end</td><td>strand</td><td>MM</td><td>target_seq</td><td>PAM</td><td>alignment</td><td>distance</td><td>position</td><td>gene name</td><td>gene id</td></tr>
<tr><td>GU305914.1</td><td>217730</td><td>217752</td><td>-</td><td>0</td><td>CGATGCGATGTTTCACGTAC</td><td>GGG</td><td>||||||||[||||||||||||]PAM</td><td>NA</td><td>Intergenic</td><td>NA</td><td>NA</td></tr>
<tr><td></td></tr>
<tr><td>T18166</td><td>TTTCACGTACGGGAGATGATAGG</td><td>1000</td></tr>
<tr><td>Oligo fwd</td><td>CACCTTTCACGTACGGGAGATGAT</td></tr>
<tr><td>Oligo rev</td><td>AAACATCATCTCCCGTACGTGAAA</td></tr>
<tr><td>Chromosome</td><td>start</td><td>end</td><td>strand</td><td>MM</td><td>target_seq</td><td>PAM</td><td>alignment</td><td>distance</td><td>position</td><td>gene name</td><td>gene id</td></tr>
<tr><td>GU305914.1</td><td>217720</td><td>217742</td><td>-</td><td>0</td><td>TTTCACGTACGGGAGATGAT</td><td>AGG</td><td>||||||||[||||||||||||]PAM</td><td>NA</td><td>Intergenic</td><td>NA</td><td>NA</td></tr>
<tr><td></td></tr>
<tr><td>T18167</td><td>CCTCGCTGATGCTTCTGATCCGG</td><td>1000</td></tr>
<tr><td>Oligo fwd</td><td>CACCCCTCGCTGATGCTTCTGATC</td></tr>
<tr><td>Oligo rev</td><td>AAACGATCAGAAGCATCAGCGAGG</td></tr>
<tr><td>Chromosome</td><td>start</td><td>end</td><td>strand</td><td>MM</td><td>target_seq</td><td>PAM</td><td>alignment</td><td>distance</td><td>position</td><td>gene name</td><td>gene id</td></tr>
<tr><td>GU305914.1</td><td>217680</td><td>217702</td><td>-</td><td>0</td><td>CCTCGCTGATGCTTCTGATC</td><td>CGG</td><td>||||||||[||||||||||||]PAM</td><td>NA</td><td>Intergenic</td><td>NA</td><td>NA</td></tr>
<tr><td></td></tr>
<tr><td>T18168</td><td>TTCTGATCCGGACATCAGATTGG</td><td>1000</td></tr>
<tr><td>Oligo fwd</td><td>CACCTTCTGATCCGGACATCAGAT</td></tr>
<tr><td>Oligo rev</td><td>AAACATCTGATGTCCGGATCAGAA</td></tr>
<tr><td>Chromosome</td><td>start</td><td>end</td><td>strand</td><td>MM</td><td>target_seq</td><td>PAM</td><td>alignment</td><td>distance</td><td>position</td><td>gene name</td><td>gene id</td></tr>
<tr><td>GU305914.1</td><td>217668</td><td>217690</td><td>-</td><td>0</td><td>TTCTGATCCGGACATCAGAT</td><td>TGG</td><td>||||||||[||||||||||||]PAM</td><td>NA</td><td>Intergenic</td><td>NA</td><td>NA</td></tr>
</table>

T18169 AGATTGGATCATACAAAACCTGG 1000
Oligo fwd CACCAGATTGGATCATACAAAACC
Oligo rev AAACGGTTTTGTATGATCCAATCT
Chromosome start end strand MM target_seq PAM alignment distance position gene name gene id
GU305914.1 217652 217674 - 0 AGATTGGATCATACAAAACC TGG ||||||||[||||||||||||]PAM NA Intergenic NA NA

T18170 CCGAGCGTATCAAAAGAGTCTGG 1000
Oligo fwd CACCCCGAGCGTATCAAAAGAGTC
Oligo rev AAACGACTCTTTTGATACGCTCGG
Chromosome start end strand MM target_seq PAM alignment distance position gene name gene id
GU305914.1 217606 217628 - 0 CCGAGCGTATCAAAAGAGTC TGG ||||||||[||||||||||||]PAM NA Intergenic NA NA

T18171 CTTGATAAATTAATACAATTCGG 1000
Oligo fwd CACCCTTGATAAATTAATACAATT
Oligo rev AAACAATTGTATTAATTTATCAAG
Chromosome start end strand MM target_seq PAM alignment distance position gene name gene id
GU305914.1 217574 217596 - 0 CTTGATAAATTAATACAATT CGG ||||||||[||||||||||||]PAM NA Intergenic NA NA

T18172 TTGATAAATTAATACAATTCGGG 1000
Oligo fwd CACCTTGATAAATTAATACAATTC
Oligo rev AAACGAATTGTATTAATTTATCAA
Chromosome start end strand MM target_seq PAM alignment distance position gene name gene id
GU305914.1 217573 217595 - 0 TTGATAAATTAATACAATTC GGG ||||||||[||||||||||||]PAM NA Intergenic NA NA

T18173 CGGGAAACCGCCGCATACCGAGG 1000
Oligo fwd CACCCGGGAAACCGCCGCATACCG
Oligo rev AAACCGGTATGCGGCGGTTTCCCG
Chromosome start end strand MM target_seq PAM alignment distance position gene name gene id
GU305914.1 217554 217576 - 0 CGGGAAACCGCCGCATACCG AGG ||||||||[||||||||||||]PAM NA Intergenic NA NA

T18174 GTAGAATTCTGTAATGTGACCGG 1000
Oligo fwd CACCGTAGAATTCTGTAATGTGAC
Oligo rev AAACGTCACATTACAGAATTCTAC
Chromosome start end strand MM target_seq PAM alignment distance position gene name gene id
GU305914.1 217532 217554 - 0 GTAGAATTCTGTAATGTGAC CGG ||||||||[||||||||||||]PAM NA Intergenic NA NA

T18175 GAATTCTGTAATGTGACCGGCGG 1000
Oligo fwd CACCGAATTCTGTAATGTGACCGG
Oligo rev AAACCCGGTCACATTACAGAATTC
Chromosome start end strand MM target_seq PAM alignment distance position gene name gene id
GU305914.1 217529 217551 - 0 GAATTCTGTAATGTGACCGG CGG ||||||||[||||||||||||]PAM NA Intergenic NA NA

T18176 TTCGCTGCACTGCGATCTCGAGG 1000
Oligo fwd CACCTTCGCTGCACTGCGATCTCG
Oligo rev AAACCGAGATCGCAGTGCAGCGAA
Chromosome start end strand MM target_seq PAM alignment distance position gene name gene id
GU305914.1 217504 217526 - 0 TTCGCTGCACTGCGATCTCG AGG ||||||||[||||||||||||]PAM NA Intergenic NA NA

T18177 CGATCTCGAGGTCGTTTCTGCGG 1000
Oligo fwd CACCCGATCTCGAGGTCGTTTCTG
Oligo rev AAACCAGAAACGACCTCGAGATCG
Chromosome start end strand MM target_seq PAM alignment distance position gene name gene id
GU305914.1 217492 217514 - 0 CGATCTCGAGGTCGTTTCTG CGG ||||||||[||||||||||||]PAM NA Intergenic NA NA

T18178 GAGGTCGTTTCTGCGGCTCACGG 1000
Oligo fwd CACCGAGGTCGTTTCTGCGGCTCA
Oligo rev AAACTGAGCCGCAGAAACGACCTC
Chromosome start end strand MM target_seq PAM alignment distance position gene name gene id
GU305914.1 217485 217507 - 0 GAGGTCGTTTCTGCGGCTCA CGG ||||||||[||||||||||||]PAM NA Intergenic NA NA

T18179 TCCCGTGTTACATTACGCGAAGG 1000
Oligo fwd CACCTCCCGTGTTACATTACGCGA
Oligo rev AAACTCGCGTAATGTAACACGGGA
Chromosome start end strand MM target_seq PAM alignment distance position gene name gene id
GU305914.1 217419 217441 - 0 TCCCGTGTTACATTACGCGA AGG ||||||||[||||||||||||]PAM NA Intergenic NA NA

T18180 TTACGCGAAGGCCAGCGTGACGG 1000
Oligo fwd CACCTTACGCGAAGGCCAGCGTGA
Oligo rev AAACTCACGCTGGCCTTCGCGTAA
Chromosome start end strand MM target_seq PAM alignment distance position gene name gene id
GU305914.1 217407 217429 - 0 TTACGCGAAGGCCAGCGTGA CGG ||||||||[||||||||||||]PAM NA Intergenic NA NA

T18181 CGCGAAGGCCAGCGTGACGGCGG 1000
Oligo fwd CACCCGCGAAGGCCAGCGTGACGG
Oligo rev AAACCCGTCACGCTGGCCTTCGCG
Chromosome start end strand MM target_seq PAM alignment distance position gene name gene id
GU305914.1 217404 217426 - 0 CGCGAAGGCCAGCGTGACGG CGG ||||||||[||||||||||||]PAM NA Intergenic NA NA

T18182 AAGGCCAGCGTGACGGCGGCTGG 1000
Oligo fwd CACCAAGGCCAGCGTGACGGCGGC
Oligo rev AAACGCCGCCGTCACGCTGGCCTT
Chromosome start end strand MM target_seq PAM alignment distance position gene name gene id
GU305914.1 217400 217422 - 0 AAGGCCAGCGTGACGGCGGC TGG ||||||||[||||||||||||]PAM NA Intergenic NA NA

T18183 GACGGCGGCTGGAAACTTCTCGG 1000
Oligo fwd CACCGACGGCGGCTGGAAACTTCT
Oligo rev AAACAGAAGTTTCCAGCCGCCGTC
Chromosome start end strand MM target_seq PAM alignment distance position gene name gene id
GU305914.1 217389 217411 - 0 GACGGCGGCTGGAAACTTCT CGG ||||||||[||||||||||||]PAM NA Intergenic NA NA

T18184 TGTAAGATCTACTCTGCTACCGG 1000
Oligo fwd CACCTGTAAGATCTACTCTGCTAC
Oligo rev AAACGTAGCAGAGTAGATCTTACA
Chromosome start end strand MM target_seq PAM alignment distance position gene name gene id
GU305914.1 217355 217377 - 0 TGTAAGATCTACTCTGCTAC CGG ||||||||[||||||||||||]PAM NA Intergenic NA NA

T18185 AGATCTACTCTGCTACCGGCTGG 1000
Oligo fwd CACCAGATCTACTCTGCTACCGGC
Oligo rev AAACGCCGGTAGCAGAGTAGATCT
Chromosome start end strand MM target_seq PAM alignment distance position gene name gene id
GU305914.1 217351 217373 - 0 AGATCTACTCTGCTACCGGC TGG ||||||||[||||||||||||]PAM NA Intergenic NA NA

T18186 CTACTCTGCTACCGGCTGGTTGG 1000
Oligo fwd CACCCTACTCTGCTACCGGCTGGT
Oligo rev AAACACCAGCCGGTAGCAGAGTAG
Chromosome start end strand MM target_seq PAM alignment distance position gene name gene id
GU305914.1 217347 217369 - 0 CTACTCTGCTACCGGCTGGT TGG ||||||||[||||||||||||]PAM NA Intergenic NA NA

T18187 GTTGGCGTCAGTGAAGTACGAGG 1000
Oligo fwd CACCGTTGGCGTCAGTGAAGTACG
Oligo rev AAACCGTACTTCACTGACGCCAAC
Chromosome start end strand MM target_seq PAM alignment distance position gene name gene id
GU305914.1 217329 217351 - 0 GTTGGCGTCAGTGAAGTACG AGG ||||||||[||||||||||||]PAM NA Intergenic NA NA

T18188 TTGGCGTCAGTGAAGTACGAGGG 1000
Oligo fwd CACCTTGGCGTCAGTGAAGTACGA
Oligo rev AAACTCGTACTTCACTGACGCCAA
Chromosome start end strand MM target_seq PAM alignment distance position gene name gene id
GU305914.1 217328 217350 - 0 TTGGCGTCAGTGAAGTACGA GGG ||||||||[||||||||||||]PAM NA Intergenic NA NA

T18189 TACGAGGGCGATCTCCAACACGG 1000
Oligo fwd CACCTACGAGGGCGATCTCCAACA
Oligo rev AAACTGTTGGAGATCGCCCTCGTA
Chromosome start end strand MM target_seq PAM alignment distance position gene name gene id
GU305914.1 217313 217335 - 0 TACGAGGGCGATCTCCAACA CGG ||||||||[||||||||||||]PAM NA Intergenic NA NA

T18190 CCGTAGACCGCTGACTTACGAGG 1000
Oligo fwd CACCCCGTAGACCGCTGACTTACG
Oligo rev AAACCGTAAGTCAGCGGTCTACGG
Chromosome start end strand MM target_seq PAM alignment distance position gene name gene id
GU305914.1 217284 217306 - 0 CCGTAGACCGCTGACTTACG AGG ||||||||[||||||||||||]PAM NA Intergenic NA NA

T18191 GACTTACGAGGACGTCGTCCCGG 1000
Oligo fwd CACCGACTTACGAGGACGTCGTCC
Oligo rev AAACGGACGACGTCCTCGTAAGTC
Chromosome start end strand MM target_seq PAM alignment distance position gene name gene id
GU305914.1 217272 217294 - 0 GACTTACGAGGACGTCGTCC CGG ||||||||[||||||||||||]PAM NA Intergenic NA NA

T18192 ACTTACGAGGACGTCGTCCCGGG 1000
Oligo fwd CACCACTTACGAGGACGTCGTCCC
Oligo rev AAACGGGACGACGTCCTCGTAAGT
Chromosome start end strand MM target_seq PAM alignment distance position gene name gene id
GU305914.1 217271 217293 - 0 ACTTACGAGGACGTCGTCCC GGG ||||||||[||||||||||||]PAM NA Intergenic NA NA

T18193 CGTCCCGGGTGTCGCGTCCGAGG 1000
Oligo fwd CACCCGTCCCGGGTGTCGCGTCCG
Oligo rev AAACCGGACGCGACACCCGGGACG
Chromosome start end strand MM target_seq PAM alignment distance position gene name gene id
GU305914.1 217257 217279 - 0 CGTCCCGGGTGTCGCGTCCG AGG ||||||||[||||||||||||]PAM NA Intergenic NA NA

T18194 CGCGTCCGAGGACGAAGTCTTGG 1000
Oligo fwd CACCCGCGTCCGAGGACGAAGTCT
Oligo rev AAACAGACTTCGTCCTCGGACGCG
Chromosome start end strand MM target_seq PAM alignment distance position gene name gene id
GU305914.1 217245 217267 - 0 CGCGTCCGAGGACGAAGTCT TGG ||||||||[||||||||||||]PAM NA Intergenic NA NA

T18195 GCCGCGAACGACCGTCAAACCGG 1000
Oligo fwd CACCGCCGCGAACGACCGTCAAAC
Oligo rev AAACGTTTGACGGTCGTTCGCGGC
Chromosome start end strand MM target_seq PAM alignment distance position gene name gene id
GU305914.1 217191 217213 - 0 GCCGCGAACGACCGTCAAAC CGG ||||||||[||||||||||||]PAM NA Intergenic NA NA

T18196 ACCGGTGAGAAACCGTACAGCGG 1000
Oligo fwd CACCACCGGTGAGAAACCGTACAG
Oligo rev AAACCTGTACGGTTTCTCACCGGT
Chromosome start end strand MM target_seq PAM alignment distance position gene name gene id
GU305914.1 217173 217195 - 0 ACCGGTGAGAAACCGTACAG CGG ||||||||[||||||||||||]PAM NA Intergenic NA NA

T18197 AGCGGCGTCCGTCTCTGTAGAGG 1000
Oligo fwd CACCAGCGGCGTCCGTCTCTGTAG
Oligo rev AAACCTACAGAGACGGACGCCGCT
Chromosome start end strand MM target_seq PAM alignment distance position gene name gene id
GU305914.1 217155 217177 - 0 AGCGGCGTCCGTCTCTGTAG AGG ||||||||[||||||||||||]PAM NA Intergenic NA NA

T18198 AGGATAATGATACAACCCTGCGG 1000
Oligo fwd CACCAGGATAATGATACAACCCTG
Oligo rev AAACCAGGGTTGTATCATTATCCT
Chromosome start end strand MM target_seq PAM alignment distance position gene name gene id

GU305914.1 217135 217157 - 0 AGGATAATGATACAACCCTG CGG ||||||||[|||||||||||||]PAM NA Intergenic NA NA

T18199 ATACAACCCTGCGGACTCCGTGG 1000
Oligo fwd CACCATACAACCCTGCGGACTCCG
Oligo rev AAACCGGAGTCCGCAGGGTTGTAT
Chromosome start end strand MM target_seq PAM alignment distance position gene name gene id
GU305914.1 217126 217148 - 0 ATACAACCCTGCGGACTCCG TGG ||||||||[|||||||||||||]PAM NA Intergenic NA NA

T18200 TCCGTGGTTAGACGTGTTAGTGG 1000
Oligo fwd CACCTCCGTGGTTAGACGTGTTAG
Oligo rev AAACCTAACACGTCTAACCACGGA
Chromosome start end strand MM target_seq PAM alignment distance position gene name gene id
GU305914.1 217110 217132 - 0 TCCGTGGTTAGACGTGTTAG TGG ||||||||[|||||||||||||]PAM NA Intergenic NA NA

T18201 GGTGTCACTCTTGTTAGTGCCGG 1000
Oligo fwd CACCGGTGTCACTCTTGTTAGTGC
Oligo rev AAACGCACTAACAAGAGTGACACC
Chromosome start end strand MM target_seq PAM alignment distance position gene name gene id
GU305914.1 217089 217111 - 0 GGTGTCACTCTTGTTAGTGC CGG ||||||||[|||||||||||||]PAM NA Intergenic NA NA

T18202 CTTGTTAGTGCCGGTGTCTGTGG 1000
Oligo fwd CACCCTTGTTAGTGCCGGTGTCTG
Oligo rev AAACCAGACACCGGCACTAACAAG
Chromosome start end strand MM target_seq PAM alignment distance position gene name gene id
GU305914.1 217080 217102 - 0 CTTGTTAGTGCCGGTGTCTG TGG ||||||||[|||||||||||||]PAM NA Intergenic NA NA

T18203 TTGTTAGTGCCGGTGTCTGTGGG 1000
Oligo fwd CACCTTGTTAGTGCCGGTGTCTGT
Oligo rev AAACACAGACACCGGCACTAACAA
Chromosome start end strand MM target_seq PAM alignment distance position gene name gene id
GU305914.1 217079 217101 - 0 TTGTTAGTGCCGGTGTCTGT GGG ||||||||[|||||||||||||]PAM NA Intergenic NA NA

T18204 CTGTGGGTGCTTATTTCTTTTGG 1000
Oligo fwd CACCCTGTGGGTGCTTATTTCTTT
Oligo rev AAACAAAGAAATAAGCACCCACAG
Chromosome start end strand MM target_seq PAM alignment distance position gene name gene id
GU305914.1 217063 217085 - 0 CTGTGGGTGCTTATTTCTTT TGG ||||||||[|||||||||||||]PAM NA Intergenic NA NA

T18205 TGGGTGCTTATTTCTTTTGGAGG 1000
Oligo fwd CACCTGGGTGCTTATTTCTTTTGG
Oligo rev AAACCCAAAAGAAATAAGCACCCA
Chromosome start end strand MM target_seq PAM alignment distance position gene name gene id
GU305914.1 217060 217082 - 0 TGGGTGCTTATTTCTTTTGG AGG ||||||||[|||||||||||||]PAM NA Intergenic NA NA

T18206 GTGCTTATTTCTTTTGGAGGTGG 1000
Oligo fwd CACCGTGCTTATTTCTTTTGGAGG
Oligo rev AAACCCTCCAAAAGAAATAAGCAC
Chromosome start end strand MM target_seq PAM alignment distance position gene name gene id
GU305914.1 217057 217079 - 0 GTGCTTATTTCTTTTGGAGG TGG ||||||||[|||||||||||||]PAM NA Intergenic NA NA

T18207 GGAGGTGGCGTGTGAAACGCAGG 1000
Oligo fwd CACCGGAGGTGGCGTGTGAAACGC
Oligo rev AAACGCGTTTCACACGCCACCTCC
Chromosome start end strand MM target_seq PAM alignment distance position gene name gene id
GU305914.1 217042 217064 - 0 GGAGGTGGCGTGTGAAACGC AGG ||||||||[|||||||||||||]PAM NA Intergenic NA NA

T18208 ATATTCAGTACCTCTTGACATGG 1000
Oligo fwd CACCATATTCAGTACCTCTTGACA
Oligo rev AAACTGTCAAGAGGTACTGAATAT
Chromosome start end strand MM target_seq PAM alignment distance position gene name gene id
GU305914.1 216982 217004 - 0 ATATTCAGTACCTCTTGACA TGG ||||||||[|||||||||||||]PAM NA Intergenic NA NA

T18209 ACTCTTGATTGTGTTTATCTCGG 1000
Oligo fwd CACCACTCTTGATTGTGTTTATCT
Oligo rev AAACAGATAAACACAATCAAGAGT
Chromosome start end strand MM target_seq PAM alignment distance position gene name gene id
GU305914.1 216937 216959 - 0 ACTCTTGATTGTGTTTATCT CGG ||||||||[|||||||||||||]PAM NA Intergenic NA NA

T18210 TGACACATATATTGTGAATATGG 1000
Oligo fwd CACCTGACACATATATTGTGAATA
Oligo rev AAACTATTCACAATATATGTGTCA
Chromosome start end strand MM target_seq PAM alignment distance position gene name gene id
GU305914.1 216880 216902 - 0 TGACACATATATTGTGAATA TGG ||||||||[|||||||||||||]PAM NA Intergenic NA NA

T18211 CTTTCAATTTACATTCACTATGG 1000
Oligo fwd CACCCTTTCAATTTACATTCACTA
Oligo rev AAACTAGTGAATGTAAATTGAAAG
Chromosome start end strand MM target_seq PAM alignment distance position gene name gene id
GU305914.1 216853 216875 - 0 CTTTCAATTTACATTCACTA TGG ||||||||[|||||||||||||]PAM NA Intergenic NA NA

T18212 TTCGAAGTTACCGTTACAAGAGG 1000
Oligo fwd CACCTTCGAAGTTACCGTTACAAG
Oligo rev AAACCTTGTAACGGTAACTTCGAA
Chromosome start end strand MM target_seq PAM alignment distance position gene name gene id
GU305914.1 216828 216850 - 0 TTCGAAGTTACCGTTACAAG AGG ||||||||[|||||||||||||]PAM NA Intergenic NA NA

T18213 GAAGTTACCGTTACAAGAGGTGG 1000
Oligo fwd CACCGAAGTTACCGTTACAAGAGG
Oligo rev AAACCCTCTTGTAACGGTAACTTC

Chromosome start end strand MM target_seq PAM alignment distance position gene name gene id
GU305914.1 216825 216847 - 0 GAAGTTACCGTTACAAGAGG TGG ||||||||[||||||||||||]PAM NA Intergenic NA NA

T18214 CGTACGATTTCTGTAGACAACGG 1000
Oligo fwd CACCCGTACGATTTCTGTAGACAA
Oligo rev AAACTTGTCTACAGAAATCGTACG
Chromosome start end strand MM target_seq PAM alignment distance position gene name gene id
GU305914.1 216792 216814 - 0 CGTACGATTTCTGTAGACAA CGG ||||||||[||||||||||||]PAM NA Intergenic NA NA

T18215 ACGGACGTCCAGTCGTCGTGTGG 1000
Oligo fwd CACCACGGACGTCCAGTCGTCGTG
Oligo rev AAACCACGACGACTGGACGTCCGT
Chromosome start end strand MM target_seq PAM alignment distance position gene name gene id
GU305914.1 216773 216795 - 0 ACGGACGTCCAGTCGTCGTG TGG ||||||||[||||||||||||]PAM NA Intergenic NA NA

T18216 CGGACGTCCAGTCGTCGTGTGGG 1000
Oligo fwd CACCCGGACGTCCAGTCGTCGTGT
Oligo rev AAACACACGACGACTGGACGTCCG
Chromosome start end strand MM target_seq PAM alignment distance position gene name gene id
GU305914.1 216772 216794 - 0 CGGACGTCCAGTCGTCGTGT GGG ||||||||[||||||||||||]PAM NA Intergenic NA NA

T18217 CCAGTCGTCGTGTGGGACGTTGG 1000
Oligo fwd CACCCCAGTCGTCGTGTGGGACGT
Oligo rev AAACACGTCCCACACGACGACTGG
Chromosome start end strand MM target_seq PAM alignment distance position gene name gene id
GU305914.1 216765 216787 - 0 CCAGTCGTCGTGTGGGACGT TGG ||||||||[||||||||||||]PAM NA Intergenic NA NA

T18218 CAAGATATGTAAAATCTGTCCGG 1000
Oligo fwd CACCCAAGATATGTAAAATCTGTC
Oligo rev AAACGACAGATTTTACATATCTTG
Chromosome start end strand MM target_seq PAM alignment distance position gene name gene id
GU305914.1 216730 216752 - 0 CAAGATATGTAAAATCTGTC CGG ||||||||[||||||||||||]PAM NA Intergenic NA NA

T18219 ACCTATTGACACAAAGTCTCTGG 1000
Oligo fwd CACCACCTATTGACACAAAGTCTC
Oligo rev AAACGAGACTTTGTGTCAATAGGT
Chromosome start end strand MM target_seq PAM alignment distance position gene name gene id
GU305914.1 216647 216669 - 0 ACCTATTGACACAAAGTCTC TGG ||||||||[||||||||||||]PAM NA Intergenic NA NA

T18220 CCTATTGACACAAAGTCTCTGGG 1000
Oligo fwd CACCCCTATTGACACAAAGTCTCT
Oligo rev AAACAGAGACTTTGTGTCAATAGG
Chromosome start end strand MM target_seq PAM alignment distance position gene name gene id
GU305914.1 216646 216668 - 0 CCTATTGACACAAAGTCTCT GGG ||||||||[||||||||||||]PAM NA Intergenic NA NA

T18221 AAAGTCTCTGGGACACACAAAGG 1000
Oligo fwd CACCAAAGTCTCTGGGACACACAA
Oligo rev AAACTTGTGTGTCCCAGAGACTTT
Chromosome start end strand MM target_seq PAM alignment distance position gene name gene id
GU305914.1 216635 216657 - 0 AAAGTCTCTGGGACACACAA AGG ||||||||[||||||||||||]PAM NA Intergenic NA NA

T18222 ACGATATGCATGTCTGTTTTTGG 1000
Oligo fwd CACCACGATATGCATGTCTGTTTT
Oligo rev AAACAAAACAGACATGCATATCGT
Chromosome start end strand MM target_seq PAM alignment distance position gene name gene id
GU305914.1 216604 216626 - 0 ACGATATGCATGTCTGTTTT TGG ||||||||[||||||||||||]PAM NA Intergenic NA NA

T18223 CGATATGCATGTCTGTTTTTGGG 1000
Oligo fwd CACCCGATATGCATGTCTGTTTTT
Oligo rev AAACAAAAACAGACATGCATATCG
Chromosome start end strand MM target_seq PAM alignment distance position gene name gene id
GU305914.1 216603 216625 - 0 CGATATGCATGTCTGTTTTT GGG ||||||||[||||||||||||]PAM NA Intergenic NA NA

T18224 TGTCTATCATACTACCGATACGG 1000
Oligo fwd CACCTGTCTATCATACTACCGATA
Oligo rev AAACTATCGGTAGTATGATAGACA
Chromosome start end strand MM target_seq PAM alignment distance position gene name gene id
GU305914.1 216562 216584 - 0 TGTCTATCATACTACCGATA CGG ||||||||[||||||||||||]PAM NA Intergenic NA NA

T18225 ATCATACTACCGATACGGTCAGG 1000
Oligo fwd CACCATCATACTACCGATACGGTC
Oligo rev AAACGACCGTATCGGTAGTATGAT
Chromosome start end strand MM target_seq PAM alignment distance position gene name gene id
GU305914.1 216557 216579 - 0 ATCATACTACCGATACGGTC AGG ||||||||[||||||||||||]PAM NA Intergenic NA NA

T18226 TCATACTACCGATACGGTCAGGG 1000
Oligo fwd CACCTCATACTACCGATACGGTCA
Oligo rev AAACTGACCGTATCGGTAGTATGA
Chromosome start end strand MM target_seq PAM alignment distance position gene name gene id
GU305914.1 216556 216578 - 0 TCATACTACCGATACGGTCA GGG ||||||||[||||||||||||]PAM NA Intergenic NA NA

T18227 ACGGTCAGGGTAGCGTATACAGG 1000
Oligo fwd CACCACGGTCAGGGTAGCGTATAC
Oligo rev AAACGTATACGCTACCCTGACCGT
Chromosome start end strand MM target_seq PAM alignment distance position gene name gene id
GU305914.1 216543 216565 - 0 ACGGTCAGGGTAGCGTATAC AGG ||||||||[||||||||||||]PAM NA Intergenic NA NA

T18228 GTAGCGTATACAGGACAGACTGG 1000
Oligo fwd CACCGTAGCGTATACAGGACAGAC

<table>
<tr><td>Oligo rev</td><td>AAACGTCTGTCCTGTATACGCTAC</td><td></td><td></td><td></td><td></td><td></td><td></td><td></td><td></td><td></td><td></td><td></td></tr>
<tr><td>Chromosome</td><td>start</td><td></td><td>end</td><td>strand</td><td>MM</td><td>target_seq</td><td>PAM</td><td>alignment</td><td>distance</td><td>position</td><td>gene name</td><td>gene id</td></tr>
<tr><td>GU305914.1</td><td></td><td>216534</td><td>216556</td><td>-</td><td>0</td><td>GTAGCGTATACAGGACAGAC</td><td>TGG</td><td>||||||||[||||||||||||]PAM</td><td>NA</td><td>Intergenic</td><td>NA</td><td>NA</td></tr>
<tr><td></td><td></td><td></td><td></td><td></td><td></td><td></td><td></td><td></td><td></td><td></td><td></td><td></td></tr>
<tr><td>T18229</td><td>ACTGGTAAGATTAATATTCAAGG</td><td></td><td>1000</td><td></td><td></td><td></td><td></td><td></td><td></td><td></td><td></td><td></td></tr>
<tr><td>Oligo fwd</td><td>CACCACTGGTAAGATTAATATTCA</td><td></td><td></td><td></td><td></td><td></td><td></td><td></td><td></td><td></td><td></td><td></td></tr>
<tr><td>Oligo rev</td><td>AAACTGAATATTAATCTTACCAGT</td><td></td><td></td><td></td><td></td><td></td><td></td><td></td><td></td><td></td><td></td><td></td></tr>
<tr><td>Chromosome</td><td>start</td><td></td><td>end</td><td>strand</td><td>MM</td><td>target_seq</td><td>PAM</td><td>alignment</td><td>distance</td><td>position</td><td>gene name</td><td>gene id</td></tr>
<tr><td>GU305914.1</td><td></td><td>216516</td><td>216538</td><td>-</td><td>0</td><td>ACTGGTAAGATTAATATTCA</td><td>AGG</td><td>||||||||[||||||||||||]PAM</td><td>NA</td><td>Intergenic</td><td>NA</td><td>NA</td></tr>
<tr><td></td><td></td><td></td><td></td><td></td><td></td><td></td><td></td><td></td><td></td><td></td><td></td><td></td></tr>
<tr><td>T18230</td><td>AAGATTAATATTCAAGGATCAGG</td><td></td><td>1000</td><td></td><td></td><td></td><td></td><td></td><td></td><td></td><td></td><td></td></tr>
<tr><td>Oligo fwd</td><td>CACCAAGATTAATATTCAAGGATC</td><td></td><td></td><td></td><td></td><td></td><td></td><td></td><td></td><td></td><td></td><td></td></tr>
<tr><td>Oligo rev</td><td>AAACGATCCTTGAATATTAATCTT</td><td></td><td></td><td></td><td></td><td></td><td></td><td></td><td></td><td></td><td></td><td></td></tr>
<tr><td>Chromosome</td><td>start</td><td></td><td>end</td><td>strand</td><td>MM</td><td>target_seq</td><td>PAM</td><td>alignment</td><td>distance</td><td>position</td><td>gene name</td><td>gene id</td></tr>
<tr><td>GU305914.1</td><td></td><td>216510</td><td>216532</td><td>-</td><td>0</td><td>AAGATTAATATTCAAGGATC</td><td>AGG</td><td>||||||||[||||||||||||]PAM</td><td>NA</td><td>Intergenic</td><td>NA</td><td>NA</td></tr>
<tr><td></td><td></td><td></td><td></td><td></td><td></td><td></td><td></td><td></td><td></td><td></td><td></td><td></td></tr>
<tr><td>T18231</td><td>ACTTCCGACGCAAAAGAAATTGG</td><td></td><td>1000</td><td></td><td></td><td></td><td></td><td></td><td></td><td></td><td></td><td></td></tr>
<tr><td>Oligo fwd</td><td>CACCACTTCCGACGCAAAAGAAAT</td><td></td><td></td><td></td><td></td><td></td><td></td><td></td><td></td><td></td><td></td><td></td></tr>
<tr><td>Oligo rev</td><td>AAACATTTCTTTTGCGTCGGAAGT</td><td></td><td></td><td></td><td></td><td></td><td></td><td></td><td></td><td></td><td></td><td></td></tr>
<tr><td>Chromosome</td><td>start</td><td></td><td>end</td><td>strand</td><td>MM</td><td>target_seq</td><td>PAM</td><td>alignment</td><td>distance</td><td>position</td><td>gene name</td><td>gene id</td></tr>
<tr><td>GU305914.1</td><td></td><td>216477</td><td>216499</td><td>-</td><td>0</td><td>ACTTCCGACGCAAAAGAAAT</td><td>TGG</td><td>||||||||[||||||||||||]PAM</td><td>NA</td><td>Intergenic</td><td>NA</td><td>NA</td></tr>
<tr><td></td><td></td><td></td><td></td><td></td><td></td><td></td><td></td><td></td><td></td><td></td><td></td><td></td></tr>
<tr><td>T18232</td><td>ATATGATAAAAAGCAACGTTCGG</td><td></td><td>1000</td><td></td><td></td><td></td><td></td><td></td><td></td><td></td><td></td><td></td></tr>
<tr><td>Oligo fwd</td><td>CACCATATGATAAAAAGCAACGTT</td><td></td><td></td><td></td><td></td><td></td><td></td><td></td><td></td><td></td><td></td><td></td></tr>
<tr><td>Oligo rev</td><td>AAACAACGTTGCTTTTTATCATAT</td><td></td><td></td><td></td><td></td><td></td><td></td><td></td><td></td><td></td><td></td><td></td></tr>
<tr><td>Chromosome</td><td>start</td><td></td><td>end</td><td>strand</td><td>MM</td><td>target_seq</td><td>PAM</td><td>alignment</td><td>distance</td><td>position</td><td>gene name</td><td>gene id</td></tr>
<tr><td>GU305914.1</td><td></td><td>216449</td><td>216471</td><td>-</td><td>0</td><td>ATATGATAAAAAGCAACGTT</td><td>CGG</td><td>||||||||[||||||||||||]PAM</td><td>NA</td><td>Intergenic</td><td>NA</td><td>NA</td></tr>
<tr><td></td><td></td><td></td><td></td><td></td><td></td><td></td><td></td><td></td><td></td><td></td><td></td><td></td></tr>
<tr><td>T18233</td><td>TATGATAAAAAGCAACGTTCGGG</td><td></td><td>1000</td><td></td><td></td><td></td><td></td><td></td><td></td><td></td><td></td><td></td></tr>
<tr><td>Oligo fwd</td><td>CACCTATGATAAAAAGCAACGTTC</td><td></td><td></td><td></td><td></td><td></td><td></td><td></td><td></td><td></td><td></td><td></td></tr>
<tr><td>Oligo rev</td><td>AAACGAACGTTGCTTTTTATCATA</td><td></td><td></td><td></td><td></td><td></td><td></td><td></td><td></td><td></td><td></td><td></td></tr>
<tr><td>Chromosome</td><td>start</td><td></td><td>end</td><td>strand</td><td>MM</td><td>target_seq</td><td>PAM</td><td>alignment</td><td>distance</td><td>position</td><td>gene name</td><td>gene id</td></tr>
<tr><td>GU305914.1</td><td></td><td>216448</td><td>216470</td><td>-</td><td>0</td><td>TATGATAAAAAGCAACGTTC</td><td>GGG</td><td>||||||||[||||||||||||]PAM</td><td>NA</td><td>Intergenic</td><td>NA</td><td>NA</td></tr>
<tr><td></td><td></td><td></td><td></td><td></td><td></td><td></td><td></td><td></td><td></td><td></td><td></td><td></td></tr>
<tr><td>T18234</td><td>TTCGGGAAATCAAGAACAGATGG</td><td></td><td>1000</td><td></td><td></td><td></td><td></td><td></td><td></td><td></td><td></td><td></td></tr>
<tr><td>Oligo fwd</td><td>CACCTTCGGGAAATCAAGAACAGA</td><td></td><td></td><td></td><td></td><td></td><td></td><td></td><td></td><td></td><td></td><td></td></tr>
<tr><td>Oligo rev</td><td>AAACTCTGTTCTTGATTTCCCGAA</td><td></td><td></td><td></td><td></td><td></td><td></td><td></td><td></td><td></td><td></td><td></td></tr>
<tr><td>Chromosome</td><td>start</td><td></td><td>end</td><td>strand</td><td>MM</td><td>target_seq</td><td>PAM</td><td>alignment</td><td>distance</td><td>position</td><td>gene name</td><td>gene id</td></tr>
<tr><td>GU305914.1</td><td></td><td>216431</td><td>216453</td><td>-</td><td>0</td><td>TTCGGGAAATCAAGAACAGA</td><td>TGG</td><td>||||||||[||||||||||||]PAM</td><td>NA</td><td>Intergenic</td><td>NA</td><td>NA</td></tr>
<tr><td></td><td></td><td></td><td></td><td></td><td></td><td></td><td></td><td></td><td></td><td></td><td></td><td></td></tr>
<tr><td>T18235</td><td>GGGAAATCAAGAACAGATGGCGG</td><td></td><td>1000</td><td></td><td></td><td></td><td></td><td></td><td></td><td></td><td></td><td></td></tr>
<tr><td>Oligo fwd</td><td>CACCGGGAAATCAAGAACAGATGG</td><td></td><td></td><td></td><td></td><td></td><td></td><td></td><td></td><td></td><td></td><td></td></tr>
<tr><td>Oligo rev</td><td>AAACCCATCTGTTCTTGATTTCCC</td><td></td><td></td><td></td><td></td><td></td><td></td><td></td><td></td><td></td><td></td><td></td></tr>
<tr><td>Chromosome</td><td>start</td><td></td><td>end</td><td>strand</td><td>MM</td><td>target_seq</td><td>PAM</td><td>alignment</td><td>distance</td><td>position</td><td>gene name</td><td>gene id</td></tr>
<tr><td>GU305914.1</td><td></td><td>216428</td><td>216450</td><td>-</td><td>0</td><td>GGGAAATCAAGAACAGATGG</td><td>CGG</td><td>||||||||[||||||||||||]PAM</td><td>NA</td><td>Intergenic</td><td>NA</td><td>NA</td></tr>
<tr><td></td><td></td><td></td><td></td><td></td><td></td><td></td><td></td><td></td><td></td><td></td><td></td><td></td></tr>
<tr><td>T18236</td><td>ACGCGACCGAAGTCGAGTTCTGG</td><td></td><td>1000</td><td></td><td></td><td></td><td></td><td></td><td></td><td></td><td></td><td></td></tr>
<tr><td>Oligo fwd</td><td>CACCACGCGACCGAAGTCGAGTTC</td><td></td><td></td><td></td><td></td><td></td><td></td><td></td><td></td><td></td><td></td><td></td></tr>
<tr><td>Oligo rev</td><td>AAACGAACTCGACTTCGGTCGCGT</td><td></td><td></td><td></td><td></td><td></td><td></td><td></td><td></td><td></td><td></td><td></td></tr>
<tr><td>Chromosome</td><td>start</td><td></td><td>end</td><td>strand</td><td>MM</td><td>target_seq</td><td>PAM</td><td>alignment</td><td>distance</td><td>position</td><td>gene name</td><td>gene id</td></tr>
<tr><td>GU305914.1</td><td></td><td>216368</td><td>216390</td><td>-</td><td>0</td><td>ACGCGACCGAAGTCGAGTTC</td><td>TGG</td><td>||||||||[||||||||||||]PAM</td><td>NA</td><td>Intergenic</td><td>NA</td><td>NA</td></tr>
<tr><td></td><td></td><td></td><td></td><td></td><td></td><td></td><td></td><td></td><td></td><td></td><td></td><td></td></tr>
<tr><td>T18237</td><td>GAGTTCTGGTACAACACGACCGG</td><td></td><td>1000</td><td></td><td></td><td></td><td></td><td></td><td></td><td></td><td></td><td></td></tr>
<tr><td>Oligo fwd</td><td>CACCGAGTTCTGGTACAACACGAC</td><td></td><td></td><td></td><td></td><td></td><td></td><td></td><td></td><td></td><td></td><td></td></tr>
<tr><td>Oligo rev</td><td>AAACGTCGTGTTGTACCAGAACTC</td><td></td><td></td><td></td><td></td><td></td><td></td><td></td><td></td><td></td><td></td><td></td></tr>
<tr><td>Chromosome</td><td>start</td><td></td><td>end</td><td>strand</td><td>MM</td><td>target_seq</td><td>PAM</td><td>alignment</td><td>distance</td><td>position</td><td>gene name</td><td>gene id</td></tr>
<tr><td>GU305914.1</td><td></td><td>216354</td><td>216376</td><td>-</td><td>0</td><td>GAGTTCTGGTACAACACGAC</td><td>CGG</td><td>||||||||[||||||||||||]PAM</td><td>NA</td><td>Intergenic</td><td>NA</td><td>NA</td></tr>
<tr><td></td><td></td><td></td><td></td><td></td><td></td><td></td><td></td><td></td><td></td><td></td><td></td><td></td></tr>
<tr><td>T18238</td><td>GACCGGCTTGACCACGTGTGTGG</td><td></td><td>1000</td><td></td><td></td><td></td><td></td><td></td><td></td><td></td><td></td><td></td></tr>
<tr><td>Oligo fwd</td><td>CACCGACCGGCTTGACCACGTGTG</td><td></td><td></td><td></td><td></td><td></td><td></td><td></td><td></td><td></td><td></td><td></td></tr>
<tr><td>Oligo rev</td><td>AAACCACACGTGGTCAAGCCGGTC</td><td></td><td></td><td></td><td></td><td></td><td></td><td></td><td></td><td></td><td></td><td></td></tr>
<tr><td>Chromosome</td><td>start</td><td></td><td>end</td><td>strand</td><td>MM</td><td>target_seq</td><td>PAM</td><td>alignment</td><td>distance</td><td>position</td><td>gene name</td><td>gene id</td></tr>
<tr><td>GU305914.1</td><td></td><td>216337</td><td>216359</td><td>-</td><td>0</td><td>GACCGGCTTGACCACGTGTG</td><td>TGG</td><td>||||||||[||||||||||||]PAM</td><td>NA</td><td>Intergenic</td><td>NA</td><td>NA</td></tr>
<tr><td></td><td></td><td></td><td></td><td></td><td></td><td></td><td></td><td></td><td></td><td></td><td></td><td></td></tr>
<tr><td>T18239</td><td>CCACGTGTGTGGTCACATCGAGG</td><td></td><td>1000</td><td></td><td></td><td></td><td></td><td></td><td></td><td></td><td></td><td></td></tr>
<tr><td>Oligo fwd</td><td>CACCCCACGTGTGTGGTCACATCG</td><td></td><td></td><td></td><td></td><td></td><td></td><td></td><td></td><td></td><td></td><td></td></tr>
<tr><td>Oligo rev</td><td>AAACCGATGTGACCACACACGTGG</td><td></td><td></td><td></td><td></td><td></td><td></td><td></td><td></td><td></td><td></td><td></td></tr>
<tr><td>Chromosome</td><td>start</td><td></td><td>end</td><td>strand</td><td>MM</td><td>target_seq</td><td>PAM</td><td>alignment</td><td>distance</td><td>position</td><td>gene name</td><td>gene id</td></tr>
<tr><td>GU305914.1</td><td></td><td>216326</td><td>216348</td><td>-</td><td>0</td><td>CCACGTGTGTGGTCACATCG</td><td>AGG</td><td>||||||||[||||||||||||]PAM</td><td>NA</td><td>Intergenic</td><td>NA</td><td>NA</td></tr>
<tr><td></td><td></td><td></td><td></td><td></td><td></td><td></td><td></td><td></td><td></td><td></td><td></td><td></td></tr>
<tr><td>T18240</td><td>GATCGTTACTGACGAATCGACGG</td><td></td><td>1000</td><td></td><td></td><td></td><td></td><td></td><td></td><td></td><td></td><td></td></tr>
<tr><td>Oligo fwd</td><td>CACCGATCGTTACTGACGAATCGA</td><td></td><td></td><td></td><td></td><td></td><td></td><td></td><td></td><td></td><td></td><td></td></tr>
<tr><td>Oligo rev</td><td>AAACTCGATTCGTCAGTAACGATC</td><td></td><td></td><td></td><td></td><td></td><td></td><td></td><td></td><td></td><td></td><td></td></tr>
<tr><td>Chromosome</td><td>start</td><td></td><td>end</td><td>strand</td><td>MM</td><td>target_seq</td><td>PAM</td><td>alignment</td><td>distance</td><td>position</td><td>gene name</td><td>gene id</td></tr>
<tr><td>GU305914.1</td><td></td><td>216253</td><td>216275</td><td>-</td><td>0</td><td>GATCGTTACTGACGAATCGA</td><td>CGG</td><td>||||||||[||||||||||||]PAM</td><td>NA</td><td>Intergenic</td><td>NA</td><td>NA</td></tr>
<tr><td>GU305914.1</td><td></td><td>4007</td><td>4029</td><td>-</td><td>4</td><td>AATCATTACTGAAGAAACGA</td><td>CGG</td><td>-|||-|||[||||-|||-|||]PAM</td><td>NA</td><td>Intergenic</td><td>NA</td><td>NA</td></tr>
<tr><td></td><td></td><td></td><td></td><td></td><td></td><td></td><td></td><td></td><td></td><td></td><td></td><td></td></tr>
<tr><td>T18241</td><td>CTGTCAGATCCTTACCGTGAAGG</td><td></td><td>1000</td><td></td><td></td><td></td><td></td><td></td><td></td><td></td><td></td><td></td></tr>
<tr><td>Oligo fwd</td><td>CACCCTGTCAGATCCTTACCGTGA</td><td></td><td></td><td></td><td></td><td></td><td></td><td></td><td></td><td></td><td></td><td></td></tr>
<tr><td>Oligo rev</td><td>AAACTCACGGTAAGGATCTGACAG</td><td></td><td></td><td></td><td></td><td></td><td></td><td></td><td></td><td></td><td></td><td></td></tr>
<tr><td>Chromosome</td><td>start</td><td></td><td>end</td><td>strand</td><td>MM</td><td>target_seq</td><td>PAM</td><td>alignment</td><td>distance</td><td>position</td><td>gene name</td><td>gene id</td></tr>
<tr><td>GU305914.1</td><td></td><td>216226</td><td>216248</td><td>-</td><td>0</td><td>CTGTCAGATCCTTACCGTGA</td><td>AGG</td><td>||||||||[||||||||||||]PAM</td><td>NA</td><td>Intergenic</td><td>NA</td><td>NA</td></tr>
<tr><td></td><td></td><td></td><td></td><td></td><td></td><td></td><td></td><td></td><td></td><td></td><td></td><td></td></tr>
<tr><td>T18242</td><td>GATCCTTACCGTGAAGGCCCCGG</td><td></td><td>1000</td><td></td><td></td><td></td><td></td><td></td><td></td><td></td><td></td><td></td></tr>
<tr><td>Oligo fwd</td><td>CACCGATCCTTACCGTGAAGGCCC</td><td></td><td></td><td></td><td></td><td></td><td></td><td></td><td></td><td></td><td></td><td></td></tr>
<tr><td>Oligo rev</td><td>AAACGGGCCTTCACGGTAAGGATC</td><td></td><td></td><td></td><td></td><td></td><td></td><td></td><td></td><td></td><td></td><td></td></tr>
<tr><td>Chromosome</td><td>start</td><td></td><td>end</td><td>strand</td><td>MM</td><td>target_seq</td><td>PAM</td><td>alignment</td><td>distance</td><td>position</td><td>gene name</td><td>gene id</td></tr>
<tr><td>GU305914.1</td><td></td><td>216220</td><td>216242</td><td>-</td><td>0</td><td>GATCCTTACCGTGAAGGCCC</td><td>CGG</td><td>||||||||[||||||||||||]PAM</td><td>NA</td><td>Intergenic</td><td>NA</td><td>NA</td></tr>
</table>

| | | | | | | | | | | | | |
|---|---|---|---|---|---|---|---|---|---|---|---|---|
| T18243 | ATCCTTACCGTGAAGGCCCCGGG | 1000 | | | | | | | | | | |
| Oligo fwd | CACCATCCTTACCGTGAAGGCCCC | | | | | | | | | | | |
| Oligo rev | AAACGGGGCCTTCACGGTAAGGAT | | | | | | | | | | | |
| Chromosome | start | | end | strand | MM | target_seq | PAM | alignment | distance | position | gene name | gene id |
| GU305914.1 | | 216219 | 216241 | - | 0 | ATCCTTACCGTGAAGGCCCC | GGG | \|\|\|\|\|\|\|\|[\|\|\|\|\|\|\|\|\|\|\|\|]PAM | NA | Intergenic | NA | NA |
| T18244 | ATGTTACGTGACTTCGTCCTTGG | 1000 | | | | | | | | | | |
| Oligo fwd | CACCATGTTACGTGACTTCGTCCT | | | | | | | | | | | |
| Oligo rev | AAACAGGACGAAGTCACGTAACAT | | | | | | | | | | | |
| Chromosome | start | | end | strand | MM | target_seq | PAM | alignment | distance | position | gene name | gene id |
| GU305914.1 | | 216181 | 216203 | - | 0 | ATGTTACGTGACTTCGTCCT | TGG | \|\|\|\|\|\|\|\|[\|\|\|\|\|\|\|\|\|\|\|\|]PAM | NA | Intergenic | NA | NA |
| GU305914.1 | | 217240 | 217262 | + | 4 | ATGCTCCAAGACTTCGTCCT | CGG | \|\|\|-\|-\|-[-\|\|\|\|\|\|\|\|\|\|\|]PAM | NA | Intergenic | NA | NA |
| T18245 | TGTTACGTGACTTCGTCCTTGGG | 1000 | | | | | | | | | | |
| Oligo fwd | CACCTGTTACGTGACTTCGTCCTT | | | | | | | | | | | |
| Oligo rev | AAACAAGGACGAAGTCACGTAACA | | | | | | | | | | | |
| Chromosome | start | | end | strand | MM | target_seq | PAM | alignment | distance | position | gene name | gene id |
| GU305914.1 | | 216180 | 216202 | - | 0 | TGTTACGTGACTTCGTCCTT | GGG | \|\|\|\|\|\|\|\|[\|\|\|\|\|\|\|\|\|\|\|\|]PAM | NA | Intergenic | NA | NA |
| T18246 | ACGTGACTTCGTCCTTGGGATGG | 1000 | | | | | | | | | | |
| Oligo fwd | CACCACGTGACTTCGTCCTTGGGA | | | | | | | | | | | |
| Oligo rev | AAACTCCCAAGGACGAAGTCACGT | | | | | | | | | | | |
| Chromosome | start | | end | strand | MM | target_seq | PAM | alignment | distance | position | gene name | gene id |
| GU305914.1 | | 216176 | 216198 | - | 0 | ACGTGACTTCGTCCTTGGGA | TGG | \|\|\|\|\|\|\|\|[\|\|\|\|\|\|\|\|\|\|\|\|]PAM | NA | Intergenic | NA | NA |
| T18247 | GACTTCGTCCTTGGGATGGAAGG | 1000 | | | | | | | | | | |
| Oligo fwd | CACCGACTTCGTCCTTGGGATGGA | | | | | | | | | | | |
| Oligo rev | AAACTCCATCCCAAGGACGAAGTC | | | | | | | | | | | |
| Chromosome | start | | end | strand | MM | target_seq | PAM | alignment | distance | position | gene name | gene id |
| GU305914.1 | | 216172 | 216194 | - | 0 | GACTTCGTCCTTGGGATGGA | AGG | \|\|\|\|\|\|\|\|[\|\|\|\|\|\|\|\|\|\|\|\|]PAM | NA | Intergenic | NA | NA |
| T18248 | ACTTCGTCCTTGGGATGGAAGGG | 1000 | | | | | | | | | | |
| Oligo fwd | CACCACTTCGTCCTTGGGATGGAA | | | | | | | | | | | |
| Oligo rev | AAACTTCCATCCCAAGGACGAAGT | | | | | | | | | | | |
| Chromosome | start | | end | strand | MM | target_seq | PAM | alignment | distance | position | gene name | gene id |
| GU305914.1 | | 216171 | 216193 | - | 0 | ACTTCGTCCTTGGGATGGAA | GGG | \|\|\|\|\|\|\|\|[\|\|\|\|\|\|\|\|\|\|\|\|]PAM | NA | Intergenic | NA | NA |
| T18249 | TCAATATCAGCTCTTCGAAATGG | 1000 | | | | | | | | | | |
| Oligo fwd | CACCTCAATATCAGCTCTTCGAAA | | | | | | | | | | | |
| Oligo rev | AAACTTTCGAAGAGCTGATATTGA | | | | | | | | | | | |
| Chromosome | start | | end | strand | MM | target_seq | PAM | alignment | distance | position | gene name | gene id |
| GU305914.1 | | 216104 | 216126 | - | 0 | TCAATATCAGCTCTTCGAAA | TGG | \|\|\|\|\|\|\|\|[\|\|\|\|\|\|\|\|\|\|\|\|]PAM | NA | Intergenic | NA | NA |
| T18250 | TATCAGCTCTTCGAAATGGACGG | 1000 | | | | | | | | | | |
| Oligo fwd | CACCTATCAGCTCTTCGAAATGGA | | | | | | | | | | | |
| Oligo rev | AAACTCCATTTCGAAGAGCTGATA | | | | | | | | | | | |
| Chromosome | start | | end | strand | MM | target_seq | PAM | alignment | distance | position | gene name | gene id |
| GU305914.1 | | 216100 | 216122 | - | 0 | TATCAGCTCTTCGAAATGGA | CGG | \|\|\|\|\|\|\|\|[\|\|\|\|\|\|\|\|\|\|\|\|]PAM | NA | Intergenic | NA | NA |
| T18251 | ATCAGCTCTTCGAAATGGACGGG | 1000 | | | | | | | | | | |
| Oligo fwd | CACCATCAGCTCTTCGAAATGGAC | | | | | | | | | | | |
| Oligo rev | AAACGTCCATTTCGAAGAGCTGAT | | | | | | | | | | | |
| Chromosome | start | | end | strand | MM | target_seq | PAM | alignment | distance | position | gene name | gene id |
| GU305914.1 | | 216099 | 216121 | - | 0 | ATCAGCTCTTCGAAATGGAC | GGG | \|\|\|\|\|\|\|\|[\|\|\|\|\|\|\|\|\|\|\|\|]PAM | NA | Intergenic | NA | NA |
| T18252 | TCAGCTCTTCGAAATGGACGGGG | 1000 | | | | | | | | | | |
| Oligo fwd | CACCTCAGCTCTTCGAAATGGACG | | | | | | | | | | | |
| Oligo rev | AAACCGTCCATTTCGAAGAGCTGA | | | | | | | | | | | |
| Chromosome | start | | end | strand | MM | target_seq | PAM | alignment | distance | position | gene name | gene id |
| GU305914.1 | | 216098 | 216120 | - | 0 | TCAGCTCTTCGAAATGGACG | GGG | \|\|\|\|\|\|\|\|[\|\|\|\|\|\|\|\|\|\|\|\|]PAM | NA | Intergenic | NA | NA |
| T18253 | AATGGACGGGGATCGTCAACTGG | 1000 | | | | | | | | | | |
| Oligo fwd | CACCAATGGACGGGGATCGTCAAC | | | | | | | | | | | |
| Oligo rev | AAACGTTGACGATCCCCGTCCATT | | | | | | | | | | | |
| Chromosome | start | | end | strand | MM | target_seq | PAM | alignment | distance | position | gene name | gene id |
| GU305914.1 | | 216086 | 216108 | - | 0 | AATGGACGGGGATCGTCAAC | TGG | \|\|\|\|\|\|\|\|[\|\|\|\|\|\|\|\|\|\|\|\|]PAM | NA | Intergenic | NA | NA |
| T18254 | ACGGGGATCGTCAACTGGAAAGG | 1000 | | | | | | | | | | |
| Oligo fwd | CACCACGGGGATCGTCAACTGGAA | | | | | | | | | | | |
| Oligo rev | AAACTTCCAGTTGACGATCCCCGT | | | | | | | | | | | |
| Chromosome | start | | end | strand | MM | target_seq | PAM | alignment | distance | position | gene name | gene id |
| GU305914.1 | | 216081 | 216103 | - | 0 | ACGGGGATCGTCAACTGGAA | AGG | \|\|\|\|\|\|\|\|[\|\|\|\|\|\|\|\|\|\|\|\|]PAM | NA | Intergenic | NA | NA |
| T18255 | CGGGGATCGTCAACTGGAAAGGG | 1000 | | | | | | | | | | |
| Oligo fwd | CACCCGGGGATCGTCAACTGGAAA | | | | | | | | | | | |
| Oligo rev | AAACTTTCCAGTTGACGATCCCCG | | | | | | | | | | | |
| Chromosome | start | | end | strand | MM | target_seq | PAM | alignment | distance | position | gene name | gene id |
| GU305914.1 | | 216080 | 216102 | - | 0 | CGGGGATCGTCAACTGGAAA | GGG | \|\|\|\|\|\|\|\|[\|\|\|\|\|\|\|\|\|\|\|\|]PAM | NA | Intergenic | NA | NA |
| T18256 | ACCGATACTCATCCTATGTGTGG | 1000 | | | | | | | | | | |
| Oligo fwd | CACCACCGATACTCATCCTATGTG | | | | | | | | | | | |
| Oligo rev | AAACCACATAGGATGAGTATCGGT | | | | | | | | | | | |
| Chromosome | start | | end | strand | MM | target_seq | PAM | alignment | distance | position | gene name | gene id |
| GU305914.1 | | 216019 | 216041 | - | 0 | ACCGATACTCATCCTATGTG | TGG | \|\|\|\|\|\|\|\|[\|\|\|\|\|\|\|\|\|\|\|\|]PAM | NA | Intergenic | NA | NA |
| T18257 | ATGTGTGGTATTTATGCGTCTGG | 1000 | | | | | | | | | | |
| Oligo fwd | CACCATGTGTGGTATTTATGCGTC | | | | | | | | | | | |
| Oligo rev | AAACGACGCATAAATACCACACAT | | | | | | | | | | | |
| Chromosome | start | | end | strand | MM | target_seq | PAM | alignment | distance | position | gene name | gene id |

GU305914.1 216004 216026 - 0 ATGTGTGGTATTTATGCGTC TGG ||||||||[||||||||||||]PAM NA Intergenic NA NA

T18258 TTTGACGACTCTAGTGACGATGG 1000
Oligo fwd CACCTTTGACGACTCTAGTGACGA
Oligo rev AAACTCGTCACTAGAGTCGTCAAA
Chromosome start end strand MM target_seq PAM alignment distance position gene name gene id
GU305914.1 215978 216000 - 0 TTTGACGACTCTAGTGACGA TGG ||||||||[||||||||||||]PAM NA Intergenic NA NA

T18259 TTGACGACTCTAGTGACGATGGG 1000
Oligo fwd CACCTTGACGACTCTAGTGACGAT
Oligo rev AAACATCGTCACTAGAGTCGTCAA
Chromosome start end strand MM target_seq PAM alignment distance position gene name gene id
GU305914.1 215977 215999 - 0 TTGACGACTCTAGTGACGAT GGG ||||||||[||||||||||||]PAM NA Intergenic NA NA

T18260 TCTAGTGACGATGGGAATTATGG 1000
Oligo fwd CACCTCTAGTGACGATGGGAATTA
Oligo rev AAACTAATTCCCATCGTCACTAGA
Chromosome start end strand MM target_seq PAM alignment distance position gene name gene id
GU305914.1 215969 215991 - 0 TCTAGTGACGATGGGAATTA TGG ||||||||[||||||||||||]PAM NA Intergenic NA NA

T18261 AGTGACGATGGGAATTATGGTGG 1000
Oligo fwd CACCAGTGACGATGGGAATTATGG
Oligo rev AAACCCATAATTCCCATCGTCACT
Chromosome start end strand MM target_seq PAM alignment distance position gene name gene id
GU305914.1 215966 215988 - 0 AGTGACGATGGGAATTATGG TGG ||||||||[||||||||||||]PAM NA Intergenic NA NA

T18262 GTGACGATGGGAATTATGGTGGG 1000
Oligo fwd CACCGTGACGATGGGAATTATGGT
Oligo rev AAACACCATAATTCCCATCGTCAC
Chromosome start end strand MM target_seq PAM alignment distance position gene name gene id
GU305914.1 215965 215987 - 0 GTGACGATGGGAATTATGGT GGG ||||||||[||||||||||||]PAM NA Intergenic NA NA

T18263 AATTATGGTGGGAACTATCTTGG 1000
Oligo fwd CACCAATTATGGTGGGAACTATCT
Oligo rev AAACAGATAGTTCCCACCATAATT
Chromosome start end strand MM target_seq PAM alignment distance position gene name gene id
GU305914.1 215954 215976 - 0 AATTATGGTGGGAACTATCT TGG ||||||||[||||||||||||]PAM NA Intergenic NA NA

T18264 GGGAACTATCTTGGTGATAATGG 1000
Oligo fwd CACCGGGAACTATCTTGGTGATAA
Oligo rev AAACTTATCACCAAGATAGTTCCC
Chromosome start end strand MM target_seq PAM alignment distance position gene name gene id
GU305914.1 215945 215967 - 0 GGGAACTATCTTGGTGATAA TGG ||||||||[||||||||||||]PAM NA Intergenic NA NA

T18265 AATGGTCACGATCTCACTGACGG 1000
Oligo fwd CACCAATGGTCACGATCTCACTGA
Oligo rev AAACTCAGTGAGATCGTGACCATT
Chromosome start end strand MM target_seq PAM alignment distance position gene name gene id
GU305914.1 215927 215949 - 0 AATGGTCACGATCTCACTGA CGG ||||||||[||||||||||||]PAM NA Intergenic NA NA

T18266 CTCACTGACGGTCGTGACTTCGG 1000
Oligo fwd CACCCTCACTGACGGTCGTGACTT
Oligo rev AAACAAGTCACGACCGTCAGTGAG
Chromosome start end strand MM target_seq PAM alignment distance position gene name gene id
GU305914.1 215915 215937 - 0 CTCACTGACGGTCGTGACTT CGG ||||||||[||||||||||||]PAM NA Intergenic NA NA

T18267 ATACCGACCGATGTAGTCCCCGG 1000
Oligo fwd CACCATACCGACCGATGTAGTCCC
Oligo rev AAACGGGACTACATCGGTCGGTAT
Chromosome start end strand MM target_seq PAM alignment distance position gene name gene id
GU305914.1 215840 215862 - 0 ATACCGACCGATGTAGTCCC CGG ||||||||[||||||||||||]PAM NA Intergenic NA NA

T18268 TTGAACTTCAATGACCCGTCTGG 1000
Oligo fwd CACCTTGAACTTCAATGACCCGTC
Oligo rev AAACGACGGGTCATTGAAGTTCAA
Chromosome start end strand MM target_seq PAM alignment distance position gene name gene id
GU305914.1 215789 215811 - 0 TTGAACTTCAATGACCCGTC TGG ||||||||[||||||||||||]PAM NA Intergenic NA NA

T18269 TTCAGTGCCAGAAAGATCTTTGG 1000
Oligo fwd CACCTTCAGTGCCAGAAAGATCTT
Oligo rev AAACAAGATCTTTCTGGCACTGAA
Chromosome start end strand MM target_seq PAM alignment distance position gene name gene id
GU305914.1 215762 215784 - 0 TTCAGTGCCAGAAAGATCTT TGG ||||||||[||||||||||||]PAM NA Intergenic NA NA

T18270 GACAACGACGATTACTGCGATGG 1000
Oligo fwd CACCGACAACGACGATTACTGCGA
Oligo rev AAACTCGCAGTAATCGTCGTTGTC
Chromosome start end strand MM target_seq PAM alignment distance position gene name gene id
GU305914.1 215568 215590 - 0 GACAACGACGATTACTGCGA TGG ||||||||[||||||||||||]PAM NA Intergenic NA NA
GU305914.1 20756 20778 + 4 GACGACGGCGATCACGGCGA GGG |||-|||-[||||-||-||||]PAM NA Intergenic NA NA

T18271 GATCTCATGACCATCCAAGACGG 1000
Oligo fwd CACCGATCTCATGACCATCCAAGA
Oligo rev AAACTCTTGGATGGTCATGAGATC
Chromosome start end strand MM target_seq PAM alignment distance position gene name gene id
GU305914.1 215466 215488 - 0 GATCTCATGACCATCCAAGA CGG ||||||||[||||||||||||]PAM NA Intergenic NA NA

T18272 CAGATCACGATGTCATGTAACGG 1000
Oligo fwd CACCCAGATCACGATGTCATGTAA

<table>
<tr><td>Oligo rev</td><td>AAACTTACATGACATCGTGATCTG</td><td></td><td></td><td></td><td></td><td></td><td></td><td></td><td></td><td></td><td></td></tr>
<tr><td>Chromosome</td><td>start</td><td>end</td><td>strand</td><td>MM</td><td>target_seq</td><td>PAM</td><td>alignment</td><td>distance</td><td>position</td><td>gene name</td><td>gene id</td></tr>
<tr><td>GU305914.1</td><td>215442</td><td>215464</td><td>-</td><td>0</td><td>CAGATCACGATGTCATGTAA</td><td>CGG</td><td>||||||||[||||||||||||]PAM</td><td>NA</td><td>Intergenic</td><td>NA</td><td>NA</td></tr>
<tr><td>T18273</td><td>TAACGGCTGCCCTAACGCTACGG</td><td>1000</td><td></td><td></td><td></td><td></td><td></td><td></td><td></td><td></td><td></td></tr>
<tr><td>Oligo fwd</td><td>CACCTAACGGCTGCCCTAACGCTA</td><td></td><td></td><td></td><td></td><td></td><td></td><td></td><td></td><td></td><td></td></tr>
<tr><td>Oligo rev</td><td>AAACTAGCGTTAGGGCAGCCGTTA</td><td></td><td></td><td></td><td></td><td></td><td></td><td></td><td></td><td></td><td></td></tr>
<tr><td>Chromosome</td><td>start</td><td>end</td><td>strand</td><td>MM</td><td>target_seq</td><td>PAM</td><td>alignment</td><td>distance</td><td>position</td><td>gene name</td><td>gene id</td></tr>
<tr><td>GU305914.1</td><td>215425</td><td>215447</td><td>-</td><td>0</td><td>TAACGGCTGCCCTAACGCTA</td><td>CGG</td><td>||||||||[||||||||||||]PAM</td><td>NA</td><td>Intergenic</td><td>NA</td><td>NA</td></tr>
<tr><td>T18274</td><td>TGCCCTAACGCTACGGATCTAGG</td><td>1000</td><td></td><td></td><td></td><td></td><td></td><td></td><td></td><td></td><td></td></tr>
<tr><td>Oligo fwd</td><td>CACCTGCCCTAACGCTACGGATCT</td><td></td><td></td><td></td><td></td><td></td><td></td><td></td><td></td><td></td><td></td></tr>
<tr><td>Oligo rev</td><td>AAACAGATCCGTAGCGTTAGGGCA</td><td></td><td></td><td></td><td></td><td></td><td></td><td></td><td></td><td></td><td></td></tr>
<tr><td>Chromosome</td><td>start</td><td>end</td><td>strand</td><td>MM</td><td>target_seq</td><td>PAM</td><td>alignment</td><td>distance</td><td>position</td><td>gene name</td><td>gene id</td></tr>
<tr><td>GU305914.1</td><td>215418</td><td>215440</td><td>-</td><td>0</td><td>TGCCCTAACGCTACGGATCT</td><td>AGG</td><td>||||||||[||||||||||||]PAM</td><td>NA</td><td>Intergenic</td><td>NA</td><td>NA</td></tr>
<tr><td>T18275</td><td>GCCCTAACGCTACGGATCTAGGG</td><td>1000</td><td></td><td></td><td></td><td></td><td></td><td></td><td></td><td></td><td></td></tr>
<tr><td>Oligo fwd</td><td>CACCGCCCTAACGCTACGGATCTA</td><td></td><td></td><td></td><td></td><td></td><td></td><td></td><td></td><td></td><td></td></tr>
<tr><td>Oligo rev</td><td>AAACTAGATCCGTAGCGTTAGGGC</td><td></td><td></td><td></td><td></td><td></td><td></td><td></td><td></td><td></td><td></td></tr>
<tr><td>Chromosome</td><td>start</td><td>end</td><td>strand</td><td>MM</td><td>target_seq</td><td>PAM</td><td>alignment</td><td>distance</td><td>position</td><td>gene name</td><td>gene id</td></tr>
<tr><td>GU305914.1</td><td>215417</td><td>215439</td><td>-</td><td>0</td><td>GCCCTAACGCTACGGATCTA</td><td>GGG</td><td>||||||||[||||||||||||]PAM</td><td>NA</td><td>Intergenic</td><td>NA</td><td>NA</td></tr>
<tr><td>T18276</td><td>TACGGATCTAGGGTCCGAACTGG</td><td>1000</td><td></td><td></td><td></td><td></td><td></td><td></td><td></td><td></td><td></td></tr>
<tr><td>Oligo fwd</td><td>CACCTACGGATCTAGGGTCCGAAC</td><td></td><td></td><td></td><td></td><td></td><td></td><td></td><td></td><td></td><td></td></tr>
<tr><td>Oligo rev</td><td>AAACGTTCGGACCCTAGATCCGTA</td><td></td><td></td><td></td><td></td><td></td><td></td><td></td><td></td><td></td><td></td></tr>
<tr><td>Chromosome</td><td>start</td><td>end</td><td>strand</td><td>MM</td><td>target_seq</td><td>PAM</td><td>alignment</td><td>distance</td><td>position</td><td>gene name</td><td>gene id</td></tr>
<tr><td>GU305914.1</td><td>215407</td><td>215429</td><td>-</td><td>0</td><td>TACGGATCTAGGGTCCGAAC</td><td>TGG</td><td>||||||||[||||||||||||]PAM</td><td>NA</td><td>Intergenic</td><td>NA</td><td>NA</td></tr>
<tr><td>T18277</td><td>GCACGTACCTGCAGACCGTGAGG</td><td>1000</td><td></td><td></td><td></td><td></td><td></td><td></td><td></td><td></td><td></td></tr>
<tr><td>Oligo fwd</td><td>CACCGCACGTACCTGCAGACCGTG</td><td></td><td></td><td></td><td></td><td></td><td></td><td></td><td></td><td></td><td></td></tr>
<tr><td>Oligo rev</td><td>AAACCACGGTCTGCAGGTACGTGC</td><td></td><td></td><td></td><td></td><td></td><td></td><td></td><td></td><td></td><td></td></tr>
<tr><td>Chromosome</td><td>start</td><td>end</td><td>strand</td><td>MM</td><td>target_seq</td><td>PAM</td><td>alignment</td><td>distance</td><td>position</td><td>gene name</td><td>gene id</td></tr>
<tr><td>GU305914.1</td><td>215366</td><td>215388</td><td>-</td><td>0</td><td>GCACGTACCTGCAGACCGTG</td><td>AGG</td><td>||||||||[||||||||||||]PAM</td><td>NA</td><td>Intergenic</td><td>NA</td><td>NA</td></tr>
<tr><td>T18278</td><td>CACGTACCTGCAGACCGTGAGGG</td><td>1000</td><td></td><td></td><td></td><td></td><td></td><td></td><td></td><td></td><td></td></tr>
<tr><td>Oligo fwd</td><td>CACCCACGTACCTGCAGACCGTGA</td><td></td><td></td><td></td><td></td><td></td><td></td><td></td><td></td><td></td><td></td></tr>
<tr><td>Oligo rev</td><td>AAACTCACGGTCTGCAGGTACGTG</td><td></td><td></td><td></td><td></td><td></td><td></td><td></td><td></td><td></td><td></td></tr>
<tr><td>Chromosome</td><td>start</td><td>end</td><td>strand</td><td>MM</td><td>target_seq</td><td>PAM</td><td>alignment</td><td>distance</td><td>position</td><td>gene name</td><td>gene id</td></tr>
<tr><td>GU305914.1</td><td>215365</td><td>215387</td><td>-</td><td>0</td><td>CACGTACCTGCAGACCGTGA</td><td>GGG</td><td>||||||||[||||||||||||]PAM</td><td>NA</td><td>Intergenic</td><td>NA</td><td>NA</td></tr>
<tr><td>GU305914.1</td><td>69370</td><td>69392</td><td>+</td><td>4</td><td>CGCGTACGTGCAGGCGGTGA</td><td>CGG</td><td>|-|||||-[|||||-|-||||]PAM</td><td>NA</td><td>Intergenic</td><td>NA</td><td>NA</td></tr>
<tr><td>T18279</td><td>GCAGACCGTGAGGGAGCTCGTGG</td><td>1000</td><td></td><td></td><td></td><td></td><td></td><td></td><td></td><td></td><td></td></tr>
<tr><td>Oligo fwd</td><td>CACCGCAGACCGTGAGGGAGCTCG</td><td></td><td></td><td></td><td></td><td></td><td></td><td></td><td></td><td></td><td></td></tr>
<tr><td>Oligo rev</td><td>AAACCGAGCTCCCTCACGGTCTGC</td><td></td><td></td><td></td><td></td><td></td><td></td><td></td><td></td><td></td><td></td></tr>
<tr><td>Chromosome</td><td>start</td><td>end</td><td>strand</td><td>MM</td><td>target_seq</td><td>PAM</td><td>alignment</td><td>distance</td><td>position</td><td>gene name</td><td>gene id</td></tr>
<tr><td>GU305914.1</td><td>215356</td><td>215378</td><td>-</td><td>0</td><td>GCAGACCGTGAGGGAGCTCG</td><td>TGG</td><td>||||||||[||||||||||||]PAM</td><td>NA</td><td>Intergenic</td><td>NA</td><td>NA</td></tr>
<tr><td>T18280</td><td>CAGACCGTGAGGGAGCTCGTGGG</td><td>1000</td><td></td><td></td><td></td><td></td><td></td><td></td><td></td><td></td><td></td></tr>
<tr><td>Oligo fwd</td><td>CACCCAGACCGTGAGGGAGCTCGT</td><td></td><td></td><td></td><td></td><td></td><td></td><td></td><td></td><td></td><td></td></tr>
<tr><td>Oligo rev</td><td>AAACACGAGCTCCCTCACGGTCTG</td><td></td><td></td><td></td><td></td><td></td><td></td><td></td><td></td><td></td><td></td></tr>
<tr><td>Chromosome</td><td>start</td><td>end</td><td>strand</td><td>MM</td><td>target_seq</td><td>PAM</td><td>alignment</td><td>distance</td><td>position</td><td>gene name</td><td>gene id</td></tr>
<tr><td>GU305914.1</td><td>215355</td><td>215377</td><td>-</td><td>0</td><td>CAGACCGTGAGGGAGCTCGT</td><td>GGG</td><td>||||||||[||||||||||||]PAM</td><td>NA</td><td>Intergenic</td><td>NA</td><td>NA</td></tr>
<tr><td>T18281</td><td>GTGAGGGAGCTCGTGGGTGACGG</td><td>1000</td><td></td><td></td><td></td><td></td><td></td><td></td><td></td><td></td><td></td></tr>
<tr><td>Oligo fwd</td><td>CACCGTGAGGGAGCTCGTGGGTGA</td><td></td><td></td><td></td><td></td><td></td><td></td><td></td><td></td><td></td><td></td></tr>
<tr><td>Oligo rev</td><td>AAACTCACCCACGAGCTCCCTCAC</td><td></td><td></td><td></td><td></td><td></td><td></td><td></td><td></td><td></td><td></td></tr>
<tr><td>Chromosome</td><td>start</td><td>end</td><td>strand</td><td>MM</td><td>target_seq</td><td>PAM</td><td>alignment</td><td>distance</td><td>position</td><td>gene name</td><td>gene id</td></tr>
<tr><td>GU305914.1</td><td>215349</td><td>215371</td><td>-</td><td>0</td><td>GTGAGGGAGCTCGTGGGTGA</td><td>CGG</td><td>||||||||[||||||||||||]PAM</td><td>NA</td><td>Intergenic</td><td>NA</td><td>NA</td></tr>
<tr><td>T18282</td><td>GCTCGTGGGTGACGGCCCTCTGG</td><td>1000</td><td></td><td></td><td></td><td></td><td></td><td></td><td></td><td></td><td></td></tr>
<tr><td>Oligo fwd</td><td>CACCGCTCGTGGGTGACGGCCCTC</td><td></td><td></td><td></td><td></td><td></td><td></td><td></td><td></td><td></td><td></td></tr>
<tr><td>Oligo rev</td><td>AAACGAGGGCCGTCACCCACGAGC</td><td></td><td></td><td></td><td></td><td></td><td></td><td></td><td></td><td></td><td></td></tr>
<tr><td>Chromosome</td><td>start</td><td>end</td><td>strand</td><td>MM</td><td>target_seq</td><td>PAM</td><td>alignment</td><td>distance</td><td>position</td><td>gene name</td><td>gene id</td></tr>
<tr><td>GU305914.1</td><td>215341</td><td>215363</td><td>-</td><td>0</td><td>GCTCGTGGGTGACGGCCCTC</td><td>TGG</td><td>||||||||[||||||||||||]PAM</td><td>NA</td><td>Intergenic</td><td>NA</td><td>NA</td></tr>
<tr><td>T18283</td><td>GGGTGACGGCCCTCTGGAATTGG</td><td>1000</td><td></td><td></td><td></td><td></td><td></td><td></td><td></td><td></td><td></td></tr>
<tr><td>Oligo fwd</td><td>CACCGGGTGACGGCCCTCTGGAAT</td><td></td><td></td><td></td><td></td><td></td><td></td><td></td><td></td><td></td><td></td></tr>
<tr><td>Oligo rev</td><td>AAACATTCCAGAGGGCCGTCACCC</td><td></td><td></td><td></td><td></td><td></td><td></td><td></td><td></td><td></td><td></td></tr>
<tr><td>Chromosome</td><td>start</td><td>end</td><td>strand</td><td>MM</td><td>target_seq</td><td>PAM</td><td>alignment</td><td>distance</td><td>position</td><td>gene name</td><td>gene id</td></tr>
<tr><td>GU305914.1</td><td>215335</td><td>215357</td><td>-</td><td>0</td><td>GGGTGACGGCCCTCTGGAAT</td><td>TGG</td><td>||||||||[||||||||||||]PAM</td><td>NA</td><td>Intergenic</td><td>NA</td><td>NA</td></tr>
<tr><td>T18284</td><td>CCTGAGACCCAACAACATGTCGG</td><td>1000</td><td></td><td></td><td></td><td></td><td></td><td></td><td></td><td></td><td></td></tr>
<tr><td>Oligo fwd</td><td>CACCCCTGAGACCCAACAACATGT</td><td></td><td></td><td></td><td></td><td></td><td></td><td></td><td></td><td></td><td></td></tr>
<tr><td>Oligo rev</td><td>AAACACATGTTGTTGGGTCTCAGG</td><td></td><td></td><td></td><td></td><td></td><td></td><td></td><td></td><td></td><td></td></tr>
<tr><td>Chromosome</td><td>start</td><td>end</td><td>strand</td><td>MM</td><td>target_seq</td><td>PAM</td><td>alignment</td><td>distance</td><td>position</td><td>gene name</td><td>gene id</td></tr>
<tr><td>GU305914.1</td><td>215299</td><td>215321</td><td>-</td><td>0</td><td>CCTGAGACCCAACAACATGT</td><td>CGG</td><td>||||||||[||||||||||||]PAM</td><td>NA</td><td>Intergenic</td><td>NA</td><td>NA</td></tr>
<tr><td>T18285</td><td>AACATGTCGGATTACTACATCGG</td><td>1000</td><td></td><td></td><td></td><td></td><td></td><td></td><td></td><td></td><td></td></tr>
<tr><td>Oligo fwd</td><td>CACCAACATGTCGGATTACTACAT</td><td></td><td></td><td></td><td></td><td></td><td></td><td></td><td></td><td></td><td></td></tr>
<tr><td>Oligo rev</td><td>AAACATGTAGTAATCCGACATGTT</td><td></td><td></td><td></td><td></td><td></td><td></td><td></td><td></td><td></td><td></td></tr>
<tr><td>Chromosome</td><td>start</td><td>end</td><td>strand</td><td>MM</td><td>target_seq</td><td>PAM</td><td>alignment</td><td>distance</td><td>position</td><td>gene name</td><td>gene id</td></tr>
<tr><td>GU305914.1</td><td>215286</td><td>215308</td><td>-</td><td>0</td><td>AACATGTCGGATTACTACAT</td><td>CGG</td><td>||||||||[||||||||||||]PAM</td><td>NA</td><td>Intergenic</td><td>NA</td><td>NA</td></tr>
<tr><td>T18286</td><td>CGGATTACTACATCGGTCTTTGG</td><td>1000</td><td></td><td></td><td></td><td></td><td></td><td></td><td></td><td></td><td></td></tr>
<tr><td>Oligo fwd</td><td>CACCCGGATTACTACATCGGTCTT</td><td></td><td></td><td></td><td></td><td></td><td></td><td></td><td></td><td></td><td></td></tr>
<tr><td>Oligo rev</td><td>AAACAAGACCGATGTAGTAATCCG</td><td></td><td></td><td></td><td></td><td></td><td></td><td></td><td></td><td></td><td></td></tr>
<tr><td>Chromosome</td><td>start</td><td>end</td><td>strand</td><td>MM</td><td>target_seq</td><td>PAM</td><td>alignment</td><td>distance</td><td>position</td><td>gene name</td><td>gene id</td></tr>
<tr><td>GU305914.1</td><td>215279</td><td>215301</td><td>-</td><td>0</td><td>CGGATTACTACATCGGTCTT</td><td>TGG</td><td>||||||||[||||||||||||]PAM</td><td>NA</td><td>Intergenic</td><td>NA</td><td>NA</td></tr>
</table>

<table>
<tr><td>T18287</td><td>AAGAATCGTACCATAACCTACGG</td><td>1000</td><td></td><td></td><td></td><td></td><td></td><td></td><td></td><td></td><td></td></tr>
<tr><td>Oligo fwd</td><td>CACCAAGAATCGTACCATAACCTA</td><td></td><td></td><td></td><td></td><td></td><td></td><td></td><td></td><td></td><td></td></tr>
<tr><td>Oligo rev</td><td>AAACTAGGTTATGGTACGATTCTT</td><td></td><td></td><td></td><td></td><td></td><td></td><td></td><td></td><td></td><td></td></tr>
<tr><td>Chromosome</td><td>start</td><td>end</td><td>strand</td><td>MM</td><td>target_seq</td><td>PAM</td><td>alignment</td><td>distance</td><td>position</td><td>gene name</td><td>gene id</td></tr>
<tr><td>GU305914.1</td><td>215223</td><td>215245</td><td>-</td><td>0</td><td>AAGAATCGTACCATAACCTA</td><td>CGG</td><td>||||||||[||||||||||||]PAM</td><td>NA</td><td>Intergenic</td><td>NA</td><td>NA</td></tr>
<tr><td>T18288</td><td>GTTCGTTGAACGATACGCTGTGG</td><td>1000</td><td></td><td></td><td></td><td></td><td></td><td></td><td></td><td></td><td></td></tr>
<tr><td>Oligo fwd</td><td>CACCGTTCGTTGAACGATACGCTG</td><td></td><td></td><td></td><td></td><td></td><td></td><td></td><td></td><td></td><td></td></tr>
<tr><td>Oligo rev</td><td>AAACCAGCGTATCGTTCAACGAAC</td><td></td><td></td><td></td><td></td><td></td><td></td><td></td><td></td><td></td><td></td></tr>
<tr><td>Chromosome</td><td>start</td><td>end</td><td>strand</td><td>MM</td><td>target_seq</td><td>PAM</td><td>alignment</td><td>distance</td><td>position</td><td>gene name</td><td>gene id</td></tr>
<tr><td>GU305914.1</td><td>215180</td><td>215202</td><td>-</td><td>0</td><td>GTTCGTTGAACGATACGCTG</td><td>TGG</td><td>||||||||[||||||||||||]PAM</td><td>NA</td><td>Intergenic</td><td>NA</td><td>NA</td></tr>
<tr><td>T18289</td><td>TTCGTTGAACGATACGCTGTGGG</td><td>1000</td><td></td><td></td><td></td><td></td><td></td><td></td><td></td><td></td><td></td></tr>
<tr><td>Oligo fwd</td><td>CACCTTCGTTGAACGATACGCTGT</td><td></td><td></td><td></td><td></td><td></td><td></td><td></td><td></td><td></td><td></td></tr>
<tr><td>Oligo rev</td><td>AAACACAGCGTATCGTTCAACGAA</td><td></td><td></td><td></td><td></td><td></td><td></td><td></td><td></td><td></td><td></td></tr>
<tr><td>Chromosome</td><td>start</td><td>end</td><td>strand</td><td>MM</td><td>target_seq</td><td>PAM</td><td>alignment</td><td>distance</td><td>position</td><td>gene name</td><td>gene id</td></tr>
<tr><td>GU305914.1</td><td>215179</td><td>215201</td><td>-</td><td>0</td><td>TTCGTTGAACGATACGCTGT</td><td>GGG</td><td>||||||||[||||||||||||]PAM</td><td>NA</td><td>Intergenic</td><td>NA</td><td>NA</td></tr>
<tr><td>T18290</td><td>GAACGATACGCTGTGGGACGCGG</td><td>1000</td><td></td><td></td><td></td><td></td><td></td><td></td><td></td><td></td><td></td></tr>
<tr><td>Oligo fwd</td><td>CACCGAACGATACGCTGTGGGACG</td><td></td><td></td><td></td><td></td><td></td><td></td><td></td><td></td><td></td><td></td></tr>
<tr><td>Oligo rev</td><td>AAACCGTCCCACAGCGTATCGTTC</td><td></td><td></td><td></td><td></td><td></td><td></td><td></td><td></td><td></td><td></td></tr>
<tr><td>Chromosome</td><td>start</td><td>end</td><td>strand</td><td>MM</td><td>target_seq</td><td>PAM</td><td>alignment</td><td>distance</td><td>position</td><td>gene name</td><td>gene id</td></tr>
<tr><td>GU305914.1</td><td>215173</td><td>215195</td><td>-</td><td>0</td><td>GAACGATACGCTGTGGGACG</td><td>CGG</td><td>||||||||[||||||||||||]PAM</td><td>NA</td><td>Intergenic</td><td>NA</td><td>NA</td></tr>
<tr><td>T18291</td><td>TGTGGGACGCGGACAACGCGAGG</td><td>1000</td><td></td><td></td><td></td><td></td><td></td><td></td><td></td><td></td><td></td></tr>
<tr><td>Oligo fwd</td><td>CACCTGTGGGACGCGGACAACGCG</td><td></td><td></td><td></td><td></td><td></td><td></td><td></td><td></td><td></td><td></td></tr>
<tr><td>Oligo rev</td><td>AAACCGCGTTGTCCGCGTCCCACA</td><td></td><td></td><td></td><td></td><td></td><td></td><td></td><td></td><td></td><td></td></tr>
<tr><td>Chromosome</td><td>start</td><td>end</td><td>strand</td><td>MM</td><td>target_seq</td><td>PAM</td><td>alignment</td><td>distance</td><td>position</td><td>gene name</td><td>gene id</td></tr>
<tr><td>GU305914.1</td><td>215162</td><td>215184</td><td>-</td><td>0</td><td>TGTGGGACGCGGACAACGCG</td><td>AGG</td><td>||||||||[||||||||||||]PAM</td><td>NA</td><td>Intergenic</td><td>NA</td><td>NA</td></tr>
<tr><td>T18292</td><td>ACGTGAGTTTCTCGAACACGCGG</td><td>1000</td><td></td><td></td><td></td><td></td><td></td><td></td><td></td><td></td><td></td></tr>
<tr><td>Oligo fwd</td><td>CACCACGTGAGTTTCTCGAACACG</td><td></td><td></td><td></td><td></td><td></td><td></td><td></td><td></td><td></td><td></td></tr>
<tr><td>Oligo rev</td><td>AAACCGTGTTCGAGAAACTCACGT</td><td></td><td></td><td></td><td></td><td></td><td></td><td></td><td></td><td></td><td></td></tr>
<tr><td>Chromosome</td><td>start</td><td>end</td><td>strand</td><td>MM</td><td>target_seq</td><td>PAM</td><td>alignment</td><td>distance</td><td>position</td><td>gene name</td><td>gene id</td></tr>
<tr><td>GU305914.1</td><td>215134</td><td>215156</td><td>-</td><td>0</td><td>ACGTGAGTTTCTCGAACACG</td><td>CGG</td><td>||||||||[||||||||||||]PAM</td><td>NA</td><td>Intergenic</td><td>NA</td><td>NA</td></tr>
<tr><td>T18293</td><td>CGGATGATGTGAGAGAGCGTTGG</td><td>1000</td><td></td><td></td><td></td><td></td><td></td><td></td><td></td><td></td><td></td></tr>
<tr><td>Oligo fwd</td><td>CACCCGGATGATGTGAGAGAGCGT</td><td></td><td></td><td></td><td></td><td></td><td></td><td></td><td></td><td></td><td></td></tr>
<tr><td>Oligo rev</td><td>AAACACGCTCTCTCACATCATCCG</td><td></td><td></td><td></td><td></td><td></td><td></td><td></td><td></td><td></td><td></td></tr>
<tr><td>Chromosome</td><td>start</td><td>end</td><td>strand</td><td>MM</td><td>target_seq</td><td>PAM</td><td>alignment</td><td>distance</td><td>position</td><td>gene name</td><td>gene id</td></tr>
<tr><td>GU305914.1</td><td>215114</td><td>215136</td><td>-</td><td>0</td><td>CGGATGATGTGAGAGAGCGT</td><td>TGG</td><td>||||||||[||||||||||||]PAM</td><td>NA</td><td>Intergenic</td><td>NA</td><td>NA</td></tr>
<tr><td>T18294</td><td>GTTGGAACGCAACGTGCCTGTGG</td><td>1000</td><td></td><td></td><td></td><td></td><td></td><td></td><td></td><td></td><td></td></tr>
<tr><td>Oligo fwd</td><td>CACCGTTGGAACGCAACGTGCCTG</td><td></td><td></td><td></td><td></td><td></td><td></td><td></td><td></td><td></td><td></td></tr>
<tr><td>Oligo rev</td><td>AAACCAGGCACGTTGCGTTCCAAC</td><td></td><td></td><td></td><td></td><td></td><td></td><td></td><td></td><td></td><td></td></tr>
<tr><td>Chromosome</td><td>start</td><td>end</td><td>strand</td><td>MM</td><td>target_seq</td><td>PAM</td><td>alignment</td><td>distance</td><td>position</td><td>gene name</td><td>gene id</td></tr>
<tr><td>GU305914.1</td><td>215096</td><td>215118</td><td>-</td><td>0</td><td>GTTGGAACGCAACGTGCCTG</td><td>TGG</td><td>||||||||[||||||||||||]PAM</td><td>NA</td><td>Intergenic</td><td>NA</td><td>NA</td></tr>
<tr><td>T18295</td><td>CAACGTGCCTGTGGAACGTCAGG</td><td>1000</td><td></td><td></td><td></td><td></td><td></td><td></td><td></td><td></td><td></td></tr>
<tr><td>Oligo fwd</td><td>CACCCAACGTGCCTGTGGAACGTC</td><td></td><td></td><td></td><td></td><td></td><td></td><td></td><td></td><td></td><td></td></tr>
<tr><td>Oligo rev</td><td>AAACGACGTTCCACAGGCACGTTG</td><td></td><td></td><td></td><td></td><td></td><td></td><td></td><td></td><td></td><td></td></tr>
<tr><td>Chromosome</td><td>start</td><td>end</td><td>strand</td><td>MM</td><td>target_seq</td><td>PAM</td><td>alignment</td><td>distance</td><td>position</td><td>gene name</td><td>gene id</td></tr>
<tr><td>GU305914.1</td><td>215087</td><td>215109</td><td>-</td><td>0</td><td>CAACGTGCCTGTGGAACGTC</td><td>AGG</td><td>||||||||[||||||||||||]PAM</td><td>NA</td><td>Intergenic</td><td>NA</td><td>NA</td></tr>
<tr><td>T18296</td><td>GTGCCTGTGGAACGTCAGGTCGG</td><td>1000</td><td></td><td></td><td></td><td></td><td></td><td></td><td></td><td></td><td></td></tr>
<tr><td>Oligo fwd</td><td>CACCGTGCCTGTGGAACGTCAGGT</td><td></td><td></td><td></td><td></td><td></td><td></td><td></td><td></td><td></td><td></td></tr>
<tr><td>Oligo rev</td><td>AAACACCTGACGTTCCACAGGCAC</td><td></td><td></td><td></td><td></td><td></td><td></td><td></td><td></td><td></td><td></td></tr>
<tr><td>Chromosome</td><td>start</td><td>end</td><td>strand</td><td>MM</td><td>target_seq</td><td>PAM</td><td>alignment</td><td>distance</td><td>position</td><td>gene name</td><td>gene id</td></tr>
<tr><td>GU305914.1</td><td>215083</td><td>215105</td><td>-</td><td>0</td><td>GTGCCTGTGGAACGTCAGGT</td><td>CGG</td><td>||||||||[||||||||||||]PAM</td><td>NA</td><td>Intergenic</td><td>NA</td><td>NA</td></tr>
<tr><td>T18297</td><td>CGAGACTTGAGATAGATTTCTGG</td><td>1000</td><td></td><td></td><td></td><td></td><td></td><td></td><td></td><td></td><td></td></tr>
<tr><td>Oligo fwd</td><td>CACCCGAGACTTGAGATAGATTTC</td><td></td><td></td><td></td><td></td><td></td><td></td><td></td><td></td><td></td><td></td></tr>
<tr><td>Oligo rev</td><td>AAACGAAATCTATCTCAAGTCTCG</td><td></td><td></td><td></td><td></td><td></td><td></td><td></td><td></td><td></td><td></td></tr>
<tr><td>Chromosome</td><td>start</td><td>end</td><td>strand</td><td>MM</td><td>target_seq</td><td>PAM</td><td>alignment</td><td>distance</td><td>position</td><td>gene name</td><td>gene id</td></tr>
<tr><td>GU305914.1</td><td>215051</td><td>215073</td><td>-</td><td>0</td><td>CGAGACTTGAGATAGATTTC</td><td>TGG</td><td>||||||||[||||||||||||]PAM</td><td>NA</td><td>Intergenic</td><td>NA</td><td>NA</td></tr>
<tr><td>T18298</td><td>GATAGATTTCTGGTACGAAGTGG</td><td>1000</td><td></td><td></td><td></td><td></td><td></td><td></td><td></td><td></td><td></td></tr>
<tr><td>Oligo fwd</td><td>CACCGATAGATTTCTGGTACGAAG</td><td></td><td></td><td></td><td></td><td></td><td></td><td></td><td></td><td></td><td></td></tr>
<tr><td>Oligo rev</td><td>AAACCTTCGTACCAGAAATCTATC</td><td></td><td></td><td></td><td></td><td></td><td></td><td></td><td></td><td></td><td></td></tr>
<tr><td>Chromosome</td><td>start</td><td>end</td><td>strand</td><td>MM</td><td>target_seq</td><td>PAM</td><td>alignment</td><td>distance</td><td>position</td><td>gene name</td><td>gene id</td></tr>
<tr><td>GU305914.1</td><td>215041</td><td>215063</td><td>-</td><td>0</td><td>GATAGATTTCTGGTACGAAG</td><td>TGG</td><td>||||||||[||||||||||||]PAM</td><td>NA</td><td>Intergenic</td><td>NA</td><td>NA</td></tr>
<tr><td>T18299</td><td>TTCTGGTACGAAGTGGAAGACGG</td><td>1000</td><td></td><td></td><td></td><td></td><td></td><td></td><td></td><td></td><td></td></tr>
<tr><td>Oligo fwd</td><td>CACCTTCTGGTACGAAGTGGAAGA</td><td></td><td></td><td></td><td></td><td></td><td></td><td></td><td></td><td></td><td></td></tr>
<tr><td>Oligo rev</td><td>AAACTCTTCCACTTCGTACCAGAA</td><td></td><td></td><td></td><td></td><td></td><td></td><td></td><td></td><td></td><td></td></tr>
<tr><td>Chromosome</td><td>start</td><td>end</td><td>strand</td><td>MM</td><td>target_seq</td><td>PAM</td><td>alignment</td><td>distance</td><td>position</td><td>gene name</td><td>gene id</td></tr>
<tr><td>GU305914.1</td><td>215034</td><td>215056</td><td>-</td><td>0</td><td>TTCTGGTACGAAGTGGAAGA</td><td>CGG</td><td>||||||||[||||||||||||]PAM</td><td>NA</td><td>Intergenic</td><td>NA</td><td>NA</td></tr>
<tr><td>T18300</td><td>GGTACGAAGTGGAAGACGGTCGG</td><td>1000</td><td></td><td></td><td></td><td></td><td></td><td></td><td></td><td></td><td></td></tr>
<tr><td>Oligo fwd</td><td>CACCGGTACGAAGTGGAAGACGGT</td><td></td><td></td><td></td><td></td><td></td><td></td><td></td><td></td><td></td><td></td></tr>
<tr><td>Oligo rev</td><td>AAACACCGTCTTCCACTTCGTACC</td><td></td><td></td><td></td><td></td><td></td><td></td><td></td><td></td><td></td><td></td></tr>
<tr><td>Chromosome</td><td>start</td><td>end</td><td>strand</td><td>MM</td><td>target_seq</td><td>PAM</td><td>alignment</td><td>distance</td><td>position</td><td>gene name</td><td>gene id</td></tr>
<tr><td>GU305914.1</td><td>215030</td><td>215052</td><td>-</td><td>0</td><td>GGTACGAAGTGGAAGACGGT</td><td>CGG</td><td>||||||||[||||||||||||]PAM</td><td>NA</td><td>Intergenic</td><td>NA</td><td>NA</td></tr>
<tr><td>T18301</td><td>GGTCGGAATCTCACAAAATGCGG</td><td>1000</td><td></td><td></td><td></td><td></td><td></td><td></td><td></td><td></td><td></td></tr>
<tr><td>Oligo fwd</td><td>CACCGGTCGGAATCTCACAAAATG</td><td></td><td></td><td></td><td></td><td></td><td></td><td></td><td></td><td></td><td></td></tr>
<tr><td>Oligo rev</td><td>AAACCATTTTGTGAGATTCCGACC</td><td></td><td></td><td></td><td></td><td></td><td></td><td></td><td></td><td></td><td></td></tr>
<tr><td>Chromosome</td><td>start</td><td>end</td><td>strand</td><td>MM</td><td>target_seq</td><td>PAM</td><td>alignment</td><td>distance</td><td>position</td><td>gene name</td><td>gene id</td></tr>
<tr><td>GU305914.1</td><td>215013</td><td>215035</td><td>-</td><td>0</td><td>GGTCGGAATCTCACAAAATG</td><td>CGG</td><td>||||||||[||||||||||||]PAM</td><td>NA</td><td>Intergenic</td><td>NA</td><td>NA</td></tr>
</table>

T18302 AAATGCGGCGTGAAGTCAGAAGG 1000
Oligo fwd CACCAAATGCGGCGTGAAGTCAGA
Oligo rev AAACTCTGACTTCACGCCGCATTT
Chromosome start end strand MM target_seq PAM alignment distance position gene name gene id
GU305914.1 214998 215020 - 0 AAATGCGGCGTGAAGTCAGA AGG ||||||||[||||||||||||]PAM NA Intergenic NA NA

T18303 GCGTGAAGTCAGAAGGACGAAGG 1000
Oligo fwd CACCGCGTGAAGTCAGAAGGACGA
Oligo rev AAACTCGTCCTTCTGACTTCACGC
Chromosome start end strand MM target_seq PAM alignment distance position gene name gene id
GU305914.1 214991 215013 - 0 GCGTGAAGTCAGAAGGACGA AGG ||||||||[||||||||||||]PAM NA Intergenic NA NA

T18304 TGAAGTCAGAAGGACGAAGGAGG 1000
Oligo fwd CACCTGAAGTCAGAAGGACGAAGG
Oligo rev AAACCCTTCGTCCTTCTGACTTCA
Chromosome start end strand MM target_seq PAM alignment distance position gene name gene id
GU305914.1 214988 215010 - 0 TGAAGTCAGAAGGACGAAGG AGG ||||||||[||||||||||||]PAM NA Intergenic NA NA

T18305 GAGGATGCACGTCGCCATCACGG 1000
Oligo fwd CACCGAGGATGCACGTCGCCATCA
Oligo rev AAACTGATGGCGACGTGCATCCTC
Chromosome start end strand MM target_seq PAM alignment distance position gene name gene id
GU305914.1 214969 214991 - 0 GAGGATGCACGTCGCCATCA CGG ||||||||[||||||||||||]PAM NA Intergenic NA NA

T18306 GTCGCCATCACGGACGAATGCGG 1000
Oligo fwd CACCGTCGCCATCACGGACGAATG
Oligo rev AAACCATTCGTCCGTGATGGCGAC
Chromosome start end strand MM target_seq PAM alignment distance position gene name gene id
GU305914.1 214959 214981 - 0 GTCGCCATCACGGACGAATG CGG ||||||||[||||||||||||]PAM NA Intergenic NA NA

T18307 ACGGACGAATGCGGTAACATAGG 1000
Oligo fwd CACCACGGACGAATGCGGTAACAT
Oligo rev AAACATGTTACCGCATTCGTCCGT
Chromosome start end strand MM target_seq PAM alignment distance position gene name gene id
GU305914.1 214950 214972 - 0 ACGGACGAATGCGGTAACAT AGG ||||||||[||||||||||||]PAM NA Intergenic NA NA

T18308 TGCGGTAACATAGGATACGCCGG 1000
Oligo fwd CACCTGCGGTAACATAGGATACGC
Oligo rev AAACGCGTATCCTATGTTACCGCA
Chromosome start end strand MM target_seq PAM alignment distance position gene name gene id
GU305914.1 214941 214963 - 0 TGCGGTAACATAGGATACGC CGG ||||||||[||||||||||||]PAM NA Intergenic NA NA

T18309 GGTAACATAGGATACGCCGGCGG 1000
Oligo fwd CACCGGTAACATAGGATACGCCGG
Oligo rev AAACCCGGCGTATCCTATGTTACC
Chromosome start end strand MM target_seq PAM alignment distance position gene name gene id
GU305914.1 214938 214960 - 0 GGTAACATAGGATACGCCGG CGG ||||||||[||||||||||||]PAM NA Intergenic NA NA

T18310 ACATAGGATACGCCGGCGGTCGG 1000
Oligo fwd CACCACATAGGATACGCCGGCGGT
Oligo rev AAACACCGCCGGCGTATCCTATGT
Chromosome start end strand MM target_seq PAM alignment distance position gene name gene id
GU305914.1 214934 214956 - 0 ACATAGGATACGCCGGCGGT CGG ||||||||[||||||||||||]PAM NA Intergenic NA NA

T18311 CGGCGGTCGGCTCAATTACACGG 1000
Oligo fwd CACCCGGCGGTCGGCTCAATTACA
Oligo rev AAACTGTAATTGAGCCGACCGCCG
Chromosome start end strand MM target_seq PAM alignment distance position gene name gene id
GU305914.1 214921 214943 - 0 CGGCGGTCGGCTCAATTACA CGG ||||||||[||||||||||||]PAM NA Intergenic NA NA

T18312 GGCGGTCGGCTCAATTACACGGG 1000
Oligo fwd CACCGGCGGTCGGCTCAATTACAC
Oligo rev AAACGTGTAATTGAGCCGACCGCC
Chromosome start end strand MM target_seq PAM alignment distance position gene name gene id
GU305914.1 214920 214942 - 0 GGCGGTCGGCTCAATTACAC GGG ||||||||[||||||||||||]PAM NA Intergenic NA NA

T18313 GGTCGGCTCAATTACACGGGAGG 1000
Oligo fwd CACCGGTCGGCTCAATTACACGGG
Oligo rev AAACCCCGTGTAATTGAGCCGACC
Chromosome start end strand MM target_seq PAM alignment distance position gene name gene id
GU305914.1 214917 214939 - 0 GGTCGGCTCAATTACACGGG AGG ||||||||[||||||||||||]PAM NA Intergenic NA NA

T18314 CGGCTCAATTACACGGGAGGCGG 1000
Oligo fwd CACCCGGCTCAATTACACGGGAGG
Oligo rev AAACCCTCCCGTGTAATTGAGCCG
Chromosome start end strand MM target_seq PAM alignment distance position gene name gene id
GU305914.1 214914 214936 - 0 CGGCTCAATTACACGGGAGG CGG ||||||||[||||||||||||]PAM NA Intergenic NA NA

T18315 CACGGGAGGCGGCTTTTATCTGG 1000
Oligo fwd CACCCACGGGAGGCGGCTTTTATC
Oligo rev AAACGATAAAAGCCGCCTCCCGTG
Chromosome start end strand MM target_seq PAM alignment distance position gene name gene id
GU305914.1 214903 214925 - 0 CACGGGAGGCGGCTTTTATC TGG ||||||||[||||||||||||]PAM NA Intergenic NA NA

T18316 CTTTTATCTGGAAGCCGTGACGG 1000
Oligo fwd CACCCTTTTATCTGGAAGCCGTGA
Oligo rev AAACTCACGGCTTCCAGATAAAAG
Chromosome start end strand MM target_seq PAM alignment distance position gene name gene id

<table>
<tr><td>GU305914.1</td><td>214891</td><td>214913</td><td>-</td><td>0</td><td>CTTTTATCTGGAAGCCGTGA</td><td>CGG</td><td>||||||||[||||||||||||]PAM</td><td>NA</td><td>Intergenic</td><td>NA</td><td>NA</td></tr>
<tr><td>T18317</td><td>GCCGTGACGGAGAGTGCACCCGG</td><td>1000</td><td></td><td></td><td></td><td></td><td></td><td></td><td></td><td></td><td></td></tr>
<tr><td>Oligo fwd</td><td>CACCGCCGTGACGGAGAGTGCACC</td><td></td><td></td><td></td><td></td><td></td><td></td><td></td><td></td><td></td><td></td></tr>
<tr><td>Oligo rev</td><td>AAACGGTGCACTCTCCGTCACGGC</td><td></td><td></td><td></td><td></td><td></td><td></td><td></td><td></td><td></td><td></td></tr>
<tr><td>Chromosome</td><td>start</td><td>end</td><td>strand</td><td>MM</td><td>target_seq</td><td>PAM</td><td>alignment</td><td>distance</td><td>position</td><td>gene name</td><td>gene id</td></tr>
<tr><td>GU305914.1</td><td>214878</td><td>214900</td><td>-</td><td>0</td><td>GCCGTGACGGAGAGTGCACC</td><td>CGG</td><td>||||||||[||||||||||||]PAM</td><td>NA</td><td>Intergenic</td><td>NA</td><td>NA</td></tr>
<tr><td>T18318</td><td>TGCACCCGGTCTCGTATGCACGG</td><td>1000</td><td></td><td></td><td></td><td></td><td></td><td></td><td></td><td></td><td></td></tr>
<tr><td>Oligo fwd</td><td>CACCTGCACCCGGTCTCGTATGCA</td><td></td><td></td><td></td><td></td><td></td><td></td><td></td><td></td><td></td><td></td></tr>
<tr><td>Oligo rev</td><td>AAACTGCATACGAGACCGGGTGCA</td><td></td><td></td><td></td><td></td><td></td><td></td><td></td><td></td><td></td><td></td></tr>
<tr><td>Chromosome</td><td>start</td><td>end</td><td>strand</td><td>MM</td><td>target_seq</td><td>PAM</td><td>alignment</td><td>distance</td><td>position</td><td>gene name</td><td>gene id</td></tr>
<tr><td>GU305914.1</td><td>214864</td><td>214886</td><td>-</td><td>0</td><td>TGCACCCGGTCTCGTATGCA</td><td>CGG</td><td>||||||||[||||||||||||]PAM</td><td>NA</td><td>Intergenic</td><td>NA</td><td>NA</td></tr>
<tr><td>T18319</td><td>TATGCACGGTGCGAACGACCTGG</td><td>1000</td><td></td><td></td><td></td><td></td><td></td><td></td><td></td><td></td><td></td></tr>
<tr><td>Oligo fwd</td><td>CACCTATGCACGGTGCGAACGACC</td><td></td><td></td><td></td><td></td><td></td><td></td><td></td><td></td><td></td><td></td></tr>
<tr><td>Oligo rev</td><td>AAACGGTCGTTCGCACCGTGCATA</td><td></td><td></td><td></td><td></td><td></td><td></td><td></td><td></td><td></td><td></td></tr>
<tr><td>Chromosome</td><td>start</td><td>end</td><td>strand</td><td>MM</td><td>target_seq</td><td>PAM</td><td>alignment</td><td>distance</td><td>position</td><td>gene name</td><td>gene id</td></tr>
<tr><td>GU305914.1</td><td>214850</td><td>214872</td><td>-</td><td>0</td><td>TATGCACGGTGCGAACGACC</td><td>TGG</td><td>||||||||[||||||||||||]PAM</td><td>NA</td><td>Intergenic</td><td>NA</td><td>NA</td></tr>
<tr><td>T18320</td><td>CACGGTGCGAACGACCTGGCTGG</td><td>1000</td><td></td><td></td><td></td><td></td><td></td><td></td><td></td><td></td><td></td></tr>
<tr><td>Oligo fwd</td><td>CACCCACGGTGCGAACGACCTGGC</td><td></td><td></td><td></td><td></td><td></td><td></td><td></td><td></td><td></td><td></td></tr>
<tr><td>Oligo rev</td><td>AAACGCCAGGTCGTTCGCACCGTG</td><td></td><td></td><td></td><td></td><td></td><td></td><td></td><td></td><td></td><td></td></tr>
<tr><td>Chromosome</td><td>start</td><td>end</td><td>strand</td><td>MM</td><td>target_seq</td><td>PAM</td><td>alignment</td><td>distance</td><td>position</td><td>gene name</td><td>gene id</td></tr>
<tr><td>GU305914.1</td><td>214846</td><td>214868</td><td>-</td><td>0</td><td>CACGGTGCGAACGACCTGGC</td><td>TGG</td><td>||||||||[||||||||||||]PAM</td><td>NA</td><td>Intergenic</td><td>NA</td><td>NA</td></tr>
<tr><td>T18321</td><td>ACGGTGCGAACGACCTGGCTGGG</td><td>1000</td><td></td><td></td><td></td><td></td><td></td><td></td><td></td><td></td><td></td></tr>
<tr><td>Oligo fwd</td><td>CACCACGGTGCGAACGACCTGGCT</td><td></td><td></td><td></td><td></td><td></td><td></td><td></td><td></td><td></td><td></td></tr>
<tr><td>Oligo rev</td><td>AAACAGCCAGGTCGTTCGCACCGT</td><td></td><td></td><td></td><td></td><td></td><td></td><td></td><td></td><td></td><td></td></tr>
<tr><td>Chromosome</td><td>start</td><td>end</td><td>strand</td><td>MM</td><td>target_seq</td><td>PAM</td><td>alignment</td><td>distance</td><td>position</td><td>gene name</td><td>gene id</td></tr>
<tr><td>GU305914.1</td><td>214845</td><td>214867</td><td>-</td><td>0</td><td>ACGGTGCGAACGACCTGGCT</td><td>GGG</td><td>||||||||[||||||||||||]PAM</td><td>NA</td><td>Intergenic</td><td>NA</td><td>NA</td></tr>
<tr><td>T18322</td><td>GCGAACGACCTGGCTGGGCTCGG</td><td>1000</td><td></td><td></td><td></td><td></td><td></td><td></td><td></td><td></td><td></td></tr>
<tr><td>Oligo fwd</td><td>CACCGCGAACGACCTGGCTGGGCT</td><td></td><td></td><td></td><td></td><td></td><td></td><td></td><td></td><td></td><td></td></tr>
<tr><td>Oligo rev</td><td>AAACAGCCCAGCCAGGTCGTTCGC</td><td></td><td></td><td></td><td></td><td></td><td></td><td></td><td></td><td></td><td></td></tr>
<tr><td>Chromosome</td><td>start</td><td>end</td><td>strand</td><td>MM</td><td>target_seq</td><td>PAM</td><td>alignment</td><td>distance</td><td>position</td><td>gene name</td><td>gene id</td></tr>
<tr><td>GU305914.1</td><td>214840</td><td>214862</td><td>-</td><td>0</td><td>GCGAACGACCTGGCTGGGCT</td><td>CGG</td><td>||||||||[||||||||||||]PAM</td><td>NA</td><td>Intergenic</td><td>NA</td><td>NA</td></tr>
<tr><td>T18323</td><td>CGACCTGGCTGGGCTCGGTGAGG</td><td>1000</td><td></td><td></td><td></td><td></td><td></td><td></td><td></td><td></td><td></td></tr>
<tr><td>Oligo fwd</td><td>CACCCGACCTGGCTGGGCTCGGTG</td><td></td><td></td><td></td><td></td><td></td><td></td><td></td><td></td><td></td><td></td></tr>
<tr><td>Oligo rev</td><td>AAACCACCGAGCCCAGCCAGGTCG</td><td></td><td></td><td></td><td></td><td></td><td></td><td></td><td></td><td></td><td></td></tr>
<tr><td>Chromosome</td><td>start</td><td>end</td><td>strand</td><td>MM</td><td>target_seq</td><td>PAM</td><td>alignment</td><td>distance</td><td>position</td><td>gene name</td><td>gene id</td></tr>
<tr><td>GU305914.1</td><td>214835</td><td>214857</td><td>-</td><td>0</td><td>CGACCTGGCTGGGCTCGGTG</td><td>AGG</td><td>||||||||[||||||||||||]PAM</td><td>NA</td><td>Intergenic</td><td>NA</td><td>NA</td></tr>
<tr><td>T18324</td><td>CTCGGTGAGGAATGCCACTTCGG</td><td>1000</td><td></td><td></td><td></td><td></td><td></td><td></td><td></td><td></td><td></td></tr>
<tr><td>Oligo fwd</td><td>CACCCTCGGTGAGGAATGCCACTT</td><td></td><td></td><td></td><td></td><td></td><td></td><td></td><td></td><td></td><td></td></tr>
<tr><td>Oligo rev</td><td>AAACAAGTGGCATTCCTCACCGAG</td><td></td><td></td><td></td><td></td><td></td><td></td><td></td><td></td><td></td><td></td></tr>
<tr><td>Chromosome</td><td>start</td><td>end</td><td>strand</td><td>MM</td><td>target_seq</td><td>PAM</td><td>alignment</td><td>distance</td><td>position</td><td>gene name</td><td>gene id</td></tr>
<tr><td>GU305914.1</td><td>214822</td><td>214844</td><td>-</td><td>0</td><td>CTCGGTGAGGAATGCCACTT</td><td>CGG</td><td>||||||||[||||||||||||]PAM</td><td>NA</td><td>Intergenic</td><td>NA</td><td>NA</td></tr>
<tr><td>T18325</td><td>TCGGTGAGGAATGCCACTTCGGG</td><td>1000</td><td></td><td></td><td></td><td></td><td></td><td></td><td></td><td></td><td></td></tr>
<tr><td>Oligo fwd</td><td>CACCTCGGTGAGGAATGCCACTTC</td><td></td><td></td><td></td><td></td><td></td><td></td><td></td><td></td><td></td><td></td></tr>
<tr><td>Oligo rev</td><td>AAACGAAGTGGCATTCCTCACCGA</td><td></td><td></td><td></td><td></td><td></td><td></td><td></td><td></td><td></td><td></td></tr>
<tr><td>Chromosome</td><td>start</td><td>end</td><td>strand</td><td>MM</td><td>target_seq</td><td>PAM</td><td>alignment</td><td>distance</td><td>position</td><td>gene name</td><td>gene id</td></tr>
<tr><td>GU305914.1</td><td>214821</td><td>214843</td><td>-</td><td>0</td><td>TCGGTGAGGAATGCCACTTC</td><td>GGG</td><td>||||||||[||||||||||||]PAM</td><td>NA</td><td>Intergenic</td><td>NA</td><td>NA</td></tr>
<tr><td>T18326</td><td>GGGACGCGTGCGTCGTAAGTCGG</td><td>1000</td><td></td><td></td><td></td><td></td><td></td><td></td><td></td><td></td><td></td></tr>
<tr><td>Oligo fwd</td><td>CACCGGGACGCGTGCGTCGTAAGT</td><td></td><td></td><td></td><td></td><td></td><td></td><td></td><td></td><td></td><td></td></tr>
<tr><td>Oligo rev</td><td>AAACACTTACGACGCACGCGTCCC</td><td></td><td></td><td></td><td></td><td></td><td></td><td></td><td></td><td></td><td></td></tr>
<tr><td>Chromosome</td><td>start</td><td>end</td><td>strand</td><td>MM</td><td>target_seq</td><td>PAM</td><td>alignment</td><td>distance</td><td>position</td><td>gene name</td><td>gene id</td></tr>
<tr><td>GU305914.1</td><td>214801</td><td>214823</td><td>-</td><td>0</td><td>GGGACGCGTGCGTCGTAAGT</td><td>CGG</td><td>||||||||[||||||||||||]PAM</td><td>NA</td><td>Intergenic</td><td>NA</td><td>NA</td></tr>
<tr><td>T18327</td><td>CGTGCGTCGTAAGTCGGTGCAGG</td><td>1000</td><td></td><td></td><td></td><td></td><td></td><td></td><td></td><td></td><td></td></tr>
<tr><td>Oligo fwd</td><td>CACCCGTGCGTCGTAAGTCGGTGC</td><td></td><td></td><td></td><td></td><td></td><td></td><td></td><td></td><td></td><td></td></tr>
<tr><td>Oligo rev</td><td>AAACGCACCGACTTACGACGCACG</td><td></td><td></td><td></td><td></td><td></td><td></td><td></td><td></td><td></td><td></td></tr>
<tr><td>Chromosome</td><td>start</td><td>end</td><td>strand</td><td>MM</td><td>target_seq</td><td>PAM</td><td>alignment</td><td>distance</td><td>position</td><td>gene name</td><td>gene id</td></tr>
<tr><td>GU305914.1</td><td>214795</td><td>214817</td><td>-</td><td>0</td><td>CGTGCGTCGTAAGTCGGTGC</td><td>AGG</td><td>||||||||[||||||||||||]PAM</td><td>NA</td><td>Intergenic</td><td>NA</td><td>NA</td></tr>
<tr><td>T18328</td><td>GTGCGTCGTAAGTCGGTGCAGGG</td><td>1000</td><td></td><td></td><td></td><td></td><td></td><td></td><td></td><td></td><td></td></tr>
<tr><td>Oligo fwd</td><td>CACCGTGCGTCGTAAGTCGGTGCA</td><td></td><td></td><td></td><td></td><td></td><td></td><td></td><td></td><td></td><td></td></tr>
<tr><td>Oligo rev</td><td>AAACTGCACCGACTTACGACGCAC</td><td></td><td></td><td></td><td></td><td></td><td></td><td></td><td></td><td></td><td></td></tr>
<tr><td>Chromosome</td><td>start</td><td>end</td><td>strand</td><td>MM</td><td>target_seq</td><td>PAM</td><td>alignment</td><td>distance</td><td>position</td><td>gene name</td><td>gene id</td></tr>
<tr><td>GU305914.1</td><td>214794</td><td>214816</td><td>-</td><td>0</td><td>GTGCGTCGTAAGTCGGTGCA</td><td>GGG</td><td>||||||||[||||||||||||]PAM</td><td>NA</td><td>Intergenic</td><td>NA</td><td>NA</td></tr>
<tr><td>T18329</td><td>ATCCCAGATGCCCGCTCAACAGG</td><td>1000</td><td></td><td></td><td></td><td></td><td></td><td></td><td></td><td></td><td></td></tr>
<tr><td>Oligo fwd</td><td>CACCATCCCAGATGCCCGCTCAAC</td><td></td><td></td><td></td><td></td><td></td><td></td><td></td><td></td><td></td><td></td></tr>
<tr><td>Oligo rev</td><td>AAACGTTGAGCGGGCATCTGGGAT</td><td></td><td></td><td></td><td></td><td></td><td></td><td></td><td></td><td></td><td></td></tr>
<tr><td>Chromosome</td><td>start</td><td>end</td><td>strand</td><td>MM</td><td>target_seq</td><td>PAM</td><td>alignment</td><td>distance</td><td>position</td><td>gene name</td><td>gene id</td></tr>
<tr><td>GU305914.1</td><td>214769</td><td>214791</td><td>-</td><td>0</td><td>ATCCCAGATGCCCGCTCAAC</td><td>AGG</td><td>||||||||[||||||||||||]PAM</td><td>NA</td><td>Intergenic</td><td>NA</td><td>NA</td></tr>
<tr><td>T18330</td><td>CCAGATGCCCGCTCAACAGGAGG</td><td>1000</td><td></td><td></td><td></td><td></td><td></td><td></td><td></td><td></td><td></td></tr>
<tr><td>Oligo fwd</td><td>CACCCCAGATGCCCGCTCAACAGG</td><td></td><td></td><td></td><td></td><td></td><td></td><td></td><td></td><td></td><td></td></tr>
<tr><td>Oligo rev</td><td>AAACCCTGTTGAGCGGGCATCTGG</td><td></td><td></td><td></td><td></td><td></td><td></td><td></td><td></td><td></td><td></td></tr>
<tr><td>Chromosome</td><td>start</td><td>end</td><td>strand</td><td>MM</td><td>target_seq</td><td>PAM</td><td>alignment</td><td>distance</td><td>position</td><td>gene name</td><td>gene id</td></tr>
<tr><td>GU305914.1</td><td>214766</td><td>214788</td><td>-</td><td>0</td><td>CCAGATGCCCGCTCAACAGG</td><td>AGG</td><td>||||||||[||||||||||||]PAM</td><td>NA</td><td>Intergenic</td><td>NA</td><td>NA</td></tr>
<tr><td>T18331</td><td>CAGATGCCCGCTCAACAGGAGGG</td><td>1000</td><td></td><td></td><td></td><td></td><td></td><td></td><td></td><td></td><td></td></tr>
<tr><td>Oligo fwd</td><td>CACCCAGATGCCCGCTCAACAGGA</td><td></td><td></td><td></td><td></td><td></td><td></td><td></td><td></td><td></td><td></td></tr>
<tr><td>Oligo rev</td><td>AAACTCCTGTTGAGCGGGCATCTG</td><td></td><td></td><td></td><td></td><td></td><td></td><td></td><td></td><td></td><td></td></tr>
</table>

| | | | | | | | | | | | |
|---|---|---|---|---|---|---|---|---|---|---|---|
| Chromosome | start | end | strand | MM | target_seq | PAM | alignment | distance | position | gene name | gene id |
| GU305914.1 | 214765 | 214787 | - | 0 | CAGATGCCCGCTCAACAGGA | GGG | \|\|\|\|\|\|\|\|[\|\|\|\|\|\|\|\|\|\|\|\|]PAM | NA | Intergenic | NA | NA |
| T18332 | GCTCAACAGGAGGGTTCGCTCGG | 1000 | | | | | | | | | |
| Oligo fwd | CACCGCTCAACAGGAGGGTTCGCT | | | | | | | | | | |
| Oligo rev | AAACAGCGAACCCTCCTGTTGAGC | | | | | | | | | | |
| Chromosome | start | end | strand | MM | target_seq | PAM | alignment | distance | position | gene name | gene id |
| GU305914.1 | 214756 | 214778 | - | 0 | GCTCAACAGGAGGGTTCGCT | CGG | \|\|\|\|\|\|\|\|[\|\|\|\|\|\|\|\|\|\|\|\|]PAM | NA | Intergenic | NA | NA |
| T18333 | CTCGGAACCATCATCCGTCAAGG | 1000 | | | | | | | | | |
| Oligo fwd | CACCCTCGGAACCATCATCCGTCA | | | | | | | | | | |
| Oligo rev | AAACTGACGGATGATGGTTCCGAG | | | | | | | | | | |
| Chromosome | start | end | strand | MM | target_seq | PAM | alignment | distance | position | gene name | gene id |
| GU305914.1 | 214738 | 214760 | - | 0 | CTCGGAACCATCATCCGTCA | AGG | \|\|\|\|\|\|\|\|[\|\|\|\|\|\|\|\|\|\|\|\|]PAM | NA | Intergenic | NA | NA |
| T18334 | GCCGAAGAATCATACCAAAGTGG | 1000 | | | | | | | | | |
| Oligo fwd | CACCGCCGAAGAATCATACCAAAG | | | | | | | | | | |
| Oligo rev | AAACCTTTGGTATGATTCTTCGGC | | | | | | | | | | |
| Chromosome | start | end | strand | MM | target_seq | PAM | alignment | distance | position | gene name | gene id |
| GU305914.1 | 214705 | 214727 | - | 0 | GCCGAAGAATCATACCAAAG | TGG | \|\|\|\|\|\|\|\|[\|\|\|\|\|\|\|\|\|\|\|\|]PAM | NA | Intergenic | NA | NA |
| T18335 | CAACGTCACGTACACCCTACTGG | 1000 | | | | | | | | | |
| Oligo fwd | CACCCAACGTCACGTACACCCTAC | | | | | | | | | | |
| Oligo rev | AAACGTAGGGTGTACGTGACGTTG | | | | | | | | | | |
| Chromosome | start | end | strand | MM | target_seq | PAM | alignment | distance | position | gene name | gene id |
| GU305914.1 | 214651 | 214673 | - | 0 | CAACGTCACGTACACCCTAC | TGG | \|\|\|\|\|\|\|\|[\|\|\|\|\|\|\|\|\|\|\|\|]PAM | NA | Intergenic | NA | NA |
| T18336 | TACTCCTCGTCGTGATGTGCTGG | 1000 | | | | | | | | | |
| Oligo fwd | CACCTACTCCTCGTCGTGATGTGC | | | | | | | | | | |
| Oligo rev | AAACGCACATCACGACGAGGAGTA | | | | | | | | | | |
| Chromosome | start | end | strand | MM | target_seq | PAM | alignment | distance | position | gene name | gene id |
| GU305914.1 | 214616 | 214638 | - | 0 | TACTCCTCGTCGTGATGTGC | TGG | \|\|\|\|\|\|\|\|[\|\|\|\|\|\|\|\|\|\|\|\|]PAM | NA | Intergenic | NA | NA |
| T18337 | TCGTGATGTGCTGGATGCTGCGG | 1000 | | | | | | | | | |
| Oligo fwd | CACCTCGTGATGTGCTGGATGCTG | | | | | | | | | | |
| Oligo rev | AAACCAGCATCCAGCACATCACGA | | | | | | | | | | |
| Chromosome | start | end | strand | MM | target_seq | PAM | alignment | distance | position | gene name | gene id |
| GU305914.1 | 214607 | 214629 | - | 0 | TCGTGATGTGCTGGATGCTG | CGG | \|\|\|\|\|\|\|\|[\|\|\|\|\|\|\|\|\|\|\|\|]PAM | NA | Intergenic | NA | NA |
| T18338 | TGCTTCGCCGCCTCCTGTGAAGG | 1000 | | | | | | | | | |
| Oligo fwd | CACCTGCTTCGCCGCCTCCTGTGA | | | | | | | | | | |
| Oligo rev | AAACTCACAGGAGGCGGCGAAGCA | | | | | | | | | | |
| Chromosome | start | end | strand | MM | target_seq | PAM | alignment | distance | position | gene name | gene id |
| GU305914.1 | 214569 | 214591 | - | 0 | TGCTTCGCCGCCTCCTGTGA | AGG | \|\|\|\|\|\|\|\|[\|\|\|\|\|\|\|\|\|\|\|\|]PAM | NA | Intergenic | NA | NA |
| T18339 | GCTTCGCCGCCTCCTGTGAAGGG | 1000 | | | | | | | | | |
| Oligo fwd | CACCGCTTCGCCGCCTCCTGTGAA | | | | | | | | | | |
| Oligo rev | AAACTTCACAGGAGGCGGCGAAGC | | | | | | | | | | |
| Chromosome | start | end | strand | MM | target_seq | PAM | alignment | distance | position | gene name | gene id |
| GU305914.1 | 214568 | 214590 | - | 0 | GCTTCGCCGCCTCCTGTGAA | GGG | \|\|\|\|\|\|\|\|[\|\|\|\|\|\|\|\|\|\|\|\|]PAM | NA | Intergenic | NA | NA |
| T18340 | CTTCGCCGCCTCCTGTGAAGGGG | 1000 | | | | | | | | | |
| Oligo fwd | CACCCTTCGCCGCCTCCTGTGAAG | | | | | | | | | | |
| Oligo rev | AAACCTTCACAGGAGGCGGCGAAG | | | | | | | | | | |
| Chromosome | start | end | strand | MM | target_seq | PAM | alignment | distance | position | gene name | gene id |
| GU305914.1 | 214567 | 214589 | - | 0 | CTTCGCCGCCTCCTGTGAAG | GGG | \|\|\|\|\|\|\|\|[\|\|\|\|\|\|\|\|\|\|\|\|]PAM | NA | Intergenic | NA | NA |
| T18341 | CCGTCTACAAATGATCACCCCGG | 1000 | | | | | | | | | |
| Oligo fwd | CACCCCGTCTACAAATGATCACCC | | | | | | | | | | |
| Oligo rev | AAACGGGTGATCATTTGTAGACGG | | | | | | | | | | |
| Chromosome | start | end | strand | MM | target_seq | PAM | alignment | distance | position | gene name | gene id |
| GU305914.1 | 214526 | 214548 | - | 0 | CCGTCTACAAATGATCACCC | CGG | \|\|\|\|\|\|\|\|[\|\|\|\|\|\|\|\|\|\|\|\|]PAM | NA | Intergenic | NA | NA |
| T18342 | GTATCGCGTCCTTTTATTGTCGG | 1000 | | | | | | | | | |
| Oligo fwd | CACCGTATCGCGTCCTTTTATTGT | | | | | | | | | | |
| Oligo rev | AAACACAATAAAAGGACGCGATAC | | | | | | | | | | |
| Chromosome | start | end | strand | MM | target_seq | PAM | alignment | distance | position | gene name | gene id |
| GU305914.1 | 214448 | 214470 | - | 0 | GTATCGCGTCCTTTTATTGT | CGG | \|\|\|\|\|\|\|\|[\|\|\|\|\|\|\|\|\|\|\|\|]PAM | NA | Intergenic | NA | NA |
| T18343 | TTATTGTCGGCGCGTGCGACCGG | 1000 | | | | | | | | | |
| Oligo fwd | CACCTTATTGTCGGCGCGTGCGAC | | | | | | | | | | |
| Oligo rev | AAACGTCGCACGCGCCGACAATAA | | | | | | | | | | |
| Chromosome | start | end | strand | MM | target_seq | PAM | alignment | distance | position | gene name | gene id |
| GU305914.1 | 214435 | 214457 | - | 0 | TTATTGTCGGCGCGTGCGAC | CGG | \|\|\|\|\|\|\|\|[\|\|\|\|\|\|\|\|\|\|\|\|]PAM | NA | Intergenic | NA | NA |
| T18344 | CACGAGTCAGATTTCTCTCGCGG | 1000 | | | | | | | | | |
| Oligo fwd | CACCCACGAGTCAGATTTCTCTCG | | | | | | | | | | |
| Oligo rev | AAACCGAGAGAAATCTGACTCGTG | | | | | | | | | | |
| Chromosome | start | end | strand | MM | target_seq | PAM | alignment | distance | position | gene name | gene id |
| GU305914.1 | 214409 | 214431 | - | 0 | CACGAGTCAGATTTCTCTCG | CGG | \|\|\|\|\|\|\|\|[\|\|\|\|\|\|\|\|\|\|\|\|]PAM | NA | Intergenic | NA | NA |
| T18345 | CAGATTTCTCTCGCGGATCGTGG | 1000 | | | | | | | | | |
| Oligo fwd | CACCCAGATTTCTCTCGCGGATCG | | | | | | | | | | |
| Oligo rev | AAACCGATCCGCGAGAGAAATCTG | | | | | | | | | | |
| Chromosome | start | end | strand | MM | target_seq | PAM | alignment | distance | position | gene name | gene id |
| GU305914.1 | 214402 | 214424 | - | 0 | CAGATTTCTCTCGCGGATCG | TGG | \|\|\|\|\|\|\|\|[\|\|\|\|\|\|\|\|\|\|\|\|]PAM | NA | Intergenic | NA | NA |
| T18346 | GGATCGTGGTCTTATATATAAGG | 1000 | | | | | | | | | |
| Oligo fwd | CACCGGATCGTGGTCTTATATATA | | | | | | | | | | |

Oligo rev AAACTATATATAAGACCACGATCC
Chromosome start end strand MM target_seq PAM alignment distance position gene name gene id
GU305914.1 214388 214410 - 0 GGATCGTGGTCTTATATATA AGG ||||||||[||||||||||||]PAM NA Intergenic NA NA

T18347 GATCGTGGTCTTATATATAAGGG 1000
Oligo fwd CACCGATCGTGGTCTTATATATAA
Oligo rev AAACTTATATATAAGACCACGATC
Chromosome start end strand MM target_seq PAM alignment distance position gene name gene id
GU305914.1 214387 214409 - 0 GATCGTGGTCTTATATATAA GGG ||||||||[||||||||||||]PAM NA Intergenic NA NA

T18348 GATGTCCTGACAGATTATCGTGG 1000
Oligo fwd CACCGATGTCCTGACAGATTATCG
Oligo rev AAACCGATAATCTGTCAGGACATC
Chromosome start end strand MM target_seq PAM alignment distance position gene name gene id
GU305914.1 214337 214359 - 0 GATGTCCTGACAGATTATCG TGG ||||||||[||||||||||||]PAM NA Intergenic NA NA

T18349 GTGATATGACTACGATTGTCAGG 1000
Oligo fwd CACCGTGATATGACTACGATTGTC
Oligo rev AAACGACAATCGTAGTCATATCAC
Chromosome start end strand MM target_seq PAM alignment distance position gene name gene id
GU305914.1 214289 214311 - 0 GTGATATGACTACGATTGTC AGG ||||||||[||||||||||||]PAM NA Intergenic NA NA

T18350 TGATATGACTACGATTGTCAGGG 1000
Oligo fwd CACCTGATATGACTACGATTGTCA
Oligo rev AAACTGACAATCGTAGTCATATCA
Chromosome start end strand MM target_seq PAM alignment distance position gene name gene id
GU305914.1 214288 214310 - 0 TGATATGACTACGATTGTCA GGG ||||||||[||||||||||||]PAM NA Intergenic NA NA

T18351 GATATGACTACGATTGTCAGGGG 1000
Oligo fwd CACCGATATGACTACGATTGTCAG
Oligo rev AAACCTGACAATCGTAGTCATATC
Chromosome start end strand MM target_seq PAM alignment distance position gene name gene id
GU305914.1 214287 214309 - 0 GATATGACTACGATTGTCAG GGG ||||||||[||||||||||||]PAM NA Intergenic NA NA

T18352 ATATGACTACGATTGTCAGGGGG 1000
Oligo fwd CACCATATGACTACGATTGTCAGG
Oligo rev AAACCCTGACAATCGTAGTCATAT
Chromosome start end strand MM target_seq PAM alignment distance position gene name gene id
GU305914.1 214286 214308 - 0 ATATGACTACGATTGTCAGG GGG ||||||||[||||||||||||]PAM NA Intergenic NA NA

T18353 TCAGGGGGACCGTAGATATCTGG 1000
Oligo fwd CACCTCAGGGGGACCGTAGATATC
Oligo rev AAACGATATCTACGGTCCCCCTGA
Chromosome start end strand MM target_seq PAM alignment distance position gene name gene id
GU305914.1 214271 214293 - 0 TCAGGGGGACCGTAGATATC TGG ||||||||[||||||||||||]PAM NA Intergenic NA NA

T18354 CAGGGGGACCGTAGATATCTGGG 1000
Oligo fwd CACCCAGGGGGACCGTAGATATCT
Oligo rev AAACAGATATCTACGGTCCCCCTG
Chromosome start end strand MM target_seq PAM alignment distance position gene name gene id
GU305914.1 214270 214292 - 0 CAGGGGGACCGTAGATATCT GGG ||||||||[||||||||||||]PAM NA Intergenic NA NA

T18355 GGACCGTAGATATCTGGGTGTGG 1000
Oligo fwd CACCGGACCGTAGATATCTGGGTG
Oligo rev AAACCACCCAGATATCTACGGTCC
Chromosome start end strand MM target_seq PAM alignment distance position gene name gene id
GU305914.1 214265 214287 - 0 GGACCGTAGATATCTGGGTG TGG ||||||||[||||||||||||]PAM NA Intergenic NA NA

T18356 TGTGGCCACTCAAACATGAAAGG 1000
Oligo fwd CACCTGTGGCCACTCAAACATGAA
Oligo rev AAACTTCATGTTTGAGTGGCCACA
Chromosome start end strand MM target_seq PAM alignment distance position gene name gene id
GU305914.1 214247 214269 - 0 TGTGGCCACTCAAACATGAA AGG ||||||||[||||||||||||]PAM NA Intergenic NA NA

T18357 AAAGGCAATATCTTGACGATAGG 1000
Oligo fwd CACCAAAGGCAATATCTTGACGAT
Oligo rev AAACATCGTCAAGATATTGCCTTT
Chromosome start end strand MM target_seq PAM alignment distance position gene name gene id
GU305914.1 214229 214251 - 0 AAAGGCAATATCTTGACGAT AGG ||||||||[||||||||||||]PAM NA Intergenic NA NA

T18358 GCCCGCCAATCACATTCACGAGG 1000
Oligo fwd CACCGCCCGCCAATCACATTCACG
Oligo rev AAACCGTGAATGTGATTGGCGGGC
Chromosome start end strand MM target_seq PAM alignment distance position gene name gene id
GU305914.1 214179 214201 - 0 GCCCGCCAATCACATTCACG AGG ||||||||[||||||||||||]PAM NA Intergenic NA NA

T18359 CCCGCCAATCACATTCACGAGGG 1000
Oligo fwd CACCCCCGCCAATCACATTCACGA
Oligo rev AAACTCGTGAATGTGATTGGCGGG
Chromosome start end strand MM target_seq PAM alignment distance position gene name gene id
GU305914.1 214178 214200 - 0 CCCGCCAATCACATTCACGA GGG ||||||||[||||||||||||]PAM NA Intergenic NA NA

T18360 CCGCCAATCACATTCACGAGGGG 1000
Oligo fwd CACCCCGCCAATCACATTCACGAG
Oligo rev AAACCTCGTGAATGTGATTGGCGG
Chromosome start end strand MM target_seq PAM alignment distance position gene name gene id
GU305914.1 214177 214199 - 0 CCGCCAATCACATTCACGAG GGG ||||||||[||||||||||||]PAM NA Intergenic NA NA

T18361 CGCCAATCACATTCACGAGGGGG 1000

<table>
<tr><td>Oligo fwd</td><td>CACCCGCCAATCACATTCACGAGG</td><td></td><td></td><td></td><td></td><td></td><td></td><td></td><td></td><td></td><td></td></tr>
<tr><td>Oligo rev</td><td>AAACCCTCGTGAATGTGATTGGCG</td><td></td><td></td><td></td><td></td><td></td><td></td><td></td><td></td><td></td><td></td></tr>
<tr><td>Chromosome</td><td>start</td><td>end</td><td>strand</td><td>MM</td><td>target_seq</td><td>PAM</td><td>alignment</td><td>distance</td><td>position</td><td>gene name</td><td>gene id</td></tr>
<tr><td>GU305914.1</td><td>214176</td><td>214198</td><td>-</td><td>0</td><td>CGCCAATCACATTCACGAGG</td><td>GGG</td><td>||||||||[||||||||||||]PAM</td><td>NA</td><td>Intergenic</td><td>NA</td><td>NA</td></tr>
<tr><td></td><td></td><td></td><td></td><td></td><td></td><td></td><td></td><td></td><td></td><td></td><td></td></tr>
<tr><td>T18362</td><td>CGAGGGGGTGCTCCGAGATACGG</td><td>1000</td><td></td><td></td><td></td><td></td><td></td><td></td><td></td><td></td><td></td></tr>
<tr><td>Oligo fwd</td><td>CACCCGAGGGGGTGCTCCGAGATA</td><td></td><td></td><td></td><td></td><td></td><td></td><td></td><td></td><td></td><td></td></tr>
<tr><td>Oligo rev</td><td>AAACTATCTCGGAGCACCCCCTCG</td><td></td><td></td><td></td><td></td><td></td><td></td><td></td><td></td><td></td><td></td></tr>
<tr><td>Chromosome</td><td>start</td><td>end</td><td>strand</td><td>MM</td><td>target_seq</td><td>PAM</td><td>alignment</td><td>distance</td><td>position</td><td>gene name</td><td>gene id</td></tr>
<tr><td>GU305914.1</td><td>214161</td><td>214183</td><td>-</td><td>0</td><td>CGAGGGGGTGCTCCGAGATA</td><td>CGG</td><td>||||||||[||||||||||||]PAM</td><td>NA</td><td>Intergenic</td><td>NA</td><td>NA</td></tr>
<tr><td></td><td></td><td></td><td></td><td></td><td></td><td></td><td></td><td></td><td></td><td></td><td></td></tr>
<tr><td>T18363</td><td>ATACGGTCTCGACCACGATGCGG</td><td>1000</td><td></td><td></td><td></td><td></td><td></td><td></td><td></td><td></td><td></td></tr>
<tr><td>Oligo fwd</td><td>CACCATACGGTCTCGACCACGATG</td><td></td><td></td><td></td><td></td><td></td><td></td><td></td><td></td><td></td><td></td></tr>
<tr><td>Oligo rev</td><td>AAACCATCGTGGTCGAGACCGTAT</td><td></td><td></td><td></td><td></td><td></td><td></td><td></td><td></td><td></td><td></td></tr>
<tr><td>Chromosome</td><td>start</td><td>end</td><td>strand</td><td>MM</td><td>target_seq</td><td>PAM</td><td>alignment</td><td>distance</td><td>position</td><td>gene name</td><td>gene id</td></tr>
<tr><td>GU305914.1</td><td>214144</td><td>214166</td><td>-</td><td>0</td><td>ATACGGTCTCGACCACGATG</td><td>CGG</td><td>||||||||[||||||||||||]PAM</td><td>NA</td><td>Intergenic</td><td>NA</td><td>NA</td></tr>
<tr><td></td><td></td><td></td><td></td><td></td><td></td><td></td><td></td><td></td><td></td><td></td><td></td></tr>
<tr><td>T18364</td><td>TACGGTCTCGACCACGATGCGGG</td><td>1000</td><td></td><td></td><td></td><td></td><td></td><td></td><td></td><td></td><td></td></tr>
<tr><td>Oligo fwd</td><td>CACCTACGGTCTCGACCACGATGC</td><td></td><td></td><td></td><td></td><td></td><td></td><td></td><td></td><td></td><td></td></tr>
<tr><td>Oligo rev</td><td>AAACGCATCGTGGTCGAGACCGTA</td><td></td><td></td><td></td><td></td><td></td><td></td><td></td><td></td><td></td><td></td></tr>
<tr><td>Chromosome</td><td>start</td><td>end</td><td>strand</td><td>MM</td><td>target_seq</td><td>PAM</td><td>alignment</td><td>distance</td><td>position</td><td>gene name</td><td>gene id</td></tr>
<tr><td>GU305914.1</td><td>214143</td><td>214165</td><td>-</td><td>0</td><td>TACGGTCTCGACCACGATGC</td><td>GGG</td><td>||||||||[||||||||||||]PAM</td><td>NA</td><td>Intergenic</td><td>NA</td><td>NA</td></tr>
<tr><td>GU305914.1</td><td>154016</td><td>154038</td><td>+</td><td>4</td><td>TACCACCTCGACCACGATTC</td><td>TGG</td><td>|||---||[||||||||||-|]PAM</td><td>NA</td><td>Intergenic</td><td>NA</td><td>NA</td></tr>
<tr><td></td><td></td><td></td><td></td><td></td><td></td><td></td><td></td><td></td><td></td><td></td><td></td></tr>
<tr><td>T18365</td><td>CCACGATGCGGGCGATGTTACGG</td><td>1000</td><td></td><td></td><td></td><td></td><td></td><td></td><td></td><td></td><td></td></tr>
<tr><td>Oligo fwd</td><td>CACCCCACGATGCGGGCGATGTTA</td><td></td><td></td><td></td><td></td><td></td><td></td><td></td><td></td><td></td><td></td></tr>
<tr><td>Oligo rev</td><td>AAACTAACATCGCCCGCATCGTGG</td><td></td><td></td><td></td><td></td><td></td><td></td><td></td><td></td><td></td><td></td></tr>
<tr><td>Chromosome</td><td>start</td><td>end</td><td>strand</td><td>MM</td><td>target_seq</td><td>PAM</td><td>alignment</td><td>distance</td><td>position</td><td>gene name</td><td>gene id</td></tr>
<tr><td>GU305914.1</td><td>214132</td><td>214154</td><td>-</td><td>0</td><td>CCACGATGCGGGCGATGTTA</td><td>CGG</td><td>||||||||[||||||||||||]PAM</td><td>NA</td><td>Intergenic</td><td>NA</td><td>NA</td></tr>
<tr><td></td><td></td><td></td><td></td><td></td><td></td><td></td><td></td><td></td><td></td><td></td><td></td></tr>
<tr><td>T18366</td><td>GATATATGCGCTAAGCTCGCTGG</td><td>1000</td><td></td><td></td><td></td><td></td><td></td><td></td><td></td><td></td><td></td></tr>
<tr><td>Oligo fwd</td><td>CACCGATATATGCGCTAAGCTCGC</td><td></td><td></td><td></td><td></td><td></td><td></td><td></td><td></td><td></td><td></td></tr>
<tr><td>Oligo rev</td><td>AAACGCGAGCTTAGCGCATATATC</td><td></td><td></td><td></td><td></td><td></td><td></td><td></td><td></td><td></td><td></td></tr>
<tr><td>Chromosome</td><td>start</td><td>end</td><td>strand</td><td>MM</td><td>target_seq</td><td>PAM</td><td>alignment</td><td>distance</td><td>position</td><td>gene name</td><td>gene id</td></tr>
<tr><td>GU305914.1</td><td>214110</td><td>214132</td><td>-</td><td>0</td><td>GATATATGCGCTAAGCTCGC</td><td>TGG</td><td>||||||||[||||||||||||]PAM</td><td>NA</td><td>Intergenic</td><td>NA</td><td>NA</td></tr>
<tr><td></td><td></td><td></td><td></td><td></td><td></td><td></td><td></td><td></td><td></td><td></td><td></td></tr>
<tr><td>T18367</td><td>ATTCCATCCCGTCGTCGCATTGG</td><td>1000</td><td></td><td></td><td></td><td></td><td></td><td></td><td></td><td></td><td></td></tr>
<tr><td>Oligo fwd</td><td>CACCATTCCATCCCGTCGTCGCAT</td><td></td><td></td><td></td><td></td><td></td><td></td><td></td><td></td><td></td><td></td></tr>
<tr><td>Oligo rev</td><td>AAACATGCGACGACGGGATGGAAT</td><td></td><td></td><td></td><td></td><td></td><td></td><td></td><td></td><td></td><td></td></tr>
<tr><td>Chromosome</td><td>start</td><td>end</td><td>strand</td><td>MM</td><td>target_seq</td><td>PAM</td><td>alignment</td><td>distance</td><td>position</td><td>gene name</td><td>gene id</td></tr>
<tr><td>GU305914.1</td><td>214077</td><td>214099</td><td>-</td><td>0</td><td>ATTCCATCCCGTCGTCGCAT</td><td>TGG</td><td>||||||||[||||||||||||]PAM</td><td>NA</td><td>Intergenic</td><td>NA</td><td>NA</td></tr>
<tr><td></td><td></td><td></td><td></td><td></td><td></td><td></td><td></td><td></td><td></td><td></td><td></td></tr>
<tr><td>T18368</td><td>TTCCATCCCGTCGTCGCATTGGG</td><td>1000</td><td></td><td></td><td></td><td></td><td></td><td></td><td></td><td></td><td></td></tr>
<tr><td>Oligo fwd</td><td>CACCTTCCATCCCGTCGTCGCATT</td><td></td><td></td><td></td><td></td><td></td><td></td><td></td><td></td><td></td><td></td></tr>
<tr><td>Oligo rev</td><td>AAACAATGCGACGACGGGATGGAA</td><td></td><td></td><td></td><td></td><td></td><td></td><td></td><td></td><td></td><td></td></tr>
<tr><td>Chromosome</td><td>start</td><td>end</td><td>strand</td><td>MM</td><td>target_seq</td><td>PAM</td><td>alignment</td><td>distance</td><td>position</td><td>gene name</td><td>gene id</td></tr>
<tr><td>GU305914.1</td><td>214076</td><td>214098</td><td>-</td><td>0</td><td>TTCCATCCCGTCGTCGCATT</td><td>GGG</td><td>||||||||[||||||||||||]PAM</td><td>NA</td><td>Intergenic</td><td>NA</td><td>NA</td></tr>
<tr><td></td><td></td><td></td><td></td><td></td><td></td><td></td><td></td><td></td><td></td><td></td><td></td></tr>
<tr><td>T18369</td><td>ATTGGGTCGTTTAGAGCTTTTGG</td><td>1000</td><td></td><td></td><td></td><td></td><td></td><td></td><td></td><td></td><td></td></tr>
<tr><td>Oligo fwd</td><td>CACCATTGGGTCGTTTAGAGCTTT</td><td></td><td></td><td></td><td></td><td></td><td></td><td></td><td></td><td></td><td></td></tr>
<tr><td>Oligo rev</td><td>AAACAAAGCTCTAAACGACCCAAT</td><td></td><td></td><td></td><td></td><td></td><td></td><td></td><td></td><td></td><td></td></tr>
<tr><td>Chromosome</td><td>start</td><td>end</td><td>strand</td><td>MM</td><td>target_seq</td><td>PAM</td><td>alignment</td><td>distance</td><td>position</td><td>gene name</td><td>gene id</td></tr>
<tr><td>GU305914.1</td><td>214059</td><td>214081</td><td>-</td><td>0</td><td>ATTGGGTCGTTTAGAGCTTT</td><td>TGG</td><td>||||||||[||||||||||||]PAM</td><td>NA</td><td>Intergenic</td><td>NA</td><td>NA</td></tr>
<tr><td></td><td></td><td></td><td></td><td></td><td></td><td></td><td></td><td></td><td></td><td></td><td></td></tr>
<tr><td>T18370</td><td>GCTTTTGGCGTTAGTCTCCTCGG</td><td>1000</td><td></td><td></td><td></td><td></td><td></td><td></td><td></td><td></td><td></td></tr>
<tr><td>Oligo fwd</td><td>CACCGCTTTTGGCGTTAGTCTCCT</td><td></td><td></td><td></td><td></td><td></td><td></td><td></td><td></td><td></td><td></td></tr>
<tr><td>Oligo rev</td><td>AAACAGGAGACTAACGCCAAAAGC</td><td></td><td></td><td></td><td></td><td></td><td></td><td></td><td></td><td></td><td></td></tr>
<tr><td>Chromosome</td><td>start</td><td>end</td><td>strand</td><td>MM</td><td>target_seq</td><td>PAM</td><td>alignment</td><td>distance</td><td>position</td><td>gene name</td><td>gene id</td></tr>
<tr><td>GU305914.1</td><td>214044</td><td>214066</td><td>-</td><td>0</td><td>GCTTTTGGCGTTAGTCTCCT</td><td>CGG</td><td>||||||||[||||||||||||]PAM</td><td>NA</td><td>Intergenic</td><td>NA</td><td>NA</td></tr>
<tr><td></td><td></td><td></td><td></td><td></td><td></td><td></td><td></td><td></td><td></td><td></td><td></td></tr>
<tr><td>T18371</td><td>TTGGCGTTAGTCTCCTCGGTAGG</td><td>1000</td><td></td><td></td><td></td><td></td><td></td><td></td><td></td><td></td><td></td></tr>
<tr><td>Oligo fwd</td><td>CACCTTGGCGTTAGTCTCCTCGGT</td><td></td><td></td><td></td><td></td><td></td><td></td><td></td><td></td><td></td><td></td></tr>
<tr><td>Oligo rev</td><td>AAACACCGAGGAGACTAACGCCAA</td><td></td><td></td><td></td><td></td><td></td><td></td><td></td><td></td><td></td><td></td></tr>
<tr><td>Chromosome</td><td>start</td><td>end</td><td>strand</td><td>MM</td><td>target_seq</td><td>PAM</td><td>alignment</td><td>distance</td><td>position</td><td>gene name</td><td>gene id</td></tr>
<tr><td>GU305914.1</td><td>214040</td><td>214062</td><td>-</td><td>0</td><td>TTGGCGTTAGTCTCCTCGGT</td><td>AGG</td><td>||||||||[||||||||||||]PAM</td><td>NA</td><td>Intergenic</td><td>NA</td><td>NA</td></tr>
<tr><td></td><td></td><td></td><td></td><td></td><td></td><td></td><td></td><td></td><td></td><td></td><td></td></tr>
<tr><td>T18372</td><td>GTTAGTCTCCTCGGTAGGAATGG</td><td>1000</td><td></td><td></td><td></td><td></td><td></td><td></td><td></td><td></td><td></td></tr>
<tr><td>Oligo fwd</td><td>CACCGTTAGTCTCCTCGGTAGGAA</td><td></td><td></td><td></td><td></td><td></td><td></td><td></td><td></td><td></td><td></td></tr>
<tr><td>Oligo rev</td><td>AAACTTCCTACCGAGGAGACTAAC</td><td></td><td></td><td></td><td></td><td></td><td></td><td></td><td></td><td></td><td></td></tr>
<tr><td>Chromosome</td><td>start</td><td>end</td><td>strand</td><td>MM</td><td>target_seq</td><td>PAM</td><td>alignment</td><td>distance</td><td>position</td><td>gene name</td><td>gene id</td></tr>
<tr><td>GU305914.1</td><td>214035</td><td>214057</td><td>-</td><td>0</td><td>GTTAGTCTCCTCGGTAGGAA</td><td>TGG</td><td>||||||||[||||||||||||]PAM</td><td>NA</td><td>Intergenic</td><td>NA</td><td>NA</td></tr>
<tr><td></td><td></td><td></td><td></td><td></td><td></td><td></td><td></td><td></td><td></td><td></td><td></td></tr>
<tr><td>T18373</td><td>GTAGGAATGGAACTTTCTGTCGG</td><td>1000</td><td></td><td></td><td></td><td></td><td></td><td></td><td></td><td></td><td></td></tr>
<tr><td>Oligo fwd</td><td>CACCGTAGGAATGGAACTTTCTGT</td><td></td><td></td><td></td><td></td><td></td><td></td><td></td><td></td><td></td><td></td></tr>
<tr><td>Oligo rev</td><td>AAACACAGAAAGTTCCATTCCTAC</td><td></td><td></td><td></td><td></td><td></td><td></td><td></td><td></td><td></td><td></td></tr>
<tr><td>Chromosome</td><td>start</td><td>end</td><td>strand</td><td>MM</td><td>target_seq</td><td>PAM</td><td>alignment</td><td>distance</td><td>position</td><td>gene name</td><td>gene id</td></tr>
<tr><td>GU305914.1</td><td>214022</td><td>214044</td><td>-</td><td>0</td><td>GTAGGAATGGAACTTTCTGT</td><td>CGG</td><td>||||||||[||||||||||||]PAM</td><td>NA</td><td>Intergenic</td><td>NA</td><td>NA</td></tr>
<tr><td></td><td></td><td></td><td></td><td></td><td></td><td></td><td></td><td></td><td></td><td></td><td></td></tr>
<tr><td>T18374</td><td>TGTCGGTGCCACTTCTACCACGG</td><td>1000</td><td></td><td></td><td></td><td></td><td></td><td></td><td></td><td></td><td></td></tr>
<tr><td>Oligo fwd</td><td>CACCTGTCGGTGCCACTTCTACCA</td><td></td><td></td><td></td><td></td><td></td><td></td><td></td><td></td><td></td><td></td></tr>
<tr><td>Oligo rev</td><td>AAACTGGTAGAAGTGGCACCGACA</td><td></td><td></td><td></td><td></td><td></td><td></td><td></td><td></td><td></td><td></td></tr>
<tr><td>Chromosome</td><td>start</td><td>end</td><td>strand</td><td>MM</td><td>target_seq</td><td>PAM</td><td>alignment</td><td>distance</td><td>position</td><td>gene name</td><td>gene id</td></tr>
<tr><td>GU305914.1</td><td>214005</td><td>214027</td><td>-</td><td>0</td><td>TGTCGGTGCCACTTCTACCA</td><td>CGG</td><td>||||||||[||||||||||||]PAM</td><td>NA</td><td>Intergenic</td><td>NA</td><td>NA</td></tr>
<tr><td></td><td></td><td></td><td></td><td></td><td></td><td></td><td></td><td></td><td></td><td></td><td></td></tr>
<tr><td>T18375</td><td>CGAGACTACCCTAGAGATGACGG</td><td>1000</td><td></td><td></td><td></td><td></td><td></td><td></td><td></td><td></td><td></td></tr>
<tr><td>Oligo fwd</td><td>CACCCGAGACTACCCTAGAGATGA</td><td></td><td></td><td></td><td></td><td></td><td></td><td></td><td></td><td></td><td></td></tr>
<tr><td>Oligo rev</td><td>AAACTCATCTCTAGGGTAGTCTCG</td><td></td><td></td><td></td><td></td><td></td><td></td><td></td><td></td><td></td><td></td></tr>
<tr><td>Chromosome</td><td>start</td><td>end</td><td>strand</td><td>MM</td><td>target_seq</td><td>PAM</td><td>alignment</td><td>distance</td><td>position</td><td>gene name</td><td>gene id</td></tr>
<tr><td>GU305914.1</td><td>213963</td><td>213985</td><td>-</td><td>0</td><td>CGAGACTACCCTAGAGATGA</td><td>CGG</td><td>||||||||[||||||||||||]PAM</td><td>NA</td><td>Intergenic</td><td>NA</td><td>NA</td></tr>
</table>

T18376 GAGACTACCCTAGAGATGACGGG 1000
Oligo fwd CACCGAGACTACCCTAGAGATGAC
Oligo rev AAACGTCATCTCTAGGGTAGTCTC
Chromosome start end strand MM target_seq PAM alignment distance position gene name gene id
GU305914.1 213962 213984 - 0 GAGACTACCCTAGAGATGAC GGG ||||||||[||||||||||||]PAM NA Intergenic NA NA

T18377 AGACTACCCTAGAGATGACGGGG 1000
Oligo fwd CACCAGACTACCCTAGAGATGACG
Oligo rev AAACCGTCATCTCTAGGGTAGTCT
Chromosome start end strand MM target_seq PAM alignment distance position gene name gene id
GU305914.1 213961 213983 - 0 AGACTACCCTAGAGATGACG GGG ||||||||[||||||||||||]PAM NA Intergenic NA NA

T18378 ACCCTAGAGATGACGGGGTCCGG 1000
Oligo fwd CACCACCCTAGAGATGACGGGGTC
Oligo rev AAACGACCCCGTCATCTCTAGGGT
Chromosome start end strand MM target_seq PAM alignment distance position gene name gene id
GU305914.1 213956 213978 - 0 ACCCTAGAGATGACGGGGTC CGG ||||||||[||||||||||||]PAM NA Intergenic NA NA

T18379 CCCTAGAGATGACGGGGTCCGGG 1000
Oligo fwd CACCCCCTAGAGATGACGGGGTCC
Oligo rev AAACGGACCCCGTCATCTCTAGGG
Chromosome start end strand MM target_seq PAM alignment distance position gene name gene id
GU305914.1 213955 213977 - 0 CCCTAGAGATGACGGGGTCC GGG ||||||||[||||||||||||]PAM NA Intergenic NA NA

T18380 CCTAGAGATGACGGGGTCCGGGG 1000
Oligo fwd CACCCCTAGAGATGACGGGGTCCG
Oligo rev AAACCGGACCCCGTCATCTCTAGG
Chromosome start end strand MM target_seq PAM alignment distance position gene name gene id
GU305914.1 213954 213976 - 0 CCTAGAGATGACGGGGTCCG GGG ||||||||[||||||||||||]PAM NA Intergenic NA NA

T18381 CCCGAGCCCACTAGCGAGCCCGG 1000
Oligo fwd CACCCCCGAGCCCACTAGCGAGCC
Oligo rev AAACGGCTCGCTAGTGGGCTCGGG
Chromosome start end strand MM target_seq PAM alignment distance position gene name gene id
GU305914.1 213902 213924 - 0 CCCGAGCCCACTAGCGAGCC CGG ||||||||[||||||||||||]PAM NA Intergenic NA NA

T18382 GTTAGATGTGTCAACTTTCGTGG 1000
Oligo fwd CACCGTTAGATGTGTCAACTTTCG
Oligo rev AAACCGAAAGTTGACACATCTAAC
Chromosome start end strand MM target_seq PAM alignment distance position gene name gene id
GU305914.1 213822 213844 - 0 GTTAGATGTGTCAACTTTCG TGG ||||||||[||||||||||||]PAM NA Intergenic NA NA

T18383 GTCAACTTTCGTGGCGTCGACGG 1000
Oligo fwd CACCGTCAACTTTCGTGGCGTCGA
Oligo rev AAACTCGACGCCACGAAAGTTGAC
Chromosome start end strand MM target_seq PAM alignment distance position gene name gene id
GU305914.1 213813 213835 - 0 GTCAACTTTCGTGGCGTCGA CGG ||||||||[||||||||||||]PAM NA Intergenic NA NA

T18384 TTTCGTGGCGTCGACGGTCGCGG 1000
Oligo fwd CACCTTTCGTGGCGTCGACGGTCG
Oligo rev AAACCGACCGTCGACGCCACGAAA
Chromosome start end strand MM target_seq PAM alignment distance position gene name gene id
GU305914.1 213807 213829 - 0 TTTCGTGGCGTCGACGGTCG CGG ||||||||[||||||||||||]PAM NA Intergenic NA NA

T18385 TACACCTCAGCCCGAGACTACGG 1000
Oligo fwd CACCTACACCTCAGCCCGAGACTA
Oligo rev AAACTAGTCTCGGGCTGAGGTGTA
Chromosome start end strand MM target_seq PAM alignment distance position gene name gene id
GU305914.1 213771 213793 - 0 TACACCTCAGCCCGAGACTA CGG ||||||||[||||||||||||]PAM NA Intergenic NA NA

T18386 GCCCGATACCTCTACCGCTGCGG 1000
Oligo fwd CACCGCCCGATACCTCTACCGCTG
Oligo rev AAACCAGCGGTAGAGGTATCGGGC
Chromosome start end strand MM target_seq PAM alignment distance position gene name gene id
GU305914.1 213747 213769 - 0 GCCCGATACCTCTACCGCTG CGG ||||||||[||||||||||||]PAM NA Intergenic NA NA

T18387 CGCTGCGGACGCGATCTCTTCGG 1000
Oligo fwd CACCCGCTGCGGACGCGATCTCTT
Oligo rev AAACAAGAGATCGCGTCCGCAGCG
Chromosome start end strand MM target_seq PAM alignment distance position gene name gene id
GU305914.1 213732 213754 - 0 CGCTGCGGACGCGATCTCTT CGG ||||||||[||||||||||||]PAM NA Intergenic NA NA

T18388 TGCGGACGCGATCTCTTCGGCGG 1000
Oligo fwd CACCTGCGGACGCGATCTCTTCGG
Oligo rev AAACCCGAAGAGATCGCGTCCGCA
Chromosome start end strand MM target_seq PAM alignment distance position gene name gene id
GU305914.1 213729 213751 - 0 TGCGGACGCGATCTCTTCGG CGG ||||||||[||||||||||||]PAM NA Intergenic NA NA

T18389 TCGGCGGCAACCCCTAGTGCAGG 1000
Oligo fwd CACCTCGGCGGCAACCCCTAGTGC
Oligo rev AAACGCACTAGGGGTTGCCGCCGA
Chromosome start end strand MM target_seq PAM alignment distance position gene name gene id
GU305914.1 213713 213735 - 0 TCGGCGGCAACCCCTAGTGC AGG ||||||||[||||||||||||]PAM NA Intergenic NA NA

T18390 TCCAAGCCCTGTTACCACGAAGG 1000
Oligo fwd CACCTCCAAGCCCTGTTACCACGA
Oligo rev AAACTCGTGGTAACAGGGCTTGGA
Chromosome start end strand MM target_seq PAM alignment distance position gene name gene id

GU305914.1 213675 213697 - 0 TCCAAGCCCTGTTACCACGA AGG ||||||||[||||||||||||]PAM NA Intergenic NA NA

T18391 CCAAGCCCTGTTACCACGAAGGG 1000
Oligo fwd CACCCCAAGCCCTGTTACCACGAA
Oligo rev AAACTTCGTGGTAACAGGGCTTGG
Chromosome start end strand MM target_seq PAM alignment distance position gene name gene id
GU305914.1 213674 213696 - 0 CCAAGCCCTGTTACCACGAA GGG ||||||||[||||||||||||]PAM NA Intergenic NA NA

T18392 GACGACCGAACCGTCGTCCACGG 1000
Oligo fwd CACCGACGACCGAACCGTCGTCCA
Oligo rev AAACTGGACGACGGTTCGGTCGTC
Chromosome start end strand MM target_seq PAM alignment distance position gene name gene id
GU305914.1 213612 213634 - 0 GACGACCGAACCGTCGTCCA CGG ||||||||[||||||||||||]PAM NA Intergenic NA NA
GU305914.1 213543 213565 - 1 GACGACCGAACCATCGTCCA CGG ||||||||[||||-|||||||]PAM NA Intergenic NA NA

T18393 GACGACCGAACCATCGTCCACGG 1000
Oligo fwd CACCGACGACCGAACCATCGTCCA
Oligo rev AAACTGGACGATGGTTCGGTCGTC
Chromosome start end strand MM target_seq PAM alignment distance position gene name gene id
GU305914.1 213543 213565 - 0 GACGACCGAACCATCGTCCA CGG ||||||||[||||||||||||]PAM NA Intergenic NA NA
GU305914.1 213612 213634 - 1 GACGACCGAACCGTCGTCCA CGG ||||||||[||||-|||||||]PAM NA Intergenic NA NA

T18394 CTCCAGCCCCAACGCCACTACGG 1000
Oligo fwd CACCCTCCAGCCCCAACGCCACTA
Oligo rev AAACTAGTGGCGTTGGGGCTGGAG
Chromosome start end strand MM target_seq PAM alignment distance position gene name gene id
GU305914.1 213459 213481 - 0 CTCCAGCCCCAACGCCACTA CGG ||||||||[||||||||||||]PAM NA Intergenic NA NA

T18395 CGCCACTACGGTGACCACGCAGG 1000
Oligo fwd CACCCGCCACTACGGTGACCACGC
Oligo rev AAACGCGTGGTCACCGTAGTGGCG
Chromosome start end strand MM target_seq PAM alignment distance position gene name gene id
GU305914.1 213447 213469 - 0 CGCCACTACGGTGACCACGC AGG ||||||||[||||||||||||]PAM NA Intergenic NA NA

T18396 GCCACTACGGTGACCACGCAGGG 1000
Oligo fwd CACCGCCACTACGGTGACCACGCA
Oligo rev AAACTGCGTGGTCACCGTAGTGGC
Chromosome start end strand MM target_seq PAM alignment distance position gene name gene id
GU305914.1 213446 213468 - 0 GCCACTACGGTGACCACGCA GGG ||||||||[||||||||||||]PAM NA Intergenic NA NA

T18397 ATACACGACTAGCATGAAGTTGG 1000
Oligo fwd CACCATACACGACTAGCATGAAGT
Oligo rev AAACACTTCATGCTAGTCGTGTAT
Chromosome start end strand MM target_seq PAM alignment distance position gene name gene id
GU305914.1 213423 213445 - 0 ATACACGACTAGCATGAAGT TGG ||||||||[||||||||||||]PAM NA Intergenic NA NA

T18398 GATGCCGATGAATAGCACGACGG 1000
Oligo fwd CACCGATGCCGATGAATAGCACGA
Oligo rev AAACTCGTGCTATTCATCGGCATC
Chromosome start end strand MM target_seq PAM alignment distance position gene name gene id
GU305914.1 213381 213403 - 0 GATGCCGATGAATAGCACGA CGG ||||||||[||||||||||||]PAM NA Intergenic NA NA

T18399 ATAGCACGACGGCCGTTTCGAGG 1000
Oligo fwd CACCATAGCACGACGGCCGTTTCG
Oligo rev AAACCGAAACGGCCGTCGTGCTAT
Chromosome start end strand MM target_seq PAM alignment distance position gene name gene id
GU305914.1 213370 213392 - 0 ATAGCACGACGGCCGTTTCG AGG ||||||||[||||||||||||]PAM NA Intergenic NA NA

T18400 GCACGACGGCCGTTTCGAGGAGG 1000
Oligo fwd CACCGCACGACGGCCGTTTCGAGG
Oligo rev AAACCCTCGAAACGGCCGTCGTGC
Chromosome start end strand MM target_seq PAM alignment distance position gene name gene id
GU305914.1 213367 213389 - 0 GCACGACGGCCGTTTCGAGG AGG ||||||||[||||||||||||]PAM NA Intergenic NA NA

T18401 GCCGTTTCGAGGAGGTCTACCGG 1000
Oligo fwd CACCGCCGTTTCGAGGAGGTCTAC
Oligo rev AAACGTAGACCTCCTCGAAACGGC
Chromosome start end strand MM target_seq PAM alignment distance position gene name gene id
GU305914.1 213359 213381 - 0 GCCGTTTCGAGGAGGTCTAC CGG ||||||||[||||||||||||]PAM NA Intergenic NA NA

T18402 CCGTTTCGAGGAGGTCTACCGGG 1000
Oligo fwd CACCCCGTTTCGAGGAGGTCTACC
Oligo rev AAACGGTAGACCTCCTCGAAACGG
Chromosome start end strand MM target_seq PAM alignment distance position gene name gene id
GU305914.1 213358 213380 - 0 CCGTTTCGAGGAGGTCTACC GGG ||||||||[||||||||||||]PAM NA Intergenic NA NA

T18403 CCACGACCTTGAGTACGACCAGG 1000
Oligo fwd CACCCCACGACCTTGAGTACGACC
Oligo rev AAACGGTCGTACTCAAGGTCGTGG
Chromosome start end strand MM target_seq PAM alignment distance position gene name gene id
GU305914.1 213307 213329 - 0 CCACGACCTTGAGTACGACC AGG ||||||||[||||||||||||]PAM NA Intergenic NA NA

T18404 AGTACGACCAGGCTGAGAACAGG 1000
Oligo fwd CACCAGTACGACCAGGCTGAGAAC
Oligo rev AAACGTTCTCAGCCTGGTCGTACT
Chromosome start end strand MM target_seq PAM alignment distance position gene name gene id
GU305914.1 213296 213318 - 0 AGTACGACCAGGCTGAGAAC AGG ||||||||[||||||||||||]PAM NA Intergenic NA NA

T18405 GTACGACCAGGCTGAGAACAGGG 1000

<table>
<tr><td>Oligo fwd</td><td>CACCGTACGACCAGGCTGAGAACA</td><td></td><td></td><td></td><td></td><td></td><td></td><td></td><td></td><td></td><td></td></tr>
<tr><td>Oligo rev</td><td>AAACTGTTCTCAGCCTGGTCGTAC</td><td></td><td></td><td></td><td></td><td></td><td></td><td></td><td></td><td></td><td></td></tr>
<tr><td>Chromosome</td><td>start</td><td>end</td><td>strand</td><td>MM</td><td>target_seq</td><td>PAM</td><td>alignment</td><td>distance</td><td>position</td><td>gene name</td><td>gene id</td></tr>
<tr><td>GU305914.1</td><td>213295</td><td>213317</td><td>-</td><td>0</td><td>GTACGACCAGGCTGAGAACA</td><td>GGG</td><td>||||||||[||||||||||||]PAM</td><td>NA</td><td>Intergenic</td><td>NA</td><td>NA</td></tr>
<tr><td></td><td></td><td></td><td></td><td></td><td></td><td></td><td></td><td></td><td></td><td></td><td></td></tr>
<tr><td>T18406</td><td>TCCGACCGCCACCTCCGACGAGG</td><td>1000</td><td></td><td></td><td></td><td></td><td></td><td></td><td></td><td></td><td></td></tr>
<tr><td>Oligo fwd</td><td>CACCTCCGACCGCCACCTCCGACG</td><td></td><td></td><td></td><td></td><td></td><td></td><td></td><td></td><td></td><td></td></tr>
<tr><td>Oligo rev</td><td>AAACCGTCGGAGGTGGCGGTCGGA</td><td></td><td></td><td></td><td></td><td></td><td></td><td></td><td></td><td></td><td></td></tr>
<tr><td>Chromosome</td><td>start</td><td>end</td><td>strand</td><td>MM</td><td>target_seq</td><td>PAM</td><td>alignment</td><td>distance</td><td>position</td><td>gene name</td><td>gene id</td></tr>
<tr><td>GU305914.1</td><td>213261</td><td>213283</td><td>-</td><td>0</td><td>TCCGACCGCCACCTCCGACG</td><td>AGG</td><td>||||||||[||||||||||||]PAM</td><td>NA</td><td>Intergenic</td><td>NA</td><td>NA</td></tr>
<tr><td></td><td></td><td></td><td></td><td></td><td></td><td></td><td></td><td></td><td></td><td></td><td></td></tr>
<tr><td>T18407</td><td>CCGACCGCCACCTCCGACGAGGG</td><td>1000</td><td></td><td></td><td></td><td></td><td></td><td></td><td></td><td></td><td></td></tr>
<tr><td>Oligo fwd</td><td>CACCCCGACCGCCACCTCCGACGA</td><td></td><td></td><td></td><td></td><td></td><td></td><td></td><td></td><td></td><td></td></tr>
<tr><td>Oligo rev</td><td>AAACTCGTCGGAGGTGGCGGTCGG</td><td></td><td></td><td></td><td></td><td></td><td></td><td></td><td></td><td></td><td></td></tr>
<tr><td>Chromosome</td><td>start</td><td>end</td><td>strand</td><td>MM</td><td>target_seq</td><td>PAM</td><td>alignment</td><td>distance</td><td>position</td><td>gene name</td><td>gene id</td></tr>
<tr><td>GU305914.1</td><td>213260</td><td>213282</td><td>-</td><td>0</td><td>CCGACCGCCACCTCCGACGA</td><td>GGG</td><td>||||||||[||||||||||||]PAM</td><td>NA</td><td>Intergenic</td><td>NA</td><td>NA</td></tr>
<tr><td></td><td></td><td></td><td></td><td></td><td></td><td></td><td></td><td></td><td></td><td></td><td></td></tr>
<tr><td>T18408</td><td>CCCTCACGACGCTGATCGACCGG</td><td>1000</td><td></td><td></td><td></td><td></td><td></td><td></td><td></td><td></td><td></td></tr>
<tr><td>Oligo fwd</td><td>CACCCCCTCACGACGCTGATCGAC</td><td></td><td></td><td></td><td></td><td></td><td></td><td></td><td></td><td></td><td></td></tr>
<tr><td>Oligo rev</td><td>AAACGTCGATCAGCGTCGTGAGGG</td><td></td><td></td><td></td><td></td><td></td><td></td><td></td><td></td><td></td><td></td></tr>
<tr><td>Chromosome</td><td>start</td><td>end</td><td>strand</td><td>MM</td><td>target_seq</td><td>PAM</td><td>alignment</td><td>distance</td><td>position</td><td>gene name</td><td>gene id</td></tr>
<tr><td>GU305914.1</td><td>213223</td><td>213245</td><td>-</td><td>0</td><td>CCCTCACGACGCTGATCGAC</td><td>CGG</td><td>||||||||[||||||||||||]PAM</td><td>NA</td><td>Intergenic</td><td>NA</td><td>NA</td></tr>
<tr><td></td><td></td><td></td><td></td><td></td><td></td><td></td><td></td><td></td><td></td><td></td><td></td></tr>
<tr><td>T18409</td><td>CCTCACGACGCTGATCGACCGGG</td><td>1000</td><td></td><td></td><td></td><td></td><td></td><td></td><td></td><td></td><td></td></tr>
<tr><td>Oligo fwd</td><td>CACCCCTCACGACGCTGATCGACC</td><td></td><td></td><td></td><td></td><td></td><td></td><td></td><td></td><td></td><td></td></tr>
<tr><td>Oligo rev</td><td>AAACGGTCGATCAGCGTCGTGAGG</td><td></td><td></td><td></td><td></td><td></td><td></td><td></td><td></td><td></td><td></td></tr>
<tr><td>Chromosome</td><td>start</td><td>end</td><td>strand</td><td>MM</td><td>target_seq</td><td>PAM</td><td>alignment</td><td>distance</td><td>position</td><td>gene name</td><td>gene id</td></tr>
<tr><td>GU305914.1</td><td>213222</td><td>213244</td><td>-</td><td>0</td><td>CCTCACGACGCTGATCGACC</td><td>GGG</td><td>||||||||[||||||||||||]PAM</td><td>NA</td><td>Intergenic</td><td>NA</td><td>NA</td></tr>
<tr><td>GU305914.1</td><td>77698</td><td>77720</td><td>+</td><td>4</td><td>CAACACGACGCTGACGGACC</td><td>TGG</td><td>|--|||||[||||||--||||]PAM</td><td>NA</td><td>Intergenic</td><td>NA</td><td>NA</td></tr>
<tr><td>GU305914.1</td><td>87687</td><td>87709</td><td>-</td><td>4</td><td>CCTGGCGACGCTGAACGACG</td><td>TGG</td><td>|||--|||[||||||-||||-]PAM</td><td>NA</td><td>Intergenic</td><td>NA</td><td>NA</td></tr>
<tr><td></td><td></td><td></td><td></td><td></td><td></td><td></td><td></td><td></td><td></td><td></td><td></td></tr>
<tr><td>T18410</td><td>GACGCTGATCGACCGGGATGAGG</td><td>1000</td><td></td><td></td><td></td><td></td><td></td><td></td><td></td><td></td><td></td></tr>
<tr><td>Oligo fwd</td><td>CACCGACGCTGATCGACCGGGATG</td><td></td><td></td><td></td><td></td><td></td><td></td><td></td><td></td><td></td><td></td></tr>
<tr><td>Oligo rev</td><td>AAACCATCCCGGTCGATCAGCGTC</td><td></td><td></td><td></td><td></td><td></td><td></td><td></td><td></td><td></td><td></td></tr>
<tr><td>Chromosome</td><td>start</td><td>end</td><td>strand</td><td>MM</td><td>target_seq</td><td>PAM</td><td>alignment</td><td>distance</td><td>position</td><td>gene name</td><td>gene id</td></tr>
<tr><td>GU305914.1</td><td>213216</td><td>213238</td><td>-</td><td>0</td><td>GACGCTGATCGACCGGGATG</td><td>AGG</td><td>||||||||[||||||||||||]PAM</td><td>NA</td><td>Intergenic</td><td>NA</td><td>NA</td></tr>
<tr><td></td><td></td><td></td><td></td><td></td><td></td><td></td><td></td><td></td><td></td><td></td><td></td></tr>
<tr><td>T18411</td><td>GATCGACCGGGATGAGGATACGG</td><td>1000</td><td></td><td></td><td></td><td></td><td></td><td></td><td></td><td></td><td></td></tr>
<tr><td>Oligo fwd</td><td>CACCGATCGACCGGGATGAGGATA</td><td></td><td></td><td></td><td></td><td></td><td></td><td></td><td></td><td></td><td></td></tr>
<tr><td>Oligo rev</td><td>AAACTATCCTCATCCCGGTCGATC</td><td></td><td></td><td></td><td></td><td></td><td></td><td></td><td></td><td></td><td></td></tr>
<tr><td>Chromosome</td><td>start</td><td>end</td><td>strand</td><td>MM</td><td>target_seq</td><td>PAM</td><td>alignment</td><td>distance</td><td>position</td><td>gene name</td><td>gene id</td></tr>
<tr><td>GU305914.1</td><td>213210</td><td>213232</td><td>-</td><td>0</td><td>GATCGACCGGGATGAGGATA</td><td>CGG</td><td>||||||||[||||||||||||]PAM</td><td>NA</td><td>Intergenic</td><td>NA</td><td>NA</td></tr>
<tr><td></td><td></td><td></td><td></td><td></td><td></td><td></td><td></td><td></td><td></td><td></td><td></td></tr>
<tr><td>T18412</td><td>GAGGATACGGAAAACCAGATAGG</td><td>1000</td><td></td><td></td><td></td><td></td><td></td><td></td><td></td><td></td><td></td></tr>
<tr><td>Oligo fwd</td><td>CACCGAGGATACGGAAAACCAGAT</td><td></td><td></td><td></td><td></td><td></td><td></td><td></td><td></td><td></td><td></td></tr>
<tr><td>Oligo rev</td><td>AAACATCTGGTTTTCCGTATCCTC</td><td></td><td></td><td></td><td></td><td></td><td></td><td></td><td></td><td></td><td></td></tr>
<tr><td>Chromosome</td><td>start</td><td>end</td><td>strand</td><td>MM</td><td>target_seq</td><td>PAM</td><td>alignment</td><td>distance</td><td>position</td><td>gene name</td><td>gene id</td></tr>
<tr><td>GU305914.1</td><td>213197</td><td>213219</td><td>-</td><td>0</td><td>GAGGATACGGAAAACCAGAT</td><td>AGG</td><td>||||||||[||||||||||||]PAM</td><td>NA</td><td>Intergenic</td><td>NA</td><td>NA</td></tr>
<tr><td></td><td></td><td></td><td></td><td></td><td></td><td></td><td></td><td></td><td></td><td></td><td></td></tr>
<tr><td>T18413</td><td>AAAACCAGATAGGCCCGAATCGG</td><td>1000</td><td></td><td></td><td></td><td></td><td></td><td></td><td></td><td></td><td></td></tr>
<tr><td>Oligo fwd</td><td>CACCAAAACCAGATAGGCCCGAAT</td><td></td><td></td><td></td><td></td><td></td><td></td><td></td><td></td><td></td><td></td></tr>
<tr><td>Oligo rev</td><td>AAACATTCGGGCCTATCTGGTTTT</td><td></td><td></td><td></td><td></td><td></td><td></td><td></td><td></td><td></td><td></td></tr>
<tr><td>Chromosome</td><td>start</td><td>end</td><td>strand</td><td>MM</td><td>target_seq</td><td>PAM</td><td>alignment</td><td>distance</td><td>position</td><td>gene name</td><td>gene id</td></tr>
<tr><td>GU305914.1</td><td>213187</td><td>213209</td><td>-</td><td>0</td><td>AAAACCAGATAGGCCCGAAT</td><td>CGG</td><td>||||||||[||||||||||||]PAM</td><td>NA</td><td>Intergenic</td><td>NA</td><td>NA</td></tr>
<tr><td></td><td></td><td></td><td></td><td></td><td></td><td></td><td></td><td></td><td></td><td></td><td></td></tr>
<tr><td>T18414</td><td>ATCACCGTTGTGATTTTATCTGG</td><td>1000</td><td></td><td></td><td></td><td></td><td></td><td></td><td></td><td></td><td></td></tr>
<tr><td>Oligo fwd</td><td>CACCATCACCGTTGTGATTTTATC</td><td></td><td></td><td></td><td></td><td></td><td></td><td></td><td></td><td></td><td></td></tr>
<tr><td>Oligo rev</td><td>AAACGATAAAATCACAACGGTGAT</td><td></td><td></td><td></td><td></td><td></td><td></td><td></td><td></td><td></td><td></td></tr>
<tr><td>Chromosome</td><td>start</td><td>end</td><td>strand</td><td>MM</td><td>target_seq</td><td>PAM</td><td>alignment</td><td>distance</td><td>position</td><td>gene name</td><td>gene id</td></tr>
<tr><td>GU305914.1</td><td>213140</td><td>213162</td><td>-</td><td>0</td><td>ATCACCGTTGTGATTTTATC</td><td>TGG</td><td>||||||||[||||||||||||]PAM</td><td>NA</td><td>Intergenic</td><td>NA</td><td>NA</td></tr>
<tr><td></td><td></td><td></td><td></td><td></td><td></td><td></td><td></td><td></td><td></td><td></td><td></td></tr>
<tr><td>T18415</td><td>TCACCGTTGTGATTTTATCTGGG</td><td>1000</td><td></td><td></td><td></td><td></td><td></td><td></td><td></td><td></td><td></td></tr>
<tr><td>Oligo fwd</td><td>CACCTCACCGTTGTGATTTTATCT</td><td></td><td></td><td></td><td></td><td></td><td></td><td></td><td></td><td></td><td></td></tr>
<tr><td>Oligo rev</td><td>AAACAGATAAAATCACAACGGTGA</td><td></td><td></td><td></td><td></td><td></td><td></td><td></td><td></td><td></td><td></td></tr>
<tr><td>Chromosome</td><td>start</td><td>end</td><td>strand</td><td>MM</td><td>target_seq</td><td>PAM</td><td>alignment</td><td>distance</td><td>position</td><td>gene name</td><td>gene id</td></tr>
<tr><td>GU305914.1</td><td>213139</td><td>213161</td><td>-</td><td>0</td><td>TCACCGTTGTGATTTTATCT</td><td>GGG</td><td>||||||||[||||||||||||]PAM</td><td>NA</td><td>Intergenic</td><td>NA</td><td>NA</td></tr>
<tr><td></td><td></td><td></td><td></td><td></td><td></td><td></td><td></td><td></td><td></td><td></td><td></td></tr>
<tr><td>T18416</td><td>GATTTTATCTGGGATCGCCGCGG</td><td>1000</td><td></td><td></td><td></td><td></td><td></td><td></td><td></td><td></td><td></td></tr>
<tr><td>Oligo fwd</td><td>CACCGATTTTATCTGGGATCGCCG</td><td></td><td></td><td></td><td></td><td></td><td></td><td></td><td></td><td></td><td></td></tr>
<tr><td>Oligo rev</td><td>AAACCGGCGATCCCAGATAAAATC</td><td></td><td></td><td></td><td></td><td></td><td></td><td></td><td></td><td></td><td></td></tr>
<tr><td>Chromosome</td><td>start</td><td>end</td><td>strand</td><td>MM</td><td>target_seq</td><td>PAM</td><td>alignment</td><td>distance</td><td>position</td><td>gene name</td><td>gene id</td></tr>
<tr><td>GU305914.1</td><td>213129</td><td>213151</td><td>-</td><td>0</td><td>GATTTTATCTGGGATCGCCG</td><td>CGG</td><td>||||||||[||||||||||||]PAM</td><td>NA</td><td>Intergenic</td><td>NA</td><td>NA</td></tr>
<tr><td></td><td></td><td></td><td></td><td></td><td></td><td></td><td></td><td></td><td></td><td></td><td></td></tr>
<tr><td>T18417</td><td>ATTTTATCTGGGATCGCCGCGGG</td><td>1000</td><td></td><td></td><td></td><td></td><td></td><td></td><td></td><td></td><td></td></tr>
<tr><td>Oligo fwd</td><td>CACCATTTTATCTGGGATCGCCGC</td><td></td><td></td><td></td><td></td><td></td><td></td><td></td><td></td><td></td><td></td></tr>
<tr><td>Oligo rev</td><td>AAACGCGGCGATCCCAGATAAAAT</td><td></td><td></td><td></td><td></td><td></td><td></td><td></td><td></td><td></td><td></td></tr>
<tr><td>Chromosome</td><td>start</td><td>end</td><td>strand</td><td>MM</td><td>target_seq</td><td>PAM</td><td>alignment</td><td>distance</td><td>position</td><td>gene name</td><td>gene id</td></tr>
<tr><td>GU305914.1</td><td>213128</td><td>213150</td><td>-</td><td>0</td><td>ATTTTATCTGGGATCGCCGC</td><td>GGG</td><td>||||||||[||||||||||||]PAM</td><td>NA</td><td>Intergenic</td><td>NA</td><td>NA</td></tr>
<tr><td></td><td></td><td></td><td></td><td></td><td></td><td></td><td></td><td></td><td></td><td></td><td></td></tr>
<tr><td>T18418</td><td>TACTCCTGATCACACACCGTTGG</td><td>1000</td><td></td><td></td><td></td><td></td><td></td><td></td><td></td><td></td><td></td></tr>
<tr><td>Oligo fwd</td><td>CACCTACTCCTGATCACACACCGT</td><td></td><td></td><td></td><td></td><td></td><td></td><td></td><td></td><td></td><td></td></tr>
<tr><td>Oligo rev</td><td>AAACACGGTGTGTGATCAGGAGTA</td><td></td><td></td><td></td><td></td><td></td><td></td><td></td><td></td><td></td><td></td></tr>
<tr><td>Chromosome</td><td>start</td><td>end</td><td>strand</td><td>MM</td><td>target_seq</td><td>PAM</td><td>alignment</td><td>distance</td><td>position</td><td>gene name</td><td>gene id</td></tr>
<tr><td>GU305914.1</td><td>213103</td><td>213125</td><td>-</td><td>0</td><td>TACTCCTGATCACACACCGT</td><td>TGG</td><td>||||||||[||||||||||||]PAM</td><td>NA</td><td>Intergenic</td><td>NA</td><td>NA</td></tr>
<tr><td></td><td></td><td></td><td></td><td></td><td></td><td></td><td></td><td></td><td></td><td></td><td></td></tr>
<tr><td>T18419</td><td>ACTCCTGATCACACACCGTTGGG</td><td>1000</td><td></td><td></td><td></td><td></td><td></td><td></td><td></td><td></td><td></td></tr>
<tr><td>Oligo fwd</td><td>CACCACTCCTGATCACACACCGTT</td><td></td><td></td><td></td><td></td><td></td><td></td><td></td><td></td><td></td><td></td></tr>
<tr><td>Oligo rev</td><td>AAACAACGGTGTGTGATCAGGAGT</td><td></td><td></td><td></td><td></td><td></td><td></td><td></td><td></td><td></td><td></td></tr>
<tr><td>Chromosome</td><td>start</td><td>end</td><td>strand</td><td>MM</td><td>target_seq</td><td>PAM</td><td>alignment</td><td>distance</td><td>position</td><td>gene name</td><td>gene id</td></tr>
</table>

GU305914.1 213102 213124 - 0 ACTCCTGATCACACACCGTT GGG ||||||||[||||||||||||]PAM NA Intergenic NA NA

T18420 ACACCGTTGGGAAGACGATAAGG 1000
Oligo fwd CACCACACCGTTGGGAAGACGATA
Oligo rev AAACTATCGTCTTCCCAACGGTGT
Chromosome start end strand MM target_seq PAM alignment distance position gene name gene id
GU305914.1 213090 213112 - 0 ACACCGTTGGGAAGACGATA AGG ||||||||[||||||||||||]PAM NA Intergenic NA NA

T18421 CACCGTTGGGAAGACGATAAGGG 1000
Oligo fwd CACCCACCGTTGGGAAGACGATAA
Oligo rev AAACTTATCGTCTTCCCAACGGTG
Chromosome start end strand MM target_seq PAM alignment distance position gene name gene id
GU305914.1 213089 213111 - 0 CACCGTTGGGAAGACGATAA GGG ||||||||[||||||||||||]PAM NA Intergenic NA NA

T18422 CGTTGGGAAGACGATAAGGGTGG 1000
Oligo fwd CACCCGTTGGGAAGACGATAAGGG
Oligo rev AAACCCCTTATCGTCTTCCCAACG
Chromosome start end strand MM target_seq PAM alignment distance position gene name gene id
GU305914.1 213086 213108 - 0 CGTTGGGAAGACGATAAGGG TGG ||||||||[||||||||||||]PAM NA Intergenic NA NA

T18423 GTTGGGAAGACGATAAGGGTGGG 1000
Oligo fwd CACCGTTGGGAAGACGATAAGGGT
Oligo rev AAACACCCTTATCGTCTTCCCAAC
Chromosome start end strand MM target_seq PAM alignment distance position gene name gene id
GU305914.1 213085 213107 - 0 GTTGGGAAGACGATAAGGGT GGG ||||||||[||||||||||||]PAM NA Intergenic NA NA

T18424 TTGGGAAGACGATAAGGGTGGGG 1000
Oligo fwd CACCTTGGGAAGACGATAAGGGTG
Oligo rev AAACCACCCTTATCGTCTTCCCAA
Chromosome start end strand MM target_seq PAM alignment distance position gene name gene id
GU305914.1 213084 213106 - 0 TTGGGAAGACGATAAGGGTG GGG ||||||||[||||||||||||]PAM NA Intergenic NA NA

T18425 GGAAGACGATAAGGGTGGGGAGG 1000
Oligo fwd CACCGGAAGACGATAAGGGTGGGG
Oligo rev AAACCCCCACCCTTATCGTCTTCC
Chromosome start end strand MM target_seq PAM alignment distance position gene name gene id
GU305914.1 213081 213103 - 0 GGAAGACGATAAGGGTGGGG AGG ||||||||[||||||||||||]PAM NA Intergenic NA NA

T18426 AGACGATAAGGGTGGGGAGGTGG 1000
Oligo fwd CACCAGACGATAAGGGTGGGGAGG
Oligo rev AAACCCTCCCCACCCTTATCGTCT
Chromosome start end strand MM target_seq PAM alignment distance position gene name gene id
GU305914.1 213078 213100 - 0 AGACGATAAGGGTGGGGAGG TGG ||||||||[||||||||||||]PAM NA Intergenic NA NA

T18427 AAGGGTGGGGAGGTGGCACTCGG 1000
Oligo fwd CACCAAGGGTGGGGAGGTGGCACT
Oligo rev AAACAGTGCCACCTCCCCACCCTT
Chromosome start end strand MM target_seq PAM alignment distance position gene name gene id
GU305914.1 213071 213093 - 0 AAGGGTGGGGAGGTGGCACT CGG ||||||||[||||||||||||]PAM NA Intergenic NA NA

T18428 AGGGTGGGGAGGTGGCACTCGGG 1000
Oligo fwd CACCAGGGTGGGGAGGTGGCACTC
Oligo rev AAACGAGTGCCACCTCCCCACCCT
Chromosome start end strand MM target_seq PAM alignment distance position gene name gene id
GU305914.1 213070 213092 - 0 AGGGTGGGGAGGTGGCACTC GGG ||||||||[||||||||||||]PAM NA Intergenic NA NA

T18429 GGGTGGGGAGGTGGCACTCGGGG 1000
Oligo fwd CACCGGGTGGGGAGGTGGCACTCG
Oligo rev AAACCGAGTGCCACCTCCCCACCC
Chromosome start end strand MM target_seq PAM alignment distance position gene name gene id
GU305914.1 213069 213091 - 0 GGGTGGGGAGGTGGCACTCG GGG ||||||||[||||||||||||]PAM NA Intergenic NA NA

T18430 GGGGAGGTGGCACTCGGGGAAGG 1000
Oligo fwd CACCGGGGAGGTGGCACTCGGGGA
Oligo rev AAACTCCCCGAGTGCCACCTCCCC
Chromosome start end strand MM target_seq PAM alignment distance position gene name gene id
GU305914.1 213065 213087 - 0 GGGGAGGTGGCACTCGGGGA AGG ||||||||[||||||||||||]PAM NA Intergenic NA NA

T18431 CAATAAAACTGCTTTATACTTGG 1000
Oligo fwd CACCCAATAAAACTGCTTTATACT
Oligo rev AAACAGTATAAAGCAGTTTTATTG
Chromosome start end strand MM target_seq PAM alignment distance position gene name gene id
GU305914.1 213010 213032 - 0 CAATAAAACTGCTTTATACT TGG ||||||||[||||||||||||]PAM NA Intergenic NA NA

T18432 ACTGCTTTATACTTGGTCTCTGG 1000
Oligo fwd CACCACTGCTTTATACTTGGTCTC
Oligo rev AAACGAGACCAAGTATAAAGCAGT
Chromosome start end strand MM target_seq PAM alignment distance position gene name gene id
GU305914.1 213003 213025 - 0 ACTGCTTTATACTTGGTCTC TGG ||||||||[||||||||||||]PAM NA Intergenic NA NA

T18433 TTTATACTTGGTCTCTGGTGCGG 1000
Oligo fwd CACCTTTATACTTGGTCTCTGGTG
Oligo rev AAACCACCAGAGACCAAGTATAAA
Chromosome start end strand MM target_seq PAM alignment distance position gene name gene id
GU305914.1 212998 213020 - 0 TTTATACTTGGTCTCTGGTG CGG ||||||||[||||||||||||]PAM NA Intergenic NA NA

T18434 TCTCTGGTGCGGTGTTTATACGG 1000
Oligo fwd CACCTCTCTGGTGCGGTGTTTATA
Oligo rev AAACTATAAACACCGCACCAGAGA

Chromosome start end strand MM target_seq PAM alignment distance position gene name gene id
GU305914.1 212987 213009 - 0 TCTCTGGTGCGGTGTTTATA CGG ||||||||[||||||||||||]PAM NA Intergenic NA NA

T18435 CTCTGGTGCGGTGTTTATACGGG 1000
Oligo fwd CACCCTCTGGTGCGGTGTTTATAC
Oligo rev AAACGTATAAACACCGCACCAGAG
Chromosome start end strand MM target_seq PAM alignment distance position gene name gene id
GU305914.1 212986 213008 - 0 CTCTGGTGCGGTGTTTATAC GGG ||||||||[||||||||||||]PAM NA Intergenic NA NA

T18436 ATACGGGCATTCGAACCATAAGG 1000
Oligo fwd CACCATACGGGCATTCGAACCATA
Oligo rev AAACTATGGTTCGAATGCCCGTAT
Chromosome start end strand MM target_seq PAM alignment distance position gene name gene id
GU305914.1 212970 212992 - 0 ATACGGGCATTCGAACCATA AGG ||||||||[||||||||||||]PAM NA Intergenic NA NA

T18437 TACGGGCATTCGAACCATAAGGG 1000
Oligo fwd CACCTACGGGCATTCGAACCATAA
Oligo rev AAACTTATGGTTCGAATGCCCGTA
Chromosome start end strand MM target_seq PAM alignment distance position gene name gene id
GU305914.1 212969 212991 - 0 TACGGGCATTCGAACCATAA GGG ||||||||[||||||||||||]PAM NA Intergenic NA NA

T18438 ACCATAAGGGCGTGATTTATCGG 1000
Oligo fwd CACCACCATAAGGGCGTGATTTAT
Oligo rev AAACATAAATCACGCCCTTATGGT
Chromosome start end strand MM target_seq PAM alignment distance position gene name gene id
GU305914.1 212956 212978 - 0 ACCATAAGGGCGTGATTTAT CGG ||||||||[||||||||||||]PAM NA Intergenic NA NA

T18439 ATCGGCGCGCTTATTTATTTTGG 1000
Oligo fwd CACCATCGGCGCGCTTATTTATTT
Oligo rev AAACAAATAAATAAGCGCGCCGAT
Chromosome start end strand MM target_seq PAM alignment distance position gene name gene id
GU305914.1 212938 212960 - 0 ATCGGCGCGCTTATTTATTT TGG ||||||||[||||||||||||]PAM NA Intergenic NA NA

T18440 TTTGGCCCCACACACAAAGCTGG 1000
Oligo fwd CACCTTTGGCCCCACACACAAAGC
Oligo rev AAACGCTTTGTGTGTGGGGCCAAA
Chromosome start end strand MM target_seq PAM alignment distance position gene name gene id
GU305914.1 212920 212942 - 0 TTTGGCCCCACACACAAAGC TGG ||||||||[||||||||||||]PAM NA Intergenic NA NA

T18441 TTGGCCCCACACACAAAGCTGGG 1000
Oligo fwd CACCTTGGCCCCACACACAAAGCT
Oligo rev AAACAGCTTTGTGTGTGGGGCCAA
Chromosome start end strand MM target_seq PAM alignment distance position gene name gene id
GU305914.1 212919 212941 - 0 TTGGCCCCACACACAAAGCT GGG ||||||||[||||||||||||]PAM NA Intergenic NA NA

T18442 CCACACACAAAGCTGGGTACCGG 1000
Oligo fwd CACCCCACACACAAAGCTGGGTAC
Oligo rev AAACGTACCCAGCTTTGTGTGTGG
Chromosome start end strand MM target_seq PAM alignment distance position gene name gene id
GU305914.1 212913 212935 - 0 CCACACACAAAGCTGGGTAC CGG ||||||||[||||||||||||]PAM NA Intergenic NA NA

T18443 CACACACAAAGCTGGGTACCGGG 1000
Oligo fwd CACCCACACACAAAGCTGGGTACC
Oligo rev AAACGGTACCCAGCTTTGTGTGTG
Chromosome start end strand MM target_seq PAM alignment distance position gene name gene id
GU305914.1 212912 212934 - 0 CACACACAAAGCTGGGTACC GGG ||||||||[||||||||||||]PAM NA Intergenic NA NA

T18444 TCCCCTTCTCCTTCTGCCGTTGG 1000
Oligo fwd CACCTCCCCTTCTCCTTCTGCCGT
Oligo rev AAACACGGCAGAAGGAGAAGGGGA
Chromosome start end strand MM target_seq PAM alignment distance position gene name gene id
GU305914.1 212878 212900 - 0 TCCCCTTCTCCTTCTGCCGT TGG ||||||||[||||||||||||]PAM NA Intergenic NA NA
GU305914.1 122037 122059 + 4 TCCCTCTCTCCTTCTGCGGA CGG ||||--||[|||||||||-|-]PAM NA Intergenic NA NA

T18445 CTGCCGTTGGTTCTCATCTCAGG 1000
Oligo fwd CACCCTGCCGTTGGTTCTCATCTC
Oligo rev AAACGAGATGAGAACCAACGGCAG
Chromosome start end strand MM target_seq PAM alignment distance position gene name gene id
GU305914.1 212865 212887 - 0 CTGCCGTTGGTTCTCATCTC AGG ||||||||[||||||||||||]PAM NA Intergenic NA NA

T18446 CCGTTGGTTCTCATCTCAGGAGG 1000
Oligo fwd CACCCCGTTGGTTCTCATCTCAGG
Oligo rev AAACCCTGAGATGAGAACCAACGG
Chromosome start end strand MM target_seq PAM alignment distance position gene name gene id
GU305914.1 212862 212884 - 0 CCGTTGGTTCTCATCTCAGG AGG ||||||||[||||||||||||]PAM NA Intergenic NA NA

T18447 CGTTGGTTCTCATCTCAGGAGGG 1000
Oligo fwd CACCCGTTGGTTCTCATCTCAGGA
Oligo rev AAACTCCTGAGATGAGAACCAACG
Chromosome start end strand MM target_seq PAM alignment distance position gene name gene id
GU305914.1 212861 212883 - 0 CGTTGGTTCTCATCTCAGGA GGG ||||||||[||||||||||||]PAM NA Intergenic NA NA

T18448 TTCTCATCTCAGGAGGGTTCTGG 1000
Oligo fwd CACCTTCTCATCTCAGGAGGGTTC
Oligo rev AAACGAACCCTCCTGAGATGAGAA
Chromosome start end strand MM target_seq PAM alignment distance position gene name gene id
GU305914.1 212855 212877 - 0 TTCTCATCTCAGGAGGGTTC TGG ||||||||[||||||||||||]PAM NA Intergenic NA NA

T18449 TCTCATCTCAGGAGGGTTCTGGG 1000

Oligo fwd CACCTCTCATCTCAGGAGGGTTCT
Oligo rev AAACAGAACCCTCCTGAGATGAGA
Chromosome start end strand MM target_seq PAM alignment distance position gene name gene id
GU305914.1 212854 212876 - 0 TCTCATCTCAGGAGGGTTCT GGG ||||||||[||||||||||||]PAM NA Intergenic NA NA

T18450 TCAGGAGGGTTCTGGGCGTGCGG 1000
Oligo fwd CACCTCAGGAGGGTTCTGGGCGTG
Oligo rev AAACCACGCCCAGAACCCTCCTGA
Chromosome start end strand MM target_seq PAM alignment distance position gene name gene id
GU305914.1 212847 212869 - 0 TCAGGAGGGTTCTGGGCGTG CGG ||||||||[||||||||||||]PAM NA Intergenic NA NA

T18451 GTTCTGGGCGTGCGGCTCAGCGG 1000
Oligo fwd CACCGTTCTGGGCGTGCGGCTCAG
Oligo rev AAACCTGAGCCGCACGCCCAGAAC
Chromosome start end strand MM target_seq PAM alignment distance position gene name gene id
GU305914.1 212839 212861 - 0 GTTCTGGGCGTGCGGCTCAG CGG ||||||||[||||||||||||]PAM NA Intergenic NA NA

T18452 CTGGGCGTGCGGCTCAGCGGAGG 1000
Oligo fwd CACCCTGGGCGTGCGGCTCAGCGG
Oligo rev AAACCCGCTGAGCCGCACGCCCAG
Chromosome start end strand MM target_seq PAM alignment distance position gene name gene id
GU305914.1 212836 212858 - 0 CTGGGCGTGCGGCTCAGCGG AGG ||||||||[||||||||||||]PAM NA Intergenic NA NA

T18453 GCGGCTCAGCGGAGGTCGTGCGG 1000
Oligo fwd CACCGCGGCTCAGCGGAGGTCGTG
Oligo rev AAACCACGACCTCCGCTGAGCCGC
Chromosome start end strand MM target_seq PAM alignment distance position gene name gene id
GU305914.1 212828 212850 - 0 GCGGCTCAGCGGAGGTCGTG CGG ||||||||[||||||||||||]PAM NA Intergenic NA NA

T18454 TCGTGCGGCCCGAAGTTAACAGG 1000
Oligo fwd CACCTCGTGCGGCCCGAAGTTAAC
Oligo rev AAACGTTAACTTCGGGCCGCACGA
Chromosome start end strand MM target_seq PAM alignment distance position gene name gene id
GU305914.1 212813 212835 - 0 TCGTGCGGCCCGAAGTTAAC AGG ||||||||[||||||||||||]PAM NA Intergenic NA NA

T18455 GCGGCCCGAAGTTAACAGGACGG 1000
Oligo fwd CACCGCGGCCCGAAGTTAACAGGA
Oligo rev AAACTCCTGTTAACTTCGGGCCGC
Chromosome start end strand MM target_seq PAM alignment distance position gene name gene id
GU305914.1 212809 212831 - 0 GCGGCCCGAAGTTAACAGGA CGG ||||||||[||||||||||||]PAM NA Intergenic NA NA

T18456 CGGCCCGAAGTTAACAGGACGGG 1000
Oligo fwd CACCCGGCCCGAAGTTAACAGGAC
Oligo rev AAACGTCCTGTTAACTTCGGGCCG
Chromosome start end strand MM target_seq PAM alignment distance position gene name gene id
GU305914.1 212808 212830 - 0 CGGCCCGAAGTTAACAGGAC GGG ||||||||[||||||||||||]PAM NA Intergenic NA NA

T18457 CGAAGTTAACAGGACGGGAACGG 1000
Oligo fwd CACCCGAAGTTAACAGGACGGGAA
Oligo rev AAACTTCCCGTCCTGTTAACTTCG
Chromosome start end strand MM target_seq PAM alignment distance position gene name gene id
GU305914.1 212803 212825 - 0 CGAAGTTAACAGGACGGGAA CGG ||||||||[||||||||||||]PAM NA Intergenic NA NA

T18458 GGGAACGGTCGATATCTGCCAGG 1000
Oligo fwd CACCGGGAACGGTCGATATCTGCC
Oligo rev AAACGGCAGATATCGACCGTTCCC
Chromosome start end strand MM target_seq PAM alignment distance position gene name gene id
GU305914.1 212788 212810 - 0 GGGAACGGTCGATATCTGCC AGG ||||||||[||||||||||||]PAM NA Intergenic NA NA

T18459 GGAACGGTCGATATCTGCCAGGG 1000
Oligo fwd CACCGGAACGGTCGATATCTGCCA
Oligo rev AAACTGGCAGATATCGACCGTTCC
Chromosome start end strand MM target_seq PAM alignment distance position gene name gene id
GU305914.1 212787 212809 - 0 GGAACGGTCGATATCTGCCA GGG ||||||||[||||||||||||]PAM NA Intergenic NA NA

T18460 CGATATCTGCCAGGGACCTATGG 1000
Oligo fwd CACCCGATATCTGCCAGGGACCTA
Oligo rev AAACTAGGTCCCTGGCAGATATCG
Chromosome start end strand MM target_seq PAM alignment distance position gene name gene id
GU305914.1 212779 212801 - 0 CGATATCTGCCAGGGACCTA TGG ||||||||[||||||||||||]PAM NA Intergenic NA NA

T18461 TCCGTCTCCAGAACATCGAGCGG 1000
Oligo fwd CACCTCCGTCTCCAGAACATCGAG
Oligo rev AAACCTCGATGTTCTGGAGACGGA
Chromosome start end strand MM target_seq PAM alignment distance position gene name gene id
GU305914.1 212745 212767 - 0 TCCGTCTCCAGAACATCGAG CGG ||||||||[||||||||||||]PAM NA Intergenic NA NA

T18462 CCGTCTCCAGAACATCGAGCGGG 1000
Oligo fwd CACCCCGTCTCCAGAACATCGAGC
Oligo rev AAACGCTCGATGTTCTGGAGACGG
Chromosome start end strand MM target_seq PAM alignment distance position gene name gene id
GU305914.1 212744 212766 - 0 CCGTCTCCAGAACATCGAGC GGG ||||||||[||||||||||||]PAM NA Intergenic NA NA

T18463 CGTCTCCAGAACATCGAGCGGGG 1000
Oligo fwd CACCCGTCTCCAGAACATCGAGCG
Oligo rev AAACCGCTCGATGTTCTGGAGACG
Chromosome start end strand MM target_seq PAM alignment distance position gene name gene id
GU305914.1 212743 212765 - 0 CGTCTCCAGAACATCGAGCG GGG ||||||||[||||||||||||]PAM NA Intergenic NA NA

<table>
<tr><td>T18464</td><td>CAGAACATCGAGCGGGGCGACGG</td><td>1000</td><td></td><td></td><td></td><td></td><td></td><td></td><td></td><td></td><td></td></tr>
<tr><td>Oligo fwd</td><td>CACCCAGAACATCGAGCGGGGCGA</td><td></td><td></td><td></td><td></td><td></td><td></td><td></td><td></td><td></td><td></td></tr>
<tr><td>Oligo rev</td><td>AAACTCGCCCCGCTCGATGTTCTG</td><td></td><td></td><td></td><td></td><td></td><td></td><td></td><td></td><td></td><td></td></tr>
<tr><td>Chromosome</td><td>start</td><td>end</td><td>strand</td><td>MM</td><td>target_seq</td><td>PAM</td><td>alignment</td><td>distance</td><td>position</td><td>gene name</td><td>gene id</td></tr>
<tr><td>GU305914.1</td><td>212737</td><td>212759</td><td>-</td><td>0</td><td>CAGAACATCGAGCGGGGCGA</td><td>CGG</td><td>||||||||[||||||||||||]PAM</td><td>NA</td><td>Intergenic</td><td>NA</td><td>NA</td></tr>
<tr><td>T18465</td><td>AGAACATCGAGCGGGGCGACGGG</td><td>1000</td><td></td><td></td><td></td><td></td><td></td><td></td><td></td><td></td><td></td></tr>
<tr><td>Oligo fwd</td><td>CACCAGAACATCGAGCGGGGCGAC</td><td></td><td></td><td></td><td></td><td></td><td></td><td></td><td></td><td></td><td></td></tr>
<tr><td>Oligo rev</td><td>AAACGTCGCCCCGCTCGATGTTCT</td><td></td><td></td><td></td><td></td><td></td><td></td><td></td><td></td><td></td><td></td></tr>
<tr><td>Chromosome</td><td>start</td><td>end</td><td>strand</td><td>MM</td><td>target_seq</td><td>PAM</td><td>alignment</td><td>distance</td><td>position</td><td>gene name</td><td>gene id</td></tr>
<tr><td>GU305914.1</td><td>212736</td><td>212758</td><td>-</td><td>0</td><td>AGAACATCGAGCGGGGCGAC</td><td>GGG</td><td>||||||||[||||||||||||]PAM</td><td>NA</td><td>Intergenic</td><td>NA</td><td>NA</td></tr>
<tr><td>T18466</td><td>CGAGCGGGGCGACGGGAGAGAGG</td><td>1000</td><td></td><td></td><td></td><td></td><td></td><td></td><td></td><td></td><td></td></tr>
<tr><td>Oligo fwd</td><td>CACCCGAGCGGGGCGACGGGAGAG</td><td></td><td></td><td></td><td></td><td></td><td></td><td></td><td></td><td></td><td></td></tr>
<tr><td>Oligo rev</td><td>AAACCTCTCCCGTCGCCCCGCTCG</td><td></td><td></td><td></td><td></td><td></td><td></td><td></td><td></td><td></td><td></td></tr>
<tr><td>Chromosome</td><td>start</td><td>end</td><td>strand</td><td>MM</td><td>target_seq</td><td>PAM</td><td>alignment</td><td>distance</td><td>position</td><td>gene name</td><td>gene id</td></tr>
<tr><td>GU305914.1</td><td>212729</td><td>212751</td><td>-</td><td>0</td><td>CGAGCGGGGCGACGGGAGAG</td><td>AGG</td><td>||||||||[||||||||||||]PAM</td><td>NA</td><td>Intergenic</td><td>NA</td><td>NA</td></tr>
<tr><td>T18467</td><td>CACCCTCTCGATTGTCAGCATGG</td><td>1000</td><td></td><td></td><td></td><td></td><td></td><td></td><td></td><td></td><td></td></tr>
<tr><td>Oligo fwd</td><td>CACCCACCCTCTCGATTGTCAGCA</td><td></td><td></td><td></td><td></td><td></td><td></td><td></td><td></td><td></td><td></td></tr>
<tr><td>Oligo rev</td><td>AAACTGCTGACAATCGAGAGGGTG</td><td></td><td></td><td></td><td></td><td></td><td></td><td></td><td></td><td></td><td></td></tr>
<tr><td>Chromosome</td><td>start</td><td>end</td><td>strand</td><td>MM</td><td>target_seq</td><td>PAM</td><td>alignment</td><td>distance</td><td>position</td><td>gene name</td><td>gene id</td></tr>
<tr><td>GU305914.1</td><td>212692</td><td>212714</td><td>-</td><td>0</td><td>CACCCTCTCGATTGTCAGCA</td><td>TGG</td><td>||||||||[||||||||||||]PAM</td><td>NA</td><td>Intergenic</td><td>NA</td><td>NA</td></tr>
<tr><td>T18468</td><td>TCGATTGTCAGCATGGAGAACGG</td><td>1000</td><td></td><td></td><td></td><td></td><td></td><td></td><td></td><td></td><td></td></tr>
<tr><td>Oligo fwd</td><td>CACCTCGATTGTCAGCATGGAGAA</td><td></td><td></td><td></td><td></td><td></td><td></td><td></td><td></td><td></td><td></td></tr>
<tr><td>Oligo rev</td><td>AAACTTCTCCATGCTGACAATCGA</td><td></td><td></td><td></td><td></td><td></td><td></td><td></td><td></td><td></td><td></td></tr>
<tr><td>Chromosome</td><td>start</td><td>end</td><td>strand</td><td>MM</td><td>target_seq</td><td>PAM</td><td>alignment</td><td>distance</td><td>position</td><td>gene name</td><td>gene id</td></tr>
<tr><td>GU305914.1</td><td>212685</td><td>212707</td><td>-</td><td>0</td><td>TCGATTGTCAGCATGGAGAA</td><td>CGG</td><td>||||||||[||||||||||||]PAM</td><td>NA</td><td>Intergenic</td><td>NA</td><td>NA</td></tr>
<tr><td>T18469</td><td>CGATTGTCAGCATGGAGAACGGG</td><td>1000</td><td></td><td></td><td></td><td></td><td></td><td></td><td></td><td></td><td></td></tr>
<tr><td>Oligo fwd</td><td>CACCCGATTGTCAGCATGGAGAAC</td><td></td><td></td><td></td><td></td><td></td><td></td><td></td><td></td><td></td><td></td></tr>
<tr><td>Oligo rev</td><td>AAACGTTCTCCATGCTGACAATCG</td><td></td><td></td><td></td><td></td><td></td><td></td><td></td><td></td><td></td><td></td></tr>
<tr><td>Chromosome</td><td>start</td><td>end</td><td>strand</td><td>MM</td><td>target_seq</td><td>PAM</td><td>alignment</td><td>distance</td><td>position</td><td>gene name</td><td>gene id</td></tr>
<tr><td>GU305914.1</td><td>212684</td><td>212706</td><td>-</td><td>0</td><td>CGATTGTCAGCATGGAGAAC</td><td>GGG</td><td>||||||||[||||||||||||]PAM</td><td>NA</td><td>Intergenic</td><td>NA</td><td>NA</td></tr>
<tr><td>T18470</td><td>GATTGTCAGCATGGAGAACGGGG</td><td>1000</td><td></td><td></td><td></td><td></td><td></td><td></td><td></td><td></td><td></td></tr>
<tr><td>Oligo fwd</td><td>CACCGATTGTCAGCATGGAGAACG</td><td></td><td></td><td></td><td></td><td></td><td></td><td></td><td></td><td></td><td></td></tr>
<tr><td>Oligo rev</td><td>AAACCGTTCTCCATGCTGACAATC</td><td></td><td></td><td></td><td></td><td></td><td></td><td></td><td></td><td></td><td></td></tr>
<tr><td>Chromosome</td><td>start</td><td>end</td><td>strand</td><td>MM</td><td>target_seq</td><td>PAM</td><td>alignment</td><td>distance</td><td>position</td><td>gene name</td><td>gene id</td></tr>
<tr><td>GU305914.1</td><td>212683</td><td>212705</td><td>-</td><td>0</td><td>GATTGTCAGCATGGAGAACG</td><td>GGG</td><td>||||||||[||||||||||||]PAM</td><td>NA</td><td>Intergenic</td><td>NA</td><td>NA</td></tr>
<tr><td>T18471</td><td>ATTGTCAGCATGGAGAACGGGGG</td><td>1000</td><td></td><td></td><td></td><td></td><td></td><td></td><td></td><td></td><td></td></tr>
<tr><td>Oligo fwd</td><td>CACCATTGTCAGCATGGAGAACGG</td><td></td><td></td><td></td><td></td><td></td><td></td><td></td><td></td><td></td><td></td></tr>
<tr><td>Oligo rev</td><td>AAACCCGTTCTCCATGCTGACAAT</td><td></td><td></td><td></td><td></td><td></td><td></td><td></td><td></td><td></td><td></td></tr>
<tr><td>Chromosome</td><td>start</td><td>end</td><td>strand</td><td>MM</td><td>target_seq</td><td>PAM</td><td>alignment</td><td>distance</td><td>position</td><td>gene name</td><td>gene id</td></tr>
<tr><td>GU305914.1</td><td>212682</td><td>212704</td><td>-</td><td>0</td><td>ATTGTCAGCATGGAGAACGG</td><td>GGG</td><td>||||||||[||||||||||||]PAM</td><td>NA</td><td>Intergenic</td><td>NA</td><td>NA</td></tr>
<tr><td>T18472</td><td>TTGTCAGCATGGAGAACGGGGGG</td><td>1000</td><td></td><td></td><td></td><td></td><td></td><td></td><td></td><td></td><td></td></tr>
<tr><td>Oligo fwd</td><td>CACCTTGTCAGCATGGAGAACGGG</td><td></td><td></td><td></td><td></td><td></td><td></td><td></td><td></td><td></td><td></td></tr>
<tr><td>Oligo rev</td><td>AAACCCCGTTCTCCATGCTGACAA</td><td></td><td></td><td></td><td></td><td></td><td></td><td></td><td></td><td></td><td></td></tr>
<tr><td>Chromosome</td><td>start</td><td>end</td><td>strand</td><td>MM</td><td>target_seq</td><td>PAM</td><td>alignment</td><td>distance</td><td>position</td><td>gene name</td><td>gene id</td></tr>
<tr><td>GU305914.1</td><td>212681</td><td>212703</td><td>-</td><td>0</td><td>TTGTCAGCATGGAGAACGGG</td><td>GGG</td><td>||||||||[||||||||||||]PAM</td><td>NA</td><td>Intergenic</td><td>NA</td><td>NA</td></tr>
<tr><td>T18473</td><td>ATGGAGAACGGGGGGAAGCCTGG</td><td>1000</td><td></td><td></td><td></td><td></td><td></td><td></td><td></td><td></td><td></td></tr>
<tr><td>Oligo fwd</td><td>CACCATGGAGAACGGGGGGAAGCC</td><td></td><td></td><td></td><td></td><td></td><td></td><td></td><td></td><td></td><td></td></tr>
<tr><td>Oligo rev</td><td>AAACGGCTTCCCCCCGTTCTCCAT</td><td></td><td></td><td></td><td></td><td></td><td></td><td></td><td></td><td></td><td></td></tr>
<tr><td>Chromosome</td><td>start</td><td>end</td><td>strand</td><td>MM</td><td>target_seq</td><td>PAM</td><td>alignment</td><td>distance</td><td>position</td><td>gene name</td><td>gene id</td></tr>
<tr><td>GU305914.1</td><td>212673</td><td>212695</td><td>-</td><td>0</td><td>ATGGAGAACGGGGGGAAGCC</td><td>TGG</td><td>||||||||[||||||||||||]PAM</td><td>NA</td><td>Intergenic</td><td>NA</td><td>NA</td></tr>
<tr><td>T18474</td><td>GGAAGCCTGGCACGTATGAGTGG</td><td>1000</td><td></td><td></td><td></td><td></td><td></td><td></td><td></td><td></td><td></td></tr>
<tr><td>Oligo fwd</td><td>CACCGGAAGCCTGGCACGTATGAG</td><td></td><td></td><td></td><td></td><td></td><td></td><td></td><td></td><td></td><td></td></tr>
<tr><td>Oligo rev</td><td>AAACCTCATACGTGCCAGGCTTCC</td><td></td><td></td><td></td><td></td><td></td><td></td><td></td><td></td><td></td><td></td></tr>
<tr><td>Chromosome</td><td>start</td><td>end</td><td>strand</td><td>MM</td><td>target_seq</td><td>PAM</td><td>alignment</td><td>distance</td><td>position</td><td>gene name</td><td>gene id</td></tr>
<tr><td>GU305914.1</td><td>212660</td><td>212682</td><td>-</td><td>0</td><td>GGAAGCCTGGCACGTATGAG</td><td>TGG</td><td>||||||||[||||||||||||]PAM</td><td>NA</td><td>Intergenic</td><td>NA</td><td>NA</td></tr>
<tr><td>T18475</td><td>CGTTTCCCGCGAACGAGTCGTGG</td><td>1000</td><td></td><td></td><td></td><td></td><td></td><td></td><td></td><td></td><td></td></tr>
<tr><td>Oligo fwd</td><td>CACCCGTTTCCCGCGAACGAGTCG</td><td></td><td></td><td></td><td></td><td></td><td></td><td></td><td></td><td></td><td></td></tr>
<tr><td>Oligo rev</td><td>AAACCGACTCGTTCGCGGGAAACG</td><td></td><td></td><td></td><td></td><td></td><td></td><td></td><td></td><td></td><td></td></tr>
<tr><td>Chromosome</td><td>start</td><td>end</td><td>strand</td><td>MM</td><td>target_seq</td><td>PAM</td><td>alignment</td><td>distance</td><td>position</td><td>gene name</td><td>gene id</td></tr>
<tr><td>GU305914.1</td><td>212636</td><td>212658</td><td>-</td><td>0</td><td>CGTTTCCCGCGAACGAGTCG</td><td>TGG</td><td>||||||||[||||||||||||]PAM</td><td>NA</td><td>Intergenic</td><td>NA</td><td>NA</td></tr>
<tr><td>T18476</td><td>GTTCGTCTCGAAGTACTACGCGG</td><td>1000</td><td></td><td></td><td></td><td></td><td></td><td></td><td></td><td></td><td></td></tr>
<tr><td>Oligo fwd</td><td>CACCGTTCGTCTCGAAGTACTACG</td><td></td><td></td><td></td><td></td><td></td><td></td><td></td><td></td><td></td><td></td></tr>
<tr><td>Oligo rev</td><td>AAACCGTAGTACTTCGAGACGAAC</td><td></td><td></td><td></td><td></td><td></td><td></td><td></td><td></td><td></td><td></td></tr>
<tr><td>Chromosome</td><td>start</td><td>end</td><td>strand</td><td>MM</td><td>target_seq</td><td>PAM</td><td>alignment</td><td>distance</td><td>position</td><td>gene name</td><td>gene id</td></tr>
<tr><td>GU305914.1</td><td>212581</td><td>212603</td><td>-</td><td>0</td><td>GTTCGTCTCGAAGTACTACG</td><td>CGG</td><td>||||||||[||||||||||||]PAM</td><td>NA</td><td>Intergenic</td><td>NA</td><td>NA</td></tr>
<tr><td>T18477</td><td>CTCGAAGTACTACGCGGATGTGG</td><td>1000</td><td></td><td></td><td></td><td></td><td></td><td></td><td></td><td></td><td></td></tr>
<tr><td>Oligo fwd</td><td>CACCCTCGAAGTACTACGCGGATG</td><td></td><td></td><td></td><td></td><td></td><td></td><td></td><td></td><td></td><td></td></tr>
<tr><td>Oligo rev</td><td>AAACCATCCGCGTAGTACTTCGAG</td><td></td><td></td><td></td><td></td><td></td><td></td><td></td><td></td><td></td><td></td></tr>
<tr><td>Chromosome</td><td>start</td><td>end</td><td>strand</td><td>MM</td><td>target_seq</td><td>PAM</td><td>alignment</td><td>distance</td><td>position</td><td>gene name</td><td>gene id</td></tr>
<tr><td>GU305914.1</td><td>212575</td><td>212597</td><td>-</td><td>0</td><td>CTCGAAGTACTACGCGGATG</td><td>TGG</td><td>||||||||[||||||||||||]PAM</td><td>NA</td><td>Intergenic</td><td>NA</td><td>NA</td></tr>
<tr><td>T18478</td><td>GCGGATGTGGTCCAGACCCCCGG</td><td>1000</td><td></td><td></td><td></td><td></td><td></td><td></td><td></td><td></td><td></td></tr>
<tr><td>Oligo fwd</td><td>CACCGCGGATGTGGTCCAGACCCC</td><td></td><td></td><td></td><td></td><td></td><td></td><td></td><td></td><td></td><td></td></tr>
<tr><td>Oligo rev</td><td>AAACGGGGTCTGGACCACATCCGC</td><td></td><td></td><td></td><td></td><td></td><td></td><td></td><td></td><td></td><td></td></tr>
<tr><td>Chromosome</td><td>start</td><td>end</td><td>strand</td><td>MM</td><td>target_seq</td><td>PAM</td><td>alignment</td><td>distance</td><td>position</td><td>gene name</td><td>gene id</td></tr>
<tr><td>GU305914.1</td><td>212562</td><td>212584</td><td>-</td><td>0</td><td>GCGGATGTGGTCCAGACCCC</td><td>CGG</td><td>||||||||[||||||||||||]PAM</td><td>NA</td><td>Intergenic</td><td>NA</td><td>NA</td></tr>
</table>

T18479 CGGATGTGGTCCAGACCCCCGGG 1000
Oligo fwd CACCCGGATGTGGTCCAGACCCCC
Oligo rev AAACGGGGGTCTGGACCACATCCG
Chromosome start end strand MM target_seq PAM alignment distance position gene name gene id
GU305914.1 212561 212583 - 0 CGGATGTGGTCCAGACCCCC GGG ||||||||[||||||||||||]PAM NA Intergenic NA NA

T18480 GGATGTGGTCCAGACCCCCGGGG 1000
Oligo fwd CACCGGATGTGGTCCAGACCCCCG
Oligo rev AAACCGGGGGTCTGGACCACATCC
Chromosome start end strand MM target_seq PAM alignment distance position gene name gene id
GU305914.1 212560 212582 - 0 GGATGTGGTCCAGACCCCCG GGG ||||||||[||||||||||||]PAM NA Intergenic NA NA

T18481 GGTCCAGACCCCCGGGGAGCTGG 1000
Oligo fwd CACCGGTCCAGACCCCCGGGGAGC
Oligo rev AAACGCTCCCCGGGGGTCTGGACC
Chromosome start end strand MM target_seq PAM alignment distance position gene name gene id
GU305914.1 212554 212576 - 0 GGTCCAGACCCCCGGGGAGC TGG ||||||||[||||||||||||]PAM NA Intergenic NA NA

T18482 GGTCGTAGAGTATAGATGTCCGG 1000
Oligo fwd CACCGGTCGTAGAGTATAGATGTC
Oligo rev AAACGACATCTATACTCTACGACC
Chromosome start end strand MM target_seq PAM alignment distance position gene name gene id
GU305914.1 212533 212555 - 0 GGTCGTAGAGTATAGATGTC CGG ||||||||[||||||||||||]PAM NA Intergenic NA NA

T18483 TCAACTGTACGATTACACATCGG 1000
Oligo fwd CACCTCAACTGTACGATTACACAT
Oligo rev AAACATGTGTAATCGTACAGTTGA
Chromosome start end strand MM target_seq PAM alignment distance position gene name gene id
GU305914.1 212501 212523 - 0 TCAACTGTACGATTACACAT CGG ||||||||[||||||||||||]PAM NA Intergenic NA NA

T18484 ACTGTACGATTACACATCGGTGG 1000
Oligo fwd CACCACTGTACGATTACACATCGG
Oligo rev AAACCCGATGTGTAATCGTACAGT
Chromosome start end strand MM target_seq PAM alignment distance position gene name gene id
GU305914.1 212498 212520 - 0 ACTGTACGATTACACATCGG TGG ||||||||[||||||||||||]PAM NA Intergenic NA NA

T18485 ACGATTACACATCGGTGGAAAGG 1000
Oligo fwd CACCACGATTACACATCGGTGGAA
Oligo rev AAACTTCCACCGATGTGTAATCGT
Chromosome start end strand MM target_seq PAM alignment distance position gene name gene id
GU305914.1 212493 212515 - 0 ACGATTACACATCGGTGGAA AGG ||||||||[||||||||||||]PAM NA Intergenic NA NA

T18486 GAGACGATCATGTCGTTCGACGG 1000
Oligo fwd CACCGAGACGATCATGTCGTTCGA
Oligo rev AAACTCGAACGACATGATCGTCTC
Chromosome start end strand MM target_seq PAM alignment distance position gene name gene id
GU305914.1 212469 212491 - 0 GAGACGATCATGTCGTTCGA CGG ||||||||[||||||||||||]PAM NA Intergenic NA NA

T18487 TGCTATACAGACGATACGATCGG 1000
Oligo fwd CACCTGCTATACAGACGATACGAT
Oligo rev AAACATCGTATCGTCTGTATAGCA
Chromosome start end strand MM target_seq PAM alignment distance position gene name gene id
GU305914.1 212446 212468 - 0 TGCTATACAGACGATACGAT CGG ||||||||[||||||||||||]PAM NA Intergenic NA NA

T18488 GACGATACGATCGGTGACATCGG 1000
Oligo fwd CACCGACGATACGATCGGTGACAT
Oligo rev AAACATGTCACCGATCGTATCGTC
Chromosome start end strand MM target_seq PAM alignment distance position gene name gene id
GU305914.1 212437 212459 - 0 GACGATACGATCGGTGACAT CGG ||||||||[||||||||||||]PAM NA Intergenic NA NA

T18489 GGAATACACGACGAAGAACGAGG 1000
Oligo fwd CACCGGAATACACGACGAAGAACG
Oligo rev AAACCGTTCTTCGTCGTGTATTCC
Chromosome start end strand MM target_seq PAM alignment distance position gene name gene id
GU305914.1 212416 212438 - 0 GGAATACACGACGAAGAACG AGG ||||||||[||||||||||||]PAM NA Intergenic NA NA
GU305914.1 68437 68459 + 4 GGAGAACACGACGACGACCG AGG |||--|||[||||||-||-||]PAM NA Intergenic NA NA

T18490 GACGAAGAACGAGGATACCTTGG 1000
Oligo fwd CACCGACGAAGAACGAGGATACCT
Oligo rev AAACAGGTATCCTCGTTCTTCGTC
Chromosome start end strand MM target_seq PAM alignment distance position gene name gene id
GU305914.1 212407 212429 - 0 GACGAAGAACGAGGATACCT TGG ||||||||[||||||||||||]PAM NA Intergenic NA NA

T18491 ACCTTGGTGAAGTATATCAGAGG 1000
Oligo fwd CACCACCTTGGTGAAGTATATCAG
Oligo rev AAACCTGATATACTTCACCAAGGT
Chromosome start end strand MM target_seq PAM alignment distance position gene name gene id
GU305914.1 212391 212413 - 0 ACCTTGGTGAAGTATATCAG AGG ||||||||[||||||||||||]PAM NA Intergenic NA NA

T18492 TCTGAACGTGACACTGCTGACGG 1000
Oligo fwd CACCTCTGAACGTGACACTGCTGA
Oligo rev AAACTCAGCAGTGTCACGTTCAGA
Chromosome start end strand MM target_seq PAM alignment distance position gene name gene id
GU305914.1 212368 212390 - 0 TCTGAACGTGACACTGCTGA CGG ||||||||[||||||||||||]PAM NA Intergenic NA NA

T18493 ACGCGAAGTCGATCGAACACCGG 1000
Oligo fwd CACCACGCGAAGTCGATCGAACAC
Oligo rev AAACGTGTTCGATCGACTTCGCGT

<table>
<tr><td>Chromosome</td><td>start</td><td>end</td><td>strand</td><td>MM</td><td>target_seq</td><td>PAM</td><td>alignment</td><td>distance</td><td>position</td><td>gene name</td><td>gene id</td></tr>
<tr><td>GU305914.1</td><td>212342</td><td>212364</td><td>-</td><td>0</td><td>ACGCGAAGTCGATCGAACAC</td><td>CGG</td><td>||||||||[||||||||||||]PAM</td><td>NA</td><td>Intergenic</td><td>NA</td><td>NA</td></tr>
<tr><td>T18494</td><td>CGAAGTCGATCGAACACCGGTGG</td><td>1000</td><td></td><td></td><td></td><td></td><td></td><td></td><td></td><td></td><td></td></tr>
<tr><td>Oligo fwd</td><td>CACCCGAAGTCGATCGAACACCGG</td><td></td><td></td><td></td><td></td><td></td><td></td><td></td><td></td><td></td><td></td></tr>
<tr><td>Oligo rev</td><td>AAACCCGGTGTTCGATCGACTTCG</td><td></td><td></td><td></td><td></td><td></td><td></td><td></td><td></td><td></td><td></td></tr>
<tr><td>Chromosome</td><td>start</td><td>end</td><td>strand</td><td>MM</td><td>target_seq</td><td>PAM</td><td>alignment</td><td>distance</td><td>position</td><td>gene name</td><td>gene id</td></tr>
<tr><td>GU305914.1</td><td>212339</td><td>212361</td><td>-</td><td>0</td><td>CGAAGTCGATCGAACACCGG</td><td>TGG</td><td>||||||||[||||||||||||]PAM</td><td>NA</td><td>Intergenic</td><td>NA</td><td>NA</td></tr>
<tr><td>T18495</td><td>TGAGATATGTAAGAAGCTCAAGG</td><td>1000</td><td></td><td></td><td></td><td></td><td></td><td></td><td></td><td></td><td></td></tr>
<tr><td>Oligo fwd</td><td>CACCTGAGATATGTAAGAAGCTCA</td><td></td><td></td><td></td><td></td><td></td><td></td><td></td><td></td><td></td><td></td></tr>
<tr><td>Oligo rev</td><td>AAACTGAGCTTCTTACATATCTCA</td><td></td><td></td><td></td><td></td><td></td><td></td><td></td><td></td><td></td><td></td></tr>
<tr><td>Chromosome</td><td>start</td><td>end</td><td>strand</td><td>MM</td><td>target_seq</td><td>PAM</td><td>alignment</td><td>distance</td><td>position</td><td>gene name</td><td>gene id</td></tr>
<tr><td>GU305914.1</td><td>212314</td><td>212336</td><td>-</td><td>0</td><td>TGAGATATGTAAGAAGCTCA</td><td>AGG</td><td>||||||||[||||||||||||]PAM</td><td>NA</td><td>Intergenic</td><td>NA</td><td>NA</td></tr>
<tr><td>T18496</td><td>GCTCAAGGATGCCGACCGTCCGG</td><td>1000</td><td></td><td></td><td></td><td></td><td></td><td></td><td></td><td></td><td></td></tr>
<tr><td>Oligo fwd</td><td>CACCGCTCAAGGATGCCGACCGTC</td><td></td><td></td><td></td><td></td><td></td><td></td><td></td><td></td><td></td><td></td></tr>
<tr><td>Oligo rev</td><td>AAACGACGGTCGGCATCCTTGAGC</td><td></td><td></td><td></td><td></td><td></td><td></td><td></td><td></td><td></td><td></td></tr>
<tr><td>Chromosome</td><td>start</td><td>end</td><td>strand</td><td>MM</td><td>target_seq</td><td>PAM</td><td>alignment</td><td>distance</td><td>position</td><td>gene name</td><td>gene id</td></tr>
<tr><td>GU305914.1</td><td>212299</td><td>212321</td><td>-</td><td>0</td><td>GCTCAAGGATGCCGACCGTC</td><td>CGG</td><td>||||||||[||||||||||||]PAM</td><td>NA</td><td>Intergenic</td><td>NA</td><td>NA</td></tr>
<tr><td>T18497</td><td>TCTAGAAGATGACATCGAGATGG</td><td>1000</td><td></td><td></td><td></td><td></td><td></td><td></td><td></td><td></td><td></td></tr>
<tr><td>Oligo fwd</td><td>CACCTCTAGAAGATGACATCGAGA</td><td></td><td></td><td></td><td></td><td></td><td></td><td></td><td></td><td></td><td></td></tr>
<tr><td>Oligo rev</td><td>AAACTCTCGATGTCATCTTCTAGA</td><td></td><td></td><td></td><td></td><td></td><td></td><td></td><td></td><td></td><td></td></tr>
<tr><td>Chromosome</td><td>start</td><td>end</td><td>strand</td><td>MM</td><td>target_seq</td><td>PAM</td><td>alignment</td><td>distance</td><td>position</td><td>gene name</td><td>gene id</td></tr>
<tr><td>GU305914.1</td><td>212245</td><td>212267</td><td>-</td><td>0</td><td>TCTAGAAGATGACATCGAGA</td><td>TGG</td><td>||||||||[||||||||||||]PAM</td><td>NA</td><td>Intergenic</td><td>NA</td><td>NA</td></tr>
<tr><td>T18498</td><td>ACAGTGTCAGATGACGACCCAGG</td><td>1000</td><td></td><td></td><td></td><td></td><td></td><td></td><td></td><td></td><td></td></tr>
<tr><td>Oligo fwd</td><td>CACCACAGTGTCAGATGACGACCC</td><td></td><td></td><td></td><td></td><td></td><td></td><td></td><td></td><td></td><td></td></tr>
<tr><td>Oligo rev</td><td>AAACGGGTCGTCATCTGACACTGT</td><td></td><td></td><td></td><td></td><td></td><td></td><td></td><td></td><td></td><td></td></tr>
<tr><td>Chromosome</td><td>start</td><td>end</td><td>strand</td><td>MM</td><td>target_seq</td><td>PAM</td><td>alignment</td><td>distance</td><td>position</td><td>gene name</td><td>gene id</td></tr>
<tr><td>GU305914.1</td><td>212215</td><td>212237</td><td>-</td><td>0</td><td>ACAGTGTCAGATGACGACCC</td><td>AGG</td><td>||||||||[||||||||||||]PAM</td><td>NA</td><td>Intergenic</td><td>NA</td><td>NA</td></tr>
<tr><td>T18499</td><td>TGAAATACCATATGACCGTGTGG</td><td>1000</td><td></td><td></td><td></td><td></td><td></td><td></td><td></td><td></td><td></td></tr>
<tr><td>Oligo fwd</td><td>CACCTGAAATACCATATGACCGTG</td><td></td><td></td><td></td><td></td><td></td><td></td><td></td><td></td><td></td><td></td></tr>
<tr><td>Oligo rev</td><td>AAACCACGGTCATATGGTATTTCA</td><td></td><td></td><td></td><td></td><td></td><td></td><td></td><td></td><td></td><td></td></tr>
<tr><td>Chromosome</td><td>start</td><td>end</td><td>strand</td><td>MM</td><td>target_seq</td><td>PAM</td><td>alignment</td><td>distance</td><td>position</td><td>gene name</td><td>gene id</td></tr>
<tr><td>GU305914.1</td><td>212186</td><td>212208</td><td>-</td><td>0</td><td>TGAAATACCATATGACCGTG</td><td>TGG</td><td>||||||||[||||||||||||]PAM</td><td>NA</td><td>Intergenic</td><td>NA</td><td>NA</td></tr>
<tr><td>T18500</td><td>CATATGACCGTGTGGAGCGCCGG</td><td>1000</td><td></td><td></td><td></td><td></td><td></td><td></td><td></td><td></td><td></td></tr>
<tr><td>Oligo fwd</td><td>CACCCATATGACCGTGTGGAGCGC</td><td></td><td></td><td></td><td></td><td></td><td></td><td></td><td></td><td></td><td></td></tr>
<tr><td>Oligo rev</td><td>AAACGCGCTCCACACGGTCATATG</td><td></td><td></td><td></td><td></td><td></td><td></td><td></td><td></td><td></td><td></td></tr>
<tr><td>Chromosome</td><td>start</td><td>end</td><td>strand</td><td>MM</td><td>target_seq</td><td>PAM</td><td>alignment</td><td>distance</td><td>position</td><td>gene name</td><td>gene id</td></tr>
<tr><td>GU305914.1</td><td>212178</td><td>212200</td><td>-</td><td>0</td><td>CATATGACCGTGTGGAGCGC</td><td>CGG</td><td>||||||||[||||||||||||]PAM</td><td>NA</td><td>Intergenic</td><td>NA</td><td>NA</td></tr>
<tr><td>T18501</td><td>TGACCGTGTGGAGCGCCGGCCGG</td><td>1000</td><td></td><td></td><td></td><td></td><td></td><td></td><td></td><td></td><td></td></tr>
<tr><td>Oligo fwd</td><td>CACCTGACCGTGTGGAGCGCCGGC</td><td></td><td></td><td></td><td></td><td></td><td></td><td></td><td></td><td></td><td></td></tr>
<tr><td>Oligo rev</td><td>AAACGCCGGCGCTCCACACGGTCA</td><td></td><td></td><td></td><td></td><td></td><td></td><td></td><td></td><td></td><td></td></tr>
<tr><td>Chromosome</td><td>start</td><td>end</td><td>strand</td><td>MM</td><td>target_seq</td><td>PAM</td><td>alignment</td><td>distance</td><td>position</td><td>gene name</td><td>gene id</td></tr>
<tr><td>GU305914.1</td><td>212174</td><td>212196</td><td>-</td><td>0</td><td>TGACCGTGTGGAGCGCCGGC</td><td>CGG</td><td>||||||||[||||||||||||]PAM</td><td>NA</td><td>Intergenic</td><td>NA</td><td>NA</td></tr>
<tr><td>T18502</td><td>GACCGTGTGGAGCGCCGGCCGGG</td><td>1000</td><td></td><td></td><td></td><td></td><td></td><td></td><td></td><td></td><td></td></tr>
<tr><td>Oligo fwd</td><td>CACCGACCGTGTGGAGCGCCGGCC</td><td></td><td></td><td></td><td></td><td></td><td></td><td></td><td></td><td></td><td></td></tr>
<tr><td>Oligo rev</td><td>AAACGGCCGGCGCTCCACACGGTC</td><td></td><td></td><td></td><td></td><td></td><td></td><td></td><td></td><td></td><td></td></tr>
<tr><td>Chromosome</td><td>start</td><td>end</td><td>strand</td><td>MM</td><td>target_seq</td><td>PAM</td><td>alignment</td><td>distance</td><td>position</td><td>gene name</td><td>gene id</td></tr>
<tr><td>GU305914.1</td><td>212173</td><td>212195</td><td>-</td><td>0</td><td>GACCGTGTGGAGCGCCGGCC</td><td>GGG</td><td>||||||||[||||||||||||]PAM</td><td>NA</td><td>Intergenic</td><td>NA</td><td>NA</td></tr>
<tr><td>T18503</td><td>CCGTGCGATAGCCTTGTCCGCGG</td><td>1000</td><td></td><td></td><td></td><td></td><td></td><td></td><td></td><td></td><td></td></tr>
<tr><td>Oligo fwd</td><td>CACCCCGTGCGATAGCCTTGTCCG</td><td></td><td></td><td></td><td></td><td></td><td></td><td></td><td></td><td></td><td></td></tr>
<tr><td>Oligo rev</td><td>AAACCGGACAAGGCTATCGCACGG</td><td></td><td></td><td></td><td></td><td></td><td></td><td></td><td></td><td></td><td></td></tr>
<tr><td>Chromosome</td><td>start</td><td>end</td><td>strand</td><td>MM</td><td>target_seq</td><td>PAM</td><td>alignment</td><td>distance</td><td>position</td><td>gene name</td><td>gene id</td></tr>
<tr><td>GU305914.1</td><td>212146</td><td>212168</td><td>-</td><td>0</td><td>CCGTGCGATAGCCTTGTCCG</td><td>CGG</td><td>||||||||[||||||||||||]PAM</td><td>NA</td><td>Intergenic</td><td>NA</td><td>NA</td></tr>
<tr><td>T18504</td><td>CTTGTCCGCGGACTACTACACGG</td><td>1000</td><td></td><td></td><td></td><td></td><td></td><td></td><td></td><td></td><td></td></tr>
<tr><td>Oligo fwd</td><td>CACCCTTGTCCGCGGACTACTACA</td><td></td><td></td><td></td><td></td><td></td><td></td><td></td><td></td><td></td><td></td></tr>
<tr><td>Oligo rev</td><td>AAACTGTAGTAGTCCGCGGACAAG</td><td></td><td></td><td></td><td></td><td></td><td></td><td></td><td></td><td></td><td></td></tr>
<tr><td>Chromosome</td><td>start</td><td>end</td><td>strand</td><td>MM</td><td>target_seq</td><td>PAM</td><td>alignment</td><td>distance</td><td>position</td><td>gene name</td><td>gene id</td></tr>
<tr><td>GU305914.1</td><td>212134</td><td>212156</td><td>-</td><td>0</td><td>CTTGTCCGCGGACTACTACA</td><td>CGG</td><td>||||||||[||||||||||||]PAM</td><td>NA</td><td>Intergenic</td><td>NA</td><td>NA</td></tr>
<tr><td>T18505</td><td>CTACTACACGGACATCGAAGTGG</td><td>1000</td><td></td><td></td><td></td><td></td><td></td><td></td><td></td><td></td><td></td></tr>
<tr><td>Oligo fwd</td><td>CACCCTACTACACGGACATCGAAG</td><td></td><td></td><td></td><td></td><td></td><td></td><td></td><td></td><td></td><td></td></tr>
<tr><td>Oligo rev</td><td>AAACCTTCGATGTCCGTGTAGTAG</td><td></td><td></td><td></td><td></td><td></td><td></td><td></td><td></td><td></td><td></td></tr>
<tr><td>Chromosome</td><td>start</td><td>end</td><td>strand</td><td>MM</td><td>target_seq</td><td>PAM</td><td>alignment</td><td>distance</td><td>position</td><td>gene name</td><td>gene id</td></tr>
<tr><td>GU305914.1</td><td>212122</td><td>212144</td><td>-</td><td>0</td><td>CTACTACACGGACATCGAAG</td><td>TGG</td><td>||||||||[||||||||||||]PAM</td><td>NA</td><td>Intergenic</td><td>NA</td><td>NA</td></tr>
<tr><td>T18506</td><td>CGAAGTGGCTTCCTATCTGCCGG</td><td>1000</td><td></td><td></td><td></td><td></td><td></td><td></td><td></td><td></td><td></td></tr>
<tr><td>Oligo fwd</td><td>CACCCGAAGTGGCTTCCTATCTGC</td><td></td><td></td><td></td><td></td><td></td><td></td><td></td><td></td><td></td><td></td></tr>
<tr><td>Oligo rev</td><td>AAACGCAGATAGGAAGCCACTTCG</td><td></td><td></td><td></td><td></td><td></td><td></td><td></td><td></td><td></td><td></td></tr>
<tr><td>Chromosome</td><td>start</td><td>end</td><td>strand</td><td>MM</td><td>target_seq</td><td>PAM</td><td>alignment</td><td>distance</td><td>position</td><td>gene name</td><td>gene id</td></tr>
<tr><td>GU305914.1</td><td>212107</td><td>212129</td><td>-</td><td>0</td><td>CGAAGTGGCTTCCTATCTGC</td><td>CGG</td><td>||||||||[||||||||||||]PAM</td><td>NA</td><td>Intergenic</td><td>NA</td><td>NA</td></tr>
<tr><td>T18507</td><td>TGCGAGATCACCTCGTCGAGCGG</td><td>1000</td><td></td><td></td><td></td><td></td><td></td><td></td><td></td><td></td><td></td></tr>
<tr><td>Oligo fwd</td><td>CACCTGCGAGATCACCTCGTCGAG</td><td></td><td></td><td></td><td></td><td></td><td></td><td></td><td></td><td></td><td></td></tr>
<tr><td>Oligo rev</td><td>AAACCTCGACGAGGTGATCTCGCA</td><td></td><td></td><td></td><td></td><td></td><td></td><td></td><td></td><td></td><td></td></tr>
<tr><td>Chromosome</td><td>start</td><td>end</td><td>strand</td><td>MM</td><td>target_seq</td><td>PAM</td><td>alignment</td><td>distance</td><td>position</td><td>gene name</td><td>gene id</td></tr>
<tr><td>GU305914.1</td><td>212055</td><td>212077</td><td>-</td><td>0</td><td>TGCGAGATCACCTCGTCGAG</td><td>CGG</td><td>||||||||[||||||||||||]PAM</td><td>NA</td><td>Intergenic</td><td>NA</td><td>NA</td></tr>
<tr><td>T18508</td><td>AGATCACCTCGTCGAGCGGCTGG</td><td>1000</td><td></td><td></td><td></td><td></td><td></td><td></td><td></td><td></td><td></td></tr>
<tr><td>Oligo fwd</td><td>CACCAGATCACCTCGTCGAGCGGC</td><td></td><td></td><td></td><td></td><td></td><td></td><td></td><td></td><td></td><td></td></tr>
</table>

<table>
<tr><td>Oligo rev</td><td>AAACGCCGCTCGACGAGGTGATCT</td><td></td><td></td><td></td><td></td><td></td><td></td><td></td><td></td><td></td><td></td></tr>
<tr><td>Chromosome</td><td>start</td><td>end</td><td>strand</td><td>MM</td><td>target_seq</td><td>PAM</td><td>alignment</td><td>distance</td><td>position</td><td>gene name</td><td>gene id</td></tr>
<tr><td>GU305914.1</td><td>212051</td><td>212073</td><td>-</td><td>0</td><td>AGATCACCTCGTCGAGCGGC</td><td>TGG</td><td>||||||||[||||||||||||]PAM</td><td>NA</td><td>Intergenic</td><td>NA</td><td>NA</td></tr>
<tr><td>T18509</td><td>CACCTCGTCGAGCGGCTGGACGG</td><td>1000</td><td></td><td></td><td></td><td></td><td></td><td></td><td></td><td></td><td></td></tr>
<tr><td>Oligo fwd</td><td>CACCCACCTCGTCGAGCGGCTGGA</td><td></td><td></td><td></td><td></td><td></td><td></td><td></td><td></td><td></td><td></td></tr>
<tr><td>Oligo rev</td><td>AAACTCCAGCCGCTCGACGAGGTG</td><td></td><td></td><td></td><td></td><td></td><td></td><td></td><td></td><td></td><td></td></tr>
<tr><td>Chromosome</td><td>start</td><td>end</td><td>strand</td><td>MM</td><td>target_seq</td><td>PAM</td><td>alignment</td><td>distance</td><td>position</td><td>gene name</td><td>gene id</td></tr>
<tr><td>GU305914.1</td><td>212047</td><td>212069</td><td>-</td><td>0</td><td>CACCTCGTCGAGCGGCTGGA</td><td>CGG</td><td>||||||||[||||||||||||]PAM</td><td>NA</td><td>Intergenic</td><td>NA</td><td>NA</td></tr>
<tr><td>T18510</td><td>GCGGCTGGACGGTCCGTCTTCGG</td><td>1000</td><td></td><td></td><td></td><td></td><td></td><td></td><td></td><td></td><td></td></tr>
<tr><td>Oligo fwd</td><td>CACCGCGGCTGGACGGTCCGTCTT</td><td></td><td></td><td></td><td></td><td></td><td></td><td></td><td></td><td></td><td></td></tr>
<tr><td>Oligo rev</td><td>AAACAAGACGGACCGTCCAGCCGC</td><td></td><td></td><td></td><td></td><td></td><td></td><td></td><td></td><td></td><td></td></tr>
<tr><td>Chromosome</td><td>start</td><td>end</td><td>strand</td><td>MM</td><td>target_seq</td><td>PAM</td><td>alignment</td><td>distance</td><td>position</td><td>gene name</td><td>gene id</td></tr>
<tr><td>GU305914.1</td><td>212036</td><td>212058</td><td>-</td><td>0</td><td>GCGGCTGGACGGTCCGTCTT</td><td>CGG</td><td>||||||||[||||||||||||]PAM</td><td>NA</td><td>Intergenic</td><td>NA</td><td>NA</td></tr>
<tr><td>T18511</td><td>GGTCCGTCTTCGGTTCTCCGAGG</td><td>1000</td><td></td><td></td><td></td><td></td><td></td><td></td><td></td><td></td><td></td></tr>
<tr><td>Oligo fwd</td><td>CACCGGTCCGTCTTCGGTTCTCCG</td><td></td><td></td><td></td><td></td><td></td><td></td><td></td><td></td><td></td><td></td></tr>
<tr><td>Oligo rev</td><td>AAACCGGAGAACCGAAGACGGACC</td><td></td><td></td><td></td><td></td><td></td><td></td><td></td><td></td><td></td><td></td></tr>
<tr><td>Chromosome</td><td>start</td><td>end</td><td>strand</td><td>MM</td><td>target_seq</td><td>PAM</td><td>alignment</td><td>distance</td><td>position</td><td>gene name</td><td>gene id</td></tr>
<tr><td>GU305914.1</td><td>212026</td><td>212048</td><td>-</td><td>0</td><td>GGTCCGTCTTCGGTTCTCCG</td><td>AGG</td><td>||||||||[||||||||||||]PAM</td><td>NA</td><td>Intergenic</td><td>NA</td><td>NA</td></tr>
<tr><td>T18512</td><td>TCTTCGGTTCTCCGAGGAGATGG</td><td>1000</td><td></td><td></td><td></td><td></td><td></td><td></td><td></td><td></td><td></td></tr>
<tr><td>Oligo fwd</td><td>CACCTCTTCGGTTCTCCGAGGAGA</td><td></td><td></td><td></td><td></td><td></td><td></td><td></td><td></td><td></td><td></td></tr>
<tr><td>Oligo rev</td><td>AAACTCTCCTCGGAGAACCGAAGA</td><td></td><td></td><td></td><td></td><td></td><td></td><td></td><td></td><td></td><td></td></tr>
<tr><td>Chromosome</td><td>start</td><td>end</td><td>strand</td><td>MM</td><td>target_seq</td><td>PAM</td><td>alignment</td><td>distance</td><td>position</td><td>gene name</td><td>gene id</td></tr>
<tr><td>GU305914.1</td><td>212020</td><td>212042</td><td>-</td><td>0</td><td>TCTTCGGTTCTCCGAGGAGA</td><td>TGG</td><td>||||||||[||||||||||||]PAM</td><td>NA</td><td>Intergenic</td><td>NA</td><td>NA</td></tr>
<tr><td>T18513</td><td>GTTCTCCGAGGAGATGGTAGCGG</td><td>1000</td><td></td><td></td><td></td><td></td><td></td><td></td><td></td><td></td><td></td></tr>
<tr><td>Oligo fwd</td><td>CACCGTTCTCCGAGGAGATGGTAG</td><td></td><td></td><td></td><td></td><td></td><td></td><td></td><td></td><td></td><td></td></tr>
<tr><td>Oligo rev</td><td>AAACCTACCATCTCCTCGGAGAAC</td><td></td><td></td><td></td><td></td><td></td><td></td><td></td><td></td><td></td><td></td></tr>
<tr><td>Chromosome</td><td>start</td><td>end</td><td>strand</td><td>MM</td><td>target_seq</td><td>PAM</td><td>alignment</td><td>distance</td><td>position</td><td>gene name</td><td>gene id</td></tr>
<tr><td>GU305914.1</td><td>212014</td><td>212036</td><td>-</td><td>0</td><td>GTTCTCCGAGGAGATGGTAG</td><td>CGG</td><td>||||||||[||||||||||||]PAM</td><td>NA</td><td>Intergenic</td><td>NA</td><td>NA</td></tr>
<tr><td>T18514</td><td>TGGTAGCGGCCTCGAAAGCGAGG</td><td>1000</td><td></td><td></td><td></td><td></td><td></td><td></td><td></td><td></td><td></td></tr>
<tr><td>Oligo fwd</td><td>CACCTGGTAGCGGCCTCGAAAGCG</td><td></td><td></td><td></td><td></td><td></td><td></td><td></td><td></td><td></td><td></td></tr>
<tr><td>Oligo rev</td><td>AAACCGCTTTCGAGGCCGCTACCA</td><td></td><td></td><td></td><td></td><td></td><td></td><td></td><td></td><td></td><td></td></tr>
<tr><td>Chromosome</td><td>start</td><td>end</td><td>strand</td><td>MM</td><td>target_seq</td><td>PAM</td><td>alignment</td><td>distance</td><td>position</td><td>gene name</td><td>gene id</td></tr>
<tr><td>GU305914.1</td><td>212000</td><td>212022</td><td>-</td><td>0</td><td>TGGTAGCGGCCTCGAAAGCG</td><td>AGG</td><td>||||||||[||||||||||||]PAM</td><td>NA</td><td>Intergenic</td><td>NA</td><td>NA</td></tr>
<tr><td>T18515</td><td>AGCGGCCTCGAAAGCGAGGCAGG</td><td>1000</td><td></td><td></td><td></td><td></td><td></td><td></td><td></td><td></td><td></td></tr>
<tr><td>Oligo fwd</td><td>CACCAGCGGCCTCGAAAGCGAGGC</td><td></td><td></td><td></td><td></td><td></td><td></td><td></td><td></td><td></td><td></td></tr>
<tr><td>Oligo rev</td><td>AAACGCCTCGCTTTCGAGGCCGCT</td><td></td><td></td><td></td><td></td><td></td><td></td><td></td><td></td><td></td><td></td></tr>
<tr><td>Chromosome</td><td>start</td><td>end</td><td>strand</td><td>MM</td><td>target_seq</td><td>PAM</td><td>alignment</td><td>distance</td><td>position</td><td>gene name</td><td>gene id</td></tr>
<tr><td>GU305914.1</td><td>211996</td><td>212018</td><td>-</td><td>0</td><td>AGCGGCCTCGAAAGCGAGGC</td><td>AGG</td><td>||||||||[||||||||||||]PAM</td><td>NA</td><td>Intergenic</td><td>NA</td><td>NA</td></tr>
<tr><td>T18516</td><td>AAGCGAGGCAGGCTCAAAAGAGG</td><td>1000</td><td></td><td></td><td></td><td></td><td></td><td></td><td></td><td></td><td></td></tr>
<tr><td>Oligo fwd</td><td>CACCAAGCGAGGCAGGCTCAAAAG</td><td></td><td></td><td></td><td></td><td></td><td></td><td></td><td></td><td></td><td></td></tr>
<tr><td>Oligo rev</td><td>AAACCTTTTGAGCCTGCCTCGCTT</td><td></td><td></td><td></td><td></td><td></td><td></td><td></td><td></td><td></td><td></td></tr>
<tr><td>Chromosome</td><td>start</td><td>end</td><td>strand</td><td>MM</td><td>target_seq</td><td>PAM</td><td>alignment</td><td>distance</td><td>position</td><td>gene name</td><td>gene id</td></tr>
<tr><td>GU305914.1</td><td>211985</td><td>212007</td><td>-</td><td>0</td><td>AAGCGAGGCAGGCTCAAAAG</td><td>AGG</td><td>||||||||[||||||||||||]PAM</td><td>NA</td><td>Intergenic</td><td>NA</td><td>NA</td></tr>
<tr><td>T18517</td><td>CTGCTCCCCGTAGAACCGCACGG</td><td>1000</td><td></td><td></td><td></td><td></td><td></td><td></td><td></td><td></td><td></td></tr>
<tr><td>Oligo fwd</td><td>CACCCTGCTCCCCGTAGAACCGCA</td><td></td><td></td><td></td><td></td><td></td><td></td><td></td><td></td><td></td><td></td></tr>
<tr><td>Oligo rev</td><td>AAACTGCGGTTCTACGGGGAGCAG</td><td></td><td></td><td></td><td></td><td></td><td></td><td></td><td></td><td></td><td></td></tr>
<tr><td>Chromosome</td><td>start</td><td>end</td><td>strand</td><td>MM</td><td>target_seq</td><td>PAM</td><td>alignment</td><td>distance</td><td>position</td><td>gene name</td><td>gene id</td></tr>
<tr><td>GU305914.1</td><td>211959</td><td>211981</td><td>-</td><td>0</td><td>CTGCTCCCCGTAGAACCGCA</td><td>CGG</td><td>||||||||[||||||||||||]PAM</td><td>NA</td><td>Intergenic</td><td>NA</td><td>NA</td></tr>
<tr><td>T18518</td><td>GGATTCATGTCCGATGAACACGG</td><td>1000</td><td></td><td></td><td></td><td></td><td></td><td></td><td></td><td></td><td></td></tr>
<tr><td>Oligo fwd</td><td>CACCGGATTCATGTCCGATGAACA</td><td></td><td></td><td></td><td></td><td></td><td></td><td></td><td></td><td></td><td></td></tr>
<tr><td>Oligo rev</td><td>AAACTGTTCATCGGACATGAATCC</td><td></td><td></td><td></td><td></td><td></td><td></td><td></td><td></td><td></td><td></td></tr>
<tr><td>Chromosome</td><td>start</td><td>end</td><td>strand</td><td>MM</td><td>target_seq</td><td>PAM</td><td>alignment</td><td>distance</td><td>position</td><td>gene name</td><td>gene id</td></tr>
<tr><td>GU305914.1</td><td>211938</td><td>211960</td><td>-</td><td>0</td><td>GGATTCATGTCCGATGAACA</td><td>CGG</td><td>||||||||[||||||||||||]PAM</td><td>NA</td><td>Intergenic</td><td>NA</td><td>NA</td></tr>
<tr><td>T18519</td><td>CATGTCCGATGAACACGGTCCGG</td><td>1000</td><td></td><td></td><td></td><td></td><td></td><td></td><td></td><td></td><td></td></tr>
<tr><td>Oligo fwd</td><td>CACCCATGTCCGATGAACACGGTC</td><td></td><td></td><td></td><td></td><td></td><td></td><td></td><td></td><td></td><td></td></tr>
<tr><td>Oligo rev</td><td>AAACGACCGTGTTCATCGGACATG</td><td></td><td></td><td></td><td></td><td></td><td></td><td></td><td></td><td></td><td></td></tr>
<tr><td>Chromosome</td><td>start</td><td>end</td><td>strand</td><td>MM</td><td>target_seq</td><td>PAM</td><td>alignment</td><td>distance</td><td>position</td><td>gene name</td><td>gene id</td></tr>
<tr><td>GU305914.1</td><td>211933</td><td>211955</td><td>-</td><td>0</td><td>CATGTCCGATGAACACGGTC</td><td>CGG</td><td>||||||||[||||||||||||]PAM</td><td>NA</td><td>Intergenic</td><td>NA</td><td>NA</td></tr>
<tr><td>T18520</td><td>ACGGTCCGGCCTTCGTCCAGAGG</td><td>1000</td><td></td><td></td><td></td><td></td><td></td><td></td><td></td><td></td><td></td></tr>
<tr><td>Oligo fwd</td><td>CACCACGGTCCGGCCTTCGTCCAG</td><td></td><td></td><td></td><td></td><td></td><td></td><td></td><td></td><td></td><td></td></tr>
<tr><td>Oligo rev</td><td>AAACCTGGACGAAGGCCGGACCGT</td><td></td><td></td><td></td><td></td><td></td><td></td><td></td><td></td><td></td><td></td></tr>
<tr><td>Chromosome</td><td>start</td><td>end</td><td>strand</td><td>MM</td><td>target_seq</td><td>PAM</td><td>alignment</td><td>distance</td><td>position</td><td>gene name</td><td>gene id</td></tr>
<tr><td>GU305914.1</td><td>211919</td><td>211941</td><td>-</td><td>0</td><td>ACGGTCCGGCCTTCGTCCAG</td><td>AGG</td><td>||||||||[||||||||||||]PAM</td><td>NA</td><td>Intergenic</td><td>NA</td><td>NA</td></tr>
<tr><td>T18521</td><td>CAACGATTCGAGACTGACCACGG</td><td>1000</td><td></td><td></td><td></td><td></td><td></td><td></td><td></td><td></td><td></td></tr>
<tr><td>Oligo fwd</td><td>CACCCAACGATTCGAGACTGACCA</td><td></td><td></td><td></td><td></td><td></td><td></td><td></td><td></td><td></td><td></td></tr>
<tr><td>Oligo rev</td><td>AAACTGGTCAGTCTCGAATCGTTG</td><td></td><td></td><td></td><td></td><td></td><td></td><td></td><td></td><td></td><td></td></tr>
<tr><td>Chromosome</td><td>start</td><td>end</td><td>strand</td><td>MM</td><td>target_seq</td><td>PAM</td><td>alignment</td><td>distance</td><td>position</td><td>gene name</td><td>gene id</td></tr>
<tr><td>GU305914.1</td><td>211891</td><td>211913</td><td>-</td><td>0</td><td>CAACGATTCGAGACTGACCA</td><td>CGG</td><td>||||||||[||||||||||||]PAM</td><td>NA</td><td>Intergenic</td><td>NA</td><td>NA</td></tr>
<tr><td>T18522</td><td>CGCATTCGTCGCGTTATTCGTGG</td><td>1000</td><td></td><td></td><td></td><td></td><td></td><td></td><td></td><td></td><td></td></tr>
<tr><td>Oligo fwd</td><td>CACCCGCATTCGTCGCGTTATTCG</td><td></td><td></td><td></td><td></td><td></td><td></td><td></td><td></td><td></td><td></td></tr>
<tr><td>Oligo rev</td><td>AAACCGAATAACGCGACGAATGCG</td><td></td><td></td><td></td><td></td><td></td><td></td><td></td><td></td><td></td><td></td></tr>
<tr><td>Chromosome</td><td>start</td><td>end</td><td>strand</td><td>MM</td><td>target_seq</td><td>PAM</td><td>alignment</td><td>distance</td><td>position</td><td>gene name</td><td>gene id</td></tr>
<tr><td>GU305914.1</td><td>211846</td><td>211868</td><td>-</td><td>0</td><td>CGCATTCGTCGCGTTATTCG</td><td>TGG</td><td>||||||||[||||||||||||]PAM</td><td>NA</td><td>Intergenic</td><td>NA</td><td>NA</td></tr>
<tr><td>T18523</td><td>GTCGCGTTATTCGTGGTCGTCGG</td><td>1000</td><td></td><td></td><td></td><td></td><td></td><td></td><td></td><td></td><td></td></tr>
</table>

<table>
<tr><td>Oligo fwd</td><td>CACCGTCGCGTTATTCGTGGTCGT</td></tr>
<tr><td>Oligo rev</td><td>AAACACGACCACGAATAACGCGAC</td></tr>
<tr><td>Chromosome</td><td>start</td><td></td><td>end</td><td>strand</td><td>MM</td><td>target_seq</td><td>PAM</td><td>alignment</td><td>distance</td><td>position</td><td>gene name</td><td>gene id</td></tr>
<tr><td>GU305914.1</td><td></td><td>211839</td><td>211861</td><td>-</td><td>0</td><td>GTCGCGTTATTCGTGGTCGT</td><td>CGG</td><td>||||||||[||||||||||||]PAM</td><td>NA</td><td>Intergenic</td><td>NA</td><td>NA</td></tr>
<tr><td></td></tr>
<tr><td>T18524</td><td>ATTCGTGGTCGTCGGACTCCTGG</td><td></td><td>1000</td></tr>
<tr><td>Oligo fwd</td><td>CACCATTCGTGGTCGTCGGACTCC</td></tr>
<tr><td>Oligo rev</td><td>AAACGGAGTCCGACGACCACGAAT</td></tr>
<tr><td>Chromosome</td><td>start</td><td></td><td>end</td><td>strand</td><td>MM</td><td>target_seq</td><td>PAM</td><td>alignment</td><td>distance</td><td>position</td><td>gene name</td><td>gene id</td></tr>
<tr><td>GU305914.1</td><td></td><td>211831</td><td>211853</td><td>-</td><td>0</td><td>ATTCGTGGTCGTCGGACTCC</td><td>TGG</td><td>||||||||[||||||||||||]PAM</td><td>NA</td><td>Intergenic</td><td>NA</td><td>NA</td></tr>
<tr><td></td></tr>
<tr><td>T18525</td><td>GTCGTCGGACTCCTGGTGCACGG</td><td></td><td>1000</td></tr>
<tr><td>Oligo fwd</td><td>CACCGTCGTCGGACTCCTGGTGCA</td></tr>
<tr><td>Oligo rev</td><td>AAACTGCACCAGGAGTCCGACGAC</td></tr>
<tr><td>Chromosome</td><td>start</td><td></td><td>end</td><td>strand</td><td>MM</td><td>target_seq</td><td>PAM</td><td>alignment</td><td>distance</td><td>position</td><td>gene name</td><td>gene id</td></tr>
<tr><td>GU305914.1</td><td></td><td>211824</td><td>211846</td><td>-</td><td>0</td><td>GTCGTCGGACTCCTGGTGCA</td><td>CGG</td><td>||||||||[||||||||||||]PAM</td><td>NA</td><td>Intergenic</td><td>NA</td><td>NA</td></tr>
<tr><td></td></tr>
<tr><td>T18526</td><td>TCGTCGGACTCCTGGTGCACGGG</td><td></td><td>1000</td></tr>
<tr><td>Oligo fwd</td><td>CACCTCGTCGGACTCCTGGTGCAC</td></tr>
<tr><td>Oligo rev</td><td>AAACGTGCACCAGGAGTCCGACGA</td></tr>
<tr><td>Chromosome</td><td>start</td><td></td><td>end</td><td>strand</td><td>MM</td><td>target_seq</td><td>PAM</td><td>alignment</td><td>distance</td><td>position</td><td>gene name</td><td>gene id</td></tr>
<tr><td>GU305914.1</td><td></td><td>211823</td><td>211845</td><td>-</td><td>0</td><td>TCGTCGGACTCCTGGTGCAC</td><td>GGG</td><td>||||||||[||||||||||||]PAM</td><td>NA</td><td>Intergenic</td><td>NA</td><td>NA</td></tr>
<tr><td></td></tr>
<tr><td>T18527</td><td>CTGGTGCACGGGATCGTCACCGG</td><td></td><td>1000</td></tr>
<tr><td>Oligo fwd</td><td>CACCCTGGTGCACGGGATCGTCAC</td></tr>
<tr><td>Oligo rev</td><td>AAACGTGACGATCCCGTGCACCAG</td></tr>
<tr><td>Chromosome</td><td>start</td><td></td><td>end</td><td>strand</td><td>MM</td><td>target_seq</td><td>PAM</td><td>alignment</td><td>distance</td><td>position</td><td>gene name</td><td>gene id</td></tr>
<tr><td>GU305914.1</td><td></td><td>211812</td><td>211834</td><td>-</td><td>0</td><td>CTGGTGCACGGGATCGTCAC</td><td>CGG</td><td>||||||||[||||||||||||]PAM</td><td>NA</td><td>Intergenic</td><td>NA</td><td>NA</td></tr>
<tr><td></td></tr>
<tr><td>T18528</td><td>TGGTGCACGGGATCGTCACCGGG</td><td></td><td>1000</td></tr>
<tr><td>Oligo fwd</td><td>CACCTGGTGCACGGGATCGTCACC</td></tr>
<tr><td>Oligo rev</td><td>AAACGGTGACGATCCCGTGCACCA</td></tr>
<tr><td>Chromosome</td><td>start</td><td></td><td>end</td><td>strand</td><td>MM</td><td>target_seq</td><td>PAM</td><td>alignment</td><td>distance</td><td>position</td><td>gene name</td><td>gene id</td></tr>
<tr><td>GU305914.1</td><td></td><td>211811</td><td>211833</td><td>-</td><td>0</td><td>TGGTGCACGGGATCGTCACC</td><td>GGG</td><td>||||||||[||||||||||||]PAM</td><td>NA</td><td>Intergenic</td><td>NA</td><td>NA</td></tr>
<tr><td></td></tr>
<tr><td>T18529</td><td>CGGGATCGTCACCGGGCGCCTGG</td><td></td><td>1000</td></tr>
<tr><td>Oligo fwd</td><td>CACCCGGGATCGTCACCGGGCGCC</td></tr>
<tr><td>Oligo rev</td><td>AAACGGCGCCCGGTGACGATCCCG</td></tr>
<tr><td>Chromosome</td><td>start</td><td></td><td>end</td><td>strand</td><td>MM</td><td>target_seq</td><td>PAM</td><td>alignment</td><td>distance</td><td>position</td><td>gene name</td><td>gene id</td></tr>
<tr><td>GU305914.1</td><td></td><td>211804</td><td>211826</td><td>-</td><td>0</td><td>CGGGATCGTCACCGGGCGCC</td><td>TGG</td><td>||||||||[||||||||||||]PAM</td><td>NA</td><td>Intergenic</td><td>NA</td><td>NA</td></tr>
<tr><td></td></tr>
<tr><td>T18530</td><td>CACCGGGCGCCTGGCTCGCATGG</td><td></td><td>1000</td></tr>
<tr><td>Oligo fwd</td><td>CACCCACCGGGCGCCTGGCTCGCA</td></tr>
<tr><td>Oligo rev</td><td>AAACTGCGAGCCAGGCGCCCGGTG</td></tr>
<tr><td>Chromosome</td><td>start</td><td></td><td>end</td><td>strand</td><td>MM</td><td>target_seq</td><td>PAM</td><td>alignment</td><td>distance</td><td>position</td><td>gene name</td><td>gene id</td></tr>
<tr><td>GU305914.1</td><td></td><td>211795</td><td>211817</td><td>-</td><td>0</td><td>CACCGGGCGCCTGGCTCGCA</td><td>TGG</td><td>||||||||[||||||||||||]PAM</td><td>NA</td><td>Intergenic</td><td>NA</td><td>NA</td></tr>
<tr><td></td></tr>
<tr><td>T18531</td><td>GCCTGGCTCGCATGGCCGAACGG</td><td></td><td>1000</td></tr>
<tr><td>Oligo fwd</td><td>CACCGCCTGGCTCGCATGGCCGAA</td></tr>
<tr><td>Oligo rev</td><td>AAACTTCGGCCATGCGAGCCAGGC</td></tr>
<tr><td>Chromosome</td><td>start</td><td></td><td>end</td><td>strand</td><td>MM</td><td>target_seq</td><td>PAM</td><td>alignment</td><td>distance</td><td>position</td><td>gene name</td><td>gene id</td></tr>
<tr><td>GU305914.1</td><td></td><td>211787</td><td>211809</td><td>-</td><td>0</td><td>GCCTGGCTCGCATGGCCGAA</td><td>CGG</td><td>||||||||[||||||||||||]PAM</td><td>NA</td><td>Intergenic</td><td>NA</td><td>NA</td></tr>
<tr><td></td></tr>
<tr><td>T18532</td><td>TTCGATCCGTTACACGCGCGAGG</td><td></td><td>1000</td></tr>
<tr><td>Oligo fwd</td><td>CACCTTCGATCCGTTACACGCGCG</td></tr>
<tr><td>Oligo rev</td><td>AAACCGCGCGTGTAACGGATCGAA</td></tr>
<tr><td>Chromosome</td><td>start</td><td></td><td>end</td><td>strand</td><td>MM</td><td>target_seq</td><td>PAM</td><td>alignment</td><td>distance</td><td>position</td><td>gene name</td><td>gene id</td></tr>
<tr><td>GU305914.1</td><td></td><td>211756</td><td>211778</td><td>-</td><td>0</td><td>TTCGATCCGTTACACGCGCG</td><td>AGG</td><td>||||||||[||||||||||||]PAM</td><td>NA</td><td>Intergenic</td><td>NA</td><td>NA</td></tr>
<tr><td></td></tr>
<tr><td>T18533</td><td>TTACACGCGCGAGGATGAAACGG</td><td></td><td>1000</td></tr>
<tr><td>Oligo fwd</td><td>CACCTTACACGCGCGAGGATGAAA</td></tr>
<tr><td>Oligo rev</td><td>AAACTTTCATCCTCGCGCGTGTAA</td></tr>
<tr><td>Chromosome</td><td>start</td><td></td><td>end</td><td>strand</td><td>MM</td><td>target_seq</td><td>PAM</td><td>alignment</td><td>distance</td><td>position</td><td>gene name</td><td>gene id</td></tr>
<tr><td>GU305914.1</td><td></td><td>211747</td><td>211769</td><td>-</td><td>0</td><td>TTACACGCGCGAGGATGAAA</td><td>CGG</td><td>||||||||[||||||||||||]PAM</td><td>NA</td><td>Intergenic</td><td>NA</td><td>NA</td></tr>
<tr><td></td></tr>
<tr><td>T18534</td><td>GGATGAAACGGTCTCGATGTTGG</td><td></td><td>1000</td></tr>
<tr><td>Oligo fwd</td><td>CACCGGATGAAACGGTCTCGATGT</td></tr>
<tr><td>Oligo rev</td><td>AAACACATCGAGACCGTTTCATCC</td></tr>
<tr><td>Chromosome</td><td>start</td><td></td><td>end</td><td>strand</td><td>MM</td><td>target_seq</td><td>PAM</td><td>alignment</td><td>distance</td><td>position</td><td>gene name</td><td>gene id</td></tr>
<tr><td>GU305914.1</td><td></td><td>211735</td><td>211757</td><td>-</td><td>0</td><td>GGATGAAACGGTCTCGATGT</td><td>TGG</td><td>||||||||[||||||||||||]PAM</td><td>NA</td><td>Intergenic</td><td>NA</td><td>NA</td></tr>
<tr><td></td></tr>
<tr><td>T18535</td><td>GATGAAACGGTCTCGATGTTGGG</td><td></td><td>1000</td></tr>
<tr><td>Oligo fwd</td><td>CACCGATGAAACGGTCTCGATGTT</td></tr>
<tr><td>Oligo rev</td><td>AAACAACATCGAGACCGTTTCATC</td></tr>
<tr><td>Chromosome</td><td>start</td><td></td><td>end</td><td>strand</td><td>MM</td><td>target_seq</td><td>PAM</td><td>alignment</td><td>distance</td><td>position</td><td>gene name</td><td>gene id</td></tr>
<tr><td>GU305914.1</td><td></td><td>211734</td><td>211756</td><td>-</td><td>0</td><td>GATGAAACGGTCTCGATGTT</td><td>GGG</td><td>||||||||[||||||||||||]PAM</td><td>NA</td><td>Intergenic</td><td>NA</td><td>NA</td></tr>
<tr><td></td></tr>
<tr><td>T18536</td><td>ATGAAACGGTCTCGATGTTGGGG</td><td></td><td>1000</td></tr>
<tr><td>Oligo fwd</td><td>CACCATGAAACGGTCTCGATGTTG</td></tr>
<tr><td>Oligo rev</td><td>AAACCAACATCGAGACCGTTTCAT</td></tr>
<tr><td>Chromosome</td><td>start</td><td></td><td>end</td><td>strand</td><td>MM</td><td>target_seq</td><td>PAM</td><td>alignment</td><td>distance</td><td>position</td><td>gene name</td><td>gene id</td></tr>
<tr><td>GU305914.1</td><td></td><td>211733</td><td>211755</td><td>-</td><td>0</td><td>ATGAAACGGTCTCGATGTTG</td><td>GGG</td><td>||||||||[||||||||||||]PAM</td><td>NA</td><td>Intergenic</td><td>NA</td><td>NA</td></tr>
<tr><td></td></tr>
<tr><td>T18537</td><td>GTTACAAGAGAACGAGTGCTCGG</td><td></td><td>1000</td></tr>
<tr><td>Oligo fwd</td><td>CACCGTTACAAGAGAACGAGTGCT</td></tr>
<tr><td>Oligo rev</td><td>AAACAGCACTCGTTCTCTTGTAAC</td></tr>
<tr><td>Chromosome</td><td>start</td><td></td><td>end</td><td>strand</td><td>MM</td><td>target_seq</td><td>PAM</td><td>alignment</td><td>distance</td><td>position</td><td>gene name</td><td>gene id</td></tr>
<tr><td>GU305914.1</td><td></td><td>211711</td><td>211733</td><td>-</td><td>0</td><td>GTTACAAGAGAACGAGTGCT</td><td>CGG</td><td>||||||||[||||||||||||]PAM</td><td>NA</td><td>Intergenic</td><td>NA</td><td>NA</td></tr>
</table>

T18538 AGAGAACGAGTGCTCGGATACGG 1000
Oligo fwd CACCAGAGAACGAGTGCTCGGATA
Oligo rev AAACTATCCGAGCACTCGTTCTCT
Chromosome start end strand MM target_seq PAM alignment distance position gene name gene id
GU305914.1 211705 211727 - 0 AGAGAACGAGTGCTCGGATA CGG ||||||||[||||||||||||]PAM NA Intergenic NA NA

T18539 GAACGAGTGCTCGGATACGGAGG 1000
Oligo fwd CACCGAACGAGTGCTCGGATACGG
Oligo rev AAACCCGTATCCGAGCACTCGTTC
Chromosome start end strand MM target_seq PAM alignment distance position gene name gene id
GU305914.1 211702 211724 - 0 GAACGAGTGCTCGGATACGG AGG ||||||||[||||||||||||]PAM NA Intergenic NA NA

T18540 CTCGGATACGGAGGAGAATGTGG 1000
Oligo fwd CACCCTCGGATACGGAGGAGAATG
Oligo rev AAACCATTCTCCTCCGTATCCGAG
Chromosome start end strand MM target_seq PAM alignment distance position gene name gene id
GU305914.1 211693 211715 - 0 CTCGGATACGGAGGAGAATG TGG ||||||||[||||||||||||]PAM NA Intergenic NA NA

T18541 ATCGACGCACGAGTCCGCCTCGG 1000
Oligo fwd CACCATCGACGCACGAGTCCGCCT
Oligo rev AAACAGGCGGACTCGTGCGTCGAT
Chromosome start end strand MM target_seq PAM alignment distance position gene name gene id
GU305914.1 211666 211688 - 0 ATCGACGCACGAGTCCGCCT CGG ||||||||[||||||||||||]PAM NA Intergenic NA NA

T18542 ACGCACGAGTCCGCCTCGGTCGG 1000
Oligo fwd CACCACGCACGAGTCCGCCTCGGT
Oligo rev AAACACCGAGGCGGACTCGTGCGT
Chromosome start end strand MM target_seq PAM alignment distance position gene name gene id
GU305914.1 211662 211684 - 0 ACGCACGAGTCCGCCTCGGT CGG ||||||||[||||||||||||]PAM NA Intergenic NA NA

T18543 GAGTCCGCCTCGGTCGGCTCAGG 1000
Oligo fwd CACCGAGTCCGCCTCGGTCGGCTC
Oligo rev AAACGAGCCGACCGAGGCGGACTC
Chromosome start end strand MM target_seq PAM alignment distance position gene name gene id
GU305914.1 211656 211678 - 0 GAGTCCGCCTCGGTCGGCTC AGG ||||||||[||||||||||||]PAM NA Intergenic NA NA

T18544 CCTCGGTCGGCTCAGGCCTTTGG 1000
Oligo fwd CACCCCTCGGTCGGCTCAGGCCTT
Oligo rev AAACAAGGCCTGAGCCGACCGAGG
Chromosome start end strand MM target_seq PAM alignment distance position gene name gene id
GU305914.1 211649 211671 - 0 CCTCGGTCGGCTCAGGCCTT TGG ||||||||[||||||||||||]PAM NA Intergenic NA NA

T18545 CTCGGTCGGCTCAGGCCTTTGGG 1000
Oligo fwd CACCCTCGGTCGGCTCAGGCCTTT
Oligo rev AAACAAAGGCCTGAGCCGACCGAG
Chromosome start end strand MM target_seq PAM alignment distance position gene name gene id
GU305914.1 211648 211670 - 0 CTCGGTCGGCTCAGGCCTTT GGG ||||||||[||||||||||||]PAM NA Intergenic NA NA

T18546 CTCAGGCCTTTGGGAAAGTCCGG 1000
Oligo fwd CACCCTCAGGCCTTTGGGAAAGTC
Oligo rev AAACGACTTTCCCAAAGGCCTGAG
Chromosome start end strand MM target_seq PAM alignment distance position gene name gene id
GU305914.1 211639 211661 - 0 CTCAGGCCTTTGGGAAAGTC CGG ||||||||[||||||||||||]PAM NA Intergenic NA NA

T18547 GGACTTTCGCTTTCGATGTTAGG 1000
Oligo fwd CACCGGACTTTCGCTTTCGATGTT
Oligo rev AAACAACATCGAAAGCGAAAGTCC
Chromosome start end strand MM target_seq PAM alignment distance position gene name gene id
GU305914.1 211618 211640 - 0 GGACTTTCGCTTTCGATGTT AGG ||||||||[||||||||||||]PAM NA Intergenic NA NA

T18548 CTTTCGATGTTAGGTAAACATGG 1000
Oligo fwd CACCCTTTCGATGTTAGGTAAACA
Oligo rev AAACTGTTTACCTAACATCGAAAG
Chromosome start end strand MM target_seq PAM alignment distance position gene name gene id
GU305914.1 211609 211631 - 0 CTTTCGATGTTAGGTAAACA TGG ||||||||[||||||||||||]PAM NA Intergenic NA NA

T18549 ACATAGTATAAAAAGAGCAGAGG 1000
Oligo fwd CACCACATAGTATAAAAAGAGCAG
Oligo rev AAACCTGCTCTTTTTATACTATGT
Chromosome start end strand MM target_seq PAM alignment distance position gene name gene id
GU305914.1 211584 211606 - 0 ACATAGTATAAAAAGAGCAG AGG ||||||||[||||||||||||]PAM NA Intergenic NA NA

T18550 GCTCACAGAGAACTCAGACGCGG 1000
Oligo fwd CACCGCTCACAGAGAACTCAGACG
Oligo rev AAACCGTCTGAGTTCTCTGTGAGC
Chromosome start end strand MM target_seq PAM alignment distance position gene name gene id
GU305914.1 211560 211582 - 0 GCTCACAGAGAACTCAGACG CGG ||||||||[||||||||||||]PAM NA Intergenic NA NA

T18551 CTCACAGAGAACTCAGACGCGGG 1000
Oligo fwd CACCCTCACAGAGAACTCAGACGC
Oligo rev AAACGCGTCTGAGTTCTCTGTGAG
Chromosome start end strand MM target_seq PAM alignment distance position gene name gene id
GU305914.1 211559 211581 - 0 CTCACAGAGAACTCAGACGC GGG ||||||||[||||||||||||]PAM NA Intergenic NA NA

T18552 ACCGCTCCACGTTTCACCGTCGG 1000
Oligo fwd CACCACCGCTCCACGTTTCACCGT
Oligo rev AAACACGGTGAAACGTGGAGCGGT
Chromosome start end strand MM target_seq PAM alignment distance position gene name gene id
GU305914.1 211502 211524 - 0 ACCGCTCCACGTTTCACCGT CGG ||||||||[||||||||||||]PAM NA Intergenic NA NA

| | | | | | | | | | | | |
|---|---|---|---|---|---|---|---|---|---|---|---|
| T18553 | ATCGCTAGCCTGTACAATGCTGG | 1000 | | | | | | | | | |
| Oligo fwd | CACCATCGCTAGCCTGTACAATGC | | | | | | | | | | |
| Oligo rev | AAACGCATTGTACAGGCTAGCGAT | | | | | | | | | | |
| Chromosome | start | end | strand | MM | target_seq | PAM | alignment | distance | position | gene name | gene id |
| GU305914.1 | 211472 | 211494 | - | 0 | ATCGCTAGCCTGTACAATGC | TGG | \|\|\|\|\|\|\|\|[\|\|\|\|\|\|\|\|\|\|\|\|]PAM | NA | Intergenic | NA | NA |
| T18554 | TCGCTAGCCTGTACAATGCTGGG | 1000 | | | | | | | | | |
| Oligo fwd | CACCTCGCTAGCCTGTACAATGCT | | | | | | | | | | |
| Oligo rev | AAACAGCATTGTACAGGCTAGCGA | | | | | | | | | | |
| Chromosome | start | end | strand | MM | target_seq | PAM | alignment | distance | position | gene name | gene id |
| GU305914.1 | 211471 | 211493 | - | 0 | TCGCTAGCCTGTACAATGCT | GGG | \|\|\|\|\|\|\|\|[\|\|\|\|\|\|\|\|\|\|\|\|]PAM | NA | Intergenic | NA | NA |
| T18555 | TATCACCTACTTGCTCCTCTCGG | 1000 | | | | | | | | | |
| Oligo fwd | CACCTATCACCTACTTGCTCCTCT | | | | | | | | | | |
| Oligo rev | AAACAGAGGAGCAAGTAGGTGATA | | | | | | | | | | |
| Chromosome | start | end | strand | MM | target_seq | PAM | alignment | distance | position | gene name | gene id |
| GU305914.1 | 211445 | 211467 | - | 0 | TATCACCTACTTGCTCCTCT | CGG | \|\|\|\|\|\|\|\|[\|\|\|\|\|\|\|\|\|\|\|\|]PAM | NA | Intergenic | NA | NA |
| T18556 | CTCTCGGTTCTCATAAACCGAGG | 1000 | | | | | | | | | |
| Oligo fwd | CACCCTCTCGGTTCTCATAAACCG | | | | | | | | | | |
| Oligo rev | AAACCGGTTTATGAGAACCGAGAG | | | | | | | | | | |
| Chromosome | start | end | strand | MM | target_seq | PAM | alignment | distance | position | gene name | gene id |
| GU305914.1 | 211429 | 211451 | - | 0 | CTCTCGGTTCTCATAAACCG | AGG | \|\|\|\|\|\|\|\|[\|\|\|\|\|\|\|\|\|\|\|\|]PAM | NA | Intergenic | NA | NA |
| T18557 | TCTCATAAACCGAGGCGAGACGG | 1000 | | | | | | | | | |
| Oligo fwd | CACCTCTCATAAACCGAGGCGAGA | | | | | | | | | | |
| Oligo rev | AAACTCTCGCCTCGGTTTATGAGA | | | | | | | | | | |
| Chromosome | start | end | strand | MM | target_seq | PAM | alignment | distance | position | gene name | gene id |
| GU305914.1 | 211421 | 211443 | - | 0 | TCTCATAAACCGAGGCGAGA | CGG | \|\|\|\|\|\|\|\|[\|\|\|\|\|\|\|\|\|\|\|\|]PAM | NA | Intergenic | NA | NA |
| T18558 | CATAAACCGAGGCGAGACGGCGG | 1000 | | | | | | | | | |
| Oligo fwd | CACCCATAAACCGAGGCGAGACGG | | | | | | | | | | |
| Oligo rev | AAACCCGTCTCGCCTCGGTTTATG | | | | | | | | | | |
| Chromosome | start | end | strand | MM | target_seq | PAM | alignment | distance | position | gene name | gene id |
| GU305914.1 | 211418 | 211440 | - | 0 | CATAAACCGAGGCGAGACGG | CGG | \|\|\|\|\|\|\|\|[\|\|\|\|\|\|\|\|\|\|\|\|]PAM | NA | Intergenic | NA | NA |
| T18559 | ATAAACCGAGGCGAGACGGCGGG | 1000 | | | | | | | | | |
| Oligo fwd | CACCATAAACCGAGGCGAGACGGC | | | | | | | | | | |
| Oligo rev | AAACGCCGTCTCGCCTCGGTTTAT | | | | | | | | | | |
| Chromosome | start | end | strand | MM | target_seq | PAM | alignment | distance | position | gene name | gene id |
| GU305914.1 | 211417 | 211439 | - | 0 | ATAAACCGAGGCGAGACGGC | GGG | \|\|\|\|\|\|\|\|[\|\|\|\|\|\|\|\|\|\|\|\|]PAM | NA | Intergenic | NA | NA |
| T18560 | GACGGCGGGCAGCAGCTATATGG | 1000 | | | | | | | | | |
| Oligo fwd | CACCGACGGCGGGCAGCAGCTATA | | | | | | | | | | |
| Oligo rev | AAACTATAGCTGCTGCCCGCCGTC | | | | | | | | | | |
| Chromosome | start | end | strand | MM | target_seq | PAM | alignment | distance | position | gene name | gene id |
| GU305914.1 | 211403 | 211425 | - | 0 | GACGGCGGGCAGCAGCTATA | TGG | \|\|\|\|\|\|\|\|[\|\|\|\|\|\|\|\|\|\|\|\|]PAM | NA | Intergenic | NA | NA |
| T18561 | TATGGACGTGCGCATATTCGAGG | 1000 | | | | | | | | | |
| Oligo fwd | CACCTATGGACGTGCGCATATTCG | | | | | | | | | | |
| Oligo rev | AAACCGAATATGCGCACGTCCATA | | | | | | | | | | |
| Chromosome | start | end | strand | MM | target_seq | PAM | alignment | distance | position | gene name | gene id |
| GU305914.1 | 211385 | 211407 | - | 0 | TATGGACGTGCGCATATTCG | AGG | \|\|\|\|\|\|\|\|[\|\|\|\|\|\|\|\|\|\|\|\|]PAM | NA | Intergenic | NA | NA |
| T18562 | TGCGCATATTCGAGGATGAGCGG | 1000 | | | | | | | | | |
| Oligo fwd | CACCTGCGCATATTCGAGGATGAG | | | | | | | | | | |
| Oligo rev | AAACCTCATCCTCGAATATGCGCA | | | | | | | | | | |
| Chromosome | start | end | strand | MM | target_seq | PAM | alignment | distance | position | gene name | gene id |
| GU305914.1 | 211377 | 211399 | - | 0 | TGCGCATATTCGAGGATGAG | CGG | \|\|\|\|\|\|\|\|[\|\|\|\|\|\|\|\|\|\|\|\|]PAM | NA | Intergenic | NA | NA |
| T18563 | GCGCATATTCGAGGATGAGCGGG | 1000 | | | | | | | | | |
| Oligo fwd | CACCGCGCATATTCGAGGATGAGC | | | | | | | | | | |
| Oligo rev | AAACGCTCATCCTCGAATATGCGC | | | | | | | | | | |
| Chromosome | start | end | strand | MM | target_seq | PAM | alignment | distance | position | gene name | gene id |
| GU305914.1 | 211376 | 211398 | - | 0 | GCGCATATTCGAGGATGAGC | GGG | \|\|\|\|\|\|\|\|[\|\|\|\|\|\|\|\|\|\|\|\|]PAM | NA | Intergenic | NA | NA |
| T18564 | CATATTCGAGGATGAGCGGGTGG | 1000 | | | | | | | | | |
| Oligo fwd | CACCCATATTCGAGGATGAGCGGG | | | | | | | | | | |
| Oligo rev | AAACCCCGCTCATCCTCGAATATG | | | | | | | | | | |
| Chromosome | start | end | strand | MM | target_seq | PAM | alignment | distance | position | gene name | gene id |
| GU305914.1 | 211373 | 211395 | - | 0 | CATATTCGAGGATGAGCGGG | TGG | \|\|\|\|\|\|\|\|[\|\|\|\|\|\|\|\|\|\|\|\|]PAM | NA | Intergenic | NA | NA |
| T18565 | GGACATCTGTCAAGACCTGACGG | 1000 | | | | | | | | | |
| Oligo fwd | CACCGGACATCTGTCAAGACCTGA | | | | | | | | | | |
| Oligo rev | AAACTCAGGTCTTGACAGATGTCC | | | | | | | | | | |
| Chromosome | start | end | strand | MM | target_seq | PAM | alignment | distance | position | gene name | gene id |
| GU305914.1 | 211352 | 211374 | - | 0 | GGACATCTGTCAAGACCTGA | CGG | \|\|\|\|\|\|\|\|[\|\|\|\|\|\|\|\|\|\|\|\|]PAM | NA | Intergenic | NA | NA |
| T18566 | ACGTTCATCTCGTACAGAGAAGG | 1000 | | | | | | | | | |
| Oligo fwd | CACCACGTTCATCTCGTACAGAGA | | | | | | | | | | |
| Oligo rev | AAACTCTCTGTACGAGATGAACGT | | | | | | | | | | |
| Chromosome | start | end | strand | MM | target_seq | PAM | alignment | distance | position | gene name | gene id |
| GU305914.1 | 211327 | 211349 | - | 0 | ACGTTCATCTCGTACAGAGA | AGG | \|\|\|\|\|\|\|\|[\|\|\|\|\|\|\|\|\|\|\|\|]PAM | NA | Intergenic | NA | NA |
| T18567 | CATCTCGTACAGAGAAGGTCCGG | 1000 | | | | | | | | | |
| Oligo fwd | CACCCATCTCGTACAGAGAAGGTC | | | | | | | | | | |
| Oligo rev | AAACGACCTTCTCTGTACGAGATG | | | | | | | | | | |
| Chromosome | start | end | strand | MM | target_seq | PAM | alignment | distance | position | gene name | gene id |

<table>
<tr><td>GU305914.1</td><td>211322</td><td>211344</td><td>-</td><td>0</td><td>CATCTCGTACAGAGAAGGTC</td><td>CGG</td><td>||||||||[||||||||||||]PAM</td><td>NA</td><td>Intergenic</td><td>NA</td><td>NA</td></tr>
<tr><td>T18568</td><td>AGCAGTCGTCTGATATCTTTCGG</td><td>1000</td><td></td><td></td><td></td><td></td><td></td><td></td><td></td><td></td><td></td></tr>
<tr><td>Oligo fwd</td><td>CACCAGCAGTCGTCTGATATCTTT</td><td></td><td></td><td></td><td></td><td></td><td></td><td></td><td></td><td></td><td></td></tr>
<tr><td>Oligo rev</td><td>AAACAAAGATATCAGACGACTGCT</td><td></td><td></td><td></td><td></td><td></td><td></td><td></td><td></td><td></td><td></td></tr>
<tr><td>Chromosome</td><td>start</td><td>end</td><td>strand</td><td>MM</td><td>target_seq</td><td>PAM</td><td>alignment</td><td>distance</td><td>position</td><td>gene name</td><td>gene id</td></tr>
<tr><td>GU305914.1</td><td>211272</td><td>211294</td><td>-</td><td>0</td><td>AGCAGTCGTCTGATATCTTT</td><td>CGG</td><td>||||||||[||||||||||||]PAM</td><td>NA</td><td>Intergenic</td><td>NA</td><td>NA</td></tr>
<tr><td>T18569</td><td>ATCTTTCGGATCGAAGCCTCCGG</td><td>1000</td><td></td><td></td><td></td><td></td><td></td><td></td><td></td><td></td><td></td></tr>
<tr><td>Oligo fwd</td><td>CACCATCTTTCGGATCGAAGCCTC</td><td></td><td></td><td></td><td></td><td></td><td></td><td></td><td></td><td></td><td></td></tr>
<tr><td>Oligo rev</td><td>AAACGAGGCTTCGATCCGAAAGAT</td><td></td><td></td><td></td><td></td><td></td><td></td><td></td><td></td><td></td><td></td></tr>
<tr><td>Chromosome</td><td>start</td><td>end</td><td>strand</td><td>MM</td><td>target_seq</td><td>PAM</td><td>alignment</td><td>distance</td><td>position</td><td>gene name</td><td>gene id</td></tr>
<tr><td>GU305914.1</td><td>211258</td><td>211280</td><td>-</td><td>0</td><td>ATCTTTCGGATCGAAGCCTC</td><td>CGG</td><td>||||||||[||||||||||||]PAM</td><td>NA</td><td>Intergenic</td><td>NA</td><td>NA</td></tr>
<tr><td>T18570</td><td>TCGGATCGAAGCCTCCGGAGAGG</td><td>1000</td><td></td><td></td><td></td><td></td><td></td><td></td><td></td><td></td><td></td></tr>
<tr><td>Oligo fwd</td><td>CACCTCGGATCGAAGCCTCCGGAG</td><td></td><td></td><td></td><td></td><td></td><td></td><td></td><td></td><td></td><td></td></tr>
<tr><td>Oligo rev</td><td>AAACCTCCGGAGGCTTCGATCCGA</td><td></td><td></td><td></td><td></td><td></td><td></td><td></td><td></td><td></td><td></td></tr>
<tr><td>Chromosome</td><td>start</td><td>end</td><td>strand</td><td>MM</td><td>target_seq</td><td>PAM</td><td>alignment</td><td>distance</td><td>position</td><td>gene name</td><td>gene id</td></tr>
<tr><td>GU305914.1</td><td>211253</td><td>211275</td><td>-</td><td>0</td><td>TCGGATCGAAGCCTCCGGAG</td><td>AGG</td><td>||||||||[||||||||||||]PAM</td><td>NA</td><td>Intergenic</td><td>NA</td><td>NA</td></tr>
<tr><td>T18571</td><td>GAGAGGTGAAACATTTTCCTTGG</td><td>1000</td><td></td><td></td><td></td><td></td><td></td><td></td><td></td><td></td><td></td></tr>
<tr><td>Oligo fwd</td><td>CACCGAGAGGTGAAACATTTTCCT</td><td></td><td></td><td></td><td></td><td></td><td></td><td></td><td></td><td></td><td></td></tr>
<tr><td>Oligo rev</td><td>AAACAGGAAAATGTTTCACCTCTC</td><td></td><td></td><td></td><td></td><td></td><td></td><td></td><td></td><td></td><td></td></tr>
<tr><td>Chromosome</td><td>start</td><td>end</td><td>strand</td><td>MM</td><td>target_seq</td><td>PAM</td><td>alignment</td><td>distance</td><td>position</td><td>gene name</td><td>gene id</td></tr>
<tr><td>GU305914.1</td><td>211236</td><td>211258</td><td>-</td><td>0</td><td>GAGAGGTGAAACATTTTCCT</td><td>TGG</td><td>||||||||[||||||||||||]PAM</td><td>NA</td><td>Intergenic</td><td>NA</td><td>NA</td></tr>
<tr><td>T18572</td><td>TTGGATGAACGTGAGCGAGCTGG</td><td>1000</td><td></td><td></td><td></td><td></td><td></td><td></td><td></td><td></td><td></td></tr>
<tr><td>Oligo fwd</td><td>CACCTTGGATGAACGTGAGCGAGC</td><td></td><td></td><td></td><td></td><td></td><td></td><td></td><td></td><td></td><td></td></tr>
<tr><td>Oligo rev</td><td>AAACGCTCGCTCACGTTCATCCAA</td><td></td><td></td><td></td><td></td><td></td><td></td><td></td><td></td><td></td><td></td></tr>
<tr><td>Chromosome</td><td>start</td><td>end</td><td>strand</td><td>MM</td><td>target_seq</td><td>PAM</td><td>alignment</td><td>distance</td><td>position</td><td>gene name</td><td>gene id</td></tr>
<tr><td>GU305914.1</td><td>211217</td><td>211239</td><td>-</td><td>0</td><td>TTGGATGAACGTGAGCGAGC</td><td>TGG</td><td>||||||||[||||||||||||]PAM</td><td>NA</td><td>Intergenic</td><td>NA</td><td>NA</td></tr>
<tr><td>T18573</td><td>GAACGTGAGCGAGCTGGCGCAGG</td><td>1000</td><td></td><td></td><td></td><td></td><td></td><td></td><td></td><td></td><td></td></tr>
<tr><td>Oligo fwd</td><td>CACCGAACGTGAGCGAGCTGGCGC</td><td></td><td></td><td></td><td></td><td></td><td></td><td></td><td></td><td></td><td></td></tr>
<tr><td>Oligo rev</td><td>AAACGCGCCAGCTCGCTCACGTTC</td><td></td><td></td><td></td><td></td><td></td><td></td><td></td><td></td><td></td><td></td></tr>
<tr><td>Chromosome</td><td>start</td><td>end</td><td>strand</td><td>MM</td><td>target_seq</td><td>PAM</td><td>alignment</td><td>distance</td><td>position</td><td>gene name</td><td>gene id</td></tr>
<tr><td>GU305914.1</td><td>211211</td><td>211233</td><td>-</td><td>0</td><td>GAACGTGAGCGAGCTGGCGC</td><td>AGG</td><td>||||||||[||||||||||||]PAM</td><td>NA</td><td>Intergenic</td><td>NA</td><td>NA</td></tr>
<tr><td>T18574</td><td>GCAGGAGAGTGCGTTCTTCGTGG</td><td>1000</td><td></td><td></td><td></td><td></td><td></td><td></td><td></td><td></td><td></td></tr>
<tr><td>Oligo fwd</td><td>CACCGCAGGAGAGTGCGTTCTTCG</td><td></td><td></td><td></td><td></td><td></td><td></td><td></td><td></td><td></td><td></td></tr>
<tr><td>Oligo rev</td><td>AAACCGAAGAACGCACTCTCCTGC</td><td></td><td></td><td></td><td></td><td></td><td></td><td></td><td></td><td></td><td></td></tr>
<tr><td>Chromosome</td><td>start</td><td>end</td><td>strand</td><td>MM</td><td>target_seq</td><td>PAM</td><td>alignment</td><td>distance</td><td>position</td><td>gene name</td><td>gene id</td></tr>
<tr><td>GU305914.1</td><td>211193</td><td>211215</td><td>-</td><td>0</td><td>GCAGGAGAGTGCGTTCTTCG</td><td>TGG</td><td>||||||||[||||||||||||]PAM</td><td>NA</td><td>Intergenic</td><td>NA</td><td>NA</td></tr>
<tr><td>T18575</td><td>GAGTGCGTTCTTCGTGGAGCAGG</td><td>1000</td><td></td><td></td><td></td><td></td><td></td><td></td><td></td><td></td><td></td></tr>
<tr><td>Oligo fwd</td><td>CACCGAGTGCGTTCTTCGTGGAGC</td><td></td><td></td><td></td><td></td><td></td><td></td><td></td><td></td><td></td><td></td></tr>
<tr><td>Oligo rev</td><td>AAACGCTCCACGAAGAACGCACTC</td><td></td><td></td><td></td><td></td><td></td><td></td><td></td><td></td><td></td><td></td></tr>
<tr><td>Chromosome</td><td>start</td><td>end</td><td>strand</td><td>MM</td><td>target_seq</td><td>PAM</td><td>alignment</td><td>distance</td><td>position</td><td>gene name</td><td>gene id</td></tr>
<tr><td>GU305914.1</td><td>211187</td><td>211209</td><td>-</td><td>0</td><td>GAGTGCGTTCTTCGTGGAGC</td><td>AGG</td><td>||||||||[||||||||||||]PAM</td><td>NA</td><td>Intergenic</td><td>NA</td><td>NA</td></tr>
<tr><td>T18576</td><td>CGTTCTTCGTGGAGCAGGAGAGG</td><td>1000</td><td></td><td></td><td></td><td></td><td></td><td></td><td></td><td></td><td></td></tr>
<tr><td>Oligo fwd</td><td>CACCCGTTCTTCGTGGAGCAGGAG</td><td></td><td></td><td></td><td></td><td></td><td></td><td></td><td></td><td></td><td></td></tr>
<tr><td>Oligo rev</td><td>AAACCTCCTGCTCCACGAAGAACG</td><td></td><td></td><td></td><td></td><td></td><td></td><td></td><td></td><td></td><td></td></tr>
<tr><td>Chromosome</td><td>start</td><td>end</td><td>strand</td><td>MM</td><td>target_seq</td><td>PAM</td><td>alignment</td><td>distance</td><td>position</td><td>gene name</td><td>gene id</td></tr>
<tr><td>GU305914.1</td><td>211182</td><td>211204</td><td>-</td><td>0</td><td>CGTTCTTCGTGGAGCAGGAG</td><td>AGG</td><td>||||||||[||||||||||||]PAM</td><td>NA</td><td>Intergenic</td><td>NA</td><td>NA</td></tr>
<tr><td>T18577</td><td>ATTATGAATGTCTTCAAAGCCGG</td><td>1000</td><td></td><td></td><td></td><td></td><td></td><td></td><td></td><td></td><td></td></tr>
<tr><td>Oligo fwd</td><td>CACCATTATGAATGTCTTCAAAGC</td><td></td><td></td><td></td><td></td><td></td><td></td><td></td><td></td><td></td><td></td></tr>
<tr><td>Oligo rev</td><td>AAACGCTTTGAAGACATTCATAAT</td><td></td><td></td><td></td><td></td><td></td><td></td><td></td><td></td><td></td><td></td></tr>
<tr><td>Chromosome</td><td>start</td><td>end</td><td>strand</td><td>MM</td><td>target_seq</td><td>PAM</td><td>alignment</td><td>distance</td><td>position</td><td>gene name</td><td>gene id</td></tr>
<tr><td>GU305914.1</td><td>211144</td><td>211166</td><td>-</td><td>0</td><td>ATTATGAATGTCTTCAAAGC</td><td>CGG</td><td>||||||||[||||||||||||]PAM</td><td>NA</td><td>Intergenic</td><td>NA</td><td>NA</td></tr>
<tr><td>T18578</td><td>TGAATGTCTTCAAAGCCGGACGG</td><td>1000</td><td></td><td></td><td></td><td></td><td></td><td></td><td></td><td></td><td></td></tr>
<tr><td>Oligo fwd</td><td>CACCTGAATGTCTTCAAAGCCGGA</td><td></td><td></td><td></td><td></td><td></td><td></td><td></td><td></td><td></td><td></td></tr>
<tr><td>Oligo rev</td><td>AAACTCCGGCTTTGAAGACATTCA</td><td></td><td></td><td></td><td></td><td></td><td></td><td></td><td></td><td></td><td></td></tr>
<tr><td>Chromosome</td><td>start</td><td>end</td><td>strand</td><td>MM</td><td>target_seq</td><td>PAM</td><td>alignment</td><td>distance</td><td>position</td><td>gene name</td><td>gene id</td></tr>
<tr><td>GU305914.1</td><td>211140</td><td>211162</td><td>-</td><td>0</td><td>TGAATGTCTTCAAAGCCGGA</td><td>CGG</td><td>||||||||[||||||||||||]PAM</td><td>NA</td><td>Intergenic</td><td>NA</td><td>NA</td></tr>
<tr><td>T18579</td><td>TGTCTTCAAAGCCGGACGGCCGG</td><td>1000</td><td></td><td></td><td></td><td></td><td></td><td></td><td></td><td></td><td></td></tr>
<tr><td>Oligo fwd</td><td>CACCTGTCTTCAAAGCCGGACGGC</td><td></td><td></td><td></td><td></td><td></td><td></td><td></td><td></td><td></td><td></td></tr>
<tr><td>Oligo rev</td><td>AAACGCCGTCCGGCTTTGAAGACA</td><td></td><td></td><td></td><td></td><td></td><td></td><td></td><td></td><td></td><td></td></tr>
<tr><td>Chromosome</td><td>start</td><td>end</td><td>strand</td><td>MM</td><td>target_seq</td><td>PAM</td><td>alignment</td><td>distance</td><td>position</td><td>gene name</td><td>gene id</td></tr>
<tr><td>GU305914.1</td><td>211136</td><td>211158</td><td>-</td><td>0</td><td>TGTCTTCAAAGCCGGACGGC</td><td>CGG</td><td>||||||||[||||||||||||]PAM</td><td>NA</td><td>Intergenic</td><td>NA</td><td>NA</td></tr>
<tr><td>T18580</td><td>ATGTACCGTACTTCAGATGATGG</td><td>1000</td><td></td><td></td><td></td><td></td><td></td><td></td><td></td><td></td><td></td></tr>
<tr><td>Oligo fwd</td><td>CACCATGTACCGTACTTCAGATGA</td><td></td><td></td><td></td><td></td><td></td><td></td><td></td><td></td><td></td><td></td></tr>
<tr><td>Oligo rev</td><td>AAACTCATCTGAAGTACGGTACAT</td><td></td><td></td><td></td><td></td><td></td><td></td><td></td><td></td><td></td><td></td></tr>
<tr><td>Chromosome</td><td>start</td><td>end</td><td>strand</td><td>MM</td><td>target_seq</td><td>PAM</td><td>alignment</td><td>distance</td><td>position</td><td>gene name</td><td>gene id</td></tr>
<tr><td>GU305914.1</td><td>211076</td><td>211098</td><td>-</td><td>0</td><td>ATGTACCGTACTTCAGATGA</td><td>TGG</td><td>||||||||[||||||||||||]PAM</td><td>NA</td><td>Intergenic</td><td>NA</td><td>NA</td></tr>
<tr><td>T18581</td><td>ACCGTACTTCAGATGATGGACGG</td><td>1000</td><td></td><td></td><td></td><td></td><td></td><td></td><td></td><td></td><td></td></tr>
<tr><td>Oligo fwd</td><td>CACCACCGTACTTCAGATGATGGA</td><td></td><td></td><td></td><td></td><td></td><td></td><td></td><td></td><td></td><td></td></tr>
<tr><td>Oligo rev</td><td>AAACTCCATCATCTGAAGTACGGT</td><td></td><td></td><td></td><td></td><td></td><td></td><td></td><td></td><td></td><td></td></tr>
<tr><td>Chromosome</td><td>start</td><td>end</td><td>strand</td><td>MM</td><td>target_seq</td><td>PAM</td><td>alignment</td><td>distance</td><td>position</td><td>gene name</td><td>gene id</td></tr>
<tr><td>GU305914.1</td><td>211072</td><td>211094</td><td>-</td><td>0</td><td>ACCGTACTTCAGATGATGGA</td><td>CGG</td><td>||||||||[||||||||||||]PAM</td><td>NA</td><td>Intergenic</td><td>NA</td><td>NA</td></tr>
<tr><td>T18582</td><td>ACACGTTACACCGTAGACAAAGG</td><td>1000</td><td></td><td></td><td></td><td></td><td></td><td></td><td></td><td></td><td></td></tr>
<tr><td>Oligo fwd</td><td>CACCACACGTTACACCGTAGACAA</td><td></td><td></td><td></td><td></td><td></td><td></td><td></td><td></td><td></td><td></td></tr>
<tr><td>Oligo rev</td><td>AAACTTGTCTACGGTGTAACGTGT</td><td></td><td></td><td></td><td></td><td></td><td></td><td></td><td></td><td></td><td></td></tr>
</table>

<table>
<tr><td>Chromosome</td><td>start</td><td>end</td><td>strand</td><td>MM</td><td>target_seq</td><td>PAM</td><td>alignment</td><td>distance</td><td>position</td><td>gene name</td><td>gene id</td></tr>
<tr><td>GU305914.1</td><td>211039</td><td>211061</td><td>-</td><td>0</td><td>ACACGTTACACCGTAGACAA</td><td>AGG</td><td>||||||||[||||||||||||]PAM</td><td>NA</td><td>Intergenic</td><td>NA</td><td>NA</td></tr>
<tr><td>T18583</td><td>GACAAAGGCGTCGAAACGCTCGG</td><td>1000</td></tr>
<tr><td>Oligo fwd</td><td>CACCGACAAAGGCGTCGAAACGCT</td></tr>
<tr><td>Oligo rev</td><td>AAACAGCGTTTCGACGCCTTTGTC</td></tr>
<tr><td>Chromosome</td><td>start</td><td>end</td><td>strand</td><td>MM</td><td>target_seq</td><td>PAM</td><td>alignment</td><td>distance</td><td>position</td><td>gene name</td><td>gene id</td></tr>
<tr><td>GU305914.1</td><td>211024</td><td>211046</td><td>-</td><td>0</td><td>GACAAAGGCGTCGAAACGCT</td><td>CGG</td><td>||||||||[||||||||||||]PAM</td><td>NA</td><td>Intergenic</td><td>NA</td><td>NA</td></tr>
<tr><td>T18584</td><td>ACAAAGGCGTCGAAACGCTCGGG</td><td>1000</td></tr>
<tr><td>Oligo fwd</td><td>CACCACAAAGGCGTCGAAACGCTC</td></tr>
<tr><td>Oligo rev</td><td>AAACGAGCGTTTCGACGCCTTTGT</td></tr>
<tr><td>Chromosome</td><td>start</td><td>end</td><td>strand</td><td>MM</td><td>target_seq</td><td>PAM</td><td>alignment</td><td>distance</td><td>position</td><td>gene name</td><td>gene id</td></tr>
<tr><td>GU305914.1</td><td>211023</td><td>211045</td><td>-</td><td>0</td><td>ACAAAGGCGTCGAAACGCTC</td><td>GGG</td><td>||||||||[||||||||||||]PAM</td><td>NA</td><td>Intergenic</td><td>NA</td><td>NA</td></tr>
<tr><td>T18585</td><td>GTACTCTCCCGACGTATCCGAGG</td><td>1000</td></tr>
<tr><td>Oligo fwd</td><td>CACCGTACTCTCCCGACGTATCCG</td></tr>
<tr><td>Oligo rev</td><td>AAACCGGATACGTCGGGAGAGTAC</td></tr>
<tr><td>Chromosome</td><td>start</td><td>end</td><td>strand</td><td>MM</td><td>target_seq</td><td>PAM</td><td>alignment</td><td>distance</td><td>position</td><td>gene name</td><td>gene id</td></tr>
<tr><td>GU305914.1</td><td>210992</td><td>211014</td><td>-</td><td>0</td><td>GTACTCTCCCGACGTATCCG</td><td>AGG</td><td>||||||||[||||||||||||]PAM</td><td>NA</td><td>Intergenic</td><td>NA</td><td>NA</td></tr>
<tr><td>T18586</td><td>GAGGATGACATCGCGCGCTACGG</td><td>1000</td></tr>
<tr><td>Oligo fwd</td><td>CACCGAGGATGACATCGCGCGCTA</td></tr>
<tr><td>Oligo rev</td><td>AAACTAGCGCGCGATGTCATCCTC</td></tr>
<tr><td>Chromosome</td><td>start</td><td>end</td><td>strand</td><td>MM</td><td>target_seq</td><td>PAM</td><td>alignment</td><td>distance</td><td>position</td><td>gene name</td><td>gene id</td></tr>
<tr><td>GU305914.1</td><td>210973</td><td>210995</td><td>-</td><td>0</td><td>GAGGATGACATCGCGCGCTA</td><td>CGG</td><td>||||||||[||||||||||||]PAM</td><td>NA</td><td>Intergenic</td><td>NA</td><td>NA</td></tr>
<tr><td>T18587</td><td>GACATCGCGCGCTACGGACAAGG</td><td>1000</td></tr>
<tr><td>Oligo fwd</td><td>CACCGACATCGCGCGCTACGGACA</td></tr>
<tr><td>Oligo rev</td><td>AAACTGTCCGTAGCGCGCGATGTC</td></tr>
<tr><td>Chromosome</td><td>start</td><td>end</td><td>strand</td><td>MM</td><td>target_seq</td><td>PAM</td><td>alignment</td><td>distance</td><td>position</td><td>gene name</td><td>gene id</td></tr>
<tr><td>GU305914.1</td><td>210967</td><td>210989</td><td>-</td><td>0</td><td>GACATCGCGCGCTACGGACA</td><td>AGG</td><td>||||||||[||||||||||||]PAM</td><td>NA</td><td>Intergenic</td><td>NA</td><td>NA</td></tr>
<tr><td>T18588</td><td>ACATCGCGCGCTACGGACAAGGG</td><td>1000</td></tr>
<tr><td>Oligo fwd</td><td>CACCACATCGCGCGCTACGGACAA</td></tr>
<tr><td>Oligo rev</td><td>AAACTTGTCCGTAGCGCGCGATGT</td></tr>
<tr><td>Chromosome</td><td>start</td><td>end</td><td>strand</td><td>MM</td><td>target_seq</td><td>PAM</td><td>alignment</td><td>distance</td><td>position</td><td>gene name</td><td>gene id</td></tr>
<tr><td>GU305914.1</td><td>210966</td><td>210988</td><td>-</td><td>0</td><td>ACATCGCGCGCTACGGACAA</td><td>GGG</td><td>||||||||[||||||||||||]PAM</td><td>NA</td><td>Intergenic</td><td>NA</td><td>NA</td></tr>
<tr><td>T18589</td><td>GCGCGCTACGGACAAGGGTCCGG</td><td>1000</td></tr>
<tr><td>Oligo fwd</td><td>CACCGCGCGCTACGGACAAGGGTC</td></tr>
<tr><td>Oligo rev</td><td>AAACGACCCTTGTCCGTAGCGCGC</td></tr>
<tr><td>Chromosome</td><td>start</td><td>end</td><td>strand</td><td>MM</td><td>target_seq</td><td>PAM</td><td>alignment</td><td>distance</td><td>position</td><td>gene name</td><td>gene id</td></tr>
<tr><td>GU305914.1</td><td>210961</td><td>210983</td><td>-</td><td>0</td><td>GCGCGCTACGGACAAGGGTC</td><td>CGG</td><td>||||||||[||||||||||||]PAM</td><td>NA</td><td>Intergenic</td><td>NA</td><td>NA</td></tr>
<tr><td>T18590</td><td>GGTCCGGAATCTCTATCTTGAGG</td><td>1000</td></tr>
<tr><td>Oligo fwd</td><td>CACCGGTCCGGAATCTCTATCTTG</td></tr>
<tr><td>Oligo rev</td><td>AAACCAAGATAGAGATTCCGGACC</td></tr>
<tr><td>Chromosome</td><td>start</td><td>end</td><td>strand</td><td>MM</td><td>target_seq</td><td>PAM</td><td>alignment</td><td>distance</td><td>position</td><td>gene name</td><td>gene id</td></tr>
<tr><td>GU305914.1</td><td>210945</td><td>210967</td><td>-</td><td>0</td><td>GGTCCGGAATCTCTATCTTG</td><td>AGG</td><td>||||||||[||||||||||||]PAM</td><td>NA</td><td>Intergenic</td><td>NA</td><td>NA</td></tr>
<tr><td>T18591</td><td>GTCCGGAATCTCTATCTTGAGGG</td><td>1000</td></tr>
<tr><td>Oligo fwd</td><td>CACCGTCCGGAATCTCTATCTTGA</td></tr>
<tr><td>Oligo rev</td><td>AAACTCAAGATAGAGATTCCGGAC</td></tr>
<tr><td>Chromosome</td><td>start</td><td>end</td><td>strand</td><td>MM</td><td>target_seq</td><td>PAM</td><td>alignment</td><td>distance</td><td>position</td><td>gene name</td><td>gene id</td></tr>
<tr><td>GU305914.1</td><td>210944</td><td>210966</td><td>-</td><td>0</td><td>GTCCGGAATCTCTATCTTGA</td><td>GGG</td><td>||||||||[||||||||||||]PAM</td><td>NA</td><td>Intergenic</td><td>NA</td><td>NA</td></tr>
<tr><td>T18592</td><td>CTGCTCTACTCCAGAAACGCTGG</td><td>1000</td></tr>
<tr><td>Oligo fwd</td><td>CACCCTGCTCTACTCCAGAAACGC</td></tr>
<tr><td>Oligo rev</td><td>AAACGCGTTTCTGGAGTAGAGCAG</td></tr>
<tr><td>Chromosome</td><td>start</td><td>end</td><td>strand</td><td>MM</td><td>target_seq</td><td>PAM</td><td>alignment</td><td>distance</td><td>position</td><td>gene name</td><td>gene id</td></tr>
<tr><td>GU305914.1</td><td>210915</td><td>210937</td><td>-</td><td>0</td><td>CTGCTCTACTCCAGAAACGC</td><td>TGG</td><td>||||||||[||||||||||||]PAM</td><td>NA</td><td>Intergenic</td><td>NA</td><td>NA</td></tr>
<tr><td>T18593</td><td>AACGCTGGACGTCCTTCTGTCGG</td><td>1000</td></tr>
<tr><td>Oligo fwd</td><td>CACCAACGCTGGACGTCCTTCTGT</td></tr>
<tr><td>Oligo rev</td><td>AAACACAGAAGGACGTCCAGCGTT</td></tr>
<tr><td>Chromosome</td><td>start</td><td>end</td><td>strand</td><td>MM</td><td>target_seq</td><td>PAM</td><td>alignment</td><td>distance</td><td>position</td><td>gene name</td><td>gene id</td></tr>
<tr><td>GU305914.1</td><td>210900</td><td>210922</td><td>-</td><td>0</td><td>AACGCTGGACGTCCTTCTGT</td><td>CGG</td><td>||||||||[||||||||||||]PAM</td><td>NA</td><td>Intergenic</td><td>NA</td><td>NA</td></tr>
<tr><td>T18594</td><td>CTGTCGGAAGATCGTCGCCATGG</td><td>1000</td></tr>
<tr><td>Oligo fwd</td><td>CACCCTGTCGGAAGATCGTCGCCA</td></tr>
<tr><td>Oligo rev</td><td>AAACTGGCGACGATCTTCCGACAG</td></tr>
<tr><td>Chromosome</td><td>start</td><td>end</td><td>strand</td><td>MM</td><td>target_seq</td><td>PAM</td><td>alignment</td><td>distance</td><td>position</td><td>gene name</td><td>gene id</td></tr>
<tr><td>GU305914.1</td><td>210884</td><td>210906</td><td>-</td><td>0</td><td>CTGTCGGAAGATCGTCGCCA</td><td>TGG</td><td>||||||||[||||||||||||]PAM</td><td>NA</td><td>Intergenic</td><td>NA</td><td>NA</td></tr>
<tr><td>T18595</td><td>TACTCGCTGTACAGTAATCGAGG</td><td>1000</td></tr>
<tr><td>Oligo fwd</td><td>CACCTACTCGCTGTACAGTAATCG</td></tr>
<tr><td>Oligo rev</td><td>AAACCGATTACTGTACAGCGAGTA</td></tr>
<tr><td>Chromosome</td><td>start</td><td>end</td><td>strand</td><td>MM</td><td>target_seq</td><td>PAM</td><td>alignment</td><td>distance</td><td>position</td><td>gene name</td><td>gene id</td></tr>
<tr><td>GU305914.1</td><td>210841</td><td>210863</td><td>-</td><td>0</td><td>TACTCGCTGTACAGTAATCG</td><td>AGG</td><td>||||||||[||||||||||||]PAM</td><td>NA</td><td>Intergenic</td><td>NA</td><td>NA</td></tr>
<tr><td>T18596</td><td>CTGTACAGTAATCGAGGCAACGG</td><td>1000</td></tr>
<tr><td>Oligo fwd</td><td>CACCCTGTACAGTAATCGAGGCAA</td></tr>
<tr><td>Oligo rev</td><td>AAACTTGCCTCGATTACTGTACAG</td></tr>
<tr><td>Chromosome</td><td>start</td><td>end</td><td>strand</td><td>MM</td><td>target_seq</td><td>PAM</td><td>alignment</td><td>distance</td><td>position</td><td>gene name</td><td>gene id</td></tr>
<tr><td>GU305914.1</td><td>210835</td><td>210857</td><td>-</td><td>0</td><td>CTGTACAGTAATCGAGGCAA</td><td>CGG</td><td>||||||||[||||||||||||]PAM</td><td>NA</td><td>Intergenic</td><td>NA</td><td>NA</td></tr>
<tr><td>T18597</td><td>GTCCTGTACGATGCGCACTCAGG</td><td>1000</td></tr>
<tr><td>Oligo fwd</td><td>CACCGTCCTGTACGATGCGCACTC</td></tr>
</table>

<table>
<tr><td>Oligo rev</td><td>AAACGAGTGCGCATCGTACAGGAC</td><td></td><td></td><td></td><td></td><td></td><td></td><td></td><td></td><td></td><td></td><td></td></tr>
<tr><td>Chromosome</td><td>start</td><td></td><td>end</td><td>strand</td><td>MM</td><td>target_seq</td><td>PAM</td><td>alignment</td><td>distance</td><td>position</td><td>gene name</td><td>gene id</td></tr>
<tr><td>GU305914.1</td><td></td><td>210806</td><td>210828</td><td>-</td><td>0</td><td>GTCCTGTACGATGCGCACTC</td><td>AGG</td><td>||||||||[||||||||||||]PAM</td><td>NA</td><td>Intergenic</td><td>NA</td><td>NA</td></tr>
<tr><td></td><td></td><td></td><td></td><td></td><td></td><td></td><td></td><td></td><td></td><td></td><td></td><td></td></tr>
<tr><td>T18598</td><td>GATGCGCACTCAGGTTCCGTTGG</td><td></td><td>1000</td><td></td><td></td><td></td><td></td><td></td><td></td><td></td><td></td><td></td></tr>
<tr><td>Oligo fwd</td><td>CACCGATGCGCACTCAGGTTCCGT</td><td></td><td></td><td></td><td></td><td></td><td></td><td></td><td></td><td></td><td></td><td></td></tr>
<tr><td>Oligo rev</td><td>AAACACGGAACCTGAGTGCGCATC</td><td></td><td></td><td></td><td></td><td></td><td></td><td></td><td></td><td></td><td></td><td></td></tr>
<tr><td>Chromosome</td><td>start</td><td></td><td>end</td><td>strand</td><td>MM</td><td>target_seq</td><td>PAM</td><td>alignment</td><td>distance</td><td>position</td><td>gene name</td><td>gene id</td></tr>
<tr><td>GU305914.1</td><td></td><td>210797</td><td>210819</td><td>-</td><td>0</td><td>GATGCGCACTCAGGTTCCGT</td><td>TGG</td><td>||||||||[||||||||||||]PAM</td><td>NA</td><td>Intergenic</td><td>NA</td><td>NA</td></tr>
<tr><td></td><td></td><td></td><td></td><td></td><td></td><td></td><td></td><td></td><td></td><td></td><td></td><td></td></tr>
<tr><td>T18599</td><td>TACAACATCAGTCTCGCGAACGG</td><td></td><td>1000</td><td></td><td></td><td></td><td></td><td></td><td></td><td></td><td></td><td></td></tr>
<tr><td>Oligo fwd</td><td>CACCTACAACATCAGTCTCGCGAA</td><td></td><td></td><td></td><td></td><td></td><td></td><td></td><td></td><td></td><td></td><td></td></tr>
<tr><td>Oligo rev</td><td>AAACTTCGCGAGACTGATGTTGTA</td><td></td><td></td><td></td><td></td><td></td><td></td><td></td><td></td><td></td><td></td><td></td></tr>
<tr><td>Chromosome</td><td>start</td><td></td><td>end</td><td>strand</td><td>MM</td><td>target_seq</td><td>PAM</td><td>alignment</td><td>distance</td><td>position</td><td>gene name</td><td>gene id</td></tr>
<tr><td>GU305914.1</td><td></td><td>210772</td><td>210794</td><td>-</td><td>0</td><td>TACAACATCAGTCTCGCGAA</td><td>CGG</td><td>||||||||[||||||||||||]PAM</td><td>NA</td><td>Intergenic</td><td>NA</td><td>NA</td></tr>
<tr><td>GU305914.1</td><td></td><td>172298</td><td>172320</td><td>-</td><td>4</td><td>TCGAACATCAGTCTCGCCAT</td><td>CGG</td><td>|--|||||[|||||||||-|-]PAM</td><td>NA</td><td>Intergenic</td><td>NA</td><td>NA</td></tr>
<tr><td></td><td></td><td></td><td></td><td></td><td></td><td></td><td></td><td></td><td></td><td></td><td></td><td></td></tr>
<tr><td>T18600</td><td>CATCAGTCTCGCGAACGGAGTGG</td><td></td><td>1000</td><td></td><td></td><td></td><td></td><td></td><td></td><td></td><td></td><td></td></tr>
<tr><td>Oligo fwd</td><td>CACCCATCAGTCTCGCGAACGGAG</td><td></td><td></td><td></td><td></td><td></td><td></td><td></td><td></td><td></td><td></td><td></td></tr>
<tr><td>Oligo rev</td><td>AAACCTCCGTTCGCGAGACTGATG</td><td></td><td></td><td></td><td></td><td></td><td></td><td></td><td></td><td></td><td></td><td></td></tr>
<tr><td>Chromosome</td><td>start</td><td></td><td>end</td><td>strand</td><td>MM</td><td>target_seq</td><td>PAM</td><td>alignment</td><td>distance</td><td>position</td><td>gene name</td><td>gene id</td></tr>
<tr><td>GU305914.1</td><td></td><td>210767</td><td>210789</td><td>-</td><td>0</td><td>CATCAGTCTCGCGAACGGAG</td><td>TGG</td><td>||||||||[||||||||||||]PAM</td><td>NA</td><td>Intergenic</td><td>NA</td><td>NA</td></tr>
<tr><td></td><td></td><td></td><td></td><td></td><td></td><td></td><td></td><td></td><td></td><td></td><td></td><td></td></tr>
<tr><td>T18601</td><td>AAGTACATGCGCATGTATTCTGG</td><td></td><td>1000</td><td></td><td></td><td></td><td></td><td></td><td></td><td></td><td></td><td></td></tr>
<tr><td>Oligo fwd</td><td>CACCAAGTACATGCGCATGTATTC</td><td></td><td></td><td></td><td></td><td></td><td></td><td></td><td></td><td></td><td></td><td></td></tr>
<tr><td>Oligo rev</td><td>AAACGAATACATGCGCATGTACTT</td><td></td><td></td><td></td><td></td><td></td><td></td><td></td><td></td><td></td><td></td><td></td></tr>
<tr><td>Chromosome</td><td>start</td><td></td><td>end</td><td>strand</td><td>MM</td><td>target_seq</td><td>PAM</td><td>alignment</td><td>distance</td><td>position</td><td>gene name</td><td>gene id</td></tr>
<tr><td>GU305914.1</td><td></td><td>210736</td><td>210758</td><td>-</td><td>0</td><td>AAGTACATGCGCATGTATTC</td><td>TGG</td><td>||||||||[||||||||||||]PAM</td><td>NA</td><td>Intergenic</td><td>NA</td><td>NA</td></tr>
<tr><td></td><td></td><td></td><td></td><td></td><td></td><td></td><td></td><td></td><td></td><td></td><td></td><td></td></tr>
<tr><td>T18602</td><td>TACATGCGCATGTATTCTGGTGG</td><td></td><td>1000</td><td></td><td></td><td></td><td></td><td></td><td></td><td></td><td></td><td></td></tr>
<tr><td>Oligo fwd</td><td>CACCTACATGCGCATGTATTCTGG</td><td></td><td></td><td></td><td></td><td></td><td></td><td></td><td></td><td></td><td></td><td></td></tr>
<tr><td>Oligo rev</td><td>AAACCCAGAATACATGCGCATGTA</td><td></td><td></td><td></td><td></td><td></td><td></td><td></td><td></td><td></td><td></td><td></td></tr>
<tr><td>Chromosome</td><td>start</td><td></td><td>end</td><td>strand</td><td>MM</td><td>target_seq</td><td>PAM</td><td>alignment</td><td>distance</td><td>position</td><td>gene name</td><td>gene id</td></tr>
<tr><td>GU305914.1</td><td></td><td>210733</td><td>210755</td><td>-</td><td>0</td><td>TACATGCGCATGTATTCTGG</td><td>TGG</td><td>||||||||[||||||||||||]PAM</td><td>NA</td><td>Intergenic</td><td>NA</td><td>NA</td></tr>
<tr><td></td><td></td><td></td><td></td><td></td><td></td><td></td><td></td><td></td><td></td><td></td><td></td><td></td></tr>
<tr><td>T18603</td><td>GTATTCTGGTGGACGATTGAAGG</td><td></td><td>1000</td><td></td><td></td><td></td><td></td><td></td><td></td><td></td><td></td><td></td></tr>
<tr><td>Oligo fwd</td><td>CACCGTATTCTGGTGGACGATTGA</td><td></td><td></td><td></td><td></td><td></td><td></td><td></td><td></td><td></td><td></td><td></td></tr>
<tr><td>Oligo rev</td><td>AAACTCAATCGTCCACCAGAATAC</td><td></td><td></td><td></td><td></td><td></td><td></td><td></td><td></td><td></td><td></td><td></td></tr>
<tr><td>Chromosome</td><td>start</td><td></td><td>end</td><td>strand</td><td>MM</td><td>target_seq</td><td>PAM</td><td>alignment</td><td>distance</td><td>position</td><td>gene name</td><td>gene id</td></tr>
<tr><td>GU305914.1</td><td></td><td>210722</td><td>210744</td><td>-</td><td>0</td><td>GTATTCTGGTGGACGATTGA</td><td>AGG</td><td>||||||||[||||||||||||]PAM</td><td>NA</td><td>Intergenic</td><td>NA</td><td>NA</td></tr>
<tr><td></td><td></td><td></td><td></td><td></td><td></td><td></td><td></td><td></td><td></td><td></td><td></td><td></td></tr>
<tr><td>T18604</td><td>TTCTGGTGGACGATTGAAGGTGG</td><td></td><td>1000</td><td></td><td></td><td></td><td></td><td></td><td></td><td></td><td></td><td></td></tr>
<tr><td>Oligo fwd</td><td>CACCTTCTGGTGGACGATTGAAGG</td><td></td><td></td><td></td><td></td><td></td><td></td><td></td><td></td><td></td><td></td><td></td></tr>
<tr><td>Oligo rev</td><td>AAACCCTTCAATCGTCCACCAGAA</td><td></td><td></td><td></td><td></td><td></td><td></td><td></td><td></td><td></td><td></td><td></td></tr>
<tr><td>Chromosome</td><td>start</td><td></td><td>end</td><td>strand</td><td>MM</td><td>target_seq</td><td>PAM</td><td>alignment</td><td>distance</td><td>position</td><td>gene name</td><td>gene id</td></tr>
<tr><td>GU305914.1</td><td></td><td>210719</td><td>210741</td><td>-</td><td>0</td><td>TTCTGGTGGACGATTGAAGG</td><td>TGG</td><td>||||||||[||||||||||||]PAM</td><td>NA</td><td>Intergenic</td><td>NA</td><td>NA</td></tr>
<tr><td></td><td></td><td></td><td></td><td></td><td></td><td></td><td></td><td></td><td></td><td></td><td></td><td></td></tr>
<tr><td>T18605</td><td>GACGATTGAAGGTGGAAGCGTGG</td><td></td><td>1000</td><td></td><td></td><td></td><td></td><td></td><td></td><td></td><td></td><td></td></tr>
<tr><td>Oligo fwd</td><td>CACCGACGATTGAAGGTGGAAGCG</td><td></td><td></td><td></td><td></td><td></td><td></td><td></td><td></td><td></td><td></td><td></td></tr>
<tr><td>Oligo rev</td><td>AAACCGCTTCCACCTTCAATCGTC</td><td></td><td></td><td></td><td></td><td></td><td></td><td></td><td></td><td></td><td></td><td></td></tr>
<tr><td>Chromosome</td><td>start</td><td></td><td>end</td><td>strand</td><td>MM</td><td>target_seq</td><td>PAM</td><td>alignment</td><td>distance</td><td>position</td><td>gene name</td><td>gene id</td></tr>
<tr><td>GU305914.1</td><td></td><td>210711</td><td>210733</td><td>-</td><td>0</td><td>GACGATTGAAGGTGGAAGCG</td><td>TGG</td><td>||||||||[||||||||||||]PAM</td><td>NA</td><td>Intergenic</td><td>NA</td><td>NA</td></tr>
<tr><td></td><td></td><td></td><td></td><td></td><td></td><td></td><td></td><td></td><td></td><td></td><td></td><td></td></tr>
<tr><td>T18606</td><td>CTCGATCTCAGAGACCTGAACGG</td><td></td><td>1000</td><td></td><td></td><td></td><td></td><td></td><td></td><td></td><td></td><td></td></tr>
<tr><td>Oligo fwd</td><td>CACCCTCGATCTCAGAGACCTGAA</td><td></td><td></td><td></td><td></td><td></td><td></td><td></td><td></td><td></td><td></td><td></td></tr>
<tr><td>Oligo rev</td><td>AAACTTCAGGTCTCTGAGATCGAG</td><td></td><td></td><td></td><td></td><td></td><td></td><td></td><td></td><td></td><td></td><td></td></tr>
<tr><td>Chromosome</td><td>start</td><td></td><td>end</td><td>strand</td><td>MM</td><td>target_seq</td><td>PAM</td><td>alignment</td><td>distance</td><td>position</td><td>gene name</td><td>gene id</td></tr>
<tr><td>GU305914.1</td><td></td><td>210688</td><td>210710</td><td>-</td><td>0</td><td>CTCGATCTCAGAGACCTGAA</td><td>CGG</td><td>||||||||[||||||||||||]PAM</td><td>NA</td><td>Intergenic</td><td>NA</td><td>NA</td></tr>
<tr><td></td><td></td><td></td><td></td><td></td><td></td><td></td><td></td><td></td><td></td><td></td><td></td><td></td></tr>
<tr><td>T18607</td><td>GTTCGTGATTTCTTCCCCGACGG</td><td></td><td>1000</td><td></td><td></td><td></td><td></td><td></td><td></td><td></td><td></td><td></td></tr>
<tr><td>Oligo fwd</td><td>CACCGTTCGTGATTTCTTCCCCGA</td><td></td><td></td><td></td><td></td><td></td><td></td><td></td><td></td><td></td><td></td><td></td></tr>
<tr><td>Oligo rev</td><td>AAACTCGGGGAAGAAATCACGAAC</td><td></td><td></td><td></td><td></td><td></td><td></td><td></td><td></td><td></td><td></td><td></td></tr>
<tr><td>Chromosome</td><td>start</td><td></td><td>end</td><td>strand</td><td>MM</td><td>target_seq</td><td>PAM</td><td>alignment</td><td>distance</td><td>position</td><td>gene name</td><td>gene id</td></tr>
<tr><td>GU305914.1</td><td></td><td>210650</td><td>210672</td><td>-</td><td>0</td><td>GTTCGTGATTTCTTCCCCGA</td><td>CGG</td><td>||||||||[||||||||||||]PAM</td><td>NA</td><td>Intergenic</td><td>NA</td><td>NA</td></tr>
<tr><td></td><td></td><td></td><td></td><td></td><td></td><td></td><td></td><td></td><td></td><td></td><td></td><td></td></tr>
<tr><td>T18608</td><td>TTCGTGATTTCTTCCCCGACGGG</td><td></td><td>1000</td><td></td><td></td><td></td><td></td><td></td><td></td><td></td><td></td><td></td></tr>
<tr><td>Oligo fwd</td><td>CACCTTCGTGATTTCTTCCCCGAC</td><td></td><td></td><td></td><td></td><td></td><td></td><td></td><td></td><td></td><td></td><td></td></tr>
<tr><td>Oligo rev</td><td>AAACGTCGGGGAAGAAATCACGAA</td><td></td><td></td><td></td><td></td><td></td><td></td><td></td><td></td><td></td><td></td><td></td></tr>
<tr><td>Chromosome</td><td>start</td><td></td><td>end</td><td>strand</td><td>MM</td><td>target_seq</td><td>PAM</td><td>alignment</td><td>distance</td><td>position</td><td>gene name</td><td>gene id</td></tr>
<tr><td>GU305914.1</td><td></td><td>210649</td><td>210671</td><td>-</td><td>0</td><td>TTCGTGATTTCTTCCCCGAC</td><td>GGG</td><td>||||||||[||||||||||||]PAM</td><td>NA</td><td>Intergenic</td><td>NA</td><td>NA</td></tr>
<tr><td></td><td></td><td></td><td></td><td></td><td></td><td></td><td></td><td></td><td></td><td></td><td></td><td></td></tr>
<tr><td>T18609</td><td>TGATTTCTTCCCCGACGGGATGG</td><td></td><td>1000</td><td></td><td></td><td></td><td></td><td></td><td></td><td></td><td></td><td></td></tr>
<tr><td>Oligo fwd</td><td>CACCTGATTTCTTCCCCGACGGGA</td><td></td><td></td><td></td><td></td><td></td><td></td><td></td><td></td><td></td><td></td><td></td></tr>
<tr><td>Oligo rev</td><td>AAACTCCCGTCGGGGAAGAAATCA</td><td></td><td></td><td></td><td></td><td></td><td></td><td></td><td></td><td></td><td></td><td></td></tr>
<tr><td>Chromosome</td><td>start</td><td></td><td>end</td><td>strand</td><td>MM</td><td>target_seq</td><td>PAM</td><td>alignment</td><td>distance</td><td>position</td><td>gene name</td><td>gene id</td></tr>
<tr><td>GU305914.1</td><td></td><td>210645</td><td>210667</td><td>-</td><td>0</td><td>TGATTTCTTCCCCGACGGGA</td><td>TGG</td><td>||||||||[||||||||||||]PAM</td><td>NA</td><td>Intergenic</td><td>NA</td><td>NA</td></tr>
<tr><td></td><td></td><td></td><td></td><td></td><td></td><td></td><td></td><td></td><td></td><td></td><td></td><td></td></tr>
<tr><td>T18610</td><td>CCCGACGGGATGGTACGCTACGG</td><td></td><td>1000</td><td></td><td></td><td></td><td></td><td></td><td></td><td></td><td></td><td></td></tr>
<tr><td>Oligo fwd</td><td>CACCCCCGACGGGATGGTACGCTA</td><td></td><td></td><td></td><td></td><td></td><td></td><td></td><td></td><td></td><td></td><td></td></tr>
<tr><td>Oligo rev</td><td>AAACTAGCGTACCATCCCGTCGGG</td><td></td><td></td><td></td><td></td><td></td><td></td><td></td><td></td><td></td><td></td><td></td></tr>
<tr><td>Chromosome</td><td>start</td><td></td><td>end</td><td>strand</td><td>MM</td><td>target_seq</td><td>PAM</td><td>alignment</td><td>distance</td><td>position</td><td>gene name</td><td>gene id</td></tr>
<tr><td>GU305914.1</td><td></td><td>210635</td><td>210657</td><td>-</td><td>0</td><td>CCCGACGGGATGGTACGCTA</td><td>CGG</td><td>||||||||[||||||||||||]PAM</td><td>NA</td><td>Intergenic</td><td>NA</td><td>NA</td></tr>
<tr><td></td><td></td><td></td><td></td><td></td><td></td><td></td><td></td><td></td><td></td><td></td><td></td><td></td></tr>
<tr><td>T18611</td><td>GAGTACCCTCAACAGAGTCCCGG</td><td></td><td>1000</td><td></td><td></td><td></td><td></td><td></td><td></td><td></td><td></td><td></td></tr>
<tr><td>Oligo fwd</td><td>CACCGAGTACCCTCAACAGAGTCC</td><td></td><td></td><td></td><td></td><td></td><td></td><td></td><td></td><td></td><td></td><td></td></tr>
<tr><td>Oligo rev</td><td>AAACGGACTCTGTTGAGGGTACTC</td><td></td><td></td><td></td><td></td><td></td><td></td><td></td><td></td><td></td><td></td><td></td></tr>
<tr><td>Chromosome</td><td>start</td><td></td><td>end</td><td>strand</td><td>MM</td><td>target_seq</td><td>PAM</td><td>alignment</td><td>distance</td><td>position</td><td>gene name</td><td>gene id</td></tr>
<tr><td>GU305914.1</td><td></td><td>210601</td><td>210623</td><td>-</td><td>0</td><td>GAGTACCCTCAACAGAGTCC</td><td>CGG</td><td>||||||||[||||||||||||]PAM</td><td>NA</td><td>Intergenic</td><td>NA</td><td>NA</td></tr>
</table>

<table>
<tr><td>T18612</td><td>ATGCTGTTGTCGTCGATCGATGG</td><td></td><td>1000</td><td></td><td></td><td></td><td></td><td></td><td></td><td></td><td></td><td></td></tr>
<tr><td>Oligo fwd</td><td>CACCATGCTGTTGTCGTCGATCGA</td><td></td><td></td><td></td><td></td><td></td><td></td><td></td><td></td><td></td><td></td><td></td></tr>
<tr><td>Oligo rev</td><td>AAACTCGATCGACGACAACAGCAT</td><td></td><td></td><td></td><td></td><td></td><td></td><td></td><td></td><td></td><td></td><td></td></tr>
<tr><td>Chromosome</td><td>start</td><td></td><td>end</td><td>strand</td><td>MM</td><td>target_seq</td><td>PAM</td><td>alignment</td><td>distance</td><td>position</td><td>gene name</td><td>gene id</td></tr>
<tr><td>GU305914.1</td><td></td><td>210577</td><td>210599</td><td>-</td><td>0</td><td>ATGCTGTTGTCGTCGATCGA</td><td>TGG</td><td>||||||||[||||||||||||]PAM</td><td>NA</td><td>Intergenic</td><td>NA</td><td>NA</td></tr>
<tr><td>T18613</td><td>TGCTGTTGTCGTCGATCGATGGG</td><td></td><td>1000</td><td></td><td></td><td></td><td></td><td></td><td></td><td></td><td></td><td></td></tr>
<tr><td>Oligo fwd</td><td>CACCTGCTGTTGTCGTCGATCGAT</td><td></td><td></td><td></td><td></td><td></td><td></td><td></td><td></td><td></td><td></td><td></td></tr>
<tr><td>Oligo rev</td><td>AAACATCGATCGACGACAACAGCA</td><td></td><td></td><td></td><td></td><td></td><td></td><td></td><td></td><td></td><td></td><td></td></tr>
<tr><td>Chromosome</td><td>start</td><td></td><td>end</td><td>strand</td><td>MM</td><td>target_seq</td><td>PAM</td><td>alignment</td><td>distance</td><td>position</td><td>gene name</td><td>gene id</td></tr>
<tr><td>GU305914.1</td><td></td><td>210576</td><td>210598</td><td>-</td><td>0</td><td>TGCTGTTGTCGTCGATCGAT</td><td>GGG</td><td>||||||||[||||||||||||]PAM</td><td>NA</td><td>Intergenic</td><td>NA</td><td>NA</td></tr>
<tr><td>T18614</td><td>TGGGCAGTTCGAGTCGTCCGCGG</td><td></td><td>1000</td><td></td><td></td><td></td><td></td><td></td><td></td><td></td><td></td><td></td></tr>
<tr><td>Oligo fwd</td><td>CACCTGGGCAGTTCGAGTCGTCCG</td><td></td><td></td><td></td><td></td><td></td><td></td><td></td><td></td><td></td><td></td><td></td></tr>
<tr><td>Oligo rev</td><td>AAACCGGACGACTCGAACTGCCCA</td><td></td><td></td><td></td><td></td><td></td><td></td><td></td><td></td><td></td><td></td><td></td></tr>
<tr><td>Chromosome</td><td>start</td><td></td><td>end</td><td>strand</td><td>MM</td><td>target_seq</td><td>PAM</td><td>alignment</td><td>distance</td><td>position</td><td>gene name</td><td>gene id</td></tr>
<tr><td>GU305914.1</td><td></td><td>210557</td><td>210579</td><td>-</td><td>0</td><td>TGGGCAGTTCGAGTCGTCCG</td><td>CGG</td><td>||||||||[||||||||||||]PAM</td><td>NA</td><td>Intergenic</td><td>NA</td><td>NA</td></tr>
<tr><td>T18615</td><td>AGTCGTCCGCGGTCGTCTCGTGG</td><td></td><td>1000</td><td></td><td></td><td></td><td></td><td></td><td></td><td></td><td></td><td></td></tr>
<tr><td>Oligo fwd</td><td>CACCAGTCGTCCGCGGTCGTCTCG</td><td></td><td></td><td></td><td></td><td></td><td></td><td></td><td></td><td></td><td></td><td></td></tr>
<tr><td>Oligo rev</td><td>AAACCGAGACGACCGCGGACGACT</td><td></td><td></td><td></td><td></td><td></td><td></td><td></td><td></td><td></td><td></td><td></td></tr>
<tr><td>Chromosome</td><td>start</td><td></td><td>end</td><td>strand</td><td>MM</td><td>target_seq</td><td>PAM</td><td>alignment</td><td>distance</td><td>position</td><td>gene name</td><td>gene id</td></tr>
<tr><td>GU305914.1</td><td></td><td>210546</td><td>210568</td><td>-</td><td>0</td><td>AGTCGTCCGCGGTCGTCTCG</td><td>TGG</td><td>||||||||[||||||||||||]PAM</td><td>NA</td><td>Intergenic</td><td>NA</td><td>NA</td></tr>
<tr><td>T18616</td><td>CCGCGGTCGTCTCGTGGCACAGG</td><td></td><td>1000</td><td></td><td></td><td></td><td></td><td></td><td></td><td></td><td></td><td></td></tr>
<tr><td>Oligo fwd</td><td>CACCCCGCGGTCGTCTCGTGGCAC</td><td></td><td></td><td></td><td></td><td></td><td></td><td></td><td></td><td></td><td></td><td></td></tr>
<tr><td>Oligo rev</td><td>AAACGTGCCACGAGACGACCGCGG</td><td></td><td></td><td></td><td></td><td></td><td></td><td></td><td></td><td></td><td></td><td></td></tr>
<tr><td>Chromosome</td><td>start</td><td></td><td>end</td><td>strand</td><td>MM</td><td>target_seq</td><td>PAM</td><td>alignment</td><td>distance</td><td>position</td><td>gene name</td><td>gene id</td></tr>
<tr><td>GU305914.1</td><td></td><td>210540</td><td>210562</td><td>-</td><td>0</td><td>CCGCGGTCGTCTCGTGGCAC</td><td>AGG</td><td>||||||||[||||||||||||]PAM</td><td>NA</td><td>Intergenic</td><td>NA</td><td>NA</td></tr>
<tr><td>T18617</td><td>CGCGGTCGTCTCGTGGCACAGGG</td><td></td><td>1000</td><td></td><td></td><td></td><td></td><td></td><td></td><td></td><td></td><td></td></tr>
<tr><td>Oligo fwd</td><td>CACCCGCGGTCGTCTCGTGGCACA</td><td></td><td></td><td></td><td></td><td></td><td></td><td></td><td></td><td></td><td></td><td></td></tr>
<tr><td>Oligo rev</td><td>AAACTGTGCCACGAGACGACCGCG</td><td></td><td></td><td></td><td></td><td></td><td></td><td></td><td></td><td></td><td></td><td></td></tr>
<tr><td>Chromosome</td><td>start</td><td></td><td>end</td><td>strand</td><td>MM</td><td>target_seq</td><td>PAM</td><td>alignment</td><td>distance</td><td>position</td><td>gene name</td><td>gene id</td></tr>
<tr><td>GU305914.1</td><td></td><td>210539</td><td>210561</td><td>-</td><td>0</td><td>CGCGGTCGTCTCGTGGCACA</td><td>GGG</td><td>||||||||[||||||||||||]PAM</td><td>NA</td><td>Intergenic</td><td>NA</td><td>NA</td></tr>
<tr><td>T18618</td><td>GCGGTCGTCTCGTGGCACAGGGG</td><td></td><td>1000</td><td></td><td></td><td></td><td></td><td></td><td></td><td></td><td></td><td></td></tr>
<tr><td>Oligo fwd</td><td>CACCGCGGTCGTCTCGTGGCACAG</td><td></td><td></td><td></td><td></td><td></td><td></td><td></td><td></td><td></td><td></td><td></td></tr>
<tr><td>Oligo rev</td><td>AAACCTGTGCCACGAGACGACCGC</td><td></td><td></td><td></td><td></td><td></td><td></td><td></td><td></td><td></td><td></td><td></td></tr>
<tr><td>Chromosome</td><td>start</td><td></td><td>end</td><td>strand</td><td>MM</td><td>target_seq</td><td>PAM</td><td>alignment</td><td>distance</td><td>position</td><td>gene name</td><td>gene id</td></tr>
<tr><td>GU305914.1</td><td></td><td>210538</td><td>210560</td><td>-</td><td>0</td><td>GCGGTCGTCTCGTGGCACAG</td><td>GGG</td><td>||||||||[||||||||||||]PAM</td><td>NA</td><td>Intergenic</td><td>NA</td><td>NA</td></tr>
<tr><td>T18619</td><td>GTCTCGTGGCACAGGGGACACGG</td><td></td><td>1000</td><td></td><td></td><td></td><td></td><td></td><td></td><td></td><td></td><td></td></tr>
<tr><td>Oligo fwd</td><td>CACCGTCTCGTGGCACAGGGGACA</td><td></td><td></td><td></td><td></td><td></td><td></td><td></td><td></td><td></td><td></td><td></td></tr>
<tr><td>Oligo rev</td><td>AAACTGTCCCCTGTGCCACGAGAC</td><td></td><td></td><td></td><td></td><td></td><td></td><td></td><td></td><td></td><td></td><td></td></tr>
<tr><td>Chromosome</td><td>start</td><td></td><td>end</td><td>strand</td><td>MM</td><td>target_seq</td><td>PAM</td><td>alignment</td><td>distance</td><td>position</td><td>gene name</td><td>gene id</td></tr>
<tr><td>GU305914.1</td><td></td><td>210532</td><td>210554</td><td>-</td><td>0</td><td>GTCTCGTGGCACAGGGGACA</td><td>CGG</td><td>||||||||[||||||||||||]PAM</td><td>NA</td><td>Intergenic</td><td>NA</td><td>NA</td></tr>
<tr><td>T18620</td><td>GCACGCTCCTCCCGTCTCCGCGG</td><td></td><td>1000</td><td></td><td></td><td></td><td></td><td></td><td></td><td></td><td></td><td></td></tr>
<tr><td>Oligo fwd</td><td>CACCGCACGCTCCTCCCGTCTCCG</td><td></td><td></td><td></td><td></td><td></td><td></td><td></td><td></td><td></td><td></td><td></td></tr>
<tr><td>Oligo rev</td><td>AAACCGGAGACGGGAGGAGCGTGC</td><td></td><td></td><td></td><td></td><td></td><td></td><td></td><td></td><td></td><td></td><td></td></tr>
<tr><td>Chromosome</td><td>start</td><td></td><td>end</td><td>strand</td><td>MM</td><td>target_seq</td><td>PAM</td><td>alignment</td><td>distance</td><td>position</td><td>gene name</td><td>gene id</td></tr>
<tr><td>GU305914.1</td><td></td><td>210503</td><td>210525</td><td>-</td><td>0</td><td>GCACGCTCCTCCCGTCTCCG</td><td>CGG</td><td>||||||||[||||||||||||]PAM</td><td>NA</td><td>Intergenic</td><td>NA</td><td>NA</td></tr>
<tr><td>T18621</td><td>GGAGTACTCCATCTTCTTCATGG</td><td></td><td>1000</td><td></td><td></td><td></td><td></td><td></td><td></td><td></td><td></td><td></td></tr>
<tr><td>Oligo fwd</td><td>CACCGGAGTACTCCATCTTCTTCA</td><td></td><td></td><td></td><td></td><td></td><td></td><td></td><td></td><td></td><td></td><td></td></tr>
<tr><td>Oligo rev</td><td>AAACTGAAGAAGATGGAGTACTCC</td><td></td><td></td><td></td><td></td><td></td><td></td><td></td><td></td><td></td><td></td><td></td></tr>
<tr><td>Chromosome</td><td>start</td><td></td><td>end</td><td>strand</td><td>MM</td><td>target_seq</td><td>PAM</td><td>alignment</td><td>distance</td><td>position</td><td>gene name</td><td>gene id</td></tr>
<tr><td>GU305914.1</td><td></td><td>210482</td><td>210504</td><td>-</td><td>0</td><td>GGAGTACTCCATCTTCTTCA</td><td>TGG</td><td>||||||||[||||||||||||]PAM</td><td>NA</td><td>Intergenic</td><td>NA</td><td>NA</td></tr>
<tr><td>T18622</td><td>CCATCTTCTTCATGGACGTGTGG</td><td></td><td>1000</td><td></td><td></td><td></td><td></td><td></td><td></td><td></td><td></td><td></td></tr>
<tr><td>Oligo fwd</td><td>CACCCCATCTTCTTCATGGACGTG</td><td></td><td></td><td></td><td></td><td></td><td></td><td></td><td></td><td></td><td></td><td></td></tr>
<tr><td>Oligo rev</td><td>AAACCACGTCCATGAAGAAGATGG</td><td></td><td></td><td></td><td></td><td></td><td></td><td></td><td></td><td></td><td></td><td></td></tr>
<tr><td>Chromosome</td><td>start</td><td></td><td>end</td><td>strand</td><td>MM</td><td>target_seq</td><td>PAM</td><td>alignment</td><td>distance</td><td>position</td><td>gene name</td><td>gene id</td></tr>
<tr><td>GU305914.1</td><td></td><td>210474</td><td>210496</td><td>-</td><td>0</td><td>CCATCTTCTTCATGGACGTG</td><td>TGG</td><td>||||||||[||||||||||||]PAM</td><td>NA</td><td>Intergenic</td><td>NA</td><td>NA</td></tr>
<tr><td>T18623</td><td>GTGTGGTCCTTGATCGCGATCGG</td><td></td><td>1000</td><td></td><td></td><td></td><td></td><td></td><td></td><td></td><td></td><td></td></tr>
<tr><td>Oligo fwd</td><td>CACCGTGTGGTCCTTGATCGCGAT</td><td></td><td></td><td></td><td></td><td></td><td></td><td></td><td></td><td></td><td></td><td></td></tr>
<tr><td>Oligo rev</td><td>AAACATCGCGATCAAGGACCACAC</td><td></td><td></td><td></td><td></td><td></td><td></td><td></td><td></td><td></td><td></td><td></td></tr>
<tr><td>Chromosome</td><td>start</td><td></td><td>end</td><td>strand</td><td>MM</td><td>target_seq</td><td>PAM</td><td>alignment</td><td>distance</td><td>position</td><td>gene name</td><td>gene id</td></tr>
<tr><td>GU305914.1</td><td></td><td>210457</td><td>210479</td><td>-</td><td>0</td><td>GTGTGGTCCTTGATCGCGAT</td><td>CGG</td><td>||||||||[||||||||||||]PAM</td><td>NA</td><td>Intergenic</td><td>NA</td><td>NA</td></tr>
<tr><td>T18624</td><td>CGTGATCGTCTTCATGTATCTGG</td><td></td><td>1000</td><td></td><td></td><td></td><td></td><td></td><td></td><td></td><td></td><td></td></tr>
<tr><td>Oligo fwd</td><td>CACCCGTGATCGTCTTCATGTATC</td><td></td><td></td><td></td><td></td><td></td><td></td><td></td><td></td><td></td><td></td><td></td></tr>
<tr><td>Oligo rev</td><td>AAACGATACATGAAGACGATCACG</td><td></td><td></td><td></td><td></td><td></td><td></td><td></td><td></td><td></td><td></td><td></td></tr>
<tr><td>Chromosome</td><td>start</td><td></td><td>end</td><td>strand</td><td>MM</td><td>target_seq</td><td>PAM</td><td>alignment</td><td>distance</td><td>position</td><td>gene name</td><td>gene id</td></tr>
<tr><td>GU305914.1</td><td></td><td>210425</td><td>210447</td><td>-</td><td>0</td><td>CGTGATCGTCTTCATGTATC</td><td>TGG</td><td>||||||||[||||||||||||]PAM</td><td>NA</td><td>Intergenic</td><td>NA</td><td>NA</td></tr>
<tr><td>T18625</td><td>TCATGTATCTGGTGAAGTTACGG</td><td></td><td>1000</td><td></td><td></td><td></td><td></td><td></td><td></td><td></td><td></td><td></td></tr>
<tr><td>Oligo fwd</td><td>CACCTCATGTATCTGGTGAAGTTA</td><td></td><td></td><td></td><td></td><td></td><td></td><td></td><td></td><td></td><td></td><td></td></tr>
<tr><td>Oligo rev</td><td>AAACTAACTTCACCAGATACATGA</td><td></td><td></td><td></td><td></td><td></td><td></td><td></td><td></td><td></td><td></td><td></td></tr>
<tr><td>Chromosome</td><td>start</td><td></td><td>end</td><td>strand</td><td>MM</td><td>target_seq</td><td>PAM</td><td>alignment</td><td>distance</td><td>position</td><td>gene name</td><td>gene id</td></tr>
<tr><td>GU305914.1</td><td></td><td>210414</td><td>210436</td><td>-</td><td>0</td><td>TCATGTATCTGGTGAAGTTA</td><td>CGG</td><td>||||||||[||||||||||||]PAM</td><td>NA</td><td>Intergenic</td><td>NA</td><td>NA</td></tr>
<tr><td>T18626</td><td>CATGTATCTGGTGAAGTTACGGG</td><td></td><td>1000</td><td></td><td></td><td></td><td></td><td></td><td></td><td></td><td></td><td></td></tr>
<tr><td>Oligo fwd</td><td>CACCCATGTATCTGGTGAAGTTAC</td><td></td><td></td><td></td><td></td><td></td><td></td><td></td><td></td><td></td><td></td><td></td></tr>
<tr><td>Oligo rev</td><td>AAACGTAACTTCACCAGATACATG</td><td></td><td></td><td></td><td></td><td></td><td></td><td></td><td></td><td></td><td></td><td></td></tr>
<tr><td>Chromosome</td><td>start</td><td></td><td>end</td><td>strand</td><td>MM</td><td>target_seq</td><td>PAM</td><td>alignment</td><td>distance</td><td>position</td><td>gene name</td><td>gene id</td></tr>
<tr><td>GU305914.1</td><td></td><td>210413</td><td>210435</td><td>-</td><td>0</td><td>CATGTATCTGGTGAAGTTAC</td><td>GGG</td><td>||||||||[||||||||||||]PAM</td><td>NA</td><td>Intergenic</td><td>NA</td><td>NA</td></tr>
</table>

<table>
<tr><td>T18627</td><td>GTATCTGGTGAAGTTACGGGTGG</td><td>1000</td><td></td><td></td><td></td><td></td><td></td><td></td><td></td><td></td><td></td></tr>
<tr><td>Oligo fwd</td><td>CACCGTATCTGGTGAAGTTACGGG</td><td></td><td></td><td></td><td></td><td></td><td></td><td></td><td></td><td></td><td></td></tr>
<tr><td>Oligo rev</td><td>AAACCCCGTAACTTCACCAGATAC</td><td></td><td></td><td></td><td></td><td></td><td></td><td></td><td></td><td></td><td></td></tr>
<tr><td>Chromosome</td><td>start</td><td>end</td><td>strand</td><td>MM</td><td>target_seq</td><td>PAM</td><td>alignment</td><td>distance</td><td>position</td><td>gene name</td><td>gene id</td></tr>
<tr><td>GU305914.1</td><td>210410</td><td>210432</td><td>-</td><td>0</td><td>GTATCTGGTGAAGTTACGGG</td><td>TGG</td><td>||||||||[||||||||||||]PAM</td><td>NA</td><td>Intergenic</td><td>NA</td><td>NA</td></tr>
<tr><td>T18628</td><td>TGGTGAAGTTACGGGTGGTGTGG</td><td>1000</td><td></td><td></td><td></td><td></td><td></td><td></td><td></td><td></td><td></td></tr>
<tr><td>Oligo fwd</td><td>CACCTGGTGAAGTTACGGGTGGTG</td><td></td><td></td><td></td><td></td><td></td><td></td><td></td><td></td><td></td><td></td></tr>
<tr><td>Oligo rev</td><td>AAACCACCACCCGTAACTTCACCA</td><td></td><td></td><td></td><td></td><td></td><td></td><td></td><td></td><td></td><td></td></tr>
<tr><td>Chromosome</td><td>start</td><td>end</td><td>strand</td><td>MM</td><td>target_seq</td><td>PAM</td><td>alignment</td><td>distance</td><td>position</td><td>gene name</td><td>gene id</td></tr>
<tr><td>GU305914.1</td><td>210405</td><td>210427</td><td>-</td><td>0</td><td>TGGTGAAGTTACGGGTGGTG</td><td>TGG</td><td>||||||||[||||||||||||]PAM</td><td>NA</td><td>Intergenic</td><td>NA</td><td>NA</td></tr>
<tr><td>T18629</td><td>TACGGGTGGTGTGGATCAATCGG</td><td>1000</td><td></td><td></td><td></td><td></td><td></td><td></td><td></td><td></td><td></td></tr>
<tr><td>Oligo fwd</td><td>CACCTACGGGTGGTGTGGATCAAT</td><td></td><td></td><td></td><td></td><td></td><td></td><td></td><td></td><td></td><td></td></tr>
<tr><td>Oligo rev</td><td>AAACATTGATCCACACCACCCGTA</td><td></td><td></td><td></td><td></td><td></td><td></td><td></td><td></td><td></td><td></td></tr>
<tr><td>Chromosome</td><td>start</td><td>end</td><td>strand</td><td>MM</td><td>target_seq</td><td>PAM</td><td>alignment</td><td>distance</td><td>position</td><td>gene name</td><td>gene id</td></tr>
<tr><td>GU305914.1</td><td>210396</td><td>210418</td><td>-</td><td>0</td><td>TACGGGTGGTGTGGATCAAT</td><td>CGG</td><td>||||||||[||||||||||||]PAM</td><td>NA</td><td>Intergenic</td><td>NA</td><td>NA</td></tr>
<tr><td>T18630</td><td>ACGGGTGGTGTGGATCAATCGGG</td><td>1000</td><td></td><td></td><td></td><td></td><td></td><td></td><td></td><td></td><td></td></tr>
<tr><td>Oligo fwd</td><td>CACCACGGGTGGTGTGGATCAATC</td><td></td><td></td><td></td><td></td><td></td><td></td><td></td><td></td><td></td><td></td></tr>
<tr><td>Oligo rev</td><td>AAACGATTGATCCACACCACCCGT</td><td></td><td></td><td></td><td></td><td></td><td></td><td></td><td></td><td></td><td></td></tr>
<tr><td>Chromosome</td><td>start</td><td>end</td><td>strand</td><td>MM</td><td>target_seq</td><td>PAM</td><td>alignment</td><td>distance</td><td>position</td><td>gene name</td><td>gene id</td></tr>
<tr><td>GU305914.1</td><td>210395</td><td>210417</td><td>-</td><td>0</td><td>ACGGGTGGTGTGGATCAATC</td><td>GGG</td><td>||||||||[||||||||||||]PAM</td><td>NA</td><td>Intergenic</td><td>NA</td><td>NA</td></tr>
<tr><td>T18631</td><td>TGGTGTGGATCAATCGGGTCTGG</td><td>1000</td><td></td><td></td><td></td><td></td><td></td><td></td><td></td><td></td><td></td></tr>
<tr><td>Oligo fwd</td><td>CACCTGGTGTGGATCAATCGGGTC</td><td></td><td></td><td></td><td></td><td></td><td></td><td></td><td></td><td></td><td></td></tr>
<tr><td>Oligo rev</td><td>AAACGACCCGATTGATCCACACCA</td><td></td><td></td><td></td><td></td><td></td><td></td><td></td><td></td><td></td><td></td></tr>
<tr><td>Chromosome</td><td>start</td><td>end</td><td>strand</td><td>MM</td><td>target_seq</td><td>PAM</td><td>alignment</td><td>distance</td><td>position</td><td>gene name</td><td>gene id</td></tr>
<tr><td>GU305914.1</td><td>210390</td><td>210412</td><td>-</td><td>0</td><td>TGGTGTGGATCAATCGGGTC</td><td>TGG</td><td>||||||||[||||||||||||]PAM</td><td>NA</td><td>Intergenic</td><td>NA</td><td>NA</td></tr>
<tr><td>T18632</td><td>ATCGGGTCTGGCCTCGTATGCGG</td><td>1000</td><td></td><td></td><td></td><td></td><td></td><td></td><td></td><td></td><td></td></tr>
<tr><td>Oligo fwd</td><td>CACCATCGGGTCTGGCCTCGTATG</td><td></td><td></td><td></td><td></td><td></td><td></td><td></td><td></td><td></td><td></td></tr>
<tr><td>Oligo rev</td><td>AAACCATACGAGGCCAGACCCGAT</td><td></td><td></td><td></td><td></td><td></td><td></td><td></td><td></td><td></td><td></td></tr>
<tr><td>Chromosome</td><td>start</td><td>end</td><td>strand</td><td>MM</td><td>target_seq</td><td>PAM</td><td>alignment</td><td>distance</td><td>position</td><td>gene name</td><td>gene id</td></tr>
<tr><td>GU305914.1</td><td>210378</td><td>210400</td><td>-</td><td>0</td><td>ATCGGGTCTGGCCTCGTATG</td><td>CGG</td><td>||||||||[||||||||||||]PAM</td><td>NA</td><td>Intergenic</td><td>NA</td><td>NA</td></tr>
<tr><td>T18633</td><td>GCCTCGTATGCGGTATCGCCTGG</td><td>1000</td><td></td><td></td><td></td><td></td><td></td><td></td><td></td><td></td><td></td></tr>
<tr><td>Oligo fwd</td><td>CACCGCCTCGTATGCGGTATCGCC</td><td></td><td></td><td></td><td></td><td></td><td></td><td></td><td></td><td></td><td></td></tr>
<tr><td>Oligo rev</td><td>AAACGGCGATACCGCATACGAGGC</td><td></td><td></td><td></td><td></td><td></td><td></td><td></td><td></td><td></td><td></td></tr>
<tr><td>Chromosome</td><td>start</td><td>end</td><td>strand</td><td>MM</td><td>target_seq</td><td>PAM</td><td>alignment</td><td>distance</td><td>position</td><td>gene name</td><td>gene id</td></tr>
<tr><td>GU305914.1</td><td>210368</td><td>210390</td><td>-</td><td>0</td><td>GCCTCGTATGCGGTATCGCC</td><td>TGG</td><td>||||||||[||||||||||||]PAM</td><td>NA</td><td>Intergenic</td><td>NA</td><td>NA</td></tr>
<tr><td>T18634</td><td>TCTACATCAACTGCCGCGTGTGG</td><td>1000</td><td></td><td></td><td></td><td></td><td></td><td></td><td></td><td></td><td></td></tr>
<tr><td>Oligo fwd</td><td>CACCTCTACATCAACTGCCGCGTG</td><td></td><td></td><td></td><td></td><td></td><td></td><td></td><td></td><td></td><td></td></tr>
<tr><td>Oligo rev</td><td>AAACCACGCGGCAGTTGATGTAGA</td><td></td><td></td><td></td><td></td><td></td><td></td><td></td><td></td><td></td><td></td></tr>
<tr><td>Chromosome</td><td>start</td><td>end</td><td>strand</td><td>MM</td><td>target_seq</td><td>PAM</td><td>alignment</td><td>distance</td><td>position</td><td>gene name</td><td>gene id</td></tr>
<tr><td>GU305914.1</td><td>210345</td><td>210367</td><td>-</td><td>0</td><td>TCTACATCAACTGCCGCGTG</td><td>TGG</td><td>||||||||[||||||||||||]PAM</td><td>NA</td><td>Intergenic</td><td>NA</td><td>NA</td></tr>
<tr><td>T18635</td><td>TGTGGTGATGTTCGTGCGACCGG</td><td>1000</td><td></td><td></td><td></td><td></td><td></td><td></td><td></td><td></td><td></td></tr>
<tr><td>Oligo fwd</td><td>CACCTGTGGTGATGTTCGTGCGAC</td><td></td><td></td><td></td><td></td><td></td><td></td><td></td><td></td><td></td><td></td></tr>
<tr><td>Oligo rev</td><td>AAACGTCGCACGAACATCACCACA</td><td></td><td></td><td></td><td></td><td></td><td></td><td></td><td></td><td></td><td></td></tr>
<tr><td>Chromosome</td><td>start</td><td>end</td><td>strand</td><td>MM</td><td>target_seq</td><td>PAM</td><td>alignment</td><td>distance</td><td>position</td><td>gene name</td><td>gene id</td></tr>
<tr><td>GU305914.1</td><td>210327</td><td>210349</td><td>-</td><td>0</td><td>TGTGGTGATGTTCGTGCGAC</td><td>CGG</td><td>||||||||[||||||||||||]PAM</td><td>NA</td><td>Intergenic</td><td>NA</td><td>NA</td></tr>
<tr><td>T18636</td><td>TGTTCGTGCGACCGGCACACTGG</td><td>1000</td><td></td><td></td><td></td><td></td><td></td><td></td><td></td><td></td><td></td></tr>
<tr><td>Oligo fwd</td><td>CACCTGTTCGTGCGACCGGCACAC</td><td></td><td></td><td></td><td></td><td></td><td></td><td></td><td></td><td></td><td></td></tr>
<tr><td>Oligo rev</td><td>AAACGTGTGCCGGTCGCACGAACA</td><td></td><td></td><td></td><td></td><td></td><td></td><td></td><td></td><td></td><td></td></tr>
<tr><td>Chromosome</td><td>start</td><td>end</td><td>strand</td><td>MM</td><td>target_seq</td><td>PAM</td><td>alignment</td><td>distance</td><td>position</td><td>gene name</td><td>gene id</td></tr>
<tr><td>GU305914.1</td><td>210319</td><td>210341</td><td>-</td><td>0</td><td>TGTTCGTGCGACCGGCACAC</td><td>TGG</td><td>||||||||[||||||||||||]PAM</td><td>NA</td><td>Intergenic</td><td>NA</td><td>NA</td></tr>
<tr><td>T18637</td><td>ACTGGACAATGTCAGATCGTCGG</td><td>1000</td><td></td><td></td><td></td><td></td><td></td><td></td><td></td><td></td><td></td></tr>
<tr><td>Oligo fwd</td><td>CACCACTGGACAATGTCAGATCGT</td><td></td><td></td><td></td><td></td><td></td><td></td><td></td><td></td><td></td><td></td></tr>
<tr><td>Oligo rev</td><td>AAACACGATCTGACATTGTCCAGT</td><td></td><td></td><td></td><td></td><td></td><td></td><td></td><td></td><td></td><td></td></tr>
<tr><td>Chromosome</td><td>start</td><td>end</td><td>strand</td><td>MM</td><td>target_seq</td><td>PAM</td><td>alignment</td><td>distance</td><td>position</td><td>gene name</td><td>gene id</td></tr>
<tr><td>GU305914.1</td><td>210301</td><td>210323</td><td>-</td><td>0</td><td>ACTGGACAATGTCAGATCGT</td><td>CGG</td><td>||||||||[||||||||||||]PAM</td><td>NA</td><td>Intergenic</td><td>NA</td><td>NA</td></tr>
<tr><td>T18638</td><td>CTGGACAATGTCAGATCGTCGGG</td><td>1000</td><td></td><td></td><td></td><td></td><td></td><td></td><td></td><td></td><td></td></tr>
<tr><td>Oligo fwd</td><td>CACCCTGGACAATGTCAGATCGTC</td><td></td><td></td><td></td><td></td><td></td><td></td><td></td><td></td><td></td><td></td></tr>
<tr><td>Oligo rev</td><td>AAACGACGATCTGACATTGTCCAG</td><td></td><td></td><td></td><td></td><td></td><td></td><td></td><td></td><td></td><td></td></tr>
<tr><td>Chromosome</td><td>start</td><td>end</td><td>strand</td><td>MM</td><td>target_seq</td><td>PAM</td><td>alignment</td><td>distance</td><td>position</td><td>gene name</td><td>gene id</td></tr>
<tr><td>GU305914.1</td><td>210300</td><td>210322</td><td>-</td><td>0</td><td>CTGGACAATGTCAGATCGTC</td><td>GGG</td><td>||||||||[||||||||||||]PAM</td><td>NA</td><td>Intergenic</td><td>NA</td><td>NA</td></tr>
<tr><td>T18639</td><td>CTCTCGCTCGAGCCGCCCGTCGG</td><td>1000</td><td></td><td></td><td></td><td></td><td></td><td></td><td></td><td></td><td></td></tr>
<tr><td>Oligo fwd</td><td>CACCCTCTCGCTCGAGCCGCCCGT</td><td></td><td></td><td></td><td></td><td></td><td></td><td></td><td></td><td></td><td></td></tr>
<tr><td>Oligo rev</td><td>AAACACGGGCGGCTCGAGCGAGAG</td><td></td><td></td><td></td><td></td><td></td><td></td><td></td><td></td><td></td><td></td></tr>
<tr><td>Chromosome</td><td>start</td><td>end</td><td>strand</td><td>MM</td><td>target_seq</td><td>PAM</td><td>alignment</td><td>distance</td><td>position</td><td>gene name</td><td>gene id</td></tr>
<tr><td>GU305914.1</td><td>210275</td><td>210297</td><td>-</td><td>0</td><td>CTCTCGCTCGAGCCGCCCGT</td><td>CGG</td><td>||||||||[||||||||||||]PAM</td><td>NA</td><td>Intergenic</td><td>NA</td><td>NA</td></tr>
<tr><td>T18640</td><td>TCTCGCTCGAGCCGCCCGTCGGG</td><td>1000</td><td></td><td></td><td></td><td></td><td></td><td></td><td></td><td></td><td></td></tr>
<tr><td>Oligo fwd</td><td>CACCTCTCGCTCGAGCCGCCCGTC</td><td></td><td></td><td></td><td></td><td></td><td></td><td></td><td></td><td></td><td></td></tr>
<tr><td>Oligo rev</td><td>AAACGACGGGCGGCTCGAGCGAGA</td><td></td><td></td><td></td><td></td><td></td><td></td><td></td><td></td><td></td><td></td></tr>
<tr><td>Chromosome</td><td>start</td><td>end</td><td>strand</td><td>MM</td><td>target_seq</td><td>PAM</td><td>alignment</td><td>distance</td><td>position</td><td>gene name</td><td>gene id</td></tr>
<tr><td>GU305914.1</td><td>210274</td><td>210296</td><td>-</td><td>0</td><td>TCTCGCTCGAGCCGCCCGTC</td><td>GGG</td><td>||||||||[||||||||||||]PAM</td><td>NA</td><td>Intergenic</td><td>NA</td><td>NA</td></tr>
<tr><td>T18641</td><td>GTCGGGCCACATACACTAACTGG</td><td>1000</td><td></td><td></td><td></td><td></td><td></td><td></td><td></td><td></td><td></td></tr>
<tr><td>Oligo fwd</td><td>CACCGTCGGGCCACATACACTAAC</td><td></td><td></td><td></td><td></td><td></td><td></td><td></td><td></td><td></td><td></td></tr>
<tr><td>Oligo rev</td><td>AAACGTTAGTGTATGTGGCCCGAC</td><td></td><td></td><td></td><td></td><td></td><td></td><td></td><td></td><td></td><td></td></tr>
<tr><td>Chromosome</td><td>start</td><td>end</td><td>strand</td><td>MM</td><td>target_seq</td><td>PAM</td><td>alignment</td><td>distance</td><td>position</td><td>gene name</td><td>gene id</td></tr>
</table>

<table>
<tr><td>GU305914.1</td><td>210257</td><td>210279</td><td>-</td><td>0</td><td>GTCGGGCCACATACACTAAC</td><td>TGG</td><td>||||||||[||||||||||||]PAM</td><td>NA</td><td>Intergenic</td><td>NA</td><td>NA</td></tr>
<tr><td>T18642</td><td>AACTGGACAATGATCAACCTCGG</td><td>1000</td><td></td><td></td><td></td><td></td><td></td><td></td><td></td><td></td><td></td></tr>
<tr><td>Oligo fwd</td><td>CACCAACTGGACAATGATCAACCT</td><td></td><td></td><td></td><td></td><td></td><td></td><td></td><td></td><td></td><td></td></tr>
<tr><td>Oligo rev</td><td>AAACAGGTTGATCATTGTCCAGTT</td><td></td><td></td><td></td><td></td><td></td><td></td><td></td><td></td><td></td><td></td></tr>
<tr><td>Chromosome</td><td>start</td><td>end</td><td>strand</td><td>MM</td><td>target_seq</td><td>PAM</td><td>alignment</td><td>distance</td><td>position</td><td>gene name</td><td>gene id</td></tr>
<tr><td>GU305914.1</td><td>210240</td><td>210262</td><td>-</td><td>0</td><td>AACTGGACAATGATCAACCT</td><td>CGG</td><td>||||||||[||||||||||||]PAM</td><td>NA</td><td>Intergenic</td><td>NA</td><td>NA</td></tr>
<tr><td>T18643</td><td>TGGACAATGATCAACCTCGGTGG</td><td>1000</td><td></td><td></td><td></td><td></td><td></td><td></td><td></td><td></td><td></td></tr>
<tr><td>Oligo fwd</td><td>CACCTGGACAATGATCAACCTCGG</td><td></td><td></td><td></td><td></td><td></td><td></td><td></td><td></td><td></td><td></td></tr>
<tr><td>Oligo rev</td><td>AAACCCGAGGTTGATCATTGTCCA</td><td></td><td></td><td></td><td></td><td></td><td></td><td></td><td></td><td></td><td></td></tr>
<tr><td>Chromosome</td><td>start</td><td>end</td><td>strand</td><td>MM</td><td>target_seq</td><td>PAM</td><td>alignment</td><td>distance</td><td>position</td><td>gene name</td><td>gene id</td></tr>
<tr><td>GU305914.1</td><td>210237</td><td>210259</td><td>-</td><td>0</td><td>TGGACAATGATCAACCTCGG</td><td>TGG</td><td>||||||||[||||||||||||]PAM</td><td>NA</td><td>Intergenic</td><td>NA</td><td>NA</td></tr>
<tr><td>T18644</td><td>GGACAATGATCAACCTCGGTGGG</td><td>1000</td><td></td><td></td><td></td><td></td><td></td><td></td><td></td><td></td><td></td></tr>
<tr><td>Oligo fwd</td><td>CACCGGACAATGATCAACCTCGGT</td><td></td><td></td><td></td><td></td><td></td><td></td><td></td><td></td><td></td><td></td></tr>
<tr><td>Oligo rev</td><td>AAACACCGAGGTTGATCATTGTCC</td><td></td><td></td><td></td><td></td><td></td><td></td><td></td><td></td><td></td><td></td></tr>
<tr><td>Chromosome</td><td>start</td><td>end</td><td>strand</td><td>MM</td><td>target_seq</td><td>PAM</td><td>alignment</td><td>distance</td><td>position</td><td>gene name</td><td>gene id</td></tr>
<tr><td>GU305914.1</td><td>210236</td><td>210258</td><td>-</td><td>0</td><td>GGACAATGATCAACCTCGGT</td><td>GGG</td><td>||||||||[||||||||||||]PAM</td><td>NA</td><td>Intergenic</td><td>NA</td><td>NA</td></tr>
<tr><td>T18645</td><td>GACAATGATCAACCTCGGTGGGG</td><td>1000</td><td></td><td></td><td></td><td></td><td></td><td></td><td></td><td></td><td></td></tr>
<tr><td>Oligo fwd</td><td>CACCGACAATGATCAACCTCGGTG</td><td></td><td></td><td></td><td></td><td></td><td></td><td></td><td></td><td></td><td></td></tr>
<tr><td>Oligo rev</td><td>AAACCACCGAGGTTGATCATTGTC</td><td></td><td></td><td></td><td></td><td></td><td></td><td></td><td></td><td></td><td></td></tr>
<tr><td>Chromosome</td><td>start</td><td>end</td><td>strand</td><td>MM</td><td>target_seq</td><td>PAM</td><td>alignment</td><td>distance</td><td>position</td><td>gene name</td><td>gene id</td></tr>
<tr><td>GU305914.1</td><td>210235</td><td>210257</td><td>-</td><td>0</td><td>GACAATGATCAACCTCGGTG</td><td>GGG</td><td>||||||||[||||||||||||]PAM</td><td>NA</td><td>Intergenic</td><td>NA</td><td>NA</td></tr>
<tr><td>T18646</td><td>ATCAACCTCGGTGGGGAATCTGG</td><td>1000</td><td></td><td></td><td></td><td></td><td></td><td></td><td></td><td></td><td></td></tr>
<tr><td>Oligo fwd</td><td>CACCATCAACCTCGGTGGGGAATC</td><td></td><td></td><td></td><td></td><td></td><td></td><td></td><td></td><td></td><td></td></tr>
<tr><td>Oligo rev</td><td>AAACGATTCCCCACCGAGGTTGAT</td><td></td><td></td><td></td><td></td><td></td><td></td><td></td><td></td><td></td><td></td></tr>
<tr><td>Chromosome</td><td>start</td><td>end</td><td>strand</td><td>MM</td><td>target_seq</td><td>PAM</td><td>alignment</td><td>distance</td><td>position</td><td>gene name</td><td>gene id</td></tr>
<tr><td>GU305914.1</td><td>210228</td><td>210250</td><td>-</td><td>0</td><td>ATCAACCTCGGTGGGGAATC</td><td>TGG</td><td>||||||||[||||||||||||]PAM</td><td>NA</td><td>Intergenic</td><td>NA</td><td>NA</td></tr>
<tr><td>T18647</td><td>TCAACCTCGGTGGGGAATCTGGG</td><td>1000</td><td></td><td></td><td></td><td></td><td></td><td></td><td></td><td></td><td></td></tr>
<tr><td>Oligo fwd</td><td>CACCTCAACCTCGGTGGGGAATCT</td><td></td><td></td><td></td><td></td><td></td><td></td><td></td><td></td><td></td><td></td></tr>
<tr><td>Oligo rev</td><td>AAACAGATTCCCCACCGAGGTTGA</td><td></td><td></td><td></td><td></td><td></td><td></td><td></td><td></td><td></td><td></td></tr>
<tr><td>Chromosome</td><td>start</td><td>end</td><td>strand</td><td>MM</td><td>target_seq</td><td>PAM</td><td>alignment</td><td>distance</td><td>position</td><td>gene name</td><td>gene id</td></tr>
<tr><td>GU305914.1</td><td>210227</td><td>210249</td><td>-</td><td>0</td><td>TCAACCTCGGTGGGGAATCT</td><td>GGG</td><td>||||||||[||||||||||||]PAM</td><td>NA</td><td>Intergenic</td><td>NA</td><td>NA</td></tr>
<tr><td>T18648</td><td>CAACCTCGGTGGGGAATCTGGGG</td><td>1000</td><td></td><td></td><td></td><td></td><td></td><td></td><td></td><td></td><td></td></tr>
<tr><td>Oligo fwd</td><td>CACCCAACCTCGGTGGGGAATCTG</td><td></td><td></td><td></td><td></td><td></td><td></td><td></td><td></td><td></td><td></td></tr>
<tr><td>Oligo rev</td><td>AAACCAGATTCCCCACCGAGGTTG</td><td></td><td></td><td></td><td></td><td></td><td></td><td></td><td></td><td></td><td></td></tr>
<tr><td>Chromosome</td><td>start</td><td>end</td><td>strand</td><td>MM</td><td>target_seq</td><td>PAM</td><td>alignment</td><td>distance</td><td>position</td><td>gene name</td><td>gene id</td></tr>
<tr><td>GU305914.1</td><td>210226</td><td>210248</td><td>-</td><td>0</td><td>CAACCTCGGTGGGGAATCTG</td><td>GGG</td><td>||||||||[||||||||||||]PAM</td><td>NA</td><td>Intergenic</td><td>NA</td><td>NA</td></tr>
<tr><td>T18649</td><td>CCTCGGTGGGGAATCTGGGGCGG</td><td>1000</td><td></td><td></td><td></td><td></td><td></td><td></td><td></td><td></td><td></td></tr>
<tr><td>Oligo fwd</td><td>CACCCCTCGGTGGGGAATCTGGGG</td><td></td><td></td><td></td><td></td><td></td><td></td><td></td><td></td><td></td><td></td></tr>
<tr><td>Oligo rev</td><td>AAACCCCCAGATTCCCCACCGAGG</td><td></td><td></td><td></td><td></td><td></td><td></td><td></td><td></td><td></td><td></td></tr>
<tr><td>Chromosome</td><td>start</td><td>end</td><td>strand</td><td>MM</td><td>target_seq</td><td>PAM</td><td>alignment</td><td>distance</td><td>position</td><td>gene name</td><td>gene id</td></tr>
<tr><td>GU305914.1</td><td>210223</td><td>210245</td><td>-</td><td>0</td><td>CCTCGGTGGGGAATCTGGGG</td><td>CGG</td><td>||||||||[||||||||||||]PAM</td><td>NA</td><td>Intergenic</td><td>NA</td><td>NA</td></tr>
<tr><td>T18650</td><td>CTCGGTGGGGAATCTGGGGCGGG</td><td>1000</td><td></td><td></td><td></td><td></td><td></td><td></td><td></td><td></td><td></td></tr>
<tr><td>Oligo fwd</td><td>CACCCTCGGTGGGGAATCTGGGGC</td><td></td><td></td><td></td><td></td><td></td><td></td><td></td><td></td><td></td><td></td></tr>
<tr><td>Oligo rev</td><td>AAACGCCCCAGATTCCCCACCGAG</td><td></td><td></td><td></td><td></td><td></td><td></td><td></td><td></td><td></td><td></td></tr>
<tr><td>Chromosome</td><td>start</td><td>end</td><td>strand</td><td>MM</td><td>target_seq</td><td>PAM</td><td>alignment</td><td>distance</td><td>position</td><td>gene name</td><td>gene id</td></tr>
<tr><td>GU305914.1</td><td>210222</td><td>210244</td><td>-</td><td>0</td><td>CTCGGTGGGGAATCTGGGGC</td><td>GGG</td><td>||||||||[||||||||||||]PAM</td><td>NA</td><td>Intergenic</td><td>NA</td><td>NA</td></tr>
<tr><td>T18651</td><td>GAATCTGGGGCGGGTTGACCTGG</td><td>1000</td><td></td><td></td><td></td><td></td><td></td><td></td><td></td><td></td><td></td></tr>
<tr><td>Oligo fwd</td><td>CACCGAATCTGGGGCGGGTTGACC</td><td></td><td></td><td></td><td></td><td></td><td></td><td></td><td></td><td></td><td></td></tr>
<tr><td>Oligo rev</td><td>AAACGGTCAACCCGCCCCAGATTC</td><td></td><td></td><td></td><td></td><td></td><td></td><td></td><td></td><td></td><td></td></tr>
<tr><td>Chromosome</td><td>start</td><td>end</td><td>strand</td><td>MM</td><td>target_seq</td><td>PAM</td><td>alignment</td><td>distance</td><td>position</td><td>gene name</td><td>gene id</td></tr>
<tr><td>GU305914.1</td><td>210213</td><td>210235</td><td>-</td><td>0</td><td>GAATCTGGGGCGGGTTGACC</td><td>TGG</td><td>||||||||[||||||||||||]PAM</td><td>NA</td><td>Intergenic</td><td>NA</td><td>NA</td></tr>
<tr><td>T18652</td><td>AATCTGGGGCGGGTTGACCTGGG</td><td>1000</td><td></td><td></td><td></td><td></td><td></td><td></td><td></td><td></td><td></td></tr>
<tr><td>Oligo fwd</td><td>CACCAATCTGGGGCGGGTTGACCT</td><td></td><td></td><td></td><td></td><td></td><td></td><td></td><td></td><td></td><td></td></tr>
<tr><td>Oligo rev</td><td>AAACAGGTCAACCCGCCCCAGATT</td><td></td><td></td><td></td><td></td><td></td><td></td><td></td><td></td><td></td><td></td></tr>
<tr><td>Chromosome</td><td>start</td><td>end</td><td>strand</td><td>MM</td><td>target_seq</td><td>PAM</td><td>alignment</td><td>distance</td><td>position</td><td>gene name</td><td>gene id</td></tr>
<tr><td>GU305914.1</td><td>210212</td><td>210234</td><td>-</td><td>0</td><td>AATCTGGGGCGGGTTGACCT</td><td>GGG</td><td>||||||||[||||||||||||]PAM</td><td>NA</td><td>Intergenic</td><td>NA</td><td>NA</td></tr>
<tr><td>T18653</td><td>GGGGCGGGTTGACCTGGGAATGG</td><td>1000</td><td></td><td></td><td></td><td></td><td></td><td></td><td></td><td></td><td></td></tr>
<tr><td>Oligo fwd</td><td>CACCGGGGCGGGTTGACCTGGGAA</td><td></td><td></td><td></td><td></td><td></td><td></td><td></td><td></td><td></td><td></td></tr>
<tr><td>Oligo rev</td><td>AAACTTCCCAGGTCAACCCGCCCC</td><td></td><td></td><td></td><td></td><td></td><td></td><td></td><td></td><td></td><td></td></tr>
<tr><td>Chromosome</td><td>start</td><td>end</td><td>strand</td><td>MM</td><td>target_seq</td><td>PAM</td><td>alignment</td><td>distance</td><td>position</td><td>gene name</td><td>gene id</td></tr>
<tr><td>GU305914.1</td><td>210207</td><td>210229</td><td>-</td><td>0</td><td>GGGGCGGGTTGACCTGGGAA</td><td>TGG</td><td>||||||||[||||||||||||]PAM</td><td>NA</td><td>Intergenic</td><td>NA</td><td>NA</td></tr>
<tr><td>T18654</td><td>GGGCGGGTTGACCTGGGAATGGG</td><td>1000</td><td></td><td></td><td></td><td></td><td></td><td></td><td></td><td></td><td></td></tr>
<tr><td>Oligo fwd</td><td>CACCGGGCGGGTTGACCTGGGAAT</td><td></td><td></td><td></td><td></td><td></td><td></td><td></td><td></td><td></td><td></td></tr>
<tr><td>Oligo rev</td><td>AAACATTCCCAGGTCAACCCGCCC</td><td></td><td></td><td></td><td></td><td></td><td></td><td></td><td></td><td></td><td></td></tr>
<tr><td>Chromosome</td><td>start</td><td>end</td><td>strand</td><td>MM</td><td>target_seq</td><td>PAM</td><td>alignment</td><td>distance</td><td>position</td><td>gene name</td><td>gene id</td></tr>
<tr><td>GU305914.1</td><td>210206</td><td>210228</td><td>-</td><td>0</td><td>GGGCGGGTTGACCTGGGAAT</td><td>GGG</td><td>||||||||[||||||||||||]PAM</td><td>NA</td><td>Intergenic</td><td>NA</td><td>NA</td></tr>
<tr><td>T18655</td><td>GTTGACCTGGGAATGGGCTCAGG</td><td>1000</td><td></td><td></td><td></td><td></td><td></td><td></td><td></td><td></td><td></td></tr>
<tr><td>Oligo fwd</td><td>CACCGTTGACCTGGGAATGGGCTC</td><td></td><td></td><td></td><td></td><td></td><td></td><td></td><td></td><td></td><td></td></tr>
<tr><td>Oligo rev</td><td>AAACGAGCCCATTCCCAGGTCAAC</td><td></td><td></td><td></td><td></td><td></td><td></td><td></td><td></td><td></td><td></td></tr>
<tr><td>Chromosome</td><td>start</td><td>end</td><td>strand</td><td>MM</td><td>target_seq</td><td>PAM</td><td>alignment</td><td>distance</td><td>position</td><td>gene name</td><td>gene id</td></tr>
<tr><td>GU305914.1</td><td>210200</td><td>210222</td><td>-</td><td>0</td><td>GTTGACCTGGGAATGGGCTC</td><td>AGG</td><td>||||||||[||||||||||||]PAM</td><td>NA</td><td>Intergenic</td><td>NA</td><td>NA</td></tr>
<tr><td>T18656</td><td>TTGACCTGGGAATGGGCTCAGGG</td><td>1000</td><td></td><td></td><td></td><td></td><td></td><td></td><td></td><td></td><td></td></tr>
<tr><td>Oligo fwd</td><td>CACCTTGACCTGGGAATGGGCTCA</td><td></td><td></td><td></td><td></td><td></td><td></td><td></td><td></td><td></td><td></td></tr>
<tr><td>Oligo rev</td><td>AAACTGAGCCCATTCCCAGGTCAA</td><td></td><td></td><td></td><td></td><td></td><td></td><td></td><td></td><td></td><td></td></tr>
</table>

| Chromosome | start | end | strand | MM | target_seq | PAM | alignment | distance | position | gene name | gene id |
|---|---|---|---|---|---|---|---|---|---|---|---|
| GU305914.1 | 210199 | 210221 | - | 0 | TTGACCTGGGAATGGCTCA | GGG | \|\|\|\|\|\|\|\|[\|\|\|\|\|\|\|\|\|\|\|\|]PAM | NA | Intergenic | NA | NA |
| T18657 | GTCAACTCTTTATTCCGCTGTGG | 1000 | | | | | | | | | |
| Oligo fwd | CACCGTCAACTCTTTATTCCGCTG | | | | | | | | | | |
| Oligo rev | AAACCAGCGGAATAAAGAGTTGAC | | | | | | | | | | |
| Chromosome | start | end | strand | MM | target_seq | PAM | alignment | distance | position | gene name | gene id |
| GU305914.1 | 210138 | 210160 | - | 0 | GTCAACTCTTTATTCCGCTG | TGG | \|\|\|\|\|\|\|\|[\|\|\|\|\|\|\|\|\|\|\|\|]PAM | NA | Intergenic | NA | NA |
| T18658 | TGAACGAACATATTCGCGCATGG | 1000 | | | | | | | | | |
| Oligo fwd | CACCTGAACGAACATATTCGCGCA | | | | | | | | | | |
| Oligo rev | AAACTGCGCGAATATGTTCGTTCA | | | | | | | | | | |
| Chromosome | start | end | strand | MM | target_seq | PAM | alignment | distance | position | gene name | gene id |
| GU305914.1 | 210107 | 210129 | - | 0 | TGAACGAACATATTCGCGCA | TGG | \|\|\|\|\|\|\|\|[\|\|\|\|\|\|\|\|\|\|\|\|]PAM | NA | Intergenic | NA | NA |
| T18659 | ACGAACATATTCGCGCATGGAGG | 1000 | | | | | | | | | |
| Oligo fwd | CACCACGAACATATTCGCGCATGG | | | | | | | | | | |
| Oligo rev | AAACCCATGCGCGAATATGTTCGT | | | | | | | | | | |
| Chromosome | start | end | strand | MM | target_seq | PAM | alignment | distance | position | gene name | gene id |
| GU305914.1 | 210104 | 210126 | - | 0 | ACGAACATATTCGCGCATGG | AGG | \|\|\|\|\|\|\|\|[\|\|\|\|\|\|\|\|\|\|\|\|]PAM | NA | Intergenic | NA | NA |
| T18660 | CGAACATATTCGCGCATGGAGGG | 1000 | | | | | | | | | |
| Oligo fwd | CACCCGAACATATTCGCGCATGGA | | | | | | | | | | |
| Oligo rev | AAACTCCATGCGCGAATATGTTCG | | | | | | | | | | |
| Chromosome | start | end | strand | MM | target_seq | PAM | alignment | distance | position | gene name | gene id |
| GU305914.1 | 210103 | 210125 | - | 0 | CGAACATATTCGCGCATGGA | GGG | \|\|\|\|\|\|\|\|[\|\|\|\|\|\|\|\|\|\|\|\|]PAM | NA | Intergenic | NA | NA |
| T18661 | CATATTCGCGCATGGAGGGACGG | 1000 | | | | | | | | | |
| Oligo fwd | CACCCATATTCGCGCATGGAGGGA | | | | | | | | | | |
| Oligo rev | AAACTCCCTCCATGCGCGAATATG | | | | | | | | | | |
| Chromosome | start | end | strand | MM | target_seq | PAM | alignment | distance | position | gene name | gene id |
| GU305914.1 | 210099 | 210121 | - | 0 | CATATTCGCGCATGGAGGGA | CGG | \|\|\|\|\|\|\|\|[\|\|\|\|\|\|\|\|\|\|\|\|]PAM | NA | Intergenic | NA | NA |
| T18662 | ATATTCGCGCATGGAGGGACGGG | 1000 | | | | | | | | | |
| Oligo fwd | CACCATATTCGCGCATGGAGGGAC | | | | | | | | | | |
| Oligo rev | AAACGTCCCTCCATGCGCGAATAT | | | | | | | | | | |
| Chromosome | start | end | strand | MM | target_seq | PAM | alignment | distance | position | gene name | gene id |
| GU305914.1 | 210098 | 210120 | - | 0 | ATATTCGCGCATGGAGGGAC | GGG | \|\|\|\|\|\|\|\|[\|\|\|\|\|\|\|\|\|\|\|\|]PAM | NA | Intergenic | NA | NA |
| T18663 | TATTCGCGCATGGAGGGACGGGG | 1000 | | | | | | | | | |
| Oligo fwd | CACCTATTCGCGCATGGAGGGACG | | | | | | | | | | |
| Oligo rev | AAACCGTCCCTCCATGCGCGAATA | | | | | | | | | | |
| Chromosome | start | end | strand | MM | target_seq | PAM | alignment | distance | position | gene name | gene id |
| GU305914.1 | 210097 | 210119 | - | 0 | TATTCGCGCATGGAGGGACG | GGG | \|\|\|\|\|\|\|\|[\|\|\|\|\|\|\|\|\|\|\|\|]PAM | NA | Intergenic | NA | NA |
| T18664 | ATTCGCGCATGGAGGGACGGGGG | 1000 | | | | | | | | | |
| Oligo fwd | CACCATTCGCGCATGGAGGGACGG | | | | | | | | | | |
| Oligo rev | AAACCCGTCCCTCCATGCGCGAAT | | | | | | | | | | |
| Chromosome | start | end | strand | MM | target_seq | PAM | alignment | distance | position | gene name | gene id |
| GU305914.1 | 210096 | 210118 | - | 0 | ATTCGCGCATGGAGGGACGG | GGG | \|\|\|\|\|\|\|\|[\|\|\|\|\|\|\|\|\|\|\|\|]PAM | NA | Intergenic | NA | NA |
| T18665 | GGAGGGACGGGGGCTCGTTGAGG | 1000 | | | | | | | | | |
| Oligo fwd | CACCGGAGGGACGGGGGCTCGTTG | | | | | | | | | | |
| Oligo rev | AAACCAACGAGCCCCCGTCCCTCC | | | | | | | | | | |
| Chromosome | start | end | strand | MM | target_seq | PAM | alignment | distance | position | gene name | gene id |
| GU305914.1 | 210086 | 210108 | - | 0 | GGAGGGACGGGGGCTCGTTG | AGG | \|\|\|\|\|\|\|\|[\|\|\|\|\|\|\|\|\|\|\|\|]PAM | NA | Intergenic | NA | NA |
| T18666 | GGGGCTCGTTGAGGTATGATTGG | 1000 | | | | | | | | | |
| Oligo fwd | CACCGGGGCTCGTTGAGGTATGAT | | | | | | | | | | |
| Oligo rev | AAACATCATACCTCAACGAGCCCC | | | | | | | | | | |
| Chromosome | start | end | strand | MM | target_seq | PAM | alignment | distance | position | gene name | gene id |
| GU305914.1 | 210077 | 210099 | - | 0 | GGGGCTCGTTGAGGTATGAT | TGG | \|\|\|\|\|\|\|\|[\|\|\|\|\|\|\|\|\|\|\|\|]PAM | NA | Intergenic | NA | NA |
| T18667 | GCGTGATACGTGCGCTCGTGTGG | 1000 | | | | | | | | | |
| Oligo fwd | CACCGCGTGATACGTGCGCTCGTG | | | | | | | | | | |
| Oligo rev | AAACCACGAGCGCACGTATCACGC | | | | | | | | | | |
| Chromosome | start | end | strand | MM | target_seq | PAM | alignment | distance | position | gene name | gene id |
| GU305914.1 | 210055 | 210077 | - | 0 | GCGTGATACGTGCGCTCGTG | TGG | \|\|\|\|\|\|\|\|[\|\|\|\|\|\|\|\|\|\|\|\|]PAM | NA | Intergenic | NA | NA |
| T18668 | TGATACGTGCGCTCGTGTGGTGG | 1000 | | | | | | | | | |
| Oligo fwd | CACCTGATACGTGCGCTCGTGTGG | | | | | | | | | | |
| Oligo rev | AAACCCACACGAGCGCACGTATCA | | | | | | | | | | |
| Chromosome | start | end | strand | MM | target_seq | PAM | alignment | distance | position | gene name | gene id |
| GU305914.1 | 210052 | 210074 | - | 0 | TGATACGTGCGCTCGTGTGG | TGG | \|\|\|\|\|\|\|\|[\|\|\|\|\|\|\|\|\|\|\|\|]PAM | NA | Intergenic | NA | NA |
| T18669 | TGCGCTCGTGTGGTGGACTTTGG | 1000 | | | | | | | | | |
| Oligo fwd | CACCTGCGCTCGTGTGGTGGACTT | | | | | | | | | | |
| Oligo rev | AAACAAGTCCACCACACGAGCGCA | | | | | | | | | | |
| Chromosome | start | end | strand | MM | target_seq | PAM | alignment | distance | position | gene name | gene id |
| GU305914.1 | 210045 | 210067 | - | 0 | TGCGCTCGTGTGGTGGACTT | TGG | \|\|\|\|\|\|\|\|[\|\|\|\|\|\|\|\|\|\|\|\|]PAM | NA | Intergenic | NA | NA |
| T18670 | AGCCTCTGTGTGTGAGTCGCCGG | 1000 | | | | | | | | | |
| Oligo fwd | CACCAGCCTCTGTGTGTGAGTCGC | | | | | | | | | | |
| Oligo rev | AAACGCGACTCACACACAGAGGCT | | | | | | | | | | |
| Chromosome | start | end | strand | MM | target_seq | PAM | alignment | distance | position | gene name | gene id |
| GU305914.1 | 210009 | 210031 | - | 0 | AGCCTCTGTGTGTGAGTCGC | CGG | \|\|\|\|\|\|\|\|[\|\|\|\|\|\|\|\|\|\|\|\|]PAM | NA | Intergenic | NA | NA |
| T18671 | GTCGCCGGTCCTCGTTCTTACGG | 1000 | | | | | | | | | |
| Oligo fwd | CACCGTCGCCGGTCCTCGTTCTTA | | | | | | | | | | |

<table>
<tr><td>Oligo rev</td><td>AAACTAAGAACGAGGACCGGCGAC</td><td></td><td></td><td></td><td></td><td></td><td></td><td></td><td></td><td></td><td></td></tr>
<tr><td>Chromosome</td><td>start</td><td>end</td><td>strand</td><td>MM</td><td>target_seq</td><td>PAM</td><td>alignment</td><td>distance</td><td>position</td><td>gene name</td><td>gene id</td></tr>
<tr><td>GU305914.1</td><td>209994</td><td>210016</td><td>-</td><td>0</td><td>GTCGCCGGTCCTCGTTCTTA</td><td>CGG</td><td>||||||||[||||||||||||]PAM</td><td>NA</td><td>Intergenic</td><td>NA</td><td>NA</td></tr>
<tr><td>T18672</td><td>CCTCGTTCTTACGGCGTCGACGG</td><td>1000</td><td></td><td></td><td></td><td></td><td></td><td></td><td></td><td></td><td></td></tr>
<tr><td>Oligo fwd</td><td>CACCCCTCGTTCTTACGGCGTCGA</td><td></td><td></td><td></td><td></td><td></td><td></td><td></td><td></td><td></td><td></td></tr>
<tr><td>Oligo rev</td><td>AAACTCGACGCCGTAAGAACGAGG</td><td></td><td></td><td></td><td></td><td></td><td></td><td></td><td></td><td></td><td></td></tr>
<tr><td>Chromosome</td><td>start</td><td>end</td><td>strand</td><td>MM</td><td>target_seq</td><td>PAM</td><td>alignment</td><td>distance</td><td>position</td><td>gene name</td><td>gene id</td></tr>
<tr><td>GU305914.1</td><td>209985</td><td>210007</td><td>-</td><td>0</td><td>CCTCGTTCTTACGGCGTCGA</td><td>CGG</td><td>||||||||[||||||||||||]PAM</td><td>NA</td><td>Intergenic</td><td>NA</td><td>NA</td></tr>
<tr><td>T18673</td><td>GTTCTTACGGCGTCGACGGCCGG</td><td>1000</td><td></td><td></td><td></td><td></td><td></td><td></td><td></td><td></td><td></td></tr>
<tr><td>Oligo fwd</td><td>CACCGTTCTTACGGCGTCGACGGC</td><td></td><td></td><td></td><td></td><td></td><td></td><td></td><td></td><td></td><td></td></tr>
<tr><td>Oligo rev</td><td>AAACGCCGTCGACGCCGTAAGAAC</td><td></td><td></td><td></td><td></td><td></td><td></td><td></td><td></td><td></td><td></td></tr>
<tr><td>Chromosome</td><td>start</td><td>end</td><td>strand</td><td>MM</td><td>target_seq</td><td>PAM</td><td>alignment</td><td>distance</td><td>position</td><td>gene name</td><td>gene id</td></tr>
<tr><td>GU305914.1</td><td>209981</td><td>210003</td><td>-</td><td>0</td><td>GTTCTTACGGCGTCGACGGC</td><td>CGG</td><td>||||||||[||||||||||||]PAM</td><td>NA</td><td>Intergenic</td><td>NA</td><td>NA</td></tr>
<tr><td>T18674</td><td>TTCTTACGGCGTCGACGGCCGGG</td><td>1000</td><td></td><td></td><td></td><td></td><td></td><td></td><td></td><td></td><td></td></tr>
<tr><td>Oligo fwd</td><td>CACCTTCTTACGGCGTCGACGGCC</td><td></td><td></td><td></td><td></td><td></td><td></td><td></td><td></td><td></td><td></td></tr>
<tr><td>Oligo rev</td><td>AAACGGCCGTCGACGCCGTAAGAA</td><td></td><td></td><td></td><td></td><td></td><td></td><td></td><td></td><td></td><td></td></tr>
<tr><td>Chromosome</td><td>start</td><td>end</td><td>strand</td><td>MM</td><td>target_seq</td><td>PAM</td><td>alignment</td><td>distance</td><td>position</td><td>gene name</td><td>gene id</td></tr>
<tr><td>GU305914.1</td><td>209980</td><td>210002</td><td>-</td><td>0</td><td>TTCTTACGGCGTCGACGGCC</td><td>GGG</td><td>||||||||[||||||||||||]PAM</td><td>NA</td><td>Intergenic</td><td>NA</td><td>NA</td></tr>
<tr><td>T18675</td><td>GACGGCCGGGCTTCATGATGCGG</td><td>1000</td><td></td><td></td><td></td><td></td><td></td><td></td><td></td><td></td><td></td></tr>
<tr><td>Oligo fwd</td><td>CACCGACGGCCGGGCTTCATGATG</td><td></td><td></td><td></td><td></td><td></td><td></td><td></td><td></td><td></td><td></td></tr>
<tr><td>Oligo rev</td><td>AAACCATCATGAAGCCCGGCCGTC</td><td></td><td></td><td></td><td></td><td></td><td></td><td></td><td></td><td></td><td></td></tr>
<tr><td>Chromosome</td><td>start</td><td>end</td><td>strand</td><td>MM</td><td>target_seq</td><td>PAM</td><td>alignment</td><td>distance</td><td>position</td><td>gene name</td><td>gene id</td></tr>
<tr><td>GU305914.1</td><td>209967</td><td>209989</td><td>-</td><td>0</td><td>GACGGCCGGGCTTCATGATG</td><td>CGG</td><td>||||||||[||||||||||||]PAM</td><td>NA</td><td>Intergenic</td><td>NA</td><td>NA</td></tr>
<tr><td>T18676</td><td>GATGCGGTCACAAAGACCGTCGG</td><td>1000</td><td></td><td></td><td></td><td></td><td></td><td></td><td></td><td></td><td></td></tr>
<tr><td>Oligo fwd</td><td>CACCGATGCGGTCACAAAGACCGT</td><td></td><td></td><td></td><td></td><td></td><td></td><td></td><td></td><td></td><td></td></tr>
<tr><td>Oligo rev</td><td>AAACACGGTCTTTGTGACCGCATC</td><td></td><td></td><td></td><td></td><td></td><td></td><td></td><td></td><td></td><td></td></tr>
<tr><td>Chromosome</td><td>start</td><td>end</td><td>strand</td><td>MM</td><td>target_seq</td><td>PAM</td><td>alignment</td><td>distance</td><td>position</td><td>gene name</td><td>gene id</td></tr>
<tr><td>GU305914.1</td><td>209951</td><td>209973</td><td>-</td><td>0</td><td>GATGCGGTCACAAAGACCGT</td><td>CGG</td><td>||||||||[||||||||||||]PAM</td><td>NA</td><td>Intergenic</td><td>NA</td><td>NA</td></tr>
<tr><td>GU305914.1</td><td>112022</td><td>112044</td><td>-</td><td>4</td><td>GATCCGCGCACAAAGACCGG</td><td>CGG</td><td>|||-||--[|||||||||||-]PAM</td><td>NA</td><td>Intergenic</td><td>NA</td><td>NA</td></tr>
<tr><td>T18677</td><td>GCAGACAGTGCCGCTGTTGACGG</td><td>1000</td><td></td><td></td><td></td><td></td><td></td><td></td><td></td><td></td><td></td></tr>
<tr><td>Oligo fwd</td><td>CACCGCAGACAGTGCCGCTGTTGA</td><td></td><td></td><td></td><td></td><td></td><td></td><td></td><td></td><td></td><td></td></tr>
<tr><td>Oligo rev</td><td>AAACTCAACAGCGGCACTGTCTGC</td><td></td><td></td><td></td><td></td><td></td><td></td><td></td><td></td><td></td><td></td></tr>
<tr><td>Chromosome</td><td>start</td><td>end</td><td>strand</td><td>MM</td><td>target_seq</td><td>PAM</td><td>alignment</td><td>distance</td><td>position</td><td>gene name</td><td>gene id</td></tr>
<tr><td>GU305914.1</td><td>209922</td><td>209944</td><td>-</td><td>0</td><td>GCAGACAGTGCCGCTGTTGA</td><td>CGG</td><td>||||||||[||||||||||||]PAM</td><td>NA</td><td>Intergenic</td><td>NA</td><td>NA</td></tr>
<tr><td>T18678</td><td>CTGTTGACGGTGTACGATAACGG</td><td>1000</td><td></td><td></td><td></td><td></td><td></td><td></td><td></td><td></td><td></td></tr>
<tr><td>Oligo fwd</td><td>CACCCTGTTGACGGTGTACGATAA</td><td></td><td></td><td></td><td></td><td></td><td></td><td></td><td></td><td></td><td></td></tr>
<tr><td>Oligo rev</td><td>AAACTTATCGTACACCGTCAACAG</td><td></td><td></td><td></td><td></td><td></td><td></td><td></td><td></td><td></td><td></td></tr>
<tr><td>Chromosome</td><td>start</td><td>end</td><td>strand</td><td>MM</td><td>target_seq</td><td>PAM</td><td>alignment</td><td>distance</td><td>position</td><td>gene name</td><td>gene id</td></tr>
<tr><td>GU305914.1</td><td>209909</td><td>209931</td><td>-</td><td>0</td><td>CTGTTGACGGTGTACGATAA</td><td>CGG</td><td>||||||||[||||||||||||]PAM</td><td>NA</td><td>Intergenic</td><td>NA</td><td>NA</td></tr>
<tr><td>T18679</td><td>TACGATAACGGCACCGTCTGTGG</td><td>1000</td><td></td><td></td><td></td><td></td><td></td><td></td><td></td><td></td><td></td></tr>
<tr><td>Oligo fwd</td><td>CACCTACGATAACGGCACCGTCTG</td><td></td><td></td><td></td><td></td><td></td><td></td><td></td><td></td><td></td><td></td></tr>
<tr><td>Oligo rev</td><td>AAACCAGACGGTGCCGTTATCGTA</td><td></td><td></td><td></td><td></td><td></td><td></td><td></td><td></td><td></td><td></td></tr>
<tr><td>Chromosome</td><td>start</td><td>end</td><td>strand</td><td>MM</td><td>target_seq</td><td>PAM</td><td>alignment</td><td>distance</td><td>position</td><td>gene name</td><td>gene id</td></tr>
<tr><td>GU305914.1</td><td>209897</td><td>209919</td><td>-</td><td>0</td><td>TACGATAACGGCACCGTCTG</td><td>TGG</td><td>||||||||[||||||||||||]PAM</td><td>NA</td><td>Intergenic</td><td>NA</td><td>NA</td></tr>
<tr><td>T18680</td><td>AACGGCACCGTCTGTGGTACAGG</td><td>1000</td><td></td><td></td><td></td><td></td><td></td><td></td><td></td><td></td><td></td></tr>
<tr><td>Oligo fwd</td><td>CACCAACGGCACCGTCTGTGGTAC</td><td></td><td></td><td></td><td></td><td></td><td></td><td></td><td></td><td></td><td></td></tr>
<tr><td>Oligo rev</td><td>AAACGTACCACAGACGGTGCCGTT</td><td></td><td></td><td></td><td></td><td></td><td></td><td></td><td></td><td></td><td></td></tr>
<tr><td>Chromosome</td><td>start</td><td>end</td><td>strand</td><td>MM</td><td>target_seq</td><td>PAM</td><td>alignment</td><td>distance</td><td>position</td><td>gene name</td><td>gene id</td></tr>
<tr><td>GU305914.1</td><td>209891</td><td>209913</td><td>-</td><td>0</td><td>AACGGCACCGTCTGTGGTAC</td><td>AGG</td><td>||||||||[||||||||||||]PAM</td><td>NA</td><td>Intergenic</td><td>NA</td><td>NA</td></tr>
<tr><td>T18681</td><td>ACGGCACCGTCTGTGGTACAGGG</td><td>1000</td><td></td><td></td><td></td><td></td><td></td><td></td><td></td><td></td><td></td></tr>
<tr><td>Oligo fwd</td><td>CACCACGGCACCGTCTGTGGTACA</td><td></td><td></td><td></td><td></td><td></td><td></td><td></td><td></td><td></td><td></td></tr>
<tr><td>Oligo rev</td><td>AAACTGTACCACAGACGGTGCCGT</td><td></td><td></td><td></td><td></td><td></td><td></td><td></td><td></td><td></td><td></td></tr>
<tr><td>Chromosome</td><td>start</td><td>end</td><td>strand</td><td>MM</td><td>target_seq</td><td>PAM</td><td>alignment</td><td>distance</td><td>position</td><td>gene name</td><td>gene id</td></tr>
<tr><td>GU305914.1</td><td>209890</td><td>209912</td><td>-</td><td>0</td><td>ACGGCACCGTCTGTGGTACA</td><td>GGG</td><td>||||||||[||||||||||||]PAM</td><td>NA</td><td>Intergenic</td><td>NA</td><td>NA</td></tr>
<tr><td>T18682</td><td>GCACCGTCTGTGGTACAGGGTGG</td><td>1000</td><td></td><td></td><td></td><td></td><td></td><td></td><td></td><td></td><td></td></tr>
<tr><td>Oligo fwd</td><td>CACCGCACCGTCTGTGGTACAGGG</td><td></td><td></td><td></td><td></td><td></td><td></td><td></td><td></td><td></td><td></td></tr>
<tr><td>Oligo rev</td><td>AAACCCCTGTACCACAGACGGTGC</td><td></td><td></td><td></td><td></td><td></td><td></td><td></td><td></td><td></td><td></td></tr>
<tr><td>Chromosome</td><td>start</td><td>end</td><td>strand</td><td>MM</td><td>target_seq</td><td>PAM</td><td>alignment</td><td>distance</td><td>position</td><td>gene name</td><td>gene id</td></tr>
<tr><td>GU305914.1</td><td>209887</td><td>209909</td><td>-</td><td>0</td><td>GCACCGTCTGTGGTACAGGG</td><td>TGG</td><td>||||||||[||||||||||||]PAM</td><td>NA</td><td>Intergenic</td><td>NA</td><td>NA</td></tr>
<tr><td>T18683</td><td>CACCGTCTGTGGTACAGGGTGGG</td><td>1000</td><td></td><td></td><td></td><td></td><td></td><td></td><td></td><td></td><td></td></tr>
<tr><td>Oligo fwd</td><td>CACCCACCGTCTGTGGTACAGGGT</td><td></td><td></td><td></td><td></td><td></td><td></td><td></td><td></td><td></td><td></td></tr>
<tr><td>Oligo rev</td><td>AAACACCCTGTACCACAGACGGTG</td><td></td><td></td><td></td><td></td><td></td><td></td><td></td><td></td><td></td><td></td></tr>
<tr><td>Chromosome</td><td>start</td><td>end</td><td>strand</td><td>MM</td><td>target_seq</td><td>PAM</td><td>alignment</td><td>distance</td><td>position</td><td>gene name</td><td>gene id</td></tr>
<tr><td>GU305914.1</td><td>209886</td><td>209908</td><td>-</td><td>0</td><td>CACCGTCTGTGGTACAGGGT</td><td>GGG</td><td>||||||||[||||||||||||]PAM</td><td>NA</td><td>Intergenic</td><td>NA</td><td>NA</td></tr>
<tr><td>T18684</td><td>AGGGTGGGTCATGTTGACGAAGG</td><td>1000</td><td></td><td></td><td></td><td></td><td></td><td></td><td></td><td></td><td></td></tr>
<tr><td>Oligo fwd</td><td>CACCAGGGTGGGTCATGTTGACGA</td><td></td><td></td><td></td><td></td><td></td><td></td><td></td><td></td><td></td><td></td></tr>
<tr><td>Oligo rev</td><td>AAACTCGTCAACATGACCCACCCT</td><td></td><td></td><td></td><td></td><td></td><td></td><td></td><td></td><td></td><td></td></tr>
<tr><td>Chromosome</td><td>start</td><td>end</td><td>strand</td><td>MM</td><td>target_seq</td><td>PAM</td><td>alignment</td><td>distance</td><td>position</td><td>gene name</td><td>gene id</td></tr>
<tr><td>GU305914.1</td><td>209871</td><td>209893</td><td>-</td><td>0</td><td>AGGGTGGGTCATGTTGACGA</td><td>AGG</td><td>||||||||[||||||||||||]PAM</td><td>NA</td><td>Intergenic</td><td>NA</td><td>NA</td></tr>
<tr><td>T18685</td><td>CCTGAATTCATTTTCATGCAAGG</td><td>1000</td><td></td><td></td><td></td><td></td><td></td><td></td><td></td><td></td><td></td></tr>
<tr><td>Oligo fwd</td><td>CACCCCTGAATTCATTTTCATGCA</td><td></td><td></td><td></td><td></td><td></td><td></td><td></td><td></td><td></td><td></td></tr>
<tr><td>Oligo rev</td><td>AAACTGCATGAAAATGAATTCAGG</td><td></td><td></td><td></td><td></td><td></td><td></td><td></td><td></td><td></td><td></td></tr>
<tr><td>Chromosome</td><td>start</td><td>end</td><td>strand</td><td>MM</td><td>target_seq</td><td>PAM</td><td>alignment</td><td>distance</td><td>position</td><td>gene name</td><td>gene id</td></tr>
<tr><td>GU305914.1</td><td>209843</td><td>209865</td><td>-</td><td>0</td><td>CCTGAATTCATTTTCATGCA</td><td>AGG</td><td>||||||||[||||||||||||]PAM</td><td>NA</td><td>Intergenic</td><td>NA</td><td>NA</td></tr>
</table>

| | | | | | | | | | | | |
|---|---|---|---|---|---|---|---|---|---|---|---|
| T18686 | CAACATCTGTCCCAAGTCCGAGG | 1000 | | | | | | | | | |
| Oligo fwd | CACCCAACATCTGTCCCAAGTCCG | | | | | | | | | | |
| Oligo rev | AAACCGGACTTGGGACAGATGTTG | | | | | | | | | | |
| Chromosome | start | end | strand | MM | target_seq | PAM | alignment | distance | position | gene name | gene id |
| GU305914.1 | 209813 | 209835 | - | 0 | CAACATCTGTCCCAAGTCCG | AGG | \|\|\|\|\|\|\|\|[\|\|\|\|\|\|\|\|\|\|\|\|]PAM | NA | Intergenic | NA | NA |
| T18687 | AGTCCGAGGACTGCTTTCCGAGG | 1000 | | | | | | | | | |
| Oligo fwd | CACCAGTCCGAGGACTGCTTTCCG | | | | | | | | | | |
| Oligo rev | AAACCGGAAAGCAGTCCTCGGACT | | | | | | | | | | |
| Chromosome | start | end | strand | MM | target_seq | PAM | alignment | distance | position | gene name | gene id |
| GU305914.1 | 209799 | 209821 | - | 0 | AGTCCGAGGACTGCTTTCCG | AGG | \|\|\|\|\|\|\|\|[\|\|\|\|\|\|\|\|\|\|\|\|]PAM | NA | Intergenic | NA | NA |
| T18688 | CGAGGACTGCTTTCCGAGGCTGG | 1000 | | | | | | | | | |
| Oligo fwd | CACCCGAGGACTGCTTTCCGAGGC | | | | | | | | | | |
| Oligo rev | AAACGCCTCGGAAAGCAGTCCTCG | | | | | | | | | | |
| Chromosome | start | end | strand | MM | target_seq | PAM | alignment | distance | position | gene name | gene id |
| GU305914.1 | 209795 | 209817 | - | 0 | CGAGGACTGCTTTCCGAGGC | TGG | \|\|\|\|\|\|\|\|[\|\|\|\|\|\|\|\|\|\|\|\|]PAM | NA | Intergenic | NA | NA |
| T18689 | CGAGGCTGGTGAGATAAGTGTGG | 1000 | | | | | | | | | |
| Oligo fwd | CACCCGAGGCTGGTGAGATAAGTG | | | | | | | | | | |
| Oligo rev | AAACCACTTATCTCACCAGCCTCG | | | | | | | | | | |
| Chromosome | start | end | strand | MM | target_seq | PAM | alignment | distance | position | gene name | gene id |
| GU305914.1 | 209781 | 209803 | - | 0 | CGAGGCTGGTGAGATAAGTG | TGG | \|\|\|\|\|\|\|\|[\|\|\|\|\|\|\|\|\|\|\|\|]PAM | NA | Intergenic | NA | NA |
| T18690 | GGTGTACGAGTGCCATCTATCGG | 1000 | | | | | | | | | |
| Oligo fwd | CACCGGTGTACGAGTGCCATCTAT | | | | | | | | | | |
| Oligo rev | AAACATAGATGGCACTCGTACACC | | | | | | | | | | |
| Chromosome | start | end | strand | MM | target_seq | PAM | alignment | distance | position | gene name | gene id |
| GU305914.1 | 209760 | 209782 | - | 0 | GGTGTACGAGTGCCATCTAT | CGG | \|\|\|\|\|\|\|\|[\|\|\|\|\|\|\|\|\|\|\|\|]PAM | NA | Intergenic | NA | NA |
| T18691 | CAAGCACAGTCTAAACGCCACGG | 1000 | | | | | | | | | |
| Oligo fwd | CACCCAAGCACAGTCTAAACGCCA | | | | | | | | | | |
| Oligo rev | AAACTGGCGTTTAGACTGTGCTTG | | | | | | | | | | |
| Chromosome | start | end | strand | MM | target_seq | PAM | alignment | distance | position | gene name | gene id |
| GU305914.1 | 209718 | 209740 | - | 0 | CAAGCACAGTCTAAACGCCA | CGG | \|\|\|\|\|\|\|\|[\|\|\|\|\|\|\|\|\|\|\|\|]PAM | NA | Intergenic | NA | NA |
| T18692 | GATGTGTTATATTATAATGTCGG | 1000 | | | | | | | | | |
| Oligo fwd | CACCGATGTGTTATATTATAATGT | | | | | | | | | | |
| Oligo rev | AAACACATTATAATATAACACATC | | | | | | | | | | |
| Chromosome | start | end | strand | MM | target_seq | PAM | alignment | distance | position | gene name | gene id |
| GU305914.1 | 209696 | 209718 | - | 0 | GATGTGTTATATTATAATGT | CGG | \|\|\|\|\|\|\|\|[\|\|\|\|\|\|\|\|\|\|\|\|]PAM | NA | Intergenic | NA | NA |
| T18693 | ATGTGTTATATTATAATGTCGGG | 1000 | | | | | | | | | |
| Oligo fwd | CACCATGTGTTATATTATAATGTC | | | | | | | | | | |
| Oligo rev | AAACGACATTATAATATAACACAT | | | | | | | | | | |
| Chromosome | start | end | strand | MM | target_seq | PAM | alignment | distance | position | gene name | gene id |
| GU305914.1 | 209695 | 209717 | - | 0 | ATGTGTTATATTATAATGTC | GGG | \|\|\|\|\|\|\|\|[\|\|\|\|\|\|\|\|\|\|\|\|]PAM | NA | Intergenic | NA | NA |
| T18694 | TCGCCTTTCGACAGTGATGATGG | 1000 | | | | | | | | | |
| Oligo fwd | CACCTCGCCTTTCGACAGTGATGA | | | | | | | | | | |
| Oligo rev | AAACTCATCACTGTCGAAAGGCGA | | | | | | | | | | |
| Chromosome | start | end | strand | MM | target_seq | PAM | alignment | distance | position | gene name | gene id |
| GU305914.1 | 209654 | 209676 | - | 0 | TCGCCTTTCGACAGTGATGA | TGG | \|\|\|\|\|\|\|\|[\|\|\|\|\|\|\|\|\|\|\|\|]PAM | NA | Intergenic | NA | NA |
| T18695 | TTCGACAGTGATGATGGTGATGG | 1000 | | | | | | | | | |
| Oligo fwd | CACCTTCGACAGTGATGATGGTGA | | | | | | | | | | |
| Oligo rev | AAACTCACCATCATCACTGTCGAA | | | | | | | | | | |
| Chromosome | start | end | strand | MM | target_seq | PAM | alignment | distance | position | gene name | gene id |
| GU305914.1 | 209648 | 209670 | - | 0 | TTCGACAGTGATGATGGTGA | TGG | \|\|\|\|\|\|\|\|[\|\|\|\|\|\|\|\|\|\|\|\|]PAM | NA | Intergenic | NA | NA |
| T18696 | TGTGAATGCATTCAGATCCATGG | 1000 | | | | | | | | | |
| Oligo fwd | CACCTGTGAATGCATTCAGATCCA | | | | | | | | | | |
| Oligo rev | AAACTGGATCTGAATGCATTCACA | | | | | | | | | | |
| Chromosome | start | end | strand | MM | target_seq | PAM | alignment | distance | position | gene name | gene id |
| GU305914.1 | 209613 | 209635 | - | 0 | TGTGAATGCATTCAGATCCA | TGG | \|\|\|\|\|\|\|\|[\|\|\|\|\|\|\|\|\|\|\|\|]PAM | NA | Intergenic | NA | NA |
| T18697 | GCATTCAGATCCATGGCTTCAGG | 1000 | | | | | | | | | |
| Oligo fwd | CACCGCATTCAGATCCATGGCTTC | | | | | | | | | | |
| Oligo rev | AAACGAAGCCATGGATCTGAATGC | | | | | | | | | | |
| Chromosome | start | end | strand | MM | target_seq | PAM | alignment | distance | position | gene name | gene id |
| GU305914.1 | 209606 | 209628 | - | 0 | GCATTCAGATCCATGGCTTC | AGG | \|\|\|\|\|\|\|\|[\|\|\|\|\|\|\|\|\|\|\|\|]PAM | NA | Intergenic | NA | NA |
| T18698 | CATTCAGATCCATGGCTTCAGGG | 1000 | | | | | | | | | |
| Oligo fwd | CACCCATTCAGATCCATGGCTTCA | | | | | | | | | | |
| Oligo rev | AAACTGAAGCCATGGATCTGAATG | | | | | | | | | | |
| Chromosome | start | end | strand | MM | target_seq | PAM | alignment | distance | position | gene name | gene id |
| GU305914.1 | 209605 | 209627 | - | 0 | CATTCAGATCCATGGCTTCA | GGG | \|\|\|\|\|\|\|\|[\|\|\|\|\|\|\|\|\|\|\|\|]PAM | NA | Intergenic | NA | NA |
| T18699 | AGATCCATGGCTTCAGGGTATGG | 1000 | | | | | | | | | |
| Oligo fwd | CACCAGATCCATGGCTTCAGGGTA | | | | | | | | | | |
| Oligo rev | AAACTACCCTGAAGCCATGGATCT | | | | | | | | | | |
| Chromosome | start | end | strand | MM | target_seq | PAM | alignment | distance | position | gene name | gene id |
| GU305914.1 | 209600 | 209622 | - | 0 | AGATCCATGGCTTCAGGGTA | TGG | \|\|\|\|\|\|\|\|[\|\|\|\|\|\|\|\|\|\|\|\|]PAM | NA | Intergenic | NA | NA |
| T18700 | GATCCATGGCTTCAGGGTATGGG | 1000 | | | | | | | | | |
| Oligo fwd | CACCGATCCATGGCTTCAGGGTAT | | | | | | | | | | |
| Oligo rev | AAACATACCCTGAAGCCATGGATC | | | | | | | | | | |
| Chromosome | start | end | strand | MM | target_seq | PAM | alignment | distance | position | gene name | gene id |
| GU305914.1 | 209599 | 209621 | - | 0 | GATCCATGGCTTCAGGGTAT | GGG | \|\|\|\|\|\|\|\|[\|\|\|\|\|\|\|\|\|\|\|\|]PAM | NA | Intergenic | NA | NA |

T18701 ATCCATGGCTTCAGGGTATGGGG 1000
Oligo fwd CACCATCCATGGCTTCAGGGTATG
Oligo rev AAACCATACCCTGAAGCCATGGAT
Chromosome start end strand MM target_seq PAM alignment distance position gene name gene id
GU305914.1 209598 209620 - 0 ATCCATGGCTTCAGGGTATG GGG ||||||||[||||||||||||]PAM NA Intergenic NA NA

T18702 TTCGACCTAACCACACACAGTGG 1000
Oligo fwd CACCTTCGACCTAACCACACACAG
Oligo rev AAACCTGTGTGTGGTTAGGTCGAA
Chromosome start end strand MM target_seq PAM alignment distance position gene name gene id
GU305914.1 209573 209595 - 0 TTCGACCTAACCACACACAG TGG ||||||||[||||||||||||]PAM NA Intergenic NA NA

T18703 GTGGTGATATTCGACGCAGATGG 1000
Oligo fwd CACCGTGGTGATATTCGACGCAGA
Oligo rev AAACTCTGCGTCGAATATCACCAC
Chromosome start end strand MM target_seq PAM alignment distance position gene name gene id
GU305914.1 209554 209576 - 0 GTGGTGATATTCGACGCAGA TGG ||||||||[||||||||||||]PAM NA Intergenic NA NA

T18704 GAGCATATGTACGCTCATCATGG 1000
Oligo fwd CACCGAGCATATGTACGCTCATCA
Oligo rev AAACTGATGAGCGTACATATGCTC
Chromosome start end strand MM target_seq PAM alignment distance position gene name gene id
GU305914.1 209529 209551 - 0 GAGCATATGTACGCTCATCA TGG ||||||||[||||||||||||]PAM NA Intergenic NA NA

T18705 TGGAGAAAGCGACCAAAGAACGG 1000
Oligo fwd CACCTGGAGAAAGCGACCAAAGAA
Oligo rev AAACTTCTTTGGTCGCTTTCTCCA
Chromosome start end strand MM target_seq PAM alignment distance position gene name gene id
GU305914.1 209509 209531 - 0 TGGAGAAAGCGACCAAAGAA CGG ||||||||[||||||||||||]PAM NA Intergenic NA NA

T18706 GAAAGCGACCAAAGAACGGTCGG 1000
Oligo fwd CACCGAAAGCGACCAAAGAACGGT
Oligo rev AAACACCGTTCTTTGGTCGCTTTC
Chromosome start end strand MM target_seq PAM alignment distance position gene name gene id
GU305914.1 209505 209527 - 0 GAAAGCGACCAAAGAACGGT CGG ||||||||[||||||||||||]PAM NA Intergenic NA NA

T18707 AGTTTCTCTAGTATGACGAACGG 1000
Oligo fwd CACCAGTTTCTCTAGTATGACGAA
Oligo rev AAACTTCGTCATACTAGAGAAACT
Chromosome start end strand MM target_seq PAM alignment distance position gene name gene id
GU305914.1 209477 209499 - 0 AGTTTCTCTAGTATGACGAA CGG ||||||||[||||||||||||]PAM NA Intergenic NA NA

T18708 CGAAGCAACAGTCTCGTCACCGG 1000
Oligo fwd CACCCGAAGCAACAGTCTCGTCAC
Oligo rev AAACGTGACGAGACTGTTGCTTCG
Chromosome start end strand MM target_seq PAM alignment distance position gene name gene id
GU305914.1 209445 209467 - 0 CGAAGCAACAGTCTCGTCAC CGG ||||||||[||||||||||||]PAM NA Intergenic NA NA

T18709 TCTCGTCACCGGTGATACTGTGG 1000
Oligo fwd CACCTCTCGTCACCGGTGATACTG
Oligo rev AAACCAGTATCACCGGTGACGAGA
Chromosome start end strand MM target_seq PAM alignment distance position gene name gene id
GU305914.1 209434 209456 - 0 TCTCGTCACCGGTGATACTG TGG ||||||||[||||||||||||]PAM NA Intergenic NA NA

T18710 ATACTGTGGATGACTATCGTCGG 1000
Oligo fwd CACCATACTGTGGATGACTATCGT
Oligo rev AAACACGATAGTCATCCACAGTAT
Chromosome start end strand MM target_seq PAM alignment distance position gene name gene id
GU305914.1 209420 209442 - 0 ATACTGTGGATGACTATCGT CGG ||||||||[||||||||||||]PAM NA Intergenic NA NA

T18711 TAACACTAATAACATCTCTTCGG 1000
Oligo fwd CACCTAACACTAATAACATCTCTT
Oligo rev AAACAAGAGATGTTATTAGTGTTA
Chromosome start end strand MM target_seq PAM alignment distance position gene name gene id
GU305914.1 209361 209383 - 0 TAACACTAATAACATCTCTT CGG ||||||||[||||||||||||]PAM NA Intergenic NA NA

T18712 TCTCCCCGAACAACTCACCGCGG 1000
Oligo fwd CACCTCTCCCCGAACAACTCACCG
Oligo rev AAACCGGTGAGTTGTTCGGGGAGA
Chromosome start end strand MM target_seq PAM alignment distance position gene name gene id
GU305914.1 209320 209342 - 0 TCTCCCCGAACAACTCACCG CGG ||||||||[||||||||||||]PAM NA Intergenic NA NA

T18713 GTGCGAGATAATATGCTCCGCGG 1000
Oligo fwd CACCGTGCGAGATAATATGCTCCG
Oligo rev AAACCGGAGCATATTATCTCGCAC
Chromosome start end strand MM target_seq PAM alignment distance position gene name gene id
GU305914.1 209298 209320 - 0 GTGCGAGATAATATGCTCCG CGG ||||||||[||||||||||||]PAM NA Intergenic NA NA

T18714 TGCGAGATAATATGCTCCGCGGG 1000
Oligo fwd CACCTGCGAGATAATATGCTCCGC
Oligo rev AAACGCGGAGCATATTATCTCGCA
Chromosome start end strand MM target_seq PAM alignment distance position gene name gene id
GU305914.1 209297 209319 - 0 TGCGAGATAATATGCTCCGC GGG ||||||||[||||||||||||]PAM NA Intergenic NA NA

T18715 AGATAATATGCTCCGCGGGCTGG 1000
Oligo fwd CACCAGATAATATGCTCCGCGGGC
Oligo rev AAACGCCCGCGGAGCATATTATCT
Chromosome start end strand MM target_seq PAM alignment distance position gene name gene id

GU305914.1 209293 209315 - 0 AGATAATATGCTCCGCGGGC TGG ||||||||[||||||||||||]PAM NA Intergenic NA NA

T18716 CGGGCTGGAACGTCACCATCCGG 1000
Oligo fwd CACCCGGGCTGGAACGTCACCATC
Oligo rev AAACGATGGTGACGTTCCAGCCCG
Chromosome start end strand MM target_seq PAM alignment distance position gene name gene id
GU305914.1 209278 209300 - 0 CGGGCTGGAACGTCACCATC CGG ||||||||[||||||||||||]PAM NA Intergenic NA NA

T18717 CATCCGGCAGCCGAACAGTTCGG 1000
Oligo fwd CACCCATCCGGCAGCCGAACAGTT
Oligo rev AAACAACTGTTCGGCTGCCGGATG
Chromosome start end strand MM target_seq PAM alignment distance position gene name gene id
GU305914.1 209262 209284 - 0 CATCCGGCAGCCGAACAGTT CGG ||||||||[||||||||||||]PAM NA Intergenic NA NA

T18718 GAACAGTTCGGCCGCCACGACGG 1000
Oligo fwd CACCGAACAGTTCGGCCGCCACGA
Oligo rev AAACTCGTGGCGGCCGAACTGTTC
Chromosome start end strand MM target_seq PAM alignment distance position gene name gene id
GU305914.1 209250 209272 - 0 GAACAGTTCGGCCGCCACGA CGG ||||||||[||||||||||||]PAM NA Intergenic NA NA

T18719 CGCCACGACGGAGAGACGTGTGG 1000
Oligo fwd CACCCGCCACGACGGAGAGACGTG
Oligo rev AAACCACGTCTCTCCGTCGTGGCG
Chromosome start end strand MM target_seq PAM alignment distance position gene name gene id
GU305914.1 209238 209260 - 0 CGCCACGACGGAGAGACGTG TGG ||||||||[||||||||||||]PAM NA Intergenic NA NA

T18720 GTGGCACTTTCTGCTGCGCCCGG 1000
Oligo fwd CACCGTGGCACTTTCTGCTGCGCC
Oligo rev AAACGGCGCAGCAGAAAGTGCCAC
Chromosome start end strand MM target_seq PAM alignment distance position gene name gene id
GU305914.1 209219 209241 - 0 GTGGCACTTTCTGCTGCGCC CGG ||||||||[||||||||||||]PAM NA Intergenic NA NA

T18721 ACCGTCCTCTCCGTCTCGAACGG 1000
Oligo fwd CACCACCGTCCTCTCCGTCTCGAA
Oligo rev AAACTTCGAGACGGAGAGGACGGT
Chromosome start end strand MM target_seq PAM alignment distance position gene name gene id
GU305914.1 209174 209196 - 0 ACCGTCCTCTCCGTCTCGAA CGG ||||||||[||||||||||||]PAM NA Intergenic NA NA

T18722 CGTCTCGAACGGCTCAGAACTGG 1000
Oligo fwd CACCCGTCTCGAACGGCTCAGAAC
Oligo rev AAACGTTCTGAGCCGTTCGAGACG
Chromosome start end strand MM target_seq PAM alignment distance position gene name gene id
GU305914.1 209163 209185 - 0 CGTCTCGAACGGCTCAGAAC TGG ||||||||[||||||||||||]PAM NA Intergenic NA NA

T18723 ATCTTCCCATATACACCAATCGG 1000
Oligo fwd CACCATCTTCCCATATACACCAAT
Oligo rev AAACATTGGTGTATATGGGAAGAT
Chromosome start end strand MM target_seq PAM alignment distance position gene name gene id
GU305914.1 209137 209159 - 0 ATCTTCCCATATACACCAAT CGG ||||||||[||||||||||||]PAM NA Intergenic NA NA

T18724 CGTCGTGAGCTACGCCGAAGAGG 1000
Oligo fwd CACCCGTCGTGAGCTACGCCGAAG
Oligo rev AAACCTTCGGCGTAGCTCACGACG
Chromosome start end strand MM target_seq PAM alignment distance position gene name gene id
GU305914.1 209109 209131 - 0 CGTCGTGAGCTACGCCGAAG AGG ||||||||[||||||||||||]PAM NA Intergenic NA NA

T18725 GAAGAGGCGACATCGCATGTCGG 1000
Oligo fwd CACCGAAGAGGCGACATCGCATGT
Oligo rev AAACACATGCGATGTCGCCTCTTC
Chromosome start end strand MM target_seq PAM alignment distance position gene name gene id
GU305914.1 209093 209115 - 0 GAAGAGGCGACATCGCATGT CGG ||||||||[||||||||||||]PAM NA Intergenic NA NA

T18726 CTATCTGCTCGCCGTTCCCGCGG 1000
Oligo fwd CACCCTATCTGCTCGCCGTTCCCG
Oligo rev AAACCGGGAACGGCGAGCAGATAG
Chromosome start end strand MM target_seq PAM alignment distance position gene name gene id
GU305914.1 209046 209068 - 0 CTATCTGCTCGCCGTTCCCG CGG ||||||||[||||||||||||]PAM NA Intergenic NA NA

T18727 ACTCGTCGTGCTATCCTCCGTGG 1000
Oligo fwd CACCACTCGTCGTGCTATCCTCCG
Oligo rev AAACCGGAGGATAGCACGACGAGT
Chromosome start end strand MM target_seq PAM alignment distance position gene name gene id
GU305914.1 209019 209041 - 0 ACTCGTCGTGCTATCCTCCG TGG ||||||||[||||||||||||]PAM NA Intergenic NA NA

T18728 CAAGATACGACTGCTGCTCATGG 1000
Oligo fwd CACCCAAGATACGACTGCTGCTCA
Oligo rev AAACTGAGCAGCAGTCGTATCTTG
Chromosome start end strand MM target_seq PAM alignment distance position gene name gene id
GU305914.1 208983 209005 - 0 CAAGATACGACTGCTGCTCA TGG ||||||||[||||||||||||]PAM NA Intergenic NA NA

T18729 TACGACTGCTGCTCATGGATAGG 1000
Oligo fwd CACCTACGACTGCTGCTCATGGAT
Oligo rev AAACATCCATGAGCAGCAGTCGTA
Chromosome start end strand MM target_seq PAM alignment distance position gene name gene id
GU305914.1 208978 209000 - 0 TACGACTGCTGCTCATGGAT AGG ||||||||[||||||||||||]PAM NA Intergenic NA NA

T18730 ACTGCTGCTCATGGATAGGACGG 1000
Oligo fwd CACCACTGCTGCTCATGGATAGGA
Oligo rev AAACTCCTATCCATGAGCAGCAGT

<table>
<tr><td>Chromosome</td><td>start</td><td>end</td><td>strand</td><td>MM</td><td>target_seq</td><td>PAM</td><td>alignment</td><td>distance</td><td>position</td><td>gene name</td><td>gene id</td></tr>
<tr><td>GU305914.1</td><td>208974</td><td>208996</td><td>-</td><td>0</td><td>ACTGCTGCTCATGGATAGGA</td><td>CGG</td><td>||||||||[||||||||||||]PAM</td><td>NA</td><td>Intergenic</td><td>NA</td><td>NA</td></tr>
<tr><td>T18731</td><td>TCATGGATAGGACGGTCACCAGG</td><td>1000</td><td></td><td></td><td></td><td></td><td></td><td></td><td></td><td></td><td></td></tr>
<tr><td>Oligo fwd</td><td>CACCTCATGGATAGGACGGTCACC</td><td></td><td></td><td></td><td></td><td></td><td></td><td></td><td></td><td></td><td></td></tr>
<tr><td>Oligo rev</td><td>AAACGGTGACCGTCCTATCCATGA</td><td></td><td></td><td></td><td></td><td></td><td></td><td></td><td></td><td></td><td></td></tr>
<tr><td>Chromosome</td><td>start</td><td>end</td><td>strand</td><td>MM</td><td>target_seq</td><td>PAM</td><td>alignment</td><td>distance</td><td>position</td><td>gene name</td><td>gene id</td></tr>
<tr><td>GU305914.1</td><td>208966</td><td>208988</td><td>-</td><td>0</td><td>TCATGGATAGGACGGTCACC</td><td>AGG</td><td>||||||||[||||||||||||]PAM</td><td>NA</td><td>Intergenic</td><td>NA</td><td>NA</td></tr>
<tr><td>T18732</td><td>AGCTTTCGATAACTGATTGTCGG</td><td>1000</td><td></td><td></td><td></td><td></td><td></td><td></td><td></td><td></td><td></td></tr>
<tr><td>Oligo fwd</td><td>CACCAGCTTTCGATAACTGATTGT</td><td></td><td></td><td></td><td></td><td></td><td></td><td></td><td></td><td></td><td></td></tr>
<tr><td>Oligo rev</td><td>AAACACAATCAGTTATCGAAAGCT</td><td></td><td></td><td></td><td></td><td></td><td></td><td></td><td></td><td></td><td></td></tr>
<tr><td>Chromosome</td><td>start</td><td>end</td><td>strand</td><td>MM</td><td>target_seq</td><td>PAM</td><td>alignment</td><td>distance</td><td>position</td><td>gene name</td><td>gene id</td></tr>
<tr><td>GU305914.1</td><td>208904</td><td>208926</td><td>-</td><td>0</td><td>AGCTTTCGATAACTGATTGT</td><td>CGG</td><td>||||||||[||||||||||||]PAM</td><td>NA</td><td>Intergenic</td><td>NA</td><td>NA</td></tr>
<tr><td>T18733</td><td>TTTCGATAACTGATTGTCGGCGG</td><td>1000</td><td></td><td></td><td></td><td></td><td></td><td></td><td></td><td></td><td></td></tr>
<tr><td>Oligo fwd</td><td>CACCTTTCGATAACTGATTGTCGG</td><td></td><td></td><td></td><td></td><td></td><td></td><td></td><td></td><td></td><td></td></tr>
<tr><td>Oligo rev</td><td>AAACCCGACAATCAGTTATCGAAA</td><td></td><td></td><td></td><td></td><td></td><td></td><td></td><td></td><td></td><td></td></tr>
<tr><td>Chromosome</td><td>start</td><td>end</td><td>strand</td><td>MM</td><td>target_seq</td><td>PAM</td><td>alignment</td><td>distance</td><td>position</td><td>gene name</td><td>gene id</td></tr>
<tr><td>GU305914.1</td><td>208901</td><td>208923</td><td>-</td><td>0</td><td>TTTCGATAACTGATTGTCGG</td><td>CGG</td><td>||||||||[||||||||||||]PAM</td><td>NA</td><td>Intergenic</td><td>NA</td><td>NA</td></tr>
<tr><td>T18734</td><td>AATAAAAATGTGTATTGTCGCGG</td><td>1000</td><td></td><td></td><td></td><td></td><td></td><td></td><td></td><td></td><td></td></tr>
<tr><td>Oligo fwd</td><td>CACCAATAAAAATGTGTATTGTCG</td><td></td><td></td><td></td><td></td><td></td><td></td><td></td><td></td><td></td><td></td></tr>
<tr><td>Oligo rev</td><td>AAACCGACAATACACATTTTTATT</td><td></td><td></td><td></td><td></td><td></td><td></td><td></td><td></td><td></td><td></td></tr>
<tr><td>Chromosome</td><td>start</td><td>end</td><td>strand</td><td>MM</td><td>target_seq</td><td>PAM</td><td>alignment</td><td>distance</td><td>position</td><td>gene name</td><td>gene id</td></tr>
<tr><td>GU305914.1</td><td>208875</td><td>208897</td><td>-</td><td>0</td><td>AATAAAAATGTGTATTGTCG</td><td>CGG</td><td>||||||||[||||||||||||]PAM</td><td>NA</td><td>Intergenic</td><td>NA</td><td>NA</td></tr>
<tr><td>T18735</td><td>ACTCTCCACGTCTGTCGTGATGG</td><td>1000</td><td></td><td></td><td></td><td></td><td></td><td></td><td></td><td></td><td></td></tr>
<tr><td>Oligo fwd</td><td>CACCACTCTCCACGTCTGTCGTGA</td><td></td><td></td><td></td><td></td><td></td><td></td><td></td><td></td><td></td><td></td></tr>
<tr><td>Oligo rev</td><td>AAACTCACGACAGACGTGGAGAGT</td><td></td><td></td><td></td><td></td><td></td><td></td><td></td><td></td><td></td><td></td></tr>
<tr><td>Chromosome</td><td>start</td><td>end</td><td>strand</td><td>MM</td><td>target_seq</td><td>PAM</td><td>alignment</td><td>distance</td><td>position</td><td>gene name</td><td>gene id</td></tr>
<tr><td>GU305914.1</td><td>208839</td><td>208861</td><td>-</td><td>0</td><td>ACTCTCCACGTCTGTCGTGA</td><td>TGG</td><td>||||||||[||||||||||||]PAM</td><td>NA</td><td>Intergenic</td><td>NA</td><td>NA</td></tr>
<tr><td>T18736</td><td>CTCCACGTCTGTCGTGATGGCGG</td><td>1000</td><td></td><td></td><td></td><td></td><td></td><td></td><td></td><td></td><td></td></tr>
<tr><td>Oligo fwd</td><td>CACCCTCCACGTCTGTCGTGATGG</td><td></td><td></td><td></td><td></td><td></td><td></td><td></td><td></td><td></td><td></td></tr>
<tr><td>Oligo rev</td><td>AAACCCATCACGACAGACGTGGAG</td><td></td><td></td><td></td><td></td><td></td><td></td><td></td><td></td><td></td><td></td></tr>
<tr><td>Chromosome</td><td>start</td><td>end</td><td>strand</td><td>MM</td><td>target_seq</td><td>PAM</td><td>alignment</td><td>distance</td><td>position</td><td>gene name</td><td>gene id</td></tr>
<tr><td>GU305914.1</td><td>208836</td><td>208858</td><td>-</td><td>0</td><td>CTCCACGTCTGTCGTGATGG</td><td>CGG</td><td>||||||||[||||||||||||]PAM</td><td>NA</td><td>Intergenic</td><td>NA</td><td>NA</td></tr>
<tr><td>T18737</td><td>CACGTCTGTCGTGATGGCGGTGG</td><td>1000</td><td></td><td></td><td></td><td></td><td></td><td></td><td></td><td></td><td></td></tr>
<tr><td>Oligo fwd</td><td>CACCCACGTCTGTCGTGATGGCGG</td><td></td><td></td><td></td><td></td><td></td><td></td><td></td><td></td><td></td><td></td></tr>
<tr><td>Oligo rev</td><td>AAACCCGCCATCACGACAGACGTG</td><td></td><td></td><td></td><td></td><td></td><td></td><td></td><td></td><td></td><td></td></tr>
<tr><td>Chromosome</td><td>start</td><td>end</td><td>strand</td><td>MM</td><td>target_seq</td><td>PAM</td><td>alignment</td><td>distance</td><td>position</td><td>gene name</td><td>gene id</td></tr>
<tr><td>GU305914.1</td><td>208833</td><td>208855</td><td>-</td><td>0</td><td>CACGTCTGTCGTGATGGCGG</td><td>TGG</td><td>||||||||[||||||||||||]PAM</td><td>NA</td><td>Intergenic</td><td>NA</td><td>NA</td></tr>
<tr><td>T18738</td><td>CGGTGGTCCGTGTGCGAACGTGG</td><td>1000</td><td></td><td></td><td></td><td></td><td></td><td></td><td></td><td></td><td></td></tr>
<tr><td>Oligo fwd</td><td>CACCCGGTGGTCCGTGTGCGAACG</td><td></td><td></td><td></td><td></td><td></td><td></td><td></td><td></td><td></td><td></td></tr>
<tr><td>Oligo rev</td><td>AAACCGTTCGCACACGGACCACCG</td><td></td><td></td><td></td><td></td><td></td><td></td><td></td><td></td><td></td><td></td></tr>
<tr><td>Chromosome</td><td>start</td><td>end</td><td>strand</td><td>MM</td><td>target_seq</td><td>PAM</td><td>alignment</td><td>distance</td><td>position</td><td>gene name</td><td>gene id</td></tr>
<tr><td>GU305914.1</td><td>208816</td><td>208838</td><td>-</td><td>0</td><td>CGGTGGTCCGTGTGCGAACG</td><td>TGG</td><td>||||||||[||||||||||||]PAM</td><td>NA</td><td>Intergenic</td><td>NA</td><td>NA</td></tr>
<tr><td>T18739</td><td>GGTGGTCCGTGTGCGAACGTGGG</td><td>1000</td><td></td><td></td><td></td><td></td><td></td><td></td><td></td><td></td><td></td></tr>
<tr><td>Oligo fwd</td><td>CACCGGTGGTCCGTGTGCGAACGT</td><td></td><td></td><td></td><td></td><td></td><td></td><td></td><td></td><td></td><td></td></tr>
<tr><td>Oligo rev</td><td>AAACACGTTCGCACACGGACCACC</td><td></td><td></td><td></td><td></td><td></td><td></td><td></td><td></td><td></td><td></td></tr>
<tr><td>Chromosome</td><td>start</td><td>end</td><td>strand</td><td>MM</td><td>target_seq</td><td>PAM</td><td>alignment</td><td>distance</td><td>position</td><td>gene name</td><td>gene id</td></tr>
<tr><td>GU305914.1</td><td>208815</td><td>208837</td><td>-</td><td>0</td><td>GGTGGTCCGTGTGCGAACGT</td><td>GGG</td><td>||||||||[||||||||||||]PAM</td><td>NA</td><td>Intergenic</td><td>NA</td><td>NA</td></tr>
<tr><td>T18740</td><td>GTGGTCCGTGTGCGAACGTGGGG</td><td>1000</td><td></td><td></td><td></td><td></td><td></td><td></td><td></td><td></td><td></td></tr>
<tr><td>Oligo fwd</td><td>CACCGTGGTCCGTGTGCGAACGTG</td><td></td><td></td><td></td><td></td><td></td><td></td><td></td><td></td><td></td><td></td></tr>
<tr><td>Oligo rev</td><td>AAACCACGTTCGCACACGGACCAC</td><td></td><td></td><td></td><td></td><td></td><td></td><td></td><td></td><td></td><td></td></tr>
<tr><td>Chromosome</td><td>start</td><td>end</td><td>strand</td><td>MM</td><td>target_seq</td><td>PAM</td><td>alignment</td><td>distance</td><td>position</td><td>gene name</td><td>gene id</td></tr>
<tr><td>GU305914.1</td><td>208814</td><td>208836</td><td>-</td><td>0</td><td>GTGGTCCGTGTGCGAACGTG</td><td>GGG</td><td>||||||||[||||||||||||]PAM</td><td>NA</td><td>Intergenic</td><td>NA</td><td>NA</td></tr>
<tr><td>T18741</td><td>TGGTCCGTGTGCGAACGTGGGGG</td><td>1000</td><td></td><td></td><td></td><td></td><td></td><td></td><td></td><td></td><td></td></tr>
<tr><td>Oligo fwd</td><td>CACCTGGTCCGTGTGCGAACGTGG</td><td></td><td></td><td></td><td></td><td></td><td></td><td></td><td></td><td></td><td></td></tr>
<tr><td>Oligo rev</td><td>AAACCCACGTTCGCACACGGACCA</td><td></td><td></td><td></td><td></td><td></td><td></td><td></td><td></td><td></td><td></td></tr>
<tr><td>Chromosome</td><td>start</td><td>end</td><td>strand</td><td>MM</td><td>target_seq</td><td>PAM</td><td>alignment</td><td>distance</td><td>position</td><td>gene name</td><td>gene id</td></tr>
<tr><td>GU305914.1</td><td>208813</td><td>208835</td><td>-</td><td>0</td><td>TGGTCCGTGTGCGAACGTGG</td><td>GGG</td><td>||||||||[||||||||||||]PAM</td><td>NA</td><td>Intergenic</td><td>NA</td><td>NA</td></tr>
<tr><td>T18742</td><td>GGTCCGTGTGCGAACGTGGGGGG</td><td>1000</td><td></td><td></td><td></td><td></td><td></td><td></td><td></td><td></td><td></td></tr>
<tr><td>Oligo fwd</td><td>CACCGGTCCGTGTGCGAACGTGGG</td><td></td><td></td><td></td><td></td><td></td><td></td><td></td><td></td><td></td><td></td></tr>
<tr><td>Oligo rev</td><td>AAACCCCACGTTCGCACACGGACC</td><td></td><td></td><td></td><td></td><td></td><td></td><td></td><td></td><td></td><td></td></tr>
<tr><td>Chromosome</td><td>start</td><td>end</td><td>strand</td><td>MM</td><td>target_seq</td><td>PAM</td><td>alignment</td><td>distance</td><td>position</td><td>gene name</td><td>gene id</td></tr>
<tr><td>GU305914.1</td><td>208812</td><td>208834</td><td>-</td><td>0</td><td>GGTCCGTGTGCGAACGTGGG</td><td>GGG</td><td>||||||||[||||||||||||]PAM</td><td>NA</td><td>Intergenic</td><td>NA</td><td>NA</td></tr>
<tr><td>T18743</td><td>GCGAACGTGGGGGGTTATCCTGG</td><td>1000</td><td></td><td></td><td></td><td></td><td></td><td></td><td></td><td></td><td></td></tr>
<tr><td>Oligo fwd</td><td>CACCGCGAACGTGGGGGGTTATCC</td><td></td><td></td><td></td><td></td><td></td><td></td><td></td><td></td><td></td><td></td></tr>
<tr><td>Oligo rev</td><td>AAACGGATAACCCCCCACGTTCGC</td><td></td><td></td><td></td><td></td><td></td><td></td><td></td><td></td><td></td><td></td></tr>
<tr><td>Chromosome</td><td>start</td><td>end</td><td>strand</td><td>MM</td><td>target_seq</td><td>PAM</td><td>alignment</td><td>distance</td><td>position</td><td>gene name</td><td>gene id</td></tr>
<tr><td>GU305914.1</td><td>208803</td><td>208825</td><td>-</td><td>0</td><td>GCGAACGTGGGGGGTTATCC</td><td>TGG</td><td>||||||||[||||||||||||]PAM</td><td>NA</td><td>Intergenic</td><td>NA</td><td>NA</td></tr>
<tr><td>T18744</td><td>CGCAGTCGCGTTAGTCTTCTCGG</td><td>1000</td><td></td><td></td><td></td><td></td><td></td><td></td><td></td><td></td><td></td></tr>
<tr><td>Oligo fwd</td><td>CACCCGCAGTCGCGTTAGTCTTCT</td><td></td><td></td><td></td><td></td><td></td><td></td><td></td><td></td><td></td><td></td></tr>
<tr><td>Oligo rev</td><td>AAACAGAAGACTAACGCGACTGCG</td><td></td><td></td><td></td><td></td><td></td><td></td><td></td><td></td><td></td><td></td></tr>
<tr><td>Chromosome</td><td>start</td><td>end</td><td>strand</td><td>MM</td><td>target_seq</td><td>PAM</td><td>alignment</td><td>distance</td><td>position</td><td>gene name</td><td>gene id</td></tr>
<tr><td>GU305914.1</td><td>208752</td><td>208774</td><td>-</td><td>0</td><td>CGCAGTCGCGTTAGTCTTCT</td><td>CGG</td><td>||||||||[||||||||||||]PAM</td><td>NA</td><td>Intergenic</td><td>NA</td><td>NA</td></tr>
<tr><td>T18745</td><td>TTTCAGCGATAGCTTCGTTGCGG</td><td>1000</td><td></td><td></td><td></td><td></td><td></td><td></td><td></td><td></td><td></td></tr>
<tr><td>Oligo fwd</td><td>CACCTTTCAGCGATAGCTTCGTTG</td><td></td><td></td><td></td><td></td><td></td><td></td><td></td><td></td><td></td><td></td></tr>
</table>

<table>
<tr><td>Oligo rev</td><td>AAACCAACGAAGCTATCGCTGAAA</td><td></td><td></td><td></td><td></td><td></td><td></td><td></td><td></td><td></td><td></td></tr>
<tr><td>Chromosome</td><td>start</td><td>end</td><td>strand</td><td>MM</td><td>target_seq</td><td>PAM</td><td>alignment</td><td>distance</td><td>position</td><td>gene name</td><td>gene id</td></tr>
<tr><td>GU305914.1</td><td>208722</td><td>208744</td><td>-</td><td>0</td><td>TTTCAGCGATAGCTTCGTTG</td><td>CGG</td><td>||||||||[||||||||||||]PAM</td><td>NA</td><td>Intergenic</td><td>NA</td><td>NA</td></tr>
<tr><td></td><td></td><td></td><td></td><td></td><td></td><td></td><td></td><td></td><td></td><td></td><td></td></tr>
<tr><td>T18746</td><td>TTCAGCGATAGCTTCGTTGCGGG</td><td>1000</td><td></td><td></td><td></td><td></td><td></td><td></td><td></td><td></td><td></td></tr>
<tr><td>Oligo fwd</td><td>CACCTTCAGCGATAGCTTCGTTGC</td><td></td><td></td><td></td><td></td><td></td><td></td><td></td><td></td><td></td><td></td></tr>
<tr><td>Oligo rev</td><td>AAACGCAACGAAGCTATCGCTGAA</td><td></td><td></td><td></td><td></td><td></td><td></td><td></td><td></td><td></td><td></td></tr>
<tr><td>Chromosome</td><td>start</td><td>end</td><td>strand</td><td>MM</td><td>target_seq</td><td>PAM</td><td>alignment</td><td>distance</td><td>position</td><td>gene name</td><td>gene id</td></tr>
<tr><td>GU305914.1</td><td>208721</td><td>208743</td><td>-</td><td>0</td><td>TTCAGCGATAGCTTCGTTGC</td><td>GGG</td><td>||||||||[||||||||||||]PAM</td><td>NA</td><td>Intergenic</td><td>NA</td><td>NA</td></tr>
<tr><td></td><td></td><td></td><td></td><td></td><td></td><td></td><td></td><td></td><td></td><td></td><td></td></tr>
<tr><td>T18747</td><td>TCAGCGATAGCTTCGTTGCGGGG</td><td>1000</td><td></td><td></td><td></td><td></td><td></td><td></td><td></td><td></td><td></td></tr>
<tr><td>Oligo fwd</td><td>CACCTCAGCGATAGCTTCGTTGCG</td><td></td><td></td><td></td><td></td><td></td><td></td><td></td><td></td><td></td><td></td></tr>
<tr><td>Oligo rev</td><td>AAACCGCAACGAAGCTATCGCTGA</td><td></td><td></td><td></td><td></td><td></td><td></td><td></td><td></td><td></td><td></td></tr>
<tr><td>Chromosome</td><td>start</td><td>end</td><td>strand</td><td>MM</td><td>target_seq</td><td>PAM</td><td>alignment</td><td>distance</td><td>position</td><td>gene name</td><td>gene id</td></tr>
<tr><td>GU305914.1</td><td>208720</td><td>208742</td><td>-</td><td>0</td><td>TCAGCGATAGCTTCGTTGCG</td><td>GGG</td><td>||||||||[||||||||||||]PAM</td><td>NA</td><td>Intergenic</td><td>NA</td><td>NA</td></tr>
<tr><td></td><td></td><td></td><td></td><td></td><td></td><td></td><td></td><td></td><td></td><td></td><td></td></tr>
<tr><td>T18748</td><td>TTTCCCCTCGTAAGAATAGACGG</td><td>1000</td><td></td><td></td><td></td><td></td><td></td><td></td><td></td><td></td><td></td></tr>
<tr><td>Oligo fwd</td><td>CACCTTTCCCCTCGTAAGAATAGA</td><td></td><td></td><td></td><td></td><td></td><td></td><td></td><td></td><td></td><td></td></tr>
<tr><td>Oligo rev</td><td>AAACTCTATTCTTACGAGGGGAAA</td><td></td><td></td><td></td><td></td><td></td><td></td><td></td><td></td><td></td><td></td></tr>
<tr><td>Chromosome</td><td>start</td><td>end</td><td>strand</td><td>MM</td><td>target_seq</td><td>PAM</td><td>alignment</td><td>distance</td><td>position</td><td>gene name</td><td>gene id</td></tr>
<tr><td>GU305914.1</td><td>208676</td><td>208698</td><td>-</td><td>0</td><td>TTTCCCCTCGTAAGAATAGA</td><td>CGG</td><td>||||||||[||||||||||||]PAM</td><td>NA</td><td>Intergenic</td><td>NA</td><td>NA</td></tr>
<tr><td></td><td></td><td></td><td></td><td></td><td></td><td></td><td></td><td></td><td></td><td></td><td></td></tr>
<tr><td>T18749</td><td>CTCGTAAGAATAGACGGTCGCGG</td><td>1000</td><td></td><td></td><td></td><td></td><td></td><td></td><td></td><td></td><td></td></tr>
<tr><td>Oligo fwd</td><td>CACCCTCGTAAGAATAGACGGTCG</td><td></td><td></td><td></td><td></td><td></td><td></td><td></td><td></td><td></td><td></td></tr>
<tr><td>Oligo rev</td><td>AAACCGACCGTCTATTCTTACGAG</td><td></td><td></td><td></td><td></td><td></td><td></td><td></td><td></td><td></td><td></td></tr>
<tr><td>Chromosome</td><td>start</td><td>end</td><td>strand</td><td>MM</td><td>target_seq</td><td>PAM</td><td>alignment</td><td>distance</td><td>position</td><td>gene name</td><td>gene id</td></tr>
<tr><td>GU305914.1</td><td>208670</td><td>208692</td><td>-</td><td>0</td><td>CTCGTAAGAATAGACGGTCG</td><td>CGG</td><td>||||||||[||||||||||||]PAM</td><td>NA</td><td>Intergenic</td><td>NA</td><td>NA</td></tr>
<tr><td></td><td></td><td></td><td></td><td></td><td></td><td></td><td></td><td></td><td></td><td></td><td></td></tr>
<tr><td>T18750</td><td>TCCCTCTGTATCCGACCTGAAGG</td><td>1000</td><td></td><td></td><td></td><td></td><td></td><td></td><td></td><td></td><td></td></tr>
<tr><td>Oligo fwd</td><td>CACCTCCCTCTGTATCCGACCTGA</td><td></td><td></td><td></td><td></td><td></td><td></td><td></td><td></td><td></td><td></td></tr>
<tr><td>Oligo rev</td><td>AAACTCAGGTCGGATACAGAGGGA</td><td></td><td></td><td></td><td></td><td></td><td></td><td></td><td></td><td></td><td></td></tr>
<tr><td>Chromosome</td><td>start</td><td>end</td><td>strand</td><td>MM</td><td>target_seq</td><td>PAM</td><td>alignment</td><td>distance</td><td>position</td><td>gene name</td><td>gene id</td></tr>
<tr><td>GU305914.1</td><td>208629</td><td>208651</td><td>-</td><td>0</td><td>TCCCTCTGTATCCGACCTGA</td><td>AGG</td><td>||||||||[||||||||||||]PAM</td><td>NA</td><td>Intergenic</td><td>NA</td><td>NA</td></tr>
<tr><td></td><td></td><td></td><td></td><td></td><td></td><td></td><td></td><td></td><td></td><td></td><td></td></tr>
<tr><td>T18751</td><td>ATCCCGAAGTACATTTTCTGTGG</td><td>1000</td><td></td><td></td><td></td><td></td><td></td><td></td><td></td><td></td><td></td></tr>
<tr><td>Oligo fwd</td><td>CACCATCCCGAAGTACATTTTCTG</td><td></td><td></td><td></td><td></td><td></td><td></td><td></td><td></td><td></td><td></td></tr>
<tr><td>Oligo rev</td><td>AAACCAGAAAATGTACTTCGGGAT</td><td></td><td></td><td></td><td></td><td></td><td></td><td></td><td></td><td></td><td></td></tr>
<tr><td>Chromosome</td><td>start</td><td>end</td><td>strand</td><td>MM</td><td>target_seq</td><td>PAM</td><td>alignment</td><td>distance</td><td>position</td><td>gene name</td><td>gene id</td></tr>
<tr><td>GU305914.1</td><td>208600</td><td>208622</td><td>-</td><td>0</td><td>ATCCCGAAGTACATTTTCTG</td><td>TGG</td><td>||||||||[||||||||||||]PAM</td><td>NA</td><td>Intergenic</td><td>NA</td><td>NA</td></tr>
<tr><td></td><td></td><td></td><td></td><td></td><td></td><td></td><td></td><td></td><td></td><td></td><td></td></tr>
<tr><td>T18752</td><td>TCCCGAAGTACATTTTCTGTGGG</td><td>1000</td><td></td><td></td><td></td><td></td><td></td><td></td><td></td><td></td><td></td></tr>
<tr><td>Oligo fwd</td><td>CACCTCCCGAAGTACATTTTCTGT</td><td></td><td></td><td></td><td></td><td></td><td></td><td></td><td></td><td></td><td></td></tr>
<tr><td>Oligo rev</td><td>AAACACAGAAAATGTACTTCGGGA</td><td></td><td></td><td></td><td></td><td></td><td></td><td></td><td></td><td></td><td></td></tr>
<tr><td>Chromosome</td><td>start</td><td>end</td><td>strand</td><td>MM</td><td>target_seq</td><td>PAM</td><td>alignment</td><td>distance</td><td>position</td><td>gene name</td><td>gene id</td></tr>
<tr><td>GU305914.1</td><td>208599</td><td>208621</td><td>-</td><td>0</td><td>TCCCGAAGTACATTTTCTGT</td><td>GGG</td><td>||||||||[||||||||||||]PAM</td><td>NA</td><td>Intergenic</td><td>NA</td><td>NA</td></tr>
<tr><td></td><td></td><td></td><td></td><td></td><td></td><td></td><td></td><td></td><td></td><td></td><td></td></tr>
<tr><td>T18753</td><td>TACATTTTCTGTGGGATCAGCGG</td><td>1000</td><td></td><td></td><td></td><td></td><td></td><td></td><td></td><td></td><td></td></tr>
<tr><td>Oligo fwd</td><td>CACCTACATTTTCTGTGGGATCAG</td><td></td><td></td><td></td><td></td><td></td><td></td><td></td><td></td><td></td><td></td></tr>
<tr><td>Oligo rev</td><td>AAACCTGATCCCACAGAAAATGTA</td><td></td><td></td><td></td><td></td><td></td><td></td><td></td><td></td><td></td><td></td></tr>
<tr><td>Chromosome</td><td>start</td><td>end</td><td>strand</td><td>MM</td><td>target_seq</td><td>PAM</td><td>alignment</td><td>distance</td><td>position</td><td>gene name</td><td>gene id</td></tr>
<tr><td>GU305914.1</td><td>208591</td><td>208613</td><td>-</td><td>0</td><td>TACATTTTCTGTGGGATCAG</td><td>CGG</td><td>||||||||[||||||||||||]PAM</td><td>NA</td><td>Intergenic</td><td>NA</td><td>NA</td></tr>
<tr><td></td><td></td><td></td><td></td><td></td><td></td><td></td><td></td><td></td><td></td><td></td><td></td></tr>
<tr><td>T18754</td><td>TCCACCTGACCGCGTTCCGTCGG</td><td>1000</td><td></td><td></td><td></td><td></td><td></td><td></td><td></td><td></td><td></td></tr>
<tr><td>Oligo fwd</td><td>CACCTCCACCTGACCGCGTTCCGT</td><td></td><td></td><td></td><td></td><td></td><td></td><td></td><td></td><td></td><td></td></tr>
<tr><td>Oligo rev</td><td>AAACACGGAACGCGGTCAGGTGGA</td><td></td><td></td><td></td><td></td><td></td><td></td><td></td><td></td><td></td><td></td></tr>
<tr><td>Chromosome</td><td>start</td><td>end</td><td>strand</td><td>MM</td><td>target_seq</td><td>PAM</td><td>alignment</td><td>distance</td><td>position</td><td>gene name</td><td>gene id</td></tr>
<tr><td>GU305914.1</td><td>208567</td><td>208589</td><td>-</td><td>0</td><td>TCCACCTGACCGCGTTCCGT</td><td>CGG</td><td>||||||||[||||||||||||]PAM</td><td>NA</td><td>Intergenic</td><td>NA</td><td>NA</td></tr>
<tr><td>GU305914.1</td><td>224707</td><td>224729</td><td>+</td><td>4</td><td>TCCATGTGGCGGCGTTCCGT</td><td>CGG</td><td>||||--||[-|-|||||||||]PAM</td><td>NA</td><td>Intergenic</td><td>NA</td><td>NA</td></tr>
<tr><td></td><td></td><td></td><td></td><td></td><td></td><td></td><td></td><td></td><td></td><td></td><td></td></tr>
<tr><td>T18755</td><td>ACCGCGTTCCGTCGGCTCCTCGG</td><td>1000</td><td></td><td></td><td></td><td></td><td></td><td></td><td></td><td></td><td></td></tr>
<tr><td>Oligo fwd</td><td>CACCACCGCGTTCCGTCGGCTCCT</td><td></td><td></td><td></td><td></td><td></td><td></td><td></td><td></td><td></td><td></td></tr>
<tr><td>Oligo rev</td><td>AAACAGGAGCCGACGGAACGCGGT</td><td></td><td></td><td></td><td></td><td></td><td></td><td></td><td></td><td></td><td></td></tr>
<tr><td>Chromosome</td><td>start</td><td>end</td><td>strand</td><td>MM</td><td>target_seq</td><td>PAM</td><td>alignment</td><td>distance</td><td>position</td><td>gene name</td><td>gene id</td></tr>
<tr><td>GU305914.1</td><td>208559</td><td>208581</td><td>-</td><td>0</td><td>ACCGCGTTCCGTCGGCTCCT</td><td>CGG</td><td>||||||||[||||||||||||]PAM</td><td>NA</td><td>Intergenic</td><td>NA</td><td>NA</td></tr>
<tr><td></td><td></td><td></td><td></td><td></td><td></td><td></td><td></td><td></td><td></td><td></td><td></td></tr>
<tr><td>T18756</td><td>TCGGCTCCTCGGTCAAAGCGCGG</td><td>1000</td><td></td><td></td><td></td><td></td><td></td><td></td><td></td><td></td><td></td></tr>
<tr><td>Oligo fwd</td><td>CACCTCGGCTCCTCGGTCAAAGCG</td><td></td><td></td><td></td><td></td><td></td><td></td><td></td><td></td><td></td><td></td></tr>
<tr><td>Oligo rev</td><td>AAACCGCTTTGACCGAGGAGCCGA</td><td></td><td></td><td></td><td></td><td></td><td></td><td></td><td></td><td></td><td></td></tr>
<tr><td>Chromosome</td><td>start</td><td>end</td><td>strand</td><td>MM</td><td>target_seq</td><td>PAM</td><td>alignment</td><td>distance</td><td>position</td><td>gene name</td><td>gene id</td></tr>
<tr><td>GU305914.1</td><td>208548</td><td>208570</td><td>-</td><td>0</td><td>TCGGCTCCTCGGTCAAAGCG</td><td>CGG</td><td>||||||||[||||||||||||]PAM</td><td>NA</td><td>Intergenic</td><td>NA</td><td>NA</td></tr>
<tr><td></td><td></td><td></td><td></td><td></td><td></td><td></td><td></td><td></td><td></td><td></td><td></td></tr>
<tr><td>T18757</td><td>GTGCGATTTCGCACCGCAGTTGG</td><td>1000</td><td></td><td></td><td></td><td></td><td></td><td></td><td></td><td></td><td></td></tr>
<tr><td>Oligo fwd</td><td>CACCGTGCGATTTCGCACCGCAGT</td><td></td><td></td><td></td><td></td><td></td><td></td><td></td><td></td><td></td><td></td></tr>
<tr><td>Oligo rev</td><td>AAACACTGCGGTGCGAAATCGCAC</td><td></td><td></td><td></td><td></td><td></td><td></td><td></td><td></td><td></td><td></td></tr>
<tr><td>Chromosome</td><td>start</td><td>end</td><td>strand</td><td>MM</td><td>target_seq</td><td>PAM</td><td>alignment</td><td>distance</td><td>position</td><td>gene name</td><td>gene id</td></tr>
<tr><td>GU305914.1</td><td>208503</td><td>208525</td><td>-</td><td>0</td><td>GTGCGATTTCGCACCGCAGT</td><td>TGG</td><td>||||||||[||||||||||||]PAM</td><td>NA</td><td>Intergenic</td><td>NA</td><td>NA</td></tr>
<tr><td></td><td></td><td></td><td></td><td></td><td></td><td></td><td></td><td></td><td></td><td></td><td></td></tr>
<tr><td>T18758</td><td>GATTTCGCACCGCAGTTGGTCGG</td><td>1000</td><td></td><td></td><td></td><td></td><td></td><td></td><td></td><td></td><td></td></tr>
<tr><td>Oligo fwd</td><td>CACCGATTTCGCACCGCAGTTGGT</td><td></td><td></td><td></td><td></td><td></td><td></td><td></td><td></td><td></td><td></td></tr>
<tr><td>Oligo rev</td><td>AAACACCAACTGCGGTGCGAAATC</td><td></td><td></td><td></td><td></td><td></td><td></td><td></td><td></td><td></td><td></td></tr>
<tr><td>Chromosome</td><td>start</td><td>end</td><td>strand</td><td>MM</td><td>target_seq</td><td>PAM</td><td>alignment</td><td>distance</td><td>position</td><td>gene name</td><td>gene id</td></tr>
<tr><td>GU305914.1</td><td>208499</td><td>208521</td><td>-</td><td>0</td><td>GATTTCGCACCGCAGTTGGT</td><td>CGG</td><td>||||||||[||||||||||||]PAM</td><td>NA</td><td>Intergenic</td><td>NA</td><td>NA</td></tr>
<tr><td></td><td></td><td></td><td></td><td></td><td></td><td></td><td></td><td></td><td></td><td></td><td></td></tr>
<tr><td>T18759</td><td>ATGCCTCGTTGTACACACCGCGG</td><td>1000</td><td></td><td></td><td></td><td></td><td></td><td></td><td></td><td></td><td></td></tr>
<tr><td>Oligo fwd</td><td>CACCATGCCTCGTTGTACACACCG</td><td></td><td></td><td></td><td></td><td></td><td></td><td></td><td></td><td></td><td></td></tr>
<tr><td>Oligo rev</td><td>AAACCGGTGTGTACAACGAGGCAT</td><td></td><td></td><td></td><td></td><td></td><td></td><td></td><td></td><td></td><td></td></tr>
<tr><td>Chromosome</td><td>start</td><td>end</td><td>strand</td><td>MM</td><td>target_seq</td><td>PAM</td><td>alignment</td><td>distance</td><td>position</td><td>gene name</td><td>gene id</td></tr>
<tr><td>GU305914.1</td><td>208476</td><td>208498</td><td>-</td><td>0</td><td>ATGCCTCGTTGTACACACCG</td><td>CGG</td><td>||||||||[||||||||||||]PAM</td><td>NA</td><td>Intergenic</td><td>NA</td><td>NA</td></tr>
</table>

<table>
<tr><td>T18760</td><td>CTCGTTGTACACACCGCGGACGG</td><td>1000</td><td></td><td></td><td></td><td></td><td></td><td></td><td></td><td></td><td></td></tr>
<tr><td>Oligo fwd</td><td>CACCCTCGTTGTACACACCGCGGA</td><td></td><td></td><td></td><td></td><td></td><td></td><td></td><td></td><td></td><td></td></tr>
<tr><td>Oligo rev</td><td>AAACTCCGCGGTGTGTACAACGAG</td><td></td><td></td><td></td><td></td><td></td><td></td><td></td><td></td><td></td><td></td></tr>
<tr><td>Chromosome</td><td>start</td><td>end</td><td>strand</td><td>MM</td><td>target_seq</td><td>PAM</td><td>alignment</td><td>distance</td><td>position</td><td>gene name</td><td>gene id</td></tr>
<tr><td>GU305914.1</td><td>208472</td><td>208494</td><td>-</td><td>0</td><td>CTCGTTGTACACACCGCGGA</td><td>CGG</td><td>||||||||[||||||||||||]PAM</td><td>NA</td><td>Intergenic</td><td>NA</td><td>NA</td></tr>
<tr><td>T18761</td><td>GGACGGTACGAACGTTAGATCGG</td><td>1000</td><td></td><td></td><td></td><td></td><td></td><td></td><td></td><td></td><td></td></tr>
<tr><td>Oligo fwd</td><td>CACCGGACGGTACGAACGTTAGAT</td><td></td><td></td><td></td><td></td><td></td><td></td><td></td><td></td><td></td><td></td></tr>
<tr><td>Oligo rev</td><td>AAACATCTAACGTTCGTACCGTCC</td><td></td><td></td><td></td><td></td><td></td><td></td><td></td><td></td><td></td><td></td></tr>
<tr><td>Chromosome</td><td>start</td><td>end</td><td>strand</td><td>MM</td><td>target_seq</td><td>PAM</td><td>alignment</td><td>distance</td><td>position</td><td>gene name</td><td>gene id</td></tr>
<tr><td>GU305914.1</td><td>208455</td><td>208477</td><td>-</td><td>0</td><td>GGACGGTACGAACGTTAGAT</td><td>CGG</td><td>||||||||[||||||||||||]PAM</td><td>NA</td><td>Intergenic</td><td>NA</td><td>NA</td></tr>
<tr><td>T18762</td><td>CGTTAGATCGGCGATCGTAACGG</td><td>1000</td><td></td><td></td><td></td><td></td><td></td><td></td><td></td><td></td><td></td></tr>
<tr><td>Oligo fwd</td><td>CACCCGTTAGATCGGCGATCGTAA</td><td></td><td></td><td></td><td></td><td></td><td></td><td></td><td></td><td></td><td></td></tr>
<tr><td>Oligo rev</td><td>AAACTTACGATCGCCGATCTAACG</td><td></td><td></td><td></td><td></td><td></td><td></td><td></td><td></td><td></td><td></td></tr>
<tr><td>Chromosome</td><td>start</td><td>end</td><td>strand</td><td>MM</td><td>target_seq</td><td>PAM</td><td>alignment</td><td>distance</td><td>position</td><td>gene name</td><td>gene id</td></tr>
<tr><td>GU305914.1</td><td>208443</td><td>208465</td><td>-</td><td>0</td><td>CGTTAGATCGGCGATCGTAA</td><td>CGG</td><td>||||||||[||||||||||||]PAM</td><td>NA</td><td>Intergenic</td><td>NA</td><td>NA</td></tr>
<tr><td>T18763</td><td>AACGGTACGAGATAAGAAGAAGG</td><td>1000</td><td></td><td></td><td></td><td></td><td></td><td></td><td></td><td></td><td></td></tr>
<tr><td>Oligo fwd</td><td>CACCAACGGTACGAGATAAGAAGA</td><td></td><td></td><td></td><td></td><td></td><td></td><td></td><td></td><td></td><td></td></tr>
<tr><td>Oligo rev</td><td>AAACTCTTCTTATCTCGTACCGTT</td><td></td><td></td><td></td><td></td><td></td><td></td><td></td><td></td><td></td><td></td></tr>
<tr><td>Chromosome</td><td>start</td><td>end</td><td>strand</td><td>MM</td><td>target_seq</td><td>PAM</td><td>alignment</td><td>distance</td><td>position</td><td>gene name</td><td>gene id</td></tr>
<tr><td>GU305914.1</td><td>208425</td><td>208447</td><td>-</td><td>0</td><td>AACGGTACGAGATAAGAAGA</td><td>AGG</td><td>||||||||[||||||||||||]PAM</td><td>NA</td><td>Intergenic</td><td>NA</td><td>NA</td></tr>
<tr><td>T18764</td><td>GGTACGAGATAAGAAGAAGGTGG</td><td>1000</td><td></td><td></td><td></td><td></td><td></td><td></td><td></td><td></td><td></td></tr>
<tr><td>Oligo fwd</td><td>CACCGGTACGAGATAAGAAGAAGG</td><td></td><td></td><td></td><td></td><td></td><td></td><td></td><td></td><td></td><td></td></tr>
<tr><td>Oligo rev</td><td>AAACCCTTCTTCTTATCTCGTACC</td><td></td><td></td><td></td><td></td><td></td><td></td><td></td><td></td><td></td><td></td></tr>
<tr><td>Chromosome</td><td>start</td><td>end</td><td>strand</td><td>MM</td><td>target_seq</td><td>PAM</td><td>alignment</td><td>distance</td><td>position</td><td>gene name</td><td>gene id</td></tr>
<tr><td>GU305914.1</td><td>208422</td><td>208444</td><td>-</td><td>0</td><td>GGTACGAGATAAGAAGAAGG</td><td>TGG</td><td>||||||||[||||||||||||]PAM</td><td>NA</td><td>Intergenic</td><td>NA</td><td>NA</td></tr>
<tr><td>T18765</td><td>TAAGAAGAAGGTGGTGATCGCGG</td><td>1000</td><td></td><td></td><td></td><td></td><td></td><td></td><td></td><td></td><td></td></tr>
<tr><td>Oligo fwd</td><td>CACCTAAGAAGAAGGTGGTGATCG</td><td></td><td></td><td></td><td></td><td></td><td></td><td></td><td></td><td></td><td></td></tr>
<tr><td>Oligo rev</td><td>AAACCGATCACCACCTTCTTCTTA</td><td></td><td></td><td></td><td></td><td></td><td></td><td></td><td></td><td></td><td></td></tr>
<tr><td>Chromosome</td><td>start</td><td>end</td><td>strand</td><td>MM</td><td>target_seq</td><td>PAM</td><td>alignment</td><td>distance</td><td>position</td><td>gene name</td><td>gene id</td></tr>
<tr><td>GU305914.1</td><td>208413</td><td>208435</td><td>-</td><td>0</td><td>TAAGAAGAAGGTGGTGATCG</td><td>CGG</td><td>||||||||[||||||||||||]PAM</td><td>NA</td><td>Intergenic</td><td>NA</td><td>NA</td></tr>
<tr><td>T18766</td><td>GGTGATCGCGGATCACTCATCGG</td><td>1000</td><td></td><td></td><td></td><td></td><td></td><td></td><td></td><td></td><td></td></tr>
<tr><td>Oligo fwd</td><td>CACCGGTGATCGCGGATCACTCAT</td><td></td><td></td><td></td><td></td><td></td><td></td><td></td><td></td><td></td><td></td></tr>
<tr><td>Oligo rev</td><td>AAACATGAGTGATCCGCGATCACC</td><td></td><td></td><td></td><td></td><td></td><td></td><td></td><td></td><td></td><td></td></tr>
<tr><td>Chromosome</td><td>start</td><td>end</td><td>strand</td><td>MM</td><td>target_seq</td><td>PAM</td><td>alignment</td><td>distance</td><td>position</td><td>gene name</td><td>gene id</td></tr>
<tr><td>GU305914.1</td><td>208401</td><td>208423</td><td>-</td><td>0</td><td>GGTGATCGCGGATCACTCAT</td><td>CGG</td><td>||||||||[||||||||||||]PAM</td><td>NA</td><td>Intergenic</td><td>NA</td><td>NA</td></tr>
<tr><td>T18767</td><td>GAGCGTGCTGCGTCGTAACGAGG</td><td>1000</td><td></td><td></td><td></td><td></td><td></td><td></td><td></td><td></td><td></td></tr>
<tr><td>Oligo fwd</td><td>CACCGAGCGTGCTGCGTCGTAACG</td><td></td><td></td><td></td><td></td><td></td><td></td><td></td><td></td><td></td><td></td></tr>
<tr><td>Oligo rev</td><td>AAACCGTTACGACGCAGCACGCTC</td><td></td><td></td><td></td><td></td><td></td><td></td><td></td><td></td><td></td><td></td></tr>
<tr><td>Chromosome</td><td>start</td><td>end</td><td>strand</td><td>MM</td><td>target_seq</td><td>PAM</td><td>alignment</td><td>distance</td><td>position</td><td>gene name</td><td>gene id</td></tr>
<tr><td>GU305914.1</td><td>208344</td><td>208366</td><td>-</td><td>0</td><td>GAGCGTGCTGCGTCGTAACG</td><td>AGG</td><td>||||||||[||||||||||||]PAM</td><td>NA</td><td>Intergenic</td><td>NA</td><td>NA</td></tr>
<tr><td>T18768</td><td>CGTAACGAGGTCTCGATTCACGG</td><td>1000</td><td></td><td></td><td></td><td></td><td></td><td></td><td></td><td></td><td></td></tr>
<tr><td>Oligo fwd</td><td>CACCCGTAACGAGGTCTCGATTCA</td><td></td><td></td><td></td><td></td><td></td><td></td><td></td><td></td><td></td><td></td></tr>
<tr><td>Oligo rev</td><td>AAACTGAATCGAGACCTCGTTACG</td><td></td><td></td><td></td><td></td><td></td><td></td><td></td><td></td><td></td><td></td></tr>
<tr><td>Chromosome</td><td>start</td><td>end</td><td>strand</td><td>MM</td><td>target_seq</td><td>PAM</td><td>alignment</td><td>distance</td><td>position</td><td>gene name</td><td>gene id</td></tr>
<tr><td>GU305914.1</td><td>208331</td><td>208353</td><td>-</td><td>0</td><td>CGTAACGAGGTCTCGATTCA</td><td>CGG</td><td>||||||||[||||||||||||]PAM</td><td>NA</td><td>Intergenic</td><td>NA</td><td>NA</td></tr>
<tr><td>T18769</td><td>AGGTCTCGATTCACGGTCGTTGG</td><td>1000</td><td></td><td></td><td></td><td></td><td></td><td></td><td></td><td></td><td></td></tr>
<tr><td>Oligo fwd</td><td>CACCAGGTCTCGATTCACGGTCGT</td><td></td><td></td><td></td><td></td><td></td><td></td><td></td><td></td><td></td><td></td></tr>
<tr><td>Oligo rev</td><td>AAACACGACCGTGAATCGAGACCT</td><td></td><td></td><td></td><td></td><td></td><td></td><td></td><td></td><td></td><td></td></tr>
<tr><td>Chromosome</td><td>start</td><td>end</td><td>strand</td><td>MM</td><td>target_seq</td><td>PAM</td><td>alignment</td><td>distance</td><td>position</td><td>gene name</td><td>gene id</td></tr>
<tr><td>GU305914.1</td><td>208324</td><td>208346</td><td>-</td><td>0</td><td>AGGTCTCGATTCACGGTCGT</td><td>TGG</td><td>||||||||[||||||||||||]PAM</td><td>NA</td><td>Intergenic</td><td>NA</td><td>NA</td></tr>
<tr><td>T18770</td><td>GCGTTTGAAGATCATGAAGCTGG</td><td>1000</td><td></td><td></td><td></td><td></td><td></td><td></td><td></td><td></td><td></td></tr>
<tr><td>Oligo fwd</td><td>CACCGCGTTTGAAGATCATGAAGC</td><td></td><td></td><td></td><td></td><td></td><td></td><td></td><td></td><td></td><td></td></tr>
<tr><td>Oligo rev</td><td>AAACGCTTCATGATCTTCAAACGC</td><td></td><td></td><td></td><td></td><td></td><td></td><td></td><td></td><td></td><td></td></tr>
<tr><td>Chromosome</td><td>start</td><td>end</td><td>strand</td><td>MM</td><td>target_seq</td><td>PAM</td><td>alignment</td><td>distance</td><td>position</td><td>gene name</td><td>gene id</td></tr>
<tr><td>GU305914.1</td><td>208293</td><td>208315</td><td>-</td><td>0</td><td>GCGTTTGAAGATCATGAAGC</td><td>TGG</td><td>||||||||[||||||||||||]PAM</td><td>NA</td><td>Intergenic</td><td>NA</td><td>NA</td></tr>
<tr><td>T18771</td><td>TTTGAAGATCATGAAGCTGGCGG</td><td>1000</td><td></td><td></td><td></td><td></td><td></td><td></td><td></td><td></td><td></td></tr>
<tr><td>Oligo fwd</td><td>CACCTTTGAAGATCATGAAGCTGG</td><td></td><td></td><td></td><td></td><td></td><td></td><td></td><td></td><td></td><td></td></tr>
<tr><td>Oligo rev</td><td>AAACCCAGCTTCATGATCTTCAAA</td><td></td><td></td><td></td><td></td><td></td><td></td><td></td><td></td><td></td><td></td></tr>
<tr><td>Chromosome</td><td>start</td><td>end</td><td>strand</td><td>MM</td><td>target_seq</td><td>PAM</td><td>alignment</td><td>distance</td><td>position</td><td>gene name</td><td>gene id</td></tr>
<tr><td>GU305914.1</td><td>208290</td><td>208312</td><td>-</td><td>0</td><td>TTTGAAGATCATGAAGCTGG</td><td>CGG</td><td>||||||||[||||||||||||]PAM</td><td>NA</td><td>Intergenic</td><td>NA</td><td>NA</td></tr>
<tr><td>T18772</td><td>GCTGGCGGCTCCCGAGCACTTGG</td><td>1000</td><td></td><td></td><td></td><td></td><td></td><td></td><td></td><td></td><td></td></tr>
<tr><td>Oligo fwd</td><td>CACCGCTGGCGGCTCCCGAGCACT</td><td></td><td></td><td></td><td></td><td></td><td></td><td></td><td></td><td></td><td></td></tr>
<tr><td>Oligo rev</td><td>AAACAGTGCTCGGGAGCCGCCAGC</td><td></td><td></td><td></td><td></td><td></td><td></td><td></td><td></td><td></td><td></td></tr>
<tr><td>Chromosome</td><td>start</td><td>end</td><td>strand</td><td>MM</td><td>target_seq</td><td>PAM</td><td>alignment</td><td>distance</td><td>position</td><td>gene name</td><td>gene id</td></tr>
<tr><td>GU305914.1</td><td>208275</td><td>208297</td><td>-</td><td>0</td><td>GCTGGCGGCTCCCGAGCACT</td><td>TGG</td><td>||||||||[||||||||||||]PAM</td><td>NA</td><td>Intergenic</td><td>NA</td><td>NA</td></tr>
<tr><td>T18773</td><td>GGCGGCTCCCGAGCACTTGGAGG</td><td>1000</td><td></td><td></td><td></td><td></td><td></td><td></td><td></td><td></td><td></td></tr>
<tr><td>Oligo fwd</td><td>CACCGGCGGCTCCCGAGCACTTGG</td><td></td><td></td><td></td><td></td><td></td><td></td><td></td><td></td><td></td><td></td></tr>
<tr><td>Oligo rev</td><td>AAACCCAAGTGCTCGGGAGCCGCC</td><td></td><td></td><td></td><td></td><td></td><td></td><td></td><td></td><td></td><td></td></tr>
<tr><td>Chromosome</td><td>start</td><td>end</td><td>strand</td><td>MM</td><td>target_seq</td><td>PAM</td><td>alignment</td><td>distance</td><td>position</td><td>gene name</td><td>gene id</td></tr>
<tr><td>GU305914.1</td><td>208272</td><td>208294</td><td>-</td><td>0</td><td>GGCGGCTCCCGAGCACTTGG</td><td>AGG</td><td>||||||||[||||||||||||]PAM</td><td>NA</td><td>Intergenic</td><td>NA</td><td>NA</td></tr>
<tr><td>T18774</td><td>GGTTAGTTTTCACGTTTCCGTGG</td><td>1000</td><td></td><td></td><td></td><td></td><td></td><td></td><td></td><td></td><td></td></tr>
<tr><td>Oligo fwd</td><td>CACCGGTTAGTTTTCACGTTTCCG</td><td></td><td></td><td></td><td></td><td></td><td></td><td></td><td></td><td></td><td></td></tr>
<tr><td>Oligo rev</td><td>AAACCGGAAACGTGAAAACTAACC</td><td></td><td></td><td></td><td></td><td></td><td></td><td></td><td></td><td></td><td></td></tr>
<tr><td>Chromosome</td><td>start</td><td>end</td><td>strand</td><td>MM</td><td>target_seq</td><td>PAM</td><td>alignment</td><td>distance</td><td>position</td><td>gene name</td><td>gene id</td></tr>
<tr><td>GU305914.1</td><td>208251</td><td>208273</td><td>-</td><td>0</td><td>GGTTAGTTTTCACGTTTCCG</td><td>TGG</td><td>||||||||[||||||||||||]PAM</td><td>NA</td><td>Intergenic</td><td>NA</td><td>NA</td></tr>
</table>

T18775 GTTAGTTTTCACGTTTCCGTGGG 1000
Oligo fwd CACCGTTAGTTTTCACGTTTCCGT
Oligo rev AAACACGGAAACGTGAAAACTAAC
Chromosome start end strand MM target_seq PAM alignment distance position gene name gene id
GU305914.1 208250 208272 - 0 GTTAGTTTTCACGTTTCCGT GGG ||||||||[||||||||||||]PAM NA Intergenic NA NA

T18776 CACGTTTCCGTGGGAAGACACGG 1000
Oligo fwd CACCCACGTTTCCGTGGGAAGACA
Oligo rev AAACTGTCTTCCCACGGAAACGTG
Chromosome start end strand MM target_seq PAM alignment distance position gene name gene id
GU305914.1 208241 208263 - 0 CACGTTTCCGTGGGAAGACA CGG ||||||||[||||||||||||]PAM NA Intergenic NA NA

T18777 ACACGGATTCTTCTGCTGTTCGG 1000
Oligo fwd CACCACACGGATTCTTCTGCTGTT
Oligo rev AAACAACAGCAGAAGAATCCGTGT
Chromosome start end strand MM target_seq PAM alignment distance position gene name gene id
GU305914.1 208224 208246 - 0 ACACGGATTCTTCTGCTGTT CGG ||||||||[||||||||||||]PAM NA Intergenic NA NA

T18778 GATTCTTCTGCTGTTCGGTGTGG 1000
Oligo fwd CACCGATTCTTCTGCTGTTCGGTG
Oligo rev AAACCACCGAACAGCAGAAGAATC
Chromosome start end strand MM target_seq PAM alignment distance position gene name gene id
GU305914.1 208219 208241 - 0 GATTCTTCTGCTGTTCGGTG TGG ||||||||[||||||||||||]PAM NA Intergenic NA NA

T18779 CTGCTGTTCGGTGTGGAGCCCGG 1000
Oligo fwd CACCCTGCTGTTCGGTGTGGAGCC
Oligo rev AAACGGCTCCACACCGAACAGCAG
Chromosome start end strand MM target_seq PAM alignment distance position gene name gene id
GU305914.1 208212 208234 - 0 CTGCTGTTCGGTGTGGAGCC CGG ||||||||[||||||||||||]PAM NA Intergenic NA NA

T18780 CCCGCTACCCTTCGACGTGACGG 1000
Oligo fwd CACCCCCGCTACCCTTCGACGTGA
Oligo rev AAACTCACGTCGAAGGGTAGCGGG
Chromosome start end strand MM target_seq PAM alignment distance position gene name gene id
GU305914.1 208188 208210 - 0 CCCGCTACCCTTCGACGTGA CGG ||||||||[||||||||||||]PAM NA Intergenic NA NA

T18781 TACCCTTCGACGTGACGGTGAGG 1000
Oligo fwd CACCTACCCTTCGACGTGACGGTG
Oligo rev AAACCACCGTCACGTCGAAGGGTA
Chromosome start end strand MM target_seq PAM alignment distance position gene name gene id
GU305914.1 208183 208205 - 0 TACCCTTCGACGTGACGGTG AGG ||||||||[||||||||||||]PAM NA Intergenic NA NA

T18782 ACCCTTCGACGTGACGGTGAGGG 1000
Oligo fwd CACCACCCTTCGACGTGACGGTGA
Oligo rev AAACTCACCGTCACGTCGAAGGGT
Chromosome start end strand MM target_seq PAM alignment distance position gene name gene id
GU305914.1 208182 208204 - 0 ACCCTTCGACGTGACGGTGA GGG ||||||||[||||||||||||]PAM NA Intergenic NA NA

T18783 CCCTTCGACGTGACGGTGAGGGG 1000
Oligo fwd CACCCCCTTCGACGTGACGGTGAG
Oligo rev AAACCTCACCGTCACGTCGAAGGG
Chromosome start end strand MM target_seq PAM alignment distance position gene name gene id
GU305914.1 208181 208203 - 0 CCCTTCGACGTGACGGTGAG GGG ||||||||[||||||||||||]PAM NA Intergenic NA NA

T18784 TTCGACGTGACGGTGAGGGGCGG 1000
Oligo fwd CACCTTCGACGTGACGGTGAGGGG
Oligo rev AAACCCCCTCACCGTCACGTCGAA
Chromosome start end strand MM target_seq PAM alignment distance position gene name gene id
GU305914.1 208178 208200 - 0 TTCGACGTGACGGTGAGGGG CGG ||||||||[||||||||||||]PAM NA Intergenic NA NA

T18785 GACGTGACGGTGAGGGGCGGTGG 1000
Oligo fwd CACCGACGTGACGGTGAGGGGCGG
Oligo rev AAACCCGCCCCTCACCGTCACGTC
Chromosome start end strand MM target_seq PAM alignment distance position gene name gene id
GU305914.1 208175 208197 - 0 GACGTGACGGTGAGGGGCGG TGG ||||||||[||||||||||||]PAM NA Intergenic NA NA

T18786 ACGTGACGGTGAGGGGCGGTGGG 1000
Oligo fwd CACCACGTGACGGTGAGGGGCGGT
Oligo rev AAACACCGCCCCTCACCGTCACGT
Chromosome start end strand MM target_seq PAM alignment distance position gene name gene id
GU305914.1 208174 208196 - 0 ACGTGACGGTGAGGGGCGGT GGG ||||||||[||||||||||||]PAM NA Intergenic NA NA

T18787 GAGGGGCGGTGGGTTGAGATCGG 1000
Oligo fwd CACCGAGGGGCGGTGGGTTGAGAT
Oligo rev AAACATCTCAACCCACCGCCCCTC
Chromosome start end strand MM target_seq PAM alignment distance position gene name gene id
GU305914.1 208164 208186 - 0 GAGGGGCGGTGGGTTGAGAT CGG ||||||||[||||||||||||]PAM NA Intergenic NA NA

T18788 TCTCTCTGAACGCGTCCCGCCGG 1000
Oligo fwd CACCTCTCTCTGAACGCGTCCCGC
Oligo rev AAACGCGGGACGCGTTCAGAGAGA
Chromosome start end strand MM target_seq PAM alignment distance position gene name gene id
GU305914.1 208141 208163 - 0 TCTCTCTGAACGCGTCCCGC CGG ||||||||[||||||||||||]PAM NA Intergenic NA NA
GU305914.1 165929 165951 - 4 CCTCTCGGATCGCGTCCTGC AGG -|||||-|[|-|||||||-||]PAM NA Intergenic NA NA

T18789 CGCGTCCCGCCGGTTCTTTTCGG 1000
Oligo fwd CACCCGCGTCCCGCCGGTTCTTTT
Oligo rev AAACAAAAGAACCGGCGGGACGCG

Chromosome start end strand MM target_seq PAM alignment distance position gene name gene id
GU305914.1 208131 208153 - 0 CGCGTCCCGCCGGTTCTTTT CGG ||||||||[||||||||||||]PAM NA Intergenic NA NA

T18790 CCGCCGGTTCTTTTCGGATACGG 1000
Oligo fwd CACCCCGCCGGTTCTTTTCGGATA
Oligo rev AAACTATCCGAAAAGAACCGGCGG
Chromosome start end strand MM target_seq PAM alignment distance position gene name gene id
GU305914.1 208125 208147 - 0 CCGCCGGTTCTTTTCGGATA CGG ||||||||[||||||||||||]PAM NA Intergenic NA NA

T18791 CCGGTTCTTTTCGGATACGGTGG 1000
Oligo fwd CACCCCGGTTCTTTTCGGATACGG
Oligo rev AAACCCGTATCCGAAAAGAACCGG
Chromosome start end strand MM target_seq PAM alignment distance position gene name gene id
GU305914.1 208122 208144 - 0 CCGGTTCTTTTCGGATACGG TGG ||||||||[||||||||||||]PAM NA Intergenic NA NA

T18792 GACCTCGAGCTTCGAGCCCTCGG 1000
Oligo fwd CACCGACCTCGAGCTTCGAGCCCT
Oligo rev AAACAGGGCTCGAAGCTCGAGGTC
Chromosome start end strand MM target_seq PAM alignment distance position gene name gene id
GU305914.1 208080 208102 - 0 GACCTCGAGCTTCGAGCCCT CGG ||||||||[||||||||||||]PAM NA Intergenic NA NA

T18793 GAGCTTCGAGCCCTCGGATCTGG 1000
Oligo fwd CACCGAGCTTCGAGCCCTCGGATC
Oligo rev AAACGATCCGAGGGCTCGAAGCTC
Chromosome start end strand MM target_seq PAM alignment distance position gene name gene id
GU305914.1 208074 208096 - 0 GAGCTTCGAGCCCTCGGATC TGG ||||||||[||||||||||||]PAM NA Intergenic NA NA

T18794 GGTCTGTGTCATTCGATCGTCGG 1000
Oligo fwd CACCGGTCTGTGTCATTCGATCGT
Oligo rev AAACACGATCGAATGACACAGACC
Chromosome start end strand MM target_seq PAM alignment distance position gene name gene id
GU305914.1 208053 208075 - 0 GGTCTGTGTCATTCGATCGT CGG ||||||||[||||||||||||]PAM NA Intergenic NA NA

T18795 CTGTGTCATTCGATCGTCGGTGG 1000
Oligo fwd CACCCTGTGTCATTCGATCGTCGG
Oligo rev AAACCCGACGATCGAATGACACAG
Chromosome start end strand MM target_seq PAM alignment distance position gene name gene id
GU305914.1 208050 208072 - 0 CTGTGTCATTCGATCGTCGG TGG ||||||||[||||||||||||]PAM NA Intergenic NA NA

T18796 TGTGTCATTCGATCGTCGGTGGG 1000
Oligo fwd CACCTGTGTCATTCGATCGTCGGT
Oligo rev AAACACCGACGATCGAATGACACA
Chromosome start end strand MM target_seq PAM alignment distance position gene name gene id
GU305914.1 208049 208071 - 0 TGTGTCATTCGATCGTCGGT GGG ||||||||[||||||||||||]PAM NA Intergenic NA NA

T18797 GTGTCATTCGATCGTCGGTGGGG 1000
Oligo fwd CACCGTGTCATTCGATCGTCGGTG
Oligo rev AAACCACCGACGATCGAATGACAC
Chromosome start end strand MM target_seq PAM alignment distance position gene name gene id
GU305914.1 208048 208070 - 0 GTGTCATTCGATCGTCGGTG GGG ||||||||[||||||||||||]PAM NA Intergenic NA NA

T18798 TCATTCGATCGTCGGTGGGGTGG 1000
Oligo fwd CACCTCATTCGATCGTCGGTGGGG
Oligo rev AAACCCCCACCGACGATCGAATGA
Chromosome start end strand MM target_seq PAM alignment distance position gene name gene id
GU305914.1 208045 208067 - 0 TCATTCGATCGTCGGTGGGG TGG ||||||||[||||||||||||]PAM NA Intergenic NA NA

T18799 TCGATCGTCGGTGGGGTGGACGG 1000
Oligo fwd CACCTCGATCGTCGGTGGGGTGGA
Oligo rev AAACTCCACCCCACCGACGATCGA
Chromosome start end strand MM target_seq PAM alignment distance position gene name gene id
GU305914.1 208041 208063 - 0 TCGATCGTCGGTGGGGTGGA CGG ||||||||[||||||||||||]PAM NA Intergenic NA NA

T18800 GACGACCCCGTCGCACGACGAGG 1000
Oligo fwd CACCGACGACCCCGTCGCACGACG
Oligo rev AAACCGTCGTGCGACGGGGTCGTC
Chromosome start end strand MM target_seq PAM alignment distance position gene name gene id
GU305914.1 208011 208033 - 0 GACGACCCCGTCGCACGACG AGG ||||||||[||||||||||||]PAM NA Intergenic NA NA

T18801 CGAGGCCAAGATCTTGAGAACGG 1000
Oligo fwd CACCCGAGGCCAAGATCTTGAGAA
Oligo rev AAACTTCTCAAGATCTTGGCCTCG
Chromosome start end strand MM target_seq PAM alignment distance position gene name gene id
GU305914.1 207993 208015 - 0 CGAGGCCAAGATCTTGAGAA CGG ||||||||[||||||||||||]PAM NA Intergenic NA NA

T18802 GAACGGCAACGACGACAGAGAGG 1000
Oligo fwd CACCGAACGGCAACGACGACAGAG
Oligo rev AAACCTCTGTCGTCGTTGCCGTTC
Chromosome start end strand MM target_seq PAM alignment distance position gene name gene id
GU305914.1 207976 207998 - 0 GAACGGCAACGACGACAGAG AGG ||||||||[||||||||||||]PAM NA Intergenic NA NA

T18803 AACGGCAACGACGACAGAGAGGG 1000
Oligo fwd CACCAACGGCAACGACGACAGAGA
Oligo rev AAACTCTCTGTCGTCGTTGCCGTT
Chromosome start end strand MM target_seq PAM alignment distance position gene name gene id
GU305914.1 207975 207997 - 0 AACGGCAACGACGACAGAGA GGG ||||||||[||||||||||||]PAM NA Intergenic NA NA

T18804 AACGACGACAGAGAGGGAGCCGG 1000
Oligo fwd CACCAACGACGACAGAGAGGGAGC

| | | | | | | | | | | | |
|---|---|---|---|---|---|---|---|---|---|---|---|
| Oligo rev | AAACGCTCCCTCTCTGTCGTCGTT | | | | | | | | | | |
| Chromosome | start | end | strand | MM | target_seq | PAM | alignment | distance | position | gene name | gene id |
| GU305914.1 | 207969 | 207991 | - | 0 | AACGACGACAGAGAGGGAGC | CGG | ||||||||[||||||||||||]PAM | NA | Intergenic | NA | NA |
| T18805 | GAGAGGGAGCCGGCCGCGTGCGG | 1000 | | | | | | | | | |
| Oligo fwd | CACCGAGAGGGAGCCGGCCGCGTG | | | | | | | | | | |
| Oligo rev | AAACCACGCGGCCGGCTCCCTCTC | | | | | | | | | | |
| Chromosome | start | end | strand | MM | target_seq | PAM | alignment | distance | position | gene name | gene id |
| GU305914.1 | 207959 | 207981 | - | 0 | GAGAGGGAGCCGGCCGCGTG | CGG | ||||||||[||||||||||||]PAM | NA | Intergenic | NA | NA |
| GU305914.1 | 197813 | 197835 | + | 4 | CCGAGGGACCCGGCCGCGCG | CGG | --||||||[-|||||||||-|]PAM | NA | Intergenic | NA | NA |
| T18806 | GGAGCCGGCCGCGTGCGGCATGG | 1000 | | | | | | | | | |
| Oligo fwd | CACCGGAGCCGGCCGCGTGCGGCA | | | | | | | | | | |
| Oligo rev | AAACTGCCGCACGCGGCCGGCTCC | | | | | | | | | | |
| Chromosome | start | end | strand | MM | target_seq | PAM | alignment | distance | position | gene name | gene id |
| GU305914.1 | 207954 | 207976 | - | 0 | GGAGCCGGCCGCGTGCGGCA | TGG | ||||||||[||||||||||||]PAM | NA | Intergenic | NA | NA |
| T18807 | GTGCGGCATGGAGCTGTCCCTGG | 1000 | | | | | | | | | |
| Oligo fwd | CACCGTGCGGCATGGAGCTGTCCC | | | | | | | | | | |
| Oligo rev | AAACGGGACAGCTCCATGCCGCAC | | | | | | | | | | |
| Chromosome | start | end | strand | MM | target_seq | PAM | alignment | distance | position | gene name | gene id |
| GU305914.1 | 207942 | 207964 | - | 0 | GTGCGGCATGGAGCTGTCCC | TGG | ||||||||[||||||||||||]PAM | NA | Intergenic | NA | NA |
| T18808 | GCTGTCCCTGGTCACCGTCGTGG | 1000 | | | | | | | | | |
| Oligo fwd | CACCGCTGTCCCTGGTCACCGTCG | | | | | | | | | | |
| Oligo rev | AAACCGACGGTGACCAGGGACAGC | | | | | | | | | | |
| Chromosome | start | end | strand | MM | target_seq | PAM | alignment | distance | position | gene name | gene id |
| GU305914.1 | 207930 | 207952 | - | 0 | GCTGTCCCTGGTCACCGTCG | TGG | ||||||||[||||||||||||]PAM | NA | Intergenic | NA | NA |
| T18809 | GGCGATCGTATCGACGTTTACGG | 1000 | | | | | | | | | |
| Oligo fwd | CACCGGCGATCGTATCGACGTTTA | | | | | | | | | | |
| Oligo rev | AAACTAAACGTCGATACGATCGCC | | | | | | | | | | |
| Chromosome | start | end | strand | MM | target_seq | PAM | alignment | distance | position | gene name | gene id |
| GU305914.1 | 207909 | 207931 | - | 0 | GGCGATCGTATCGACGTTTA | CGG | ||||||||[||||||||||||]PAM | NA | Intergenic | NA | NA |
| T18810 | ATCGACGTTTACGGTGTTGAAGG | 1000 | | | | | | | | | |
| Oligo fwd | CACCATCGACGTTTACGGTGTTGA | | | | | | | | | | |
| Oligo rev | AAACTCAACACCGTAAACGTCGAT | | | | | | | | | | |
| Chromosome | start | end | strand | MM | target_seq | PAM | alignment | distance | position | gene name | gene id |
| GU305914.1 | 207900 | 207922 | - | 0 | ATCGACGTTTACGGTGTTGA | AGG | ||||||||[||||||||||||]PAM | NA | Intergenic | NA | NA |
| T18811 | TCGACGTTTACGGTGTTGAAGGG | 1000 | | | | | | | | | |
| Oligo fwd | CACCTCGACGTTTACGGTGTTGAA | | | | | | | | | | |
| Oligo rev | AAACTTCAACACCGTAAACGTCGA | | | | | | | | | | |
| Chromosome | start | end | strand | MM | target_seq | PAM | alignment | distance | position | gene name | gene id |
| GU305914.1 | 207899 | 207921 | - | 0 | TCGACGTTTACGGTGTTGAA | GGG | ||||||||[||||||||||||]PAM | NA | Intergenic | NA | NA |
| T18812 | CGACGTTTACGGTGTTGAAGGGG | 1000 | | | | | | | | | |
| Oligo fwd | CACCCGACGTTTACGGTGTTGAAG | | | | | | | | | | |
| Oligo rev | AAACCTTCAACACCGTAAACGTCG | | | | | | | | | | |
| Chromosome | start | end | strand | MM | target_seq | PAM | alignment | distance | position | gene name | gene id |
| GU305914.1 | 207898 | 207920 | - | 0 | CGACGTTTACGGTGTTGAAG | GGG | ||||||||[||||||||||||]PAM | NA | Intergenic | NA | NA |
| T18813 | GACGTTTACGGTGTTGAAGGGGG | 1000 | | | | | | | | | |
| Oligo fwd | CACCGACGTTTACGGTGTTGAAGG | | | | | | | | | | |
| Oligo rev | AAACCCTTCAACACCGTAAACGTC | | | | | | | | | | |
| Chromosome | start | end | strand | MM | target_seq | PAM | alignment | distance | position | gene name | gene id |
| GU305914.1 | 207897 | 207919 | - | 0 | GACGTTTACGGTGTTGAAGG | GGG | ||||||||[||||||||||||]PAM | NA | Intergenic | NA | NA |
| T18814 | TGTGTGCCTATGTGTTTCTCAGG | 1000 | | | | | | | | | |
| Oligo fwd | CACCTGTGTGCCTATGTGTTTCTC | | | | | | | | | | |
| Oligo rev | AAACGAGAAACACATAGGCACACA | | | | | | | | | | |
| Chromosome | start | end | strand | MM | target_seq | PAM | alignment | distance | position | gene name | gene id |
| GU305914.1 | 207871 | 207893 | - | 0 | TGTGTGCCTATGTGTTTCTC | AGG | ||||||||[||||||||||||]PAM | NA | Intergenic | NA | NA |
| T18815 | GTTTCTCAGGCGCGACGACGCGG | 1000 | | | | | | | | | |
| Oligo fwd | CACCGTTTCTCAGGCGCGACGACG | | | | | | | | | | |
| Oligo rev | AAACCGTCGTCGCGCCTGAGAAAC | | | | | | | | | | |
| Chromosome | start | end | strand | MM | target_seq | PAM | alignment | distance | position | gene name | gene id |
| GU305914.1 | 207858 | 207880 | - | 0 | GTTTCTCAGGCGCGACGACG | CGG | ||||||||[||||||||||||]PAM | NA | Intergenic | NA | NA |
| T18816 | GCGCGACGACGCGGAGCGCGCGG | 1000 | | | | | | | | | |
| Oligo fwd | CACCGCGCGACGACGCGGAGCGCG | | | | | | | | | | |
| Oligo rev | AAACCGCGCTCCGCGTCGTCGCGC | | | | | | | | | | |
| Chromosome | start | end | strand | MM | target_seq | PAM | alignment | distance | position | gene name | gene id |
| GU305914.1 | 207849 | 207871 | - | 0 | GCGCGACGACGCGGAGCGCG | CGG | ||||||||[||||||||||||]PAM | NA | Intergenic | NA | NA |
| T18817 | ACGACGCGGAGCGCGCGGCGAGG | 1000 | | | | | | | | | |
| Oligo fwd | CACCACGACGCGGAGCGCGCGGCG | | | | | | | | | | |
| Oligo rev | AAACCGCCGCGCGCTCCGCGTCGT | | | | | | | | | | |
| Chromosome | start | end | strand | MM | target_seq | PAM | alignment | distance | position | gene name | gene id |
| GU305914.1 | 207844 | 207866 | - | 0 | ACGACGCGGAGCGCGCGGCG | AGG | ||||||||[||||||||||||]PAM | NA | Intergenic | NA | NA |
| T18818 | ACGCGGAGCGCGCGGCGAGGAGG | 1000 | | | | | | | | | |
| Oligo fwd | CACCACGCGGAGCGCGCGGCGAGG | | | | | | | | | | |
| Oligo rev | AAACCCTCGCCGCGCGCTCCGCGT | | | | | | | | | | |
| Chromosome | start | end | strand | MM | target_seq | PAM | alignment | distance | position | gene name | gene id |
| GU305914.1 | 207841 | 207863 | - | 0 | ACGCGGAGCGCGCGGCGAGG | AGG | ||||||||[||||||||||||]PAM | NA | Intergenic | NA | NA |

T18819 CCGCTATCGCGCCGCGAGTCTGG 1000
Oligo fwd CACCCCGCTATCGCGCCGCGAGTC
Oligo rev AAACGACTCGCGGCGCGATAGCGG
Chromosome start end strand MM target_seq PAM alignment distance position gene name gene id
GU305914.1 207810 207832 - 0 CCGCTATCGCGCCGCGAGTC TGG ||||||||[||||||||||||]PAM NA Intergenic NA NA

T18820 TGGACGAGCTGACTCCGACGCGG 1000
Oligo fwd CACCTGGACGAGCTGACTCCGACG
Oligo rev AAACCGTCGGAGTCAGCTCGTCCA
Chromosome start end strand MM target_seq PAM alignment distance position gene name gene id
GU305914.1 207790 207812 - 0 TGGACGAGCTGACTCCGACG CGG ||||||||[||||||||||||]PAM NA Intergenic NA NA

T18821 ATCAGCGCACGCTATGTCTTCGG 1000
Oligo fwd CACCATCAGCGCACGCTATGTCTT
Oligo rev AAACAAGACATAGCGTGCGCTGAT
Chromosome start end strand MM target_seq PAM alignment distance position gene name gene id
GU305914.1 207758 207780 - 0 ATCAGCGCACGCTATGTCTT CGG ||||||||[||||||||||||]PAM NA Intergenic NA NA

T18822 TATGTCTTCGGCAGACCGCCTGG 1000
Oligo fwd CACCTATGTCTTCGGCAGACCGCC
Oligo rev AAACGGCGGTCTGCCGAAGACATA
Chromosome start end strand MM target_seq PAM alignment distance position gene name gene id
GU305914.1 207746 207768 - 0 TATGTCTTCGGCAGACCGCC TGG ||||||||[||||||||||||]PAM NA Intergenic NA NA

T18823 ACCGCCTGGCGAGTCGAGCACGG 1000
Oligo fwd CACCACCGCCTGGCGAGTCGAGCA
Oligo rev AAACTGCTCGACTCGCCAGGCGGT
Chromosome start end strand MM target_seq PAM alignment distance position gene name gene id
GU305914.1 207732 207754 - 0 ACCGCCTGGCGAGTCGAGCA CGG ||||||||[||||||||||||]PAM NA Intergenic NA NA

T18824 AGCACGGACGTGTGATCGACCGG 1000
Oligo fwd CACCAGCACGGACGTGTGATCGAC
Oligo rev AAACGTCGATCACACGTCCGTGCT
Chromosome start end strand MM target_seq PAM alignment distance position gene name gene id
GU305914.1 207716 207738 - 0 AGCACGGACGTGTGATCGAC CGG ||||||||[||||||||||||]PAM NA Intergenic NA NA

T18825 TCGACCGGTCGATTGACACGAGG 1000
Oligo fwd CACCTCGACCGGTCGATTGACACG
Oligo rev AAACCGTGTCAATCGACCGGTCGA
Chromosome start end strand MM target_seq PAM alignment distance position gene name gene id
GU305914.1 207701 207723 - 0 TCGACCGGTCGATTGACACG AGG ||||||||[||||||||||||]PAM NA Intergenic NA NA

T18826 TTGACACGAGGAAGCATCACCGG 1000
Oligo fwd CACCTTGACACGAGGAAGCATCAC
Oligo rev AAACGTGATGCTTCCTCGTGTCAA
Chromosome start end strand MM target_seq PAM alignment distance position gene name gene id
GU305914.1 207689 207711 - 0 TTGACACGAGGAAGCATCAC CGG ||||||||[||||||||||||]PAM NA Intergenic NA NA

T18827 CTTAGACTTTGTATATAAAGCGG 1000
Oligo fwd CACCCTTAGACTTTGTATATAAAG
Oligo rev AAACCTTTATATACAAAGTCTAAG
Chromosome start end strand MM target_seq PAM alignment distance position gene name gene id
GU305914.1 207644 207666 - 0 CTTAGACTTTGTATATAAAG CGG ||||||||[||||||||||||]PAM NA Intergenic NA NA

T18828 TTTGTATATAAAGCGGTTTTTGG 1000
Oligo fwd CACCTTTGTATATAAAGCGGTTTT
Oligo rev AAACAAAACCGCTTTATATACAAA
Chromosome start end strand MM target_seq PAM alignment distance position gene name gene id
GU305914.1 207637 207659 - 0 TTTGTATATAAAGCGGTTTT TGG ||||||||[||||||||||||]PAM NA Intergenic NA NA

T18829 TCCACCTCAGACCTTTCAGATGG 1000
Oligo fwd CACCTCCACCTCAGACCTTTCAGA
Oligo rev AAACTCTGAAAGGTCTGAGGTGGA
Chromosome start end strand MM target_seq PAM alignment distance position gene name gene id
GU305914.1 207607 207629 - 0 TCCACCTCAGACCTTTCAGA TGG ||||||||[||||||||||||]PAM NA Intergenic NA NA

T18830 TGGCTCATGAGTGATCACCCCGG 1000
Oligo fwd CACCTGGCTCATGAGTGATCACCC
Oligo rev AAACGGGTGATCACTCATGAGCCA
Chromosome start end strand MM target_seq PAM alignment distance position gene name gene id
GU305914.1 207587 207609 - 0 TGGCTCATGAGTGATCACCC CGG ||||||||[||||||||||||]PAM NA Intergenic NA NA

T18831 CTCATGAGTGATCACCCCGGAGG 1000
Oligo fwd CACCCTCATGAGTGATCACCCCGG
Oligo rev AAACCCGGGGTGATCACTCATGAG
Chromosome start end strand MM target_seq PAM alignment distance position gene name gene id
GU305914.1 207584 207606 - 0 CTCATGAGTGATCACCCCGG AGG ||||||||[||||||||||||]PAM NA Intergenic NA NA

T18832 CGGAGGATGCGTCTTCTTGATGG 1000
Oligo fwd CACCCGGAGGATGCGTCTTCTTGA
Oligo rev AAACTCAAGAAGACGCATCCTCCG
Chromosome start end strand MM target_seq PAM alignment distance position gene name gene id
GU305914.1 207567 207589 - 0 CGGAGGATGCGTCTTCTTGA TGG ||||||||[||||||||||||]PAM NA Intergenic NA NA

T18833 AGGATGCGTCTTCTTGATGGTGG 1000
Oligo fwd CACCAGGATGCGTCTTCTTGATGG
Oligo rev AAACCCATCAAGAAGACGCATCCT
Chromosome start end strand MM target_seq PAM alignment distance position gene name gene id
GU305914.1 207564 207586 - 0 AGGATGCGTCTTCTTGATGG TGG ||||||||[||||||||||||]PAM NA Intergenic NA NA

T18834 GGATGCGTCTTCTTGATGGTGGG 1000
Oligo fwd CACCGGATGCGTCTTCTTGATGGT
Oligo rev AAACACCATCAAGAAGACGCATCC
Chromosome start end strand MM target_seq PAM alignment distance position gene name gene id
GU305914.1 207563 207585 - 0 GGATGCGTCTTCTTGATGGT GGG ||||||||[||||||||||||]PAM NA Intergenic NA NA

T18835 TCTTGATGGTGGGTACCGTGCGG 1000
Oligo fwd CACCTCTTGATGGTGGGTACCGTG
Oligo rev AAACCACGGTACCCACCATCAAGA
Chromosome start end strand MM target_seq PAM alignment distance position gene name gene id
GU305914.1 207553 207575 - 0 TCTTGATGGTGGGTACCGTG CGG ||||||||[||||||||||||]PAM NA Intergenic NA NA

T18836 GTGGGTACCGTGCGGCTCTGTGG 1000
Oligo fwd CACCGTGGGTACCGTGCGGCTCTG
Oligo rev AAACCAGAGCCGCACGGTACCCAC
Chromosome start end strand MM target_seq PAM alignment distance position gene name gene id
GU305914.1 207545 207567 - 0 GTGGGTACCGTGCGGCTCTG TGG ||||||||[||||||||||||]PAM NA Intergenic NA NA

T18837 TGGGTACCGTGCGGCTCTGTGGG 1000
Oligo fwd CACCTGGGTACCGTGCGGCTCTGT
Oligo rev AAACACAGAGCCGCACGGTACCCA
Chromosome start end strand MM target_seq PAM alignment distance position gene name gene id
GU305914.1 207544 207566 - 0 TGGGTACCGTGCGGCTCTGT GGG ||||||||[||||||||||||]PAM NA Intergenic NA NA

T18838 GGGTACCGTGCGGCTCTGTGGGG 1000
Oligo fwd CACCGGGTACCGTGCGGCTCTGTG
Oligo rev AAACCACAGAGCCGCACGGTACCC
Chromosome start end strand MM target_seq PAM alignment distance position gene name gene id
GU305914.1 207543 207565 - 0 GGGTACCGTGCGGCTCTGTG GGG ||||||||[||||||||||||]PAM NA Intergenic NA NA

T18839 GGTACCGTGCGGCTCTGTGGGGG 1000
Oligo fwd CACCGGTACCGTGCGGCTCTGTGG
Oligo rev AAACCCACAGAGCCGCACGGTACC
Chromosome start end strand MM target_seq PAM alignment distance position gene name gene id
GU305914.1 207542 207564 - 0 GGTACCGTGCGGCTCTGTGG GGG ||||||||[||||||||||||]PAM NA Intergenic NA NA

T18840 ACCGTGCGGCTCTGTGGGGGTGG 1000
Oligo fwd CACCACCGTGCGGCTCTGTGGGGG
Oligo rev AAACCCCCCACAGAGCCGCACGGT
Chromosome start end strand MM target_seq PAM alignment distance position gene name gene id
GU305914.1 207539 207561 - 0 ACCGTGCGGCTCTGTGGGGG TGG ||||||||[||||||||||||]PAM NA Intergenic NA NA

T18841 TGCGGCTCTGTGGGGGTGGTTGG 1000
Oligo fwd CACCTGCGGCTCTGTGGGGGTGGT
Oligo rev AAACACCACCCCCACAGAGCCGCA
Chromosome start end strand MM target_seq PAM alignment distance position gene name gene id
GU305914.1 207535 207557 - 0 TGCGGCTCTGTGGGGGTGGT TGG ||||||||[||||||||||||]PAM NA Intergenic NA NA

T18842 CTCGTGTACGTCACCGACGACGG 1000
Oligo fwd CACCCTCGTGTACGTCACCGACGA
Oligo rev AAACTCGTCGGTGACGTACACGAG
Chromosome start end strand MM target_seq PAM alignment distance position gene name gene id
GU305914.1 207503 207525 - 0 CTCGTGTACGTCACCGACGA CGG ||||||||[||||||||||||]PAM NA Intergenic NA NA

T18843 ACGTCACCGACGACGGAGTCTGG 1000
Oligo fwd CACCACGTCACCGACGACGGAGTC
Oligo rev AAACGACTCCGTCGTCGGTGACGT
Chromosome start end strand MM target_seq PAM alignment distance position gene name gene id
GU305914.1 207496 207518 - 0 ACGTCACCGACGACGGAGTC TGG ||||||||[||||||||||||]PAM NA Intergenic NA NA

T18844 CGTCACCGACGACGGAGTCTGGG 1000
Oligo fwd CACCCGTCACCGACGACGGAGTCT
Oligo rev AAACAGACTCCGTCGTCGGTGACG
Chromosome start end strand MM target_seq PAM alignment distance position gene name gene id
GU305914.1 207495 207517 - 0 CGTCACCGACGACGGAGTCT GGG ||||||||[||||||||||||]PAM NA Intergenic NA NA

T18845 ACCGACGACGGAGTCTGGGACGG 1000
Oligo fwd CACCACCGACGACGGAGTCTGGGA
Oligo rev AAACTCCCAGACTCCGTCGTCGGT
Chromosome start end strand MM target_seq PAM alignment distance position gene name gene id
GU305914.1 207491 207513 - 0 ACCGACGACGGAGTCTGGGA CGG ||||||||[||||||||||||]PAM NA Intergenic NA NA

T18846 GGAGTCTGGGACGGTGTCGAAGG 1000
Oligo fwd CACCGGAGTCTGGGACGGTGTCGA
Oligo rev AAACTCGACACCGTCCCAGACTCC
Chromosome start end strand MM target_seq PAM alignment distance position gene name gene id
GU305914.1 207482 207504 - 0 GGAGTCTGGGACGGTGTCGA AGG ||||||||[||||||||||||]PAM NA Intergenic NA NA

T18847 GTCTGGGACGGTGTCGAAGGTGG 1000
Oligo fwd CACCGTCTGGGACGGTGTCGAAGG
Oligo rev AAACCCTTCGACACCGTCCCAGAC
Chromosome start end strand MM target_seq PAM alignment distance position gene name gene id
GU305914.1 207479 207501 - 0 GTCTGGGACGGTGTCGAAGG TGG ||||||||[||||||||||||]PAM NA Intergenic NA NA

T18848 TGGGACGGTGTCGAAGGTGGCGG 1000
Oligo fwd CACCTGGGACGGTGTCGAAGGTGG
Oligo rev AAACCCACCTTCGACACCGTCCCA
Chromosome start end strand MM target_seq PAM alignment distance position gene name gene id

<table>
<tr><td>GU305914.1</td><td>207476</td><td>207498</td><td>-</td><td>0</td><td>TGGGACGGTGTCGAAGGTGG</td><td>CGG</td><td>||||||||[||||||||||||]PAM</td><td>NA</td><td>Intergenic</td><td>NA</td><td>NA</td></tr>
<tr><td></td><td></td><td></td><td></td><td></td><td></td><td></td><td></td><td></td><td></td><td></td><td></td></tr>
<tr><td>T18849</td><td>TAAATTCAGCGAGAAGAATTCGG</td><td>1000</td><td></td><td></td><td></td><td></td><td></td><td></td><td></td><td></td><td></td></tr>
<tr><td>Oligo fwd</td><td>CACCTAAATTCAGCGAGAAGAATT</td><td></td><td></td><td></td><td></td><td></td><td></td><td></td><td></td><td></td><td></td></tr>
<tr><td>Oligo rev</td><td>AAACAATTCTTCTCGCTGAATTTA</td><td></td><td></td><td></td><td></td><td></td><td></td><td></td><td></td><td></td><td></td></tr>
<tr><td>Chromosome</td><td>start</td><td>end</td><td>strand</td><td>MM</td><td>target_seq</td><td>PAM</td><td>alignment</td><td>distance</td><td>position</td><td>gene name</td><td>gene id</td></tr>
<tr><td>GU305914.1</td><td>207453</td><td>207475</td><td>-</td><td>0</td><td>TAAATTCAGCGAGAAGAATT</td><td>CGG</td><td>||||||||[||||||||||||]PAM</td><td>NA</td><td>Intergenic</td><td>NA</td><td>NA</td></tr>
<tr><td></td><td></td><td></td><td></td><td></td><td></td><td></td><td></td><td></td><td></td><td></td><td></td></tr>
<tr><td>T18850</td><td>ATTCAGCGAGAAGAATTCGGAGG</td><td>1000</td><td></td><td></td><td></td><td></td><td></td><td></td><td></td><td></td><td></td></tr>
<tr><td>Oligo fwd</td><td>CACCATTCAGCGAGAAGAATTCGG</td><td></td><td></td><td></td><td></td><td></td><td></td><td></td><td></td><td></td><td></td></tr>
<tr><td>Oligo rev</td><td>AAACCCGAATTCTTCTCGCTGAAT</td><td></td><td></td><td></td><td></td><td></td><td></td><td></td><td></td><td></td><td></td></tr>
<tr><td>Chromosome</td><td>start</td><td>end</td><td>strand</td><td>MM</td><td>target_seq</td><td>PAM</td><td>alignment</td><td>distance</td><td>position</td><td>gene name</td><td>gene id</td></tr>
<tr><td>GU305914.1</td><td>207450</td><td>207472</td><td>-</td><td>0</td><td>ATTCAGCGAGAAGAATTCGG</td><td>AGG</td><td>||||||||[||||||||||||]PAM</td><td>NA</td><td>Intergenic</td><td>NA</td><td>NA</td></tr>
<tr><td></td><td></td><td></td><td></td><td></td><td></td><td></td><td></td><td></td><td></td><td></td><td></td></tr>
<tr><td>T18851</td><td>GAGGTCTCTGAAGAACGAGACGG</td><td>1000</td><td></td><td></td><td></td><td></td><td></td><td></td><td></td><td></td><td></td></tr>
<tr><td>Oligo fwd</td><td>CACCGAGGTCTCTGAAGAACGAGA</td><td></td><td></td><td></td><td></td><td></td><td></td><td></td><td></td><td></td><td></td></tr>
<tr><td>Oligo rev</td><td>AAACTCTCGTTCTTCAGAGACCTC</td><td></td><td></td><td></td><td></td><td></td><td></td><td></td><td></td><td></td><td></td></tr>
<tr><td>Chromosome</td><td>start</td><td>end</td><td>strand</td><td>MM</td><td>target_seq</td><td>PAM</td><td>alignment</td><td>distance</td><td>position</td><td>gene name</td><td>gene id</td></tr>
<tr><td>GU305914.1</td><td>207431</td><td>207453</td><td>-</td><td>0</td><td>GAGGTCTCTGAAGAACGAGA</td><td>CGG</td><td>||||||||[||||||||||||]PAM</td><td>NA</td><td>Intergenic</td><td>NA</td><td>NA</td></tr>
<tr><td></td><td></td><td></td><td></td><td></td><td></td><td></td><td></td><td></td><td></td><td></td><td></td></tr>
<tr><td>T18852</td><td>GAACGAGACGGTCATCCTGCTGG</td><td>1000</td><td></td><td></td><td></td><td></td><td></td><td></td><td></td><td></td><td></td></tr>
<tr><td>Oligo fwd</td><td>CACCGAACGAGACGGTCATCCTGC</td><td></td><td></td><td></td><td></td><td></td><td></td><td></td><td></td><td></td><td></td></tr>
<tr><td>Oligo rev</td><td>AAACGCAGGATGACCGTCTCGTTC</td><td></td><td></td><td></td><td></td><td></td><td></td><td></td><td></td><td></td><td></td></tr>
<tr><td>Chromosome</td><td>start</td><td>end</td><td>strand</td><td>MM</td><td>target_seq</td><td>PAM</td><td>alignment</td><td>distance</td><td>position</td><td>gene name</td><td>gene id</td></tr>
<tr><td>GU305914.1</td><td>207419</td><td>207441</td><td>-</td><td>0</td><td>GAACGAGACGGTCATCCTGC</td><td>TGG</td><td>||||||||[||||||||||||]PAM</td><td>NA</td><td>Intergenic</td><td>NA</td><td>NA</td></tr>
<tr><td></td><td></td><td></td><td></td><td></td><td></td><td></td><td></td><td></td><td></td><td></td><td></td></tr>
<tr><td>T18853</td><td>AACGAGACGGTCATCCTGCTGGG</td><td>1000</td><td></td><td></td><td></td><td></td><td></td><td></td><td></td><td></td><td></td></tr>
<tr><td>Oligo fwd</td><td>CACCAACGAGACGGTCATCCTGCT</td><td></td><td></td><td></td><td></td><td></td><td></td><td></td><td></td><td></td><td></td></tr>
<tr><td>Oligo rev</td><td>AAACAGCAGGATGACCGTCTCGTT</td><td></td><td></td><td></td><td></td><td></td><td></td><td></td><td></td><td></td><td></td></tr>
<tr><td>Chromosome</td><td>start</td><td>end</td><td>strand</td><td>MM</td><td>target_seq</td><td>PAM</td><td>alignment</td><td>distance</td><td>position</td><td>gene name</td><td>gene id</td></tr>
<tr><td>GU305914.1</td><td>207418</td><td>207440</td><td>-</td><td>0</td><td>AACGAGACGGTCATCCTGCT</td><td>GGG</td><td>||||||||[||||||||||||]PAM</td><td>NA</td><td>Intergenic</td><td>NA</td><td>NA</td></tr>
<tr><td></td><td></td><td></td><td></td><td></td><td></td><td></td><td></td><td></td><td></td><td></td><td></td></tr>
<tr><td>T18854</td><td>TTCACATACAGACTGATTTGTGG</td><td>1000</td><td></td><td></td><td></td><td></td><td></td><td></td><td></td><td></td><td></td></tr>
<tr><td>Oligo fwd</td><td>CACCTTCACATACAGACTGATTTG</td><td></td><td></td><td></td><td></td><td></td><td></td><td></td><td></td><td></td><td></td></tr>
<tr><td>Oligo rev</td><td>AAACCAAATCAGTCTGTATGTGAA</td><td></td><td></td><td></td><td></td><td></td><td></td><td></td><td></td><td></td><td></td></tr>
<tr><td>Chromosome</td><td>start</td><td>end</td><td>strand</td><td>MM</td><td>target_seq</td><td>PAM</td><td>alignment</td><td>distance</td><td>position</td><td>gene name</td><td>gene id</td></tr>
<tr><td>GU305914.1</td><td>207353</td><td>207375</td><td>-</td><td>0</td><td>TTCACATACAGACTGATTTG</td><td>TGG</td><td>||||||||[||||||||||||]PAM</td><td>NA</td><td>Intergenic</td><td>NA</td><td>NA</td></tr>
<tr><td></td><td></td><td></td><td></td><td></td><td></td><td></td><td></td><td></td><td></td><td></td><td></td></tr>
<tr><td>T18855</td><td>GAATAAGATGCAACGAAGACCGG</td><td>1000</td><td></td><td></td><td></td><td></td><td></td><td></td><td></td><td></td><td></td></tr>
<tr><td>Oligo fwd</td><td>CACCGAATAAGATGCAACGAAGAC</td><td></td><td></td><td></td><td></td><td></td><td></td><td></td><td></td><td></td><td></td></tr>
<tr><td>Oligo rev</td><td>AAACGTCTTCGTTGCATCTTATTC</td><td></td><td></td><td></td><td></td><td></td><td></td><td></td><td></td><td></td><td></td></tr>
<tr><td>Chromosome</td><td>start</td><td>end</td><td>strand</td><td>MM</td><td>target_seq</td><td>PAM</td><td>alignment</td><td>distance</td><td>position</td><td>gene name</td><td>gene id</td></tr>
<tr><td>GU305914.1</td><td>207331</td><td>207353</td><td>-</td><td>0</td><td>GAATAAGATGCAACGAAGAC</td><td>CGG</td><td>||||||||[||||||||||||]PAM</td><td>NA</td><td>Intergenic</td><td>NA</td><td>NA</td></tr>
<tr><td></td><td></td><td></td><td></td><td></td><td></td><td></td><td></td><td></td><td></td><td></td><td></td></tr>
<tr><td>T18856</td><td>AATAAGATGCAACGAAGACCGGG</td><td>1000</td><td></td><td></td><td></td><td></td><td></td><td></td><td></td><td></td><td></td></tr>
<tr><td>Oligo fwd</td><td>CACCAATAAGATGCAACGAAGACC</td><td></td><td></td><td></td><td></td><td></td><td></td><td></td><td></td><td></td><td></td></tr>
<tr><td>Oligo rev</td><td>AAACGGTCTTCGTTGCATCTTATT</td><td></td><td></td><td></td><td></td><td></td><td></td><td></td><td></td><td></td><td></td></tr>
<tr><td>Chromosome</td><td>start</td><td>end</td><td>strand</td><td>MM</td><td>target_seq</td><td>PAM</td><td>alignment</td><td>distance</td><td>position</td><td>gene name</td><td>gene id</td></tr>
<tr><td>GU305914.1</td><td>207330</td><td>207352</td><td>-</td><td>0</td><td>AATAAGATGCAACGAAGACC</td><td>GGG</td><td>||||||||[||||||||||||]PAM</td><td>NA</td><td>Intergenic</td><td>NA</td><td>NA</td></tr>
<tr><td></td><td></td><td></td><td></td><td></td><td></td><td></td><td></td><td></td><td></td><td></td><td></td></tr>
<tr><td>T18857</td><td>GATGCAACGAAGACCGGGATCGG</td><td>1000</td><td></td><td></td><td></td><td></td><td></td><td></td><td></td><td></td><td></td></tr>
<tr><td>Oligo fwd</td><td>CACCGATGCAACGAAGACCGGGAT</td><td></td><td></td><td></td><td></td><td></td><td></td><td></td><td></td><td></td><td></td></tr>
<tr><td>Oligo rev</td><td>AAACATCCCGGTCTTCGTTGCATC</td><td></td><td></td><td></td><td></td><td></td><td></td><td></td><td></td><td></td><td></td></tr>
<tr><td>Chromosome</td><td>start</td><td>end</td><td>strand</td><td>MM</td><td>target_seq</td><td>PAM</td><td>alignment</td><td>distance</td><td>position</td><td>gene name</td><td>gene id</td></tr>
<tr><td>GU305914.1</td><td>207325</td><td>207347</td><td>-</td><td>0</td><td>GATGCAACGAAGACCGGGAT</td><td>CGG</td><td>||||||||[||||||||||||]PAM</td><td>NA</td><td>Intergenic</td><td>NA</td><td>NA</td></tr>
<tr><td></td><td></td><td></td><td></td><td></td><td></td><td></td><td></td><td></td><td></td><td></td><td></td></tr>
<tr><td>T18858</td><td>GGAAACAAAAAGTTATCTGAAGG</td><td>1000</td><td></td><td></td><td></td><td></td><td></td><td></td><td></td><td></td><td></td></tr>
<tr><td>Oligo fwd</td><td>CACCGGAAACAAAAAGTTATCTGA</td><td></td><td></td><td></td><td></td><td></td><td></td><td></td><td></td><td></td><td></td></tr>
<tr><td>Oligo rev</td><td>AAACTCAGATAACTTTTTGTTTCC</td><td></td><td></td><td></td><td></td><td></td><td></td><td></td><td></td><td></td><td></td></tr>
<tr><td>Chromosome</td><td>start</td><td>end</td><td>strand</td><td>MM</td><td>target_seq</td><td>PAM</td><td>alignment</td><td>distance</td><td>position</td><td>gene name</td><td>gene id</td></tr>
<tr><td>GU305914.1</td><td>207304</td><td>207326</td><td>-</td><td>0</td><td>GGAAACAAAAAGTTATCTGA</td><td>AGG</td><td>||||||||[||||||||||||]PAM</td><td>NA</td><td>Intergenic</td><td>NA</td><td>NA</td></tr>
<tr><td></td><td></td><td></td><td></td><td></td><td></td><td></td><td></td><td></td><td></td><td></td><td></td></tr>
<tr><td>T18859</td><td>GAAACAAAAAGTTATCTGAAGGG</td><td>1000</td><td></td><td></td><td></td><td></td><td></td><td></td><td></td><td></td><td></td></tr>
<tr><td>Oligo fwd</td><td>CACCGAAACAAAAAGTTATCTGAA</td><td></td><td></td><td></td><td></td><td></td><td></td><td></td><td></td><td></td><td></td></tr>
<tr><td>Oligo rev</td><td>AAACTTCAGATAACTTTTTGTTTC</td><td></td><td></td><td></td><td></td><td></td><td></td><td></td><td></td><td></td><td></td></tr>
<tr><td>Chromosome</td><td>start</td><td>end</td><td>strand</td><td>MM</td><td>target_seq</td><td>PAM</td><td>alignment</td><td>distance</td><td>position</td><td>gene name</td><td>gene id</td></tr>
<tr><td>GU305914.1</td><td>207303</td><td>207325</td><td>-</td><td>0</td><td>GAAACAAAAAGTTATCTGAA</td><td>GGG</td><td>||||||||[||||||||||||]PAM</td><td>NA</td><td>Intergenic</td><td>NA</td><td>NA</td></tr>
<tr><td></td><td></td><td></td><td></td><td></td><td></td><td></td><td></td><td></td><td></td><td></td><td></td></tr>
<tr><td>T18860</td><td>CTACCTGACGTTCAGAGTGAAGG</td><td>1000</td><td></td><td></td><td></td><td></td><td></td><td></td><td></td><td></td><td></td></tr>
<tr><td>Oligo fwd</td><td>CACCCTACCTGACGTTCAGAGTGA</td><td></td><td></td><td></td><td></td><td></td><td></td><td></td><td></td><td></td><td></td></tr>
<tr><td>Oligo rev</td><td>AAACTCACTCTGAACGTCAGGTAG</td><td></td><td></td><td></td><td></td><td></td><td></td><td></td><td></td><td></td><td></td></tr>
<tr><td>Chromosome</td><td>start</td><td>end</td><td>strand</td><td>MM</td><td>target_seq</td><td>PAM</td><td>alignment</td><td>distance</td><td>position</td><td>gene name</td><td>gene id</td></tr>
<tr><td>GU305914.1</td><td>207277</td><td>207299</td><td>-</td><td>0</td><td>CTACCTGACGTTCAGAGTGA</td><td>AGG</td><td>||||||||[||||||||||||]PAM</td><td>NA</td><td>Intergenic</td><td>NA</td><td>NA</td></tr>
<tr><td></td><td></td><td></td><td></td><td></td><td></td><td></td><td></td><td></td><td></td><td></td><td></td></tr>
<tr><td>T18861</td><td>TGACGTTCAGAGTGAAGGATCGG</td><td>1000</td><td></td><td></td><td></td><td></td><td></td><td></td><td></td><td></td><td></td></tr>
<tr><td>Oligo fwd</td><td>CACCTGACGTTCAGAGTGAAGGAT</td><td></td><td></td><td></td><td></td><td></td><td></td><td></td><td></td><td></td><td></td></tr>
<tr><td>Oligo rev</td><td>AAACATCCTTCACTCTGAACGTCA</td><td></td><td></td><td></td><td></td><td></td><td></td><td></td><td></td><td></td><td></td></tr>
<tr><td>Chromosome</td><td>start</td><td>end</td><td>strand</td><td>MM</td><td>target_seq</td><td>PAM</td><td>alignment</td><td>distance</td><td>position</td><td>gene name</td><td>gene id</td></tr>
<tr><td>GU305914.1</td><td>207272</td><td>207294</td><td>-</td><td>0</td><td>TGACGTTCAGAGTGAAGGAT</td><td>CGG</td><td>||||||||[||||||||||||]PAM</td><td>NA</td><td>Intergenic</td><td>NA</td><td>NA</td></tr>
<tr><td></td><td></td><td></td><td></td><td></td><td></td><td></td><td></td><td></td><td></td><td></td><td></td></tr>
<tr><td>T18862</td><td>AGTGAAGGATCGGAGAGACGCGG</td><td>1000</td><td></td><td></td><td></td><td></td><td></td><td></td><td></td><td></td><td></td></tr>
<tr><td>Oligo fwd</td><td>CACCAGTGAAGGATCGGAGAGACG</td><td></td><td></td><td></td><td></td><td></td><td></td><td></td><td></td><td></td><td></td></tr>
<tr><td>Oligo rev</td><td>AAACCGTCTCTCCGATCCTTCACT</td><td></td><td></td><td></td><td></td><td></td><td></td><td></td><td></td><td></td><td></td></tr>
<tr><td>Chromosome</td><td>start</td><td>end</td><td>strand</td><td>MM</td><td>target_seq</td><td>PAM</td><td>alignment</td><td>distance</td><td>position</td><td>gene name</td><td>gene id</td></tr>
<tr><td>GU305914.1</td><td>207262</td><td>207284</td><td>-</td><td>0</td><td>AGTGAAGGATCGGAGAGACG</td><td>CGG</td><td>||||||||[||||||||||||]PAM</td><td>NA</td><td>Intergenic</td><td>NA</td><td>NA</td></tr>
<tr><td></td><td></td><td></td><td></td><td></td><td></td><td></td><td></td><td></td><td></td><td></td><td></td></tr>
<tr><td>T18863</td><td>TCGGAGAGACGCGGACCGAGAGG</td><td>1000</td><td></td><td></td><td></td><td></td><td></td><td></td><td></td><td></td><td></td></tr>
<tr><td>Oligo fwd</td><td>CACCTCGGAGAGACGCGGACCGAG</td><td></td><td></td><td></td><td></td><td></td><td></td><td></td><td></td><td></td><td></td></tr>
<tr><td>Oligo rev</td><td>AAACCTCGGTCCGCGTCTCTCCGA</td><td></td><td></td><td></td><td></td><td></td><td></td><td></td><td></td><td></td><td></td></tr>
</table>

Chromosome start end strand MM target_seq PAM alignment distance position gene name gene id
GU305914.1 207253 207275 - 0 TCGGAGAGACGCGGACCGAG AGG ||||||||[||||||||||||]PAM NA Intergenic NA NA

T18864 GAGAGACGCGGACCGAGAGGAGG 1000
Oligo fwd CACCGAGAGACGCGGACCGAGAGG
Oligo rev AAACCCTCTCGGTCCGCGTCTCTC
Chromosome start end strand MM target_seq PAM alignment distance position gene name gene id
GU305914.1 207250 207272 - 0 GAGAGACGCGGACCGAGAGG AGG ||||||||[||||||||||||]PAM NA Intergenic NA NA

T18865 CGCGGACCGAGAGGAGGACGAGG 1000
Oligo fwd CACCCGCGGACCGAGAGGAGGACG
Oligo rev AAACCGTCCTCCTCTCGGTCCGCG
Chromosome start end strand MM target_seq PAM alignment distance position gene name gene id
GU305914.1 207244 207266 - 0 CGCGGACCGAGAGGAGGACG AGG ||||||||[||||||||||||]PAM NA Intergenic NA NA

T18866 AGGAGGACGAGGTACCGAAAAGG 1000
Oligo fwd CACCAGGAGGACGAGGTACCGAAA
Oligo rev AAACTTTCGGTACCTCGTCCTCCT
Chromosome start end strand MM target_seq PAM alignment distance position gene name gene id
GU305914.1 207233 207255 - 0 AGGAGGACGAGGTACCGAAA AGG ||||||||[||||||||||||]PAM NA Intergenic NA NA

T18867 GGACGAGGTACCGAAAAGGCCGG 1000
Oligo fwd CACCGGACGAGGTACCGAAAAGGC
Oligo rev AAACGCCTTTTCGGTACCTCGTCC
Chromosome start end strand MM target_seq PAM alignment distance position gene name gene id
GU305914.1 207229 207251 - 0 GGACGAGGTACCGAAAAGGC CGG ||||||||[||||||||||||]PAM NA Intergenic NA NA

T18868 AGGTACCGAAAAGGCCGGACTGG 1000
Oligo fwd CACCAGGTACCGAAAAGGCCGGAC
Oligo rev AAACGTCCGGCCTTTTCGGTACCT
Chromosome start end strand MM target_seq PAM alignment distance position gene name gene id
GU305914.1 207224 207246 - 0 AGGTACCGAAAAGGCCGGAC TGG ||||||||[||||||||||||]PAM NA Intergenic NA NA

T18869 CCACCGCGATAGTCTTGAAATGG 1000
Oligo fwd CACCCCACCGCGATAGTCTTGAAA
Oligo rev AAACTTTCAAGACTATCGCGGTGG
Chromosome start end strand MM target_seq PAM alignment distance position gene name gene id
GU305914.1 207200 207222 - 0 CCACCGCGATAGTCTTGAAA TGG ||||||||[||||||||||||]PAM NA Intergenic NA NA

T18870 TCTACTTTGTGATCGCCTTTCGG 1000
Oligo fwd CACCTCTACTTTGTGATCGCCTTT
Oligo rev AAACAAAGGCGATCACAAAGTAGA
Chromosome start end strand MM target_seq PAM alignment distance position gene name gene id
GU305914.1 207155 207177 - 0 TCTACTTTGTGATCGCCTTT CGG ||||||||[||||||||||||]PAM NA Intergenic NA NA

T18871 CTTTGTGATCGCCTTTCGGCTGG 1000
Oligo fwd CACCCTTTGTGATCGCCTTTCGGC
Oligo rev AAACGCCGAAAGGCGATCACAAAG
Chromosome start end strand MM target_seq PAM alignment distance position gene name gene id
GU305914.1 207151 207173 - 0 CTTTGTGATCGCCTTTCGGC TGG ||||||||[||||||||||||]PAM NA Intergenic NA NA

T18872 GCTGGTGTATCTGACCGTGCAGG 1000
Oligo fwd CACCGCTGGTGTATCTGACCGTGC
Oligo rev AAACGCACGGTCAGATACACCAGC
Chromosome start end strand MM target_seq PAM alignment distance position gene name gene id
GU305914.1 207133 207155 - 0 GCTGGTGTATCTGACCGTGC AGG ||||||||[||||||||||||]PAM NA Intergenic NA NA

T18873 TATCTGACCGTGCAGGCGCACGG 1000
Oligo fwd CACCTATCTGACCGTGCAGGCGCA
Oligo rev AAACTGCGCCTGCACGGTCAGATA
Chromosome start end strand MM target_seq PAM alignment distance position gene name gene id
GU305914.1 207126 207148 - 0 TATCTGACCGTGCAGGCGCA CGG ||||||||[||||||||||||]PAM NA Intergenic NA NA

T18874 CCGTGCAGGCGCACGGACTCTGG 1000
Oligo fwd CACCCCGTGCAGGCGCACGGACTC
Oligo rev AAACGAGTCCGTGCGCCTGCACGG
Chromosome start end strand MM target_seq PAM alignment distance position gene name gene id
GU305914.1 207119 207141 - 0 CCGTGCAGGCGCACGGACTC TGG ||||||||[||||||||||||]PAM NA Intergenic NA NA

T18875 GACTCTGGACGCGACACGAGCGG 1000
Oligo fwd CACCGACTCTGGACGCGACACGAG
Oligo rev AAACCTCGTGTCGCGTCCAGAGTC
Chromosome start end strand MM target_seq PAM alignment distance position gene name gene id
GU305914.1 207104 207126 - 0 GACTCTGGACGCGACACGAG CGG ||||||||[||||||||||||]PAM NA Intergenic NA NA

T18876 CGCGACACGAGCGGCATCAGAGG 1000
Oligo fwd CACCCGCGACACGAGCGGCATCAG
Oligo rev AAACCTGATGCCGCTCGTGTCGCG
Chromosome start end strand MM target_seq PAM alignment distance position gene name gene id
GU305914.1 207095 207117 - 0 CGCGACACGAGCGGCATCAG AGG ||||||||[||||||||||||]PAM NA Intergenic NA NA

T18877 GCGACACGAGCGGCATCAGAGGG 1000
Oligo fwd CACCGCGACACGAGCGGCATCAGA
Oligo rev AAACTCTGATGCCGCTCGTGTCGC
Chromosome start end strand MM target_seq PAM alignment distance position gene name gene id
GU305914.1 207094 207116 - 0 GCGACACGAGCGGCATCAGA GGG ||||||||[||||||||||||]PAM NA Intergenic NA NA

T18878 CGACACGAGCGGCATCAGAGGGG 1000
Oligo fwd CACCCGACACGAGCGGCATCAGAG

Oligo rev AAACCTCTGATGCCGCTCGTGTCG
Chromosome start end strand MM target_seq PAM alignment distance position gene name gene id
GU305914.1 207093 207115 - 0 CGACACGAGCGGCATCAGAG GGG ||||||||[||||||||||||]PAM NA Intergenic NA NA

T18879 GACACGAGCGGCATCAGAGGGGG 1000
Oligo fwd CACCGACACGAGCGGCATCAGAGG
Oligo rev AAACCCTCTGATGCCGCTCGTGTC
Chromosome start end strand MM target_seq PAM alignment distance position gene name gene id
GU305914.1 207092 207114 - 0 GACACGAGCGGCATCAGAGG GGG ||||||||[||||||||||||]PAM NA Intergenic NA NA

T18880 GCATCAGAGGGGGACCGACGCGG 1000
Oligo fwd CACCGCATCAGAGGGGGACCGACG
Oligo rev AAACCGTCGGTCCCCCTCTGATGC
Chromosome start end strand MM target_seq PAM alignment distance position gene name gene id
GU305914.1 207082 207104 - 0 GCATCAGAGGGGGACCGACG CGG ||||||||[||||||||||||]PAM NA Intergenic NA NA

T18881 GGGGACCGACGCGGTACGACTGG 1000
Oligo fwd CACCGGGGACCGACGCGGTACGAC
Oligo rev AAACGTCGTACCGCGTCGGTCCCC
Chromosome start end strand MM target_seq PAM alignment distance position gene name gene id
GU305914.1 207073 207095 - 0 GGGGACCGACGCGGTACGAC TGG ||||||||[||||||||||||]PAM NA Intergenic NA NA

T18882 ACGACTGGCGCCCGTAGCCGCGG 1000
Oligo fwd CACCACGACTGGCGCCCGTAGCCG
Oligo rev AAACCGGCTACGGGCGCCAGTCGT
Chromosome start end strand MM target_seq PAM alignment distance position gene name gene id
GU305914.1 207058 207080 - 0 ACGACTGGCGCCCGTAGCCG CGG ||||||||[||||||||||||]PAM NA Intergenic NA NA

T18883 GTAGCCGCGGCCTTCGTTATCGG 1000
Oligo fwd CACCGTAGCCGCGGCCTTCGTTAT
Oligo rev AAACATAACGAAGGCCGCGGCTAC
Chromosome start end strand MM target_seq PAM alignment distance position gene name gene id
GU305914.1 207045 207067 - 0 GTAGCCGCGGCCTTCGTTAT CGG ||||||||[||||||||||||]PAM NA Intergenic NA NA

T18884 GTGCATGCATATTCCCTCCGTGG 1000
Oligo fwd CACCGTGCATGCATATTCCCTCCG
Oligo rev AAACCGGAGGGAATATGCATGCAC
Chromosome start end strand MM target_seq PAM alignment distance position gene name gene id
GU305914.1 207012 207034 - 0 GTGCATGCATATTCCCTCCG TGG ||||||||[||||||||||||]PAM NA Intergenic NA NA

T18885 TGCATGCATATTCCCTCCGTGGG 1000
Oligo fwd CACCTGCATGCATATTCCCTCCGT
Oligo rev AAACACGGAGGGAATATGCATGCA
Chromosome start end strand MM target_seq PAM alignment distance position gene name gene id
GU305914.1 207011 207033 - 0 TGCATGCATATTCCCTCCGT GGG ||||||||[||||||||||||]PAM NA Intergenic NA NA

T18886 GATAGCTAACAGATATGTAGCGG 1000
Oligo fwd CACCGATAGCTAACAGATATGTAG
Oligo rev AAACCTACATATCTGTTAGCTATC
Chromosome start end strand MM target_seq PAM alignment distance position gene name gene id
GU305914.1 206989 207011 - 0 GATAGCTAACAGATATGTAG CGG ||||||||[||||||||||||]PAM NA Intergenic NA NA

T18887 AGATATGTAGCGGAACCCTGCGG 1000
Oligo fwd CACCAGATATGTAGCGGAACCCTG
Oligo rev AAACCAGGGTTCCGCTACATATCT
Chromosome start end strand MM target_seq PAM alignment distance position gene name gene id
GU305914.1 206979 207001 - 0 AGATATGTAGCGGAACCCTG CGG ||||||||[||||||||||||]PAM NA Intergenic NA NA

T18888 ATAAAAAAATTCAACCTTTCCGG 1000
Oligo fwd CACCATAAAAAAATTCAACCTTTC
Oligo rev AAACGAAAGGTTGAATTTTTTTAT
Chromosome start end strand MM target_seq PAM alignment distance position gene name gene id
GU305914.1 206944 206966 - 0 ATAAAAAAATTCAACCTTTC CGG ||||||||[||||||||||||]PAM NA Intergenic NA NA

T18889 GTCTCTATCCGTATCGCGCGCGG 1000
Oligo fwd CACCGTCTCTATCCGTATCGCGCG
Oligo rev AAACCGCGCGATACGGATAGAGAC
Chromosome start end strand MM target_seq PAM alignment distance position gene name gene id
GU305914.1 206886 206908 - 0 GTCTCTATCCGTATCGCGCG CGG ||||||||[||||||||||||]PAM NA Intergenic NA NA

T18890 GCGTCCGCGCTATCGCCGTCTGG 1000
Oligo fwd CACCGCGTCCGCGCTATCGCCGTC
Oligo rev AAACGACGGCGATAGCGCGGACGC
Chromosome start end strand MM target_seq PAM alignment distance position gene name gene id
GU305914.1 206832 206854 - 0 GCGTCCGCGCTATCGCCGTC TGG ||||||||[||||||||||||]PAM NA Intergenic NA NA

T18891 CGCTATCGCCGTCTGGTGCGTGG 1000
Oligo fwd CACCCGCTATCGCCGTCTGGTGCG
Oligo rev AAACCGCACCAGACGGCGATAGCG
Chromosome start end strand MM target_seq PAM alignment distance position gene name gene id
GU305914.1 206825 206847 - 0 CGCTATCGCCGTCTGGTGCG TGG ||||||||[||||||||||||]PAM NA Intergenic NA NA
GU305914.1 61386 61408 - 4 GGCTCTCGCCGTCGAGTGCG AGG -|||-|||[|||||--|||||]PAM NA Intergenic NA NA

T18892 TCTGGTGCGTGGTCTCGCTTTGG 1000
Oligo fwd CACCTCTGGTGCGTGGTCTCGCTT
Oligo rev AAACAAGCGAGACCACGCACCAGA
Chromosome start end strand MM target_seq PAM alignment distance position gene name gene id
GU305914.1 206814 206836 - 0 TCTGGTGCGTGGTCTCGCTT TGG ||||||||[||||||||||||]PAM NA Intergenic NA NA

| | | | | | | | | | | | |
|---|---|---|---|---|---|---|---|---|---|---|---|
| T18893 | CTGGTGCGTGGTCTCGCTTTGGG | 1000 | | | | | | | | | |
| Oligo fwd | CACCCTGGTGCGTGGTCTCGCTTT | | | | | | | | | | |
| Oligo rev | AAACAAAGCGAGACCACGCACCAG | | | | | | | | | | |
| Chromosome | start | end | strand | MM | target_seq | PAM | alignment | distance | position | gene name | gene id |
| GU305914.1 | 206813 | 206835 | - | 0 | CTGGTGCGTGGTCTCGCTTT | GGG | \|\|\|\|\|\|\|\|[\|\|\|\|\|\|\|\|\|\|\|\|]PAM | NA | Intergenic | NA | NA |
| T18894 | GTGCGTGGTCTCGCTTTGGGTGG | 1000 | | | | | | | | | |
| Oligo fwd | CACCGTGCGTGGTCTCGCTTTGGG | | | | | | | | | | |
| Oligo rev | AAACCCCAAAGCGAGACCACGCAC | | | | | | | | | | |
| Chromosome | start | end | strand | MM | target_seq | PAM | alignment | distance | position | gene name | gene id |
| GU305914.1 | 206810 | 206832 | - | 0 | GTGCGTGGTCTCGCTTTGGG | TGG | \|\|\|\|\|\|\|\|[\|\|\|\|\|\|\|\|\|\|\|\|]PAM | NA | Intergenic | NA | NA |
| T18895 | TGCGTGGTCTCGCTTTGGGTGGG | 1000 | | | | | | | | | |
| Oligo fwd | CACCTGCGTGGTCTCGCTTTGGGT | | | | | | | | | | |
| Oligo rev | AAACACCCAAAGCGAGACCACGCA | | | | | | | | | | |
| Chromosome | start | end | strand | MM | target_seq | PAM | alignment | distance | position | gene name | gene id |
| GU305914.1 | 206809 | 206831 | - | 0 | TGCGTGGTCTCGCTTTGGGT | GGG | \|\|\|\|\|\|\|\|[\|\|\|\|\|\|\|\|\|\|\|\|]PAM | NA | Intergenic | NA | NA |
| T18896 | GGGCTACGTGACGAACGCCCAGG | 1000 | | | | | | | | | |
| Oligo fwd | CACCGGGCTACGTGACGAACGCCC | | | | | | | | | | |
| Oligo rev | AAACGGGCGTTCGTCACGTAGCCC | | | | | | | | | | |
| Chromosome | start | end | strand | MM | target_seq | PAM | alignment | distance | position | gene name | gene id |
| GU305914.1 | 206789 | 206811 | - | 0 | GGGCTACGTGACGAACGCCC | AGG | \|\|\|\|\|\|\|\|[\|\|\|\|\|\|\|\|\|\|\|\|]PAM | NA | Intergenic | NA | NA |
| T18897 | TCTCTTTCGTCGTACGCAGCAGG | 1000 | | | | | | | | | |
| Oligo fwd | CACCTCTCTTTCGTCGTACGCAGC | | | | | | | | | | |
| Oligo rev | AAACGCTGCGTACGACGAAAGAGA | | | | | | | | | | |
| Chromosome | start | end | strand | MM | target_seq | PAM | alignment | distance | position | gene name | gene id |
| GU305914.1 | 206763 | 206785 | - | 0 | TCTCTTTCGTCGTACGCAGC | AGG | \|\|\|\|\|\|\|\|[\|\|\|\|\|\|\|\|\|\|\|\|]PAM | NA | Intergenic | NA | NA |
| T18898 | CTATAGCGTCTCTGAACTCGCGG | 1000 | | | | | | | | | |
| Oligo fwd | CACCCTATAGCGTCTCTGAACTCG | | | | | | | | | | |
| Oligo rev | AAACCGAGTTCAGAGACGCTATAG | | | | | | | | | | |
| Chromosome | start | end | strand | MM | target_seq | PAM | alignment | distance | position | gene name | gene id |
| GU305914.1 | 206685 | 206707 | - | 0 | CTATAGCGTCTCTGAACTCG | CGG | \|\|\|\|\|\|\|\|[\|\|\|\|\|\|\|\|\|\|\|\|]PAM | NA | Intergenic | NA | NA |
| T18899 | GCGTCTCTGAACTCGCGGTCAGG | 1000 | | | | | | | | | |
| Oligo fwd | CACCGCGTCTCTGAACTCGCGGTC | | | | | | | | | | |
| Oligo rev | AAACGACCGCGAGTTCAGAGACGC | | | | | | | | | | |
| Chromosome | start | end | strand | MM | target_seq | PAM | alignment | distance | position | gene name | gene id |
| GU305914.1 | 206680 | 206702 | - | 0 | GCGTCTCTGAACTCGCGGTC | AGG | \|\|\|\|\|\|\|\|[\|\|\|\|\|\|\|\|\|\|\|\|]PAM | NA | Intergenic | NA | NA |
| T18900 | TCTGAACTCGCGGTCAGGAAAGG | 1000 | | | | | | | | | |
| Oligo fwd | CACCTCTGAACTCGCGGTCAGGAA | | | | | | | | | | |
| Oligo rev | AAACTTCCTGACCGCGAGTTCAGA | | | | | | | | | | |
| Chromosome | start | end | strand | MM | target_seq | PAM | alignment | distance | position | gene name | gene id |
| GU305914.1 | 206675 | 206697 | - | 0 | TCTGAACTCGCGGTCAGGAA | AGG | \|\|\|\|\|\|\|\|[\|\|\|\|\|\|\|\|\|\|\|\|]PAM | NA | Intergenic | NA | NA |
| T18901 | GCGGTCAGGAAAGGATACGACGG | 1000 | | | | | | | | | |
| Oligo fwd | CACCGCGGTCAGGAAAGGATACGA | | | | | | | | | | |
| Oligo rev | AAACTCGTATCCTTTCCTGACCGC | | | | | | | | | | |
| Chromosome | start | end | strand | MM | target_seq | PAM | alignment | distance | position | gene name | gene id |
| GU305914.1 | 206666 | 206688 | - | 0 | GCGGTCAGGAAAGGATACGA | CGG | \|\|\|\|\|\|\|\|[\|\|\|\|\|\|\|\|\|\|\|\|]PAM | NA | Intergenic | NA | NA |
| T18902 | TGATACGAATGCGTTTGATCCGG | 1000 | | | | | | | | | |
| Oligo fwd | CACCTGATACGAATGCGTTTGATC | | | | | | | | | | |
| Oligo rev | AAACGATCAAACGCATTCGTATCA | | | | | | | | | | |
| Chromosome | start | end | strand | MM | target_seq | PAM | alignment | distance | position | gene name | gene id |
| GU305914.1 | 206624 | 206646 | - | 0 | TGATACGAATGCGTTTGATC | CGG | \|\|\|\|\|\|\|\|[\|\|\|\|\|\|\|\|\|\|\|\|]PAM | NA | Intergenic | NA | NA |
| T18903 | ATCCGGTTCTCCGTTTACATAGG | 1000 | | | | | | | | | |
| Oligo fwd | CACCATCCGGTTCTCCGTTTACAT | | | | | | | | | | |
| Oligo rev | AAACATGTAAACGGAGAACCGGAT | | | | | | | | | | |
| Chromosome | start | end | strand | MM | target_seq | PAM | alignment | distance | position | gene name | gene id |
| GU305914.1 | 206607 | 206629 | - | 0 | ATCCGGTTCTCCGTTTACAT | AGG | \|\|\|\|\|\|\|\|[\|\|\|\|\|\|\|\|\|\|\|\|]PAM | NA | Intergenic | NA | NA |
| T18904 | TCCGGTTCTCCGTTTACATAGGG | 1000 | | | | | | | | | |
| Oligo fwd | CACCTCCGGTTCTCCGTTTACATA | | | | | | | | | | |
| Oligo rev | AAACTATGTAAACGGAGAACCGGA | | | | | | | | | | |
| Chromosome | start | end | strand | MM | target_seq | PAM | alignment | distance | position | gene name | gene id |
| GU305914.1 | 206606 | 206628 | - | 0 | TCCGGTTCTCCGTTTACATA | GGG | \|\|\|\|\|\|\|\|[\|\|\|\|\|\|\|\|\|\|\|\|]PAM | NA | Intergenic | NA | NA |
| T18905 | AAGACTCTACACACAAGCTTCGG | 1000 | | | | | | | | | |
| Oligo fwd | CACCAAGACTCTACACACAAGCTT | | | | | | | | | | |
| Oligo rev | AAACAAGCTTGTGTGTAGAGTCTT | | | | | | | | | | |
| Chromosome | start | end | strand | MM | target_seq | PAM | alignment | distance | position | gene name | gene id |
| GU305914.1 | 206561 | 206583 | - | 0 | AAGACTCTACACACAAGCTT | CGG | \|\|\|\|\|\|\|\|[\|\|\|\|\|\|\|\|\|\|\|\|]PAM | NA | Intergenic | NA | NA |
| T18906 | TCAGCACAGCACGAATGTATCGG | 1000 | | | | | | | | | |
| Oligo fwd | CACCTCAGCACAGCACGAATGTAT | | | | | | | | | | |
| Oligo rev | AAACATACATTCGTGCTGTGCTGA | | | | | | | | | | |
| Chromosome | start | end | strand | MM | target_seq | PAM | alignment | distance | position | gene name | gene id |
| GU305914.1 | 206536 | 206558 | - | 0 | TCAGCACAGCACGAATGTAT | CGG | \|\|\|\|\|\|\|\|[\|\|\|\|\|\|\|\|\|\|\|\|]PAM | NA | Intergenic | NA | NA |
| T18907 | ATTTCTGTGATCTCGATAGCCGG | 1000 | | | | | | | | | |
| Oligo fwd | CACCATTTCTGTGATCTCGATAGC | | | | | | | | | | |
| Oligo rev | AAACGCTATCGAGATCACAGAAAT | | | | | | | | | | |
| Chromosome | start | end | strand | MM | target_seq | PAM | alignment | distance | position | gene name | gene id |
| GU305914.1 | 206458 | 206480 | - | 0 | ATTTCTGTGATCTCGATAGC | CGG | \|\|\|\|\|\|\|\|[\|\|\|\|\|\|\|\|\|\|\|\|]PAM | NA | Intergenic | NA | NA |

<table>
<tr><td>T18908</td><td>TCGATAGCCGGCGTTACAGATGG</td><td>1000</td><td></td><td></td><td></td><td></td><td></td><td></td><td></td><td></td><td></td></tr>
<tr><td>Oligo fwd</td><td>CACCTCGATAGCCGGCGTTACAGA</td><td></td><td></td><td></td><td></td><td></td><td></td><td></td><td></td><td></td><td></td></tr>
<tr><td>Oligo rev</td><td>AAACTCTGTAACGCCGGCTATCGA</td><td></td><td></td><td></td><td></td><td></td><td></td><td></td><td></td><td></td><td></td></tr>
<tr><td>Chromosome</td><td>start</td><td>end</td><td>strand</td><td>MM</td><td>target_seq</td><td>PAM</td><td>alignment</td><td>distance</td><td>position</td><td>gene name</td><td>gene id</td></tr>
<tr><td>GU305914.1</td><td>206446</td><td>206468</td><td>-</td><td>0</td><td>TCGATAGCCGGCGTTACAGA</td><td>TGG</td><td>||||||||[||||||||||||]PAM</td><td>NA</td><td>Intergenic</td><td>NA</td><td>NA</td></tr>
<tr><td>T18909</td><td>TTTGAGAAGAACCCTGAAAACGG</td><td>1000</td><td></td><td></td><td></td><td></td><td></td><td></td><td></td><td></td><td></td></tr>
<tr><td>Oligo fwd</td><td>CACCTTTGAGAAGAACCCTGAAAA</td><td></td><td></td><td></td><td></td><td></td><td></td><td></td><td></td><td></td><td></td></tr>
<tr><td>Oligo rev</td><td>AAACTTTTCAGGGTTCTTCTCAAA</td><td></td><td></td><td></td><td></td><td></td><td></td><td></td><td></td><td></td><td></td></tr>
<tr><td>Chromosome</td><td>start</td><td>end</td><td>strand</td><td>MM</td><td>target_seq</td><td>PAM</td><td>alignment</td><td>distance</td><td>position</td><td>gene name</td><td>gene id</td></tr>
<tr><td>GU305914.1</td><td>206404</td><td>206426</td><td>-</td><td>0</td><td>TTTGAGAAGAACCCTGAAAA</td><td>CGG</td><td>||||||||[||||||||||||]PAM</td><td>NA</td><td>Intergenic</td><td>NA</td><td>NA</td></tr>
<tr><td>T18910</td><td>AAGAACCCTGAAAACGGAGACGG</td><td>1000</td><td></td><td></td><td></td><td></td><td></td><td></td><td></td><td></td><td></td></tr>
<tr><td>Oligo fwd</td><td>CACCAAGAACCCTGAAAACGGAGA</td><td></td><td></td><td></td><td></td><td></td><td></td><td></td><td></td><td></td><td></td></tr>
<tr><td>Oligo rev</td><td>AAACTCTCCGTTTTCAGGGTTCTT</td><td></td><td></td><td></td><td></td><td></td><td></td><td></td><td></td><td></td><td></td></tr>
<tr><td>Chromosome</td><td>start</td><td>end</td><td>strand</td><td>MM</td><td>target_seq</td><td>PAM</td><td>alignment</td><td>distance</td><td>position</td><td>gene name</td><td>gene id</td></tr>
<tr><td>GU305914.1</td><td>206398</td><td>206420</td><td>-</td><td>0</td><td>AAGAACCCTGAAAACGGAGA</td><td>CGG</td><td>||||||||[||||||||||||]PAM</td><td>NA</td><td>Intergenic</td><td>NA</td><td>NA</td></tr>
<tr><td>T18911</td><td>GATGTGATAAACGTGTTAACTGG</td><td>1000</td><td></td><td></td><td></td><td></td><td></td><td></td><td></td><td></td><td></td></tr>
<tr><td>Oligo fwd</td><td>CACCGATGTGATAAACGTGTTAAC</td><td></td><td></td><td></td><td></td><td></td><td></td><td></td><td></td><td></td><td></td></tr>
<tr><td>Oligo rev</td><td>AAACGTTAACACGTTTATCACATC</td><td></td><td></td><td></td><td></td><td></td><td></td><td></td><td></td><td></td><td></td></tr>
<tr><td>Chromosome</td><td>start</td><td>end</td><td>strand</td><td>MM</td><td>target_seq</td><td>PAM</td><td>alignment</td><td>distance</td><td>position</td><td>gene name</td><td>gene id</td></tr>
<tr><td>GU305914.1</td><td>206362</td><td>206384</td><td>-</td><td>0</td><td>GATGTGATAAACGTGTTAAC</td><td>TGG</td><td>||||||||[||||||||||||]PAM</td><td>NA</td><td>Intergenic</td><td>NA</td><td>NA</td></tr>
<tr><td>T18912</td><td>ATTATTCTACAGTTTACAGCCGG</td><td>1000</td><td></td><td></td><td></td><td></td><td></td><td></td><td></td><td></td><td></td></tr>
<tr><td>Oligo fwd</td><td>CACCATTATTCTACAGTTTACAGC</td><td></td><td></td><td></td><td></td><td></td><td></td><td></td><td></td><td></td><td></td></tr>
<tr><td>Oligo rev</td><td>AAACGCTGTAAACTGTAGAATAAT</td><td></td><td></td><td></td><td></td><td></td><td></td><td></td><td></td><td></td><td></td></tr>
<tr><td>Chromosome</td><td>start</td><td>end</td><td>strand</td><td>MM</td><td>target_seq</td><td>PAM</td><td>alignment</td><td>distance</td><td>position</td><td>gene name</td><td>gene id</td></tr>
<tr><td>GU305914.1</td><td>206322</td><td>206344</td><td>-</td><td>0</td><td>ATTATTCTACAGTTTACAGC</td><td>CGG</td><td>||||||||[||||||||||||]PAM</td><td>NA</td><td>Intergenic</td><td>NA</td><td>NA</td></tr>
<tr><td>T18913</td><td>ATTCTCAATAATTGTGTAGTTGG</td><td>1000</td><td></td><td></td><td></td><td></td><td></td><td></td><td></td><td></td><td></td></tr>
<tr><td>Oligo fwd</td><td>CACCATTCTCAATAATTGTGTAGT</td><td></td><td></td><td></td><td></td><td></td><td></td><td></td><td></td><td></td><td></td></tr>
<tr><td>Oligo rev</td><td>AAACACTACACAATTATTGAGAAT</td><td></td><td></td><td></td><td></td><td></td><td></td><td></td><td></td><td></td><td></td></tr>
<tr><td>Chromosome</td><td>start</td><td>end</td><td>strand</td><td>MM</td><td>target_seq</td><td>PAM</td><td>alignment</td><td>distance</td><td>position</td><td>gene name</td><td>gene id</td></tr>
<tr><td>GU305914.1</td><td>206278</td><td>206300</td><td>-</td><td>0</td><td>ATTCTCAATAATTGTGTAGT</td><td>TGG</td><td>||||||||[||||||||||||]PAM</td><td>NA</td><td>Intergenic</td><td>NA</td><td>NA</td></tr>
<tr><td>T18914</td><td>TTCTCAATAATTGTGTAGTTGGG</td><td>1000</td><td></td><td></td><td></td><td></td><td></td><td></td><td></td><td></td><td></td></tr>
<tr><td>Oligo fwd</td><td>CACCTTCTCAATAATTGTGTAGTT</td><td></td><td></td><td></td><td></td><td></td><td></td><td></td><td></td><td></td><td></td></tr>
<tr><td>Oligo rev</td><td>AAACAACTACACAATTATTGAGAA</td><td></td><td></td><td></td><td></td><td></td><td></td><td></td><td></td><td></td><td></td></tr>
<tr><td>Chromosome</td><td>start</td><td>end</td><td>strand</td><td>MM</td><td>target_seq</td><td>PAM</td><td>alignment</td><td>distance</td><td>position</td><td>gene name</td><td>gene id</td></tr>
<tr><td>GU305914.1</td><td>206277</td><td>206299</td><td>-</td><td>0</td><td>TTCTCAATAATTGTGTAGTT</td><td>GGG</td><td>||||||||[||||||||||||]PAM</td><td>NA</td><td>Intergenic</td><td>NA</td><td>NA</td></tr>
<tr><td>T18915</td><td>GATACTAGACACTAAGTTTTCGG</td><td>1000</td><td></td><td></td><td></td><td></td><td></td><td></td><td></td><td></td><td></td></tr>
<tr><td>Oligo fwd</td><td>CACCGATACTAGACACTAAGTTTT</td><td></td><td></td><td></td><td></td><td></td><td></td><td></td><td></td><td></td><td></td></tr>
<tr><td>Oligo rev</td><td>AAACAAAACTTAGTGTCTAGTATC</td><td></td><td></td><td></td><td></td><td></td><td></td><td></td><td></td><td></td><td></td></tr>
<tr><td>Chromosome</td><td>start</td><td>end</td><td>strand</td><td>MM</td><td>target_seq</td><td>PAM</td><td>alignment</td><td>distance</td><td>position</td><td>gene name</td><td>gene id</td></tr>
<tr><td>GU305914.1</td><td>206255</td><td>206277</td><td>-</td><td>0</td><td>GATACTAGACACTAAGTTTT</td><td>CGG</td><td>||||||||[||||||||||||]PAM</td><td>NA</td><td>Intergenic</td><td>NA</td><td>NA</td></tr>
<tr><td>T18916</td><td>CTAAGTTTTCGGCATATAAACGG</td><td>1000</td><td></td><td></td><td></td><td></td><td></td><td></td><td></td><td></td><td></td></tr>
<tr><td>Oligo fwd</td><td>CACCCTAAGTTTTCGGCATATAAA</td><td></td><td></td><td></td><td></td><td></td><td></td><td></td><td></td><td></td><td></td></tr>
<tr><td>Oligo rev</td><td>AAACTTTATATGCCGAAAACTTAG</td><td></td><td></td><td></td><td></td><td></td><td></td><td></td><td></td><td></td><td></td></tr>
<tr><td>Chromosome</td><td>start</td><td>end</td><td>strand</td><td>MM</td><td>target_seq</td><td>PAM</td><td>alignment</td><td>distance</td><td>position</td><td>gene name</td><td>gene id</td></tr>
<tr><td>GU305914.1</td><td>206244</td><td>206266</td><td>-</td><td>0</td><td>CTAAGTTTTCGGCATATAAA</td><td>CGG</td><td>||||||||[||||||||||||]PAM</td><td>NA</td><td>Intergenic</td><td>NA</td><td>NA</td></tr>
<tr><td>T18917</td><td>TAAGTTTTCGGCATATAAACGGG</td><td>1000</td><td></td><td></td><td></td><td></td><td></td><td></td><td></td><td></td><td></td></tr>
<tr><td>Oligo fwd</td><td>CACCTAAGTTTTCGGCATATAAAC</td><td></td><td></td><td></td><td></td><td></td><td></td><td></td><td></td><td></td><td></td></tr>
<tr><td>Oligo rev</td><td>AAACGTTTATATGCCGAAAACTTA</td><td></td><td></td><td></td><td></td><td></td><td></td><td></td><td></td><td></td><td></td></tr>
<tr><td>Chromosome</td><td>start</td><td>end</td><td>strand</td><td>MM</td><td>target_seq</td><td>PAM</td><td>alignment</td><td>distance</td><td>position</td><td>gene name</td><td>gene id</td></tr>
<tr><td>GU305914.1</td><td>206243</td><td>206265</td><td>-</td><td>0</td><td>TAAGTTTTCGGCATATAAAC</td><td>GGG</td><td>||||||||[||||||||||||]PAM</td><td>NA</td><td>Intergenic</td><td>NA</td><td>NA</td></tr>
<tr><td>T18918</td><td>GATGATTCATTTGACGTGTACGG</td><td>1000</td><td></td><td></td><td></td><td></td><td></td><td></td><td></td><td></td><td></td></tr>
<tr><td>Oligo fwd</td><td>CACCGATGATTCATTTGACGTGTA</td><td></td><td></td><td></td><td></td><td></td><td></td><td></td><td></td><td></td><td></td></tr>
<tr><td>Oligo rev</td><td>AAACTACACGTCAAATGAATCATC</td><td></td><td></td><td></td><td></td><td></td><td></td><td></td><td></td><td></td><td></td></tr>
<tr><td>Chromosome</td><td>start</td><td>end</td><td>strand</td><td>MM</td><td>target_seq</td><td>PAM</td><td>alignment</td><td>distance</td><td>position</td><td>gene name</td><td>gene id</td></tr>
<tr><td>GU305914.1</td><td>206216</td><td>206238</td><td>-</td><td>0</td><td>GATGATTCATTTGACGTGTA</td><td>CGG</td><td>||||||||[||||||||||||]PAM</td><td>NA</td><td>Intergenic</td><td>NA</td><td>NA</td></tr>
<tr><td>T18919</td><td>AATGTATGTTTCACTGTAAATGG</td><td>1000</td><td></td><td></td><td></td><td></td><td></td><td></td><td></td><td></td><td></td></tr>
<tr><td>Oligo fwd</td><td>CACCAATGTATGTTTCACTGTAAA</td><td></td><td></td><td></td><td></td><td></td><td></td><td></td><td></td><td></td><td></td></tr>
<tr><td>Oligo rev</td><td>AAACTTTACAGTGAAACATACATT</td><td></td><td></td><td></td><td></td><td></td><td></td><td></td><td></td><td></td><td></td></tr>
<tr><td>Chromosome</td><td>start</td><td>end</td><td>strand</td><td>MM</td><td>target_seq</td><td>PAM</td><td>alignment</td><td>distance</td><td>position</td><td>gene name</td><td>gene id</td></tr>
<tr><td>GU305914.1</td><td>206167</td><td>206189</td><td>-</td><td>0</td><td>AATGTATGTTTCACTGTAAA</td><td>TGG</td><td>||||||||[||||||||||||]PAM</td><td>NA</td><td>Intergenic</td><td>NA</td><td>NA</td></tr>
<tr><td>T18920</td><td>GTATGTTTCACTGTAAATGGTGG</td><td>1000</td><td></td><td></td><td></td><td></td><td></td><td></td><td></td><td></td><td></td></tr>
<tr><td>Oligo fwd</td><td>CACCGTATGTTTCACTGTAAATGG</td><td></td><td></td><td></td><td></td><td></td><td></td><td></td><td></td><td></td><td></td></tr>
<tr><td>Oligo rev</td><td>AAACCCATTTACAGTGAAACATAC</td><td></td><td></td><td></td><td></td><td></td><td></td><td></td><td></td><td></td><td></td></tr>
<tr><td>Chromosome</td><td>start</td><td>end</td><td>strand</td><td>MM</td><td>target_seq</td><td>PAM</td><td>alignment</td><td>distance</td><td>position</td><td>gene name</td><td>gene id</td></tr>
<tr><td>GU305914.1</td><td>206164</td><td>206186</td><td>-</td><td>0</td><td>GTATGTTTCACTGTAAATGG</td><td>TGG</td><td>||||||||[||||||||||||]PAM</td><td>NA</td><td>Intergenic</td><td>NA</td><td>NA</td></tr>
<tr><td>T18921</td><td>TATGTTTCACTGTAAATGGTGGG</td><td>1000</td><td></td><td></td><td></td><td></td><td></td><td></td><td></td><td></td><td></td></tr>
<tr><td>Oligo fwd</td><td>CACCTATGTTTCACTGTAAATGGT</td><td></td><td></td><td></td><td></td><td></td><td></td><td></td><td></td><td></td><td></td></tr>
<tr><td>Oligo rev</td><td>AAACACCATTTACAGTGAAACATA</td><td></td><td></td><td></td><td></td><td></td><td></td><td></td><td></td><td></td><td></td></tr>
<tr><td>Chromosome</td><td>start</td><td>end</td><td>strand</td><td>MM</td><td>target_seq</td><td>PAM</td><td>alignment</td><td>distance</td><td>position</td><td>gene name</td><td>gene id</td></tr>
<tr><td>GU305914.1</td><td>206163</td><td>206185</td><td>-</td><td>0</td><td>TATGTTTCACTGTAAATGGT</td><td>GGG</td><td>||||||||[||||||||||||]PAM</td><td>NA</td><td>Intergenic</td><td>NA</td><td>NA</td></tr>
<tr><td>T18922</td><td>GCGACGCACGAGCAGACAACAGG</td><td>1000</td><td></td><td></td><td></td><td></td><td></td><td></td><td></td><td></td><td></td></tr>
<tr><td>Oligo fwd</td><td>CACCGCGACGCACGAGCAGACAAC</td><td></td><td></td><td></td><td></td><td></td><td></td><td></td><td></td><td></td><td></td></tr>
<tr><td>Oligo rev</td><td>AAACGTTGTCTGCTCGTGCGTCGC</td><td></td><td></td><td></td><td></td><td></td><td></td><td></td><td></td><td></td><td></td></tr>
<tr><td>Chromosome</td><td>start</td><td>end</td><td>strand</td><td>MM</td><td>target_seq</td><td>PAM</td><td>alignment</td><td>distance</td><td>position</td><td>gene name</td><td>gene id</td></tr>
</table>

GU305914.1 206139 206161 - 0 GCGACGCACGAGCAGACAAC AGG ||||||||[||||||||||||]PAM NA Intergenic NA NA

T18923 CGACGCACGAGCAGACAACAGGG 1000
Oligo fwd CACCCGACGCACGAGCAGACAACA
Oligo rev AAACTGTTGTCTGCTCGTGCGTCG
Chromosome start end strand MM target_seq PAM alignment distance position gene name gene id
GU305914.1 206138 206160 - 0 CGACGCACGAGCAGACAACA GGG ||||||||[||||||||||||]PAM NA Intergenic NA NA

T18924 AGGGCACAACCGACATCATCCGG 1000
Oligo fwd CACCAGGGCACAACCGACATCATC
Oligo rev AAACGATGATGTCGGTTGTGCCCT
Chromosome start end strand MM target_seq PAM alignment distance position gene name gene id
GU305914.1 206119 206141 - 0 AGGGCACAACCGACATCATC CGG ||||||||[||||||||||||]PAM NA Intergenic NA NA

T18925 GGGCACAACCGACATCATCCGGG 1000
Oligo fwd CACCGGGCACAACCGACATCATCC
Oligo rev AAACGGATGATGTCGGTTGTGCCC
Chromosome start end strand MM target_seq PAM alignment distance position gene name gene id
GU305914.1 206118 206140 - 0 GGGCACAACCGACATCATCC GGG ||||||||[||||||||||||]PAM NA Intergenic NA NA

T18926 GGCACAACCGACATCATCCGGGG 1000
Oligo fwd CACCGGCACAACCGACATCATCCG
Oligo rev AAACCGGATGATGTCGGTTGTGCC
Chromosome start end strand MM target_seq PAM alignment distance position gene name gene id
GU305914.1 206117 206139 - 0 GGCACAACCGACATCATCCG GGG ||||||||[||||||||||||]PAM NA Intergenic NA NA

T18927 ACAACCGACATCATCCGGGGAGG 1000
Oligo fwd CACCACAACCGACATCATCCGGGG
Oligo rev AAACCCCCGGATGATGTCGGTTGT
Chromosome start end strand MM target_seq PAM alignment distance position gene name gene id
GU305914.1 206114 206136 - 0 ACAACCGACATCATCCGGGG AGG ||||||||[||||||||||||]PAM NA Intergenic NA NA

T18928 ACGCAGCAAATACCTAAGTTTGG 1000
Oligo fwd CACCACGCAGCAAATACCTAAGTT
Oligo rev AAACAACTTAGGTATTTGCTGCGT
Chromosome start end strand MM target_seq PAM alignment distance position gene name gene id
GU305914.1 206047 206069 - 0 ACGCAGCAAATACCTAAGTT TGG ||||||||[||||||||||||]PAM NA Intergenic NA NA

T18929 AGTTTGGTCATAATCCCGTATGG 1000
Oligo fwd CACCAGTTTGGTCATAATCCCGTA
Oligo rev AAACTACGGGATTATGACCAAACT
Chromosome start end strand MM target_seq PAM alignment distance position gene name gene id
GU305914.1 206031 206053 - 0 AGTTTGGTCATAATCCCGTA TGG ||||||||[||||||||||||]PAM NA Intergenic NA NA

T18930 AAGTTGTGATGTTGAATCGAAGG 1000
Oligo fwd CACCAAGTTGTGATGTTGAATCGA
Oligo rev AAACTCGATTCAACATCACAACTT
Chromosome start end strand MM target_seq PAM alignment distance position gene name gene id
GU305914.1 206006 206028 - 0 AAGTTGTGATGTTGAATCGA AGG ||||||||[||||||||||||]PAM NA Intergenic NA NA

T18931 TGTGATGTTGAATCGAAGGTCGG 1000
Oligo fwd CACCTGTGATGTTGAATCGAAGGT
Oligo rev AAACACCTTCGATTCAACATCACA
Chromosome start end strand MM target_seq PAM alignment distance position gene name gene id
GU305914.1 206002 206024 - 0 TGTGATGTTGAATCGAAGGT CGG ||||||||[||||||||||||]PAM NA Intergenic NA NA

T18932 GTGATGTTGAATCGAAGGTCGGG 1000
Oligo fwd CACCGTGATGTTGAATCGAAGGTC
Oligo rev AAACGACCTTCGATTCAACATCAC
Chromosome start end strand MM target_seq PAM alignment distance position gene name gene id
GU305914.1 206001 206023 - 0 GTGATGTTGAATCGAAGGTC GGG ||||||||[||||||||||||]PAM NA Intergenic NA NA

T18933 ATGTTGAATCGAAGGTCGGGTGG 1000
Oligo fwd CACCATGTTGAATCGAAGGTCGGG
Oligo rev AAACCCCGACCTTCGATTCAACAT
Chromosome start end strand MM target_seq PAM alignment distance position gene name gene id
GU305914.1 205998 206020 - 0 ATGTTGAATCGAAGGTCGGG TGG ||||||||[||||||||||||]PAM NA Intergenic NA NA
GU305914.1 115 137 + 4 CTGTTGCATCGATGGTCGTG AGG -|||||-|[||||-|||||-|]PAM NA Intergenic NA NA

T18934 GCGTGTCACAGATTGCCACGCGG 1000
Oligo fwd CACCGCGTGTCACAGATTGCCACG
Oligo rev AAACCGTGGCAATCTGTGACACGC
Chromosome start end strand MM target_seq PAM alignment distance position gene name gene id
GU305914.1 205890 205912 - 0 GCGTGTCACAGATTGCCACG CGG ||||||||[||||||||||||]PAM NA Intergenic NA NA

T18935 GATCGAAACCTTTCGATCCGAGG 1000
Oligo fwd CACCGATCGAAACCTTTCGATCCG
Oligo rev AAACCGGATCGAAAGGTTTCGATC
Chromosome start end strand MM target_seq PAM alignment distance position gene name gene id
GU305914.1 205845 205867 - 0 GATCGAAACCTTTCGATCCG AGG ||||||||[||||||||||||]PAM NA Intergenic NA NA

T18936 ATCGAAACCTTTCGATCCGAGGG 1000
Oligo fwd CACCATCGAAACCTTTCGATCCGA
Oligo rev AAACTCGGATCGAAAGGTTTCGAT
Chromosome start end strand MM target_seq PAM alignment distance position gene name gene id
GU305914.1 205844 205866 - 0 ATCGAAACCTTTCGATCCGA GGG ||||||||[||||||||||||]PAM NA Intergenic NA NA

T18937 GAAACCTTTCGATCCGAGGGCGG 1000
Oligo fwd CACCGAAACCTTTCGATCCGAGGG

<table>
<tr><td>Oligo rev</td><td>AAACCCCTCGGATCGAAAGGTTTC</td></tr>
<tr><td>Chromosome</td><td>start</td><td></td><td>end</td><td>strand</td><td>MM</td><td>target_seq</td><td>PAM</td><td>alignment</td><td>distance</td><td>position</td><td>gene name</td><td>gene id</td></tr>
<tr><td>GU305914.1</td><td></td><td>205841</td><td>205863</td><td>-</td><td>0</td><td>GAAACCTTTCGATCCGAGGG</td><td>CGG</td><td>||||||||[||||||||||||]PAM</td><td>NA</td><td>Intergenic</td><td>NA</td><td>NA</td></tr>
<tr><td></td></tr>
<tr><td>T18938</td><td>CGATCCGAGGGCGGTCATCGTGG</td><td></td><td>1000</td></tr>
<tr><td>Oligo fwd</td><td>CACCCGATCCGAGGGCGGTCATCG</td></tr>
<tr><td>Oligo rev</td><td>AAACCGATGACCGCCCTCGGATCG</td></tr>
<tr><td>Chromosome</td><td>start</td><td></td><td>end</td><td>strand</td><td>MM</td><td>target_seq</td><td>PAM</td><td>alignment</td><td>distance</td><td>position</td><td>gene name</td><td>gene id</td></tr>
<tr><td>GU305914.1</td><td></td><td>205832</td><td>205854</td><td>-</td><td>0</td><td>CGATCCGAGGGCGGTCATCG</td><td>TGG</td><td>||||||||[||||||||||||]PAM</td><td>NA</td><td>Intergenic</td><td>NA</td><td>NA</td></tr>
<tr><td></td></tr>
<tr><td>T18939</td><td>GATCCGAGGGCGGTCATCGTGGG</td><td></td><td>1000</td></tr>
<tr><td>Oligo fwd</td><td>CACCGATCCGAGGGCGGTCATCGT</td></tr>
<tr><td>Oligo rev</td><td>AAACACGATGACCGCCCTCGGATC</td></tr>
<tr><td>Chromosome</td><td>start</td><td></td><td>end</td><td>strand</td><td>MM</td><td>target_seq</td><td>PAM</td><td>alignment</td><td>distance</td><td>position</td><td>gene name</td><td>gene id</td></tr>
<tr><td>GU305914.1</td><td></td><td>205831</td><td>205853</td><td>-</td><td>0</td><td>GATCCGAGGGCGGTCATCGT</td><td>GGG</td><td>||||||||[||||||||||||]PAM</td><td>NA</td><td>Intergenic</td><td>NA</td><td>NA</td></tr>
<tr><td></td></tr>
<tr><td>T18940</td><td>GTCATCGTGGGCATTATCGCAGG</td><td></td><td>1000</td></tr>
<tr><td>Oligo fwd</td><td>CACCGTCATCGTGGGCATTATCGC</td></tr>
<tr><td>Oligo rev</td><td>AAACGCGATAATGCCCACGATGAC</td></tr>
<tr><td>Chromosome</td><td>start</td><td></td><td>end</td><td>strand</td><td>MM</td><td>target_seq</td><td>PAM</td><td>alignment</td><td>distance</td><td>position</td><td>gene name</td><td>gene id</td></tr>
<tr><td>GU305914.1</td><td></td><td>205819</td><td>205841</td><td>-</td><td>0</td><td>GTCATCGTGGGCATTATCGC</td><td>AGG</td><td>||||||||[||||||||||||]PAM</td><td>NA</td><td>Intergenic</td><td>NA</td><td>NA</td></tr>
<tr><td></td></tr>
<tr><td>T18941</td><td>GTTATTTATTCTCACTGTACTGG</td><td></td><td>1000</td></tr>
<tr><td>Oligo fwd</td><td>CACCGTTATTTATTCTCACTGTAC</td></tr>
<tr><td>Oligo rev</td><td>AAACGTACAGTGAGAATAAATAAC</td></tr>
<tr><td>Chromosome</td><td>start</td><td></td><td>end</td><td>strand</td><td>MM</td><td>target_seq</td><td>PAM</td><td>alignment</td><td>distance</td><td>position</td><td>gene name</td><td>gene id</td></tr>
<tr><td>GU305914.1</td><td></td><td>205793</td><td>205815</td><td>-</td><td>0</td><td>GTTATTTATTCTCACTGTAC</td><td>TGG</td><td>||||||||[||||||||||||]PAM</td><td>NA</td><td>Intergenic</td><td>NA</td><td>NA</td></tr>
<tr><td></td></tr>
<tr><td>T18942</td><td>GTGTGCGTGTAAGAGTCAGATGG</td><td></td><td>1000</td></tr>
<tr><td>Oligo fwd</td><td>CACCGTGTGCGTGTAAGAGTCAGA</td></tr>
<tr><td>Oligo rev</td><td>AAACTCTGACTCTTACACGCACAC</td></tr>
<tr><td>Chromosome</td><td>start</td><td></td><td>end</td><td>strand</td><td>MM</td><td>target_seq</td><td>PAM</td><td>alignment</td><td>distance</td><td>position</td><td>gene name</td><td>gene id</td></tr>
<tr><td>GU305914.1</td><td></td><td>205732</td><td>205754</td><td>-</td><td>0</td><td>GTGTGCGTGTAAGAGTCAGA</td><td>TGG</td><td>||||||||[||||||||||||]PAM</td><td>NA</td><td>Intergenic</td><td>NA</td><td>NA</td></tr>
<tr><td></td></tr>
<tr><td>T18943</td><td>ATGAACTGCGTCAAATTGAAAGG</td><td></td><td>1000</td></tr>
<tr><td>Oligo fwd</td><td>CACCATGAACTGCGTCAAATTGAA</td></tr>
<tr><td>Oligo rev</td><td>AAACTTCAATTTGACGCAGTTCAT</td></tr>
<tr><td>Chromosome</td><td>start</td><td></td><td>end</td><td>strand</td><td>MM</td><td>target_seq</td><td>PAM</td><td>alignment</td><td>distance</td><td>position</td><td>gene name</td><td>gene id</td></tr>
<tr><td>GU305914.1</td><td></td><td>205674</td><td>205696</td><td>-</td><td>0</td><td>ATGAACTGCGTCAAATTGAA</td><td>AGG</td><td>||||||||[||||||||||||]PAM</td><td>NA</td><td>Intergenic</td><td>NA</td><td>NA</td></tr>
<tr><td></td></tr>
<tr><td>T18944</td><td>TGAACTGCGTCAAATTGAAAGGG</td><td></td><td>1000</td></tr>
<tr><td>Oligo fwd</td><td>CACCTGAACTGCGTCAAATTGAAA</td></tr>
<tr><td>Oligo rev</td><td>AAACTTTCAATTTGACGCAGTTCA</td></tr>
<tr><td>Chromosome</td><td>start</td><td></td><td>end</td><td>strand</td><td>MM</td><td>target_seq</td><td>PAM</td><td>alignment</td><td>distance</td><td>position</td><td>gene name</td><td>gene id</td></tr>
<tr><td>GU305914.1</td><td></td><td>205673</td><td>205695</td><td>-</td><td>0</td><td>TGAACTGCGTCAAATTGAAA</td><td>GGG</td><td>||||||||[||||||||||||]PAM</td><td>NA</td><td>Intergenic</td><td>NA</td><td>NA</td></tr>
<tr><td></td></tr>
<tr><td>T18945</td><td>GGTTATGTAAACGAGCTCATCGG</td><td></td><td>1000</td></tr>
<tr><td>Oligo fwd</td><td>CACCGGTTATGTAAACGAGCTCAT</td></tr>
<tr><td>Oligo rev</td><td>AAACATGAGCTCGTTTACATAACC</td></tr>
<tr><td>Chromosome</td><td>start</td><td></td><td>end</td><td>strand</td><td>MM</td><td>target_seq</td><td>PAM</td><td>alignment</td><td>distance</td><td>position</td><td>gene name</td><td>gene id</td></tr>
<tr><td>GU305914.1</td><td></td><td>205652</td><td>205674</td><td>-</td><td>0</td><td>GGTTATGTAAACGAGCTCAT</td><td>CGG</td><td>||||||||[||||||||||||]PAM</td><td>NA</td><td>Intergenic</td><td>NA</td><td>NA</td></tr>
<tr><td></td></tr>
<tr><td>T18946</td><td>GTTATGTAAACGAGCTCATCGGG</td><td></td><td>1000</td></tr>
<tr><td>Oligo fwd</td><td>CACCGTTATGTAAACGAGCTCATC</td></tr>
<tr><td>Oligo rev</td><td>AAACGATGAGCTCGTTTACATAAC</td></tr>
<tr><td>Chromosome</td><td>start</td><td></td><td>end</td><td>strand</td><td>MM</td><td>target_seq</td><td>PAM</td><td>alignment</td><td>distance</td><td>position</td><td>gene name</td><td>gene id</td></tr>
<tr><td>GU305914.1</td><td></td><td>205651</td><td>205673</td><td>-</td><td>0</td><td>GTTATGTAAACGAGCTCATC</td><td>GGG</td><td>||||||||[||||||||||||]PAM</td><td>NA</td><td>Intergenic</td><td>NA</td><td>NA</td></tr>
<tr><td></td></tr>
<tr><td>T18947</td><td>TTAAAACCGCACACAGATGTAGG</td><td></td><td>1000</td></tr>
<tr><td>Oligo fwd</td><td>CACCTTAAAACCGCACACAGATGT</td></tr>
<tr><td>Oligo rev</td><td>AAACACATCTGTGTGCGGTTTTAA</td></tr>
<tr><td>Chromosome</td><td>start</td><td></td><td>end</td><td>strand</td><td>MM</td><td>target_seq</td><td>PAM</td><td>alignment</td><td>distance</td><td>position</td><td>gene name</td><td>gene id</td></tr>
<tr><td>GU305914.1</td><td></td><td>205628</td><td>205650</td><td>-</td><td>0</td><td>TTAAAACCGCACACAGATGT</td><td>AGG</td><td>||||||||[||||||||||||]PAM</td><td>NA</td><td>Intergenic</td><td>NA</td><td>NA</td></tr>
<tr><td></td></tr>
<tr><td>T18948</td><td>TAAAACCGCACACAGATGTAGGG</td><td></td><td>1000</td></tr>
<tr><td>Oligo fwd</td><td>CACCTAAAACCGCACACAGATGTA</td></tr>
<tr><td>Oligo rev</td><td>AAACTACATCTGTGTGCGGTTTTA</td></tr>
<tr><td>Chromosome</td><td>start</td><td></td><td>end</td><td>strand</td><td>MM</td><td>target_seq</td><td>PAM</td><td>alignment</td><td>distance</td><td>position</td><td>gene name</td><td>gene id</td></tr>
<tr><td>GU305914.1</td><td></td><td>205627</td><td>205649</td><td>-</td><td>0</td><td>TAAAACCGCACACAGATGTA</td><td>GGG</td><td>||||||||[||||||||||||]PAM</td><td>NA</td><td>Intergenic</td><td>NA</td><td>NA</td></tr>
<tr><td></td></tr>
<tr><td>T18949</td><td>AAAACCGCACACAGATGTAGGGG</td><td></td><td>1000</td></tr>
<tr><td>Oligo fwd</td><td>CACCAAAACCGCACACAGATGTAG</td></tr>
<tr><td>Oligo rev</td><td>AAACCTACATCTGTGTGCGGTTTT</td></tr>
<tr><td>Chromosome</td><td>start</td><td></td><td>end</td><td>strand</td><td>MM</td><td>target_seq</td><td>PAM</td><td>alignment</td><td>distance</td><td>position</td><td>gene name</td><td>gene id</td></tr>
<tr><td>GU305914.1</td><td></td><td>205626</td><td>205648</td><td>-</td><td>0</td><td>AAAACCGCACACAGATGTAG</td><td>GGG</td><td>||||||||[||||||||||||]PAM</td><td>NA</td><td>Intergenic</td><td>NA</td><td>NA</td></tr>
<tr><td></td></tr>
<tr><td>T18950</td><td>AGATGTAGGGGCAGACTCTGAGG</td><td></td><td>1000</td></tr>
<tr><td>Oligo fwd</td><td>CACCAGATGTAGGGGCAGACTCTG</td></tr>
<tr><td>Oligo rev</td><td>AAACCAGAGTCTGCCCCTACATCT</td></tr>
<tr><td>Chromosome</td><td>start</td><td></td><td>end</td><td>strand</td><td>MM</td><td>target_seq</td><td>PAM</td><td>alignment</td><td>distance</td><td>position</td><td>gene name</td><td>gene id</td></tr>
<tr><td>GU305914.1</td><td></td><td>205614</td><td>205636</td><td>-</td><td>0</td><td>AGATGTAGGGGCAGACTCTG</td><td>AGG</td><td>||||||||[||||||||||||]PAM</td><td>NA</td><td>Intergenic</td><td>NA</td><td>NA</td></tr>
<tr><td></td></tr>
<tr><td>T18951</td><td>TAGGGGCAGACTCTGAGGACCGG</td><td></td><td>1000</td></tr>
<tr><td>Oligo fwd</td><td>CACCTAGGGGCAGACTCTGAGGAC</td></tr>
<tr><td>Oligo rev</td><td>AAACGTCCTCAGAGTCTGCCCCTA</td></tr>
<tr><td>Chromosome</td><td>start</td><td></td><td>end</td><td>strand</td><td>MM</td><td>target_seq</td><td>PAM</td><td>alignment</td><td>distance</td><td>position</td><td>gene name</td><td>gene id</td></tr>
<tr><td>GU305914.1</td><td></td><td>205609</td><td>205631</td><td>-</td><td>0</td><td>TAGGGGCAGACTCTGAGGAC</td><td>CGG</td><td>||||||||[||||||||||||]PAM</td><td>NA</td><td>Intergenic</td><td>NA</td><td>NA</td></tr>
<tr><td></td></tr>
<tr><td>T18952</td><td>CCGGTGTTTCAACTCCGCGATGG</td><td></td><td>1000</td></tr>
</table>

| | | | | | | | | | | | |
|---|---|---|---|---|---|---|---|---|---|---|---|
| Oligo fwd | CACCCCGGTGTTTCAACTCCGCGA | | | | | | | | | | |
| Oligo rev | AAACTCGCGGAGTTGAAACACCGG | | | | | | | | | | |
| Chromosome | start | end | strand | MM | target_seq | PAM | alignment | distance | position | gene name | gene id |
| GU305914.1 | 205590 | 205612 | - | 0 | CCGGTGTTTCAACTCCGCGA | TGG | \|\|\|\|\|\|\|\|[\|\|\|\|\|\|\|\|\|\|\|\|]PAM | NA | Intergenic | NA | NA |
| T18953 | TCAACTCCGCGATGGACCGTCGG | 1000 | | | | | | | | | |
| Oligo fwd | CACCTCAACTCCGCGATGGACCGT | | | | | | | | | | |
| Oligo rev | AAACACGGTCCATCGCGGAGTTGA | | | | | | | | | | |
| Chromosome | start | end | strand | MM | target_seq | PAM | alignment | distance | position | gene name | gene id |
| GU305914.1 | 205582 | 205604 | - | 0 | TCAACTCCGCGATGGACCGT | CGG | \|\|\|\|\|\|\|\|[\|\|\|\|\|\|\|\|\|\|\|\|]PAM | NA | Intergenic | NA | NA |
| T18954 | CAACTCCGCGATGGACCGTCGGG | 1000 | | | | | | | | | |
| Oligo fwd | CACCCAACTCCGCGATGGACCGTC | | | | | | | | | | |
| Oligo rev | AAACGACGGTCCATCGCGGAGTTG | | | | | | | | | | |
| Chromosome | start | end | strand | MM | target_seq | PAM | alignment | distance | position | gene name | gene id |
| GU305914.1 | 205581 | 205603 | - | 0 | CAACTCCGCGATGGACCGTC | GGG | \|\|\|\|\|\|\|\|[\|\|\|\|\|\|\|\|\|\|\|\|]PAM | NA | Intergenic | NA | NA |
| T18955 | CTCCGCGATGGACCGTCGGGTGG | 1000 | | | | | | | | | |
| Oligo fwd | CACCCTCCGCGATGGACCGTCGGG | | | | | | | | | | |
| Oligo rev | AAACCCCGACGGTCCATCGCGGAG | | | | | | | | | | |
| Chromosome | start | end | strand | MM | target_seq | PAM | alignment | distance | position | gene name | gene id |
| GU305914.1 | 205578 | 205600 | - | 0 | CTCCGCGATGGACCGTCGGG | TGG | \|\|\|\|\|\|\|\|[\|\|\|\|\|\|\|\|\|\|\|\|]PAM | NA | Intergenic | NA | NA |
| T18956 | CTCATACCTAATGTGCGCCTTGG | 1000 | | | | | | | | | |
| Oligo fwd | CACCCTCATACCTAATGTGCGCCT | | | | | | | | | | |
| Oligo rev | AAACAGGCGCACATTAGGTATGAG | | | | | | | | | | |
| Chromosome | start | end | strand | MM | target_seq | PAM | alignment | distance | position | gene name | gene id |
| GU305914.1 | 205554 | 205576 | - | 0 | CTCATACCTAATGTGCGCCT | TGG | \|\|\|\|\|\|\|\|[\|\|\|\|\|\|\|\|\|\|\|\|]PAM | NA | Intergenic | NA | NA |
| T18957 | ATACCTAATGTGCGCCTTGGCGG | 1000 | | | | | | | | | |
| Oligo fwd | CACCATACCTAATGTGCGCCTTGG | | | | | | | | | | |
| Oligo rev | AAACCCAAGGCGCACATTAGGTAT | | | | | | | | | | |
| Chromosome | start | end | strand | MM | target_seq | PAM | alignment | distance | position | gene name | gene id |
| GU305914.1 | 205551 | 205573 | - | 0 | ATACCTAATGTGCGCCTTGG | CGG | \|\|\|\|\|\|\|\|[\|\|\|\|\|\|\|\|\|\|\|\|]PAM | NA | Intergenic | NA | NA |
| T18958 | AGCTACGATCAACGTACCCCCGG | 1000 | | | | | | | | | |
| Oligo fwd | CACCAGCTACGATCAACGTACCCC | | | | | | | | | | |
| Oligo rev | AAACGGGGTACGTTGATCGTAGCT | | | | | | | | | | |
| Chromosome | start | end | strand | MM | target_seq | PAM | alignment | distance | position | gene name | gene id |
| GU305914.1 | 205517 | 205539 | - | 0 | AGCTACGATCAACGTACCCC | CGG | \|\|\|\|\|\|\|\|[\|\|\|\|\|\|\|\|\|\|\|\|]PAM | NA | Intergenic | NA | NA |
| T18959 | GTAGATCCCGAATGTGCTCGCGG | 1000 | | | | | | | | | |
| Oligo fwd | CACCGTAGATCCCGAATGTGCTCG | | | | | | | | | | |
| Oligo rev | AAACCGAGCACATTCGGGATCTAC | | | | | | | | | | |
| Chromosome | start | end | strand | MM | target_seq | PAM | alignment | distance | position | gene name | gene id |
| GU305914.1 | 205490 | 205512 | - | 0 | GTAGATCCCGAATGTGCTCG | CGG | \|\|\|\|\|\|\|\|[\|\|\|\|\|\|\|\|\|\|\|\|]PAM | NA | Intergenic | NA | NA |
| T18960 | TAGATCCCGAATGTGCTCGCGGG | 1000 | | | | | | | | | |
| Oligo fwd | CACCTAGATCCCGAATGTGCTCGC | | | | | | | | | | |
| Oligo rev | AAACGCGAGCACATTCGGGATCTA | | | | | | | | | | |
| Chromosome | start | end | strand | MM | target_seq | PAM | alignment | distance | position | gene name | gene id |
| GU305914.1 | 205489 | 205511 | - | 0 | TAGATCCCGAATGTGCTCGC | GGG | \|\|\|\|\|\|\|\|[\|\|\|\|\|\|\|\|\|\|\|\|]PAM | NA | Intergenic | NA | NA |
| T18961 | CGACCGCGTGAACCTGACACTGG | 1000 | | | | | | | | | |
| Oligo fwd | CACCCGACCGCGTGAACCTGACAC | | | | | | | | | | |
| Oligo rev | AAACGTGTCAGGTTCACGCGGTCG | | | | | | | | | | |
| Chromosome | start | end | strand | MM | target_seq | PAM | alignment | distance | position | gene name | gene id |
| GU305914.1 | 205449 | 205471 | - | 0 | CGACCGCGTGAACCTGACAC | TGG | \|\|\|\|\|\|\|\|[\|\|\|\|\|\|\|\|\|\|\|\|]PAM | NA | Intergenic | NA | NA |
| T18962 | CTTCCTAGCTACGATCGCGTTGG | 1000 | | | | | | | | | |
| Oligo fwd | CACCCTTCCTAGCTACGATCGCGT | | | | | | | | | | |
| Oligo rev | AAACACGCGATCGTAGCTAGGAAG | | | | | | | | | | |
| Chromosome | start | end | strand | MM | target_seq | PAM | alignment | distance | position | gene name | gene id |
| GU305914.1 | 205392 | 205414 | - | 0 | CTTCCTAGCTACGATCGCGT | TGG | \|\|\|\|\|\|\|\|[\|\|\|\|\|\|\|\|\|\|\|\|]PAM | NA | Intergenic | NA | NA |
| T18963 | TGACACTACCAGTTTTCTGACGG | 1000 | | | | | | | | | |
| Oligo fwd | CACCTGACACTACCAGTTTTCTGA | | | | | | | | | | |
| Oligo rev | AAACTCAGAAAACTGGTAGTGTCA | | | | | | | | | | |
| Chromosome | start | end | strand | MM | target_seq | PAM | alignment | distance | position | gene name | gene id |
| GU305914.1 | 205368 | 205390 | - | 0 | TGACACTACCAGTTTTCTGA | CGG | \|\|\|\|\|\|\|\|[\|\|\|\|\|\|\|\|\|\|\|\|]PAM | NA | Intergenic | NA | NA |
| T18964 | CCAGTTTTCTGACGGTCAAATGG | 1000 | | | | | | | | | |
| Oligo fwd | CACCCCAGTTTTCTGACGGTCAAA | | | | | | | | | | |
| Oligo rev | AAACTTTGACCGTCAGAAAACTGG | | | | | | | | | | |
| Chromosome | start | end | strand | MM | target_seq | PAM | alignment | distance | position | gene name | gene id |
| GU305914.1 | 205360 | 205382 | - | 0 | CCAGTTTTCTGACGGTCAAA | TGG | \|\|\|\|\|\|\|\|[\|\|\|\|\|\|\|\|\|\|\|\|]PAM | NA | Intergenic | NA | NA |
| T18965 | AATGGAACAATACTCCGAGATGG | 1000 | | | | | | | | | |
| Oligo fwd | CACCAATGGAACAATACTCCGAGA | | | | | | | | | | |
| Oligo rev | AAACTCTCGGAGTATTGTTCCATT | | | | | | | | | | |
| Chromosome | start | end | strand | MM | target_seq | PAM | alignment | distance | position | gene name | gene id |
| GU305914.1 | 205342 | 205364 | - | 0 | AATGGAACAATACTCCGAGA | TGG | \|\|\|\|\|\|\|\|[\|\|\|\|\|\|\|\|\|\|\|\|]PAM | NA | Intergenic | NA | NA |
| T18966 | ATGGAACAATACTCCGAGATGGG | 1000 | | | | | | | | | |
| Oligo fwd | CACCATGGAACAATACTCCGAGAT | | | | | | | | | | |
| Oligo rev | AAACATCTCGGAGTATTGTTCCAT | | | | | | | | | | |
| Chromosome | start | end | strand | MM | target_seq | PAM | alignment | distance | position | gene name | gene id |
| GU305914.1 | 205341 | 205363 | - | 0 | ATGGAACAATACTCCGAGAT | GGG | \|\|\|\|\|\|\|\|[\|\|\|\|\|\|\|\|\|\|\|\|]PAM | NA | Intergenic | NA | NA |

T18967 GAGATGGGATCATACTGATGAGG 1000
Oligo fwd CACCGAGATGGGATCATACTGATG
Oligo rev AAACCATCAGTATGATCCCATCTC
Chromosome start end strand MM target_seq PAM alignment distance position gene name gene id
GU305914.1 205326 205348 - 0 GAGATGGGATCATACTGATG AGG ||||||||[||||||||||||]PAM NA Intergenic NA NA

T18968 AGATGGGATCATACTGATGAGGG 1000
Oligo fwd CACCAGATGGGATCATACTGATGA
Oligo rev AAACTCATCAGTATGATCCCATCT
Chromosome start end strand MM target_seq PAM alignment distance position gene name gene id
GU305914.1 205325 205347 - 0 AGATGGGATCATACTGATGA GGG ||||||||[||||||||||||]PAM NA Intergenic NA NA

T18969 GGAACCGTCACTGACAGTTACGG 1000
Oligo fwd CACCGGAACCGTCACTGACAGTTA
Oligo rev AAACTAACTGTCAGTGACGGTTCC
Chromosome start end strand MM target_seq PAM alignment distance position gene name gene id
GU305914.1 205304 205326 - 0 GGAACCGTCACTGACAGTTA CGG ||||||||[||||||||||||]PAM NA Intergenic NA NA

T18970 CCGTCACTGACAGTTACGGATGG 1000
Oligo fwd CACCCCGTCACTGACAGTTACGGA
Oligo rev AAACTCCGTAACTGTCAGTGACGG
Chromosome start end strand MM target_seq PAM alignment distance position gene name gene id
GU305914.1 205300 205322 - 0 CCGTCACTGACAGTTACGGA TGG ||||||||[||||||||||||]PAM NA Intergenic NA NA

T18971 CGTCACTGACAGTTACGGATGGG 1000
Oligo fwd CACCCGTCACTGACAGTTACGGAT
Oligo rev AAACATCCGTAACTGTCAGTGACG
Chromosome start end strand MM target_seq PAM alignment distance position gene name gene id
GU305914.1 205299 205321 - 0 CGTCACTGACAGTTACGGAT GGG ||||||||[||||||||||||]PAM NA Intergenic NA NA

T18972 CCAGCGAGAATTTCTCAAAACGG 1000
Oligo fwd CACCCCAGCGAGAATTTCTCAAAA
Oligo rev AAACTTTTGAGAAATTCTCGCTGG
Chromosome start end strand MM target_seq PAM alignment distance position gene name gene id
GU305914.1 205233 205255 - 0 CCAGCGAGAATTTCTCAAAA CGG ||||||||[||||||||||||]PAM NA Intergenic NA NA

T18973 GAGAATTTCTCAAAACGGTTCGG 1000
Oligo fwd CACCGAGAATTTCTCAAAACGGTT
Oligo rev AAACAACCGTTTTGAGAAATTCTC
Chromosome start end strand MM target_seq PAM alignment distance position gene name gene id
GU305914.1 205228 205250 - 0 GAGAATTTCTCAAAACGGTT CGG ||||||||[||||||||||||]PAM NA Intergenic NA NA

T18974 AGAATTTCTCAAAACGGTTCGGG 1000
Oligo fwd CACCAGAATTTCTCAAAACGGTTC
Oligo rev AAACGAACCGTTTTGAGAAATTCT
Chromosome start end strand MM target_seq PAM alignment distance position gene name gene id
GU305914.1 205227 205249 - 0 AGAATTTCTCAAAACGGTTC GGG ||||||||[||||||||||||]PAM NA Intergenic NA NA

T18975 ATTTCTCAAAACGGTTCGGGCGG 1000
Oligo fwd CACCATTTCTCAAAACGGTTCGGG
Oligo rev AAACCCCGAACCGTTTTGAGAAAT
Chromosome start end strand MM target_seq PAM alignment distance position gene name gene id
GU305914.1 205224 205246 - 0 ATTTCTCAAAACGGTTCGGG CGG ||||||||[||||||||||||]PAM NA Intergenic NA NA

T18976 AACGGTTCGGGCGGTTCTGATGG 1000
Oligo fwd CACCAACGGTTCGGGCGGTTCTGA
Oligo rev AAACTCAGAACCGCCCGAACCGTT
Chromosome start end strand MM target_seq PAM alignment distance position gene name gene id
GU305914.1 205215 205237 - 0 AACGGTTCGGGCGGTTCTGA TGG ||||||||[||||||||||||]PAM NA Intergenic NA NA

T18977 TTCTGATGGAACTGAGATCACGG 1000
Oligo fwd CACCTTCTGATGGAACTGAGATCA
Oligo rev AAACTGATCTCAGTTCCATCAGAA
Chromosome start end strand MM target_seq PAM alignment distance position gene name gene id
GU305914.1 205201 205223 - 0 TTCTGATGGAACTGAGATCA CGG ||||||||[||||||||||||]PAM NA Intergenic NA NA

T18978 ATGGAACTGAGATCACGGTGTGG 1000
Oligo fwd CACCATGGAACTGAGATCACGGTG
Oligo rev AAACCACCGTGATCTCAGTTCCAT
Chromosome start end strand MM target_seq PAM alignment distance position gene name gene id
GU305914.1 205196 205218 - 0 ATGGAACTGAGATCACGGTG TGG ||||||||[||||||||||||]PAM NA Intergenic NA NA

T18979 GTACCTTCAACGCTACATTCTGG 1000
Oligo fwd CACCGTACCTTCAACGCTACATTC
Oligo rev AAACGAATGTAGCGTTGAAGGTAC
Chromosome start end strand MM target_seq PAM alignment distance position gene name gene id
GU305914.1 205159 205181 - 0 GTACCTTCAACGCTACATTC TGG ||||||||[||||||||||||]PAM NA Intergenic NA NA

T18980 TTCAACGCTACATTCTGGTACGG 1000
Oligo fwd CACCTTCAACGCTACATTCTGGTA
Oligo rev AAACTACCAGAATGTAGCGTTGAA
Chromosome start end strand MM target_seq PAM alignment distance position gene name gene id
GU305914.1 205154 205176 - 0 TTCAACGCTACATTCTGGTA CGG ||||||||[||||||||||||]PAM NA Intergenic NA NA

T18981 TCAACGCTACATTCTGGTACGGG 1000
Oligo fwd CACCTCAACGCTACATTCTGGTAC
Oligo rev AAACGTACCAGAATGTAGCGTTGA
Chromosome start end strand MM target_seq PAM alignment distance position gene name gene id
GU305914.1 205153 205175 - 0 TCAACGCTACATTCTGGTAC GGG ||||||||[||||||||||||]PAM NA Intergenic NA NA

T18982 CGTTTTGTTGAAATATAATCCGG 1000
Oligo fwd CACCCGTTTTGTTGAAATATAATC
Oligo rev AAACGATTATATTTCAACAAAACG
Chromosome start end strand MM target_seq PAM alignment distance position gene name gene id
GU305914.1 205101 205123 - 0 CGTTTTGTTGAAATATAATC CGG ||||||||[||||||||||||]PAM NA Intergenic NA NA

T18983 TATAATCCGGCTTTTGTGAATGG 1000
Oligo fwd CACCTATAATCCGGCTTTTGTGAA
Oligo rev AAACTTCACAAAAGCCGGATTATA
Chromosome start end strand MM target_seq PAM alignment distance position gene name gene id
GU305914.1 205088 205110 - 0 TATAATCCGGCTTTTGTGAA TGG ||||||||[||||||||||||]PAM NA Intergenic NA NA

T18984 ATAATCCGGCTTTTGTGAATGGG 1000
Oligo fwd CACCATAATCCGGCTTTTGTGAAT
Oligo rev AAACATTCACAAAAGCCGGATTAT
Chromosome start end strand MM target_seq PAM alignment distance position gene name gene id
GU305914.1 205087 205109 - 0 ATAATCCGGCTTTTGTGAAT GGG ||||||||[||||||||||||]PAM NA Intergenic NA NA

T18985 AAAATTGAAGAACTACGCTGAGG 1000
Oligo fwd CACCAAAATTGAAGAACTACGCTG
Oligo rev AAACCAGCGTAGTTCTTCAATTTT
Chromosome start end strand MM target_seq PAM alignment distance position gene name gene id
GU305914.1 205029 205051 - 0 AAAATTGAAGAACTACGCTG AGG ||||||||[||||||||||||]PAM NA Intergenic NA NA

T18986 AAATTGAAGAACTACGCTGAGGG 1000
Oligo fwd CACCAAATTGAAGAACTACGCTGA
Oligo rev AAACTCAGCGTAGTTCTTCAATTT
Chromosome start end strand MM target_seq PAM alignment distance position gene name gene id
GU305914.1 205028 205050 - 0 AAATTGAAGAACTACGCTGA GGG ||||||||[||||||||||||]PAM NA Intergenic NA NA

T18987 TGAAGAACTACGCTGAGGGACGG 1000
Oligo fwd CACCTGAAGAACTACGCTGAGGGA
Oligo rev AAACTCCCTCAGCGTAGTTCTTCA
Chromosome start end strand MM target_seq PAM alignment distance position gene name gene id
GU305914.1 205024 205046 - 0 TGAAGAACTACGCTGAGGGA CGG ||||||||[||||||||||||]PAM NA Intergenic NA NA

T18988 GAAGAACTACGCTGAGGGACGGG 1000
Oligo fwd CACCGAAGAACTACGCTGAGGGAC
Oligo rev AAACGTCCCTCAGCGTAGTTCTTC
Chromosome start end strand MM target_seq PAM alignment distance position gene name gene id
GU305914.1 205023 205045 - 0 GAAGAACTACGCTGAGGGAC GGG ||||||||[||||||||||||]PAM NA Intergenic NA NA

T18989 CACCGATCTACTGAAAAACAAGG 1000
Oligo fwd CACCCACCGATCTACTGAAAAACA
Oligo rev AAACTGTTTTTCAGTAGATCGGTG
Chromosome start end strand MM target_seq PAM alignment distance position gene name gene id
GU305914.1 204999 205021 - 0 CACCGATCTACTGAAAAACA AGG ||||||||[||||||||||||]PAM NA Intergenic NA NA

T18990 AGGTCACAAGTCTCTACCCCAGG 1000
Oligo fwd CACCAGGTCACAAGTCTCTACCCC
Oligo rev AAACGGGGTAGAGACTTGTGACCT
Chromosome start end strand MM target_seq PAM alignment distance position gene name gene id
GU305914.1 204979 205001 - 0 AGGTCACAAGTCTCTACCCC AGG ||||||||[||||||||||||]PAM NA Intergenic NA NA

T18991 TCACAAGTCTCTACCCCAGGTGG 1000
Oligo fwd CACCTCACAAGTCTCTACCCCAGG
Oligo rev AAACCCTGGGGTAGAGACTTGTGA
Chromosome start end strand MM target_seq PAM alignment distance position gene name gene id
GU305914.1 204976 204998 - 0 TCACAAGTCTCTACCCCAGG TGG ||||||||[||||||||||||]PAM NA Intergenic NA NA

T18992 TACAAAGCCGCTGCCAAACTCGG 1000
Oligo fwd CACCTACAAAGCCGCTGCCAAACT
Oligo rev AAACAGTTTGGCAGCGGCTTTGTA
Chromosome start end strand MM target_seq PAM alignment distance position gene name gene id
GU305914.1 204944 204966 - 0 TACAAAGCCGCTGCCAAACT CGG ||||||||[||||||||||||]PAM NA Intergenic NA NA

T18993 AAAGCCGCTGCCAAACTCGGCGG 1000
Oligo fwd CACCAAAGCCGCTGCCAAACTCGG
Oligo rev AAACCCGAGTTTGGCAGCGGCTTT
Chromosome start end strand MM target_seq PAM alignment distance position gene name gene id
GU305914.1 204941 204963 - 0 AAAGCCGCTGCCAAACTCGG CGG ||||||||[||||||||||||]PAM NA Intergenic NA NA

T18994 GAACAACACAACGATCTGCGAGG 1000
Oligo fwd CACCGAACAACACAACGATCTGCG
Oligo rev AAACCGCAGATCGTTGTGTTGTTC
Chromosome start end strand MM target_seq PAM alignment distance position gene name gene id
GU305914.1 204885 204907 - 0 GAACAACACAACGATCTGCG AGG ||||||||[||||||||||||]PAM NA Intergenic NA NA

T18995 ACACAACGATCTGCGAGGTGTGG 1000
Oligo fwd CACCACACAACGATCTGCGAGGTG
Oligo rev AAACCACCTCGCAGATCGTTGTGT
Chromosome start end strand MM target_seq PAM alignment distance position gene name gene id
GU305914.1 204880 204902 - 0 ACACAACGATCTGCGAGGTG TGG ||||||||[||||||||||||]PAM NA Intergenic NA NA

T18996 CTATTTTTCTATAGAGATAGAGG 1000
Oligo fwd CACCCTATTTTTCTATAGAGATAG
Oligo rev AAACCTATCTCTATAGAAAAATAG
Chromosome start end strand MM target_seq PAM alignment distance position gene name gene id

GU305914.1 204843 204865 - 0 CTATTTTTCTATAGAGATAG AGG ||||||||[||||||||||||]PAM NA Intergenic NA NA

T18997 TCTATAGAGATAGAGGTCGCCGG 1000
Oligo fwd CACCTCTATAGAGATAGAGGTCGC
Oligo rev AAACGCGACCTCTATCTCTATAGA
Chromosome start end strand MM target_seq PAM alignment distance position gene name gene id
GU305914.1 204836 204858 - 0 TCTATAGAGATAGAGGTCGC CGG ||||||||[||||||||||||]PAM NA Intergenic NA NA

T18998 GCCGGACTCCCTCCCGTAAAAGG 1000
Oligo fwd CACCGCCGGACTCCCTCCCGTAAA
Oligo rev AAACTTTACGGGAGGGAGTCCGGC
Chromosome start end strand MM target_seq PAM alignment distance position gene name gene id
GU305914.1 204818 204840 - 0 GCCGGACTCCCTCCCGTAAA AGG ||||||||[||||||||||||]PAM NA Intergenic NA NA

T18999 AAGGCGAAGTATCCAAGAGCTGG 1000
Oligo fwd CACCAAGGCGAAGTATCCAAGAGC
Oligo rev AAACGCTCTTGGATACTTCGCCTT
Chromosome start end strand MM target_seq PAM alignment distance position gene name gene id
GU305914.1 204799 204821 - 0 AAGGCGAAGTATCCAAGAGC TGG ||||||||[||||||||||||]PAM NA Intergenic NA NA

T19000 CTAAGAGCATCTGTAAGATCCGG 1000
Oligo fwd CACCCTAAGAGCATCTGTAAGATC
Oligo rev AAACGATCTTACAGATGCTCTTAG
Chromosome start end strand MM target_seq PAM alignment distance position gene name gene id
GU305914.1 204727 204749 - 0 CTAAGAGCATCTGTAAGATC CGG ||||||||[||||||||||||]PAM NA Intergenic NA NA

T19001 CATCTGTAAGATCCGGTCTTCGG 1000
Oligo fwd CACCCATCTGTAAGATCCGGTCTT
Oligo rev AAACAAGACCGGATCTTACAGATG
Chromosome start end strand MM target_seq PAM alignment distance position gene name gene id
GU305914.1 204720 204742 - 0 CATCTGTAAGATCCGGTCTT CGG ||||||||[||||||||||||]PAM NA Intergenic NA NA

T19002 AAGATCCGGTCTTCGGTTCGCGG 1000
Oligo fwd CACCAAGATCCGGTCTTCGGTTCG
Oligo rev AAACCGAACCGAAGACCGGATCTT
Chromosome start end strand MM target_seq PAM alignment distance position gene name gene id
GU305914.1 204713 204735 - 0 AAGATCCGGTCTTCGGTTCG CGG ||||||||[||||||||||||]PAM NA Intergenic NA NA

T19003 TTCGGTTCGCGGAGCTTGTAAGG 1000
Oligo fwd CACCTTCGGTTCGCGGAGCTTGTA
Oligo rev AAACTACAAGCTCCGCGAACCGAA
Chromosome start end strand MM target_seq PAM alignment distance position gene name gene id
GU305914.1 204702 204724 - 0 TTCGGTTCGCGGAGCTTGTA AGG ||||||||[||||||||||||]PAM NA Intergenic NA NA

T19004 CACGACGTCCCCGACTACCACGG 1000
Oligo fwd CACCCACGACGTCCCCGACTACCA
Oligo rev AAACTGGTAGTCGGGGACGTCGTG
Chromosome start end strand MM target_seq PAM alignment distance position gene name gene id
GU305914.1 204645 204667 - 0 CACGACGTCCCCGACTACCA CGG ||||||||[||||||||||||]PAM NA Intergenic NA NA

T19005 CTCTTGACCGTCAGTACTCTAGG 1000
Oligo fwd CACCCTCTTGACCGTCAGTACTCT
Oligo rev AAACAGAGTACTGACGGTCAAGAG
Chromosome start end strand MM target_seq PAM alignment distance position gene name gene id
GU305914.1 204539 204561 - 0 CTCTTGACCGTCAGTACTCT AGG ||||||||[||||||||||||]PAM NA Intergenic NA NA

T19006 TCAGTACTCTAGGACCGACAAGG 1000
Oligo fwd CACCTCAGTACTCTAGGACCGACA
Oligo rev AAACTGTCGGTCCTAGAGTACTGA
Chromosome start end strand MM target_seq PAM alignment distance position gene name gene id
GU305914.1 204529 204551 - 0 TCAGTACTCTAGGACCGACA AGG ||||||||[||||||||||||]PAM NA Intergenic NA NA

T19007 AAGGCCGACTATCATCCCGACGG 1000
Oligo fwd CACCAAGGCCGACTATCATCCCGA
Oligo rev AAACTCGGGATGATAGTCGGCCTT
Chromosome start end strand MM target_seq PAM alignment distance position gene name gene id
GU305914.1 204510 204532 - 0 AAGGCCGACTATCATCCCGA CGG ||||||||[||||||||||||]PAM NA Intergenic NA NA

T19008 TTGACATGATATCGAGTGTCAGG 1000
Oligo fwd CACCTTGACATGATATCGAGTGTC
Oligo rev AAACGACACTCGATATCATGTCAA
Chromosome start end strand MM target_seq PAM alignment distance position gene name gene id
GU305914.1 204472 204494 - 0 TTGACATGATATCGAGTGTC AGG ||||||||[||||||||||||]PAM NA Intergenic NA NA

T19009 TACCAAGACCGCGACCGTTGAGG 1000
Oligo fwd CACCTACCAAGACCGCGACCGTTG
Oligo rev AAACCAACGGTCGCGGTCTTGGTA
Chromosome start end strand MM target_seq PAM alignment distance position gene name gene id
GU305914.1 204441 204463 - 0 TACCAAGACCGCGACCGTTG AGG ||||||||[||||||||||||]PAM NA Intergenic NA NA

T19010 TGAGGTCTCTACGAACATCGAGG 1000
Oligo fwd CACCTGAGGTCTCTACGAACATCG
Oligo rev AAACCGATGTTCGTAGAGACCTCA
Chromosome start end strand MM target_seq PAM alignment distance position gene name gene id
GU305914.1 204423 204445 - 0 TGAGGTCTCTACGAACATCG AGG ||||||||[||||||||||||]PAM NA Intergenic NA NA

T19011 CTCTACGAACATCGAGGTCTCGG 1000
Oligo fwd CACCCTCTACGAACATCGAGGTCT
Oligo rev AAACAGACCTCGATGTTCGTAGAG

<table>
<tr><td>Chromosome</td><td>start</td><td>end</td><td>strand</td><td>MM</td><td>target_seq</td><td>PAM</td><td>alignment</td><td>distance</td><td>position</td><td>gene name</td><td>gene id</td></tr>
<tr><td>GU305914.1</td><td>204417</td><td>204439</td><td>-</td><td>0</td><td>CTCTACGAACATCGAGGTCT</td><td>CGG</td><td>||||||||[||||||||||||]PAM</td><td>NA</td><td>Intergenic</td><td>NA</td><td>NA</td></tr>
<tr><td></td><td></td><td></td><td></td><td></td><td></td><td></td><td></td><td></td><td></td><td></td><td></td></tr>
<tr><td>T19012</td><td>CTCGGAAACCACGTCCTCCTCGG</td><td>1000</td><td></td><td></td><td></td><td></td><td></td><td></td><td></td><td></td><td></td></tr>
<tr><td>Oligo fwd</td><td>CACCCTCGGAAACCACGTCCTCCT</td><td></td><td></td><td></td><td></td><td></td><td></td><td></td><td></td><td></td><td></td></tr>
<tr><td>Oligo rev</td><td>AAACAGGAGGACGTGGTTTCCGAG</td><td></td><td></td><td></td><td></td><td></td><td></td><td></td><td></td><td></td><td></td></tr>
<tr><td>Chromosome</td><td>start</td><td>end</td><td>strand</td><td>MM</td><td>target_seq</td><td>PAM</td><td>alignment</td><td>distance</td><td>position</td><td>gene name</td><td>gene id</td></tr>
<tr><td>GU305914.1</td><td>204399</td><td>204421</td><td>-</td><td>0</td><td>CTCGGAAACCACGTCCTCCT</td><td>CGG</td><td>||||||||[||||||||||||]PAM</td><td>NA</td><td>Intergenic</td><td>NA</td><td>NA</td></tr>
<tr><td></td><td></td><td></td><td></td><td></td><td></td><td></td><td></td><td></td><td></td><td></td><td></td></tr>
<tr><td>T19013</td><td>TTTAATGATCGATGTCTTGACGG</td><td>1000</td><td></td><td></td><td></td><td></td><td></td><td></td><td></td><td></td><td></td></tr>
<tr><td>Oligo fwd</td><td>CACCTTTAATGATCGATGTCTTGA</td><td></td><td></td><td></td><td></td><td></td><td></td><td></td><td></td><td></td><td></td></tr>
<tr><td>Oligo rev</td><td>AAACTCAAGACATCGATCATTAAA</td><td></td><td></td><td></td><td></td><td></td><td></td><td></td><td></td><td></td><td></td></tr>
<tr><td>Chromosome</td><td>start</td><td>end</td><td>strand</td><td>MM</td><td>target_seq</td><td>PAM</td><td>alignment</td><td>distance</td><td>position</td><td>gene name</td><td>gene id</td></tr>
<tr><td>GU305914.1</td><td>204366</td><td>204388</td><td>-</td><td>0</td><td>TTTAATGATCGATGTCTTGA</td><td>CGG</td><td>||||||||[||||||||||||]PAM</td><td>NA</td><td>Intergenic</td><td>NA</td><td>NA</td></tr>
<tr><td></td><td></td><td></td><td></td><td></td><td></td><td></td><td></td><td></td><td></td><td></td><td></td></tr>
<tr><td>T19014</td><td>TTAATGATCGATGTCTTGACGGG</td><td>1000</td><td></td><td></td><td></td><td></td><td></td><td></td><td></td><td></td><td></td></tr>
<tr><td>Oligo fwd</td><td>CACCTTAATGATCGATGTCTTGAC</td><td></td><td></td><td></td><td></td><td></td><td></td><td></td><td></td><td></td><td></td></tr>
<tr><td>Oligo rev</td><td>AAACGTCAAGACATCGATCATTAA</td><td></td><td></td><td></td><td></td><td></td><td></td><td></td><td></td><td></td><td></td></tr>
<tr><td>Chromosome</td><td>start</td><td>end</td><td>strand</td><td>MM</td><td>target_seq</td><td>PAM</td><td>alignment</td><td>distance</td><td>position</td><td>gene name</td><td>gene id</td></tr>
<tr><td>GU305914.1</td><td>204365</td><td>204387</td><td>-</td><td>0</td><td>TTAATGATCGATGTCTTGAC</td><td>GGG</td><td>||||||||[||||||||||||]PAM</td><td>NA</td><td>Intergenic</td><td>NA</td><td>NA</td></tr>
<tr><td></td><td></td><td></td><td></td><td></td><td></td><td></td><td></td><td></td><td></td><td></td><td></td></tr>
<tr><td>T19015</td><td>GTCTTGACGGGTATCGTAACTGG</td><td>1000</td><td></td><td></td><td></td><td></td><td></td><td></td><td></td><td></td><td></td></tr>
<tr><td>Oligo fwd</td><td>CACCGTCTTGACGGGTATCGTAAC</td><td></td><td></td><td></td><td></td><td></td><td></td><td></td><td></td><td></td><td></td></tr>
<tr><td>Oligo rev</td><td>AAACGTTACGATACCCGTCAAGAC</td><td></td><td></td><td></td><td></td><td></td><td></td><td></td><td></td><td></td><td></td></tr>
<tr><td>Chromosome</td><td>start</td><td>end</td><td>strand</td><td>MM</td><td>target_seq</td><td>PAM</td><td>alignment</td><td>distance</td><td>position</td><td>gene name</td><td>gene id</td></tr>
<tr><td>GU305914.1</td><td>204353</td><td>204375</td><td>-</td><td>0</td><td>GTCTTGACGGGTATCGTAAC</td><td>TGG</td><td>||||||||[||||||||||||]PAM</td><td>NA</td><td>Intergenic</td><td>NA</td><td>NA</td></tr>
<tr><td></td><td></td><td></td><td></td><td></td><td></td><td></td><td></td><td></td><td></td><td></td><td></td></tr>
<tr><td>T19016</td><td>GGTATCGTAACTGGCGAGCAAGG</td><td>1000</td><td></td><td></td><td></td><td></td><td></td><td></td><td></td><td></td><td></td></tr>
<tr><td>Oligo fwd</td><td>CACCGGTATCGTAACTGGCGAGCA</td><td></td><td></td><td></td><td></td><td></td><td></td><td></td><td></td><td></td><td></td></tr>
<tr><td>Oligo rev</td><td>AAACTGCTCGCCAGTTACGATACC</td><td></td><td></td><td></td><td></td><td></td><td></td><td></td><td></td><td></td><td></td></tr>
<tr><td>Chromosome</td><td>start</td><td>end</td><td>strand</td><td>MM</td><td>target_seq</td><td>PAM</td><td>alignment</td><td>distance</td><td>position</td><td>gene name</td><td>gene id</td></tr>
<tr><td>GU305914.1</td><td>204344</td><td>204366</td><td>-</td><td>0</td><td>GGTATCGTAACTGGCGAGCA</td><td>AGG</td><td>||||||||[||||||||||||]PAM</td><td>NA</td><td>Intergenic</td><td>NA</td><td>NA</td></tr>
<tr><td></td><td></td><td></td><td></td><td></td><td></td><td></td><td></td><td></td><td></td><td></td><td></td></tr>
<tr><td>T19017</td><td>CGAGCAAGGAAAATATTACTCGG</td><td>1000</td><td></td><td></td><td></td><td></td><td></td><td></td><td></td><td></td><td></td></tr>
<tr><td>Oligo fwd</td><td>CACCCGAGCAAGGAAAATATTACT</td><td></td><td></td><td></td><td></td><td></td><td></td><td></td><td></td><td></td><td></td></tr>
<tr><td>Oligo rev</td><td>AAACAGTAATATTTTCCTTGCTCG</td><td></td><td></td><td></td><td></td><td></td><td></td><td></td><td></td><td></td><td></td></tr>
<tr><td>Chromosome</td><td>start</td><td>end</td><td>strand</td><td>MM</td><td>target_seq</td><td>PAM</td><td>alignment</td><td>distance</td><td>position</td><td>gene name</td><td>gene id</td></tr>
<tr><td>GU305914.1</td><td>204330</td><td>204352</td><td>-</td><td>0</td><td>CGAGCAAGGAAAATATTACT</td><td>CGG</td><td>||||||||[||||||||||||]PAM</td><td>NA</td><td>Intergenic</td><td>NA</td><td>NA</td></tr>
<tr><td></td><td></td><td></td><td></td><td></td><td></td><td></td><td></td><td></td><td></td><td></td><td></td></tr>
<tr><td>T19018</td><td>GAGCAAGGAAAATATTACTCGGG</td><td>1000</td><td></td><td></td><td></td><td></td><td></td><td></td><td></td><td></td><td></td></tr>
<tr><td>Oligo fwd</td><td>CACCGAGCAAGGAAAATATTACTC</td><td></td><td></td><td></td><td></td><td></td><td></td><td></td><td></td><td></td><td></td></tr>
<tr><td>Oligo rev</td><td>AAACGAGTAATATTTTCCTTGCTC</td><td></td><td></td><td></td><td></td><td></td><td></td><td></td><td></td><td></td><td></td></tr>
<tr><td>Chromosome</td><td>start</td><td>end</td><td>strand</td><td>MM</td><td>target_seq</td><td>PAM</td><td>alignment</td><td>distance</td><td>position</td><td>gene name</td><td>gene id</td></tr>
<tr><td>GU305914.1</td><td>204329</td><td>204351</td><td>-</td><td>0</td><td>GAGCAAGGAAAATATTACTC</td><td>GGG</td><td>||||||||[||||||||||||]PAM</td><td>NA</td><td>Intergenic</td><td>NA</td><td>NA</td></tr>
<tr><td></td><td></td><td></td><td></td><td></td><td></td><td></td><td></td><td></td><td></td><td></td><td></td></tr>
<tr><td>T19019</td><td>AAAATATTACTCGGGCTATTCGG</td><td>1000</td><td></td><td></td><td></td><td></td><td></td><td></td><td></td><td></td><td></td></tr>
<tr><td>Oligo fwd</td><td>CACCAAAATATTACTCGGGCTATT</td><td></td><td></td><td></td><td></td><td></td><td></td><td></td><td></td><td></td><td></td></tr>
<tr><td>Oligo rev</td><td>AAACAATAGCCCGAGTAATATTTT</td><td></td><td></td><td></td><td></td><td></td><td></td><td></td><td></td><td></td><td></td></tr>
<tr><td>Chromosome</td><td>start</td><td>end</td><td>strand</td><td>MM</td><td>target_seq</td><td>PAM</td><td>alignment</td><td>distance</td><td>position</td><td>gene name</td><td>gene id</td></tr>
<tr><td>GU305914.1</td><td>204321</td><td>204343</td><td>-</td><td>0</td><td>AAAATATTACTCGGGCTATT</td><td>CGG</td><td>||||||||[||||||||||||]PAM</td><td>NA</td><td>Intergenic</td><td>NA</td><td>NA</td></tr>
<tr><td></td><td></td><td></td><td></td><td></td><td></td><td></td><td></td><td></td><td></td><td></td><td></td></tr>
<tr><td>T19020</td><td>TTTAGTCGCCATGTTCGTCGCGG</td><td>1000</td><td></td><td></td><td></td><td></td><td></td><td></td><td></td><td></td><td></td></tr>
<tr><td>Oligo fwd</td><td>CACCTTTAGTCGCCATGTTCGTCG</td><td></td><td></td><td></td><td></td><td></td><td></td><td></td><td></td><td></td><td></td></tr>
<tr><td>Oligo rev</td><td>AAACCGACGAACATGGCGACTAAA</td><td></td><td></td><td></td><td></td><td></td><td></td><td></td><td></td><td></td><td></td></tr>
<tr><td>Chromosome</td><td>start</td><td>end</td><td>strand</td><td>MM</td><td>target_seq</td><td>PAM</td><td>alignment</td><td>distance</td><td>position</td><td>gene name</td><td>gene id</td></tr>
<tr><td>GU305914.1</td><td>204285</td><td>204307</td><td>-</td><td>0</td><td>TTTAGTCGCCATGTTCGTCG</td><td>CGG</td><td>||||||||[||||||||||||]PAM</td><td>NA</td><td>Intergenic</td><td>NA</td><td>NA</td></tr>
<tr><td></td><td></td><td></td><td></td><td></td><td></td><td></td><td></td><td></td><td></td><td></td><td></td></tr>
<tr><td>T19021</td><td>CGCCATGTTCGTCGCGGTAACGG</td><td>1000</td><td></td><td></td><td></td><td></td><td></td><td></td><td></td><td></td><td></td></tr>
<tr><td>Oligo fwd</td><td>CACCCGCCATGTTCGTCGCGGTAA</td><td></td><td></td><td></td><td></td><td></td><td></td><td></td><td></td><td></td><td></td></tr>
<tr><td>Oligo rev</td><td>AAACTTACCGCGACGAACATGGCG</td><td></td><td></td><td></td><td></td><td></td><td></td><td></td><td></td><td></td><td></td></tr>
<tr><td>Chromosome</td><td>start</td><td>end</td><td>strand</td><td>MM</td><td>target_seq</td><td>PAM</td><td>alignment</td><td>distance</td><td>position</td><td>gene name</td><td>gene id</td></tr>
<tr><td>GU305914.1</td><td>204279</td><td>204301</td><td>-</td><td>0</td><td>CGCCATGTTCGTCGCGGTAA</td><td>CGG</td><td>||||||||[||||||||||||]PAM</td><td>NA</td><td>Intergenic</td><td>NA</td><td>NA</td></tr>
<tr><td></td><td></td><td></td><td></td><td></td><td></td><td></td><td></td><td></td><td></td><td></td><td></td></tr>
<tr><td>T19022</td><td>GGCCCTGTGCATCTGTGCAATGG</td><td>1000</td><td></td><td></td><td></td><td></td><td></td><td></td><td></td><td></td><td></td></tr>
<tr><td>Oligo fwd</td><td>CACCGGCCCTGTGCATCTGTGCAA</td><td></td><td></td><td></td><td></td><td></td><td></td><td></td><td></td><td></td><td></td></tr>
<tr><td>Oligo rev</td><td>AAACTTGCACAGATGCACAGGGCC</td><td></td><td></td><td></td><td></td><td></td><td></td><td></td><td></td><td></td><td></td></tr>
<tr><td>Chromosome</td><td>start</td><td>end</td><td>strand</td><td>MM</td><td>target_seq</td><td>PAM</td><td>alignment</td><td>distance</td><td>position</td><td>gene name</td><td>gene id</td></tr>
<tr><td>GU305914.1</td><td>204258</td><td>204280</td><td>-</td><td>0</td><td>GGCCCTGTGCATCTGTGCAA</td><td>TGG</td><td>||||||||[||||||||||||]PAM</td><td>NA</td><td>Intergenic</td><td>NA</td><td>NA</td></tr>
<tr><td></td><td></td><td></td><td></td><td></td><td></td><td></td><td></td><td></td><td></td><td></td><td></td></tr>
<tr><td>T19023</td><td>GAGCCGCACAAAACTACGTTCGG</td><td>1000</td><td></td><td></td><td></td><td></td><td></td><td></td><td></td><td></td><td></td></tr>
<tr><td>Oligo fwd</td><td>CACCGAGCCGCACAAAACTACGTT</td><td></td><td></td><td></td><td></td><td></td><td></td><td></td><td></td><td></td><td></td></tr>
<tr><td>Oligo rev</td><td>AAACAACGTAGTTTTGTGCGGCTC</td><td></td><td></td><td></td><td></td><td></td><td></td><td></td><td></td><td></td><td></td></tr>
<tr><td>Chromosome</td><td>start</td><td>end</td><td>strand</td><td>MM</td><td>target_seq</td><td>PAM</td><td>alignment</td><td>distance</td><td>position</td><td>gene name</td><td>gene id</td></tr>
<tr><td>GU305914.1</td><td>204210</td><td>204232</td><td>-</td><td>0</td><td>GAGCCGCACAAAACTACGTT</td><td>CGG</td><td>||||||||[||||||||||||]PAM</td><td>NA</td><td>Intergenic</td><td>NA</td><td>NA</td></tr>
<tr><td></td><td></td><td></td><td></td><td></td><td></td><td></td><td></td><td></td><td></td><td></td><td></td></tr>
<tr><td>T19024</td><td>AGCCGCACAAAACTACGTTCGGG</td><td>1000</td><td></td><td></td><td></td><td></td><td></td><td></td><td></td><td></td><td></td></tr>
<tr><td>Oligo fwd</td><td>CACCAGCCGCACAAAACTACGTTC</td><td></td><td></td><td></td><td></td><td></td><td></td><td></td><td></td><td></td><td></td></tr>
<tr><td>Oligo rev</td><td>AAACGAACGTAGTTTTGTGCGGCT</td><td></td><td></td><td></td><td></td><td></td><td></td><td></td><td></td><td></td><td></td></tr>
<tr><td>Chromosome</td><td>start</td><td>end</td><td>strand</td><td>MM</td><td>target_seq</td><td>PAM</td><td>alignment</td><td>distance</td><td>position</td><td>gene name</td><td>gene id</td></tr>
<tr><td>GU305914.1</td><td>204209</td><td>204231</td><td>-</td><td>0</td><td>AGCCGCACAAAACTACGTTC</td><td>GGG</td><td>||||||||[||||||||||||]PAM</td><td>NA</td><td>Intergenic</td><td>NA</td><td>NA</td></tr>
<tr><td></td><td></td><td></td><td></td><td></td><td></td><td></td><td></td><td></td><td></td><td></td><td></td></tr>
<tr><td>T19025</td><td>GGGAGAACTCGACATCGAAAAGG</td><td>1000</td><td></td><td></td><td></td><td></td><td></td><td></td><td></td><td></td><td></td></tr>
<tr><td>Oligo fwd</td><td>CACCGGGAGAACTCGACATCGAAA</td><td></td><td></td><td></td><td></td><td></td><td></td><td></td><td></td><td></td><td></td></tr>
<tr><td>Oligo rev</td><td>AAACTTTCGATGTCGAGTTCTCCC</td><td></td><td></td><td></td><td></td><td></td><td></td><td></td><td></td><td></td><td></td></tr>
<tr><td>Chromosome</td><td>start</td><td>end</td><td>strand</td><td>MM</td><td>target_seq</td><td>PAM</td><td>alignment</td><td>distance</td><td>position</td><td>gene name</td><td>gene id</td></tr>
<tr><td>GU305914.1</td><td>204189</td><td>204211</td><td>-</td><td>0</td><td>GGGAGAACTCGACATCGAAA</td><td>AGG</td><td>||||||||[||||||||||||]PAM</td><td>NA</td><td>Intergenic</td><td>NA</td><td>NA</td></tr>
<tr><td></td><td></td><td></td><td></td><td></td><td></td><td></td><td></td><td></td><td></td><td></td><td></td></tr>
<tr><td>T19026</td><td>TAAAAACAGATTGTCTACTGTGG</td><td>1000</td><td></td><td></td><td></td><td></td><td></td><td></td><td></td><td></td><td></td></tr>
<tr><td>Oligo fwd</td><td>CACCTAAAAACAGATTGTCTACTG</td><td></td><td></td><td></td><td></td><td></td><td></td><td></td><td></td><td></td><td></td></tr>
</table>

<table>
<tr><td>Oligo rev</td><td>AAACCAGTAGACAATCTGTTTTTA</td><td></td><td></td><td></td><td></td><td></td><td></td><td></td><td></td><td></td><td></td></tr>
<tr><td>Chromosome</td><td>start</td><td>end</td><td>strand</td><td>MM</td><td>target_seq</td><td>PAM</td><td>alignment</td><td>distance</td><td>position</td><td>gene name</td><td>gene id</td></tr>
<tr><td>GU305914.1</td><td>204159</td><td>204181</td><td>-</td><td>0</td><td>TAAAAACAGATTGTCTACTG</td><td>TGG</td><td>||||||||[||||||||||||]PAM</td><td>NA</td><td>Intergenic</td><td>NA</td><td>NA</td></tr>
<tr><td>T19027</td><td>GATTGTCTACTGTGGTGTCTTGG</td><td>1000</td><td></td><td></td><td></td><td></td><td></td><td></td><td></td><td></td><td></td></tr>
<tr><td>Oligo fwd</td><td>CACCGATTGTCTACTGTGGTGTCT</td><td></td><td></td><td></td><td></td><td></td><td></td><td></td><td></td><td></td><td></td></tr>
<tr><td>Oligo rev</td><td>AAACAGACACCACAGTAGACAATC</td><td></td><td></td><td></td><td></td><td></td><td></td><td></td><td></td><td></td><td></td></tr>
<tr><td>Chromosome</td><td>start</td><td>end</td><td>strand</td><td>MM</td><td>target_seq</td><td>PAM</td><td>alignment</td><td>distance</td><td>position</td><td>gene name</td><td>gene id</td></tr>
<tr><td>GU305914.1</td><td>204151</td><td>204173</td><td>-</td><td>0</td><td>GATTGTCTACTGTGGTGTCT</td><td>TGG</td><td>||||||||[||||||||||||]PAM</td><td>NA</td><td>Intergenic</td><td>NA</td><td>NA</td></tr>
<tr><td>T19028</td><td>GTCTTGGTATAAGACGATAGAGG</td><td>1000</td><td></td><td></td><td></td><td></td><td></td><td></td><td></td><td></td><td></td></tr>
<tr><td>Oligo fwd</td><td>CACCGTCTTGGTATAAGACGATAG</td><td></td><td></td><td></td><td></td><td></td><td></td><td></td><td></td><td></td><td></td></tr>
<tr><td>Oligo rev</td><td>AAACCTATCGTCTTATACCAAGAC</td><td></td><td></td><td></td><td></td><td></td><td></td><td></td><td></td><td></td><td></td></tr>
<tr><td>Chromosome</td><td>start</td><td>end</td><td>strand</td><td>MM</td><td>target_seq</td><td>PAM</td><td>alignment</td><td>distance</td><td>position</td><td>gene name</td><td>gene id</td></tr>
<tr><td>GU305914.1</td><td>204135</td><td>204157</td><td>-</td><td>0</td><td>GTCTTGGTATAAGACGATAG</td><td>AGG</td><td>||||||||[||||||||||||]PAM</td><td>NA</td><td>Intergenic</td><td>NA</td><td>NA</td></tr>
<tr><td>T19029</td><td>TCTGTCGTGAGACAATAAACAGG</td><td>1000</td><td></td><td></td><td></td><td></td><td></td><td></td><td></td><td></td><td></td></tr>
<tr><td>Oligo fwd</td><td>CACCTCTGTCGTGAGACAATAAAC</td><td></td><td></td><td></td><td></td><td></td><td></td><td></td><td></td><td></td><td></td></tr>
<tr><td>Oligo rev</td><td>AAACGTTTATTGTCTCACGACAGA</td><td></td><td></td><td></td><td></td><td></td><td></td><td></td><td></td><td></td><td></td></tr>
<tr><td>Chromosome</td><td>start</td><td>end</td><td>strand</td><td>MM</td><td>target_seq</td><td>PAM</td><td>alignment</td><td>distance</td><td>position</td><td>gene name</td><td>gene id</td></tr>
<tr><td>GU305914.1</td><td>204088</td><td>204110</td><td>-</td><td>0</td><td>TCTGTCGTGAGACAATAAAC</td><td>AGG</td><td>||||||||[||||||||||||]PAM</td><td>NA</td><td>Intergenic</td><td>NA</td><td>NA</td></tr>
<tr><td>T19030</td><td>CATGAATGTGAGATGTCGCATGG</td><td>1000</td><td></td><td></td><td></td><td></td><td></td><td></td><td></td><td></td><td></td></tr>
<tr><td>Oligo fwd</td><td>CACCCATGAATGTGAGATGTCGCA</td><td></td><td></td><td></td><td></td><td></td><td></td><td></td><td></td><td></td><td></td></tr>
<tr><td>Oligo rev</td><td>AAACTGCGACATCTCACATTCATG</td><td></td><td></td><td></td><td></td><td></td><td></td><td></td><td></td><td></td><td></td></tr>
<tr><td>Chromosome</td><td>start</td><td>end</td><td>strand</td><td>MM</td><td>target_seq</td><td>PAM</td><td>alignment</td><td>distance</td><td>position</td><td>gene name</td><td>gene id</td></tr>
<tr><td>GU305914.1</td><td>204019</td><td>204041</td><td>-</td><td>0</td><td>CATGAATGTGAGATGTCGCA</td><td>TGG</td><td>||||||||[||||||||||||]PAM</td><td>NA</td><td>Intergenic</td><td>NA</td><td>NA</td></tr>
<tr><td>T19031</td><td>ATGTGTGATGTGTAATTATGAGG</td><td>1000</td><td></td><td></td><td></td><td></td><td></td><td></td><td></td><td></td><td></td></tr>
<tr><td>Oligo fwd</td><td>CACCATGTGTGATGTGTAATTATG</td><td></td><td></td><td></td><td></td><td></td><td></td><td></td><td></td><td></td><td></td></tr>
<tr><td>Oligo rev</td><td>AAACCATAATTACACATCACACAT</td><td></td><td></td><td></td><td></td><td></td><td></td><td></td><td></td><td></td><td></td></tr>
<tr><td>Chromosome</td><td>start</td><td>end</td><td>strand</td><td>MM</td><td>target_seq</td><td>PAM</td><td>alignment</td><td>distance</td><td>position</td><td>gene name</td><td>gene id</td></tr>
<tr><td>GU305914.1</td><td>203989</td><td>204011</td><td>-</td><td>0</td><td>ATGTGTGATGTGTAATTATG</td><td>AGG</td><td>||||||||[||||||||||||]PAM</td><td>NA</td><td>Intergenic</td><td>NA</td><td>NA</td></tr>
<tr><td>T19032</td><td>TGTGTGATGTGTAATTATGAGGG</td><td>1000</td><td></td><td></td><td></td><td></td><td></td><td></td><td></td><td></td><td></td></tr>
<tr><td>Oligo fwd</td><td>CACCTGTGTGATGTGTAATTATGA</td><td></td><td></td><td></td><td></td><td></td><td></td><td></td><td></td><td></td><td></td></tr>
<tr><td>Oligo rev</td><td>AAACTCATAATTACACATCACACA</td><td></td><td></td><td></td><td></td><td></td><td></td><td></td><td></td><td></td><td></td></tr>
<tr><td>Chromosome</td><td>start</td><td>end</td><td>strand</td><td>MM</td><td>target_seq</td><td>PAM</td><td>alignment</td><td>distance</td><td>position</td><td>gene name</td><td>gene id</td></tr>
<tr><td>GU305914.1</td><td>203988</td><td>204010</td><td>-</td><td>0</td><td>TGTGTGATGTGTAATTATGA</td><td>GGG</td><td>||||||||[||||||||||||]PAM</td><td>NA</td><td>Intergenic</td><td>NA</td><td>NA</td></tr>
<tr><td>T19033</td><td>ATGTGTAATTATGAGGGCTCTGG</td><td>1000</td><td></td><td></td><td></td><td></td><td></td><td></td><td></td><td></td><td></td></tr>
<tr><td>Oligo fwd</td><td>CACCATGTGTAATTATGAGGGCTC</td><td></td><td></td><td></td><td></td><td></td><td></td><td></td><td></td><td></td><td></td></tr>
<tr><td>Oligo rev</td><td>AAACGAGCCCTCATAATTACACAT</td><td></td><td></td><td></td><td></td><td></td><td></td><td></td><td></td><td></td><td></td></tr>
<tr><td>Chromosome</td><td>start</td><td>end</td><td>strand</td><td>MM</td><td>target_seq</td><td>PAM</td><td>alignment</td><td>distance</td><td>position</td><td>gene name</td><td>gene id</td></tr>
<tr><td>GU305914.1</td><td>203982</td><td>204004</td><td>-</td><td>0</td><td>ATGTGTAATTATGAGGGCTC</td><td>TGG</td><td>||||||||[||||||||||||]PAM</td><td>NA</td><td>Intergenic</td><td>NA</td><td>NA</td></tr>
<tr><td>T19034</td><td>GGCTCTGGCGCTGATATGCGTGG</td><td>1000</td><td></td><td></td><td></td><td></td><td></td><td></td><td></td><td></td><td></td></tr>
<tr><td>Oligo fwd</td><td>CACCGGCTCTGGCGCTGATATGCG</td><td></td><td></td><td></td><td></td><td></td><td></td><td></td><td></td><td></td><td></td></tr>
<tr><td>Oligo rev</td><td>AAACCGCATATCAGCGCCAGAGCC</td><td></td><td></td><td></td><td></td><td></td><td></td><td></td><td></td><td></td><td></td></tr>
<tr><td>Chromosome</td><td>start</td><td>end</td><td>strand</td><td>MM</td><td>target_seq</td><td>PAM</td><td>alignment</td><td>distance</td><td>position</td><td>gene name</td><td>gene id</td></tr>
<tr><td>GU305914.1</td><td>203967</td><td>203989</td><td>-</td><td>0</td><td>GGCTCTGGCGCTGATATGCG</td><td>TGG</td><td>||||||||[||||||||||||]PAM</td><td>NA</td><td>Intergenic</td><td>NA</td><td>NA</td></tr>
<tr><td>T19035</td><td>TGGCGCTGATATGCGTGGTGTGG</td><td>1000</td><td></td><td></td><td></td><td></td><td></td><td></td><td></td><td></td><td></td></tr>
<tr><td>Oligo fwd</td><td>CACCTGGCGCTGATATGCGTGGTG</td><td></td><td></td><td></td><td></td><td></td><td></td><td></td><td></td><td></td><td></td></tr>
<tr><td>Oligo rev</td><td>AAACCACCACGCATATCAGCGCCA</td><td></td><td></td><td></td><td></td><td></td><td></td><td></td><td></td><td></td><td></td></tr>
<tr><td>Chromosome</td><td>start</td><td>end</td><td>strand</td><td>MM</td><td>target_seq</td><td>PAM</td><td>alignment</td><td>distance</td><td>position</td><td>gene name</td><td>gene id</td></tr>
<tr><td>GU305914.1</td><td>203962</td><td>203984</td><td>-</td><td>0</td><td>TGGCGCTGATATGCGTGGTG</td><td>TGG</td><td>||||||||[||||||||||||]PAM</td><td>NA</td><td>Intergenic</td><td>NA</td><td>NA</td></tr>
<tr><td>T19036</td><td>GGCGCTGATATGCGTGGTGTGGG</td><td>1000</td><td></td><td></td><td></td><td></td><td></td><td></td><td></td><td></td><td></td></tr>
<tr><td>Oligo fwd</td><td>CACCGGCGCTGATATGCGTGGTGT</td><td></td><td></td><td></td><td></td><td></td><td></td><td></td><td></td><td></td><td></td></tr>
<tr><td>Oligo rev</td><td>AAACACACCACGCATATCAGCGCC</td><td></td><td></td><td></td><td></td><td></td><td></td><td></td><td></td><td></td><td></td></tr>
<tr><td>Chromosome</td><td>start</td><td>end</td><td>strand</td><td>MM</td><td>target_seq</td><td>PAM</td><td>alignment</td><td>distance</td><td>position</td><td>gene name</td><td>gene id</td></tr>
<tr><td>GU305914.1</td><td>203961</td><td>203983</td><td>-</td><td>0</td><td>GGCGCTGATATGCGTGGTGT</td><td>GGG</td><td>||||||||[||||||||||||]PAM</td><td>NA</td><td>Intergenic</td><td>NA</td><td>NA</td></tr>
<tr><td>T19037</td><td>TATGCGTGGTGTGGGTGTGTTGG</td><td>1000</td><td></td><td></td><td></td><td></td><td></td><td></td><td></td><td></td><td></td></tr>
<tr><td>Oligo fwd</td><td>CACCTATGCGTGGTGTGGGTGTGT</td><td></td><td></td><td></td><td></td><td></td><td></td><td></td><td></td><td></td><td></td></tr>
<tr><td>Oligo rev</td><td>AAACACACACCCACACCACGCATA</td><td></td><td></td><td></td><td></td><td></td><td></td><td></td><td></td><td></td><td></td></tr>
<tr><td>Chromosome</td><td>start</td><td>end</td><td>strand</td><td>MM</td><td>target_seq</td><td>PAM</td><td>alignment</td><td>distance</td><td>position</td><td>gene name</td><td>gene id</td></tr>
<tr><td>GU305914.1</td><td>203953</td><td>203975</td><td>-</td><td>0</td><td>TATGCGTGGTGTGGGTGTGT</td><td>TGG</td><td>||||||||[||||||||||||]PAM</td><td>NA</td><td>Intergenic</td><td>NA</td><td>NA</td></tr>
<tr><td>T19038</td><td>CGAGAGAGTTGCGCTCGCCACGG</td><td>1000</td><td></td><td></td><td></td><td></td><td></td><td></td><td></td><td></td><td></td></tr>
<tr><td>Oligo fwd</td><td>CACCCGAGAGAGTTGCGCTCGCCA</td><td></td><td></td><td></td><td></td><td></td><td></td><td></td><td></td><td></td><td></td></tr>
<tr><td>Oligo rev</td><td>AAACTGGCGAGCGCAACTCTCTCG</td><td></td><td></td><td></td><td></td><td></td><td></td><td></td><td></td><td></td><td></td></tr>
<tr><td>Chromosome</td><td>start</td><td>end</td><td>strand</td><td>MM</td><td>target_seq</td><td>PAM</td><td>alignment</td><td>distance</td><td>position</td><td>gene name</td><td>gene id</td></tr>
<tr><td>GU305914.1</td><td>203930</td><td>203952</td><td>-</td><td>0</td><td>CGAGAGAGTTGCGCTCGCCA</td><td>CGG</td><td>||||||||[||||||||||||]PAM</td><td>NA</td><td>Intergenic</td><td>NA</td><td>NA</td></tr>
<tr><td>T19039</td><td>GAGAGAGTTGCGCTCGCCACGGG</td><td>1000</td><td></td><td></td><td></td><td></td><td></td><td></td><td></td><td></td><td></td></tr>
<tr><td>Oligo fwd</td><td>CACCGAGAGAGTTGCGCTCGCCAC</td><td></td><td></td><td></td><td></td><td></td><td></td><td></td><td></td><td></td><td></td></tr>
<tr><td>Oligo rev</td><td>AAACGTGGCGAGCGCAACTCTCTC</td><td></td><td></td><td></td><td></td><td></td><td></td><td></td><td></td><td></td><td></td></tr>
<tr><td>Chromosome</td><td>start</td><td>end</td><td>strand</td><td>MM</td><td>target_seq</td><td>PAM</td><td>alignment</td><td>distance</td><td>position</td><td>gene name</td><td>gene id</td></tr>
<tr><td>GU305914.1</td><td>203929</td><td>203951</td><td>-</td><td>0</td><td>GAGAGAGTTGCGCTCGCCAC</td><td>GGG</td><td>||||||||[||||||||||||]PAM</td><td>NA</td><td>Intergenic</td><td>NA</td><td>NA</td></tr>
<tr><td>T19040</td><td>TTGCGCTCGCCACGGGACAGAGG</td><td>1000</td><td></td><td></td><td></td><td></td><td></td><td></td><td></td><td></td><td></td></tr>
<tr><td>Oligo fwd</td><td>CACCTTGCGCTCGCCACGGGACAG</td><td></td><td></td><td></td><td></td><td></td><td></td><td></td><td></td><td></td><td></td></tr>
<tr><td>Oligo rev</td><td>AAACCTGTCCCGTGGCGAGCGCAA</td><td></td><td></td><td></td><td></td><td></td><td></td><td></td><td></td><td></td><td></td></tr>
<tr><td>Chromosome</td><td>start</td><td>end</td><td>strand</td><td>MM</td><td>target_seq</td><td>PAM</td><td>alignment</td><td>distance</td><td>position</td><td>gene name</td><td>gene id</td></tr>
<tr><td>GU305914.1</td><td>203922</td><td>203944</td><td>-</td><td>0</td><td>TTGCGCTCGCCACGGGACAG</td><td>AGG</td><td>||||||||[||||||||||||]PAM</td><td>NA</td><td>Intergenic</td><td>NA</td><td>NA</td></tr>
<tr><td>T19041</td><td>ACAGAGGACTCGTCCGAGTCCGG</td><td>1000</td><td></td><td></td><td></td><td></td><td></td><td></td><td></td><td></td><td></td></tr>
</table>

<table>
<tr><td>Oligo fwd</td><td>CACCACAGAGGACTCGTCCGAGTC</td><td></td><td></td><td></td><td></td><td></td><td></td><td></td><td></td><td></td><td></td><td></td></tr>
<tr><td>Oligo rev</td><td>AAACGACTCGGACGAGTCCTCTGT</td><td></td><td></td><td></td><td></td><td></td><td></td><td></td><td></td><td></td><td></td><td></td></tr>
<tr><td>Chromosome</td><td>start</td><td></td><td>end</td><td>strand</td><td>MM</td><td>target_seq</td><td>PAM</td><td>alignment</td><td>distance</td><td>position</td><td>gene name</td><td>gene id</td></tr>
<tr><td>GU305914.1</td><td></td><td>203906</td><td>203928</td><td>-</td><td>0</td><td>ACAGAGGACTCGTCCGAGTC</td><td>CGG</td><td>||||||||[||||||||||||]PAM</td><td>NA</td><td>Intergenic</td><td>NA</td><td>NA</td></tr>
<tr><td>T19042</td><td>ACTCGTCCGAGTCCGGACTGCGG</td><td></td><td>1000</td><td></td><td></td><td></td><td></td><td></td><td></td><td></td><td></td><td></td></tr>
<tr><td>Oligo fwd</td><td>CACCACTCGTCCGAGTCCGGACTG</td><td></td><td></td><td></td><td></td><td></td><td></td><td></td><td></td><td></td><td></td><td></td></tr>
<tr><td>Oligo rev</td><td>AAACCAGTCCGGACTCGGACGAGT</td><td></td><td></td><td></td><td></td><td></td><td></td><td></td><td></td><td></td><td></td><td></td></tr>
<tr><td>Chromosome</td><td>start</td><td></td><td>end</td><td>strand</td><td>MM</td><td>target_seq</td><td>PAM</td><td>alignment</td><td>distance</td><td>position</td><td>gene name</td><td>gene id</td></tr>
<tr><td>GU305914.1</td><td></td><td>203899</td><td>203921</td><td>-</td><td>0</td><td>ACTCGTCCGAGTCCGGACTG</td><td>CGG</td><td>||||||||[||||||||||||]PAM</td><td>NA</td><td>Intergenic</td><td>NA</td><td>NA</td></tr>
<tr><td>T19043</td><td>ACTGCGGTACGCATACACGTTGG</td><td></td><td>1000</td><td></td><td></td><td></td><td></td><td></td><td></td><td></td><td></td><td></td></tr>
<tr><td>Oligo fwd</td><td>CACCACTGCGGTACGCATACACGT</td><td></td><td></td><td></td><td></td><td></td><td></td><td></td><td></td><td></td><td></td><td></td></tr>
<tr><td>Oligo rev</td><td>AAACACGTGTATGCGTACCGCAGT</td><td></td><td></td><td></td><td></td><td></td><td></td><td></td><td></td><td></td><td></td><td></td></tr>
<tr><td>Chromosome</td><td>start</td><td></td><td>end</td><td>strand</td><td>MM</td><td>target_seq</td><td>PAM</td><td>alignment</td><td>distance</td><td>position</td><td>gene name</td><td>gene id</td></tr>
<tr><td>GU305914.1</td><td></td><td>203883</td><td>203905</td><td>-</td><td>0</td><td>ACTGCGGTACGCATACACGT</td><td>TGG</td><td>||||||||[||||||||||||]PAM</td><td>NA</td><td>Intergenic</td><td>NA</td><td>NA</td></tr>
<tr><td>T19044</td><td>GCGGTACGCATACACGTTGGTGG</td><td></td><td>1000</td><td></td><td></td><td></td><td></td><td></td><td></td><td></td><td></td><td></td></tr>
<tr><td>Oligo fwd</td><td>CACCGCGGTACGCATACACGTTGG</td><td></td><td></td><td></td><td></td><td></td><td></td><td></td><td></td><td></td><td></td><td></td></tr>
<tr><td>Oligo rev</td><td>AAACCCAACGTGTATGCGTACCGC</td><td></td><td></td><td></td><td></td><td></td><td></td><td></td><td></td><td></td><td></td><td></td></tr>
<tr><td>Chromosome</td><td>start</td><td></td><td>end</td><td>strand</td><td>MM</td><td>target_seq</td><td>PAM</td><td>alignment</td><td>distance</td><td>position</td><td>gene name</td><td>gene id</td></tr>
<tr><td>GU305914.1</td><td></td><td>203880</td><td>203902</td><td>-</td><td>0</td><td>GCGGTACGCATACACGTTGG</td><td>TGG</td><td>||||||||[||||||||||||]PAM</td><td>NA</td><td>Intergenic</td><td>NA</td><td>NA</td></tr>
<tr><td>T19045</td><td>GCATACACGTTGGTGGTCGACGG</td><td></td><td>1000</td><td></td><td></td><td></td><td></td><td></td><td></td><td></td><td></td><td></td></tr>
<tr><td>Oligo fwd</td><td>CACCGCATACACGTTGGTGGTCGA</td><td></td><td></td><td></td><td></td><td></td><td></td><td></td><td></td><td></td><td></td><td></td></tr>
<tr><td>Oligo rev</td><td>AAACTCGACCACCAACGTGTATGC</td><td></td><td></td><td></td><td></td><td></td><td></td><td></td><td></td><td></td><td></td><td></td></tr>
<tr><td>Chromosome</td><td>start</td><td></td><td>end</td><td>strand</td><td>MM</td><td>target_seq</td><td>PAM</td><td>alignment</td><td>distance</td><td>position</td><td>gene name</td><td>gene id</td></tr>
<tr><td>GU305914.1</td><td></td><td>203873</td><td>203895</td><td>-</td><td>0</td><td>GCATACACGTTGGTGGTCGA</td><td>CGG</td><td>||||||||[||||||||||||]PAM</td><td>NA</td><td>Intergenic</td><td>NA</td><td>NA</td></tr>
<tr><td>T19046</td><td>GCAAATACGCGCCGTTGTTTTGG</td><td></td><td>1000</td><td></td><td></td><td></td><td></td><td></td><td></td><td></td><td></td><td></td></tr>
<tr><td>Oligo fwd</td><td>CACCGCAAATACGCGCCGTTGTTT</td><td></td><td></td><td></td><td></td><td></td><td></td><td></td><td></td><td></td><td></td><td></td></tr>
<tr><td>Oligo rev</td><td>AAACAAACAACGGCGCGTATTTGC</td><td></td><td></td><td></td><td></td><td></td><td></td><td></td><td></td><td></td><td></td><td></td></tr>
<tr><td>Chromosome</td><td>start</td><td></td><td>end</td><td>strand</td><td>MM</td><td>target_seq</td><td>PAM</td><td>alignment</td><td>distance</td><td>position</td><td>gene name</td><td>gene id</td></tr>
<tr><td>GU305914.1</td><td></td><td>203846</td><td>203868</td><td>-</td><td>0</td><td>GCAAATACGCGCCGTTGTTT</td><td>TGG</td><td>||||||||[||||||||||||]PAM</td><td>NA</td><td>Intergenic</td><td>NA</td><td>NA</td></tr>
<tr><td>T19047</td><td>CAAATACGCGCCGTTGTTTTGGG</td><td></td><td>1000</td><td></td><td></td><td></td><td></td><td></td><td></td><td></td><td></td><td></td></tr>
<tr><td>Oligo fwd</td><td>CACCCAAATACGCGCCGTTGTTTT</td><td></td><td></td><td></td><td></td><td></td><td></td><td></td><td></td><td></td><td></td><td></td></tr>
<tr><td>Oligo rev</td><td>AAACAAAACAACGGCGCGTATTTG</td><td></td><td></td><td></td><td></td><td></td><td></td><td></td><td></td><td></td><td></td><td></td></tr>
<tr><td>Chromosome</td><td>start</td><td></td><td>end</td><td>strand</td><td>MM</td><td>target_seq</td><td>PAM</td><td>alignment</td><td>distance</td><td>position</td><td>gene name</td><td>gene id</td></tr>
<tr><td>GU305914.1</td><td></td><td>203845</td><td>203867</td><td>-</td><td>0</td><td>CAAATACGCGCCGTTGTTTT</td><td>GGG</td><td>||||||||[||||||||||||]PAM</td><td>NA</td><td>Intergenic</td><td>NA</td><td>NA</td></tr>
<tr><td>T19048</td><td>ACGCGCCGTTGTTTTGGGACCGG</td><td></td><td>1000</td><td></td><td></td><td></td><td></td><td></td><td></td><td></td><td></td><td></td></tr>
<tr><td>Oligo fwd</td><td>CACCACGCGCCGTTGTTTTGGGAC</td><td></td><td></td><td></td><td></td><td></td><td></td><td></td><td></td><td></td><td></td><td></td></tr>
<tr><td>Oligo rev</td><td>AAACGTCCCAAAACAACGGCGCGT</td><td></td><td></td><td></td><td></td><td></td><td></td><td></td><td></td><td></td><td></td><td></td></tr>
<tr><td>Chromosome</td><td>start</td><td></td><td>end</td><td>strand</td><td>MM</td><td>target_seq</td><td>PAM</td><td>alignment</td><td>distance</td><td>position</td><td>gene name</td><td>gene id</td></tr>
<tr><td>GU305914.1</td><td></td><td>203840</td><td>203862</td><td>-</td><td>0</td><td>ACGCGCCGTTGTTTTGGGAC</td><td>CGG</td><td>||||||||[||||||||||||]PAM</td><td>NA</td><td>Intergenic</td><td>NA</td><td>NA</td></tr>
<tr><td>T19049</td><td>TTTGGGACCGGACACGTCGACGG</td><td></td><td>1000</td><td></td><td></td><td></td><td></td><td></td><td></td><td></td><td></td><td></td></tr>
<tr><td>Oligo fwd</td><td>CACCTTTGGGACCGGACACGTCGA</td><td></td><td></td><td></td><td></td><td></td><td></td><td></td><td></td><td></td><td></td><td></td></tr>
<tr><td>Oligo rev</td><td>AAACTCGACGTGTCCGGTCCCAAA</td><td></td><td></td><td></td><td></td><td></td><td></td><td></td><td></td><td></td><td></td><td></td></tr>
<tr><td>Chromosome</td><td>start</td><td></td><td>end</td><td>strand</td><td>MM</td><td>target_seq</td><td>PAM</td><td>alignment</td><td>distance</td><td>position</td><td>gene name</td><td>gene id</td></tr>
<tr><td>GU305914.1</td><td></td><td>203828</td><td>203850</td><td>-</td><td>0</td><td>TTTGGGACCGGACACGTCGA</td><td>CGG</td><td>||||||||[||||||||||||]PAM</td><td>NA</td><td>Intergenic</td><td>NA</td><td>NA</td></tr>
<tr><td>T19050</td><td>TTGGGACCGGACACGTCGACGGG</td><td></td><td>1000</td><td></td><td></td><td></td><td></td><td></td><td></td><td></td><td></td><td></td></tr>
<tr><td>Oligo fwd</td><td>CACCTTGGGACCGGACACGTCGAC</td><td></td><td></td><td></td><td></td><td></td><td></td><td></td><td></td><td></td><td></td><td></td></tr>
<tr><td>Oligo rev</td><td>AAACGTCGACGTGTCCGGTCCCAA</td><td></td><td></td><td></td><td></td><td></td><td></td><td></td><td></td><td></td><td></td><td></td></tr>
<tr><td>Chromosome</td><td>start</td><td></td><td>end</td><td>strand</td><td>MM</td><td>target_seq</td><td>PAM</td><td>alignment</td><td>distance</td><td>position</td><td>gene name</td><td>gene id</td></tr>
<tr><td>GU305914.1</td><td></td><td>203827</td><td>203849</td><td>-</td><td>0</td><td>TTGGGACCGGACACGTCGAC</td><td>GGG</td><td>||||||||[||||||||||||]PAM</td><td>NA</td><td>Intergenic</td><td>NA</td><td>NA</td></tr>
<tr><td>T19051</td><td>TGGGACCGGACACGTCGACGGGG</td><td></td><td>1000</td><td></td><td></td><td></td><td></td><td></td><td></td><td></td><td></td><td></td></tr>
<tr><td>Oligo fwd</td><td>CACCTGGGACCGGACACGTCGACG</td><td></td><td></td><td></td><td></td><td></td><td></td><td></td><td></td><td></td><td></td><td></td></tr>
<tr><td>Oligo rev</td><td>AAACCGTCGACGTGTCCGGTCCCA</td><td></td><td></td><td></td><td></td><td></td><td></td><td></td><td></td><td></td><td></td><td></td></tr>
<tr><td>Chromosome</td><td>start</td><td></td><td>end</td><td>strand</td><td>MM</td><td>target_seq</td><td>PAM</td><td>alignment</td><td>distance</td><td>position</td><td>gene name</td><td>gene id</td></tr>
<tr><td>GU305914.1</td><td></td><td>203826</td><td>203848</td><td>-</td><td>0</td><td>TGGGACCGGACACGTCGACG</td><td>GGG</td><td>||||||||[||||||||||||]PAM</td><td>NA</td><td>Intergenic</td><td>NA</td><td>NA</td></tr>
<tr><td>T19052</td><td>CGTCGACGGGGAAGCCTTTGTGG</td><td></td><td>1000</td><td></td><td></td><td></td><td></td><td></td><td></td><td></td><td></td><td></td></tr>
<tr><td>Oligo fwd</td><td>CACCCGTCGACGGGGAAGCCTTTG</td><td></td><td></td><td></td><td></td><td></td><td></td><td></td><td></td><td></td><td></td><td></td></tr>
<tr><td>Oligo rev</td><td>AAACCAAAGGCTTCCCCGTCGACG</td><td></td><td></td><td></td><td></td><td></td><td></td><td></td><td></td><td></td><td></td><td></td></tr>
<tr><td>Chromosome</td><td>start</td><td></td><td>end</td><td>strand</td><td>MM</td><td>target_seq</td><td>PAM</td><td>alignment</td><td>distance</td><td>position</td><td>gene name</td><td>gene id</td></tr>
<tr><td>GU305914.1</td><td></td><td>203814</td><td>203836</td><td>-</td><td>0</td><td>CGTCGACGGGGAAGCCTTTG</td><td>TGG</td><td>||||||||[||||||||||||]PAM</td><td>NA</td><td>Intergenic</td><td>NA</td><td>NA</td></tr>
<tr><td>T19053</td><td>GTCGACGGGGAAGCCTTTGTGGG</td><td></td><td>1000</td><td></td><td></td><td></td><td></td><td></td><td></td><td></td><td></td><td></td></tr>
<tr><td>Oligo fwd</td><td>CACCGTCGACGGGGAAGCCTTTGT</td><td></td><td></td><td></td><td></td><td></td><td></td><td></td><td></td><td></td><td></td><td></td></tr>
<tr><td>Oligo rev</td><td>AAACACAAAGGCTTCCCCGTCGAC</td><td></td><td></td><td></td><td></td><td></td><td></td><td></td><td></td><td></td><td></td><td></td></tr>
<tr><td>Chromosome</td><td>start</td><td></td><td>end</td><td>strand</td><td>MM</td><td>target_seq</td><td>PAM</td><td>alignment</td><td>distance</td><td>position</td><td>gene name</td><td>gene id</td></tr>
<tr><td>GU305914.1</td><td></td><td>203813</td><td>203835</td><td>-</td><td>0</td><td>GTCGACGGGGAAGCCTTTGT</td><td>GGG</td><td>||||||||[||||||||||||]PAM</td><td>NA</td><td>Intergenic</td><td>NA</td><td>NA</td></tr>
<tr><td>T19054</td><td>TACTCCAATAACAAGACACACGG</td><td></td><td>1000</td><td></td><td></td><td></td><td></td><td></td><td></td><td></td><td></td><td></td></tr>
<tr><td>Oligo fwd</td><td>CACCTACTCCAATAACAAGACACA</td><td></td><td></td><td></td><td></td><td></td><td></td><td></td><td></td><td></td><td></td><td></td></tr>
<tr><td>Oligo rev</td><td>AAACTGTGTCTTGTTATTGGAGTA</td><td></td><td></td><td></td><td></td><td></td><td></td><td></td><td></td><td></td><td></td><td></td></tr>
<tr><td>Chromosome</td><td>start</td><td></td><td>end</td><td>strand</td><td>MM</td><td>target_seq</td><td>PAM</td><td>alignment</td><td>distance</td><td>position</td><td>gene name</td><td>gene id</td></tr>
<tr><td>GU305914.1</td><td></td><td>203789</td><td>203811</td><td>-</td><td>0</td><td>TACTCCAATAACAAGACACA</td><td>CGG</td><td>||||||||[||||||||||||]PAM</td><td>NA</td><td>Intergenic</td><td>NA</td><td>NA</td></tr>
<tr><td>T19055</td><td>ACTCCAATAACAAGACACACGGG</td><td></td><td>1000</td><td></td><td></td><td></td><td></td><td></td><td></td><td></td><td></td><td></td></tr>
<tr><td>Oligo fwd</td><td>CACCACTCCAATAACAAGACACAC</td><td></td><td></td><td></td><td></td><td></td><td></td><td></td><td></td><td></td><td></td><td></td></tr>
<tr><td>Oligo rev</td><td>AAACGTGTGTCTTGTTATTGGAGT</td><td></td><td></td><td></td><td></td><td></td><td></td><td></td><td></td><td></td><td></td><td></td></tr>
<tr><td>Chromosome</td><td>start</td><td></td><td>end</td><td>strand</td><td>MM</td><td>target_seq</td><td>PAM</td><td>alignment</td><td>distance</td><td>position</td><td>gene name</td><td>gene id</td></tr>
<tr><td>GU305914.1</td><td></td><td>203788</td><td>203810</td><td>-</td><td>0</td><td>ACTCCAATAACAAGACACAC</td><td>GGG</td><td>||||||||[||||||||||||]PAM</td><td>NA</td><td>Intergenic</td><td>NA</td><td>NA</td></tr>
</table>

T19056 AATAACAAGACACACGGGATCGG 1000
Oligo fwd CACCAATAACAAGACACACGGGAT
Oligo rev AAACATCCCGTGTGTCTTGTTATT
Chromosome start end strand MM target_seq PAM alignment distance position gene name gene id
GU305914.1 203783 203805 - 0 AATAACAAGACACACGGGAT CGG ||||||||[||||||||||||]PAM NA Intergenic NA NA

T19057 AGACACACGGGATCGGTAGATGG 1000
Oligo fwd CACCAGACACACGGGATCGGTAGA
Oligo rev AAACTCTACCGATCCCGTGTGTCT
Chromosome start end strand MM target_seq PAM alignment distance position gene name gene id
GU305914.1 203776 203798 - 0 AGACACACGGGATCGGTAGA TGG ||||||||[||||||||||||]PAM NA Intergenic NA NA

T19058 GACACACGGGATCGGTAGATGGG 1000
Oligo fwd CACCGACACACGGGATCGGTAGAT
Oligo rev AAACATCTACCGATCCCGTGTGTC
Chromosome start end strand MM target_seq PAM alignment distance position gene name gene id
GU305914.1 203775 203797 - 0 GACACACGGGATCGGTAGAT GGG ||||||||[||||||||||||]PAM NA Intergenic NA NA

T19059 GGTGAACGCATCTCACGTAGAGG 1000
Oligo fwd CACCGGTGAACGCATCTCACGTAG
Oligo rev AAACCTACGTGAGATGCGTTCACC
Chromosome start end strand MM target_seq PAM alignment distance position gene name gene id
GU305914.1 203754 203776 - 0 GGTGAACGCATCTCACGTAG AGG ||||||||[||||||||||||]PAM NA Intergenic NA NA

T19060 AAGAGAACAAAGAGTTCGTGAGG 1000
Oligo fwd CACCAAGAGAACAAAGAGTTCGTG
Oligo rev AAACCACGAACTCTTTGTTCTCTT
Chromosome start end strand MM target_seq PAM alignment distance position gene name gene id
GU305914.1 203731 203753 - 0 AAGAGAACAAAGAGTTCGTG AGG ||||||||[||||||||||||]PAM NA Intergenic NA NA

T19061 AGAGTTCGTGAGGCAGTGTAAGG 1000
Oligo fwd CACCAGAGTTCGTGAGGCAGTGTA
Oligo rev AAACTACACTGCCTCACGAACTCT
Chromosome start end strand MM target_seq PAM alignment distance position gene name gene id
GU305914.1 203721 203743 - 0 AGAGTTCGTGAGGCAGTGTA AGG ||||||||[||||||||||||]PAM NA Intergenic NA NA

T19062 TAAAATGCAAAACAACTCAAAGG 1000
Oligo fwd CACCTAAAATGCAAAACAACTCAA
Oligo rev AAACTTGAGTTGTTTTGCATTTTA
Chromosome start end strand MM target_seq PAM alignment distance position gene name gene id
GU305914.1 203679 203701 - 0 TAAAATGCAAAACAACTCAA AGG ||||||||[||||||||||||]PAM NA Intergenic NA NA

T19063 CAAAACAACTCAAAGGTGATCGG 1000
Oligo fwd CACCCAAAACAACTCAAAGGTGAT
Oligo rev AAACATCACCTTTGAGTTGTTTTG
Chromosome start end strand MM target_seq PAM alignment distance position gene name gene id
GU305914.1 203672 203694 - 0 CAAAACAACTCAAAGGTGAT CGG ||||||||[||||||||||||]PAM NA Intergenic NA NA

T19064 AAGACCGTACAACTAGACGTCGG 1000
Oligo fwd CACCAAGACCGTACAACTAGACGT
Oligo rev AAACACGTCTAGTTGTACGGTCTT
Chromosome start end strand MM target_seq PAM alignment distance position gene name gene id
GU305914.1 203645 203667 - 0 AAGACCGTACAACTAGACGT CGG ||||||||[||||||||||||]PAM NA Intergenic NA NA

T19065 GAGAAACATTACGCGTATGACGG 1000
Oligo fwd CACCGAGAAACATTACGCGTATGA
Oligo rev AAACTCATACGCGTAATGTTTCTC
Chromosome start end strand MM target_seq PAM alignment distance position gene name gene id
GU305914.1 203606 203628 - 0 GAGAAACATTACGCGTATGA CGG ||||||||[||||||||||||]PAM NA Intergenic NA NA

T19066 GTATGACGGCAACGAAACCGAGG 1000
Oligo fwd CACCGTATGACGGCAACGAAACCG
Oligo rev AAACCGGTTTCGTTGCCGTCATAC
Chromosome start end strand MM target_seq PAM alignment distance position gene name gene id
GU305914.1 203592 203614 - 0 GTATGACGGCAACGAAACCG AGG ||||||||[||||||||||||]PAM NA Intergenic NA NA

T19067 CAACGAAACCGAGGATGATACGG 1000
Oligo fwd CACCCAACGAAACCGAGGATGATA
Oligo rev AAACTATCATCCTCGGTTTCGTTG
Chromosome start end strand MM target_seq PAM alignment distance position gene name gene id
GU305914.1 203583 203605 - 0 CAACGAAACCGAGGATGATA CGG ||||||||[||||||||||||]PAM NA Intergenic NA NA

T19068 AGAAAAAACTAACTGAATATAGG 1000
Oligo fwd CACCAGAAAAAACTAACTGAATAT
Oligo rev AAACATATTCAGTTAGTTTTTTCT
Chromosome start end strand MM target_seq PAM alignment distance position gene name gene id
GU305914.1 203527 203549 - 0 AGAAAAAACTAACTGAATAT AGG ||||||||[||||||||||||]PAM NA Intergenic NA NA

T19069 GTCCGCCGTATCACCACAGCTGG 1000
Oligo fwd CACCGTCCGCCGTATCACCACAGC
Oligo rev AAACGCTGTGGTGATACGGCGGAC
Chromosome start end strand MM target_seq PAM alignment distance position gene name gene id
GU305914.1 203490 203512 - 0 GTCCGCCGTATCACCACAGC TGG ||||||||[||||||||||||]PAM NA Intergenic NA NA

T19070 CACAGCTGGAAGTCGAGAGACGG 1000
Oligo fwd CACCCACAGCTGGAAGTCGAGAGA
Oligo rev AAACTCTCTCGACTTCCAGCTGTG
Chromosome start end strand MM target_seq PAM alignment distance position gene name gene id
GU305914.1 203476 203498 - 0 CACAGCTGGAAGTCGAGAGA CGG ||||||||[||||||||||||]PAM NA Intergenic NA NA

T19071 GGAAGTCGAGAGACGGTCCTCGG 1000
Oligo fwd CACCGGAAGTCGAGAGACGGTCCT
Oligo rev AAACAGGACCGTCTCTCGACTTCC
Chromosome start end strand MM target_seq PAM alignment distance position gene name gene id
GU305914.1 203469 203491 - 0 GGAAGTCGAGAGACGGTCCT CGG ||||||||[||||||||||||]PAM NA Intergenic NA NA

T19072 GAAGTCGAGAGACGGTCCTCGGG 1000
Oligo fwd CACCGAAGTCGAGAGACGGTCCTC
Oligo rev AAACGAGGACCGTCTCTCGACTTC
Chromosome start end strand MM target_seq PAM alignment distance position gene name gene id
GU305914.1 203468 203490 - 0 GAAGTCGAGAGACGGTCCTC GGG ||||||||[||||||||||||]PAM NA Intergenic NA NA

T19073 GAGACGGTCCTCGGGTCGCGAGG 1000
Oligo fwd CACCGAGACGGTCCTCGGGTCGCG
Oligo rev AAACCGCGACCCGAGGACCGTCTC
Chromosome start end strand MM target_seq PAM alignment distance position gene name gene id
GU305914.1 203460 203482 - 0 GAGACGGTCCTCGGGTCGCG AGG ||||||||[||||||||||||]PAM NA Intergenic NA NA

T19074 AGACGGTCCTCGGGTCGCGAGGG 1000
Oligo fwd CACCAGACGGTCCTCGGGTCGCGA
Oligo rev AAACTCGCGACCCGAGGACCGTCT
Chromosome start end strand MM target_seq PAM alignment distance position gene name gene id
GU305914.1 203459 203481 - 0 AGACGGTCCTCGGGTCGCGA GGG ||||||||[||||||||||||]PAM NA Intergenic NA NA

T19075 CGGTCCTCGGGTCGCGAGGGCGG 1000
Oligo fwd CACCCGGTCCTCGGGTCGCGAGGG
Oligo rev AAACCCCTCGCGACCCGAGGACCG
Chromosome start end strand MM target_seq PAM alignment distance position gene name gene id
GU305914.1 203456 203478 - 0 CGGTCCTCGGGTCGCGAGGG CGG ||||||||[||||||||||||]PAM NA Intergenic NA NA

T19076 GCGAGGGCGGCATGAGACTACGG 1000
Oligo fwd CACCGCGAGGGCGGCATGAGACTA
Oligo rev AAACTAGTCTCATGCCGCCCTCGC
Chromosome start end strand MM target_seq PAM alignment distance position gene name gene id
GU305914.1 203443 203465 - 0 GCGAGGGCGGCATGAGACTA CGG ||||||||[||||||||||||]PAM NA Intergenic NA NA

T19077 TGAGACTACGGTGTTTCGCTAGG 1000
Oligo fwd CACCTGAGACTACGGTGTTTCGCT
Oligo rev AAACAGCGAAACACCGTAGTCTCA
Chromosome start end strand MM target_seq PAM alignment distance position gene name gene id
GU305914.1 203431 203453 - 0 TGAGACTACGGTGTTTCGCT AGG ||||||||[||||||||||||]PAM NA Intergenic NA NA

T19078 GAGACTACGGTGTTTCGCTAGGG 1000
Oligo fwd CACCGAGACTACGGTGTTTCGCTA
Oligo rev AAACTAGCGAAACACCGTAGTCTC
Chromosome start end strand MM target_seq PAM alignment distance position gene name gene id
GU305914.1 203430 203452 - 0 GAGACTACGGTGTTTCGCTA GGG ||||||||[||||||||||||]PAM NA Intergenic NA NA

T19079 ATCCCGCCGATTTAGAGATTCGG 1000
Oligo fwd CACCATCCCGCCGATTTAGAGATT
Oligo rev AAACAATCTCTAAATCGGCGGGAT
Chromosome start end strand MM target_seq PAM alignment distance position gene name gene id
GU305914.1 203401 203423 - 0 ATCCCGCCGATTTAGAGATT CGG ||||||||[||||||||||||]PAM NA Intergenic NA NA

T19080 CCGCCGATTTAGAGATTCGGTGG 1000
Oligo fwd CACCCCGCCGATTTAGAGATTCGG
Oligo rev AAACCCGAATCTCTAAATCGGCGG
Chromosome start end strand MM target_seq PAM alignment distance position gene name gene id
GU305914.1 203398 203420 - 0 CCGCCGATTTAGAGATTCGG TGG ||||||||[||||||||||||]PAM NA Intergenic NA NA

T19081 CCGATTTAGAGATTCGGTGGTGG 1000
Oligo fwd CACCCCGATTTAGAGATTCGGTGG
Oligo rev AAACCCACCGAATCTCTAAATCGG
Chromosome start end strand MM target_seq PAM alignment distance position gene name gene id
GU305914.1 203395 203417 - 0 CCGATTTAGAGATTCGGTGG TGG ||||||||[||||||||||||]PAM NA Intergenic NA NA

T19082 ATTCGGTGGTGGAAAGATGATGG 1000
Oligo fwd CACCATTCGGTGGTGGAAAGATGA
Oligo rev AAACTCATCTTTCCACCACCGAAT
Chromosome start end strand MM target_seq PAM alignment distance position gene name gene id
GU305914.1 203384 203406 - 0 ATTCGGTGGTGGAAAGATGA TGG ||||||||[||||||||||||]PAM NA Intergenic NA NA

T19083 CGGTGGTGGAAAGATGATGGCGG 1000
Oligo fwd CACCCGGTGGTGGAAAGATGATGG
Oligo rev AAACCCATCATCTTTCCACCACCG
Chromosome start end strand MM target_seq PAM alignment distance position gene name gene id
GU305914.1 203381 203403 - 0 CGGTGGTGGAAAGATGATGG CGG ||||||||[||||||||||||]PAM NA Intergenic NA NA

T19084 TGGTGGAAAGATGATGGCGGTGG 1000
Oligo fwd CACCTGGTGGAAAGATGATGGCGG
Oligo rev AAACCCGCCATCATCTTTCCACCA
Chromosome start end strand MM target_seq PAM alignment distance position gene name gene id
GU305914.1 203378 203400 - 0 TGGTGGAAAGATGATGGCGG TGG ||||||||[||||||||||||]PAM NA Intergenic NA NA

T19085 TGGAAAGATGATGGCGGTGGTGG 1000
Oligo fwd CACCTGGAAAGATGATGGCGGTGG
Oligo rev AAACCCACCGCCATCATCTTTCCA
Chromosome start end strand MM target_seq PAM alignment distance position gene name gene id

GU305914.1 203375 203397 - 0 TGGAAAGATGATGGCGGTGG TGG ||||||||[||||||||||||]PAM NA Intergenic NA NA

T19086 CACCACGACCCGCTACCGTCCGG 1000
Oligo fwd CACCCACCACGACCCGCTACCGTC
Oligo rev AAACGACGGTAGCGGGTCGTGGTG
Chromosome start end strand MM target_seq PAM alignment distance position gene name gene id
GU305914.1 203327 203349 - 0 CACCACGACCCGCTACCGTC CGG ||||||||[||||||||||||]PAM NA Intergenic NA NA

T19087 GACCCGCTACCGTCCGGTAACGG 1000
Oligo fwd CACCGACCCGCTACCGTCCGGTAA
Oligo rev AAACTTACCGGACGGTAGCGGGTC
Chromosome start end strand MM target_seq PAM alignment distance position gene name gene id
GU305914.1 203321 203343 - 0 GACCCGCTACCGTCCGGTAA CGG ||||||||[||||||||||||]PAM NA Intergenic NA NA

T19088 ACCCGCTACCGTCCGGTAACGGG 1000
Oligo fwd CACCACCCGCTACCGTCCGGTAAC
Oligo rev AAACGTTACCGGACGGTAGCGGGT
Chromosome start end strand MM target_seq PAM alignment distance position gene name gene id
GU305914.1 203320 203342 - 0 ACCCGCTACCGTCCGGTAAC GGG ||||||||[||||||||||||]PAM NA Intergenic NA NA

T19089 AAAACATATCGACGTCTATGTGG 1000
Oligo fwd CACCAAAACATATCGACGTCTATG
Oligo rev AAACCATAGACGTCGATATGTTTT
Chromosome start end strand MM target_seq PAM alignment distance position gene name gene id
GU305914.1 203289 203311 - 0 AAAACATATCGACGTCTATG TGG ||||||||[||||||||||||]PAM NA Intergenic NA NA

T19090 CATATCGACGTCTATGTGGACGG 1000
Oligo fwd CACCCATATCGACGTCTATGTGGA
Oligo rev AAACTCCACATAGACGTCGATATG
Chromosome start end strand MM target_seq PAM alignment distance position gene name gene id
GU305914.1 203285 203307 - 0 CATATCGACGTCTATGTGGA CGG ||||||||[||||||||||||]PAM NA Intergenic NA NA

T19091 ATCGACGTCTATGTGGACGGAGG 1000
Oligo fwd CACCATCGACGTCTATGTGGACGG
Oligo rev AAACCCGTCCACATAGACGTCGAT
Chromosome start end strand MM target_seq PAM alignment distance position gene name gene id
GU305914.1 203282 203304 - 0 ATCGACGTCTATGTGGACGG AGG ||||||||[||||||||||||]PAM NA Intergenic NA NA

T19092 CGTCTATGTGGACGGAGGCTTGG 1000
Oligo fwd CACCCGTCTATGTGGACGGAGGCT
Oligo rev AAACAGCCTCCGTCCACATAGACG
Chromosome start end strand MM target_seq PAM alignment distance position gene name gene id
GU305914.1 203277 203299 - 0 CGTCTATGTGGACGGAGGCT TGG ||||||||[||||||||||||]PAM NA Intergenic NA NA

T19093 CGTGTACTCGTGTCGAGTCAAGG 1000
Oligo fwd CACCCGTGTACTCGTGTCGAGTCA
Oligo rev AAACTGACTCGACACGAGTACACG
Chromosome start end strand MM target_seq PAM alignment distance position gene name gene id
GU305914.1 203250 203272 - 0 CGTGTACTCGTGTCGAGTCA AGG ||||||||[||||||||||||]PAM NA Intergenic NA NA

T19094 GTGTACTCGTGTCGAGTCAAGGG 1000
Oligo fwd CACCGTGTACTCGTGTCGAGTCAA
Oligo rev AAACTTGACTCGACACGAGTACAC
Chromosome start end strand MM target_seq PAM alignment distance position gene name gene id
GU305914.1 203249 203271 - 0 GTGTACTCGTGTCGAGTCAA GGG ||||||||[||||||||||||]PAM NA Intergenic NA NA

T19095 CGAGTCAAGGGTATCGCGACCGG 1000
Oligo fwd CACCCGAGTCAAGGGTATCGCGAC
Oligo rev AAACGTCGCGATACCCTTGACTCG
Chromosome start end strand MM target_seq PAM alignment distance position gene name gene id
GU305914.1 203237 203259 - 0 CGAGTCAAGGGTATCGCGAC CGG ||||||||[||||||||||||]PAM NA Intergenic NA NA

T19096 GAGTCAAGGGTATCGCGACCGGG 1000
Oligo fwd CACCGAGTCAAGGGTATCGCGACC
Oligo rev AAACGGTCGCGATACCCTTGACTC
Chromosome start end strand MM target_seq PAM alignment distance position gene name gene id
GU305914.1 203236 203258 - 0 GAGTCAAGGGTATCGCGACC GGG ||||||||[||||||||||||]PAM NA Intergenic NA NA

T19097 GGCTCGAACTGCAGATCGTGAGG 1000
Oligo fwd CACCGGCTCGAACTGCAGATCGTG
Oligo rev AAACCACGATCTGCAGTTCGAGCC
Chromosome start end strand MM target_seq PAM alignment distance position gene name gene id
GU305914.1 203215 203237 - 0 GGCTCGAACTGCAGATCGTG AGG ||||||||[||||||||||||]PAM NA Intergenic NA NA
GU305914.1 113328 113350 - 4 GGATCGAGCTGCTGATCCTG AGG ||-||||-[||||-||||-||]PAM NA Intergenic NA NA

T19098 TCGAACTGCAGATCGTGAGGTGG 1000
Oligo fwd CACCTCGAACTGCAGATCGTGAGG
Oligo rev AAACCCTCACGATCTGCAGTTCGA
Chromosome start end strand MM target_seq PAM alignment distance position gene name gene id
GU305914.1 203212 203234 - 0 TCGAACTGCAGATCGTGAGG TGG ||||||||[||||||||||||]PAM NA Intergenic NA NA

T19099 ACTGCAGATCGTGAGGTGGAAGG 1000
Oligo fwd CACCACTGCAGATCGTGAGGTGGA
Oligo rev AAACTCCACCTCACGATCTGCAGT
Chromosome start end strand MM target_seq PAM alignment distance position gene name gene id
GU305914.1 203208 203230 - 0 ACTGCAGATCGTGAGGTGGA AGG ||||||||[||||||||||||]PAM NA Intergenic NA NA

T19100 CTGCAGATCGTGAGGTGGAAGGG 1000
Oligo fwd CACCCTGCAGATCGTGAGGTGGAA

Oligo rev AAACTTCCACCTCACGATCTGCAG
Chromosome start end strand MM target_seq PAM alignment distance position gene name gene id
GU305914.1 203207 203229 - 0 CTGCAGATCGTGAGGTGGAA GGG ||||||||[||||||||||||]PAM NA Intergenic NA NA

T19101 AGGTGGAAGGGCTACGCGAGAGG 1000
Oligo fwd CACCAGGTGGAAGGGCTACGCGAG
Oligo rev AAACCTCGCGTAGCCCTTCCACCT
Chromosome start end strand MM target_seq PAM alignment distance position gene name gene id
GU305914.1 203195 203217 - 0 AGGTGGAAGGGCTACGCGAG AGG ||||||||[||||||||||||]PAM NA Intergenic NA NA

T19102 GAAGGGCTACGCGAGAGGCGCGG 1000
Oligo fwd CACCGAAGGGCTACGCGAGAGGCG
Oligo rev AAACCGCCTCTCGCGTAGCCCTTC
Chromosome start end strand MM target_seq PAM alignment distance position gene name gene id
GU305914.1 203190 203212 - 0 GAAGGGCTACGCGAGAGGCG CGG ||||||||[||||||||||||]PAM NA Intergenic NA NA

T19103 AAGGGCTACGCGAGAGGCGCGGG 1000
Oligo fwd CACCAAGGGCTACGCGAGAGGCGC
Oligo rev AAACGCGCCTCTCGCGTAGCCCTT
Chromosome start end strand MM target_seq PAM alignment distance position gene name gene id
GU305914.1 203189 203211 - 0 AAGGGCTACGCGAGAGGCGC GGG ||||||||[||||||||||||]PAM NA Intergenic NA NA

T19104 AGGGCTACGCGAGAGGCGCGGGG 1000
Oligo fwd CACCAGGGCTACGCGAGAGGCGCG
Oligo rev AAACCGCGCCTCTCGCGTAGCCCT
Chromosome start end strand MM target_seq PAM alignment distance position gene name gene id
GU305914.1 203188 203210 - 0 AGGGCTACGCGAGAGGCGCG GGG ||||||||[||||||||||||]PAM NA Intergenic NA NA

T19105 CGCGAGAGGCGCGGGGAACTCGG 1000
Oligo fwd CACCCGCGAGAGGCGCGGGGAACT
Oligo rev AAACAGTTCCCCGCGCCTCTCGCG
Chromosome start end strand MM target_seq PAM alignment distance position gene name gene id
GU305914.1 203181 203203 - 0 CGCGAGAGGCGCGGGGAACT CGG ||||||||[||||||||||||]PAM NA Intergenic NA NA

T19106 GGGGAACTCGGTCGTCCTGTTGG 1000
Oligo fwd CACCGGGGAACTCGGTCGTCCTGT
Oligo rev AAACACAGGACGACCGAGTTCCCC
Chromosome start end strand MM target_seq PAM alignment distance position gene name gene id
GU305914.1 203169 203191 - 0 GGGGAACTCGGTCGTCCTGT TGG ||||||||[||||||||||||]PAM NA Intergenic NA NA

T19107 GTTCATACCCGCGAGCGTCATGG 1000
Oligo fwd CACCGTTCATACCCGCGAGCGTCA
Oligo rev AAACTGACGCTCGCGGGTATGAAC
Chromosome start end strand MM target_seq PAM alignment distance position gene name gene id
GU305914.1 203142 203164 - 0 GTTCATACCCGCGAGCGTCA TGG ||||||||[||||||||||||]PAM NA Intergenic NA NA

T19108 CATACCCGCGAGCGTCATGGCGG 1000
Oligo fwd CACCCATACCCGCGAGCGTCATGG
Oligo rev AAACCCATGACGCTCGCGGGTATG
Chromosome start end strand MM target_seq PAM alignment distance position gene name gene id
GU305914.1 203139 203161 - 0 CATACCCGCGAGCGTCATGG CGG ||||||||[||||||||||||]PAM NA Intergenic NA NA

T19109 ACCCGCGAGCGTCATGGCGGCGG 1000
Oligo fwd CACCACCCGCGAGCGTCATGGCGG
Oligo rev AAACCCGCCATGACGCTCGCGGGT
Chromosome start end strand MM target_seq PAM alignment distance position gene name gene id
GU305914.1 203136 203158 - 0 ACCCGCGAGCGTCATGGCGG CGG ||||||||[||||||||||||]PAM NA Intergenic NA NA

T19110 GAGCGTCATGGCGGCGGTCGTGG 1000
Oligo fwd CACCGAGCGTCATGGCGGCGGTCG
Oligo rev AAACCGACCGCCGCCATGACGCTC
Chromosome start end strand MM target_seq PAM alignment distance position gene name gene id
GU305914.1 203130 203152 - 0 GAGCGTCATGGCGGCGGTCG TGG ||||||||[||||||||||||]PAM NA Intergenic NA NA
GU305914.1 107509 107531 - 4 GAGCGTTCTGGCGTCGGTAG TGG ||||||--[|||||-||||-|]PAM NA Intergenic NA NA

T19111 GTCATGGCGGCGGTCGTGGTCGG 1000
Oligo fwd CACCGTCATGGCGGCGGTCGTGGT
Oligo rev AAACACCACGACCGCCGCCATGAC
Chromosome start end strand MM target_seq PAM alignment distance position gene name gene id
GU305914.1 203126 203148 - 0 GTCATGGCGGCGGTCGTGGT CGG ||||||||[||||||||||||]PAM NA Intergenic NA NA

T19112 GTTGATACGAAAGAAGAGCAAGG 1000
Oligo fwd CACCGTTGATACGAAAGAAGAGCA
Oligo rev AAACTGCTCTTCTTTCGTATCAAC
Chromosome start end strand MM target_seq PAM alignment distance position gene name gene id
GU305914.1 203097 203119 - 0 GTTGATACGAAAGAAGAGCA AGG ||||||||[||||||||||||]PAM NA Intergenic NA NA

T19113 GAAAGAAGAGCAAGGAACAGAGG 1000
Oligo fwd CACCGAAAGAAGAGCAAGGAACAG
Oligo rev AAACCTGTTCCTTGCTCTTCTTTC
Chromosome start end strand MM target_seq PAM alignment distance position gene name gene id
GU305914.1 203089 203111 - 0 GAAAGAAGAGCAAGGAACAG AGG ||||||||[||||||||||||]PAM NA Intergenic NA NA

T19114 AGGAACAGAGGAAGACGCGTAGG 1000
Oligo fwd CACCAGGAACAGAGGAAGACGCGT
Oligo rev AAACACGCGTCTTCCTCTGTTCCT
Chromosome start end strand MM target_seq PAM alignment distance position gene name gene id
GU305914.1 203077 203099 - 0 AGGAACAGAGGAAGACGCGT AGG ||||||||[||||||||||||]PAM NA Intergenic NA NA

<table>
<tr><td>T19115</td><td>AACAGAGGAAGACGCGTAGGAGG</td><td>1000</td><td></td><td></td><td></td><td></td><td></td><td></td><td></td><td></td><td></td></tr>
<tr><td>Oligo fwd</td><td>CACCAACAGAGGAAGACGCGTAGG</td><td></td><td></td><td></td><td></td><td></td><td></td><td></td><td></td><td></td><td></td></tr>
<tr><td>Oligo rev</td><td>AAACCCTACGCGTCTTCCTCTGTT</td><td></td><td></td><td></td><td></td><td></td><td></td><td></td><td></td><td></td><td></td></tr>
<tr><td>Chromosome</td><td>start</td><td>end</td><td>strand</td><td>MM</td><td>target_seq</td><td>PAM</td><td>alignment</td><td>distance</td><td>position</td><td>gene name</td><td>gene id</td></tr>
<tr><td>GU305914.1</td><td>203074</td><td>203096</td><td>-</td><td>0</td><td>AACAGAGGAAGACGCGTAGG</td><td>AGG</td><td>||||||||[||||||||||||]PAM</td><td>NA</td><td>Intergenic</td><td>NA</td><td>NA</td></tr>
<tr><td>T19116</td><td>AGGAAGACGCGTAGGAGGTTCGG</td><td>1000</td><td></td><td></td><td></td><td></td><td></td><td></td><td></td><td></td><td></td></tr>
<tr><td>Oligo fwd</td><td>CACCAGGAAGACGCGTAGGAGGTT</td><td></td><td></td><td></td><td></td><td></td><td></td><td></td><td></td><td></td><td></td></tr>
<tr><td>Oligo rev</td><td>AAACAACCTCCTACGCGTCTTCCT</td><td></td><td></td><td></td><td></td><td></td><td></td><td></td><td></td><td></td><td></td></tr>
<tr><td>Chromosome</td><td>start</td><td>end</td><td>strand</td><td>MM</td><td>target_seq</td><td>PAM</td><td>alignment</td><td>distance</td><td>position</td><td>gene name</td><td>gene id</td></tr>
<tr><td>GU305914.1</td><td>203069</td><td>203091</td><td>-</td><td>0</td><td>AGGAAGACGCGTAGGAGGTT</td><td>CGG</td><td>||||||||[||||||||||||]PAM</td><td>NA</td><td>Intergenic</td><td>NA</td><td>NA</td></tr>
<tr><td>T19117</td><td>AGACGCGTAGGAGGTTCGGACGG</td><td>1000</td><td></td><td></td><td></td><td></td><td></td><td></td><td></td><td></td><td></td></tr>
<tr><td>Oligo fwd</td><td>CACCAGACGCGTAGGAGGTTCGGA</td><td></td><td></td><td></td><td></td><td></td><td></td><td></td><td></td><td></td><td></td></tr>
<tr><td>Oligo rev</td><td>AAACTCCGAACCTCCTACGCGTCT</td><td></td><td></td><td></td><td></td><td></td><td></td><td></td><td></td><td></td><td></td></tr>
<tr><td>Chromosome</td><td>start</td><td>end</td><td>strand</td><td>MM</td><td>target_seq</td><td>PAM</td><td>alignment</td><td>distance</td><td>position</td><td>gene name</td><td>gene id</td></tr>
<tr><td>GU305914.1</td><td>203065</td><td>203087</td><td>-</td><td>0</td><td>AGACGCGTAGGAGGTTCGGA</td><td>CGG</td><td>||||||||[||||||||||||]PAM</td><td>NA</td><td>Intergenic</td><td>NA</td><td>NA</td></tr>
<tr><td>T19118</td><td>TAGGAGGTTCGGACGGCGCTCGG</td><td>1000</td><td></td><td></td><td></td><td></td><td></td><td></td><td></td><td></td><td></td></tr>
<tr><td>Oligo fwd</td><td>CACCTAGGAGGTTCGGACGGCGCT</td><td></td><td></td><td></td><td></td><td></td><td></td><td></td><td></td><td></td><td></td></tr>
<tr><td>Oligo rev</td><td>AAACAGCGCCGTCCGAACCTCCTA</td><td></td><td></td><td></td><td></td><td></td><td></td><td></td><td></td><td></td><td></td></tr>
<tr><td>Chromosome</td><td>start</td><td>end</td><td>strand</td><td>MM</td><td>target_seq</td><td>PAM</td><td>alignment</td><td>distance</td><td>position</td><td>gene name</td><td>gene id</td></tr>
<tr><td>GU305914.1</td><td>203058</td><td>203080</td><td>-</td><td>0</td><td>TAGGAGGTTCGGACGGCGCT</td><td>CGG</td><td>||||||||[||||||||||||]PAM</td><td>NA</td><td>Intergenic</td><td>NA</td><td>NA</td></tr>
<tr><td>T19119</td><td>AGGAGGTTCGGACGGCGCTCGGG</td><td>1000</td><td></td><td></td><td></td><td></td><td></td><td></td><td></td><td></td><td></td></tr>
<tr><td>Oligo fwd</td><td>CACCAGGAGGTTCGGACGGCGCTC</td><td></td><td></td><td></td><td></td><td></td><td></td><td></td><td></td><td></td><td></td></tr>
<tr><td>Oligo rev</td><td>AAACGAGCGCCGTCCGAACCTCCT</td><td></td><td></td><td></td><td></td><td></td><td></td><td></td><td></td><td></td><td></td></tr>
<tr><td>Chromosome</td><td>start</td><td>end</td><td>strand</td><td>MM</td><td>target_seq</td><td>PAM</td><td>alignment</td><td>distance</td><td>position</td><td>gene name</td><td>gene id</td></tr>
<tr><td>GU305914.1</td><td>203057</td><td>203079</td><td>-</td><td>0</td><td>AGGAGGTTCGGACGGCGCTC</td><td>GGG</td><td>||||||||[||||||||||||]PAM</td><td>NA</td><td>Intergenic</td><td>NA</td><td>NA</td></tr>
<tr><td>GU305914.1</td><td>169598</td><td>169620</td><td>-</td><td>4</td><td>AGAAGTTTCGAACGGCGCGC</td><td>GGG</td><td>||-||-||[||-|||||||-|]PAM</td><td>NA</td><td>Intergenic</td><td>NA</td><td>NA</td></tr>
<tr><td>T19120</td><td>GCTCGGGACACGAGCCGAAACGG</td><td>1000</td><td></td><td></td><td></td><td></td><td></td><td></td><td></td><td></td><td></td></tr>
<tr><td>Oligo fwd</td><td>CACCGCTCGGGACACGAGCCGAAA</td><td></td><td></td><td></td><td></td><td></td><td></td><td></td><td></td><td></td><td></td></tr>
<tr><td>Oligo rev</td><td>AAACTTTCGGCTCGTGTCCCGAGC</td><td></td><td></td><td></td><td></td><td></td><td></td><td></td><td></td><td></td><td></td></tr>
<tr><td>Chromosome</td><td>start</td><td>end</td><td>strand</td><td>MM</td><td>target_seq</td><td>PAM</td><td>alignment</td><td>distance</td><td>position</td><td>gene name</td><td>gene id</td></tr>
<tr><td>GU305914.1</td><td>203041</td><td>203063</td><td>-</td><td>0</td><td>GCTCGGGACACGAGCCGAAA</td><td>CGG</td><td>||||||||[||||||||||||]PAM</td><td>NA</td><td>Intergenic</td><td>NA</td><td>NA</td></tr>
<tr><td>T19121</td><td>GCCGAAACGGCCGTCCTACCAGG</td><td>1000</td><td></td><td></td><td></td><td></td><td></td><td></td><td></td><td></td><td></td></tr>
<tr><td>Oligo fwd</td><td>CACCGCCGAAACGGCCGTCCTACC</td><td></td><td></td><td></td><td></td><td></td><td></td><td></td><td></td><td></td><td></td></tr>
<tr><td>Oligo rev</td><td>AAACGGTAGGACGGCCGTTTCGGC</td><td></td><td></td><td></td><td></td><td></td><td></td><td></td><td></td><td></td><td></td></tr>
<tr><td>Chromosome</td><td>start</td><td>end</td><td>strand</td><td>MM</td><td>target_seq</td><td>PAM</td><td>alignment</td><td>distance</td><td>position</td><td>gene name</td><td>gene id</td></tr>
<tr><td>GU305914.1</td><td>203028</td><td>203050</td><td>-</td><td>0</td><td>GCCGAAACGGCCGTCCTACC</td><td>AGG</td><td>||||||||[||||||||||||]PAM</td><td>NA</td><td>Intergenic</td><td>NA</td><td>NA</td></tr>
<tr><td>T19122</td><td>CTACCAGGTGAAGAGACGCGCGG</td><td>1000</td><td></td><td></td><td></td><td></td><td></td><td></td><td></td><td></td><td></td></tr>
<tr><td>Oligo fwd</td><td>CACCCTACCAGGTGAAGAGACGCG</td><td></td><td></td><td></td><td></td><td></td><td></td><td></td><td></td><td></td><td></td></tr>
<tr><td>Oligo rev</td><td>AAACCGCGTCTCTTCACCTGGTAG</td><td></td><td></td><td></td><td></td><td></td><td></td><td></td><td></td><td></td><td></td></tr>
<tr><td>Chromosome</td><td>start</td><td>end</td><td>strand</td><td>MM</td><td>target_seq</td><td>PAM</td><td>alignment</td><td>distance</td><td>position</td><td>gene name</td><td>gene id</td></tr>
<tr><td>GU305914.1</td><td>203013</td><td>203035</td><td>-</td><td>0</td><td>CTACCAGGTGAAGAGACGCG</td><td>CGG</td><td>||||||||[||||||||||||]PAM</td><td>NA</td><td>Intergenic</td><td>NA</td><td>NA</td></tr>
<tr><td>T19123</td><td>GCGACCTACCGATGACCATCTGG</td><td>1000</td><td></td><td></td><td></td><td></td><td></td><td></td><td></td><td></td><td></td></tr>
<tr><td>Oligo fwd</td><td>CACCGCGACCTACCGATGACCATC</td><td></td><td></td><td></td><td></td><td></td><td></td><td></td><td></td><td></td><td></td></tr>
<tr><td>Oligo rev</td><td>AAACGATGGTCATCGGTAGGTCGC</td><td></td><td></td><td></td><td></td><td></td><td></td><td></td><td></td><td></td><td></td></tr>
<tr><td>Chromosome</td><td>start</td><td>end</td><td>strand</td><td>MM</td><td>target_seq</td><td>PAM</td><td>alignment</td><td>distance</td><td>position</td><td>gene name</td><td>gene id</td></tr>
<tr><td>GU305914.1</td><td>202981</td><td>203003</td><td>-</td><td>0</td><td>GCGACCTACCGATGACCATC</td><td>TGG</td><td>||||||||[||||||||||||]PAM</td><td>NA</td><td>Intergenic</td><td>NA</td><td>NA</td></tr>
<tr><td>T19124</td><td>CCGATGACCATCTGGTTCCGCGG</td><td>1000</td><td></td><td></td><td></td><td></td><td></td><td></td><td></td><td></td><td></td></tr>
<tr><td>Oligo fwd</td><td>CACCCCGATGACCATCTGGTTCCG</td><td></td><td></td><td></td><td></td><td></td><td></td><td></td><td></td><td></td><td></td></tr>
<tr><td>Oligo rev</td><td>AAACCGGAACCAGATGGTCATCGG</td><td></td><td></td><td></td><td></td><td></td><td></td><td></td><td></td><td></td><td></td></tr>
<tr><td>Chromosome</td><td>start</td><td>end</td><td>strand</td><td>MM</td><td>target_seq</td><td>PAM</td><td>alignment</td><td>distance</td><td>position</td><td>gene name</td><td>gene id</td></tr>
<tr><td>GU305914.1</td><td>202973</td><td>202995</td><td>-</td><td>0</td><td>CCGATGACCATCTGGTTCCG</td><td>CGG</td><td>||||||||[||||||||||||]PAM</td><td>NA</td><td>Intergenic</td><td>NA</td><td>NA</td></tr>
<tr><td>T19125</td><td>CGACAACGTGATGAGCACTCAGG</td><td>1000</td><td></td><td></td><td></td><td></td><td></td><td></td><td></td><td></td><td></td></tr>
<tr><td>Oligo fwd</td><td>CACCCGACAACGTGATGAGCACTC</td><td></td><td></td><td></td><td></td><td></td><td></td><td></td><td></td><td></td><td></td></tr>
<tr><td>Oligo rev</td><td>AAACGAGTGCTCATCACGTTGTCG</td><td></td><td></td><td></td><td></td><td></td><td></td><td></td><td></td><td></td><td></td></tr>
<tr><td>Chromosome</td><td>start</td><td>end</td><td>strand</td><td>MM</td><td>target_seq</td><td>PAM</td><td>alignment</td><td>distance</td><td>position</td><td>gene name</td><td>gene id</td></tr>
<tr><td>GU305914.1</td><td>202950</td><td>202972</td><td>-</td><td>0</td><td>CGACAACGTGATGAGCACTC</td><td>AGG</td><td>||||||||[||||||||||||]PAM</td><td>NA</td><td>Intergenic</td><td>NA</td><td>NA</td></tr>
<tr><td>T19126</td><td>CGTGATGAGCACTCAGGTCGAGG</td><td>1000</td><td></td><td></td><td></td><td></td><td></td><td></td><td></td><td></td><td></td></tr>
<tr><td>Oligo fwd</td><td>CACCCGTGATGAGCACTCAGGTCG</td><td></td><td></td><td></td><td></td><td></td><td></td><td></td><td></td><td></td><td></td></tr>
<tr><td>Oligo rev</td><td>AAACCGACCTGAGTGCTCATCACG</td><td></td><td></td><td></td><td></td><td></td><td></td><td></td><td></td><td></td><td></td></tr>
<tr><td>Chromosome</td><td>start</td><td>end</td><td>strand</td><td>MM</td><td>target_seq</td><td>PAM</td><td>alignment</td><td>distance</td><td>position</td><td>gene name</td><td>gene id</td></tr>
<tr><td>GU305914.1</td><td>202944</td><td>202966</td><td>-</td><td>0</td><td>CGTGATGAGCACTCAGGTCG</td><td>AGG</td><td>||||||||[||||||||||||]PAM</td><td>NA</td><td>Intergenic</td><td>NA</td><td>NA</td></tr>
<tr><td>T19127</td><td>CACTCAGGTCGAGGCCTGTCCGG</td><td>1000</td><td></td><td></td><td></td><td></td><td></td><td></td><td></td><td></td><td></td></tr>
<tr><td>Oligo fwd</td><td>CACCCACTCAGGTCGAGGCCTGTC</td><td></td><td></td><td></td><td></td><td></td><td></td><td></td><td></td><td></td><td></td></tr>
<tr><td>Oligo rev</td><td>AAACGACAGGCCTCGACCTGAGTG</td><td></td><td></td><td></td><td></td><td></td><td></td><td></td><td></td><td></td><td></td></tr>
<tr><td>Chromosome</td><td>start</td><td>end</td><td>strand</td><td>MM</td><td>target_seq</td><td>PAM</td><td>alignment</td><td>distance</td><td>position</td><td>gene name</td><td>gene id</td></tr>
<tr><td>GU305914.1</td><td>202935</td><td>202957</td><td>-</td><td>0</td><td>CACTCAGGTCGAGGCCTGTC</td><td>CGG</td><td>||||||||[||||||||||||]PAM</td><td>NA</td><td>Intergenic</td><td>NA</td><td>NA</td></tr>
<tr><td>T19128</td><td>GATGTCCGCCCGCGAGTTGTCGG</td><td>1000</td><td></td><td></td><td></td><td></td><td></td><td></td><td></td><td></td><td></td></tr>
<tr><td>Oligo fwd</td><td>CACCGATGTCCGCCCGCGAGTTGT</td><td></td><td></td><td></td><td></td><td></td><td></td><td></td><td></td><td></td><td></td></tr>
<tr><td>Oligo rev</td><td>AAACACAACTCGCGGGCGGACATC</td><td></td><td></td><td></td><td></td><td></td><td></td><td></td><td></td><td></td><td></td></tr>
<tr><td>Chromosome</td><td>start</td><td>end</td><td>strand</td><td>MM</td><td>target_seq</td><td>PAM</td><td>alignment</td><td>distance</td><td>position</td><td>gene name</td><td>gene id</td></tr>
<tr><td>GU305914.1</td><td>202899</td><td>202921</td><td>-</td><td>0</td><td>GATGTCCGCCCGCGAGTTGT</td><td>CGG</td><td>||||||||[||||||||||||]PAM</td><td>NA</td><td>Intergenic</td><td>NA</td><td>NA</td></tr>
<tr><td>T19129</td><td>CCCGCGAGTTGTCGGACGCCTGG</td><td>1000</td><td></td><td></td><td></td><td></td><td></td><td></td><td></td><td></td><td></td></tr>
<tr><td>Oligo fwd</td><td>CACCCCCGCGAGTTGTCGGACGCC</td><td></td><td></td><td></td><td></td><td></td><td></td><td></td><td></td><td></td><td></td></tr>
<tr><td>Oligo rev</td><td>AAACGGCGTCCGACAACTCGCGGG</td><td></td><td></td><td></td><td></td><td></td><td></td><td></td><td></td><td></td><td></td></tr>
<tr><td>Chromosome</td><td>start</td><td>end</td><td>strand</td><td>MM</td><td>target_seq</td><td>PAM</td><td>alignment</td><td>distance</td><td>position</td><td>gene name</td><td>gene id</td></tr>
</table>

GU305914.1 202891 202913 - 0 CCCGCGAGTTGTCGGACGCC TGG ||||||||[||||||||||||]PAM NA Intergenic NA NA

T19130 TTGTCGGACGCCTGGTCGAACGG 1000
Oligo fwd CACCTTGTCGGACGCCTGGTCGAA
Oligo rev AAACTTCGACCAGGCGTCCGACAA
Chromosome start end strand MM target_seq PAM alignment distance position gene name gene id
GU305914.1 202883 202905 - 0 TTGTCGGACGCCTGGTCGAA CGG ||||||||[||||||||||||]PAM NA Intergenic NA NA

T19131 TGTCGGACGCCTGGTCGAACGGG 1000
Oligo fwd CACCTGTCGGACGCCTGGTCGAAC
Oligo rev AAACGTTCGACCAGGCGTCCGACA
Chromosome start end strand MM target_seq PAM alignment distance position gene name gene id
GU305914.1 202882 202904 - 0 TGTCGGACGCCTGGTCGAAC GGG ||||||||[||||||||||||]PAM NA Intergenic NA NA

T19132 GTCGGACGCCTGGTCGAACGGGG 1000
Oligo fwd CACCGTCGGACGCCTGGTCGAACG
Oligo rev AAACCGTTCGACCAGGCGTCCGAC
Chromosome start end strand MM target_seq PAM alignment distance position gene name gene id
GU305914.1 202881 202903 - 0 GTCGGACGCCTGGTCGAACG GGG ||||||||[||||||||||||]PAM NA Intergenic NA NA

T19133 GACGCCTGGTCGAACGGGGACGG 1000
Oligo fwd CACCGACGCCTGGTCGAACGGGGA
Oligo rev AAACTCCCCGTTCGACCAGGCGTC
Chromosome start end strand MM target_seq PAM alignment distance position gene name gene id
GU305914.1 202877 202899 - 0 GACGCCTGGTCGAACGGGGA CGG ||||||||[||||||||||||]PAM NA Intergenic NA NA

T19134 CGGGGACGGTCCGATCGTCACGG 1000
Oligo fwd CACCCGGGGACGGTCCGATCGTCA
Oligo rev AAACTGACGATCGGACCGTCCCCG
Chromosome start end strand MM target_seq PAM alignment distance position gene name gene id
GU305914.1 202863 202885 - 0 CGGGGACGGTCCGATCGTCA CGG ||||||||[||||||||||||]PAM NA Intergenic NA NA

T19135 CGGTCCGATCGTCACGGTCCCGG 1000
Oligo fwd CACCCGGTCCGATCGTCACGGTCC
Oligo rev AAACGGACCGTGACGATCGGACCG
Chromosome start end strand MM target_seq PAM alignment distance position gene name gene id
GU305914.1 202857 202879 - 0 CGGTCCGATCGTCACGGTCC CGG ||||||||[||||||||||||]PAM NA Intergenic NA NA

T19136 GAGCGCGCCTCTGATCTGTCTGG 1000
Oligo fwd CACCGAGCGCGCCTCTGATCTGTC
Oligo rev AAACGACAGATCAGAGGCGCGCTC
Chromosome start end strand MM target_seq PAM alignment distance position gene name gene id
GU305914.1 202822 202844 - 0 GAGCGCGCCTCTGATCTGTC TGG ||||||||[||||||||||||]PAM NA Intergenic NA NA

T19137 AGCGCGCCTCTGATCTGTCTGGG 1000
Oligo fwd CACCAGCGCGCCTCTGATCTGTCT
Oligo rev AAACAGACAGATCAGAGGCGCGCT
Chromosome start end strand MM target_seq PAM alignment distance position gene name gene id
GU305914.1 202821 202843 - 0 AGCGCGCCTCTGATCTGTCT GGG ||||||||[||||||||||||]PAM NA Intergenic NA NA

T19138 GCCTCTGATCTGTCTGGGTGTGG 1000
Oligo fwd CACCGCCTCTGATCTGTCTGGGTG
Oligo rev AAACCACCCAGACAGATCAGAGGC
Chromosome start end strand MM target_seq PAM alignment distance position gene name gene id
GU305914.1 202816 202838 - 0 GCCTCTGATCTGTCTGGGTG TGG ||||||||[||||||||||||]PAM NA Intergenic NA NA

T19139 TCTGTCTGGGTGTGGTCACCCGG 1000
Oligo fwd CACCTCTGTCTGGGTGTGGTCACC
Oligo rev AAACGGTGACCACACCCAGACAGA
Chromosome start end strand MM target_seq PAM alignment distance position gene name gene id
GU305914.1 202808 202830 - 0 TCTGTCTGGGTGTGGTCACC CGG ||||||||[||||||||||||]PAM NA Intergenic NA NA

T19140 TCTGGGTGTGGTCACCCGGATGG 1000
Oligo fwd CACCTCTGGGTGTGGTCACCCGGA
Oligo rev AAACTCCGGGTGACCACACCCAGA
Chromosome start end strand MM target_seq PAM alignment distance position gene name gene id
GU305914.1 202804 202826 - 0 TCTGGGTGTGGTCACCCGGA TGG ||||||||[||||||||||||]PAM NA Intergenic NA NA

T19141 TCACCCGGATGGTGTAAAAGAGG 1000
Oligo fwd CACCTCACCCGGATGGTGTAAAAG
Oligo rev AAACCTTTTACACCATCCGGGTGA
Chromosome start end strand MM target_seq PAM alignment distance position gene name gene id
GU305914.1 202793 202815 - 0 TCACCCGGATGGTGTAAAAG AGG ||||||||[||||||||||||]PAM NA Intergenic NA NA

T19142 CACCCGGATGGTGTAAAAGAGGG 1000
Oligo fwd CACCCACCCGGATGGTGTAAAAGA
Oligo rev AAACTCTTTTACACCATCCGGGTG
Chromosome start end strand MM target_seq PAM alignment distance position gene name gene id
GU305914.1 202792 202814 - 0 CACCCGGATGGTGTAAAAGA GGG ||||||||[||||||||||||]PAM NA Intergenic NA NA

T19143 GGGCCGCGTAACTCCAGAAACGG 1000
Oligo fwd CACCGGGCCGCGTAACTCCAGAAA
Oligo rev AAACTTTCTGGAGTTACGCGGCCC
Chromosome start end strand MM target_seq PAM alignment distance position gene name gene id
GU305914.1 202772 202794 - 0 GGGCCGCGTAACTCCAGAAA CGG ||||||||[||||||||||||]PAM NA Intergenic NA NA

T19144 TCTCTGTTTCCACAGCAGAGAGG 1000
Oligo fwd CACCTCTCTGTTTCCACAGCAGAG
Oligo rev AAACCTCTGCTGTGGAAACAGAGA

| | | | | | | | | | | | |
|---|---|---|---|---|---|---|---|---|---|---|---|
| Chromosome | start | end | strand | MM | target_seq | PAM | alignment | distance | position | gene name | gene id |
| GU305914.1 | 202741 | 202763 | - | 0 | TCTCTGTTTCCACAGCAGAG | AGG | \|\|\|\|\|\|\|\|[\|\|\|\|\|\|\|\|\|\|\|\|]PAM | NA | Intergenic | NA | NA |
| T19145 | CTGTTTCCACAGCAGAGAGGTGG | 1000 | | | | | | | | | |
| Oligo fwd | CACCCTGTTTCCACAGCAGAGAGG | | | | | | | | | | |
| Oligo rev | AAACCCTCTCTGCTGTGGAAACAG | | | | | | | | | | |
| Chromosome | start | end | strand | MM | target_seq | PAM | alignment | distance | position | gene name | gene id |
| GU305914.1 | 202738 | 202760 | - | 0 | CTGTTTCCACAGCAGAGAGG | TGG | \|\|\|\|\|\|\|\|[\|\|\|\|\|\|\|\|\|\|\|\|]PAM | NA | Intergenic | NA | NA |
| T19146 | CAGCAGAGAGGTGGTTGCCTCGG | 1000 | | | | | | | | | |
| Oligo fwd | CACCCAGCAGAGAGGTGGTTGCCT | | | | | | | | | | |
| Oligo rev | AAACAGGCAACCACCTCTCTGCTG | | | | | | | | | | |
| Chromosome | start | end | strand | MM | target_seq | PAM | alignment | distance | position | gene name | gene id |
| GU305914.1 | 202729 | 202751 | - | 0 | CAGCAGAGAGGTGGTTGCCT | CGG | \|\|\|\|\|\|\|\|[\|\|\|\|\|\|\|\|\|\|\|\|]PAM | NA | Intergenic | NA | NA |
| T19147 | CCGTCTCGTGCGCGATGTCTTGG | 1000 | | | | | | | | | |
| Oligo fwd | CACCCCGTCTCGTGCGCGATGTCT | | | | | | | | | | |
| Oligo rev | AAACAGACATCGCGCACGAGACGG | | | | | | | | | | |
| Chromosome | start | end | strand | MM | target_seq | PAM | alignment | distance | position | gene name | gene id |
| GU305914.1 | 202686 | 202708 | - | 0 | CCGTCTCGTGCGCGATGTCT | TGG | \|\|\|\|\|\|\|\|[\|\|\|\|\|\|\|\|\|\|\|\|]PAM | NA | Intergenic | NA | NA |
| T19148 | CGTCTCGTGCGCGATGTCTTGGG | 1000 | | | | | | | | | |
| Oligo fwd | CACCCGTCTCGTGCGCGATGTCTT | | | | | | | | | | |
| Oligo rev | AAACAAGACATCGCGCACGAGACG | | | | | | | | | | |
| Chromosome | start | end | strand | MM | target_seq | PAM | alignment | distance | position | gene name | gene id |
| GU305914.1 | 202685 | 202707 | - | 0 | CGTCTCGTGCGCGATGTCTT | GGG | \|\|\|\|\|\|\|\|[\|\|\|\|\|\|\|\|\|\|\|\|]PAM | NA | Intergenic | NA | NA |
| T19149 | TGCGCGATGTCTTGGGTGACCGG | 1000 | | | | | | | | | |
| Oligo fwd | CACCTGCGCGATGTCTTGGGTGAC | | | | | | | | | | |
| Oligo rev | AAACGTCACCCAAGACATCGCGCA | | | | | | | | | | |
| Chromosome | start | end | strand | MM | target_seq | PAM | alignment | distance | position | gene name | gene id |
| GU305914.1 | 202678 | 202700 | - | 0 | TGCGCGATGTCTTGGGTGAC | CGG | \|\|\|\|\|\|\|\|[\|\|\|\|\|\|\|\|\|\|\|\|]PAM | NA | Intergenic | NA | NA |
| GU305914.1 | 18000 | 18022 | + | 4 | CGCGCGACCTCTTGGGTGTC | GGG | -\|\|\|\|\|\|-[-\|\|\|\|\|\|\|\|\|-\|]PAM | NA | Intergenic | NA | NA |
| T19150 | GTCTTGGGTGACCGGAGATCCGG | 1000 | | | | | | | | | |
| Oligo fwd | CACCGTCTTGGGTGACCGGAGATC | | | | | | | | | | |
| Oligo rev | AAACGATCTCCGGTCACCCAAGAC | | | | | | | | | | |
| Chromosome | start | end | strand | MM | target_seq | PAM | alignment | distance | position | gene name | gene id |
| GU305914.1 | 202670 | 202692 | - | 0 | GTCTTGGGTGACCGGAGATC | CGG | \|\|\|\|\|\|\|\|[\|\|\|\|\|\|\|\|\|\|\|\|]PAM | NA | Intergenic | NA | NA |
| T19151 | GGGTGACCGGAGATCCGGCCTGG | 1000 | | | | | | | | | |
| Oligo fwd | CACCGGGTGACCGGAGATCCGGCC | | | | | | | | | | |
| Oligo rev | AAACGGCCGGATCTCCGGTCACCC | | | | | | | | | | |
| Chromosome | start | end | strand | MM | target_seq | PAM | alignment | distance | position | gene name | gene id |
| GU305914.1 | 202665 | 202687 | - | 0 | GGGTGACCGGAGATCCGGCC | TGG | \|\|\|\|\|\|\|\|[\|\|\|\|\|\|\|\|\|\|\|\|]PAM | NA | Intergenic | NA | NA |
| GU305914.1 | 149409 | 149431 | + | 4 | GGGGGGCCGGAGACCTGGCC | TGG | \|\|\|-\|-\|\|[\|\|\|\|\|-\|-\|\|\|\|]PAM | NA | Intergenic | NA | NA |
| T19152 | TGACCGGAGATCCGGCCTGGAGG | 1000 | | | | | | | | | |
| Oligo fwd | CACCTGACCGGAGATCCGGCCTGG | | | | | | | | | | |
| Oligo rev | AAACCCAGGCCGGATCTCCGGTCA | | | | | | | | | | |
| Chromosome | start | end | strand | MM | target_seq | PAM | alignment | distance | position | gene name | gene id |
| GU305914.1 | 202662 | 202684 | - | 0 | TGACCGGAGATCCGGCCTGG | AGG | \|\|\|\|\|\|\|\|[\|\|\|\|\|\|\|\|\|\|\|\|]PAM | NA | Intergenic | NA | NA |
| T19153 | CGGAGATCCGGCCTGGAGGCCGG | 1000 | | | | | | | | | |
| Oligo fwd | CACCCGGAGATCCGGCCTGGAGGC | | | | | | | | | | |
| Oligo rev | AAACGCCTCCAGGCCGGATCTCCG | | | | | | | | | | |
| Chromosome | start | end | strand | MM | target_seq | PAM | alignment | distance | position | gene name | gene id |
| GU305914.1 | 202658 | 202680 | - | 0 | CGGAGATCCGGCCTGGAGGC | CGG | \|\|\|\|\|\|\|\|[\|\|\|\|\|\|\|\|\|\|\|\|]PAM | NA | Intergenic | NA | NA |
| T19154 | AGATCCGGCCTGGAGGCCGGAGG | 1000 | | | | | | | | | |
| Oligo fwd | CACCAGATCCGGCCTGGAGGCCGG | | | | | | | | | | |
| Oligo rev | AAACCCGGCCTCCAGGCCGGATCT | | | | | | | | | | |
| Chromosome | start | end | strand | MM | target_seq | PAM | alignment | distance | position | gene name | gene id |
| GU305914.1 | 202655 | 202677 | - | 0 | AGATCCGGCCTGGAGGCCGG | AGG | \|\|\|\|\|\|\|\|[\|\|\|\|\|\|\|\|\|\|\|\|]PAM | NA | Intergenic | NA | NA |
| T19155 | GCCGGAGGCTCTGATCCGCCCGG | 1000 | | | | | | | | | |
| Oligo fwd | CACCGCCGGAGGCTCTGATCCGCC | | | | | | | | | | |
| Oligo rev | AAACGGCGGATCAGAGCCTCCGGC | | | | | | | | | | |
| Chromosome | start | end | strand | MM | target_seq | PAM | alignment | distance | position | gene name | gene id |
| GU305914.1 | 202640 | 202662 | - | 0 | GCCGGAGGCTCTGATCCGCC | CGG | \|\|\|\|\|\|\|\|[\|\|\|\|\|\|\|\|\|\|\|\|]PAM | NA | Intergenic | NA | NA |
| T19156 | GGAGGCTCTGATCCGCCCGGCGG | 1000 | | | | | | | | | |
| Oligo fwd | CACCGGAGGCTCTGATCCGCCCGG | | | | | | | | | | |
| Oligo rev | AAACCCGGGCGGATCAGAGCCTCC | | | | | | | | | | |
| Chromosome | start | end | strand | MM | target_seq | PAM | alignment | distance | position | gene name | gene id |
| GU305914.1 | 202637 | 202659 | - | 0 | GGAGGCTCTGATCCGCCCGG | CGG | \|\|\|\|\|\|\|\|[\|\|\|\|\|\|\|\|\|\|\|\|]PAM | NA | Intergenic | NA | NA |
| T19157 | TTTTTTCCTCACGCAGACGTCGG | 1000 | | | | | | | | | |
| Oligo fwd | CACCTTTTTTCCTCACGCAGACGT | | | | | | | | | | |
| Oligo rev | AAACACGTCTGCGTGAGGAAAAAA | | | | | | | | | | |
| Chromosome | start | end | strand | MM | target_seq | PAM | alignment | distance | position | gene name | gene id |
| GU305914.1 | 202583 | 202605 | - | 0 | TTTTTTCCTCACGCAGACGT | CGG | \|\|\|\|\|\|\|\|[\|\|\|\|\|\|\|\|\|\|\|\|]PAM | NA | Intergenic | NA | NA |
| T19158 | CTCTGAAAAAGACAGTCAAGCGG | 1000 | | | | | | | | | |
| Oligo fwd | CACCCTCTGAAAAAGACAGTCAAG | | | | | | | | | | |
| Oligo rev | AAACCTTGACTGTCTTTTTCAGAG | | | | | | | | | | |
| Chromosome | start | end | strand | MM | target_seq | PAM | alignment | distance | position | gene name | gene id |
| GU305914.1 | 202554 | 202576 | - | 0 | CTCTGAAAAAGACAGTCAAG | CGG | \|\|\|\|\|\|\|\|[\|\|\|\|\|\|\|\|\|\|\|\|]PAM | NA | Intergenic | NA | NA |

<table>
<tr><td>T19159</td><td>TCTGAAAAAGACAGTCAAGCGGG</td><td>1000</td><td></td><td></td><td></td><td></td><td></td><td></td><td></td><td></td><td></td></tr>
<tr><td>Oligo fwd</td><td>CACCTCTGAAAAAGACAGTCAAGC</td><td></td><td></td><td></td><td></td><td></td><td></td><td></td><td></td><td></td><td></td></tr>
<tr><td>Oligo rev</td><td>AAACGCTTGACTGTCTTTTTCAGA</td><td></td><td></td><td></td><td></td><td></td><td></td><td></td><td></td><td></td><td></td></tr>
<tr><td>Chromosome</td><td>start</td><td>end</td><td>strand</td><td>MM</td><td>target_seq</td><td>PAM</td><td>alignment</td><td>distance</td><td>position</td><td>gene name</td><td>gene id</td></tr>
<tr><td>GU305914.1</td><td>202553</td><td>202575</td><td>-</td><td>0</td><td>TCTGAAAAAGACAGTCAAGC</td><td>GGG</td><td>||||||||[||||||||||||]PAM</td><td>NA</td><td>Intergenic</td><td>NA</td><td>NA</td></tr>
<tr><td>T19160</td><td>AAGACAGTCAAGCGGGAGAGCGG</td><td>1000</td><td></td><td></td><td></td><td></td><td></td><td></td><td></td><td></td><td></td></tr>
<tr><td>Oligo fwd</td><td>CACCAAGACAGTCAAGCGGGAGAG</td><td></td><td></td><td></td><td></td><td></td><td></td><td></td><td></td><td></td><td></td></tr>
<tr><td>Oligo rev</td><td>AAACCTCTCCCGCTTGACTGTCTT</td><td></td><td></td><td></td><td></td><td></td><td></td><td></td><td></td><td></td><td></td></tr>
<tr><td>Chromosome</td><td>start</td><td>end</td><td>strand</td><td>MM</td><td>target_seq</td><td>PAM</td><td>alignment</td><td>distance</td><td>position</td><td>gene name</td><td>gene id</td></tr>
<tr><td>GU305914.1</td><td>202546</td><td>202568</td><td>-</td><td>0</td><td>AAGACAGTCAAGCGGGAGAG</td><td>CGG</td><td>||||||||[||||||||||||]PAM</td><td>NA</td><td>Intergenic</td><td>NA</td><td>NA</td></tr>
<tr><td>T19161</td><td>AGCGGCACCAAGCTCTGTGTCGG</td><td>1000</td><td></td><td></td><td></td><td></td><td></td><td></td><td></td><td></td><td></td></tr>
<tr><td>Oligo fwd</td><td>CACCAGCGGCACCAAGCTCTGTGT</td><td></td><td></td><td></td><td></td><td></td><td></td><td></td><td></td><td></td><td></td></tr>
<tr><td>Oligo rev</td><td>AAACACACAGAGCTTGGTGCCGCT</td><td></td><td></td><td></td><td></td><td></td><td></td><td></td><td></td><td></td><td></td></tr>
<tr><td>Chromosome</td><td>start</td><td>end</td><td>strand</td><td>MM</td><td>target_seq</td><td>PAM</td><td>alignment</td><td>distance</td><td>position</td><td>gene name</td><td>gene id</td></tr>
<tr><td>GU305914.1</td><td>202528</td><td>202550</td><td>-</td><td>0</td><td>AGCGGCACCAAGCTCTGTGT</td><td>CGG</td><td>||||||||[||||||||||||]PAM</td><td>NA</td><td>Intergenic</td><td>NA</td><td>NA</td></tr>
<tr><td>T19162</td><td>GCGGCACCAAGCTCTGTGTCGGG</td><td>1000</td><td></td><td></td><td></td><td></td><td></td><td></td><td></td><td></td><td></td></tr>
<tr><td>Oligo fwd</td><td>CACCGCGGCACCAAGCTCTGTGTC</td><td></td><td></td><td></td><td></td><td></td><td></td><td></td><td></td><td></td><td></td></tr>
<tr><td>Oligo rev</td><td>AAACGACACAGAGCTTGGTGCCGC</td><td></td><td></td><td></td><td></td><td></td><td></td><td></td><td></td><td></td><td></td></tr>
<tr><td>Chromosome</td><td>start</td><td>end</td><td>strand</td><td>MM</td><td>target_seq</td><td>PAM</td><td>alignment</td><td>distance</td><td>position</td><td>gene name</td><td>gene id</td></tr>
<tr><td>GU305914.1</td><td>202527</td><td>202549</td><td>-</td><td>0</td><td>GCGGCACCAAGCTCTGTGTC</td><td>GGG</td><td>||||||||[||||||||||||]PAM</td><td>NA</td><td>Intergenic</td><td>NA</td><td>NA</td></tr>
<tr><td>T19163</td><td>GCTCTGTGTCGGGTACCCCCCGG</td><td>1000</td><td></td><td></td><td></td><td></td><td></td><td></td><td></td><td></td><td></td></tr>
<tr><td>Oligo fwd</td><td>CACCGCTCTGTGTCGGGTACCCCC</td><td></td><td></td><td></td><td></td><td></td><td></td><td></td><td></td><td></td><td></td></tr>
<tr><td>Oligo rev</td><td>AAACGGGGGTACCCGACACAGAGC</td><td></td><td></td><td></td><td></td><td></td><td></td><td></td><td></td><td></td><td></td></tr>
<tr><td>Chromosome</td><td>start</td><td>end</td><td>strand</td><td>MM</td><td>target_seq</td><td>PAM</td><td>alignment</td><td>distance</td><td>position</td><td>gene name</td><td>gene id</td></tr>
<tr><td>GU305914.1</td><td>202517</td><td>202539</td><td>-</td><td>0</td><td>GCTCTGTGTCGGGTACCCCC</td><td>CGG</td><td>||||||||[||||||||||||]PAM</td><td>NA</td><td>Intergenic</td><td>NA</td><td>NA</td></tr>
<tr><td>T19164</td><td>CTCTGTGTCGGGTACCCCCCGGG</td><td>1000</td><td></td><td></td><td></td><td></td><td></td><td></td><td></td><td></td><td></td></tr>
<tr><td>Oligo fwd</td><td>CACCCTCTGTGTCGGGTACCCCCC</td><td></td><td></td><td></td><td></td><td></td><td></td><td></td><td></td><td></td><td></td></tr>
<tr><td>Oligo rev</td><td>AAACGGGGGGTACCCGACACAGAG</td><td></td><td></td><td></td><td></td><td></td><td></td><td></td><td></td><td></td><td></td></tr>
<tr><td>Chromosome</td><td>start</td><td>end</td><td>strand</td><td>MM</td><td>target_seq</td><td>PAM</td><td>alignment</td><td>distance</td><td>position</td><td>gene name</td><td>gene id</td></tr>
<tr><td>GU305914.1</td><td>202516</td><td>202538</td><td>-</td><td>0</td><td>CTCTGTGTCGGGTACCCCCC</td><td>GGG</td><td>||||||||[||||||||||||]PAM</td><td>NA</td><td>Intergenic</td><td>NA</td><td>NA</td></tr>
<tr><td>T19165</td><td>TCTGTGTCGGGTACCCCCCGGGG</td><td>1000</td><td></td><td></td><td></td><td></td><td></td><td></td><td></td><td></td><td></td></tr>
<tr><td>Oligo fwd</td><td>CACCTCTGTGTCGGGTACCCCCCG</td><td></td><td></td><td></td><td></td><td></td><td></td><td></td><td></td><td></td><td></td></tr>
<tr><td>Oligo rev</td><td>AAACCGGGGGGTACCCGACACAGA</td><td></td><td></td><td></td><td></td><td></td><td></td><td></td><td></td><td></td><td></td></tr>
<tr><td>Chromosome</td><td>start</td><td>end</td><td>strand</td><td>MM</td><td>target_seq</td><td>PAM</td><td>alignment</td><td>distance</td><td>position</td><td>gene name</td><td>gene id</td></tr>
<tr><td>GU305914.1</td><td>202515</td><td>202537</td><td>-</td><td>0</td><td>TCTGTGTCGGGTACCCCCCG</td><td>GGG</td><td>||||||||[||||||||||||]PAM</td><td>NA</td><td>Intergenic</td><td>NA</td><td>NA</td></tr>
<tr><td>T19166</td><td>GTGTCGGGTACCCCCCGGGGTGG</td><td>1000</td><td></td><td></td><td></td><td></td><td></td><td></td><td></td><td></td><td></td></tr>
<tr><td>Oligo fwd</td><td>CACCGTGTCGGGTACCCCCCGGGG</td><td></td><td></td><td></td><td></td><td></td><td></td><td></td><td></td><td></td><td></td></tr>
<tr><td>Oligo rev</td><td>AAACCCCCGGGGGGTACCCGACAC</td><td></td><td></td><td></td><td></td><td></td><td></td><td></td><td></td><td></td><td></td></tr>
<tr><td>Chromosome</td><td>start</td><td>end</td><td>strand</td><td>MM</td><td>target_seq</td><td>PAM</td><td>alignment</td><td>distance</td><td>position</td><td>gene name</td><td>gene id</td></tr>
<tr><td>GU305914.1</td><td>202512</td><td>202534</td><td>-</td><td>0</td><td>GTGTCGGGTACCCCCCGGGG</td><td>TGG</td><td>||||||||[||||||||||||]PAM</td><td>NA</td><td>Intergenic</td><td>NA</td><td>NA</td></tr>
<tr><td>T19167</td><td>TCGGGTACCCCCCGGGGTGGTGG</td><td>1000</td><td></td><td></td><td></td><td></td><td></td><td></td><td></td><td></td><td></td></tr>
<tr><td>Oligo fwd</td><td>CACCTCGGGTACCCCCCGGGGTGG</td><td></td><td></td><td></td><td></td><td></td><td></td><td></td><td></td><td></td><td></td></tr>
<tr><td>Oligo rev</td><td>AAACCCACCCCGGGGGGTACCCGA</td><td></td><td></td><td></td><td></td><td></td><td></td><td></td><td></td><td></td><td></td></tr>
<tr><td>Chromosome</td><td>start</td><td>end</td><td>strand</td><td>MM</td><td>target_seq</td><td>PAM</td><td>alignment</td><td>distance</td><td>position</td><td>gene name</td><td>gene id</td></tr>
<tr><td>GU305914.1</td><td>202509</td><td>202531</td><td>-</td><td>0</td><td>TCGGGTACCCCCCGGGGTGG</td><td>TGG</td><td>||||||||[||||||||||||]PAM</td><td>NA</td><td>Intergenic</td><td>NA</td><td>NA</td></tr>
<tr><td>T19168</td><td>GTACCCCCCGGGGTGGTGGCTGG</td><td>1000</td><td></td><td></td><td></td><td></td><td></td><td></td><td></td><td></td><td></td></tr>
<tr><td>Oligo fwd</td><td>CACCGTACCCCCCGGGGTGGTGGC</td><td></td><td></td><td></td><td></td><td></td><td></td><td></td><td></td><td></td><td></td></tr>
<tr><td>Oligo rev</td><td>AAACGCCACCACCCCGGGGGGTAC</td><td></td><td></td><td></td><td></td><td></td><td></td><td></td><td></td><td></td><td></td></tr>
<tr><td>Chromosome</td><td>start</td><td>end</td><td>strand</td><td>MM</td><td>target_seq</td><td>PAM</td><td>alignment</td><td>distance</td><td>position</td><td>gene name</td><td>gene id</td></tr>
<tr><td>GU305914.1</td><td>202505</td><td>202527</td><td>-</td><td>0</td><td>GTACCCCCCGGGGTGGTGGC</td><td>TGG</td><td>||||||||[||||||||||||]PAM</td><td>NA</td><td>Intergenic</td><td>NA</td><td>NA</td></tr>
<tr><td>T19169</td><td>CCCGGGGTGGTGGCTGGCCGTGG</td><td>1000</td><td></td><td></td><td></td><td></td><td></td><td></td><td></td><td></td><td></td></tr>
<tr><td>Oligo fwd</td><td>CACCCCCGGGGTGGTGGCTGGCCG</td><td></td><td></td><td></td><td></td><td></td><td></td><td></td><td></td><td></td><td></td></tr>
<tr><td>Oligo rev</td><td>AAACCGGCCAGCCACCACCCCGGG</td><td></td><td></td><td></td><td></td><td></td><td></td><td></td><td></td><td></td><td></td></tr>
<tr><td>Chromosome</td><td>start</td><td>end</td><td>strand</td><td>MM</td><td>target_seq</td><td>PAM</td><td>alignment</td><td>distance</td><td>position</td><td>gene name</td><td>gene id</td></tr>
<tr><td>GU305914.1</td><td>202499</td><td>202521</td><td>-</td><td>0</td><td>CCCGGGGTGGTGGCTGGCCG</td><td>TGG</td><td>||||||||[||||||||||||]PAM</td><td>NA</td><td>Intergenic</td><td>NA</td><td>NA</td></tr>
<tr><td>T19170</td><td>GGTGGCTGGCCGTGGTGCCGCGG</td><td>1000</td><td></td><td></td><td></td><td></td><td></td><td></td><td></td><td></td><td></td></tr>
<tr><td>Oligo fwd</td><td>CACCGGTGGCTGGCCGTGGTGCCG</td><td></td><td></td><td></td><td></td><td></td><td></td><td></td><td></td><td></td><td></td></tr>
<tr><td>Oligo rev</td><td>AAACCGGCACCACGGCCAGCCACC</td><td></td><td></td><td></td><td></td><td></td><td></td><td></td><td></td><td></td><td></td></tr>
<tr><td>Chromosome</td><td>start</td><td>end</td><td>strand</td><td>MM</td><td>target_seq</td><td>PAM</td><td>alignment</td><td>distance</td><td>position</td><td>gene name</td><td>gene id</td></tr>
<tr><td>GU305914.1</td><td>202491</td><td>202513</td><td>-</td><td>0</td><td>GGTGGCTGGCCGTGGTGCCG</td><td>CGG</td><td>||||||||[||||||||||||]PAM</td><td>NA</td><td>Intergenic</td><td>NA</td><td>NA</td></tr>
<tr><td>GU305914.1</td><td>168889</td><td>168911</td><td>+</td><td>4</td><td>GGCTGCTGGCGCTGGTGCCG</td><td>CGG</td><td>||--||||[||--||||||||]PAM</td><td>NA</td><td>Intergenic</td><td>NA</td><td>NA</td></tr>
<tr><td>T19171</td><td>GTGGCTGGCCGTGGTGCCGCGGG</td><td>1000</td><td></td><td></td><td></td><td></td><td></td><td></td><td></td><td></td><td></td></tr>
<tr><td>Oligo fwd</td><td>CACCGTGGCTGGCCGTGGTGCCGC</td><td></td><td></td><td></td><td></td><td></td><td></td><td></td><td></td><td></td><td></td></tr>
<tr><td>Oligo rev</td><td>AAACGCGGCACCACGGCCAGCCAC</td><td></td><td></td><td></td><td></td><td></td><td></td><td></td><td></td><td></td><td></td></tr>
<tr><td>Chromosome</td><td>start</td><td>end</td><td>strand</td><td>MM</td><td>target_seq</td><td>PAM</td><td>alignment</td><td>distance</td><td>position</td><td>gene name</td><td>gene id</td></tr>
<tr><td>GU305914.1</td><td>202490</td><td>202512</td><td>-</td><td>0</td><td>GTGGCTGGCCGTGGTGCCGC</td><td>GGG</td><td>||||||||[||||||||||||]PAM</td><td>NA</td><td>Intergenic</td><td>NA</td><td>NA</td></tr>
<tr><td>GU305914.1</td><td>168890</td><td>168912</td><td>+</td><td>4</td><td>GCTGCTGGCGCTGGTGCCGC</td><td>GGG</td><td>|--|||||[|--|||||||||]PAM</td><td>NA</td><td>Intergenic</td><td>NA</td><td>NA</td></tr>
<tr><td>T19172</td><td>TGGCTGGCCGTGGTGCCGCGGGG</td><td>1000</td><td></td><td></td><td></td><td></td><td></td><td></td><td></td><td></td><td></td></tr>
<tr><td>Oligo fwd</td><td>CACCTGGCTGGCCGTGGTGCCGCG</td><td></td><td></td><td></td><td></td><td></td><td></td><td></td><td></td><td></td><td></td></tr>
<tr><td>Oligo rev</td><td>AAACCGCGGCACCACGGCCAGCCA</td><td></td><td></td><td></td><td></td><td></td><td></td><td></td><td></td><td></td><td></td></tr>
<tr><td>Chromosome</td><td>start</td><td>end</td><td>strand</td><td>MM</td><td>target_seq</td><td>PAM</td><td>alignment</td><td>distance</td><td>position</td><td>gene name</td><td>gene id</td></tr>
<tr><td>GU305914.1</td><td>202489</td><td>202511</td><td>-</td><td>0</td><td>TGGCTGGCCGTGGTGCCGCG</td><td>GGG</td><td>||||||||[||||||||||||]PAM</td><td>NA</td><td>Intergenic</td><td>NA</td><td>NA</td></tr>
<tr><td>T19173</td><td>CGTGGTGCCGCGGGGCGAACTGG</td><td>1000</td><td></td><td></td><td></td><td></td><td></td><td></td><td></td><td></td><td></td></tr>
<tr><td>Oligo fwd</td><td>CACCCGTGGTGCCGCGGGGCGAAC</td><td></td><td></td><td></td><td></td><td></td><td></td><td></td><td></td><td></td><td></td></tr>
<tr><td>Oligo rev</td><td>AAACGTTCGCCCCGCGGCACCACG</td><td></td><td></td><td></td><td></td><td></td><td></td><td></td><td></td><td></td><td></td></tr>
</table>

<table>
<tr><td>Chromosome</td><td>start</td><td>end</td><td>strand</td><td>MM</td><td>target_seq</td><td>PAM</td><td>alignment</td><td>distance</td><td>position</td><td>gene name</td><td>gene id</td></tr>
<tr><td>GU305914.1</td><td>202481</td><td>202503</td><td>-</td><td>0</td><td>CGTGGTGCCGCGGGGCGAAC</td><td>TGG</td><td>||||||||[||||||||||||]PAM</td><td>NA</td><td>Intergenic</td><td>NA</td><td>NA</td></tr>
<tr><td>T19174</td><td>GCGGGGCGAACTGGTCGAGATGG</td><td>1000</td><td></td><td></td><td></td><td></td><td></td><td></td><td></td><td></td><td></td></tr>
<tr><td>Oligo fwd</td><td>CACCGCGGGGCGAACTGGTCGAGA</td><td></td><td></td><td></td><td></td><td></td><td></td><td></td><td></td><td></td><td></td></tr>
<tr><td>Oligo rev</td><td>AAACTCTCGACCAGTTCGCCCCGC</td><td></td><td></td><td></td><td></td><td></td><td></td><td></td><td></td><td></td><td></td></tr>
<tr><td>Chromosome</td><td>start</td><td>end</td><td>strand</td><td>MM</td><td>target_seq</td><td>PAM</td><td>alignment</td><td>distance</td><td>position</td><td>gene name</td><td>gene id</td></tr>
<tr><td>GU305914.1</td><td>202472</td><td>202494</td><td>-</td><td>0</td><td>GCGGGGCGAACTGGTCGAGA</td><td>TGG</td><td>||||||||[||||||||||||]PAM</td><td>NA</td><td>Intergenic</td><td>NA</td><td>NA</td></tr>
<tr><td>T19175</td><td>GGCGAACTGGTCGAGATGGACGG</td><td>1000</td><td></td><td></td><td></td><td></td><td></td><td></td><td></td><td></td><td></td></tr>
<tr><td>Oligo fwd</td><td>CACCGGCGAACTGGTCGAGATGGA</td><td></td><td></td><td></td><td></td><td></td><td></td><td></td><td></td><td></td><td></td></tr>
<tr><td>Oligo rev</td><td>AAACTCCATCTCGACCAGTTCGCC</td><td></td><td></td><td></td><td></td><td></td><td></td><td></td><td></td><td></td><td></td></tr>
<tr><td>Chromosome</td><td>start</td><td>end</td><td>strand</td><td>MM</td><td>target_seq</td><td>PAM</td><td>alignment</td><td>distance</td><td>position</td><td>gene name</td><td>gene id</td></tr>
<tr><td>GU305914.1</td><td>202468</td><td>202490</td><td>-</td><td>0</td><td>GGCGAACTGGTCGAGATGGA</td><td>CGG</td><td>||||||||[||||||||||||]PAM</td><td>NA</td><td>Intergenic</td><td>NA</td><td>NA</td></tr>
<tr><td>T19176</td><td>AACTGGTCGAGATGGACGGCCGG</td><td>1000</td><td></td><td></td><td></td><td></td><td></td><td></td><td></td><td></td><td></td></tr>
<tr><td>Oligo fwd</td><td>CACCAACTGGTCGAGATGGACGGC</td><td></td><td></td><td></td><td></td><td></td><td></td><td></td><td></td><td></td><td></td></tr>
<tr><td>Oligo rev</td><td>AAACGCCGTCCATCTCGACCAGTT</td><td></td><td></td><td></td><td></td><td></td><td></td><td></td><td></td><td></td><td></td></tr>
<tr><td>Chromosome</td><td>start</td><td>end</td><td>strand</td><td>MM</td><td>target_seq</td><td>PAM</td><td>alignment</td><td>distance</td><td>position</td><td>gene name</td><td>gene id</td></tr>
<tr><td>GU305914.1</td><td>202464</td><td>202486</td><td>-</td><td>0</td><td>AACTGGTCGAGATGGACGGC</td><td>CGG</td><td>||||||||[||||||||||||]PAM</td><td>NA</td><td>Intergenic</td><td>NA</td><td>NA</td></tr>
<tr><td>T19177</td><td>ACTGGTCGAGATGGACGGCCGGG</td><td>1000</td><td></td><td></td><td></td><td></td><td></td><td></td><td></td><td></td><td></td></tr>
<tr><td>Oligo fwd</td><td>CACCACTGGTCGAGATGGACGGCC</td><td></td><td></td><td></td><td></td><td></td><td></td><td></td><td></td><td></td><td></td></tr>
<tr><td>Oligo rev</td><td>AAACGGCCGTCCATCTCGACCAGT</td><td></td><td></td><td></td><td></td><td></td><td></td><td></td><td></td><td></td><td></td></tr>
<tr><td>Chromosome</td><td>start</td><td>end</td><td>strand</td><td>MM</td><td>target_seq</td><td>PAM</td><td>alignment</td><td>distance</td><td>position</td><td>gene name</td><td>gene id</td></tr>
<tr><td>GU305914.1</td><td>202463</td><td>202485</td><td>-</td><td>0</td><td>ACTGGTCGAGATGGACGGCC</td><td>GGG</td><td>||||||||[||||||||||||]PAM</td><td>NA</td><td>Intergenic</td><td>NA</td><td>NA</td></tr>
<tr><td>T19178</td><td>CCTCCGTCCGATCCTGCCGACGG</td><td>1000</td><td></td><td></td><td></td><td></td><td></td><td></td><td></td><td></td><td></td></tr>
<tr><td>Oligo fwd</td><td>CACCCCTCCGTCCGATCCTGCCGA</td><td></td><td></td><td></td><td></td><td></td><td></td><td></td><td></td><td></td><td></td></tr>
<tr><td>Oligo rev</td><td>AAACTCGGCAGGATCGGACGGAGG</td><td></td><td></td><td></td><td></td><td></td><td></td><td></td><td></td><td></td><td></td></tr>
<tr><td>Chromosome</td><td>start</td><td>end</td><td>strand</td><td>MM</td><td>target_seq</td><td>PAM</td><td>alignment</td><td>distance</td><td>position</td><td>gene name</td><td>gene id</td></tr>
<tr><td>GU305914.1</td><td>202439</td><td>202461</td><td>-</td><td>0</td><td>CCTCCGTCCGATCCTGCCGA</td><td>CGG</td><td>||||||||[||||||||||||]PAM</td><td>NA</td><td>Intergenic</td><td>NA</td><td>NA</td></tr>
<tr><td>T19179</td><td>CTCCGTCCGATCCTGCCGACGGG</td><td>1000</td><td></td><td></td><td></td><td></td><td></td><td></td><td></td><td></td><td></td></tr>
<tr><td>Oligo fwd</td><td>CACCCTCCGTCCGATCCTGCCGAC</td><td></td><td></td><td></td><td></td><td></td><td></td><td></td><td></td><td></td><td></td></tr>
<tr><td>Oligo rev</td><td>AAACGTCGGCAGGATCGGACGGAG</td><td></td><td></td><td></td><td></td><td></td><td></td><td></td><td></td><td></td><td></td></tr>
<tr><td>Chromosome</td><td>start</td><td>end</td><td>strand</td><td>MM</td><td>target_seq</td><td>PAM</td><td>alignment</td><td>distance</td><td>position</td><td>gene name</td><td>gene id</td></tr>
<tr><td>GU305914.1</td><td>202438</td><td>202460</td><td>-</td><td>0</td><td>CTCCGTCCGATCCTGCCGAC</td><td>GGG</td><td>||||||||[||||||||||||]PAM</td><td>NA</td><td>Intergenic</td><td>NA</td><td>NA</td></tr>
<tr><td>T19180</td><td>CGATCCTGCCGACGGGCGAGTGG</td><td>1000</td><td></td><td></td><td></td><td></td><td></td><td></td><td></td><td></td><td></td></tr>
<tr><td>Oligo fwd</td><td>CACCCGATCCTGCCGACGGGCGAG</td><td></td><td></td><td></td><td></td><td></td><td></td><td></td><td></td><td></td><td></td></tr>
<tr><td>Oligo rev</td><td>AAACCTCGCCCGTCGGCAGGATCG</td><td></td><td></td><td></td><td></td><td></td><td></td><td></td><td></td><td></td><td></td></tr>
<tr><td>Chromosome</td><td>start</td><td>end</td><td>strand</td><td>MM</td><td>target_seq</td><td>PAM</td><td>alignment</td><td>distance</td><td>position</td><td>gene name</td><td>gene id</td></tr>
<tr><td>GU305914.1</td><td>202431</td><td>202453</td><td>-</td><td>0</td><td>CGATCCTGCCGACGGGCGAG</td><td>TGG</td><td>||||||||[||||||||||||]PAM</td><td>NA</td><td>Intergenic</td><td>NA</td><td>NA</td></tr>
<tr><td>T19181</td><td>GATCCTGCCGACGGGCGAGTGGG</td><td>1000</td><td></td><td></td><td></td><td></td><td></td><td></td><td></td><td></td><td></td></tr>
<tr><td>Oligo fwd</td><td>CACCGATCCTGCCGACGGGCGAGT</td><td></td><td></td><td></td><td></td><td></td><td></td><td></td><td></td><td></td><td></td></tr>
<tr><td>Oligo rev</td><td>AAACACTCGCCCGTCGGCAGGATC</td><td></td><td></td><td></td><td></td><td></td><td></td><td></td><td></td><td></td><td></td></tr>
<tr><td>Chromosome</td><td>start</td><td>end</td><td>strand</td><td>MM</td><td>target_seq</td><td>PAM</td><td>alignment</td><td>distance</td><td>position</td><td>gene name</td><td>gene id</td></tr>
<tr><td>GU305914.1</td><td>202430</td><td>202452</td><td>-</td><td>0</td><td>GATCCTGCCGACGGGCGAGT</td><td>GGG</td><td>||||||||[||||||||||||]PAM</td><td>NA</td><td>Intergenic</td><td>NA</td><td>NA</td></tr>
<tr><td>T19182</td><td>CCTGCCGACGGGCGAGTGGGTGG</td><td>1000</td><td></td><td></td><td></td><td></td><td></td><td></td><td></td><td></td><td></td></tr>
<tr><td>Oligo fwd</td><td>CACCCCTGCCGACGGGCGAGTGGG</td><td></td><td></td><td></td><td></td><td></td><td></td><td></td><td></td><td></td><td></td></tr>
<tr><td>Oligo rev</td><td>AAACCCCACTCGCCCGTCGGCAGG</td><td></td><td></td><td></td><td></td><td></td><td></td><td></td><td></td><td></td><td></td></tr>
<tr><td>Chromosome</td><td>start</td><td>end</td><td>strand</td><td>MM</td><td>target_seq</td><td>PAM</td><td>alignment</td><td>distance</td><td>position</td><td>gene name</td><td>gene id</td></tr>
<tr><td>GU305914.1</td><td>202427</td><td>202449</td><td>-</td><td>0</td><td>CCTGCCGACGGGCGAGTGGG</td><td>TGG</td><td>||||||||[||||||||||||]PAM</td><td>NA</td><td>Intergenic</td><td>NA</td><td>NA</td></tr>
<tr><td>T19183</td><td>GGGCGAGTGGGTGGTCGTGTTGG</td><td>1000</td><td></td><td></td><td></td><td></td><td></td><td></td><td></td><td></td><td></td></tr>
<tr><td>Oligo fwd</td><td>CACCGGGCGAGTGGGTGGTCGTGT</td><td></td><td></td><td></td><td></td><td></td><td></td><td></td><td></td><td></td><td></td></tr>
<tr><td>Oligo rev</td><td>AAACACACGACCACCCACTCGCCC</td><td></td><td></td><td></td><td></td><td></td><td></td><td></td><td></td><td></td><td></td></tr>
<tr><td>Chromosome</td><td>start</td><td>end</td><td>strand</td><td>MM</td><td>target_seq</td><td>PAM</td><td>alignment</td><td>distance</td><td>position</td><td>gene name</td><td>gene id</td></tr>
<tr><td>GU305914.1</td><td>202418</td><td>202440</td><td>-</td><td>0</td><td>GGGCGAGTGGGTGGTCGTGT</td><td>TGG</td><td>||||||||[||||||||||||]PAM</td><td>NA</td><td>Intergenic</td><td>NA</td><td>NA</td></tr>
<tr><td>T19184</td><td>GGCGAGTGGGTGGTCGTGTTGGG</td><td>1000</td><td></td><td></td><td></td><td></td><td></td><td></td><td></td><td></td><td></td></tr>
<tr><td>Oligo fwd</td><td>CACCGGCGAGTGGGTGGTCGTGTT</td><td></td><td></td><td></td><td></td><td></td><td></td><td></td><td></td><td></td><td></td></tr>
<tr><td>Oligo rev</td><td>AAACAACACGACCACCCACTCGCC</td><td></td><td></td><td></td><td></td><td></td><td></td><td></td><td></td><td></td><td></td></tr>
<tr><td>Chromosome</td><td>start</td><td>end</td><td>strand</td><td>MM</td><td>target_seq</td><td>PAM</td><td>alignment</td><td>distance</td><td>position</td><td>gene name</td><td>gene id</td></tr>
<tr><td>GU305914.1</td><td>202417</td><td>202439</td><td>-</td><td>0</td><td>GGCGAGTGGGTGGTCGTGTT</td><td>GGG</td><td>||||||||[||||||||||||]PAM</td><td>NA</td><td>Intergenic</td><td>NA</td><td>NA</td></tr>
<tr><td>T19185</td><td>GCGAGTGGGTGGTCGTGTTGGGG</td><td>1000</td><td></td><td></td><td></td><td></td><td></td><td></td><td></td><td></td><td></td></tr>
<tr><td>Oligo fwd</td><td>CACCGCGAGTGGGTGGTCGTGTTG</td><td></td><td></td><td></td><td></td><td></td><td></td><td></td><td></td><td></td><td></td></tr>
<tr><td>Oligo rev</td><td>AAACCAACACGACCACCCACTCGC</td><td></td><td></td><td></td><td></td><td></td><td></td><td></td><td></td><td></td><td></td></tr>
<tr><td>Chromosome</td><td>start</td><td>end</td><td>strand</td><td>MM</td><td>target_seq</td><td>PAM</td><td>alignment</td><td>distance</td><td>position</td><td>gene name</td><td>gene id</td></tr>
<tr><td>GU305914.1</td><td>202416</td><td>202438</td><td>-</td><td>0</td><td>GCGAGTGGGTGGTCGTGTTG</td><td>GGG</td><td>||||||||[||||||||||||]PAM</td><td>NA</td><td>Intergenic</td><td>NA</td><td>NA</td></tr>
<tr><td>T19186</td><td>CGAGTGGGTGGTCGTGTTGGGGG</td><td>1000</td><td></td><td></td><td></td><td></td><td></td><td></td><td></td><td></td><td></td></tr>
<tr><td>Oligo fwd</td><td>CACCCGAGTGGGTGGTCGTGTTGG</td><td></td><td></td><td></td><td></td><td></td><td></td><td></td><td></td><td></td><td></td></tr>
<tr><td>Oligo rev</td><td>AAACCCAACACGACCACCCACTCG</td><td></td><td></td><td></td><td></td><td></td><td></td><td></td><td></td><td></td><td></td></tr>
<tr><td>Chromosome</td><td>start</td><td>end</td><td>strand</td><td>MM</td><td>target_seq</td><td>PAM</td><td>alignment</td><td>distance</td><td>position</td><td>gene name</td><td>gene id</td></tr>
<tr><td>GU305914.1</td><td>202415</td><td>202437</td><td>-</td><td>0</td><td>CGAGTGGGTGGTCGTGTTGG</td><td>GGG</td><td>||||||||[||||||||||||]PAM</td><td>NA</td><td>Intergenic</td><td>NA</td><td>NA</td></tr>
<tr><td>T19187</td><td>TGGTCGTGTTGGGGGAGATCAGG</td><td>1000</td><td></td><td></td><td></td><td></td><td></td><td></td><td></td><td></td><td></td></tr>
<tr><td>Oligo fwd</td><td>CACCTGGTCGTGTTGGGGGAGATC</td><td></td><td></td><td></td><td></td><td></td><td></td><td></td><td></td><td></td><td></td></tr>
<tr><td>Oligo rev</td><td>AAACGATCTCCCCCAACACGACCA</td><td></td><td></td><td></td><td></td><td></td><td></td><td></td><td></td><td></td><td></td></tr>
<tr><td>Chromosome</td><td>start</td><td>end</td><td>strand</td><td>MM</td><td>target_seq</td><td>PAM</td><td>alignment</td><td>distance</td><td>position</td><td>gene name</td><td>gene id</td></tr>
<tr><td>GU305914.1</td><td>202407</td><td>202429</td><td>-</td><td>0</td><td>TGGTCGTGTTGGGGGAGATC</td><td>AGG</td><td>||||||||[||||||||||||]PAM</td><td>NA</td><td>Intergenic</td><td>NA</td><td>NA</td></tr>
<tr><td>T19188</td><td>GGGGGAGATCAGGTCGCCGATGG</td><td>1000</td><td></td><td></td><td></td><td></td><td></td><td></td><td></td><td></td><td></td></tr>
<tr><td>Oligo fwd</td><td>CACCGGGGGAGATCAGGTCGCCGA</td><td></td><td></td><td></td><td></td><td></td><td></td><td></td><td></td><td></td><td></td></tr>
</table>

| | | | | | | | | | | | |
|---|---|---|---|---|---|---|---|---|---|---|---|
| Oligo rev | AAACTCGGCGACCTGATCTCCCCC | | | | | | | | | | |
| Chromosome | start | end | strand | MM | target_seq | PAM | alignment | distance | position | gene name | gene id |
| GU305914.1 | 202397 | 202419 | - | 0 | GGGGGAGATCAGGTCGCCGA | TGG | \|\|\|\|\|\|\|\|[\|\|\|\|\|\|\|\|\|\|\|\|]PAM | NA | Intergenic | NA | NA |
| T19189 | CACGTCCCTGTACCTCGCCATGG | 1000 | | | | | | | | | |
| Oligo fwd | CACCCACGTCCCTGTACCTCGCCA | | | | | | | | | | |
| Oligo rev | AAACTGGCGAGGTACAGGGACGTG | | | | | | | | | | |
| Chromosome | start | end | strand | MM | target_seq | PAM | alignment | distance | position | gene name | gene id |
| GU305914.1 | 202367 | 202389 | - | 0 | CACGTCCCTGTACCTCGCCA | TGG | \|\|\|\|\|\|\|\|[\|\|\|\|\|\|\|\|\|\|\|\|]PAM | NA | Intergenic | NA | NA |
| T19190 | ACGTCCCTGTACCTCGCCATGGG | 1000 | | | | | | | | | |
| Oligo fwd | CACCACGTCCCTGTACCTCGCCAT | | | | | | | | | | |
| Oligo rev | AAACATGGCGAGGTACAGGGACGT | | | | | | | | | | |
| Chromosome | start | end | strand | MM | target_seq | PAM | alignment | distance | position | gene name | gene id |
| GU305914.1 | 202366 | 202388 | - | 0 | ACGTCCCTGTACCTCGCCAT | GGG | \|\|\|\|\|\|\|\|[\|\|\|\|\|\|\|\|\|\|\|\|]PAM | NA | Intergenic | NA | NA |
| T19191 | CCTGTACCTCGCCATGGGCAAGG | 1000 | | | | | | | | | |
| Oligo fwd | CACCCCTGTACCTCGCCATGGGCA | | | | | | | | | | |
| Oligo rev | AAACTGCCCATGGCGAGGTACAGG | | | | | | | | | | |
| Chromosome | start | end | strand | MM | target_seq | PAM | alignment | distance | position | gene name | gene id |
| GU305914.1 | 202361 | 202383 | - | 0 | CCTGTACCTCGCCATGGGCA | AGG | \|\|\|\|\|\|\|\|[\|\|\|\|\|\|\|\|\|\|\|\|]PAM | NA | Intergenic | NA | NA |
| T19192 | TCGCCATGGGCAAGGAGTCGAGG | 1000 | | | | | | | | | |
| Oligo fwd | CACCTCGCCATGGGCAAGGAGTCG | | | | | | | | | | |
| Oligo rev | AAACCGACTCCTTGCCCATGGCGA | | | | | | | | | | |
| Chromosome | start | end | strand | MM | target_seq | PAM | alignment | distance | position | gene name | gene id |
| GU305914.1 | 202353 | 202375 | - | 0 | TCGCCATGGGCAAGGAGTCG | AGG | \|\|\|\|\|\|\|\|[\|\|\|\|\|\|\|\|\|\|\|\|]PAM | NA | Intergenic | NA | NA |
| T19193 | GATGTTCGTCTACAGCGCCGAGG | 1000 | | | | | | | | | |
| Oligo fwd | CACCGATGTTCGTCTACAGCGCCG | | | | | | | | | | |
| Oligo rev | AAACCGGCGCTGTAGACGAACATC | | | | | | | | | | |
| Chromosome | start | end | strand | MM | target_seq | PAM | alignment | distance | position | gene name | gene id |
| GU305914.1 | 202331 | 202353 | - | 0 | GATGTTCGTCTACAGCGCCG | AGG | \|\|\|\|\|\|\|\|[\|\|\|\|\|\|\|\|\|\|\|\|]PAM | NA | Intergenic | NA | NA |
| T19194 | GTTCGTCTACAGCGCCGAGGAGG | 1000 | | | | | | | | | |
| Oligo fwd | CACCGTTCGTCTACAGCGCCGAGG | | | | | | | | | | |
| Oligo rev | AAACCCTCGGCGCTGTAGACGAAC | | | | | | | | | | |
| Chromosome | start | end | strand | MM | target_seq | PAM | alignment | distance | position | gene name | gene id |
| GU305914.1 | 202328 | 202350 | - | 0 | GTTCGTCTACAGCGCCGAGG | AGG | \|\|\|\|\|\|\|\|[\|\|\|\|\|\|\|\|\|\|\|\|]PAM | NA | Intergenic | NA | NA |
| T19195 | CGAGGAGGACGCCATCGCGTTGG | 1000 | | | | | | | | | |
| Oligo fwd | CACCCGAGGAGGACGCCATCGCGT | | | | | | | | | | |
| Oligo rev | AAACACGCGATGGCGTCCTCCTCG | | | | | | | | | | |
| Chromosome | start | end | strand | MM | target_seq | PAM | alignment | distance | position | gene name | gene id |
| GU305914.1 | 202313 | 202335 | - | 0 | CGAGGAGGACGCCATCGCGT | TGG | \|\|\|\|\|\|\|\|[\|\|\|\|\|\|\|\|\|\|\|\|]PAM | NA | Intergenic | NA | NA |
| T19196 | GGAGGACGCCATCGCGTTGGCGG | 1000 | | | | | | | | | |
| Oligo fwd | CACCGGAGGACGCCATCGCGTTGG | | | | | | | | | | |
| Oligo rev | AAACCCAACGCGATGGCGTCCTCC | | | | | | | | | | |
| Chromosome | start | end | strand | MM | target_seq | PAM | alignment | distance | position | gene name | gene id |
| GU305914.1 | 202310 | 202332 | - | 0 | GGAGGACGCCATCGCGTTGG | CGG | \|\|\|\|\|\|\|\|[\|\|\|\|\|\|\|\|\|\|\|\|]PAM | NA | Intergenic | NA | NA |
| T19197 | ACGACCTCGACGAGTTCTCGCGG | 1000 | | | | | | | | | |
| Oligo fwd | CACCACGACCTCGACGAGTTCTCG | | | | | | | | | | |
| Oligo rev | AAACCGAGAACTCGTCGAGGTCGT | | | | | | | | | | |
| Chromosome | start | end | strand | MM | target_seq | PAM | alignment | distance | position | gene name | gene id |
| GU305914.1 | 202284 | 202306 | - | 0 | ACGACCTCGACGAGTTCTCG | CGG | \|\|\|\|\|\|\|\|[\|\|\|\|\|\|\|\|\|\|\|\|]PAM | NA | Intergenic | NA | NA |
| T19198 | CTCGACGAGTTCTCGCGGATAGG | 1000 | | | | | | | | | |
| Oligo fwd | CACCCTCGACGAGTTCTCGCGGAT | | | | | | | | | | |
| Oligo rev | AAACATCCGCGAGAACTCGTCGAG | | | | | | | | | | |
| Chromosome | start | end | strand | MM | target_seq | PAM | alignment | distance | position | gene name | gene id |
| GU305914.1 | 202279 | 202301 | - | 0 | CTCGACGAGTTCTCGCGGAT | AGG | \|\|\|\|\|\|\|\|[\|\|\|\|\|\|\|\|\|\|\|\|]PAM | NA | Intergenic | NA | NA |
| T19199 | CTCGCGGATAGGTCTGCTGACGG | 1000 | | | | | | | | | |
| Oligo fwd | CACCCTCGCGGATAGGTCTGCTGA | | | | | | | | | | |
| Oligo rev | AAACTCAGCAGACCTATCCGCGAG | | | | | | | | | | |
| Chromosome | start | end | strand | MM | target_seq | PAM | alignment | distance | position | gene name | gene id |
| GU305914.1 | 202268 | 202290 | - | 0 | CTCGCGGATAGGTCTGCTGA | CGG | \|\|\|\|\|\|\|\|[\|\|\|\|\|\|\|\|\|\|\|\|]PAM | NA | Intergenic | NA | NA |
| T19200 | GCGGATAGGTCTGCTGACGGTGG | 1000 | | | | | | | | | |
| Oligo fwd | CACCGCGGATAGGTCTGCTGACGG | | | | | | | | | | |
| Oligo rev | AAACCCGTCAGCAGACCTATCCGC | | | | | | | | | | |
| Chromosome | start | end | strand | MM | target_seq | PAM | alignment | distance | position | gene name | gene id |
| GU305914.1 | 202265 | 202287 | - | 0 | GCGGATAGGTCTGCTGACGG | TGG | \|\|\|\|\|\|\|\|[\|\|\|\|\|\|\|\|\|\|\|\|]PAM | NA | Intergenic | NA | NA |
| T19201 | TGACGGTGGAGTTCATCTACAGG | 1000 | | | | | | | | | |
| Oligo fwd | CACCTGACGGTGGAGTTCATCTAC | | | | | | | | | | |
| Oligo rev | AAACGTAGATGAACTCCACCGTCA | | | | | | | | | | |
| Chromosome | start | end | strand | MM | target_seq | PAM | alignment | distance | position | gene name | gene id |
| GU305914.1 | 202251 | 202273 | - | 0 | TGACGGTGGAGTTCATCTAC | AGG | \|\|\|\|\|\|\|\|[\|\|\|\|\|\|\|\|\|\|\|\|]PAM | NA | Intergenic | NA | NA |
| T19202 | GACGGTGGAGTTCATCTACAGGG | 1000 | | | | | | | | | |
| Oligo fwd | CACCGACGGTGGAGTTCATCTACA | | | | | | | | | | |
| Oligo rev | AAACTGTAGATGAACTCCACCGTC | | | | | | | | | | |
| Chromosome | start | end | strand | MM | target_seq | PAM | alignment | distance | position | gene name | gene id |
| GU305914.1 | 202250 | 202272 | - | 0 | GACGGTGGAGTTCATCTACA | GGG | \|\|\|\|\|\|\|\|[\|\|\|\|\|\|\|\|\|\|\|\|]PAM | NA | Intergenic | NA | NA |
| T19203 | GATACCGTTGAAGAAAGACCCGG | 1000 | | | | | | | | | |

<table>
<tr><td>Oligo fwd</td><td>CACCGATACCGTTGAAGAAAGACC</td><td></td><td></td><td></td><td></td><td></td><td></td><td></td><td></td><td></td><td></td></tr>
<tr><td>Oligo rev</td><td>AAACGGTCTTTCTTCAACGGTATC</td><td></td><td></td><td></td><td></td><td></td><td></td><td></td><td></td><td></td><td></td></tr>
<tr><td>Chromosome</td><td>start</td><td>end</td><td>strand</td><td>MM</td><td>target_seq</td><td>PAM</td><td>alignment</td><td>distance</td><td>position</td><td>gene name</td><td>gene id</td></tr>
<tr><td>GU305914.1</td><td>202217</td><td>202239</td><td>-</td><td>0</td><td>GATACCGTTGAAGAAAGACC</td><td>CGG</td><td>||||||||[||||||||||||]PAM</td><td>NA</td><td>Intergenic</td><td>NA</td><td>NA</td></tr>
<tr><td>T19204</td><td>CGTTGAAGAAAGACCCGGTCTGG</td><td>1000</td><td></td><td></td><td></td><td></td><td></td><td></td><td></td><td></td><td></td></tr>
<tr><td>Oligo fwd</td><td>CACCCGTTGAAGAAAGACCCGGTC</td><td></td><td></td><td></td><td></td><td></td><td></td><td></td><td></td><td></td><td></td></tr>
<tr><td>Oligo rev</td><td>AAACGACCGGGTCTTTCTTCAACG</td><td></td><td></td><td></td><td></td><td></td><td></td><td></td><td></td><td></td><td></td></tr>
<tr><td>Chromosome</td><td>start</td><td>end</td><td>strand</td><td>MM</td><td>target_seq</td><td>PAM</td><td>alignment</td><td>distance</td><td>position</td><td>gene name</td><td>gene id</td></tr>
<tr><td>GU305914.1</td><td>202212</td><td>202234</td><td>-</td><td>0</td><td>CGTTGAAGAAAGACCCGGTC</td><td>TGG</td><td>||||||||[||||||||||||]PAM</td><td>NA</td><td>Intergenic</td><td>NA</td><td>NA</td></tr>
<tr><td>T19205</td><td>GTTGAAGAAAGACCCGGTCTGGG</td><td>1000</td><td></td><td></td><td></td><td></td><td></td><td></td><td></td><td></td><td></td></tr>
<tr><td>Oligo fwd</td><td>CACCGTTGAAGAAAGACCCGGTCT</td><td></td><td></td><td></td><td></td><td></td><td></td><td></td><td></td><td></td><td></td></tr>
<tr><td>Oligo rev</td><td>AAACAGACCGGGTCTTTCTTCAAC</td><td></td><td></td><td></td><td></td><td></td><td></td><td></td><td></td><td></td><td></td></tr>
<tr><td>Chromosome</td><td>start</td><td>end</td><td>strand</td><td>MM</td><td>target_seq</td><td>PAM</td><td>alignment</td><td>distance</td><td>position</td><td>gene name</td><td>gene id</td></tr>
<tr><td>GU305914.1</td><td>202211</td><td>202233</td><td>-</td><td>0</td><td>GTTGAAGAAAGACCCGGTCT</td><td>GGG</td><td>||||||||[||||||||||||]PAM</td><td>NA</td><td>Intergenic</td><td>NA</td><td>NA</td></tr>
<tr><td>T19206</td><td>GACCCGGTCTGGGACGCCATCGG</td><td>1000</td><td></td><td></td><td></td><td></td><td></td><td></td><td></td><td></td><td></td></tr>
<tr><td>Oligo fwd</td><td>CACCGACCCGGTCTGGGACGCCAT</td><td></td><td></td><td></td><td></td><td></td><td></td><td></td><td></td><td></td><td></td></tr>
<tr><td>Oligo rev</td><td>AAACATGGCGTCCCAGACCGGGTC</td><td></td><td></td><td></td><td></td><td></td><td></td><td></td><td></td><td></td><td></td></tr>
<tr><td>Chromosome</td><td>start</td><td>end</td><td>strand</td><td>MM</td><td>target_seq</td><td>PAM</td><td>alignment</td><td>distance</td><td>position</td><td>gene name</td><td>gene id</td></tr>
<tr><td>GU305914.1</td><td>202201</td><td>202223</td><td>-</td><td>0</td><td>GACCCGGTCTGGGACGCCAT</td><td>CGG</td><td>||||||||[||||||||||||]PAM</td><td>NA</td><td>Intergenic</td><td>NA</td><td>NA</td></tr>
<tr><td>T19207</td><td>GCCATCGGCCTCTGCGTGACCGG</td><td>1000</td><td></td><td></td><td></td><td></td><td></td><td></td><td></td><td></td><td></td></tr>
<tr><td>Oligo fwd</td><td>CACCGCCATCGGCCTCTGCGTGAC</td><td></td><td></td><td></td><td></td><td></td><td></td><td></td><td></td><td></td><td></td></tr>
<tr><td>Oligo rev</td><td>AAACGTCACGCAGAGGCCGATGGC</td><td></td><td></td><td></td><td></td><td></td><td></td><td></td><td></td><td></td><td></td></tr>
<tr><td>Chromosome</td><td>start</td><td>end</td><td>strand</td><td>MM</td><td>target_seq</td><td>PAM</td><td>alignment</td><td>distance</td><td>position</td><td>gene name</td><td>gene id</td></tr>
<tr><td>GU305914.1</td><td>202186</td><td>202208</td><td>-</td><td>0</td><td>GCCATCGGCCTCTGCGTGAC</td><td>CGG</td><td>||||||||[||||||||||||]PAM</td><td>NA</td><td>Intergenic</td><td>NA</td><td>NA</td></tr>
<tr><td>T19208</td><td>TCTGTCCACGTACCGCAACTCGG</td><td>1000</td><td></td><td></td><td></td><td></td><td></td><td></td><td></td><td></td><td></td></tr>
<tr><td>Oligo fwd</td><td>CACCTCTGTCCACGTACCGCAACT</td><td></td><td></td><td></td><td></td><td></td><td></td><td></td><td></td><td></td><td></td></tr>
<tr><td>Oligo rev</td><td>AAACAGTTGCGGTACGTGGACAGA</td><td></td><td></td><td></td><td></td><td></td><td></td><td></td><td></td><td></td><td></td></tr>
<tr><td>Chromosome</td><td>start</td><td>end</td><td>strand</td><td>MM</td><td>target_seq</td><td>PAM</td><td>alignment</td><td>distance</td><td>position</td><td>gene name</td><td>gene id</td></tr>
<tr><td>GU305914.1</td><td>202145</td><td>202167</td><td>-</td><td>0</td><td>TCTGTCCACGTACCGCAACT</td><td>CGG</td><td>||||||||[||||||||||||]PAM</td><td>NA</td><td>Intergenic</td><td>NA</td><td>NA</td></tr>
<tr><td>T19209</td><td>GCAACTCGGTGATCGAGCTGCGG</td><td>1000</td><td></td><td></td><td></td><td></td><td></td><td></td><td></td><td></td><td></td></tr>
<tr><td>Oligo fwd</td><td>CACCGCAACTCGGTGATCGAGCTG</td><td></td><td></td><td></td><td></td><td></td><td></td><td></td><td></td><td></td><td></td></tr>
<tr><td>Oligo rev</td><td>AAACCAGCTCGATCACCGAGTTGC</td><td></td><td></td><td></td><td></td><td></td><td></td><td></td><td></td><td></td><td></td></tr>
<tr><td>Chromosome</td><td>start</td><td>end</td><td>strand</td><td>MM</td><td>target_seq</td><td>PAM</td><td>alignment</td><td>distance</td><td>position</td><td>gene name</td><td>gene id</td></tr>
<tr><td>GU305914.1</td><td>202131</td><td>202153</td><td>-</td><td>0</td><td>GCAACTCGGTGATCGAGCTG</td><td>CGG</td><td>||||||||[||||||||||||]PAM</td><td>NA</td><td>Intergenic</td><td>NA</td><td>NA</td></tr>
<tr><td>T19210</td><td>GTGATCGAGCTGCGGACGCCCGG</td><td>1000</td><td></td><td></td><td></td><td></td><td></td><td></td><td></td><td></td><td></td></tr>
<tr><td>Oligo fwd</td><td>CACCGTGATCGAGCTGCGGACGCC</td><td></td><td></td><td></td><td></td><td></td><td></td><td></td><td></td><td></td><td></td></tr>
<tr><td>Oligo rev</td><td>AAACGGCGTCCGCAGCTCGATCAC</td><td></td><td></td><td></td><td></td><td></td><td></td><td></td><td></td><td></td><td></td></tr>
<tr><td>Chromosome</td><td>start</td><td>end</td><td>strand</td><td>MM</td><td>target_seq</td><td>PAM</td><td>alignment</td><td>distance</td><td>position</td><td>gene name</td><td>gene id</td></tr>
<tr><td>GU305914.1</td><td>202123</td><td>202145</td><td>-</td><td>0</td><td>GTGATCGAGCTGCGGACGCC</td><td>CGG</td><td>||||||||[||||||||||||]PAM</td><td>NA</td><td>Intergenic</td><td>NA</td><td>NA</td></tr>
<tr><td>T19211</td><td>GCTGCGGACGCCCGGCAAGCAGG</td><td>1000</td><td></td><td></td><td></td><td></td><td></td><td></td><td></td><td></td><td></td></tr>
<tr><td>Oligo fwd</td><td>CACCGCTGCGGACGCCCGGCAAGC</td><td></td><td></td><td></td><td></td><td></td><td></td><td></td><td></td><td></td><td></td></tr>
<tr><td>Oligo rev</td><td>AAACGCTTGCCGGGCGTCCGCAGC</td><td></td><td></td><td></td><td></td><td></td><td></td><td></td><td></td><td></td><td></td></tr>
<tr><td>Chromosome</td><td>start</td><td>end</td><td>strand</td><td>MM</td><td>target_seq</td><td>PAM</td><td>alignment</td><td>distance</td><td>position</td><td>gene name</td><td>gene id</td></tr>
<tr><td>GU305914.1</td><td>202115</td><td>202137</td><td>-</td><td>0</td><td>GCTGCGGACGCCCGGCAAGC</td><td>AGG</td><td>||||||||[||||||||||||]PAM</td><td>NA</td><td>Intergenic</td><td>NA</td><td>NA</td></tr>
<tr><td>T19212</td><td>GACCAACCCCCTGATCCTACTGG</td><td>1000</td><td></td><td></td><td></td><td></td><td></td><td></td><td></td><td></td><td></td></tr>
<tr><td>Oligo fwd</td><td>CACCGACCAACCCCCTGATCCTAC</td><td></td><td></td><td></td><td></td><td></td><td></td><td></td><td></td><td></td><td></td></tr>
<tr><td>Oligo rev</td><td>AAACGTAGGATCAGGGGGTTGGTC</td><td></td><td></td><td></td><td></td><td></td><td></td><td></td><td></td><td></td><td></td></tr>
<tr><td>Chromosome</td><td>start</td><td>end</td><td>strand</td><td>MM</td><td>target_seq</td><td>PAM</td><td>alignment</td><td>distance</td><td>position</td><td>gene name</td><td>gene id</td></tr>
<tr><td>GU305914.1</td><td>202091</td><td>202113</td><td>-</td><td>0</td><td>GACCAACCCCCTGATCCTAC</td><td>TGG</td><td>||||||||[||||||||||||]PAM</td><td>NA</td><td>Intergenic</td><td>NA</td><td>NA</td></tr>
<tr><td>T19213</td><td>ACCCCCTGATCCTACTGGACCGG</td><td>1000</td><td></td><td></td><td></td><td></td><td></td><td></td><td></td><td></td><td></td></tr>
<tr><td>Oligo fwd</td><td>CACCACCCCCTGATCCTACTGGAC</td><td></td><td></td><td></td><td></td><td></td><td></td><td></td><td></td><td></td><td></td></tr>
<tr><td>Oligo rev</td><td>AAACGTCCAGTAGGATCAGGGGGT</td><td></td><td></td><td></td><td></td><td></td><td></td><td></td><td></td><td></td><td></td></tr>
<tr><td>Chromosome</td><td>start</td><td>end</td><td>strand</td><td>MM</td><td>target_seq</td><td>PAM</td><td>alignment</td><td>distance</td><td>position</td><td>gene name</td><td>gene id</td></tr>
<tr><td>GU305914.1</td><td>202086</td><td>202108</td><td>-</td><td>0</td><td>ACCCCCTGATCCTACTGGAC</td><td>CGG</td><td>||||||||[||||||||||||]PAM</td><td>NA</td><td>Intergenic</td><td>NA</td><td>NA</td></tr>
<tr><td>T19214</td><td>TCGAACTCTGCTGCAAGTACTGG</td><td>1000</td><td></td><td></td><td></td><td></td><td></td><td></td><td></td><td></td><td></td></tr>
<tr><td>Oligo fwd</td><td>CACCTCGAACTCTGCTGCAAGTAC</td><td></td><td></td><td></td><td></td><td></td><td></td><td></td><td></td><td></td><td></td></tr>
<tr><td>Oligo rev</td><td>AAACGTACTTGCAGCAGAGTTCGA</td><td></td><td></td><td></td><td></td><td></td><td></td><td></td><td></td><td></td><td></td></tr>
<tr><td>Chromosome</td><td>start</td><td>end</td><td>strand</td><td>MM</td><td>target_seq</td><td>PAM</td><td>alignment</td><td>distance</td><td>position</td><td>gene name</td><td>gene id</td></tr>
<tr><td>GU305914.1</td><td>202062</td><td>202084</td><td>-</td><td>0</td><td>TCGAACTCTGCTGCAAGTAC</td><td>TGG</td><td>||||||||[||||||||||||]PAM</td><td>NA</td><td>Intergenic</td><td>NA</td><td>NA</td></tr>
<tr><td>T19215</td><td>CTGCAAGTACTGGCCGCTCGTGG</td><td>1000</td><td></td><td></td><td></td><td></td><td></td><td></td><td></td><td></td><td></td></tr>
<tr><td>Oligo fwd</td><td>CACCCTGCAAGTACTGGCCGCTCG</td><td></td><td></td><td></td><td></td><td></td><td></td><td></td><td></td><td></td><td></td></tr>
<tr><td>Oligo rev</td><td>AAACCGAGCGGCCAGTACTTGCAG</td><td></td><td></td><td></td><td></td><td></td><td></td><td></td><td></td><td></td><td></td></tr>
<tr><td>Chromosome</td><td>start</td><td>end</td><td>strand</td><td>MM</td><td>target_seq</td><td>PAM</td><td>alignment</td><td>distance</td><td>position</td><td>gene name</td><td>gene id</td></tr>
<tr><td>GU305914.1</td><td>202052</td><td>202074</td><td>-</td><td>0</td><td>CTGCAAGTACTGGCCGCTCG</td><td>TGG</td><td>||||||||[||||||||||||]PAM</td><td>NA</td><td>Intergenic</td><td>NA</td><td>NA</td></tr>
<tr><td>T19216</td><td>GTACTGGCCGCTCGTGGCGATGG</td><td>1000</td><td></td><td></td><td></td><td></td><td></td><td></td><td></td><td></td><td></td></tr>
<tr><td>Oligo fwd</td><td>CACCGTACTGGCCGCTCGTGGCGA</td><td></td><td></td><td></td><td></td><td></td><td></td><td></td><td></td><td></td><td></td></tr>
<tr><td>Oligo rev</td><td>AAACTCGCCACGAGCGGCCAGTAC</td><td></td><td></td><td></td><td></td><td></td><td></td><td></td><td></td><td></td><td></td></tr>
<tr><td>Chromosome</td><td>start</td><td>end</td><td>strand</td><td>MM</td><td>target_seq</td><td>PAM</td><td>alignment</td><td>distance</td><td>position</td><td>gene name</td><td>gene id</td></tr>
<tr><td>GU305914.1</td><td>202046</td><td>202068</td><td>-</td><td>0</td><td>GTACTGGCCGCTCGTGGCGA</td><td>TGG</td><td>||||||||[||||||||||||]PAM</td><td>NA</td><td>Intergenic</td><td>NA</td><td>NA</td></tr>
<tr><td>T19217</td><td>TCGTGGCGATGGACAAATCTCGG</td><td>1000</td><td></td><td></td><td></td><td></td><td></td><td></td><td></td><td></td><td></td></tr>
<tr><td>Oligo fwd</td><td>CACCTCGTGGCGATGGACAAATCT</td><td></td><td></td><td></td><td></td><td></td><td></td><td></td><td></td><td></td><td></td></tr>
<tr><td>Oligo rev</td><td>AAACAGATTTGTCCATCGCCACGA</td><td></td><td></td><td></td><td></td><td></td><td></td><td></td><td></td><td></td><td></td></tr>
<tr><td>Chromosome</td><td>start</td><td>end</td><td>strand</td><td>MM</td><td>target_seq</td><td>PAM</td><td>alignment</td><td>distance</td><td>position</td><td>gene name</td><td>gene id</td></tr>
<tr><td>GU305914.1</td><td>202035</td><td>202057</td><td>-</td><td>0</td><td>TCGTGGCGATGGACAAATCT</td><td>CGG</td><td>||||||||[||||||||||||]PAM</td><td>NA</td><td>Intergenic</td><td>NA</td><td>NA</td></tr>
</table>

T19218 ATGGACAAATCTCGGATCGACGG 1000
Oligo fwd CACCATGGACAAATCTCGGATCGA
Oligo rev AAACTCGATCCGAGATTTGTCCAT
Chromosome start end strand MM target_seq PAM alignment distance position gene name gene id
GU305914.1 202027 202049 - 0 ATGGACAAATCTCGGATCGA CGG ||||||||[||||||||||||]PAM NA Intergenic NA NA

T19219 CTCGAAAGATGTCCTCGCGCTGG 1000
Oligo fwd CACCCTCGAAAGATGTCCTCGCGC
Oligo rev AAACGCGCGAGGACATCTTTCGAG
Chromosome start end strand MM target_seq PAM alignment distance position gene name gene id
GU305914.1 201987 202009 - 0 CTCGAAAGATGTCCTCGCGC TGG ||||||||[||||||||||||]PAM NA Intergenic NA NA

T19220 TCCTCGCGCTGGTACACGCTCGG 1000
Oligo fwd CACCTCCTCGCGCTGGTACACGCT
Oligo rev AAACAGCGTGTACCAGCGCGAGGA
Chromosome start end strand MM target_seq PAM alignment distance position gene name gene id
GU305914.1 201976 201998 - 0 TCCTCGCGCTGGTACACGCT CGG ||||||||[||||||||||||]PAM NA Intergenic NA NA

T19221 TGGTACACGCTCGGACTCGTCGG 1000
Oligo fwd CACCTGGTACACGCTCGGACTCGT
Oligo rev AAACACGAGTCCGAGCGTGTACCA
Chromosome start end strand MM target_seq PAM alignment distance position gene name gene id
GU305914.1 201967 201989 - 0 TGGTACACGCTCGGACTCGT CGG ||||||||[||||||||||||]PAM NA Intergenic NA NA

T19222 TCGGACTCGTCGGCGTCTACCGG 1000
Oligo fwd CACCTCGGACTCGTCGGCGTCTAC
Oligo rev AAACGTAGACGCCGACGAGTCCGA
Chromosome start end strand MM target_seq PAM alignment distance position gene name gene id
GU305914.1 201957 201979 - 0 TCGGACTCGTCGGCGTCTAC CGG ||||||||[||||||||||||]PAM NA Intergenic NA NA

T19223 CGGACTCGTCGGCGTCTACCGGG 1000
Oligo fwd CACCCGGACTCGTCGGCGTCTACC
Oligo rev AAACGGTAGACGCCGACGAGTCCG
Chromosome start end strand MM target_seq PAM alignment distance position gene name gene id
GU305914.1 201956 201978 - 0 CGGACTCGTCGGCGTCTACC GGG ||||||||[||||||||||||]PAM NA Intergenic NA NA

T19224 GTCGGCGTCTACCGGGACACCGG 1000
Oligo fwd CACCGTCGGCGTCTACCGGGACAC
Oligo rev AAACGTGTCCCGGTAGACGCCGAC
Chromosome start end strand MM target_seq PAM alignment distance position gene name gene id
GU305914.1 201949 201971 - 0 GTCGGCGTCTACCGGGACAC CGG ||||||||[||||||||||||]PAM NA Intergenic NA NA

T19225 TCGGCGTCTACCGGGACACCGGG 1000
Oligo fwd CACCTCGGCGTCTACCGGGACACC
Oligo rev AAACGGTGTCCCGGTAGACGCCGA
Chromosome start end strand MM target_seq PAM alignment distance position gene name gene id
GU305914.1 201948 201970 - 0 TCGGCGTCTACCGGGACACC GGG ||||||||[||||||||||||]PAM NA Intergenic NA NA

T19226 CGGCGTCTACCGGGACACCGGGG 1000
Oligo fwd CACCCGGCGTCTACCGGGACACCG
Oligo rev AAACCGGTGTCCCGGTAGACGCCG
Chromosome start end strand MM target_seq PAM alignment distance position gene name gene id
GU305914.1 201947 201969 - 0 CGGCGTCTACCGGGACACCG GGG ||||||||[||||||||||||]PAM NA Intergenic NA NA

T19227 CGGGACACCGGGGTGTTCCACGG 1000
Oligo fwd CACCCGGGACACCGGGGTGTTCCA
Oligo rev AAACTGGAACACCCCGGTGTCCCG
Chromosome start end strand MM target_seq PAM alignment distance position gene name gene id
GU305914.1 201937 201959 - 0 CGGGACACCGGGGTGTTCCA CGG ||||||||[||||||||||||]PAM NA Intergenic NA NA

T19228 GGGACACCGGGGTGTTCCACGGG 1000
Oligo fwd CACCGGGACACCGGGGTGTTCCAC
Oligo rev AAACGTGGAACACCCCGGTGTCCC
Chromosome start end strand MM target_seq PAM alignment distance position gene name gene id
GU305914.1 201936 201958 - 0 GGGACACCGGGGTGTTCCAC GGG ||||||||[||||||||||||]PAM NA Intergenic NA NA

T19229 GGACACCGGGGTGTTCCACGGGG 1000
Oligo fwd CACCGGACACCGGGGTGTTCCACG
Oligo rev AAACCGTGGAACACCCCGGTGTCC
Chromosome start end strand MM target_seq PAM alignment distance position gene name gene id
GU305914.1 201935 201957 - 0 GGACACCGGGGTGTTCCACG GGG ||||||||[||||||||||||]PAM NA Intergenic NA NA

T19230 GTTCCACGGGGCGCAGCTCCTGG 1000
Oligo fwd CACCGTTCCACGGGGCGCAGCTCC
Oligo rev AAACGGAGCTGCGCCCCGTGGAAC
Chromosome start end strand MM target_seq PAM alignment distance position gene name gene id
GU305914.1 201923 201945 - 0 GTTCCACGGGGCGCAGCTCC TGG ||||||||[||||||||||||]PAM NA Intergenic NA NA

T19231 GCTCCTGGTCTTCGACGACTCGG 1000
Oligo fwd CACCGCTCCTGGTCTTCGACGACT
Oligo rev AAACAGTCGTCGAAGACCAGGAGC
Chromosome start end strand MM target_seq PAM alignment distance position gene name gene id
GU305914.1 201908 201930 - 0 GCTCCTGGTCTTCGACGACT CGG ||||||||[||||||||||||]PAM NA Intergenic NA NA

T19232 CTCCTGGTCTTCGACGACTCGGG 1000
Oligo fwd CACCCTCCTGGTCTTCGACGACTC
Oligo rev AAACGAGTCGTCGAAGACCAGGAG
Chromosome start end strand MM target_seq PAM alignment distance position gene name gene id
GU305914.1 201907 201929 - 0 CTCCTGGTCTTCGACGACTC GGG ||||||||[||||||||||||]PAM NA Intergenic NA NA

T19233 TTCTATCTATCGCTGATCAGCGG 1000
Oligo fwd CACCTTCTATCTATCGCTGATCAG
Oligo rev AAACCTGATCAGCGATAGATAGAA
Chromosome start end strand MM target_seq PAM alignment distance position gene name gene id
GU305914.1 201877 201899 - 0 TTCTATCTATCGCTGATCAG CGG ||||||||[||||||||||||]PAM NA Intergenic NA NA

T19234 CGCTGATCAGCGGCGAGTTGTGG 1000
Oligo fwd CACCCGCTGATCAGCGGCGAGTTG
Oligo rev AAACCAACTCGCCGCTGATCAGCG
Chromosome start end strand MM target_seq PAM alignment distance position gene name gene id
GU305914.1 201867 201889 - 0 CGCTGATCAGCGGCGAGTTG TGG ||||||||[||||||||||||]PAM NA Intergenic NA NA

T19235 CAGCGGCGAGTTGTGGCGTCTGG 1000
Oligo fwd CACCCAGCGGCGAGTTGTGGCGTC
Oligo rev AAACGACGCCACAACTCGCCGCTG
Chromosome start end strand MM target_seq PAM alignment distance position gene name gene id
GU305914.1 201860 201882 - 0 CAGCGGCGAGTTGTGGCGTC TGG ||||||||[||||||||||||]PAM NA Intergenic NA NA

T19236 GCGTCTGGCCGACAGCGCCGCGG 1000
Oligo fwd CACCGCGTCTGGCCGACAGCGCCG
Oligo rev AAACCGGCGCTGTCGGCCAGACGC
Chromosome start end strand MM target_seq PAM alignment distance position gene name gene id
GU305914.1 201845 201867 - 0 GCGTCTGGCCGACAGCGCCG CGG ||||||||[||||||||||||]PAM NA Intergenic NA NA

T19237 CGCCGCGGAGCTCCGTCGCATGG 1000
Oligo fwd CACCCGCCGCGGAGCTCCGTCGCA
Oligo rev AAACTGCGACGGAGCTCCGCGGCG
Chromosome start end strand MM target_seq PAM alignment distance position gene name gene id
GU305914.1 201830 201852 - 0 CGCCGCGGAGCTCCGTCGCA TGG ||||||||[||||||||||||]PAM NA Intergenic NA NA

T19238 GCCGCGGAGCTCCGTCGCATGGG 1000
Oligo fwd CACCGCCGCGGAGCTCCGTCGCAT
Oligo rev AAACATGCGACGGAGCTCCGCGGC
Chromosome start end strand MM target_seq PAM alignment distance position gene name gene id
GU305914.1 201829 201851 - 0 GCCGCGGAGCTCCGTCGCAT GGG ||||||||[||||||||||||]PAM NA Intergenic NA NA

T19239 ACTGCTGAAGATCTTCACCGCGG 1000
Oligo fwd CACCACTGCTGAAGATCTTCACCG
Oligo rev AAACCGGTGAAGATCTTCAGCAGT
Chromosome start end strand MM target_seq PAM alignment distance position gene name gene id
GU305914.1 201806 201828 - 0 ACTGCTGAAGATCTTCACCG CGG ||||||||[||||||||||||]PAM NA Intergenic NA NA

T19240 CTGCTGAAGATCTTCACCGCGGG 1000
Oligo fwd CACCCTGCTGAAGATCTTCACCGC
Oligo rev AAACGCGGTGAAGATCTTCAGCAG
Chromosome start end strand MM target_seq PAM alignment distance position gene name gene id
GU305914.1 201805 201827 - 0 CTGCTGAAGATCTTCACCGC GGG ||||||||[||||||||||||]PAM NA Intergenic NA NA

T19241 AGATCTTCACCGCGGGCCGTCGG 1000
Oligo fwd CACCAGATCTTCACCGCGGGCCGT
Oligo rev AAACACGGCCCGCGGTGAAGATCT
Chromosome start end strand MM target_seq PAM alignment distance position gene name gene id
GU305914.1 201798 201820 - 0 AGATCTTCACCGCGGGCCGT CGG ||||||||[||||||||||||]PAM NA Intergenic NA NA

T19242 GATCTTCACCGCGGGCCGTCGGG 1000
Oligo fwd CACCGATCTTCACCGCGGGCCGTC
Oligo rev AAACGACGGCCCGCGGTGAAGATC
Chromosome start end strand MM target_seq PAM alignment distance position gene name gene id
GU305914.1 201797 201819 - 0 GATCTTCACCGCGGGCCGTC GGG ||||||||[||||||||||||]PAM NA Intergenic NA NA

T19243 CCGCGGGCCGTCGGGTCGACCGG 1000
Oligo fwd CACCCCGCGGGCCGTCGGGTCGAC
Oligo rev AAACGTCGACCCGACGGCCCGCGG
Chromosome start end strand MM target_seq PAM alignment distance position gene name gene id
GU305914.1 201789 201811 - 0 CCGCGGGCCGTCGGGTCGAC CGG ||||||||[||||||||||||]PAM NA Intergenic NA NA

T19244 CGCGGGCCGTCGGGTCGACCGGG 1000
Oligo fwd CACCCGCGGGCCGTCGGGTCGACC
Oligo rev AAACGGTCGACCCGACGGCCCGCG
Chromosome start end strand MM target_seq PAM alignment distance position gene name gene id
GU305914.1 201788 201810 - 0 CGCGGGCCGTCGGGTCGACC GGG ||||||||[||||||||||||]PAM NA Intergenic NA NA

T19245 TCGGGTCGACCGGGATCTCGCGG 1000
Oligo fwd CACCTCGGGTCGACCGGGATCTCG
Oligo rev AAACCGAGATCCCGGTCGACCCGA
Chromosome start end strand MM target_seq PAM alignment distance position gene name gene id
GU305914.1 201779 201801 - 0 TCGGGTCGACCGGGATCTCG CGG ||||||||[||||||||||||]PAM NA Intergenic NA NA

T19246 CGGGTCGACCGGGATCTCGCGGG 1000
Oligo fwd CACCCGGGTCGACCGGGATCTCGC
Oligo rev AAACGCGAGATCCCGGTCGACCCG
Chromosome start end strand MM target_seq PAM alignment distance position gene name gene id
GU305914.1 201778 201800 - 0 CGGGTCGACCGGGATCTCGC GGG ||||||||[||||||||||||]PAM NA Intergenic NA NA

T19247 GGGTCGACCGGGATCTCGCGGGG 1000
Oligo fwd CACCGGGTCGACCGGGATCTCGCG
Oligo rev AAACCGCGAGATCCCGGTCGACCC
Chromosome start end strand MM target_seq PAM alignment distance position gene name gene id

<table>
<tr><td>GU305914.1</td><td>201777</td><td>201799</td><td>-</td><td>0</td><td>GGGTCGACCGGGATCTCGCG</td><td>GGG</td><td>||||||||[||||||||||||]PAM</td><td>NA</td><td>Intergenic</td><td>NA</td><td>NA</td></tr>
<tr><td></td><td></td><td></td><td></td><td></td><td></td><td></td><td></td><td></td><td></td><td></td><td></td></tr>
<tr><td>T19248</td><td>GGGATCTCGCGGGGAGCGTCCGG</td><td>1000</td><td></td><td></td><td></td><td></td><td></td><td></td><td></td><td></td><td></td></tr>
<tr><td>Oligo fwd</td><td>CACCGGGATCTCGCGGGGAGCGTC</td><td></td><td></td><td></td><td></td><td></td><td></td><td></td><td></td><td></td><td></td></tr>
<tr><td>Oligo rev</td><td>AAACGACGCTCCCCGCGAGATCCC</td><td></td><td></td><td></td><td></td><td></td><td></td><td></td><td></td><td></td><td></td></tr>
<tr><td>Chromosome</td><td>start</td><td>end</td><td>strand</td><td>MM</td><td>target_seq</td><td>PAM</td><td>alignment</td><td>distance</td><td>position</td><td>gene name</td><td>gene id</td></tr>
<tr><td>GU305914.1</td><td>201768</td><td>201790</td><td>-</td><td>0</td><td>GGGATCTCGCGGGGAGCGTC</td><td>CGG</td><td>||||||||[||||||||||||]PAM</td><td>NA</td><td>Intergenic</td><td>NA</td><td>NA</td></tr>
<tr><td></td><td></td><td></td><td></td><td></td><td></td><td></td><td></td><td></td><td></td><td></td><td></td></tr>
<tr><td>T19249</td><td>GCTCGAGCCCCCACCCGACCAGG</td><td>1000</td><td></td><td></td><td></td><td></td><td></td><td></td><td></td><td></td><td></td></tr>
<tr><td>Oligo fwd</td><td>CACCGCTCGAGCCCCCACCCGACC</td><td></td><td></td><td></td><td></td><td></td><td></td><td></td><td></td><td></td><td></td></tr>
<tr><td>Oligo rev</td><td>AAACGGTCGGGTGGGGGCTCGAGC</td><td></td><td></td><td></td><td></td><td></td><td></td><td></td><td></td><td></td><td></td></tr>
<tr><td>Chromosome</td><td>start</td><td>end</td><td>strand</td><td>MM</td><td>target_seq</td><td>PAM</td><td>alignment</td><td>distance</td><td>position</td><td>gene name</td><td>gene id</td></tr>
<tr><td>GU305914.1</td><td>201746</td><td>201768</td><td>-</td><td>0</td><td>GCTCGAGCCCCCACCCGACC</td><td>AGG</td><td>||||||||[||||||||||||]PAM</td><td>NA</td><td>Intergenic</td><td>NA</td><td>NA</td></tr>
<tr><td></td><td></td><td></td><td></td><td></td><td></td><td></td><td></td><td></td><td></td><td></td><td></td></tr>
<tr><td>T19250</td><td>CCCCACCCGACCAGGAGCTGTGG</td><td>1000</td><td></td><td></td><td></td><td></td><td></td><td></td><td></td><td></td><td></td></tr>
<tr><td>Oligo fwd</td><td>CACCCCCCACCCGACCAGGAGCTG</td><td></td><td></td><td></td><td></td><td></td><td></td><td></td><td></td><td></td><td></td></tr>
<tr><td>Oligo rev</td><td>AAACCAGCTCCTGGTCGGGTGGGG</td><td></td><td></td><td></td><td></td><td></td><td></td><td></td><td></td><td></td><td></td></tr>
<tr><td>Chromosome</td><td>start</td><td>end</td><td>strand</td><td>MM</td><td>target_seq</td><td>PAM</td><td>alignment</td><td>distance</td><td>position</td><td>gene name</td><td>gene id</td></tr>
<tr><td>GU305914.1</td><td>201738</td><td>201760</td><td>-</td><td>0</td><td>CCCCACCCGACCAGGAGCTG</td><td>TGG</td><td>||||||||[||||||||||||]PAM</td><td>NA</td><td>Intergenic</td><td>NA</td><td>NA</td></tr>
<tr><td></td><td></td><td></td><td></td><td></td><td></td><td></td><td></td><td></td><td></td><td></td><td></td></tr>
<tr><td>T19251</td><td>GACGCATCTGTGCGTCGACGTGG</td><td>1000</td><td></td><td></td><td></td><td></td><td></td><td></td><td></td><td></td><td></td></tr>
<tr><td>Oligo fwd</td><td>CACCGACGCATCTGTGCGTCGACG</td><td></td><td></td><td></td><td></td><td></td><td></td><td></td><td></td><td></td><td></td></tr>
<tr><td>Oligo rev</td><td>AAACCGTCGACGCACAGATGCGTC</td><td></td><td></td><td></td><td></td><td></td><td></td><td></td><td></td><td></td><td></td></tr>
<tr><td>Chromosome</td><td>start</td><td>end</td><td>strand</td><td>MM</td><td>target_seq</td><td>PAM</td><td>alignment</td><td>distance</td><td>position</td><td>gene name</td><td>gene id</td></tr>
<tr><td>GU305914.1</td><td>201695</td><td>201717</td><td>-</td><td>0</td><td>GACGCATCTGTGCGTCGACG</td><td>TGG</td><td>||||||||[||||||||||||]PAM</td><td>NA</td><td>Intergenic</td><td>NA</td><td>NA</td></tr>
<tr><td></td><td></td><td></td><td></td><td></td><td></td><td></td><td></td><td></td><td></td><td></td><td></td></tr>
<tr><td>T19252</td><td>TGCGTCGACGTGGCACCCCCCGG</td><td>1000</td><td></td><td></td><td></td><td></td><td></td><td></td><td></td><td></td><td></td></tr>
<tr><td>Oligo fwd</td><td>CACCTGCGTCGACGTGGCACCCCC</td><td></td><td></td><td></td><td></td><td></td><td></td><td></td><td></td><td></td><td></td></tr>
<tr><td>Oligo rev</td><td>AAACGGGGGTGCCACGTCGACGCA</td><td></td><td></td><td></td><td></td><td></td><td></td><td></td><td></td><td></td><td></td></tr>
<tr><td>Chromosome</td><td>start</td><td>end</td><td>strand</td><td>MM</td><td>target_seq</td><td>PAM</td><td>alignment</td><td>distance</td><td>position</td><td>gene name</td><td>gene id</td></tr>
<tr><td>GU305914.1</td><td>201685</td><td>201707</td><td>-</td><td>0</td><td>TGCGTCGACGTGGCACCCCC</td><td>CGG</td><td>||||||||[||||||||||||]PAM</td><td>NA</td><td>Intergenic</td><td>NA</td><td>NA</td></tr>
<tr><td></td><td></td><td></td><td></td><td></td><td></td><td></td><td></td><td></td><td></td><td></td><td></td></tr>
<tr><td>T19253</td><td>CGACGTGGCACCCCCCGGCGAGG</td><td>1000</td><td></td><td></td><td></td><td></td><td></td><td></td><td></td><td></td><td></td></tr>
<tr><td>Oligo fwd</td><td>CACCCGACGTGGCACCCCCCGGCG</td><td></td><td></td><td></td><td></td><td></td><td></td><td></td><td></td><td></td><td></td></tr>
<tr><td>Oligo rev</td><td>AAACCGCCGGGGGGTGCCACGTCG</td><td></td><td></td><td></td><td></td><td></td><td></td><td></td><td></td><td></td><td></td></tr>
<tr><td>Chromosome</td><td>start</td><td>end</td><td>strand</td><td>MM</td><td>target_seq</td><td>PAM</td><td>alignment</td><td>distance</td><td>position</td><td>gene name</td><td>gene id</td></tr>
<tr><td>GU305914.1</td><td>201680</td><td>201702</td><td>-</td><td>0</td><td>CGACGTGGCACCCCCCGGCG</td><td>AGG</td><td>||||||||[||||||||||||]PAM</td><td>NA</td><td>Intergenic</td><td>NA</td><td>NA</td></tr>
<tr><td>GU305914.1</td><td>76803</td><td>76825</td><td>+</td><td>4</td><td>CGAGGTTGCACTCCTCGGCG</td><td>GGG</td><td>|||-||-|[|||-||-|||||]PAM</td><td>NA</td><td>Intergenic</td><td>NA</td><td>NA</td></tr>
<tr><td></td><td></td><td></td><td></td><td></td><td></td><td></td><td></td><td></td><td></td><td></td><td></td></tr>
<tr><td>T19254</td><td>AGCTCGCGCAGCAGTTCGCGTGG</td><td>1000</td><td></td><td></td><td></td><td></td><td></td><td></td><td></td><td></td><td></td></tr>
<tr><td>Oligo fwd</td><td>CACCAGCTCGCGCAGCAGTTCGCG</td><td></td><td></td><td></td><td></td><td></td><td></td><td></td><td></td><td></td><td></td></tr>
<tr><td>Oligo rev</td><td>AAACCGCGAACTGCTGCGCGAGCT</td><td></td><td></td><td></td><td></td><td></td><td></td><td></td><td></td><td></td><td></td></tr>
<tr><td>Chromosome</td><td>start</td><td>end</td><td>strand</td><td>MM</td><td>target_seq</td><td>PAM</td><td>alignment</td><td>distance</td><td>position</td><td>gene name</td><td>gene id</td></tr>
<tr><td>GU305914.1</td><td>201654</td><td>201676</td><td>-</td><td>0</td><td>AGCTCGCGCAGCAGTTCGCG</td><td>TGG</td><td>||||||||[||||||||||||]PAM</td><td>NA</td><td>Intergenic</td><td>NA</td><td>NA</td></tr>
<tr><td></td><td></td><td></td><td></td><td></td><td></td><td></td><td></td><td></td><td></td><td></td><td></td></tr>
<tr><td>T19255</td><td>TTCGCGTGGATCTCACGTCCCGG</td><td>1000</td><td></td><td></td><td></td><td></td><td></td><td></td><td></td><td></td><td></td></tr>
<tr><td>Oligo fwd</td><td>CACCTTCGCGTGGATCTCACGTCC</td><td></td><td></td><td></td><td></td><td></td><td></td><td></td><td></td><td></td><td></td></tr>
<tr><td>Oligo rev</td><td>AAACGGACGTGAGATCCACGCGAA</td><td></td><td></td><td></td><td></td><td></td><td></td><td></td><td></td><td></td><td></td></tr>
<tr><td>Chromosome</td><td>start</td><td>end</td><td>strand</td><td>MM</td><td>target_seq</td><td>PAM</td><td>alignment</td><td>distance</td><td>position</td><td>gene name</td><td>gene id</td></tr>
<tr><td>GU305914.1</td><td>201640</td><td>201662</td><td>-</td><td>0</td><td>TTCGCGTGGATCTCACGTCC</td><td>CGG</td><td>||||||||[||||||||||||]PAM</td><td>NA</td><td>Intergenic</td><td>NA</td><td>NA</td></tr>
<tr><td></td><td></td><td></td><td></td><td></td><td></td><td></td><td></td><td></td><td></td><td></td><td></td></tr>
<tr><td>T19256</td><td>CGTGGATCTCACGTCCCGGCAGG</td><td>1000</td><td></td><td></td><td></td><td></td><td></td><td></td><td></td><td></td><td></td></tr>
<tr><td>Oligo fwd</td><td>CACCCGTGGATCTCACGTCCCGGC</td><td></td><td></td><td></td><td></td><td></td><td></td><td></td><td></td><td></td><td></td></tr>
<tr><td>Oligo rev</td><td>AAACGCCGGGACGTGAGATCCACG</td><td></td><td></td><td></td><td></td><td></td><td></td><td></td><td></td><td></td><td></td></tr>
<tr><td>Chromosome</td><td>start</td><td>end</td><td>strand</td><td>MM</td><td>target_seq</td><td>PAM</td><td>alignment</td><td>distance</td><td>position</td><td>gene name</td><td>gene id</td></tr>
<tr><td>GU305914.1</td><td>201636</td><td>201658</td><td>-</td><td>0</td><td>CGTGGATCTCACGTCCCGGC</td><td>AGG</td><td>||||||||[||||||||||||]PAM</td><td>NA</td><td>Intergenic</td><td>NA</td><td>NA</td></tr>
<tr><td></td><td></td><td></td><td></td><td></td><td></td><td></td><td></td><td></td><td></td><td></td><td></td></tr>
<tr><td>T19257</td><td>GTGGATCTCACGTCCCGGCAGGG</td><td>1000</td><td></td><td></td><td></td><td></td><td></td><td></td><td></td><td></td><td></td></tr>
<tr><td>Oligo fwd</td><td>CACCGTGGATCTCACGTCCCGGCA</td><td></td><td></td><td></td><td></td><td></td><td></td><td></td><td></td><td></td><td></td></tr>
<tr><td>Oligo rev</td><td>AAACTGCCGGGACGTGAGATCCAC</td><td></td><td></td><td></td><td></td><td></td><td></td><td></td><td></td><td></td><td></td></tr>
<tr><td>Chromosome</td><td>start</td><td>end</td><td>strand</td><td>MM</td><td>target_seq</td><td>PAM</td><td>alignment</td><td>distance</td><td>position</td><td>gene name</td><td>gene id</td></tr>
<tr><td>GU305914.1</td><td>201635</td><td>201657</td><td>-</td><td>0</td><td>GTGGATCTCACGTCCCGGCA</td><td>GGG</td><td>||||||||[||||||||||||]PAM</td><td>NA</td><td>Intergenic</td><td>NA</td><td>NA</td></tr>
<tr><td></td><td></td><td></td><td></td><td></td><td></td><td></td><td></td><td></td><td></td><td></td><td></td></tr>
<tr><td>T19258</td><td>GATCTCACGTCCCGGCAGGGCGG</td><td>1000</td><td></td><td></td><td></td><td></td><td></td><td></td><td></td><td></td><td></td></tr>
<tr><td>Oligo fwd</td><td>CACCGATCTCACGTCCCGGCAGGG</td><td></td><td></td><td></td><td></td><td></td><td></td><td></td><td></td><td></td><td></td></tr>
<tr><td>Oligo rev</td><td>AAACCCCTGCCGGGACGTGAGATC</td><td></td><td></td><td></td><td></td><td></td><td></td><td></td><td></td><td></td><td></td></tr>
<tr><td>Chromosome</td><td>start</td><td>end</td><td>strand</td><td>MM</td><td>target_seq</td><td>PAM</td><td>alignment</td><td>distance</td><td>position</td><td>gene name</td><td>gene id</td></tr>
<tr><td>GU305914.1</td><td>201632</td><td>201654</td><td>-</td><td>0</td><td>GATCTCACGTCCCGGCAGGG</td><td>CGG</td><td>||||||||[||||||||||||]PAM</td><td>NA</td><td>Intergenic</td><td>NA</td><td>NA</td></tr>
<tr><td></td><td></td><td></td><td></td><td></td><td></td><td></td><td></td><td></td><td></td><td></td><td></td></tr>
<tr><td>T19259</td><td>CAGGGCGGAGATCCTGTCCGAGG</td><td>1000</td><td></td><td></td><td></td><td></td><td></td><td></td><td></td><td></td><td></td></tr>
<tr><td>Oligo fwd</td><td>CACCCAGGGCGGAGATCCTGTCCG</td><td></td><td></td><td></td><td></td><td></td><td></td><td></td><td></td><td></td><td></td></tr>
<tr><td>Oligo rev</td><td>AAACCGGACAGGATCTCCGCCCTG</td><td></td><td></td><td></td><td></td><td></td><td></td><td></td><td></td><td></td><td></td></tr>
<tr><td>Chromosome</td><td>start</td><td>end</td><td>strand</td><td>MM</td><td>target_seq</td><td>PAM</td><td>alignment</td><td>distance</td><td>position</td><td>gene name</td><td>gene id</td></tr>
<tr><td>GU305914.1</td><td>201617</td><td>201639</td><td>-</td><td>0</td><td>CAGGGCGGAGATCCTGTCCG</td><td>AGG</td><td>||||||||[||||||||||||]PAM</td><td>NA</td><td>Intergenic</td><td>NA</td><td>NA</td></tr>
<tr><td></td><td></td><td></td><td></td><td></td><td></td><td></td><td></td><td></td><td></td><td></td><td></td></tr>
<tr><td>T19260</td><td>GGCGGAGATCCTGTCCGAGGAGG</td><td>1000</td><td></td><td></td><td></td><td></td><td></td><td></td><td></td><td></td><td></td></tr>
<tr><td>Oligo fwd</td><td>CACCGGCGGAGATCCTGTCCGAGG</td><td></td><td></td><td></td><td></td><td></td><td></td><td></td><td></td><td></td><td></td></tr>
<tr><td>Oligo rev</td><td>AAACCCTCGGACAGGATCTCCGCC</td><td></td><td></td><td></td><td></td><td></td><td></td><td></td><td></td><td></td><td></td></tr>
<tr><td>Chromosome</td><td>start</td><td>end</td><td>strand</td><td>MM</td><td>target_seq</td><td>PAM</td><td>alignment</td><td>distance</td><td>position</td><td>gene name</td><td>gene id</td></tr>
<tr><td>GU305914.1</td><td>201614</td><td>201636</td><td>-</td><td>0</td><td>GGCGGAGATCCTGTCCGAGG</td><td>AGG</td><td>||||||||[||||||||||||]PAM</td><td>NA</td><td>Intergenic</td><td>NA</td><td>NA</td></tr>
<tr><td></td><td></td><td></td><td></td><td></td><td></td><td></td><td></td><td></td><td></td><td></td><td></td></tr>
<tr><td>T19261</td><td>CCTGTCCGAGGAGGAGACCGCGG</td><td>1000</td><td></td><td></td><td></td><td></td><td></td><td></td><td></td><td></td><td></td></tr>
<tr><td>Oligo fwd</td><td>CACCCCTGTCCGAGGAGGAGACCG</td><td></td><td></td><td></td><td></td><td></td><td></td><td></td><td></td><td></td><td></td></tr>
<tr><td>Oligo rev</td><td>AAACCGGTCTCCTCCTCGGACAGG</td><td></td><td></td><td></td><td></td><td></td><td></td><td></td><td></td><td></td><td></td></tr>
<tr><td>Chromosome</td><td>start</td><td>end</td><td>strand</td><td>MM</td><td>target_seq</td><td>PAM</td><td>alignment</td><td>distance</td><td>position</td><td>gene name</td><td>gene id</td></tr>
<tr><td>GU305914.1</td><td>201605</td><td>201627</td><td>-</td><td>0</td><td>CCTGTCCGAGGAGGAGACCG</td><td>CGG</td><td>||||||||[||||||||||||]PAM</td><td>NA</td><td>Intergenic</td><td>NA</td><td>NA</td></tr>
<tr><td></td><td></td><td></td><td></td><td></td><td></td><td></td><td></td><td></td><td></td><td></td><td></td></tr>
<tr><td>T19262</td><td>GTCCGAGGAGGAGACCGCGGTGG</td><td>1000</td><td></td><td></td><td></td><td></td><td></td><td></td><td></td><td></td><td></td></tr>
<tr><td>Oligo fwd</td><td>CACCGTCCGAGGAGGAGACCGCGG</td><td></td><td></td><td></td><td></td><td></td><td></td><td></td><td></td><td></td><td></td></tr>
</table>

| | | | | | | | | | | | |
|---|---|---|---|---|---|---|---|---|---|---|---|
| Oligo rev | AAACCCGCGGTCTCCTCCTCGGAC | | | | | | | | | | |
| Chromosome | start | end | strand | MM | target_seq | PAM | alignment | distance | position | gene name | gene id |
| GU305914.1 | 201602 | 201624 | - | 0 | GTCCGAGGAGGAGACCGCGG | TGG | \|\|\|\|\|\|\|\|[\|\|\|\|\|\|\|\|\|\|\|\|]PAM | NA | Intergenic | NA | NA |
| GU305914.1 | 99490 | 99512 | - | 4 | TTACGAGGAGGAGAGGGCGG | TGG | -\|-\|\|\|\|\|[\|\|\|\|\|\|--\|\|\|\|]PAM | NA | Intergenic | NA | NA |
| GU305914.1 | 165098 | 165120 | + | 3 | TTCCGAGGAGGAAACCGCGA | TGG | -\|\|\|\|\|\|\|[\|\|\|\|-\|\|\|\|\|\|-]PAM | NA | Intergenic | NA | NA |
| T19263 | CGAGGAGGAGACCGCGGTGGCGG | 1000 | | | | | | | | | |
| Oligo fwd | CACCCGAGGAGGAGACCGCGGTGG | | | | | | | | | | |
| Oligo rev | AAACCCACCGCGGTCTCCTCCTCG | | | | | | | | | | |
| Chromosome | start | end | strand | MM | target_seq | PAM | alignment | distance | position | gene name | gene id |
| GU305914.1 | 201599 | 201621 | - | 0 | CGAGGAGGAGACCGCGGTGG | CGG | \|\|\|\|\|\|\|\|[\|\|\|\|\|\|\|\|\|\|\|\|]PAM | NA | Intergenic | NA | NA |
| GU305914.1 | 114670 | 114692 | - | 4 | CGATGGTGAGACGGCGGTGG | TGG | \|\|\|-\|--\|[\|\|\|\|-\|\|\|\|\|\|\|]PAM | NA | Intergenic | NA | NA |
| T19264 | ACGCCTTCCTGCGCCTGTCGAGG | 1000 | | | | | | | | | |
| Oligo fwd | CACCACGCCTTCCTGCGCCTGTCG | | | | | | | | | | |
| Oligo rev | AAACCGACAGGCGCAGGAAGGCGT | | | | | | | | | | |
| Chromosome | start | end | strand | MM | target_seq | PAM | alignment | distance | position | gene name | gene id |
| GU305914.1 | 201576 | 201598 | - | 0 | ACGCCTTCCTGCGCCTGTCG | AGG | \|\|\|\|\|\|\|\|[\|\|\|\|\|\|\|\|\|\|\|\|]PAM | NA | Intergenic | NA | NA |
| T19265 | CCTGCGCCTGTCGAGGCTCAAGG | 1000 | | | | | | | | | |
| Oligo fwd | CACCCCTGCGCCTGTCGAGGCTCA | | | | | | | | | | |
| Oligo rev | AAACTGAGCCTCGACAGGCGCAGG | | | | | | | | | | |
| Chromosome | start | end | strand | MM | target_seq | PAM | alignment | distance | position | gene name | gene id |
| GU305914.1 | 201569 | 201591 | - | 0 | CCTGCGCCTGTCGAGGCTCA | AGG | \|\|\|\|\|\|\|\|[\|\|\|\|\|\|\|\|\|\|\|\|]PAM | NA | Intergenic | NA | NA |
| T19266 | CCTGTCGAGGCTCAAGGTGACGG | 1000 | | | | | | | | | |
| Oligo fwd | CACCCCTGTCGAGGCTCAAGGTGA | | | | | | | | | | |
| Oligo rev | AAACTCACCTTGAGCCTCGACAGG | | | | | | | | | | |
| Chromosome | start | end | strand | MM | target_seq | PAM | alignment | distance | position | gene name | gene id |
| GU305914.1 | 201563 | 201585 | - | 0 | CCTGTCGAGGCTCAAGGTGA | CGG | \|\|\|\|\|\|\|\|[\|\|\|\|\|\|\|\|\|\|\|\|]PAM | NA | Intergenic | NA | NA |
| T19267 | GAGGCTCAAGGTGACGGTGACGG | 1000 | | | | | | | | | |
| Oligo fwd | CACCGAGGCTCAAGGTGACGGTGA | | | | | | | | | | |
| Oligo rev | AAACTCACCGTCACCTTGAGCCTC | | | | | | | | | | |
| Chromosome | start | end | strand | MM | target_seq | PAM | alignment | distance | position | gene name | gene id |
| GU305914.1 | 201557 | 201579 | - | 0 | GAGGCTCAAGGTGACGGTGA | CGG | \|\|\|\|\|\|\|\|[\|\|\|\|\|\|\|\|\|\|\|\|]PAM | NA | Intergenic | NA | NA |
| T19268 | GCTCAAGGTGACGGTGACGGAGG | 1000 | | | | | | | | | |
| Oligo fwd | CACCGCTCAAGGTGACGGTGACGG | | | | | | | | | | |
| Oligo rev | AAACCCGTCACCGTCACCTTGAGC | | | | | | | | | | |
| Chromosome | start | end | strand | MM | target_seq | PAM | alignment | distance | position | gene name | gene id |
| GU305914.1 | 201554 | 201576 | - | 0 | GCTCAAGGTGACGGTGACGG | AGG | \|\|\|\|\|\|\|\|[\|\|\|\|\|\|\|\|\|\|\|\|]PAM | NA | Intergenic | NA | NA |
| T19269 | TGACGGAGGAGAACAGCTCGCGG | 1000 | | | | | | | | | |
| Oligo fwd | CACCTGACGGAGGAGAACAGCTCG | | | | | | | | | | |
| Oligo rev | AAACCGAGCTGTTCTCCTCCGTCA | | | | | | | | | | |
| Chromosome | start | end | strand | MM | target_seq | PAM | alignment | distance | position | gene name | gene id |
| GU305914.1 | 201540 | 201562 | - | 0 | TGACGGAGGAGAACAGCTCG | CGG | \|\|\|\|\|\|\|\|[\|\|\|\|\|\|\|\|\|\|\|\|]PAM | NA | Intergenic | NA | NA |
| T19270 | GACGGAGGAGAACAGCTCGCGGG | 1000 | | | | | | | | | |
| Oligo fwd | CACCGACGGAGGAGAACAGCTCGC | | | | | | | | | | |
| Oligo rev | AAACGCGAGCTGTTCTCCTCCGTC | | | | | | | | | | |
| Chromosome | start | end | strand | MM | target_seq | PAM | alignment | distance | position | gene name | gene id |
| GU305914.1 | 201539 | 201561 | - | 0 | GACGGAGGAGAACAGCTCGC | GGG | \|\|\|\|\|\|\|\|[\|\|\|\|\|\|\|\|\|\|\|\|]PAM | NA | Intergenic | NA | NA |
| GU305914.1 | 77470 | 77492 | + | 4 | GATGGTGGAGAACATCACGC | TGG | \|\|-\|\|-\|\|[\|\|\|\|\|\|-\|-\|\|\|]PAM | NA | Intergenic | NA | NA |
| T19271 | TCGCCGCGCCGCCCGAGCCGCGG | 1000 | | | | | | | | | |
| Oligo fwd | CACCTCGCCGCGCCGCCCGAGCCG | | | | | | | | | | |
| Oligo rev | AAACCGGCTCGGGCGGCGCGGCGA | | | | | | | | | | |
| Chromosome | start | end | strand | MM | target_seq | PAM | alignment | distance | position | gene name | gene id |
| GU305914.1 | 201501 | 201523 | - | 0 | TCGCCGCGCCGCCCGAGCCG | CGG | \|\|\|\|\|\|\|\|[\|\|\|\|\|\|\|\|\|\|\|\|]PAM | NA | Intergenic | NA | NA |
| T19272 | CGAGCCGCGGACGCGCCGCTTGG | 1000 | | | | | | | | | |
| Oligo fwd | CACCCGAGCCGCGGACGCGCCGCT | | | | | | | | | | |
| Oligo rev | AAACAGCGGCGCGTCCGCGGCTCG | | | | | | | | | | |
| Chromosome | start | end | strand | MM | target_seq | PAM | alignment | distance | position | gene name | gene id |
| GU305914.1 | 201488 | 201510 | - | 0 | CGAGCCGCGGACGCGCCGCT | TGG | \|\|\|\|\|\|\|\|[\|\|\|\|\|\|\|\|\|\|\|\|]PAM | NA | Intergenic | NA | NA |
| T19273 | GCGGACGCGCCGCTTGGTGTCGG | 1000 | | | | | | | | | |
| Oligo fwd | CACCGCGGACGCGCCGCTTGGTGT | | | | | | | | | | |
| Oligo rev | AAACACACCAAGCGGCGCGTCCGC | | | | | | | | | | |
| Chromosome | start | end | strand | MM | target_seq | PAM | alignment | distance | position | gene name | gene id |
| GU305914.1 | 201482 | 201504 | - | 0 | GCGGACGCGCCGCTTGGTGT | CGG | \|\|\|\|\|\|\|\|[\|\|\|\|\|\|\|\|\|\|\|\|]PAM | NA | Intergenic | NA | NA |
| T19274 | GTCGGCGCTGCTGCGTGACGAGG | 1000 | | | | | | | | | |
| Oligo fwd | CACCGTCGGCGCTGCTGCGTGACG | | | | | | | | | | |
| Oligo rev | AAACCGTCACGCAGCAGCGCCGAC | | | | | | | | | | |
| Chromosome | start | end | strand | MM | target_seq | PAM | alignment | distance | position | gene name | gene id |
| GU305914.1 | 201464 | 201486 | - | 0 | GTCGGCGCTGCTGCGTGACG | AGG | \|\|\|\|\|\|\|\|[\|\|\|\|\|\|\|\|\|\|\|\|]PAM | NA | Intergenic | NA | NA |
| GU305914.1 | 124826 | 124848 | - | 4 | GGCCGCGCTGGTGCGGGACG | TGG | \|-\|-\|\|\|\|[\|\|-\|\|\|\|-\|\|\|\|]PAM | NA | Intergenic | NA | NA |
| T19275 | TGCTGCGTGACGAGGACTGCTGG | 1000 | | | | | | | | | |
| Oligo fwd | CACCTGCTGCGTGACGAGGACTGC | | | | | | | | | | |
| Oligo rev | AAACGCAGTCCTCGTCACGCAGCA | | | | | | | | | | |
| Chromosome | start | end | strand | MM | target_seq | PAM | alignment | distance | position | gene name | gene id |
| GU305914.1 | 201456 | 201478 | - | 0 | TGCTGCGTGACGAGGACTGC | TGG | \|\|\|\|\|\|\|\|[\|\|\|\|\|\|\|\|\|\|\|\|]PAM | NA | Intergenic | NA | NA |
| T19276 | GGACTGCTGGCGCCCCGAGTCGG | 1000 | | | | | | | | | |
| Oligo fwd | CACCGGACTGCTGGCGCCCCGAGT | | | | | | | | | | |

| | | | | | | | | | | | |
|---|---|---|---|---|---|---|---|---|---|---|---|
| Oligo rev | AAACACTCGGGGCGCCAGCAGTCC | | | | | | | | | | |
| Chromosome | start | end | strand | MM | target_seq | PAM | alignment | distance | position | gene name | gene id |
| GU305914.1 | 201443 | 201465 | - | 0 | GGACTGCTGGCGCCCCGAGT | CGG | \|\|\|\|\|\|\|\|[\|\|\|\|\|\|\|\|\|\|\|\|]PAM | NA | Intergenic | NA | NA |
| T19277 | CTGCTGGCGCCCCGAGTCGGCGG | 1000 | | | | | | | | | |
| Oligo fwd | CACCCTGCTGGCGCCCCGAGTCGG | | | | | | | | | | |
| Oligo rev | AAACCCGACTCGGGGCGCCAGCAG | | | | | | | | | | |
| Chromosome | start | end | strand | MM | target_seq | PAM | alignment | distance | position | gene name | gene id |
| GU305914.1 | 201440 | 201462 | - | 0 | CTGCTGGCGCCCCGAGTCGG | CGG | \|\|\|\|\|\|\|\|[\|\|\|\|\|\|\|\|\|\|\|\|]PAM | NA | Intergenic | NA | NA |
| T19278 | GCGCCCCGAGTCGGCGGACAAGG | 1000 | | | | | | | | | |
| Oligo fwd | CACCGCGCCCCGAGTCGGCGGACA | | | | | | | | | | |
| Oligo rev | AAACTGTCCGCCGACTCGGGGCGC | | | | | | | | | | |
| Chromosome | start | end | strand | MM | target_seq | PAM | alignment | distance | position | gene name | gene id |
| GU305914.1 | 201434 | 201456 | - | 0 | GCGCCCCGAGTCGGCGGACA | AGG | \|\|\|\|\|\|\|\|[\|\|\|\|\|\|\|\|\|\|\|\|]PAM | NA | Intergenic | NA | NA |
| T19279 | CCCCGAGTCGGCGGACAAGGCGG | 1000 | | | | | | | | | |
| Oligo fwd | CACCCCCCGAGTCGGCGGACAAGG | | | | | | | | | | |
| Oligo rev | AAACCCTTGTCCGCCGACTCGGGG | | | | | | | | | | |
| Chromosome | start | end | strand | MM | target_seq | PAM | alignment | distance | position | gene name | gene id |
| GU305914.1 | 201431 | 201453 | - | 0 | CCCCGAGTCGGCGGACAAGG | CGG | \|\|\|\|\|\|\|\|[\|\|\|\|\|\|\|\|\|\|\|\|]PAM | NA | Intergenic | NA | NA |
| T19280 | ACAAGGCGGACATCGTGCCTCGG | 1000 | | | | | | | | | |
| Oligo fwd | CACCACAAGGCGGACATCGTGCCT | | | | | | | | | | |
| Oligo rev | AAACAGGCACGATGTCCGCCTTGT | | | | | | | | | | |
| Chromosome | start | end | strand | MM | target_seq | PAM | alignment | distance | position | gene name | gene id |
| GU305914.1 | 201417 | 201439 | - | 0 | ACAAGGCGGACATCGTGCCT | CGG | \|\|\|\|\|\|\|\|[\|\|\|\|\|\|\|\|\|\|\|\|]PAM | NA | Intergenic | NA | NA |
| T19281 | GGACATCGTGCCTCGGTTCAAGG | 1000 | | | | | | | | | |
| Oligo fwd | CACCGGACATCGTGCCTCGGTTCA | | | | | | | | | | |
| Oligo rev | AAACTGAACCGAGGCACGATGTCC | | | | | | | | | | |
| Chromosome | start | end | strand | MM | target_seq | PAM | alignment | distance | position | gene name | gene id |
| GU305914.1 | 201410 | 201432 | - | 0 | GGACATCGTGCCTCGGTTCA | AGG | \|\|\|\|\|\|\|\|[\|\|\|\|\|\|\|\|\|\|\|\|]PAM | NA | Intergenic | NA | NA |
| T19282 | GAAGCCGCCGCCTCCGTCGCCGG | 1000 | | | | | | | | | |
| Oligo fwd | CACCGAAGCCGCCGCCTCCGTCGC | | | | | | | | | | |
| Oligo rev | AAACGCGACGGAGGCGGCGGCTTC | | | | | | | | | | |
| Chromosome | start | end | strand | MM | target_seq | PAM | alignment | distance | position | gene name | gene id |
| GU305914.1 | 201386 | 201408 | - | 0 | GAAGCCGCCGCCTCCGTCGC | CGG | \|\|\|\|\|\|\|\|[\|\|\|\|\|\|\|\|\|\|\|\|]PAM | NA | Intergenic | NA | NA |
| GU305914.1 | 86843 | 86865 | + | 3 | GAAGCTGCCGCCCCCGCCGC | GGG | \|\|\|\|\|-\|\|[\|\|\|\|-\|\|\|-\|\|\|]PAM | NA | Intergenic | NA | NA |
| T19283 | GCCGCCGCCTCCGTCGCCGGAGG | 1000 | | | | | | | | | |
| Oligo fwd | CACCGCCGCCGCCTCCGTCGCCGG | | | | | | | | | | |
| Oligo rev | AAACCCGGCGACGGAGGCGGCGGC | | | | | | | | | | |
| Chromosome | start | end | strand | MM | target_seq | PAM | alignment | distance | position | gene name | gene id |
| GU305914.1 | 201383 | 201405 | - | 0 | GCCGCCGCCTCCGTCGCCGG | AGG | \|\|\|\|\|\|\|\|[\|\|\|\|\|\|\|\|\|\|\|\|]PAM | NA | Intergenic | NA | NA |
| T19284 | GCCGCCTCCGTCGCCGGAGGCGG | 1000 | | | | | | | | | |
| Oligo fwd | CACCGCCGCCTCCGTCGCCGGAGG | | | | | | | | | | |
| Oligo rev | AAACCCTCCGGCGACGGAGGCGGC | | | | | | | | | | |
| Chromosome | start | end | strand | MM | target_seq | PAM | alignment | distance | position | gene name | gene id |
| GU305914.1 | 201380 | 201402 | - | 0 | GCCGCCTCCGTCGCCGGAGG | CGG | \|\|\|\|\|\|\|\|[\|\|\|\|\|\|\|\|\|\|\|\|]PAM | NA | Intergenic | NA | NA |
| GU305914.1 | 201379 | 201401 | + | 3 | CCCGCCTCCGGCGACGGAGG | CGG | -\|\|\|\|\|\|\|[\|\|-\|\|-\|\|\|\|\|\|]PAM | NA | Intergenic | NA | NA |
| T19285 | CCGCCTCCGTCGCCGGAGGCGGG | 1000 | | | | | | | | | |
| Oligo fwd | CACCCCGCCTCCGTCGCCGGAGGC | | | | | | | | | | |
| Oligo rev | AAACGCCTCCGGCGACGGAGGCGG | | | | | | | | | | |
| Chromosome | start | end | strand | MM | target_seq | PAM | alignment | distance | position | gene name | gene id |
| GU305914.1 | 201379 | 201401 | - | 0 | CCGCCTCCGTCGCCGGAGGC | GGG | \|\|\|\|\|\|\|\|[\|\|\|\|\|\|\|\|\|\|\|\|]PAM | NA | Intergenic | NA | NA |
| T19286 | CGCCTCCGTCGCCGGAGGCGGGG | 1000 | | | | | | | | | |
| Oligo fwd | CACCCGCCTCCGTCGCCGGAGGCG | | | | | | | | | | |
| Oligo rev | AAACCGCCTCCGGCGACGGAGGCG | | | | | | | | | | |
| Chromosome | start | end | strand | MM | target_seq | PAM | alignment | distance | position | gene name | gene id |
| GU305914.1 | 201378 | 201400 | - | 0 | CGCCTCCGTCGCCGGAGGCG | GGG | \|\|\|\|\|\|\|\|[\|\|\|\|\|\|\|\|\|\|\|\|]PAM | NA | Intergenic | NA | NA |
| T19287 | GGAGGCGGGGCGCTACGACACGG | 1000 | | | | | | | | | |
| Oligo fwd | CACCGGAGGCGGGGCGCTACGACA | | | | | | | | | | |
| Oligo rev | AAACTGTCGTAGCGCCCCGCCTCC | | | | | | | | | | |
| Chromosome | start | end | strand | MM | target_seq | PAM | alignment | distance | position | gene name | gene id |
| GU305914.1 | 201365 | 201387 | - | 0 | GGAGGCGGGGCGCTACGACA | CGG | \|\|\|\|\|\|\|\|[\|\|\|\|\|\|\|\|\|\|\|\|]PAM | NA | Intergenic | NA | NA |
| GU305914.1 | 69196 | 69218 | + | 4 | GGACGCGCGGCGCGCCGACA | AGG | \|\|\|-\|\|\|-[\|\|\|\|\|--\|\|\|\|\|]PAM | NA | Intergenic | NA | NA |
| T19288 | CTACGACACGGATCAGTCTCCGG | 1000 | | | | | | | | | |
| Oligo fwd | CACCCTACGACACGGATCAGTCTC | | | | | | | | | | |
| Oligo rev | AAACGAGACTGATCCGTGTCGTAG | | | | | | | | | | |
| Chromosome | start | end | strand | MM | target_seq | PAM | alignment | distance | position | gene name | gene id |
| GU305914.1 | 201353 | 201375 | - | 0 | CTACGACACGGATCAGTCTC | CGG | \|\|\|\|\|\|\|\|[\|\|\|\|\|\|\|\|\|\|\|\|]PAM | NA | Intergenic | NA | NA |
| T19289 | CGACACGGATCAGTCTCCGGCGG | 1000 | | | | | | | | | |
| Oligo fwd | CACCCGACACGGATCAGTCTCCGG | | | | | | | | | | |
| Oligo rev | AAACCCGGAGACTGATCCGTGTCG | | | | | | | | | | |
| Chromosome | start | end | strand | MM | target_seq | PAM | alignment | distance | position | gene name | gene id |
| GU305914.1 | 201350 | 201372 | - | 0 | CGACACGGATCAGTCTCCGG | CGG | \|\|\|\|\|\|\|\|[\|\|\|\|\|\|\|\|\|\|\|\|]PAM | NA | Intergenic | NA | NA |
| T19290 | CACGGATCAGTCTCCGGCGGCGG | 1000 | | | | | | | | | |
| Oligo fwd | CACCCACGGATCAGTCTCCGGCGG | | | | | | | | | | |
| Oligo rev | AAACCCGCCGGAGACTGATCCGTG | | | | | | | | | | |
| Chromosome | start | end | strand | MM | target_seq | PAM | alignment | distance | position | gene name | gene id |

GU305914.1 201347 201369 - 0 CACGGATCAGTCTCCGGCGG CGG ||||||||[||||||||||||]PAM NA Intergenic NA NA

T19291 GGATCAGTCTCCGGCGGCGGTGG 1000
Oligo fwd CACCGGATCAGTCTCCGGCGGCGG
Oligo rev AAACCCGCCGCCGGAGACTGATCC
Chromosome start end strand MM target_seq PAM alignment distance position gene name gene id
GU305914.1 201344 201366 - 0 GGATCAGTCTCCGGCGGCGG TGG ||||||||[||||||||||||]PAM NA Intergenic NA NA
GU305914.1 149488 149510 - 4 GGAGCAGGCGACGGCGGCGG CGG |||-|||-[|--|||||||||]PAM NA Intergenic NA NA

T19292 AGTCTCCGGCGGCGGTGGCGAGG 1000
Oligo fwd CACCAGTCTCCGGCGGCGGTGGCG
Oligo rev AAACCGCCACCGCCGCCGGAGACT
Chromosome start end strand MM target_seq PAM alignment distance position gene name gene id
GU305914.1 201339 201361 - 0 AGTCTCCGGCGGCGGTGGCG AGG ||||||||[||||||||||||]PAM NA Intergenic NA NA

T19293 GGCGAGGCGCCGCGCGCACGCGG 1000
Oligo fwd CACCGGCGAGGCGCCGCGCGCACG
Oligo rev AAACCGTGCGCGCGGCGCCTCGCC
Chromosome start end strand MM target_seq PAM alignment distance position gene name gene id
GU305914.1 201323 201345 - 0 GGCGAGGCGCCGCGCGCACG CGG ||||||||[||||||||||||]PAM NA Intergenic NA NA

T19294 CACGCGGCCTCGAGTCAGACCGG 1000
Oligo fwd CACCCACGCGGCCTCGAGTCAGAC
Oligo rev AAACGTCTGACTCGAGGCCGCGTG
Chromosome start end strand MM target_seq PAM alignment distance position gene name gene id
GU305914.1 201307 201329 - 0 CACGCGGCCTCGAGTCAGAC CGG ||||||||[||||||||||||]PAM NA Intergenic NA NA

T19295 CCGTCACCACGCACCGATCCTGG 1000
Oligo fwd CACCCCGTCACCACGCACCGATCC
Oligo rev AAACGGATCGGTGCGTGGTGACGG
Chromosome start end strand MM target_seq PAM alignment distance position gene name gene id
GU305914.1 201275 201297 - 0 CCGTCACCACGCACCGATCC TGG ||||||||[||||||||||||]PAM NA Intergenic NA NA

T19296 GCCGCGCTCCCTGCGCTCCGAGG 1000
Oligo fwd CACCGCCGCGCTCCCTGCGCTCCG
Oligo rev AAACCGGAGCGCAGGGAGCGCGGC
Chromosome start end strand MM target_seq PAM alignment distance position gene name gene id
GU305914.1 201248 201270 - 0 GCCGCGCTCCCTGCGCTCCG AGG ||||||||[||||||||||||]PAM NA Intergenic NA NA

T19297 GCGCTCCCTGCGCTCCGAGGCGG 1000
Oligo fwd CACCGCGCTCCCTGCGCTCCGAGG
Oligo rev AAACCCTCGGAGCGCAGGGAGCGC
Chromosome start end strand MM target_seq PAM alignment distance position gene name gene id
GU305914.1 201245 201267 - 0 GCGCTCCCTGCGCTCCGAGG CGG ||||||||[||||||||||||]PAM NA Intergenic NA NA

T19298 CTCCCTGCGCTCCGAGGCGGAGG 1000
Oligo fwd CACCCTCCCTGCGCTCCGAGGCGG
Oligo rev AAACCCGCCTCGGAGCGCAGGGAG
Chromosome start end strand MM target_seq PAM alignment distance position gene name gene id
GU305914.1 201242 201264 - 0 CTCCCTGCGCTCCGAGGCGG AGG ||||||||[||||||||||||]PAM NA Intergenic NA NA

T19299 TCCCTGCGCTCCGAGGCGGAGGG 1000
Oligo fwd CACCTCCCTGCGCTCCGAGGCGGA
Oligo rev AAACTCCGCCTCGGAGCGCAGGGA
Chromosome start end strand MM target_seq PAM alignment distance position gene name gene id
GU305914.1 201241 201263 - 0 TCCCTGCGCTCCGAGGCGGA GGG ||||||||[||||||||||||]PAM NA Intergenic NA NA

T19300 GTCACGAATGCCTCCGAGAGCGG 1000
Oligo fwd CACCGTCACGAATGCCTCCGAGAG
Oligo rev AAACCTCTCGGAGGCATTCGTGAC
Chromosome start end strand MM target_seq PAM alignment distance position gene name gene id
GU305914.1 201213 201235 - 0 GTCACGAATGCCTCCGAGAG CGG ||||||||[||||||||||||]PAM NA Intergenic NA NA

T19301 CCGTCGACGCCGAACTCCACCGG 1000
Oligo fwd CACCCCGTCGACGCCGAACTCCAC
Oligo rev AAACGTGGAGTTCGGCGTCGACGG
Chromosome start end strand MM target_seq PAM alignment distance position gene name gene id
GU305914.1 201148 201170 - 0 CCGTCGACGCCGAACTCCAC CGG ||||||||[||||||||||||]PAM NA Intergenic NA NA

T19302 CGTCGACGCCGAACTCCACCGGG 1000
Oligo fwd CACCCGTCGACGCCGAACTCCACC
Oligo rev AAACGGTGGAGTTCGGCGTCGACG
Chromosome start end strand MM target_seq PAM alignment distance position gene name gene id
GU305914.1 201147 201169 - 0 CGTCGACGCCGAACTCCACC GGG ||||||||[||||||||||||]PAM NA Intergenic NA NA

T19303 ACGCGTGAGCCGCCTCGTCGAGG 1000
Oligo fwd CACCACGCGTGAGCCGCCTCGTCG
Oligo rev AAACCGACGAGGCGGCTCACGCGT
Chromosome start end strand MM target_seq PAM alignment distance position gene name gene id
GU305914.1 201048 201070 - 0 ACGCGTGAGCCGCCTCGTCG AGG ||||||||[||||||||||||]PAM NA Intergenic NA NA

T19304 CGCCTCGTCGAGGTCCACGACGG 1000
Oligo fwd CACCCGCCTCGTCGAGGTCCACGA
Oligo rev AAACTCGTGGACCTCGACGAGGCG
Chromosome start end strand MM target_seq PAM alignment distance position gene name gene id
GU305914.1 201038 201060 - 0 CGCCTCGTCGAGGTCCACGA CGG ||||||||[||||||||||||]PAM NA Intergenic NA NA

T19305 GCCTCGTCGAGGTCCACGACGGG 1000
Oligo fwd CACCGCCTCGTCGAGGTCCACGAC

Oligo rev AAACGTCGTGGACCTCGACGAGGC
Chromosome start end strand MM target_seq PAM alignment distance position gene name gene id
GU305914.1 201037 201059 - 0 GCCTCGTCGAGGTCCACGAC GGG ||||||||[||||||||||||]PAM NA Intergenic NA NA

T19306 CCTCGTCGAGGTCCACGACGGGG 1000
Oligo fwd CACCCCTCGTCGAGGTCCACGACG
Oligo rev AAACCGTCGTGGACCTCGACGAGG
Chromosome start end strand MM target_seq PAM alignment distance position gene name gene id
GU305914.1 201036 201058 - 0 CCTCGTCGAGGTCCACGACG GGG ||||||||[||||||||||||]PAM NA Intergenic NA NA

T19307 CGTCGAGGTCCACGACGGGGCGG 1000
Oligo fwd CACCCGTCGAGGTCCACGACGGGG
Oligo rev AAACCCCCGTCGTGGACCTCGACG
Chromosome start end strand MM target_seq PAM alignment distance position gene name gene id
GU305914.1 201033 201055 - 0 CGTCGAGGTCCACGACGGGG CGG ||||||||[||||||||||||]PAM NA Intergenic NA NA
GU305914.1 142266 142288 + 4 CGTCGTCGTCGACGACGGTG GGG |||||--|[||-|||||||-|]PAM NA Intergenic NA NA

T19308 ACCCGCCCCTCTCTCACACTTGG 1000
Oligo fwd CACCACCCGCCCCTCTCTCACACT
Oligo rev AAACAGTGTGAGAGAGGGGCGGGT
Chromosome start end strand MM target_seq PAM alignment distance position gene name gene id
GU305914.1 200976 200998 - 0 ACCCGCCCCTCTCTCACACT TGG ||||||||[||||||||||||]PAM NA Intergenic NA NA

T19309 CCCGCCCCTCTCTCACACTTGGG 1000
Oligo fwd CACCCCCGCCCCTCTCTCACACTT
Oligo rev AAACAAGTGTGAGAGAGGGGCGGG
Chromosome start end strand MM target_seq PAM alignment distance position gene name gene id
GU305914.1 200975 200997 - 0 CCCGCCCCTCTCTCACACTT GGG ||||||||[||||||||||||]PAM NA Intergenic NA NA

T19310 CGCCTCGAGCGCGACGCCATCGG 1000
Oligo fwd CACCCGCCTCGAGCGCGACGCCAT
Oligo rev AAACATGGCGTCGCGCTCGAGGCG
Chromosome start end strand MM target_seq PAM alignment distance position gene name gene id
GU305914.1 200935 200957 - 0 CGCCTCGAGCGCGACGCCAT CGG ||||||||[||||||||||||]PAM NA Intergenic NA NA

T19311 AGCGCGACGCCATCGGTATTCGG 1000
Oligo fwd CACCAGCGCGACGCCATCGGTATT
Oligo rev AAACAATACCGATGGCGTCGCGCT
Chromosome start end strand MM target_seq PAM alignment distance position gene name gene id
GU305914.1 200928 200950 - 0 AGCGCGACGCCATCGGTATT CGG ||||||||[||||||||||||]PAM NA Intergenic NA NA

T19312 ATTCGGAGTGTGTGAGTGATAGG 1000
Oligo fwd CACCATTCGGAGTGTGTGAGTGAT
Oligo rev AAACATCACTCACACACTCCGAAT
Chromosome start end strand MM target_seq PAM alignment distance position gene name gene id
GU305914.1 200911 200933 - 0 ATTCGGAGTGTGTGAGTGAT AGG ||||||||[||||||||||||]PAM NA Intergenic NA NA

T19313 GCCGCCCGTCGCCCTCGCCATGG 1000
Oligo fwd CACCGCCGCCCGTCGCCCTCGCCA
Oligo rev AAACTGGCGAGGGCGACGGGCGGC
Chromosome start end strand MM target_seq PAM alignment distance position gene name gene id
GU305914.1 200845 200867 - 0 GCCGCCCGTCGCCCTCGCCA TGG ||||||||[||||||||||||]PAM NA Intergenic NA NA
GU305914.1 200191 200213 + 4 GCGGCCGGTCGCCCTCCCCG GGG ||-|||-|[||||||||-||-]PAM NA Intergenic NA NA

T19314 CGCCATGGACGCCCTGTGCGCGG 1000
Oligo fwd CACCCGCCATGGACGCCCTGTGCG
Oligo rev AAACCGCACAGGGCGTCCATGGCG
Chromosome start end strand MM target_seq PAM alignment distance position gene name gene id
GU305914.1 200830 200852 - 0 CGCCATGGACGCCCTGTGCG CGG ||||||||[||||||||||||]PAM NA Intergenic NA NA

T19315 CCTGTGCGCGGCGATCCGATCGG 1000
Oligo fwd CACCCCTGTGCGCGGCGATCCGAT
Oligo rev AAACATCGGATCGCCGCGCACAGG
Chromosome start end strand MM target_seq PAM alignment distance position gene name gene id
GU305914.1 200818 200840 - 0 CCTGTGCGCGGCGATCCGAT CGG ||||||||[||||||||||||]PAM NA Intergenic NA NA

T19316 CGGATCACTTCAGAGACCAGAGG 1000
Oligo fwd CACCCGGATCACTTCAGAGACCAG
Oligo rev AAACCTGGTCTCTGAAGTGATCCG
Chromosome start end strand MM target_seq PAM alignment distance position gene name gene id
GU305914.1 200798 200820 - 0 CGGATCACTTCAGAGACCAG AGG ||||||||[||||||||||||]PAM NA Intergenic NA NA

T19317 TTCAGAGACCAGAGGCCTGTCGG 1000
Oligo fwd CACCTTCAGAGACCAGAGGCCTGT
Oligo rev AAACACAGGCCTCTGGTCTCTGAA
Chromosome start end strand MM target_seq PAM alignment distance position gene name gene id
GU305914.1 200790 200812 - 0 TTCAGAGACCAGAGGCCTGT CGG ||||||||[||||||||||||]PAM NA Intergenic NA NA

T19318 ACCAGAGGCCTGTCGGCATCAGG 1000
Oligo fwd CACCACCAGAGGCCTGTCGGCATC
Oligo rev AAACGATGCCGACAGGCCTCTGGT
Chromosome start end strand MM target_seq PAM alignment distance position gene name gene id
GU305914.1 200783 200805 - 0 ACCAGAGGCCTGTCGGCATC AGG ||||||||[||||||||||||]PAM NA Intergenic NA NA

T19319 GCCGCGCTCCCGCTATCTGACGG 1000
Oligo fwd CACCGCCGCGCTCCCGCTATCTGA
Oligo rev AAACTCAGATAGCGGGAGCGCGGC
Chromosome start end strand MM target_seq PAM alignment distance position gene name gene id
GU305914.1 200761 200783 - 0 GCCGCGCTCCCGCTATCTGA CGG ||||||||[||||||||||||]PAM NA Intergenic NA NA

T19320 GCGCTCCCGCTATCTGACGGCGG 1000
Oligo fwd CACCGCGCTCCCGCTATCTGACGG
Oligo rev AAACCCGTCAGATAGCGGGAGCGC
Chromosome start end strand MM target_seq PAM alignment distance position gene name gene id
GU305914.1 200758 200780 - 0 GCGCTCCCGCTATCTGACGG CGG ||||||||[||||||||||||]PAM NA Intergenic NA NA

T19321 CCGCTATCTGACGGCGGCCGCGG 1000
Oligo fwd CACCCCGCTATCTGACGGCGGCCG
Oligo rev AAACCGGCCGCCGTCAGATAGCGG
Chromosome start end strand MM target_seq PAM alignment distance position gene name gene id
GU305914.1 200752 200774 - 0 CCGCTATCTGACGGCGGCCG CGG ||||||||[||||||||||||]PAM NA Intergenic NA NA

T19322 GACGGCGGCCGCGGTGACCGCGG 1000
Oligo fwd CACCGACGGCGGCCGCGGTGACCG
Oligo rev AAACCGGTCACCGCGGCCGCCGTC
Chromosome start end strand MM target_seq PAM alignment distance position gene name gene id
GU305914.1 200743 200765 - 0 GACGGCGGCCGCGGTGACCG CGG ||||||||[||||||||||||]PAM NA Intergenic NA NA

T19323 CGCGGTGACCGCGGTCCTGCAGG 1000
Oligo fwd CACCCGCGGTGACCGCGGTCCTGC
Oligo rev AAACGCAGGACCGCGGTCACCGCG
Chromosome start end strand MM target_seq PAM alignment distance position gene name gene id
GU305914.1 200734 200756 - 0 CGCGGTGACCGCGGTCCTGC AGG ||||||||[||||||||||||]PAM NA Intergenic NA NA
GU305914.1 157973 157995 - 4 CGCCGCGACCGCGGTCCCGA CGG |||-|-||[|||||||||-|-]PAM NA Intergenic NA NA

T19324 CGCGGTCCTGCAGGACTTCGCGG 1000
Oligo fwd CACCCGCGGTCCTGCAGGACTTCG
Oligo rev AAACCGAAGTCCTGCAGGACCGCG
Chromosome start end strand MM target_seq PAM alignment distance position gene name gene id
GU305914.1 200725 200747 - 0 CGCGGTCCTGCAGGACTTCG CGG ||||||||[||||||||||||]PAM NA Intergenic NA NA

T19325 GGACTTCGCGGACATCTTCCTGG 1000
Oligo fwd CACCGGACTTCGCGGACATCTTCC
Oligo rev AAACGGAAGATGTCCGCGAAGTCC
Chromosome start end strand MM target_seq PAM alignment distance position gene name gene id
GU305914.1 200713 200735 - 0 GGACTTCGCGGACATCTTCC TGG ||||||||[||||||||||||]PAM NA Intergenic NA NA

T19326 CCACACGTCCGTCGAGAACGTGG 1000
Oligo fwd CACCCCACACGTCCGTCGAGAACG
Oligo rev AAACCGTTCTCGACGGACGTGTGG
Chromosome start end strand MM target_seq PAM alignment distance position gene name gene id
GU305914.1 200680 200702 - 0 CCACACGTCCGTCGAGAACG TGG ||||||||[||||||||||||]PAM NA Intergenic NA NA

T19327 CGTCCGTCGAGAACGTGGTCAGG 1000
Oligo fwd CACCCGTCCGTCGAGAACGTGGTC
Oligo rev AAACGACCACGTTCTCGACGGACG
Chromosome start end strand MM target_seq PAM alignment distance position gene name gene id
GU305914.1 200675 200697 - 0 CGTCCGTCGAGAACGTGGTC AGG ||||||||[||||||||||||]PAM NA Intergenic NA NA

T19328 AACGTGGTCAGGAAAGAGTCCGG 1000
Oligo fwd CACCAACGTGGTCAGGAAAGAGTC
Oligo rev AAACGACTCTTTCCTGACCACGTT
Chromosome start end strand MM target_seq PAM alignment distance position gene name gene id
GU305914.1 200664 200686 - 0 AACGTGGTCAGGAAAGAGTC CGG ||||||||[||||||||||||]PAM NA Intergenic NA NA

T19329 TGGTCAGGAAAGAGTCCGGCAGG 1000
Oligo fwd CACCTGGTCAGGAAAGAGTCCGGC
Oligo rev AAACGCCGGACTCTTTCCTGACCA
Chromosome start end strand MM target_seq PAM alignment distance position gene name gene id
GU305914.1 200660 200682 - 0 TGGTCAGGAAAGAGTCCGGC AGG ||||||||[||||||||||||]PAM NA Intergenic NA NA

T19330 TCCGGCAGGTTGCTGAGCCTCGG 1000
Oligo fwd CACCTCCGGCAGGTTGCTGAGCCT
Oligo rev AAACAGGCTCAGCAACCTGCCGGA
Chromosome start end strand MM target_seq PAM alignment distance position gene name gene id
GU305914.1 200646 200668 - 0 TCCGGCAGGTTGCTGAGCCT CGG ||||||||[||||||||||||]PAM NA Intergenic NA NA

T19331 GCTGAGCCTCGGCGCCCCGACGG 1000
Oligo fwd CACCGCTGAGCCTCGGCGCCCCGA
Oligo rev AAACTCGGGGCGCCGAGGCTCAGC
Chromosome start end strand MM target_seq PAM alignment distance position gene name gene id
GU305914.1 200635 200657 - 0 GCTGAGCCTCGGCGCCCCGA CGG ||||||||[||||||||||||]PAM NA Intergenic NA NA

T19332 CTGAGCCTCGGCGCCCCGACGGG 1000
Oligo fwd CACCCTGAGCCTCGGCGCCCCGAC
Oligo rev AAACGTCGGGGCGCCGAGGCTCAG
Chromosome start end strand MM target_seq PAM alignment distance position gene name gene id
GU305914.1 200634 200656 - 0 CTGAGCCTCGGCGCCCCGAC GGG ||||||||[||||||||||||]PAM NA Intergenic NA NA

T19333 GCCTCGGCGCCCCGACGGGCTGG 1000
Oligo fwd CACCGCCTCGGCGCCCCGACGGGC
Oligo rev AAACGCCCGTCGGGGCGCCGAGGC
Chromosome start end strand MM target_seq PAM alignment distance position gene name gene id
GU305914.1 200630 200652 - 0 GCCTCGGCGCCCCGACGGGC TGG ||||||||[||||||||||||]PAM NA Intergenic NA NA

T19334 CTGCCTGCGTTCCGCCGATCAGG 1000
Oligo fwd CACCCTGCCTGCGTTCCGCCGATC
Oligo rev AAACGATCGGCGGAACGCAGGCAG

Chromosome start end strand MM target_seq PAM alignment distance position gene name gene id
GU305914.1 200602 200624 - 0 CTGCCTGCGTTCCGCCGATC AGG ||||||||[||||||||||||]PAM NA Intergenic NA NA

T19335 GATCAGGTCGAGAGCTTCCCCGG 1000
Oligo fwd CACCGATCAGGTCGAGAGCTTCCC
Oligo rev AAACGGGAAGCTCTCGACCTGATC
Chromosome start end strand MM target_seq PAM alignment distance position gene name gene id
GU305914.1 200586 200608 - 0 GATCAGGTCGAGAGCTTCCC CGG ||||||||[||||||||||||]PAM NA Intergenic NA NA

T19336 GCTTCCCCGGCTATCCCGTCTGG 1000
Oligo fwd CACCGCTTCCCCGGCTATCCCGTC
Oligo rev AAACGACGGGATAGCCGGGGAAGC
Chromosome start end strand MM target_seq PAM alignment distance position gene name gene id
GU305914.1 200573 200595 - 0 GCTTCCCCGGCTATCCCGTC TGG ||||||||[||||||||||||]PAM NA Intergenic NA NA

T19337 CTTCCCCGGCTATCCCGTCTGGG 1000
Oligo fwd CACCCTTCCCCGGCTATCCCGTCT
Oligo rev AAACAGACGGGATAGCCGGGGAAG
Chromosome start end strand MM target_seq PAM alignment distance position gene name gene id
GU305914.1 200572 200594 - 0 CTTCCCCGGCTATCCCGTCT GGG ||||||||[||||||||||||]PAM NA Intergenic NA NA

T19338 TATCCCGTCTGGGAGACGTACGG 1000
Oligo fwd CACCTATCCCGTCTGGGAGACGTA
Oligo rev AAACTACGTCTCCCAGACGGGATA
Chromosome start end strand MM target_seq PAM alignment distance position gene name gene id
GU305914.1 200562 200584 - 0 TATCCCGTCTGGGAGACGTA CGG ||||||||[||||||||||||]PAM NA Intergenic NA NA

T19339 ATCCCGTCTGGGAGACGTACGGG 1000
Oligo fwd CACCATCCCGTCTGGGAGACGTAC
Oligo rev AAACGTACGTCTCCCAGACGGGAT
Chromosome start end strand MM target_seq PAM alignment distance position gene name gene id
GU305914.1 200561 200583 - 0 ATCCCGTCTGGGAGACGTAC GGG ||||||||[||||||||||||]PAM NA Intergenic NA NA

T19340 TCCCGTCTGGGAGACGTACGGGG 1000
Oligo fwd CACCTCCCGTCTGGGAGACGTACG
Oligo rev AAACCGTACGTCTCCCAGACGGGA
Chromosome start end strand MM target_seq PAM alignment distance position gene name gene id
GU305914.1 200560 200582 - 0 TCCCGTCTGGGAGACGTACG GGG ||||||||[||||||||||||]PAM NA Intergenic NA NA

T19341 CCCGTCTGGGAGACGTACGGGGG 1000
Oligo fwd CACCCCCGTCTGGGAGACGTACGG
Oligo rev AAACCCGTACGTCTCCCAGACGGG
Chromosome start end strand MM target_seq PAM alignment distance position gene name gene id
GU305914.1 200559 200581 - 0 CCCGTCTGGGAGACGTACGG GGG ||||||||[||||||||||||]PAM NA Intergenic NA NA

T19342 GGAGACGTACGGGGGCGCTCCGG 1000
Oligo fwd CACCGGAGACGTACGGGGGCGCTC
Oligo rev AAACGAGCGCCCCCGTACGTCTCC
Chromosome start end strand MM target_seq PAM alignment distance position gene name gene id
GU305914.1 200551 200573 - 0 GGAGACGTACGGGGGCGCTC CGG ||||||||[||||||||||||]PAM NA Intergenic NA NA

T19343 GACGTACGGGGGCGCTCCGGCGG 1000
Oligo fwd CACCGACGTACGGGGGCGCTCCGG
Oligo rev AAACCCGGAGCGCCCCCGTACGTC
Chromosome start end strand MM target_seq PAM alignment distance position gene name gene id
GU305914.1 200548 200570 - 0 GACGTACGGGGGCGCTCCGG CGG ||||||||[||||||||||||]PAM NA Intergenic NA NA

T19344 ACGTACGGGGGCGCTCCGGCGGG 1000
Oligo fwd CACCACGTACGGGGGCGCTCCGGC
Oligo rev AAACGCCGGAGCGCCCCCGTACGT
Chromosome start end strand MM target_seq PAM alignment distance position gene name gene id
GU305914.1 200547 200569 - 0 ACGTACGGGGGCGCTCCGGC GGG ||||||||[||||||||||||]PAM NA Intergenic NA NA

T19345 GCGGGCCGTTACACGCTGCTCGG 1000
Oligo fwd CACCGCGGGCCGTTACACGCTGCT
Oligo rev AAACAGCAGCGTGTAACGGCCCGC
Chromosome start end strand MM target_seq PAM alignment distance position gene name gene id
GU305914.1 200529 200551 - 0 GCGGGCCGTTACACGCTGCT CGG ||||||||[||||||||||||]PAM NA Intergenic NA NA

T19346 CCGTTACACGCTGCTCGGCCAGG 1000
Oligo fwd CACCCCGTTACACGCTGCTCGGCC
Oligo rev AAACGGCCGAGCAGCGTGTAACGG
Chromosome start end strand MM target_seq PAM alignment distance position gene name gene id
GU305914.1 200524 200546 - 0 CCGTTACACGCTGCTCGGCC AGG ||||||||[||||||||||||]PAM NA Intergenic NA NA

T19347 CGCTGCTCGGCCAGGTCGTCCGG 1000
Oligo fwd CACCCGCTGCTCGGCCAGGTCGTC
Oligo rev AAACGACGACCTGGCCGAGCAGCG
Chromosome start end strand MM target_seq PAM alignment distance position gene name gene id
GU305914.1 200516 200538 - 0 CGCTGCTCGGCCAGGTCGTC CGG ||||||||[||||||||||||]PAM NA Intergenic NA NA

T19348 CAGGTCGTCCGGAGATTCCGAGG 1000
Oligo fwd CACCCAGGTCGTCCGGAGATTCCG
Oligo rev AAACCGGAATCTCCGGACGACCTG
Chromosome start end strand MM target_seq PAM alignment distance position gene name gene id
GU305914.1 200505 200527 - 0 CAGGTCGTCCGGAGATTCCG AGG ||||||||[||||||||||||]PAM NA Intergenic NA NA

T19349 CCGAGGCGTCGTTCCGACCGAGG 1000
Oligo fwd CACCCCGAGGCGTCGTTCCGACCG

<table>
<tr><td>Oligo rev</td><td>AAACCGGTCGGAACGACGCCTCGG</td><td></td><td></td><td></td><td></td><td></td><td></td><td></td><td></td><td></td><td></td><td></td></tr>
<tr><td>Chromosome</td><td>start</td><td></td><td>end</td><td>strand</td><td>MM</td><td>target_seq</td><td>PAM</td><td>alignment</td><td>distance</td><td>position</td><td>gene name</td><td>gene id</td></tr>
<tr><td>GU305914.1</td><td></td><td>200488</td><td>200510</td><td>-</td><td>0</td><td>CCGAGGCGTCGTTCCGACCG</td><td>AGG</td><td>||||||||[||||||||||||]PAM</td><td>NA</td><td>Intergenic</td><td>NA</td><td>NA</td></tr>
<tr><td></td><td></td><td></td><td></td><td></td><td></td><td></td><td></td><td></td><td></td><td></td><td></td><td></td></tr>
<tr><td>T19350</td><td>TTCCGACCGAGGAGCTGAGATGG</td><td></td><td>1000</td><td></td><td></td><td></td><td></td><td></td><td></td><td></td><td></td><td></td></tr>
<tr><td>Oligo fwd</td><td>CACCTTCCGACCGAGGAGCTGAGA</td><td></td><td></td><td></td><td></td><td></td><td></td><td></td><td></td><td></td><td></td><td></td></tr>
<tr><td>Oligo rev</td><td>AAACTCTCAGCTCCTCGGTCGGAA</td><td></td><td></td><td></td><td></td><td></td><td></td><td></td><td></td><td></td><td></td><td></td></tr>
<tr><td>Chromosome</td><td>start</td><td></td><td>end</td><td>strand</td><td>MM</td><td>target_seq</td><td>PAM</td><td>alignment</td><td>distance</td><td>position</td><td>gene name</td><td>gene id</td></tr>
<tr><td>GU305914.1</td><td></td><td>200477</td><td>200499</td><td>-</td><td>0</td><td>TTCCGACCGAGGAGCTGAGA</td><td>TGG</td><td>||||||||[||||||||||||]PAM</td><td>NA</td><td>Intergenic</td><td>NA</td><td>NA</td></tr>
<tr><td></td><td></td><td></td><td></td><td></td><td></td><td></td><td></td><td></td><td></td><td></td><td></td><td></td></tr>
<tr><td>T19351</td><td>GCTGAGATGGTTCGTGTACCTGG</td><td></td><td>1000</td><td></td><td></td><td></td><td></td><td></td><td></td><td></td><td></td><td></td></tr>
<tr><td>Oligo fwd</td><td>CACCGCTGAGATGGTTCGTGTACC</td><td></td><td></td><td></td><td></td><td></td><td></td><td></td><td></td><td></td><td></td><td></td></tr>
<tr><td>Oligo rev</td><td>AAACGGTACACGAACCATCTCAGC</td><td></td><td></td><td></td><td></td><td></td><td></td><td></td><td></td><td></td><td></td><td></td></tr>
<tr><td>Chromosome</td><td>start</td><td></td><td>end</td><td>strand</td><td>MM</td><td>target_seq</td><td>PAM</td><td>alignment</td><td>distance</td><td>position</td><td>gene name</td><td>gene id</td></tr>
<tr><td>GU305914.1</td><td></td><td>200464</td><td>200486</td><td>-</td><td>0</td><td>GCTGAGATGGTTCGTGTACC</td><td>TGG</td><td>||||||||[||||||||||||]PAM</td><td>NA</td><td>Intergenic</td><td>NA</td><td>NA</td></tr>
<tr><td></td><td></td><td></td><td></td><td></td><td></td><td></td><td></td><td></td><td></td><td></td><td></td><td></td></tr>
<tr><td>T19352</td><td>CTGAGATGGTTCGTGTACCTGGG</td><td></td><td>1000</td><td></td><td></td><td></td><td></td><td></td><td></td><td></td><td></td><td></td></tr>
<tr><td>Oligo fwd</td><td>CACCCTGAGATGGTTCGTGTACCT</td><td></td><td></td><td></td><td></td><td></td><td></td><td></td><td></td><td></td><td></td><td></td></tr>
<tr><td>Oligo rev</td><td>AAACAGGTACACGAACCATCTCAG</td><td></td><td></td><td></td><td></td><td></td><td></td><td></td><td></td><td></td><td></td><td></td></tr>
<tr><td>Chromosome</td><td>start</td><td></td><td>end</td><td>strand</td><td>MM</td><td>target_seq</td><td>PAM</td><td>alignment</td><td>distance</td><td>position</td><td>gene name</td><td>gene id</td></tr>
<tr><td>GU305914.1</td><td></td><td>200463</td><td>200485</td><td>-</td><td>0</td><td>CTGAGATGGTTCGTGTACCT</td><td>GGG</td><td>||||||||[||||||||||||]PAM</td><td>NA</td><td>Intergenic</td><td>NA</td><td>NA</td></tr>
<tr><td></td><td></td><td></td><td></td><td></td><td></td><td></td><td></td><td></td><td></td><td></td><td></td><td></td></tr>
<tr><td>T19353</td><td>GTTCGTGTACCTGGGTCCCCAGG</td><td></td><td>1000</td><td></td><td></td><td></td><td></td><td></td><td></td><td></td><td></td><td></td></tr>
<tr><td>Oligo fwd</td><td>CACCGTTCGTGTACCTGGGTCCCC</td><td></td><td></td><td></td><td></td><td></td><td></td><td></td><td></td><td></td><td></td><td></td></tr>
<tr><td>Oligo rev</td><td>AAACGGGGACCCAGGTACACGAAC</td><td></td><td></td><td></td><td></td><td></td><td></td><td></td><td></td><td></td><td></td><td></td></tr>
<tr><td>Chromosome</td><td>start</td><td></td><td>end</td><td>strand</td><td>MM</td><td>target_seq</td><td>PAM</td><td>alignment</td><td>distance</td><td>position</td><td>gene name</td><td>gene id</td></tr>
<tr><td>GU305914.1</td><td></td><td>200455</td><td>200477</td><td>-</td><td>0</td><td>GTTCGTGTACCTGGGTCCCC</td><td>AGG</td><td>||||||||[||||||||||||]PAM</td><td>NA</td><td>Intergenic</td><td>NA</td><td>NA</td></tr>
<tr><td></td><td></td><td></td><td></td><td></td><td></td><td></td><td></td><td></td><td></td><td></td><td></td><td></td></tr>
<tr><td>T19354</td><td>TTCGTGTACCTGGGTCCCCAGGG</td><td></td><td>1000</td><td></td><td></td><td></td><td></td><td></td><td></td><td></td><td></td><td></td></tr>
<tr><td>Oligo fwd</td><td>CACCTTCGTGTACCTGGGTCCCCA</td><td></td><td></td><td></td><td></td><td></td><td></td><td></td><td></td><td></td><td></td><td></td></tr>
<tr><td>Oligo rev</td><td>AAACTGGGGACCCAGGTACACGAA</td><td></td><td></td><td></td><td></td><td></td><td></td><td></td><td></td><td></td><td></td><td></td></tr>
<tr><td>Chromosome</td><td>start</td><td></td><td>end</td><td>strand</td><td>MM</td><td>target_seq</td><td>PAM</td><td>alignment</td><td>distance</td><td>position</td><td>gene name</td><td>gene id</td></tr>
<tr><td>GU305914.1</td><td></td><td>200454</td><td>200476</td><td>-</td><td>0</td><td>TTCGTGTACCTGGGTCCCCA</td><td>GGG</td><td>||||||||[||||||||||||]PAM</td><td>NA</td><td>Intergenic</td><td>NA</td><td>NA</td></tr>
<tr><td></td><td></td><td></td><td></td><td></td><td></td><td></td><td></td><td></td><td></td><td></td><td></td><td></td></tr>
<tr><td>T19355</td><td>GGGCCAGCTGCTCTGTCACCAGG</td><td></td><td>1000</td><td></td><td></td><td></td><td></td><td></td><td></td><td></td><td></td><td></td></tr>
<tr><td>Oligo fwd</td><td>CACCGGGCCAGCTGCTCTGTCACC</td><td></td><td></td><td></td><td></td><td></td><td></td><td></td><td></td><td></td><td></td><td></td></tr>
<tr><td>Oligo rev</td><td>AAACGGTGACAGAGCAGCTGGCCC</td><td></td><td></td><td></td><td></td><td></td><td></td><td></td><td></td><td></td><td></td><td></td></tr>
<tr><td>Chromosome</td><td>start</td><td></td><td>end</td><td>strand</td><td>MM</td><td>target_seq</td><td>PAM</td><td>alignment</td><td>distance</td><td>position</td><td>gene name</td><td>gene id</td></tr>
<tr><td>GU305914.1</td><td></td><td>200434</td><td>200456</td><td>-</td><td>0</td><td>GGGCCAGCTGCTCTGTCACC</td><td>AGG</td><td>||||||||[||||||||||||]PAM</td><td>NA</td><td>Intergenic</td><td>NA</td><td>NA</td></tr>
<tr><td></td><td></td><td></td><td></td><td></td><td></td><td></td><td></td><td></td><td></td><td></td><td></td><td></td></tr>
<tr><td>T19356</td><td>AGCTGCTCTGTCACCAGGAGCGG</td><td></td><td>1000</td><td></td><td></td><td></td><td></td><td></td><td></td><td></td><td></td><td></td></tr>
<tr><td>Oligo fwd</td><td>CACCAGCTGCTCTGTCACCAGGAG</td><td></td><td></td><td></td><td></td><td></td><td></td><td></td><td></td><td></td><td></td><td></td></tr>
<tr><td>Oligo rev</td><td>AAACCTCCTGGTGACAGAGCAGCT</td><td></td><td></td><td></td><td></td><td></td><td></td><td></td><td></td><td></td><td></td><td></td></tr>
<tr><td>Chromosome</td><td>start</td><td></td><td>end</td><td>strand</td><td>MM</td><td>target_seq</td><td>PAM</td><td>alignment</td><td>distance</td><td>position</td><td>gene name</td><td>gene id</td></tr>
<tr><td>GU305914.1</td><td></td><td>200429</td><td>200451</td><td>-</td><td>0</td><td>AGCTGCTCTGTCACCAGGAG</td><td>CGG</td><td>||||||||[||||||||||||]PAM</td><td>NA</td><td>Intergenic</td><td>NA</td><td>NA</td></tr>
<tr><td></td><td></td><td></td><td></td><td></td><td></td><td></td><td></td><td></td><td></td><td></td><td></td><td></td></tr>
<tr><td>T19357</td><td>GAGTCTGAGCATCGACGAGCTGG</td><td></td><td>1000</td><td></td><td></td><td></td><td></td><td></td><td></td><td></td><td></td><td></td></tr>
<tr><td>Oligo fwd</td><td>CACCGAGTCTGAGCATCGACGAGC</td><td></td><td></td><td></td><td></td><td></td><td></td><td></td><td></td><td></td><td></td><td></td></tr>
<tr><td>Oligo rev</td><td>AAACGCTCGTCGATGCTCAGACTC</td><td></td><td></td><td></td><td></td><td></td><td></td><td></td><td></td><td></td><td></td><td></td></tr>
<tr><td>Chromosome</td><td>start</td><td></td><td>end</td><td>strand</td><td>MM</td><td>target_seq</td><td>PAM</td><td>alignment</td><td>distance</td><td>position</td><td>gene name</td><td>gene id</td></tr>
<tr><td>GU305914.1</td><td></td><td>200386</td><td>200408</td><td>-</td><td>0</td><td>GAGTCTGAGCATCGACGAGC</td><td>TGG</td><td>||||||||[||||||||||||]PAM</td><td>NA</td><td>Intergenic</td><td>NA</td><td>NA</td></tr>
<tr><td></td><td></td><td></td><td></td><td></td><td></td><td></td><td></td><td></td><td></td><td></td><td></td><td></td></tr>
<tr><td>T19358</td><td>TGAGCATCGACGAGCTGGCGAGG</td><td></td><td>1000</td><td></td><td></td><td></td><td></td><td></td><td></td><td></td><td></td><td></td></tr>
<tr><td>Oligo fwd</td><td>CACCTGAGCATCGACGAGCTGGCG</td><td></td><td></td><td></td><td></td><td></td><td></td><td></td><td></td><td></td><td></td><td></td></tr>
<tr><td>Oligo rev</td><td>AAACCGCCAGCTCGTCGATGCTCA</td><td></td><td></td><td></td><td></td><td></td><td></td><td></td><td></td><td></td><td></td><td></td></tr>
<tr><td>Chromosome</td><td>start</td><td></td><td>end</td><td>strand</td><td>MM</td><td>target_seq</td><td>PAM</td><td>alignment</td><td>distance</td><td>position</td><td>gene name</td><td>gene id</td></tr>
<tr><td>GU305914.1</td><td></td><td>200381</td><td>200403</td><td>-</td><td>0</td><td>TGAGCATCGACGAGCTGGCG</td><td>AGG</td><td>||||||||[||||||||||||]PAM</td><td>NA</td><td>Intergenic</td><td>NA</td><td>NA</td></tr>
<tr><td></td><td></td><td></td><td></td><td></td><td></td><td></td><td></td><td></td><td></td><td></td><td></td><td></td></tr>
<tr><td>T19359</td><td>ATCGACGAGCTGGCGAGGAGAGG</td><td></td><td>1000</td><td></td><td></td><td></td><td></td><td></td><td></td><td></td><td></td><td></td></tr>
<tr><td>Oligo fwd</td><td>CACCATCGACGAGCTGGCGAGGAG</td><td></td><td></td><td></td><td></td><td></td><td></td><td></td><td></td><td></td><td></td><td></td></tr>
<tr><td>Oligo rev</td><td>AAACCTCCTCGCCAGCTCGTCGAT</td><td></td><td></td><td></td><td></td><td></td><td></td><td></td><td></td><td></td><td></td><td></td></tr>
<tr><td>Chromosome</td><td>start</td><td></td><td>end</td><td>strand</td><td>MM</td><td>target_seq</td><td>PAM</td><td>alignment</td><td>distance</td><td>position</td><td>gene name</td><td>gene id</td></tr>
<tr><td>GU305914.1</td><td></td><td>200376</td><td>200398</td><td>-</td><td>0</td><td>ATCGACGAGCTGGCGAGGAG</td><td>AGG</td><td>||||||||[||||||||||||]PAM</td><td>NA</td><td>Intergenic</td><td>NA</td><td>NA</td></tr>
<tr><td>GU305914.1</td><td></td><td>195248</td><td>195270</td><td>-</td><td>3</td><td>ATCGACCAGCTGGCGAGACG</td><td>GGG</td><td>||||||-|[|||||||||--|]PAM</td><td>NA</td><td>Intergenic</td><td>NA</td><td>NA</td></tr>
<tr><td></td><td></td><td></td><td></td><td></td><td></td><td></td><td></td><td></td><td></td><td></td><td></td><td></td></tr>
<tr><td>T19360</td><td>CGAGCTGGCGAGGAGAGGCCTGG</td><td></td><td>1000</td><td></td><td></td><td></td><td></td><td></td><td></td><td></td><td></td><td></td></tr>
<tr><td>Oligo fwd</td><td>CACCCGAGCTGGCGAGGAGAGGCC</td><td></td><td></td><td></td><td></td><td></td><td></td><td></td><td></td><td></td><td></td><td></td></tr>
<tr><td>Oligo rev</td><td>AAACGGCCTCTCCTCGCCAGCTCG</td><td></td><td></td><td></td><td></td><td></td><td></td><td></td><td></td><td></td><td></td><td></td></tr>
<tr><td>Chromosome</td><td>start</td><td></td><td>end</td><td>strand</td><td>MM</td><td>target_seq</td><td>PAM</td><td>alignment</td><td>distance</td><td>position</td><td>gene name</td><td>gene id</td></tr>
<tr><td>GU305914.1</td><td></td><td>200371</td><td>200393</td><td>-</td><td>0</td><td>CGAGCTGGCGAGGAGAGGCC</td><td>TGG</td><td>||||||||[||||||||||||]PAM</td><td>NA</td><td>Intergenic</td><td>NA</td><td>NA</td></tr>
<tr><td></td><td></td><td></td><td></td><td></td><td></td><td></td><td></td><td></td><td></td><td></td><td></td><td></td></tr>
<tr><td>T19361</td><td>GAACTTCGAACCCCTGTACGCGG</td><td></td><td>1000</td><td></td><td></td><td></td><td></td><td></td><td></td><td></td><td></td><td></td></tr>
<tr><td>Oligo fwd</td><td>CACCGAACTTCGAACCCCTGTACG</td><td></td><td></td><td></td><td></td><td></td><td></td><td></td><td></td><td></td><td></td><td></td></tr>
<tr><td>Oligo rev</td><td>AAACCGTACAGGGGTTCGAAGTTC</td><td></td><td></td><td></td><td></td><td></td><td></td><td></td><td></td><td></td><td></td><td></td></tr>
<tr><td>Chromosome</td><td>start</td><td></td><td>end</td><td>strand</td><td>MM</td><td>target_seq</td><td>PAM</td><td>alignment</td><td>distance</td><td>position</td><td>gene name</td><td>gene id</td></tr>
<tr><td>GU305914.1</td><td></td><td>200347</td><td>200369</td><td>-</td><td>0</td><td>GAACTTCGAACCCCTGTACG</td><td>CGG</td><td>||||||||[||||||||||||]PAM</td><td>NA</td><td>Intergenic</td><td>NA</td><td>NA</td></tr>
<tr><td></td><td></td><td></td><td></td><td></td><td></td><td></td><td></td><td></td><td></td><td></td><td></td><td></td></tr>
<tr><td>T19362</td><td>TCGAACCCCTGTACGCGGAACGG</td><td></td><td>1000</td><td></td><td></td><td></td><td></td><td></td><td></td><td></td><td></td><td></td></tr>
<tr><td>Oligo fwd</td><td>CACCTCGAACCCCTGTACGCGGAA</td><td></td><td></td><td></td><td></td><td></td><td></td><td></td><td></td><td></td><td></td><td></td></tr>
<tr><td>Oligo rev</td><td>AAACTTCCGCGTACAGGGGTTCGA</td><td></td><td></td><td></td><td></td><td></td><td></td><td></td><td></td><td></td><td></td><td></td></tr>
<tr><td>Chromosome</td><td>start</td><td></td><td>end</td><td>strand</td><td>MM</td><td>target_seq</td><td>PAM</td><td>alignment</td><td>distance</td><td>position</td><td>gene name</td><td>gene id</td></tr>
<tr><td>GU305914.1</td><td></td><td>200342</td><td>200364</td><td>-</td><td>0</td><td>TCGAACCCCTGTACGCGGAA</td><td>CGG</td><td>||||||||[||||||||||||]PAM</td><td>NA</td><td>Intergenic</td><td>NA</td><td>NA</td></tr>
<tr><td></td><td></td><td></td><td></td><td></td><td></td><td></td><td></td><td></td><td></td><td></td><td></td><td></td></tr>
<tr><td>T19363</td><td>GCACCTGCCCCTGACGACCGCGG</td><td></td><td>1000</td><td></td><td></td><td></td><td></td><td></td><td></td><td></td><td></td><td></td></tr>
<tr><td>Oligo fwd</td><td>CACCGCACCTGCCCCTGACGACCG</td><td></td><td></td><td></td><td></td><td></td><td></td><td></td><td></td><td></td><td></td><td></td></tr>
<tr><td>Oligo rev</td><td>AAACCGGTCGTCAGGGGCAGGTGC</td><td></td><td></td><td></td><td></td><td></td><td></td><td></td><td></td><td></td><td></td><td></td></tr>
<tr><td>Chromosome</td><td>start</td><td></td><td>end</td><td>strand</td><td>MM</td><td>target_seq</td><td>PAM</td><td>alignment</td><td>distance</td><td>position</td><td>gene name</td><td>gene id</td></tr>
<tr><td>GU305914.1</td><td></td><td>200320</td><td>200342</td><td>-</td><td>0</td><td>GCACCTGCCCCTGACGACCG</td><td>CGG</td><td>||||||||[||||||||||||]PAM</td><td>NA</td><td>Intergenic</td><td>NA</td><td>NA</td></tr>
</table>

T19364 GACCGCGGTGCCCGTCAAGCTGG 1000
Oligo fwd CACCGACCGCGGTGCCCGTCAAGC
Oligo rev AAACGCTTGACGGGCACCGCGGTC
Chromosome start end strand MM target_seq PAM alignment distance position gene name gene id
GU305914.1 200305 200327 - 0 GACCGCGGTGCCCGTCAAGC TGG ||||||||[||||||||||||]PAM NA Intergenic NA NA

T19365 CGCGGTGCCCGTCAAGCTGGTGG 1000
Oligo fwd CACCCGCGGTGCCCGTCAAGCTGG
Oligo rev AAACCCAGCTTGACGGGCACCGCG
Chromosome start end strand MM target_seq PAM alignment distance position gene name gene id
GU305914.1 200302 200324 - 0 CGCGGTGCCCGTCAAGCTGG TGG ||||||||[||||||||||||]PAM NA Intergenic NA NA

T19366 GGTGCCCGTCAAGCTGGTGGCGG 1000
Oligo fwd CACCGGTGCCCGTCAAGCTGGTGG
Oligo rev AAACCCACCAGCTTGACGGGCACC
Chromosome start end strand MM target_seq PAM alignment distance position gene name gene id
GU305914.1 200299 200321 - 0 GGTGCCCGTCAAGCTGGTGG CGG ||||||||[||||||||||||]PAM NA Intergenic NA NA

T19367 CAAGCTGGTGGCGGAGTTCTCGG 1000
Oligo fwd CACCCAAGCTGGTGGCGGAGTTCT
Oligo rev AAACAGAACTCCGCCACCAGCTTG
Chromosome start end strand MM target_seq PAM alignment distance position gene name gene id
GU305914.1 200290 200312 - 0 CAAGCTGGTGGCGGAGTTCT CGG ||||||||[||||||||||||]PAM NA Intergenic NA NA

T19368 TCGGCGCTCGATCCCAGAGACGG 1000
Oligo fwd CACCTCGGCGCTCGATCCCAGAGA
Oligo rev AAACTCTCTGGGATCGAGCGCCGA
Chromosome start end strand MM target_seq PAM alignment distance position gene name gene id
GU305914.1 200271 200293 - 0 TCGGCGCTCGATCCCAGAGA CGG ||||||||[||||||||||||]PAM NA Intergenic NA NA

T19369 CGGCGCTCGATCCCAGAGACGGG 1000
Oligo fwd CACCCGGCGCTCGATCCCAGAGAC
Oligo rev AAACGTCTCTGGGATCGAGCGCCG
Chromosome start end strand MM target_seq PAM alignment distance position gene name gene id
GU305914.1 200270 200292 - 0 CGGCGCTCGATCCCAGAGAC GGG ||||||||[||||||||||||]PAM NA Intergenic NA NA

T19370 TCCCAGAGACGGGACCGCCGTGG 1000
Oligo fwd CACCTCCCAGAGACGGGACCGCCG
Oligo rev AAACCGGCGGTCCCGTCTCTGGGA
Chromosome start end strand MM target_seq PAM alignment distance position gene name gene id
GU305914.1 200260 200282 - 0 TCCCAGAGACGGGACCGCCG TGG ||||||||[||||||||||||]PAM NA Intergenic NA NA

T19371 CAGAGACGGGACCGCCGTGGCGG 1000
Oligo fwd CACCCAGAGACGGGACCGCCGTGG
Oligo rev AAACCCACGGCGGTCCCGTCTCTG
Chromosome start end strand MM target_seq PAM alignment distance position gene name gene id
GU305914.1 200257 200279 - 0 CAGAGACGGGACCGCCGTGG CGG ||||||||[||||||||||||]PAM NA Intergenic NA NA

T19372 GGAGCGCGCGCTGATGTGTCAGG 1000
Oligo fwd CACCGGAGCGCGCGCTGATGTGTC
Oligo rev AAACGACACATCAGCGCGCGCTCC
Chromosome start end strand MM target_seq PAM alignment distance position gene name gene id
GU305914.1 200236 200258 - 0 GGAGCGCGCGCTGATGTGTC AGG ||||||||[||||||||||||]PAM NA Intergenic NA NA

T19373 GAGCGCGCGCTGATGTGTCAGGG 1000
Oligo fwd CACCGAGCGCGCGCTGATGTGTCA
Oligo rev AAACTGACACATCAGCGCGCGCTC
Chromosome start end strand MM target_seq PAM alignment distance position gene name gene id
GU305914.1 200235 200257 - 0 GAGCGCGCGCTGATGTGTCA GGG ||||||||[||||||||||||]PAM NA Intergenic NA NA

T19374 GCGCGCTGATGTGTCAGGGACGG 1000
Oligo fwd CACCGCGCGCTGATGTGTCAGGGA
Oligo rev AAACTCCCTGACACATCAGCGCGC
Chromosome start end strand MM target_seq PAM alignment distance position gene name gene id
GU305914.1 200231 200253 - 0 GCGCGCTGATGTGTCAGGGA CGG ||||||||[||||||||||||]PAM NA Intergenic NA NA

T19375 ACGATCCTGCTACACACCCCCGG 1000
Oligo fwd CACCACGATCCTGCTACACACCCC
Oligo rev AAACGGGGTGTGTAGCAGGATCGT
Chromosome start end strand MM target_seq PAM alignment distance position gene name gene id
GU305914.1 200208 200230 - 0 ACGATCCTGCTACACACCCC CGG ||||||||[||||||||||||]PAM NA Intergenic NA NA

T19376 CGATCCTGCTACACACCCCCGGG 1000
Oligo fwd CACCCGATCCTGCTACACACCCCC
Oligo rev AAACGGGGGTGTGTAGCAGGATCG
Chromosome start end strand MM target_seq PAM alignment distance position gene name gene id
GU305914.1 200207 200229 - 0 CGATCCTGCTACACACCCCC GGG ||||||||[||||||||||||]PAM NA Intergenic NA NA

T19377 GATCCTGCTACACACCCCCGGGG 1000
Oligo fwd CACCGATCCTGCTACACACCCCCG
Oligo rev AAACCGGGGGTGTGTAGCAGGATC
Chromosome start end strand MM target_seq PAM alignment distance position gene name gene id
GU305914.1 200206 200228 - 0 GATCCTGCTACACACCCCCG GGG ||||||||[||||||||||||]PAM NA Intergenic NA NA

T19378 CCTGCTACACACCCCCGGGGAGG 1000
Oligo fwd CACCCCTGCTACACACCCCCGGGG
Oligo rev AAACCCCCGGGGGTGTGTAGCAGG
Chromosome start end strand MM target_seq PAM alignment distance position gene name gene id
GU305914.1 200203 200225 - 0 CCTGCTACACACCCCCGGGG AGG ||||||||[||||||||||||]PAM NA Intergenic NA NA

T19379 CTGCTACACACCCCCGGGGAGGG 1000
Oligo fwd CACCCTGCTACACACCCCCGGGGA
Oligo rev AAACTCCCCGGGGGTGTGTAGCAG
Chromosome start end strand MM target_seq PAM alignment distance position gene name gene id
GU305914.1 200202 200224 - 0 CTGCTACACACCCCCGGGGA GGG ||||||||[||||||||||||]PAM NA Intergenic NA NA

T19380 ACACCCCCGGGGAGGGCGACCGG 1000
Oligo fwd CACCACACCCCCGGGGAGGGCGAC
Oligo rev AAACGTCGCCCTCCCCGGGGGTGT
Chromosome start end strand MM target_seq PAM alignment distance position gene name gene id
GU305914.1 200195 200217 - 0 ACACCCCCGGGGAGGGCGAC CGG ||||||||[||||||||||||]PAM NA Intergenic NA NA

T19381 GACCGGCCGCTCGTGCTGTGCGG 1000
Oligo fwd CACCGACCGGCCGCTCGTGCTGTG
Oligo rev AAACCACAGCACGAGCGGCCGGTC
Chromosome start end strand MM target_seq PAM alignment distance position gene name gene id
GU305914.1 200178 200200 - 0 GACCGGCCGCTCGTGCTGTG CGG ||||||||[||||||||||||]PAM NA Intergenic NA NA

T19382 GCGGCACCGACGAGTGCCTGCGG 1000
Oligo fwd CACCGCGGCACCGACGAGTGCCTG
Oligo rev AAACCAGGCACTCGTCGGTGCCGC
Chromosome start end strand MM target_seq PAM alignment distance position gene name gene id
GU305914.1 200159 200181 - 0 GCGGCACCGACGAGTGCCTG CGG ||||||||[||||||||||||]PAM NA Intergenic NA NA

T19383 GCACCGACGAGTGCCTGCGGCGG 1000
Oligo fwd CACCGCACCGACGAGTGCCTGCGG
Oligo rev AAACCCGCAGGCACTCGTCGGTGC
Chromosome start end strand MM target_seq PAM alignment distance position gene name gene id
GU305914.1 200156 200178 - 0 GCACCGACGAGTGCCTGCGG CGG ||||||||[||||||||||||]PAM NA Intergenic NA NA

T19384 CCGACGAGTGCCTGCGGCGGTGG 1000
Oligo fwd CACCCCGACGAGTGCCTGCGGCGG
Oligo rev AAACCCGCCGCAGGCACTCGTCGG
Chromosome start end strand MM target_seq PAM alignment distance position gene name gene id
GU305914.1 200153 200175 - 0 CCGACGAGTGCCTGCGGCGG TGG ||||||||[||||||||||||]PAM NA Intergenic NA NA

T19385 ACGAGTGCCTGCGGCGGTGGTGG 1000
Oligo fwd CACCACGAGTGCCTGCGGCGGTGG
Oligo rev AAACCCACCGCCGCAGGCACTCGT
Chromosome start end strand MM target_seq PAM alignment distance position gene name gene id
GU305914.1 200150 200172 - 0 ACGAGTGCCTGCGGCGGTGG TGG ||||||||[||||||||||||]PAM NA Intergenic NA NA
GU305914.1 48150 48172 + 4 ACCTGTGGCGGCGGCGGTGG CGG ||--|||-[|-||||||||||]PAM NA Intergenic NA NA

T19386 GGCGGTGGTGGCCGTTCTGCAGG 1000
Oligo fwd CACCGGCGGTGGTGGCCGTTCTGC
Oligo rev AAACGCAGAACGGCCACCACCGCC
Chromosome start end strand MM target_seq PAM alignment distance position gene name gene id
GU305914.1 200138 200160 - 0 GGCGGTGGTGGCCGTTCTGC AGG ||||||||[||||||||||||]PAM NA Intergenic NA NA

T19387 CCGTTCTGCAGGATGTCCGACGG 1000
Oligo fwd CACCCCGTTCTGCAGGATGTCCGA
Oligo rev AAACTCGGACATCCTGCAGAACGG
Chromosome start end strand MM target_seq PAM alignment distance position gene name gene id
GU305914.1 200127 200149 - 0 CCGTTCTGCAGGATGTCCGA CGG ||||||||[||||||||||||]PAM NA Intergenic NA NA

T19388 GAAACCTCAAGACGAGCCACTGG 1000
Oligo fwd CACCGAAACCTCAAGACGAGCCAC
Oligo rev AAACGTGGCTCGTCTTGAGGTTTC
Chromosome start end strand MM target_seq PAM alignment distance position gene name gene id
GU305914.1 200072 200094 - 0 GAAACCTCAAGACGAGCCAC TGG ||||||||[||||||||||||]PAM NA Intergenic NA NA

T19389 ACGAGCCACTGGAAGCACTTCGG 1000
Oligo fwd CACCACGAGCCACTGGAAGCACTT
Oligo rev AAACAAGTGCTTCCAGTGGCTCGT
Chromosome start end strand MM target_seq PAM alignment distance position gene name gene id
GU305914.1 200061 200083 - 0 ACGAGCCACTGGAAGCACTT CGG ||||||||[||||||||||||]PAM NA Intergenic NA NA

T19390 CGAGCCACTGGAAGCACTTCGGG 1000
Oligo fwd CACCCGAGCCACTGGAAGCACTTC
Oligo rev AAACGAAGTGCTTCCAGTGGCTCG
Chromosome start end strand MM target_seq PAM alignment distance position gene name gene id
GU305914.1 200060 200082 - 0 CGAGCCACTGGAAGCACTTC GGG ||||||||[||||||||||||]PAM NA Intergenic NA NA

T19391 GAGCCACTGGAAGCACTTCGGGG 1000
Oligo fwd CACCGAGCCACTGGAAGCACTTCG
Oligo rev AAACCGAAGTGCTTCCAGTGGCTC
Chromosome start end strand MM target_seq PAM alignment distance position gene name gene id
GU305914.1 200059 200081 - 0 GAGCCACTGGAAGCACTTCG GGG ||||||||[||||||||||||]PAM NA Intergenic NA NA

T19392 CTGGAAGCACTTCGGGGTCGTGG 1000
Oligo fwd CACCCTGGAAGCACTTCGGGGTCG
Oligo rev AAACCGACCCCGAAGTGCTTCCAG
Chromosome start end strand MM target_seq PAM alignment distance position gene name gene id
GU305914.1 200053 200075 - 0 CTGGAAGCACTTCGGGGTCG TGG ||||||||[||||||||||||]PAM NA Intergenic NA NA

T19393 TGGAAGCACTTCGGGGTCGTGGG 1000
Oligo fwd CACCTGGAAGCACTTCGGGGTCGT
Oligo rev AAACACGACCCCGAAGTGCTTCCA

| | | | | | | | | | | | |
|---|---|---|---|---|---|---|---|---|---|---|---|
| Chromosome | start | end | strand | MM | target_seq | PAM | alignment | distance | position | gene name | gene id |
| GU305914.1 | 200052 | 200074 | - | 0 | TGGAAGCACTTCGGGGTCGT | GGG | \|\|\|\|\|\|\|\|[\|\|\|\|\|\|\|\|\|\|\|\|]PAM | NA | Intergenic | NA | NA |
| T19394 | ACTTCGGGGTCGTGGGCCTGCGG | 1000 | | | | | | | | | |
| Oligo fwd | CACCACTTCGGGGTCGTGGGCCTG | | | | | | | | | | |
| Oligo rev | AAACCAGGCCCACGACCCCGAAGT | | | | | | | | | | |
| Chromosome | start | end | strand | MM | target_seq | PAM | alignment | distance | position | gene name | gene id |
| GU305914.1 | 200045 | 200067 | - | 0 | ACTTCGGGGTCGTGGGCCTG | CGG | \|\|\|\|\|\|\|\|[\|\|\|\|\|\|\|\|\|\|\|\|]PAM | NA | Intergenic | NA | NA |
| T19395 | CTTCGGGGTCGTGGGCCTGCGGG | 1000 | | | | | | | | | |
| Oligo fwd | CACCCTTCGGGGTCGTGGGCCTGC | | | | | | | | | | |
| Oligo rev | AAACGCAGGCCCACGACCCCGAAG | | | | | | | | | | |
| Chromosome | start | end | strand | MM | target_seq | PAM | alignment | distance | position | gene name | gene id |
| GU305914.1 | 200044 | 200066 | - | 0 | CTTCGGGGTCGTGGGCCTGC | GGG | \|\|\|\|\|\|\|\|[\|\|\|\|\|\|\|\|\|\|\|\|]PAM | NA | Intergenic | NA | NA |
| T19396 | GGGGTCGTGGGCCTGCGGGTCGG | 1000 | | | | | | | | | |
| Oligo fwd | CACCGGGGTCGTGGGCCTGCGGGT | | | | | | | | | | |
| Oligo rev | AAACACCCGCAGGCCCACGACCCC | | | | | | | | | | |
| Chromosome | start | end | strand | MM | target_seq | PAM | alignment | distance | position | gene name | gene id |
| GU305914.1 | 200040 | 200062 | - | 0 | GGGGTCGTGGGCCTGCGGGT | CGG | \|\|\|\|\|\|\|\|[\|\|\|\|\|\|\|\|\|\|\|\|]PAM | NA | Intergenic | NA | NA |
| T19397 | GGGTCGTGGGCCTGCGGGTCGGG | 1000 | | | | | | | | | |
| Oligo fwd | CACCGGGTCGTGGGCCTGCGGGTC | | | | | | | | | | |
| Oligo rev | AAACGACCCGCAGGCCCACGACCC | | | | | | | | | | |
| Chromosome | start | end | strand | MM | target_seq | PAM | alignment | distance | position | gene name | gene id |
| GU305914.1 | 200039 | 200061 | - | 0 | GGGTCGTGGGCCTGCGGGTC | GGG | \|\|\|\|\|\|\|\|[\|\|\|\|\|\|\|\|\|\|\|\|]PAM | NA | Intergenic | NA | NA |
| T19398 | CGTGGGCCTGCGGGTCGGGCAGG | 1000 | | | | | | | | | |
| Oligo fwd | CACCCGTGGGCCTGCGGGTCGGGC | | | | | | | | | | |
| Oligo rev | AAACGCCCGACCCGCAGGCCCACG | | | | | | | | | | |
| Chromosome | start | end | strand | MM | target_seq | PAM | alignment | distance | position | gene name | gene id |
| GU305914.1 | 200035 | 200057 | - | 0 | CGTGGGCCTGCGGGTCGGGC | AGG | \|\|\|\|\|\|\|\|[\|\|\|\|\|\|\|\|\|\|\|\|]PAM | NA | Intergenic | NA | NA |
| T19399 | GGATCCTTTTCGCGTCGAGTCGG | 1000 | | | | | | | | | |
| Oligo fwd | CACCGGATCCTTTTCGCGTCGAGT | | | | | | | | | | |
| Oligo rev | AAACACTCGACGCGAAAAGGATCC | | | | | | | | | | |
| Chromosome | start | end | strand | MM | target_seq | PAM | alignment | distance | position | gene name | gene id |
| GU305914.1 | 200014 | 200036 | - | 0 | GGATCCTTTTCGCGTCGAGT | CGG | \|\|\|\|\|\|\|\|[\|\|\|\|\|\|\|\|\|\|\|\|]PAM | NA | Intergenic | NA | NA |
| T19400 | GAGTCGGTGCTGAGCGTGATCGG | 1000 | | | | | | | | | |
| Oligo fwd | CACCGAGTCGGTGCTGAGCGTGAT | | | | | | | | | | |
| Oligo rev | AAACATCACGCTCAGCACCGACTC | | | | | | | | | | |
| Chromosome | start | end | strand | MM | target_seq | PAM | alignment | distance | position | gene name | gene id |
| GU305914.1 | 199998 | 200020 | - | 0 | GAGTCGGTGCTGAGCGTGAT | CGG | \|\|\|\|\|\|\|\|[\|\|\|\|\|\|\|\|\|\|\|\|]PAM | NA | Intergenic | NA | NA |
| T19401 | GTGCTGAGCGTGATCGGAAACGG | 1000 | | | | | | | | | |
| Oligo fwd | CACCGTGCTGAGCGTGATCGGAAA | | | | | | | | | | |
| Oligo rev | AAACTTTCCGATCACGCTCAGCAC | | | | | | | | | | |
| Chromosome | start | end | strand | MM | target_seq | PAM | alignment | distance | position | gene name | gene id |
| GU305914.1 | 199992 | 200014 | - | 0 | GTGCTGAGCGTGATCGGAAA | CGG | \|\|\|\|\|\|\|\|[\|\|\|\|\|\|\|\|\|\|\|\|]PAM | NA | Intergenic | NA | NA |
| T19402 | TGCTGAGCGTGATCGGAAACGGG | 1000 | | | | | | | | | |
| Oligo fwd | CACCTGCTGAGCGTGATCGGAAAC | | | | | | | | | | |
| Oligo rev | AAACGTTTCCGATCACGCTCAGCA | | | | | | | | | | |
| Chromosome | start | end | strand | MM | target_seq | PAM | alignment | distance | position | gene name | gene id |
| GU305914.1 | 199991 | 200013 | - | 0 | TGCTGAGCGTGATCGGAAAC | GGG | \|\|\|\|\|\|\|\|[\|\|\|\|\|\|\|\|\|\|\|\|]PAM | NA | Intergenic | NA | NA |
| T19403 | GCTGAGCGTGATCGGAAACGGGG | 1000 | | | | | | | | | |
| Oligo fwd | CACCGCTGAGCGTGATCGGAAACG | | | | | | | | | | |
| Oligo rev | AAACCGTTTCCGATCACGCTCAGC | | | | | | | | | | |
| Chromosome | start | end | strand | MM | target_seq | PAM | alignment | distance | position | gene name | gene id |
| GU305914.1 | 199990 | 200012 | - | 0 | GCTGAGCGTGATCGGAAACG | GGG | \|\|\|\|\|\|\|\|[\|\|\|\|\|\|\|\|\|\|\|\|]PAM | NA | Intergenic | NA | NA |
| T19404 | AAACGGGGCGATCTTCCACACGG | 1000 | | | | | | | | | |
| Oligo fwd | CACCAAACGGGGCGATCTTCCACA | | | | | | | | | | |
| Oligo rev | AAACTGTGGAAGATCGCCCCGTTT | | | | | | | | | | |
| Chromosome | start | end | strand | MM | target_seq | PAM | alignment | distance | position | gene name | gene id |
| GU305914.1 | 199975 | 199997 | - | 0 | AAACGGGGCGATCTTCCACA | CGG | \|\|\|\|\|\|\|\|[\|\|\|\|\|\|\|\|\|\|\|\|]PAM | NA | Intergenic | NA | NA |
| T19405 | GGCGATCTTCCACACGGACCCGG | 1000 | | | | | | | | | |
| Oligo fwd | CACCGGCGATCTTCCACACGGACC | | | | | | | | | | |
| Oligo rev | AAACGGTCCGTGTGGAAGATCGCC | | | | | | | | | | |
| Chromosome | start | end | strand | MM | target_seq | PAM | alignment | distance | position | gene name | gene id |
| GU305914.1 | 199969 | 199991 | - | 0 | GGCGATCTTCCACACGGACC | CGG | \|\|\|\|\|\|\|\|[\|\|\|\|\|\|\|\|\|\|\|\|]PAM | NA | Intergenic | NA | NA |
| T19406 | CTTCCACACGGACCCGGACATGG | 1000 | | | | | | | | | |
| Oligo fwd | CACCCTTCCACACGGACCCGGACA | | | | | | | | | | |
| Oligo rev | AAACTGTCCGGGTCCGTGTGGAAG | | | | | | | | | | |
| Chromosome | start | end | strand | MM | target_seq | PAM | alignment | distance | position | gene name | gene id |
| GU305914.1 | 199963 | 199985 | - | 0 | CTTCCACACGGACCCGGACA | TGG | \|\|\|\|\|\|\|\|[\|\|\|\|\|\|\|\|\|\|\|\|]PAM | NA | Intergenic | NA | NA |
| T19407 | ACCCGGACATGGACGTGACCTGG | 1000 | | | | | | | | | |
| Oligo fwd | CACCACCCGGACATGGACGTGACC | | | | | | | | | | |
| Oligo rev | AAACGGTCACGTCCATGTCCGGGT | | | | | | | | | | |
| Chromosome | start | end | strand | MM | target_seq | PAM | alignment | distance | position | gene name | gene id |
| GU305914.1 | 199952 | 199974 | - | 0 | ACCCGGACATGGACGTGACC | TGG | \|\|\|\|\|\|\|\|[\|\|\|\|\|\|\|\|\|\|\|\|]PAM | NA | Intergenic | NA | NA |
| T19408 | CGGACATGGACGTGACCTGGCGG | 1000 | | | | | | | | | |
| Oligo fwd | CACCCGGACATGGACGTGACCTGG | | | | | | | | | | |

<table>
<tr><td>Oligo rev</td><td>AAACCCAGGTCACGTCCATGTCCG</td><td></td><td></td><td></td><td></td><td></td><td></td><td></td><td></td><td></td><td></td></tr>
<tr><td>Chromosome</td><td>start</td><td>end</td><td>strand</td><td>MM</td><td>target_seq</td><td>PAM</td><td>alignment</td><td>distance</td><td>position</td><td>gene name</td><td>gene id</td></tr>
<tr><td>GU305914.1</td><td>199949</td><td>199971</td><td>-</td><td>0</td><td>CGGACATGGACGTGACCTGG</td><td>CGG</td><td>||||||||[||||||||||||]PAM</td><td>NA</td><td>Intergenic</td><td>NA</td><td>NA</td></tr>
<tr><td>GU305914.1</td><td>128550</td><td>128572</td><td>+</td><td>4</td><td>CGGACCACGACGTGATCTGG</td><td>AGG</td><td>|||||---[|||||||-||||]PAM</td><td>NA</td><td>Intergenic</td><td>NA</td><td>NA</td></tr>
<tr><td></td><td></td><td></td><td></td><td></td><td></td><td></td><td></td><td></td><td></td><td></td><td></td></tr>
<tr><td>T19409</td><td>GGACGTGACCTGGCGGATCGCGG</td><td>1000</td><td></td><td></td><td></td><td></td><td></td><td></td><td></td><td></td><td></td></tr>
<tr><td>Oligo fwd</td><td>CACCGGACGTGACCTGGCGGATCG</td><td></td><td></td><td></td><td></td><td></td><td></td><td></td><td></td><td></td><td></td></tr>
<tr><td>Oligo rev</td><td>AAACCGATCCGCCAGGTCACGTCC</td><td></td><td></td><td></td><td></td><td></td><td></td><td></td><td></td><td></td><td></td></tr>
<tr><td>Chromosome</td><td>start</td><td>end</td><td>strand</td><td>MM</td><td>target_seq</td><td>PAM</td><td>alignment</td><td>distance</td><td>position</td><td>gene name</td><td>gene id</td></tr>
<tr><td>GU305914.1</td><td>199942</td><td>199964</td><td>-</td><td>0</td><td>GGACGTGACCTGGCGGATCG</td><td>CGG</td><td>||||||||[||||||||||||]PAM</td><td>NA</td><td>Intergenic</td><td>NA</td><td>NA</td></tr>
<tr><td></td><td></td><td></td><td></td><td></td><td></td><td></td><td></td><td></td><td></td><td></td><td></td></tr>
<tr><td>T19410</td><td>CTGGCGGATCGCGGACAACGTGG</td><td>1000</td><td></td><td></td><td></td><td></td><td></td><td></td><td></td><td></td><td></td></tr>
<tr><td>Oligo fwd</td><td>CACCCTGGCGGATCGCGGACAACG</td><td></td><td></td><td></td><td></td><td></td><td></td><td></td><td></td><td></td><td></td></tr>
<tr><td>Oligo rev</td><td>AAACCGTTGTCCGCGATCCGCCAG</td><td></td><td></td><td></td><td></td><td></td><td></td><td></td><td></td><td></td><td></td></tr>
<tr><td>Chromosome</td><td>start</td><td>end</td><td>strand</td><td>MM</td><td>target_seq</td><td>PAM</td><td>alignment</td><td>distance</td><td>position</td><td>gene name</td><td>gene id</td></tr>
<tr><td>GU305914.1</td><td>199933</td><td>199955</td><td>-</td><td>0</td><td>CTGGCGGATCGCGGACAACG</td><td>TGG</td><td>||||||||[||||||||||||]PAM</td><td>NA</td><td>Intergenic</td><td>NA</td><td>NA</td></tr>
<tr><td></td><td></td><td></td><td></td><td></td><td></td><td></td><td></td><td></td><td></td><td></td><td></td></tr>
<tr><td>T19411</td><td>CGTGGACGAGCTGTTTCGTATGG</td><td>1000</td><td></td><td></td><td></td><td></td><td></td><td></td><td></td><td></td><td></td></tr>
<tr><td>Oligo fwd</td><td>CACCCGTGGACGAGCTGTTTCGTA</td><td></td><td></td><td></td><td></td><td></td><td></td><td></td><td></td><td></td><td></td></tr>
<tr><td>Oligo rev</td><td>AAACTACGAAACAGCTCGTCCACG</td><td></td><td></td><td></td><td></td><td></td><td></td><td></td><td></td><td></td><td></td></tr>
<tr><td>Chromosome</td><td>start</td><td>end</td><td>strand</td><td>MM</td><td>target_seq</td><td>PAM</td><td>alignment</td><td>distance</td><td>position</td><td>gene name</td><td>gene id</td></tr>
<tr><td>GU305914.1</td><td>199915</td><td>199937</td><td>-</td><td>0</td><td>CGTGGACGAGCTGTTTCGTA</td><td>TGG</td><td>||||||||[||||||||||||]PAM</td><td>NA</td><td>Intergenic</td><td>NA</td><td>NA</td></tr>
<tr><td></td><td></td><td></td><td></td><td></td><td></td><td></td><td></td><td></td><td></td><td></td><td></td></tr>
<tr><td>T19412</td><td>GTGGACGAGCTGTTTCGTATGGG</td><td>1000</td><td></td><td></td><td></td><td></td><td></td><td></td><td></td><td></td><td></td></tr>
<tr><td>Oligo fwd</td><td>CACCGTGGACGAGCTGTTTCGTAT</td><td></td><td></td><td></td><td></td><td></td><td></td><td></td><td></td><td></td><td></td></tr>
<tr><td>Oligo rev</td><td>AAACATACGAAACAGCTCGTCCAC</td><td></td><td></td><td></td><td></td><td></td><td></td><td></td><td></td><td></td><td></td></tr>
<tr><td>Chromosome</td><td>start</td><td>end</td><td>strand</td><td>MM</td><td>target_seq</td><td>PAM</td><td>alignment</td><td>distance</td><td>position</td><td>gene name</td><td>gene id</td></tr>
<tr><td>GU305914.1</td><td>199914</td><td>199936</td><td>-</td><td>0</td><td>GTGGACGAGCTGTTTCGTAT</td><td>GGG</td><td>||||||||[||||||||||||]PAM</td><td>NA</td><td>Intergenic</td><td>NA</td><td>NA</td></tr>
<tr><td></td><td></td><td></td><td></td><td></td><td></td><td></td><td></td><td></td><td></td><td></td><td></td></tr>
<tr><td>T19413</td><td>CCAAGCTGTACCTGCCGAAGCGG</td><td>1000</td><td></td><td></td><td></td><td></td><td></td><td></td><td></td><td></td><td></td></tr>
<tr><td>Oligo fwd</td><td>CACCCCAAGCTGTACCTGCCGAAG</td><td></td><td></td><td></td><td></td><td></td><td></td><td></td><td></td><td></td><td></td></tr>
<tr><td>Oligo rev</td><td>AAACCTTCGGCAGGTACAGCTTGG</td><td></td><td></td><td></td><td></td><td></td><td></td><td></td><td></td><td></td><td></td></tr>
<tr><td>Chromosome</td><td>start</td><td>end</td><td>strand</td><td>MM</td><td>target_seq</td><td>PAM</td><td>alignment</td><td>distance</td><td>position</td><td>gene name</td><td>gene id</td></tr>
<tr><td>GU305914.1</td><td>199886</td><td>199908</td><td>-</td><td>0</td><td>CCAAGCTGTACCTGCCGAAG</td><td>CGG</td><td>||||||||[||||||||||||]PAM</td><td>NA</td><td>Intergenic</td><td>NA</td><td>NA</td></tr>
<tr><td></td><td></td><td></td><td></td><td></td><td></td><td></td><td></td><td></td><td></td><td></td><td></td></tr>
<tr><td>T19414</td><td>GTACCTGCCGAAGCGGCGTCTGG</td><td>1000</td><td></td><td></td><td></td><td></td><td></td><td></td><td></td><td></td><td></td></tr>
<tr><td>Oligo fwd</td><td>CACCGTACCTGCCGAAGCGGCGTC</td><td></td><td></td><td></td><td></td><td></td><td></td><td></td><td></td><td></td><td></td></tr>
<tr><td>Oligo rev</td><td>AAACGACGCCGCTTCGGCAGGTAC</td><td></td><td></td><td></td><td></td><td></td><td></td><td></td><td></td><td></td><td></td></tr>
<tr><td>Chromosome</td><td>start</td><td>end</td><td>strand</td><td>MM</td><td>target_seq</td><td>PAM</td><td>alignment</td><td>distance</td><td>position</td><td>gene name</td><td>gene id</td></tr>
<tr><td>GU305914.1</td><td>199879</td><td>199901</td><td>-</td><td>0</td><td>GTACCTGCCGAAGCGGCGTC</td><td>TGG</td><td>||||||||[||||||||||||]PAM</td><td>NA</td><td>Intergenic</td><td>NA</td><td>NA</td></tr>
<tr><td></td><td></td><td></td><td></td><td></td><td></td><td></td><td></td><td></td><td></td><td></td><td></td></tr>
<tr><td>T19415</td><td>CCGAAGCGGCGTCTGGACCGCGG</td><td>1000</td><td></td><td></td><td></td><td></td><td></td><td></td><td></td><td></td><td></td></tr>
<tr><td>Oligo fwd</td><td>CACCCCGAAGCGGCGTCTGGACCG</td><td></td><td></td><td></td><td></td><td></td><td></td><td></td><td></td><td></td><td></td></tr>
<tr><td>Oligo rev</td><td>AAACCGGTCCAGACGCCGCTTCGG</td><td></td><td></td><td></td><td></td><td></td><td></td><td></td><td></td><td></td><td></td></tr>
<tr><td>Chromosome</td><td>start</td><td>end</td><td>strand</td><td>MM</td><td>target_seq</td><td>PAM</td><td>alignment</td><td>distance</td><td>position</td><td>gene name</td><td>gene id</td></tr>
<tr><td>GU305914.1</td><td>199872</td><td>199894</td><td>-</td><td>0</td><td>CCGAAGCGGCGTCTGGACCG</td><td>CGG</td><td>||||||||[||||||||||||]PAM</td><td>NA</td><td>Intergenic</td><td>NA</td><td>NA</td></tr>
<tr><td></td><td></td><td></td><td></td><td></td><td></td><td></td><td></td><td></td><td></td><td></td><td></td></tr>
<tr><td>T19416</td><td>GCGTCTGGACCGCGGCACGATGG</td><td>1000</td><td></td><td></td><td></td><td></td><td></td><td></td><td></td><td></td><td></td></tr>
<tr><td>Oligo fwd</td><td>CACCGCGTCTGGACCGCGGCACGA</td><td></td><td></td><td></td><td></td><td></td><td></td><td></td><td></td><td></td><td></td></tr>
<tr><td>Oligo rev</td><td>AAACTCGTGCCGCGGTCCAGACGC</td><td></td><td></td><td></td><td></td><td></td><td></td><td></td><td></td><td></td><td></td></tr>
<tr><td>Chromosome</td><td>start</td><td>end</td><td>strand</td><td>MM</td><td>target_seq</td><td>PAM</td><td>alignment</td><td>distance</td><td>position</td><td>gene name</td><td>gene id</td></tr>
<tr><td>GU305914.1</td><td>199864</td><td>199886</td><td>-</td><td>0</td><td>GCGTCTGGACCGCGGCACGA</td><td>TGG</td><td>||||||||[||||||||||||]PAM</td><td>NA</td><td>Intergenic</td><td>NA</td><td>NA</td></tr>
<tr><td></td><td></td><td></td><td></td><td></td><td></td><td></td><td></td><td></td><td></td><td></td><td></td></tr>
<tr><td>T19417</td><td>CGTCTGGACCGCGGCACGATGGG</td><td>1000</td><td></td><td></td><td></td><td></td><td></td><td></td><td></td><td></td><td></td></tr>
<tr><td>Oligo fwd</td><td>CACCCGTCTGGACCGCGGCACGAT</td><td></td><td></td><td></td><td></td><td></td><td></td><td></td><td></td><td></td><td></td></tr>
<tr><td>Oligo rev</td><td>AAACATCGTGCCGCGGTCCAGACG</td><td></td><td></td><td></td><td></td><td></td><td></td><td></td><td></td><td></td><td></td></tr>
<tr><td>Chromosome</td><td>start</td><td>end</td><td>strand</td><td>MM</td><td>target_seq</td><td>PAM</td><td>alignment</td><td>distance</td><td>position</td><td>gene name</td><td>gene id</td></tr>
<tr><td>GU305914.1</td><td>199863</td><td>199885</td><td>-</td><td>0</td><td>CGTCTGGACCGCGGCACGAT</td><td>GGG</td><td>||||||||[||||||||||||]PAM</td><td>NA</td><td>Intergenic</td><td>NA</td><td>NA</td></tr>
<tr><td></td><td></td><td></td><td></td><td></td><td></td><td></td><td></td><td></td><td></td><td></td><td></td></tr>
<tr><td>T19418</td><td>GTCTGGACCGCGGCACGATGGGG</td><td>1000</td><td></td><td></td><td></td><td></td><td></td><td></td><td></td><td></td><td></td></tr>
<tr><td>Oligo fwd</td><td>CACCGTCTGGACCGCGGCACGATG</td><td></td><td></td><td></td><td></td><td></td><td></td><td></td><td></td><td></td><td></td></tr>
<tr><td>Oligo rev</td><td>AAACCATCGTGCCGCGGTCCAGAC</td><td></td><td></td><td></td><td></td><td></td><td></td><td></td><td></td><td></td><td></td></tr>
<tr><td>Chromosome</td><td>start</td><td>end</td><td>strand</td><td>MM</td><td>target_seq</td><td>PAM</td><td>alignment</td><td>distance</td><td>position</td><td>gene name</td><td>gene id</td></tr>
<tr><td>GU305914.1</td><td>199862</td><td>199884</td><td>-</td><td>0</td><td>GTCTGGACCGCGGCACGATG</td><td>GGG</td><td>||||||||[||||||||||||]PAM</td><td>NA</td><td>Intergenic</td><td>NA</td><td>NA</td></tr>
<tr><td></td><td></td><td></td><td></td><td></td><td></td><td></td><td></td><td></td><td></td><td></td><td></td></tr>
<tr><td>T19419</td><td>GCGGCACGATGGGGCGCGCCCGG</td><td>1000</td><td></td><td></td><td></td><td></td><td></td><td></td><td></td><td></td><td></td></tr>
<tr><td>Oligo fwd</td><td>CACCGCGGCACGATGGGGCGCGCC</td><td></td><td></td><td></td><td></td><td></td><td></td><td></td><td></td><td></td><td></td></tr>
<tr><td>Oligo rev</td><td>AAACGGCGCGCCCCATCGTGCCGC</td><td></td><td></td><td></td><td></td><td></td><td></td><td></td><td></td><td></td><td></td></tr>
<tr><td>Chromosome</td><td>start</td><td>end</td><td>strand</td><td>MM</td><td>target_seq</td><td>PAM</td><td>alignment</td><td>distance</td><td>position</td><td>gene name</td><td>gene id</td></tr>
<tr><td>GU305914.1</td><td>199853</td><td>199875</td><td>-</td><td>0</td><td>GCGGCACGATGGGGCGCGCC</td><td>CGG</td><td>||||||||[||||||||||||]PAM</td><td>NA</td><td>Intergenic</td><td>NA</td><td>NA</td></tr>
<tr><td></td><td></td><td></td><td></td><td></td><td></td><td></td><td></td><td></td><td></td><td></td><td></td></tr>
<tr><td>T19420</td><td>CACGATGGGGCGCGCCCGGCTGG</td><td>1000</td><td></td><td></td><td></td><td></td><td></td><td></td><td></td><td></td><td></td></tr>
<tr><td>Oligo fwd</td><td>CACCCACGATGGGGCGCGCCCGGC</td><td></td><td></td><td></td><td></td><td></td><td></td><td></td><td></td><td></td><td></td></tr>
<tr><td>Oligo rev</td><td>AAACGCCGGGCGCGCCCCATCGTG</td><td></td><td></td><td></td><td></td><td></td><td></td><td></td><td></td><td></td><td></td></tr>
<tr><td>Chromosome</td><td>start</td><td>end</td><td>strand</td><td>MM</td><td>target_seq</td><td>PAM</td><td>alignment</td><td>distance</td><td>position</td><td>gene name</td><td>gene id</td></tr>
<tr><td>GU305914.1</td><td>199849</td><td>199871</td><td>-</td><td>0</td><td>CACGATGGGGCGCGCCCGGC</td><td>TGG</td><td>||||||||[||||||||||||]PAM</td><td>NA</td><td>Intergenic</td><td>NA</td><td>NA</td></tr>
<tr><td></td><td></td><td></td><td></td><td></td><td></td><td></td><td></td><td></td><td></td><td></td><td></td></tr>
<tr><td>T19421</td><td>GATGGGGCGCGCCCGGCTGGAGG</td><td>1000</td><td></td><td></td><td></td><td></td><td></td><td></td><td></td><td></td><td></td></tr>
<tr><td>Oligo fwd</td><td>CACCGATGGGGCGCGCCCGGCTGG</td><td></td><td></td><td></td><td></td><td></td><td></td><td></td><td></td><td></td><td></td></tr>
<tr><td>Oligo rev</td><td>AAACCCAGCCGGGCGCGCCCCATC</td><td></td><td></td><td></td><td></td><td></td><td></td><td></td><td></td><td></td><td></td></tr>
<tr><td>Chromosome</td><td>start</td><td>end</td><td>strand</td><td>MM</td><td>target_seq</td><td>PAM</td><td>alignment</td><td>distance</td><td>position</td><td>gene name</td><td>gene id</td></tr>
<tr><td>GU305914.1</td><td>199846</td><td>199868</td><td>-</td><td>0</td><td>GATGGGGCGCGCCCGGCTGG</td><td>AGG</td><td>||||||||[||||||||||||]PAM</td><td>NA</td><td>Intergenic</td><td>NA</td><td>NA</td></tr>
<tr><td></td><td></td><td></td><td></td><td></td><td></td><td></td><td></td><td></td><td></td><td></td><td></td></tr>
<tr><td>T19422</td><td>CGCGCCCGGCTGGAGGCGCCCGG</td><td>1000</td><td></td><td></td><td></td><td></td><td></td><td></td><td></td><td></td><td></td></tr>
<tr><td>Oligo fwd</td><td>CACCCGCGCCCGGCTGGAGGCGCC</td><td></td><td></td><td></td><td></td><td></td><td></td><td></td><td></td><td></td><td></td></tr>
<tr><td>Oligo rev</td><td>AAACGGCGCCTCCAGCCGGGCGCG</td><td></td><td></td><td></td><td></td><td></td><td></td><td></td><td></td><td></td><td></td></tr>
<tr><td>Chromosome</td><td>start</td><td>end</td><td>strand</td><td>MM</td><td>target_seq</td><td>PAM</td><td>alignment</td><td>distance</td><td>position</td><td>gene name</td><td>gene id</td></tr>
<tr><td>GU305914.1</td><td>199839</td><td>199861</td><td>-</td><td>0</td><td>CGCGCCCGGCTGGAGGCGCC</td><td>CGG</td><td>||||||||[||||||||||||]PAM</td><td>NA</td><td>Intergenic</td><td>NA</td><td>NA</td></tr>
</table>

T19423 GGCGCCCGGCTGCGCGCACGTGG 1000
Oligo fwd CACCGGCGCCCGGCTGCGCGCACG
Oligo rev AAACCGTGCGCGCAGCCGGGCGCC
Chromosome start end strand MM target_seq PAM alignment distance position gene name gene id
GU305914.1 199825 199847 - 0 GGCGCCCGGCTGCGCGCACG TGG ||||||||[||||||||||||]PAM NA Intergenic NA NA
GU305914.1 83300 83322 - 4 TCCTCCCGGCGGCGCGCACG GGG --|-||||[||-|||||||||]PAM NA Intergenic NA NA

T19424 GCCCGGCTGCGCGCACGTGGCGG 1000
Oligo fwd CACCGCCCGGCTGCGCGCACGTGG
Oligo rev AAACCCACGTGCGCGCAGCCGGGC
Chromosome start end strand MM target_seq PAM alignment distance position gene name gene id
GU305914.1 199822 199844 - 0 GCCCGGCTGCGCGCACGTGG CGG ||||||||[||||||||||||]PAM NA Intergenic NA NA

T19425 CGCACGTGGCGGCCGCCGCGTGG 1000
Oligo fwd CACCCGCACGTGGCGGCCGCCGCG
Oligo rev AAACCGCGGCGGCCGCCACGTGCG
Chromosome start end strand MM target_seq PAM alignment distance position gene name gene id
GU305914.1 199811 199833 - 0 CGCACGTGGCGGCCGCCGCG TGG ||||||||[||||||||||||]PAM NA Intergenic NA NA
GU305914.1 81264 81286 + 4 CGATCGCGGCGGCCGCCTCG TGG ||--||-|[|||||||||-||]PAM NA Intergenic NA NA

T19426 GCACGTGGCGGCCGCCGCGTGGG 1000
Oligo fwd CACCGCACGTGGCGGCCGCCGCGT
Oligo rev AAACACGCGGCGGCCGCCACGTGC
Chromosome start end strand MM target_seq PAM alignment distance position gene name gene id
GU305914.1 199810 199832 - 0 GCACGTGGCGGCCGCCGCGT GGG ||||||||[||||||||||||]PAM NA Intergenic NA NA
GU305914.1 110387 110409 - 4 GCGCGTCGCGGCCGCCGCAG AGG ||-|||-|[||||||||||--]PAM NA Intergenic NA NA

T19427 GAAGAGACGTACGTGATCTTCGG 1000
Oligo fwd CACCGAAGAGACGTACGTGATCTT
Oligo rev AAACAAGATCACGTACGTCTCTTC
Chromosome start end strand MM target_seq PAM alignment distance position gene name gene id
GU305914.1 199788 199810 - 0 GAAGAGACGTACGTGATCTT CGG ||||||||[||||||||||||]PAM NA Intergenic NA NA

T19428 CCGCGACTTCGCGTCCCAGCTGG 1000
Oligo fwd CACCCCGCGACTTCGCGTCCCAGC
Oligo rev AAACGCTGGGACGCGAAGTCGCGG
Chromosome start end strand MM target_seq PAM alignment distance position gene name gene id
GU305914.1 199753 199775 - 0 CCGCGACTTCGCGTCCCAGC TGG ||||||||[||||||||||||]PAM NA Intergenic NA NA

T19429 CGCGACTTCGCGTCCCAGCTGGG 1000
Oligo fwd CACCCGCGACTTCGCGTCCCAGCT
Oligo rev AAACAGCTGGGACGCGAAGTCGCG
Chromosome start end strand MM target_seq PAM alignment distance position gene name gene id
GU305914.1 199752 199774 - 0 CGCGACTTCGCGTCCCAGCT GGG ||||||||[||||||||||||]PAM NA Intergenic NA NA

T19430 GCGACTTCGCGTCCCAGCTGGGG 1000
Oligo fwd CACCGCGACTTCGCGTCCCAGCTG
Oligo rev AAACCAGCTGGGACGCGAAGTCGC
Chromosome start end strand MM target_seq PAM alignment distance position gene name gene id
GU305914.1 199751 199773 - 0 GCGACTTCGCGTCCCAGCTG GGG ||||||||[||||||||||||]PAM NA Intergenic NA NA

T19431 ACTTCGCGTCCCAGCTGGGGTGG 1000
Oligo fwd CACCACTTCGCGTCCCAGCTGGGG
Oligo rev AAACCCCCAGCTGGGACGCGAAGT
Chromosome start end strand MM target_seq PAM alignment distance position gene name gene id
GU305914.1 199748 199770 - 0 ACTTCGCGTCCCAGCTGGGG TGG ||||||||[||||||||||||]PAM NA Intergenic NA NA

T19432 CTGGGGTGGCTGATGAGAGAAGG 1000
Oligo fwd CACCCTGGGGTGGCTGATGAGAGA
Oligo rev AAACTCTCTCATCAGCCACCCCAG
Chromosome start end strand MM target_seq PAM alignment distance position gene name gene id
GU305914.1 199734 199756 - 0 CTGGGGTGGCTGATGAGAGA AGG ||||||||[||||||||||||]PAM NA Intergenic NA NA

T19433 GGTGGCTGATGAGAGAAGGTCGG 1000
Oligo fwd CACCGGTGGCTGATGAGAGAAGGT
Oligo rev AAACACCTTCTCTCATCAGCCACC
Chromosome start end strand MM target_seq PAM alignment distance position gene name gene id
GU305914.1 199730 199752 - 0 GGTGGCTGATGAGAGAAGGT CGG ||||||||[||||||||||||]PAM NA Intergenic NA NA

T19434 GGTCGGTTCGACGAGTCGAGCGG 1000
Oligo fwd CACCGGTCGGTTCGACGAGTCGAG
Oligo rev AAACCTCGACTCGTCGAACCGACC
Chromosome start end strand MM target_seq PAM alignment distance position gene name gene id
GU305914.1 199713 199735 - 0 GGTCGGTTCGACGAGTCGAG CGG ||||||||[||||||||||||]PAM NA Intergenic NA NA

T19435 TCGACGAGTCGAGCGGCACCTGG 1000
Oligo fwd CACCTCGACGAGTCGAGCGGCACC
Oligo rev AAACGGTGCCGCTCGACTCGTCGA
Chromosome start end strand MM target_seq PAM alignment distance position gene name gene id
GU305914.1 199706 199728 - 0 TCGACGAGTCGAGCGGCACC TGG ||||||||[||||||||||||]PAM NA Intergenic NA NA

T19436 CGACGAGTCGAGCGGCACCTGGG 1000
Oligo fwd CACCCGACGAGTCGAGCGGCACCT
Oligo rev AAACAGGTGCCGCTCGACTCGTCG
Chromosome start end strand MM target_seq PAM alignment distance position gene name gene id
GU305914.1 199705 199727 - 0 CGACGAGTCGAGCGGCACCT GGG ||||||||[||||||||||||]PAM NA Intergenic NA NA

T19437 GAGCGGCACCTGGGACGAGATGG 1000
Oligo fwd CACCGAGCGGCACCTGGGACGAGA

| | | | | | | | | | | | |
|---|---|---|---|---|---|---|---|---|---|---|---|
| Oligo rev | AAACTCTCGTCCCAGGTGCCGCTC | | | | | | | | | | |
| Chromosome | start | end | strand | MM | target_seq | PAM | alignment | distance | position | gene name | gene id |
| GU305914.1 | 199696 | 199718 | - | 0 | GAGCGGCACCTGGGACGAGA | TGG | \|\|\|\|\|\|\|\|[\|\|\|\|\|\|\|\|\|\|\|\|]PAM | NA | Intergenic | NA | NA |
| T19438 | ACGAGATGGACGACGCCATGAGG | 1000 | | | | | | | | | |
| Oligo fwd | CACCACGAGATGGACGACGCCATG | | | | | | | | | | |
| Oligo rev | AAACCATGGCGTCGTCCATCTCGT | | | | | | | | | | |
| Chromosome | start | end | strand | MM | target_seq | PAM | alignment | distance | position | gene name | gene id |
| GU305914.1 | 199682 | 199704 | - | 0 | ACGAGATGGACGACGCCATG | AGG | \|\|\|\|\|\|\|\|[\|\|\|\|\|\|\|\|\|\|\|\|]PAM | NA | Intergenic | NA | NA |
| T19439 | CATGAGGAAGACGTTCGACGCGG | 1000 | | | | | | | | | |
| Oligo fwd | CACCCATGAGGAAGACGTTCGACG | | | | | | | | | | |
| Oligo rev | AAACCGTCGAACGTCTTCCTCATG | | | | | | | | | | |
| Chromosome | start | end | strand | MM | target_seq | PAM | alignment | distance | position | gene name | gene id |
| GU305914.1 | 199666 | 199688 | - | 0 | CATGAGGAAGACGTTCGACG | CGG | \|\|\|\|\|\|\|\|[\|\|\|\|\|\|\|\|\|\|\|\|]PAM | NA | Intergenic | NA | NA |
| T19440 | GACGTTCGACGCGGAGAGCAAGG | 1000 | | | | | | | | | |
| Oligo fwd | CACCGACGTTCGACGCGGAGAGCA | | | | | | | | | | |
| Oligo rev | AAACTGCTCTCCGCGTCGAACGTC | | | | | | | | | | |
| Chromosome | start | end | strand | MM | target_seq | PAM | alignment | distance | position | gene name | gene id |
| GU305914.1 | 199657 | 199679 | - | 0 | GACGTTCGACGCGGAGAGCA | AGG | \|\|\|\|\|\|\|\|[\|\|\|\|\|\|\|\|\|\|\|\|]PAM | NA | Intergenic | NA | NA |
| T19441 | CGACGCGGAGAGCAAGGAGCTGG | 1000 | | | | | | | | | |
| Oligo fwd | CACCCGACGCGGAGAGCAAGGAGC | | | | | | | | | | |
| Oligo rev | AAACGCTCCTTGCTCTCCGCGTCG | | | | | | | | | | |
| Chromosome | start | end | strand | MM | target_seq | PAM | alignment | distance | position | gene name | gene id |
| GU305914.1 | 199651 | 199673 | - | 0 | CGACGCGGAGAGCAAGGAGC | TGG | \|\|\|\|\|\|\|\|[\|\|\|\|\|\|\|\|\|\|\|\|]PAM | NA | Intergenic | NA | NA |
| T19442 | GCTGGTCCGTCGCGACGTCGAGG | 1000 | | | | | | | | | |
| Oligo fwd | CACCGCTGGTCCGTCGCGACGTCG | | | | | | | | | | |
| Oligo rev | AAACCGACGTCGCGACGGACCAGC | | | | | | | | | | |
| Chromosome | start | end | strand | MM | target_seq | PAM | alignment | distance | position | gene name | gene id |
| GU305914.1 | 199633 | 199655 | - | 0 | GCTGGTCCGTCGCGACGTCG | AGG | \|\|\|\|\|\|\|\|[\|\|\|\|\|\|\|\|\|\|\|\|]PAM | NA | Intergenic | NA | NA |
| GU305914.1 | 113610 | 113632 | + | 3 | GCGGCTCCCTCGCGACGTCG | TGG | \|\|-\|-\|\|\|[-\|\|\|\|\|\|\|\|\|\|\|]PAM | NA | Intergenic | NA | NA |
| T19443 | GGTCCGTCGCGACGTCGAGGTGG | 1000 | | | | | | | | | |
| Oligo fwd | CACCGGTCCGTCGCGACGTCGAGG | | | | | | | | | | |
| Oligo rev | AAACCCTCGACGTCGCGACGGACC | | | | | | | | | | |
| Chromosome | start | end | strand | MM | target_seq | PAM | alignment | distance | position | gene name | gene id |
| GU305914.1 | 199630 | 199652 | - | 0 | GGTCCGTCGCGACGTCGAGG | TGG | \|\|\|\|\|\|\|\|[\|\|\|\|\|\|\|\|\|\|\|\|]PAM | NA | Intergenic | NA | NA |
| T19444 | CCGTCGCGACGTCGAGGTGGCGG | 1000 | | | | | | | | | |
| Oligo fwd | CACCCCGTCGCGACGTCGAGGTGG | | | | | | | | | | |
| Oligo rev | AAACCCACCTCGACGTCGCGACGG | | | | | | | | | | |
| Chromosome | start | end | strand | MM | target_seq | PAM | alignment | distance | position | gene name | gene id |
| GU305914.1 | 199627 | 199649 | - | 0 | CCGTCGCGACGTCGAGGTGG | CGG | \|\|\|\|\|\|\|\|[\|\|\|\|\|\|\|\|\|\|\|\|]PAM | NA | Intergenic | NA | NA |
| T19445 | GGCGGACGCGTTGAACGACGTGG | 1000 | | | | | | | | | |
| Oligo fwd | CACCGGCGGACGCGTTGAACGACG | | | | | | | | | | |
| Oligo rev | AAACCGTCGTTCAACGCGTCCGCC | | | | | | | | | | |
| Chromosome | start | end | strand | MM | target_seq | PAM | alignment | distance | position | gene name | gene id |
| GU305914.1 | 199609 | 199631 | - | 0 | GGCGGACGCGTTGAACGACG | TGG | \|\|\|\|\|\|\|\|[\|\|\|\|\|\|\|\|\|\|\|\|]PAM | NA | Intergenic | NA | NA |
| T19446 | GACGTGGACGCCACCTTCAGAGG | 1000 | | | | | | | | | |
| Oligo fwd | CACCGACGTGGACGCCACCTTCAG | | | | | | | | | | |
| Oligo rev | AAACCTGAAGGTGGCGTCCACGTC | | | | | | | | | | |
| Chromosome | start | end | strand | MM | target_seq | PAM | alignment | distance | position | gene name | gene id |
| GU305914.1 | 199593 | 199615 | - | 0 | GACGTGGACGCCACCTTCAG | AGG | \|\|\|\|\|\|\|\|[\|\|\|\|\|\|\|\|\|\|\|\|]PAM | NA | Intergenic | NA | NA |
| T19447 | CCACCTTCAGAGGCTGCCCTCGG | 1000 | | | | | | | | | |
| Oligo fwd | CACCCCACCTTCAGAGGCTGCCCT | | | | | | | | | | |
| Oligo rev | AAACAGGGCAGCCTCTGAAGGTGG | | | | | | | | | | |
| Chromosome | start | end | strand | MM | target_seq | PAM | alignment | distance | position | gene name | gene id |
| GU305914.1 | 199583 | 199605 | - | 0 | CCACCTTCAGAGGCTGCCCT | CGG | \|\|\|\|\|\|\|\|[\|\|\|\|\|\|\|\|\|\|\|\|]PAM | NA | Intergenic | NA | NA |
| T19448 | GAGGCTGCCCTCGGAATCCGAGG | 1000 | | | | | | | | | |
| Oligo fwd | CACCGAGGCTGCCCTCGGAATCCG | | | | | | | | | | |
| Oligo rev | AAACCGGATTCCGAGGGCAGCCTC | | | | | | | | | | |
| Chromosome | start | end | strand | MM | target_seq | PAM | alignment | distance | position | gene name | gene id |
| GU305914.1 | 199574 | 199596 | - | 0 | GAGGCTGCCCTCGGAATCCG | AGG | \|\|\|\|\|\|\|\|[\|\|\|\|\|\|\|\|\|\|\|\|]PAM | NA | Intergenic | NA | NA |
| T19449 | CTGCCCTCGGAATCCGAGGTCGG | 1000 | | | | | | | | | |
| Oligo fwd | CACCCTGCCCTCGGAATCCGAGGT | | | | | | | | | | |
| Oligo rev | AAACACCTCGGATTCCGAGGGCAG | | | | | | | | | | |
| Chromosome | start | end | strand | MM | target_seq | PAM | alignment | distance | position | gene name | gene id |
| GU305914.1 | 199570 | 199592 | - | 0 | CTGCCCTCGGAATCCGAGGT | CGG | \|\|\|\|\|\|\|\|[\|\|\|\|\|\|\|\|\|\|\|\|]PAM | NA | Intergenic | NA | NA |
| T19450 | CCCTCGGAATCCGAGGTCGGCGG | 1000 | | | | | | | | | |
| Oligo fwd | CACCCCCTCGGAATCCGAGGTCGG | | | | | | | | | | |
| Oligo rev | AAACCCGACCTCGGATTCCGAGGG | | | | | | | | | | |
| Chromosome | start | end | strand | MM | target_seq | PAM | alignment | distance | position | gene name | gene id |
| GU305914.1 | 199567 | 199589 | - | 0 | CCCTCGGAATCCGAGGTCGG | CGG | \|\|\|\|\|\|\|\|[\|\|\|\|\|\|\|\|\|\|\|\|]PAM | NA | Intergenic | NA | NA |
| T19451 | GAATCCGAGGTCGGCGGACGCGG | 1000 | | | | | | | | | |
| Oligo fwd | CACCGAATCCGAGGTCGGCGGACG | | | | | | | | | | |
| Oligo rev | AAACCGTCCGCCGACCTCGGATTC | | | | | | | | | | |
| Chromosome | start | end | strand | MM | target_seq | PAM | alignment | distance | position | gene name | gene id |
| GU305914.1 | 199561 | 199583 | - | 0 | GAATCCGAGGTCGGCGGACG | CGG | \|\|\|\|\|\|\|\|[\|\|\|\|\|\|\|\|\|\|\|\|]PAM | NA | Intergenic | NA | NA |

<table>
<tr><td>T19452</td><td>ACGCCGACGATGACGACTGACGG</td><td>1000</td><td></td><td></td><td></td><td></td><td></td><td></td><td></td><td></td><td></td></tr>
<tr><td>Oligo fwd</td><td>CACCACGCCGACGATGACGACTGA</td><td></td><td></td><td></td><td></td><td></td><td></td><td></td><td></td><td></td><td></td></tr>
<tr><td>Oligo rev</td><td>AAACTCAGTCGTCATCGTCGGCGT</td><td></td><td></td><td></td><td></td><td></td><td></td><td></td><td></td><td></td><td></td></tr>
<tr><td>Chromosome</td><td>start</td><td>end</td><td>strand</td><td>MM</td><td>target_seq</td><td>PAM</td><td>alignment</td><td>distance</td><td>position</td><td>gene name</td><td>gene id</td></tr>
<tr><td>GU305914.1</td><td>199538</td><td>199560</td><td>-</td><td>0</td><td>ACGCCGACGATGACGACTGA</td><td>CGG</td><td>||||||||[||||||||||||]PAM</td><td>NA</td><td>Intergenic</td><td>NA</td><td>NA</td></tr>
<tr><td>T19453</td><td>CGACGATGACGACTGACGGACGG</td><td>1000</td><td></td><td></td><td></td><td></td><td></td><td></td><td></td><td></td><td></td></tr>
<tr><td>Oligo fwd</td><td>CACCCGACGATGACGACTGACGGA</td><td></td><td></td><td></td><td></td><td></td><td></td><td></td><td></td><td></td><td></td></tr>
<tr><td>Oligo rev</td><td>AAACTCCGTCAGTCGTCATCGTCG</td><td></td><td></td><td></td><td></td><td></td><td></td><td></td><td></td><td></td><td></td></tr>
<tr><td>Chromosome</td><td>start</td><td>end</td><td>strand</td><td>MM</td><td>target_seq</td><td>PAM</td><td>alignment</td><td>distance</td><td>position</td><td>gene name</td><td>gene id</td></tr>
<tr><td>GU305914.1</td><td>199534</td><td>199556</td><td>-</td><td>0</td><td>CGACGATGACGACTGACGGA</td><td>CGG</td><td>||||||||[||||||||||||]PAM</td><td>NA</td><td>Intergenic</td><td>NA</td><td>NA</td></tr>
<tr><td>T19454</td><td>GATGACGACTGACGGACGGACGG</td><td>1000</td><td></td><td></td><td></td><td></td><td></td><td></td><td></td><td></td><td></td></tr>
<tr><td>Oligo fwd</td><td>CACCGATGACGACTGACGGACGGA</td><td></td><td></td><td></td><td></td><td></td><td></td><td></td><td></td><td></td><td></td></tr>
<tr><td>Oligo rev</td><td>AAACTCCGTCCGTCAGTCGTCATC</td><td></td><td></td><td></td><td></td><td></td><td></td><td></td><td></td><td></td><td></td></tr>
<tr><td>Chromosome</td><td>start</td><td>end</td><td>strand</td><td>MM</td><td>target_seq</td><td>PAM</td><td>alignment</td><td>distance</td><td>position</td><td>gene name</td><td>gene id</td></tr>
<tr><td>GU305914.1</td><td>199530</td><td>199552</td><td>-</td><td>0</td><td>GATGACGACTGACGGACGGA</td><td>CGG</td><td>||||||||[||||||||||||]PAM</td><td>NA</td><td>Intergenic</td><td>NA</td><td>NA</td></tr>
<tr><td>T19455</td><td>CCATCTTCTCAGATCACTTCTGG</td><td>1000</td><td></td><td></td><td></td><td></td><td></td><td></td><td></td><td></td><td></td></tr>
<tr><td>Oligo fwd</td><td>CACCCCATCTTCTCAGATCACTTC</td><td></td><td></td><td></td><td></td><td></td><td></td><td></td><td></td><td></td><td></td></tr>
<tr><td>Oligo rev</td><td>AAACGAAGTGATCTGAGAAGATGG</td><td></td><td></td><td></td><td></td><td></td><td></td><td></td><td></td><td></td><td></td></tr>
<tr><td>Chromosome</td><td>start</td><td>end</td><td>strand</td><td>MM</td><td>target_seq</td><td>PAM</td><td>alignment</td><td>distance</td><td>position</td><td>gene name</td><td>gene id</td></tr>
<tr><td>GU305914.1</td><td>199437</td><td>199459</td><td>-</td><td>0</td><td>CCATCTTCTCAGATCACTTC</td><td>TGG</td><td>||||||||[||||||||||||]PAM</td><td>NA</td><td>Intergenic</td><td>NA</td><td>NA</td></tr>
<tr><td>T19456</td><td>TCTCAGATCACTTCTGGCTCTGG</td><td>1000</td><td></td><td></td><td></td><td></td><td></td><td></td><td></td><td></td><td></td></tr>
<tr><td>Oligo fwd</td><td>CACCTCTCAGATCACTTCTGGCTC</td><td></td><td></td><td></td><td></td><td></td><td></td><td></td><td></td><td></td><td></td></tr>
<tr><td>Oligo rev</td><td>AAACGAGCCAGAAGTGATCTGAGA</td><td></td><td></td><td></td><td></td><td></td><td></td><td></td><td></td><td></td><td></td></tr>
<tr><td>Chromosome</td><td>start</td><td>end</td><td>strand</td><td>MM</td><td>target_seq</td><td>PAM</td><td>alignment</td><td>distance</td><td>position</td><td>gene name</td><td>gene id</td></tr>
<tr><td>GU305914.1</td><td>199431</td><td>199453</td><td>-</td><td>0</td><td>TCTCAGATCACTTCTGGCTC</td><td>TGG</td><td>||||||||[||||||||||||]PAM</td><td>NA</td><td>Intergenic</td><td>NA</td><td>NA</td></tr>
<tr><td>T19457</td><td>TCGTCTGTCATTCATCCTCTCGG</td><td>1000</td><td></td><td></td><td></td><td></td><td></td><td></td><td></td><td></td><td></td></tr>
<tr><td>Oligo fwd</td><td>CACCTCGTCTGTCATTCATCCTCT</td><td></td><td></td><td></td><td></td><td></td><td></td><td></td><td></td><td></td><td></td></tr>
<tr><td>Oligo rev</td><td>AAACAGAGGATGAATGACAGACGA</td><td></td><td></td><td></td><td></td><td></td><td></td><td></td><td></td><td></td><td></td></tr>
<tr><td>Chromosome</td><td>start</td><td>end</td><td>strand</td><td>MM</td><td>target_seq</td><td>PAM</td><td>alignment</td><td>distance</td><td>position</td><td>gene name</td><td>gene id</td></tr>
<tr><td>GU305914.1</td><td>199390</td><td>199412</td><td>-</td><td>0</td><td>TCGTCTGTCATTCATCCTCT</td><td>CGG</td><td>||||||||[||||||||||||]PAM</td><td>NA</td><td>Intergenic</td><td>NA</td><td>NA</td></tr>
<tr><td>T19458</td><td>CGTCTGTCATTCATCCTCTCGGG</td><td>1000</td><td></td><td></td><td></td><td></td><td></td><td></td><td></td><td></td><td></td></tr>
<tr><td>Oligo fwd</td><td>CACCCGTCTGTCATTCATCCTCTC</td><td></td><td></td><td></td><td></td><td></td><td></td><td></td><td></td><td></td><td></td></tr>
<tr><td>Oligo rev</td><td>AAACGAGAGGATGAATGACAGACG</td><td></td><td></td><td></td><td></td><td></td><td></td><td></td><td></td><td></td><td></td></tr>
<tr><td>Chromosome</td><td>start</td><td>end</td><td>strand</td><td>MM</td><td>target_seq</td><td>PAM</td><td>alignment</td><td>distance</td><td>position</td><td>gene name</td><td>gene id</td></tr>
<tr><td>GU305914.1</td><td>199389</td><td>199411</td><td>-</td><td>0</td><td>CGTCTGTCATTCATCCTCTC</td><td>GGG</td><td>||||||||[||||||||||||]PAM</td><td>NA</td><td>Intergenic</td><td>NA</td><td>NA</td></tr>
<tr><td>T19459</td><td>TTCATCCTCTCGGGTGTCCTAGG</td><td>1000</td><td></td><td></td><td></td><td></td><td></td><td></td><td></td><td></td><td></td></tr>
<tr><td>Oligo fwd</td><td>CACCTTCATCCTCTCGGGTGTCCT</td><td></td><td></td><td></td><td></td><td></td><td></td><td></td><td></td><td></td><td></td></tr>
<tr><td>Oligo rev</td><td>AAACAGGACACCCGAGAGGATGAA</td><td></td><td></td><td></td><td></td><td></td><td></td><td></td><td></td><td></td><td></td></tr>
<tr><td>Chromosome</td><td>start</td><td>end</td><td>strand</td><td>MM</td><td>target_seq</td><td>PAM</td><td>alignment</td><td>distance</td><td>position</td><td>gene name</td><td>gene id</td></tr>
<tr><td>GU305914.1</td><td>199380</td><td>199402</td><td>-</td><td>0</td><td>TTCATCCTCTCGGGTGTCCT</td><td>AGG</td><td>||||||||[||||||||||||]PAM</td><td>NA</td><td>Intergenic</td><td>NA</td><td>NA</td></tr>
<tr><td>T19460</td><td>CTCTCGGGTGTCCTAGGCAGAGG</td><td>1000</td><td></td><td></td><td></td><td></td><td></td><td></td><td></td><td></td><td></td></tr>
<tr><td>Oligo fwd</td><td>CACCCTCTCGGGTGTCCTAGGCAG</td><td></td><td></td><td></td><td></td><td></td><td></td><td></td><td></td><td></td><td></td></tr>
<tr><td>Oligo rev</td><td>AAACCTGCCTAGGACACCCGAGAG</td><td></td><td></td><td></td><td></td><td></td><td></td><td></td><td></td><td></td><td></td></tr>
<tr><td>Chromosome</td><td>start</td><td>end</td><td>strand</td><td>MM</td><td>target_seq</td><td>PAM</td><td>alignment</td><td>distance</td><td>position</td><td>gene name</td><td>gene id</td></tr>
<tr><td>GU305914.1</td><td>199374</td><td>199396</td><td>-</td><td>0</td><td>CTCTCGGGTGTCCTAGGCAG</td><td>AGG</td><td>||||||||[||||||||||||]PAM</td><td>NA</td><td>Intergenic</td><td>NA</td><td>NA</td></tr>
<tr><td>T19461</td><td>GGGTGTCCTAGGCAGAGGAGAGG</td><td>1000</td><td></td><td></td><td></td><td></td><td></td><td></td><td></td><td></td><td></td></tr>
<tr><td>Oligo fwd</td><td>CACCGGGTGTCCTAGGCAGAGGAG</td><td></td><td></td><td></td><td></td><td></td><td></td><td></td><td></td><td></td><td></td></tr>
<tr><td>Oligo rev</td><td>AAACCTCCTCTGCCTAGGACACCC</td><td></td><td></td><td></td><td></td><td></td><td></td><td></td><td></td><td></td><td></td></tr>
<tr><td>Chromosome</td><td>start</td><td>end</td><td>strand</td><td>MM</td><td>target_seq</td><td>PAM</td><td>alignment</td><td>distance</td><td>position</td><td>gene name</td><td>gene id</td></tr>
<tr><td>GU305914.1</td><td>199369</td><td>199391</td><td>-</td><td>0</td><td>GGGTGTCCTAGGCAGAGGAG</td><td>AGG</td><td>||||||||[||||||||||||]PAM</td><td>NA</td><td>Intergenic</td><td>NA</td><td>NA</td></tr>
<tr><td>T19462</td><td>GTCCTAGGCAGAGGAGAGGAAGG</td><td>1000</td><td></td><td></td><td></td><td></td><td></td><td></td><td></td><td></td><td></td></tr>
<tr><td>Oligo fwd</td><td>CACCGTCCTAGGCAGAGGAGAGGA</td><td></td><td></td><td></td><td></td><td></td><td></td><td></td><td></td><td></td><td></td></tr>
<tr><td>Oligo rev</td><td>AAACTCCTCTCCTCTGCCTAGGAC</td><td></td><td></td><td></td><td></td><td></td><td></td><td></td><td></td><td></td><td></td></tr>
<tr><td>Chromosome</td><td>start</td><td>end</td><td>strand</td><td>MM</td><td>target_seq</td><td>PAM</td><td>alignment</td><td>distance</td><td>position</td><td>gene name</td><td>gene id</td></tr>
<tr><td>GU305914.1</td><td>199365</td><td>199387</td><td>-</td><td>0</td><td>GTCCTAGGCAGAGGAGAGGA</td><td>AGG</td><td>||||||||[||||||||||||]PAM</td><td>NA</td><td>Intergenic</td><td>NA</td><td>NA</td></tr>
<tr><td>T19463</td><td>TCCTAGGCAGAGGAGAGGAAGGG</td><td>1000</td><td></td><td></td><td></td><td></td><td></td><td></td><td></td><td></td><td></td></tr>
<tr><td>Oligo fwd</td><td>CACCTCCTAGGCAGAGGAGAGGAA</td><td></td><td></td><td></td><td></td><td></td><td></td><td></td><td></td><td></td><td></td></tr>
<tr><td>Oligo rev</td><td>AAACTTCCTCTCCTCTGCCTAGGA</td><td></td><td></td><td></td><td></td><td></td><td></td><td></td><td></td><td></td><td></td></tr>
<tr><td>Chromosome</td><td>start</td><td>end</td><td>strand</td><td>MM</td><td>target_seq</td><td>PAM</td><td>alignment</td><td>distance</td><td>position</td><td>gene name</td><td>gene id</td></tr>
<tr><td>GU305914.1</td><td>199364</td><td>199386</td><td>-</td><td>0</td><td>TCCTAGGCAGAGGAGAGGAA</td><td>GGG</td><td>||||||||[||||||||||||]PAM</td><td>NA</td><td>Intergenic</td><td>NA</td><td>NA</td></tr>
<tr><td>T19464</td><td>TAGGCAGAGGAGAGGAAGGGTGG</td><td>1000</td><td></td><td></td><td></td><td></td><td></td><td></td><td></td><td></td><td></td></tr>
<tr><td>Oligo fwd</td><td>CACCTAGGCAGAGGAGAGGAAGGG</td><td></td><td></td><td></td><td></td><td></td><td></td><td></td><td></td><td></td><td></td></tr>
<tr><td>Oligo rev</td><td>AAACCCCTTCCTCTCCTCTGCCTA</td><td></td><td></td><td></td><td></td><td></td><td></td><td></td><td></td><td></td><td></td></tr>
<tr><td>Chromosome</td><td>start</td><td>end</td><td>strand</td><td>MM</td><td>target_seq</td><td>PAM</td><td>alignment</td><td>distance</td><td>position</td><td>gene name</td><td>gene id</td></tr>
<tr><td>GU305914.1</td><td>199361</td><td>199383</td><td>-</td><td>0</td><td>TAGGCAGAGGAGAGGAAGGG</td><td>TGG</td><td>||||||||[||||||||||||]PAM</td><td>NA</td><td>Intergenic</td><td>NA</td><td>NA</td></tr>
<tr><td>T19465</td><td>CAGAGGAGAGGAAGGGTGGTTGG</td><td>1000</td><td></td><td></td><td></td><td></td><td></td><td></td><td></td><td></td><td></td></tr>
<tr><td>Oligo fwd</td><td>CACCCAGAGGAGAGGAAGGGTGGT</td><td></td><td></td><td></td><td></td><td></td><td></td><td></td><td></td><td></td><td></td></tr>
<tr><td>Oligo rev</td><td>AAACACCACCCTTCCTCTCCTCTG</td><td></td><td></td><td></td><td></td><td></td><td></td><td></td><td></td><td></td><td></td></tr>
<tr><td>Chromosome</td><td>start</td><td>end</td><td>strand</td><td>MM</td><td>target_seq</td><td>PAM</td><td>alignment</td><td>distance</td><td>position</td><td>gene name</td><td>gene id</td></tr>
<tr><td>GU305914.1</td><td>199357</td><td>199379</td><td>-</td><td>0</td><td>CAGAGGAGAGGAAGGGTGGT</td><td>TGG</td><td>||||||||[||||||||||||]PAM</td><td>NA</td><td>Intergenic</td><td>NA</td><td>NA</td></tr>
<tr><td>T19466</td><td>TGGTTGGTGTGTGTGTGCGCCGG</td><td>1000</td><td></td><td></td><td></td><td></td><td></td><td></td><td></td><td></td><td></td></tr>
<tr><td>Oligo fwd</td><td>CACCTGGTTGGTGTGTGTGTGCGC</td><td></td><td></td><td></td><td></td><td></td><td></td><td></td><td></td><td></td><td></td></tr>
<tr><td>Oligo rev</td><td>AAACGCGCACACACACACCAACCA</td><td></td><td></td><td></td><td></td><td></td><td></td><td></td><td></td><td></td><td></td></tr>
<tr><td>Chromosome</td><td>start</td><td>end</td><td>strand</td><td>MM</td><td>target_seq</td><td>PAM</td><td>alignment</td><td>distance</td><td>position</td><td>gene name</td><td>gene id</td></tr>
<tr><td>GU305914.1</td><td>199341</td><td>199363</td><td>-</td><td>0</td><td>TGGTTGGTGTGTGTGTGCGC</td><td>CGG</td><td>||||||||[||||||||||||]PAM</td><td>NA</td><td>Intergenic</td><td>NA</td><td>NA</td></tr>
</table>

T19467 GGTTGGTGTGTGTGTGCGCCGGG 1000
Oligo fwd CACCGGTTGGTGTGTGTGTGCGCC
Oligo rev AAACGGCGCACACACACACCAACC
Chromosome start end strand MM target_seq PAM alignment distance position gene name gene id
GU305914.1 199340 199362 - 0 GGTTGGTGTGTGTGTGCGCC GGG ||||||||[||||||||||||]PAM NA Intergenic NA NA

T19468 GTTGGTGTGTGTGTGCGCCGGGG 1000
Oligo fwd CACCGTTGGTGTGTGTGTGCGCCG
Oligo rev AAACCGGCGCACACACACACCAAC
Chromosome start end strand MM target_seq PAM alignment distance position gene name gene id
GU305914.1 199339 199361 - 0 GTTGGTGTGTGTGTGCGCCG GGG ||||||||[||||||||||||]PAM NA Intergenic NA NA
GU305914.1 1179 1201 + 4 GTTGTTGCTCGTGTGCGCCG CGG ||||-||-[--||||||||||]PAM NA Intergenic NA NA

T19469 GGTGTGTGTGTGCGCCGGGGTGG 1000
Oligo fwd CACCGGTGTGTGTGTGCGCCGGGG
Oligo rev AAACCCCCGGCGCACACACACACC
Chromosome start end strand MM target_seq PAM alignment distance position gene name gene id
GU305914.1 199336 199358 - 0 GGTGTGTGTGTGCGCCGGGG TGG ||||||||[||||||||||||]PAM NA Intergenic NA NA

T19470 TGTGCGCCGGGGTGGCGAGATGG 1000
Oligo fwd CACCTGTGCGCCGGGGTGGCGAGA
Oligo rev AAACTCTCGCCACCCCGGCGCACA
Chromosome start end strand MM target_seq PAM alignment distance position gene name gene id
GU305914.1 199328 199350 - 0 TGTGCGCCGGGGTGGCGAGA TGG ||||||||[||||||||||||]PAM NA Intergenic NA NA

T19471 GCCGGGGTGGCGAGATGGAGAGG 1000
Oligo fwd CACCGCCGGGGTGGCGAGATGGAG
Oligo rev AAACCTCCATCTCGCCACCCCGGC
Chromosome start end strand MM target_seq PAM alignment distance position gene name gene id
GU305914.1 199323 199345 - 0 GCCGGGGTGGCGAGATGGAG AGG ||||||||[||||||||||||]PAM NA Intergenic NA NA

T19472 GGGTGGCGAGATGGAGAGGCCGG 1000
Oligo fwd CACCGGGTGGCGAGATGGAGAGGC
Oligo rev AAACGCCTCTCCATCTCGCCACCC
Chromosome start end strand MM target_seq PAM alignment distance position gene name gene id
GU305914.1 199319 199341 - 0 GGGTGGCGAGATGGAGAGGC CGG ||||||||[||||||||||||]PAM NA Intergenic NA NA
GU305914.1 152275 152297 - 4 GGCGTGCGAGATGAAGAGGC GGG ||---|||[|||||-||||||]PAM NA Intergenic NA NA

T19473 CGAGATGGAGAGGCCGGTGACGG 1000
Oligo fwd CACCCGAGATGGAGAGGCCGGTGA
Oligo rev AAACTCACCGGCCTCTCCATCTCG
Chromosome start end strand MM target_seq PAM alignment distance position gene name gene id
GU305914.1 199313 199335 - 0 CGAGATGGAGAGGCCGGTGA CGG ||||||||[||||||||||||]PAM NA Intergenic NA NA

T19474 ACGCCATCTCCGCGTACAAGCGG 1000
Oligo fwd CACCACGCCATCTCCGCGTACAAG
Oligo rev AAACCTTGTACGCGGAGATGGCGT
Chromosome start end strand MM target_seq PAM alignment distance position gene name gene id
GU305914.1 199287 199309 - 0 ACGCCATCTCCGCGTACAAG CGG ||||||||[||||||||||||]PAM NA Intergenic NA NA

T19475 CGCCATCTCCGCGTACAAGCGGG 1000
Oligo fwd CACCCGCCATCTCCGCGTACAAGC
Oligo rev AAACGCTTGTACGCGGAGATGGCG
Chromosome start end strand MM target_seq PAM alignment distance position gene name gene id
GU305914.1 199286 199308 - 0 CGCCATCTCCGCGTACAAGC GGG ||||||||[||||||||||||]PAM NA Intergenic NA NA

T19476 GGGTGATAGACGCCTTCTCGCGG 1000
Oligo fwd CACCGGGTGATAGACGCCTTCTCG
Oligo rev AAACCGAGAAGGCGTCTATCACCC
Chromosome start end strand MM target_seq PAM alignment distance position gene name gene id
GU305914.1 199266 199288 - 0 GGGTGATAGACGCCTTCTCG CGG ||||||||[||||||||||||]PAM NA Intergenic NA NA

T19477 GATAGACGCCTTCTCGCGGCTGG 1000
Oligo fwd CACCGATAGACGCCTTCTCGCGGC
Oligo rev AAACGCCGCGAGAAGGCGTCTATC
Chromosome start end strand MM target_seq PAM alignment distance position gene name gene id
GU305914.1 199262 199284 - 0 GATAGACGCCTTCTCGCGGC TGG ||||||||[||||||||||||]PAM NA Intergenic NA NA
GU305914.1 197901 197923 + 4 CATAGACTCCGTCTCGTGGC AGG -||||||-[||-|||||-|||]PAM NA Intergenic NA NA

T19478 AGACGCCTTCTCGCGGCTGGTGG 1000
Oligo fwd CACCAGACGCCTTCTCGCGGCTGG
Oligo rev AAACCCAGCCGCGAGAAGGCGTCT
Chromosome start end strand MM target_seq PAM alignment distance position gene name gene id
GU305914.1 199259 199281 - 0 AGACGCCTTCTCGCGGCTGG TGG ||||||||[||||||||||||]PAM NA Intergenic NA NA

T19479 CTTCTCGCGGCTGGTGGCCACGG 1000
Oligo fwd CACCCTTCTCGCGGCTGGTGGCCA
Oligo rev AAACTGGCCACCAGCCGCGAGAAG
Chromosome start end strand MM target_seq PAM alignment distance position gene name gene id
GU305914.1 199253 199275 - 0 CTTCTCGCGGCTGGTGGCCA CGG ||||||||[||||||||||||]PAM NA Intergenic NA NA

T19480 TTCTCGCGGCTGGTGGCCACGGG 1000
Oligo fwd CACCTTCTCGCGGCTGGTGGCCAC
Oligo rev AAACGTGGCCACCAGCCGCGAGAA
Chromosome start end strand MM target_seq PAM alignment distance position gene name gene id
GU305914.1 199252 199274 - 0 TTCTCGCGGCTGGTGGCCAC GGG ||||||||[||||||||||||]PAM NA Intergenic NA NA

T19481 TCGCGGCTGGTGGCCACGGGCGG 1000

<table>
<tr><td>Oligo fwd</td><td>CACCTCGCGGCTGGTGGCCACGGG</td><td></td><td></td><td></td><td></td><td></td><td></td><td></td><td></td><td></td><td></td></tr>
<tr><td>Oligo rev</td><td>AAACCCCGTGGCCACCAGCCGCGA</td><td></td><td></td><td></td><td></td><td></td><td></td><td></td><td></td><td></td><td></td></tr>
<tr><td>Chromosome</td><td>start</td><td>end</td><td>strand</td><td>MM</td><td>target_seq</td><td>PAM</td><td>alignment</td><td>distance</td><td>position</td><td>gene name</td><td>gene id</td></tr>
<tr><td>GU305914.1</td><td>199249</td><td>199271</td><td>-</td><td>0</td><td>TCGCGGCTGGTGGCCACGGG</td><td>CGG</td><td>||||||||[||||||||||||]PAM</td><td>NA</td><td>Intergenic</td><td>NA</td><td>NA</td></tr>
<tr><td>T19482</td><td>CGGGCGGCGATCAGAACGCCCGG</td><td>1000</td><td></td><td></td><td></td><td></td><td></td><td></td><td></td><td></td><td></td></tr>
<tr><td>Oligo fwd</td><td>CACCCGGGCGGCGATCAGAACGCC</td><td></td><td></td><td></td><td></td><td></td><td></td><td></td><td></td><td></td><td></td></tr>
<tr><td>Oligo rev</td><td>AAACGGCGTTCTGATCGCCGCCCG</td><td></td><td></td><td></td><td></td><td></td><td></td><td></td><td></td><td></td><td></td></tr>
<tr><td>Chromosome</td><td>start</td><td>end</td><td>strand</td><td>MM</td><td>target_seq</td><td>PAM</td><td>alignment</td><td>distance</td><td>position</td><td>gene name</td><td>gene id</td></tr>
<tr><td>GU305914.1</td><td>199233</td><td>199255</td><td>-</td><td>0</td><td>CGGGCGGCGATCAGAACGCC</td><td>CGG</td><td>||||||||[||||||||||||]PAM</td><td>NA</td><td>Intergenic</td><td>NA</td><td>NA</td></tr>
<tr><td>T19483</td><td>GCGGCGATCAGAACGCCCGGCGG</td><td>1000</td><td></td><td></td><td></td><td></td><td></td><td></td><td></td><td></td><td></td></tr>
<tr><td>Oligo fwd</td><td>CACCGCGGCGATCAGAACGCCCGG</td><td></td><td></td><td></td><td></td><td></td><td></td><td></td><td></td><td></td><td></td></tr>
<tr><td>Oligo rev</td><td>AAACCCGGGCGTTCTGATCGCCGC</td><td></td><td></td><td></td><td></td><td></td><td></td><td></td><td></td><td></td><td></td></tr>
<tr><td>Chromosome</td><td>start</td><td>end</td><td>strand</td><td>MM</td><td>target_seq</td><td>PAM</td><td>alignment</td><td>distance</td><td>position</td><td>gene name</td><td>gene id</td></tr>
<tr><td>GU305914.1</td><td>199230</td><td>199252</td><td>-</td><td>0</td><td>GCGGCGATCAGAACGCCCGG</td><td>CGG</td><td>||||||||[||||||||||||]PAM</td><td>NA</td><td>Intergenic</td><td>NA</td><td>NA</td></tr>
<tr><td>T19484</td><td>CCCGGCGGCGCGCGATCGAGAGG</td><td>1000</td><td></td><td></td><td></td><td></td><td></td><td></td><td></td><td></td><td></td></tr>
<tr><td>Oligo fwd</td><td>CACCCCCGGCGGCGCGCGATCGAG</td><td></td><td></td><td></td><td></td><td></td><td></td><td></td><td></td><td></td><td></td></tr>
<tr><td>Oligo rev</td><td>AAACCTCGATCGCGCGCCGCCGGG</td><td></td><td></td><td></td><td></td><td></td><td></td><td></td><td></td><td></td><td></td></tr>
<tr><td>Chromosome</td><td>start</td><td>end</td><td>strand</td><td>MM</td><td>target_seq</td><td>PAM</td><td>alignment</td><td>distance</td><td>position</td><td>gene name</td><td>gene id</td></tr>
<tr><td>GU305914.1</td><td>199215</td><td>199237</td><td>-</td><td>0</td><td>CCCGGCGGCGCGCGATCGAG</td><td>AGG</td><td>||||||||[||||||||||||]PAM</td><td>NA</td><td>Intergenic</td><td>NA</td><td>NA</td></tr>
<tr><td>T19485</td><td>GCGGCGCGCGATCGAGAGGCTGG</td><td>1000</td><td></td><td></td><td></td><td></td><td></td><td></td><td></td><td></td><td></td></tr>
<tr><td>Oligo fwd</td><td>CACCGCGGCGCGCGATCGAGAGGC</td><td></td><td></td><td></td><td></td><td></td><td></td><td></td><td></td><td></td><td></td></tr>
<tr><td>Oligo rev</td><td>AAACGCCTCTCGATCGCGCGCCGC</td><td></td><td></td><td></td><td></td><td></td><td></td><td></td><td></td><td></td><td></td></tr>
<tr><td>Chromosome</td><td>start</td><td>end</td><td>strand</td><td>MM</td><td>target_seq</td><td>PAM</td><td>alignment</td><td>distance</td><td>position</td><td>gene name</td><td>gene id</td></tr>
<tr><td>GU305914.1</td><td>199211</td><td>199233</td><td>-</td><td>0</td><td>GCGGCGCGCGATCGAGAGGC</td><td>TGG</td><td>||||||||[||||||||||||]PAM</td><td>NA</td><td>Intergenic</td><td>NA</td><td>NA</td></tr>
<tr><td>T19486</td><td>CGCGCGATCGAGAGGCTGGACGG</td><td>1000</td><td></td><td></td><td></td><td></td><td></td><td></td><td></td><td></td><td></td></tr>
<tr><td>Oligo fwd</td><td>CACCCGCGCGATCGAGAGGCTGGA</td><td></td><td></td><td></td><td></td><td></td><td></td><td></td><td></td><td></td><td></td></tr>
<tr><td>Oligo rev</td><td>AAACTCCAGCCTCTCGATCGCGCG</td><td></td><td></td><td></td><td></td><td></td><td></td><td></td><td></td><td></td><td></td></tr>
<tr><td>Chromosome</td><td>start</td><td>end</td><td>strand</td><td>MM</td><td>target_seq</td><td>PAM</td><td>alignment</td><td>distance</td><td>position</td><td>gene name</td><td>gene id</td></tr>
<tr><td>GU305914.1</td><td>199207</td><td>199229</td><td>-</td><td>0</td><td>CGCGCGATCGAGAGGCTGGA</td><td>CGG</td><td>||||||||[||||||||||||]PAM</td><td>NA</td><td>Intergenic</td><td>NA</td><td>NA</td></tr>
<tr><td>T19487</td><td>GCGCGATCGAGAGGCTGGACGGG</td><td>1000</td><td></td><td></td><td></td><td></td><td></td><td></td><td></td><td></td><td></td></tr>
<tr><td>Oligo fwd</td><td>CACCGCGCGATCGAGAGGCTGGAC</td><td></td><td></td><td></td><td></td><td></td><td></td><td></td><td></td><td></td><td></td></tr>
<tr><td>Oligo rev</td><td>AAACGTCCAGCCTCTCGATCGCGC</td><td></td><td></td><td></td><td></td><td></td><td></td><td></td><td></td><td></td><td></td></tr>
<tr><td>Chromosome</td><td>start</td><td>end</td><td>strand</td><td>MM</td><td>target_seq</td><td>PAM</td><td>alignment</td><td>distance</td><td>position</td><td>gene name</td><td>gene id</td></tr>
<tr><td>GU305914.1</td><td>199206</td><td>199228</td><td>-</td><td>0</td><td>GCGCGATCGAGAGGCTGGAC</td><td>GGG</td><td>||||||||[||||||||||||]PAM</td><td>NA</td><td>Intergenic</td><td>NA</td><td>NA</td></tr>
<tr><td>T19488</td><td>GATCGAGAGGCTGGACGGGAAGG</td><td>1000</td><td></td><td></td><td></td><td></td><td></td><td></td><td></td><td></td><td></td></tr>
<tr><td>Oligo fwd</td><td>CACCGATCGAGAGGCTGGACGGGA</td><td></td><td></td><td></td><td></td><td></td><td></td><td></td><td></td><td></td><td></td></tr>
<tr><td>Oligo rev</td><td>AAACTCCCGTCCAGCCTCTCGATC</td><td></td><td></td><td></td><td></td><td></td><td></td><td></td><td></td><td></td><td></td></tr>
<tr><td>Chromosome</td><td>start</td><td>end</td><td>strand</td><td>MM</td><td>target_seq</td><td>PAM</td><td>alignment</td><td>distance</td><td>position</td><td>gene name</td><td>gene id</td></tr>
<tr><td>GU305914.1</td><td>199202</td><td>199224</td><td>-</td><td>0</td><td>GATCGAGAGGCTGGACGGGA</td><td>AGG</td><td>||||||||[||||||||||||]PAM</td><td>NA</td><td>Intergenic</td><td>NA</td><td>NA</td></tr>
<tr><td>T19489</td><td>GGACGGGAAGGTGATCCCCCTGG</td><td>1000</td><td></td><td></td><td></td><td></td><td></td><td></td><td></td><td></td><td></td></tr>
<tr><td>Oligo fwd</td><td>CACCGGACGGGAAGGTGATCCCCC</td><td></td><td></td><td></td><td></td><td></td><td></td><td></td><td></td><td></td><td></td></tr>
<tr><td>Oligo rev</td><td>AAACGGGGGATCACCTTCCCGTCC</td><td></td><td></td><td></td><td></td><td></td><td></td><td></td><td></td><td></td><td></td></tr>
<tr><td>Chromosome</td><td>start</td><td>end</td><td>strand</td><td>MM</td><td>target_seq</td><td>PAM</td><td>alignment</td><td>distance</td><td>position</td><td>gene name</td><td>gene id</td></tr>
<tr><td>GU305914.1</td><td>199190</td><td>199212</td><td>-</td><td>0</td><td>GGACGGGAAGGTGATCCCCC</td><td>TGG</td><td>||||||||[||||||||||||]PAM</td><td>NA</td><td>Intergenic</td><td>NA</td><td>NA</td></tr>
<tr><td>T19490</td><td>GACGGGAAGGTGATCCCCCTGGG</td><td>1000</td><td></td><td></td><td></td><td></td><td></td><td></td><td></td><td></td><td></td></tr>
<tr><td>Oligo fwd</td><td>CACCGACGGGAAGGTGATCCCCCT</td><td></td><td></td><td></td><td></td><td></td><td></td><td></td><td></td><td></td><td></td></tr>
<tr><td>Oligo rev</td><td>AAACAGGGGGATCACCTTCCCGTC</td><td></td><td></td><td></td><td></td><td></td><td></td><td></td><td></td><td></td><td></td></tr>
<tr><td>Chromosome</td><td>start</td><td>end</td><td>strand</td><td>MM</td><td>target_seq</td><td>PAM</td><td>alignment</td><td>distance</td><td>position</td><td>gene name</td><td>gene id</td></tr>
<tr><td>GU305914.1</td><td>199189</td><td>199211</td><td>-</td><td>0</td><td>GACGGGAAGGTGATCCCCCT</td><td>GGG</td><td>||||||||[||||||||||||]PAM</td><td>NA</td><td>Intergenic</td><td>NA</td><td>NA</td></tr>
<tr><td>T19491</td><td>ACGGGAAGGTGATCCCCCTGGGG</td><td>1000</td><td></td><td></td><td></td><td></td><td></td><td></td><td></td><td></td><td></td></tr>
<tr><td>Oligo fwd</td><td>CACCACGGGAAGGTGATCCCCCTG</td><td></td><td></td><td></td><td></td><td></td><td></td><td></td><td></td><td></td><td></td></tr>
<tr><td>Oligo rev</td><td>AAACCAGGGGGATCACCTTCCCGT</td><td></td><td></td><td></td><td></td><td></td><td></td><td></td><td></td><td></td><td></td></tr>
<tr><td>Chromosome</td><td>start</td><td>end</td><td>strand</td><td>MM</td><td>target_seq</td><td>PAM</td><td>alignment</td><td>distance</td><td>position</td><td>gene name</td><td>gene id</td></tr>
<tr><td>GU305914.1</td><td>199188</td><td>199210</td><td>-</td><td>0</td><td>ACGGGAAGGTGATCCCCCTG</td><td>GGG</td><td>||||||||[||||||||||||]PAM</td><td>NA</td><td>Intergenic</td><td>NA</td><td>NA</td></tr>
<tr><td>T19492</td><td>GATCCCCCTGGGGTTTCCGCAGG</td><td>1000</td><td></td><td></td><td></td><td></td><td></td><td></td><td></td><td></td><td></td></tr>
<tr><td>Oligo fwd</td><td>CACCGATCCCCCTGGGGTTTCCGC</td><td></td><td></td><td></td><td></td><td></td><td></td><td></td><td></td><td></td><td></td></tr>
<tr><td>Oligo rev</td><td>AAACGCGGAAACCCCAGGGGGATC</td><td></td><td></td><td></td><td></td><td></td><td></td><td></td><td></td><td></td><td></td></tr>
<tr><td>Chromosome</td><td>start</td><td>end</td><td>strand</td><td>MM</td><td>target_seq</td><td>PAM</td><td>alignment</td><td>distance</td><td>position</td><td>gene name</td><td>gene id</td></tr>
<tr><td>GU305914.1</td><td>199178</td><td>199200</td><td>-</td><td>0</td><td>GATCCCCCTGGGGTTTCCGC</td><td>AGG</td><td>||||||||[||||||||||||]PAM</td><td>NA</td><td>Intergenic</td><td>NA</td><td>NA</td></tr>
<tr><td>T19493</td><td>CCCTGGGGTTTCCGCAGGACTGG</td><td>1000</td><td></td><td></td><td></td><td></td><td></td><td></td><td></td><td></td><td></td></tr>
<tr><td>Oligo fwd</td><td>CACCCCCTGGGGTTTCCGCAGGAC</td><td></td><td></td><td></td><td></td><td></td><td></td><td></td><td></td><td></td><td></td></tr>
<tr><td>Oligo rev</td><td>AAACGTCCTGCGGAAACCCCAGGG</td><td></td><td></td><td></td><td></td><td></td><td></td><td></td><td></td><td></td><td></td></tr>
<tr><td>Chromosome</td><td>start</td><td>end</td><td>strand</td><td>MM</td><td>target_seq</td><td>PAM</td><td>alignment</td><td>distance</td><td>position</td><td>gene name</td><td>gene id</td></tr>
<tr><td>GU305914.1</td><td>199173</td><td>199195</td><td>-</td><td>0</td><td>CCCTGGGGTTTCCGCAGGAC</td><td>TGG</td><td>||||||||[||||||||||||]PAM</td><td>NA</td><td>Intergenic</td><td>NA</td><td>NA</td></tr>
<tr><td>T19494</td><td>GTTTCCGCAGGACTGGTACGTGG</td><td>1000</td><td></td><td></td><td></td><td></td><td></td><td></td><td></td><td></td><td></td></tr>
<tr><td>Oligo fwd</td><td>CACCGTTTCCGCAGGACTGGTACG</td><td></td><td></td><td></td><td></td><td></td><td></td><td></td><td></td><td></td><td></td></tr>
<tr><td>Oligo rev</td><td>AAACCGTACCAGTCCTGCGGAAAC</td><td></td><td></td><td></td><td></td><td></td><td></td><td></td><td></td><td></td><td></td></tr>
<tr><td>Chromosome</td><td>start</td><td>end</td><td>strand</td><td>MM</td><td>target_seq</td><td>PAM</td><td>alignment</td><td>distance</td><td>position</td><td>gene name</td><td>gene id</td></tr>
<tr><td>GU305914.1</td><td>199166</td><td>199188</td><td>-</td><td>0</td><td>GTTTCCGCAGGACTGGTACG</td><td>TGG</td><td>||||||||[||||||||||||]PAM</td><td>NA</td><td>Intergenic</td><td>NA</td><td>NA</td></tr>
<tr><td>T19495</td><td>CAGGACTGGTACGTGGTGTGCGG</td><td>1000</td><td></td><td></td><td></td><td></td><td></td><td></td><td></td><td></td><td></td></tr>
<tr><td>Oligo fwd</td><td>CACCCAGGACTGGTACGTGGTGTG</td><td></td><td></td><td></td><td></td><td></td><td></td><td></td><td></td><td></td><td></td></tr>
<tr><td>Oligo rev</td><td>AAACCACACCACGTACCAGTCCTG</td><td></td><td></td><td></td><td></td><td></td><td></td><td></td><td></td><td></td><td></td></tr>
<tr><td>Chromosome</td><td>start</td><td>end</td><td>strand</td><td>MM</td><td>target_seq</td><td>PAM</td><td>alignment</td><td>distance</td><td>position</td><td>gene name</td><td>gene id</td></tr>
<tr><td>GU305914.1</td><td>199159</td><td>199181</td><td>-</td><td>0</td><td>CAGGACTGGTACGTGGTGTG</td><td>CGG</td><td>||||||||[||||||||||||]PAM</td><td>NA</td><td>Intergenic</td><td>NA</td><td>NA</td></tr>
</table>

<table>
<tr><td>T19496</td><td>CTGGTACGTGGTGTGCGGATCGG</td><td>1000</td><td></td><td></td><td></td><td></td><td></td><td></td><td></td><td></td><td></td></tr>
<tr><td>Oligo fwd</td><td>CACCCTGGTACGTGGTGTGCGGAT</td><td></td><td></td><td></td><td></td><td></td><td></td><td></td><td></td><td></td><td></td></tr>
<tr><td>Oligo rev</td><td>AAACATCCGCACACCACGTACCAG</td><td></td><td></td><td></td><td></td><td></td><td></td><td></td><td></td><td></td><td></td></tr>
<tr><td>Chromosome</td><td>start</td><td>end</td><td>strand</td><td>MM</td><td>target_seq</td><td>PAM</td><td>alignment</td><td>distance</td><td>position</td><td>gene name</td><td>gene id</td></tr>
<tr><td>GU305914.1</td><td>199154</td><td>199176</td><td>-</td><td>0</td><td>CTGGTACGTGGTGTGCGGAT</td><td>CGG</td><td>||||||||[||||||||||||]PAM</td><td>NA</td><td>Intergenic</td><td>NA</td><td>NA</td></tr>
<tr><td>T19497</td><td>TGGTACGTGGTGTGCGGATCGGG</td><td>1000</td><td></td><td></td><td></td><td></td><td></td><td></td><td></td><td></td><td></td></tr>
<tr><td>Oligo fwd</td><td>CACCTGGTACGTGGTGTGCGGATC</td><td></td><td></td><td></td><td></td><td></td><td></td><td></td><td></td><td></td><td></td></tr>
<tr><td>Oligo rev</td><td>AAACGATCCGCACACCACGTACCA</td><td></td><td></td><td></td><td></td><td></td><td></td><td></td><td></td><td></td><td></td></tr>
<tr><td>Chromosome</td><td>start</td><td>end</td><td>strand</td><td>MM</td><td>target_seq</td><td>PAM</td><td>alignment</td><td>distance</td><td>position</td><td>gene name</td><td>gene id</td></tr>
<tr><td>GU305914.1</td><td>199153</td><td>199175</td><td>-</td><td>0</td><td>TGGTACGTGGTGTGCGGATC</td><td>GGG</td><td>||||||||[||||||||||||]PAM</td><td>NA</td><td>Intergenic</td><td>NA</td><td>NA</td></tr>
<tr><td>T19498</td><td>GCGGATCGGGCAAGAGTTGCTGG</td><td>1000</td><td></td><td></td><td></td><td></td><td></td><td></td><td></td><td></td><td></td></tr>
<tr><td>Oligo fwd</td><td>CACCGCGGATCGGGCAAGAGTTGC</td><td></td><td></td><td></td><td></td><td></td><td></td><td></td><td></td><td></td><td></td></tr>
<tr><td>Oligo rev</td><td>AAACGCAACTCTTGCCCGATCCGC</td><td></td><td></td><td></td><td></td><td></td><td></td><td></td><td></td><td></td><td></td></tr>
<tr><td>Chromosome</td><td>start</td><td>end</td><td>strand</td><td>MM</td><td>target_seq</td><td>PAM</td><td>alignment</td><td>distance</td><td>position</td><td>gene name</td><td>gene id</td></tr>
<tr><td>GU305914.1</td><td>199140</td><td>199162</td><td>-</td><td>0</td><td>GCGGATCGGGCAAGAGTTGC</td><td>TGG</td><td>||||||||[||||||||||||]PAM</td><td>NA</td><td>Intergenic</td><td>NA</td><td>NA</td></tr>
<tr><td>T19499</td><td>GAGTTGCTGGTCCGTTCTGACGG</td><td>1000</td><td></td><td></td><td></td><td></td><td></td><td></td><td></td><td></td><td></td></tr>
<tr><td>Oligo fwd</td><td>CACCGAGTTGCTGGTCCGTTCTGA</td><td></td><td></td><td></td><td></td><td></td><td></td><td></td><td></td><td></td><td></td></tr>
<tr><td>Oligo rev</td><td>AAACTCAGAACGGACCAGCAACTC</td><td></td><td></td><td></td><td></td><td></td><td></td><td></td><td></td><td></td><td></td></tr>
<tr><td>Chromosome</td><td>start</td><td>end</td><td>strand</td><td>MM</td><td>target_seq</td><td>PAM</td><td>alignment</td><td>distance</td><td>position</td><td>gene name</td><td>gene id</td></tr>
<tr><td>GU305914.1</td><td>199127</td><td>199149</td><td>-</td><td>0</td><td>GAGTTGCTGGTCCGTTCTGA</td><td>CGG</td><td>||||||||[||||||||||||]PAM</td><td>NA</td><td>Intergenic</td><td>NA</td><td>NA</td></tr>
<tr><td>T19500</td><td>GCTGGTCCGTTCTGACGGACAGG</td><td>1000</td><td></td><td></td><td></td><td></td><td></td><td></td><td></td><td></td><td></td></tr>
<tr><td>Oligo fwd</td><td>CACCGCTGGTCCGTTCTGACGGAC</td><td></td><td></td><td></td><td></td><td></td><td></td><td></td><td></td><td></td><td></td></tr>
<tr><td>Oligo rev</td><td>AAACGTCCGTCAGAACGGACCAGC</td><td></td><td></td><td></td><td></td><td></td><td></td><td></td><td></td><td></td><td></td></tr>
<tr><td>Chromosome</td><td>start</td><td>end</td><td>strand</td><td>MM</td><td>target_seq</td><td>PAM</td><td>alignment</td><td>distance</td><td>position</td><td>gene name</td><td>gene id</td></tr>
<tr><td>GU305914.1</td><td>199122</td><td>199144</td><td>-</td><td>0</td><td>GCTGGTCCGTTCTGACGGAC</td><td>AGG</td><td>||||||||[||||||||||||]PAM</td><td>NA</td><td>Intergenic</td><td>NA</td><td>NA</td></tr>
<tr><td>T19501</td><td>CTGGTCCGTTCTGACGGACAGGG</td><td>1000</td><td></td><td></td><td></td><td></td><td></td><td></td><td></td><td></td><td></td></tr>
<tr><td>Oligo fwd</td><td>CACCCTGGTCCGTTCTGACGGACA</td><td></td><td></td><td></td><td></td><td></td><td></td><td></td><td></td><td></td><td></td></tr>
<tr><td>Oligo rev</td><td>AAACTGTCCGTCAGAACGGACCAG</td><td></td><td></td><td></td><td></td><td></td><td></td><td></td><td></td><td></td><td></td></tr>
<tr><td>Chromosome</td><td>start</td><td>end</td><td>strand</td><td>MM</td><td>target_seq</td><td>PAM</td><td>alignment</td><td>distance</td><td>position</td><td>gene name</td><td>gene id</td></tr>
<tr><td>GU305914.1</td><td>199121</td><td>199143</td><td>-</td><td>0</td><td>CTGGTCCGTTCTGACGGACA</td><td>GGG</td><td>||||||||[||||||||||||]PAM</td><td>NA</td><td>Intergenic</td><td>NA</td><td>NA</td></tr>
<tr><td>T19502</td><td>CGCGCTCCTGTGTTGCGAAGAGG</td><td>1000</td><td></td><td></td><td></td><td></td><td></td><td></td><td></td><td></td><td></td></tr>
<tr><td>Oligo fwd</td><td>CACCCGCGCTCCTGTGTTGCGAAG</td><td></td><td></td><td></td><td></td><td></td><td></td><td></td><td></td><td></td><td></td></tr>
<tr><td>Oligo rev</td><td>AAACCTTCGCAACACAGGAGCGCG</td><td></td><td></td><td></td><td></td><td></td><td></td><td></td><td></td><td></td><td></td></tr>
<tr><td>Chromosome</td><td>start</td><td>end</td><td>strand</td><td>MM</td><td>target_seq</td><td>PAM</td><td>alignment</td><td>distance</td><td>position</td><td>gene name</td><td>gene id</td></tr>
<tr><td>GU305914.1</td><td>199094</td><td>199116</td><td>-</td><td>0</td><td>CGCGCTCCTGTGTTGCGAAG</td><td>AGG</td><td>||||||||[||||||||||||]PAM</td><td>NA</td><td>Intergenic</td><td>NA</td><td>NA</td></tr>
<tr><td>T19503</td><td>CGAAGAGGCCCTGATCCCGTTGG</td><td>1000</td><td></td><td></td><td></td><td></td><td></td><td></td><td></td><td></td><td></td></tr>
<tr><td>Oligo fwd</td><td>CACCCGAAGAGGCCCTGATCCCGT</td><td></td><td></td><td></td><td></td><td></td><td></td><td></td><td></td><td></td><td></td></tr>
<tr><td>Oligo rev</td><td>AAACACGGGATCAGGGCCTCTTCG</td><td></td><td></td><td></td><td></td><td></td><td></td><td></td><td></td><td></td><td></td></tr>
<tr><td>Chromosome</td><td>start</td><td>end</td><td>strand</td><td>MM</td><td>target_seq</td><td>PAM</td><td>alignment</td><td>distance</td><td>position</td><td>gene name</td><td>gene id</td></tr>
<tr><td>GU305914.1</td><td>199079</td><td>199101</td><td>-</td><td>0</td><td>CGAAGAGGCCCTGATCCCGT</td><td>TGG</td><td>||||||||[||||||||||||]PAM</td><td>NA</td><td>Intergenic</td><td>NA</td><td>NA</td></tr>
<tr><td>T19504</td><td>GAAGAGGCCCTGATCCCGTTGGG</td><td>1000</td><td></td><td></td><td></td><td></td><td></td><td></td><td></td><td></td><td></td></tr>
<tr><td>Oligo fwd</td><td>CACCGAAGAGGCCCTGATCCCGTT</td><td></td><td></td><td></td><td></td><td></td><td></td><td></td><td></td><td></td><td></td></tr>
<tr><td>Oligo rev</td><td>AAACAACGGGATCAGGGCCTCTTC</td><td></td><td></td><td></td><td></td><td></td><td></td><td></td><td></td><td></td><td></td></tr>
<tr><td>Chromosome</td><td>start</td><td>end</td><td>strand</td><td>MM</td><td>target_seq</td><td>PAM</td><td>alignment</td><td>distance</td><td>position</td><td>gene name</td><td>gene id</td></tr>
<tr><td>GU305914.1</td><td>199078</td><td>199100</td><td>-</td><td>0</td><td>GAAGAGGCCCTGATCCCGTT</td><td>GGG</td><td>||||||||[||||||||||||]PAM</td><td>NA</td><td>Intergenic</td><td>NA</td><td>NA</td></tr>
<tr><td>T19505</td><td>CTGATCCCGTTGGGTCTGTGCGG</td><td>1000</td><td></td><td></td><td></td><td></td><td></td><td></td><td></td><td></td><td></td></tr>
<tr><td>Oligo fwd</td><td>CACCCTGATCCCGTTGGGTCTGTG</td><td></td><td></td><td></td><td></td><td></td><td></td><td></td><td></td><td></td><td></td></tr>
<tr><td>Oligo rev</td><td>AAACCACAGACCCAACGGGATCAG</td><td></td><td></td><td></td><td></td><td></td><td></td><td></td><td></td><td></td><td></td></tr>
<tr><td>Chromosome</td><td>start</td><td>end</td><td>strand</td><td>MM</td><td>target_seq</td><td>PAM</td><td>alignment</td><td>distance</td><td>position</td><td>gene name</td><td>gene id</td></tr>
<tr><td>GU305914.1</td><td>199069</td><td>199091</td><td>-</td><td>0</td><td>CTGATCCCGTTGGGTCTGTG</td><td>CGG</td><td>||||||||[||||||||||||]PAM</td><td>NA</td><td>Intergenic</td><td>NA</td><td>NA</td></tr>
<tr><td>T19506</td><td>TCCCGTTGGGTCTGTGCGGACGG</td><td>1000</td><td></td><td></td><td></td><td></td><td></td><td></td><td></td><td></td><td></td></tr>
<tr><td>Oligo fwd</td><td>CACCTCCCGTTGGGTCTGTGCGGA</td><td></td><td></td><td></td><td></td><td></td><td></td><td></td><td></td><td></td><td></td></tr>
<tr><td>Oligo rev</td><td>AAACTCCGCACAGACCCAACGGGA</td><td></td><td></td><td></td><td></td><td></td><td></td><td></td><td></td><td></td><td></td></tr>
<tr><td>Chromosome</td><td>start</td><td>end</td><td>strand</td><td>MM</td><td>target_seq</td><td>PAM</td><td>alignment</td><td>distance</td><td>position</td><td>gene name</td><td>gene id</td></tr>
<tr><td>GU305914.1</td><td>199065</td><td>199087</td><td>-</td><td>0</td><td>TCCCGTTGGGTCTGTGCGGA</td><td>CGG</td><td>||||||||[||||||||||||]PAM</td><td>NA</td><td>Intergenic</td><td>NA</td><td>NA</td></tr>
<tr><td>T19507</td><td>TCCGACCCGTACGACACGCGAGG</td><td>1000</td><td></td><td></td><td></td><td></td><td></td><td></td><td></td><td></td><td></td></tr>
<tr><td>Oligo fwd</td><td>CACCTCCGACCCGTACGACACGCG</td><td></td><td></td><td></td><td></td><td></td><td></td><td></td><td></td><td></td><td></td></tr>
<tr><td>Oligo rev</td><td>AAACCGCGTGTCGTACGGGTCGGA</td><td></td><td></td><td></td><td></td><td></td><td></td><td></td><td></td><td></td><td></td></tr>
<tr><td>Chromosome</td><td>start</td><td>end</td><td>strand</td><td>MM</td><td>target_seq</td><td>PAM</td><td>alignment</td><td>distance</td><td>position</td><td>gene name</td><td>gene id</td></tr>
<tr><td>GU305914.1</td><td>199039</td><td>199061</td><td>-</td><td>0</td><td>TCCGACCCGTACGACACGCG</td><td>AGG</td><td>||||||||[||||||||||||]PAM</td><td>NA</td><td>Intergenic</td><td>NA</td><td>NA</td></tr>
<tr><td>T19508</td><td>CCGACCCGTACGACACGCGAGGG</td><td>1000</td><td></td><td></td><td></td><td></td><td></td><td></td><td></td><td></td><td></td></tr>
<tr><td>Oligo fwd</td><td>CACCCCGACCCGTACGACACGCGA</td><td></td><td></td><td></td><td></td><td></td><td></td><td></td><td></td><td></td><td></td></tr>
<tr><td>Oligo rev</td><td>AAACTCGCGTGTCGTACGGGTCGG</td><td></td><td></td><td></td><td></td><td></td><td></td><td></td><td></td><td></td><td></td></tr>
<tr><td>Chromosome</td><td>start</td><td>end</td><td>strand</td><td>MM</td><td>target_seq</td><td>PAM</td><td>alignment</td><td>distance</td><td>position</td><td>gene name</td><td>gene id</td></tr>
<tr><td>GU305914.1</td><td>199038</td><td>199060</td><td>-</td><td>0</td><td>CCGACCCGTACGACACGCGA</td><td>GGG</td><td>||||||||[||||||||||||]PAM</td><td>NA</td><td>Intergenic</td><td>NA</td><td>NA</td></tr>
<tr><td>T19509</td><td>TACGACACGCGAGGGTTGCGAGG</td><td>1000</td><td></td><td></td><td></td><td></td><td></td><td></td><td></td><td></td><td></td></tr>
<tr><td>Oligo fwd</td><td>CACCTACGACACGCGAGGGTTGCG</td><td></td><td></td><td></td><td></td><td></td><td></td><td></td><td></td><td></td><td></td></tr>
<tr><td>Oligo rev</td><td>AAACCGCAACCCTCGCGTGTCGTA</td><td></td><td></td><td></td><td></td><td></td><td></td><td></td><td></td><td></td><td></td></tr>
<tr><td>Chromosome</td><td>start</td><td>end</td><td>strand</td><td>MM</td><td>target_seq</td><td>PAM</td><td>alignment</td><td>distance</td><td>position</td><td>gene name</td><td>gene id</td></tr>
<tr><td>GU305914.1</td><td>199030</td><td>199052</td><td>-</td><td>0</td><td>TACGACACGCGAGGGTTGCG</td><td>AGG</td><td>||||||||[||||||||||||]PAM</td><td>NA</td><td>Intergenic</td><td>NA</td><td>NA</td></tr>
<tr><td>T19510</td><td>ACGCGAGGGTTGCGAGGATACGG</td><td>1000</td><td></td><td></td><td></td><td></td><td></td><td></td><td></td><td></td><td></td></tr>
<tr><td>Oligo fwd</td><td>CACCACGCGAGGGTTGCGAGGATA</td><td></td><td></td><td></td><td></td><td></td><td></td><td></td><td></td><td></td><td></td></tr>
<tr><td>Oligo rev</td><td>AAACTATCCTCGCAACCCTCGCGT</td><td></td><td></td><td></td><td></td><td></td><td></td><td></td><td></td><td></td><td></td></tr>
<tr><td>Chromosome</td><td>start</td><td>end</td><td>strand</td><td>MM</td><td>target_seq</td><td>PAM</td><td>alignment</td><td>distance</td><td>position</td><td>gene name</td><td>gene id</td></tr>
<tr><td>GU305914.1</td><td>199024</td><td>199046</td><td>-</td><td>0</td><td>ACGCGAGGGTTGCGAGGATA</td><td>CGG</td><td>||||||||[||||||||||||]PAM</td><td>NA</td><td>Intergenic</td><td>NA</td><td>NA</td></tr>
</table>

| | | | | | | | | | | | |
|---|---|---|---|---|---|---|---|---|---|---|---|
| T19511 | CGCGAGGGTTGCGAGGATACGGG | 1000 | | | | | | | | | |
| Oligo fwd | CACCCGCGAGGGTTGCGAGGATAC | | | | | | | | | | |
| Oligo rev | AAACGTATCCTCGCAACCCTCGCG | | | | | | | | | | |
| Chromosome | start | end | strand | MM | target_seq | PAM | alignment | distance | position | gene name | gene id |
| GU305914.1 | 199023 | 199045 | - | 0 | CGCGAGGGTTGCGAGGATAC | GGG | ||||||||[||||||||||||]PAM | NA | Intergenic | NA | NA |
| T19512 | GCGAGGGTTGCGAGGATACGGGG | 1000 | | | | | | | | | |
| Oligo fwd | CACCGCGAGGGTTGCGAGGATACG | | | | | | | | | | |
| Oligo rev | AAACCGTATCCTCGCAACCCTCGC | | | | | | | | | | |
| Chromosome | start | end | strand | MM | target_seq | PAM | alignment | distance | position | gene name | gene id |
| GU305914.1 | 199022 | 199044 | - | 0 | GCGAGGGTTGCGAGGATACG | GGG | ||||||||[||||||||||||]PAM | NA | Intergenic | NA | NA |
| GU305914.1 | 26392 | 26414 | + | 4 | GCGACGGCGGCGGGGATACG | GGG | ||||-||-[-|||-|||||||]PAM | NA | Intergenic | NA | NA |
| T19513 | CGGGGTGTCCCTCGTGCTGTCGG | 1000 | | | | | | | | | |
| Oligo fwd | CACCCGGGGTGTCCCTCGTGCTGT | | | | | | | | | | |
| Oligo rev | AAACACAGCACGAGGGACACCCCG | | | | | | | | | | |
| Chromosome | start | end | strand | MM | target_seq | PAM | alignment | distance | position | gene name | gene id |
| GU305914.1 | 199004 | 199026 | - | 0 | CGGGGTGTCCCTCGTGCTGT | CGG | ||||||||[||||||||||||]PAM | NA | Intergenic | NA | NA |
| T19514 | GGGGTGTCCCTCGTGCTGTCGGG | 1000 | | | | | | | | | |
| Oligo fwd | CACCGGGGTGTCCCTCGTGCTGTC | | | | | | | | | | |
| Oligo rev | AAACGACAGCACGAGGGACACCCC | | | | | | | | | | |
| Chromosome | start | end | strand | MM | target_seq | PAM | alignment | distance | position | gene name | gene id |
| GU305914.1 | 199003 | 199025 | - | 0 | GGGGTGTCCCTCGTGCTGTC | GGG | ||||||||[||||||||||||]PAM | NA | Intergenic | NA | NA |
| T19515 | CTCGTGCTGTCGGGCGAGTACGG | 1000 | | | | | | | | | |
| Oligo fwd | CACCCTCGTGCTGTCGGGCGAGTA | | | | | | | | | | |
| Oligo rev | AAACTACTCGCCCGACAGCACGAG | | | | | | | | | | |
| Chromosome | start | end | strand | MM | target_seq | PAM | alignment | distance | position | gene name | gene id |
| GU305914.1 | 198994 | 199016 | - | 0 | CTCGTGCTGTCGGGCGAGTA | CGG | ||||||||[||||||||||||]PAM | NA | Intergenic | NA | NA |
| T19516 | TGCTGTCGGGCGAGTACGGACGG | 1000 | | | | | | | | | |
| Oligo fwd | CACCTGCTGTCGGGCGAGTACGGA | | | | | | | | | | |
| Oligo rev | AAACTCCGTACTCGCCCGACAGCA | | | | | | | | | | |
| Chromosome | start | end | strand | MM | target_seq | PAM | alignment | distance | position | gene name | gene id |
| GU305914.1 | 198990 | 199012 | - | 0 | TGCTGTCGGGCGAGTACGGA | CGG | ||||||||[||||||||||||]PAM | NA | Intergenic | NA | NA |
| T19517 | GCTGTCGGGCGAGTACGGACGGG | 1000 | | | | | | | | | |
| Oligo fwd | CACCGCTGTCGGGCGAGTACGGAC | | | | | | | | | | |
| Oligo rev | AAACGTCCGTACTCGCCCGACAGC | | | | | | | | | | |
| Chromosome | start | end | strand | MM | target_seq | PAM | alignment | distance | position | gene name | gene id |
| GU305914.1 | 198989 | 199011 | - | 0 | GCTGTCGGGCGAGTACGGAC | GGG | ||||||||[||||||||||||]PAM | NA | Intergenic | NA | NA |
| T19518 | CGGACGGGTGTACGTGTACACGG | 1000 | | | | | | | | | |
| Oligo fwd | CACCCGGACGGGTGTACGTGTACA | | | | | | | | | | |
| Oligo rev | AAACTGTACACGTACACCCGTCCG | | | | | | | | | | |
| Chromosome | start | end | strand | MM | target_seq | PAM | alignment | distance | position | gene name | gene id |
| GU305914.1 | 198974 | 198996 | - | 0 | CGGACGGGTGTACGTGTACA | CGG | ||||||||[||||||||||||]PAM | NA | Intergenic | NA | NA |
| T19519 | GGTGTACGTGTACACGGCGTCGG | 1000 | | | | | | | | | |
| Oligo fwd | CACCGGTGTACGTGTACACGGCGT | | | | | | | | | | |
| Oligo rev | AAACACGCCGTGTACACGTACACC | | | | | | | | | | |
| Chromosome | start | end | strand | MM | target_seq | PAM | alignment | distance | position | gene name | gene id |
| GU305914.1 | 198968 | 198990 | - | 0 | GGTGTACGTGTACACGGCGT | CGG | ||||||||[||||||||||||]PAM | NA | Intergenic | NA | NA |
| T19520 | GTCGGACGACACGCTGCACCTGG | 1000 | | | | | | | | | |
| Oligo fwd | CACCGTCGGACGACACGCTGCACC | | | | | | | | | | |
| Oligo rev | AAACGGTGCAGCGTGTCGTCCGAC | | | | | | | | | | |
| Chromosome | start | end | strand | MM | target_seq | PAM | alignment | distance | position | gene name | gene id |
| GU305914.1 | 198950 | 198972 | - | 0 | GTCGGACGACACGCTGCACC | TGG | ||||||||[||||||||||||]PAM | NA | Intergenic | NA | NA |
| T19521 | GGACGACACGCTGCACCTGGTGG | 1000 | | | | | | | | | |
| Oligo fwd | CACCGGACGACACGCTGCACCTGG | | | | | | | | | | |
| Oligo rev | AAACCCAGGTGCAGCGTGTCGTCC | | | | | | | | | | |
| Chromosome | start | end | strand | MM | target_seq | PAM | alignment | distance | position | gene name | gene id |
| GU305914.1 | 198947 | 198969 | - | 0 | GGACGACACGCTGCACCTGG | TGG | ||||||||[||||||||||||]PAM | NA | Intergenic | NA | NA |
| T19522 | GCACCTGGTGGCGCGCAACCTGG | 1000 | | | | | | | | | |
| Oligo fwd | CACCGCACCTGGTGGCGCGCAACC | | | | | | | | | | |
| Oligo rev | AAACGGTTGCGCGCCACCAGGTGC | | | | | | | | | | |
| Chromosome | start | end | strand | MM | target_seq | PAM | alignment | distance | position | gene name | gene id |
| GU305914.1 | 198935 | 198957 | - | 0 | GCACCTGGTGGCGCGCAACC | TGG | ||||||||[||||||||||||]PAM | NA | Intergenic | NA | NA |
| GU305914.1 | 125114 | 125136 | - | 4 | GCACATCGAGGAGCGCAACC | CGG | ||||-|-|[-||-||||||||]PAM | NA | Intergenic | NA | NA |
| T19523 | CCTGGTGGCGCGCAACCTGGAGG | 1000 | | | | | | | | | |
| Oligo fwd | CACCCCTGGTGGCGCGCAACCTGG | | | | | | | | | | |
| Oligo rev | AAACCCAGGTTGCGCGCCACCAGG | | | | | | | | | | |
| Chromosome | start | end | strand | MM | target_seq | PAM | alignment | distance | position | gene name | gene id |
| GU305914.1 | 198932 | 198954 | - | 0 | CCTGGTGGCGCGCAACCTGG | AGG | ||||||||[||||||||||||]PAM | NA | Intergenic | NA | NA |
| T19524 | GCGCGCAACCTGGAGGAGCTCGG | 1000 | | | | | | | | | |
| Oligo fwd | CACCGCGCGCAACCTGGAGGAGCT | | | | | | | | | | |
| Oligo rev | AAACAGCTCCTCCAGGTTGCGCGC | | | | | | | | | | |
| Chromosome | start | end | strand | MM | target_seq | PAM | alignment | distance | position | gene name | gene id |
| GU305914.1 | 198925 | 198947 | - | 0 | GCGCGCAACCTGGAGGAGCT | CGG | ||||||||[||||||||||||]PAM | NA | Intergenic | NA | NA |
| T19525 | CTGGAGGAGCTCGGCCGTTTCGG | 1000 | | | | | | | | | |
| Oligo fwd | CACCCTGGAGGAGCTCGGCCGTTT | | | | | | | | | | |

<table>
<tr><td>Oligo rev</td><td>AAACAAACGGCCGAGCTCCTCCAG</td><td></td><td></td><td></td><td></td><td></td><td></td><td></td><td></td><td></td><td></td></tr>
<tr><td>Chromosome</td><td>start</td><td>end</td><td>strand</td><td>MM</td><td>target_seq</td><td>PAM</td><td>alignment</td><td>distance</td><td>position</td><td>gene name</td><td>gene id</td></tr>
<tr><td>GU305914.1</td><td>198916</td><td>198938</td><td>-</td><td>0</td><td>CTGGAGGAGCTCGGCCGTTT</td><td>CGG</td><td>||||||||[||||||||||||]PAM</td><td>NA</td><td>Intergenic</td><td>NA</td><td>NA</td></tr>
<tr><td>T19526</td><td>TGGAGGAGCTCGGCCGTTTCGGG</td><td>1000</td><td></td><td></td><td></td><td></td><td></td><td></td><td></td><td></td><td></td></tr>
<tr><td>Oligo fwd</td><td>CACCTGGAGGAGCTCGGCCGTTTC</td><td></td><td></td><td></td><td></td><td></td><td></td><td></td><td></td><td></td><td></td></tr>
<tr><td>Oligo rev</td><td>AAACGAAACGGCCGAGCTCCTCCA</td><td></td><td></td><td></td><td></td><td></td><td></td><td></td><td></td><td></td><td></td></tr>
<tr><td>Chromosome</td><td>start</td><td>end</td><td>strand</td><td>MM</td><td>target_seq</td><td>PAM</td><td>alignment</td><td>distance</td><td>position</td><td>gene name</td><td>gene id</td></tr>
<tr><td>GU305914.1</td><td>198915</td><td>198937</td><td>-</td><td>0</td><td>TGGAGGAGCTCGGCCGTTTC</td><td>GGG</td><td>||||||||[||||||||||||]PAM</td><td>NA</td><td>Intergenic</td><td>NA</td><td>NA</td></tr>
<tr><td>T19527</td><td>CCTACAAAGAAGACATGCAGCGG</td><td>1000</td><td></td><td></td><td></td><td></td><td></td><td></td><td></td><td></td><td></td></tr>
<tr><td>Oligo fwd</td><td>CACCCCTACAAAGAAGACATGCAG</td><td></td><td></td><td></td><td></td><td></td><td></td><td></td><td></td><td></td><td></td></tr>
<tr><td>Oligo rev</td><td>AAACCTGCATGTCTTCTTTGTAGG</td><td></td><td></td><td></td><td></td><td></td><td></td><td></td><td></td><td></td><td></td></tr>
<tr><td>Chromosome</td><td>start</td><td>end</td><td>strand</td><td>MM</td><td>target_seq</td><td>PAM</td><td>alignment</td><td>distance</td><td>position</td><td>gene name</td><td>gene id</td></tr>
<tr><td>GU305914.1</td><td>198873</td><td>198895</td><td>-</td><td>0</td><td>CCTACAAAGAAGACATGCAG</td><td>CGG</td><td>||||||||[||||||||||||]PAM</td><td>NA</td><td>Intergenic</td><td>NA</td><td>NA</td></tr>
<tr><td>T19528</td><td>GCAGCGGTCCGTGCCGCGCGCGG</td><td>1000</td><td></td><td></td><td></td><td></td><td></td><td></td><td></td><td></td><td></td></tr>
<tr><td>Oligo fwd</td><td>CACCGCAGCGGTCCGTGCCGCGCG</td><td></td><td></td><td></td><td></td><td></td><td></td><td></td><td></td><td></td><td></td></tr>
<tr><td>Oligo rev</td><td>AAACCGCGCGGCACGGACCGCTGC</td><td></td><td></td><td></td><td></td><td></td><td></td><td></td><td></td><td></td><td></td></tr>
<tr><td>Chromosome</td><td>start</td><td>end</td><td>strand</td><td>MM</td><td>target_seq</td><td>PAM</td><td>alignment</td><td>distance</td><td>position</td><td>gene name</td><td>gene id</td></tr>
<tr><td>GU305914.1</td><td>198857</td><td>198879</td><td>-</td><td>0</td><td>GCAGCGGTCCGTGCCGCGCG</td><td>CGG</td><td>||||||||[||||||||||||]PAM</td><td>NA</td><td>Intergenic</td><td>NA</td><td>NA</td></tr>
<tr><td>GU305914.1</td><td>100777</td><td>100799</td><td>-</td><td>4</td><td>GGACCGGTACGTGCCGCGGG</td><td>CGG</td><td>|-|-||||[-|||||||||-|]PAM</td><td>NA</td><td>Intergenic</td><td>NA</td><td>NA</td></tr>
<tr><td>T19529</td><td>GTCCGTGCCGCGCGCGGACCTGG</td><td>1000</td><td></td><td></td><td></td><td></td><td></td><td></td><td></td><td></td><td></td></tr>
<tr><td>Oligo fwd</td><td>CACCGTCCGTGCCGCGCGCGGACC</td><td></td><td></td><td></td><td></td><td></td><td></td><td></td><td></td><td></td><td></td></tr>
<tr><td>Oligo rev</td><td>AAACGGTCCGCGCGCGGCACGGAC</td><td></td><td></td><td></td><td></td><td></td><td></td><td></td><td></td><td></td><td></td></tr>
<tr><td>Chromosome</td><td>start</td><td>end</td><td>strand</td><td>MM</td><td>target_seq</td><td>PAM</td><td>alignment</td><td>distance</td><td>position</td><td>gene name</td><td>gene id</td></tr>
<tr><td>GU305914.1</td><td>198851</td><td>198873</td><td>-</td><td>0</td><td>GTCCGTGCCGCGCGCGGACC</td><td>TGG</td><td>||||||||[||||||||||||]PAM</td><td>NA</td><td>Intergenic</td><td>NA</td><td>NA</td></tr>
<tr><td>T19530</td><td>TGCCGCGCGCGGACCTGGTGCGG</td><td>1000</td><td></td><td></td><td></td><td></td><td></td><td></td><td></td><td></td><td></td></tr>
<tr><td>Oligo fwd</td><td>CACCTGCCGCGCGCGGACCTGGTG</td><td></td><td></td><td></td><td></td><td></td><td></td><td></td><td></td><td></td><td></td></tr>
<tr><td>Oligo rev</td><td>AAACCACCAGGTCCGCGCGCGGCA</td><td></td><td></td><td></td><td></td><td></td><td></td><td></td><td></td><td></td><td></td></tr>
<tr><td>Chromosome</td><td>start</td><td>end</td><td>strand</td><td>MM</td><td>target_seq</td><td>PAM</td><td>alignment</td><td>distance</td><td>position</td><td>gene name</td><td>gene id</td></tr>
<tr><td>GU305914.1</td><td>198846</td><td>198868</td><td>-</td><td>0</td><td>TGCCGCGCGCGGACCTGGTG</td><td>CGG</td><td>||||||||[||||||||||||]PAM</td><td>NA</td><td>Intergenic</td><td>NA</td><td>NA</td></tr>
<tr><td>T19531</td><td>GCGCGGACCTGGTGCGGCTGAGG</td><td>1000</td><td></td><td></td><td></td><td></td><td></td><td></td><td></td><td></td><td></td></tr>
<tr><td>Oligo fwd</td><td>CACCGCGCGGACCTGGTGCGGCTG</td><td></td><td></td><td></td><td></td><td></td><td></td><td></td><td></td><td></td><td></td></tr>
<tr><td>Oligo rev</td><td>AAACCAGCCGCACCAGGTCCGCGC</td><td></td><td></td><td></td><td></td><td></td><td></td><td></td><td></td><td></td><td></td></tr>
<tr><td>Chromosome</td><td>start</td><td>end</td><td>strand</td><td>MM</td><td>target_seq</td><td>PAM</td><td>alignment</td><td>distance</td><td>position</td><td>gene name</td><td>gene id</td></tr>
<tr><td>GU305914.1</td><td>198840</td><td>198862</td><td>-</td><td>0</td><td>GCGCGGACCTGGTGCGGCTG</td><td>AGG</td><td>||||||||[||||||||||||]PAM</td><td>NA</td><td>Intergenic</td><td>NA</td><td>NA</td></tr>
<tr><td>T19532</td><td>TCGCCTCGTGCGACGTGCCGCGG</td><td>1000</td><td></td><td></td><td></td><td></td><td></td><td></td><td></td><td></td><td></td></tr>
<tr><td>Oligo fwd</td><td>CACCTCGCCTCGTGCGACGTGCCG</td><td></td><td></td><td></td><td></td><td></td><td></td><td></td><td></td><td></td><td></td></tr>
<tr><td>Oligo rev</td><td>AAACCGGCACGTCGCACGAGGCGA</td><td></td><td></td><td></td><td></td><td></td><td></td><td></td><td></td><td></td><td></td></tr>
<tr><td>Chromosome</td><td>start</td><td>end</td><td>strand</td><td>MM</td><td>target_seq</td><td>PAM</td><td>alignment</td><td>distance</td><td>position</td><td>gene name</td><td>gene id</td></tr>
<tr><td>GU305914.1</td><td>198813</td><td>198835</td><td>-</td><td>0</td><td>TCGCCTCGTGCGACGTGCCG</td><td>CGG</td><td>||||||||[||||||||||||]PAM</td><td>NA</td><td>Intergenic</td><td>NA</td><td>NA</td></tr>
<tr><td>T19533</td><td>CGCCTCGTGCGACGTGCCGCGGG</td><td>1000</td><td></td><td></td><td></td><td></td><td></td><td></td><td></td><td></td><td></td></tr>
<tr><td>Oligo fwd</td><td>CACCCGCCTCGTGCGACGTGCCGC</td><td></td><td></td><td></td><td></td><td></td><td></td><td></td><td></td><td></td><td></td></tr>
<tr><td>Oligo rev</td><td>AAACGCGGCACGTCGCACGAGGCG</td><td></td><td></td><td></td><td></td><td></td><td></td><td></td><td></td><td></td><td></td></tr>
<tr><td>Chromosome</td><td>start</td><td>end</td><td>strand</td><td>MM</td><td>target_seq</td><td>PAM</td><td>alignment</td><td>distance</td><td>position</td><td>gene name</td><td>gene id</td></tr>
<tr><td>GU305914.1</td><td>198812</td><td>198834</td><td>-</td><td>0</td><td>CGCCTCGTGCGACGTGCCGC</td><td>GGG</td><td>||||||||[||||||||||||]PAM</td><td>NA</td><td>Intergenic</td><td>NA</td><td>NA</td></tr>
<tr><td>GU305914.1</td><td>38096</td><td>38118</td><td>+</td><td>4</td><td>CGACCCGTCCAACGTGCCGC</td><td>CGG</td><td>||-|-|||[-|-|||||||||]PAM</td><td>NA</td><td>Intergenic</td><td>NA</td><td>NA</td></tr>
<tr><td>T19534</td><td>GCCTCGTGCGACGTGCCGCGGGG</td><td>1000</td><td></td><td></td><td></td><td></td><td></td><td></td><td></td><td></td><td></td></tr>
<tr><td>Oligo fwd</td><td>CACCGCCTCGTGCGACGTGCCGCG</td><td></td><td></td><td></td><td></td><td></td><td></td><td></td><td></td><td></td><td></td></tr>
<tr><td>Oligo rev</td><td>AAACCGCGGCACGTCGCACGAGGC</td><td></td><td></td><td></td><td></td><td></td><td></td><td></td><td></td><td></td><td></td></tr>
<tr><td>Chromosome</td><td>start</td><td>end</td><td>strand</td><td>MM</td><td>target_seq</td><td>PAM</td><td>alignment</td><td>distance</td><td>position</td><td>gene name</td><td>gene id</td></tr>
<tr><td>GU305914.1</td><td>198811</td><td>198833</td><td>-</td><td>0</td><td>GCCTCGTGCGACGTGCCGCG</td><td>GGG</td><td>||||||||[||||||||||||]PAM</td><td>NA</td><td>Intergenic</td><td>NA</td><td>NA</td></tr>
<tr><td>T19535</td><td>GACGTGCCGCGGGGCTACCCCGG</td><td>1000</td><td></td><td></td><td></td><td></td><td></td><td></td><td></td><td></td><td></td></tr>
<tr><td>Oligo fwd</td><td>CACCGACGTGCCGCGGGGCTACCC</td><td></td><td></td><td></td><td></td><td></td><td></td><td></td><td></td><td></td><td></td></tr>
<tr><td>Oligo rev</td><td>AAACGGGTAGCCCCGCGGCACGTC</td><td></td><td></td><td></td><td></td><td></td><td></td><td></td><td></td><td></td><td></td></tr>
<tr><td>Chromosome</td><td>start</td><td>end</td><td>strand</td><td>MM</td><td>target_seq</td><td>PAM</td><td>alignment</td><td>distance</td><td>position</td><td>gene name</td><td>gene id</td></tr>
<tr><td>GU305914.1</td><td>198802</td><td>198824</td><td>-</td><td>0</td><td>GACGTGCCGCGGGGCTACCC</td><td>CGG</td><td>||||||||[||||||||||||]PAM</td><td>NA</td><td>Intergenic</td><td>NA</td><td>NA</td></tr>
<tr><td>T19536</td><td>GCCGCGGGGCTACCCCGGCATGG</td><td>1000</td><td></td><td></td><td></td><td></td><td></td><td></td><td></td><td></td><td></td></tr>
<tr><td>Oligo fwd</td><td>CACCGCCGCGGGGCTACCCCGGCA</td><td></td><td></td><td></td><td></td><td></td><td></td><td></td><td></td><td></td><td></td></tr>
<tr><td>Oligo rev</td><td>AAACTGCCGGGGTAGCCCCGCGGC</td><td></td><td></td><td></td><td></td><td></td><td></td><td></td><td></td><td></td><td></td></tr>
<tr><td>Chromosome</td><td>start</td><td>end</td><td>strand</td><td>MM</td><td>target_seq</td><td>PAM</td><td>alignment</td><td>distance</td><td>position</td><td>gene name</td><td>gene id</td></tr>
<tr><td>GU305914.1</td><td>198797</td><td>198819</td><td>-</td><td>0</td><td>GCCGCGGGGCTACCCCGGCA</td><td>TGG</td><td>||||||||[||||||||||||]PAM</td><td>NA</td><td>Intergenic</td><td>NA</td><td>NA</td></tr>
<tr><td>T19537</td><td>CTACCCCGGCATGGAGCTGCTGG</td><td>1000</td><td></td><td></td><td></td><td></td><td></td><td></td><td></td><td></td><td></td></tr>
<tr><td>Oligo fwd</td><td>CACCCTACCCCGGCATGGAGCTGC</td><td></td><td></td><td></td><td></td><td></td><td></td><td></td><td></td><td></td><td></td></tr>
<tr><td>Oligo rev</td><td>AAACGCAGCTCCATGCCGGGGTAG</td><td></td><td></td><td></td><td></td><td></td><td></td><td></td><td></td><td></td><td></td></tr>
<tr><td>Chromosome</td><td>start</td><td>end</td><td>strand</td><td>MM</td><td>target_seq</td><td>PAM</td><td>alignment</td><td>distance</td><td>position</td><td>gene name</td><td>gene id</td></tr>
<tr><td>GU305914.1</td><td>198788</td><td>198810</td><td>-</td><td>0</td><td>CTACCCCGGCATGGAGCTGC</td><td>TGG</td><td>||||||||[||||||||||||]PAM</td><td>NA</td><td>Intergenic</td><td>NA</td><td>NA</td></tr>
<tr><td>T19538</td><td>CCCCGGCATGGAGCTGCTGGTGG</td><td>1000</td><td></td><td></td><td></td><td></td><td></td><td></td><td></td><td></td><td></td></tr>
<tr><td>Oligo fwd</td><td>CACCCCCCGGCATGGAGCTGCTGG</td><td></td><td></td><td></td><td></td><td></td><td></td><td></td><td></td><td></td><td></td></tr>
<tr><td>Oligo rev</td><td>AAACCCAGCAGCTCCATGCCGGGG</td><td></td><td></td><td></td><td></td><td></td><td></td><td></td><td></td><td></td><td></td></tr>
<tr><td>Chromosome</td><td>start</td><td>end</td><td>strand</td><td>MM</td><td>target_seq</td><td>PAM</td><td>alignment</td><td>distance</td><td>position</td><td>gene name</td><td>gene id</td></tr>
<tr><td>GU305914.1</td><td>198785</td><td>198807</td><td>-</td><td>0</td><td>CCCCGGCATGGAGCTGCTGG</td><td>TGG</td><td>||||||||[||||||||||||]PAM</td><td>NA</td><td>Intergenic</td><td>NA</td><td>NA</td></tr>
<tr><td>T19539</td><td>GGTGTTCTCCGACCCGAACGCGG</td><td>1000</td><td></td><td></td><td></td><td></td><td></td><td></td><td></td><td></td><td></td></tr>
<tr><td>Oligo fwd</td><td>CACCGGTGTTCTCCGACCCGAACG</td><td></td><td></td><td></td><td></td><td></td><td></td><td></td><td></td><td></td><td></td></tr>
<tr><td>Oligo rev</td><td>AAACCGTTCGGGTCGGAGAACACC</td><td></td><td></td><td></td><td></td><td></td><td></td><td></td><td></td><td></td><td></td></tr>
<tr><td>Chromosome</td><td>start</td><td>end</td><td>strand</td><td>MM</td><td>target_seq</td><td>PAM</td><td>alignment</td><td>distance</td><td>position</td><td>gene name</td><td>gene id</td></tr>
<tr><td>GU305914.1</td><td>198764</td><td>198786</td><td>-</td><td>0</td><td>GGTGTTCTCCGACCCGAACG</td><td>CGG</td><td>||||||||[||||||||||||]PAM</td><td>NA</td><td>Intergenic</td><td>NA</td><td>NA</td></tr>
</table>

| | | | | | | | | | | | |
|---|---|---|---|---|---|---|---|---|---|---|---|
| GU305914.1 | 96577 | 96599 | - | 4 | GGAGTTCGCCGAGCCGATCG | AGG | \|\|-\|\|\|\|-[\|\|\|\|-\|\|\|\|-\|\|]PAM | NA | Intergenic | NA | NA |
| T19540 | CTCCGACCCGAACGCGGACGCGG | 1000 | | | | | | | | | |
| Oligo fwd | CACCCTCCGACCCGAACGCGGACG | | | | | | | | | | |
| Oligo rev | AAACCGTCCGCGTTCGGGTCGGAG | | | | | | | | | | |
| Chromosome | start | end | strand | MM | target_seq | PAM | alignment | distance | position | gene name | gene id |
| GU305914.1 | 198758 | 198780 | - | 0 | CTCCGACCCGAACGCGGACG | CGG | \|\|\|\|\|\|\|\|[\|\|\|\|\|\|\|\|\|\|\|\|]PAM | NA | Intergenic | NA | NA |
| T19541 | GAACGCGGACGCGGCGACGTCGG | 1000 | | | | | | | | | |
| Oligo fwd | CACCGAACGCGGACGCGGCGACGT | | | | | | | | | | |
| Oligo rev | AAACACGTCGCCGCGTCCGCGTTC | | | | | | | | | | |
| Chromosome | start | end | strand | MM | target_seq | PAM | alignment | distance | position | gene name | gene id |
| GU305914.1 | 198749 | 198771 | - | 0 | GAACGCGGACGCGGCGACGT | CGG | \|\|\|\|\|\|\|\|[\|\|\|\|\|\|\|\|\|\|\|\|]PAM | NA | Intergenic | NA | NA |
| GU305914.1 | 110086 | 110108 | - | 4 | CGACGCGGCCGCGGCGACGC | CGG | --\|\|\|\|\|\|[-\|\|\|\|\|\|\|\|\|\|-]PAM | NA | Intergenic | NA | NA |
| T19542 | CGCGGACGCGGCGACGTCGGTGG | 1000 | | | | | | | | | |
| Oligo fwd | CACCCGCGGACGCGGCGACGTCGG | | | | | | | | | | |
| Oligo rev | AAACCCGACGTCGCCGCGTCCGCG | | | | | | | | | | |
| Chromosome | start | end | strand | MM | target_seq | PAM | alignment | distance | position | gene name | gene id |
| GU305914.1 | 198746 | 198768 | - | 0 | CGCGGACGCGGCGACGTCGG | TGG | \|\|\|\|\|\|\|\|[\|\|\|\|\|\|\|\|\|\|\|\|]PAM | NA | Intergenic | NA | NA |
| GU305914.1 | 128803 | 128825 | + | 4 | CGCAGCGGCAGCGACGTCGG | CGG | \|\|\|-\|--\|[\|-\|\|\|\|\|\|\|\|\|\|]PAM | NA | Intergenic | NA | NA |
| GU305914.1 | 110083 | 110105 | - | 2 | CGCGGCCGCGGCGACGCCGG | TGG | \|\|\|\|\|-\|\|[\|\|\|\|\|\|\|\|-\|\|\|]PAM | NA | Intergenic | NA | NA |
| T19543 | GCGGACGCGGCGACGTCGGTGGG | 1000 | | | | | | | | | |
| Oligo fwd | CACCGCGGACGCGGCGACGTCGGT | | | | | | | | | | |
| Oligo rev | AAACACCGACGTCGCCGCGTCCGC | | | | | | | | | | |
| Chromosome | start | end | strand | MM | target_seq | PAM | alignment | distance | position | gene name | gene id |
| GU305914.1 | 198745 | 198767 | - | 0 | GCGGACGCGGCGACGTCGGT | GGG | \|\|\|\|\|\|\|\|[\|\|\|\|\|\|\|\|\|\|\|\|]PAM | NA | Intergenic | NA | NA |
| T19544 | GGTGGGCTACACTCGCTTCGTGG | 1000 | | | | | | | | | |
| Oligo fwd | CACCGGTGGGCTACACTCGCTTCG | | | | | | | | | | |
| Oligo rev | AAACCGAAGCGAGTGTAGCCCACC | | | | | | | | | | |
| Chromosome | start | end | strand | MM | target_seq | PAM | alignment | distance | position | gene name | gene id |
| GU305914.1 | 198728 | 198750 | - | 0 | GGTGGGCTACACTCGCTTCG | TGG | \|\|\|\|\|\|\|\|[\|\|\|\|\|\|\|\|\|\|\|\|]PAM | NA | Intergenic | NA | NA |
| T19545 | GGCTACACTCGCTTCGTGGACGG | 1000 | | | | | | | | | |
| Oligo fwd | CACCGGCTACACTCGCTTCGTGGA | | | | | | | | | | |
| Oligo rev | AAACTCCACGAAGCGAGTGTAGCC | | | | | | | | | | |
| Chromosome | start | end | strand | MM | target_seq | PAM | alignment | distance | position | gene name | gene id |
| GU305914.1 | 198724 | 198746 | - | 0 | GGCTACACTCGCTTCGTGGA | CGG | \|\|\|\|\|\|\|\|[\|\|\|\|\|\|\|\|\|\|\|\|]PAM | NA | Intergenic | NA | NA |
| T19546 | TCGTGGACGGCAAGTTCGTGCGG | 1000 | | | | | | | | | |
| Oligo fwd | CACCTCGTGGACGGCAAGTTCGTG | | | | | | | | | | |
| Oligo rev | AAACCACGAACTTGCCGTCCACGA | | | | | | | | | | |
| Chromosome | start | end | strand | MM | target_seq | PAM | alignment | distance | position | gene name | gene id |
| GU305914.1 | 198711 | 198733 | - | 0 | TCGTGGACGGCAAGTTCGTG | CGG | \|\|\|\|\|\|\|\|[\|\|\|\|\|\|\|\|\|\|\|\|]PAM | NA | Intergenic | NA | NA |
| T19547 | GTTCGTGCGGCTGACGACGCCGG | 1000 | | | | | | | | | |
| Oligo fwd | CACCGTTCGTGCGGCTGACGACGC | | | | | | | | | | |
| Oligo rev | AAACGCGTCGTCAGCCGCACGAAC | | | | | | | | | | |
| Chromosome | start | end | strand | MM | target_seq | PAM | alignment | distance | position | gene name | gene id |
| GU305914.1 | 198698 | 198720 | - | 0 | GTTCGTGCGGCTGACGACGC | CGG | \|\|\|\|\|\|\|\|[\|\|\|\|\|\|\|\|\|\|\|\|]PAM | NA | Intergenic | NA | NA |
| T19548 | TTCGTGCGGCTGACGACGCCGGG | 1000 | | | | | | | | | |
| Oligo fwd | CACCTTCGTGCGGCTGACGACGCC | | | | | | | | | | |
| Oligo rev | AAACGGCGTCGTCAGCCGCACGAA | | | | | | | | | | |
| Chromosome | start | end | strand | MM | target_seq | PAM | alignment | distance | position | gene name | gene id |
| GU305914.1 | 198697 | 198719 | - | 0 | TTCGTGCGGCTGACGACGCC | GGG | \|\|\|\|\|\|\|\|[\|\|\|\|\|\|\|\|\|\|\|\|]PAM | NA | Intergenic | NA | NA |
| T19549 | CGGCTGACGACGCCGGGCTACGG | 1000 | | | | | | | | | |
| Oligo fwd | CACCCGGCTGACGACGCCGGGCTA | | | | | | | | | | |
| Oligo rev | AAACTAGCCCGGCGTCGTCAGCCG | | | | | | | | | | |
| Chromosome | start | end | strand | MM | target_seq | PAM | alignment | distance | position | gene name | gene id |
| GU305914.1 | 198691 | 198713 | - | 0 | CGGCTGACGACGCCGGGCTA | CGG | \|\|\|\|\|\|\|\|[\|\|\|\|\|\|\|\|\|\|\|\|]PAM | NA | Intergenic | NA | NA |
| T19550 | GGCTGACGACGCCGGGCTACGGG | 1000 | | | | | | | | | |
| Oligo fwd | CACCGGCTGACGACGCCGGGCTAC | | | | | | | | | | |
| Oligo rev | AAACGTAGCCCGGCGTCGTCAGCC | | | | | | | | | | |
| Chromosome | start | end | strand | MM | target_seq | PAM | alignment | distance | position | gene name | gene id |
| GU305914.1 | 198690 | 198712 | - | 0 | GGCTGACGACGCCGGGCTAC | GGG | \|\|\|\|\|\|\|\|[\|\|\|\|\|\|\|\|\|\|\|\|]PAM | NA | Intergenic | NA | NA |
| T19551 | GACGCCGGGCTACGGGACGTCGG | 1000 | | | | | | | | | |
| Oligo fwd | CACCGACGCCGGGCTACGGGACGT | | | | | | | | | | |
| Oligo rev | AAACACGTCCCGTAGCCCGGCGTC | | | | | | | | | | |
| Chromosome | start | end | strand | MM | target_seq | PAM | alignment | distance | position | gene name | gene id |
| GU305914.1 | 198683 | 198705 | - | 0 | GACGCCGGGCTACGGGACGT | CGG | \|\|\|\|\|\|\|\|[\|\|\|\|\|\|\|\|\|\|\|\|]PAM | NA | Intergenic | NA | NA |
| T19552 | CGGGACGTCGGCGCTGTTCCTGG | 1000 | | | | | | | | | |
| Oligo fwd | CACCCGGGACGTCGGCGCTGTTCC | | | | | | | | | | |
| Oligo rev | AAACGGAACAGCGCCGACGTCCCG | | | | | | | | | | |
| Chromosome | start | end | strand | MM | target_seq | PAM | alignment | distance | position | gene name | gene id |
| GU305914.1 | 198671 | 198693 | - | 0 | CGGGACGTCGGCGCTGTTCC | TGG | \|\|\|\|\|\|\|\|[\|\|\|\|\|\|\|\|\|\|\|\|]PAM | NA | Intergenic | NA | NA |
| T19553 | GTCGGCGCTGTTCCTGGCGAAGG | 1000 | | | | | | | | | |
| Oligo fwd | CACCGTCGGCGCTGTTCCTGGCGA | | | | | | | | | | |
| Oligo rev | AAACTCGCCAGGAACAGCGCCGAC | | | | | | | | | | |
| Chromosome | start | end | strand | MM | target_seq | PAM | alignment | distance | position | gene name | gene id |
| GU305914.1 | 198665 | 198687 | - | 0 | GTCGGCGCTGTTCCTGGCGA | AGG | \|\|\|\|\|\|\|\|[\|\|\|\|\|\|\|\|\|\|\|\|]PAM | NA | Intergenic | NA | NA |

<table>
<tr><td>T19554</td><td>CTGTTCCTGGCGAAGGACCCCGG</td><td>1000</td><td></td><td></td><td></td><td></td><td></td><td></td><td></td><td></td><td></td></tr>
<tr><td>Oligo fwd</td><td>CACCCTGTTCCTGGCGAAGGACCC</td><td></td><td></td><td></td><td></td><td></td><td></td><td></td><td></td><td></td><td></td></tr>
<tr><td>Oligo rev</td><td>AAACGGGTCCTTCGCCAGGAACAG</td><td></td><td></td><td></td><td></td><td></td><td></td><td></td><td></td><td></td><td></td></tr>
<tr><td>Chromosome</td><td>start</td><td>end</td><td>strand</td><td>MM</td><td>target_seq</td><td>PAM</td><td>alignment</td><td>distance</td><td>position</td><td>gene name</td><td>gene id</td></tr>
<tr><td>GU305914.1</td><td>198658</td><td>198680</td><td>-</td><td>0</td><td>CTGTTCCTGGCGAAGGACCC</td><td>CGG</td><td>||||||||[||||||||||||]PAM</td><td>NA</td><td>Intergenic</td><td>NA</td><td>NA</td></tr>
<tr><td>T19555</td><td>CGAAGGACCCCGGCGAGCTGAGG</td><td>1000</td><td></td><td></td><td></td><td></td><td></td><td></td><td></td><td></td><td></td></tr>
<tr><td>Oligo fwd</td><td>CACCCGAAGGACCCCGGCGAGCTG</td><td></td><td></td><td></td><td></td><td></td><td></td><td></td><td></td><td></td><td></td></tr>
<tr><td>Oligo rev</td><td>AAACCAGCTCGCCGGGGTCCTTCG</td><td></td><td></td><td></td><td></td><td></td><td></td><td></td><td></td><td></td><td></td></tr>
<tr><td>Chromosome</td><td>start</td><td>end</td><td>strand</td><td>MM</td><td>target_seq</td><td>PAM</td><td>alignment</td><td>distance</td><td>position</td><td>gene name</td><td>gene id</td></tr>
<tr><td>GU305914.1</td><td>198648</td><td>198670</td><td>-</td><td>0</td><td>CGAAGGACCCCGGCGAGCTG</td><td>AGG</td><td>||||||||[||||||||||||]PAM</td><td>NA</td><td>Intergenic</td><td>NA</td><td>NA</td></tr>
<tr><td>T19556</td><td>GAAGGACCCCGGCGAGCTGAGGG</td><td>1000</td><td></td><td></td><td></td><td></td><td></td><td></td><td></td><td></td><td></td></tr>
<tr><td>Oligo fwd</td><td>CACCGAAGGACCCCGGCGAGCTGA</td><td></td><td></td><td></td><td></td><td></td><td></td><td></td><td></td><td></td><td></td></tr>
<tr><td>Oligo rev</td><td>AAACTCAGCTCGCCGGGGTCCTTC</td><td></td><td></td><td></td><td></td><td></td><td></td><td></td><td></td><td></td><td></td></tr>
<tr><td>Chromosome</td><td>start</td><td>end</td><td>strand</td><td>MM</td><td>target_seq</td><td>PAM</td><td>alignment</td><td>distance</td><td>position</td><td>gene name</td><td>gene id</td></tr>
<tr><td>GU305914.1</td><td>198647</td><td>198669</td><td>-</td><td>0</td><td>GAAGGACCCCGGCGAGCTGA</td><td>GGG</td><td>||||||||[||||||||||||]PAM</td><td>NA</td><td>Intergenic</td><td>NA</td><td>NA</td></tr>
<tr><td>T19557</td><td>AAGGACCCCGGCGAGCTGAGGGG</td><td>1000</td><td></td><td></td><td></td><td></td><td></td><td></td><td></td><td></td><td></td></tr>
<tr><td>Oligo fwd</td><td>CACCAAGGACCCCGGCGAGCTGAG</td><td></td><td></td><td></td><td></td><td></td><td></td><td></td><td></td><td></td><td></td></tr>
<tr><td>Oligo rev</td><td>AAACCTCAGCTCGCCGGGGTCCTT</td><td></td><td></td><td></td><td></td><td></td><td></td><td></td><td></td><td></td><td></td></tr>
<tr><td>Chromosome</td><td>start</td><td>end</td><td>strand</td><td>MM</td><td>target_seq</td><td>PAM</td><td>alignment</td><td>distance</td><td>position</td><td>gene name</td><td>gene id</td></tr>
<tr><td>GU305914.1</td><td>198646</td><td>198668</td><td>-</td><td>0</td><td>AAGGACCCCGGCGAGCTGAG</td><td>GGG</td><td>||||||||[||||||||||||]PAM</td><td>NA</td><td>Intergenic</td><td>NA</td><td>NA</td></tr>
<tr><td>T19558</td><td>AGGACCCCGGCGAGCTGAGGGGG</td><td>1000</td><td></td><td></td><td></td><td></td><td></td><td></td><td></td><td></td><td></td></tr>
<tr><td>Oligo fwd</td><td>CACCAGGACCCCGGCGAGCTGAGG</td><td></td><td></td><td></td><td></td><td></td><td></td><td></td><td></td><td></td><td></td></tr>
<tr><td>Oligo rev</td><td>AAACCCTCAGCTCGCCGGGGTCCT</td><td></td><td></td><td></td><td></td><td></td><td></td><td></td><td></td><td></td><td></td></tr>
<tr><td>Chromosome</td><td>start</td><td>end</td><td>strand</td><td>MM</td><td>target_seq</td><td>PAM</td><td>alignment</td><td>distance</td><td>position</td><td>gene name</td><td>gene id</td></tr>
<tr><td>GU305914.1</td><td>198645</td><td>198667</td><td>-</td><td>0</td><td>AGGACCCCGGCGAGCTGAGG</td><td>GGG</td><td>||||||||[||||||||||||]PAM</td><td>NA</td><td>Intergenic</td><td>NA</td><td>NA</td></tr>
<tr><td>T19559</td><td>CCGGCGAGCTGAGGGGGATCTGG</td><td>1000</td><td></td><td></td><td></td><td></td><td></td><td></td><td></td><td></td><td></td></tr>
<tr><td>Oligo fwd</td><td>CACCCCGGCGAGCTGAGGGGGATC</td><td></td><td></td><td></td><td></td><td></td><td></td><td></td><td></td><td></td><td></td></tr>
<tr><td>Oligo rev</td><td>AAACGATCCCCCTCAGCTCGCCGG</td><td></td><td></td><td></td><td></td><td></td><td></td><td></td><td></td><td></td><td></td></tr>
<tr><td>Chromosome</td><td>start</td><td>end</td><td>strand</td><td>MM</td><td>target_seq</td><td>PAM</td><td>alignment</td><td>distance</td><td>position</td><td>gene name</td><td>gene id</td></tr>
<tr><td>GU305914.1</td><td>198639</td><td>198661</td><td>-</td><td>0</td><td>CCGGCGAGCTGAGGGGGATC</td><td>TGG</td><td>||||||||[||||||||||||]PAM</td><td>NA</td><td>Intergenic</td><td>NA</td><td>NA</td></tr>
<tr><td>T19560</td><td>AGGGGGATCTGGCCGTTCCGCGG</td><td>1000</td><td></td><td></td><td></td><td></td><td></td><td></td><td></td><td></td><td></td></tr>
<tr><td>Oligo fwd</td><td>CACCAGGGGGATCTGGCCGTTCCG</td><td></td><td></td><td></td><td></td><td></td><td></td><td></td><td></td><td></td><td></td></tr>
<tr><td>Oligo rev</td><td>AAACCGGAACGGCCAGATCCCCCT</td><td></td><td></td><td></td><td></td><td></td><td></td><td></td><td></td><td></td><td></td></tr>
<tr><td>Chromosome</td><td>start</td><td>end</td><td>strand</td><td>MM</td><td>target_seq</td><td>PAM</td><td>alignment</td><td>distance</td><td>position</td><td>gene name</td><td>gene id</td></tr>
<tr><td>GU305914.1</td><td>198628</td><td>198650</td><td>-</td><td>0</td><td>AGGGGGATCTGGCCGTTCCG</td><td>CGG</td><td>||||||||[||||||||||||]PAM</td><td>NA</td><td>Intergenic</td><td>NA</td><td>NA</td></tr>
<tr><td>T19561</td><td>TGTGCGACCCCGAGTTCAAGCGG</td><td>1000</td><td></td><td></td><td></td><td></td><td></td><td></td><td></td><td></td><td></td></tr>
<tr><td>Oligo fwd</td><td>CACCTGTGCGACCCCGAGTTCAAG</td><td></td><td></td><td></td><td></td><td></td><td></td><td></td><td></td><td></td><td></td></tr>
<tr><td>Oligo rev</td><td>AAACCTTGAACTCGGGGTCGCACA</td><td></td><td></td><td></td><td></td><td></td><td></td><td></td><td></td><td></td><td></td></tr>
<tr><td>Chromosome</td><td>start</td><td>end</td><td>strand</td><td>MM</td><td>target_seq</td><td>PAM</td><td>alignment</td><td>distance</td><td>position</td><td>gene name</td><td>gene id</td></tr>
<tr><td>GU305914.1</td><td>198603</td><td>198625</td><td>-</td><td>0</td><td>TGTGCGACCCCGAGTTCAAG</td><td>CGG</td><td>||||||||[||||||||||||]PAM</td><td>NA</td><td>Intergenic</td><td>NA</td><td>NA</td></tr>
<tr><td>T19562</td><td>ACCCCGAGTTCAAGCGGTGCTGG</td><td>1000</td><td></td><td></td><td></td><td></td><td></td><td></td><td></td><td></td><td></td></tr>
<tr><td>Oligo fwd</td><td>CACCACCCCGAGTTCAAGCGGTGC</td><td></td><td></td><td></td><td></td><td></td><td></td><td></td><td></td><td></td><td></td></tr>
<tr><td>Oligo rev</td><td>AAACGCACCGCTTGAACTCGGGGT</td><td></td><td></td><td></td><td></td><td></td><td></td><td></td><td></td><td></td><td></td></tr>
<tr><td>Chromosome</td><td>start</td><td>end</td><td>strand</td><td>MM</td><td>target_seq</td><td>PAM</td><td>alignment</td><td>distance</td><td>position</td><td>gene name</td><td>gene id</td></tr>
<tr><td>GU305914.1</td><td>198597</td><td>198619</td><td>-</td><td>0</td><td>ACCCCGAGTTCAAGCGGTGC</td><td>TGG</td><td>||||||||[||||||||||||]PAM</td><td>NA</td><td>Intergenic</td><td>NA</td><td>NA</td></tr>
<tr><td>T19563</td><td>GTTCAAGCGGTGCTGGAACGCGG</td><td>1000</td><td></td><td></td><td></td><td></td><td></td><td></td><td></td><td></td><td></td></tr>
<tr><td>Oligo fwd</td><td>CACCGTTCAAGCGGTGCTGGAACG</td><td></td><td></td><td></td><td></td><td></td><td></td><td></td><td></td><td></td><td></td></tr>
<tr><td>Oligo rev</td><td>AAACCGTTCCAGCACCGCTTGAAC</td><td></td><td></td><td></td><td></td><td></td><td></td><td></td><td></td><td></td><td></td></tr>
<tr><td>Chromosome</td><td>start</td><td>end</td><td>strand</td><td>MM</td><td>target_seq</td><td>PAM</td><td>alignment</td><td>distance</td><td>position</td><td>gene name</td><td>gene id</td></tr>
<tr><td>GU305914.1</td><td>198590</td><td>198612</td><td>-</td><td>0</td><td>GTTCAAGCGGTGCTGGAACG</td><td>CGG</td><td>||||||||[||||||||||||]PAM</td><td>NA</td><td>Intergenic</td><td>NA</td><td>NA</td></tr>
<tr><td>T19564</td><td>GCGGTGCTGGAACGCGGTGTCGG</td><td>1000</td><td></td><td></td><td></td><td></td><td></td><td></td><td></td><td></td><td></td></tr>
<tr><td>Oligo fwd</td><td>CACCGCGGTGCTGGAACGCGGTGT</td><td></td><td></td><td></td><td></td><td></td><td></td><td></td><td></td><td></td><td></td></tr>
<tr><td>Oligo rev</td><td>AAACACACCGCGTTCCAGCACCGC</td><td></td><td></td><td></td><td></td><td></td><td></td><td></td><td></td><td></td><td></td></tr>
<tr><td>Chromosome</td><td>start</td><td>end</td><td>strand</td><td>MM</td><td>target_seq</td><td>PAM</td><td>alignment</td><td>distance</td><td>position</td><td>gene name</td><td>gene id</td></tr>
<tr><td>GU305914.1</td><td>198584</td><td>198606</td><td>-</td><td>0</td><td>GCGGTGCTGGAACGCGGTGT</td><td>CGG</td><td>||||||||[||||||||||||]PAM</td><td>NA</td><td>Intergenic</td><td>NA</td><td>NA</td></tr>
<tr><td>T19565</td><td>CGGACGCGTTGCACTGCTCGTGG</td><td>1000</td><td></td><td></td><td></td><td></td><td></td><td></td><td></td><td></td><td></td></tr>
<tr><td>Oligo fwd</td><td>CACCCGGACGCGTTGCACTGCTCG</td><td></td><td></td><td></td><td></td><td></td><td></td><td></td><td></td><td></td><td></td></tr>
<tr><td>Oligo rev</td><td>AAACCGAGCAGTGCAACGCGTCCG</td><td></td><td></td><td></td><td></td><td></td><td></td><td></td><td></td><td></td><td></td></tr>
<tr><td>Chromosome</td><td>start</td><td>end</td><td>strand</td><td>MM</td><td>target_seq</td><td>PAM</td><td>alignment</td><td>distance</td><td>position</td><td>gene name</td><td>gene id</td></tr>
<tr><td>GU305914.1</td><td>198564</td><td>198586</td><td>-</td><td>0</td><td>CGGACGCGTTGCACTGCTCG</td><td>TGG</td><td>||||||||[||||||||||||]PAM</td><td>NA</td><td>Intergenic</td><td>NA</td><td>NA</td></tr>
<tr><td>T19566</td><td>CACTGCTCGTGGTCGTTGATCGG</td><td>1000</td><td></td><td></td><td></td><td></td><td></td><td></td><td></td><td></td><td></td></tr>
<tr><td>Oligo fwd</td><td>CACCCACTGCTCGTGGTCGTTGAT</td><td></td><td></td><td></td><td></td><td></td><td></td><td></td><td></td><td></td><td></td></tr>
<tr><td>Oligo rev</td><td>AAACATCAACGACCACGAGCAGTG</td><td></td><td></td><td></td><td></td><td></td><td></td><td></td><td></td><td></td><td></td></tr>
<tr><td>Chromosome</td><td>start</td><td>end</td><td>strand</td><td>MM</td><td>target_seq</td><td>PAM</td><td>alignment</td><td>distance</td><td>position</td><td>gene name</td><td>gene id</td></tr>
<tr><td>GU305914.1</td><td>198553</td><td>198575</td><td>-</td><td>0</td><td>CACTGCTCGTGGTCGTTGAT</td><td>CGG</td><td>||||||||[||||||||||||]PAM</td><td>NA</td><td>Intergenic</td><td>NA</td><td>NA</td></tr>
<tr><td>T19567</td><td>ACTGCTCGTGGTCGTTGATCGGG</td><td>1000</td><td></td><td></td><td></td><td></td><td></td><td></td><td></td><td></td><td></td></tr>
<tr><td>Oligo fwd</td><td>CACCACTGCTCGTGGTCGTTGATC</td><td></td><td></td><td></td><td></td><td></td><td></td><td></td><td></td><td></td><td></td></tr>
<tr><td>Oligo rev</td><td>AAACGATCAACGACCACGAGCAGT</td><td></td><td></td><td></td><td></td><td></td><td></td><td></td><td></td><td></td><td></td></tr>
<tr><td>Chromosome</td><td>start</td><td>end</td><td>strand</td><td>MM</td><td>target_seq</td><td>PAM</td><td>alignment</td><td>distance</td><td>position</td><td>gene name</td><td>gene id</td></tr>
<tr><td>GU305914.1</td><td>198552</td><td>198574</td><td>-</td><td>0</td><td>ACTGCTCGTGGTCGTTGATC</td><td>GGG</td><td>||||||||[||||||||||||]PAM</td><td>NA</td><td>Intergenic</td><td>NA</td><td>NA</td></tr>
<tr><td>GU305914.1</td><td>8066</td><td>8088</td><td>-</td><td>3</td><td>ACTGACCGTGGTCGTTGGTC</td><td>TGG</td><td>||||--||[|||||||||-||]PAM</td><td>NA</td><td>Intergenic</td><td>NA</td><td>NA</td></tr>
<tr><td>T19568</td><td>CTGCTCGTGGTCGTTGATCGGGG</td><td>1000</td><td></td><td></td><td></td><td></td><td></td><td></td><td></td><td></td><td></td></tr>
<tr><td>Oligo fwd</td><td>CACCCTGCTCGTGGTCGTTGATCG</td><td></td><td></td><td></td><td></td><td></td><td></td><td></td><td></td><td></td><td></td></tr>
<tr><td>Oligo rev</td><td>AAACCGATCAACGACCACGAGCAG</td><td></td><td></td><td></td><td></td><td></td><td></td><td></td><td></td><td></td><td></td></tr>
<tr><td>Chromosome</td><td>start</td><td>end</td><td>strand</td><td>MM</td><td>target_seq</td><td>PAM</td><td>alignment</td><td>distance</td><td>position</td><td>gene name</td><td>gene id</td></tr>
</table>

| | | | | | | | | | | | |
|---|---|---|---|---|---|---|---|---|---|---|---|
| GU305914.1 | 198551 | 198573 | - | 0 | CTGCTCGTGGTCGTTGATCG | GGG | \|\|\|\|\|\|\|\|[\|\|\|\|\|\|\|\|\|\|\|\|]PAM | NA | Intergenic | NA | NA |
| T19569 | GTGGTCGTTGATCGGGGTGCTGG | 1000 | | | | | | | | | |
| Oligo fwd | CACCGTGGTCGTTGATCGGGGTGC | | | | | | | | | | |
| Oligo rev | AAACGCACCCCGATCAACGACCAC | | | | | | | | | | |
| Chromosome | start | end | strand | MM | target_seq | PAM | alignment | distance | position | gene name | gene id |
| GU305914.1 | 198545 | 198567 | - | 0 | GTGGTCGTTGATCGGGGTGC | TGG | \|\|\|\|\|\|\|\|[\|\|\|\|\|\|\|\|\|\|\|\|]PAM | NA | Intergenic | NA | NA |
| T19570 | TGGTCGTTGATCGGGGTGCTGGG | 1000 | | | | | | | | | |
| Oligo fwd | CACCTGGTCGTTGATCGGGGTGCT | | | | | | | | | | |
| Oligo rev | AAACAGCACCCCGATCAACGACCA | | | | | | | | | | |
| Chromosome | start | end | strand | MM | target_seq | PAM | alignment | distance | position | gene name | gene id |
| GU305914.1 | 198544 | 198566 | - | 0 | TGGTCGTTGATCGGGGTGCT | GGG | \|\|\|\|\|\|\|\|[\|\|\|\|\|\|\|\|\|\|\|\|]PAM | NA | Intergenic | NA | NA |
| T19571 | TGATCGGGGTGCTGGGCTACAGG | 1000 | | | | | | | | | |
| Oligo fwd | CACCTGATCGGGGTGCTGGGCTAC | | | | | | | | | | |
| Oligo rev | AAACGTAGCCCAGCACCCCGATCA | | | | | | | | | | |
| Chromosome | start | end | strand | MM | target_seq | PAM | alignment | distance | position | gene name | gene id |
| GU305914.1 | 198537 | 198559 | - | 0 | TGATCGGGGTGCTGGGCTAC | AGG | \|\|\|\|\|\|\|\|[\|\|\|\|\|\|\|\|\|\|\|\|]PAM | NA | Intergenic | NA | NA |
| T19572 | CGGGGTGCTGGGCTACAGGCCGG | 1000 | | | | | | | | | |
| Oligo fwd | CACCCGGGGTGCTGGGCTACAGGC | | | | | | | | | | |
| Oligo rev | AAACGCCTGTAGCCCAGCACCCCG | | | | | | | | | | |
| Chromosome | start | end | strand | MM | target_seq | PAM | alignment | distance | position | gene name | gene id |
| GU305914.1 | 198533 | 198555 | - | 0 | CGGGGTGCTGGGCTACAGGC | CGG | \|\|\|\|\|\|\|\|[\|\|\|\|\|\|\|\|\|\|\|\|]PAM | NA | Intergenic | NA | NA |
| T19573 | GGGGTGCTGGGCTACAGGCCGGG | 1000 | | | | | | | | | |
| Oligo fwd | CACCGGGGTGCTGGGCTACAGGCC | | | | | | | | | | |
| Oligo rev | AAACGGCCTGTAGCCCAGCACCCC | | | | | | | | | | |
| Chromosome | start | end | strand | MM | target_seq | PAM | alignment | distance | position | gene name | gene id |
| GU305914.1 | 198532 | 198554 | - | 0 | GGGGTGCTGGGCTACAGGCC | GGG | \|\|\|\|\|\|\|\|[\|\|\|\|\|\|\|\|\|\|\|\|]PAM | NA | Intergenic | NA | NA |
| T19574 | GGGTGCTGGGCTACAGGCCGGGG | 1000 | | | | | | | | | |
| Oligo fwd | CACCGGGTGCTGGGCTACAGGCCG | | | | | | | | | | |
| Oligo rev | AAACCGGCCTGTAGCCCAGCACCC | | | | | | | | | | |
| Chromosome | start | end | strand | MM | target_seq | PAM | alignment | distance | position | gene name | gene id |
| GU305914.1 | 198531 | 198553 | - | 0 | GGGTGCTGGGCTACAGGCCG | GGG | \|\|\|\|\|\|\|\|[\|\|\|\|\|\|\|\|\|\|\|\|]PAM | NA | Intergenic | NA | NA |
| T19575 | GGTGCTGGGCTACAGGCCGGGGG | 1000 | | | | | | | | | |
| Oligo fwd | CACCGGTGCTGGGCTACAGGCCGG | | | | | | | | | | |
| Oligo rev | AAACCCGGCCTGTAGCCCAGCACC | | | | | | | | | | |
| Chromosome | start | end | strand | MM | target_seq | PAM | alignment | distance | position | gene name | gene id |
| GU305914.1 | 198530 | 198552 | - | 0 | GGTGCTGGGCTACAGGCCGG | GGG | \|\|\|\|\|\|\|\|[\|\|\|\|\|\|\|\|\|\|\|\|]PAM | NA | Intergenic | NA | NA |
| T19576 | GTGCTGGGCTACAGGCCGGGGGG | 1000 | | | | | | | | | |
| Oligo fwd | CACCGTGCTGGGCTACAGGCCGGG | | | | | | | | | | |
| Oligo rev | AAACCCCGGCCTGTAGCCCAGCAC | | | | | | | | | | |
| Chromosome | start | end | strand | MM | target_seq | PAM | alignment | distance | position | gene name | gene id |
| GU305914.1 | 198529 | 198551 | - | 0 | GTGCTGGGCTACAGGCCGGG | GGG | \|\|\|\|\|\|\|\|[\|\|\|\|\|\|\|\|\|\|\|\|]PAM | NA | Intergenic | NA | NA |
| T19577 | CTACAGGCCGGGGGGCCTGCCGG | 1000 | | | | | | | | | |
| Oligo fwd | CACCCTACAGGCCGGGGGGCCTGC | | | | | | | | | | |
| Oligo rev | AAACGCAGGCCCCCCGGCCTGTAG | | | | | | | | | | |
| Chromosome | start | end | strand | MM | target_seq | PAM | alignment | distance | position | gene name | gene id |
| GU305914.1 | 198521 | 198543 | - | 0 | CTACAGGCCGGGGGGCCTGC | CGG | \|\|\|\|\|\|\|\|[\|\|\|\|\|\|\|\|\|\|\|\|]PAM | NA | Intergenic | NA | NA |
| T19578 | GCCGGGGGGCCTGCCGGACGCGG | 1000 | | | | | | | | | |
| Oligo fwd | CACCGCCGGGGGGCCTGCCGGACG | | | | | | | | | | |
| Oligo rev | AAACCGTCCGGCAGGCCCCCCGGC | | | | | | | | | | |
| Chromosome | start | end | strand | MM | target_seq | PAM | alignment | distance | position | gene name | gene id |
| GU305914.1 | 198515 | 198537 | - | 0 | GCCGGGGGGCCTGCCGGACG | CGG | \|\|\|\|\|\|\|\|[\|\|\|\|\|\|\|\|\|\|\|\|]PAM | NA | Intergenic | NA | NA |
| T19579 | GCCTGCCGGACGCGGAGTTCAGG | 1000 | | | | | | | | | |
| Oligo fwd | CACCGCCTGCCGGACGCGGAGTTC | | | | | | | | | | |
| Oligo rev | AAACGAACTCCGCGTCCGGCAGGC | | | | | | | | | | |
| Chromosome | start | end | strand | MM | target_seq | PAM | alignment | distance | position | gene name | gene id |
| GU305914.1 | 198507 | 198529 | - | 0 | GCCTGCCGGACGCGGAGTTC | AGG | \|\|\|\|\|\|\|\|[\|\|\|\|\|\|\|\|\|\|\|\|]PAM | NA | Intergenic | NA | NA |
| T19580 | CCTGCCGGACGCGGAGTTCAGGG | 1000 | | | | | | | | | |
| Oligo fwd | CACCCCTGCCGGACGCGGAGTTCA | | | | | | | | | | |
| Oligo rev | AAACTGAACTCCGCGTCCGGCAGG | | | | | | | | | | |
| Chromosome | start | end | strand | MM | target_seq | PAM | alignment | distance | position | gene name | gene id |
| GU305914.1 | 198506 | 198528 | - | 0 | CCTGCCGGACGCGGAGTTCA | GGG | \|\|\|\|\|\|\|\|[\|\|\|\|\|\|\|\|\|\|\|\|]PAM | NA | Intergenic | NA | NA |
| T19581 | CAGGGCCGACCACGTGATCGTGG | 1000 | | | | | | | | | |
| Oligo fwd | CACCCAGGGCCGACCACGTGATCG | | | | | | | | | | |
| Oligo rev | AAACCGATCACGTGGTCGGCCCTG | | | | | | | | | | |
| Chromosome | start | end | strand | MM | target_seq | PAM | alignment | distance | position | gene name | gene id |
| GU305914.1 | 198488 | 198510 | - | 0 | CAGGGCCGACCACGTGATCG | TGG | \|\|\|\|\|\|\|\|[\|\|\|\|\|\|\|\|\|\|\|\|]PAM | NA | Intergenic | NA | NA |
| T19582 | GTGATCGTGGCCGACAAGTTCGG | 1000 | | | | | | | | | |
| Oligo fwd | CACCGTGATCGTGGCCGACAAGTT | | | | | | | | | | |
| Oligo rev | AAACAACTTGTCGGCCACGATCAC | | | | | | | | | | |
| Chromosome | start | end | strand | MM | target_seq | PAM | alignment | distance | position | gene name | gene id |
| GU305914.1 | 198475 | 198497 | - | 0 | GTGATCGTGGCCGACAAGTT | CGG | \|\|\|\|\|\|\|\|[\|\|\|\|\|\|\|\|\|\|\|\|]PAM | NA | Intergenic | NA | NA |
| T19583 | TGATCGTGGCCGACAAGTTCGGG | 1000 | | | | | | | | | |
| Oligo fwd | CACCTGATCGTGGCCGACAAGTTC | | | | | | | | | | |
| Oligo rev | AAACGAACTTGTCGGCCACGATCA | | | | | | | | | | |

Chromosome start end strand MM target_seq PAM alignment distance position gene name gene id
GU305914.1 198474 198496 - 0 TGATCGTGGCCGACAAGTTC GGG ||||||||[||||||||||||]PAM NA Intergenic NA NA

T19584 GATCGTGGCCGACAAGTTCGGGG 1000
Oligo fwd CACCGATCGTGGCCGACAAGTTCG
Oligo rev AAACCGAACTTGTCGGCCACGATC
Chromosome start end strand MM target_seq PAM alignment distance position gene name gene id
GU305914.1 198473 198495 - 0 GATCGTGGCCGACAAGTTCG GGG ||||||||[||||||||||||]PAM NA Intergenic NA NA

T19585 CGGGGCCGTGTACAGCCTGCTGG 1000
Oligo fwd CACCCGGGGCCGTGTACAGCCTGC
Oligo rev AAACGCAGGCTGTACACGGCCCCG
Chromosome start end strand MM target_seq PAM alignment distance position gene name gene id
GU305914.1 198455 198477 - 0 CGGGGCCGTGTACAGCCTGC TGG ||||||||[||||||||||||]PAM NA Intergenic NA NA

T19586 CCTGCTGGTGTACTCGAGAGCGG 1000
Oligo fwd CACCCCTGCTGGTGTACTCGAGAG
Oligo rev AAACCTCTCGAGTACACCAGCAGG
Chromosome start end strand MM target_seq PAM alignment distance position gene name gene id
GU305914.1 198440 198462 - 0 CCTGCTGGTGTACTCGAGAG CGG ||||||||[||||||||||||]PAM NA Intergenic NA NA

T19587 CTGCTGGTGTACTCGAGAGCGGG 1000
Oligo fwd CACCCTGCTGGTGTACTCGAGAGC
Oligo rev AAACGCTCTCGAGTACACCAGCAG
Chromosome start end strand MM target_seq PAM alignment distance position gene name gene id
GU305914.1 198439 198461 - 0 CTGCTGGTGTACTCGAGAGC GGG ||||||||[||||||||||||]PAM NA Intergenic NA NA

T19588 ACTCGAGAGCGGGCCTGACGCGG 1000
Oligo fwd CACCACTCGAGAGCGGGCCTGACG
Oligo rev AAACCGTCAGGCCCGCTCTCGAGT
Chromosome start end strand MM target_seq PAM alignment distance position gene name gene id
GU305914.1 198429 198451 - 0 ACTCGAGAGCGGGCCTGACG CGG ||||||||[||||||||||||]PAM NA Intergenic NA NA

T19589 GCGGCGCGTCGCCGACAGCGTGG 1000
Oligo fwd CACCGCGGCGCGTCGCCGACAGCG
Oligo rev AAACCGCTGTCGGCGACGCGCCGC
Chromosome start end strand MM target_seq PAM alignment distance position gene name gene id
GU305914.1 198410 198432 - 0 GCGGCGCGTCGCCGACAGCG TGG ||||||||[||||||||||||]PAM NA Intergenic NA NA

T19590 CGTGGCCGAATTCTTCCAGATGG 1000
Oligo fwd CACCCGTGGCCGAATTCTTCCAGA
Oligo rev AAACTCTGGAAGAATTCGGCCACG
Chromosome start end strand MM target_seq PAM alignment distance position gene name gene id
GU305914.1 198392 198414 - 0 CGTGGCCGAATTCTTCCAGA TGG ||||||||[||||||||||||]PAM NA Intergenic NA NA

T19591 GTGGCCGAATTCTTCCAGATGGG 1000
Oligo fwd CACCGTGGCCGAATTCTTCCAGAT
Oligo rev AAACATCTGGAAGAATTCGGCCAC
Chromosome start end strand MM target_seq PAM alignment distance position gene name gene id
GU305914.1 198391 198413 - 0 GTGGCCGAATTCTTCCAGAT GGG ||||||||[||||||||||||]PAM NA Intergenic NA NA

T19592 TCCAGATGGGCATGCTGAAGAGG 1000
Oligo fwd CACCTCCAGATGGGCATGCTGAAG
Oligo rev AAACCTTCAGCATGCCCATCTGGA
Chromosome start end strand MM target_seq PAM alignment distance position gene name gene id
GU305914.1 198378 198400 - 0 TCCAGATGGGCATGCTGAAG AGG ||||||||[||||||||||||]PAM NA Intergenic NA NA

T19593 CCAGATGGGCATGCTGAAGAGGG 1000
Oligo fwd CACCCCAGATGGGCATGCTGAAGA
Oligo rev AAACTCTTCAGCATGCCCATCTGG
Chromosome start end strand MM target_seq PAM alignment distance position gene name gene id
GU305914.1 198377 198399 - 0 CCAGATGGGCATGCTGAAGA GGG ||||||||[||||||||||||]PAM NA Intergenic NA NA

T19594 CAGACAGAGAGAGCGCCGCAAGG 1000
Oligo fwd CACCCAGACAGAGAGAGCGCCGCA
Oligo rev AAACTGCGGCGCTCTCTCTGTCTG
Chromosome start end strand MM target_seq PAM alignment distance position gene name gene id
GU305914.1 198332 198354 - 0 CAGACAGAGAGAGCGCCGCA AGG ||||||||[||||||||||||]PAM NA Intergenic NA NA

T19595 AGAGCGCCGCAAGGAGCGTCTGG 1000
Oligo fwd CACCAGAGCGCCGCAAGGAGCGTC
Oligo rev AAACGACGCTCCTTGCGGCGCTCT
Chromosome start end strand MM target_seq PAM alignment distance position gene name gene id
GU305914.1 198323 198345 - 0 AGAGCGCCGCAAGGAGCGTC TGG ||||||||[||||||||||||]PAM NA Intergenic NA NA

T19596 AGCGTCTGGAGAGCATGCCCAGG 1000
Oligo fwd CACCAGCGTCTGGAGAGCATGCCC
Oligo rev AAACGGGCATGCTCTCCAGACGCT
Chromosome start end strand MM target_seq PAM alignment distance position gene name gene id
GU305914.1 198309 198331 - 0 AGCGTCTGGAGAGCATGCCC AGG ||||||||[||||||||||||]PAM NA Intergenic NA NA

T19597 GCATGCCCAGGTGCCCGCACTGG 1000
Oligo fwd CACCGCATGCCCAGGTGCCCGCAC
Oligo rev AAACGTGCGGGCACCTGGGCATGC
Chromosome start end strand MM target_seq PAM alignment distance position gene name gene id
GU305914.1 198297 198319 - 0 GCATGCCCAGGTGCCCGCAC TGG ||||||||[||||||||||||]PAM NA Intergenic NA NA

T19598 GCAGAAGTTCAAGTACATCGAGG 1000
Oligo fwd CACCGCAGAAGTTCAAGTACATCG

| | | | | | | | | | | | |
|---|---|---|---|---|---|---|---|---|---|---|---|
| Oligo rev | AAACCGATGTACTTGAACTTCTGC | | | | | | | | | | |
| Chromosome | start | end | strand | MM | target_seq | PAM | alignment | distance | position | gene name | gene id |
| GU305914.1 | 198269 | 198291 | - | 0 | GCAGAAGTTCAAGTACATCG | AGG | \|\|\|\|\|\|\|\|[\|\|\|\|\|\|\|\|\|\|\|\|]PAM | NA | Intergenic | NA | NA |
| T19599 | CGACTCGCCGACCGACGATGAGG | 1000 | | | | | | | | | |
| Oligo fwd | CACCCGACTCGCCGACCGACGATG | | | | | | | | | | |
| Oligo rev | AAACCATCGTCGGTCGGCGAGTCG | | | | | | | | | | |
| Chromosome | start | end | strand | MM | target_seq | PAM | alignment | distance | position | gene name | gene id |
| GU305914.1 | 198227 | 198249 | - | 0 | CGACTCGCCGACCGACGATG | AGG | \|\|\|\|\|\|\|\|[\|\|\|\|\|\|\|\|\|\|\|\|]PAM | NA | Intergenic | NA | NA |
| T19600 | CTACGTGCCCGCCGATTGCATGG | 1000 | | | | | | | | | |
| Oligo fwd | CACCCTACGTGCCCGCCGATTGCA | | | | | | | | | | |
| Oligo rev | AAACTGCAATCGGCGGGCACGTAG | | | | | | | | | | |
| Chromosome | start | end | strand | MM | target_seq | PAM | alignment | distance | position | gene name | gene id |
| GU305914.1 | 198203 | 198225 | - | 0 | CTACGTGCCCGCCGATTGCA | TGG | \|\|\|\|\|\|\|\|[\|\|\|\|\|\|\|\|\|\|\|\|]PAM | NA | Intergenic | NA | NA |
| T19601 | GCATGGCCGAGCACTACGCCTGG | 1000 | | | | | | | | | |
| Oligo fwd | CACCGCATGGCCGAGCACTACGCC | | | | | | | | | | |
| Oligo rev | AAACGGCGTAGTGCTCGGCCATGC | | | | | | | | | | |
| Chromosome | start | end | strand | MM | target_seq | PAM | alignment | distance | position | gene name | gene id |
| GU305914.1 | 198186 | 198208 | - | 0 | GCATGGCCGAGCACTACGCC | TGG | \|\|\|\|\|\|\|\|[\|\|\|\|\|\|\|\|\|\|\|\|]PAM | NA | Intergenic | NA | NA |
| T19602 | GGCCGAGCACTACGCCTGGCTGG | 1000 | | | | | | | | | |
| Oligo fwd | CACCGGCCGAGCACTACGCCTGGC | | | | | | | | | | |
| Oligo rev | AAACGCCAGGCGTAGTGCTCGGCC | | | | | | | | | | |
| Chromosome | start | end | strand | MM | target_seq | PAM | alignment | distance | position | gene name | gene id |
| GU305914.1 | 198182 | 198204 | - | 0 | GGCCGAGCACTACGCCTGGC | TGG | \|\|\|\|\|\|\|\|[\|\|\|\|\|\|\|\|\|\|\|\|]PAM | NA | Intergenic | NA | NA |
| T19603 | GCTGGTGCAGTCCACCGTCAAGG | 1000 | | | | | | | | | |
| Oligo fwd | CACCGCTGGTGCAGTCCACCGTCA | | | | | | | | | | |
| Oligo rev | AAACTGACGGTGGACTGCACCAGC | | | | | | | | | | |
| Chromosome | start | end | strand | MM | target_seq | PAM | alignment | distance | position | gene name | gene id |
| GU305914.1 | 198164 | 198186 | - | 0 | GCTGGTGCAGTCCACCGTCA | AGG | \|\|\|\|\|\|\|\|[\|\|\|\|\|\|\|\|\|\|\|\|]PAM | NA | Intergenic | NA | NA |
| T19604 | AGGTGCACTCCTGCCGCCCCTGG | 1000 | | | | | | | | | |
| Oligo fwd | CACCAGGTGCACTCCTGCCGCCCC | | | | | | | | | | |
| Oligo rev | AAACGGGGCGGCAGGAGTGCACCT | | | | | | | | | | |
| Chromosome | start | end | strand | MM | target_seq | PAM | alignment | distance | position | gene name | gene id |
| GU305914.1 | 198144 | 198166 | - | 0 | AGGTGCACTCCTGCCGCCCC | TGG | \|\|\|\|\|\|\|\|[\|\|\|\|\|\|\|\|\|\|\|\|]PAM | NA | Intergenic | NA | NA |
| T19605 | GGTGCACTCCTGCCGCCCCTGGG | 1000 | | | | | | | | | |
| Oligo fwd | CACCGGTGCACTCCTGCCGCCCCT | | | | | | | | | | |
| Oligo rev | AAACAGGGGCGGCAGGAGTGCACC | | | | | | | | | | |
| Chromosome | start | end | strand | MM | target_seq | PAM | alignment | distance | position | gene name | gene id |
| GU305914.1 | 198143 | 198165 | - | 0 | GGTGCACTCCTGCCGCCCCT | GGG | \|\|\|\|\|\|\|\|[\|\|\|\|\|\|\|\|\|\|\|\|]PAM | NA | Intergenic | NA | NA |
| T19606 | CTCCTGCCGCCCCTGGGACCTGG | 1000 | | | | | | | | | |
| Oligo fwd | CACCCTCCTGCCGCCCCTGGGACC | | | | | | | | | | |
| Oligo rev | AAACGGTCCCAGGGGCGGCAGGAG | | | | | | | | | | |
| Chromosome | start | end | strand | MM | target_seq | PAM | alignment | distance | position | gene name | gene id |
| GU305914.1 | 198137 | 198159 | - | 0 | CTCCTGCCGCCCCTGGGACC | TGG | \|\|\|\|\|\|\|\|[\|\|\|\|\|\|\|\|\|\|\|\|]PAM | NA | Intergenic | NA | NA |
| T19607 | GGACCTGGACGACCCCAACGTGG | 1000 | | | | | | | | | |
| Oligo fwd | CACCGGACCTGGACGACCCCAACG | | | | | | | | | | |
| Oligo rev | AAACCGTTGGGGTCGTCCAGGTCC | | | | | | | | | | |
| Chromosome | start | end | strand | MM | target_seq | PAM | alignment | distance | position | gene name | gene id |
| GU305914.1 | 198122 | 198144 | - | 0 | GGACCTGGACGACCCCAACG | TGG | \|\|\|\|\|\|\|\|[\|\|\|\|\|\|\|\|\|\|\|\|]PAM | NA | Intergenic | NA | NA |
| T19608 | ACGACCCCAACGTGGCGCCGCGG | 1000 | | | | | | | | | |
| Oligo fwd | CACCACGACCCCAACGTGGCGCCG | | | | | | | | | | |
| Oligo rev | AAACCGGCGCCACGTTGGGGTCGT | | | | | | | | | | |
| Chromosome | start | end | strand | MM | target_seq | PAM | alignment | distance | position | gene name | gene id |
| GU305914.1 | 198114 | 198136 | - | 0 | ACGACCCCAACGTGGCGCCG | CGG | \|\|\|\|\|\|\|\|[\|\|\|\|\|\|\|\|\|\|\|\|]PAM | NA | Intergenic | NA | NA |
| T19609 | CGACCCCAACGTGGCGCCGCGGG | 1000 | | | | | | | | | |
| Oligo fwd | CACCCGACCCCAACGTGGCGCCGC | | | | | | | | | | |
| Oligo rev | AAACGCGGCGCCACGTTGGGGTCG | | | | | | | | | | |
| Chromosome | start | end | strand | MM | target_seq | PAM | alignment | distance | position | gene name | gene id |
| GU305914.1 | 198113 | 198135 | - | 0 | CGACCCCAACGTGGCGCCGC | GGG | \|\|\|\|\|\|\|\|[\|\|\|\|\|\|\|\|\|\|\|\|]PAM | NA | Intergenic | NA | NA |
| T19610 | CAACGTGGCGCCGCGGGCGCTGG | 1000 | | | | | | | | | |
| Oligo fwd | CACCCAACGTGGCGCCGCGGGCGC | | | | | | | | | | |
| Oligo rev | AAACGCGCCCGCGGCGCCACGTTG | | | | | | | | | | |
| Chromosome | start | end | strand | MM | target_seq | PAM | alignment | distance | position | gene name | gene id |
| GU305914.1 | 198107 | 198129 | - | 0 | CAACGTGGCGCCGCGGGCGC | TGG | \|\|\|\|\|\|\|\|[\|\|\|\|\|\|\|\|\|\|\|\|]PAM | NA | Intergenic | NA | NA |
| T19611 | GCGGGCGCTGGTGACCAACGCGG | 1000 | | | | | | | | | |
| Oligo fwd | CACCGCGGGCGCTGGTGACCAACG | | | | | | | | | | |
| Oligo rev | AAACCGTTGGTCACCAGCGCCCGC | | | | | | | | | | |
| Chromosome | start | end | strand | MM | target_seq | PAM | alignment | distance | position | gene name | gene id |
| GU305914.1 | 198095 | 198117 | - | 0 | GCGGGCGCTGGTGACCAACG | CGG | \|\|\|\|\|\|\|\|[\|\|\|\|\|\|\|\|\|\|\|\|]PAM | NA | Intergenic | NA | NA |
| T19612 | CTGGTGACCAACGCGGACACCGG | 1000 | | | | | | | | | |
| Oligo fwd | CACCCTGGTGACCAACGCGGACAC | | | | | | | | | | |
| Oligo rev | AAACGTGTCCGCGTTGGTCACCAG | | | | | | | | | | |
| Chromosome | start | end | strand | MM | target_seq | PAM | alignment | distance | position | gene name | gene id |
| GU305914.1 | 198088 | 198110 | - | 0 | CTGGTGACCAACGCGGACAC | CGG | \|\|\|\|\|\|\|\|[\|\|\|\|\|\|\|\|\|\|\|\|]PAM | NA | Intergenic | NA | NA |
| T19613 | GCGATCCGATGAAGACCTACTGG | 1000 | | | | | | | | | |

| | | | | | | | | | | | |
|---|---|---|---|---|---|---|---|---|---|---|---|
| Oligo fwd | CACCGCGATCCGATGAAGACCTAC | | | | | | | | | | |
| Oligo rev | AAACGTAGGTCTTCATCGGATCGC | | | | | | | | | | |
| Chromosome | start | end | strand | MM | target_seq | PAM | alignment | distance | position | gene name | gene id |
| GU305914.1 | 198066 | 198088 | - | 0 | GCGATCCGATGAAGACCTAC | TGG | \|\|\|\|\|\|\|\|[\|\|\|\|\|\|\|\|\|\|\|\|]PAM | NA | Intergenic | NA | NA |
| T19614 | CGATCCGATGAAGACCTACTGGG | 1000 | | | | | | | | | |
| Oligo fwd | CACCCGATCCGATGAAGACCTACT | | | | | | | | | | |
| Oligo rev | AAACAGTAGGTCTTCATCGGATCG | | | | | | | | | | |
| Chromosome | start | end | strand | MM | target_seq | PAM | alignment | distance | position | gene name | gene id |
| GU305914.1 | 198065 | 198087 | - | 0 | CGATCCGATGAAGACCTACT | GGG | \|\|\|\|\|\|\|\|[\|\|\|\|\|\|\|\|\|\|\|\|]PAM | NA | Intergenic | NA | NA |
| GU305914.1 | 179284 | 179306 | + | 4 | TGATCTGAGGAAGACCTGCT | GGG | -\|\|\|\|-\|\|[-\|\|\|\|\|\|\|\|-\|\|]PAM | NA | Intergenic | NA | NA |
| T19615 | GAAGACCTACTGGGACCGCCTGG | 1000 | | | | | | | | | |
| Oligo fwd | CACCGAAGACCTACTGGGACCGCC | | | | | | | | | | |
| Oligo rev | AAACGGCGGTCCCAGTAGGTCTTC | | | | | | | | | | |
| Chromosome | start | end | strand | MM | target_seq | PAM | alignment | distance | position | gene name | gene id |
| GU305914.1 | 198056 | 198078 | - | 0 | GAAGACCTACTGGGACCGCC | TGG | \|\|\|\|\|\|\|\|[\|\|\|\|\|\|\|\|\|\|\|\|]PAM | NA | Intergenic | NA | NA |
| T19616 | CTACTGGGACCGCCTGGAGTCGG | 1000 | | | | | | | | | |
| Oligo fwd | CACCCTACTGGGACCGCCTGGAGT | | | | | | | | | | |
| Oligo rev | AAACACTCCAGGCGGTCCCAGTAG | | | | | | | | | | |
| Chromosome | start | end | strand | MM | target_seq | PAM | alignment | distance | position | gene name | gene id |
| GU305914.1 | 198050 | 198072 | - | 0 | CTACTGGGACCGCCTGGAGT | CGG | \|\|\|\|\|\|\|\|[\|\|\|\|\|\|\|\|\|\|\|\|]PAM | NA | Intergenic | NA | NA |
| T19617 | CCTGGAGTCGGCGATCGCCAAGG | 1000 | | | | | | | | | |
| Oligo fwd | CACCCCTGGAGTCGGCGATCGCCA | | | | | | | | | | |
| Oligo rev | AAACTGGCGATCGCCGACTCCAGG | | | | | | | | | | |
| Chromosome | start | end | strand | MM | target_seq | PAM | alignment | distance | position | gene name | gene id |
| GU305914.1 | 198038 | 198060 | - | 0 | CCTGGAGTCGGCGATCGCCA | AGG | \|\|\|\|\|\|\|\|[\|\|\|\|\|\|\|\|\|\|\|\|]PAM | NA | Intergenic | NA | NA |
| T19618 | GATCGCCAAGGTGAAGCTCACGG | 1000 | | | | | | | | | |
| Oligo fwd | CACCGATCGCCAAGGTGAAGCTCA | | | | | | | | | | |
| Oligo rev | AAACTGAGCTTCACCTTGGCGATC | | | | | | | | | | |
| Chromosome | start | end | strand | MM | target_seq | PAM | alignment | distance | position | gene name | gene id |
| GU305914.1 | 198026 | 198048 | - | 0 | GATCGCCAAGGTGAAGCTCA | CGG | \|\|\|\|\|\|\|\|[\|\|\|\|\|\|\|\|\|\|\|\|]PAM | NA | Intergenic | NA | NA |
| T19619 | CGCCAAGGTGAAGCTCACGGCGG | 1000 | | | | | | | | | |
| Oligo fwd | CACCCGCCAAGGTGAAGCTCACGG | | | | | | | | | | |
| Oligo rev | AAACCCGTGAGCTTCACCTTGGCG | | | | | | | | | | |
| Chromosome | start | end | strand | MM | target_seq | PAM | alignment | distance | position | gene name | gene id |
| GU305914.1 | 198023 | 198045 | - | 0 | CGCCAAGGTGAAGCTCACGG | CGG | \|\|\|\|\|\|\|\|[\|\|\|\|\|\|\|\|\|\|\|\|]PAM | NA | Intergenic | NA | NA |
| T19620 | GAAGCTCACGGCGGCCGCGACGG | 1000 | | | | | | | | | |
| Oligo fwd | CACCGAAGCTCACGGCGGCCGCGA | | | | | | | | | | |
| Oligo rev | AAACTCGCGGCCGCCGTGAGCTTC | | | | | | | | | | |
| Chromosome | start | end | strand | MM | target_seq | PAM | alignment | distance | position | gene name | gene id |
| GU305914.1 | 198014 | 198036 | - | 0 | GAAGCTCACGGCGGCCGCGA | CGG | \|\|\|\|\|\|\|\|[\|\|\|\|\|\|\|\|\|\|\|\|]PAM | NA | Intergenic | NA | NA |
| T19621 | CACGGCGGCCGCGACGGCGTCGG | 1000 | | | | | | | | | |
| Oligo fwd | CACCCACGGCGGCCGCGACGGCGT | | | | | | | | | | |
| Oligo rev | AAACACGCCGTCGCGGCCGCCGTG | | | | | | | | | | |
| Chromosome | start | end | strand | MM | target_seq | PAM | alignment | distance | position | gene name | gene id |
| GU305914.1 | 198008 | 198030 | - | 0 | CACGGCGGCCGCGACGGCGT | CGG | \|\|\|\|\|\|\|\|[\|\|\|\|\|\|\|\|\|\|\|\|]PAM | NA | Intergenic | NA | NA |
| GU305914.1 | 113912 | 113934 | - | 4 | GGCGGCAGCGGCGACGGCGT | CGG | --\|\|\|\|-\|[\|-\|\|\|\|\|\|\|\|\|\|]PAM | NA | Intergenic | NA | NA |
| T19622 | GCGTCGGACGTCGAGTCGAGCGG | 1000 | | | | | | | | | |
| Oligo fwd | CACCGCGTCGGACGTCGAGTCGAG | | | | | | | | | | |
| Oligo rev | AAACCTCGACTCGACGTCCGACGC | | | | | | | | | | |
| Chromosome | start | end | strand | MM | target_seq | PAM | alignment | distance | position | gene name | gene id |
| GU305914.1 | 197992 | 198014 | - | 0 | GCGTCGGACGTCGAGTCGAG | CGG | \|\|\|\|\|\|\|\|[\|\|\|\|\|\|\|\|\|\|\|\|]PAM | NA | Intergenic | NA | NA |
| T19623 | TCGGACGTCGAGTCGAGCGGCGG | 1000 | | | | | | | | | |
| Oligo fwd | CACCTCGGACGTCGAGTCGAGCGG | | | | | | | | | | |
| Oligo rev | AAACCCGCTCGACTCGACGTCCGA | | | | | | | | | | |
| Chromosome | start | end | strand | MM | target_seq | PAM | alignment | distance | position | gene name | gene id |
| GU305914.1 | 197989 | 198011 | - | 0 | TCGGACGTCGAGTCGAGCGG | CGG | \|\|\|\|\|\|\|\|[\|\|\|\|\|\|\|\|\|\|\|\|]PAM | NA | Intergenic | NA | NA |
| GU305914.1 | 77765 | 77787 | + | 4 | TCGGAGATCGAGCCGAGCGT | GGG | \|\|\|\|\|--\|[\|\|\|\|-\|\|\|\|\|\|-]PAM | NA | Intergenic | NA | NA |
| T19624 | ACGTCGAGTCGAGCGGCGGTCGG | 1000 | | | | | | | | | |
| Oligo fwd | CACCACGTCGAGTCGAGCGGCGGT | | | | | | | | | | |
| Oligo rev | AAACACCGCCGCTCGACTCGACGT | | | | | | | | | | |
| Chromosome | start | end | strand | MM | target_seq | PAM | alignment | distance | position | gene name | gene id |
| GU305914.1 | 197985 | 198007 | - | 0 | ACGTCGAGTCGAGCGGCGGT | CGG | \|\|\|\|\|\|\|\|[\|\|\|\|\|\|\|\|\|\|\|\|]PAM | NA | Intergenic | NA | NA |
| T19625 | GAGTCGAGCGGCGGTCGGTACGG | 1000 | | | | | | | | | |
| Oligo fwd | CACCGAGTCGAGCGGCGGTCGGTA | | | | | | | | | | |
| Oligo rev | AAACTACCGACCGCCGCTCGACTC | | | | | | | | | | |
| Chromosome | start | end | strand | MM | target_seq | PAM | alignment | distance | position | gene name | gene id |
| GU305914.1 | 197980 | 198002 | - | 0 | GAGTCGAGCGGCGGTCGGTA | CGG | \|\|\|\|\|\|\|\|[\|\|\|\|\|\|\|\|\|\|\|\|]PAM | NA | Intergenic | NA | NA |
| T19626 | GAGCGGCGGTCGGTACGGCGAGG | 1000 | | | | | | | | | |
| Oligo fwd | CACCGAGCGGCGGTCGGTACGGCG | | | | | | | | | | |
| Oligo rev | AAACCGCCGTACCGACCGCCGCTC | | | | | | | | | | |
| Chromosome | start | end | strand | MM | target_seq | PAM | alignment | distance | position | gene name | gene id |
| GU305914.1 | 197975 | 197997 | - | 0 | GAGCGGCGGTCGGTACGGCG | AGG | \|\|\|\|\|\|\|\|[\|\|\|\|\|\|\|\|\|\|\|\|]PAM | NA | Intergenic | NA | NA |
| T19627 | AGCGGCGGTCGGTACGGCGAGGG | 1000 | | | | | | | | | |
| Oligo fwd | CACCAGCGGCGGTCGGTACGGCGA | | | | | | | | | | |
| Oligo rev | AAACTCGCCGTACCGACCGCCGCT | | | | | | | | | | |

| | | | | | | | | | | | |
|---|---|---|---|---|---|---|---|---|---|---|---|
| Chromosome | start | end | strand | MM | target_seq | PAM | alignment | distance | position | gene name | gene id |
| GU305914.1 | 197974 | 197996 | - | 0 | AGCGGCGGTCGGTACGGCGA | GGG | \|\|\|\|\|\|\|\|[\|\|\|\|\|\|\|\|\|\|\|\|]PAM | NA | Intergenic | NA | NA |
| T19628 | CTACGTGTACAAGCTGAGCCCGG | 1000 | | | | | | | | | |
| Oligo fwd | CACCCTACGTGTACAAGCTGAGCC | | | | | | | | | | |
| Oligo rev | AAACGGCTCAGCTTGTACACGTAG | | | | | | | | | | |
| Chromosome | start | end | strand | MM | target_seq | PAM | alignment | distance | position | gene name | gene id |
| GU305914.1 | 197948 | 197970 | - | 0 | CTACGTGTACAAGCTGAGCC | CGG | \|\|\|\|\|\|\|\|[\|\|\|\|\|\|\|\|\|\|\|\|]PAM | NA | Intergenic | NA | NA |
| T19629 | CGCGTTGACCTGCCACGAGACGG | 1000 | | | | | | | | | |
| Oligo fwd | CACCCGCGTTGACCTGCCACGAGA | | | | | | | | | | |
| Oligo rev | AAACTCTCGTGGCAGGTCAACGCG | | | | | | | | | | |
| Chromosome | start | end | strand | MM | target_seq | PAM | alignment | distance | position | gene name | gene id |
| GU305914.1 | 197909 | 197931 | - | 0 | CGCGTTGACCTGCCACGAGA | CGG | \|\|\|\|\|\|\|\|[\|\|\|\|\|\|\|\|\|\|\|\|]PAM | NA | Intergenic | NA | NA |
| T19630 | CGAGACGGAGTCTATGACGATGG | 1000 | | | | | | | | | |
| Oligo fwd | CACCCGAGACGGAGTCTATGACGA | | | | | | | | | | |
| Oligo rev | AAACTCGTCATAGACTCCGTCTCG | | | | | | | | | | |
| Chromosome | start | end | strand | MM | target_seq | PAM | alignment | distance | position | gene name | gene id |
| GU305914.1 | 197894 | 197916 | - | 0 | CGAGACGGAGTCTATGACGA | TGG | \|\|\|\|\|\|\|\|[\|\|\|\|\|\|\|\|\|\|\|\|]PAM | NA | Intergenic | NA | NA |
| GU305914.1 | 191891 | 191913 | - | 4 | GGAGAGGGCGTCTACGACGA | CGG | -\|\|\|\|-\|\|[-\|\|\|\|\|-\|\|\|\|\|]PAM | NA | Intergenic | NA | NA |
| T19631 | GACGGAGTCTATGACGATGGAGG | 1000 | | | | | | | | | |
| Oligo fwd | CACCGACGGAGTCTATGACGATGG | | | | | | | | | | |
| Oligo rev | AAACCCATCGTCATAGACTCCGTC | | | | | | | | | | |
| Chromosome | start | end | strand | MM | target_seq | PAM | alignment | distance | position | gene name | gene id |
| GU305914.1 | 197891 | 197913 | - | 0 | GACGGAGTCTATGACGATGG | AGG | \|\|\|\|\|\|\|\|[\|\|\|\|\|\|\|\|\|\|\|\|]PAM | NA | Intergenic | NA | NA |
| T19632 | GGAGTCTATGACGATGGAGGAGG | 1000 | | | | | | | | | |
| Oligo fwd | CACCGGAGTCTATGACGATGGAGG | | | | | | | | | | |
| Oligo rev | AAACCCTCCATCGTCATAGACTCC | | | | | | | | | | |
| Chromosome | start | end | strand | MM | target_seq | PAM | alignment | distance | position | gene name | gene id |
| GU305914.1 | 197888 | 197910 | - | 0 | GGAGTCTATGACGATGGAGG | AGG | \|\|\|\|\|\|\|\|[\|\|\|\|\|\|\|\|\|\|\|\|]PAM | NA | Intergenic | NA | NA |
| T19633 | AGGTGTACGCGTGCCGCCGACGG | 1000 | | | | | | | | | |
| Oligo fwd | CACCAGGTGTACGCGTGCCGCCGA | | | | | | | | | | |
| Oligo rev | AAACTCGGCGGCACGCGTACACCT | | | | | | | | | | |
| Chromosome | start | end | strand | MM | target_seq | PAM | alignment | distance | position | gene name | gene id |
| GU305914.1 | 197868 | 197890 | - | 0 | AGGTGTACGCGTGCCGCCGA | CGG | \|\|\|\|\|\|\|\|[\|\|\|\|\|\|\|\|\|\|\|\|]PAM | NA | Intergenic | NA | NA |
| T19634 | TCGTCCCCTTCTACCGCGCGCGG | 1000 | | | | | | | | | |
| Oligo fwd | CACCTCGTCCCCTTCTACCGCGCG | | | | | | | | | | |
| Oligo rev | AAACCGCGCGGTAGAAGGGGACGA | | | | | | | | | | |
| Chromosome | start | end | strand | MM | target_seq | PAM | alignment | distance | position | gene name | gene id |
| GU305914.1 | 197826 | 197848 | - | 0 | TCGTCCCCTTCTACCGCGCG | CGG | \|\|\|\|\|\|\|\|[\|\|\|\|\|\|\|\|\|\|\|\|]PAM | NA | Intergenic | NA | NA |
| T19635 | CCCCTTCTACCGCGCGCGGCCGG | 1000 | | | | | | | | | |
| Oligo fwd | CACCCCCCTTCTACCGCGCGCGGC | | | | | | | | | | |
| Oligo rev | AAACGCCGCGCGCGGTAGAAGGGG | | | | | | | | | | |
| Chromosome | start | end | strand | MM | target_seq | PAM | alignment | distance | position | gene name | gene id |
| GU305914.1 | 197822 | 197844 | - | 0 | CCCCTTCTACCGCGCGCGGC | CGG | \|\|\|\|\|\|\|\|[\|\|\|\|\|\|\|\|\|\|\|\|]PAM | NA | Intergenic | NA | NA |
| T19636 | CCCTTCTACCGCGCGCGGCCGGG | 1000 | | | | | | | | | |
| Oligo fwd | CACCCCCTTCTACCGCGCGCGGCC | | | | | | | | | | |
| Oligo rev | AAACGGCCGCGCGCGGTAGAAGGG | | | | | | | | | | |
| Chromosome | start | end | strand | MM | target_seq | PAM | alignment | distance | position | gene name | gene id |
| GU305914.1 | 197821 | 197843 | - | 0 | CCCTTCTACCGCGCGCGGCC | GGG | \|\|\|\|\|\|\|\|[\|\|\|\|\|\|\|\|\|\|\|\|]PAM | NA | Intergenic | NA | NA |
| T19637 | CCGCGCGCGGCCGGGTCCCTCGG | 1000 | | | | | | | | | |
| Oligo fwd | CACCCCGCGCGCGGCCGGGTCCCT | | | | | | | | | | |
| Oligo rev | AAACAGGGACCCGGCCGCGCGCGG | | | | | | | | | | |
| Chromosome | start | end | strand | MM | target_seq | PAM | alignment | distance | position | gene name | gene id |
| GU305914.1 | 197813 | 197835 | - | 0 | CCGCGCGCGGCCGGGTCCCT | CGG | \|\|\|\|\|\|\|\|[\|\|\|\|\|\|\|\|\|\|\|\|]PAM | NA | Intergenic | NA | NA |
| T19638 | ACCCCCCCCCACCCCACCAGCGG | 1000 | | | | | | | | | |
| Oligo fwd | CACCACCCCCCCCCACCCCACCAG | | | | | | | | | | |
| Oligo rev | AAACCTGGTGGGGTGGGGGGGGGT | | | | | | | | | | |
| Chromosome | start | end | strand | MM | target_seq | PAM | alignment | distance | position | gene name | gene id |
| GU305914.1 | 197766 | 197788 | - | 0 | ACCCCCCCCCACCCCACCAG | CGG | \|\|\|\|\|\|\|\|[\|\|\|\|\|\|\|\|\|\|\|\|]PAM | NA | Intergenic | NA | NA |
| T19639 | GCATCAGATCTAATAACCCTCGG | 1000 | | | | | | | | | |
| Oligo fwd | CACCGCATCAGATCTAATAACCCT | | | | | | | | | | |
| Oligo rev | AAACAGGGTTATTAGATCTGATGC | | | | | | | | | | |
| Chromosome | start | end | strand | MM | target_seq | PAM | alignment | distance | position | gene name | gene id |
| GU305914.1 | 197714 | 197736 | - | 0 | GCATCAGATCTAATAACCCT | CGG | \|\|\|\|\|\|\|\|[\|\|\|\|\|\|\|\|\|\|\|\|]PAM | NA | Intergenic | NA | NA |
| T19640 | TAATAACCCTCGGCCGTCACCGG | 1000 | | | | | | | | | |
| Oligo fwd | CACCTAATAACCCTCGGCCGTCAC | | | | | | | | | | |
| Oligo rev | AAACGTGACGGCCGAGGGTTATTA | | | | | | | | | | |
| Chromosome | start | end | strand | MM | target_seq | PAM | alignment | distance | position | gene name | gene id |
| GU305914.1 | 197704 | 197726 | - | 0 | TAATAACCCTCGGCCGTCAC | CGG | \|\|\|\|\|\|\|\|[\|\|\|\|\|\|\|\|\|\|\|\|]PAM | NA | Intergenic | NA | NA |
| T19641 | AATAACCCTCGGCCGTCACCGGG | 1000 | | | | | | | | | |
| Oligo fwd | CACCAATAACCCTCGGCCGTCACC | | | | | | | | | | |
| Oligo rev | AAACGGTGACGGCCGAGGGTTATT | | | | | | | | | | |
| Chromosome | start | end | strand | MM | target_seq | PAM | alignment | distance | position | gene name | gene id |
| GU305914.1 | 197703 | 197725 | - | 0 | AATAACCCTCGGCCGTCACC | GGG | \|\|\|\|\|\|\|\|[\|\|\|\|\|\|\|\|\|\|\|\|]PAM | NA | Intergenic | NA | NA |
| T19642 | ACTGCCTGTCAACATCGCCACGG | 1000 | | | | | | | | | |

<table>
<tr><td>Oligo fwd</td><td>CACCACTGCCTGTCAACATCGCCA</td><td></td><td></td><td></td><td></td><td></td><td></td><td></td><td></td><td></td><td></td></tr>
<tr><td>Oligo rev</td><td>AAACTGGCGATGTTGACAGGCAGT</td><td></td><td></td><td></td><td></td><td></td><td></td><td></td><td></td><td></td><td></td></tr>
<tr><td>Chromosome</td><td>start</td><td>end</td><td>strand</td><td>MM</td><td>target_seq</td><td>PAM</td><td>alignment</td><td>distance</td><td>position</td><td>gene name</td><td>gene id</td></tr>
<tr><td>GU305914.1</td><td>197680</td><td>197702</td><td>-</td><td>0</td><td>ACTGCCTGTCAACATCGCCA</td><td>CGG</td><td>||||||||[||||||||||||]PAM</td><td>NA</td><td>Intergenic</td><td>NA</td><td>NA</td></tr>
<tr><td>T19643</td><td>GTCAACATCGCCACGGCCCGCGG</td><td>1000</td><td></td><td></td><td></td><td></td><td></td><td></td><td></td><td></td><td></td></tr>
<tr><td>Oligo fwd</td><td>CACCGTCAACATCGCCACGGCCCG</td><td></td><td></td><td></td><td></td><td></td><td></td><td></td><td></td><td></td><td></td></tr>
<tr><td>Oligo rev</td><td>AAACCGGGCCGTGGCGATGTTGAC</td><td></td><td></td><td></td><td></td><td></td><td></td><td></td><td></td><td></td><td></td></tr>
<tr><td>Chromosome</td><td>start</td><td>end</td><td>strand</td><td>MM</td><td>target_seq</td><td>PAM</td><td>alignment</td><td>distance</td><td>position</td><td>gene name</td><td>gene id</td></tr>
<tr><td>GU305914.1</td><td>197673</td><td>197695</td><td>-</td><td>0</td><td>GTCAACATCGCCACGGCCCG</td><td>CGG</td><td>||||||||[||||||||||||]PAM</td><td>NA</td><td>Intergenic</td><td>NA</td><td>NA</td></tr>
<tr><td>T19644</td><td>AACATCGCCACGGCCCGCGGCGG</td><td>1000</td><td></td><td></td><td></td><td></td><td></td><td></td><td></td><td></td><td></td></tr>
<tr><td>Oligo fwd</td><td>CACCAACATCGCCACGGCCCGCGG</td><td></td><td></td><td></td><td></td><td></td><td></td><td></td><td></td><td></td><td></td></tr>
<tr><td>Oligo rev</td><td>AAACCCGCGGGCCGTGGCGATGTT</td><td></td><td></td><td></td><td></td><td></td><td></td><td></td><td></td><td></td><td></td></tr>
<tr><td>Chromosome</td><td>start</td><td>end</td><td>strand</td><td>MM</td><td>target_seq</td><td>PAM</td><td>alignment</td><td>distance</td><td>position</td><td>gene name</td><td>gene id</td></tr>
<tr><td>GU305914.1</td><td>197670</td><td>197692</td><td>-</td><td>0</td><td>AACATCGCCACGGCCCGCGG</td><td>CGG</td><td>||||||||[||||||||||||]PAM</td><td>NA</td><td>Intergenic</td><td>NA</td><td>NA</td></tr>
<tr><td>T19645</td><td>CACGGCCCGCGGCGGAGAATCGG</td><td>1000</td><td></td><td></td><td></td><td></td><td></td><td></td><td></td><td></td><td></td></tr>
<tr><td>Oligo fwd</td><td>CACCCACGGCCCGCGGCGGAGAAT</td><td></td><td></td><td></td><td></td><td></td><td></td><td></td><td></td><td></td><td></td></tr>
<tr><td>Oligo rev</td><td>AAACATTCTCCGCCGCGGGCCGTG</td><td></td><td></td><td></td><td></td><td></td><td></td><td></td><td></td><td></td><td></td></tr>
<tr><td>Chromosome</td><td>start</td><td>end</td><td>strand</td><td>MM</td><td>target_seq</td><td>PAM</td><td>alignment</td><td>distance</td><td>position</td><td>gene name</td><td>gene id</td></tr>
<tr><td>GU305914.1</td><td>197662</td><td>197684</td><td>-</td><td>0</td><td>CACGGCCCGCGGCGGAGAAT</td><td>CGG</td><td>||||||||[||||||||||||]PAM</td><td>NA</td><td>Intergenic</td><td>NA</td><td>NA</td></tr>
<tr><td>T19646</td><td>ATTATCACCATCATCTCATCCGG</td><td>1000</td><td></td><td></td><td></td><td></td><td></td><td></td><td></td><td></td><td></td></tr>
<tr><td>Oligo fwd</td><td>CACCATTATCACCATCATCTCATC</td><td></td><td></td><td></td><td></td><td></td><td></td><td></td><td></td><td></td><td></td></tr>
<tr><td>Oligo rev</td><td>AAACGATGAGATGATGGTGATAAT</td><td></td><td></td><td></td><td></td><td></td><td></td><td></td><td></td><td></td><td></td></tr>
<tr><td>Chromosome</td><td>start</td><td>end</td><td>strand</td><td>MM</td><td>target_seq</td><td>PAM</td><td>alignment</td><td>distance</td><td>position</td><td>gene name</td><td>gene id</td></tr>
<tr><td>GU305914.1</td><td>197621</td><td>197643</td><td>-</td><td>0</td><td>ATTATCACCATCATCTCATC</td><td>CGG</td><td>||||||||[||||||||||||]PAM</td><td>NA</td><td>Intergenic</td><td>NA</td><td>NA</td></tr>
<tr><td>T19647</td><td>CATCATCTCATCCGGCGAGATGG</td><td>1000</td><td></td><td></td><td></td><td></td><td></td><td></td><td></td><td></td><td></td></tr>
<tr><td>Oligo fwd</td><td>CACCCATCATCTCATCCGGCGAGA</td><td></td><td></td><td></td><td></td><td></td><td></td><td></td><td></td><td></td><td></td></tr>
<tr><td>Oligo rev</td><td>AAACTCTCGCCGGATGAGATGATG</td><td></td><td></td><td></td><td></td><td></td><td></td><td></td><td></td><td></td><td></td></tr>
<tr><td>Chromosome</td><td>start</td><td>end</td><td>strand</td><td>MM</td><td>target_seq</td><td>PAM</td><td>alignment</td><td>distance</td><td>position</td><td>gene name</td><td>gene id</td></tr>
<tr><td>GU305914.1</td><td>197613</td><td>197635</td><td>-</td><td>0</td><td>CATCATCTCATCCGGCGAGA</td><td>TGG</td><td>||||||||[||||||||||||]PAM</td><td>NA</td><td>Intergenic</td><td>NA</td><td>NA</td></tr>
<tr><td>T19648</td><td>GCGAGATGGACAGCACACTGTGG</td><td>1000</td><td></td><td></td><td></td><td></td><td></td><td></td><td></td><td></td><td></td></tr>
<tr><td>Oligo fwd</td><td>CACCGCGAGATGGACAGCACACTG</td><td></td><td></td><td></td><td></td><td></td><td></td><td></td><td></td><td></td><td></td></tr>
<tr><td>Oligo rev</td><td>AAACCAGTGTGCTGTCCATCTCGC</td><td></td><td></td><td></td><td></td><td></td><td></td><td></td><td></td><td></td><td></td></tr>
<tr><td>Chromosome</td><td>start</td><td>end</td><td>strand</td><td>MM</td><td>target_seq</td><td>PAM</td><td>alignment</td><td>distance</td><td>position</td><td>gene name</td><td>gene id</td></tr>
<tr><td>GU305914.1</td><td>197599</td><td>197621</td><td>-</td><td>0</td><td>GCGAGATGGACAGCACACTG</td><td>TGG</td><td>||||||||[||||||||||||]PAM</td><td>NA</td><td>Intergenic</td><td>NA</td><td>NA</td></tr>
<tr><td>T19649</td><td>CGAGATGGACAGCACACTGTGGG</td><td>1000</td><td></td><td></td><td></td><td></td><td></td><td></td><td></td><td></td><td></td></tr>
<tr><td>Oligo fwd</td><td>CACCCGAGATGGACAGCACACTGT</td><td></td><td></td><td></td><td></td><td></td><td></td><td></td><td></td><td></td><td></td></tr>
<tr><td>Oligo rev</td><td>AAACACAGTGTGCTGTCCATCTCG</td><td></td><td></td><td></td><td></td><td></td><td></td><td></td><td></td><td></td><td></td></tr>
<tr><td>Chromosome</td><td>start</td><td>end</td><td>strand</td><td>MM</td><td>target_seq</td><td>PAM</td><td>alignment</td><td>distance</td><td>position</td><td>gene name</td><td>gene id</td></tr>
<tr><td>GU305914.1</td><td>197598</td><td>197620</td><td>-</td><td>0</td><td>CGAGATGGACAGCACACTGT</td><td>GGG</td><td>||||||||[||||||||||||]PAM</td><td>NA</td><td>Intergenic</td><td>NA</td><td>NA</td></tr>
<tr><td>T19650</td><td>GCTCCCCGTGACCGATGATCCGG</td><td>1000</td><td></td><td></td><td></td><td></td><td></td><td></td><td></td><td></td><td></td></tr>
<tr><td>Oligo fwd</td><td>CACCGCTCCCCGTGACCGATGATC</td><td></td><td></td><td></td><td></td><td></td><td></td><td></td><td></td><td></td><td></td></tr>
<tr><td>Oligo rev</td><td>AAACGATCATCGGTCACGGGGAGC</td><td></td><td></td><td></td><td></td><td></td><td></td><td></td><td></td><td></td><td></td></tr>
<tr><td>Chromosome</td><td>start</td><td>end</td><td>strand</td><td>MM</td><td>target_seq</td><td>PAM</td><td>alignment</td><td>distance</td><td>position</td><td>gene name</td><td>gene id</td></tr>
<tr><td>GU305914.1</td><td>197571</td><td>197593</td><td>-</td><td>0</td><td>GCTCCCCGTGACCGATGATC</td><td>CGG</td><td>||||||||[||||||||||||]PAM</td><td>NA</td><td>Intergenic</td><td>NA</td><td>NA</td></tr>
<tr><td>T19651</td><td>CCGATGATCCGGCACCCTATCGG</td><td>1000</td><td></td><td></td><td></td><td></td><td></td><td></td><td></td><td></td><td></td></tr>
<tr><td>Oligo fwd</td><td>CACCCCGATGATCCGGCACCCTAT</td><td></td><td></td><td></td><td></td><td></td><td></td><td></td><td></td><td></td><td></td></tr>
<tr><td>Oligo rev</td><td>AAACATAGGGTGCCGGATCATCGG</td><td></td><td></td><td></td><td></td><td></td><td></td><td></td><td></td><td></td><td></td></tr>
<tr><td>Chromosome</td><td>start</td><td>end</td><td>strand</td><td>MM</td><td>target_seq</td><td>PAM</td><td>alignment</td><td>distance</td><td>position</td><td>gene name</td><td>gene id</td></tr>
<tr><td>GU305914.1</td><td>197560</td><td>197582</td><td>-</td><td>0</td><td>CCGATGATCCGGCACCCTAT</td><td>CGG</td><td>||||||||[||||||||||||]PAM</td><td>NA</td><td>Intergenic</td><td>NA</td><td>NA</td></tr>
<tr><td>T19652</td><td>TACCGCCACCAGTCGAACCGCGG</td><td>1000</td><td></td><td></td><td></td><td></td><td></td><td></td><td></td><td></td><td></td></tr>
<tr><td>Oligo fwd</td><td>CACCTACCGCCACCAGTCGAACCG</td><td></td><td></td><td></td><td></td><td></td><td></td><td></td><td></td><td></td><td></td></tr>
<tr><td>Oligo rev</td><td>AAACCGGTTCGACTGGTGGCGGTA</td><td></td><td></td><td></td><td></td><td></td><td></td><td></td><td></td><td></td><td></td></tr>
<tr><td>Chromosome</td><td>start</td><td>end</td><td>strand</td><td>MM</td><td>target_seq</td><td>PAM</td><td>alignment</td><td>distance</td><td>position</td><td>gene name</td><td>gene id</td></tr>
<tr><td>GU305914.1</td><td>197486</td><td>197508</td><td>-</td><td>0</td><td>TACCGCCACCAGTCGAACCG</td><td>CGG</td><td>||||||||[||||||||||||]PAM</td><td>NA</td><td>Intergenic</td><td>NA</td><td>NA</td></tr>
<tr><td>T19653</td><td>ACCGCCACCAGTCGAACCGCGGG</td><td>1000</td><td></td><td></td><td></td><td></td><td></td><td></td><td></td><td></td><td></td></tr>
<tr><td>Oligo fwd</td><td>CACCACCGCCACCAGTCGAACCGC</td><td></td><td></td><td></td><td></td><td></td><td></td><td></td><td></td><td></td><td></td></tr>
<tr><td>Oligo rev</td><td>AAACGCGGTTCGACTGGTGGCGGT</td><td></td><td></td><td></td><td></td><td></td><td></td><td></td><td></td><td></td><td></td></tr>
<tr><td>Chromosome</td><td>start</td><td>end</td><td>strand</td><td>MM</td><td>target_seq</td><td>PAM</td><td>alignment</td><td>distance</td><td>position</td><td>gene name</td><td>gene id</td></tr>
<tr><td>GU305914.1</td><td>197485</td><td>197507</td><td>-</td><td>0</td><td>ACCGCCACCAGTCGAACCGC</td><td>GGG</td><td>||||||||[||||||||||||]PAM</td><td>NA</td><td>Intergenic</td><td>NA</td><td>NA</td></tr>
<tr><td>T19654</td><td>CCGCCACCAGTCGAACCGCGGGG</td><td>1000</td><td></td><td></td><td></td><td></td><td></td><td></td><td></td><td></td><td></td></tr>
<tr><td>Oligo fwd</td><td>CACCCCGCCACCAGTCGAACCGCG</td><td></td><td></td><td></td><td></td><td></td><td></td><td></td><td></td><td></td><td></td></tr>
<tr><td>Oligo rev</td><td>AAACCGCGGTTCGACTGGTGGCGG</td><td></td><td></td><td></td><td></td><td></td><td></td><td></td><td></td><td></td><td></td></tr>
<tr><td>Chromosome</td><td>start</td><td>end</td><td>strand</td><td>MM</td><td>target_seq</td><td>PAM</td><td>alignment</td><td>distance</td><td>position</td><td>gene name</td><td>gene id</td></tr>
<tr><td>GU305914.1</td><td>197484</td><td>197506</td><td>-</td><td>0</td><td>CCGCCACCAGTCGAACCGCG</td><td>GGG</td><td>||||||||[||||||||||||]PAM</td><td>NA</td><td>Intergenic</td><td>NA</td><td>NA</td></tr>
<tr><td>T19655</td><td>CCAGTCGAACCGCGGGGAGCTGG</td><td>1000</td><td></td><td></td><td></td><td></td><td></td><td></td><td></td><td></td><td></td></tr>
<tr><td>Oligo fwd</td><td>CACCCCAGTCGAACCGCGGGGAGC</td><td></td><td></td><td></td><td></td><td></td><td></td><td></td><td></td><td></td><td></td></tr>
<tr><td>Oligo rev</td><td>AAACGCTCCCCGCGGTTCGACTGG</td><td></td><td></td><td></td><td></td><td></td><td></td><td></td><td></td><td></td><td></td></tr>
<tr><td>Chromosome</td><td>start</td><td>end</td><td>strand</td><td>MM</td><td>target_seq</td><td>PAM</td><td>alignment</td><td>distance</td><td>position</td><td>gene name</td><td>gene id</td></tr>
<tr><td>GU305914.1</td><td>197478</td><td>197500</td><td>-</td><td>0</td><td>CCAGTCGAACCGCGGGGAGC</td><td>TGG</td><td>||||||||[||||||||||||]PAM</td><td>NA</td><td>Intergenic</td><td>NA</td><td>NA</td></tr>
<tr><td>T19656</td><td>CCGCGGGGAGCTGGCGTCCCTGG</td><td>1000</td><td></td><td></td><td></td><td></td><td></td><td></td><td></td><td></td><td></td></tr>
<tr><td>Oligo fwd</td><td>CACCCCGCGGGGAGCTGGCGTCCC</td><td></td><td></td><td></td><td></td><td></td><td></td><td></td><td></td><td></td><td></td></tr>
<tr><td>Oligo rev</td><td>AAACGGGACGCCAGCTCCCCGCGG</td><td></td><td></td><td></td><td></td><td></td><td></td><td></td><td></td><td></td><td></td></tr>
<tr><td>Chromosome</td><td>start</td><td>end</td><td>strand</td><td>MM</td><td>target_seq</td><td>PAM</td><td>alignment</td><td>distance</td><td>position</td><td>gene name</td><td>gene id</td></tr>
<tr><td>GU305914.1</td><td>197469</td><td>197491</td><td>-</td><td>0</td><td>CCGCGGGGAGCTGGCGTCCC</td><td>TGG</td><td>||||||||[||||||||||||]PAM</td><td>NA</td><td>Intergenic</td><td>NA</td><td>NA</td></tr>
<tr><td>GU305914.1</td><td>18892</td><td>18914</td><td>-</td><td>4</td><td>CGGGGGGGAGCGGGCGTGCC</td><td>GGG</td><td>|-|-||||[|||-|||||-||]PAM</td><td>NA</td><td>Intergenic</td><td>NA</td><td>NA</td></tr>
</table>

| | | | | | | | | | | | |
|---|---|---|---|---|---|---|---|---|---|---|---|
| GU305914.1 | 68698 | 68720 | + | 4 | CAGCGAGCAGCTGGCGTCGC | TGG | \|-\|\|\|-\|-[\|\|\|\|\|\|\|\|\|\|-\|]PAM | NA | Intergenic | NA | NA |
| T19657 | AGCTGGCGTCCCTGGTCGAGAGG | 1000 | | | | | | | | | |
| Oligo fwd | CACCAGCTGGCGTCCCTGGTCGAG | | | | | | | | | | |
| Oligo rev | AAACCTCGACCAGGGACGCCAGCT | | | | | | | | | | |
| Chromosome | start | end | strand | MM | target_seq | PAM | alignment | distance | position | gene name | gene id |
| GU305914.1 | 197461 | 197483 | - | 0 | AGCTGGCGTCCCTGGTCGAG | AGG | \|\|\|\|\|\|\|\|[\|\|\|\|\|\|\|\|\|\|\|\|]PAM | NA | Intergenic | NA | NA |
| T19658 | TCCCTGGTCGAGAGGCGCGCCGG | 1000 | | | | | | | | | |
| Oligo fwd | CACCTCCCTGGTCGAGAGGCGCGC | | | | | | | | | | |
| Oligo rev | AAACGCGCGCCTCTCGACCAGGGA | | | | | | | | | | |
| Chromosome | start | end | strand | MM | target_seq | PAM | alignment | distance | position | gene name | gene id |
| GU305914.1 | 197453 | 197475 | - | 0 | TCCCTGGTCGAGAGGCGCGC | CGG | \|\|\|\|\|\|\|\|[\|\|\|\|\|\|\|\|\|\|\|\|]PAM | NA | Intergenic | NA | NA |
| T19659 | CGCCGGCGAGCGTCTGCCGCTGG | 1000 | | | | | | | | | |
| Oligo fwd | CACCCGCCGGCGAGCGTCTGCCGC | | | | | | | | | | |
| Oligo rev | AAACGCGGCAGACGCTCGCCGGCG | | | | | | | | | | |
| Chromosome | start | end | strand | MM | target_seq | PAM | alignment | distance | position | gene name | gene id |
| GU305914.1 | 197436 | 197458 | - | 0 | CGCCGGCGAGCGTCTGCCGC | TGG | \|\|\|\|\|\|\|\|[\|\|\|\|\|\|\|\|\|\|\|\|]PAM | NA | Intergenic | NA | NA |
| GU305914.1 | 107656 | 107678 | + | 4 | CGTGGTCGGGCGTCTGCCGC | AGG | \|\|--\|-\|\|[-\|\|\|\|\|\|\|\|\|\|\|]PAM | NA | Intergenic | NA | NA |
| T19660 | GCCGGCGAGCGTCTGCCGCTGGG | 1000 | | | | | | | | | |
| Oligo fwd | CACCGCCGGCGAGCGTCTGCCGCT | | | | | | | | | | |
| Oligo rev | AAACAGCGGCAGACGCTCGCCGGC | | | | | | | | | | |
| Chromosome | start | end | strand | MM | target_seq | PAM | alignment | distance | position | gene name | gene id |
| GU305914.1 | 197435 | 197457 | - | 0 | GCCGGCGAGCGTCTGCCGCT | GGG | \|\|\|\|\|\|\|\|[\|\|\|\|\|\|\|\|\|\|\|\|]PAM | NA | Intergenic | NA | NA |
| T19661 | CGCTGGGCATCCCCCACAACTGG | 1000 | | | | | | | | | |
| Oligo fwd | CACCCGCTGGGCATCCCCCACAAC | | | | | | | | | | |
| Oligo rev | AAACGTTGTGGGGGATGCCCAGCG | | | | | | | | | | |
| Chromosome | start | end | strand | MM | target_seq | PAM | alignment | distance | position | gene name | gene id |
| GU305914.1 | 197419 | 197441 | - | 0 | CGCTGGGCATCCCCCACAAC | TGG | \|\|\|\|\|\|\|\|[\|\|\|\|\|\|\|\|\|\|\|\|]PAM | NA | Intergenic | NA | NA |
| T19662 | CATCCCCCACAACTGGTTCGTGG | 1000 | | | | | | | | | |
| Oligo fwd | CACCCATCCCCCACAACTGGTTCG | | | | | | | | | | |
| Oligo rev | AAACCGAACCAGTTGTGGGGGATG | | | | | | | | | | |
| Chromosome | start | end | strand | MM | target_seq | PAM | alignment | distance | position | gene name | gene id |
| GU305914.1 | 197412 | 197434 | - | 0 | CATCCCCCACAACTGGTTCG | TGG | \|\|\|\|\|\|\|\|[\|\|\|\|\|\|\|\|\|\|\|\|]PAM | NA | Intergenic | NA | NA |
| T19663 | GACCCCCGCGAGCCGCTACGAGG | 1000 | | | | | | | | | |
| Oligo fwd | CACCGACCCCCGCGAGCCGCTACG | | | | | | | | | | |
| Oligo rev | AAACCGTAGCGGCTCGCGGGGGTC | | | | | | | | | | |
| Chromosome | start | end | strand | MM | target_seq | PAM | alignment | distance | position | gene name | gene id |
| GU305914.1 | 197385 | 197407 | - | 0 | GACCCCCGCGAGCCGCTACG | AGG | \|\|\|\|\|\|\|\|[\|\|\|\|\|\|\|\|\|\|\|\|]PAM | NA | Intergenic | NA | NA |
| T19664 | AGCCGCTACGAGGAACTGAAAGG | 1000 | | | | | | | | | |
| Oligo fwd | CACCAGCCGCTACGAGGAACTGAA | | | | | | | | | | |
| Oligo rev | AAACTTCAGTTCCTCGTAGCGGCT | | | | | | | | | | |
| Chromosome | start | end | strand | MM | target_seq | PAM | alignment | distance | position | gene name | gene id |
| GU305914.1 | 197375 | 197397 | - | 0 | AGCCGCTACGAGGAACTGAA | AGG | \|\|\|\|\|\|\|\|[\|\|\|\|\|\|\|\|\|\|\|\|]PAM | NA | Intergenic | NA | NA |
| T19665 | CTGAAAGGCGTCGACATCAACGG | 1000 | | | | | | | | | |
| Oligo fwd | CACCCTGAAAGGCGTCGACATCAA | | | | | | | | | | |
| Oligo rev | AAACTTGATGTCGACGCCTTTCAG | | | | | | | | | | |
| Chromosome | start | end | strand | MM | target_seq | PAM | alignment | distance | position | gene name | gene id |
| GU305914.1 | 197360 | 197382 | - | 0 | CTGAAAGGCGTCGACATCAA | CGG | \|\|\|\|\|\|\|\|[\|\|\|\|\|\|\|\|\|\|\|\|]PAM | NA | Intergenic | NA | NA |
| T19666 | GTGTTGCGACGAACCCCTGACGG | 1000 | | | | | | | | | |
| Oligo fwd | CACCGTGTTGCGACGAACCCCTGA | | | | | | | | | | |
| Oligo rev | AAACTCAGGGGTTCGTCGCAACAC | | | | | | | | | | |
| Chromosome | start | end | strand | MM | target_seq | PAM | alignment | distance | position | gene name | gene id |
| GU305914.1 | 197331 | 197353 | - | 0 | GTGTTGCGACGAACCCCTGA | CGG | \|\|\|\|\|\|\|\|[\|\|\|\|\|\|\|\|\|\|\|\|]PAM | NA | Intergenic | NA | NA |
| T19667 | GACGAACCCCTGACGGTCGTCGG | 1000 | | | | | | | | | |
| Oligo fwd | CACCGACGAACCCCTGACGGTCGT | | | | | | | | | | |
| Oligo rev | AAACACGACCGTCAGGGGTTCGTC | | | | | | | | | | |
| Chromosome | start | end | strand | MM | target_seq | PAM | alignment | distance | position | gene name | gene id |
| GU305914.1 | 197324 | 197346 | - | 0 | GACGAACCCCTGACGGTCGT | CGG | \|\|\|\|\|\|\|\|[\|\|\|\|\|\|\|\|\|\|\|\|]PAM | NA | Intergenic | NA | NA |
| T19668 | AACCCCTGACGGTCGTCGGTCGG | 1000 | | | | | | | | | |
| Oligo fwd | CACCAACCCCTGACGGTCGTCGGT | | | | | | | | | | |
| Oligo rev | AAACACCGACGACCGTCAGGGGTT | | | | | | | | | | |
| Chromosome | start | end | strand | MM | target_seq | PAM | alignment | distance | position | gene name | gene id |
| GU305914.1 | 197320 | 197342 | - | 0 | AACCCCTGACGGTCGTCGGT | CGG | \|\|\|\|\|\|\|\|[\|\|\|\|\|\|\|\|\|\|\|\|]PAM | NA | Intergenic | NA | NA |
| T19669 | ATCGTCATCCGCAACAACGACGG | 1000 | | | | | | | | | |
| Oligo fwd | CACCATCGTCATCCGCAACAACGA | | | | | | | | | | |
| Oligo rev | AAACTCGTTGTTGCGGATGACGAT | | | | | | | | | | |
| Chromosome | start | end | strand | MM | target_seq | PAM | alignment | distance | position | gene name | gene id |
| GU305914.1 | 197297 | 197319 | - | 0 | ATCGTCATCCGCAACAACGA | CGG | \|\|\|\|\|\|\|\|[\|\|\|\|\|\|\|\|\|\|\|\|]PAM | NA | Intergenic | NA | NA |
| T19670 | CCGCAACAACGACGGATACGAGG | 1000 | | | | | | | | | |
| Oligo fwd | CACCCCGCAACAACGACGGATACG | | | | | | | | | | |
| Oligo rev | AAACCGTATCCGTCGTTGTTGCGG | | | | | | | | | | |
| Chromosome | start | end | strand | MM | target_seq | PAM | alignment | distance | position | gene name | gene id |
| GU305914.1 | 197289 | 197311 | - | 0 | CCGCAACAACGACGGATACG | AGG | \|\|\|\|\|\|\|\|[\|\|\|\|\|\|\|\|\|\|\|\|]PAM | NA | Intergenic | NA | NA |
| T19671 | CAACGACGGATACGAGGATGCGG | 1000 | | | | | | | | | |
| Oligo fwd | CACCCAACGACGGATACGAGGATG | | | | | | | | | | |

| | | | | | | | | | | | | |
|---|---|---|---|---|---|---|---|---|---|---|---|---|
| Oligo rev | AAACCATCCTCGTATCCGTCGTTG | | | | | | | | | | | |
| Chromosome | start | | end | strand | MM | target_seq | PAM | alignment | distance | position | gene name | gene id |
| GU305914.1 | | 197283 | 197305 | - | 0 | CAACGACGGATACGAGGATG | CGG | \|\|\|\|\|\|\|\|[\|\|\|\|\|\|\|\|\|\|\|\|]PAM | NA | Intergenic | NA | NA |
| T19672 | AACGACGGATACGAGGATGCGGG | | 1000 | | | | | | | | | |
| Oligo fwd | CACCAACGACGGATACGAGGATGC | | | | | | | | | | | |
| Oligo rev | AAACGCATCCTCGTATCCGTCGTT | | | | | | | | | | | |
| Chromosome | start | | end | strand | MM | target_seq | PAM | alignment | distance | position | gene name | gene id |
| GU305914.1 | | 197282 | 197304 | - | 0 | AACGACGGATACGAGGATGC | GGG | \|\|\|\|\|\|\|\|[\|\|\|\|\|\|\|\|\|\|\|\|]PAM | NA | Intergenic | NA | NA |
| T19673 | TGCGGGCGCCGCCTTGTGTCTGG | | 1000 | | | | | | | | | |
| Oligo fwd | CACCTGCGGGCGCCGCCTTGTGTC | | | | | | | | | | | |
| Oligo rev | AAACGACACAAGGCGGCGCCCGCA | | | | | | | | | | | |
| Chromosome | start | | end | strand | MM | target_seq | PAM | alignment | distance | position | gene name | gene id |
| GU305914.1 | | 197265 | 197287 | - | 0 | TGCGGGCGCCGCCTTGTGTC | TGG | \|\|\|\|\|\|\|\|[\|\|\|\|\|\|\|\|\|\|\|\|]PAM | NA | Intergenic | NA | NA |
| T19674 | GCGGGCGCCGCCTTGTGTCTGGG | | 1000 | | | | | | | | | |
| Oligo fwd | CACCGCGGGCGCCGCCTTGTGTCT | | | | | | | | | | | |
| Oligo rev | AAACAGACACAAGGCGGCGCCCGC | | | | | | | | | | | |
| Chromosome | start | | end | strand | MM | target_seq | PAM | alignment | distance | position | gene name | gene id |
| GU305914.1 | | 197264 | 197286 | - | 0 | GCGGGCGCCGCCTTGTGTCT | GGG | \|\|\|\|\|\|\|\|[\|\|\|\|\|\|\|\|\|\|\|\|]PAM | NA | Intergenic | NA | NA |
| T19675 | CCGCCTTGTGTCTGGGAGCGTGG | | 1000 | | | | | | | | | |
| Oligo fwd | CACCCCGCCTTGTGTCTGGGAGCG | | | | | | | | | | | |
| Oligo rev | AAACCGCTCCCAGACACAAGGCGG | | | | | | | | | | | |
| Chromosome | start | | end | strand | MM | target_seq | PAM | alignment | distance | position | gene name | gene id |
| GU305914.1 | | 197257 | 197279 | - | 0 | CCGCCTTGTGTCTGGGAGCG | TGG | \|\|\|\|\|\|\|\|[\|\|\|\|\|\|\|\|\|\|\|\|]PAM | NA | Intergenic | NA | NA |
| T19676 | TGTGTCTGGGAGCGTGGACCCGG | | 1000 | | | | | | | | | |
| Oligo fwd | CACCTGTGTCTGGGAGCGTGGACC | | | | | | | | | | | |
| Oligo rev | AAACGGTCCACGCTCCCAGACACA | | | | | | | | | | | |
| Chromosome | start | | end | strand | MM | target_seq | PAM | alignment | distance | position | gene name | gene id |
| GU305914.1 | | 197251 | 197273 | - | 0 | TGTGTCTGGGAGCGTGGACC | CGG | \|\|\|\|\|\|\|\|[\|\|\|\|\|\|\|\|\|\|\|\|]PAM | NA | Intergenic | NA | NA |
| T19677 | GTGTCTGGGAGCGTGGACCCGGG | | 1000 | | | | | | | | | |
| Oligo fwd | CACCGTGTCTGGGAGCGTGGACCC | | | | | | | | | | | |
| Oligo rev | AAACGGGTCCACGCTCCCAGACAC | | | | | | | | | | | |
| Chromosome | start | | end | strand | MM | target_seq | PAM | alignment | distance | position | gene name | gene id |
| GU305914.1 | | 197250 | 197272 | - | 0 | GTGTCTGGGAGCGTGGACCC | GGG | \|\|\|\|\|\|\|\|[\|\|\|\|\|\|\|\|\|\|\|\|]PAM | NA | Intergenic | NA | NA |
| T19678 | GGTCTTCGTCTACGAGATTCAGG | | 1000 | | | | | | | | | |
| Oligo fwd | CACCGGTCTTCGTCTACGAGATTC | | | | | | | | | | | |
| Oligo rev | AAACGAATCTCGTAGACGAAGACC | | | | | | | | | | | |
| Chromosome | start | | end | strand | MM | target_seq | PAM | alignment | distance | position | gene name | gene id |
| GU305914.1 | | 197229 | 197251 | - | 0 | GGTCTTCGTCTACGAGATTC | AGG | \|\|\|\|\|\|\|\|[\|\|\|\|\|\|\|\|\|\|\|\|]PAM | NA | Intergenic | NA | NA |
| T19679 | CTTCGTCTACGAGATTCAGGAGG | | 1000 | | | | | | | | | |
| Oligo fwd | CACCCTTCGTCTACGAGATTCAGG | | | | | | | | | | | |
| Oligo rev | AAACCCTGAATCTCGTAGACGAAG | | | | | | | | | | | |
| Chromosome | start | | end | strand | MM | target_seq | PAM | alignment | distance | position | gene name | gene id |
| GU305914.1 | | 197226 | 197248 | - | 0 | CTTCGTCTACGAGATTCAGG | AGG | \|\|\|\|\|\|\|\|[\|\|\|\|\|\|\|\|\|\|\|\|]PAM | NA | Intergenic | NA | NA |
| T19680 | TCAGGAGGACGCCATGATCCTGG | | 1000 | | | | | | | | | |
| Oligo fwd | CACCTCAGGAGGACGCCATGATCC | | | | | | | | | | | |
| Oligo rev | AAACGGATCATGGCGTCCTCCTGA | | | | | | | | | | | |
| Chromosome | start | | end | strand | MM | target_seq | PAM | alignment | distance | position | gene name | gene id |
| GU305914.1 | | 197211 | 197233 | - | 0 | TCAGGAGGACGCCATGATCC | TGG | \|\|\|\|\|\|\|\|[\|\|\|\|\|\|\|\|\|\|\|\|]PAM | NA | Intergenic | NA | NA |
| T19681 | GGAGGACGCCATGATCCTGGTGG | | 1000 | | | | | | | | | |
| Oligo fwd | CACCGGAGGACGCCATGATCCTGG | | | | | | | | | | | |
| Oligo rev | AAACCCAGGATCATGGCGTCCTCC | | | | | | | | | | | |
| Chromosome | start | | end | strand | MM | target_seq | PAM | alignment | distance | position | gene name | gene id |
| GU305914.1 | | 197208 | 197230 | - | 0 | GGAGGACGCCATGATCCTGG | TGG | \|\|\|\|\|\|\|\|[\|\|\|\|\|\|\|\|\|\|\|\|]PAM | NA | Intergenic | NA | NA |
| T19682 | CGCCATGATCCTGGTGGCCCCGG | | 1000 | | | | | | | | | |
| Oligo fwd | CACCCGCCATGATCCTGGTGGCCC | | | | | | | | | | | |
| Oligo rev | AAACGGGCCACCAGGATCATGGCG | | | | | | | | | | | |
| Chromosome | start | | end | strand | MM | target_seq | PAM | alignment | distance | position | gene name | gene id |
| GU305914.1 | | 197202 | 197224 | - | 0 | CGCCATGATCCTGGTGGCCC | CGG | \|\|\|\|\|\|\|\|[\|\|\|\|\|\|\|\|\|\|\|\|]PAM | NA | Intergenic | NA | NA |
| T19683 | GATCCTGGTGGCCCCGGACCTGG | | 1000 | | | | | | | | | |
| Oligo fwd | CACCGATCCTGGTGGCCCCGGACC | | | | | | | | | | | |
| Oligo rev | AAACGGTCCGGGGCCACCAGGATC | | | | | | | | | | | |
| Chromosome | start | | end | strand | MM | target_seq | PAM | alignment | distance | position | gene name | gene id |
| GU305914.1 | | 197196 | 197218 | - | 0 | GATCCTGGTGGCCCCGGACC | TGG | \|\|\|\|\|\|\|\|[\|\|\|\|\|\|\|\|\|\|\|\|]PAM | NA | Intergenic | NA | NA |
| GU305914.1 | | 124715 | 124737 | - | 4 | GATCCGCGCGGCCCTGGACC | CGG | \|\|\|\|\|--\|[-\|\|\|\|\|-\|\|\|\|\|]PAM | NA | Intergenic | NA | NA |
| T19684 | GGCCCCGGACCTGGATAAACTGG | | 1000 | | | | | | | | | |
| Oligo fwd | CACCGGCCCCGGACCTGGATAAAC | | | | | | | | | | | |
| Oligo rev | AAACGTTTATCCAGGTCCGGGGCC | | | | | | | | | | | |
| Chromosome | start | | end | strand | MM | target_seq | PAM | alignment | distance | position | gene name | gene id |
| GU305914.1 | | 197187 | 197209 | - | 0 | GGCCCCGGACCTGGATAAAC | TGG | \|\|\|\|\|\|\|\|[\|\|\|\|\|\|\|\|\|\|\|\|]PAM | NA | Intergenic | NA | NA |
| T19685 | CTGGATAAACTGGCGCGTTTCGG | | 1000 | | | | | | | | | |
| Oligo fwd | CACCCTGGATAAACTGGCGCGTTT | | | | | | | | | | | |
| Oligo rev | AAACAAACGCGCCAGTTTATCCAG | | | | | | | | | | | |
| Chromosome | start | | end | strand | MM | target_seq | PAM | alignment | distance | position | gene name | gene id |
| GU305914.1 | | 197177 | 197199 | - | 0 | CTGGATAAACTGGCGCGTTT | CGG | \|\|\|\|\|\|\|\|[\|\|\|\|\|\|\|\|\|\|\|\|]PAM | NA | Intergenic | NA | NA |

<table>
<tr><td>T19686</td><td>CCGTCGCCCGCACCTGCCGCAGG</td><td>1000</td><td></td><td></td><td></td><td></td><td></td><td></td><td></td><td></td><td></td></tr>
<tr><td>Oligo fwd</td><td>CACCCCGTCGCCCGCACCTGCCGC</td><td></td><td></td><td></td><td></td><td></td><td></td><td></td><td></td><td></td><td></td></tr>
<tr><td>Oligo rev</td><td>AAACGCGGCAGGTGCGGGCGACGG</td><td></td><td></td><td></td><td></td><td></td><td></td><td></td><td></td><td></td><td></td></tr>
<tr><td>Chromosome</td><td>start</td><td>end</td><td>strand</td><td>MM</td><td>target_seq</td><td>PAM</td><td>alignment</td><td>distance</td><td>position</td><td>gene name</td><td>gene id</td></tr>
<tr><td>GU305914.1</td><td>197130</td><td>197152</td><td>-</td><td>0</td><td>CCGTCGCCCGCACCTGCCGC</td><td>AGG</td><td>||||||||[||||||||||||]PAM</td><td>NA</td><td>Intergenic</td><td>NA</td><td>NA</td></tr>
<tr><td>T19687</td><td>GACCCCCCACCGCCTCGTCGTGG</td><td>1000</td><td></td><td></td><td></td><td></td><td></td><td></td><td></td><td></td><td></td></tr>
<tr><td>Oligo fwd</td><td>CACCGACCCCCCACCGCCTCGTCG</td><td></td><td></td><td></td><td></td><td></td><td></td><td></td><td></td><td></td><td></td></tr>
<tr><td>Oligo rev</td><td>AAACCGACGAGGCGGTGGGGGGTC</td><td></td><td></td><td></td><td></td><td></td><td></td><td></td><td></td><td></td><td></td></tr>
<tr><td>Chromosome</td><td>start</td><td>end</td><td>strand</td><td>MM</td><td>target_seq</td><td>PAM</td><td>alignment</td><td>distance</td><td>position</td><td>gene name</td><td>gene id</td></tr>
<tr><td>GU305914.1</td><td>197100</td><td>197122</td><td>-</td><td>0</td><td>GACCCCCCACCGCCTCGTCG</td><td>TGG</td><td>||||||||[||||||||||||]PAM</td><td>NA</td><td>Intergenic</td><td>NA</td><td>NA</td></tr>
<tr><td>T19688</td><td>ACCCCCCACCGCCTCGTCGTGGG</td><td>1000</td><td></td><td></td><td></td><td></td><td></td><td></td><td></td><td></td><td></td></tr>
<tr><td>Oligo fwd</td><td>CACCACCCCCCACCGCCTCGTCGT</td><td></td><td></td><td></td><td></td><td></td><td></td><td></td><td></td><td></td><td></td></tr>
<tr><td>Oligo rev</td><td>AAACACGACGAGGCGGTGGGGGGT</td><td></td><td></td><td></td><td></td><td></td><td></td><td></td><td></td><td></td><td></td></tr>
<tr><td>Chromosome</td><td>start</td><td>end</td><td>strand</td><td>MM</td><td>target_seq</td><td>PAM</td><td>alignment</td><td>distance</td><td>position</td><td>gene name</td><td>gene id</td></tr>
<tr><td>GU305914.1</td><td>197099</td><td>197121</td><td>-</td><td>0</td><td>ACCCCCCACCGCCTCGTCGT</td><td>GGG</td><td>||||||||[||||||||||||]PAM</td><td>NA</td><td>Intergenic</td><td>NA</td><td>NA</td></tr>
<tr><td>T19689</td><td>CCCCCCACCGCCTCGTCGTGGGG</td><td>1000</td><td></td><td></td><td></td><td></td><td></td><td></td><td></td><td></td><td></td></tr>
<tr><td>Oligo fwd</td><td>CACCCCCCCCACCGCCTCGTCGTG</td><td></td><td></td><td></td><td></td><td></td><td></td><td></td><td></td><td></td><td></td></tr>
<tr><td>Oligo rev</td><td>AAACCACGACGAGGCGGTGGGGGG</td><td></td><td></td><td></td><td></td><td></td><td></td><td></td><td></td><td></td><td></td></tr>
<tr><td>Chromosome</td><td>start</td><td>end</td><td>strand</td><td>MM</td><td>target_seq</td><td>PAM</td><td>alignment</td><td>distance</td><td>position</td><td>gene name</td><td>gene id</td></tr>
<tr><td>GU305914.1</td><td>197098</td><td>197120</td><td>-</td><td>0</td><td>CCCCCCACCGCCTCGTCGTG</td><td>GGG</td><td>||||||||[||||||||||||]PAM</td><td>NA</td><td>Intergenic</td><td>NA</td><td>NA</td></tr>
<tr><td>T19690</td><td>CGTGGGGCTGATCATGTGCCAGG</td><td>1000</td><td></td><td></td><td></td><td></td><td></td><td></td><td></td><td></td><td></td></tr>
<tr><td>Oligo fwd</td><td>CACCCGTGGGGCTGATCATGTGCC</td><td></td><td></td><td></td><td></td><td></td><td></td><td></td><td></td><td></td><td></td></tr>
<tr><td>Oligo rev</td><td>AAACGGCACATGATCAGCCCCACG</td><td></td><td></td><td></td><td></td><td></td><td></td><td></td><td></td><td></td><td></td></tr>
<tr><td>Chromosome</td><td>start</td><td>end</td><td>strand</td><td>MM</td><td>target_seq</td><td>PAM</td><td>alignment</td><td>distance</td><td>position</td><td>gene name</td><td>gene id</td></tr>
<tr><td>GU305914.1</td><td>197082</td><td>197104</td><td>-</td><td>0</td><td>CGTGGGGCTGATCATGTGCC</td><td>AGG</td><td>||||||||[||||||||||||]PAM</td><td>NA</td><td>Intergenic</td><td>NA</td><td>NA</td></tr>
<tr><td>T19691</td><td>TGTGCCAGGACGATCTCGACCGG</td><td>1000</td><td></td><td></td><td></td><td></td><td></td><td></td><td></td><td></td><td></td></tr>
<tr><td>Oligo fwd</td><td>CACCTGTGCCAGGACGATCTCGAC</td><td></td><td></td><td></td><td></td><td></td><td></td><td></td><td></td><td></td><td></td></tr>
<tr><td>Oligo rev</td><td>AAACGTCGAGATCGTCCTGGCACA</td><td></td><td></td><td></td><td></td><td></td><td></td><td></td><td></td><td></td><td></td></tr>
<tr><td>Chromosome</td><td>start</td><td>end</td><td>strand</td><td>MM</td><td>target_seq</td><td>PAM</td><td>alignment</td><td>distance</td><td>position</td><td>gene name</td><td>gene id</td></tr>
<tr><td>GU305914.1</td><td>197068</td><td>197090</td><td>-</td><td>0</td><td>TGTGCCAGGACGATCTCGAC</td><td>CGG</td><td>||||||||[||||||||||||]PAM</td><td>NA</td><td>Intergenic</td><td>NA</td><td>NA</td></tr>
<tr><td>T19692</td><td>GTGCCAGGACGATCTCGACCGGG</td><td>1000</td><td></td><td></td><td></td><td></td><td></td><td></td><td></td><td></td><td></td></tr>
<tr><td>Oligo fwd</td><td>CACCGTGCCAGGACGATCTCGACC</td><td></td><td></td><td></td><td></td><td></td><td></td><td></td><td></td><td></td><td></td></tr>
<tr><td>Oligo rev</td><td>AAACGGTCGAGATCGTCCTGGCAC</td><td></td><td></td><td></td><td></td><td></td><td></td><td></td><td></td><td></td><td></td></tr>
<tr><td>Chromosome</td><td>start</td><td>end</td><td>strand</td><td>MM</td><td>target_seq</td><td>PAM</td><td>alignment</td><td>distance</td><td>position</td><td>gene name</td><td>gene id</td></tr>
<tr><td>GU305914.1</td><td>197067</td><td>197089</td><td>-</td><td>0</td><td>GTGCCAGGACGATCTCGACC</td><td>GGG</td><td>||||||||[||||||||||||]PAM</td><td>NA</td><td>Intergenic</td><td>NA</td><td>NA</td></tr>
<tr><td>T19693</td><td>CCGGGTGTCCGATTACTGCCAGG</td><td>1000</td><td></td><td></td><td></td><td></td><td></td><td></td><td></td><td></td><td></td></tr>
<tr><td>Oligo fwd</td><td>CACCCCGGGTGTCCGATTACTGCC</td><td></td><td></td><td></td><td></td><td></td><td></td><td></td><td></td><td></td><td></td></tr>
<tr><td>Oligo rev</td><td>AAACGGCAGTAATCGGACACCCGG</td><td></td><td></td><td></td><td></td><td></td><td></td><td></td><td></td><td></td><td></td></tr>
<tr><td>Chromosome</td><td>start</td><td>end</td><td>strand</td><td>MM</td><td>target_seq</td><td>PAM</td><td>alignment</td><td>distance</td><td>position</td><td>gene name</td><td>gene id</td></tr>
<tr><td>GU305914.1</td><td>197049</td><td>197071</td><td>-</td><td>0</td><td>CCGGGTGTCCGATTACTGCC</td><td>AGG</td><td>||||||||[||||||||||||]PAM</td><td>NA</td><td>Intergenic</td><td>NA</td><td>NA</td></tr>
<tr><td>T19694</td><td>GATTACTGCCAGGAGAACAGCGG</td><td>1000</td><td></td><td></td><td></td><td></td><td></td><td></td><td></td><td></td><td></td></tr>
<tr><td>Oligo fwd</td><td>CACCGATTACTGCCAGGAGAACAG</td><td></td><td></td><td></td><td></td><td></td><td></td><td></td><td></td><td></td><td></td></tr>
<tr><td>Oligo rev</td><td>AAACCTGTTCTCCTGGCAGTAATC</td><td></td><td></td><td></td><td></td><td></td><td></td><td></td><td></td><td></td><td></td></tr>
<tr><td>Chromosome</td><td>start</td><td>end</td><td>strand</td><td>MM</td><td>target_seq</td><td>PAM</td><td>alignment</td><td>distance</td><td>position</td><td>gene name</td><td>gene id</td></tr>
<tr><td>GU305914.1</td><td>197039</td><td>197061</td><td>-</td><td>0</td><td>GATTACTGCCAGGAGAACAG</td><td>CGG</td><td>||||||||[||||||||||||]PAM</td><td>NA</td><td>Intergenic</td><td>NA</td><td>NA</td></tr>
<tr><td>T19695</td><td>ATTACTGCCAGGAGAACAGCGGG</td><td>1000</td><td></td><td></td><td></td><td></td><td></td><td></td><td></td><td></td><td></td></tr>
<tr><td>Oligo fwd</td><td>CACCATTACTGCCAGGAGAACAGC</td><td></td><td></td><td></td><td></td><td></td><td></td><td></td><td></td><td></td><td></td></tr>
<tr><td>Oligo rev</td><td>AAACGCTGTTCTCCTGGCAGTAAT</td><td></td><td></td><td></td><td></td><td></td><td></td><td></td><td></td><td></td><td></td></tr>
<tr><td>Chromosome</td><td>start</td><td>end</td><td>strand</td><td>MM</td><td>target_seq</td><td>PAM</td><td>alignment</td><td>distance</td><td>position</td><td>gene name</td><td>gene id</td></tr>
<tr><td>GU305914.1</td><td>197038</td><td>197060</td><td>-</td><td>0</td><td>ATTACTGCCAGGAGAACAGC</td><td>GGG</td><td>||||||||[||||||||||||]PAM</td><td>NA</td><td>Intergenic</td><td>NA</td><td>NA</td></tr>
<tr><td>T19696</td><td>GACGTCGCGCTGTACACCCCCGG</td><td>1000</td><td></td><td></td><td></td><td></td><td></td><td></td><td></td><td></td><td></td></tr>
<tr><td>Oligo fwd</td><td>CACCGACGTCGCGCTGTACACCCC</td><td></td><td></td><td></td><td></td><td></td><td></td><td></td><td></td><td></td><td></td></tr>
<tr><td>Oligo rev</td><td>AAACGGGGTGTACAGCGCGACGTC</td><td></td><td></td><td></td><td></td><td></td><td></td><td></td><td></td><td></td><td></td></tr>
<tr><td>Chromosome</td><td>start</td><td>end</td><td>strand</td><td>MM</td><td>target_seq</td><td>PAM</td><td>alignment</td><td>distance</td><td>position</td><td>gene name</td><td>gene id</td></tr>
<tr><td>GU305914.1</td><td>197012</td><td>197034</td><td>-</td><td>0</td><td>GACGTCGCGCTGTACACCCC</td><td>CGG</td><td>||||||||[||||||||||||]PAM</td><td>NA</td><td>Intergenic</td><td>NA</td><td>NA</td></tr>
<tr><td>T19697</td><td>GTACCAGCCGATGAAACTGCTGG</td><td>1000</td><td></td><td></td><td></td><td></td><td></td><td></td><td></td><td></td><td></td></tr>
<tr><td>Oligo fwd</td><td>CACCGTACCAGCCGATGAAACTGC</td><td></td><td></td><td></td><td></td><td></td><td></td><td></td><td></td><td></td><td></td></tr>
<tr><td>Oligo rev</td><td>AAACGCAGTTTCATCGGCTGGTAC</td><td></td><td></td><td></td><td></td><td></td><td></td><td></td><td></td><td></td><td></td></tr>
<tr><td>Chromosome</td><td>start</td><td>end</td><td>strand</td><td>MM</td><td>target_seq</td><td>PAM</td><td>alignment</td><td>distance</td><td>position</td><td>gene name</td><td>gene id</td></tr>
<tr><td>GU305914.1</td><td>196983</td><td>197005</td><td>-</td><td>0</td><td>GTACCAGCCGATGAAACTGC</td><td>TGG</td><td>||||||||[||||||||||||]PAM</td><td>NA</td><td>Intergenic</td><td>NA</td><td>NA</td></tr>
<tr><td>T19698</td><td>TACCAGCCGATGAAACTGCTGGG</td><td>1000</td><td></td><td></td><td></td><td></td><td></td><td></td><td></td><td></td><td></td></tr>
<tr><td>Oligo fwd</td><td>CACCTACCAGCCGATGAAACTGCT</td><td></td><td></td><td></td><td></td><td></td><td></td><td></td><td></td><td></td><td></td></tr>
<tr><td>Oligo rev</td><td>AAACAGCAGTTTCATCGGCTGGTA</td><td></td><td></td><td></td><td></td><td></td><td></td><td></td><td></td><td></td><td></td></tr>
<tr><td>Chromosome</td><td>start</td><td>end</td><td>strand</td><td>MM</td><td>target_seq</td><td>PAM</td><td>alignment</td><td>distance</td><td>position</td><td>gene name</td><td>gene id</td></tr>
<tr><td>GU305914.1</td><td>196982</td><td>197004</td><td>-</td><td>0</td><td>TACCAGCCGATGAAACTGCT</td><td>GGG</td><td>||||||||[||||||||||||]PAM</td><td>NA</td><td>Intergenic</td><td>NA</td><td>NA</td></tr>
<tr><td>T19699</td><td>CAGCCGATGAAACTGCTGGGTGG</td><td>1000</td><td></td><td></td><td></td><td></td><td></td><td></td><td></td><td></td><td></td></tr>
<tr><td>Oligo fwd</td><td>CACCCAGCCGATGAAACTGCTGGG</td><td></td><td></td><td></td><td></td><td></td><td></td><td></td><td></td><td></td><td></td></tr>
<tr><td>Oligo rev</td><td>AAACCCCAGCAGTTTCATCGGCTG</td><td></td><td></td><td></td><td></td><td></td><td></td><td></td><td></td><td></td><td></td></tr>
<tr><td>Chromosome</td><td>start</td><td>end</td><td>strand</td><td>MM</td><td>target_seq</td><td>PAM</td><td>alignment</td><td>distance</td><td>position</td><td>gene name</td><td>gene id</td></tr>
<tr><td>GU305914.1</td><td>196979</td><td>197001</td><td>-</td><td>0</td><td>CAGCCGATGAAACTGCTGGG</td><td>TGG</td><td>||||||||[||||||||||||]PAM</td><td>NA</td><td>Intergenic</td><td>NA</td><td>NA</td></tr>
<tr><td>T19700</td><td>TGAGAGACGCCGCGCGCTACTGG</td><td>1000</td><td></td><td></td><td></td><td></td><td></td><td></td><td></td><td></td><td></td></tr>
<tr><td>Oligo fwd</td><td>CACCTGAGAGACGCCGCGCGCTAC</td><td></td><td></td><td></td><td></td><td></td><td></td><td></td><td></td><td></td><td></td></tr>
<tr><td>Oligo rev</td><td>AAACGTAGCGCGCGGCGTCTCTCA</td><td></td><td></td><td></td><td></td><td></td><td></td><td></td><td></td><td></td><td></td></tr>
<tr><td>Chromosome</td><td>start</td><td>end</td><td>strand</td><td>MM</td><td>target_seq</td><td>PAM</td><td>alignment</td><td>distance</td><td>position</td><td>gene name</td><td>gene id</td></tr>
<tr><td>GU305914.1</td><td>196954</td><td>196976</td><td>-</td><td>0</td><td>TGAGAGACGCCGCGCGCTAC</td><td>TGG</td><td>||||||||[||||||||||||]PAM</td><td>NA</td><td>Intergenic</td><td>NA</td><td>NA</td></tr>
</table>

| | | | | | | | | | | | |
|---|---|---|---|---|---|---|---|---|---|---|---|
| T19701 | CGCCGCGCGCTACTGGCCCCTGG | 1000 | | | | | | | | | |
| Oligo fwd | CACCCGCCGCGCGCTACTGGCCCC | | | | | | | | | | |
| Oligo rev | AAACGGGGCCAGTAGCGCGCGGCG | | | | | | | | | | |
| Chromosome | start | end | strand | MM | target_seq | PAM | alignment | distance | position | gene name | gene id |
| GU305914.1 | 196947 | 196969 | - | 0 | CGCCGCGCGCTACTGGCCCC | TGG | \|\|\|\|\|\|\|\|[\|\|\|\|\|\|\|\|\|\|\|\|]PAM | NA | Intergenic | NA | NA |
| T19702 | CATGAACCCGTCGAACCTGAAGG | 1000 | | | | | | | | | |
| Oligo fwd | CACCCATGAACCCGTCGAACCTGA | | | | | | | | | | |
| Oligo rev | AAACTCAGGTTCGACGGGTTCATG | | | | | | | | | | |
| Chromosome | start | end | strand | MM | target_seq | PAM | alignment | distance | position | gene name | gene id |
| GU305914.1 | 196920 | 196942 | - | 0 | CATGAACCCGTCGAACCTGA | AGG | \|\|\|\|\|\|\|\|[\|\|\|\|\|\|\|\|\|\|\|\|]PAM | NA | Intergenic | NA | NA |
| T19703 | GTCGAACCTGAAGGCCTGCCTGG | 1000 | | | | | | | | | |
| Oligo fwd | CACCGTCGAACCTGAAGGCCTGCC | | | | | | | | | | |
| Oligo rev | AAACGGCAGGCCTTCAGGTTCGAC | | | | | | | | | | |
| Chromosome | start | end | strand | MM | target_seq | PAM | alignment | distance | position | gene name | gene id |
| GU305914.1 | 196911 | 196933 | - | 0 | GTCGAACCTGAAGGCCTGCC | TGG | \|\|\|\|\|\|\|\|[\|\|\|\|\|\|\|\|\|\|\|\|]PAM | NA | Intergenic | NA | NA |
| GU305914.1 | 131952 | 131974 | - | 4 | GTCGTACATCAAGGCCTCCC | AGG | \|\|\|\|-\|\|-[\|-\|\|\|\|\|\|\|-\|\|]PAM | NA | Intergenic | NA | NA |
| T19704 | GGCCTGCCTGGACGAGATCACGG | 1000 | | | | | | | | | |
| Oligo fwd | CACCGGCCTGCCTGGACGAGATCA | | | | | | | | | | |
| Oligo rev | AAACTGATCTCGTCCAGGCAGGCC | | | | | | | | | | |
| Chromosome | start | end | strand | MM | target_seq | PAM | alignment | distance | position | gene name | gene id |
| GU305914.1 | 196899 | 196921 | - | 0 | GGCCTGCCTGGACGAGATCA | CGG | \|\|\|\|\|\|\|\|[\|\|\|\|\|\|\|\|\|\|\|\|]PAM | NA | Intergenic | NA | NA |
| T19705 | GCCTGCCTGGACGAGATCACGGG | 1000 | | | | | | | | | |
| Oligo fwd | CACCGCCTGCCTGGACGAGATCAC | | | | | | | | | | |
| Oligo rev | AAACGTGATCTCGTCCAGGCAGGC | | | | | | | | | | |
| Chromosome | start | end | strand | MM | target_seq | PAM | alignment | distance | position | gene name | gene id |
| GU305914.1 | 196898 | 196920 | - | 0 | GCCTGCCTGGACGAGATCAC | GGG | \|\|\|\|\|\|\|\|[\|\|\|\|\|\|\|\|\|\|\|\|]PAM | NA | Intergenic | NA | NA |
| T19706 | CCTGCCTGGACGAGATCACGGGG | 1000 | | | | | | | | | |
| Oligo fwd | CACCCCTGCCTGGACGAGATCACG | | | | | | | | | | |
| Oligo rev | AAACCGTGATCTCGTCCAGGCAGG | | | | | | | | | | |
| Chromosome | start | end | strand | MM | target_seq | PAM | alignment | distance | position | gene name | gene id |
| GU305914.1 | 196897 | 196919 | - | 0 | CCTGCCTGGACGAGATCACG | GGG | \|\|\|\|\|\|\|\|[\|\|\|\|\|\|\|\|\|\|\|\|]PAM | NA | Intergenic | NA | NA |
| GU305914.1 | 83661 | 83683 | - | 3 | TCAGCCTGGACGAGATCATG | CGG | -\|-\|\|\|\|\|[\|\|\|\|\|\|\|\|\|\|-\|]PAM | NA | Intergenic | NA | NA |
| T19707 | CGGGGCGACTGTGCTGCTTCTGG | 1000 | | | | | | | | | |
| Oligo fwd | CACCCGGGGCGACTGTGCTGCTTC | | | | | | | | | | |
| Oligo rev | AAACGAAGCAGCACAGTCGCCCCG | | | | | | | | | | |
| Chromosome | start | end | strand | MM | target_seq | PAM | alignment | distance | position | gene name | gene id |
| GU305914.1 | 196879 | 196901 | - | 0 | CGGGGCGACTGTGCTGCTTC | TGG | \|\|\|\|\|\|\|\|[\|\|\|\|\|\|\|\|\|\|\|\|]PAM | NA | Intergenic | NA | NA |
| T19708 | CTGCTTCTGGCACGCGTTCGCGG | 1000 | | | | | | | | | |
| Oligo fwd | CACCCTGCTTCTGGCACGCGTTCG | | | | | | | | | | |
| Oligo rev | AAACCGAACGCGTGCCAGAAGCAG | | | | | | | | | | |
| Chromosome | start | end | strand | MM | target_seq | PAM | alignment | distance | position | gene name | gene id |
| GU305914.1 | 196866 | 196888 | - | 0 | CTGCTTCTGGCACGCGTTCG | CGG | \|\|\|\|\|\|\|\|[\|\|\|\|\|\|\|\|\|\|\|\|]PAM | NA | Intergenic | NA | NA |
| T19709 | CTGGCACGCGTTCGCGGCCGTGG | 1000 | | | | | | | | | |
| Oligo fwd | CACCCTGGCACGCGTTCGCGGCCG | | | | | | | | | | |
| Oligo rev | AAACCGGCCGCGAACGCGTGCCAG | | | | | | | | | | |
| Chromosome | start | end | strand | MM | target_seq | PAM | alignment | distance | position | gene name | gene id |
| GU305914.1 | 196860 | 196882 | - | 0 | CTGGCACGCGTTCGCGGCCG | TGG | \|\|\|\|\|\|\|\|[\|\|\|\|\|\|\|\|\|\|\|\|]PAM | NA | Intergenic | NA | NA |
| T19710 | TGGCACGCGTTCGCGGCCGTGGG | 1000 | | | | | | | | | |
| Oligo fwd | CACCTGGCACGCGTTCGCGGCCGT | | | | | | | | | | |
| Oligo rev | AAACACGGCCGCGAACGCGTGCCA | | | | | | | | | | |
| Chromosome | start | end | strand | MM | target_seq | PAM | alignment | distance | position | gene name | gene id |
| GU305914.1 | 196859 | 196881 | - | 0 | TGGCACGCGTTCGCGGCCGT | GGG | \|\|\|\|\|\|\|\|[\|\|\|\|\|\|\|\|\|\|\|\|]PAM | NA | Intergenic | NA | NA |
| T19711 | GGCCGTGGGCGCCTACGCGCCGG | 1000 | | | | | | | | | |
| Oligo fwd | CACCGGCCGTGGGCGCCTACGCGC | | | | | | | | | | |
| Oligo rev | AAACGCGCGTAGGCGCCCACGGCC | | | | | | | | | | |
| Chromosome | start | end | strand | MM | target_seq | PAM | alignment | distance | position | gene name | gene id |
| GU305914.1 | 196845 | 196867 | - | 0 | GGCCGTGGGCGCCTACGCGC | CGG | \|\|\|\|\|\|\|\|[\|\|\|\|\|\|\|\|\|\|\|\|]PAM | NA | Intergenic | NA | NA |
| T19712 | GTGGGCGCCTACGCGCCGGCAGG | 1000 | | | | | | | | | |
| Oligo fwd | CACCGTGGGCGCCTACGCGCCGGC | | | | | | | | | | |
| Oligo rev | AAACGCCGGCGCGTAGGCGCCCAC | | | | | | | | | | |
| Chromosome | start | end | strand | MM | target_seq | PAM | alignment | distance | position | gene name | gene id |
| GU305914.1 | 196841 | 196863 | - | 0 | GTGGGCGCCTACGCGCCGGC | AGG | \|\|\|\|\|\|\|\|[\|\|\|\|\|\|\|\|\|\|\|\|]PAM | NA | Intergenic | NA | NA |
| T19713 | CTGCTCGTCATCGACACGTTCGG | 1000 | | | | | | | | | |
| Oligo fwd | CACCCTGCTCGTCATCGACACGTT | | | | | | | | | | |
| Oligo rev | AAACAACGTGTCGATGACGAGCAG | | | | | | | | | | |
| Chromosome | start | end | strand | MM | target_seq | PAM | alignment | distance | position | gene name | gene id |
| GU305914.1 | 196799 | 196821 | - | 0 | CTGCTCGTCATCGACACGTT | CGG | \|\|\|\|\|\|\|\|[\|\|\|\|\|\|\|\|\|\|\|\|]PAM | NA | Intergenic | NA | NA |
| GU305914.1 | 119171 | 119193 | + | 4 | CTGTTCGGCATCCACACGCT | CGG | \|\|\|-\|\|\|-[\|\|\|\|-\|\|\|\|\|-\|]PAM | NA | Intergenic | NA | NA |
| T19714 | GTTCGGCGCGATCTACACGCTGG | 1000 | | | | | | | | | |
| Oligo fwd | CACCGTTCGGCGCGATCTACACGC | | | | | | | | | | |
| Oligo rev | AAACGCGTGTAGATCGCGCCGAAC | | | | | | | | | | |
| Chromosome | start | end | strand | MM | target_seq | PAM | alignment | distance | position | gene name | gene id |
| GU305914.1 | 196782 | 196804 | - | 0 | GTTCGGCGCGATCTACACGC | TGG | \|\|\|\|\|\|\|\|[\|\|\|\|\|\|\|\|\|\|\|\|]PAM | NA | Intergenic | NA | NA |
| T19715 | GCGCGAGAAGTTCTACCGCGTGG | 1000 | | | | | | | | | |

<table>
<tr><td>Oligo fwd</td><td>CACCGCGCGAGAAGTTCTACCGCG</td><td></td><td></td><td></td><td></td><td></td><td></td><td></td><td></td><td></td><td></td></tr>
<tr><td>Oligo rev</td><td>AAACCGCGGTAGAACTTCTCGCGC</td><td></td><td></td><td></td><td></td><td></td><td></td><td></td><td></td><td></td><td></td></tr>
<tr><td>Chromosome</td><td>start</td><td>end</td><td>strand</td><td>MM</td><td>target_seq</td><td>PAM</td><td>alignment</td><td>distance</td><td>position</td><td>gene name</td><td>gene id</td></tr>
<tr><td>GU305914.1</td><td>196752</td><td>196774</td><td>-</td><td>0</td><td>GCGCGAGAAGTTCTACCGCG</td><td>TGG</td><td>||||||||[||||||||||||]PAM</td><td>NA</td><td>Intergenic</td><td>NA</td><td>NA</td></tr>
<tr><td>T19716</td><td>CGAGAAGTTCTACCGCGTGGCGG</td><td>1000</td><td></td><td></td><td></td><td></td><td></td><td></td><td></td><td></td><td></td></tr>
<tr><td>Oligo fwd</td><td>CACCCGAGAAGTTCTACCGCGTGG</td><td></td><td></td><td></td><td></td><td></td><td></td><td></td><td></td><td></td><td></td></tr>
<tr><td>Oligo rev</td><td>AAACCCACGCGGTAGAACTTCTCG</td><td></td><td></td><td></td><td></td><td></td><td></td><td></td><td></td><td></td><td></td></tr>
<tr><td>Chromosome</td><td>start</td><td>end</td><td>strand</td><td>MM</td><td>target_seq</td><td>PAM</td><td>alignment</td><td>distance</td><td>position</td><td>gene name</td><td>gene id</td></tr>
<tr><td>GU305914.1</td><td>196749</td><td>196771</td><td>-</td><td>0</td><td>CGAGAAGTTCTACCGCGTGG</td><td>CGG</td><td>||||||||[||||||||||||]PAM</td><td>NA</td><td>Intergenic</td><td>NA</td><td>NA</td></tr>
<tr><td>T19717</td><td>AAGTTCTACCGCGTGGCGGACGG</td><td>1000</td><td></td><td></td><td></td><td></td><td></td><td></td><td></td><td></td><td></td></tr>
<tr><td>Oligo fwd</td><td>CACCAAGTTCTACCGCGTGGCGGA</td><td></td><td></td><td></td><td></td><td></td><td></td><td></td><td></td><td></td><td></td></tr>
<tr><td>Oligo rev</td><td>AAACTCCGCCACGCGGTAGAACTT</td><td></td><td></td><td></td><td></td><td></td><td></td><td></td><td></td><td></td><td></td></tr>
<tr><td>Chromosome</td><td>start</td><td>end</td><td>strand</td><td>MM</td><td>target_seq</td><td>PAM</td><td>alignment</td><td>distance</td><td>position</td><td>gene name</td><td>gene id</td></tr>
<tr><td>GU305914.1</td><td>196745</td><td>196767</td><td>-</td><td>0</td><td>AAGTTCTACCGCGTGGCGGA</td><td>CGG</td><td>||||||||[||||||||||||]PAM</td><td>NA</td><td>Intergenic</td><td>NA</td><td>NA</td></tr>
<tr><td>T19718</td><td>ACGGCATCACGATGCTGCTGCGG</td><td>1000</td><td></td><td></td><td></td><td></td><td></td><td></td><td></td><td></td><td></td></tr>
<tr><td>Oligo fwd</td><td>CACCACGGCATCACGATGCTGCTG</td><td></td><td></td><td></td><td></td><td></td><td></td><td></td><td></td><td></td><td></td></tr>
<tr><td>Oligo rev</td><td>AAACCAGCAGCATCGTGATGCCGT</td><td></td><td></td><td></td><td></td><td></td><td></td><td></td><td></td><td></td><td></td></tr>
<tr><td>Chromosome</td><td>start</td><td>end</td><td>strand</td><td>MM</td><td>target_seq</td><td>PAM</td><td>alignment</td><td>distance</td><td>position</td><td>gene name</td><td>gene id</td></tr>
<tr><td>GU305914.1</td><td>196726</td><td>196748</td><td>-</td><td>0</td><td>ACGGCATCACGATGCTGCTG</td><td>CGG</td><td>||||||||[||||||||||||]PAM</td><td>NA</td><td>Intergenic</td><td>NA</td><td>NA</td></tr>
<tr><td>T19719</td><td>CGGCATCACGATGCTGCTGCGGG</td><td>1000</td><td></td><td></td><td></td><td></td><td></td><td></td><td></td><td></td><td></td></tr>
<tr><td>Oligo fwd</td><td>CACCCGGCATCACGATGCTGCTGC</td><td></td><td></td><td></td><td></td><td></td><td></td><td></td><td></td><td></td><td></td></tr>
<tr><td>Oligo rev</td><td>AAACGCAGCAGCATCGTGATGCCG</td><td></td><td></td><td></td><td></td><td></td><td></td><td></td><td></td><td></td><td></td></tr>
<tr><td>Chromosome</td><td>start</td><td>end</td><td>strand</td><td>MM</td><td>target_seq</td><td>PAM</td><td>alignment</td><td>distance</td><td>position</td><td>gene name</td><td>gene id</td></tr>
<tr><td>GU305914.1</td><td>196725</td><td>196747</td><td>-</td><td>0</td><td>CGGCATCACGATGCTGCTGC</td><td>GGG</td><td>||||||||[||||||||||||]PAM</td><td>NA</td><td>Intergenic</td><td>NA</td><td>NA</td></tr>
<tr><td>T19720</td><td>ATCACGATGCTGCTGCGGGCCGG</td><td>1000</td><td></td><td></td><td></td><td></td><td></td><td></td><td></td><td></td><td></td></tr>
<tr><td>Oligo fwd</td><td>CACCATCACGATGCTGCTGCGGGC</td><td></td><td></td><td></td><td></td><td></td><td></td><td></td><td></td><td></td><td></td></tr>
<tr><td>Oligo rev</td><td>AAACGCCCGCAGCAGCATCGTGAT</td><td></td><td></td><td></td><td></td><td></td><td></td><td></td><td></td><td></td><td></td></tr>
<tr><td>Chromosome</td><td>start</td><td>end</td><td>strand</td><td>MM</td><td>target_seq</td><td>PAM</td><td>alignment</td><td>distance</td><td>position</td><td>gene name</td><td>gene id</td></tr>
<tr><td>GU305914.1</td><td>196721</td><td>196743</td><td>-</td><td>0</td><td>ATCACGATGCTGCTGCGGGC</td><td>CGG</td><td>||||||||[||||||||||||]PAM</td><td>NA</td><td>Intergenic</td><td>NA</td><td>NA</td></tr>
<tr><td>T19721</td><td>GATGCTGCTGCGGGCCGGCATGG</td><td>1000</td><td></td><td></td><td></td><td></td><td></td><td></td><td></td><td></td><td></td></tr>
<tr><td>Oligo fwd</td><td>CACCGATGCTGCTGCGGGCCGGCA</td><td></td><td></td><td></td><td></td><td></td><td></td><td></td><td></td><td></td><td></td></tr>
<tr><td>Oligo rev</td><td>AAACTGCCGGCCCGCAGCAGCATC</td><td></td><td></td><td></td><td></td><td></td><td></td><td></td><td></td><td></td><td></td></tr>
<tr><td>Chromosome</td><td>start</td><td>end</td><td>strand</td><td>MM</td><td>target_seq</td><td>PAM</td><td>alignment</td><td>distance</td><td>position</td><td>gene name</td><td>gene id</td></tr>
<tr><td>GU305914.1</td><td>196716</td><td>196738</td><td>-</td><td>0</td><td>GATGCTGCTGCGGGCCGGCA</td><td>TGG</td><td>||||||||[||||||||||||]PAM</td><td>NA</td><td>Intergenic</td><td>NA</td><td>NA</td></tr>
<tr><td>T19722</td><td>GCTGCGGGCCGGCATGGCCAAGG</td><td>1000</td><td></td><td></td><td></td><td></td><td></td><td></td><td></td><td></td><td></td></tr>
<tr><td>Oligo fwd</td><td>CACCGCTGCGGGCCGGCATGGCCA</td><td></td><td></td><td></td><td></td><td></td><td></td><td></td><td></td><td></td><td></td></tr>
<tr><td>Oligo rev</td><td>AAACTGGCCATGCCGGCCCGCAGC</td><td></td><td></td><td></td><td></td><td></td><td></td><td></td><td></td><td></td><td></td></tr>
<tr><td>Chromosome</td><td>start</td><td>end</td><td>strand</td><td>MM</td><td>target_seq</td><td>PAM</td><td>alignment</td><td>distance</td><td>position</td><td>gene name</td><td>gene id</td></tr>
<tr><td>GU305914.1</td><td>196710</td><td>196732</td><td>-</td><td>0</td><td>GCTGCGGGCCGGCATGGCCA</td><td>AGG</td><td>||||||||[||||||||||||]PAM</td><td>NA</td><td>Intergenic</td><td>NA</td><td>NA</td></tr>
<tr><td>T19723</td><td>ATGGCCAAGGCCATCGCGTTCGG</td><td>1000</td><td></td><td></td><td></td><td></td><td></td><td></td><td></td><td></td><td></td></tr>
<tr><td>Oligo fwd</td><td>CACCATGGCCAAGGCCATCGCGTT</td><td></td><td></td><td></td><td></td><td></td><td></td><td></td><td></td><td></td><td></td></tr>
<tr><td>Oligo rev</td><td>AAACAACGCGATGGCCTTGGCCAT</td><td></td><td></td><td></td><td></td><td></td><td></td><td></td><td></td><td></td><td></td></tr>
<tr><td>Chromosome</td><td>start</td><td>end</td><td>strand</td><td>MM</td><td>target_seq</td><td>PAM</td><td>alignment</td><td>distance</td><td>position</td><td>gene name</td><td>gene id</td></tr>
<tr><td>GU305914.1</td><td>196697</td><td>196719</td><td>-</td><td>0</td><td>ATGGCCAAGGCCATCGCGTT</td><td>CGG</td><td>||||||||[||||||||||||]PAM</td><td>NA</td><td>Intergenic</td><td>NA</td><td>NA</td></tr>
<tr><td>T19724</td><td>TGGCCAAGGCCATCGCGTTCGGG</td><td>1000</td><td></td><td></td><td></td><td></td><td></td><td></td><td></td><td></td><td></td></tr>
<tr><td>Oligo fwd</td><td>CACCTGGCCAAGGCCATCGCGTTC</td><td></td><td></td><td></td><td></td><td></td><td></td><td></td><td></td><td></td><td></td></tr>
<tr><td>Oligo rev</td><td>AAACGAACGCGATGGCCTTGGCCA</td><td></td><td></td><td></td><td></td><td></td><td></td><td></td><td></td><td></td><td></td></tr>
<tr><td>Chromosome</td><td>start</td><td>end</td><td>strand</td><td>MM</td><td>target_seq</td><td>PAM</td><td>alignment</td><td>distance</td><td>position</td><td>gene name</td><td>gene id</td></tr>
<tr><td>GU305914.1</td><td>196696</td><td>196718</td><td>-</td><td>0</td><td>TGGCCAAGGCCATCGCGTTC</td><td>GGG</td><td>||||||||[||||||||||||]PAM</td><td>NA</td><td>Intergenic</td><td>NA</td><td>NA</td></tr>
<tr><td>T19725</td><td>GGCCAAGGCCATCGCGTTCGGGG</td><td>1000</td><td></td><td></td><td></td><td></td><td></td><td></td><td></td><td></td><td></td></tr>
<tr><td>Oligo fwd</td><td>CACCGGCCAAGGCCATCGCGTTCG</td><td></td><td></td><td></td><td></td><td></td><td></td><td></td><td></td><td></td><td></td></tr>
<tr><td>Oligo rev</td><td>AAACCGAACGCGATGGCCTTGGCC</td><td></td><td></td><td></td><td></td><td></td><td></td><td></td><td></td><td></td><td></td></tr>
<tr><td>Chromosome</td><td>start</td><td>end</td><td>strand</td><td>MM</td><td>target_seq</td><td>PAM</td><td>alignment</td><td>distance</td><td>position</td><td>gene name</td><td>gene id</td></tr>
<tr><td>GU305914.1</td><td>196695</td><td>196717</td><td>-</td><td>0</td><td>GGCCAAGGCCATCGCGTTCG</td><td>GGG</td><td>||||||||[||||||||||||]PAM</td><td>NA</td><td>Intergenic</td><td>NA</td><td>NA</td></tr>
<tr><td>T19726</td><td>CGGGGCGCGCTTCGACCGTCCGG</td><td>1000</td><td></td><td></td><td></td><td></td><td></td><td></td><td></td><td></td><td></td></tr>
<tr><td>Oligo fwd</td><td>CACCCGGGGCGCGCTTCGACCGTC</td><td></td><td></td><td></td><td></td><td></td><td></td><td></td><td></td><td></td><td></td></tr>
<tr><td>Oligo rev</td><td>AAACGACGGTCGAAGCGCGCCCCG</td><td></td><td></td><td></td><td></td><td></td><td></td><td></td><td></td><td></td><td></td></tr>
<tr><td>Chromosome</td><td>start</td><td>end</td><td>strand</td><td>MM</td><td>target_seq</td><td>PAM</td><td>alignment</td><td>distance</td><td>position</td><td>gene name</td><td>gene id</td></tr>
<tr><td>GU305914.1</td><td>196677</td><td>196699</td><td>-</td><td>0</td><td>CGGGGCGCGCTTCGACCGTC</td><td>CGG</td><td>||||||||[||||||||||||]PAM</td><td>NA</td><td>Intergenic</td><td>NA</td><td>NA</td></tr>
<tr><td>T19727</td><td>CGCTTCGACCGTCCGGCGCGCGG</td><td>1000</td><td></td><td></td><td></td><td></td><td></td><td></td><td></td><td></td><td></td></tr>
<tr><td>Oligo fwd</td><td>CACCCGCTTCGACCGTCCGGCGCG</td><td></td><td></td><td></td><td></td><td></td><td></td><td></td><td></td><td></td><td></td></tr>
<tr><td>Oligo rev</td><td>AAACCGCGCCGGACGGTCGAAGCG</td><td></td><td></td><td></td><td></td><td></td><td></td><td></td><td></td><td></td><td></td></tr>
<tr><td>Chromosome</td><td>start</td><td>end</td><td>strand</td><td>MM</td><td>target_seq</td><td>PAM</td><td>alignment</td><td>distance</td><td>position</td><td>gene name</td><td>gene id</td></tr>
<tr><td>GU305914.1</td><td>196670</td><td>196692</td><td>-</td><td>0</td><td>CGCTTCGACCGTCCGGCGCG</td><td>CGG</td><td>||||||||[||||||||||||]PAM</td><td>NA</td><td>Intergenic</td><td>NA</td><td>NA</td></tr>
<tr><td>T19728</td><td>GCTTCGACCGTCCGGCGCGCGGG</td><td>1000</td><td></td><td></td><td></td><td></td><td></td><td></td><td></td><td></td><td></td></tr>
<tr><td>Oligo fwd</td><td>CACCGCTTCGACCGTCCGGCGCGC</td><td></td><td></td><td></td><td></td><td></td><td></td><td></td><td></td><td></td><td></td></tr>
<tr><td>Oligo rev</td><td>AAACGCGCGCCGGACGGTCGAAGC</td><td></td><td></td><td></td><td></td><td></td><td></td><td></td><td></td><td></td><td></td></tr>
<tr><td>Chromosome</td><td>start</td><td>end</td><td>strand</td><td>MM</td><td>target_seq</td><td>PAM</td><td>alignment</td><td>distance</td><td>position</td><td>gene name</td><td>gene id</td></tr>
<tr><td>GU305914.1</td><td>196669</td><td>196691</td><td>-</td><td>0</td><td>GCTTCGACCGTCCGGCGCGC</td><td>GGG</td><td>||||||||[||||||||||||]PAM</td><td>NA</td><td>Intergenic</td><td>NA</td><td>NA</td></tr>
<tr><td>T19729</td><td>CTTCGACCGTCCGGCGCGCGGGG</td><td>1000</td><td></td><td></td><td></td><td></td><td></td><td></td><td></td><td></td><td></td></tr>
<tr><td>Oligo fwd</td><td>CACCCTTCGACCGTCCGGCGCGCG</td><td></td><td></td><td></td><td></td><td></td><td></td><td></td><td></td><td></td><td></td></tr>
<tr><td>Oligo rev</td><td>AAACCGCGCGCCGGACGGTCGAAG</td><td></td><td></td><td></td><td></td><td></td><td></td><td></td><td></td><td></td><td></td></tr>
<tr><td>Chromosome</td><td>start</td><td>end</td><td>strand</td><td>MM</td><td>target_seq</td><td>PAM</td><td>alignment</td><td>distance</td><td>position</td><td>gene name</td><td>gene id</td></tr>
<tr><td>GU305914.1</td><td>196668</td><td>196690</td><td>-</td><td>0</td><td>CTTCGACCGTCCGGCGCGCG</td><td>GGG</td><td>||||||||[||||||||||||]PAM</td><td>NA</td><td>Intergenic</td><td>NA</td><td>NA</td></tr>
</table>

<table>
<tr><td>T19730</td><td>CGACCGTCCGGCGCGCGGGGAGG</td><td>1000</td><td></td><td></td><td></td><td></td><td></td><td></td><td></td><td></td><td></td></tr>
<tr><td>Oligo fwd</td><td>CACCCGACCGTCCGGCGCGCGGGG</td><td></td><td></td><td></td><td></td><td></td><td></td><td></td><td></td><td></td><td></td></tr>
<tr><td>Oligo rev</td><td>AAACCCCCGCGCGCCGGACGGTCG</td><td></td><td></td><td></td><td></td><td></td><td></td><td></td><td></td><td></td><td></td></tr>
<tr><td>Chromosome</td><td>start</td><td>end</td><td>strand</td><td>MM</td><td>target_seq</td><td>PAM</td><td>alignment</td><td>distance</td><td>position</td><td>gene name</td><td>gene id</td></tr>
<tr><td>GU305914.1</td><td>196665</td><td>196687</td><td>-</td><td>0</td><td>CGACCGTCCGGCGCGCGGGG</td><td>AGG</td><td>||||||||[||||||||||||]PAM</td><td>NA</td><td>Intergenic</td><td>NA</td><td>NA</td></tr>
<tr><td>T19731</td><td>GACCGTCCGGCGCGCGGGGAGGG</td><td>1000</td><td></td><td></td><td></td><td></td><td></td><td></td><td></td><td></td><td></td></tr>
<tr><td>Oligo fwd</td><td>CACCGACCGTCCGGCGCGCGGGGA</td><td></td><td></td><td></td><td></td><td></td><td></td><td></td><td></td><td></td><td></td></tr>
<tr><td>Oligo rev</td><td>AAACTCCCCGCGCGCCGGACGGTC</td><td></td><td></td><td></td><td></td><td></td><td></td><td></td><td></td><td></td><td></td></tr>
<tr><td>Chromosome</td><td>start</td><td>end</td><td>strand</td><td>MM</td><td>target_seq</td><td>PAM</td><td>alignment</td><td>distance</td><td>position</td><td>gene name</td><td>gene id</td></tr>
<tr><td>GU305914.1</td><td>196664</td><td>196686</td><td>-</td><td>0</td><td>GACCGTCCGGCGCGCGGGGA</td><td>GGG</td><td>||||||||[||||||||||||]PAM</td><td>NA</td><td>Intergenic</td><td>NA</td><td>NA</td></tr>
<tr><td>T19732</td><td>CCTGCCCGATCACCGCAAGTCGG</td><td>1000</td><td></td><td></td><td></td><td></td><td></td><td></td><td></td><td></td><td></td></tr>
<tr><td>Oligo fwd</td><td>CACCCCTGCCCGATCACCGCAAGT</td><td></td><td></td><td></td><td></td><td></td><td></td><td></td><td></td><td></td><td></td></tr>
<tr><td>Oligo rev</td><td>AAACACTTGCGGTGATCGGGCAGG</td><td></td><td></td><td></td><td></td><td></td><td></td><td></td><td></td><td></td><td></td></tr>
<tr><td>Chromosome</td><td>start</td><td>end</td><td>strand</td><td>MM</td><td>target_seq</td><td>PAM</td><td>alignment</td><td>distance</td><td>position</td><td>gene name</td><td>gene id</td></tr>
<tr><td>GU305914.1</td><td>196611</td><td>196633</td><td>-</td><td>0</td><td>CCTGCCCGATCACCGCAAGT</td><td>CGG</td><td>||||||||[||||||||||||]PAM</td><td>NA</td><td>Intergenic</td><td>NA</td><td>NA</td></tr>
<tr><td>T19733</td><td>CCGCAAGTCGGAGCGCAGCGAGG</td><td>1000</td><td></td><td></td><td></td><td></td><td></td><td></td><td></td><td></td><td></td></tr>
<tr><td>Oligo fwd</td><td>CACCCCGCAAGTCGGAGCGCAGCG</td><td></td><td></td><td></td><td></td><td></td><td></td><td></td><td></td><td></td><td></td></tr>
<tr><td>Oligo rev</td><td>AAACCGCTGCGCTCCGACTTGCGG</td><td></td><td></td><td></td><td></td><td></td><td></td><td></td><td></td><td></td><td></td></tr>
<tr><td>Chromosome</td><td>start</td><td>end</td><td>strand</td><td>MM</td><td>target_seq</td><td>PAM</td><td>alignment</td><td>distance</td><td>position</td><td>gene name</td><td>gene id</td></tr>
<tr><td>GU305914.1</td><td>196599</td><td>196621</td><td>-</td><td>0</td><td>CCGCAAGTCGGAGCGCAGCG</td><td>AGG</td><td>||||||||[||||||||||||]PAM</td><td>NA</td><td>Intergenic</td><td>NA</td><td>NA</td></tr>
<tr><td>T19734</td><td>ATTACGCCAACGAACACGAGTGG</td><td>1000</td><td></td><td></td><td></td><td></td><td></td><td></td><td></td><td></td><td></td></tr>
<tr><td>Oligo fwd</td><td>CACCATTACGCCAACGAACACGAG</td><td></td><td></td><td></td><td></td><td></td><td></td><td></td><td></td><td></td><td></td></tr>
<tr><td>Oligo rev</td><td>AAACCTCGTGTTCGTTGGCGTAAT</td><td></td><td></td><td></td><td></td><td></td><td></td><td></td><td></td><td></td><td></td></tr>
<tr><td>Chromosome</td><td>start</td><td>end</td><td>strand</td><td>MM</td><td>target_seq</td><td>PAM</td><td>alignment</td><td>distance</td><td>position</td><td>gene name</td><td>gene id</td></tr>
<tr><td>GU305914.1</td><td>196573</td><td>196595</td><td>-</td><td>0</td><td>ATTACGCCAACGAACACGAG</td><td>TGG</td><td>||||||||[||||||||||||]PAM</td><td>NA</td><td>Intergenic</td><td>NA</td><td>NA</td></tr>
<tr><td>T19735</td><td>AACACGAGTGGCTGTGTCGCCGG</td><td>1000</td><td></td><td></td><td></td><td></td><td></td><td></td><td></td><td></td><td></td></tr>
<tr><td>Oligo fwd</td><td>CACCAACACGAGTGGCTGTGTCGC</td><td></td><td></td><td></td><td></td><td></td><td></td><td></td><td></td><td></td><td></td></tr>
<tr><td>Oligo rev</td><td>AAACGCGACACAGCCACTCGTGTT</td><td></td><td></td><td></td><td></td><td></td><td></td><td></td><td></td><td></td><td></td></tr>
<tr><td>Chromosome</td><td>start</td><td>end</td><td>strand</td><td>MM</td><td>target_seq</td><td>PAM</td><td>alignment</td><td>distance</td><td>position</td><td>gene name</td><td>gene id</td></tr>
<tr><td>GU305914.1</td><td>196561</td><td>196583</td><td>-</td><td>0</td><td>AACACGAGTGGCTGTGTCGC</td><td>CGG</td><td>||||||||[||||||||||||]PAM</td><td>NA</td><td>Intergenic</td><td>NA</td><td>NA</td></tr>
<tr><td>T19736</td><td>ACACGAGTGGCTGTGTCGCCGGG</td><td>1000</td><td></td><td></td><td></td><td></td><td></td><td></td><td></td><td></td><td></td></tr>
<tr><td>Oligo fwd</td><td>CACCACACGAGTGGCTGTGTCGCC</td><td></td><td></td><td></td><td></td><td></td><td></td><td></td><td></td><td></td><td></td></tr>
<tr><td>Oligo rev</td><td>AAACGGCGACACAGCCACTCGTGT</td><td></td><td></td><td></td><td></td><td></td><td></td><td></td><td></td><td></td><td></td></tr>
<tr><td>Chromosome</td><td>start</td><td>end</td><td>strand</td><td>MM</td><td>target_seq</td><td>PAM</td><td>alignment</td><td>distance</td><td>position</td><td>gene name</td><td>gene id</td></tr>
<tr><td>GU305914.1</td><td>196560</td><td>196582</td><td>-</td><td>0</td><td>ACACGAGTGGCTGTGTCGCC</td><td>GGG</td><td>||||||||[||||||||||||]PAM</td><td>NA</td><td>Intergenic</td><td>NA</td><td>NA</td></tr>
<tr><td>T19737</td><td>TGTGTCGCCGGGACCGCTTTCGG</td><td>1000</td><td></td><td></td><td></td><td></td><td></td><td></td><td></td><td></td><td></td></tr>
<tr><td>Oligo fwd</td><td>CACCTGTGTCGCCGGGACCGCTTT</td><td></td><td></td><td></td><td></td><td></td><td></td><td></td><td></td><td></td><td></td></tr>
<tr><td>Oligo rev</td><td>AAACAAAGCGGTCCCGGCGACACA</td><td></td><td></td><td></td><td></td><td></td><td></td><td></td><td></td><td></td><td></td></tr>
<tr><td>Chromosome</td><td>start</td><td>end</td><td>strand</td><td>MM</td><td>target_seq</td><td>PAM</td><td>alignment</td><td>distance</td><td>position</td><td>gene name</td><td>gene id</td></tr>
<tr><td>GU305914.1</td><td>196549</td><td>196571</td><td>-</td><td>0</td><td>TGTGTCGCCGGGACCGCTTT</td><td>CGG</td><td>||||||||[||||||||||||]PAM</td><td>NA</td><td>Intergenic</td><td>NA</td><td>NA</td></tr>
<tr><td>T19738</td><td>ACCGCTTTCGGCCCGACATGCGG</td><td>1000</td><td></td><td></td><td></td><td></td><td></td><td></td><td></td><td></td><td></td></tr>
<tr><td>Oligo fwd</td><td>CACCACCGCTTTCGGCCCGACATG</td><td></td><td></td><td></td><td></td><td></td><td></td><td></td><td></td><td></td><td></td></tr>
<tr><td>Oligo rev</td><td>AAACCATGTCGGGCCGAAAGCGGT</td><td></td><td></td><td></td><td></td><td></td><td></td><td></td><td></td><td></td><td></td></tr>
<tr><td>Chromosome</td><td>start</td><td>end</td><td>strand</td><td>MM</td><td>target_seq</td><td>PAM</td><td>alignment</td><td>distance</td><td>position</td><td>gene name</td><td>gene id</td></tr>
<tr><td>GU305914.1</td><td>196537</td><td>196559</td><td>-</td><td>0</td><td>ACCGCTTTCGGCCCGACATG</td><td>CGG</td><td>||||||||[||||||||||||]PAM</td><td>NA</td><td>Intergenic</td><td>NA</td><td>NA</td></tr>
<tr><td>T19739</td><td>TTCGGCCCGACATGCGGACGTGG</td><td>1000</td><td></td><td></td><td></td><td></td><td></td><td></td><td></td><td></td><td></td></tr>
<tr><td>Oligo fwd</td><td>CACCTTCGGCCCGACATGCGGACG</td><td></td><td></td><td></td><td></td><td></td><td></td><td></td><td></td><td></td><td></td></tr>
<tr><td>Oligo rev</td><td>AAACCGTCCGCATGTCGGGCCGAA</td><td></td><td></td><td></td><td></td><td></td><td></td><td></td><td></td><td></td><td></td></tr>
<tr><td>Chromosome</td><td>start</td><td>end</td><td>strand</td><td>MM</td><td>target_seq</td><td>PAM</td><td>alignment</td><td>distance</td><td>position</td><td>gene name</td><td>gene id</td></tr>
<tr><td>GU305914.1</td><td>196531</td><td>196553</td><td>-</td><td>0</td><td>TTCGGCCCGACATGCGGACG</td><td>TGG</td><td>||||||||[||||||||||||]PAM</td><td>NA</td><td>Intergenic</td><td>NA</td><td>NA</td></tr>
<tr><td>T19740</td><td>TCGGCCCGACATGCGGACGTGGG</td><td>1000</td><td></td><td></td><td></td><td></td><td></td><td></td><td></td><td></td><td></td></tr>
<tr><td>Oligo fwd</td><td>CACCTCGGCCCGACATGCGGACGT</td><td></td><td></td><td></td><td></td><td></td><td></td><td></td><td></td><td></td><td></td></tr>
<tr><td>Oligo rev</td><td>AAACACGTCCGCATGTCGGGCCGA</td><td></td><td></td><td></td><td></td><td></td><td></td><td></td><td></td><td></td><td></td></tr>
<tr><td>Chromosome</td><td>start</td><td>end</td><td>strand</td><td>MM</td><td>target_seq</td><td>PAM</td><td>alignment</td><td>distance</td><td>position</td><td>gene name</td><td>gene id</td></tr>
<tr><td>GU305914.1</td><td>196530</td><td>196552</td><td>-</td><td>0</td><td>TCGGCCCGACATGCGGACGT</td><td>GGG</td><td>||||||||[||||||||||||]PAM</td><td>NA</td><td>Intergenic</td><td>NA</td><td>NA</td></tr>
<tr><td>T19741</td><td>CATGCGGACGTGGGACGACGCGG</td><td>1000</td><td></td><td></td><td></td><td></td><td></td><td></td><td></td><td></td><td></td></tr>
<tr><td>Oligo fwd</td><td>CACCCATGCGGACGTGGGACGACG</td><td></td><td></td><td></td><td></td><td></td><td></td><td></td><td></td><td></td><td></td></tr>
<tr><td>Oligo rev</td><td>AAACCGTCGTCCCACGTCCGCATG</td><td></td><td></td><td></td><td></td><td></td><td></td><td></td><td></td><td></td><td></td></tr>
<tr><td>Chromosome</td><td>start</td><td>end</td><td>strand</td><td>MM</td><td>target_seq</td><td>PAM</td><td>alignment</td><td>distance</td><td>position</td><td>gene name</td><td>gene id</td></tr>
<tr><td>GU305914.1</td><td>196521</td><td>196543</td><td>-</td><td>0</td><td>CATGCGGACGTGGGACGACG</td><td>CGG</td><td>||||||||[||||||||||||]PAM</td><td>NA</td><td>Intergenic</td><td>NA</td><td>NA</td></tr>
<tr><td>T19742</td><td>GTGGGACGACGCGGACAAGCTGG</td><td>1000</td><td></td><td></td><td></td><td></td><td></td><td></td><td></td><td></td><td></td></tr>
<tr><td>Oligo fwd</td><td>CACCGTGGGACGACGCGGACAAGC</td><td></td><td></td><td></td><td></td><td></td><td></td><td></td><td></td><td></td><td></td></tr>
<tr><td>Oligo rev</td><td>AAACGCTTGTCCGCGTCGTCCCAC</td><td></td><td></td><td></td><td></td><td></td><td></td><td></td><td></td><td></td><td></td></tr>
<tr><td>Chromosome</td><td>start</td><td>end</td><td>strand</td><td>MM</td><td>target_seq</td><td>PAM</td><td>alignment</td><td>distance</td><td>position</td><td>gene name</td><td>gene id</td></tr>
<tr><td>GU305914.1</td><td>196512</td><td>196534</td><td>-</td><td>0</td><td>GTGGGACGACGCGGACAAGC</td><td>TGG</td><td>||||||||[||||||||||||]PAM</td><td>NA</td><td>Intergenic</td><td>NA</td><td>NA</td></tr>
<tr><td>T19743</td><td>CGTGCGCGCCATGAAGAGAAAGG</td><td>1000</td><td></td><td></td><td></td><td></td><td></td><td></td><td></td><td></td><td></td></tr>
<tr><td>Oligo fwd</td><td>CACCCGTGCGCGCCATGAAGAGAA</td><td></td><td></td><td></td><td></td><td></td><td></td><td></td><td></td><td></td><td></td></tr>
<tr><td>Oligo rev</td><td>AAACTTCTCTTCATGGCGCGCACG</td><td></td><td></td><td></td><td></td><td></td><td></td><td></td><td></td><td></td><td></td></tr>
<tr><td>Chromosome</td><td>start</td><td>end</td><td>strand</td><td>MM</td><td>target_seq</td><td>PAM</td><td>alignment</td><td>distance</td><td>position</td><td>gene name</td><td>gene id</td></tr>
<tr><td>GU305914.1</td><td>196476</td><td>196498</td><td>-</td><td>0</td><td>CGTGCGCGCCATGAAGAGAA</td><td>AGG</td><td>||||||||[||||||||||||]PAM</td><td>NA</td><td>Intergenic</td><td>NA</td><td>NA</td></tr>
<tr><td>T19744</td><td>CGCGCCATGAAGAGAAAGGCCGG</td><td>1000</td><td></td><td></td><td></td><td></td><td></td><td></td><td></td><td></td><td></td></tr>
<tr><td>Oligo fwd</td><td>CACCCGCGCCATGAAGAGAAAGGC</td><td></td><td></td><td></td><td></td><td></td><td></td><td></td><td></td><td></td><td></td></tr>
<tr><td>Oligo rev</td><td>AAACGCCTTTCTCTTCATGGCGCG</td><td></td><td></td><td></td><td></td><td></td><td></td><td></td><td></td><td></td><td></td></tr>
<tr><td>Chromosome</td><td>start</td><td>end</td><td>strand</td><td>MM</td><td>target_seq</td><td>PAM</td><td>alignment</td><td>distance</td><td>position</td><td>gene name</td><td>gene id</td></tr>
<tr><td>GU305914.1</td><td>196472</td><td>196494</td><td>-</td><td>0</td><td>CGCGCCATGAAGAGAAAGGC</td><td>CGG</td><td>||||||||[||||||||||||]PAM</td><td>NA</td><td>Intergenic</td><td>NA</td><td>NA</td></tr>
</table>

T19745 GCGCCATGAAGAGAAAGGCCGGG 1000
Oligo fwd CACCGCGCCATGAAGAGAAAGGCC
Oligo rev AAACGGCCTTTCTCTTCATGGCGC
Chromosome start end strand MM target_seq PAM alignment distance position gene name gene id
GU305914.1 196471 196493 - 0 GCGCCATGAAGAGAAAGGCC GGG ||||||||[||||||||||||]PAM NA Intergenic NA NA

T19746 CGCCATGAAGAGAAAGGCCGGGG 1000
Oligo fwd CACCCGCCATGAAGAGAAAGGCCG
Oligo rev AAACCGGCCTTTCTCTTCATGGCG
Chromosome start end strand MM target_seq PAM alignment distance position gene name gene id
GU305914.1 196470 196492 - 0 CGCCATGAAGAGAAAGGCCG GGG ||||||||[||||||||||||]PAM NA Intergenic NA NA

T19747 CGGGGAGATGTACACGAACGAGG 1000
Oligo fwd CACCCGGGGAGATGTACACGAACG
Oligo rev AAACCGTTCGTGTACATCTCCCCG
Chromosome start end strand MM target_seq PAM alignment distance position gene name gene id
GU305914.1 196452 196474 - 0 CGGGGAGATGTACACGAACG AGG ||||||||[||||||||||||]PAM NA Intergenic NA NA

T19748 GGAGATGTACACGAACGAGGAGG 1000
Oligo fwd CACCGGAGATGTACACGAACGAGG
Oligo rev AAACCCTCGTTCGTGTACATCTCC
Chromosome start end strand MM target_seq PAM alignment distance position gene name gene id
GU305914.1 196449 196471 - 0 GGAGATGTACACGAACGAGG AGG ||||||||[||||||||||||]PAM NA Intergenic NA NA

T19749 GATGTACACGAACGAGGAGGAGG 1000
Oligo fwd CACCGATGTACACGAACGAGGAGG
Oligo rev AAACCCTCCTCGTTCGTGTACATC
Chromosome start end strand MM target_seq PAM alignment distance position gene name gene id
GU305914.1 196446 196468 - 0 GATGTACACGAACGAGGAGG AGG ||||||||[||||||||||||]PAM NA Intergenic NA NA

T19750 ACACGAACGAGGAGGAGGACTGG 1000
Oligo fwd CACCACACGAACGAGGAGGAGGAC
Oligo rev AAACGTCCTCCTCCTCGTTCGTGT
Chromosome start end strand MM target_seq PAM alignment distance position gene name gene id
GU305914.1 196441 196463 - 0 ACACGAACGAGGAGGAGGAC TGG ||||||||[||||||||||||]PAM NA Intergenic NA NA
GU305914.1 160681 160703 + 4 AAACCAACGAAGAGGAGGAG AGG |-||-|||[||-||||||||-]PAM NA Intergenic NA NA

T19751 GAACGAGGAGGAGGACTGGACGG 1000
Oligo fwd CACCGAACGAGGAGGAGGACTGGA
Oligo rev AAACTCCAGTCCTCCTCCTCGTTC
Chromosome start end strand MM target_seq PAM alignment distance position gene name gene id
GU305914.1 196437 196459 - 0 GAACGAGGAGGAGGACTGGA CGG ||||||||[||||||||||||]PAM NA Intergenic NA NA

T19752 ACTGGACGGATACCAACGAGTGG 1000
Oligo fwd CACCACTGGACGGATACCAACGAG
Oligo rev AAACCTCGTTGGTATCCGTCCAGT
Chromosome start end strand MM target_seq PAM alignment distance position gene name gene id
GU305914.1 196423 196445 - 0 ACTGGACGGATACCAACGAG TGG ||||||||[||||||||||||]PAM NA Intergenic NA NA

T19753 CTGGACGGATACCAACGAGTGGG 1000
Oligo fwd CACCCTGGACGGATACCAACGAGT
Oligo rev AAACACTCGTTGGTATCCGTCCAG
Chromosome start end strand MM target_seq PAM alignment distance position gene name gene id
GU305914.1 196422 196444 - 0 CTGGACGGATACCAACGAGT GGG ||||||||[||||||||||||]PAM NA Intergenic NA NA

T19754 GGATACCAACGAGTGGGAACAGG 1000
Oligo fwd CACCGGATACCAACGAGTGGGAAC
Oligo rev AAACGTTCCCACTCGTTGGTATCC
Chromosome start end strand MM target_seq PAM alignment distance position gene name gene id
GU305914.1 196416 196438 - 0 GGATACCAACGAGTGGGAAC AGG ||||||||[||||||||||||]PAM NA Intergenic NA NA

T19755 TACCAACGAGTGGGAACAGGAGG 1000
Oligo fwd CACCTACCAACGAGTGGGAACAGG
Oligo rev AAACCCTGTTCCCACTCGTTGGTA
Chromosome start end strand MM target_seq PAM alignment distance position gene name gene id
GU305914.1 196413 196435 - 0 TACCAACGAGTGGGAACAGG AGG ||||||||[||||||||||||]PAM NA Intergenic NA NA

T19756 AACAGGAGGACGACGACAACAGG 1000
Oligo fwd CACCAACAGGAGGACGACGACAAC
Oligo rev AAACGTTGTCGTCGTCCTCCTGTT
Chromosome start end strand MM target_seq PAM alignment distance position gene name gene id
GU305914.1 196399 196421 - 0 AACAGGAGGACGACGACAAC AGG ||||||||[||||||||||||]PAM NA Intergenic NA NA

T19757 ACAGGAGGACGACGACAACAGGG 1000
Oligo fwd CACCACAGGAGGACGACGACAACA
Oligo rev AAACTGTTGTCGTCGTCCTCCTGT
Chromosome start end strand MM target_seq PAM alignment distance position gene name gene id
GU305914.1 196398 196420 - 0 ACAGGAGGACGACGACAACA GGG ||||||||[||||||||||||]PAM NA Intergenic NA NA
GU305914.1 32834 32856 - 4 CTACGAGGACGACGACAACG AGG --|-||||[|||||||||||-]PAM NA Intergenic NA NA

T19758 CAGGAGGACGACGACAACAGGGG 1000
Oligo fwd CACCCAGGAGGACGACGACAACAG
Oligo rev AAACCTGTTGTCGTCGTCCTCCTG
Chromosome start end strand MM target_seq PAM alignment distance position gene name gene id
GU305914.1 196397 196419 - 0 CAGGAGGACGACGACAACAG GGG ||||||||[||||||||||||]PAM NA Intergenic NA NA

T19759 GACGACAACAGGGGATTCGACGG 1000
Oligo fwd CACCGACGACAACAGGGGATTCGA

<table>
<tr><td>Oligo rev</td><td>AAACTCGAATCCCCTGTTGTCGTC</td><td></td><td></td><td></td><td></td><td></td><td></td><td></td><td></td><td></td><td></td></tr>
<tr><td>Chromosome</td><td>start</td><td>end</td><td>strand</td><td>MM</td><td>target_seq</td><td>PAM</td><td>alignment</td><td>distance</td><td>position</td><td>gene name</td><td>gene id</td></tr>
<tr><td>GU305914.1</td><td>196388</td><td>196410</td><td>-</td><td>0</td><td>GACGACAACAGGGGATTCGA</td><td>CGG</td><td>||||||||[||||||||||||]PAM</td><td>NA</td><td>Intergenic</td><td>NA</td><td>NA</td></tr>
<tr><td></td><td></td><td></td><td></td><td></td><td></td><td></td><td></td><td></td><td></td><td></td><td></td></tr>
<tr><td>T19760</td><td>GGGATTCGACGGACACGCGATGG</td><td>1000</td><td></td><td></td><td></td><td></td><td></td><td></td><td></td><td></td><td></td></tr>
<tr><td>Oligo fwd</td><td>CACCGGGATTCGACGGACACGCGA</td><td></td><td></td><td></td><td></td><td></td><td></td><td></td><td></td><td></td><td></td></tr>
<tr><td>Oligo rev</td><td>AAACTCGCGTGTCCGTCGAATCCC</td><td></td><td></td><td></td><td></td><td></td><td></td><td></td><td></td><td></td><td></td></tr>
<tr><td>Chromosome</td><td>start</td><td>end</td><td>strand</td><td>MM</td><td>target_seq</td><td>PAM</td><td>alignment</td><td>distance</td><td>position</td><td>gene name</td><td>gene id</td></tr>
<tr><td>GU305914.1</td><td>196377</td><td>196399</td><td>-</td><td>0</td><td>GGGATTCGACGGACACGCGA</td><td>TGG</td><td>||||||||[||||||||||||]PAM</td><td>NA</td><td>Intergenic</td><td>NA</td><td>NA</td></tr>
<tr><td></td><td></td><td></td><td></td><td></td><td></td><td></td><td></td><td></td><td></td><td></td><td></td></tr>
<tr><td>T19761</td><td>CGGACACGCGATGGACATCACGG</td><td>1000</td><td></td><td></td><td></td><td></td><td></td><td></td><td></td><td></td><td></td></tr>
<tr><td>Oligo fwd</td><td>CACCCGGACACGCGATGGACATCA</td><td></td><td></td><td></td><td></td><td></td><td></td><td></td><td></td><td></td><td></td></tr>
<tr><td>Oligo rev</td><td>AAACTGATGTCCATCGCGTGTCCG</td><td></td><td></td><td></td><td></td><td></td><td></td><td></td><td></td><td></td><td></td></tr>
<tr><td>Chromosome</td><td>start</td><td>end</td><td>strand</td><td>MM</td><td>target_seq</td><td>PAM</td><td>alignment</td><td>distance</td><td>position</td><td>gene name</td><td>gene id</td></tr>
<tr><td>GU305914.1</td><td>196368</td><td>196390</td><td>-</td><td>0</td><td>CGGACACGCGATGGACATCA</td><td>CGG</td><td>||||||||[||||||||||||]PAM</td><td>NA</td><td>Intergenic</td><td>NA</td><td>NA</td></tr>
<tr><td></td><td></td><td></td><td></td><td></td><td></td><td></td><td></td><td></td><td></td><td></td><td></td></tr>
<tr><td>T19762</td><td>GGACACGCGATGGACATCACGGG</td><td>1000</td><td></td><td></td><td></td><td></td><td></td><td></td><td></td><td></td><td></td></tr>
<tr><td>Oligo fwd</td><td>CACCGGACACGCGATGGACATCAC</td><td></td><td></td><td></td><td></td><td></td><td></td><td></td><td></td><td></td><td></td></tr>
<tr><td>Oligo rev</td><td>AAACGTGATGTCCATCGCGTGTCC</td><td></td><td></td><td></td><td></td><td></td><td></td><td></td><td></td><td></td><td></td></tr>
<tr><td>Chromosome</td><td>start</td><td>end</td><td>strand</td><td>MM</td><td>target_seq</td><td>PAM</td><td>alignment</td><td>distance</td><td>position</td><td>gene name</td><td>gene id</td></tr>
<tr><td>GU305914.1</td><td>196367</td><td>196389</td><td>-</td><td>0</td><td>GGACACGCGATGGACATCAC</td><td>GGG</td><td>||||||||[||||||||||||]PAM</td><td>NA</td><td>Intergenic</td><td>NA</td><td>NA</td></tr>
<tr><td></td><td></td><td></td><td></td><td></td><td></td><td></td><td></td><td></td><td></td><td></td><td></td></tr>
<tr><td>T19763</td><td>GACATCACGGGCCCCGACGATGG</td><td>1000</td><td></td><td></td><td></td><td></td><td></td><td></td><td></td><td></td><td></td></tr>
<tr><td>Oligo fwd</td><td>CACCGACATCACGGGCCCCGACGA</td><td></td><td></td><td></td><td></td><td></td><td></td><td></td><td></td><td></td><td></td></tr>
<tr><td>Oligo rev</td><td>AAACTCGTCGGGGCCCGTGATGTC</td><td></td><td></td><td></td><td></td><td></td><td></td><td></td><td></td><td></td><td></td></tr>
<tr><td>Chromosome</td><td>start</td><td>end</td><td>strand</td><td>MM</td><td>target_seq</td><td>PAM</td><td>alignment</td><td>distance</td><td>position</td><td>gene name</td><td>gene id</td></tr>
<tr><td>GU305914.1</td><td>196355</td><td>196377</td><td>-</td><td>0</td><td>GACATCACGGGCCCCGACGA</td><td>TGG</td><td>||||||||[||||||||||||]PAM</td><td>NA</td><td>Intergenic</td><td>NA</td><td>NA</td></tr>
<tr><td></td><td></td><td></td><td></td><td></td><td></td><td></td><td></td><td></td><td></td><td></td><td></td></tr>
<tr><td>T19764</td><td>ATGGCTCGCGCTCATGCACGTGG</td><td>1000</td><td></td><td></td><td></td><td></td><td></td><td></td><td></td><td></td><td></td></tr>
<tr><td>Oligo fwd</td><td>CACCATGGCTCGCGCTCATGCACG</td><td></td><td></td><td></td><td></td><td></td><td></td><td></td><td></td><td></td><td></td></tr>
<tr><td>Oligo rev</td><td>AAACCGTGCATGAGCGCGAGCCAT</td><td></td><td></td><td></td><td></td><td></td><td></td><td></td><td></td><td></td><td></td></tr>
<tr><td>Chromosome</td><td>start</td><td>end</td><td>strand</td><td>MM</td><td>target_seq</td><td>PAM</td><td>alignment</td><td>distance</td><td>position</td><td>gene name</td><td>gene id</td></tr>
<tr><td>GU305914.1</td><td>196336</td><td>196358</td><td>-</td><td>0</td><td>ATGGCTCGCGCTCATGCACG</td><td>TGG</td><td>||||||||[||||||||||||]PAM</td><td>NA</td><td>Intergenic</td><td>NA</td><td>NA</td></tr>
<tr><td></td><td></td><td></td><td></td><td></td><td></td><td></td><td></td><td></td><td></td><td></td><td></td></tr>
<tr><td>T19765</td><td>GCTCGCGCTCATGCACGTGGTGG</td><td>1000</td><td></td><td></td><td></td><td></td><td></td><td></td><td></td><td></td><td></td></tr>
<tr><td>Oligo fwd</td><td>CACCGCTCGCGCTCATGCACGTGG</td><td></td><td></td><td></td><td></td><td></td><td></td><td></td><td></td><td></td><td></td></tr>
<tr><td>Oligo rev</td><td>AAACCCACGTGCATGAGCGCGAGC</td><td></td><td></td><td></td><td></td><td></td><td></td><td></td><td></td><td></td><td></td></tr>
<tr><td>Chromosome</td><td>start</td><td>end</td><td>strand</td><td>MM</td><td>target_seq</td><td>PAM</td><td>alignment</td><td>distance</td><td>position</td><td>gene name</td><td>gene id</td></tr>
<tr><td>GU305914.1</td><td>196333</td><td>196355</td><td>-</td><td>0</td><td>GCTCGCGCTCATGCACGTGG</td><td>TGG</td><td>||||||||[||||||||||||]PAM</td><td>NA</td><td>Intergenic</td><td>NA</td><td>NA</td></tr>
<tr><td></td><td></td><td></td><td></td><td></td><td></td><td></td><td></td><td></td><td></td><td></td><td></td></tr>
<tr><td>T19766</td><td>GAGCGTGCTAACGACACGCCCGG</td><td>1000</td><td></td><td></td><td></td><td></td><td></td><td></td><td></td><td></td><td></td></tr>
<tr><td>Oligo fwd</td><td>CACCGAGCGTGCTAACGACACGCC</td><td></td><td></td><td></td><td></td><td></td><td></td><td></td><td></td><td></td><td></td></tr>
<tr><td>Oligo rev</td><td>AAACGGCGTGTCGTTAGCACGCTC</td><td></td><td></td><td></td><td></td><td></td><td></td><td></td><td></td><td></td><td></td></tr>
<tr><td>Chromosome</td><td>start</td><td>end</td><td>strand</td><td>MM</td><td>target_seq</td><td>PAM</td><td>alignment</td><td>distance</td><td>position</td><td>gene name</td><td>gene id</td></tr>
<tr><td>GU305914.1</td><td>196305</td><td>196327</td><td>-</td><td>0</td><td>GAGCGTGCTAACGACACGCC</td><td>CGG</td><td>||||||||[||||||||||||]PAM</td><td>NA</td><td>Intergenic</td><td>NA</td><td>NA</td></tr>
<tr><td></td><td></td><td></td><td></td><td></td><td></td><td></td><td></td><td></td><td></td><td></td><td></td></tr>
<tr><td>T19767</td><td>GCGTTACATGTCAGAGAAAAAGG</td><td>1000</td><td></td><td></td><td></td><td></td><td></td><td></td><td></td><td></td><td></td></tr>
<tr><td>Oligo fwd</td><td>CACCGCGTTACATGTCAGAGAAAA</td><td></td><td></td><td></td><td></td><td></td><td></td><td></td><td></td><td></td><td></td></tr>
<tr><td>Oligo rev</td><td>AAACTTTTCTCTGACATGTAACGC</td><td></td><td></td><td></td><td></td><td></td><td></td><td></td><td></td><td></td><td></td></tr>
<tr><td>Chromosome</td><td>start</td><td>end</td><td>strand</td><td>MM</td><td>target_seq</td><td>PAM</td><td>alignment</td><td>distance</td><td>position</td><td>gene name</td><td>gene id</td></tr>
<tr><td>GU305914.1</td><td>196254</td><td>196276</td><td>-</td><td>0</td><td>GCGTTACATGTCAGAGAAAA</td><td>AGG</td><td>||||||||[||||||||||||]PAM</td><td>NA</td><td>Intergenic</td><td>NA</td><td>NA</td></tr>
<tr><td></td><td></td><td></td><td></td><td></td><td></td><td></td><td></td><td></td><td></td><td></td><td></td></tr>
<tr><td>T19768</td><td>GAAAAAGGTCACGTTTCAAGAGG</td><td>1000</td><td></td><td></td><td></td><td></td><td></td><td></td><td></td><td></td><td></td></tr>
<tr><td>Oligo fwd</td><td>CACCGAAAAAGGTCACGTTTCAAG</td><td></td><td></td><td></td><td></td><td></td><td></td><td></td><td></td><td></td><td></td></tr>
<tr><td>Oligo rev</td><td>AAACCTTGAAACGTGACCTTTTTC</td><td></td><td></td><td></td><td></td><td></td><td></td><td></td><td></td><td></td><td></td></tr>
<tr><td>Chromosome</td><td>start</td><td>end</td><td>strand</td><td>MM</td><td>target_seq</td><td>PAM</td><td>alignment</td><td>distance</td><td>position</td><td>gene name</td><td>gene id</td></tr>
<tr><td>GU305914.1</td><td>196239</td><td>196261</td><td>-</td><td>0</td><td>GAAAAAGGTCACGTTTCAAG</td><td>AGG</td><td>||||||||[||||||||||||]PAM</td><td>NA</td><td>Intergenic</td><td>NA</td><td>NA</td></tr>
<tr><td></td><td></td><td></td><td></td><td></td><td></td><td></td><td></td><td></td><td></td><td></td><td></td></tr>
<tr><td>T19769</td><td>AAAGGTCACGTTTCAAGAGGTGG</td><td>1000</td><td></td><td></td><td></td><td></td><td></td><td></td><td></td><td></td><td></td></tr>
<tr><td>Oligo fwd</td><td>CACCAAAGGTCACGTTTCAAGAGG</td><td></td><td></td><td></td><td></td><td></td><td></td><td></td><td></td><td></td><td></td></tr>
<tr><td>Oligo rev</td><td>AAACCCTCTTGAAACGTGACCTTT</td><td></td><td></td><td></td><td></td><td></td><td></td><td></td><td></td><td></td><td></td></tr>
<tr><td>Chromosome</td><td>start</td><td>end</td><td>strand</td><td>MM</td><td>target_seq</td><td>PAM</td><td>alignment</td><td>distance</td><td>position</td><td>gene name</td><td>gene id</td></tr>
<tr><td>GU305914.1</td><td>196236</td><td>196258</td><td>-</td><td>0</td><td>AAAGGTCACGTTTCAAGAGG</td><td>TGG</td><td>||||||||[||||||||||||]PAM</td><td>NA</td><td>Intergenic</td><td>NA</td><td>NA</td></tr>
<tr><td></td><td></td><td></td><td></td><td></td><td></td><td></td><td></td><td></td><td></td><td></td><td></td></tr>
<tr><td>T19770</td><td>TTCAAGAGGTGGCGACCAGAAGG</td><td>1000</td><td></td><td></td><td></td><td></td><td></td><td></td><td></td><td></td><td></td></tr>
<tr><td>Oligo fwd</td><td>CACCTTCAAGAGGTGGCGACCAGA</td><td></td><td></td><td></td><td></td><td></td><td></td><td></td><td></td><td></td><td></td></tr>
<tr><td>Oligo rev</td><td>AAACTCTGGTCGCCACCTCTTGAA</td><td></td><td></td><td></td><td></td><td></td><td></td><td></td><td></td><td></td><td></td></tr>
<tr><td>Chromosome</td><td>start</td><td>end</td><td>strand</td><td>MM</td><td>target_seq</td><td>PAM</td><td>alignment</td><td>distance</td><td>position</td><td>gene name</td><td>gene id</td></tr>
<tr><td>GU305914.1</td><td>196225</td><td>196247</td><td>-</td><td>0</td><td>TTCAAGAGGTGGCGACCAGA</td><td>AGG</td><td>||||||||[||||||||||||]PAM</td><td>NA</td><td>Intergenic</td><td>NA</td><td>NA</td></tr>
<tr><td></td><td></td><td></td><td></td><td></td><td></td><td></td><td></td><td></td><td></td><td></td><td></td></tr>
<tr><td>T19771</td><td>GGTGGCGACCAGAAGGCTCTTGG</td><td>1000</td><td></td><td></td><td></td><td></td><td></td><td></td><td></td><td></td><td></td></tr>
<tr><td>Oligo fwd</td><td>CACCGGTGGCGACCAGAAGGCTCT</td><td></td><td></td><td></td><td></td><td></td><td></td><td></td><td></td><td></td><td></td></tr>
<tr><td>Oligo rev</td><td>AAACAGAGCCTTCTGGTCGCCACC</td><td></td><td></td><td></td><td></td><td></td><td></td><td></td><td></td><td></td><td></td></tr>
<tr><td>Chromosome</td><td>start</td><td>end</td><td>strand</td><td>MM</td><td>target_seq</td><td>PAM</td><td>alignment</td><td>distance</td><td>position</td><td>gene name</td><td>gene id</td></tr>
<tr><td>GU305914.1</td><td>196218</td><td>196240</td><td>-</td><td>0</td><td>GGTGGCGACCAGAAGGCTCT</td><td>TGG</td><td>||||||||[||||||||||||]PAM</td><td>NA</td><td>Intergenic</td><td>NA</td><td>NA</td></tr>
<tr><td></td><td></td><td></td><td></td><td></td><td></td><td></td><td></td><td></td><td></td><td></td><td></td></tr>
<tr><td>T19772</td><td>CGACCAGAAGGCTCTTGGAGAGG</td><td>1000</td><td></td><td></td><td></td><td></td><td></td><td></td><td></td><td></td><td></td></tr>
<tr><td>Oligo fwd</td><td>CACCCGACCAGAAGGCTCTTGGAG</td><td></td><td></td><td></td><td></td><td></td><td></td><td></td><td></td><td></td><td></td></tr>
<tr><td>Oligo rev</td><td>AAACCTCCAAGAGCCTTCTGGTCG</td><td></td><td></td><td></td><td></td><td></td><td></td><td></td><td></td><td></td><td></td></tr>
<tr><td>Chromosome</td><td>start</td><td>end</td><td>strand</td><td>MM</td><td>target_seq</td><td>PAM</td><td>alignment</td><td>distance</td><td>position</td><td>gene name</td><td>gene id</td></tr>
<tr><td>GU305914.1</td><td>196213</td><td>196235</td><td>-</td><td>0</td><td>CGACCAGAAGGCTCTTGGAG</td><td>AGG</td><td>||||||||[||||||||||||]PAM</td><td>NA</td><td>Intergenic</td><td>NA</td><td>NA</td></tr>
<tr><td></td><td></td><td></td><td></td><td></td><td></td><td></td><td></td><td></td><td></td><td></td><td></td></tr>
<tr><td>T19773</td><td>CTCTGCAGACCGTGAGTTGACGG</td><td>1000</td><td></td><td></td><td></td><td></td><td></td><td></td><td></td><td></td><td></td></tr>
<tr><td>Oligo fwd</td><td>CACCCTCTGCAGACCGTGAGTTGA</td><td></td><td></td><td></td><td></td><td></td><td></td><td></td><td></td><td></td><td></td></tr>
<tr><td>Oligo rev</td><td>AAACTCAACTCACGGTCTGCAGAG</td><td></td><td></td><td></td><td></td><td></td><td></td><td></td><td></td><td></td><td></td></tr>
<tr><td>Chromosome</td><td>start</td><td>end</td><td>strand</td><td>MM</td><td>target_seq</td><td>PAM</td><td>alignment</td><td>distance</td><td>position</td><td>gene name</td><td>gene id</td></tr>
<tr><td>GU305914.1</td><td>196154</td><td>196176</td><td>-</td><td>0</td><td>CTCTGCAGACCGTGAGTTGA</td><td>CGG</td><td>||||||||[||||||||||||]PAM</td><td>NA</td><td>Intergenic</td><td>NA</td><td>NA</td></tr>
<tr><td>GU305914.1</td><td>194051</td><td>194073</td><td>+</td><td>3</td><td>CTCTGCCGACCGAGAGATGA</td><td>TGG</td><td>||||||-|[||||-|||-|||]PAM</td><td>NA</td><td>Intergenic</td><td>NA</td><td>NA</td></tr>
</table>

T19774 AGACCGTGAGTTGACGGCGCCGG 1000
Oligo fwd CACCAGACCGTGAGTTGACGGCGC
Oligo rev AAACGCGCCGTCAACTCACGGTCT
Chromosome start end strand MM target_seq PAM alignment distance position gene name gene id
GU305914.1 196148 196170 - 0 AGACCGTGAGTTGACGGCGC CGG ||||||||[||||||||||||]PAM NA Intergenic NA NA

T19775 TTGACGGCGCCGGCGCCAGACGG 1000
Oligo fwd CACCTTGACGGCGCCGGCGCCAGA
Oligo rev AAACTCTGGCGCCGGCGCCGTCAA
Chromosome start end strand MM target_seq PAM alignment distance position gene name gene id
GU305914.1 196138 196160 - 0 TTGACGGCGCCGGCGCCAGA CGG ||||||||[||||||||||||]PAM NA Intergenic NA NA

T19776 GGAGCAGACAGAGAGAGAGAAGG 1000
Oligo fwd CACCGGAGCAGACAGAGAGAGAGA
Oligo rev AAACTCTCTCTCTCTGTCTGCTCC
Chromosome start end strand MM target_seq PAM alignment distance position gene name gene id
GU305914.1 196117 196139 - 0 GGAGCAGACAGAGAGAGAGA AGG ||||||||[||||||||||||]PAM NA Intergenic NA NA

T19777 GACAGAGAGAGAGAAGGATGTGG 1000
Oligo fwd CACCGACAGAGAGAGAGAAGGATG
Oligo rev AAACCATCCTTCTCTCTCTCTGTC
Chromosome start end strand MM target_seq PAM alignment distance position gene name gene id
GU305914.1 196111 196133 - 0 GACAGAGAGAGAGAAGGATG TGG ||||||||[||||||||||||]PAM NA Intergenic NA NA

T19778 AGAAGGATGTGGTCTCTCCGCGG 1000
Oligo fwd CACCAGAAGGATGTGGTCTCTCCG
Oligo rev AAACCGGAGAGACCACATCCTTCT
Chromosome start end strand MM target_seq PAM alignment distance position gene name gene id
GU305914.1 196100 196122 - 0 AGAAGGATGTGGTCTCTCCG CGG ||||||||[||||||||||||]PAM NA Intergenic NA NA

T19779 AGACGCGACCTGCGTTTCGATGG 1000
Oligo fwd CACCAGACGCGACCTGCGTTTCGA
Oligo rev AAACTCGAAACGCAGGTCGCGTCT
Chromosome start end strand MM target_seq PAM alignment distance position gene name gene id
GU305914.1 196070 196092 - 0 AGACGCGACCTGCGTTTCGA TGG ||||||||[||||||||||||]PAM NA Intergenic NA NA

T19780 CGACCTGCGTTTCGATGGTCAGG 1000
Oligo fwd CACCCGACCTGCGTTTCGATGGTC
Oligo rev AAACGACCATCGAAACGCAGGTCG
Chromosome start end strand MM target_seq PAM alignment distance position gene name gene id
GU305914.1 196065 196087 - 0 CGACCTGCGTTTCGATGGTC AGG ||||||||[||||||||||||]PAM NA Intergenic NA NA

T19781 TCGATGGTCAGGACGACGAGTGG 1000
Oligo fwd CACCTCGATGGTCAGGACGACGAG
Oligo rev AAACCTCGTCGTCCTGACCATCGA
Chromosome start end strand MM target_seq PAM alignment distance position gene name gene id
GU305914.1 196054 196076 - 0 TCGATGGTCAGGACGACGAG TGG ||||||||[||||||||||||]PAM NA Intergenic NA NA

T19782 TCAGGACGACGAGTGGTGTTTGG 1000
Oligo fwd CACCTCAGGACGACGAGTGGTGTT
Oligo rev AAACAACACCACTCGTCGTCCTGA
Chromosome start end strand MM target_seq PAM alignment distance position gene name gene id
GU305914.1 196047 196069 - 0 TCAGGACGACGAGTGGTGTT TGG ||||||||[||||||||||||]PAM NA Intergenic NA NA

T19783 CGAGTGGTGTTTGGCGTACGAGG 1000
Oligo fwd CACCCGAGTGGTGTTTGGCGTACG
Oligo rev AAACCGTACGCCAAACACCACTCG
Chromosome start end strand MM target_seq PAM alignment distance position gene name gene id
GU305914.1 196038 196060 - 0 CGAGTGGTGTTTGGCGTACG AGG ||||||||[||||||||||||]PAM NA Intergenic NA NA

T19784 TTTGGCGTACGAGGATTGCATGG 1000
Oligo fwd CACCTTTGGCGTACGAGGATTGCA
Oligo rev AAACTGCAATCCTCGTACGCCAAA
Chromosome start end strand MM target_seq PAM alignment distance position gene name gene id
GU305914.1 196029 196051 - 0 TTTGGCGTACGAGGATTGCA TGG ||||||||[||||||||||||]PAM NA Intergenic NA NA

T19785 GATTGCATGGACCAGAAGTTCGG 1000
Oligo fwd CACCGATTGCATGGACCAGAAGTT
Oligo rev AAACAACTTCTGGTCCATGCAATC
Chromosome start end strand MM target_seq PAM alignment distance position gene name gene id
GU305914.1 196016 196038 - 0 GATTGCATGGACCAGAAGTT CGG ||||||||[||||||||||||]PAM NA Intergenic NA NA

T19786 GACCAGAAGTTCGGAGAATACGG 1000
Oligo fwd CACCGACCAGAAGTTCGGAGAATA
Oligo rev AAACTATTCTCCGAACTTCTGGTC
Chromosome start end strand MM target_seq PAM alignment distance position gene name gene id
GU305914.1 196007 196029 - 0 GACCAGAAGTTCGGAGAATA CGG ||||||||[||||||||||||]PAM NA Intergenic NA NA

T19787 CAGAAGTTCGGAGAATACGGCGG 1000
Oligo fwd CACCCAGAAGTTCGGAGAATACGG
Oligo rev AAACCCGTATTCTCCGAACTTCTG
Chromosome start end strand MM target_seq PAM alignment distance position gene name gene id
GU305914.1 196004 196026 - 0 CAGAAGTTCGGAGAATACGG CGG ||||||||[||||||||||||]PAM NA Intergenic NA NA

T19788 GTTCGGAGAATACGGCGGCGTGG 1000
Oligo fwd CACCGTTCGGAGAATACGGCGGCG
Oligo rev AAACCGCCGCCGTATTCTCCGAAC
Chromosome start end strand MM target_seq PAM alignment distance position gene name gene id
GU305914.1 195999 196021 - 0 GTTCGGAGAATACGGCGGCG TGG ||||||||[||||||||||||]PAM NA Intergenic NA NA

<table>
<tr><td>T19789</td><td>AATACGGCGGCGTGGACGAGAGG</td><td>1000</td><td></td><td></td><td></td><td></td><td></td><td></td><td></td><td></td><td></td></tr>
<tr><td>Oligo fwd</td><td>CACCAATACGGCGGCGTGGACGAG</td><td></td><td></td><td></td><td></td><td></td><td></td><td></td><td></td><td></td><td></td></tr>
<tr><td>Oligo rev</td><td>AAACCTCGTCCACGCCGCCGTATT</td><td></td><td></td><td></td><td></td><td></td><td></td><td></td><td></td><td></td><td></td></tr>
<tr><td>Chromosome</td><td>start</td><td>end</td><td>strand</td><td>MM</td><td>target_seq</td><td>PAM</td><td>alignment</td><td>distance</td><td>position</td><td>gene name</td><td>gene id</td></tr>
<tr><td>GU305914.1</td><td>195991</td><td>196013</td><td>-</td><td>0</td><td>AATACGGCGGCGTGGACGAG</td><td>AGG</td><td>||||||||[||||||||||||]PAM</td><td>NA</td><td>Intergenic</td><td>NA</td><td>NA</td></tr>
<tr><td>T19790</td><td>AGAGGAGCCTTCCCCCGATCAGG</td><td>1000</td><td></td><td></td><td></td><td></td><td></td><td></td><td></td><td></td><td></td></tr>
<tr><td>Oligo fwd</td><td>CACCAGAGGAGCCTTCCCCCGATC</td><td></td><td></td><td></td><td></td><td></td><td></td><td></td><td></td><td></td><td></td></tr>
<tr><td>Oligo rev</td><td>AAACGATCGGGGGAAGGCTCCTCT</td><td></td><td></td><td></td><td></td><td></td><td></td><td></td><td></td><td></td><td></td></tr>
<tr><td>Chromosome</td><td>start</td><td>end</td><td>strand</td><td>MM</td><td>target_seq</td><td>PAM</td><td>alignment</td><td>distance</td><td>position</td><td>gene name</td><td>gene id</td></tr>
<tr><td>GU305914.1</td><td>195973</td><td>195995</td><td>-</td><td>0</td><td>AGAGGAGCCTTCCCCCGATC</td><td>AGG</td><td>||||||||[||||||||||||]PAM</td><td>NA</td><td>Intergenic</td><td>NA</td><td>NA</td></tr>
<tr><td>T19791</td><td>ATAAATGACCAGTTTTCCTTCGG</td><td>1000</td><td></td><td></td><td></td><td></td><td></td><td></td><td></td><td></td><td></td></tr>
<tr><td>Oligo fwd</td><td>CACCATAAATGACCAGTTTTCCTT</td><td></td><td></td><td></td><td></td><td></td><td></td><td></td><td></td><td></td><td></td></tr>
<tr><td>Oligo rev</td><td>AAACAAGGAAAACTGGTCATTTAT</td><td></td><td></td><td></td><td></td><td></td><td></td><td></td><td></td><td></td><td></td></tr>
<tr><td>Chromosome</td><td>start</td><td>end</td><td>strand</td><td>MM</td><td>target_seq</td><td>PAM</td><td>alignment</td><td>distance</td><td>position</td><td>gene name</td><td>gene id</td></tr>
<tr><td>GU305914.1</td><td>195923</td><td>195945</td><td>-</td><td>0</td><td>ATAAATGACCAGTTTTCCTT</td><td>CGG</td><td>||||||||[||||||||||||]PAM</td><td>NA</td><td>Intergenic</td><td>NA</td><td>NA</td></tr>
<tr><td>T19792</td><td>GTTTTCCTTCGGCCTGCCATTGG</td><td>1000</td><td></td><td></td><td></td><td></td><td></td><td></td><td></td><td></td><td></td></tr>
<tr><td>Oligo fwd</td><td>CACCGTTTTCCTTCGGCCTGCCAT</td><td></td><td></td><td></td><td></td><td></td><td></td><td></td><td></td><td></td><td></td></tr>
<tr><td>Oligo rev</td><td>AAACATGGCAGGCCGAAGGAAAAC</td><td></td><td></td><td></td><td></td><td></td><td></td><td></td><td></td><td></td><td></td></tr>
<tr><td>Chromosome</td><td>start</td><td>end</td><td>strand</td><td>MM</td><td>target_seq</td><td>PAM</td><td>alignment</td><td>distance</td><td>position</td><td>gene name</td><td>gene id</td></tr>
<tr><td>GU305914.1</td><td>195912</td><td>195934</td><td>-</td><td>0</td><td>GTTTTCCTTCGGCCTGCCAT</td><td>TGG</td><td>||||||||[||||||||||||]PAM</td><td>NA</td><td>Intergenic</td><td>NA</td><td>NA</td></tr>
<tr><td>T19793</td><td>CGGCCTGCCATTGGACTCCACGG</td><td>1000</td><td></td><td></td><td></td><td></td><td></td><td></td><td></td><td></td><td></td></tr>
<tr><td>Oligo fwd</td><td>CACCCGGCCTGCCATTGGACTCCA</td><td></td><td></td><td></td><td></td><td></td><td></td><td></td><td></td><td></td><td></td></tr>
<tr><td>Oligo rev</td><td>AAACTGGAGTCCAATGGCAGGCCG</td><td></td><td></td><td></td><td></td><td></td><td></td><td></td><td></td><td></td><td></td></tr>
<tr><td>Chromosome</td><td>start</td><td>end</td><td>strand</td><td>MM</td><td>target_seq</td><td>PAM</td><td>alignment</td><td>distance</td><td>position</td><td>gene name</td><td>gene id</td></tr>
<tr><td>GU305914.1</td><td>195903</td><td>195925</td><td>-</td><td>0</td><td>CGGCCTGCCATTGGACTCCA</td><td>CGG</td><td>||||||||[||||||||||||]PAM</td><td>NA</td><td>Intergenic</td><td>NA</td><td>NA</td></tr>
<tr><td>T19794</td><td>ATTGGACTCCACGGATCGCACGG</td><td>1000</td><td></td><td></td><td></td><td></td><td></td><td></td><td></td><td></td><td></td></tr>
<tr><td>Oligo fwd</td><td>CACCATTGGACTCCACGGATCGCA</td><td></td><td></td><td></td><td></td><td></td><td></td><td></td><td></td><td></td><td></td></tr>
<tr><td>Oligo rev</td><td>AAACTGCGATCCGTGGAGTCCAAT</td><td></td><td></td><td></td><td></td><td></td><td></td><td></td><td></td><td></td><td></td></tr>
<tr><td>Chromosome</td><td>start</td><td>end</td><td>strand</td><td>MM</td><td>target_seq</td><td>PAM</td><td>alignment</td><td>distance</td><td>position</td><td>gene name</td><td>gene id</td></tr>
<tr><td>GU305914.1</td><td>195894</td><td>195916</td><td>-</td><td>0</td><td>ATTGGACTCCACGGATCGCA</td><td>CGG</td><td>||||||||[||||||||||||]PAM</td><td>NA</td><td>Intergenic</td><td>NA</td><td>NA</td></tr>
<tr><td>T19795</td><td>GACTCCACGGATCGCACGGCCGG</td><td>1000</td><td></td><td></td><td></td><td></td><td></td><td></td><td></td><td></td><td></td></tr>
<tr><td>Oligo fwd</td><td>CACCGACTCCACGGATCGCACGGC</td><td></td><td></td><td></td><td></td><td></td><td></td><td></td><td></td><td></td><td></td></tr>
<tr><td>Oligo rev</td><td>AAACGCCGTGCGATCCGTGGAGTC</td><td></td><td></td><td></td><td></td><td></td><td></td><td></td><td></td><td></td><td></td></tr>
<tr><td>Chromosome</td><td>start</td><td>end</td><td>strand</td><td>MM</td><td>target_seq</td><td>PAM</td><td>alignment</td><td>distance</td><td>position</td><td>gene name</td><td>gene id</td></tr>
<tr><td>GU305914.1</td><td>195890</td><td>195912</td><td>-</td><td>0</td><td>GACTCCACGGATCGCACGGC</td><td>CGG</td><td>||||||||[||||||||||||]PAM</td><td>NA</td><td>Intergenic</td><td>NA</td><td>NA</td></tr>
<tr><td>T19796</td><td>ACTCCACGGATCGCACGGCCGGG</td><td>1000</td><td></td><td></td><td></td><td></td><td></td><td></td><td></td><td></td><td></td></tr>
<tr><td>Oligo fwd</td><td>CACCACTCCACGGATCGCACGGCC</td><td></td><td></td><td></td><td></td><td></td><td></td><td></td><td></td><td></td><td></td></tr>
<tr><td>Oligo rev</td><td>AAACGGCCGTGCGATCCGTGGAGT</td><td></td><td></td><td></td><td></td><td></td><td></td><td></td><td></td><td></td><td></td></tr>
<tr><td>Chromosome</td><td>start</td><td>end</td><td>strand</td><td>MM</td><td>target_seq</td><td>PAM</td><td>alignment</td><td>distance</td><td>position</td><td>gene name</td><td>gene id</td></tr>
<tr><td>GU305914.1</td><td>195889</td><td>195911</td><td>-</td><td>0</td><td>ACTCCACGGATCGCACGGCC</td><td>GGG</td><td>||||||||[||||||||||||]PAM</td><td>NA</td><td>Intergenic</td><td>NA</td><td>NA</td></tr>
<tr><td>T19797</td><td>CTCCACGGATCGCACGGCCGGGG</td><td>1000</td><td></td><td></td><td></td><td></td><td></td><td></td><td></td><td></td><td></td></tr>
<tr><td>Oligo fwd</td><td>CACCCTCCACGGATCGCACGGCCG</td><td></td><td></td><td></td><td></td><td></td><td></td><td></td><td></td><td></td><td></td></tr>
<tr><td>Oligo rev</td><td>AAACCGGCCGTGCGATCCGTGGAG</td><td></td><td></td><td></td><td></td><td></td><td></td><td></td><td></td><td></td><td></td></tr>
<tr><td>Chromosome</td><td>start</td><td>end</td><td>strand</td><td>MM</td><td>target_seq</td><td>PAM</td><td>alignment</td><td>distance</td><td>position</td><td>gene name</td><td>gene id</td></tr>
<tr><td>GU305914.1</td><td>195888</td><td>195910</td><td>-</td><td>0</td><td>CTCCACGGATCGCACGGCCG</td><td>GGG</td><td>||||||||[||||||||||||]PAM</td><td>NA</td><td>Intergenic</td><td>NA</td><td>NA</td></tr>
<tr><td>GU305914.1</td><td>65339</td><td>65361</td><td>+</td><td>3</td><td>CGCCACGTACCGCACGGCCG</td><td>CGG</td><td>|-|||||-[|-||||||||||]PAM</td><td>NA</td><td>Intergenic</td><td>NA</td><td>NA</td></tr>
<tr><td>T19798</td><td>TCCACGGATCGCACGGCCGGGGG</td><td>1000</td><td></td><td></td><td></td><td></td><td></td><td></td><td></td><td></td><td></td></tr>
<tr><td>Oligo fwd</td><td>CACCTCCACGGATCGCACGGCCGG</td><td></td><td></td><td></td><td></td><td></td><td></td><td></td><td></td><td></td><td></td></tr>
<tr><td>Oligo rev</td><td>AAACCCGGCCGTGCGATCCGTGGA</td><td></td><td></td><td></td><td></td><td></td><td></td><td></td><td></td><td></td><td></td></tr>
<tr><td>Chromosome</td><td>start</td><td>end</td><td>strand</td><td>MM</td><td>target_seq</td><td>PAM</td><td>alignment</td><td>distance</td><td>position</td><td>gene name</td><td>gene id</td></tr>
<tr><td>GU305914.1</td><td>195887</td><td>195909</td><td>-</td><td>0</td><td>TCCACGGATCGCACGGCCGG</td><td>GGG</td><td>||||||||[||||||||||||]PAM</td><td>NA</td><td>Intergenic</td><td>NA</td><td>NA</td></tr>
<tr><td>T19799</td><td>GCCGGGGGACGCGAATCCTGCGG</td><td>1000</td><td></td><td></td><td></td><td></td><td></td><td></td><td></td><td></td><td></td></tr>
<tr><td>Oligo fwd</td><td>CACCGCCGGGGGACGCGAATCCTG</td><td></td><td></td><td></td><td></td><td></td><td></td><td></td><td></td><td></td><td></td></tr>
<tr><td>Oligo rev</td><td>AAACCAGGATTCGCGTCCCCCGGC</td><td></td><td></td><td></td><td></td><td></td><td></td><td></td><td></td><td></td><td></td></tr>
<tr><td>Chromosome</td><td>start</td><td>end</td><td>strand</td><td>MM</td><td>target_seq</td><td>PAM</td><td>alignment</td><td>distance</td><td>position</td><td>gene name</td><td>gene id</td></tr>
<tr><td>GU305914.1</td><td>195872</td><td>195894</td><td>-</td><td>0</td><td>GCCGGGGGACGCGAATCCTG</td><td>CGG</td><td>||||||||[||||||||||||]PAM</td><td>NA</td><td>Intergenic</td><td>NA</td><td>NA</td></tr>
<tr><td>T19800</td><td>ACGCGAATCCTGCGGCGCGATGG</td><td>1000</td><td></td><td></td><td></td><td></td><td></td><td></td><td></td><td></td><td></td></tr>
<tr><td>Oligo fwd</td><td>CACCACGCGAATCCTGCGGCGCGA</td><td></td><td></td><td></td><td></td><td></td><td></td><td></td><td></td><td></td><td></td></tr>
<tr><td>Oligo rev</td><td>AAACTCGCGCCGCAGGATTCGCGT</td><td></td><td></td><td></td><td></td><td></td><td></td><td></td><td></td><td></td><td></td></tr>
<tr><td>Chromosome</td><td>start</td><td>end</td><td>strand</td><td>MM</td><td>target_seq</td><td>PAM</td><td>alignment</td><td>distance</td><td>position</td><td>gene name</td><td>gene id</td></tr>
<tr><td>GU305914.1</td><td>195864</td><td>195886</td><td>-</td><td>0</td><td>ACGCGAATCCTGCGGCGCGA</td><td>TGG</td><td>||||||||[||||||||||||]PAM</td><td>NA</td><td>Intergenic</td><td>NA</td><td>NA</td></tr>
<tr><td>T19801</td><td>CGAATCCTGCGGCGCGATGGCGG</td><td>1000</td><td></td><td></td><td></td><td></td><td></td><td></td><td></td><td></td><td></td></tr>
<tr><td>Oligo fwd</td><td>CACCCGAATCCTGCGGCGCGATGG</td><td></td><td></td><td></td><td></td><td></td><td></td><td></td><td></td><td></td><td></td></tr>
<tr><td>Oligo rev</td><td>AAACCCATCGCGCCGCAGGATTCG</td><td></td><td></td><td></td><td></td><td></td><td></td><td></td><td></td><td></td><td></td></tr>
<tr><td>Chromosome</td><td>start</td><td>end</td><td>strand</td><td>MM</td><td>target_seq</td><td>PAM</td><td>alignment</td><td>distance</td><td>position</td><td>gene name</td><td>gene id</td></tr>
<tr><td>GU305914.1</td><td>195861</td><td>195883</td><td>-</td><td>0</td><td>CGAATCCTGCGGCGCGATGG</td><td>CGG</td><td>||||||||[||||||||||||]PAM</td><td>NA</td><td>Intergenic</td><td>NA</td><td>NA</td></tr>
<tr><td>T19802</td><td>GTTCATAGATCGCGCCGCGCTGG</td><td>1000</td><td></td><td></td><td></td><td></td><td></td><td></td><td></td><td></td><td></td></tr>
<tr><td>Oligo fwd</td><td>CACCGTTCATAGATCGCGCCGCGC</td><td></td><td></td><td></td><td></td><td></td><td></td><td></td><td></td><td></td><td></td></tr>
<tr><td>Oligo rev</td><td>AAACGCGCGGCGCGATCTATGAAC</td><td></td><td></td><td></td><td></td><td></td><td></td><td></td><td></td><td></td><td></td></tr>
<tr><td>Chromosome</td><td>start</td><td>end</td><td>strand</td><td>MM</td><td>target_seq</td><td>PAM</td><td>alignment</td><td>distance</td><td>position</td><td>gene name</td><td>gene id</td></tr>
<tr><td>GU305914.1</td><td>195828</td><td>195850</td><td>-</td><td>0</td><td>GTTCATAGATCGCGCCGCGC</td><td>TGG</td><td>||||||||[||||||||||||]PAM</td><td>NA</td><td>Intergenic</td><td>NA</td><td>NA</td></tr>
<tr><td>T19803</td><td>ATAGATCGCGCCGCGCTGGACGG</td><td>1000</td><td></td><td></td><td></td><td></td><td></td><td></td><td></td><td></td><td></td></tr>
<tr><td>Oligo fwd</td><td>CACCATAGATCGCGCCGCGCTGGA</td><td></td><td></td><td></td><td></td><td></td><td></td><td></td><td></td><td></td><td></td></tr>
<tr><td>Oligo rev</td><td>AAACTCCAGCGCGGCGCGATCTAT</td><td></td><td></td><td></td><td></td><td></td><td></td><td></td><td></td><td></td><td></td></tr>
</table>

| | | | | | | | | | | | |
|---|---|---|---|---|---|---|---|---|---|---|---|
| Chromosome | start | end | strand | MM | target_seq | PAM | alignment | distance | position | gene name | gene id |
| GU305914.1 | 195824 | 195846 | - | 0 | ATAGATCGCGCCGCGCTGGA | CGG | \|\|\|\|\|\|\|\|[\|\|\|\|\|\|\|\|\|\|\|\|]PAM | NA | Intergenic | NA | NA |
| T19804 | TAGATCGCGCCGCGCTGGACGGG | 1000 | | | | | | | | | |
| Oligo fwd | CACCTAGATCGCGCCGCGCTGGAC | | | | | | | | | | |
| Oligo rev | AAACGTCCAGCGCGGCGCGATCTA | | | | | | | | | | |
| Chromosome | start | end | strand | MM | target_seq | PAM | alignment | distance | position | gene name | gene id |
| GU305914.1 | 195823 | 195845 | - | 0 | TAGATCGCGCCGCGCTGGAC | GGG | \|\|\|\|\|\|\|\|[\|\|\|\|\|\|\|\|\|\|\|\|]PAM | NA | Intergenic | NA | NA |
| T19805 | CGCGCCGCGCTGGACGGGACCGG | 1000 | | | | | | | | | |
| Oligo fwd | CACCCGCGCCGCGCTGGACGGGAC | | | | | | | | | | |
| Oligo rev | AAACGTCCCGTCCAGCGCGGCGCG | | | | | | | | | | |
| Chromosome | start | end | strand | MM | target_seq | PAM | alignment | distance | position | gene name | gene id |
| GU305914.1 | 195818 | 195840 | - | 0 | CGCGCCGCGCTGGACGGGAC | CGG | \|\|\|\|\|\|\|\|[\|\|\|\|\|\|\|\|\|\|\|\|]PAM | NA | Intergenic | NA | NA |
| T19806 | GCGCCGCGCTGGACGGGACCGGG | 1000 | | | | | | | | | |
| Oligo fwd | CACCGCGCCGCGCTGGACGGGACC | | | | | | | | | | |
| Oligo rev | AAACGGTCCCGTCCAGCGCGGCGC | | | | | | | | | | |
| Chromosome | start | end | strand | MM | target_seq | PAM | alignment | distance | position | gene name | gene id |
| GU305914.1 | 195817 | 195839 | - | 0 | GCGCCGCGCTGGACGGGACC | GGG | \|\|\|\|\|\|\|\|[\|\|\|\|\|\|\|\|\|\|\|\|]PAM | NA | Intergenic | NA | NA |
| T19807 | CGGGACCGGGCGCTCTGACGCGG | 1000 | | | | | | | | | |
| Oligo fwd | CACCCGGGACCGGGCGCTCTGACG | | | | | | | | | | |
| Oligo rev | AAACCGTCAGAGCGCCCGGTCCCG | | | | | | | | | | |
| Chromosome | start | end | strand | MM | target_seq | PAM | alignment | distance | position | gene name | gene id |
| GU305914.1 | 195804 | 195826 | - | 0 | CGGGACCGGGCGCTCTGACG | CGG | \|\|\|\|\|\|\|\|[\|\|\|\|\|\|\|\|\|\|\|\|]PAM | NA | Intergenic | NA | NA |
| T19808 | GGGACCGGGCGCTCTGACGCGGG | 1000 | | | | | | | | | |
| Oligo fwd | CACCGGGACCGGGCGCTCTGACGC | | | | | | | | | | |
| Oligo rev | AAACGCGTCAGAGCGCCCGGTCCC | | | | | | | | | | |
| Chromosome | start | end | strand | MM | target_seq | PAM | alignment | distance | position | gene name | gene id |
| GU305914.1 | 195803 | 195825 | - | 0 | GGGACCGGGCGCTCTGACGC | GGG | \|\|\|\|\|\|\|\|[\|\|\|\|\|\|\|\|\|\|\|\|]PAM | NA | Intergenic | NA | NA |
| T19809 | GAGCGACTCAGACTTCGACATGG | 1000 | | | | | | | | | |
| Oligo fwd | CACCGAGCGACTCAGACTTCGACA | | | | | | | | | | |
| Oligo rev | AAACTGTCGAAGTCTGAGTCGCTC | | | | | | | | | | |
| Chromosome | start | end | strand | MM | target_seq | PAM | alignment | distance | position | gene name | gene id |
| GU305914.1 | 195765 | 195787 | - | 0 | GAGCGACTCAGACTTCGACA | TGG | \|\|\|\|\|\|\|\|[\|\|\|\|\|\|\|\|\|\|\|\|]PAM | NA | Intergenic | NA | NA |
| GU305914.1 | 72778 | 72800 | + | 3 | GAGCGACCCGGACTTCGTCA | AGG | \|\|\|\|\|\|\|-[\|-\|\|\|\|\|\|\|-\|\|]PAM | NA | Intergenic | NA | NA |
| T19810 | CTCAGACTTCGACATGGAGTCGG | 1000 | | | | | | | | | |
| Oligo fwd | CACCCTCAGACTTCGACATGGAGT | | | | | | | | | | |
| Oligo rev | AAACACTCCATGTCGAAGTCTGAG | | | | | | | | | | |
| Chromosome | start | end | strand | MM | target_seq | PAM | alignment | distance | position | gene name | gene id |
| GU305914.1 | 195759 | 195781 | - | 0 | CTCAGACTTCGACATGGAGT | CGG | \|\|\|\|\|\|\|\|[\|\|\|\|\|\|\|\|\|\|\|\|]PAM | NA | Intergenic | NA | NA |
| T19811 | CTTCGACATGGAGTCGGACTCGG | 1000 | | | | | | | | | |
| Oligo fwd | CACCCTTCGACATGGAGTCGGACT | | | | | | | | | | |
| Oligo rev | AAACAGTCCGACTCCATGTCGAAG | | | | | | | | | | |
| Chromosome | start | end | strand | MM | target_seq | PAM | alignment | distance | position | gene name | gene id |
| GU305914.1 | 195753 | 195775 | - | 0 | CTTCGACATGGAGTCGGACT | CGG | \|\|\|\|\|\|\|\|[\|\|\|\|\|\|\|\|\|\|\|\|]PAM | NA | Intergenic | NA | NA |
| T19812 | GACTCGGACCTGTCCGACGACGG | 1000 | | | | | | | | | |
| Oligo fwd | CACCGACTCGGACCTGTCCGACGA | | | | | | | | | | |
| Oligo rev | AAACTCGTCGGACAGGTCCGAGTC | | | | | | | | | | |
| Chromosome | start | end | strand | MM | target_seq | PAM | alignment | distance | position | gene name | gene id |
| GU305914.1 | 195737 | 195759 | - | 0 | GACTCGGACCTGTCCGACGA | CGG | \|\|\|\|\|\|\|\|[\|\|\|\|\|\|\|\|\|\|\|\|]PAM | NA | Intergenic | NA | NA |
| T19813 | CCTGTCCGACGACGGCGACGCGG | 1000 | | | | | | | | | |
| Oligo fwd | CACCCCTGTCCGACGACGGCGACG | | | | | | | | | | |
| Oligo rev | AAACCGTCGCCGTCGTCGGACAGG | | | | | | | | | | |
| Chromosome | start | end | strand | MM | target_seq | PAM | alignment | distance | position | gene name | gene id |
| GU305914.1 | 195729 | 195751 | - | 0 | CCTGTCCGACGACGGCGACG | CGG | \|\|\|\|\|\|\|\|[\|\|\|\|\|\|\|\|\|\|\|\|]PAM | NA | Intergenic | NA | NA |
| T19814 | GTCCGACGACGGCGACGCGGCGG | 1000 | | | | | | | | | |
| Oligo fwd | CACCGTCCGACGACGGCGACGCGG | | | | | | | | | | |
| Oligo rev | AAACCCGCGTCGCCGTCGTCGGAC | | | | | | | | | | |
| Chromosome | start | end | strand | MM | target_seq | PAM | alignment | distance | position | gene name | gene id |
| GU305914.1 | 195726 | 195748 | - | 0 | GTCCGACGACGGCGACGCGG | CGG | \|\|\|\|\|\|\|\|[\|\|\|\|\|\|\|\|\|\|\|\|]PAM | NA | Intergenic | NA | NA |
| T19815 | TCCGACGACGGCGACGCGGCGGG | 1000 | | | | | | | | | |
| Oligo fwd | CACCTCCGACGACGGCGACGCGGC | | | | | | | | | | |
| Oligo rev | AAACGCCGCGTCGCCGTCGTCGGA | | | | | | | | | | |
| Chromosome | start | end | strand | MM | target_seq | PAM | alignment | distance | position | gene name | gene id |
| GU305914.1 | 195725 | 195747 | - | 0 | TCCGACGACGGCGACGCGGC | GGG | \|\|\|\|\|\|\|\|[\|\|\|\|\|\|\|\|\|\|\|\|]PAM | NA | Intergenic | NA | NA |
| T19816 | GCGGCGGGAGCGTACGTGTACGG | 1000 | | | | | | | | | |
| Oligo fwd | CACCGCGGCGGGAGCGTACGTGTA | | | | | | | | | | |
| Oligo rev | AAACTACACGTACGCTCCCGCCGC | | | | | | | | | | |
| Chromosome | start | end | strand | MM | target_seq | PAM | alignment | distance | position | gene name | gene id |
| GU305914.1 | 195710 | 195732 | - | 0 | GCGGCGGGAGCGTACGTGTA | CGG | \|\|\|\|\|\|\|\|[\|\|\|\|\|\|\|\|\|\|\|\|]PAM | NA | Intergenic | NA | NA |
| T19817 | CGTACGTGTACGGCTCGCCGTGG | 1000 | | | | | | | | | |
| Oligo fwd | CACCCGTACGTGTACGGCTCGCCG | | | | | | | | | | |
| Oligo rev | AAACCGGCGAGCCGTACACGTACG | | | | | | | | | | |
| Chromosome | start | end | strand | MM | target_seq | PAM | alignment | distance | position | gene name | gene id |
| GU305914.1 | 195700 | 195722 | - | 0 | CGTACGTGTACGGCTCGCCG | TGG | \|\|\|\|\|\|\|\|[\|\|\|\|\|\|\|\|\|\|\|\|]PAM | NA | Intergenic | NA | NA |
| T19818 | CGGCTCGCCGTGGAAGACCTCGG | 1000 | | | | | | | | | |

| | | | | | | | | | | | |
|---|---|---|---|---|---|---|---|---|---|---|---|
| Oligo fwd | CACCCGGCTCGCCGTGGAAGACCT | | | | | | | | | | |
| Oligo rev | AAACAGGTCTTCCACGGCGAGCCG | | | | | | | | | | |
| Chromosome | start | end | strand | MM | target_seq | PAM | alignment | distance | position | gene name | gene id |
| GU305914.1 | 195690 | 195712 | - | 0 | CGGCTCGCCGTGGAAGACCT | CGG | \|\|\|\|\|\|\|\|[\|\|\|\|\|\|\|\|\|\|\|\|]PAM | NA | Intergenic | NA | NA |
| T19819 | GTGGAAGACCTCGGCCGTCACGG | 1000 | | | | | | | | | |
| Oligo fwd | CACCGTGGAAGACCTCGGCCGTCA | | | | | | | | | | |
| Oligo rev | AAACTGACGGCCGAGGTCTTCCAC | | | | | | | | | | |
| Chromosome | start | end | strand | MM | target_seq | PAM | alignment | distance | position | gene name | gene id |
| GU305914.1 | 195681 | 195703 | - | 0 | GTGGAAGACCTCGGCCGTCA | CGG | \|\|\|\|\|\|\|\|[\|\|\|\|\|\|\|\|\|\|\|\|]PAM | NA | Intergenic | NA | NA |
| T19820 | CCTCGGCCGTCACGGTGTCCAGG | 1000 | | | | | | | | | |
| Oligo fwd | CACCCCTCGGCCGTCACGGTGTCC | | | | | | | | | | |
| Oligo rev | AAACGGACACCGTGACGGCCGAGG | | | | | | | | | | |
| Chromosome | start | end | strand | MM | target_seq | PAM | alignment | distance | position | gene name | gene id |
| GU305914.1 | 195673 | 195695 | - | 0 | CCTCGGCCGTCACGGTGTCC | AGG | \|\|\|\|\|\|\|\|[\|\|\|\|\|\|\|\|\|\|\|\|]PAM | NA | Intergenic | NA | NA |
| T19821 | GGCCGTCACGGTGTCCAGGAAGG | 1000 | | | | | | | | | |
| Oligo fwd | CACCGGCCGTCACGGTGTCCAGGA | | | | | | | | | | |
| Oligo rev | AAACTCCTGGACACCGTGACGGCC | | | | | | | | | | |
| Chromosome | start | end | strand | MM | target_seq | PAM | alignment | distance | position | gene name | gene id |
| GU305914.1 | 195669 | 195691 | - | 0 | GGCCGTCACGGTGTCCAGGA | AGG | \|\|\|\|\|\|\|\|[\|\|\|\|\|\|\|\|\|\|\|\|]PAM | NA | Intergenic | NA | NA |
| T19822 | GAAATACGAGAAGAAGAAGCTGG | 1000 | | | | | | | | | |
| Oligo fwd | CACCGAAATACGAGAAGAAGAAGC | | | | | | | | | | |
| Oligo rev | AAACGCTTCTTCTTCTCGTATTTC | | | | | | | | | | |
| Chromosome | start | end | strand | MM | target_seq | PAM | alignment | distance | position | gene name | gene id |
| GU305914.1 | 195639 | 195661 | - | 0 | GAAATACGAGAAGAAGAAGC | TGG | \|\|\|\|\|\|\|\|[\|\|\|\|\|\|\|\|\|\|\|\|]PAM | NA | Intergenic | NA | NA |
| T19823 | CGACTTCAAGAGCTTCCACCTGG | 1000 | | | | | | | | | |
| Oligo fwd | CACCCGACTTCAAGAGCTTCCACC | | | | | | | | | | |
| Oligo rev | AAACGGTGGAAGCTCTTGAAGTCG | | | | | | | | | | |
| Chromosome | start | end | strand | MM | target_seq | PAM | alignment | distance | position | gene name | gene id |
| GU305914.1 | 195588 | 195610 | - | 0 | CGACTTCAAGAGCTTCCACC | TGG | \|\|\|\|\|\|\|\|[\|\|\|\|\|\|\|\|\|\|\|\|]PAM | NA | Intergenic | NA | NA |
| T19824 | CCTGGCCCTCAGTTTCCCGCCGG | 1000 | | | | | | | | | |
| Oligo fwd | CACCCCTGGCCCTCAGTTTCCCGC | | | | | | | | | | |
| Oligo rev | AAACGCGGGAAACTGAGGGCCAGG | | | | | | | | | | |
| Chromosome | start | end | strand | MM | target_seq | PAM | alignment | distance | position | gene name | gene id |
| GU305914.1 | 195570 | 195592 | - | 0 | CCTGGCCCTCAGTTTCCCGC | CGG | \|\|\|\|\|\|\|\|[\|\|\|\|\|\|\|\|\|\|\|\|]PAM | NA | Intergenic | NA | NA |
| T19825 | CTGGCCCTCAGTTTCCCGCCGGG | 1000 | | | | | | | | | |
| Oligo fwd | CACCCTGGCCCTCAGTTTCCCGCC | | | | | | | | | | |
| Oligo rev | AAACGGCGGGAAACTGAGGGCCAG | | | | | | | | | | |
| Chromosome | start | end | strand | MM | target_seq | PAM | alignment | distance | position | gene name | gene id |
| GU305914.1 | 195569 | 195591 | - | 0 | CTGGCCCTCAGTTTCCCGCC | GGG | \|\|\|\|\|\|\|\|[\|\|\|\|\|\|\|\|\|\|\|\|]PAM | NA | Intergenic | NA | NA |
| T19826 | TTTCCCGCCGGGCCAGATCGCGG | 1000 | | | | | | | | | |
| Oligo fwd | CACCTTTCCCGCCGGGCCAGATCG | | | | | | | | | | |
| Oligo rev | AAACCGATCTGGCCCGGCGGGAAA | | | | | | | | | | |
| Chromosome | start | end | strand | MM | target_seq | PAM | alignment | distance | position | gene name | gene id |
| GU305914.1 | 195558 | 195580 | - | 0 | TTTCCCGCCGGGCCAGATCG | CGG | \|\|\|\|\|\|\|\|[\|\|\|\|\|\|\|\|\|\|\|\|]PAM | NA | Intergenic | NA | NA |
| T19827 | GCCGGGCCAGATCGCGGCCGCGG | 1000 | | | | | | | | | |
| Oligo fwd | CACCGCCGGGCCAGATCGCGGCCG | | | | | | | | | | |
| Oligo rev | AAACCGGCCGCGATCTGGCCCGGC | | | | | | | | | | |
| Chromosome | start | end | strand | MM | target_seq | PAM | alignment | distance | position | gene name | gene id |
| GU305914.1 | 195552 | 195574 | - | 0 | GCCGGGCCAGATCGCGGCCG | CGG | \|\|\|\|\|\|\|\|[\|\|\|\|\|\|\|\|\|\|\|\|]PAM | NA | Intergenic | NA | NA |
| T19828 | GGGCCAGATCGCGGCCGCGGTGG | 1000 | | | | | | | | | |
| Oligo fwd | CACCGGGCCAGATCGCGGCCGCGG | | | | | | | | | | |
| Oligo rev | AAACCCGCGGCCGCGATCTGGCCC | | | | | | | | | | |
| Chromosome | start | end | strand | MM | target_seq | PAM | alignment | distance | position | gene name | gene id |
| GU305914.1 | 195549 | 195571 | - | 0 | GGGCCAGATCGCGGCCGCGG | TGG | \|\|\|\|\|\|\|\|[\|\|\|\|\|\|\|\|\|\|\|\|]PAM | NA | Intergenic | NA | NA |
| T19829 | GCCGCGGTGGTGCACTACCTCGG | 1000 | | | | | | | | | |
| Oligo fwd | CACCGCCGCGGTGGTGCACTACCT | | | | | | | | | | |
| Oligo rev | AAACAGGTAGTGCACCACCGCGGC | | | | | | | | | | |
| Chromosome | start | end | strand | MM | target_seq | PAM | alignment | distance | position | gene name | gene id |
| GU305914.1 | 195536 | 195558 | - | 0 | GCCGCGGTGGTGCACTACCT | CGG | \|\|\|\|\|\|\|\|[\|\|\|\|\|\|\|\|\|\|\|\|]PAM | NA | Intergenic | NA | NA |
| GU305914.1 | 101073 | 101095 | - | 4 | GCGGCGGAGGAGAACTACCT | GGG | \|\|-\|\|\|\|-[\|\|-\|-\|\|\|\|\|\|\|]PAM | NA | Intergenic | NA | NA |
| T19830 | GCCCCTCGTGCTCCGCTACCCGG | 1000 | | | | | | | | | |
| Oligo fwd | CACCGCCCCTCGTGCTCCGCTACC | | | | | | | | | | |
| Oligo rev | AAACGGTAGCGGAGCACGAGGGGC | | | | | | | | | | |
| Chromosome | start | end | strand | MM | target_seq | PAM | alignment | distance | position | gene name | gene id |
| GU305914.1 | 195510 | 195532 | - | 0 | GCCCCTCGTGCTCCGCTACC | CGG | \|\|\|\|\|\|\|\|[\|\|\|\|\|\|\|\|\|\|\|\|]PAM | NA | Intergenic | NA | NA |
| T19831 | TGCTCCGCTACCCGGAGAACTGG | 1000 | | | | | | | | | |
| Oligo fwd | CACCTGCTCCGCTACCCGGAGAAC | | | | | | | | | | |
| Oligo rev | AAACGTTCTCCGGGTAGCGGAGCA | | | | | | | | | | |
| Chromosome | start | end | strand | MM | target_seq | PAM | alignment | distance | position | gene name | gene id |
| GU305914.1 | 195502 | 195524 | - | 0 | TGCTCCGCTACCCGGAGAAC | TGG | \|\|\|\|\|\|\|\|[\|\|\|\|\|\|\|\|\|\|\|\|]PAM | NA | Intergenic | NA | NA |
| T19832 | TCCGAGATCCGCGTGCTGCGCGG | 1000 | | | | | | | | | |
| Oligo fwd | CACCTCCGAGATCCGCGTGCTGCG | | | | | | | | | | |
| Oligo rev | AAACCGCAGCACGCGGATCTCGGA | | | | | | | | | | |
| Chromosome | start | end | strand | MM | target_seq | PAM | alignment | distance | position | gene name | gene id |
| GU305914.1 | 195458 | 195480 | - | 0 | TCCGAGATCCGCGTGCTGCG | CGG | \|\|\|\|\|\|\|\|[\|\|\|\|\|\|\|\|\|\|\|\|]PAM | NA | Intergenic | NA | NA |

T19833 GATCCGCGTGCTGCGCGGCGTGG 1000
Oligo fwd CACCGATCCGCGTGCTGCGCGGCG
Oligo rev AAACCGCCGCGCAGCACGCGGATC
Chromosome start end strand MM target_seq PAM alignment distance position gene name gene id
GU305914.1 195453 195475 - 0 GATCCGCGTGCTGCGCGGCG TGG ||||||||[||||||||||||]PAM NA Intergenic NA NA

T19834 CTGTCCAAGTATCTGTGCTGCGG 1000
Oligo fwd CACCCTGTCCAAGTATCTGTGCTG
Oligo rev AAACCAGCACAGATACTTGGACAG
Chromosome start end strand MM target_seq PAM alignment distance position gene name gene id
GU305914.1 195428 195450 - 0 CTGTCCAAGTATCTGTGCTG CGG ||||||||[||||||||||||]PAM NA Intergenic NA NA

T19835 CTGTGCTGCGGCGACACCGCCGG 1000
Oligo fwd CACCCTGTGCTGCGGCGACACCGC
Oligo rev AAACGCGGTGTCGCCGCAGCACAG
Chromosome start end strand MM target_seq PAM alignment distance position gene name gene id
GU305914.1 195416 195438 - 0 CTGTGCTGCGGCGACACCGC CGG ||||||||[||||||||||||]PAM NA Intergenic NA NA
GU305914.1 56596 56618 + 4 CAGAGTGGCGGCGACACCGC AGG |-|-|--|[||||||||||||]PAM NA Intergenic NA NA

T19836 CGCCGGCGAGCTGACGCCCCTGG 1000
Oligo fwd CACCCGCCGGCGAGCTGACGCCCC
Oligo rev AAACGGGGCGTCAGCTCGCCGGCG
Chromosome start end strand MM target_seq PAM alignment distance position gene name gene id
GU305914.1 195399 195421 - 0 CGCCGGCGAGCTGACGCCCC TGG ||||||||[||||||||||||]PAM NA Intergenic NA NA

T19837 GCCGGCGAGCTGACGCCCCTGGG 1000
Oligo fwd CACCGCCGGCGAGCTGACGCCCCT
Oligo rev AAACAGGGGCGTCAGCTCGCCGGC
Chromosome start end strand MM target_seq PAM alignment distance position gene name gene id
GU305914.1 195398 195420 - 0 GCCGGCGAGCTGACGCCCCT GGG ||||||||[||||||||||||]PAM NA Intergenic NA NA

T19838 CCGGCGAGCTGACGCCCCTGGGG 1000
Oligo fwd CACCCCGGCGAGCTGACGCCCCTG
Oligo rev AAACCAGGGGCGTCAGCTCGCCGG
Chromosome start end strand MM target_seq PAM alignment distance position gene name gene id
GU305914.1 195397 195419 - 0 CCGGCGAGCTGACGCCCCTG GGG ||||||||[||||||||||||]PAM NA Intergenic NA NA

T19839 CGGCGAGCTGACGCCCCTGGGGG 1000
Oligo fwd CACCCGGCGAGCTGACGCCCCTGG
Oligo rev AAACCCAGGGGCGTCAGCTCGCCG
Chromosome start end strand MM target_seq PAM alignment distance position gene name gene id
GU305914.1 195396 195418 - 0 CGGCGAGCTGACGCCCCTGG GGG ||||||||[||||||||||||]PAM NA Intergenic NA NA

T19840 CAGAGAGACGCCGTGCGCCCTGG 1000
Oligo fwd CACCCAGAGAGACGCCGTGCGCCC
Oligo rev AAACGGGCGCACGGCGTCTCTCTG
Chromosome start end strand MM target_seq PAM alignment distance position gene name gene id
GU305914.1 195342 195364 - 0 CAGAGAGACGCCGTGCGCCC TGG ||||||||[||||||||||||]PAM NA Intergenic NA NA

T19841 GACGCCGTGCGCCCTGGTCATGG 1000
Oligo fwd CACCGACGCCGTGCGCCCTGGTCA
Oligo rev AAACTGACCAGGGCGCACGGCGTC
Chromosome start end strand MM target_seq PAM alignment distance position gene name gene id
GU305914.1 195336 195358 - 0 GACGCCGTGCGCCCTGGTCA TGG ||||||||[||||||||||||]PAM NA Intergenic NA NA

T19842 GCCCTGGTCATGGACGAGCGCGG 1000
Oligo fwd CACCGCCCTGGTCATGGACGAGCG
Oligo rev AAACCGCTCGTCCATGACCAGGGC
Chromosome start end strand MM target_seq PAM alignment distance position gene name gene id
GU305914.1 195326 195348 - 0 GCCCTGGTCATGGACGAGCG CGG ||||||||[||||||||||||]PAM NA Intergenic NA NA
GU305914.1 141738 141760 + 4 GTCAGGGTCCTGGACGAGCG CGG |-|--|||[|-||||||||||]PAM NA Intergenic NA NA

T19843 CTGTACGACGCCGAGTCCGACGG 1000
Oligo fwd CACCCTGTACGACGCCGAGTCCGA
Oligo rev AAACTCGGACTCGGCGTCGTACAG
Chromosome start end strand MM target_seq PAM alignment distance position gene name gene id
GU305914.1 195293 195315 - 0 CTGTACGACGCCGAGTCCGA CGG ||||||||[||||||||||||]PAM NA Intergenic NA NA

T19844 TGTACGACGCCGAGTCCGACGGG 1000
Oligo fwd CACCTGTACGACGCCGAGTCCGAC
Oligo rev AAACGTCGGACTCGGCGTCGTACA
Chromosome start end strand MM target_seq PAM alignment distance position gene name gene id
GU305914.1 195292 195314 - 0 TGTACGACGCCGAGTCCGAC GGG ||||||||[||||||||||||]PAM NA Intergenic NA NA

T19845 GTCCGACGGGCTGTACTACGCGG 1000
Oligo fwd CACCGTCCGACGGGCTGTACTACG
Oligo rev AAACCGTAGTACAGCCCGTCGGAC
Chromosome start end strand MM target_seq PAM alignment distance position gene name gene id
GU305914.1 195279 195301 - 0 GTCCGACGGGCTGTACTACG CGG ||||||||[||||||||||||]PAM NA Intergenic NA NA

T19846 ACGGGCTGTACTACGCGGCCAGG 1000
Oligo fwd CACCACGGGCTGTACTACGCGGCC
Oligo rev AAACGGCCGCGTAGTACAGCCCGT
Chromosome start end strand MM target_seq PAM alignment distance position gene name gene id
GU305914.1 195274 195296 - 0 ACGGGCTGTACTACGCGGCC AGG ||||||||[||||||||||||]PAM NA Intergenic NA NA

T19847 GGCCAGGAACATCGACCAGCTGG 1000
Oligo fwd CACCGGCCAGGAACATCGACCAGC

<table>
<tr><td>Oligo rev</td><td>AAACGCTGGTCGATGTTCCTGGCC</td><td></td><td></td><td></td><td></td><td></td><td></td><td></td><td></td><td></td><td></td><td></td></tr>
<tr><td>Chromosome</td><td>start</td><td></td><td>end</td><td>strand</td><td>MM</td><td>target_seq</td><td>PAM</td><td>alignment</td><td>distance</td><td>position</td><td>gene name</td><td>gene id</td></tr>
<tr><td>GU305914.1</td><td></td><td>195258</td><td>195280</td><td>-</td><td>0</td><td>GGCCAGGAACATCGACCAGC</td><td>TGG</td><td>||||||||[||||||||||||]PAM</td><td>NA</td><td>Intergenic</td><td>NA</td><td>NA</td></tr>
<tr><td></td><td></td><td></td><td></td><td></td><td></td><td></td><td></td><td></td><td></td><td></td><td></td><td></td></tr>
<tr><td>T19848</td><td>ACATCGACCAGCTGGCGAGACGG</td><td></td><td>1000</td><td></td><td></td><td></td><td></td><td></td><td></td><td></td><td></td><td></td></tr>
<tr><td>Oligo fwd</td><td>CACCACATCGACCAGCTGGCGAGA</td><td></td><td></td><td></td><td></td><td></td><td></td><td></td><td></td><td></td><td></td><td></td></tr>
<tr><td>Oligo rev</td><td>AAACTCTCGCCAGCTGGTCGATGT</td><td></td><td></td><td></td><td></td><td></td><td></td><td></td><td></td><td></td><td></td><td></td></tr>
<tr><td>Chromosome</td><td>start</td><td></td><td>end</td><td>strand</td><td>MM</td><td>target_seq</td><td>PAM</td><td>alignment</td><td>distance</td><td>position</td><td>gene name</td><td>gene id</td></tr>
<tr><td>GU305914.1</td><td></td><td>195250</td><td>195272</td><td>-</td><td>0</td><td>ACATCGACCAGCTGGCGAGA</td><td>CGG</td><td>||||||||[||||||||||||]PAM</td><td>NA</td><td>Intergenic</td><td>NA</td><td>NA</td></tr>
<tr><td></td><td></td><td></td><td></td><td></td><td></td><td></td><td></td><td></td><td></td><td></td><td></td><td></td></tr>
<tr><td>T19849</td><td>CATCGACCAGCTGGCGAGACGGG</td><td></td><td>1000</td><td></td><td></td><td></td><td></td><td></td><td></td><td></td><td></td><td></td></tr>
<tr><td>Oligo fwd</td><td>CACCCATCGACCAGCTGGCGAGAC</td><td></td><td></td><td></td><td></td><td></td><td></td><td></td><td></td><td></td><td></td><td></td></tr>
<tr><td>Oligo rev</td><td>AAACGTCTCGCCAGCTGGTCGATG</td><td></td><td></td><td></td><td></td><td></td><td></td><td></td><td></td><td></td><td></td><td></td></tr>
<tr><td>Chromosome</td><td>start</td><td></td><td>end</td><td>strand</td><td>MM</td><td>target_seq</td><td>PAM</td><td>alignment</td><td>distance</td><td>position</td><td>gene name</td><td>gene id</td></tr>
<tr><td>GU305914.1</td><td></td><td>195249</td><td>195271</td><td>-</td><td>0</td><td>CATCGACCAGCTGGCGAGAC</td><td>GGG</td><td>||||||||[||||||||||||]PAM</td><td>NA</td><td>Intergenic</td><td>NA</td><td>NA</td></tr>
<tr><td></td><td></td><td></td><td></td><td></td><td></td><td></td><td></td><td></td><td></td><td></td><td></td><td></td></tr>
<tr><td>T19850</td><td>ATCGACCAGCTGGCGAGACGGGG</td><td></td><td>1000</td><td></td><td></td><td></td><td></td><td></td><td></td><td></td><td></td><td></td></tr>
<tr><td>Oligo fwd</td><td>CACCATCGACCAGCTGGCGAGACG</td><td></td><td></td><td></td><td></td><td></td><td></td><td></td><td></td><td></td><td></td><td></td></tr>
<tr><td>Oligo rev</td><td>AAACCGTCTCGCCAGCTGGTCGAT</td><td></td><td></td><td></td><td></td><td></td><td></td><td></td><td></td><td></td><td></td><td></td></tr>
<tr><td>Chromosome</td><td>start</td><td></td><td>end</td><td>strand</td><td>MM</td><td>target_seq</td><td>PAM</td><td>alignment</td><td>distance</td><td>position</td><td>gene name</td><td>gene id</td></tr>
<tr><td>GU305914.1</td><td></td><td>195248</td><td>195270</td><td>-</td><td>0</td><td>ATCGACCAGCTGGCGAGACG</td><td>GGG</td><td>||||||||[||||||||||||]PAM</td><td>NA</td><td>Intergenic</td><td>NA</td><td>NA</td></tr>
<tr><td>GU305914.1</td><td></td><td>200376</td><td>200398</td><td>-</td><td>3</td><td>ATCGACGAGCTGGCGAGGAG</td><td>AGG</td><td>||||||-|[|||||||||--|]PAM</td><td>NA</td><td>Intergenic</td><td>NA</td><td>NA</td></tr>
<tr><td></td><td></td><td></td><td></td><td></td><td></td><td></td><td></td><td></td><td></td><td></td><td></td><td></td></tr>
<tr><td>T19851</td><td>TCGACCAGCTGGCGAGACGGGGG</td><td></td><td>1000</td><td></td><td></td><td></td><td></td><td></td><td></td><td></td><td></td><td></td></tr>
<tr><td>Oligo fwd</td><td>CACCTCGACCAGCTGGCGAGACGG</td><td></td><td></td><td></td><td></td><td></td><td></td><td></td><td></td><td></td><td></td><td></td></tr>
<tr><td>Oligo rev</td><td>AAACCCGTCTCGCCAGCTGGTCGA</td><td></td><td></td><td></td><td></td><td></td><td></td><td></td><td></td><td></td><td></td><td></td></tr>
<tr><td>Chromosome</td><td>start</td><td></td><td>end</td><td>strand</td><td>MM</td><td>target_seq</td><td>PAM</td><td>alignment</td><td>distance</td><td>position</td><td>gene name</td><td>gene id</td></tr>
<tr><td>GU305914.1</td><td></td><td>195247</td><td>195269</td><td>-</td><td>0</td><td>TCGACCAGCTGGCGAGACGG</td><td>GGG</td><td>||||||||[||||||||||||]PAM</td><td>NA</td><td>Intergenic</td><td>NA</td><td>NA</td></tr>
<tr><td></td><td></td><td></td><td></td><td></td><td></td><td></td><td></td><td></td><td></td><td></td><td></td><td></td></tr>
<tr><td>T19852</td><td>GGGGCTGTCTCTGTGCGAGCCGG</td><td></td><td>1000</td><td></td><td></td><td></td><td></td><td></td><td></td><td></td><td></td><td></td></tr>
<tr><td>Oligo fwd</td><td>CACCGGGGCTGTCTCTGTGCGAGC</td><td></td><td></td><td></td><td></td><td></td><td></td><td></td><td></td><td></td><td></td><td></td></tr>
<tr><td>Oligo rev</td><td>AAACGCTCGCACAGAGACAGCCCC</td><td></td><td></td><td></td><td></td><td></td><td></td><td></td><td></td><td></td><td></td><td></td></tr>
<tr><td>Chromosome</td><td>start</td><td></td><td>end</td><td>strand</td><td>MM</td><td>target_seq</td><td>PAM</td><td>alignment</td><td>distance</td><td>position</td><td>gene name</td><td>gene id</td></tr>
<tr><td>GU305914.1</td><td></td><td>195228</td><td>195250</td><td>-</td><td>0</td><td>GGGGCTGTCTCTGTGCGAGC</td><td>CGG</td><td>||||||||[||||||||||||]PAM</td><td>NA</td><td>Intergenic</td><td>NA</td><td>NA</td></tr>
<tr><td></td><td></td><td></td><td></td><td></td><td></td><td></td><td></td><td></td><td></td><td></td><td></td><td></td></tr>
<tr><td>T19853</td><td>TGCGAGCCGGTGTATCGAGACGG</td><td></td><td>1000</td><td></td><td></td><td></td><td></td><td></td><td></td><td></td><td></td><td></td></tr>
<tr><td>Oligo fwd</td><td>CACCTGCGAGCCGGTGTATCGAGA</td><td></td><td></td><td></td><td></td><td></td><td></td><td></td><td></td><td></td><td></td><td></td></tr>
<tr><td>Oligo rev</td><td>AAACTCTCGATACACCGGCTCGCA</td><td></td><td></td><td></td><td></td><td></td><td></td><td></td><td></td><td></td><td></td><td></td></tr>
<tr><td>Chromosome</td><td>start</td><td></td><td>end</td><td>strand</td><td>MM</td><td>target_seq</td><td>PAM</td><td>alignment</td><td>distance</td><td>position</td><td>gene name</td><td>gene id</td></tr>
<tr><td>GU305914.1</td><td></td><td>195215</td><td>195237</td><td>-</td><td>0</td><td>TGCGAGCCGGTGTATCGAGA</td><td>CGG</td><td>||||||||[||||||||||||]PAM</td><td>NA</td><td>Intergenic</td><td>NA</td><td>NA</td></tr>
<tr><td></td><td></td><td></td><td></td><td></td><td></td><td></td><td></td><td></td><td></td><td></td><td></td><td></td></tr>
<tr><td>T19854</td><td>GAGCCGGTGTATCGAGACGGCGG</td><td></td><td>1000</td><td></td><td></td><td></td><td></td><td></td><td></td><td></td><td></td><td></td></tr>
<tr><td>Oligo fwd</td><td>CACCGAGCCGGTGTATCGAGACGG</td><td></td><td></td><td></td><td></td><td></td><td></td><td></td><td></td><td></td><td></td><td></td></tr>
<tr><td>Oligo rev</td><td>AAACCCGTCTCGATACACCGGCTC</td><td></td><td></td><td></td><td></td><td></td><td></td><td></td><td></td><td></td><td></td><td></td></tr>
<tr><td>Chromosome</td><td>start</td><td></td><td>end</td><td>strand</td><td>MM</td><td>target_seq</td><td>PAM</td><td>alignment</td><td>distance</td><td>position</td><td>gene name</td><td>gene id</td></tr>
<tr><td>GU305914.1</td><td></td><td>195212</td><td>195234</td><td>-</td><td>0</td><td>GAGCCGGTGTATCGAGACGG</td><td>CGG</td><td>||||||||[||||||||||||]PAM</td><td>NA</td><td>Intergenic</td><td>NA</td><td>NA</td></tr>
<tr><td>GU305914.1</td><td></td><td>21081</td><td>21103</td><td>-</td><td>4</td><td>GACACGGTGTGGCGAGACGG</td><td>GGG</td><td>||--||||[||--||||||||]PAM</td><td>NA</td><td>Intergenic</td><td>NA</td><td>NA</td></tr>
<tr><td></td><td></td><td></td><td></td><td></td><td></td><td></td><td></td><td></td><td></td><td></td><td></td><td></td></tr>
<tr><td>T19855</td><td>GTATCGAGACGGCGGCGCCGTGG</td><td></td><td>1000</td><td></td><td></td><td></td><td></td><td></td><td></td><td></td><td></td><td></td></tr>
<tr><td>Oligo fwd</td><td>CACCGTATCGAGACGGCGGCGCCG</td><td></td><td></td><td></td><td></td><td></td><td></td><td></td><td></td><td></td><td></td><td></td></tr>
<tr><td>Oligo rev</td><td>AAACCGGCGCCGCCGTCTCGATAC</td><td></td><td></td><td></td><td></td><td></td><td></td><td></td><td></td><td></td><td></td><td></td></tr>
<tr><td>Chromosome</td><td>start</td><td></td><td>end</td><td>strand</td><td>MM</td><td>target_seq</td><td>PAM</td><td>alignment</td><td>distance</td><td>position</td><td>gene name</td><td>gene id</td></tr>
<tr><td>GU305914.1</td><td></td><td>195204</td><td>195226</td><td>-</td><td>0</td><td>GTATCGAGACGGCGGCGCCG</td><td>TGG</td><td>||||||||[||||||||||||]PAM</td><td>NA</td><td>Intergenic</td><td>NA</td><td>NA</td></tr>
<tr><td>GU305914.1</td><td></td><td>37262</td><td>37284</td><td>-</td><td>4</td><td>GCATGGAGACGGCGGCGTCT</td><td>GGG</td><td>|-||-|||[|||||||||-|-]PAM</td><td>NA</td><td>Intergenic</td><td>NA</td><td>NA</td></tr>
<tr><td></td><td></td><td></td><td></td><td></td><td></td><td></td><td></td><td></td><td></td><td></td><td></td><td></td></tr>
<tr><td>T19856</td><td>GATGCCGAAGCCCAAGACCCTGG</td><td></td><td>1000</td><td></td><td></td><td></td><td></td><td></td><td></td><td></td><td></td><td></td></tr>
<tr><td>Oligo fwd</td><td>CACCGATGCCGAAGCCCAAGACCC</td><td></td><td></td><td></td><td></td><td></td><td></td><td></td><td></td><td></td><td></td><td></td></tr>
<tr><td>Oligo rev</td><td>AAACGGGTCTTGGGCTTCGGCATC</td><td></td><td></td><td></td><td></td><td></td><td></td><td></td><td></td><td></td><td></td><td></td></tr>
<tr><td>Chromosome</td><td>start</td><td></td><td>end</td><td>strand</td><td>MM</td><td>target_seq</td><td>PAM</td><td>alignment</td><td>distance</td><td>position</td><td>gene name</td><td>gene id</td></tr>
<tr><td>GU305914.1</td><td></td><td>195177</td><td>195199</td><td>-</td><td>0</td><td>GATGCCGAAGCCCAAGACCC</td><td>TGG</td><td>||||||||[||||||||||||]PAM</td><td>NA</td><td>Intergenic</td><td>NA</td><td>NA</td></tr>
<tr><td></td><td></td><td></td><td></td><td></td><td></td><td></td><td></td><td></td><td></td><td></td><td></td><td></td></tr>
<tr><td>T19857</td><td>CGAAGCCCAAGACCCTGGTCAGG</td><td></td><td>1000</td><td></td><td></td><td></td><td></td><td></td><td></td><td></td><td></td><td></td></tr>
<tr><td>Oligo fwd</td><td>CACCCGAAGCCCAAGACCCTGGTC</td><td></td><td></td><td></td><td></td><td></td><td></td><td></td><td></td><td></td><td></td><td></td></tr>
<tr><td>Oligo rev</td><td>AAACGACCAGGGTCTTGGGCTTCG</td><td></td><td></td><td></td><td></td><td></td><td></td><td></td><td></td><td></td><td></td><td></td></tr>
<tr><td>Chromosome</td><td>start</td><td></td><td>end</td><td>strand</td><td>MM</td><td>target_seq</td><td>PAM</td><td>alignment</td><td>distance</td><td>position</td><td>gene name</td><td>gene id</td></tr>
<tr><td>GU305914.1</td><td></td><td>195172</td><td>195194</td><td>-</td><td>0</td><td>CGAAGCCCAAGACCCTGGTC</td><td>AGG</td><td>||||||||[||||||||||||]PAM</td><td>NA</td><td>Intergenic</td><td>NA</td><td>NA</td></tr>
<tr><td></td><td></td><td></td><td></td><td></td><td></td><td></td><td></td><td></td><td></td><td></td><td></td><td></td></tr>
<tr><td>T19858</td><td>GAAGATCGTGTCCGCCGCCGTGG</td><td></td><td>1000</td><td></td><td></td><td></td><td></td><td></td><td></td><td></td><td></td><td></td></tr>
<tr><td>Oligo fwd</td><td>CACCGAAGATCGTGTCCGCCGCCG</td><td></td><td></td><td></td><td></td><td></td><td></td><td></td><td></td><td></td><td></td><td></td></tr>
<tr><td>Oligo rev</td><td>AAACCGGCGGCGGACACGATCTTC</td><td></td><td></td><td></td><td></td><td></td><td></td><td></td><td></td><td></td><td></td><td></td></tr>
<tr><td>Chromosome</td><td>start</td><td></td><td>end</td><td>strand</td><td>MM</td><td>target_seq</td><td>PAM</td><td>alignment</td><td>distance</td><td>position</td><td>gene name</td><td>gene id</td></tr>
<tr><td>GU305914.1</td><td></td><td>195150</td><td>195172</td><td>-</td><td>0</td><td>GAAGATCGTGTCCGCCGCCG</td><td>TGG</td><td>||||||||[||||||||||||]PAM</td><td>NA</td><td>Intergenic</td><td>NA</td><td>NA</td></tr>
<tr><td></td><td></td><td></td><td></td><td></td><td></td><td></td><td></td><td></td><td></td><td></td><td></td><td></td></tr>
<tr><td>T19859</td><td>ATCGTGTCCGCCGCCGTGGTCGG</td><td></td><td>1000</td><td></td><td></td><td></td><td></td><td></td><td></td><td></td><td></td><td></td></tr>
<tr><td>Oligo fwd</td><td>CACCATCGTGTCCGCCGCCGTGGT</td><td></td><td></td><td></td><td></td><td></td><td></td><td></td><td></td><td></td><td></td><td></td></tr>
<tr><td>Oligo rev</td><td>AAACACCACGGCGGCGGACACGAT</td><td></td><td></td><td></td><td></td><td></td><td></td><td></td><td></td><td></td><td></td><td></td></tr>
<tr><td>Chromosome</td><td>start</td><td></td><td>end</td><td>strand</td><td>MM</td><td>target_seq</td><td>PAM</td><td>alignment</td><td>distance</td><td>position</td><td>gene name</td><td>gene id</td></tr>
<tr><td>GU305914.1</td><td></td><td>195146</td><td>195168</td><td>-</td><td>0</td><td>ATCGTGTCCGCCGCCGTGGT</td><td>CGG</td><td>||||||||[||||||||||||]PAM</td><td>NA</td><td>Intergenic</td><td>NA</td><td>NA</td></tr>
<tr><td></td><td></td><td></td><td></td><td></td><td></td><td></td><td></td><td></td><td></td><td></td><td></td><td></td></tr>
<tr><td>T19860</td><td>GTCCGCCGCCGTGGTCGGCCTGG</td><td></td><td>1000</td><td></td><td></td><td></td><td></td><td></td><td></td><td></td><td></td><td></td></tr>
<tr><td>Oligo fwd</td><td>CACCGTCCGCCGCCGTGGTCGGCC</td><td></td><td></td><td></td><td></td><td></td><td></td><td></td><td></td><td></td><td></td><td></td></tr>
<tr><td>Oligo rev</td><td>AAACGGCCGACCACGGCGGCGGAC</td><td></td><td></td><td></td><td></td><td></td><td></td><td></td><td></td><td></td><td></td><td></td></tr>
<tr><td>Chromosome</td><td>start</td><td></td><td>end</td><td>strand</td><td>MM</td><td>target_seq</td><td>PAM</td><td>alignment</td><td>distance</td><td>position</td><td>gene name</td><td>gene id</td></tr>
<tr><td>GU305914.1</td><td></td><td>195141</td><td>195163</td><td>-</td><td>0</td><td>GTCCGCCGCCGTGGTCGGCC</td><td>TGG</td><td>||||||||[||||||||||||]PAM</td><td>NA</td><td>Intergenic</td><td>NA</td><td>NA</td></tr>
<tr><td></td><td></td><td></td><td></td><td></td><td></td><td></td><td></td><td></td><td></td><td></td><td></td><td></td></tr>
<tr><td>T19861</td><td>CGTGGTCGGCCTGGAGAACGTGG</td><td></td><td>1000</td><td></td><td></td><td></td><td></td><td></td><td></td><td></td><td></td><td></td></tr>
<tr><td>Oligo fwd</td><td>CACCCGTGGTCGGCCTGGAGAACG</td><td></td><td></td><td></td><td></td><td></td><td></td><td></td><td></td><td></td><td></td><td></td></tr>
<tr><td>Oligo rev</td><td>AAACCGTTCTCCAGGCCGACCACG</td><td></td><td></td><td></td><td></td><td></td><td></td><td></td><td></td><td></td><td></td><td></td></tr>
<tr><td>Chromosome</td><td>start</td><td></td><td>end</td><td>strand</td><td>MM</td><td>target_seq</td><td>PAM</td><td>alignment</td><td>distance</td><td>position</td><td>gene name</td><td>gene id</td></tr>
</table>

GU305914.1 195132 195154 - 0 CGTGGTCGGCCTGGAGAACG TGG ||||||||[||||||||||||]PAM NA Intergenic NA NA

T19862 GGTCGGCCTGGAGAACGTGGCGG 1000
Oligo fwd CACCGGTCGGCCTGGAGAACGTGG
Oligo rev AAACCCACGTTCTCCAGGCCGACC
Chromosome start end strand MM target_seq PAM alignment distance position gene name gene id
GU305914.1 195129 195151 - 0 GGTCGGCCTGGAGAACGTGG CGG ||||||||[||||||||||||]PAM NA Intergenic NA NA

T19863 GAACGTGGCGGCCACCGCCACGG 1000
Oligo fwd CACCGAACGTGGCGGCCACCGCCA
Oligo rev AAACTGGCGGTGGCCGCCACGTTC
Chromosome start end strand MM target_seq PAM alignment distance position gene name gene id
GU305914.1 195117 195139 - 0 GAACGTGGCGGCCACCGCCA CGG ||||||||[||||||||||||]PAM NA Intergenic NA NA

T19864 CGGCCACCGCCACGGCGTTCAGG 1000
Oligo fwd CACCCGGCCACCGCCACGGCGTTC
Oligo rev AAACGAACGCCGTGGCGGTGGCCG
Chromosome start end strand MM target_seq PAM alignment distance position gene name gene id
GU305914.1 195109 195131 - 0 CGGCCACCGCCACGGCGTTC AGG ||||||||[||||||||||||]PAM NA Intergenic NA NA
GU305914.1 130320 130342 + 4 CAGCCGCCGCCGCGGCGGTC CGG |-|||-||[|||-|||||-||]PAM NA Intergenic NA NA

T19865 GGCCACCGCCACGGCGTTCAGGG 1000
Oligo fwd CACCGGCCACCGCCACGGCGTTCA
Oligo rev AAACTGAACGCCGTGGCGGTGGCC
Chromosome start end strand MM target_seq PAM alignment distance position gene name gene id
GU305914.1 195108 195130 - 0 GGCCACCGCCACGGCGTTCA GGG ||||||||[||||||||||||]PAM NA Intergenic NA NA

T19866 GCCACCGCCACGGCGTTCAGGGG 1000
Oligo fwd CACCGCCACCGCCACGGCGTTCAG
Oligo rev AAACCTGAACGCCGTGGCGGTGGC
Chromosome start end strand MM target_seq PAM alignment distance position gene name gene id
GU305914.1 195107 195129 - 0 GCCACCGCCACGGCGTTCAG GGG ||||||||[||||||||||||]PAM NA Intergenic NA NA
GU305914.1 100233 100255 + 4 GCCGCCCCGCGGCGTTCTG CGG |||-||-|[|-||||||||-|]PAM NA Intergenic NA NA

T19867 GACGATCGCCCTGCGCGACCCGG 1000
Oligo fwd CACCGACGATCGCCCTGCGCGACC
Oligo rev AAACGGTCGCGCAGGGCGATCGTC
Chromosome start end strand MM target_seq PAM alignment distance position gene name gene id
GU305914.1 195081 195103 - 0 GACGATCGCCCTGCGCGACC CGG ||||||||[||||||||||||]PAM NA Intergenic NA NA

T19868 CGCCCTGCGCGACCCGGTCTCGG 1000
Oligo fwd CACCCGCCCTGCGCGACCCGGTCT
Oligo rev AAACAGACCGGGTCGCGCAGGGCG
Chromosome start end strand MM target_seq PAM alignment distance position gene name gene id
GU305914.1 195075 195097 - 0 CGCCCTGCGCGACCCGGTCT CGG ||||||||[||||||||||||]PAM NA Intergenic NA NA

T19869 GCCCTGCGCGACCCGGTCTCGGG 1000
Oligo fwd CACCGCCCTGCGCGACCCGGTCTC
Oligo rev AAACGAGACCGGGTCGCGCAGGGC
Chromosome start end strand MM target_seq PAM alignment distance position gene name gene id
GU305914.1 195074 195096 - 0 GCCCTGCGCGACCCGGTCTC GGG ||||||||[||||||||||||]PAM NA Intergenic NA NA

T19870 TGCGCGACCCGGTCTCGGGACGG 1000
Oligo fwd CACCTGCGCGACCCGGTCTCGGGA
Oligo rev AAACTCCCGAGACCGGGTCGCGCA
Chromosome start end strand MM target_seq PAM alignment distance position gene name gene id
GU305914.1 195070 195092 - 0 TGCGCGACCCGGTCTCGGGA CGG ||||||||[||||||||||||]PAM NA Intergenic NA NA

T19871 GGGACGGCGCGAGACGTTTCAGG 1000
Oligo fwd CACCGGGACGGCGCGAGACGTTTC
Oligo rev AAACGAAACGTCTCGCGCCGTCCC
Chromosome start end strand MM target_seq PAM alignment distance position gene name gene id
GU305914.1 195054 195076 - 0 GGGACGGCGCGAGACGTTTC AGG ||||||||[||||||||||||]PAM NA Intergenic NA NA

T19872 CGCGAGACGTTTCAGGTGTTCGG 1000
Oligo fwd CACCCGCGAGACGTTTCAGGTGTT
Oligo rev AAACAACACCTGAAACGTCTCGCG
Chromosome start end strand MM target_seq PAM alignment distance position gene name gene id
GU305914.1 195047 195069 - 0 CGCGAGACGTTTCAGGTGTT CGG ||||||||[||||||||||||]PAM NA Intergenic NA NA

T19873 GCGAGACGTTTCAGGTGTTCGGG 1000
Oligo fwd CACCGCGAGACGTTTCAGGTGTTC
Oligo rev AAACGAACACCTGAAACGTCTCGC
Chromosome start end strand MM target_seq PAM alignment distance position gene name gene id
GU305914.1 195046 195068 - 0 GCGAGACGTTTCAGGTGTTC GGG ||||||||[||||||||||||]PAM NA Intergenic NA NA

T19874 CGAGACGTTTCAGGTGTTCGGGG 1000
Oligo fwd CACCCGAGACGTTTCAGGTGTTCG
Oligo rev AAACCGAACACCTGAAACGTCTCG
Chromosome start end strand MM target_seq PAM alignment distance position gene name gene id
GU305914.1 195045 195067 - 0 CGAGACGTTTCAGGTGTTCG GGG ||||||||[||||||||||||]PAM NA Intergenic NA NA

T19875 TCGGGGCGTCAGACCTGAAGAGG 1000
Oligo fwd CACCTCGGGGCGTCAGACCTGAAG
Oligo rev AAACCTTCAGGTCTGACGCCCCGA
Chromosome start end strand MM target_seq PAM alignment distance position gene name gene id
GU305914.1 195028 195050 - 0 TCGGGGCGTCAGACCTGAAG AGG ||||||||[||||||||||||]PAM NA Intergenic NA NA

T19876 AGAGGAAGCCGCCCTTCTCGCGG 1000

| | | | | | | | | | | | |
|---|---|---|---|---|---|---|---|---|---|---|---|
| Oligo fwd | CACCAGAGGAAGCCGCCCTTCTCG | | | | | | | | | | |
| Oligo rev | AAACCGAGAAGGGCGGCTTCCTCT | | | | | | | | | | |
| Chromosome | start | end | strand | MM | target_seq | PAM | alignment | distance | position | gene name | gene id |
| GU305914.1 | 195010 | 195032 | - | 0 | AGAGGAAGCCGCCCTTCTCG | CGG | \|\|\|\|\|\|\|\|[\|\|\|\|\|\|\|\|\|\|\|\|]PAM | NA | Intergenic | NA | NA |
| T19877 | GAAGCCGCCCTTCTCGCGGATGG | 1000 | | | | | | | | | |
| Oligo fwd | CACCGAAGCCGCCCTTCTCGCGGA | | | | | | | | | | |
| Oligo rev | AAACTCCGCGAGAAGGGCGGCTTC | | | | | | | | | | |
| Chromosome | start | end | strand | MM | target_seq | PAM | alignment | distance | position | gene name | gene id |
| GU305914.1 | 195006 | 195028 | - | 0 | GAAGCCGCCCTTCTCGCGGA | TGG | \|\|\|\|\|\|\|\|[\|\|\|\|\|\|\|\|\|\|\|\|]PAM | NA | Intergenic | NA | NA |
| T19878 | ACACGACGTACACACTCGTGCGG | 1000 | | | | | | | | | |
| Oligo fwd | CACCACACGACGTACACACTCGTG | | | | | | | | | | |
| Oligo rev | AAACCACGAGTGTGTACGTCGTGT | | | | | | | | | | |
| Chromosome | start | end | strand | MM | target_seq | PAM | alignment | distance | position | gene name | gene id |
| GU305914.1 | 194980 | 195002 | - | 0 | ACACGACGTACACACTCGTG | CGG | \|\|\|\|\|\|\|\|[\|\|\|\|\|\|\|\|\|\|\|\|]PAM | NA | Intergenic | NA | NA |
| T19879 | CACGACGTACACACTCGTGCGGG | 1000 | | | | | | | | | |
| Oligo fwd | CACCCACGACGTACACACTCGTGC | | | | | | | | | | |
| Oligo rev | AAACGCACGAGTGTGTACGTCGTG | | | | | | | | | | |
| Chromosome | start | end | strand | MM | target_seq | PAM | alignment | distance | position | gene name | gene id |
| GU305914.1 | 194979 | 195001 | - | 0 | CACGACGTACACACTCGTGC | GGG | \|\|\|\|\|\|\|\|[\|\|\|\|\|\|\|\|\|\|\|\|]PAM | NA | Intergenic | NA | NA |
| T19880 | TGCGGGAGTACATCACCTTTCGG | 1000 | | | | | | | | | |
| Oligo fwd | CACCTGCGGGAGTACATCACCTTT | | | | | | | | | | |
| Oligo rev | AAACAAAGGTGATGTACTCCCGCA | | | | | | | | | | |
| Chromosome | start | end | strand | MM | target_seq | PAM | alignment | distance | position | gene name | gene id |
| GU305914.1 | 194962 | 194984 | - | 0 | TGCGGGAGTACATCACCTTT | CGG | \|\|\|\|\|\|\|\|[\|\|\|\|\|\|\|\|\|\|\|\|]PAM | NA | Intergenic | NA | NA |
| T19881 | GGAGTACATCACCTTTCGGCTGG | 1000 | | | | | | | | | |
| Oligo fwd | CACCGGAGTACATCACCTTTCGGC | | | | | | | | | | |
| Oligo rev | AAACGCCGAAAGGTGATGTACTCC | | | | | | | | | | |
| Chromosome | start | end | strand | MM | target_seq | PAM | alignment | distance | position | gene name | gene id |
| GU305914.1 | 194958 | 194980 | - | 0 | GGAGTACATCACCTTTCGGC | TGG | \|\|\|\|\|\|\|\|[\|\|\|\|\|\|\|\|\|\|\|\|]PAM | NA | Intergenic | NA | NA |
| T19882 | CATCACCTTTCGGCTGGCCGAGG | 1000 | | | | | | | | | |
| Oligo fwd | CACCCATCACCTTTCGGCTGGCCG | | | | | | | | | | |
| Oligo rev | AAACCGGCCAGCCGAAAGGTGATG | | | | | | | | | | |
| Chromosome | start | end | strand | MM | target_seq | PAM | alignment | distance | position | gene name | gene id |
| GU305914.1 | 194952 | 194974 | - | 0 | CATCACCTTTCGGCTGGCCG | AGG | \|\|\|\|\|\|\|\|[\|\|\|\|\|\|\|\|\|\|\|\|]PAM | NA | Intergenic | NA | NA |
| T19883 | CCTTTCGGCTGGCCGAGGCGTGG | 1000 | | | | | | | | | |
| Oligo fwd | CACCCCTTTCGGCTGGCCGAGGCG | | | | | | | | | | |
| Oligo rev | AAACCGCCTCGGCCAGCCGAAAGG | | | | | | | | | | |
| Chromosome | start | end | strand | MM | target_seq | PAM | alignment | distance | position | gene name | gene id |
| GU305914.1 | 194947 | 194969 | - | 0 | CCTTTCGGCTGGCCGAGGCG | TGG | \|\|\|\|\|\|\|\|[\|\|\|\|\|\|\|\|\|\|\|\|]PAM | NA | Intergenic | NA | NA |
| T19884 | GCCGAGGCGTGGACCGTCATCGG | 1000 | | | | | | | | | |
| Oligo fwd | CACCGCCGAGGCGTGGACCGTCAT | | | | | | | | | | |
| Oligo rev | AAACATGACGGTCCACGCCTCGGC | | | | | | | | | | |
| Chromosome | start | end | strand | MM | target_seq | PAM | alignment | distance | position | gene name | gene id |
| GU305914.1 | 194936 | 194958 | - | 0 | GCCGAGGCGTGGACCGTCAT | CGG | \|\|\|\|\|\|\|\|[\|\|\|\|\|\|\|\|\|\|\|\|]PAM | NA | Intergenic | NA | NA |
| T19885 | GTGGACCGTCATCGGCGCCGTGG | 1000 | | | | | | | | | |
| Oligo fwd | CACCGTGGACCGTCATCGGCGCCG | | | | | | | | | | |
| Oligo rev | AAACCGGCGCCGATGACGGTCCAC | | | | | | | | | | |
| Chromosome | start | end | strand | MM | target_seq | PAM | alignment | distance | position | gene name | gene id |
| GU305914.1 | 194928 | 194950 | - | 0 | GTGGACCGTCATCGGCGCCG | TGG | \|\|\|\|\|\|\|\|[\|\|\|\|\|\|\|\|\|\|\|\|]PAM | NA | Intergenic | NA | NA |
| T19886 | TGGACCGTCATCGGCGCCGTGGG | 1000 | | | | | | | | | |
| Oligo fwd | CACCTGGACCGTCATCGGCGCCGT | | | | | | | | | | |
| Oligo rev | AAACACGGCGCCGATGACGGTCCA | | | | | | | | | | |
| Chromosome | start | end | strand | MM | target_seq | PAM | alignment | distance | position | gene name | gene id |
| GU305914.1 | 194927 | 194949 | - | 0 | TGGACCGTCATCGGCGCCGT | GGG | \|\|\|\|\|\|\|\|[\|\|\|\|\|\|\|\|\|\|\|\|]PAM | NA | Intergenic | NA | NA |
| GU305914.1 | 149874 | 149896 | + | 3 | TCCACCGTCACCGGCGCCGT | CGG | \|--\|\|\|\|\|[\|\|-\|\|\|\|\|\|\|\|\|]PAM | NA | Intergenic | NA | NA |
| T19887 | TCGGCGCCGTGGGAGAGTACCGG | 1000 | | | | | | | | | |
| Oligo fwd | CACCTCGGCGCCGTGGGAGAGTAC | | | | | | | | | | |
| Oligo rev | AAACGTACTCTCCCACGGCGCCGA | | | | | | | | | | |
| Chromosome | start | end | strand | MM | target_seq | PAM | alignment | distance | position | gene name | gene id |
| GU305914.1 | 194917 | 194939 | - | 0 | TCGGCGCCGTGGGAGAGTAC | CGG | \|\|\|\|\|\|\|\|[\|\|\|\|\|\|\|\|\|\|\|\|]PAM | NA | Intergenic | NA | NA |
| T19888 | CGGCGCCGTGGGAGAGTACCGGG | 1000 | | | | | | | | | |
| Oligo fwd | CACCCGGCGCCGTGGGAGAGTACC | | | | | | | | | | |
| Oligo rev | AAACGGTACTCTCCCACGGCGCCG | | | | | | | | | | |
| Chromosome | start | end | strand | MM | target_seq | PAM | alignment | distance | position | gene name | gene id |
| GU305914.1 | 194916 | 194938 | - | 0 | CGGCGCCGTGGGAGAGTACC | GGG | \|\|\|\|\|\|\|\|[\|\|\|\|\|\|\|\|\|\|\|\|]PAM | NA | Intergenic | NA | NA |
| T19889 | GTGGGAGAGTACCGGGACGACGG | 1000 | | | | | | | | | |
| Oligo fwd | CACCGTGGGAGAGTACCGGGACGA | | | | | | | | | | |
| Oligo rev | AAACTCGTCCCGGTACTCTCCCAC | | | | | | | | | | |
| Chromosome | start | end | strand | MM | target_seq | PAM | alignment | distance | position | gene name | gene id |
| GU305914.1 | 194909 | 194931 | - | 0 | GTGGGAGAGTACCGGGACGA | CGG | \|\|\|\|\|\|\|\|[\|\|\|\|\|\|\|\|\|\|\|\|]PAM | NA | Intergenic | NA | NA |
| T19890 | GGACGACGGCTTCGTCTTCGAGG | 1000 | | | | | | | | | |
| Oligo fwd | CACCGGACGACGGCTTCGTCTTCG | | | | | | | | | | |
| Oligo rev | AAACCGAAGACGAAGCCGTCGTCC | | | | | | | | | | |
| Chromosome | start | end | strand | MM | target_seq | PAM | alignment | distance | position | gene name | gene id |
| GU305914.1 | 194895 | 194917 | - | 0 | GGACGACGGCTTCGTCTTCG | AGG | \|\|\|\|\|\|\|\|[\|\|\|\|\|\|\|\|\|\|\|\|]PAM | NA | Intergenic | NA | NA |

<table>
<tr><td>GU305914.1</td><td>152579</td><td>152601</td><td>+</td><td>4</td><td>CGATGACGGCTTCGACCTCG</td><td>AGG</td><td>-||-||||[||||||-|-|||]PAM</td><td>NA</td><td>Intergenic</td><td>NA</td><td>NA</td></tr>
<tr><td></td><td></td><td></td><td></td><td></td><td></td><td></td><td></td><td></td><td></td><td></td><td></td></tr>
<tr><td>T19891</td><td>CGAGGTGTCCACCGTCGTGCTGG</td><td>1000</td><td></td><td></td><td></td><td></td><td></td><td></td><td></td><td></td><td></td></tr>
<tr><td>Oligo fwd</td><td>CACCCGAGGTGTCCACCGTCGTGC</td><td></td><td></td><td></td><td></td><td></td><td></td><td></td><td></td><td></td><td></td></tr>
<tr><td>Oligo rev</td><td>AAACGCACGACGGTGGACACCTCG</td><td></td><td></td><td></td><td></td><td></td><td></td><td></td><td></td><td></td><td></td></tr>
<tr><td>Chromosome</td><td>start</td><td>end</td><td>strand</td><td>MM</td><td>target_seq</td><td>PAM</td><td>alignment</td><td>distance</td><td>position</td><td>gene name</td><td>gene id</td></tr>
<tr><td>GU305914.1</td><td>194877</td><td>194899</td><td>-</td><td>0</td><td>CGAGGTGTCCACCGTCGTGC</td><td>TGG</td><td>||||||||[||||||||||||]PAM</td><td>NA</td><td>Intergenic</td><td>NA</td><td>NA</td></tr>
<tr><td></td><td></td><td></td><td></td><td></td><td></td><td></td><td></td><td></td><td></td><td></td><td></td></tr>
<tr><td>T19892</td><td>GTGTCCACCGTCGTGCTGGTCGG</td><td>1000</td><td></td><td></td><td></td><td></td><td></td><td></td><td></td><td></td><td></td></tr>
<tr><td>Oligo fwd</td><td>CACCGTGTCCACCGTCGTGCTGGT</td><td></td><td></td><td></td><td></td><td></td><td></td><td></td><td></td><td></td><td></td></tr>
<tr><td>Oligo rev</td><td>AAACACCAGCACGACGGTGGACAC</td><td></td><td></td><td></td><td></td><td></td><td></td><td></td><td></td><td></td><td></td></tr>
<tr><td>Chromosome</td><td>start</td><td>end</td><td>strand</td><td>MM</td><td>target_seq</td><td>PAM</td><td>alignment</td><td>distance</td><td>position</td><td>gene name</td><td>gene id</td></tr>
<tr><td>GU305914.1</td><td>194873</td><td>194895</td><td>-</td><td>0</td><td>GTGTCCACCGTCGTGCTGGT</td><td>CGG</td><td>||||||||[||||||||||||]PAM</td><td>NA</td><td>Intergenic</td><td>NA</td><td>NA</td></tr>
<tr><td></td><td></td><td></td><td></td><td></td><td></td><td></td><td></td><td></td><td></td><td></td><td></td></tr>
<tr><td>T19893</td><td>GTCGTGCTGGTCGGAGCCCGCGG</td><td>1000</td><td></td><td></td><td></td><td></td><td></td><td></td><td></td><td></td><td></td></tr>
<tr><td>Oligo fwd</td><td>CACCGTCGTGCTGGTCGGAGCCCG</td><td></td><td></td><td></td><td></td><td></td><td></td><td></td><td></td><td></td><td></td></tr>
<tr><td>Oligo rev</td><td>AAACCGGGCTCCGACCAGCACGAC</td><td></td><td></td><td></td><td></td><td></td><td></td><td></td><td></td><td></td><td></td></tr>
<tr><td>Chromosome</td><td>start</td><td>end</td><td>strand</td><td>MM</td><td>target_seq</td><td>PAM</td><td>alignment</td><td>distance</td><td>position</td><td>gene name</td><td>gene id</td></tr>
<tr><td>GU305914.1</td><td>194864</td><td>194886</td><td>-</td><td>0</td><td>GTCGTGCTGGTCGGAGCCCG</td><td>CGG</td><td>||||||||[||||||||||||]PAM</td><td>NA</td><td>Intergenic</td><td>NA</td><td>NA</td></tr>
<tr><td></td><td></td><td></td><td></td><td></td><td></td><td></td><td></td><td></td><td></td><td></td><td></td></tr>
<tr><td>T19894</td><td>GCTGGTCGGAGCCCGCGGAACGG</td><td>1000</td><td></td><td></td><td></td><td></td><td></td><td></td><td></td><td></td><td></td></tr>
<tr><td>Oligo fwd</td><td>CACCGCTGGTCGGAGCCCGCGGAA</td><td></td><td></td><td></td><td></td><td></td><td></td><td></td><td></td><td></td><td></td></tr>
<tr><td>Oligo rev</td><td>AAACTTCCGCGGGCTCCGACCAGC</td><td></td><td></td><td></td><td></td><td></td><td></td><td></td><td></td><td></td><td></td></tr>
<tr><td>Chromosome</td><td>start</td><td>end</td><td>strand</td><td>MM</td><td>target_seq</td><td>PAM</td><td>alignment</td><td>distance</td><td>position</td><td>gene name</td><td>gene id</td></tr>
<tr><td>GU305914.1</td><td>194859</td><td>194881</td><td>-</td><td>0</td><td>GCTGGTCGGAGCCCGCGGAA</td><td>CGG</td><td>||||||||[||||||||||||]PAM</td><td>NA</td><td>Intergenic</td><td>NA</td><td>NA</td></tr>
<tr><td></td><td></td><td></td><td></td><td></td><td></td><td></td><td></td><td></td><td></td><td></td><td></td></tr>
<tr><td>T19895</td><td>GGAGCCCGCGGAACGGTGTACGG</td><td>1000</td><td></td><td></td><td></td><td></td><td></td><td></td><td></td><td></td><td></td></tr>
<tr><td>Oligo fwd</td><td>CACCGGAGCCCGCGGAACGGTGTA</td><td></td><td></td><td></td><td></td><td></td><td></td><td></td><td></td><td></td><td></td></tr>
<tr><td>Oligo rev</td><td>AAACTACACCGTTCCGCGGGCTCC</td><td></td><td></td><td></td><td></td><td></td><td></td><td></td><td></td><td></td><td></td></tr>
<tr><td>Chromosome</td><td>start</td><td>end</td><td>strand</td><td>MM</td><td>target_seq</td><td>PAM</td><td>alignment</td><td>distance</td><td>position</td><td>gene name</td><td>gene id</td></tr>
<tr><td>GU305914.1</td><td>194852</td><td>194874</td><td>-</td><td>0</td><td>GGAGCCCGCGGAACGGTGTA</td><td>CGG</td><td>||||||||[||||||||||||]PAM</td><td>NA</td><td>Intergenic</td><td>NA</td><td>NA</td></tr>
<tr><td></td><td></td><td></td><td></td><td></td><td></td><td></td><td></td><td></td><td></td><td></td><td></td></tr>
<tr><td>T19896</td><td>TGCTGAGTAACGACGTCTTCAGG</td><td>1000</td><td></td><td></td><td></td><td></td><td></td><td></td><td></td><td></td><td></td></tr>
<tr><td>Oligo fwd</td><td>CACCTGCTGAGTAACGACGTCTTC</td><td></td><td></td><td></td><td></td><td></td><td></td><td></td><td></td><td></td><td></td></tr>
<tr><td>Oligo rev</td><td>AAACGAAGACGTCGTTACTCAGCA</td><td></td><td></td><td></td><td></td><td></td><td></td><td></td><td></td><td></td><td></td></tr>
<tr><td>Chromosome</td><td>start</td><td>end</td><td>strand</td><td>MM</td><td>target_seq</td><td>PAM</td><td>alignment</td><td>distance</td><td>position</td><td>gene name</td><td>gene id</td></tr>
<tr><td>GU305914.1</td><td>194821</td><td>194843</td><td>-</td><td>0</td><td>TGCTGAGTAACGACGTCTTC</td><td>AGG</td><td>||||||||[||||||||||||]PAM</td><td>NA</td><td>Intergenic</td><td>NA</td><td>NA</td></tr>
<tr><td></td><td></td><td></td><td></td><td></td><td></td><td></td><td></td><td></td><td></td><td></td><td></td></tr>
<tr><td>T19897</td><td>CGACGTCTTCAGGATCGCCGAGG</td><td>1000</td><td></td><td></td><td></td><td></td><td></td><td></td><td></td><td></td><td></td></tr>
<tr><td>Oligo fwd</td><td>CACCCGACGTCTTCAGGATCGCCG</td><td></td><td></td><td></td><td></td><td></td><td></td><td></td><td></td><td></td><td></td></tr>
<tr><td>Oligo rev</td><td>AAACCGGCGATCCTGAAGACGTCG</td><td></td><td></td><td></td><td></td><td></td><td></td><td></td><td></td><td></td><td></td></tr>
<tr><td>Chromosome</td><td>start</td><td>end</td><td>strand</td><td>MM</td><td>target_seq</td><td>PAM</td><td>alignment</td><td>distance</td><td>position</td><td>gene name</td><td>gene id</td></tr>
<tr><td>GU305914.1</td><td>194811</td><td>194833</td><td>-</td><td>0</td><td>CGACGTCTTCAGGATCGCCG</td><td>AGG</td><td>||||||||[||||||||||||]PAM</td><td>NA</td><td>Intergenic</td><td>NA</td><td>NA</td></tr>
<tr><td></td><td></td><td></td><td></td><td></td><td></td><td></td><td></td><td></td><td></td><td></td><td></td></tr>
<tr><td>T19898</td><td>CAGGATCGCCGAGGACATCTCGG</td><td>1000</td><td></td><td></td><td></td><td></td><td></td><td></td><td></td><td></td><td></td></tr>
<tr><td>Oligo fwd</td><td>CACCCAGGATCGCCGAGGACATCT</td><td></td><td></td><td></td><td></td><td></td><td></td><td></td><td></td><td></td><td></td></tr>
<tr><td>Oligo rev</td><td>AAACAGATGTCCTCGGCGATCCTG</td><td></td><td></td><td></td><td></td><td></td><td></td><td></td><td></td><td></td><td></td></tr>
<tr><td>Chromosome</td><td>start</td><td>end</td><td>strand</td><td>MM</td><td>target_seq</td><td>PAM</td><td>alignment</td><td>distance</td><td>position</td><td>gene name</td><td>gene id</td></tr>
<tr><td>GU305914.1</td><td>194802</td><td>194824</td><td>-</td><td>0</td><td>CAGGATCGCCGAGGACATCT</td><td>CGG</td><td>||||||||[||||||||||||]PAM</td><td>NA</td><td>Intergenic</td><td>NA</td><td>NA</td></tr>
<tr><td></td><td></td><td></td><td></td><td></td><td></td><td></td><td></td><td></td><td></td><td></td><td></td></tr>
<tr><td>T19899</td><td>ATCTCGGTCTTCTTCAAGAGAGG</td><td>1000</td><td></td><td></td><td></td><td></td><td></td><td></td><td></td><td></td><td></td></tr>
<tr><td>Oligo fwd</td><td>CACCATCTCGGTCTTCTTCAAGAG</td><td></td><td></td><td></td><td></td><td></td><td></td><td></td><td></td><td></td><td></td></tr>
<tr><td>Oligo rev</td><td>AAACCTCTTGAAGAAGACCGAGAT</td><td></td><td></td><td></td><td></td><td></td><td></td><td></td><td></td><td></td><td></td></tr>
<tr><td>Chromosome</td><td>start</td><td>end</td><td>strand</td><td>MM</td><td>target_seq</td><td>PAM</td><td>alignment</td><td>distance</td><td>position</td><td>gene name</td><td>gene id</td></tr>
<tr><td>GU305914.1</td><td>194786</td><td>194808</td><td>-</td><td>0</td><td>ATCTCGGTCTTCTTCAAGAG</td><td>AGG</td><td>||||||||[||||||||||||]PAM</td><td>NA</td><td>Intergenic</td><td>NA</td><td>NA</td></tr>
<tr><td></td><td></td><td></td><td></td><td></td><td></td><td></td><td></td><td></td><td></td><td></td><td></td></tr>
<tr><td>T19900</td><td>TCGGTCTTCTTCAAGAGAGGAGG</td><td>1000</td><td></td><td></td><td></td><td></td><td></td><td></td><td></td><td></td><td></td></tr>
<tr><td>Oligo fwd</td><td>CACCTCGGTCTTCTTCAAGAGAGG</td><td></td><td></td><td></td><td></td><td></td><td></td><td></td><td></td><td></td><td></td></tr>
<tr><td>Oligo rev</td><td>AAACCCTCTCTTGAAGAAGACCGA</td><td></td><td></td><td></td><td></td><td></td><td></td><td></td><td></td><td></td><td></td></tr>
<tr><td>Chromosome</td><td>start</td><td>end</td><td>strand</td><td>MM</td><td>target_seq</td><td>PAM</td><td>alignment</td><td>distance</td><td>position</td><td>gene name</td><td>gene id</td></tr>
<tr><td>GU305914.1</td><td>194783</td><td>194805</td><td>-</td><td>0</td><td>TCGGTCTTCTTCAAGAGAGG</td><td>AGG</td><td>||||||||[||||||||||||]PAM</td><td>NA</td><td>Intergenic</td><td>NA</td><td>NA</td></tr>
<tr><td></td><td></td><td></td><td></td><td></td><td></td><td></td><td></td><td></td><td></td><td></td><td></td></tr>
<tr><td>T19901</td><td>TTCAAGAGAGGAGGCGTGAGCGG</td><td>1000</td><td></td><td></td><td></td><td></td><td></td><td></td><td></td><td></td><td></td></tr>
<tr><td>Oligo fwd</td><td>CACCTTCAAGAGAGGAGGCGTGAG</td><td></td><td></td><td></td><td></td><td></td><td></td><td></td><td></td><td></td><td></td></tr>
<tr><td>Oligo rev</td><td>AAACCTCACGCCTCCTCTCTTGAA</td><td></td><td></td><td></td><td></td><td></td><td></td><td></td><td></td><td></td><td></td></tr>
<tr><td>Chromosome</td><td>start</td><td>end</td><td>strand</td><td>MM</td><td>target_seq</td><td>PAM</td><td>alignment</td><td>distance</td><td>position</td><td>gene name</td><td>gene id</td></tr>
<tr><td>GU305914.1</td><td>194774</td><td>194796</td><td>-</td><td>0</td><td>TTCAAGAGAGGAGGCGTGAG</td><td>CGG</td><td>||||||||[||||||||||||]PAM</td><td>NA</td><td>Intergenic</td><td>NA</td><td>NA</td></tr>
<tr><td></td><td></td><td></td><td></td><td></td><td></td><td></td><td></td><td></td><td></td><td></td><td></td></tr>
<tr><td>T19902</td><td>AAGAGAGGAGGCGTGAGCGGCGG</td><td>1000</td><td></td><td></td><td></td><td></td><td></td><td></td><td></td><td></td><td></td></tr>
<tr><td>Oligo fwd</td><td>CACCAAGAGAGGAGGCGTGAGCGG</td><td></td><td></td><td></td><td></td><td></td><td></td><td></td><td></td><td></td><td></td></tr>
<tr><td>Oligo rev</td><td>AAACCCGCTCACGCCTCCTCTCTT</td><td></td><td></td><td></td><td></td><td></td><td></td><td></td><td></td><td></td><td></td></tr>
<tr><td>Chromosome</td><td>start</td><td>end</td><td>strand</td><td>MM</td><td>target_seq</td><td>PAM</td><td>alignment</td><td>distance</td><td>position</td><td>gene name</td><td>gene id</td></tr>
<tr><td>GU305914.1</td><td>194771</td><td>194793</td><td>-</td><td>0</td><td>AAGAGAGGAGGCGTGAGCGG</td><td>CGG</td><td>||||||||[||||||||||||]PAM</td><td>NA</td><td>Intergenic</td><td>NA</td><td>NA</td></tr>
<tr><td></td><td></td><td></td><td></td><td></td><td></td><td></td><td></td><td></td><td></td><td></td><td></td></tr>
<tr><td>T19903</td><td>GTGCCCAACCGCTTCGACCGAGG</td><td>1000</td><td></td><td></td><td></td><td></td><td></td><td></td><td></td><td></td><td></td></tr>
<tr><td>Oligo fwd</td><td>CACCGTGCCCAACCGCTTCGACCG</td><td></td><td></td><td></td><td></td><td></td><td></td><td></td><td></td><td></td><td></td></tr>
<tr><td>Oligo rev</td><td>AAACCGGTCGAAGCGGTTGGGCAC</td><td></td><td></td><td></td><td></td><td></td><td></td><td></td><td></td><td></td><td></td></tr>
<tr><td>Chromosome</td><td>start</td><td>end</td><td>strand</td><td>MM</td><td>target_seq</td><td>PAM</td><td>alignment</td><td>distance</td><td>position</td><td>gene name</td><td>gene id</td></tr>
<tr><td>GU305914.1</td><td>194741</td><td>194763</td><td>-</td><td>0</td><td>GTGCCCAACCGCTTCGACCG</td><td>AGG</td><td>||||||||[||||||||||||]PAM</td><td>NA</td><td>Intergenic</td><td>NA</td><td>NA</td></tr>
<tr><td></td><td></td><td></td><td></td><td></td><td></td><td></td><td></td><td></td><td></td><td></td><td></td></tr>
<tr><td>T19904</td><td>ACCGCTTCGACCGAGGCGCCCGG</td><td>1000</td><td></td><td></td><td></td><td></td><td></td><td></td><td></td><td></td><td></td></tr>
<tr><td>Oligo fwd</td><td>CACCACCGCTTCGACCGAGGCGCC</td><td></td><td></td><td></td><td></td><td></td><td></td><td></td><td></td><td></td><td></td></tr>
<tr><td>Oligo rev</td><td>AAACGGCGCCTCGGTCGAAGCGGT</td><td></td><td></td><td></td><td></td><td></td><td></td><td></td><td></td><td></td><td></td></tr>
<tr><td>Chromosome</td><td>start</td><td>end</td><td>strand</td><td>MM</td><td>target_seq</td><td>PAM</td><td>alignment</td><td>distance</td><td>position</td><td>gene name</td><td>gene id</td></tr>
<tr><td>GU305914.1</td><td>194734</td><td>194756</td><td>-</td><td>0</td><td>ACCGCTTCGACCGAGGCGCC</td><td>CGG</td><td>||||||||[||||||||||||]PAM</td><td>NA</td><td>Intergenic</td><td>NA</td><td>NA</td></tr>
<tr><td></td><td></td><td></td><td></td><td></td><td></td><td></td><td></td><td></td><td></td><td></td><td></td></tr>
<tr><td>T19905</td><td>CCGCTTCGACCGAGGCGCCCGGG</td><td>1000</td><td></td><td></td><td></td><td></td><td></td><td></td><td></td><td></td><td></td></tr>
<tr><td>Oligo fwd</td><td>CACCCCGCTTCGACCGAGGCGCCC</td><td></td><td></td><td></td><td></td><td></td><td></td><td></td><td></td><td></td><td></td></tr>
<tr><td>Oligo rev</td><td>AAACGGGCGCCTCGGTCGAAGCGG</td><td></td><td></td><td></td><td></td><td></td><td></td><td></td><td></td><td></td><td></td></tr>
</table>

<table>
<tr><td>Chromosome</td><td>start</td><td>end</td><td>strand</td><td>MM</td><td>target_seq</td><td>PAM</td><td>alignment</td><td>distance</td><td>position</td><td>gene name</td><td>gene id</td></tr>
<tr><td>GU305914.1</td><td>194733</td><td>194755</td><td>-</td><td>0</td><td>CCGCTTCGACCGAGGCGCCC</td><td>GGG</td><td>||||||||[||||||||||||]PAM</td><td>NA</td><td>Intergenic</td><td>NA</td><td>NA</td></tr>
<tr><td></td><td></td><td></td><td></td><td></td><td></td><td></td><td></td><td></td><td></td><td></td><td></td></tr>
<tr><td>T19906</td><td>CGCTTCGACCGAGGCGCCCGGGG</td><td>1000</td><td></td><td></td><td></td><td></td><td></td><td></td><td></td><td></td><td></td></tr>
<tr><td>Oligo fwd</td><td>CACCCGCTTCGACCGAGGCGCCCG</td><td></td><td></td><td></td><td></td><td></td><td></td><td></td><td></td><td></td><td></td></tr>
<tr><td>Oligo rev</td><td>AAACCGGGCGCCTCGGTCGAAGCG</td><td></td><td></td><td></td><td></td><td></td><td></td><td></td><td></td><td></td><td></td></tr>
<tr><td>Chromosome</td><td>start</td><td>end</td><td>strand</td><td>MM</td><td>target_seq</td><td>PAM</td><td>alignment</td><td>distance</td><td>position</td><td>gene name</td><td>gene id</td></tr>
<tr><td>GU305914.1</td><td>194732</td><td>194754</td><td>-</td><td>0</td><td>CGCTTCGACCGAGGCGCCCG</td><td>GGG</td><td>||||||||[||||||||||||]PAM</td><td>NA</td><td>Intergenic</td><td>NA</td><td>NA</td></tr>
<tr><td></td><td></td><td></td><td></td><td></td><td></td><td></td><td></td><td></td><td></td><td></td><td></td></tr>
<tr><td>T19907</td><td>CGCCCGGGGAGAACTCCGTCTGG</td><td>1000</td><td></td><td></td><td></td><td></td><td></td><td></td><td></td><td></td><td></td></tr>
<tr><td>Oligo fwd</td><td>CACCCGCCCGGGGAGAACTCCGTC</td><td></td><td></td><td></td><td></td><td></td><td></td><td></td><td></td><td></td><td></td></tr>
<tr><td>Oligo rev</td><td>AAACGACGGAGTTCTCCCCGGGCG</td><td></td><td></td><td></td><td></td><td></td><td></td><td></td><td></td><td></td><td></td></tr>
<tr><td>Chromosome</td><td>start</td><td>end</td><td>strand</td><td>MM</td><td>target_seq</td><td>PAM</td><td>alignment</td><td>distance</td><td>position</td><td>gene name</td><td>gene id</td></tr>
<tr><td>GU305914.1</td><td>194718</td><td>194740</td><td>-</td><td>0</td><td>CGCCCGGGGAGAACTCCGTC</td><td>TGG</td><td>||||||||[||||||||||||]PAM</td><td>NA</td><td>Intergenic</td><td>NA</td><td>NA</td></tr>
<tr><td></td><td></td><td></td><td></td><td></td><td></td><td></td><td></td><td></td><td></td><td></td><td></td></tr>
<tr><td>T19908</td><td>GCCCGCCGCTCTGTCCCCACAGG</td><td>1000</td><td></td><td></td><td></td><td></td><td></td><td></td><td></td><td></td><td></td></tr>
<tr><td>Oligo fwd</td><td>CACCGCCCGCCGCTCTGTCCCCAC</td><td></td><td></td><td></td><td></td><td></td><td></td><td></td><td></td><td></td><td></td></tr>
<tr><td>Oligo rev</td><td>AAACGTGGGGACAGAGCGGCGGGC</td><td></td><td></td><td></td><td></td><td></td><td></td><td></td><td></td><td></td><td></td></tr>
<tr><td>Chromosome</td><td>start</td><td>end</td><td>strand</td><td>MM</td><td>target_seq</td><td>PAM</td><td>alignment</td><td>distance</td><td>position</td><td>gene name</td><td>gene id</td></tr>
<tr><td>GU305914.1</td><td>194692</td><td>194714</td><td>-</td><td>0</td><td>GCCCGCCGCTCTGTCCCCAC</td><td>AGG</td><td>||||||||[||||||||||||]PAM</td><td>NA</td><td>Intergenic</td><td>NA</td><td>NA</td></tr>
<tr><td></td><td></td><td></td><td></td><td></td><td></td><td></td><td></td><td></td><td></td><td></td><td></td></tr>
<tr><td>T19909</td><td>CCCGCCGCTCTGTCCCCACAGGG</td><td>1000</td><td></td><td></td><td></td><td></td><td></td><td></td><td></td><td></td><td></td></tr>
<tr><td>Oligo fwd</td><td>CACCCCCGCCGCTCTGTCCCCACA</td><td></td><td></td><td></td><td></td><td></td><td></td><td></td><td></td><td></td><td></td></tr>
<tr><td>Oligo rev</td><td>AAACTGTGGGGACAGAGCGGCGGG</td><td></td><td></td><td></td><td></td><td></td><td></td><td></td><td></td><td></td><td></td></tr>
<tr><td>Chromosome</td><td>start</td><td>end</td><td>strand</td><td>MM</td><td>target_seq</td><td>PAM</td><td>alignment</td><td>distance</td><td>position</td><td>gene name</td><td>gene id</td></tr>
<tr><td>GU305914.1</td><td>194691</td><td>194713</td><td>-</td><td>0</td><td>CCCGCCGCTCTGTCCCCACA</td><td>GGG</td><td>||||||||[||||||||||||]PAM</td><td>NA</td><td>Intergenic</td><td>NA</td><td>NA</td></tr>
<tr><td></td><td></td><td></td><td></td><td></td><td></td><td></td><td></td><td></td><td></td><td></td><td></td></tr>
<tr><td>T19910</td><td>GTCCCCACAGGGACGAAGACCGG</td><td>1000</td><td></td><td></td><td></td><td></td><td></td><td></td><td></td><td></td><td></td></tr>
<tr><td>Oligo fwd</td><td>CACCGTCCCCACAGGGACGAAGAC</td><td></td><td></td><td></td><td></td><td></td><td></td><td></td><td></td><td></td><td></td></tr>
<tr><td>Oligo rev</td><td>AAACGTCTTCGTCCCTGTGGGGAC</td><td></td><td></td><td></td><td></td><td></td><td></td><td></td><td></td><td></td><td></td></tr>
<tr><td>Chromosome</td><td>start</td><td>end</td><td>strand</td><td>MM</td><td>target_seq</td><td>PAM</td><td>alignment</td><td>distance</td><td>position</td><td>gene name</td><td>gene id</td></tr>
<tr><td>GU305914.1</td><td>194680</td><td>194702</td><td>-</td><td>0</td><td>GTCCCCACAGGGACGAAGAC</td><td>CGG</td><td>||||||||[||||||||||||]PAM</td><td>NA</td><td>Intergenic</td><td>NA</td><td>NA</td></tr>
<tr><td></td><td></td><td></td><td></td><td></td><td></td><td></td><td></td><td></td><td></td><td></td><td></td></tr>
<tr><td>T19911</td><td>CGCCACGTCGCTCGCGAGCCAGG</td><td>1000</td><td></td><td></td><td></td><td></td><td></td><td></td><td></td><td></td><td></td></tr>
<tr><td>Oligo fwd</td><td>CACCCGCCACGTCGCTCGCGAGCC</td><td></td><td></td><td></td><td></td><td></td><td></td><td></td><td></td><td></td><td></td></tr>
<tr><td>Oligo rev</td><td>AAACGGCTCGCGAGCGACGTGGCG</td><td></td><td></td><td></td><td></td><td></td><td></td><td></td><td></td><td></td><td></td></tr>
<tr><td>Chromosome</td><td>start</td><td>end</td><td>strand</td><td>MM</td><td>target_seq</td><td>PAM</td><td>alignment</td><td>distance</td><td>position</td><td>gene name</td><td>gene id</td></tr>
<tr><td>GU305914.1</td><td>194649</td><td>194671</td><td>-</td><td>0</td><td>CGCCACGTCGCTCGCGAGCC</td><td>AGG</td><td>||||||||[||||||||||||]PAM</td><td>NA</td><td>Intergenic</td><td>NA</td><td>NA</td></tr>
<tr><td></td><td></td><td></td><td></td><td></td><td></td><td></td><td></td><td></td><td></td><td></td><td></td></tr>
<tr><td>T19912</td><td>GGAGATCGCGCGTCGCGACCTGG</td><td>1000</td><td></td><td></td><td></td><td></td><td></td><td></td><td></td><td></td><td></td></tr>
<tr><td>Oligo fwd</td><td>CACCGGAGATCGCGCGTCGCGACC</td><td></td><td></td><td></td><td></td><td></td><td></td><td></td><td></td><td></td><td></td></tr>
<tr><td>Oligo rev</td><td>AAACGGTCGCGACGCGCGATCTCC</td><td></td><td></td><td></td><td></td><td></td><td></td><td></td><td></td><td></td><td></td></tr>
<tr><td>Chromosome</td><td>start</td><td>end</td><td>strand</td><td>MM</td><td>target_seq</td><td>PAM</td><td>alignment</td><td>distance</td><td>position</td><td>gene name</td><td>gene id</td></tr>
<tr><td>GU305914.1</td><td>194628</td><td>194650</td><td>-</td><td>0</td><td>GGAGATCGCGCGTCGCGACC</td><td>TGG</td><td>||||||||[||||||||||||]PAM</td><td>NA</td><td>Intergenic</td><td>NA</td><td>NA</td></tr>
<tr><td></td><td></td><td></td><td></td><td></td><td></td><td></td><td></td><td></td><td></td><td></td><td></td></tr>
<tr><td>T19913</td><td>CGCGTCGCGACCTGGAAGACTGG</td><td>1000</td><td></td><td></td><td></td><td></td><td></td><td></td><td></td><td></td><td></td></tr>
<tr><td>Oligo fwd</td><td>CACCCGCGTCGCGACCTGGAAGAC</td><td></td><td></td><td></td><td></td><td></td><td></td><td></td><td></td><td></td><td></td></tr>
<tr><td>Oligo rev</td><td>AAACGTCTTCCAGGTCGCGACGCG</td><td></td><td></td><td></td><td></td><td></td><td></td><td></td><td></td><td></td><td></td></tr>
<tr><td>Chromosome</td><td>start</td><td>end</td><td>strand</td><td>MM</td><td>target_seq</td><td>PAM</td><td>alignment</td><td>distance</td><td>position</td><td>gene name</td><td>gene id</td></tr>
<tr><td>GU305914.1</td><td>194620</td><td>194642</td><td>-</td><td>0</td><td>CGCGTCGCGACCTGGAAGAC</td><td>TGG</td><td>||||||||[||||||||||||]PAM</td><td>NA</td><td>Intergenic</td><td>NA</td><td>NA</td></tr>
<tr><td></td><td></td><td></td><td></td><td></td><td></td><td></td><td></td><td></td><td></td><td></td><td></td></tr>
<tr><td>T19914</td><td>ACCTGGAAGACTGGTACCGCTGG</td><td>1000</td><td></td><td></td><td></td><td></td><td></td><td></td><td></td><td></td><td></td></tr>
<tr><td>Oligo fwd</td><td>CACCACCTGGAAGACTGGTACCGC</td><td></td><td></td><td></td><td></td><td></td><td></td><td></td><td></td><td></td><td></td></tr>
<tr><td>Oligo rev</td><td>AAACGCGGTACCAGTCTTCCAGGT</td><td></td><td></td><td></td><td></td><td></td><td></td><td></td><td></td><td></td><td></td></tr>
<tr><td>Chromosome</td><td>start</td><td>end</td><td>strand</td><td>MM</td><td>target_seq</td><td>PAM</td><td>alignment</td><td>distance</td><td>position</td><td>gene name</td><td>gene id</td></tr>
<tr><td>GU305914.1</td><td>194611</td><td>194633</td><td>-</td><td>0</td><td>ACCTGGAAGACTGGTACCGC</td><td>TGG</td><td>||||||||[||||||||||||]PAM</td><td>NA</td><td>Intergenic</td><td>NA</td><td>NA</td></tr>
<tr><td></td><td></td><td></td><td></td><td></td><td></td><td></td><td></td><td></td><td></td><td></td><td></td></tr>
<tr><td>T19915</td><td>GACTGGTACCGCTGGCGACTCGG</td><td>1000</td><td></td><td></td><td></td><td></td><td></td><td></td><td></td><td></td><td></td></tr>
<tr><td>Oligo fwd</td><td>CACCGACTGGTACCGCTGGCGACT</td><td></td><td></td><td></td><td></td><td></td><td></td><td></td><td></td><td></td><td></td></tr>
<tr><td>Oligo rev</td><td>AAACAGTCGCCAGCGGTACCAGTC</td><td></td><td></td><td></td><td></td><td></td><td></td><td></td><td></td><td></td><td></td></tr>
<tr><td>Chromosome</td><td>start</td><td>end</td><td>strand</td><td>MM</td><td>target_seq</td><td>PAM</td><td>alignment</td><td>distance</td><td>position</td><td>gene name</td><td>gene id</td></tr>
<tr><td>GU305914.1</td><td>194603</td><td>194625</td><td>-</td><td>0</td><td>GACTGGTACCGCTGGCGACT</td><td>CGG</td><td>||||||||[||||||||||||]PAM</td><td>NA</td><td>Intergenic</td><td>NA</td><td>NA</td></tr>
<tr><td></td><td></td><td></td><td></td><td></td><td></td><td></td><td></td><td></td><td></td><td></td><td></td></tr>
<tr><td>T19916</td><td>GGCGACTCGGCGCCAGATCGCGG</td><td>1000</td><td></td><td></td><td></td><td></td><td></td><td></td><td></td><td></td><td></td></tr>
<tr><td>Oligo fwd</td><td>CACCGGCGACTCGGCGCCAGATCG</td><td></td><td></td><td></td><td></td><td></td><td></td><td></td><td></td><td></td><td></td></tr>
<tr><td>Oligo rev</td><td>AAACCGATCTGGCGCCGAGTCGCC</td><td></td><td></td><td></td><td></td><td></td><td></td><td></td><td></td><td></td><td></td></tr>
<tr><td>Chromosome</td><td>start</td><td>end</td><td>strand</td><td>MM</td><td>target_seq</td><td>PAM</td><td>alignment</td><td>distance</td><td>position</td><td>gene name</td><td>gene id</td></tr>
<tr><td>GU305914.1</td><td>194590</td><td>194612</td><td>-</td><td>0</td><td>GGCGACTCGGCGCCAGATCG</td><td>CGG</td><td>||||||||[||||||||||||]PAM</td><td>NA</td><td>Intergenic</td><td>NA</td><td>NA</td></tr>
<tr><td></td><td></td><td></td><td></td><td></td><td></td><td></td><td></td><td></td><td></td><td></td><td></td></tr>
<tr><td>T19917</td><td>CGGCGCCAGATCGCGGTTGCAGG</td><td>1000</td><td></td><td></td><td></td><td></td><td></td><td></td><td></td><td></td><td></td></tr>
<tr><td>Oligo fwd</td><td>CACCCGGCGCCAGATCGCGGTTGC</td><td></td><td></td><td></td><td></td><td></td><td></td><td></td><td></td><td></td><td></td></tr>
<tr><td>Oligo rev</td><td>AAACGCAACCGCGATCTGGCGCCG</td><td></td><td></td><td></td><td></td><td></td><td></td><td></td><td></td><td></td><td></td></tr>
<tr><td>Chromosome</td><td>start</td><td>end</td><td>strand</td><td>MM</td><td>target_seq</td><td>PAM</td><td>alignment</td><td>distance</td><td>position</td><td>gene name</td><td>gene id</td></tr>
<tr><td>GU305914.1</td><td>194583</td><td>194605</td><td>-</td><td>0</td><td>CGGCGCCAGATCGCGGTTGC</td><td>AGG</td><td>||||||||[||||||||||||]PAM</td><td>NA</td><td>Intergenic</td><td>NA</td><td>NA</td></tr>
<tr><td></td><td></td><td></td><td></td><td></td><td></td><td></td><td></td><td></td><td></td><td></td><td></td></tr>
<tr><td>T19918</td><td>GCCAGATCGCGGTTGCAGGACGG</td><td>1000</td><td></td><td></td><td></td><td></td><td></td><td></td><td></td><td></td><td></td></tr>
<tr><td>Oligo fwd</td><td>CACCGCCAGATCGCGGTTGCAGGA</td><td></td><td></td><td></td><td></td><td></td><td></td><td></td><td></td><td></td><td></td></tr>
<tr><td>Oligo rev</td><td>AAACTCCTGCAACCGCGATCTGGC</td><td></td><td></td><td></td><td></td><td></td><td></td><td></td><td></td><td></td><td></td></tr>
<tr><td>Chromosome</td><td>start</td><td>end</td><td>strand</td><td>MM</td><td>target_seq</td><td>PAM</td><td>alignment</td><td>distance</td><td>position</td><td>gene name</td><td>gene id</td></tr>
<tr><td>GU305914.1</td><td>194579</td><td>194601</td><td>-</td><td>0</td><td>GCCAGATCGCGGTTGCAGGA</td><td>CGG</td><td>||||||||[||||||||||||]PAM</td><td>NA</td><td>Intergenic</td><td>NA</td><td>NA</td></tr>
<tr><td></td><td></td><td></td><td></td><td></td><td></td><td></td><td></td><td></td><td></td><td></td><td></td></tr>
<tr><td>T19919</td><td>ATCGCGGTTGCAGGACGGCGTGG</td><td>1000</td><td></td><td></td><td></td><td></td><td></td><td></td><td></td><td></td><td></td></tr>
<tr><td>Oligo fwd</td><td>CACCATCGCGGTTGCAGGACGGCG</td><td></td><td></td><td></td><td></td><td></td><td></td><td></td><td></td><td></td><td></td></tr>
<tr><td>Oligo rev</td><td>AAACCGCCGTCCTGCAACCGCGAT</td><td></td><td></td><td></td><td></td><td></td><td></td><td></td><td></td><td></td><td></td></tr>
<tr><td>Chromosome</td><td>start</td><td>end</td><td>strand</td><td>MM</td><td>target_seq</td><td>PAM</td><td>alignment</td><td>distance</td><td>position</td><td>gene name</td><td>gene id</td></tr>
<tr><td>GU305914.1</td><td>194574</td><td>194596</td><td>-</td><td>0</td><td>ATCGCGGTTGCAGGACGGCG</td><td>TGG</td><td>||||||||[||||||||||||]PAM</td><td>NA</td><td>Intergenic</td><td>NA</td><td>NA</td></tr>
<tr><td></td><td></td><td></td><td></td><td></td><td></td><td></td><td></td><td></td><td></td><td></td><td></td></tr>
<tr><td>T19920</td><td>GGCCCTGTCGAACGTGAGCGAGG</td><td>1000</td><td></td><td></td><td></td><td></td><td></td><td></td><td></td><td></td><td></td></tr>
<tr><td>Oligo fwd</td><td>CACCGGCCCTGTCGAACGTGAGCG</td><td></td><td></td><td></td><td></td><td></td><td></td><td></td><td></td><td></td><td></td></tr>
</table>

```
Oligo rev     AAACCGCTCACGTTCGACAGGGCC
Chromosome    start                       end     strand  MM  target_seq            PAM  alignment                     distance  position    gene name  gene id
GU305914.1                      194553    194575  -        0  GGCCCTGTCGAACGTGAGCG  AGG  ||||||||[||||||||||||]PAM     NA        Intergenic  NA         NA

T19921        CGAACGTGAGCGAGGCGAAGAGG     1000
Oligo fwd     CACCCGAACGTGAGCGAGGCGAAG
Oligo rev     AAACCTTCGCCTCGCTCACGTTCG
Chromosome    start                       end     strand  MM  target_seq            PAM  alignment                     distance  position    gene name  gene id
GU305914.1                      194545    194567  -        0  CGAACGTGAGCGAGGCGAAG  AGG  ||||||||[||||||||||||]PAM     NA        Intergenic  NA         NA

T19922        CAGCTGACCCATCCCGCCAAAGG     1000
Oligo fwd     CACCCAGCTGACCCATCCCGCCAA
Oligo rev     AAACTTGGCGGGATGGGTCAGCTG
Chromosome    start                       end     strand  MM  target_seq            PAM  alignment                     distance  position    gene name  gene id
GU305914.1                      194522    194544  -        0  CAGCTGACCCATCCCGCCAA  AGG  ||||||||[||||||||||||]PAM     NA        Intergenic  NA         NA

T19923        AGCTGACCCATCCCGCCAAAGGG     1000
Oligo fwd     CACCAGCTGACCCATCCCGCCAAA
Oligo rev     AAACTTTGGCGGGATGGGTCAGCT
Chromosome    start                       end     strand  MM  target_seq            PAM  alignment                     distance  position    gene name  gene id
GU305914.1                      194521    194543  -        0  AGCTGACCCATCCCGCCAAA  GGG  ||||||||[||||||||||||]PAM     NA        Intergenic  NA         NA

T19924        GATCCCCGTCTCCATCGTGACGG     1000
Oligo fwd     CACCGATCCCCGTCTCCATCGTGA
Oligo rev     AAACTCACGATGGAGACGGGGATC
Chromosome    start                       end     strand  MM  target_seq            PAM  alignment                     distance  position    gene name  gene id
GU305914.1                      194499    194521  -        0  GATCCCCGTCTCCATCGTGA  CGG  ||||||||[||||||||||||]PAM     NA        Intergenic  NA         NA

T19925        CCCCGTCTCCATCGTGACGGCGG     1000
Oligo fwd     CACCCCCCGTCTCCATCGTGACGG
Oligo rev     AAACCCGTCACGATGGAGACGGGG
Chromosome    start                       end     strand  MM  target_seq            PAM  alignment                     distance  position    gene name  gene id
GU305914.1                      194496    194518  -        0  CCCCGTCTCCATCGTGACGG  CGG  ||||||||[||||||||||||]PAM     NA        Intergenic  NA         NA
GU305914.1                       19512     19534  -        4  CGCCGTCCCCATCGCGACGA  TGG  |-|||||-[||||||-||||-]PAM     NA        Intergenic  NA         NA

T19926        CGGCGGAGTCTTACCACCCCCGG     1000
Oligo fwd     CACCCGGCGGAGTCTTACCACCCC
Oligo rev     AAACGGGGTGGTAAGACTCCGCCG
Chromosome    start                       end     strand  MM  target_seq            PAM  alignment                     distance  position    gene name  gene id
GU305914.1                      194479    194501  -        0  CGGCGGAGTCTTACCACCCC  CGG  ||||||||[||||||||||||]PAM     NA        Intergenic  NA         NA

T19927        GGCGGAGTCTTACCACCCCCGGG     1000
Oligo fwd     CACCGGCGGAGTCTTACCACCCCC
Oligo rev     AAACGGGGGTGGTAAGACTCCGCC
Chromosome    start                       end     strand  MM  target_seq            PAM  alignment                     distance  position    gene name  gene id
GU305914.1                      194478    194500  -        0  GGCGGAGTCTTACCACCCCC  GGG  ||||||||[||||||||||||]PAM     NA        Intergenic  NA         NA

T19928        GCGGAGTCTTACCACCCCCGGGG     1000
Oligo fwd     CACCGCGGAGTCTTACCACCCCCG
Oligo rev     AAACCGGGGGTGGTAAGACTCCGC
Chromosome    start                       end     strand  MM  target_seq            PAM  alignment                     distance  position    gene name  gene id
GU305914.1                      194477    194499  -        0  GCGGAGTCTTACCACCCCCG  GGG  ||||||||[||||||||||||]PAM     NA        Intergenic  NA         NA

T19929        CGGAGTCTTACCACCCCCGGGGG     1000
Oligo fwd     CACCCGGAGTCTTACCACCCCCGG
Oligo rev     AAACCCGGGGGTGGTAAGACTCCG
Chromosome    start                       end     strand  MM  target_seq            PAM  alignment                     distance  position    gene name  gene id
GU305914.1                      194476    194498  -        0  CGGAGTCTTACCACCCCCGG  GGG  ||||||||[||||||||||||]PAM     NA        Intergenic  NA         NA

T19930        GGAGTCTTACCACCCCCGGGGGG     1000
Oligo fwd     CACCGGAGTCTTACCACCCCCGGG
Oligo rev     AAACCCCGGGGGTGGTAAGACTCC
Chromosome    start                       end     strand  MM  target_seq            PAM  alignment                     distance  position    gene name  gene id
GU305914.1                      194475    194497  -        0  GGAGTCTTACCACCCCCGGG  GGG  ||||||||[||||||||||||]PAM     NA        Intergenic  NA         NA

T19931        CCACCCCCGGGGGGATCCCGAGG     1000
Oligo fwd     CACCCCACCCCCGGGGGGATCCCG
Oligo rev     AAACCGGGATCCCCCCGGGGGTGG
Chromosome    start                       end     strand  MM  target_seq            PAM  alignment                     distance  position    gene name  gene id
GU305914.1                      194466    194488  -        0  CCACCCCCGGGGGGATCCCG  AGG  ||||||||[||||||||||||]PAM     NA        Intergenic  NA         NA

T19932        GGAAGCCTTCGAATCCCTGTCGG     1000
Oligo fwd     CACCGGAAGCCTTCGAATCCCTGT
Oligo rev     AAACACAGGGATTCGAAGGCTTCC
Chromosome    start                       end     strand  MM  target_seq            PAM  alignment                     distance  position    gene name  gene id
GU305914.1                      194445    194467  -        0  GGAAGCCTTCGAATCCCTGT  CGG  ||||||||[||||||||||||]PAM     NA        Intergenic  NA         NA

T19933        GAAGCCTTCGAATCCCTGTCGGG     1000
Oligo fwd     CACCGAAGCCTTCGAATCCCTGTC
Oligo rev     AAACGACAGGGATTCGAAGGCTTC
Chromosome    start                       end     strand  MM  target_seq            PAM  alignment                     distance  position    gene name  gene id
GU305914.1                      194444    194466  -        0  GAAGCCTTCGAATCCCTGTC  GGG  ||||||||[||||||||||||]PAM     NA        Intergenic  NA         NA

T19934        AATCCCTGTCGGGACAGACGAGG     1000
Oligo fwd     CACCAATCCCTGTCGGGACAGACG
Oligo rev     AAACCGTCTGTCCCGACAGGGATT
Chromosome    start                       end     strand  MM  target_seq            PAM  alignment                     distance  position    gene name  gene id
GU305914.1                      194434    194456  -        0  AATCCCTGTCGGGACAGACG  AGG  ||||||||[||||||||||||]PAM     NA        Intergenic  NA         NA
```

<table>
<tr><td>T19935</td><td>CGAGGCCGTACCGATACCCGCGG</td><td>1000</td><td></td><td></td><td></td><td></td><td></td><td></td><td></td><td></td><td></td></tr>
<tr><td>Oligo fwd</td><td>CACCCGAGGCCGTACCGATACCCG</td><td></td><td></td><td></td><td></td><td></td><td></td><td></td><td></td><td></td><td></td></tr>
<tr><td>Oligo rev</td><td>AAACCGGGTATCGGTACGGCCTCG</td><td></td><td></td><td></td><td></td><td></td><td></td><td></td><td></td><td></td><td></td></tr>
<tr><td>Chromosome</td><td>start</td><td>end</td><td>strand</td><td>MM</td><td>target_seq</td><td>PAM</td><td>alignment</td><td>distance</td><td>position</td><td>gene name</td><td>gene id</td></tr>
<tr><td>GU305914.1</td><td>194416</td><td>194438</td><td>-</td><td>0</td><td>CGAGGCCGTACCGATACCCG</td><td>CGG</td><td>||||||||[||||||||||||]PAM</td><td>NA</td><td>Intergenic</td><td>NA</td><td>NA</td></tr>
<tr><td></td><td></td><td></td><td></td><td></td><td></td><td></td><td></td><td></td><td></td><td></td><td></td></tr>
<tr><td>T19936</td><td>GAGGCCGTACCGATACCCGCGGG</td><td>1000</td><td></td><td></td><td></td><td></td><td></td><td></td><td></td><td></td><td></td></tr>
<tr><td>Oligo fwd</td><td>CACCGAGGCCGTACCGATACCCGC</td><td></td><td></td><td></td><td></td><td></td><td></td><td></td><td></td><td></td><td></td></tr>
<tr><td>Oligo rev</td><td>AAACGCGGGTATCGGTACGGCCTC</td><td></td><td></td><td></td><td></td><td></td><td></td><td></td><td></td><td></td><td></td></tr>
<tr><td>Chromosome</td><td>start</td><td>end</td><td>strand</td><td>MM</td><td>target_seq</td><td>PAM</td><td>alignment</td><td>distance</td><td>position</td><td>gene name</td><td>gene id</td></tr>
<tr><td>GU305914.1</td><td>194415</td><td>194437</td><td>-</td><td>0</td><td>GAGGCCGTACCGATACCCGC</td><td>GGG</td><td>||||||||[||||||||||||]PAM</td><td>NA</td><td>Intergenic</td><td>NA</td><td>NA</td></tr>
<tr><td></td><td></td><td></td><td></td><td></td><td></td><td></td><td></td><td></td><td></td><td></td><td></td></tr>
<tr><td>T19937</td><td>TCGTCGTTACGTCCGCGCCATGG</td><td>1000</td><td></td><td></td><td></td><td></td><td></td><td></td><td></td><td></td><td></td></tr>
<tr><td>Oligo fwd</td><td>CACCTCGTCGTTACGTCCGCGCCA</td><td></td><td></td><td></td><td></td><td></td><td></td><td></td><td></td><td></td><td></td></tr>
<tr><td>Oligo rev</td><td>AAACTGGCGCGGACGTAACGACGA</td><td></td><td></td><td></td><td></td><td></td><td></td><td></td><td></td><td></td><td></td></tr>
<tr><td>Chromosome</td><td>start</td><td>end</td><td>strand</td><td>MM</td><td>target_seq</td><td>PAM</td><td>alignment</td><td>distance</td><td>position</td><td>gene name</td><td>gene id</td></tr>
<tr><td>GU305914.1</td><td>194352</td><td>194374</td><td>-</td><td>0</td><td>TCGTCGTTACGTCCGCGCCA</td><td>TGG</td><td>||||||||[||||||||||||]PAM</td><td>NA</td><td>Intergenic</td><td>NA</td><td>NA</td></tr>
<tr><td></td><td></td><td></td><td></td><td></td><td></td><td></td><td></td><td></td><td></td><td></td><td></td></tr>
<tr><td>T19938</td><td>CGCGCCATGGACATGTTCGACGG</td><td>1000</td><td></td><td></td><td></td><td></td><td></td><td></td><td></td><td></td><td></td></tr>
<tr><td>Oligo fwd</td><td>CACCCGCGCCATGGACATGTTCGA</td><td></td><td></td><td></td><td></td><td></td><td></td><td></td><td></td><td></td><td></td></tr>
<tr><td>Oligo rev</td><td>AAACTCGAACATGTCCATGGCGCG</td><td></td><td></td><td></td><td></td><td></td><td></td><td></td><td></td><td></td><td></td></tr>
<tr><td>Chromosome</td><td>start</td><td>end</td><td>strand</td><td>MM</td><td>target_seq</td><td>PAM</td><td>alignment</td><td>distance</td><td>position</td><td>gene name</td><td>gene id</td></tr>
<tr><td>GU305914.1</td><td>194339</td><td>194361</td><td>-</td><td>0</td><td>CGCGCCATGGACATGTTCGA</td><td>CGG</td><td>||||||||[||||||||||||]PAM</td><td>NA</td><td>Intergenic</td><td>NA</td><td>NA</td></tr>
<tr><td></td><td></td><td></td><td></td><td></td><td></td><td></td><td></td><td></td><td></td><td></td><td></td></tr>
<tr><td>T19939</td><td>GCCATGGACATGTTCGACGGCGG</td><td>1000</td><td></td><td></td><td></td><td></td><td></td><td></td><td></td><td></td><td></td></tr>
<tr><td>Oligo fwd</td><td>CACCGCCATGGACATGTTCGACGG</td><td></td><td></td><td></td><td></td><td></td><td></td><td></td><td></td><td></td><td></td></tr>
<tr><td>Oligo rev</td><td>AAACCCGTCGAACATGTCCATGGC</td><td></td><td></td><td></td><td></td><td></td><td></td><td></td><td></td><td></td><td></td></tr>
<tr><td>Chromosome</td><td>start</td><td>end</td><td>strand</td><td>MM</td><td>target_seq</td><td>PAM</td><td>alignment</td><td>distance</td><td>position</td><td>gene name</td><td>gene id</td></tr>
<tr><td>GU305914.1</td><td>194336</td><td>194358</td><td>-</td><td>0</td><td>GCCATGGACATGTTCGACGG</td><td>CGG</td><td>||||||||[||||||||||||]PAM</td><td>NA</td><td>Intergenic</td><td>NA</td><td>NA</td></tr>
<tr><td>GU305914.1</td><td>163494</td><td>163516</td><td>+</td><td>4</td><td>GTCATGAACATGTTCCACCG</td><td>CGG</td><td>|-||||-|[|||||||-||-|]PAM</td><td>NA</td><td>Intergenic</td><td>NA</td><td>NA</td></tr>
<tr><td></td><td></td><td></td><td></td><td></td><td></td><td></td><td></td><td></td><td></td><td></td><td></td></tr>
<tr><td>T19940</td><td>GGACATGTTCGACGGCGGCAAGG</td><td>1000</td><td></td><td></td><td></td><td></td><td></td><td></td><td></td><td></td><td></td></tr>
<tr><td>Oligo fwd</td><td>CACCGGACATGTTCGACGGCGGCA</td><td></td><td></td><td></td><td></td><td></td><td></td><td></td><td></td><td></td><td></td></tr>
<tr><td>Oligo rev</td><td>AAACTGCCGCCGTCGAACATGTCC</td><td></td><td></td><td></td><td></td><td></td><td></td><td></td><td></td><td></td><td></td></tr>
<tr><td>Chromosome</td><td>start</td><td>end</td><td>strand</td><td>MM</td><td>target_seq</td><td>PAM</td><td>alignment</td><td>distance</td><td>position</td><td>gene name</td><td>gene id</td></tr>
<tr><td>GU305914.1</td><td>194331</td><td>194353</td><td>-</td><td>0</td><td>GGACATGTTCGACGGCGGCA</td><td>AGG</td><td>||||||||[||||||||||||]PAM</td><td>NA</td><td>Intergenic</td><td>NA</td><td>NA</td></tr>
<tr><td></td><td></td><td></td><td></td><td></td><td></td><td></td><td></td><td></td><td></td><td></td><td></td></tr>
<tr><td>T19941</td><td>AGCGCGAGATTATGTTCATCAGG</td><td>1000</td><td></td><td></td><td></td><td></td><td></td><td></td><td></td><td></td><td></td></tr>
<tr><td>Oligo fwd</td><td>CACCAGCGCGAGATTATGTTCATC</td><td></td><td></td><td></td><td></td><td></td><td></td><td></td><td></td><td></td><td></td></tr>
<tr><td>Oligo rev</td><td>AAACGATGAACATAATCTCGCGCT</td><td></td><td></td><td></td><td></td><td></td><td></td><td></td><td></td><td></td><td></td></tr>
<tr><td>Chromosome</td><td>start</td><td>end</td><td>strand</td><td>MM</td><td>target_seq</td><td>PAM</td><td>alignment</td><td>distance</td><td>position</td><td>gene name</td><td>gene id</td></tr>
<tr><td>GU305914.1</td><td>194299</td><td>194321</td><td>-</td><td>0</td><td>AGCGCGAGATTATGTTCATC</td><td>AGG</td><td>||||||||[||||||||||||]PAM</td><td>NA</td><td>Intergenic</td><td>NA</td><td>NA</td></tr>
<tr><td></td><td></td><td></td><td></td><td></td><td></td><td></td><td></td><td></td><td></td><td></td><td></td></tr>
<tr><td>T19942</td><td>GCGCGAGATTATGTTCATCAGGG</td><td>1000</td><td></td><td></td><td></td><td></td><td></td><td></td><td></td><td></td><td></td></tr>
<tr><td>Oligo fwd</td><td>CACCGCGCGAGATTATGTTCATCA</td><td></td><td></td><td></td><td></td><td></td><td></td><td></td><td></td><td></td><td></td></tr>
<tr><td>Oligo rev</td><td>AAACTGATGAACATAATCTCGCGC</td><td></td><td></td><td></td><td></td><td></td><td></td><td></td><td></td><td></td><td></td></tr>
<tr><td>Chromosome</td><td>start</td><td>end</td><td>strand</td><td>MM</td><td>target_seq</td><td>PAM</td><td>alignment</td><td>distance</td><td>position</td><td>gene name</td><td>gene id</td></tr>
<tr><td>GU305914.1</td><td>194298</td><td>194320</td><td>-</td><td>0</td><td>GCGCGAGATTATGTTCATCA</td><td>GGG</td><td>||||||||[||||||||||||]PAM</td><td>NA</td><td>Intergenic</td><td>NA</td><td>NA</td></tr>
<tr><td></td><td></td><td></td><td></td><td></td><td></td><td></td><td></td><td></td><td></td><td></td><td></td></tr>
<tr><td>T19943</td><td>TTATGTTCATCAGGGCCGAGAGG</td><td>1000</td><td></td><td></td><td></td><td></td><td></td><td></td><td></td><td></td><td></td></tr>
<tr><td>Oligo fwd</td><td>CACCTTATGTTCATCAGGGCCGAG</td><td></td><td></td><td></td><td></td><td></td><td></td><td></td><td></td><td></td><td></td></tr>
<tr><td>Oligo rev</td><td>AAACCTCGGCCCTGATGAACATAA</td><td></td><td></td><td></td><td></td><td></td><td></td><td></td><td></td><td></td><td></td></tr>
<tr><td>Chromosome</td><td>start</td><td>end</td><td>strand</td><td>MM</td><td>target_seq</td><td>PAM</td><td>alignment</td><td>distance</td><td>position</td><td>gene name</td><td>gene id</td></tr>
<tr><td>GU305914.1</td><td>194290</td><td>194312</td><td>-</td><td>0</td><td>TTATGTTCATCAGGGCCGAG</td><td>AGG</td><td>||||||||[||||||||||||]PAM</td><td>NA</td><td>Intergenic</td><td>NA</td><td>NA</td></tr>
<tr><td></td><td></td><td></td><td></td><td></td><td></td><td></td><td></td><td></td><td></td><td></td><td></td></tr>
<tr><td>T19944</td><td>TATGTTCATCAGGGCCGAGAGGG</td><td>1000</td><td></td><td></td><td></td><td></td><td></td><td></td><td></td><td></td><td></td></tr>
<tr><td>Oligo fwd</td><td>CACCTATGTTCATCAGGGCCGAGA</td><td></td><td></td><td></td><td></td><td></td><td></td><td></td><td></td><td></td><td></td></tr>
<tr><td>Oligo rev</td><td>AAACTCTCGGCCCTGATGAACATA</td><td></td><td></td><td></td><td></td><td></td><td></td><td></td><td></td><td></td><td></td></tr>
<tr><td>Chromosome</td><td>start</td><td>end</td><td>strand</td><td>MM</td><td>target_seq</td><td>PAM</td><td>alignment</td><td>distance</td><td>position</td><td>gene name</td><td>gene id</td></tr>
<tr><td>GU305914.1</td><td>194289</td><td>194311</td><td>-</td><td>0</td><td>TATGTTCATCAGGGCCGAGA</td><td>GGG</td><td>||||||||[||||||||||||]PAM</td><td>NA</td><td>Intergenic</td><td>NA</td><td>NA</td></tr>
<tr><td></td><td></td><td></td><td></td><td></td><td></td><td></td><td></td><td></td><td></td><td></td><td></td></tr>
<tr><td>T19945</td><td>TTCATCAGGGCCGAGAGGGTCGG</td><td>1000</td><td></td><td></td><td></td><td></td><td></td><td></td><td></td><td></td><td></td></tr>
<tr><td>Oligo fwd</td><td>CACCTTCATCAGGGCCGAGAGGGT</td><td></td><td></td><td></td><td></td><td></td><td></td><td></td><td></td><td></td><td></td></tr>
<tr><td>Oligo rev</td><td>AAACACCCTCTCGGCCCTGATGAA</td><td></td><td></td><td></td><td></td><td></td><td></td><td></td><td></td><td></td><td></td></tr>
<tr><td>Chromosome</td><td>start</td><td>end</td><td>strand</td><td>MM</td><td>target_seq</td><td>PAM</td><td>alignment</td><td>distance</td><td>position</td><td>gene name</td><td>gene id</td></tr>
<tr><td>GU305914.1</td><td>194285</td><td>194307</td><td>-</td><td>0</td><td>TTCATCAGGGCCGAGAGGGT</td><td>CGG</td><td>||||||||[||||||||||||]PAM</td><td>NA</td><td>Intergenic</td><td>NA</td><td>NA</td></tr>
<tr><td></td><td></td><td></td><td></td><td></td><td></td><td></td><td></td><td></td><td></td><td></td><td></td></tr>
<tr><td>T19946</td><td>GGTCGGCTCGCTCCAGAAGACGG</td><td>1000</td><td></td><td></td><td></td><td></td><td></td><td></td><td></td><td></td><td></td></tr>
<tr><td>Oligo fwd</td><td>CACCGGTCGGCTCGCTCCAGAAGA</td><td></td><td></td><td></td><td></td><td></td><td></td><td></td><td></td><td></td><td></td></tr>
<tr><td>Oligo rev</td><td>AAACTCTTCTGGAGCGAGCCGACC</td><td></td><td></td><td></td><td></td><td></td><td></td><td></td><td></td><td></td><td></td></tr>
<tr><td>Chromosome</td><td>start</td><td>end</td><td>strand</td><td>MM</td><td>target_seq</td><td>PAM</td><td>alignment</td><td>distance</td><td>position</td><td>gene name</td><td>gene id</td></tr>
<tr><td>GU305914.1</td><td>194268</td><td>194290</td><td>-</td><td>0</td><td>GGTCGGCTCGCTCCAGAAGA</td><td>CGG</td><td>||||||||[||||||||||||]PAM</td><td>NA</td><td>Intergenic</td><td>NA</td><td>NA</td></tr>
<tr><td></td><td></td><td></td><td></td><td></td><td></td><td></td><td></td><td></td><td></td><td></td><td></td></tr>
<tr><td>T19947</td><td>GTCGGCTCGCTCCAGAAGACGGG</td><td>1000</td><td></td><td></td><td></td><td></td><td></td><td></td><td></td><td></td><td></td></tr>
<tr><td>Oligo fwd</td><td>CACCGTCGGCTCGCTCCAGAAGAC</td><td></td><td></td><td></td><td></td><td></td><td></td><td></td><td></td><td></td><td></td></tr>
<tr><td>Oligo rev</td><td>AAACGTCTTCTGGAGCGAGCCGAC</td><td></td><td></td><td></td><td></td><td></td><td></td><td></td><td></td><td></td><td></td></tr>
<tr><td>Chromosome</td><td>start</td><td>end</td><td>strand</td><td>MM</td><td>target_seq</td><td>PAM</td><td>alignment</td><td>distance</td><td>position</td><td>gene name</td><td>gene id</td></tr>
<tr><td>GU305914.1</td><td>194267</td><td>194289</td><td>-</td><td>0</td><td>GTCGGCTCGCTCCAGAAGAC</td><td>GGG</td><td>||||||||[||||||||||||]PAM</td><td>NA</td><td>Intergenic</td><td>NA</td><td>NA</td></tr>
<tr><td></td><td></td><td></td><td></td><td></td><td></td><td></td><td></td><td></td><td></td><td></td><td></td></tr>
<tr><td>T19948</td><td>CCCATTGAAACTGCCACCTCTGG</td><td>1000</td><td></td><td></td><td></td><td></td><td></td><td></td><td></td><td></td><td></td></tr>
<tr><td>Oligo fwd</td><td>CACCCCCATTGAAACTGCCACCTC</td><td></td><td></td><td></td><td></td><td></td><td></td><td></td><td></td><td></td><td></td></tr>
<tr><td>Oligo rev</td><td>AAACGAGGTGGCAGTTTCAATGGG</td><td></td><td></td><td></td><td></td><td></td><td></td><td></td><td></td><td></td><td></td></tr>
<tr><td>Chromosome</td><td>start</td><td>end</td><td>strand</td><td>MM</td><td>target_seq</td><td>PAM</td><td>alignment</td><td>distance</td><td>position</td><td>gene name</td><td>gene id</td></tr>
<tr><td>GU305914.1</td><td>194241</td><td>194263</td><td>-</td><td>0</td><td>CCCATTGAAACTGCCACCTC</td><td>TGG</td><td>||||||||[||||||||||||]PAM</td><td>NA</td><td>Intergenic</td><td>NA</td><td>NA</td></tr>
<tr><td></td><td></td><td></td><td></td><td></td><td></td><td></td><td></td><td></td><td></td><td></td><td></td></tr>
<tr><td>T19949</td><td>ACCTCTGGTCCGCGCGACTCAGG</td><td>1000</td><td></td><td></td><td></td><td></td><td></td><td></td><td></td><td></td><td></td></tr>
<tr><td>Oligo fwd</td><td>CACCACCTCTGGTCCGCGCGACTC</td><td></td><td></td><td></td><td></td><td></td><td></td><td></td><td></td><td></td><td></td></tr>
<tr><td>Oligo rev</td><td>AAACGAGTCGCGCGGACCAGAGGT</td><td></td><td></td><td></td><td></td><td></td><td></td><td></td><td></td><td></td><td></td></tr>
<tr><td>Chromosome</td><td>start</td><td>end</td><td>strand</td><td>MM</td><td>target_seq</td><td>PAM</td><td>alignment</td><td>distance</td><td>position</td><td>gene name</td><td>gene id</td></tr>
</table>

| | | | | | | | | | | | |
|---|---|---|---|---|---|---|---|---|---|---|---|
| GU305914.1 | 194226 | 194248 | - | 0 | ACCTCTGGTCCGCGCGACTC | AGG | \|\|\|\|\|\|\|\|[\|\|\|\|\|\|\|\|\|\|\|\|]PAM | NA | Intergenic | NA | NA |
| T19950 | TCTGGTCCGCGCGACTCAGGCGG | 1000 | | | | | | | | | |
| Oligo fwd | CACCTCTGGTCCGCGCGACTCAGG | | | | | | | | | | |
| Oligo rev | AAACCCTGAGTCGCGCGGACCAGA | | | | | | | | | | |
| Chromosome | start | end | strand | MM | target_seq | PAM | alignment | distance | position | gene name | gene id |
| GU305914.1 | 194223 | 194245 | - | 0 | TCTGGTCCGCGCGACTCAGG | CGG | \|\|\|\|\|\|\|\|[\|\|\|\|\|\|\|\|\|\|\|\|]PAM | NA | Intergenic | NA | NA |
| T19951 | CTGGTCCGCGCGACTCAGGCGGG | 1000 | | | | | | | | | |
| Oligo fwd | CACCCTGGTCCGCGCGACTCAGGC | | | | | | | | | | |
| Oligo rev | AAACGCCTGAGTCGCGCGGACCAG | | | | | | | | | | |
| Chromosome | start | end | strand | MM | target_seq | PAM | alignment | distance | position | gene name | gene id |
| GU305914.1 | 194222 | 194244 | - | 0 | CTGGTCCGCGCGACTCAGGC | GGG | \|\|\|\|\|\|\|\|[\|\|\|\|\|\|\|\|\|\|\|\|]PAM | NA | Intergenic | NA | NA |
| T19952 | GGGATCGACGAGCAACAACATGG | 1000 | | | | | | | | | |
| Oligo fwd | CACCGGGATCGACGAGCAACAACA | | | | | | | | | | |
| Oligo rev | AAACTGTTGTTGCTCGTCGATCCC | | | | | | | | | | |
| Chromosome | start | end | strand | MM | target_seq | PAM | alignment | distance | position | gene name | gene id |
| GU305914.1 | 194202 | 194224 | - | 0 | GGGATCGACGAGCAACAACA | TGG | \|\|\|\|\|\|\|\|[\|\|\|\|\|\|\|\|\|\|\|\|]PAM | NA | Intergenic | NA | NA |
| T19953 | ACGAGCAACAACATGGAAACCGG | 1000 | | | | | | | | | |
| Oligo fwd | CACCACGAGCAACAACATGGAAAC | | | | | | | | | | |
| Oligo rev | AAACGTTTCCATGTTGTTGCTCGT | | | | | | | | | | |
| Chromosome | start | end | strand | MM | target_seq | PAM | alignment | distance | position | gene name | gene id |
| GU305914.1 | 194195 | 194217 | - | 0 | ACGAGCAACAACATGGAAAC | CGG | \|\|\|\|\|\|\|\|[\|\|\|\|\|\|\|\|\|\|\|\|]PAM | NA | Intergenic | NA | NA |
| T19954 | GGCTCCGATCGCTAGTCCAGCGG | 1000 | | | | | | | | | |
| Oligo fwd | CACCGGCTCCGATCGCTAGTCCAG | | | | | | | | | | |
| Oligo rev | AAACCTGGACTAGCGATCGGAGCC | | | | | | | | | | |
| Chromosome | start | end | strand | MM | target_seq | PAM | alignment | distance | position | gene name | gene id |
| GU305914.1 | 194174 | 194196 | - | 0 | GGCTCCGATCGCTAGTCCAG | CGG | \|\|\|\|\|\|\|\|[\|\|\|\|\|\|\|\|\|\|\|\|]PAM | NA | Intergenic | NA | NA |
| T19955 | GGCGCTGTGTCAGCCGCTCGCGG | 1000 | | | | | | | | | |
| Oligo fwd | CACCGGCGCTGTGTCAGCCGCTCG | | | | | | | | | | |
| Oligo rev | AAACCGAGCGGCTGACACAGCGCC | | | | | | | | | | |
| Chromosome | start | end | strand | MM | target_seq | PAM | alignment | distance | position | gene name | gene id |
| GU305914.1 | 194153 | 194175 | - | 0 | GGCGCTGTGTCAGCCGCTCG | CGG | \|\|\|\|\|\|\|\|[\|\|\|\|\|\|\|\|\|\|\|\|]PAM | NA | Intergenic | NA | NA |
| T19956 | GCGCTGTGTCAGCCGCTCGCGGG | 1000 | | | | | | | | | |
| Oligo fwd | CACCGCGCTGTGTCAGCCGCTCGC | | | | | | | | | | |
| Oligo rev | AAACGCGAGCGGCTGACACAGCGC | | | | | | | | | | |
| Chromosome | start | end | strand | MM | target_seq | PAM | alignment | distance | position | gene name | gene id |
| GU305914.1 | 194152 | 194174 | - | 0 | GCGCTGTGTCAGCCGCTCGC | GGG | \|\|\|\|\|\|\|\|[\|\|\|\|\|\|\|\|\|\|\|\|]PAM | NA | Intergenic | NA | NA |
| T19957 | CGCTGTGTCAGCCGCTCGCGGGG | 1000 | | | | | | | | | |
| Oligo fwd | CACCCGCTGTGTCAGCCGCTCGCG | | | | | | | | | | |
| Oligo rev | AAACCGCGAGCGGCTGACACAGCG | | | | | | | | | | |
| Chromosome | start | end | strand | MM | target_seq | PAM | alignment | distance | position | gene name | gene id |
| GU305914.1 | 194151 | 194173 | - | 0 | CGCTGTGTCAGCCGCTCGCG | GGG | \|\|\|\|\|\|\|\|[\|\|\|\|\|\|\|\|\|\|\|\|]PAM | NA | Intergenic | NA | NA |
| T19958 | GCTGTGTCAGCCGCTCGCGGGGG | 1000 | | | | | | | | | |
| Oligo fwd | CACCGCTGTGTCAGCCGCTCGCGG | | | | | | | | | | |
| Oligo rev | AAACCCGCGAGCGGCTGACACAGC | | | | | | | | | | |
| Chromosome | start | end | strand | MM | target_seq | PAM | alignment | distance | position | gene name | gene id |
| GU305914.1 | 194150 | 194172 | - | 0 | GCTGTGTCAGCCGCTCGCGG | GGG | \|\|\|\|\|\|\|\|[\|\|\|\|\|\|\|\|\|\|\|\|]PAM | NA | Intergenic | NA | NA |
| T19959 | CTGTGTCAGCCGCTCGCGGGGGG | 1000 | | | | | | | | | |
| Oligo fwd | CACCCTGTGTCAGCCGCTCGCGGG | | | | | | | | | | |
| Oligo rev | AAACCCCGCGAGCGGCTGACACAG | | | | | | | | | | |
| Chromosome | start | end | strand | MM | target_seq | PAM | alignment | distance | position | gene name | gene id |
| GU305914.1 | 194149 | 194171 | - | 0 | CTGTGTCAGCCGCTCGCGGG | GGG | \|\|\|\|\|\|\|\|[\|\|\|\|\|\|\|\|\|\|\|\|]PAM | NA | Intergenic | NA | NA |
| T19960 | TGTGTCAGCCGCTCGCGGGGGGG | 1000 | | | | | | | | | |
| Oligo fwd | CACCTGTGTCAGCCGCTCGCGGGG | | | | | | | | | | |
| Oligo rev | AAACCCCCGCGAGCGGCTGACACA | | | | | | | | | | |
| Chromosome | start | end | strand | MM | target_seq | PAM | alignment | distance | position | gene name | gene id |
| GU305914.1 | 194148 | 194170 | - | 0 | TGTGTCAGCCGCTCGCGGGG | GGG | \|\|\|\|\|\|\|\|[\|\|\|\|\|\|\|\|\|\|\|\|]PAM | NA | Intergenic | NA | NA |
| T19961 | CTCAAGTCGCCATCATCTCTCGG | 1000 | | | | | | | | | |
| Oligo fwd | CACCCTCAAGTCGCCATCATCTCT | | | | | | | | | | |
| Oligo rev | AAACAGAGATGATGGCGACTTGAG | | | | | | | | | | |
| Chromosome | start | end | strand | MM | target_seq | PAM | alignment | distance | position | gene name | gene id |
| GU305914.1 | 194060 | 194082 | - | 0 | CTCAAGTCGCCATCATCTCT | CGG | \|\|\|\|\|\|\|\|[\|\|\|\|\|\|\|\|\|\|\|\|]PAM | NA | Intergenic | NA | NA |
| T19962 | AGTCGCCATCATCTCTCGGTCGG | 1000 | | | | | | | | | |
| Oligo fwd | CACCAGTCGCCATCATCTCTCGGT | | | | | | | | | | |
| Oligo rev | AAACACCGAGAGATGATGGCGACT | | | | | | | | | | |
| Chromosome | start | end | strand | MM | target_seq | PAM | alignment | distance | position | gene name | gene id |
| GU305914.1 | 194056 | 194078 | - | 0 | AGTCGCCATCATCTCTCGGT | CGG | \|\|\|\|\|\|\|\|[\|\|\|\|\|\|\|\|\|\|\|\|]PAM | NA | Intergenic | NA | NA |
| T19963 | TCTCTCGGTCGGCAGAGCCGAGG | 1000 | | | | | | | | | |
| Oligo fwd | CACCTCTCTCGGTCGGCAGAGCCG | | | | | | | | | | |
| Oligo rev | AAACCGGCTCTGCCGACCGAGAGA | | | | | | | | | | |
| Chromosome | start | end | strand | MM | target_seq | PAM | alignment | distance | position | gene name | gene id |
| GU305914.1 | 194045 | 194067 | - | 0 | TCTCTCGGTCGGCAGAGCCG | AGG | \|\|\|\|\|\|\|\|[\|\|\|\|\|\|\|\|\|\|\|\|]PAM | NA | Intergenic | NA | NA |
| T19964 | GGTCGGCAGAGCCGAGGCGATGG | 1000 | | | | | | | | | |
| Oligo fwd | CACCGGTCGGCAGAGCCGAGGCGA | | | | | | | | | | |
| Oligo rev | AAACTCGCCTCGGCTCTGCCGACC | | | | | | | | | | |

<table>
<tr><td>Chromosome</td><td>start</td><td>end</td><td>strand</td><td>MM</td><td>target_seq</td><td>PAM</td><td>alignment</td><td>distance</td><td>position</td><td>gene name</td><td>gene id</td></tr>
<tr><td>GU305914.1</td><td>194039</td><td>194061</td><td>-</td><td>0</td><td>GGTCGGCAGAGCCGAGGCGA</td><td>TGG</td><td>||||||||[||||||||||||]PAM</td><td>NA</td><td>Intergenic</td><td>NA</td><td>NA</td></tr>
<tr><td></td><td></td><td></td><td></td><td></td><td></td><td></td><td></td><td></td><td></td><td></td><td></td></tr>
<tr><td>T19965</td><td>GATCGCACTCTGCCTCCTCGCGG</td><td>1000</td><td></td><td></td><td></td><td></td><td></td><td></td><td></td><td></td><td></td></tr>
<tr><td>Oligo fwd</td><td>CACCGATCGCACTCTGCCTCCTCG</td><td></td><td></td><td></td><td></td><td></td><td></td><td></td><td></td><td></td><td></td></tr>
<tr><td>Oligo rev</td><td>AAACCGAGGAGGCAGAGTGCGATC</td><td></td><td></td><td></td><td></td><td></td><td></td><td></td><td></td><td></td><td></td></tr>
<tr><td>Chromosome</td><td>start</td><td>end</td><td>strand</td><td>MM</td><td>target_seq</td><td>PAM</td><td>alignment</td><td>distance</td><td>position</td><td>gene name</td><td>gene id</td></tr>
<tr><td>GU305914.1</td><td>194003</td><td>194025</td><td>-</td><td>0</td><td>GATCGCACTCTGCCTCCTCG</td><td>CGG</td><td>||||||||[||||||||||||]PAM</td><td>NA</td><td>Intergenic</td><td>NA</td><td>NA</td></tr>
<tr><td></td><td></td><td></td><td></td><td></td><td></td><td></td><td></td><td></td><td></td><td></td><td></td></tr>
<tr><td>T19966</td><td>CTGCCTCCTCGCGGCCGTGACGG</td><td>1000</td><td></td><td></td><td></td><td></td><td></td><td></td><td></td><td></td><td></td></tr>
<tr><td>Oligo fwd</td><td>CACCCTGCCTCCTCGCGGCCGTGA</td><td></td><td></td><td></td><td></td><td></td><td></td><td></td><td></td><td></td><td></td></tr>
<tr><td>Oligo rev</td><td>AAACTCACGGCCGCGAGGAGGCAG</td><td></td><td></td><td></td><td></td><td></td><td></td><td></td><td></td><td></td><td></td></tr>
<tr><td>Chromosome</td><td>start</td><td>end</td><td>strand</td><td>MM</td><td>target_seq</td><td>PAM</td><td>alignment</td><td>distance</td><td>position</td><td>gene name</td><td>gene id</td></tr>
<tr><td>GU305914.1</td><td>193994</td><td>194016</td><td>-</td><td>0</td><td>CTGCCTCCTCGCGGCCGTGA</td><td>CGG</td><td>||||||||[||||||||||||]PAM</td><td>NA</td><td>Intergenic</td><td>NA</td><td>NA</td></tr>
<tr><td></td><td></td><td></td><td></td><td></td><td></td><td></td><td></td><td></td><td></td><td></td><td></td></tr>
<tr><td>T19967</td><td>ATCAATTACCTGCGTGCCAGTGG</td><td>1000</td><td></td><td></td><td></td><td></td><td></td><td></td><td></td><td></td><td></td></tr>
<tr><td>Oligo fwd</td><td>CACCATCAATTACCTGCGTGCCAG</td><td></td><td></td><td></td><td></td><td></td><td></td><td></td><td></td><td></td><td></td></tr>
<tr><td>Oligo rev</td><td>AAACCTGGCACGCAGGTAATTGAT</td><td></td><td></td><td></td><td></td><td></td><td></td><td></td><td></td><td></td><td></td></tr>
<tr><td>Chromosome</td><td>start</td><td>end</td><td>strand</td><td>MM</td><td>target_seq</td><td>PAM</td><td>alignment</td><td>distance</td><td>position</td><td>gene name</td><td>gene id</td></tr>
<tr><td>GU305914.1</td><td>193968</td><td>193990</td><td>-</td><td>0</td><td>ATCAATTACCTGCGTGCCAG</td><td>TGG</td><td>||||||||[||||||||||||]PAM</td><td>NA</td><td>Intergenic</td><td>NA</td><td>NA</td></tr>
<tr><td></td><td></td><td></td><td></td><td></td><td></td><td></td><td></td><td></td><td></td><td></td><td></td></tr>
<tr><td>T19968</td><td>GTTCAATATCTCCTGCGACCTGG</td><td>1000</td><td></td><td></td><td></td><td></td><td></td><td></td><td></td><td></td><td></td></tr>
<tr><td>Oligo fwd</td><td>CACCGTTCAATATCTCCTGCGACC</td><td></td><td></td><td></td><td></td><td></td><td></td><td></td><td></td><td></td><td></td></tr>
<tr><td>Oligo rev</td><td>AAACGGTCGCAGGAGATATTGAAC</td><td></td><td></td><td></td><td></td><td></td><td></td><td></td><td></td><td></td><td></td></tr>
<tr><td>Chromosome</td><td>start</td><td>end</td><td>strand</td><td>MM</td><td>target_seq</td><td>PAM</td><td>alignment</td><td>distance</td><td>position</td><td>gene name</td><td>gene id</td></tr>
<tr><td>GU305914.1</td><td>193919</td><td>193941</td><td>-</td><td>0</td><td>GTTCAATATCTCCTGCGACC</td><td>TGG</td><td>||||||||[||||||||||||]PAM</td><td>NA</td><td>Intergenic</td><td>NA</td><td>NA</td></tr>
<tr><td></td><td></td><td></td><td></td><td></td><td></td><td></td><td></td><td></td><td></td><td></td><td></td></tr>
<tr><td>T19969</td><td>ATCTCCTGCGACCTGGTGAACGG</td><td>1000</td><td></td><td></td><td></td><td></td><td></td><td></td><td></td><td></td><td></td></tr>
<tr><td>Oligo fwd</td><td>CACCATCTCCTGCGACCTGGTGAA</td><td></td><td></td><td></td><td></td><td></td><td></td><td></td><td></td><td></td><td></td></tr>
<tr><td>Oligo rev</td><td>AAACTTCACCAGGTCGCAGGAGAT</td><td></td><td></td><td></td><td></td><td></td><td></td><td></td><td></td><td></td><td></td></tr>
<tr><td>Chromosome</td><td>start</td><td>end</td><td>strand</td><td>MM</td><td>target_seq</td><td>PAM</td><td>alignment</td><td>distance</td><td>position</td><td>gene name</td><td>gene id</td></tr>
<tr><td>GU305914.1</td><td>193912</td><td>193934</td><td>-</td><td>0</td><td>ATCTCCTGCGACCTGGTGAA</td><td>CGG</td><td>||||||||[||||||||||||]PAM</td><td>NA</td><td>Intergenic</td><td>NA</td><td>NA</td></tr>
<tr><td></td><td></td><td></td><td></td><td></td><td></td><td></td><td></td><td></td><td></td><td></td><td></td></tr>
<tr><td>T19970</td><td>CGACCTGGTGAACGGCTCCACGG</td><td>1000</td><td></td><td></td><td></td><td></td><td></td><td></td><td></td><td></td><td></td></tr>
<tr><td>Oligo fwd</td><td>CACCCGACCTGGTGAACGGCTCCA</td><td></td><td></td><td></td><td></td><td></td><td></td><td></td><td></td><td></td><td></td></tr>
<tr><td>Oligo rev</td><td>AAACTGGAGCCGTTCACCAGGTCG</td><td></td><td></td><td></td><td></td><td></td><td></td><td></td><td></td><td></td><td></td></tr>
<tr><td>Chromosome</td><td>start</td><td>end</td><td>strand</td><td>MM</td><td>target_seq</td><td>PAM</td><td>alignment</td><td>distance</td><td>position</td><td>gene name</td><td>gene id</td></tr>
<tr><td>GU305914.1</td><td>193904</td><td>193926</td><td>-</td><td>0</td><td>CGACCTGGTGAACGGCTCCA</td><td>CGG</td><td>||||||||[||||||||||||]PAM</td><td>NA</td><td>Intergenic</td><td>NA</td><td>NA</td></tr>
<tr><td></td><td></td><td></td><td></td><td></td><td></td><td></td><td></td><td></td><td></td><td></td><td></td></tr>
<tr><td>T19971</td><td>GGTGAACGGCTCCACGGTCAAGG</td><td>1000</td><td></td><td></td><td></td><td></td><td></td><td></td><td></td><td></td><td></td></tr>
<tr><td>Oligo fwd</td><td>CACCGGTGAACGGCTCCACGGTCA</td><td></td><td></td><td></td><td></td><td></td><td></td><td></td><td></td><td></td><td></td></tr>
<tr><td>Oligo rev</td><td>AAACTGACCGTGGAGCCGTTCACC</td><td></td><td></td><td></td><td></td><td></td><td></td><td></td><td></td><td></td><td></td></tr>
<tr><td>Chromosome</td><td>start</td><td>end</td><td>strand</td><td>MM</td><td>target_seq</td><td>PAM</td><td>alignment</td><td>distance</td><td>position</td><td>gene name</td><td>gene id</td></tr>
<tr><td>GU305914.1</td><td>193898</td><td>193920</td><td>-</td><td>0</td><td>GGTGAACGGCTCCACGGTCA</td><td>AGG</td><td>||||||||[||||||||||||]PAM</td><td>NA</td><td>Intergenic</td><td>NA</td><td>NA</td></tr>
<tr><td></td><td></td><td></td><td></td><td></td><td></td><td></td><td></td><td></td><td></td><td></td><td></td></tr>
<tr><td>T19972</td><td>ACGGCTCCACGGTCAAGGTCTGG</td><td>1000</td><td></td><td></td><td></td><td></td><td></td><td></td><td></td><td></td><td></td></tr>
<tr><td>Oligo fwd</td><td>CACCACGGCTCCACGGTCAAGGTC</td><td></td><td></td><td></td><td></td><td></td><td></td><td></td><td></td><td></td><td></td></tr>
<tr><td>Oligo rev</td><td>AAACGACCTTGACCGTGGAGCCGT</td><td></td><td></td><td></td><td></td><td></td><td></td><td></td><td></td><td></td><td></td></tr>
<tr><td>Chromosome</td><td>start</td><td>end</td><td>strand</td><td>MM</td><td>target_seq</td><td>PAM</td><td>alignment</td><td>distance</td><td>position</td><td>gene name</td><td>gene id</td></tr>
<tr><td>GU305914.1</td><td>193893</td><td>193915</td><td>-</td><td>0</td><td>ACGGCTCCACGGTCAAGGTC</td><td>TGG</td><td>||||||||[||||||||||||]PAM</td><td>NA</td><td>Intergenic</td><td>NA</td><td>NA</td></tr>
<tr><td></td><td></td><td></td><td></td><td></td><td></td><td></td><td></td><td></td><td></td><td></td><td></td></tr>
<tr><td>T19973</td><td>AAGGTCTGGCGAGACCTCTGCGG</td><td>1000</td><td></td><td></td><td></td><td></td><td></td><td></td><td></td><td></td><td></td></tr>
<tr><td>Oligo fwd</td><td>CACCAAGGTCTGGCGAGACCTCTG</td><td></td><td></td><td></td><td></td><td></td><td></td><td></td><td></td><td></td><td></td></tr>
<tr><td>Oligo rev</td><td>AAACCAGAGGTCTCGCCAGACCTT</td><td></td><td></td><td></td><td></td><td></td><td></td><td></td><td></td><td></td><td></td></tr>
<tr><td>Chromosome</td><td>start</td><td>end</td><td>strand</td><td>MM</td><td>target_seq</td><td>PAM</td><td>alignment</td><td>distance</td><td>position</td><td>gene name</td><td>gene id</td></tr>
<tr><td>GU305914.1</td><td>193879</td><td>193901</td><td>-</td><td>0</td><td>AAGGTCTGGCGAGACCTCTG</td><td>CGG</td><td>||||||||[||||||||||||]PAM</td><td>NA</td><td>Intergenic</td><td>NA</td><td>NA</td></tr>
<tr><td></td><td></td><td></td><td></td><td></td><td></td><td></td><td></td><td></td><td></td><td></td><td></td></tr>
<tr><td>T19974</td><td>TGGCGAGACCTCTGCGGCGAAGG</td><td>1000</td><td></td><td></td><td></td><td></td><td></td><td></td><td></td><td></td><td></td></tr>
<tr><td>Oligo fwd</td><td>CACCTGGCGAGACCTCTGCGGCGA</td><td></td><td></td><td></td><td></td><td></td><td></td><td></td><td></td><td></td><td></td></tr>
<tr><td>Oligo rev</td><td>AAACTCGCCGCAGAGGTCTCGCCA</td><td></td><td></td><td></td><td></td><td></td><td></td><td></td><td></td><td></td><td></td></tr>
<tr><td>Chromosome</td><td>start</td><td>end</td><td>strand</td><td>MM</td><td>target_seq</td><td>PAM</td><td>alignment</td><td>distance</td><td>position</td><td>gene name</td><td>gene id</td></tr>
<tr><td>GU305914.1</td><td>193873</td><td>193895</td><td>-</td><td>0</td><td>TGGCGAGACCTCTGCGGCGA</td><td>AGG</td><td>||||||||[||||||||||||]PAM</td><td>NA</td><td>Intergenic</td><td>NA</td><td>NA</td></tr>
<tr><td></td><td></td><td></td><td></td><td></td><td></td><td></td><td></td><td></td><td></td><td></td><td></td></tr>
<tr><td>T19975</td><td>CGGCGAAGGCAATGCTACCGAGG</td><td>1000</td><td></td><td></td><td></td><td></td><td></td><td></td><td></td><td></td><td></td></tr>
<tr><td>Oligo fwd</td><td>CACCCGGCGAAGGCAATGCTACCG</td><td></td><td></td><td></td><td></td><td></td><td></td><td></td><td></td><td></td><td></td></tr>
<tr><td>Oligo rev</td><td>AAACCGGTAGCATTGCCTTCGCCG</td><td></td><td></td><td></td><td></td><td></td><td></td><td></td><td></td><td></td><td></td></tr>
<tr><td>Chromosome</td><td>start</td><td>end</td><td>strand</td><td>MM</td><td>target_seq</td><td>PAM</td><td>alignment</td><td>distance</td><td>position</td><td>gene name</td><td>gene id</td></tr>
<tr><td>GU305914.1</td><td>193859</td><td>193881</td><td>-</td><td>0</td><td>CGGCGAAGGCAATGCTACCG</td><td>AGG</td><td>||||||||[||||||||||||]PAM</td><td>NA</td><td>Intergenic</td><td>NA</td><td>NA</td></tr>
<tr><td></td><td></td><td></td><td></td><td></td><td></td><td></td><td></td><td></td><td></td><td></td><td></td></tr>
<tr><td>T19976</td><td>GGCGAAGGCAATGCTACCGAGGG</td><td>1000</td><td></td><td></td><td></td><td></td><td></td><td></td><td></td><td></td><td></td></tr>
<tr><td>Oligo fwd</td><td>CACCGGCGAAGGCAATGCTACCGA</td><td></td><td></td><td></td><td></td><td></td><td></td><td></td><td></td><td></td><td></td></tr>
<tr><td>Oligo rev</td><td>AAACTCGGTAGCATTGCCTTCGCC</td><td></td><td></td><td></td><td></td><td></td><td></td><td></td><td></td><td></td><td></td></tr>
<tr><td>Chromosome</td><td>start</td><td>end</td><td>strand</td><td>MM</td><td>target_seq</td><td>PAM</td><td>alignment</td><td>distance</td><td>position</td><td>gene name</td><td>gene id</td></tr>
<tr><td>GU305914.1</td><td>193858</td><td>193880</td><td>-</td><td>0</td><td>GGCGAAGGCAATGCTACCGA</td><td>GGG</td><td>||||||||[||||||||||||]PAM</td><td>NA</td><td>Intergenic</td><td>NA</td><td>NA</td></tr>
<tr><td></td><td></td><td></td><td></td><td></td><td></td><td></td><td></td><td></td><td></td><td></td><td></td></tr>
<tr><td>T19977</td><td>AAGGCAATGCTACCGAGGGATGG</td><td>1000</td><td></td><td></td><td></td><td></td><td></td><td></td><td></td><td></td><td></td></tr>
<tr><td>Oligo fwd</td><td>CACCAAGGCAATGCTACCGAGGGA</td><td></td><td></td><td></td><td></td><td></td><td></td><td></td><td></td><td></td><td></td></tr>
<tr><td>Oligo rev</td><td>AAACTCCCTCGGTAGCATTGCCTT</td><td></td><td></td><td></td><td></td><td></td><td></td><td></td><td></td><td></td><td></td></tr>
<tr><td>Chromosome</td><td>start</td><td>end</td><td>strand</td><td>MM</td><td>target_seq</td><td>PAM</td><td>alignment</td><td>distance</td><td>position</td><td>gene name</td><td>gene id</td></tr>
<tr><td>GU305914.1</td><td>193854</td><td>193876</td><td>-</td><td>0</td><td>AAGGCAATGCTACCGAGGGA</td><td>TGG</td><td>||||||||[||||||||||||]PAM</td><td>NA</td><td>Intergenic</td><td>NA</td><td>NA</td></tr>
<tr><td></td><td></td><td></td><td></td><td></td><td></td><td></td><td></td><td></td><td></td><td></td><td></td></tr>
<tr><td>T19978</td><td>AATGCTACCGAGGGATGGATCGG</td><td>1000</td><td></td><td></td><td></td><td></td><td></td><td></td><td></td><td></td><td></td></tr>
<tr><td>Oligo fwd</td><td>CACCAATGCTACCGAGGGATGGAT</td><td></td><td></td><td></td><td></td><td></td><td></td><td></td><td></td><td></td><td></td></tr>
<tr><td>Oligo rev</td><td>AAACATCCATCCCTCGGTAGCATT</td><td></td><td></td><td></td><td></td><td></td><td></td><td></td><td></td><td></td><td></td></tr>
<tr><td>Chromosome</td><td>start</td><td>end</td><td>strand</td><td>MM</td><td>target_seq</td><td>PAM</td><td>alignment</td><td>distance</td><td>position</td><td>gene name</td><td>gene id</td></tr>
<tr><td>GU305914.1</td><td>193849</td><td>193871</td><td>-</td><td>0</td><td>AATGCTACCGAGGGATGGAT</td><td>CGG</td><td>||||||||[||||||||||||]PAM</td><td>NA</td><td>Intergenic</td><td>NA</td><td>NA</td></tr>
<tr><td></td><td></td><td></td><td></td><td></td><td></td><td></td><td></td><td></td><td></td><td></td><td></td></tr>
<tr><td>T19979</td><td>ATGCTACCGAGGGATGGATCGGG</td><td>1000</td><td></td><td></td><td></td><td></td><td></td><td></td><td></td><td></td><td></td></tr>
<tr><td>Oligo fwd</td><td>CACCATGCTACCGAGGGATGGATC</td><td></td><td></td><td></td><td></td><td></td><td></td><td></td><td></td><td></td><td></td></tr>
</table>

Oligo rev AAACGATCCATCCCTCGGTAGCAT
Chromosome start end strand MM target_seq PAM alignment distance position gene name gene id
GU305914.1 193848 193870 - 0 ATGCTACCGAGGGATGGATC GGG ||||||||[||||||||||||]PAM NA Intergenic NA NA

T19980 TACCGAGGGATGGATCGGGATGG 1000
Oligo fwd CACCTACCGAGGGATGGATCGGGA
Oligo rev AAACTCCCGATCCATCCCTCGGTA
Chromosome start end strand MM target_seq PAM alignment distance position gene name gene id
GU305914.1 193844 193866 - 0 TACCGAGGGATGGATCGGGA TGG ||||||||[||||||||||||]PAM NA Intergenic NA NA

T19981 AGGGATGGATCGGGATGGCCCGG 1000
Oligo fwd CACCAGGGATGGATCGGGATGGCC
Oligo rev AAACGGCCATCCCGATCCATCCCT
Chromosome start end strand MM target_seq PAM alignment distance position gene name gene id
GU305914.1 193839 193861 - 0 AGGGATGGATCGGGATGGCC CGG ||||||||[||||||||||||]PAM NA Intergenic NA NA

T19982 TGGATCGGGATGGCCCGGCGCGG 1000
Oligo fwd CACCTGGATCGGGATGGCCCGGCG
Oligo rev AAACCGCCGGGCCATCCCGATCCA
Chromosome start end strand MM target_seq PAM alignment distance position gene name gene id
GU305914.1 193834 193856 - 0 TGGATCGGGATGGCCCGGCG CGG ||||||||[||||||||||||]PAM NA Intergenic NA NA

T19983 ACAAATTACATTTCTCCGCCAGG 1000
Oligo fwd CACCACAAATTACATTTCTCCGCC
Oligo rev AAACGGCGGAGAAATGTAATTTGT
Chromosome start end strand MM target_seq PAM alignment distance position gene name gene id
GU305914.1 193809 193831 - 0 ACAAATTACATTTCTCCGCC AGG ||||||||[||||||||||||]PAM NA Intergenic NA NA

T19984 AATTACATTTCTCCGCCAGGTGG 1000
Oligo fwd CACCAATTACATTTCTCCGCCAGG
Oligo rev AAACCCTGGCGGAGAAATGTAATT
Chromosome start end strand MM target_seq PAM alignment distance position gene name gene id
GU305914.1 193806 193828 - 0 AATTACATTTCTCCGCCAGG TGG ||||||||[||||||||||||]PAM NA Intergenic NA NA

T19985 AACTACTCGACCGCCGACATCGG 1000
Oligo fwd CACCAACTACTCGACCGCCGACAT
Oligo rev AAACATGTCGGCGGTCGAGTAGTT
Chromosome start end strand MM target_seq PAM alignment distance position gene name gene id
GU305914.1 193783 193805 - 0 AACTACTCGACCGCCGACAT CGG ||||||||[||||||||||||]PAM NA Intergenic NA NA

T19986 ACTACTCGACCGCCGACATCGGG 1000
Oligo fwd CACCACTACTCGACCGCCGACATC
Oligo rev AAACGATGTCGGCGGTCGAGTAGT
Chromosome start end strand MM target_seq PAM alignment distance position gene name gene id
GU305914.1 193782 193804 - 0 ACTACTCGACCGCCGACATC GGG ||||||||[||||||||||||]PAM NA Intergenic NA NA

T19987 CTACTCGACCGCCGACATCGGGG 1000
Oligo fwd CACCCTACTCGACCGCCGACATCG
Oligo rev AAACCGATGTCGGCGGTCGAGTAG
Chromosome start end strand MM target_seq PAM alignment distance position gene name gene id
GU305914.1 193781 193803 - 0 CTACTCGACCGCCGACATCG GGG ||||||||[||||||||||||]PAM NA Intergenic NA NA

T19988 TCACCCCACCGATGACGATCTGG 1000
Oligo fwd CACCTCACCCCACCGATGACGATC
Oligo rev AAACGATCGTCATCGGTGGGGTGA
Chromosome start end strand MM target_seq PAM alignment distance position gene name gene id
GU305914.1 193704 193726 - 0 TCACCCCACCGATGACGATC TGG ||||||||[||||||||||||]PAM NA Intergenic NA NA

T19989 CACCCCACCGATGACGATCTGGG 1000
Oligo fwd CACCCACCCCACCGATGACGATCT
Oligo rev AAACAGATCGTCATCGGTGGGGTG
Chromosome start end strand MM target_seq PAM alignment distance position gene name gene id
GU305914.1 193703 193725 - 0 CACCCCACCGATGACGATCT GGG ||||||||[||||||||||||]PAM NA Intergenic NA NA

T19990 ACCCCACCGATGACGATCTGGGG 1000
Oligo fwd CACCACCCCACCGATGACGATCTG
Oligo rev AAACCAGATCGTCATCGGTGGGGT
Chromosome start end strand MM target_seq PAM alignment distance position gene name gene id
GU305914.1 193702 193724 - 0 ACCCCACCGATGACGATCTG GGG ||||||||[||||||||||||]PAM NA Intergenic NA NA

T19991 CGATGACGATCTGGGGCATGAGG 1000
Oligo fwd CACCCGATGACGATCTGGGGCATG
Oligo rev AAACCATGCCCCAGATCGTCATCG
Chromosome start end strand MM target_seq PAM alignment distance position gene name gene id
GU305914.1 193695 193717 - 0 CGATGACGATCTGGGGCATG AGG ||||||||[||||||||||||]PAM NA Intergenic NA NA

T19992 GACGATCTGGGGCATGAGGCAGG 1000
Oligo fwd CACCGACGATCTGGGGCATGAGGC
Oligo rev AAACGCCTCATGCCCCAGATCGTC
Chromosome start end strand MM target_seq PAM alignment distance position gene name gene id
GU305914.1 193691 193713 - 0 GACGATCTGGGGCATGAGGC AGG ||||||||[||||||||||||]PAM NA Intergenic NA NA

T19993 ACGATCTGGGGCATGAGGCAGGG 1000
Oligo fwd CACCACGATCTGGGGCATGAGGCA
Oligo rev AAACTGCCTCATGCCCCAGATCGT
Chromosome start end strand MM target_seq PAM alignment distance position gene name gene id
GU305914.1 193690 193712 - 0 ACGATCTGGGGCATGAGGCA GGG ||||||||[||||||||||||]PAM NA Intergenic NA NA

T19994 ATCTGGGGCATGAGGCAGGGCGG 1000

Oligo fwd CACCATCTGGGGCATGAGGCAGGG
Oligo rev AAACCCCTGCCTCATGCCCCAGAT
Chromosome start end strand MM target_seq PAM alignment distance position gene name gene id
GU305914.1 193687 193709 - 0 ATCTGGGGCATGAGGCAGGG CGG ||||||||[||||||||||||]PAM NA Intergenic NA NA

T19995 CTTTGTCCTCTGCACCGTGACGG 1000
Oligo fwd CACCCTTTGTCCTCTGCACCGTGA
Oligo rev AAACTCACGGTGCAGAGGACAAAG
Chromosome start end strand MM target_seq PAM alignment distance position gene name gene id
GU305914.1 193652 193674 - 0 CTTTGTCCTCTGCACCGTGA CGG ||||||||[||||||||||||]PAM NA Intergenic NA NA

T19996 TTTGTCCTCTGCACCGTGACGGG 1000
Oligo fwd CACCTTTGTCCTCTGCACCGTGAC
Oligo rev AAACGTCACGGTGCAGAGGACAAA
Chromosome start end strand MM target_seq PAM alignment distance position gene name gene id
GU305914.1 193651 193673 - 0 TTTGTCCTCTGCACCGTGAC GGG ||||||||[||||||||||||]PAM NA Intergenic NA NA

T19997 CGTGACGGGCATCTTCTCGAAGG 1000
Oligo fwd CACCCGTGACGGGCATCTTCTCGA
Oligo rev AAACTCGAGAAGATGCCCGTCACG
Chromosome start end strand MM target_seq PAM alignment distance position gene name gene id
GU305914.1 193637 193659 - 0 CGTGACGGGCATCTTCTCGA AGG ||||||||[||||||||||||]PAM NA Intergenic NA NA

T19998 GTGACGGGCATCTTCTCGAAGGG 1000
Oligo fwd CACCGTGACGGGCATCTTCTCGAA
Oligo rev AAACTTCGAGAAGATGCCCGTCAC
Chromosome start end strand MM target_seq PAM alignment distance position gene name gene id
GU305914.1 193636 193658 - 0 GTGACGGGCATCTTCTCGAA GGG ||||||||[||||||||||||]PAM NA Intergenic NA NA

T19999 GAAGGGTGAATTCACACTCGAGG 1000
Oligo fwd CACCGAAGGGTGAATTCACACTCG
Oligo rev AAACCGAGTGTGAATTCACCCTTC
Chromosome start end strand MM target_seq PAM alignment distance position gene name gene id
GU305914.1 193619 193641 - 0 GAAGGGTGAATTCACACTCG AGG ||||||||[||||||||||||]PAM NA Intergenic NA NA

T20000 GAATTCACACTCGAGGCGAACGG 1000
Oligo fwd CACCGAATTCACACTCGAGGCGAA
Oligo rev AAACTTCGCCTCGAGTGTGAATTC
Chromosome start end strand MM target_seq PAM alignment distance position gene name gene id
GU305914.1 193612 193634 - 0 GAATTCACACTCGAGGCGAA CGG ||||||||[||||||||||||]PAM NA Intergenic NA NA

T20001 ACTCGAGGCGAACGGCGCGACGG 1000
Oligo fwd CACCACTCGAGGCGAACGGCGCGA
Oligo rev AAACTCGCGCCGTTCGCCTCGAGT
Chromosome start end strand MM target_seq PAM alignment distance position gene name gene id
GU305914.1 193604 193626 - 0 ACTCGAGGCGAACGGCGCGA CGG ||||||||[||||||||||||]PAM NA Intergenic NA NA

T20002 TCTGTCAACTTCACAAAACCCGG 1000
Oligo fwd CACCTCTGTCAACTTCACAAAACC
Oligo rev AAACGGTTTTGTGAAGTTGACAGA
Chromosome start end strand MM target_seq PAM alignment distance position gene name gene id
GU305914.1 193576 193598 - 0 TCTGTCAACTTCACAAAACC CGG ||||||||[||||||||||||]PAM NA Intergenic NA NA
GU305914.1 205082 205104 + 4 TCTGTCCCATTCACAAAAGC CGG ||||||--[-|||||||||-|]PAM NA Intergenic NA NA

T20003 CTAGACCTCTCACATCGCAACGG 1000
Oligo fwd CACCCTAGACCTCTCACATCGCAA
Oligo rev AAACTTGCGATGTGAGAGGTCTAG
Chromosome start end strand MM target_seq PAM alignment distance position gene name gene id
GU305914.1 193522 193544 - 0 CTAGACCTCTCACATCGCAA CGG ||||||||[||||||||||||]PAM NA Intergenic NA NA

T20004 CGATTTCTCGTCTCGATCAGAGG 1000
Oligo fwd CACCCGATTTCTCGTCTCGATCAG
Oligo rev AAACCTGATCGAGACGAGAAATCG
Chromosome start end strand MM target_seq PAM alignment distance position gene name gene id
GU305914.1 193492 193514 - 0 CGATTTCTCGTCTCGATCAG AGG ||||||||[||||||||||||]PAM NA Intergenic NA NA

T20005 GATTTCTCGTCTCGATCAGAGGG 1000
Oligo fwd CACCGATTTCTCGTCTCGATCAGA
Oligo rev AAACTCTGATCGAGACGAGAAATC
Chromosome start end strand MM target_seq PAM alignment distance position gene name gene id
GU305914.1 193491 193513 - 0 GATTTCTCGTCTCGATCAGA GGG ||||||||[||||||||||||]PAM NA Intergenic NA NA

T20006 ATTTCTCGTCTCGATCAGAGGGG 1000
Oligo fwd CACCATTTCTCGTCTCGATCAGAG
Oligo rev AAACCTCTGATCGAGACGAGAAAT
Chromosome start end strand MM target_seq PAM alignment distance position gene name gene id
GU305914.1 193490 193512 - 0 ATTTCTCGTCTCGATCAGAG GGG ||||||||[||||||||||||]PAM NA Intergenic NA NA

T20007 TTTCTCGTCTCGATCAGAGGGGG 1000
Oligo fwd CACCTTTCTCGTCTCGATCAGAGG
Oligo rev AAACCCTCTGATCGAGACGAGAAA
Chromosome start end strand MM target_seq PAM alignment distance position gene name gene id
GU305914.1 193489 193511 - 0 TTTCTCGTCTCGATCAGAGG GGG ||||||||[||||||||||||]PAM NA Intergenic NA NA

T20008 TTCTCGTCTCGATCAGAGGGGGG 1000
Oligo fwd CACCTTCTCGTCTCGATCAGAGGG
Oligo rev AAACCCCTCTGATCGAGACGAGAA
Chromosome start end strand MM target_seq PAM alignment distance position gene name gene id
GU305914.1 193488 193510 - 0 TTCTCGTCTCGATCAGAGGG GGG ||||||||[||||||||||||]PAM NA Intergenic NA NA

```
T20009          TCTCTCGTGCGTTCCTTTTGCGG              1000
Oligo fwd       CACCTCTCTCGTGCGTTCCTTTTG
Oligo rev       AAACCAAAAGGAACGCACGAGAGA
Chromosome      start          end     strand  MM  target_seq              PAM  alignment                    distance  position    gene name  gene id
GU305914.1             193427  193449  -        0  TCTCTCGTGCGTTCCTTTTG    CGG  ||||||||[||||||||||||]PAM    NA        Intergenic  NA         NA

T20010          GGTCAGCAAGCCTCTGATGTTGG              1000
Oligo fwd       CACCGGTCAGCAAGCCTCTGATGT
Oligo rev       AAACACATCAGAGGCTTGCTGACC
Chromosome      start          end     strand  MM  target_seq              PAM  alignment                    distance  position    gene name  gene id
GU305914.1             193406  193428  -        0  GGTCAGCAAGCCTCTGATGT    TGG  ||||||||[||||||||||||]PAM    NA        Intergenic  NA         NA

T20011          ATACAGACAGACCATGCCATCGG              1000
Oligo fwd       CACCATACAGACAGACCATGCCAT
Oligo rev       AAACATGGCATGGTCTGTCTGTAT
Chromosome      start          end     strand  MM  target_seq              PAM  alignment                    distance  position    gene name  gene id
GU305914.1             193382  193404  -        0  ATACAGACAGACCATGCCAT    CGG  ||||||||[||||||||||||]PAM    NA        Intergenic  NA         NA

T20012          TACAGACAGACCATGCCATCGGG              1000
Oligo fwd       CACCTACAGACAGACCATGCCATC
Oligo rev       AAACGATGGCATGGTCTGTCTGTA
Chromosome      start          end     strand  MM  target_seq              PAM  alignment                    distance  position    gene name  gene id
GU305914.1             193381  193403  -        0  TACAGACAGACCATGCCATC    GGG  ||||||||[||||||||||||]PAM    NA        Intergenic  NA         NA

T20013          CCATGCCATCGGGAGTGAAAAGG              1000
Oligo fwd       CACCCCATGCCATCGGGAGTGAAA
Oligo rev       AAACTTTCACTCCCGATGGCATGG
Chromosome      start          end     strand  MM  target_seq              PAM  alignment                    distance  position    gene name  gene id
GU305914.1             193371  193393  -        0  CCATGCCATCGGGAGTGAAA    AGG  ||||||||[||||||||||||]PAM    NA        Intergenic  NA         NA

T20014          GGGAGTGAAAAGGTACACCAAGG              1000
Oligo fwd       CACCGGGAGTGAAAAGGTACACCA
Oligo rev       AAACTGGTGTACCTTTTCACTCCC
Chromosome      start          end     strand  MM  target_seq              PAM  alignment                    distance  position    gene name  gene id
GU305914.1             193361  193383  -        0  GGGAGTGAAAAGGTACACCA    AGG  ||||||||[||||||||||||]PAM    NA        Intergenic  NA         NA

T20015          AGGTACACCAAGGACTCCACCGG              1000
Oligo fwd       CACCAGGTACACCAAGGACTCCAC
Oligo rev       AAACGTGGAGTCCTTGGTGTACCT
Chromosome      start          end     strand  MM  target_seq              PAM  alignment                    distance  position    gene name  gene id
GU305914.1             193351  193373  -        0  AGGTACACCAAGGACTCCAC    CGG  ||||||||[||||||||||||]PAM    NA        Intergenic  NA         NA

T20016          GGTACACCAAGGACTCCACCGGG              1000
Oligo fwd       CACCGGTACACCAAGGACTCCACC
Oligo rev       AAACGGTGGAGTCCTTGGTGTACC
Chromosome      start          end     strand  MM  target_seq              PAM  alignment                    distance  position    gene name  gene id
GU305914.1             193350  193372  -        0  GGTACACCAAGGACTCCACC    GGG  ||||||||[||||||||||||]PAM    NA        Intergenic  NA         NA

T20017          AGCTCGATGATTACCATAACTGG              1000
Oligo fwd       CACCAGCTCGATGATTACCATAAC
Oligo rev       AAACGTTATGGTAATCATCGAGCT
Chromosome      start          end     strand  MM  target_seq              PAM  alignment                    distance  position    gene name  gene id
GU305914.1             193299  193321  -        0  AGCTCGATGATTACCATAAC    TGG  ||||||||[||||||||||||]PAM    NA        Intergenic  NA         NA

T20018          GCTCGATGATTACCATAACTGGG              1000
Oligo fwd       CACCGCTCGATGATTACCATAACT
Oligo rev       AAACAGTTATGGTAATCATCGAGC
Chromosome      start          end     strand  MM  target_seq              PAM  alignment                    distance  position    gene name  gene id
GU305914.1             193298  193320  -        0  GCTCGATGATTACCATAACT    GGG  ||||||||[||||||||||||]PAM    NA        Intergenic  NA         NA

T20019          GTACATGTTCTACATAGCCGCGG              1000
Oligo fwd       CACCGTACATGTTCTACATAGCCG
Oligo rev       AAACCGGCTATGTAGAACATGTAC
Chromosome      start          end     strand  MM  target_seq              PAM  alignment                    distance  position    gene name  gene id
GU305914.1             193193  193215  -        0  GTACATGTTCTACATAGCCG    CGG  ||||||||[||||||||||||]PAM    NA        Intergenic  NA         NA

T20020          CATGTTCTACATAGCCGCGGTGG              1000
Oligo fwd       CACCCATGTTCTACATAGCCGCGG
Oligo rev       AAACCCGCGGCTATGTAGAACATG
Chromosome      start          end     strand  MM  target_seq              PAM  alignment                    distance  position    gene name  gene id
GU305914.1             193190  193212  -        0  CATGTTCTACATAGCCGCGG    TGG  ||||||||[||||||||||||]PAM    NA        Intergenic  NA         NA

T20021          AGCCGCGGTGGAAAAGCCGCTGG              1000
Oligo fwd       CACCAGCCGCGGTGGAAAAGCCGC
Oligo rev       AAACGCGGCTTTTCCACCGCGGCT
Chromosome      start          end     strand  MM  target_seq              PAM  alignment                    distance  position    gene name  gene id
GU305914.1             193178  193200  -        0  AGCCGCGGTGGAAAAGCCGC    TGG  ||||||||[||||||||||||]PAM    NA        Intergenic  NA         NA

T20022          GGTGGAAAAGCCGCTGGAATCGG              1000
Oligo fwd       CACCGGTGGAAAAGCCGCTGGAAT
Oligo rev       AAACATTCCAGCGGCTTTTCCACC
Chromosome      start          end     strand  MM  target_seq              PAM  alignment                    distance  position    gene name  gene id
GU305914.1             193172  193194  -        0  GGTGGAAAAGCCGCTGGAAT    CGG  ||||||||[||||||||||||]PAM    NA        Intergenic  NA         NA

T20023          TGCTGTACGACGTCGACACATGG              1000
Oligo fwd       CACCTGCTGTACGACGTCGACACA
Oligo rev       AAACTGTGTCGACGTCGTACAGCA
Chromosome      start          end     strand  MM  target_seq              PAM  alignment                    distance  position    gene name  gene id
```

<table>
<tr><td>GU305914.1</td><td>193143</td><td>193165</td><td>-</td><td>0</td><td>TGCTGTACGACGTCGACACA</td><td>TGG</td><td>||||||||[||||||||||||]PAM</td><td>NA</td><td>Intergenic</td><td>NA</td><td>NA</td></tr>
<tr><td></td><td></td><td></td><td></td><td></td><td></td><td></td><td></td><td></td><td></td><td></td><td></td></tr>
<tr><td>T20024</td><td>AACTACGCTTTCACCTCCAGTGG</td><td>1000</td><td></td><td></td><td></td><td></td><td></td><td></td><td></td><td></td><td></td></tr>
<tr><td>Oligo fwd</td><td>CACCAACTACGCTTTCACCTCCAG</td><td></td><td></td><td></td><td></td><td></td><td></td><td></td><td></td><td></td><td></td></tr>
<tr><td>Oligo rev</td><td>AAACCTGGAGGTGAAAGCGTAGTT</td><td></td><td></td><td></td><td></td><td></td><td></td><td></td><td></td><td></td><td></td></tr>
<tr><td>Chromosome</td><td>start</td><td>end</td><td>strand</td><td>MM</td><td>target_seq</td><td>PAM</td><td>alignment</td><td>distance</td><td>position</td><td>gene name</td><td>gene id</td></tr>
<tr><td>GU305914.1</td><td>193120</td><td>193142</td><td>-</td><td>0</td><td>AACTACGCTTTCACCTCCAG</td><td>TGG</td><td>||||||||[||||||||||||]PAM</td><td>NA</td><td>Intergenic</td><td>NA</td><td>NA</td></tr>
<tr><td></td><td></td><td></td><td></td><td></td><td></td><td></td><td></td><td></td><td></td><td></td><td></td></tr>
<tr><td>T20025</td><td>CATACAAACATCTAAAAAGACGG</td><td>1000</td><td></td><td></td><td></td><td></td><td></td><td></td><td></td><td></td><td></td></tr>
<tr><td>Oligo fwd</td><td>CACCCATACAAACATCTAAAAAGA</td><td></td><td></td><td></td><td></td><td></td><td></td><td></td><td></td><td></td><td></td></tr>
<tr><td>Oligo rev</td><td>AAACTCTTTTTAGATGTTTGTATG</td><td></td><td></td><td></td><td></td><td></td><td></td><td></td><td></td><td></td><td></td></tr>
<tr><td>Chromosome</td><td>start</td><td>end</td><td>strand</td><td>MM</td><td>target_seq</td><td>PAM</td><td>alignment</td><td>distance</td><td>position</td><td>gene name</td><td>gene id</td></tr>
<tr><td>GU305914.1</td><td>193091</td><td>193113</td><td>-</td><td>0</td><td>CATACAAACATCTAAAAAGA</td><td>CGG</td><td>||||||||[||||||||||||]PAM</td><td>NA</td><td>Intergenic</td><td>NA</td><td>NA</td></tr>
<tr><td></td><td></td><td></td><td></td><td></td><td></td><td></td><td></td><td></td><td></td><td></td><td></td></tr>
<tr><td>T20026</td><td>ATACAAACATCTAAAAAGACGGG</td><td>1000</td><td></td><td></td><td></td><td></td><td></td><td></td><td></td><td></td><td></td></tr>
<tr><td>Oligo fwd</td><td>CACCATACAAACATCTAAAAAGAC</td><td></td><td></td><td></td><td></td><td></td><td></td><td></td><td></td><td></td><td></td></tr>
<tr><td>Oligo rev</td><td>AAACGTCTTTTTAGATGTTTGTAT</td><td></td><td></td><td></td><td></td><td></td><td></td><td></td><td></td><td></td><td></td></tr>
<tr><td>Chromosome</td><td>start</td><td>end</td><td>strand</td><td>MM</td><td>target_seq</td><td>PAM</td><td>alignment</td><td>distance</td><td>position</td><td>gene name</td><td>gene id</td></tr>
<tr><td>GU305914.1</td><td>193090</td><td>193112</td><td>-</td><td>0</td><td>ATACAAACATCTAAAAAGAC</td><td>GGG</td><td>||||||||[||||||||||||]PAM</td><td>NA</td><td>Intergenic</td><td>NA</td><td>NA</td></tr>
<tr><td></td><td></td><td></td><td></td><td></td><td></td><td></td><td></td><td></td><td></td><td></td><td></td></tr>
<tr><td>T20027</td><td>ACATCTAAAAAGACGGGTGTAGG</td><td>1000</td><td></td><td></td><td></td><td></td><td></td><td></td><td></td><td></td><td></td></tr>
<tr><td>Oligo fwd</td><td>CACCACATCTAAAAAGACGGGTGT</td><td></td><td></td><td></td><td></td><td></td><td></td><td></td><td></td><td></td><td></td></tr>
<tr><td>Oligo rev</td><td>AAACACACCCGTCTTTTTAGATGT</td><td></td><td></td><td></td><td></td><td></td><td></td><td></td><td></td><td></td><td></td></tr>
<tr><td>Chromosome</td><td>start</td><td>end</td><td>strand</td><td>MM</td><td>target_seq</td><td>PAM</td><td>alignment</td><td>distance</td><td>position</td><td>gene name</td><td>gene id</td></tr>
<tr><td>GU305914.1</td><td>193084</td><td>193106</td><td>-</td><td>0</td><td>ACATCTAAAAAGACGGGTGT</td><td>AGG</td><td>||||||||[||||||||||||]PAM</td><td>NA</td><td>Intergenic</td><td>NA</td><td>NA</td></tr>
<tr><td></td><td></td><td></td><td></td><td></td><td></td><td></td><td></td><td></td><td></td><td></td><td></td></tr>
<tr><td>T20028</td><td>GACGGGTGTAGGCCTCCCCCCGG</td><td>1000</td><td></td><td></td><td></td><td></td><td></td><td></td><td></td><td></td><td></td></tr>
<tr><td>Oligo fwd</td><td>CACCGACGGGTGTAGGCCTCCCCC</td><td></td><td></td><td></td><td></td><td></td><td></td><td></td><td></td><td></td><td></td></tr>
<tr><td>Oligo rev</td><td>AAACGGGGGAGGCCTACACCCGTC</td><td></td><td></td><td></td><td></td><td></td><td></td><td></td><td></td><td></td><td></td></tr>
<tr><td>Chromosome</td><td>start</td><td>end</td><td>strand</td><td>MM</td><td>target_seq</td><td>PAM</td><td>alignment</td><td>distance</td><td>position</td><td>gene name</td><td>gene id</td></tr>
<tr><td>GU305914.1</td><td>193073</td><td>193095</td><td>-</td><td>0</td><td>GACGGGTGTAGGCCTCCCCC</td><td>CGG</td><td>||||||||[||||||||||||]PAM</td><td>NA</td><td>Intergenic</td><td>NA</td><td>NA</td></tr>
<tr><td></td><td></td><td></td><td></td><td></td><td></td><td></td><td></td><td></td><td></td><td></td><td></td></tr>
<tr><td>T20029</td><td>ACGGGTGTAGGCCTCCCCCCGGG</td><td>1000</td><td></td><td></td><td></td><td></td><td></td><td></td><td></td><td></td><td></td></tr>
<tr><td>Oligo fwd</td><td>CACCACGGGTGTAGGCCTCCCCCC</td><td></td><td></td><td></td><td></td><td></td><td></td><td></td><td></td><td></td><td></td></tr>
<tr><td>Oligo rev</td><td>AAACGGGGGGAGGCCTACACCCGT</td><td></td><td></td><td></td><td></td><td></td><td></td><td></td><td></td><td></td><td></td></tr>
<tr><td>Chromosome</td><td>start</td><td>end</td><td>strand</td><td>MM</td><td>target_seq</td><td>PAM</td><td>alignment</td><td>distance</td><td>position</td><td>gene name</td><td>gene id</td></tr>
<tr><td>GU305914.1</td><td>193072</td><td>193094</td><td>-</td><td>0</td><td>ACGGGTGTAGGCCTCCCCCC</td><td>GGG</td><td>||||||||[||||||||||||]PAM</td><td>NA</td><td>Intergenic</td><td>NA</td><td>NA</td></tr>
<tr><td></td><td></td><td></td><td></td><td></td><td></td><td></td><td></td><td></td><td></td><td></td><td></td></tr>
<tr><td>T20030</td><td>TGTATCTTCGAGAGAGAGAGTGG</td><td>1000</td><td></td><td></td><td></td><td></td><td></td><td></td><td></td><td></td><td></td></tr>
<tr><td>Oligo fwd</td><td>CACCTGTATCTTCGAGAGAGAGAG</td><td></td><td></td><td></td><td></td><td></td><td></td><td></td><td></td><td></td><td></td></tr>
<tr><td>Oligo rev</td><td>AAACCTCTCTCTCTCGAAGATACA</td><td></td><td></td><td></td><td></td><td></td><td></td><td></td><td></td><td></td><td></td></tr>
<tr><td>Chromosome</td><td>start</td><td>end</td><td>strand</td><td>MM</td><td>target_seq</td><td>PAM</td><td>alignment</td><td>distance</td><td>position</td><td>gene name</td><td>gene id</td></tr>
<tr><td>GU305914.1</td><td>193039</td><td>193061</td><td>-</td><td>0</td><td>TGTATCTTCGAGAGAGAGAG</td><td>TGG</td><td>||||||||[||||||||||||]PAM</td><td>NA</td><td>Intergenic</td><td>NA</td><td>NA</td></tr>
<tr><td></td><td></td><td></td><td></td><td></td><td></td><td></td><td></td><td></td><td></td><td></td><td></td></tr>
<tr><td>T20031</td><td>AGAGAGAGTGGATCGTTATACGG</td><td>1000</td><td></td><td></td><td></td><td></td><td></td><td></td><td></td><td></td><td></td></tr>
<tr><td>Oligo fwd</td><td>CACCAGAGAGAGTGGATCGTTATA</td><td></td><td></td><td></td><td></td><td></td><td></td><td></td><td></td><td></td><td></td></tr>
<tr><td>Oligo rev</td><td>AAACTATAACGATCCACTCTCTCT</td><td></td><td></td><td></td><td></td><td></td><td></td><td></td><td></td><td></td><td></td></tr>
<tr><td>Chromosome</td><td>start</td><td>end</td><td>strand</td><td>MM</td><td>target_seq</td><td>PAM</td><td>alignment</td><td>distance</td><td>position</td><td>gene name</td><td>gene id</td></tr>
<tr><td>GU305914.1</td><td>193027</td><td>193049</td><td>-</td><td>0</td><td>AGAGAGAGTGGATCGTTATA</td><td>CGG</td><td>||||||||[||||||||||||]PAM</td><td>NA</td><td>Intergenic</td><td>NA</td><td>NA</td></tr>
<tr><td></td><td></td><td></td><td></td><td></td><td></td><td></td><td></td><td></td><td></td><td></td><td></td></tr>
<tr><td>T20032</td><td>AGAGTGGATCGTTATACGGCCGG</td><td>1000</td><td></td><td></td><td></td><td></td><td></td><td></td><td></td><td></td><td></td></tr>
<tr><td>Oligo fwd</td><td>CACCAGAGTGGATCGTTATACGGC</td><td></td><td></td><td></td><td></td><td></td><td></td><td></td><td></td><td></td><td></td></tr>
<tr><td>Oligo rev</td><td>AAACGCCGTATAACGATCCACTCT</td><td></td><td></td><td></td><td></td><td></td><td></td><td></td><td></td><td></td><td></td></tr>
<tr><td>Chromosome</td><td>start</td><td>end</td><td>strand</td><td>MM</td><td>target_seq</td><td>PAM</td><td>alignment</td><td>distance</td><td>position</td><td>gene name</td><td>gene id</td></tr>
<tr><td>GU305914.1</td><td>193023</td><td>193045</td><td>-</td><td>0</td><td>AGAGTGGATCGTTATACGGC</td><td>CGG</td><td>||||||||[||||||||||||]PAM</td><td>NA</td><td>Intergenic</td><td>NA</td><td>NA</td></tr>
<tr><td></td><td></td><td></td><td></td><td></td><td></td><td></td><td></td><td></td><td></td><td></td><td></td></tr>
<tr><td>T20033</td><td>GTTATACGGCCGGTCGCTGATGG</td><td>1000</td><td></td><td></td><td></td><td></td><td></td><td></td><td></td><td></td><td></td></tr>
<tr><td>Oligo fwd</td><td>CACCGTTATACGGCCGGTCGCTGA</td><td></td><td></td><td></td><td></td><td></td><td></td><td></td><td></td><td></td><td></td></tr>
<tr><td>Oligo rev</td><td>AAACTCAGCGACCGGCCGTATAAC</td><td></td><td></td><td></td><td></td><td></td><td></td><td></td><td></td><td></td><td></td></tr>
<tr><td>Chromosome</td><td>start</td><td>end</td><td>strand</td><td>MM</td><td>target_seq</td><td>PAM</td><td>alignment</td><td>distance</td><td>position</td><td>gene name</td><td>gene id</td></tr>
<tr><td>GU305914.1</td><td>193013</td><td>193035</td><td>-</td><td>0</td><td>GTTATACGGCCGGTCGCTGA</td><td>TGG</td><td>||||||||[||||||||||||]PAM</td><td>NA</td><td>Intergenic</td><td>NA</td><td>NA</td></tr>
<tr><td></td><td></td><td></td><td></td><td></td><td></td><td></td><td></td><td></td><td></td><td></td><td></td></tr>
<tr><td>T20034</td><td>GGCCGGTCGCTGATGGTACACGG</td><td>1000</td><td></td><td></td><td></td><td></td><td></td><td></td><td></td><td></td><td></td></tr>
<tr><td>Oligo fwd</td><td>CACCGGCCGGTCGCTGATGGTACA</td><td></td><td></td><td></td><td></td><td></td><td></td><td></td><td></td><td></td><td></td></tr>
<tr><td>Oligo rev</td><td>AAACTGTACCATCAGCGACCGGCC</td><td></td><td></td><td></td><td></td><td></td><td></td><td></td><td></td><td></td><td></td></tr>
<tr><td>Chromosome</td><td>start</td><td>end</td><td>strand</td><td>MM</td><td>target_seq</td><td>PAM</td><td>alignment</td><td>distance</td><td>position</td><td>gene name</td><td>gene id</td></tr>
<tr><td>GU305914.1</td><td>193006</td><td>193028</td><td>-</td><td>0</td><td>GGCCGGTCGCTGATGGTACA</td><td>CGG</td><td>||||||||[||||||||||||]PAM</td><td>NA</td><td>Intergenic</td><td>NA</td><td>NA</td></tr>
<tr><td></td><td></td><td></td><td></td><td></td><td></td><td></td><td></td><td></td><td></td><td></td><td></td></tr>
<tr><td>T20035</td><td>ACACGGCAGCATAACGATCGAGG</td><td>1000</td><td></td><td></td><td></td><td></td><td></td><td></td><td></td><td></td><td></td></tr>
<tr><td>Oligo fwd</td><td>CACCACACGGCAGCATAACGATCG</td><td></td><td></td><td></td><td></td><td></td><td></td><td></td><td></td><td></td><td></td></tr>
<tr><td>Oligo rev</td><td>AAACCGATCGTTATGCTGCCGTGT</td><td></td><td></td><td></td><td></td><td></td><td></td><td></td><td></td><td></td><td></td></tr>
<tr><td>Chromosome</td><td>start</td><td>end</td><td>strand</td><td>MM</td><td>target_seq</td><td>PAM</td><td>alignment</td><td>distance</td><td>position</td><td>gene name</td><td>gene id</td></tr>
<tr><td>GU305914.1</td><td>192989</td><td>193011</td><td>-</td><td>0</td><td>ACACGGCAGCATAACGATCG</td><td>AGG</td><td>||||||||[||||||||||||]PAM</td><td>NA</td><td>Intergenic</td><td>NA</td><td>NA</td></tr>
<tr><td></td><td></td><td></td><td></td><td></td><td></td><td></td><td></td><td></td><td></td><td></td><td></td></tr>
<tr><td>T20036</td><td>AATAACGTGTACCTTCGACGCGG</td><td>1000</td><td></td><td></td><td></td><td></td><td></td><td></td><td></td><td></td><td></td></tr>
<tr><td>Oligo fwd</td><td>CACCAATAACGTGTACCTTCGACG</td><td></td><td></td><td></td><td></td><td></td><td></td><td></td><td></td><td></td><td></td></tr>
<tr><td>Oligo rev</td><td>AAACCGTCGAAGGTACACGTTATT</td><td></td><td></td><td></td><td></td><td></td><td></td><td></td><td></td><td></td><td></td></tr>
<tr><td>Chromosome</td><td>start</td><td>end</td><td>strand</td><td>MM</td><td>target_seq</td><td>PAM</td><td>alignment</td><td>distance</td><td>position</td><td>gene name</td><td>gene id</td></tr>
<tr><td>GU305914.1</td><td>192926</td><td>192948</td><td>-</td><td>0</td><td>AATAACGTGTACCTTCGACG</td><td>CGG</td><td>||||||||[||||||||||||]PAM</td><td>NA</td><td>Intergenic</td><td>NA</td><td>NA</td></tr>
<tr><td></td><td></td><td></td><td></td><td></td><td></td><td></td><td></td><td></td><td></td><td></td><td></td></tr>
<tr><td>T20037</td><td>ACCTTCGACGCGGTGTCCGACGG</td><td>1000</td><td></td><td></td><td></td><td></td><td></td><td></td><td></td><td></td><td></td></tr>
<tr><td>Oligo fwd</td><td>CACCACCTTCGACGCGGTGTCCGA</td><td></td><td></td><td></td><td></td><td></td><td></td><td></td><td></td><td></td><td></td></tr>
<tr><td>Oligo rev</td><td>AAACTCGGACACCGCGTCGAAGGT</td><td></td><td></td><td></td><td></td><td></td><td></td><td></td><td></td><td></td><td></td></tr>
<tr><td>Chromosome</td><td>start</td><td>end</td><td>strand</td><td>MM</td><td>target_seq</td><td>PAM</td><td>alignment</td><td>distance</td><td>position</td><td>gene name</td><td>gene id</td></tr>
<tr><td>GU305914.1</td><td>192916</td><td>192938</td><td>-</td><td>0</td><td>ACCTTCGACGCGGTGTCCGA</td><td>CGG</td><td>||||||||[||||||||||||]PAM</td><td>NA</td><td>Intergenic</td><td>NA</td><td>NA</td></tr>
<tr><td></td><td></td><td></td><td></td><td></td><td></td><td></td><td></td><td></td><td></td><td></td><td></td></tr>
<tr><td>T20038</td><td>GTCCGACGGCGCGTTCACGCTGG</td><td>1000</td><td></td><td></td><td></td><td></td><td></td><td></td><td></td><td></td><td></td></tr>
<tr><td>Oligo fwd</td><td>CACCGTCCGACGGCGCGTTCACGC</td><td></td><td></td><td></td><td></td><td></td><td></td><td></td><td></td><td></td><td></td></tr>
<tr><td>Oligo rev</td><td>AAACGCGTGAACGCGCCGTCGGAC</td><td></td><td></td><td></td><td></td><td></td><td></td><td></td><td></td><td></td><td></td></tr>
</table>

<table>
<tr><td>Chromosome</td><td>start</td><td>end</td><td>strand</td><td>MM</td><td>target_seq</td><td>PAM</td><td>alignment</td><td>distance</td><td>position</td><td>gene name</td><td>gene id</td></tr>
<tr><td>GU305914.1</td><td>192902</td><td>192924</td><td>-</td><td>0</td><td>GTCCGACGGCGCGTTCACGC</td><td>TGG</td><td>||||||||[||||||||||||]PAM</td><td>NA</td><td>Intergenic</td><td>NA</td><td>NA</td></tr>
<tr><td>GU305914.1</td><td>65678</td><td>65700</td><td>+</td><td>4</td><td>GGACGACATCGCGTTCACGC</td><td>TGG</td><td>|--||||-[-|||||||||||]PAM</td><td>NA</td><td>Intergenic</td><td>NA</td><td>NA</td></tr>
<tr><td>GU305914.1</td><td>77638</td><td>77660</td><td>+</td><td>4</td><td>GACCGAGGGCTCGTTCGCGC</td><td>GGG</td><td>|-||||-|[||-|||||-|||]PAM</td><td>NA</td><td>Intergenic</td><td>NA</td><td>NA</td></tr>
<tr><td></td><td></td><td></td><td></td><td></td><td></td><td></td><td></td><td></td><td></td><td></td><td></td></tr>
<tr><td>T20039</td><td>CGCGTTCACGCTGGAGCGCGTGG</td><td>1000</td><td></td><td></td><td></td><td></td><td></td><td></td><td></td><td></td><td></td></tr>
<tr><td>Oligo fwd</td><td>CACCCGCGTTCACGCTGGAGCGCG</td><td></td><td></td><td></td><td></td><td></td><td></td><td></td><td></td><td></td><td></td></tr>
<tr><td>Oligo rev</td><td>AAACCGCGCTCCAGCGTGAACGCG</td><td></td><td></td><td></td><td></td><td></td><td></td><td></td><td></td><td></td><td></td></tr>
<tr><td>Chromosome</td><td>start</td><td>end</td><td>strand</td><td>MM</td><td>target_seq</td><td>PAM</td><td>alignment</td><td>distance</td><td>position</td><td>gene name</td><td>gene id</td></tr>
<tr><td>GU305914.1</td><td>192893</td><td>192915</td><td>-</td><td>0</td><td>CGCGTTCACGCTGGAGCGCG</td><td>TGG</td><td>||||||||[||||||||||||]PAM</td><td>NA</td><td>Intergenic</td><td>NA</td><td>NA</td></tr>
<tr><td></td><td></td><td></td><td></td><td></td><td></td><td></td><td></td><td></td><td></td><td></td><td></td></tr>
<tr><td>T20040</td><td>ACGCTGGAGCGCGTGGAAGACGG</td><td>1000</td><td></td><td></td><td></td><td></td><td></td><td></td><td></td><td></td><td></td></tr>
<tr><td>Oligo fwd</td><td>CACCACGCTGGAGCGCGTGGAAGA</td><td></td><td></td><td></td><td></td><td></td><td></td><td></td><td></td><td></td><td></td></tr>
<tr><td>Oligo rev</td><td>AAACTCTTCCACGCGCTCCAGCGT</td><td></td><td></td><td></td><td></td><td></td><td></td><td></td><td></td><td></td><td></td></tr>
<tr><td>Chromosome</td><td>start</td><td>end</td><td>strand</td><td>MM</td><td>target_seq</td><td>PAM</td><td>alignment</td><td>distance</td><td>position</td><td>gene name</td><td>gene id</td></tr>
<tr><td>GU305914.1</td><td>192886</td><td>192908</td><td>-</td><td>0</td><td>ACGCTGGAGCGCGTGGAAGA</td><td>CGG</td><td>||||||||[||||||||||||]PAM</td><td>NA</td><td>Intergenic</td><td>NA</td><td>NA</td></tr>
<tr><td></td><td></td><td></td><td></td><td></td><td></td><td></td><td></td><td></td><td></td><td></td><td></td></tr>
<tr><td>T20041</td><td>GCGCGTGGAAGACGGACAGCCGG</td><td>1000</td><td></td><td></td><td></td><td></td><td></td><td></td><td></td><td></td><td></td></tr>
<tr><td>Oligo fwd</td><td>CACCGCGCGTGGAAGACGGACAGC</td><td></td><td></td><td></td><td></td><td></td><td></td><td></td><td></td><td></td><td></td></tr>
<tr><td>Oligo rev</td><td>AAACGCTGTCCGTCTTCCACGCGC</td><td></td><td></td><td></td><td></td><td></td><td></td><td></td><td></td><td></td><td></td></tr>
<tr><td>Chromosome</td><td>start</td><td>end</td><td>strand</td><td>MM</td><td>target_seq</td><td>PAM</td><td>alignment</td><td>distance</td><td>position</td><td>gene name</td><td>gene id</td></tr>
<tr><td>GU305914.1</td><td>192878</td><td>192900</td><td>-</td><td>0</td><td>GCGCGTGGAAGACGGACAGC</td><td>CGG</td><td>||||||||[||||||||||||]PAM</td><td>NA</td><td>Intergenic</td><td>NA</td><td>NA</td></tr>
<tr><td></td><td></td><td></td><td></td><td></td><td></td><td></td><td></td><td></td><td></td><td></td><td></td></tr>
<tr><td>T20042</td><td>CCGGTGTATCAGAGCCGTGTCGG</td><td>1000</td><td></td><td></td><td></td><td></td><td></td><td></td><td></td><td></td><td></td></tr>
<tr><td>Oligo fwd</td><td>CACCCCGGTGTATCAGAGCCGTGT</td><td></td><td></td><td></td><td></td><td></td><td></td><td></td><td></td><td></td><td></td></tr>
<tr><td>Oligo rev</td><td>AAACACACGGCTCTGATACACCGG</td><td></td><td></td><td></td><td></td><td></td><td></td><td></td><td></td><td></td><td></td></tr>
<tr><td>Chromosome</td><td>start</td><td>end</td><td>strand</td><td>MM</td><td>target_seq</td><td>PAM</td><td>alignment</td><td>distance</td><td>position</td><td>gene name</td><td>gene id</td></tr>
<tr><td>GU305914.1</td><td>192859</td><td>192881</td><td>-</td><td>0</td><td>CCGGTGTATCAGAGCCGTGT</td><td>CGG</td><td>||||||||[||||||||||||]PAM</td><td>NA</td><td>Intergenic</td><td>NA</td><td>NA</td></tr>
<tr><td></td><td></td><td></td><td></td><td></td><td></td><td></td><td></td><td></td><td></td><td></td><td></td></tr>
<tr><td>T20043</td><td>GACGACTCCATCTCGATCAACGG</td><td>1000</td><td></td><td></td><td></td><td></td><td></td><td></td><td></td><td></td><td></td></tr>
<tr><td>Oligo fwd</td><td>CACCGACGACTCCATCTCGATCAA</td><td></td><td></td><td></td><td></td><td></td><td></td><td></td><td></td><td></td><td></td></tr>
<tr><td>Oligo rev</td><td>AAACTTGATCGAGATGGAGTCGTC</td><td></td><td></td><td></td><td></td><td></td><td></td><td></td><td></td><td></td><td></td></tr>
<tr><td>Chromosome</td><td>start</td><td>end</td><td>strand</td><td>MM</td><td>target_seq</td><td>PAM</td><td>alignment</td><td>distance</td><td>position</td><td>gene name</td><td>gene id</td></tr>
<tr><td>GU305914.1</td><td>192814</td><td>192836</td><td>-</td><td>0</td><td>GACGACTCCATCTCGATCAA</td><td>CGG</td><td>||||||||[||||||||||||]PAM</td><td>NA</td><td>Intergenic</td><td>NA</td><td>NA</td></tr>
<tr><td></td><td></td><td></td><td></td><td></td><td></td><td></td><td></td><td></td><td></td><td></td><td></td></tr>
<tr><td>T20044</td><td>CCATCTCGATCAACGGCTCGCGG</td><td>1000</td><td></td><td></td><td></td><td></td><td></td><td></td><td></td><td></td><td></td></tr>
<tr><td>Oligo fwd</td><td>CACCCCATCTCGATCAACGGCTCG</td><td></td><td></td><td></td><td></td><td></td><td></td><td></td><td></td><td></td><td></td></tr>
<tr><td>Oligo rev</td><td>AAACCGAGCCGTTGATCGAGATGG</td><td></td><td></td><td></td><td></td><td></td><td></td><td></td><td></td><td></td><td></td></tr>
<tr><td>Chromosome</td><td>start</td><td>end</td><td>strand</td><td>MM</td><td>target_seq</td><td>PAM</td><td>alignment</td><td>distance</td><td>position</td><td>gene name</td><td>gene id</td></tr>
<tr><td>GU305914.1</td><td>192807</td><td>192829</td><td>-</td><td>0</td><td>CCATCTCGATCAACGGCTCG</td><td>CGG</td><td>||||||||[||||||||||||]PAM</td><td>NA</td><td>Intergenic</td><td>NA</td><td>NA</td></tr>
<tr><td></td><td></td><td></td><td></td><td></td><td></td><td></td><td></td><td></td><td></td><td></td><td></td></tr>
<tr><td>T20045</td><td>CAACGGCTCGCGGCTGTCCGAGG</td><td>1000</td><td></td><td></td><td></td><td></td><td></td><td></td><td></td><td></td><td></td></tr>
<tr><td>Oligo fwd</td><td>CACCCAACGGCTCGCGGCTGTCCG</td><td></td><td></td><td></td><td></td><td></td><td></td><td></td><td></td><td></td><td></td></tr>
<tr><td>Oligo rev</td><td>AAACCGGACAGCCGCGAGCCGTTG</td><td></td><td></td><td></td><td></td><td></td><td></td><td></td><td></td><td></td><td></td></tr>
<tr><td>Chromosome</td><td>start</td><td>end</td><td>strand</td><td>MM</td><td>target_seq</td><td>PAM</td><td>alignment</td><td>distance</td><td>position</td><td>gene name</td><td>gene id</td></tr>
<tr><td>GU305914.1</td><td>192797</td><td>192819</td><td>-</td><td>0</td><td>CAACGGCTCGCGGCTGTCCG</td><td>AGG</td><td>||||||||[||||||||||||]PAM</td><td>NA</td><td>Intergenic</td><td>NA</td><td>NA</td></tr>
<tr><td></td><td></td><td></td><td></td><td></td><td></td><td></td><td></td><td></td><td></td><td></td><td></td></tr>
<tr><td>T20046</td><td>AACGGCTCGCGGCTGTCCGAGGG</td><td>1000</td><td></td><td></td><td></td><td></td><td></td><td></td><td></td><td></td><td></td></tr>
<tr><td>Oligo fwd</td><td>CACCAACGGCTCGCGGCTGTCCGA</td><td></td><td></td><td></td><td></td><td></td><td></td><td></td><td></td><td></td><td></td></tr>
<tr><td>Oligo rev</td><td>AAACTCGGACAGCCGCGAGCCGTT</td><td></td><td></td><td></td><td></td><td></td><td></td><td></td><td></td><td></td><td></td></tr>
<tr><td>Chromosome</td><td>start</td><td>end</td><td>strand</td><td>MM</td><td>target_seq</td><td>PAM</td><td>alignment</td><td>distance</td><td>position</td><td>gene name</td><td>gene id</td></tr>
<tr><td>GU305914.1</td><td>192796</td><td>192818</td><td>-</td><td>0</td><td>AACGGCTCGCGGCTGTCCGA</td><td>GGG</td><td>||||||||[||||||||||||]PAM</td><td>NA</td><td>Intergenic</td><td>NA</td><td>NA</td></tr>
<tr><td></td><td></td><td></td><td></td><td></td><td></td><td></td><td></td><td></td><td></td><td></td><td></td></tr>
<tr><td>T20047</td><td>GCTGTCCGAGGGCAAGATCTCGG</td><td>1000</td><td></td><td></td><td></td><td></td><td></td><td></td><td></td><td></td><td></td></tr>
<tr><td>Oligo fwd</td><td>CACCGCTGTCCGAGGGCAAGATCT</td><td></td><td></td><td></td><td></td><td></td><td></td><td></td><td></td><td></td><td></td></tr>
<tr><td>Oligo rev</td><td>AAACAGATCTTGCCCTCGGACAGC</td><td></td><td></td><td></td><td></td><td></td><td></td><td></td><td></td><td></td><td></td></tr>
<tr><td>Chromosome</td><td>start</td><td>end</td><td>strand</td><td>MM</td><td>target_seq</td><td>PAM</td><td>alignment</td><td>distance</td><td>position</td><td>gene name</td><td>gene id</td></tr>
<tr><td>GU305914.1</td><td>192785</td><td>192807</td><td>-</td><td>0</td><td>GCTGTCCGAGGGCAAGATCT</td><td>CGG</td><td>||||||||[||||||||||||]PAM</td><td>NA</td><td>Intergenic</td><td>NA</td><td>NA</td></tr>
<tr><td></td><td></td><td></td><td></td><td></td><td></td><td></td><td></td><td></td><td></td><td></td><td></td></tr>
<tr><td>T20048</td><td>GTGTACCTGAGAGAAGACGATGG</td><td>1000</td><td></td><td></td><td></td><td></td><td></td><td></td><td></td><td></td><td></td></tr>
<tr><td>Oligo fwd</td><td>CACCGTGTACCTGAGAGAAGACGA</td><td></td><td></td><td></td><td></td><td></td><td></td><td></td><td></td><td></td><td></td></tr>
<tr><td>Oligo rev</td><td>AAACTCGTCTTCTCTCAGGTACAC</td><td></td><td></td><td></td><td></td><td></td><td></td><td></td><td></td><td></td><td></td></tr>
<tr><td>Chromosome</td><td>start</td><td>end</td><td>strand</td><td>MM</td><td>target_seq</td><td>PAM</td><td>alignment</td><td>distance</td><td>position</td><td>gene name</td><td>gene id</td></tr>
<tr><td>GU305914.1</td><td>192757</td><td>192779</td><td>-</td><td>0</td><td>GTGTACCTGAGAGAAGACGA</td><td>TGG</td><td>||||||||[||||||||||||]PAM</td><td>NA</td><td>Intergenic</td><td>NA</td><td>NA</td></tr>
<tr><td></td><td></td><td></td><td></td><td></td><td></td><td></td><td></td><td></td><td></td><td></td><td></td></tr>
<tr><td>T20049</td><td>AGAGAAGACGATGGCGTGATCGG</td><td>1000</td><td></td><td></td><td></td><td></td><td></td><td></td><td></td><td></td><td></td></tr>
<tr><td>Oligo fwd</td><td>CACCAGAGAAGACGATGGCGTGAT</td><td></td><td></td><td></td><td></td><td></td><td></td><td></td><td></td><td></td><td></td></tr>
<tr><td>Oligo rev</td><td>AAACATCACGCCATCGTCTTCTCT</td><td></td><td></td><td></td><td></td><td></td><td></td><td></td><td></td><td></td><td></td></tr>
<tr><td>Chromosome</td><td>start</td><td>end</td><td>strand</td><td>MM</td><td>target_seq</td><td>PAM</td><td>alignment</td><td>distance</td><td>position</td><td>gene name</td><td>gene id</td></tr>
<tr><td>GU305914.1</td><td>192748</td><td>192770</td><td>-</td><td>0</td><td>AGAGAAGACGATGGCGTGAT</td><td>CGG</td><td>||||||||[||||||||||||]PAM</td><td>NA</td><td>Intergenic</td><td>NA</td><td>NA</td></tr>
<tr><td></td><td></td><td></td><td></td><td></td><td></td><td></td><td></td><td></td><td></td><td></td><td></td></tr>
<tr><td>T20050</td><td>AAGACGATGGCGTGATCGGCCGG</td><td>1000</td><td></td><td></td><td></td><td></td><td></td><td></td><td></td><td></td><td></td></tr>
<tr><td>Oligo fwd</td><td>CACCAAGACGATGGCGTGATCGGC</td><td></td><td></td><td></td><td></td><td></td><td></td><td></td><td></td><td></td><td></td></tr>
<tr><td>Oligo rev</td><td>AAACGCCGATCACGCCATCGTCTT</td><td></td><td></td><td></td><td></td><td></td><td></td><td></td><td></td><td></td><td></td></tr>
<tr><td>Chromosome</td><td>start</td><td>end</td><td>strand</td><td>MM</td><td>target_seq</td><td>PAM</td><td>alignment</td><td>distance</td><td>position</td><td>gene name</td><td>gene id</td></tr>
<tr><td>GU305914.1</td><td>192744</td><td>192766</td><td>-</td><td>0</td><td>AAGACGATGGCGTGATCGGC</td><td>CGG</td><td>||||||||[||||||||||||]PAM</td><td>NA</td><td>Intergenic</td><td>NA</td><td>NA</td></tr>
<tr><td></td><td></td><td></td><td></td><td></td><td></td><td></td><td></td><td></td><td></td><td></td><td></td></tr>
<tr><td>T20051</td><td>ATGGCGTGATCGGCCGGTTTAGG</td><td>1000</td><td></td><td></td><td></td><td></td><td></td><td></td><td></td><td></td><td></td></tr>
<tr><td>Oligo fwd</td><td>CACCATGGCGTGATCGGCCGGTTT</td><td></td><td></td><td></td><td></td><td></td><td></td><td></td><td></td><td></td><td></td></tr>
<tr><td>Oligo rev</td><td>AAACAAACCGGCCGATCACGCCAT</td><td></td><td></td><td></td><td></td><td></td><td></td><td></td><td></td><td></td><td></td></tr>
<tr><td>Chromosome</td><td>start</td><td>end</td><td>strand</td><td>MM</td><td>target_seq</td><td>PAM</td><td>alignment</td><td>distance</td><td>position</td><td>gene name</td><td>gene id</td></tr>
<tr><td>GU305914.1</td><td>192738</td><td>192760</td><td>-</td><td>0</td><td>ATGGCGTGATCGGCCGGTTT</td><td>AGG</td><td>||||||||[||||||||||||]PAM</td><td>NA</td><td>Intergenic</td><td>NA</td><td>NA</td></tr>
<tr><td></td><td></td><td></td><td></td><td></td><td></td><td></td><td></td><td></td><td></td><td></td><td></td></tr>
<tr><td>T20052</td><td>GTTTAGGTGCCGCGTCCGCTCGG</td><td>1000</td><td></td><td></td><td></td><td></td><td></td><td></td><td></td><td></td><td></td></tr>
<tr><td>Oligo fwd</td><td>CACCGTTTAGGTGCCGCGTCCGCT</td><td></td><td></td><td></td><td></td><td></td><td></td><td></td><td></td><td></td><td></td></tr>
<tr><td>Oligo rev</td><td>AAACAGCGGACGCGGCACCTAAAC</td><td></td><td></td><td></td><td></td><td></td><td></td><td></td><td></td><td></td><td></td></tr>
<tr><td>Chromosome</td><td>start</td><td>end</td><td>strand</td><td>MM</td><td>target_seq</td><td>PAM</td><td>alignment</td><td>distance</td><td>position</td><td>gene name</td><td>gene id</td></tr>
<tr><td>GU305914.1</td><td>192722</td><td>192744</td><td>-</td><td>0</td><td>GTTTAGGTGCCGCGTCCGCT</td><td>CGG</td><td>||||||||[||||||||||||]PAM</td><td>NA</td><td>Intergenic</td><td>NA</td><td>NA</td></tr>
</table>

<table>
<tr><td>T20053</td><td>TTTAGGTGCCGCGTCCGCTCGGG</td><td>1000</td><td></td><td></td><td></td><td></td><td></td><td></td><td></td><td></td><td></td></tr>
<tr><td>Oligo fwd</td><td>CACCTTTAGGTGCCGCGTCCGCTC</td><td></td><td></td><td></td><td></td><td></td><td></td><td></td><td></td><td></td><td></td></tr>
<tr><td>Oligo rev</td><td>AAACGAGCGGACGCGGCACCTAAA</td><td></td><td></td><td></td><td></td><td></td><td></td><td></td><td></td><td></td><td></td></tr>
<tr><td>Chromosome</td><td>start</td><td>end</td><td>strand</td><td>MM</td><td>target_seq</td><td>PAM</td><td>alignment</td><td>distance</td><td>position</td><td>gene name</td><td>gene id</td></tr>
<tr><td>GU305914.1</td><td>192721</td><td>192743</td><td>-</td><td>0</td><td>TTTAGGTGCCGCGTCCGCTC</td><td>GGG</td><td>||||||||[||||||||||||]PAM</td><td>NA</td><td>Intergenic</td><td>NA</td><td>NA</td></tr>
<tr><td></td><td></td><td></td><td></td><td></td><td></td><td></td><td></td><td></td><td></td><td></td><td></td></tr>
<tr><td>T20054</td><td>CCGCTCGGGATGCTTGAATCTGG</td><td>1000</td><td></td><td></td><td></td><td></td><td></td><td></td><td></td><td></td><td></td></tr>
<tr><td>Oligo fwd</td><td>CACCCCGCTCGGGATGCTTGAATC</td><td></td><td></td><td></td><td></td><td></td><td></td><td></td><td></td><td></td><td></td></tr>
<tr><td>Oligo rev</td><td>AAACGATTCAAGCATCCCGAGCGG</td><td></td><td></td><td></td><td></td><td></td><td></td><td></td><td></td><td></td><td></td></tr>
<tr><td>Chromosome</td><td>start</td><td>end</td><td>strand</td><td>MM</td><td>target_seq</td><td>PAM</td><td>alignment</td><td>distance</td><td>position</td><td>gene name</td><td>gene id</td></tr>
<tr><td>GU305914.1</td><td>192707</td><td>192729</td><td>-</td><td>0</td><td>CCGCTCGGGATGCTTGAATC</td><td>TGG</td><td>||||||||[||||||||||||]PAM</td><td>NA</td><td>Intergenic</td><td>NA</td><td>NA</td></tr>
<tr><td></td><td></td><td></td><td></td><td></td><td></td><td></td><td></td><td></td><td></td><td></td><td></td></tr>
<tr><td>T20055</td><td>TCACCGTGACGAAGCCCACCAGG</td><td>1000</td><td></td><td></td><td></td><td></td><td></td><td></td><td></td><td></td><td></td></tr>
<tr><td>Oligo fwd</td><td>CACCTCACCGTGACGAAGCCCACC</td><td></td><td></td><td></td><td></td><td></td><td></td><td></td><td></td><td></td><td></td></tr>
<tr><td>Oligo rev</td><td>AAACGGTGGGCTTCGTCACGGTGA</td><td></td><td></td><td></td><td></td><td></td><td></td><td></td><td></td><td></td><td></td></tr>
<tr><td>Chromosome</td><td>start</td><td>end</td><td>strand</td><td>MM</td><td>target_seq</td><td>PAM</td><td>alignment</td><td>distance</td><td>position</td><td>gene name</td><td>gene id</td></tr>
<tr><td>GU305914.1</td><td>192672</td><td>192694</td><td>-</td><td>0</td><td>TCACCGTGACGAAGCCCACC</td><td>AGG</td><td>||||||||[||||||||||||]PAM</td><td>NA</td><td>Intergenic</td><td>NA</td><td>NA</td></tr>
<tr><td>GU305914.1</td><td>189034</td><td>189056</td><td>+</td><td>4</td><td>GCACGGCGGCGAAGCCCACC</td><td>AGG</td><td>-|||-|-|[-|||||||||||]PAM</td><td>NA</td><td>Intergenic</td><td>NA</td><td>NA</td></tr>
<tr><td></td><td></td><td></td><td></td><td></td><td></td><td></td><td></td><td></td><td></td><td></td><td></td></tr>
<tr><td>T20056</td><td>CGTGACGAAGCCCACCAGGATGG</td><td>1000</td><td></td><td></td><td></td><td></td><td></td><td></td><td></td><td></td><td></td></tr>
<tr><td>Oligo fwd</td><td>CACCCGTGACGAAGCCCACCAGGA</td><td></td><td></td><td></td><td></td><td></td><td></td><td></td><td></td><td></td><td></td></tr>
<tr><td>Oligo rev</td><td>AAACTCCTGGTGGGCTTCGTCACG</td><td></td><td></td><td></td><td></td><td></td><td></td><td></td><td></td><td></td><td></td></tr>
<tr><td>Chromosome</td><td>start</td><td>end</td><td>strand</td><td>MM</td><td>target_seq</td><td>PAM</td><td>alignment</td><td>distance</td><td>position</td><td>gene name</td><td>gene id</td></tr>
<tr><td>GU305914.1</td><td>192668</td><td>192690</td><td>-</td><td>0</td><td>CGTGACGAAGCCCACCAGGA</td><td>TGG</td><td>||||||||[||||||||||||]PAM</td><td>NA</td><td>Intergenic</td><td>NA</td><td>NA</td></tr>
<tr><td></td><td></td><td></td><td></td><td></td><td></td><td></td><td></td><td></td><td></td><td></td><td></td></tr>
<tr><td>T20057</td><td>GGAATACGCCCTCGTGACAGAGG</td><td>1000</td><td></td><td></td><td></td><td></td><td></td><td></td><td></td><td></td><td></td></tr>
<tr><td>Oligo fwd</td><td>CACCGGAATACGCCCTCGTGACAG</td><td></td><td></td><td></td><td></td><td></td><td></td><td></td><td></td><td></td><td></td></tr>
<tr><td>Oligo rev</td><td>AAACCTGTCACGAGGGCGTATTCC</td><td></td><td></td><td></td><td></td><td></td><td></td><td></td><td></td><td></td><td></td></tr>
<tr><td>Chromosome</td><td>start</td><td>end</td><td>strand</td><td>MM</td><td>target_seq</td><td>PAM</td><td>alignment</td><td>distance</td><td>position</td><td>gene name</td><td>gene id</td></tr>
<tr><td>GU305914.1</td><td>192647</td><td>192669</td><td>-</td><td>0</td><td>GGAATACGCCCTCGTGACAG</td><td>AGG</td><td>||||||||[||||||||||||]PAM</td><td>NA</td><td>Intergenic</td><td>NA</td><td>NA</td></tr>
<tr><td></td><td></td><td></td><td></td><td></td><td></td><td></td><td></td><td></td><td></td><td></td><td></td></tr>
<tr><td>T20058</td><td>ACTAGAAGAGATCAGAATAAAGG</td><td>1000</td><td></td><td></td><td></td><td></td><td></td><td></td><td></td><td></td><td></td></tr>
<tr><td>Oligo fwd</td><td>CACCACTAGAAGAGATCAGAATAA</td><td></td><td></td><td></td><td></td><td></td><td></td><td></td><td></td><td></td><td></td></tr>
<tr><td>Oligo rev</td><td>AAACTTATTCTGATCTCTTCTAGT</td><td></td><td></td><td></td><td></td><td></td><td></td><td></td><td></td><td></td><td></td></tr>
<tr><td>Chromosome</td><td>start</td><td>end</td><td>strand</td><td>MM</td><td>target_seq</td><td>PAM</td><td>alignment</td><td>distance</td><td>position</td><td>gene name</td><td>gene id</td></tr>
<tr><td>GU305914.1</td><td>192545</td><td>192567</td><td>-</td><td>0</td><td>ACTAGAAGAGATCAGAATAA</td><td>AGG</td><td>||||||||[||||||||||||]PAM</td><td>NA</td><td>Intergenic</td><td>NA</td><td>NA</td></tr>
<tr><td></td><td></td><td></td><td></td><td></td><td></td><td></td><td></td><td></td><td></td><td></td><td></td></tr>
<tr><td>T20059</td><td>GAGATCAGAATAAAGGTCGACGG</td><td>1000</td><td></td><td></td><td></td><td></td><td></td><td></td><td></td><td></td><td></td></tr>
<tr><td>Oligo fwd</td><td>CACCGAGATCAGAATAAAGGTCGA</td><td></td><td></td><td></td><td></td><td></td><td></td><td></td><td></td><td></td><td></td></tr>
<tr><td>Oligo rev</td><td>AAACTCGACCTTTATTCTGATCTC</td><td></td><td></td><td></td><td></td><td></td><td></td><td></td><td></td><td></td><td></td></tr>
<tr><td>Chromosome</td><td>start</td><td>end</td><td>strand</td><td>MM</td><td>target_seq</td><td>PAM</td><td>alignment</td><td>distance</td><td>position</td><td>gene name</td><td>gene id</td></tr>
<tr><td>GU305914.1</td><td>192538</td><td>192560</td><td>-</td><td>0</td><td>GAGATCAGAATAAAGGTCGA</td><td>CGG</td><td>||||||||[||||||||||||]PAM</td><td>NA</td><td>Intergenic</td><td>NA</td><td>NA</td></tr>
<tr><td></td><td></td><td></td><td></td><td></td><td></td><td></td><td></td><td></td><td></td><td></td><td></td></tr>
<tr><td>T20060</td><td>CGACGACGATGACTTTCTGTTGG</td><td>1000</td><td></td><td></td><td></td><td></td><td></td><td></td><td></td><td></td><td></td></tr>
<tr><td>Oligo fwd</td><td>CACCCGACGACGATGACTTTCTGT</td><td></td><td></td><td></td><td></td><td></td><td></td><td></td><td></td><td></td><td></td></tr>
<tr><td>Oligo rev</td><td>AAACACAGAAAGTCATCGTCGTCG</td><td></td><td></td><td></td><td></td><td></td><td></td><td></td><td></td><td></td><td></td></tr>
<tr><td>Chromosome</td><td>start</td><td>end</td><td>strand</td><td>MM</td><td>target_seq</td><td>PAM</td><td>alignment</td><td>distance</td><td>position</td><td>gene name</td><td>gene id</td></tr>
<tr><td>GU305914.1</td><td>192512</td><td>192534</td><td>-</td><td>0</td><td>CGACGACGATGACTTTCTGT</td><td>TGG</td><td>||||||||[||||||||||||]PAM</td><td>NA</td><td>Intergenic</td><td>NA</td><td>NA</td></tr>
<tr><td></td><td></td><td></td><td></td><td></td><td></td><td></td><td></td><td></td><td></td><td></td><td></td></tr>
<tr><td>T20061</td><td>TCTATTCGCTCGCGTCAGCGCGG</td><td>1000</td><td></td><td></td><td></td><td></td><td></td><td></td><td></td><td></td><td></td></tr>
<tr><td>Oligo fwd</td><td>CACCTCTATTCGCTCGCGTCAGCG</td><td></td><td></td><td></td><td></td><td></td><td></td><td></td><td></td><td></td><td></td></tr>
<tr><td>Oligo rev</td><td>AAACCGCTGACGCGAGCGAATAGA</td><td></td><td></td><td></td><td></td><td></td><td></td><td></td><td></td><td></td><td></td></tr>
<tr><td>Chromosome</td><td>start</td><td>end</td><td>strand</td><td>MM</td><td>target_seq</td><td>PAM</td><td>alignment</td><td>distance</td><td>position</td><td>gene name</td><td>gene id</td></tr>
<tr><td>GU305914.1</td><td>192452</td><td>192474</td><td>-</td><td>0</td><td>TCTATTCGCTCGCGTCAGCG</td><td>CGG</td><td>||||||||[||||||||||||]PAM</td><td>NA</td><td>Intergenic</td><td>NA</td><td>NA</td></tr>
<tr><td></td><td></td><td></td><td></td><td></td><td></td><td></td><td></td><td></td><td></td><td></td><td></td></tr>
<tr><td>T20062</td><td>TTCGCTCGCGTCAGCGCGGAAGG</td><td>1000</td><td></td><td></td><td></td><td></td><td></td><td></td><td></td><td></td><td></td></tr>
<tr><td>Oligo fwd</td><td>CACCTTCGCTCGCGTCAGCGCGGA</td><td></td><td></td><td></td><td></td><td></td><td></td><td></td><td></td><td></td><td></td></tr>
<tr><td>Oligo rev</td><td>AAACTCCGCGCTGACGCGAGCGAA</td><td></td><td></td><td></td><td></td><td></td><td></td><td></td><td></td><td></td><td></td></tr>
<tr><td>Chromosome</td><td>start</td><td>end</td><td>strand</td><td>MM</td><td>target_seq</td><td>PAM</td><td>alignment</td><td>distance</td><td>position</td><td>gene name</td><td>gene id</td></tr>
<tr><td>GU305914.1</td><td>192448</td><td>192470</td><td>-</td><td>0</td><td>TTCGCTCGCGTCAGCGCGGA</td><td>AGG</td><td>||||||||[||||||||||||]PAM</td><td>NA</td><td>Intergenic</td><td>NA</td><td>NA</td></tr>
<tr><td></td><td></td><td></td><td></td><td></td><td></td><td></td><td></td><td></td><td></td><td></td><td></td></tr>
<tr><td>T20063</td><td>CGTCAGCGCGGAAGGCCGTCTGG</td><td>1000</td><td></td><td></td><td></td><td></td><td></td><td></td><td></td><td></td><td></td></tr>
<tr><td>Oligo fwd</td><td>CACCCGTCAGCGCGGAAGGCCGTC</td><td></td><td></td><td></td><td></td><td></td><td></td><td></td><td></td><td></td><td></td></tr>
<tr><td>Oligo rev</td><td>AAACGACGGCCTTCCGCGCTGACG</td><td></td><td></td><td></td><td></td><td></td><td></td><td></td><td></td><td></td><td></td></tr>
<tr><td>Chromosome</td><td>start</td><td>end</td><td>strand</td><td>MM</td><td>target_seq</td><td>PAM</td><td>alignment</td><td>distance</td><td>position</td><td>gene name</td><td>gene id</td></tr>
<tr><td>GU305914.1</td><td>192440</td><td>192462</td><td>-</td><td>0</td><td>CGTCAGCGCGGAAGGCCGTC</td><td>TGG</td><td>||||||||[||||||||||||]PAM</td><td>NA</td><td>Intergenic</td><td>NA</td><td>NA</td></tr>
<tr><td></td><td></td><td></td><td></td><td></td><td></td><td></td><td></td><td></td><td></td><td></td><td></td></tr>
<tr><td>T20064</td><td>CAGCGCGGAAGGCCGTCTGGCGG</td><td>1000</td><td></td><td></td><td></td><td></td><td></td><td></td><td></td><td></td><td></td></tr>
<tr><td>Oligo fwd</td><td>CACCCAGCGCGGAAGGCCGTCTGG</td><td></td><td></td><td></td><td></td><td></td><td></td><td></td><td></td><td></td><td></td></tr>
<tr><td>Oligo rev</td><td>AAACCCAGACGGCCTTCCGCGCTG</td><td></td><td></td><td></td><td></td><td></td><td></td><td></td><td></td><td></td><td></td></tr>
<tr><td>Chromosome</td><td>start</td><td>end</td><td>strand</td><td>MM</td><td>target_seq</td><td>PAM</td><td>alignment</td><td>distance</td><td>position</td><td>gene name</td><td>gene id</td></tr>
<tr><td>GU305914.1</td><td>192437</td><td>192459</td><td>-</td><td>0</td><td>CAGCGCGGAAGGCCGTCTGG</td><td>CGG</td><td>||||||||[||||||||||||]PAM</td><td>NA</td><td>Intergenic</td><td>NA</td><td>NA</td></tr>
<tr><td></td><td></td><td></td><td></td><td></td><td></td><td></td><td></td><td></td><td></td><td></td><td></td></tr>
<tr><td>T20065</td><td>CCTAGCCGCGATCCTCGTGTTGG</td><td>1000</td><td></td><td></td><td></td><td></td><td></td><td></td><td></td><td></td><td></td></tr>
<tr><td>Oligo fwd</td><td>CACCCCTAGCCGCGATCCTCGTGT</td><td></td><td></td><td></td><td></td><td></td><td></td><td></td><td></td><td></td><td></td></tr>
<tr><td>Oligo rev</td><td>AAACACACGAGGATCGCGGCTAGG</td><td></td><td></td><td></td><td></td><td></td><td></td><td></td><td></td><td></td><td></td></tr>
<tr><td>Chromosome</td><td>start</td><td>end</td><td>strand</td><td>MM</td><td>target_seq</td><td>PAM</td><td>alignment</td><td>distance</td><td>position</td><td>gene name</td><td>gene id</td></tr>
<tr><td>GU305914.1</td><td>192398</td><td>192420</td><td>-</td><td>0</td><td>CCTAGCCGCGATCCTCGTGT</td><td>TGG</td><td>||||||||[||||||||||||]PAM</td><td>NA</td><td>Intergenic</td><td>NA</td><td>NA</td></tr>
<tr><td></td><td></td><td></td><td></td><td></td><td></td><td></td><td></td><td></td><td></td><td></td><td></td></tr>
<tr><td>T20066</td><td>TGTTGGTCCACATCTCGTTCTGG</td><td>1000</td><td></td><td></td><td></td><td></td><td></td><td></td><td></td><td></td><td></td></tr>
<tr><td>Oligo fwd</td><td>CACCTGTTGGTCCACATCTCGTTC</td><td></td><td></td><td></td><td></td><td></td><td></td><td></td><td></td><td></td><td></td></tr>
<tr><td>Oligo rev</td><td>AAACGAACGAGATGTGGACCAACA</td><td></td><td></td><td></td><td></td><td></td><td></td><td></td><td></td><td></td><td></td></tr>
<tr><td>Chromosome</td><td>start</td><td>end</td><td>strand</td><td>MM</td><td>target_seq</td><td>PAM</td><td>alignment</td><td>distance</td><td>position</td><td>gene name</td><td>gene id</td></tr>
<tr><td>GU305914.1</td><td>192381</td><td>192403</td><td>-</td><td>0</td><td>TGTTGGTCCACATCTCGTTC</td><td>TGG</td><td>||||||||[||||||||||||]PAM</td><td>NA</td><td>Intergenic</td><td>NA</td><td>NA</td></tr>
<tr><td></td><td></td><td></td><td></td><td></td><td></td><td></td><td></td><td></td><td></td><td></td><td></td></tr>
<tr><td>T20067</td><td>GTCGCCGCCGTGTAACCTGCCGG</td><td>1000</td><td></td><td></td><td></td><td></td><td></td><td></td><td></td><td></td><td></td></tr>
<tr><td>Oligo fwd</td><td>CACCGTCGCCGCCGTGTAACCTGC</td><td></td><td></td><td></td><td></td><td></td><td></td><td></td><td></td><td></td><td></td></tr>
<tr><td>Oligo rev</td><td>AAACGCAGGTTACACGGCGGCGAC</td><td></td><td></td><td></td><td></td><td></td><td></td><td></td><td></td><td></td><td></td></tr>
<tr><td>Chromosome</td><td>start</td><td>end</td><td>strand</td><td>MM</td><td>target_seq</td><td>PAM</td><td>alignment</td><td>distance</td><td>position</td><td>gene name</td><td>gene id</td></tr>
</table>

GU305914.1 192353 192375 - 0 GTCGCCGCCGTGTAACCTGC CGG ||||||||[||||||||||||]PAM NA Intergenic NA NA

T20068 TCGCCGCCGTGTAACCTGCCGGG 1000
Oligo fwd CACCTCGCCGCCGTGTAACCTGCC
Oligo rev AAACGGCAGGTTACACGGCGGCGA
Chromosome start end strand MM target_seq PAM alignment distance position gene name gene id
GU305914.1 192352 192374 - 0 TCGCCGCCGTGTAACCTGCC GGG ||||||||[||||||||||||]PAM NA Intergenic NA NA

T20069 AAAAGCTCTCAAACTAGACACGG 1000
Oligo fwd CACCAAAAGCTCTCAAACTAGACA
Oligo rev AAACTGTCTAGTTTGAGAGCTTTT
Chromosome start end strand MM target_seq PAM alignment distance position gene name gene id
GU305914.1 192255 192277 - 0 AAAAGCTCTCAAACTAGACA CGG ||||||||[||||||||||||]PAM NA Intergenic NA NA

T20070 CTTCATTACTCTGTTATTTTCGG 1000
Oligo fwd CACCCTTCATTACTCTGTTATTTT
Oligo rev AAACAAAATAACAGAGTAATGAAG
Chromosome start end strand MM target_seq PAM alignment distance position gene name gene id
GU305914.1 192229 192251 - 0 CTTCATTACTCTGTTATTTT CGG ||||||||[||||||||||||]PAM NA Intergenic NA NA

T20071 CTGTTATTTTCGGTCTCCGACGG 1000
Oligo fwd CACCCTGTTATTTTCGGTCTCCGA
Oligo rev AAACTCGGAGACCGAAAATAACAG
Chromosome start end strand MM target_seq PAM alignment distance position gene name gene id
GU305914.1 192219 192241 - 0 CTGTTATTTTCGGTCTCCGA CGG ||||||||[||||||||||||]PAM NA Intergenic NA NA

T20072 TGTTATTTTCGGTCTCCGACGGG 1000
Oligo fwd CACCTGTTATTTTCGGTCTCCGAC
Oligo rev AAACGTCGGAGACCGAAAATAACA
Chromosome start end strand MM target_seq PAM alignment distance position gene name gene id
GU305914.1 192218 192240 - 0 TGTTATTTTCGGTCTCCGAC GGG ||||||||[||||||||||||]PAM NA Intergenic NA NA

T20073 TTTCGGTCTCCGACGGGTCACGG 1000
Oligo fwd CACCTTTCGGTCTCCGACGGGTCA
Oligo rev AAACTGACCCGTCGGAGACCGAAA
Chromosome start end strand MM target_seq PAM alignment distance position gene name gene id
GU305914.1 192212 192234 - 0 TTTCGGTCTCCGACGGGTCA CGG ||||||||[||||||||||||]PAM NA Intergenic NA NA

T20074 CTCTTGCCTCTCTCTCCATGTGG 1000
Oligo fwd CACCCTCTTGCCTCTCTCTCCATG
Oligo rev AAACCATGGAGAGAGAGGCAAGAG
Chromosome start end strand MM target_seq PAM alignment distance position gene name gene id
GU305914.1 192187 192209 - 0 CTCTTGCCTCTCTCTCCATG TGG ||||||||[||||||||||||]PAM NA Intergenic NA NA

T20075 CTCTCTCCATGTGGCGATACTGG 1000
Oligo fwd CACCCTCTCTCCATGTGGCGATAC
Oligo rev AAACGTATCGCCACATGGAGAGAG
Chromosome start end strand MM target_seq PAM alignment distance position gene name gene id
GU305914.1 192178 192200 - 0 CTCTCTCCATGTGGCGATAC TGG ||||||||[||||||||||||]PAM NA Intergenic NA NA

T20076 CCATGTGGCGATACTGGTATCGG 1000
Oligo fwd CACCCCATGTGGCGATACTGGTAT
Oligo rev AAACATACCAGTATCGCCACATGG
Chromosome start end strand MM target_seq PAM alignment distance position gene name gene id
GU305914.1 192172 192194 - 0 CCATGTGGCGATACTGGTAT CGG ||||||||[||||||||||||]PAM NA Intergenic NA NA

T20077 CATGTGGCGATACTGGTATCGGG 1000
Oligo fwd CACCCATGTGGCGATACTGGTATC
Oligo rev AAACGATACCAGTATCGCCACATG
Chromosome start end strand MM target_seq PAM alignment distance position gene name gene id
GU305914.1 192171 192193 - 0 CATGTGGCGATACTGGTATC GGG ||||||||[||||||||||||]PAM NA Intergenic NA NA

T20078 ATGTGGCGATACTGGTATCGGGG 1000
Oligo fwd CACCATGTGGCGATACTGGTATCG
Oligo rev AAACCGATACCAGTATCGCCACAT
Chromosome start end strand MM target_seq PAM alignment distance position gene name gene id
GU305914.1 192170 192192 - 0 ATGTGGCGATACTGGTATCG GGG ||||||||[||||||||||||]PAM NA Intergenic NA NA

T20079 CTGGTATCGGGGCGCCATCGCGG 1000
Oligo fwd CACCCTGGTATCGGGGCGCCATCG
Oligo rev AAACCGATGGCGCCCCGATACCAG
Chromosome start end strand MM target_seq PAM alignment distance position gene name gene id
GU305914.1 192159 192181 - 0 CTGGTATCGGGGCGCCATCG CGG ||||||||[||||||||||||]PAM NA Intergenic NA NA

T20080 CGCCATCGCGGCGATAGTCGCGG 1000
Oligo fwd CACCCGCCATCGCGGCGATAGTCG
Oligo rev AAACCGACTATCGCCGCGATGGCG
Chromosome start end strand MM target_seq PAM alignment distance position gene name gene id
GU305914.1 192147 192169 - 0 CGCCATCGCGGCGATAGTCG CGG ||||||||[||||||||||||]PAM NA Intergenic NA NA

T20081 AGTCGCGGCCGTCACCACGACGG 1000
Oligo fwd CACCAGTCGCGGCCGTCACCACGA
Oligo rev AAACTCGTGGTGACGGCCGCGACT
Chromosome start end strand MM target_seq PAM alignment distance position gene name gene id
GU305914.1 192132 192154 - 0 AGTCGCGGCCGTCACCACGA CGG ||||||||[||||||||||||]PAM NA Intergenic NA NA
GU305914.1 144918 144940 + 4 GGTCACGAGCGTCACCACGA AGG -|||-||-[-|||||||||||]PAM NA Intergenic NA NA

T20082 CACGACGGCACTGATAGCCCCGG 1000
Oligo fwd CACCCACGACGGCACTGATAGCCC

Oligo rev AAACGGGCTATCAGTGCCGTCGTG
Chromosome start end strand MM target_seq PAM alignment distance position gene name gene id
GU305914.1 192117 192139 - 0 CACGACGGCACTGATAGCCC CGG ||||||||[||||||||||||]PAM NA Intergenic NA NA

T20083 ACGACGGCACTGATAGCCCCGGG 1000
Oligo fwd CACCACGACGGCACTGATAGCCCC
Oligo rev AAACGGGGCTATCAGTGCCGTCGT
Chromosome start end strand MM target_seq PAM alignment distance position gene name gene id
GU305914.1 192116 192138 - 0 ACGACGGCACTGATAGCCCC GGG ||||||||[||||||||||||]PAM NA Intergenic NA NA

T20084 CGACGGCACTGATAGCCCCGGGG 1000
Oligo fwd CACCCGACGGCACTGATAGCCCCG
Oligo rev AAACCGGGGCTATCAGTGCCGTCG
Chromosome start end strand MM target_seq PAM alignment distance position gene name gene id
GU305914.1 192115 192137 - 0 CGACGGCACTGATAGCCCCG GGG ||||||||[||||||||||||]PAM NA Intergenic NA NA

T20085 GACGGCACTGATAGCCCCGGGGG 1000
Oligo fwd CACCGACGGCACTGATAGCCCCGG
Oligo rev AAACCCGGGGCTATCAGTGCCGTC
Chromosome start end strand MM target_seq PAM alignment distance position gene name gene id
GU305914.1 192114 192136 - 0 GACGGCACTGATAGCCCCGG GGG ||||||||[||||||||||||]PAM NA Intergenic NA NA

T20086 ACGGCACTGATAGCCCCGGGGGG 1000
Oligo fwd CACCACGGCACTGATAGCCCCGGG
Oligo rev AAACCCCGGGGCTATCAGTGCCGT
Chromosome start end strand MM target_seq PAM alignment distance position gene name gene id
GU305914.1 192113 192135 - 0 ACGGCACTGATAGCCCCGGG GGG ||||||||[||||||||||||]PAM NA Intergenic NA NA

T20087 CGGCACTGATAGCCCCGGGGGGG 1000
Oligo fwd CACCCGGCACTGATAGCCCCGGGG
Oligo rev AAACCCCCGGGGCTATCAGTGCCG
Chromosome start end strand MM target_seq PAM alignment distance position gene name gene id
GU305914.1 192112 192134 - 0 CGGCACTGATAGCCCCGGGG GGG ||||||||[||||||||||||]PAM NA Intergenic NA NA

T20088 CACTGATAGCCCCGGGGGGGTGG 1000
Oligo fwd CACCCACTGATAGCCCCGGGGGGG
Oligo rev AAACCCCCCCCGGGGCTATCAGTG
Chromosome start end strand MM target_seq PAM alignment distance position gene name gene id
GU305914.1 192109 192131 - 0 CACTGATAGCCCCGGGGGGG TGG ||||||||[||||||||||||]PAM NA Intergenic NA NA

T20089 ACTGATAGCCCCGGGGGGGTGGG 1000
Oligo fwd CACCACTGATAGCCCCGGGGGGGT
Oligo rev AAACACCCCCCCGGGGCTATCAGT
Chromosome start end strand MM target_seq PAM alignment distance position gene name gene id
GU305914.1 192108 192130 - 0 ACTGATAGCCCCGGGGGGGT GGG ||||||||[||||||||||||]PAM NA Intergenic NA NA

T20090 CCGGGGGGGTGGGATCCCTCCGG 1000
Oligo fwd CACCCCGGGGGGGTGGGATCCCTC
Oligo rev AAACGAGGGATCCCACCCCCCCGG
Chromosome start end strand MM target_seq PAM alignment distance position gene name gene id
GU305914.1 192098 192120 - 0 CCGGGGGGGTGGGATCCCTC CGG ||||||||[||||||||||||]PAM NA Intergenic NA NA

T20091 TCGAACAGCCCGAGTTCGTGAGG 1000
Oligo fwd CACCTCGAACAGCCCGAGTTCGTG
Oligo rev AAACCACGAACTCGGGCTGTTCGA
Chromosome start end strand MM target_seq PAM alignment distance position gene name gene id
GU305914.1 192067 192089 - 0 TCGAACAGCCCGAGTTCGTG AGG ||||||||[||||||||||||]PAM NA Intergenic NA NA

T20092 CTGACGACCCGATTCTCCCGCGG 1000
Oligo fwd CACCCTGACGACCCGATTCTCCCG
Oligo rev AAACCGGGAGAATCGGGTCGTCAG
Chromosome start end strand MM target_seq PAM alignment distance position gene name gene id
GU305914.1 192005 192027 - 0 CTGACGACCCGATTCTCCCG CGG ||||||||[||||||||||||]PAM NA Intergenic NA NA

T20093 TGACGACCCGATTCTCCCGCGGG 1000
Oligo fwd CACCTGACGACCCGATTCTCCCGC
Oligo rev AAACGCGGGAGAATCGGGTCGTCA
Chromosome start end strand MM target_seq PAM alignment distance position gene name gene id
GU305914.1 192004 192026 - 0 TGACGACCCGATTCTCCCGC GGG ||||||||[||||||||||||]PAM NA Intergenic NA NA

T20094 GACGACCCGATTCTCCCGCGGGG 1000
Oligo fwd CACCGACGACCCGATTCTCCCGCG
Oligo rev AAACCGCGGGAGAATCGGGTCGTC
Chromosome start end strand MM target_seq PAM alignment distance position gene name gene id
GU305914.1 192003 192025 - 0 GACGACCCGATTCTCCCGCG GGG ||||||||[||||||||||||]PAM NA Intergenic NA NA

T20095 GACCCGATTCTCCCGCGGGGTGG 1000
Oligo fwd CACCGACCCGATTCTCCCGCGGGG
Oligo rev AAACCCCCGCGGGAGAATCGGGTC
Chromosome start end strand MM target_seq PAM alignment distance position gene name gene id
GU305914.1 192000 192022 - 0 GACCCGATTCTCCCGCGGGG TGG ||||||||[||||||||||||]PAM NA Intergenic NA NA

T20096 CCGATTCTCCCGCGGGGTGGTGG 1000
Oligo fwd CACCCCGATTCTCCCGCGGGGTGG
Oligo rev AAACCCACCCCGCGGGAGAATCGG
Chromosome start end strand MM target_seq PAM alignment distance position gene name gene id
GU305914.1 191997 192019 - 0 CCGATTCTCCCGCGGGGTGG TGG ||||||||[||||||||||||]PAM NA Intergenic NA NA

T20097 CGATTCTCCCGCGGGGTGGTGGG 1000

Oligo fwd CACCCGATTCTCCCGCGGGGTGGT
Oligo rev AAACACCACCCCGCGGGAGAATCG
Chromosome start end strand MM target_seq PAM alignment distance position gene name gene id
GU305914.1 191996 192018 - 0 CGATTCTCCCGCGGGGTGGT GGG ||||||||[||||||||||||]PAM NA Intergenic NA NA

T20098 TCTCCCGCGGGGTGGTGGGCTGG 1000
Oligo fwd CACCTCTCCCGCGGGGTGGTGGGC
Oligo rev AAACGCCCACCACCCCGCGGGAGA
Chromosome start end strand MM target_seq PAM alignment distance position gene name gene id
GU305914.1 191992 192014 - 0 TCTCCCGCGGGGTGGTGGGC TGG ||||||||[||||||||||||]PAM NA Intergenic NA NA

T20099 CTCCCGCGGGGTGGTGGGCTGGG 1000
Oligo fwd CACCCTCCCGCGGGGTGGTGGGCT
Oligo rev AAACAGCCCACCACCCCGCGGGAG
Chromosome start end strand MM target_seq PAM alignment distance position gene name gene id
GU305914.1 191991 192013 - 0 CTCCCGCGGGGTGGTGGGCT GGG ||||||||[||||||||||||]PAM NA Intergenic NA NA

T20100 GGTGGGCTGGGTCAAATGTTCGG 1000
Oligo fwd CACCGGTGGGCTGGGTCAAATGTT
Oligo rev AAACAACATTTGACCCAGCCCACC
Chromosome start end strand MM target_seq PAM alignment distance position gene name gene id
GU305914.1 191979 192001 - 0 GGTGGGCTGGGTCAAATGTT CGG ||||||||[||||||||||||]PAM NA Intergenic NA NA

T20101 GTGGGCTGGGTCAAATGTTCGGG 1000
Oligo fwd CACCGTGGGCTGGGTCAAATGTTC
Oligo rev AAACGAACATTTGACCCAGCCCAC
Chromosome start end strand MM target_seq PAM alignment distance position gene name gene id
GU305914.1 191978 192000 - 0 GTGGGCTGGGTCAAATGTTC GGG ||||||||[||||||||||||]PAM NA Intergenic NA NA

T20102 GGCTGGGTCAAATGTTCGGGCGG 1000
Oligo fwd CACCGGCTGGGTCAAATGTTCGGG
Oligo rev AAACCCCGAACATTTGACCCAGCC
Chromosome start end strand MM target_seq PAM alignment distance position gene name gene id
GU305914.1 191975 191997 - 0 GGCTGGGTCAAATGTTCGGG CGG ||||||||[||||||||||||]PAM NA Intergenic NA NA

T20103 AGAAGAGATGTCGTTCGATGTGG 1000
Oligo fwd CACCAGAAGAGATGTCGTTCGATG
Oligo rev AAACCATCGAACGACATCTCTTCT
Chromosome start end strand MM target_seq PAM alignment distance position gene name gene id
GU305914.1 191944 191966 - 0 AGAAGAGATGTCGTTCGATG TGG ||||||||[||||||||||||]PAM NA Intergenic NA NA

T20104 GAAGAGATGTCGTTCGATGTGGG 1000
Oligo fwd CACCGAAGAGATGTCGTTCGATGT
Oligo rev AAACACATCGAACGACATCTCTTC
Chromosome start end strand MM target_seq PAM alignment distance position gene name gene id
GU305914.1 191943 191965 - 0 GAAGAGATGTCGTTCGATGT GGG ||||||||[||||||||||||]PAM NA Intergenic NA NA

T20105 GAGATGTCGTTCGATGTGGGTGG 1000
Oligo fwd CACCGAGATGTCGTTCGATGTGGG
Oligo rev AAACCCCACATCGAACGACATCTC
Chromosome start end strand MM target_seq PAM alignment distance position gene name gene id
GU305914.1 191940 191962 - 0 GAGATGTCGTTCGATGTGGG TGG ||||||||[||||||||||||]PAM NA Intergenic NA NA

T20106 TCGATGTGGGTGGAATACGTCGG 1000
Oligo fwd CACCTCGATGTGGGTGGAATACGT
Oligo rev AAACACGTATTCCACCCACATCGA
Chromosome start end strand MM target_seq PAM alignment distance position gene name gene id
GU305914.1 191930 191952 - 0 TCGATGTGGGTGGAATACGT CGG ||||||||[||||||||||||]PAM NA Intergenic NA NA

T20107 CGATGTGGGTGGAATACGTCGGG 1000
Oligo fwd CACCCGATGTGGGTGGAATACGTC
Oligo rev AAACGACGTATTCCACCCACATCG
Chromosome start end strand MM target_seq PAM alignment distance position gene name gene id
GU305914.1 191929 191951 - 0 CGATGTGGGTGGAATACGTC GGG ||||||||[||||||||||||]PAM NA Intergenic NA NA

T20108 GATGTGGGTGGAATACGTCGGGG 1000
Oligo fwd CACCGATGTGGGTGGAATACGTCG
Oligo rev AAACCGACGTATTCCACCCACATC
Chromosome start end strand MM target_seq PAM alignment distance position gene name gene id
GU305914.1 191928 191950 - 0 GATGTGGGTGGAATACGTCG GGG ||||||||[||||||||||||]PAM NA Intergenic NA NA

T20109 GGTGGAATACGTCGGGGACACGG 1000
Oligo fwd CACCGGTGGAATACGTCGGGGACA
Oligo rev AAACTGTCCCCGACGTATTCCACC
Chromosome start end strand MM target_seq PAM alignment distance position gene name gene id
GU305914.1 191922 191944 - 0 GGTGGAATACGTCGGGGACA CGG ||||||||[||||||||||||]PAM NA Intergenic NA NA

T20110 CGTCGGGGACACGGACTTCGCGG 1000
Oligo fwd CACCCGTCGGGGACACGGACTTCG
Oligo rev AAACCGAAGTCCGTGTCCCCGACG
Chromosome start end strand MM target_seq PAM alignment distance position gene name gene id
GU305914.1 191913 191935 - 0 CGTCGGGGACACGGACTTCG CGG ||||||||[||||||||||||]PAM NA Intergenic NA NA

T20111 GTCGGGGACACGGACTTCGCGGG 1000
Oligo fwd CACCGTCGGGGACACGGACTTCGC
Oligo rev AAACGCGAAGTCCGTGTCCCCGAC
Chromosome start end strand MM target_seq PAM alignment distance position gene name gene id
GU305914.1 191912 191934 - 0 GTCGGGGACACGGACTTCGC GGG ||||||||[||||||||||||]PAM NA Intergenic NA NA

<table>
<tr><td>T20112</td><td>GGACACGGACTTCGCGGGAGAGG</td><td>1000</td><td></td><td></td><td></td><td></td><td></td><td></td><td></td><td></td><td></td></tr>
<tr><td>Oligo fwd</td><td>CACCGGACACGGACTTCGCGGGAG</td><td></td><td></td><td></td><td></td><td></td><td></td><td></td><td></td><td></td><td></td></tr>
<tr><td>Oligo rev</td><td>AAACCTCCCGCGAAGTCCGTGTCC</td><td></td><td></td><td></td><td></td><td></td><td></td><td></td><td></td><td></td><td></td></tr>
<tr><td>Chromosome</td><td>start</td><td>end</td><td>strand</td><td>MM</td><td>target_seq</td><td>PAM</td><td>alignment</td><td>distance</td><td>position</td><td>gene name</td><td>gene id</td></tr>
<tr><td>GU305914.1</td><td>191907</td><td>191929</td><td>-</td><td>0</td><td>GGACACGGACTTCGCGGGAG</td><td>AGG</td><td>||||||||[||||||||||||]PAM</td><td>NA</td><td>Intergenic</td><td>NA</td><td>NA</td></tr>
<tr><td></td><td></td><td></td><td></td><td></td><td></td><td></td><td></td><td></td><td></td><td></td><td></td></tr>
<tr><td>T20113</td><td>GACACGGACTTCGCGGGAGAGGG</td><td>1000</td><td></td><td></td><td></td><td></td><td></td><td></td><td></td><td></td><td></td></tr>
<tr><td>Oligo fwd</td><td>CACCGACACGGACTTCGCGGGAGA</td><td></td><td></td><td></td><td></td><td></td><td></td><td></td><td></td><td></td><td></td></tr>
<tr><td>Oligo rev</td><td>AAACTCTCCCGCGAAGTCCGTGTC</td><td></td><td></td><td></td><td></td><td></td><td></td><td></td><td></td><td></td><td></td></tr>
<tr><td>Chromosome</td><td>start</td><td>end</td><td>strand</td><td>MM</td><td>target_seq</td><td>PAM</td><td>alignment</td><td>distance</td><td>position</td><td>gene name</td><td>gene id</td></tr>
<tr><td>GU305914.1</td><td>191906</td><td>191928</td><td>-</td><td>0</td><td>GACACGGACTTCGCGGGAGA</td><td>GGG</td><td>||||||||[||||||||||||]PAM</td><td>NA</td><td>Intergenic</td><td>NA</td><td>NA</td></tr>
<tr><td></td><td></td><td></td><td></td><td></td><td></td><td></td><td></td><td></td><td></td><td></td><td></td></tr>
<tr><td>T20114</td><td>GGAGAGGGCGTCTACGACGACGG</td><td>1000</td><td></td><td></td><td></td><td></td><td></td><td></td><td></td><td></td><td></td></tr>
<tr><td>Oligo fwd</td><td>CACCGGAGAGGGCGTCTACGACGA</td><td></td><td></td><td></td><td></td><td></td><td></td><td></td><td></td><td></td><td></td></tr>
<tr><td>Oligo rev</td><td>AAACTCGTCGTAGACGCCCTCTCC</td><td></td><td></td><td></td><td></td><td></td><td></td><td></td><td></td><td></td><td></td></tr>
<tr><td>Chromosome</td><td>start</td><td>end</td><td>strand</td><td>MM</td><td>target_seq</td><td>PAM</td><td>alignment</td><td>distance</td><td>position</td><td>gene name</td><td>gene id</td></tr>
<tr><td>GU305914.1</td><td>191891</td><td>191913</td><td>-</td><td>0</td><td>GGAGAGGGCGTCTACGACGA</td><td>CGG</td><td>||||||||[||||||||||||]PAM</td><td>NA</td><td>Intergenic</td><td>NA</td><td>NA</td></tr>
<tr><td>GU305914.1</td><td>197894</td><td>197916</td><td>-</td><td>4</td><td>CGAGACGGAGTCTATGACGA</td><td>TGG</td><td>-||||-||[-|||||-|||||]PAM</td><td>NA</td><td>Intergenic</td><td>NA</td><td>NA</td></tr>
<tr><td></td><td></td><td></td><td></td><td></td><td></td><td></td><td></td><td></td><td></td><td></td><td></td></tr>
<tr><td>T20115</td><td>CGCCGATCACCGCACGCGTGCGG</td><td>1000</td><td></td><td></td><td></td><td></td><td></td><td></td><td></td><td></td><td></td></tr>
<tr><td>Oligo fwd</td><td>CACCCGCCGATCACCGCACGCGTG</td><td></td><td></td><td></td><td></td><td></td><td></td><td></td><td></td><td></td><td></td></tr>
<tr><td>Oligo rev</td><td>AAACCACGCGTGCGGTGATCGGCG</td><td></td><td></td><td></td><td></td><td></td><td></td><td></td><td></td><td></td><td></td></tr>
<tr><td>Chromosome</td><td>start</td><td>end</td><td>strand</td><td>MM</td><td>target_seq</td><td>PAM</td><td>alignment</td><td>distance</td><td>position</td><td>gene name</td><td>gene id</td></tr>
<tr><td>GU305914.1</td><td>191860</td><td>191882</td><td>-</td><td>0</td><td>CGCCGATCACCGCACGCGTG</td><td>CGG</td><td>||||||||[||||||||||||]PAM</td><td>NA</td><td>Intergenic</td><td>NA</td><td>NA</td></tr>
<tr><td></td><td></td><td></td><td></td><td></td><td></td><td></td><td></td><td></td><td></td><td></td><td></td></tr>
<tr><td>T20116</td><td>GAAGAGTGAGCCGCCCTTGCTGG</td><td>1000</td><td></td><td></td><td></td><td></td><td></td><td></td><td></td><td></td><td></td></tr>
<tr><td>Oligo fwd</td><td>CACCGAAGAGTGAGCCGCCCTTGC</td><td></td><td></td><td></td><td></td><td></td><td></td><td></td><td></td><td></td><td></td></tr>
<tr><td>Oligo rev</td><td>AAACGCAAGGGCGGCTCACTCTTC</td><td></td><td></td><td></td><td></td><td></td><td></td><td></td><td></td><td></td><td></td></tr>
<tr><td>Chromosome</td><td>start</td><td>end</td><td>strand</td><td>MM</td><td>target_seq</td><td>PAM</td><td>alignment</td><td>distance</td><td>position</td><td>gene name</td><td>gene id</td></tr>
<tr><td>GU305914.1</td><td>191838</td><td>191860</td><td>-</td><td>0</td><td>GAAGAGTGAGCCGCCCTTGC</td><td>TGG</td><td>||||||||[||||||||||||]PAM</td><td>NA</td><td>Intergenic</td><td>NA</td><td>NA</td></tr>
<tr><td></td><td></td><td></td><td></td><td></td><td></td><td></td><td></td><td></td><td></td><td></td><td></td></tr>
<tr><td>T20117</td><td>TGAGCCGCCCTTGCTGGACGTGG</td><td>1000</td><td></td><td></td><td></td><td></td><td></td><td></td><td></td><td></td><td></td></tr>
<tr><td>Oligo fwd</td><td>CACCTGAGCCGCCCTTGCTGGACG</td><td></td><td></td><td></td><td></td><td></td><td></td><td></td><td></td><td></td><td></td></tr>
<tr><td>Oligo rev</td><td>AAACCGTCCAGCAAGGGCGGCTCA</td><td></td><td></td><td></td><td></td><td></td><td></td><td></td><td></td><td></td><td></td></tr>
<tr><td>Chromosome</td><td>start</td><td>end</td><td>strand</td><td>MM</td><td>target_seq</td><td>PAM</td><td>alignment</td><td>distance</td><td>position</td><td>gene name</td><td>gene id</td></tr>
<tr><td>GU305914.1</td><td>191832</td><td>191854</td><td>-</td><td>0</td><td>TGAGCCGCCCTTGCTGGACG</td><td>TGG</td><td>||||||||[||||||||||||]PAM</td><td>NA</td><td>Intergenic</td><td>NA</td><td>NA</td></tr>
<tr><td></td><td></td><td></td><td></td><td></td><td></td><td></td><td></td><td></td><td></td><td></td><td></td></tr>
<tr><td>T20118</td><td>CCCTTGCTGGACGTGGACCTCGG</td><td>1000</td><td></td><td></td><td></td><td></td><td></td><td></td><td></td><td></td><td></td></tr>
<tr><td>Oligo fwd</td><td>CACCCCCTTGCTGGACGTGGACCT</td><td></td><td></td><td></td><td></td><td></td><td></td><td></td><td></td><td></td><td></td></tr>
<tr><td>Oligo rev</td><td>AAACAGGTCCACGTCCAGCAAGGG</td><td></td><td></td><td></td><td></td><td></td><td></td><td></td><td></td><td></td><td></td></tr>
<tr><td>Chromosome</td><td>start</td><td>end</td><td>strand</td><td>MM</td><td>target_seq</td><td>PAM</td><td>alignment</td><td>distance</td><td>position</td><td>gene name</td><td>gene id</td></tr>
<tr><td>GU305914.1</td><td>191825</td><td>191847</td><td>-</td><td>0</td><td>CCCTTGCTGGACGTGGACCT</td><td>CGG</td><td>||||||||[||||||||||||]PAM</td><td>NA</td><td>Intergenic</td><td>NA</td><td>NA</td></tr>
<tr><td></td><td></td><td></td><td></td><td></td><td></td><td></td><td></td><td></td><td></td><td></td><td></td></tr>
<tr><td>T20119</td><td>CCTTGCTGGACGTGGACCTCGGG</td><td>1000</td><td></td><td></td><td></td><td></td><td></td><td></td><td></td><td></td><td></td></tr>
<tr><td>Oligo fwd</td><td>CACCCCTTGCTGGACGTGGACCTC</td><td></td><td></td><td></td><td></td><td></td><td></td><td></td><td></td><td></td><td></td></tr>
<tr><td>Oligo rev</td><td>AAACGAGGTCCACGTCCAGCAAGG</td><td></td><td></td><td></td><td></td><td></td><td></td><td></td><td></td><td></td><td></td></tr>
<tr><td>Chromosome</td><td>start</td><td>end</td><td>strand</td><td>MM</td><td>target_seq</td><td>PAM</td><td>alignment</td><td>distance</td><td>position</td><td>gene name</td><td>gene id</td></tr>
<tr><td>GU305914.1</td><td>191824</td><td>191846</td><td>-</td><td>0</td><td>CCTTGCTGGACGTGGACCTC</td><td>GGG</td><td>||||||||[||||||||||||]PAM</td><td>NA</td><td>Intergenic</td><td>NA</td><td>NA</td></tr>
<tr><td></td><td></td><td></td><td></td><td></td><td></td><td></td><td></td><td></td><td></td><td></td><td></td></tr>
<tr><td>T20120</td><td>CGTGGACCTCGGGATCGAGCTGG</td><td>1000</td><td></td><td></td><td></td><td></td><td></td><td></td><td></td><td></td><td></td></tr>
<tr><td>Oligo fwd</td><td>CACCCGTGGACCTCGGGATCGAGC</td><td></td><td></td><td></td><td></td><td></td><td></td><td></td><td></td><td></td><td></td></tr>
<tr><td>Oligo rev</td><td>AAACGCTCGATCCCGAGGTCCACG</td><td></td><td></td><td></td><td></td><td></td><td></td><td></td><td></td><td></td><td></td></tr>
<tr><td>Chromosome</td><td>start</td><td>end</td><td>strand</td><td>MM</td><td>target_seq</td><td>PAM</td><td>alignment</td><td>distance</td><td>position</td><td>gene name</td><td>gene id</td></tr>
<tr><td>GU305914.1</td><td>191814</td><td>191836</td><td>-</td><td>0</td><td>CGTGGACCTCGGGATCGAGC</td><td>TGG</td><td>||||||||[||||||||||||]PAM</td><td>NA</td><td>Intergenic</td><td>NA</td><td>NA</td></tr>
<tr><td></td><td></td><td></td><td></td><td></td><td></td><td></td><td></td><td></td><td></td><td></td><td></td></tr>
<tr><td>T20121</td><td>TCGGGATCGAGCTGGAGCCGAGG</td><td>1000</td><td></td><td></td><td></td><td></td><td></td><td></td><td></td><td></td><td></td></tr>
<tr><td>Oligo fwd</td><td>CACCTCGGGATCGAGCTGGAGCCG</td><td></td><td></td><td></td><td></td><td></td><td></td><td></td><td></td><td></td><td></td></tr>
<tr><td>Oligo rev</td><td>AAACCGGCTCCAGCTCGATCCCGA</td><td></td><td></td><td></td><td></td><td></td><td></td><td></td><td></td><td></td><td></td></tr>
<tr><td>Chromosome</td><td>start</td><td>end</td><td>strand</td><td>MM</td><td>target_seq</td><td>PAM</td><td>alignment</td><td>distance</td><td>position</td><td>gene name</td><td>gene id</td></tr>
<tr><td>GU305914.1</td><td>191806</td><td>191828</td><td>-</td><td>0</td><td>TCGGGATCGAGCTGGAGCCG</td><td>AGG</td><td>||||||||[||||||||||||]PAM</td><td>NA</td><td>Intergenic</td><td>NA</td><td>NA</td></tr>
<tr><td></td><td></td><td></td><td></td><td></td><td></td><td></td><td></td><td></td><td></td><td></td><td></td></tr>
<tr><td>T20122</td><td>GGAGCCGAGGCACGAGAACCTGG</td><td>1000</td><td></td><td></td><td></td><td></td><td></td><td></td><td></td><td></td><td></td></tr>
<tr><td>Oligo fwd</td><td>CACCGGAGCCGAGGCACGAGAACC</td><td></td><td></td><td></td><td></td><td></td><td></td><td></td><td></td><td></td><td></td></tr>
<tr><td>Oligo rev</td><td>AAACGGTTCTCGTGCCTCGGCTCC</td><td></td><td></td><td></td><td></td><td></td><td></td><td></td><td></td><td></td><td></td></tr>
<tr><td>Chromosome</td><td>start</td><td>end</td><td>strand</td><td>MM</td><td>target_seq</td><td>PAM</td><td>alignment</td><td>distance</td><td>position</td><td>gene name</td><td>gene id</td></tr>
<tr><td>GU305914.1</td><td>191793</td><td>191815</td><td>-</td><td>0</td><td>GGAGCCGAGGCACGAGAACC</td><td>TGG</td><td>||||||||[||||||||||||]PAM</td><td>NA</td><td>Intergenic</td><td>NA</td><td>NA</td></tr>
<tr><td></td><td></td><td></td><td></td><td></td><td></td><td></td><td></td><td></td><td></td><td></td><td></td></tr>
<tr><td>T20123</td><td>GAGCCGAGGCACGAGAACCTGGG</td><td>1000</td><td></td><td></td><td></td><td></td><td></td><td></td><td></td><td></td><td></td></tr>
<tr><td>Oligo fwd</td><td>CACCGAGCCGAGGCACGAGAACCT</td><td></td><td></td><td></td><td></td><td></td><td></td><td></td><td></td><td></td><td></td></tr>
<tr><td>Oligo rev</td><td>AAACAGGTTCTCGTGCCTCGGCTC</td><td></td><td></td><td></td><td></td><td></td><td></td><td></td><td></td><td></td><td></td></tr>
<tr><td>Chromosome</td><td>start</td><td>end</td><td>strand</td><td>MM</td><td>target_seq</td><td>PAM</td><td>alignment</td><td>distance</td><td>position</td><td>gene name</td><td>gene id</td></tr>
<tr><td>GU305914.1</td><td>191792</td><td>191814</td><td>-</td><td>0</td><td>GAGCCGAGGCACGAGAACCT</td><td>GGG</td><td>||||||||[||||||||||||]PAM</td><td>NA</td><td>Intergenic</td><td>NA</td><td>NA</td></tr>
<tr><td></td><td></td><td></td><td></td><td></td><td></td><td></td><td></td><td></td><td></td><td></td><td></td></tr>
<tr><td>T20124</td><td>AGCCGAGGCACGAGAACCTGGGG</td><td>1000</td><td></td><td></td><td></td><td></td><td></td><td></td><td></td><td></td><td></td></tr>
<tr><td>Oligo fwd</td><td>CACCAGCCGAGGCACGAGAACCTG</td><td></td><td></td><td></td><td></td><td></td><td></td><td></td><td></td><td></td><td></td></tr>
<tr><td>Oligo rev</td><td>AAACCAGGTTCTCGTGCCTCGGCT</td><td></td><td></td><td></td><td></td><td></td><td></td><td></td><td></td><td></td><td></td></tr>
<tr><td>Chromosome</td><td>start</td><td>end</td><td>strand</td><td>MM</td><td>target_seq</td><td>PAM</td><td>alignment</td><td>distance</td><td>position</td><td>gene name</td><td>gene id</td></tr>
<tr><td>GU305914.1</td><td>191791</td><td>191813</td><td>-</td><td>0</td><td>AGCCGAGGCACGAGAACCTG</td><td>GGG</td><td>||||||||[||||||||||||]PAM</td><td>NA</td><td>Intergenic</td><td>NA</td><td>NA</td></tr>
<tr><td></td><td></td><td></td><td></td><td></td><td></td><td></td><td></td><td></td><td></td><td></td><td></td></tr>
<tr><td>T20125</td><td>GCCGAGGCACGAGAACCTGGGGG</td><td>1000</td><td></td><td></td><td></td><td></td><td></td><td></td><td></td><td></td><td></td></tr>
<tr><td>Oligo fwd</td><td>CACCGCCGAGGCACGAGAACCTGG</td><td></td><td></td><td></td><td></td><td></td><td></td><td></td><td></td><td></td><td></td></tr>
<tr><td>Oligo rev</td><td>AAACCCAGGTTCTCGTGCCTCGGC</td><td></td><td></td><td></td><td></td><td></td><td></td><td></td><td></td><td></td><td></td></tr>
<tr><td>Chromosome</td><td>start</td><td>end</td><td>strand</td><td>MM</td><td>target_seq</td><td>PAM</td><td>alignment</td><td>distance</td><td>position</td><td>gene name</td><td>gene id</td></tr>
<tr><td>GU305914.1</td><td>191790</td><td>191812</td><td>-</td><td>0</td><td>GCCGAGGCACGAGAACCTGG</td><td>GGG</td><td>||||||||[||||||||||||]PAM</td><td>NA</td><td>Intergenic</td><td>NA</td><td>NA</td></tr>
<tr><td></td><td></td><td></td><td></td><td></td><td></td><td></td><td></td><td></td><td></td><td></td><td></td></tr>
<tr><td>T20126</td><td>CCGAGGCACGAGAACCTGGGGGG</td><td>1000</td><td></td><td></td><td></td><td></td><td></td><td></td><td></td><td></td><td></td></tr>
<tr><td>Oligo fwd</td><td>CACCCCGAGGCACGAGAACCTGGG</td><td></td><td></td><td></td><td></td><td></td><td></td><td></td><td></td><td></td><td></td></tr>
<tr><td>Oligo rev</td><td>AAACCCCAGGTTCTCGTGCCTCGG</td><td></td><td></td><td></td><td></td><td></td><td></td><td></td><td></td><td></td><td></td></tr>
<tr><td>Chromosome</td><td>start</td><td>end</td><td>strand</td><td>MM</td><td>target_seq</td><td>PAM</td><td>alignment</td><td>distance</td><td>position</td><td>gene name</td><td>gene id</td></tr>
</table>

<table>
<tr><td>GU305914.1</td><td>191789</td><td>191811</td><td>-</td><td>0</td><td>CCGAGGCACGAGAACCTGGG</td><td>GGG</td><td>||||||||[||||||||||||]PAM</td><td>NA</td><td>Intergenic</td><td>NA</td><td>NA</td></tr>
<tr><td></td></tr>
<tr><td>T20127</td><td>CCTGGGGGGACACGCGTGCGTGG</td><td>1000</td></tr>
<tr><td>Oligo fwd</td><td>CACCCCTGGGGGGACACGCGTGCG</td></tr>
<tr><td>Oligo rev</td><td>AAACCGCACGCGTGTCCCCCCAGG</td></tr>
<tr><td>Chromosome</td><td>start</td><td>end</td><td>strand</td><td>MM</td><td>target_seq</td><td>PAM</td><td>alignment</td><td>distance</td><td>position</td><td>gene name</td><td>gene id</td></tr>
<tr><td>GU305914.1</td><td>191775</td><td>191797</td><td>-</td><td>0</td><td>CCTGGGGGGACACGCGTGCG</td><td>TGG</td><td>||||||||[||||||||||||]PAM</td><td>NA</td><td>Intergenic</td><td>NA</td><td>NA</td></tr>
<tr><td></td></tr>
<tr><td>T20128</td><td>GGGGGGACACGCGTGCGTGGTGG</td><td>1000</td></tr>
<tr><td>Oligo fwd</td><td>CACCGGGGGGACACGCGTGCGTGG</td></tr>
<tr><td>Oligo rev</td><td>AAACCCACGCACGCGTGTCCCCCC</td></tr>
<tr><td>Chromosome</td><td>start</td><td>end</td><td>strand</td><td>MM</td><td>target_seq</td><td>PAM</td><td>alignment</td><td>distance</td><td>position</td><td>gene name</td><td>gene id</td></tr>
<tr><td>GU305914.1</td><td>191772</td><td>191794</td><td>-</td><td>0</td><td>GGGGGGACACGCGTGCGTGG</td><td>TGG</td><td>||||||||[||||||||||||]PAM</td><td>NA</td><td>Intergenic</td><td>NA</td><td>NA</td></tr>
<tr><td></td></tr>
<tr><td>T20129</td><td>TGCGTGGTGGTGCCGCACCTCGG</td><td>1000</td></tr>
<tr><td>Oligo fwd</td><td>CACCTGCGTGGTGGTGCCGCACCT</td></tr>
<tr><td>Oligo rev</td><td>AAACAGGTGCGGCACCACCACGCA</td></tr>
<tr><td>Chromosome</td><td>start</td><td>end</td><td>strand</td><td>MM</td><td>target_seq</td><td>PAM</td><td>alignment</td><td>distance</td><td>position</td><td>gene name</td><td>gene id</td></tr>
<tr><td>GU305914.1</td><td>191759</td><td>191781</td><td>-</td><td>0</td><td>TGCGTGGTGGTGCCGCACCT</td><td>CGG</td><td>||||||||[||||||||||||]PAM</td><td>NA</td><td>Intergenic</td><td>NA</td><td>NA</td></tr>
<tr><td></td></tr>
<tr><td>T20130</td><td>GGTGCCGCACCTCGGCGTCGAGG</td><td>1000</td></tr>
<tr><td>Oligo fwd</td><td>CACCGGTGCCGCACCTCGGCGTCG</td></tr>
<tr><td>Oligo rev</td><td>AAACCGACGCCGAGGTGCGGCACC</td></tr>
<tr><td>Chromosome</td><td>start</td><td>end</td><td>strand</td><td>MM</td><td>target_seq</td><td>PAM</td><td>alignment</td><td>distance</td><td>position</td><td>gene name</td><td>gene id</td></tr>
<tr><td>GU305914.1</td><td>191751</td><td>191773</td><td>-</td><td>0</td><td>GGTGCCGCACCTCGGCGTCG</td><td>AGG</td><td>||||||||[||||||||||||]PAM</td><td>NA</td><td>Intergenic</td><td>NA</td><td>NA</td></tr>
<tr><td></td></tr>
<tr><td>T20131</td><td>CCTCGGCGTCGAGGACTCCTCGG</td><td>1000</td></tr>
<tr><td>Oligo fwd</td><td>CACCCCTCGGCGTCGAGGACTCCT</td></tr>
<tr><td>Oligo rev</td><td>AAACAGGAGTCCTCGACGCCGAGG</td></tr>
<tr><td>Chromosome</td><td>start</td><td>end</td><td>strand</td><td>MM</td><td>target_seq</td><td>PAM</td><td>alignment</td><td>distance</td><td>position</td><td>gene name</td><td>gene id</td></tr>
<tr><td>GU305914.1</td><td>191742</td><td>191764</td><td>-</td><td>0</td><td>CCTCGGCGTCGAGGACTCCT</td><td>CGG</td><td>||||||||[||||||||||||]PAM</td><td>NA</td><td>Intergenic</td><td>NA</td><td>NA</td></tr>
<tr><td></td></tr>
<tr><td>T20132</td><td>GCGCGCGACCTTCATACTCACGG</td><td>1000</td></tr>
<tr><td>Oligo fwd</td><td>CACCGCGCGCGACCTTCATACTCA</td></tr>
<tr><td>Oligo rev</td><td>AAACTGAGTATGAAGGTCGCGCGC</td></tr>
<tr><td>Chromosome</td><td>start</td><td>end</td><td>strand</td><td>MM</td><td>target_seq</td><td>PAM</td><td>alignment</td><td>distance</td><td>position</td><td>gene name</td><td>gene id</td></tr>
<tr><td>GU305914.1</td><td>191700</td><td>191722</td><td>-</td><td>0</td><td>GCGCGCGACCTTCATACTCA</td><td>CGG</td><td>||||||||[||||||||||||]PAM</td><td>NA</td><td>Intergenic</td><td>NA</td><td>NA</td></tr>
<tr><td></td></tr>
<tr><td>T20133</td><td>CGACCTTCATACTCACGGCGAGG</td><td>1000</td></tr>
<tr><td>Oligo fwd</td><td>CACCCGACCTTCATACTCACGGCG</td></tr>
<tr><td>Oligo rev</td><td>AAACCGCCGTGAGTATGAAGGTCG</td></tr>
<tr><td>Chromosome</td><td>start</td><td>end</td><td>strand</td><td>MM</td><td>target_seq</td><td>PAM</td><td>alignment</td><td>distance</td><td>position</td><td>gene name</td><td>gene id</td></tr>
<tr><td>GU305914.1</td><td>191695</td><td>191717</td><td>-</td><td>0</td><td>CGACCTTCATACTCACGGCG</td><td>AGG</td><td>||||||||[||||||||||||]PAM</td><td>NA</td><td>Intergenic</td><td>NA</td><td>NA</td></tr>
<tr><td></td></tr>
<tr><td>T20134</td><td>GACCTTCATACTCACGGCGAGGG</td><td>1000</td></tr>
<tr><td>Oligo fwd</td><td>CACCGACCTTCATACTCACGGCGA</td></tr>
<tr><td>Oligo rev</td><td>AAACTCGCCGTGAGTATGAAGGTC</td></tr>
<tr><td>Chromosome</td><td>start</td><td>end</td><td>strand</td><td>MM</td><td>target_seq</td><td>PAM</td><td>alignment</td><td>distance</td><td>position</td><td>gene name</td><td>gene id</td></tr>
<tr><td>GU305914.1</td><td>191694</td><td>191716</td><td>-</td><td>0</td><td>GACCTTCATACTCACGGCGA</td><td>GGG</td><td>||||||||[||||||||||||]PAM</td><td>NA</td><td>Intergenic</td><td>NA</td><td>NA</td></tr>
<tr><td></td></tr>
<tr><td>T20135</td><td>CGTGAAAGACTCCATCCTGTCGG</td><td>1000</td></tr>
<tr><td>Oligo fwd</td><td>CACCCGTGAAAGACTCCATCCTGT</td></tr>
<tr><td>Oligo rev</td><td>AAACACAGGATGGAGTCTTTCACG</td></tr>
<tr><td>Chromosome</td><td>start</td><td>end</td><td>strand</td><td>MM</td><td>target_seq</td><td>PAM</td><td>alignment</td><td>distance</td><td>position</td><td>gene name</td><td>gene id</td></tr>
<tr><td>GU305914.1</td><td>191655</td><td>191677</td><td>-</td><td>0</td><td>CGTGAAAGACTCCATCCTGT</td><td>CGG</td><td>||||||||[||||||||||||]PAM</td><td>NA</td><td>Intergenic</td><td>NA</td><td>NA</td></tr>
<tr><td></td></tr>
<tr><td>T20136</td><td>ATTCCACATGTCGAGCGCGATGG</td><td>1000</td></tr>
<tr><td>Oligo fwd</td><td>CACCATTCCACATGTCGAGCGCGA</td></tr>
<tr><td>Oligo rev</td><td>AAACTCGCGCTCGACATGTGGAAT</td></tr>
<tr><td>Chromosome</td><td>start</td><td>end</td><td>strand</td><td>MM</td><td>target_seq</td><td>PAM</td><td>alignment</td><td>distance</td><td>position</td><td>gene name</td><td>gene id</td></tr>
<tr><td>GU305914.1</td><td>191628</td><td>191650</td><td>-</td><td>0</td><td>ATTCCACATGTCGAGCGCGA</td><td>TGG</td><td>||||||||[||||||||||||]PAM</td><td>NA</td><td>Intergenic</td><td>NA</td><td>NA</td></tr>
<tr><td></td></tr>
<tr><td>T20137</td><td>GAGCGCGATGGTGTCACGCGCGG</td><td>1000</td></tr>
<tr><td>Oligo fwd</td><td>CACCGAGCGCGATGGTGTCACGCG</td></tr>
<tr><td>Oligo rev</td><td>AAACCGCGTGACACCATCGCGCTC</td></tr>
<tr><td>Chromosome</td><td>start</td><td>end</td><td>strand</td><td>MM</td><td>target_seq</td><td>PAM</td><td>alignment</td><td>distance</td><td>position</td><td>gene name</td><td>gene id</td></tr>
<tr><td>GU305914.1</td><td>191616</td><td>191638</td><td>-</td><td>0</td><td>GAGCGCGATGGTGTCACGCG</td><td>CGG</td><td>||||||||[||||||||||||]PAM</td><td>NA</td><td>Intergenic</td><td>NA</td><td>NA</td></tr>
<tr><td></td></tr>
<tr><td>T20138</td><td>GCGATGGTGTCACGCGCGGCCGG</td><td>1000</td></tr>
<tr><td>Oligo fwd</td><td>CACCGCGATGGTGTCACGCGCGGC</td></tr>
<tr><td>Oligo rev</td><td>AAACGCCGCGCGTGACACCATCGC</td></tr>
<tr><td>Chromosome</td><td>start</td><td>end</td><td>strand</td><td>MM</td><td>target_seq</td><td>PAM</td><td>alignment</td><td>distance</td><td>position</td><td>gene name</td><td>gene id</td></tr>
<tr><td>GU305914.1</td><td>191612</td><td>191634</td><td>-</td><td>0</td><td>GCGATGGTGTCACGCGCGGC</td><td>CGG</td><td>||||||||[||||||||||||]PAM</td><td>NA</td><td>Intergenic</td><td>NA</td><td>NA</td></tr>
<tr><td></td></tr>
<tr><td>T20139</td><td>CATCCTGCTCCGCCTCAGAGAGG</td><td>1000</td></tr>
<tr><td>Oligo fwd</td><td>CACCCATCCTGCTCCGCCTCAGAG</td></tr>
<tr><td>Oligo rev</td><td>AAACCTCTGAGGCGGAGCAGGATG</td></tr>
<tr><td>Chromosome</td><td>start</td><td>end</td><td>strand</td><td>MM</td><td>target_seq</td><td>PAM</td><td>alignment</td><td>distance</td><td>position</td><td>gene name</td><td>gene id</td></tr>
<tr><td>GU305914.1</td><td>191586</td><td>191608</td><td>-</td><td>0</td><td>CATCCTGCTCCGCCTCAGAG</td><td>AGG</td><td>||||||||[||||||||||||]PAM</td><td>NA</td><td>Intergenic</td><td>NA</td><td>NA</td></tr>
<tr><td></td></tr>
<tr><td>T20140</td><td>ATCCTGCTCCGCCTCAGAGAGGG</td><td>1000</td></tr>
<tr><td>Oligo fwd</td><td>CACCATCCTGCTCCGCCTCAGAGA</td></tr>
<tr><td>Oligo rev</td><td>AAACTCTCTGAGGCGGAGCAGGAT</td></tr>
<tr><td>Chromosome</td><td>start</td><td>end</td><td>strand</td><td>MM</td><td>target_seq</td><td>PAM</td><td>alignment</td><td>distance</td><td>position</td><td>gene name</td><td>gene id</td></tr>
<tr><td>GU305914.1</td><td>191585</td><td>191607</td><td>-</td><td>0</td><td>ATCCTGCTCCGCCTCAGAGA</td><td>GGG</td><td>||||||||[||||||||||||]PAM</td><td>NA</td><td>Intergenic</td><td>NA</td><td>NA</td></tr>
<tr><td></td></tr>
<tr><td>T20141</td><td>CCGAAGTGCACTTCGAGACGAGG</td><td>1000</td></tr>
<tr><td>Oligo fwd</td><td>CACCCCGAAGTGCACTTCGAGACG</td></tr>
<tr><td>Oligo rev</td><td>AAACCGTCTCGAAGTGCACTTCGG</td></tr>
</table>

<table>
<tr><td>Chromosome</td><td>start</td><td>end</td><td>strand</td><td>MM</td><td>target_seq</td><td>PAM</td><td>alignment</td><td>distance</td><td>position</td><td>gene name</td><td>gene id</td></tr>
<tr><td>GU305914.1</td><td>191560</td><td>191582</td><td>-</td><td>0</td><td>CCGAAGTGCACTTCGAGACG</td><td>AGG</td><td>||||||||[||||||||||||]PAM</td><td>NA</td><td>Intergenic</td><td>NA</td><td>NA</td></tr>
<tr><td>T20142</td><td>AGTGCACTTCGAGACGAGGCTGG</td><td>1000</td></tr>
<tr><td>Oligo fwd</td><td>CACCAGTGCACTTCGAGACGAGGC</td></tr>
<tr><td>Oligo rev</td><td>AAACGCCTCGTCTCGAAGTGCACT</td></tr>
<tr><td>Chromosome</td><td>start</td><td>end</td><td>strand</td><td>MM</td><td>target_seq</td><td>PAM</td><td>alignment</td><td>distance</td><td>position</td><td>gene name</td><td>gene id</td></tr>
<tr><td>GU305914.1</td><td>191556</td><td>191578</td><td>-</td><td>0</td><td>AGTGCACTTCGAGACGAGGC</td><td>TGG</td><td>||||||||[||||||||||||]PAM</td><td>NA</td><td>Intergenic</td><td>NA</td><td>NA</td></tr>
<tr><td>T20143</td><td>GAGGCTGGATCACATCAAGTCGG</td><td>1000</td></tr>
<tr><td>Oligo fwd</td><td>CACCGAGGCTGGATCACATCAAGT</td></tr>
<tr><td>Oligo rev</td><td>AAACACTTGATGTGATCCAGCCTC</td></tr>
<tr><td>Chromosome</td><td>start</td><td>end</td><td>strand</td><td>MM</td><td>target_seq</td><td>PAM</td><td>alignment</td><td>distance</td><td>position</td><td>gene name</td><td>gene id</td></tr>
<tr><td>GU305914.1</td><td>191541</td><td>191563</td><td>-</td><td>0</td><td>GAGGCTGGATCACATCAAGT</td><td>CGG</td><td>||||||||[||||||||||||]PAM</td><td>NA</td><td>Intergenic</td><td>NA</td><td>NA</td></tr>
<tr><td>T20144</td><td>AAGTCGGCGATCAGCAACCCCGG</td><td>1000</td></tr>
<tr><td>Oligo fwd</td><td>CACCAAGTCGGCGATCAGCAACCC</td></tr>
<tr><td>Oligo rev</td><td>AAACGGGTTGCTGATCGCCGACTT</td></tr>
<tr><td>Chromosome</td><td>start</td><td>end</td><td>strand</td><td>MM</td><td>target_seq</td><td>PAM</td><td>alignment</td><td>distance</td><td>position</td><td>gene name</td><td>gene id</td></tr>
<tr><td>GU305914.1</td><td>191525</td><td>191547</td><td>-</td><td>0</td><td>AAGTCGGCGATCAGCAACCC</td><td>CGG</td><td>||||||||[||||||||||||]PAM</td><td>NA</td><td>Intergenic</td><td>NA</td><td>NA</td></tr>
<tr><td>T20145</td><td>AGTCGGCGATCAGCAACCCCGGG</td><td>1000</td></tr>
<tr><td>Oligo fwd</td><td>CACCAGTCGGCGATCAGCAACCCC</td></tr>
<tr><td>Oligo rev</td><td>AAACGGGGTTGCTGATCGCCGACT</td></tr>
<tr><td>Chromosome</td><td>start</td><td>end</td><td>strand</td><td>MM</td><td>target_seq</td><td>PAM</td><td>alignment</td><td>distance</td><td>position</td><td>gene name</td><td>gene id</td></tr>
<tr><td>GU305914.1</td><td>191524</td><td>191546</td><td>-</td><td>0</td><td>AGTCGGCGATCAGCAACCCC</td><td>GGG</td><td>||||||||[||||||||||||]PAM</td><td>NA</td><td>Intergenic</td><td>NA</td><td>NA</td></tr>
<tr><td>T20146</td><td>GCTGAACTTCTCGAAAACGACGG</td><td>1000</td></tr>
<tr><td>Oligo fwd</td><td>CACCGCTGAACTTCTCGAAAACGA</td></tr>
<tr><td>Oligo rev</td><td>AAACTCGTTTTCGAGAAGTTCAGC</td></tr>
<tr><td>Chromosome</td><td>start</td><td>end</td><td>strand</td><td>MM</td><td>target_seq</td><td>PAM</td><td>alignment</td><td>distance</td><td>position</td><td>gene name</td><td>gene id</td></tr>
<tr><td>GU305914.1</td><td>191502</td><td>191524</td><td>-</td><td>0</td><td>GCTGAACTTCTCGAAAACGA</td><td>CGG</td><td>||||||||[||||||||||||]PAM</td><td>NA</td><td>Intergenic</td><td>NA</td><td>NA</td></tr>
<tr><td>T20147</td><td>CTTCTCGAAAACGACGGAGAAGG</td><td>1000</td></tr>
<tr><td>Oligo fwd</td><td>CACCCTTCTCGAAAACGACGGAGA</td></tr>
<tr><td>Oligo rev</td><td>AAACTCTCCGTCGTTTTCGAGAAG</td></tr>
<tr><td>Chromosome</td><td>start</td><td>end</td><td>strand</td><td>MM</td><td>target_seq</td><td>PAM</td><td>alignment</td><td>distance</td><td>position</td><td>gene name</td><td>gene id</td></tr>
<tr><td>GU305914.1</td><td>191496</td><td>191518</td><td>-</td><td>0</td><td>CTTCTCGAAAACGACGGAGA</td><td>AGG</td><td>||||||||[||||||||||||]PAM</td><td>NA</td><td>Intergenic</td><td>NA</td><td>NA</td></tr>
<tr><td>T20148</td><td>CGTCACGACGTTCGCGCTGACGG</td><td>1000</td></tr>
<tr><td>Oligo fwd</td><td>CACCCGTCACGACGTTCGCGCTGA</td></tr>
<tr><td>Oligo rev</td><td>AAACTCAGCGCGAACGTCGTGACG</td></tr>
<tr><td>Chromosome</td><td>start</td><td>end</td><td>strand</td><td>MM</td><td>target_seq</td><td>PAM</td><td>alignment</td><td>distance</td><td>position</td><td>gene name</td><td>gene id</td></tr>
<tr><td>GU305914.1</td><td>191469</td><td>191491</td><td>-</td><td>0</td><td>CGTCACGACGTTCGCGCTGA</td><td>CGG</td><td>||||||||[||||||||||||]PAM</td><td>NA</td><td>Intergenic</td><td>NA</td><td>NA</td></tr>
<tr><td>T20149</td><td>CAAGATCGACCCGTTCGATCTGG</td><td>1000</td></tr>
<tr><td>Oligo fwd</td><td>CACCCAAGATCGACCCGTTCGATC</td></tr>
<tr><td>Oligo rev</td><td>AAACGATCGAACGGGTCGATCTTG</td></tr>
<tr><td>Chromosome</td><td>start</td><td>end</td><td>strand</td><td>MM</td><td>target_seq</td><td>PAM</td><td>alignment</td><td>distance</td><td>position</td><td>gene name</td><td>gene id</td></tr>
<tr><td>GU305914.1</td><td>191445</td><td>191467</td><td>-</td><td>0</td><td>CAAGATCGACCCGTTCGATC</td><td>TGG</td><td>||||||||[||||||||||||]PAM</td><td>NA</td><td>Intergenic</td><td>NA</td><td>NA</td></tr>
<tr><td>T20150</td><td>AAGATCGACCCGTTCGATCTGGG</td><td>1000</td></tr>
<tr><td>Oligo fwd</td><td>CACCAAGATCGACCCGTTCGATCT</td></tr>
<tr><td>Oligo rev</td><td>AAACAGATCGAACGGGTCGATCTT</td></tr>
<tr><td>Chromosome</td><td>start</td><td>end</td><td>strand</td><td>MM</td><td>target_seq</td><td>PAM</td><td>alignment</td><td>distance</td><td>position</td><td>gene name</td><td>gene id</td></tr>
<tr><td>GU305914.1</td><td>191444</td><td>191466</td><td>-</td><td>0</td><td>AAGATCGACCCGTTCGATCT</td><td>GGG</td><td>||||||||[||||||||||||]PAM</td><td>NA</td><td>Intergenic</td><td>NA</td><td>NA</td></tr>
<tr><td>T20151</td><td>GTTCGATCTGGGTTTCAGCATGG</td><td>1000</td></tr>
<tr><td>Oligo fwd</td><td>CACCGTTCGATCTGGGTTTCAGCA</td></tr>
<tr><td>Oligo rev</td><td>AAACTGCTGAAACCCAGATCGAAC</td></tr>
<tr><td>Chromosome</td><td>start</td><td>end</td><td>strand</td><td>MM</td><td>target_seq</td><td>PAM</td><td>alignment</td><td>distance</td><td>position</td><td>gene name</td><td>gene id</td></tr>
<tr><td>GU305914.1</td><td>191433</td><td>191455</td><td>-</td><td>0</td><td>GTTCGATCTGGGTTTCAGCA</td><td>TGG</td><td>||||||||[||||||||||||]PAM</td><td>NA</td><td>Intergenic</td><td>NA</td><td>NA</td></tr>
<tr><td>T20152</td><td>CGATCTGGGTTTCAGCATGGAGG</td><td>1000</td></tr>
<tr><td>Oligo fwd</td><td>CACCCGATCTGGGTTTCAGCATGG</td></tr>
<tr><td>Oligo rev</td><td>AAACCCATGCTGAAACCCAGATCG</td></tr>
<tr><td>Chromosome</td><td>start</td><td>end</td><td>strand</td><td>MM</td><td>target_seq</td><td>PAM</td><td>alignment</td><td>distance</td><td>position</td><td>gene name</td><td>gene id</td></tr>
<tr><td>GU305914.1</td><td>191430</td><td>191452</td><td>-</td><td>0</td><td>CGATCTGGGTTTCAGCATGG</td><td>AGG</td><td>||||||||[||||||||||||]PAM</td><td>NA</td><td>Intergenic</td><td>NA</td><td>NA</td></tr>
<tr><td>T20153</td><td>TTCAGCATGGAGGTCGACGACGG</td><td>1000</td></tr>
<tr><td>Oligo fwd</td><td>CACCTTCAGCATGGAGGTCGACGA</td></tr>
<tr><td>Oligo rev</td><td>AAACTCGTCGACCTCCATGCTGAA</td></tr>
<tr><td>Chromosome</td><td>start</td><td>end</td><td>strand</td><td>MM</td><td>target_seq</td><td>PAM</td><td>alignment</td><td>distance</td><td>position</td><td>gene name</td><td>gene id</td></tr>
<tr><td>GU305914.1</td><td>191420</td><td>191442</td><td>-</td><td>0</td><td>TTCAGCATGGAGGTCGACGA</td><td>CGG</td><td>||||||||[||||||||||||]PAM</td><td>NA</td><td>Intergenic</td><td>NA</td><td>NA</td></tr>
<tr><td>T20154</td><td>CATGGAGGTCGACGACGGTCAGG</td><td>1000</td></tr>
<tr><td>Oligo fwd</td><td>CACCCATGGAGGTCGACGACGGTC</td></tr>
<tr><td>Oligo rev</td><td>AAACGACCGTCGTCGACCTCCATG</td></tr>
<tr><td>Chromosome</td><td>start</td><td>end</td><td>strand</td><td>MM</td><td>target_seq</td><td>PAM</td><td>alignment</td><td>distance</td><td>position</td><td>gene name</td><td>gene id</td></tr>
<tr><td>GU305914.1</td><td>191415</td><td>191437</td><td>-</td><td>0</td><td>CATGGAGGTCGACGACGGTC</td><td>AGG</td><td>||||||||[||||||||||||]PAM</td><td>NA</td><td>Intergenic</td><td>NA</td><td>NA</td></tr>
<tr><td>T20155</td><td>ATGGAGGTCGACGACGGTCAGGG</td><td>1000</td></tr>
<tr><td>Oligo fwd</td><td>CACCATGGAGGTCGACGACGGTCA</td></tr>
<tr><td>Oligo rev</td><td>AAACTGACCGTCGTCGACCTCCAT</td></tr>
<tr><td>Chromosome</td><td>start</td><td>end</td><td>strand</td><td>MM</td><td>target_seq</td><td>PAM</td><td>alignment</td><td>distance</td><td>position</td><td>gene name</td><td>gene id</td></tr>
<tr><td>GU305914.1</td><td>191414</td><td>191436</td><td>-</td><td>0</td><td>ATGGAGGTCGACGACGGTCA</td><td>GGG</td><td>||||||||[||||||||||||]PAM</td><td>NA</td><td>Intergenic</td><td>NA</td><td>NA</td></tr>
<tr><td>T20156</td><td>GAGGTCGACGACGGTCAGGGCGG</td><td>1000</td></tr>
<tr><td>Oligo fwd</td><td>CACCGAGGTCGACGACGGTCAGGG</td></tr>
</table>

Oligo rev AAACCCCTGACCGTCGTCGACCTC
Chromosome start end strand MM target_seq PAM alignment distance position gene name gene id
GU305914.1 191411 191433 - 0 GAGGTCGACGACGGTCAGGG CGG ||||||||[||||||||||||]PAM NA Intergenic NA NA

T20157 GACGGTCAGGGCGGTTTCGTCGG 1000
Oligo fwd CACCGACGGTCAGGGCGGTTTCGT
Oligo rev AAACACGAAACCGCCCTGACCGTC
Chromosome start end strand MM target_seq PAM alignment distance position gene name gene id
GU305914.1 191402 191424 - 0 GACGGTCAGGGCGGTTTCGT CGG ||||||||[||||||||||||]PAM NA Intergenic NA NA

T20158 GTCAGGGCGGTTTCGTCGGCAGG 1000
Oligo fwd CACCGTCAGGGCGGTTTCGTCGGC
Oligo rev AAACGCCGACGAAACCGCCCTGAC
Chromosome start end strand MM target_seq PAM alignment distance position gene name gene id
GU305914.1 191398 191420 - 0 GTCAGGGCGGTTTCGTCGGC AGG ||||||||[||||||||||||]PAM NA Intergenic NA NA

T20159 GGGCGGTTTCGTCGGCAGGTCGG 1000
Oligo fwd CACCGGGCGGTTTCGTCGGCAGGT
Oligo rev AAACACCTGCCGACGAAACCGCCC
Chromosome start end strand MM target_seq PAM alignment distance position gene name gene id
GU305914.1 191394 191416 - 0 GGGCGGTTTCGTCGGCAGGT CGG ||||||||[||||||||||||]PAM NA Intergenic NA NA

T20160 GGCGGTTTCGTCGGCAGGTCGGG 1000
Oligo fwd CACCGGCGGTTTCGTCGGCAGGTC
Oligo rev AAACGACCTGCCGACGAAACCGCC
Chromosome start end strand MM target_seq PAM alignment distance position gene name gene id
GU305914.1 191393 191415 - 0 GGCGGTTTCGTCGGCAGGTC GGG ||||||||[||||||||||||]PAM NA Intergenic NA NA

T20161 GGGTGTCGAGCTCGAGCGACCGG 1000
Oligo fwd CACCGGGTGTCGAGCTCGAGCGAC
Oligo rev AAACGTCGCTCGAGCTCGACACCC
Chromosome start end strand MM target_seq PAM alignment distance position gene name gene id
GU305914.1 191373 191395 - 0 GGGTGTCGAGCTCGAGCGAC CGG ||||||||[||||||||||||]PAM NA Intergenic NA NA

T20162 TCGAGCTCGAGCGACCGGCGCGG 1000
Oligo fwd CACCTCGAGCTCGAGCGACCGGCG
Oligo rev AAACCGCCGGTCGCTCGAGCTCGA
Chromosome start end strand MM target_seq PAM alignment distance position gene name gene id
GU305914.1 191368 191390 - 0 TCGAGCTCGAGCGACCGGCG CGG ||||||||[||||||||||||]PAM NA Intergenic NA NA

T20163 CGAGCTCGAGCGACCGGCGCGGG 1000
Oligo fwd CACCCGAGCTCGAGCGACCGGCGC
Oligo rev AAACGCGCCGGTCGCTCGAGCTCG
Chromosome start end strand MM target_seq PAM alignment distance position gene name gene id
GU305914.1 191367 191389 - 0 CGAGCTCGAGCGACCGGCGC GGG ||||||||[||||||||||||]PAM NA Intergenic NA NA

T20164 GAGCTCGAGCGACCGGCGCGGGG 1000
Oligo fwd CACCGAGCTCGAGCGACCGGCGCG
Oligo rev AAACCGCGCCGGTCGCTCGAGCTC
Chromosome start end strand MM target_seq PAM alignment distance position gene name gene id
GU305914.1 191366 191388 - 0 GAGCTCGAGCGACCGGCGCG GGG ||||||||[||||||||||||]PAM NA Intergenic NA NA

T20165 AGCTCGAGCGACCGGCGCGGGGG 1000
Oligo fwd CACCAGCTCGAGCGACCGGCGCGG
Oligo rev AAACCCGCGCCGGTCGCTCGAGCT
Chromosome start end strand MM target_seq PAM alignment distance position gene name gene id
GU305914.1 191365 191387 - 0 AGCTCGAGCGACCGGCGCGG GGG ||||||||[||||||||||||]PAM NA Intergenic NA NA

T20166 AGCGACCGGCGCGGGGGTTCTGG 1000
Oligo fwd CACCAGCGACCGGCGCGGGGGTTC
Oligo rev AAACGAACCCCCGCGCCGGTCGCT
Chromosome start end strand MM target_seq PAM alignment distance position gene name gene id
GU305914.1 191359 191381 - 0 AGCGACCGGCGCGGGGGTTC TGG ||||||||[||||||||||||]PAM NA Intergenic NA NA

T20167 GACCGGCGCGGGGGTTCTGGTGG 1000
Oligo fwd CACCGACCGGCGCGGGGGTTCTGG
Oligo rev AAACCCAGAACCCCCGCGCCGGTC
Chromosome start end strand MM target_seq PAM alignment distance position gene name gene id
GU305914.1 191356 191378 - 0 GACCGGCGCGGGGGTTCTGG TGG ||||||||[||||||||||||]PAM NA Intergenic NA NA

T20168 GGCGCGGGGGTTCTGGTGGAAGG 1000
Oligo fwd CACCGGCGCGGGGGTTCTGGTGGA
Oligo rev AAACTCCACCAGAACCCCCGCGCC
Chromosome start end strand MM target_seq PAM alignment distance position gene name gene id
GU305914.1 191352 191374 - 0 GGCGCGGGGGTTCTGGTGGA AGG ||||||||[||||||||||||]PAM NA Intergenic NA NA

T20169 GATCGTCGCGACCGTGCTACTGG 1000
Oligo fwd CACCGATCGTCGCGACCGTGCTAC
Oligo rev AAACGTAGCACGGTCGCGACGATC
Chromosome start end strand MM target_seq PAM alignment distance position gene name gene id
GU305914.1 191313 191335 - 0 GATCGTCGCGACCGTGCTAC TGG ||||||||[||||||||||||]PAM NA Intergenic NA NA
GU305914.1 213883 213905 + 4 GGTCGTAGCGAACGTGGTAC CGG |-||||-|[|||-||||-|||]PAM NA Intergenic NA NA

T20170 GACCGTGCTACTGGCCCTGCTGG 1000
Oligo fwd CACCGACCGTGCTACTGGCCCTGC
Oligo rev AAACGCAGGGCCAGTAGCACGGTC
Chromosome start end strand MM target_seq PAM alignment distance position gene name gene id
GU305914.1 191304 191326 - 0 GACCGTGCTACTGGCCCTGC TGG ||||||||[||||||||||||]PAM NA Intergenic NA NA

T20171 ACCGTGCTACTGGCCCTGCTGGG 1000
Oligo fwd CACCACCGTGCTACTGGCCCTGCT
Oligo rev AAACAGCAGGGCCAGTAGCACGGT
Chromosome start end strand MM target_seq PAM alignment distance position gene name gene id
GU305914.1 191303 191325 - 0 ACCGTGCTACTGGCCCTGCT GGG ||||||||[||||||||||||]PAM NA Intergenic NA NA

T20172 GTACTTCGTGACCCCGCACACGG 1000
Oligo fwd CACCGTACTTCGTGACCCCGCACA
Oligo rev AAACTGTGCGGGGTCACGAAGTAC
Chromosome start end strand MM target_seq PAM alignment distance position gene name gene id
GU305914.1 191274 191296 - 0 GTACTTCGTGACCCCGCACA CGG ||||||||[||||||||||||]PAM NA Intergenic NA NA

T20173 CGTGACCCCGCACACGGAGAAGG 1000
Oligo fwd CACCCGTGACCCCGCACACGGAGA
Oligo rev AAACTCTCCGTGTGCGGGGTCACG
Chromosome start end strand MM target_seq PAM alignment distance position gene name gene id
GU305914.1 191268 191290 - 0 CGTGACCCCGCACACGGAGA AGG ||||||||[||||||||||||]PAM NA Intergenic NA NA

T20174 CGGAGAAGGCATTCATCGCCTGG 1000
Oligo fwd CACCCGGAGAAGGCATTCATCGCC
Oligo rev AAACGGCGATGAATGCCTTCTCCG
Chromosome start end strand MM target_seq PAM alignment distance position gene name gene id
GU305914.1 191254 191276 - 0 CGGAGAAGGCATTCATCGCC TGG ||||||||[||||||||||||]PAM NA Intergenic NA NA

T20175 GCCTGGTGCGCGATGTTCATCGG 1000
Oligo fwd CACCGCCTGGTGCGCGATGTTCAT
Oligo rev AAACATGAACATCGCGCACCAGGC
Chromosome start end strand MM target_seq PAM alignment distance position gene name gene id
GU305914.1 191237 191259 - 0 GCCTGGTGCGCGATGTTCAT CGG ||||||||[||||||||||||]PAM NA Intergenic NA NA

T20176 GGTGCGCGATGTTCATCGGATGG 1000
Oligo fwd CACCGGTGCGCGATGTTCATCGGA
Oligo rev AAACTCCGATGAACATCGCGCACC
Chromosome start end strand MM target_seq PAM alignment distance position gene name gene id
GU305914.1 191233 191255 - 0 GGTGCGCGATGTTCATCGGA TGG ||||||||[||||||||||||]PAM NA Intergenic NA NA

T20177 GTGCGCGATGTTCATCGGATGGG 1000
Oligo fwd CACCGTGCGCGATGTTCATCGGAT
Oligo rev AAACATCCGATGAACATCGCGCAC
Chromosome start end strand MM target_seq PAM alignment distance position gene name gene id
GU305914.1 191232 191254 - 0 GTGCGCGATGTTCATCGGAT GGG ||||||||[||||||||||||]PAM NA Intergenic NA NA

T20178 TTCATCGGATGGGTGCTCCGAGG 1000
Oligo fwd CACCTTCATCGGATGGGTGCTCCG
Oligo rev AAACCGGAGCACCCATCCGATGAA
Chromosome start end strand MM target_seq PAM alignment distance position gene name gene id
GU305914.1 191222 191244 - 0 TTCATCGGATGGGTGCTCCG AGG ||||||||[||||||||||||]PAM NA Intergenic NA NA

T20179 AAGATTTACAAGAACTGTGACGG 1000
Oligo fwd CACCAAGATTTACAAGAACTGTGA
Oligo rev AAACTCACAGTTCTTGTAAATCTT
Chromosome start end strand MM target_seq PAM alignment distance position gene name gene id
GU305914.1 191185 191207 - 0 AAGATTTACAAGAACTGTGA CGG ||||||||[||||||||||||]PAM NA Intergenic NA NA

T20180 AGATTTACAAGAACTGTGACGGG 1000
Oligo fwd CACCAGATTTACAAGAACTGTGAC
Oligo rev AAACGTCACAGTTCTTGTAAATCT
Chromosome start end strand MM target_seq PAM alignment distance position gene name gene id
GU305914.1 191184 191206 - 0 AGATTTACAAGAACTGTGAC GGG ||||||||[||||||||||||]PAM NA Intergenic NA NA

T20181 GATTTACAAGAACTGTGACGGGG 1000
Oligo fwd CACCGATTTACAAGAACTGTGACG
Oligo rev AAACCGTCACAGTTCTTGTAAATC
Chromosome start end strand MM target_seq PAM alignment distance position gene name gene id
GU305914.1 191183 191205 - 0 GATTTACAAGAACTGTGACG GGG ||||||||[||||||||||||]PAM NA Intergenic NA NA

T20182 CAATAAAATGTTTTTACCGACGG 1000
Oligo fwd CACCCAATAAAATGTTTTTACCGA
Oligo rev AAACTCGGTAAAAACATTTTATTG
Chromosome start end strand MM target_seq PAM alignment distance position gene name gene id
GU305914.1 191156 191178 - 0 CAATAAAATGTTTTTACCGA CGG ||||||||[||||||||||||]PAM NA Intergenic NA NA

T20183 TAAAATGTTTTTACCGACGGCGG 1000
Oligo fwd CACCTAAAATGTTTTTACCGACGG
Oligo rev AAACCCGTCGGTAAAAACATTTTA
Chromosome start end strand MM target_seq PAM alignment distance position gene name gene id
GU305914.1 191153 191175 - 0 TAAAATGTTTTTACCGACGG CGG ||||||||[||||||||||||]PAM NA Intergenic NA NA

T20184 AATGTTTTTACCGACGGCGGCGG 1000
Oligo fwd CACCAATGTTTTTACCGACGGCGG
Oligo rev AAACCCGCCGTCGGTAAAAACATT
Chromosome start end strand MM target_seq PAM alignment distance position gene name gene id
GU305914.1 191150 191172 - 0 AATGTTTTTACCGACGGCGG CGG ||||||||[||||||||||||]PAM NA Intergenic NA NA

T20185 TTTTACCGACGGCGGCGGCGTGG 1000
Oligo fwd CACCTTTTACCGACGGCGGCGGCG
Oligo rev AAACCGCCGCCGCCGTCGGTAAAA
Chromosome start end strand MM target_seq PAM alignment distance position gene name gene id
GU305914.1 191145 191167 - 0 TTTTACCGACGGCGGCGGCG TGG ||||||||[||||||||||||]PAM NA Intergenic NA NA

T20186 GCGTGGTTATTTCTCGCGCACGG 1000
Oligo fwd CACCGCGTGGTTATTTCTCGCGCA
Oligo rev AAACTGCGCGAGAAATAACCACGC
Chromosome start end strand MM target_seq PAM alignment distance position gene name gene id
GU305914.1 191128 191150 - 0 GCGTGGTTATTTCTCGCGCA CGG ||||||||[||||||||||||]PAM NA Intergenic NA NA

T20187 ACTACCGTCTCGCAGACGCGAGG 1000
Oligo fwd CACCACTACCGTCTCGCAGACGCG
Oligo rev AAACCGCGTCTGCGAGACGGTAGT
Chromosome start end strand MM target_seq PAM alignment distance position gene name gene id
GU305914.1 191086 191108 - 0 ACTACCGTCTCGCAGACGCG AGG ||||||||[||||||||||||]PAM NA Intergenic NA NA

T20188 GTAACAAAACCTACATCGAGTGG 1000
Oligo fwd CACCGTAACAAAACCTACATCGAG
Oligo rev AAACCTCGATGTAGGTTTTGTTAC
Chromosome start end strand MM target_seq PAM alignment distance position gene name gene id
GU305914.1 191049 191071 - 0 GTAACAAAACCTACATCGAG TGG ||||||||[||||||||||||]PAM NA Intergenic NA NA

T20189 GGTGCTTTCGTACGAACGAAAGG 1000
Oligo fwd CACCGGTGCTTTCGTACGAACGAA
Oligo rev AAACTTCGTTCGTACGAAAGCACC
Chromosome start end strand MM target_seq PAM alignment distance position gene name gene id
GU305914.1 191028 191050 - 0 GGTGCTTTCGTACGAACGAA AGG ||||||||[||||||||||||]PAM NA Intergenic NA NA

T20190 TCGCTTCCCTAAACGTAACGTGG 1000
Oligo fwd CACCTCGCTTCCCTAAACGTAACG
Oligo rev AAACCGTTACGTTTAGGGAAGCGA
Chromosome start end strand MM target_seq PAM alignment distance position gene name gene id
GU305914.1 191004 191026 - 0 TCGCTTCCCTAAACGTAACG TGG ||||||||[||||||||||||]PAM NA Intergenic NA NA

T20191 CCCTAAACGTAACGTGGAACAGG 1000
Oligo fwd CACCCCCTAAACGTAACGTGGAAC
Oligo rev AAACGTTCCACGTTACGTTTAGGG
Chromosome start end strand MM target_seq PAM alignment distance position gene name gene id
GU305914.1 190998 191020 - 0 CCCTAAACGTAACGTGGAAC AGG ||||||||[||||||||||||]PAM NA Intergenic NA NA

T20192 AGATCCTCTTCGAGACCTCGTGG 1000
Oligo fwd CACCAGATCCTCTTCGAGACCTCG
Oligo rev AAACCGAGGTCTCGAAGAGGATCT
Chromosome start end strand MM target_seq PAM alignment distance position gene name gene id
GU305914.1 190959 190981 - 0 AGATCCTCTTCGAGACCTCG TGG ||||||||[||||||||||||]PAM NA Intergenic NA NA

T20193 GAGATGCGTCGCACGCAAGTTGG 1000
Oligo fwd CACCGAGATGCGTCGCACGCAAGT
Oligo rev AAACACTTGCGTGCGACGCATCTC
Chromosome start end strand MM target_seq PAM alignment distance position gene name gene id
GU305914.1 190919 190941 - 0 GAGATGCGTCGCACGCAAGT TGG ||||||||[||||||||||||]PAM NA Intergenic NA NA

T20194 TGCGTCGCACGCAAGTTGGTCGG 1000
Oligo fwd CACCTGCGTCGCACGCAAGTTGGT
Oligo rev AAACACCAACTTGCGTGCGACGCA
Chromosome start end strand MM target_seq PAM alignment distance position gene name gene id
GU305914.1 190915 190937 - 0 TGCGTCGCACGCAAGTTGGT CGG ||||||||[||||||||||||]PAM NA Intergenic NA NA

T20195 GCGTCGCACGCAAGTTGGTCGGG 1000
Oligo fwd CACCGCGTCGCACGCAAGTTGGTC
Oligo rev AAACGACCAACTTGCGTGCGACGC
Chromosome start end strand MM target_seq PAM alignment distance position gene name gene id
GU305914.1 190914 190936 - 0 GCGTCGCACGCAAGTTGGTC GGG ||||||||[||||||||||||]PAM NA Intergenic NA NA

T20196 CGTCGCACGCAAGTTGGTCGGGG 1000
Oligo fwd CACCCGTCGCACGCAAGTTGGTCG
Oligo rev AAACCGACCAACTTGCGTGCGACG
Chromosome start end strand MM target_seq PAM alignment distance position gene name gene id
GU305914.1 190913 190935 - 0 CGTCGCACGCAAGTTGGTCG GGG ||||||||[||||||||||||]PAM NA Intergenic NA NA

T20197 ACGCAAGTTGGTCGGGGCCGAGG 1000
Oligo fwd CACCACGCAAGTTGGTCGGGGCCG
Oligo rev AAACCGGCCCCGACCAACTTGCGT
Chromosome start end strand MM target_seq PAM alignment distance position gene name gene id
GU305914.1 190907 190929 - 0 ACGCAAGTTGGTCGGGGCCG AGG ||||||||[||||||||||||]PAM NA Intergenic NA NA

T20198 TGATCCTGACAACAAAAAGATGG 1000
Oligo fwd CACCTGATCCTGACAACAAAAAGA
Oligo rev AAACTCTTTTTGTTGTCAGGATCA
Chromosome start end strand MM target_seq PAM alignment distance position gene name gene id
GU305914.1 190859 190881 - 0 TGATCCTGACAACAAAAAGA TGG ||||||||[||||||||||||]PAM NA Intergenic NA NA

T20199 AGAGTTGCGTGTGAAATTTGAGG 1000
Oligo fwd CACCAGAGTTGCGTGTGAAATTTG
Oligo rev AAACCAAATTTCACACGCAACTCT
Chromosome start end strand MM target_seq PAM alignment distance position gene name gene id
GU305914.1 190835 190857 - 0 AGAGTTGCGTGTGAAATTTG AGG ||||||||[||||||||||||]PAM NA Intergenic NA NA

T20200 GAGTTGCGTGTGAAATTTGAGGG 1000
Oligo fwd CACCGAGTTGCGTGTGAAATTTGA
Oligo rev AAACTCAAATTTCACACGCAACTC
Chromosome start end strand MM target_seq PAM alignment distance position gene name gene id

<table>
<tr><td>GU305914.1</td><td>190834</td><td>190856</td><td>-</td><td>0</td><td>GAGTTGCGTGTGAAATTTGA</td><td>GGG</td><td>||||||||[||||||||||||]PAM</td><td>NA</td><td>Intergenic</td><td>NA</td><td>NA</td></tr>
<tr><td></td><td></td><td></td><td></td><td></td><td></td><td></td><td></td><td></td><td></td><td></td><td></td></tr>
<tr><td>T20201</td><td>AGTTGCGTGTGAAATTTGAGGGG</td><td>1000</td><td></td><td></td><td></td><td></td><td></td><td></td><td></td><td></td><td></td></tr>
<tr><td>Oligo fwd</td><td>CACCAGTTGCGTGTGAAATTTGAG</td><td></td><td></td><td></td><td></td><td></td><td></td><td></td><td></td><td></td><td></td></tr>
<tr><td>Oligo rev</td><td>AAACCTCAAATTTCACACGCAACT</td><td></td><td></td><td></td><td></td><td></td><td></td><td></td><td></td><td></td><td></td></tr>
<tr><td>Chromosome</td><td>start</td><td>end</td><td>strand</td><td>MM</td><td>target_seq</td><td>PAM</td><td>alignment</td><td>distance</td><td>position</td><td>gene name</td><td>gene id</td></tr>
<tr><td>GU305914.1</td><td>190833</td><td>190855</td><td>-</td><td>0</td><td>AGTTGCGTGTGAAATTTGAG</td><td>GGG</td><td>||||||||[||||||||||||]PAM</td><td>NA</td><td>Intergenic</td><td>NA</td><td>NA</td></tr>
<tr><td></td><td></td><td></td><td></td><td></td><td></td><td></td><td></td><td></td><td></td><td></td><td></td></tr>
<tr><td>T20202</td><td>GACCATAAACCACGATTCAACGG</td><td>1000</td><td></td><td></td><td></td><td></td><td></td><td></td><td></td><td></td><td></td></tr>
<tr><td>Oligo fwd</td><td>CACCGACCATAAACCACGATTCAA</td><td></td><td></td><td></td><td></td><td></td><td></td><td></td><td></td><td></td><td></td></tr>
<tr><td>Oligo rev</td><td>AAACTTGAATCGTGGTTTATGGTC</td><td></td><td></td><td></td><td></td><td></td><td></td><td></td><td></td><td></td><td></td></tr>
<tr><td>Chromosome</td><td>start</td><td>end</td><td>strand</td><td>MM</td><td>target_seq</td><td>PAM</td><td>alignment</td><td>distance</td><td>position</td><td>gene name</td><td>gene id</td></tr>
<tr><td>GU305914.1</td><td>190811</td><td>190833</td><td>-</td><td>0</td><td>GACCATAAACCACGATTCAA</td><td>CGG</td><td>||||||||[||||||||||||]PAM</td><td>NA</td><td>Intergenic</td><td>NA</td><td>NA</td></tr>
<tr><td></td><td></td><td></td><td></td><td></td><td></td><td></td><td></td><td></td><td></td><td></td><td></td></tr>
<tr><td>T20203</td><td>ACCATAAACCACGATTCAACGGG</td><td>1000</td><td></td><td></td><td></td><td></td><td></td><td></td><td></td><td></td><td></td></tr>
<tr><td>Oligo fwd</td><td>CACCACCATAAACCACGATTCAAC</td><td></td><td></td><td></td><td></td><td></td><td></td><td></td><td></td><td></td><td></td></tr>
<tr><td>Oligo rev</td><td>AAACGTTGAATCGTGGTTTATGGT</td><td></td><td></td><td></td><td></td><td></td><td></td><td></td><td></td><td></td><td></td></tr>
<tr><td>Chromosome</td><td>start</td><td>end</td><td>strand</td><td>MM</td><td>target_seq</td><td>PAM</td><td>alignment</td><td>distance</td><td>position</td><td>gene name</td><td>gene id</td></tr>
<tr><td>GU305914.1</td><td>190810</td><td>190832</td><td>-</td><td>0</td><td>ACCATAAACCACGATTCAAC</td><td>GGG</td><td>||||||||[||||||||||||]PAM</td><td>NA</td><td>Intergenic</td><td>NA</td><td>NA</td></tr>
<tr><td></td><td></td><td></td><td></td><td></td><td></td><td></td><td></td><td></td><td></td><td></td><td></td></tr>
<tr><td>T20204</td><td>CCATAAACCACGATTCAACGGGG</td><td>1000</td><td></td><td></td><td></td><td></td><td></td><td></td><td></td><td></td><td></td></tr>
<tr><td>Oligo fwd</td><td>CACCCCATAAACCACGATTCAACG</td><td></td><td></td><td></td><td></td><td></td><td></td><td></td><td></td><td></td><td></td></tr>
<tr><td>Oligo rev</td><td>AAACCGTTGAATCGTGGTTTATGG</td><td></td><td></td><td></td><td></td><td></td><td></td><td></td><td></td><td></td><td></td></tr>
<tr><td>Chromosome</td><td>start</td><td>end</td><td>strand</td><td>MM</td><td>target_seq</td><td>PAM</td><td>alignment</td><td>distance</td><td>position</td><td>gene name</td><td>gene id</td></tr>
<tr><td>GU305914.1</td><td>190809</td><td>190831</td><td>-</td><td>0</td><td>CCATAAACCACGATTCAACG</td><td>GGG</td><td>||||||||[||||||||||||]PAM</td><td>NA</td><td>Intergenic</td><td>NA</td><td>NA</td></tr>
<tr><td></td><td></td><td></td><td></td><td></td><td></td><td></td><td></td><td></td><td></td><td></td><td></td></tr>
<tr><td>T20205</td><td>TCCTACATCAACACACAATCAGG</td><td>1000</td><td></td><td></td><td></td><td></td><td></td><td></td><td></td><td></td><td></td></tr>
<tr><td>Oligo fwd</td><td>CACCTCCTACATCAACACACAATC</td><td></td><td></td><td></td><td></td><td></td><td></td><td></td><td></td><td></td><td></td></tr>
<tr><td>Oligo rev</td><td>AAACGATTGTGTGTTGATGTAGGA</td><td></td><td></td><td></td><td></td><td></td><td></td><td></td><td></td><td></td><td></td></tr>
<tr><td>Chromosome</td><td>start</td><td>end</td><td>strand</td><td>MM</td><td>target_seq</td><td>PAM</td><td>alignment</td><td>distance</td><td>position</td><td>gene name</td><td>gene id</td></tr>
<tr><td>GU305914.1</td><td>190777</td><td>190799</td><td>-</td><td>0</td><td>TCCTACATCAACACACAATC</td><td>AGG</td><td>||||||||[||||||||||||]PAM</td><td>NA</td><td>Intergenic</td><td>NA</td><td>NA</td></tr>
<tr><td></td><td></td><td></td><td></td><td></td><td></td><td></td><td></td><td></td><td></td><td></td><td></td></tr>
<tr><td>T20206</td><td>TCAACACACAATCAGGTAGATGG</td><td>1000</td><td></td><td></td><td></td><td></td><td></td><td></td><td></td><td></td><td></td></tr>
<tr><td>Oligo fwd</td><td>CACCTCAACACACAATCAGGTAGA</td><td></td><td></td><td></td><td></td><td></td><td></td><td></td><td></td><td></td><td></td></tr>
<tr><td>Oligo rev</td><td>AAACTCTACCTGATTGTGTGTTGA</td><td></td><td></td><td></td><td></td><td></td><td></td><td></td><td></td><td></td><td></td></tr>
<tr><td>Chromosome</td><td>start</td><td>end</td><td>strand</td><td>MM</td><td>target_seq</td><td>PAM</td><td>alignment</td><td>distance</td><td>position</td><td>gene name</td><td>gene id</td></tr>
<tr><td>GU305914.1</td><td>190770</td><td>190792</td><td>-</td><td>0</td><td>TCAACACACAATCAGGTAGA</td><td>TGG</td><td>||||||||[||||||||||||]PAM</td><td>NA</td><td>Intergenic</td><td>NA</td><td>NA</td></tr>
<tr><td></td><td></td><td></td><td></td><td></td><td></td><td></td><td></td><td></td><td></td><td></td><td></td></tr>
<tr><td>T20207</td><td>CACACAATCAGGTAGATGGTCGG</td><td>1000</td><td></td><td></td><td></td><td></td><td></td><td></td><td></td><td></td><td></td></tr>
<tr><td>Oligo fwd</td><td>CACCCACACAATCAGGTAGATGGT</td><td></td><td></td><td></td><td></td><td></td><td></td><td></td><td></td><td></td><td></td></tr>
<tr><td>Oligo rev</td><td>AAACACCATCTACCTGATTGTGTG</td><td></td><td></td><td></td><td></td><td></td><td></td><td></td><td></td><td></td><td></td></tr>
<tr><td>Chromosome</td><td>start</td><td>end</td><td>strand</td><td>MM</td><td>target_seq</td><td>PAM</td><td>alignment</td><td>distance</td><td>position</td><td>gene name</td><td>gene id</td></tr>
<tr><td>GU305914.1</td><td>190766</td><td>190788</td><td>-</td><td>0</td><td>CACACAATCAGGTAGATGGT</td><td>CGG</td><td>||||||||[||||||||||||]PAM</td><td>NA</td><td>Intergenic</td><td>NA</td><td>NA</td></tr>
<tr><td></td><td></td><td></td><td></td><td></td><td></td><td></td><td></td><td></td><td></td><td></td><td></td></tr>
<tr><td>T20208</td><td>ACAATCAGGTAGATGGTCGGTGG</td><td>1000</td><td></td><td></td><td></td><td></td><td></td><td></td><td></td><td></td><td></td></tr>
<tr><td>Oligo fwd</td><td>CACCACAATCAGGTAGATGGTCGG</td><td></td><td></td><td></td><td></td><td></td><td></td><td></td><td></td><td></td><td></td></tr>
<tr><td>Oligo rev</td><td>AAACCCGACCATCTACCTGATTGT</td><td></td><td></td><td></td><td></td><td></td><td></td><td></td><td></td><td></td><td></td></tr>
<tr><td>Chromosome</td><td>start</td><td>end</td><td>strand</td><td>MM</td><td>target_seq</td><td>PAM</td><td>alignment</td><td>distance</td><td>position</td><td>gene name</td><td>gene id</td></tr>
<tr><td>GU305914.1</td><td>190763</td><td>190785</td><td>-</td><td>0</td><td>ACAATCAGGTAGATGGTCGG</td><td>TGG</td><td>||||||||[||||||||||||]PAM</td><td>NA</td><td>Intergenic</td><td>NA</td><td>NA</td></tr>
<tr><td></td><td></td><td></td><td></td><td></td><td></td><td></td><td></td><td></td><td></td><td></td><td></td></tr>
<tr><td>T20209</td><td>AGGTAGATGGTCGGTGGAGAAGG</td><td>1000</td><td></td><td></td><td></td><td></td><td></td><td></td><td></td><td></td><td></td></tr>
<tr><td>Oligo fwd</td><td>CACCAGGTAGATGGTCGGTGGAGA</td><td></td><td></td><td></td><td></td><td></td><td></td><td></td><td></td><td></td><td></td></tr>
<tr><td>Oligo rev</td><td>AAACTCTCCACCGACCATCTACCT</td><td></td><td></td><td></td><td></td><td></td><td></td><td></td><td></td><td></td><td></td></tr>
<tr><td>Chromosome</td><td>start</td><td>end</td><td>strand</td><td>MM</td><td>target_seq</td><td>PAM</td><td>alignment</td><td>distance</td><td>position</td><td>gene name</td><td>gene id</td></tr>
<tr><td>GU305914.1</td><td>190757</td><td>190779</td><td>-</td><td>0</td><td>AGGTAGATGGTCGGTGGAGA</td><td>AGG</td><td>||||||||[||||||||||||]PAM</td><td>NA</td><td>Intergenic</td><td>NA</td><td>NA</td></tr>
<tr><td></td><td></td><td></td><td></td><td></td><td></td><td></td><td></td><td></td><td></td><td></td><td></td></tr>
<tr><td>T20210</td><td>GTCGGTGGAGAAGGTTATGAAGG</td><td>1000</td><td></td><td></td><td></td><td></td><td></td><td></td><td></td><td></td><td></td></tr>
<tr><td>Oligo fwd</td><td>CACCGTCGGTGGAGAAGGTTATGA</td><td></td><td></td><td></td><td></td><td></td><td></td><td></td><td></td><td></td><td></td></tr>
<tr><td>Oligo rev</td><td>AAACTCATAACCTTCTCCACCGAC</td><td></td><td></td><td></td><td></td><td></td><td></td><td></td><td></td><td></td><td></td></tr>
<tr><td>Chromosome</td><td>start</td><td>end</td><td>strand</td><td>MM</td><td>target_seq</td><td>PAM</td><td>alignment</td><td>distance</td><td>position</td><td>gene name</td><td>gene id</td></tr>
<tr><td>GU305914.1</td><td>190748</td><td>190770</td><td>-</td><td>0</td><td>GTCGGTGGAGAAGGTTATGA</td><td>AGG</td><td>||||||||[||||||||||||]PAM</td><td>NA</td><td>Intergenic</td><td>NA</td><td>NA</td></tr>
<tr><td></td><td></td><td></td><td></td><td></td><td></td><td></td><td></td><td></td><td></td><td></td><td></td></tr>
<tr><td>T20211</td><td>GTGTGAATTGAAAGTCAGTGAGG</td><td>1000</td><td></td><td></td><td></td><td></td><td></td><td></td><td></td><td></td><td></td></tr>
<tr><td>Oligo fwd</td><td>CACCGTGTGAATTGAAAGTCAGTG</td><td></td><td></td><td></td><td></td><td></td><td></td><td></td><td></td><td></td><td></td></tr>
<tr><td>Oligo rev</td><td>AAACCACTGACTTTCAATTCACAC</td><td></td><td></td><td></td><td></td><td></td><td></td><td></td><td></td><td></td><td></td></tr>
<tr><td>Chromosome</td><td>start</td><td>end</td><td>strand</td><td>MM</td><td>target_seq</td><td>PAM</td><td>alignment</td><td>distance</td><td>position</td><td>gene name</td><td>gene id</td></tr>
<tr><td>GU305914.1</td><td>190709</td><td>190731</td><td>-</td><td>0</td><td>GTGTGAATTGAAAGTCAGTG</td><td>AGG</td><td>||||||||[||||||||||||]PAM</td><td>NA</td><td>Intergenic</td><td>NA</td><td>NA</td></tr>
<tr><td></td><td></td><td></td><td></td><td></td><td></td><td></td><td></td><td></td><td></td><td></td><td></td></tr>
<tr><td>T20212</td><td>AGATACAGCTGCGAGCCTGAAGG</td><td>1000</td><td></td><td></td><td></td><td></td><td></td><td></td><td></td><td></td><td></td></tr>
<tr><td>Oligo fwd</td><td>CACCAGATACAGCTGCGAGCCTGA</td><td></td><td></td><td></td><td></td><td></td><td></td><td></td><td></td><td></td><td></td></tr>
<tr><td>Oligo rev</td><td>AAACTCAGGCTCGCAGCTGTATCT</td><td></td><td></td><td></td><td></td><td></td><td></td><td></td><td></td><td></td><td></td></tr>
<tr><td>Chromosome</td><td>start</td><td>end</td><td>strand</td><td>MM</td><td>target_seq</td><td>PAM</td><td>alignment</td><td>distance</td><td>position</td><td>gene name</td><td>gene id</td></tr>
<tr><td>GU305914.1</td><td>190678</td><td>190700</td><td>-</td><td>0</td><td>AGATACAGCTGCGAGCCTGA</td><td>AGG</td><td>||||||||[||||||||||||]PAM</td><td>NA</td><td>Intergenic</td><td>NA</td><td>NA</td></tr>
<tr><td></td><td></td><td></td><td></td><td></td><td></td><td></td><td></td><td></td><td></td><td></td><td></td></tr>
<tr><td>T20213</td><td>AGCTGCGAGCCTGAAGGCTACGG</td><td>1000</td><td></td><td></td><td></td><td></td><td></td><td></td><td></td><td></td><td></td></tr>
<tr><td>Oligo fwd</td><td>CACCAGCTGCGAGCCTGAAGGCTA</td><td></td><td></td><td></td><td></td><td></td><td></td><td></td><td></td><td></td><td></td></tr>
<tr><td>Oligo rev</td><td>AAACTAGCCTTCAGGCTCGCAGCT</td><td></td><td></td><td></td><td></td><td></td><td></td><td></td><td></td><td></td><td></td></tr>
<tr><td>Chromosome</td><td>start</td><td>end</td><td>strand</td><td>MM</td><td>target_seq</td><td>PAM</td><td>alignment</td><td>distance</td><td>position</td><td>gene name</td><td>gene id</td></tr>
<tr><td>GU305914.1</td><td>190672</td><td>190694</td><td>-</td><td>0</td><td>AGCTGCGAGCCTGAAGGCTA</td><td>CGG</td><td>||||||||[||||||||||||]PAM</td><td>NA</td><td>Intergenic</td><td>NA</td><td>NA</td></tr>
<tr><td></td><td></td><td></td><td></td><td></td><td></td><td></td><td></td><td></td><td></td><td></td><td></td></tr>
<tr><td>T20214</td><td>GCTGCGAGCCTGAAGGCTACGGG</td><td>1000</td><td></td><td></td><td></td><td></td><td></td><td></td><td></td><td></td><td></td></tr>
<tr><td>Oligo fwd</td><td>CACCGCTGCGAGCCTGAAGGCTAC</td><td></td><td></td><td></td><td></td><td></td><td></td><td></td><td></td><td></td><td></td></tr>
<tr><td>Oligo rev</td><td>AAACGTAGCCTTCAGGCTCGCAGC</td><td></td><td></td><td></td><td></td><td></td><td></td><td></td><td></td><td></td><td></td></tr>
<tr><td>Chromosome</td><td>start</td><td>end</td><td>strand</td><td>MM</td><td>target_seq</td><td>PAM</td><td>alignment</td><td>distance</td><td>position</td><td>gene name</td><td>gene id</td></tr>
<tr><td>GU305914.1</td><td>190671</td><td>190693</td><td>-</td><td>0</td><td>GCTGCGAGCCTGAAGGCTAC</td><td>GGG</td><td>||||||||[||||||||||||]PAM</td><td>NA</td><td>Intergenic</td><td>NA</td><td>NA</td></tr>
<tr><td></td><td></td><td></td><td></td><td></td><td></td><td></td><td></td><td></td><td></td><td></td><td></td></tr>
<tr><td>T20215</td><td>GCGAGCCTGAAGGCTACGGGTGG</td><td>1000</td><td></td><td></td><td></td><td></td><td></td><td></td><td></td><td></td><td></td></tr>
<tr><td>Oligo fwd</td><td>CACCGCGAGCCTGAAGGCTACGGG</td><td></td><td></td><td></td><td></td><td></td><td></td><td></td><td></td><td></td><td></td></tr>
<tr><td>Oligo rev</td><td>AAACCCCGTAGCCTTCAGGCTCGC</td><td></td><td></td><td></td><td></td><td></td><td></td><td></td><td></td><td></td><td></td></tr>
</table>

<table>
<tr><td>Chromosome</td><td>start</td><td>end</td><td>strand</td><td>MM</td><td>target_seq</td><td>PAM</td><td>alignment</td><td>distance</td><td>position</td><td>gene name</td><td>gene id</td></tr>
<tr><td>GU305914.1</td><td>190668</td><td>190690</td><td>-</td><td>0</td><td>GCGAGCCTGAAGGCTACGGG</td><td>TGG</td><td>||||||||[||||||||||||]PAM</td><td>NA</td><td>Intergenic</td><td>NA</td><td>NA</td></tr>
<tr><td>T20216</td><td>GGCTACGGGTGGCTGACCAAAGG</td><td>1000</td><td></td><td></td><td></td><td></td><td></td><td></td><td></td><td></td><td></td></tr>
<tr><td>Oligo fwd</td><td>CACCGGCTACGGGTGGCTGACCAA</td><td></td><td></td><td></td><td></td><td></td><td></td><td></td><td></td><td></td><td></td></tr>
<tr><td>Oligo rev</td><td>AAACTTGGTCAGCCACCCGTAGCC</td><td></td><td></td><td></td><td></td><td></td><td></td><td></td><td></td><td></td><td></td></tr>
<tr><td>Chromosome</td><td>start</td><td>end</td><td>strand</td><td>MM</td><td>target_seq</td><td>PAM</td><td>alignment</td><td>distance</td><td>position</td><td>gene name</td><td>gene id</td></tr>
<tr><td>GU305914.1</td><td>190657</td><td>190679</td><td>-</td><td>0</td><td>GGCTACGGGTGGCTGACCAA</td><td>AGG</td><td>||||||||[||||||||||||]PAM</td><td>NA</td><td>Intergenic</td><td>NA</td><td>NA</td></tr>
<tr><td>T20217</td><td>ACCGCGACCGTTCGTGTGAATGG</td><td>1000</td><td></td><td></td><td></td><td></td><td></td><td></td><td></td><td></td><td></td></tr>
<tr><td>Oligo fwd</td><td>CACCACCGCGACCGTTCGTGTGAA</td><td></td><td></td><td></td><td></td><td></td><td></td><td></td><td></td><td></td><td></td></tr>
<tr><td>Oligo rev</td><td>AAACTTCACACGAACGGTCGCGGT</td><td></td><td></td><td></td><td></td><td></td><td></td><td></td><td></td><td></td><td></td></tr>
<tr><td>Chromosome</td><td>start</td><td>end</td><td>strand</td><td>MM</td><td>target_seq</td><td>PAM</td><td>alignment</td><td>distance</td><td>position</td><td>gene name</td><td>gene id</td></tr>
<tr><td>GU305914.1</td><td>190627</td><td>190649</td><td>-</td><td>0</td><td>ACCGCGACCGTTCGTGTGAA</td><td>TGG</td><td>||||||||[||||||||||||]PAM</td><td>NA</td><td>Intergenic</td><td>NA</td><td>NA</td></tr>
<tr><td>T20218</td><td>AGCATCCACCTACCAGAGCCAGG</td><td>1000</td><td></td><td></td><td></td><td></td><td></td><td></td><td></td><td></td><td></td></tr>
<tr><td>Oligo fwd</td><td>CACCAGCATCCACCTACCAGAGCC</td><td></td><td></td><td></td><td></td><td></td><td></td><td></td><td></td><td></td><td></td></tr>
<tr><td>Oligo rev</td><td>AAACGGCTCTGGTAGGTGGATGCT</td><td></td><td></td><td></td><td></td><td></td><td></td><td></td><td></td><td></td><td></td></tr>
<tr><td>Chromosome</td><td>start</td><td>end</td><td>strand</td><td>MM</td><td>target_seq</td><td>PAM</td><td>alignment</td><td>distance</td><td>position</td><td>gene name</td><td>gene id</td></tr>
<tr><td>GU305914.1</td><td>190594</td><td>190616</td><td>-</td><td>0</td><td>AGCATCCACCTACCAGAGCC</td><td>AGG</td><td>||||||||[||||||||||||]PAM</td><td>NA</td><td>Intergenic</td><td>NA</td><td>NA</td></tr>
<tr><td>T20219</td><td>GCATCCACCTACCAGAGCCAGGG</td><td>1000</td><td></td><td></td><td></td><td></td><td></td><td></td><td></td><td></td><td></td></tr>
<tr><td>Oligo fwd</td><td>CACCGCATCCACCTACCAGAGCCA</td><td></td><td></td><td></td><td></td><td></td><td></td><td></td><td></td><td></td><td></td></tr>
<tr><td>Oligo rev</td><td>AAACTGGCTCTGGTAGGTGGATGC</td><td></td><td></td><td></td><td></td><td></td><td></td><td></td><td></td><td></td><td></td></tr>
<tr><td>Chromosome</td><td>start</td><td>end</td><td>strand</td><td>MM</td><td>target_seq</td><td>PAM</td><td>alignment</td><td>distance</td><td>position</td><td>gene name</td><td>gene id</td></tr>
<tr><td>GU305914.1</td><td>190593</td><td>190615</td><td>-</td><td>0</td><td>GCATCCACCTACCAGAGCCA</td><td>GGG</td><td>||||||||[||||||||||||]PAM</td><td>NA</td><td>Intergenic</td><td>NA</td><td>NA</td></tr>
<tr><td>T20220</td><td>CATCCACCTACCAGAGCCAGGGG</td><td>1000</td><td></td><td></td><td></td><td></td><td></td><td></td><td></td><td></td><td></td></tr>
<tr><td>Oligo fwd</td><td>CACCCATCCACCTACCAGAGCCAG</td><td></td><td></td><td></td><td></td><td></td><td></td><td></td><td></td><td></td><td></td></tr>
<tr><td>Oligo rev</td><td>AAACCTGGCTCTGGTAGGTGGATG</td><td></td><td></td><td></td><td></td><td></td><td></td><td></td><td></td><td></td><td></td></tr>
<tr><td>Chromosome</td><td>start</td><td>end</td><td>strand</td><td>MM</td><td>target_seq</td><td>PAM</td><td>alignment</td><td>distance</td><td>position</td><td>gene name</td><td>gene id</td></tr>
<tr><td>GU305914.1</td><td>190592</td><td>190614</td><td>-</td><td>0</td><td>CATCCACCTACCAGAGCCAG</td><td>GGG</td><td>||||||||[||||||||||||]PAM</td><td>NA</td><td>Intergenic</td><td>NA</td><td>NA</td></tr>
<tr><td>T20221</td><td>ATCCACCTACCAGAGCCAGGGGG</td><td>1000</td><td></td><td></td><td></td><td></td><td></td><td></td><td></td><td></td><td></td></tr>
<tr><td>Oligo fwd</td><td>CACCATCCACCTACCAGAGCCAGG</td><td></td><td></td><td></td><td></td><td></td><td></td><td></td><td></td><td></td><td></td></tr>
<tr><td>Oligo rev</td><td>AAACCCTGGCTCTGGTAGGTGGAT</td><td></td><td></td><td></td><td></td><td></td><td></td><td></td><td></td><td></td><td></td></tr>
<tr><td>Chromosome</td><td>start</td><td>end</td><td>strand</td><td>MM</td><td>target_seq</td><td>PAM</td><td>alignment</td><td>distance</td><td>position</td><td>gene name</td><td>gene id</td></tr>
<tr><td>GU305914.1</td><td>190591</td><td>190613</td><td>-</td><td>0</td><td>ATCCACCTACCAGAGCCAGG</td><td>GGG</td><td>||||||||[||||||||||||]PAM</td><td>NA</td><td>Intergenic</td><td>NA</td><td>NA</td></tr>
<tr><td>T20222</td><td>GCGCGTATATCGAGTACGAACGG</td><td>1000</td><td></td><td></td><td></td><td></td><td></td><td></td><td></td><td></td><td></td></tr>
<tr><td>Oligo fwd</td><td>CACCGCGCGTATATCGAGTACGAA</td><td></td><td></td><td></td><td></td><td></td><td></td><td></td><td></td><td></td><td></td></tr>
<tr><td>Oligo rev</td><td>AAACTTCGTACTCGATATACGCGC</td><td></td><td></td><td></td><td></td><td></td><td></td><td></td><td></td><td></td><td></td></tr>
<tr><td>Chromosome</td><td>start</td><td>end</td><td>strand</td><td>MM</td><td>target_seq</td><td>PAM</td><td>alignment</td><td>distance</td><td>position</td><td>gene name</td><td>gene id</td></tr>
<tr><td>GU305914.1</td><td>190569</td><td>190591</td><td>-</td><td>0</td><td>GCGCGTATATCGAGTACGAA</td><td>CGG</td><td>||||||||[||||||||||||]PAM</td><td>NA</td><td>Intergenic</td><td>NA</td><td>NA</td></tr>
<tr><td>T20223</td><td>CGCGTATATCGAGTACGAACGGG</td><td>1000</td><td></td><td></td><td></td><td></td><td></td><td></td><td></td><td></td><td></td></tr>
<tr><td>Oligo fwd</td><td>CACCCGCGTATATCGAGTACGAAC</td><td></td><td></td><td></td><td></td><td></td><td></td><td></td><td></td><td></td><td></td></tr>
<tr><td>Oligo rev</td><td>AAACGTTCGTACTCGATATACGCG</td><td></td><td></td><td></td><td></td><td></td><td></td><td></td><td></td><td></td><td></td></tr>
<tr><td>Chromosome</td><td>start</td><td>end</td><td>strand</td><td>MM</td><td>target_seq</td><td>PAM</td><td>alignment</td><td>distance</td><td>position</td><td>gene name</td><td>gene id</td></tr>
<tr><td>GU305914.1</td><td>190568</td><td>190590</td><td>-</td><td>0</td><td>CGCGTATATCGAGTACGAAC</td><td>GGG</td><td>||||||||[||||||||||||]PAM</td><td>NA</td><td>Intergenic</td><td>NA</td><td>NA</td></tr>
<tr><td>T20224</td><td>GCGTATATCGAGTACGAACGGGG</td><td>1000</td><td></td><td></td><td></td><td></td><td></td><td></td><td></td><td></td><td></td></tr>
<tr><td>Oligo fwd</td><td>CACCGCGTATATCGAGTACGAACG</td><td></td><td></td><td></td><td></td><td></td><td></td><td></td><td></td><td></td><td></td></tr>
<tr><td>Oligo rev</td><td>AAACCGTTCGTACTCGATATACGC</td><td></td><td></td><td></td><td></td><td></td><td></td><td></td><td></td><td></td><td></td></tr>
<tr><td>Chromosome</td><td>start</td><td>end</td><td>strand</td><td>MM</td><td>target_seq</td><td>PAM</td><td>alignment</td><td>distance</td><td>position</td><td>gene name</td><td>gene id</td></tr>
<tr><td>GU305914.1</td><td>190567</td><td>190589</td><td>-</td><td>0</td><td>GCGTATATCGAGTACGAACG</td><td>GGG</td><td>||||||||[||||||||||||]PAM</td><td>NA</td><td>Intergenic</td><td>NA</td><td>NA</td></tr>
<tr><td>T20225</td><td>CGTATATCGAGTACGAACGGGGG</td><td>1000</td><td></td><td></td><td></td><td></td><td></td><td></td><td></td><td></td><td></td></tr>
<tr><td>Oligo fwd</td><td>CACCCGTATATCGAGTACGAACGG</td><td></td><td></td><td></td><td></td><td></td><td></td><td></td><td></td><td></td><td></td></tr>
<tr><td>Oligo rev</td><td>AAACCCGTTCGTACTCGATATACG</td><td></td><td></td><td></td><td></td><td></td><td></td><td></td><td></td><td></td><td></td></tr>
<tr><td>Chromosome</td><td>start</td><td>end</td><td>strand</td><td>MM</td><td>target_seq</td><td>PAM</td><td>alignment</td><td>distance</td><td>position</td><td>gene name</td><td>gene id</td></tr>
<tr><td>GU305914.1</td><td>190566</td><td>190588</td><td>-</td><td>0</td><td>CGTATATCGAGTACGAACGG</td><td>GGG</td><td>||||||||[||||||||||||]PAM</td><td>NA</td><td>Intergenic</td><td>NA</td><td>NA</td></tr>
<tr><td>T20226</td><td>CGAACGGGGGAGTTTCCTATCGG</td><td>1000</td><td></td><td></td><td></td><td></td><td></td><td></td><td></td><td></td><td></td></tr>
<tr><td>Oligo fwd</td><td>CACCCGAACGGGGGAGTTTCCTAT</td><td></td><td></td><td></td><td></td><td></td><td></td><td></td><td></td><td></td><td></td></tr>
<tr><td>Oligo rev</td><td>AAACATAGGAAACTCCCCCGTTCG</td><td></td><td></td><td></td><td></td><td></td><td></td><td></td><td></td><td></td><td></td></tr>
<tr><td>Chromosome</td><td>start</td><td>end</td><td>strand</td><td>MM</td><td>target_seq</td><td>PAM</td><td>alignment</td><td>distance</td><td>position</td><td>gene name</td><td>gene id</td></tr>
<tr><td>GU305914.1</td><td>190553</td><td>190575</td><td>-</td><td>0</td><td>CGAACGGGGGAGTTTCCTAT</td><td>CGG</td><td>||||||||[||||||||||||]PAM</td><td>NA</td><td>Intergenic</td><td>NA</td><td>NA</td></tr>
<tr><td>T20227</td><td>GAACGGGGGAGTTTCCTATCGGG</td><td>1000</td><td></td><td></td><td></td><td></td><td></td><td></td><td></td><td></td><td></td></tr>
<tr><td>Oligo fwd</td><td>CACCGAACGGGGGAGTTTCCTATC</td><td></td><td></td><td></td><td></td><td></td><td></td><td></td><td></td><td></td><td></td></tr>
<tr><td>Oligo rev</td><td>AAACGATAGGAAACTCCCCCGTTC</td><td></td><td></td><td></td><td></td><td></td><td></td><td></td><td></td><td></td><td></td></tr>
<tr><td>Chromosome</td><td>start</td><td>end</td><td>strand</td><td>MM</td><td>target_seq</td><td>PAM</td><td>alignment</td><td>distance</td><td>position</td><td>gene name</td><td>gene id</td></tr>
<tr><td>GU305914.1</td><td>190552</td><td>190574</td><td>-</td><td>0</td><td>GAACGGGGGAGTTTCCTATC</td><td>GGG</td><td>||||||||[||||||||||||]PAM</td><td>NA</td><td>Intergenic</td><td>NA</td><td>NA</td></tr>
<tr><td>T20228</td><td>AGTTTCCTATCGGGTTACTTAGG</td><td>1000</td><td></td><td></td><td></td><td></td><td></td><td></td><td></td><td></td><td></td></tr>
<tr><td>Oligo fwd</td><td>CACCAGTTTCCTATCGGGTTACTT</td><td></td><td></td><td></td><td></td><td></td><td></td><td></td><td></td><td></td><td></td></tr>
<tr><td>Oligo rev</td><td>AAACAAGTAACCCGATAGGAAACT</td><td></td><td></td><td></td><td></td><td></td><td></td><td></td><td></td><td></td><td></td></tr>
<tr><td>Chromosome</td><td>start</td><td>end</td><td>strand</td><td>MM</td><td>target_seq</td><td>PAM</td><td>alignment</td><td>distance</td><td>position</td><td>gene name</td><td>gene id</td></tr>
<tr><td>GU305914.1</td><td>190543</td><td>190565</td><td>-</td><td>0</td><td>AGTTTCCTATCGGGTTACTT</td><td>AGG</td><td>||||||||[||||||||||||]PAM</td><td>NA</td><td>Intergenic</td><td>NA</td><td>NA</td></tr>
<tr><td>T20229</td><td>CTATCGGGTTACTTAGGACCCGG</td><td>1000</td><td></td><td></td><td></td><td></td><td></td><td></td><td></td><td></td><td></td></tr>
<tr><td>Oligo fwd</td><td>CACCCTATCGGGTTACTTAGGACC</td><td></td><td></td><td></td><td></td><td></td><td></td><td></td><td></td><td></td><td></td></tr>
<tr><td>Oligo rev</td><td>AAACGGTCCTAAGTAACCCGATAG</td><td></td><td></td><td></td><td></td><td></td><td></td><td></td><td></td><td></td><td></td></tr>
<tr><td>Chromosome</td><td>start</td><td>end</td><td>strand</td><td>MM</td><td>target_seq</td><td>PAM</td><td>alignment</td><td>distance</td><td>position</td><td>gene name</td><td>gene id</td></tr>
<tr><td>GU305914.1</td><td>190537</td><td>190559</td><td>-</td><td>0</td><td>CTATCGGGTTACTTAGGACC</td><td>CGG</td><td>||||||||[||||||||||||]PAM</td><td>NA</td><td>Intergenic</td><td>NA</td><td>NA</td></tr>
<tr><td>T20230</td><td>TCGGGTTACTTAGGACCCGGCGG</td><td>1000</td><td></td><td></td><td></td><td></td><td></td><td></td><td></td><td></td><td></td></tr>
<tr><td>Oligo fwd</td><td>CACCTCGGGTTACTTAGGACCCGG</td><td></td><td></td><td></td><td></td><td></td><td></td><td></td><td></td><td></td><td></td></tr>
</table>

Oligo rev AAACCCGGGTCCTAAGTAACCCGA
Chromosome start end strand MM target_seq PAM alignment distance position gene name gene id
GU305914.1 190534 190556 - 0 TCGGGTTACTTAGGACCCGG CGG ||||||||[||||||||||||]PAM NA Intergenic NA NA

T20231 CGGGTTACTTAGGACCCGGCGGG 1000
Oligo fwd CACCCGGGTTACTTAGGACCCGGC
Oligo rev AAACGCCGGGTCCTAAGTAACCCG
Chromosome start end strand MM target_seq PAM alignment distance position gene name gene id
GU305914.1 190533 190555 - 0 CGGGTTACTTAGGACCCGGC GGG ||||||||[||||||||||||]PAM NA Intergenic NA NA

T20232 TGCTGCAACCGTGTCCACCGAGG 1000
Oligo fwd CACCTGCTGCAACCGTGTCCACCG
Oligo rev AAACCGGTGGACACGGTTGCAGCA
Chromosome start end strand MM target_seq PAM alignment distance position gene name gene id
GU305914.1 190491 190513 - 0 TGCTGCAACCGTGTCCACCG AGG ||||||||[||||||||||||]PAM NA Intergenic NA NA

T20233 GCTGCAACCGTGTCCACCGAGGG 1000
Oligo fwd CACCGCTGCAACCGTGTCCACCGA
Oligo rev AAACTCGGTGGACACGGTTGCAGC
Chromosome start end strand MM target_seq PAM alignment distance position gene name gene id
GU305914.1 190490 190512 - 0 GCTGCAACCGTGTCCACCGA GGG ||||||||[||||||||||||]PAM NA Intergenic NA NA

T20234 GTATCAGAGTTTTGTAATTCCGG 1000
Oligo fwd CACCGTATCAGAGTTTTGTAATTC
Oligo rev AAACGAATTACAAAACTCTGATAC
Chromosome start end strand MM target_seq PAM alignment distance position gene name gene id
GU305914.1 190427 190449 - 0 GTATCAGAGTTTTGTAATTC CGG ||||||||[||||||||||||]PAM NA Intergenic NA NA

T20235 GTTTTGTAATTCCGGAGCGCCGG 1000
Oligo fwd CACCGTTTTGTAATTCCGGAGCGC
Oligo rev AAACGCGCTCCGGAATTACAAAAC
Chromosome start end strand MM target_seq PAM alignment distance position gene name gene id
GU305914.1 190419 190441 - 0 GTTTTGTAATTCCGGAGCGC CGG ||||||||[||||||||||||]PAM NA Intergenic NA NA

T20236 TTTTGTAATTCCGGAGCGCCGGG 1000
Oligo fwd CACCTTTTGTAATTCCGGAGCGCC
Oligo rev AAACGGCGCTCCGGAATTACAAAA
Chromosome start end strand MM target_seq PAM alignment distance position gene name gene id
GU305914.1 190418 190440 - 0 TTTTGTAATTCCGGAGCGCC GGG ||||||||[||||||||||||]PAM NA Intergenic NA NA

T20237 TTTGTAATTCCGGAGCGCCGGGG 1000
Oligo fwd CACCTTTGTAATTCCGGAGCGCCG
Oligo rev AAACCGGCGCTCCGGAATTACAAA
Chromosome start end strand MM target_seq PAM alignment distance position gene name gene id
GU305914.1 190417 190439 - 0 TTTGTAATTCCGGAGCGCCG GGG ||||||||[||||||||||||]PAM NA Intergenic NA NA

T20238 TTGTAATTCCGGAGCGCCGGGGG 1000
Oligo fwd CACCTTGTAATTCCGGAGCGCCGG
Oligo rev AAACCCGGCGCTCCGGAATTACAA
Chromosome start end strand MM target_seq PAM alignment distance position gene name gene id
GU305914.1 190416 190438 - 0 TTGTAATTCCGGAGCGCCGG GGG ||||||||[||||||||||||]PAM NA Intergenic NA NA

T20239 TGTAATTCCGGAGCGCCGGGGGG 1000
Oligo fwd CACCTGTAATTCCGGAGCGCCGGG
Oligo rev AAACCCCGGCGCTCCGGAATTACA
Chromosome start end strand MM target_seq PAM alignment distance position gene name gene id
GU305914.1 190415 190437 - 0 TGTAATTCCGGAGCGCCGGG GGG ||||||||[||||||||||||]PAM NA Intergenic NA NA

T20240 ATAAAAACTTCGCGATCCTCCGG 1000
Oligo fwd CACCATAAAAACTTCGCGATCCTC
Oligo rev AAACGAGGATCGCGAAGTTTTTAT
Chromosome start end strand MM target_seq PAM alignment distance position gene name gene id
GU305914.1 190391 190413 - 0 ATAAAAACTTCGCGATCCTC CGG ||||||||[||||||||||||]PAM NA Intergenic NA NA

T20241 TTCGCGATCCTCCGGTGCCGCGG 1000
Oligo fwd CACCTTCGCGATCCTCCGGTGCCG
Oligo rev AAACCGGCACCGGAGGATCGCGAA
Chromosome start end strand MM target_seq PAM alignment distance position gene name gene id
GU305914.1 190383 190405 - 0 TTCGCGATCCTCCGGTGCCG CGG ||||||||[||||||||||||]PAM NA Intergenic NA NA

T20242 CCGGTGCCGCGGCTCATTGTCGG 1000
Oligo fwd CACCCCGGTGCCGCGGCTCATTGT
Oligo rev AAACACAATGAGCCGCGGCACCGG
Chromosome start end strand MM target_seq PAM alignment distance position gene name gene id
GU305914.1 190372 190394 - 0 CCGGTGCCGCGGCTCATTGT CGG ||||||||[||||||||||||]PAM NA Intergenic NA NA

T20243 AACAGAACTCTCTGTGAGAGAGG 1000
Oligo fwd CACCAACAGAACTCTCTGTGAGAG
Oligo rev AAACCTCTCACAGAGAGTTCTGTT
Chromosome start end strand MM target_seq PAM alignment distance position gene name gene id
GU305914.1 190345 190367 - 0 AACAGAACTCTCTGTGAGAG AGG ||||||||[||||||||||||]PAM NA Intergenic NA NA

T20244 ACAGAACTCTCTGTGAGAGAGGG 1000
Oligo fwd CACCACAGAACTCTCTGTGAGAGA
Oligo rev AAACTCTCTCACAGAGAGTTCTGT
Chromosome start end strand MM target_seq PAM alignment distance position gene name gene id
GU305914.1 190344 190366 - 0 ACAGAACTCTCTGTGAGAGA GGG ||||||||[||||||||||||]PAM NA Intergenic NA NA

T20245 CAGAACTCTCTGTGAGAGAGGGG 1000

| | | | | | | | | | | | |
|---|---|---|---|---|---|---|---|---|---|---|---|
| Oligo fwd | CACCCAGAACTCTCTGTGAGAGAG | | | | | | | | | | |
| Oligo rev | AAACCTCTCTCACAGAGAGTTCTG | | | | | | | | | | |
| Chromosome | start | end | strand | MM | target_seq | PAM | alignment | distance | position | gene name | gene id |
| GU305914.1 | 190343 | 190365 | - | 0 | CAGAACTCTCTGTGAGAGAG | GGG | \|\|\|\|\|\|\|\|[\|\|\|\|\|\|\|\|\|\|\|\|]PAM | NA | Intergenic | NA | NA |
| T20246 | CTCTCTGTGAGAGAGGGGACAGG | 1000 | | | | | | | | | |
| Oligo fwd | CACCCTCTCTGTGAGAGAGGGGAC | | | | | | | | | | |
| Oligo rev | AAACGTCCCCTCTCTCACAGAGAG | | | | | | | | | | |
| Chromosome | start | end | strand | MM | target_seq | PAM | alignment | distance | position | gene name | gene id |
| GU305914.1 | 190338 | 190360 | - | 0 | CTCTCTGTGAGAGAGGGGAC | AGG | \|\|\|\|\|\|\|\|[\|\|\|\|\|\|\|\|\|\|\|\|]PAM | NA | Intergenic | NA | NA |
| T20247 | GAGAGGGGACAGGAGAATCGCGG | 1000 | | | | | | | | | |
| Oligo fwd | CACCGAGAGGGGACAGGAGAATCG | | | | | | | | | | |
| Oligo rev | AAACCGATTCTCCTGTCCCCTCTC | | | | | | | | | | |
| Chromosome | start | end | strand | MM | target_seq | PAM | alignment | distance | position | gene name | gene id |
| GU305914.1 | 190328 | 190350 | - | 0 | GAGAGGGGACAGGAGAATCG | CGG | \|\|\|\|\|\|\|\|[\|\|\|\|\|\|\|\|\|\|\|\|]PAM | NA | Intergenic | NA | NA |
| T20248 | GGGACAGGAGAATCGCGGCGAGG | 1000 | | | | | | | | | |
| Oligo fwd | CACCGGGACAGGAGAATCGCGGCG | | | | | | | | | | |
| Oligo rev | AAACCGCCGCGATTCTCCTGTCCC | | | | | | | | | | |
| Chromosome | start | end | strand | MM | target_seq | PAM | alignment | distance | position | gene name | gene id |
| GU305914.1 | 190323 | 190345 | - | 0 | GGGACAGGAGAATCGCGGCG | AGG | \|\|\|\|\|\|\|\|[\|\|\|\|\|\|\|\|\|\|\|\|]PAM | NA | Intergenic | NA | NA |
| T20249 | AGGAGAATCGCGGCGAGGCATGG | 1000 | | | | | | | | | |
| Oligo fwd | CACCAGGAGAATCGCGGCGAGGCA | | | | | | | | | | |
| Oligo rev | AAACTGCCTCGCCGCGATTCTCCT | | | | | | | | | | |
| Chromosome | start | end | strand | MM | target_seq | PAM | alignment | distance | position | gene name | gene id |
| GU305914.1 | 190318 | 190340 | - | 0 | AGGAGAATCGCGGCGAGGCA | TGG | \|\|\|\|\|\|\|\|[\|\|\|\|\|\|\|\|\|\|\|\|]PAM | NA | Intergenic | NA | NA |
| T20250 | GACCTGCGCCGCCCTGACTTCGG | 1000 | | | | | | | | | |
| Oligo fwd | CACCGACCTGCGCCGCCCTGACTT | | | | | | | | | | |
| Oligo rev | AAACAAGTCAGGGCGGCGCAGGTC | | | | | | | | | | |
| Chromosome | start | end | strand | MM | target_seq | PAM | alignment | distance | position | gene name | gene id |
| GU305914.1 | 190234 | 190256 | - | 0 | GACCTGCGCCGCCCTGACTT | CGG | \|\|\|\|\|\|\|\|[\|\|\|\|\|\|\|\|\|\|\|\|]PAM | NA | Intergenic | NA | NA |
| T20251 | CGCCCTGACTTCGGTGTTGTTGG | 1000 | | | | | | | | | |
| Oligo fwd | CACCCGCCCTGACTTCGGTGTTGT | | | | | | | | | | |
| Oligo rev | AAACACAACACCGAAGTCAGGGCG | | | | | | | | | | |
| Chromosome | start | end | strand | MM | target_seq | PAM | alignment | distance | position | gene name | gene id |
| GU305914.1 | 190225 | 190247 | - | 0 | CGCCCTGACTTCGGTGTTGT | TGG | \|\|\|\|\|\|\|\|[\|\|\|\|\|\|\|\|\|\|\|\|]PAM | NA | Intergenic | NA | NA |
| T20252 | CCTGACTTCGGTGTTGTTGGCGG | 1000 | | | | | | | | | |
| Oligo fwd | CACCCCTGACTTCGGTGTTGTTGG | | | | | | | | | | |
| Oligo rev | AAACCCAACAACACCGAAGTCAGG | | | | | | | | | | |
| Chromosome | start | end | strand | MM | target_seq | PAM | alignment | distance | position | gene name | gene id |
| GU305914.1 | 190222 | 190244 | - | 0 | CCTGACTTCGGTGTTGTTGG | CGG | \|\|\|\|\|\|\|\|[\|\|\|\|\|\|\|\|\|\|\|\|]PAM | NA | Intergenic | NA | NA |
| T20253 | GTTGTTGGCGGCCGCTATATTGG | 1000 | | | | | | | | | |
| Oligo fwd | CACCGTTGTTGGCGGCCGCTATAT | | | | | | | | | | |
| Oligo rev | AAACATATAGCGGCCGCCAACAAC | | | | | | | | | | |
| Chromosome | start | end | strand | MM | target_seq | PAM | alignment | distance | position | gene name | gene id |
| GU305914.1 | 190210 | 190232 | - | 0 | GTTGTTGGCGGCCGCTATAT | TGG | \|\|\|\|\|\|\|\|[\|\|\|\|\|\|\|\|\|\|\|\|]PAM | NA | Intergenic | NA | NA |
| T20254 | GTTGGCGGCCGCTATATTGGTGG | 1000 | | | | | | | | | |
| Oligo fwd | CACCGTTGGCGGCCGCTATATTGG | | | | | | | | | | |
| Oligo rev | AAACCCAATATAGCGGCCGCCAAC | | | | | | | | | | |
| Chromosome | start | end | strand | MM | target_seq | PAM | alignment | distance | position | gene name | gene id |
| GU305914.1 | 190207 | 190229 | - | 0 | GTTGGCGGCCGCTATATTGG | TGG | \|\|\|\|\|\|\|\|[\|\|\|\|\|\|\|\|\|\|\|\|]PAM | NA | Intergenic | NA | NA |
| GU305914.1 | 165116 | 165138 | + | 4 | GATGGCCGCCGCTATGTTAG | AGG | \|-\|\|\|\|-\|[\|\|\|\|\|\|\|-\|\|-\|]PAM | NA | Intergenic | NA | NA |
| T20255 | GGCGGCCGCTATATTGGTGGTGG | 1000 | | | | | | | | | |
| Oligo fwd | CACCGGCGGCCGCTATATTGGTGG | | | | | | | | | | |
| Oligo rev | AAACCCACCAATATAGCGGCCGCC | | | | | | | | | | |
| Chromosome | start | end | strand | MM | target_seq | PAM | alignment | distance | position | gene name | gene id |
| GU305914.1 | 190204 | 190226 | - | 0 | GGCGGCCGCTATATTGGTGG | TGG | \|\|\|\|\|\|\|\|[\|\|\|\|\|\|\|\|\|\|\|\|]PAM | NA | Intergenic | NA | NA |
| T20256 | CGCTCGTGAGACGCTGCCGACGG | 1000 | | | | | | | | | |
| Oligo fwd | CACCCGCTCGTGAGACGCTGCCGA | | | | | | | | | | |
| Oligo rev | AAACTCGGCAGCGTCTCACGAGCG | | | | | | | | | | |
| Chromosome | start | end | strand | MM | target_seq | PAM | alignment | distance | position | gene name | gene id |
| GU305914.1 | 190172 | 190194 | - | 0 | CGCTCGTGAGACGCTGCCGA | CGG | \|\|\|\|\|\|\|\|[\|\|\|\|\|\|\|\|\|\|\|\|]PAM | NA | Intergenic | NA | NA |
| T20257 | CGACGGCAGCAAACCCCTCGCGG | 1000 | | | | | | | | | |
| Oligo fwd | CACCCGACGGCAGCAAACCCCTCG | | | | | | | | | | |
| Oligo rev | AAACCGAGGGGTTTGCTGCCGTCG | | | | | | | | | | |
| Chromosome | start | end | strand | MM | target_seq | PAM | alignment | distance | position | gene name | gene id |
| GU305914.1 | 190155 | 190177 | - | 0 | CGACGGCAGCAAACCCCTCG | CGG | \|\|\|\|\|\|\|\|[\|\|\|\|\|\|\|\|\|\|\|\|]PAM | NA | Intergenic | NA | NA |
| T20258 | AACCCCTCGCGGCGACGAGCAGG | 1000 | | | | | | | | | |
| Oligo fwd | CACCAACCCCTCGCGGCGACGAGC | | | | | | | | | | |
| Oligo rev | AAACGCTCGTCGCCGCGAGGGGTT | | | | | | | | | | |
| Chromosome | start | end | strand | MM | target_seq | PAM | alignment | distance | position | gene name | gene id |
| GU305914.1 | 190144 | 190166 | - | 0 | AACCCCTCGCGGCGACGAGC | AGG | \|\|\|\|\|\|\|\|[\|\|\|\|\|\|\|\|\|\|\|\|]PAM | NA | Intergenic | NA | NA |
| T20259 | GATCGAATGTAGCGTCTATTTGG | 1000 | | | | | | | | | |
| Oligo fwd | CACCGATCGAATGTAGCGTCTATT | | | | | | | | | | |
| Oligo rev | AAACAATAGACGCTACATTCGATC | | | | | | | | | | |
| Chromosome | start | end | strand | MM | target_seq | PAM | alignment | distance | position | gene name | gene id |
| GU305914.1 | 190117 | 190139 | - | 0 | GATCGAATGTAGCGTCTATT | TGG | \|\|\|\|\|\|\|\|[\|\|\|\|\|\|\|\|\|\|\|\|]PAM | NA | Intergenic | NA | NA |

T20260 GTCTATTTGGTACAGAACAACGG 1000
Oligo fwd CACCGTCTATTTGGTACAGAACAA
Oligo rev AAACTTGTTCTGTACCAAATAGAC
Chromosome start end strand MM target_seq PAM alignment distance position gene name gene id
GU305914.1 190104 190126 - 0 GTCTATTTGGTACAGAACAA CGG ||||||||[||||||||||||]PAM NA Intergenic NA NA

T20261 TCTATTTGGTACAGAACAACGGG 1000
Oligo fwd CACCTCTATTTGGTACAGAACAAC
Oligo rev AAACGTTGTTCTGTACCAAATAGA
Chromosome start end strand MM target_seq PAM alignment distance position gene name gene id
GU305914.1 190103 190125 - 0 TCTATTTGGTACAGAACAAC GGG ||||||||[||||||||||||]PAM NA Intergenic NA NA

T20262 CTATTTGGTACAGAACAACGGGG 1000
Oligo fwd CACCCTATTTGGTACAGAACAACG
Oligo rev AAACCGTTGTTCTGTACCAAATAG
Chromosome start end strand MM target_seq PAM alignment distance position gene name gene id
GU305914.1 190102 190124 - 0 CTATTTGGTACAGAACAACG GGG ||||||||[||||||||||||]PAM NA Intergenic NA NA

T20263 TTTGGTACAGAACAACGGGGTGG 1000
Oligo fwd CACCTTTGGTACAGAACAACGGGG
Oligo rev AAACCCCCGTTGTTCTGTACCAAA
Chromosome start end strand MM target_seq PAM alignment distance position gene name gene id
GU305914.1 190099 190121 - 0 TTTGGTACAGAACAACGGGG TGG ||||||||[||||||||||||]PAM NA Intergenic NA NA

T20264 GGACGACGACGACATCCCCCCGG 1000
Oligo fwd CACCGGACGACGACGACATCCCCC
Oligo rev AAACGGGGGATGTCGTCGTCGTCC
Chromosome start end strand MM target_seq PAM alignment distance position gene name gene id
GU305914.1 190078 190100 - 0 GGACGACGACGACATCCCCC CGG ||||||||[||||||||||||]PAM NA Intergenic NA NA

T20265 GACGACGACGACATCCCCCCGGG 1000
Oligo fwd CACCGACGACGACGACATCCCCCC
Oligo rev AAACGGGGGGATGTCGTCGTCGTC
Chromosome start end strand MM target_seq PAM alignment distance position gene name gene id
GU305914.1 190077 190099 - 0 GACGACGACGACATCCCCCC GGG ||||||||[||||||||||||]PAM NA Intergenic NA NA

T20266 ACGACGACGACATCCCCCCGGGG 1000
Oligo fwd CACCACGACGACGACATCCCCCCG
Oligo rev AAACCGGGGGGATGTCGTCGTCGT
Chromosome start end strand MM target_seq PAM alignment distance position gene name gene id
GU305914.1 190076 190098 - 0 ACGACGACGACATCCCCCCG GGG ||||||||[||||||||||||]PAM NA Intergenic NA NA

T20267 CGACATCCCCCCGGGGTTTCTGG 1000
Oligo fwd CACCCGACATCCCCCCGGGGTTTC
Oligo rev AAACGAAACCCCGGGGGGATGTCG
Chromosome start end strand MM target_seq PAM alignment distance position gene name gene id
GU305914.1 190069 190091 - 0 CGACATCCCCCCGGGGTTTC TGG ||||||||[||||||||||||]PAM NA Intergenic NA NA

T20268 GACATCCCCCCGGGGTTTCTGGG 1000
Oligo fwd CACCGACATCCCCCCGGGGTTTCT
Oligo rev AAACAGAAACCCCGGGGGGATGTC
Chromosome start end strand MM target_seq PAM alignment distance position gene name gene id
GU305914.1 190068 190090 - 0 GACATCCCCCCGGGGTTTCT GGG ||||||||[||||||||||||]PAM NA Intergenic NA NA

T20269 ACATCCCCCCGGGGTTTCTGGGG 1000
Oligo fwd CACCACATCCCCCCGGGGTTTCTG
Oligo rev AAACCAGAAACCCCGGGGGGATGT
Chromosome start end strand MM target_seq PAM alignment distance position gene name gene id
GU305914.1 190067 190089 - 0 ACATCCCCCCGGGGTTTCTG GGG ||||||||[||||||||||||]PAM NA Intergenic NA NA

T20270 GGGTTTCTGGGGCGCCCCGCCGG 1000
Oligo fwd CACCGGGTTTCTGGGGCGCCCCGC
Oligo rev AAACGCGGGGCGCCCCAGAAACCC
Chromosome start end strand MM target_seq PAM alignment distance position gene name gene id
GU305914.1 190056 190078 - 0 GGGTTTCTGGGGCGCCCCGC CGG ||||||||[||||||||||||]PAM NA Intergenic NA NA

T20271 GGTTTCTGGGGCGCCCCGCCGGG 1000
Oligo fwd CACCGGTTTCTGGGGCGCCCCGCC
Oligo rev AAACGGCGGGGCGCCCCAGAAACC
Chromosome start end strand MM target_seq PAM alignment distance position gene name gene id
GU305914.1 190055 190077 - 0 GGTTTCTGGGGCGCCCCGCC GGG ||||||||[||||||||||||]PAM NA Intergenic NA NA

T20272 GAGCCCGCACTCCTTCCCGCTGG 1000
Oligo fwd CACCGAGCCCGCACTCCTTCCCGC
Oligo rev AAACGCGGGAAGGAGTGCGGGCTC
Chromosome start end strand MM target_seq PAM alignment distance position gene name gene id
GU305914.1 190021 190043 - 0 GAGCCCGCACTCCTTCCCGC TGG ||||||||[||||||||||||]PAM NA Intergenic NA NA

T20273 TTACGAGAGTATCGATTGAACGG 1000
Oligo fwd CACCTTACGAGAGTATCGATTGAA
Oligo rev AAACTTCAATCGATACTCTCGTAA
Chromosome start end strand MM target_seq PAM alignment distance position gene name gene id
GU305914.1 189991 190013 - 0 TTACGAGAGTATCGATTGAA CGG ||||||||[||||||||||||]PAM NA Intergenic NA NA

T20274 CGACTGCCATCCCCCAAACCCGG 1000
Oligo fwd CACCCGACTGCCATCCCCCAAACC
Oligo rev AAACGGTTTGGGGGATGGCAGTCG
Chromosome start end strand MM target_seq PAM alignment distance position gene name gene id

| | | | | | | | | | | | |
|---|---|---|---|---|---|---|---|---|---|---|---|
| GU305914.1 | | 189955 | 189977 | - | 0 | CGACTGCCATCCCCCAAACC | CGG | \|\|\|\|\|\|\|\|[\|\|\|\|\|\|\|\|\|\|\|\|]PAM | NA | Intergenic | NA | NA |
| T20275 | TACACACTTTCGCACCCGACAGG | | 1000 | | | | | | | | | |
| Oligo fwd | CACCTACACACTTTCGCACCCGAC | | | | | | | | | | | |
| Oligo rev | AAACGTCGGGTGCGAAAGTGTGTA | | | | | | | | | | | |
| Chromosome | start | | end | strand | MM | target_seq | PAM | alignment | distance | position | gene name | gene id |
| GU305914.1 | | 189900 | 189922 | - | 0 | TACACACTTTCGCACCCGAC | AGG | \|\|\|\|\|\|\|\|[\|\|\|\|\|\|\|\|\|\|\|\|]PAM | NA | Intergenic | NA | NA |
| T20276 | GATTTCTCTCTTTTTCGCGCTGG | | 1000 | | | | | | | | | |
| Oligo fwd | CACCGATTTCTCTCTTTTTCGCGC | | | | | | | | | | | |
| Oligo rev | AAACGCGCGAAAAAGAGAGAAATC | | | | | | | | | | | |
| Chromosome | start | | end | strand | MM | target_seq | PAM | alignment | distance | position | gene name | gene id |
| GU305914.1 | | 189850 | 189872 | - | 0 | GATTTCTCTCTTTTTCGCGC | TGG | \|\|\|\|\|\|\|\|[\|\|\|\|\|\|\|\|\|\|\|\|]PAM | NA | Intergenic | NA | NA |
| T20277 | ATTTCTCTCTTTTTCGCGCTGGG | | 1000 | | | | | | | | | |
| Oligo fwd | CACCATTTCTCTCTTTTTCGCGCT | | | | | | | | | | | |
| Oligo rev | AAACAGCGCGAAAAAGAGAGAAAT | | | | | | | | | | | |
| Chromosome | start | | end | strand | MM | target_seq | PAM | alignment | distance | position | gene name | gene id |
| GU305914.1 | | 189849 | 189871 | - | 0 | ATTTCTCTCTTTTTCGCGCT | GGG | \|\|\|\|\|\|\|\|[\|\|\|\|\|\|\|\|\|\|\|\|]PAM | NA | Intergenic | NA | NA |
| T20278 | TTTCTCTCTTTTTCGCGCTGGGG | | 1000 | | | | | | | | | |
| Oligo fwd | CACCTTTCTCTCTTTTTCGCGCTG | | | | | | | | | | | |
| Oligo rev | AAACCAGCGCGAAAAAGAGAGAAA | | | | | | | | | | | |
| Chromosome | start | | end | strand | MM | target_seq | PAM | alignment | distance | position | gene name | gene id |
| GU305914.1 | | 189848 | 189870 | - | 0 | TTTCTCTCTTTTTCGCGCTG | GGG | \|\|\|\|\|\|\|\|[\|\|\|\|\|\|\|\|\|\|\|\|]PAM | NA | Intergenic | NA | NA |
| T20279 | TCTTTTTCGCGCTGGGGTCCAGG | | 1000 | | | | | | | | | |
| Oligo fwd | CACCTCTTTTTCGCGCTGGGGTCC | | | | | | | | | | | |
| Oligo rev | AAACGGACCCCAGCGCGAAAAAGA | | | | | | | | | | | |
| Chromosome | start | | end | strand | MM | target_seq | PAM | alignment | distance | position | gene name | gene id |
| GU305914.1 | | 189842 | 189864 | - | 0 | TCTTTTTCGCGCTGGGGTCC | AGG | \|\|\|\|\|\|\|\|[\|\|\|\|\|\|\|\|\|\|\|\|]PAM | NA | Intergenic | NA | NA |
| T20280 | GCTGGGGTCCAGGCCGCTCGCGG | | 1000 | | | | | | | | | |
| Oligo fwd | CACCGCTGGGGTCCAGGCCGCTCG | | | | | | | | | | | |
| Oligo rev | AAACCGAGCGGCCTGGACCCCAGC | | | | | | | | | | | |
| Chromosome | start | | end | strand | MM | target_seq | PAM | alignment | distance | position | gene name | gene id |
| GU305914.1 | | 189832 | 189854 | - | 0 | GCTGGGGTCCAGGCCGCTCG | CGG | \|\|\|\|\|\|\|\|[\|\|\|\|\|\|\|\|\|\|\|\|]PAM | NA | Intergenic | NA | NA |
| T20281 | CTGGGGTCCAGGCCGCTCGCGGG | | 1000 | | | | | | | | | |
| Oligo fwd | CACCCTGGGGTCCAGGCCGCTCGC | | | | | | | | | | | |
| Oligo rev | AAACGCGAGCGGCCTGGACCCCAG | | | | | | | | | | | |
| Chromosome | start | | end | strand | MM | target_seq | PAM | alignment | distance | position | gene name | gene id |
| GU305914.1 | | 189831 | 189853 | - | 0 | CTGGGGTCCAGGCCGCTCGC | GGG | \|\|\|\|\|\|\|\|[\|\|\|\|\|\|\|\|\|\|\|\|]PAM | NA | Intergenic | NA | NA |
| T20282 | GCTCGCGGGCTATAGCAATTTGG | | 1000 | | | | | | | | | |
| Oligo fwd | CACCGCTCGCGGGCTATAGCAATT | | | | | | | | | | | |
| Oligo rev | AAACAATTGCTATAGCCCGCGAGC | | | | | | | | | | | |
| Chromosome | start | | end | strand | MM | target_seq | PAM | alignment | distance | position | gene name | gene id |
| GU305914.1 | | 189817 | 189839 | - | 0 | GCTCGCGGGCTATAGCAATT | TGG | \|\|\|\|\|\|\|\|[\|\|\|\|\|\|\|\|\|\|\|\|]PAM | NA | Intergenic | NA | NA |
| T20283 | GCTATAGCAATTTGGCAGTGTGG | | 1000 | | | | | | | | | |
| Oligo fwd | CACCGCTATAGCAATTTGGCAGTG | | | | | | | | | | | |
| Oligo rev | AAACCACTGCCAAATTGCTATAGC | | | | | | | | | | | |
| Chromosome | start | | end | strand | MM | target_seq | PAM | alignment | distance | position | gene name | gene id |
| GU305914.1 | | 189809 | 189831 | - | 0 | GCTATAGCAATTTGGCAGTG | TGG | \|\|\|\|\|\|\|\|[\|\|\|\|\|\|\|\|\|\|\|\|]PAM | NA | Intergenic | NA | NA |
| T20284 | TGGTACTCCGATTATGAAAACGG | | 1000 | | | | | | | | | |
| Oligo fwd | CACCTGGTACTCCGATTATGAAAA | | | | | | | | | | | |
| Oligo rev | AAACTTTTCATAATCGGAGTACCA | | | | | | | | | | | |
| Chromosome | start | | end | strand | MM | target_seq | PAM | alignment | distance | position | gene name | gene id |
| GU305914.1 | | 189789 | 189811 | - | 0 | TGGTACTCCGATTATGAAAA | CGG | \|\|\|\|\|\|\|\|[\|\|\|\|\|\|\|\|\|\|\|\|]PAM | NA | Intergenic | NA | NA |
| T20285 | TCCCGAGAAAATCAAATCCTCGG | | 1000 | | | | | | | | | |
| Oligo fwd | CACCTCCCGAGAAAATCAAATCCT | | | | | | | | | | | |
| Oligo rev | AAACAGGATTTGATTTTCTCGGGA | | | | | | | | | | | |
| Chromosome | start | | end | strand | MM | target_seq | PAM | alignment | distance | position | gene name | gene id |
| GU305914.1 | | 189697 | 189719 | - | 0 | TCCCGAGAAAATCAAATCCT | CGG | \|\|\|\|\|\|\|\|[\|\|\|\|\|\|\|\|\|\|\|\|]PAM | NA | Intergenic | NA | NA |
| T20286 | GAAAATCAAATCCTCGGCGACGG | | 1000 | | | | | | | | | |
| Oligo fwd | CACCGAAAATCAAATCCTCGGCGA | | | | | | | | | | | |
| Oligo rev | AAACTCGCCGAGGATTTGATTTTC | | | | | | | | | | | |
| Chromosome | start | | end | strand | MM | target_seq | PAM | alignment | distance | position | gene name | gene id |
| GU305914.1 | | 189691 | 189713 | - | 0 | GAAAATCAAATCCTCGGCGA | CGG | \|\|\|\|\|\|\|\|[\|\|\|\|\|\|\|\|\|\|\|\|]PAM | NA | Intergenic | NA | NA |
| T20287 | TCGGCGACGGTGACGCCGAACGG | | 1000 | | | | | | | | | |
| Oligo fwd | CACCTCGGCGACGGTGACGCCGAA | | | | | | | | | | | |
| Oligo rev | AAACTTCGGCGTCACCGTCGCCGA | | | | | | | | | | | |
| Chromosome | start | | end | strand | MM | target_seq | PAM | alignment | distance | position | gene name | gene id |
| GU305914.1 | | 189678 | 189700 | - | 0 | TCGGCGACGGTGACGCCGAA | CGG | \|\|\|\|\|\|\|\|[\|\|\|\|\|\|\|\|\|\|\|\|]PAM | NA | Intergenic | NA | NA |
| T20288 | CAACTACCTGACCACCAAGTTGG | | 1000 | | | | | | | | | |
| Oligo fwd | CACCCAACTACCTGACCACCAAGT | | | | | | | | | | | |
| Oligo rev | AAACACTTGGTGGTCAGGTAGTTG | | | | | | | | | | | |
| Chromosome | start | | end | strand | MM | target_seq | PAM | alignment | distance | position | gene name | gene id |
| GU305914.1 | | 189637 | 189659 | - | 0 | CAACTACCTGACCACCAAGT | TGG | \|\|\|\|\|\|\|\|[\|\|\|\|\|\|\|\|\|\|\|\|]PAM | NA | Intergenic | NA | NA |
| T20289 | CTACCTGACCACCAAGTTGGTGG | | 1000 | | | | | | | | | |
| Oligo fwd | CACCCTACCTGACCACCAAGTTGG | | | | | | | | | | | |
| Oligo rev | AAACCCAACTTGGTGGTCAGGTAG | | | | | | | | | | | |

Chromosome start end strand MM target_seq PAM alignment distance position gene name gene id
GU305914.1 189634 189656 - 0 CTACCTGACCACCAAGTTGG TGG ||||||||[||||||||||||]PAM NA Intergenic NA NA

T20290 CCTGACCACCAAGTTGGTGGAGG 1000
Oligo fwd CACCCCTGACCACCAAGTTGGTGG
Oligo rev AAACCCACCAACTTGGTGGTCAGG
Chromosome start end strand MM target_seq PAM alignment distance position gene name gene id
GU305914.1 189631 189653 - 0 CCTGACCACCAAGTTGGTGG AGG ||||||||[||||||||||||]PAM NA Intergenic NA NA

T20291 TTGGTGGAGGATAGTTCGACCGG 1000
Oligo fwd CACCTTGGTGGAGGATAGTTCGAC
Oligo rev AAACGTCGAACTATCCTCCACCAA
Chromosome start end strand MM target_seq PAM alignment distance position gene name gene id
GU305914.1 189618 189640 - 0 TTGGTGGAGGATAGTTCGAC CGG ||||||||[||||||||||||]PAM NA Intergenic NA NA

T20292 AGGATAGTTCGACCGGCAAGCGG 1000
Oligo fwd CACCAGGATAGTTCGACCGGCAAG
Oligo rev AAACCTTGCCGGTCGAACTATCCT
Chromosome start end strand MM target_seq PAM alignment distance position gene name gene id
GU305914.1 189611 189633 - 0 AGGATAGTTCGACCGGCAAG CGG ||||||||[||||||||||||]PAM NA Intergenic NA NA

T20293 CGACCGGCAAGCGGCATCGCCGG 1000
Oligo fwd CACCCGACCGGCAAGCGGCATCGC
Oligo rev AAACGCGATGCCGCTTGCCGGTCG
Chromosome start end strand MM target_seq PAM alignment distance position gene name gene id
GU305914.1 189602 189624 - 0 CGACCGGCAAGCGGCATCGC CGG ||||||||[||||||||||||]PAM NA Intergenic NA NA

T20294 CCGGCTGTCGTTCAGAGACCCGG 1000
Oligo fwd CACCCCGGCTGTCGTTCAGAGACC
Oligo rev AAACGGTCTCTGAACGACAGCCGG
Chromosome start end strand MM target_seq PAM alignment distance position gene name gene id
GU305914.1 189583 189605 - 0 CCGGCTGTCGTTCAGAGACC CGG ||||||||[||||||||||||]PAM NA Intergenic NA NA

T20295 GTGTCTTTACTGCGAGAGACCGG 1000
Oligo fwd CACCGTGTCTTTACTGCGAGAGAC
Oligo rev AAACGTCTCTCGCAGTAAAGACAC
Chromosome start end strand MM target_seq PAM alignment distance position gene name gene id
GU305914.1 189541 189563 - 0 GTGTCTTTACTGCGAGAGAC CGG ||||||||[||||||||||||]PAM NA Intergenic NA NA

T20296 TGTCTTTACTGCGAGAGACCGGG 1000
Oligo fwd CACCTGTCTTTACTGCGAGAGACC
Oligo rev AAACGGTCTCTCGCAGTAAAGACA
Chromosome start end strand MM target_seq PAM alignment distance position gene name gene id
GU305914.1 189540 189562 - 0 TGTCTTTACTGCGAGAGACC GGG ||||||||[||||||||||||]PAM NA Intergenic NA NA

T20297 CGAGAGACCGGGCGATTCGATGG 1000
Oligo fwd CACCCGAGAGACCGGGCGATTCGA
Oligo rev AAACTCGAATCGCCCGGTCTCTCG
Chromosome start end strand MM target_seq PAM alignment distance position gene name gene id
GU305914.1 189529 189551 - 0 CGAGAGACCGGGCGATTCGA TGG ||||||||[||||||||||||]PAM NA Intergenic NA NA

T20298 ACCGGGCGATTCGATGGATATGG 1000
Oligo fwd CACCACCGGGCGATTCGATGGATA
Oligo rev AAACTATCCATCGAATCGCCCGGT
Chromosome start end strand MM target_seq PAM alignment distance position gene name gene id
GU305914.1 189523 189545 - 0 ACCGGGCGATTCGATGGATA TGG ||||||||[||||||||||||]PAM NA Intergenic NA NA

T20299 GGAGAGATGTCTGATGTGCAAGG 1000
Oligo fwd CACCGGAGAGATGTCTGATGTGCA
Oligo rev AAACTGCACATCAGACATCTCTCC
Chromosome start end strand MM target_seq PAM alignment distance position gene name gene id
GU305914.1 189502 189524 - 0 GGAGAGATGTCTGATGTGCA AGG ||||||||[||||||||||||]PAM NA Intergenic NA NA

T20300 GATGTGCAAGGACACGACAATGG 1000
Oligo fwd CACCGATGTGCAAGGACACGACAA
Oligo rev AAACTTGTCGTGTCCTTGCACATC
Chromosome start end strand MM target_seq PAM alignment distance position gene name gene id
GU305914.1 189490 189512 - 0 GATGTGCAAGGACACGACAA TGG ||||||||[||||||||||||]PAM NA Intergenic NA NA

T20301 GTGCAAGGACACGACAATGGCGG 1000
Oligo fwd CACCGTGCAAGGACACGACAATGG
Oligo rev AAACCCATTGTCGTGTCCTTGCAC
Chromosome start end strand MM target_seq PAM alignment distance position gene name gene id
GU305914.1 189487 189509 - 0 GTGCAAGGACACGACAATGG CGG ||||||||[||||||||||||]PAM NA Intergenic NA NA

T20302 GACAATGGCGGTCTTACTAAAGG 1000
Oligo fwd CACCGACAATGGCGGTCTTACTAA
Oligo rev AAACTTAGTAAGACCGCCATTGTC
Chromosome start end strand MM target_seq PAM alignment distance position gene name gene id
GU305914.1 189475 189497 - 0 GACAATGGCGGTCTTACTAA AGG ||||||||[||||||||||||]PAM NA Intergenic NA NA

T20303 AGGATTTACACACGCCCAAACGG 1000
Oligo fwd CACCAGGATTTACACACGCCCAAA
Oligo rev AAACTTTGGGCGTGTGTAAATCCT
Chromosome start end strand MM target_seq PAM alignment distance position gene name gene id
GU305914.1 189455 189477 - 0 AGGATTTACACACGCCCAAA CGG ||||||||[||||||||||||]PAM NA Intergenic NA NA

T20304 GCGCGATTTTTCGCCTATGTTGG 1000
Oligo fwd CACCGCGCGATTTTTCGCCTATGT

<table>
<tr><td>Oligo rev</td><td>AAACACATAGGCGAAAAATCGCGC</td><td></td><td></td><td></td><td></td><td></td><td></td><td></td><td></td><td></td><td></td><td></td></tr>
<tr><td>Chromosome</td><td>start</td><td></td><td>end</td><td>strand</td><td>MM</td><td>target_seq</td><td>PAM</td><td>alignment</td><td>distance</td><td>position</td><td>gene name</td><td>gene id</td></tr>
<tr><td>GU305914.1</td><td></td><td>189433</td><td>189455</td><td>-</td><td>0</td><td>GCGCGATTTTTCGCCTATGT</td><td>TGG</td><td>||||||||[||||||||||||]PAM</td><td>NA</td><td>Intergenic</td><td>NA</td><td>NA</td></tr>
<tr><td>T20305</td><td>ATTTTTCGCCTATGTTGGTCCGG</td><td></td><td>1000</td><td></td><td></td><td></td><td></td><td></td><td></td><td></td><td></td><td></td></tr>
<tr><td>Oligo fwd</td><td>CACCATTTTTCGCCTATGTTGGTC</td><td></td><td></td><td></td><td></td><td></td><td></td><td></td><td></td><td></td><td></td><td></td></tr>
<tr><td>Oligo rev</td><td>AAACGACCAACATAGGCGAAAAAT</td><td></td><td></td><td></td><td></td><td></td><td></td><td></td><td></td><td></td><td></td><td></td></tr>
<tr><td>Chromosome</td><td>start</td><td></td><td>end</td><td>strand</td><td>MM</td><td>target_seq</td><td>PAM</td><td>alignment</td><td>distance</td><td>position</td><td>gene name</td><td>gene id</td></tr>
<tr><td>GU305914.1</td><td></td><td>189428</td><td>189450</td><td>-</td><td>0</td><td>ATTTTTCGCCTATGTTGGTC</td><td>CGG</td><td>||||||||[||||||||||||]PAM</td><td>NA</td><td>Intergenic</td><td>NA</td><td>NA</td></tr>
<tr><td>T20306</td><td>TTTTTCGCCTATGTTGGTCCGGG</td><td></td><td>1000</td><td></td><td></td><td></td><td></td><td></td><td></td><td></td><td></td><td></td></tr>
<tr><td>Oligo fwd</td><td>CACCTTTTTCGCCTATGTTGGTCC</td><td></td><td></td><td></td><td></td><td></td><td></td><td></td><td></td><td></td><td></td><td></td></tr>
<tr><td>Oligo rev</td><td>AAACGGACCAACATAGGCGAAAAA</td><td></td><td></td><td></td><td></td><td></td><td></td><td></td><td></td><td></td><td></td><td></td></tr>
<tr><td>Chromosome</td><td>start</td><td></td><td>end</td><td>strand</td><td>MM</td><td>target_seq</td><td>PAM</td><td>alignment</td><td>distance</td><td>position</td><td>gene name</td><td>gene id</td></tr>
<tr><td>GU305914.1</td><td></td><td>189427</td><td>189449</td><td>-</td><td>0</td><td>TTTTTCGCCTATGTTGGTCC</td><td>GGG</td><td>||||||||[||||||||||||]PAM</td><td>NA</td><td>Intergenic</td><td>NA</td><td>NA</td></tr>
<tr><td>T20307</td><td>GCCTATGTTGGTCCGGGCCACGG</td><td></td><td>1000</td><td></td><td></td><td></td><td></td><td></td><td></td><td></td><td></td><td></td></tr>
<tr><td>Oligo fwd</td><td>CACCGCCTATGTTGGTCCGGGCCA</td><td></td><td></td><td></td><td></td><td></td><td></td><td></td><td></td><td></td><td></td><td></td></tr>
<tr><td>Oligo rev</td><td>AAACTGGCCCGGACCAACATAGGC</td><td></td><td></td><td></td><td></td><td></td><td></td><td></td><td></td><td></td><td></td><td></td></tr>
<tr><td>Chromosome</td><td>start</td><td></td><td>end</td><td>strand</td><td>MM</td><td>target_seq</td><td>PAM</td><td>alignment</td><td>distance</td><td>position</td><td>gene name</td><td>gene id</td></tr>
<tr><td>GU305914.1</td><td></td><td>189421</td><td>189443</td><td>-</td><td>0</td><td>GCCTATGTTGGTCCGGGCCA</td><td>CGG</td><td>||||||||[||||||||||||]PAM</td><td>NA</td><td>Intergenic</td><td>NA</td><td>NA</td></tr>
<tr><td>T20308</td><td>TATGTTGGTCCGGGCCACGGTGG</td><td></td><td>1000</td><td></td><td></td><td></td><td></td><td></td><td></td><td></td><td></td><td></td></tr>
<tr><td>Oligo fwd</td><td>CACCTATGTTGGTCCGGGCCACGG</td><td></td><td></td><td></td><td></td><td></td><td></td><td></td><td></td><td></td><td></td><td></td></tr>
<tr><td>Oligo rev</td><td>AAACCCGTGGCCCGGACCAACATA</td><td></td><td></td><td></td><td></td><td></td><td></td><td></td><td></td><td></td><td></td><td></td></tr>
<tr><td>Chromosome</td><td>start</td><td></td><td>end</td><td>strand</td><td>MM</td><td>target_seq</td><td>PAM</td><td>alignment</td><td>distance</td><td>position</td><td>gene name</td><td>gene id</td></tr>
<tr><td>GU305914.1</td><td></td><td>189418</td><td>189440</td><td>-</td><td>0</td><td>TATGTTGGTCCGGGCCACGG</td><td>TGG</td><td>||||||||[||||||||||||]PAM</td><td>NA</td><td>Intergenic</td><td>NA</td><td>NA</td></tr>
<tr><td>T20309</td><td>CGGTGGAGACCGTCAATCACCGG</td><td></td><td>1000</td><td></td><td></td><td></td><td></td><td></td><td></td><td></td><td></td><td></td></tr>
<tr><td>Oligo fwd</td><td>CACCCGGTGGAGACCGTCAATCAC</td><td></td><td></td><td></td><td></td><td></td><td></td><td></td><td></td><td></td><td></td><td></td></tr>
<tr><td>Oligo rev</td><td>AAACGTGATTGACGGTCTCCACCG</td><td></td><td></td><td></td><td></td><td></td><td></td><td></td><td></td><td></td><td></td><td></td></tr>
<tr><td>Chromosome</td><td>start</td><td></td><td>end</td><td>strand</td><td>MM</td><td>target_seq</td><td>PAM</td><td>alignment</td><td>distance</td><td>position</td><td>gene name</td><td>gene id</td></tr>
<tr><td>GU305914.1</td><td></td><td>189401</td><td>189423</td><td>-</td><td>0</td><td>CGGTGGAGACCGTCAATCAC</td><td>CGG</td><td>||||||||[||||||||||||]PAM</td><td>NA</td><td>Intergenic</td><td>NA</td><td>NA</td></tr>
<tr><td>T20310</td><td>ACCGGCCTTATTTCACGTGCCGG</td><td></td><td>1000</td><td></td><td></td><td></td><td></td><td></td><td></td><td></td><td></td><td></td></tr>
<tr><td>Oligo fwd</td><td>CACCACCGGCCTTATTTCACGTGC</td><td></td><td></td><td></td><td></td><td></td><td></td><td></td><td></td><td></td><td></td><td></td></tr>
<tr><td>Oligo rev</td><td>AAACGCACGTGAAATAAGGCCGGT</td><td></td><td></td><td></td><td></td><td></td><td></td><td></td><td></td><td></td><td></td><td></td></tr>
<tr><td>Chromosome</td><td>start</td><td></td><td>end</td><td>strand</td><td>MM</td><td>target_seq</td><td>PAM</td><td>alignment</td><td>distance</td><td>position</td><td>gene name</td><td>gene id</td></tr>
<tr><td>GU305914.1</td><td></td><td>189383</td><td>189405</td><td>-</td><td>0</td><td>ACCGGCCTTATTTCACGTGC</td><td>CGG</td><td>||||||||[||||||||||||]PAM</td><td>NA</td><td>Intergenic</td><td>NA</td><td>NA</td></tr>
<tr><td>T20311</td><td>GTGCCGGTTTCGCGTAACTCCGG</td><td></td><td>1000</td><td></td><td></td><td></td><td></td><td></td><td></td><td></td><td></td><td></td></tr>
<tr><td>Oligo fwd</td><td>CACCGTGCCGGTTTCGCGTAACTC</td><td></td><td></td><td></td><td></td><td></td><td></td><td></td><td></td><td></td><td></td><td></td></tr>
<tr><td>Oligo rev</td><td>AAACGAGTTACGCGAAACCGGCAC</td><td></td><td></td><td></td><td></td><td></td><td></td><td></td><td></td><td></td><td></td><td></td></tr>
<tr><td>Chromosome</td><td>start</td><td></td><td>end</td><td>strand</td><td>MM</td><td>target_seq</td><td>PAM</td><td>alignment</td><td>distance</td><td>position</td><td>gene name</td><td>gene id</td></tr>
<tr><td>GU305914.1</td><td></td><td>189367</td><td>189389</td><td>-</td><td>0</td><td>GTGCCGGTTTCGCGTAACTC</td><td>CGG</td><td>||||||||[||||||||||||]PAM</td><td>NA</td><td>Intergenic</td><td>NA</td><td>NA</td></tr>
<tr><td>T20312</td><td>GGACACCCAGAAGATCGAGACGG</td><td></td><td>1000</td><td></td><td></td><td></td><td></td><td></td><td></td><td></td><td></td><td></td></tr>
<tr><td>Oligo fwd</td><td>CACCGGACACCCAGAAGATCGAGA</td><td></td><td></td><td></td><td></td><td></td><td></td><td></td><td></td><td></td><td></td><td></td></tr>
<tr><td>Oligo rev</td><td>AAACTCTCGATCTTCTGGGTGTCC</td><td></td><td></td><td></td><td></td><td></td><td></td><td></td><td></td><td></td><td></td><td></td></tr>
<tr><td>Chromosome</td><td>start</td><td></td><td>end</td><td>strand</td><td>MM</td><td>target_seq</td><td>PAM</td><td>alignment</td><td>distance</td><td>position</td><td>gene name</td><td>gene id</td></tr>
<tr><td>GU305914.1</td><td></td><td>189346</td><td>189368</td><td>-</td><td>0</td><td>GGACACCCAGAAGATCGAGA</td><td>CGG</td><td>||||||||[||||||||||||]PAM</td><td>NA</td><td>Intergenic</td><td>NA</td><td>NA</td></tr>
<tr><td>T20313</td><td>CCCAGAAGATCGAGACGGCGTGG</td><td></td><td>1000</td><td></td><td></td><td></td><td></td><td></td><td></td><td></td><td></td><td></td></tr>
<tr><td>Oligo fwd</td><td>CACCCCCAGAAGATCGAGACGGCG</td><td></td><td></td><td></td><td></td><td></td><td></td><td></td><td></td><td></td><td></td><td></td></tr>
<tr><td>Oligo rev</td><td>AAACCGCCGTCTCGATCTTCTGGG</td><td></td><td></td><td></td><td></td><td></td><td></td><td></td><td></td><td></td><td></td><td></td></tr>
<tr><td>Chromosome</td><td>start</td><td></td><td>end</td><td>strand</td><td>MM</td><td>target_seq</td><td>PAM</td><td>alignment</td><td>distance</td><td>position</td><td>gene name</td><td>gene id</td></tr>
<tr><td>GU305914.1</td><td></td><td>189341</td><td>189363</td><td>-</td><td>0</td><td>CCCAGAAGATCGAGACGGCG</td><td>TGG</td><td>||||||||[||||||||||||]PAM</td><td>NA</td><td>Intergenic</td><td>NA</td><td>NA</td></tr>
<tr><td>T20314</td><td>TTGAACGACGACTGCACCTTCGG</td><td></td><td>1000</td><td></td><td></td><td></td><td></td><td></td><td></td><td></td><td></td><td></td></tr>
<tr><td>Oligo fwd</td><td>CACCTTGAACGACGACTGCACCTT</td><td></td><td></td><td></td><td></td><td></td><td></td><td></td><td></td><td></td><td></td><td></td></tr>
<tr><td>Oligo rev</td><td>AAACAAGGTGCAGTCGTCGTTCAA</td><td></td><td></td><td></td><td></td><td></td><td></td><td></td><td></td><td></td><td></td><td></td></tr>
<tr><td>Chromosome</td><td>start</td><td></td><td>end</td><td>strand</td><td>MM</td><td>target_seq</td><td>PAM</td><td>alignment</td><td>distance</td><td>position</td><td>gene name</td><td>gene id</td></tr>
<tr><td>GU305914.1</td><td></td><td>189318</td><td>189340</td><td>-</td><td>0</td><td>TTGAACGACGACTGCACCTT</td><td>CGG</td><td>||||||||[||||||||||||]PAM</td><td>NA</td><td>Intergenic</td><td>NA</td><td>NA</td></tr>
<tr><td>T20315</td><td>CAAAACGAAAGATCCGATCGAGG</td><td></td><td>1000</td><td></td><td></td><td></td><td></td><td></td><td></td><td></td><td></td><td></td></tr>
<tr><td>Oligo fwd</td><td>CACCCAAAACGAAAGATCCGATCG</td><td></td><td></td><td></td><td></td><td></td><td></td><td></td><td></td><td></td><td></td><td></td></tr>
<tr><td>Oligo rev</td><td>AAACCGATCGGATCTTTCGTTTTG</td><td></td><td></td><td></td><td></td><td></td><td></td><td></td><td></td><td></td><td></td><td></td></tr>
<tr><td>Chromosome</td><td>start</td><td></td><td>end</td><td>strand</td><td>MM</td><td>target_seq</td><td>PAM</td><td>alignment</td><td>distance</td><td>position</td><td>gene name</td><td>gene id</td></tr>
<tr><td>GU305914.1</td><td></td><td>189292</td><td>189314</td><td>-</td><td>0</td><td>CAAAACGAAAGATCCGATCG</td><td>AGG</td><td>||||||||[||||||||||||]PAM</td><td>NA</td><td>Intergenic</td><td>NA</td><td>NA</td></tr>
<tr><td>T20316</td><td>GATCGAGGCCTGCGCCAAAACGG</td><td></td><td>1000</td><td></td><td></td><td></td><td></td><td></td><td></td><td></td><td></td><td></td></tr>
<tr><td>Oligo fwd</td><td>CACCGATCGAGGCCTGCGCCAAAA</td><td></td><td></td><td></td><td></td><td></td><td></td><td></td><td></td><td></td><td></td><td></td></tr>
<tr><td>Oligo rev</td><td>AAACTTTTGGCGCAGGCCTCGATC</td><td></td><td></td><td></td><td></td><td></td><td></td><td></td><td></td><td></td><td></td><td></td></tr>
<tr><td>Chromosome</td><td>start</td><td></td><td>end</td><td>strand</td><td>MM</td><td>target_seq</td><td>PAM</td><td>alignment</td><td>distance</td><td>position</td><td>gene name</td><td>gene id</td></tr>
<tr><td>GU305914.1</td><td></td><td>189277</td><td>189299</td><td>-</td><td>0</td><td>GATCGAGGCCTGCGCCAAAA</td><td>CGG</td><td>||||||||[||||||||||||]PAM</td><td>NA</td><td>Intergenic</td><td>NA</td><td>NA</td></tr>
<tr><td>T20317</td><td>CTACAAGAAATCCATACGCGAGG</td><td></td><td>1000</td><td></td><td></td><td></td><td></td><td></td><td></td><td></td><td></td><td></td></tr>
<tr><td>Oligo fwd</td><td>CACCCTACAAGAAATCCATACGCG</td><td></td><td></td><td></td><td></td><td></td><td></td><td></td><td></td><td></td><td></td><td></td></tr>
<tr><td>Oligo rev</td><td>AAACCGCGTATGGATTTCTTGTAG</td><td></td><td></td><td></td><td></td><td></td><td></td><td></td><td></td><td></td><td></td><td></td></tr>
<tr><td>Chromosome</td><td>start</td><td></td><td>end</td><td>strand</td><td>MM</td><td>target_seq</td><td>PAM</td><td>alignment</td><td>distance</td><td>position</td><td>gene name</td><td>gene id</td></tr>
<tr><td>GU305914.1</td><td></td><td>189250</td><td>189272</td><td>-</td><td>0</td><td>CTACAAGAAATCCATACGCG</td><td>AGG</td><td>||||||||[||||||||||||]PAM</td><td>NA</td><td>Intergenic</td><td>NA</td><td>NA</td></tr>
<tr><td>T20318</td><td>TACAAGAAATCCATACGCGAGGG</td><td></td><td>1000</td><td></td><td></td><td></td><td></td><td></td><td></td><td></td><td></td><td></td></tr>
<tr><td>Oligo fwd</td><td>CACCTACAAGAAATCCATACGCGA</td><td></td><td></td><td></td><td></td><td></td><td></td><td></td><td></td><td></td><td></td><td></td></tr>
<tr><td>Oligo rev</td><td>AAACTCGCGTATGGATTTCTTGTA</td><td></td><td></td><td></td><td></td><td></td><td></td><td></td><td></td><td></td><td></td><td></td></tr>
<tr><td>Chromosome</td><td>start</td><td></td><td>end</td><td>strand</td><td>MM</td><td>target_seq</td><td>PAM</td><td>alignment</td><td>distance</td><td>position</td><td>gene name</td><td>gene id</td></tr>
<tr><td>GU305914.1</td><td></td><td>189249</td><td>189271</td><td>-</td><td>0</td><td>TACAAGAAATCCATACGCGA</td><td>GGG</td><td>||||||||[||||||||||||]PAM</td><td>NA</td><td>Intergenic</td><td>NA</td><td>NA</td></tr>
<tr><td>T20319</td><td>ATACGTGACCATGAGCGCCAAGG</td><td></td><td>1000</td><td></td><td></td><td></td><td></td><td></td><td></td><td></td><td></td><td></td></tr>
</table>

| | | | | | | | | | | | |
|---|---|---|---|---|---|---|---|---|---|---|---|
| Oligo fwd | CACCATACGTGACCATGAGCGCCA | | | | | | | | | | |
| Oligo rev | AAACTGGCGCTCATGGTCACGTAT | | | | | | | | | | |
| Chromosome | start | end | strand | MM | target_seq | PAM | alignment | distance | position | gene name | gene id |
| GU305914.1 | 189223 | 189245 | - | 0 | ATACGTGACCATGAGCGCCA | AGG | \|\|\|\|\|\|\|\|[\|\|\|\|\|\|\|\|\|\|\|\|]PAM | NA | Intergenic | NA | NA |
| T20320 | ATGAGCGCCAAGGTGAAACCCGG | 1000 | | | | | | | | | |
| Oligo fwd | CACCATGAGCGCCAAGGTGAAACC | | | | | | | | | | |
| Oligo rev | AAACGGTTTCACCTTGGCGCTCAT | | | | | | | | | | |
| Chromosome | start | end | strand | MM | target_seq | PAM | alignment | distance | position | gene name | gene id |
| GU305914.1 | 189213 | 189235 | - | 0 | ATGAGCGCCAAGGTGAAACC | CGG | \|\|\|\|\|\|\|\|[\|\|\|\|\|\|\|\|\|\|\|\|]PAM | NA | Intergenic | NA | NA |
| T20321 | CCGCGTATGCCTGCTGAAGACGG | 1000 | | | | | | | | | |
| Oligo fwd | CACCCCGCGTATGCCTGCTGAAGA | | | | | | | | | | |
| Oligo rev | AAACTCTTCAGCAGGCATACGCGG | | | | | | | | | | |
| Chromosome | start | end | strand | MM | target_seq | PAM | alignment | distance | position | gene name | gene id |
| GU305914.1 | 189175 | 189197 | - | 0 | CCGCGTATGCCTGCTGAAGA | CGG | \|\|\|\|\|\|\|\|[\|\|\|\|\|\|\|\|\|\|\|\|]PAM | NA | Intergenic | NA | NA |
| T20322 | GGACAACAAGTACGTGCTGATGG | 1000 | | | | | | | | | |
| Oligo fwd | CACCGGACAACAAGTACGTGCTGA | | | | | | | | | | |
| Oligo rev | AAACTCAGCACGTACTTGTTGTCC | | | | | | | | | | |
| Chromosome | start | end | strand | MM | target_seq | PAM | alignment | distance | position | gene name | gene id |
| GU305914.1 | 189154 | 189176 | - | 0 | GGACAACAAGTACGTGCTGA | TGG | \|\|\|\|\|\|\|\|[\|\|\|\|\|\|\|\|\|\|\|\|]PAM | NA | Intergenic | NA | NA |
| T20323 | GATGGCGCGTCCATGCCTGACGG | 1000 | | | | | | | | | |
| Oligo fwd | CACCGATGGCGCGTCCATGCCTGA | | | | | | | | | | |
| Oligo rev | AAACTCAGGCATGGACGCGCCATC | | | | | | | | | | |
| Chromosome | start | end | strand | MM | target_seq | PAM | alignment | distance | position | gene name | gene id |
| GU305914.1 | 189136 | 189158 | - | 0 | GATGGCGCGTCCATGCCTGA | CGG | \|\|\|\|\|\|\|\|[\|\|\|\|\|\|\|\|\|\|\|\|]PAM | NA | Intergenic | NA | NA |
| T20324 | GTCCATGCCTGACGGACAAGCGG | 1000 | | | | | | | | | |
| Oligo fwd | CACCGTCCATGCCTGACGGACAAG | | | | | | | | | | |
| Oligo rev | AAACCTTGTCCGTCAGGCATGGAC | | | | | | | | | | |
| Chromosome | start | end | strand | MM | target_seq | PAM | alignment | distance | position | gene name | gene id |
| GU305914.1 | 189128 | 189150 | - | 0 | GTCCATGCCTGACGGACAAG | CGG | \|\|\|\|\|\|\|\|[\|\|\|\|\|\|\|\|\|\|\|\|]PAM | NA | Intergenic | NA | NA |
| T20325 | CCTGACGGACAAGCGGCACCAGG | 1000 | | | | | | | | | |
| Oligo fwd | CACCCCTGACGGACAAGCGGCACC | | | | | | | | | | |
| Oligo rev | AAACGGTGCCGCTTGTCCGTCAGG | | | | | | | | | | |
| Chromosome | start | end | strand | MM | target_seq | PAM | alignment | distance | position | gene name | gene id |
| GU305914.1 | 189121 | 189143 | - | 0 | CCTGACGGACAAGCGGCACC | AGG | \|\|\|\|\|\|\|\|[\|\|\|\|\|\|\|\|\|\|\|\|]PAM | NA | Intergenic | NA | NA |
| T20326 | CGTGCGCATCGTGCACGCGTTGG | 1000 | | | | | | | | | |
| Oligo fwd | CACCCGTGCGCATCGTGCACGCGT | | | | | | | | | | |
| Oligo rev | AAACACGCGTGCACGATGCGCACG | | | | | | | | | | |
| Chromosome | start | end | strand | MM | target_seq | PAM | alignment | distance | position | gene name | gene id |
| GU305914.1 | 189085 | 189107 | - | 0 | CGTGCGCATCGTGCACGCGT | TGG | \|\|\|\|\|\|\|\|[\|\|\|\|\|\|\|\|\|\|\|\|]PAM | NA | Intergenic | NA | NA |
| T20327 | ACGCGTTGGTGATCAGCGTGTGG | 1000 | | | | | | | | | |
| Oligo fwd | CACCACGCGTTGGTGATCAGCGTG | | | | | | | | | | |
| Oligo rev | AAACCACGCTGATCACCAACGCGT | | | | | | | | | | |
| Chromosome | start | end | strand | MM | target_seq | PAM | alignment | distance | position | gene name | gene id |
| GU305914.1 | 189071 | 189093 | - | 0 | ACGCGTTGGTGATCAGCGTG | TGG | \|\|\|\|\|\|\|\|[\|\|\|\|\|\|\|\|\|\|\|\|]PAM | NA | Intergenic | NA | NA |
| T20328 | CGCGTTGGTGATCAGCGTGTGGG | 1000 | | | | | | | | | |
| Oligo fwd | CACCCGCGTTGGTGATCAGCGTGT | | | | | | | | | | |
| Oligo rev | AAACACACGCTGATCACCAACGCG | | | | | | | | | | |
| Chromosome | start | end | strand | MM | target_seq | PAM | alignment | distance | position | gene name | gene id |
| GU305914.1 | 189070 | 189092 | - | 0 | CGCGTTGGTGATCAGCGTGT | GGG | \|\|\|\|\|\|\|\|[\|\|\|\|\|\|\|\|\|\|\|\|]PAM | NA | Intergenic | NA | NA |
| T20329 | GTTGGTGATCAGCGTGTGGGCGG | 1000 | | | | | | | | | |
| Oligo fwd | CACCGTTGGTGATCAGCGTGTGGG | | | | | | | | | | |
| Oligo rev | AAACCCCACACGCTGATCACCAAC | | | | | | | | | | |
| Chromosome | start | end | strand | MM | target_seq | PAM | alignment | distance | position | gene name | gene id |
| GU305914.1 | 189067 | 189089 | - | 0 | GTTGGTGATCAGCGTGTGGG | CGG | \|\|\|\|\|\|\|\|[\|\|\|\|\|\|\|\|\|\|\|\|]PAM | NA | Intergenic | NA | NA |
| T20330 | GTGGGCGGTCGCGATCGCCCTGG | 1000 | | | | | | | | | |
| Oligo fwd | CACCGTGGGCGGTCGCGATCGCCC | | | | | | | | | | |
| Oligo rev | AAACGGGCGATCGCGACCGCCCAC | | | | | | | | | | |
| Chromosome | start | end | strand | MM | target_seq | PAM | alignment | distance | position | gene name | gene id |
| GU305914.1 | 189052 | 189074 | - | 0 | GTGGGCGGTCGCGATCGCCC | TGG | \|\|\|\|\|\|\|\|[\|\|\|\|\|\|\|\|\|\|\|\|]PAM | NA | Intergenic | NA | NA |
| T20331 | GGCGGTCGCGATCGCCCTGGTGG | 1000 | | | | | | | | | |
| Oligo fwd | CACCGGCGGTCGCGATCGCCCTGG | | | | | | | | | | |
| Oligo rev | AAACCCAGGGCGATCGCGACCGCC | | | | | | | | | | |
| Chromosome | start | end | strand | MM | target_seq | PAM | alignment | distance | position | gene name | gene id |
| GU305914.1 | 189049 | 189071 | - | 0 | GGCGGTCGCGATCGCCCTGG | TGG | \|\|\|\|\|\|\|\|[\|\|\|\|\|\|\|\|\|\|\|\|]PAM | NA | Intergenic | NA | NA |
| GU305914.1 | 127383 | 127405 | + | 4 | GGCGGCGGCGATCGCCCTCT | CGG | \|\|\|\|\|--\|[\|\|\|\|\|\|\|\|\|\|--]PAM | NA | Intergenic | NA | NA |
| T20332 | GCGGTCGCGATCGCCCTGGTGGG | 1000 | | | | | | | | | |
| Oligo fwd | CACCGCGGTCGCGATCGCCCTGGT | | | | | | | | | | |
| Oligo rev | AAACACCAGGGCGATCGCGACCGC | | | | | | | | | | |
| Chromosome | start | end | strand | MM | target_seq | PAM | alignment | distance | position | gene name | gene id |
| GU305914.1 | 189048 | 189070 | - | 0 | GCGGTCGCGATCGCCCTGGT | GGG | \|\|\|\|\|\|\|\|[\|\|\|\|\|\|\|\|\|\|\|\|]PAM | NA | Intergenic | NA | NA |
| T20333 | CGTGCGCCTGCTATTAAGCCTGG | 1000 | | | | | | | | | |
| Oligo fwd | CACCCGTGCGCCTGCTATTAAGCC | | | | | | | | | | |
| Oligo rev | AAACGGCTTAATAGCAGGCGCACG | | | | | | | | | | |
| Chromosome | start | end | strand | MM | target_seq | PAM | alignment | distance | position | gene name | gene id |
| GU305914.1 | 189016 | 189038 | - | 0 | CGTGCGCCTGCTATTAAGCC | TGG | \|\|\|\|\|\|\|\|[\|\|\|\|\|\|\|\|\|\|\|\|]PAM | NA | Intergenic | NA | NA |

T20334 TAAGCCTGGACAGCGTCGCCCGG 1000
Oligo fwd CACCTAAGCCTGGACAGCGTCGCC
Oligo rev AAACGGCGACGCTGTCCAGGCTTA
Chromosome start end strand MM target_seq PAM alignment distance position gene name gene id
GU305914.1 189002 189024 - 0 TAAGCCTGGACAGCGTCGCC CGG ||||||||[||||||||||||]PAM NA Intergenic NA NA

T20335 CCTGGACAGCGTCGCCCGGCTGG 1000
Oligo fwd CACCCCTGGACAGCGTCGCCCGGC
Oligo rev AAACGCCGGGCGACGCTGTCCAGG
Chromosome start end strand MM target_seq PAM alignment distance position gene name gene id
GU305914.1 188998 189020 - 0 CCTGGACAGCGTCGCCCGGC TGG ||||||||[||||||||||||]PAM NA Intergenic NA NA

T20336 GCTGGTGAAAAAAATGTTCGCGG 1000
Oligo fwd CACCGCTGGTGAAAAAAATGTTCG
Oligo rev AAACCGAACATTTTTTTCACCAGC
Chromosome start end strand MM target_seq PAM alignment distance position gene name gene id
GU305914.1 188980 189002 - 0 GCTGGTGAAAAAAATGTTCG CGG ||||||||[||||||||||||]PAM NA Intergenic NA NA

T20337 ACTGCCGTTACCGTTACCGCTGG 1000
Oligo fwd CACCACTGCCGTTACCGTTACCGC
Oligo rev AAACGCGGTAACGGTAACGGCAGT
Chromosome start end strand MM target_seq PAM alignment distance position gene name gene id
GU305914.1 188950 188972 - 0 ACTGCCGTTACCGTTACCGC TGG ||||||||[||||||||||||]PAM NA Intergenic NA NA

T20338 GTTACCGTTACCGCTGGTCGTGG 1000
Oligo fwd CACCGTTACCGTTACCGCTGGTCG
Oligo rev AAACCGACCAGCGGTAACGGTAAC
Chromosome start end strand MM target_seq PAM alignment distance position gene name gene id
GU305914.1 188944 188966 - 0 GTTACCGTTACCGCTGGTCG TGG ||||||||[||||||||||||]PAM NA Intergenic NA NA

T20339 CGTTACCGCTGGTCGTGGTGTGG 1000
Oligo fwd CACCCGTTACCGCTGGTCGTGGTG
Oligo rev AAACCACCACGACCAGCGGTAACG
Chromosome start end strand MM target_seq PAM alignment distance position gene name gene id
GU305914.1 188939 188961 - 0 CGTTACCGCTGGTCGTGGTG TGG ||||||||[||||||||||||]PAM NA Intergenic NA NA

T20340 GTGTGGTACGTGCTCGCCGCCGG 1000
Oligo fwd CACCGTGTGGTACGTGCTCGCCGC
Oligo rev AAACGCGGCGAGCACGTACCACAC
Chromosome start end strand MM target_seq PAM alignment distance position gene name gene id
GU305914.1 188922 188944 - 0 GTGTGGTACGTGCTCGCCGC CGG ||||||||[||||||||||||]PAM NA Intergenic NA NA

T20341 CGTGCTCGCCGCCGGCGACACGG 1000
Oligo fwd CACCCGTGCTCGCCGCCGGCGACA
Oligo rev AAACTGTCGCCGGCGGCGAGCACG
Chromosome start end strand MM target_seq PAM alignment distance position gene name gene id
GU305914.1 188914 188936 - 0 CGTGCTCGCCGCCGGCGACA CGG ||||||||[||||||||||||]PAM NA Intergenic NA NA

T20342 CGGCGACACGGTCGCCGTGACGG 1000
Oligo fwd CACCCGGCGACACGGTCGCCGTGA
Oligo rev AAACTCACGGCGACCGTGTCGCCG
Chromosome start end strand MM target_seq PAM alignment distance position gene name gene id
GU305914.1 188902 188924 - 0 CGGCGACACGGTCGCCGTGA CGG ||||||||[||||||||||||]PAM NA Intergenic NA NA
GU305914.1 169252 169274 - 4 CTCGGACACGCTCGCCGTGA GGG |---||||[||-|||||||||]PAM NA Intergenic NA NA

T20343 GTCGCCGTGACGGACCGAGACGG 1000
Oligo fwd CACCGTCGCCGTGACGGACCGAGA
Oligo rev AAACTCTCGGTCCGTCACGGCGAC
Chromosome start end strand MM target_seq PAM alignment distance position gene name gene id
GU305914.1 188892 188914 - 0 GTCGCCGTGACGGACCGAGA CGG ||||||||[||||||||||||]PAM NA Intergenic NA NA

T20344 TCGCCGTGACGGACCGAGACGGG 1000
Oligo fwd CACCTCGCCGTGACGGACCGAGAC
Oligo rev AAACGTCTCGGTCCGTCACGGCGA
Chromosome start end strand MM target_seq PAM alignment distance position gene name gene id
GU305914.1 188891 188913 - 0 TCGCCGTGACGGACCGAGAC GGG ||||||||[||||||||||||]PAM NA Intergenic NA NA

T20345 CGCCGTGACGGACCGAGACGGGG 1000
Oligo fwd CACCCGCCGTGACGGACCGAGACG
Oligo rev AAACCGTCTCGGTCCGTCACGGCG
Chromosome start end strand MM target_seq PAM alignment distance position gene name gene id
GU305914.1 188890 188912 - 0 CGCCGTGACGGACCGAGACG GGG ||||||||[||||||||||||]PAM NA Intergenic NA NA

T20346 GACGGACCGAGACGGGGTGCTGG 1000
Oligo fwd CACCGACGGACCGAGACGGGGTGC
Oligo rev AAACGCACCCCGTCTCGGTCCGTC
Chromosome start end strand MM target_seq PAM alignment distance position gene name gene id
GU305914.1 188884 188906 - 0 GACGGACCGAGACGGGGTGC TGG ||||||||[||||||||||||]PAM NA Intergenic NA NA

T20347 CCGAGACGGGGTGCTGGCCGTGG 1000
Oligo fwd CACCCCGAGACGGGGTGCTGGCCG
Oligo rev AAACCGGCCAGCACCCCGTCTCGG
Chromosome start end strand MM target_seq PAM alignment distance position gene name gene id
GU305914.1 188878 188900 - 0 CCGAGACGGGGTGCTGGCCG TGG ||||||||[||||||||||||]PAM NA Intergenic NA NA

T20348 GTGGAGAGCGTCCTGCTCTCCGG 1000
Oligo fwd CACCGTGGAGAGCGTCCTGCTCTC
Oligo rev AAACGAGAGCAGGACGCTCTCCAC

| | | | | | | | | | | | |
|---|---|---|---|---|---|---|---|---|---|---|---|
| Chromosome | start | end | strand | MM | target_seq | PAM | alignment | distance | position | gene name | gene id |
| GU305914.1 | 188859 | 188881 | - | 0 | GTGGAGAGCGTCCTGCTCTC | CGG | ||||||||[||||||||||||]PAM | NA | Intergenic | NA | NA |
| T20349 | TGGAGAGCGTCCTGCTCTCCGGG | 1000 | | | | | | | | | |
| Oligo fwd | CACCTGGAGAGCGTCCTGCTCTCC | | | | | | | | | | |
| Oligo rev | AAACGGAGAGCAGGACGCTCTCCA | | | | | | | | | | |
| Chromosome | start | end | strand | MM | target_seq | PAM | alignment | distance | position | gene name | gene id |
| GU305914.1 | 188858 | 188880 | - | 0 | TGGAGAGCGTCCTGCTCTCC | GGG | ||||||||[||||||||||||]PAM | NA | Intergenic | NA | NA |
| T20350 | CCTGCTCTCCGGGCTCGAGCTGG | 1000 | | | | | | | | | |
| Oligo fwd | CACCCCTGCTCTCCGGGCTCGAGC | | | | | | | | | | |
| Oligo rev | AAACGCTCGAGCCCGGAGAGCAGG | | | | | | | | | | |
| Chromosome | start | end | strand | MM | target_seq | PAM | alignment | distance | position | gene name | gene id |
| GU305914.1 | 188848 | 188870 | - | 0 | CCTGCTCTCCGGGCTCGAGC | TGG | ||||||||[||||||||||||]PAM | NA | Intergenic | NA | NA |
| GU305914.1 | 93316 | 93338 | + | 4 | CCCGCCCTCCCAGCTCGAGC | TGG | ||-||-||[||--||||||||]PAM | NA | Intergenic | NA | NA |
| T20351 | CTGCTCTCCGGGCTCGAGCTGGG | 1000 | | | | | | | | | |
| Oligo fwd | CACCCTGCTCTCCGGGCTCGAGCT | | | | | | | | | | |
| Oligo rev | AAACAGCTCGAGCCCGGAGAGCAG | | | | | | | | | | |
| Chromosome | start | end | strand | MM | target_seq | PAM | alignment | distance | position | gene name | gene id |
| GU305914.1 | 188847 | 188869 | - | 0 | CTGCTCTCCGGGCTCGAGCT | GGG | ||||||||[||||||||||||]PAM | NA | Intergenic | NA | NA |
| T20352 | TGCTCTCCGGGCTCGAGCTGGGG | 1000 | | | | | | | | | |
| Oligo fwd | CACCTGCTCTCCGGGCTCGAGCTG | | | | | | | | | | |
| Oligo rev | AAACCAGCTCGAGCCCGGAGAGCA | | | | | | | | | | |
| Chromosome | start | end | strand | MM | target_seq | PAM | alignment | distance | position | gene name | gene id |
| GU305914.1 | 188846 | 188868 | - | 0 | TGCTCTCCGGGCTCGAGCTG | GGG | ||||||||[||||||||||||]PAM | NA | Intergenic | NA | NA |
| T20353 | GAACGCGTCCCAAATGAAACTGG | 1000 | | | | | | | | | |
| Oligo fwd | CACCGAACGCGTCCCAAATGAAAC | | | | | | | | | | |
| Oligo rev | AAACGTTTCATTTGGGACGCGTTC | | | | | | | | | | |
| Chromosome | start | end | strand | MM | target_seq | PAM | alignment | distance | position | gene name | gene id |
| GU305914.1 | 188806 | 188828 | - | 0 | GAACGCGTCCCAAATGAAAC | TGG | ||||||||[||||||||||||]PAM | NA | Intergenic | NA | NA |
| T20354 | GTCCCAAATGAAACTGGCGCTGG | 1000 | | | | | | | | | |
| Oligo fwd | CACCGTCCCAAATGAAACTGGCGC | | | | | | | | | | |
| Oligo rev | AAACGCGCCAGTTTCATTTGGGAC | | | | | | | | | | |
| Chromosome | start | end | strand | MM | target_seq | PAM | alignment | distance | position | gene name | gene id |
| GU305914.1 | 188800 | 188822 | - | 0 | GTCCCAAATGAAACTGGCGC | TGG | ||||||||[||||||||||||]PAM | NA | Intergenic | NA | NA |
| T20355 | AAATGAAACTGGCGCTGGAGAGG | 1000 | | | | | | | | | |
| Oligo fwd | CACCAAATGAAACTGGCGCTGGAG | | | | | | | | | | |
| Oligo rev | AAACCTCCAGCGCCAGTTTCATTT | | | | | | | | | | |
| Chromosome | start | end | strand | MM | target_seq | PAM | alignment | distance | position | gene name | gene id |
| GU305914.1 | 188795 | 188817 | - | 0 | AAATGAAACTGGCGCTGGAG | AGG | ||||||||[||||||||||||]PAM | NA | Intergenic | NA | NA |
| T20356 | GAGGTAGCTCTTCCGAGAAACGG | 1000 | | | | | | | | | |
| Oligo fwd | CACCGAGGTAGCTCTTCCGAGAAA | | | | | | | | | | |
| Oligo rev | AAACTTTCTCGGAAGAGCTACCTC | | | | | | | | | | |
| Chromosome | start | end | strand | MM | target_seq | PAM | alignment | distance | position | gene name | gene id |
| GU305914.1 | 188776 | 188798 | - | 0 | GAGGTAGCTCTTCCGAGAAA | CGG | ||||||||[||||||||||||]PAM | NA | Intergenic | NA | NA |
| T20357 | TAGCTCTTCCGAGAAACGGTCGG | 1000 | | | | | | | | | |
| Oligo fwd | CACCTAGCTCTTCCGAGAAACGGT | | | | | | | | | | |
| Oligo rev | AAACACCGTTTCTCGGAAGAGCTA | | | | | | | | | | |
| Chromosome | start | end | strand | MM | target_seq | PAM | alignment | distance | position | gene name | gene id |
| GU305914.1 | 188772 | 188794 | - | 0 | TAGCTCTTCCGAGAAACGGT | CGG | ||||||||[||||||||||||]PAM | NA | Intergenic | NA | NA |
| T20358 | AGCTCTTCCGAGAAACGGTCGGG | 1000 | | | | | | | | | |
| Oligo fwd | CACCAGCTCTTCCGAGAAACGGTC | | | | | | | | | | |
| Oligo rev | AAACGACCGTTTCTCGGAAGAGCT | | | | | | | | | | |
| Chromosome | start | end | strand | MM | target_seq | PAM | alignment | distance | position | gene name | gene id |
| GU305914.1 | 188771 | 188793 | - | 0 | AGCTCTTCCGAGAAACGGTC | GGG | ||||||||[||||||||||||]PAM | NA | Intergenic | NA | NA |
| T20359 | GAAACGGTCGGGTTCCGAGAAGG | 1000 | | | | | | | | | |
| Oligo fwd | CACCGAAACGGTCGGGTTCCGAGA | | | | | | | | | | |
| Oligo rev | AAACTCTCGGAACCCGACCGTTTC | | | | | | | | | | |
| Chromosome | start | end | strand | MM | target_seq | PAM | alignment | distance | position | gene name | gene id |
| GU305914.1 | 188760 | 188782 | - | 0 | GAAACGGTCGGGTTCCGAGA | AGG | ||||||||[||||||||||||]PAM | NA | Intergenic | NA | NA |
| T20360 | AAACGGTCGGGTTCCGAGAAGGG | 1000 | | | | | | | | | |
| Oligo fwd | CACCAAACGGTCGGGTTCCGAGAA | | | | | | | | | | |
| Oligo rev | AAACTTCTCGGAACCCGACCGTTT | | | | | | | | | | |
| Chromosome | start | end | strand | MM | target_seq | PAM | alignment | distance | position | gene name | gene id |
| GU305914.1 | 188759 | 188781 | - | 0 | AAACGGTCGGGTTCCGAGAA | GGG | ||||||||[||||||||||||]PAM | NA | Intergenic | NA | NA |
| T20361 | CTCTCTGACTAATCACCCCCCGG | 1000 | | | | | | | | | |
| Oligo fwd | CACCCTCTCTGACTAATCACCCCC | | | | | | | | | | |
| Oligo rev | AAACGGGGGTGATTAGTCAGAGAG | | | | | | | | | | |
| Chromosome | start | end | strand | MM | target_seq | PAM | alignment | distance | position | gene name | gene id |
| GU305914.1 | 188732 | 188754 | - | 0 | CTCTCTGACTAATCACCCCC | CGG | ||||||||[||||||||||||]PAM | NA | Intergenic | NA | NA |
| T20362 | TGTGCCCGTTTTTGTCTTTCAGG | 1000 | | | | | | | | | |
| Oligo fwd | CACCTGTGCCCGTTTTTGTCTTTC | | | | | | | | | | |
| Oligo rev | AAACGAAAGACAAAAACGGGCACA | | | | | | | | | | |
| Chromosome | start | end | strand | MM | target_seq | PAM | alignment | distance | position | gene name | gene id |
| GU305914.1 | 188701 | 188723 | - | 0 | TGTGCCCGTTTTTGTCTTTC | AGG | ||||||||[||||||||||||]PAM | NA | Intergenic | NA | NA |
| T20363 | CCCGTTTTTGTCTTTCAGGCTGG | 1000 | | | | | | | | | |

| | | | | | | | | | | | | |
|---|---|---|---|---|---|---|---|---|---|---|---|---|
| Oligo fwd | CACCCCCGTTTTTGTCTTTCAGGC | | | | | | | | | | | |
| Oligo rev | AAACGCCTGAAAGACAAAAACGGG | | | | | | | | | | | |
| Chromosome | start | | end | strand | MM | target_seq | PAM | alignment | distance | position | gene name | gene id |
| GU305914.1 | | 188697 | 188719 | - | 0 | CCCGTTTTTGTCTTTCAGGC | TGG | \|\|\|\|\|\|\|\|[\|\|\|\|\|\|\|\|\|\|\|\|]PAM | NA | Intergenic | NA | NA |
| T20364 | TTGTCTTTCAGGCTGGATATAGG | | 1000 | | | | | | | | | |
| Oligo fwd | CACCTTGTCTTTCAGGCTGGATAT | | | | | | | | | | | |
| Oligo rev | AAACATATCCAGCCTGAAAGACAA | | | | | | | | | | | |
| Chromosome | start | | end | strand | MM | target_seq | PAM | alignment | distance | position | gene name | gene id |
| GU305914.1 | | 188690 | 188712 | - | 0 | TTGTCTTTCAGGCTGGATAT | AGG | \|\|\|\|\|\|\|\|[\|\|\|\|\|\|\|\|\|\|\|\|]PAM | NA | Intergenic | NA | NA |
| T20365 | TCTTTCAGGCTGGATATAGGAGG | | 1000 | | | | | | | | | |
| Oligo fwd | CACCTCTTTCAGGCTGGATATAGG | | | | | | | | | | | |
| Oligo rev | AAACCCTATATCCAGCCTGAAAGA | | | | | | | | | | | |
| Chromosome | start | | end | strand | MM | target_seq | PAM | alignment | distance | position | gene name | gene id |
| GU305914.1 | | 188687 | 188709 | - | 0 | TCTTTCAGGCTGGATATAGG | AGG | \|\|\|\|\|\|\|\|[\|\|\|\|\|\|\|\|\|\|\|\|]PAM | NA | Intergenic | NA | NA |
| T20366 | TCTTAAGACTATCGATGTGTTGG | | 1000 | | | | | | | | | |
| Oligo fwd | CACCTCTTAAGACTATCGATGTGT | | | | | | | | | | | |
| Oligo rev | AAACACACATCGATAGTCTTAAGA | | | | | | | | | | | |
| Chromosome | start | | end | strand | MM | target_seq | PAM | alignment | distance | position | gene name | gene id |
| GU305914.1 | | 188661 | 188683 | - | 0 | TCTTAAGACTATCGATGTGT | TGG | \|\|\|\|\|\|\|\|[\|\|\|\|\|\|\|\|\|\|\|\|]PAM | NA | Intergenic | NA | NA |
| T20367 | AAGACTATCGATGTGTTGGTCGG | | 1000 | | | | | | | | | |
| Oligo fwd | CACCAAGACTATCGATGTGTTGGT | | | | | | | | | | | |
| Oligo rev | AAACACCAACACATCGATAGTCTT | | | | | | | | | | | |
| Chromosome | start | | end | strand | MM | target_seq | PAM | alignment | distance | position | gene name | gene id |
| GU305914.1 | | 188657 | 188679 | - | 0 | AAGACTATCGATGTGTTGGT | CGG | \|\|\|\|\|\|\|\|[\|\|\|\|\|\|\|\|\|\|\|\|]PAM | NA | Intergenic | NA | NA |
| T20368 | TGGTCGGCTCATCTGACGCACGG | | 1000 | | | | | | | | | |
| Oligo fwd | CACCTGGTCGGCTCATCTGACGCA | | | | | | | | | | | |
| Oligo rev | AAACTGCGTCAGATGAGCCGACCA | | | | | | | | | | | |
| Chromosome | start | | end | strand | MM | target_seq | PAM | alignment | distance | position | gene name | gene id |
| GU305914.1 | | 188641 | 188663 | - | 0 | TGGTCGGCTCATCTGACGCA | CGG | \|\|\|\|\|\|\|\|[\|\|\|\|\|\|\|\|\|\|\|\|]PAM | NA | Intergenic | NA | NA |
| T20369 | GGTCGGCTCATCTGACGCACGGG | | 1000 | | | | | | | | | |
| Oligo fwd | CACCGGTCGGCTCATCTGACGCAC | | | | | | | | | | | |
| Oligo rev | AAACGTGCGTCAGATGAGCCGACC | | | | | | | | | | | |
| Chromosome | start | | end | strand | MM | target_seq | PAM | alignment | distance | position | gene name | gene id |
| GU305914.1 | | 188640 | 188662 | - | 0 | GGTCGGCTCATCTGACGCAC | GGG | \|\|\|\|\|\|\|\|[\|\|\|\|\|\|\|\|\|\|\|\|]PAM | NA | Intergenic | NA | NA |
| T20370 | CGGCTCATCTGACGCACGGGAGG | | 1000 | | | | | | | | | |
| Oligo fwd | CACCCGGCTCATCTGACGCACGGG | | | | | | | | | | | |
| Oligo rev | AAACCCCGTGCGTCAGATGAGCCG | | | | | | | | | | | |
| Chromosome | start | | end | strand | MM | target_seq | PAM | alignment | distance | position | gene name | gene id |
| GU305914.1 | | 188637 | 188659 | - | 0 | CGGCTCATCTGACGCACGGG | AGG | \|\|\|\|\|\|\|\|[\|\|\|\|\|\|\|\|\|\|\|\|]PAM | NA | Intergenic | NA | NA |
| T20371 | TGACGCACGGGAGGTGAGACAGG | | 1000 | | | | | | | | | |
| Oligo fwd | CACCTGACGCACGGGAGGTGAGAC | | | | | | | | | | | |
| Oligo rev | AAACGTCTCACCTCCCGTGCGTCA | | | | | | | | | | | |
| Chromosome | start | | end | strand | MM | target_seq | PAM | alignment | distance | position | gene name | gene id |
| GU305914.1 | | 188628 | 188650 | - | 0 | TGACGCACGGGAGGTGAGAC | AGG | \|\|\|\|\|\|\|\|[\|\|\|\|\|\|\|\|\|\|\|\|]PAM | NA | Intergenic | NA | NA |
| T20372 | GACGCACGGGAGGTGAGACAGGG | | 1000 | | | | | | | | | |
| Oligo fwd | CACCGACGCACGGGAGGTGAGACA | | | | | | | | | | | |
| Oligo rev | AAACTGTCTCACCTCCCGTGCGTC | | | | | | | | | | | |
| Chromosome | start | | end | strand | MM | target_seq | PAM | alignment | distance | position | gene name | gene id |
| GU305914.1 | | 188627 | 188649 | - | 0 | GACGCACGGGAGGTGAGACA | GGG | \|\|\|\|\|\|\|\|[\|\|\|\|\|\|\|\|\|\|\|\|]PAM | NA | Intergenic | NA | NA |
| T20373 | TGAGACAGGGAAGAGAAAACAGG | | 1000 | | | | | | | | | |
| Oligo fwd | CACCTGAGACAGGGAAGAGAAAAC | | | | | | | | | | | |
| Oligo rev | AAACGTTTTCTCTTCCCTGTCTCA | | | | | | | | | | | |
| Chromosome | start | | end | strand | MM | target_seq | PAM | alignment | distance | position | gene name | gene id |
| GU305914.1 | | 188614 | 188636 | - | 0 | TGAGACAGGGAAGAGAAAAC | AGG | \|\|\|\|\|\|\|\|[\|\|\|\|\|\|\|\|\|\|\|\|]PAM | NA | Intergenic | NA | NA |
| T20374 | GACCGTCGTACGCCCCGTGCTGG | | 1000 | | | | | | | | | |
| Oligo fwd | CACCGACCGTCGTACGCCCCGTGC | | | | | | | | | | | |
| Oligo rev | AAACGCACGGGGCGTACGACGGTC | | | | | | | | | | | |
| Chromosome | start | | end | strand | MM | target_seq | PAM | alignment | distance | position | gene name | gene id |
| GU305914.1 | | 188592 | 188614 | - | 0 | GACCGTCGTACGCCCCGTGC | TGG | \|\|\|\|\|\|\|\|[\|\|\|\|\|\|\|\|\|\|\|\|]PAM | NA | Intergenic | NA | NA |
| T20375 | ACCGTCGTACGCCCCGTGCTGGG | | 1000 | | | | | | | | | |
| Oligo fwd | CACCACCGTCGTACGCCCCGTGCT | | | | | | | | | | | |
| Oligo rev | AAACAGCACGGGGCGTACGACGGT | | | | | | | | | | | |
| Chromosome | start | | end | strand | MM | target_seq | PAM | alignment | distance | position | gene name | gene id |
| GU305914.1 | | 188591 | 188613 | - | 0 | ACCGTCGTACGCCCCGTGCT | GGG | \|\|\|\|\|\|\|\|[\|\|\|\|\|\|\|\|\|\|\|\|]PAM | NA | Intergenic | NA | NA |
| T20376 | TGCTGGGAAGTTACAACGCGAGG | | 1000 | | | | | | | | | |
| Oligo fwd | CACCTGCTGGGAAGTTACAACGCG | | | | | | | | | | | |
| Oligo rev | AAACCGCGTTGTAACTTCCCAGCA | | | | | | | | | | | |
| Chromosome | start | | end | strand | MM | target_seq | PAM | alignment | distance | position | gene name | gene id |
| GU305914.1 | | 188575 | 188597 | - | 0 | TGCTGGGAAGTTACAACGCG | AGG | \|\|\|\|\|\|\|\|[\|\|\|\|\|\|\|\|\|\|\|\|]PAM | NA | Intergenic | NA | NA |
| T20377 | GCTGGGAAGTTACAACGCGAGGG | | 1000 | | | | | | | | | |
| Oligo fwd | CACCGCTGGGAAGTTACAACGCGA | | | | | | | | | | | |
| Oligo rev | AAACTCGCGTTGTAACTTCCCAGC | | | | | | | | | | | |
| Chromosome | start | | end | strand | MM | target_seq | PAM | alignment | distance | position | gene name | gene id |
| GU305914.1 | | 188574 | 188596 | - | 0 | GCTGGGAAGTTACAACGCGA | GGG | \|\|\|\|\|\|\|\|[\|\|\|\|\|\|\|\|\|\|\|\|]PAM | NA | Intergenic | NA | NA |

<table>
<tr><td>T20378</td><td>GGGAAGTTACAACGCGAGGGTGG</td><td>1000</td><td></td><td></td><td></td><td></td><td></td><td></td><td></td><td></td><td></td></tr>
<tr><td>Oligo fwd</td><td>CACCGGGAAGTTACAACGCGAGGG</td><td></td><td></td><td></td><td></td><td></td><td></td><td></td><td></td><td></td><td></td></tr>
<tr><td>Oligo rev</td><td>AAACCCCTCGCGTTGTAACTTCCC</td><td></td><td></td><td></td><td></td><td></td><td></td><td></td><td></td><td></td><td></td></tr>
<tr><td>Chromosome</td><td>start</td><td>end</td><td>strand</td><td>MM</td><td>target_seq</td><td>PAM</td><td>alignment</td><td>distance</td><td>position</td><td>gene name</td><td>gene id</td></tr>
<tr><td>GU305914.1</td><td>188571</td><td>188593</td><td>-</td><td>0</td><td>GGGAAGTTACAACGCGAGGG</td><td>TGG</td><td>||||||||[||||||||||||]PAM</td><td>NA</td><td>Intergenic</td><td>NA</td><td>NA</td></tr>
<tr><td>GU305914.1</td><td>229909</td><td>229931</td><td>+</td><td>4</td><td>CGGACGTTACGCCGCGAGGG</td><td>CGG</td><td>-|||-|||[||--||||||||]PAM</td><td>NA</td><td>Intergenic</td><td>NA</td><td>NA</td></tr>
<tr><td>T20379</td><td>CTGTCCGCCTCGACACAACCCGG</td><td>1000</td><td></td><td></td><td></td><td></td><td></td><td></td><td></td><td></td><td></td></tr>
<tr><td>Oligo fwd</td><td>CACCCTGTCCGCCTCGACACAACC</td><td></td><td></td><td></td><td></td><td></td><td></td><td></td><td></td><td></td><td></td></tr>
<tr><td>Oligo rev</td><td>AAACGGTTGTGTCGAGGCGGACAG</td><td></td><td></td><td></td><td></td><td></td><td></td><td></td><td></td><td></td><td></td></tr>
<tr><td>Chromosome</td><td>start</td><td>end</td><td>strand</td><td>MM</td><td>target_seq</td><td>PAM</td><td>alignment</td><td>distance</td><td>position</td><td>gene name</td><td>gene id</td></tr>
<tr><td>GU305914.1</td><td>188531</td><td>188553</td><td>-</td><td>0</td><td>CTGTCCGCCTCGACACAACC</td><td>CGG</td><td>||||||||[||||||||||||]PAM</td><td>NA</td><td>Intergenic</td><td>NA</td><td>NA</td></tr>
<tr><td>T20380</td><td>TGTCCGCCTCGACACAACCCGGG</td><td>1000</td><td></td><td></td><td></td><td></td><td></td><td></td><td></td><td></td><td></td></tr>
<tr><td>Oligo fwd</td><td>CACCTGTCCGCCTCGACACAACCC</td><td></td><td></td><td></td><td></td><td></td><td></td><td></td><td></td><td></td><td></td></tr>
<tr><td>Oligo rev</td><td>AAACGGGTTGTGTCGAGGCGGACA</td><td></td><td></td><td></td><td></td><td></td><td></td><td></td><td></td><td></td><td></td></tr>
<tr><td>Chromosome</td><td>start</td><td>end</td><td>strand</td><td>MM</td><td>target_seq</td><td>PAM</td><td>alignment</td><td>distance</td><td>position</td><td>gene name</td><td>gene id</td></tr>
<tr><td>GU305914.1</td><td>188530</td><td>188552</td><td>-</td><td>0</td><td>TGTCCGCCTCGACACAACCC</td><td>GGG</td><td>||||||||[||||||||||||]PAM</td><td>NA</td><td>Intergenic</td><td>NA</td><td>NA</td></tr>
<tr><td>T20381</td><td>CTCGACACAACCCGGGAAAAAGG</td><td>1000</td><td></td><td></td><td></td><td></td><td></td><td></td><td></td><td></td><td></td></tr>
<tr><td>Oligo fwd</td><td>CACCCTCGACACAACCCGGGAAAA</td><td></td><td></td><td></td><td></td><td></td><td></td><td></td><td></td><td></td><td></td></tr>
<tr><td>Oligo rev</td><td>AAACTTTTCCCGGGTTGTGTCGAG</td><td></td><td></td><td></td><td></td><td></td><td></td><td></td><td></td><td></td><td></td></tr>
<tr><td>Chromosome</td><td>start</td><td>end</td><td>strand</td><td>MM</td><td>target_seq</td><td>PAM</td><td>alignment</td><td>distance</td><td>position</td><td>gene name</td><td>gene id</td></tr>
<tr><td>GU305914.1</td><td>188523</td><td>188545</td><td>-</td><td>0</td><td>CTCGACACAACCCGGGAAAA</td><td>AGG</td><td>||||||||[||||||||||||]PAM</td><td>NA</td><td>Intergenic</td><td>NA</td><td>NA</td></tr>
<tr><td>T20382</td><td>TAGAGTATTCATCAGCGTGCAGG</td><td>1000</td><td></td><td></td><td></td><td></td><td></td><td></td><td></td><td></td><td></td></tr>
<tr><td>Oligo fwd</td><td>CACCTAGAGTATTCATCAGCGTGC</td><td></td><td></td><td></td><td></td><td></td><td></td><td></td><td></td><td></td><td></td></tr>
<tr><td>Oligo rev</td><td>AAACGCACGCTGATGAATACTCTA</td><td></td><td></td><td></td><td></td><td></td><td></td><td></td><td></td><td></td><td></td></tr>
<tr><td>Chromosome</td><td>start</td><td>end</td><td>strand</td><td>MM</td><td>target_seq</td><td>PAM</td><td>alignment</td><td>distance</td><td>position</td><td>gene name</td><td>gene id</td></tr>
<tr><td>GU305914.1</td><td>188490</td><td>188512</td><td>-</td><td>0</td><td>TAGAGTATTCATCAGCGTGC</td><td>AGG</td><td>||||||||[||||||||||||]PAM</td><td>NA</td><td>Intergenic</td><td>NA</td><td>NA</td></tr>
<tr><td>T20383</td><td>AGGATGACGTGTGACGATGAAGG</td><td>1000</td><td></td><td></td><td></td><td></td><td></td><td></td><td></td><td></td><td></td></tr>
<tr><td>Oligo fwd</td><td>CACCAGGATGACGTGTGACGATGA</td><td></td><td></td><td></td><td></td><td></td><td></td><td></td><td></td><td></td><td></td></tr>
<tr><td>Oligo rev</td><td>AAACTCATCGTCACACGTCATCCT</td><td></td><td></td><td></td><td></td><td></td><td></td><td></td><td></td><td></td><td></td></tr>
<tr><td>Chromosome</td><td>start</td><td>end</td><td>strand</td><td>MM</td><td>target_seq</td><td>PAM</td><td>alignment</td><td>distance</td><td>position</td><td>gene name</td><td>gene id</td></tr>
<tr><td>GU305914.1</td><td>188470</td><td>188492</td><td>-</td><td>0</td><td>AGGATGACGTGTGACGATGA</td><td>AGG</td><td>||||||||[||||||||||||]PAM</td><td>NA</td><td>Intergenic</td><td>NA</td><td>NA</td></tr>
<tr><td>GU305914.1</td><td>44627</td><td>44649</td><td>-</td><td>4</td><td>AGGACGAGGAGTGATGATGA</td><td>AGG</td><td>||||-||-[|-||||-|||||]PAM</td><td>NA</td><td>Intergenic</td><td>NA</td><td>NA</td></tr>
<tr><td>T20384</td><td>GAAGGTGTCGTTACTGCCTATGG</td><td>1000</td><td></td><td></td><td></td><td></td><td></td><td></td><td></td><td></td><td></td></tr>
<tr><td>Oligo fwd</td><td>CACCGAAGGTGTCGTTACTGCCTA</td><td></td><td></td><td></td><td></td><td></td><td></td><td></td><td></td><td></td><td></td></tr>
<tr><td>Oligo rev</td><td>AAACTAGGCAGTAACGACACCTTC</td><td></td><td></td><td></td><td></td><td></td><td></td><td></td><td></td><td></td><td></td></tr>
<tr><td>Chromosome</td><td>start</td><td>end</td><td>strand</td><td>MM</td><td>target_seq</td><td>PAM</td><td>alignment</td><td>distance</td><td>position</td><td>gene name</td><td>gene id</td></tr>
<tr><td>GU305914.1</td><td>188452</td><td>188474</td><td>-</td><td>0</td><td>GAAGGTGTCGTTACTGCCTA</td><td>TGG</td><td>||||||||[||||||||||||]PAM</td><td>NA</td><td>Intergenic</td><td>NA</td><td>NA</td></tr>
<tr><td>T20385</td><td>AAGGTGTCGTTACTGCCTATGGG</td><td>1000</td><td></td><td></td><td></td><td></td><td></td><td></td><td></td><td></td><td></td></tr>
<tr><td>Oligo fwd</td><td>CACCAAGGTGTCGTTACTGCCTAT</td><td></td><td></td><td></td><td></td><td></td><td></td><td></td><td></td><td></td><td></td></tr>
<tr><td>Oligo rev</td><td>AAACATAGGCAGTAACGACACCTT</td><td></td><td></td><td></td><td></td><td></td><td></td><td></td><td></td><td></td><td></td></tr>
<tr><td>Chromosome</td><td>start</td><td>end</td><td>strand</td><td>MM</td><td>target_seq</td><td>PAM</td><td>alignment</td><td>distance</td><td>position</td><td>gene name</td><td>gene id</td></tr>
<tr><td>GU305914.1</td><td>188451</td><td>188473</td><td>-</td><td>0</td><td>AAGGTGTCGTTACTGCCTAT</td><td>GGG</td><td>||||||||[||||||||||||]PAM</td><td>NA</td><td>Intergenic</td><td>NA</td><td>NA</td></tr>
<tr><td>T20386</td><td>CGTTACTGCCTATGGGAAGCTGG</td><td>1000</td><td></td><td></td><td></td><td></td><td></td><td></td><td></td><td></td><td></td></tr>
<tr><td>Oligo fwd</td><td>CACCCGTTACTGCCTATGGGAAGC</td><td></td><td></td><td></td><td></td><td></td><td></td><td></td><td></td><td></td><td></td></tr>
<tr><td>Oligo rev</td><td>AAACGCTTCCCATAGGCAGTAACG</td><td></td><td></td><td></td><td></td><td></td><td></td><td></td><td></td><td></td><td></td></tr>
<tr><td>Chromosome</td><td>start</td><td>end</td><td>strand</td><td>MM</td><td>target_seq</td><td>PAM</td><td>alignment</td><td>distance</td><td>position</td><td>gene name</td><td>gene id</td></tr>
<tr><td>GU305914.1</td><td>188444</td><td>188466</td><td>-</td><td>0</td><td>CGTTACTGCCTATGGGAAGC</td><td>TGG</td><td>||||||||[||||||||||||]PAM</td><td>NA</td><td>Intergenic</td><td>NA</td><td>NA</td></tr>
<tr><td>T20387</td><td>ATGGGAAGCTGGAGACCCCACGG</td><td>1000</td><td></td><td></td><td></td><td></td><td></td><td></td><td></td><td></td><td></td></tr>
<tr><td>Oligo fwd</td><td>CACCATGGGAAGCTGGAGACCCCA</td><td></td><td></td><td></td><td></td><td></td><td></td><td></td><td></td><td></td><td></td></tr>
<tr><td>Oligo rev</td><td>AAACTGGGGTCTCCAGCTTCCCAT</td><td></td><td></td><td></td><td></td><td></td><td></td><td></td><td></td><td></td><td></td></tr>
<tr><td>Chromosome</td><td>start</td><td>end</td><td>strand</td><td>MM</td><td>target_seq</td><td>PAM</td><td>alignment</td><td>distance</td><td>position</td><td>gene name</td><td>gene id</td></tr>
<tr><td>GU305914.1</td><td>188433</td><td>188455</td><td>-</td><td>0</td><td>ATGGGAAGCTGGAGACCCCA</td><td>CGG</td><td>||||||||[||||||||||||]PAM</td><td>NA</td><td>Intergenic</td><td>NA</td><td>NA</td></tr>
<tr><td>T20388</td><td>CAATAAAATCTCATCTGAAATGG</td><td>1000</td><td></td><td></td><td></td><td></td><td></td><td></td><td></td><td></td><td></td></tr>
<tr><td>Oligo fwd</td><td>CACCCAATAAAATCTCATCTGAAA</td><td></td><td></td><td></td><td></td><td></td><td></td><td></td><td></td><td></td><td></td></tr>
<tr><td>Oligo rev</td><td>AAACTTTCAGATGAGATTTTATTG</td><td></td><td></td><td></td><td></td><td></td><td></td><td></td><td></td><td></td><td></td></tr>
<tr><td>Chromosome</td><td>start</td><td>end</td><td>strand</td><td>MM</td><td>target_seq</td><td>PAM</td><td>alignment</td><td>distance</td><td>position</td><td>gene name</td><td>gene id</td></tr>
<tr><td>GU305914.1</td><td>188368</td><td>188390</td><td>-</td><td>0</td><td>CAATAAAATCTCATCTGAAA</td><td>TGG</td><td>||||||||[||||||||||||]PAM</td><td>NA</td><td>Intergenic</td><td>NA</td><td>NA</td></tr>
<tr><td>T20389</td><td>AATAAAATCTCATCTGAAATGGG</td><td>1000</td><td></td><td></td><td></td><td></td><td></td><td></td><td></td><td></td><td></td></tr>
<tr><td>Oligo fwd</td><td>CACCAATAAAATCTCATCTGAAAT</td><td></td><td></td><td></td><td></td><td></td><td></td><td></td><td></td><td></td><td></td></tr>
<tr><td>Oligo rev</td><td>AAACATTTCAGATGAGATTTTATT</td><td></td><td></td><td></td><td></td><td></td><td></td><td></td><td></td><td></td><td></td></tr>
<tr><td>Chromosome</td><td>start</td><td>end</td><td>strand</td><td>MM</td><td>target_seq</td><td>PAM</td><td>alignment</td><td>distance</td><td>position</td><td>gene name</td><td>gene id</td></tr>
<tr><td>GU305914.1</td><td>188367</td><td>188389</td><td>-</td><td>0</td><td>AATAAAATCTCATCTGAAAT</td><td>GGG</td><td>||||||||[||||||||||||]PAM</td><td>NA</td><td>Intergenic</td><td>NA</td><td>NA</td></tr>
<tr><td>T20390</td><td>CCTCGTCTGTTGCGTCGTGTTGG</td><td>1000</td><td></td><td></td><td></td><td></td><td></td><td></td><td></td><td></td><td></td></tr>
<tr><td>Oligo fwd</td><td>CACCCCTCGTCTGTTGCGTCGTGT</td><td></td><td></td><td></td><td></td><td></td><td></td><td></td><td></td><td></td><td></td></tr>
<tr><td>Oligo rev</td><td>AAACACACGACGCAACAGACGAGG</td><td></td><td></td><td></td><td></td><td></td><td></td><td></td><td></td><td></td><td></td></tr>
<tr><td>Chromosome</td><td>start</td><td>end</td><td>strand</td><td>MM</td><td>target_seq</td><td>PAM</td><td>alignment</td><td>distance</td><td>position</td><td>gene name</td><td>gene id</td></tr>
<tr><td>GU305914.1</td><td>188326</td><td>188348</td><td>-</td><td>0</td><td>CCTCGTCTGTTGCGTCGTGT</td><td>TGG</td><td>||||||||[||||||||||||]PAM</td><td>NA</td><td>Intergenic</td><td>NA</td><td>NA</td></tr>
<tr><td>T20391</td><td>GTTGGTCGTGTCTACCGTCGCGG</td><td>1000</td><td></td><td></td><td></td><td></td><td></td><td></td><td></td><td></td><td></td></tr>
<tr><td>Oligo fwd</td><td>CACCGTTGGTCGTGTCTACCGTCG</td><td></td><td></td><td></td><td></td><td></td><td></td><td></td><td></td><td></td><td></td></tr>
<tr><td>Oligo rev</td><td>AAACCGACGGTAGACACGACCAAC</td><td></td><td></td><td></td><td></td><td></td><td></td><td></td><td></td><td></td><td></td></tr>
<tr><td>Chromosome</td><td>start</td><td>end</td><td>strand</td><td>MM</td><td>target_seq</td><td>PAM</td><td>alignment</td><td>distance</td><td>position</td><td>gene name</td><td>gene id</td></tr>
<tr><td>GU305914.1</td><td>188308</td><td>188330</td><td>-</td><td>0</td><td>GTTGGTCGTGTCTACCGTCG</td><td>CGG</td><td>||||||||[||||||||||||]PAM</td><td>NA</td><td>Intergenic</td><td>NA</td><td>NA</td></tr>
<tr><td>T20392</td><td>TCCGCGAGCCATGTTGCGCGCGG</td><td>1000</td><td></td><td></td><td></td><td></td><td></td><td></td><td></td><td></td><td></td></tr>
<tr><td>Oligo fwd</td><td>CACCTCCGCGAGCCATGTTGCGCG</td><td></td><td></td><td></td><td></td><td></td><td></td><td></td><td></td><td></td><td></td></tr>
<tr><td>Oligo rev</td><td>AAACCGCGCAACATGGCTCGCGGA</td><td></td><td></td><td></td><td></td><td></td><td></td><td></td><td></td><td></td><td></td></tr>
</table>

Chromosome start end strand MM target_seq PAM alignment distance position gene name gene id
GU305914.1 188282 188304 - 0 TCCGCGAGCCATGTTGCGCG CGG ||||||||[||||||||||||]PAM NA Intergenic NA NA

T20393 GCACCCTCTCCCGCTGTACGCGG 1000
Oligo fwd CACCGCACCCTCTCCCGCTGTACG
Oligo rev AAACCGTACAGCGGGAGAGGGTGC
Chromosome start end strand MM target_seq PAM alignment distance position gene name gene id
GU305914.1 188251 188273 - 0 GCACCCTCTCCCGCTGTACG CGG ||||||||[||||||||||||]PAM NA Intergenic NA NA

T20394 GAATACACTAACACTTCGTGCGG 1000
Oligo fwd CACCGAATACACTAACACTTCGTG
Oligo rev AAACCACGAAGTGTTAGTGTATTC
Chromosome start end strand MM target_seq PAM alignment distance position gene name gene id
GU305914.1 188217 188239 - 0 GAATACACTAACACTTCGTG CGG ||||||||[||||||||||||]PAM NA Intergenic NA NA

T20395 TAACACTTCGTGCGGACGAGAGG 1000
Oligo fwd CACCTAACACTTCGTGCGGACGAG
Oligo rev AAACCTCGTCCGCACGAAGTGTTA
Chromosome start end strand MM target_seq PAM alignment distance position gene name gene id
GU305914.1 188209 188231 - 0 TAACACTTCGTGCGGACGAG AGG ||||||||[||||||||||||]PAM NA Intergenic NA NA

T20396 CACTTCGTGCGGACGAGAGGTGG 1000
Oligo fwd CACCCACTTCGTGCGGACGAGAGG
Oligo rev AAACCCTCTCGTCCGCACGAAGTG
Chromosome start end strand MM target_seq PAM alignment distance position gene name gene id
GU305914.1 188206 188228 - 0 CACTTCGTGCGGACGAGAGG TGG ||||||||[||||||||||||]PAM NA Intergenic NA NA

T20397 GTGGTTTTCACTACCTTCTCTGG 1000
Oligo fwd CACCGTGGTTTTCACTACCTTCTC
Oligo rev AAACGAGAAGGTAGTGAAAACCAC
Chromosome start end strand MM target_seq PAM alignment distance position gene name gene id
GU305914.1 188187 188209 - 0 GTGGTTTTCACTACCTTCTC TGG ||||||||[||||||||||||]PAM NA Intergenic NA NA

T20398 TGGTTTTCACTACCTTCTCTGGG 1000
Oligo fwd CACCTGGTTTTCACTACCTTCTCT
Oligo rev AAACAGAGAAGGTAGTGAAAACCA
Chromosome start end strand MM target_seq PAM alignment distance position gene name gene id
GU305914.1 188186 188208 - 0 TGGTTTTCACTACCTTCTCT GGG ||||||||[||||||||||||]PAM NA Intergenic NA NA

T20399 TCACTACCTTCTCTGGGATGAGG 1000
Oligo fwd CACCTCACTACCTTCTCTGGGATG
Oligo rev AAACCATCCCAGAGAAGGTAGTGA
Chromosome start end strand MM target_seq PAM alignment distance position gene name gene id
GU305914.1 188180 188202 - 0 TCACTACCTTCTCTGGGATG AGG ||||||||[||||||||||||]PAM NA Intergenic NA NA

T20400 CACTACCTTCTCTGGGATGAGGG 1000
Oligo fwd CACCCACTACCTTCTCTGGGATGA
Oligo rev AAACTCATCCCAGAGAAGGTAGTG
Chromosome start end strand MM target_seq PAM alignment distance position gene name gene id
GU305914.1 188179 188201 - 0 CACTACCTTCTCTGGGATGA GGG ||||||||[||||||||||||]PAM NA Intergenic NA NA

T20401 GGATGAGGGTGTGCGCTAAGCGG 1000
Oligo fwd CACCGGATGAGGGTGTGCGCTAAG
Oligo rev AAACCTTAGCGCACACCCTCATCC
Chromosome start end strand MM target_seq PAM alignment distance position gene name gene id
GU305914.1 188165 188187 - 0 GGATGAGGGTGTGCGCTAAG CGG ||||||||[||||||||||||]PAM NA Intergenic NA NA

T20402 GATGAGGGTGTGCGCTAAGCGGG 1000
Oligo fwd CACCGATGAGGGTGTGCGCTAAGC
Oligo rev AAACGCTTAGCGCACACCCTCATC
Chromosome start end strand MM target_seq PAM alignment distance position gene name gene id
GU305914.1 188164 188186 - 0 GATGAGGGTGTGCGCTAAGC GGG ||||||||[||||||||||||]PAM NA Intergenic NA NA

T20403 GGGTGTGCGCTAAGCGGGCCTGG 1000
Oligo fwd CACCGGGTGTGCGCTAAGCGGGCC
Oligo rev AAACGGCCCGCTTAGCGCACACCC
Chromosome start end strand MM target_seq PAM alignment distance position gene name gene id
GU305914.1 188159 188181 - 0 GGGTGTGCGCTAAGCGGGCC TGG ||||||||[||||||||||||]PAM NA Intergenic NA NA

T20404 TGTGCGCTAAGCGGGCCTGGTGG 1000
Oligo fwd CACCTGTGCGCTAAGCGGGCCTGG
Oligo rev AAACCCAGGCCCGCTTAGCGCACA
Chromosome start end strand MM target_seq PAM alignment distance position gene name gene id
GU305914.1 188156 188178 - 0 TGTGCGCTAAGCGGGCCTGG TGG ||||||||[||||||||||||]PAM NA Intergenic NA NA

T20405 CGCTAAGCGGGCCTGGTGGTCGG 1000
Oligo fwd CACCCGCTAAGCGGGCCTGGTGGT
Oligo rev AAACACCACCAGGCCCGCTTAGCG
Chromosome start end strand MM target_seq PAM alignment distance position gene name gene id
GU305914.1 188152 188174 - 0 CGCTAAGCGGGCCTGGTGGT CGG ||||||||[||||||||||||]PAM NA Intergenic NA NA

T20406 GTCGGACAGACTCTTGTGTCTGG 1000
Oligo fwd CACCGTCGGACAGACTCTTGTGTC
Oligo rev AAACGACACAAGAGTCTGTCCGAC
Chromosome start end strand MM target_seq PAM alignment distance position gene name gene id
GU305914.1 188134 188156 - 0 GTCGGACAGACTCTTGTGTC TGG ||||||||[||||||||||||]PAM NA Intergenic NA NA

T20407 ACAGACTCTTGTGTCTGGTCAGG 1000
Oligo fwd CACCACAGACTCTTGTGTCTGGTC

Oligo rev AAACGACCAGACACAAGAGTCTGT
Chromosome start end strand MM target_seq PAM alignment distance position gene name gene id
GU305914.1 188129 188151 - 0 ACAGACTCTTGTGTCTGGTC AGG ||||||||[||||||||||||]PAM NA Intergenic NA NA
GU305914.1 113738 113760 + 4 CCGGACTCTTCTGTCTGGGC CGG -|-|||||[||-|||||||-|]PAM NA Intergenic NA NA

T20408 GGTGAGTGTTTTGATCTGAAAGG 1000
Oligo fwd CACCGGTGAGTGTTTTGATCTGAA
Oligo rev AAACTTCAGATCAAAACACTCACC
Chromosome start end strand MM target_seq PAM alignment distance position gene name gene id
GU305914.1 188108 188130 - 0 GGTGAGTGTTTTGATCTGAA AGG ||||||||[||||||||||||]PAM NA Intergenic NA NA

T20409 GTGTTTTGATCTGAAAGGAGAGG 1000
Oligo fwd CACCGTGTTTTGATCTGAAAGGAG
Oligo rev AAACCTCCTTTCAGATCAAAACAC
Chromosome start end strand MM target_seq PAM alignment distance position gene name gene id
GU305914.1 188103 188125 - 0 GTGTTTTGATCTGAAAGGAG AGG ||||||||[||||||||||||]PAM NA Intergenic NA NA

T20410 TGTTTTGATCTGAAAGGAGAGGG 1000
Oligo fwd CACCTGTTTTGATCTGAAAGGAGA
Oligo rev AAACTCTCCTTTCAGATCAAAACA
Chromosome start end strand MM target_seq PAM alignment distance position gene name gene id
GU305914.1 188102 188124 - 0 TGTTTTGATCTGAAAGGAGA GGG ||||||||[||||||||||||]PAM NA Intergenic NA NA

T20411 TGATCTGAAAGGAGAGGGATCGG 1000
Oligo fwd CACCTGATCTGAAAGGAGAGGGAT
Oligo rev AAACATCCCTCTCCTTTCAGATCA
Chromosome start end strand MM target_seq PAM alignment distance position gene name gene id
GU305914.1 188097 188119 - 0 TGATCTGAAAGGAGAGGGAT CGG ||||||||[||||||||||||]PAM NA Intergenic NA NA

T20412 TCTGAAAGGAGAGGGATCGGCGG 1000
Oligo fwd CACCTCTGAAAGGAGAGGGATCGG
Oligo rev AAACCCGATCCCTCTCCTTTCAGA
Chromosome start end strand MM target_seq PAM alignment distance position gene name gene id
GU305914.1 188094 188116 - 0 TCTGAAAGGAGAGGGATCGG CGG ||||||||[||||||||||||]PAM NA Intergenic NA NA

T20413 CTGAAAGGAGAGGGATCGGCGGG 1000
Oligo fwd CACCCTGAAAGGAGAGGGATCGGC
Oligo rev AAACGCCGATCCCTCTCCTTTCAG
Chromosome start end strand MM target_seq PAM alignment distance position gene name gene id
GU305914.1 188093 188115 - 0 CTGAAAGGAGAGGGATCGGC GGG ||||||||[||||||||||||]PAM NA Intergenic NA NA

T20414 TGAAAGGAGAGGGATCGGCGGGG 1000
Oligo fwd CACCTGAAAGGAGAGGGATCGGCG
Oligo rev AAACCGCCGATCCCTCTCCTTTCA
Chromosome start end strand MM target_seq PAM alignment distance position gene name gene id
GU305914.1 188092 188114 - 0 TGAAAGGAGAGGGATCGGCG GGG ||||||||[||||||||||||]PAM NA Intergenic NA NA

T20415 GAAAGGAGAGGGATCGGCGGGGG 1000
Oligo fwd CACCGAAAGGAGAGGGATCGGCGG
Oligo rev AAACCCGCCGATCCCTCTCCTTTC
Chromosome start end strand MM target_seq PAM alignment distance position gene name gene id
GU305914.1 188091 188113 - 0 GAAAGGAGAGGGATCGGCGG GGG ||||||||[||||||||||||]PAM NA Intergenic NA NA

T20416 AAAGGAGAGGGATCGGCGGGGGG 1000
Oligo fwd CACCAAAGGAGAGGGATCGGCGGG
Oligo rev AAACCCCGCCGATCCCTCTCCTTT
Chromosome start end strand MM target_seq PAM alignment distance position gene name gene id
GU305914.1 188090 188112 - 0 AAAGGAGAGGGATCGGCGGG GGG ||||||||[||||||||||||]PAM NA Intergenic NA NA

T20417 GGATCGGCGGGGGGCGCGATAGG 1000
Oligo fwd CACCGGATCGGCGGGGGGCGCGAT
Oligo rev AAACATCGCGCCCCCCGCCGATCC
Chromosome start end strand MM target_seq PAM alignment distance position gene name gene id
GU305914.1 188081 188103 - 0 GGATCGGCGGGGGGCGCGAT AGG ||||||||[||||||||||||]PAM NA Intergenic NA NA

T20418 GATCGGCGGGGGGCGCGATAGGG 1000
Oligo fwd CACCGATCGGCGGGGGGCGCGATA
Oligo rev AAACTATCGCGCCCCCCGCCGATC
Chromosome start end strand MM target_seq PAM alignment distance position gene name gene id
GU305914.1 188080 188102 - 0 GATCGGCGGGGGGCGCGATA GGG ||||||||[||||||||||||]PAM NA Intergenic NA NA

T20419 ATCGGCGGGGGGCGCGATAGGGG 1000
Oligo fwd CACCATCGGCGGGGGGCGCGATAG
Oligo rev AAACCTATCGCGCCCCCCGCCGAT
Chromosome start end strand MM target_seq PAM alignment distance position gene name gene id
GU305914.1 188079 188101 - 0 ATCGGCGGGGGGCGCGATAG GGG ||||||||[||||||||||||]PAM NA Intergenic NA NA
GU305914.1 61922 61944 - 4 TTCCGCGCGGGGCGCGACAG CGG -||-|||-[|||||||||-||]PAM NA Intergenic NA NA

T20420 TCGGCGGGGGGCGCGATAGGGGG 1000
Oligo fwd CACCTCGGCGGGGGGCGCGATAGG
Oligo rev AAACCCTATCGCGCCCCCCGCCGA
Chromosome start end strand MM target_seq PAM alignment distance position gene name gene id
GU305914.1 188078 188100 - 0 TCGGCGGGGGGCGCGATAGG GGG ||||||||[||||||||||||]PAM NA Intergenic NA NA
GU305914.1 61921 61943 - 4 TCCGCGCGGGGCGCGACAGC GGG ||-|||-|[||||||||-||-]PAM NA Intergenic NA NA

T20421 GTCTGTGTGTATATAGATCTCGG 1000
Oligo fwd CACCGTCTGTGTGTATATAGATCT
Oligo rev AAACAGATCTATATACACACAGAC
Chromosome start end strand MM target_seq PAM alignment distance position gene name gene id

<table>
<tr><td>GU305914.1</td><td></td><td>188041</td><td>188063</td><td>-</td><td>0</td><td>GTCTGTGTGTATATAGATCT</td><td>CGG</td><td>||||||||[||||||||||||]PAM</td><td>NA</td><td>Intergenic</td><td>NA</td><td>NA</td></tr>
<tr><td>T20422</td><td>CTGCGTGTCCTGCTTATCTTTGG</td><td></td><td>1000</td><td></td><td></td><td></td><td></td><td></td><td></td><td></td><td></td><td></td></tr>
<tr><td>Oligo fwd</td><td>CACCCTGCGTGTCCTGCTTATCTT</td><td></td><td></td><td></td><td></td><td></td><td></td><td></td><td></td><td></td><td></td><td></td></tr>
<tr><td>Oligo rev</td><td>AAACAAGATAAGCAGGACACGCAG</td><td></td><td></td><td></td><td></td><td></td><td></td><td></td><td></td><td></td><td></td><td></td></tr>
<tr><td>Chromosome</td><td>start</td><td></td><td>end</td><td>strand</td><td>MM</td><td>target_seq</td><td>PAM</td><td>alignment</td><td>distance</td><td>position</td><td>gene name</td><td>gene id</td></tr>
<tr><td>GU305914.1</td><td></td><td>188006</td><td>188028</td><td>-</td><td>0</td><td>CTGCGTGTCCTGCTTATCTT</td><td>TGG</td><td>||||||||[||||||||||||]PAM</td><td>NA</td><td>Intergenic</td><td>NA</td><td>NA</td></tr>
<tr><td>T20423</td><td>AAGCTTTCAACACCACCATCAGG</td><td></td><td>1000</td><td></td><td></td><td></td><td></td><td></td><td></td><td></td><td></td><td></td></tr>
<tr><td>Oligo fwd</td><td>CACCAAGCTTTCAACACCACCATC</td><td></td><td></td><td></td><td></td><td></td><td></td><td></td><td></td><td></td><td></td><td></td></tr>
<tr><td>Oligo rev</td><td>AAACGATGGTGGTGTTGAAAGCTT</td><td></td><td></td><td></td><td></td><td></td><td></td><td></td><td></td><td></td><td></td><td></td></tr>
<tr><td>Chromosome</td><td>start</td><td></td><td>end</td><td>strand</td><td>MM</td><td>target_seq</td><td>PAM</td><td>alignment</td><td>distance</td><td>position</td><td>gene name</td><td>gene id</td></tr>
<tr><td>GU305914.1</td><td></td><td>187972</td><td>187994</td><td>-</td><td>0</td><td>AAGCTTTCAACACCACCATC</td><td>AGG</td><td>||||||||[||||||||||||]PAM</td><td>NA</td><td>Intergenic</td><td>NA</td><td>NA</td></tr>
<tr><td>T20424</td><td>CACCATCAGGCTCGTGCCCGTGG</td><td></td><td>1000</td><td></td><td></td><td></td><td></td><td></td><td></td><td></td><td></td><td></td></tr>
<tr><td>Oligo fwd</td><td>CACCCACCATCAGGCTCGTGCCCG</td><td></td><td></td><td></td><td></td><td></td><td></td><td></td><td></td><td></td><td></td><td></td></tr>
<tr><td>Oligo rev</td><td>AAACCGGGCACGAGCCTGATGGTG</td><td></td><td></td><td></td><td></td><td></td><td></td><td></td><td></td><td></td><td></td><td></td></tr>
<tr><td>Chromosome</td><td>start</td><td></td><td>end</td><td>strand</td><td>MM</td><td>target_seq</td><td>PAM</td><td>alignment</td><td>distance</td><td>position</td><td>gene name</td><td>gene id</td></tr>
<tr><td>GU305914.1</td><td></td><td>187959</td><td>187981</td><td>-</td><td>0</td><td>CACCATCAGGCTCGTGCCCG</td><td>TGG</td><td>||||||||[||||||||||||]PAM</td><td>NA</td><td>Intergenic</td><td>NA</td><td>NA</td></tr>
<tr><td>T20425</td><td>CCCCGCCGACATGATGTACGTGG</td><td></td><td>1000</td><td></td><td></td><td></td><td></td><td></td><td></td><td></td><td></td><td></td></tr>
<tr><td>Oligo fwd</td><td>CACCCCCCGCCGACATGATGTACG</td><td></td><td></td><td></td><td></td><td></td><td></td><td></td><td></td><td></td><td></td><td></td></tr>
<tr><td>Oligo rev</td><td>AAACCGTACATCATGTCGGCGGGG</td><td></td><td></td><td></td><td></td><td></td><td></td><td></td><td></td><td></td><td></td><td></td></tr>
<tr><td>Chromosome</td><td>start</td><td></td><td>end</td><td>strand</td><td>MM</td><td>target_seq</td><td>PAM</td><td>alignment</td><td>distance</td><td>position</td><td>gene name</td><td>gene id</td></tr>
<tr><td>GU305914.1</td><td></td><td>187935</td><td>187957</td><td>-</td><td>0</td><td>CCCCGCCGACATGATGTACG</td><td>TGG</td><td>||||||||[||||||||||||]PAM</td><td>NA</td><td>Intergenic</td><td>NA</td><td>NA</td></tr>
<tr><td>T20426</td><td>GTACGTGGCCGATGTGATGACGG</td><td></td><td>1000</td><td></td><td></td><td></td><td></td><td></td><td></td><td></td><td></td><td></td></tr>
<tr><td>Oligo fwd</td><td>CACCGTACGTGGCCGATGTGATGA</td><td></td><td></td><td></td><td></td><td></td><td></td><td></td><td></td><td></td><td></td><td></td></tr>
<tr><td>Oligo rev</td><td>AAACTCATCACATCGGCCACGTAC</td><td></td><td></td><td></td><td></td><td></td><td></td><td></td><td></td><td></td><td></td><td></td></tr>
<tr><td>Chromosome</td><td>start</td><td></td><td>end</td><td>strand</td><td>MM</td><td>target_seq</td><td>PAM</td><td>alignment</td><td>distance</td><td>position</td><td>gene name</td><td>gene id</td></tr>
<tr><td>GU305914.1</td><td></td><td>187920</td><td>187942</td><td>-</td><td>0</td><td>GTACGTGGCCGATGTGATGA</td><td>CGG</td><td>||||||||[||||||||||||]PAM</td><td>NA</td><td>Intergenic</td><td>NA</td><td>NA</td></tr>
<tr><td>T20427</td><td>GACGGAATCGCCACGCATCACGG</td><td></td><td>1000</td><td></td><td></td><td></td><td></td><td></td><td></td><td></td><td></td><td></td></tr>
<tr><td>Oligo fwd</td><td>CACCGACGGAATCGCCACGCATCA</td><td></td><td></td><td></td><td></td><td></td><td></td><td></td><td></td><td></td><td></td><td></td></tr>
<tr><td>Oligo rev</td><td>AAACTGATGCGTGGCGATTCCGTC</td><td></td><td></td><td></td><td></td><td></td><td></td><td></td><td></td><td></td><td></td><td></td></tr>
<tr><td>Chromosome</td><td>start</td><td></td><td>end</td><td>strand</td><td>MM</td><td>target_seq</td><td>PAM</td><td>alignment</td><td>distance</td><td>position</td><td>gene name</td><td>gene id</td></tr>
<tr><td>GU305914.1</td><td></td><td>187902</td><td>187924</td><td>-</td><td>0</td><td>GACGGAATCGCCACGCATCA</td><td>CGG</td><td>||||||||[||||||||||||]PAM</td><td>NA</td><td>Intergenic</td><td>NA</td><td>NA</td></tr>
<tr><td>T20428</td><td>GGAATCGCCACGCATCACGGTGG</td><td></td><td>1000</td><td></td><td></td><td></td><td></td><td></td><td></td><td></td><td></td><td></td></tr>
<tr><td>Oligo fwd</td><td>CACCGGAATCGCCACGCATCACGG</td><td></td><td></td><td></td><td></td><td></td><td></td><td></td><td></td><td></td><td></td><td></td></tr>
<tr><td>Oligo rev</td><td>AAACCCGTGATGCGTGGCGATTCC</td><td></td><td></td><td></td><td></td><td></td><td></td><td></td><td></td><td></td><td></td><td></td></tr>
<tr><td>Chromosome</td><td>start</td><td></td><td>end</td><td>strand</td><td>MM</td><td>target_seq</td><td>PAM</td><td>alignment</td><td>distance</td><td>position</td><td>gene name</td><td>gene id</td></tr>
<tr><td>GU305914.1</td><td></td><td>187899</td><td>187921</td><td>-</td><td>0</td><td>GGAATCGCCACGCATCACGG</td><td>TGG</td><td>||||||||[||||||||||||]PAM</td><td>NA</td><td>Intergenic</td><td>NA</td><td>NA</td></tr>
<tr><td>T20429</td><td>GAATCGCCACGCATCACGGTGGG</td><td></td><td>1000</td><td></td><td></td><td></td><td></td><td></td><td></td><td></td><td></td><td></td></tr>
<tr><td>Oligo fwd</td><td>CACCGAATCGCCACGCATCACGGT</td><td></td><td></td><td></td><td></td><td></td><td></td><td></td><td></td><td></td><td></td><td></td></tr>
<tr><td>Oligo rev</td><td>AAACACCGTGATGCGTGGCGATTC</td><td></td><td></td><td></td><td></td><td></td><td></td><td></td><td></td><td></td><td></td><td></td></tr>
<tr><td>Chromosome</td><td>start</td><td></td><td>end</td><td>strand</td><td>MM</td><td>target_seq</td><td>PAM</td><td>alignment</td><td>distance</td><td>position</td><td>gene name</td><td>gene id</td></tr>
<tr><td>GU305914.1</td><td></td><td>187898</td><td>187920</td><td>-</td><td>0</td><td>GAATCGCCACGCATCACGGT</td><td>GGG</td><td>||||||||[||||||||||||]PAM</td><td>NA</td><td>Intergenic</td><td>NA</td><td>NA</td></tr>
<tr><td>T20430</td><td>GGTGGGCAAGTACCCCTACGAGG</td><td></td><td>1000</td><td></td><td></td><td></td><td></td><td></td><td></td><td></td><td></td><td></td></tr>
<tr><td>Oligo fwd</td><td>CACCGGTGGGCAAGTACCCCTACG</td><td></td><td></td><td></td><td></td><td></td><td></td><td></td><td></td><td></td><td></td><td></td></tr>
<tr><td>Oligo rev</td><td>AAACCGTAGGGGTACTTGCCCACC</td><td></td><td></td><td></td><td></td><td></td><td></td><td></td><td></td><td></td><td></td><td></td></tr>
<tr><td>Chromosome</td><td>start</td><td></td><td>end</td><td>strand</td><td>MM</td><td>target_seq</td><td>PAM</td><td>alignment</td><td>distance</td><td>position</td><td>gene name</td><td>gene id</td></tr>
<tr><td>GU305914.1</td><td></td><td>187881</td><td>187903</td><td>-</td><td>0</td><td>GGTGGGCAAGTACCCCTACG</td><td>AGG</td><td>||||||||[||||||||||||]PAM</td><td>NA</td><td>Intergenic</td><td>NA</td><td>NA</td></tr>
<tr><td>T20431</td><td>CAAGTACCCCTACGAGGTGAAGG</td><td></td><td>1000</td><td></td><td></td><td></td><td></td><td></td><td></td><td></td><td></td><td></td></tr>
<tr><td>Oligo fwd</td><td>CACCCAAGTACCCCTACGAGGTGA</td><td></td><td></td><td></td><td></td><td></td><td></td><td></td><td></td><td></td><td></td><td></td></tr>
<tr><td>Oligo rev</td><td>AAACTCACCTCGTAGGGGTACTTG</td><td></td><td></td><td></td><td></td><td></td><td></td><td></td><td></td><td></td><td></td><td></td></tr>
<tr><td>Chromosome</td><td>start</td><td></td><td>end</td><td>strand</td><td>MM</td><td>target_seq</td><td>PAM</td><td>alignment</td><td>distance</td><td>position</td><td>gene name</td><td>gene id</td></tr>
<tr><td>GU305914.1</td><td></td><td>187875</td><td>187897</td><td>-</td><td>0</td><td>CAAGTACCCCTACGAGGTGA</td><td>AGG</td><td>||||||||[||||||||||||]PAM</td><td>NA</td><td>Intergenic</td><td>NA</td><td>NA</td></tr>
<tr><td>T20432</td><td>TACCCCTACGAGGTGAAGGACGG</td><td></td><td>1000</td><td></td><td></td><td></td><td></td><td></td><td></td><td></td><td></td><td></td></tr>
<tr><td>Oligo fwd</td><td>CACCTACCCCTACGAGGTGAAGGA</td><td></td><td></td><td></td><td></td><td></td><td></td><td></td><td></td><td></td><td></td><td></td></tr>
<tr><td>Oligo rev</td><td>AAACTCCTTCACCTCGTAGGGGTA</td><td></td><td></td><td></td><td></td><td></td><td></td><td></td><td></td><td></td><td></td><td></td></tr>
<tr><td>Chromosome</td><td>start</td><td></td><td>end</td><td>strand</td><td>MM</td><td>target_seq</td><td>PAM</td><td>alignment</td><td>distance</td><td>position</td><td>gene name</td><td>gene id</td></tr>
<tr><td>GU305914.1</td><td></td><td>187871</td><td>187893</td><td>-</td><td>0</td><td>TACCCCTACGAGGTGAAGGA</td><td>CGG</td><td>||||||||[||||||||||||]PAM</td><td>NA</td><td>Intergenic</td><td>NA</td><td>NA</td></tr>
<tr><td>T20433</td><td>AGGTGAAGGACGGTGAAAACTGG</td><td></td><td>1000</td><td></td><td></td><td></td><td></td><td></td><td></td><td></td><td></td><td></td></tr>
<tr><td>Oligo fwd</td><td>CACCAGGTGAAGGACGGTGAAAAC</td><td></td><td></td><td></td><td></td><td></td><td></td><td></td><td></td><td></td><td></td><td></td></tr>
<tr><td>Oligo rev</td><td>AAACGTTTTCACCGTCCTTCACCT</td><td></td><td></td><td></td><td></td><td></td><td></td><td></td><td></td><td></td><td></td><td></td></tr>
<tr><td>Chromosome</td><td>start</td><td></td><td>end</td><td>strand</td><td>MM</td><td>target_seq</td><td>PAM</td><td>alignment</td><td>distance</td><td>position</td><td>gene name</td><td>gene id</td></tr>
<tr><td>GU305914.1</td><td></td><td>187861</td><td>187883</td><td>-</td><td>0</td><td>AGGTGAAGGACGGTGAAAAC</td><td>TGG</td><td>||||||||[||||||||||||]PAM</td><td>NA</td><td>Intergenic</td><td>NA</td><td>NA</td></tr>
<tr><td>T20434</td><td>AGTGTGTCTGTGAGATCACCTGG</td><td></td><td>1000</td><td></td><td></td><td></td><td></td><td></td><td></td><td></td><td></td><td></td></tr>
<tr><td>Oligo fwd</td><td>CACCAGTGTGTCTGTGAGATCACC</td><td></td><td></td><td></td><td></td><td></td><td></td><td></td><td></td><td></td><td></td><td></td></tr>
<tr><td>Oligo rev</td><td>AAACGGTGATCTCACAGACACACT</td><td></td><td></td><td></td><td></td><td></td><td></td><td></td><td></td><td></td><td></td><td></td></tr>
<tr><td>Chromosome</td><td>start</td><td></td><td>end</td><td>strand</td><td>MM</td><td>target_seq</td><td>PAM</td><td>alignment</td><td>distance</td><td>position</td><td>gene name</td><td>gene id</td></tr>
<tr><td>GU305914.1</td><td></td><td>187834</td><td>187856</td><td>-</td><td>0</td><td>AGTGTGTCTGTGAGATCACC</td><td>TGG</td><td>||||||||[||||||||||||]PAM</td><td>NA</td><td>Intergenic</td><td>NA</td><td>NA</td></tr>
<tr><td>T20435</td><td>GTGTGTCTGTGAGATCACCTGGG</td><td></td><td>1000</td><td></td><td></td><td></td><td></td><td></td><td></td><td></td><td></td><td></td></tr>
<tr><td>Oligo fwd</td><td>CACCGTGTGTCTGTGAGATCACCT</td><td></td><td></td><td></td><td></td><td></td><td></td><td></td><td></td><td></td><td></td><td></td></tr>
<tr><td>Oligo rev</td><td>AAACAGGTGATCTCACAGACACAC</td><td></td><td></td><td></td><td></td><td></td><td></td><td></td><td></td><td></td><td></td><td></td></tr>
<tr><td>Chromosome</td><td>start</td><td></td><td>end</td><td>strand</td><td>MM</td><td>target_seq</td><td>PAM</td><td>alignment</td><td>distance</td><td>position</td><td>gene name</td><td>gene id</td></tr>
<tr><td>GU305914.1</td><td></td><td>187833</td><td>187855</td><td>-</td><td>0</td><td>GTGTGTCTGTGAGATCACCT</td><td>GGG</td><td>||||||||[||||||||||||]PAM</td><td>NA</td><td>Intergenic</td><td>NA</td><td>NA</td></tr>
<tr><td>T20436</td><td>GTCTGTGAGATCACCTGGGTCGG</td><td></td><td>1000</td><td></td><td></td><td></td><td></td><td></td><td></td><td></td><td></td><td></td></tr>
<tr><td>Oligo fwd</td><td>CACCGTCTGTGAGATCACCTGGGT</td><td></td><td></td><td></td><td></td><td></td><td></td><td></td><td></td><td></td><td></td><td></td></tr>
<tr><td>Oligo rev</td><td>AAACACCCAGGTGATCTCACAGAC</td><td></td><td></td><td></td><td></td><td></td><td></td><td></td><td></td><td></td><td></td><td></td></tr>
</table>

| | | | | | | | | | | | |
|---|---|---|---|---|---|---|---|---|---|---|---|
| Chromosome | start | end | strand | MM | target_seq | PAM | alignment | distance | position | gene name | gene id |
| GU305914.1 | 187829 | 187851 | - | 0 | GTCTGTGAGATCACCTGGGT | CGG | \|\|\|\|\|\|\|\|[\|\|\|\|\|\|\|\|\|\|\|\|]PAM | NA | Intergenic | NA | NA |
| T20437 | TCTGTGAGATCACCTGGGTCGGG | 1000 | | | | | | | | | |
| Oligo fwd | CACCTCTGTGAGATCACCTGGGTC | | | | | | | | | | |
| Oligo rev | AAACGACCCAGGTGATCTCACAGA | | | | | | | | | | |
| Chromosome | start | end | strand | MM | target_seq | PAM | alignment | distance | position | gene name | gene id |
| GU305914.1 | 187828 | 187850 | - | 0 | TCTGTGAGATCACCTGGGTC | GGG | \|\|\|\|\|\|\|\|[\|\|\|\|\|\|\|\|\|\|\|\|]PAM | NA | Intergenic | NA | NA |
| T20438 | GAGATCACCTGGGTCGGGAACGG | 1000 | | | | | | | | | |
| Oligo fwd | CACCGAGATCACCTGGGTCGGGAA | | | | | | | | | | |
| Oligo rev | AAACTTCCCGACCCAGGTGATCTC | | | | | | | | | | |
| Chromosome | start | end | strand | MM | target_seq | PAM | alignment | distance | position | gene name | gene id |
| GU305914.1 | 187823 | 187845 | - | 0 | GAGATCACCTGGGTCGGGAA | CGG | \|\|\|\|\|\|\|\|[\|\|\|\|\|\|\|\|\|\|\|\|]PAM | NA | Intergenic | NA | NA |
| T20439 | GGGTCGGGAACGGTACGAACTGG | 1000 | | | | | | | | | |
| Oligo fwd | CACCGGGTCGGGAACGGTACGAAC | | | | | | | | | | |
| Oligo rev | AAACGTTCGTACCGTTCCCGACCC | | | | | | | | | | |
| Chromosome | start | end | strand | MM | target_seq | PAM | alignment | distance | position | gene name | gene id |
| GU305914.1 | 187813 | 187835 | - | 0 | GGGTCGGGAACGGTACGAAC | TGG | \|\|\|\|\|\|\|\|[\|\|\|\|\|\|\|\|\|\|\|\|]PAM | NA | Intergenic | NA | NA |
| T20440 | AACGGTACGAACTGGCCGCTCGG | 1000 | | | | | | | | | |
| Oligo fwd | CACCAACGGTACGAACTGGCCGCT | | | | | | | | | | |
| Oligo rev | AAACAGCGGCCAGTTCGTACCGTT | | | | | | | | | | |
| Chromosome | start | end | strand | MM | target_seq | PAM | alignment | distance | position | gene name | gene id |
| GU305914.1 | 187805 | 187827 | - | 0 | AACGGTACGAACTGGCCGCT | CGG | \|\|\|\|\|\|\|\|[\|\|\|\|\|\|\|\|\|\|\|\|]PAM | NA | Intergenic | NA | NA |
| T20441 | CATACATCCGTTCAAACCCACGG | 1000 | | | | | | | | | |
| Oligo fwd | CACCCATACATCCGTTCAAACCCA | | | | | | | | | | |
| Oligo rev | AAACTGGGTTTGAACGGATGTATG | | | | | | | | | | |
| Chromosome | start | end | strand | MM | target_seq | PAM | alignment | distance | position | gene name | gene id |
| GU305914.1 | 187782 | 187804 | - | 0 | CATACATCCGTTCAAACCCA | CGG | \|\|\|\|\|\|\|\|[\|\|\|\|\|\|\|\|\|\|\|\|]PAM | NA | Intergenic | NA | NA |
| T20442 | ATCCGTTCAAACCCACGGTGTGG | 1000 | | | | | | | | | |
| Oligo fwd | CACCATCCGTTCAAACCCACGGTG | | | | | | | | | | |
| Oligo rev | AAACCACCGTGGGTTTGAACGGAT | | | | | | | | | | |
| Chromosome | start | end | strand | MM | target_seq | PAM | alignment | distance | position | gene name | gene id |
| GU305914.1 | 187777 | 187799 | - | 0 | ATCCGTTCAAACCCACGGTG | TGG | \|\|\|\|\|\|\|\|[\|\|\|\|\|\|\|\|\|\|\|\|]PAM | NA | Intergenic | NA | NA |
| T20443 | ACGTTGCAGATGACGTTCTCCGG | 1000 | | | | | | | | | |
| Oligo fwd | CACCACGTTGCAGATGACGTTCTC | | | | | | | | | | |
| Oligo rev | AAACGAGAACGTCATCTGCAACGT | | | | | | | | | | |
| Chromosome | start | end | strand | MM | target_seq | PAM | alignment | distance | position | gene name | gene id |
| GU305914.1 | 187745 | 187767 | - | 0 | ACGTTGCAGATGACGTTCTC | CGG | \|\|\|\|\|\|\|\|[\|\|\|\|\|\|\|\|\|\|\|\|]PAM | NA | Intergenic | NA | NA |
| T20444 | GCAGATGACGTTCTCCGGCAAGG | 1000 | | | | | | | | | |
| Oligo fwd | CACCGCAGATGACGTTCTCCGGCA | | | | | | | | | | |
| Oligo rev | AAACTGCCGGAGAACGTCATCTGC | | | | | | | | | | |
| Chromosome | start | end | strand | MM | target_seq | PAM | alignment | distance | position | gene name | gene id |
| GU305914.1 | 187740 | 187762 | - | 0 | GCAGATGACGTTCTCCGGCA | AGG | \|\|\|\|\|\|\|\|[\|\|\|\|\|\|\|\|\|\|\|\|]PAM | NA | Intergenic | NA | NA |
| T20445 | GCTCGCACGTGTCCGAGAGACGG | 1000 | | | | | | | | | |
| Oligo fwd | CACCGCTCGCACGTGTCCGAGAGA | | | | | | | | | | |
| Oligo rev | AAACTCTCTCGGACACGTGCGAGC | | | | | | | | | | |
| Chromosome | start | end | strand | MM | target_seq | PAM | alignment | distance | position | gene name | gene id |
| GU305914.1 | 187687 | 187709 | - | 0 | GCTCGCACGTGTCCGAGAGA | CGG | \|\|\|\|\|\|\|\|[\|\|\|\|\|\|\|\|\|\|\|\|]PAM | NA | Intergenic | NA | NA |
| T20446 | CGCACGTGTCCGAGAGACGGTGG | 1000 | | | | | | | | | |
| Oligo fwd | CACCCGCACGTGTCCGAGAGACGG | | | | | | | | | | |
| Oligo rev | AAACCCGTCTCTCGGACACGTGCG | | | | | | | | | | |
| Chromosome | start | end | strand | MM | target_seq | PAM | alignment | distance | position | gene name | gene id |
| GU305914.1 | 187684 | 187706 | - | 0 | CGCACGTGTCCGAGAGACGG | TGG | \|\|\|\|\|\|\|\|[\|\|\|\|\|\|\|\|\|\|\|\|]PAM | NA | Intergenic | NA | NA |
| T20447 | GTCCGAGAGACGGTGGACCGTGG | 1000 | | | | | | | | | |
| Oligo fwd | CACCGTCCGAGAGACGGTGGACCG | | | | | | | | | | |
| Oligo rev | AAACCGGTCCACCGTCTCTCGGAC | | | | | | | | | | |
| Chromosome | start | end | strand | MM | target_seq | PAM | alignment | distance | position | gene name | gene id |
| GU305914.1 | 187677 | 187699 | - | 0 | GTCCGAGAGACGGTGGACCG | TGG | \|\|\|\|\|\|\|\|[\|\|\|\|\|\|\|\|\|\|\|\|]PAM | NA | Intergenic | NA | NA |
| T20448 | CGTAGCAACTTTATTATCCTCGG | 1000 | | | | | | | | | |
| Oligo fwd | CACCCGTAGCAACTTTATTATCCT | | | | | | | | | | |
| Oligo rev | AAACAGGATAATAAAGTTGCTACG | | | | | | | | | | |
| Chromosome | start | end | strand | MM | target_seq | PAM | alignment | distance | position | gene name | gene id |
| GU305914.1 | 187620 | 187642 | - | 0 | CGTAGCAACTTTATTATCCT | CGG | \|\|\|\|\|\|\|\|[\|\|\|\|\|\|\|\|\|\|\|\|]PAM | NA | Intergenic | NA | NA |
| T20449 | TCCTCGGCCTCCCCCCCTACCGG | 1000 | | | | | | | | | |
| Oligo fwd | CACCTCCTCGGCCTCCCCCCCTAC | | | | | | | | | | |
| Oligo rev | AAACGTAGGGGGGGAGGCCGAGGA | | | | | | | | | | |
| Chromosome | start | end | strand | MM | target_seq | PAM | alignment | distance | position | gene name | gene id |
| GU305914.1 | 187604 | 187626 | - | 0 | TCCTCGGCCTCCCCCCCTAC | CGG | \|\|\|\|\|\|\|\|[\|\|\|\|\|\|\|\|\|\|\|\|]PAM | NA | Intergenic | NA | NA |
| T20450 | AAAAACTTTATTAACCCCCACGG | 1000 | | | | | | | | | |
| Oligo fwd | CACCAAAAACTTTATTAACCCCCA | | | | | | | | | | |
| Oligo rev | AAACTGGGGGTTAATAAAGTTTTT | | | | | | | | | | |
| Chromosome | start | end | strand | MM | target_seq | PAM | alignment | distance | position | gene name | gene id |
| GU305914.1 | 187575 | 187597 | - | 0 | AAAAACTTTATTAACCCCCA | CGG | \|\|\|\|\|\|\|\|[\|\|\|\|\|\|\|\|\|\|\|\|]PAM | NA | Intergenic | NA | NA |
| T20451 | AACTTTATTAACCCCCACGGCGG | 1000 | | | | | | | | | |
| Oligo fwd | CACCAACTTTATTAACCCCCACGG | | | | | | | | | | |

Oligo rev AAACCCGTGGGGGTTAATAAAGTT
Chromosome start end strand MM target_seq PAM alignment distance position gene name gene id
GU305914.1 187572 187594 - 0 AACTTTATTAACCCCCACGG CGG ||||||||[||||||||||||]PAM NA Intergenic NA NA

T20452 AGAACGCATCCGAGACACTCAGG 1000
Oligo fwd CACCAGAACGCATCCGAGACACTC
Oligo rev AAACGAGTGTCTCGGATGCGTTCT
Chromosome start end strand MM target_seq PAM alignment distance position gene name gene id
GU305914.1 187534 187556 - 0 AGAACGCATCCGAGACACTC AGG ||||||||[||||||||||||]PAM NA Intergenic NA NA

T20453 CTACTGCATGCCCCTCACATCGG 1000
Oligo fwd CACCCTACTGCATGCCCCTCACAT
Oligo rev AAACATGTGAGGGGCATGCAGTAG
Chromosome start end strand MM target_seq PAM alignment distance position gene name gene id
GU305914.1 187503 187525 - 0 CTACTGCATGCCCCTCACAT CGG ||||||||[||||||||||||]PAM NA Intergenic NA NA

T20454 TACTGCATGCCCCTCACATCGGG 1000
Oligo fwd CACCTACTGCATGCCCCTCACATC
Oligo rev AAACGATGTGAGGGGCATGCAGTA
Chromosome start end strand MM target_seq PAM alignment distance position gene name gene id
GU305914.1 187502 187524 - 0 TACTGCATGCCCCTCACATC GGG ||||||||[||||||||||||]PAM NA Intergenic NA NA

T20455 ACTGCATGCCCCTCACATCGGGG 1000
Oligo fwd CACCACTGCATGCCCCTCACATCG
Oligo rev AAACCGATGTGAGGGGCATGCAGT
Chromosome start end strand MM target_seq PAM alignment distance position gene name gene id
GU305914.1 187501 187523 - 0 ACTGCATGCCCCTCACATCG GGG ||||||||[||||||||||||]PAM NA Intergenic NA NA

T20456 CTGCATGCCCCTCACATCGGGGG 1000
Oligo fwd CACCCTGCATGCCCCTCACATCGG
Oligo rev AAACCCGATGTGAGGGGCATGCAG
Chromosome start end strand MM target_seq PAM alignment distance position gene name gene id
GU305914.1 187500 187522 - 0 CTGCATGCCCCTCACATCGG GGG ||||||||[||||||||||||]PAM NA Intergenic NA NA

T20457 CACATCGGGGGAACCCCCTCCGG 1000
Oligo fwd CACCCACATCGGGGGAACCCCCTC
Oligo rev AAACGAGGGGGTTCCCCCGATGTG
Chromosome start end strand MM target_seq PAM alignment distance position gene name gene id
GU305914.1 187488 187510 - 0 CACATCGGGGGAACCCCCTC CGG ||||||||[||||||||||||]PAM NA Intergenic NA NA

T20458 ACATCGGGGGAACCCCCTCCGGG 1000
Oligo fwd CACCACATCGGGGGAACCCCCTCC
Oligo rev AAACGGAGGGGGTTCCCCCGATGT
Chromosome start end strand MM target_seq PAM alignment distance position gene name gene id
GU305914.1 187487 187509 - 0 ACATCGGGGGAACCCCCTCC GGG ||||||||[||||||||||||]PAM NA Intergenic NA NA

T20459 CATCGGGGGAACCCCCTCCGGGG 1000
Oligo fwd CACCCATCGGGGGAACCCCCTCCG
Oligo rev AAACCGGAGGGGGTTCCCCCGATG
Chromosome start end strand MM target_seq PAM alignment distance position gene name gene id
GU305914.1 187486 187508 - 0 CATCGGGGGAACCCCCTCCG GGG ||||||||[||||||||||||]PAM NA Intergenic NA NA

T20460 CTCCGGGGCGAGTCTTTTACAGG 1000
Oligo fwd CACCCTCCGGGGCGAGTCTTTTAC
Oligo rev AAACGTAAAAGACTCGCCCCGGAG
Chromosome start end strand MM target_seq PAM alignment distance position gene name gene id
GU305914.1 187471 187493 - 0 CTCCGGGGCGAGTCTTTTAC AGG ||||||||[||||||||||||]PAM NA Intergenic NA NA

T20461 TCACTGAATCTTCTTCCTGACGG 1000
Oligo fwd CACCTCACTGAATCTTCTTCCTGA
Oligo rev AAACTCAGGAAGAAGATTCAGTGA
Chromosome start end strand MM target_seq PAM alignment distance position gene name gene id
GU305914.1 187377 187399 - 0 TCACTGAATCTTCTTCCTGA CGG ||||||||[||||||||||||]PAM NA Intergenic NA NA

T20462 TCTTCTTCCTGACGGTCACCAGG 1000
Oligo fwd CACCTCTTCTTCCTGACGGTCACC
Oligo rev AAACGGTGACCGTCAGGAAGAAGA
Chromosome start end strand MM target_seq PAM alignment distance position gene name gene id
GU305914.1 187369 187391 - 0 TCTTCTTCCTGACGGTCACC AGG ||||||||[||||||||||||]PAM NA Intergenic NA NA

T20463 CCTGACGGTCACCAGGTCTCTGG 1000
Oligo fwd CACCCCTGACGGTCACCAGGTCTC
Oligo rev AAACGAGACCTGGTGACCGTCAGG
Chromosome start end strand MM target_seq PAM alignment distance position gene name gene id
GU305914.1 187362 187384 - 0 CCTGACGGTCACCAGGTCTC TGG ||||||||[||||||||||||]PAM NA Intergenic NA NA

T20464 TGGTCAAGAGAAAGAACCCGTGG 1000
Oligo fwd CACCTGGTCAAGAGAAAGAACCCG
Oligo rev AAACCGGGTTCTTTCTCTTGACCA
Chromosome start end strand MM target_seq PAM alignment distance position gene name gene id
GU305914.1 187342 187364 - 0 TGGTCAAGAGAAAGAACCCG TGG ||||||||[||||||||||||]PAM NA Intergenic NA NA

T20465 GAGAAAGAACCCGTGGTTCTTGG 1000
Oligo fwd CACCGAGAAAGAACCCGTGGTTCT
Oligo rev AAACAGAACCACGGGTTCTTTCTC
Chromosome start end strand MM target_seq PAM alignment distance position gene name gene id
GU305914.1 187335 187357 - 0 GAGAAAGAACCCGTGGTTCT TGG ||||||||[||||||||||||]PAM NA Intergenic NA NA

T20466 GTGGTTCTTGGTCCCAAACGAGG 1000

<table>
<tr><td>Oligo fwd</td><td>CACCGTGGTTCTTGGTCCCAAACG</td><td></td><td></td><td></td><td></td><td></td><td></td><td></td><td></td><td></td><td></td></tr>
<tr><td>Oligo rev</td><td>AAACCGTTTGGGACCAAGAACCAC</td><td></td><td></td><td></td><td></td><td></td><td></td><td></td><td></td><td></td><td></td></tr>
<tr><td>Chromosome</td><td>start</td><td>end</td><td>strand</td><td>MM</td><td>target_seq</td><td>PAM</td><td>alignment</td><td>distance</td><td>position</td><td>gene name</td><td>gene id</td></tr>
<tr><td>GU305914.1</td><td>187323</td><td>187345</td><td>-</td><td>0</td><td>GTGGTTCTTGGTCCCAAACG</td><td>AGG</td><td>||||||||[||||||||||||]PAM</td><td>NA</td><td>Intergenic</td><td>NA</td><td>NA</td></tr>
<tr><td>T20467</td><td>AAACGAGGTCTCCTCGTACTCGG</td><td>1000</td><td></td><td></td><td></td><td></td><td></td><td></td><td></td><td></td><td></td></tr>
<tr><td>Oligo fwd</td><td>CACCAAACGAGGTCTCCTCGTACT</td><td></td><td></td><td></td><td></td><td></td><td></td><td></td><td></td><td></td><td></td></tr>
<tr><td>Oligo rev</td><td>AAACAGTACGAGGAGACCTCGTTT</td><td></td><td></td><td></td><td></td><td></td><td></td><td></td><td></td><td></td><td></td></tr>
<tr><td>Chromosome</td><td>start</td><td>end</td><td>strand</td><td>MM</td><td>target_seq</td><td>PAM</td><td>alignment</td><td>distance</td><td>position</td><td>gene name</td><td>gene id</td></tr>
<tr><td>GU305914.1</td><td>187308</td><td>187330</td><td>-</td><td>0</td><td>AAACGAGGTCTCCTCGTACT</td><td>CGG</td><td>||||||||[||||||||||||]PAM</td><td>NA</td><td>Intergenic</td><td>NA</td><td>NA</td></tr>
<tr><td>T20468</td><td>GACTATCTCCCTTAACCATCTGG</td><td>1000</td><td></td><td></td><td></td><td></td><td></td><td></td><td></td><td></td><td></td></tr>
<tr><td>Oligo fwd</td><td>CACCGACTATCTCCCTTAACCATC</td><td></td><td></td><td></td><td></td><td></td><td></td><td></td><td></td><td></td><td></td></tr>
<tr><td>Oligo rev</td><td>AAACGATGGTTAAGGGAGATAGTC</td><td></td><td></td><td></td><td></td><td></td><td></td><td></td><td></td><td></td><td></td></tr>
<tr><td>Chromosome</td><td>start</td><td>end</td><td>strand</td><td>MM</td><td>target_seq</td><td>PAM</td><td>alignment</td><td>distance</td><td>position</td><td>gene name</td><td>gene id</td></tr>
<tr><td>GU305914.1</td><td>187279</td><td>187301</td><td>-</td><td>0</td><td>GACTATCTCCCTTAACCATC</td><td>TGG</td><td>||||||||[||||||||||||]PAM</td><td>NA</td><td>Intergenic</td><td>NA</td><td>NA</td></tr>
<tr><td>T20469</td><td>GGAAATTGAAATAAGCCCTCCGG</td><td>1000</td><td></td><td></td><td></td><td></td><td></td><td></td><td></td><td></td><td></td></tr>
<tr><td>Oligo fwd</td><td>CACCGGAAATTGAAATAAGCCCTC</td><td></td><td></td><td></td><td></td><td></td><td></td><td></td><td></td><td></td><td></td></tr>
<tr><td>Oligo rev</td><td>AAACGAGGGCTTATTTCAATTTCC</td><td></td><td></td><td></td><td></td><td></td><td></td><td></td><td></td><td></td><td></td></tr>
<tr><td>Chromosome</td><td>start</td><td>end</td><td>strand</td><td>MM</td><td>target_seq</td><td>PAM</td><td>alignment</td><td>distance</td><td>position</td><td>gene name</td><td>gene id</td></tr>
<tr><td>GU305914.1</td><td>187258</td><td>187280</td><td>-</td><td>0</td><td>GGAAATTGAAATAAGCCCTC</td><td>CGG</td><td>||||||||[||||||||||||]PAM</td><td>NA</td><td>Intergenic</td><td>NA</td><td>NA</td></tr>
<tr><td>T20470</td><td>GAAATTGAAATAAGCCCTCCGGG</td><td>1000</td><td></td><td></td><td></td><td></td><td></td><td></td><td></td><td></td><td></td></tr>
<tr><td>Oligo fwd</td><td>CACCGAAATTGAAATAAGCCCTCC</td><td></td><td></td><td></td><td></td><td></td><td></td><td></td><td></td><td></td><td></td></tr>
<tr><td>Oligo rev</td><td>AAACGGAGGGCTTATTTCAATTTC</td><td></td><td></td><td></td><td></td><td></td><td></td><td></td><td></td><td></td><td></td></tr>
<tr><td>Chromosome</td><td>start</td><td>end</td><td>strand</td><td>MM</td><td>target_seq</td><td>PAM</td><td>alignment</td><td>distance</td><td>position</td><td>gene name</td><td>gene id</td></tr>
<tr><td>GU305914.1</td><td>187257</td><td>187279</td><td>-</td><td>0</td><td>GAAATTGAAATAAGCCCTCC</td><td>GGG</td><td>||||||||[||||||||||||]PAM</td><td>NA</td><td>Intergenic</td><td>NA</td><td>NA</td></tr>
<tr><td>T20471</td><td>GGGTGCCCATCCGCATGAACTGG</td><td>1000</td><td></td><td></td><td></td><td></td><td></td><td></td><td></td><td></td><td></td></tr>
<tr><td>Oligo fwd</td><td>CACCGGGTGCCCATCCGCATGAAC</td><td></td><td></td><td></td><td></td><td></td><td></td><td></td><td></td><td></td><td></td></tr>
<tr><td>Oligo rev</td><td>AAACGTTCATGCGGATGGGCACCC</td><td></td><td></td><td></td><td></td><td></td><td></td><td></td><td></td><td></td><td></td></tr>
<tr><td>Chromosome</td><td>start</td><td>end</td><td>strand</td><td>MM</td><td>target_seq</td><td>PAM</td><td>alignment</td><td>distance</td><td>position</td><td>gene name</td><td>gene id</td></tr>
<tr><td>GU305914.1</td><td>187237</td><td>187259</td><td>-</td><td>0</td><td>GGGTGCCCATCCGCATGAAC</td><td>TGG</td><td>||||||||[||||||||||||]PAM</td><td>NA</td><td>Intergenic</td><td>NA</td><td>NA</td></tr>
<tr><td>T20472</td><td>ATCTAACATGTGCCCGTAGATGG</td><td>1000</td><td></td><td></td><td></td><td></td><td></td><td></td><td></td><td></td><td></td></tr>
<tr><td>Oligo fwd</td><td>CACCATCTAACATGTGCCCGTAGA</td><td></td><td></td><td></td><td></td><td></td><td></td><td></td><td></td><td></td><td></td></tr>
<tr><td>Oligo rev</td><td>AAACTCTACGGGCACATGTTAGAT</td><td></td><td></td><td></td><td></td><td></td><td></td><td></td><td></td><td></td><td></td></tr>
<tr><td>Chromosome</td><td>start</td><td>end</td><td>strand</td><td>MM</td><td>target_seq</td><td>PAM</td><td>alignment</td><td>distance</td><td>position</td><td>gene name</td><td>gene id</td></tr>
<tr><td>GU305914.1</td><td>187179</td><td>187201</td><td>-</td><td>0</td><td>ATCTAACATGTGCCCGTAGA</td><td>TGG</td><td>||||||||[||||||||||||]PAM</td><td>NA</td><td>Intergenic</td><td>NA</td><td>NA</td></tr>
<tr><td>T20473</td><td>GTGCCCGTAGATGGCCCCCCCGG</td><td>1000</td><td></td><td></td><td></td><td></td><td></td><td></td><td></td><td></td><td></td></tr>
<tr><td>Oligo fwd</td><td>CACCGTGCCCGTAGATGGCCCCCC</td><td></td><td></td><td></td><td></td><td></td><td></td><td></td><td></td><td></td><td></td></tr>
<tr><td>Oligo rev</td><td>AAACGGGGGGCCATCTACGGGCAC</td><td></td><td></td><td></td><td></td><td></td><td></td><td></td><td></td><td></td><td></td></tr>
<tr><td>Chromosome</td><td>start</td><td>end</td><td>strand</td><td>MM</td><td>target_seq</td><td>PAM</td><td>alignment</td><td>distance</td><td>position</td><td>gene name</td><td>gene id</td></tr>
<tr><td>GU305914.1</td><td>187170</td><td>187192</td><td>-</td><td>0</td><td>GTGCCCGTAGATGGCCCCCC</td><td>CGG</td><td>||||||||[||||||||||||]PAM</td><td>NA</td><td>Intergenic</td><td>NA</td><td>NA</td></tr>
<tr><td>T20474</td><td>CATGATAAACATCTCACACTCGG</td><td>1000</td><td></td><td></td><td></td><td></td><td></td><td></td><td></td><td></td><td></td></tr>
<tr><td>Oligo fwd</td><td>CACCCATGATAAACATCTCACACT</td><td></td><td></td><td></td><td></td><td></td><td></td><td></td><td></td><td></td><td></td></tr>
<tr><td>Oligo rev</td><td>AAACAGTGTGAGATGTTTATCATG</td><td></td><td></td><td></td><td></td><td></td><td></td><td></td><td></td><td></td><td></td></tr>
<tr><td>Chromosome</td><td>start</td><td>end</td><td>strand</td><td>MM</td><td>target_seq</td><td>PAM</td><td>alignment</td><td>distance</td><td>position</td><td>gene name</td><td>gene id</td></tr>
<tr><td>GU305914.1</td><td>187140</td><td>187162</td><td>-</td><td>0</td><td>CATGATAAACATCTCACACT</td><td>CGG</td><td>||||||||[||||||||||||]PAM</td><td>NA</td><td>Intergenic</td><td>NA</td><td>NA</td></tr>
<tr><td>T20475</td><td>ATGATAAACATCTCACACTCGGG</td><td>1000</td><td></td><td></td><td></td><td></td><td></td><td></td><td></td><td></td><td></td></tr>
<tr><td>Oligo fwd</td><td>CACCATGATAAACATCTCACACTC</td><td></td><td></td><td></td><td></td><td></td><td></td><td></td><td></td><td></td><td></td></tr>
<tr><td>Oligo rev</td><td>AAACGAGTGTGAGATGTTTATCAT</td><td></td><td></td><td></td><td></td><td></td><td></td><td></td><td></td><td></td><td></td></tr>
<tr><td>Chromosome</td><td>start</td><td>end</td><td>strand</td><td>MM</td><td>target_seq</td><td>PAM</td><td>alignment</td><td>distance</td><td>position</td><td>gene name</td><td>gene id</td></tr>
<tr><td>GU305914.1</td><td>187139</td><td>187161</td><td>-</td><td>0</td><td>ATGATAAACATCTCACACTC</td><td>GGG</td><td>||||||||[||||||||||||]PAM</td><td>NA</td><td>Intergenic</td><td>NA</td><td>NA</td></tr>
<tr><td>T20476</td><td>CATCTCACACTCGGGATCCTCGG</td><td>1000</td><td></td><td></td><td></td><td></td><td></td><td></td><td></td><td></td><td></td></tr>
<tr><td>Oligo fwd</td><td>CACCCATCTCACACTCGGGATCCT</td><td></td><td></td><td></td><td></td><td></td><td></td><td></td><td></td><td></td><td></td></tr>
<tr><td>Oligo rev</td><td>AAACAGGATCCCGAGTGTGAGATG</td><td></td><td></td><td></td><td></td><td></td><td></td><td></td><td></td><td></td><td></td></tr>
<tr><td>Chromosome</td><td>start</td><td>end</td><td>strand</td><td>MM</td><td>target_seq</td><td>PAM</td><td>alignment</td><td>distance</td><td>position</td><td>gene name</td><td>gene id</td></tr>
<tr><td>GU305914.1</td><td>187131</td><td>187153</td><td>-</td><td>0</td><td>CATCTCACACTCGGGATCCT</td><td>CGG</td><td>||||||||[||||||||||||]PAM</td><td>NA</td><td>Intergenic</td><td>NA</td><td>NA</td></tr>
<tr><td>T20477</td><td>ATCTCACACTCGGGATCCTCGGG</td><td>1000</td><td></td><td></td><td></td><td></td><td></td><td></td><td></td><td></td><td></td></tr>
<tr><td>Oligo fwd</td><td>CACCATCTCACACTCGGGATCCTC</td><td></td><td></td><td></td><td></td><td></td><td></td><td></td><td></td><td></td><td></td></tr>
<tr><td>Oligo rev</td><td>AAACGAGGATCCCGAGTGTGAGAT</td><td></td><td></td><td></td><td></td><td></td><td></td><td></td><td></td><td></td><td></td></tr>
<tr><td>Chromosome</td><td>start</td><td>end</td><td>strand</td><td>MM</td><td>target_seq</td><td>PAM</td><td>alignment</td><td>distance</td><td>position</td><td>gene name</td><td>gene id</td></tr>
<tr><td>GU305914.1</td><td>187130</td><td>187152</td><td>-</td><td>0</td><td>ATCTCACACTCGGGATCCTC</td><td>GGG</td><td>||||||||[||||||||||||]PAM</td><td>NA</td><td>Intergenic</td><td>NA</td><td>NA</td></tr>
<tr><td>T20478</td><td>TCTCACACTCGGGATCCTCGGGG</td><td>1000</td><td></td><td></td><td></td><td></td><td></td><td></td><td></td><td></td><td></td></tr>
<tr><td>Oligo fwd</td><td>CACCTCTCACACTCGGGATCCTCG</td><td></td><td></td><td></td><td></td><td></td><td></td><td></td><td></td><td></td><td></td></tr>
<tr><td>Oligo rev</td><td>AAACCGAGGATCCCGAGTGTGAGA</td><td></td><td></td><td></td><td></td><td></td><td></td><td></td><td></td><td></td><td></td></tr>
<tr><td>Chromosome</td><td>start</td><td>end</td><td>strand</td><td>MM</td><td>target_seq</td><td>PAM</td><td>alignment</td><td>distance</td><td>position</td><td>gene name</td><td>gene id</td></tr>
<tr><td>GU305914.1</td><td>187129</td><td>187151</td><td>-</td><td>0</td><td>TCTCACACTCGGGATCCTCG</td><td>GGG</td><td>||||||||[||||||||||||]PAM</td><td>NA</td><td>Intergenic</td><td>NA</td><td>NA</td></tr>
<tr><td>T20479</td><td>CTCACACTCGGGATCCTCGGGGG</td><td>1000</td><td></td><td></td><td></td><td></td><td></td><td></td><td></td><td></td><td></td></tr>
<tr><td>Oligo fwd</td><td>CACCCTCACACTCGGGATCCTCGG</td><td></td><td></td><td></td><td></td><td></td><td></td><td></td><td></td><td></td><td></td></tr>
<tr><td>Oligo rev</td><td>AAACCCGAGGATCCCGAGTGTGAG</td><td></td><td></td><td></td><td></td><td></td><td></td><td></td><td></td><td></td><td></td></tr>
<tr><td>Chromosome</td><td>start</td><td>end</td><td>strand</td><td>MM</td><td>target_seq</td><td>PAM</td><td>alignment</td><td>distance</td><td>position</td><td>gene name</td><td>gene id</td></tr>
<tr><td>GU305914.1</td><td>187128</td><td>187150</td><td>-</td><td>0</td><td>CTCACACTCGGGATCCTCGG</td><td>GGG</td><td>||||||||[||||||||||||]PAM</td><td>NA</td><td>Intergenic</td><td>NA</td><td>NA</td></tr>
<tr><td>T20480</td><td>ACTCGGGATCCTCGGGGGAACGG</td><td>1000</td><td></td><td></td><td></td><td></td><td></td><td></td><td></td><td></td><td></td></tr>
<tr><td>Oligo fwd</td><td>CACCACTCGGGATCCTCGGGGGAA</td><td></td><td></td><td></td><td></td><td></td><td></td><td></td><td></td><td></td><td></td></tr>
<tr><td>Oligo rev</td><td>AAACTTCCCCCGAGGATCCCGAGT</td><td></td><td></td><td></td><td></td><td></td><td></td><td></td><td></td><td></td><td></td></tr>
<tr><td>Chromosome</td><td>start</td><td>end</td><td>strand</td><td>MM</td><td>target_seq</td><td>PAM</td><td>alignment</td><td>distance</td><td>position</td><td>gene name</td><td>gene id</td></tr>
<tr><td>GU305914.1</td><td>187123</td><td>187145</td><td>-</td><td>0</td><td>ACTCGGGATCCTCGGGGGAA</td><td>CGG</td><td>||||||||[||||||||||||]PAM</td><td>NA</td><td>Intergenic</td><td>NA</td><td>NA</td></tr>
</table>

<table>
<tr><td>T20481</td><td>CTCGGGATCCTCGGGGGAACGGG</td><td>1000</td><td colspan="9"></td></tr>
<tr><td>Oligo fwd</td><td>CACCCTCGGGATCCTCGGGGGAAC</td><td colspan="10"></td></tr>
<tr><td>Oligo rev</td><td>AAACGTTCCCCCGAGGATCCCGAG</td><td colspan="10"></td></tr>
<tr><td>Chromosome</td><td>start</td><td>end</td><td>strand</td><td>MM</td><td>target_seq</td><td>PAM</td><td>alignment</td><td>distance</td><td>position</td><td>gene name</td><td>gene id</td></tr>
<tr><td>GU305914.1</td><td>187122</td><td>187144</td><td>-</td><td>0</td><td>CTCGGGATCCTCGGGGGAAC</td><td>GGG</td><td>||||||||[||||||||||||]PAM</td><td>NA</td><td>Intergenic</td><td>NA</td><td>NA</td></tr>
<tr><td>T20482</td><td>GGAACGGGTCACCCTACCGATGG</td><td>1000</td><td colspan="9"></td></tr>
<tr><td>Oligo fwd</td><td>CACCGGAACGGGTCACCCTACCGA</td><td colspan="10"></td></tr>
<tr><td>Oligo rev</td><td>AAACTCGGTAGGGTGACCCGTTCC</td><td colspan="10"></td></tr>
<tr><td>Chromosome</td><td>start</td><td>end</td><td>strand</td><td>MM</td><td>target_seq</td><td>PAM</td><td>alignment</td><td>distance</td><td>position</td><td>gene name</td><td>gene id</td></tr>
<tr><td>GU305914.1</td><td>187107</td><td>187129</td><td>-</td><td>0</td><td>GGAACGGGTCACCCTACCGA</td><td>TGG</td><td>||||||||[||||||||||||]PAM</td><td>NA</td><td>Intergenic</td><td>NA</td><td>NA</td></tr>
<tr><td>T20483</td><td>GAACGGGTCACCCTACCGATGGG</td><td>1000</td><td colspan="9"></td></tr>
<tr><td>Oligo fwd</td><td>CACCGAACGGGTCACCCTACCGAT</td><td colspan="10"></td></tr>
<tr><td>Oligo rev</td><td>AAACATCGGTAGGGTGACCCGTTC</td><td colspan="10"></td></tr>
<tr><td>Chromosome</td><td>start</td><td>end</td><td>strand</td><td>MM</td><td>target_seq</td><td>PAM</td><td>alignment</td><td>distance</td><td>position</td><td>gene name</td><td>gene id</td></tr>
<tr><td>GU305914.1</td><td>187106</td><td>187128</td><td>-</td><td>0</td><td>GAACGGGTCACCCTACCGAT</td><td>GGG</td><td>||||||||[||||||||||||]PAM</td><td>NA</td><td>Intergenic</td><td>NA</td><td>NA</td></tr>
<tr><td>T20484</td><td>ACCTATGTCCAAAGAGTTCCAGG</td><td>1000</td><td colspan="9"></td></tr>
<tr><td>Oligo fwd</td><td>CACCACCTATGTCCAAAGAGTTCC</td><td colspan="10"></td></tr>
<tr><td>Oligo rev</td><td>AAACGGAACTCTTTGGACATAGGT</td><td colspan="10"></td></tr>
<tr><td>Chromosome</td><td>start</td><td>end</td><td>strand</td><td>MM</td><td>target_seq</td><td>PAM</td><td>alignment</td><td>distance</td><td>position</td><td>gene name</td><td>gene id</td></tr>
<tr><td>GU305914.1</td><td>187078</td><td>187100</td><td>-</td><td>0</td><td>ACCTATGTCCAAAGAGTTCC</td><td>AGG</td><td>||||||||[||||||||||||]PAM</td><td>NA</td><td>Intergenic</td><td>NA</td><td>NA</td></tr>
<tr><td>T20485</td><td>CAAAGAGTTCCAGGTTCACGTGG</td><td>1000</td><td colspan="9"></td></tr>
<tr><td>Oligo fwd</td><td>CACCCAAAGAGTTCCAGGTTCACG</td><td colspan="10"></td></tr>
<tr><td>Oligo rev</td><td>AAACCGTGAACCTGGAACTCTTTG</td><td colspan="10"></td></tr>
<tr><td>Chromosome</td><td>start</td><td>end</td><td>strand</td><td>MM</td><td>target_seq</td><td>PAM</td><td>alignment</td><td>distance</td><td>position</td><td>gene name</td><td>gene id</td></tr>
<tr><td>GU305914.1</td><td>187069</td><td>187091</td><td>-</td><td>0</td><td>CAAAGAGTTCCAGGTTCACG</td><td>TGG</td><td>||||||||[||||||||||||]PAM</td><td>NA</td><td>Intergenic</td><td>NA</td><td>NA</td></tr>
<tr><td>T20486</td><td>AGAGTTCCAGGTTCACGTGGCGG</td><td>1000</td><td colspan="9"></td></tr>
<tr><td>Oligo fwd</td><td>CACCAGAGTTCCAGGTTCACGTGG</td><td colspan="10"></td></tr>
<tr><td>Oligo rev</td><td>AAACCCACGTGAACCTGGAACTCT</td><td colspan="10"></td></tr>
<tr><td>Chromosome</td><td>start</td><td>end</td><td>strand</td><td>MM</td><td>target_seq</td><td>PAM</td><td>alignment</td><td>distance</td><td>position</td><td>gene name</td><td>gene id</td></tr>
<tr><td>GU305914.1</td><td>187066</td><td>187088</td><td>-</td><td>0</td><td>AGAGTTCCAGGTTCACGTGG</td><td>CGG</td><td>||||||||[||||||||||||]PAM</td><td>NA</td><td>Intergenic</td><td>NA</td><td>NA</td></tr>
<tr><td>T20487</td><td>CAGGTTCACGTGGCGGACGAAGG</td><td>1000</td><td colspan="9"></td></tr>
<tr><td>Oligo fwd</td><td>CACCCAGGTTCACGTGGCGGACGA</td><td colspan="10"></td></tr>
<tr><td>Oligo rev</td><td>AAACTCGTCCGCCACGTGAACCTG</td><td colspan="10"></td></tr>
<tr><td>Chromosome</td><td>start</td><td>end</td><td>strand</td><td>MM</td><td>target_seq</td><td>PAM</td><td>alignment</td><td>distance</td><td>position</td><td>gene name</td><td>gene id</td></tr>
<tr><td>GU305914.1</td><td>187059</td><td>187081</td><td>-</td><td>0</td><td>CAGGTTCACGTGGCGGACGA</td><td>AGG</td><td>||||||||[||||||||||||]PAM</td><td>NA</td><td>Intergenic</td><td>NA</td><td>NA</td></tr>
<tr><td>T20488</td><td>AGGTTCACGTGGCGGACGAAGGG</td><td>1000</td><td colspan="9"></td></tr>
<tr><td>Oligo fwd</td><td>CACCAGGTTCACGTGGCGGACGAA</td><td colspan="10"></td></tr>
<tr><td>Oligo rev</td><td>AAACTTCGTCCGCCACGTGAACCT</td><td colspan="10"></td></tr>
<tr><td>Chromosome</td><td>start</td><td>end</td><td>strand</td><td>MM</td><td>target_seq</td><td>PAM</td><td>alignment</td><td>distance</td><td>position</td><td>gene name</td><td>gene id</td></tr>
<tr><td>GU305914.1</td><td>187058</td><td>187080</td><td>-</td><td>0</td><td>AGGTTCACGTGGCGGACGAA</td><td>GGG</td><td>||||||||[||||||||||||]PAM</td><td>NA</td><td>Intergenic</td><td>NA</td><td>NA</td></tr>
<tr><td>T20489</td><td>GGTTCACGTGGCGGACGAAGGGG</td><td>1000</td><td colspan="9"></td></tr>
<tr><td>Oligo fwd</td><td>CACCGGTTCACGTGGCGGACGAAG</td><td colspan="10"></td></tr>
<tr><td>Oligo rev</td><td>AAACCTTCGTCCGCCACGTGAACC</td><td colspan="10"></td></tr>
<tr><td>Chromosome</td><td>start</td><td>end</td><td>strand</td><td>MM</td><td>target_seq</td><td>PAM</td><td>alignment</td><td>distance</td><td>position</td><td>gene name</td><td>gene id</td></tr>
<tr><td>GU305914.1</td><td>187057</td><td>187079</td><td>-</td><td>0</td><td>GGTTCACGTGGCGGACGAAG</td><td>GGG</td><td>||||||||[||||||||||||]PAM</td><td>NA</td><td>Intergenic</td><td>NA</td><td>NA</td></tr>
<tr><td>T20490</td><td>GTTCACGTGGCGGACGAAGGGGG</td><td>1000</td><td colspan="9"></td></tr>
<tr><td>Oligo fwd</td><td>CACCGTTCACGTGGCGGACGAAGG</td><td colspan="10"></td></tr>
<tr><td>Oligo rev</td><td>AAACCCTTCGTCCGCCACGTGAAC</td><td colspan="10"></td></tr>
<tr><td>Chromosome</td><td>start</td><td>end</td><td>strand</td><td>MM</td><td>target_seq</td><td>PAM</td><td>alignment</td><td>distance</td><td>position</td><td>gene name</td><td>gene id</td></tr>
<tr><td>GU305914.1</td><td>187056</td><td>187078</td><td>-</td><td>0</td><td>GTTCACGTGGCGGACGAAGG</td><td>GGG</td><td>||||||||[||||||||||||]PAM</td><td>NA</td><td>Intergenic</td><td>NA</td><td>NA</td></tr>
<tr><td>T20491</td><td>TTCACGTGGCGGACGAAGGGGGG</td><td>1000</td><td colspan="9"></td></tr>
<tr><td>Oligo fwd</td><td>CACCTTCACGTGGCGGACGAAGGG</td><td colspan="10"></td></tr>
<tr><td>Oligo rev</td><td>AAACCCCTTCGTCCGCCACGTGAA</td><td colspan="10"></td></tr>
<tr><td>Chromosome</td><td>start</td><td>end</td><td>strand</td><td>MM</td><td>target_seq</td><td>PAM</td><td>alignment</td><td>distance</td><td>position</td><td>gene name</td><td>gene id</td></tr>
<tr><td>GU305914.1</td><td>187055</td><td>187077</td><td>-</td><td>0</td><td>TTCACGTGGCGGACGAAGGG</td><td>GGG</td><td>||||||||[||||||||||||]PAM</td><td>NA</td><td>Intergenic</td><td>NA</td><td>NA</td></tr>
<tr><td>T20492</td><td>TCACGTGGCGGACGAAGGGGGGG</td><td>1000</td><td colspan="9"></td></tr>
<tr><td>Oligo fwd</td><td>CACCTCACGTGGCGGACGAAGGGG</td><td colspan="10"></td></tr>
<tr><td>Oligo rev</td><td>AAACCCCCTTCGTCCGCCACGTGA</td><td colspan="10"></td></tr>
<tr><td>Chromosome</td><td>start</td><td>end</td><td>strand</td><td>MM</td><td>target_seq</td><td>PAM</td><td>alignment</td><td>distance</td><td>position</td><td>gene name</td><td>gene id</td></tr>
<tr><td>GU305914.1</td><td>187054</td><td>187076</td><td>-</td><td>0</td><td>TCACGTGGCGGACGAAGGGG</td><td>GGG</td><td>||||||||[||||||||||||]PAM</td><td>NA</td><td>Intergenic</td><td>NA</td><td>NA</td></tr>
<tr><td>T20493</td><td>CGGACGAAGGGGGGGATGTCCGG</td><td>1000</td><td colspan="9"></td></tr>
<tr><td>Oligo fwd</td><td>CACCCGGACGAAGGGGGGGATGTC</td><td colspan="10"></td></tr>
<tr><td>Oligo rev</td><td>AAACGACATCCCCCCCTTCGTCCG</td><td colspan="10"></td></tr>
<tr><td>Chromosome</td><td>start</td><td>end</td><td>strand</td><td>MM</td><td>target_seq</td><td>PAM</td><td>alignment</td><td>distance</td><td>position</td><td>gene name</td><td>gene id</td></tr>
<tr><td>GU305914.1</td><td>187046</td><td>187068</td><td>-</td><td>0</td><td>CGGACGAAGGGGGGGATGTC</td><td>CGG</td><td>||||||||[||||||||||||]PAM</td><td>NA</td><td>Intergenic</td><td>NA</td><td>NA</td></tr>
<tr><td>T20494</td><td>GAAGGGGGGGATGTCCGGCGTGG</td><td>1000</td><td colspan="9"></td></tr>
<tr><td>Oligo fwd</td><td>CACCGAAGGGGGGGATGTCCGGCG</td><td colspan="10"></td></tr>
<tr><td>Oligo rev</td><td>AAACCGCCGGACATCCCCCCCTTC</td><td colspan="10"></td></tr>
<tr><td>Chromosome</td><td>start</td><td>end</td><td>strand</td><td>MM</td><td>target_seq</td><td>PAM</td><td>alignment</td><td>distance</td><td>position</td><td>gene name</td><td>gene id</td></tr>
<tr><td>GU305914.1</td><td>187041</td><td>187063</td><td>-</td><td>0</td><td>GAAGGGGGGGATGTCCGGCG</td><td>TGG</td><td>||||||||[||||||||||||]PAM</td><td>NA</td><td>Intergenic</td><td>NA</td><td>NA</td></tr>
<tr><td>T20495</td><td>CACCCTGTCCTTCAGCATGCCGG</td><td>1000</td><td colspan="9"></td></tr>
<tr><td>Oligo fwd</td><td>CACCCACCCTGTCCTTCAGCATGC</td><td colspan="10"></td></tr>
<tr><td>Oligo rev</td><td>AAACGCATGCTGAAGGACAGGGTG</td><td colspan="10"></td></tr>
<tr><td>Chromosome</td><td>start</td><td>end</td><td>strand</td><td>MM</td><td>target_seq</td><td>PAM</td><td>alignment</td><td>distance</td><td>position</td><td>gene name</td><td>gene id</td></tr>
<tr><td>GU305914.1</td><td>187011</td><td>187033</td><td>-</td><td>0</td><td>CACCCTGTCCTTCAGCATGC</td><td>CGG</td><td>||||||||[||||||||||||]PAM</td><td>NA</td><td>Intergenic</td><td>NA</td><td>NA</td></tr>
</table>

T20496 ACCCTGTCCTTCAGCATGCCGGG 1000
Oligo fwd CACCACCCTGTCCTTCAGCATGCC
Oligo rev AAACGGCATGCTGAAGGACAGGGT
Chromosome start end strand MM target_seq PAM alignment distance position gene name gene id
GU305914.1 187010 187032 - 0 ACCCTGTCCTTCAGCATGCC GGG ||||||||[||||||||||||]PAM NA Intergenic NA NA

T20497 CCCTGTCCTTCAGCATGCCGGGG 1000
Oligo fwd CACCCCCTGTCCTTCAGCATGCCG
Oligo rev AAACCGGCATGCTGAAGGACAGGG
Chromosome start end strand MM target_seq PAM alignment distance position gene name gene id
GU305914.1 187009 187031 - 0 CCCTGTCCTTCAGCATGCCG GGG ||||||||[||||||||||||]PAM NA Intergenic NA NA

T20498 GCATGCCGGGGCAACCGTTCAGG 1000
Oligo fwd CACCGCATGCCGGGGCAACCGTTC
Oligo rev AAACGAACGGTTGCCCCGGCATGC
Chromosome start end strand MM target_seq PAM alignment distance position gene name gene id
GU305914.1 186997 187019 - 0 GCATGCCGGGGCAACCGTTC AGG ||||||||[||||||||||||]PAM NA Intergenic NA NA

T20499 GGGGCAACCGTTCAGGTACCAGG 1000
Oligo fwd CACCGGGGCAACCGTTCAGGTACC
Oligo rev AAACGGTACCTGAACGGTTGCCCC
Chromosome start end strand MM target_seq PAM alignment distance position gene name gene id
GU305914.1 186990 187012 - 0 GGGGCAACCGTTCAGGTACC AGG ||||||||[||||||||||||]PAM NA Intergenic NA NA

T20500 CCTTGATCTTACAGATCTGCTGG 1000
Oligo fwd CACCCCTTGATCTTACAGATCTGC
Oligo rev AAACGCAGATCTGTAAGATCAAGG
Chromosome start end strand MM target_seq PAM alignment distance position gene name gene id
GU305914.1 186961 186983 - 0 CCTTGATCTTACAGATCTGC TGG ||||||||[||||||||||||]PAM NA Intergenic NA NA

T20501 TGATCTTACAGATCTGCTGGTGG 1000
Oligo fwd CACCTGATCTTACAGATCTGCTGG
Oligo rev AAACCCAGCAGATCTGTAAGATCA
Chromosome start end strand MM target_seq PAM alignment distance position gene name gene id
GU305914.1 186958 186980 - 0 TGATCTTACAGATCTGCTGG TGG ||||||||[||||||||||||]PAM NA Intergenic NA NA

T20502 GGAGCGCCTTCCCGCCGAGATGG 1000
Oligo fwd CACCGGAGCGCCTTCCCGCCGAGA
Oligo rev AAACTCTCGGCGGGAAGGCGCTCC
Chromosome start end strand MM target_seq PAM alignment distance position gene name gene id
GU305914.1 186937 186959 - 0 GGAGCGCCTTCCCGCCGAGA TGG ||||||||[||||||||||||]PAM NA Intergenic NA NA

T20503 GAGCGCCTTCCCGCCGAGATGGG 1000
Oligo fwd CACCGAGCGCCTTCCCGCCGAGAT
Oligo rev AAACATCTCGGCGGGAAGGCGCTC
Chromosome start end strand MM target_seq PAM alignment distance position gene name gene id
GU305914.1 186936 186958 - 0 GAGCGCCTTCCCGCCGAGAT GGG ||||||||[||||||||||||]PAM NA Intergenic NA NA

T20504 AGCGCCTTCCCGCCGAGATGGGG 1000
Oligo fwd CACCAGCGCCTTCCCGCCGAGATG
Oligo rev AAACCATCTCGGCGGGAAGGCGCT
Chromosome start end strand MM target_seq PAM alignment distance position gene name gene id
GU305914.1 186935 186957 - 0 AGCGCCTTCCCGCCGAGATG GGG ||||||||[||||||||||||]PAM NA Intergenic NA NA

T20505 GAACACGATGTTCTTGAGCACGG 1000
Oligo fwd CACCGAACACGATGTTCTTGAGCA
Oligo rev AAACTGCTCAAGAACATCGTGTTC
Chromosome start end strand MM target_seq PAM alignment distance position gene name gene id
GU305914.1 186909 186931 - 0 GAACACGATGTTCTTGAGCA CGG ||||||||[||||||||||||]PAM NA Intergenic NA NA

T20506 AACACGATGTTCTTGAGCACGGG 1000
Oligo fwd CACCAACACGATGTTCTTGAGCAC
Oligo rev AAACGTGCTCAAGAACATCGTGTT
Chromosome start end strand MM target_seq PAM alignment distance position gene name gene id
GU305914.1 186908 186930 - 0 AACACGATGTTCTTGAGCAC GGG ||||||||[||||||||||||]PAM NA Intergenic NA NA

T20507 ACACGATGTTCTTGAGCACGGGG 1000
Oligo fwd CACCACACGATGTTCTTGAGCACG
Oligo rev AAACCGTGCTCAAGAACATCGTGT
Chromosome start end strand MM target_seq PAM alignment distance position gene name gene id
GU305914.1 186907 186929 - 0 ACACGATGTTCTTGAGCACG GGG ||||||||[||||||||||||]PAM NA Intergenic NA NA

T20508 GTTCTTGAGCACGGGGTCCTCGG 1000
Oligo fwd CACCGTTCTTGAGCACGGGGTCCT
Oligo rev AAACAGGACCCCGTGCTCAAGAAC
Chromosome start end strand MM target_seq PAM alignment distance position gene name gene id
GU305914.1 186900 186922 - 0 GTTCTTGAGCACGGGGTCCT CGG ||||||||[||||||||||||]PAM NA Intergenic NA NA

T20509 TTCTTGAGCACGGGGTCCTCGGG 1000
Oligo fwd CACCTTCTTGAGCACGGGGTCCTC
Oligo rev AAACGAGGACCCCGTGCTCAAGAA
Chromosome start end strand MM target_seq PAM alignment distance position gene name gene id
GU305914.1 186899 186921 - 0 TTCTTGAGCACGGGGTCCTC GGG ||||||||[||||||||||||]PAM NA Intergenic NA NA

T20510 GGGTCCTCGGGAGTAGCGTACGG 1000
Oligo fwd CACCGGGTCCTCGGGAGTAGCGTA
Oligo rev AAACTACGCTACTCCCGAGGACCC
Chromosome start end strand MM target_seq PAM alignment distance position gene name gene id

| | | | | | | | | | | | |
|---|---|---|---|---|---|---|---|---|---|---|---|
| GU305914.1 | 186887 | 186909 | - | 0 | GGGTCCTCGGGAGTAGCGTA | CGG | \|\|\|\|\|\|\|\|[\|\|\|\|\|\|\|\|\|\|\|\|]PAM | NA | Intergenic | NA | NA |
| T20511 | GTACGGAGCGCCGCCGTCCTCGG | 1000 | | | | | | | | | |
| Oligo fwd | CACCGTACGGAGCGCCGCCGTCCT | | | | | | | | | | |
| Oligo rev | AAACAGGACGGCGGCGCTCCGTAC | | | | | | | | | | |
| Chromosome | start | end | strand | MM | target_seq | PAM | alignment | distance | position | gene name | gene id |
| GU305914.1 | 186870 | 186892 | - | 0 | GTACGGAGCGCCGCCGTCCT | CGG | \|\|\|\|\|\|\|\|[\|\|\|\|\|\|\|\|\|\|\|\|]PAM | NA | Intergenic | NA | NA |
| T20512 | GCAGAGTCCTTCCCTACCGAAGG | 1000 | | | | | | | | | |
| Oligo fwd | CACCGCAGAGTCCTTCCCTACCGA | | | | | | | | | | |
| Oligo rev | AAACTCGGTAGGGAAGGACTCTGC | | | | | | | | | | |
| Chromosome | start | end | strand | MM | target_seq | PAM | alignment | distance | position | gene name | gene id |
| GU305914.1 | 186828 | 186850 | - | 0 | GCAGAGTCCTTCCCTACCGA | AGG | \|\|\|\|\|\|\|\|[\|\|\|\|\|\|\|\|\|\|\|\|]PAM | NA | Intergenic | NA | NA |
| T20513 | CTTCCCTACCGAAGGACTGCAGG | 1000 | | | | | | | | | |
| Oligo fwd | CACCCTTCCCTACCGAAGGACTGC | | | | | | | | | | |
| Oligo rev | AAACGCAGTCCTTCGGTAGGGAAG | | | | | | | | | | |
| Chromosome | start | end | strand | MM | target_seq | PAM | alignment | distance | position | gene name | gene id |
| GU305914.1 | 186820 | 186842 | - | 0 | CTTCCCTACCGAAGGACTGC | AGG | \|\|\|\|\|\|\|\|[\|\|\|\|\|\|\|\|\|\|\|\|]PAM | NA | Intergenic | NA | NA |
| T20514 | TTCCCTACCGAAGGACTGCAGGG | 1000 | | | | | | | | | |
| Oligo fwd | CACCTTCCCTACCGAAGGACTGCA | | | | | | | | | | |
| Oligo rev | AAACTGCAGTCCTTCGGTAGGGAA | | | | | | | | | | |
| Chromosome | start | end | strand | MM | target_seq | PAM | alignment | distance | position | gene name | gene id |
| GU305914.1 | 186819 | 186841 | - | 0 | TTCCCTACCGAAGGACTGCA | GGG | \|\|\|\|\|\|\|\|[\|\|\|\|\|\|\|\|\|\|\|\|]PAM | NA | Intergenic | NA | NA |
| T20515 | ACCGAAGGACTGCAGGGACTCGG | 1000 | | | | | | | | | |
| Oligo fwd | CACCACCGAAGGACTGCAGGGACT | | | | | | | | | | |
| Oligo rev | AAACAGTCCCTGCAGTCCTTCGGT | | | | | | | | | | |
| Chromosome | start | end | strand | MM | target_seq | PAM | alignment | distance | position | gene name | gene id |
| GU305914.1 | 186813 | 186835 | - | 0 | ACCGAAGGACTGCAGGGACT | CGG | \|\|\|\|\|\|\|\|[\|\|\|\|\|\|\|\|\|\|\|\|]PAM | NA | Intergenic | NA | NA |
| T20516 | GAAGGACTGCAGGGACTCGGCGG | 1000 | | | | | | | | | |
| Oligo fwd | CACCGAAGGACTGCAGGGACTCGG | | | | | | | | | | |
| Oligo rev | AAACCCGAGTCCCTGCAGTCCTTC | | | | | | | | | | |
| Chromosome | start | end | strand | MM | target_seq | PAM | alignment | distance | position | gene name | gene id |
| GU305914.1 | 186810 | 186832 | - | 0 | GAAGGACTGCAGGGACTCGG | CGG | \|\|\|\|\|\|\|\|[\|\|\|\|\|\|\|\|\|\|\|\|]PAM | NA | Intergenic | NA | NA |
| T20517 | ACTGCAGGGACTCGGCGGCCAGG | 1000 | | | | | | | | | |
| Oligo fwd | CACCACTGCAGGGACTCGGCGGCC | | | | | | | | | | |
| Oligo rev | AAACGGCCGCCGAGTCCCTGCAGT | | | | | | | | | | |
| Chromosome | start | end | strand | MM | target_seq | PAM | alignment | distance | position | gene name | gene id |
| GU305914.1 | 186805 | 186827 | - | 0 | ACTGCAGGGACTCGGCGGCC | AGG | \|\|\|\|\|\|\|\|[\|\|\|\|\|\|\|\|\|\|\|\|]PAM | NA | Intergenic | NA | NA |
| T20518 | GGCGGCCAGGAAGAGTCGATCGG | 1000 | | | | | | | | | |
| Oligo fwd | CACCGGCGGCCAGGAAGAGTCGAT | | | | | | | | | | |
| Oligo rev | AAACATCGACTCTTCCTGGCCGCC | | | | | | | | | | |
| Chromosome | start | end | strand | MM | target_seq | PAM | alignment | distance | position | gene name | gene id |
| GU305914.1 | 186792 | 186814 | - | 0 | GGCGGCCAGGAAGAGTCGAT | CGG | \|\|\|\|\|\|\|\|[\|\|\|\|\|\|\|\|\|\|\|\|]PAM | NA | Intergenic | NA | NA |
| T20519 | AGTCGATCGGCGTCCCGCTCCGG | 1000 | | | | | | | | | |
| Oligo fwd | CACCAGTCGATCGGCGTCCCGCTC | | | | | | | | | | |
| Oligo rev | AAACGAGCGGGACGCCGATCGACT | | | | | | | | | | |
| Chromosome | start | end | strand | MM | target_seq | PAM | alignment | distance | position | gene name | gene id |
| GU305914.1 | 186779 | 186801 | - | 0 | AGTCGATCGGCGTCCCGCTC | CGG | \|\|\|\|\|\|\|\|[\|\|\|\|\|\|\|\|\|\|\|\|]PAM | NA | Intergenic | NA | NA |
| T20520 | GTCCCGCTCCGGATCGTAGTCGG | 1000 | | | | | | | | | |
| Oligo fwd | CACCGTCCCGCTCCGGATCGTAGT | | | | | | | | | | |
| Oligo rev | AAACACTACGATCCGGAGCGGGAC | | | | | | | | | | |
| Chromosome | start | end | strand | MM | target_seq | PAM | alignment | distance | position | gene name | gene id |
| GU305914.1 | 186768 | 186790 | - | 0 | GTCCCGCTCCGGATCGTAGT | CGG | \|\|\|\|\|\|\|\|[\|\|\|\|\|\|\|\|\|\|\|\|]PAM | NA | Intergenic | NA | NA |
| T20521 | TCCCGCTCCGGATCGTAGTCGGG | 1000 | | | | | | | | | |
| Oligo fwd | CACCTCCCGCTCCGGATCGTAGTC | | | | | | | | | | |
| Oligo rev | AAACGACTACGATCCGGAGCGGGA | | | | | | | | | | |
| Chromosome | start | end | strand | MM | target_seq | PAM | alignment | distance | position | gene name | gene id |
| GU305914.1 | 186767 | 186789 | - | 0 | TCCCGCTCCGGATCGTAGTC | GGG | \|\|\|\|\|\|\|\|[\|\|\|\|\|\|\|\|\|\|\|\|]PAM | NA | Intergenic | NA | NA |
| T20522 | ATCGTAGTCGGGACTCCAGACGG | 1000 | | | | | | | | | |
| Oligo fwd | CACCATCGTAGTCGGGACTCCAGA | | | | | | | | | | |
| Oligo rev | AAACTCTGGAGTCCCGACTACGAT | | | | | | | | | | |
| Chromosome | start | end | strand | MM | target_seq | PAM | alignment | distance | position | gene name | gene id |
| GU305914.1 | 186756 | 186778 | - | 0 | ATCGTAGTCGGGACTCCAGA | CGG | \|\|\|\|\|\|\|\|[\|\|\|\|\|\|\|\|\|\|\|\|]PAM | NA | Intergenic | NA | NA |
| T20523 | TCGTAGTCGGGACTCCAGACGGG | 1000 | | | | | | | | | |
| Oligo fwd | CACCTCGTAGTCGGGACTCCAGAC | | | | | | | | | | |
| Oligo rev | AAACGTCTGGAGTCCCGACTACGA | | | | | | | | | | |
| Chromosome | start | end | strand | MM | target_seq | PAM | alignment | distance | position | gene name | gene id |
| GU305914.1 | 186755 | 186777 | - | 0 | TCGTAGTCGGGACTCCAGAC | GGG | \|\|\|\|\|\|\|\|[\|\|\|\|\|\|\|\|\|\|\|\|]PAM | NA | Intergenic | NA | NA |
| T20524 | ACTCCAGACGGGCCTGCCTCTGG | 1000 | | | | | | | | | |
| Oligo fwd | CACCACTCCAGACGGGCCTGCCTC | | | | | | | | | | |
| Oligo rev | AAACGAGGCAGGCCCGTCTGGAGT | | | | | | | | | | |
| Chromosome | start | end | strand | MM | target_seq | PAM | alignment | distance | position | gene name | gene id |
| GU305914.1 | 186744 | 186766 | - | 0 | ACTCCAGACGGGCCTGCCTC | TGG | \|\|\|\|\|\|\|\|[\|\|\|\|\|\|\|\|\|\|\|\|]PAM | NA | Intergenic | NA | NA |
| T20525 | AGACGGGCCTGCCTCTGGCGTGG | 1000 | | | | | | | | | |
| Oligo fwd | CACCAGACGGGCCTGCCTCTGGCG | | | | | | | | | | |
| Oligo rev | AAACCGCCAGAGGCAGGCCCGTCT | | | | | | | | | | |

Chromosome start end strand MM target_seq PAM alignment distance position gene name gene id
GU305914.1 186739 186761 - 0 AGACGGGCCTGCCTCTGGCG TGG ||||||||[||||||||||||]PAM NA Intergenic NA NA

T20526 CGGGCCTGCCTCTGGCGTGGAGG 1000
Oligo fwd CACCCGGGCCTGCCTCTGGCGTGG
Oligo rev AAACCCACGCCAGAGGCAGGCCCG
Chromosome start end strand MM target_seq PAM alignment distance position gene name gene id
GU305914.1 186736 186758 - 0 CGGGCCTGCCTCTGGCGTGG AGG ||||||||[||||||||||||]PAM NA Intergenic NA NA

T20527 GCCTCTGGCGTGGAGGAAGATGG 1000
Oligo fwd CACCGCCTCTGGCGTGGAGGAAGA
Oligo rev AAACTCTTCCTCCACGCCAGAGGC
Chromosome start end strand MM target_seq PAM alignment distance position gene name gene id
GU305914.1 186729 186751 - 0 GCCTCTGGCGTGGAGGAAGA TGG ||||||||[||||||||||||]PAM NA Intergenic NA NA
GU305914.1 53426 53448 - 3 GCATCTGGCGTGGTGGAGGA CGG ||-|||||[|||||-|||-||]PAM NA Intergenic NA NA

T20528 GGCGTGGAGGAAGATGGTCCCGG 1000
Oligo fwd CACCGGCGTGGAGGAAGATGGTCC
Oligo rev AAACGGACCATCTTCCTCCACGCC
Chromosome start end strand MM target_seq PAM alignment distance position gene name gene id
GU305914.1 186723 186745 - 0 GGCGTGGAGGAAGATGGTCC CGG ||||||||[||||||||||||]PAM NA Intergenic NA NA

T20529 GGAGGAAGATGGTCCCGGTGAGG 1000
Oligo fwd CACCGGAGGAAGATGGTCCCGGTG
Oligo rev AAACCACCGGGACCATCTTCCTCC
Chromosome start end strand MM target_seq PAM alignment distance position gene name gene id
GU305914.1 186718 186740 - 0 GGAGGAAGATGGTCCCGGTG AGG ||||||||[||||||||||||]PAM NA Intergenic NA NA

T20530 TCCCGGTGAGGCCCATCAAGAGG 1000
Oligo fwd CACCTCCCGGTGAGGCCCATCAAG
Oligo rev AAACCTTGATGGGCCTCACCGGGA
Chromosome start end strand MM target_seq PAM alignment distance position gene name gene id
GU305914.1 186706 186728 - 0 TCCCGGTGAGGCCCATCAAG AGG ||||||||[||||||||||||]PAM NA Intergenic NA NA

T20531 GGTGAGGCCCATCAAGAGGACGG 1000
Oligo fwd CACCGGTGAGGCCCATCAAGAGGA
Oligo rev AAACTCCTCTTGATGGGCCTCACC
Chromosome start end strand MM target_seq PAM alignment distance position gene name gene id
GU305914.1 186702 186724 - 0 GGTGAGGCCCATCAAGAGGA CGG ||||||||[||||||||||||]PAM NA Intergenic NA NA

T20532 GTGAGGCCCATCAAGAGGACGGG 1000
Oligo fwd CACCGTGAGGCCCATCAAGAGGAC
Oligo rev AAACGTCCTCTTGATGGGCCTCAC
Chromosome start end strand MM target_seq PAM alignment distance position gene name gene id
GU305914.1 186701 186723 - 0 GTGAGGCCCATCAAGAGGAC GGG ||||||||[||||||||||||]PAM NA Intergenic NA NA

T20533 TGAGGCCCATCAAGAGGACGGGG 1000
Oligo fwd CACCTGAGGCCCATCAAGAGGACG
Oligo rev AAACCGTCCTCTTGATGGGCCTCA
Chromosome start end strand MM target_seq PAM alignment distance position gene name gene id
GU305914.1 186700 186722 - 0 TGAGGCCCATCAAGAGGACG GGG ||||||||[||||||||||||]PAM NA Intergenic NA NA

T20534 CCATCAAGAGGACGGGGTGCTGG 1000
Oligo fwd CACCCCATCAAGAGGACGGGGTGC
Oligo rev AAACGCACCCCGTCCTCTTGATGG
Chromosome start end strand MM target_seq PAM alignment distance position gene name gene id
GU305914.1 186694 186716 - 0 CCATCAAGAGGACGGGGTGC TGG ||||||||[||||||||||||]PAM NA Intergenic NA NA

T20535 CGGGGTGCTGGTCGCTGTAGAGG 1000
Oligo fwd CACCCGGGGTGCTGGTCGCTGTAG
Oligo rev AAACCTACAGCGACCAGCACCCCG
Chromosome start end strand MM target_seq PAM alignment distance position gene name gene id
GU305914.1 186682 186704 - 0 CGGGGTGCTGGTCGCTGTAG AGG ||||||||[||||||||||||]PAM NA Intergenic NA NA

T20536 GGGGTGCTGGTCGCTGTAGAGGG 1000
Oligo fwd CACCGGGGTGCTGGTCGCTGTAGA
Oligo rev AAACTCTACAGCGACCAGCACCCC
Chromosome start end strand MM target_seq PAM alignment distance position gene name gene id
GU305914.1 186681 186703 - 0 GGGGTGCTGGTCGCTGTAGA GGG ||||||||[||||||||||||]PAM NA Intergenic NA NA

T20537 GTCGCTGTAGAGGGACTGTCCGG 1000
Oligo fwd CACCGTCGCTGTAGAGGGACTGTC
Oligo rev AAACGACAGTCCCTCTACAGCGAC
Chromosome start end strand MM target_seq PAM alignment distance position gene name gene id
GU305914.1 186672 186694 - 0 GTCGCTGTAGAGGGACTGTC CGG ||||||||[||||||||||||]PAM NA Intergenic NA NA

T20538 TCGCTGTAGAGGGACTGTCCGGG 1000
Oligo fwd CACCTCGCTGTAGAGGGACTGTCC
Oligo rev AAACGGACAGTCCCTCTACAGCGA
Chromosome start end strand MM target_seq PAM alignment distance position gene name gene id
GU305914.1 186671 186693 - 0 TCGCTGTAGAGGGACTGTCC GGG ||||||||[||||||||||||]PAM NA Intergenic NA NA

T20539 AGAGGGACTGTCCGGGTCCGAGG 1000
Oligo fwd CACCAGAGGGACTGTCCGGGTCCG
Oligo rev AAACCGGACCCGGACAGTCCCTCT
Chromosome start end strand MM target_seq PAM alignment distance position gene name gene id
GU305914.1 186664 186686 - 0 AGAGGGACTGTCCGGGTCCG AGG ||||||||[||||||||||||]PAM NA Intergenic NA NA

T20540 GGATCACGCCGAGAACCTCGAGG 1000

<table>
<tr><td>Oligo fwd</td><td>CACCGGATCACGCCGAGAACCTCG</td><td></td><td></td><td></td><td></td><td></td><td></td><td></td><td></td><td></td><td></td></tr>
<tr><td>Oligo rev</td><td>AAACCGAGGTTCTCGGCGTGATCC</td><td></td><td></td><td></td><td></td><td></td><td></td><td></td><td></td><td></td><td></td></tr>
<tr><td>Chromosome</td><td>start</td><td>end</td><td>strand</td><td>MM</td><td>target_seq</td><td>PAM</td><td>alignment</td><td>distance</td><td>position</td><td>gene name</td><td>gene id</td></tr>
<tr><td>GU305914.1</td><td>186643</td><td>186665</td><td>-</td><td>0</td><td>GGATCACGCCGAGAACCTCG</td><td>AGG</td><td>||||||||[||||||||||||]PAM</td><td>NA</td><td>Intergenic</td><td>NA</td><td>NA</td></tr>
<tr><td>T20541</td><td>GATCACGCCGAGAACCTCGAGGG</td><td>1000</td><td></td><td></td><td></td><td></td><td></td><td></td><td></td><td></td><td></td></tr>
<tr><td>Oligo fwd</td><td>CACCGATCACGCCGAGAACCTCGA</td><td></td><td></td><td></td><td></td><td></td><td></td><td></td><td></td><td></td><td></td></tr>
<tr><td>Oligo rev</td><td>AAACTCGAGGTTCTCGGCGTGATC</td><td></td><td></td><td></td><td></td><td></td><td></td><td></td><td></td><td></td><td></td></tr>
<tr><td>Chromosome</td><td>start</td><td>end</td><td>strand</td><td>MM</td><td>target_seq</td><td>PAM</td><td>alignment</td><td>distance</td><td>position</td><td>gene name</td><td>gene id</td></tr>
<tr><td>GU305914.1</td><td>186642</td><td>186664</td><td>-</td><td>0</td><td>GATCACGCCGAGAACCTCGA</td><td>GGG</td><td>||||||||[||||||||||||]PAM</td><td>NA</td><td>Intergenic</td><td>NA</td><td>NA</td></tr>
<tr><td>GU305914.1</td><td>147800</td><td>147822</td><td>-</td><td>4</td><td>GGTCACCCTGAGCACCTCGA</td><td>AGG</td><td>|-||||-|[-|||-|||||||]PAM</td><td>NA</td><td>Intergenic</td><td>NA</td><td>NA</td></tr>
<tr><td>T20542</td><td>ATCACGCCGAGAACCTCGAGGGG</td><td>1000</td><td></td><td></td><td></td><td></td><td></td><td></td><td></td><td></td><td></td></tr>
<tr><td>Oligo fwd</td><td>CACCATCACGCCGAGAACCTCGAG</td><td></td><td></td><td></td><td></td><td></td><td></td><td></td><td></td><td></td><td></td></tr>
<tr><td>Oligo rev</td><td>AAACCTCGAGGTTCTCGGCGTGAT</td><td></td><td></td><td></td><td></td><td></td><td></td><td></td><td></td><td></td><td></td></tr>
<tr><td>Chromosome</td><td>start</td><td>end</td><td>strand</td><td>MM</td><td>target_seq</td><td>PAM</td><td>alignment</td><td>distance</td><td>position</td><td>gene name</td><td>gene id</td></tr>
<tr><td>GU305914.1</td><td>186641</td><td>186663</td><td>-</td><td>0</td><td>ATCACGCCGAGAACCTCGAG</td><td>GGG</td><td>||||||||[||||||||||||]PAM</td><td>NA</td><td>Intergenic</td><td>NA</td><td>NA</td></tr>
<tr><td>GU305914.1</td><td>152200</td><td>152222</td><td>+</td><td>4</td><td>AGCACGTCGAGTACCTCGAC</td><td>AGG</td><td>|-||||-|[|||-|||||||-]PAM</td><td>NA</td><td>Intergenic</td><td>NA</td><td>NA</td></tr>
<tr><td>T20543</td><td>CTCGAGGGGACCGTTGCACATGG</td><td>1000</td><td></td><td></td><td></td><td></td><td></td><td></td><td></td><td></td><td></td></tr>
<tr><td>Oligo fwd</td><td>CACCCTCGAGGGGACCGTTGCACA</td><td></td><td></td><td></td><td></td><td></td><td></td><td></td><td></td><td></td><td></td></tr>
<tr><td>Oligo rev</td><td>AAACTGTGCAACGGTCCCCTCGAG</td><td></td><td></td><td></td><td></td><td></td><td></td><td></td><td></td><td></td><td></td></tr>
<tr><td>Chromosome</td><td>start</td><td>end</td><td>strand</td><td>MM</td><td>target_seq</td><td>PAM</td><td>alignment</td><td>distance</td><td>position</td><td>gene name</td><td>gene id</td></tr>
<tr><td>GU305914.1</td><td>186627</td><td>186649</td><td>-</td><td>0</td><td>CTCGAGGGGACCGTTGCACA</td><td>TGG</td><td>||||||||[||||||||||||]PAM</td><td>NA</td><td>Intergenic</td><td>NA</td><td>NA</td></tr>
<tr><td>T20544</td><td>TCGAGGGGACCGTTGCACATGGG</td><td>1000</td><td></td><td></td><td></td><td></td><td></td><td></td><td></td><td></td><td></td></tr>
<tr><td>Oligo fwd</td><td>CACCTCGAGGGGACCGTTGCACAT</td><td></td><td></td><td></td><td></td><td></td><td></td><td></td><td></td><td></td><td></td></tr>
<tr><td>Oligo rev</td><td>AAACATGTGCAACGGTCCCCTCGA</td><td></td><td></td><td></td><td></td><td></td><td></td><td></td><td></td><td></td><td></td></tr>
<tr><td>Chromosome</td><td>start</td><td>end</td><td>strand</td><td>MM</td><td>target_seq</td><td>PAM</td><td>alignment</td><td>distance</td><td>position</td><td>gene name</td><td>gene id</td></tr>
<tr><td>GU305914.1</td><td>186626</td><td>186648</td><td>-</td><td>0</td><td>TCGAGGGGACCGTTGCACAT</td><td>GGG</td><td>||||||||[||||||||||||]PAM</td><td>NA</td><td>Intergenic</td><td>NA</td><td>NA</td></tr>
<tr><td>T20545</td><td>CGAGGGGACCGTTGCACATGGGG</td><td>1000</td><td></td><td></td><td></td><td></td><td></td><td></td><td></td><td></td><td></td></tr>
<tr><td>Oligo fwd</td><td>CACCCGAGGGGACCGTTGCACATG</td><td></td><td></td><td></td><td></td><td></td><td></td><td></td><td></td><td></td><td></td></tr>
<tr><td>Oligo rev</td><td>AAACCATGTGCAACGGTCCCCTCG</td><td></td><td></td><td></td><td></td><td></td><td></td><td></td><td></td><td></td><td></td></tr>
<tr><td>Chromosome</td><td>start</td><td>end</td><td>strand</td><td>MM</td><td>target_seq</td><td>PAM</td><td>alignment</td><td>distance</td><td>position</td><td>gene name</td><td>gene id</td></tr>
<tr><td>GU305914.1</td><td>186625</td><td>186647</td><td>-</td><td>0</td><td>CGAGGGGACCGTTGCACATG</td><td>GGG</td><td>||||||||[||||||||||||]PAM</td><td>NA</td><td>Intergenic</td><td>NA</td><td>NA</td></tr>
<tr><td>T20546</td><td>CCAGCTTTCTGCCCGACGTCAGG</td><td>1000</td><td></td><td></td><td></td><td></td><td></td><td></td><td></td><td></td><td></td></tr>
<tr><td>Oligo fwd</td><td>CACCCCAGCTTTCTGCCCGACGTC</td><td></td><td></td><td></td><td></td><td></td><td></td><td></td><td></td><td></td><td></td></tr>
<tr><td>Oligo rev</td><td>AAACGACGTCGGGCAGAAAGCTGG</td><td></td><td></td><td></td><td></td><td></td><td></td><td></td><td></td><td></td><td></td></tr>
<tr><td>Chromosome</td><td>start</td><td>end</td><td>strand</td><td>MM</td><td>target_seq</td><td>PAM</td><td>alignment</td><td>distance</td><td>position</td><td>gene name</td><td>gene id</td></tr>
<tr><td>GU305914.1</td><td>186565</td><td>186587</td><td>-</td><td>0</td><td>CCAGCTTTCTGCCCGACGTC</td><td>AGG</td><td>||||||||[||||||||||||]PAM</td><td>NA</td><td>Intergenic</td><td>NA</td><td>NA</td></tr>
<tr><td>T20547</td><td>TTCTGCCCGACGTCAGGCGAAGG</td><td>1000</td><td></td><td></td><td></td><td></td><td></td><td></td><td></td><td></td><td></td></tr>
<tr><td>Oligo fwd</td><td>CACCTTCTGCCCGACGTCAGGCGA</td><td></td><td></td><td></td><td></td><td></td><td></td><td></td><td></td><td></td><td></td></tr>
<tr><td>Oligo rev</td><td>AAACTCGCCTGACGTCGGGCAGAA</td><td></td><td></td><td></td><td></td><td></td><td></td><td></td><td></td><td></td><td></td></tr>
<tr><td>Chromosome</td><td>start</td><td>end</td><td>strand</td><td>MM</td><td>target_seq</td><td>PAM</td><td>alignment</td><td>distance</td><td>position</td><td>gene name</td><td>gene id</td></tr>
<tr><td>GU305914.1</td><td>186559</td><td>186581</td><td>-</td><td>0</td><td>TTCTGCCCGACGTCAGGCGA</td><td>AGG</td><td>||||||||[||||||||||||]PAM</td><td>NA</td><td>Intergenic</td><td>NA</td><td>NA</td></tr>
<tr><td>T20548</td><td>GTCAGGCGAAGGTACCTGTTCGG</td><td>1000</td><td></td><td></td><td></td><td></td><td></td><td></td><td></td><td></td><td></td></tr>
<tr><td>Oligo fwd</td><td>CACCGTCAGGCGAAGGTACCTGTT</td><td></td><td></td><td></td><td></td><td></td><td></td><td></td><td></td><td></td><td></td></tr>
<tr><td>Oligo rev</td><td>AAACAACAGGTACCTTCGCCTGAC</td><td></td><td></td><td></td><td></td><td></td><td></td><td></td><td></td><td></td><td></td></tr>
<tr><td>Chromosome</td><td>start</td><td>end</td><td>strand</td><td>MM</td><td>target_seq</td><td>PAM</td><td>alignment</td><td>distance</td><td>position</td><td>gene name</td><td>gene id</td></tr>
<tr><td>GU305914.1</td><td>186548</td><td>186570</td><td>-</td><td>0</td><td>GTCAGGCGAAGGTACCTGTT</td><td>CGG</td><td>||||||||[||||||||||||]PAM</td><td>NA</td><td>Intergenic</td><td>NA</td><td>NA</td></tr>
<tr><td>T20549</td><td>TCAGGCGAAGGTACCTGTTCGGG</td><td>1000</td><td></td><td></td><td></td><td></td><td></td><td></td><td></td><td></td><td></td></tr>
<tr><td>Oligo fwd</td><td>CACCTCAGGCGAAGGTACCTGTTC</td><td></td><td></td><td></td><td></td><td></td><td></td><td></td><td></td><td></td><td></td></tr>
<tr><td>Oligo rev</td><td>AAACGAACAGGTACCTTCGCCTGA</td><td></td><td></td><td></td><td></td><td></td><td></td><td></td><td></td><td></td><td></td></tr>
<tr><td>Chromosome</td><td>start</td><td>end</td><td>strand</td><td>MM</td><td>target_seq</td><td>PAM</td><td>alignment</td><td>distance</td><td>position</td><td>gene name</td><td>gene id</td></tr>
<tr><td>GU305914.1</td><td>186547</td><td>186569</td><td>-</td><td>0</td><td>TCAGGCGAAGGTACCTGTTC</td><td>GGG</td><td>||||||||[||||||||||||]PAM</td><td>NA</td><td>Intergenic</td><td>NA</td><td>NA</td></tr>
<tr><td>T20550</td><td>CAGGCGAAGGTACCTGTTCGGGG</td><td>1000</td><td></td><td></td><td></td><td></td><td></td><td></td><td></td><td></td><td></td></tr>
<tr><td>Oligo fwd</td><td>CACCCAGGCGAAGGTACCTGTTCG</td><td></td><td></td><td></td><td></td><td></td><td></td><td></td><td></td><td></td><td></td></tr>
<tr><td>Oligo rev</td><td>AAACCGAACAGGTACCTTCGCCTG</td><td></td><td></td><td></td><td></td><td></td><td></td><td></td><td></td><td></td><td></td></tr>
<tr><td>Chromosome</td><td>start</td><td>end</td><td>strand</td><td>MM</td><td>target_seq</td><td>PAM</td><td>alignment</td><td>distance</td><td>position</td><td>gene name</td><td>gene id</td></tr>
<tr><td>GU305914.1</td><td>186546</td><td>186568</td><td>-</td><td>0</td><td>CAGGCGAAGGTACCTGTTCG</td><td>GGG</td><td>||||||||[||||||||||||]PAM</td><td>NA</td><td>Intergenic</td><td>NA</td><td>NA</td></tr>
<tr><td>T20551</td><td>AGGCGAAGGTACCTGTTCGGGGG</td><td>1000</td><td></td><td></td><td></td><td></td><td></td><td></td><td></td><td></td><td></td></tr>
<tr><td>Oligo fwd</td><td>CACCAGGCGAAGGTACCTGTTCGG</td><td></td><td></td><td></td><td></td><td></td><td></td><td></td><td></td><td></td><td></td></tr>
<tr><td>Oligo rev</td><td>AAACCCGAACAGGTACCTTCGCCT</td><td></td><td></td><td></td><td></td><td></td><td></td><td></td><td></td><td></td><td></td></tr>
<tr><td>Chromosome</td><td>start</td><td>end</td><td>strand</td><td>MM</td><td>target_seq</td><td>PAM</td><td>alignment</td><td>distance</td><td>position</td><td>gene name</td><td>gene id</td></tr>
<tr><td>GU305914.1</td><td>186545</td><td>186567</td><td>-</td><td>0</td><td>AGGCGAAGGTACCTGTTCGG</td><td>GGG</td><td>||||||||[||||||||||||]PAM</td><td>NA</td><td>Intergenic</td><td>NA</td><td>NA</td></tr>
<tr><td>T20552</td><td>AAGGTACCTGTTCGGGGGCCAGG</td><td>1000</td><td></td><td></td><td></td><td></td><td></td><td></td><td></td><td></td><td></td></tr>
<tr><td>Oligo fwd</td><td>CACCAAGGTACCTGTTCGGGGGCC</td><td></td><td></td><td></td><td></td><td></td><td></td><td></td><td></td><td></td><td></td></tr>
<tr><td>Oligo rev</td><td>AAACGGCCCCCGAACAGGTACCTT</td><td></td><td></td><td></td><td></td><td></td><td></td><td></td><td></td><td></td><td></td></tr>
<tr><td>Chromosome</td><td>start</td><td>end</td><td>strand</td><td>MM</td><td>target_seq</td><td>PAM</td><td>alignment</td><td>distance</td><td>position</td><td>gene name</td><td>gene id</td></tr>
<tr><td>GU305914.1</td><td>186540</td><td>186562</td><td>-</td><td>0</td><td>AAGGTACCTGTTCGGGGGCC</td><td>AGG</td><td>||||||||[||||||||||||]PAM</td><td>NA</td><td>Intergenic</td><td>NA</td><td>NA</td></tr>
<tr><td>T20553</td><td>ACCTGTTCGGGGGCCAGGCCAGG</td><td>1000</td><td></td><td></td><td></td><td></td><td></td><td></td><td></td><td></td><td></td></tr>
<tr><td>Oligo fwd</td><td>CACCACCTGTTCGGGGGCCAGGCC</td><td></td><td></td><td></td><td></td><td></td><td></td><td></td><td></td><td></td><td></td></tr>
<tr><td>Oligo rev</td><td>AAACGGCCTGGCCCCCGAACAGGT</td><td></td><td></td><td></td><td></td><td></td><td></td><td></td><td></td><td></td><td></td></tr>
<tr><td>Chromosome</td><td>start</td><td>end</td><td>strand</td><td>MM</td><td>target_seq</td><td>PAM</td><td>alignment</td><td>distance</td><td>position</td><td>gene name</td><td>gene id</td></tr>
<tr><td>GU305914.1</td><td>186535</td><td>186557</td><td>-</td><td>0</td><td>ACCTGTTCGGGGGCCAGGCC</td><td>AGG</td><td>||||||||[||||||||||||]PAM</td><td>NA</td><td>Intergenic</td><td>NA</td><td>NA</td></tr>
<tr><td>T20554</td><td>CCTGTTCGGGGGCCAGGCCAGGG</td><td>1000</td><td></td><td></td><td></td><td></td><td></td><td></td><td></td><td></td><td></td></tr>
<tr><td>Oligo fwd</td><td>CACCCCTGTTCGGGGGCCAGGCCA</td><td></td><td></td><td></td><td></td><td></td><td></td><td></td><td></td><td></td><td></td></tr>
<tr><td>Oligo rev</td><td>AAACTGGCCTGGCCCCCGAACAGG</td><td></td><td></td><td></td><td></td><td></td><td></td><td></td><td></td><td></td><td></td></tr>
<tr><td>Chromosome</td><td>start</td><td>end</td><td>strand</td><td>MM</td><td>target_seq</td><td>PAM</td><td>alignment</td><td>distance</td><td>position</td><td>gene name</td><td>gene id</td></tr>
</table>

GU305914.1 186534 186556 - 0 CCTGTTCGGGGGCCAGGCCA GGG ||||||||[||||||||||||]PAM NA Intergenic NA NA

T20555 CTGTTCGGGGGCCAGGCCAGGGG 1000
Oligo fwd CACCCTGTTCGGGGGCCAGGCCAG
Oligo rev AAACCTGGCCTGGCCCCCGAACAG
Chromosome start end strand MM target_seq PAM alignment distance position gene name gene id
GU305914.1 186533 186555 - 0 CTGTTCGGGGGCCAGGCCAG GGG ||||||||[||||||||||||]PAM NA Intergenic NA NA

T20556 GGGGCAGCATCGCTCCATCGTGG 1000
Oligo fwd CACCGGGGCAGCATCGCTCCATCG
Oligo rev AAACCGATGGAGCGATGCTGCCCC
Chromosome start end strand MM target_seq PAM alignment distance position gene name gene id
GU305914.1 186514 186536 - 0 GGGGCAGCATCGCTCCATCG TGG ||||||||[||||||||||||]PAM NA Intergenic NA NA

T20557 ATCGTGGTCCTCCAAGAACTTGG 1000
Oligo fwd CACCATCGTGGTCCTCCAAGAACT
Oligo rev AAACAGTTCTTGGAGGACCACGAT
Chromosome start end strand MM target_seq PAM alignment distance position gene name gene id
GU305914.1 186498 186520 - 0 ATCGTGGTCCTCCAAGAACT TGG ||||||||[||||||||||||]PAM NA Intergenic NA NA

T20558 CCTCCAAGAACTTGGTCAGCTGG 1000
Oligo fwd CACCCCTCCAAGAACTTGGTCAGC
Oligo rev AAACGCTGACCAAGTTCTTGGAGG
Chromosome start end strand MM target_seq PAM alignment distance position gene name gene id
GU305914.1 186490 186512 - 0 CCTCCAAGAACTTGGTCAGC TGG ||||||||[||||||||||||]PAM NA Intergenic NA NA

T20559 CAAGAACTTGGTCAGCTGGCCGG 1000
Oligo fwd CACCCAAGAACTTGGTCAGCTGGC
Oligo rev AAACGCCAGCTGACCAAGTTCTTG
Chromosome start end strand MM target_seq PAM alignment distance position gene name gene id
GU305914.1 186486 186508 - 0 CAAGAACTTGGTCAGCTGGC CGG ||||||||[||||||||||||]PAM NA Intergenic NA NA

T20560 GGTCAGCTGGCCGGACAGAGCGG 1000
Oligo fwd CACCGGTCAGCTGGCCGGACAGAG
Oligo rev AAACCTCTGTCCGGCCAGCTGACC
Chromosome start end strand MM target_seq PAM alignment distance position gene name gene id
GU305914.1 186477 186499 - 0 GGTCAGCTGGCCGGACAGAG CGG ||||||||[||||||||||||]PAM NA Intergenic NA NA

T20561 CGGACAGAGCGGAGACGCCGTGG 1000
Oligo fwd CACCCGGACAGAGCGGAGACGCCG
Oligo rev AAACCGGCGTCTCCGCTCTGTCCG
Chromosome start end strand MM target_seq PAM alignment distance position gene name gene id
GU305914.1 186466 186488 - 0 CGGACAGAGCGGAGACGCCG TGG ||||||||[||||||||||||]PAM NA Intergenic NA NA

T20562 AGCGGAGACGCCGTGGAGTTCGG 1000
Oligo fwd CACCAGCGGAGACGCCGTGGAGTT
Oligo rev AAACAACTCCACGGCGTCTCCGCT
Chromosome start end strand MM target_seq PAM alignment distance position gene name gene id
GU305914.1 186459 186481 - 0 AGCGGAGACGCCGTGGAGTT CGG ||||||||[||||||||||||]PAM NA Intergenic NA NA

T20563 GGAGACGCCGTGGAGTTCGGTGG 1000
Oligo fwd CACCGGAGACGCCGTGGAGTTCGG
Oligo rev AAACCCGAACTCCACGGCGTCTCC
Chromosome start end strand MM target_seq PAM alignment distance position gene name gene id
GU305914.1 186456 186478 - 0 GGAGACGCCGTGGAGTTCGG TGG ||||||||[||||||||||||]PAM NA Intergenic NA NA

T20564 GACGCCGTGGAGTTCGGTGGCGG 1000
Oligo fwd CACCGACGCCGTGGAGTTCGGTGG
Oligo rev AAACCCACCGAACTCCACGGCGTC
Chromosome start end strand MM target_seq PAM alignment distance position gene name gene id
GU305914.1 186453 186475 - 0 GACGCCGTGGAGTTCGGTGG CGG ||||||||[||||||||||||]PAM NA Intergenic NA NA

T20565 GAGTTCGGTGGCGGCCTCCACGG 1000
Oligo fwd CACCGAGTTCGGTGGCGGCCTCCA
Oligo rev AAACTGGAGGCCGCCACCGAACTC
Chromosome start end strand MM target_seq PAM alignment distance position gene name gene id
GU305914.1 186444 186466 - 0 GAGTTCGGTGGCGGCCTCCA CGG ||||||||[||||||||||||]PAM NA Intergenic NA NA

T20566 CGGCCTCCACGGCCACCCTCAGG 1000
Oligo fwd CACCCGGCCTCCACGGCCACCCTC
Oligo rev AAACGAGGGTGGCCGTGGAGGCCG
Chromosome start end strand MM target_seq PAM alignment distance position gene name gene id
GU305914.1 186433 186455 - 0 CGGCCTCCACGGCCACCCTC AGG ||||||||[||||||||||||]PAM NA Intergenic NA NA

T20567 GGCCTCCACGGCCACCCTCAGGG 1000
Oligo fwd CACCGGCCTCCACGGCCACCCTCA
Oligo rev AAACTGAGGGTGGCCGTGGAGGCC
Chromosome start end strand MM target_seq PAM alignment distance position gene name gene id
GU305914.1 186432 186454 - 0 GGCCTCCACGGCCACCCTCA GGG ||||||||[||||||||||||]PAM NA Intergenic NA NA

T20568 CCTCAGGGTTCTTCTGTATTCGG 1000
Oligo fwd CACCCCTCAGGGTTCTTCTGTATT
Oligo rev AAACAATACAGAAGAACCCTGAGG
Chromosome start end strand MM target_seq PAM alignment distance position gene name gene id
GU305914.1 186417 186439 - 0 CCTCAGGGTTCTTCTGTATT CGG ||||||||[||||||||||||]PAM NA Intergenic NA NA

T20569 CTCAGGGTTCTTCTGTATTCGGG 1000
Oligo fwd CACCCTCAGGGTTCTTCTGTATTC
Oligo rev AAACGAATACAGAAGAACCCTGAG

<table>
<tr><td>Chromosome</td><td>start</td><td>end</td><td>strand</td><td>MM</td><td>target_seq</td><td>PAM</td><td>alignment</td><td>distance</td><td>position</td><td>gene name</td><td>gene id</td></tr>
<tr><td>GU305914.1</td><td>186416</td><td>186438</td><td>-</td><td>0</td><td>CTCAGGGTTCTTCTGTATTC</td><td>GGG</td><td>||||||||[||||||||||||]PAM</td><td>NA</td><td>Intergenic</td><td>NA</td><td>NA</td></tr>
<tr><td></td><td></td><td></td><td></td><td></td><td></td><td></td><td></td><td></td><td></td><td></td><td></td></tr>
<tr><td>T20570</td><td>TCAGGGTTCTTCTGTATTCGGGG</td><td>1000</td><td></td><td></td><td></td><td></td><td></td><td></td><td></td><td></td><td></td></tr>
<tr><td>Oligo fwd</td><td>CACCTCAGGGTTCTTCTGTATTCG</td><td></td><td></td><td></td><td></td><td></td><td></td><td></td><td></td><td></td><td></td></tr>
<tr><td>Oligo rev</td><td>AAACCGAATACAGAAGAACCCTGA</td><td></td><td></td><td></td><td></td><td></td><td></td><td></td><td></td><td></td><td></td></tr>
<tr><td>Chromosome</td><td>start</td><td>end</td><td>strand</td><td>MM</td><td>target_seq</td><td>PAM</td><td>alignment</td><td>distance</td><td>position</td><td>gene name</td><td>gene id</td></tr>
<tr><td>GU305914.1</td><td>186415</td><td>186437</td><td>-</td><td>0</td><td>TCAGGGTTCTTCTGTATTCG</td><td>GGG</td><td>||||||||[||||||||||||]PAM</td><td>NA</td><td>Intergenic</td><td>NA</td><td>NA</td></tr>
<tr><td></td><td></td><td></td><td></td><td></td><td></td><td></td><td></td><td></td><td></td><td></td><td></td></tr>
<tr><td>T20571</td><td>AAGCTCGTGAGACATTCGACCGG</td><td>1000</td><td></td><td></td><td></td><td></td><td></td><td></td><td></td><td></td><td></td></tr>
<tr><td>Oligo fwd</td><td>CACCAAGCTCGTGAGACATTCGAC</td><td></td><td></td><td></td><td></td><td></td><td></td><td></td><td></td><td></td><td></td></tr>
<tr><td>Oligo rev</td><td>AAACGTCGAATGTCTCACGAGCTT</td><td></td><td></td><td></td><td></td><td></td><td></td><td></td><td></td><td></td><td></td></tr>
<tr><td>Chromosome</td><td>start</td><td>end</td><td>strand</td><td>MM</td><td>target_seq</td><td>PAM</td><td>alignment</td><td>distance</td><td>position</td><td>gene name</td><td>gene id</td></tr>
<tr><td>GU305914.1</td><td>186392</td><td>186414</td><td>-</td><td>0</td><td>AAGCTCGTGAGACATTCGAC</td><td>CGG</td><td>||||||||[||||||||||||]PAM</td><td>NA</td><td>Intergenic</td><td>NA</td><td>NA</td></tr>
<tr><td></td><td></td><td></td><td></td><td></td><td></td><td></td><td></td><td></td><td></td><td></td><td></td></tr>
<tr><td>T20572</td><td>CTGCATAAAGATATCGTCCACGG</td><td>1000</td><td></td><td></td><td></td><td></td><td></td><td></td><td></td><td></td><td></td></tr>
<tr><td>Oligo fwd</td><td>CACCCTGCATAAAGATATCGTCCA</td><td></td><td></td><td></td><td></td><td></td><td></td><td></td><td></td><td></td><td></td></tr>
<tr><td>Oligo rev</td><td>AAACTGGACGATATCTTTATGCAG</td><td></td><td></td><td></td><td></td><td></td><td></td><td></td><td></td><td></td><td></td></tr>
<tr><td>Chromosome</td><td>start</td><td>end</td><td>strand</td><td>MM</td><td>target_seq</td><td>PAM</td><td>alignment</td><td>distance</td><td>position</td><td>gene name</td><td>gene id</td></tr>
<tr><td>GU305914.1</td><td>186336</td><td>186358</td><td>-</td><td>0</td><td>CTGCATAAAGATATCGTCCA</td><td>CGG</td><td>||||||||[||||||||||||]PAM</td><td>NA</td><td>Intergenic</td><td>NA</td><td>NA</td></tr>
<tr><td></td><td></td><td></td><td></td><td></td><td></td><td></td><td></td><td></td><td></td><td></td><td></td></tr>
<tr><td>T20573</td><td>AAAGATATCGTCCACGGTCGCGG</td><td>1000</td><td></td><td></td><td></td><td></td><td></td><td></td><td></td><td></td><td></td></tr>
<tr><td>Oligo fwd</td><td>CACCAAAGATATCGTCCACGGTCG</td><td></td><td></td><td></td><td></td><td></td><td></td><td></td><td></td><td></td><td></td></tr>
<tr><td>Oligo rev</td><td>AAACCGACCGTGGACGATATCTTT</td><td></td><td></td><td></td><td></td><td></td><td></td><td></td><td></td><td></td><td></td></tr>
<tr><td>Chromosome</td><td>start</td><td>end</td><td>strand</td><td>MM</td><td>target_seq</td><td>PAM</td><td>alignment</td><td>distance</td><td>position</td><td>gene name</td><td>gene id</td></tr>
<tr><td>GU305914.1</td><td>186330</td><td>186352</td><td>-</td><td>0</td><td>AAAGATATCGTCCACGGTCG</td><td>CGG</td><td>||||||||[||||||||||||]PAM</td><td>NA</td><td>Intergenic</td><td>NA</td><td>NA</td></tr>
<tr><td></td><td></td><td></td><td></td><td></td><td></td><td></td><td></td><td></td><td></td><td></td><td></td></tr>
<tr><td>T20574</td><td>AAGATATCGTCCACGGTCGCGGG</td><td>1000</td><td></td><td></td><td></td><td></td><td></td><td></td><td></td><td></td><td></td></tr>
<tr><td>Oligo fwd</td><td>CACCAAGATATCGTCCACGGTCGC</td><td></td><td></td><td></td><td></td><td></td><td></td><td></td><td></td><td></td><td></td></tr>
<tr><td>Oligo rev</td><td>AAACGCGACCGTGGACGATATCTT</td><td></td><td></td><td></td><td></td><td></td><td></td><td></td><td></td><td></td><td></td></tr>
<tr><td>Chromosome</td><td>start</td><td>end</td><td>strand</td><td>MM</td><td>target_seq</td><td>PAM</td><td>alignment</td><td>distance</td><td>position</td><td>gene name</td><td>gene id</td></tr>
<tr><td>GU305914.1</td><td>186329</td><td>186351</td><td>-</td><td>0</td><td>AAGATATCGTCCACGGTCGC</td><td>GGG</td><td>||||||||[||||||||||||]PAM</td><td>NA</td><td>Intergenic</td><td>NA</td><td>NA</td></tr>
<tr><td></td><td></td><td></td><td></td><td></td><td></td><td></td><td></td><td></td><td></td><td></td><td></td></tr>
<tr><td>T20575</td><td>AGATATCGTCCACGGTCGCGGGG</td><td>1000</td><td></td><td></td><td></td><td></td><td></td><td></td><td></td><td></td><td></td></tr>
<tr><td>Oligo fwd</td><td>CACCAGATATCGTCCACGGTCGCG</td><td></td><td></td><td></td><td></td><td></td><td></td><td></td><td></td><td></td><td></td></tr>
<tr><td>Oligo rev</td><td>AAACCGCGACCGTGGACGATATCT</td><td></td><td></td><td></td><td></td><td></td><td></td><td></td><td></td><td></td><td></td></tr>
<tr><td>Chromosome</td><td>start</td><td>end</td><td>strand</td><td>MM</td><td>target_seq</td><td>PAM</td><td>alignment</td><td>distance</td><td>position</td><td>gene name</td><td>gene id</td></tr>
<tr><td>GU305914.1</td><td>186328</td><td>186350</td><td>-</td><td>0</td><td>AGATATCGTCCACGGTCGCG</td><td>GGG</td><td>||||||||[||||||||||||]PAM</td><td>NA</td><td>Intergenic</td><td>NA</td><td>NA</td></tr>
<tr><td></td><td></td><td></td><td></td><td></td><td></td><td></td><td></td><td></td><td></td><td></td><td></td></tr>
<tr><td>T20576</td><td>TCGCGGGGTCCCTGTGCTCCAGG</td><td>1000</td><td></td><td></td><td></td><td></td><td></td><td></td><td></td><td></td><td></td></tr>
<tr><td>Oligo fwd</td><td>CACCTCGCGGGGTCCCTGTGCTCC</td><td></td><td></td><td></td><td></td><td></td><td></td><td></td><td></td><td></td><td></td></tr>
<tr><td>Oligo rev</td><td>AAACGGAGCACAGGGACCCCGCGA</td><td></td><td></td><td></td><td></td><td></td><td></td><td></td><td></td><td></td><td></td></tr>
<tr><td>Chromosome</td><td>start</td><td>end</td><td>strand</td><td>MM</td><td>target_seq</td><td>PAM</td><td>alignment</td><td>distance</td><td>position</td><td>gene name</td><td>gene id</td></tr>
<tr><td>GU305914.1</td><td>186313</td><td>186335</td><td>-</td><td>0</td><td>TCGCGGGGTCCCTGTGCTCC</td><td>AGG</td><td>||||||||[||||||||||||]PAM</td><td>NA</td><td>Intergenic</td><td>NA</td><td>NA</td></tr>
<tr><td></td><td></td><td></td><td></td><td></td><td></td><td></td><td></td><td></td><td></td><td></td><td></td></tr>
<tr><td>T20577</td><td>CCAGGATACTTACAACCCCCCGG</td><td>1000</td><td></td><td></td><td></td><td></td><td></td><td></td><td></td><td></td><td></td></tr>
<tr><td>Oligo fwd</td><td>CACCCCAGGATACTTACAACCCCC</td><td></td><td></td><td></td><td></td><td></td><td></td><td></td><td></td><td></td><td></td></tr>
<tr><td>Oligo rev</td><td>AAACGGGGGTTGTAAGTATCCTGG</td><td></td><td></td><td></td><td></td><td></td><td></td><td></td><td></td><td></td><td></td></tr>
<tr><td>Chromosome</td><td>start</td><td>end</td><td>strand</td><td>MM</td><td>target_seq</td><td>PAM</td><td>alignment</td><td>distance</td><td>position</td><td>gene name</td><td>gene id</td></tr>
<tr><td>GU305914.1</td><td>186295</td><td>186317</td><td>-</td><td>0</td><td>CCAGGATACTTACAACCCCC</td><td>CGG</td><td>||||||||[||||||||||||]PAM</td><td>NA</td><td>Intergenic</td><td>NA</td><td>NA</td></tr>
<tr><td></td><td></td><td></td><td></td><td></td><td></td><td></td><td></td><td></td><td></td><td></td><td></td></tr>
<tr><td>T20578</td><td>CAGGATACTTACAACCCCCCGGG</td><td>1000</td><td></td><td></td><td></td><td></td><td></td><td></td><td></td><td></td><td></td></tr>
<tr><td>Oligo fwd</td><td>CACCCAGGATACTTACAACCCCCC</td><td></td><td></td><td></td><td></td><td></td><td></td><td></td><td></td><td></td><td></td></tr>
<tr><td>Oligo rev</td><td>AAACGGGGGGTTGTAAGTATCCTG</td><td></td><td></td><td></td><td></td><td></td><td></td><td></td><td></td><td></td><td></td></tr>
<tr><td>Chromosome</td><td>start</td><td>end</td><td>strand</td><td>MM</td><td>target_seq</td><td>PAM</td><td>alignment</td><td>distance</td><td>position</td><td>gene name</td><td>gene id</td></tr>
<tr><td>GU305914.1</td><td>186294</td><td>186316</td><td>-</td><td>0</td><td>CAGGATACTTACAACCCCCC</td><td>GGG</td><td>||||||||[||||||||||||]PAM</td><td>NA</td><td>Intergenic</td><td>NA</td><td>NA</td></tr>
<tr><td></td><td></td><td></td><td></td><td></td><td></td><td></td><td></td><td></td><td></td><td></td><td></td></tr>
<tr><td>T20579</td><td>CGAAGTGCAATCCAGCTTGAAGG</td><td>1000</td><td></td><td></td><td></td><td></td><td></td><td></td><td></td><td></td><td></td></tr>
<tr><td>Oligo fwd</td><td>CACCCGAAGTGCAATCCAGCTTGA</td><td></td><td></td><td></td><td></td><td></td><td></td><td></td><td></td><td></td><td></td></tr>
<tr><td>Oligo rev</td><td>AAACTCAAGCTGGATTGCACTTCG</td><td></td><td></td><td></td><td></td><td></td><td></td><td></td><td></td><td></td><td></td></tr>
<tr><td>Chromosome</td><td>start</td><td>end</td><td>strand</td><td>MM</td><td>target_seq</td><td>PAM</td><td>alignment</td><td>distance</td><td>position</td><td>gene name</td><td>gene id</td></tr>
<tr><td>GU305914.1</td><td>186264</td><td>186286</td><td>-</td><td>0</td><td>CGAAGTGCAATCCAGCTTGA</td><td>AGG</td><td>||||||||[||||||||||||]PAM</td><td>NA</td><td>Intergenic</td><td>NA</td><td>NA</td></tr>
<tr><td></td><td></td><td></td><td></td><td></td><td></td><td></td><td></td><td></td><td></td><td></td><td></td></tr>
<tr><td>T20580</td><td>GCTTGAAGGTCGCCATGCAGTGG</td><td>1000</td><td></td><td></td><td></td><td></td><td></td><td></td><td></td><td></td><td></td></tr>
<tr><td>Oligo fwd</td><td>CACCGCTTGAAGGTCGCCATGCAG</td><td></td><td></td><td></td><td></td><td></td><td></td><td></td><td></td><td></td><td></td></tr>
<tr><td>Oligo rev</td><td>AAACCTGCATGGCGACCTTCAAGC</td><td></td><td></td><td></td><td></td><td></td><td></td><td></td><td></td><td></td><td></td></tr>
<tr><td>Chromosome</td><td>start</td><td>end</td><td>strand</td><td>MM</td><td>target_seq</td><td>PAM</td><td>alignment</td><td>distance</td><td>position</td><td>gene name</td><td>gene id</td></tr>
<tr><td>GU305914.1</td><td>186250</td><td>186272</td><td>-</td><td>0</td><td>GCTTGAAGGTCGCCATGCAG</td><td>TGG</td><td>||||||||[||||||||||||]PAM</td><td>NA</td><td>Intergenic</td><td>NA</td><td>NA</td></tr>
<tr><td></td><td></td><td></td><td></td><td></td><td></td><td></td><td></td><td></td><td></td><td></td><td></td></tr>
<tr><td>T20581</td><td>AGTGGTAGTTGTGATCGCCGTGG</td><td>1000</td><td></td><td></td><td></td><td></td><td></td><td></td><td></td><td></td><td></td></tr>
<tr><td>Oligo fwd</td><td>CACCAGTGGTAGTTGTGATCGCCG</td><td></td><td></td><td></td><td></td><td></td><td></td><td></td><td></td><td></td><td></td></tr>
<tr><td>Oligo rev</td><td>AAACCGGCGATCACAACTACCACT</td><td></td><td></td><td></td><td></td><td></td><td></td><td></td><td></td><td></td><td></td></tr>
<tr><td>Chromosome</td><td>start</td><td>end</td><td>strand</td><td>MM</td><td>target_seq</td><td>PAM</td><td>alignment</td><td>distance</td><td>position</td><td>gene name</td><td>gene id</td></tr>
<tr><td>GU305914.1</td><td>186232</td><td>186254</td><td>-</td><td>0</td><td>AGTGGTAGTTGTGATCGCCG</td><td>TGG</td><td>||||||||[||||||||||||]PAM</td><td>NA</td><td>Intergenic</td><td>NA</td><td>NA</td></tr>
<tr><td></td><td></td><td></td><td></td><td></td><td></td><td></td><td></td><td></td><td></td><td></td><td></td></tr>
<tr><td>T20582</td><td>GGTAGTTGTGATCGCCGTGGAGG</td><td>1000</td><td></td><td></td><td></td><td></td><td></td><td></td><td></td><td></td><td></td></tr>
<tr><td>Oligo fwd</td><td>CACCGGTAGTTGTGATCGCCGTGG</td><td></td><td></td><td></td><td></td><td></td><td></td><td></td><td></td><td></td><td></td></tr>
<tr><td>Oligo rev</td><td>AAACCCACGGCGATCACAACTACC</td><td></td><td></td><td></td><td></td><td></td><td></td><td></td><td></td><td></td><td></td></tr>
<tr><td>Chromosome</td><td>start</td><td>end</td><td>strand</td><td>MM</td><td>target_seq</td><td>PAM</td><td>alignment</td><td>distance</td><td>position</td><td>gene name</td><td>gene id</td></tr>
<tr><td>GU305914.1</td><td>186229</td><td>186251</td><td>-</td><td>0</td><td>GGTAGTTGTGATCGCCGTGG</td><td>AGG</td><td>||||||||[||||||||||||]PAM</td><td>NA</td><td>Intergenic</td><td>NA</td><td>NA</td></tr>
<tr><td></td><td></td><td></td><td></td><td></td><td></td><td></td><td></td><td></td><td></td><td></td><td></td></tr>
<tr><td>T20583</td><td>GTAGTTGTGATCGCCGTGGAGGG</td><td>1000</td><td></td><td></td><td></td><td></td><td></td><td></td><td></td><td></td><td></td></tr>
<tr><td>Oligo fwd</td><td>CACCGTAGTTGTGATCGCCGTGGA</td><td></td><td></td><td></td><td></td><td></td><td></td><td></td><td></td><td></td><td></td></tr>
<tr><td>Oligo rev</td><td>AAACTCCACGGCGATCACAACTAC</td><td></td><td></td><td></td><td></td><td></td><td></td><td></td><td></td><td></td><td></td></tr>
<tr><td>Chromosome</td><td>start</td><td>end</td><td>strand</td><td>MM</td><td>target_seq</td><td>PAM</td><td>alignment</td><td>distance</td><td>position</td><td>gene name</td><td>gene id</td></tr>
<tr><td>GU305914.1</td><td>186228</td><td>186250</td><td>-</td><td>0</td><td>GTAGTTGTGATCGCCGTGGA</td><td>GGG</td><td>||||||||[||||||||||||]PAM</td><td>NA</td><td>Intergenic</td><td>NA</td><td>NA</td></tr>
<tr><td></td><td></td><td></td><td></td><td></td><td></td><td></td><td></td><td></td><td></td><td></td><td></td></tr>
<tr><td>T20584</td><td>GAGGGTCGCCACCGAGTCTGAGG</td><td>1000</td><td></td><td></td><td></td><td></td><td></td><td></td><td></td><td></td><td></td></tr>
<tr><td>Oligo fwd</td><td>CACCGAGGGTCGCCACCGAGTCTG</td><td></td><td></td><td></td><td></td><td></td><td></td><td></td><td></td><td></td><td></td></tr>
</table>

<table>
<tr><td>Oligo rev</td><td>AAACCAGACTCGGTGGCGACCCTC</td><td></td><td></td><td></td><td></td><td></td><td></td><td></td><td></td><td></td><td></td></tr>
<tr><td>Chromosome</td><td>start</td><td>end</td><td>strand</td><td>MM</td><td>target_seq</td><td>PAM</td><td>alignment</td><td>distance</td><td>position</td><td>gene name</td><td>gene id</td></tr>
<tr><td>GU305914.1</td><td>186210</td><td>186232</td><td>-</td><td>0</td><td>GAGGGTCGCCACCGAGTCTG</td><td>AGG</td><td>||||||||[||||||||||||]PAM</td><td>NA</td><td>Intergenic</td><td>NA</td><td>NA</td></tr>
<tr><td>T20585</td><td>GGTCGCCACCGAGTCTGAGGAGG</td><td>1000</td><td></td><td></td><td></td><td></td><td></td><td></td><td></td><td></td><td></td></tr>
<tr><td>Oligo fwd</td><td>CACCGGTCGCCACCGAGTCTGAGG</td><td></td><td></td><td></td><td></td><td></td><td></td><td></td><td></td><td></td><td></td></tr>
<tr><td>Oligo rev</td><td>AAACCCTCAGACTCGGTGGCGACC</td><td></td><td></td><td></td><td></td><td></td><td></td><td></td><td></td><td></td><td></td></tr>
<tr><td>Chromosome</td><td>start</td><td>end</td><td>strand</td><td>MM</td><td>target_seq</td><td>PAM</td><td>alignment</td><td>distance</td><td>position</td><td>gene name</td><td>gene id</td></tr>
<tr><td>GU305914.1</td><td>186207</td><td>186229</td><td>-</td><td>0</td><td>GGTCGCCACCGAGTCTGAGG</td><td>AGG</td><td>||||||||[||||||||||||]PAM</td><td>NA</td><td>Intergenic</td><td>NA</td><td>NA</td></tr>
<tr><td>T20586</td><td>AGTCGCGTCCCTCACGCCCCCGG</td><td>1000</td><td></td><td></td><td></td><td></td><td></td><td></td><td></td><td></td><td></td></tr>
<tr><td>Oligo fwd</td><td>CACCAGTCGCGTCCCTCACGCCCC</td><td></td><td></td><td></td><td></td><td></td><td></td><td></td><td></td><td></td><td></td></tr>
<tr><td>Oligo rev</td><td>AAACGGGGCGTGAGGGACGCGACT</td><td></td><td></td><td></td><td></td><td></td><td></td><td></td><td></td><td></td><td></td></tr>
<tr><td>Chromosome</td><td>start</td><td>end</td><td>strand</td><td>MM</td><td>target_seq</td><td>PAM</td><td>alignment</td><td>distance</td><td>position</td><td>gene name</td><td>gene id</td></tr>
<tr><td>GU305914.1</td><td>186136</td><td>186158</td><td>-</td><td>0</td><td>AGTCGCGTCCCTCACGCCCC</td><td>CGG</td><td>||||||||[||||||||||||]PAM</td><td>NA</td><td>Intergenic</td><td>NA</td><td>NA</td></tr>
<tr><td>T20587</td><td>GCTGCAATCAGAAAGAAGAATGG</td><td>1000</td><td></td><td></td><td></td><td></td><td></td><td></td><td></td><td></td><td></td></tr>
<tr><td>Oligo fwd</td><td>CACCGCTGCAATCAGAAAGAAGAA</td><td></td><td></td><td></td><td></td><td></td><td></td><td></td><td></td><td></td><td></td></tr>
<tr><td>Oligo rev</td><td>AAACTTCTTCTTTCTGATTGCAGC</td><td></td><td></td><td></td><td></td><td></td><td></td><td></td><td></td><td></td><td></td></tr>
<tr><td>Chromosome</td><td>start</td><td>end</td><td>strand</td><td>MM</td><td>target_seq</td><td>PAM</td><td>alignment</td><td>distance</td><td>position</td><td>gene name</td><td>gene id</td></tr>
<tr><td>GU305914.1</td><td>186072</td><td>186094</td><td>-</td><td>0</td><td>GCTGCAATCAGAAAGAAGAA</td><td>TGG</td><td>||||||||[||||||||||||]PAM</td><td>NA</td><td>Intergenic</td><td>NA</td><td>NA</td></tr>
<tr><td>T20588</td><td>CGAGCGAGCGAGCGAACGACCGG</td><td>1000</td><td></td><td></td><td></td><td></td><td></td><td></td><td></td><td></td><td></td></tr>
<tr><td>Oligo fwd</td><td>CACCCGAGCGAGCGAGCGAACGAC</td><td></td><td></td><td></td><td></td><td></td><td></td><td></td><td></td><td></td><td></td></tr>
<tr><td>Oligo rev</td><td>AAACGTCGTTCGCTCGCTCGCTCG</td><td></td><td></td><td></td><td></td><td></td><td></td><td></td><td></td><td></td><td></td></tr>
<tr><td>Chromosome</td><td>start</td><td>end</td><td>strand</td><td>MM</td><td>target_seq</td><td>PAM</td><td>alignment</td><td>distance</td><td>position</td><td>gene name</td><td>gene id</td></tr>
<tr><td>GU305914.1</td><td>186047</td><td>186069</td><td>-</td><td>0</td><td>CGAGCGAGCGAGCGAACGAC</td><td>CGG</td><td>||||||||[||||||||||||]PAM</td><td>NA</td><td>Intergenic</td><td>NA</td><td>NA</td></tr>
<tr><td>T20589</td><td>GAGCGAGCGAGCGAACGACCGGG</td><td>1000</td><td></td><td></td><td></td><td></td><td></td><td></td><td></td><td></td><td></td></tr>
<tr><td>Oligo fwd</td><td>CACCGAGCGAGCGAGCGAACGACC</td><td></td><td></td><td></td><td></td><td></td><td></td><td></td><td></td><td></td><td></td></tr>
<tr><td>Oligo rev</td><td>AAACGGTCGTTCGCTCGCTCGCTC</td><td></td><td></td><td></td><td></td><td></td><td></td><td></td><td></td><td></td><td></td></tr>
<tr><td>Chromosome</td><td>start</td><td>end</td><td>strand</td><td>MM</td><td>target_seq</td><td>PAM</td><td>alignment</td><td>distance</td><td>position</td><td>gene name</td><td>gene id</td></tr>
<tr><td>GU305914.1</td><td>186046</td><td>186068</td><td>-</td><td>0</td><td>GAGCGAGCGAGCGAACGACC</td><td>GGG</td><td>||||||||[||||||||||||]PAM</td><td>NA</td><td>Intergenic</td><td>NA</td><td>NA</td></tr>
<tr><td>T20590</td><td>CGAGCGAGCGAACGACCGGGCGG</td><td>1000</td><td></td><td></td><td></td><td></td><td></td><td></td><td></td><td></td><td></td></tr>
<tr><td>Oligo fwd</td><td>CACCCGAGCGAGCGAACGACCGGG</td><td></td><td></td><td></td><td></td><td></td><td></td><td></td><td></td><td></td><td></td></tr>
<tr><td>Oligo rev</td><td>AAACCCCGGTCGTTCGCTCGCTCG</td><td></td><td></td><td></td><td></td><td></td><td></td><td></td><td></td><td></td><td></td></tr>
<tr><td>Chromosome</td><td>start</td><td>end</td><td>strand</td><td>MM</td><td>target_seq</td><td>PAM</td><td>alignment</td><td>distance</td><td>position</td><td>gene name</td><td>gene id</td></tr>
<tr><td>GU305914.1</td><td>186043</td><td>186065</td><td>-</td><td>0</td><td>CGAGCGAGCGAACGACCGGG</td><td>CGG</td><td>||||||||[||||||||||||]PAM</td><td>NA</td><td>Intergenic</td><td>NA</td><td>NA</td></tr>
<tr><td>T20591</td><td>GAGCGAGCGAACGACCGGGCGGG</td><td>1000</td><td></td><td></td><td></td><td></td><td></td><td></td><td></td><td></td><td></td></tr>
<tr><td>Oligo fwd</td><td>CACCGAGCGAGCGAACGACCGGGC</td><td></td><td></td><td></td><td></td><td></td><td></td><td></td><td></td><td></td><td></td></tr>
<tr><td>Oligo rev</td><td>AAACGCCCGGTCGTTCGCTCGCTC</td><td></td><td></td><td></td><td></td><td></td><td></td><td></td><td></td><td></td><td></td></tr>
<tr><td>Chromosome</td><td>start</td><td>end</td><td>strand</td><td>MM</td><td>target_seq</td><td>PAM</td><td>alignment</td><td>distance</td><td>position</td><td>gene name</td><td>gene id</td></tr>
<tr><td>GU305914.1</td><td>186042</td><td>186064</td><td>-</td><td>0</td><td>GAGCGAGCGAACGACCGGGC</td><td>GGG</td><td>||||||||[||||||||||||]PAM</td><td>NA</td><td>Intergenic</td><td>NA</td><td>NA</td></tr>
<tr><td>T20592</td><td>CGAGCGAACGACCGGGCGGGCGG</td><td>1000</td><td></td><td></td><td></td><td></td><td></td><td></td><td></td><td></td><td></td></tr>
<tr><td>Oligo fwd</td><td>CACCCGAGCGAACGACCGGGCGGG</td><td></td><td></td><td></td><td></td><td></td><td></td><td></td><td></td><td></td><td></td></tr>
<tr><td>Oligo rev</td><td>AAACCCCGCCCGGTCGTTCGCTCG</td><td></td><td></td><td></td><td></td><td></td><td></td><td></td><td></td><td></td><td></td></tr>
<tr><td>Chromosome</td><td>start</td><td>end</td><td>strand</td><td>MM</td><td>target_seq</td><td>PAM</td><td>alignment</td><td>distance</td><td>position</td><td>gene name</td><td>gene id</td></tr>
<tr><td>GU305914.1</td><td>186039</td><td>186061</td><td>-</td><td>0</td><td>CGAGCGAACGACCGGGCGGG</td><td>CGG</td><td>||||||||[||||||||||||]PAM</td><td>NA</td><td>Intergenic</td><td>NA</td><td>NA</td></tr>
<tr><td>T20593</td><td>GAGCGAACGACCGGGCGGGCGGG</td><td>1000</td><td></td><td></td><td></td><td></td><td></td><td></td><td></td><td></td><td></td></tr>
<tr><td>Oligo fwd</td><td>CACCGAGCGAACGACCGGGCGGGC</td><td></td><td></td><td></td><td></td><td></td><td></td><td></td><td></td><td></td><td></td></tr>
<tr><td>Oligo rev</td><td>AAACGCCCGCCCGGTCGTTCGCTC</td><td></td><td></td><td></td><td></td><td></td><td></td><td></td><td></td><td></td><td></td></tr>
<tr><td>Chromosome</td><td>start</td><td>end</td><td>strand</td><td>MM</td><td>target_seq</td><td>PAM</td><td>alignment</td><td>distance</td><td>position</td><td>gene name</td><td>gene id</td></tr>
<tr><td>GU305914.1</td><td>186038</td><td>186060</td><td>-</td><td>0</td><td>GAGCGAACGACCGGGCGGGC</td><td>GGG</td><td>||||||||[||||||||||||]PAM</td><td>NA</td><td>Intergenic</td><td>NA</td><td>NA</td></tr>
<tr><td>T20594</td><td>TCCCTTCACCAAGAAATGACCGG</td><td>1000</td><td></td><td></td><td></td><td></td><td></td><td></td><td></td><td></td><td></td></tr>
<tr><td>Oligo fwd</td><td>CACCTCCCTTCACCAAGAAATGAC</td><td></td><td></td><td></td><td></td><td></td><td></td><td></td><td></td><td></td><td></td></tr>
<tr><td>Oligo rev</td><td>AAACGTCATTTCTTGGTGAAGGGA</td><td></td><td></td><td></td><td></td><td></td><td></td><td></td><td></td><td></td><td></td></tr>
<tr><td>Chromosome</td><td>start</td><td>end</td><td>strand</td><td>MM</td><td>target_seq</td><td>PAM</td><td>alignment</td><td>distance</td><td>position</td><td>gene name</td><td>gene id</td></tr>
<tr><td>GU305914.1</td><td>186003</td><td>186025</td><td>-</td><td>0</td><td>TCCCTTCACCAAGAAATGAC</td><td>CGG</td><td>||||||||[||||||||||||]PAM</td><td>NA</td><td>Intergenic</td><td>NA</td><td>NA</td></tr>
<tr><td>T20595</td><td>CGTCTTACCTGCTCCGTGCTCGG</td><td>1000</td><td></td><td></td><td></td><td></td><td></td><td></td><td></td><td></td><td></td></tr>
<tr><td>Oligo fwd</td><td>CACCCGTCTTACCTGCTCCGTGCT</td><td></td><td></td><td></td><td></td><td></td><td></td><td></td><td></td><td></td><td></td></tr>
<tr><td>Oligo rev</td><td>AAACAGCACGGAGCAGGTAAGACG</td><td></td><td></td><td></td><td></td><td></td><td></td><td></td><td></td><td></td><td></td></tr>
<tr><td>Chromosome</td><td>start</td><td>end</td><td>strand</td><td>MM</td><td>target_seq</td><td>PAM</td><td>alignment</td><td>distance</td><td>position</td><td>gene name</td><td>gene id</td></tr>
<tr><td>GU305914.1</td><td>185952</td><td>185974</td><td>-</td><td>0</td><td>CGTCTTACCTGCTCCGTGCT</td><td>CGG</td><td>||||||||[||||||||||||]PAM</td><td>NA</td><td>Intergenic</td><td>NA</td><td>NA</td></tr>
<tr><td>T20596</td><td>TTATCTCACCTGTGCAGCCTAGG</td><td>1000</td><td></td><td></td><td></td><td></td><td></td><td></td><td></td><td></td><td></td></tr>
<tr><td>Oligo fwd</td><td>CACCTTATCTCACCTGTGCAGCCT</td><td></td><td></td><td></td><td></td><td></td><td></td><td></td><td></td><td></td><td></td></tr>
<tr><td>Oligo rev</td><td>AAACAGGCTGCACAGGTGAGATAA</td><td></td><td></td><td></td><td></td><td></td><td></td><td></td><td></td><td></td><td></td></tr>
<tr><td>Chromosome</td><td>start</td><td>end</td><td>strand</td><td>MM</td><td>target_seq</td><td>PAM</td><td>alignment</td><td>distance</td><td>position</td><td>gene name</td><td>gene id</td></tr>
<tr><td>GU305914.1</td><td>185912</td><td>185934</td><td>-</td><td>0</td><td>TTATCTCACCTGTGCAGCCT</td><td>AGG</td><td>||||||||[||||||||||||]PAM</td><td>NA</td><td>Intergenic</td><td>NA</td><td>NA</td></tr>
<tr><td>T20597</td><td>TCCCACAATGCTATTCTGTCAGG</td><td>1000</td><td></td><td></td><td></td><td></td><td></td><td></td><td></td><td></td><td></td></tr>
<tr><td>Oligo fwd</td><td>CACCTCCCACAATGCTATTCTGTC</td><td></td><td></td><td></td><td></td><td></td><td></td><td></td><td></td><td></td><td></td></tr>
<tr><td>Oligo rev</td><td>AAACGACAGAATAGCATTGTGGGA</td><td></td><td></td><td></td><td></td><td></td><td></td><td></td><td></td><td></td><td></td></tr>
<tr><td>Chromosome</td><td>start</td><td>end</td><td>strand</td><td>MM</td><td>target_seq</td><td>PAM</td><td>alignment</td><td>distance</td><td>position</td><td>gene name</td><td>gene id</td></tr>
<tr><td>GU305914.1</td><td>185875</td><td>185897</td><td>-</td><td>0</td><td>TCCCACAATGCTATTCTGTC</td><td>AGG</td><td>||||||||[||||||||||||]PAM</td><td>NA</td><td>Intergenic</td><td>NA</td><td>NA</td></tr>
<tr><td>T20598</td><td>AACTATTGTATAAGATCCGCAGG</td><td>1000</td><td></td><td></td><td></td><td></td><td></td><td></td><td></td><td></td><td></td></tr>
<tr><td>Oligo fwd</td><td>CACCAACTATTGTATAAGATCCGC</td><td></td><td></td><td></td><td></td><td></td><td></td><td></td><td></td><td></td><td></td></tr>
<tr><td>Oligo rev</td><td>AAACGCGGATCTTATACAATAGTT</td><td></td><td></td><td></td><td></td><td></td><td></td><td></td><td></td><td></td><td></td></tr>
<tr><td>Chromosome</td><td>start</td><td>end</td><td>strand</td><td>MM</td><td>target_seq</td><td>PAM</td><td>alignment</td><td>distance</td><td>position</td><td>gene name</td><td>gene id</td></tr>
<tr><td>GU305914.1</td><td>185845</td><td>185867</td><td>-</td><td>0</td><td>AACTATTGTATAAGATCCGC</td><td>AGG</td><td>||||||||[||||||||||||]PAM</td><td>NA</td><td>Intergenic</td><td>NA</td><td>NA</td></tr>
<tr><td>T20599</td><td>ACTATTGTATAAGATCCGCAGGG</td><td>1000</td><td></td><td></td><td></td><td></td><td></td><td></td><td></td><td></td><td></td></tr>
</table>

Oligo fwd CACCACTATTGTATAAGATCCGCA
Oligo rev AAACTGCGGATCTTATACAATAGT
Chromosome start end strand MM target_seq PAM alignment distance position gene name gene id
GU305914.1 185844 185866 - 0 ACTATTGTATAAGATCCGCA GGG ||||||||[||||||||||||]PAM NA Intergenic NA NA

T20600 CTGTACTCACTAGTAACAACAGG 1000
Oligo fwd CACCCTGTACTCACTAGTAACAAC
Oligo rev AAACGTTGTTACTAGTGAGTACAG
Chromosome start end strand MM target_seq PAM alignment distance position gene name gene id
GU305914.1 185805 185827 - 0 CTGTACTCACTAGTAACAAC AGG ||||||||[||||||||||||]PAM NA Intergenic NA NA

T20601 CAGGTCAGCAGCACCTGTCCCGG 1000
Oligo fwd CACCCAGGTCAGCAGCACCTGTCC
Oligo rev AAACGGACAGGTGCTGCTGACCTG
Chromosome start end strand MM target_seq PAM alignment distance position gene name gene id
GU305914.1 185786 185808 - 0 CAGGTCAGCAGCACCTGTCC CGG ||||||||[||||||||||||]PAM NA Intergenic NA NA

T20602 CCGGTGTCCTCATCTCCCCGAGG 1000
Oligo fwd CACCCCGGTGTCCTCATCTCCCCG
Oligo rev AAACCGGGGAGATGAGGACACCGG
Chromosome start end strand MM target_seq PAM alignment distance position gene name gene id
GU305914.1 185767 185789 - 0 CCGGTGTCCTCATCTCCCCG AGG ||||||||[||||||||||||]PAM NA Intergenic NA NA

T20603 TCCCCGAGGAAGTCCTCTAGCGG 1000
Oligo fwd CACCTCCCCGAGGAAGTCCTCTAG
Oligo rev AAACCTAGAGGACTTCCTCGGGGA
Chromosome start end strand MM target_seq PAM alignment distance position gene name gene id
GU305914.1 185753 185775 - 0 TCCCCGAGGAAGTCCTCTAG CGG ||||||||[||||||||||||]PAM NA Intergenic NA NA

T20604 GGCGTCAGTCTGAAGAACAAAGG 1000
Oligo fwd CACCGGCGTCAGTCTGAAGAACAA
Oligo rev AAACTTGTTCTTCAGACTGACGCC
Chromosome start end strand MM target_seq PAM alignment distance position gene name gene id
GU305914.1 185732 185754 - 0 GGCGTCAGTCTGAAGAACAA AGG ||||||||[||||||||||||]PAM NA Intergenic NA NA

T20605 AGGTATCTCAGTAGCGAGTCCGG 1000
Oligo fwd CACCAGGTATCTCAGTAGCGAGTC
Oligo rev AAACGACTCGCTACTGAGATACCT
Chromosome start end strand MM target_seq PAM alignment distance position gene name gene id
GU305914.1 185712 185734 - 0 AGGTATCTCAGTAGCGAGTC CGG ||||||||[||||||||||||]PAM NA Intergenic NA NA

T20606 CTCAGTAGCGAGTCCGGAGCAGG 1000
Oligo fwd CACCCTCAGTAGCGAGTCCGGAGC
Oligo rev AAACGCTCCGGACTCGCTACTGAG
Chromosome start end strand MM target_seq PAM alignment distance position gene name gene id
GU305914.1 185706 185728 - 0 CTCAGTAGCGAGTCCGGAGC AGG ||||||||[||||||||||||]PAM NA Intergenic NA NA

T20607 TCAGTAGCGAGTCCGGAGCAGGG 1000
Oligo fwd CACCTCAGTAGCGAGTCCGGAGCA
Oligo rev AAACTGCTCCGGACTCGCTACTGA
Chromosome start end strand MM target_seq PAM alignment distance position gene name gene id
GU305914.1 185705 185727 - 0 TCAGTAGCGAGTCCGGAGCA GGG ||||||||[||||||||||||]PAM NA Intergenic NA NA

T20608 CAGTAGCGAGTCCGGAGCAGGGG 1000
Oligo fwd CACCCAGTAGCGAGTCCGGAGCAG
Oligo rev AAACCTGCTCCGGACTCGCTACTG
Chromosome start end strand MM target_seq PAM alignment distance position gene name gene id
GU305914.1 185704 185726 - 0 CAGTAGCGAGTCCGGAGCAG GGG ||||||||[||||||||||||]PAM NA Intergenic NA NA

T20609 GCAGGGGCGTAGCTGACCTATGG 1000
Oligo fwd CACCGCAGGGGCGTAGCTGACCTA
Oligo rev AAACTAGGTCAGCTACGCCCCTGC
Chromosome start end strand MM target_seq PAM alignment distance position gene name gene id
GU305914.1 185688 185710 - 0 GCAGGGGCGTAGCTGACCTA TGG ||||||||[||||||||||||]PAM NA Intergenic NA NA

T20610 CAAGAGAGATCAGTCTCGAAAGG 1000
Oligo fwd CACCCAAGAGAGATCAGTCTCGAA
Oligo rev AAACTTCGAGACTGATCTCTCTTG
Chromosome start end strand MM target_seq PAM alignment distance position gene name gene id
GU305914.1 185661 185683 - 0 CAAGAGAGATCAGTCTCGAA AGG ||||||||[||||||||||||]PAM NA Intergenic NA NA

T20611 ACTCGAGTCATTCAAAGATCAGG 1000
Oligo fwd CACCACTCGAGTCATTCAAAGATC
Oligo rev AAACGATCTTTGAATGACTCGAGT
Chromosome start end strand MM target_seq PAM alignment distance position gene name gene id
GU305914.1 185627 185649 - 0 ACTCGAGTCATTCAAAGATC AGG ||||||||[||||||||||||]PAM NA Intergenic NA NA

T20612 TTCAAAGATCAGGAAGACTGCGG 1000
Oligo fwd CACCTTCAAAGATCAGGAAGACTG
Oligo rev AAACCAGTCTTCCTGATCTTTGAA
Chromosome start end strand MM target_seq PAM alignment distance position gene name gene id
GU305914.1 185617 185639 - 0 TTCAAAGATCAGGAAGACTG CGG ||||||||[||||||||||||]PAM NA Intergenic NA NA

T20613 TGTCCGCACGTCAATCTACCTGG 1000
Oligo fwd CACCTGTCCGCACGTCAATCTACC
Oligo rev AAACGGTAGATTGACGTGCGGACA
Chromosome start end strand MM target_seq PAM alignment distance position gene name gene id
GU305914.1 185584 185606 - 0 TGTCCGCACGTCAATCTACC TGG ||||||||[||||||||||||]PAM NA Intergenic NA NA

| | | | | | | | | | | | |
|---|---|---|---|---|---|---|---|---|---|---|---|
| T20614 | CACGTCAATCTACCTGGATCAGG | 1000 | | | | | | | | | |
| Oligo fwd | CACCCACGTCAATCTACCTGGATC | | | | | | | | | | |
| Oligo rev | AAACGATCCAGGTAGATTGACGTG | | | | | | | | | | |
| Chromosome | start | end | strand | MM | target_seq | PAM | alignment | distance | position | gene name | gene id |
| GU305914.1 | 185578 | 185600 | - | 0 | CACGTCAATCTACCTGGATC | AGG | \|\|\|\|\|\|\|\|[\|\|\|\|\|\|\|\|\|\|\|\|]PAM | NA | Intergenic | NA | NA |
| T20615 | ATCTACCTGGATCAGGAAGAAGG | 1000 | | | | | | | | | |
| Oligo fwd | CACCATCTACCTGGATCAGGAAGA | | | | | | | | | | |
| Oligo rev | AAACTCTTCCTGATCCAGGTAGAT | | | | | | | | | | |
| Chromosome | start | end | strand | MM | target_seq | PAM | alignment | distance | position | gene name | gene id |
| GU305914.1 | 185571 | 185593 | - | 0 | ATCTACCTGGATCAGGAAGA | AGG | \|\|\|\|\|\|\|\|[\|\|\|\|\|\|\|\|\|\|\|\|]PAM | NA | Intergenic | NA | NA |
| T20616 | GGATCAGGAAGAAGGAAGCCCGG | 1000 | | | | | | | | | |
| Oligo fwd | CACCGGATCAGGAAGAAGGAAGCC | | | | | | | | | | |
| Oligo rev | AAACGGCTTCCTTCTTCCTGATCC | | | | | | | | | | |
| Chromosome | start | end | strand | MM | target_seq | PAM | alignment | distance | position | gene name | gene id |
| GU305914.1 | 185563 | 185585 | - | 0 | GGATCAGGAAGAAGGAAGCC | CGG | \|\|\|\|\|\|\|\|[\|\|\|\|\|\|\|\|\|\|\|\|]PAM | NA | Intergenic | NA | NA |
| T20617 | GATCAGGAAGAAGGAAGCCCGGG | 1000 | | | | | | | | | |
| Oligo fwd | CACCGATCAGGAAGAAGGAAGCCC | | | | | | | | | | |
| Oligo rev | AAACGGGCTTCCTTCTTCCTGATC | | | | | | | | | | |
| Chromosome | start | end | strand | MM | target_seq | PAM | alignment | distance | position | gene name | gene id |
| GU305914.1 | 185562 | 185584 | - | 0 | GATCAGGAAGAAGGAAGCCC | GGG | \|\|\|\|\|\|\|\|[\|\|\|\|\|\|\|\|\|\|\|\|]PAM | NA | Intergenic | NA | NA |
| T20618 | GGAAGAAGGAAGCCCGGGCCAGG | 1000 | | | | | | | | | |
| Oligo fwd | CACCGGAAGAAGGAAGCCCGGGCC | | | | | | | | | | |
| Oligo rev | AAACGGCCCGGGCTTCCTTCTTCC | | | | | | | | | | |
| Chromosome | start | end | strand | MM | target_seq | PAM | alignment | distance | position | gene name | gene id |
| GU305914.1 | 185557 | 185579 | - | 0 | GGAAGAAGGAAGCCCGGGCC | AGG | \|\|\|\|\|\|\|\|[\|\|\|\|\|\|\|\|\|\|\|\|]PAM | NA | Intergenic | NA | NA |
| T20619 | GAAGAAGGAAGCCCGGGCCAGGG | 1000 | | | | | | | | | |
| Oligo fwd | CACCGAAGAAGGAAGCCCGGGCCA | | | | | | | | | | |
| Oligo rev | AAACTGGCCCGGGCTTCCTTCTTC | | | | | | | | | | |
| Chromosome | start | end | strand | MM | target_seq | PAM | alignment | distance | position | gene name | gene id |
| GU305914.1 | 185556 | 185578 | - | 0 | GAAGAAGGAAGCCCGGGCCA | GGG | \|\|\|\|\|\|\|\|[\|\|\|\|\|\|\|\|\|\|\|\|]PAM | NA | Intergenic | NA | NA |
| T20620 | GCCCGGGCCAGGGTCGTCCGAGG | 1000 | | | | | | | | | |
| Oligo fwd | CACCGCCCGGGCCAGGGTCGTCCG | | | | | | | | | | |
| Oligo rev | AAACCGGACGACCCTGGCCCGGGC | | | | | | | | | | |
| Chromosome | start | end | strand | MM | target_seq | PAM | alignment | distance | position | gene name | gene id |
| GU305914.1 | 185546 | 185568 | - | 0 | GCCCGGGCCAGGGTCGTCCG | AGG | \|\|\|\|\|\|\|\|[\|\|\|\|\|\|\|\|\|\|\|\|]PAM | NA | Intergenic | NA | NA |
| T20621 | GGCCAGGGTCGTCCGAGGCCTGG | 1000 | | | | | | | | | |
| Oligo fwd | CACCGGCCAGGGTCGTCCGAGGCC | | | | | | | | | | |
| Oligo rev | AAACGGCCTCGGACGACCCTGGCC | | | | | | | | | | |
| Chromosome | start | end | strand | MM | target_seq | PAM | alignment | distance | position | gene name | gene id |
| GU305914.1 | 185541 | 185563 | - | 0 | GGCCAGGGTCGTCCGAGGCC | TGG | \|\|\|\|\|\|\|\|[\|\|\|\|\|\|\|\|\|\|\|\|]PAM | NA | Intergenic | NA | NA |
| T20622 | GCCAGGGTCGTCCGAGGCCTGGG | 1000 | | | | | | | | | |
| Oligo fwd | CACCGCCAGGGTCGTCCGAGGCCT | | | | | | | | | | |
| Oligo rev | AAACAGGCCTCGGACGACCCTGGC | | | | | | | | | | |
| Chromosome | start | end | strand | MM | target_seq | PAM | alignment | distance | position | gene name | gene id |
| GU305914.1 | 185540 | 185562 | - | 0 | GCCAGGGTCGTCCGAGGCCT | GGG | \|\|\|\|\|\|\|\|[\|\|\|\|\|\|\|\|\|\|\|\|]PAM | NA | Intergenic | NA | NA |
| T20623 | ACCGTCCACAATGAACAAGAAGG | 1000 | | | | | | | | | |
| Oligo fwd | CACCACCGTCCACAATGAACAAGA | | | | | | | | | | |
| Oligo rev | AAACTCTTGTTCATTGTGGACGGT | | | | | | | | | | |
| Chromosome | start | end | strand | MM | target_seq | PAM | alignment | distance | position | gene name | gene id |
| GU305914.1 | 185511 | 185533 | - | 0 | ACCGTCCACAATGAACAAGA | AGG | \|\|\|\|\|\|\|\|[\|\|\|\|\|\|\|\|\|\|\|\|]PAM | NA | Intergenic | NA | NA |
| T20624 | CCAGACAATGCACCTCAGTGAGG | 1000 | | | | | | | | | |
| Oligo fwd | CACCCCAGACAATGCACCTCAGTG | | | | | | | | | | |
| Oligo rev | AAACCACTGAGGTGCATTGTCTGG | | | | | | | | | | |
| Chromosome | start | end | strand | MM | target_seq | PAM | alignment | distance | position | gene name | gene id |
| GU305914.1 | 185476 | 185498 | - | 0 | CCAGACAATGCACCTCAGTG | AGG | \|\|\|\|\|\|\|\|[\|\|\|\|\|\|\|\|\|\|\|\|]PAM | NA | Intergenic | NA | NA |
| T20625 | CAATGCACCTCAGTGAGGAACGG | 1000 | | | | | | | | | |
| Oligo fwd | CACCCAATGCACCTCAGTGAGGAA | | | | | | | | | | |
| Oligo rev | AAACTTCCTCACTGAGGTGCATTG | | | | | | | | | | |
| Chromosome | start | end | strand | MM | target_seq | PAM | alignment | distance | position | gene name | gene id |
| GU305914.1 | 185471 | 185493 | - | 0 | CAATGCACCTCAGTGAGGAA | CGG | \|\|\|\|\|\|\|\|[\|\|\|\|\|\|\|\|\|\|\|\|]PAM | NA | Intergenic | NA | NA |
| T20626 | TCAGTGAGGAACGGTGTTTGTGG | 1000 | | | | | | | | | |
| Oligo fwd | CACCTCAGTGAGGAACGGTGTTTG | | | | | | | | | | |
| Oligo rev | AAACCAAACACCGTTCCTCACTGA | | | | | | | | | | |
| Chromosome | start | end | strand | MM | target_seq | PAM | alignment | distance | position | gene name | gene id |
| GU305914.1 | 185462 | 185484 | - | 0 | TCAGTGAGGAACGGTGTTTG | TGG | \|\|\|\|\|\|\|\|[\|\|\|\|\|\|\|\|\|\|\|\|]PAM | NA | Intergenic | NA | NA |
| T20627 | AAGCGTGTGTCCTTCCAAAGAGG | 1000 | | | | | | | | | |
| Oligo fwd | CACCAAGCGTGTGTCCTTCCAAAG | | | | | | | | | | |
| Oligo rev | AAACCTTTGGAAGGACACACGCTT | | | | | | | | | | |
| Chromosome | start | end | strand | MM | target_seq | PAM | alignment | distance | position | gene name | gene id |
| GU305914.1 | 185439 | 185461 | - | 0 | AAGCGTGTGTCCTTCCAAAG | AGG | \|\|\|\|\|\|\|\|[\|\|\|\|\|\|\|\|\|\|\|\|]PAM | NA | Intergenic | NA | NA |
| T20628 | TATCAATTGAGCTCTGTCAGTGG | 1000 | | | | | | | | | |
| Oligo fwd | CACCTATCAATTGAGCTCTGTCAG | | | | | | | | | | |
| Oligo rev | AAACCTGACAGAGCTCAATTGATA | | | | | | | | | | |
| Chromosome | start | end | strand | MM | target_seq | PAM | alignment | distance | position | gene name | gene id |
| GU305914.1 | 185408 | 185430 | - | 0 | TATCAATTGAGCTCTGTCAG | TGG | \|\|\|\|\|\|\|\|[\|\|\|\|\|\|\|\|\|\|\|\|]PAM | NA | Intergenic | NA | NA |

T20629 ATCAATTGAGCTCTGTCAGTGGG 1000
Oligo fwd CACCATCAATTGAGCTCTGTCAGT
Oligo rev AAACACTGACAGAGCTCAATTGAT
Chromosome start end strand MM target_seq PAM alignment distance position gene name gene id
GU305914.1 185407 185429 - 0 ATCAATTGAGCTCTGTCAGT GGG ||||||||[||||||||||||]PAM NA Intergenic NA NA

T20630 AATTGAGCTCTGTCAGTGGGCGG 1000
Oligo fwd CACCAATTGAGCTCTGTCAGTGGG
Oligo rev AAACCCCACTGACAGAGCTCAATT
Chromosome start end strand MM target_seq PAM alignment distance position gene name gene id
GU305914.1 185404 185426 - 0 AATTGAGCTCTGTCAGTGGG CGG ||||||||[||||||||||||]PAM NA Intergenic NA NA

T20631 ATTGAGCTCTGTCAGTGGGCGGG 1000
Oligo fwd CACCATTGAGCTCTGTCAGTGGGC
Oligo rev AAACGCCCACTGACAGAGCTCAAT
Chromosome start end strand MM target_seq PAM alignment distance position gene name gene id
GU305914.1 185403 185425 - 0 ATTGAGCTCTGTCAGTGGGC GGG ||||||||[||||||||||||]PAM NA Intergenic NA NA

T20632 TTGAGCTCTGTCAGTGGGCGGGG 1000
Oligo fwd CACCTTGAGCTCTGTCAGTGGGCG
Oligo rev AAACCGCCCACTGACAGAGCTCAA
Chromosome start end strand MM target_seq PAM alignment distance position gene name gene id
GU305914.1 185402 185424 - 0 TTGAGCTCTGTCAGTGGGCG GGG ||||||||[||||||||||||]PAM NA Intergenic NA NA

T20633 TGAGCTCTGTCAGTGGGCGGGGG 1000
Oligo fwd CACCTGAGCTCTGTCAGTGGGCGG
Oligo rev AAACCCGCCCACTGACAGAGCTCA
Chromosome start end strand MM target_seq PAM alignment distance position gene name gene id
GU305914.1 185401 185423 - 0 TGAGCTCTGTCAGTGGGCGG GGG ||||||||[||||||||||||]PAM NA Intergenic NA NA

T20634 TCTGTCAGTGGGCGGGGGTCTGG 1000
Oligo fwd CACCTCTGTCAGTGGGCGGGGGTC
Oligo rev AAACGACCCCCGCCCACTGACAGA
Chromosome start end strand MM target_seq PAM alignment distance position gene name gene id
GU305914.1 185396 185418 - 0 TCTGTCAGTGGGCGGGGGTC TGG ||||||||[||||||||||||]PAM NA Intergenic NA NA

T20635 GGGGTCTGGCAGAGATGAGCAGG 1000
Oligo fwd CACCGGGGTCTGGCAGAGATGAGC
Oligo rev AAACGCTCATCTCTGCCAGACCCC
Chromosome start end strand MM target_seq PAM alignment distance position gene name gene id
GU305914.1 185382 185404 - 0 GGGGTCTGGCAGAGATGAGC AGG ||||||||[||||||||||||]PAM NA Intergenic NA NA

T20636 TTTTAGAGCTGCTTTTCAACAGG 1000
Oligo fwd CACCTTTTAGAGCTGCTTTTCAAC
Oligo rev AAACGTTGAAAAGCAGCTCTAAAA
Chromosome start end strand MM target_seq PAM alignment distance position gene name gene id
GU305914.1 185353 185375 - 0 TTTTAGAGCTGCTTTTCAAC AGG ||||||||[||||||||||||]PAM NA Intergenic NA NA

T20637 TTTTCAACAGGAGTGTCTCTTGG 1000
Oligo fwd CACCTTTTCAACAGGAGTGTCTCT
Oligo rev AAACAGAGACACTCCTGTTGAAAA
Chromosome start end strand MM target_seq PAM alignment distance position gene name gene id
GU305914.1 185341 185363 - 0 TTTTCAACAGGAGTGTCTCT TGG ||||||||[||||||||||||]PAM NA Intergenic NA NA

T20638 GGAGTGTCTCTTGGCAGTCTCGG 1000
Oligo fwd CACCGGAGTGTCTCTTGGCAGTCT
Oligo rev AAACAGACTGCCAAGAGACACTCC
Chromosome start end strand MM target_seq PAM alignment distance position gene name gene id
GU305914.1 185332 185354 - 0 GGAGTGTCTCTTGGCAGTCT CGG ||||||||[||||||||||||]PAM NA Intergenic NA NA

T20639 CTCTTGGCAGTCTCGGACACTGG 1000
Oligo fwd CACCCTCTTGGCAGTCTCGGACAC
Oligo rev AAACGTGTCCGAGACTGCCAAGAG
Chromosome start end strand MM target_seq PAM alignment distance position gene name gene id
GU305914.1 185325 185347 - 0 CTCTTGGCAGTCTCGGACAC TGG ||||||||[||||||||||||]PAM NA Intergenic NA NA

T20640 TCTTGGCAGTCTCGGACACTGGG 1000
Oligo fwd CACCTCTTGGCAGTCTCGGACACT
Oligo rev AAACAGTGTCCGAGACTGCCAAGA
Chromosome start end strand MM target_seq PAM alignment distance position gene name gene id
GU305914.1 185324 185346 - 0 TCTTGGCAGTCTCGGACACT GGG ||||||||[||||||||||||]PAM NA Intergenic NA NA

T20641 TGGCAGTCTCGGACACTGGGAGG 1000
Oligo fwd CACCTGGCAGTCTCGGACACTGGG
Oligo rev AAACCCCAGTGTCCGAGACTGCCA
Chromosome start end strand MM target_seq PAM alignment distance position gene name gene id
GU305914.1 185321 185343 - 0 TGGCAGTCTCGGACACTGGG AGG ||||||||[||||||||||||]PAM NA Intergenic NA NA

T20642 CTCGGACACTGGGAGGATCTCGG 1000
Oligo fwd CACCCTCGGACACTGGGAGGATCT
Oligo rev AAACAGATCCTCCCAGTGTCCGAG
Chromosome start end strand MM target_seq PAM alignment distance position gene name gene id
GU305914.1 185314 185336 - 0 CTCGGACACTGGGAGGATCT CGG ||||||||[||||||||||||]PAM NA Intergenic NA NA

T20643 GGATCTCGGACCCCTTCCCCTGG 1000
Oligo fwd CACCGGATCTCGGACCCCTTCCCC
Oligo rev AAACGGGGAAGGGGTCCGAGATCC
Chromosome start end strand MM target_seq PAM alignment distance position gene name gene id

GU305914.1 185300 185322 - 0 GGATCTCGGACCCCTTCCCC TGG ||||||||[||||||||||||]PAM NA Intergenic NA NA

T20644 GATCTCGGACCCCTTCCCCTGGG 1000
Oligo fwd CACCGATCTCGGACCCCTTCCCCT
Oligo rev AAACAGGGGAAGGGGTCCGAGATC
Chromosome start end strand MM target_seq PAM alignment distance position gene name gene id
GU305914.1 185299 185321 - 0 GATCTCGGACCCCTTCCCCT GGG ||||||||[||||||||||||]PAM NA Intergenic NA NA

T20645 TTTCCCAACTGTTCAGTGTCAGG 1000
Oligo fwd CACCTTTCCCAACTGTTCAGTGTC
Oligo rev AAACGACACTGAACAGTTGGGAAA
Chromosome start end strand MM target_seq PAM alignment distance position gene name gene id
GU305914.1 185256 185278 - 0 TTTCCCAACTGTTCAGTGTC AGG ||||||||[||||||||||||]PAM NA Intergenic NA NA

T20646 AGTGTCAGGACACGCAGCTCTGG 1000
Oligo fwd CACCAGTGTCAGGACACGCAGCTC
Oligo rev AAACGAGCTGCGTGTCCTGACACT
Chromosome start end strand MM target_seq PAM alignment distance position gene name gene id
GU305914.1 185242 185264 - 0 AGTGTCAGGACACGCAGCTC TGG ||||||||[||||||||||||]PAM NA Intergenic NA NA

T20647 GAGCTCTGCAGACCGTCTCCCGG 1000
Oligo fwd CACCGAGCTCTGCAGACCGTCTCC
Oligo rev AAACGGAGACGGTCTGCAGAGCTC
Chromosome start end strand MM target_seq PAM alignment distance position gene name gene id
GU305914.1 185220 185242 - 0 GAGCTCTGCAGACCGTCTCC CGG ||||||||[||||||||||||]PAM NA Intergenic NA NA

T20648 CCGGCCGCTCCCGAGCCACCCGG 1000
Oligo fwd CACCCCGGCCGCTCCCGAGCCACC
Oligo rev AAACGGTGGCTCGGGAGCGGCCGG
Chromosome start end strand MM target_seq PAM alignment distance position gene name gene id
GU305914.1 185201 185223 - 0 CCGGCCGCTCCCGAGCCACC CGG ||||||||[||||||||||||]PAM NA Intergenic NA NA

T20649 CGGCCGCTCCCGAGCCACCCGGG 1000
Oligo fwd CACCCGGCCGCTCCCGAGCCACCC
Oligo rev AAACGGGTGGCTCGGGAGCGGCCG
Chromosome start end strand MM target_seq PAM alignment distance position gene name gene id
GU305914.1 185200 185222 - 0 CGGCCGCTCCCGAGCCACCC GGG ||||||||[||||||||||||]PAM NA Intergenic NA NA

T20650 TCCCGATCCTCTCTCGTTCCCGG 1000
Oligo fwd CACCTCCCGATCCTCTCTCGTTCC
Oligo rev AAACGGAACGAGAGAGGATCGGGA
Chromosome start end strand MM target_seq PAM alignment distance position gene name gene id
GU305914.1 185170 185192 - 0 TCCCGATCCTCTCTCGTTCC CGG ||||||||[||||||||||||]PAM NA Intergenic NA NA

T20651 GCCCGCTCGACTCGATCCCTCGG 1000
Oligo fwd CACCGCCCGCTCGACTCGATCCCT
Oligo rev AAACAGGGATCGAGTCGAGCGGGC
Chromosome start end strand MM target_seq PAM alignment distance position gene name gene id
GU305914.1 185138 185160 - 0 GCCCGCTCGACTCGATCCCT CGG ||||||||[||||||||||||]PAM NA Intergenic NA NA

T20652 CCCTCGGCCGAACCCAGCAACGG 1000
Oligo fwd CACCCCCTCGGCCGAACCCAGCAA
Oligo rev AAACTTGCTGGGTTCGGCCGAGGG
Chromosome start end strand MM target_seq PAM alignment distance position gene name gene id
GU305914.1 185122 185144 - 0 CCCTCGGCCGAACCCAGCAA CGG ||||||||[||||||||||||]PAM NA Intergenic NA NA

T20653 CCCAGCAACGGAGACTACGCTGG 1000
Oligo fwd CACCCCCAGCAACGGAGACTACGC
Oligo rev AAACGCGTAGTCTCCGTTGCTGGG
Chromosome start end strand MM target_seq PAM alignment distance position gene name gene id
GU305914.1 185110 185132 - 0 CCCAGCAACGGAGACTACGC TGG ||||||||[||||||||||||]PAM NA Intergenic NA NA

T20654 GGAGACTACGCTGGCCCTCCCGG 1000
Oligo fwd CACCGGAGACTACGCTGGCCCTCC
Oligo rev AAACGGAGGGCCAGCGTAGTCTCC
Chromosome start end strand MM target_seq PAM alignment distance position gene name gene id
GU305914.1 185101 185123 - 0 GGAGACTACGCTGGCCCTCC CGG ||||||||[||||||||||||]PAM NA Intergenic NA NA

T20655 GAGACTACGCTGGCCCTCCCGGG 1000
Oligo fwd CACCGAGACTACGCTGGCCCTCCC
Oligo rev AAACGGGAGGGCCAGCGTAGTCTC
Chromosome start end strand MM target_seq PAM alignment distance position gene name gene id
GU305914.1 185100 185122 - 0 GAGACTACGCTGGCCCTCCC GGG ||||||||[||||||||||||]PAM NA Intergenic NA NA

T20656 ATGATCCGAACAACGCGTTTCGG 1000
Oligo fwd CACCATGATCCGAACAACGCGTTT
Oligo rev AAACAAACGCGTTGTTCGGATCAT
Chromosome start end strand MM target_seq PAM alignment distance position gene name gene id
GU305914.1 185077 185099 - 0 ATGATCCGAACAACGCGTTT CGG ||||||||[||||||||||||]PAM NA Intergenic NA NA

T20657 TCCGAACAACGCGTTTCGGCAGG 1000
Oligo fwd CACCTCCGAACAACGCGTTTCGGC
Oligo rev AAACGCCGAAACGCGTTGTTCGGA
Chromosome start end strand MM target_seq PAM alignment distance position gene name gene id
GU305914.1 185073 185095 - 0 TCCGAACAACGCGTTTCGGC AGG ||||||||[||||||||||||]PAM NA Intergenic NA NA

T20658 TTTCTAGCCCTTTTTCTGCTCGG 1000
Oligo fwd CACCTTTCTAGCCCTTTTTCTGCT
Oligo rev AAACAGCAGAAAAAGGGCTAGAAA

| Chromosome | start | end | strand | MM | target_seq | PAM | alignment | distance | position | gene name | gene id |
|---|---|---|---|---|---|---|---|---|---|---|---|
| GU305914.1 | 185040 | 185062 | - | 0 | TTTCTAGCCCTTTTTCTGCT | CGG | \|\|\|\|\|\|\|\|[\|\|\|\|\|\|\|\|\|\|\|\|]PAM | NA | Intergenic | NA | NA |
| T20659 | TGCTCGGAATCCGAGCCGAGCGG | 1000 | | | | | | | | | |
| Oligo fwd | CACCTGCTCGGAATCCGAGCCGAG | | | | | | | | | | |
| Oligo rev | AAACCTCGGCTCGGATTCCGAGCA | | | | | | | | | | |
| Chromosome | start | end | strand | MM | target_seq | PAM | alignment | distance | position | gene name | gene id |
| GU305914.1 | 185024 | 185046 | - | 0 | TGCTCGGAATCCGAGCCGAG | CGG | \|\|\|\|\|\|\|\|[\|\|\|\|\|\|\|\|\|\|\|\|]PAM | NA | Intergenic | NA | NA |
| T20660 | AGAAAACGAAAGTAAAAACCCGG | 1000 | | | | | | | | | |
| Oligo fwd | CACCAGAAAACGAAAGTAAAAACC | | | | | | | | | | |
| Oligo rev | AAACGGTTTTTACTTTCGTTTTCT | | | | | | | | | | |
| Chromosome | start | end | strand | MM | target_seq | PAM | alignment | distance | position | gene name | gene id |
| GU305914.1 | 184995 | 185017 | - | 0 | AGAAAACGAAAGTAAAAACC | CGG | \|\|\|\|\|\|\|\|[\|\|\|\|\|\|\|\|\|\|\|\|]PAM | NA | Intergenic | NA | NA |
| T20661 | GAAAACGAAAGTAAAAACCCGGG | 1000 | | | | | | | | | |
| Oligo fwd | CACCGAAAACGAAAGTAAAAACCC | | | | | | | | | | |
| Oligo rev | AAACGGGTTTTTACTTTCGTTTTC | | | | | | | | | | |
| Chromosome | start | end | strand | MM | target_seq | PAM | alignment | distance | position | gene name | gene id |
| GU305914.1 | 184994 | 185016 | - | 0 | GAAAACGAAAGTAAAAACCC | GGG | \|\|\|\|\|\|\|\|[\|\|\|\|\|\|\|\|\|\|\|\|]PAM | NA | Intergenic | NA | NA |
| T20662 | AACGAAAGTAAAAACCCGGGCGG | 1000 | | | | | | | | | |
| Oligo fwd | CACCAACGAAAGTAAAAACCCGGG | | | | | | | | | | |
| Oligo rev | AAACCCCGGGTTTTTACTTTCGTT | | | | | | | | | | |
| Chromosome | start | end | strand | MM | target_seq | PAM | alignment | distance | position | gene name | gene id |
| GU305914.1 | 184991 | 185013 | - | 0 | AACGAAAGTAAAAACCCGGG | CGG | \|\|\|\|\|\|\|\|[\|\|\|\|\|\|\|\|\|\|\|\|]PAM | NA | Intergenic | NA | NA |
| T20663 | GTAAAAACCCGGGCGGTCCGCGG | 1000 | | | | | | | | | |
| Oligo fwd | CACCGTAAAAACCCGGGCGGTCCG | | | | | | | | | | |
| Oligo rev | AAACCGGACCGCCCGGGTTTTTAC | | | | | | | | | | |
| Chromosome | start | end | strand | MM | target_seq | PAM | alignment | distance | position | gene name | gene id |
| GU305914.1 | 184984 | 185006 | - | 0 | GTAAAAACCCGGGCGGTCCG | CGG | \|\|\|\|\|\|\|\|[\|\|\|\|\|\|\|\|\|\|\|\|]PAM | NA | Intergenic | NA | NA |
| GU305914.1 | 135443 | 135465 | - | 4 | GAAGAAACCCGGGCCGTCGG | CGG | \|-\|-\|\|\|\|[\|\|\|\|\|\|-\|\|\|-\|]PAM | NA | Intergenic | NA | NA |
| T20664 | TCGCTCGAATAAAATTATAGAGG | 1000 | | | | | | | | | |
| Oligo fwd | CACCTCGCTCGAATAAAATTATAG | | | | | | | | | | |
| Oligo rev | AAACCTATAATTTTATTCGAGCGA | | | | | | | | | | |
| Chromosome | start | end | strand | MM | target_seq | PAM | alignment | distance | position | gene name | gene id |
| GU305914.1 | 184954 | 184976 | - | 0 | TCGCTCGAATAAAATTATAG | AGG | \|\|\|\|\|\|\|\|[\|\|\|\|\|\|\|\|\|\|\|\|]PAM | NA | Intergenic | NA | NA |
| T20665 | CGCTCGAATAAAATTATAGAGGG | 1000 | | | | | | | | | |
| Oligo fwd | CACCCGCTCGAATAAAATTATAGA | | | | | | | | | | |
| Oligo rev | AAACTCTATAATTTTATTCGAGCG | | | | | | | | | | |
| Chromosome | start | end | strand | MM | target_seq | PAM | alignment | distance | position | gene name | gene id |
| GU305914.1 | 184953 | 184975 | - | 0 | CGCTCGAATAAAATTATAGA | GGG | \|\|\|\|\|\|\|\|[\|\|\|\|\|\|\|\|\|\|\|\|]PAM | NA | Intergenic | NA | NA |
| T20666 | GCTCGAATAAAATTATAGAGGGG | 1000 | | | | | | | | | |
| Oligo fwd | CACCGCTCGAATAAAATTATAGAG | | | | | | | | | | |
| Oligo rev | AAACCTCTATAATTTTATTCGAGC | | | | | | | | | | |
| Chromosome | start | end | strand | MM | target_seq | PAM | alignment | distance | position | gene name | gene id |
| GU305914.1 | 184952 | 184974 | - | 0 | GCTCGAATAAAATTATAGAG | GGG | \|\|\|\|\|\|\|\|[\|\|\|\|\|\|\|\|\|\|\|\|]PAM | NA | Intergenic | NA | NA |
| T20667 | CTAAAATTTTTCCTGCGTCCTGG | 1000 | | | | | | | | | |
| Oligo fwd | CACCCTAAAATTTTTCCTGCGTCC | | | | | | | | | | |
| Oligo rev | AAACGGACGCAGGAAAAATTTTAG | | | | | | | | | | |
| Chromosome | start | end | strand | MM | target_seq | PAM | alignment | distance | position | gene name | gene id |
| GU305914.1 | 184927 | 184949 | - | 0 | CTAAAATTTTTCCTGCGTCC | TGG | \|\|\|\|\|\|\|\|[\|\|\|\|\|\|\|\|\|\|\|\|]PAM | NA | Intergenic | NA | NA |
| T20668 | TAAAATTTTTCCTGCGTCCTGGG | 1000 | | | | | | | | | |
| Oligo fwd | CACCTAAAATTTTTCCTGCGTCCT | | | | | | | | | | |
| Oligo rev | AAACAGGACGCAGGAAAAATTTTA | | | | | | | | | | |
| Chromosome | start | end | strand | MM | target_seq | PAM | alignment | distance | position | gene name | gene id |
| GU305914.1 | 184926 | 184948 | - | 0 | TAAAATTTTTCCTGCGTCCT | GGG | \|\|\|\|\|\|\|\|[\|\|\|\|\|\|\|\|\|\|\|\|]PAM | NA | Intergenic | NA | NA |
| T20669 | AAAATTTTTCCTGCGTCCTGGGG | 1000 | | | | | | | | | |
| Oligo fwd | CACCAAAATTTTTCCTGCGTCCTG | | | | | | | | | | |
| Oligo rev | AAACCAGGACGCAGGAAAAATTTT | | | | | | | | | | |
| Chromosome | start | end | strand | MM | target_seq | PAM | alignment | distance | position | gene name | gene id |
| GU305914.1 | 184925 | 184947 | - | 0 | AAAATTTTTCCTGCGTCCTG | GGG | \|\|\|\|\|\|\|\|[\|\|\|\|\|\|\|\|\|\|\|\|]PAM | NA | Intergenic | NA | NA |
| T20670 | AAATTTTTCCTGCGTCCTGGGGG | 1000 | | | | | | | | | |
| Oligo fwd | CACCAAATTTTTCCTGCGTCCTGG | | | | | | | | | | |
| Oligo rev | AAACCCAGGACGCAGGAAAAATTT | | | | | | | | | | |
| Chromosome | start | end | strand | MM | target_seq | PAM | alignment | distance | position | gene name | gene id |
| GU305914.1 | 184924 | 184946 | - | 0 | AAATTTTTCCTGCGTCCTGG | GGG | \|\|\|\|\|\|\|\|[\|\|\|\|\|\|\|\|\|\|\|\|]PAM | NA | Intergenic | NA | NA |
| T20671 | AATTTTTCCTGCGTCCTGGGGGG | 1000 | | | | | | | | | |
| Oligo fwd | CACCAATTTTTCCTGCGTCCTGGG | | | | | | | | | | |
| Oligo rev | AAACCCCAGGACGCAGGAAAAATT | | | | | | | | | | |
| Chromosome | start | end | strand | MM | target_seq | PAM | alignment | distance | position | gene name | gene id |
| GU305914.1 | 184923 | 184945 | - | 0 | AATTTTTCCTGCGTCCTGGG | GGG | \|\|\|\|\|\|\|\|[\|\|\|\|\|\|\|\|\|\|\|\|]PAM | NA | Intergenic | NA | NA |
| T20672 | GCGTCCTGGGGGGCATCTATAGG | 1000 | | | | | | | | | |
| Oligo fwd | CACCGCGTCCTGGGGGGCATCTAT | | | | | | | | | | |
| Oligo rev | AAACATAGATGCCCCCCAGGACGC | | | | | | | | | | |
| Chromosome | start | end | strand | MM | target_seq | PAM | alignment | distance | position | gene name | gene id |
| GU305914.1 | 184913 | 184935 | - | 0 | GCGTCCTGGGGGGCATCTAT | AGG | \|\|\|\|\|\|\|\|[\|\|\|\|\|\|\|\|\|\|\|\|]PAM | NA | Intergenic | NA | NA |
| GU305914.1 | 45870 | 45892 | + | 4 | GCGTGCGGGTGGGCATGTAT | CGG | \|\|\|\|-\|-\|[\|-\|\|\|\|\|\|-\|\|\|]PAM | NA | Intergenic | NA | NA |

T20673 CGTCCTGGGGGGCATCTATAGGG 1000
Oligo fwd CACCCGTCCTGGGGGGCATCTATA
Oligo rev AAACTATAGATGCCCCCCAGGACG
Chromosome start end strand MM target_seq PAM alignment distance position gene name gene id
GU305914.1 184912 184934 - 0 CGTCCTGGGGGGCATCTATA GGG ||||||||[||||||||||||]PAM NA Intergenic NA NA

T20674 GGGGGCATCTATAGGGTCTAAGG 1000
Oligo fwd CACCGGGGGCATCTATAGGGTCTA
Oligo rev AAACTAGACCCTATAGATGCCCCC
Chromosome start end strand MM target_seq PAM alignment distance position gene name gene id
GU305914.1 184905 184927 - 0 GGGGGCATCTATAGGGTCTA AGG ||||||||[||||||||||||]PAM NA Intergenic NA NA

T20675 AGGGTCTAAGGCACATACCCCGG 1000
Oligo fwd CACCAGGGTCTAAGGCACATACCC
Oligo rev AAACGGGTATGTGCCTTAGACCCT
Chromosome start end strand MM target_seq PAM alignment distance position gene name gene id
GU305914.1 184893 184915 - 0 AGGGTCTAAGGCACATACCC CGG ||||||||[||||||||||||]PAM NA Intergenic NA NA

T20676 GGGTCTAAGGCACATACCCCGGG 1000
Oligo fwd CACCGGGTCTAAGGCACATACCCC
Oligo rev AAACGGGGTATGTGCCTTAGACCC
Chromosome start end strand MM target_seq PAM alignment distance position gene name gene id
GU305914.1 184892 184914 - 0 GGGTCTAAGGCACATACCCC GGG ||||||||[||||||||||||]PAM NA Intergenic NA NA

T20677 GGTCTAAGGCACATACCCCGGGG 1000
Oligo fwd CACCGGTCTAAGGCACATACCCCG
Oligo rev AAACCGGGGTATGTGCCTTAGACC
Chromosome start end strand MM target_seq PAM alignment distance position gene name gene id
GU305914.1 184891 184913 - 0 GGTCTAAGGCACATACCCCG GGG ||||||||[||||||||||||]PAM NA Intergenic NA NA

T20678 GTCTAAGGCACATACCCCGGGGG 1000
Oligo fwd CACCGTCTAAGGCACATACCCCGG
Oligo rev AAACCCGGGGTATGTGCCTTAGAC
Chromosome start end strand MM target_seq PAM alignment distance position gene name gene id
GU305914.1 184890 184912 - 0 GTCTAAGGCACATACCCCGG GGG ||||||||[||||||||||||]PAM NA Intergenic NA NA

T20679 TAAGGCACATACCCCGGGGGTGG 1000
Oligo fwd CACCTAAGGCACATACCCCGGGGG
Oligo rev AAACCCCCCGGGGTATGTGCCTTA
Chromosome start end strand MM target_seq PAM alignment distance position gene name gene id
GU305914.1 184887 184909 - 0 TAAGGCACATACCCCGGGGG TGG ||||||||[||||||||||||]PAM NA Intergenic NA NA

T20680 AAGGCACATACCCCGGGGGTGGG 1000
Oligo fwd CACCAAGGCACATACCCCGGGGGT
Oligo rev AAACACCCCCGGGGTATGTGCCTT
Chromosome start end strand MM target_seq PAM alignment distance position gene name gene id
GU305914.1 184886 184908 - 0 AAGGCACATACCCCGGGGGT GGG ||||||||[||||||||||||]PAM NA Intergenic NA NA

T20681 CCCGGGGGTGGGTTGCGTTTCGG 1000
Oligo fwd CACCCCCGGGGGTGGGTTGCGTTT
Oligo rev AAACAAACGCAACCCACCCCCGGG
Chromosome start end strand MM target_seq PAM alignment distance position gene name gene id
GU305914.1 184875 184897 - 0 CCCGGGGGTGGGTTGCGTTT CGG ||||||||[||||||||||||]PAM NA Intergenic NA NA

T20682 GTGGGTTGCGTTTCGGAAGCAGG 1000
Oligo fwd CACCGTGGGTTGCGTTTCGGAAGC
Oligo rev AAACGCTTCCGAAACGCAACCCAC
Chromosome start end strand MM target_seq PAM alignment distance position gene name gene id
GU305914.1 184868 184890 - 0 GTGGGTTGCGTTTCGGAAGC AGG ||||||||[||||||||||||]PAM NA Intergenic NA NA

T20683 GCGTTTCGGAAGCAGGATCGAGG 1000
Oligo fwd CACCGCGTTTCGGAAGCAGGATCG
Oligo rev AAACCGATCCTGCTTCCGAAACGC
Chromosome start end strand MM target_seq PAM alignment distance position gene name gene id
GU305914.1 184861 184883 - 0 GCGTTTCGGAAGCAGGATCG AGG ||||||||[||||||||||||]PAM NA Intergenic NA NA

T20684 CGTTTCGGAAGCAGGATCGAGGG 1000
Oligo fwd CACCCGTTTCGGAAGCAGGATCGA
Oligo rev AAACTCGATCCTGCTTCCGAAACG
Chromosome start end strand MM target_seq PAM alignment distance position gene name gene id
GU305914.1 184860 184882 - 0 CGTTTCGGAAGCAGGATCGA GGG ||||||||[||||||||||||]PAM NA Intergenic NA NA

T20685 GAAAAAGCGATCGAAAGTAGCGG 1000
Oligo fwd CACCGAAAAAGCGATCGAAAGTAG
Oligo rev AAACCTACTTTCGATCGCTTTTTC
Chromosome start end strand MM target_seq PAM alignment distance position gene name gene id
GU305914.1 184835 184857 - 0 GAAAAAGCGATCGAAAGTAG CGG ||||||||[||||||||||||]PAM NA Intergenic NA NA

T20686 TATAGCCCTGACAGCTGTTACGG 1000
Oligo fwd CACCTATAGCCCTGACAGCTGTTA
Oligo rev AAACTAACAGCTGTCAGGGCTATA
Chromosome start end strand MM target_seq PAM alignment distance position gene name gene id
GU305914.1 184801 184823 - 0 TATAGCCCTGACAGCTGTTA CGG ||||||||[||||||||||||]PAM NA Intergenic NA NA

T20687 ATAGCCCTGACAGCTGTTACGGG 1000
Oligo fwd CACCATAGCCCTGACAGCTGTTAC
Oligo rev AAACGTAACAGCTGTCAGGGCTAT
Chromosome start end strand MM target_seq PAM alignment distance position gene name gene id
GU305914.1 184800 184822 - 0 ATAGCCCTGACAGCTGTTAC GGG ||||||||[||||||||||||]PAM NA Intergenic NA NA

<table>
<tr><td>T20688</td><td>GCCCTGACAGCTGTTACGGGCGG</td><td>1000</td><td></td><td></td><td></td><td></td><td></td><td></td><td></td><td></td><td></td></tr>
<tr><td>Oligo fwd</td><td>CACCGCCCTGACAGCTGTTACGGG</td><td></td><td></td><td></td><td></td><td></td><td></td><td></td><td></td><td></td><td></td></tr>
<tr><td>Oligo rev</td><td>AAACCCCGTAACAGCTGTCAGGGC</td><td></td><td></td><td></td><td></td><td></td><td></td><td></td><td></td><td></td><td></td></tr>
<tr><td>Chromosome</td><td>start</td><td>end</td><td>strand</td><td>MM</td><td>target_seq</td><td>PAM</td><td>alignment</td><td>distance</td><td>position</td><td>gene name</td><td>gene id</td></tr>
<tr><td>GU305914.1</td><td>184797</td><td>184819</td><td>-</td><td>0</td><td>GCCCTGACAGCTGTTACGGG</td><td>CGG</td><td>||||||||[||||||||||||]PAM</td><td>NA</td><td>Intergenic</td><td>NA</td><td>NA</td></tr>
<tr><td>T20689</td><td>TAAAAATACAAGCAGCCGCTAGG</td><td>1000</td><td></td><td></td><td></td><td></td><td></td><td></td><td></td><td></td><td></td></tr>
<tr><td>Oligo fwd</td><td>CACCTAAAAATACAAGCAGCCGCT</td><td></td><td></td><td></td><td></td><td></td><td></td><td></td><td></td><td></td><td></td></tr>
<tr><td>Oligo rev</td><td>AAACAGCGGCTGCTTGTATTTTTA</td><td></td><td></td><td></td><td></td><td></td><td></td><td></td><td></td><td></td><td></td></tr>
<tr><td>Chromosome</td><td>start</td><td>end</td><td>strand</td><td>MM</td><td>target_seq</td><td>PAM</td><td>alignment</td><td>distance</td><td>position</td><td>gene name</td><td>gene id</td></tr>
<tr><td>GU305914.1</td><td>184752</td><td>184774</td><td>-</td><td>0</td><td>TAAAAATACAAGCAGCCGCT</td><td>AGG</td><td>||||||||[||||||||||||]PAM</td><td>NA</td><td>Intergenic</td><td>NA</td><td>NA</td></tr>
<tr><td>T20690</td><td>AAAAATACAAGCAGCCGCTAGGG</td><td>1000</td><td></td><td></td><td></td><td></td><td></td><td></td><td></td><td></td><td></td></tr>
<tr><td>Oligo fwd</td><td>CACCAAAAATACAAGCAGCCGCTA</td><td></td><td></td><td></td><td></td><td></td><td></td><td></td><td></td><td></td><td></td></tr>
<tr><td>Oligo rev</td><td>AAACTAGCGGCTGCTTGTATTTTT</td><td></td><td></td><td></td><td></td><td></td><td></td><td></td><td></td><td></td><td></td></tr>
<tr><td>Chromosome</td><td>start</td><td>end</td><td>strand</td><td>MM</td><td>target_seq</td><td>PAM</td><td>alignment</td><td>distance</td><td>position</td><td>gene name</td><td>gene id</td></tr>
<tr><td>GU305914.1</td><td>184751</td><td>184773</td><td>-</td><td>0</td><td>AAAAATACAAGCAGCCGCTA</td><td>GGG</td><td>||||||||[||||||||||||]PAM</td><td>NA</td><td>Intergenic</td><td>NA</td><td>NA</td></tr>
<tr><td>T20691</td><td>AAAATACAAGCAGCCGCTAGGGG</td><td>1000</td><td></td><td></td><td></td><td></td><td></td><td></td><td></td><td></td><td></td></tr>
<tr><td>Oligo fwd</td><td>CACCAAAATACAAGCAGCCGCTAG</td><td></td><td></td><td></td><td></td><td></td><td></td><td></td><td></td><td></td><td></td></tr>
<tr><td>Oligo rev</td><td>AAACCTAGCGGCTGCTTGTATTTT</td><td></td><td></td><td></td><td></td><td></td><td></td><td></td><td></td><td></td><td></td></tr>
<tr><td>Chromosome</td><td>start</td><td>end</td><td>strand</td><td>MM</td><td>target_seq</td><td>PAM</td><td>alignment</td><td>distance</td><td>position</td><td>gene name</td><td>gene id</td></tr>
<tr><td>GU305914.1</td><td>184750</td><td>184772</td><td>-</td><td>0</td><td>AAAATACAAGCAGCCGCTAG</td><td>GGG</td><td>||||||||[||||||||||||]PAM</td><td>NA</td><td>Intergenic</td><td>NA</td><td>NA</td></tr>
<tr><td>T20692</td><td>TAGGGGTCGCCCACCTATCCAGG</td><td>1000</td><td></td><td></td><td></td><td></td><td></td><td></td><td></td><td></td><td></td></tr>
<tr><td>Oligo fwd</td><td>CACCTAGGGGTCGCCCACCTATCC</td><td></td><td></td><td></td><td></td><td></td><td></td><td></td><td></td><td></td><td></td></tr>
<tr><td>Oligo rev</td><td>AAACGGATAGGTGGGCGACCCCTA</td><td></td><td></td><td></td><td></td><td></td><td></td><td></td><td></td><td></td><td></td></tr>
<tr><td>Chromosome</td><td>start</td><td>end</td><td>strand</td><td>MM</td><td>target_seq</td><td>PAM</td><td>alignment</td><td>distance</td><td>position</td><td>gene name</td><td>gene id</td></tr>
<tr><td>GU305914.1</td><td>184733</td><td>184755</td><td>-</td><td>0</td><td>TAGGGGTCGCCCACCTATCC</td><td>AGG</td><td>||||||||[||||||||||||]PAM</td><td>NA</td><td>Intergenic</td><td>NA</td><td>NA</td></tr>
<tr><td>T20693</td><td>ACCTATCCAGGCACGTACCCTGG</td><td>1000</td><td></td><td></td><td></td><td></td><td></td><td></td><td></td><td></td><td></td></tr>
<tr><td>Oligo fwd</td><td>CACCACCTATCCAGGCACGTACCC</td><td></td><td></td><td></td><td></td><td></td><td></td><td></td><td></td><td></td><td></td></tr>
<tr><td>Oligo rev</td><td>AAACGGGTACGTGCCTGGATAGGT</td><td></td><td></td><td></td><td></td><td></td><td></td><td></td><td></td><td></td><td></td></tr>
<tr><td>Chromosome</td><td>start</td><td>end</td><td>strand</td><td>MM</td><td>target_seq</td><td>PAM</td><td>alignment</td><td>distance</td><td>position</td><td>gene name</td><td>gene id</td></tr>
<tr><td>GU305914.1</td><td>184721</td><td>184743</td><td>-</td><td>0</td><td>ACCTATCCAGGCACGTACCC</td><td>TGG</td><td>||||||||[||||||||||||]PAM</td><td>NA</td><td>Intergenic</td><td>NA</td><td>NA</td></tr>
<tr><td>T20694</td><td>CCTATCCAGGCACGTACCCTGGG</td><td>1000</td><td></td><td></td><td></td><td></td><td></td><td></td><td></td><td></td><td></td></tr>
<tr><td>Oligo fwd</td><td>CACCCCTATCCAGGCACGTACCCT</td><td></td><td></td><td></td><td></td><td></td><td></td><td></td><td></td><td></td><td></td></tr>
<tr><td>Oligo rev</td><td>AAACAGGGTACGTGCCTGGATAGG</td><td></td><td></td><td></td><td></td><td></td><td></td><td></td><td></td><td></td><td></td></tr>
<tr><td>Chromosome</td><td>start</td><td>end</td><td>strand</td><td>MM</td><td>target_seq</td><td>PAM</td><td>alignment</td><td>distance</td><td>position</td><td>gene name</td><td>gene id</td></tr>
<tr><td>GU305914.1</td><td>184720</td><td>184742</td><td>-</td><td>0</td><td>CCTATCCAGGCACGTACCCT</td><td>GGG</td><td>||||||||[||||||||||||]PAM</td><td>NA</td><td>Intergenic</td><td>NA</td><td>NA</td></tr>
<tr><td>T20695</td><td>CTATCCAGGCACGTACCCTGGGG</td><td>1000</td><td></td><td></td><td></td><td></td><td></td><td></td><td></td><td></td><td></td></tr>
<tr><td>Oligo fwd</td><td>CACCCTATCCAGGCACGTACCCTG</td><td></td><td></td><td></td><td></td><td></td><td></td><td></td><td></td><td></td><td></td></tr>
<tr><td>Oligo rev</td><td>AAACCAGGGTACGTGCCTGGATAG</td><td></td><td></td><td></td><td></td><td></td><td></td><td></td><td></td><td></td><td></td></tr>
<tr><td>Chromosome</td><td>start</td><td>end</td><td>strand</td><td>MM</td><td>target_seq</td><td>PAM</td><td>alignment</td><td>distance</td><td>position</td><td>gene name</td><td>gene id</td></tr>
<tr><td>GU305914.1</td><td>184719</td><td>184741</td><td>-</td><td>0</td><td>CTATCCAGGCACGTACCCTG</td><td>GGG</td><td>||||||||[||||||||||||]PAM</td><td>NA</td><td>Intergenic</td><td>NA</td><td>NA</td></tr>
<tr><td>T20696</td><td>GGCACGTACCCTGGGGTCTAAGG</td><td>1000</td><td></td><td></td><td></td><td></td><td></td><td></td><td></td><td></td><td></td></tr>
<tr><td>Oligo fwd</td><td>CACCGGCACGTACCCTGGGGTCTA</td><td></td><td></td><td></td><td></td><td></td><td></td><td></td><td></td><td></td><td></td></tr>
<tr><td>Oligo rev</td><td>AAACTAGACCCCAGGGTACGTGCC</td><td></td><td></td><td></td><td></td><td></td><td></td><td></td><td></td><td></td><td></td></tr>
<tr><td>Chromosome</td><td>start</td><td>end</td><td>strand</td><td>MM</td><td>target_seq</td><td>PAM</td><td>alignment</td><td>distance</td><td>position</td><td>gene name</td><td>gene id</td></tr>
<tr><td>GU305914.1</td><td>184712</td><td>184734</td><td>-</td><td>0</td><td>GGCACGTACCCTGGGGTCTA</td><td>AGG</td><td>||||||||[||||||||||||]PAM</td><td>NA</td><td>Intergenic</td><td>NA</td><td>NA</td></tr>
<tr><td>T20697</td><td>GGGGTCTAAGGCACAGATCATGG</td><td>1000</td><td></td><td></td><td></td><td></td><td></td><td></td><td></td><td></td><td></td></tr>
<tr><td>Oligo fwd</td><td>CACCGGGGTCTAAGGCACAGATCA</td><td></td><td></td><td></td><td></td><td></td><td></td><td></td><td></td><td></td><td></td></tr>
<tr><td>Oligo rev</td><td>AAACTGATCTGTGCCTTAGACCCC</td><td></td><td></td><td></td><td></td><td></td><td></td><td></td><td></td><td></td><td></td></tr>
<tr><td>Chromosome</td><td>start</td><td>end</td><td>strand</td><td>MM</td><td>target_seq</td><td>PAM</td><td>alignment</td><td>distance</td><td>position</td><td>gene name</td><td>gene id</td></tr>
<tr><td>GU305914.1</td><td>184700</td><td>184722</td><td>-</td><td>0</td><td>GGGGTCTAAGGCACAGATCA</td><td>TGG</td><td>||||||||[||||||||||||]PAM</td><td>NA</td><td>Intergenic</td><td>NA</td><td>NA</td></tr>
<tr><td>T20698</td><td>GGGTCTAAGGCACAGATCATGGG</td><td>1000</td><td></td><td></td><td></td><td></td><td></td><td></td><td></td><td></td><td></td></tr>
<tr><td>Oligo fwd</td><td>CACCGGGTCTAAGGCACAGATCAT</td><td></td><td></td><td></td><td></td><td></td><td></td><td></td><td></td><td></td><td></td></tr>
<tr><td>Oligo rev</td><td>AAACATGATCTGTGCCTTAGACCC</td><td></td><td></td><td></td><td></td><td></td><td></td><td></td><td></td><td></td><td></td></tr>
<tr><td>Chromosome</td><td>start</td><td>end</td><td>strand</td><td>MM</td><td>target_seq</td><td>PAM</td><td>alignment</td><td>distance</td><td>position</td><td>gene name</td><td>gene id</td></tr>
<tr><td>GU305914.1</td><td>184699</td><td>184721</td><td>-</td><td>0</td><td>GGGTCTAAGGCACAGATCAT</td><td>GGG</td><td>||||||||[||||||||||||]PAM</td><td>NA</td><td>Intergenic</td><td>NA</td><td>NA</td></tr>
<tr><td>T20699</td><td>GGTCTAAGGCACAGATCATGGGG</td><td>1000</td><td></td><td></td><td></td><td></td><td></td><td></td><td></td><td></td><td></td></tr>
<tr><td>Oligo fwd</td><td>CACCGGTCTAAGGCACAGATCATG</td><td></td><td></td><td></td><td></td><td></td><td></td><td></td><td></td><td></td><td></td></tr>
<tr><td>Oligo rev</td><td>AAACCATGATCTGTGCCTTAGACC</td><td></td><td></td><td></td><td></td><td></td><td></td><td></td><td></td><td></td><td></td></tr>
<tr><td>Chromosome</td><td>start</td><td>end</td><td>strand</td><td>MM</td><td>target_seq</td><td>PAM</td><td>alignment</td><td>distance</td><td>position</td><td>gene name</td><td>gene id</td></tr>
<tr><td>GU305914.1</td><td>184698</td><td>184720</td><td>-</td><td>0</td><td>GGTCTAAGGCACAGATCATG</td><td>GGG</td><td>||||||||[||||||||||||]PAM</td><td>NA</td><td>Intergenic</td><td>NA</td><td>NA</td></tr>
<tr><td>T20700</td><td>GTCTAAGGCACAGATCATGGGGG</td><td>1000</td><td></td><td></td><td></td><td></td><td></td><td></td><td></td><td></td><td></td></tr>
<tr><td>Oligo fwd</td><td>CACCGTCTAAGGCACAGATCATGG</td><td></td><td></td><td></td><td></td><td></td><td></td><td></td><td></td><td></td><td></td></tr>
<tr><td>Oligo rev</td><td>AAACCCATGATCTGTGCCTTAGAC</td><td></td><td></td><td></td><td></td><td></td><td></td><td></td><td></td><td></td><td></td></tr>
<tr><td>Chromosome</td><td>start</td><td>end</td><td>strand</td><td>MM</td><td>target_seq</td><td>PAM</td><td>alignment</td><td>distance</td><td>position</td><td>gene name</td><td>gene id</td></tr>
<tr><td>GU305914.1</td><td>184697</td><td>184719</td><td>-</td><td>0</td><td>GTCTAAGGCACAGATCATGG</td><td>GGG</td><td>||||||||[||||||||||||]PAM</td><td>NA</td><td>Intergenic</td><td>NA</td><td>NA</td></tr>
<tr><td>T20701</td><td>AAGGCACAGATCATGGGGGTCGG</td><td>1000</td><td></td><td></td><td></td><td></td><td></td><td></td><td></td><td></td><td></td></tr>
<tr><td>Oligo fwd</td><td>CACCAAGGCACAGATCATGGGGGT</td><td></td><td></td><td></td><td></td><td></td><td></td><td></td><td></td><td></td><td></td></tr>
<tr><td>Oligo rev</td><td>AAACACCCCCATGATCTGTGCCTT</td><td></td><td></td><td></td><td></td><td></td><td></td><td></td><td></td><td></td><td></td></tr>
<tr><td>Chromosome</td><td>start</td><td>end</td><td>strand</td><td>MM</td><td>target_seq</td><td>PAM</td><td>alignment</td><td>distance</td><td>position</td><td>gene name</td><td>gene id</td></tr>
<tr><td>GU305914.1</td><td>184693</td><td>184715</td><td>-</td><td>0</td><td>AAGGCACAGATCATGGGGGT</td><td>CGG</td><td>||||||||[||||||||||||]PAM</td><td>NA</td><td>Intergenic</td><td>NA</td><td>NA</td></tr>
<tr><td>T20702</td><td>AGCTATAGAAATGAGACCCCCGG</td><td>1000</td><td></td><td></td><td></td><td></td><td></td><td></td><td></td><td></td><td></td></tr>
<tr><td>Oligo fwd</td><td>CACCAGCTATAGAAATGAGACCCC</td><td></td><td></td><td></td><td></td><td></td><td></td><td></td><td></td><td></td><td></td></tr>
<tr><td>Oligo rev</td><td>AAACGGGGTCTCATTTCTATAGCT</td><td></td><td></td><td></td><td></td><td></td><td></td><td></td><td></td><td></td><td></td></tr>
<tr><td>Chromosome</td><td>start</td><td>end</td><td>strand</td><td>MM</td><td>target_seq</td><td>PAM</td><td>alignment</td><td>distance</td><td>position</td><td>gene name</td><td>gene id</td></tr>
</table>

| | | | | | | | | | | | |
|---|---|---|---|---|---|---|---|---|---|---|---|
| GU305914.1 | 184661 | 184683 | - | 0 | AGCTATAGAAATGAGACCCC | CGG | \|\|\|\|\|\|\|\|[\|\|\|\|\|\|\|\|\|\|\|\|]PAM | NA | Intergenic | NA | NA |
| T20703 | GCTATAGAAATGAGACCCCCGGG | 1000 | | | | | | | | | |
| Oligo fwd | CACCGCTATAGAAATGAGACCCCC | | | | | | | | | | |
| Oligo rev | AAACGGGGGTCTCATTTCTATAGC | | | | | | | | | | |
| Chromosome | start | end | strand | MM | target_seq | PAM | alignment | distance | position | gene name | gene id |
| GU305914.1 | 184660 | 184682 | - | 0 | GCTATAGAAATGAGACCCCC | GGG | \|\|\|\|\|\|\|\|[\|\|\|\|\|\|\|\|\|\|\|\|]PAM | NA | Intergenic | NA | NA |
| T20704 | CCGGGAGCGATCGAAAACCGCGG | 1000 | | | | | | | | | |
| Oligo fwd | CACCCCGGGAGCGATCGAAAACCG | | | | | | | | | | |
| Oligo rev | AAACCGGTTTTCGATCGCTCCCGG | | | | | | | | | | |
| Chromosome | start | end | strand | MM | target_seq | PAM | alignment | distance | position | gene name | gene id |
| GU305914.1 | 184642 | 184664 | - | 0 | CCGGGAGCGATCGAAAACCG | CGG | \|\|\|\|\|\|\|\|[\|\|\|\|\|\|\|\|\|\|\|\|]PAM | NA | Intergenic | NA | NA |
| T20705 | TAGAACCGAAAGTACCGCTATGG | 1000 | | | | | | | | | |
| Oligo fwd | CACCTAGAACCGAAAGTACCGCTA | | | | | | | | | | |
| Oligo rev | AAACTAGCGGTACTTTCGGTTCTA | | | | | | | | | | |
| Chromosome | start | end | strand | MM | target_seq | PAM | alignment | distance | position | gene name | gene id |
| GU305914.1 | 184606 | 184628 | - | 0 | TAGAACCGAAAGTACCGCTA | TGG | \|\|\|\|\|\|\|\|[\|\|\|\|\|\|\|\|\|\|\|\|]PAM | NA | Intergenic | NA | NA |
| T20706 | AGAACCGAAAGTACCGCTATGGG | 1000 | | | | | | | | | |
| Oligo fwd | CACCAGAACCGAAAGTACCGCTAT | | | | | | | | | | |
| Oligo rev | AAACATAGCGGTACTTTCGGTTCT | | | | | | | | | | |
| Chromosome | start | end | strand | MM | target_seq | PAM | alignment | distance | position | gene name | gene id |
| GU305914.1 | 184605 | 184627 | - | 0 | AGAACCGAAAGTACCGCTAT | GGG | \|\|\|\|\|\|\|\|[\|\|\|\|\|\|\|\|\|\|\|\|]PAM | NA | Intergenic | NA | NA |
| T20707 | ACCGAAAGTACCGCTATGGGCGG | 1000 | | | | | | | | | |
| Oligo fwd | CACCACCGAAAGTACCGCTATGGG | | | | | | | | | | |
| Oligo rev | AAACCCCATAGCGGTACTTTCGGT | | | | | | | | | | |
| Chromosome | start | end | strand | MM | target_seq | PAM | alignment | distance | position | gene name | gene id |
| GU305914.1 | 184602 | 184624 | - | 0 | ACCGAAAGTACCGCTATGGG | CGG | \|\|\|\|\|\|\|\|[\|\|\|\|\|\|\|\|\|\|\|\|]PAM | NA | Intergenic | NA | NA |
| T20708 | CCGAAAGTACCGCTATGGGCGGG | 1000 | | | | | | | | | |
| Oligo fwd | CACCCCGAAAGTACCGCTATGGGC | | | | | | | | | | |
| Oligo rev | AAACGCCCATAGCGGTACTTTCGG | | | | | | | | | | |
| Chromosome | start | end | strand | MM | target_seq | PAM | alignment | distance | position | gene name | gene id |
| GU305914.1 | 184601 | 184623 | - | 0 | CCGAAAGTACCGCTATGGGC | GGG | \|\|\|\|\|\|\|\|[\|\|\|\|\|\|\|\|\|\|\|\|]PAM | NA | Intergenic | NA | NA |
| T20709 | CCGCTATGGGCGGGCGCCAATGG | 1000 | | | | | | | | | |
| Oligo fwd | CACCCCGCTATGGGCGGGCGCCAA | | | | | | | | | | |
| Oligo rev | AAACTTGGCGCCCGCCCATAGCGG | | | | | | | | | | |
| Chromosome | start | end | strand | MM | target_seq | PAM | alignment | distance | position | gene name | gene id |
| GU305914.1 | 184592 | 184614 | - | 0 | CCGCTATGGGCGGGCGCCAA | TGG | \|\|\|\|\|\|\|\|[\|\|\|\|\|\|\|\|\|\|\|\|]PAM | NA | Intergenic | NA | NA |
| T20710 | ATGGGCGGGCGCCAATGGCGTGG | 1000 | | | | | | | | | |
| Oligo fwd | CACCATGGGCGGGCGCCAATGGCG | | | | | | | | | | |
| Oligo rev | AAACCGCCATTGGCGCCCGCCCAT | | | | | | | | | | |
| Chromosome | start | end | strand | MM | target_seq | PAM | alignment | distance | position | gene name | gene id |
| GU305914.1 | 184587 | 184609 | - | 0 | ATGGGCGGGCGCCAATGGCG | TGG | \|\|\|\|\|\|\|\|[\|\|\|\|\|\|\|\|\|\|\|\|]PAM | NA | Intergenic | NA | NA |
| T20711 | GCGGGCGCCAATGGCGTGGTAGG | 1000 | | | | | | | | | |
| Oligo fwd | CACCGCGGGCGCCAATGGCGTGGT | | | | | | | | | | |
| Oligo rev | AAACACCACGCCATTGGCGCCCGC | | | | | | | | | | |
| Chromosome | start | end | strand | MM | target_seq | PAM | alignment | distance | position | gene name | gene id |
| GU305914.1 | 184583 | 184605 | - | 0 | GCGGGCGCCAATGGCGTGGT | AGG | \|\|\|\|\|\|\|\|[\|\|\|\|\|\|\|\|\|\|\|\|]PAM | NA | Intergenic | NA | NA |
| T20712 | CGGGCGCCAATGGCGTGGTAGGG | 1000 | | | | | | | | | |
| Oligo fwd | CACCCGGGCGCCAATGGCGTGGTA | | | | | | | | | | |
| Oligo rev | AAACTACCACGCCATTGGCGCCCG | | | | | | | | | | |
| Chromosome | start | end | strand | MM | target_seq | PAM | alignment | distance | position | gene name | gene id |
| GU305914.1 | 184582 | 184604 | - | 0 | CGGGCGCCAATGGCGTGGTA | GGG | \|\|\|\|\|\|\|\|[\|\|\|\|\|\|\|\|\|\|\|\|]PAM | NA | Intergenic | NA | NA |
| T20713 | GGGCGCCAATGGCGTGGTAGGGG | 1000 | | | | | | | | | |
| Oligo fwd | CACCGGGCGCCAATGGCGTGGTAG | | | | | | | | | | |
| Oligo rev | AAACCTACCACGCCATTGGCGCCC | | | | | | | | | | |
| Chromosome | start | end | strand | MM | target_seq | PAM | alignment | distance | position | gene name | gene id |
| GU305914.1 | 184581 | 184603 | - | 0 | GGGCGCCAATGGCGTGGTAG | GGG | \|\|\|\|\|\|\|\|[\|\|\|\|\|\|\|\|\|\|\|\|]PAM | NA | Intergenic | NA | NA |
| T20714 | GGCGCCAATGGCGTGGTAGGGGG | 1000 | | | | | | | | | |
| Oligo fwd | CACCGGCGCCAATGGCGTGGTAGG | | | | | | | | | | |
| Oligo rev | AAACCCTACCACGCCATTGGCGCC | | | | | | | | | | |
| Chromosome | start | end | strand | MM | target_seq | PAM | alignment | distance | position | gene name | gene id |
| GU305914.1 | 184580 | 184602 | - | 0 | GGCGCCAATGGCGTGGTAGG | GGG | \|\|\|\|\|\|\|\|[\|\|\|\|\|\|\|\|\|\|\|\|]PAM | NA | Intergenic | NA | NA |
| T20715 | ATTACGTGACGCAATATGTATGG | 1000 | | | | | | | | | |
| Oligo fwd | CACCATTACGTGACGCAATATGTA | | | | | | | | | | |
| Oligo rev | AAACTACATATTGCGTCACGTAAT | | | | | | | | | | |
| Chromosome | start | end | strand | MM | target_seq | PAM | alignment | distance | position | gene name | gene id |
| GU305914.1 | 184551 | 184573 | - | 0 | ATTACGTGACGCAATATGTA | TGG | \|\|\|\|\|\|\|\|[\|\|\|\|\|\|\|\|\|\|\|\|]PAM | NA | Intergenic | NA | NA |
| T20716 | TTACGTGACGCAATATGTATGGG | 1000 | | | | | | | | | |
| Oligo fwd | CACCTTACGTGACGCAATATGTAT | | | | | | | | | | |
| Oligo rev | AAACATACATATTGCGTCACGTAA | | | | | | | | | | |
| Chromosome | start | end | strand | MM | target_seq | PAM | alignment | distance | position | gene name | gene id |
| GU305914.1 | 184550 | 184572 | - | 0 | TTACGTGACGCAATATGTAT | GGG | \|\|\|\|\|\|\|\|[\|\|\|\|\|\|\|\|\|\|\|\|]PAM | NA | Intergenic | NA | NA |
| T20717 | GTGACGCAATATGTATGGGTAGG | 1000 | | | | | | | | | |
| Oligo fwd | CACCGTGACGCAATATGTATGGGT | | | | | | | | | | |
| Oligo rev | AAACACCCATACATATTGCGTCAC | | | | | | | | | | |

Chromosome start end strand MM target_seq PAM alignment distance position gene name gene id
GU305914.1 184546 184568 - 0 GTGACGCAATATGTATGGGT AGG ||||||||[||||||||||||]PAM NA Intergenic NA NA

T20718 TGACGCAATATGTATGGGTAGGG 1000
Oligo fwd CACCTGACGCAATATGTATGGGTA
Oligo rev AAACTACCCATACATATTGCGTCA
Chromosome start end strand MM target_seq PAM alignment distance position gene name gene id
GU305914.1 184545 184567 - 0 TGACGCAATATGTATGGGTA GGG ||||||||[||||||||||||]PAM NA Intergenic NA NA

T20719 GACGCAATATGTATGGGTAGGGG 1000
Oligo fwd CACCGACGCAATATGTATGGGTAG
Oligo rev AAACCTACCCATACATATTGCGTC
Chromosome start end strand MM target_seq PAM alignment distance position gene name gene id
GU305914.1 184544 184566 - 0 GACGCAATATGTATGGGTAG GGG ||||||||[||||||||||||]PAM NA Intergenic NA NA

T20720 ACGCAATATGTATGGGTAGGGGG 1000
Oligo fwd CACCACGCAATATGTATGGGTAGG
Oligo rev AAACCCTACCCATACATATTGCGT
Chromosome start end strand MM target_seq PAM alignment distance position gene name gene id
GU305914.1 184543 184565 - 0 ACGCAATATGTATGGGTAGG GGG ||||||||[||||||||||||]PAM NA Intergenic NA NA

T20721 CCCCACCCATATTAAAGTACCGG 1000
Oligo fwd CACCCCCCACCCATATTAAAGTAC
Oligo rev AAACGTACTTTAATATGGGTGGGG
Chromosome start end strand MM target_seq PAM alignment distance position gene name gene id
GU305914.1 184489 184511 - 0 CCCCACCCATATTAAAGTAC CGG ||||||||[||||||||||||]PAM NA Intergenic NA NA

T20722 CCCACCCATATTAAAGTACCGGG 1000
Oligo fwd CACCCCCACCCATATTAAAGTACC
Oligo rev AAACGGTACTTTAATATGGGTGGG
Chromosome start end strand MM target_seq PAM alignment distance position gene name gene id
GU305914.1 184488 184510 - 0 CCCACCCATATTAAAGTACC GGG ||||||||[||||||||||||]PAM NA Intergenic NA NA

T20723 GAGCCCAATGCAACCTTACCCGG 1000
Oligo fwd CACCGAGCCCAATGCAACCTTACC
Oligo rev AAACGGTAAGGTTGCATTGGGCTC
Chromosome start end strand MM target_seq PAM alignment distance position gene name gene id
GU305914.1 184444 184466 - 0 GAGCCCAATGCAACCTTACC CGG ||||||||[||||||||||||]PAM NA Intergenic NA NA

T20724 CAATGCAACCTTACCCGGCCTGG 1000
Oligo fwd CACCCAATGCAACCTTACCCGGCC
Oligo rev AAACGGCCGGGTAAGGTTGCATTG
Chromosome start end strand MM target_seq PAM alignment distance position gene name gene id
GU305914.1 184439 184461 - 0 CAATGCAACCTTACCCGGCC TGG ||||||||[||||||||||||]PAM NA Intergenic NA NA

T20725 AATGCAACCTTACCCGGCCTGGG 1000
Oligo fwd CACCAATGCAACCTTACCCGGCCT
Oligo rev AAACAGGCCGGGTAAGGTTGCATT
Chromosome start end strand MM target_seq PAM alignment distance position gene name gene id
GU305914.1 184438 184460 - 0 AATGCAACCTTACCCGGCCT GGG ||||||||[||||||||||||]PAM NA Intergenic NA NA

T20726 ATGCAACCTTACCCGGCCTGGGG 1000
Oligo fwd CACCATGCAACCTTACCCGGCCTG
Oligo rev AAACCAGGCCGGGTAAGGTTGCAT
Chromosome start end strand MM target_seq PAM alignment distance position gene name gene id
GU305914.1 184437 184459 - 0 ATGCAACCTTACCCGGCCTG GGG ||||||||[||||||||||||]PAM NA Intergenic NA NA

T20727 TGCAACCTTACCCGGCCTGGGGG 1000
Oligo fwd CACCTGCAACCTTACCCGGCCTGG
Oligo rev AAACCCAGGCCGGGTAAGGTTGCA
Chromosome start end strand MM target_seq PAM alignment distance position gene name gene id
GU305914.1 184436 184458 - 0 TGCAACCTTACCCGGCCTGG GGG ||||||||[||||||||||||]PAM NA Intergenic NA NA

T20728 GCAACCTTACCCGGCCTGGGGGG 1000
Oligo fwd CACCGCAACCTTACCCGGCCTGGG
Oligo rev AAACCCCAGGCCGGGTAAGGTTGC
Chromosome start end strand MM target_seq PAM alignment distance position gene name gene id
GU305914.1 184435 184457 - 0 GCAACCTTACCCGGCCTGGG GGG ||||||||[||||||||||||]PAM NA Intergenic NA NA

T20729 ACCCGCTCGTTAACGTAGATAGG 1000
Oligo fwd CACCACCCGCTCGTTAACGTAGAT
Oligo rev AAACATCTACGTTAACGAGCGGGT
Chromosome start end strand MM target_seq PAM alignment distance position gene name gene id
GU305914.1 184404 184426 - 0 ACCCGCTCGTTAACGTAGAT AGG ||||||||[||||||||||||]PAM NA Intergenic NA NA

T20730 CCCGCTCGTTAACGTAGATAGGG 1000
Oligo fwd CACCCCCGCTCGTTAACGTAGATA
Oligo rev AAACTATCTACGTTAACGAGCGGG
Chromosome start end strand MM target_seq PAM alignment distance position gene name gene id
GU305914.1 184403 184425 - 0 CCCGCTCGTTAACGTAGATA GGG ||||||||[||||||||||||]PAM NA Intergenic NA NA

T20731 GATAGGGTCGTATCGAGCCGCGG 1000
Oligo fwd CACCGATAGGGTCGTATCGAGCCG
Oligo rev AAACCGGCTCGATACGACCCTATC
Chromosome start end strand MM target_seq PAM alignment distance position gene name gene id
GU305914.1 184387 184409 - 0 GATAGGGTCGTATCGAGCCG CGG ||||||||[||||||||||||]PAM NA Intergenic NA NA

T20732 AGCCGCGGTCCGTCCTGCTCTGG 1000
Oligo fwd CACCAGCCGCGGTCCGTCCTGCTC

Oligo rev AAACGAGCAGGACGGACCGCGGCT
Chromosome start end strand MM target_seq PAM alignment distance position gene name gene id
GU305914.1 184372 184394 - 0 AGCCGCGGTCCGTCCTGCTC TGG ||||||||[||||||||||||]PAM NA Intergenic NA NA

T20733 GCCGCGGTCCGTCCTGCTCTGGG 1000
Oligo fwd CACCGCCGCGGTCCGTCCTGCTCT
Oligo rev AAACAGAGCAGGACGGACCGCGGC
Chromosome start end strand MM target_seq PAM alignment distance position gene name gene id
GU305914.1 184371 184393 - 0 GCCGCGGTCCGTCCTGCTCT GGG ||||||||[||||||||||||]PAM NA Intergenic NA NA

T20734 CGTCCTGCTCTGGGCTCGAATGG 1000
Oligo fwd CACCCGTCCTGCTCTGGGCTCGAA
Oligo rev AAACTTCGAGCCCAGAGCAGGACG
Chromosome start end strand MM target_seq PAM alignment distance position gene name gene id
GU305914.1 184362 184384 - 0 CGTCCTGCTCTGGGCTCGAA TGG ||||||||[||||||||||||]PAM NA Intergenic NA NA

T20735 TGCTCTGGGCTCGAATGGCATGG 1000
Oligo fwd CACCTGCTCTGGGCTCGAATGGCA
Oligo rev AAACTGCCATTCGAGCCCAGAGCA
Chromosome start end strand MM target_seq PAM alignment distance position gene name gene id
GU305914.1 184357 184379 - 0 TGCTCTGGGCTCGAATGGCA TGG ||||||||[||||||||||||]PAM NA Intergenic NA NA

T20736 GCTCTGGGCTCGAATGGCATGGG 1000
Oligo fwd CACCGCTCTGGGCTCGAATGGCAT
Oligo rev AAACATGCCATTCGAGCCCAGAGC
Chromosome start end strand MM target_seq PAM alignment distance position gene name gene id
GU305914.1 184356 184378 - 0 GCTCTGGGCTCGAATGGCAT GGG ||||||||[||||||||||||]PAM NA Intergenic NA NA

T20737 CTCTGGGCTCGAATGGCATGGGG 1000
Oligo fwd CACCCTCTGGGCTCGAATGGCATG
Oligo rev AAACCATGCCATTCGAGCCCAGAG
Chromosome start end strand MM target_seq PAM alignment distance position gene name gene id
GU305914.1 184355 184377 - 0 CTCTGGGCTCGAATGGCATG GGG ||||||||[||||||||||||]PAM NA Intergenic NA NA

T20738 TCTGGGCTCGAATGGCATGGGGG 1000
Oligo fwd CACCTCTGGGCTCGAATGGCATGG
Oligo rev AAACCCATGCCATTCGAGCCCAGA
Chromosome start end strand MM target_seq PAM alignment distance position gene name gene id
GU305914.1 184354 184376 - 0 TCTGGGCTCGAATGGCATGG GGG ||||||||[||||||||||||]PAM NA Intergenic NA NA

T20739 CATGGGGGACAGCTTTTATATGG 1000
Oligo fwd CACCCATGGGGGACAGCTTTTATA
Oligo rev AAACTATAAAAGCTGTCCCCCATG
Chromosome start end strand MM target_seq PAM alignment distance position gene name gene id
GU305914.1 184339 184361 - 0 CATGGGGGACAGCTTTTATA TGG ||||||||[||||||||||||]PAM NA Intergenic NA NA

T20740 CGCCCCCTATGTGAGTAATACGG 1000
Oligo fwd CACCCGCCCCCTATGTGAGTAATA
Oligo rev AAACTATTACTCACATAGGGGGCG
Chromosome start end strand MM target_seq PAM alignment distance position gene name gene id
GU305914.1 184236 184258 - 0 CGCCCCCTATGTGAGTAATA CGG ||||||||[||||||||||||]PAM NA Intergenic NA NA

T20741 GCCCCCTATGTGAGTAATACGGG 1000
Oligo fwd CACCGCCCCCTATGTGAGTAATAC
Oligo rev AAACGTATTACTCACATAGGGGGC
Chromosome start end strand MM target_seq PAM alignment distance position gene name gene id
GU305914.1 184235 184257 - 0 GCCCCCTATGTGAGTAATAC GGG ||||||||[||||||||||||]PAM NA Intergenic NA NA

T20742 CCCCCTATGTGAGTAATACGGGG 1000
Oligo fwd CACCCCCCCTATGTGAGTAATACG
Oligo rev AAACCGTATTACTCACATAGGGGG
Chromosome start end strand MM target_seq PAM alignment distance position gene name gene id
GU305914.1 184234 184256 - 0 CCCCCTATGTGAGTAATACG GGG ||||||||[||||||||||||]PAM NA Intergenic NA NA

T20743 TTTTTACCCAATTTCCCAAGCGG 1000
Oligo fwd CACCTTTTTACCCAATTTCCCAAG
Oligo rev AAACCTTGGGAAATTGGGTAAAAA
Chromosome start end strand MM target_seq PAM alignment distance position gene name gene id
GU305914.1 184209 184231 - 0 TTTTTACCCAATTTCCCAAG CGG ||||||||[||||||||||||]PAM NA Intergenic NA NA

T20744 AGCCCCCTAATACACTCATATGG 1000
Oligo fwd CACCAGCCCCCTAATACACTCATA
Oligo rev AAACTATGAGTGTATTAGGGGGCT
Chromosome start end strand MM target_seq PAM alignment distance position gene name gene id
GU305914.1 184184 184206 - 0 AGCCCCCTAATACACTCATA TGG ||||||||[||||||||||||]PAM NA Intergenic NA NA

T20745 ATATGGCATATGAATCAGCACGG 1000
Oligo fwd CACCATATGGCATATGAATCAGCA
Oligo rev AAACTGCTGATTCATATGCCATAT
Chromosome start end strand MM target_seq PAM alignment distance position gene name gene id
GU305914.1 184167 184189 - 0 ATATGGCATATGAATCAGCA CGG ||||||||[||||||||||||]PAM NA Intergenic NA NA

T20746 AGCACGGTCATGCACTCTAATGG 1000
Oligo fwd CACCAGCACGGTCATGCACTCTAA
Oligo rev AAACTTAGAGTGCATGACCGTGCT
Chromosome start end strand MM target_seq PAM alignment distance position gene name gene id
GU305914.1 184151 184173 - 0 AGCACGGTCATGCACTCTAA TGG ||||||||[||||||||||||]PAM NA Intergenic NA NA

T20747 ACGGTCATGCACTCTAATGGCGG 1000

<table>
<tr><td>Oligo fwd</td><td>CACCACGGTCATGCACTCTAATGG</td><td></td><td></td><td></td><td></td><td></td><td></td><td></td><td></td><td></td><td></td></tr>
<tr><td>Oligo rev</td><td>AAACCCATTAGAGTGCATGACCGT</td><td></td><td></td><td></td><td></td><td></td><td></td><td></td><td></td><td></td><td></td></tr>
<tr><td>Chromosome</td><td>start</td><td>end</td><td>strand</td><td>MM</td><td>target_seq</td><td>PAM</td><td>alignment</td><td>distance</td><td>position</td><td>gene name</td><td>gene id</td></tr>
<tr><td>GU305914.1</td><td>184148</td><td>184170</td><td>-</td><td>0</td><td>ACGGTCATGCACTCTAATGG</td><td>CGG</td><td>||||||||[||||||||||||]PAM</td><td>NA</td><td>Intergenic</td><td>NA</td><td>NA</td></tr>
<tr><td>T20748</td><td>GCACTCTAATGGCGGCCCATAGG</td><td>1000</td><td></td><td></td><td></td><td></td><td></td><td></td><td></td><td></td><td></td></tr>
<tr><td>Oligo fwd</td><td>CACCGCACTCTAATGGCGGCCCAT</td><td></td><td></td><td></td><td></td><td></td><td></td><td></td><td></td><td></td><td></td></tr>
<tr><td>Oligo rev</td><td>AAACATGGGCCGCCATTAGAGTGC</td><td></td><td></td><td></td><td></td><td></td><td></td><td></td><td></td><td></td><td></td></tr>
<tr><td>Chromosome</td><td>start</td><td>end</td><td>strand</td><td>MM</td><td>target_seq</td><td>PAM</td><td>alignment</td><td>distance</td><td>position</td><td>gene name</td><td>gene id</td></tr>
<tr><td>GU305914.1</td><td>184140</td><td>184162</td><td>-</td><td>0</td><td>GCACTCTAATGGCGGCCCAT</td><td>AGG</td><td>||||||||[||||||||||||]PAM</td><td>NA</td><td>Intergenic</td><td>NA</td><td>NA</td></tr>
<tr><td>T20749</td><td>CACTCTAATGGCGGCCCATAGGG</td><td>1000</td><td></td><td></td><td></td><td></td><td></td><td></td><td></td><td></td><td></td></tr>
<tr><td>Oligo fwd</td><td>CACCCACTCTAATGGCGGCCCATA</td><td></td><td></td><td></td><td></td><td></td><td></td><td></td><td></td><td></td><td></td></tr>
<tr><td>Oligo rev</td><td>AAACTATGGGCCGCCATTAGAGTG</td><td></td><td></td><td></td><td></td><td></td><td></td><td></td><td></td><td></td><td></td></tr>
<tr><td>Chromosome</td><td>start</td><td>end</td><td>strand</td><td>MM</td><td>target_seq</td><td>PAM</td><td>alignment</td><td>distance</td><td>position</td><td>gene name</td><td>gene id</td></tr>
<tr><td>GU305914.1</td><td>184139</td><td>184161</td><td>-</td><td>0</td><td>CACTCTAATGGCGGCCCATA</td><td>GGG</td><td>||||||||[||||||||||||]PAM</td><td>NA</td><td>Intergenic</td><td>NA</td><td>NA</td></tr>
<tr><td>T20750</td><td>CCCATAGGGACTTTCCACATAGG</td><td>1000</td><td></td><td></td><td></td><td></td><td></td><td></td><td></td><td></td><td></td></tr>
<tr><td>Oligo fwd</td><td>CACCCCCATAGGGACTTTCCACAT</td><td></td><td></td><td></td><td></td><td></td><td></td><td></td><td></td><td></td><td></td></tr>
<tr><td>Oligo rev</td><td>AAACATGTGGAAAGTCCCTATGGG</td><td></td><td></td><td></td><td></td><td></td><td></td><td></td><td></td><td></td><td></td></tr>
<tr><td>Chromosome</td><td>start</td><td>end</td><td>strand</td><td>MM</td><td>target_seq</td><td>PAM</td><td>alignment</td><td>distance</td><td>position</td><td>gene name</td><td>gene id</td></tr>
<tr><td>GU305914.1</td><td>184125</td><td>184147</td><td>-</td><td>0</td><td>CCCATAGGGACTTTCCACAT</td><td>AGG</td><td>||||||||[||||||||||||]PAM</td><td>NA</td><td>Intergenic</td><td>NA</td><td>NA</td></tr>
<tr><td>T20751</td><td>CCATAGGGACTTTCCACATAGGG</td><td>1000</td><td></td><td></td><td></td><td></td><td></td><td></td><td></td><td></td><td></td></tr>
<tr><td>Oligo fwd</td><td>CACCCCATAGGGACTTTCCACATA</td><td></td><td></td><td></td><td></td><td></td><td></td><td></td><td></td><td></td><td></td></tr>
<tr><td>Oligo rev</td><td>AAACTATGTGGAAAGTCCCTATGG</td><td></td><td></td><td></td><td></td><td></td><td></td><td></td><td></td><td></td><td></td></tr>
<tr><td>Chromosome</td><td>start</td><td>end</td><td>strand</td><td>MM</td><td>target_seq</td><td>PAM</td><td>alignment</td><td>distance</td><td>position</td><td>gene name</td><td>gene id</td></tr>
<tr><td>GU305914.1</td><td>184124</td><td>184146</td><td>-</td><td>0</td><td>CCATAGGGACTTTCCACATA</td><td>GGG</td><td>||||||||[||||||||||||]PAM</td><td>NA</td><td>Intergenic</td><td>NA</td><td>NA</td></tr>
<tr><td>T20752</td><td>CATAGGGACTTTCCACATAGGGG</td><td>1000</td><td></td><td></td><td></td><td></td><td></td><td></td><td></td><td></td><td></td></tr>
<tr><td>Oligo fwd</td><td>CACCCATAGGGACTTTCCACATAG</td><td></td><td></td><td></td><td></td><td></td><td></td><td></td><td></td><td></td><td></td></tr>
<tr><td>Oligo rev</td><td>AAACCTATGTGGAAAGTCCCTATG</td><td></td><td></td><td></td><td></td><td></td><td></td><td></td><td></td><td></td><td></td></tr>
<tr><td>Chromosome</td><td>start</td><td>end</td><td>strand</td><td>MM</td><td>target_seq</td><td>PAM</td><td>alignment</td><td>distance</td><td>position</td><td>gene name</td><td>gene id</td></tr>
<tr><td>GU305914.1</td><td>184123</td><td>184145</td><td>-</td><td>0</td><td>CATAGGGACTTTCCACATAG</td><td>GGG</td><td>||||||||[||||||||||||]PAM</td><td>NA</td><td>Intergenic</td><td>NA</td><td>NA</td></tr>
<tr><td>T20753</td><td>ATAGGGACTTTCCACATAGGGGG</td><td>1000</td><td></td><td></td><td></td><td></td><td></td><td></td><td></td><td></td><td></td></tr>
<tr><td>Oligo fwd</td><td>CACCATAGGGACTTTCCACATAGG</td><td></td><td></td><td></td><td></td><td></td><td></td><td></td><td></td><td></td><td></td></tr>
<tr><td>Oligo rev</td><td>AAACCCTATGTGGAAAGTCCCTAT</td><td></td><td></td><td></td><td></td><td></td><td></td><td></td><td></td><td></td><td></td></tr>
<tr><td>Chromosome</td><td>start</td><td>end</td><td>strand</td><td>MM</td><td>target_seq</td><td>PAM</td><td>alignment</td><td>distance</td><td>position</td><td>gene name</td><td>gene id</td></tr>
<tr><td>GU305914.1</td><td>184122</td><td>184144</td><td>-</td><td>0</td><td>ATAGGGACTTTCCACATAGG</td><td>GGG</td><td>||||||||[||||||||||||]PAM</td><td>NA</td><td>Intergenic</td><td>NA</td><td>NA</td></tr>
<tr><td>T20754</td><td>CGTTCACCATTTCCCAGCATAGG</td><td>1000</td><td></td><td></td><td></td><td></td><td></td><td></td><td></td><td></td><td></td></tr>
<tr><td>Oligo fwd</td><td>CACCCGTTCACCATTTCCCAGCAT</td><td></td><td></td><td></td><td></td><td></td><td></td><td></td><td></td><td></td><td></td></tr>
<tr><td>Oligo rev</td><td>AAACATGCTGGGAAATGGTGAACG</td><td></td><td></td><td></td><td></td><td></td><td></td><td></td><td></td><td></td><td></td></tr>
<tr><td>Chromosome</td><td>start</td><td>end</td><td>strand</td><td>MM</td><td>target_seq</td><td>PAM</td><td>alignment</td><td>distance</td><td>position</td><td>gene name</td><td>gene id</td></tr>
<tr><td>GU305914.1</td><td>184099</td><td>184121</td><td>-</td><td>0</td><td>CGTTCACCATTTCCCAGCAT</td><td>AGG</td><td>||||||||[||||||||||||]PAM</td><td>NA</td><td>Intergenic</td><td>NA</td><td>NA</td></tr>
<tr><td>T20755</td><td>GTTCACCATTTCCCAGCATAGGG</td><td>1000</td><td></td><td></td><td></td><td></td><td></td><td></td><td></td><td></td><td></td></tr>
<tr><td>Oligo fwd</td><td>CACCGTTCACCATTTCCCAGCATA</td><td></td><td></td><td></td><td></td><td></td><td></td><td></td><td></td><td></td><td></td></tr>
<tr><td>Oligo rev</td><td>AAACTATGCTGGGAAATGGTGAAC</td><td></td><td></td><td></td><td></td><td></td><td></td><td></td><td></td><td></td><td></td></tr>
<tr><td>Chromosome</td><td>start</td><td>end</td><td>strand</td><td>MM</td><td>target_seq</td><td>PAM</td><td>alignment</td><td>distance</td><td>position</td><td>gene name</td><td>gene id</td></tr>
<tr><td>GU305914.1</td><td>184098</td><td>184120</td><td>-</td><td>0</td><td>GTTCACCATTTCCCAGCATA</td><td>GGG</td><td>||||||||[||||||||||||]PAM</td><td>NA</td><td>Intergenic</td><td>NA</td><td>NA</td></tr>
<tr><td>T20756</td><td>TTCACCATTTCCCAGCATAGGGG</td><td>1000</td><td></td><td></td><td></td><td></td><td></td><td></td><td></td><td></td><td></td></tr>
<tr><td>Oligo fwd</td><td>CACCTTCACCATTTCCCAGCATAG</td><td></td><td></td><td></td><td></td><td></td><td></td><td></td><td></td><td></td><td></td></tr>
<tr><td>Oligo rev</td><td>AAACCTATGCTGGGAAATGGTGAA</td><td></td><td></td><td></td><td></td><td></td><td></td><td></td><td></td><td></td><td></td></tr>
<tr><td>Chromosome</td><td>start</td><td>end</td><td>strand</td><td>MM</td><td>target_seq</td><td>PAM</td><td>alignment</td><td>distance</td><td>position</td><td>gene name</td><td>gene id</td></tr>
<tr><td>GU305914.1</td><td>184097</td><td>184119</td><td>-</td><td>0</td><td>TTCACCATTTCCCAGCATAG</td><td>GGG</td><td>||||||||[||||||||||||]PAM</td><td>NA</td><td>Intergenic</td><td>NA</td><td>NA</td></tr>
<tr><td>T20757</td><td>ACCATTTCCCAGCATAGGGGTGG</td><td>1000</td><td></td><td></td><td></td><td></td><td></td><td></td><td></td><td></td><td></td></tr>
<tr><td>Oligo fwd</td><td>CACCACCATTTCCCAGCATAGGGG</td><td></td><td></td><td></td><td></td><td></td><td></td><td></td><td></td><td></td><td></td></tr>
<tr><td>Oligo rev</td><td>AAACCCCCTATGCTGGGAAATGGT</td><td></td><td></td><td></td><td></td><td></td><td></td><td></td><td></td><td></td><td></td></tr>
<tr><td>Chromosome</td><td>start</td><td>end</td><td>strand</td><td>MM</td><td>target_seq</td><td>PAM</td><td>alignment</td><td>distance</td><td>position</td><td>gene name</td><td>gene id</td></tr>
<tr><td>GU305914.1</td><td>184094</td><td>184116</td><td>-</td><td>0</td><td>ACCATTTCCCAGCATAGGGG</td><td>TGG</td><td>||||||||[||||||||||||]PAM</td><td>NA</td><td>Intergenic</td><td>NA</td><td>NA</td></tr>
<tr><td>T20758</td><td>GCATAGGGGTGGTGACTCAATGG</td><td>1000</td><td></td><td></td><td></td><td></td><td></td><td></td><td></td><td></td><td></td></tr>
<tr><td>Oligo fwd</td><td>CACCGCATAGGGGTGGTGACTCAA</td><td></td><td></td><td></td><td></td><td></td><td></td><td></td><td></td><td></td><td></td></tr>
<tr><td>Oligo rev</td><td>AAACTTGAGTCACCACCCCTATGC</td><td></td><td></td><td></td><td></td><td></td><td></td><td></td><td></td><td></td><td></td></tr>
<tr><td>Chromosome</td><td>start</td><td>end</td><td>strand</td><td>MM</td><td>target_seq</td><td>PAM</td><td>alignment</td><td>distance</td><td>position</td><td>gene name</td><td>gene id</td></tr>
<tr><td>GU305914.1</td><td>184083</td><td>184105</td><td>-</td><td>0</td><td>GCATAGGGGTGGTGACTCAA</td><td>TGG</td><td>||||||||[||||||||||||]PAM</td><td>NA</td><td>Intergenic</td><td>NA</td><td>NA</td></tr>
<tr><td>T20759</td><td>TGGCCTTTACCCAAGTACATTGG</td><td>1000</td><td></td><td></td><td></td><td></td><td></td><td></td><td></td><td></td><td></td></tr>
<tr><td>Oligo fwd</td><td>CACCTGGCCTTTACCCAAGTACAT</td><td></td><td></td><td></td><td></td><td></td><td></td><td></td><td></td><td></td><td></td></tr>
<tr><td>Oligo rev</td><td>AAACATGTACTTGGGTAAAGGCCA</td><td></td><td></td><td></td><td></td><td></td><td></td><td></td><td></td><td></td><td></td></tr>
<tr><td>Chromosome</td><td>start</td><td>end</td><td>strand</td><td>MM</td><td>target_seq</td><td>PAM</td><td>alignment</td><td>distance</td><td>position</td><td>gene name</td><td>gene id</td></tr>
<tr><td>GU305914.1</td><td>184063</td><td>184085</td><td>-</td><td>0</td><td>TGGCCTTTACCCAAGTACAT</td><td>TGG</td><td>||||||||[||||||||||||]PAM</td><td>NA</td><td>Intergenic</td><td>NA</td><td>NA</td></tr>
<tr><td>GU305914.1</td><td>183969</td><td>183991</td><td>-</td><td>3</td><td>TGGGTTTTGCCCAAGTACAT</td><td>TGG</td><td>|||--|||[-|||||||||||]PAM</td><td>NA</td><td>Intergenic</td><td>NA</td><td>NA</td></tr>
<tr><td>T20760</td><td>GGCCTTTACCCAAGTACATTGGG</td><td>1000</td><td></td><td></td><td></td><td></td><td></td><td></td><td></td><td></td><td></td></tr>
<tr><td>Oligo fwd</td><td>CACCGGCCTTTACCCAAGTACATT</td><td></td><td></td><td></td><td></td><td></td><td></td><td></td><td></td><td></td><td></td></tr>
<tr><td>Oligo rev</td><td>AAACAATGTACTTGGGTAAAGGCC</td><td></td><td></td><td></td><td></td><td></td><td></td><td></td><td></td><td></td><td></td></tr>
<tr><td>Chromosome</td><td>start</td><td>end</td><td>strand</td><td>MM</td><td>target_seq</td><td>PAM</td><td>alignment</td><td>distance</td><td>position</td><td>gene name</td><td>gene id</td></tr>
<tr><td>GU305914.1</td><td>184062</td><td>184084</td><td>-</td><td>0</td><td>GGCCTTTACCCAAGTACATT</td><td>GGG</td><td>||||||||[||||||||||||]PAM</td><td>NA</td><td>Intergenic</td><td>NA</td><td>NA</td></tr>
<tr><td>GU305914.1</td><td>183968</td><td>183990</td><td>-</td><td>3</td><td>GGGTTTTGCCCAAGTACATT</td><td>GGG</td><td>||--|||-[||||||||||||]PAM</td><td>NA</td><td>Intergenic</td><td>NA</td><td>NA</td></tr>
<tr><td>T20761</td><td>ACCCAAGTACATTGGGTCAATGG</td><td>1000</td><td></td><td></td><td></td><td></td><td></td><td></td><td></td><td></td><td></td></tr>
<tr><td>Oligo fwd</td><td>CACCACCCAAGTACATTGGGTCAA</td><td></td><td></td><td></td><td></td><td></td><td></td><td></td><td></td><td></td><td></td></tr>
<tr><td>Oligo rev</td><td>AAACTTGACCCAATGTACTTGGGT</td><td></td><td></td><td></td><td></td><td></td><td></td><td></td><td></td><td></td><td></td></tr>
<tr><td>Chromosome</td><td>start</td><td>end</td><td>strand</td><td>MM</td><td>target_seq</td><td>PAM</td><td>alignment</td><td>distance</td><td>position</td><td>gene name</td><td>gene id</td></tr>
</table>

<table>
<tr><td>GU305914.1</td><td></td><td>184055</td><td>184077</td><td>-</td><td>0</td><td>ACCCAAGTACATTGGGTCAA</td><td>TGG</td><td>||||||||[||||||||||||]PAM</td><td>NA</td><td>Intergenic</td><td>NA</td><td>NA</td></tr>
<tr><td>GU305914.1</td><td></td><td>183961</td><td>183983</td><td>-</td><td>1</td><td>GCCCAAGTACATTGGGTCAA</td><td>TGG</td><td>-|||||||[||||||||||||]PAM</td><td>NA</td><td>Intergenic</td><td>NA</td><td>NA</td></tr>
<tr><td></td><td></td><td></td><td></td><td></td><td></td><td></td><td></td><td></td><td></td><td></td><td></td><td></td></tr>
<tr><td>T20762</td><td>CCCAAGTACATTGGGTCAATGGG</td><td></td><td>1000</td><td></td><td></td><td></td><td></td><td></td><td></td><td></td><td></td><td></td></tr>
<tr><td>Oligo fwd</td><td>CACCCCCAAGTACATTGGGTCAAT</td><td></td><td></td><td></td><td></td><td></td><td></td><td></td><td></td><td></td><td></td><td></td></tr>
<tr><td>Oligo rev</td><td>AAACATTGACCCAATGTACTTGGG</td><td></td><td></td><td></td><td></td><td></td><td></td><td></td><td></td><td></td><td></td><td></td></tr>
<tr><td>Chromosome</td><td>start</td><td></td><td>end</td><td>strand</td><td>MM</td><td>target_seq</td><td>PAM</td><td>alignment</td><td>distance</td><td>position</td><td>gene name</td><td>gene id</td></tr>
<tr><td>GU305914.1</td><td></td><td>184054</td><td>184076</td><td>-</td><td>0</td><td>CCCAAGTACATTGGGTCAAT</td><td>GGG</td><td>||||||||[||||||||||||]PAM</td><td>NA</td><td>Intergenic</td><td>NA</td><td>NA</td></tr>
<tr><td>GU305914.1</td><td></td><td>183960</td><td>183982</td><td>-</td><td>0</td><td>CCCAAGTACATTGGGTCAAT</td><td>GGG</td><td>||||||||[||||||||||||]PAM</td><td>NA</td><td>Intergenic</td><td>NA</td><td>NA</td></tr>
<tr><td>GU305914.1</td><td></td><td>183565</td><td>183587</td><td>-</td><td>2</td><td>GCCAAGTACATTGAGTCAAT</td><td>AGG</td><td>-|||||||[|||||-||||||]PAM</td><td>NA</td><td>Intergenic</td><td>NA</td><td>NA</td></tr>
<tr><td>GU305914.1</td><td></td><td>183380</td><td>183402</td><td>-</td><td>3</td><td>GCCAAGTACACTGAGTCAAT</td><td>AGG</td><td>-|||||||[||-||-||||||]PAM</td><td>NA</td><td>Intergenic</td><td>NA</td><td>NA</td></tr>
<tr><td>GU305914.1</td><td></td><td>183751</td><td>183773</td><td>-</td><td>3</td><td>GCCAAGTACACTGCGTCAAT</td><td>AGG</td><td>-|||||||[||-||-||||||]PAM</td><td>NA</td><td>Intergenic</td><td>NA</td><td>NA</td></tr>
<tr><td>GU305914.1</td><td></td><td>183524</td><td>183546</td><td>-</td><td>4</td><td>GCCCAGTACATAAGGTCAAT</td><td>GGG</td><td>-||-||||[|||--|||||||]PAM</td><td>NA</td><td>Intergenic</td><td>NA</td><td>NA</td></tr>
<tr><td>GU305914.1</td><td></td><td>183710</td><td>183732</td><td>-</td><td>4</td><td>GCCCAGTACATAAGGTCAAT</td><td>AGG</td><td>-||-||||[|||--|||||||]PAM</td><td>NA</td><td>Intergenic</td><td>NA</td><td>NA</td></tr>
<tr><td>GU305914.1</td><td></td><td>183431</td><td>183453</td><td>-</td><td>4</td><td>GCCCAGTACATAAGGTCAAT</td><td>AGG</td><td>-||-||||[|||--|||||||]PAM</td><td>NA</td><td>Intergenic</td><td>NA</td><td>NA</td></tr>
<tr><td>GU305914.1</td><td></td><td>183617</td><td>183639</td><td>-</td><td>4</td><td>GCCCAGTACATAAGGTCAAT</td><td>AGG</td><td>-||-||||[|||--|||||||]PAM</td><td>NA</td><td>Intergenic</td><td>NA</td><td>NA</td></tr>
<tr><td>GU305914.1</td><td></td><td>183803</td><td>183825</td><td>-</td><td>4</td><td>GCCCAGTACATAAGGTCAAT</td><td>AGG</td><td>-||-||||[|||--|||||||]PAM</td><td>NA</td><td>Intergenic</td><td>NA</td><td>NA</td></tr>
<tr><td></td><td></td><td></td><td></td><td></td><td></td><td></td><td></td><td></td><td></td><td></td><td></td><td></td></tr>
<tr><td>T20763</td><td>AAGTACATTGGGTCAATGGGAGG</td><td></td><td>1000</td><td></td><td></td><td></td><td></td><td></td><td></td><td></td><td></td><td></td></tr>
<tr><td>Oligo fwd</td><td>CACCAAGTACATTGGGTCAATGGG</td><td></td><td></td><td></td><td></td><td></td><td></td><td></td><td></td><td></td><td></td><td></td></tr>
<tr><td>Oligo rev</td><td>AAACCCCATTGACCCAATGTACTT</td><td></td><td></td><td></td><td></td><td></td><td></td><td></td><td></td><td></td><td></td><td></td></tr>
<tr><td>Chromosome</td><td>start</td><td></td><td>end</td><td>strand</td><td>MM</td><td>target_seq</td><td>PAM</td><td>alignment</td><td>distance</td><td>position</td><td>gene name</td><td>gene id</td></tr>
<tr><td>GU305914.1</td><td></td><td>183957</td><td>183979</td><td>-</td><td>0</td><td>AAGTACATTGGGTCAATGGG</td><td>AGG</td><td>||||||||[||||||||||||]PAM</td><td>NA</td><td>Intergenic</td><td>NA</td><td>NA</td></tr>
<tr><td>GU305914.1</td><td></td><td>184051</td><td>184073</td><td>-</td><td>0</td><td>AAGTACATTGGGTCAATGGG</td><td>AGG</td><td>||||||||[||||||||||||]PAM</td><td>NA</td><td>Intergenic</td><td>NA</td><td>NA</td></tr>
<tr><td>GU305914.1</td><td></td><td>183521</td><td>183543</td><td>-</td><td>3</td><td>CAGTACATAAGGTCAATGGG</td><td>AGG</td><td>-|||||||[--||||||||||]PAM</td><td>NA</td><td>Intergenic</td><td>NA</td><td>NA</td></tr>
<tr><td></td><td></td><td></td><td></td><td></td><td></td><td></td><td></td><td></td><td></td><td></td><td></td><td></td></tr>
<tr><td>T20764</td><td>GTCAATGGGAGGTAAGCCAATGG</td><td></td><td>1000</td><td></td><td></td><td></td><td></td><td></td><td></td><td></td><td></td><td></td></tr>
<tr><td>Oligo fwd</td><td>CACCGTCAATGGGAGGTAAGCCAA</td><td></td><td></td><td></td><td></td><td></td><td></td><td></td><td></td><td></td><td></td><td></td></tr>
<tr><td>Oligo rev</td><td>AAACTTGGCTTACCTCCCATTGAC</td><td></td><td></td><td></td><td></td><td></td><td></td><td></td><td></td><td></td><td></td><td></td></tr>
<tr><td>Chromosome</td><td>start</td><td></td><td>end</td><td>strand</td><td>MM</td><td>target_seq</td><td>PAM</td><td>alignment</td><td>distance</td><td>position</td><td>gene name</td><td>gene id</td></tr>
<tr><td>GU305914.1</td><td></td><td>184040</td><td>184062</td><td>-</td><td>0</td><td>GTCAATGGGAGGTAAGCCAA</td><td>TGG</td><td>||||||||[||||||||||||]PAM</td><td>NA</td><td>Intergenic</td><td>NA</td><td>NA</td></tr>
<tr><td>GU305914.1</td><td></td><td>183510</td><td>183532</td><td>-</td><td>0</td><td>GTCAATGGGAGGTAAGCCAA</td><td>TGG</td><td>||||||||[||||||||||||]PAM</td><td>NA</td><td>Intergenic</td><td>NA</td><td>NA</td></tr>
<tr><td>GU305914.1</td><td></td><td>183946</td><td>183968</td><td>-</td><td>1</td><td>GTCAATGGGAGGTGAGCCAA</td><td>TGG</td><td>||||||||[|||||-||||||]PAM</td><td>NA</td><td>Intergenic</td><td>NA</td><td>NA</td></tr>
<tr><td></td><td></td><td></td><td></td><td></td><td></td><td></td><td></td><td></td><td></td><td></td><td></td><td></td></tr>
<tr><td>T20765</td><td>TCAATGGGAGGTAAGCCAATGGG</td><td></td><td>1000</td><td></td><td></td><td></td><td></td><td></td><td></td><td></td><td></td><td></td></tr>
<tr><td>Oligo fwd</td><td>CACCTCAATGGGAGGTAAGCCAAT</td><td></td><td></td><td></td><td></td><td></td><td></td><td></td><td></td><td></td><td></td><td></td></tr>
<tr><td>Oligo rev</td><td>AAACATTGGCTTACCTCCCATTGA</td><td></td><td></td><td></td><td></td><td></td><td></td><td></td><td></td><td></td><td></td><td></td></tr>
<tr><td>Chromosome</td><td>start</td><td></td><td>end</td><td>strand</td><td>MM</td><td>target_seq</td><td>PAM</td><td>alignment</td><td>distance</td><td>position</td><td>gene name</td><td>gene id</td></tr>
<tr><td>GU305914.1</td><td></td><td>184039</td><td>184061</td><td>-</td><td>0</td><td>TCAATGGGAGGTAAGCCAAT</td><td>GGG</td><td>||||||||[||||||||||||]PAM</td><td>NA</td><td>Intergenic</td><td>NA</td><td>NA</td></tr>
<tr><td>GU305914.1</td><td></td><td>183509</td><td>183531</td><td>-</td><td>0</td><td>TCAATGGGAGGTAAGCCAAT</td><td>GGG</td><td>||||||||[||||||||||||]PAM</td><td>NA</td><td>Intergenic</td><td>NA</td><td>NA</td></tr>
<tr><td>GU305914.1</td><td></td><td>183945</td><td>183967</td><td>-</td><td>1</td><td>TCAATGGGAGGTGAGCCAAT</td><td>GGG</td><td>||||||||[||||-|||||||]PAM</td><td>NA</td><td>Intergenic</td><td>NA</td><td>NA</td></tr>
<tr><td></td><td></td><td></td><td></td><td></td><td></td><td></td><td></td><td></td><td></td><td></td><td></td><td></td></tr>
<tr><td>T20766</td><td>CAATGGGTTTTTCCCATTACTGG</td><td></td><td>1000</td><td></td><td></td><td></td><td></td><td></td><td></td><td></td><td></td><td></td></tr>
<tr><td>Oligo fwd</td><td>CACCCAATGGGTTTTTCCCATTAC</td><td></td><td></td><td></td><td></td><td></td><td></td><td></td><td></td><td></td><td></td><td></td></tr>
<tr><td>Oligo rev</td><td>AAACGTAATGGGAAAAACCCATTG</td><td></td><td></td><td></td><td></td><td></td><td></td><td></td><td></td><td></td><td></td><td></td></tr>
<tr><td>Chromosome</td><td>start</td><td></td><td>end</td><td>strand</td><td>MM</td><td>target_seq</td><td>PAM</td><td>alignment</td><td>distance</td><td>position</td><td>gene name</td><td>gene id</td></tr>
<tr><td>GU305914.1</td><td></td><td>184023</td><td>184045</td><td>-</td><td>0</td><td>CAATGGGTTTTTCCCATTAC</td><td>TGG</td><td>||||||||[||||||||||||]PAM</td><td>NA</td><td>Intergenic</td><td>NA</td><td>NA</td></tr>
<tr><td>GU305914.1</td><td></td><td>183493</td><td>183515</td><td>-</td><td>0</td><td>CAATGGGTTTTTCCCATTAC</td><td>TGG</td><td>||||||||[||||||||||||]PAM</td><td>NA</td><td>Intergenic</td><td>NA</td><td>NA</td></tr>
<tr><td>GU305914.1</td><td></td><td>183308</td><td>183330</td><td>-</td><td>1</td><td>CAATGGGTTTTTCCCATTAT</td><td>TGG</td><td>||||||||[|||||||||||-]PAM</td><td>NA</td><td>Intergenic</td><td>NA</td><td>NA</td></tr>
<tr><td></td><td></td><td></td><td></td><td></td><td></td><td></td><td></td><td></td><td></td><td></td><td></td><td></td></tr>
<tr><td>T20767</td><td>GGCAAGCACACTGAGTCAAATGG</td><td></td><td>1000</td><td></td><td></td><td></td><td></td><td></td><td></td><td></td><td></td><td></td></tr>
<tr><td>Oligo fwd</td><td>CACCGGCAAGCACACTGAGTCAAA</td><td></td><td></td><td></td><td></td><td></td><td></td><td></td><td></td><td></td><td></td><td></td></tr>
<tr><td>Oligo rev</td><td>AAACTTTGACTCAGTGTGCTTGCC</td><td></td><td></td><td></td><td></td><td></td><td></td><td></td><td></td><td></td><td></td><td></td></tr>
<tr><td>Chromosome</td><td>start</td><td></td><td>end</td><td>strand</td><td>MM</td><td>target_seq</td><td>PAM</td><td>alignment</td><td>distance</td><td>position</td><td>gene name</td><td>gene id</td></tr>
<tr><td>GU305914.1</td><td></td><td>184002</td><td>184024</td><td>-</td><td>0</td><td>GGCAAGCACACTGAGTCAAA</td><td>TGG</td><td>||||||||[||||||||||||]PAM</td><td>NA</td><td>Intergenic</td><td>NA</td><td>NA</td></tr>
<tr><td>GU305914.1</td><td></td><td>183380</td><td>183402</td><td>-</td><td>3</td><td>GCCAAGTACACTGAGTCAAT</td><td>AGG</td><td>|-||||-|[|||||||||||-]PAM</td><td>NA</td><td>Intergenic</td><td>NA</td><td>NA</td></tr>
<tr><td>GU305914.1</td><td></td><td>183565</td><td>183587</td><td>-</td><td>4</td><td>GCCAAGTACATTGAGTCAAT</td><td>AGG</td><td>|-||||-|[||-||||||||-]PAM</td><td>NA</td><td>Intergenic</td><td>NA</td><td>NA</td></tr>
<tr><td>GU305914.1</td><td></td><td>183751</td><td>183773</td><td>-</td><td>4</td><td>GCCAAGTACACTGCGTCAAT</td><td>AGG</td><td>|-||||-|[|||||-|||||-]PAM</td><td>NA</td><td>Intergenic</td><td>NA</td><td>NA</td></tr>
<tr><td>GU305914.1</td><td></td><td>183658</td><td>183680</td><td>-</td><td>4</td><td>GCCAAGTACACTGACTCAAT</td><td>AGG</td><td>|-||||-|[||||||-||||-]PAM</td><td>NA</td><td>Intergenic</td><td>NA</td><td>NA</td></tr>
<tr><td>GU305914.1</td><td></td><td>183908</td><td>183930</td><td>-</td><td>4</td><td>GCCAAGTACACTGACTCAAT</td><td>AGG</td><td>|-||||-|[||||||-||||-]PAM</td><td>NA</td><td>Intergenic</td><td>NA</td><td>NA</td></tr>
<tr><td></td><td></td><td></td><td></td><td></td><td></td><td></td><td></td><td></td><td></td><td></td><td></td><td></td></tr>
<tr><td>T20768</td><td>GCAAGCACACTGAGTCAAATGGG</td><td></td><td>1000</td><td></td><td></td><td></td><td></td><td></td><td></td><td></td><td></td><td></td></tr>
<tr><td>Oligo fwd</td><td>CACCGCAAGCACACTGAGTCAAAT</td><td></td><td></td><td></td><td></td><td></td><td></td><td></td><td></td><td></td><td></td><td></td></tr>
<tr><td>Oligo rev</td><td>AAACATTTGACTCAGTGTGCTTGC</td><td></td><td></td><td></td><td></td><td></td><td></td><td></td><td></td><td></td><td></td><td></td></tr>
<tr><td>Chromosome</td><td>start</td><td></td><td>end</td><td>strand</td><td>MM</td><td>target_seq</td><td>PAM</td><td>alignment</td><td>distance</td><td>position</td><td>gene name</td><td>gene id</td></tr>
<tr><td>GU305914.1</td><td></td><td>184001</td><td>184023</td><td>-</td><td>0</td><td>GCAAGCACACTGAGTCAAAT</td><td>GGG</td><td>||||||||[||||||||||||]PAM</td><td>NA</td><td>Intergenic</td><td>NA</td><td>NA</td></tr>
<tr><td>GU305914.1</td><td></td><td>183379</td><td>183401</td><td>-</td><td>4</td><td>CCAAGTACACTGAGTCAATA</td><td>GGG</td><td>-||||-||[||||||||||--]PAM</td><td>NA</td><td>Intergenic</td><td>NA</td><td>NA</td></tr>
<tr><td></td><td></td><td></td><td></td><td></td><td></td><td></td><td></td><td></td><td></td><td></td><td></td><td></td></tr>
<tr><td>T20769</td><td>AGTCAAATGGGACTTTCCACTGG</td><td></td><td>1000</td><td></td><td></td><td></td><td></td><td></td><td></td><td></td><td></td><td></td></tr>
<tr><td>Oligo fwd</td><td>CACCAGTCAAATGGGACTTTCCAC</td><td></td><td></td><td></td><td></td><td></td><td></td><td></td><td></td><td></td><td></td><td></td></tr>
<tr><td>Oligo rev</td><td>AAACGTGGAAAGTCCCATTTGACT</td><td></td><td></td><td></td><td></td><td></td><td></td><td></td><td></td><td></td><td></td><td></td></tr>
<tr><td>Chromosome</td><td>start</td><td></td><td>end</td><td>strand</td><td>MM</td><td>target_seq</td><td>PAM</td><td>alignment</td><td>distance</td><td>position</td><td>gene name</td><td>gene id</td></tr>
<tr><td>GU305914.1</td><td></td><td>183989</td><td>184011</td><td>-</td><td>0</td><td>AGTCAAATGGGACTTTCCAC</td><td>TGG</td><td>||||||||[||||||||||||]PAM</td><td>NA</td><td>Intergenic</td><td>NA</td><td>NA</td></tr>
<tr><td>GU305914.1</td><td></td><td>183552</td><td>183574</td><td>-</td><td>3</td><td>AGTCAATAGGGACTTTCCAA</td><td>TGG</td><td>||||||--[|||||||||||-]PAM</td><td>NA</td><td>Intergenic</td><td>NA</td><td>NA</td></tr>
<tr><td>GU305914.1</td><td></td><td>183367</td><td>183389</td><td>-</td><td>3</td><td>AGTCAATAGGGACTTTCCAT</td><td>TGG</td><td>||||||--[|||||||||||-]PAM</td><td>NA</td><td>Intergenic</td><td>NA</td><td>NA</td></tr>
<tr><td>GU305914.1</td><td></td><td>183738</td><td>183760</td><td>-</td><td>4</td><td>CGTCAATAGGGACTTTCCAT</td><td>TGG</td><td>-|||||--[|||||||||||-]PAM</td><td>NA</td><td>Intergenic</td><td>NA</td><td>NA</td></tr>
<tr><td>GU305914.1</td><td></td><td>183895</td><td>183917</td><td>-</td><td>4</td><td>ACTCAATAGGGACTTTCCAA</td><td>TGG</td><td>|-||||--[|||||||||||-]PAM</td><td>NA</td><td>Intergenic</td><td>NA</td><td>NA</td></tr>
<tr><td>GU305914.1</td><td></td><td>183645</td><td>183667</td><td>-</td><td>4</td><td>ACTCAATAGGGACTTTCCAT</td><td>TGG</td><td>|-||||--[|||||||||||-]PAM</td><td>NA</td><td>Intergenic</td><td>NA</td><td>NA</td></tr>
<tr><td>GU305914.1</td><td></td><td>183459</td><td>183481</td><td>-</td><td>4</td><td>AGTCATTAGGGACTTTCCAA</td><td>TGG</td><td>|||||---[|||||||||||-]PAM</td><td>NA</td><td>Intergenic</td><td>NA</td><td>NA</td></tr>
<tr><td></td><td></td><td></td><td></td><td></td><td></td><td></td><td></td><td></td><td></td><td></td><td></td><td></td></tr>
<tr><td>T20770</td><td>GTCAAATGGGACTTTCCACTGGG</td><td></td><td>1000</td><td></td><td></td><td></td><td></td><td></td><td></td><td></td><td></td><td></td></tr>
<tr><td>Oligo fwd</td><td>CACCGTCAAATGGGACTTTCCACT</td><td></td><td></td><td></td><td></td><td></td><td></td><td></td><td></td><td></td><td></td><td></td></tr>
<tr><td>Oligo rev</td><td>AAACAGTGGAAAGTCCCATTTGAC</td><td></td><td></td><td></td><td></td><td></td><td></td><td></td><td></td><td></td><td></td><td></td></tr>
<tr><td>Chromosome</td><td>start</td><td></td><td>end</td><td>strand</td><td>MM</td><td>target_seq</td><td>PAM</td><td>alignment</td><td>distance</td><td>position</td><td>gene name</td><td>gene id</td></tr>
<tr><td>GU305914.1</td><td></td><td>183988</td><td>184010</td><td>-</td><td>0</td><td>GTCAAATGGGACTTTCCACT</td><td>GGG</td><td>||||||||[||||||||||||]PAM</td><td>NA</td><td>Intergenic</td><td>NA</td><td>NA</td></tr>
<tr><td>GU305914.1</td><td></td><td>183366</td><td>183388</td><td>-</td><td>3</td><td>GTCAATAGGGACTTTCCATT</td><td>GGG</td><td>|||||--|[||||||||||-|]PAM</td><td>NA</td><td>Intergenic</td><td>NA</td><td>NA</td></tr>
<tr><td>GU305914.1</td><td></td><td>183737</td><td>183759</td><td>-</td><td>3</td><td>GTCAATAGGGACTTTCCATT</td><td>GGG</td><td>|||||--|[||||||||||-|]PAM</td><td>NA</td><td>Intergenic</td><td>NA</td><td>NA</td></tr>
<tr><td>GU305914.1</td><td></td><td>183551</td><td>183573</td><td>-</td><td>3</td><td>GTCAATAGGGACTTTCCAAT</td><td>GGG</td><td>|||||--|[||||||||||-|]PAM</td><td>NA</td><td>Intergenic</td><td>NA</td><td>NA</td></tr>
<tr><td>GU305914.1</td><td></td><td>183644</td><td>183666</td><td>-</td><td>4</td><td>CTCAATAGGGACTTTCCATT</td><td>GGG</td><td>-||||--|[||||||||||-|]PAM</td><td>NA</td><td>Intergenic</td><td>NA</td><td>NA</td></tr>
</table>

<table>
<tr><td>GU305914.1</td><td>183894</td><td>183916</td><td>-</td><td>4</td><td>CTCAATAGGGACTTTCCAAT</td><td>GGG</td><td>-||||--|[||||||||||-|]PAM</td><td>NA</td><td>Intergenic</td><td>NA</td><td>NA</td></tr>
<tr><td>GU305914.1</td><td>183458</td><td>183480</td><td>-</td><td>4</td><td>GTCATTAGGGACTTTCCAAT</td><td>GGG</td><td>||||---|[||||||||||-|]PAM</td><td>NA</td><td>Intergenic</td><td>NA</td><td>NA</td></tr>
<tr><td></td><td></td><td></td><td></td><td></td><td></td><td></td><td></td><td></td><td></td><td></td><td></td></tr>
<tr><td>T20771</td><td>TGGGTTTTGCCCAAGTACATTGG</td><td>1000</td><td></td><td></td><td></td><td></td><td></td><td></td><td></td><td></td><td></td></tr>
<tr><td>Oligo fwd</td><td>CACCTGGGTTTTGCCCAAGTACAT</td><td></td><td></td><td></td><td></td><td></td><td></td><td></td><td></td><td></td><td></td></tr>
<tr><td>Oligo rev</td><td>AAACATGTACTTGGGCAAAACCCA</td><td></td><td></td><td></td><td></td><td></td><td></td><td></td><td></td><td></td><td></td></tr>
<tr><td>Chromosome</td><td>start</td><td>end</td><td>strand</td><td>MM</td><td>target_seq</td><td>PAM</td><td>alignment</td><td>distance</td><td>position</td><td>gene name</td><td>gene id</td></tr>
<tr><td>GU305914.1</td><td>183969</td><td>183991</td><td>-</td><td>0</td><td>TGGGTTTTGCCCAAGTACAT</td><td>TGG</td><td>||||||||[||||||||||||]PAM</td><td>NA</td><td>Intergenic</td><td>NA</td><td>NA</td></tr>
<tr><td>GU305914.1</td><td>184063</td><td>184085</td><td>-</td><td>3</td><td>TGGCCTTTACCCAAGTACAT</td><td>TGG</td><td>|||--|||[-|||||||||||]PAM</td><td>NA</td><td>Intergenic</td><td>NA</td><td>NA</td></tr>
<tr><td></td><td></td><td></td><td></td><td></td><td></td><td></td><td></td><td></td><td></td><td></td><td></td></tr>
<tr><td>T20772</td><td>GGGTTTTGCCCAAGTACATTGGG</td><td>1000</td><td></td><td></td><td></td><td></td><td></td><td></td><td></td><td></td><td></td></tr>
<tr><td>Oligo fwd</td><td>CACCGGGTTTTGCCCAAGTACATT</td><td></td><td></td><td></td><td></td><td></td><td></td><td></td><td></td><td></td><td></td></tr>
<tr><td>Oligo rev</td><td>AAACAATGTACTTGGGCAAAACCC</td><td></td><td></td><td></td><td></td><td></td><td></td><td></td><td></td><td></td><td></td></tr>
<tr><td>Chromosome</td><td>start</td><td>end</td><td>strand</td><td>MM</td><td>target_seq</td><td>PAM</td><td>alignment</td><td>distance</td><td>position</td><td>gene name</td><td>gene id</td></tr>
<tr><td>GU305914.1</td><td>183968</td><td>183990</td><td>-</td><td>0</td><td>GGGTTTTGCCCAAGTACATT</td><td>GGG</td><td>||||||||[||||||||||||]PAM</td><td>NA</td><td>Intergenic</td><td>NA</td><td>NA</td></tr>
<tr><td>GU305914.1</td><td>184062</td><td>184084</td><td>-</td><td>3</td><td>GGCCTTTACCCAAGTACATT</td><td>GGG</td><td>||--|||-[||||||||||||]PAM</td><td>NA</td><td>Intergenic</td><td>NA</td><td>NA</td></tr>
<tr><td></td><td></td><td></td><td></td><td></td><td></td><td></td><td></td><td></td><td></td><td></td><td></td></tr>
<tr><td>T20773</td><td>GCCCAAGTACATTGGGTCAATGG</td><td>1000</td><td></td><td></td><td></td><td></td><td></td><td></td><td></td><td></td><td></td></tr>
<tr><td>Oligo fwd</td><td>CACCGCCCAAGTACATTGGGTCAA</td><td></td><td></td><td></td><td></td><td></td><td></td><td></td><td></td><td></td><td></td></tr>
<tr><td>Oligo rev</td><td>AAACTTGACCCAATGTACTTGGGC</td><td></td><td></td><td></td><td></td><td></td><td></td><td></td><td></td><td></td><td></td></tr>
<tr><td>Chromosome</td><td>start</td><td>end</td><td>strand</td><td>MM</td><td>target_seq</td><td>PAM</td><td>alignment</td><td>distance</td><td>position</td><td>gene name</td><td>gene id</td></tr>
<tr><td>GU305914.1</td><td>183961</td><td>183983</td><td>-</td><td>0</td><td>GCCCAAGTACATTGGGTCAA</td><td>TGG</td><td>||||||||[||||||||||||]PAM</td><td>NA</td><td>Intergenic</td><td>NA</td><td>NA</td></tr>
<tr><td>GU305914.1</td><td>184055</td><td>184077</td><td>-</td><td>1</td><td>ACCCAAGTACATTGGGTCAA</td><td>TGG</td><td>-|||||||[||||||||||||]PAM</td><td>NA</td><td>Intergenic</td><td>NA</td><td>NA</td></tr>
<tr><td></td><td></td><td></td><td></td><td></td><td></td><td></td><td></td><td></td><td></td><td></td><td></td></tr>
<tr><td>T20774</td><td>CCCAAGTACATTGGGTCAATGGG</td><td>1000</td><td></td><td></td><td></td><td></td><td></td><td></td><td></td><td></td><td></td></tr>
<tr><td>Oligo fwd</td><td>CACCCCCAAGTACATTGGGTCAAT</td><td></td><td></td><td></td><td></td><td></td><td></td><td></td><td></td><td></td><td></td></tr>
<tr><td>Oligo rev</td><td>AAACATTGACCCAATGTACTTGGG</td><td></td><td></td><td></td><td></td><td></td><td></td><td></td><td></td><td></td><td></td></tr>
<tr><td>Chromosome</td><td>start</td><td>end</td><td>strand</td><td>MM</td><td>target_seq</td><td>PAM</td><td>alignment</td><td>distance</td><td>position</td><td>gene name</td><td>gene id</td></tr>
<tr><td>GU305914.1</td><td>184054</td><td>184076</td><td>-</td><td>0</td><td>CCCAAGTACATTGGGTCAAT</td><td>GGG</td><td>||||||||[||||||||||||]PAM</td><td>NA</td><td>Intergenic</td><td>NA</td><td>NA</td></tr>
<tr><td>GU305914.1</td><td>183960</td><td>183982</td><td>-</td><td>0</td><td>CCCAAGTACATTGGGTCAAT</td><td>GGG</td><td>||||||||[||||||||||||]PAM</td><td>NA</td><td>Intergenic</td><td>NA</td><td>NA</td></tr>
<tr><td>GU305914.1</td><td>183565</td><td>183587</td><td>-</td><td>2</td><td>GCCAAGTACATTGAGTCAAT</td><td>AGG</td><td>-|||||||[|||||-||||||]PAM</td><td>NA</td><td>Intergenic</td><td>NA</td><td>NA</td></tr>
<tr><td>GU305914.1</td><td>183380</td><td>183402</td><td>-</td><td>3</td><td>GCCAAGTACACTGAGTCAAT</td><td>AGG</td><td>-|||||||[||-||-||||||]PAM</td><td>NA</td><td>Intergenic</td><td>NA</td><td>NA</td></tr>
<tr><td>GU305914.1</td><td>183751</td><td>183773</td><td>-</td><td>3</td><td>GCCAAGTACACTGCGTCAAT</td><td>AGG</td><td>-|||||||[||-||-||||||]PAM</td><td>NA</td><td>Intergenic</td><td>NA</td><td>NA</td></tr>
<tr><td>GU305914.1</td><td>183524</td><td>183546</td><td>-</td><td>4</td><td>GCCCAGTACATAAGGTCAAT</td><td>GGG</td><td>-||-||||[|||--|||||||]PAM</td><td>NA</td><td>Intergenic</td><td>NA</td><td>NA</td></tr>
<tr><td>GU305914.1</td><td>183710</td><td>183732</td><td>-</td><td>4</td><td>GCCCAGTACATAAGGTCAAT</td><td>AGG</td><td>-||-||||[|||--|||||||]PAM</td><td>NA</td><td>Intergenic</td><td>NA</td><td>NA</td></tr>
<tr><td>GU305914.1</td><td>183431</td><td>183453</td><td>-</td><td>4</td><td>GCCCAGTACATAAGGTCAAT</td><td>AGG</td><td>-||-||||[|||--|||||||]PAM</td><td>NA</td><td>Intergenic</td><td>NA</td><td>NA</td></tr>
<tr><td>GU305914.1</td><td>183617</td><td>183639</td><td>-</td><td>4</td><td>GCCCAGTACATAAGGTCAAT</td><td>AGG</td><td>-||-||||[|||--|||||||]PAM</td><td>NA</td><td>Intergenic</td><td>NA</td><td>NA</td></tr>
<tr><td>GU305914.1</td><td>183803</td><td>183825</td><td>-</td><td>4</td><td>GCCCAGTACATAAGGTCAAT</td><td>AGG</td><td>-||-||||[|||--|||||||]PAM</td><td>NA</td><td>Intergenic</td><td>NA</td><td>NA</td></tr>
<tr><td></td><td></td><td></td><td></td><td></td><td></td><td></td><td></td><td></td><td></td><td></td><td></td></tr>
<tr><td>T20775</td><td>AAGTACATTGGGTCAATGGGAGG</td><td>1000</td><td></td><td></td><td></td><td></td><td></td><td></td><td></td><td></td><td></td></tr>
<tr><td>Oligo fwd</td><td>CACCAAGTACATTGGGTCAATGGG</td><td></td><td></td><td></td><td></td><td></td><td></td><td></td><td></td><td></td><td></td></tr>
<tr><td>Oligo rev</td><td>AAACCCCATTGACCCAATGTACTT</td><td></td><td></td><td></td><td></td><td></td><td></td><td></td><td></td><td></td><td></td></tr>
<tr><td>Chromosome</td><td>start</td><td>end</td><td>strand</td><td>MM</td><td>target_seq</td><td>PAM</td><td>alignment</td><td>distance</td><td>position</td><td>gene name</td><td>gene id</td></tr>
<tr><td>GU305914.1</td><td>183957</td><td>183979</td><td>-</td><td>0</td><td>AAGTACATTGGGTCAATGGG</td><td>AGG</td><td>||||||||[||||||||||||]PAM</td><td>NA</td><td>Intergenic</td><td>NA</td><td>NA</td></tr>
<tr><td>GU305914.1</td><td>184051</td><td>184073</td><td>-</td><td>0</td><td>AAGTACATTGGGTCAATGGG</td><td>AGG</td><td>||||||||[||||||||||||]PAM</td><td>NA</td><td>Intergenic</td><td>NA</td><td>NA</td></tr>
<tr><td>GU305914.1</td><td>183521</td><td>183543</td><td>-</td><td>3</td><td>CAGTACATAAGGTCAATGGG</td><td>AGG</td><td>-|||||||[--||||||||||]PAM</td><td>NA</td><td>Intergenic</td><td>NA</td><td>NA</td></tr>
<tr><td></td><td></td><td></td><td></td><td></td><td></td><td></td><td></td><td></td><td></td><td></td><td></td></tr>
<tr><td>T20776</td><td>GTCAATGGGAGGTGAGCCAATGG</td><td>1000</td><td></td><td></td><td></td><td></td><td></td><td></td><td></td><td></td><td></td></tr>
<tr><td>Oligo fwd</td><td>CACCGTCAATGGGAGGTGAGCCAA</td><td></td><td></td><td></td><td></td><td></td><td></td><td></td><td></td><td></td><td></td></tr>
<tr><td>Oligo rev</td><td>AAACTTGGCTCACCTCCCATTGAC</td><td></td><td></td><td></td><td></td><td></td><td></td><td></td><td></td><td></td><td></td></tr>
<tr><td>Chromosome</td><td>start</td><td>end</td><td>strand</td><td>MM</td><td>target_seq</td><td>PAM</td><td>alignment</td><td>distance</td><td>position</td><td>gene name</td><td>gene id</td></tr>
<tr><td>GU305914.1</td><td>183946</td><td>183968</td><td>-</td><td>0</td><td>GTCAATGGGAGGTGAGCCAA</td><td>TGG</td><td>||||||||[||||||||||||]PAM</td><td>NA</td><td>Intergenic</td><td>NA</td><td>NA</td></tr>
<tr><td>GU305914.1</td><td>184040</td><td>184062</td><td>-</td><td>1</td><td>GTCAATGGGAGGTAAGCCAA</td><td>TGG</td><td>||||||||[|||||-||||||]PAM</td><td>NA</td><td>Intergenic</td><td>NA</td><td>NA</td></tr>
<tr><td>GU305914.1</td><td>183510</td><td>183532</td><td>-</td><td>1</td><td>GTCAATGGGAGGTAAGCCAA</td><td>TGG</td><td>||||||||[|||||-||||||]PAM</td><td>NA</td><td>Intergenic</td><td>NA</td><td>NA</td></tr>
<tr><td>GU305914.1</td><td>183325</td><td>183347</td><td>-</td><td>3</td><td>GTCAATAGGGGGTGAGTCAA</td><td>TGG</td><td>||||||-|[|-||||||-|||]PAM</td><td>NA</td><td>Intergenic</td><td>NA</td><td>NA</td></tr>
<tr><td></td><td></td><td></td><td></td><td></td><td></td><td></td><td></td><td></td><td></td><td></td><td></td></tr>
<tr><td>T20777</td><td>TCAATGGGAGGTGAGCCAATGGG</td><td>1000</td><td></td><td></td><td></td><td></td><td></td><td></td><td></td><td></td><td></td></tr>
<tr><td>Oligo fwd</td><td>CACCTCAATGGGAGGTGAGCCAAT</td><td></td><td></td><td></td><td></td><td></td><td></td><td></td><td></td><td></td><td></td></tr>
<tr><td>Oligo rev</td><td>AAACATTGGCTCACCTCCCATTGA</td><td></td><td></td><td></td><td></td><td></td><td></td><td></td><td></td><td></td><td></td></tr>
<tr><td>Chromosome</td><td>start</td><td>end</td><td>strand</td><td>MM</td><td>target_seq</td><td>PAM</td><td>alignment</td><td>distance</td><td>position</td><td>gene name</td><td>gene id</td></tr>
<tr><td>GU305914.1</td><td>183945</td><td>183967</td><td>-</td><td>0</td><td>TCAATGGGAGGTGAGCCAAT</td><td>GGG</td><td>||||||||[||||||||||||]PAM</td><td>NA</td><td>Intergenic</td><td>NA</td><td>NA</td></tr>
<tr><td>GU305914.1</td><td>184039</td><td>184061</td><td>-</td><td>1</td><td>TCAATGGGAGGTAAGCCAAT</td><td>GGG</td><td>||||||||[||||-|||||||]PAM</td><td>NA</td><td>Intergenic</td><td>NA</td><td>NA</td></tr>
<tr><td>GU305914.1</td><td>183509</td><td>183531</td><td>-</td><td>1</td><td>TCAATGGGAGGTAAGCCAAT</td><td>GGG</td><td>||||||||[||||-|||||||]PAM</td><td>NA</td><td>Intergenic</td><td>NA</td><td>NA</td></tr>
<tr><td>GU305914.1</td><td>183324</td><td>183346</td><td>-</td><td>3</td><td>TCAATAGGGGGTGAGTCAAT</td><td>GGG</td><td>|||||-||[-||||||-||||]PAM</td><td>NA</td><td>Intergenic</td><td>NA</td><td>NA</td></tr>
<tr><td>GU305914.1</td><td>116270</td><td>116292</td><td>+</td><td>4</td><td>TCAAGGAGAGTTGAGCGAAT</td><td>TGG</td><td>||||-|-|[||-|||||-|||]PAM</td><td>NA</td><td>Intergenic</td><td>NA</td><td>NA</td></tr>
<tr><td></td><td></td><td></td><td></td><td></td><td></td><td></td><td></td><td></td><td></td><td></td><td></td></tr>
<tr><td>T20778</td><td>GCCAAGTACACTGACTCAATAGG</td><td>1000</td><td></td><td></td><td></td><td></td><td></td><td></td><td></td><td></td><td></td></tr>
<tr><td>Oligo fwd</td><td>CACCGCCAAGTACACTGACTCAAT</td><td></td><td></td><td></td><td></td><td></td><td></td><td></td><td></td><td></td><td></td></tr>
<tr><td>Oligo rev</td><td>AAACATTGAGTCAGTGTACTTGGC</td><td></td><td></td><td></td><td></td><td></td><td></td><td></td><td></td><td></td><td></td></tr>
<tr><td>Chromosome</td><td>start</td><td>end</td><td>strand</td><td>MM</td><td>target_seq</td><td>PAM</td><td>alignment</td><td>distance</td><td>position</td><td>gene name</td><td>gene id</td></tr>
<tr><td>GU305914.1</td><td>183908</td><td>183930</td><td>-</td><td>0</td><td>GCCAAGTACACTGACTCAAT</td><td>AGG</td><td>||||||||[||||||||||||]PAM</td><td>NA</td><td>Intergenic</td><td>NA</td><td>NA</td></tr>
<tr><td>GU305914.1</td><td>183658</td><td>183680</td><td>-</td><td>0</td><td>GCCAAGTACACTGACTCAAT</td><td>AGG</td><td>||||||||[||||||||||||]PAM</td><td>NA</td><td>Intergenic</td><td>NA</td><td>NA</td></tr>
<tr><td>GU305914.1</td><td>183380</td><td>183402</td><td>-</td><td>1</td><td>GCCAAGTACACTGAGTCAAT</td><td>AGG</td><td>||||||||[||||||-|||||]PAM</td><td>NA</td><td>Intergenic</td><td>NA</td><td>NA</td></tr>
<tr><td>GU305914.1</td><td>183565</td><td>183587</td><td>-</td><td>2</td><td>GCCAAGTACATTGAGTCAAT</td><td>AGG</td><td>||||||||[||-|||-|||||]PAM</td><td>NA</td><td>Intergenic</td><td>NA</td><td>NA</td></tr>
<tr><td>GU305914.1</td><td>183751</td><td>183773</td><td>-</td><td>2</td><td>GCCAAGTACACTGCGTCAAT</td><td>AGG</td><td>||||||||[|||||--|||||]PAM</td><td>NA</td><td>Intergenic</td><td>NA</td><td>NA</td></tr>
<tr><td>GU305914.1</td><td>184002</td><td>184024</td><td>-</td><td>4</td><td>GGCAAGCACACTGAGTCAAA</td><td>TGG</td><td>|-||||-|[||||||-||||-]PAM</td><td>NA</td><td>Intergenic</td><td>NA</td><td>NA</td></tr>
<tr><td></td><td></td><td></td><td></td><td></td><td></td><td></td><td></td><td></td><td></td><td></td><td></td></tr>
<tr><td>T20779</td><td>CCAAGTACACTGACTCAATAGGG</td><td>1000</td><td></td><td></td><td></td><td></td><td></td><td></td><td></td><td></td><td></td></tr>
<tr><td>Oligo fwd</td><td>CACCCCAAGTACACTGACTCAATA</td><td></td><td></td><td></td><td></td><td></td><td></td><td></td><td></td><td></td><td></td></tr>
<tr><td>Oligo rev</td><td>AAACTATTGAGTCAGTGTACTTGG</td><td></td><td></td><td></td><td></td><td></td><td></td><td></td><td></td><td></td><td></td></tr>
<tr><td>Chromosome</td><td>start</td><td>end</td><td>strand</td><td>MM</td><td>target_seq</td><td>PAM</td><td>alignment</td><td>distance</td><td>position</td><td>gene name</td><td>gene id</td></tr>
<tr><td>GU305914.1</td><td>183907</td><td>183929</td><td>-</td><td>0</td><td>CCAAGTACACTGACTCAATA</td><td>GGG</td><td>||||||||[||||||||||||]PAM</td><td>NA</td><td>Intergenic</td><td>NA</td><td>NA</td></tr>
<tr><td>GU305914.1</td><td>183657</td><td>183679</td><td>-</td><td>0</td><td>CCAAGTACACTGACTCAATA</td><td>GGG</td><td>||||||||[||||||||||||]PAM</td><td>NA</td><td>Intergenic</td><td>NA</td><td>NA</td></tr>
<tr><td>GU305914.1</td><td>183379</td><td>183401</td><td>-</td><td>1</td><td>CCAAGTACACTGAGTCAATA</td><td>GGG</td><td>||||||||[|||||-||||||]PAM</td><td>NA</td><td>Intergenic</td><td>NA</td><td>NA</td></tr>
<tr><td>GU305914.1</td><td>183564</td><td>183586</td><td>-</td><td>2</td><td>CCAAGTACATTGAGTCAATA</td><td>GGG</td><td>||||||||[|-|||-||||||]PAM</td><td>NA</td><td>Intergenic</td><td>NA</td><td>NA</td></tr>
<tr><td>GU305914.1</td><td>183750</td><td>183772</td><td>-</td><td>2</td><td>CCAAGTACACTGCGTCAATA</td><td>GGG</td><td>||||||||[||||--||||||]PAM</td><td>NA</td><td>Intergenic</td><td>NA</td><td>NA</td></tr>
<tr><td></td><td></td><td></td><td></td><td></td><td></td><td></td><td></td><td></td><td></td><td></td><td></td></tr>
<tr><td>T20780</td><td>ACTCAATAGGGACTTTCCAATGG</td><td>1000</td><td></td><td></td><td></td><td></td><td></td><td></td><td></td><td></td><td></td></tr>
<tr><td>Oligo fwd</td><td>CACCACTCAATAGGGACTTTCCAA</td><td></td><td></td><td></td><td></td><td></td><td></td><td></td><td></td><td></td><td></td></tr>
</table>

<table>
<tr><td>Oligo rev</td><td>AAACTTGGAAAGTCCCTATTGAGT</td><td></td><td></td><td></td><td></td><td></td><td></td><td></td><td></td><td></td><td></td></tr>
<tr><td>Chromosome</td><td>start</td><td>end</td><td>strand</td><td>MM</td><td>target_seq</td><td>PAM</td><td>alignment</td><td>distance</td><td>position</td><td>gene name</td><td>gene id</td></tr>
<tr><td>GU305914.1</td><td>183895</td><td>183917</td><td>-</td><td>0</td><td>ACTCAATAGGGACTTTCCAA</td><td>TGG</td><td>||||||||[||||||||||||]PAM</td><td>NA</td><td>Intergenic</td><td>NA</td><td>NA</td></tr>
<tr><td>GU305914.1</td><td>183552</td><td>183574</td><td>-</td><td>1</td><td>AGTCAATAGGGACTTTCCAA</td><td>TGG</td><td>|-||||||[||||||||||||]PAM</td><td>NA</td><td>Intergenic</td><td>NA</td><td>NA</td></tr>
<tr><td>GU305914.1</td><td>183459</td><td>183481</td><td>-</td><td>2</td><td>AGTCATTAGGGACTTTCCAA</td><td>TGG</td><td>|-|||-||[||||||||||||]PAM</td><td>NA</td><td>Intergenic</td><td>NA</td><td>NA</td></tr>
<tr><td>GU305914.1</td><td>183645</td><td>183667</td><td>-</td><td>1</td><td>ACTCAATAGGGACTTTCCAT</td><td>TGG</td><td>||||||||[|||||||||||-]PAM</td><td>NA</td><td>Intergenic</td><td>NA</td><td>NA</td></tr>
<tr><td>GU305914.1</td><td>183367</td><td>183389</td><td>-</td><td>2</td><td>AGTCAATAGGGACTTTCCAT</td><td>TGG</td><td>|-||||||[|||||||||||-]PAM</td><td>NA</td><td>Intergenic</td><td>NA</td><td>NA</td></tr>
<tr><td>GU305914.1</td><td>183738</td><td>183760</td><td>-</td><td>3</td><td>CGTCAATAGGGACTTTCCAT</td><td>TGG</td><td>--||||||[|||||||||||-]PAM</td><td>NA</td><td>Intergenic</td><td>NA</td><td>NA</td></tr>
<tr><td>GU305914.1</td><td>183989</td><td>184011</td><td>-</td><td>4</td><td>AGTCAAATGGGACTTTCCAC</td><td>TGG</td><td>|-||||--[|||||||||||-]PAM</td><td>NA</td><td>Intergenic</td><td>NA</td><td>NA</td></tr>
<tr><td></td><td></td><td></td><td></td><td></td><td></td><td></td><td></td><td></td><td></td><td></td><td></td></tr>
<tr><td>T20781</td><td>CTCAATAGGGACTTTCCAATGGG</td><td>1000</td><td></td><td></td><td></td><td></td><td></td><td></td><td></td><td></td><td></td></tr>
<tr><td>Oligo fwd</td><td>CACCCTCAATAGGGACTTTCCAAT</td><td></td><td></td><td></td><td></td><td></td><td></td><td></td><td></td><td></td><td></td></tr>
<tr><td>Oligo rev</td><td>AAACATTGGAAAGTCCCTATTGAG</td><td></td><td></td><td></td><td></td><td></td><td></td><td></td><td></td><td></td><td></td></tr>
<tr><td>Chromosome</td><td>start</td><td>end</td><td>strand</td><td>MM</td><td>target_seq</td><td>PAM</td><td>alignment</td><td>distance</td><td>position</td><td>gene name</td><td>gene id</td></tr>
<tr><td>GU305914.1</td><td>183894</td><td>183916</td><td>-</td><td>0</td><td>CTCAATAGGGACTTTCCAAT</td><td>GGG</td><td>||||||||[||||||||||||]PAM</td><td>NA</td><td>Intergenic</td><td>NA</td><td>NA</td></tr>
<tr><td>GU305914.1</td><td>183551</td><td>183573</td><td>-</td><td>1</td><td>GTCAATAGGGACTTTCCAAT</td><td>GGG</td><td>-|||||||[||||||||||||]PAM</td><td>NA</td><td>Intergenic</td><td>NA</td><td>NA</td></tr>
<tr><td>GU305914.1</td><td>183458</td><td>183480</td><td>-</td><td>2</td><td>GTCATTAGGGACTTTCCAAT</td><td>GGG</td><td>-|||-|||[||||||||||||]PAM</td><td>NA</td><td>Intergenic</td><td>NA</td><td>NA</td></tr>
<tr><td>GU305914.1</td><td>183644</td><td>183666</td><td>-</td><td>1</td><td>CTCAATAGGGACTTTCCATT</td><td>GGG</td><td>||||||||[||||||||||-|]PAM</td><td>NA</td><td>Intergenic</td><td>NA</td><td>NA</td></tr>
<tr><td>GU305914.1</td><td>183737</td><td>183759</td><td>-</td><td>2</td><td>GTCAATAGGGACTTTCCATT</td><td>GGG</td><td>-|||||||[||||||||||-|]PAM</td><td>NA</td><td>Intergenic</td><td>NA</td><td>NA</td></tr>
<tr><td>GU305914.1</td><td>183366</td><td>183388</td><td>-</td><td>2</td><td>GTCAATAGGGACTTTCCATT</td><td>GGG</td><td>-|||||||[||||||||||-|]PAM</td><td>NA</td><td>Intergenic</td><td>NA</td><td>NA</td></tr>
<tr><td>GU305914.1</td><td>183988</td><td>184010</td><td>-</td><td>4</td><td>GTCAAATGGGACTTTCCACT</td><td>GGG</td><td>-||||--|[||||||||||-|]PAM</td><td>NA</td><td>Intergenic</td><td>NA</td><td>NA</td></tr>
<tr><td></td><td></td><td></td><td></td><td></td><td></td><td></td><td></td><td></td><td></td><td></td><td></td></tr>
<tr><td>T20782</td><td>CCAATGGGTTTTTCCATTGTTGG</td><td>1000</td><td></td><td></td><td></td><td></td><td></td><td></td><td></td><td></td><td></td></tr>
<tr><td>Oligo fwd</td><td>CACCCCAATGGGTTTTTCCATTGT</td><td></td><td></td><td></td><td></td><td></td><td></td><td></td><td></td><td></td><td></td></tr>
<tr><td>Oligo rev</td><td>AAACACAATGGAAAAACCCATTGG</td><td></td><td></td><td></td><td></td><td></td><td></td><td></td><td></td><td></td><td></td></tr>
<tr><td>Chromosome</td><td>start</td><td>end</td><td>strand</td><td>MM</td><td>target_seq</td><td>PAM</td><td>alignment</td><td>distance</td><td>position</td><td>gene name</td><td>gene id</td></tr>
<tr><td>GU305914.1</td><td>183879</td><td>183901</td><td>-</td><td>0</td><td>CCAATGGGTTTTTCCATTGT</td><td>TGG</td><td>||||||||[||||||||||||]PAM</td><td>NA</td><td>Intergenic</td><td>NA</td><td>NA</td></tr>
<tr><td></td><td></td><td></td><td></td><td></td><td></td><td></td><td></td><td></td><td></td><td></td><td></td></tr>
<tr><td>T20783</td><td>CCATTGTTGGCAAGCATATAAGG</td><td>1000</td><td></td><td></td><td></td><td></td><td></td><td></td><td></td><td></td><td></td></tr>
<tr><td>Oligo fwd</td><td>CACCCCATTGTTGGCAAGCATATA</td><td></td><td></td><td></td><td></td><td></td><td></td><td></td><td></td><td></td><td></td></tr>
<tr><td>Oligo rev</td><td>AAACTATATGCTTGCCAACAATGG</td><td></td><td></td><td></td><td></td><td></td><td></td><td></td><td></td><td></td><td></td></tr>
<tr><td>Chromosome</td><td>start</td><td>end</td><td>strand</td><td>MM</td><td>target_seq</td><td>PAM</td><td>alignment</td><td>distance</td><td>position</td><td>gene name</td><td>gene id</td></tr>
<tr><td>GU305914.1</td><td>183866</td><td>183888</td><td>-</td><td>0</td><td>CCATTGTTGGCAAGCATATA</td><td>AGG</td><td>||||||||[||||||||||||]PAM</td><td>NA</td><td>Intergenic</td><td>NA</td><td>NA</td></tr>
<tr><td></td><td></td><td></td><td></td><td></td><td></td><td></td><td></td><td></td><td></td><td></td><td></td></tr>
<tr><td>T20784</td><td>GCAAGCATATAAGGTCAATGTGG</td><td>1000</td><td></td><td></td><td></td><td></td><td></td><td></td><td></td><td></td><td></td></tr>
<tr><td>Oligo fwd</td><td>CACCGCAAGCATATAAGGTCAATG</td><td></td><td></td><td></td><td></td><td></td><td></td><td></td><td></td><td></td><td></td></tr>
<tr><td>Oligo rev</td><td>AAACCATTGACCTTATATGCTTGC</td><td></td><td></td><td></td><td></td><td></td><td></td><td></td><td></td><td></td><td></td></tr>
<tr><td>Chromosome</td><td>start</td><td>end</td><td>strand</td><td>MM</td><td>target_seq</td><td>PAM</td><td>alignment</td><td>distance</td><td>position</td><td>gene name</td><td>gene id</td></tr>
<tr><td>GU305914.1</td><td>183857</td><td>183879</td><td>-</td><td>0</td><td>GCAAGCATATAAGGTCAATG</td><td>TGG</td><td>||||||||[||||||||||||]PAM</td><td>NA</td><td>Intergenic</td><td>NA</td><td>NA</td></tr>
<tr><td>GU305914.1</td><td>183286</td><td>183308</td><td>-</td><td>4</td><td>GCACGTACATAAGGTCAATA</td><td>GGG</td><td>|||-|-|-[|||||||||||-]PAM</td><td>NA</td><td>Intergenic</td><td>NA</td><td>NA</td></tr>
<tr><td></td><td></td><td></td><td></td><td></td><td></td><td></td><td></td><td></td><td></td><td></td><td></td></tr>
<tr><td>T20785</td><td>CAAGCATATAAGGTCAATGTGGG</td><td>1000</td><td></td><td></td><td></td><td></td><td></td><td></td><td></td><td></td><td></td></tr>
<tr><td>Oligo fwd</td><td>CACCCAAGCATATAAGGTCAATGT</td><td></td><td></td><td></td><td></td><td></td><td></td><td></td><td></td><td></td><td></td></tr>
<tr><td>Oligo rev</td><td>AAACACATTGACCTTATATGCTTG</td><td></td><td></td><td></td><td></td><td></td><td></td><td></td><td></td><td></td><td></td></tr>
<tr><td>Chromosome</td><td>start</td><td>end</td><td>strand</td><td>MM</td><td>target_seq</td><td>PAM</td><td>alignment</td><td>distance</td><td>position</td><td>gene name</td><td>gene id</td></tr>
<tr><td>GU305914.1</td><td>183856</td><td>183878</td><td>-</td><td>0</td><td>CAAGCATATAAGGTCAATGT</td><td>GGG</td><td>||||||||[||||||||||||]PAM</td><td>NA</td><td>Intergenic</td><td>NA</td><td>NA</td></tr>
<tr><td></td><td></td><td></td><td></td><td></td><td></td><td></td><td></td><td></td><td></td><td></td><td></td></tr>
<tr><td>T20786</td><td>GTCAATGTGGGTGAGTCAATAGG</td><td>1000</td><td></td><td></td><td></td><td></td><td></td><td></td><td></td><td></td><td></td></tr>
<tr><td>Oligo fwd</td><td>CACCGTCAATGTGGGTGAGTCAAT</td><td></td><td></td><td></td><td></td><td></td><td></td><td></td><td></td><td></td><td></td></tr>
<tr><td>Oligo rev</td><td>AAACATTGACTCACCCACATTGAC</td><td></td><td></td><td></td><td></td><td></td><td></td><td></td><td></td><td></td><td></td></tr>
<tr><td>Chromosome</td><td>start</td><td>end</td><td>strand</td><td>MM</td><td>target_seq</td><td>PAM</td><td>alignment</td><td>distance</td><td>position</td><td>gene name</td><td>gene id</td></tr>
<tr><td>GU305914.1</td><td>183844</td><td>183866</td><td>-</td><td>0</td><td>GTCAATGTGGGTGAGTCAAT</td><td>AGG</td><td>||||||||[||||||||||||]PAM</td><td>NA</td><td>Intergenic</td><td>NA</td><td>NA</td></tr>
<tr><td>GU305914.1</td><td>183273</td><td>183295</td><td>-</td><td>3</td><td>GTCAATAGGGGTGAGTCATT</td><td>GGG</td><td>||||||--[||||||||||-|]PAM</td><td>NA</td><td>Intergenic</td><td>NA</td><td>NA</td></tr>
<tr><td>GU305914.1</td><td>183417</td><td>183439</td><td>-</td><td>4</td><td>GTCAATAGGGGTGAATCAAC</td><td>AGG</td><td>||||||--[||||||-||||-]PAM</td><td>NA</td><td>Intergenic</td><td>NA</td><td>NA</td></tr>
<tr><td></td><td></td><td></td><td></td><td></td><td></td><td></td><td></td><td></td><td></td><td></td><td></td></tr>
<tr><td>T20787</td><td>TCAATGTGGGTGAGTCAATAGGG</td><td>1000</td><td></td><td></td><td></td><td></td><td></td><td></td><td></td><td></td><td></td></tr>
<tr><td>Oligo fwd</td><td>CACCTCAATGTGGGTGAGTCAATA</td><td></td><td></td><td></td><td></td><td></td><td></td><td></td><td></td><td></td><td></td></tr>
<tr><td>Oligo rev</td><td>AAACTATTGACTCACCCACATTGA</td><td></td><td></td><td></td><td></td><td></td><td></td><td></td><td></td><td></td><td></td></tr>
<tr><td>Chromosome</td><td>start</td><td>end</td><td>strand</td><td>MM</td><td>target_seq</td><td>PAM</td><td>alignment</td><td>distance</td><td>position</td><td>gene name</td><td>gene id</td></tr>
<tr><td>GU305914.1</td><td>183843</td><td>183865</td><td>-</td><td>0</td><td>TCAATGTGGGTGAGTCAATA</td><td>GGG</td><td>||||||||[||||||||||||]PAM</td><td>NA</td><td>Intergenic</td><td>NA</td><td>NA</td></tr>
<tr><td></td><td></td><td></td><td></td><td></td><td></td><td></td><td></td><td></td><td></td><td></td><td></td></tr>
<tr><td>T20788</td><td>TGTATTCTGCCCAGTACATAAGG</td><td>1000</td><td></td><td></td><td></td><td></td><td></td><td></td><td></td><td></td><td></td></tr>
<tr><td>Oligo fwd</td><td>CACCTGTATTCTGCCCAGTACATA</td><td></td><td></td><td></td><td></td><td></td><td></td><td></td><td></td><td></td><td></td></tr>
<tr><td>Oligo rev</td><td>AAACTATGTACTGGGCAGAATACA</td><td></td><td></td><td></td><td></td><td></td><td></td><td></td><td></td><td></td><td></td></tr>
<tr><td>Chromosome</td><td>start</td><td>end</td><td>strand</td><td>MM</td><td>target_seq</td><td>PAM</td><td>alignment</td><td>distance</td><td>position</td><td>gene name</td><td>gene id</td></tr>
<tr><td>GU305914.1</td><td>183811</td><td>183833</td><td>-</td><td>0</td><td>TGTATTCTGCCCAGTACATA</td><td>AGG</td><td>||||||||[||||||||||||]PAM</td><td>NA</td><td>Intergenic</td><td>NA</td><td>NA</td></tr>
<tr><td>GU305914.1</td><td>183532</td><td>183554</td><td>-</td><td>3</td><td>TGGGTTTTGCCCAGTACATA</td><td>AGG</td><td>||--||-|[||||||||||||]PAM</td><td>NA</td><td>Intergenic</td><td>NA</td><td>NA</td></tr>
<tr><td>GU305914.1</td><td>183439</td><td>183461</td><td>-</td><td>3</td><td>TGGGTTTTGCCCAGTACATA</td><td>AGG</td><td>||--||-|[||||||||||||]PAM</td><td>NA</td><td>Intergenic</td><td>NA</td><td>NA</td></tr>
<tr><td>GU305914.1</td><td>183625</td><td>183647</td><td>-</td><td>3</td><td>TGGGTTTTGCCCAGTACATA</td><td>AGG</td><td>||--||-|[||||||||||||]PAM</td><td>NA</td><td>Intergenic</td><td>NA</td><td>NA</td></tr>
<tr><td>GU305914.1</td><td>183718</td><td>183740</td><td>-</td><td>3</td><td>TGGGTTTTGCCCAGTACATA</td><td>AGG</td><td>||--||-|[||||||||||||]PAM</td><td>NA</td><td>Intergenic</td><td>NA</td><td>NA</td></tr>
<tr><td>GU305914.1</td><td>183347</td><td>183369</td><td>-</td><td>4</td><td>TGGGTTTTGCCCAGTACAAA</td><td>AGG</td><td>||--||-|[||||||||||-|]PAM</td><td>NA</td><td>Intergenic</td><td>NA</td><td>NA</td></tr>
<tr><td></td><td></td><td></td><td></td><td></td><td></td><td></td><td></td><td></td><td></td><td></td><td></td></tr>
<tr><td>T20789</td><td>GCCCAGTACATAAGGTCAATAGG</td><td>1000</td><td></td><td></td><td></td><td></td><td></td><td></td><td></td><td></td><td></td></tr>
<tr><td>Oligo fwd</td><td>CACCGCCCAGTACATAAGGTCAAT</td><td></td><td></td><td></td><td></td><td></td><td></td><td></td><td></td><td></td><td></td></tr>
<tr><td>Oligo rev</td><td>AAACATTGACCTTATGTACTGGGC</td><td></td><td></td><td></td><td></td><td></td><td></td><td></td><td></td><td></td><td></td></tr>
<tr><td>Chromosome</td><td>start</td><td>end</td><td>strand</td><td>MM</td><td>target_seq</td><td>PAM</td><td>alignment</td><td>distance</td><td>position</td><td>gene name</td><td>gene id</td></tr>
<tr><td>GU305914.1</td><td>183524</td><td>183546</td><td>-</td><td>0</td><td>GCCCAGTACATAAGGTCAAT</td><td>GGG</td><td>||||||||[||||||||||||]PAM</td><td>NA</td><td>Intergenic</td><td>NA</td><td>NA</td></tr>
<tr><td>GU305914.1</td><td>183617</td><td>183639</td><td>-</td><td>0</td><td>GCCCAGTACATAAGGTCAAT</td><td>AGG</td><td>||||||||[||||||||||||]PAM</td><td>NA</td><td>Intergenic</td><td>NA</td><td>NA</td></tr>
<tr><td>GU305914.1</td><td>183803</td><td>183825</td><td>-</td><td>0</td><td>GCCCAGTACATAAGGTCAAT</td><td>AGG</td><td>||||||||[||||||||||||]PAM</td><td>NA</td><td>Intergenic</td><td>NA</td><td>NA</td></tr>
<tr><td>GU305914.1</td><td>183710</td><td>183732</td><td>-</td><td>0</td><td>GCCCAGTACATAAGGTCAAT</td><td>AGG</td><td>||||||||[||||||||||||]PAM</td><td>NA</td><td>Intergenic</td><td>NA</td><td>NA</td></tr>
<tr><td>GU305914.1</td><td>183431</td><td>183453</td><td>-</td><td>0</td><td>GCCCAGTACATAAGGTCAAT</td><td>AGG</td><td>||||||||[||||||||||||]PAM</td><td>NA</td><td>Intergenic</td><td>NA</td><td>NA</td></tr>
<tr><td>GU305914.1</td><td>183287</td><td>183309</td><td>-</td><td>3</td><td>GGCACGTACATAAGGTCAAT</td><td>AGG</td><td>|-|--|||[||||||||||||]PAM</td><td>NA</td><td>Intergenic</td><td>NA</td><td>NA</td></tr>
<tr><td>GU305914.1</td><td>183339</td><td>183361</td><td>-</td><td>1</td><td>GCCCAGTACAAAAGGTCAAT</td><td>AGG</td><td>||||||||[||-|||||||||]PAM</td><td>NA</td><td>Intergenic</td><td>NA</td><td>NA</td></tr>
<tr><td>GU305914.1</td><td>184054</td><td>184076</td><td>-</td><td>4</td><td>CCCAAGTACATTGGGTCAAT</td><td>GGG</td><td>-||-||||[|||--|||||||]PAM</td><td>NA</td><td>Intergenic</td><td>NA</td><td>NA</td></tr>
<tr><td>GU305914.1</td><td>183960</td><td>183982</td><td>-</td><td>4</td><td>CCCAAGTACATTGGGTCAAT</td><td>GGG</td><td>-||-||||[|||--|||||||]PAM</td><td>NA</td><td>Intergenic</td><td>NA</td><td>NA</td></tr>
<tr><td></td><td></td><td></td><td></td><td></td><td></td><td></td><td></td><td></td><td></td><td></td><td></td></tr>
<tr><td>T20790</td><td>CCCAGTACATAAGGTCAATAGGG</td><td>1000</td><td></td><td></td><td></td><td></td><td></td><td></td><td></td><td></td><td></td></tr>
<tr><td>Oligo fwd</td><td>CACCCCCAGTACATAAGGTCAATA</td><td></td><td></td><td></td><td></td><td></td><td></td><td></td><td></td><td></td><td></td></tr>
<tr><td>Oligo rev</td><td>AAACTATTGACCTTATGTACTGGG</td><td></td><td></td><td></td><td></td><td></td><td></td><td></td><td></td><td></td><td></td></tr>
</table>

<table>
<tr><td>Chromosome</td><td>start</td><td>end</td><td>strand</td><td>MM</td><td>target_seq</td><td>PAM</td><td>alignment</td><td>distance</td><td>position</td><td>gene name</td><td>gene id</td></tr>
<tr><td>GU305914.1</td><td>183709</td><td>183731</td><td>-</td><td>0</td><td>CCCAGTACATAAGGTCAATA</td><td>GGG</td><td>||||||||[||||||||||||]PAM</td><td>NA</td><td>Intergenic</td><td>NA</td><td>NA</td></tr>
<tr><td>GU305914.1</td><td>183430</td><td>183452</td><td>-</td><td>0</td><td>CCCAGTACATAAGGTCAATA</td><td>GGG</td><td>||||||||[||||||||||||]PAM</td><td>NA</td><td>Intergenic</td><td>NA</td><td>NA</td></tr>
<tr><td>GU305914.1</td><td>183616</td><td>183638</td><td>-</td><td>0</td><td>CCCAGTACATAAGGTCAATA</td><td>GGG</td><td>||||||||[||||||||||||]PAM</td><td>NA</td><td>Intergenic</td><td>NA</td><td>NA</td></tr>
<tr><td>GU305914.1</td><td>183802</td><td>183824</td><td>-</td><td>0</td><td>CCCAGTACATAAGGTCAATA</td><td>GGG</td><td>||||||||[||||||||||||]PAM</td><td>NA</td><td>Intergenic</td><td>NA</td><td>NA</td></tr>
<tr><td>GU305914.1</td><td>183286</td><td>183308</td><td>-</td><td>3</td><td>GCACGTACATAAGGTCAATA</td><td>GGG</td><td>-|--||||[||||||||||||]PAM</td><td>NA</td><td>Intergenic</td><td>NA</td><td>NA</td></tr>
<tr><td>GU305914.1</td><td>183338</td><td>183360</td><td>-</td><td>1</td><td>CCCAGTACAAAAGGTCAATA</td><td>GGG</td><td>||||||||[|-||||||||||]PAM</td><td>NA</td><td>Intergenic</td><td>NA</td><td>NA</td></tr>
<tr><td>T20791</td><td>CCAGTACATAAGGTCAATAGGGG</td><td>1000</td><td></td><td></td><td></td><td></td><td></td><td></td><td></td><td></td><td></td></tr>
<tr><td>Oligo fwd</td><td>CACCCCAGTACATAAGGTCAATAG</td><td></td><td></td><td></td><td></td><td></td><td></td><td></td><td></td><td></td><td></td></tr>
<tr><td>Oligo rev</td><td>AAACCTATTGACCTTATGTACTGG</td><td></td><td></td><td></td><td></td><td></td><td></td><td></td><td></td><td></td><td></td></tr>
<tr><td>Chromosome</td><td>start</td><td>end</td><td>strand</td><td>MM</td><td>target_seq</td><td>PAM</td><td>alignment</td><td>distance</td><td>position</td><td>gene name</td><td>gene id</td></tr>
<tr><td>GU305914.1</td><td>183708</td><td>183730</td><td>-</td><td>0</td><td>CCAGTACATAAGGTCAATAG</td><td>GGG</td><td>||||||||[||||||||||||]PAM</td><td>NA</td><td>Intergenic</td><td>NA</td><td>NA</td></tr>
<tr><td>GU305914.1</td><td>183429</td><td>183451</td><td>-</td><td>0</td><td>CCAGTACATAAGGTCAATAG</td><td>GGG</td><td>||||||||[||||||||||||]PAM</td><td>NA</td><td>Intergenic</td><td>NA</td><td>NA</td></tr>
<tr><td>GU305914.1</td><td>183615</td><td>183637</td><td>-</td><td>0</td><td>CCAGTACATAAGGTCAATAG</td><td>GGG</td><td>||||||||[||||||||||||]PAM</td><td>NA</td><td>Intergenic</td><td>NA</td><td>NA</td></tr>
<tr><td>GU305914.1</td><td>183801</td><td>183823</td><td>-</td><td>0</td><td>CCAGTACATAAGGTCAATAG</td><td>GGG</td><td>||||||||[||||||||||||]PAM</td><td>NA</td><td>Intergenic</td><td>NA</td><td>NA</td></tr>
<tr><td>GU305914.1</td><td>183285</td><td>183307</td><td>-</td><td>2</td><td>CACGTACATAAGGTCAATAG</td><td>GGG</td><td>|--|||||[||||||||||||]PAM</td><td>NA</td><td>Intergenic</td><td>NA</td><td>NA</td></tr>
<tr><td>GU305914.1</td><td>183337</td><td>183359</td><td>-</td><td>1</td><td>CCAGTACAAAAGGTCAATAG</td><td>GGG</td><td>||||||||[-|||||||||||]PAM</td><td>NA</td><td>Intergenic</td><td>NA</td><td>NA</td></tr>
<tr><td>T20792</td><td>CAGTACATAAGGTCAATAGGGGG</td><td>1000</td><td></td><td></td><td></td><td></td><td></td><td></td><td></td><td></td><td></td></tr>
<tr><td>Oligo fwd</td><td>CACCCAGTACATAAGGTCAATAGG</td><td></td><td></td><td></td><td></td><td></td><td></td><td></td><td></td><td></td><td></td></tr>
<tr><td>Oligo rev</td><td>AAACCCTATTGACCTTATGTACTG</td><td></td><td></td><td></td><td></td><td></td><td></td><td></td><td></td><td></td><td></td></tr>
<tr><td>Chromosome</td><td>start</td><td>end</td><td>strand</td><td>MM</td><td>target_seq</td><td>PAM</td><td>alignment</td><td>distance</td><td>position</td><td>gene name</td><td>gene id</td></tr>
<tr><td>GU305914.1</td><td>183800</td><td>183822</td><td>-</td><td>0</td><td>CAGTACATAAGGTCAATAGG</td><td>GGG</td><td>||||||||[||||||||||||]PAM</td><td>NA</td><td>Intergenic</td><td>NA</td><td>NA</td></tr>
<tr><td>GU305914.1</td><td>183614</td><td>183636</td><td>-</td><td>0</td><td>CAGTACATAAGGTCAATAGG</td><td>GGG</td><td>||||||||[||||||||||||]PAM</td><td>NA</td><td>Intergenic</td><td>NA</td><td>NA</td></tr>
<tr><td>GU305914.1</td><td>183336</td><td>183358</td><td>-</td><td>1</td><td>CAGTACAAAAGGTCAATAGG</td><td>GGG</td><td>|||||||-[||||||||||||]PAM</td><td>NA</td><td>Intergenic</td><td>NA</td><td>NA</td></tr>
<tr><td>GU305914.1</td><td>183521</td><td>183543</td><td>-</td><td>1</td><td>CAGTACATAAGGTCAATGGG</td><td>AGG</td><td>||||||||[|||||||||-||]PAM</td><td>NA</td><td>Intergenic</td><td>NA</td><td>NA</td></tr>
<tr><td>T20793</td><td>TCAATAGGGGGTGAATCAACAGG</td><td>1000</td><td></td><td></td><td></td><td></td><td></td><td></td><td></td><td></td><td></td></tr>
<tr><td>Oligo fwd</td><td>CACCTCAATAGGGGGTGAATCAAC</td><td></td><td></td><td></td><td></td><td></td><td></td><td></td><td></td><td></td><td></td></tr>
<tr><td>Oligo rev</td><td>AAACGTTGATTCACCCCCTATTGA</td><td></td><td></td><td></td><td></td><td></td><td></td><td></td><td></td><td></td><td></td></tr>
<tr><td>Chromosome</td><td>start</td><td>end</td><td>strand</td><td>MM</td><td>target_seq</td><td>PAM</td><td>alignment</td><td>distance</td><td>position</td><td>gene name</td><td>gene id</td></tr>
<tr><td>GU305914.1</td><td>183788</td><td>183810</td><td>-</td><td>0</td><td>TCAATAGGGGGTGAATCAAC</td><td>AGG</td><td>||||||||[||||||||||||]PAM</td><td>NA</td><td>Intergenic</td><td>NA</td><td>NA</td></tr>
<tr><td>GU305914.1</td><td>183602</td><td>183624</td><td>-</td><td>1</td><td>TCAATAGGGGGTGAGTCAAC</td><td>AGG</td><td>||||||||[|||||-||||||]PAM</td><td>NA</td><td>Intergenic</td><td>NA</td><td>NA</td></tr>
<tr><td>GU305914.1</td><td>183324</td><td>183346</td><td>-</td><td>2</td><td>TCAATAGGGGGTGAGTCAAT</td><td>GGG</td><td>||||||||[|||||-|||||-]PAM</td><td>NA</td><td>Intergenic</td><td>NA</td><td>NA</td></tr>
<tr><td>T20794</td><td>AATCAACAGGAAAGTCCCATTGG</td><td>1000</td><td></td><td></td><td></td><td></td><td></td><td></td><td></td><td></td><td></td></tr>
<tr><td>Oligo fwd</td><td>CACCAATCAACAGGAAAGTCCCAT</td><td></td><td></td><td></td><td></td><td></td><td></td><td></td><td></td><td></td><td></td></tr>
<tr><td>Oligo rev</td><td>AAACATGGGACTTTCCTGTTGATT</td><td></td><td></td><td></td><td></td><td></td><td></td><td></td><td></td><td></td><td></td></tr>
<tr><td>Chromosome</td><td>start</td><td>end</td><td>strand</td><td>MM</td><td>target_seq</td><td>PAM</td><td>alignment</td><td>distance</td><td>position</td><td>gene name</td><td>gene id</td></tr>
<tr><td>GU305914.1</td><td>183775</td><td>183797</td><td>-</td><td>0</td><td>AATCAACAGGAAAGTCCCAT</td><td>TGG</td><td>||||||||[||||||||||||]PAM</td><td>NA</td><td>Intergenic</td><td>NA</td><td>NA</td></tr>
<tr><td>GU305914.1</td><td>183404</td><td>183426</td><td>-</td><td>0</td><td>AATCAACAGGAAAGTCCCAT</td><td>TGG</td><td>||||||||[||||||||||||]PAM</td><td>NA</td><td>Intergenic</td><td>NA</td><td>NA</td></tr>
<tr><td>GU305914.1</td><td>183589</td><td>183611</td><td>-</td><td>1</td><td>AGTCAACAGGAAAGTCCCAT</td><td>TGG</td><td>|-||||||[||||||||||||]PAM</td><td>NA</td><td>Intergenic</td><td>NA</td><td>NA</td></tr>
<tr><td>T20795</td><td>GCCAAGTACACTGCGTCAATAGG</td><td>1000</td><td></td><td></td><td></td><td></td><td></td><td></td><td></td><td></td><td></td></tr>
<tr><td>Oligo fwd</td><td>CACCGCCAAGTACACTGCGTCAAT</td><td></td><td></td><td></td><td></td><td></td><td></td><td></td><td></td><td></td><td></td></tr>
<tr><td>Oligo rev</td><td>AAACATTGACGCAGTGTACTTGGC</td><td></td><td></td><td></td><td></td><td></td><td></td><td></td><td></td><td></td><td></td></tr>
<tr><td>Chromosome</td><td>start</td><td>end</td><td>strand</td><td>MM</td><td>target_seq</td><td>PAM</td><td>alignment</td><td>distance</td><td>position</td><td>gene name</td><td>gene id</td></tr>
<tr><td>GU305914.1</td><td>183751</td><td>183773</td><td>-</td><td>0</td><td>GCCAAGTACACTGCGTCAAT</td><td>AGG</td><td>||||||||[||||||||||||]PAM</td><td>NA</td><td>Intergenic</td><td>NA</td><td>NA</td></tr>
<tr><td>GU305914.1</td><td>183380</td><td>183402</td><td>-</td><td>1</td><td>GCCAAGTACACTGAGTCAAT</td><td>AGG</td><td>||||||||[|||||-||||||]PAM</td><td>NA</td><td>Intergenic</td><td>NA</td><td>NA</td></tr>
<tr><td>GU305914.1</td><td>183565</td><td>183587</td><td>-</td><td>2</td><td>GCCAAGTACATTGAGTCAAT</td><td>AGG</td><td>||||||||[||-||-||||||]PAM</td><td>NA</td><td>Intergenic</td><td>NA</td><td>NA</td></tr>
<tr><td>GU305914.1</td><td>183960</td><td>183982</td><td>-</td><td>3</td><td>CCCAAGTACATTGGGTCAAT</td><td>GGG</td><td>-|||||||[||-||-||||||]PAM</td><td>NA</td><td>Intergenic</td><td>NA</td><td>NA</td></tr>
<tr><td>GU305914.1</td><td>184054</td><td>184076</td><td>-</td><td>3</td><td>CCCAAGTACATTGGGTCAAT</td><td>GGG</td><td>-|||||||[||-||-||||||]PAM</td><td>NA</td><td>Intergenic</td><td>NA</td><td>NA</td></tr>
<tr><td>GU305914.1</td><td>183658</td><td>183680</td><td>-</td><td>2</td><td>GCCAAGTACACTGACTCAAT</td><td>AGG</td><td>||||||||[|||||--|||||]PAM</td><td>NA</td><td>Intergenic</td><td>NA</td><td>NA</td></tr>
<tr><td>GU305914.1</td><td>183908</td><td>183930</td><td>-</td><td>2</td><td>GCCAAGTACACTGACTCAAT</td><td>AGG</td><td>||||||||[|||||--|||||]PAM</td><td>NA</td><td>Intergenic</td><td>NA</td><td>NA</td></tr>
<tr><td>GU305914.1</td><td>184002</td><td>184024</td><td>-</td><td>4</td><td>GGCAAGCACACTGAGTCAAA</td><td>TGG</td><td>|-||||-|[|||||-|||||-]PAM</td><td>NA</td><td>Intergenic</td><td>NA</td><td>NA</td></tr>
<tr><td>T20796</td><td>CCAAGTACACTGCGTCAATAGGG</td><td>1000</td><td></td><td></td><td></td><td></td><td></td><td></td><td></td><td></td><td></td></tr>
<tr><td>Oligo fwd</td><td>CACCCCAAGTACACTGCGTCAATA</td><td></td><td></td><td></td><td></td><td></td><td></td><td></td><td></td><td></td><td></td></tr>
<tr><td>Oligo rev</td><td>AAACTATTGACGCAGTGTACTTGG</td><td></td><td></td><td></td><td></td><td></td><td></td><td></td><td></td><td></td><td></td></tr>
<tr><td>Chromosome</td><td>start</td><td>end</td><td>strand</td><td>MM</td><td>target_seq</td><td>PAM</td><td>alignment</td><td>distance</td><td>position</td><td>gene name</td><td>gene id</td></tr>
<tr><td>GU305914.1</td><td>183750</td><td>183772</td><td>-</td><td>0</td><td>CCAAGTACACTGCGTCAATA</td><td>GGG</td><td>||||||||[||||||||||||]PAM</td><td>NA</td><td>Intergenic</td><td>NA</td><td>NA</td></tr>
<tr><td>GU305914.1</td><td>183379</td><td>183401</td><td>-</td><td>1</td><td>CCAAGTACACTGAGTCAATA</td><td>GGG</td><td>||||||||[||||-|||||||]PAM</td><td>NA</td><td>Intergenic</td><td>NA</td><td>NA</td></tr>
<tr><td>GU305914.1</td><td>183564</td><td>183586</td><td>-</td><td>2</td><td>CCAAGTACATTGAGTCAATA</td><td>GGG</td><td>||||||||[|-||-|||||||]PAM</td><td>NA</td><td>Intergenic</td><td>NA</td><td>NA</td></tr>
<tr><td>GU305914.1</td><td>183657</td><td>183679</td><td>-</td><td>2</td><td>CCAAGTACACTGACTCAATA</td><td>GGG</td><td>||||||||[||||--||||||]PAM</td><td>NA</td><td>Intergenic</td><td>NA</td><td>NA</td></tr>
<tr><td>GU305914.1</td><td>183907</td><td>183929</td><td>-</td><td>2</td><td>CCAAGTACACTGACTCAATA</td><td>GGG</td><td>||||||||[||||--||||||]PAM</td><td>NA</td><td>Intergenic</td><td>NA</td><td>NA</td></tr>
<tr><td>T20797</td><td>CGTCAATAGGGACTTTCCATTGG</td><td>1000</td><td></td><td></td><td></td><td></td><td></td><td></td><td></td><td></td><td></td></tr>
<tr><td>Oligo fwd</td><td>CACCCGTCAATAGGGACTTTCCAT</td><td></td><td></td><td></td><td></td><td></td><td></td><td></td><td></td><td></td><td></td></tr>
<tr><td>Oligo rev</td><td>AAACATGGAAAGTCCCTATTGACG</td><td></td><td></td><td></td><td></td><td></td><td></td><td></td><td></td><td></td><td></td></tr>
<tr><td>Chromosome</td><td>start</td><td>end</td><td>strand</td><td>MM</td><td>target_seq</td><td>PAM</td><td>alignment</td><td>distance</td><td>position</td><td>gene name</td><td>gene id</td></tr>
<tr><td>GU305914.1</td><td>183738</td><td>183760</td><td>-</td><td>0</td><td>CGTCAATAGGGACTTTCCAT</td><td>TGG</td><td>||||||||[||||||||||||]PAM</td><td>NA</td><td>Intergenic</td><td>NA</td><td>NA</td></tr>
<tr><td>GU305914.1</td><td>183367</td><td>183389</td><td>-</td><td>1</td><td>AGTCAATAGGGACTTTCCAT</td><td>TGG</td><td>-|||||||[||||||||||||]PAM</td><td>NA</td><td>Intergenic</td><td>NA</td><td>NA</td></tr>
<tr><td>GU305914.1</td><td>183645</td><td>183667</td><td>-</td><td>2</td><td>ACTCAATAGGGACTTTCCAT</td><td>TGG</td><td>--||||||[||||||||||||]PAM</td><td>NA</td><td>Intergenic</td><td>NA</td><td>NA</td></tr>
<tr><td>GU305914.1</td><td>183552</td><td>183574</td><td>-</td><td>2</td><td>AGTCAATAGGGACTTTCCAA</td><td>TGG</td><td>-|||||||[|||||||||||-]PAM</td><td>NA</td><td>Intergenic</td><td>NA</td><td>NA</td></tr>
<tr><td>GU305914.1</td><td>183895</td><td>183917</td><td>-</td><td>3</td><td>ACTCAATAGGGACTTTCCAA</td><td>TGG</td><td>--||||||[|||||||||||-]PAM</td><td>NA</td><td>Intergenic</td><td>NA</td><td>NA</td></tr>
<tr><td>GU305914.1</td><td>183459</td><td>183481</td><td>-</td><td>3</td><td>AGTCATTAGGGACTTTCCAA</td><td>TGG</td><td>-||||-||[|||||||||||-]PAM</td><td>NA</td><td>Intergenic</td><td>NA</td><td>NA</td></tr>
<tr><td>GU305914.1</td><td>183989</td><td>184011</td><td>-</td><td>4</td><td>AGTCAAATGGGACTTTCCAC</td><td>TGG</td><td>-|||||--[|||||||||||-]PAM</td><td>NA</td><td>Intergenic</td><td>NA</td><td>NA</td></tr>
<tr><td>T20798</td><td>GTCAATAGGGACTTTCCATTGGG</td><td>1000</td><td></td><td></td><td></td><td></td><td></td><td></td><td></td><td></td><td></td></tr>
<tr><td>Oligo fwd</td><td>CACCGTCAATAGGGACTTTCCATT</td><td></td><td></td><td></td><td></td><td></td><td></td><td></td><td></td><td></td><td></td></tr>
<tr><td>Oligo rev</td><td>AAACAATGGAAAGTCCCTATTGAC</td><td></td><td></td><td></td><td></td><td></td><td></td><td></td><td></td><td></td><td></td></tr>
<tr><td>Chromosome</td><td>start</td><td>end</td><td>strand</td><td>MM</td><td>target_seq</td><td>PAM</td><td>alignment</td><td>distance</td><td>position</td><td>gene name</td><td>gene id</td></tr>
<tr><td>GU305914.1</td><td>183737</td><td>183759</td><td>-</td><td>0</td><td>GTCAATAGGGACTTTCCATT</td><td>GGG</td><td>||||||||[||||||||||||]PAM</td><td>NA</td><td>Intergenic</td><td>NA</td><td>NA</td></tr>
<tr><td>GU305914.1</td><td>183366</td><td>183388</td><td>-</td><td>0</td><td>GTCAATAGGGACTTTCCATT</td><td>GGG</td><td>||||||||[||||||||||||]PAM</td><td>NA</td><td>Intergenic</td><td>NA</td><td>NA</td></tr>
<tr><td>GU305914.1</td><td>183644</td><td>183666</td><td>-</td><td>1</td><td>CTCAATAGGGACTTTCCATT</td><td>GGG</td><td>-|||||||[||||||||||||]PAM</td><td>NA</td><td>Intergenic</td><td>NA</td><td>NA</td></tr>
<tr><td>GU305914.1</td><td>183551</td><td>183573</td><td>-</td><td>1</td><td>GTCAATAGGGACTTTCCAAT</td><td>GGG</td><td>||||||||[||||||||||-|]PAM</td><td>NA</td><td>Intergenic</td><td>NA</td><td>NA</td></tr>
<tr><td>GU305914.1</td><td>183894</td><td>183916</td><td>-</td><td>2</td><td>CTCAATAGGGACTTTCCAAT</td><td>GGG</td><td>-|||||||[||||||||||-|]PAM</td><td>NA</td><td>Intergenic</td><td>NA</td><td>NA</td></tr>
<tr><td>GU305914.1</td><td>183458</td><td>183480</td><td>-</td><td>2</td><td>GTCATTAGGGACTTTCCAAT</td><td>GGG</td><td>||||-|||[||||||||||-|]PAM</td><td>NA</td><td>Intergenic</td><td>NA</td><td>NA</td></tr>
</table>

| | | | | | | | | | | | |
|---|---|---|---|---|---|---|---|---|---|---|---|
| GU305914.1 | 183988 | 184010 | - | 3 | GTCAAATGGGACTTTCCACT | GGG | \|\|\|\|\|--\|[\|\|\|\|\|\|\|\|\|\|-\|]PAM | NA | Intergenic | NA | NA |

T20799 TGGGTTTTGCCCAGTACATAAGG 1000
Oligo fwd CACCTGGGTTTTGCCCAGTACATA
Oligo rev AAACTATGTACTGGGCAAAACCCA

| Chromosome | start | end | strand | MM | target_seq | PAM | alignment | distance | position | gene name | gene id |
|---|---|---|---|---|---|---|---|---|---|---|---|
| GU305914.1 | 183439 | 183461 | - | 0 | TGGGTTTTGCCCAGTACATA | AGG | \|\|\|\|\|\|\|\|[\|\|\|\|\|\|\|\|\|\|\|\|]PAM | NA | Intergenic | NA | NA |
| GU305914.1 | 183625 | 183647 | - | 0 | TGGGTTTTGCCCAGTACATA | AGG | \|\|\|\|\|\|\|\|[\|\|\|\|\|\|\|\|\|\|\|\|]PAM | NA | Intergenic | NA | NA |
| GU305914.1 | 183718 | 183740 | - | 0 | TGGGTTTTGCCCAGTACATA | AGG | \|\|\|\|\|\|\|\|[\|\|\|\|\|\|\|\|\|\|\|\|]PAM | NA | Intergenic | NA | NA |
| GU305914.1 | 183532 | 183554 | - | 0 | TGGGTTTTGCCCAGTACATA | AGG | \|\|\|\|\|\|\|\|[\|\|\|\|\|\|\|\|\|\|\|\|]PAM | NA | Intergenic | NA | NA |
| GU305914.1 | 183811 | 183833 | - | 3 | TGTATTCTGCCCAGTACATA | AGG | \|\|--\|\|-\|[\|\|\|\|\|\|\|\|\|\|\|\|]PAM | NA | Intergenic | NA | NA |
| GU305914.1 | 183347 | 183369 | - | 1 | TGGGTTTTGCCCAGTACAAA | AGG | \|\|\|\|\|\|\|\|[\|\|\|\|\|\|\|\|\|\|-\|]PAM | NA | Intergenic | NA | NA |

T20800 GCCCAGTACATAAGGTCAATAGG 1000
Oligo fwd CACCGCCCAGTACATAAGGTCAAT
Oligo rev AAACATTGACCTTATGTACTGGGC

| Chromosome | start | end | strand | MM | target_seq | PAM | alignment | distance | position | gene name | gene id |
|---|---|---|---|---|---|---|---|---|---|---|---|
| GU305914.1 | 183524 | 183546 | - | 0 | GCCCAGTACATAAGGTCAAT | GGG | \|\|\|\|\|\|\|\|[\|\|\|\|\|\|\|\|\|\|\|\|]PAM | NA | Intergenic | NA | NA |
| GU305914.1 | 183617 | 183639 | - | 0 | GCCCAGTACATAAGGTCAAT | AGG | \|\|\|\|\|\|\|\|[\|\|\|\|\|\|\|\|\|\|\|\|]PAM | NA | Intergenic | NA | NA |
| GU305914.1 | 183803 | 183825 | - | 0 | GCCCAGTACATAAGGTCAAT | AGG | \|\|\|\|\|\|\|\|[\|\|\|\|\|\|\|\|\|\|\|\|]PAM | NA | Intergenic | NA | NA |
| GU305914.1 | 183710 | 183732 | - | 0 | GCCCAGTACATAAGGTCAAT | AGG | \|\|\|\|\|\|\|\|[\|\|\|\|\|\|\|\|\|\|\|\|]PAM | NA | Intergenic | NA | NA |
| GU305914.1 | 183431 | 183453 | - | 0 | GCCCAGTACATAAGGTCAAT | AGG | \|\|\|\|\|\|\|\|[\|\|\|\|\|\|\|\|\|\|\|\|]PAM | NA | Intergenic | NA | NA |
| GU305914.1 | 183287 | 183309 | - | 3 | GGCACGTACATAAGGTCAAT | AGG | \|-\|--\|\|\|[\|\|\|\|\|\|\|\|\|\|\|\|]PAM | NA | Intergenic | NA | NA |
| GU305914.1 | 183339 | 183361 | - | 1 | GCCCAGTACAAAAGGTCAAT | AGG | \|\|\|\|\|\|\|\|[\|\|-\|\|\|\|\|\|\|\|\|]PAM | NA | Intergenic | NA | NA |
| GU305914.1 | 184054 | 184076 | - | 4 | CCCAAGTACATTGGGTCAAT | GGG | -\|\|-\|\|\|\|[\|\|\|--\|\|\|\|\|\|\|]PAM | NA | Intergenic | NA | NA |
| GU305914.1 | 183960 | 183982 | - | 4 | CCCAAGTACATTGGGTCAAT | GGG | -\|\|-\|\|\|\|[\|\|\|--\|\|\|\|\|\|\|]PAM | NA | Intergenic | NA | NA |

T20801 CCCAGTACATAAGGTCAATAGGG 1000
Oligo fwd CACCCCCAGTACATAAGGTCAATA
Oligo rev AAACTATTGACCTTATGTACTGGG

| Chromosome | start | end | strand | MM | target_seq | PAM | alignment | distance | position | gene name | gene id |
|---|---|---|---|---|---|---|---|---|---|---|---|
| GU305914.1 | 183709 | 183731 | - | 0 | CCCAGTACATAAGGTCAATA | GGG | \|\|\|\|\|\|\|\|[\|\|\|\|\|\|\|\|\|\|\|\|]PAM | NA | Intergenic | NA | NA |
| GU305914.1 | 183430 | 183452 | - | 0 | CCCAGTACATAAGGTCAATA | GGG | \|\|\|\|\|\|\|\|[\|\|\|\|\|\|\|\|\|\|\|\|]PAM | NA | Intergenic | NA | NA |
| GU305914.1 | 183616 | 183638 | - | 0 | CCCAGTACATAAGGTCAATA | GGG | \|\|\|\|\|\|\|\|[\|\|\|\|\|\|\|\|\|\|\|\|]PAM | NA | Intergenic | NA | NA |
| GU305914.1 | 183802 | 183824 | - | 0 | CCCAGTACATAAGGTCAATA | GGG | \|\|\|\|\|\|\|\|[\|\|\|\|\|\|\|\|\|\|\|\|]PAM | NA | Intergenic | NA | NA |
| GU305914.1 | 183286 | 183308 | - | 3 | GCACGTACATAAGGTCAATA | GGG | -\|--\|\|\|\|[\|\|\|\|\|\|\|\|\|\|\|\|]PAM | NA | Intergenic | NA | NA |
| GU305914.1 | 183338 | 183360 | - | 1 | CCCAGTACAAAAGGTCAATA | GGG | \|\|\|\|\|\|\|\|[\|-\|\|\|\|\|\|\|\|\|\|]PAM | NA | Intergenic | NA | NA |

T20802 CCAGTACATAAGGTCAATAGGGG 1000
Oligo fwd CACCCCAGTACATAAGGTCAATAG
Oligo rev AAACCTATTGACCTTATGTACTGG

| Chromosome | start | end | strand | MM | target_seq | PAM | alignment | distance | position | gene name | gene id |
|---|---|---|---|---|---|---|---|---|---|---|---|
| GU305914.1 | 183708 | 183730 | - | 0 | CCAGTACATAAGGTCAATAG | GGG | \|\|\|\|\|\|\|\|[\|\|\|\|\|\|\|\|\|\|\|\|]PAM | NA | Intergenic | NA | NA |
| GU305914.1 | 183429 | 183451 | - | 0 | CCAGTACATAAGGTCAATAG | GGG | \|\|\|\|\|\|\|\|[\|\|\|\|\|\|\|\|\|\|\|\|]PAM | NA | Intergenic | NA | NA |
| GU305914.1 | 183615 | 183637 | - | 0 | CCAGTACATAAGGTCAATAG | GGG | \|\|\|\|\|\|\|\|[\|\|\|\|\|\|\|\|\|\|\|\|]PAM | NA | Intergenic | NA | NA |
| GU305914.1 | 183801 | 183823 | - | 0 | CCAGTACATAAGGTCAATAG | GGG | \|\|\|\|\|\|\|\|[\|\|\|\|\|\|\|\|\|\|\|\|]PAM | NA | Intergenic | NA | NA |
| GU305914.1 | 183285 | 183307 | - | 2 | CACGTACATAAGGTCAATAG | GGG | \|--\|\|\|\|\|[\|\|\|\|\|\|\|\|\|\|\|\|]PAM | NA | Intergenic | NA | NA |
| GU305914.1 | 183337 | 183359 | - | 1 | CCAGTACAAAAGGTCAATAG | GGG | \|\|\|\|\|\|\|\|[-\|\|\|\|\|\|\|\|\|\|\|]PAM | NA | Intergenic | NA | NA |

T20803 GTCAATAGGGGATGAGTCAATGG 1000
Oligo fwd CACCGTCAATAGGGGATGAGTCAA
Oligo rev AAACTTGACTCATCCCCTATTGAC

| Chromosome | start | end | strand | MM | target_seq | PAM | alignment | distance | position | gene name | gene id |
|---|---|---|---|---|---|---|---|---|---|---|---|
| GU305914.1 | 183696 | 183718 | - | 0 | GTCAATAGGGGATGAGTCAA | TGG | \|\|\|\|\|\|\|\|[\|\|\|\|\|\|\|\|\|\|\|\|]PAM | NA | Intergenic | NA | NA |
| GU305914.1 | 183325 | 183347 | - | 1 | GTCAATAGGGGGTGAGTCAA | TGG | \|\|\|\|\|\|\|\|[\|\|\|-\|\|\|\|\|\|\|\|]PAM | NA | Intergenic | NA | NA |

T20804 TCAATAGGGGATGAGTCAATGGG 1000
Oligo fwd CACCTCAATAGGGGATGAGTCAAT
Oligo rev AAACATTGACTCATCCCCTATTGA

| Chromosome | start | end | strand | MM | target_seq | PAM | alignment | distance | position | gene name | gene id |
|---|---|---|---|---|---|---|---|---|---|---|---|
| GU305914.1 | 183695 | 183717 | - | 0 | TCAATAGGGGATGAGTCAAT | GGG | \|\|\|\|\|\|\|\|[\|\|\|\|\|\|\|\|\|\|\|\|]PAM | NA | Intergenic | NA | NA |
| GU305914.1 | 183324 | 183346 | - | 1 | TCAATAGGGGGTGAGTCAAT | GGG | \|\|\|\|\|\|\|\|[\|\|-\|\|\|\|\|\|\|\|\|]PAM | NA | Intergenic | NA | NA |
| GU305914.1 | 183602 | 183624 | - | 2 | TCAATAGGGGGTGAGTCAAC | AGG | \|\|\|\|\|\|\|\|[\|\|-\|\|\|\|\|\|\|\|-]PAM | NA | Intergenic | NA | NA |

T20805 AGTCAATGGGAAAAACCCATTGG 1000
Oligo fwd CACCAGTCAATGGGAAAAACCCAT
Oligo rev AAACATGGGTTTTTCCCATTGACT

| Chromosome | start | end | strand | MM | target_seq | PAM | alignment | distance | position | gene name | gene id |
|---|---|---|---|---|---|---|---|---|---|---|---|
| GU305914.1 | 183682 | 183704 | - | 0 | AGTCAATGGGAAAAACCCAT | TGG | \|\|\|\|\|\|\|\|[\|\|\|\|\|\|\|\|\|\|\|\|]PAM | NA | Intergenic | NA | NA |
| GU305914.1 | 184024 | 184046 | + | 4 | CAGTAATGGGAAAAACCCAT | TGG | ----\|\|\|\|[\|\|\|\|\|\|\|\|\|\|\|\|]PAM | NA | Intergenic | NA | NA |
| GU305914.1 | 183494 | 183516 | + | 4 | CAGTAATGGGAAAAACCCAT | TGG | ----\|\|\|\|[\|\|\|\|\|\|\|\|\|\|\|\|]PAM | NA | Intergenic | NA | NA |
| GU305914.1 | 183589 | 183611 | - | 4 | AGTCAACAGGAAAGTCCCAT | TGG | \|\|\|\|\|\|--[\|\|\|\|\|--\|\|\|\|\|]PAM | NA | Intergenic | NA | NA |

T20806 GCCAAGTACACTGACTCAATAGG 1000
Oligo fwd CACCGCCAAGTACACTGACTCAAT
Oligo rev AAACATTGAGTCAGTGTACTTGGC

| Chromosome | start | end | strand | MM | target_seq | PAM | alignment | distance | position | gene name | gene id |
|---|---|---|---|---|---|---|---|---|---|---|---|
| GU305914.1 | 183908 | 183930 | - | 0 | GCCAAGTACACTGACTCAAT | AGG | \|\|\|\|\|\|\|\|[\|\|\|\|\|\|\|\|\|\|\|\|]PAM | NA | Intergenic | NA | NA |
| GU305914.1 | 183658 | 183680 | - | 0 | GCCAAGTACACTGACTCAAT | AGG | \|\|\|\|\|\|\|\|[\|\|\|\|\|\|\|\|\|\|\|\|]PAM | NA | Intergenic | NA | NA |
| GU305914.1 | 183380 | 183402 | - | 1 | GCCAAGTACACTGAGTCAAT | AGG | \|\|\|\|\|\|\|\|[\|\|\|\|\|\|-\|\|\|\|\|]PAM | NA | Intergenic | NA | NA |
| GU305914.1 | 183565 | 183587 | - | 2 | GCCAAGTACATTGAGTCAAT | AGG | \|\|\|\|\|\|\|\|[\|\|-\|\|\|-\|\|\|\|\|]PAM | NA | Intergenic | NA | NA |
| GU305914.1 | 183751 | 183773 | - | 2 | GCCAAGTACACTGCGTCAAT | AGG | \|\|\|\|\|\|\|\|[\|\|\|\|\|--\|\|\|\|\|]PAM | NA | Intergenic | NA | NA |
| GU305914.1 | 184002 | 184024 | - | 4 | GGCAAGCACACTGAGTCAAA | TGG | \|-\|\|\|\|-\|[\|\|\|\|\|\|-\|\|\|\|-]PAM | NA | Intergenic | NA | NA |

T20807 CCAAGTACACTGACTCAATAGGG 1000
Oligo fwd CACCCCAAGTACACTGACTCAATA
Oligo rev AAACTATTGAGTCAGTGTACTTGG

| Chromosome | start | end | strand | MM | target_seq | PAM | alignment | distance | position | gene name | gene id |
|---|---|---|---|---|---|---|---|---|---|---|---|
| GU305914.1 | 183907 | 183929 | - | 0 | CCAAGTACACTGACTCAATA | GGG | \|\|\|\|\|\|\|\|[\|\|\|\|\|\|\|\|\|\|\|\|]PAM | NA | Intergenic | NA | NA |

| | | | | | | | | | | | |
|---|---|---|---|---|---|---|---|---|---|---|---|
| GU305914.1 | 183657 | 183679 | - | 0 | CCAAGTACACTGACTCAATA | GGG | \|\|\|\|\|\|\|\|[\|\|\|\|\|\|\|\|\|\|\|\|]PAM | NA | Intergenic | NA | NA |
| GU305914.1 | 183379 | 183401 | - | 1 | CCAAGTACACTGAGTCAATA | GGG | \|\|\|\|\|\|\|\|[\|\|\|\|\|-\|\|\|\|\|\|]PAM | NA | Intergenic | NA | NA |
| GU305914.1 | 183564 | 183586 | - | 2 | CCAAGTACATTGAGTCAATA | GGG | \|\|\|\|\|\|\|\|[\|-\|\|\|-\|\|\|\|\|\|]PAM | NA | Intergenic | NA | NA |
| GU305914.1 | 183750 | 183772 | - | 2 | CCAAGTACACTGCGTCAATA | GGG | \|\|\|\|\|\|\|\|[\|\|\|\|--\|\|\|\|\|\|]PAM | NA | Intergenic | NA | NA |
| T20808 | ACTCAATAGGGACTTTCCATTGG | 1000 | | | | | | | | | |
| Oligo fwd | CACCACTCAATAGGGACTTTCCAT | | | | | | | | | | |
| Oligo rev | AAACATGGAAAGTCCCTATTGAGT | | | | | | | | | | |
| Chromosome | start | end | strand | MM | target_seq | PAM | alignment | distance | position | gene name | gene id |
| GU305914.1 | 183645 | 183667 | - | 0 | ACTCAATAGGGACTTTCCAT | TGG | \|\|\|\|\|\|\|\|[\|\|\|\|\|\|\|\|\|\|\|\|]PAM | NA | Intergenic | NA | NA |
| GU305914.1 | 183367 | 183389 | - | 1 | AGTCAATAGGGACTTTCCAT | TGG | \|-\|\|\|\|\|\|[\|\|\|\|\|\|\|\|\|\|\|\|]PAM | NA | Intergenic | NA | NA |
| GU305914.1 | 183738 | 183760 | - | 2 | CGTCAATAGGGACTTTCCAT | TGG | --\|\|\|\|\|\|[\|\|\|\|\|\|\|\|\|\|\|\|]PAM | NA | Intergenic | NA | NA |
| GU305914.1 | 183895 | 183917 | - | 1 | ACTCAATAGGGACTTTCCAA | TGG | \|\|\|\|\|\|\|\|[\|\|\|\|\|\|\|\|\|\|\|-]PAM | NA | Intergenic | NA | NA |
| GU305914.1 | 183552 | 183574 | - | 2 | AGTCAATAGGGACTTTCCAA | TGG | \|-\|\|\|\|\|\|[\|\|\|\|\|\|\|\|\|\|\|-]PAM | NA | Intergenic | NA | NA |
| GU305914.1 | 183459 | 183481 | - | 3 | AGTCATTAGGGACTTTCCAA | TGG | \|-\|\|\|-\|\|[\|\|\|\|\|\|\|\|\|\|\|-]PAM | NA | Intergenic | NA | NA |
| GU305914.1 | 183989 | 184011 | - | 4 | AGTCAAATGGGACTTTCCAC | TGG | \|-\|\|\|\|--[\|\|\|\|\|\|\|\|\|\|\|-]PAM | NA | Intergenic | NA | NA |
| T20809 | CTCAATAGGGACTTTCCATTGGG | 1000 | | | | | | | | | |
| Oligo fwd | CACCCTCAATAGGGACTTTCCATT | | | | | | | | | | |
| Oligo rev | AAACAATGGAAAGTCCCTATTGAG | | | | | | | | | | |
| Chromosome | start | end | strand | MM | target_seq | PAM | alignment | distance | position | gene name | gene id |
| GU305914.1 | 183644 | 183666 | - | 0 | CTCAATAGGGACTTTCCATT | GGG | \|\|\|\|\|\|\|\|[\|\|\|\|\|\|\|\|\|\|\|\|]PAM | NA | Intergenic | NA | NA |
| GU305914.1 | 183737 | 183759 | - | 1 | GTCAATAGGGACTTTCCATT | GGG | -\|\|\|\|\|\|\|[\|\|\|\|\|\|\|\|\|\|\|\|]PAM | NA | Intergenic | NA | NA |
| GU305914.1 | 183366 | 183388 | - | 1 | GTCAATAGGGACTTTCCATT | GGG | -\|\|\|\|\|\|\|[\|\|\|\|\|\|\|\|\|\|\|\|]PAM | NA | Intergenic | NA | NA |
| GU305914.1 | 183894 | 183916 | - | 1 | CTCAATAGGGACTTTCCAAT | GGG | \|\|\|\|\|\|\|\|[\|\|\|\|\|\|\|\|\|\|-\|]PAM | NA | Intergenic | NA | NA |
| GU305914.1 | 183551 | 183573 | - | 2 | GTCAATAGGGACTTTCCAAT | GGG | -\|\|\|\|\|\|\|[\|\|\|\|\|\|\|\|\|\|-\|]PAM | NA | Intergenic | NA | NA |
| GU305914.1 | 183458 | 183480 | - | 3 | GTCATTAGGGACTTTCCAAT | GGG | -\|\|\|-\|\|\|[\|\|\|\|\|\|\|\|\|\|-\|]PAM | NA | Intergenic | NA | NA |
| GU305914.1 | 183988 | 184010 | - | 4 | GTCAAATGGGACTTTCCACT | GGG | -\|\|\|\|--\|[\|\|\|\|\|\|\|\|\|\|-\|]PAM | NA | Intergenic | NA | NA |
| T20810 | TGGGTTTTGCCCAGTACATAAGG | 1000 | | | | | | | | | |
| Oligo fwd | CACCTGGGTTTTGCCCAGTACATA | | | | | | | | | | |
| Oligo rev | AAACTATGTACTGGGCAAAACCCA | | | | | | | | | | |
| Chromosome | start | end | strand | MM | target_seq | PAM | alignment | distance | position | gene name | gene id |
| GU305914.1 | 183439 | 183461 | - | 0 | TGGGTTTTGCCCAGTACATA | AGG | \|\|\|\|\|\|\|\|[\|\|\|\|\|\|\|\|\|\|\|\|]PAM | NA | Intergenic | NA | NA |
| GU305914.1 | 183625 | 183647 | - | 0 | TGGGTTTTGCCCAGTACATA | AGG | \|\|\|\|\|\|\|\|[\|\|\|\|\|\|\|\|\|\|\|\|]PAM | NA | Intergenic | NA | NA |
| GU305914.1 | 183718 | 183740 | - | 0 | TGGGTTTTGCCCAGTACATA | AGG | \|\|\|\|\|\|\|\|[\|\|\|\|\|\|\|\|\|\|\|\|]PAM | NA | Intergenic | NA | NA |
| GU305914.1 | 183532 | 183554 | - | 0 | TGGGTTTTGCCCAGTACATA | AGG | \|\|\|\|\|\|\|\|[\|\|\|\|\|\|\|\|\|\|\|\|]PAM | NA | Intergenic | NA | NA |
| GU305914.1 | 183811 | 183833 | - | 3 | TGTATTCTGCCCAGTACATA | AGG | \|\|--\|\|-\|[\|\|\|\|\|\|\|\|\|\|\|\|]PAM | NA | Intergenic | NA | NA |
| GU305914.1 | 183347 | 183369 | - | 1 | TGGGTTTTGCCCAGTACAAA | AGG | \|\|\|\|\|\|\|\|[\|\|\|\|\|\|\|\|\|\|-\|]PAM | NA | Intergenic | NA | NA |
| T20811 | GCCCAGTACATAAGGTCAATAGG | 1000 | | | | | | | | | |
| Oligo fwd | CACCGCCCAGTACATAAGGTCAAT | | | | | | | | | | |
| Oligo rev | AAACATTGACCTTATGTACTGGGC | | | | | | | | | | |
| Chromosome | start | end | strand | MM | target_seq | PAM | alignment | distance | position | gene name | gene id |
| GU305914.1 | 183524 | 183546 | - | 0 | GCCCAGTACATAAGGTCAAT | GGG | \|\|\|\|\|\|\|\|[\|\|\|\|\|\|\|\|\|\|\|\|]PAM | NA | Intergenic | NA | NA |
| GU305914.1 | 183617 | 183639 | - | 0 | GCCCAGTACATAAGGTCAAT | AGG | \|\|\|\|\|\|\|\|[\|\|\|\|\|\|\|\|\|\|\|\|]PAM | NA | Intergenic | NA | NA |
| GU305914.1 | 183803 | 183825 | - | 0 | GCCCAGTACATAAGGTCAAT | AGG | \|\|\|\|\|\|\|\|[\|\|\|\|\|\|\|\|\|\|\|\|]PAM | NA | Intergenic | NA | NA |
| GU305914.1 | 183710 | 183732 | - | 0 | GCCCAGTACATAAGGTCAAT | AGG | \|\|\|\|\|\|\|\|[\|\|\|\|\|\|\|\|\|\|\|\|]PAM | NA | Intergenic | NA | NA |
| GU305914.1 | 183431 | 183453 | - | 0 | GCCCAGTACATAAGGTCAAT | AGG | \|\|\|\|\|\|\|\|[\|\|\|\|\|\|\|\|\|\|\|\|]PAM | NA | Intergenic | NA | NA |
| GU305914.1 | 183287 | 183309 | - | 3 | GGCACGTACATAAGGTCAAT | AGG | \|-\|--\|\|\|[\|\|\|\|\|\|\|\|\|\|\|\|]PAM | NA | Intergenic | NA | NA |
| GU305914.1 | 183339 | 183361 | - | 1 | GCCCAGTACAAAAGGTCAAT | AGG | \|\|\|\|\|\|\|\|[\|\|-\|\|\|\|\|\|\|\|\|]PAM | NA | Intergenic | NA | NA |
| GU305914.1 | 184054 | 184076 | - | 4 | CCCAAGTACATTGGGTCAAT | GGG | -\|\|-\|\|\|\|[\|\|\|--\|\|\|\|\|\|\|]PAM | NA | Intergenic | NA | NA |
| GU305914.1 | 183960 | 183982 | - | 4 | CCCAAGTACATTGGGTCAAT | GGG | -\|\|-\|\|\|\|[\|\|\|--\|\|\|\|\|\|\|]PAM | NA | Intergenic | NA | NA |
| T20812 | CCCAGTACATAAGGTCAATAGGG | 1000 | | | | | | | | | |
| Oligo fwd | CACCCCCAGTACATAAGGTCAATA | | | | | | | | | | |
| Oligo rev | AAACTATTGACCTTATGTACTGGG | | | | | | | | | | |
| Chromosome | start | end | strand | MM | target_seq | PAM | alignment | distance | position | gene name | gene id |
| GU305914.1 | 183709 | 183731 | - | 0 | CCCAGTACATAAGGTCAATA | GGG | \|\|\|\|\|\|\|\|[\|\|\|\|\|\|\|\|\|\|\|\|]PAM | NA | Intergenic | NA | NA |
| GU305914.1 | 183430 | 183452 | - | 0 | CCCAGTACATAAGGTCAATA | GGG | \|\|\|\|\|\|\|\|[\|\|\|\|\|\|\|\|\|\|\|\|]PAM | NA | Intergenic | NA | NA |
| GU305914.1 | 183616 | 183638 | - | 0 | CCCAGTACATAAGGTCAATA | GGG | \|\|\|\|\|\|\|\|[\|\|\|\|\|\|\|\|\|\|\|\|]PAM | NA | Intergenic | NA | NA |
| GU305914.1 | 183802 | 183824 | - | 0 | CCCAGTACATAAGGTCAATA | GGG | \|\|\|\|\|\|\|\|[\|\|\|\|\|\|\|\|\|\|\|\|]PAM | NA | Intergenic | NA | NA |
| GU305914.1 | 183286 | 183308 | - | 3 | GCACGTACATAAGGTCAATA | GGG | -\|--\|\|\|\|[\|\|\|\|\|\|\|\|\|\|\|\|]PAM | NA | Intergenic | NA | NA |
| GU305914.1 | 183338 | 183360 | - | 1 | CCCAGTACAAAAGGTCAATA | GGG | \|\|\|\|\|\|\|\|[\|-\|\|\|\|\|\|\|\|\|\|]PAM | NA | Intergenic | NA | NA |
| T20813 | CCAGTACATAAGGTCAATAGGGG | 1000 | | | | | | | | | |
| Oligo fwd | CACCCCAGTACATAAGGTCAATAG | | | | | | | | | | |
| Oligo rev | AAACCTATTGACCTTATGTACTGG | | | | | | | | | | |
| Chromosome | start | end | strand | MM | target_seq | PAM | alignment | distance | position | gene name | gene id |
| GU305914.1 | 183708 | 183730 | - | 0 | CCAGTACATAAGGTCAATAG | GGG | \|\|\|\|\|\|\|\|[\|\|\|\|\|\|\|\|\|\|\|\|]PAM | NA | Intergenic | NA | NA |
| GU305914.1 | 183429 | 183451 | - | 0 | CCAGTACATAAGGTCAATAG | GGG | \|\|\|\|\|\|\|\|[\|\|\|\|\|\|\|\|\|\|\|\|]PAM | NA | Intergenic | NA | NA |
| GU305914.1 | 183615 | 183637 | - | 0 | CCAGTACATAAGGTCAATAG | GGG | \|\|\|\|\|\|\|\|[\|\|\|\|\|\|\|\|\|\|\|\|]PAM | NA | Intergenic | NA | NA |
| GU305914.1 | 183801 | 183823 | - | 0 | CCAGTACATAAGGTCAATAG | GGG | \|\|\|\|\|\|\|\|[\|\|\|\|\|\|\|\|\|\|\|\|]PAM | NA | Intergenic | NA | NA |
| GU305914.1 | 183285 | 183307 | - | 2 | CACGTACATAAGGTCAATAG | GGG | \|--\|\|\|\|\|[\|\|\|\|\|\|\|\|\|\|\|\|]PAM | NA | Intergenic | NA | NA |
| GU305914.1 | 183337 | 183359 | - | 1 | CCAGTACAAAAGGTCAATAG | GGG | \|\|\|\|\|\|\|\|[-\|\|\|\|\|\|\|\|\|\|\|]PAM | NA | Intergenic | NA | NA |
| T20814 | CAGTACATAAGGTCAATAGGGGG | 1000 | | | | | | | | | |
| Oligo fwd | CACCCAGTACATAAGGTCAATAGG | | | | | | | | | | |
| Oligo rev | AAACCCTATTGACCTTATGTACTG | | | | | | | | | | |
| Chromosome | start | end | strand | MM | target_seq | PAM | alignment | distance | position | gene name | gene id |
| GU305914.1 | 183800 | 183822 | - | 0 | CAGTACATAAGGTCAATAGG | GGG | \|\|\|\|\|\|\|\|[\|\|\|\|\|\|\|\|\|\|\|\|]PAM | NA | Intergenic | NA | NA |
| GU305914.1 | 183614 | 183636 | - | 0 | CAGTACATAAGGTCAATAGG | GGG | \|\|\|\|\|\|\|\|[\|\|\|\|\|\|\|\|\|\|\|\|]PAM | NA | Intergenic | NA | NA |
| GU305914.1 | 183336 | 183358 | - | 1 | CAGTACAAAAGGTCAATAGG | GGG | \|\|\|\|\|\|\|-[\|\|\|\|\|\|\|\|\|\|\|\|]PAM | NA | Intergenic | NA | NA |
| GU305914.1 | 183521 | 183543 | - | 1 | CAGTACATAAGGTCAATGGG | AGG | \|\|\|\|\|\|\|\|[\|\|\|\|\|\|\|\|\|-\|\|]PAM | NA | Intergenic | NA | NA |
| T20815 | TCAATAGGGGGTGAGTCAACAGG | 1000 | | | | | | | | | |
| Oligo fwd | CACCTCAATAGGGGGTGAGTCAAC | | | | | | | | | | |
| Oligo rev | AAACGTTGACTCACCCCCTATTGA | | | | | | | | | | |
| Chromosome | start | end | strand | MM | target_seq | PAM | alignment | distance | position | gene name | gene id |

<table>
<tr><td>GU305914.1</td><td>183602</td><td>183624</td><td>-</td><td>0</td><td>TCAATAGGGGGTGAGTCAAC</td><td>AGG</td><td>||||||||[||||||||||||]PAM</td><td>NA</td><td>Intergenic</td><td>NA</td><td>NA</td></tr>
<tr><td>GU305914.1</td><td>183788</td><td>183810</td><td>-</td><td>1</td><td>TCAATAGGGGGTGAATCAAC</td><td>AGG</td><td>||||||||[||||||-|||||]PAM</td><td>NA</td><td>Intergenic</td><td>NA</td><td>NA</td></tr>
<tr><td>GU305914.1</td><td>183324</td><td>183346</td><td>-</td><td>1</td><td>TCAATAGGGGGTGAGTCAAT</td><td>GGG</td><td>||||||||[|||||||||||-]PAM</td><td>NA</td><td>Intergenic</td><td>NA</td><td>NA</td></tr>
<tr><td>GU305914.1</td><td>183695</td><td>183717</td><td>-</td><td>2</td><td>TCAATAGGGGATGAGTCAAT</td><td>GGG</td><td>||||||||[||-||||||||-]PAM</td><td>NA</td><td>Intergenic</td><td>NA</td><td>NA</td></tr>
<tr><td></td><td></td><td></td><td></td><td></td><td></td><td></td><td></td><td></td><td></td><td></td><td></td></tr>
<tr><td>T20816</td><td>AGTCAACAGGAAAGTCCCATTGG</td><td>1000</td><td></td><td></td><td></td><td></td><td></td><td></td><td></td><td></td><td></td></tr>
<tr><td>Oligo fwd</td><td>CACCAGTCAACAGGAAAGTCCCAT</td><td></td><td></td><td></td><td></td><td></td><td></td><td></td><td></td><td></td><td></td></tr>
<tr><td>Oligo rev</td><td>AAACATGGGACTTTCCTGTTGACT</td><td></td><td></td><td></td><td></td><td></td><td></td><td></td><td></td><td></td><td></td></tr>
<tr><td>Chromosome</td><td>start</td><td>end</td><td>strand</td><td>MM</td><td>target_seq</td><td>PAM</td><td>alignment</td><td>distance</td><td>position</td><td>gene name</td><td>gene id</td></tr>
<tr><td>GU305914.1</td><td>183589</td><td>183611</td><td>-</td><td>0</td><td>AGTCAACAGGAAAGTCCCAT</td><td>TGG</td><td>||||||||[||||||||||||]PAM</td><td>NA</td><td>Intergenic</td><td>NA</td><td>NA</td></tr>
<tr><td>GU305914.1</td><td>183404</td><td>183426</td><td>-</td><td>1</td><td>AATCAACAGGAAAGTCCCAT</td><td>TGG</td><td>|-||||||[||||||||||||]PAM</td><td>NA</td><td>Intergenic</td><td>NA</td><td>NA</td></tr>
<tr><td>GU305914.1</td><td>183775</td><td>183797</td><td>-</td><td>1</td><td>AATCAACAGGAAAGTCCCAT</td><td>TGG</td><td>|-||||||[||||||||||||]PAM</td><td>NA</td><td>Intergenic</td><td>NA</td><td>NA</td></tr>
<tr><td>GU305914.1</td><td>183682</td><td>183704</td><td>-</td><td>4</td><td>AGTCAATGGGAAAAACCCAT</td><td>TGG</td><td>||||||--[|||||--|||||]PAM</td><td>NA</td><td>Intergenic</td><td>NA</td><td>NA</td></tr>
<tr><td></td><td></td><td></td><td></td><td></td><td></td><td></td><td></td><td></td><td></td><td></td><td></td></tr>
<tr><td>T20817</td><td>GCCAAGTACATTGAGTCAATAGG</td><td>1000</td><td></td><td></td><td></td><td></td><td></td><td></td><td></td><td></td><td></td></tr>
<tr><td>Oligo fwd</td><td>CACCGCCAAGTACATTGAGTCAAT</td><td></td><td></td><td></td><td></td><td></td><td></td><td></td><td></td><td></td><td></td></tr>
<tr><td>Oligo rev</td><td>AAACATTGACTCAATGTACTTGGC</td><td></td><td></td><td></td><td></td><td></td><td></td><td></td><td></td><td></td><td></td></tr>
<tr><td>Chromosome</td><td>start</td><td>end</td><td>strand</td><td>MM</td><td>target_seq</td><td>PAM</td><td>alignment</td><td>distance</td><td>position</td><td>gene name</td><td>gene id</td></tr>
<tr><td>GU305914.1</td><td>183565</td><td>183587</td><td>-</td><td>0</td><td>GCCAAGTACATTGAGTCAAT</td><td>AGG</td><td>||||||||[||||||||||||]PAM</td><td>NA</td><td>Intergenic</td><td>NA</td><td>NA</td></tr>
<tr><td>GU305914.1</td><td>183380</td><td>183402</td><td>-</td><td>1</td><td>GCCAAGTACACTGAGTCAAT</td><td>AGG</td><td>||||||||[||-|||||||||]PAM</td><td>NA</td><td>Intergenic</td><td>NA</td><td>NA</td></tr>
<tr><td>GU305914.1</td><td>184054</td><td>184076</td><td>-</td><td>2</td><td>CCCAAGTACATTGGGTCAAT</td><td>GGG</td><td>-|||||||[|||||-||||||]PAM</td><td>NA</td><td>Intergenic</td><td>NA</td><td>NA</td></tr>
<tr><td>GU305914.1</td><td>183960</td><td>183982</td><td>-</td><td>2</td><td>CCCAAGTACATTGGGTCAAT</td><td>GGG</td><td>-|||||||[|||||-||||||]PAM</td><td>NA</td><td>Intergenic</td><td>NA</td><td>NA</td></tr>
<tr><td>GU305914.1</td><td>183751</td><td>183773</td><td>-</td><td>2</td><td>GCCAAGTACACTGCGTCAAT</td><td>AGG</td><td>||||||||[||-||-||||||]PAM</td><td>NA</td><td>Intergenic</td><td>NA</td><td>NA</td></tr>
<tr><td>GU305914.1</td><td>183908</td><td>183930</td><td>-</td><td>2</td><td>GCCAAGTACACTGACTCAAT</td><td>AGG</td><td>||||||||[||-|||-|||||]PAM</td><td>NA</td><td>Intergenic</td><td>NA</td><td>NA</td></tr>
<tr><td>GU305914.1</td><td>183658</td><td>183680</td><td>-</td><td>2</td><td>GCCAAGTACACTGACTCAAT</td><td>AGG</td><td>||||||||[||-|||-|||||]PAM</td><td>NA</td><td>Intergenic</td><td>NA</td><td>NA</td></tr>
<tr><td>GU305914.1</td><td>184002</td><td>184024</td><td>-</td><td>4</td><td>GGCAAGCACACTGAGTCAAA</td><td>TGG</td><td>|-||||-|[||-||||||||-]PAM</td><td>NA</td><td>Intergenic</td><td>NA</td><td>NA</td></tr>
<tr><td></td><td></td><td></td><td></td><td></td><td></td><td></td><td></td><td></td><td></td><td></td><td></td></tr>
<tr><td>T20818</td><td>CCAAGTACATTGAGTCAATAGGG</td><td>1000</td><td></td><td></td><td></td><td></td><td></td><td></td><td></td><td></td><td></td></tr>
<tr><td>Oligo fwd</td><td>CACCCCAAGTACATTGAGTCAATA</td><td></td><td></td><td></td><td></td><td></td><td></td><td></td><td></td><td></td><td></td></tr>
<tr><td>Oligo rev</td><td>AAACTATTGACTCAATGTACTTGG</td><td></td><td></td><td></td><td></td><td></td><td></td><td></td><td></td><td></td><td></td></tr>
<tr><td>Chromosome</td><td>start</td><td>end</td><td>strand</td><td>MM</td><td>target_seq</td><td>PAM</td><td>alignment</td><td>distance</td><td>position</td><td>gene name</td><td>gene id</td></tr>
<tr><td>GU305914.1</td><td>183564</td><td>183586</td><td>-</td><td>0</td><td>CCAAGTACATTGAGTCAATA</td><td>GGG</td><td>||||||||[||||||||||||]PAM</td><td>NA</td><td>Intergenic</td><td>NA</td><td>NA</td></tr>
<tr><td>GU305914.1</td><td>183379</td><td>183401</td><td>-</td><td>1</td><td>CCAAGTACACTGAGTCAATA</td><td>GGG</td><td>||||||||[|-||||||||||]PAM</td><td>NA</td><td>Intergenic</td><td>NA</td><td>NA</td></tr>
<tr><td>GU305914.1</td><td>183750</td><td>183772</td><td>-</td><td>2</td><td>CCAAGTACACTGCGTCAATA</td><td>GGG</td><td>||||||||[|-||-|||||||]PAM</td><td>NA</td><td>Intergenic</td><td>NA</td><td>NA</td></tr>
<tr><td>GU305914.1</td><td>183657</td><td>183679</td><td>-</td><td>2</td><td>CCAAGTACACTGACTCAATA</td><td>GGG</td><td>||||||||[|-|||-||||||]PAM</td><td>NA</td><td>Intergenic</td><td>NA</td><td>NA</td></tr>
<tr><td>GU305914.1</td><td>183907</td><td>183929</td><td>-</td><td>2</td><td>CCAAGTACACTGACTCAATA</td><td>GGG</td><td>||||||||[|-|||-||||||]PAM</td><td>NA</td><td>Intergenic</td><td>NA</td><td>NA</td></tr>
<tr><td></td><td></td><td></td><td></td><td></td><td></td><td></td><td></td><td></td><td></td><td></td><td></td></tr>
<tr><td>T20819</td><td>AGTCAATAGGGACTTTCCAATGG</td><td>1000</td><td></td><td></td><td></td><td></td><td></td><td></td><td></td><td></td><td></td></tr>
<tr><td>Oligo fwd</td><td>CACCAGTCAATAGGGACTTTCCAA</td><td></td><td></td><td></td><td></td><td></td><td></td><td></td><td></td><td></td><td></td></tr>
<tr><td>Oligo rev</td><td>AAACTTGGAAAGTCCCTATTGACT</td><td></td><td></td><td></td><td></td><td></td><td></td><td></td><td></td><td></td><td></td></tr>
<tr><td>Chromosome</td><td>start</td><td>end</td><td>strand</td><td>MM</td><td>target_seq</td><td>PAM</td><td>alignment</td><td>distance</td><td>position</td><td>gene name</td><td>gene id</td></tr>
<tr><td>GU305914.1</td><td>183552</td><td>183574</td><td>-</td><td>0</td><td>AGTCAATAGGGACTTTCCAA</td><td>TGG</td><td>||||||||[||||||||||||]PAM</td><td>NA</td><td>Intergenic</td><td>NA</td><td>NA</td></tr>
<tr><td>GU305914.1</td><td>183895</td><td>183917</td><td>-</td><td>1</td><td>ACTCAATAGGGACTTTCCAA</td><td>TGG</td><td>|-||||||[||||||||||||]PAM</td><td>NA</td><td>Intergenic</td><td>NA</td><td>NA</td></tr>
<tr><td>GU305914.1</td><td>183459</td><td>183481</td><td>-</td><td>1</td><td>AGTCATTAGGGACTTTCCAA</td><td>TGG</td><td>|||||-||[||||||||||||]PAM</td><td>NA</td><td>Intergenic</td><td>NA</td><td>NA</td></tr>
<tr><td>GU305914.1</td><td>183367</td><td>183389</td><td>-</td><td>1</td><td>AGTCAATAGGGACTTTCCAT</td><td>TGG</td><td>||||||||[|||||||||||-]PAM</td><td>NA</td><td>Intergenic</td><td>NA</td><td>NA</td></tr>
<tr><td>GU305914.1</td><td>183738</td><td>183760</td><td>-</td><td>2</td><td>CGTCAATAGGGACTTTCCAT</td><td>TGG</td><td>-|||||||[|||||||||||-]PAM</td><td>NA</td><td>Intergenic</td><td>NA</td><td>NA</td></tr>
<tr><td>GU305914.1</td><td>183645</td><td>183667</td><td>-</td><td>2</td><td>ACTCAATAGGGACTTTCCAT</td><td>TGG</td><td>|-||||||[|||||||||||-]PAM</td><td>NA</td><td>Intergenic</td><td>NA</td><td>NA</td></tr>
<tr><td>GU305914.1</td><td>183989</td><td>184011</td><td>-</td><td>3</td><td>AGTCAAATGGGACTTTCCAC</td><td>TGG</td><td>||||||--[|||||||||||-]PAM</td><td>NA</td><td>Intergenic</td><td>NA</td><td>NA</td></tr>
<tr><td></td><td></td><td></td><td></td><td></td><td></td><td></td><td></td><td></td><td></td><td></td><td></td></tr>
<tr><td>T20820</td><td>GTCAATAGGGACTTTCCAATGGG</td><td>1000</td><td></td><td></td><td></td><td></td><td></td><td></td><td></td><td></td><td></td></tr>
<tr><td>Oligo fwd</td><td>CACCGTCAATAGGGACTTTCCAAT</td><td></td><td></td><td></td><td></td><td></td><td></td><td></td><td></td><td></td><td></td></tr>
<tr><td>Oligo rev</td><td>AAACATTGGAAAGTCCCTATTGAC</td><td></td><td></td><td></td><td></td><td></td><td></td><td></td><td></td><td></td><td></td></tr>
<tr><td>Chromosome</td><td>start</td><td>end</td><td>strand</td><td>MM</td><td>target_seq</td><td>PAM</td><td>alignment</td><td>distance</td><td>position</td><td>gene name</td><td>gene id</td></tr>
<tr><td>GU305914.1</td><td>183551</td><td>183573</td><td>-</td><td>0</td><td>GTCAATAGGGACTTTCCAAT</td><td>GGG</td><td>||||||||[||||||||||||]PAM</td><td>NA</td><td>Intergenic</td><td>NA</td><td>NA</td></tr>
<tr><td>GU305914.1</td><td>183894</td><td>183916</td><td>-</td><td>1</td><td>CTCAATAGGGACTTTCCAAT</td><td>GGG</td><td>-|||||||[||||||||||||]PAM</td><td>NA</td><td>Intergenic</td><td>NA</td><td>NA</td></tr>
<tr><td>GU305914.1</td><td>183458</td><td>183480</td><td>-</td><td>1</td><td>GTCATTAGGGACTTTCCAAT</td><td>GGG</td><td>||||-|||[||||||||||||]PAM</td><td>NA</td><td>Intergenic</td><td>NA</td><td>NA</td></tr>
<tr><td>GU305914.1</td><td>183366</td><td>183388</td><td>-</td><td>1</td><td>GTCAATAGGGACTTTCCATT</td><td>GGG</td><td>||||||||[||||||||||-|]PAM</td><td>NA</td><td>Intergenic</td><td>NA</td><td>NA</td></tr>
<tr><td>GU305914.1</td><td>183737</td><td>183759</td><td>-</td><td>1</td><td>GTCAATAGGGACTTTCCATT</td><td>GGG</td><td>||||||||[||||||||||-|]PAM</td><td>NA</td><td>Intergenic</td><td>NA</td><td>NA</td></tr>
<tr><td>GU305914.1</td><td>183644</td><td>183666</td><td>-</td><td>2</td><td>CTCAATAGGGACTTTCCATT</td><td>GGG</td><td>-|||||||[||||||||||-|]PAM</td><td>NA</td><td>Intergenic</td><td>NA</td><td>NA</td></tr>
<tr><td>GU305914.1</td><td>183988</td><td>184010</td><td>-</td><td>3</td><td>GTCAAATGGGACTTTCCACT</td><td>GGG</td><td>|||||--|[||||||||||-|]PAM</td><td>NA</td><td>Intergenic</td><td>NA</td><td>NA</td></tr>
<tr><td></td><td></td><td></td><td></td><td></td><td></td><td></td><td></td><td></td><td></td><td></td><td></td></tr>
<tr><td>T20821</td><td>TGGGTTTTGCCCAGTACATAAGG</td><td>1000</td><td></td><td></td><td></td><td></td><td></td><td></td><td></td><td></td><td></td></tr>
<tr><td>Oligo fwd</td><td>CACCTGGGTTTTGCCCAGTACATA</td><td></td><td></td><td></td><td></td><td></td><td></td><td></td><td></td><td></td><td></td></tr>
<tr><td>Oligo rev</td><td>AAACTATGTACTGGGCAAAACCCA</td><td></td><td></td><td></td><td></td><td></td><td></td><td></td><td></td><td></td><td></td></tr>
<tr><td>Chromosome</td><td>start</td><td>end</td><td>strand</td><td>MM</td><td>target_seq</td><td>PAM</td><td>alignment</td><td>distance</td><td>position</td><td>gene name</td><td>gene id</td></tr>
<tr><td>GU305914.1</td><td>183439</td><td>183461</td><td>-</td><td>0</td><td>TGGGTTTTGCCCAGTACATA</td><td>AGG</td><td>||||||||[||||||||||||]PAM</td><td>NA</td><td>Intergenic</td><td>NA</td><td>NA</td></tr>
<tr><td>GU305914.1</td><td>183625</td><td>183647</td><td>-</td><td>0</td><td>TGGGTTTTGCCCAGTACATA</td><td>AGG</td><td>||||||||[||||||||||||]PAM</td><td>NA</td><td>Intergenic</td><td>NA</td><td>NA</td></tr>
<tr><td>GU305914.1</td><td>183718</td><td>183740</td><td>-</td><td>0</td><td>TGGGTTTTGCCCAGTACATA</td><td>AGG</td><td>||||||||[||||||||||||]PAM</td><td>NA</td><td>Intergenic</td><td>NA</td><td>NA</td></tr>
<tr><td>GU305914.1</td><td>183532</td><td>183554</td><td>-</td><td>0</td><td>TGGGTTTTGCCCAGTACATA</td><td>AGG</td><td>||||||||[||||||||||||]PAM</td><td>NA</td><td>Intergenic</td><td>NA</td><td>NA</td></tr>
<tr><td>GU305914.1</td><td>183811</td><td>183833</td><td>-</td><td>3</td><td>TGTATTCTGCCCAGTACATA</td><td>AGG</td><td>||--||-|[||||||||||||]PAM</td><td>NA</td><td>Intergenic</td><td>NA</td><td>NA</td></tr>
<tr><td>GU305914.1</td><td>183347</td><td>183369</td><td>-</td><td>1</td><td>TGGGTTTTGCCCAGTACAAA</td><td>AGG</td><td>||||||||[||||||||||-|]PAM</td><td>NA</td><td>Intergenic</td><td>NA</td><td>NA</td></tr>
<tr><td></td><td></td><td></td><td></td><td></td><td></td><td></td><td></td><td></td><td></td><td></td><td></td></tr>
<tr><td>T20822</td><td>TGCCCAGTACATAAGGTCAATGG</td><td>1000</td><td></td><td></td><td></td><td></td><td></td><td></td><td></td><td></td><td></td></tr>
<tr><td>Oligo fwd</td><td>CACCTGCCCAGTACATAAGGTCAA</td><td></td><td></td><td></td><td></td><td></td><td></td><td></td><td></td><td></td><td></td></tr>
<tr><td>Oligo rev</td><td>AAACTTGACCTTATGTACTGGGCA</td><td></td><td></td><td></td><td></td><td></td><td></td><td></td><td></td><td></td><td></td></tr>
<tr><td>Chromosome</td><td>start</td><td>end</td><td>strand</td><td>MM</td><td>target_seq</td><td>PAM</td><td>alignment</td><td>distance</td><td>position</td><td>gene name</td><td>gene id</td></tr>
<tr><td>GU305914.1</td><td>183525</td><td>183547</td><td>-</td><td>0</td><td>TGCCCAGTACATAAGGTCAA</td><td>TGG</td><td>||||||||[||||||||||||]PAM</td><td>NA</td><td>Intergenic</td><td>NA</td><td>NA</td></tr>
<tr><td></td><td></td><td></td><td></td><td></td><td></td><td></td><td></td><td></td><td></td><td></td><td></td></tr>
<tr><td>T20823</td><td>GCCCAGTACATAAGGTCAATGGG</td><td>1000</td><td></td><td></td><td></td><td></td><td></td><td></td><td></td><td></td><td></td></tr>
<tr><td>Oligo fwd</td><td>CACCGCCCAGTACATAAGGTCAAT</td><td></td><td></td><td></td><td></td><td></td><td></td><td></td><td></td><td></td><td></td></tr>
<tr><td>Oligo rev</td><td>AAACATTGACCTTATGTACTGGGC</td><td></td><td></td><td></td><td></td><td></td><td></td><td></td><td></td><td></td><td></td></tr>
<tr><td>Chromosome</td><td>start</td><td>end</td><td>strand</td><td>MM</td><td>target_seq</td><td>PAM</td><td>alignment</td><td>distance</td><td>position</td><td>gene name</td><td>gene id</td></tr>
<tr><td>GU305914.1</td><td>183524</td><td>183546</td><td>-</td><td>0</td><td>GCCCAGTACATAAGGTCAAT</td><td>GGG</td><td>||||||||[||||||||||||]PAM</td><td>NA</td><td>Intergenic</td><td>NA</td><td>NA</td></tr>
<tr><td>GU305914.1</td><td>183617</td><td>183639</td><td>-</td><td>0</td><td>GCCCAGTACATAAGGTCAAT</td><td>AGG</td><td>||||||||[||||||||||||]PAM</td><td>NA</td><td>Intergenic</td><td>NA</td><td>NA</td></tr>
<tr><td>GU305914.1</td><td>183803</td><td>183825</td><td>-</td><td>0</td><td>GCCCAGTACATAAGGTCAAT</td><td>AGG</td><td>||||||||[||||||||||||]PAM</td><td>NA</td><td>Intergenic</td><td>NA</td><td>NA</td></tr>
<tr><td>GU305914.1</td><td>183710</td><td>183732</td><td>-</td><td>0</td><td>GCCCAGTACATAAGGTCAAT</td><td>AGG</td><td>||||||||[||||||||||||]PAM</td><td>NA</td><td>Intergenic</td><td>NA</td><td>NA</td></tr>
<tr><td>GU305914.1</td><td>183431</td><td>183453</td><td>-</td><td>0</td><td>GCCCAGTACATAAGGTCAAT</td><td>AGG</td><td>||||||||[||||||||||||]PAM</td><td>NA</td><td>Intergenic</td><td>NA</td><td>NA</td></tr>
<tr><td>GU305914.1</td><td>183287</td><td>183309</td><td>-</td><td>3</td><td>GGCACGTACATAAGGTCAAT</td><td>AGG</td><td>|-|--|||[||||||||||||]PAM</td><td>NA</td><td>Intergenic</td><td>NA</td><td>NA</td></tr>
<tr><td>GU305914.1</td><td>183339</td><td>183361</td><td>-</td><td>1</td><td>GCCCAGTACAAAAGGTCAAT</td><td>AGG</td><td>||||||||[||-|||||||||]PAM</td><td>NA</td><td>Intergenic</td><td>NA</td><td>NA</td></tr>
</table>

<table>
<tr><td>GU305914.1</td><td>184054</td><td>184076</td><td>-</td><td>4</td><td>CCCAAGTACATTGGGTCAAT</td><td>GGG</td><td>-||-||||[|||--|||||||]PAM</td><td>NA</td><td>Intergenic</td><td>NA</td><td>NA</td></tr>
<tr><td>GU305914.1</td><td>183960</td><td>183982</td><td>-</td><td>4</td><td>CCCAAGTACATTGGGTCAAT</td><td>GGG</td><td>-||-||||[|||--|||||||]PAM</td><td>NA</td><td>Intergenic</td><td>NA</td><td>NA</td></tr>
<tr><td></td><td></td><td></td><td></td><td></td><td></td><td></td><td></td><td></td><td></td><td></td><td></td></tr>
<tr><td>T20824</td><td>CAGTACATAAGGTCAATGGGAGG</td><td>1000</td><td></td><td></td><td></td><td></td><td></td><td></td><td></td><td></td><td></td></tr>
<tr><td>Oligo fwd</td><td>CACCCAGTACATAAGGTCAATGGG</td><td></td><td></td><td></td><td></td><td></td><td></td><td></td><td></td><td></td><td></td></tr>
<tr><td>Oligo rev</td><td>AAACCCCATTGACCTTATGTACTG</td><td></td><td></td><td></td><td></td><td></td><td></td><td></td><td></td><td></td><td></td></tr>
<tr><td>Chromosome</td><td>start</td><td>end</td><td>strand</td><td>MM</td><td>target_seq</td><td>PAM</td><td>alignment</td><td>distance</td><td>position</td><td>gene name</td><td>gene id</td></tr>
<tr><td>GU305914.1</td><td>183521</td><td>183543</td><td>-</td><td>0</td><td>CAGTACATAAGGTCAATGGG</td><td>AGG</td><td>||||||||[||||||||||||]PAM</td><td>NA</td><td>Intergenic</td><td>NA</td><td>NA</td></tr>
<tr><td>GU305914.1</td><td>183957</td><td>183979</td><td>-</td><td>3</td><td>AAGTACATTGGGTCAATGGG</td><td>AGG</td><td>-|||||||[--||||||||||]PAM</td><td>NA</td><td>Intergenic</td><td>NA</td><td>NA</td></tr>
<tr><td>GU305914.1</td><td>184051</td><td>184073</td><td>-</td><td>3</td><td>AAGTACATTGGGTCAATGGG</td><td>AGG</td><td>-|||||||[--||||||||||]PAM</td><td>NA</td><td>Intergenic</td><td>NA</td><td>NA</td></tr>
<tr><td>GU305914.1</td><td>183614</td><td>183636</td><td>-</td><td>1</td><td>CAGTACATAAGGTCAATAGG</td><td>GGG</td><td>||||||||[|||||||||-||]PAM</td><td>NA</td><td>Intergenic</td><td>NA</td><td>NA</td></tr>
<tr><td>GU305914.1</td><td>183800</td><td>183822</td><td>-</td><td>1</td><td>CAGTACATAAGGTCAATAGG</td><td>GGG</td><td>||||||||[|||||||||-||]PAM</td><td>NA</td><td>Intergenic</td><td>NA</td><td>NA</td></tr>
<tr><td>GU305914.1</td><td>183336</td><td>183358</td><td>-</td><td>2</td><td>CAGTACAAAAGGTCAATAGG</td><td>GGG</td><td>|||||||-[|||||||||-||]PAM</td><td>NA</td><td>Intergenic</td><td>NA</td><td>NA</td></tr>
<tr><td></td><td></td><td></td><td></td><td></td><td></td><td></td><td></td><td></td><td></td><td></td><td></td></tr>
<tr><td>T20825</td><td>GTCAATGGGAGGTAAGCCAATGG</td><td>1000</td><td></td><td></td><td></td><td></td><td></td><td></td><td></td><td></td><td></td></tr>
<tr><td>Oligo fwd</td><td>CACCGTCAATGGGAGGTAAGCCAA</td><td></td><td></td><td></td><td></td><td></td><td></td><td></td><td></td><td></td><td></td></tr>
<tr><td>Oligo rev</td><td>AAACTTGGCTTACCTCCCATTGAC</td><td></td><td></td><td></td><td></td><td></td><td></td><td></td><td></td><td></td><td></td></tr>
<tr><td>Chromosome</td><td>start</td><td>end</td><td>strand</td><td>MM</td><td>target_seq</td><td>PAM</td><td>alignment</td><td>distance</td><td>position</td><td>gene name</td><td>gene id</td></tr>
<tr><td>GU305914.1</td><td>184040</td><td>184062</td><td>-</td><td>0</td><td>GTCAATGGGAGGTAAGCCAA</td><td>TGG</td><td>||||||||[||||||||||||]PAM</td><td>NA</td><td>Intergenic</td><td>NA</td><td>NA</td></tr>
<tr><td>GU305914.1</td><td>183510</td><td>183532</td><td>-</td><td>0</td><td>GTCAATGGGAGGTAAGCCAA</td><td>TGG</td><td>||||||||[||||||||||||]PAM</td><td>NA</td><td>Intergenic</td><td>NA</td><td>NA</td></tr>
<tr><td>GU305914.1</td><td>183946</td><td>183968</td><td>-</td><td>1</td><td>GTCAATGGGAGGTGAGCCAA</td><td>TGG</td><td>||||||||[|||||-||||||]PAM</td><td>NA</td><td>Intergenic</td><td>NA</td><td>NA</td></tr>
<tr><td></td><td></td><td></td><td></td><td></td><td></td><td></td><td></td><td></td><td></td><td></td><td></td></tr>
<tr><td>T20826</td><td>TCAATGGGAGGTAAGCCAATGGG</td><td>1000</td><td></td><td></td><td></td><td></td><td></td><td></td><td></td><td></td><td></td></tr>
<tr><td>Oligo fwd</td><td>CACCTCAATGGGAGGTAAGCCAAT</td><td></td><td></td><td></td><td></td><td></td><td></td><td></td><td></td><td></td><td></td></tr>
<tr><td>Oligo rev</td><td>AAACATTGGCTTACCTCCCATTGA</td><td></td><td></td><td></td><td></td><td></td><td></td><td></td><td></td><td></td><td></td></tr>
<tr><td>Chromosome</td><td>start</td><td>end</td><td>strand</td><td>MM</td><td>target_seq</td><td>PAM</td><td>alignment</td><td>distance</td><td>position</td><td>gene name</td><td>gene id</td></tr>
<tr><td>GU305914.1</td><td>184039</td><td>184061</td><td>-</td><td>0</td><td>TCAATGGGAGGTAAGCCAAT</td><td>GGG</td><td>||||||||[||||||||||||]PAM</td><td>NA</td><td>Intergenic</td><td>NA</td><td>NA</td></tr>
<tr><td>GU305914.1</td><td>183509</td><td>183531</td><td>-</td><td>0</td><td>TCAATGGGAGGTAAGCCAAT</td><td>GGG</td><td>||||||||[||||||||||||]PAM</td><td>NA</td><td>Intergenic</td><td>NA</td><td>NA</td></tr>
<tr><td>GU305914.1</td><td>183945</td><td>183967</td><td>-</td><td>1</td><td>TCAATGGGAGGTGAGCCAAT</td><td>GGG</td><td>||||||||[||||-|||||||]PAM</td><td>NA</td><td>Intergenic</td><td>NA</td><td>NA</td></tr>
<tr><td></td><td></td><td></td><td></td><td></td><td></td><td></td><td></td><td></td><td></td><td></td><td></td></tr>
<tr><td>T20827</td><td>CAATGGGTTTTTCCCATTACTGG</td><td>1000</td><td></td><td></td><td></td><td></td><td></td><td></td><td></td><td></td><td></td></tr>
<tr><td>Oligo fwd</td><td>CACCCAATGGGTTTTTCCCATTAC</td><td></td><td></td><td></td><td></td><td></td><td></td><td></td><td></td><td></td><td></td></tr>
<tr><td>Oligo rev</td><td>AAACGTAATGGGAAAAACCCATTG</td><td></td><td></td><td></td><td></td><td></td><td></td><td></td><td></td><td></td><td></td></tr>
<tr><td>Chromosome</td><td>start</td><td>end</td><td>strand</td><td>MM</td><td>target_seq</td><td>PAM</td><td>alignment</td><td>distance</td><td>position</td><td>gene name</td><td>gene id</td></tr>
<tr><td>GU305914.1</td><td>184023</td><td>184045</td><td>-</td><td>0</td><td>CAATGGGTTTTTCCCATTAC</td><td>TGG</td><td>||||||||[||||||||||||]PAM</td><td>NA</td><td>Intergenic</td><td>NA</td><td>NA</td></tr>
<tr><td>GU305914.1</td><td>183493</td><td>183515</td><td>-</td><td>0</td><td>CAATGGGTTTTTCCCATTAC</td><td>TGG</td><td>||||||||[||||||||||||]PAM</td><td>NA</td><td>Intergenic</td><td>NA</td><td>NA</td></tr>
<tr><td>GU305914.1</td><td>183308</td><td>183330</td><td>-</td><td>1</td><td>CAATGGGTTTTTCCCATTAT</td><td>TGG</td><td>||||||||[|||||||||||-]PAM</td><td>NA</td><td>Intergenic</td><td>NA</td><td>NA</td></tr>
<tr><td></td><td></td><td></td><td></td><td></td><td></td><td></td><td></td><td></td><td></td><td></td><td></td></tr>
<tr><td>T20828</td><td>GGCACGTATACTGAGTCATTAGG</td><td>1000</td><td></td><td></td><td></td><td></td><td></td><td></td><td></td><td></td><td></td></tr>
<tr><td>Oligo fwd</td><td>CACCGGCACGTATACTGAGTCATT</td><td></td><td></td><td></td><td></td><td></td><td></td><td></td><td></td><td></td><td></td></tr>
<tr><td>Oligo rev</td><td>AAACAATGACTCAGTATACGTGCC</td><td></td><td></td><td></td><td></td><td></td><td></td><td></td><td></td><td></td><td></td></tr>
<tr><td>Chromosome</td><td>start</td><td>end</td><td>strand</td><td>MM</td><td>target_seq</td><td>PAM</td><td>alignment</td><td>distance</td><td>position</td><td>gene name</td><td>gene id</td></tr>
<tr><td>GU305914.1</td><td>183472</td><td>183494</td><td>-</td><td>0</td><td>GGCACGTATACTGAGTCATT</td><td>AGG</td><td>||||||||[||||||||||||]PAM</td><td>NA</td><td>Intergenic</td><td>NA</td><td>NA</td></tr>
<tr><td>GU305914.1</td><td>183380</td><td>183402</td><td>-</td><td>4</td><td>GCCAAGTACACTGAGTCAAT</td><td>AGG</td><td>|-||-|||[-|||||||||-|]PAM</td><td>NA</td><td>Intergenic</td><td>NA</td><td>NA</td></tr>
<tr><td></td><td></td><td></td><td></td><td></td><td></td><td></td><td></td><td></td><td></td><td></td><td></td></tr>
<tr><td>T20829</td><td>GCACGTATACTGAGTCATTAGGG</td><td>1000</td><td></td><td></td><td></td><td></td><td></td><td></td><td></td><td></td><td></td></tr>
<tr><td>Oligo fwd</td><td>CACCGCACGTATACTGAGTCATTA</td><td></td><td></td><td></td><td></td><td></td><td></td><td></td><td></td><td></td><td></td></tr>
<tr><td>Oligo rev</td><td>AAACTAATGACTCAGTATACGTGC</td><td></td><td></td><td></td><td></td><td></td><td></td><td></td><td></td><td></td><td></td></tr>
<tr><td>Chromosome</td><td>start</td><td>end</td><td>strand</td><td>MM</td><td>target_seq</td><td>PAM</td><td>alignment</td><td>distance</td><td>position</td><td>gene name</td><td>gene id</td></tr>
<tr><td>GU305914.1</td><td>183471</td><td>183493</td><td>-</td><td>0</td><td>GCACGTATACTGAGTCATTA</td><td>GGG</td><td>||||||||[||||||||||||]PAM</td><td>NA</td><td>Intergenic</td><td>NA</td><td>NA</td></tr>
<tr><td>GU305914.1</td><td>183379</td><td>183401</td><td>-</td><td>4</td><td>CCAAGTACACTGAGTCAATA</td><td>GGG</td><td>-||-|||-[|||||||||-||]PAM</td><td>NA</td><td>Intergenic</td><td>NA</td><td>NA</td></tr>
<tr><td></td><td></td><td></td><td></td><td></td><td></td><td></td><td></td><td></td><td></td><td></td><td></td></tr>
<tr><td>T20830</td><td>AGTCATTAGGGACTTTCCAATGG</td><td>1000</td><td></td><td></td><td></td><td></td><td></td><td></td><td></td><td></td><td></td></tr>
<tr><td>Oligo fwd</td><td>CACCAGTCATTAGGGACTTTCCAA</td><td></td><td></td><td></td><td></td><td></td><td></td><td></td><td></td><td></td><td></td></tr>
<tr><td>Oligo rev</td><td>AAACTTGGAAAGTCCCTAATGACT</td><td></td><td></td><td></td><td></td><td></td><td></td><td></td><td></td><td></td><td></td></tr>
<tr><td>Chromosome</td><td>start</td><td>end</td><td>strand</td><td>MM</td><td>target_seq</td><td>PAM</td><td>alignment</td><td>distance</td><td>position</td><td>gene name</td><td>gene id</td></tr>
<tr><td>GU305914.1</td><td>183459</td><td>183481</td><td>-</td><td>0</td><td>AGTCATTAGGGACTTTCCAA</td><td>TGG</td><td>||||||||[||||||||||||]PAM</td><td>NA</td><td>Intergenic</td><td>NA</td><td>NA</td></tr>
<tr><td>GU305914.1</td><td>183552</td><td>183574</td><td>-</td><td>1</td><td>AGTCAATAGGGACTTTCCAA</td><td>TGG</td><td>|||||-||[||||||||||||]PAM</td><td>NA</td><td>Intergenic</td><td>NA</td><td>NA</td></tr>
<tr><td>GU305914.1</td><td>183895</td><td>183917</td><td>-</td><td>2</td><td>ACTCAATAGGGACTTTCCAA</td><td>TGG</td><td>|-|||-||[||||||||||||]PAM</td><td>NA</td><td>Intergenic</td><td>NA</td><td>NA</td></tr>
<tr><td>GU305914.1</td><td>183367</td><td>183389</td><td>-</td><td>2</td><td>AGTCAATAGGGACTTTCCAT</td><td>TGG</td><td>|||||-||[|||||||||||-]PAM</td><td>NA</td><td>Intergenic</td><td>NA</td><td>NA</td></tr>
<tr><td>GU305914.1</td><td>183738</td><td>183760</td><td>-</td><td>3</td><td>CGTCAATAGGGACTTTCCAT</td><td>TGG</td><td>-||||-||[|||||||||||-]PAM</td><td>NA</td><td>Intergenic</td><td>NA</td><td>NA</td></tr>
<tr><td>GU305914.1</td><td>183645</td><td>183667</td><td>-</td><td>3</td><td>ACTCAATAGGGACTTTCCAT</td><td>TGG</td><td>|-|||-||[|||||||||||-]PAM</td><td>NA</td><td>Intergenic</td><td>NA</td><td>NA</td></tr>
<tr><td>GU305914.1</td><td>183989</td><td>184011</td><td>-</td><td>4</td><td>AGTCAAATGGGACTTTCCAC</td><td>TGG</td><td>|||||---[|||||||||||-]PAM</td><td>NA</td><td>Intergenic</td><td>NA</td><td>NA</td></tr>
<tr><td></td><td></td><td></td><td></td><td></td><td></td><td></td><td></td><td></td><td></td><td></td><td></td></tr>
<tr><td>T20831</td><td>GTCATTAGGGACTTTCCAATGGG</td><td>1000</td><td></td><td></td><td></td><td></td><td></td><td></td><td></td><td></td><td></td></tr>
<tr><td>Oligo fwd</td><td>CACCGTCATTAGGGACTTTCCAAT</td><td></td><td></td><td></td><td></td><td></td><td></td><td></td><td></td><td></td><td></td></tr>
<tr><td>Oligo rev</td><td>AAACATTGGAAAGTCCCTAATGAC</td><td></td><td></td><td></td><td></td><td></td><td></td><td></td><td></td><td></td><td></td></tr>
<tr><td>Chromosome</td><td>start</td><td>end</td><td>strand</td><td>MM</td><td>target_seq</td><td>PAM</td><td>alignment</td><td>distance</td><td>position</td><td>gene name</td><td>gene id</td></tr>
<tr><td>GU305914.1</td><td>183458</td><td>183480</td><td>-</td><td>0</td><td>GTCATTAGGGACTTTCCAAT</td><td>GGG</td><td>||||||||[||||||||||||]PAM</td><td>NA</td><td>Intergenic</td><td>NA</td><td>NA</td></tr>
<tr><td>GU305914.1</td><td>183551</td><td>183573</td><td>-</td><td>1</td><td>GTCAATAGGGACTTTCCAAT</td><td>GGG</td><td>||||-|||[||||||||||||]PAM</td><td>NA</td><td>Intergenic</td><td>NA</td><td>NA</td></tr>
<tr><td>GU305914.1</td><td>183894</td><td>183916</td><td>-</td><td>2</td><td>CTCAATAGGGACTTTCCAAT</td><td>GGG</td><td>-|||-|||[||||||||||||]PAM</td><td>NA</td><td>Intergenic</td><td>NA</td><td>NA</td></tr>
<tr><td>GU305914.1</td><td>183737</td><td>183759</td><td>-</td><td>2</td><td>GTCAATAGGGACTTTCCATT</td><td>GGG</td><td>||||-|||[||||||||||-|]PAM</td><td>NA</td><td>Intergenic</td><td>NA</td><td>NA</td></tr>
<tr><td>GU305914.1</td><td>183366</td><td>183388</td><td>-</td><td>2</td><td>GTCAATAGGGACTTTCCATT</td><td>GGG</td><td>||||-|||[||||||||||-|]PAM</td><td>NA</td><td>Intergenic</td><td>NA</td><td>NA</td></tr>
<tr><td>GU305914.1</td><td>183644</td><td>183666</td><td>-</td><td>3</td><td>CTCAATAGGGACTTTCCATT</td><td>GGG</td><td>-|||-|||[||||||||||-|]PAM</td><td>NA</td><td>Intergenic</td><td>NA</td><td>NA</td></tr>
<tr><td>GU305914.1</td><td>183988</td><td>184010</td><td>-</td><td>4</td><td>GTCAAATGGGACTTTCCACT</td><td>GGG</td><td>||||---|[||||||||||-|]PAM</td><td>NA</td><td>Intergenic</td><td>NA</td><td>NA</td></tr>
<tr><td></td><td></td><td></td><td></td><td></td><td></td><td></td><td></td><td></td><td></td><td></td><td></td></tr>
<tr><td>T20832</td><td>TGGGTTTTGCCCAGTACATAAGG</td><td>1000</td><td></td><td></td><td></td><td></td><td></td><td></td><td></td><td></td><td></td></tr>
<tr><td>Oligo fwd</td><td>CACCTGGGTTTTGCCCAGTACATA</td><td></td><td></td><td></td><td></td><td></td><td></td><td></td><td></td><td></td><td></td></tr>
<tr><td>Oligo rev</td><td>AAACTATGTACTGGGCAAAACCCA</td><td></td><td></td><td></td><td></td><td></td><td></td><td></td><td></td><td></td><td></td></tr>
<tr><td>Chromosome</td><td>start</td><td>end</td><td>strand</td><td>MM</td><td>target_seq</td><td>PAM</td><td>alignment</td><td>distance</td><td>position</td><td>gene name</td><td>gene id</td></tr>
<tr><td>GU305914.1</td><td>183439</td><td>183461</td><td>-</td><td>0</td><td>TGGGTTTTGCCCAGTACATA</td><td>AGG</td><td>||||||||[||||||||||||]PAM</td><td>NA</td><td>Intergenic</td><td>NA</td><td>NA</td></tr>
<tr><td>GU305914.1</td><td>183625</td><td>183647</td><td>-</td><td>0</td><td>TGGGTTTTGCCCAGTACATA</td><td>AGG</td><td>||||||||[||||||||||||]PAM</td><td>NA</td><td>Intergenic</td><td>NA</td><td>NA</td></tr>
<tr><td>GU305914.1</td><td>183718</td><td>183740</td><td>-</td><td>0</td><td>TGGGTTTTGCCCAGTACATA</td><td>AGG</td><td>||||||||[||||||||||||]PAM</td><td>NA</td><td>Intergenic</td><td>NA</td><td>NA</td></tr>
<tr><td>GU305914.1</td><td>183532</td><td>183554</td><td>-</td><td>0</td><td>TGGGTTTTGCCCAGTACATA</td><td>AGG</td><td>||||||||[||||||||||||]PAM</td><td>NA</td><td>Intergenic</td><td>NA</td><td>NA</td></tr>
<tr><td>GU305914.1</td><td>183811</td><td>183833</td><td>-</td><td>3</td><td>TGTATTCTGCCCAGTACATA</td><td>AGG</td><td>||--||-|[||||||||||||]PAM</td><td>NA</td><td>Intergenic</td><td>NA</td><td>NA</td></tr>
<tr><td>GU305914.1</td><td>183347</td><td>183369</td><td>-</td><td>1</td><td>TGGGTTTTGCCCAGTACAAA</td><td>AGG</td><td>||||||||[||||||||||-|]PAM</td><td>NA</td><td>Intergenic</td><td>NA</td><td>NA</td></tr>
<tr><td></td><td></td><td></td><td></td><td></td><td></td><td></td><td></td><td></td><td></td><td></td><td></td></tr>
<tr><td>T20833</td><td>GCCCAGTACATAAGGTCAATAGG</td><td>1000</td><td></td><td></td><td></td><td></td><td></td><td></td><td></td><td></td><td></td></tr>
<tr><td>Oligo fwd</td><td>CACCGCCCAGTACATAAGGTCAAT</td><td></td><td></td><td></td><td></td><td></td><td></td><td></td><td></td><td></td><td></td></tr>
</table>

<table>
<tr><td>Oligo rev</td><td>AAACATTGACCTTATGTACTGGGC</td><td></td><td></td><td></td><td></td><td></td><td></td><td></td><td></td><td></td><td></td></tr>
<tr><td>Chromosome</td><td>start</td><td>end</td><td>strand</td><td>MM</td><td>target_seq</td><td>PAM</td><td>alignment</td><td>distance</td><td>position</td><td>gene name</td><td>gene id</td></tr>
<tr><td>GU305914.1</td><td>183524</td><td>183546</td><td>-</td><td>0</td><td>GCCCAGTACATAAGGTCAAT</td><td>GGG</td><td>||||||||[||||||||||||]PAM</td><td>NA</td><td>Intergenic</td><td>NA</td><td>NA</td></tr>
<tr><td>GU305914.1</td><td>183617</td><td>183639</td><td>-</td><td>0</td><td>GCCCAGTACATAAGGTCAAT</td><td>AGG</td><td>||||||||[||||||||||||]PAM</td><td>NA</td><td>Intergenic</td><td>NA</td><td>NA</td></tr>
<tr><td>GU305914.1</td><td>183803</td><td>183825</td><td>-</td><td>0</td><td>GCCCAGTACATAAGGTCAAT</td><td>AGG</td><td>||||||||[||||||||||||]PAM</td><td>NA</td><td>Intergenic</td><td>NA</td><td>NA</td></tr>
<tr><td>GU305914.1</td><td>183710</td><td>183732</td><td>-</td><td>0</td><td>GCCCAGTACATAAGGTCAAT</td><td>AGG</td><td>||||||||[||||||||||||]PAM</td><td>NA</td><td>Intergenic</td><td>NA</td><td>NA</td></tr>
<tr><td>GU305914.1</td><td>183431</td><td>183453</td><td>-</td><td>0</td><td>GCCCAGTACATAAGGTCAAT</td><td>AGG</td><td>||||||||[||||||||||||]PAM</td><td>NA</td><td>Intergenic</td><td>NA</td><td>NA</td></tr>
<tr><td>GU305914.1</td><td>183287</td><td>183309</td><td>-</td><td>3</td><td>GGCACGTACATAAGGTCAAT</td><td>AGG</td><td>|-|--|||[||||||||||||]PAM</td><td>NA</td><td>Intergenic</td><td>NA</td><td>NA</td></tr>
<tr><td>GU305914.1</td><td>183339</td><td>183361</td><td>-</td><td>1</td><td>GCCCAGTACAAAAGGTCAAT</td><td>AGG</td><td>||||||||[||-|||||||||]PAM</td><td>NA</td><td>Intergenic</td><td>NA</td><td>NA</td></tr>
<tr><td>GU305914.1</td><td>184054</td><td>184076</td><td>-</td><td>4</td><td>CCCAAGTACATTGGGTCAAT</td><td>GGG</td><td>-||-||||[|||--|||||||]PAM</td><td>NA</td><td>Intergenic</td><td>NA</td><td>NA</td></tr>
<tr><td>GU305914.1</td><td>183960</td><td>183982</td><td>-</td><td>4</td><td>CCCAAGTACATTGGGTCAAT</td><td>GGG</td><td>-||-||||[|||--|||||||]PAM</td><td>NA</td><td>Intergenic</td><td>NA</td><td>NA</td></tr>
<tr><td></td><td></td><td></td><td></td><td></td><td></td><td></td><td></td><td></td><td></td><td></td><td></td></tr>
<tr><td>T20834</td><td>CCCAGTACATAAGGTCAATAGGG</td><td>1000</td><td></td><td></td><td></td><td></td><td></td><td></td><td></td><td></td><td></td></tr>
<tr><td>Oligo fwd</td><td>CACCCCCAGTACATAAGGTCAATA</td><td></td><td></td><td></td><td></td><td></td><td></td><td></td><td></td><td></td><td></td></tr>
<tr><td>Oligo rev</td><td>AAACTATTGACCTTATGTACTGGG</td><td></td><td></td><td></td><td></td><td></td><td></td><td></td><td></td><td></td><td></td></tr>
<tr><td>Chromosome</td><td>start</td><td>end</td><td>strand</td><td>MM</td><td>target_seq</td><td>PAM</td><td>alignment</td><td>distance</td><td>position</td><td>gene name</td><td>gene id</td></tr>
<tr><td>GU305914.1</td><td>183709</td><td>183731</td><td>-</td><td>0</td><td>CCCAGTACATAAGGTCAATA</td><td>GGG</td><td>||||||||[||||||||||||]PAM</td><td>NA</td><td>Intergenic</td><td>NA</td><td>NA</td></tr>
<tr><td>GU305914.1</td><td>183430</td><td>183452</td><td>-</td><td>0</td><td>CCCAGTACATAAGGTCAATA</td><td>GGG</td><td>||||||||[||||||||||||]PAM</td><td>NA</td><td>Intergenic</td><td>NA</td><td>NA</td></tr>
<tr><td>GU305914.1</td><td>183616</td><td>183638</td><td>-</td><td>0</td><td>CCCAGTACATAAGGTCAATA</td><td>GGG</td><td>||||||||[||||||||||||]PAM</td><td>NA</td><td>Intergenic</td><td>NA</td><td>NA</td></tr>
<tr><td>GU305914.1</td><td>183802</td><td>183824</td><td>-</td><td>0</td><td>CCCAGTACATAAGGTCAATA</td><td>GGG</td><td>||||||||[||||||||||||]PAM</td><td>NA</td><td>Intergenic</td><td>NA</td><td>NA</td></tr>
<tr><td>GU305914.1</td><td>183286</td><td>183308</td><td>-</td><td>3</td><td>GCACGTACATAAGGTCAATA</td><td>GGG</td><td>-|--||||[||||||||||||]PAM</td><td>NA</td><td>Intergenic</td><td>NA</td><td>NA</td></tr>
<tr><td>GU305914.1</td><td>183338</td><td>183360</td><td>-</td><td>1</td><td>CCCAGTACAAAAGGTCAATA</td><td>GGG</td><td>||||||||[|-||||||||||]PAM</td><td>NA</td><td>Intergenic</td><td>NA</td><td>NA</td></tr>
<tr><td></td><td></td><td></td><td></td><td></td><td></td><td></td><td></td><td></td><td></td><td></td><td></td></tr>
<tr><td>T20835</td><td>CCAGTACATAAGGTCAATAGGGG</td><td>1000</td><td></td><td></td><td></td><td></td><td></td><td></td><td></td><td></td><td></td></tr>
<tr><td>Oligo fwd</td><td>CACCCCAGTACATAAGGTCAATAG</td><td></td><td></td><td></td><td></td><td></td><td></td><td></td><td></td><td></td><td></td></tr>
<tr><td>Oligo rev</td><td>AAACCTATTGACCTTATGTACTGG</td><td></td><td></td><td></td><td></td><td></td><td></td><td></td><td></td><td></td><td></td></tr>
<tr><td>Chromosome</td><td>start</td><td>end</td><td>strand</td><td>MM</td><td>target_seq</td><td>PAM</td><td>alignment</td><td>distance</td><td>position</td><td>gene name</td><td>gene id</td></tr>
<tr><td>GU305914.1</td><td>183708</td><td>183730</td><td>-</td><td>0</td><td>CCAGTACATAAGGTCAATAG</td><td>GGG</td><td>||||||||[||||||||||||]PAM</td><td>NA</td><td>Intergenic</td><td>NA</td><td>NA</td></tr>
<tr><td>GU305914.1</td><td>183429</td><td>183451</td><td>-</td><td>0</td><td>CCAGTACATAAGGTCAATAG</td><td>GGG</td><td>||||||||[||||||||||||]PAM</td><td>NA</td><td>Intergenic</td><td>NA</td><td>NA</td></tr>
<tr><td>GU305914.1</td><td>183615</td><td>183637</td><td>-</td><td>0</td><td>CCAGTACATAAGGTCAATAG</td><td>GGG</td><td>||||||||[||||||||||||]PAM</td><td>NA</td><td>Intergenic</td><td>NA</td><td>NA</td></tr>
<tr><td>GU305914.1</td><td>183801</td><td>183823</td><td>-</td><td>0</td><td>CCAGTACATAAGGTCAATAG</td><td>GGG</td><td>||||||||[||||||||||||]PAM</td><td>NA</td><td>Intergenic</td><td>NA</td><td>NA</td></tr>
<tr><td>GU305914.1</td><td>183285</td><td>183307</td><td>-</td><td>2</td><td>CACGTACATAAGGTCAATAG</td><td>GGG</td><td>|--|||||[||||||||||||]PAM</td><td>NA</td><td>Intergenic</td><td>NA</td><td>NA</td></tr>
<tr><td>GU305914.1</td><td>183337</td><td>183359</td><td>-</td><td>1</td><td>CCAGTACAAAAGGTCAATAG</td><td>GGG</td><td>||||||||[-|||||||||||]PAM</td><td>NA</td><td>Intergenic</td><td>NA</td><td>NA</td></tr>
<tr><td></td><td></td><td></td><td></td><td></td><td></td><td></td><td></td><td></td><td></td><td></td><td></td></tr>
<tr><td>T20836</td><td>GTCAATAGGGGTGAATCAACAGG</td><td>1000</td><td></td><td></td><td></td><td></td><td></td><td></td><td></td><td></td><td></td></tr>
<tr><td>Oligo fwd</td><td>CACCGTCAATAGGGGTGAATCAAC</td><td></td><td></td><td></td><td></td><td></td><td></td><td></td><td></td><td></td><td></td></tr>
<tr><td>Oligo rev</td><td>AAACGTTGATTCACCCCTATTGAC</td><td></td><td></td><td></td><td></td><td></td><td></td><td></td><td></td><td></td><td></td></tr>
<tr><td>Chromosome</td><td>start</td><td>end</td><td>strand</td><td>MM</td><td>target_seq</td><td>PAM</td><td>alignment</td><td>distance</td><td>position</td><td>gene name</td><td>gene id</td></tr>
<tr><td>GU305914.1</td><td>183417</td><td>183439</td><td>-</td><td>0</td><td>GTCAATAGGGGTGAATCAAC</td><td>AGG</td><td>||||||||[||||||||||||]PAM</td><td>NA</td><td>Intergenic</td><td>NA</td><td>NA</td></tr>
<tr><td>GU305914.1</td><td>183844</td><td>183866</td><td>-</td><td>4</td><td>GTCAATGTGGGTGAGTCAAT</td><td>AGG</td><td>||||||--[||||||-||||-]PAM</td><td>NA</td><td>Intergenic</td><td>NA</td><td>NA</td></tr>
<tr><td></td><td></td><td></td><td></td><td></td><td></td><td></td><td></td><td></td><td></td><td></td><td></td></tr>
<tr><td>T20837</td><td>AATCAACAGGAAAGTCCCATTGG</td><td>1000</td><td></td><td></td><td></td><td></td><td></td><td></td><td></td><td></td><td></td></tr>
<tr><td>Oligo fwd</td><td>CACCAATCAACAGGAAAGTCCCAT</td><td></td><td></td><td></td><td></td><td></td><td></td><td></td><td></td><td></td><td></td></tr>
<tr><td>Oligo rev</td><td>AAACATGGGACTTTCCTGTTGATT</td><td></td><td></td><td></td><td></td><td></td><td></td><td></td><td></td><td></td><td></td></tr>
<tr><td>Chromosome</td><td>start</td><td>end</td><td>strand</td><td>MM</td><td>target_seq</td><td>PAM</td><td>alignment</td><td>distance</td><td>position</td><td>gene name</td><td>gene id</td></tr>
<tr><td>GU305914.1</td><td>183775</td><td>183797</td><td>-</td><td>0</td><td>AATCAACAGGAAAGTCCCAT</td><td>TGG</td><td>||||||||[||||||||||||]PAM</td><td>NA</td><td>Intergenic</td><td>NA</td><td>NA</td></tr>
<tr><td>GU305914.1</td><td>183404</td><td>183426</td><td>-</td><td>0</td><td>AATCAACAGGAAAGTCCCAT</td><td>TGG</td><td>||||||||[||||||||||||]PAM</td><td>NA</td><td>Intergenic</td><td>NA</td><td>NA</td></tr>
<tr><td>GU305914.1</td><td>183589</td><td>183611</td><td>-</td><td>1</td><td>AGTCAACAGGAAAGTCCCAT</td><td>TGG</td><td>|-||||||[||||||||||||]PAM</td><td>NA</td><td>Intergenic</td><td>NA</td><td>NA</td></tr>
<tr><td></td><td></td><td></td><td></td><td></td><td></td><td></td><td></td><td></td><td></td><td></td><td></td></tr>
<tr><td>T20838</td><td>GCCAAGTACACTGAGTCAATAGG</td><td>1000</td><td></td><td></td><td></td><td></td><td></td><td></td><td></td><td></td><td></td></tr>
<tr><td>Oligo fwd</td><td>CACCGCCAAGTACACTGAGTCAAT</td><td></td><td></td><td></td><td></td><td></td><td></td><td></td><td></td><td></td><td></td></tr>
<tr><td>Oligo rev</td><td>AAACATTGACTCAGTGTACTTGGC</td><td></td><td></td><td></td><td></td><td></td><td></td><td></td><td></td><td></td><td></td></tr>
<tr><td>Chromosome</td><td>start</td><td>end</td><td>strand</td><td>MM</td><td>target_seq</td><td>PAM</td><td>alignment</td><td>distance</td><td>position</td><td>gene name</td><td>gene id</td></tr>
<tr><td>GU305914.1</td><td>183380</td><td>183402</td><td>-</td><td>0</td><td>GCCAAGTACACTGAGTCAAT</td><td>AGG</td><td>||||||||[||||||||||||]PAM</td><td>NA</td><td>Intergenic</td><td>NA</td><td>NA</td></tr>
<tr><td>GU305914.1</td><td>183565</td><td>183587</td><td>-</td><td>1</td><td>GCCAAGTACATTGAGTCAAT</td><td>AGG</td><td>||||||||[||-|||||||||]PAM</td><td>NA</td><td>Intergenic</td><td>NA</td><td>NA</td></tr>
<tr><td>GU305914.1</td><td>183751</td><td>183773</td><td>-</td><td>1</td><td>GCCAAGTACACTGCGTCAAT</td><td>AGG</td><td>||||||||[|||||-||||||]PAM</td><td>NA</td><td>Intergenic</td><td>NA</td><td>NA</td></tr>
<tr><td>GU305914.1</td><td>183908</td><td>183930</td><td>-</td><td>1</td><td>GCCAAGTACACTGACTCAAT</td><td>AGG</td><td>||||||||[||||||-|||||]PAM</td><td>NA</td><td>Intergenic</td><td>NA</td><td>NA</td></tr>
<tr><td>GU305914.1</td><td>183658</td><td>183680</td><td>-</td><td>1</td><td>GCCAAGTACACTGACTCAAT</td><td>AGG</td><td>||||||||[||||||-|||||]PAM</td><td>NA</td><td>Intergenic</td><td>NA</td><td>NA</td></tr>
<tr><td>GU305914.1</td><td>184054</td><td>184076</td><td>-</td><td>3</td><td>CCCAAGTACATTGGGTCAAT</td><td>GGG</td><td>-|||||||[||-||-||||||]PAM</td><td>NA</td><td>Intergenic</td><td>NA</td><td>NA</td></tr>
<tr><td>GU305914.1</td><td>183960</td><td>183982</td><td>-</td><td>3</td><td>CCCAAGTACATTGGGTCAAT</td><td>GGG</td><td>-|||||||[||-||-||||||]PAM</td><td>NA</td><td>Intergenic</td><td>NA</td><td>NA</td></tr>
<tr><td>GU305914.1</td><td>183472</td><td>183494</td><td>-</td><td>4</td><td>GGCACGTATACTGAGTCATT</td><td>AGG</td><td>|-||-|||[-|||||||||-|]PAM</td><td>NA</td><td>Intergenic</td><td>NA</td><td>NA</td></tr>
<tr><td>GU305914.1</td><td>184002</td><td>184024</td><td>-</td><td>3</td><td>GGCAAGCACACTGAGTCAAA</td><td>TGG</td><td>|-||||-|[|||||||||||-]PAM</td><td>NA</td><td>Intergenic</td><td>NA</td><td>NA</td></tr>
<tr><td></td><td></td><td></td><td></td><td></td><td></td><td></td><td></td><td></td><td></td><td></td><td></td></tr>
<tr><td>T20839</td><td>CCAAGTACACTGAGTCAATAGGG</td><td>1000</td><td></td><td></td><td></td><td></td><td></td><td></td><td></td><td></td><td></td></tr>
<tr><td>Oligo fwd</td><td>CACCCCAAGTACACTGAGTCAATA</td><td></td><td></td><td></td><td></td><td></td><td></td><td></td><td></td><td></td><td></td></tr>
<tr><td>Oligo rev</td><td>AAACTATTGACTCAGTGTACTTGG</td><td></td><td></td><td></td><td></td><td></td><td></td><td></td><td></td><td></td><td></td></tr>
<tr><td>Chromosome</td><td>start</td><td>end</td><td>strand</td><td>MM</td><td>target_seq</td><td>PAM</td><td>alignment</td><td>distance</td><td>position</td><td>gene name</td><td>gene id</td></tr>
<tr><td>GU305914.1</td><td>183379</td><td>183401</td><td>-</td><td>0</td><td>CCAAGTACACTGAGTCAATA</td><td>GGG</td><td>||||||||[||||||||||||]PAM</td><td>NA</td><td>Intergenic</td><td>NA</td><td>NA</td></tr>
<tr><td>GU305914.1</td><td>183564</td><td>183586</td><td>-</td><td>1</td><td>CCAAGTACATTGAGTCAATA</td><td>GGG</td><td>||||||||[|-||||||||||]PAM</td><td>NA</td><td>Intergenic</td><td>NA</td><td>NA</td></tr>
<tr><td>GU305914.1</td><td>183750</td><td>183772</td><td>-</td><td>1</td><td>CCAAGTACACTGCGTCAATA</td><td>GGG</td><td>||||||||[||||-|||||||]PAM</td><td>NA</td><td>Intergenic</td><td>NA</td><td>NA</td></tr>
<tr><td>GU305914.1</td><td>183907</td><td>183929</td><td>-</td><td>1</td><td>CCAAGTACACTGACTCAATA</td><td>GGG</td><td>||||||||[|||||-||||||]PAM</td><td>NA</td><td>Intergenic</td><td>NA</td><td>NA</td></tr>
<tr><td>GU305914.1</td><td>183657</td><td>183679</td><td>-</td><td>1</td><td>CCAAGTACACTGACTCAATA</td><td>GGG</td><td>||||||||[|||||-||||||]PAM</td><td>NA</td><td>Intergenic</td><td>NA</td><td>NA</td></tr>
<tr><td>GU305914.1</td><td>183471</td><td>183493</td><td>-</td><td>4</td><td>GCACGTATACTGAGTCATTA</td><td>GGG</td><td>-||-|||-[|||||||||-||]PAM</td><td>NA</td><td>Intergenic</td><td>NA</td><td>NA</td></tr>
<tr><td>GU305914.1</td><td>184001</td><td>184023</td><td>-</td><td>4</td><td>GCAAGCACACTGAGTCAAAT</td><td>GGG</td><td>-||||-||[||||||||||--]PAM</td><td>NA</td><td>Intergenic</td><td>NA</td><td>NA</td></tr>
<tr><td></td><td></td><td></td><td></td><td></td><td></td><td></td><td></td><td></td><td></td><td></td><td></td></tr>
<tr><td>T20840</td><td>AGTCAATAGGGACTTTCCATTGG</td><td>1000</td><td></td><td></td><td></td><td></td><td></td><td></td><td></td><td></td><td></td></tr>
<tr><td>Oligo fwd</td><td>CACCAGTCAATAGGGACTTTCCAT</td><td></td><td></td><td></td><td></td><td></td><td></td><td></td><td></td><td></td><td></td></tr>
<tr><td>Oligo rev</td><td>AAACATGGAAAGTCCCTATTGACT</td><td></td><td></td><td></td><td></td><td></td><td></td><td></td><td></td><td></td><td></td></tr>
<tr><td>Chromosome</td><td>start</td><td>end</td><td>strand</td><td>MM</td><td>target_seq</td><td>PAM</td><td>alignment</td><td>distance</td><td>position</td><td>gene name</td><td>gene id</td></tr>
<tr><td>GU305914.1</td><td>183367</td><td>183389</td><td>-</td><td>0</td><td>AGTCAATAGGGACTTTCCAT</td><td>TGG</td><td>||||||||[||||||||||||]PAM</td><td>NA</td><td>Intergenic</td><td>NA</td><td>NA</td></tr>
<tr><td>GU305914.1</td><td>183738</td><td>183760</td><td>-</td><td>1</td><td>CGTCAATAGGGACTTTCCAT</td><td>TGG</td><td>-|||||||[||||||||||||]PAM</td><td>NA</td><td>Intergenic</td><td>NA</td><td>NA</td></tr>
<tr><td>GU305914.1</td><td>183645</td><td>183667</td><td>-</td><td>1</td><td>ACTCAATAGGGACTTTCCAT</td><td>TGG</td><td>|-||||||[||||||||||||]PAM</td><td>NA</td><td>Intergenic</td><td>NA</td><td>NA</td></tr>
<tr><td>GU305914.1</td><td>183552</td><td>183574</td><td>-</td><td>1</td><td>AGTCAATAGGGACTTTCCAA</td><td>TGG</td><td>||||||||[|||||||||||-]PAM</td><td>NA</td><td>Intergenic</td><td>NA</td><td>NA</td></tr>
<tr><td>GU305914.1</td><td>183895</td><td>183917</td><td>-</td><td>2</td><td>ACTCAATAGGGACTTTCCAA</td><td>TGG</td><td>|-||||||[|||||||||||-]PAM</td><td>NA</td><td>Intergenic</td><td>NA</td><td>NA</td></tr>
<tr><td>GU305914.1</td><td>183459</td><td>183481</td><td>-</td><td>2</td><td>AGTCATTAGGGACTTTCCAA</td><td>TGG</td><td>|||||-||[|||||||||||-]PAM</td><td>NA</td><td>Intergenic</td><td>NA</td><td>NA</td></tr>
<tr><td>GU305914.1</td><td>183989</td><td>184011</td><td>-</td><td>3</td><td>AGTCAAATGGGACTTTCCAC</td><td>TGG</td><td>||||||--[|||||||||||-]PAM</td><td>NA</td><td>Intergenic</td><td>NA</td><td>NA</td></tr>
<tr><td></td><td></td><td></td><td></td><td></td><td></td><td></td><td></td><td></td><td></td><td></td><td></td></tr>
<tr><td>T20841</td><td>GTCAATAGGGACTTTCCATTGGG</td><td>1000</td><td></td><td></td><td></td><td></td><td></td><td></td><td></td><td></td><td></td></tr>
<tr><td>Oligo fwd</td><td>CACCGTCAATAGGGACTTTCCATT</td><td></td><td></td><td></td><td></td><td></td><td></td><td></td><td></td><td></td><td></td></tr>
</table>

| | | | | | | | | | | | |
|---|---|---|---|---|---|---|---|---|---|---|---|
| Oligo rev | AAACAATGGAAAGTCCCTATTGAC | | | | | | | | | | |
| Chromosome | start | end | strand | MM | target_seq | PAM | alignment | distance | position | gene name | gene id |
| GU305914.1 | 183737 | 183759 | - | 0 | GTCAATAGGGACTTTCCATT | GGG | \|\|\|\|\|\|\|\|[\|\|\|\|\|\|\|\|\|\|\|\|]PAM | NA | Intergenic | NA | NA |
| GU305914.1 | 183366 | 183388 | - | 0 | GTCAATAGGGACTTTCCATT | GGG | \|\|\|\|\|\|\|\|[\|\|\|\|\|\|\|\|\|\|\|\|]PAM | NA | Intergenic | NA | NA |
| GU305914.1 | 183644 | 183666 | - | 1 | CTCAATAGGGACTTTCCATT | GGG | -\|\|\|\|\|\|\|[\|\|\|\|\|\|\|\|\|\|\|\|]PAM | NA | Intergenic | NA | NA |
| GU305914.1 | 183551 | 183573 | - | 1 | GTCAATAGGGACTTTCCAAT | GGG | \|\|\|\|\|\|\|\|[\|\|\|\|\|\|\|\|\|\|-\|]PAM | NA | Intergenic | NA | NA |
| GU305914.1 | 183894 | 183916 | - | 2 | CTCAATAGGGACTTTCCAAT | GGG | -\|\|\|\|\|\|\|[\|\|\|\|\|\|\|\|\|\|-\|]PAM | NA | Intergenic | NA | NA |
| GU305914.1 | 183458 | 183480 | - | 2 | GTCATTAGGGACTTTCCAAT | GGG | \|\|\|\|-\|\|\|[\|\|\|\|\|\|\|\|\|\|-\|]PAM | NA | Intergenic | NA | NA |
| GU305914.1 | 183988 | 184010 | - | 3 | GTCAAATGGGACTTTCCACT | GGG | \|\|\|\|\|--\|[\|\|\|\|\|\|\|\|\|\|-\|]PAM | NA | Intergenic | NA | NA |
| T20842 | TGGGTTTTGCCCAGTACAAAAGG | 1000 | | | | | | | | | |
| Oligo fwd | CACCTGGGTTTTGCCCAGTACAAA | | | | | | | | | | |
| Oligo rev | AAACTTTGTACTGGGCAAAACCCA | | | | | | | | | | |
| Chromosome | start | end | strand | MM | target_seq | PAM | alignment | distance | position | gene name | gene id |
| GU305914.1 | 183347 | 183369 | - | 0 | TGGGTTTTGCCCAGTACAAA | AGG | \|\|\|\|\|\|\|\|[\|\|\|\|\|\|\|\|\|\|\|\|]PAM | NA | Intergenic | NA | NA |
| GU305914.1 | 183718 | 183740 | - | 1 | TGGGTTTTGCCCAGTACATA | AGG | \|\|\|\|\|\|\|\|[\|\|\|\|\|\|\|\|\|\|-\|]PAM | NA | Intergenic | NA | NA |
| GU305914.1 | 183532 | 183554 | - | 1 | TGGGTTTTGCCCAGTACATA | AGG | \|\|\|\|\|\|\|\|[\|\|\|\|\|\|\|\|\|\|-\|]PAM | NA | Intergenic | NA | NA |
| GU305914.1 | 183439 | 183461 | - | 1 | TGGGTTTTGCCCAGTACATA | AGG | \|\|\|\|\|\|\|\|[\|\|\|\|\|\|\|\|\|\|-\|]PAM | NA | Intergenic | NA | NA |
| GU305914.1 | 183625 | 183647 | - | 1 | TGGGTTTTGCCCAGTACATA | AGG | \|\|\|\|\|\|\|\|[\|\|\|\|\|\|\|\|\|\|-\|]PAM | NA | Intergenic | NA | NA |
| GU305914.1 | 183811 | 183833 | - | 4 | TGTATTCTGCCCAGTACATA | AGG | \|\|--\|\|-\|[\|\|\|\|\|\|\|\|\|\|-\|]PAM | NA | Intergenic | NA | NA |
| T20843 | GCCCAGTACAAAAGGTCAATAGG | 1000 | | | | | | | | | |
| Oligo fwd | CACCGCCCAGTACAAAAGGTCAAT | | | | | | | | | | |
| Oligo rev | AAACATTGACCTTTTGTACTGGGC | | | | | | | | | | |
| Chromosome | start | end | strand | MM | target_seq | PAM | alignment | distance | position | gene name | gene id |
| GU305914.1 | 183339 | 183361 | - | 0 | GCCCAGTACAAAAGGTCAAT | AGG | \|\|\|\|\|\|\|\|[\|\|\|\|\|\|\|\|\|\|\|\|]PAM | NA | Intergenic | NA | NA |
| GU305914.1 | 183524 | 183546 | - | 1 | GCCCAGTACATAAGGTCAAT | GGG | \|\|\|\|\|\|\|\|[\|\|-\|\|\|\|\|\|\|\|\|]PAM | NA | Intergenic | NA | NA |
| GU305914.1 | 183617 | 183639 | - | 1 | GCCCAGTACATAAGGTCAAT | AGG | \|\|\|\|\|\|\|\|[\|\|-\|\|\|\|\|\|\|\|\|]PAM | NA | Intergenic | NA | NA |
| GU305914.1 | 183803 | 183825 | - | 1 | GCCCAGTACATAAGGTCAAT | AGG | \|\|\|\|\|\|\|\|[\|\|-\|\|\|\|\|\|\|\|\|]PAM | NA | Intergenic | NA | NA |
| GU305914.1 | 183710 | 183732 | - | 1 | GCCCAGTACATAAGGTCAAT | AGG | \|\|\|\|\|\|\|\|[\|\|-\|\|\|\|\|\|\|\|\|]PAM | NA | Intergenic | NA | NA |
| GU305914.1 | 183431 | 183453 | - | 1 | GCCCAGTACATAAGGTCAAT | AGG | \|\|\|\|\|\|\|\|[\|\|-\|\|\|\|\|\|\|\|\|]PAM | NA | Intergenic | NA | NA |
| GU305914.1 | 183287 | 183309 | - | 4 | GGCACGTACATAAGGTCAAT | AGG | \|-\|--\|\|\|[\|\|-\|\|\|\|\|\|\|\|\|]PAM | NA | Intergenic | NA | NA |
| T20844 | CCCAGTACAAAAGGTCAATAGGG | 1000 | | | | | | | | | |
| Oligo fwd | CACCCCCAGTACAAAAGGTCAATA | | | | | | | | | | |
| Oligo rev | AAACTATTGACCTTTTGTACTGGG | | | | | | | | | | |
| Chromosome | start | end | strand | MM | target_seq | PAM | alignment | distance | position | gene name | gene id |
| GU305914.1 | 183338 | 183360 | - | 0 | CCCAGTACAAAAGGTCAATA | GGG | \|\|\|\|\|\|\|\|[\|\|\|\|\|\|\|\|\|\|\|\|]PAM | NA | Intergenic | NA | NA |
| GU305914.1 | 183802 | 183824 | - | 1 | CCCAGTACATAAGGTCAATA | GGG | \|\|\|\|\|\|\|\|[\|-\|\|\|\|\|\|\|\|\|\|]PAM | NA | Intergenic | NA | NA |
| GU305914.1 | 183709 | 183731 | - | 1 | CCCAGTACATAAGGTCAATA | GGG | \|\|\|\|\|\|\|\|[\|-\|\|\|\|\|\|\|\|\|\|]PAM | NA | Intergenic | NA | NA |
| GU305914.1 | 183430 | 183452 | - | 1 | CCCAGTACATAAGGTCAATA | GGG | \|\|\|\|\|\|\|\|[\|-\|\|\|\|\|\|\|\|\|\|]PAM | NA | Intergenic | NA | NA |
| GU305914.1 | 183616 | 183638 | - | 1 | CCCAGTACATAAGGTCAATA | GGG | \|\|\|\|\|\|\|\|[\|-\|\|\|\|\|\|\|\|\|\|]PAM | NA | Intergenic | NA | NA |
| GU305914.1 | 183286 | 183308 | - | 4 | GCACGTACATAAGGTCAATA | GGG | -\|--\|\|\|\|[\|-\|\|\|\|\|\|\|\|\|\|]PAM | NA | Intergenic | NA | NA |
| T20845 | CCAGTACAAAAGGTCAATAGGGG | 1000 | | | | | | | | | |
| Oligo fwd | CACCCCAGTACAAAAGGTCAATAG | | | | | | | | | | |
| Oligo rev | AAACCTATTGACCTTTTGTACTGG | | | | | | | | | | |
| Chromosome | start | end | strand | MM | target_seq | PAM | alignment | distance | position | gene name | gene id |
| GU305914.1 | 183337 | 183359 | - | 0 | CCAGTACAAAAGGTCAATAG | GGG | \|\|\|\|\|\|\|\|[\|\|\|\|\|\|\|\|\|\|\|\|]PAM | NA | Intergenic | NA | NA |
| GU305914.1 | 183801 | 183823 | - | 1 | CCAGTACATAAGGTCAATAG | GGG | \|\|\|\|\|\|\|\|[-\|\|\|\|\|\|\|\|\|\|\|]PAM | NA | Intergenic | NA | NA |
| GU305914.1 | 183708 | 183730 | - | 1 | CCAGTACATAAGGTCAATAG | GGG | \|\|\|\|\|\|\|\|[-\|\|\|\|\|\|\|\|\|\|\|]PAM | NA | Intergenic | NA | NA |
| GU305914.1 | 183429 | 183451 | - | 1 | CCAGTACATAAGGTCAATAG | GGG | \|\|\|\|\|\|\|\|[-\|\|\|\|\|\|\|\|\|\|\|]PAM | NA | Intergenic | NA | NA |
| GU305914.1 | 183615 | 183637 | - | 1 | CCAGTACATAAGGTCAATAG | GGG | \|\|\|\|\|\|\|\|[-\|\|\|\|\|\|\|\|\|\|\|]PAM | NA | Intergenic | NA | NA |
| GU305914.1 | 183285 | 183307 | - | 3 | CACGTACATAAGGTCAATAG | GGG | \|--\|\|\|\|\|[-\|\|\|\|\|\|\|\|\|\|\|]PAM | NA | Intergenic | NA | NA |
| T20846 | CAGTACAAAAGGTCAATAGGGGG | 1000 | | | | | | | | | |
| Oligo fwd | CACCCAGTACAAAAGGTCAATAGG | | | | | | | | | | |
| Oligo rev | AAACCCTATTGACCTTTTGTACTG | | | | | | | | | | |
| Chromosome | start | end | strand | MM | target_seq | PAM | alignment | distance | position | gene name | gene id |
| GU305914.1 | 183336 | 183358 | - | 0 | CAGTACAAAAGGTCAATAGG | GGG | \|\|\|\|\|\|\|\|[\|\|\|\|\|\|\|\|\|\|\|\|]PAM | NA | Intergenic | NA | NA |
| GU305914.1 | 183614 | 183636 | - | 1 | CAGTACATAAGGTCAATAGG | GGG | \|\|\|\|\|\|\|-[\|\|\|\|\|\|\|\|\|\|\|\|]PAM | NA | Intergenic | NA | NA |
| GU305914.1 | 183800 | 183822 | - | 1 | CAGTACATAAGGTCAATAGG | GGG | \|\|\|\|\|\|\|-[\|\|\|\|\|\|\|\|\|\|\|\|]PAM | NA | Intergenic | NA | NA |
| GU305914.1 | 183521 | 183543 | - | 2 | CAGTACATAAGGTCAATGGG | AGG | \|\|\|\|\|\|\|-[\|\|\|\|\|\|\|\|\|-\|\|]PAM | NA | Intergenic | NA | NA |
| T20847 | GTCAATAGGGGGTGAGTCAATGG | 1000 | | | | | | | | | |
| Oligo fwd | CACCGTCAATAGGGGGTGAGTCAA | | | | | | | | | | |
| Oligo rev | AAACTTGACTCACCCCCTATTGAC | | | | | | | | | | |
| Chromosome | start | end | strand | MM | target_seq | PAM | alignment | distance | position | gene name | gene id |
| GU305914.1 | 183325 | 183347 | - | 0 | GTCAATAGGGGGTGAGTCAA | TGG | \|\|\|\|\|\|\|\|[\|\|\|\|\|\|\|\|\|\|\|\|]PAM | NA | Intergenic | NA | NA |
| GU305914.1 | 183696 | 183718 | - | 1 | GTCAATAGGGGATGAGTCAA | TGG | \|\|\|\|\|\|\|\|[\|\|\|-\|\|\|\|\|\|\|\|]PAM | NA | Intergenic | NA | NA |
| GU305914.1 | 183946 | 183968 | - | 3 | GTCAATGGGAGGTGAGCCAA | TGG | \|\|\|\|\|\|-\|[\|-\|\|\|\|\|-\|\|\|]PAM | NA | Intergenic | NA | NA |
| T20848 | TCAATAGGGGGTGAGTCAATGGG | 1000 | | | | | | | | | |
| Oligo fwd | CACCTCAATAGGGGGTGAGTCAAT | | | | | | | | | | |
| Oligo rev | AAACATTGACTCACCCCCTATTGA | | | | | | | | | | |
| Chromosome | start | end | strand | MM | target_seq | PAM | alignment | distance | position | gene name | gene id |
| GU305914.1 | 183324 | 183346 | - | 0 | TCAATAGGGGGTGAGTCAAT | GGG | \|\|\|\|\|\|\|\|[\|\|\|\|\|\|\|\|\|\|\|\|]PAM | NA | Intergenic | NA | NA |
| GU305914.1 | 183695 | 183717 | - | 1 | TCAATAGGGGATGAGTCAAT | GGG | \|\|\|\|\|\|\|\|[\|\|-\|\|\|\|\|\|\|\|\|]PAM | NA | Intergenic | NA | NA |
| GU305914.1 | 183945 | 183967 | - | 3 | TCAATGGGAGGTGAGCCAAT | GGG | \|\|\|\|\|-\|\|[-\|\|\|\|\|-\|\|\|\|]PAM | NA | Intergenic | NA | NA |
| GU305914.1 | 183602 | 183624 | - | 1 | TCAATAGGGGGTGAGTCAAC | AGG | \|\|\|\|\|\|\|\|[\|\|\|\|\|\|\|\|\|\|\|-]PAM | NA | Intergenic | NA | NA |
| GU305914.1 | 183788 | 183810 | - | 2 | TCAATAGGGGGTGAATCAAC | AGG | \|\|\|\|\|\|\|\|[\|\|\|\|\|\|-\|\|\|\|-]PAM | NA | Intergenic | NA | NA |
| T20849 | CAATGGGTTTTTCCCATTATTGG | 1000 | | | | | | | | | |
| Oligo fwd | CACCCAATGGGTTTTTCCCATTAT | | | | | | | | | | |
| Oligo rev | AAACATAATGGGAAAAACCCATTG | | | | | | | | | | |
| Chromosome | start | end | strand | MM | target_seq | PAM | alignment | distance | position | gene name | gene id |
| GU305914.1 | 183308 | 183330 | - | 0 | CAATGGGTTTTTCCCATTAT | TGG | \|\|\|\|\|\|\|\|[\|\|\|\|\|\|\|\|\|\|\|\|]PAM | NA | Intergenic | NA | NA |
| GU305914.1 | 184023 | 184045 | - | 1 | CAATGGGTTTTTCCCATTAC | TGG | \|\|\|\|\|\|\|\|[\|\|\|\|\|\|\|\|\|\|\|-]PAM | NA | Intergenic | NA | NA |
| GU305914.1 | 183493 | 183515 | - | 1 | CAATGGGTTTTTCCCATTAC | TGG | \|\|\|\|\|\|\|\|[\|\|\|\|\|\|\|\|\|\|\|-]PAM | NA | Intergenic | NA | NA |

| | | | | | | | | | | | |
|---|---|---|---|---|---|---|---|---|---|---|---|
| T20850 | CCATTATTGGCACGTACATAAGG | 1000 | | | | | | | | | |
| Oligo fwd | CACCCCATTATTGGCACGTACATA | | | | | | | | | | |
| Oligo rev | AAACTATGTACGTGCCAATAATGG | | | | | | | | | | |
| Chromosome | start | end | strand | MM | target_seq | PAM | alignment | distance | position | gene name | gene id |
| GU305914.1 | 183295 | 183317 | - | 0 | CCATTATTGGCACGTACATA | AGG | \|\|\|\|\|\|\|\|[\|\|\|\|\|\|\|\|\|\|\|\|]PAM | NA | Intergenic | NA | NA |

| | | | | | | | | | | | |
|---|---|---|---|---|---|---|---|---|---|---|---|
| T20851 | GGCACGTACATAAGGTCAATAGG | 1000 | | | | | | | | | |
| Oligo fwd | CACCGGCACGTACATAAGGTCAAT | | | | | | | | | | |
| Oligo rev | AAACATTGACCTTATGTACGTGCC | | | | | | | | | | |
| Chromosome | start | end | strand | MM | target_seq | PAM | alignment | distance | position | gene name | gene id |
| GU305914.1 | 183287 | 183309 | - | 0 | GGCACGTACATAAGGTCAAT | AGG | \|\|\|\|\|\|\|\|[\|\|\|\|\|\|\|\|\|\|\|\|]PAM | NA | Intergenic | NA | NA |
| GU305914.1 | 183617 | 183639 | - | 3 | GCCCAGTACATAAGGTCAAT | AGG | \|-\|--\|\|\|[\|\|\|\|\|\|\|\|\|\|\|\|]PAM | NA | Intergenic | NA | NA |
| GU305914.1 | 183803 | 183825 | - | 3 | GCCCAGTACATAAGGTCAAT | AGG | \|-\|--\|\|\|[\|\|\|\|\|\|\|\|\|\|\|\|]PAM | NA | Intergenic | NA | NA |
| GU305914.1 | 183710 | 183732 | - | 3 | GCCCAGTACATAAGGTCAAT | AGG | \|-\|--\|\|\|[\|\|\|\|\|\|\|\|\|\|\|\|]PAM | NA | Intergenic | NA | NA |
| GU305914.1 | 183431 | 183453 | - | 3 | GCCCAGTACATAAGGTCAAT | AGG | \|-\|--\|\|\|[\|\|\|\|\|\|\|\|\|\|\|\|]PAM | NA | Intergenic | NA | NA |
| GU305914.1 | 183524 | 183546 | - | 3 | GCCCAGTACATAAGGTCAAT | GGG | \|-\|--\|\|\|[\|\|\|\|\|\|\|\|\|\|\|\|]PAM | NA | Intergenic | NA | NA |
| GU305914.1 | 183339 | 183361 | - | 4 | GCCCAGTACAAAAGGTCAAT | AGG | \|-\|--\|\|\|[\|\|-\|\|\|\|\|\|\|\|\|]PAM | NA | Intergenic | NA | NA |

| | | | | | | | | | | | |
|---|---|---|---|---|---|---|---|---|---|---|---|
| T20852 | GCACGTACATAAGGTCAATAGGG | 1000 | | | | | | | | | |
| Oligo fwd | CACCGCACGTACATAAGGTCAATA | | | | | | | | | | |
| Oligo rev | AAACTATTGACCTTATGTACGTGC | | | | | | | | | | |
| Chromosome | start | end | strand | MM | target_seq | PAM | alignment | distance | position | gene name | gene id |
| GU305914.1 | 183286 | 183308 | - | 0 | GCACGTACATAAGGTCAATA | GGG | \|\|\|\|\|\|\|\|[\|\|\|\|\|\|\|\|\|\|\|\|]PAM | NA | Intergenic | NA | NA |
| GU305914.1 | 183616 | 183638 | - | 3 | CCCAGTACATAAGGTCAATA | GGG | -\|--\|\|\|\|[\|\|\|\|\|\|\|\|\|\|\|\|]PAM | NA | Intergenic | NA | NA |
| GU305914.1 | 183802 | 183824 | - | 3 | CCCAGTACATAAGGTCAATA | GGG | -\|--\|\|\|\|[\|\|\|\|\|\|\|\|\|\|\|\|]PAM | NA | Intergenic | NA | NA |
| GU305914.1 | 183709 | 183731 | - | 3 | CCCAGTACATAAGGTCAATA | GGG | -\|--\|\|\|\|[\|\|\|\|\|\|\|\|\|\|\|\|]PAM | NA | Intergenic | NA | NA |
| GU305914.1 | 183430 | 183452 | - | 3 | CCCAGTACATAAGGTCAATA | GGG | -\|--\|\|\|\|[\|\|\|\|\|\|\|\|\|\|\|\|]PAM | NA | Intergenic | NA | NA |
| GU305914.1 | 183338 | 183360 | - | 4 | CCCAGTACAAAAGGTCAATA | GGG | -\|--\|\|\|\|[\|-\|\|\|\|\|\|\|\|\|\|]PAM | NA | Intergenic | NA | NA |
| GU305914.1 | 183857 | 183879 | - | 4 | GCAAGCATATAAGGTCAATG | TGG | \|\|\|-\|-\|-[\|\|\|\|\|\|\|\|\|\|\|-]PAM | NA | Intergenic | NA | NA |

| | | | | | | | | | | | |
|---|---|---|---|---|---|---|---|---|---|---|---|
| T20853 | CACGTACATAAGGTCAATAGGGG | 1000 | | | | | | | | | |
| Oligo fwd | CACCCACGTACATAAGGTCAATAG | | | | | | | | | | |
| Oligo rev | AAACCTATTGACCTTATGTACGTG | | | | | | | | | | |
| Chromosome | start | end | strand | MM | target_seq | PAM | alignment | distance | position | gene name | gene id |
| GU305914.1 | 183285 | 183307 | - | 0 | CACGTACATAAGGTCAATAG | GGG | \|\|\|\|\|\|\|\|[\|\|\|\|\|\|\|\|\|\|\|\|]PAM | NA | Intergenic | NA | NA |
| GU305914.1 | 183708 | 183730 | - | 2 | CCAGTACATAAGGTCAATAG | GGG | \|--\|\|\|\|\|[\|\|\|\|\|\|\|\|\|\|\|\|]PAM | NA | Intergenic | NA | NA |
| GU305914.1 | 183429 | 183451 | - | 2 | CCAGTACATAAGGTCAATAG | GGG | \|--\|\|\|\|\|[\|\|\|\|\|\|\|\|\|\|\|\|]PAM | NA | Intergenic | NA | NA |
| GU305914.1 | 183615 | 183637 | - | 2 | CCAGTACATAAGGTCAATAG | GGG | \|--\|\|\|\|\|[\|\|\|\|\|\|\|\|\|\|\|\|]PAM | NA | Intergenic | NA | NA |
| GU305914.1 | 183801 | 183823 | - | 2 | CCAGTACATAAGGTCAATAG | GGG | \|--\|\|\|\|\|[\|\|\|\|\|\|\|\|\|\|\|\|]PAM | NA | Intergenic | NA | NA |
| GU305914.1 | 183337 | 183359 | - | 3 | CCAGTACAAAAGGTCAATAG | GGG | \|--\|\|\|\|\|[-\|\|\|\|\|\|\|\|\|\|\|]PAM | NA | Intergenic | NA | NA |

| | | | | | | | | | | | |
|---|---|---|---|---|---|---|---|---|---|---|---|
| T20854 | GGTCAATAGGGGTGAGTCATTGG | 1000 | | | | | | | | | |
| Oligo fwd | CACCGGTCAATAGGGGTGAGTCAT | | | | | | | | | | |
| Oligo rev | AAACATGACTCACCCCTATTGACC | | | | | | | | | | |
| Chromosome | start | end | strand | MM | target_seq | PAM | alignment | distance | position | gene name | gene id |
| GU305914.1 | 183274 | 183296 | - | 0 | GGTCAATAGGGGTGAGTCAT | TGG | \|\|\|\|\|\|\|\|[\|\|\|\|\|\|\|\|\|\|\|\|]PAM | NA | Intergenic | NA | NA |

| | | | | | | | | | | | |
|---|---|---|---|---|---|---|---|---|---|---|---|
| T20855 | GTCAATAGGGGTGAGTCATTGGG | 1000 | | | | | | | | | |
| Oligo fwd | CACCGTCAATAGGGGTGAGTCATT | | | | | | | | | | |
| Oligo rev | AAACAATGACTCACCCCTATTGAC | | | | | | | | | | |
| Chromosome | start | end | strand | MM | target_seq | PAM | alignment | distance | position | gene name | gene id |
| GU305914.1 | 183273 | 183295 | - | 0 | GTCAATAGGGGTGAGTCATT | GGG | \|\|\|\|\|\|\|\|[\|\|\|\|\|\|\|\|\|\|\|\|]PAM | NA | Intergenic | NA | NA |
| GU305914.1 | 183844 | 183866 | - | 3 | GTCAATGTGGGTGAGTCAAT | AGG | \|\|\|\|\|\|--[\|\|\|\|\|\|\|\|\|\|-\|]PAM | NA | Intergenic | NA | NA |

| | | | | | | | | | | | |
|---|---|---|---|---|---|---|---|---|---|---|---|
| T20856 | TTTCCCACCATTGACGTCAATGG | 1000 | | | | | | | | | |
| Oligo fwd | CACCTTTCCCACCATTGACGTCAA | | | | | | | | | | |
| Oligo rev | AAACTTGACGTCAATGGTGGGAAA | | | | | | | | | | |
| Chromosome | start | end | strand | MM | target_seq | PAM | alignment | distance | position | gene name | gene id |
| GU305914.1 | 183216 | 183238 | - | 0 | TTTCCCACCATTGACGTCAA | TGG | \|\|\|\|\|\|\|\|[\|\|\|\|\|\|\|\|\|\|\|\|]PAM | NA | Intergenic | NA | NA |

| | | | | | | | | | | | |
|---|---|---|---|---|---|---|---|---|---|---|---|
| T20857 | TTCCCACCATTGACGTCAATGGG | 1000 | | | | | | | | | |
| Oligo fwd | CACCTTCCCACCATTGACGTCAAT | | | | | | | | | | |
| Oligo rev | AAACATTGACGTCAATGGTGGGAA | | | | | | | | | | |
| Chromosome | start | end | strand | MM | target_seq | PAM | alignment | distance | position | gene name | gene id |
| GU305914.1 | 183215 | 183237 | - | 0 | TTCCCACCATTGACGTCAAT | GGG | \|\|\|\|\|\|\|\|[\|\|\|\|\|\|\|\|\|\|\|\|]PAM | NA | Intergenic | NA | NA |

| | | | | | | | | | | | |
|---|---|---|---|---|---|---|---|---|---|---|---|
| T20858 | AATGCAACGTGACCTTTAAACGG | 1000 | | | | | | | | | |
| Oligo fwd | CACCAATGCAACGTGACCTTTAAA | | | | | | | | | | |
| Oligo rev | AAACTTTAAAGGTCACGTTGCATT | | | | | | | | | | |
| Chromosome | start | end | strand | MM | target_seq | PAM | alignment | distance | position | gene name | gene id |
| GU305914.1 | 183181 | 183203 | - | 0 | AATGCAACGTGACCTTTAAA | CGG | \|\|\|\|\|\|\|\|[\|\|\|\|\|\|\|\|\|\|\|\|]PAM | NA | Intergenic | NA | NA |

| | | | | | | | | | | | |
|---|---|---|---|---|---|---|---|---|---|---|---|
| T20859 | ACTTTCCCATAGCTGATTAATGG | 1000 | | | | | | | | | |
| Oligo fwd | CACCACTTTCCCATAGCTGATTAA | | | | | | | | | | |
| Oligo rev | AAACTTAATCAGCTATGGGAAAGT | | | | | | | | | | |
| Chromosome | start | end | strand | MM | target_seq | PAM | alignment | distance | position | gene name | gene id |
| GU305914.1 | 183157 | 183179 | - | 0 | ACTTTCCCATAGCTGATTAA | TGG | \|\|\|\|\|\|\|\|[\|\|\|\|\|\|\|\|\|\|\|\|]PAM | NA | Intergenic | NA | NA |

| | | | | | | | | | | | |
|---|---|---|---|---|---|---|---|---|---|---|---|
| T20860 | CTTTCCCATAGCTGATTAATGGG | 1000 | | | | | | | | | |
| Oligo fwd | CACCCTTTCCCATAGCTGATTAAT | | | | | | | | | | |
| Oligo rev | AAACATTAATCAGCTATGGGAAAG | | | | | | | | | | |
| Chromosome | start | end | strand | MM | target_seq | PAM | alignment | distance | position | gene name | gene id |
| GU305914.1 | 183156 | 183178 | - | 0 | CTTTCCCATAGCTGATTAAT | GGG | \|\|\|\|\|\|\|\|[\|\|\|\|\|\|\|\|\|\|\|\|]PAM | NA | Intergenic | NA | NA |

| | | | | | | | | | | | |
|---|---|---|---|---|---|---|---|---|---|---|---|
| T20861 | CTCGAGCCAATACACGTCAATGG | 1000 | | | | | | | | | |
| Oligo fwd | CACCCTCGAGCCAATACACGTCAA | | | | | | | | | | |
| Oligo rev | AAACTTGACGTGTATTGGCTCGAG | | | | | | | | | | |
| Chromosome | start | end | strand | MM | target_seq | PAM | alignment | distance | position | gene name | gene id |

GU305914.1 183122 183144 - 0 CTCGAGCCAATACACGTCAA TGG ||||||||[||||||||||||]PAM NA Intergenic NA NA

T20862 TCGAGCCAATACACGTCAATGGG 1000
Oligo fwd CACCTCGAGCCAATACACGTCAAT
Oligo rev AAACATTGACGTGTATTGGCTCGA
Chromosome start end strand MM target_seq PAM alignment distance position gene name gene id
GU305914.1 183121 183143 - 0 TCGAGCCAATACACGTCAAT GGG ||||||||[||||||||||||]PAM NA Intergenic NA NA

T20863 ACACGTCAATGGGAAGTGAAAGG 1000
Oligo fwd CACCACACGTCAATGGGAAGTGAA
Oligo rev AAACTTCACTTCCCATTGACGTGT
Chromosome start end strand MM target_seq PAM alignment distance position gene name gene id
GU305914.1 183111 183133 - 0 ACACGTCAATGGGAAGTGAA AGG ||||||||[||||||||||||]PAM NA Intergenic NA NA

T20864 CACGTCAATGGGAAGTGAAAGGG 1000
Oligo fwd CACCCACGTCAATGGGAAGTGAAA
Oligo rev AAACTTTCACTTCCCATTGACGTG
Chromosome start end strand MM target_seq PAM alignment distance position gene name gene id
GU305914.1 183110 183132 - 0 CACGTCAATGGGAAGTGAAA GGG ||||||||[||||||||||||]PAM NA Intergenic NA NA

T20865 GCCAAAACGTAACACCGCCCCGG 1000
Oligo fwd CACCGCCAAAACGTAACACCGCCC
Oligo rev AAACGGGCGGTGTTACGTTTTGGC
Chromosome start end strand MM target_seq PAM alignment distance position gene name gene id
GU305914.1 183085 183107 - 0 GCCAAAACGTAACACCGCCC CGG ||||||||[||||||||||||]PAM NA Intergenic NA NA

T20866 ACACCGCCCCGGTTTTCCCCTGG 1000
Oligo fwd CACCACACCGCCCCGGTTTTCCCC
Oligo rev AAACGGGGAAAACCGGGGCGGTGT
Chromosome start end strand MM target_seq PAM alignment distance position gene name gene id
GU305914.1 183074 183096 - 0 ACACCGCCCCGGTTTTCCCC TGG ||||||||[||||||||||||]PAM NA Intergenic NA NA

T20867 TTCCCCTGGAAATTCCATATTGG 1000
Oligo fwd CACCTTCCCCTGGAAATTCCATAT
Oligo rev AAACATATGGAATTTCCAGGGGAA
Chromosome start end strand MM target_seq PAM alignment distance position gene name gene id
GU305914.1 183060 183082 - 0 TTCCCCTGGAAATTCCATAT TGG ||||||||[||||||||||||]PAM NA Intergenic NA NA

T20868 CATATTGGCACGCATTCTATTGG 1000
Oligo fwd CACCCATATTGGCACGCATTCTAT
Oligo rev AAACATAGAATGCGTGCCAATATG
Chromosome start end strand MM target_seq PAM alignment distance position gene name gene id
GU305914.1 183045 183067 - 0 CATATTGGCACGCATTCTAT TGG ||||||||[||||||||||||]PAM NA Intergenic NA NA

T20869 TGGCTGAGCTGCGTTCTACGTGG 1000
Oligo fwd CACCTGGCTGAGCTGCGTTCTACG
Oligo rev AAACCGTAGAACGCAGCTCAGCCA
Chromosome start end strand MM target_seq PAM alignment distance position gene name gene id
GU305914.1 183025 183047 - 0 TGGCTGAGCTGCGTTCTACG TGG ||||||||[||||||||||||]PAM NA Intergenic NA NA

T20870 GGCTGAGCTGCGTTCTACGTGGG 1000
Oligo fwd CACCGGCTGAGCTGCGTTCTACGT
Oligo rev AAACACGTAGAACGCAGCTCAGCC
Chromosome start end strand MM target_seq PAM alignment distance position gene name gene id
GU305914.1 183024 183046 - 0 GGCTGAGCTGCGTTCTACGT GGG ||||||||[||||||||||||]PAM NA Intergenic NA NA

T20871 GCGTTCTACGTGGGTATAAGAGG 1000
Oligo fwd CACCGCGTTCTACGTGGGTATAAG
Oligo rev AAACCTTATACCCACGTAGAACGC
Chromosome start end strand MM target_seq PAM alignment distance position gene name gene id
GU305914.1 183015 183037 - 0 GCGTTCTACGTGGGTATAAG AGG ||||||||[||||||||||||]PAM NA Intergenic NA NA

T20872 ATAAGAGGCGCGACCAGCGTCGG 1000
Oligo fwd CACCATAAGAGGCGCGACCAGCGT
Oligo rev AAACACGCTGGTCGCGCCTCTTAT
Chromosome start end strand MM target_seq PAM alignment distance position gene name gene id
GU305914.1 183000 183022 - 0 ATAAGAGGCGCGACCAGCGT CGG ||||||||[||||||||||||]PAM NA Intergenic NA NA

T20873 GTCGGTACCGTCGCAGTCTTCGG 1000
Oligo fwd CACCGTCGGTACCGTCGCAGTCTT
Oligo rev AAACAAGACTGCGACGGTACCGAC
Chromosome start end strand MM target_seq PAM alignment distance position gene name gene id
GU305914.1 182982 183004 - 0 GTCGGTACCGTCGCAGTCTT CGG ||||||||[||||||||||||]PAM NA Intergenic NA NA

T20874 CTCGCTGCAGTATCCTGCCCAGG 1000
Oligo fwd CACCCTCGCTGCAGTATCCTGCCC
Oligo rev AAACGGGCAGGATACTGCAGCGAG
Chromosome start end strand MM target_seq PAM alignment distance position gene name gene id
GU305914.1 182933 182955 - 0 CTCGCTGCAGTATCCTGCCC AGG ||||||||[||||||||||||]PAM NA Intergenic NA NA

T20875 CAGGAAAGAGCGACCTTCCCCGG 1000
Oligo fwd CACCCAGGAAAGAGCGACCTTCCC
Oligo rev AAACGGGAAGGTCGCTCTTTCCTG
Chromosome start end strand MM target_seq PAM alignment distance position gene name gene id
GU305914.1 182914 182936 - 0 CAGGAAAGAGCGACCTTCCC CGG ||||||||[||||||||||||]PAM NA Intergenic NA NA

T20876 CCCCGGCTACTGAAGCGCATCGG 1000
Oligo fwd CACCCCCCGGCTACTGAAGCGCAT
Oligo rev AAACATGCGCTTCAGTAGCCGGGG

Chromosome start end strand MM target_seq PAM alignment distance position gene name gene id
GU305914.1 182897 182919 - 0 CCCCGGCTACTGAAGCGCAT CGG ||||||||[||||||||||||]PAM NA Intergenic NA NA

T20877 TGAAGCGCATCGGACCCGCTCGG 1000
Oligo fwd CACCTGAAGCGCATCGGACCCGCT
Oligo rev AAACAGCGGGTCCGATGCGCTTCA
Chromosome start end strand MM target_seq PAM alignment distance position gene name gene id
GU305914.1 182887 182909 - 0 TGAAGCGCATCGGACCCGCT CGG ||||||||[||||||||||||]PAM NA Intergenic NA NA

T20878 TCGGACCCGCTCGGTTCTATTGG 1000
Oligo fwd CACCTCGGACCCGCTCGGTTCTAT
Oligo rev AAACATAGAACCGAGCGGGTCCGA
Chromosome start end strand MM target_seq PAM alignment distance position gene name gene id
GU305914.1 182878 182900 - 0 TCGGACCCGCTCGGTTCTAT TGG ||||||||[||||||||||||]PAM NA Intergenic NA NA

T20879 CGGACCCGCTCGGTTCTATTGGG 1000
Oligo fwd CACCCGGACCCGCTCGGTTCTATT
Oligo rev AAACAATAGAACCGAGCGGGTCCG
Chromosome start end strand MM target_seq PAM alignment distance position gene name gene id
GU305914.1 182877 182899 - 0 CGGACCCGCTCGGTTCTATT GGG ||||||||[||||||||||||]PAM NA Intergenic NA NA

T20880 ACCACGCCCATCCGCCATATTGG 1000
Oligo fwd CACCACCACGCCCATCCGCCATAT
Oligo rev AAACATATGGCGGATGGGCGTGGT
Chromosome start end strand MM target_seq PAM alignment distance position gene name gene id
GU305914.1 182854 182876 - 0 ACCACGCCCATCCGCCATAT TGG ||||||||[||||||||||||]PAM NA Intergenic NA NA

T20881 TATTGGAACAGCCGCCATATTGG 1000
Oligo fwd CACCTATTGGAACAGCCGCCATAT
Oligo rev AAACATATGGCGGCTGTTCCAATA
Chromosome start end strand MM target_seq PAM alignment distance position gene name gene id
GU305914.1 182837 182859 - 0 TATTGGAACAGCCGCCATAT TGG ||||||||[||||||||||||]PAM NA Intergenic NA NA

T20882 GGTAACGCCGCCATCTTGATTGG 1000
Oligo fwd CACCGGTAACGCCGCCATCTTGAT
Oligo rev AAACATCAAGATGGCGGCGTTACC
Chromosome start end strand MM target_seq PAM alignment distance position gene name gene id
GU305914.1 182816 182838 - 0 GGTAACGCCGCCATCTTGAT TGG ||||||||[||||||||||||]PAM NA Intergenic NA NA

T20883 GTAACGCCGCCATCTTGATTGGG 1000
Oligo fwd CACCGTAACGCCGCCATCTTGATT
Oligo rev AAACAATCAAGATGGCGGCGTTAC
Chromosome start end strand MM target_seq PAM alignment distance position gene name gene id
GU305914.1 182815 182837 - 0 GTAACGCCGCCATCTTGATT GGG ||||||||[||||||||||||]PAM NA Intergenic NA NA

T20884 ATTGGGACCACCGCCCCTATTGG 1000
Oligo fwd CACCATTGGGACCACCGCCCCTAT
Oligo rev AAACATAGGGGCGGTGGTCCCAAT
Chromosome start end strand MM target_seq PAM alignment distance position gene name gene id
GU305914.1 182798 182820 - 0 ATTGGGACCACCGCCCCTAT TGG ||||||||[||||||||||||]PAM NA Intergenic NA NA

T20885 TTGGGACCACCGCCCCTATTGGG 1000
Oligo fwd CACCTTGGGACCACCGCCCCTATT
Oligo rev AAACAATAGGGGCGGTGGTCCCAA
Chromosome start end strand MM target_seq PAM alignment distance position gene name gene id
GU305914.1 182797 182819 - 0 TTGGGACCACCGCCCCTATT GGG ||||||||[||||||||||||]PAM NA Intergenic NA NA

T20886 TATTGGGACCACCGCCCAATTGG 1000
Oligo fwd CACCTATTGGGACCACCGCCCAAT
Oligo rev AAACATTGGGCGGTGGTCCCAATA
Chromosome start end strand MM target_seq PAM alignment distance position gene name gene id
GU305914.1 182781 182803 - 0 TATTGGGACCACCGCCCAAT TGG ||||||||[||||||||||||]PAM NA Intergenic NA NA

T20887 ATTGGGACCACCGCCCAATTGGG 1000
Oligo fwd CACCATTGGGACCACCGCCCAATT
Oligo rev AAACAATTGGGCGGTGGTCCCAAT
Chromosome start end strand MM target_seq PAM alignment distance position gene name gene id
GU305914.1 182780 182802 - 0 ATTGGGACCACCGCCCAATT GGG ||||||||[||||||||||||]PAM NA Intergenic NA NA

T20888 CACCGCCCACCCAGCGACCCAGG 1000
Oligo fwd CACCCACCGCCCACCCAGCGACCC
Oligo rev AAACGGGTCGCTGGGTGGGCGGTG
Chromosome start end strand MM target_seq PAM alignment distance position gene name gene id
GU305914.1 182755 182777 - 0 CACCGCCCACCCAGCGACCC AGG ||||||||[||||||||||||]PAM NA Intergenic NA NA

T20889 ACCCAGCGACCCAGGCCCAGAGG 1000
Oligo fwd CACCACCCAGCGACCCAGGCCCAG
Oligo rev AAACCTGGGCCTGGGTCGCTGGGT
Chromosome start end strand MM target_seq PAM alignment distance position gene name gene id
GU305914.1 182747 182769 - 0 ACCCAGCGACCCAGGCCCAG AGG ||||||||[||||||||||||]PAM NA Intergenic NA NA

T20890 GGCCCAGAGGCTACGTCACCAGG 1000
Oligo fwd CACCGGCCCAGAGGCTACGTCACC
Oligo rev AAACGGTGACGTAGCCTCTGGGCC
Chromosome start end strand MM target_seq PAM alignment distance position gene name gene id
GU305914.1 182734 182756 - 0 GGCCCAGAGGCTACGTCACC AGG ||||||||[||||||||||||]PAM NA Intergenic NA NA

T20891 GGCTACGTCACCAGGACCCCCGG 1000
Oligo fwd CACCGGCTACGTCACCAGGACCCC

Oligo rev AAACGGGGTCCTGGTGACGTAGCC
Chromosome start end strand MM target_seq PAM alignment distance position gene name gene id
GU305914.1 182726 182748 - 0 GGCTACGTCACCAGGACCCC CGG ||||||||[||||||||||||]PAM NA Intergenic NA NA

T20892 GGCTGCCCAGCGCCCCCTCGAGG 1000
Oligo fwd CACCGGCTGCCCAGCGCCCCCTCG
Oligo rev AAACCGAGGGGGCGCTGGGCAGCC
Chromosome start end strand MM target_seq PAM alignment distance position gene name gene id
GU305914.1 182705 182727 - 0 GGCTGCCCAGCGCCCCCTCG AGG ||||||||[||||||||||||]PAM NA Intergenic NA NA

T20893 CACACAATGTAAGTAGACCAAGG 1000
Oligo fwd CACCCACACAATGTAAGTAGACCA
Oligo rev AAACTGGTCTACTTACATTGTGTG
Chromosome start end strand MM target_seq PAM alignment distance position gene name gene id
GU305914.1 182681 182703 - 0 CACACAATGTAAGTAGACCA AGG ||||||||[||||||||||||]PAM NA Intergenic NA NA

T20894 AGTAGACCAAGGCCTTTTATTGG 1000
Oligo fwd CACCAGTAGACCAAGGCCTTTTAT
Oligo rev AAACATAAAAGGCCTTGGTCTACT
Chromosome start end strand MM target_seq PAM alignment distance position gene name gene id
GU305914.1 182670 182692 - 0 AGTAGACCAAGGCCTTTTAT TGG ||||||||[||||||||||||]PAM NA Intergenic NA NA

T20895 CCAAGGCCTTTTATTGGCTGCGG 1000
Oligo fwd CACCCCAAGGCCTTTTATTGGCTG
Oligo rev AAACCAGCCAATAAAAGGCCTTGG
Chromosome start end strand MM target_seq PAM alignment distance position gene name gene id
GU305914.1 182664 182686 - 0 CCAAGGCCTTTTATTGGCTG CGG ||||||||[||||||||||||]PAM NA Intergenic NA NA

T20896 TTTTATTGGCTGCGGTGTTAAGG 1000
Oligo fwd CACCTTTTATTGGCTGCGGTGTTA
Oligo rev AAACTAACACCGCAGCCAATAAAA
Chromosome start end strand MM target_seq PAM alignment distance position gene name gene id
GU305914.1 182656 182678 - 0 TTTTATTGGCTGCGGTGTTA AGG ||||||||[||||||||||||]PAM NA Intergenic NA NA

T20897 CTATCTATCTACTATCCACCTGG 1000
Oligo fwd CACCCTATCTATCTACTATCCACC
Oligo rev AAACGGTGGATAGTAGATAGATAG
Chromosome start end strand MM target_seq PAM alignment distance position gene name gene id
GU305914.1 182615 182637 - 0 CTATCTATCTACTATCCACC TGG ||||||||[||||||||||||]PAM NA Intergenic NA NA

T20898 CTCTAGCTCCATACTGCGTACGG 1000
Oligo fwd CACCCTCTAGCTCCATACTGCGTA
Oligo rev AAACTACGCAGTATGGAGCTAGAG
Chromosome start end strand MM target_seq PAM alignment distance position gene name gene id
GU305914.1 182590 182612 - 0 CTCTAGCTCCATACTGCGTA CGG ||||||||[||||||||||||]PAM NA Intergenic NA NA

T20899 CGTACGGTGTTTGTTAGCTTAGG 1000
Oligo fwd CACCCGTACGGTGTTTGTTAGCTT
Oligo rev AAACAAGCTAACAAACACCGTACG
Chromosome start end strand MM target_seq PAM alignment distance position gene name gene id
GU305914.1 182574 182596 - 0 CGTACGGTGTTTGTTAGCTT AGG ||||||||[||||||||||||]PAM NA Intergenic NA NA

T20900 GTACGGTGTTTGTTAGCTTAGGG 1000
Oligo fwd CACCGTACGGTGTTTGTTAGCTTA
Oligo rev AAACTAAGCTAACAAACACCGTAC
Chromosome start end strand MM target_seq PAM alignment distance position gene name gene id
GU305914.1 182573 182595 - 0 GTACGGTGTTTGTTAGCTTA GGG ||||||||[||||||||||||]PAM NA Intergenic NA NA

T20901 GGTGATAGCAAGTAGAACGCTGG 1000
Oligo fwd CACCGGTGATAGCAAGTAGAACGC
Oligo rev AAACGCGTTCTACTTGCTATCACC
Chromosome start end strand MM target_seq PAM alignment distance position gene name gene id
GU305914.1 182552 182574 - 0 GGTGATAGCAAGTAGAACGC TGG ||||||||[||||||||||||]PAM NA Intergenic NA NA

T20902 AGCAAGTAGAACGCTGGTGATGG 1000
Oligo fwd CACCAGCAAGTAGAACGCTGGTGA
Oligo rev AAACTCACCAGCGTTCTACTTGCT
Chromosome start end strand MM target_seq PAM alignment distance position gene name gene id
GU305914.1 182546 182568 - 0 AGCAAGTAGAACGCTGGTGA TGG ||||||||[||||||||||||]PAM NA Intergenic NA NA

T20903 GCTCTCTATCAGTTTAGCGCTGG 1000
Oligo fwd CACCGCTCTCTATCAGTTTAGCGC
Oligo rev AAACGCGCTAAACTGATAGAGAGC
Chromosome start end strand MM target_seq PAM alignment distance position gene name gene id
GU305914.1 182521 182543 - 0 GCTCTCTATCAGTTTAGCGC TGG ||||||||[||||||||||||]PAM NA Intergenic NA NA

T20904 TATCAGTTTAGCGCTGGTGATGG 1000
Oligo fwd CACCTATCAGTTTAGCGCTGGTGA
Oligo rev AAACTCACCAGCGCTAAACTGATA
Chromosome start end strand MM target_seq PAM alignment distance position gene name gene id
GU305914.1 182515 182537 - 0 TATCAGTTTAGCGCTGGTGA TGG ||||||||[||||||||||||]PAM NA Intergenic NA NA

T20905 CAGTTTAGCGCTGGTGATGGTGG 1000
Oligo fwd CACCCAGTTTAGCGCTGGTGATGG
Oligo rev AAACCCATCACCAGCGCTAAACTG
Chromosome start end strand MM target_seq PAM alignment distance position gene name gene id
GU305914.1 182512 182534 - 0 CAGTTTAGCGCTGGTGATGG TGG ||||||||[||||||||||||]PAM NA Intergenic NA NA

T20906 CATTTCTTATATATATATGCTGG 1000

Oligo fwd CACCCATTTCTTATATATATATGC
Oligo rev AAACGCATATATATATAAGAAATG
Chromosome start end strand MM target_seq PAM alignment distance position gene name gene id
GU305914.1 182479 182501 - 0 CATTTCTTATATATATATGC TGG ||||||||[||||||||||||]PAM NA Intergenic NA NA

T20907 TTATATATATATGCTGGTGATGG 1000
Oligo fwd CACCTTATATATATATGCTGGTGA
Oligo rev AAACTCACCAGCATATATATATAA
Chromosome start end strand MM target_seq PAM alignment distance position gene name gene id
GU305914.1 182473 182495 - 0 TTATATATATATGCTGGTGA TGG ||||||||[||||||||||||]PAM NA Intergenic NA NA

T20908 TGGTGATGGTCCCTCTTCCATGG 1000
Oligo fwd CACCTGGTGATGGTCCCTCTTCCA
Oligo rev AAACTGGAAGAGGGACCATCACCA
Chromosome start end strand MM target_seq PAM alignment distance position gene name gene id
GU305914.1 182459 182481 - 0 TGGTGATGGTCCCTCTTCCA TGG ||||||||[||||||||||||]PAM NA Intergenic NA NA

T20909 TGGTCCCTCTTCCATGGCCGCGG 1000
Oligo fwd CACCTGGTCCCTCTTCCATGGCCG
Oligo rev AAACCGGCCATGGAAGAGGGACCA
Chromosome start end strand MM target_seq PAM alignment distance position gene name gene id
GU305914.1 182453 182475 - 0 TGGTCCCTCTTCCATGGCCG CGG ||||||||[||||||||||||]PAM NA Intergenic NA NA

T20910 TCCATGGCCGCGGTGTGCTTCGG 1000
Oligo fwd CACCTCCATGGCCGCGGTGTGCTT
Oligo rev AAACAAGCACACCGCGGCCATGGA
Chromosome start end strand MM target_seq PAM alignment distance position gene name gene id
GU305914.1 182443 182465 - 0 TCCATGGCCGCGGTGTGCTT CGG ||||||||[||||||||||||]PAM NA Intergenic NA NA
GU305914.1 100155 100177 + 4 ACCATCGCCGCGGCGTGCTC GGG -||||-||[|||||-|||||-]PAM NA Intergenic NA NA

T20911 GCCGCGGTGTGCTTCGGTGCCGG 1000
Oligo fwd CACCGCCGCGGTGTGCTTCGGTGC
Oligo rev AAACGCACCGAAGCACACCGCGGC
Chromosome start end strand MM target_seq PAM alignment distance position gene name gene id
GU305914.1 182437 182459 - 0 GCCGCGGTGTGCTTCGGTGC CGG ||||||||[||||||||||||]PAM NA Intergenic NA NA

T20912 CCGCGGTGTGCTTCGGTGCCGGG 1000
Oligo fwd CACCCCGCGGTGTGCTTCGGTGCC
Oligo rev AAACGGCACCGAAGCACACCGCGG
Chromosome start end strand MM target_seq PAM alignment distance position gene name gene id
GU305914.1 182436 182458 - 0 CCGCGGTGTGCTTCGGTGCC GGG ||||||||[||||||||||||]PAM NA Intergenic NA NA

T20913 CGGTGCCGGGTACCGTAACGTGG 1000
Oligo fwd CACCCGGTGCCGGGTACCGTAACG
Oligo rev AAACCGTTACGGTACCCGGCACCG
Chromosome start end strand MM target_seq PAM alignment distance position gene name gene id
GU305914.1 182423 182445 - 0 CGGTGCCGGGTACCGTAACG TGG ||||||||[||||||||||||]PAM NA Intergenic NA NA

T20914 CCGTAACGTGGTACAGAAAAAGG 1000
Oligo fwd CACCCCGTAACGTGGTACAGAAAA
Oligo rev AAACTTTTCTGTACCACGTTACGG
Chromosome start end strand MM target_seq PAM alignment distance position gene name gene id
GU305914.1 182411 182433 - 0 CCGTAACGTGGTACAGAAAA AGG ||||||||[||||||||||||]PAM NA Intergenic NA NA

T20915 ACACGTACGCGTCTCTCCCCAGG 1000
Oligo fwd CACCACACGTACGCGTCTCTCCCC
Oligo rev AAACGGGGAGAGACGCGTACGTGT
Chromosome start end strand MM target_seq PAM alignment distance position gene name gene id
GU305914.1 182384 182406 - 0 ACACGTACGCGTCTCTCCCC AGG ||||||||[||||||||||||]PAM NA Intergenic NA NA
GU305914.1 191775 191797 + 4 CCACGCACGCGTGTCCCCCC AGG -||||-||[||||-||-||||]PAM NA Intergenic NA NA

T20916 ATGCTATGTGTCTGTAGCACCGG 1000
Oligo fwd CACCATGCTATGTGTCTGTAGCAC
Oligo rev AAACGTGCTACAGACACATAGCAT
Chromosome start end strand MM target_seq PAM alignment distance position gene name gene id
GU305914.1 182361 182383 - 0 ATGCTATGTGTCTGTAGCAC CGG ||||||||[||||||||||||]PAM NA Intergenic NA NA

T20917 CTATGTGTCTGTAGCACCGGCGG 1000
Oligo fwd CACCCTATGTGTCTGTAGCACCGG
Oligo rev AAACCCGGTGCTACAGACACATAG
Chromosome start end strand MM target_seq PAM alignment distance position gene name gene id
GU305914.1 182358 182380 - 0 CTATGTGTCTGTAGCACCGG CGG ||||||||[||||||||||||]PAM NA Intergenic NA NA

T20918 ACACCAGATATTACAAGCGACGG 1000
Oligo fwd CACCACACCAGATATTACAAGCGA
Oligo rev AAACTCGCTTGTAATATCTGGTGT
Chromosome start end strand MM target_seq PAM alignment distance position gene name gene id
GU305914.1 182271 182293 - 0 ACACCAGATATTACAAGCGA CGG ||||||||[||||||||||||]PAM NA Intergenic NA NA

T20919 ATACAGAATATTGTACTCAGAGG 1000
Oligo fwd CACCATACAGAATATTGTACTCAG
Oligo rev AAACCTGAGTACAATATTCTGTAT
Chromosome start end strand MM target_seq PAM alignment distance position gene name gene id
GU305914.1 182219 182241 - 0 ATACAGAATATTGTACTCAG AGG ||||||||[||||||||||||]PAM NA Intergenic NA NA

T20920 CTCAGAGGCTGCAGCCATCTTGG 1000
Oligo fwd CACCCTCAGAGGCTGCAGCCATCT
Oligo rev AAACAGATGGCTGCAGCCTCTGAG
Chromosome start end strand MM target_seq PAM alignment distance position gene name gene id

<table>
<tr><td>GU305914.1</td><td>182204</td><td>182226</td><td>-</td><td>0</td><td>CTCAGAGGCTGCAGCCATCT</td><td>TGG</td><td>||||||||[||||||||||||]PAM</td><td>NA</td><td>Intergenic</td><td>NA</td><td>NA</td></tr>
<tr><td>T20921</td><td>AGAGGCTGCAGCCATCTTGGTGG</td><td>1000</td><td></td><td></td><td></td><td></td><td></td><td></td><td></td><td></td><td></td></tr>
<tr><td>Oligo fwd</td><td>CACCAGAGGCTGCAGCCATCTTGG</td><td></td><td></td><td></td><td></td><td></td><td></td><td></td><td></td><td></td><td></td></tr>
<tr><td>Oligo rev</td><td>AAACCCAAGATGGCTGCAGCCTCT</td><td></td><td></td><td></td><td></td><td></td><td></td><td></td><td></td><td></td><td></td></tr>
<tr><td>Chromosome</td><td>start</td><td>end</td><td>strand</td><td>MM</td><td>target_seq</td><td>PAM</td><td>alignment</td><td>distance</td><td>position</td><td>gene name</td><td>gene id</td></tr>
<tr><td>GU305914.1</td><td>182201</td><td>182223</td><td>-</td><td>0</td><td>AGAGGCTGCAGCCATCTTGG</td><td>TGG</td><td>||||||||[||||||||||||]PAM</td><td>NA</td><td>Intergenic</td><td>NA</td><td>NA</td></tr>
<tr><td>T20922</td><td>GCCATCTTGGTGGCCTGCTGTGG</td><td>1000</td><td></td><td></td><td></td><td></td><td></td><td></td><td></td><td></td><td></td></tr>
<tr><td>Oligo fwd</td><td>CACCGCCATCTTGGTGGCCTGCTG</td><td></td><td></td><td></td><td></td><td></td><td></td><td></td><td></td><td></td><td></td></tr>
<tr><td>Oligo rev</td><td>AAACCAGCAGGCCACCAAGATGGC</td><td></td><td></td><td></td><td></td><td></td><td></td><td></td><td></td><td></td><td></td></tr>
<tr><td>Chromosome</td><td>start</td><td>end</td><td>strand</td><td>MM</td><td>target_seq</td><td>PAM</td><td>alignment</td><td>distance</td><td>position</td><td>gene name</td><td>gene id</td></tr>
<tr><td>GU305914.1</td><td>182191</td><td>182213</td><td>-</td><td>0</td><td>GCCATCTTGGTGGCCTGCTG</td><td>TGG</td><td>||||||||[||||||||||||]PAM</td><td>NA</td><td>Intergenic</td><td>NA</td><td>NA</td></tr>
<tr><td>T20923</td><td>GCTGTGGTGCTCTTTTCCCGTGG</td><td>1000</td><td></td><td></td><td></td><td></td><td></td><td></td><td></td><td></td><td></td></tr>
<tr><td>Oligo fwd</td><td>CACCGCTGTGGTGCTCTTTTCCCG</td><td></td><td></td><td></td><td></td><td></td><td></td><td></td><td></td><td></td><td></td></tr>
<tr><td>Oligo rev</td><td>AAACCGGGAAAAGAGCACCACAGC</td><td></td><td></td><td></td><td></td><td></td><td></td><td></td><td></td><td></td><td></td></tr>
<tr><td>Chromosome</td><td>start</td><td>end</td><td>strand</td><td>MM</td><td>target_seq</td><td>PAM</td><td>alignment</td><td>distance</td><td>position</td><td>gene name</td><td>gene id</td></tr>
<tr><td>GU305914.1</td><td>182175</td><td>182197</td><td>-</td><td>0</td><td>GCTGTGGTGCTCTTTTCCCG</td><td>TGG</td><td>||||||||[||||||||||||]PAM</td><td>NA</td><td>Intergenic</td><td>NA</td><td>NA</td></tr>
<tr><td>T20924</td><td>GTGGTGCTCTTTTCCCGTGGAGG</td><td>1000</td><td></td><td></td><td></td><td></td><td></td><td></td><td></td><td></td><td></td></tr>
<tr><td>Oligo fwd</td><td>CACCGTGGTGCTCTTTTCCCGTGG</td><td></td><td></td><td></td><td></td><td></td><td></td><td></td><td></td><td></td><td></td></tr>
<tr><td>Oligo rev</td><td>AAACCCACGGGAAAAGAGCACCAC</td><td></td><td></td><td></td><td></td><td></td><td></td><td></td><td></td><td></td><td></td></tr>
<tr><td>Chromosome</td><td>start</td><td>end</td><td>strand</td><td>MM</td><td>target_seq</td><td>PAM</td><td>alignment</td><td>distance</td><td>position</td><td>gene name</td><td>gene id</td></tr>
<tr><td>GU305914.1</td><td>182172</td><td>182194</td><td>-</td><td>0</td><td>GTGGTGCTCTTTTCCCGTGG</td><td>AGG</td><td>||||||||[||||||||||||]PAM</td><td>NA</td><td>Intergenic</td><td>NA</td><td>NA</td></tr>
<tr><td>T20925</td><td>TTGCCCCGTCCTGAAAACCTAGG</td><td>1000</td><td></td><td></td><td></td><td></td><td></td><td></td><td></td><td></td><td></td></tr>
<tr><td>Oligo fwd</td><td>CACCTTGCCCCGTCCTGAAAACCT</td><td></td><td></td><td></td><td></td><td></td><td></td><td></td><td></td><td></td><td></td></tr>
<tr><td>Oligo rev</td><td>AAACAGGTTTTCAGGACGGGGCAA</td><td></td><td></td><td></td><td></td><td></td><td></td><td></td><td></td><td></td><td></td></tr>
<tr><td>Chromosome</td><td>start</td><td>end</td><td>strand</td><td>MM</td><td>target_seq</td><td>PAM</td><td>alignment</td><td>distance</td><td>position</td><td>gene name</td><td>gene id</td></tr>
<tr><td>GU305914.1</td><td>182102</td><td>182124</td><td>-</td><td>0</td><td>TTGCCCCGTCCTGAAAACCT</td><td>AGG</td><td>||||||||[||||||||||||]PAM</td><td>NA</td><td>Intergenic</td><td>NA</td><td>NA</td></tr>
<tr><td>T20926</td><td>CCTAGGCCCTGACAGAAAAAAGG</td><td>1000</td><td></td><td></td><td></td><td></td><td></td><td></td><td></td><td></td><td></td></tr>
<tr><td>Oligo fwd</td><td>CACCCCTAGGCCCTGACAGAAAAA</td><td></td><td></td><td></td><td></td><td></td><td></td><td></td><td></td><td></td><td></td></tr>
<tr><td>Oligo rev</td><td>AAACTTTTTCTGTCAGGGCCTAGG</td><td></td><td></td><td></td><td></td><td></td><td></td><td></td><td></td><td></td><td></td></tr>
<tr><td>Chromosome</td><td>start</td><td>end</td><td>strand</td><td>MM</td><td>target_seq</td><td>PAM</td><td>alignment</td><td>distance</td><td>position</td><td>gene name</td><td>gene id</td></tr>
<tr><td>GU305914.1</td><td>182085</td><td>182107</td><td>-</td><td>0</td><td>CCTAGGCCCTGACAGAAAAA</td><td>AGG</td><td>||||||||[||||||||||||]PAM</td><td>NA</td><td>Intergenic</td><td>NA</td><td>NA</td></tr>
<tr><td>T20927</td><td>CTAGGCCCTGACAGAAAAAAGGG</td><td>1000</td><td></td><td></td><td></td><td></td><td></td><td></td><td></td><td></td><td></td></tr>
<tr><td>Oligo fwd</td><td>CACCCTAGGCCCTGACAGAAAAAA</td><td></td><td></td><td></td><td></td><td></td><td></td><td></td><td></td><td></td><td></td></tr>
<tr><td>Oligo rev</td><td>AAACTTTTTTCTGTCAGGGCCTAG</td><td></td><td></td><td></td><td></td><td></td><td></td><td></td><td></td><td></td><td></td></tr>
<tr><td>Chromosome</td><td>start</td><td>end</td><td>strand</td><td>MM</td><td>target_seq</td><td>PAM</td><td>alignment</td><td>distance</td><td>position</td><td>gene name</td><td>gene id</td></tr>
<tr><td>GU305914.1</td><td>182084</td><td>182106</td><td>-</td><td>0</td><td>CTAGGCCCTGACAGAAAAAA</td><td>GGG</td><td>||||||||[||||||||||||]PAM</td><td>NA</td><td>Intergenic</td><td>NA</td><td>NA</td></tr>
<tr><td>T20928</td><td>GACAGAAAAAAGGGAGAGTTAGG</td><td>1000</td><td></td><td></td><td></td><td></td><td></td><td></td><td></td><td></td><td></td></tr>
<tr><td>Oligo fwd</td><td>CACCGACAGAAAAAAGGGAGAGTT</td><td></td><td></td><td></td><td></td><td></td><td></td><td></td><td></td><td></td><td></td></tr>
<tr><td>Oligo rev</td><td>AAACAACTCTCCCTTTTTTCTGTC</td><td></td><td></td><td></td><td></td><td></td><td></td><td></td><td></td><td></td><td></td></tr>
<tr><td>Chromosome</td><td>start</td><td>end</td><td>strand</td><td>MM</td><td>target_seq</td><td>PAM</td><td>alignment</td><td>distance</td><td>position</td><td>gene name</td><td>gene id</td></tr>
<tr><td>GU305914.1</td><td>182075</td><td>182097</td><td>-</td><td>0</td><td>GACAGAAAAAAGGGAGAGTT</td><td>AGG</td><td>||||||||[||||||||||||]PAM</td><td>NA</td><td>Intergenic</td><td>NA</td><td>NA</td></tr>
<tr><td>T20929</td><td>AAAAAAGGGAGAGTTAGGAGAGG</td><td>1000</td><td></td><td></td><td></td><td></td><td></td><td></td><td></td><td></td><td></td></tr>
<tr><td>Oligo fwd</td><td>CACCAAAAAAGGGAGAGTTAGGAG</td><td></td><td></td><td></td><td></td><td></td><td></td><td></td><td></td><td></td><td></td></tr>
<tr><td>Oligo rev</td><td>AAACCTCCTAACTCTCCCTTTTTT</td><td></td><td></td><td></td><td></td><td></td><td></td><td></td><td></td><td></td><td></td></tr>
<tr><td>Chromosome</td><td>start</td><td>end</td><td>strand</td><td>MM</td><td>target_seq</td><td>PAM</td><td>alignment</td><td>distance</td><td>position</td><td>gene name</td><td>gene id</td></tr>
<tr><td>GU305914.1</td><td>182070</td><td>182092</td><td>-</td><td>0</td><td>AAAAAAGGGAGAGTTAGGAG</td><td>AGG</td><td>||||||||[||||||||||||]PAM</td><td>NA</td><td>Intergenic</td><td>NA</td><td>NA</td></tr>
<tr><td>T20930</td><td>AAAAAGGGAGAGTTAGGAGAGGG</td><td>1000</td><td></td><td></td><td></td><td></td><td></td><td></td><td></td><td></td><td></td></tr>
<tr><td>Oligo fwd</td><td>CACCAAAAAGGGAGAGTTAGGAGA</td><td></td><td></td><td></td><td></td><td></td><td></td><td></td><td></td><td></td><td></td></tr>
<tr><td>Oligo rev</td><td>AAACTCTCCTAACTCTCCCTTTTT</td><td></td><td></td><td></td><td></td><td></td><td></td><td></td><td></td><td></td><td></td></tr>
<tr><td>Chromosome</td><td>start</td><td>end</td><td>strand</td><td>MM</td><td>target_seq</td><td>PAM</td><td>alignment</td><td>distance</td><td>position</td><td>gene name</td><td>gene id</td></tr>
<tr><td>GU305914.1</td><td>182069</td><td>182091</td><td>-</td><td>0</td><td>AAAAAGGGAGAGTTAGGAGA</td><td>GGG</td><td>||||||||[||||||||||||]PAM</td><td>NA</td><td>Intergenic</td><td>NA</td><td>NA</td></tr>
<tr><td>T20931</td><td>AGAGTTAGGAGAGGGCACAGAGG</td><td>1000</td><td></td><td></td><td></td><td></td><td></td><td></td><td></td><td></td><td></td></tr>
<tr><td>Oligo fwd</td><td>CACCAGAGTTAGGAGAGGGCACAG</td><td></td><td></td><td></td><td></td><td></td><td></td><td></td><td></td><td></td><td></td></tr>
<tr><td>Oligo rev</td><td>AAACCTGTGCCCTCTCCTAACTCT</td><td></td><td></td><td></td><td></td><td></td><td></td><td></td><td></td><td></td><td></td></tr>
<tr><td>Chromosome</td><td>start</td><td>end</td><td>strand</td><td>MM</td><td>target_seq</td><td>PAM</td><td>alignment</td><td>distance</td><td>position</td><td>gene name</td><td>gene id</td></tr>
<tr><td>GU305914.1</td><td>182061</td><td>182083</td><td>-</td><td>0</td><td>AGAGTTAGGAGAGGGCACAG</td><td>AGG</td><td>||||||||[||||||||||||]PAM</td><td>NA</td><td>Intergenic</td><td>NA</td><td>NA</td></tr>
<tr><td>T20932</td><td>AGAGGATTCTGTCTGTGTCAAGG</td><td>1000</td><td></td><td></td><td></td><td></td><td></td><td></td><td></td><td></td><td></td></tr>
<tr><td>Oligo fwd</td><td>CACCAGAGGATTCTGTCTGTGTCA</td><td></td><td></td><td></td><td></td><td></td><td></td><td></td><td></td><td></td><td></td></tr>
<tr><td>Oligo rev</td><td>AAACTGACACAGACAGAATCCTCT</td><td></td><td></td><td></td><td></td><td></td><td></td><td></td><td></td><td></td><td></td></tr>
<tr><td>Chromosome</td><td>start</td><td>end</td><td>strand</td><td>MM</td><td>target_seq</td><td>PAM</td><td>alignment</td><td>distance</td><td>position</td><td>gene name</td><td>gene id</td></tr>
<tr><td>GU305914.1</td><td>182043</td><td>182065</td><td>-</td><td>0</td><td>AGAGGATTCTGTCTGTGTCA</td><td>AGG</td><td>||||||||[||||||||||||]PAM</td><td>NA</td><td>Intergenic</td><td>NA</td><td>NA</td></tr>
<tr><td>T20933</td><td>GAGGATTCTGTCTGTGTCAAGGG</td><td>1000</td><td></td><td></td><td></td><td></td><td></td><td></td><td></td><td></td><td></td></tr>
<tr><td>Oligo fwd</td><td>CACCGAGGATTCTGTCTGTGTCAA</td><td></td><td></td><td></td><td></td><td></td><td></td><td></td><td></td><td></td><td></td></tr>
<tr><td>Oligo rev</td><td>AAACTTGACACAGACAGAATCCTC</td><td></td><td></td><td></td><td></td><td></td><td></td><td></td><td></td><td></td><td></td></tr>
<tr><td>Chromosome</td><td>start</td><td>end</td><td>strand</td><td>MM</td><td>target_seq</td><td>PAM</td><td>alignment</td><td>distance</td><td>position</td><td>gene name</td><td>gene id</td></tr>
<tr><td>GU305914.1</td><td>182042</td><td>182064</td><td>-</td><td>0</td><td>GAGGATTCTGTCTGTGTCAA</td><td>GGG</td><td>||||||||[||||||||||||]PAM</td><td>NA</td><td>Intergenic</td><td>NA</td><td>NA</td></tr>
<tr><td>T20934</td><td>AGGATTCTGTCTGTGTCAAGGGG</td><td>1000</td><td></td><td></td><td></td><td></td><td></td><td></td><td></td><td></td><td></td></tr>
<tr><td>Oligo fwd</td><td>CACCAGGATTCTGTCTGTGTCAAG</td><td></td><td></td><td></td><td></td><td></td><td></td><td></td><td></td><td></td><td></td></tr>
<tr><td>Oligo rev</td><td>AAACCTTGACACAGACAGAATCCT</td><td></td><td></td><td></td><td></td><td></td><td></td><td></td><td></td><td></td><td></td></tr>
<tr><td>Chromosome</td><td>start</td><td>end</td><td>strand</td><td>MM</td><td>target_seq</td><td>PAM</td><td>alignment</td><td>distance</td><td>position</td><td>gene name</td><td>gene id</td></tr>
<tr><td>GU305914.1</td><td>182041</td><td>182063</td><td>-</td><td>0</td><td>AGGATTCTGTCTGTGTCAAG</td><td>GGG</td><td>||||||||[||||||||||||]PAM</td><td>NA</td><td>Intergenic</td><td>NA</td><td>NA</td></tr>
<tr><td>T20935</td><td>GGATTCTGTCTGTGTCAAGGGGG</td><td>1000</td><td></td><td></td><td></td><td></td><td></td><td></td><td></td><td></td><td></td></tr>
<tr><td>Oligo fwd</td><td>CACCGGATTCTGTCTGTGTCAAGG</td><td></td><td></td><td></td><td></td><td></td><td></td><td></td><td></td><td></td><td></td></tr>
<tr><td>Oligo rev</td><td>AAACCCTTGACACAGACAGAATCC</td><td></td><td></td><td></td><td></td><td></td><td></td><td></td><td></td><td></td><td></td></tr>
</table>

<table>
<tr><td>Chromosome</td><td>start</td><td>end</td><td>strand</td><td>MM</td><td>target_seq</td><td>PAM</td><td>alignment</td><td>distance</td><td>position</td><td>gene name</td><td>gene id</td></tr>
<tr><td>GU305914.1</td><td>182040</td><td>182062</td><td>-</td><td>0</td><td>GGATTCTGTCTGTGTCAAGG</td><td>GGG</td><td>||||||||[||||||||||||]PAM</td><td>NA</td><td>Intergenic</td><td>NA</td><td>NA</td></tr>
<tr><td>T20936</td><td>GTTTTTTATCAGCAACTGACAGG</td><td>1000</td><td></td><td></td><td></td><td></td><td></td><td></td><td></td><td></td><td></td></tr>
<tr><td>Oligo fwd</td><td>CACCGTTTTTTATCAGCAACTGAC</td><td></td><td></td><td></td><td></td><td></td><td></td><td></td><td></td><td></td><td></td></tr>
<tr><td>Oligo rev</td><td>AAACGTCAGTTGCTGATAAAAAAC</td><td></td><td></td><td></td><td></td><td></td><td></td><td></td><td></td><td></td><td></td></tr>
<tr><td>Chromosome</td><td>start</td><td>end</td><td>strand</td><td>MM</td><td>target_seq</td><td>PAM</td><td>alignment</td><td>distance</td><td>position</td><td>gene name</td><td>gene id</td></tr>
<tr><td>GU305914.1</td><td>181976</td><td>181998</td><td>-</td><td>0</td><td>GTTTTTTATCAGCAACTGAC</td><td>AGG</td><td>||||||||[||||||||||||]PAM</td><td>NA</td><td>Intergenic</td><td>NA</td><td>NA</td></tr>
<tr><td>T20937</td><td>TGTATACTTTAGATATCTTCTGG</td><td>1000</td><td></td><td></td><td></td><td></td><td></td><td></td><td></td><td></td><td></td></tr>
<tr><td>Oligo fwd</td><td>CACCTGTATACTTTAGATATCTTC</td><td></td><td></td><td></td><td></td><td></td><td></td><td></td><td></td><td></td><td></td></tr>
<tr><td>Oligo rev</td><td>AAACGAAGATATCTAAAGTATACA</td><td></td><td></td><td></td><td></td><td></td><td></td><td></td><td></td><td></td><td></td></tr>
<tr><td>Chromosome</td><td>start</td><td>end</td><td>strand</td><td>MM</td><td>target_seq</td><td>PAM</td><td>alignment</td><td>distance</td><td>position</td><td>gene name</td><td>gene id</td></tr>
<tr><td>GU305914.1</td><td>181915</td><td>181937</td><td>-</td><td>0</td><td>TGTATACTTTAGATATCTTC</td><td>TGG</td><td>||||||||[||||||||||||]PAM</td><td>NA</td><td>Intergenic</td><td>NA</td><td>NA</td></tr>
<tr><td>T20938</td><td>TAGATATCTTCTGGTCTCTGTGG</td><td>1000</td><td></td><td></td><td></td><td></td><td></td><td></td><td></td><td></td><td></td></tr>
<tr><td>Oligo fwd</td><td>CACCTAGATATCTTCTGGTCTCTG</td><td></td><td></td><td></td><td></td><td></td><td></td><td></td><td></td><td></td><td></td></tr>
<tr><td>Oligo rev</td><td>AAACCAGAGACCAGAAGATATCTA</td><td></td><td></td><td></td><td></td><td></td><td></td><td></td><td></td><td></td><td></td></tr>
<tr><td>Chromosome</td><td>start</td><td>end</td><td>strand</td><td>MM</td><td>target_seq</td><td>PAM</td><td>alignment</td><td>distance</td><td>position</td><td>gene name</td><td>gene id</td></tr>
<tr><td>GU305914.1</td><td>181906</td><td>181928</td><td>-</td><td>0</td><td>TAGATATCTTCTGGTCTCTG</td><td>TGG</td><td>||||||||[||||||||||||]PAM</td><td>NA</td><td>Intergenic</td><td>NA</td><td>NA</td></tr>
<tr><td>T20939</td><td>ATTATATTTTTTTAGAGAGATGG</td><td>1000</td><td></td><td></td><td></td><td></td><td></td><td></td><td></td><td></td><td></td></tr>
<tr><td>Oligo fwd</td><td>CACCATTATATTTTTTTAGAGAGA</td><td></td><td></td><td></td><td></td><td></td><td></td><td></td><td></td><td></td><td></td></tr>
<tr><td>Oligo rev</td><td>AAACTCTCTCTAAAAAAATATAAT</td><td></td><td></td><td></td><td></td><td></td><td></td><td></td><td></td><td></td><td></td></tr>
<tr><td>Chromosome</td><td>start</td><td>end</td><td>strand</td><td>MM</td><td>target_seq</td><td>PAM</td><td>alignment</td><td>distance</td><td>position</td><td>gene name</td><td>gene id</td></tr>
<tr><td>GU305914.1</td><td>181863</td><td>181885</td><td>-</td><td>0</td><td>ATTATATTTTTTTAGAGAGA</td><td>TGG</td><td>||||||||[||||||||||||]PAM</td><td>NA</td><td>Intergenic</td><td>NA</td><td>NA</td></tr>
<tr><td>T20940</td><td>CATGATCATGATCGCCGATCAGG</td><td>1000</td><td></td><td></td><td></td><td></td><td></td><td></td><td></td><td></td><td></td></tr>
<tr><td>Oligo fwd</td><td>CACCCATGATCATGATCGCCGATC</td><td></td><td></td><td></td><td></td><td></td><td></td><td></td><td></td><td></td><td></td></tr>
<tr><td>Oligo rev</td><td>AAACGATCGGCGATCATGATCATG</td><td></td><td></td><td></td><td></td><td></td><td></td><td></td><td></td><td></td><td></td></tr>
<tr><td>Chromosome</td><td>start</td><td>end</td><td>strand</td><td>MM</td><td>target_seq</td><td>PAM</td><td>alignment</td><td>distance</td><td>position</td><td>gene name</td><td>gene id</td></tr>
<tr><td>GU305914.1</td><td>181818</td><td>181840</td><td>-</td><td>0</td><td>CATGATCATGATCGCCGATC</td><td>AGG</td><td>||||||||[||||||||||||]PAM</td><td>NA</td><td>Intergenic</td><td>NA</td><td>NA</td></tr>
<tr><td>T20941</td><td>CAGGCATCCGTCAACGCCCACGG</td><td>1000</td><td></td><td></td><td></td><td></td><td></td><td></td><td></td><td></td><td></td></tr>
<tr><td>Oligo fwd</td><td>CACCCAGGCATCCGTCAACGCCCA</td><td></td><td></td><td></td><td></td><td></td><td></td><td></td><td></td><td></td><td></td></tr>
<tr><td>Oligo rev</td><td>AAACTGGGCGTTGACGGATGCCTG</td><td></td><td></td><td></td><td></td><td></td><td></td><td></td><td></td><td></td><td></td></tr>
<tr><td>Chromosome</td><td>start</td><td>end</td><td>strand</td><td>MM</td><td>target_seq</td><td>PAM</td><td>alignment</td><td>distance</td><td>position</td><td>gene name</td><td>gene id</td></tr>
<tr><td>GU305914.1</td><td>181799</td><td>181821</td><td>-</td><td>0</td><td>CAGGCATCCGTCAACGCCCA</td><td>CGG</td><td>||||||||[||||||||||||]PAM</td><td>NA</td><td>Intergenic</td><td>NA</td><td>NA</td></tr>
<tr><td>T20942</td><td>CAACGCCCACGGCCGTCACTTGG</td><td>1000</td><td></td><td></td><td></td><td></td><td></td><td></td><td></td><td></td><td></td></tr>
<tr><td>Oligo fwd</td><td>CACCCAACGCCCACGGCCGTCACT</td><td></td><td></td><td></td><td></td><td></td><td></td><td></td><td></td><td></td><td></td></tr>
<tr><td>Oligo rev</td><td>AAACAGTGACGGCCGTGGGCGTTG</td><td></td><td></td><td></td><td></td><td></td><td></td><td></td><td></td><td></td><td></td></tr>
<tr><td>Chromosome</td><td>start</td><td>end</td><td>strand</td><td>MM</td><td>target_seq</td><td>PAM</td><td>alignment</td><td>distance</td><td>position</td><td>gene name</td><td>gene id</td></tr>
<tr><td>GU305914.1</td><td>181788</td><td>181810</td><td>-</td><td>0</td><td>CAACGCCCACGGCCGTCACT</td><td>TGG</td><td>||||||||[||||||||||||]PAM</td><td>NA</td><td>Intergenic</td><td>NA</td><td>NA</td></tr>
<tr><td>T20943</td><td>TTTACATAACCCCTGACTCATGG</td><td>1000</td><td></td><td></td><td></td><td></td><td></td><td></td><td></td><td></td><td></td></tr>
<tr><td>Oligo fwd</td><td>CACCTTTACATAACCCCTGACTCA</td><td></td><td></td><td></td><td></td><td></td><td></td><td></td><td></td><td></td><td></td></tr>
<tr><td>Oligo rev</td><td>AAACTGAGTCAGGGGTTATGTAAA</td><td></td><td></td><td></td><td></td><td></td><td></td><td></td><td></td><td></td><td></td></tr>
<tr><td>Chromosome</td><td>start</td><td>end</td><td>strand</td><td>MM</td><td>target_seq</td><td>PAM</td><td>alignment</td><td>distance</td><td>position</td><td>gene name</td><td>gene id</td></tr>
<tr><td>GU305914.1</td><td>181686</td><td>181708</td><td>-</td><td>0</td><td>TTTACATAACCCCTGACTCA</td><td>TGG</td><td>||||||||[||||||||||||]PAM</td><td>NA</td><td>Intergenic</td><td>NA</td><td>NA</td></tr>
<tr><td>T20944</td><td>CTTCCCACTCTTGTTCTCTCAGG</td><td>1000</td><td></td><td></td><td></td><td></td><td></td><td></td><td></td><td></td><td></td></tr>
<tr><td>Oligo fwd</td><td>CACCCTTCCCACTCTTGTTCTCTC</td><td></td><td></td><td></td><td></td><td></td><td></td><td></td><td></td><td></td><td></td></tr>
<tr><td>Oligo rev</td><td>AAACGAGAGAACAAGAGTGGGAAG</td><td></td><td></td><td></td><td></td><td></td><td></td><td></td><td></td><td></td><td></td></tr>
<tr><td>Chromosome</td><td>start</td><td>end</td><td>strand</td><td>MM</td><td>target_seq</td><td>PAM</td><td>alignment</td><td>distance</td><td>position</td><td>gene name</td><td>gene id</td></tr>
<tr><td>GU305914.1</td><td>181662</td><td>181684</td><td>-</td><td>0</td><td>CTTCCCACTCTTGTTCTCTC</td><td>AGG</td><td>||||||||[||||||||||||]PAM</td><td>NA</td><td>Intergenic</td><td>NA</td><td>NA</td></tr>
<tr><td>T20945</td><td>TTCCCACTCTTGTTCTCTCAGGG</td><td>1000</td><td></td><td></td><td></td><td></td><td></td><td></td><td></td><td></td><td></td></tr>
<tr><td>Oligo fwd</td><td>CACCTTCCCACTCTTGTTCTCTCA</td><td></td><td></td><td></td><td></td><td></td><td></td><td></td><td></td><td></td><td></td></tr>
<tr><td>Oligo rev</td><td>AAACTGAGAGAACAAGAGTGGGAA</td><td></td><td></td><td></td><td></td><td></td><td></td><td></td><td></td><td></td><td></td></tr>
<tr><td>Chromosome</td><td>start</td><td>end</td><td>strand</td><td>MM</td><td>target_seq</td><td>PAM</td><td>alignment</td><td>distance</td><td>position</td><td>gene name</td><td>gene id</td></tr>
<tr><td>GU305914.1</td><td>181661</td><td>181683</td><td>-</td><td>0</td><td>TTCCCACTCTTGTTCTCTCA</td><td>GGG</td><td>||||||||[||||||||||||]PAM</td><td>NA</td><td>Intergenic</td><td>NA</td><td>NA</td></tr>
<tr><td>T20946</td><td>GCCAACAAGATCCTCGAGTCTGG</td><td>1000</td><td></td><td></td><td></td><td></td><td></td><td></td><td></td><td></td><td></td></tr>
<tr><td>Oligo fwd</td><td>CACCGCCAACAAGATCCTCGAGTC</td><td></td><td></td><td></td><td></td><td></td><td></td><td></td><td></td><td></td><td></td></tr>
<tr><td>Oligo rev</td><td>AAACGACTCGAGGATCTTGTTGGC</td><td></td><td></td><td></td><td></td><td></td><td></td><td></td><td></td><td></td><td></td></tr>
<tr><td>Chromosome</td><td>start</td><td>end</td><td>strand</td><td>MM</td><td>target_seq</td><td>PAM</td><td>alignment</td><td>distance</td><td>position</td><td>gene name</td><td>gene id</td></tr>
<tr><td>GU305914.1</td><td>181624</td><td>181646</td><td>-</td><td>0</td><td>GCCAACAAGATCCTCGAGTC</td><td>TGG</td><td>||||||||[||||||||||||]PAM</td><td>NA</td><td>Intergenic</td><td>NA</td><td>NA</td></tr>
<tr><td>T20947</td><td>CGTCCGCTGTGACCTGACTCTGG</td><td>1000</td><td></td><td></td><td></td><td></td><td></td><td></td><td></td><td></td><td></td></tr>
<tr><td>Oligo fwd</td><td>CACCCGTCCGCTGTGACCTGACTC</td><td></td><td></td><td></td><td></td><td></td><td></td><td></td><td></td><td></td><td></td></tr>
<tr><td>Oligo rev</td><td>AAACGAGTCAGGTCACAGCGGACG</td><td></td><td></td><td></td><td></td><td></td><td></td><td></td><td></td><td></td><td></td></tr>
<tr><td>Chromosome</td><td>start</td><td>end</td><td>strand</td><td>MM</td><td>target_seq</td><td>PAM</td><td>alignment</td><td>distance</td><td>position</td><td>gene name</td><td>gene id</td></tr>
<tr><td>GU305914.1</td><td>181592</td><td>181614</td><td>-</td><td>0</td><td>CGTCCGCTGTGACCTGACTC</td><td>TGG</td><td>||||||||[||||||||||||]PAM</td><td>NA</td><td>Intergenic</td><td>NA</td><td>NA</td></tr>
<tr><td>T20948</td><td>CCGCTGTGACCTGACTCTGGAGG</td><td>1000</td><td></td><td></td><td></td><td></td><td></td><td></td><td></td><td></td><td></td></tr>
<tr><td>Oligo fwd</td><td>CACCCCGCTGTGACCTGACTCTGG</td><td></td><td></td><td></td><td></td><td></td><td></td><td></td><td></td><td></td><td></td></tr>
<tr><td>Oligo rev</td><td>AAACCCAGAGTCAGGTCACAGCGG</td><td></td><td></td><td></td><td></td><td></td><td></td><td></td><td></td><td></td><td></td></tr>
<tr><td>Chromosome</td><td>start</td><td>end</td><td>strand</td><td>MM</td><td>target_seq</td><td>PAM</td><td>alignment</td><td>distance</td><td>position</td><td>gene name</td><td>gene id</td></tr>
<tr><td>GU305914.1</td><td>181589</td><td>181611</td><td>-</td><td>0</td><td>CCGCTGTGACCTGACTCTGG</td><td>AGG</td><td>||||||||[||||||||||||]PAM</td><td>NA</td><td>Intergenic</td><td>NA</td><td>NA</td></tr>
<tr><td>T20949</td><td>CCTGACTCTGGAGGACATGTTGG</td><td>1000</td><td></td><td></td><td></td><td></td><td></td><td></td><td></td><td></td><td></td></tr>
<tr><td>Oligo fwd</td><td>CACCCCTGACTCTGGAGGACATGT</td><td></td><td></td><td></td><td></td><td></td><td></td><td></td><td></td><td></td><td></td></tr>
<tr><td>Oligo rev</td><td>AAACACATGTCCTCCAGAGTCAGG</td><td></td><td></td><td></td><td></td><td></td><td></td><td></td><td></td><td></td><td></td></tr>
<tr><td>Chromosome</td><td>start</td><td>end</td><td>strand</td><td>MM</td><td>target_seq</td><td>PAM</td><td>alignment</td><td>distance</td><td>position</td><td>gene name</td><td>gene id</td></tr>
<tr><td>GU305914.1</td><td>181580</td><td>181602</td><td>-</td><td>0</td><td>CCTGACTCTGGAGGACATGT</td><td>TGG</td><td>||||||||[||||||||||||]PAM</td><td>NA</td><td>Intergenic</td><td>NA</td><td>NA</td></tr>
<tr><td>T20950</td><td>CTGACTCTGGAGGACATGTTGGG</td><td>1000</td><td></td><td></td><td></td><td></td><td></td><td></td><td></td><td></td><td></td></tr>
<tr><td>Oligo fwd</td><td>CACCCTGACTCTGGAGGACATGTT</td><td></td><td></td><td></td><td></td><td></td><td></td><td></td><td></td><td></td><td></td></tr>
</table>

Oligo rev AAACAACATGTCCTCCAGAGTCAG
Chromosome start end strand MM target_seq PAM alignment distance position gene name gene id
GU305914.1 181579 181601 - 0 CTGACTCTGGAGGACATGTT GGG ||||||||[||||||||||||]PAM NA Intergenic NA NA

T20951 TGAGTATGATGATCCCACTGAGG 1000
Oligo fwd CACCTGAGTATGATGATCCCACTG
Oligo rev AAACCAGTGGGATCATCATACTCA
Chromosome start end strand MM target_seq PAM alignment distance position gene name gene id
GU305914.1 181550 181572 - 0 TGAGTATGATGATCCCACTG AGG ||||||||[||||||||||||]PAM NA Intergenic NA NA

T20952 TGAGGAAGAGAAGATCCTCATGG 1000
Oligo fwd CACCTGAGGAAGAGAAGATCCTCA
Oligo rev AAACTGAGGATCTTCTCTTCCTCA
Chromosome start end strand MM target_seq PAM alignment distance position gene name gene id
GU305914.1 181532 181554 - 0 TGAGGAAGAGAAGATCCTCA TGG ||||||||[||||||||||||]PAM NA Intergenic NA NA

T20953 GGACCGCATCGCTGACCACGTGG 1000
Oligo fwd CACCGGACCGCATCGCTGACCACG
Oligo rev AAACCGTGGTCAGCGATGCGGTCC
Chromosome start end strand MM target_seq PAM alignment distance position gene name gene id
GU305914.1 181511 181533 - 0 GGACCGCATCGCTGACCACG TGG ||||||||[||||||||||||]PAM NA Intergenic NA NA

T20954 GACCGCATCGCTGACCACGTGGG 1000
Oligo fwd CACCGACCGCATCGCTGACCACGT
Oligo rev AAACACGTGGTCAGCGATGCGGTC
Chromosome start end strand MM target_seq PAM alignment distance position gene name gene id
GU305914.1 181510 181532 - 0 GACCGCATCGCTGACCACGT GGG ||||||||[||||||||||||]PAM NA Intergenic NA NA

T20955 ACCGCATCGCTGACCACGTGGGG 1000
Oligo fwd CACCACCGCATCGCTGACCACGTG
Oligo rev AAACCACGTGGTCAGCGATGCGGT
Chromosome start end strand MM target_seq PAM alignment distance position gene name gene id
GU305914.1 181509 181531 - 0 ACCGCATCGCTGACCACGTG GGG ||||||||[||||||||||||]PAM NA Intergenic NA NA

T20956 GGGGAATGATAACAGCGACATGG 1000
Oligo fwd CACCGGGGAATGATAACAGCGACA
Oligo rev AAACTGTCGCTGTTATCATTCCCC
Chromosome start end strand MM target_seq PAM alignment distance position gene name gene id
GU305914.1 181490 181512 - 0 GGGGAATGATAACAGCGACA TGG ||||||||[||||||||||||]PAM NA Intergenic NA NA

T20957 ATGGCCATCAAGCATGCTGCAGG 1000
Oligo fwd CACCATGGCCATCAAGCATGCTGC
Oligo rev AAACGCAGCATGCTTGATGGCCAT
Chromosome start end strand MM target_seq PAM alignment distance position gene name gene id
GU305914.1 181471 181493 - 0 ATGGCCATCAAGCATGCTGC AGG ||||||||[||||||||||||]PAM NA Intergenic NA NA

T20958 ACCACTCTTATATTCCAGTGAGG 1000
Oligo fwd CACCACCACTCTTATATTCCAGTG
Oligo rev AAACCACTGGAATATAAGAGTGGT
Chromosome start end strand MM target_seq PAM alignment distance position gene name gene id
GU305914.1 181345 181367 - 0 ACCACTCTTATATTCCAGTG AGG ||||||||[||||||||||||]PAM NA Intergenic NA NA

T20959 AGACAACGCAAGATGATATACGG 1000
Oligo fwd CACCAGACAACGCAAGATGATATA
Oligo rev AAACTATATCATCTTGCGTTGTCT
Chromosome start end strand MM target_seq PAM alignment distance position gene name gene id
GU305914.1 181202 181224 - 0 AGACAACGCAAGATGATATA CGG ||||||||[||||||||||||]PAM NA Intergenic NA NA

T20960 GAGTTTGATAATGTCCAGCTAGG 1000
Oligo fwd CACCGAGTTTGATAATGTCCAGCT
Oligo rev AAACAGCTGGACATTATCAAACTC
Chromosome start end strand MM target_seq PAM alignment distance position gene name gene id
GU305914.1 181166 181188 - 0 GAGTTTGATAATGTCCAGCT AGG ||||||||[||||||||||||]PAM NA Intergenic NA NA

T20961 AGTTTGATAATGTCCAGCTAGGG 1000
Oligo fwd CACCAGTTTGATAATGTCCAGCTA
Oligo rev AAACTAGCTGGACATTATCAAACT
Chromosome start end strand MM target_seq PAM alignment distance position gene name gene id
GU305914.1 181165 181187 - 0 AGTTTGATAATGTCCAGCTA GGG ||||||||[||||||||||||]PAM NA Intergenic NA NA

T20962 TTGATAATGTCCAGCTAGGGAGG 1000
Oligo fwd CACCTTGATAATGTCCAGCTAGGG
Oligo rev AAACCCCTAGCTGGACATTATCAA
Chromosome start end strand MM target_seq PAM alignment distance position gene name gene id
GU305914.1 181162 181184 - 0 TTGATAATGTCCAGCTAGGG AGG ||||||||[||||||||||||]PAM NA Intergenic NA NA

T20963 CACTTCATGCCCACTAATCTAGG 1000
Oligo fwd CACCCACTTCATGCCCACTAATCT
Oligo rev AAACAGATTAGTGGGCATGAAGTG
Chromosome start end strand MM target_seq PAM alignment distance position gene name gene id
GU305914.1 181118 181140 - 0 CACTTCATGCCCACTAATCT AGG ||||||||[||||||||||||]PAM NA Intergenic NA NA

T20964 ACTTCATGCCCACTAATCTAGGG 1000
Oligo fwd CACCACTTCATGCCCACTAATCTA
Oligo rev AAACTAGATTAGTGGGCATGAAGT
Chromosome start end strand MM target_seq PAM alignment distance position gene name gene id
GU305914.1 181117 181139 - 0 ACTTCATGCCCACTAATCTA GGG ||||||||[||||||||||||]PAM NA Intergenic NA NA

T20965 GGCCCTCAGAAAAGAGAGTCTGG 1000

<table>
<tr><td>Oligo fwd</td><td>CACCGGCCCTCAGAAAAGAGAGTC</td><td></td><td></td><td></td><td></td><td></td><td></td><td></td><td></td><td></td><td></td><td></td></tr>
<tr><td>Oligo rev</td><td>AAACGACTCTCTTTTCTGAGGGCC</td><td></td><td></td><td></td><td></td><td></td><td></td><td></td><td></td><td></td><td></td><td></td></tr>
<tr><td>Chromosome</td><td>start</td><td></td><td>end</td><td>strand</td><td>MM</td><td>target_seq</td><td>PAM</td><td>alignment</td><td>distance</td><td>position</td><td>gene name</td><td>gene id</td></tr>
<tr><td>GU305914.1</td><td></td><td>181096</td><td>181118</td><td>-</td><td>0</td><td>GGCCCTCAGAAAAGAGAGTC</td><td>TGG</td><td>||||||||[||||||||||||]PAM</td><td>NA</td><td>Intergenic</td><td>NA</td><td>NA</td></tr>
<tr><td>T20966</td><td>AGAGTCTGGATGAGCTACGTAGG</td><td></td><td>1000</td><td></td><td></td><td></td><td></td><td></td><td></td><td></td><td></td><td></td></tr>
<tr><td>Oligo fwd</td><td>CACCAGAGTCTGGATGAGCTACGT</td><td></td><td></td><td></td><td></td><td></td><td></td><td></td><td></td><td></td><td></td><td></td></tr>
<tr><td>Oligo rev</td><td>AAACACGTAGCTCATCCAGACTCT</td><td></td><td></td><td></td><td></td><td></td><td></td><td></td><td></td><td></td><td></td><td></td></tr>
<tr><td>Chromosome</td><td>start</td><td></td><td>end</td><td>strand</td><td>MM</td><td>target_seq</td><td>PAM</td><td>alignment</td><td>distance</td><td>position</td><td>gene name</td><td>gene id</td></tr>
<tr><td>GU305914.1</td><td></td><td>181082</td><td>181104</td><td>-</td><td>0</td><td>AGAGTCTGGATGAGCTACGT</td><td>AGG</td><td>||||||||[||||||||||||]PAM</td><td>NA</td><td>Intergenic</td><td>NA</td><td>NA</td></tr>
<tr><td>GU305914.1</td><td></td><td>166887</td><td>166909</td><td>-</td><td>4</td><td>AGCGACTGGATGAGCTTCTT</td><td>GGG</td><td>||-|-|||[||||||||-|-|]PAM</td><td>NA</td><td>Intergenic</td><td>NA</td><td>NA</td></tr>
<tr><td>T20967</td><td>CTGGATGAGCTACGTAGGTGAGG</td><td></td><td>1000</td><td></td><td></td><td></td><td></td><td></td><td></td><td></td><td></td><td></td></tr>
<tr><td>Oligo fwd</td><td>CACCCTGGATGAGCTACGTAGGTG</td><td></td><td></td><td></td><td></td><td></td><td></td><td></td><td></td><td></td><td></td><td></td></tr>
<tr><td>Oligo rev</td><td>AAACCACCTACGTAGCTCATCCAG</td><td></td><td></td><td></td><td></td><td></td><td></td><td></td><td></td><td></td><td></td><td></td></tr>
<tr><td>Chromosome</td><td>start</td><td></td><td>end</td><td>strand</td><td>MM</td><td>target_seq</td><td>PAM</td><td>alignment</td><td>distance</td><td>position</td><td>gene name</td><td>gene id</td></tr>
<tr><td>GU305914.1</td><td></td><td>181077</td><td>181099</td><td>-</td><td>0</td><td>CTGGATGAGCTACGTAGGTG</td><td>AGG</td><td>||||||||[||||||||||||]PAM</td><td>NA</td><td>Intergenic</td><td>NA</td><td>NA</td></tr>
<tr><td>T20968</td><td>CTACGTAGGTGAGGCCATAGTGG</td><td></td><td>1000</td><td></td><td></td><td></td><td></td><td></td><td></td><td></td><td></td><td></td></tr>
<tr><td>Oligo fwd</td><td>CACCCTACGTAGGTGAGGCCATAG</td><td></td><td></td><td></td><td></td><td></td><td></td><td></td><td></td><td></td><td></td><td></td></tr>
<tr><td>Oligo rev</td><td>AAACCTATGGCCTCACCTACGTAG</td><td></td><td></td><td></td><td></td><td></td><td></td><td></td><td></td><td></td><td></td><td></td></tr>
<tr><td>Chromosome</td><td>start</td><td></td><td>end</td><td>strand</td><td>MM</td><td>target_seq</td><td>PAM</td><td>alignment</td><td>distance</td><td>position</td><td>gene name</td><td>gene id</td></tr>
<tr><td>GU305914.1</td><td></td><td>181068</td><td>181090</td><td>-</td><td>0</td><td>CTACGTAGGTGAGGCCATAG</td><td>TGG</td><td>||||||||[||||||||||||]PAM</td><td>NA</td><td>Intergenic</td><td>NA</td><td>NA</td></tr>
<tr><td>T20969</td><td>CACTAGATGAGCGTGCCGCATGG</td><td></td><td>1000</td><td></td><td></td><td></td><td></td><td></td><td></td><td></td><td></td><td></td></tr>
<tr><td>Oligo fwd</td><td>CACCCACTAGATGAGCGTGCCGCA</td><td></td><td></td><td></td><td></td><td></td><td></td><td></td><td></td><td></td><td></td><td></td></tr>
<tr><td>Oligo rev</td><td>AAACTGCGGCACGCTCATCTAGTG</td><td></td><td></td><td></td><td></td><td></td><td></td><td></td><td></td><td></td><td></td><td></td></tr>
<tr><td>Chromosome</td><td>start</td><td></td><td>end</td><td>strand</td><td>MM</td><td>target_seq</td><td>PAM</td><td>alignment</td><td>distance</td><td>position</td><td>gene name</td><td>gene id</td></tr>
<tr><td>GU305914.1</td><td></td><td>181024</td><td>181046</td><td>-</td><td>0</td><td>CACTAGATGAGCGTGCCGCA</td><td>TGG</td><td>||||||||[||||||||||||]PAM</td><td>NA</td><td>Intergenic</td><td>NA</td><td>NA</td></tr>
<tr><td>T20970</td><td>ACTAGATGAGCGTGCCGCATGGG</td><td></td><td>1000</td><td></td><td></td><td></td><td></td><td></td><td></td><td></td><td></td><td></td></tr>
<tr><td>Oligo fwd</td><td>CACCACTAGATGAGCGTGCCGCAT</td><td></td><td></td><td></td><td></td><td></td><td></td><td></td><td></td><td></td><td></td><td></td></tr>
<tr><td>Oligo rev</td><td>AAACATGCGGCACGCTCATCTAGT</td><td></td><td></td><td></td><td></td><td></td><td></td><td></td><td></td><td></td><td></td><td></td></tr>
<tr><td>Chromosome</td><td>start</td><td></td><td>end</td><td>strand</td><td>MM</td><td>target_seq</td><td>PAM</td><td>alignment</td><td>distance</td><td>position</td><td>gene name</td><td>gene id</td></tr>
<tr><td>GU305914.1</td><td></td><td>181023</td><td>181045</td><td>-</td><td>0</td><td>ACTAGATGAGCGTGCCGCAT</td><td>GGG</td><td>||||||||[||||||||||||]PAM</td><td>NA</td><td>Intergenic</td><td>NA</td><td>NA</td></tr>
<tr><td>T20971</td><td>AAGACTGATTGTTCATTGCCTGG</td><td></td><td>1000</td><td></td><td></td><td></td><td></td><td></td><td></td><td></td><td></td><td></td></tr>
<tr><td>Oligo fwd</td><td>CACCAAGACTGATTGTTCATTGCC</td><td></td><td></td><td></td><td></td><td></td><td></td><td></td><td></td><td></td><td></td><td></td></tr>
<tr><td>Oligo rev</td><td>AAACGGCAATGAACAATCAGTCTT</td><td></td><td></td><td></td><td></td><td></td><td></td><td></td><td></td><td></td><td></td><td></td></tr>
<tr><td>Chromosome</td><td>start</td><td></td><td>end</td><td>strand</td><td>MM</td><td>target_seq</td><td>PAM</td><td>alignment</td><td>distance</td><td>position</td><td>gene name</td><td>gene id</td></tr>
<tr><td>GU305914.1</td><td></td><td>180998</td><td>181020</td><td>-</td><td>0</td><td>AAGACTGATTGTTCATTGCC</td><td>TGG</td><td>||||||||[||||||||||||]PAM</td><td>NA</td><td>Intergenic</td><td>NA</td><td>NA</td></tr>
<tr><td>T20972</td><td>AGACTGATTGTTCATTGCCTGGG</td><td></td><td>1000</td><td></td><td></td><td></td><td></td><td></td><td></td><td></td><td></td><td></td></tr>
<tr><td>Oligo fwd</td><td>CACCAGACTGATTGTTCATTGCCT</td><td></td><td></td><td></td><td></td><td></td><td></td><td></td><td></td><td></td><td></td><td></td></tr>
<tr><td>Oligo rev</td><td>AAACAGGCAATGAACAATCAGTCT</td><td></td><td></td><td></td><td></td><td></td><td></td><td></td><td></td><td></td><td></td><td></td></tr>
<tr><td>Chromosome</td><td>start</td><td></td><td>end</td><td>strand</td><td>MM</td><td>target_seq</td><td>PAM</td><td>alignment</td><td>distance</td><td>position</td><td>gene name</td><td>gene id</td></tr>
<tr><td>GU305914.1</td><td></td><td>180997</td><td>181019</td><td>-</td><td>0</td><td>AGACTGATTGTTCATTGCCT</td><td>GGG</td><td>||||||||[||||||||||||]PAM</td><td>NA</td><td>Intergenic</td><td>NA</td><td>NA</td></tr>
<tr><td>T20973</td><td>GACTGATTGTTCATTGCCTGGGG</td><td></td><td>1000</td><td></td><td></td><td></td><td></td><td></td><td></td><td></td><td></td><td></td></tr>
<tr><td>Oligo fwd</td><td>CACCGACTGATTGTTCATTGCCTG</td><td></td><td></td><td></td><td></td><td></td><td></td><td></td><td></td><td></td><td></td><td></td></tr>
<tr><td>Oligo rev</td><td>AAACCAGGCAATGAACAATCAGTC</td><td></td><td></td><td></td><td></td><td></td><td></td><td></td><td></td><td></td><td></td><td></td></tr>
<tr><td>Chromosome</td><td>start</td><td></td><td>end</td><td>strand</td><td>MM</td><td>target_seq</td><td>PAM</td><td>alignment</td><td>distance</td><td>position</td><td>gene name</td><td>gene id</td></tr>
<tr><td>GU305914.1</td><td></td><td>180996</td><td>181018</td><td>-</td><td>0</td><td>GACTGATTGTTCATTGCCTG</td><td>GGG</td><td>||||||||[||||||||||||]PAM</td><td>NA</td><td>Intergenic</td><td>NA</td><td>NA</td></tr>
<tr><td>T20974</td><td>TAAGTCAGAGATGTGTGTCCTGG</td><td></td><td>1000</td><td></td><td></td><td></td><td></td><td></td><td></td><td></td><td></td><td></td></tr>
<tr><td>Oligo fwd</td><td>CACCTAAGTCAGAGATGTGTGTCC</td><td></td><td></td><td></td><td></td><td></td><td></td><td></td><td></td><td></td><td></td><td></td></tr>
<tr><td>Oligo rev</td><td>AAACGGACACACATCTCTGACTTA</td><td></td><td></td><td></td><td></td><td></td><td></td><td></td><td></td><td></td><td></td><td></td></tr>
<tr><td>Chromosome</td><td>start</td><td></td><td>end</td><td>strand</td><td>MM</td><td>target_seq</td><td>PAM</td><td>alignment</td><td>distance</td><td>position</td><td>gene name</td><td>gene id</td></tr>
<tr><td>GU305914.1</td><td></td><td>180969</td><td>180991</td><td>-</td><td>0</td><td>TAAGTCAGAGATGTGTGTCC</td><td>TGG</td><td>||||||||[||||||||||||]PAM</td><td>NA</td><td>Intergenic</td><td>NA</td><td>NA</td></tr>
<tr><td>T20975</td><td>TCAGAGATGTGTGTCCTGGTTGG</td><td></td><td>1000</td><td></td><td></td><td></td><td></td><td></td><td></td><td></td><td></td><td></td></tr>
<tr><td>Oligo fwd</td><td>CACCTCAGAGATGTGTGTCCTGGT</td><td></td><td></td><td></td><td></td><td></td><td></td><td></td><td></td><td></td><td></td><td></td></tr>
<tr><td>Oligo rev</td><td>AAACACCAGGACACACATCTCTGA</td><td></td><td></td><td></td><td></td><td></td><td></td><td></td><td></td><td></td><td></td><td></td></tr>
<tr><td>Chromosome</td><td>start</td><td></td><td>end</td><td>strand</td><td>MM</td><td>target_seq</td><td>PAM</td><td>alignment</td><td>distance</td><td>position</td><td>gene name</td><td>gene id</td></tr>
<tr><td>GU305914.1</td><td></td><td>180965</td><td>180987</td><td>-</td><td>0</td><td>TCAGAGATGTGTGTCCTGGT</td><td>TGG</td><td>||||||||[||||||||||||]PAM</td><td>NA</td><td>Intergenic</td><td>NA</td><td>NA</td></tr>
<tr><td>T20976</td><td>CAGAGATGTGTGTCCTGGTTGGG</td><td></td><td>1000</td><td></td><td></td><td></td><td></td><td></td><td></td><td></td><td></td><td></td></tr>
<tr><td>Oligo fwd</td><td>CACCCAGAGATGTGTGTCCTGGTT</td><td></td><td></td><td></td><td></td><td></td><td></td><td></td><td></td><td></td><td></td><td></td></tr>
<tr><td>Oligo rev</td><td>AAACAACCAGGACACACATCTCTG</td><td></td><td></td><td></td><td></td><td></td><td></td><td></td><td></td><td></td><td></td><td></td></tr>
<tr><td>Chromosome</td><td>start</td><td></td><td>end</td><td>strand</td><td>MM</td><td>target_seq</td><td>PAM</td><td>alignment</td><td>distance</td><td>position</td><td>gene name</td><td>gene id</td></tr>
<tr><td>GU305914.1</td><td></td><td>180964</td><td>180986</td><td>-</td><td>0</td><td>CAGAGATGTGTGTCCTGGTT</td><td>GGG</td><td>||||||||[||||||||||||]PAM</td><td>NA</td><td>Intergenic</td><td>NA</td><td>NA</td></tr>
<tr><td>T20977</td><td>AGAGATGTGTGTCCTGGTTGGGG</td><td></td><td>1000</td><td></td><td></td><td></td><td></td><td></td><td></td><td></td><td></td><td></td></tr>
<tr><td>Oligo fwd</td><td>CACCAGAGATGTGTGTCCTGGTTG</td><td></td><td></td><td></td><td></td><td></td><td></td><td></td><td></td><td></td><td></td><td></td></tr>
<tr><td>Oligo rev</td><td>AAACCAACCAGGACACACATCTCT</td><td></td><td></td><td></td><td></td><td></td><td></td><td></td><td></td><td></td><td></td><td></td></tr>
<tr><td>Chromosome</td><td>start</td><td></td><td>end</td><td>strand</td><td>MM</td><td>target_seq</td><td>PAM</td><td>alignment</td><td>distance</td><td>position</td><td>gene name</td><td>gene id</td></tr>
<tr><td>GU305914.1</td><td></td><td>180963</td><td>180985</td><td>-</td><td>0</td><td>AGAGATGTGTGTCCTGGTTG</td><td>GGG</td><td>||||||||[||||||||||||]PAM</td><td>NA</td><td>Intergenic</td><td>NA</td><td>NA</td></tr>
<tr><td>T20978</td><td>GTGTCCTGGTTGGGGCCATCAGG</td><td></td><td>1000</td><td></td><td></td><td></td><td></td><td></td><td></td><td></td><td></td><td></td></tr>
<tr><td>Oligo fwd</td><td>CACCGTGTCCTGGTTGGGGCCATC</td><td></td><td></td><td></td><td></td><td></td><td></td><td></td><td></td><td></td><td></td><td></td></tr>
<tr><td>Oligo rev</td><td>AAACGATGGCCCCAACCAGGACAC</td><td></td><td></td><td></td><td></td><td></td><td></td><td></td><td></td><td></td><td></td><td></td></tr>
<tr><td>Chromosome</td><td>start</td><td></td><td>end</td><td>strand</td><td>MM</td><td>target_seq</td><td>PAM</td><td>alignment</td><td>distance</td><td>position</td><td>gene name</td><td>gene id</td></tr>
<tr><td>GU305914.1</td><td></td><td>180955</td><td>180977</td><td>-</td><td>0</td><td>GTGTCCTGGTTGGGGCCATC</td><td>AGG</td><td>||||||||[||||||||||||]PAM</td><td>NA</td><td>Intergenic</td><td>NA</td><td>NA</td></tr>
<tr><td>T20979</td><td>TCCTGGTTGGGGCCATCAGGAGG</td><td></td><td>1000</td><td></td><td></td><td></td><td></td><td></td><td></td><td></td><td></td><td></td></tr>
<tr><td>Oligo fwd</td><td>CACCTCCTGGTTGGGGCCATCAGG</td><td></td><td></td><td></td><td></td><td></td><td></td><td></td><td></td><td></td><td></td><td></td></tr>
<tr><td>Oligo rev</td><td>AAACCCTGATGGCCCCAACCAGGA</td><td></td><td></td><td></td><td></td><td></td><td></td><td></td><td></td><td></td><td></td><td></td></tr>
<tr><td>Chromosome</td><td>start</td><td></td><td>end</td><td>strand</td><td>MM</td><td>target_seq</td><td>PAM</td><td>alignment</td><td>distance</td><td>position</td><td>gene name</td><td>gene id</td></tr>
<tr><td>GU305914.1</td><td></td><td>180952</td><td>180974</td><td>-</td><td>0</td><td>TCCTGGTTGGGGCCATCAGG</td><td>AGG</td><td>||||||||[||||||||||||]PAM</td><td>NA</td><td>Intergenic</td><td>NA</td><td>NA</td></tr>
</table>

T20980 CTGTTACACCAAGCCTTTCCTGG 1000
Oligo fwd CACCCTGTTACACCAAGCCTTTCC
Oligo rev AAACGGAAAGGCTTGGTGTAACAG
Chromosome start end strand MM target_seq PAM alignment distance position gene name gene id
GU305914.1 180915 180937 - 0 CTGTTACACCAAGCCTTTCC TGG ||||||||[||||||||||||]PAM NA Intergenic NA NA

T20981 CCAAGCCTTTCCTGGATGCCAGG 1000
Oligo fwd CACCCCAAGCCTTTCCTGGATGCC
Oligo rev AAACGGCATCCAGGAAAGGCTTGG
Chromosome start end strand MM target_seq PAM alignment distance position gene name gene id
GU305914.1 180907 180929 - 0 CCAAGCCTTTCCTGGATGCC AGG ||||||||[||||||||||||]PAM NA Intergenic NA NA

T20982 CCTGGATGCCAGGTCTCAGCTGG 1000
Oligo fwd CACCCCTGGATGCCAGGTCTCAGC
Oligo rev AAACGCTGAGACCTGGCATCCAGG
Chromosome start end strand MM target_seq PAM alignment distance position gene name gene id
GU305914.1 180897 180919 - 0 CCTGGATGCCAGGTCTCAGC TGG ||||||||[||||||||||||]PAM NA Intergenic NA NA

T20983 CCAGGTCTCAGCTGGCAGTATGG 1000
Oligo fwd CACCCCAGGTCTCAGCTGGCAGTA
Oligo rev AAACTACTGCCAGCTGAGACCTGG
Chromosome start end strand MM target_seq PAM alignment distance position gene name gene id
GU305914.1 180889 180911 - 0 CCAGGTCTCAGCTGGCAGTA TGG ||||||||[||||||||||||]PAM NA Intergenic NA NA

T20984 GGCAGTATGGCAGCAGATGAAGG 1000
Oligo fwd CACCGGCAGTATGGCAGCAGATGA
Oligo rev AAACTCATCTGCTGCCATACTGCC
Chromosome start end strand MM target_seq PAM alignment distance position gene name gene id
GU305914.1 180876 180898 - 0 GGCAGTATGGCAGCAGATGA AGG ||||||||[||||||||||||]PAM NA Intergenic NA NA

T20985 TGAGAGCGTATCCACACATGTGG 1000
Oligo fwd CACCTGAGAGCGTATCCACACATG
Oligo rev AAACCATGTGTGGATACGCTCTCA
Chromosome start end strand MM target_seq PAM alignment distance position gene name gene id
GU305914.1 180843 180865 - 0 TGAGAGCGTATCCACACATG TGG ||||||||[||||||||||||]PAM NA Intergenic NA NA

T20986 CGTATCCACACATGTGGTAGTGG 1000
Oligo fwd CACCCGTATCCACACATGTGGTAG
Oligo rev AAACCTACCACATGTGTGGATACG
Chromosome start end strand MM target_seq PAM alignment distance position gene name gene id
GU305914.1 180837 180859 - 0 CGTATCCACACATGTGGTAG TGG ||||||||[||||||||||||]PAM NA Intergenic NA NA

T20987 ATCCACACATGTGGTAGTGGTGG 1000
Oligo fwd CACCATCCACACATGTGGTAGTGG
Oligo rev AAACCCACTACCACATGTGTGGAT
Chromosome start end strand MM target_seq PAM alignment distance position gene name gene id
GU305914.1 180834 180856 - 0 ATCCACACATGTGGTAGTGG TGG ||||||||[||||||||||||]PAM NA Intergenic NA NA

T20988 TGGTGGAAGCTTTGAAGCTCAGG 1000
Oligo fwd CACCTGGTGGAAGCTTTGAAGCTC
Oligo rev AAACGAGCTTCAAAGCTTCCACCA
Chromosome start end strand MM target_seq PAM alignment distance position gene name gene id
GU305914.1 180817 180839 - 0 TGGTGGAAGCTTTGAAGCTC AGG ||||||||[||||||||||||]PAM NA Intergenic NA NA

T20989 GGTGGAAGCTTTGAAGCTCAGGG 1000
Oligo fwd CACCGGTGGAAGCTTTGAAGCTCA
Oligo rev AAACTGAGCTTCAAAGCTTCCACC
Chromosome start end strand MM target_seq PAM alignment distance position gene name gene id
GU305914.1 180816 180838 - 0 GGTGGAAGCTTTGAAGCTCA GGG ||||||||[||||||||||||]PAM NA Intergenic NA NA

T20990 TTTGAAGCTCAGGGAGAATCTGG 1000
Oligo fwd CACCTTTGAAGCTCAGGGAGAATC
Oligo rev AAACGATTCTCCCTGAGCTTCAAA
Chromosome start end strand MM target_seq PAM alignment distance position gene name gene id
GU305914.1 180807 180829 - 0 TTTGAAGCTCAGGGAGAATC TGG ||||||||[||||||||||||]PAM NA Intergenic NA NA

T20991 GCTCAGGGAGAATCTGGCTAAGG 1000
Oligo fwd CACCGCTCAGGGAGAATCTGGCTA
Oligo rev AAACTAGCCAGATTCTCCCTGAGC
Chromosome start end strand MM target_seq PAM alignment distance position gene name gene id
GU305914.1 180801 180823 - 0 GCTCAGGGAGAATCTGGCTA AGG ||||||||[||||||||||||]PAM NA Intergenic NA NA

T20992 GAATCTGGCTAAGGCAGTGCAGG 1000
Oligo fwd CACCGAATCTGGCTAAGGCAGTGC
Oligo rev AAACGCACTGCCTTAGCCAGATTC
Chromosome start end strand MM target_seq PAM alignment distance position gene name gene id
GU305914.1 180792 180814 - 0 GAATCTGGCTAAGGCAGTGC AGG ||||||||[||||||||||||]PAM NA Intergenic NA NA

T20993 ATGAGAGACACCAATATCACAGG 1000
Oligo fwd CACCATGAGAGACACCAATATCAC
Oligo rev AAACGTGATATTGGTGTCTCTCAT
Chromosome start end strand MM target_seq PAM alignment distance position gene name gene id
GU305914.1 180757 180779 - 0 ATGAGAGACACCAATATCAC AGG ||||||||[||||||||||||]PAM NA Intergenic NA NA

T20994 TGAGAGACACCAATATCACAGGG 1000
Oligo fwd CACCTGAGAGACACCAATATCACA
Oligo rev AAACTGTGATATTGGTGTCTCTCA
Chromosome start end strand MM target_seq PAM alignment distance position gene name gene id

GU305914.1 180756 180778 - 0 TGAGAGACACCAATATCACA GGG ||||||||[||||||||||||]PAM NA Intergenic NA NA

T20995 TCTAGAGACCACCTGTATGCTGG 1000
Oligo fwd CACCTCTAGAGACCACCTGTATGC
Oligo rev AAACGCATACAGGTGGTCTCTAGA
Chromosome start end strand MM target_seq PAM alignment distance position gene name gene id
GU305914.1 180699 180721 - 0 TCTAGAGACCACCTGTATGC TGG ||||||||[||||||||||||]PAM NA Intergenic NA NA

T20996 ACCACCTGTATGCTGGCCAGAGG 1000
Oligo fwd CACCACCACCTGTATGCTGGCCAG
Oligo rev AAACCTGGCCAGCATACAGGTGGT
Chromosome start end strand MM target_seq PAM alignment distance position gene name gene id
GU305914.1 180692 180714 - 0 ACCACCTGTATGCTGGCCAG AGG ||||||||[||||||||||||]PAM NA Intergenic NA NA

T20997 GCTGGCCAGAGGCAGAACCTTGG 1000
Oligo fwd CACCGCTGGCCAGAGGCAGAACCT
Oligo rev AAACAGGTTCTGCCTCTGGCCAGC
Chromosome start end strand MM target_seq PAM alignment distance position gene name gene id
GU305914.1 180681 180703 - 0 GCTGGCCAGAGGCAGAACCT TGG ||||||||[||||||||||||]PAM NA Intergenic NA NA

T20998 TGGCTACCTTAGCTGATCTGAGG 1000
Oligo fwd CACCTGGCTACCTTAGCTGATCTG
Oligo rev AAACCAGATCAGCTAAGGTAGCCA
Chromosome start end strand MM target_seq PAM alignment distance position gene name gene id
GU305914.1 180661 180683 - 0 TGGCTACCTTAGCTGATCTG AGG ||||||||[||||||||||||]PAM NA Intergenic NA NA

T20999 TTGATAGCTTCACCATGCCTAGG 1000
Oligo fwd CACCTTGATAGCTTCACCATGCCT
Oligo rev AAACAGGCATGGTGAAGCTATCAA
Chromosome start end strand MM target_seq PAM alignment distance position gene name gene id
GU305914.1 180589 180611 - 0 TTGATAGCTTCACCATGCCT AGG ||||||||[||||||||||||]PAM NA Intergenic NA NA

T21000 TTCACCATGCCTAGGATCAGAGG 1000
Oligo fwd CACCTTCACCATGCCTAGGATCAG
Oligo rev AAACCTGATCCTAGGCATGGTGAA
Chromosome start end strand MM target_seq PAM alignment distance position gene name gene id
GU305914.1 180581 180603 - 0 TTCACCATGCCTAGGATCAG AGG ||||||||[||||||||||||]PAM NA Intergenic NA NA

T21001 GAGGTGCTATGAAACAAGCAAGG 1000
Oligo fwd CACCGAGGTGCTATGAAACAAGCA
Oligo rev AAACTGCTTGTTTCATAGCACCTC
Chromosome start end strand MM target_seq PAM alignment distance position gene name gene id
GU305914.1 180562 180584 - 0 GAGGTGCTATGAAACAAGCA AGG ||||||||[||||||||||||]PAM NA Intergenic NA NA

T21002 AGCAAGGTGTATGAGTTATGTGG 1000
Oligo fwd CACCAGCAAGGTGTATGAGTTATG
Oligo rev AAACCATAACTCATACACCTTGCT
Chromosome start end strand MM target_seq PAM alignment distance position gene name gene id
GU305914.1 180546 180568 - 0 AGCAAGGTGTATGAGTTATG TGG ||||||||[||||||||||||]PAM NA Intergenic NA NA

T21003 GGTGTATGAGTTATGTGGAGAGG 1000
Oligo fwd CACCGGTGTATGAGTTATGTGGAG
Oligo rev AAACCTCCACATAACTCATACACC
Chromosome start end strand MM target_seq PAM alignment distance position gene name gene id
GU305914.1 180541 180563 - 0 GGTGTATGAGTTATGTGGAG AGG ||||||||[||||||||||||]PAM NA Intergenic NA NA

T21004 TGTGGAGAGGACCATCTCTCTGG 1000
Oligo fwd CACCTGTGGAGAGGACCATCTCTC
Oligo rev AAACGAGAGATGGTCCTCTCCACA
Chromosome start end strand MM target_seq PAM alignment distance position gene name gene id
GU305914.1 180528 180550 - 0 TGTGGAGAGGACCATCTCTC TGG ||||||||[||||||||||||]PAM NA Intergenic NA NA

T21005 TAGCTGATAGAGTCCATAACAGG 1000
Oligo fwd CACCTAGCTGATAGAGTCCATAAC
Oligo rev AAACGTTATGGACTCTATCAGCTA
Chromosome start end strand MM target_seq PAM alignment distance position gene name gene id
GU305914.1 180490 180512 - 0 TAGCTGATAGAGTCCATAAC AGG ||||||||[||||||||||||]PAM NA Intergenic NA NA

T21006 CAGGAGTGCACCATCACCCCAGG 1000
Oligo fwd CACCCAGGAGTGCACCATCACCCC
Oligo rev AAACGGGGTGATGGTGCACTCCTG
Chromosome start end strand MM target_seq PAM alignment distance position gene name gene id
GU305914.1 180471 180493 - 0 CAGGAGTGCACCATCACCCC AGG ||||||||[||||||||||||]PAM NA Intergenic NA NA

T21007 AGGAGTGCACCATCACCCCAGGG 1000
Oligo fwd CACCAGGAGTGCACCATCACCCCA
Oligo rev AAACTGGGGTGATGGTGCACTCCT
Chromosome start end strand MM target_seq PAM alignment distance position gene name gene id
GU305914.1 180470 180492 - 0 AGGAGTGCACCATCACCCCA GGG ||||||||[||||||||||||]PAM NA Intergenic NA NA

T21008 AGTGATAGAAGAACAGCAGCAGG 1000
Oligo fwd CACCAGTGATAGAAGAACAGCAGC
Oligo rev AAACGCTGCTGTTCTTCTATCACT
Chromosome start end strand MM target_seq PAM alignment distance position gene name gene id
GU305914.1 180447 180469 - 0 AGTGATAGAAGAACAGCAGC AGG ||||||||[||||||||||||]PAM NA Intergenic NA NA

T21009 ATAGAAGAACAGCAGCAGGCAGG 1000
Oligo fwd CACCATAGAAGAACAGCAGCAGGC
Oligo rev AAACGCCTGCTGCTGTTCTTCTAT

<table>
<tr><td>Chromosome</td><td>start</td><td>end</td><td>strand</td><td>MM</td><td>target_seq</td><td>PAM</td><td>alignment</td><td>distance</td><td>position</td><td>gene name</td><td>gene id</td></tr>
<tr><td>GU305914.1</td><td>180443</td><td>180465</td><td>-</td><td>0</td><td>ATAGAAGAACAGCAGCAGGC</td><td>AGG</td><td>||||||||[||||||||||||]PAM</td><td>NA</td><td>Intergenic</td><td>NA</td><td>NA</td></tr>
<tr><td></td><td></td><td></td><td></td><td></td><td></td><td></td><td></td><td></td><td></td><td></td><td></td></tr>
<tr><td>T21010</td><td>TAGAAGAACAGCAGCAGGCAGGG</td><td>1000</td><td></td><td></td><td></td><td></td><td></td><td></td><td></td><td></td><td></td></tr>
<tr><td>Oligo fwd</td><td>CACCTAGAAGAACAGCAGCAGGCA</td><td></td><td></td><td></td><td></td><td></td><td></td><td></td><td></td><td></td><td></td></tr>
<tr><td>Oligo rev</td><td>AAACTGCCTGCTGCTGTTCTTCTA</td><td></td><td></td><td></td><td></td><td></td><td></td><td></td><td></td><td></td><td></td></tr>
<tr><td>Chromosome</td><td>start</td><td>end</td><td>strand</td><td>MM</td><td>target_seq</td><td>PAM</td><td>alignment</td><td>distance</td><td>position</td><td>gene name</td><td>gene id</td></tr>
<tr><td>GU305914.1</td><td>180442</td><td>180464</td><td>-</td><td>0</td><td>TAGAAGAACAGCAGCAGGCA</td><td>GGG</td><td>||||||||[||||||||||||]PAM</td><td>NA</td><td>Intergenic</td><td>NA</td><td>NA</td></tr>
<tr><td></td><td></td><td></td><td></td><td></td><td></td><td></td><td></td><td></td><td></td><td></td><td></td></tr>
<tr><td>T21011</td><td>AGAAGAACAGCAGCAGGCAGGGG</td><td>1000</td><td></td><td></td><td></td><td></td><td></td><td></td><td></td><td></td><td></td></tr>
<tr><td>Oligo fwd</td><td>CACCAGAAGAACAGCAGCAGGCAG</td><td></td><td></td><td></td><td></td><td></td><td></td><td></td><td></td><td></td><td></td></tr>
<tr><td>Oligo rev</td><td>AAACCTGCCTGCTGCTGTTCTTCT</td><td></td><td></td><td></td><td></td><td></td><td></td><td></td><td></td><td></td><td></td></tr>
<tr><td>Chromosome</td><td>start</td><td>end</td><td>strand</td><td>MM</td><td>target_seq</td><td>PAM</td><td>alignment</td><td>distance</td><td>position</td><td>gene name</td><td>gene id</td></tr>
<tr><td>GU305914.1</td><td>180441</td><td>180463</td><td>-</td><td>0</td><td>AGAAGAACAGCAGCAGGCAG</td><td>GGG</td><td>||||||||[||||||||||||]PAM</td><td>NA</td><td>Intergenic</td><td>NA</td><td>NA</td></tr>
<tr><td></td><td></td><td></td><td></td><td></td><td></td><td></td><td></td><td></td><td></td><td></td><td></td></tr>
<tr><td>T21012</td><td>AGAACAGCAGCAGGCAGGGGAGG</td><td>1000</td><td></td><td></td><td></td><td></td><td></td><td></td><td></td><td></td><td></td></tr>
<tr><td>Oligo fwd</td><td>CACCAGAACAGCAGCAGGCAGGGG</td><td></td><td></td><td></td><td></td><td></td><td></td><td></td><td></td><td></td><td></td></tr>
<tr><td>Oligo rev</td><td>AAACCCCCTGCCTGCTGCTGTTCT</td><td></td><td></td><td></td><td></td><td></td><td></td><td></td><td></td><td></td><td></td></tr>
<tr><td>Chromosome</td><td>start</td><td>end</td><td>strand</td><td>MM</td><td>target_seq</td><td>PAM</td><td>alignment</td><td>distance</td><td>position</td><td>gene name</td><td>gene id</td></tr>
<tr><td>GU305914.1</td><td>180438</td><td>180460</td><td>-</td><td>0</td><td>AGAACAGCAGCAGGCAGGGG</td><td>AGG</td><td>||||||||[||||||||||||]PAM</td><td>NA</td><td>Intergenic</td><td>NA</td><td>NA</td></tr>
<tr><td></td><td></td><td></td><td></td><td></td><td></td><td></td><td></td><td></td><td></td><td></td><td></td></tr>
<tr><td>T21013</td><td>GGAAGAACAGCAACAGCAGCAGG</td><td>1000</td><td></td><td></td><td></td><td></td><td></td><td></td><td></td><td></td><td></td></tr>
<tr><td>Oligo fwd</td><td>CACCGGAAGAACAGCAACAGCAGC</td><td></td><td></td><td></td><td></td><td></td><td></td><td></td><td></td><td></td><td></td></tr>
<tr><td>Oligo rev</td><td>AAACGCTGCTGTTGCTGTTCTTCC</td><td></td><td></td><td></td><td></td><td></td><td></td><td></td><td></td><td></td><td></td></tr>
<tr><td>Chromosome</td><td>start</td><td>end</td><td>strand</td><td>MM</td><td>target_seq</td><td>PAM</td><td>alignment</td><td>distance</td><td>position</td><td>gene name</td><td>gene id</td></tr>
<tr><td>GU305914.1</td><td>180417</td><td>180439</td><td>-</td><td>0</td><td>GGAAGAACAGCAACAGCAGC</td><td>AGG</td><td>||||||||[||||||||||||]PAM</td><td>NA</td><td>Intergenic</td><td>NA</td><td>NA</td></tr>
<tr><td></td><td></td><td></td><td></td><td></td><td></td><td></td><td></td><td></td><td></td><td></td><td></td></tr>
<tr><td>T21014</td><td>TGATCCTGAGATGCCCCCTCTGG</td><td>1000</td><td></td><td></td><td></td><td></td><td></td><td></td><td></td><td></td><td></td></tr>
<tr><td>Oligo fwd</td><td>CACCTGATCCTGAGATGCCCCCTC</td><td></td><td></td><td></td><td></td><td></td><td></td><td></td><td></td><td></td><td></td></tr>
<tr><td>Oligo rev</td><td>AAACGAGGGGGCATCTCAGGATCA</td><td></td><td></td><td></td><td></td><td></td><td></td><td></td><td></td><td></td><td></td></tr>
<tr><td>Chromosome</td><td>start</td><td>end</td><td>strand</td><td>MM</td><td>target_seq</td><td>PAM</td><td>alignment</td><td>distance</td><td>position</td><td>gene name</td><td>gene id</td></tr>
<tr><td>GU305914.1</td><td>180384</td><td>180406</td><td>-</td><td>0</td><td>TGATCCTGAGATGCCCCCTC</td><td>TGG</td><td>||||||||[||||||||||||]PAM</td><td>NA</td><td>Intergenic</td><td>NA</td><td>NA</td></tr>
<tr><td></td><td></td><td></td><td></td><td></td><td></td><td></td><td></td><td></td><td></td><td></td><td></td></tr>
<tr><td>T21015</td><td>GATGCCCCCTCTGGAAAGAGAGG</td><td>1000</td><td></td><td></td><td></td><td></td><td></td><td></td><td></td><td></td><td></td></tr>
<tr><td>Oligo fwd</td><td>CACCGATGCCCCCTCTGGAAAGAG</td><td></td><td></td><td></td><td></td><td></td><td></td><td></td><td></td><td></td><td></td></tr>
<tr><td>Oligo rev</td><td>AAACCTCTTTCCAGAGGGGGCATC</td><td></td><td></td><td></td><td></td><td></td><td></td><td></td><td></td><td></td><td></td></tr>
<tr><td>Chromosome</td><td>start</td><td>end</td><td>strand</td><td>MM</td><td>target_seq</td><td>PAM</td><td>alignment</td><td>distance</td><td>position</td><td>gene name</td><td>gene id</td></tr>
<tr><td>GU305914.1</td><td>180375</td><td>180397</td><td>-</td><td>0</td><td>GATGCCCCCTCTGGAAAGAG</td><td>AGG</td><td>||||||||[||||||||||||]PAM</td><td>NA</td><td>Intergenic</td><td>NA</td><td>NA</td></tr>
<tr><td></td><td></td><td></td><td></td><td></td><td></td><td></td><td></td><td></td><td></td><td></td><td></td></tr>
<tr><td>T21016</td><td>TCTGGAAAGAGAGGAAGAGCAGG</td><td>1000</td><td></td><td></td><td></td><td></td><td></td><td></td><td></td><td></td><td></td></tr>
<tr><td>Oligo fwd</td><td>CACCTCTGGAAAGAGAGGAAGAGC</td><td></td><td></td><td></td><td></td><td></td><td></td><td></td><td></td><td></td><td></td></tr>
<tr><td>Oligo rev</td><td>AAACGCTCTTCCTCTCTTTCCAGA</td><td></td><td></td><td></td><td></td><td></td><td></td><td></td><td></td><td></td><td></td></tr>
<tr><td>Chromosome</td><td>start</td><td>end</td><td>strand</td><td>MM</td><td>target_seq</td><td>PAM</td><td>alignment</td><td>distance</td><td>position</td><td>gene name</td><td>gene id</td></tr>
<tr><td>GU305914.1</td><td>180366</td><td>180388</td><td>-</td><td>0</td><td>TCTGGAAAGAGAGGAAGAGC</td><td>AGG</td><td>||||||||[||||||||||||]PAM</td><td>NA</td><td>Intergenic</td><td>NA</td><td>NA</td></tr>
<tr><td></td><td></td><td></td><td></td><td></td><td></td><td></td><td></td><td></td><td></td><td></td><td></td></tr>
<tr><td>T21017</td><td>GGAAAGAGAGGAAGAGCAGGAGG</td><td>1000</td><td></td><td></td><td></td><td></td><td></td><td></td><td></td><td></td><td></td></tr>
<tr><td>Oligo fwd</td><td>CACCGGAAAGAGAGGAAGAGCAGG</td><td></td><td></td><td></td><td></td><td></td><td></td><td></td><td></td><td></td><td></td></tr>
<tr><td>Oligo rev</td><td>AAACCCTGCTCTTCCTCTCTTTCC</td><td></td><td></td><td></td><td></td><td></td><td></td><td></td><td></td><td></td><td></td></tr>
<tr><td>Chromosome</td><td>start</td><td>end</td><td>strand</td><td>MM</td><td>target_seq</td><td>PAM</td><td>alignment</td><td>distance</td><td>position</td><td>gene name</td><td>gene id</td></tr>
<tr><td>GU305914.1</td><td>180363</td><td>180385</td><td>-</td><td>0</td><td>GGAAAGAGAGGAAGAGCAGG</td><td>AGG</td><td>||||||||[||||||||||||]PAM</td><td>NA</td><td>Intergenic</td><td>NA</td><td>NA</td></tr>
<tr><td></td><td></td><td></td><td></td><td></td><td></td><td></td><td></td><td></td><td></td><td></td><td></td></tr>
<tr><td>T21018</td><td>GGAAGAGCAGGAGGATGAGCAGG</td><td>1000</td><td></td><td></td><td></td><td></td><td></td><td></td><td></td><td></td><td></td></tr>
<tr><td>Oligo fwd</td><td>CACCGGAAGAGCAGGAGGATGAGC</td><td></td><td></td><td></td><td></td><td></td><td></td><td></td><td></td><td></td><td></td></tr>
<tr><td>Oligo rev</td><td>AAACGCTCATCCTCCTGCTCTTCC</td><td></td><td></td><td></td><td></td><td></td><td></td><td></td><td></td><td></td><td></td></tr>
<tr><td>Chromosome</td><td>start</td><td>end</td><td>strand</td><td>MM</td><td>target_seq</td><td>PAM</td><td>alignment</td><td>distance</td><td>position</td><td>gene name</td><td>gene id</td></tr>
<tr><td>GU305914.1</td><td>180354</td><td>180376</td><td>-</td><td>0</td><td>GGAAGAGCAGGAGGATGAGC</td><td>AGG</td><td>||||||||[||||||||||||]PAM</td><td>NA</td><td>Intergenic</td><td>NA</td><td>NA</td></tr>
<tr><td></td><td></td><td></td><td></td><td></td><td></td><td></td><td></td><td></td><td></td><td></td><td></td></tr>
<tr><td>T21019</td><td>GCAGGAGGATGAGCAGGTAGAGG</td><td>1000</td><td></td><td></td><td></td><td></td><td></td><td></td><td></td><td></td><td></td></tr>
<tr><td>Oligo fwd</td><td>CACCGCAGGAGGATGAGCAGGTAG</td><td></td><td></td><td></td><td></td><td></td><td></td><td></td><td></td><td></td><td></td></tr>
<tr><td>Oligo rev</td><td>AAACCTACCTGCTCATCCTCCTGC</td><td></td><td></td><td></td><td></td><td></td><td></td><td></td><td></td><td></td><td></td></tr>
<tr><td>Chromosome</td><td>start</td><td>end</td><td>strand</td><td>MM</td><td>target_seq</td><td>PAM</td><td>alignment</td><td>distance</td><td>position</td><td>gene name</td><td>gene id</td></tr>
<tr><td>GU305914.1</td><td>180348</td><td>180370</td><td>-</td><td>0</td><td>GCAGGAGGATGAGCAGGTAG</td><td>AGG</td><td>||||||||[||||||||||||]PAM</td><td>NA</td><td>Intergenic</td><td>NA</td><td>NA</td></tr>
<tr><td></td><td></td><td></td><td></td><td></td><td></td><td></td><td></td><td></td><td></td><td></td><td></td></tr>
<tr><td>T21020</td><td>GGAGGATGAGCAGGTAGAGGAGG</td><td>1000</td><td></td><td></td><td></td><td></td><td></td><td></td><td></td><td></td><td></td></tr>
<tr><td>Oligo fwd</td><td>CACCGGAGGATGAGCAGGTAGAGG</td><td></td><td></td><td></td><td></td><td></td><td></td><td></td><td></td><td></td><td></td></tr>
<tr><td>Oligo rev</td><td>AAACCCTCTACCTGCTCATCCTCC</td><td></td><td></td><td></td><td></td><td></td><td></td><td></td><td></td><td></td><td></td></tr>
<tr><td>Chromosome</td><td>start</td><td>end</td><td>strand</td><td>MM</td><td>target_seq</td><td>PAM</td><td>alignment</td><td>distance</td><td>position</td><td>gene name</td><td>gene id</td></tr>
<tr><td>GU305914.1</td><td>180345</td><td>180367</td><td>-</td><td>0</td><td>GGAGGATGAGCAGGTAGAGG</td><td>AGG</td><td>||||||||[||||||||||||]PAM</td><td>NA</td><td>Intergenic</td><td>NA</td><td>NA</td></tr>
<tr><td></td><td></td><td></td><td></td><td></td><td></td><td></td><td></td><td></td><td></td><td></td><td></td></tr>
<tr><td>T21021</td><td>GGAGGAGCCCCCTGCAGATGAGG</td><td>1000</td><td></td><td></td><td></td><td></td><td></td><td></td><td></td><td></td><td></td></tr>
<tr><td>Oligo fwd</td><td>CACCGGAGGAGCCCCCTGCAGATG</td><td></td><td></td><td></td><td></td><td></td><td></td><td></td><td></td><td></td><td></td></tr>
<tr><td>Oligo rev</td><td>AAACCATCTGCAGGGGGCTCCTCC</td><td></td><td></td><td></td><td></td><td></td><td></td><td></td><td></td><td></td><td></td></tr>
<tr><td>Chromosome</td><td>start</td><td>end</td><td>strand</td><td>MM</td><td>target_seq</td><td>PAM</td><td>alignment</td><td>distance</td><td>position</td><td>gene name</td><td>gene id</td></tr>
<tr><td>GU305914.1</td><td>180327</td><td>180349</td><td>-</td><td>0</td><td>GGAGGAGCCCCCTGCAGATG</td><td>AGG</td><td>||||||||[||||||||||||]PAM</td><td>NA</td><td>Intergenic</td><td>NA</td><td>NA</td></tr>
<tr><td></td><td></td><td></td><td></td><td></td><td></td><td></td><td></td><td></td><td></td><td></td><td></td></tr>
<tr><td>T21022</td><td>GGAGCCCCCTGCAGATGAGGAGG</td><td>1000</td><td></td><td></td><td></td><td></td><td></td><td></td><td></td><td></td><td></td></tr>
<tr><td>Oligo fwd</td><td>CACCGGAGCCCCCTGCAGATGAGG</td><td></td><td></td><td></td><td></td><td></td><td></td><td></td><td></td><td></td><td></td></tr>
<tr><td>Oligo rev</td><td>AAACCCTCATCTGCAGGGGGCTCC</td><td></td><td></td><td></td><td></td><td></td><td></td><td></td><td></td><td></td><td></td></tr>
<tr><td>Chromosome</td><td>start</td><td>end</td><td>strand</td><td>MM</td><td>target_seq</td><td>PAM</td><td>alignment</td><td>distance</td><td>position</td><td>gene name</td><td>gene id</td></tr>
<tr><td>GU305914.1</td><td>180324</td><td>180346</td><td>-</td><td>0</td><td>GGAGCCCCCTGCAGATGAGG</td><td>AGG</td><td>||||||||[||||||||||||]PAM</td><td>NA</td><td>Intergenic</td><td>NA</td><td>NA</td></tr>
<tr><td></td><td></td><td></td><td></td><td></td><td></td><td></td><td></td><td></td><td></td><td></td><td></td></tr>
<tr><td>T21023</td><td>CCCCCTGCAGATGAGGAGGAAGG</td><td>1000</td><td></td><td></td><td></td><td></td><td></td><td></td><td></td><td></td><td></td></tr>
<tr><td>Oligo fwd</td><td>CACCCCCCCTGCAGATGAGGAGGA</td><td></td><td></td><td></td><td></td><td></td><td></td><td></td><td></td><td></td><td></td></tr>
<tr><td>Oligo rev</td><td>AAACTCCTCCTCATCTGCAGGGGG</td><td></td><td></td><td></td><td></td><td></td><td></td><td></td><td></td><td></td><td></td></tr>
<tr><td>Chromosome</td><td>start</td><td>end</td><td>strand</td><td>MM</td><td>target_seq</td><td>PAM</td><td>alignment</td><td>distance</td><td>position</td><td>gene name</td><td>gene id</td></tr>
<tr><td>GU305914.1</td><td>180320</td><td>180342</td><td>-</td><td>0</td><td>CCCCCTGCAGATGAGGAGGA</td><td>AGG</td><td>||||||||[||||||||||||]PAM</td><td>NA</td><td>Intergenic</td><td>NA</td><td>NA</td></tr>
<tr><td>GU305914.1</td><td>38884</td><td>38906</td><td>-</td><td>4</td><td>CCCGCGGCAGATGTGGAGGC</td><td>TGG</td><td>|||-|-||[|||||-|||||-]PAM</td><td>NA</td><td>Intergenic</td><td>NA</td><td>NA</td></tr>
<tr><td></td><td></td><td></td><td></td><td></td><td></td><td></td><td></td><td></td><td></td><td></td><td></td></tr>
<tr><td>T21024</td><td>CCTGCAGATGAGGAGGAAGGAGG</td><td>1000</td><td></td><td></td><td></td><td></td><td></td><td></td><td></td><td></td><td></td></tr>
</table>

<table>
<tr><td>Oligo fwd</td><td>CACCCCTGCAGATGAGGAGGAAGG</td></tr>
<tr><td>Oligo rev</td><td>AAACCCTTCCTCCTCATCTGCAGG</td></tr>
<tr><td>Chromosome</td><td>start</td><td>end</td><td>strand</td><td>MM</td><td>target_seq</td><td>PAM</td><td>alignment</td><td>distance</td><td>position</td><td>gene name</td><td>gene id</td></tr>
<tr><td>GU305914.1</td><td>180317</td><td>180339</td><td>-</td><td>0</td><td>CCTGCAGATGAGGAGGAAGG</td><td>AGG</td><td>||||||||[||||||||||||]PAM</td><td>NA</td><td>Intergenic</td><td>NA</td><td>NA</td></tr>
<tr><td>T21025</td><td>GAGGAGGAAGGAGGAGCTGTTGG</td><td>1000</td></tr>
<tr><td>Oligo fwd</td><td>CACCGAGGAGGAAGGAGGAGCTGT</td></tr>
<tr><td>Oligo rev</td><td>AAACACAGCTCCTCCTTCCTCCTC</td></tr>
<tr><td>Chromosome</td><td>start</td><td>end</td><td>strand</td><td>MM</td><td>target_seq</td><td>PAM</td><td>alignment</td><td>distance</td><td>position</td><td>gene name</td><td>gene id</td></tr>
<tr><td>GU305914.1</td><td>180308</td><td>180330</td><td>-</td><td>0</td><td>GAGGAGGAAGGAGGAGCTGT</td><td>TGG</td><td>||||||||[||||||||||||]PAM</td><td>NA</td><td>Intergenic</td><td>NA</td><td>NA</td></tr>
<tr><td>T21026</td><td>GAGGAAGGAGGAGCTGTTGGTGG</td><td>1000</td></tr>
<tr><td>Oligo fwd</td><td>CACCGAGGAAGGAGGAGCTGTTGG</td></tr>
<tr><td>Oligo rev</td><td>AAACCCAACAGCTCCTCCTTCCTC</td></tr>
<tr><td>Chromosome</td><td>start</td><td>end</td><td>strand</td><td>MM</td><td>target_seq</td><td>PAM</td><td>alignment</td><td>distance</td><td>position</td><td>gene name</td><td>gene id</td></tr>
<tr><td>GU305914.1</td><td>180305</td><td>180327</td><td>-</td><td>0</td><td>GAGGAAGGAGGAGCTGTTGG</td><td>TGG</td><td>||||||||[||||||||||||]PAM</td><td>NA</td><td>Intergenic</td><td>NA</td><td>NA</td></tr>
<tr><td>T21027</td><td>TGTTGGTGGTGTCACTCAAGAGG</td><td>1000</td></tr>
<tr><td>Oligo fwd</td><td>CACCTGTTGGTGGTGTCACTCAAG</td></tr>
<tr><td>Oligo rev</td><td>AAACCTTGAGTGACACCACCAACA</td></tr>
<tr><td>Chromosome</td><td>start</td><td>end</td><td>strand</td><td>MM</td><td>target_seq</td><td>PAM</td><td>alignment</td><td>distance</td><td>position</td><td>gene name</td><td>gene id</td></tr>
<tr><td>GU305914.1</td><td>180291</td><td>180313</td><td>-</td><td>0</td><td>TGTTGGTGGTGTCACTCAAG</td><td>AGG</td><td>||||||||[||||||||||||]PAM</td><td>NA</td><td>Intergenic</td><td>NA</td><td>NA</td></tr>
<tr><td>T21028</td><td>GTCACTCAAGAGGAACCTGCTGG</td><td>1000</td></tr>
<tr><td>Oligo fwd</td><td>CACCGTCACTCAAGAGGAACCTGC</td></tr>
<tr><td>Oligo rev</td><td>AAACGCAGGTTCCTCTTGAGTGAC</td></tr>
<tr><td>Chromosome</td><td>start</td><td>end</td><td>strand</td><td>MM</td><td>target_seq</td><td>PAM</td><td>alignment</td><td>distance</td><td>position</td><td>gene name</td><td>gene id</td></tr>
<tr><td>GU305914.1</td><td>180281</td><td>180303</td><td>-</td><td>0</td><td>GTCACTCAAGAGGAACCTGC</td><td>TGG</td><td>||||||||[||||||||||||]PAM</td><td>NA</td><td>Intergenic</td><td>NA</td><td>NA</td></tr>
<tr><td>T21029</td><td>ACCTGCTGGTGAAGCCACTGAGG</td><td>1000</td></tr>
<tr><td>Oligo fwd</td><td>CACCACCTGCTGGTGAAGCCACTG</td></tr>
<tr><td>Oligo rev</td><td>AAACCAGTGGCTTCACCAGCAGGT</td></tr>
<tr><td>Chromosome</td><td>start</td><td>end</td><td>strand</td><td>MM</td><td>target_seq</td><td>PAM</td><td>alignment</td><td>distance</td><td>position</td><td>gene name</td><td>gene id</td></tr>
<tr><td>GU305914.1</td><td>180267</td><td>180289</td><td>-</td><td>0</td><td>ACCTGCTGGTGAAGCCACTG</td><td>AGG</td><td>||||||||[||||||||||||]PAM</td><td>NA</td><td>Intergenic</td><td>NA</td><td>NA</td></tr>
<tr><td>T21030</td><td>GAAGAAGATGAGTCCCAGCCTGG</td><td>1000</td></tr>
<tr><td>Oligo fwd</td><td>CACCGAAGAAGATGAGTCCCAGCC</td></tr>
<tr><td>Oligo rev</td><td>AAACGGCTGGGACTCATCTTCTTC</td></tr>
<tr><td>Chromosome</td><td>start</td><td>end</td><td>strand</td><td>MM</td><td>target_seq</td><td>PAM</td><td>alignment</td><td>distance</td><td>position</td><td>gene name</td><td>gene id</td></tr>
<tr><td>GU305914.1</td><td>180239</td><td>180261</td><td>-</td><td>0</td><td>GAAGAAGATGAGTCCCAGCC</td><td>TGG</td><td>||||||||[||||||||||||]PAM</td><td>NA</td><td>Intergenic</td><td>NA</td><td>NA</td></tr>
<tr><td>T21031</td><td>GCCTGGACCAAGTGACAACCAGG</td><td>1000</td></tr>
<tr><td>Oligo fwd</td><td>CACCGCCTGGACCAAGTGACAACC</td></tr>
<tr><td>Oligo rev</td><td>AAACGGTTGTCACTTGGTCCAGGC</td></tr>
<tr><td>Chromosome</td><td>start</td><td>end</td><td>strand</td><td>MM</td><td>target_seq</td><td>PAM</td><td>alignment</td><td>distance</td><td>position</td><td>gene name</td><td>gene id</td></tr>
<tr><td>GU305914.1</td><td>180222</td><td>180244</td><td>-</td><td>0</td><td>GCCTGGACCAAGTGACAACC</td><td>AGG</td><td>||||||||[||||||||||||]PAM</td><td>NA</td><td>Intergenic</td><td>NA</td><td>NA</td></tr>
<tr><td>T21032</td><td>TGGACCAAGTGACAACCAGGTGG</td><td>1000</td></tr>
<tr><td>Oligo fwd</td><td>CACCTGGACCAAGTGACAACCAGG</td></tr>
<tr><td>Oligo rev</td><td>AAACCCTGGTTGTCACTTGGTCCA</td></tr>
<tr><td>Chromosome</td><td>start</td><td>end</td><td>strand</td><td>MM</td><td>target_seq</td><td>PAM</td><td>alignment</td><td>distance</td><td>position</td><td>gene name</td><td>gene id</td></tr>
<tr><td>GU305914.1</td><td>180219</td><td>180241</td><td>-</td><td>0</td><td>TGGACCAAGTGACAACCAGG</td><td>TGG</td><td>||||||||[||||||||||||]PAM</td><td>NA</td><td>Intergenic</td><td>NA</td><td>NA</td></tr>
<tr><td>T21033</td><td>AGAGACACCAACACCTGCTGAGG</td><td>1000</td></tr>
<tr><td>Oligo fwd</td><td>CACCAGAGACACCAACACCTGCTG</td></tr>
<tr><td>Oligo rev</td><td>AAACCAGCAGGTGTTGGTGTCTCT</td></tr>
<tr><td>Chromosome</td><td>start</td><td>end</td><td>strand</td><td>MM</td><td>target_seq</td><td>PAM</td><td>alignment</td><td>distance</td><td>position</td><td>gene name</td><td>gene id</td></tr>
<tr><td>GU305914.1</td><td>180183</td><td>180205</td><td>-</td><td>0</td><td>AGAGACACCAACACCTGCTG</td><td>AGG</td><td>||||||||[||||||||||||]PAM</td><td>NA</td><td>Intergenic</td><td>NA</td><td>NA</td></tr>
<tr><td>T21034</td><td>ACCAACACCTGCTGAGGATGAGG</td><td>1000</td></tr>
<tr><td>Oligo fwd</td><td>CACCACCAACACCTGCTGAGGATG</td></tr>
<tr><td>Oligo rev</td><td>AAACCATCCTCAGCAGGTGTTGGT</td></tr>
<tr><td>Chromosome</td><td>start</td><td>end</td><td>strand</td><td>MM</td><td>target_seq</td><td>PAM</td><td>alignment</td><td>distance</td><td>position</td><td>gene name</td><td>gene id</td></tr>
<tr><td>GU305914.1</td><td>180177</td><td>180199</td><td>-</td><td>0</td><td>ACCAACACCTGCTGAGGATG</td><td>AGG</td><td>||||||||[||||||||||||]PAM</td><td>NA</td><td>Intergenic</td><td>NA</td><td>NA</td></tr>
<tr><td>T21035</td><td>GAGACCCAAAGTGCTGATGAAGG</td><td>1000</td></tr>
<tr><td>Oligo fwd</td><td>CACCGAGACCCAAAGTGCTGATGA</td></tr>
<tr><td>Oligo rev</td><td>AAACTCATCAGCACTTTGGGTCTC</td></tr>
<tr><td>Chromosome</td><td>start</td><td>end</td><td>strand</td><td>MM</td><td>target_seq</td><td>PAM</td><td>alignment</td><td>distance</td><td>position</td><td>gene name</td><td>gene id</td></tr>
<tr><td>GU305914.1</td><td>180155</td><td>180177</td><td>-</td><td>0</td><td>GAGACCCAAAGTGCTGATGA</td><td>AGG</td><td>||||||||[||||||||||||]PAM</td><td>NA</td><td>Intergenic</td><td>NA</td><td>NA</td></tr>
<tr><td>T21036</td><td>TGCTGATGAAGGAGAATCTCAGG</td><td>1000</td></tr>
<tr><td>Oligo fwd</td><td>CACCTGCTGATGAAGGAGAATCTC</td></tr>
<tr><td>Oligo rev</td><td>AAACGAGATTCTCCTTCATCAGCA</td></tr>
<tr><td>Chromosome</td><td>start</td><td>end</td><td>strand</td><td>MM</td><td>target_seq</td><td>PAM</td><td>alignment</td><td>distance</td><td>position</td><td>gene name</td><td>gene id</td></tr>
<tr><td>GU305914.1</td><td>180144</td><td>180166</td><td>-</td><td>0</td><td>TGCTGATGAAGGAGAATCTC</td><td>AGG</td><td>||||||||[||||||||||||]PAM</td><td>NA</td><td>Intergenic</td><td>NA</td><td>NA</td></tr>
<tr><td>T21037</td><td>TGAAGGAGAATCTCAGGAGCTGG</td><td>1000</td></tr>
<tr><td>Oligo fwd</td><td>CACCTGAAGGAGAATCTCAGGAGC</td></tr>
<tr><td>Oligo rev</td><td>AAACGCTCCTGAGATTCTCCTTCA</td></tr>
<tr><td>Chromosome</td><td>start</td><td>end</td><td>strand</td><td>MM</td><td>target_seq</td><td>PAM</td><td>alignment</td><td>distance</td><td>position</td><td>gene name</td><td>gene id</td></tr>
<tr><td>GU305914.1</td><td>180138</td><td>180160</td><td>-</td><td>0</td><td>TGAAGGAGAATCTCAGGAGC</td><td>TGG</td><td>||||||||[||||||||||||]PAM</td><td>NA</td><td>Intergenic</td><td>NA</td><td>NA</td></tr>
<tr><td>T21038</td><td>GGAGAATCTCAGGAGCTGGAAGG</td><td>1000</td></tr>
<tr><td>Oligo fwd</td><td>CACCGGAGAATCTCAGGAGCTGGA</td></tr>
<tr><td>Oligo rev</td><td>AAACTCCAGCTCCTGAGATTCTCC</td></tr>
<tr><td>Chromosome</td><td>start</td><td>end</td><td>strand</td><td>MM</td><td>target_seq</td><td>PAM</td><td>alignment</td><td>distance</td><td>position</td><td>gene name</td><td>gene id</td></tr>
<tr><td>GU305914.1</td><td>180134</td><td>180156</td><td>-</td><td>0</td><td>GGAGAATCTCAGGAGCTGGA</td><td>AGG</td><td>||||||||[||||||||||||]PAM</td><td>NA</td><td>Intergenic</td><td>NA</td><td>NA</td></tr>
</table>

| | | | | | | | | | | | | |
|---|---|---|---|---|---|---|---|---|---|---|---|---|
| T21039 | TGATTCTGAAGATGATGATGAGG | | 1000 | | | | | | | | | |
| Oligo fwd | CACCTGATTCTGAAGATGATGATG | | | | | | | | | | | |
| Oligo rev | AAACCATCATCATCTTCAGAATCA | | | | | | | | | | | |
| Chromosome | start | | end | strand | MM | target_seq | PAM | alignment | distance | position | gene name | gene id |
| GU305914.1 | | 180051 | 180073 | - | 0 | TGATTCTGAAGATGATGATG | AGG | ||||||||[||||||||||||]PAM | NA | Intergenic | NA | NA |
| GU305914.1 | | 111710 | 111732 | - | 4 | AGAGCCTGGAGATGATGATG | AGG | -||--|||[-|||||||||||]PAM | NA | Intergenic | NA | NA |
| GU305914.1 | | 168589 | 168611 | + | 4 | TGAATATGAGGATGATGAAG | AGG | |||-|-||[|-||||||||-|]PAM | NA | Intergenic | NA | NA |
| T21040 | TGAGGTGACCCGCATCCCAGTGG | | 1000 | | | | | | | | | |
| Oligo fwd | CACCTGAGGTGACCCGCATCCCAG | | | | | | | | | | | |
| Oligo rev | AAACCTGGGATGCGGGTCACCTCA | | | | | | | | | | | |
| Chromosome | start | | end | strand | MM | target_seq | PAM | alignment | distance | position | gene name | gene id |
| GU305914.1 | | 180033 | 180055 | - | 0 | TGAGGTGACCCGCATCCCAG | TGG | ||||||||[||||||||||||]PAM | NA | Intergenic | NA | NA |
| T21041 | GAGGTGACCCGCATCCCAGTGGG | | 1000 | | | | | | | | | |
| Oligo fwd | CACCGAGGTGACCCGCATCCCAGT | | | | | | | | | | | |
| Oligo rev | AAACACTGGGATGCGGGTCACCTC | | | | | | | | | | | |
| Chromosome | start | | end | strand | MM | target_seq | PAM | alignment | distance | position | gene name | gene id |
| GU305914.1 | | 180032 | 180054 | - | 0 | GAGGTGACCCGCATCCCAGT | GGG | ||||||||[||||||||||||]PAM | NA | Intergenic | NA | NA |
| T21042 | TACGACCAGATCTGCAACAGCGG | | 1000 | | | | | | | | | |
| Oligo fwd | CACCTACGACCAGATCTGCAACAG | | | | | | | | | | | |
| Oligo rev | AAACCTGTTGCAGATCTGGTCGTA | | | | | | | | | | | |
| Chromosome | start | | end | strand | MM | target_seq | PAM | alignment | distance | position | gene name | gene id |
| GU305914.1 | | 179976 | 179998 | - | 0 | TACGACCAGATCTGCAACAG | CGG | ||||||||[||||||||||||]PAM | NA | Intergenic | NA | NA |
| T21043 | GCAACAGCGGCAGCTTCTTCTGG | | 1000 | | | | | | | | | |
| Oligo fwd | CACCGCAACAGCGGCAGCTTCTTC | | | | | | | | | | | |
| Oligo rev | AAACGAAGAAGCTGCCGCTGTTGC | | | | | | | | | | | |
| Chromosome | start | | end | strand | MM | target_seq | PAM | alignment | distance | position | gene name | gene id |
| GU305914.1 | | 179963 | 179985 | - | 0 | GCAACAGCGGCAGCTTCTTC | TGG | ||||||||[||||||||||||]PAM | NA | Intergenic | NA | NA |
| T21044 | CTCCTCGCCCAGCTCTCAAGAGG | | 1000 | | | | | | | | | |
| Oligo fwd | CACCCTCCTCGCCCAGCTCTCAAG | | | | | | | | | | | |
| Oligo rev | AAACCTTGAGAGCTGGGCGAGGAG | | | | | | | | | | | |
| Chromosome | start | | end | strand | MM | target_seq | PAM | alignment | distance | position | gene name | gene id |
| GU305914.1 | | 179935 | 179957 | - | 0 | CTCCTCGCCCAGCTCTCAAG | AGG | ||||||||[||||||||||||]PAM | NA | Intergenic | NA | NA |
| T21045 | TCTCAAGAGGCAGTACGCCATGG | | 1000 | | | | | | | | | |
| Oligo fwd | CACCTCTCAAGAGGCAGTACGCCA | | | | | | | | | | | |
| Oligo rev | AAACTGGCGTACTGCCTCTTGAGA | | | | | | | | | | | |
| Chromosome | start | | end | strand | MM | target_seq | PAM | alignment | distance | position | gene name | gene id |
| GU305914.1 | | 179922 | 179944 | - | 0 | TCTCAAGAGGCAGTACGCCA | TGG | ||||||||[||||||||||||]PAM | NA | Intergenic | NA | NA |
| T21046 | CAGCTCCCCACCCATCACCATGG | | 1000 | | | | | | | | | |
| Oligo fwd | CACCCAGCTCCCCACCCATCACCA | | | | | | | | | | | |
| Oligo rev | AAACTGGTGATGGGTGGGGAGCTG | | | | | | | | | | | |
| Chromosome | start | | end | strand | MM | target_seq | PAM | alignment | distance | position | gene name | gene id |
| GU305914.1 | | 179826 | 179848 | - | 0 | CAGCTCCCCACCCATCACCA | TGG | ||||||||[||||||||||||]PAM | NA | Intergenic | NA | NA |
| T21047 | AATGTTTTTTTACTAACACTTGG | | 1000 | | | | | | | | | |
| Oligo fwd | CACCAATGTTTTTTTACTAACACT | | | | | | | | | | | |
| Oligo rev | AAACAGTGTTAGTAAAAAAACATT | | | | | | | | | | | |
| Chromosome | start | | end | strand | MM | target_seq | PAM | alignment | distance | position | gene name | gene id |
| GU305914.1 | | 179801 | 179823 | - | 0 | AATGTTTTTTTACTAACACT | TGG | ||||||||[||||||||||||]PAM | NA | Intergenic | NA | NA |
| GU305914.1 | | 179769 | 179791 | - | 3 | GATACTTTTTTACTAACACT | TGG | -||--|||[||||||||||||]PAM | NA | Intergenic | NA | NA |
| T21048 | TACTAACACTTGGACAATGATGG | | 1000 | | | | | | | | | |
| Oligo fwd | CACCTACTAACACTTGGACAATGA | | | | | | | | | | | |
| Oligo rev | AAACTCATTGTCCAAGTGTTAGTA | | | | | | | | | | | |
| Chromosome | start | | end | strand | MM | target_seq | PAM | alignment | distance | position | gene name | gene id |
| GU305914.1 | | 179791 | 179813 | - | 0 | TACTAACACTTGGACAATGA | TGG | ||||||||[||||||||||||]PAM | NA | Intergenic | NA | NA |
| T21049 | GATACTTTTTTACTAACACTTGG | | 1000 | | | | | | | | | |
| Oligo fwd | CACCGATACTTTTTTACTAACACT | | | | | | | | | | | |
| Oligo rev | AAACAGTGTTAGTAAAAAAGTATC | | | | | | | | | | | |
| Chromosome | start | | end | strand | MM | target_seq | PAM | alignment | distance | position | gene name | gene id |
| GU305914.1 | | 179769 | 179791 | - | 0 | GATACTTTTTTACTAACACT | TGG | ||||||||[||||||||||||]PAM | NA | Intergenic | NA | NA |
| GU305914.1 | | 179801 | 179823 | - | 3 | AATGTTTTTTTACTAACACT | TGG | -||--|||[||||||||||||]PAM | NA | Intergenic | NA | NA |
| T21050 | AAGACTTCTTTAACTCTTTATGG | | 1000 | | | | | | | | | |
| Oligo fwd | CACCAAGACTTCTTTAACTCTTTA | | | | | | | | | | | |
| Oligo rev | AAACTAAAGAGTTAAAGAAGTCTT | | | | | | | | | | | |
| Chromosome | start | | end | strand | MM | target_seq | PAM | alignment | distance | position | gene name | gene id |
| GU305914.1 | | 179625 | 179647 | - | 0 | AAGACTTCTTTAACTCTTTA | TGG | ||||||||[||||||||||||]PAM | NA | Intergenic | NA | NA |
| T21051 | TTTAACTCTTTATGGTTCACAGG | | 1000 | | | | | | | | | |
| Oligo fwd | CACCTTTAACTCTTTATGGTTCAC | | | | | | | | | | | |
| Oligo rev | AAACGTGAACCATAAAGAGTTAAA | | | | | | | | | | | |
| Chromosome | start | | end | strand | MM | target_seq | PAM | alignment | distance | position | gene name | gene id |
| GU305914.1 | | 179617 | 179639 | - | 0 | TTTAACTCTTTATGGTTCAC | AGG | ||||||||[||||||||||||]PAM | NA | Intergenic | NA | NA |
| T21052 | CATAGATATTGTTACAGCAGCGG | | 1000 | | | | | | | | | |
| Oligo fwd | CACCCATAGATATTGTTACAGCAG | | | | | | | | | | | |
| Oligo rev | AAACCTGCTGTAACAATATCTATG | | | | | | | | | | | |
| Chromosome | start | | end | strand | MM | target_seq | PAM | alignment | distance | position | gene name | gene id |
| GU305914.1 | | 179582 | 179604 | - | 0 | CATAGATATTGTTACAGCAG | CGG | ||||||||[||||||||||||]PAM | NA | Intergenic | NA | NA |
| T21053 | ACAGCAGCGGCCTCCATGTCAGG | | 1000 | | | | | | | | | |

| | | | | | | | | | | | |
|---|---|---|---|---|---|---|---|---|---|---|---|
| Oligo fwd | CACCACAGCAGCGGCCTCCATGTC | | | | | | | | | | |
| Oligo rev | AAACGACATGGAGGCCGCTGCTGT | | | | | | | | | | |
| Chromosome | start | end | strand | MM | target_seq | PAM | alignment | distance | position | gene name | gene id |
| GU305914.1 | 179569 | 179591 | - | 0 | ACAGCAGCGGCCTCCATGTC | AGG | \|\|\|\|\|\|\|\|[\|\|\|\|\|\|\|\|\|\|\|\|]PAM | NA | Intergenic | NA | NA |
| T21054 | ACATAAGCCCTCTGAGAGTGAGG | 1000 | | | | | | | | | |
| Oligo fwd | CACCACATAAGCCCTCTGAGAGTG | | | | | | | | | | |
| Oligo rev | AAACCACTCTCAGAGGGCTTATGT | | | | | | | | | | |
| Chromosome | start | end | strand | MM | target_seq | PAM | alignment | distance | position | gene name | gene id |
| GU305914.1 | 179411 | 179433 | - | 0 | ACATAAGCCCTCTGAGAGTG | AGG | \|\|\|\|\|\|\|\|[\|\|\|\|\|\|\|\|\|\|\|\|]PAM | NA | Intergenic | NA | NA |
| T21055 | CTCTGAGAGTGAGGCAGATGAGG | 1000 | | | | | | | | | |
| Oligo fwd | CACCCTCTGAGAGTGAGGCAGATG | | | | | | | | | | |
| Oligo rev | AAACCATCTGCCTCACTCTCAGAG | | | | | | | | | | |
| Chromosome | start | end | strand | MM | target_seq | PAM | alignment | distance | position | gene name | gene id |
| GU305914.1 | 179402 | 179424 | - | 0 | CTCTGAGAGTGAGGCAGATG | AGG | \|\|\|\|\|\|\|\|[\|\|\|\|\|\|\|\|\|\|\|\|]PAM | NA | Intergenic | NA | NA |
| T21056 | CTTTATTTTTCAGCAGCCTCAGG | 1000 | | | | | | | | | |
| Oligo fwd | CACCCTTTATTTTTCAGCAGCCTC | | | | | | | | | | |
| Oligo rev | AAACGAGGCTGCTGAAAAATAAAG | | | | | | | | | | |
| Chromosome | start | end | strand | MM | target_seq | PAM | alignment | distance | position | gene name | gene id |
| GU305914.1 | 179378 | 179400 | - | 0 | CTTTATTTTTCAGCAGCCTC | AGG | \|\|\|\|\|\|\|\|[\|\|\|\|\|\|\|\|\|\|\|\|]PAM | NA | Intergenic | NA | NA |
| T21057 | TATTTTTCAGCAGCCTCAGGAGG | 1000 | | | | | | | | | |
| Oligo fwd | CACCTATTTTTCAGCAGCCTCAGG | | | | | | | | | | |
| Oligo rev | AAACCCTGAGGCTGCTGAAAAATA | | | | | | | | | | |
| Chromosome | start | end | strand | MM | target_seq | PAM | alignment | distance | position | gene name | gene id |
| GU305914.1 | 179375 | 179397 | - | 0 | TATTTTTCAGCAGCCTCAGG | AGG | \|\|\|\|\|\|\|\|[\|\|\|\|\|\|\|\|\|\|\|\|]PAM | NA | Intergenic | NA | NA |
| T21058 | TTTTCAGCAGCCTCAGGAGGAGG | 1000 | | | | | | | | | |
| Oligo fwd | CACCTTTTCAGCAGCCTCAGGAGG | | | | | | | | | | |
| Oligo rev | AAACCCTCCTGAGGCTGCTGAAAA | | | | | | | | | | |
| Chromosome | start | end | strand | MM | target_seq | PAM | alignment | distance | position | gene name | gene id |
| GU305914.1 | 179372 | 179394 | - | 0 | TTTTCAGCAGCCTCAGGAGG | AGG | \|\|\|\|\|\|\|\|[\|\|\|\|\|\|\|\|\|\|\|\|]PAM | NA | Intergenic | NA | NA |
| T21059 | GCCTCAGGAGGAGGAGTATCAGG | 1000 | | | | | | | | | |
| Oligo fwd | CACCGCCTCAGGAGGAGGAGTATC | | | | | | | | | | |
| Oligo rev | AAACGATACTCCTCCTCCTGAGGC | | | | | | | | | | |
| Chromosome | start | end | strand | MM | target_seq | PAM | alignment | distance | position | gene name | gene id |
| GU305914.1 | 179363 | 179385 | - | 0 | GCCTCAGGAGGAGGAGTATC | AGG | \|\|\|\|\|\|\|\|[\|\|\|\|\|\|\|\|\|\|\|\|]PAM | NA | Intergenic | NA | NA |
| T21060 | TCAGGAGGAGGAGTATCAGGAGG | 1000 | | | | | | | | | |
| Oligo fwd | CACCTCAGGAGGAGGAGTATCAGG | | | | | | | | | | |
| Oligo rev | AAACCCTGATACTCCTCCTCCTGA | | | | | | | | | | |
| Chromosome | start | end | strand | MM | target_seq | PAM | alignment | distance | position | gene name | gene id |
| GU305914.1 | 179360 | 179382 | - | 0 | TCAGGAGGAGGAGTATCAGG | AGG | \|\|\|\|\|\|\|\|[\|\|\|\|\|\|\|\|\|\|\|\|]PAM | NA | Intergenic | NA | NA |
| T21061 | GGAGGAGTATCAGGAGGAGCAGG | 1000 | | | | | | | | | |
| Oligo fwd | CACCGGAGGAGTATCAGGAGGAGC | | | | | | | | | | |
| Oligo rev | AAACGCTCCTCCTGATACTCCTCC | | | | | | | | | | |
| Chromosome | start | end | strand | MM | target_seq | PAM | alignment | distance | position | gene name | gene id |
| GU305914.1 | 179354 | 179376 | - | 0 | GGAGGAGTATCAGGAGGAGC | AGG | \|\|\|\|\|\|\|\|[\|\|\|\|\|\|\|\|\|\|\|\|]PAM | NA | Intergenic | NA | NA |
| GU305914.1 | 152954 | 152976 | + | 4 | CGATGAGTATGAGGAGGATC | TGG | -\|\|-\|\|\|\|[\|\|-\|\|\|\|\|\|\|-\|]PAM | NA | Intergenic | NA | NA |
| T21062 | TCAGGAGGAGCAGGAGCCCCAGG | 1000 | | | | | | | | | |
| Oligo fwd | CACCTCAGGAGGAGCAGGAGCCCC | | | | | | | | | | |
| Oligo rev | AAACGGGGCTCCTGCTCCTCCTGA | | | | | | | | | | |
| Chromosome | start | end | strand | MM | target_seq | PAM | alignment | distance | position | gene name | gene id |
| GU305914.1 | 179345 | 179367 | - | 0 | TCAGGAGGAGCAGGAGCCCC | AGG | \|\|\|\|\|\|\|\|[\|\|\|\|\|\|\|\|\|\|\|\|]PAM | NA | Intergenic | NA | NA |
| T21063 | CTCCTCAACACTTTGATGAGAGG | 1000 | | | | | | | | | |
| Oligo fwd | CACCCTCCTCAACACTTTGATGAG | | | | | | | | | | |
| Oligo rev | AAACCTCATCAAAGTGTTGAGGAG | | | | | | | | | | |
| Chromosome | start | end | strand | MM | target_seq | PAM | alignment | distance | position | gene name | gene id |
| GU305914.1 | 179310 | 179332 | - | 0 | CTCCTCAACACTTTGATGAG | AGG | \|\|\|\|\|\|\|\|[\|\|\|\|\|\|\|\|\|\|\|\|]PAM | NA | Intergenic | NA | NA |
| T21064 | TTGATGAGAGGAGCCCCAGCAGG | 1000 | | | | | | | | | |
| Oligo fwd | CACCTTGATGAGAGGAGCCCCAGC | | | | | | | | | | |
| Oligo rev | AAACGCTGGGGCTCCTCTCATCAA | | | | | | | | | | |
| Chromosome | start | end | strand | MM | target_seq | PAM | alignment | distance | position | gene name | gene id |
| GU305914.1 | 179298 | 179320 | - | 0 | TTGATGAGAGGAGCCCCAGC | AGG | \|\|\|\|\|\|\|\|[\|\|\|\|\|\|\|\|\|\|\|\|]PAM | NA | Intergenic | NA | NA |
| T21065 | AGCAGGTCTTCCTCAGATCATGG | 1000 | | | | | | | | | |
| Oligo fwd | CACCAGCAGGTCTTCCTCAGATCA | | | | | | | | | | |
| Oligo rev | AAACTGATCTGAGGAAGACCTGCT | | | | | | | | | | |
| Chromosome | start | end | strand | MM | target_seq | PAM | alignment | distance | position | gene name | gene id |
| GU305914.1 | 179281 | 179303 | - | 0 | AGCAGGTCTTCCTCAGATCA | TGG | \|\|\|\|\|\|\|\|[\|\|\|\|\|\|\|\|\|\|\|\|]PAM | NA | Intergenic | NA | NA |
| T21066 | AGGTCTTCCTCAGATCATGGTGG | 1000 | | | | | | | | | |
| Oligo fwd | CACCAGGTCTTCCTCAGATCATGG | | | | | | | | | | |
| Oligo rev | AAACCCATGATCTGAGGAAGACCT | | | | | | | | | | |
| Chromosome | start | end | strand | MM | target_seq | PAM | alignment | distance | position | gene name | gene id |
| GU305914.1 | 179278 | 179300 | - | 0 | AGGTCTTCCTCAGATCATGG | TGG | \|\|\|\|\|\|\|\|[\|\|\|\|\|\|\|\|\|\|\|\|]PAM | NA | Intergenic | NA | NA |
| T21067 | GGTCTTCCTCAGATCATGGTGGG | 1000 | | | | | | | | | |
| Oligo fwd | CACCGGTCTTCCTCAGATCATGGT | | | | | | | | | | |
| Oligo rev | AAACACCATGATCTGAGGAAGACC | | | | | | | | | | |
| Chromosome | start | end | strand | MM | target_seq | PAM | alignment | distance | position | gene name | gene id |
| GU305914.1 | 179277 | 179299 | - | 0 | GGTCTTCCTCAGATCATGGT | GGG | \|\|\|\|\|\|\|\|[\|\|\|\|\|\|\|\|\|\|\|\|]PAM | NA | Intergenic | NA | NA |

T21068 TCAGATCATGGTGGGCACAGTGG 1000
Oligo fwd CACCTCAGATCATGGTGGGCACAG
Oligo rev AAACCTGTGCCCACCATGATCTGA
Chromosome start end strand MM target_seq PAM alignment distance position gene name gene id
GU305914.1 179269 179291 - 0 TCAGATCATGGTGGGCACAG TGG ||||||||[||||||||||||]PAM NA Intergenic NA NA

T21069 CAGATCATGGTGGGCACAGTGGG 1000
Oligo fwd CACCCAGATCATGGTGGGCACAGT
Oligo rev AAACACTGTGCCCACCATGATCTG
Chromosome start end strand MM target_seq PAM alignment distance position gene name gene id
GU305914.1 179268 179290 - 0 CAGATCATGGTGGGCACAGT GGG ||||||||[||||||||||||]PAM NA Intergenic NA NA

T21070 AGATCATGGTGGGCACAGTGGGG 1000
Oligo fwd CACCAGATCATGGTGGGCACAGTG
Oligo rev AAACCACTGTGCCCACCATGATCT
Chromosome start end strand MM target_seq PAM alignment distance position gene name gene id
GU305914.1 179267 179289 - 0 AGATCATGGTGGGCACAGTG GGG ||||||||[||||||||||||]PAM NA Intergenic NA NA

T21071 GATCATGGTGGGCACAGTGGGGG 1000
Oligo fwd CACCGATCATGGTGGGCACAGTGG
Oligo rev AAACCCACTGTGCCCACCATGATC
Chromosome start end strand MM target_seq PAM alignment distance position gene name gene id
GU305914.1 179266 179288 - 0 GATCATGGTGGGCACAGTGG GGG ||||||||[||||||||||||]PAM NA Intergenic NA NA

T21072 AGTGGGGGAAACAGTAGAGATGG 1000
Oligo fwd CACCAGTGGGGGAAACAGTAGAGA
Oligo rev AAACTCTCTACTGTTTCCCCCACT
Chromosome start end strand MM target_seq PAM alignment distance position gene name gene id
GU305914.1 179251 179273 - 0 AGTGGGGGAAACAGTAGAGA TGG ||||||||[||||||||||||]PAM NA Intergenic NA NA

T21073 GGACAGTTCTATCTGTCTAGTGG 1000
Oligo fwd CACCGGACAGTTCTATCTGTCTAG
Oligo rev AAACCTAGACAGATAGAACTGTCC
Chromosome start end strand MM target_seq PAM alignment distance position gene name gene id
GU305914.1 179230 179252 - 0 GGACAGTTCTATCTGTCTAG TGG ||||||||[||||||||||||]PAM NA Intergenic NA NA

T21074 CTATCTGTCTAGTGGTTCTGAGG 1000
Oligo fwd CACCCTATCTGTCTAGTGGTTCTG
Oligo rev AAACCAGAACCACTAGACAGATAG
Chromosome start end strand MM target_seq PAM alignment distance position gene name gene id
GU305914.1 179222 179244 - 0 CTATCTGTCTAGTGGTTCTG AGG ||||||||[||||||||||||]PAM NA Intergenic NA NA

T21075 TTCTGAGGATGAAGATGATGAGG 1000
Oligo fwd CACCTTCTGAGGATGAAGATGATG
Oligo rev AAACCATCATCTTCATCCTCAGAA
Chromosome start end strand MM target_seq PAM alignment distance position gene name gene id
GU305914.1 179207 179229 - 0 TTCTGAGGATGAAGATGATG AGG ||||||||[||||||||||||]PAM NA Intergenic NA NA

T21076 AAGATGATGAGGATGATGAGCGG 1000
Oligo fwd CACCAAGATGATGAGGATGATGAG
Oligo rev AAACCTCATCATCCTCATCATCTT
Chromosome start end strand MM target_seq PAM alignment distance position gene name gene id
GU305914.1 179196 179218 - 0 AAGATGATGAGGATGATGAG CGG ||||||||[||||||||||||]PAM NA Intergenic NA NA

T21077 AGATGATGAGGATGATGAGCGGG 1000
Oligo fwd CACCAGATGATGAGGATGATGAGC
Oligo rev AAACGCTCATCATCCTCATCATCT
Chromosome start end strand MM target_seq PAM alignment distance position gene name gene id
GU305914.1 179195 179217 - 0 AGATGATGAGGATGATGAGC GGG ||||||||[||||||||||||]PAM NA Intergenic NA NA

T21078 GATGATGAGCGGGTAGATAGTGG 1000
Oligo fwd CACCGATGATGAGCGGGTAGATAG
Oligo rev AAACCTATCTACCCGCTCATCATC
Chromosome start end strand MM target_seq PAM alignment distance position gene name gene id
GU305914.1 179185 179207 - 0 GATGATGAGCGGGTAGATAG TGG ||||||||[||||||||||||]PAM NA Intergenic NA NA

T21079 AGCGGGTAGATAGTGGCTACAGG 1000
Oligo fwd CACCAGCGGGTAGATAGTGGCTAC
Oligo rev AAACGTAGCCACTATCTACCCGCT
Chromosome start end strand MM target_seq PAM alignment distance position gene name gene id
GU305914.1 179178 179200 - 0 AGCGGGTAGATAGTGGCTAC AGG ||||||||[||||||||||||]PAM NA Intergenic NA NA

T21080 GCGGGTAGATAGTGGCTACAGGG 1000
Oligo fwd CACCGCGGGTAGATAGTGGCTACA
Oligo rev AAACTGTAGCCACTATCTACCCGC
Chromosome start end strand MM target_seq PAM alignment distance position gene name gene id
GU305914.1 179177 179199 - 0 GCGGGTAGATAGTGGCTACA GGG ||||||||[||||||||||||]PAM NA Intergenic NA NA

T21081 CGGGTAGATAGTGGCTACAGGGG 1000
Oligo fwd CACCCGGGTAGATAGTGGCTACAG
Oligo rev AAACCTGTAGCCACTATCTACCCG
Chromosome start end strand MM target_seq PAM alignment distance position gene name gene id
GU305914.1 179176 179198 - 0 CGGGTAGATAGTGGCTACAG GGG ||||||||[||||||||||||]PAM NA Intergenic NA NA

T21082 GGGTAGATAGTGGCTACAGGGGG 1000
Oligo fwd CACCGGGTAGATAGTGGCTACAGG
Oligo rev AAACCCTGTAGCCACTATCTACCC
Chromosome start end strand MM target_seq PAM alignment distance position gene name gene id

GU305914.1 179175 179197 - 0 GGGTAGATAGTGGCTACAGG GGG ||||||||[||||||||||||]PAM NA Intergenic NA NA

T21083 GTGGCTACAGGGGGAGCAGTAGG 1000
Oligo fwd CACCGTGGCTACAGGGGGAGCAGT
Oligo rev AAACACTGCTCCCCCTGTAGCCAC
Chromosome start end strand MM target_seq PAM alignment distance position gene name gene id
GU305914.1 179166 179188 - 0 GTGGCTACAGGGGGAGCAGT AGG ||||||||[||||||||||||]PAM NA Intergenic NA NA

T21084 GTAGGAGTGAAGACAGCAGCAGG 1000
Oligo fwd CACCGTAGGAGTGAAGACAGCAGC
Oligo rev AAACGCTGCTGTCTTCACTCCTAC
Chromosome start end strand MM target_seq PAM alignment distance position gene name gene id
GU305914.1 179148 179170 - 0 GTAGGAGTGAAGACAGCAGC AGG ||||||||[||||||||||||]PAM NA Intergenic NA NA

T21085 GTGAAGACAGCAGCAGGTACAGG 1000
Oligo fwd CACCGTGAAGACAGCAGCAGGTAC
Oligo rev AAACGTACCTGCTGCTGTCTTCAC
Chromosome start end strand MM target_seq PAM alignment distance position gene name gene id
GU305914.1 179142 179164 - 0 GTGAAGACAGCAGCAGGTAC AGG ||||||||[||||||||||||]PAM NA Intergenic NA NA

T21086 AGCAGCAGGTACAGGCCACACGG 1000
Oligo fwd CACCAGCAGCAGGTACAGGCCACA
Oligo rev AAACTGTGGCCTGTACCTGCTGCT
Chromosome start end strand MM target_seq PAM alignment distance position gene name gene id
GU305914.1 179134 179156 - 0 AGCAGCAGGTACAGGCCACA CGG ||||||||[||||||||||||]PAM NA Intergenic NA NA

T21087 AGCAGGTACAGGCCACACGGAGG 1000
Oligo fwd CACCAGCAGGTACAGGCCACACGG
Oligo rev AAACCCGTGTGGCCTGTACCTGCT
Chromosome start end strand MM target_seq PAM alignment distance position gene name gene id
GU305914.1 179131 179153 - 0 AGCAGGTACAGGCCACACGG AGG ||||||||[||||||||||||]PAM NA Intergenic NA NA

T21088 ACGGAGGCTCCCACTCATCAAGG 1000
Oligo fwd CACCACGGAGGCTCCCACTCATCA
Oligo rev AAACTGATGAGTGGGAGCCTCCGT
Chromosome start end strand MM target_seq PAM alignment distance position gene name gene id
GU305914.1 179115 179137 - 0 ACGGAGGCTCCCACTCATCA AGG ||||||||[||||||||||||]PAM NA Intergenic NA NA

T21089 AGGTCCTCCATTAAATCATCAGG 1000
Oligo fwd CACCAGGTCCTCCATTAAATCATC
Oligo rev AAACGATGATTTAATGGAGGACCT
Chromosome start end strand MM target_seq PAM alignment distance position gene name gene id
GU305914.1 179095 179117 - 0 AGGTCCTCCATTAAATCATC AGG ||||||||[||||||||||||]PAM NA Intergenic NA NA

T21090 GGTCCTCCATTAAATCATCAGGG 1000
Oligo fwd CACCGGTCCTCCATTAAATCATCA
Oligo rev AAACTGATGATTTAATGGAGGACC
Chromosome start end strand MM target_seq PAM alignment distance position gene name gene id
GU305914.1 179094 179116 - 0 GGTCCTCCATTAAATCATCA GGG ||||||||[||||||||||||]PAM NA Intergenic NA NA

T21091 TCCATTAAATCATCAGGGAGCGG 1000
Oligo fwd CACCTCCATTAAATCATCAGGGAG
Oligo rev AAACCTCCCTGATGATTTAATGGA
Chromosome start end strand MM target_seq PAM alignment distance position gene name gene id
GU305914.1 179089 179111 - 0 TCCATTAAATCATCAGGGAG CGG ||||||||[||||||||||||]PAM NA Intergenic NA NA

T21092 CCATTAAATCATCAGGGAGCGGG 1000
Oligo fwd CACCCCATTAAATCATCAGGGAGC
Oligo rev AAACGCTCCCTGATGATTTAATGG
Chromosome start end strand MM target_seq PAM alignment distance position gene name gene id
GU305914.1 179088 179110 - 0 CCATTAAATCATCAGGGAGC GGG ||||||||[||||||||||||]PAM NA Intergenic NA NA

T21093 CCTCTAGACACCACCACAAGAGG 1000
Oligo fwd CACCCCTCTAGACACCACCACAAG
Oligo rev AAACCTTGTGGTGGTGTCTAGAGG
Chromosome start end strand MM target_seq PAM alignment distance position gene name gene id
GU305914.1 179064 179086 - 0 CCTCTAGACACCACCACAAG AGG ||||||||[||||||||||||]PAM NA Intergenic NA NA

T21094 TAGACACCACCACAAGAGGAAGG 1000
Oligo fwd CACCTAGACACCACCACAAGAGGA
Oligo rev AAACTCCTCTTGTGGTGGTGTCTA
Chromosome start end strand MM target_seq PAM alignment distance position gene name gene id
GU305914.1 179060 179082 - 0 TAGACACCACCACAAGAGGA AGG ||||||||[||||||||||||]PAM NA Intergenic NA NA

T21095 ACACCACCACAAGAGGAAGGCGG 1000
Oligo fwd CACCACACCACCACAAGAGGAAGG
Oligo rev AAACCCTTCCTCTTGTGGTGGTGT
Chromosome start end strand MM target_seq PAM alignment distance position gene name gene id
GU305914.1 179057 179079 - 0 ACACCACCACAAGAGGAAGG CGG ||||||||[||||||||||||]PAM NA Intergenic NA NA

T21096 TCACACCTCCCTCTGCGAAGAGG 1000
Oligo fwd CACCTCACACCTCCCTCTGCGAAG
Oligo rev AAACCTTCGCAGAGGGAGGTGTGA
Chromosome start end strand MM target_seq PAM alignment distance position gene name gene id
GU305914.1 179013 179035 - 0 TCACACCTCCCTCTGCGAAG AGG ||||||||[||||||||||||]PAM NA Intergenic NA NA

T21097 ATATGAGTGTCCTGTGCGTGTGG 1000
Oligo fwd CACCATATGAGTGTCCTGTGCGTG
Oligo rev AAACCACGCACAGGACACTCATAT

<table>
<tr><td>Chromosome</td><td>start</td><td>end</td><td>strand</td><td>MM</td><td>target_seq</td><td>PAM</td><td>alignment</td><td>distance</td><td>position</td><td>gene name</td><td>gene id</td></tr>
<tr><td>GU305914.1</td><td>178964</td><td>178986</td><td>-</td><td>0</td><td>ATATGAGTGTCCTGTGCGTG</td><td>TGG</td><td>||||||||[||||||||||||]PAM</td><td>NA</td><td>Intergenic</td><td>NA</td><td>NA</td></tr>
<tr><td></td></tr>
<tr><td>T21098</td><td>CAGTGATGATGCCCCTACCATGG</td><td>1000</td></tr>
<tr><td>Oligo fwd</td><td>CACCCAGTGATGATGCCCCTACCA</td></tr>
<tr><td>Oligo rev</td><td>AAACTGGTAGGGGCATCATCACTG</td></tr>
<tr><td>Chromosome</td><td>start</td><td>end</td><td>strand</td><td>MM</td><td>target_seq</td><td>PAM</td><td>alignment</td><td>distance</td><td>position</td><td>gene name</td><td>gene id</td></tr>
<tr><td>GU305914.1</td><td>178931</td><td>178953</td><td>-</td><td>0</td><td>CAGTGATGATGCCCCTACCA</td><td>TGG</td><td>||||||||[||||||||||||]PAM</td><td>NA</td><td>Intergenic</td><td>NA</td><td>NA</td></tr>
<tr><td></td></tr>
<tr><td>T21099</td><td>ATGCCCCTACCATGGTACAGAGG</td><td>1000</td></tr>
<tr><td>Oligo fwd</td><td>CACCATGCCCCTACCATGGTACAG</td></tr>
<tr><td>Oligo rev</td><td>AAACCTGTACCATGGTAGGGGCAT</td></tr>
<tr><td>Chromosome</td><td>start</td><td>end</td><td>strand</td><td>MM</td><td>target_seq</td><td>PAM</td><td>alignment</td><td>distance</td><td>position</td><td>gene name</td><td>gene id</td></tr>
<tr><td>GU305914.1</td><td>178923</td><td>178945</td><td>-</td><td>0</td><td>ATGCCCCTACCATGGTACAG</td><td>AGG</td><td>||||||||[||||||||||||]PAM</td><td>NA</td><td>Intergenic</td><td>NA</td><td>NA</td></tr>
<tr><td></td></tr>
<tr><td>T21100</td><td>TGCCCCTACCATGGTACAGAGGG</td><td>1000</td></tr>
<tr><td>Oligo fwd</td><td>CACCTGCCCCTACCATGGTACAGA</td></tr>
<tr><td>Oligo rev</td><td>AAACTCTGTACCATGGTAGGGGCA</td></tr>
<tr><td>Chromosome</td><td>start</td><td>end</td><td>strand</td><td>MM</td><td>target_seq</td><td>PAM</td><td>alignment</td><td>distance</td><td>position</td><td>gene name</td><td>gene id</td></tr>
<tr><td>GU305914.1</td><td>178922</td><td>178944</td><td>-</td><td>0</td><td>TGCCCCTACCATGGTACAGA</td><td>GGG</td><td>||||||||[||||||||||||]PAM</td><td>NA</td><td>Intergenic</td><td>NA</td><td>NA</td></tr>
<tr><td></td></tr>
<tr><td>T21101</td><td>GCCCCTACCATGGTACAGAGGGG</td><td>1000</td></tr>
<tr><td>Oligo fwd</td><td>CACCGCCCCTACCATGGTACAGAG</td></tr>
<tr><td>Oligo rev</td><td>AAACCTCTGTACCATGGTAGGGGC</td></tr>
<tr><td>Chromosome</td><td>start</td><td>end</td><td>strand</td><td>MM</td><td>target_seq</td><td>PAM</td><td>alignment</td><td>distance</td><td>position</td><td>gene name</td><td>gene id</td></tr>
<tr><td>GU305914.1</td><td>178921</td><td>178943</td><td>-</td><td>0</td><td>GCCCCTACCATGGTACAGAG</td><td>GGG</td><td>||||||||[||||||||||||]PAM</td><td>NA</td><td>Intergenic</td><td>NA</td><td>NA</td></tr>
<tr><td></td></tr>
<tr><td>T21102</td><td>CCCCTACCATGGTACAGAGGGGG</td><td>1000</td></tr>
<tr><td>Oligo fwd</td><td>CACCCCCCTACCATGGTACAGAGG</td></tr>
<tr><td>Oligo rev</td><td>AAACCCTCTGTACCATGGTAGGGG</td></tr>
<tr><td>Chromosome</td><td>start</td><td>end</td><td>strand</td><td>MM</td><td>target_seq</td><td>PAM</td><td>alignment</td><td>distance</td><td>position</td><td>gene name</td><td>gene id</td></tr>
<tr><td>GU305914.1</td><td>178920</td><td>178942</td><td>-</td><td>0</td><td>CCCCTACCATGGTACAGAGG</td><td>GGG</td><td>||||||||[||||||||||||]PAM</td><td>NA</td><td>Intergenic</td><td>NA</td><td>NA</td></tr>
<tr><td></td></tr>
<tr><td>T21103</td><td>CCCTGAAGATCAAGTCCTTCAGG</td><td>1000</td></tr>
<tr><td>Oligo fwd</td><td>CACCCCCTGAAGATCAAGTCCTTC</td></tr>
<tr><td>Oligo rev</td><td>AAACGAAGGACTTGATCTTCAGGG</td></tr>
<tr><td>Chromosome</td><td>start</td><td>end</td><td>strand</td><td>MM</td><td>target_seq</td><td>PAM</td><td>alignment</td><td>distance</td><td>position</td><td>gene name</td><td>gene id</td></tr>
<tr><td>GU305914.1</td><td>178896</td><td>178918</td><td>-</td><td>0</td><td>CCCTGAAGATCAAGTCCTTC</td><td>AGG</td><td>||||||||[||||||||||||]PAM</td><td>NA</td><td>Intergenic</td><td>NA</td><td>NA</td></tr>
<tr><td></td></tr>
<tr><td>T21104</td><td>TCCTTCAGGCCCCCTAGCTCTGG</td><td>1000</td></tr>
<tr><td>Oligo fwd</td><td>CACCTCCTTCAGGCCCCCTAGCTC</td></tr>
<tr><td>Oligo rev</td><td>AAACGAGCTAGGGGGCCTGAAGGA</td></tr>
<tr><td>Chromosome</td><td>start</td><td>end</td><td>strand</td><td>MM</td><td>target_seq</td><td>PAM</td><td>alignment</td><td>distance</td><td>position</td><td>gene name</td><td>gene id</td></tr>
<tr><td>GU305914.1</td><td>178882</td><td>178904</td><td>-</td><td>0</td><td>TCCTTCAGGCCCCCTAGCTC</td><td>TGG</td><td>||||||||[||||||||||||]PAM</td><td>NA</td><td>Intergenic</td><td>NA</td><td>NA</td></tr>
<tr><td></td></tr>
<tr><td>T21105</td><td>AACAAACACAGCAGCAGCTCTGG</td><td>1000</td></tr>
<tr><td>Oligo fwd</td><td>CACCAACAAACACAGCAGCAGCTC</td></tr>
<tr><td>Oligo rev</td><td>AAACGAGCTGCTGCTGTGTTTGTT</td></tr>
<tr><td>Chromosome</td><td>start</td><td>end</td><td>strand</td><td>MM</td><td>target_seq</td><td>PAM</td><td>alignment</td><td>distance</td><td>position</td><td>gene name</td><td>gene id</td></tr>
<tr><td>GU305914.1</td><td>178849</td><td>178871</td><td>-</td><td>0</td><td>AACAAACACAGCAGCAGCTC</td><td>TGG</td><td>||||||||[||||||||||||]PAM</td><td>NA</td><td>Intergenic</td><td>NA</td><td>NA</td></tr>
<tr><td></td></tr>
<tr><td>T21106</td><td>AAACACAGCAGCAGCTCTGGTGG</td><td>1000</td></tr>
<tr><td>Oligo fwd</td><td>CACCAAACACAGCAGCAGCTCTGG</td></tr>
<tr><td>Oligo rev</td><td>AAACCCAGAGCTGCTGCTGTGTTT</td></tr>
<tr><td>Chromosome</td><td>start</td><td>end</td><td>strand</td><td>MM</td><td>target_seq</td><td>PAM</td><td>alignment</td><td>distance</td><td>position</td><td>gene name</td><td>gene id</td></tr>
<tr><td>GU305914.1</td><td>178846</td><td>178868</td><td>-</td><td>0</td><td>AAACACAGCAGCAGCTCTGG</td><td>TGG</td><td>||||||||[||||||||||||]PAM</td><td>NA</td><td>Intergenic</td><td>NA</td><td>NA</td></tr>
<tr><td></td></tr>
<tr><td>T21107</td><td>AACACAGCAGCAGCTCTGGTGGG</td><td>1000</td></tr>
<tr><td>Oligo fwd</td><td>CACCAACACAGCAGCAGCTCTGGT</td></tr>
<tr><td>Oligo rev</td><td>AAACACCAGAGCTGCTGCTGTGTT</td></tr>
<tr><td>Chromosome</td><td>start</td><td>end</td><td>strand</td><td>MM</td><td>target_seq</td><td>PAM</td><td>alignment</td><td>distance</td><td>position</td><td>gene name</td><td>gene id</td></tr>
<tr><td>GU305914.1</td><td>178845</td><td>178867</td><td>-</td><td>0</td><td>AACACAGCAGCAGCTCTGGT</td><td>GGG</td><td>||||||||[||||||||||||]PAM</td><td>NA</td><td>Intergenic</td><td>NA</td><td>NA</td></tr>
<tr><td></td></tr>
<tr><td>T21108</td><td>CAAGAAGCAACAGCAGCAGCAGG</td><td>1000</td></tr>
<tr><td>Oligo fwd</td><td>CACCCAAGAAGCAACAGCAGCAGC</td></tr>
<tr><td>Oligo rev</td><td>AAACGCTGCTGCTGTTGCTTCTTG</td></tr>
<tr><td>Chromosome</td><td>start</td><td>end</td><td>strand</td><td>MM</td><td>target_seq</td><td>PAM</td><td>alignment</td><td>distance</td><td>position</td><td>gene name</td><td>gene id</td></tr>
<tr><td>GU305914.1</td><td>178805</td><td>178827</td><td>-</td><td>0</td><td>CAAGAAGCAACAGCAGCAGC</td><td>AGG</td><td>||||||||[||||||||||||]PAM</td><td>NA</td><td>Intergenic</td><td>NA</td><td>NA</td></tr>
<tr><td></td></tr>
<tr><td>T21109</td><td>AAGAAGCCTGTGATGAGTAGTGG</td><td>1000</td></tr>
<tr><td>Oligo fwd</td><td>CACCAAGAAGCCTGTGATGAGTAG</td></tr>
<tr><td>Oligo rev</td><td>AAACCTACTCATCACAGGCTTCTT</td></tr>
<tr><td>Chromosome</td><td>start</td><td>end</td><td>strand</td><td>MM</td><td>target_seq</td><td>PAM</td><td>alignment</td><td>distance</td><td>position</td><td>gene name</td><td>gene id</td></tr>
<tr><td>GU305914.1</td><td>178774</td><td>178796</td><td>-</td><td>0</td><td>AAGAAGCCTGTGATGAGTAG</td><td>TGG</td><td>||||||||[||||||||||||]PAM</td><td>NA</td><td>Intergenic</td><td>NA</td><td>NA</td></tr>
<tr><td></td></tr>
<tr><td>T21110</td><td>AGAAGCCTGTGATGAGTAGTGGG</td><td>1000</td></tr>
<tr><td>Oligo fwd</td><td>CACCAGAAGCCTGTGATGAGTAGT</td></tr>
<tr><td>Oligo rev</td><td>AAACACTACTCATCACAGGCTTCT</td></tr>
<tr><td>Chromosome</td><td>start</td><td>end</td><td>strand</td><td>MM</td><td>target_seq</td><td>PAM</td><td>alignment</td><td>distance</td><td>position</td><td>gene name</td><td>gene id</td></tr>
<tr><td>GU305914.1</td><td>178773</td><td>178795</td><td>-</td><td>0</td><td>AGAAGCCTGTGATGAGTAGT</td><td>GGG</td><td>||||||||[||||||||||||]PAM</td><td>NA</td><td>Intergenic</td><td>NA</td><td>NA</td></tr>
<tr><td></td></tr>
<tr><td>T21111</td><td>GAGTGATAAGATCAGAGACATGG</td><td>1000</td></tr>
<tr><td>Oligo fwd</td><td>CACCGAGTGATAAGATCAGAGACA</td></tr>
<tr><td>Oligo rev</td><td>AAACTGTCTCTGATCTTATCACTC</td></tr>
<tr><td>Chromosome</td><td>start</td><td>end</td><td>strand</td><td>MM</td><td>target_seq</td><td>PAM</td><td>alignment</td><td>distance</td><td>position</td><td>gene name</td><td>gene id</td></tr>
<tr><td>GU305914.1</td><td>178751</td><td>178773</td><td>-</td><td>0</td><td>GAGTGATAAGATCAGAGACA</td><td>TGG</td><td>||||||||[||||||||||||]PAM</td><td>NA</td><td>Intergenic</td><td>NA</td><td>NA</td></tr>
<tr><td></td></tr>
<tr><td>T21112</td><td>AGATCAGAGACATGGTAGACAGG</td><td>1000</td></tr>
<tr><td>Oligo fwd</td><td>CACCAGATCAGAGACATGGTAGAC</td></tr>
</table>

<table>
<tr><td>Oligo rev</td><td>AAACGTCTACCATGTCTCTGATCT</td><td></td><td></td><td></td><td></td><td></td><td></td><td></td><td></td><td></td><td></td></tr>
<tr><td>Chromosome</td><td>start</td><td>end</td><td>strand</td><td>MM</td><td>target_seq</td><td>PAM</td><td>alignment</td><td>distance</td><td>position</td><td>gene name</td><td>gene id</td></tr>
<tr><td>GU305914.1</td><td>178743</td><td>178765</td><td>-</td><td>0</td><td>AGATCAGAGACATGGTAGAC</td><td>AGG</td><td>||||||||[||||||||||||]PAM</td><td>NA</td><td>Intergenic</td><td>NA</td><td>NA</td></tr>
<tr><td>T21113</td><td>GACATGGTAGACAGGACAGCCGG</td><td>1000</td><td></td><td></td><td></td><td></td><td></td><td></td><td></td><td></td><td></td></tr>
<tr><td>Oligo fwd</td><td>CACCGACATGGTAGACAGGACAGC</td><td></td><td></td><td></td><td></td><td></td><td></td><td></td><td></td><td></td><td></td></tr>
<tr><td>Oligo rev</td><td>AAACGCTGTCCTGTCTACCATGTC</td><td></td><td></td><td></td><td></td><td></td><td></td><td></td><td></td><td></td><td></td></tr>
<tr><td>Chromosome</td><td>start</td><td>end</td><td>strand</td><td>MM</td><td>target_seq</td><td>PAM</td><td>alignment</td><td>distance</td><td>position</td><td>gene name</td><td>gene id</td></tr>
<tr><td>GU305914.1</td><td>178735</td><td>178757</td><td>-</td><td>0</td><td>GACATGGTAGACAGGACAGC</td><td>CGG</td><td>||||||||[||||||||||||]PAM</td><td>NA</td><td>Intergenic</td><td>NA</td><td>NA</td></tr>
<tr><td>T21114</td><td>ATGGTAGACAGGACAGCCGGAGG</td><td>1000</td><td></td><td></td><td></td><td></td><td></td><td></td><td></td><td></td><td></td></tr>
<tr><td>Oligo fwd</td><td>CACCATGGTAGACAGGACAGCCGG</td><td></td><td></td><td></td><td></td><td></td><td></td><td></td><td></td><td></td><td></td></tr>
<tr><td>Oligo rev</td><td>AAACCCGGCTGTCCTGTCTACCAT</td><td></td><td></td><td></td><td></td><td></td><td></td><td></td><td></td><td></td><td></td></tr>
<tr><td>Chromosome</td><td>start</td><td>end</td><td>strand</td><td>MM</td><td>target_seq</td><td>PAM</td><td>alignment</td><td>distance</td><td>position</td><td>gene name</td><td>gene id</td></tr>
<tr><td>GU305914.1</td><td>178732</td><td>178754</td><td>-</td><td>0</td><td>ATGGTAGACAGGACAGCCGG</td><td>AGG</td><td>||||||||[||||||||||||]PAM</td><td>NA</td><td>Intergenic</td><td>NA</td><td>NA</td></tr>
<tr><td>T21115</td><td>CAGGACAGCCGGAGGCTACGTGG</td><td>1000</td><td></td><td></td><td></td><td></td><td></td><td></td><td></td><td></td><td></td></tr>
<tr><td>Oligo fwd</td><td>CACCCAGGACAGCCGGAGGCTACG</td><td></td><td></td><td></td><td></td><td></td><td></td><td></td><td></td><td></td><td></td></tr>
<tr><td>Oligo rev</td><td>AAACCGTAGCCTCCGGCTGTCCTG</td><td></td><td></td><td></td><td></td><td></td><td></td><td></td><td></td><td></td><td></td></tr>
<tr><td>Chromosome</td><td>start</td><td>end</td><td>strand</td><td>MM</td><td>target_seq</td><td>PAM</td><td>alignment</td><td>distance</td><td>position</td><td>gene name</td><td>gene id</td></tr>
<tr><td>GU305914.1</td><td>178724</td><td>178746</td><td>-</td><td>0</td><td>CAGGACAGCCGGAGGCTACG</td><td>TGG</td><td>||||||||[||||||||||||]PAM</td><td>NA</td><td>Intergenic</td><td>NA</td><td>NA</td></tr>
<tr><td>T21116</td><td>CCAACGCCCACAAGAAGTGCAGG</td><td>1000</td><td></td><td></td><td></td><td></td><td></td><td></td><td></td><td></td><td></td></tr>
<tr><td>Oligo fwd</td><td>CACCCCAACGCCCACAAGAAGTGC</td><td></td><td></td><td></td><td></td><td></td><td></td><td></td><td></td><td></td><td></td></tr>
<tr><td>Oligo rev</td><td>AAACGCACTTCTTGTGGGCGTTGG</td><td></td><td></td><td></td><td></td><td></td><td></td><td></td><td></td><td></td><td></td></tr>
<tr><td>Chromosome</td><td>start</td><td>end</td><td>strand</td><td>MM</td><td>target_seq</td><td>PAM</td><td>alignment</td><td>distance</td><td>position</td><td>gene name</td><td>gene id</td></tr>
<tr><td>GU305914.1</td><td>178698</td><td>178720</td><td>-</td><td>0</td><td>CCAACGCCCACAAGAAGTGC</td><td>AGG</td><td>||||||||[||||||||||||]PAM</td><td>NA</td><td>Intergenic</td><td>NA</td><td>NA</td></tr>
<tr><td>T21117</td><td>CAACGCCCACAAGAAGTGCAGGG</td><td>1000</td><td></td><td></td><td></td><td></td><td></td><td></td><td></td><td></td><td></td></tr>
<tr><td>Oligo fwd</td><td>CACCCAACGCCCACAAGAAGTGCA</td><td></td><td></td><td></td><td></td><td></td><td></td><td></td><td></td><td></td><td></td></tr>
<tr><td>Oligo rev</td><td>AAACTGCACTTCTTGTGGGCGTTG</td><td></td><td></td><td></td><td></td><td></td><td></td><td></td><td></td><td></td><td></td></tr>
<tr><td>Chromosome</td><td>start</td><td>end</td><td>strand</td><td>MM</td><td>target_seq</td><td>PAM</td><td>alignment</td><td>distance</td><td>position</td><td>gene name</td><td>gene id</td></tr>
<tr><td>GU305914.1</td><td>178697</td><td>178719</td><td>-</td><td>0</td><td>CAACGCCCACAAGAAGTGCA</td><td>GGG</td><td>||||||||[||||||||||||]PAM</td><td>NA</td><td>Intergenic</td><td>NA</td><td>NA</td></tr>
<tr><td>T21118</td><td>AGTGCAGGGAAGATAAGTCCAGG</td><td>1000</td><td></td><td></td><td></td><td></td><td></td><td></td><td></td><td></td><td></td></tr>
<tr><td>Oligo fwd</td><td>CACCAGTGCAGGGAAGATAAGTCC</td><td></td><td></td><td></td><td></td><td></td><td></td><td></td><td></td><td></td><td></td></tr>
<tr><td>Oligo rev</td><td>AAACGGACTTATCTTCCCTGCACT</td><td></td><td></td><td></td><td></td><td></td><td></td><td></td><td></td><td></td><td></td></tr>
<tr><td>Chromosome</td><td>start</td><td>end</td><td>strand</td><td>MM</td><td>target_seq</td><td>PAM</td><td>alignment</td><td>distance</td><td>position</td><td>gene name</td><td>gene id</td></tr>
<tr><td>GU305914.1</td><td>178683</td><td>178705</td><td>-</td><td>0</td><td>AGTGCAGGGAAGATAAGTCC</td><td>AGG</td><td>||||||||[||||||||||||]PAM</td><td>NA</td><td>Intergenic</td><td>NA</td><td>NA</td></tr>
<tr><td>T21119</td><td>AGTCCAGGAAGTACCCAGCCAGG</td><td>1000</td><td></td><td></td><td></td><td></td><td></td><td></td><td></td><td></td><td></td></tr>
<tr><td>Oligo fwd</td><td>CACCAGTCCAGGAAGTACCCAGCC</td><td></td><td></td><td></td><td></td><td></td><td></td><td></td><td></td><td></td><td></td></tr>
<tr><td>Oligo rev</td><td>AAACGGCTGGGTACTTCCTGGACT</td><td></td><td></td><td></td><td></td><td></td><td></td><td></td><td></td><td></td><td></td></tr>
<tr><td>Chromosome</td><td>start</td><td>end</td><td>strand</td><td>MM</td><td>target_seq</td><td>PAM</td><td>alignment</td><td>distance</td><td>position</td><td>gene name</td><td>gene id</td></tr>
<tr><td>GU305914.1</td><td>178668</td><td>178690</td><td>-</td><td>0</td><td>AGTCCAGGAAGTACCCAGCC</td><td>AGG</td><td>||||||||[||||||||||||]PAM</td><td>NA</td><td>Intergenic</td><td>NA</td><td>NA</td></tr>
<tr><td>T21120</td><td>GTCCAGGAAGTACCCAGCCAGGG</td><td>1000</td><td></td><td></td><td></td><td></td><td></td><td></td><td></td><td></td><td></td></tr>
<tr><td>Oligo fwd</td><td>CACCGTCCAGGAAGTACCCAGCCA</td><td></td><td></td><td></td><td></td><td></td><td></td><td></td><td></td><td></td><td></td></tr>
<tr><td>Oligo rev</td><td>AAACTGGCTGGGTACTTCCTGGAC</td><td></td><td></td><td></td><td></td><td></td><td></td><td></td><td></td><td></td><td></td></tr>
<tr><td>Chromosome</td><td>start</td><td>end</td><td>strand</td><td>MM</td><td>target_seq</td><td>PAM</td><td>alignment</td><td>distance</td><td>position</td><td>gene name</td><td>gene id</td></tr>
<tr><td>GU305914.1</td><td>178667</td><td>178689</td><td>-</td><td>0</td><td>GTCCAGGAAGTACCCAGCCA</td><td>GGG</td><td>||||||||[||||||||||||]PAM</td><td>NA</td><td>Intergenic</td><td>NA</td><td>NA</td></tr>
<tr><td>T21121</td><td>GAAGTACCCAGCCAGGGCCTTGG</td><td>1000</td><td></td><td></td><td></td><td></td><td></td><td></td><td></td><td></td><td></td></tr>
<tr><td>Oligo fwd</td><td>CACCGAAGTACCCAGCCAGGGCCT</td><td></td><td></td><td></td><td></td><td></td><td></td><td></td><td></td><td></td><td></td></tr>
<tr><td>Oligo rev</td><td>AAACAGGCCCTGGCTGGGTACTTC</td><td></td><td></td><td></td><td></td><td></td><td></td><td></td><td></td><td></td><td></td></tr>
<tr><td>Chromosome</td><td>start</td><td>end</td><td>strand</td><td>MM</td><td>target_seq</td><td>PAM</td><td>alignment</td><td>distance</td><td>position</td><td>gene name</td><td>gene id</td></tr>
<tr><td>GU305914.1</td><td>178661</td><td>178683</td><td>-</td><td>0</td><td>GAAGTACCCAGCCAGGGCCT</td><td>TGG</td><td>||||||||[||||||||||||]PAM</td><td>NA</td><td>Intergenic</td><td>NA</td><td>NA</td></tr>
<tr><td>T21122</td><td>TCAGTCCCCACAGTATCTACTGG</td><td>1000</td><td></td><td></td><td></td><td></td><td></td><td></td><td></td><td></td><td></td></tr>
<tr><td>Oligo fwd</td><td>CACCTCAGTCCCCACAGTATCTAC</td><td></td><td></td><td></td><td></td><td></td><td></td><td></td><td></td><td></td><td></td></tr>
<tr><td>Oligo rev</td><td>AAACGTAGATACTGTGGGGACTGA</td><td></td><td></td><td></td><td></td><td></td><td></td><td></td><td></td><td></td><td></td></tr>
<tr><td>Chromosome</td><td>start</td><td>end</td><td>strand</td><td>MM</td><td>target_seq</td><td>PAM</td><td>alignment</td><td>distance</td><td>position</td><td>gene name</td><td>gene id</td></tr>
<tr><td>GU305914.1</td><td>178613</td><td>178635</td><td>-</td><td>0</td><td>TCAGTCCCCACAGTATCTAC</td><td>TGG</td><td>||||||||[||||||||||||]PAM</td><td>NA</td><td>Intergenic</td><td>NA</td><td>NA</td></tr>
<tr><td>T21123</td><td>CAGTCCCCACAGTATCTACTGGG</td><td>1000</td><td></td><td></td><td></td><td></td><td></td><td></td><td></td><td></td><td></td></tr>
<tr><td>Oligo fwd</td><td>CACCCAGTCCCCACAGTATCTACT</td><td></td><td></td><td></td><td></td><td></td><td></td><td></td><td></td><td></td><td></td></tr>
<tr><td>Oligo rev</td><td>AAACAGTAGATACTGTGGGGACTG</td><td></td><td></td><td></td><td></td><td></td><td></td><td></td><td></td><td></td><td></td></tr>
<tr><td>Chromosome</td><td>start</td><td>end</td><td>strand</td><td>MM</td><td>target_seq</td><td>PAM</td><td>alignment</td><td>distance</td><td>position</td><td>gene name</td><td>gene id</td></tr>
<tr><td>GU305914.1</td><td>178612</td><td>178634</td><td>-</td><td>0</td><td>CAGTCCCCACAGTATCTACT</td><td>GGG</td><td>||||||||[||||||||||||]PAM</td><td>NA</td><td>Intergenic</td><td>NA</td><td>NA</td></tr>
<tr><td>T21124</td><td>CCCACAGTATCTACTGGGTAAGG</td><td>1000</td><td></td><td></td><td></td><td></td><td></td><td></td><td></td><td></td><td></td></tr>
<tr><td>Oligo fwd</td><td>CACCCCCACAGTATCTACTGGGTA</td><td></td><td></td><td></td><td></td><td></td><td></td><td></td><td></td><td></td><td></td></tr>
<tr><td>Oligo rev</td><td>AAACTACCCAGTAGATACTGTGGG</td><td></td><td></td><td></td><td></td><td></td><td></td><td></td><td></td><td></td><td></td></tr>
<tr><td>Chromosome</td><td>start</td><td>end</td><td>strand</td><td>MM</td><td>target_seq</td><td>PAM</td><td>alignment</td><td>distance</td><td>position</td><td>gene name</td><td>gene id</td></tr>
<tr><td>GU305914.1</td><td>178607</td><td>178629</td><td>-</td><td>0</td><td>CCCACAGTATCTACTGGGTA</td><td>AGG</td><td>||||||||[||||||||||||]PAM</td><td>NA</td><td>Intergenic</td><td>NA</td><td>NA</td></tr>
<tr><td>T21125</td><td>TAAGGCCATCCAGTTCTGTAAGG</td><td>1000</td><td></td><td></td><td></td><td></td><td></td><td></td><td></td><td></td><td></td></tr>
<tr><td>Oligo fwd</td><td>CACCTAAGGCCATCCAGTTCTGTA</td><td></td><td></td><td></td><td></td><td></td><td></td><td></td><td></td><td></td><td></td></tr>
<tr><td>Oligo rev</td><td>AAACTACAGAACTGGATGGCCTTA</td><td></td><td></td><td></td><td></td><td></td><td></td><td></td><td></td><td></td><td></td></tr>
<tr><td>Chromosome</td><td>start</td><td>end</td><td>strand</td><td>MM</td><td>target_seq</td><td>PAM</td><td>alignment</td><td>distance</td><td>position</td><td>gene name</td><td>gene id</td></tr>
<tr><td>GU305914.1</td><td>178589</td><td>178611</td><td>-</td><td>0</td><td>TAAGGCCATCCAGTTCTGTA</td><td>AGG</td><td>||||||||[||||||||||||]PAM</td><td>NA</td><td>Intergenic</td><td>NA</td><td>NA</td></tr>
<tr><td>T21126</td><td>GGCCATCCAGTTCTGTAAGGAGG</td><td>1000</td><td></td><td></td><td></td><td></td><td></td><td></td><td></td><td></td><td></td></tr>
<tr><td>Oligo fwd</td><td>CACCGGCCATCCAGTTCTGTAAGG</td><td></td><td></td><td></td><td></td><td></td><td></td><td></td><td></td><td></td><td></td></tr>
<tr><td>Oligo rev</td><td>AAACCCTTACAGAACTGGATGGCC</td><td></td><td></td><td></td><td></td><td></td><td></td><td></td><td></td><td></td><td></td></tr>
<tr><td>Chromosome</td><td>start</td><td>end</td><td>strand</td><td>MM</td><td>target_seq</td><td>PAM</td><td>alignment</td><td>distance</td><td>position</td><td>gene name</td><td>gene id</td></tr>
<tr><td>GU305914.1</td><td>178586</td><td>178608</td><td>-</td><td>0</td><td>GGCCATCCAGTTCTGTAAGG</td><td>AGG</td><td>||||||||[||||||||||||]PAM</td><td>NA</td><td>Intergenic</td><td>NA</td><td>NA</td></tr>
<tr><td>T21127</td><td>TCATCATGCTGTTCTACACCAGG</td><td>1000</td><td></td><td></td><td></td><td></td><td></td><td></td><td></td><td></td><td></td></tr>
</table>

<table>
<tr><td>Oligo fwd</td><td>CACCTCATCATGCTGTTCTACACC</td></tr>
<tr><td>Oligo rev</td><td>AAACGGTGTAGAACAGCATGATGA</td></tr>
<tr><td>Chromosome</td><td>start</td><td>end</td><td>strand</td><td>MM</td><td>target_seq</td><td>PAM</td><td>alignment</td><td>distance</td><td>position</td><td>gene name</td><td>gene id</td></tr>
<tr><td>GU305914.1</td><td>178545</td><td>178567</td><td>-</td><td>0</td><td>TCATCATGCTGTTCTACACC</td><td>AGG</td><td>||||||||[||||||||||||]PAM</td><td>NA</td><td>Intergenic</td><td>NA</td><td>NA</td></tr>
<tr><td></td></tr>
<tr><td>T21128</td><td>GCTGTTCTACACCAGGAGTCAGG</td><td>1000</td></tr>
<tr><td>Oligo fwd</td><td>CACCGCTGTTCTACACCAGGAGTC</td></tr>
<tr><td>Oligo rev</td><td>AAACGACTCCTGGTGTAGAACAGC</td></tr>
<tr><td>Chromosome</td><td>start</td><td>end</td><td>strand</td><td>MM</td><td>target_seq</td><td>PAM</td><td>alignment</td><td>distance</td><td>position</td><td>gene name</td><td>gene id</td></tr>
<tr><td>GU305914.1</td><td>178538</td><td>178560</td><td>-</td><td>0</td><td>GCTGTTCTACACCAGGAGTC</td><td>AGG</td><td>||||||||[||||||||||||]PAM</td><td>NA</td><td>Intergenic</td><td>NA</td><td>NA</td></tr>
<tr><td></td></tr>
<tr><td>T21129</td><td>ACACCAGGAGTCAGGATGTGAGG</td><td>1000</td></tr>
<tr><td>Oligo fwd</td><td>CACCACACCAGGAGTCAGGATGTG</td></tr>
<tr><td>Oligo rev</td><td>AAACCACATCCTGACTCCTGGTGT</td></tr>
<tr><td>Chromosome</td><td>start</td><td>end</td><td>strand</td><td>MM</td><td>target_seq</td><td>PAM</td><td>alignment</td><td>distance</td><td>position</td><td>gene name</td><td>gene id</td></tr>
<tr><td>GU305914.1</td><td>178530</td><td>178552</td><td>-</td><td>0</td><td>ACACCAGGAGTCAGGATGTG</td><td>AGG</td><td>||||||||[||||||||||||]PAM</td><td>NA</td><td>Intergenic</td><td>NA</td><td>NA</td></tr>
<tr><td></td></tr>
<tr><td>T21130</td><td>CAGGAGTCAGGATGTGAGGAAGG</td><td>1000</td></tr>
<tr><td>Oligo fwd</td><td>CACCCAGGAGTCAGGATGTGAGGA</td></tr>
<tr><td>Oligo rev</td><td>AAACTCCTCACATCCTGACTCCTG</td></tr>
<tr><td>Chromosome</td><td>start</td><td>end</td><td>strand</td><td>MM</td><td>target_seq</td><td>PAM</td><td>alignment</td><td>distance</td><td>position</td><td>gene name</td><td>gene id</td></tr>
<tr><td>GU305914.1</td><td>178526</td><td>178548</td><td>-</td><td>0</td><td>CAGGAGTCAGGATGTGAGGA</td><td>AGG</td><td>||||||||[||||||||||||]PAM</td><td>NA</td><td>Intergenic</td><td>NA</td><td>NA</td></tr>
<tr><td></td></tr>
<tr><td>T21131</td><td>TCAGGATGTGAGGAAGGCAGTGG</td><td>1000</td></tr>
<tr><td>Oligo fwd</td><td>CACCTCAGGATGTGAGGAAGGCAG</td></tr>
<tr><td>Oligo rev</td><td>AAACCTGCCTTCCTCACATCCTGA</td></tr>
<tr><td>Chromosome</td><td>start</td><td>end</td><td>strand</td><td>MM</td><td>target_seq</td><td>PAM</td><td>alignment</td><td>distance</td><td>position</td><td>gene name</td><td>gene id</td></tr>
<tr><td>GU305914.1</td><td>178520</td><td>178542</td><td>-</td><td>0</td><td>TCAGGATGTGAGGAAGGCAG</td><td>TGG</td><td>||||||||[||||||||||||]PAM</td><td>NA</td><td>Intergenic</td><td>NA</td><td>NA</td></tr>
<tr><td></td></tr>
<tr><td>T21132</td><td>GGATGAGACACGCGCCCGCATGG</td><td>1000</td></tr>
<tr><td>Oligo fwd</td><td>CACCGGATGAGACACGCGCCCGCA</td></tr>
<tr><td>Oligo rev</td><td>AAACTGCGGGCGCGTGTCTCATCC</td></tr>
<tr><td>Chromosome</td><td>start</td><td>end</td><td>strand</td><td>MM</td><td>target_seq</td><td>PAM</td><td>alignment</td><td>distance</td><td>position</td><td>gene name</td><td>gene id</td></tr>
<tr><td>GU305914.1</td><td>178499</td><td>178521</td><td>-</td><td>0</td><td>GGATGAGACACGCGCCCGCA</td><td>TGG</td><td>||||||||[||||||||||||]PAM</td><td>NA</td><td>Intergenic</td><td>NA</td><td>NA</td></tr>
<tr><td></td></tr>
<tr><td>T21133</td><td>GATGAGACACGCGCCCGCATGGG</td><td>1000</td></tr>
<tr><td>Oligo fwd</td><td>CACCGATGAGACACGCGCCCGCAT</td></tr>
<tr><td>Oligo rev</td><td>AAACATGCGGGCGCGTGTCTCATC</td></tr>
<tr><td>Chromosome</td><td>start</td><td>end</td><td>strand</td><td>MM</td><td>target_seq</td><td>PAM</td><td>alignment</td><td>distance</td><td>position</td><td>gene name</td><td>gene id</td></tr>
<tr><td>GU305914.1</td><td>178498</td><td>178520</td><td>-</td><td>0</td><td>GATGAGACACGCGCCCGCAT</td><td>GGG</td><td>||||||||[||||||||||||]PAM</td><td>NA</td><td>Intergenic</td><td>NA</td><td>NA</td></tr>
<tr><td></td></tr>
<tr><td>T21134</td><td>ATGAGACACGCGCCCGCATGGGG</td><td>1000</td></tr>
<tr><td>Oligo fwd</td><td>CACCATGAGACACGCGCCCGCATG</td></tr>
<tr><td>Oligo rev</td><td>AAACCATGCGGGCGCGTGTCTCAT</td></tr>
<tr><td>Chromosome</td><td>start</td><td>end</td><td>strand</td><td>MM</td><td>target_seq</td><td>PAM</td><td>alignment</td><td>distance</td><td>position</td><td>gene name</td><td>gene id</td></tr>
<tr><td>GU305914.1</td><td>178497</td><td>178519</td><td>-</td><td>0</td><td>ATGAGACACGCGCCCGCATG</td><td>GGG</td><td>||||||||[||||||||||||]PAM</td><td>NA</td><td>Intergenic</td><td>NA</td><td>NA</td></tr>
<tr><td></td></tr>
<tr><td>T21135</td><td>CCAAGCCCATTAACCACAGCAGG</td><td>1000</td></tr>
<tr><td>Oligo fwd</td><td>CACCCCAAGCCCATTAACCACAGC</td></tr>
<tr><td>Oligo rev</td><td>AAACGCTGTGGTTAATGGGCTTGG</td></tr>
<tr><td>Chromosome</td><td>start</td><td>end</td><td>strand</td><td>MM</td><td>target_seq</td><td>PAM</td><td>alignment</td><td>distance</td><td>position</td><td>gene name</td><td>gene id</td></tr>
<tr><td>GU305914.1</td><td>178428</td><td>178450</td><td>-</td><td>0</td><td>CCAAGCCCATTAACCACAGC</td><td>AGG</td><td>||||||||[||||||||||||]PAM</td><td>NA</td><td>Intergenic</td><td>NA</td><td>NA</td></tr>
<tr><td></td></tr>
<tr><td>T21136</td><td>CAAGCCCATTAACCACAGCAGGG</td><td>1000</td></tr>
<tr><td>Oligo fwd</td><td>CACCCAAGCCCATTAACCACAGCA</td></tr>
<tr><td>Oligo rev</td><td>AAACTGCTGTGGTTAATGGGCTTG</td></tr>
<tr><td>Chromosome</td><td>start</td><td>end</td><td>strand</td><td>MM</td><td>target_seq</td><td>PAM</td><td>alignment</td><td>distance</td><td>position</td><td>gene name</td><td>gene id</td></tr>
<tr><td>GU305914.1</td><td>178427</td><td>178449</td><td>-</td><td>0</td><td>CAAGCCCATTAACCACAGCA</td><td>GGG</td><td>||||||||[||||||||||||]PAM</td><td>NA</td><td>Intergenic</td><td>NA</td><td>NA</td></tr>
<tr><td></td></tr>
<tr><td>T21137</td><td>GACCATAGATAGAACCTCAGCGG</td><td>1000</td></tr>
<tr><td>Oligo fwd</td><td>CACCGACCATAGATAGAACCTCAG</td></tr>
<tr><td>Oligo rev</td><td>AAACCTGAGGTTCTATCTATGGTC</td></tr>
<tr><td>Chromosome</td><td>start</td><td>end</td><td>strand</td><td>MM</td><td>target_seq</td><td>PAM</td><td>alignment</td><td>distance</td><td>position</td><td>gene name</td><td>gene id</td></tr>
<tr><td>GU305914.1</td><td>178403</td><td>178425</td><td>-</td><td>0</td><td>GACCATAGATAGAACCTCAG</td><td>CGG</td><td>||||||||[||||||||||||]PAM</td><td>NA</td><td>Intergenic</td><td>NA</td><td>NA</td></tr>
<tr><td></td></tr>
<tr><td>T21138</td><td>ACCTCAGCGGCCTGCACCGCAGG</td><td>1000</td></tr>
<tr><td>Oligo fwd</td><td>CACCACCTCAGCGGCCTGCACCGC</td></tr>
<tr><td>Oligo rev</td><td>AAACGCGGTGCAGGCCGCTGAGGT</td></tr>
<tr><td>Chromosome</td><td>start</td><td>end</td><td>strand</td><td>MM</td><td>target_seq</td><td>PAM</td><td>alignment</td><td>distance</td><td>position</td><td>gene name</td><td>gene id</td></tr>
<tr><td>GU305914.1</td><td>178390</td><td>178412</td><td>-</td><td>0</td><td>ACCTCAGCGGCCTGCACCGC</td><td>AGG</td><td>||||||||[||||||||||||]PAM</td><td>NA</td><td>Intergenic</td><td>NA</td><td>NA</td></tr>
<tr><td></td></tr>
<tr><td>T21139</td><td>GGCCTGCACCGCAGGTACTCAGG</td><td>1000</td></tr>
<tr><td>Oligo fwd</td><td>CACCGGCCTGCACCGCAGGTACTC</td></tr>
<tr><td>Oligo rev</td><td>AAACGAGTACCTGCGGTGCAGGCC</td></tr>
<tr><td>Chromosome</td><td>start</td><td>end</td><td>strand</td><td>MM</td><td>target_seq</td><td>PAM</td><td>alignment</td><td>distance</td><td>position</td><td>gene name</td><td>gene id</td></tr>
<tr><td>GU305914.1</td><td>178382</td><td>178404</td><td>-</td><td>0</td><td>GGCCTGCACCGCAGGTACTC</td><td>AGG</td><td>||||||||[||||||||||||]PAM</td><td>NA</td><td>Intergenic</td><td>NA</td><td>NA</td></tr>
<tr><td></td></tr>
<tr><td>T21140</td><td>CCGCAGGTACTCAGGCAGTGTGG</td><td>1000</td></tr>
<tr><td>Oligo fwd</td><td>CACCCCGCAGGTACTCAGGCAGTG</td></tr>
<tr><td>Oligo rev</td><td>AAACCACTGCCTGAGTACCTGCGG</td></tr>
<tr><td>Chromosome</td><td>start</td><td>end</td><td>strand</td><td>MM</td><td>target_seq</td><td>PAM</td><td>alignment</td><td>distance</td><td>position</td><td>gene name</td><td>gene id</td></tr>
<tr><td>GU305914.1</td><td>178374</td><td>178396</td><td>-</td><td>0</td><td>CCGCAGGTACTCAGGCAGTG</td><td>TGG</td><td>||||||||[||||||||||||]PAM</td><td>NA</td><td>Intergenic</td><td>NA</td><td>NA</td></tr>
<tr><td></td></tr>
<tr><td>T21141</td><td>CGCAGGTACTCAGGCAGTGTGGG</td><td>1000</td></tr>
<tr><td>Oligo fwd</td><td>CACCCGCAGGTACTCAGGCAGTGT</td></tr>
<tr><td>Oligo rev</td><td>AAACACACTGCCTGAGTACCTGCG</td></tr>
<tr><td>Chromosome</td><td>start</td><td>end</td><td>strand</td><td>MM</td><td>target_seq</td><td>PAM</td><td>alignment</td><td>distance</td><td>position</td><td>gene name</td><td>gene id</td></tr>
<tr><td>GU305914.1</td><td>178373</td><td>178395</td><td>-</td><td>0</td><td>CGCAGGTACTCAGGCAGTGT</td><td>GGG</td><td>||||||||[||||||||||||]PAM</td><td>NA</td><td>Intergenic</td><td>NA</td><td>NA</td></tr>
</table>

<table>
<tr><td>T21142</td><td>TACTCAGGCAGTGTGGGACATGG</td><td>1000</td><td></td><td></td><td></td><td></td><td></td><td></td><td></td><td></td><td></td></tr>
<tr><td>Oligo fwd</td><td>CACCTACTCAGGCAGTGTGGGACA</td><td></td><td></td><td></td><td></td><td></td><td></td><td></td><td></td><td></td><td></td></tr>
<tr><td>Oligo rev</td><td>AAACTGTCCCACACTGCCTGAGTA</td><td></td><td></td><td></td><td></td><td></td><td></td><td></td><td></td><td></td><td></td></tr>
<tr><td>Chromosome</td><td>start</td><td>end</td><td>strand</td><td>MM</td><td>target_seq</td><td>PAM</td><td>alignment</td><td>distance</td><td>position</td><td>gene name</td><td>gene id</td></tr>
<tr><td>GU305914.1</td><td>178367</td><td>178389</td><td>-</td><td>0</td><td>TACTCAGGCAGTGTGGGACA</td><td>TGG</td><td>||||||||[||||||||||||]PAM</td><td>NA</td><td>Intergenic</td><td>NA</td><td>NA</td></tr>
<tr><td>T21143</td><td>TCAGGCAGTGTGGGACATGGAGG</td><td>1000</td><td></td><td></td><td></td><td></td><td></td><td></td><td></td><td></td><td></td></tr>
<tr><td>Oligo fwd</td><td>CACCTCAGGCAGTGTGGGACATGG</td><td></td><td></td><td></td><td></td><td></td><td></td><td></td><td></td><td></td><td></td></tr>
<tr><td>Oligo rev</td><td>AAACCCATGTCCCACACTGCCTGA</td><td></td><td></td><td></td><td></td><td></td><td></td><td></td><td></td><td></td><td></td></tr>
<tr><td>Chromosome</td><td>start</td><td>end</td><td>strand</td><td>MM</td><td>target_seq</td><td>PAM</td><td>alignment</td><td>distance</td><td>position</td><td>gene name</td><td>gene id</td></tr>
<tr><td>GU305914.1</td><td>178364</td><td>178386</td><td>-</td><td>0</td><td>TCAGGCAGTGTGGGACATGG</td><td>AGG</td><td>||||||||[||||||||||||]PAM</td><td>NA</td><td>Intergenic</td><td>NA</td><td>NA</td></tr>
<tr><td>T21144</td><td>TGGGACATGGAGGAGAGACGAGG</td><td>1000</td><td></td><td></td><td></td><td></td><td></td><td></td><td></td><td></td><td></td></tr>
<tr><td>Oligo fwd</td><td>CACCTGGGACATGGAGGAGAGACG</td><td></td><td></td><td></td><td></td><td></td><td></td><td></td><td></td><td></td><td></td></tr>
<tr><td>Oligo rev</td><td>AAACCGTCTCTCCTCCATGTCCCA</td><td></td><td></td><td></td><td></td><td></td><td></td><td></td><td></td><td></td><td></td></tr>
<tr><td>Chromosome</td><td>start</td><td>end</td><td>strand</td><td>MM</td><td>target_seq</td><td>PAM</td><td>alignment</td><td>distance</td><td>position</td><td>gene name</td><td>gene id</td></tr>
<tr><td>GU305914.1</td><td>178354</td><td>178376</td><td>-</td><td>0</td><td>TGGGACATGGAGGAGAGACG</td><td>AGG</td><td>||||||||[||||||||||||]PAM</td><td>NA</td><td>Intergenic</td><td>NA</td><td>NA</td></tr>
<tr><td>T21145</td><td>GAGGTCAGAAGTGTGTGCCCAGG</td><td>1000</td><td></td><td></td><td></td><td></td><td></td><td></td><td></td><td></td><td></td></tr>
<tr><td>Oligo fwd</td><td>CACCGAGGTCAGAAGTGTGTGCCC</td><td></td><td></td><td></td><td></td><td></td><td></td><td></td><td></td><td></td><td></td></tr>
<tr><td>Oligo rev</td><td>AAACGGGCACACACTTCTGACCTC</td><td></td><td></td><td></td><td></td><td></td><td></td><td></td><td></td><td></td><td></td></tr>
<tr><td>Chromosome</td><td>start</td><td>end</td><td>strand</td><td>MM</td><td>target_seq</td><td>PAM</td><td>alignment</td><td>distance</td><td>position</td><td>gene name</td><td>gene id</td></tr>
<tr><td>GU305914.1</td><td>178335</td><td>178357</td><td>-</td><td>0</td><td>GAGGTCAGAAGTGTGTGCCC</td><td>AGG</td><td>||||||||[||||||||||||]PAM</td><td>NA</td><td>Intergenic</td><td>NA</td><td>NA</td></tr>
<tr><td>T21146</td><td>TGCCCAGGACCTCAGACTACAGG</td><td>1000</td><td></td><td></td><td></td><td></td><td></td><td></td><td></td><td></td><td></td></tr>
<tr><td>Oligo fwd</td><td>CACCTGCCCAGGACCTCAGACTAC</td><td></td><td></td><td></td><td></td><td></td><td></td><td></td><td></td><td></td><td></td></tr>
<tr><td>Oligo rev</td><td>AAACGTAGTCTGAGGTCCTGGGCA</td><td></td><td></td><td></td><td></td><td></td><td></td><td></td><td></td><td></td><td></td></tr>
<tr><td>Chromosome</td><td>start</td><td>end</td><td>strand</td><td>MM</td><td>target_seq</td><td>PAM</td><td>alignment</td><td>distance</td><td>position</td><td>gene name</td><td>gene id</td></tr>
<tr><td>GU305914.1</td><td>178320</td><td>178342</td><td>-</td><td>0</td><td>TGCCCAGGACCTCAGACTAC</td><td>AGG</td><td>||||||||[||||||||||||]PAM</td><td>NA</td><td>Intergenic</td><td>NA</td><td>NA</td></tr>
<tr><td>T21147</td><td>CTACAGGAGCATGATAATTCAGG</td><td>1000</td><td></td><td></td><td></td><td></td><td></td><td></td><td></td><td></td><td></td></tr>
<tr><td>Oligo fwd</td><td>CACCCTACAGGAGCATGATAATTC</td><td></td><td></td><td></td><td></td><td></td><td></td><td></td><td></td><td></td><td></td></tr>
<tr><td>Oligo rev</td><td>AAACGAATTATCATGCTCCTGTAG</td><td></td><td></td><td></td><td></td><td></td><td></td><td></td><td></td><td></td><td></td></tr>
<tr><td>Chromosome</td><td>start</td><td>end</td><td>strand</td><td>MM</td><td>target_seq</td><td>PAM</td><td>alignment</td><td>distance</td><td>position</td><td>gene name</td><td>gene id</td></tr>
<tr><td>GU305914.1</td><td>178304</td><td>178326</td><td>-</td><td>0</td><td>CTACAGGAGCATGATAATTC</td><td>AGG</td><td>||||||||[||||||||||||]PAM</td><td>NA</td><td>Intergenic</td><td>NA</td><td>NA</td></tr>
<tr><td>T21148</td><td>AGCCAACCCTCCAGACTTCTTGG</td><td>1000</td><td></td><td></td><td></td><td></td><td></td><td></td><td></td><td></td><td></td></tr>
<tr><td>Oligo fwd</td><td>CACCAGCCAACCCTCCAGACTTCT</td><td></td><td></td><td></td><td></td><td></td><td></td><td></td><td></td><td></td><td></td></tr>
<tr><td>Oligo rev</td><td>AAACAGAAGTCTGGAGGGTTGGCT</td><td></td><td></td><td></td><td></td><td></td><td></td><td></td><td></td><td></td><td></td></tr>
<tr><td>Chromosome</td><td>start</td><td>end</td><td>strand</td><td>MM</td><td>target_seq</td><td>PAM</td><td>alignment</td><td>distance</td><td>position</td><td>gene name</td><td>gene id</td></tr>
<tr><td>GU305914.1</td><td>178280</td><td>178302</td><td>-</td><td>0</td><td>AGCCAACCCTCCAGACTTCT</td><td>TGG</td><td>||||||||[||||||||||||]PAM</td><td>NA</td><td>Intergenic</td><td>NA</td><td>NA</td></tr>
<tr><td>T21149</td><td>GCCAACCCTCCAGACTTCTTGGG</td><td>1000</td><td></td><td></td><td></td><td></td><td></td><td></td><td></td><td></td><td></td></tr>
<tr><td>Oligo fwd</td><td>CACCGCCAACCCTCCAGACTTCTT</td><td></td><td></td><td></td><td></td><td></td><td></td><td></td><td></td><td></td><td></td></tr>
<tr><td>Oligo rev</td><td>AAACAAGAAGTCTGGAGGGTTGGC</td><td></td><td></td><td></td><td></td><td></td><td></td><td></td><td></td><td></td><td></td></tr>
<tr><td>Chromosome</td><td>start</td><td>end</td><td>strand</td><td>MM</td><td>target_seq</td><td>PAM</td><td>alignment</td><td>distance</td><td>position</td><td>gene name</td><td>gene id</td></tr>
<tr><td>GU305914.1</td><td>178279</td><td>178301</td><td>-</td><td>0</td><td>GCCAACCCTCCAGACTTCTT</td><td>GGG</td><td>||||||||[||||||||||||]PAM</td><td>NA</td><td>Intergenic</td><td>NA</td><td>NA</td></tr>
<tr><td>T21150</td><td>CCCTCCAGACTTCTTGGGAGCGG</td><td>1000</td><td></td><td></td><td></td><td></td><td></td><td></td><td></td><td></td><td></td></tr>
<tr><td>Oligo fwd</td><td>CACCCCCTCCAGACTTCTTGGGAG</td><td></td><td></td><td></td><td></td><td></td><td></td><td></td><td></td><td></td><td></td></tr>
<tr><td>Oligo rev</td><td>AAACCTCCCAAGAAGTCTGGAGGG</td><td></td><td></td><td></td><td></td><td></td><td></td><td></td><td></td><td></td><td></td></tr>
<tr><td>Chromosome</td><td>start</td><td>end</td><td>strand</td><td>MM</td><td>target_seq</td><td>PAM</td><td>alignment</td><td>distance</td><td>position</td><td>gene name</td><td>gene id</td></tr>
<tr><td>GU305914.1</td><td>178274</td><td>178296</td><td>-</td><td>0</td><td>CCCTCCAGACTTCTTGGGAG</td><td>CGG</td><td>||||||||[||||||||||||]PAM</td><td>NA</td><td>Intergenic</td><td>NA</td><td>NA</td></tr>
<tr><td>T21151</td><td>GACCTGTCTGCATTTGAGTCAGG</td><td>1000</td><td></td><td></td><td></td><td></td><td></td><td></td><td></td><td></td><td></td></tr>
<tr><td>Oligo fwd</td><td>CACCGACCTGTCTGCATTTGAGTC</td><td></td><td></td><td></td><td></td><td></td><td></td><td></td><td></td><td></td><td></td></tr>
<tr><td>Oligo rev</td><td>AAACGACTCAAATGCAGACAGGTC</td><td></td><td></td><td></td><td></td><td></td><td></td><td></td><td></td><td></td><td></td></tr>
<tr><td>Chromosome</td><td>start</td><td>end</td><td>strand</td><td>MM</td><td>target_seq</td><td>PAM</td><td>alignment</td><td>distance</td><td>position</td><td>gene name</td><td>gene id</td></tr>
<tr><td>GU305914.1</td><td>178247</td><td>178269</td><td>-</td><td>0</td><td>GACCTGTCTGCATTTGAGTC</td><td>AGG</td><td>||||||||[||||||||||||]PAM</td><td>NA</td><td>Intergenic</td><td>NA</td><td>NA</td></tr>
<tr><td>T21152</td><td>GAGTCAGGTCTTCCCCAAGCAGG</td><td>1000</td><td></td><td></td><td></td><td></td><td></td><td></td><td></td><td></td><td></td></tr>
<tr><td>Oligo fwd</td><td>CACCGAGTCAGGTCTTCCCCAAGC</td><td></td><td></td><td></td><td></td><td></td><td></td><td></td><td></td><td></td><td></td></tr>
<tr><td>Oligo rev</td><td>AAACGCTTGGGGAAGACCTGACTC</td><td></td><td></td><td></td><td></td><td></td><td></td><td></td><td></td><td></td><td></td></tr>
<tr><td>Chromosome</td><td>start</td><td>end</td><td>strand</td><td>MM</td><td>target_seq</td><td>PAM</td><td>alignment</td><td>distance</td><td>position</td><td>gene name</td><td>gene id</td></tr>
<tr><td>GU305914.1</td><td>178232</td><td>178254</td><td>-</td><td>0</td><td>GAGTCAGGTCTTCCCCAAGC</td><td>AGG</td><td>||||||||[||||||||||||]PAM</td><td>NA</td><td>Intergenic</td><td>NA</td><td>NA</td></tr>
<tr><td>T21153</td><td>TCCCCAAGCAGGTGTGCATGAGG</td><td>1000</td><td></td><td></td><td></td><td></td><td></td><td></td><td></td><td></td><td></td></tr>
<tr><td>Oligo fwd</td><td>CACCTCCCCAAGCAGGTGTGCATG</td><td></td><td></td><td></td><td></td><td></td><td></td><td></td><td></td><td></td><td></td></tr>
<tr><td>Oligo rev</td><td>AAACCATGCACACCTGCTTGGGGA</td><td></td><td></td><td></td><td></td><td></td><td></td><td></td><td></td><td></td><td></td></tr>
<tr><td>Chromosome</td><td>start</td><td>end</td><td>strand</td><td>MM</td><td>target_seq</td><td>PAM</td><td>alignment</td><td>distance</td><td>position</td><td>gene name</td><td>gene id</td></tr>
<tr><td>GU305914.1</td><td>178221</td><td>178243</td><td>-</td><td>0</td><td>TCCCCAAGCAGGTGTGCATG</td><td>AGG</td><td>||||||||[||||||||||||]PAM</td><td>NA</td><td>Intergenic</td><td>NA</td><td>NA</td></tr>
<tr><td>T21154</td><td>ATGAGGCTCTGCAGCATAACAGG</td><td>1000</td><td></td><td></td><td></td><td></td><td></td><td></td><td></td><td></td><td></td></tr>
<tr><td>Oligo fwd</td><td>CACCATGAGGCTCTGCAGCATAAC</td><td></td><td></td><td></td><td></td><td></td><td></td><td></td><td></td><td></td><td></td></tr>
<tr><td>Oligo rev</td><td>AAACGTTATGCTGCAGAGCCTCAT</td><td></td><td></td><td></td><td></td><td></td><td></td><td></td><td></td><td></td><td></td></tr>
<tr><td>Chromosome</td><td>start</td><td>end</td><td>strand</td><td>MM</td><td>target_seq</td><td>PAM</td><td>alignment</td><td>distance</td><td>position</td><td>gene name</td><td>gene id</td></tr>
<tr><td>GU305914.1</td><td>178204</td><td>178226</td><td>-</td><td>0</td><td>ATGAGGCTCTGCAGCATAAC</td><td>AGG</td><td>||||||||[||||||||||||]PAM</td><td>NA</td><td>Intergenic</td><td>NA</td><td>NA</td></tr>
<tr><td>T21155</td><td>AGGCTCTGCAGCATAACAGGCGG</td><td>1000</td><td></td><td></td><td></td><td></td><td></td><td></td><td></td><td></td><td></td></tr>
<tr><td>Oligo fwd</td><td>CACCAGGCTCTGCAGCATAACAGG</td><td></td><td></td><td></td><td></td><td></td><td></td><td></td><td></td><td></td><td></td></tr>
<tr><td>Oligo rev</td><td>AAACCCTGTTATGCTGCAGAGCCT</td><td></td><td></td><td></td><td></td><td></td><td></td><td></td><td></td><td></td><td></td></tr>
<tr><td>Chromosome</td><td>start</td><td>end</td><td>strand</td><td>MM</td><td>target_seq</td><td>PAM</td><td>alignment</td><td>distance</td><td>position</td><td>gene name</td><td>gene id</td></tr>
<tr><td>GU305914.1</td><td>178201</td><td>178223</td><td>-</td><td>0</td><td>AGGCTCTGCAGCATAACAGG</td><td>CGG</td><td>||||||||[||||||||||||]PAM</td><td>NA</td><td>Intergenic</td><td>NA</td><td>NA</td></tr>
<tr><td>T21156</td><td>GGCTCTGCAGCATAACAGGCGGG</td><td>1000</td><td></td><td></td><td></td><td></td><td></td><td></td><td></td><td></td><td></td></tr>
<tr><td>Oligo fwd</td><td>CACCGGCTCTGCAGCATAACAGGC</td><td></td><td></td><td></td><td></td><td></td><td></td><td></td><td></td><td></td><td></td></tr>
<tr><td>Oligo rev</td><td>AAACGCCTGTTATGCTGCAGAGCC</td><td></td><td></td><td></td><td></td><td></td><td></td><td></td><td></td><td></td><td></td></tr>
<tr><td>Chromosome</td><td>start</td><td>end</td><td>strand</td><td>MM</td><td>target_seq</td><td>PAM</td><td>alignment</td><td>distance</td><td>position</td><td>gene name</td><td>gene id</td></tr>
<tr><td>GU305914.1</td><td>178200</td><td>178222</td><td>-</td><td>0</td><td>GGCTCTGCAGCATAACAGGC</td><td>GGG</td><td>||||||||[||||||||||||]PAM</td><td>NA</td><td>Intergenic</td><td>NA</td><td>NA</td></tr>
</table>

T21157 GAATCCTCTTCCCATCTATGAGG 1000
Oligo fwd CACCGAATCCTCTTCCCATCTATG
Oligo rev AAACCATAGATGGGAAGAGGATTC
Chromosome start end strand MM target_seq PAM alignment distance position gene name gene id
GU305914.1 178175 178197 - 0 GAATCCTCTTCCCATCTATG AGG ||||||||[||||||||||||]PAM NA Intergenic NA NA

T21158 TTATGTGAACGCCCAGTTTGAGG 1000
Oligo fwd CACCTTATGTGAACGCCCAGTTTG
Oligo rev AAACCAAACTGGGCGTTCACATAA
Chromosome start end strand MM target_seq PAM alignment distance position gene name gene id
GU305914.1 178139 178161 - 0 TTATGTGAACGCCCAGTTTG AGG ||||||||[||||||||||||]PAM NA Intergenic NA NA

T21159 AGATGATATAAGCCATCATGAGG 1000
Oligo fwd CACCAGATGATATAAGCCATCATG
Oligo rev AAACCATGATGGCTTATATCATCT
Chromosome start end strand MM target_seq PAM alignment distance position gene name gene id
GU305914.1 178115 178137 - 0 AGATGATATAAGCCATCATG AGG ||||||||[||||||||||||]PAM NA Intergenic NA NA

T21160 AGCCATCATGAGGATGAGTCTGG 1000
Oligo fwd CACCAGCCATCATGAGGATGAGTC
Oligo rev AAACGACTCATCCTCATGATGGCT
Chromosome start end strand MM target_seq PAM alignment distance position gene name gene id
GU305914.1 178105 178127 - 0 AGCCATCATGAGGATGAGTC TGG ||||||||[||||||||||||]PAM NA Intergenic NA NA
GU305914.1 48374 48396 + 4 ATCGATGATGAGGATGAGCC GGG |-|-||-|[||||||||||-|]PAM NA Intergenic NA NA

T21161 GCCATCATGAGGATGAGTCTGGG 1000
Oligo fwd CACCGCCATCATGAGGATGAGTCT
Oligo rev AAACAGACTCATCCTCATGATGGC
Chromosome start end strand MM target_seq PAM alignment distance position gene name gene id
GU305914.1 178104 178126 - 0 GCCATCATGAGGATGAGTCT GGG ||||||||[||||||||||||]PAM NA Intergenic NA NA

T21162 CCATCATGAGGATGAGTCTGGGG 1000
Oligo fwd CACCCCATCATGAGGATGAGTCTG
Oligo rev AAACCAGACTCATCCTCATGATGG
Chromosome start end strand MM target_seq PAM alignment distance position gene name gene id
GU305914.1 178103 178125 - 0 CCATCATGAGGATGAGTCTG GGG ||||||||[||||||||||||]PAM NA Intergenic NA NA

T21163 ACTGCGAGTGATCTGTACTGTGG 1000
Oligo fwd CACCACTGCGAGTGATCTGTACTG
Oligo rev AAACCAGTACAGATCACTCGCAGT
Chromosome start end strand MM target_seq PAM alignment distance position gene name gene id
GU305914.1 178068 178090 - 0 ACTGCGAGTGATCTGTACTG TGG ||||||||[||||||||||||]PAM NA Intergenic NA NA

T21164 AGTGATCTGTACTGTGGACTTGG 1000
Oligo fwd CACCAGTGATCTGTACTGTGGACT
Oligo rev AAACAGTCCACAGTACAGATCACT
Chromosome start end strand MM target_seq PAM alignment distance position gene name gene id
GU305914.1 178062 178084 - 0 AGTGATCTGTACTGTGGACT TGG ||||||||[||||||||||||]PAM NA Intergenic NA NA

T21165 AGTAGACACCGTCCTCCTTTTGG 1000
Oligo fwd CACCAGTAGACACCGTCCTCCTTT
Oligo rev AAACAAAGGAGGACGGTGTCTACT
Chromosome start end strand MM target_seq PAM alignment distance position gene name gene id
GU305914.1 177842 177864 - 0 AGTAGACACCGTCCTCCTTT TGG ||||||||[||||||||||||]PAM NA Intergenic NA NA

T21166 AGACACCGTCCTCCTTTTGGCGG 1000
Oligo fwd CACCAGACACCGTCCTCCTTTTGG
Oligo rev AAACCCAAAAGGAGGACGGTGTCT
Chromosome start end strand MM target_seq PAM alignment distance position gene name gene id
GU305914.1 177839 177861 - 0 AGACACCGTCCTCCTTTTGG CGG ||||||||[||||||||||||]PAM NA Intergenic NA NA

T21167 CACCGTCCTCCTTTTGGCGGTGG 1000
Oligo fwd CACCCACCGTCCTCCTTTTGGCGG
Oligo rev AAACCCGCCAAAAGGAGGACGGTG
Chromosome start end strand MM target_seq PAM alignment distance position gene name gene id
GU305914.1 177836 177858 - 0 CACCGTCCTCCTTTTGGCGG TGG ||||||||[||||||||||||]PAM NA Intergenic NA NA

T21168 ACCGTCCTCCTTTTGGCGGTGGG 1000
Oligo fwd CACCACCGTCCTCCTTTTGGCGGT
Oligo rev AAACACCGCCAAAAGGAGGACGGT
Chromosome start end strand MM target_seq PAM alignment distance position gene name gene id
GU305914.1 177835 177857 - 0 ACCGTCCTCCTTTTGGCGGT GGG ||||||||[||||||||||||]PAM NA Intergenic NA NA

T21169 CCGTCCTCCTTTTGGCGGTGGGG 1000
Oligo fwd CACCCCGTCCTCCTTTTGGCGGTG
Oligo rev AAACCACCGCCAAAAGGAGGACGG
Chromosome start end strand MM target_seq PAM alignment distance position gene name gene id
GU305914.1 177834 177856 - 0 CCGTCCTCCTTTTGGCGGTG GGG ||||||||[||||||||||||]PAM NA Intergenic NA NA

T21170 TCCTTTTGGCGGTGGGGTGTTGG 1000
Oligo fwd CACCTCCTTTTGGCGGTGGGGTGT
Oligo rev AAACACACCCCACCGCCAAAAGGA
Chromosome start end strand MM target_seq PAM alignment distance position gene name gene id
GU305914.1 177828 177850 - 0 TCCTTTTGGCGGTGGGGTGT TGG ||||||||[||||||||||||]PAM NA Intergenic NA NA

T21171 CCTTTTGGCGGTGGGGTGTTGGG 1000
Oligo fwd CACCCCTTTTGGCGGTGGGGTGTT
Oligo rev AAACAACACCCCACCGCCAAAAGG

Chromosome start end strand MM target_seq PAM alignment distance position gene name gene id
GU305914.1 177827 177849 - 0 CCTTTTGGCGGTGGGGTGTT GGG ||||||||[||||||||||||]PAM NA Intergenic NA NA

T21172 GGTGGGGTGTTGGGCGCTGTTGG 1000
Oligo fwd CACCGGTGGGGTGTTGGGCGCTGT
Oligo rev AAACACAGCGCCCAACACCCCACC
Chromosome start end strand MM target_seq PAM alignment distance position gene name gene id
GU305914.1 177818 177840 - 0 GGTGGGGTGTTGGGCGCTGT TGG ||||||||[||||||||||||]PAM NA Intergenic NA NA

T21173 TGTTGGGCGCTGTTGGCGCGCGG 1000
Oligo fwd CACCTGTTGGGCGCTGTTGGCGCG
Oligo rev AAACCGCGCCAACAGCGCCCAACA
Chromosome start end strand MM target_seq PAM alignment distance position gene name gene id
GU305914.1 177811 177833 - 0 TGTTGGGCGCTGTTGGCGCG CGG ||||||||[||||||||||||]PAM NA Intergenic NA NA

T21174 TGGGCGCTGTTGGCGCGCGGTGG 1000
Oligo fwd CACCTGGGCGCTGTTGGCGCGCGG
Oligo rev AAACCCGCGCGCCAACAGCGCCCA
Chromosome start end strand MM target_seq PAM alignment distance position gene name gene id
GU305914.1 177808 177830 - 0 TGGGCGCTGTTGGCGCGCGG TGG ||||||||[||||||||||||]PAM NA Intergenic NA NA

T21175 GCGCTGTTGGCGCGCGGTGGCGG 1000
Oligo fwd CACCGCGCTGTTGGCGCGCGGTGG
Oligo rev AAACCCACCGCGCGCCAACAGCGC
Chromosome start end strand MM target_seq PAM alignment distance position gene name gene id
GU305914.1 177805 177827 - 0 GCGCTGTTGGCGCGCGGTGG CGG ||||||||[||||||||||||]PAM NA Intergenic NA NA

T21176 CTGTTGGCGCGCGGTGGCGGTGG 1000
Oligo fwd CACCCTGTTGGCGCGCGGTGGCGG
Oligo rev AAACCCGCCACCGCGCGCCAACAG
Chromosome start end strand MM target_seq PAM alignment distance position gene name gene id
GU305914.1 177802 177824 - 0 CTGTTGGCGCGCGGTGGCGG TGG ||||||||[||||||||||||]PAM NA Intergenic NA NA

T21177 CCGCTCAAGTGACTAACACCTGG 1000
Oligo fwd CACCCCGCTCAAGTGACTAACACC
Oligo rev AAACGGTGTTAGTCACTTGAGCGG
Chromosome start end strand MM target_seq PAM alignment distance position gene name gene id
GU305914.1 177768 177790 - 0 CCGCTCAAGTGACTAACACC TGG ||||||||[||||||||||||]PAM NA Intergenic NA NA

T21178 TGGTGCCTCTTCATTGAAGAAGG 1000
Oligo fwd CACCTGGTGCCTCTTCATTGAAGA
Oligo rev AAACTCTTCAATGAAGAGGCACCA
Chromosome start end strand MM target_seq PAM alignment distance position gene name gene id
GU305914.1 177748 177770 - 0 TGGTGCCTCTTCATTGAAGA AGG ||||||||[||||||||||||]PAM NA Intergenic NA NA

T21179 TGCCTCTTCATTGAAGAAGGAGG 1000
Oligo fwd CACCTGCCTCTTCATTGAAGAAGG
Oligo rev AAACCCTTCTTCAATGAAGAGGCA
Chromosome start end strand MM target_seq PAM alignment distance position gene name gene id
GU305914.1 177745 177767 - 0 TGCCTCTTCATTGAAGAAGG AGG ||||||||[||||||||||||]PAM NA Intergenic NA NA

T21180 AGAAGGAGGCACTACTTTTGTGG 1000
Oligo fwd CACCAGAAGGAGGCACTACTTTTG
Oligo rev AAACCAAAAGTAGTGCCTCCTTCT
Chromosome start end strand MM target_seq PAM alignment distance position gene name gene id
GU305914.1 177731 177753 - 0 AGAAGGAGGCACTACTTTTG TGG ||||||||[||||||||||||]PAM NA Intergenic NA NA

T21181 GAAGGAGGCACTACTTTTGTGGG 1000
Oligo fwd CACCGAAGGAGGCACTACTTTTGT
Oligo rev AAACACAAAAGTAGTGCCTCCTTC
Chromosome start end strand MM target_seq PAM alignment distance position gene name gene id
GU305914.1 177730 177752 - 0 GAAGGAGGCACTACTTTTGT GGG ||||||||[||||||||||||]PAM NA Intergenic NA NA

T21182 AAGGAGGCACTACTTTTGTGGGG 1000
Oligo fwd CACCAAGGAGGCACTACTTTTGTG
Oligo rev AAACCACAAAAGTAGTGCCTCCTT
Chromosome start end strand MM target_seq PAM alignment distance position gene name gene id
GU305914.1 177729 177751 - 0 AAGGAGGCACTACTTTTGTG GGG ||||||||[||||||||||||]PAM NA Intergenic NA NA

T21183 AGGAGGCACTACTTTTGTGGGGG 1000
Oligo fwd CACCAGGAGGCACTACTTTTGTGG
Oligo rev AAACCCACAAAAGTAGTGCCTCCT
Chromosome start end strand MM target_seq PAM alignment distance position gene name gene id
GU305914.1 177728 177750 - 0 AGGAGGCACTACTTTTGTGG GGG ||||||||[||||||||||||]PAM NA Intergenic NA NA

T21184 AGGCACTACTTTTGTGGGGGAGG 1000
Oligo fwd CACCAGGCACTACTTTTGTGGGGG
Oligo rev AAACCCCCCACAAAAGTAGTGCCT
Chromosome start end strand MM target_seq PAM alignment distance position gene name gene id
GU305914.1 177725 177747 - 0 AGGCACTACTTTTGTGGGGG AGG ||||||||[||||||||||||]PAM NA Intergenic NA NA

T21185 CACTACTTTTGTGGGGGAGGAGG 1000
Oligo fwd CACCCACTACTTTTGTGGGGGAGG
Oligo rev AAACCCTCCCCCACAAAAGTAGTG
Chromosome start end strand MM target_seq PAM alignment distance position gene name gene id
GU305914.1 177722 177744 - 0 CACTACTTTTGTGGGGGAGG AGG ||||||||[||||||||||||]PAM NA Intergenic NA NA

T21186 ACTACTTTTGTGGGGGAGGAGGG 1000
Oligo fwd CACCACTACTTTTGTGGGGGAGGA

| | | | | | | | | | | | |
|---|---|---|---|---|---|---|---|---|---|---|---|
| Oligo rev | AAACTCCTCCCCCACAAAAGTAGT | | | | | | | | | | |
| Chromosome | start | end | strand | MM | target_seq | PAM | alignment | distance | position | gene name | gene id |
| GU305914.1 | 177721 | 177743 | - | 0 | ACTACTTTTGTGGGGGAGGA | GGG | \|\|\|\|\|\|\|\|[\|\|\|\|\|\|\|\|\|\|\|\|]PAM | NA | Intergenic | NA | NA |
| T21187 | TTTGTGGGGGAGGAGGGAACAGG | 1000 | | | | | | | | | |
| Oligo fwd | CACCTTTGTGGGGGAGGAGGGAAC | | | | | | | | | | |
| Oligo rev | AAACGTTCCCTCCTCCCCCACAAA | | | | | | | | | | |
| Chromosome | start | end | strand | MM | target_seq | PAM | alignment | distance | position | gene name | gene id |
| GU305914.1 | 177715 | 177737 | - | 0 | TTTGTGGGGGAGGAGGGAAC | AGG | \|\|\|\|\|\|\|\|[\|\|\|\|\|\|\|\|\|\|\|\|]PAM | NA | Intergenic | NA | NA |
| T21188 | CGTGATATATCAGTCAGCGTCGG | 1000 | | | | | | | | | |
| Oligo fwd | CACCCGTGATATATCAGTCAGCGT | | | | | | | | | | |
| Oligo rev | AAACACGCTGACTGATATATCACG | | | | | | | | | | |
| Chromosome | start | end | strand | MM | target_seq | PAM | alignment | distance | position | gene name | gene id |
| GU305914.1 | 177679 | 177701 | - | 0 | CGTGATATATCAGTCAGCGT | CGG | \|\|\|\|\|\|\|\|[\|\|\|\|\|\|\|\|\|\|\|\|]PAM | NA | Intergenic | NA | NA |
| T21189 | GTGATATATCAGTCAGCGTCGGG | 1000 | | | | | | | | | |
| Oligo fwd | CACCGTGATATATCAGTCAGCGTC | | | | | | | | | | |
| Oligo rev | AAACGACGCTGACTGATATATCAC | | | | | | | | | | |
| Chromosome | start | end | strand | MM | target_seq | PAM | alignment | distance | position | gene name | gene id |
| GU305914.1 | 177678 | 177700 | - | 0 | GTGATATATCAGTCAGCGTC | GGG | \|\|\|\|\|\|\|\|[\|\|\|\|\|\|\|\|\|\|\|\|]PAM | NA | Intergenic | NA | NA |
| T21190 | TCAGTCAGCGTCGGGAACACCGG | 1000 | | | | | | | | | |
| Oligo fwd | CACCTCAGTCAGCGTCGGGAACAC | | | | | | | | | | |
| Oligo rev | AAACGTGTTCCCGACGCTGACTGA | | | | | | | | | | |
| Chromosome | start | end | strand | MM | target_seq | PAM | alignment | distance | position | gene name | gene id |
| GU305914.1 | 177670 | 177692 | - | 0 | TCAGTCAGCGTCGGGAACAC | CGG | \|\|\|\|\|\|\|\|[\|\|\|\|\|\|\|\|\|\|\|\|]PAM | NA | Intergenic | NA | NA |
| T21191 | ACGTGCGAGACCGCCGAGAACGG | 1000 | | | | | | | | | |
| Oligo fwd | CACCACGTGCGAGACCGCCGAGAA | | | | | | | | | | |
| Oligo rev | AAACTTCTCGGCGGTCTCGCACGT | | | | | | | | | | |
| Chromosome | start | end | strand | MM | target_seq | PAM | alignment | distance | position | gene name | gene id |
| GU305914.1 | 177622 | 177644 | - | 0 | ACGTGCGAGACCGCCGAGAA | CGG | \|\|\|\|\|\|\|\|[\|\|\|\|\|\|\|\|\|\|\|\|]PAM | NA | Intergenic | NA | NA |
| T21192 | CTCATCGTCAACTATGAGAACGG | 1000 | | | | | | | | | |
| Oligo fwd | CACCCTCATCGTCAACTATGAGAA | | | | | | | | | | |
| Oligo rev | AAACTTCTCATAGTTGACGATGAG | | | | | | | | | | |
| Chromosome | start | end | strand | MM | target_seq | PAM | alignment | distance | position | gene name | gene id |
| GU305914.1 | 177595 | 177617 | - | 0 | CTCATCGTCAACTATGAGAA | CGG | \|\|\|\|\|\|\|\|[\|\|\|\|\|\|\|\|\|\|\|\|]PAM | NA | Intergenic | NA | NA |
| T21193 | GAACGGTACTCTGTTCTGTGAGG | 1000 | | | | | | | | | |
| Oligo fwd | CACCGAACGGTACTCTGTTCTGTG | | | | | | | | | | |
| Oligo rev | AAACCACAGAACAGAGTACCGTTC | | | | | | | | | | |
| Chromosome | start | end | strand | MM | target_seq | PAM | alignment | distance | position | gene name | gene id |
| GU305914.1 | 177578 | 177600 | - | 0 | GAACGGTACTCTGTTCTGTG | AGG | \|\|\|\|\|\|\|\|[\|\|\|\|\|\|\|\|\|\|\|\|]PAM | NA | Intergenic | NA | NA |
| T21194 | GGTACTCTGTTCTGTGAGGCCGG | 1000 | | | | | | | | | |
| Oligo fwd | CACCGGTACTCTGTTCTGTGAGGC | | | | | | | | | | |
| Oligo rev | AAACGCCTCACAGAACAGAGTACC | | | | | | | | | | |
| Chromosome | start | end | strand | MM | target_seq | PAM | alignment | distance | position | gene name | gene id |
| GU305914.1 | 177574 | 177596 | - | 0 | GGTACTCTGTTCTGTGAGGC | CGG | \|\|\|\|\|\|\|\|[\|\|\|\|\|\|\|\|\|\|\|\|]PAM | NA | Intergenic | NA | NA |
| T21195 | GTACTCTGTTCTGTGAGGCCGGG | 1000 | | | | | | | | | |
| Oligo fwd | CACCGTACTCTGTTCTGTGAGGCC | | | | | | | | | | |
| Oligo rev | AAACGGCCTCACAGAACAGAGTAC | | | | | | | | | | |
| Chromosome | start | end | strand | MM | target_seq | PAM | alignment | distance | position | gene name | gene id |
| GU305914.1 | 177573 | 177595 | - | 0 | GTACTCTGTTCTGTGAGGCC | GGG | \|\|\|\|\|\|\|\|[\|\|\|\|\|\|\|\|\|\|\|\|]PAM | NA | Intergenic | NA | NA |
| T21196 | ACGTATGTTATTCCCCAGTCAGG | 1000 | | | | | | | | | |
| Oligo fwd | CACCACGTATGTTATTCCCCAGTC | | | | | | | | | | |
| Oligo rev | AAACGACTGGGGAATAACATACGT | | | | | | | | | | |
| Chromosome | start | end | strand | MM | target_seq | PAM | alignment | distance | position | gene name | gene id |
| GU305914.1 | 177524 | 177546 | - | 0 | ACGTATGTTATTCCCCAGTC | AGG | \|\|\|\|\|\|\|\|[\|\|\|\|\|\|\|\|\|\|\|\|]PAM | NA | Intergenic | NA | NA |
| T21197 | CGTATGTTATTCCCCAGTCAGGG | 1000 | | | | | | | | | |
| Oligo fwd | CACCCGTATGTTATTCCCCAGTCA | | | | | | | | | | |
| Oligo rev | AAACTGACTGGGGAATAACATACG | | | | | | | | | | |
| Chromosome | start | end | strand | MM | target_seq | PAM | alignment | distance | position | gene name | gene id |
| GU305914.1 | 177523 | 177545 | - | 0 | CGTATGTTATTCCCCAGTCA | GGG | \|\|\|\|\|\|\|\|[\|\|\|\|\|\|\|\|\|\|\|\|]PAM | NA | Intergenic | NA | NA |
| T21198 | GTTATTCCCCAGTCAGGGAGAGG | 1000 | | | | | | | | | |
| Oligo fwd | CACCGTTATTCCCCAGTCAGGGAG | | | | | | | | | | |
| Oligo rev | AAACCTCCCTGACTGGGGAATAAC | | | | | | | | | | |
| Chromosome | start | end | strand | MM | target_seq | PAM | alignment | distance | position | gene name | gene id |
| GU305914.1 | 177518 | 177540 | - | 0 | GTTATTCCCCAGTCAGGGAG | AGG | \|\|\|\|\|\|\|\|[\|\|\|\|\|\|\|\|\|\|\|\|]PAM | NA | Intergenic | NA | NA |
| T21199 | GCTCTCTTCTGTGACAAGAACGG | 1000 | | | | | | | | | |
| Oligo fwd | CACCGCTCTCTTCTGTGACAAGAA | | | | | | | | | | |
| Oligo rev | AAACTTCTTGTCACAGAAGAGAGC | | | | | | | | | | |
| Chromosome | start | end | strand | MM | target_seq | PAM | alignment | distance | position | gene name | gene id |
| GU305914.1 | 177496 | 177518 | - | 0 | GCTCTCTTCTGTGACAAGAA | CGG | \|\|\|\|\|\|\|\|[\|\|\|\|\|\|\|\|\|\|\|\|]PAM | NA | Intergenic | NA | NA |
| T21200 | GCACCGTCAGCACGCTCGAGTGG | 1000 | | | | | | | | | |
| Oligo fwd | CACCGCACCGTCAGCACGCTCGAG | | | | | | | | | | |
| Oligo rev | AAACCTCGAGCGTGCTGACGGTGC | | | | | | | | | | |
| Chromosome | start | end | strand | MM | target_seq | PAM | alignment | distance | position | gene name | gene id |
| GU305914.1 | 177474 | 177496 | - | 0 | GCACCGTCAGCACGCTCGAG | TGG | \|\|\|\|\|\|\|\|[\|\|\|\|\|\|\|\|\|\|\|\|]PAM | NA | Intergenic | NA | NA |
| GU305914.1 | 88751 | 88773 | + | 4 | GCGCCGCCAGCACCCGCGAG | CGG | \|\|-\|\|\|-\|[\|\|\|\|\|-\|-\|\|\|\|]PAM | NA | Intergenic | NA | NA |

<table>
<tr><td>T21201</td><td>GAGTGGTTCAACATAGACTCTGG</td><td>1000</td><td></td><td></td><td></td><td></td><td></td><td></td><td></td><td></td><td></td></tr>
<tr><td>Oligo fwd</td><td>CACCGAGTGGTTCAACATAGACTC</td><td></td><td></td><td></td><td></td><td></td><td></td><td></td><td></td><td></td><td></td></tr>
<tr><td>Oligo rev</td><td>AAACGAGTCTATGTTGAACCACTC</td><td></td><td></td><td></td><td></td><td></td><td></td><td></td><td></td><td></td><td></td></tr>
<tr><td>Chromosome</td><td>start</td><td>end</td><td>strand</td><td>MM</td><td>target_seq</td><td>PAM</td><td>alignment</td><td>distance</td><td>position</td><td>gene name</td><td>gene id</td></tr>
<tr><td>GU305914.1</td><td>177457</td><td>177479</td><td>-</td><td>0</td><td>GAGTGGTTCAACATAGACTC</td><td>TGG</td><td>||||||||[||||||||||||]PAM</td><td>NA</td><td>Intergenic</td><td>NA</td><td>NA</td></tr>
<tr><td>T21202</td><td>GTTCAACATAGACTCTGGCCAGG</td><td>1000</td><td></td><td></td><td></td><td></td><td></td><td></td><td></td><td></td><td></td></tr>
<tr><td>Oligo fwd</td><td>CACCGTTCAACATAGACTCTGGCC</td><td></td><td></td><td></td><td></td><td></td><td></td><td></td><td></td><td></td><td></td></tr>
<tr><td>Oligo rev</td><td>AAACGGCCAGAGTCTATGTTGAAC</td><td></td><td></td><td></td><td></td><td></td><td></td><td></td><td></td><td></td><td></td></tr>
<tr><td>Chromosome</td><td>start</td><td>end</td><td>strand</td><td>MM</td><td>target_seq</td><td>PAM</td><td>alignment</td><td>distance</td><td>position</td><td>gene name</td><td>gene id</td></tr>
<tr><td>GU305914.1</td><td>177452</td><td>177474</td><td>-</td><td>0</td><td>GTTCAACATAGACTCTGGCC</td><td>AGG</td><td>||||||||[||||||||||||]PAM</td><td>NA</td><td>Intergenic</td><td>NA</td><td>NA</td></tr>
<tr><td>T21203</td><td>CCAGGACGCTCCTCCGATCGAGG</td><td>1000</td><td></td><td></td><td></td><td></td><td></td><td></td><td></td><td></td><td></td></tr>
<tr><td>Oligo fwd</td><td>CACCCCAGGACGCTCCTCCGATCG</td><td></td><td></td><td></td><td></td><td></td><td></td><td></td><td></td><td></td><td></td></tr>
<tr><td>Oligo rev</td><td>AAACCGATCGGAGGAGCGTCCTGG</td><td></td><td></td><td></td><td></td><td></td><td></td><td></td><td></td><td></td><td></td></tr>
<tr><td>Chromosome</td><td>start</td><td>end</td><td>strand</td><td>MM</td><td>target_seq</td><td>PAM</td><td>alignment</td><td>distance</td><td>position</td><td>gene name</td><td>gene id</td></tr>
<tr><td>GU305914.1</td><td>177434</td><td>177456</td><td>-</td><td>0</td><td>CCAGGACGCTCCTCCGATCG</td><td>AGG</td><td>||||||||[||||||||||||]PAM</td><td>NA</td><td>Intergenic</td><td>NA</td><td>NA</td></tr>
<tr><td>T21204</td><td>TCGAGGCGCAGTTCCTCCCCAGG</td><td>1000</td><td></td><td></td><td></td><td></td><td></td><td></td><td></td><td></td><td></td></tr>
<tr><td>Oligo fwd</td><td>CACCTCGAGGCGCAGTTCCTCCCC</td><td></td><td></td><td></td><td></td><td></td><td></td><td></td><td></td><td></td><td></td></tr>
<tr><td>Oligo rev</td><td>AAACGGGGAGGAACTGCGCCTCGA</td><td></td><td></td><td></td><td></td><td></td><td></td><td></td><td></td><td></td><td></td></tr>
<tr><td>Chromosome</td><td>start</td><td>end</td><td>strand</td><td>MM</td><td>target_seq</td><td>PAM</td><td>alignment</td><td>distance</td><td>position</td><td>gene name</td><td>gene id</td></tr>
<tr><td>GU305914.1</td><td>177417</td><td>177439</td><td>-</td><td>0</td><td>TCGAGGCGCAGTTCCTCCCC</td><td>AGG</td><td>||||||||[||||||||||||]PAM</td><td>NA</td><td>Intergenic</td><td>NA</td><td>NA</td></tr>
<tr><td>T21205</td><td>CTTCTTACAGACAGACACCGAGG</td><td>1000</td><td></td><td></td><td></td><td></td><td></td><td></td><td></td><td></td><td></td></tr>
<tr><td>Oligo fwd</td><td>CACCCTTCTTACAGACAGACACCG</td><td></td><td></td><td></td><td></td><td></td><td></td><td></td><td></td><td></td><td></td></tr>
<tr><td>Oligo rev</td><td>AAACCGGTGTCTGTCTGTAAGAAG</td><td></td><td></td><td></td><td></td><td></td><td></td><td></td><td></td><td></td><td></td></tr>
<tr><td>Chromosome</td><td>start</td><td>end</td><td>strand</td><td>MM</td><td>target_seq</td><td>PAM</td><td>alignment</td><td>distance</td><td>position</td><td>gene name</td><td>gene id</td></tr>
<tr><td>GU305914.1</td><td>177386</td><td>177408</td><td>-</td><td>0</td><td>CTTCTTACAGACAGACACCG</td><td>AGG</td><td>||||||||[||||||||||||]PAM</td><td>NA</td><td>Intergenic</td><td>NA</td><td>NA</td></tr>
<tr><td>T21206</td><td>TTCTTACAGACAGACACCGAGGG</td><td>1000</td><td></td><td></td><td></td><td></td><td></td><td></td><td></td><td></td><td></td></tr>
<tr><td>Oligo fwd</td><td>CACCTTCTTACAGACAGACACCGA</td><td></td><td></td><td></td><td></td><td></td><td></td><td></td><td></td><td></td><td></td></tr>
<tr><td>Oligo rev</td><td>AAACTCGGTGTCTGTCTGTAAGAA</td><td></td><td></td><td></td><td></td><td></td><td></td><td></td><td></td><td></td><td></td></tr>
<tr><td>Chromosome</td><td>start</td><td>end</td><td>strand</td><td>MM</td><td>target_seq</td><td>PAM</td><td>alignment</td><td>distance</td><td>position</td><td>gene name</td><td>gene id</td></tr>
<tr><td>GU305914.1</td><td>177385</td><td>177407</td><td>-</td><td>0</td><td>TTCTTACAGACAGACACCGA</td><td>GGG</td><td>||||||||[||||||||||||]PAM</td><td>NA</td><td>Intergenic</td><td>NA</td><td>NA</td></tr>
<tr><td>T21207</td><td>GACACTCTGCTGTGTGTCTACGG</td><td>1000</td><td></td><td></td><td></td><td></td><td></td><td></td><td></td><td></td><td></td></tr>
<tr><td>Oligo fwd</td><td>CACCGACACTCTGCTGTGTGTCTA</td><td></td><td></td><td></td><td></td><td></td><td></td><td></td><td></td><td></td><td></td></tr>
<tr><td>Oligo rev</td><td>AAACTAGACACACAGCAGAGTGTC</td><td></td><td></td><td></td><td></td><td></td><td></td><td></td><td></td><td></td><td></td></tr>
<tr><td>Chromosome</td><td>start</td><td>end</td><td>strand</td><td>MM</td><td>target_seq</td><td>PAM</td><td>alignment</td><td>distance</td><td>position</td><td>gene name</td><td>gene id</td></tr>
<tr><td>GU305914.1</td><td>177328</td><td>177350</td><td>-</td><td>0</td><td>GACACTCTGCTGTGTGTCTA</td><td>CGG</td><td>||||||||[||||||||||||]PAM</td><td>NA</td><td>Intergenic</td><td>NA</td><td>NA</td></tr>
<tr><td>T21208</td><td>TGTCTACGGTGTCCTCGATCCGG</td><td>1000</td><td></td><td></td><td></td><td></td><td></td><td></td><td></td><td></td><td></td></tr>
<tr><td>Oligo fwd</td><td>CACCTGTCTACGGTGTCCTCGATC</td><td></td><td></td><td></td><td></td><td></td><td></td><td></td><td></td><td></td><td></td></tr>
<tr><td>Oligo rev</td><td>AAACGATCGAGGACACCGTAGACA</td><td></td><td></td><td></td><td></td><td></td><td></td><td></td><td></td><td></td><td></td></tr>
<tr><td>Chromosome</td><td>start</td><td>end</td><td>strand</td><td>MM</td><td>target_seq</td><td>PAM</td><td>alignment</td><td>distance</td><td>position</td><td>gene name</td><td>gene id</td></tr>
<tr><td>GU305914.1</td><td>177314</td><td>177336</td><td>-</td><td>0</td><td>TGTCTACGGTGTCCTCGATC</td><td>CGG</td><td>||||||||[||||||||||||]PAM</td><td>NA</td><td>Intergenic</td><td>NA</td><td>NA</td></tr>
<tr><td>T21209</td><td>CTACGGTGTCCTCGATCCGGAGG</td><td>1000</td><td></td><td></td><td></td><td></td><td></td><td></td><td></td><td></td><td></td></tr>
<tr><td>Oligo fwd</td><td>CACCCTACGGTGTCCTCGATCCGG</td><td></td><td></td><td></td><td></td><td></td><td></td><td></td><td></td><td></td><td></td></tr>
<tr><td>Oligo rev</td><td>AAACCCGGATCGAGGACACCGTAG</td><td></td><td></td><td></td><td></td><td></td><td></td><td></td><td></td><td></td><td></td></tr>
<tr><td>Chromosome</td><td>start</td><td>end</td><td>strand</td><td>MM</td><td>target_seq</td><td>PAM</td><td>alignment</td><td>distance</td><td>position</td><td>gene name</td><td>gene id</td></tr>
<tr><td>GU305914.1</td><td>177311</td><td>177333</td><td>-</td><td>0</td><td>CTACGGTGTCCTCGATCCGG</td><td>AGG</td><td>||||||||[||||||||||||]PAM</td><td>NA</td><td>Intergenic</td><td>NA</td><td>NA</td></tr>
<tr><td>T21210</td><td>CGGAGGAGTTCGCTGACACGCGG</td><td>1000</td><td></td><td></td><td></td><td></td><td></td><td></td><td></td><td></td><td></td></tr>
<tr><td>Oligo fwd</td><td>CACCCGGAGGAGTTCGCTGACACG</td><td></td><td></td><td></td><td></td><td></td><td></td><td></td><td></td><td></td><td></td></tr>
<tr><td>Oligo rev</td><td>AAACCGTGTCAGCGAACTCCTCCG</td><td></td><td></td><td></td><td></td><td></td><td></td><td></td><td></td><td></td><td></td></tr>
<tr><td>Chromosome</td><td>start</td><td>end</td><td>strand</td><td>MM</td><td>target_seq</td><td>PAM</td><td>alignment</td><td>distance</td><td>position</td><td>gene name</td><td>gene id</td></tr>
<tr><td>GU305914.1</td><td>177294</td><td>177316</td><td>-</td><td>0</td><td>CGGAGGAGTTCGCTGACACG</td><td>CGG</td><td>||||||||[||||||||||||]PAM</td><td>NA</td><td>Intergenic</td><td>NA</td><td>NA</td></tr>
<tr><td>T21211</td><td>GGAGGAGTTCGCTGACACGCGGG</td><td>1000</td><td></td><td></td><td></td><td></td><td></td><td></td><td></td><td></td><td></td></tr>
<tr><td>Oligo fwd</td><td>CACCGGAGGAGTTCGCTGACACGC</td><td></td><td></td><td></td><td></td><td></td><td></td><td></td><td></td><td></td><td></td></tr>
<tr><td>Oligo rev</td><td>AAACGCGTGTCAGCGAACTCCTCC</td><td></td><td></td><td></td><td></td><td></td><td></td><td></td><td></td><td></td><td></td></tr>
<tr><td>Chromosome</td><td>start</td><td>end</td><td>strand</td><td>MM</td><td>target_seq</td><td>PAM</td><td>alignment</td><td>distance</td><td>position</td><td>gene name</td><td>gene id</td></tr>
<tr><td>GU305914.1</td><td>177293</td><td>177315</td><td>-</td><td>0</td><td>GGAGGAGTTCGCTGACACGC</td><td>GGG</td><td>||||||||[||||||||||||]PAM</td><td>NA</td><td>Intergenic</td><td>NA</td><td>NA</td></tr>
<tr><td>T21212</td><td>GGGATATATACAGCGCCCTGAGG</td><td>1000</td><td></td><td></td><td></td><td></td><td></td><td></td><td></td><td></td><td></td></tr>
<tr><td>Oligo fwd</td><td>CACCGGGATATATACAGCGCCCTG</td><td></td><td></td><td></td><td></td><td></td><td></td><td></td><td></td><td></td><td></td></tr>
<tr><td>Oligo rev</td><td>AAACCAGGGCGCTGTATATATCCC</td><td></td><td></td><td></td><td></td><td></td><td></td><td></td><td></td><td></td><td></td></tr>
<tr><td>Chromosome</td><td>start</td><td>end</td><td>strand</td><td>MM</td><td>target_seq</td><td>PAM</td><td>alignment</td><td>distance</td><td>position</td><td>gene name</td><td>gene id</td></tr>
<tr><td>GU305914.1</td><td>177273</td><td>177295</td><td>-</td><td>0</td><td>GGGATATATACAGCGCCCTG</td><td>AGG</td><td>||||||||[||||||||||||]PAM</td><td>NA</td><td>Intergenic</td><td>NA</td><td>NA</td></tr>
<tr><td>T21213</td><td>AGCGCCCTGAGGAGCGTCGCTGG</td><td>1000</td><td></td><td></td><td></td><td></td><td></td><td></td><td></td><td></td><td></td></tr>
<tr><td>Oligo fwd</td><td>CACCAGCGCCCTGAGGAGCGTCGC</td><td></td><td></td><td></td><td></td><td></td><td></td><td></td><td></td><td></td><td></td></tr>
<tr><td>Oligo rev</td><td>AAACGCGACGCTCCTCAGGGCGCT</td><td></td><td></td><td></td><td></td><td></td><td></td><td></td><td></td><td></td><td></td></tr>
<tr><td>Chromosome</td><td>start</td><td>end</td><td>strand</td><td>MM</td><td>target_seq</td><td>PAM</td><td>alignment</td><td>distance</td><td>position</td><td>gene name</td><td>gene id</td></tr>
<tr><td>GU305914.1</td><td>177262</td><td>177284</td><td>-</td><td>0</td><td>AGCGCCCTGAGGAGCGTCGC</td><td>TGG</td><td>||||||||[||||||||||||]PAM</td><td>NA</td><td>Intergenic</td><td>NA</td><td>NA</td></tr>
<tr><td>T21214</td><td>CCCTGAGGAGCGTCGCTGGTCGG</td><td>1000</td><td></td><td></td><td></td><td></td><td></td><td></td><td></td><td></td><td></td></tr>
<tr><td>Oligo fwd</td><td>CACCCCCTGAGGAGCGTCGCTGGT</td><td></td><td></td><td></td><td></td><td></td><td></td><td></td><td></td><td></td><td></td></tr>
<tr><td>Oligo rev</td><td>AAACACCAGCGACGCTCCTCAGGG</td><td></td><td></td><td></td><td></td><td></td><td></td><td></td><td></td><td></td><td></td></tr>
<tr><td>Chromosome</td><td>start</td><td>end</td><td>strand</td><td>MM</td><td>target_seq</td><td>PAM</td><td>alignment</td><td>distance</td><td>position</td><td>gene name</td><td>gene id</td></tr>
<tr><td>GU305914.1</td><td>177258</td><td>177280</td><td>-</td><td>0</td><td>CCCTGAGGAGCGTCGCTGGT</td><td>CGG</td><td>||||||||[||||||||||||]PAM</td><td>NA</td><td>Intergenic</td><td>NA</td><td>NA</td></tr>
<tr><td>T21215</td><td>CCTGAGGAGCGTCGCTGGTCGGG</td><td>1000</td><td></td><td></td><td></td><td></td><td></td><td></td><td></td><td></td><td></td></tr>
<tr><td>Oligo fwd</td><td>CACCCCTGAGGAGCGTCGCTGGTC</td><td></td><td></td><td></td><td></td><td></td><td></td><td></td><td></td><td></td><td></td></tr>
<tr><td>Oligo rev</td><td>AAACGACCAGCGACGCTCCTCAGG</td><td></td><td></td><td></td><td></td><td></td><td></td><td></td><td></td><td></td><td></td></tr>
<tr><td>Chromosome</td><td>start</td><td>end</td><td>strand</td><td>MM</td><td>target_seq</td><td>PAM</td><td>alignment</td><td>distance</td><td>position</td><td>gene name</td><td>gene id</td></tr>
<tr><td>GU305914.1</td><td>177257</td><td>177279</td><td>-</td><td>0</td><td>CCTGAGGAGCGTCGCTGGTC</td><td>GGG</td><td>||||||||[||||||||||||]PAM</td><td>NA</td><td>Intergenic</td><td>NA</td><td>NA</td></tr>
</table>

<table>
<tr><td>T21216</td><td>CTGAGGAGCGTCGCTGGTCGGGG</td><td>1000</td><td></td><td></td><td></td><td></td><td></td><td></td><td></td><td></td><td></td></tr>
<tr><td>Oligo fwd</td><td>CACCCTGAGGAGCGTCGCTGGTCG</td><td></td><td></td><td></td><td></td><td></td><td></td><td></td><td></td><td></td><td></td></tr>
<tr><td>Oligo rev</td><td>AAACCGACCAGCGACGCTCCTCAG</td><td></td><td></td><td></td><td></td><td></td><td></td><td></td><td></td><td></td><td></td></tr>
<tr><td>Chromosome</td><td>start</td><td>end</td><td>strand</td><td>MM</td><td>target_seq</td><td>PAM</td><td>alignment</td><td>distance</td><td>position</td><td>gene name</td><td>gene id</td></tr>
<tr><td>GU305914.1</td><td>177256</td><td>177278</td><td>-</td><td>0</td><td>CTGAGGAGCGTCGCTGGTCG</td><td>GGG</td><td>||||||||[||||||||||||]PAM</td><td>NA</td><td>Intergenic</td><td>NA</td><td>NA</td></tr>
<tr><td>T21217</td><td>TGAGGAGCGTCGCTGGTCGGGGG</td><td>1000</td><td></td><td></td><td></td><td></td><td></td><td></td><td></td><td></td><td></td></tr>
<tr><td>Oligo fwd</td><td>CACCTGAGGAGCGTCGCTGGTCGG</td><td></td><td></td><td></td><td></td><td></td><td></td><td></td><td></td><td></td><td></td></tr>
<tr><td>Oligo rev</td><td>AAACCCGACCAGCGACGCTCCTCA</td><td></td><td></td><td></td><td></td><td></td><td></td><td></td><td></td><td></td><td></td></tr>
<tr><td>Chromosome</td><td>start</td><td>end</td><td>strand</td><td>MM</td><td>target_seq</td><td>PAM</td><td>alignment</td><td>distance</td><td>position</td><td>gene name</td><td>gene id</td></tr>
<tr><td>GU305914.1</td><td>177255</td><td>177277</td><td>-</td><td>0</td><td>TGAGGAGCGTCGCTGGTCGG</td><td>GGG</td><td>||||||||[||||||||||||]PAM</td><td>NA</td><td>Intergenic</td><td>NA</td><td>NA</td></tr>
<tr><td>T21218</td><td>CAGAGACCGACGCCACGCTATGG</td><td>1000</td><td></td><td></td><td></td><td></td><td></td><td></td><td></td><td></td><td></td></tr>
<tr><td>Oligo fwd</td><td>CACCCAGAGACCGACGCCACGCTA</td><td></td><td></td><td></td><td></td><td></td><td></td><td></td><td></td><td></td><td></td></tr>
<tr><td>Oligo rev</td><td>AAACTAGCGTGGCGTCGGTCTCTG</td><td></td><td></td><td></td><td></td><td></td><td></td><td></td><td></td><td></td><td></td></tr>
<tr><td>Chromosome</td><td>start</td><td>end</td><td>strand</td><td>MM</td><td>target_seq</td><td>PAM</td><td>alignment</td><td>distance</td><td>position</td><td>gene name</td><td>gene id</td></tr>
<tr><td>GU305914.1</td><td>177220</td><td>177242</td><td>-</td><td>0</td><td>CAGAGACCGACGCCACGCTA</td><td>TGG</td><td>||||||||[||||||||||||]PAM</td><td>NA</td><td>Intergenic</td><td>NA</td><td>NA</td></tr>
<tr><td>T21219</td><td>CGCTATGGCGCACCAACCTCAGG</td><td>1000</td><td></td><td></td><td></td><td></td><td></td><td></td><td></td><td></td><td></td></tr>
<tr><td>Oligo fwd</td><td>CACCCGCTATGGCGCACCAACCTC</td><td></td><td></td><td></td><td></td><td></td><td></td><td></td><td></td><td></td><td></td></tr>
<tr><td>Oligo rev</td><td>AAACGAGGTTGGTGCGCCATAGCG</td><td></td><td></td><td></td><td></td><td></td><td></td><td></td><td></td><td></td><td></td></tr>
<tr><td>Chromosome</td><td>start</td><td>end</td><td>strand</td><td>MM</td><td>target_seq</td><td>PAM</td><td>alignment</td><td>distance</td><td>position</td><td>gene name</td><td>gene id</td></tr>
<tr><td>GU305914.1</td><td>177205</td><td>177227</td><td>-</td><td>0</td><td>CGCTATGGCGCACCAACCTC</td><td>AGG</td><td>||||||||[||||||||||||]PAM</td><td>NA</td><td>Intergenic</td><td>NA</td><td>NA</td></tr>
<tr><td>T21220</td><td>CTCCCTTCCCAGAGCCTCGACGG</td><td>1000</td><td></td><td></td><td></td><td></td><td></td><td></td><td></td><td></td><td></td></tr>
<tr><td>Oligo fwd</td><td>CACCCTCCCTTCCCAGAGCCTCGA</td><td></td><td></td><td></td><td></td><td></td><td></td><td></td><td></td><td></td><td></td></tr>
<tr><td>Oligo rev</td><td>AAACTCGAGGCTCTGGGAAGGGAG</td><td></td><td></td><td></td><td></td><td></td><td></td><td></td><td></td><td></td><td></td></tr>
<tr><td>Chromosome</td><td>start</td><td>end</td><td>strand</td><td>MM</td><td>target_seq</td><td>PAM</td><td>alignment</td><td>distance</td><td>position</td><td>gene name</td><td>gene id</td></tr>
<tr><td>GU305914.1</td><td>177161</td><td>177183</td><td>-</td><td>0</td><td>CTCCCTTCCCAGAGCCTCGA</td><td>CGG</td><td>||||||||[||||||||||||]PAM</td><td>NA</td><td>Intergenic</td><td>NA</td><td>NA</td></tr>
<tr><td>T21221</td><td>TCCCTTCCCAGAGCCTCGACGGG</td><td>1000</td><td></td><td></td><td></td><td></td><td></td><td></td><td></td><td></td><td></td></tr>
<tr><td>Oligo fwd</td><td>CACCTCCCTTCCCAGAGCCTCGAC</td><td></td><td></td><td></td><td></td><td></td><td></td><td></td><td></td><td></td><td></td></tr>
<tr><td>Oligo rev</td><td>AAACGTCGAGGCTCTGGGAAGGGA</td><td></td><td></td><td></td><td></td><td></td><td></td><td></td><td></td><td></td><td></td></tr>
<tr><td>Chromosome</td><td>start</td><td>end</td><td>strand</td><td>MM</td><td>target_seq</td><td>PAM</td><td>alignment</td><td>distance</td><td>position</td><td>gene name</td><td>gene id</td></tr>
<tr><td>GU305914.1</td><td>177160</td><td>177182</td><td>-</td><td>0</td><td>TCCCTTCCCAGAGCCTCGAC</td><td>GGG</td><td>||||||||[||||||||||||]PAM</td><td>NA</td><td>Intergenic</td><td>NA</td><td>NA</td></tr>
<tr><td>T21222</td><td>ACAAGAATTCTCCCTTTCATCGG</td><td>1000</td><td></td><td></td><td></td><td></td><td></td><td></td><td></td><td></td><td></td></tr>
<tr><td>Oligo fwd</td><td>CACCACAAGAATTCTCCCTTTCAT</td><td></td><td></td><td></td><td></td><td></td><td></td><td></td><td></td><td></td><td></td></tr>
<tr><td>Oligo rev</td><td>AAACATGAAAGGGAGAATTCTTGT</td><td></td><td></td><td></td><td></td><td></td><td></td><td></td><td></td><td></td><td></td></tr>
<tr><td>Chromosome</td><td>start</td><td>end</td><td>strand</td><td>MM</td><td>target_seq</td><td>PAM</td><td>alignment</td><td>distance</td><td>position</td><td>gene name</td><td>gene id</td></tr>
<tr><td>GU305914.1</td><td>177095</td><td>177117</td><td>-</td><td>0</td><td>ACAAGAATTCTCCCTTTCAT</td><td>CGG</td><td>||||||||[||||||||||||]PAM</td><td>NA</td><td>Intergenic</td><td>NA</td><td>NA</td></tr>
<tr><td>T21223</td><td>AATTCTCCCTTTCATCGGTCTGG</td><td>1000</td><td></td><td></td><td></td><td></td><td></td><td></td><td></td><td></td><td></td></tr>
<tr><td>Oligo fwd</td><td>CACCAATTCTCCCTTTCATCGGTC</td><td></td><td></td><td></td><td></td><td></td><td></td><td></td><td></td><td></td><td></td></tr>
<tr><td>Oligo rev</td><td>AAACGACCGATGAAAGGGAGAATT</td><td></td><td></td><td></td><td></td><td></td><td></td><td></td><td></td><td></td><td></td></tr>
<tr><td>Chromosome</td><td>start</td><td>end</td><td>strand</td><td>MM</td><td>target_seq</td><td>PAM</td><td>alignment</td><td>distance</td><td>position</td><td>gene name</td><td>gene id</td></tr>
<tr><td>GU305914.1</td><td>177090</td><td>177112</td><td>-</td><td>0</td><td>AATTCTCCCTTTCATCGGTC</td><td>TGG</td><td>||||||||[||||||||||||]PAM</td><td>NA</td><td>Intergenic</td><td>NA</td><td>NA</td></tr>
<tr><td>T21224</td><td>ACGCGCCTCGCCTCAAACCTTGG</td><td>1000</td><td></td><td></td><td></td><td></td><td></td><td></td><td></td><td></td><td></td></tr>
<tr><td>Oligo fwd</td><td>CACCACGCGCCTCGCCTCAAACCT</td><td></td><td></td><td></td><td></td><td></td><td></td><td></td><td></td><td></td><td></td></tr>
<tr><td>Oligo rev</td><td>AAACAGGTTTGAGGCGAGGCGCGT</td><td></td><td></td><td></td><td></td><td></td><td></td><td></td><td></td><td></td><td></td></tr>
<tr><td>Chromosome</td><td>start</td><td>end</td><td>strand</td><td>MM</td><td>target_seq</td><td>PAM</td><td>alignment</td><td>distance</td><td>position</td><td>gene name</td><td>gene id</td></tr>
<tr><td>GU305914.1</td><td>177064</td><td>177086</td><td>-</td><td>0</td><td>ACGCGCCTCGCCTCAAACCT</td><td>TGG</td><td>||||||||[||||||||||||]PAM</td><td>NA</td><td>Intergenic</td><td>NA</td><td>NA</td></tr>
<tr><td>T21225</td><td>CGCGCCTCGCCTCAAACCTTGGG</td><td>1000</td><td></td><td></td><td></td><td></td><td></td><td></td><td></td><td></td><td></td></tr>
<tr><td>Oligo fwd</td><td>CACCCGCGCCTCGCCTCAAACCTT</td><td></td><td></td><td></td><td></td><td></td><td></td><td></td><td></td><td></td><td></td></tr>
<tr><td>Oligo rev</td><td>AAACAAGGTTTGAGGCGAGGCGCG</td><td></td><td></td><td></td><td></td><td></td><td></td><td></td><td></td><td></td><td></td></tr>
<tr><td>Chromosome</td><td>start</td><td>end</td><td>strand</td><td>MM</td><td>target_seq</td><td>PAM</td><td>alignment</td><td>distance</td><td>position</td><td>gene name</td><td>gene id</td></tr>
<tr><td>GU305914.1</td><td>177063</td><td>177085</td><td>-</td><td>0</td><td>CGCGCCTCGCCTCAAACCTT</td><td>GGG</td><td>||||||||[||||||||||||]PAM</td><td>NA</td><td>Intergenic</td><td>NA</td><td>NA</td></tr>
<tr><td>T21226</td><td>GCGCCTCGCCTCAAACCTTGGGG</td><td>1000</td><td></td><td></td><td></td><td></td><td></td><td></td><td></td><td></td><td></td></tr>
<tr><td>Oligo fwd</td><td>CACCGCGCCTCGCCTCAAACCTTG</td><td></td><td></td><td></td><td></td><td></td><td></td><td></td><td></td><td></td><td></td></tr>
<tr><td>Oligo rev</td><td>AAACCAAGGTTTGAGGCGAGGCGC</td><td></td><td></td><td></td><td></td><td></td><td></td><td></td><td></td><td></td><td></td></tr>
<tr><td>Chromosome</td><td>start</td><td>end</td><td>strand</td><td>MM</td><td>target_seq</td><td>PAM</td><td>alignment</td><td>distance</td><td>position</td><td>gene name</td><td>gene id</td></tr>
<tr><td>GU305914.1</td><td>177062</td><td>177084</td><td>-</td><td>0</td><td>GCGCCTCGCCTCAAACCTTG</td><td>GGG</td><td>||||||||[||||||||||||]PAM</td><td>NA</td><td>Intergenic</td><td>NA</td><td>NA</td></tr>
<tr><td>T21227</td><td>GCCTCAAACCTTGGGGCGCGAGG</td><td>1000</td><td></td><td></td><td></td><td></td><td></td><td></td><td></td><td></td><td></td></tr>
<tr><td>Oligo fwd</td><td>CACCGCCTCAAACCTTGGGGCGCG</td><td></td><td></td><td></td><td></td><td></td><td></td><td></td><td></td><td></td><td></td></tr>
<tr><td>Oligo rev</td><td>AAACCGCGCCCCAAGGTTTGAGGC</td><td></td><td></td><td></td><td></td><td></td><td></td><td></td><td></td><td></td><td></td></tr>
<tr><td>Chromosome</td><td>start</td><td>end</td><td>strand</td><td>MM</td><td>target_seq</td><td>PAM</td><td>alignment</td><td>distance</td><td>position</td><td>gene name</td><td>gene id</td></tr>
<tr><td>GU305914.1</td><td>177055</td><td>177077</td><td>-</td><td>0</td><td>GCCTCAAACCTTGGGGCGCG</td><td>AGG</td><td>||||||||[||||||||||||]PAM</td><td>NA</td><td>Intergenic</td><td>NA</td><td>NA</td></tr>
<tr><td>T21228</td><td>ACTGATTCGCCACAACACCGCGG</td><td>1000</td><td></td><td></td><td></td><td></td><td></td><td></td><td></td><td></td><td></td></tr>
<tr><td>Oligo fwd</td><td>CACCACTGATTCGCCACAACACCG</td><td></td><td></td><td></td><td></td><td></td><td></td><td></td><td></td><td></td><td></td></tr>
<tr><td>Oligo rev</td><td>AAACCGGTGTTGTGGCGAATCAGT</td><td></td><td></td><td></td><td></td><td></td><td></td><td></td><td></td><td></td><td></td></tr>
<tr><td>Chromosome</td><td>start</td><td>end</td><td>strand</td><td>MM</td><td>target_seq</td><td>PAM</td><td>alignment</td><td>distance</td><td>position</td><td>gene name</td><td>gene id</td></tr>
<tr><td>GU305914.1</td><td>177019</td><td>177041</td><td>-</td><td>0</td><td>ACTGATTCGCCACAACACCG</td><td>CGG</td><td>||||||||[||||||||||||]PAM</td><td>NA</td><td>Intergenic</td><td>NA</td><td>NA</td></tr>
<tr><td>T21229</td><td>GTTCTCCAACCCGCTACGCGCGG</td><td>1000</td><td></td><td></td><td></td><td></td><td></td><td></td><td></td><td></td><td></td></tr>
<tr><td>Oligo fwd</td><td>CACCGTTCTCCAACCCGCTACGCG</td><td></td><td></td><td></td><td></td><td></td><td></td><td></td><td></td><td></td><td></td></tr>
<tr><td>Oligo rev</td><td>AAACCGCGTAGCGGGTTGGAGAAC</td><td></td><td></td><td></td><td></td><td></td><td></td><td></td><td></td><td></td><td></td></tr>
<tr><td>Chromosome</td><td>start</td><td>end</td><td>strand</td><td>MM</td><td>target_seq</td><td>PAM</td><td>alignment</td><td>distance</td><td>position</td><td>gene name</td><td>gene id</td></tr>
<tr><td>GU305914.1</td><td>176980</td><td>177002</td><td>-</td><td>0</td><td>GTTCTCCAACCCGCTACGCG</td><td>CGG</td><td>||||||||[||||||||||||]PAM</td><td>NA</td><td>Intergenic</td><td>NA</td><td>NA</td></tr>
<tr><td>T21230</td><td>TTCTCCAACCCGCTACGCGCGGG</td><td>1000</td><td></td><td></td><td></td><td></td><td></td><td></td><td></td><td></td><td></td></tr>
<tr><td>Oligo fwd</td><td>CACCTTCTCCAACCCGCTACGCGC</td><td></td><td></td><td></td><td></td><td></td><td></td><td></td><td></td><td></td><td></td></tr>
<tr><td>Oligo rev</td><td>AAACGCGCGTAGCGGGTTGGAGAA</td><td></td><td></td><td></td><td></td><td></td><td></td><td></td><td></td><td></td><td></td></tr>
<tr><td>Chromosome</td><td>start</td><td>end</td><td>strand</td><td>MM</td><td>target_seq</td><td>PAM</td><td>alignment</td><td>distance</td><td>position</td><td>gene name</td><td>gene id</td></tr>
</table>

<table>
<tr><td>GU305914.1</td><td>176979</td><td>177001</td><td>-</td><td>0</td><td>TTCTCCAACCCGCTACGCGC</td><td>GGG</td><td>||||||||[||||||||||||]PAM</td><td>NA</td><td>Intergenic</td><td>NA</td><td>NA</td></tr>
<tr><td>T21231</td><td>CAACCCGCTACGCGCGGGACCGG</td><td>1000</td><td></td><td></td><td></td><td></td><td></td><td></td><td></td><td></td><td></td></tr>
<tr><td>Oligo fwd</td><td>CACCCAACCCGCTACGCGCGGGAC</td><td></td><td></td><td></td><td></td><td></td><td></td><td></td><td></td><td></td><td></td></tr>
<tr><td>Oligo rev</td><td>AAACGTCCCGCGCGTAGCGGGTTG</td><td></td><td></td><td></td><td></td><td></td><td></td><td></td><td></td><td></td><td></td></tr>
<tr><td>Chromosome</td><td>start</td><td>end</td><td>strand</td><td>MM</td><td>target_seq</td><td>PAM</td><td>alignment</td><td>distance</td><td>position</td><td>gene name</td><td>gene id</td></tr>
<tr><td>GU305914.1</td><td>176974</td><td>176996</td><td>-</td><td>0</td><td>CAACCCGCTACGCGCGGGAC</td><td>CGG</td><td>||||||||[||||||||||||]PAM</td><td>NA</td><td>Intergenic</td><td>NA</td><td>NA</td></tr>
<tr><td>T21232</td><td>CGCTACGCGCGGGACCGGTGAGG</td><td>1000</td><td></td><td></td><td></td><td></td><td></td><td></td><td></td><td></td><td></td></tr>
<tr><td>Oligo fwd</td><td>CACCCGCTACGCGCGGGACCGGTG</td><td></td><td></td><td></td><td></td><td></td><td></td><td></td><td></td><td></td><td></td></tr>
<tr><td>Oligo rev</td><td>AAACCACCGGTCCCGCGCGTAGCG</td><td></td><td></td><td></td><td></td><td></td><td></td><td></td><td></td><td></td><td></td></tr>
<tr><td>Chromosome</td><td>start</td><td>end</td><td>strand</td><td>MM</td><td>target_seq</td><td>PAM</td><td>alignment</td><td>distance</td><td>position</td><td>gene name</td><td>gene id</td></tr>
<tr><td>GU305914.1</td><td>176969</td><td>176991</td><td>-</td><td>0</td><td>CGCTACGCGCGGGACCGGTG</td><td>AGG</td><td>||||||||[||||||||||||]PAM</td><td>NA</td><td>Intergenic</td><td>NA</td><td>NA</td></tr>
<tr><td>T21233</td><td>CTACAGACAGAGATCGTACGTGG</td><td>1000</td><td></td><td></td><td></td><td></td><td></td><td></td><td></td><td></td><td></td></tr>
<tr><td>Oligo fwd</td><td>CACCCTACAGACAGAGATCGTACG</td><td></td><td></td><td></td><td></td><td></td><td></td><td></td><td></td><td></td><td></td></tr>
<tr><td>Oligo rev</td><td>AAACCGTACGATCTCTGTCTGTAG</td><td></td><td></td><td></td><td></td><td></td><td></td><td></td><td></td><td></td><td></td></tr>
<tr><td>Chromosome</td><td>start</td><td>end</td><td>strand</td><td>MM</td><td>target_seq</td><td>PAM</td><td>alignment</td><td>distance</td><td>position</td><td>gene name</td><td>gene id</td></tr>
<tr><td>GU305914.1</td><td>176923</td><td>176945</td><td>-</td><td>0</td><td>CTACAGACAGAGATCGTACG</td><td>TGG</td><td>||||||||[||||||||||||]PAM</td><td>NA</td><td>Intergenic</td><td>NA</td><td>NA</td></tr>
<tr><td>T21234</td><td>CAGACAGAGATCGTACGTGGAGG</td><td>1000</td><td></td><td></td><td></td><td></td><td></td><td></td><td></td><td></td><td></td></tr>
<tr><td>Oligo fwd</td><td>CACCCAGACAGAGATCGTACGTGG</td><td></td><td></td><td></td><td></td><td></td><td></td><td></td><td></td><td></td><td></td></tr>
<tr><td>Oligo rev</td><td>AAACCCACGTACGATCTCTGTCTG</td><td></td><td></td><td></td><td></td><td></td><td></td><td></td><td></td><td></td><td></td></tr>
<tr><td>Chromosome</td><td>start</td><td>end</td><td>strand</td><td>MM</td><td>target_seq</td><td>PAM</td><td>alignment</td><td>distance</td><td>position</td><td>gene name</td><td>gene id</td></tr>
<tr><td>GU305914.1</td><td>176920</td><td>176942</td><td>-</td><td>0</td><td>CAGACAGAGATCGTACGTGG</td><td>AGG</td><td>||||||||[||||||||||||]PAM</td><td>NA</td><td>Intergenic</td><td>NA</td><td>NA</td></tr>
<tr><td>T21235</td><td>GAGATCGTACGTGGAGGCGCCGG</td><td>1000</td><td></td><td></td><td></td><td></td><td></td><td></td><td></td><td></td><td></td></tr>
<tr><td>Oligo fwd</td><td>CACCGAGATCGTACGTGGAGGCGC</td><td></td><td></td><td></td><td></td><td></td><td></td><td></td><td></td><td></td><td></td></tr>
<tr><td>Oligo rev</td><td>AAACGCGCCTCCACGTACGATCTC</td><td></td><td></td><td></td><td></td><td></td><td></td><td></td><td></td><td></td><td></td></tr>
<tr><td>Chromosome</td><td>start</td><td>end</td><td>strand</td><td>MM</td><td>target_seq</td><td>PAM</td><td>alignment</td><td>distance</td><td>position</td><td>gene name</td><td>gene id</td></tr>
<tr><td>GU305914.1</td><td>176914</td><td>176936</td><td>-</td><td>0</td><td>GAGATCGTACGTGGAGGCGC</td><td>CGG</td><td>||||||||[||||||||||||]PAM</td><td>NA</td><td>Intergenic</td><td>NA</td><td>NA</td></tr>
<tr><td>T21236</td><td>AGATCGTACGTGGAGGCGCCGGG</td><td>1000</td><td></td><td></td><td></td><td></td><td></td><td></td><td></td><td></td><td></td></tr>
<tr><td>Oligo fwd</td><td>CACCAGATCGTACGTGGAGGCGCC</td><td></td><td></td><td></td><td></td><td></td><td></td><td></td><td></td><td></td><td></td></tr>
<tr><td>Oligo rev</td><td>AAACGGCGCCTCCACGTACGATCT</td><td></td><td></td><td></td><td></td><td></td><td></td><td></td><td></td><td></td><td></td></tr>
<tr><td>Chromosome</td><td>start</td><td>end</td><td>strand</td><td>MM</td><td>target_seq</td><td>PAM</td><td>alignment</td><td>distance</td><td>position</td><td>gene name</td><td>gene id</td></tr>
<tr><td>GU305914.1</td><td>176913</td><td>176935</td><td>-</td><td>0</td><td>AGATCGTACGTGGAGGCGCC</td><td>GGG</td><td>||||||||[||||||||||||]PAM</td><td>NA</td><td>Intergenic</td><td>NA</td><td>NA</td></tr>
<tr><td>T21237</td><td>ACACTTCCGCTATCCCAGCGTGG</td><td>1000</td><td></td><td></td><td></td><td></td><td></td><td></td><td></td><td></td><td></td></tr>
<tr><td>Oligo fwd</td><td>CACCACACTTCCGCTATCCCAGCG</td><td></td><td></td><td></td><td></td><td></td><td></td><td></td><td></td><td></td><td></td></tr>
<tr><td>Oligo rev</td><td>AAACCGCTGGGATAGCGGAAGTGT</td><td></td><td></td><td></td><td></td><td></td><td></td><td></td><td></td><td></td><td></td></tr>
<tr><td>Chromosome</td><td>start</td><td>end</td><td>strand</td><td>MM</td><td>target_seq</td><td>PAM</td><td>alignment</td><td>distance</td><td>position</td><td>gene name</td><td>gene id</td></tr>
<tr><td>GU305914.1</td><td>176857</td><td>176879</td><td>-</td><td>0</td><td>ACACTTCCGCTATCCCAGCG</td><td>TGG</td><td>||||||||[||||||||||||]PAM</td><td>NA</td><td>Intergenic</td><td>NA</td><td>NA</td></tr>
<tr><td>T21238</td><td>CACTTCCGCTATCCCAGCGTGGG</td><td>1000</td><td></td><td></td><td></td><td></td><td></td><td></td><td></td><td></td><td></td></tr>
<tr><td>Oligo fwd</td><td>CACCCACTTCCGCTATCCCAGCGT</td><td></td><td></td><td></td><td></td><td></td><td></td><td></td><td></td><td></td><td></td></tr>
<tr><td>Oligo rev</td><td>AAACACGCTGGGATAGCGGAAGTG</td><td></td><td></td><td></td><td></td><td></td><td></td><td></td><td></td><td></td><td></td></tr>
<tr><td>Chromosome</td><td>start</td><td>end</td><td>strand</td><td>MM</td><td>target_seq</td><td>PAM</td><td>alignment</td><td>distance</td><td>position</td><td>gene name</td><td>gene id</td></tr>
<tr><td>GU305914.1</td><td>176856</td><td>176878</td><td>-</td><td>0</td><td>CACTTCCGCTATCCCAGCGT</td><td>GGG</td><td>||||||||[||||||||||||]PAM</td><td>NA</td><td>Intergenic</td><td>NA</td><td>NA</td></tr>
<tr><td>T21239</td><td>CTATCCCAGCGTGGGAATGATGG</td><td>1000</td><td></td><td></td><td></td><td></td><td></td><td></td><td></td><td></td><td></td></tr>
<tr><td>Oligo fwd</td><td>CACCCTATCCCAGCGTGGGAATGA</td><td></td><td></td><td></td><td></td><td></td><td></td><td></td><td></td><td></td><td></td></tr>
<tr><td>Oligo rev</td><td>AAACTCATTCCCACGCTGGGATAG</td><td></td><td></td><td></td><td></td><td></td><td></td><td></td><td></td><td></td><td></td></tr>
<tr><td>Chromosome</td><td>start</td><td>end</td><td>strand</td><td>MM</td><td>target_seq</td><td>PAM</td><td>alignment</td><td>distance</td><td>position</td><td>gene name</td><td>gene id</td></tr>
<tr><td>GU305914.1</td><td>176848</td><td>176870</td><td>-</td><td>0</td><td>CTATCCCAGCGTGGGAATGA</td><td>TGG</td><td>||||||||[||||||||||||]PAM</td><td>NA</td><td>Intergenic</td><td>NA</td><td>NA</td></tr>
<tr><td>T21240</td><td>GGCTCACCAGCCTCCGTTCTCGG</td><td>1000</td><td></td><td></td><td></td><td></td><td></td><td></td><td></td><td></td><td></td></tr>
<tr><td>Oligo fwd</td><td>CACCGGCTCACCAGCCTCCGTTCT</td><td></td><td></td><td></td><td></td><td></td><td></td><td></td><td></td><td></td><td></td></tr>
<tr><td>Oligo rev</td><td>AAACAGAACGGAGGCTGGTGAGCC</td><td></td><td></td><td></td><td></td><td></td><td></td><td></td><td></td><td></td><td></td></tr>
<tr><td>Chromosome</td><td>start</td><td>end</td><td>strand</td><td>MM</td><td>target_seq</td><td>PAM</td><td>alignment</td><td>distance</td><td>position</td><td>gene name</td><td>gene id</td></tr>
<tr><td>GU305914.1</td><td>176827</td><td>176849</td><td>-</td><td>0</td><td>GGCTCACCAGCCTCCGTTCT</td><td>CGG</td><td>||||||||[||||||||||||]PAM</td><td>NA</td><td>Intergenic</td><td>NA</td><td>NA</td></tr>
<tr><td>T21241</td><td>GCTCACCAGCCTCCGTTCTCGGG</td><td>1000</td><td></td><td></td><td></td><td></td><td></td><td></td><td></td><td></td><td></td></tr>
<tr><td>Oligo fwd</td><td>CACCGCTCACCAGCCTCCGTTCTC</td><td></td><td></td><td></td><td></td><td></td><td></td><td></td><td></td><td></td><td></td></tr>
<tr><td>Oligo rev</td><td>AAACGAGAACGGAGGCTGGTGAGC</td><td></td><td></td><td></td><td></td><td></td><td></td><td></td><td></td><td></td><td></td></tr>
<tr><td>Chromosome</td><td>start</td><td>end</td><td>strand</td><td>MM</td><td>target_seq</td><td>PAM</td><td>alignment</td><td>distance</td><td>position</td><td>gene name</td><td>gene id</td></tr>
<tr><td>GU305914.1</td><td>176826</td><td>176848</td><td>-</td><td>0</td><td>GCTCACCAGCCTCCGTTCTC</td><td>GGG</td><td>||||||||[||||||||||||]PAM</td><td>NA</td><td>Intergenic</td><td>NA</td><td>NA</td></tr>
<tr><td>T21242</td><td>CAACAACACCTCAGATCCTAAGG</td><td>1000</td><td></td><td></td><td></td><td></td><td></td><td></td><td></td><td></td><td></td></tr>
<tr><td>Oligo fwd</td><td>CACCCAACAACACCTCAGATCCTA</td><td></td><td></td><td></td><td></td><td></td><td></td><td></td><td></td><td></td><td></td></tr>
<tr><td>Oligo rev</td><td>AAACTAGGATCTGAGGTGTTGTTG</td><td></td><td></td><td></td><td></td><td></td><td></td><td></td><td></td><td></td><td></td></tr>
<tr><td>Chromosome</td><td>start</td><td>end</td><td>strand</td><td>MM</td><td>target_seq</td><td>PAM</td><td>alignment</td><td>distance</td><td>position</td><td>gene name</td><td>gene id</td></tr>
<tr><td>GU305914.1</td><td>176762</td><td>176784</td><td>-</td><td>0</td><td>CAACAACACCTCAGATCCTA</td><td>AGG</td><td>||||||||[||||||||||||]PAM</td><td>NA</td><td>Intergenic</td><td>NA</td><td>NA</td></tr>
<tr><td>T21243</td><td>ACCTCAGATCCTAAGGTTCCTGG</td><td>1000</td><td></td><td></td><td></td><td></td><td></td><td></td><td></td><td></td><td></td></tr>
<tr><td>Oligo fwd</td><td>CACCACCTCAGATCCTAAGGTTCC</td><td></td><td></td><td></td><td></td><td></td><td></td><td></td><td></td><td></td><td></td></tr>
<tr><td>Oligo rev</td><td>AAACGGAACCTTAGGATCTGAGGT</td><td></td><td></td><td></td><td></td><td></td><td></td><td></td><td></td><td></td><td></td></tr>
<tr><td>Chromosome</td><td>start</td><td>end</td><td>strand</td><td>MM</td><td>target_seq</td><td>PAM</td><td>alignment</td><td>distance</td><td>position</td><td>gene name</td><td>gene id</td></tr>
<tr><td>GU305914.1</td><td>176755</td><td>176777</td><td>-</td><td>0</td><td>ACCTCAGATCCTAAGGTTCC</td><td>TGG</td><td>||||||||[||||||||||||]PAM</td><td>NA</td><td>Intergenic</td><td>NA</td><td>NA</td></tr>
<tr><td>T21244</td><td>CCTCAGATCCTAAGGTTCCTGGG</td><td>1000</td><td></td><td></td><td></td><td></td><td></td><td></td><td></td><td></td><td></td></tr>
<tr><td>Oligo fwd</td><td>CACCCCTCAGATCCTAAGGTTCCT</td><td></td><td></td><td></td><td></td><td></td><td></td><td></td><td></td><td></td><td></td></tr>
<tr><td>Oligo rev</td><td>AAACAGGAACCTTAGGATCTGAGG</td><td></td><td></td><td></td><td></td><td></td><td></td><td></td><td></td><td></td><td></td></tr>
<tr><td>Chromosome</td><td>start</td><td>end</td><td>strand</td><td>MM</td><td>target_seq</td><td>PAM</td><td>alignment</td><td>distance</td><td>position</td><td>gene name</td><td>gene id</td></tr>
<tr><td>GU305914.1</td><td>176754</td><td>176776</td><td>-</td><td>0</td><td>CCTCAGATCCTAAGGTTCCT</td><td>GGG</td><td>||||||||[||||||||||||]PAM</td><td>NA</td><td>Intergenic</td><td>NA</td><td>NA</td></tr>
<tr><td>T21245</td><td>CATGTTCTACATCCCCAGAATGG</td><td>1000</td><td></td><td></td><td></td><td></td><td></td><td></td><td></td><td></td><td></td></tr>
<tr><td>Oligo fwd</td><td>CACCCATGTTCTACATCCCCAGAA</td><td></td><td></td><td></td><td></td><td></td><td></td><td></td><td></td><td></td><td></td></tr>
<tr><td>Oligo rev</td><td>AAACTTCTGGGGATGTAGAACATG</td><td></td><td></td><td></td><td></td><td></td><td></td><td></td><td></td><td></td><td></td></tr>
</table>

Chromosome start end strand MM target_seq PAM alignment distance position gene name gene id
GU305914.1 176710 176732 - 0 CATGTTCTACATCCCCAGAA TGG ||||||||[||||||||||||]PAM NA Intergenic NA NA

T21246 CTACATCCCCAGAATGGCACCGG 1000
Oligo fwd CACCCTACATCCCCAGAATGGCAC
Oligo rev AAACGTGCCATTCTGGGGATGTAG
Chromosome start end strand MM target_seq PAM alignment distance position gene name gene id
GU305914.1 176704 176726 - 0 CTACATCCCCAGAATGGCAC CGG ||||||||[||||||||||||]PAM NA Intergenic NA NA

T21247 TACATCCCCAGAATGGCACCGGG 1000
Oligo fwd CACCTACATCCCCAGAATGGCACC
Oligo rev AAACGGTGCCATTCTGGGGATGTA
Chromosome start end strand MM target_seq PAM alignment distance position gene name gene id
GU305914.1 176703 176725 - 0 TACATCCCCAGAATGGCACC GGG ||||||||[||||||||||||]PAM NA Intergenic NA NA

T21248 CCCCAGAATGGCACCGGGCGCGG 1000
Oligo fwd CACCCCCCAGAATGGCACCGGGCG
Oligo rev AAACCGCCCGGTGCCATTCTGGGG
Chromosome start end strand MM target_seq PAM alignment distance position gene name gene id
GU305914.1 176698 176720 - 0 CCCCAGAATGGCACCGGGCG CGG ||||||||[||||||||||||]PAM NA Intergenic NA NA

T21249 GCGCGGCGCCCTATCATCCGTGG 1000
Oligo fwd CACCGCGCGGCGCCCTATCATCCG
Oligo rev AAACCGGATGATAGGGCGCCGCGC
Chromosome start end strand MM target_seq PAM alignment distance position gene name gene id
GU305914.1 176681 176703 - 0 GCGCGGCGCCCTATCATCCG TGG ||||||||[||||||||||||]PAM NA Intergenic NA NA

T21250 TCATCCGTGGTGCGAGCAGATGG 1000
Oligo fwd CACCTCATCCGTGGTGCGAGCAGA
Oligo rev AAACTCTGCTCGCACCACGGATGA
Chromosome start end strand MM target_seq PAM alignment distance position gene name gene id
GU305914.1 176668 176690 - 0 TCATCCGTGGTGCGAGCAGA TGG ||||||||[||||||||||||]PAM NA Intergenic NA NA

T21251 TCCGTGGTGCGAGCAGATGGTGG 1000
Oligo fwd CACCTCCGTGGTGCGAGCAGATGG
Oligo rev AAACCCATCTGCTCGCACCACGGA
Chromosome start end strand MM target_seq PAM alignment distance position gene name gene id
GU305914.1 176665 176687 - 0 TCCGTGGTGCGAGCAGATGG TGG ||||||||[||||||||||||]PAM NA Intergenic NA NA

T21252 ATGGTGGCGCCTCGTCCGCAAGG 1000
Oligo fwd CACCATGGTGGCGCCTCGTCCGCA
Oligo rev AAACTGCGGACGAGGCGCCACCAT
Chromosome start end strand MM target_seq PAM alignment distance position gene name gene id
GU305914.1 176649 176671 - 0 ATGGTGGCGCCTCGTCCGCA AGG ||||||||[||||||||||||]PAM NA Intergenic NA NA

T21253 ATCATCAGATTTTGCAAGAGAGG 1000
Oligo fwd CACCATCATCAGATTTTGCAAGAG
Oligo rev AAACCTCTTGCAAAATCTGATGAT
Chromosome start end strand MM target_seq PAM alignment distance position gene name gene id
GU305914.1 176609 176631 - 0 ATCATCAGATTTTGCAAGAG AGG ||||||||[||||||||||||]PAM NA Intergenic NA NA

T21254 TCATCAGATTTTGCAAGAGAGGG 1000
Oligo fwd CACCTCATCAGATTTTGCAAGAGA
Oligo rev AAACTCTCTTGCAAAATCTGATGA
Chromosome start end strand MM target_seq PAM alignment distance position gene name gene id
GU305914.1 176608 176630 - 0 TCATCAGATTTTGCAAGAGA GGG ||||||||[||||||||||||]PAM NA Intergenic NA NA

T21255 CAACTAGAGCAGCGCATGCAAGG 1000
Oligo fwd CACCCAACTAGAGCAGCGCATGCA
Oligo rev AAACTGCATGCGCTGCTCTAGTTG
Chromosome start end strand MM target_seq PAM alignment distance position gene name gene id
GU305914.1 176559 176581 - 0 CAACTAGAGCAGCGCATGCA AGG ||||||||[||||||||||||]PAM NA Intergenic NA NA

T21256 AGCAGCGCATGCAAGGACTCAGG 1000
Oligo fwd CACCAGCAGCGCATGCAAGGACTC
Oligo rev AAACGAGTCCTTGCATGCGCTGCT
Chromosome start end strand MM target_seq PAM alignment distance position gene name gene id
GU305914.1 176552 176574 - 0 AGCAGCGCATGCAAGGACTC AGG ||||||||[||||||||||||]PAM NA Intergenic NA NA

T21257 GCGCATGCAAGGACTCAGGCTGG 1000
Oligo fwd CACCGCGCATGCAAGGACTCAGGC
Oligo rev AAACGCCTGAGTCCTTGCATGCGC
Chromosome start end strand MM target_seq PAM alignment distance position gene name gene id
GU305914.1 176548 176570 - 0 GCGCATGCAAGGACTCAGGC TGG ||||||||[||||||||||||]PAM NA Intergenic NA NA

T21258 CATGCAAGGACTCAGGCTGGAGG 1000
Oligo fwd CACCCATGCAAGGACTCAGGCTGG
Oligo rev AAACCCAGCCTGAGTCCTTGCATG
Chromosome start end strand MM target_seq PAM alignment distance position gene name gene id
GU305914.1 176545 176567 - 0 CATGCAAGGACTCAGGCTGG AGG ||||||||[||||||||||||]PAM NA Intergenic NA NA

T21259 GGAGGATCCGCGAGCCCAGCAGG 1000
Oligo fwd CACCGGAGGATCCGCGAGCCCAGC
Oligo rev AAACGCTGGGCTCGCGGATCCTCC
Chromosome start end strand MM target_seq PAM alignment distance position gene name gene id
GU305914.1 176527 176549 - 0 GGAGGATCCGCGAGCCCAGC AGG ||||||||[||||||||||||]PAM NA Intergenic NA NA

T21260 GCGAGCCCAGCAGGTCTGCGCGG 1000
Oligo fwd CACCGCGAGCCCAGCAGGTCTGCG

Oligo rev AAACCGCAGACCTGCTGGGCTCGC
Chromosome start end strand MM target_seq PAM alignment distance position gene name gene id
GU305914.1 176518 176540 - 0 GCGAGCCCAGCAGGTCTGCG CGG ||||||||[||||||||||||]PAM NA Intergenic NA NA

T21261 CTGCGCGGTCCGCCGAGAGCCGG 1000
Oligo fwd CACCCTGCGCGGTCCGCCGAGAGC
Oligo rev AAACGCTCTCGGCGGACCGCGCAG
Chromosome start end strand MM target_seq PAM alignment distance position gene name gene id
GU305914.1 176503 176525 - 0 CTGCGCGGTCCGCCGAGAGC CGG ||||||||[||||||||||||]PAM NA Intergenic NA NA

T21262 GTCGCCATCGTCTCAGCAGACGG 1000
Oligo fwd CACCGTCGCCATCGTCTCAGCAGA
Oligo rev AAACTCTGCTGAGACGATGGCGAC
Chromosome start end strand MM target_seq PAM alignment distance position gene name gene id
GU305914.1 176443 176465 - 0 GTCGCCATCGTCTCAGCAGA CGG ||||||||[||||||||||||]PAM NA Intergenic NA NA

T21263 GACGCTCCTCTGTCCGAAGAAGG 1000
Oligo fwd CACCGACGCTCCTCTGTCCGAAGA
Oligo rev AAACTCTTCGGACAGAGGAGCGTC
Chromosome start end strand MM target_seq PAM alignment distance position gene name gene id
GU305914.1 176421 176443 - 0 GACGCTCCTCTGTCCGAAGA AGG ||||||||[||||||||||||]PAM NA Intergenic NA NA

T21264 CCGAAGAAGGAGACGACGCGAGG 1000
Oligo fwd CACCCCGAAGAAGGAGACGACGCG
Oligo rev AAACCGCGTCGTCTCCTTCTTCGG
Chromosome start end strand MM target_seq PAM alignment distance position gene name gene id
GU305914.1 176408 176430 - 0 CCGAAGAAGGAGACGACGCG AGG ||||||||[||||||||||||]PAM NA Intergenic NA NA

T21265 CGACGCGAGGAGCGCCCGTCAGG 1000
Oligo fwd CACCCGACGCGAGGAGCGCCCGTC
Oligo rev AAACGACGGGCGCTCCTCGCGTCG
Chromosome start end strand MM target_seq PAM alignment distance position gene name gene id
GU305914.1 176395 176417 - 0 CGACGCGAGGAGCGCCCGTC AGG ||||||||[||||||||||||]PAM NA Intergenic NA NA

T21266 GCCCGTCAGGATACCGCTTGCGG 1000
Oligo fwd CACCGCCCGTCAGGATACCGCTTG
Oligo rev AAACCAAGCGGTATCCTGACGGGC
Chromosome start end strand MM target_seq PAM alignment distance position gene name gene id
GU305914.1 176382 176404 - 0 GCCCGTCAGGATACCGCTTG CGG ||||||||[||||||||||||]PAM NA Intergenic NA NA

T21267 CGTCAGGATACCGCTTGCGGCGG 1000
Oligo fwd CACCCGTCAGGATACCGCTTGCGG
Oligo rev AAACCCGCAAGCGGTATCCTGACG
Chromosome start end strand MM target_seq PAM alignment distance position gene name gene id
GU305914.1 176379 176401 - 0 CGTCAGGATACCGCTTGCGG CGG ||||||||[||||||||||||]PAM NA Intergenic NA NA

T21268 CCGCTTGCGGCGGCTTCTCGAGG 1000
Oligo fwd CACCCCGCTTGCGGCGGCTTCTCG
Oligo rev AAACCGAGAAGCCGCCGCAAGCGG
Chromosome start end strand MM target_seq PAM alignment distance position gene name gene id
GU305914.1 176369 176391 - 0 CCGCTTGCGGCGGCTTCTCG AGG ||||||||[||||||||||||]PAM NA Intergenic NA NA

T21269 ACAACAACATCAAATTCCACCGG 1000
Oligo fwd CACCACAACAACATCAAATTCCAC
Oligo rev AAACGTGGAATTTGATGTTGTTGT
Chromosome start end strand MM target_seq PAM alignment distance position gene name gene id
GU305914.1 176338 176360 - 0 ACAACAACATCAAATTCCAC CGG ||||||||[||||||||||||]PAM NA Intergenic NA NA

T21270 CAACAACATCAAATTCCACCGGG 1000
Oligo fwd CACCCAACAACATCAAATTCCACC
Oligo rev AAACGGTGGAATTTGATGTTGTTG
Chromosome start end strand MM target_seq PAM alignment distance position gene name gene id
GU305914.1 176337 176359 - 0 CAACAACATCAAATTCCACC GGG ||||||||[||||||||||||]PAM NA Intergenic NA NA

T21271 CAGTTACGTCGAAAGAAGTACGG 1000
Oligo fwd CACCCAGTTACGTCGAAAGAAGTA
Oligo rev AAACTACTTCTTTCGACGTAACTG
Chromosome start end strand MM target_seq PAM alignment distance position gene name gene id
GU305914.1 176307 176329 - 0 CAGTTACGTCGAAAGAAGTA CGG ||||||||[||||||||||||]PAM NA Intergenic NA NA

T21272 CAATCAGTTTCAGAAGCGAACGG 1000
Oligo fwd CACCCAATCAGTTTCAGAAGCGAA
Oligo rev AAACTTCGCTTCTGAAACTGATTG
Chromosome start end strand MM target_seq PAM alignment distance position gene name gene id
GU305914.1 176281 176303 - 0 CAATCAGTTTCAGAAGCGAA CGG ||||||||[||||||||||||]PAM NA Intergenic NA NA

T21273 AATCAGTTTCAGAAGCGAACGGG 1000
Oligo fwd CACCAATCAGTTTCAGAAGCGAAC
Oligo rev AAACGTTCGCTTCTGAAACTGATT
Chromosome start end strand MM target_seq PAM alignment distance position gene name gene id
GU305914.1 176280 176302 - 0 AATCAGTTTCAGAAGCGAAC GGG ||||||||[||||||||||||]PAM NA Intergenic NA NA

T21274 ATCAACCCCTACCAGCGTGCAGG 1000
Oligo fwd CACCATCAACCCCTACCAGCGTGC
Oligo rev AAACGCACGCTGGTAGGGGTTGAT
Chromosome start end strand MM target_seq PAM alignment distance position gene name gene id
GU305914.1 176252 176274 - 0 ATCAACCCCTACCAGCGTGC AGG ||||||||[||||||||||||]PAM NA Intergenic NA NA

T21275 AAAGACAATCTACCTCCGTACGG 1000

<table>
<tr><td>Oligo fwd</td><td>CACCAAAGACAATCTACCTCCGTA</td><td></td><td></td><td></td><td></td><td></td><td></td><td></td><td></td><td></td><td></td></tr>
<tr><td>Oligo rev</td><td>AAACTACGGAGGTAGATTGTCTTT</td><td></td><td></td><td></td><td></td><td></td><td></td><td></td><td></td><td></td><td></td></tr>
<tr><td>Chromosome</td><td>start</td><td>end</td><td>strand</td><td>MM</td><td>target_seq</td><td>PAM</td><td>alignment</td><td>distance</td><td>position</td><td>gene name</td><td>gene id</td></tr>
<tr><td>GU305914.1</td><td>176229</td><td>176251</td><td>-</td><td>0</td><td>AAAGACAATCTACCTCCGTA</td><td>CGG</td><td>||||||||[||||||||||||]PAM</td><td>NA</td><td>Intergenic</td><td>NA</td><td>NA</td></tr>
<tr><td></td><td></td><td></td><td></td><td></td><td></td><td></td><td></td><td></td><td></td><td></td><td></td></tr>
<tr><td>T21276</td><td>AAGACAATCTACCTCCGTACGGG</td><td>1000</td><td></td><td></td><td></td><td></td><td></td><td></td><td></td><td></td><td></td></tr>
<tr><td>Oligo fwd</td><td>CACCAAGACAATCTACCTCCGTAC</td><td></td><td></td><td></td><td></td><td></td><td></td><td></td><td></td><td></td><td></td></tr>
<tr><td>Oligo rev</td><td>AAACGTACGGAGGTAGATTGTCTT</td><td></td><td></td><td></td><td></td><td></td><td></td><td></td><td></td><td></td><td></td></tr>
<tr><td>Chromosome</td><td>start</td><td>end</td><td>strand</td><td>MM</td><td>target_seq</td><td>PAM</td><td>alignment</td><td>distance</td><td>position</td><td>gene name</td><td>gene id</td></tr>
<tr><td>GU305914.1</td><td>176228</td><td>176250</td><td>-</td><td>0</td><td>AAGACAATCTACCTCCGTAC</td><td>GGG</td><td>||||||||[||||||||||||]PAM</td><td>NA</td><td>Intergenic</td><td>NA</td><td>NA</td></tr>
<tr><td></td><td></td><td></td><td></td><td></td><td></td><td></td><td></td><td></td><td></td><td></td><td></td></tr>
<tr><td>T21277</td><td>TCCGTACGGGTCGTCGCAACCGG</td><td>1000</td><td></td><td></td><td></td><td></td><td></td><td></td><td></td><td></td><td></td></tr>
<tr><td>Oligo fwd</td><td>CACCTCCGTACGGGTCGTCGCAAC</td><td></td><td></td><td></td><td></td><td></td><td></td><td></td><td></td><td></td><td></td></tr>
<tr><td>Oligo rev</td><td>AAACGTTGCGACGACCCGTACGGA</td><td></td><td></td><td></td><td></td><td></td><td></td><td></td><td></td><td></td><td></td></tr>
<tr><td>Chromosome</td><td>start</td><td>end</td><td>strand</td><td>MM</td><td>target_seq</td><td>PAM</td><td>alignment</td><td>distance</td><td>position</td><td>gene name</td><td>gene id</td></tr>
<tr><td>GU305914.1</td><td>176215</td><td>176237</td><td>-</td><td>0</td><td>TCCGTACGGGTCGTCGCAAC</td><td>CGG</td><td>||||||||[||||||||||||]PAM</td><td>NA</td><td>Intergenic</td><td>NA</td><td>NA</td></tr>
<tr><td></td><td></td><td></td><td></td><td></td><td></td><td></td><td></td><td></td><td></td><td></td><td></td></tr>
<tr><td>T21278</td><td>CAGTGTTAACAGTTCAGCAGCGG</td><td>1000</td><td></td><td></td><td></td><td></td><td></td><td></td><td></td><td></td><td></td></tr>
<tr><td>Oligo fwd</td><td>CACCCAGTGTTAACAGTTCAGCAG</td><td></td><td></td><td></td><td></td><td></td><td></td><td></td><td></td><td></td><td></td></tr>
<tr><td>Oligo rev</td><td>AAACCTGCTGAACTGTTAACACTG</td><td></td><td></td><td></td><td></td><td></td><td></td><td></td><td></td><td></td><td></td></tr>
<tr><td>Chromosome</td><td>start</td><td>end</td><td>strand</td><td>MM</td><td>target_seq</td><td>PAM</td><td>alignment</td><td>distance</td><td>position</td><td>gene name</td><td>gene id</td></tr>
<tr><td>GU305914.1</td><td>176192</td><td>176214</td><td>-</td><td>0</td><td>CAGTGTTAACAGTTCAGCAG</td><td>CGG</td><td>||||||||[||||||||||||]PAM</td><td>NA</td><td>Intergenic</td><td>NA</td><td>NA</td></tr>
<tr><td></td><td></td><td></td><td></td><td></td><td></td><td></td><td></td><td></td><td></td><td></td><td></td></tr>
<tr><td>T21279</td><td>AGTGTTAACAGTTCAGCAGCGGG</td><td>1000</td><td></td><td></td><td></td><td></td><td></td><td></td><td></td><td></td><td></td></tr>
<tr><td>Oligo fwd</td><td>CACCAGTGTTAACAGTTCAGCAGC</td><td></td><td></td><td></td><td></td><td></td><td></td><td></td><td></td><td></td><td></td></tr>
<tr><td>Oligo rev</td><td>AAACGCTGCTGAACTGTTAACACT</td><td></td><td></td><td></td><td></td><td></td><td></td><td></td><td></td><td></td><td></td></tr>
<tr><td>Chromosome</td><td>start</td><td>end</td><td>strand</td><td>MM</td><td>target_seq</td><td>PAM</td><td>alignment</td><td>distance</td><td>position</td><td>gene name</td><td>gene id</td></tr>
<tr><td>GU305914.1</td><td>176191</td><td>176213</td><td>-</td><td>0</td><td>AGTGTTAACAGTTCAGCAGC</td><td>GGG</td><td>||||||||[||||||||||||]PAM</td><td>NA</td><td>Intergenic</td><td>NA</td><td>NA</td></tr>
<tr><td></td><td></td><td></td><td></td><td></td><td></td><td></td><td></td><td></td><td></td><td></td><td></td></tr>
<tr><td>T21280</td><td>GTGTTAACAGTTCAGCAGCGGGG</td><td>1000</td><td></td><td></td><td></td><td></td><td></td><td></td><td></td><td></td><td></td></tr>
<tr><td>Oligo fwd</td><td>CACCGTGTTAACAGTTCAGCAGCG</td><td></td><td></td><td></td><td></td><td></td><td></td><td></td><td></td><td></td><td></td></tr>
<tr><td>Oligo rev</td><td>AAACCGCTGCTGAACTGTTAACAC</td><td></td><td></td><td></td><td></td><td></td><td></td><td></td><td></td><td></td><td></td></tr>
<tr><td>Chromosome</td><td>start</td><td>end</td><td>strand</td><td>MM</td><td>target_seq</td><td>PAM</td><td>alignment</td><td>distance</td><td>position</td><td>gene name</td><td>gene id</td></tr>
<tr><td>GU305914.1</td><td>176190</td><td>176212</td><td>-</td><td>0</td><td>GTGTTAACAGTTCAGCAGCG</td><td>GGG</td><td>||||||||[||||||||||||]PAM</td><td>NA</td><td>Intergenic</td><td>NA</td><td>NA</td></tr>
<tr><td></td><td></td><td></td><td></td><td></td><td></td><td></td><td></td><td></td><td></td><td></td><td></td></tr>
<tr><td>T21281</td><td>AGTTCAGCAGCGGGGACCAATGG</td><td>1000</td><td></td><td></td><td></td><td></td><td></td><td></td><td></td><td></td><td></td></tr>
<tr><td>Oligo fwd</td><td>CACCAGTTCAGCAGCGGGGACCAA</td><td></td><td></td><td></td><td></td><td></td><td></td><td></td><td></td><td></td><td></td></tr>
<tr><td>Oligo rev</td><td>AAACTTGGTCCCCGCTGCTGAACT</td><td></td><td></td><td></td><td></td><td></td><td></td><td></td><td></td><td></td><td></td></tr>
<tr><td>Chromosome</td><td>start</td><td>end</td><td>strand</td><td>MM</td><td>target_seq</td><td>PAM</td><td>alignment</td><td>distance</td><td>position</td><td>gene name</td><td>gene id</td></tr>
<tr><td>GU305914.1</td><td>176182</td><td>176204</td><td>-</td><td>0</td><td>AGTTCAGCAGCGGGGACCAA</td><td>TGG</td><td>||||||||[||||||||||||]PAM</td><td>NA</td><td>Intergenic</td><td>NA</td><td>NA</td></tr>
<tr><td></td><td></td><td></td><td></td><td></td><td></td><td></td><td></td><td></td><td></td><td></td><td></td></tr>
<tr><td>T21282</td><td>GCAGCGGGGACCAATGGAACAGG</td><td>1000</td><td></td><td></td><td></td><td></td><td></td><td></td><td></td><td></td><td></td></tr>
<tr><td>Oligo fwd</td><td>CACCGCAGCGGGGACCAATGGAAC</td><td></td><td></td><td></td><td></td><td></td><td></td><td></td><td></td><td></td><td></td></tr>
<tr><td>Oligo rev</td><td>AAACGTTCCATTGGTCCCCGCTGC</td><td></td><td></td><td></td><td></td><td></td><td></td><td></td><td></td><td></td><td></td></tr>
<tr><td>Chromosome</td><td>start</td><td>end</td><td>strand</td><td>MM</td><td>target_seq</td><td>PAM</td><td>alignment</td><td>distance</td><td>position</td><td>gene name</td><td>gene id</td></tr>
<tr><td>GU305914.1</td><td>176176</td><td>176198</td><td>-</td><td>0</td><td>GCAGCGGGGACCAATGGAAC</td><td>AGG</td><td>||||||||[||||||||||||]PAM</td><td>NA</td><td>Intergenic</td><td>NA</td><td>NA</td></tr>
<tr><td></td><td></td><td></td><td></td><td></td><td></td><td></td><td></td><td></td><td></td><td></td><td></td></tr>
<tr><td>T21283</td><td>ACAGGTGCCACAACCGCATCCGG</td><td>1000</td><td></td><td></td><td></td><td></td><td></td><td></td><td></td><td></td><td></td></tr>
<tr><td>Oligo fwd</td><td>CACCACAGGTGCCACAACCGCATC</td><td></td><td></td><td></td><td></td><td></td><td></td><td></td><td></td><td></td><td></td></tr>
<tr><td>Oligo rev</td><td>AAACGATGCGGTTGTGGCACCTGT</td><td></td><td></td><td></td><td></td><td></td><td></td><td></td><td></td><td></td><td></td></tr>
<tr><td>Chromosome</td><td>start</td><td>end</td><td>strand</td><td>MM</td><td>target_seq</td><td>PAM</td><td>alignment</td><td>distance</td><td>position</td><td>gene name</td><td>gene id</td></tr>
<tr><td>GU305914.1</td><td>176158</td><td>176180</td><td>-</td><td>0</td><td>ACAGGTGCCACAACCGCATC</td><td>CGG</td><td>||||||||[||||||||||||]PAM</td><td>NA</td><td>Intergenic</td><td>NA</td><td>NA</td></tr>
<tr><td></td><td></td><td></td><td></td><td></td><td></td><td></td><td></td><td></td><td></td><td></td><td></td></tr>
<tr><td>T21284</td><td>ATCCGGAACAGTGAGCTCGAAGG</td><td>1000</td><td></td><td></td><td></td><td></td><td></td><td></td><td></td><td></td><td></td></tr>
<tr><td>Oligo fwd</td><td>CACCATCCGGAACAGTGAGCTCGA</td><td></td><td></td><td></td><td></td><td></td><td></td><td></td><td></td><td></td><td></td></tr>
<tr><td>Oligo rev</td><td>AAACTCGAGCTCACTGTTCCGGAT</td><td></td><td></td><td></td><td></td><td></td><td></td><td></td><td></td><td></td><td></td></tr>
<tr><td>Chromosome</td><td>start</td><td>end</td><td>strand</td><td>MM</td><td>target_seq</td><td>PAM</td><td>alignment</td><td>distance</td><td>position</td><td>gene name</td><td>gene id</td></tr>
<tr><td>GU305914.1</td><td>176141</td><td>176163</td><td>-</td><td>0</td><td>ATCCGGAACAGTGAGCTCGA</td><td>AGG</td><td>||||||||[||||||||||||]PAM</td><td>NA</td><td>Intergenic</td><td>NA</td><td>NA</td></tr>
<tr><td></td><td></td><td></td><td></td><td></td><td></td><td></td><td></td><td></td><td></td><td></td><td></td></tr>
<tr><td>T21285</td><td>AACCAGCAGAAAGAGCGATGCGG</td><td>1000</td><td></td><td></td><td></td><td></td><td></td><td></td><td></td><td></td><td></td></tr>
<tr><td>Oligo fwd</td><td>CACCAACCAGCAGAAAGAGCGATG</td><td></td><td></td><td></td><td></td><td></td><td></td><td></td><td></td><td></td><td></td></tr>
<tr><td>Oligo rev</td><td>AAACCATCGCTCTTTCTGCTGGTT</td><td></td><td></td><td></td><td></td><td></td><td></td><td></td><td></td><td></td><td></td></tr>
<tr><td>Chromosome</td><td>start</td><td>end</td><td>strand</td><td>MM</td><td>target_seq</td><td>PAM</td><td>alignment</td><td>distance</td><td>position</td><td>gene name</td><td>gene id</td></tr>
<tr><td>GU305914.1</td><td>176117</td><td>176139</td><td>-</td><td>0</td><td>AACCAGCAGAAAGAGCGATG</td><td>CGG</td><td>||||||||[||||||||||||]PAM</td><td>NA</td><td>Intergenic</td><td>NA</td><td>NA</td></tr>
<tr><td></td><td></td><td></td><td></td><td></td><td></td><td></td><td></td><td></td><td></td><td></td><td></td></tr>
<tr><td>T21286</td><td>AGCAGAAAGAGCGATGCGGTAGG</td><td>1000</td><td></td><td></td><td></td><td></td><td></td><td></td><td></td><td></td><td></td></tr>
<tr><td>Oligo fwd</td><td>CACCAGCAGAAAGAGCGATGCGGT</td><td></td><td></td><td></td><td></td><td></td><td></td><td></td><td></td><td></td><td></td></tr>
<tr><td>Oligo rev</td><td>AAACACCGCATCGCTCTTTCTGCT</td><td></td><td></td><td></td><td></td><td></td><td></td><td></td><td></td><td></td><td></td></tr>
<tr><td>Chromosome</td><td>start</td><td>end</td><td>strand</td><td>MM</td><td>target_seq</td><td>PAM</td><td>alignment</td><td>distance</td><td>position</td><td>gene name</td><td>gene id</td></tr>
<tr><td>GU305914.1</td><td>176113</td><td>176135</td><td>-</td><td>0</td><td>AGCAGAAAGAGCGATGCGGT</td><td>AGG</td><td>||||||||[||||||||||||]PAM</td><td>NA</td><td>Intergenic</td><td>NA</td><td>NA</td></tr>
<tr><td></td><td></td><td></td><td></td><td></td><td></td><td></td><td></td><td></td><td></td><td></td><td></td></tr>
<tr><td>T21287</td><td>GCAGAAAGAGCGATGCGGTAGGG</td><td>1000</td><td></td><td></td><td></td><td></td><td></td><td></td><td></td><td></td><td></td></tr>
<tr><td>Oligo fwd</td><td>CACCGCAGAAAGAGCGATGCGGTA</td><td></td><td></td><td></td><td></td><td></td><td></td><td></td><td></td><td></td><td></td></tr>
<tr><td>Oligo rev</td><td>AAACTACCGCATCGCTCTTTCTGC</td><td></td><td></td><td></td><td></td><td></td><td></td><td></td><td></td><td></td><td></td></tr>
<tr><td>Chromosome</td><td>start</td><td>end</td><td>strand</td><td>MM</td><td>target_seq</td><td>PAM</td><td>alignment</td><td>distance</td><td>position</td><td>gene name</td><td>gene id</td></tr>
<tr><td>GU305914.1</td><td>176112</td><td>176134</td><td>-</td><td>0</td><td>GCAGAAAGAGCGATGCGGTA</td><td>GGG</td><td>||||||||[||||||||||||]PAM</td><td>NA</td><td>Intergenic</td><td>NA</td><td>NA</td></tr>
<tr><td></td><td></td><td></td><td></td><td></td><td></td><td></td><td></td><td></td><td></td><td></td><td></td></tr>
<tr><td>T21288</td><td>GAGCGATGCGGTAGGGACATCGG</td><td>1000</td><td></td><td></td><td></td><td></td><td></td><td></td><td></td><td></td><td></td></tr>
<tr><td>Oligo fwd</td><td>CACCGAGCGATGCGGTAGGGACAT</td><td></td><td></td><td></td><td></td><td></td><td></td><td></td><td></td><td></td><td></td></tr>
<tr><td>Oligo rev</td><td>AAACATGTCCCTACCGCATCGCTC</td><td></td><td></td><td></td><td></td><td></td><td></td><td></td><td></td><td></td><td></td></tr>
<tr><td>Chromosome</td><td>start</td><td>end</td><td>strand</td><td>MM</td><td>target_seq</td><td>PAM</td><td>alignment</td><td>distance</td><td>position</td><td>gene name</td><td>gene id</td></tr>
<tr><td>GU305914.1</td><td>176105</td><td>176127</td><td>-</td><td>0</td><td>GAGCGATGCGGTAGGGACAT</td><td>CGG</td><td>||||||||[||||||||||||]PAM</td><td>NA</td><td>Intergenic</td><td>NA</td><td>NA</td></tr>
<tr><td></td><td></td><td></td><td></td><td></td><td></td><td></td><td></td><td></td><td></td><td></td><td></td></tr>
<tr><td>T21289</td><td>CGAAGTCATCCTCGCCACCGAGG</td><td>1000</td><td></td><td></td><td></td><td></td><td></td><td></td><td></td><td></td><td></td></tr>
<tr><td>Oligo fwd</td><td>CACCCGAAGTCATCCTCGCCACCG</td><td></td><td></td><td></td><td></td><td></td><td></td><td></td><td></td><td></td><td></td></tr>
<tr><td>Oligo rev</td><td>AAACCGGTGGCGAGGATGACTTCG</td><td></td><td></td><td></td><td></td><td></td><td></td><td></td><td></td><td></td><td></td></tr>
<tr><td>Chromosome</td><td>start</td><td>end</td><td>strand</td><td>MM</td><td>target_seq</td><td>PAM</td><td>alignment</td><td>distance</td><td>position</td><td>gene name</td><td>gene id</td></tr>
<tr><td>GU305914.1</td><td>176082</td><td>176104</td><td>-</td><td>0</td><td>CGAAGTCATCCTCGCCACCG</td><td>AGG</td><td>||||||||[||||||||||||]PAM</td><td>NA</td><td>Intergenic</td><td>NA</td><td>NA</td></tr>
</table>

<table>
<tr><td>T21290</td><td>CTCGCCACCGAGGTCTCATTCGG</td><td>1000</td><td></td><td></td><td></td><td></td><td></td><td></td><td></td><td></td><td></td></tr>
<tr><td>Oligo fwd</td><td>CACCCTCGCCACCGAGGTCTCATT</td><td></td><td></td><td></td><td></td><td></td><td></td><td></td><td></td><td></td><td></td></tr>
<tr><td>Oligo rev</td><td>AAACAATGAGACCTCGGTGGCGAG</td><td></td><td></td><td></td><td></td><td></td><td></td><td></td><td></td><td></td><td></td></tr>
<tr><td>Chromosome</td><td>start</td><td>end</td><td>strand</td><td>MM</td><td>target_seq</td><td>PAM</td><td>alignment</td><td>distance</td><td>position</td><td>gene name</td><td>gene id</td></tr>
<tr><td>GU305914.1</td><td>176072</td><td>176094</td><td>-</td><td>0</td><td>CTCGCCACCGAGGTCTCATT</td><td>CGG</td><td>||||||||[||||||||||||]PAM</td><td>NA</td><td>Intergenic</td><td>NA</td><td>NA</td></tr>
<tr><td>T21291</td><td>GTCTCATTCGGCGACCGTTCAGG</td><td>1000</td><td></td><td></td><td></td><td></td><td></td><td></td><td></td><td></td><td></td></tr>
<tr><td>Oligo fwd</td><td>CACCGTCTCATTCGGCGACCGTTC</td><td></td><td></td><td></td><td></td><td></td><td></td><td></td><td></td><td></td><td></td></tr>
<tr><td>Oligo rev</td><td>AAACGAACGGTCGCCGAATGAGAC</td><td></td><td></td><td></td><td></td><td></td><td></td><td></td><td></td><td></td><td></td></tr>
<tr><td>Chromosome</td><td>start</td><td>end</td><td>strand</td><td>MM</td><td>target_seq</td><td>PAM</td><td>alignment</td><td>distance</td><td>position</td><td>gene name</td><td>gene id</td></tr>
<tr><td>GU305914.1</td><td>176060</td><td>176082</td><td>-</td><td>0</td><td>GTCTCATTCGGCGACCGTTC</td><td>AGG</td><td>||||||||[||||||||||||]PAM</td><td>NA</td><td>Intergenic</td><td>NA</td><td>NA</td></tr>
<tr><td>T21292</td><td>TCTCACTCGAACACACCTATAGG</td><td>1000</td><td></td><td></td><td></td><td></td><td></td><td></td><td></td><td></td><td></td></tr>
<tr><td>Oligo fwd</td><td>CACCTCTCACTCGAACACACCTAT</td><td></td><td></td><td></td><td></td><td></td><td></td><td></td><td></td><td></td><td></td></tr>
<tr><td>Oligo rev</td><td>AAACATAGGTGTGTTCGAGTGAGA</td><td></td><td></td><td></td><td></td><td></td><td></td><td></td><td></td><td></td><td></td></tr>
<tr><td>Chromosome</td><td>start</td><td>end</td><td>strand</td><td>MM</td><td>target_seq</td><td>PAM</td><td>alignment</td><td>distance</td><td>position</td><td>gene name</td><td>gene id</td></tr>
<tr><td>GU305914.1</td><td>176035</td><td>176057</td><td>-</td><td>0</td><td>TCTCACTCGAACACACCTAT</td><td>AGG</td><td>||||||||[||||||||||||]PAM</td><td>NA</td><td>Intergenic</td><td>NA</td><td>NA</td></tr>
<tr><td>T21293</td><td>CTCACTCGAACACACCTATAGGG</td><td>1000</td><td></td><td></td><td></td><td></td><td></td><td></td><td></td><td></td><td></td></tr>
<tr><td>Oligo fwd</td><td>CACCCTCACTCGAACACACCTATA</td><td></td><td></td><td></td><td></td><td></td><td></td><td></td><td></td><td></td><td></td></tr>
<tr><td>Oligo rev</td><td>AAACTATAGGTGTGTTCGAGTGAG</td><td></td><td></td><td></td><td></td><td></td><td></td><td></td><td></td><td></td><td></td></tr>
<tr><td>Chromosome</td><td>start</td><td>end</td><td>strand</td><td>MM</td><td>target_seq</td><td>PAM</td><td>alignment</td><td>distance</td><td>position</td><td>gene name</td><td>gene id</td></tr>
<tr><td>GU305914.1</td><td>176034</td><td>176056</td><td>-</td><td>0</td><td>CTCACTCGAACACACCTATA</td><td>GGG</td><td>||||||||[||||||||||||]PAM</td><td>NA</td><td>Intergenic</td><td>NA</td><td>NA</td></tr>
<tr><td>T21294</td><td>TCGCACGCAGACTATAAGCGTGG</td><td>1000</td><td></td><td></td><td></td><td></td><td></td><td></td><td></td><td></td><td></td></tr>
<tr><td>Oligo fwd</td><td>CACCTCGCACGCAGACTATAAGCG</td><td></td><td></td><td></td><td></td><td></td><td></td><td></td><td></td><td></td><td></td></tr>
<tr><td>Oligo rev</td><td>AAACCGCTTATAGTCTGCGTGCGA</td><td></td><td></td><td></td><td></td><td></td><td></td><td></td><td></td><td></td><td></td></tr>
<tr><td>Chromosome</td><td>start</td><td>end</td><td>strand</td><td>MM</td><td>target_seq</td><td>PAM</td><td>alignment</td><td>distance</td><td>position</td><td>gene name</td><td>gene id</td></tr>
<tr><td>GU305914.1</td><td>176000</td><td>176022</td><td>-</td><td>0</td><td>TCGCACGCAGACTATAAGCG</td><td>TGG</td><td>||||||||[||||||||||||]PAM</td><td>NA</td><td>Intergenic</td><td>NA</td><td>NA</td></tr>
<tr><td>T21295</td><td>CGTGGTCGACCTTCTCCCCGTGG</td><td>1000</td><td></td><td></td><td></td><td></td><td></td><td></td><td></td><td></td><td></td></tr>
<tr><td>Oligo fwd</td><td>CACCCGTGGTCGACCTTCTCCCCG</td><td></td><td></td><td></td><td></td><td></td><td></td><td></td><td></td><td></td><td></td></tr>
<tr><td>Oligo rev</td><td>AAACCGGGGAGAAGGTCGACCACG</td><td></td><td></td><td></td><td></td><td></td><td></td><td></td><td></td><td></td><td></td></tr>
<tr><td>Chromosome</td><td>start</td><td>end</td><td>strand</td><td>MM</td><td>target_seq</td><td>PAM</td><td>alignment</td><td>distance</td><td>position</td><td>gene name</td><td>gene id</td></tr>
<tr><td>GU305914.1</td><td>175982</td><td>176004</td><td>-</td><td>0</td><td>CGTGGTCGACCTTCTCCCCG</td><td>TGG</td><td>||||||||[||||||||||||]PAM</td><td>NA</td><td>Intergenic</td><td>NA</td><td>NA</td></tr>
<tr><td>T21296</td><td>GACCTTCTCCCCGTGGTCTCAGG</td><td>1000</td><td></td><td></td><td></td><td></td><td></td><td></td><td></td><td></td><td></td></tr>
<tr><td>Oligo fwd</td><td>CACCGACCTTCTCCCCGTGGTCTC</td><td></td><td></td><td></td><td></td><td></td><td></td><td></td><td></td><td></td><td></td></tr>
<tr><td>Oligo rev</td><td>AAACGAGACCACGGGGAGAAGGTC</td><td></td><td></td><td></td><td></td><td></td><td></td><td></td><td></td><td></td><td></td></tr>
<tr><td>Chromosome</td><td>start</td><td>end</td><td>strand</td><td>MM</td><td>target_seq</td><td>PAM</td><td>alignment</td><td>distance</td><td>position</td><td>gene name</td><td>gene id</td></tr>
<tr><td>GU305914.1</td><td>175975</td><td>175997</td><td>-</td><td>0</td><td>GACCTTCTCCCCGTGGTCTC</td><td>AGG</td><td>||||||||[||||||||||||]PAM</td><td>NA</td><td>Intergenic</td><td>NA</td><td>NA</td></tr>
<tr><td>T21297</td><td>CCCCGTGGTCTCAGGTGAACCGG</td><td>1000</td><td></td><td></td><td></td><td></td><td></td><td></td><td></td><td></td><td></td></tr>
<tr><td>Oligo fwd</td><td>CACCCCCCGTGGTCTCAGGTGAAC</td><td></td><td></td><td></td><td></td><td></td><td></td><td></td><td></td><td></td><td></td></tr>
<tr><td>Oligo rev</td><td>AAACGTTCACCTGAGACCACGGGG</td><td></td><td></td><td></td><td></td><td></td><td></td><td></td><td></td><td></td><td></td></tr>
<tr><td>Chromosome</td><td>start</td><td>end</td><td>strand</td><td>MM</td><td>target_seq</td><td>PAM</td><td>alignment</td><td>distance</td><td>position</td><td>gene name</td><td>gene id</td></tr>
<tr><td>GU305914.1</td><td>175967</td><td>175989</td><td>-</td><td>0</td><td>CCCCGTGGTCTCAGGTGAAC</td><td>CGG</td><td>||||||||[||||||||||||]PAM</td><td>NA</td><td>Intergenic</td><td>NA</td><td>NA</td></tr>
<tr><td>T21298</td><td>TACCCGTTGTCTGTCACCGACGG</td><td>1000</td><td></td><td></td><td></td><td></td><td></td><td></td><td></td><td></td><td></td></tr>
<tr><td>Oligo fwd</td><td>CACCTACCCGTTGTCTGTCACCGA</td><td></td><td></td><td></td><td></td><td></td><td></td><td></td><td></td><td></td><td></td></tr>
<tr><td>Oligo rev</td><td>AAACTCGGTGACAGACAACGGGTA</td><td></td><td></td><td></td><td></td><td></td><td></td><td></td><td></td><td></td><td></td></tr>
<tr><td>Chromosome</td><td>start</td><td>end</td><td>strand</td><td>MM</td><td>target_seq</td><td>PAM</td><td>alignment</td><td>distance</td><td>position</td><td>gene name</td><td>gene id</td></tr>
<tr><td>GU305914.1</td><td>175927</td><td>175949</td><td>-</td><td>0</td><td>TACCCGTTGTCTGTCACCGA</td><td>CGG</td><td>||||||||[||||||||||||]PAM</td><td>NA</td><td>Intergenic</td><td>NA</td><td>NA</td></tr>
<tr><td>T21299</td><td>GTTGTCTGTCACCGACGGACAGG</td><td>1000</td><td></td><td></td><td></td><td></td><td></td><td></td><td></td><td></td><td></td></tr>
<tr><td>Oligo fwd</td><td>CACCGTTGTCTGTCACCGACGGAC</td><td></td><td></td><td></td><td></td><td></td><td></td><td></td><td></td><td></td><td></td></tr>
<tr><td>Oligo rev</td><td>AAACGTCCGTCGGTGACAGACAAC</td><td></td><td></td><td></td><td></td><td></td><td></td><td></td><td></td><td></td><td></td></tr>
<tr><td>Chromosome</td><td>start</td><td>end</td><td>strand</td><td>MM</td><td>target_seq</td><td>PAM</td><td>alignment</td><td>distance</td><td>position</td><td>gene name</td><td>gene id</td></tr>
<tr><td>GU305914.1</td><td>175922</td><td>175944</td><td>-</td><td>0</td><td>GTTGTCTGTCACCGACGGAC</td><td>AGG</td><td>||||||||[||||||||||||]PAM</td><td>NA</td><td>Intergenic</td><td>NA</td><td>NA</td></tr>
<tr><td>T21300</td><td>CGGACAGGTAGTAACGCCTCAGG</td><td>1000</td><td></td><td></td><td></td><td></td><td></td><td></td><td></td><td></td><td></td></tr>
<tr><td>Oligo fwd</td><td>CACCCGGACAGGTAGTAACGCCTC</td><td></td><td></td><td></td><td></td><td></td><td></td><td></td><td></td><td></td><td></td></tr>
<tr><td>Oligo rev</td><td>AAACGAGGCGTTACTACCTGTCCG</td><td></td><td></td><td></td><td></td><td></td><td></td><td></td><td></td><td></td><td></td></tr>
<tr><td>Chromosome</td><td>start</td><td>end</td><td>strand</td><td>MM</td><td>target_seq</td><td>PAM</td><td>alignment</td><td>distance</td><td>position</td><td>gene name</td><td>gene id</td></tr>
<tr><td>GU305914.1</td><td>175907</td><td>175929</td><td>-</td><td>0</td><td>CGGACAGGTAGTAACGCCTC</td><td>AGG</td><td>||||||||[||||||||||||]PAM</td><td>NA</td><td>Intergenic</td><td>NA</td><td>NA</td></tr>
<tr><td>T21301</td><td>ACAGGTAGTAACGCCTCAGGTGG</td><td>1000</td><td></td><td></td><td></td><td></td><td></td><td></td><td></td><td></td><td></td></tr>
<tr><td>Oligo fwd</td><td>CACCACAGGTAGTAACGCCTCAGG</td><td></td><td></td><td></td><td></td><td></td><td></td><td></td><td></td><td></td><td></td></tr>
<tr><td>Oligo rev</td><td>AAACCCTGAGGCGTTACTACCTGT</td><td></td><td></td><td></td><td></td><td></td><td></td><td></td><td></td><td></td><td></td></tr>
<tr><td>Chromosome</td><td>start</td><td>end</td><td>strand</td><td>MM</td><td>target_seq</td><td>PAM</td><td>alignment</td><td>distance</td><td>position</td><td>gene name</td><td>gene id</td></tr>
<tr><td>GU305914.1</td><td>175904</td><td>175926</td><td>-</td><td>0</td><td>ACAGGTAGTAACGCCTCAGG</td><td>TGG</td><td>||||||||[||||||||||||]PAM</td><td>NA</td><td>Intergenic</td><td>NA</td><td>NA</td></tr>
<tr><td>T21302</td><td>CAGGTAGTAACGCCTCAGGTGGG</td><td>1000</td><td></td><td></td><td></td><td></td><td></td><td></td><td></td><td></td><td></td></tr>
<tr><td>Oligo fwd</td><td>CACCCAGGTAGTAACGCCTCAGGT</td><td></td><td></td><td></td><td></td><td></td><td></td><td></td><td></td><td></td><td></td></tr>
<tr><td>Oligo rev</td><td>AAACACCTGAGGCGTTACTACCTG</td><td></td><td></td><td></td><td></td><td></td><td></td><td></td><td></td><td></td><td></td></tr>
<tr><td>Chromosome</td><td>start</td><td>end</td><td>strand</td><td>MM</td><td>target_seq</td><td>PAM</td><td>alignment</td><td>distance</td><td>position</td><td>gene name</td><td>gene id</td></tr>
<tr><td>GU305914.1</td><td>175903</td><td>175925</td><td>-</td><td>0</td><td>CAGGTAGTAACGCCTCAGGT</td><td>GGG</td><td>||||||||[||||||||||||]PAM</td><td>NA</td><td>Intergenic</td><td>NA</td><td>NA</td></tr>
<tr><td>T21303</td><td>AGGTAGTAACGCCTCAGGTGGGG</td><td>1000</td><td></td><td></td><td></td><td></td><td></td><td></td><td></td><td></td><td></td></tr>
<tr><td>Oligo fwd</td><td>CACCAGGTAGTAACGCCTCAGGTG</td><td></td><td></td><td></td><td></td><td></td><td></td><td></td><td></td><td></td><td></td></tr>
<tr><td>Oligo rev</td><td>AAACCACCTGAGGCGTTACTACCT</td><td></td><td></td><td></td><td></td><td></td><td></td><td></td><td></td><td></td><td></td></tr>
<tr><td>Chromosome</td><td>start</td><td>end</td><td>strand</td><td>MM</td><td>target_seq</td><td>PAM</td><td>alignment</td><td>distance</td><td>position</td><td>gene name</td><td>gene id</td></tr>
<tr><td>GU305914.1</td><td>175902</td><td>175924</td><td>-</td><td>0</td><td>AGGTAGTAACGCCTCAGGTG</td><td>GGG</td><td>||||||||[||||||||||||]PAM</td><td>NA</td><td>Intergenic</td><td>NA</td><td>NA</td></tr>
<tr><td>T21304</td><td>TAACGCCTCAGGTGGGGACGTGG</td><td>1000</td><td></td><td></td><td></td><td></td><td></td><td></td><td></td><td></td><td></td></tr>
<tr><td>Oligo fwd</td><td>CACCTAACGCCTCAGGTGGGGACG</td><td></td><td></td><td></td><td></td><td></td><td></td><td></td><td></td><td></td><td></td></tr>
<tr><td>Oligo rev</td><td>AAACCGTCCCCACCTGAGGCGTTA</td><td></td><td></td><td></td><td></td><td></td><td></td><td></td><td></td><td></td><td></td></tr>
<tr><td>Chromosome</td><td>start</td><td>end</td><td>strand</td><td>MM</td><td>target_seq</td><td>PAM</td><td>alignment</td><td>distance</td><td>position</td><td>gene name</td><td>gene id</td></tr>
<tr><td>GU305914.1</td><td>175896</td><td>175918</td><td>-</td><td>0</td><td>TAACGCCTCAGGTGGGGACG</td><td>TGG</td><td>||||||||[||||||||||||]PAM</td><td>NA</td><td>Intergenic</td><td>NA</td><td>NA</td></tr>
</table>

<table>
<tr><td>T21305</td><td>GTTTGTGACGCACCCACTCCAGG</td><td>1000</td><td></td><td></td><td></td><td></td><td></td><td></td><td></td><td></td><td></td></tr>
<tr><td>Oligo fwd</td><td>CACCGTTTGTGACGCACCCACTCC</td><td></td><td></td><td></td><td></td><td></td><td></td><td></td><td></td><td></td><td></td></tr>
<tr><td>Oligo rev</td><td>AAACGGAGTGGGTGCGTCACAAAC</td><td></td><td></td><td></td><td></td><td></td><td></td><td></td><td></td><td></td><td></td></tr>
<tr><td>Chromosome</td><td>start</td><td>end</td><td>strand</td><td>MM</td><td>target_seq</td><td>PAM</td><td>alignment</td><td>distance</td><td>position</td><td>gene name</td><td>gene id</td></tr>
<tr><td>GU305914.1</td><td>175874</td><td>175896</td><td>-</td><td>0</td><td>GTTTGTGACGCACCCACTCC</td><td>AGG</td><td>||||||||[||||||||||||]PAM</td><td>NA</td><td>Intergenic</td><td>NA</td><td>NA</td></tr>
<tr><td>T21306</td><td>GTGCTATTAATTAGCCTCGCAGG</td><td>1000</td><td></td><td></td><td></td><td></td><td></td><td></td><td></td><td></td><td></td></tr>
<tr><td>Oligo fwd</td><td>CACCGTGCTATTAATTAGCCTCGC</td><td></td><td></td><td></td><td></td><td></td><td></td><td></td><td></td><td></td><td></td></tr>
<tr><td>Oligo rev</td><td>AAACGCGAGGCTAATTAATAGCAC</td><td></td><td></td><td></td><td></td><td></td><td></td><td></td><td></td><td></td><td></td></tr>
<tr><td>Chromosome</td><td>start</td><td>end</td><td>strand</td><td>MM</td><td>target_seq</td><td>PAM</td><td>alignment</td><td>distance</td><td>position</td><td>gene name</td><td>gene id</td></tr>
<tr><td>GU305914.1</td><td>175852</td><td>175874</td><td>-</td><td>0</td><td>GTGCTATTAATTAGCCTCGC</td><td>AGG</td><td>||||||||[||||||||||||]PAM</td><td>NA</td><td>Intergenic</td><td>NA</td><td>NA</td></tr>
<tr><td>T21307</td><td>ATTAATTAGCCTCGCAGGTGCGG</td><td>1000</td><td></td><td></td><td></td><td></td><td></td><td></td><td></td><td></td><td></td></tr>
<tr><td>Oligo fwd</td><td>CACCATTAATTAGCCTCGCAGGTG</td><td></td><td></td><td></td><td></td><td></td><td></td><td></td><td></td><td></td><td></td></tr>
<tr><td>Oligo rev</td><td>AAACCACCTGCGAGGCTAATTAAT</td><td></td><td></td><td></td><td></td><td></td><td></td><td></td><td></td><td></td><td></td></tr>
<tr><td>Chromosome</td><td>start</td><td>end</td><td>strand</td><td>MM</td><td>target_seq</td><td>PAM</td><td>alignment</td><td>distance</td><td>position</td><td>gene name</td><td>gene id</td></tr>
<tr><td>GU305914.1</td><td>175847</td><td>175869</td><td>-</td><td>0</td><td>ATTAATTAGCCTCGCAGGTG</td><td>CGG</td><td>||||||||[||||||||||||]PAM</td><td>NA</td><td>Intergenic</td><td>NA</td><td>NA</td></tr>
<tr><td>T21308</td><td>CCTCGCAGGTGCGGTGTTGTTGG</td><td>1000</td><td></td><td></td><td></td><td></td><td></td><td></td><td></td><td></td><td></td></tr>
<tr><td>Oligo fwd</td><td>CACCCCTCGCAGGTGCGGTGTTGT</td><td></td><td></td><td></td><td></td><td></td><td></td><td></td><td></td><td></td><td></td></tr>
<tr><td>Oligo rev</td><td>AAACACAACACCGCACCTGCGAGG</td><td></td><td></td><td></td><td></td><td></td><td></td><td></td><td></td><td></td><td></td></tr>
<tr><td>Chromosome</td><td>start</td><td>end</td><td>strand</td><td>MM</td><td>target_seq</td><td>PAM</td><td>alignment</td><td>distance</td><td>position</td><td>gene name</td><td>gene id</td></tr>
<tr><td>GU305914.1</td><td>175838</td><td>175860</td><td>-</td><td>0</td><td>CCTCGCAGGTGCGGTGTTGT</td><td>TGG</td><td>||||||||[||||||||||||]PAM</td><td>NA</td><td>Intergenic</td><td>NA</td><td>NA</td></tr>
<tr><td>T21309</td><td>CAGGTGCGGTGTTGTTGGTGTGG</td><td>1000</td><td></td><td></td><td></td><td></td><td></td><td></td><td></td><td></td><td></td></tr>
<tr><td>Oligo fwd</td><td>CACCCAGGTGCGGTGTTGTTGGTG</td><td></td><td></td><td></td><td></td><td></td><td></td><td></td><td></td><td></td><td></td></tr>
<tr><td>Oligo rev</td><td>AAACCACCAACAACACCGCACCTG</td><td></td><td></td><td></td><td></td><td></td><td></td><td></td><td></td><td></td><td></td></tr>
<tr><td>Chromosome</td><td>start</td><td>end</td><td>strand</td><td>MM</td><td>target_seq</td><td>PAM</td><td>alignment</td><td>distance</td><td>position</td><td>gene name</td><td>gene id</td></tr>
<tr><td>GU305914.1</td><td>175833</td><td>175855</td><td>-</td><td>0</td><td>CAGGTGCGGTGTTGTTGGTG</td><td>TGG</td><td>||||||||[||||||||||||]PAM</td><td>NA</td><td>Intergenic</td><td>NA</td><td>NA</td></tr>
<tr><td>T21310</td><td>GGTGTGGATACTGACATCGTTGG</td><td>1000</td><td></td><td></td><td></td><td></td><td></td><td></td><td></td><td></td><td></td></tr>
<tr><td>Oligo fwd</td><td>CACCGGTGTGGATACTGACATCGT</td><td></td><td></td><td></td><td></td><td></td><td></td><td></td><td></td><td></td><td></td></tr>
<tr><td>Oligo rev</td><td>AAACACGATGTCAGTATCCACACC</td><td></td><td></td><td></td><td></td><td></td><td></td><td></td><td></td><td></td><td></td></tr>
<tr><td>Chromosome</td><td>start</td><td>end</td><td>strand</td><td>MM</td><td>target_seq</td><td>PAM</td><td>alignment</td><td>distance</td><td>position</td><td>gene name</td><td>gene id</td></tr>
<tr><td>GU305914.1</td><td>175817</td><td>175839</td><td>-</td><td>0</td><td>GGTGTGGATACTGACATCGT</td><td>TGG</td><td>||||||||[||||||||||||]PAM</td><td>NA</td><td>Intergenic</td><td>NA</td><td>NA</td></tr>
<tr><td>T21311</td><td>GTGGATACTGACATCGTTGGCGG</td><td>1000</td><td></td><td></td><td></td><td></td><td></td><td></td><td></td><td></td><td></td></tr>
<tr><td>Oligo fwd</td><td>CACCGTGGATACTGACATCGTTGG</td><td></td><td></td><td></td><td></td><td></td><td></td><td></td><td></td><td></td><td></td></tr>
<tr><td>Oligo rev</td><td>AAACCCAACGATGTCAGTATCCAC</td><td></td><td></td><td></td><td></td><td></td><td></td><td></td><td></td><td></td><td></td></tr>
<tr><td>Chromosome</td><td>start</td><td>end</td><td>strand</td><td>MM</td><td>target_seq</td><td>PAM</td><td>alignment</td><td>distance</td><td>position</td><td>gene name</td><td>gene id</td></tr>
<tr><td>GU305914.1</td><td>175814</td><td>175836</td><td>-</td><td>0</td><td>GTGGATACTGACATCGTTGG</td><td>CGG</td><td>||||||||[||||||||||||]PAM</td><td>NA</td><td>Intergenic</td><td>NA</td><td>NA</td></tr>
<tr><td>T21312</td><td>CGTTGGCGGCGTTACTCTTTCGG</td><td>1000</td><td></td><td></td><td></td><td></td><td></td><td></td><td></td><td></td><td></td></tr>
<tr><td>Oligo fwd</td><td>CACCCGTTGGCGGCGTTACTCTTT</td><td></td><td></td><td></td><td></td><td></td><td></td><td></td><td></td><td></td><td></td></tr>
<tr><td>Oligo rev</td><td>AAACAAAGAGTAACGCCGCCAACG</td><td></td><td></td><td></td><td></td><td></td><td></td><td></td><td></td><td></td><td></td></tr>
<tr><td>Chromosome</td><td>start</td><td>end</td><td>strand</td><td>MM</td><td>target_seq</td><td>PAM</td><td>alignment</td><td>distance</td><td>position</td><td>gene name</td><td>gene id</td></tr>
<tr><td>GU305914.1</td><td>175800</td><td>175822</td><td>-</td><td>0</td><td>CGTTGGCGGCGTTACTCTTT</td><td>CGG</td><td>||||||||[||||||||||||]PAM</td><td>NA</td><td>Intergenic</td><td>NA</td><td>NA</td></tr>
<tr><td>T21313</td><td>TGGCGGCGTTACTCTTTCGGCGG</td><td>1000</td><td></td><td></td><td></td><td></td><td></td><td></td><td></td><td></td><td></td></tr>
<tr><td>Oligo fwd</td><td>CACCTGGCGGCGTTACTCTTTCGG</td><td></td><td></td><td></td><td></td><td></td><td></td><td></td><td></td><td></td><td></td></tr>
<tr><td>Oligo rev</td><td>AAACCCGAAAGAGTAACGCCGCCA</td><td></td><td></td><td></td><td></td><td></td><td></td><td></td><td></td><td></td><td></td></tr>
<tr><td>Chromosome</td><td>start</td><td>end</td><td>strand</td><td>MM</td><td>target_seq</td><td>PAM</td><td>alignment</td><td>distance</td><td>position</td><td>gene name</td><td>gene id</td></tr>
<tr><td>GU305914.1</td><td>175797</td><td>175819</td><td>-</td><td>0</td><td>TGGCGGCGTTACTCTTTCGG</td><td>CGG</td><td>||||||||[||||||||||||]PAM</td><td>NA</td><td>Intergenic</td><td>NA</td><td>NA</td></tr>
<tr><td>T21314</td><td>GGACGCGTTCTGTCTACTGACGG</td><td>1000</td><td></td><td></td><td></td><td></td><td></td><td></td><td></td><td></td><td></td></tr>
<tr><td>Oligo fwd</td><td>CACCGGACGCGTTCTGTCTACTGA</td><td></td><td></td><td></td><td></td><td></td><td></td><td></td><td></td><td></td><td></td></tr>
<tr><td>Oligo rev</td><td>AAACTCAGTAGACAGAACGCGTCC</td><td></td><td></td><td></td><td></td><td></td><td></td><td></td><td></td><td></td><td></td></tr>
<tr><td>Chromosome</td><td>start</td><td>end</td><td>strand</td><td>MM</td><td>target_seq</td><td>PAM</td><td>alignment</td><td>distance</td><td>position</td><td>gene name</td><td>gene id</td></tr>
<tr><td>GU305914.1</td><td>175776</td><td>175798</td><td>-</td><td>0</td><td>GGACGCGTTCTGTCTACTGA</td><td>CGG</td><td>||||||||[||||||||||||]PAM</td><td>NA</td><td>Intergenic</td><td>NA</td><td>NA</td></tr>
<tr><td>T21315</td><td>TTCTGTCTACTGACGGTCCGTGG</td><td>1000</td><td></td><td></td><td></td><td></td><td></td><td></td><td></td><td></td><td></td></tr>
<tr><td>Oligo fwd</td><td>CACCTTCTGTCTACTGACGGTCCG</td><td></td><td></td><td></td><td></td><td></td><td></td><td></td><td></td><td></td><td></td></tr>
<tr><td>Oligo rev</td><td>AAACCGGACCGTCAGTAGACAGAA</td><td></td><td></td><td></td><td></td><td></td><td></td><td></td><td></td><td></td><td></td></tr>
<tr><td>Chromosome</td><td>start</td><td>end</td><td>strand</td><td>MM</td><td>target_seq</td><td>PAM</td><td>alignment</td><td>distance</td><td>position</td><td>gene name</td><td>gene id</td></tr>
<tr><td>GU305914.1</td><td>175769</td><td>175791</td><td>-</td><td>0</td><td>TTCTGTCTACTGACGGTCCG</td><td>TGG</td><td>||||||||[||||||||||||]PAM</td><td>NA</td><td>Intergenic</td><td>NA</td><td>NA</td></tr>
<tr><td>T21316</td><td>TACTGACGGTCCGTGGACGATGG</td><td>1000</td><td></td><td></td><td></td><td></td><td></td><td></td><td></td><td></td><td></td></tr>
<tr><td>Oligo fwd</td><td>CACCTACTGACGGTCCGTGGACGA</td><td></td><td></td><td></td><td></td><td></td><td></td><td></td><td></td><td></td><td></td></tr>
<tr><td>Oligo rev</td><td>AAACTCGTCCACGGACCGTCAGTA</td><td></td><td></td><td></td><td></td><td></td><td></td><td></td><td></td><td></td><td></td></tr>
<tr><td>Chromosome</td><td>start</td><td>end</td><td>strand</td><td>MM</td><td>target_seq</td><td>PAM</td><td>alignment</td><td>distance</td><td>position</td><td>gene name</td><td>gene id</td></tr>
<tr><td>GU305914.1</td><td>175762</td><td>175784</td><td>-</td><td>0</td><td>TACTGACGGTCCGTGGACGA</td><td>TGG</td><td>||||||||[||||||||||||]PAM</td><td>NA</td><td>Intergenic</td><td>NA</td><td>NA</td></tr>
<tr><td>T21317</td><td>ACTGACGGTCCGTGGACGATGGG</td><td>1000</td><td></td><td></td><td></td><td></td><td></td><td></td><td></td><td></td><td></td></tr>
<tr><td>Oligo fwd</td><td>CACCACTGACGGTCCGTGGACGAT</td><td></td><td></td><td></td><td></td><td></td><td></td><td></td><td></td><td></td><td></td></tr>
<tr><td>Oligo rev</td><td>AAACATCGTCCACGGACCGTCAGT</td><td></td><td></td><td></td><td></td><td></td><td></td><td></td><td></td><td></td><td></td></tr>
<tr><td>Chromosome</td><td>start</td><td>end</td><td>strand</td><td>MM</td><td>target_seq</td><td>PAM</td><td>alignment</td><td>distance</td><td>position</td><td>gene name</td><td>gene id</td></tr>
<tr><td>GU305914.1</td><td>175761</td><td>175783</td><td>-</td><td>0</td><td>ACTGACGGTCCGTGGACGAT</td><td>GGG</td><td>||||||||[||||||||||||]PAM</td><td>NA</td><td>Intergenic</td><td>NA</td><td>NA</td></tr>
<tr><td>T21318</td><td>CTGACGGTCCGTGGACGATGGGG</td><td>1000</td><td></td><td></td><td></td><td></td><td></td><td></td><td></td><td></td><td></td></tr>
<tr><td>Oligo fwd</td><td>CACCCTGACGGTCCGTGGACGATG</td><td></td><td></td><td></td><td></td><td></td><td></td><td></td><td></td><td></td><td></td></tr>
<tr><td>Oligo rev</td><td>AAACCATCGTCCACGGACCGTCAG</td><td></td><td></td><td></td><td></td><td></td><td></td><td></td><td></td><td></td><td></td></tr>
<tr><td>Chromosome</td><td>start</td><td>end</td><td>strand</td><td>MM</td><td>target_seq</td><td>PAM</td><td>alignment</td><td>distance</td><td>position</td><td>gene name</td><td>gene id</td></tr>
<tr><td>GU305914.1</td><td>175760</td><td>175782</td><td>-</td><td>0</td><td>CTGACGGTCCGTGGACGATG</td><td>GGG</td><td>||||||||[||||||||||||]PAM</td><td>NA</td><td>Intergenic</td><td>NA</td><td>NA</td></tr>
<tr><td>T21319</td><td>GCGTGCAATGTAAGATCGACTGG</td><td>1000</td><td></td><td></td><td></td><td></td><td></td><td></td><td></td><td></td><td></td></tr>
<tr><td>Oligo fwd</td><td>CACCGCGTGCAATGTAAGATCGAC</td><td></td><td></td><td></td><td></td><td></td><td></td><td></td><td></td><td></td><td></td></tr>
<tr><td>Oligo rev</td><td>AAACGTCGATCTTACATTGCACGC</td><td></td><td></td><td></td><td></td><td></td><td></td><td></td><td></td><td></td><td></td></tr>
<tr><td>Chromosome</td><td>start</td><td>end</td><td>strand</td><td>MM</td><td>target_seq</td><td>PAM</td><td>alignment</td><td>distance</td><td>position</td><td>gene name</td><td>gene id</td></tr>
</table>

GU305914.1 175682 175704 - 0 GCGTGCAATGTAAGATCGAC TGG ||||||||[||||||||||||]PAM NA Intergenic NA NA

T21320 ACTGGAAAGTGACCTCGACCTGG 1000
Oligo fwd CACCACTGGAAAGTGACCTCGACC
Oligo rev AAACGGTCGAGGTCACTTTCCAGT
Chromosome start end strand MM target_seq PAM alignment distance position gene name gene id
GU305914.1 175664 175686 - 0 ACTGGAAAGTGACCTCGACC TGG ||||||||[||||||||||||]PAM NA Intergenic NA NA

T21321 ACGACTAACATGACCTGTATAGG 1000
Oligo fwd CACCACGACTAACATGACCTGTAT
Oligo rev AAACATACAGGTCATGTTAGTCGT
Chromosome start end strand MM target_seq PAM alignment distance position gene name gene id
GU305914.1 175638 175660 - 0 ACGACTAACATGACCTGTAT AGG ||||||||[||||||||||||]PAM NA Intergenic NA NA

T21322 AACATGACCTGTATAGGCCACGG 1000
Oligo fwd CACCAACATGACCTGTATAGGCCA
Oligo rev AAACTGGCCTATACAGGTCATGTT
Chromosome start end strand MM target_seq PAM alignment distance position gene name gene id
GU305914.1 175632 175654 - 0 AACATGACCTGTATAGGCCA CGG ||||||||[||||||||||||]PAM NA Intergenic NA NA

T21323 TGTATAGGCCACGGTGATACAGG 1000
Oligo fwd CACCTGTATAGGCCACGGTGATAC
Oligo rev AAACGTATCACCGTGGCCTATACA
Chromosome start end strand MM target_seq PAM alignment distance position gene name gene id
GU305914.1 175623 175645 - 0 TGTATAGGCCACGGTGATAC AGG ||||||||[||||||||||||]PAM NA Intergenic NA NA

T21324 GCCACGGTGATACAGGATCGTGG 1000
Oligo fwd CACCGCCACGGTGATACAGGATCG
Oligo rev AAACCGATCCTGTATCACCGTGGC
Chromosome start end strand MM target_seq PAM alignment distance position gene name gene id
GU305914.1 175616 175638 - 0 GCCACGGTGATACAGGATCG TGG ||||||||[||||||||||||]PAM NA Intergenic NA NA

T21325 AACAAGACAGACTATCTGATCGG 1000
Oligo fwd CACCAACAAGACAGACTATCTGAT
Oligo rev AAACATCAGATAGTCTGTCTTGTT
Chromosome start end strand MM target_seq PAM alignment distance position gene name gene id
GU305914.1 175581 175603 - 0 AACAAGACAGACTATCTGAT CGG ||||||||[||||||||||||]PAM NA Intergenic NA NA

T21326 ACAAGACAGACTATCTGATCGGG 1000
Oligo fwd CACCACAAGACAGACTATCTGATC
Oligo rev AAACGATCAGATAGTCTGTCTTGT
Chromosome start end strand MM target_seq PAM alignment distance position gene name gene id
GU305914.1 175580 175602 - 0 ACAAGACAGACTATCTGATC GGG ||||||||[||||||||||||]PAM NA Intergenic NA NA

T21327 TCGAAACACGACCGTCGTCATGG 1000
Oligo fwd CACCTCGAAACACGACCGTCGTCA
Oligo rev AAACTGACGACGGTCGTGTTTCGA
Chromosome start end strand MM target_seq PAM alignment distance position gene name gene id
GU305914.1 175552 175574 - 0 TCGAAACACGACCGTCGTCA TGG ||||||||[||||||||||||]PAM NA Intergenic NA NA

T21328 ACCGTCGTCATGGAACCTTACGG 1000
Oligo fwd CACCACCGTCGTCATGGAACCTTA
Oligo rev AAACTAAGGTTCCATGACGACGGT
Chromosome start end strand MM target_seq PAM alignment distance position gene name gene id
GU305914.1 175542 175564 - 0 ACCGTCGTCATGGAACCTTA CGG ||||||||[||||||||||||]PAM NA Intergenic NA NA

T21329 ATGGAACCTTACGGTCTCGACGG 1000
Oligo fwd CACCATGGAACCTTACGGTCTCGA
Oligo rev AAACTCGAGACCGTAAGGTTCCAT
Chromosome start end strand MM target_seq PAM alignment distance position gene name gene id
GU305914.1 175533 175555 - 0 ATGGAACCTTACGGTCTCGA CGG ||||||||[||||||||||||]PAM NA Intergenic NA NA

T21330 GAACCTTACGGTCTCGACGGAGG 1000
Oligo fwd CACCGAACCTTACGGTCTCGACGG
Oligo rev AAACCCGTCGAGACCGTAAGGTTC
Chromosome start end strand MM target_seq PAM alignment distance position gene name gene id
GU305914.1 175530 175552 - 0 GAACCTTACGGTCTCGACGG AGG ||||||||[||||||||||||]PAM NA Intergenic NA NA

T21331 TACGGTCTCGACGGAGGATACGG 1000
Oligo fwd CACCTACGGTCTCGACGGAGGATA
Oligo rev AAACTATCCTCCGTCGAGACCGTA
Chromosome start end strand MM target_seq PAM alignment distance position gene name gene id
GU305914.1 175524 175546 - 0 TACGGTCTCGACGGAGGATA CGG ||||||||[||||||||||||]PAM NA Intergenic NA NA

T21332 GTATGTGAAAGTATCGAAATCGG 1000
Oligo fwd CACCGTATGTGAAAGTATCGAAAT
Oligo rev AAACATTTCGATACTTTCACATAC
Chromosome start end strand MM target_seq PAM alignment distance position gene name gene id
GU305914.1 175498 175520 - 0 GTATGTGAAAGTATCGAAAT CGG ||||||||[||||||||||||]PAM NA Intergenic NA NA

T21333 TATGTGAAAGTATCGAAATCGGG 1000
Oligo fwd CACCTATGTGAAAGTATCGAAATC
Oligo rev AAACGATTTCGATACTTTCACATA
Chromosome start end strand MM target_seq PAM alignment distance position gene name gene id
GU305914.1 175497 175519 - 0 TATGTGAAAGTATCGAAATC GGG ||||||||[||||||||||||]PAM NA Intergenic NA NA

T21334 ACTATGAACACCACACGACACGG 1000
Oligo fwd CACCACTATGAACACCACACGACA
Oligo rev AAACTGTCGTGTGGTGTTCATAGT

<table>
<tr><td>Chromosome</td><td>start</td><td>end</td><td>strand</td><td>MM</td><td>target_seq</td><td>PAM</td><td>alignment</td><td>distance</td><td>position</td><td>gene name</td><td>gene id</td></tr>
<tr><td>GU305914.1</td><td>175455</td><td>175477</td><td>-</td><td>0</td><td>ACTATGAACACCACACGACA</td><td>CGG</td><td>||||||||[||||||||||||]PAM</td><td>NA</td><td>Intergenic</td><td>NA</td><td>NA</td></tr>
<tr><td></td><td></td><td></td><td></td><td></td><td></td><td></td><td></td><td></td><td></td><td></td><td></td></tr>
<tr><td>T21335</td><td>CTATGAACACCACACGACACGGG</td><td>1000</td><td></td><td></td><td></td><td></td><td></td><td></td><td></td><td></td><td></td></tr>
<tr><td>Oligo fwd</td><td>CACCCTATGAACACCACACGACAC</td><td></td><td></td><td></td><td></td><td></td><td></td><td></td><td></td><td></td><td></td></tr>
<tr><td>Oligo rev</td><td>AAACGTGTCGTGTGGTGTTCATAG</td><td></td><td></td><td></td><td></td><td></td><td></td><td></td><td></td><td></td><td></td></tr>
<tr><td>Chromosome</td><td>start</td><td>end</td><td>strand</td><td>MM</td><td>target_seq</td><td>PAM</td><td>alignment</td><td>distance</td><td>position</td><td>gene name</td><td>gene id</td></tr>
<tr><td>GU305914.1</td><td>175454</td><td>175476</td><td>-</td><td>0</td><td>CTATGAACACCACACGACAC</td><td>GGG</td><td>||||||||[||||||||||||]PAM</td><td>NA</td><td>Intergenic</td><td>NA</td><td>NA</td></tr>
<tr><td></td><td></td><td></td><td></td><td></td><td></td><td></td><td></td><td></td><td></td><td></td><td></td></tr>
<tr><td>T21336</td><td>TATGAACACCACACGACACGGGG</td><td>1000</td><td></td><td></td><td></td><td></td><td></td><td></td><td></td><td></td><td></td></tr>
<tr><td>Oligo fwd</td><td>CACCTATGAACACCACACGACACG</td><td></td><td></td><td></td><td></td><td></td><td></td><td></td><td></td><td></td><td></td></tr>
<tr><td>Oligo rev</td><td>AAACCGTGTCGTGTGGTGTTCATA</td><td></td><td></td><td></td><td></td><td></td><td></td><td></td><td></td><td></td><td></td></tr>
<tr><td>Chromosome</td><td>start</td><td>end</td><td>strand</td><td>MM</td><td>target_seq</td><td>PAM</td><td>alignment</td><td>distance</td><td>position</td><td>gene name</td><td>gene id</td></tr>
<tr><td>GU305914.1</td><td>175453</td><td>175475</td><td>-</td><td>0</td><td>TATGAACACCACACGACACG</td><td>GGG</td><td>||||||||[||||||||||||]PAM</td><td>NA</td><td>Intergenic</td><td>NA</td><td>NA</td></tr>
<tr><td></td><td></td><td></td><td></td><td></td><td></td><td></td><td></td><td></td><td></td><td></td><td></td></tr>
<tr><td>T21337</td><td>ATGAACACCACACGACACGGGGG</td><td>1000</td><td></td><td></td><td></td><td></td><td></td><td></td><td></td><td></td><td></td></tr>
<tr><td>Oligo fwd</td><td>CACCATGAACACCACACGACACGG</td><td></td><td></td><td></td><td></td><td></td><td></td><td></td><td></td><td></td><td></td></tr>
<tr><td>Oligo rev</td><td>AAACCCGTGTCGTGTGGTGTTCAT</td><td></td><td></td><td></td><td></td><td></td><td></td><td></td><td></td><td></td><td></td></tr>
<tr><td>Chromosome</td><td>start</td><td>end</td><td>strand</td><td>MM</td><td>target_seq</td><td>PAM</td><td>alignment</td><td>distance</td><td>position</td><td>gene name</td><td>gene id</td></tr>
<tr><td>GU305914.1</td><td>175452</td><td>175474</td><td>-</td><td>0</td><td>ATGAACACCACACGACACGG</td><td>GGG</td><td>||||||||[||||||||||||]PAM</td><td>NA</td><td>Intergenic</td><td>NA</td><td>NA</td></tr>
<tr><td></td><td></td><td></td><td></td><td></td><td></td><td></td><td></td><td></td><td></td><td></td><td></td></tr>
<tr><td>T21338</td><td>GACACGGGGGTTACAAGTGTAGG</td><td>1000</td><td></td><td></td><td></td><td></td><td></td><td></td><td></td><td></td><td></td></tr>
<tr><td>Oligo fwd</td><td>CACCGACACGGGGGTTACAAGTGT</td><td></td><td></td><td></td><td></td><td></td><td></td><td></td><td></td><td></td><td></td></tr>
<tr><td>Oligo rev</td><td>AAACACACTTGTAACCCCCGTGTC</td><td></td><td></td><td></td><td></td><td></td><td></td><td></td><td></td><td></td><td></td></tr>
<tr><td>Chromosome</td><td>start</td><td>end</td><td>strand</td><td>MM</td><td>target_seq</td><td>PAM</td><td>alignment</td><td>distance</td><td>position</td><td>gene name</td><td>gene id</td></tr>
<tr><td>GU305914.1</td><td>175439</td><td>175461</td><td>-</td><td>0</td><td>GACACGGGGGTTACAAGTGT</td><td>AGG</td><td>||||||||[||||||||||||]PAM</td><td>NA</td><td>Intergenic</td><td>NA</td><td>NA</td></tr>
<tr><td></td><td></td><td></td><td></td><td></td><td></td><td></td><td></td><td></td><td></td><td></td><td></td></tr>
<tr><td>T21339</td><td>GGGGGTTACAAGTGTAGGATAGG</td><td>1000</td><td></td><td></td><td></td><td></td><td></td><td></td><td></td><td></td><td></td></tr>
<tr><td>Oligo fwd</td><td>CACCGGGGGTTACAAGTGTAGGAT</td><td></td><td></td><td></td><td></td><td></td><td></td><td></td><td></td><td></td><td></td></tr>
<tr><td>Oligo rev</td><td>AAACATCCTACACTTGTAACCCCC</td><td></td><td></td><td></td><td></td><td></td><td></td><td></td><td></td><td></td><td></td></tr>
<tr><td>Chromosome</td><td>start</td><td>end</td><td>strand</td><td>MM</td><td>target_seq</td><td>PAM</td><td>alignment</td><td>distance</td><td>position</td><td>gene name</td><td>gene id</td></tr>
<tr><td>GU305914.1</td><td>175434</td><td>175456</td><td>-</td><td>0</td><td>GGGGGTTACAAGTGTAGGAT</td><td>AGG</td><td>||||||||[||||||||||||]PAM</td><td>NA</td><td>Intergenic</td><td>NA</td><td>NA</td></tr>
<tr><td></td><td></td><td></td><td></td><td></td><td></td><td></td><td></td><td></td><td></td><td></td><td></td></tr>
<tr><td>T21340</td><td>TCTGACAGTGAGTTCAACGACGG</td><td>1000</td><td></td><td></td><td></td><td></td><td></td><td></td><td></td><td></td><td></td></tr>
<tr><td>Oligo fwd</td><td>CACCTCTGACAGTGAGTTCAACGA</td><td></td><td></td><td></td><td></td><td></td><td></td><td></td><td></td><td></td><td></td></tr>
<tr><td>Oligo rev</td><td>AAACTCGTTGAACTCACTGTCAGA</td><td></td><td></td><td></td><td></td><td></td><td></td><td></td><td></td><td></td><td></td></tr>
<tr><td>Chromosome</td><td>start</td><td>end</td><td>strand</td><td>MM</td><td>target_seq</td><td>PAM</td><td>alignment</td><td>distance</td><td>position</td><td>gene name</td><td>gene id</td></tr>
<tr><td>GU305914.1</td><td>175378</td><td>175400</td><td>-</td><td>0</td><td>TCTGACAGTGAGTTCAACGA</td><td>CGG</td><td>||||||||[||||||||||||]PAM</td><td>NA</td><td>Intergenic</td><td>NA</td><td>NA</td></tr>
<tr><td></td><td></td><td></td><td></td><td></td><td></td><td></td><td></td><td></td><td></td><td></td><td></td></tr>
<tr><td>T21341</td><td>CTGACAGTGAGTTCAACGACGGG</td><td>1000</td><td></td><td></td><td></td><td></td><td></td><td></td><td></td><td></td><td></td></tr>
<tr><td>Oligo fwd</td><td>CACCCTGACAGTGAGTTCAACGAC</td><td></td><td></td><td></td><td></td><td></td><td></td><td></td><td></td><td></td><td></td></tr>
<tr><td>Oligo rev</td><td>AAACGTCGTTGAACTCACTGTCAG</td><td></td><td></td><td></td><td></td><td></td><td></td><td></td><td></td><td></td><td></td></tr>
<tr><td>Chromosome</td><td>start</td><td>end</td><td>strand</td><td>MM</td><td>target_seq</td><td>PAM</td><td>alignment</td><td>distance</td><td>position</td><td>gene name</td><td>gene id</td></tr>
<tr><td>GU305914.1</td><td>175377</td><td>175399</td><td>-</td><td>0</td><td>CTGACAGTGAGTTCAACGAC</td><td>GGG</td><td>||||||||[||||||||||||]PAM</td><td>NA</td><td>Intergenic</td><td>NA</td><td>NA</td></tr>
<tr><td></td><td></td><td></td><td></td><td></td><td></td><td></td><td></td><td></td><td></td><td></td><td></td></tr>
<tr><td>T21342</td><td>AGTGAGTTCAACGACGGGTCTGG</td><td>1000</td><td></td><td></td><td></td><td></td><td></td><td></td><td></td><td></td><td></td></tr>
<tr><td>Oligo fwd</td><td>CACCAGTGAGTTCAACGACGGGTC</td><td></td><td></td><td></td><td></td><td></td><td></td><td></td><td></td><td></td><td></td></tr>
<tr><td>Oligo rev</td><td>AAACGACCCGTCGTTGAACTCACT</td><td></td><td></td><td></td><td></td><td></td><td></td><td></td><td></td><td></td><td></td></tr>
<tr><td>Chromosome</td><td>start</td><td>end</td><td>strand</td><td>MM</td><td>target_seq</td><td>PAM</td><td>alignment</td><td>distance</td><td>position</td><td>gene name</td><td>gene id</td></tr>
<tr><td>GU305914.1</td><td>175372</td><td>175394</td><td>-</td><td>0</td><td>AGTGAGTTCAACGACGGGTC</td><td>TGG</td><td>||||||||[||||||||||||]PAM</td><td>NA</td><td>Intergenic</td><td>NA</td><td>NA</td></tr>
<tr><td></td><td></td><td></td><td></td><td></td><td></td><td></td><td></td><td></td><td></td><td></td><td></td></tr>
<tr><td>T21343</td><td>GTGAGTTCAACGACGGGTCTGGG</td><td>1000</td><td></td><td></td><td></td><td></td><td></td><td></td><td></td><td></td><td></td></tr>
<tr><td>Oligo fwd</td><td>CACCGTGAGTTCAACGACGGGTCT</td><td></td><td></td><td></td><td></td><td></td><td></td><td></td><td></td><td></td><td></td></tr>
<tr><td>Oligo rev</td><td>AAACAGACCCGTCGTTGAACTCAC</td><td></td><td></td><td></td><td></td><td></td><td></td><td></td><td></td><td></td><td></td></tr>
<tr><td>Chromosome</td><td>start</td><td>end</td><td>strand</td><td>MM</td><td>target_seq</td><td>PAM</td><td>alignment</td><td>distance</td><td>position</td><td>gene name</td><td>gene id</td></tr>
<tr><td>GU305914.1</td><td>175371</td><td>175393</td><td>-</td><td>0</td><td>GTGAGTTCAACGACGGGTCT</td><td>GGG</td><td>||||||||[||||||||||||]PAM</td><td>NA</td><td>Intergenic</td><td>NA</td><td>NA</td></tr>
<tr><td></td><td></td><td></td><td></td><td></td><td></td><td></td><td></td><td></td><td></td><td></td><td></td></tr>
<tr><td>T21344</td><td>GGGTCTGGGCTCTATTCAGATGG</td><td>1000</td><td></td><td></td><td></td><td></td><td></td><td></td><td></td><td></td><td></td></tr>
<tr><td>Oligo fwd</td><td>CACCGGGTCTGGGCTCTATTCAGA</td><td></td><td></td><td></td><td></td><td></td><td></td><td></td><td></td><td></td><td></td></tr>
<tr><td>Oligo rev</td><td>AAACTCTGAATAGAGCCCAGACCC</td><td></td><td></td><td></td><td></td><td></td><td></td><td></td><td></td><td></td><td></td></tr>
<tr><td>Chromosome</td><td>start</td><td>end</td><td>strand</td><td>MM</td><td>target_seq</td><td>PAM</td><td>alignment</td><td>distance</td><td>position</td><td>gene name</td><td>gene id</td></tr>
<tr><td>GU305914.1</td><td>175357</td><td>175379</td><td>-</td><td>0</td><td>GGGTCTGGGCTCTATTCAGA</td><td>TGG</td><td>||||||||[||||||||||||]PAM</td><td>NA</td><td>Intergenic</td><td>NA</td><td>NA</td></tr>
<tr><td></td><td></td><td></td><td></td><td></td><td></td><td></td><td></td><td></td><td></td><td></td><td></td></tr>
<tr><td>T21345</td><td>GCTCTATTCAGATGGACTGTAGG</td><td>1000</td><td></td><td></td><td></td><td></td><td></td><td></td><td></td><td></td><td></td></tr>
<tr><td>Oligo fwd</td><td>CACCGCTCTATTCAGATGGACTGT</td><td></td><td></td><td></td><td></td><td></td><td></td><td></td><td></td><td></td><td></td></tr>
<tr><td>Oligo rev</td><td>AAACACAGTCCATCTGAATAGAGC</td><td></td><td></td><td></td><td></td><td></td><td></td><td></td><td></td><td></td><td></td></tr>
<tr><td>Chromosome</td><td>start</td><td>end</td><td>strand</td><td>MM</td><td>target_seq</td><td>PAM</td><td>alignment</td><td>distance</td><td>position</td><td>gene name</td><td>gene id</td></tr>
<tr><td>GU305914.1</td><td>175349</td><td>175371</td><td>-</td><td>0</td><td>GCTCTATTCAGATGGACTGT</td><td>AGG</td><td>||||||||[||||||||||||]PAM</td><td>NA</td><td>Intergenic</td><td>NA</td><td>NA</td></tr>
<tr><td></td><td></td><td></td><td></td><td></td><td></td><td></td><td></td><td></td><td></td><td></td><td></td></tr>
<tr><td>T21346</td><td>CTCCACCATACACCATCTTATGG</td><td>1000</td><td></td><td></td><td></td><td></td><td></td><td></td><td></td><td></td><td></td></tr>
<tr><td>Oligo fwd</td><td>CACCCTCCACCATACACCATCTTA</td><td></td><td></td><td></td><td></td><td></td><td></td><td></td><td></td><td></td><td></td></tr>
<tr><td>Oligo rev</td><td>AAACTAAGATGGTGTATGGTGGAG</td><td></td><td></td><td></td><td></td><td></td><td></td><td></td><td></td><td></td><td></td></tr>
<tr><td>Chromosome</td><td>start</td><td>end</td><td>strand</td><td>MM</td><td>target_seq</td><td>PAM</td><td>alignment</td><td>distance</td><td>position</td><td>gene name</td><td>gene id</td></tr>
<tr><td>GU305914.1</td><td>175313</td><td>175335</td><td>-</td><td>0</td><td>CTCCACCATACACCATCTTA</td><td>TGG</td><td>||||||||[||||||||||||]PAM</td><td>NA</td><td>Intergenic</td><td>NA</td><td>NA</td></tr>
<tr><td></td><td></td><td></td><td></td><td></td><td></td><td></td><td></td><td></td><td></td><td></td><td></td></tr>
<tr><td>T21347</td><td>TGGATGCTAAACTCGACGCTCGG</td><td>1000</td><td></td><td></td><td></td><td></td><td></td><td></td><td></td><td></td><td></td></tr>
<tr><td>Oligo fwd</td><td>CACCTGGATGCTAAACTCGACGCT</td><td></td><td></td><td></td><td></td><td></td><td></td><td></td><td></td><td></td><td></td></tr>
<tr><td>Oligo rev</td><td>AAACAGCGTCGAGTTTAGCATCCA</td><td></td><td></td><td></td><td></td><td></td><td></td><td></td><td></td><td></td><td></td></tr>
<tr><td>Chromosome</td><td>start</td><td>end</td><td>strand</td><td>MM</td><td>target_seq</td><td>PAM</td><td>alignment</td><td>distance</td><td>position</td><td>gene name</td><td>gene id</td></tr>
<tr><td>GU305914.1</td><td>175293</td><td>175315</td><td>-</td><td>0</td><td>TGGATGCTAAACTCGACGCT</td><td>CGG</td><td>||||||||[||||||||||||]PAM</td><td>NA</td><td>Intergenic</td><td>NA</td><td>NA</td></tr>
<tr><td></td><td></td><td></td><td></td><td></td><td></td><td></td><td></td><td></td><td></td><td></td><td></td></tr>
<tr><td>T21348</td><td>ACGTTCTACAATTTACGCAAAGG</td><td>1000</td><td></td><td></td><td></td><td></td><td></td><td></td><td></td><td></td><td></td></tr>
<tr><td>Oligo fwd</td><td>CACCACGTTCTACAATTTACGCAA</td><td></td><td></td><td></td><td></td><td></td><td></td><td></td><td></td><td></td><td></td></tr>
<tr><td>Oligo rev</td><td>AAACTTGCGTAAATTGTAGAACGT</td><td></td><td></td><td></td><td></td><td></td><td></td><td></td><td></td><td></td><td></td></tr>
<tr><td>Chromosome</td><td>start</td><td>end</td><td>strand</td><td>MM</td><td>target_seq</td><td>PAM</td><td>alignment</td><td>distance</td><td>position</td><td>gene name</td><td>gene id</td></tr>
<tr><td>GU305914.1</td><td>175267</td><td>175289</td><td>-</td><td>0</td><td>ACGTTCTACAATTTACGCAA</td><td>AGG</td><td>||||||||[||||||||||||]PAM</td><td>NA</td><td>Intergenic</td><td>NA</td><td>NA</td></tr>
<tr><td></td><td></td><td></td><td></td><td></td><td></td><td></td><td></td><td></td><td></td><td></td><td></td></tr>
<tr><td>T21349</td><td>TTACGCAAAGGATAACGTTACGG</td><td>1000</td><td></td><td></td><td></td><td></td><td></td><td></td><td></td><td></td><td></td></tr>
<tr><td>Oligo fwd</td><td>CACCTTACGCAAAGGATAACGTTA</td><td></td><td></td><td></td><td></td><td></td><td></td><td></td><td></td><td></td><td></td></tr>
</table>

<table>
<tr><td>Oligo rev</td><td>AAACTAACGTTATCCTTTGCGTAA</td><td></td><td></td><td></td><td></td><td></td><td></td><td></td><td></td><td></td><td></td></tr>
<tr><td>Chromosome</td><td>start</td><td>end</td><td>strand</td><td>MM</td><td>target_seq</td><td>PAM</td><td>alignment</td><td>distance</td><td>position</td><td>gene name</td><td>gene id</td></tr>
<tr><td>GU305914.1</td><td>175255</td><td>175277</td><td>-</td><td>0</td><td>TTACGCAAAGGATAACGTTA</td><td>CGG</td><td>||||||||[||||||||||||]PAM</td><td>NA</td><td>Intergenic</td><td>NA</td><td>NA</td></tr>
<tr><td>T21350</td><td>GGATAACGTTACGGTCAAAATGG</td><td>1000</td><td></td><td></td><td></td><td></td><td></td><td></td><td></td><td></td><td></td></tr>
<tr><td>Oligo fwd</td><td>CACCGGATAACGTTACGGTCAAAA</td><td></td><td></td><td></td><td></td><td></td><td></td><td></td><td></td><td></td><td></td></tr>
<tr><td>Oligo rev</td><td>AAACTTTTGACCGTAACGTTATCC</td><td></td><td></td><td></td><td></td><td></td><td></td><td></td><td></td><td></td><td></td></tr>
<tr><td>Chromosome</td><td>start</td><td>end</td><td>strand</td><td>MM</td><td>target_seq</td><td>PAM</td><td>alignment</td><td>distance</td><td>position</td><td>gene name</td><td>gene id</td></tr>
<tr><td>GU305914.1</td><td>175246</td><td>175268</td><td>-</td><td>0</td><td>GGATAACGTTACGGTCAAAA</td><td>TGG</td><td>||||||||[||||||||||||]PAM</td><td>NA</td><td>Intergenic</td><td>NA</td><td>NA</td></tr>
<tr><td>T21351</td><td>GTTACGGTCAAAATGGAGAACGG</td><td>1000</td><td></td><td></td><td></td><td></td><td></td><td></td><td></td><td></td><td></td></tr>
<tr><td>Oligo fwd</td><td>CACCGTTACGGTCAAAATGGAGAA</td><td></td><td></td><td></td><td></td><td></td><td></td><td></td><td></td><td></td><td></td></tr>
<tr><td>Oligo rev</td><td>AAACTTCTCCATTTTGACCGTAAC</td><td></td><td></td><td></td><td></td><td></td><td></td><td></td><td></td><td></td><td></td></tr>
<tr><td>Chromosome</td><td>start</td><td>end</td><td>strand</td><td>MM</td><td>target_seq</td><td>PAM</td><td>alignment</td><td>distance</td><td>position</td><td>gene name</td><td>gene id</td></tr>
<tr><td>GU305914.1</td><td>175239</td><td>175261</td><td>-</td><td>0</td><td>GTTACGGTCAAAATGGAGAA</td><td>CGG</td><td>||||||||[||||||||||||]PAM</td><td>NA</td><td>Intergenic</td><td>NA</td><td>NA</td></tr>
<tr><td>T21352</td><td>GTCAAAATGGAGAACGGTACCGG</td><td>1000</td><td></td><td></td><td></td><td></td><td></td><td></td><td></td><td></td><td></td></tr>
<tr><td>Oligo fwd</td><td>CACCGTCAAAATGGAGAACGGTAC</td><td></td><td></td><td></td><td></td><td></td><td></td><td></td><td></td><td></td><td></td></tr>
<tr><td>Oligo rev</td><td>AAACGTACCGTTCTCCATTTTGAC</td><td></td><td></td><td></td><td></td><td></td><td></td><td></td><td></td><td></td><td></td></tr>
<tr><td>Chromosome</td><td>start</td><td>end</td><td>strand</td><td>MM</td><td>target_seq</td><td>PAM</td><td>alignment</td><td>distance</td><td>position</td><td>gene name</td><td>gene id</td></tr>
<tr><td>GU305914.1</td><td>175233</td><td>175255</td><td>-</td><td>0</td><td>GTCAAAATGGAGAACGGTAC</td><td>CGG</td><td>||||||||[||||||||||||]PAM</td><td>NA</td><td>Intergenic</td><td>NA</td><td>NA</td></tr>
<tr><td>T21353</td><td>GAGAACGGTACCGGATTTTTCGG</td><td>1000</td><td></td><td></td><td></td><td></td><td></td><td></td><td></td><td></td><td></td></tr>
<tr><td>Oligo fwd</td><td>CACCGAGAACGGTACCGGATTTTT</td><td></td><td></td><td></td><td></td><td></td><td></td><td></td><td></td><td></td><td></td></tr>
<tr><td>Oligo rev</td><td>AAACAAAAATCCGGTACCGTTCTC</td><td></td><td></td><td></td><td></td><td></td><td></td><td></td><td></td><td></td><td></td></tr>
<tr><td>Chromosome</td><td>start</td><td>end</td><td>strand</td><td>MM</td><td>target_seq</td><td>PAM</td><td>alignment</td><td>distance</td><td>position</td><td>gene name</td><td>gene id</td></tr>
<tr><td>GU305914.1</td><td>175224</td><td>175246</td><td>-</td><td>0</td><td>GAGAACGGTACCGGATTTTT</td><td>CGG</td><td>||||||||[||||||||||||]PAM</td><td>NA</td><td>Intergenic</td><td>NA</td><td>NA</td></tr>
<tr><td>T21354</td><td>TTTTTCGGTACAGATCTGTTCGG</td><td>1000</td><td></td><td></td><td></td><td></td><td></td><td></td><td></td><td></td><td></td></tr>
<tr><td>Oligo fwd</td><td>CACCTTTTTCGGTACAGATCTGTT</td><td></td><td></td><td></td><td></td><td></td><td></td><td></td><td></td><td></td><td></td></tr>
<tr><td>Oligo rev</td><td>AAACAACAGATCTGTACCGAAAAA</td><td></td><td></td><td></td><td></td><td></td><td></td><td></td><td></td><td></td><td></td></tr>
<tr><td>Chromosome</td><td>start</td><td>end</td><td>strand</td><td>MM</td><td>target_seq</td><td>PAM</td><td>alignment</td><td>distance</td><td>position</td><td>gene name</td><td>gene id</td></tr>
<tr><td>GU305914.1</td><td>175209</td><td>175231</td><td>-</td><td>0</td><td>TTTTTCGGTACAGATCTGTT</td><td>CGG</td><td>||||||||[||||||||||||]PAM</td><td>NA</td><td>Intergenic</td><td>NA</td><td>NA</td></tr>
<tr><td>T21355</td><td>TTTTCGGTACAGATCTGTTCGGG</td><td>1000</td><td></td><td></td><td></td><td></td><td></td><td></td><td></td><td></td><td></td></tr>
<tr><td>Oligo fwd</td><td>CACCTTTTCGGTACAGATCTGTTC</td><td></td><td></td><td></td><td></td><td></td><td></td><td></td><td></td><td></td><td></td></tr>
<tr><td>Oligo rev</td><td>AAACGAACAGATCTGTACCGAAAA</td><td></td><td></td><td></td><td></td><td></td><td></td><td></td><td></td><td></td><td></td></tr>
<tr><td>Chromosome</td><td>start</td><td>end</td><td>strand</td><td>MM</td><td>target_seq</td><td>PAM</td><td>alignment</td><td>distance</td><td>position</td><td>gene name</td><td>gene id</td></tr>
<tr><td>GU305914.1</td><td>175208</td><td>175230</td><td>-</td><td>0</td><td>TTTTCGGTACAGATCTGTTC</td><td>GGG</td><td>||||||||[||||||||||||]PAM</td><td>NA</td><td>Intergenic</td><td>NA</td><td>NA</td></tr>
<tr><td>T21356</td><td>TTTCGGTACAGATCTGTTCGGGG</td><td>1000</td><td></td><td></td><td></td><td></td><td></td><td></td><td></td><td></td><td></td></tr>
<tr><td>Oligo fwd</td><td>CACCTTTCGGTACAGATCTGTTCG</td><td></td><td></td><td></td><td></td><td></td><td></td><td></td><td></td><td></td><td></td></tr>
<tr><td>Oligo rev</td><td>AAACCGAACAGATCTGTACCGAAA</td><td></td><td></td><td></td><td></td><td></td><td></td><td></td><td></td><td></td><td></td></tr>
<tr><td>Chromosome</td><td>start</td><td>end</td><td>strand</td><td>MM</td><td>target_seq</td><td>PAM</td><td>alignment</td><td>distance</td><td>position</td><td>gene name</td><td>gene id</td></tr>
<tr><td>GU305914.1</td><td>175207</td><td>175229</td><td>-</td><td>0</td><td>TTTCGGTACAGATCTGTTCG</td><td>GGG</td><td>||||||||[||||||||||||]PAM</td><td>NA</td><td>Intergenic</td><td>NA</td><td>NA</td></tr>
<tr><td>T21357</td><td>ACAGATCTGTTCGGGGTGAACGG</td><td>1000</td><td></td><td></td><td></td><td></td><td></td><td></td><td></td><td></td><td></td></tr>
<tr><td>Oligo fwd</td><td>CACCACAGATCTGTTCGGGGTGAA</td><td></td><td></td><td></td><td></td><td></td><td></td><td></td><td></td><td></td><td></td></tr>
<tr><td>Oligo rev</td><td>AAACTTCACCCCGAACAGATCTGT</td><td></td><td></td><td></td><td></td><td></td><td></td><td></td><td></td><td></td><td></td></tr>
<tr><td>Chromosome</td><td>start</td><td>end</td><td>strand</td><td>MM</td><td>target_seq</td><td>PAM</td><td>alignment</td><td>distance</td><td>position</td><td>gene name</td><td>gene id</td></tr>
<tr><td>GU305914.1</td><td>175200</td><td>175222</td><td>-</td><td>0</td><td>ACAGATCTGTTCGGGGTGAA</td><td>CGG</td><td>||||||||[||||||||||||]PAM</td><td>NA</td><td>Intergenic</td><td>NA</td><td>NA</td></tr>
<tr><td>T21358</td><td>GATCTGTTCGGGGTGAACGGTGG</td><td>1000</td><td></td><td></td><td></td><td></td><td></td><td></td><td></td><td></td><td></td></tr>
<tr><td>Oligo fwd</td><td>CACCGATCTGTTCGGGGTGAACGG</td><td></td><td></td><td></td><td></td><td></td><td></td><td></td><td></td><td></td><td></td></tr>
<tr><td>Oligo rev</td><td>AAACCCGTTCACCCCGAACAGATC</td><td></td><td></td><td></td><td></td><td></td><td></td><td></td><td></td><td></td><td></td></tr>
<tr><td>Chromosome</td><td>start</td><td>end</td><td>strand</td><td>MM</td><td>target_seq</td><td>PAM</td><td>alignment</td><td>distance</td><td>position</td><td>gene name</td><td>gene id</td></tr>
<tr><td>GU305914.1</td><td>175197</td><td>175219</td><td>-</td><td>0</td><td>GATCTGTTCGGGGTGAACGG</td><td>TGG</td><td>||||||||[||||||||||||]PAM</td><td>NA</td><td>Intergenic</td><td>NA</td><td>NA</td></tr>
<tr><td>T21359</td><td>TTCGGGGTGAACGGTGGAGTAGG</td><td>1000</td><td></td><td></td><td></td><td></td><td></td><td></td><td></td><td></td><td></td></tr>
<tr><td>Oligo fwd</td><td>CACCTTCGGGGTGAACGGTGGAGT</td><td></td><td></td><td></td><td></td><td></td><td></td><td></td><td></td><td></td><td></td></tr>
<tr><td>Oligo rev</td><td>AAACACTCCACCGTTCACCCCGAA</td><td></td><td></td><td></td><td></td><td></td><td></td><td></td><td></td><td></td><td></td></tr>
<tr><td>Chromosome</td><td>start</td><td>end</td><td>strand</td><td>MM</td><td>target_seq</td><td>PAM</td><td>alignment</td><td>distance</td><td>position</td><td>gene name</td><td>gene id</td></tr>
<tr><td>GU305914.1</td><td>175191</td><td>175213</td><td>-</td><td>0</td><td>TTCGGGGTGAACGGTGGAGT</td><td>AGG</td><td>||||||||[||||||||||||]PAM</td><td>NA</td><td>Intergenic</td><td>NA</td><td>NA</td></tr>
<tr><td>T21360</td><td>TAGGTTCGTATTACAGTCCGAGG</td><td>1000</td><td></td><td></td><td></td><td></td><td></td><td></td><td></td><td></td><td></td></tr>
<tr><td>Oligo fwd</td><td>CACCTAGGTTCGTATTACAGTCCG</td><td></td><td></td><td></td><td></td><td></td><td></td><td></td><td></td><td></td><td></td></tr>
<tr><td>Oligo rev</td><td>AAACCGGACTGTAATACGAACCTA</td><td></td><td></td><td></td><td></td><td></td><td></td><td></td><td></td><td></td><td></td></tr>
<tr><td>Chromosome</td><td>start</td><td>end</td><td>strand</td><td>MM</td><td>target_seq</td><td>PAM</td><td>alignment</td><td>distance</td><td>position</td><td>gene name</td><td>gene id</td></tr>
<tr><td>GU305914.1</td><td>175172</td><td>175194</td><td>-</td><td>0</td><td>TAGGTTCGTATTACAGTCCG</td><td>AGG</td><td>||||||||[||||||||||||]PAM</td><td>NA</td><td>Intergenic</td><td>NA</td><td>NA</td></tr>
<tr><td>T21361</td><td>TCCGAGGAGACCACCTCAACCGG</td><td>1000</td><td></td><td></td><td></td><td></td><td></td><td></td><td></td><td></td><td></td></tr>
<tr><td>Oligo fwd</td><td>CACCTCCGAGGAGACCACCTCAAC</td><td></td><td></td><td></td><td></td><td></td><td></td><td></td><td></td><td></td><td></td></tr>
<tr><td>Oligo rev</td><td>AAACGTTGAGGTGGTCTCCTCGGA</td><td></td><td></td><td></td><td></td><td></td><td></td><td></td><td></td><td></td><td></td></tr>
<tr><td>Chromosome</td><td>start</td><td>end</td><td>strand</td><td>MM</td><td>target_seq</td><td>PAM</td><td>alignment</td><td>distance</td><td>position</td><td>gene name</td><td>gene id</td></tr>
<tr><td>GU305914.1</td><td>175156</td><td>175178</td><td>-</td><td>0</td><td>TCCGAGGAGACCACCTCAAC</td><td>CGG</td><td>||||||||[||||||||||||]PAM</td><td>NA</td><td>Intergenic</td><td>NA</td><td>NA</td></tr>
<tr><td>T21362</td><td>GTATCGAAAAACGATTCGTACGG</td><td>1000</td><td></td><td></td><td></td><td></td><td></td><td></td><td></td><td></td><td></td></tr>
<tr><td>Oligo fwd</td><td>CACCGTATCGAAAAACGATTCGTA</td><td></td><td></td><td></td><td></td><td></td><td></td><td></td><td></td><td></td><td></td></tr>
<tr><td>Oligo rev</td><td>AAACTACGAATCGTTTTTCGATAC</td><td></td><td></td><td></td><td></td><td></td><td></td><td></td><td></td><td></td><td></td></tr>
<tr><td>Chromosome</td><td>start</td><td>end</td><td>strand</td><td>MM</td><td>target_seq</td><td>PAM</td><td>alignment</td><td>distance</td><td>position</td><td>gene name</td><td>gene id</td></tr>
<tr><td>GU305914.1</td><td>175116</td><td>175138</td><td>-</td><td>0</td><td>GTATCGAAAAACGATTCGTA</td><td>CGG</td><td>||||||||[||||||||||||]PAM</td><td>NA</td><td>Intergenic</td><td>NA</td><td>NA</td></tr>
<tr><td>T21363</td><td>TAGTGCCACCAGATGCACGACGG</td><td>1000</td><td></td><td></td><td></td><td></td><td></td><td></td><td></td><td></td><td></td></tr>
<tr><td>Oligo fwd</td><td>CACCTAGTGCCACCAGATGCACGA</td><td></td><td></td><td></td><td></td><td></td><td></td><td></td><td></td><td></td><td></td></tr>
<tr><td>Oligo rev</td><td>AAACTCGTGCATCTGGTGGCACTA</td><td></td><td></td><td></td><td></td><td></td><td></td><td></td><td></td><td></td><td></td></tr>
<tr><td>Chromosome</td><td>start</td><td>end</td><td>strand</td><td>MM</td><td>target_seq</td><td>PAM</td><td>alignment</td><td>distance</td><td>position</td><td>gene name</td><td>gene id</td></tr>
<tr><td>GU305914.1</td><td>175093</td><td>175115</td><td>-</td><td>0</td><td>TAGTGCCACCAGATGCACGA</td><td>CGG</td><td>||||||||[||||||||||||]PAM</td><td>NA</td><td>Intergenic</td><td>NA</td><td>NA</td></tr>
<tr><td>T21364</td><td>AGTGCCACCAGATGCACGACGGG</td><td>1000</td><td></td><td></td><td></td><td></td><td></td><td></td><td></td><td></td><td></td></tr>
</table>

<table>
<tr><td>Oligo fwd</td><td>CACCAGTGCCACCAGATGCACGAC</td><td></td><td></td><td></td><td></td><td></td><td></td><td></td><td></td><td></td><td></td></tr>
<tr><td>Oligo rev</td><td>AAACGTCGTGCATCTGGTGGCACT</td><td></td><td></td><td></td><td></td><td></td><td></td><td></td><td></td><td></td><td></td></tr>
<tr><td>Chromosome</td><td>start</td><td>end</td><td>strand</td><td>MM</td><td>target_seq</td><td>PAM</td><td>alignment</td><td>distance</td><td>position</td><td>gene name</td><td>gene id</td></tr>
<tr><td>GU305914.1</td><td>175092</td><td>175114</td><td>-</td><td>0</td><td>AGTGCCACCAGATGCACGAC</td><td>GGG</td><td>||||||||[||||||||||||]PAM</td><td>NA</td><td>Intergenic</td><td>NA</td><td>NA</td></tr>
<tr><td>T21365</td><td>GTGCCACCAGATGCACGACGGGG</td><td>1000</td><td></td><td></td><td></td><td></td><td></td><td></td><td></td><td></td><td></td></tr>
<tr><td>Oligo fwd</td><td>CACCGTGCCACCAGATGCACGACG</td><td></td><td></td><td></td><td></td><td></td><td></td><td></td><td></td><td></td><td></td></tr>
<tr><td>Oligo rev</td><td>AAACCGTCGTGCATCTGGTGGCAC</td><td></td><td></td><td></td><td></td><td></td><td></td><td></td><td></td><td></td><td></td></tr>
<tr><td>Chromosome</td><td>start</td><td>end</td><td>strand</td><td>MM</td><td>target_seq</td><td>PAM</td><td>alignment</td><td>distance</td><td>position</td><td>gene name</td><td>gene id</td></tr>
<tr><td>GU305914.1</td><td>175091</td><td>175113</td><td>-</td><td>0</td><td>GTGCCACCAGATGCACGACG</td><td>GGG</td><td>||||||||[||||||||||||]PAM</td><td>NA</td><td>Intergenic</td><td>NA</td><td>NA</td></tr>
<tr><td>T21366</td><td>CACGACGGGGATGAAAGACATGG</td><td>1000</td><td></td><td></td><td></td><td></td><td></td><td></td><td></td><td></td><td></td></tr>
<tr><td>Oligo fwd</td><td>CACCCACGACGGGGATGAAAGACA</td><td></td><td></td><td></td><td></td><td></td><td></td><td></td><td></td><td></td><td></td></tr>
<tr><td>Oligo rev</td><td>AAACTGTCTTTCATCCCCGTCGTG</td><td></td><td></td><td></td><td></td><td></td><td></td><td></td><td></td><td></td><td></td></tr>
<tr><td>Chromosome</td><td>start</td><td>end</td><td>strand</td><td>MM</td><td>target_seq</td><td>PAM</td><td>alignment</td><td>distance</td><td>position</td><td>gene name</td><td>gene id</td></tr>
<tr><td>GU305914.1</td><td>175078</td><td>175100</td><td>-</td><td>0</td><td>CACGACGGGGATGAAAGACA</td><td>TGG</td><td>||||||||[||||||||||||]PAM</td><td>NA</td><td>Intergenic</td><td>NA</td><td>NA</td></tr>
<tr><td>T21367</td><td>ACGACGGGGATGAAAGACATGGG</td><td>1000</td><td></td><td></td><td></td><td></td><td></td><td></td><td></td><td></td><td></td></tr>
<tr><td>Oligo fwd</td><td>CACCACGACGGGGATGAAAGACAT</td><td></td><td></td><td></td><td></td><td></td><td></td><td></td><td></td><td></td><td></td></tr>
<tr><td>Oligo rev</td><td>AAACATGTCTTTCATCCCCGTCGT</td><td></td><td></td><td></td><td></td><td></td><td></td><td></td><td></td><td></td><td></td></tr>
<tr><td>Chromosome</td><td>start</td><td>end</td><td>strand</td><td>MM</td><td>target_seq</td><td>PAM</td><td>alignment</td><td>distance</td><td>position</td><td>gene name</td><td>gene id</td></tr>
<tr><td>GU305914.1</td><td>175077</td><td>175099</td><td>-</td><td>0</td><td>ACGACGGGGATGAAAGACAT</td><td>GGG</td><td>||||||||[||||||||||||]PAM</td><td>NA</td><td>Intergenic</td><td>NA</td><td>NA</td></tr>
<tr><td>T21368</td><td>TCGAAAACGAAGCAATCTCCAGG</td><td>1000</td><td></td><td></td><td></td><td></td><td></td><td></td><td></td><td></td><td></td></tr>
<tr><td>Oligo fwd</td><td>CACCTCGAAAACGAAGCAATCTCC</td><td></td><td></td><td></td><td></td><td></td><td></td><td></td><td></td><td></td><td></td></tr>
<tr><td>Oligo rev</td><td>AAACGGAGATTGCTTCGTTTTCGA</td><td></td><td></td><td></td><td></td><td></td><td></td><td></td><td></td><td></td><td></td></tr>
<tr><td>Chromosome</td><td>start</td><td>end</td><td>strand</td><td>MM</td><td>target_seq</td><td>PAM</td><td>alignment</td><td>distance</td><td>position</td><td>gene name</td><td>gene id</td></tr>
<tr><td>GU305914.1</td><td>175050</td><td>175072</td><td>-</td><td>0</td><td>TCGAAAACGAAGCAATCTCC</td><td>AGG</td><td>||||||||[||||||||||||]PAM</td><td>NA</td><td>Intergenic</td><td>NA</td><td>NA</td></tr>
<tr><td>T21369</td><td>CGAAAACGAAGCAATCTCCAGGG</td><td>1000</td><td></td><td></td><td></td><td></td><td></td><td></td><td></td><td></td><td></td></tr>
<tr><td>Oligo fwd</td><td>CACCCGAAAACGAAGCAATCTCCA</td><td></td><td></td><td></td><td></td><td></td><td></td><td></td><td></td><td></td><td></td></tr>
<tr><td>Oligo rev</td><td>AAACTGGAGATTGCTTCGTTTTCG</td><td></td><td></td><td></td><td></td><td></td><td></td><td></td><td></td><td></td><td></td></tr>
<tr><td>Chromosome</td><td>start</td><td>end</td><td>strand</td><td>MM</td><td>target_seq</td><td>PAM</td><td>alignment</td><td>distance</td><td>position</td><td>gene name</td><td>gene id</td></tr>
<tr><td>GU305914.1</td><td>175049</td><td>175071</td><td>-</td><td>0</td><td>CGAAAACGAAGCAATCTCCA</td><td>GGG</td><td>||||||||[||||||||||||]PAM</td><td>NA</td><td>Intergenic</td><td>NA</td><td>NA</td></tr>
<tr><td>T21370</td><td>TAGCGAGTTTATGAGAGCCACGG</td><td>1000</td><td></td><td></td><td></td><td></td><td></td><td></td><td></td><td></td><td></td></tr>
<tr><td>Oligo fwd</td><td>CACCTAGCGAGTTTATGAGAGCCA</td><td></td><td></td><td></td><td></td><td></td><td></td><td></td><td></td><td></td><td></td></tr>
<tr><td>Oligo rev</td><td>AAACTGGCTCTCATAAACTCGCTA</td><td></td><td></td><td></td><td></td><td></td><td></td><td></td><td></td><td></td><td></td></tr>
<tr><td>Chromosome</td><td>start</td><td>end</td><td>strand</td><td>MM</td><td>target_seq</td><td>PAM</td><td>alignment</td><td>distance</td><td>position</td><td>gene name</td><td>gene id</td></tr>
<tr><td>GU305914.1</td><td>175006</td><td>175028</td><td>-</td><td>0</td><td>TAGCGAGTTTATGAGAGCCA</td><td>CGG</td><td>||||||||[||||||||||||]PAM</td><td>NA</td><td>Intergenic</td><td>NA</td><td>NA</td></tr>
<tr><td>T21371</td><td>AGCGAGTTTATGAGAGCCACGGG</td><td>1000</td><td></td><td></td><td></td><td></td><td></td><td></td><td></td><td></td><td></td></tr>
<tr><td>Oligo fwd</td><td>CACCAGCGAGTTTATGAGAGCCAC</td><td></td><td></td><td></td><td></td><td></td><td></td><td></td><td></td><td></td><td></td></tr>
<tr><td>Oligo rev</td><td>AAACGTGGCTCTCATAAACTCGCT</td><td></td><td></td><td></td><td></td><td></td><td></td><td></td><td></td><td></td><td></td></tr>
<tr><td>Chromosome</td><td>start</td><td>end</td><td>strand</td><td>MM</td><td>target_seq</td><td>PAM</td><td>alignment</td><td>distance</td><td>position</td><td>gene name</td><td>gene id</td></tr>
<tr><td>GU305914.1</td><td>175005</td><td>175027</td><td>-</td><td>0</td><td>AGCGAGTTTATGAGAGCCAC</td><td>GGG</td><td>||||||||[||||||||||||]PAM</td><td>NA</td><td>Intergenic</td><td>NA</td><td>NA</td></tr>
<tr><td>GU305914.1</td><td>82097</td><td>82119</td><td>-</td><td>4</td><td>ATCGAGTGTATGAGCGGCAC</td><td>GGG</td><td>|-|||||-[||||||-|-|||]PAM</td><td>NA</td><td>Intergenic</td><td>NA</td><td>NA</td></tr>
<tr><td>T21372</td><td>GCCACGGGCGAGAGACCGAGCGG</td><td>1000</td><td></td><td></td><td></td><td></td><td></td><td></td><td></td><td></td><td></td></tr>
<tr><td>Oligo fwd</td><td>CACCGCCACGGGCGAGAGACCGAG</td><td></td><td></td><td></td><td></td><td></td><td></td><td></td><td></td><td></td><td></td></tr>
<tr><td>Oligo rev</td><td>AAACCTCGGTCTCTCGCCCGTGGC</td><td></td><td></td><td></td><td></td><td></td><td></td><td></td><td></td><td></td><td></td></tr>
<tr><td>Chromosome</td><td>start</td><td>end</td><td>strand</td><td>MM</td><td>target_seq</td><td>PAM</td><td>alignment</td><td>distance</td><td>position</td><td>gene name</td><td>gene id</td></tr>
<tr><td>GU305914.1</td><td>174990</td><td>175012</td><td>-</td><td>0</td><td>GCCACGGGCGAGAGACCGAG</td><td>CGG</td><td>||||||||[||||||||||||]PAM</td><td>NA</td><td>Intergenic</td><td>NA</td><td>NA</td></tr>
<tr><td>T21373</td><td>ACGGGCGAGAGACCGAGCGGCGG</td><td>1000</td><td></td><td></td><td></td><td></td><td></td><td></td><td></td><td></td><td></td></tr>
<tr><td>Oligo fwd</td><td>CACCACGGGCGAGAGACCGAGCGG</td><td></td><td></td><td></td><td></td><td></td><td></td><td></td><td></td><td></td><td></td></tr>
<tr><td>Oligo rev</td><td>AAACCCGCTCGGTCTCTCGCCCGT</td><td></td><td></td><td></td><td></td><td></td><td></td><td></td><td></td><td></td><td></td></tr>
<tr><td>Chromosome</td><td>start</td><td>end</td><td>strand</td><td>MM</td><td>target_seq</td><td>PAM</td><td>alignment</td><td>distance</td><td>position</td><td>gene name</td><td>gene id</td></tr>
<tr><td>GU305914.1</td><td>174987</td><td>175009</td><td>-</td><td>0</td><td>ACGGGCGAGAGACCGAGCGG</td><td>CGG</td><td>||||||||[||||||||||||]PAM</td><td>NA</td><td>Intergenic</td><td>NA</td><td>NA</td></tr>
<tr><td>T21374</td><td>GAGAGACCGAGCGGCGGAAACGG</td><td>1000</td><td></td><td></td><td></td><td></td><td></td><td></td><td></td><td></td><td></td></tr>
<tr><td>Oligo fwd</td><td>CACCGAGAGACCGAGCGGCGGAAA</td><td></td><td></td><td></td><td></td><td></td><td></td><td></td><td></td><td></td><td></td></tr>
<tr><td>Oligo rev</td><td>AAACTTTCCGCCGCTCGGTCTCTC</td><td></td><td></td><td></td><td></td><td></td><td></td><td></td><td></td><td></td><td></td></tr>
<tr><td>Chromosome</td><td>start</td><td>end</td><td>strand</td><td>MM</td><td>target_seq</td><td>PAM</td><td>alignment</td><td>distance</td><td>position</td><td>gene name</td><td>gene id</td></tr>
<tr><td>GU305914.1</td><td>174981</td><td>175003</td><td>-</td><td>0</td><td>GAGAGACCGAGCGGCGGAAA</td><td>CGG</td><td>||||||||[||||||||||||]PAM</td><td>NA</td><td>Intergenic</td><td>NA</td><td>NA</td></tr>
<tr><td>T21375</td><td>CCGAGCGGCGGAAACGGACACGG</td><td>1000</td><td></td><td></td><td></td><td></td><td></td><td></td><td></td><td></td><td></td></tr>
<tr><td>Oligo fwd</td><td>CACCCCGAGCGGCGGAAACGGACA</td><td></td><td></td><td></td><td></td><td></td><td></td><td></td><td></td><td></td><td></td></tr>
<tr><td>Oligo rev</td><td>AAACTGTCCGTTTCCGCCGCTCGG</td><td></td><td></td><td></td><td></td><td></td><td></td><td></td><td></td><td></td><td></td></tr>
<tr><td>Chromosome</td><td>start</td><td>end</td><td>strand</td><td>MM</td><td>target_seq</td><td>PAM</td><td>alignment</td><td>distance</td><td>position</td><td>gene name</td><td>gene id</td></tr>
<tr><td>GU305914.1</td><td>174975</td><td>174997</td><td>-</td><td>0</td><td>CCGAGCGGCGGAAACGGACA</td><td>CGG</td><td>||||||||[||||||||||||]PAM</td><td>NA</td><td>Intergenic</td><td>NA</td><td>NA</td></tr>
<tr><td>GU305914.1</td><td>119818</td><td>119840</td><td>-</td><td>4</td><td>CCCAGCCGCGGAAACAGACT</td><td>CGG</td><td>||-|||-|[|||||||-|||-]PAM</td><td>NA</td><td>Intergenic</td><td>NA</td><td>NA</td></tr>
<tr><td>T21376</td><td>GGAAACGGACACGGACCCGACGG</td><td>1000</td><td></td><td></td><td></td><td></td><td></td><td></td><td></td><td></td><td></td></tr>
<tr><td>Oligo fwd</td><td>CACCGGAAACGGACACGGACCCGA</td><td></td><td></td><td></td><td></td><td></td><td></td><td></td><td></td><td></td><td></td></tr>
<tr><td>Oligo rev</td><td>AAACTCGGGTCCGTGTCCGTTTCC</td><td></td><td></td><td></td><td></td><td></td><td></td><td></td><td></td><td></td><td></td></tr>
<tr><td>Chromosome</td><td>start</td><td>end</td><td>strand</td><td>MM</td><td>target_seq</td><td>PAM</td><td>alignment</td><td>distance</td><td>position</td><td>gene name</td><td>gene id</td></tr>
<tr><td>GU305914.1</td><td>174966</td><td>174988</td><td>-</td><td>0</td><td>GGAAACGGACACGGACCCGA</td><td>CGG</td><td>||||||||[||||||||||||]PAM</td><td>NA</td><td>Intergenic</td><td>NA</td><td>NA</td></tr>
<tr><td>T21377</td><td>CACGGACCCGACGGAGAACGAGG</td><td>1000</td><td></td><td></td><td></td><td></td><td></td><td></td><td></td><td></td><td></td></tr>
<tr><td>Oligo fwd</td><td>CACCCACGGACCCGACGGAGAACG</td><td></td><td></td><td></td><td></td><td></td><td></td><td></td><td></td><td></td><td></td></tr>
<tr><td>Oligo rev</td><td>AAACCGTTCTCCGTCGGGTCCGTG</td><td></td><td></td><td></td><td></td><td></td><td></td><td></td><td></td><td></td><td></td></tr>
<tr><td>Chromosome</td><td>start</td><td>end</td><td>strand</td><td>MM</td><td>target_seq</td><td>PAM</td><td>alignment</td><td>distance</td><td>position</td><td>gene name</td><td>gene id</td></tr>
<tr><td>GU305914.1</td><td>174957</td><td>174979</td><td>-</td><td>0</td><td>CACGGACCCGACGGAGAACG</td><td>AGG</td><td>||||||||[||||||||||||]PAM</td><td>NA</td><td>Intergenic</td><td>NA</td><td>NA</td></tr>
<tr><td>T21378</td><td>CCCGACGGAGAACGAGGTCCCGG</td><td>1000</td><td></td><td></td><td></td><td></td><td></td><td></td><td></td><td></td><td></td></tr>
<tr><td>Oligo fwd</td><td>CACCCCCGACGGAGAACGAGGTCC</td><td></td><td></td><td></td><td></td><td></td><td></td><td></td><td></td><td></td><td></td></tr>
<tr><td>Oligo rev</td><td>AAACGGACCTCGTTCTCCGTCGGG</td><td></td><td></td><td></td><td></td><td></td><td></td><td></td><td></td><td></td><td></td></tr>
<tr><td>Chromosome</td><td>start</td><td>end</td><td>strand</td><td>MM</td><td>target_seq</td><td>PAM</td><td>alignment</td><td>distance</td><td>position</td><td>gene name</td><td>gene id</td></tr>
</table>

<table>
<tr><td>GU305914.1</td><td>174951</td><td>174973</td><td>-</td><td>0</td><td>CCCGACGGAGAACGAGGTCC</td><td>CGG</td><td>||||||||[||||||||||||]PAM</td><td>NA</td><td>Intergenic</td><td>NA</td><td>NA</td></tr>
<tr><td>T21379</td><td>CGGCGACGACCCCAGTCCTCCGG</td><td>1000</td><td></td><td></td><td></td><td></td><td></td><td></td><td></td><td></td><td></td></tr>
<tr><td>Oligo fwd</td><td>CACCCGGCGACGACCCCAGTCCTC</td><td></td><td></td><td></td><td></td><td></td><td></td><td></td><td></td><td></td><td></td></tr>
<tr><td>Oligo rev</td><td>AAACGAGGACTGGGGTCGTCGCCG</td><td></td><td></td><td></td><td></td><td></td><td></td><td></td><td></td><td></td><td></td></tr>
<tr><td>Chromosome</td><td>start</td><td>end</td><td>strand</td><td>MM</td><td>target_seq</td><td>PAM</td><td>alignment</td><td>distance</td><td>position</td><td>gene name</td><td>gene id</td></tr>
<tr><td>GU305914.1</td><td>174931</td><td>174953</td><td>-</td><td>0</td><td>CGGCGACGACCCCAGTCCTC</td><td>CGG</td><td>||||||||[||||||||||||]PAM</td><td>NA</td><td>Intergenic</td><td>NA</td><td>NA</td></tr>
<tr><td>T21380</td><td>TCCGGCACCCGCGCAACCGACGG</td><td>1000</td><td></td><td></td><td></td><td></td><td></td><td></td><td></td><td></td><td></td></tr>
<tr><td>Oligo fwd</td><td>CACCTCCGGCACCCGCGCAACCGA</td><td></td><td></td><td></td><td></td><td></td><td></td><td></td><td></td><td></td><td></td></tr>
<tr><td>Oligo rev</td><td>AAACTCGGTTGCGCGGGTGCCGGA</td><td></td><td></td><td></td><td></td><td></td><td></td><td></td><td></td><td></td><td></td></tr>
<tr><td>Chromosome</td><td>start</td><td>end</td><td>strand</td><td>MM</td><td>target_seq</td><td>PAM</td><td>alignment</td><td>distance</td><td>position</td><td>gene name</td><td>gene id</td></tr>
<tr><td>GU305914.1</td><td>174913</td><td>174935</td><td>-</td><td>0</td><td>TCCGGCACCCGCGCAACCGA</td><td>CGG</td><td>||||||||[||||||||||||]PAM</td><td>NA</td><td>Intergenic</td><td>NA</td><td>NA</td></tr>
<tr><td>T21381</td><td>CCGGCACCCGCGCAACCGACGGG</td><td>1000</td><td></td><td></td><td></td><td></td><td></td><td></td><td></td><td></td><td></td></tr>
<tr><td>Oligo fwd</td><td>CACCCCGGCACCCGCGCAACCGAC</td><td></td><td></td><td></td><td></td><td></td><td></td><td></td><td></td><td></td><td></td></tr>
<tr><td>Oligo rev</td><td>AAACGTCGGTTGCGCGGGTGCCGG</td><td></td><td></td><td></td><td></td><td></td><td></td><td></td><td></td><td></td><td></td></tr>
<tr><td>Chromosome</td><td>start</td><td>end</td><td>strand</td><td>MM</td><td>target_seq</td><td>PAM</td><td>alignment</td><td>distance</td><td>position</td><td>gene name</td><td>gene id</td></tr>
<tr><td>GU305914.1</td><td>174912</td><td>174934</td><td>-</td><td>0</td><td>CCGGCACCCGCGCAACCGAC</td><td>GGG</td><td>||||||||[||||||||||||]PAM</td><td>NA</td><td>Intergenic</td><td>NA</td><td>NA</td></tr>
<tr><td>T21382</td><td>GACGGGAGTGTTTCTCGTCGTGG</td><td>1000</td><td></td><td></td><td></td><td></td><td></td><td></td><td></td><td></td><td></td></tr>
<tr><td>Oligo fwd</td><td>CACCGACGGGAGTGTTTCTCGTCG</td><td></td><td></td><td></td><td></td><td></td><td></td><td></td><td></td><td></td><td></td></tr>
<tr><td>Oligo rev</td><td>AAACCGACGAGAAACACTCCCGTC</td><td></td><td></td><td></td><td></td><td></td><td></td><td></td><td></td><td></td><td></td></tr>
<tr><td>Chromosome</td><td>start</td><td>end</td><td>strand</td><td>MM</td><td>target_seq</td><td>PAM</td><td>alignment</td><td>distance</td><td>position</td><td>gene name</td><td>gene id</td></tr>
<tr><td>GU305914.1</td><td>174895</td><td>174917</td><td>-</td><td>0</td><td>GACGGGAGTGTTTCTCGTCG</td><td>TGG</td><td>||||||||[||||||||||||]PAM</td><td>NA</td><td>Intergenic</td><td>NA</td><td>NA</td></tr>
<tr><td>T21383</td><td>GGGAGTGTTTCTCGTCGTGGTGG</td><td>1000</td><td></td><td></td><td></td><td></td><td></td><td></td><td></td><td></td><td></td></tr>
<tr><td>Oligo fwd</td><td>CACCGGGAGTGTTTCTCGTCGTGG</td><td></td><td></td><td></td><td></td><td></td><td></td><td></td><td></td><td></td><td></td></tr>
<tr><td>Oligo rev</td><td>AAACCCACGACGAGAAACACTCCC</td><td></td><td></td><td></td><td></td><td></td><td></td><td></td><td></td><td></td><td></td></tr>
<tr><td>Chromosome</td><td>start</td><td>end</td><td>strand</td><td>MM</td><td>target_seq</td><td>PAM</td><td>alignment</td><td>distance</td><td>position</td><td>gene name</td><td>gene id</td></tr>
<tr><td>GU305914.1</td><td>174892</td><td>174914</td><td>-</td><td>0</td><td>GGGAGTGTTTCTCGTCGTGG</td><td>TGG</td><td>||||||||[||||||||||||]PAM</td><td>NA</td><td>Intergenic</td><td>NA</td><td>NA</td></tr>
<tr><td>T21384</td><td>AGTGTTTCTCGTCGTGGTGGTGG</td><td>1000</td><td></td><td></td><td></td><td></td><td></td><td></td><td></td><td></td><td></td></tr>
<tr><td>Oligo fwd</td><td>CACCAGTGTTTCTCGTCGTGGTGG</td><td></td><td></td><td></td><td></td><td></td><td></td><td></td><td></td><td></td><td></td></tr>
<tr><td>Oligo rev</td><td>AAACCCACCACGACGAGAAACACT</td><td></td><td></td><td></td><td></td><td></td><td></td><td></td><td></td><td></td><td></td></tr>
<tr><td>Chromosome</td><td>start</td><td>end</td><td>strand</td><td>MM</td><td>target_seq</td><td>PAM</td><td>alignment</td><td>distance</td><td>position</td><td>gene name</td><td>gene id</td></tr>
<tr><td>GU305914.1</td><td>174889</td><td>174911</td><td>-</td><td>0</td><td>AGTGTTTCTCGTCGTGGTGG</td><td>TGG</td><td>||||||||[||||||||||||]PAM</td><td>NA</td><td>Intergenic</td><td>NA</td><td>NA</td></tr>
<tr><td>T21385</td><td>GGTGGTGGCGTGCTTCCTAGCGG</td><td>1000</td><td></td><td></td><td></td><td></td><td></td><td></td><td></td><td></td><td></td></tr>
<tr><td>Oligo fwd</td><td>CACCGGTGGTGGCGTGCTTCCTAG</td><td></td><td></td><td></td><td></td><td></td><td></td><td></td><td></td><td></td><td></td></tr>
<tr><td>Oligo rev</td><td>AAACCTAGGAAGCACGCCACCACC</td><td></td><td></td><td></td><td></td><td></td><td></td><td></td><td></td><td></td><td></td></tr>
<tr><td>Chromosome</td><td>start</td><td>end</td><td>strand</td><td>MM</td><td>target_seq</td><td>PAM</td><td>alignment</td><td>distance</td><td>position</td><td>gene name</td><td>gene id</td></tr>
<tr><td>GU305914.1</td><td>174874</td><td>174896</td><td>-</td><td>0</td><td>GGTGGTGGCGTGCTTCCTAG</td><td>CGG</td><td>||||||||[||||||||||||]PAM</td><td>NA</td><td>Intergenic</td><td>NA</td><td>NA</td></tr>
<tr><td>T21386</td><td>GTGGTGGCGTGCTTCCTAGCGGG</td><td>1000</td><td></td><td></td><td></td><td></td><td></td><td></td><td></td><td></td><td></td></tr>
<tr><td>Oligo fwd</td><td>CACCGTGGTGGCGTGCTTCCTAGC</td><td></td><td></td><td></td><td></td><td></td><td></td><td></td><td></td><td></td><td></td></tr>
<tr><td>Oligo rev</td><td>AAACGCTAGGAAGCACGCCACCAC</td><td></td><td></td><td></td><td></td><td></td><td></td><td></td><td></td><td></td><td></td></tr>
<tr><td>Chromosome</td><td>start</td><td>end</td><td>strand</td><td>MM</td><td>target_seq</td><td>PAM</td><td>alignment</td><td>distance</td><td>position</td><td>gene name</td><td>gene id</td></tr>
<tr><td>GU305914.1</td><td>174873</td><td>174895</td><td>-</td><td>0</td><td>GTGGTGGCGTGCTTCCTAGC</td><td>GGG</td><td>||||||||[||||||||||||]PAM</td><td>NA</td><td>Intergenic</td><td>NA</td><td>NA</td></tr>
<tr><td>T21387</td><td>TGGTGGCGTGCTTCCTAGCGGGG</td><td>1000</td><td></td><td></td><td></td><td></td><td></td><td></td><td></td><td></td><td></td></tr>
<tr><td>Oligo fwd</td><td>CACCTGGTGGCGTGCTTCCTAGCG</td><td></td><td></td><td></td><td></td><td></td><td></td><td></td><td></td><td></td><td></td></tr>
<tr><td>Oligo rev</td><td>AAACCGCTAGGAAGCACGCCACCA</td><td></td><td></td><td></td><td></td><td></td><td></td><td></td><td></td><td></td><td></td></tr>
<tr><td>Chromosome</td><td>start</td><td>end</td><td>strand</td><td>MM</td><td>target_seq</td><td>PAM</td><td>alignment</td><td>distance</td><td>position</td><td>gene name</td><td>gene id</td></tr>
<tr><td>GU305914.1</td><td>174872</td><td>174894</td><td>-</td><td>0</td><td>TGGTGGCGTGCTTCCTAGCG</td><td>GGG</td><td>||||||||[||||||||||||]PAM</td><td>NA</td><td>Intergenic</td><td>NA</td><td>NA</td></tr>
<tr><td>T21388</td><td>GGTGGCGTGCTTCCTAGCGGGGG</td><td>1000</td><td></td><td></td><td></td><td></td><td></td><td></td><td></td><td></td><td></td></tr>
<tr><td>Oligo fwd</td><td>CACCGGTGGCGTGCTTCCTAGCGG</td><td></td><td></td><td></td><td></td><td></td><td></td><td></td><td></td><td></td><td></td></tr>
<tr><td>Oligo rev</td><td>AAACCCGCTAGGAAGCACGCCACC</td><td></td><td></td><td></td><td></td><td></td><td></td><td></td><td></td><td></td><td></td></tr>
<tr><td>Chromosome</td><td>start</td><td>end</td><td>strand</td><td>MM</td><td>target_seq</td><td>PAM</td><td>alignment</td><td>distance</td><td>position</td><td>gene name</td><td>gene id</td></tr>
<tr><td>GU305914.1</td><td>174871</td><td>174893</td><td>-</td><td>0</td><td>GGTGGCGTGCTTCCTAGCGG</td><td>GGG</td><td>||||||||[||||||||||||]PAM</td><td>NA</td><td>Intergenic</td><td>NA</td><td>NA</td></tr>
<tr><td>T21389</td><td>ATATTCACGATTCACCACAAAGG</td><td>1000</td><td></td><td></td><td></td><td></td><td></td><td></td><td></td><td></td><td></td></tr>
<tr><td>Oligo fwd</td><td>CACCATATTCACGATTCACCACAA</td><td></td><td></td><td></td><td></td><td></td><td></td><td></td><td></td><td></td><td></td></tr>
<tr><td>Oligo rev</td><td>AAACTTGTGGTGAATCGTGAATAT</td><td></td><td></td><td></td><td></td><td></td><td></td><td></td><td></td><td></td><td></td></tr>
<tr><td>Chromosome</td><td>start</td><td>end</td><td>strand</td><td>MM</td><td>target_seq</td><td>PAM</td><td>alignment</td><td>distance</td><td>position</td><td>gene name</td><td>gene id</td></tr>
<tr><td>GU305914.1</td><td>174837</td><td>174859</td><td>-</td><td>0</td><td>ATATTCACGATTCACCACAA</td><td>AGG</td><td>||||||||[||||||||||||]PAM</td><td>NA</td><td>Intergenic</td><td>NA</td><td>NA</td></tr>
<tr><td>T21390</td><td>AAACACGCGCGCCTATCTCGCGG</td><td>1000</td><td></td><td></td><td></td><td></td><td></td><td></td><td></td><td></td><td></td></tr>
<tr><td>Oligo fwd</td><td>CACCAAACACGCGCGCCTATCTCG</td><td></td><td></td><td></td><td></td><td></td><td></td><td></td><td></td><td></td><td></td></tr>
<tr><td>Oligo rev</td><td>AAACCGAGATAGGCGCGCGTGTTT</td><td></td><td></td><td></td><td></td><td></td><td></td><td></td><td></td><td></td><td></td></tr>
<tr><td>Chromosome</td><td>start</td><td>end</td><td>strand</td><td>MM</td><td>target_seq</td><td>PAM</td><td>alignment</td><td>distance</td><td>position</td><td>gene name</td><td>gene id</td></tr>
<tr><td>GU305914.1</td><td>174813</td><td>174835</td><td>-</td><td>0</td><td>AAACACGCGCGCCTATCTCG</td><td>CGG</td><td>||||||||[||||||||||||]PAM</td><td>NA</td><td>Intergenic</td><td>NA</td><td>NA</td></tr>
<tr><td>T21391</td><td>TCGCGGTCCTTCCGATATGAAGG</td><td>1000</td><td></td><td></td><td></td><td></td><td></td><td></td><td></td><td></td><td></td></tr>
<tr><td>Oligo fwd</td><td>CACCTCGCGGTCCTTCCGATATGA</td><td></td><td></td><td></td><td></td><td></td><td></td><td></td><td></td><td></td><td></td></tr>
<tr><td>Oligo rev</td><td>AAACTCATATCGGAAGGACCGCGA</td><td></td><td></td><td></td><td></td><td></td><td></td><td></td><td></td><td></td><td></td></tr>
<tr><td>Chromosome</td><td>start</td><td>end</td><td>strand</td><td>MM</td><td>target_seq</td><td>PAM</td><td>alignment</td><td>distance</td><td>position</td><td>gene name</td><td>gene id</td></tr>
<tr><td>GU305914.1</td><td>174796</td><td>174818</td><td>-</td><td>0</td><td>TCGCGGTCCTTCCGATATGA</td><td>AGG</td><td>||||||||[||||||||||||]PAM</td><td>NA</td><td>Intergenic</td><td>NA</td><td>NA</td></tr>
<tr><td>T21392</td><td>AGGTATATAAACCGTGCAACAGG</td><td>1000</td><td></td><td></td><td></td><td></td><td></td><td></td><td></td><td></td><td></td></tr>
<tr><td>Oligo fwd</td><td>CACCAGGTATATAAACCGTGCAAC</td><td></td><td></td><td></td><td></td><td></td><td></td><td></td><td></td><td></td><td></td></tr>
<tr><td>Oligo rev</td><td>AAACGTTGCACGGTTTATATACCT</td><td></td><td></td><td></td><td></td><td></td><td></td><td></td><td></td><td></td><td></td></tr>
<tr><td>Chromosome</td><td>start</td><td>end</td><td>strand</td><td>MM</td><td>target_seq</td><td>PAM</td><td>alignment</td><td>distance</td><td>position</td><td>gene name</td><td>gene id</td></tr>
<tr><td>GU305914.1</td><td>174776</td><td>174798</td><td>-</td><td>0</td><td>AGGTATATAAACCGTGCAAC</td><td>AGG</td><td>||||||||[||||||||||||]PAM</td><td>NA</td><td>Intergenic</td><td>NA</td><td>NA</td></tr>
<tr><td>T21393</td><td>GCAACAGGACCGAGAGTCAAAGG</td><td>1000</td><td></td><td></td><td></td><td></td><td></td><td></td><td></td><td></td><td></td></tr>
<tr><td>Oligo fwd</td><td>CACCGCAACAGGACCGAGAGTCAA</td><td></td><td></td><td></td><td></td><td></td><td></td><td></td><td></td><td></td><td></td></tr>
<tr><td>Oligo rev</td><td>AAACTTGACTCTCGGTCCTGTTGC</td><td></td><td></td><td></td><td></td><td></td><td></td><td></td><td></td><td></td><td></td></tr>
</table>

<table>
<tr><td>Chromosome</td><td>start</td><td>end</td><td>strand</td><td>MM</td><td>target_seq</td><td>PAM</td><td>alignment</td><td>distance</td><td>position</td><td>gene name</td><td>gene id</td></tr>
<tr><td>GU305914.1</td><td>174761</td><td>174783</td><td>-</td><td>0</td><td>GCAACAGGACCGAGAGTCAA</td><td>AGG</td><td>||||||||[||||||||||||]PAM</td><td>NA</td><td>Intergenic</td><td>NA</td><td>NA</td></tr>
<tr><td></td><td></td><td></td><td></td><td></td><td></td><td></td><td></td><td></td><td></td><td></td><td></td></tr>
<tr><td>T21394</td><td>ACCACTACCACCATCTCCTCCGG</td><td>1000</td><td></td><td></td><td></td><td></td><td></td><td></td><td></td><td></td><td></td></tr>
<tr><td>Oligo fwd</td><td>CACCACCACTACCACCATCTCCTC</td><td></td><td></td><td></td><td></td><td></td><td></td><td></td><td></td><td></td><td></td></tr>
<tr><td>Oligo rev</td><td>AAACGAGGAGATGGTGGTAGTGGT</td><td></td><td></td><td></td><td></td><td></td><td></td><td></td><td></td><td></td><td></td></tr>
<tr><td>Chromosome</td><td>start</td><td>end</td><td>strand</td><td>MM</td><td>target_seq</td><td>PAM</td><td>alignment</td><td>distance</td><td>position</td><td>gene name</td><td>gene id</td></tr>
<tr><td>GU305914.1</td><td>174637</td><td>174659</td><td>-</td><td>0</td><td>ACCACTACCACCATCTCCTC</td><td>CGG</td><td>||||||||[||||||||||||]PAM</td><td>NA</td><td>Intergenic</td><td>NA</td><td>NA</td></tr>
<tr><td></td><td></td><td></td><td></td><td></td><td></td><td></td><td></td><td></td><td></td><td></td><td></td></tr>
<tr><td>T21395</td><td>ACTACCACCATCTCCTCCGGCGG</td><td>1000</td><td></td><td></td><td></td><td></td><td></td><td></td><td></td><td></td><td></td></tr>
<tr><td>Oligo fwd</td><td>CACCACTACCACCATCTCCTCCGG</td><td></td><td></td><td></td><td></td><td></td><td></td><td></td><td></td><td></td><td></td></tr>
<tr><td>Oligo rev</td><td>AAACCCGGAGGAGATGGTGGTAGT</td><td></td><td></td><td></td><td></td><td></td><td></td><td></td><td></td><td></td><td></td></tr>
<tr><td>Chromosome</td><td>start</td><td>end</td><td>strand</td><td>MM</td><td>target_seq</td><td>PAM</td><td>alignment</td><td>distance</td><td>position</td><td>gene name</td><td>gene id</td></tr>
<tr><td>GU305914.1</td><td>174634</td><td>174656</td><td>-</td><td>0</td><td>ACTACCACCATCTCCTCCGG</td><td>CGG</td><td>||||||||[||||||||||||]PAM</td><td>NA</td><td>Intergenic</td><td>NA</td><td>NA</td></tr>
<tr><td></td><td></td><td></td><td></td><td></td><td></td><td></td><td></td><td></td><td></td><td></td><td></td></tr>
<tr><td>T21396</td><td>TATCTCCCTGATCGCATTCGCGG</td><td>1000</td><td></td><td></td><td></td><td></td><td></td><td></td><td></td><td></td><td></td></tr>
<tr><td>Oligo fwd</td><td>CACCTATCTCCCTGATCGCATTCG</td><td></td><td></td><td></td><td></td><td></td><td></td><td></td><td></td><td></td><td></td></tr>
<tr><td>Oligo rev</td><td>AAACCGAATGCGATCAGGGAGATA</td><td></td><td></td><td></td><td></td><td></td><td></td><td></td><td></td><td></td><td></td></tr>
<tr><td>Chromosome</td><td>start</td><td>end</td><td>strand</td><td>MM</td><td>target_seq</td><td>PAM</td><td>alignment</td><td>distance</td><td>position</td><td>gene name</td><td>gene id</td></tr>
<tr><td>GU305914.1</td><td>174599</td><td>174621</td><td>-</td><td>0</td><td>TATCTCCCTGATCGCATTCG</td><td>CGG</td><td>||||||||[||||||||||||]PAM</td><td>NA</td><td>Intergenic</td><td>NA</td><td>NA</td></tr>
<tr><td></td><td></td><td></td><td></td><td></td><td></td><td></td><td></td><td></td><td></td><td></td><td></td></tr>
<tr><td>T21397</td><td>ATCTCCCTGATCGCATTCGCGGG</td><td>1000</td><td></td><td></td><td></td><td></td><td></td><td></td><td></td><td></td><td></td></tr>
<tr><td>Oligo fwd</td><td>CACCATCTCCCTGATCGCATTCGC</td><td></td><td></td><td></td><td></td><td></td><td></td><td></td><td></td><td></td><td></td></tr>
<tr><td>Oligo rev</td><td>AAACGCGAATGCGATCAGGGAGAT</td><td></td><td></td><td></td><td></td><td></td><td></td><td></td><td></td><td></td><td></td></tr>
<tr><td>Chromosome</td><td>start</td><td>end</td><td>strand</td><td>MM</td><td>target_seq</td><td>PAM</td><td>alignment</td><td>distance</td><td>position</td><td>gene name</td><td>gene id</td></tr>
<tr><td>GU305914.1</td><td>174598</td><td>174620</td><td>-</td><td>0</td><td>ATCTCCCTGATCGCATTCGC</td><td>GGG</td><td>||||||||[||||||||||||]PAM</td><td>NA</td><td>Intergenic</td><td>NA</td><td>NA</td></tr>
<tr><td></td><td></td><td></td><td></td><td></td><td></td><td></td><td></td><td></td><td></td><td></td><td></td></tr>
<tr><td>T21398</td><td>TCTCCCTGATCGCATTCGCGGGG</td><td>1000</td><td></td><td></td><td></td><td></td><td></td><td></td><td></td><td></td><td></td></tr>
<tr><td>Oligo fwd</td><td>CACCTCTCCCTGATCGCATTCGCG</td><td></td><td></td><td></td><td></td><td></td><td></td><td></td><td></td><td></td><td></td></tr>
<tr><td>Oligo rev</td><td>AAACCGCGAATGCGATCAGGGAGA</td><td></td><td></td><td></td><td></td><td></td><td></td><td></td><td></td><td></td><td></td></tr>
<tr><td>Chromosome</td><td>start</td><td>end</td><td>strand</td><td>MM</td><td>target_seq</td><td>PAM</td><td>alignment</td><td>distance</td><td>position</td><td>gene name</td><td>gene id</td></tr>
<tr><td>GU305914.1</td><td>174597</td><td>174619</td><td>-</td><td>0</td><td>TCTCCCTGATCGCATTCGCG</td><td>GGG</td><td>||||||||[||||||||||||]PAM</td><td>NA</td><td>Intergenic</td><td>NA</td><td>NA</td></tr>
<tr><td></td><td></td><td></td><td></td><td></td><td></td><td></td><td></td><td></td><td></td><td></td><td></td></tr>
<tr><td>T21399</td><td>GGGGACCCTGATCATCCTACTGG</td><td>1000</td><td></td><td></td><td></td><td></td><td></td><td></td><td></td><td></td><td></td></tr>
<tr><td>Oligo fwd</td><td>CACCGGGGACCCTGATCATCCTAC</td><td></td><td></td><td></td><td></td><td></td><td></td><td></td><td></td><td></td><td></td></tr>
<tr><td>Oligo rev</td><td>AAACGTAGGATGATCAGGGTCCCC</td><td></td><td></td><td></td><td></td><td></td><td></td><td></td><td></td><td></td><td></td></tr>
<tr><td>Chromosome</td><td>start</td><td>end</td><td>strand</td><td>MM</td><td>target_seq</td><td>PAM</td><td>alignment</td><td>distance</td><td>position</td><td>gene name</td><td>gene id</td></tr>
<tr><td>GU305914.1</td><td>174578</td><td>174600</td><td>-</td><td>0</td><td>GGGGACCCTGATCATCCTAC</td><td>TGG</td><td>||||||||[||||||||||||]PAM</td><td>NA</td><td>Intergenic</td><td>NA</td><td>NA</td></tr>
<tr><td></td><td></td><td></td><td></td><td></td><td></td><td></td><td></td><td></td><td></td><td></td><td></td></tr>
<tr><td>T21400</td><td>TGTTCGCTGTCGCGTATAAATGG</td><td>1000</td><td></td><td></td><td></td><td></td><td></td><td></td><td></td><td></td><td></td></tr>
<tr><td>Oligo fwd</td><td>CACCTGTTCGCTGTCGCGTATAAA</td><td></td><td></td><td></td><td></td><td></td><td></td><td></td><td></td><td></td><td></td></tr>
<tr><td>Oligo rev</td><td>AAACTTTATACGCGACAGCGAACA</td><td></td><td></td><td></td><td></td><td></td><td></td><td></td><td></td><td></td><td></td></tr>
<tr><td>Chromosome</td><td>start</td><td>end</td><td>strand</td><td>MM</td><td>target_seq</td><td>PAM</td><td>alignment</td><td>distance</td><td>position</td><td>gene name</td><td>gene id</td></tr>
<tr><td>GU305914.1</td><td>174546</td><td>174568</td><td>-</td><td>0</td><td>TGTTCGCTGTCGCGTATAAA</td><td>TGG</td><td>||||||||[||||||||||||]PAM</td><td>NA</td><td>Intergenic</td><td>NA</td><td>NA</td></tr>
<tr><td></td><td></td><td></td><td></td><td></td><td></td><td></td><td></td><td></td><td></td><td></td><td></td></tr>
<tr><td>T21401</td><td>TCGCGTATAAATGGTCCAAGAGG</td><td>1000</td><td></td><td></td><td></td><td></td><td></td><td></td><td></td><td></td><td></td></tr>
<tr><td>Oligo fwd</td><td>CACCTCGCGTATAAATGGTCCAAG</td><td></td><td></td><td></td><td></td><td></td><td></td><td></td><td></td><td></td><td></td></tr>
<tr><td>Oligo rev</td><td>AAACCTTGGACCATTTATACGCGA</td><td></td><td></td><td></td><td></td><td></td><td></td><td></td><td></td><td></td><td></td></tr>
<tr><td>Chromosome</td><td>start</td><td>end</td><td>strand</td><td>MM</td><td>target_seq</td><td>PAM</td><td>alignment</td><td>distance</td><td>position</td><td>gene name</td><td>gene id</td></tr>
<tr><td>GU305914.1</td><td>174537</td><td>174559</td><td>-</td><td>0</td><td>TCGCGTATAAATGGTCCAAG</td><td>AGG</td><td>||||||||[||||||||||||]PAM</td><td>NA</td><td>Intergenic</td><td>NA</td><td>NA</td></tr>
<tr><td></td><td></td><td></td><td></td><td></td><td></td><td></td><td></td><td></td><td></td><td></td><td></td></tr>
<tr><td>T21402</td><td>GAATTCGTTACCACACAAGCTGG</td><td>1000</td><td></td><td></td><td></td><td></td><td></td><td></td><td></td><td></td><td></td></tr>
<tr><td>Oligo fwd</td><td>CACCGAATTCGTTACCACACAAGC</td><td></td><td></td><td></td><td></td><td></td><td></td><td></td><td></td><td></td><td></td></tr>
<tr><td>Oligo rev</td><td>AAACGCTTGTGTGGTAACGAATTC</td><td></td><td></td><td></td><td></td><td></td><td></td><td></td><td></td><td></td><td></td></tr>
<tr><td>Chromosome</td><td>start</td><td>end</td><td>strand</td><td>MM</td><td>target_seq</td><td>PAM</td><td>alignment</td><td>distance</td><td>position</td><td>gene name</td><td>gene id</td></tr>
<tr><td>GU305914.1</td><td>174515</td><td>174537</td><td>-</td><td>0</td><td>GAATTCGTTACCACACAAGC</td><td>TGG</td><td>||||||||[||||||||||||]PAM</td><td>NA</td><td>Intergenic</td><td>NA</td><td>NA</td></tr>
<tr><td></td><td></td><td></td><td></td><td></td><td></td><td></td><td></td><td></td><td></td><td></td><td></td></tr>
<tr><td>T21403</td><td>CGTCCCGCGTGTTCCCCTCGAGG</td><td>1000</td><td></td><td></td><td></td><td></td><td></td><td></td><td></td><td></td><td></td></tr>
<tr><td>Oligo fwd</td><td>CACCCGTCCCGCGTGTTCCCCTCG</td><td></td><td></td><td></td><td></td><td></td><td></td><td></td><td></td><td></td><td></td></tr>
<tr><td>Oligo rev</td><td>AAACCGAGGGGAACACGCGGGACG</td><td></td><td></td><td></td><td></td><td></td><td></td><td></td><td></td><td></td><td></td></tr>
<tr><td>Chromosome</td><td>start</td><td>end</td><td>strand</td><td>MM</td><td>target_seq</td><td>PAM</td><td>alignment</td><td>distance</td><td>position</td><td>gene name</td><td>gene id</td></tr>
<tr><td>GU305914.1</td><td>174462</td><td>174484</td><td>-</td><td>0</td><td>CGTCCCGCGTGTTCCCCTCG</td><td>AGG</td><td>||||||||[||||||||||||]PAM</td><td>NA</td><td>Intergenic</td><td>NA</td><td>NA</td></tr>
<tr><td></td><td></td><td></td><td></td><td></td><td></td><td></td><td></td><td></td><td></td><td></td><td></td></tr>
<tr><td>T21404</td><td>GTCCCGCGTGTTCCCCTCGAGGG</td><td>1000</td><td></td><td></td><td></td><td></td><td></td><td></td><td></td><td></td><td></td></tr>
<tr><td>Oligo fwd</td><td>CACCGTCCCGCGTGTTCCCCTCGA</td><td></td><td></td><td></td><td></td><td></td><td></td><td></td><td></td><td></td><td></td></tr>
<tr><td>Oligo rev</td><td>AAACTCGAGGGGAACACGCGGGAC</td><td></td><td></td><td></td><td></td><td></td><td></td><td></td><td></td><td></td><td></td></tr>
<tr><td>Chromosome</td><td>start</td><td>end</td><td>strand</td><td>MM</td><td>target_seq</td><td>PAM</td><td>alignment</td><td>distance</td><td>position</td><td>gene name</td><td>gene id</td></tr>
<tr><td>GU305914.1</td><td>174461</td><td>174483</td><td>-</td><td>0</td><td>GTCCCGCGTGTTCCCCTCGA</td><td>GGG</td><td>||||||||[||||||||||||]PAM</td><td>NA</td><td>Intergenic</td><td>NA</td><td>NA</td></tr>
<tr><td></td><td></td><td></td><td></td><td></td><td></td><td></td><td></td><td></td><td></td><td></td><td></td></tr>
<tr><td>T21405</td><td>TCCCGCGTGTTCCCCTCGAGGGG</td><td>1000</td><td></td><td></td><td></td><td></td><td></td><td></td><td></td><td></td><td></td></tr>
<tr><td>Oligo fwd</td><td>CACCTCCCGCGTGTTCCCCTCGAG</td><td></td><td></td><td></td><td></td><td></td><td></td><td></td><td></td><td></td><td></td></tr>
<tr><td>Oligo rev</td><td>AAACCTCGAGGGGAACACGCGGGA</td><td></td><td></td><td></td><td></td><td></td><td></td><td></td><td></td><td></td><td></td></tr>
<tr><td>Chromosome</td><td>start</td><td>end</td><td>strand</td><td>MM</td><td>target_seq</td><td>PAM</td><td>alignment</td><td>distance</td><td>position</td><td>gene name</td><td>gene id</td></tr>
<tr><td>GU305914.1</td><td>174460</td><td>174482</td><td>-</td><td>0</td><td>TCCCGCGTGTTCCCCTCGAG</td><td>GGG</td><td>||||||||[||||||||||||]PAM</td><td>NA</td><td>Intergenic</td><td>NA</td><td>NA</td></tr>
<tr><td></td><td></td><td></td><td></td><td></td><td></td><td></td><td></td><td></td><td></td><td></td><td></td></tr>
<tr><td>T21406</td><td>AAGACCCCGCTGCCCCGCTAAGG</td><td>1000</td><td></td><td></td><td></td><td></td><td></td><td></td><td></td><td></td><td></td></tr>
<tr><td>Oligo fwd</td><td>CACCAAGACCCCGCTGCCCCGCTA</td><td></td><td></td><td></td><td></td><td></td><td></td><td></td><td></td><td></td><td></td></tr>
<tr><td>Oligo rev</td><td>AAACTAGCGGGGCAGCGGGGTCTT</td><td></td><td></td><td></td><td></td><td></td><td></td><td></td><td></td><td></td><td></td></tr>
<tr><td>Chromosome</td><td>start</td><td>end</td><td>strand</td><td>MM</td><td>target_seq</td><td>PAM</td><td>alignment</td><td>distance</td><td>position</td><td>gene name</td><td>gene id</td></tr>
<tr><td>GU305914.1</td><td>174397</td><td>174419</td><td>-</td><td>0</td><td>AAGACCCCGCTGCCCCGCTA</td><td>AGG</td><td>||||||||[||||||||||||]PAM</td><td>NA</td><td>Intergenic</td><td>NA</td><td>NA</td></tr>
<tr><td></td><td></td><td></td><td></td><td></td><td></td><td></td><td></td><td></td><td></td><td></td><td></td></tr>
<tr><td>T21407</td><td>TCCGACGTTTCTACCACACATGG</td><td>1000</td><td></td><td></td><td></td><td></td><td></td><td></td><td></td><td></td><td></td></tr>
<tr><td>Oligo fwd</td><td>CACCTCCGACGTTTCTACCACACA</td><td></td><td></td><td></td><td></td><td></td><td></td><td></td><td></td><td></td><td></td></tr>
<tr><td>Oligo rev</td><td>AAACTGTGTGGTAGAAACGTCGGA</td><td></td><td></td><td></td><td></td><td></td><td></td><td></td><td></td><td></td><td></td></tr>
<tr><td>Chromosome</td><td>start</td><td>end</td><td>strand</td><td>MM</td><td>target_seq</td><td>PAM</td><td>alignment</td><td>distance</td><td>position</td><td>gene name</td><td>gene id</td></tr>
<tr><td>GU305914.1</td><td>174351</td><td>174373</td><td>-</td><td>0</td><td>TCCGACGTTTCTACCACACA</td><td>TGG</td><td>||||||||[||||||||||||]PAM</td><td>NA</td><td>Intergenic</td><td>NA</td><td>NA</td></tr>
<tr><td></td><td></td><td></td><td></td><td></td><td></td><td></td><td></td><td></td><td></td><td></td><td></td></tr>
<tr><td>T21408</td><td>GTTTCTACCACACATGGTCTCGG</td><td>1000</td><td></td><td></td><td></td><td></td><td></td><td></td><td></td><td></td><td></td></tr>
<tr><td>Oligo fwd</td><td>CACCGTTTCTACCACACATGGTCT</td><td></td><td></td><td></td><td></td><td></td><td></td><td></td><td></td><td></td><td></td></tr>
</table>

<table>
<tr><td>Oligo rev</td><td>AAACAGACCATGTGTGGTAGAAAC</td><td></td><td></td><td></td><td></td><td></td><td></td><td></td><td></td><td></td><td></td></tr>
<tr><td>Chromosome</td><td>start</td><td>end</td><td>strand</td><td>MM</td><td>target_seq</td><td>PAM</td><td>alignment</td><td>distance</td><td>position</td><td>gene name</td><td>gene id</td></tr>
<tr><td>GU305914.1</td><td>174345</td><td>174367</td><td>-</td><td>0</td><td>GTTTCTACCACACATGGTCT</td><td>CGG</td><td>||||||||[||||||||||||]PAM</td><td>NA</td><td>Intergenic</td><td>NA</td><td>NA</td></tr>
<tr><td>T21409</td><td>TTTCTACCACACATGGTCTCGGG</td><td>1000</td><td></td><td></td><td></td><td></td><td></td><td></td><td></td><td></td><td></td></tr>
<tr><td>Oligo fwd</td><td>CACCTTTCTACCACACATGGTCTC</td><td></td><td></td><td></td><td></td><td></td><td></td><td></td><td></td><td></td><td></td></tr>
<tr><td>Oligo rev</td><td>AAACGAGACCATGTGTGGTAGAAA</td><td></td><td></td><td></td><td></td><td></td><td></td><td></td><td></td><td></td><td></td></tr>
<tr><td>Chromosome</td><td>start</td><td>end</td><td>strand</td><td>MM</td><td>target_seq</td><td>PAM</td><td>alignment</td><td>distance</td><td>position</td><td>gene name</td><td>gene id</td></tr>
<tr><td>GU305914.1</td><td>174344</td><td>174366</td><td>-</td><td>0</td><td>TTTCTACCACACATGGTCTC</td><td>GGG</td><td>||||||||[||||||||||||]PAM</td><td>NA</td><td>Intergenic</td><td>NA</td><td>NA</td></tr>
<tr><td>T21410</td><td>GATACATTGACAGCTCCCTCTGG</td><td>1000</td><td></td><td></td><td></td><td></td><td></td><td></td><td></td><td></td><td></td></tr>
<tr><td>Oligo fwd</td><td>CACCGATACATTGACAGCTCCCTC</td><td></td><td></td><td></td><td></td><td></td><td></td><td></td><td></td><td></td><td></td></tr>
<tr><td>Oligo rev</td><td>AAACGAGGGAGCTGTCAATGTATC</td><td></td><td></td><td></td><td></td><td></td><td></td><td></td><td></td><td></td><td></td></tr>
<tr><td>Chromosome</td><td>start</td><td>end</td><td>strand</td><td>MM</td><td>target_seq</td><td>PAM</td><td>alignment</td><td>distance</td><td>position</td><td>gene name</td><td>gene id</td></tr>
<tr><td>GU305914.1</td><td>174322</td><td>174344</td><td>-</td><td>0</td><td>GATACATTGACAGCTCCCTC</td><td>TGG</td><td>||||||||[||||||||||||]PAM</td><td>NA</td><td>Intergenic</td><td>NA</td><td>NA</td></tr>
<tr><td>T21411</td><td>CCCTCTGGTCACCCATCGTCCGG</td><td>1000</td><td></td><td></td><td></td><td></td><td></td><td></td><td></td><td></td><td></td></tr>
<tr><td>Oligo fwd</td><td>CACCCCCTCTGGTCACCCATCGTC</td><td></td><td></td><td></td><td></td><td></td><td></td><td></td><td></td><td></td><td></td></tr>
<tr><td>Oligo rev</td><td>AAACGACGATGGGTGACCAGAGGG</td><td></td><td></td><td></td><td></td><td></td><td></td><td></td><td></td><td></td><td></td></tr>
<tr><td>Chromosome</td><td>start</td><td>end</td><td>strand</td><td>MM</td><td>target_seq</td><td>PAM</td><td>alignment</td><td>distance</td><td>position</td><td>gene name</td><td>gene id</td></tr>
<tr><td>GU305914.1</td><td>174307</td><td>174329</td><td>-</td><td>0</td><td>CCCTCTGGTCACCCATCGTC</td><td>CGG</td><td>||||||||[||||||||||||]PAM</td><td>NA</td><td>Intergenic</td><td>NA</td><td>NA</td></tr>
<tr><td>T21412</td><td>CCGGTTCGTCGTCATGCTGCCGG</td><td>1000</td><td></td><td></td><td></td><td></td><td></td><td></td><td></td><td></td><td></td></tr>
<tr><td>Oligo fwd</td><td>CACCCCGGTTCGTCGTCATGCTGC</td><td></td><td></td><td></td><td></td><td></td><td></td><td></td><td></td><td></td><td></td></tr>
<tr><td>Oligo rev</td><td>AAACGCAGCATGACGACGAACCGG</td><td></td><td></td><td></td><td></td><td></td><td></td><td></td><td></td><td></td><td></td></tr>
<tr><td>Chromosome</td><td>start</td><td>end</td><td>strand</td><td>MM</td><td>target_seq</td><td>PAM</td><td>alignment</td><td>distance</td><td>position</td><td>gene name</td><td>gene id</td></tr>
<tr><td>GU305914.1</td><td>174288</td><td>174310</td><td>-</td><td>0</td><td>CCGGTTCGTCGTCATGCTGC</td><td>CGG</td><td>||||||||[||||||||||||]PAM</td><td>NA</td><td>Intergenic</td><td>NA</td><td>NA</td></tr>
<tr><td>T21413</td><td>CGCCGTCGCGATCATCGCGTTGG</td><td>1000</td><td></td><td></td><td></td><td></td><td></td><td></td><td></td><td></td><td></td></tr>
<tr><td>Oligo fwd</td><td>CACCCGCCGTCGCGATCATCGCGT</td><td></td><td></td><td></td><td></td><td></td><td></td><td></td><td></td><td></td><td></td></tr>
<tr><td>Oligo rev</td><td>AAACACGCGATGATCGCGACGGCG</td><td></td><td></td><td></td><td></td><td></td><td></td><td></td><td></td><td></td><td></td></tr>
<tr><td>Chromosome</td><td>start</td><td>end</td><td>strand</td><td>MM</td><td>target_seq</td><td>PAM</td><td>alignment</td><td>distance</td><td>position</td><td>gene name</td><td>gene id</td></tr>
<tr><td>GU305914.1</td><td>174246</td><td>174268</td><td>-</td><td>0</td><td>CGCCGTCGCGATCATCGCGT</td><td>TGG</td><td>||||||||[||||||||||||]PAM</td><td>NA</td><td>Intergenic</td><td>NA</td><td>NA</td></tr>
<tr><td>GU305914.1</td><td>174244</td><td>174266</td><td>+</td><td>4</td><td>CGCCAACGCGATGATCGCGA</td><td>CGG</td><td>||||--||[||||-||||||-]PAM</td><td>NA</td><td>Intergenic</td><td>NA</td><td>NA</td></tr>
<tr><td>T21414</td><td>CGTCGCGATCATCGCGTTGGCGG</td><td>1000</td><td></td><td></td><td></td><td></td><td></td><td></td><td></td><td></td><td></td></tr>
<tr><td>Oligo fwd</td><td>CACCCGTCGCGATCATCGCGTTGG</td><td></td><td></td><td></td><td></td><td></td><td></td><td></td><td></td><td></td><td></td></tr>
<tr><td>Oligo rev</td><td>AAACCCAACGCGATGATCGCGACG</td><td></td><td></td><td></td><td></td><td></td><td></td><td></td><td></td><td></td><td></td></tr>
<tr><td>Chromosome</td><td>start</td><td>end</td><td>strand</td><td>MM</td><td>target_seq</td><td>PAM</td><td>alignment</td><td>distance</td><td>position</td><td>gene name</td><td>gene id</td></tr>
<tr><td>GU305914.1</td><td>174243</td><td>174265</td><td>-</td><td>0</td><td>CGTCGCGATCATCGCGTTGG</td><td>CGG</td><td>||||||||[||||||||||||]PAM</td><td>NA</td><td>Intergenic</td><td>NA</td><td>NA</td></tr>
<tr><td>T21415</td><td>TCGCGTTGGCGGTGATCTTTCGG</td><td>1000</td><td></td><td></td><td></td><td></td><td></td><td></td><td></td><td></td><td></td></tr>
<tr><td>Oligo fwd</td><td>CACCTCGCGTTGGCGGTGATCTTT</td><td></td><td></td><td></td><td></td><td></td><td></td><td></td><td></td><td></td><td></td></tr>
<tr><td>Oligo rev</td><td>AAACAAAGATCACCGCCAACGCGA</td><td></td><td></td><td></td><td></td><td></td><td></td><td></td><td></td><td></td><td></td></tr>
<tr><td>Chromosome</td><td>start</td><td>end</td><td>strand</td><td>MM</td><td>target_seq</td><td>PAM</td><td>alignment</td><td>distance</td><td>position</td><td>gene name</td><td>gene id</td></tr>
<tr><td>GU305914.1</td><td>174232</td><td>174254</td><td>-</td><td>0</td><td>TCGCGTTGGCGGTGATCTTT</td><td>CGG</td><td>||||||||[||||||||||||]PAM</td><td>NA</td><td>Intergenic</td><td>NA</td><td>NA</td></tr>
<tr><td>T21416</td><td>CGGTGATCTTTCGGATCGTGCGG</td><td>1000</td><td></td><td></td><td></td><td></td><td></td><td></td><td></td><td></td><td></td></tr>
<tr><td>Oligo fwd</td><td>CACCCGGTGATCTTTCGGATCGTG</td><td></td><td></td><td></td><td></td><td></td><td></td><td></td><td></td><td></td><td></td></tr>
<tr><td>Oligo rev</td><td>AAACCACGATCCGAAAGATCACCG</td><td></td><td></td><td></td><td></td><td></td><td></td><td></td><td></td><td></td><td></td></tr>
<tr><td>Chromosome</td><td>start</td><td>end</td><td>strand</td><td>MM</td><td>target_seq</td><td>PAM</td><td>alignment</td><td>distance</td><td>position</td><td>gene name</td><td>gene id</td></tr>
<tr><td>GU305914.1</td><td>174223</td><td>174245</td><td>-</td><td>0</td><td>CGGTGATCTTTCGGATCGTG</td><td>CGG</td><td>||||||||[||||||||||||]PAM</td><td>NA</td><td>Intergenic</td><td>NA</td><td>NA</td></tr>
<tr><td>T21417</td><td>CGTGCGGATGAAGATCCGCGCGG</td><td>1000</td><td></td><td></td><td></td><td></td><td></td><td></td><td></td><td></td><td></td></tr>
<tr><td>Oligo fwd</td><td>CACCCGTGCGGATGAAGATCCGCG</td><td></td><td></td><td></td><td></td><td></td><td></td><td></td><td></td><td></td><td></td></tr>
<tr><td>Oligo rev</td><td>AAACCGCGGATCTTCATCCGCACG</td><td></td><td></td><td></td><td></td><td></td><td></td><td></td><td></td><td></td><td></td></tr>
<tr><td>Chromosome</td><td>start</td><td>end</td><td>strand</td><td>MM</td><td>target_seq</td><td>PAM</td><td>alignment</td><td>distance</td><td>position</td><td>gene name</td><td>gene id</td></tr>
<tr><td>GU305914.1</td><td>174207</td><td>174229</td><td>-</td><td>0</td><td>CGTGCGGATGAAGATCCGCG</td><td>CGG</td><td>||||||||[||||||||||||]PAM</td><td>NA</td><td>Intergenic</td><td>NA</td><td>NA</td></tr>
<tr><td>T21418</td><td>TGAAGATCCGCGCGGTGTATAGG</td><td>1000</td><td></td><td></td><td></td><td></td><td></td><td></td><td></td><td></td><td></td></tr>
<tr><td>Oligo fwd</td><td>CACCTGAAGATCCGCGCGGTGTAT</td><td></td><td></td><td></td><td></td><td></td><td></td><td></td><td></td><td></td><td></td></tr>
<tr><td>Oligo rev</td><td>AAACATACACCGCGCGGATCTTCA</td><td></td><td></td><td></td><td></td><td></td><td></td><td></td><td></td><td></td><td></td></tr>
<tr><td>Chromosome</td><td>start</td><td>end</td><td>strand</td><td>MM</td><td>target_seq</td><td>PAM</td><td>alignment</td><td>distance</td><td>position</td><td>gene name</td><td>gene id</td></tr>
<tr><td>GU305914.1</td><td>174199</td><td>174221</td><td>-</td><td>0</td><td>TGAAGATCCGCGCGGTGTAT</td><td>AGG</td><td>||||||||[||||||||||||]PAM</td><td>NA</td><td>Intergenic</td><td>NA</td><td>NA</td></tr>
<tr><td>T21419</td><td>ATAGGATCTACCGAGTGTTGAGG</td><td>1000</td><td></td><td></td><td></td><td></td><td></td><td></td><td></td><td></td><td></td></tr>
<tr><td>Oligo fwd</td><td>CACCATAGGATCTACCGAGTGTTG</td><td></td><td></td><td></td><td></td><td></td><td></td><td></td><td></td><td></td><td></td></tr>
<tr><td>Oligo rev</td><td>AAACCAACACTCGGTAGATCCTAT</td><td></td><td></td><td></td><td></td><td></td><td></td><td></td><td></td><td></td><td></td></tr>
<tr><td>Chromosome</td><td>start</td><td>end</td><td>strand</td><td>MM</td><td>target_seq</td><td>PAM</td><td>alignment</td><td>distance</td><td>position</td><td>gene name</td><td>gene id</td></tr>
<tr><td>GU305914.1</td><td>174181</td><td>174203</td><td>-</td><td>0</td><td>ATAGGATCTACCGAGTGTTG</td><td>AGG</td><td>||||||||[||||||||||||]PAM</td><td>NA</td><td>Intergenic</td><td>NA</td><td>NA</td></tr>
<tr><td>T21420</td><td>GGATCTACCGAGTGTTGAGGCGG</td><td>1000</td><td></td><td></td><td></td><td></td><td></td><td></td><td></td><td></td><td></td></tr>
<tr><td>Oligo fwd</td><td>CACCGGATCTACCGAGTGTTGAGG</td><td></td><td></td><td></td><td></td><td></td><td></td><td></td><td></td><td></td><td></td></tr>
<tr><td>Oligo rev</td><td>AAACCCTCAACACTCGGTAGATCC</td><td></td><td></td><td></td><td></td><td></td><td></td><td></td><td></td><td></td><td></td></tr>
<tr><td>Chromosome</td><td>start</td><td>end</td><td>strand</td><td>MM</td><td>target_seq</td><td>PAM</td><td>alignment</td><td>distance</td><td>position</td><td>gene name</td><td>gene id</td></tr>
<tr><td>GU305914.1</td><td>174178</td><td>174200</td><td>-</td><td>0</td><td>GGATCTACCGAGTGTTGAGG</td><td>CGG</td><td>||||||||[||||||||||||]PAM</td><td>NA</td><td>Intergenic</td><td>NA</td><td>NA</td></tr>
<tr><td>T21421</td><td>TCTACCGAGTGTTGAGGCGGCGG</td><td>1000</td><td></td><td></td><td></td><td></td><td></td><td></td><td></td><td></td><td></td></tr>
<tr><td>Oligo fwd</td><td>CACCTCTACCGAGTGTTGAGGCGG</td><td></td><td></td><td></td><td></td><td></td><td></td><td></td><td></td><td></td><td></td></tr>
<tr><td>Oligo rev</td><td>AAACCCGCCTCAACACTCGGTAGA</td><td></td><td></td><td></td><td></td><td></td><td></td><td></td><td></td><td></td><td></td></tr>
<tr><td>Chromosome</td><td>start</td><td>end</td><td>strand</td><td>MM</td><td>target_seq</td><td>PAM</td><td>alignment</td><td>distance</td><td>position</td><td>gene name</td><td>gene id</td></tr>
<tr><td>GU305914.1</td><td>174175</td><td>174197</td><td>-</td><td>0</td><td>TCTACCGAGTGTTGAGGCGG</td><td>CGG</td><td>||||||||[||||||||||||]PAM</td><td>NA</td><td>Intergenic</td><td>NA</td><td>NA</td></tr>
<tr><td>T21422</td><td>ACCGAGTGTTGAGGCGGCGGCGG</td><td>1000</td><td></td><td></td><td></td><td></td><td></td><td></td><td></td><td></td><td></td></tr>
<tr><td>Oligo fwd</td><td>CACCACCGAGTGTTGAGGCGGCGG</td><td></td><td></td><td></td><td></td><td></td><td></td><td></td><td></td><td></td><td></td></tr>
<tr><td>Oligo rev</td><td>AAACCCGCCGCCTCAACACTCGGT</td><td></td><td></td><td></td><td></td><td></td><td></td><td></td><td></td><td></td><td></td></tr>
<tr><td>Chromosome</td><td>start</td><td>end</td><td>strand</td><td>MM</td><td>target_seq</td><td>PAM</td><td>alignment</td><td>distance</td><td>position</td><td>gene name</td><td>gene id</td></tr>
<tr><td>GU305914.1</td><td>174172</td><td>174194</td><td>-</td><td>0</td><td>ACCGAGTGTTGAGGCGGCGG</td><td>CGG</td><td>||||||||[||||||||||||]PAM</td><td>NA</td><td>Intergenic</td><td>NA</td><td>NA</td></tr>
</table>

T21423 GCGGCGGCGGCCCACACACACGG 1000
Oligo fwd CACCGCGGCGGCGGCCCACACACA
Oligo rev AAACTGTGTGTGGGCCGCCGCCGC
Chromosome start end strand MM target_seq PAM alignment distance position gene name gene id
GU305914.1 174159 174181 - 0 GCGGCGGCGGCCCACACACA CGG ||||||||[||||||||||||]PAM NA Intergenic NA NA

T21424 GGCCCACACACACGGCGTGACGG 1000
Oligo fwd CACCGGCCCACACACACGGCGTGA
Oligo rev AAACTCACGCCGTGTGTGTGGGCC
Chromosome start end strand MM target_seq PAM alignment distance position gene name gene id
GU305914.1 174151 174173 - 0 GGCCCACACACACGGCGTGA CGG ||||||||[||||||||||||]PAM NA Intergenic NA NA

T21425 CCGATTTCCGAGAAACCCATAGG 1000
Oligo fwd CACCCCGATTTCCGAGAAACCCAT
Oligo rev AAACATGGGTTTCTCGGAAATCGG
Chromosome start end strand MM target_seq PAM alignment distance position gene name gene id
GU305914.1 174095 174117 - 0 CCGATTTCCGAGAAACCCAT AGG ||||||||[||||||||||||]PAM NA Intergenic NA NA

T21426 CGAGAAACCCATAGGATCCTAGG 1000
Oligo fwd CACCCGAGAAACCCATAGGATCCT
Oligo rev AAACAGGATCCTATGGGTTTCTCG
Chromosome start end strand MM target_seq PAM alignment distance position gene name gene id
GU305914.1 174087 174109 - 0 CGAGAAACCCATAGGATCCT AGG ||||||||[||||||||||||]PAM NA Intergenic NA NA

T21427 GGATCCTAGGCAGTTTCCCTCGG 1000
Oligo fwd CACCGGATCCTAGGCAGTTTCCCT
Oligo rev AAACAGGGAAACTGCCTAGGATCC
Chromosome start end strand MM target_seq PAM alignment distance position gene name gene id
GU305914.1 174074 174096 - 0 GGATCCTAGGCAGTTTCCCT CGG ||||||||[||||||||||||]PAM NA Intergenic NA NA

T21428 GGCAGTTTCCCTCGGTCGACCGG 1000
Oligo fwd CACCGGCAGTTTCCCTCGGTCGAC
Oligo rev AAACGTCGACCGAGGGAAACTGCC
Chromosome start end strand MM target_seq PAM alignment distance position gene name gene id
GU305914.1 174066 174088 - 0 GGCAGTTTCCCTCGGTCGAC CGG ||||||||[||||||||||||]PAM NA Intergenic NA NA

T21429 GACCGGATCGTCGAAAAACTAGG 1000
Oligo fwd CACCGACCGGATCGTCGAAAAACT
Oligo rev AAACAGTTTTTCGACGATCCGGTC
Chromosome start end strand MM target_seq PAM alignment distance position gene name gene id
GU305914.1 174049 174071 - 0 GACCGGATCGTCGAAAAACT AGG ||||||||[||||||||||||]PAM NA Intergenic NA NA

T21430 TCGAAAAACTAGGCTATTTAAGG 1000
Oligo fwd CACCTCGAAAAACTAGGCTATTTA
Oligo rev AAACTAAATAGCCTAGTTTTTCGA
Chromosome start end strand MM target_seq PAM alignment distance position gene name gene id
GU305914.1 174039 174061 - 0 TCGAAAAACTAGGCTATTTA AGG ||||||||[||||||||||||]PAM NA Intergenic NA NA

T21431 CTATTTAAGGAGCCGTCATTCGG 1000
Oligo fwd CACCCTATTTAAGGAGCCGTCATT
Oligo rev AAACAATGACGGCTCCTTAAATAG
Chromosome start end strand MM target_seq PAM alignment distance position gene name gene id
GU305914.1 174026 174048 - 0 CTATTTAAGGAGCCGTCATT CGG ||||||||[||||||||||||]PAM NA Intergenic NA NA

T21432 GGAGCCGTCATTCGGACGAAAGG 1000
Oligo fwd CACCGGAGCCGTCATTCGGACGAA
Oligo rev AAACTTCGTCCGAATGACGGCTCC
Chromosome start end strand MM target_seq PAM alignment distance position gene name gene id
GU305914.1 174018 174040 - 0 GGAGCCGTCATTCGGACGAA AGG ||||||||[||||||||||||]PAM NA Intergenic NA NA

T21433 GAGCCGTCATTCGGACGAAAGGG 1000
Oligo fwd CACCGAGCCGTCATTCGGACGAAA
Oligo rev AAACTTTCGTCCGAATGACGGCTC
Chromosome start end strand MM target_seq PAM alignment distance position gene name gene id
GU305914.1 174017 174039 - 0 GAGCCGTCATTCGGACGAAA GGG ||||||||[||||||||||||]PAM NA Intergenic NA NA

T21434 GGTTACCCCCACTCGTCGACAGG 1000
Oligo fwd CACCGGTTACCCCCACTCGTCGAC
Oligo rev AAACGTCGACGAGTGGGGGTAACC
Chromosome start end strand MM target_seq PAM alignment distance position gene name gene id
GU305914.1 173996 174018 - 0 GGTTACCCCCACTCGTCGAC AGG ||||||||[||||||||||||]PAM NA Intergenic NA NA

T21435 GGATTATGTCATCGAGATCTCGG 1000
Oligo fwd CACCGGATTATGTCATCGAGATCT
Oligo rev AAACAGATCTCGATGACATAATCC
Chromosome start end strand MM target_seq PAM alignment distance position gene name gene id
GU305914.1 173975 173997 - 0 GGATTATGTCATCGAGATCT CGG ||||||||[||||||||||||]PAM NA Intergenic NA NA

T21436 ATCGAGATCTCGGCATCATCCGG 1000
Oligo fwd CACCATCGAGATCTCGGCATCATC
Oligo rev AAACGATGATGCCGAGATCTCGAT
Chromosome start end strand MM target_seq PAM alignment distance position gene name gene id
GU305914.1 173965 173987 - 0 ATCGAGATCTCGGCATCATC CGG ||||||||[||||||||||||]PAM NA Intergenic NA NA

T21437 TGAGACCATCCGACACAATTCGG 1000
Oligo fwd CACCTGAGACCATCCGACACAATT
Oligo rev AAACAATTGTGTCGGATGGTCTCA
Chromosome start end strand MM target_seq PAM alignment distance position gene name gene id
GU305914.1 173921 173943 - 0 TGAGACCATCCGACACAATT CGG ||||||||[||||||||||||]PAM NA Intergenic NA NA

T21438 ACCATCCGACACAATTCGGTCGG 1000
Oligo fwd CACCACCATCCGACACAATTCGGT
Oligo rev AAACACCGAATTGTGTCGGATGGT
Chromosome start end strand MM target_seq PAM alignment distance position gene name gene id
GU305914.1 173917 173939 - 0 ACCATCCGACACAATTCGGT CGG ||||||||[||||||||||||]PAM NA Intergenic NA NA

T21439 ACACAATTCGGTCGGACGCGCGG 1000
Oligo fwd CACCACACAATTCGGTCGGACGCG
Oligo rev AAACCGCGTCCGACCGAATTGTGT
Chromosome start end strand MM target_seq PAM alignment distance position gene name gene id
GU305914.1 173909 173931 - 0 ACACAATTCGGTCGGACGCG CGG ||||||||[||||||||||||]PAM NA Intergenic NA NA

T21440 GGTCGGACGCGCGGCGACATAGG 1000
Oligo fwd CACCGGTCGGACGCGCGGCGACAT
Oligo rev AAACATGTCGCCGCGCGTCCGACC
Chromosome start end strand MM target_seq PAM alignment distance position gene name gene id
GU305914.1 173900 173922 - 0 GGTCGGACGCGCGGCGACAT AGG ||||||||[||||||||||||]PAM NA Intergenic NA NA

T21441 CGGACGCGCGGCGACATAGGAGG 1000
Oligo fwd CACCCGGACGCGCGGCGACATAGG
Oligo rev AAACCCTATGTCGCCGCGCGTCCG
Chromosome start end strand MM target_seq PAM alignment distance position gene name gene id
GU305914.1 173897 173919 - 0 CGGACGCGCGGCGACATAGG AGG ||||||||[||||||||||||]PAM NA Intergenic NA NA

T21442 ACGCGCGGCGACATAGGAGGTGG 1000
Oligo fwd CACCACGCGCGGCGACATAGGAGG
Oligo rev AAACCCTCCTATGTCGCCGCGCGT
Chromosome start end strand MM target_seq PAM alignment distance position gene name gene id
GU305914.1 173894 173916 - 0 ACGCGCGGCGACATAGGAGG TGG ||||||||[||||||||||||]PAM NA Intergenic NA NA

T21443 GGCGACATAGGAGGTGGACGAGG 1000
Oligo fwd CACCGGCGACATAGGAGGTGGACG
Oligo rev AAACCGTCCACCTCCTATGTCGCC
Chromosome start end strand MM target_seq PAM alignment distance position gene name gene id
GU305914.1 173888 173910 - 0 GGCGACATAGGAGGTGGACG AGG ||||||||[||||||||||||]PAM NA Intergenic NA NA

T21444 CGACCTCCCCGCCGAAGATACGG 1000
Oligo fwd CACCCGACCTCCCCGCCGAAGATA
Oligo rev AAACTATCTTCGGCGGGGAGGTCG
Chromosome start end strand MM target_seq PAM alignment distance position gene name gene id
GU305914.1 173824 173846 - 0 CGACCTCCCCGCCGAAGATA CGG ||||||||[||||||||||||]PAM NA Intergenic NA NA

T21445 CCGCCGAAGATACGGTGATGCGG 1000
Oligo fwd CACCCCGCCGAAGATACGGTGATG
Oligo rev AAACCATCACCGTATCTTCGGCGG
Chromosome start end strand MM target_seq PAM alignment distance position gene name gene id
GU305914.1 173816 173838 - 0 CCGCCGAAGATACGGTGATG CGG ||||||||[||||||||||||]PAM NA Intergenic NA NA

T21446 CGCCGAAGATACGGTGATGCGGG 1000
Oligo fwd CACCCGCCGAAGATACGGTGATGC
Oligo rev AAACGCATCACCGTATCTTCGGCG
Chromosome start end strand MM target_seq PAM alignment distance position gene name gene id
GU305914.1 173815 173837 - 0 CGCCGAAGATACGGTGATGC GGG ||||||||[||||||||||||]PAM NA Intergenic NA NA

T21447 GATGCGGGTCGCTATACCGTTGG 1000
Oligo fwd CACCGATGCGGGTCGCTATACCGT
Oligo rev AAACACGGTATAGCGACCCGCATC
Chromosome start end strand MM target_seq PAM alignment distance position gene name gene id
GU305914.1 173800 173822 - 0 GATGCGGGTCGCTATACCGT TGG ||||||||[||||||||||||]PAM NA Intergenic NA NA

T21448 TATACCGTTGGTCTTCTGTATGG 1000
Oligo fwd CACCTATACCGTTGGTCTTCTGTA
Oligo rev AAACTACAGAAGACCAACGGTATA
Chromosome start end strand MM target_seq PAM alignment distance position gene name gene id
GU305914.1 173788 173810 - 0 TATACCGTTGGTCTTCTGTA TGG ||||||||[||||||||||||]PAM NA Intergenic NA NA

T21449 CGTTGGTCTTCTGTATGGTGTGG 1000
Oligo fwd CACCCGTTGGTCTTCTGTATGGTG
Oligo rev AAACCACCATACAGAAGACCAACG
Chromosome start end strand MM target_seq PAM alignment distance position gene name gene id
GU305914.1 173783 173805 - 0 CGTTGGTCTTCTGTATGGTG TGG ||||||||[||||||||||||]PAM NA Intergenic NA NA

T21450 CTACCTCCTGCATCCCGCTTCGG 1000
Oligo fwd CACCCTACCTCCTGCATCCCGCTT
Oligo rev AAACAAGCGGGATGCAGGAGGTAG
Chromosome start end strand MM target_seq PAM alignment distance position gene name gene id
GU305914.1 173755 173777 - 0 CTACCTCCTGCATCCCGCTT CGG ||||||||[||||||||||||]PAM NA Intergenic NA NA

T21451 CCTGCATCCCGCTTCGGTGCTGG 1000
Oligo fwd CACCCCTGCATCCCGCTTCGGTGC
Oligo rev AAACGCACCGAAGCGGGATGCAGG
Chromosome start end strand MM target_seq PAM alignment distance position gene name gene id
GU305914.1 173749 173771 - 0 CCTGCATCCCGCTTCGGTGC TGG ||||||||[||||||||||||]PAM NA Intergenic NA NA

T21452 CTGGCGTCGAGCGCCGCCACCGG 1000
Oligo fwd CACCCTGGCGTCGAGCGCCGCCAC
Oligo rev AAACGTGGCGGCGCTCGACGCCAG
Chromosome start end strand MM target_seq PAM alignment distance position gene name gene id

| | | | | | | | | | | | |
|---|---|---|---|---|---|---|---|---|---|---|---|
| GU305914.1 | 173730 | 173752 | - | 0 | CTGGCGTCGAGCGCCGCCAC | CGG | \|\|\|\|\|\|\|\|[\|\|\|\|\|\|\|\|\|\|\|\|]PAM | NA | Intergenic | NA | NA |
| T21453 | CCGACTAGCGTTCTCCTCGACGG | 1000 | | | | | | | | | |
| Oligo fwd | CACCCCGACTAGCGTTCTCCTCGA | | | | | | | | | | |
| Oligo rev | AAACTCGAGGAGAACGCTAGTCGG | | | | | | | | | | |
| Chromosome | start | end | strand | MM | target_seq | PAM | alignment | distance | position | gene name | gene id |
| GU305914.1 | 173542 | 173564 | - | 0 | CCGACTAGCGTTCTCCTCGA | CGG | \|\|\|\|\|\|\|\|[\|\|\|\|\|\|\|\|\|\|\|\|]PAM | NA | Intergenic | NA | NA |
| T21454 | AAAAGAAGAGAACAACCTCATGG | 1000 | | | | | | | | | |
| Oligo fwd | CACCAAAAGAAGAGAACAACCTCA | | | | | | | | | | |
| Oligo rev | AAACTGAGGTTGTTCTCTTCTTTT | | | | | | | | | | |
| Chromosome | start | end | strand | MM | target_seq | PAM | alignment | distance | position | gene name | gene id |
| GU305914.1 | 173487 | 173509 | - | 0 | AAAAGAAGAGAACAACCTCA | TGG | \|\|\|\|\|\|\|\|[\|\|\|\|\|\|\|\|\|\|\|\|]PAM | NA | Intergenic | NA | NA |
| T21455 | CGCTATTTTCAACCTCGTCATGG | 1000 | | | | | | | | | |
| Oligo fwd | CACCCGCTATTTTCAACCTCGTCA | | | | | | | | | | |
| Oligo rev | AAACTGACGAGGTTGAAAATAGCG | | | | | | | | | | |
| Chromosome | start | end | strand | MM | target_seq | PAM | alignment | distance | position | gene name | gene id |
| GU305914.1 | 173412 | 173434 | - | 0 | CGCTATTTTCAACCTCGTCA | TGG | \|\|\|\|\|\|\|\|[\|\|\|\|\|\|\|\|\|\|\|\|]PAM | NA | Intergenic | NA | NA |
| T21456 | TCTACCGATCTCCTCACGTACGG | 1000 | | | | | | | | | |
| Oligo fwd | CACCTCTACCGATCTCCTCACGTA | | | | | | | | | | |
| Oligo rev | AAACTACGTGAGGAGATCGGTAGA | | | | | | | | | | |
| Chromosome | start | end | strand | MM | target_seq | PAM | alignment | distance | position | gene name | gene id |
| GU305914.1 | 173381 | 173403 | - | 0 | TCTACCGATCTCCTCACGTA | CGG | \|\|\|\|\|\|\|\|[\|\|\|\|\|\|\|\|\|\|\|\|]PAM | NA | Intergenic | NA | NA |
| T21457 | TCGTGCATACTGCTAGCGCTCGG | 1000 | | | | | | | | | |
| Oligo fwd | CACCTCGTGCATACTGCTAGCGCT | | | | | | | | | | |
| Oligo rev | AAACAGCGCTAGCAGTATGCACGA | | | | | | | | | | |
| Chromosome | start | end | strand | MM | target_seq | PAM | alignment | distance | position | gene name | gene id |
| GU305914.1 | 173351 | 173373 | - | 0 | TCGTGCATACTGCTAGCGCT | CGG | \|\|\|\|\|\|\|\|[\|\|\|\|\|\|\|\|\|\|\|\|]PAM | NA | Intergenic | NA | NA |
| T21458 | CTAGCGCTCGGATTCCTCTCCGG | 1000 | | | | | | | | | |
| Oligo fwd | CACCCTAGCGCTCGGATTCCTCTC | | | | | | | | | | |
| Oligo rev | AAACGAGAGGAATCCGAGCGCTAG | | | | | | | | | | |
| Chromosome | start | end | strand | MM | target_seq | PAM | alignment | distance | position | gene name | gene id |
| GU305914.1 | 173339 | 173361 | - | 0 | CTAGCGCTCGGATTCCTCTC | CGG | \|\|\|\|\|\|\|\|[\|\|\|\|\|\|\|\|\|\|\|\|]PAM | NA | Intergenic | NA | NA |
| T21459 | TAGCGCTCGGATTCCTCTCCGGG | 1000 | | | | | | | | | |
| Oligo fwd | CACCTAGCGCTCGGATTCCTCTCC | | | | | | | | | | |
| Oligo rev | AAACGGAGAGGAATCCGAGCGCTA | | | | | | | | | | |
| Chromosome | start | end | strand | MM | target_seq | PAM | alignment | distance | position | gene name | gene id |
| GU305914.1 | 173338 | 173360 | - | 0 | TAGCGCTCGGATTCCTCTCC | GGG | \|\|\|\|\|\|\|\|[\|\|\|\|\|\|\|\|\|\|\|\|]PAM | NA | Intergenic | NA | NA |
| T21460 | ATTCCTCTCCGGGATCCTGATGG | 1000 | | | | | | | | | |
| Oligo fwd | CACCATTCCTCTCCGGGATCCTGA | | | | | | | | | | |
| Oligo rev | AAACTCAGGATCCCGGAGAGGAAT | | | | | | | | | | |
| Chromosome | start | end | strand | MM | target_seq | PAM | alignment | distance | position | gene name | gene id |
| GU305914.1 | 173328 | 173350 | - | 0 | ATTCCTCTCCGGGATCCTGA | TGG | \|\|\|\|\|\|\|\|[\|\|\|\|\|\|\|\|\|\|\|\|]PAM | NA | Intergenic | NA | NA |
| T21461 | CTCCGGGATCCTGATGGCCGTGG | 1000 | | | | | | | | | |
| Oligo fwd | CACCCTCCGGGATCCTGATGGCCG | | | | | | | | | | |
| Oligo rev | AAACCGGCCATCAGGATCCCGGAG | | | | | | | | | | |
| Chromosome | start | end | strand | MM | target_seq | PAM | alignment | distance | position | gene name | gene id |
| GU305914.1 | 173322 | 173344 | - | 0 | CTCCGGGATCCTGATGGCCG | TGG | \|\|\|\|\|\|\|\|[\|\|\|\|\|\|\|\|\|\|\|\|]PAM | NA | Intergenic | NA | NA |
| GU305914.1 | 127684 | 127706 | + | 4 | CTTCAGGACGCTGATGGCCG | CGG | \|\|-\|-\|\|\|[--\|\|\|\|\|\|\|\|\|\|]PAM | NA | Intergenic | NA | NA |
| T21462 | GCACAAGCTAAAAAAGCTTCTGG | 1000 | | | | | | | | | |
| Oligo fwd | CACCGCACAAGCTAAAAAAGCTTC | | | | | | | | | | |
| Oligo rev | AAACGAAGCTTTTTTAGCTTGTGC | | | | | | | | | | |
| Chromosome | start | end | strand | MM | target_seq | PAM | alignment | distance | position | gene name | gene id |
| GU305914.1 | 173265 | 173287 | - | 0 | GCACAAGCTAAAAAAGCTTC | TGG | \|\|\|\|\|\|\|\|[\|\|\|\|\|\|\|\|\|\|\|\|]PAM | NA | Intergenic | NA | NA |
| T21463 | CACAAGCTAAAAAAGCTTCTGGG | 1000 | | | | | | | | | |
| Oligo fwd | CACCCACAAGCTAAAAAAGCTTCT | | | | | | | | | | |
| Oligo rev | AAACAGAAGCTTTTTTAGCTTGTG | | | | | | | | | | |
| Chromosome | start | end | strand | MM | target_seq | PAM | alignment | distance | position | gene name | gene id |
| GU305914.1 | 173264 | 173286 | - | 0 | CACAAGCTAAAAAAGCTTCT | GGG | \|\|\|\|\|\|\|\|[\|\|\|\|\|\|\|\|\|\|\|\|]PAM | NA | Intergenic | NA | NA |
| T21464 | AATAAGAAAAGCAACGATCACGG | 1000 | | | | | | | | | |
| Oligo fwd | CACCAATAAGAAAAGCAACGATCA | | | | | | | | | | |
| Oligo rev | AAACTGATCGTTGCTTTTCTTATT | | | | | | | | | | |
| Chromosome | start | end | strand | MM | target_seq | PAM | alignment | distance | position | gene name | gene id |
| GU305914.1 | 173174 | 173196 | - | 0 | AATAAGAAAAGCAACGATCA | CGG | \|\|\|\|\|\|\|\|[\|\|\|\|\|\|\|\|\|\|\|\|]PAM | NA | Intergenic | NA | NA |
| T21465 | AAAAGCAACGATCACGGCGACGG | 1000 | | | | | | | | | |
| Oligo fwd | CACCAAAAGCAACGATCACGGCGA | | | | | | | | | | |
| Oligo rev | AAACTCGCCGTGATCGTTGCTTTT | | | | | | | | | | |
| Chromosome | start | end | strand | MM | target_seq | PAM | alignment | distance | position | gene name | gene id |
| GU305914.1 | 173168 | 173190 | - | 0 | AAAAGCAACGATCACGGCGA | CGG | \|\|\|\|\|\|\|\|[\|\|\|\|\|\|\|\|\|\|\|\|]PAM | NA | Intergenic | NA | NA |
| T21466 | CAACGATCACGGCGACGGCACGG | 1000 | | | | | | | | | |
| Oligo fwd | CACCCAACGATCACGGCGACGGCA | | | | | | | | | | |
| Oligo rev | AAACTGCCGTCGCCGTGATCGTTG | | | | | | | | | | |
| Chromosome | start | end | strand | MM | target_seq | PAM | alignment | distance | position | gene name | gene id |
| GU305914.1 | 173163 | 173185 | - | 0 | CAACGATCACGGCGACGGCA | CGG | \|\|\|\|\|\|\|\|[\|\|\|\|\|\|\|\|\|\|\|\|]PAM | NA | Intergenic | NA | NA |
| GU305914.1 | 140225 | 140247 | - | 4 | CCACGCTCATGGCGACGGCG | GGG | \|-\|\|\|-\|\|[\|-\|\|\|\|\|\|\|\|\|-]PAM | NA | Intergenic | NA | NA |
| T21467 | CGACGGCACGGAATCTGCCGAGG | 1000 | | | | | | | | | |

<table>
<tr><td>Oligo fwd</td><td>CACCCGACGGCACGGAATCTGCCG</td><td></td><td></td><td></td><td></td><td></td><td></td><td></td><td></td><td></td><td></td></tr>
<tr><td>Oligo rev</td><td>AAACCGGCAGATTCCGTGCCGTCG</td><td></td><td></td><td></td><td></td><td></td><td></td><td></td><td></td><td></td><td></td></tr>
<tr><td>Chromosome</td><td>start</td><td>end</td><td>strand</td><td>MM</td><td>target_seq</td><td>PAM</td><td>alignment</td><td>distance</td><td>position</td><td>gene name</td><td>gene id</td></tr>
<tr><td>GU305914.1</td><td>173151</td><td>173173</td><td>-</td><td>0</td><td>CGACGGCACGGAATCTGCCG</td><td>AGG</td><td>||||||||[||||||||||||]PAM</td><td>NA</td><td>Intergenic</td><td>NA</td><td>NA</td></tr>
<tr><td></td><td></td><td></td><td></td><td></td><td></td><td></td><td></td><td></td><td></td><td></td><td></td></tr>
<tr><td>T21468</td><td>TGTACATCCCAACGAAGTGACGG</td><td>1000</td><td></td><td></td><td></td><td></td><td></td><td></td><td></td><td></td><td></td></tr>
<tr><td>Oligo fwd</td><td>CACCTGTACATCCCAACGAAGTGA</td><td></td><td></td><td></td><td></td><td></td><td></td><td></td><td></td><td></td><td></td></tr>
<tr><td>Oligo rev</td><td>AAACTCACTTCGTTGGGATGTACA</td><td></td><td></td><td></td><td></td><td></td><td></td><td></td><td></td><td></td><td></td></tr>
<tr><td>Chromosome</td><td>start</td><td>end</td><td>strand</td><td>MM</td><td>target_seq</td><td>PAM</td><td>alignment</td><td>distance</td><td>position</td><td>gene name</td><td>gene id</td></tr>
<tr><td>GU305914.1</td><td>173119</td><td>173141</td><td>-</td><td>0</td><td>TGTACATCCCAACGAAGTGA</td><td>CGG</td><td>||||||||[||||||||||||]PAM</td><td>NA</td><td>Intergenic</td><td>NA</td><td>NA</td></tr>
<tr><td></td><td></td><td></td><td></td><td></td><td></td><td></td><td></td><td></td><td></td><td></td><td></td></tr>
<tr><td>T21469</td><td>ACGGAACCCACTGAGTCTACTGG</td><td>1000</td><td></td><td></td><td></td><td></td><td></td><td></td><td></td><td></td><td></td></tr>
<tr><td>Oligo fwd</td><td>CACCACGGAACCCACTGAGTCTAC</td><td></td><td></td><td></td><td></td><td></td><td></td><td></td><td></td><td></td><td></td></tr>
<tr><td>Oligo rev</td><td>AAACGTAGACTCAGTGGGTTCCGT</td><td></td><td></td><td></td><td></td><td></td><td></td><td></td><td></td><td></td><td></td></tr>
<tr><td>Chromosome</td><td>start</td><td>end</td><td>strand</td><td>MM</td><td>target_seq</td><td>PAM</td><td>alignment</td><td>distance</td><td>position</td><td>gene name</td><td>gene id</td></tr>
<tr><td>GU305914.1</td><td>173100</td><td>173122</td><td>-</td><td>0</td><td>ACGGAACCCACTGAGTCTAC</td><td>TGG</td><td>||||||||[||||||||||||]PAM</td><td>NA</td><td>Intergenic</td><td>NA</td><td>NA</td></tr>
<tr><td></td><td></td><td></td><td></td><td></td><td></td><td></td><td></td><td></td><td></td><td></td><td></td></tr>
<tr><td>T21470</td><td>GAGTCTACTGGCTTAACGCATGG</td><td>1000</td><td></td><td></td><td></td><td></td><td></td><td></td><td></td><td></td><td></td></tr>
<tr><td>Oligo fwd</td><td>CACCGAGTCTACTGGCTTAACGCA</td><td></td><td></td><td></td><td></td><td></td><td></td><td></td><td></td><td></td><td></td></tr>
<tr><td>Oligo rev</td><td>AAACTGCGTTAAGCCAGTAGACTC</td><td></td><td></td><td></td><td></td><td></td><td></td><td></td><td></td><td></td><td></td></tr>
<tr><td>Chromosome</td><td>start</td><td>end</td><td>strand</td><td>MM</td><td>target_seq</td><td>PAM</td><td>alignment</td><td>distance</td><td>position</td><td>gene name</td><td>gene id</td></tr>
<tr><td>GU305914.1</td><td>173088</td><td>173110</td><td>-</td><td>0</td><td>GAGTCTACTGGCTTAACGCA</td><td>TGG</td><td>||||||||[||||||||||||]PAM</td><td>NA</td><td>Intergenic</td><td>NA</td><td>NA</td></tr>
<tr><td></td><td></td><td></td><td></td><td></td><td></td><td></td><td></td><td></td><td></td><td></td><td></td></tr>
<tr><td>T21471</td><td>GCTTTTTTTATTTTGCTCCACGG</td><td>1000</td><td></td><td></td><td></td><td></td><td></td><td></td><td></td><td></td><td></td></tr>
<tr><td>Oligo fwd</td><td>CACCGCTTTTTTTATTTTGCTCCA</td><td></td><td></td><td></td><td></td><td></td><td></td><td></td><td></td><td></td><td></td></tr>
<tr><td>Oligo rev</td><td>AAACTGGAGCAAAATAAAAAAAGC</td><td></td><td></td><td></td><td></td><td></td><td></td><td></td><td></td><td></td><td></td></tr>
<tr><td>Chromosome</td><td>start</td><td>end</td><td>strand</td><td>MM</td><td>target_seq</td><td>PAM</td><td>alignment</td><td>distance</td><td>position</td><td>gene name</td><td>gene id</td></tr>
<tr><td>GU305914.1</td><td>172927</td><td>172949</td><td>-</td><td>0</td><td>GCTTTTTTTATTTTGCTCCA</td><td>CGG</td><td>||||||||[||||||||||||]PAM</td><td>NA</td><td>Intergenic</td><td>NA</td><td>NA</td></tr>
<tr><td></td><td></td><td></td><td></td><td></td><td></td><td></td><td></td><td></td><td></td><td></td><td></td></tr>
<tr><td>T21472</td><td>TGATTTGCTACTCCATCTCGTGG</td><td>1000</td><td></td><td></td><td></td><td></td><td></td><td></td><td></td><td></td><td></td></tr>
<tr><td>Oligo fwd</td><td>CACCTGATTTGCTACTCCATCTCG</td><td></td><td></td><td></td><td></td><td></td><td></td><td></td><td></td><td></td><td></td></tr>
<tr><td>Oligo rev</td><td>AAACCGAGATGGAGTAGCAAATCA</td><td></td><td></td><td></td><td></td><td></td><td></td><td></td><td></td><td></td><td></td></tr>
<tr><td>Chromosome</td><td>start</td><td>end</td><td>strand</td><td>MM</td><td>target_seq</td><td>PAM</td><td>alignment</td><td>distance</td><td>position</td><td>gene name</td><td>gene id</td></tr>
<tr><td>GU305914.1</td><td>172881</td><td>172903</td><td>-</td><td>0</td><td>TGATTTGCTACTCCATCTCG</td><td>TGG</td><td>||||||||[||||||||||||]PAM</td><td>NA</td><td>Intergenic</td><td>NA</td><td>NA</td></tr>
<tr><td></td><td></td><td></td><td></td><td></td><td></td><td></td><td></td><td></td><td></td><td></td><td></td></tr>
<tr><td>T21473</td><td>TGTGTCTCTTTTCTTAGCGTCGG</td><td>1000</td><td></td><td></td><td></td><td></td><td></td><td></td><td></td><td></td><td></td></tr>
<tr><td>Oligo fwd</td><td>CACCTGTGTCTCTTTTCTTAGCGT</td><td></td><td></td><td></td><td></td><td></td><td></td><td></td><td></td><td></td><td></td></tr>
<tr><td>Oligo rev</td><td>AAACACGCTAAGAAAAGAGACACA</td><td></td><td></td><td></td><td></td><td></td><td></td><td></td><td></td><td></td><td></td></tr>
<tr><td>Chromosome</td><td>start</td><td>end</td><td>strand</td><td>MM</td><td>target_seq</td><td>PAM</td><td>alignment</td><td>distance</td><td>position</td><td>gene name</td><td>gene id</td></tr>
<tr><td>GU305914.1</td><td>172856</td><td>172878</td><td>-</td><td>0</td><td>TGTGTCTCTTTTCTTAGCGT</td><td>CGG</td><td>||||||||[||||||||||||]PAM</td><td>NA</td><td>Intergenic</td><td>NA</td><td>NA</td></tr>
<tr><td></td><td></td><td></td><td></td><td></td><td></td><td></td><td></td><td></td><td></td><td></td><td></td></tr>
<tr><td>T21474</td><td>GTCTCTTTTCTTAGCGTCGGCGG</td><td>1000</td><td></td><td></td><td></td><td></td><td></td><td></td><td></td><td></td><td></td></tr>
<tr><td>Oligo fwd</td><td>CACCGTCTCTTTTCTTAGCGTCGG</td><td></td><td></td><td></td><td></td><td></td><td></td><td></td><td></td><td></td><td></td></tr>
<tr><td>Oligo rev</td><td>AAACCCGACGCTAAGAAAAGAGAC</td><td></td><td></td><td></td><td></td><td></td><td></td><td></td><td></td><td></td><td></td></tr>
<tr><td>Chromosome</td><td>start</td><td>end</td><td>strand</td><td>MM</td><td>target_seq</td><td>PAM</td><td>alignment</td><td>distance</td><td>position</td><td>gene name</td><td>gene id</td></tr>
<tr><td>GU305914.1</td><td>172853</td><td>172875</td><td>-</td><td>0</td><td>GTCTCTTTTCTTAGCGTCGG</td><td>CGG</td><td>||||||||[||||||||||||]PAM</td><td>NA</td><td>Intergenic</td><td>NA</td><td>NA</td></tr>
<tr><td></td><td></td><td></td><td></td><td></td><td></td><td></td><td></td><td></td><td></td><td></td><td></td></tr>
<tr><td>T21475</td><td>TCTCTTTTCTTAGCGTCGGCGGG</td><td>1000</td><td></td><td></td><td></td><td></td><td></td><td></td><td></td><td></td><td></td></tr>
<tr><td>Oligo fwd</td><td>CACCTCTCTTTTCTTAGCGTCGGC</td><td></td><td></td><td></td><td></td><td></td><td></td><td></td><td></td><td></td><td></td></tr>
<tr><td>Oligo rev</td><td>AAACGCCGACGCTAAGAAAAGAGA</td><td></td><td></td><td></td><td></td><td></td><td></td><td></td><td></td><td></td><td></td></tr>
<tr><td>Chromosome</td><td>start</td><td>end</td><td>strand</td><td>MM</td><td>target_seq</td><td>PAM</td><td>alignment</td><td>distance</td><td>position</td><td>gene name</td><td>gene id</td></tr>
<tr><td>GU305914.1</td><td>172852</td><td>172874</td><td>-</td><td>0</td><td>TCTCTTTTCTTAGCGTCGGC</td><td>GGG</td><td>||||||||[||||||||||||]PAM</td><td>NA</td><td>Intergenic</td><td>NA</td><td>NA</td></tr>
<tr><td></td><td></td><td></td><td></td><td></td><td></td><td></td><td></td><td></td><td></td><td></td><td></td></tr>
<tr><td>T21476</td><td>TGCGCATCAATTGCACCGTCCGG</td><td>1000</td><td></td><td></td><td></td><td></td><td></td><td></td><td></td><td></td><td></td></tr>
<tr><td>Oligo fwd</td><td>CACCTGCGCATCAATTGCACCGTC</td><td></td><td></td><td></td><td></td><td></td><td></td><td></td><td></td><td></td><td></td></tr>
<tr><td>Oligo rev</td><td>AAACGACGGTGCAATTGATGCGCA</td><td></td><td></td><td></td><td></td><td></td><td></td><td></td><td></td><td></td><td></td></tr>
<tr><td>Chromosome</td><td>start</td><td>end</td><td>strand</td><td>MM</td><td>target_seq</td><td>PAM</td><td>alignment</td><td>distance</td><td>position</td><td>gene name</td><td>gene id</td></tr>
<tr><td>GU305914.1</td><td>172780</td><td>172802</td><td>-</td><td>0</td><td>TGCGCATCAATTGCACCGTC</td><td>CGG</td><td>||||||||[||||||||||||]PAM</td><td>NA</td><td>Intergenic</td><td>NA</td><td>NA</td></tr>
<tr><td></td><td></td><td></td><td></td><td></td><td></td><td></td><td></td><td></td><td></td><td></td><td></td></tr>
<tr><td>T21477</td><td>CAATTGCACCGTCCGGTGCGAGG</td><td>1000</td><td></td><td></td><td></td><td></td><td></td><td></td><td></td><td></td><td></td></tr>
<tr><td>Oligo fwd</td><td>CACCCAATTGCACCGTCCGGTGCG</td><td></td><td></td><td></td><td></td><td></td><td></td><td></td><td></td><td></td><td></td></tr>
<tr><td>Oligo rev</td><td>AAACCGCACCGGACGGTGCAATTG</td><td></td><td></td><td></td><td></td><td></td><td></td><td></td><td></td><td></td><td></td></tr>
<tr><td>Chromosome</td><td>start</td><td>end</td><td>strand</td><td>MM</td><td>target_seq</td><td>PAM</td><td>alignment</td><td>distance</td><td>position</td><td>gene name</td><td>gene id</td></tr>
<tr><td>GU305914.1</td><td>172773</td><td>172795</td><td>-</td><td>0</td><td>CAATTGCACCGTCCGGTGCG</td><td>AGG</td><td>||||||||[||||||||||||]PAM</td><td>NA</td><td>Intergenic</td><td>NA</td><td>NA</td></tr>
<tr><td></td><td></td><td></td><td></td><td></td><td></td><td></td><td></td><td></td><td></td><td></td><td></td></tr>
<tr><td>T21478</td><td>AGCTCTCTGAGATACACTTTCGG</td><td>1000</td><td></td><td></td><td></td><td></td><td></td><td></td><td></td><td></td><td></td></tr>
<tr><td>Oligo fwd</td><td>CACCAGCTCTCTGAGATACACTTT</td><td></td><td></td><td></td><td></td><td></td><td></td><td></td><td></td><td></td><td></td></tr>
<tr><td>Oligo rev</td><td>AAACAAAGTGTATCTCAGAGAGCT</td><td></td><td></td><td></td><td></td><td></td><td></td><td></td><td></td><td></td><td></td></tr>
<tr><td>Chromosome</td><td>start</td><td>end</td><td>strand</td><td>MM</td><td>target_seq</td><td>PAM</td><td>alignment</td><td>distance</td><td>position</td><td>gene name</td><td>gene id</td></tr>
<tr><td>GU305914.1</td><td>172744</td><td>172766</td><td>-</td><td>0</td><td>AGCTCTCTGAGATACACTTT</td><td>CGG</td><td>||||||||[||||||||||||]PAM</td><td>NA</td><td>Intergenic</td><td>NA</td><td>NA</td></tr>
<tr><td></td><td></td><td></td><td></td><td></td><td></td><td></td><td></td><td></td><td></td><td></td><td></td></tr>
<tr><td>T21479</td><td>GCTCTCTGAGATACACTTTCGGG</td><td>1000</td><td></td><td></td><td></td><td></td><td></td><td></td><td></td><td></td><td></td></tr>
<tr><td>Oligo fwd</td><td>CACCGCTCTCTGAGATACACTTTC</td><td></td><td></td><td></td><td></td><td></td><td></td><td></td><td></td><td></td><td></td></tr>
<tr><td>Oligo rev</td><td>AAACGAAAGTGTATCTCAGAGAGC</td><td></td><td></td><td></td><td></td><td></td><td></td><td></td><td></td><td></td><td></td></tr>
<tr><td>Chromosome</td><td>start</td><td>end</td><td>strand</td><td>MM</td><td>target_seq</td><td>PAM</td><td>alignment</td><td>distance</td><td>position</td><td>gene name</td><td>gene id</td></tr>
<tr><td>GU305914.1</td><td>172743</td><td>172765</td><td>-</td><td>0</td><td>GCTCTCTGAGATACACTTTC</td><td>GGG</td><td>||||||||[||||||||||||]PAM</td><td>NA</td><td>Intergenic</td><td>NA</td><td>NA</td></tr>
<tr><td></td><td></td><td></td><td></td><td></td><td></td><td></td><td></td><td></td><td></td><td></td><td></td></tr>
<tr><td>T21480</td><td>GAGATACACTTTCGGGACTATGG</td><td>1000</td><td></td><td></td><td></td><td></td><td></td><td></td><td></td><td></td><td></td></tr>
<tr><td>Oligo fwd</td><td>CACCGAGATACACTTTCGGGACTA</td><td></td><td></td><td></td><td></td><td></td><td></td><td></td><td></td><td></td><td></td></tr>
<tr><td>Oligo rev</td><td>AAACTAGTCCCGAAAGTGTATCTC</td><td></td><td></td><td></td><td></td><td></td><td></td><td></td><td></td><td></td><td></td></tr>
<tr><td>Chromosome</td><td>start</td><td>end</td><td>strand</td><td>MM</td><td>target_seq</td><td>PAM</td><td>alignment</td><td>distance</td><td>position</td><td>gene name</td><td>gene id</td></tr>
<tr><td>GU305914.1</td><td>172736</td><td>172758</td><td>-</td><td>0</td><td>GAGATACACTTTCGGGACTA</td><td>TGG</td><td>||||||||[||||||||||||]PAM</td><td>NA</td><td>Intergenic</td><td>NA</td><td>NA</td></tr>
<tr><td></td><td></td><td></td><td></td><td></td><td></td><td></td><td></td><td></td><td></td><td></td><td></td></tr>
<tr><td>T21481</td><td>CTTTCGGGACTATGGATCGACGG</td><td>1000</td><td></td><td></td><td></td><td></td><td></td><td></td><td></td><td></td><td></td></tr>
<tr><td>Oligo fwd</td><td>CACCCTTTCGGGACTATGGATCGA</td><td></td><td></td><td></td><td></td><td></td><td></td><td></td><td></td><td></td><td></td></tr>
<tr><td>Oligo rev</td><td>AAACTCGATCCATAGTCCCGAAAG</td><td></td><td></td><td></td><td></td><td></td><td></td><td></td><td></td><td></td><td></td></tr>
<tr><td>Chromosome</td><td>start</td><td>end</td><td>strand</td><td>MM</td><td>target_seq</td><td>PAM</td><td>alignment</td><td>distance</td><td>position</td><td>gene name</td><td>gene id</td></tr>
<tr><td>GU305914.1</td><td>172728</td><td>172750</td><td>-</td><td>0</td><td>CTTTCGGGACTATGGATCGA</td><td>CGG</td><td>||||||||[||||||||||||]PAM</td><td>NA</td><td>Intergenic</td><td>NA</td><td>NA</td></tr>
</table>

<table>
<tr><td>T21482</td><td>GGACTATGGATCGACGGACTTGG</td><td>1000</td><td colspan="9"></td></tr>
<tr><td>Oligo fwd</td><td>CACCGGACTATGGATCGACGGACT</td><td colspan="10"></td></tr>
<tr><td>Oligo rev</td><td>AAACAGTCCGTCGATCCATAGTCC</td><td colspan="10"></td></tr>
<tr><td>Chromosome</td><td>start</td><td>end</td><td>strand</td><td>MM</td><td>target_seq</td><td>PAM</td><td>alignment</td><td>distance</td><td>position</td><td>gene name</td><td>gene id</td></tr>
<tr><td>GU305914.1</td><td>172722</td><td>172744</td><td>-</td><td>0</td><td>GGACTATGGATCGACGGACT</td><td>TGG</td><td>||||||||[||||||||||||]PAM</td><td>NA</td><td>Intergenic</td><td>NA</td><td>NA</td></tr>
</table>

<table>
<tr><td>T21483</td><td>TGGTAGATTTCGATAACCCGAGG</td><td>1000</td><td colspan="9"></td></tr>
<tr><td>Oligo fwd</td><td>CACCTGGTAGATTTCGATAACCCG</td><td colspan="10"></td></tr>
<tr><td>Oligo rev</td><td>AAACCGGGTTATCGAAATCTACCA</td><td colspan="10"></td></tr>
<tr><td>Chromosome</td><td>start</td><td>end</td><td>strand</td><td>MM</td><td>target_seq</td><td>PAM</td><td>alignment</td><td>distance</td><td>position</td><td>gene name</td><td>gene id</td></tr>
<tr><td>GU305914.1</td><td>172702</td><td>172724</td><td>-</td><td>0</td><td>TGGTAGATTTCGATAACCCG</td><td>AGG</td><td>||||||||[||||||||||||]PAM</td><td>NA</td><td>Intergenic</td><td>NA</td><td>NA</td></tr>
</table>

<table>
<tr><td>T21484</td><td>TCGATAACCCGAGGCTGCCGAGG</td><td>1000</td><td colspan="9"></td></tr>
<tr><td>Oligo fwd</td><td>CACCTCGATAACCCGAGGCTGCCG</td><td colspan="10"></td></tr>
<tr><td>Oligo rev</td><td>AAACCGGCAGCCTCGGGTTATCGA</td><td colspan="10"></td></tr>
<tr><td>Chromosome</td><td>start</td><td>end</td><td>strand</td><td>MM</td><td>target_seq</td><td>PAM</td><td>alignment</td><td>distance</td><td>position</td><td>gene name</td><td>gene id</td></tr>
<tr><td>GU305914.1</td><td>172693</td><td>172715</td><td>-</td><td>0</td><td>TCGATAACCCGAGGCTGCCG</td><td>AGG</td><td>||||||||[||||||||||||]PAM</td><td>NA</td><td>Intergenic</td><td>NA</td><td>NA</td></tr>
</table>

<table>
<tr><td>T21485</td><td>CCCGAGGCTGCCGAGGCACAAGG</td><td>1000</td><td colspan="9"></td></tr>
<tr><td>Oligo fwd</td><td>CACCCCCGAGGCTGCCGAGGCACA</td><td colspan="10"></td></tr>
<tr><td>Oligo rev</td><td>AAACTGTGCCTCGGCAGCCTCGGG</td><td colspan="10"></td></tr>
<tr><td>Chromosome</td><td>start</td><td>end</td><td>strand</td><td>MM</td><td>target_seq</td><td>PAM</td><td>alignment</td><td>distance</td><td>position</td><td>gene name</td><td>gene id</td></tr>
<tr><td>GU305914.1</td><td>172686</td><td>172708</td><td>-</td><td>0</td><td>CCCGAGGCTGCCGAGGCACA</td><td>AGG</td><td>||||||||[||||||||||||]PAM</td><td>NA</td><td>Intergenic</td><td>NA</td><td>NA</td></tr>
</table>

<table>
<tr><td>T21486</td><td>GAGTTATACGATCTCCGCCACGG</td><td>1000</td><td colspan="9"></td></tr>
<tr><td>Oligo fwd</td><td>CACCGAGTTATACGATCTCCGCCA</td><td colspan="10"></td></tr>
<tr><td>Oligo rev</td><td>AAACTGGCGGAGATCGTATAACTC</td><td colspan="10"></td></tr>
<tr><td>Chromosome</td><td>start</td><td>end</td><td>strand</td><td>MM</td><td>target_seq</td><td>PAM</td><td>alignment</td><td>distance</td><td>position</td><td>gene name</td><td>gene id</td></tr>
<tr><td>GU305914.1</td><td>172659</td><td>172681</td><td>-</td><td>0</td><td>GAGTTATACGATCTCCGCCA</td><td>CGG</td><td>||||||||[||||||||||||]PAM</td><td>NA</td><td>Intergenic</td><td>NA</td><td>NA</td></tr>
</table>

<table>
<tr><td>T21487</td><td>AGTTATACGATCTCCGCCACGGG</td><td>1000</td><td colspan="9"></td></tr>
<tr><td>Oligo fwd</td><td>CACCAGTTATACGATCTCCGCCAC</td><td colspan="10"></td></tr>
<tr><td>Oligo rev</td><td>AAACGTGGCGGAGATCGTATAACT</td><td colspan="10"></td></tr>
<tr><td>Chromosome</td><td>start</td><td>end</td><td>strand</td><td>MM</td><td>target_seq</td><td>PAM</td><td>alignment</td><td>distance</td><td>position</td><td>gene name</td><td>gene id</td></tr>
<tr><td>GU305914.1</td><td>172658</td><td>172680</td><td>-</td><td>0</td><td>AGTTATACGATCTCCGCCAC</td><td>GGG</td><td>||||||||[||||||||||||]PAM</td><td>NA</td><td>Intergenic</td><td>NA</td><td>NA</td></tr>
</table>

<table>
<tr><td>T21488</td><td>TCTCCGCCACGGGTTCGTTCTGG</td><td>1000</td><td colspan="9"></td></tr>
<tr><td>Oligo fwd</td><td>CACCTCTCCGCCACGGGTTCGTTC</td><td colspan="10"></td></tr>
<tr><td>Oligo rev</td><td>AAACGAACGAACCCGTGGCGGAGA</td><td colspan="10"></td></tr>
<tr><td>Chromosome</td><td>start</td><td>end</td><td>strand</td><td>MM</td><td>target_seq</td><td>PAM</td><td>alignment</td><td>distance</td><td>position</td><td>gene name</td><td>gene id</td></tr>
<tr><td>GU305914.1</td><td>172648</td><td>172670</td><td>-</td><td>0</td><td>TCTCCGCCACGGGTTCGTTC</td><td>TGG</td><td>||||||||[||||||||||||]PAM</td><td>NA</td><td>Intergenic</td><td>NA</td><td>NA</td></tr>
</table>

<table>
<tr><td>T21489</td><td>CTCCGCCACGGGTTCGTTCTGGG</td><td>1000</td><td colspan="9"></td></tr>
<tr><td>Oligo fwd</td><td>CACCCTCCGCCACGGGTTCGTTCT</td><td colspan="10"></td></tr>
<tr><td>Oligo rev</td><td>AAACAGAACGAACCCGTGGCGGAG</td><td colspan="10"></td></tr>
<tr><td>Chromosome</td><td>start</td><td>end</td><td>strand</td><td>MM</td><td>target_seq</td><td>PAM</td><td>alignment</td><td>distance</td><td>position</td><td>gene name</td><td>gene id</td></tr>
<tr><td>GU305914.1</td><td>172647</td><td>172669</td><td>-</td><td>0</td><td>CTCCGCCACGGGTTCGTTCT</td><td>GGG</td><td>||||||||[||||||||||||]PAM</td><td>NA</td><td>Intergenic</td><td>NA</td><td>NA</td></tr>
</table>

<table>
<tr><td>T21490</td><td>TCCGCCACGGGTTCGTTCTGGGG</td><td>1000</td><td colspan="9"></td></tr>
<tr><td>Oligo fwd</td><td>CACCTCCGCCACGGGTTCGTTCTG</td><td colspan="10"></td></tr>
<tr><td>Oligo rev</td><td>AAACCAGAACGAACCCGTGGCGGA</td><td colspan="10"></td></tr>
<tr><td>Chromosome</td><td>start</td><td>end</td><td>strand</td><td>MM</td><td>target_seq</td><td>PAM</td><td>alignment</td><td>distance</td><td>position</td><td>gene name</td><td>gene id</td></tr>
<tr><td>GU305914.1</td><td>172646</td><td>172668</td><td>-</td><td>0</td><td>TCCGCCACGGGTTCGTTCTG</td><td>GGG</td><td>||||||||[||||||||||||]PAM</td><td>NA</td><td>Intergenic</td><td>NA</td><td>NA</td></tr>
</table>

<table>
<tr><td>T21491</td><td>GGGAGAGATGATCCACTGTCAGG</td><td>1000</td><td colspan="9"></td></tr>
<tr><td>Oligo fwd</td><td>CACCGGGAGAGATGATCCACTGTC</td><td colspan="10"></td></tr>
<tr><td>Oligo rev</td><td>AAACGACAGTGGATCATCTCTCCC</td><td colspan="10"></td></tr>
<tr><td>Chromosome</td><td>start</td><td>end</td><td>strand</td><td>MM</td><td>target_seq</td><td>PAM</td><td>alignment</td><td>distance</td><td>position</td><td>gene name</td><td>gene id</td></tr>
<tr><td>GU305914.1</td><td>172626</td><td>172648</td><td>-</td><td>0</td><td>GGGAGAGATGATCCACTGTC</td><td>AGG</td><td>||||||||[||||||||||||]PAM</td><td>NA</td><td>Intergenic</td><td>NA</td><td>NA</td></tr>
</table>

<table>
<tr><td>T21492</td><td>AGATGATCCACTGTCAGGTGAGG</td><td>1000</td><td colspan="9"></td></tr>
<tr><td>Oligo fwd</td><td>CACCAGATGATCCACTGTCAGGTG</td><td colspan="10"></td></tr>
<tr><td>Oligo rev</td><td>AAACCACCTGACAGTGGATCATCT</td><td colspan="10"></td></tr>
<tr><td>Chromosome</td><td>start</td><td>end</td><td>strand</td><td>MM</td><td>target_seq</td><td>PAM</td><td>alignment</td><td>distance</td><td>position</td><td>gene name</td><td>gene id</td></tr>
<tr><td>GU305914.1</td><td>172621</td><td>172643</td><td>-</td><td>0</td><td>AGATGATCCACTGTCAGGTG</td><td>AGG</td><td>||||||||[||||||||||||]PAM</td><td>NA</td><td>Intergenic</td><td>NA</td><td>NA</td></tr>
</table>

<table>
<tr><td>T21493</td><td>GATGATCCACTGTCAGGTGAGGG</td><td>1000</td><td colspan="9"></td></tr>
<tr><td>Oligo fwd</td><td>CACCGATGATCCACTGTCAGGTGA</td><td colspan="10"></td></tr>
<tr><td>Oligo rev</td><td>AAACTCACCTGACAGTGGATCATC</td><td colspan="10"></td></tr>
<tr><td>Chromosome</td><td>start</td><td>end</td><td>strand</td><td>MM</td><td>target_seq</td><td>PAM</td><td>alignment</td><td>distance</td><td>position</td><td>gene name</td><td>gene id</td></tr>
<tr><td>GU305914.1</td><td>172620</td><td>172642</td><td>-</td><td>0</td><td>GATGATCCACTGTCAGGTGA</td><td>GGG</td><td>||||||||[||||||||||||]PAM</td><td>NA</td><td>Intergenic</td><td>NA</td><td>NA</td></tr>
</table>

<table>
<tr><td>T21494</td><td>ACCGTGTACTTTGTCTGTAACGG</td><td>1000</td><td colspan="9"></td></tr>
<tr><td>Oligo fwd</td><td>CACCACCGTGTACTTTGTCTGTAA</td><td colspan="10"></td></tr>
<tr><td>Oligo rev</td><td>AAACTTACAGACAAAGTACACGGT</td><td colspan="10"></td></tr>
<tr><td>Chromosome</td><td>start</td><td>end</td><td>strand</td><td>MM</td><td>target_seq</td><td>PAM</td><td>alignment</td><td>distance</td><td>position</td><td>gene name</td><td>gene id</td></tr>
<tr><td>GU305914.1</td><td>172559</td><td>172581</td><td>-</td><td>0</td><td>ACCGTGTACTTTGTCTGTAA</td><td>CGG</td><td>||||||||[||||||||||||]PAM</td><td>NA</td><td>Intergenic</td><td>NA</td><td>NA</td></tr>
</table>

<table>
<tr><td>T21495</td><td>TCTGTAACGGCACGAGCGCCAGG</td><td>1000</td><td colspan="9"></td></tr>
<tr><td>Oligo fwd</td><td>CACCTCTGTAACGGCACGAGCGCC</td><td colspan="10"></td></tr>
<tr><td>Oligo rev</td><td>AAACGGCGCTCGTGCCGTTACAGA</td><td colspan="10"></td></tr>
<tr><td>Chromosome</td><td>start</td><td>end</td><td>strand</td><td>MM</td><td>target_seq</td><td>PAM</td><td>alignment</td><td>distance</td><td>position</td><td>gene name</td><td>gene id</td></tr>
<tr><td>GU305914.1</td><td>172546</td><td>172568</td><td>-</td><td>0</td><td>TCTGTAACGGCACGAGCGCC</td><td>AGG</td><td>||||||||[||||||||||||]PAM</td><td>NA</td><td>Intergenic</td><td>NA</td><td>NA</td></tr>
</table>

<table>
<tr><td>T21496</td><td>GTAACGGCACGAGCGCCAGGTGG</td><td>1000</td><td colspan="9"></td></tr>
<tr><td>Oligo fwd</td><td>CACCGTAACGGCACGAGCGCCAGG</td><td colspan="10"></td></tr>
<tr><td>Oligo rev</td><td>AAACCCTGGCGCTCGTGCCGTTAC</td><td colspan="10"></td></tr>
<tr><td>Chromosome</td><td>start</td><td>end</td><td>strand</td><td>MM</td><td>target_seq</td><td>PAM</td><td>alignment</td><td>distance</td><td>position</td><td>gene name</td><td>gene id</td></tr>
<tr><td>GU305914.1</td><td>172543</td><td>172565</td><td>-</td><td>0</td><td>GTAACGGCACGAGCGCCAGG</td><td>TGG</td><td>||||||||[||||||||||||]PAM</td><td>NA</td><td>Intergenic</td><td>NA</td><td>NA</td></tr>
</table>

<table>
<tr><td>T21497</td><td>CGCCAGGTGGCCGCCTTACGAGG</td><td>1000</td></tr>
<tr><td>Oligo fwd</td><td>CACCCGCCAGGTGGCCGCCTTACG</td></tr>
<tr><td>Oligo rev</td><td>AAACCGTAAGGCGGCCACCTGGCG</td></tr>
<tr><td>Chromosome</td><td>start</td><td>end</td><td>strand</td><td>MM</td><td>target_seq</td><td>PAM</td><td>alignment</td><td>distance</td><td>position</td><td>gene name</td><td>gene id</td></tr>
<tr><td>GU305914.1</td><td>172530</td><td>172552</td><td>-</td><td>0</td><td>CGCCAGGTGGCCGCCTTACG</td><td>AGG</td><td>||||||||[||||||||||||]PAM</td><td>NA</td><td>Intergenic</td><td>NA</td><td>NA</td></tr>
<tr><td>T21498</td><td>GCCAGGTGGCCGCCTTACGAGGG</td><td>1000</td></tr>
<tr><td>Oligo fwd</td><td>CACCGCCAGGTGGCCGCCTTACGA</td></tr>
<tr><td>Oligo rev</td><td>AAACTCGTAAGGCGGCCACCTGGC</td></tr>
<tr><td>Chromosome</td><td>start</td><td>end</td><td>strand</td><td>MM</td><td>target_seq</td><td>PAM</td><td>alignment</td><td>distance</td><td>position</td><td>gene name</td><td>gene id</td></tr>
<tr><td>GU305914.1</td><td>172529</td><td>172551</td><td>-</td><td>0</td><td>GCCAGGTGGCCGCCTTACGA</td><td>GGG</td><td>||||||||[||||||||||||]PAM</td><td>NA</td><td>Intergenic</td><td>NA</td><td>NA</td></tr>
<tr><td>T21499</td><td>CCAGGTGGCCGCCTTACGAGGGG</td><td>1000</td></tr>
<tr><td>Oligo fwd</td><td>CACCCCAGGTGGCCGCCTTACGAG</td></tr>
<tr><td>Oligo rev</td><td>AAACCTCGTAAGGCGGCCACCTGG</td></tr>
<tr><td>Chromosome</td><td>start</td><td>end</td><td>strand</td><td>MM</td><td>target_seq</td><td>PAM</td><td>alignment</td><td>distance</td><td>position</td><td>gene name</td><td>gene id</td></tr>
<tr><td>GU305914.1</td><td>172528</td><td>172550</td><td>-</td><td>0</td><td>CCAGGTGGCCGCCTTACGAG</td><td>GGG</td><td>||||||||[||||||||||||]PAM</td><td>NA</td><td>Intergenic</td><td>NA</td><td>NA</td></tr>
<tr><td>T21500</td><td>TTACGAGGGGCCGTTCGTGACGG</td><td>1000</td></tr>
<tr><td>Oligo fwd</td><td>CACCTTACGAGGGGCCGTTCGTGA</td></tr>
<tr><td>Oligo rev</td><td>AAACTCACGAACGGCCCCTCGTAA</td></tr>
<tr><td>Chromosome</td><td>start</td><td>end</td><td>strand</td><td>MM</td><td>target_seq</td><td>PAM</td><td>alignment</td><td>distance</td><td>position</td><td>gene name</td><td>gene id</td></tr>
<tr><td>GU305914.1</td><td>172515</td><td>172537</td><td>-</td><td>0</td><td>TTACGAGGGGCCGTTCGTGA</td><td>CGG</td><td>||||||||[||||||||||||]PAM</td><td>NA</td><td>Intergenic</td><td>NA</td><td>NA</td></tr>
<tr><td>T21501</td><td>TACGAGGGGCCGTTCGTGACGGG</td><td>1000</td></tr>
<tr><td>Oligo fwd</td><td>CACCTACGAGGGGCCGTTCGTGAC</td></tr>
<tr><td>Oligo rev</td><td>AAACGTCACGAACGGCCCCTCGTA</td></tr>
<tr><td>Chromosome</td><td>start</td><td>end</td><td>strand</td><td>MM</td><td>target_seq</td><td>PAM</td><td>alignment</td><td>distance</td><td>position</td><td>gene name</td><td>gene id</td></tr>
<tr><td>GU305914.1</td><td>172514</td><td>172536</td><td>-</td><td>0</td><td>TACGAGGGGCCGTTCGTGAC</td><td>GGG</td><td>||||||||[||||||||||||]PAM</td><td>NA</td><td>Intergenic</td><td>NA</td><td>NA</td></tr>
<tr><td>T21502</td><td>GAGGGGCCGTTCGTGACGGGAGG</td><td>1000</td></tr>
<tr><td>Oligo fwd</td><td>CACCGAGGGGCCGTTCGTGACGGG</td></tr>
<tr><td>Oligo rev</td><td>AAACCCCGTCACGAACGGCCCCTC</td></tr>
<tr><td>Chromosome</td><td>start</td><td>end</td><td>strand</td><td>MM</td><td>target_seq</td><td>PAM</td><td>alignment</td><td>distance</td><td>position</td><td>gene name</td><td>gene id</td></tr>
<tr><td>GU305914.1</td><td>172511</td><td>172533</td><td>-</td><td>0</td><td>GAGGGGCCGTTCGTGACGGG</td><td>AGG</td><td>||||||||[||||||||||||]PAM</td><td>NA</td><td>Intergenic</td><td>NA</td><td>NA</td></tr>
<tr><td>T21503</td><td>TTCGTGACGGGAGGACCCGACGG</td><td>1000</td></tr>
<tr><td>Oligo fwd</td><td>CACCTTCGTGACGGGAGGACCCGA</td></tr>
<tr><td>Oligo rev</td><td>AAACTCGGGTCCTCCCGTCACGAA</td></tr>
<tr><td>Chromosome</td><td>start</td><td>end</td><td>strand</td><td>MM</td><td>target_seq</td><td>PAM</td><td>alignment</td><td>distance</td><td>position</td><td>gene name</td><td>gene id</td></tr>
<tr><td>GU305914.1</td><td>172502</td><td>172524</td><td>-</td><td>0</td><td>TTCGTGACGGGAGGACCCGA</td><td>CGG</td><td>||||||||[||||||||||||]PAM</td><td>NA</td><td>Intergenic</td><td>NA</td><td>NA</td></tr>
<tr><td>T21504</td><td>TCGTGACGGGAGGACCCGACGGG</td><td>1000</td></tr>
<tr><td>Oligo fwd</td><td>CACCTCGTGACGGGAGGACCCGAC</td></tr>
<tr><td>Oligo rev</td><td>AAACGTCGGGTCCTCCCGTCACGA</td></tr>
<tr><td>Chromosome</td><td>start</td><td>end</td><td>strand</td><td>MM</td><td>target_seq</td><td>PAM</td><td>alignment</td><td>distance</td><td>position</td><td>gene name</td><td>gene id</td></tr>
<tr><td>GU305914.1</td><td>172501</td><td>172523</td><td>-</td><td>0</td><td>TCGTGACGGGAGGACCCGAC</td><td>GGG</td><td>||||||||[||||||||||||]PAM</td><td>NA</td><td>Intergenic</td><td>NA</td><td>NA</td></tr>
<tr><td>T21505</td><td>CGTGACGGGAGGACCCGACGGGG</td><td>1000</td></tr>
<tr><td>Oligo fwd</td><td>CACCCGTGACGGGAGGACCCGACG</td></tr>
<tr><td>Oligo rev</td><td>AAACCGTCGGGTCCTCCCGTCACG</td></tr>
<tr><td>Chromosome</td><td>start</td><td>end</td><td>strand</td><td>MM</td><td>target_seq</td><td>PAM</td><td>alignment</td><td>distance</td><td>position</td><td>gene name</td><td>gene id</td></tr>
<tr><td>GU305914.1</td><td>172500</td><td>172522</td><td>-</td><td>0</td><td>CGTGACGGGAGGACCCGACG</td><td>GGG</td><td>||||||||[||||||||||||]PAM</td><td>NA</td><td>Intergenic</td><td>NA</td><td>NA</td></tr>
<tr><td>T21506</td><td>CGACGAGATCAGCGTCCCTCCGG</td><td>1000</td></tr>
<tr><td>Oligo fwd</td><td>CACCCGACGAGATCAGCGTCCCTC</td></tr>
<tr><td>Oligo rev</td><td>AAACGAGGGACGCTGATCTCGTCG</td></tr>
<tr><td>Chromosome</td><td>start</td><td>end</td><td>strand</td><td>MM</td><td>target_seq</td><td>PAM</td><td>alignment</td><td>distance</td><td>position</td><td>gene name</td><td>gene id</td></tr>
<tr><td>GU305914.1</td><td>172476</td><td>172498</td><td>-</td><td>0</td><td>CGACGAGATCAGCGTCCCTC</td><td>CGG</td><td>||||||||[||||||||||||]PAM</td><td>NA</td><td>Intergenic</td><td>NA</td><td>NA</td></tr>
<tr><td>T21507</td><td>CCTCCGGAGACGACCGACGACGG</td><td>1000</td></tr>
<tr><td>Oligo fwd</td><td>CACCCCTCCGGAGACGACCGACGA</td></tr>
<tr><td>Oligo rev</td><td>AAACTCGTCGGTCGTCTCCGGAGG</td></tr>
<tr><td>Chromosome</td><td>start</td><td>end</td><td>strand</td><td>MM</td><td>target_seq</td><td>PAM</td><td>alignment</td><td>distance</td><td>position</td><td>gene name</td><td>gene id</td></tr>
<tr><td>GU305914.1</td><td>172460</td><td>172482</td><td>-</td><td>0</td><td>CCTCCGGAGACGACCGACGA</td><td>CGG</td><td>||||||||[||||||||||||]PAM</td><td>NA</td><td>Intergenic</td><td>NA</td><td>NA</td></tr>
<tr><td>T21508</td><td>TCGCCCAGACTTCTGAAAAGCGG</td><td>1000</td></tr>
<tr><td>Oligo fwd</td><td>CACCTCGCCCAGACTTCTGAAAAG</td></tr>
<tr><td>Oligo rev</td><td>AAACCTTTTCAGAAGTCTGGGCGA</td></tr>
<tr><td>Chromosome</td><td>start</td><td>end</td><td>strand</td><td>MM</td><td>target_seq</td><td>PAM</td><td>alignment</td><td>distance</td><td>position</td><td>gene name</td><td>gene id</td></tr>
<tr><td>GU305914.1</td><td>172436</td><td>172458</td><td>-</td><td>0</td><td>TCGCCCAGACTTCTGAAAAG</td><td>CGG</td><td>||||||||[||||||||||||]PAM</td><td>NA</td><td>Intergenic</td><td>NA</td><td>NA</td></tr>
<tr><td>T21509</td><td>CCAGACTTCTGAAAAGCGGACGG</td><td>1000</td></tr>
<tr><td>Oligo fwd</td><td>CACCCCAGACTTCTGAAAAGCGGA</td></tr>
<tr><td>Oligo rev</td><td>AAACTCCGCTTTTCAGAAGTCTGG</td></tr>
<tr><td>Chromosome</td><td>start</td><td>end</td><td>strand</td><td>MM</td><td>target_seq</td><td>PAM</td><td>alignment</td><td>distance</td><td>position</td><td>gene name</td><td>gene id</td></tr>
<tr><td>GU305914.1</td><td>172432</td><td>172454</td><td>-</td><td>0</td><td>CCAGACTTCTGAAAAGCGGA</td><td>CGG</td><td>||||||||[||||||||||||]PAM</td><td>NA</td><td>Intergenic</td><td>NA</td><td>NA</td></tr>
<tr><td>T21510</td><td>CAGACTTCTGAAAAGCGGACGGG</td><td>1000</td></tr>
<tr><td>Oligo fwd</td><td>CACCCAGACTTCTGAAAAGCGGAC</td></tr>
<tr><td>Oligo rev</td><td>AAACGTCCGCTTTTCAGAAGTCTG</td></tr>
<tr><td>Chromosome</td><td>start</td><td>end</td><td>strand</td><td>MM</td><td>target_seq</td><td>PAM</td><td>alignment</td><td>distance</td><td>position</td><td>gene name</td><td>gene id</td></tr>
<tr><td>GU305914.1</td><td>172431</td><td>172453</td><td>-</td><td>0</td><td>CAGACTTCTGAAAAGCGGAC</td><td>GGG</td><td>||||||||[||||||||||||]PAM</td><td>NA</td><td>Intergenic</td><td>NA</td><td>NA</td></tr>
<tr><td>T21511</td><td>AGACTTCTGAAAAGCGGACGGGG</td><td>1000</td></tr>
<tr><td>Oligo fwd</td><td>CACCAGACTTCTGAAAAGCGGACG</td></tr>
<tr><td>Oligo rev</td><td>AAACCGTCCGCTTTTCAGAAGTCT</td></tr>
<tr><td>Chromosome</td><td>start</td><td>end</td><td>strand</td><td>MM</td><td>target_seq</td><td>PAM</td><td>alignment</td><td>distance</td><td>position</td><td>gene name</td><td>gene id</td></tr>
</table>

| | | | | | | | | | | | |
|---|---|---|---|---|---|---|---|---|---|---|---|
| GU305914.1 | 172430 | 172452 | - | 0 | AGACTTCTGAAAAGCGGACG | GGG | \|\|\|\|\|\|\|\|[\|\|\|\|\|\|\|\|\|\|\|\|]PAM | NA | Intergenic | NA | NA |
| T21512 | GGAAAGAAATGCGTCGAGTACGG | 1000 | | | | | | | | | |
| Oligo fwd | CACCGGAAAGAAATGCGTCGAGTA | | | | | | | | | | |
| Oligo rev | AAACTACTCGACGCATTTCTTTCC | | | | | | | | | | |
| Chromosome | start | end | strand | MM | target_seq | PAM | alignment | distance | position | gene name | gene id |
| GU305914.1 | 172409 | 172431 | - | 0 | GGAAAGAAATGCGTCGAGTA | CGG | \|\|\|\|\|\|\|\|[\|\|\|\|\|\|\|\|\|\|\|\|]PAM | NA | Intergenic | NA | NA |
| T21513 | GAAAGAAATGCGTCGAGTACGGG | 1000 | | | | | | | | | |
| Oligo fwd | CACCGAAAGAAATGCGTCGAGTAC | | | | | | | | | | |
| Oligo rev | AAACGTACTCGACGCATTTCTTTC | | | | | | | | | | |
| Chromosome | start | end | strand | MM | target_seq | PAM | alignment | distance | position | gene name | gene id |
| GU305914.1 | 172408 | 172430 | - | 0 | GAAAGAAATGCGTCGAGTAC | GGG | \|\|\|\|\|\|\|\|[\|\|\|\|\|\|\|\|\|\|\|\|]PAM | NA | Intergenic | NA | NA |
| T21514 | AAAGAAATGCGTCGAGTACGGGG | 1000 | | | | | | | | | |
| Oligo fwd | CACCAAAGAAATGCGTCGAGTACG | | | | | | | | | | |
| Oligo rev | AAACCGTACTCGACGCATTTCTTT | | | | | | | | | | |
| Chromosome | start | end | strand | MM | target_seq | PAM | alignment | distance | position | gene name | gene id |
| GU305914.1 | 172407 | 172429 | - | 0 | AAAGAAATGCGTCGAGTACG | GGG | \|\|\|\|\|\|\|\|[\|\|\|\|\|\|\|\|\|\|\|\|]PAM | NA | Intergenic | NA | NA |
| T21515 | ACGGGGAACTGCATAAGCTGCGG | 1000 | | | | | | | | | |
| Oligo fwd | CACCACGGGGAACTGCATAAGCTG | | | | | | | | | | |
| Oligo rev | AAACCAGCTTATGCAGTTCCCCGT | | | | | | | | | | |
| Chromosome | start | end | strand | MM | target_seq | PAM | alignment | distance | position | gene name | gene id |
| GU305914.1 | 172390 | 172412 | - | 0 | ACGGGGAACTGCATAAGCTG | CGG | \|\|\|\|\|\|\|\|[\|\|\|\|\|\|\|\|\|\|\|\|]PAM | NA | Intergenic | NA | NA |
| T21516 | GGGAACTGCATAAGCTGCGGAGG | 1000 | | | | | | | | | |
| Oligo fwd | CACCGGGAACTGCATAAGCTGCGG | | | | | | | | | | |
| Oligo rev | AAACCCGCAGCTTATGCAGTTCCC | | | | | | | | | | |
| Chromosome | start | end | strand | MM | target_seq | PAM | alignment | distance | position | gene name | gene id |
| GU305914.1 | 172387 | 172409 | - | 0 | GGGAACTGCATAAGCTGCGG | AGG | \|\|\|\|\|\|\|\|[\|\|\|\|\|\|\|\|\|\|\|\|]PAM | NA | Intergenic | NA | NA |
| T21517 | TACGTCCCCCGCTCGAACCGCGG | 1000 | | | | | | | | | |
| Oligo fwd | CACCTACGTCCCCCGCTCGAACCG | | | | | | | | | | |
| Oligo rev | AAACCGGTTCGAGCGGGGGACGTA | | | | | | | | | | |
| Chromosome | start | end | strand | MM | target_seq | PAM | alignment | distance | position | gene name | gene id |
| GU305914.1 | 172356 | 172378 | - | 0 | TACGTCCCCCGCTCGAACCG | CGG | \|\|\|\|\|\|\|\|[\|\|\|\|\|\|\|\|\|\|\|\|]PAM | NA | Intergenic | NA | NA |
| T21518 | GTCCCCCGCTCGAACCGCGGCGG | 1000 | | | | | | | | | |
| Oligo fwd | CACCGTCCCCCGCTCGAACCGCGG | | | | | | | | | | |
| Oligo rev | AAACCCGCGGTTCGAGCGGGGGAC | | | | | | | | | | |
| Chromosome | start | end | strand | MM | target_seq | PAM | alignment | distance | position | gene name | gene id |
| GU305914.1 | 172353 | 172375 | - | 0 | GTCCCCCGCTCGAACCGCGG | CGG | \|\|\|\|\|\|\|\|[\|\|\|\|\|\|\|\|\|\|\|\|]PAM | NA | Intergenic | NA | NA |
| T21519 | CGCTCGAACCGCGGCGGCCACGG | 1000 | | | | | | | | | |
| Oligo fwd | CACCCGCTCGAACCGCGGCGGCCA | | | | | | | | | | |
| Oligo rev | AAACTGGCCGCCGCGGTTCGAGCG | | | | | | | | | | |
| Chromosome | start | end | strand | MM | target_seq | PAM | alignment | distance | position | gene name | gene id |
| GU305914.1 | 172347 | 172369 | - | 0 | CGCTCGAACCGCGGCGGCCA | CGG | \|\|\|\|\|\|\|\|[\|\|\|\|\|\|\|\|\|\|\|\|]PAM | NA | Intergenic | NA | NA |
| T21520 | TCGAACCGCGGCGGCCACGGAGG | 1000 | | | | | | | | | |
| Oligo fwd | CACCTCGAACCGCGGCGGCCACGG | | | | | | | | | | |
| Oligo rev | AAACCCGTGGCCGCCGCGGTTCGA | | | | | | | | | | |
| Chromosome | start | end | strand | MM | target_seq | PAM | alignment | distance | position | gene name | gene id |
| GU305914.1 | 172344 | 172366 | - | 0 | TCGAACCGCGGCGGCCACGG | AGG | \|\|\|\|\|\|\|\|[\|\|\|\|\|\|\|\|\|\|\|\|]PAM | NA | Intergenic | NA | NA |
| T21521 | GGCCACGGAGGACCGCGACGAGG | 1000 | | | | | | | | | |
| Oligo fwd | CACCGGCCACGGAGGACCGCGACG | | | | | | | | | | |
| Oligo rev | AAACCGTCGCGGTCCTCCGTGGCC | | | | | | | | | | |
| Chromosome | start | end | strand | MM | target_seq | PAM | alignment | distance | position | gene name | gene id |
| GU305914.1 | 172332 | 172354 | - | 0 | GGCCACGGAGGACCGCGACG | AGG | \|\|\|\|\|\|\|\|[\|\|\|\|\|\|\|\|\|\|\|\|]PAM | NA | Intergenic | NA | NA |
| GU305914.1 | 88679 | 88701 | - | 4 | GCTCATGGAGGGCCGCGACG | AGG | \|--\|\|-\|\|[\|\|\|-\|\|\|\|\|\|\|\|]PAM | NA | Intergenic | NA | NA |
| T21522 | TCGAACATCAGTCTCGCCATCGG | 1000 | | | | | | | | | |
| Oligo fwd | CACCTCGAACATCAGTCTCGCCAT | | | | | | | | | | |
| Oligo rev | AAACATGGCGAGACTGATGTTCGA | | | | | | | | | | |
| Chromosome | start | end | strand | MM | target_seq | PAM | alignment | distance | position | gene name | gene id |
| GU305914.1 | 172298 | 172320 | - | 0 | TCGAACATCAGTCTCGCCAT | CGG | \|\|\|\|\|\|\|\|[\|\|\|\|\|\|\|\|\|\|\|\|]PAM | NA | Intergenic | NA | NA |
| GU305914.1 | 210772 | 210794 | - | 4 | TACAACATCAGTCTCGCGAA | CGG | \|--\|\|\|\|\|[\|\|\|\|\|\|\|\|\|-\|-]PAM | NA | Intergenic | NA | NA |
| T21523 | TCCTCGCGACGATCTGCATGTGG | 1000 | | | | | | | | | |
| Oligo fwd | CACCTCCTCGCGACGATCTGCATG | | | | | | | | | | |
| Oligo rev | AAACCATGCAGATCGTCGCGAGGA | | | | | | | | | | |
| Chromosome | start | end | strand | MM | target_seq | PAM | alignment | distance | position | gene name | gene id |
| GU305914.1 | 172252 | 172274 | - | 0 | TCCTCGCGACGATCTGCATG | TGG | \|\|\|\|\|\|\|\|[\|\|\|\|\|\|\|\|\|\|\|\|]PAM | NA | Intergenic | NA | NA |
| T21524 | GTCCGCGATCGTATCGATCTCGG | 1000 | | | | | | | | | |
| Oligo fwd | CACCGTCCGCGATCGTATCGATCT | | | | | | | | | | |
| Oligo rev | AAACAGATCGATACGATCGCGGAC | | | | | | | | | | |
| Chromosome | start | end | strand | MM | target_seq | PAM | alignment | distance | position | gene name | gene id |
| GU305914.1 | 172198 | 172220 | - | 0 | GTCCGCGATCGTATCGATCT | CGG | \|\|\|\|\|\|\|\|[\|\|\|\|\|\|\|\|\|\|\|\|]PAM | NA | Intergenic | NA | NA |
| T21525 | CGCGATCGTATCGATCTCGGTGG | 1000 | | | | | | | | | |
| Oligo fwd | CACCCGCGATCGTATCGATCTCGG | | | | | | | | | | |
| Oligo rev | AAACCCGAGATCGATACGATCGCG | | | | | | | | | | |
| Chromosome | start | end | strand | MM | target_seq | PAM | alignment | distance | position | gene name | gene id |
| GU305914.1 | 172195 | 172217 | - | 0 | CGCGATCGTATCGATCTCGG | TGG | \|\|\|\|\|\|\|\|[\|\|\|\|\|\|\|\|\|\|\|\|]PAM | NA | Intergenic | NA | NA |
| T21526 | TCTCGGTGGTTCCTCCGAGAAGG | 1000 | | | | | | | | | |

<table>
<tr><td>Oligo fwd</td><td>CACCTCTCGGTGGTTCCTCCGAGA</td><td></td><td></td><td></td><td></td><td></td><td></td><td></td><td></td><td></td><td></td><td></td></tr>
<tr><td>Oligo rev</td><td>AAACTCTCGGAGGAACCACCGAGA</td><td></td><td></td><td></td><td></td><td></td><td></td><td></td><td></td><td></td><td></td><td></td></tr>
<tr><td>Chromosome</td><td>start</td><td></td><td>end</td><td>strand</td><td>MM</td><td>target_seq</td><td>PAM</td><td>alignment</td><td>distance</td><td>position</td><td>gene name</td><td>gene id</td></tr>
<tr><td>GU305914.1</td><td></td><td>172181</td><td>172203</td><td>-</td><td>0</td><td>TCTCGGTGGTTCCTCCGAGA</td><td>AGG</td><td>||||||||[||||||||||||]PAM</td><td>NA</td><td>Intergenic</td><td>NA</td><td>NA</td></tr>
<tr><td></td><td></td><td></td><td></td><td></td><td></td><td></td><td></td><td></td><td></td><td></td><td></td><td></td></tr>
<tr><td>T21527</td><td>CGACTCTGATGATACTATGACGG</td><td></td><td>1000</td><td></td><td></td><td></td><td></td><td></td><td></td><td></td><td></td><td></td></tr>
<tr><td>Oligo fwd</td><td>CACCCGACTCTGATGATACTATGA</td><td></td><td></td><td></td><td></td><td></td><td></td><td></td><td></td><td></td><td></td><td></td></tr>
<tr><td>Oligo rev</td><td>AAACTCATAGTATCATCAGAGTCG</td><td></td><td></td><td></td><td></td><td></td><td></td><td></td><td></td><td></td><td></td><td></td></tr>
<tr><td>Chromosome</td><td>start</td><td></td><td>end</td><td>strand</td><td>MM</td><td>target_seq</td><td>PAM</td><td>alignment</td><td>distance</td><td>position</td><td>gene name</td><td>gene id</td></tr>
<tr><td>GU305914.1</td><td></td><td>172142</td><td>172164</td><td>-</td><td>0</td><td>CGACTCTGATGATACTATGA</td><td>CGG</td><td>||||||||[||||||||||||]PAM</td><td>NA</td><td>Intergenic</td><td>NA</td><td>NA</td></tr>
<tr><td></td><td></td><td></td><td></td><td></td><td></td><td></td><td></td><td></td><td></td><td></td><td></td><td></td></tr>
<tr><td>T21528</td><td>CGCCCGTCTCGTGTTTCAGAAGG</td><td></td><td>1000</td><td></td><td></td><td></td><td></td><td></td><td></td><td></td><td></td><td></td></tr>
<tr><td>Oligo fwd</td><td>CACCCGCCCGTCTCGTGTTTCAGA</td><td></td><td></td><td></td><td></td><td></td><td></td><td></td><td></td><td></td><td></td><td></td></tr>
<tr><td>Oligo rev</td><td>AAACTCTGAAACACGAGACGGGCG</td><td></td><td></td><td></td><td></td><td></td><td></td><td></td><td></td><td></td><td></td><td></td></tr>
<tr><td>Chromosome</td><td>start</td><td></td><td>end</td><td>strand</td><td>MM</td><td>target_seq</td><td>PAM</td><td>alignment</td><td>distance</td><td>position</td><td>gene name</td><td>gene id</td></tr>
<tr><td>GU305914.1</td><td></td><td>172061</td><td>172083</td><td>-</td><td>0</td><td>CGCCCGTCTCGTGTTTCAGA</td><td>AGG</td><td>||||||||[||||||||||||]PAM</td><td>NA</td><td>Intergenic</td><td>NA</td><td>NA</td></tr>
<tr><td></td><td></td><td></td><td></td><td></td><td></td><td></td><td></td><td></td><td></td><td></td><td></td><td></td></tr>
<tr><td>T21529</td><td>CGTGTTTCAGAAGGCTAGATCGG</td><td></td><td>1000</td><td></td><td></td><td></td><td></td><td></td><td></td><td></td><td></td><td></td></tr>
<tr><td>Oligo fwd</td><td>CACCCGTGTTTCAGAAGGCTAGAT</td><td></td><td></td><td></td><td></td><td></td><td></td><td></td><td></td><td></td><td></td><td></td></tr>
<tr><td>Oligo rev</td><td>AAACATCTAGCCTTCTGAAACACG</td><td></td><td></td><td></td><td></td><td></td><td></td><td></td><td></td><td></td><td></td><td></td></tr>
<tr><td>Chromosome</td><td>start</td><td></td><td>end</td><td>strand</td><td>MM</td><td>target_seq</td><td>PAM</td><td>alignment</td><td>distance</td><td>position</td><td>gene name</td><td>gene id</td></tr>
<tr><td>GU305914.1</td><td></td><td>172052</td><td>172074</td><td>-</td><td>0</td><td>CGTGTTTCAGAAGGCTAGAT</td><td>CGG</td><td>||||||||[||||||||||||]PAM</td><td>NA</td><td>Intergenic</td><td>NA</td><td>NA</td></tr>
<tr><td></td><td></td><td></td><td></td><td></td><td></td><td></td><td></td><td></td><td></td><td></td><td></td><td></td></tr>
<tr><td>T21530</td><td>TTTCAGAAGGCTAGATCGGCAGG</td><td></td><td>1000</td><td></td><td></td><td></td><td></td><td></td><td></td><td></td><td></td><td></td></tr>
<tr><td>Oligo fwd</td><td>CACCTTTCAGAAGGCTAGATCGGC</td><td></td><td></td><td></td><td></td><td></td><td></td><td></td><td></td><td></td><td></td><td></td></tr>
<tr><td>Oligo rev</td><td>AAACGCCGATCTAGCCTTCTGAAA</td><td></td><td></td><td></td><td></td><td></td><td></td><td></td><td></td><td></td><td></td><td></td></tr>
<tr><td>Chromosome</td><td>start</td><td></td><td>end</td><td>strand</td><td>MM</td><td>target_seq</td><td>PAM</td><td>alignment</td><td>distance</td><td>position</td><td>gene name</td><td>gene id</td></tr>
<tr><td>GU305914.1</td><td></td><td>172048</td><td>172070</td><td>-</td><td>0</td><td>TTTCAGAAGGCTAGATCGGC</td><td>AGG</td><td>||||||||[||||||||||||]PAM</td><td>NA</td><td>Intergenic</td><td>NA</td><td>NA</td></tr>
<tr><td></td><td></td><td></td><td></td><td></td><td></td><td></td><td></td><td></td><td></td><td></td><td></td><td></td></tr>
<tr><td>T21531</td><td>TCGGCAGGTCATGTTACGTTTGG</td><td></td><td>1000</td><td></td><td></td><td></td><td></td><td></td><td></td><td></td><td></td><td></td></tr>
<tr><td>Oligo fwd</td><td>CACCTCGGCAGGTCATGTTACGTT</td><td></td><td></td><td></td><td></td><td></td><td></td><td></td><td></td><td></td><td></td><td></td></tr>
<tr><td>Oligo rev</td><td>AAACAACGTAACATGACCTGCCGA</td><td></td><td></td><td></td><td></td><td></td><td></td><td></td><td></td><td></td><td></td><td></td></tr>
<tr><td>Chromosome</td><td>start</td><td></td><td>end</td><td>strand</td><td>MM</td><td>target_seq</td><td>PAM</td><td>alignment</td><td>distance</td><td>position</td><td>gene name</td><td>gene id</td></tr>
<tr><td>GU305914.1</td><td></td><td>172033</td><td>172055</td><td>-</td><td>0</td><td>TCGGCAGGTCATGTTACGTT</td><td>TGG</td><td>||||||||[||||||||||||]PAM</td><td>NA</td><td>Intergenic</td><td>NA</td><td>NA</td></tr>
<tr><td></td><td></td><td></td><td></td><td></td><td></td><td></td><td></td><td></td><td></td><td></td><td></td><td></td></tr>
<tr><td>T21532</td><td>GTTACGTTTGGTTCTGTTCGCGG</td><td></td><td>1000</td><td></td><td></td><td></td><td></td><td></td><td></td><td></td><td></td><td></td></tr>
<tr><td>Oligo fwd</td><td>CACCGTTACGTTTGGTTCTGTTCG</td><td></td><td></td><td></td><td></td><td></td><td></td><td></td><td></td><td></td><td></td><td></td></tr>
<tr><td>Oligo rev</td><td>AAACCGAACAGAACCAAACGTAAC</td><td></td><td></td><td></td><td></td><td></td><td></td><td></td><td></td><td></td><td></td><td></td></tr>
<tr><td>Chromosome</td><td>start</td><td></td><td>end</td><td>strand</td><td>MM</td><td>target_seq</td><td>PAM</td><td>alignment</td><td>distance</td><td>position</td><td>gene name</td><td>gene id</td></tr>
<tr><td>GU305914.1</td><td></td><td>172021</td><td>172043</td><td>-</td><td>0</td><td>GTTACGTTTGGTTCTGTTCG</td><td>CGG</td><td>||||||||[||||||||||||]PAM</td><td>NA</td><td>Intergenic</td><td>NA</td><td>NA</td></tr>
<tr><td>GU305914.1</td><td></td><td>31984</td><td>32006</td><td>-</td><td>4</td><td>GTTGCGTCTGGATCTGATCG</td><td>AGG</td><td>|||-|||-[|||-||||-|||]PAM</td><td>NA</td><td>Intergenic</td><td>NA</td><td>NA</td></tr>
<tr><td></td><td></td><td></td><td></td><td></td><td></td><td></td><td></td><td></td><td></td><td></td><td></td><td></td></tr>
<tr><td>T21533</td><td>GCCGCATTTCACACAGCGCACGG</td><td></td><td>1000</td><td></td><td></td><td></td><td></td><td></td><td></td><td></td><td></td><td></td></tr>
<tr><td>Oligo fwd</td><td>CACCGCCGCATTTCACACAGCGCA</td><td></td><td></td><td></td><td></td><td></td><td></td><td></td><td></td><td></td><td></td><td></td></tr>
<tr><td>Oligo rev</td><td>AAACTGCGCTGTGTGAAATGCGGC</td><td></td><td></td><td></td><td></td><td></td><td></td><td></td><td></td><td></td><td></td><td></td></tr>
<tr><td>Chromosome</td><td>start</td><td></td><td>end</td><td>strand</td><td>MM</td><td>target_seq</td><td>PAM</td><td>alignment</td><td>distance</td><td>position</td><td>gene name</td><td>gene id</td></tr>
<tr><td>GU305914.1</td><td></td><td>171999</td><td>172021</td><td>-</td><td>0</td><td>GCCGCATTTCACACAGCGCA</td><td>CGG</td><td>||||||||[||||||||||||]PAM</td><td>NA</td><td>Intergenic</td><td>NA</td><td>NA</td></tr>
<tr><td></td><td></td><td></td><td></td><td></td><td></td><td></td><td></td><td></td><td></td><td></td><td></td><td></td></tr>
<tr><td>T21534</td><td>CCGCATTTCACACAGCGCACGGG</td><td></td><td>1000</td><td></td><td></td><td></td><td></td><td></td><td></td><td></td><td></td><td></td></tr>
<tr><td>Oligo fwd</td><td>CACCCCGCATTTCACACAGCGCAC</td><td></td><td></td><td></td><td></td><td></td><td></td><td></td><td></td><td></td><td></td><td></td></tr>
<tr><td>Oligo rev</td><td>AAACGTGCGCTGTGTGAAATGCGG</td><td></td><td></td><td></td><td></td><td></td><td></td><td></td><td></td><td></td><td></td><td></td></tr>
<tr><td>Chromosome</td><td>start</td><td></td><td>end</td><td>strand</td><td>MM</td><td>target_seq</td><td>PAM</td><td>alignment</td><td>distance</td><td>position</td><td>gene name</td><td>gene id</td></tr>
<tr><td>GU305914.1</td><td></td><td>171998</td><td>172020</td><td>-</td><td>0</td><td>CCGCATTTCACACAGCGCAC</td><td>GGG</td><td>||||||||[||||||||||||]PAM</td><td>NA</td><td>Intergenic</td><td>NA</td><td>NA</td></tr>
<tr><td></td><td></td><td></td><td></td><td></td><td></td><td></td><td></td><td></td><td></td><td></td><td></td><td></td></tr>
<tr><td>T21535</td><td>CGCATTTCACACAGCGCACGGGG</td><td></td><td>1000</td><td></td><td></td><td></td><td></td><td></td><td></td><td></td><td></td><td></td></tr>
<tr><td>Oligo fwd</td><td>CACCCGCATTTCACACAGCGCACG</td><td></td><td></td><td></td><td></td><td></td><td></td><td></td><td></td><td></td><td></td><td></td></tr>
<tr><td>Oligo rev</td><td>AAACCGTGCGCTGTGTGAAATGCG</td><td></td><td></td><td></td><td></td><td></td><td></td><td></td><td></td><td></td><td></td><td></td></tr>
<tr><td>Chromosome</td><td>start</td><td></td><td>end</td><td>strand</td><td>MM</td><td>target_seq</td><td>PAM</td><td>alignment</td><td>distance</td><td>position</td><td>gene name</td><td>gene id</td></tr>
<tr><td>GU305914.1</td><td></td><td>171997</td><td>172019</td><td>-</td><td>0</td><td>CGCATTTCACACAGCGCACG</td><td>GGG</td><td>||||||||[||||||||||||]PAM</td><td>NA</td><td>Intergenic</td><td>NA</td><td>NA</td></tr>
<tr><td></td><td></td><td></td><td></td><td></td><td></td><td></td><td></td><td></td><td></td><td></td><td></td><td></td></tr>
<tr><td>T21536</td><td>TTCCGTGTCGTGTCACGATGCGG</td><td></td><td>1000</td><td></td><td></td><td></td><td></td><td></td><td></td><td></td><td></td><td></td></tr>
<tr><td>Oligo fwd</td><td>CACCTTCCGTGTCGTGTCACGATG</td><td></td><td></td><td></td><td></td><td></td><td></td><td></td><td></td><td></td><td></td><td></td></tr>
<tr><td>Oligo rev</td><td>AAACCATCGTGACACGACACGGAA</td><td></td><td></td><td></td><td></td><td></td><td></td><td></td><td></td><td></td><td></td><td></td></tr>
<tr><td>Chromosome</td><td>start</td><td></td><td>end</td><td>strand</td><td>MM</td><td>target_seq</td><td>PAM</td><td>alignment</td><td>distance</td><td>position</td><td>gene name</td><td>gene id</td></tr>
<tr><td>GU305914.1</td><td></td><td>171970</td><td>171992</td><td>-</td><td>0</td><td>TTCCGTGTCGTGTCACGATG</td><td>CGG</td><td>||||||||[||||||||||||]PAM</td><td>NA</td><td>Intergenic</td><td>NA</td><td>NA</td></tr>
<tr><td></td><td></td><td></td><td></td><td></td><td></td><td></td><td></td><td></td><td></td><td></td><td></td><td></td></tr>
<tr><td>T21537</td><td>GTCACGATGCGGAGACGTACTGG</td><td></td><td>1000</td><td></td><td></td><td></td><td></td><td></td><td></td><td></td><td></td><td></td></tr>
<tr><td>Oligo fwd</td><td>CACCGTCACGATGCGGAGACGTAC</td><td></td><td></td><td></td><td></td><td></td><td></td><td></td><td></td><td></td><td></td><td></td></tr>
<tr><td>Oligo rev</td><td>AAACGTACGTCTCCGCATCGTGAC</td><td></td><td></td><td></td><td></td><td></td><td></td><td></td><td></td><td></td><td></td><td></td></tr>
<tr><td>Chromosome</td><td>start</td><td></td><td>end</td><td>strand</td><td>MM</td><td>target_seq</td><td>PAM</td><td>alignment</td><td>distance</td><td>position</td><td>gene name</td><td>gene id</td></tr>
<tr><td>GU305914.1</td><td></td><td>171959</td><td>171981</td><td>-</td><td>0</td><td>GTCACGATGCGGAGACGTAC</td><td>TGG</td><td>||||||||[||||||||||||]PAM</td><td>NA</td><td>Intergenic</td><td>NA</td><td>NA</td></tr>
<tr><td></td><td></td><td></td><td></td><td></td><td></td><td></td><td></td><td></td><td></td><td></td><td></td><td></td></tr>
<tr><td>T21538</td><td>GCGGAGACGTACTGGAACTATGG</td><td></td><td>1000</td><td></td><td></td><td></td><td></td><td></td><td></td><td></td><td></td><td></td></tr>
<tr><td>Oligo fwd</td><td>CACCGCGGAGACGTACTGGAACTA</td><td></td><td></td><td></td><td></td><td></td><td></td><td></td><td></td><td></td><td></td><td></td></tr>
<tr><td>Oligo rev</td><td>AAACTAGTTCCAGTACGTCTCCGC</td><td></td><td></td><td></td><td></td><td></td><td></td><td></td><td></td><td></td><td></td><td></td></tr>
<tr><td>Chromosome</td><td>start</td><td></td><td>end</td><td>strand</td><td>MM</td><td>target_seq</td><td>PAM</td><td>alignment</td><td>distance</td><td>position</td><td>gene name</td><td>gene id</td></tr>
<tr><td>GU305914.1</td><td></td><td>171951</td><td>171973</td><td>-</td><td>0</td><td>GCGGAGACGTACTGGAACTA</td><td>TGG</td><td>||||||||[||||||||||||]PAM</td><td>NA</td><td>Intergenic</td><td>NA</td><td>NA</td></tr>
<tr><td></td><td></td><td></td><td></td><td></td><td></td><td></td><td></td><td></td><td></td><td></td><td></td><td></td></tr>
<tr><td>T21539</td><td>CGGAGACGTACTGGAACTATGGG</td><td></td><td>1000</td><td></td><td></td><td></td><td></td><td></td><td></td><td></td><td></td><td></td></tr>
<tr><td>Oligo fwd</td><td>CACCCGGAGACGTACTGGAACTAT</td><td></td><td></td><td></td><td></td><td></td><td></td><td></td><td></td><td></td><td></td><td></td></tr>
<tr><td>Oligo rev</td><td>AAACATAGTTCCAGTACGTCTCCG</td><td></td><td></td><td></td><td></td><td></td><td></td><td></td><td></td><td></td><td></td><td></td></tr>
<tr><td>Chromosome</td><td>start</td><td></td><td>end</td><td>strand</td><td>MM</td><td>target_seq</td><td>PAM</td><td>alignment</td><td>distance</td><td>position</td><td>gene name</td><td>gene id</td></tr>
<tr><td>GU305914.1</td><td></td><td>171950</td><td>171972</td><td>-</td><td>0</td><td>CGGAGACGTACTGGAACTAT</td><td>GGG</td><td>||||||||[||||||||||||]PAM</td><td>NA</td><td>Intergenic</td><td>NA</td><td>NA</td></tr>
<tr><td></td><td></td><td></td><td></td><td></td><td></td><td></td><td></td><td></td><td></td><td></td><td></td><td></td></tr>
<tr><td>T21540</td><td>GGAGACGTACTGGAACTATGGGG</td><td></td><td>1000</td><td></td><td></td><td></td><td></td><td></td><td></td><td></td><td></td><td></td></tr>
<tr><td>Oligo fwd</td><td>CACCGGAGACGTACTGGAACTATG</td><td></td><td></td><td></td><td></td><td></td><td></td><td></td><td></td><td></td><td></td><td></td></tr>
<tr><td>Oligo rev</td><td>AAACCATAGTTCCAGTACGTCTCC</td><td></td><td></td><td></td><td></td><td></td><td></td><td></td><td></td><td></td><td></td><td></td></tr>
<tr><td>Chromosome</td><td>start</td><td></td><td>end</td><td>strand</td><td>MM</td><td>target_seq</td><td>PAM</td><td>alignment</td><td>distance</td><td>position</td><td>gene name</td><td>gene id</td></tr>
<tr><td>GU305914.1</td><td></td><td>171949</td><td>171971</td><td>-</td><td>0</td><td>GGAGACGTACTGGAACTATG</td><td>GGG</td><td>||||||||[||||||||||||]PAM</td><td>NA</td><td>Intergenic</td><td>NA</td><td>NA</td></tr>
</table>

T21541 GTGACGCTATACCCTAAGAACGG 1000
Oligo fwd CACCGTGACGCTATACCCTAAGAA
Oligo rev AAACTTCTTAGGGTATAGCGTCAC
Chromosome start end strand MM target_seq PAM alignment distance position gene name gene id
GU305914.1 171909 171931 - 0 GTGACGCTATACCCTAAGAA CGG ||||||||[||||||||||||]PAM NA Intergenic NA NA

T21542 ACGCTATACCCTAAGAACGGCGG 1000
Oligo fwd CACCACGCTATACCCTAAGAACGG
Oligo rev AAACCCGTTCTTAGGGTATAGCGT
Chromosome start end strand MM target_seq PAM alignment distance position gene name gene id
GU305914.1 171906 171928 - 0 ACGCTATACCCTAAGAACGG CGG ||||||||[||||||||||||]PAM NA Intergenic NA NA

T21543 TGTTCGCCGTCGATTGTTTCTGG 1000
Oligo fwd CACCTGTTCGCCGTCGATTGTTTC
Oligo rev AAACGAAACAATCGACGGCGAACA
Chromosome start end strand MM target_seq PAM alignment distance position gene name gene id
GU305914.1 171881 171903 - 0 TGTTCGCCGTCGATTGTTTC TGG ||||||||[||||||||||||]PAM NA Intergenic NA NA

T21544 AAGCTCACGAACACAAACCGTGG 1000
Oligo fwd CACCAAGCTCACGAACACAAACCG
Oligo rev AAACCGGTTTGTGTTCGTGAGCTT
Chromosome start end strand MM target_seq PAM alignment distance position gene name gene id
GU305914.1 171857 171879 - 0 AAGCTCACGAACACAAACCG TGG ||||||||[||||||||||||]PAM NA Intergenic NA NA

T21545 AACCGTGGCACAAGATCTCGAGG 1000
Oligo fwd CACCAACCGTGGCACAAGATCTCG
Oligo rev AAACCGAGATCTTGTGCCACGGTT
Chromosome start end strand MM target_seq PAM alignment distance position gene name gene id
GU305914.1 171842 171864 - 0 AACCGTGGCACAAGATCTCG AGG ||||||||[||||||||||||]PAM NA Intergenic NA NA

T21546 GTGGCACAAGATCTCGAGGCTGG 1000
Oligo fwd CACCGTGGCACAAGATCTCGAGGC
Oligo rev AAACGCCTCGAGATCTTGTGCCAC
Chromosome start end strand MM target_seq PAM alignment distance position gene name gene id
GU305914.1 171838 171860 - 0 GTGGCACAAGATCTCGAGGC TGG ||||||||[||||||||||||]PAM NA Intergenic NA NA

T21547 ATCTCGAGGCTGGACGCCTACGG 1000
Oligo fwd CACCATCTCGAGGCTGGACGCCTA
Oligo rev AAACTAGGCGTCCAGCCTCGAGAT
Chromosome start end strand MM target_seq PAM alignment distance position gene name gene id
GU305914.1 171828 171850 - 0 ATCTCGAGGCTGGACGCCTA CGG ||||||||[||||||||||||]PAM NA Intergenic NA NA

T21548 CGAGGCTGGACGCCTACGGATGG 1000
Oligo fwd CACCCGAGGCTGGACGCCTACGGA
Oligo rev AAACTCCGTAGGCGTCCAGCCTCG
Chromosome start end strand MM target_seq PAM alignment distance position gene name gene id
GU305914.1 171824 171846 - 0 CGAGGCTGGACGCCTACGGA TGG ||||||||[||||||||||||]PAM NA Intergenic NA NA

T21549 GTAGTTAAGCTAACCGCGTACGG 1000
Oligo fwd CACCGTAGTTAAGCTAACCGCGTA
Oligo rev AAACTACGCGGTTAGCTTAACTAC
Chromosome start end strand MM target_seq PAM alignment distance position gene name gene id
GU305914.1 171724 171746 - 0 GTAGTTAAGCTAACCGCGTA CGG ||||||||[||||||||||||]PAM NA Intergenic NA NA

T21550 TAGTTAAGCTAACCGCGTACGGG 1000
Oligo fwd CACCTAGTTAAGCTAACCGCGTAC
Oligo rev AAACGTACGCGGTTAGCTTAACTA
Chromosome start end strand MM target_seq PAM alignment distance position gene name gene id
GU305914.1 171723 171745 - 0 TAGTTAAGCTAACCGCGTAC GGG ||||||||[||||||||||||]PAM NA Intergenic NA NA

T21551 AGTTAAGCTAACCGCGTACGGGG 1000
Oligo fwd CACCAGTTAAGCTAACCGCGTACG
Oligo rev AAACCGTACGCGGTTAGCTTAACT
Chromosome start end strand MM target_seq PAM alignment distance position gene name gene id
GU305914.1 171722 171744 - 0 AGTTAAGCTAACCGCGTACG GGG ||||||||[||||||||||||]PAM NA Intergenic NA NA

T21552 TAGTCACAGCGCCGTCGCTCCGG 1000
Oligo fwd CACCTAGTCACAGCGCCGTCGCTC
Oligo rev AAACGAGCGACGGCGCTGTGACTA
Chromosome start end strand MM target_seq PAM alignment distance position gene name gene id
GU305914.1 171613 171635 - 0 TAGTCACAGCGCCGTCGCTC CGG ||||||||[||||||||||||]PAM NA Intergenic NA NA

T21553 AGTCACAGCGCCGTCGCTCCGGG 1000
Oligo fwd CACCAGTCACAGCGCCGTCGCTCC
Oligo rev AAACGGAGCGACGGCGCTGTGACT
Chromosome start end strand MM target_seq PAM alignment distance position gene name gene id
GU305914.1 171612 171634 - 0 AGTCACAGCGCCGTCGCTCC GGG ||||||||[||||||||||||]PAM NA Intergenic NA NA

T21554 GTCATTTTTACCACATCGCCAGG 1000
Oligo fwd CACCGTCATTTTTACCACATCGCC
Oligo rev AAACGGCGATGTGGTAAAAATGAC
Chromosome start end strand MM target_seq PAM alignment distance position gene name gene id
GU305914.1 171572 171594 - 0 GTCATTTTTACCACATCGCC AGG ||||||||[||||||||||||]PAM NA Intergenic NA NA

T21555 CACATCGCCAGGACTAGAGACGG 1000
Oligo fwd CACCCACATCGCCAGGACTAGAGA
Oligo rev AAACTCTCTAGTCCTGGCGATGTG
Chromosome start end strand MM target_seq PAM alignment distance position gene name gene id

<table>
<tr><td>GU305914.1</td><td>171561</td><td>171583</td><td>-</td><td>0</td><td>CACATCGCCAGGACTAGAGA</td><td>CGG</td><td>||||||||[||||||||||||]PAM</td><td>NA</td><td>Intergenic</td><td>NA</td><td>NA</td></tr>
<tr><td>T21556</td><td>ATCGCCAGGACTAGAGACGGCGG</td><td>1000</td><td></td><td></td><td></td><td></td><td></td><td></td><td></td><td></td><td></td></tr>
<tr><td>Oligo fwd</td><td>CACCATCGCCAGGACTAGAGACGG</td><td></td><td></td><td></td><td></td><td></td><td></td><td></td><td></td><td></td><td></td></tr>
<tr><td>Oligo rev</td><td>AAACCCGTCTCTAGTCCTGGCGAT</td><td></td><td></td><td></td><td></td><td></td><td></td><td></td><td></td><td></td><td></td></tr>
<tr><td>Chromosome</td><td>start</td><td>end</td><td>strand</td><td>MM</td><td>target_seq</td><td>PAM</td><td>alignment</td><td>distance</td><td>position</td><td>gene name</td><td>gene id</td></tr>
<tr><td>GU305914.1</td><td>171558</td><td>171580</td><td>-</td><td>0</td><td>ATCGCCAGGACTAGAGACGG</td><td>CGG</td><td>||||||||[||||||||||||]PAM</td><td>NA</td><td>Intergenic</td><td>NA</td><td>NA</td></tr>
<tr><td>T21557</td><td>TCGCCAGGACTAGAGACGGCGGG</td><td>1000</td><td></td><td></td><td></td><td></td><td></td><td></td><td></td><td></td><td></td></tr>
<tr><td>Oligo fwd</td><td>CACCTCGCCAGGACTAGAGACGGC</td><td></td><td></td><td></td><td></td><td></td><td></td><td></td><td></td><td></td><td></td></tr>
<tr><td>Oligo rev</td><td>AAACGCCGTCTCTAGTCCTGGCGA</td><td></td><td></td><td></td><td></td><td></td><td></td><td></td><td></td><td></td><td></td></tr>
<tr><td>Chromosome</td><td>start</td><td>end</td><td>strand</td><td>MM</td><td>target_seq</td><td>PAM</td><td>alignment</td><td>distance</td><td>position</td><td>gene name</td><td>gene id</td></tr>
<tr><td>GU305914.1</td><td>171557</td><td>171579</td><td>-</td><td>0</td><td>TCGCCAGGACTAGAGACGGC</td><td>GGG</td><td>||||||||[||||||||||||]PAM</td><td>NA</td><td>Intergenic</td><td>NA</td><td>NA</td></tr>
<tr><td>T21558</td><td>CTAGAGACGGCGGGCCCGATTGG</td><td>1000</td><td></td><td></td><td></td><td></td><td></td><td></td><td></td><td></td><td></td></tr>
<tr><td>Oligo fwd</td><td>CACCCTAGAGACGGCGGGCCCGAT</td><td></td><td></td><td></td><td></td><td></td><td></td><td></td><td></td><td></td><td></td></tr>
<tr><td>Oligo rev</td><td>AAACATCGGGCCCGCCGTCTCTAG</td><td></td><td></td><td></td><td></td><td></td><td></td><td></td><td></td><td></td><td></td></tr>
<tr><td>Chromosome</td><td>start</td><td>end</td><td>strand</td><td>MM</td><td>target_seq</td><td>PAM</td><td>alignment</td><td>distance</td><td>position</td><td>gene name</td><td>gene id</td></tr>
<tr><td>GU305914.1</td><td>171548</td><td>171570</td><td>-</td><td>0</td><td>CTAGAGACGGCGGGCCCGAT</td><td>TGG</td><td>||||||||[||||||||||||]PAM</td><td>NA</td><td>Intergenic</td><td>NA</td><td>NA</td></tr>
<tr><td>T21559</td><td>CGGCGGGCCCGATTGGAAAGAGG</td><td>1000</td><td></td><td></td><td></td><td></td><td></td><td></td><td></td><td></td><td></td></tr>
<tr><td>Oligo fwd</td><td>CACCCGGCGGGCCCGATTGGAAAG</td><td></td><td></td><td></td><td></td><td></td><td></td><td></td><td></td><td></td><td></td></tr>
<tr><td>Oligo rev</td><td>AAACCTTTCCAATCGGGCCCGCCG</td><td></td><td></td><td></td><td></td><td></td><td></td><td></td><td></td><td></td><td></td></tr>
<tr><td>Chromosome</td><td>start</td><td>end</td><td>strand</td><td>MM</td><td>target_seq</td><td>PAM</td><td>alignment</td><td>distance</td><td>position</td><td>gene name</td><td>gene id</td></tr>
<tr><td>GU305914.1</td><td>171541</td><td>171563</td><td>-</td><td>0</td><td>CGGCGGGCCCGATTGGAAAG</td><td>AGG</td><td>||||||||[||||||||||||]PAM</td><td>NA</td><td>Intergenic</td><td>NA</td><td>NA</td></tr>
<tr><td>T21560</td><td>CCGATTGGAAAGAGGCGACGTGG</td><td>1000</td><td></td><td></td><td></td><td></td><td></td><td></td><td></td><td></td><td></td></tr>
<tr><td>Oligo fwd</td><td>CACCCCGATTGGAAAGAGGCGACG</td><td></td><td></td><td></td><td></td><td></td><td></td><td></td><td></td><td></td><td></td></tr>
<tr><td>Oligo rev</td><td>AAACCGTCGCCTCTTTCCAATCGG</td><td></td><td></td><td></td><td></td><td></td><td></td><td></td><td></td><td></td><td></td></tr>
<tr><td>Chromosome</td><td>start</td><td>end</td><td>strand</td><td>MM</td><td>target_seq</td><td>PAM</td><td>alignment</td><td>distance</td><td>position</td><td>gene name</td><td>gene id</td></tr>
<tr><td>GU305914.1</td><td>171533</td><td>171555</td><td>-</td><td>0</td><td>CCGATTGGAAAGAGGCGACG</td><td>TGG</td><td>||||||||[||||||||||||]PAM</td><td>NA</td><td>Intergenic</td><td>NA</td><td>NA</td></tr>
<tr><td>T21561</td><td>TTGGAAAGAGGCGACGTGGAAGG</td><td>1000</td><td></td><td></td><td></td><td></td><td></td><td></td><td></td><td></td><td></td></tr>
<tr><td>Oligo fwd</td><td>CACCTTGGAAAGAGGCGACGTGGA</td><td></td><td></td><td></td><td></td><td></td><td></td><td></td><td></td><td></td><td></td></tr>
<tr><td>Oligo rev</td><td>AAACTCCACGTCGCCTCTTTCCAA</td><td></td><td></td><td></td><td></td><td></td><td></td><td></td><td></td><td></td><td></td></tr>
<tr><td>Chromosome</td><td>start</td><td>end</td><td>strand</td><td>MM</td><td>target_seq</td><td>PAM</td><td>alignment</td><td>distance</td><td>position</td><td>gene name</td><td>gene id</td></tr>
<tr><td>GU305914.1</td><td>171529</td><td>171551</td><td>-</td><td>0</td><td>TTGGAAAGAGGCGACGTGGA</td><td>AGG</td><td>||||||||[||||||||||||]PAM</td><td>NA</td><td>Intergenic</td><td>NA</td><td>NA</td></tr>
<tr><td>T21562</td><td>AGAGGCGACGTGGAAGGTCACGG</td><td>1000</td><td></td><td></td><td></td><td></td><td></td><td></td><td></td><td></td><td></td></tr>
<tr><td>Oligo fwd</td><td>CACCAGAGGCGACGTGGAAGGTCA</td><td></td><td></td><td></td><td></td><td></td><td></td><td></td><td></td><td></td><td></td></tr>
<tr><td>Oligo rev</td><td>AAACTGACCTTCCACGTCGCCTCT</td><td></td><td></td><td></td><td></td><td></td><td></td><td></td><td></td><td></td><td></td></tr>
<tr><td>Chromosome</td><td>start</td><td>end</td><td>strand</td><td>MM</td><td>target_seq</td><td>PAM</td><td>alignment</td><td>distance</td><td>position</td><td>gene name</td><td>gene id</td></tr>
<tr><td>GU305914.1</td><td>171523</td><td>171545</td><td>-</td><td>0</td><td>AGAGGCGACGTGGAAGGTCA</td><td>CGG</td><td>||||||||[||||||||||||]PAM</td><td>NA</td><td>Intergenic</td><td>NA</td><td>NA</td></tr>
<tr><td>T21563</td><td>CCGCTCATCTCTCTGACGACCGG</td><td>1000</td><td></td><td></td><td></td><td></td><td></td><td></td><td></td><td></td><td></td></tr>
<tr><td>Oligo fwd</td><td>CACCCCGCTCATCTCTCTGACGAC</td><td></td><td></td><td></td><td></td><td></td><td></td><td></td><td></td><td></td><td></td></tr>
<tr><td>Oligo rev</td><td>AAACGTCGTCAGAGAGATGAGCGG</td><td></td><td></td><td></td><td></td><td></td><td></td><td></td><td></td><td></td><td></td></tr>
<tr><td>Chromosome</td><td>start</td><td>end</td><td>strand</td><td>MM</td><td>target_seq</td><td>PAM</td><td>alignment</td><td>distance</td><td>position</td><td>gene name</td><td>gene id</td></tr>
<tr><td>GU305914.1</td><td>171495</td><td>171517</td><td>-</td><td>0</td><td>CCGCTCATCTCTCTGACGAC</td><td>CGG</td><td>||||||||[||||||||||||]PAM</td><td>NA</td><td>Intergenic</td><td>NA</td><td>NA</td></tr>
<tr><td>T21564</td><td>CGCTCATCTCTCTGACGACCGGG</td><td>1000</td><td></td><td></td><td></td><td></td><td></td><td></td><td></td><td></td><td></td></tr>
<tr><td>Oligo fwd</td><td>CACCCGCTCATCTCTCTGACGACC</td><td></td><td></td><td></td><td></td><td></td><td></td><td></td><td></td><td></td><td></td></tr>
<tr><td>Oligo rev</td><td>AAACGGTCGTCAGAGAGATGAGCG</td><td></td><td></td><td></td><td></td><td></td><td></td><td></td><td></td><td></td><td></td></tr>
<tr><td>Chromosome</td><td>start</td><td>end</td><td>strand</td><td>MM</td><td>target_seq</td><td>PAM</td><td>alignment</td><td>distance</td><td>position</td><td>gene name</td><td>gene id</td></tr>
<tr><td>GU305914.1</td><td>171494</td><td>171516</td><td>-</td><td>0</td><td>CGCTCATCTCTCTGACGACC</td><td>GGG</td><td>||||||||[||||||||||||]PAM</td><td>NA</td><td>Intergenic</td><td>NA</td><td>NA</td></tr>
<tr><td>T21565</td><td>CTCTCTGACGACCGGGTTTCAGG</td><td>1000</td><td></td><td></td><td></td><td></td><td></td><td></td><td></td><td></td><td></td></tr>
<tr><td>Oligo fwd</td><td>CACCCTCTCTGACGACCGGGTTTC</td><td></td><td></td><td></td><td></td><td></td><td></td><td></td><td></td><td></td><td></td></tr>
<tr><td>Oligo rev</td><td>AAACGAAACCCGGTCGTCAGAGAG</td><td></td><td></td><td></td><td></td><td></td><td></td><td></td><td></td><td></td><td></td></tr>
<tr><td>Chromosome</td><td>start</td><td>end</td><td>strand</td><td>MM</td><td>target_seq</td><td>PAM</td><td>alignment</td><td>distance</td><td>position</td><td>gene name</td><td>gene id</td></tr>
<tr><td>GU305914.1</td><td>171487</td><td>171509</td><td>-</td><td>0</td><td>CTCTCTGACGACCGGGTTTC</td><td>AGG</td><td>||||||||[||||||||||||]PAM</td><td>NA</td><td>Intergenic</td><td>NA</td><td>NA</td></tr>
<tr><td>T21566</td><td>GACGACCGGGTTTCAGGTCAAGG</td><td>1000</td><td></td><td></td><td></td><td></td><td></td><td></td><td></td><td></td><td></td></tr>
<tr><td>Oligo fwd</td><td>CACCGACGACCGGGTTTCAGGTCA</td><td></td><td></td><td></td><td></td><td></td><td></td><td></td><td></td><td></td><td></td></tr>
<tr><td>Oligo rev</td><td>AAACTGACCTGAAACCCGGTCGTC</td><td></td><td></td><td></td><td></td><td></td><td></td><td></td><td></td><td></td><td></td></tr>
<tr><td>Chromosome</td><td>start</td><td>end</td><td>strand</td><td>MM</td><td>target_seq</td><td>PAM</td><td>alignment</td><td>distance</td><td>position</td><td>gene name</td><td>gene id</td></tr>
<tr><td>GU305914.1</td><td>171481</td><td>171503</td><td>-</td><td>0</td><td>GACGACCGGGTTTCAGGTCA</td><td>AGG</td><td>||||||||[||||||||||||]PAM</td><td>NA</td><td>Intergenic</td><td>NA</td><td>NA</td></tr>
<tr><td>T21567</td><td>TGCGAGTCACGTACAAAAACAGG</td><td>1000</td><td></td><td></td><td></td><td></td><td></td><td></td><td></td><td></td><td></td></tr>
<tr><td>Oligo fwd</td><td>CACCTGCGAGTCACGTACAAAAAC</td><td></td><td></td><td></td><td></td><td></td><td></td><td></td><td></td><td></td><td></td></tr>
<tr><td>Oligo rev</td><td>AAACGTTTTTGTACGTGACTCGCA</td><td></td><td></td><td></td><td></td><td></td><td></td><td></td><td></td><td></td><td></td></tr>
<tr><td>Chromosome</td><td>start</td><td>end</td><td>strand</td><td>MM</td><td>target_seq</td><td>PAM</td><td>alignment</td><td>distance</td><td>position</td><td>gene name</td><td>gene id</td></tr>
<tr><td>GU305914.1</td><td>171449</td><td>171471</td><td>-</td><td>0</td><td>TGCGAGTCACGTACAAAAAC</td><td>AGG</td><td>||||||||[||||||||||||]PAM</td><td>NA</td><td>Intergenic</td><td>NA</td><td>NA</td></tr>
<tr><td>T21568</td><td>AGTCACGTACAAAAACAGGACGG</td><td>1000</td><td></td><td></td><td></td><td></td><td></td><td></td><td></td><td></td><td></td></tr>
<tr><td>Oligo fwd</td><td>CACCAGTCACGTACAAAAACAGGA</td><td></td><td></td><td></td><td></td><td></td><td></td><td></td><td></td><td></td><td></td></tr>
<tr><td>Oligo rev</td><td>AAACTCCTGTTTTTGTACGTGACT</td><td></td><td></td><td></td><td></td><td></td><td></td><td></td><td></td><td></td><td></td></tr>
<tr><td>Chromosome</td><td>start</td><td>end</td><td>strand</td><td>MM</td><td>target_seq</td><td>PAM</td><td>alignment</td><td>distance</td><td>position</td><td>gene name</td><td>gene id</td></tr>
<tr><td>GU305914.1</td><td>171445</td><td>171467</td><td>-</td><td>0</td><td>AGTCACGTACAAAAACAGGA</td><td>CGG</td><td>||||||||[||||||||||||]PAM</td><td>NA</td><td>Intergenic</td><td>NA</td><td>NA</td></tr>
<tr><td>T21569</td><td>TCCGAACCCCATGTCGATCTCGG</td><td>1000</td><td></td><td></td><td></td><td></td><td></td><td></td><td></td><td></td><td></td></tr>
<tr><td>Oligo fwd</td><td>CACCTCCGAACCCCATGTCGATCT</td><td></td><td></td><td></td><td></td><td></td><td></td><td></td><td></td><td></td><td></td></tr>
<tr><td>Oligo rev</td><td>AAACAGATCGACATGGGGTTCGGA</td><td></td><td></td><td></td><td></td><td></td><td></td><td></td><td></td><td></td><td></td></tr>
<tr><td>Chromosome</td><td>start</td><td>end</td><td>strand</td><td>MM</td><td>target_seq</td><td>PAM</td><td>alignment</td><td>distance</td><td>position</td><td>gene name</td><td>gene id</td></tr>
<tr><td>GU305914.1</td><td>171397</td><td>171419</td><td>-</td><td>0</td><td>TCCGAACCCCATGTCGATCT</td><td>CGG</td><td>||||||||[||||||||||||]PAM</td><td>NA</td><td>Intergenic</td><td>NA</td><td>NA</td></tr>
<tr><td>T21570</td><td>ACGTCGATCCAGAACGAGAACGG</td><td>1000</td><td></td><td></td><td></td><td></td><td></td><td></td><td></td><td></td><td></td></tr>
<tr><td>Oligo fwd</td><td>CACCACGTCGATCCAGAACGAGAA</td><td></td><td></td><td></td><td></td><td></td><td></td><td></td><td></td><td></td><td></td></tr>
<tr><td>Oligo rev</td><td>AAACTTCTCGTTCTGGATCGACGT</td><td></td><td></td><td></td><td></td><td></td><td></td><td></td><td></td><td></td><td></td></tr>
</table>

<table>
<tr><td>Chromosome</td><td>start</td><td>end</td><td>strand</td><td>MM</td><td>target_seq</td><td>PAM</td><td>alignment</td><td>distance</td><td>position</td><td>gene name</td><td>gene id</td></tr>
<tr><td>GU305914.1</td><td>171333</td><td>171355</td><td>-</td><td>0</td><td>ACGTCGATCCAGAACGAGAA</td><td>CGG</td><td>||||||||[||||||||||||]PAM</td><td>NA</td><td>Intergenic</td><td>NA</td><td>NA</td></tr>
<tr><td>T21571</td><td>TGATGACGTTCAGTGTCCCGTGG</td><td>1000</td><td></td><td></td><td></td><td></td><td></td><td></td><td></td><td></td><td></td></tr>
<tr><td>Oligo fwd</td><td>CACCTGATGACGTTCAGTGTCCCG</td><td></td><td></td><td></td><td></td><td></td><td></td><td></td><td></td><td></td><td></td></tr>
<tr><td>Oligo rev</td><td>AAACCGGGACACTGAACGTCATCA</td><td></td><td></td><td></td><td></td><td></td><td></td><td></td><td></td><td></td><td></td></tr>
<tr><td>Chromosome</td><td>start</td><td>end</td><td>strand</td><td>MM</td><td>target_seq</td><td>PAM</td><td>alignment</td><td>distance</td><td>position</td><td>gene name</td><td>gene id</td></tr>
<tr><td>GU305914.1</td><td>171296</td><td>171318</td><td>-</td><td>0</td><td>TGATGACGTTCAGTGTCCCG</td><td>TGG</td><td>||||||||[||||||||||||]PAM</td><td>NA</td><td>Intergenic</td><td>NA</td><td>NA</td></tr>
<tr><td>T21572</td><td>GATGACGTTCAGTGTCCCGTGGG</td><td>1000</td><td></td><td></td><td></td><td></td><td></td><td></td><td></td><td></td><td></td></tr>
<tr><td>Oligo fwd</td><td>CACCGATGACGTTCAGTGTCCCGT</td><td></td><td></td><td></td><td></td><td></td><td></td><td></td><td></td><td></td><td></td></tr>
<tr><td>Oligo rev</td><td>AAACACGGGACACTGAACGTCATC</td><td></td><td></td><td></td><td></td><td></td><td></td><td></td><td></td><td></td><td></td></tr>
<tr><td>Chromosome</td><td>start</td><td>end</td><td>strand</td><td>MM</td><td>target_seq</td><td>PAM</td><td>alignment</td><td>distance</td><td>position</td><td>gene name</td><td>gene id</td></tr>
<tr><td>GU305914.1</td><td>171295</td><td>171317</td><td>-</td><td>0</td><td>GATGACGTTCAGTGTCCCGT</td><td>GGG</td><td>||||||||[||||||||||||]PAM</td><td>NA</td><td>Intergenic</td><td>NA</td><td>NA</td></tr>
<tr><td>T21573</td><td>TGTCCCGTGGGCAGACAGTCAGG</td><td>1000</td><td></td><td></td><td></td><td></td><td></td><td></td><td></td><td></td><td></td></tr>
<tr><td>Oligo fwd</td><td>CACCTGTCCCGTGGGCAGACAGTC</td><td></td><td></td><td></td><td></td><td></td><td></td><td></td><td></td><td></td><td></td></tr>
<tr><td>Oligo rev</td><td>AAACGACTGTCTGCCCACGGGACA</td><td></td><td></td><td></td><td></td><td></td><td></td><td></td><td></td><td></td><td></td></tr>
<tr><td>Chromosome</td><td>start</td><td>end</td><td>strand</td><td>MM</td><td>target_seq</td><td>PAM</td><td>alignment</td><td>distance</td><td>position</td><td>gene name</td><td>gene id</td></tr>
<tr><td>GU305914.1</td><td>171283</td><td>171305</td><td>-</td><td>0</td><td>TGTCCCGTGGGCAGACAGTC</td><td>AGG</td><td>||||||||[||||||||||||]PAM</td><td>NA</td><td>Intergenic</td><td>NA</td><td>NA</td></tr>
<tr><td>T21574</td><td>GTCCCGTGGGCAGACAGTCAGGG</td><td>1000</td><td></td><td></td><td></td><td></td><td></td><td></td><td></td><td></td><td></td></tr>
<tr><td>Oligo fwd</td><td>CACCGTCCCGTGGGCAGACAGTCA</td><td></td><td></td><td></td><td></td><td></td><td></td><td></td><td></td><td></td><td></td></tr>
<tr><td>Oligo rev</td><td>AAACTGACTGTCTGCCCACGGGAC</td><td></td><td></td><td></td><td></td><td></td><td></td><td></td><td></td><td></td><td></td></tr>
<tr><td>Chromosome</td><td>start</td><td>end</td><td>strand</td><td>MM</td><td>target_seq</td><td>PAM</td><td>alignment</td><td>distance</td><td>position</td><td>gene name</td><td>gene id</td></tr>
<tr><td>GU305914.1</td><td>171282</td><td>171304</td><td>-</td><td>0</td><td>GTCCCGTGGGCAGACAGTCA</td><td>GGG</td><td>||||||||[||||||||||||]PAM</td><td>NA</td><td>Intergenic</td><td>NA</td><td>NA</td></tr>
<tr><td>T21575</td><td>TCCCGTGGGCAGACAGTCAGGGG</td><td>1000</td><td></td><td></td><td></td><td></td><td></td><td></td><td></td><td></td><td></td></tr>
<tr><td>Oligo fwd</td><td>CACCTCCCGTGGGCAGACAGTCAG</td><td></td><td></td><td></td><td></td><td></td><td></td><td></td><td></td><td></td><td></td></tr>
<tr><td>Oligo rev</td><td>AAACCTGACTGTCTGCCCACGGGA</td><td></td><td></td><td></td><td></td><td></td><td></td><td></td><td></td><td></td><td></td></tr>
<tr><td>Chromosome</td><td>start</td><td>end</td><td>strand</td><td>MM</td><td>target_seq</td><td>PAM</td><td>alignment</td><td>distance</td><td>position</td><td>gene name</td><td>gene id</td></tr>
<tr><td>GU305914.1</td><td>171281</td><td>171303</td><td>-</td><td>0</td><td>TCCCGTGGGCAGACAGTCAG</td><td>GGG</td><td>||||||||[||||||||||||]PAM</td><td>NA</td><td>Intergenic</td><td>NA</td><td>NA</td></tr>
<tr><td>T21576</td><td>GAACGCGACCGTAGACGTCATGG</td><td>1000</td><td></td><td></td><td></td><td></td><td></td><td></td><td></td><td></td><td></td></tr>
<tr><td>Oligo fwd</td><td>CACCGAACGCGACCGTAGACGTCA</td><td></td><td></td><td></td><td></td><td></td><td></td><td></td><td></td><td></td><td></td></tr>
<tr><td>Oligo rev</td><td>AAACTGACGTCTACGGTCGCGTTC</td><td></td><td></td><td></td><td></td><td></td><td></td><td></td><td></td><td></td><td></td></tr>
<tr><td>Chromosome</td><td>start</td><td>end</td><td>strand</td><td>MM</td><td>target_seq</td><td>PAM</td><td>alignment</td><td>distance</td><td>position</td><td>gene name</td><td>gene id</td></tr>
<tr><td>GU305914.1</td><td>171259</td><td>171281</td><td>-</td><td>0</td><td>GAACGCGACCGTAGACGTCA</td><td>TGG</td><td>||||||||[||||||||||||]PAM</td><td>NA</td><td>Intergenic</td><td>NA</td><td>NA</td></tr>
<tr><td>T21577</td><td>TGCCGCCCTGACCGTCCTGCTGG</td><td>1000</td><td></td><td></td><td></td><td></td><td></td><td></td><td></td><td></td><td></td></tr>
<tr><td>Oligo fwd</td><td>CACCTGCCGCCCTGACCGTCCTGC</td><td></td><td></td><td></td><td></td><td></td><td></td><td></td><td></td><td></td><td></td></tr>
<tr><td>Oligo rev</td><td>AAACGCAGGACGGTCAGGGCGGCA</td><td></td><td></td><td></td><td></td><td></td><td></td><td></td><td></td><td></td><td></td></tr>
<tr><td>Chromosome</td><td>start</td><td>end</td><td>strand</td><td>MM</td><td>target_seq</td><td>PAM</td><td>alignment</td><td>distance</td><td>position</td><td>gene name</td><td>gene id</td></tr>
<tr><td>GU305914.1</td><td>171178</td><td>171200</td><td>-</td><td>0</td><td>TGCCGCCCTGACCGTCCTGC</td><td>TGG</td><td>||||||||[||||||||||||]PAM</td><td>NA</td><td>Intergenic</td><td>NA</td><td>NA</td></tr>
<tr><td>T21578</td><td>CCTGACCGTCCTGCTGGTCTTGG</td><td>1000</td><td></td><td></td><td></td><td></td><td></td><td></td><td></td><td></td><td></td></tr>
<tr><td>Oligo fwd</td><td>CACCCCTGACCGTCCTGCTGGTCT</td><td></td><td></td><td></td><td></td><td></td><td></td><td></td><td></td><td></td><td></td></tr>
<tr><td>Oligo rev</td><td>AAACAGACCAGCAGGACGGTCAGG</td><td></td><td></td><td></td><td></td><td></td><td></td><td></td><td></td><td></td><td></td></tr>
<tr><td>Chromosome</td><td>start</td><td>end</td><td>strand</td><td>MM</td><td>target_seq</td><td>PAM</td><td>alignment</td><td>distance</td><td>position</td><td>gene name</td><td>gene id</td></tr>
<tr><td>GU305914.1</td><td>171172</td><td>171194</td><td>-</td><td>0</td><td>CCTGACCGTCCTGCTGGTCT</td><td>TGG</td><td>||||||||[||||||||||||]PAM</td><td>NA</td><td>Intergenic</td><td>NA</td><td>NA</td></tr>
<tr><td>T21579</td><td>CGTACATCGTTCATCGTCATCGG</td><td>1000</td><td></td><td></td><td></td><td></td><td></td><td></td><td></td><td></td><td></td></tr>
<tr><td>Oligo fwd</td><td>CACCCGTACATCGTTCATCGTCAT</td><td></td><td></td><td></td><td></td><td></td><td></td><td></td><td></td><td></td><td></td></tr>
<tr><td>Oligo rev</td><td>AAACATGACGATGAACGATGTACG</td><td></td><td></td><td></td><td></td><td></td><td></td><td></td><td></td><td></td><td></td></tr>
<tr><td>Chromosome</td><td>start</td><td>end</td><td>strand</td><td>MM</td><td>target_seq</td><td>PAM</td><td>alignment</td><td>distance</td><td>position</td><td>gene name</td><td>gene id</td></tr>
<tr><td>GU305914.1</td><td>171131</td><td>171153</td><td>-</td><td>0</td><td>CGTACATCGTTCATCGTCAT</td><td>CGG</td><td>||||||||[||||||||||||]PAM</td><td>NA</td><td>Intergenic</td><td>NA</td><td>NA</td></tr>
<tr><td>T21580</td><td>GTACATCGTTCATCGTCATCGGG</td><td>1000</td><td></td><td></td><td></td><td></td><td></td><td></td><td></td><td></td><td></td></tr>
<tr><td>Oligo fwd</td><td>CACCGTACATCGTTCATCGTCATC</td><td></td><td></td><td></td><td></td><td></td><td></td><td></td><td></td><td></td><td></td></tr>
<tr><td>Oligo rev</td><td>AAACGATGACGATGAACGATGTAC</td><td></td><td></td><td></td><td></td><td></td><td></td><td></td><td></td><td></td><td></td></tr>
<tr><td>Chromosome</td><td>start</td><td>end</td><td>strand</td><td>MM</td><td>target_seq</td><td>PAM</td><td>alignment</td><td>distance</td><td>position</td><td>gene name</td><td>gene id</td></tr>
<tr><td>GU305914.1</td><td>171130</td><td>171152</td><td>-</td><td>0</td><td>GTACATCGTTCATCGTCATC</td><td>GGG</td><td>||||||||[||||||||||||]PAM</td><td>NA</td><td>Intergenic</td><td>NA</td><td>NA</td></tr>
<tr><td>T21581</td><td>TACATCGTTCATCGTCATCGGGG</td><td>1000</td><td></td><td></td><td></td><td></td><td></td><td></td><td></td><td></td><td></td></tr>
<tr><td>Oligo fwd</td><td>CACCTACATCGTTCATCGTCATCG</td><td></td><td></td><td></td><td></td><td></td><td></td><td></td><td></td><td></td><td></td></tr>
<tr><td>Oligo rev</td><td>AAACCGATGACGATGAACGATGTA</td><td></td><td></td><td></td><td></td><td></td><td></td><td></td><td></td><td></td><td></td></tr>
<tr><td>Chromosome</td><td>start</td><td>end</td><td>strand</td><td>MM</td><td>target_seq</td><td>PAM</td><td>alignment</td><td>distance</td><td>position</td><td>gene name</td><td>gene id</td></tr>
<tr><td>GU305914.1</td><td>171129</td><td>171151</td><td>-</td><td>0</td><td>TACATCGTTCATCGTCATCG</td><td>GGG</td><td>||||||||[||||||||||||]PAM</td><td>NA</td><td>Intergenic</td><td>NA</td><td>NA</td></tr>
<tr><td>T21582</td><td>ACATCGTTCATCGTCATCGGGGG</td><td>1000</td><td></td><td></td><td></td><td></td><td></td><td></td><td></td><td></td><td></td></tr>
<tr><td>Oligo fwd</td><td>CACCACATCGTTCATCGTCATCGG</td><td></td><td></td><td></td><td></td><td></td><td></td><td></td><td></td><td></td><td></td></tr>
<tr><td>Oligo rev</td><td>AAACCCGATGACGATGAACGATGT</td><td></td><td></td><td></td><td></td><td></td><td></td><td></td><td></td><td></td><td></td></tr>
<tr><td>Chromosome</td><td>start</td><td>end</td><td>strand</td><td>MM</td><td>target_seq</td><td>PAM</td><td>alignment</td><td>distance</td><td>position</td><td>gene name</td><td>gene id</td></tr>
<tr><td>GU305914.1</td><td>171128</td><td>171150</td><td>-</td><td>0</td><td>ACATCGTTCATCGTCATCGG</td><td>GGG</td><td>||||||||[||||||||||||]PAM</td><td>NA</td><td>Intergenic</td><td>NA</td><td>NA</td></tr>
<tr><td>T21583</td><td>TCGTCATCGGGGGACGTTGAAGG</td><td>1000</td><td></td><td></td><td></td><td></td><td></td><td></td><td></td><td></td><td></td></tr>
<tr><td>Oligo fwd</td><td>CACCTCGTCATCGGGGGACGTTGA</td><td></td><td></td><td></td><td></td><td></td><td></td><td></td><td></td><td></td><td></td></tr>
<tr><td>Oligo rev</td><td>AAACTCAACGTCCCCCGATGACGA</td><td></td><td></td><td></td><td></td><td></td><td></td><td></td><td></td><td></td><td></td></tr>
<tr><td>Chromosome</td><td>start</td><td>end</td><td>strand</td><td>MM</td><td>target_seq</td><td>PAM</td><td>alignment</td><td>distance</td><td>position</td><td>gene name</td><td>gene id</td></tr>
<tr><td>GU305914.1</td><td>171118</td><td>171140</td><td>-</td><td>0</td><td>TCGTCATCGGGGGACGTTGA</td><td>AGG</td><td>||||||||[||||||||||||]PAM</td><td>NA</td><td>Intergenic</td><td>NA</td><td>NA</td></tr>
<tr><td>T21584</td><td>TCGCGCGAACGATGATGTTTTGG</td><td>1000</td><td></td><td></td><td></td><td></td><td></td><td></td><td></td><td></td><td></td></tr>
<tr><td>Oligo fwd</td><td>CACCTCGCGCGAACGATGATGTTT</td><td></td><td></td><td></td><td></td><td></td><td></td><td></td><td></td><td></td><td></td></tr>
<tr><td>Oligo rev</td><td>AAACAAACATCATCGTTCGCGCGA</td><td></td><td></td><td></td><td></td><td></td><td></td><td></td><td></td><td></td><td></td></tr>
<tr><td>Chromosome</td><td>start</td><td>end</td><td>strand</td><td>MM</td><td>target_seq</td><td>PAM</td><td>alignment</td><td>distance</td><td>position</td><td>gene name</td><td>gene id</td></tr>
<tr><td>GU305914.1</td><td>170999</td><td>171021</td><td>-</td><td>0</td><td>TCGCGCGAACGATGATGTTT</td><td>TGG</td><td>||||||||[||||||||||||]PAM</td><td>NA</td><td>Intergenic</td><td>NA</td><td>NA</td></tr>
<tr><td>T21585</td><td>CGAACGATGATGTTTTGGAGCGG</td><td>1000</td><td></td><td></td><td></td><td></td><td></td><td></td><td></td><td></td><td></td></tr>
<tr><td>Oligo fwd</td><td>CACCCGAACGATGATGTTTTGGAG</td><td></td><td></td><td></td><td></td><td></td><td></td><td></td><td></td><td></td><td></td></tr>
</table>

Oligo rev AAACCTCCAAAACATCATCGTTCG
Chromosome start end strand MM target_seq PAM alignment distance position gene name gene id
GU305914.1 170994 171016 - 0 CGAACGATGATGTTTTGGAG CGG ||||||||[||||||||||||]PAM NA Intergenic NA NA

T21586 TTGGAGCGGCAATCGCGCCGCGG 1000
Oligo fwd CACCTTGGAGCGGCAATCGCGCCG
Oligo rev AAACCGGCGCGATTGCCGCTCCAA
Chromosome start end strand MM target_seq PAM alignment distance position gene name gene id
GU305914.1 170980 171002 - 0 TTGGAGCGGCAATCGCGCCG CGG ||||||||[||||||||||||]PAM NA Intergenic NA NA

T21587 TGGAGCGGCAATCGCGCCGCGGG 1000
Oligo fwd CACCTGGAGCGGCAATCGCGCCGC
Oligo rev AAACGCGGCGCGATTGCCGCTCCA
Chromosome start end strand MM target_seq PAM alignment distance position gene name gene id
GU305914.1 170979 171001 - 0 TGGAGCGGCAATCGCGCCGC GGG ||||||||[||||||||||||]PAM NA Intergenic NA NA

T21588 AGCGGCAATCGCGCCGCGGGCGG 1000
Oligo fwd CACCAGCGGCAATCGCGCCGCGGG
Oligo rev AAACCCCGCGGCGCGATTGCCGCT
Chromosome start end strand MM target_seq PAM alignment distance position gene name gene id
GU305914.1 170976 170998 - 0 AGCGGCAATCGCGCCGCGGG CGG ||||||||[||||||||||||]PAM NA Intergenic NA NA

T21589 GATTTCCAGCTCCGAGTGCGTGG 1000
Oligo fwd CACCGATTTCCAGCTCCGAGTGCG
Oligo rev AAACCGCACTCGGAGCTGGAAATC
Chromosome start end strand MM target_seq PAM alignment distance position gene name gene id
GU305914.1 170948 170970 - 0 GATTTCCAGCTCCGAGTGCG TGG ||||||||[||||||||||||]PAM NA Intergenic NA NA

T21590 GTCGACCATCACGTACGCTAAGG 1000
Oligo fwd CACCGTCGACCATCACGTACGCTA
Oligo rev AAACTAGCGTACGTGATGGTCGAC
Chromosome start end strand MM target_seq PAM alignment distance position gene name gene id
GU305914.1 170926 170948 - 0 GTCGACCATCACGTACGCTA AGG ||||||||[||||||||||||]PAM NA Intergenic NA NA

T21591 TCACGTACGCTAAGGTGTCTCGG 1000
Oligo fwd CACCTCACGTACGCTAAGGTGTCT
Oligo rev AAACAGACACCTTAGCGTACGTGA
Chromosome start end strand MM target_seq PAM alignment distance position gene name gene id
GU305914.1 170918 170940 - 0 TCACGTACGCTAAGGTGTCT CGG ||||||||[||||||||||||]PAM NA Intergenic NA NA

T21592 CGCTAAGGTGTCTCGGCCTCTGG 1000
Oligo fwd CACCCGCTAAGGTGTCTCGGCCTC
Oligo rev AAACGAGGCCGAGACACCTTAGCG
Chromosome start end strand MM target_seq PAM alignment distance position gene name gene id
GU305914.1 170911 170933 - 0 CGCTAAGGTGTCTCGGCCTC TGG ||||||||[||||||||||||]PAM NA Intergenic NA NA

T21593 TCTGGTCACGTCCCTGACGCCGG 1000
Oligo fwd CACCTCTGGTCACGTCCCTGACGC
Oligo rev AAACGCGTCAGGGACGTGACCAGA
Chromosome start end strand MM target_seq PAM alignment distance position gene name gene id
GU305914.1 170893 170915 - 0 TCTGGTCACGTCCCTGACGC CGG ||||||||[||||||||||||]PAM NA Intergenic NA NA

T21594 CGAACGAGACCACGTTCGCCAGG 1000
Oligo fwd CACCCGAACGAGACCACGTTCGCC
Oligo rev AAACGGCGAACGTGGTCTCGTTCG
Chromosome start end strand MM target_seq PAM alignment distance position gene name gene id
GU305914.1 170834 170856 - 0 CGAACGAGACCACGTTCGCC AGG ||||||||[||||||||||||]PAM NA Intergenic NA NA

T21595 CACGTTCGCCAGGATACCTCCGG 1000
Oligo fwd CACCCACGTTCGCCAGGATACCTC
Oligo rev AAACGAGGTATCCTGGCGAACGTG
Chromosome start end strand MM target_seq PAM alignment distance position gene name gene id
GU305914.1 170824 170846 - 0 CACGTTCGCCAGGATACCTC CGG ||||||||[||||||||||||]PAM NA Intergenic NA NA

T21596 ACGTTCGCCAGGATACCTCCGGG 1000
Oligo fwd CACCACGTTCGCCAGGATACCTCC
Oligo rev AAACGGAGGTATCCTGGCGAACGT
Chromosome start end strand MM target_seq PAM alignment distance position gene name gene id
GU305914.1 170823 170845 - 0 ACGTTCGCCAGGATACCTCC GGG ||||||||[||||||||||||]PAM NA Intergenic NA NA

T21597 TACCTCCGGGAGAATCTCCCTGG 1000
Oligo fwd CACCTACCTCCGGGAGAATCTCCC
Oligo rev AAACGGGAGATTCTCCCGGAGGTA
Chromosome start end strand MM target_seq PAM alignment distance position gene name gene id
GU305914.1 170810 170832 - 0 TACCTCCGGGAGAATCTCCC TGG ||||||||[||||||||||||]PAM NA Intergenic NA NA

T21598 ACCTCCGGGAGAATCTCCCTGGG 1000
Oligo fwd CACCACCTCCGGGAGAATCTCCCT
Oligo rev AAACAGGGAGATTCTCCCGGAGGT
Chromosome start end strand MM target_seq PAM alignment distance position gene name gene id
GU305914.1 170809 170831 - 0 ACCTCCGGGAGAATCTCCCT GGG ||||||||[||||||||||||]PAM NA Intergenic NA NA

T21599 GAGACTCGAGTTCGTATCATAGG 1000
Oligo fwd CACCGAGACTCGAGTTCGTATCAT
Oligo rev AAACATGATACGAACTCGAGTCTC
Chromosome start end strand MM target_seq PAM alignment distance position gene name gene id
GU305914.1 170787 170809 - 0 GAGACTCGAGTTCGTATCAT AGG ||||||||[||||||||||||]PAM NA Intergenic NA NA

T21600 AGACTCGAGTTCGTATCATAGGG 1000

<table>
<tr><td>Oligo fwd</td><td>CACCAGACTCGAGTTCGTATCATA</td><td></td><td></td><td></td><td></td><td></td><td></td><td></td><td></td><td></td><td></td><td></td></tr>
<tr><td>Oligo rev</td><td>AAACTATGATACGAACTCGAGTCT</td><td></td><td></td><td></td><td></td><td></td><td></td><td></td><td></td><td></td><td></td><td></td></tr>
<tr><td>Chromosome</td><td>start</td><td></td><td>end</td><td>strand</td><td>MM</td><td>target_seq</td><td>PAM</td><td>alignment</td><td>distance</td><td>position</td><td>gene name</td><td>gene id</td></tr>
<tr><td>GU305914.1</td><td></td><td>170786</td><td>170808</td><td>-</td><td>0</td><td>AGACTCGAGTTCGTATCATA</td><td>GGG</td><td>||||||||[||||||||||||]PAM</td><td>NA</td><td>Intergenic</td><td>NA</td><td>NA</td></tr>
<tr><td>T21601</td><td>CAGACGAAGAGCTCGCCGAGCGG</td><td></td><td>1000</td><td></td><td></td><td></td><td></td><td></td><td></td><td></td><td></td><td></td></tr>
<tr><td>Oligo fwd</td><td>CACCCAGACGAAGAGCTCGCCGAG</td><td></td><td></td><td></td><td></td><td></td><td></td><td></td><td></td><td></td><td></td><td></td></tr>
<tr><td>Oligo rev</td><td>AAACCTCGGCGAGCTCTTCGTCTG</td><td></td><td></td><td></td><td></td><td></td><td></td><td></td><td></td><td></td><td></td><td></td></tr>
<tr><td>Chromosome</td><td>start</td><td></td><td>end</td><td>strand</td><td>MM</td><td>target_seq</td><td>PAM</td><td>alignment</td><td>distance</td><td>position</td><td>gene name</td><td>gene id</td></tr>
<tr><td>GU305914.1</td><td></td><td>170753</td><td>170775</td><td>-</td><td>0</td><td>CAGACGAAGAGCTCGCCGAG</td><td>CGG</td><td>||||||||[||||||||||||]PAM</td><td>NA</td><td>Intergenic</td><td>NA</td><td>NA</td></tr>
<tr><td>T21602</td><td>AGACGAAGAGCTCGCCGAGCGGG</td><td></td><td>1000</td><td></td><td></td><td></td><td></td><td></td><td></td><td></td><td></td><td></td></tr>
<tr><td>Oligo fwd</td><td>CACCAGACGAAGAGCTCGCCGAGC</td><td></td><td></td><td></td><td></td><td></td><td></td><td></td><td></td><td></td><td></td><td></td></tr>
<tr><td>Oligo rev</td><td>AAACGCTCGGCGAGCTCTTCGTCT</td><td></td><td></td><td></td><td></td><td></td><td></td><td></td><td></td><td></td><td></td><td></td></tr>
<tr><td>Chromosome</td><td>start</td><td></td><td>end</td><td>strand</td><td>MM</td><td>target_seq</td><td>PAM</td><td>alignment</td><td>distance</td><td>position</td><td>gene name</td><td>gene id</td></tr>
<tr><td>GU305914.1</td><td></td><td>170752</td><td>170774</td><td>-</td><td>0</td><td>AGACGAAGAGCTCGCCGAGC</td><td>GGG</td><td>||||||||[||||||||||||]PAM</td><td>NA</td><td>Intergenic</td><td>NA</td><td>NA</td></tr>
<tr><td>T21603</td><td>AGAGCTCGCCGAGCGGGCTGCGG</td><td></td><td>1000</td><td></td><td></td><td></td><td></td><td></td><td></td><td></td><td></td><td></td></tr>
<tr><td>Oligo fwd</td><td>CACCAGAGCTCGCCGAGCGGGCTG</td><td></td><td></td><td></td><td></td><td></td><td></td><td></td><td></td><td></td><td></td><td></td></tr>
<tr><td>Oligo rev</td><td>AAACCAGCCCGCTCGGCGAGCTCT</td><td></td><td></td><td></td><td></td><td></td><td></td><td></td><td></td><td></td><td></td><td></td></tr>
<tr><td>Chromosome</td><td>start</td><td></td><td>end</td><td>strand</td><td>MM</td><td>target_seq</td><td>PAM</td><td>alignment</td><td>distance</td><td>position</td><td>gene name</td><td>gene id</td></tr>
<tr><td>GU305914.1</td><td></td><td>170746</td><td>170768</td><td>-</td><td>0</td><td>AGAGCTCGCCGAGCGGGCTG</td><td>CGG</td><td>||||||||[||||||||||||]PAM</td><td>NA</td><td>Intergenic</td><td>NA</td><td>NA</td></tr>
<tr><td>T21604</td><td>GCTCGCCGAGCGGGCTGCGGCGG</td><td></td><td>1000</td><td></td><td></td><td></td><td></td><td></td><td></td><td></td><td></td><td></td></tr>
<tr><td>Oligo fwd</td><td>CACCGCTCGCCGAGCGGGCTGCGG</td><td></td><td></td><td></td><td></td><td></td><td></td><td></td><td></td><td></td><td></td><td></td></tr>
<tr><td>Oligo rev</td><td>AAACCCGCAGCCCGCTCGGCGAGC</td><td></td><td></td><td></td><td></td><td></td><td></td><td></td><td></td><td></td><td></td><td></td></tr>
<tr><td>Chromosome</td><td>start</td><td></td><td>end</td><td>strand</td><td>MM</td><td>target_seq</td><td>PAM</td><td>alignment</td><td>distance</td><td>position</td><td>gene name</td><td>gene id</td></tr>
<tr><td>GU305914.1</td><td></td><td>170743</td><td>170765</td><td>-</td><td>0</td><td>GCTCGCCGAGCGGGCTGCGG</td><td>CGG</td><td>||||||||[||||||||||||]PAM</td><td>NA</td><td>Intergenic</td><td>NA</td><td>NA</td></tr>
<tr><td>T21605</td><td>GCGGGCTGCGGCGGCAGCGAAGG</td><td></td><td>1000</td><td></td><td></td><td></td><td></td><td></td><td></td><td></td><td></td><td></td></tr>
<tr><td>Oligo fwd</td><td>CACCGCGGGCTGCGGCGGCAGCGA</td><td></td><td></td><td></td><td></td><td></td><td></td><td></td><td></td><td></td><td></td><td></td></tr>
<tr><td>Oligo rev</td><td>AAACTCGCTGCCGCCGCAGCCCGC</td><td></td><td></td><td></td><td></td><td></td><td></td><td></td><td></td><td></td><td></td><td></td></tr>
<tr><td>Chromosome</td><td>start</td><td></td><td>end</td><td>strand</td><td>MM</td><td>target_seq</td><td>PAM</td><td>alignment</td><td>distance</td><td>position</td><td>gene name</td><td>gene id</td></tr>
<tr><td>GU305914.1</td><td></td><td>170734</td><td>170756</td><td>-</td><td>0</td><td>GCGGGCTGCGGCGGCAGCGA</td><td>AGG</td><td>||||||||[||||||||||||]PAM</td><td>NA</td><td>Intergenic</td><td>NA</td><td>NA</td></tr>
<tr><td>T21606</td><td>GGCGGCAGCGAAGGCGCCTCCGG</td><td></td><td>1000</td><td></td><td></td><td></td><td></td><td></td><td></td><td></td><td></td><td></td></tr>
<tr><td>Oligo fwd</td><td>CACCGGCGGCAGCGAAGGCGCCTC</td><td></td><td></td><td></td><td></td><td></td><td></td><td></td><td></td><td></td><td></td><td></td></tr>
<tr><td>Oligo rev</td><td>AAACGAGGCGCCTTCGCTGCCGCC</td><td></td><td></td><td></td><td></td><td></td><td></td><td></td><td></td><td></td><td></td><td></td></tr>
<tr><td>Chromosome</td><td>start</td><td></td><td>end</td><td>strand</td><td>MM</td><td>target_seq</td><td>PAM</td><td>alignment</td><td>distance</td><td>position</td><td>gene name</td><td>gene id</td></tr>
<tr><td>GU305914.1</td><td></td><td>170725</td><td>170747</td><td>-</td><td>0</td><td>GGCGGCAGCGAAGGCGCCTC</td><td>CGG</td><td>||||||||[||||||||||||]PAM</td><td>NA</td><td>Intergenic</td><td>NA</td><td>NA</td></tr>
<tr><td>T21607</td><td>GGCAGCGAAGGCGCCTCCGGCGG</td><td></td><td>1000</td><td></td><td></td><td></td><td></td><td></td><td></td><td></td><td></td><td></td></tr>
<tr><td>Oligo fwd</td><td>CACCGGCAGCGAAGGCGCCTCCGG</td><td></td><td></td><td></td><td></td><td></td><td></td><td></td><td></td><td></td><td></td><td></td></tr>
<tr><td>Oligo rev</td><td>AAACCCGGAGGCGCCTTCGCTGCC</td><td></td><td></td><td></td><td></td><td></td><td></td><td></td><td></td><td></td><td></td><td></td></tr>
<tr><td>Chromosome</td><td>start</td><td></td><td>end</td><td>strand</td><td>MM</td><td>target_seq</td><td>PAM</td><td>alignment</td><td>distance</td><td>position</td><td>gene name</td><td>gene id</td></tr>
<tr><td>GU305914.1</td><td></td><td>170722</td><td>170744</td><td>-</td><td>0</td><td>GGCAGCGAAGGCGCCTCCGG</td><td>CGG</td><td>||||||||[||||||||||||]PAM</td><td>NA</td><td>Intergenic</td><td>NA</td><td>NA</td></tr>
<tr><td>T21608</td><td>ACCTCTGCCTTTCATGCCCACGG</td><td></td><td>1000</td><td></td><td></td><td></td><td></td><td></td><td></td><td></td><td></td><td></td></tr>
<tr><td>Oligo fwd</td><td>CACCACCTCTGCCTTTCATGCCCA</td><td></td><td></td><td></td><td></td><td></td><td></td><td></td><td></td><td></td><td></td><td></td></tr>
<tr><td>Oligo rev</td><td>AAACTGGGCATGAAAGGCAGAGGT</td><td></td><td></td><td></td><td></td><td></td><td></td><td></td><td></td><td></td><td></td><td></td></tr>
<tr><td>Chromosome</td><td>start</td><td></td><td>end</td><td>strand</td><td>MM</td><td>target_seq</td><td>PAM</td><td>alignment</td><td>distance</td><td>position</td><td>gene name</td><td>gene id</td></tr>
<tr><td>GU305914.1</td><td></td><td>170638</td><td>170660</td><td>-</td><td>0</td><td>ACCTCTGCCTTTCATGCCCA</td><td>CGG</td><td>||||||||[||||||||||||]PAM</td><td>NA</td><td>Intergenic</td><td>NA</td><td>NA</td></tr>
<tr><td>T21609</td><td>CGACATCCCGACCACGTTCGCGG</td><td></td><td>1000</td><td></td><td></td><td></td><td></td><td></td><td></td><td></td><td></td><td></td></tr>
<tr><td>Oligo fwd</td><td>CACCCGACATCCCGACCACGTTCG</td><td></td><td></td><td></td><td></td><td></td><td></td><td></td><td></td><td></td><td></td><td></td></tr>
<tr><td>Oligo rev</td><td>AAACCGAACGTGGTCGGGATGTCG</td><td></td><td></td><td></td><td></td><td></td><td></td><td></td><td></td><td></td><td></td><td></td></tr>
<tr><td>Chromosome</td><td>start</td><td></td><td>end</td><td>strand</td><td>MM</td><td>target_seq</td><td>PAM</td><td>alignment</td><td>distance</td><td>position</td><td>gene name</td><td>gene id</td></tr>
<tr><td>GU305914.1</td><td></td><td>170614</td><td>170636</td><td>-</td><td>0</td><td>CGACATCCCGACCACGTTCG</td><td>CGG</td><td>||||||||[||||||||||||]PAM</td><td>NA</td><td>Intergenic</td><td>NA</td><td>NA</td></tr>
<tr><td>T21610</td><td>CCCGACCACGTTCGCGGCAGCGG</td><td></td><td>1000</td><td></td><td></td><td></td><td></td><td></td><td></td><td></td><td></td><td></td></tr>
<tr><td>Oligo fwd</td><td>CACCCCCGACCACGTTCGCGGCAG</td><td></td><td></td><td></td><td></td><td></td><td></td><td></td><td></td><td></td><td></td><td></td></tr>
<tr><td>Oligo rev</td><td>AAACCTGCCGCGAACGTGGTCGGG</td><td></td><td></td><td></td><td></td><td></td><td></td><td></td><td></td><td></td><td></td><td></td></tr>
<tr><td>Chromosome</td><td>start</td><td></td><td>end</td><td>strand</td><td>MM</td><td>target_seq</td><td>PAM</td><td>alignment</td><td>distance</td><td>position</td><td>gene name</td><td>gene id</td></tr>
<tr><td>GU305914.1</td><td></td><td>170608</td><td>170630</td><td>-</td><td>0</td><td>CCCGACCACGTTCGCGGCAG</td><td>CGG</td><td>||||||||[||||||||||||]PAM</td><td>NA</td><td>Intergenic</td><td>NA</td><td>NA</td></tr>
<tr><td>T21611</td><td>GCTGCCGATGTTTACCGACCCGG</td><td></td><td>1000</td><td></td><td></td><td></td><td></td><td></td><td></td><td></td><td></td><td></td></tr>
<tr><td>Oligo fwd</td><td>CACCGCTGCCGATGTTTACCGACC</td><td></td><td></td><td></td><td></td><td></td><td></td><td></td><td></td><td></td><td></td><td></td></tr>
<tr><td>Oligo rev</td><td>AAACGGTCGGTAAACATCGGCAGC</td><td></td><td></td><td></td><td></td><td></td><td></td><td></td><td></td><td></td><td></td><td></td></tr>
<tr><td>Chromosome</td><td>start</td><td></td><td>end</td><td>strand</td><td>MM</td><td>target_seq</td><td>PAM</td><td>alignment</td><td>distance</td><td>position</td><td>gene name</td><td>gene id</td></tr>
<tr><td>GU305914.1</td><td></td><td>170578</td><td>170600</td><td>-</td><td>0</td><td>GCTGCCGATGTTTACCGACC</td><td>CGG</td><td>||||||||[||||||||||||]PAM</td><td>NA</td><td>Intergenic</td><td>NA</td><td>NA</td></tr>
<tr><td>T21612</td><td>GCCGATGTTTACCGACCCGGCGG</td><td></td><td>1000</td><td></td><td></td><td></td><td></td><td></td><td></td><td></td><td></td><td></td></tr>
<tr><td>Oligo fwd</td><td>CACCGCCGATGTTTACCGACCCGG</td><td></td><td></td><td></td><td></td><td></td><td></td><td></td><td></td><td></td><td></td><td></td></tr>
<tr><td>Oligo rev</td><td>AAACCCGGGTCGGTAAACATCGGC</td><td></td><td></td><td></td><td></td><td></td><td></td><td></td><td></td><td></td><td></td><td></td></tr>
<tr><td>Chromosome</td><td>start</td><td></td><td>end</td><td>strand</td><td>MM</td><td>target_seq</td><td>PAM</td><td>alignment</td><td>distance</td><td>position</td><td>gene name</td><td>gene id</td></tr>
<tr><td>GU305914.1</td><td></td><td>170575</td><td>170597</td><td>-</td><td>0</td><td>GCCGATGTTTACCGACCCGG</td><td>CGG</td><td>||||||||[||||||||||||]PAM</td><td>NA</td><td>Intergenic</td><td>NA</td><td>NA</td></tr>
<tr><td>T21613</td><td>GTTTACCGACCCGGCGGCGATGG</td><td></td><td>1000</td><td></td><td></td><td></td><td></td><td></td><td></td><td></td><td></td><td></td></tr>
<tr><td>Oligo fwd</td><td>CACCGTTTACCGACCCGGCGGCGA</td><td></td><td></td><td></td><td></td><td></td><td></td><td></td><td></td><td></td><td></td><td></td></tr>
<tr><td>Oligo rev</td><td>AAACTCGCCGCCGGGTCGGTAAAC</td><td></td><td></td><td></td><td></td><td></td><td></td><td></td><td></td><td></td><td></td><td></td></tr>
<tr><td>Chromosome</td><td>start</td><td></td><td>end</td><td>strand</td><td>MM</td><td>target_seq</td><td>PAM</td><td>alignment</td><td>distance</td><td>position</td><td>gene name</td><td>gene id</td></tr>
<tr><td>GU305914.1</td><td></td><td>170569</td><td>170591</td><td>-</td><td>0</td><td>GTTTACCGACCCGGCGGCGA</td><td>TGG</td><td>||||||||[||||||||||||]PAM</td><td>NA</td><td>Intergenic</td><td>NA</td><td>NA</td></tr>
<tr><td>T21614</td><td>CGGCGATGGTGCCGACCGCGTGG</td><td></td><td>1000</td><td></td><td></td><td></td><td></td><td></td><td></td><td></td><td></td><td></td></tr>
<tr><td>Oligo fwd</td><td>CACCCGGCGATGGTGCCGACCGCG</td><td></td><td></td><td></td><td></td><td></td><td></td><td></td><td></td><td></td><td></td><td></td></tr>
<tr><td>Oligo rev</td><td>AAACCGCGGTCGGCACCATCGCCG</td><td></td><td></td><td></td><td></td><td></td><td></td><td></td><td></td><td></td><td></td><td></td></tr>
<tr><td>Chromosome</td><td>start</td><td></td><td>end</td><td>strand</td><td>MM</td><td>target_seq</td><td>PAM</td><td>alignment</td><td>distance</td><td>position</td><td>gene name</td><td>gene id</td></tr>
<tr><td>GU305914.1</td><td></td><td>170555</td><td>170577</td><td>-</td><td>0</td><td>CGGCGATGGTGCCGACCGCG</td><td>TGG</td><td>||||||||[||||||||||||]PAM</td><td>NA</td><td>Intergenic</td><td>NA</td><td>NA</td></tr>
</table>

| | | | | | | | | | | | | |
|---|---|---|---|---|---|---|---|---|---|---|---|---|
| T21615 | GGTGCCGACCGCGTGGCCGATGG | 1000 | | | | | | | | | | |
| Oligo fwd | CACCGGTGCCGACCGCGTGGCCGA | | | | | | | | | | | |
| Oligo rev | AAACTCGGCCACGCGGTCGGCACC | | | | | | | | | | | |
| Chromosome | start | | end | strand | MM | target_seq | PAM | alignment | distance | position | gene name | gene id |
| GU305914.1 | 170548 | | 170570 | - | 0 | GGTGCCGACCGCGTGGCCGA | TGG | \|\|\|\|\|\|\|\|[\|\|\|\|\|\|\|\|\|\|\|\|]PAM | NA | Intergenic | NA | NA |

| | | | | | | | | | | | | |
|---|---|---|---|---|---|---|---|---|---|---|---|---|
| T21616 | GCCGACCGCGTGGCCGATGGTGG | 1000 | | | | | | | | | | |
| Oligo fwd | CACCGCCGACCGCGTGGCCGATGG | | | | | | | | | | | |
| Oligo rev | AAACCCATCGGCCACGCGGTCGGC | | | | | | | | | | | |
| Chromosome | start | | end | strand | MM | target_seq | PAM | alignment | distance | position | gene name | gene id |
| GU305914.1 | 170545 | | 170567 | - | 0 | GCCGACCGCGTGGCCGATGG | TGG | \|\|\|\|\|\|\|\|[\|\|\|\|\|\|\|\|\|\|\|\|]PAM | NA | Intergenic | NA | NA |

| | | | | | | | | | | | | |
|---|---|---|---|---|---|---|---|---|---|---|---|---|
| T21617 | GACCGCGTGGCCGATGGTGGCGG | 1000 | | | | | | | | | | |
| Oligo fwd | CACCGACCGCGTGGCCGATGGTGG | | | | | | | | | | | |
| Oligo rev | AAACCCACCATCGGCCACGCGGTC | | | | | | | | | | | |
| Chromosome | start | | end | strand | MM | target_seq | PAM | alignment | distance | position | gene name | gene id |
| GU305914.1 | 170542 | | 170564 | - | 0 | GACCGCGTGGCCGATGGTGG | CGG | \|\|\|\|\|\|\|\|[\|\|\|\|\|\|\|\|\|\|\|\|]PAM | NA | Intergenic | NA | NA |

| | | | | | | | | | | | | |
|---|---|---|---|---|---|---|---|---|---|---|---|---|
| T21618 | CGCGTGGCCGATGGTGGCGGCGG | 1000 | | | | | | | | | | |
| Oligo fwd | CACCCGCGTGGCCGATGGTGGCGG | | | | | | | | | | | |
| Oligo rev | AAACCCGCCACCATCGGCCACGCG | | | | | | | | | | | |
| Chromosome | start | | end | strand | MM | target_seq | PAM | alignment | distance | position | gene name | gene id |
| GU305914.1 | 170539 | | 170561 | - | 0 | CGCGTGGCCGATGGTGGCGG | CGG | \|\|\|\|\|\|\|\|[\|\|\|\|\|\|\|\|\|\|\|\|]PAM | NA | Intergenic | NA | NA |

| | | | | | | | | | | | | |
|---|---|---|---|---|---|---|---|---|---|---|---|---|
| T21619 | GTGGCCGATGGTGGCGGCGGCGG | 1000 | | | | | | | | | | |
| Oligo fwd | CACCGTGGCCGATGGTGGCGGCGG | | | | | | | | | | | |
| Oligo rev | AAACCCGCCGCCACCATCGGCCAC | | | | | | | | | | | |
| Chromosome | start | | end | strand | MM | target_seq | PAM | alignment | distance | position | gene name | gene id |
| GU305914.1 | 170536 | | 170558 | - | 0 | GTGGCCGATGGTGGCGGCGG | CGG | \|\|\|\|\|\|\|\|[\|\|\|\|\|\|\|\|\|\|\|\|]PAM | NA | Intergenic | NA | NA |
| GU305914.1 | 29994 | | 30016 | + | 4 | GGGACCGGTGCTGGCGGCGG | CGG | \|-\|-\|\|\|-[\|\|-\|\|\|\|\|\|\|\|\|]PAM | NA | Intergenic | NA | NA |
| GU305914.1 | 58129 | | 58151 | - | 4 | GCGGGCGGTGGCGGCGGCGG | CGG | \|-\|\|-\|\|-[\|\|\|-\|\|\|\|\|\|\|\|]PAM | NA | Intergenic | NA | NA |
| GU305914.1 | 24872 | | 24894 | + | 4 | GCGCCCGATGTTGGCGGCGA | CGG | \|-\|-\|\|\|\|[\|\|-\|\|\|\|\|\|\|\|-]PAM | NA | Intergenic | NA | NA |

| | | | | | | | | | | | | |
|---|---|---|---|---|---|---|---|---|---|---|---|---|
| T21620 | GATGGTGGCGGCGGCGGCCACGG | 1000 | | | | | | | | | | |
| Oligo fwd | CACCGATGGTGGCGGCGGCGGCCA | | | | | | | | | | | |
| Oligo rev | AAACTGGCCGCCGCCGCCACCATC | | | | | | | | | | | |
| Chromosome | start | | end | strand | MM | target_seq | PAM | alignment | distance | position | gene name | gene id |
| GU305914.1 | 170530 | | 170552 | - | 0 | GATGGTGGCGGCGGCGGCCA | CGG | \|\|\|\|\|\|\|\|[\|\|\|\|\|\|\|\|\|\|\|\|]PAM | NA | Intergenic | NA | NA |
| GU305914.1 | 154404 | | 154426 | + | 4 | GGCGGTGGCGGCGGCGGCGG | CGG | \|--\|\|\|\|\|[\|\|\|\|\|\|\|\|\|\|--]PAM | NA | Intergenic | NA | NA |
| GU305914.1 | 58126 | | 58148 | - | 4 | GGCGGTGGCGGCGGCGGCGG | CGG | \|--\|\|\|\|\|[\|\|\|\|\|\|\|\|\|\|--]PAM | NA | Intergenic | NA | NA |
| GU305914.1 | 30000 | | 30022 | + | 4 | GGTGCTGGCGGCGGCGGCGG | CGG | \|-\|\|-\|\|\|[\|\|\|\|\|\|\|\|\|\|--]PAM | NA | Intergenic | NA | NA |
| GU305914.1 | 154407 | | 154429 | + | 4 | GGTGGCGGCGGCGGCGGCGG | CGG | \|-\|\|\|-\|\|[\|\|\|\|\|\|\|\|\|\|--]PAM | NA | Intergenic | NA | NA |
| GU305914.1 | 58123 | | 58145 | - | 4 | GGTGGCGGCGGCGGCGGCGG | AGG | \|-\|\|\|-\|\|[\|\|\|\|\|\|\|\|\|\|--]PAM | NA | Intergenic | NA | NA |

| | | | | | | | | | | | | |
|---|---|---|---|---|---|---|---|---|---|---|---|---|
| T21621 | GGCGGCCACGGCCACGATACCGG | 1000 | | | | | | | | | | |
| Oligo fwd | CACCGGCGGCCACGGCCACGATAC | | | | | | | | | | | |
| Oligo rev | AAACGTATCGTGGCCGTGGCCGCC | | | | | | | | | | | |
| Chromosome | start | | end | strand | MM | target_seq | PAM | alignment | distance | position | gene name | gene id |
| GU305914.1 | 170518 | | 170540 | - | 0 | GGCGGCCACGGCCACGATAC | CGG | \|\|\|\|\|\|\|\|[\|\|\|\|\|\|\|\|\|\|\|\|]PAM | NA | Intergenic | NA | NA |

| | | | | | | | | | | | | |
|---|---|---|---|---|---|---|---|---|---|---|---|---|
| T21622 | GCGGCCACGGCCACGATACCGGG | 1000 | | | | | | | | | | |
| Oligo fwd | CACCGCGGCCACGGCCACGATACC | | | | | | | | | | | |
| Oligo rev | AAACGGTATCGTGGCCGTGGCCGC | | | | | | | | | | | |
| Chromosome | start | | end | strand | MM | target_seq | PAM | alignment | distance | position | gene name | gene id |
| GU305914.1 | 170517 | | 170539 | - | 0 | GCGGCCACGGCCACGATACC | GGG | \|\|\|\|\|\|\|\|[\|\|\|\|\|\|\|\|\|\|\|\|]PAM | NA | Intergenic | NA | NA |

| | | | | | | | | | | | | |
|---|---|---|---|---|---|---|---|---|---|---|---|---|
| T21623 | CGGCCACGGCCACGATACCGGGG | 1000 | | | | | | | | | | |
| Oligo fwd | CACCCGGCCACGGCCACGATACCG | | | | | | | | | | | |
| Oligo rev | AAACCGGTATCGTGGCCGTGGCCG | | | | | | | | | | | |
| Chromosome | start | | end | strand | MM | target_seq | PAM | alignment | distance | position | gene name | gene id |
| GU305914.1 | 170516 | | 170538 | - | 0 | CGGCCACGGCCACGATACCG | GGG | \|\|\|\|\|\|\|\|[\|\|\|\|\|\|\|\|\|\|\|\|]PAM | NA | Intergenic | NA | NA |
| GU305914.1 | 142229 | | 142251 | + | 4 | CGGCGATGACGACGATACCG | CGG | \|\|\|\|-\|-\|[-\|-\|\|\|\|\|\|\|\|\|]PAM | NA | Intergenic | NA | NA |

| | | | | | | | | | | | | |
|---|---|---|---|---|---|---|---|---|---|---|---|---|
| T21624 | ACCCAAACTCCCGATGATAGAGG | 1000 | | | | | | | | | | |
| Oligo fwd | CACCACCCAAACTCCCGATGATAG | | | | | | | | | | | |
| Oligo rev | AAACCTATCATCGGGAGTTTGGGT | | | | | | | | | | | |
| Chromosome | start | | end | strand | MM | target_seq | PAM | alignment | distance | position | gene name | gene id |
| GU305914.1 | 170479 | | 170501 | - | 0 | ACCCAAACTCCCGATGATAG | AGG | \|\|\|\|\|\|\|\|[\|\|\|\|\|\|\|\|\|\|\|\|]PAM | NA | Intergenic | NA | NA |

| | | | | | | | | | | | | |
|---|---|---|---|---|---|---|---|---|---|---|---|---|
| T21625 | CCCATCTCGATACCCGATCCCGG | 1000 | | | | | | | | | | |
| Oligo fwd | CACCCCCATCTCGATACCCGATCC | | | | | | | | | | | |
| Oligo rev | AAACGGATCGGGTATCGAGATGGG | | | | | | | | | | | |
| Chromosome | start | | end | strand | MM | target_seq | PAM | alignment | distance | position | gene name | gene id |
| GU305914.1 | 170397 | | 170419 | - | 0 | CCCATCTCGATACCCGATCC | CGG | \|\|\|\|\|\|\|\|[\|\|\|\|\|\|\|\|\|\|\|\|]PAM | NA | Intergenic | NA | NA |

| | | | | | | | | | | | | |
|---|---|---|---|---|---|---|---|---|---|---|---|---|
| T21626 | TCTCGATACCCGATCCCGGCAGG | 1000 | | | | | | | | | | |
| Oligo fwd | CACCTCTCGATACCCGATCCCGGC | | | | | | | | | | | |
| Oligo rev | AAACGCCGGGATCGGGTATCGAGA | | | | | | | | | | | |
| Chromosome | start | | end | strand | MM | target_seq | PAM | alignment | distance | position | gene name | gene id |
| GU305914.1 | 170393 | | 170415 | - | 0 | TCTCGATACCCGATCCCGGC | AGG | \|\|\|\|\|\|\|\|[\|\|\|\|\|\|\|\|\|\|\|\|]PAM | NA | Intergenic | NA | NA |

| | | | | | | | | | | | | |
|---|---|---|---|---|---|---|---|---|---|---|---|---|
| T21627 | CACCGAGCCCATCGCGTTCCCGG | 1000 | | | | | | | | | | |
| Oligo fwd | CACCCACCGAGCCCATCGCGTTCC | | | | | | | | | | | |
| Oligo rev | AAACGGAACGCGATGGGCTCGGTG | | | | | | | | | | | |
| Chromosome | start | | end | strand | MM | target_seq | PAM | alignment | distance | position | gene name | gene id |
| GU305914.1 | 170368 | | 170390 | - | 0 | CACCGAGCCCATCGCGTTCC | CGG | \|\|\|\|\|\|\|\|[\|\|\|\|\|\|\|\|\|\|\|\|]PAM | NA | Intergenic | NA | NA |

| | | |
|---|---|---|
| T21628 | GTCCCCGAATCCGAACCCGTCGG | 1000 |
| Oligo fwd | CACCGTCCCCGAATCCGAACCCGT | |

| | | | | | | | | | | | | |
|---|---|---|---|---|---|---|---|---|---|---|---|---|
| Oligo rev | AAACACGGGTTCGGATTCGGGGAC | | | | | | | | | | | |
| Chromosome | start | | end | strand | MM | target_seq | PAM | alignment | distance | position | gene name | gene id |
| GU305914.1 | | 170244 | 170266 | - | 0 | GTCCCCGAATCCGAACCCGT | CGG | ||||||||[||||||||||||]PAM | NA | Intergenic | NA | NA |
| T21629 | CACCGTCCCCGAACCCGTCGCGG | | 1000 | | | | | | | | | |
| Oligo fwd | CACCCACCGTCCCCGAACCCGTCG | | | | | | | | | | | |
| Oligo rev | AAACCGACGGGTTCGGGGACGGTG | | | | | | | | | | | |
| Chromosome | start | | end | strand | MM | target_seq | PAM | alignment | distance | position | gene name | gene id |
| GU305914.1 | | 170125 | 170147 | - | 0 | CACCGTCCCCGAACCCGTCG | CGG | ||||||||[||||||||||||]PAM | NA | Intergenic | NA | NA |
| T21630 | TCCCGCTCCCAAACCCGTCGCGG | | 1000 | | | | | | | | | |
| Oligo fwd | CACCTCCCGCTCCCAAACCCGTCG | | | | | | | | | | | |
| Oligo rev | AAACCGACGGGTTTGGGAGCGGGA | | | | | | | | | | | |
| Chromosome | start | | end | strand | MM | target_seq | PAM | alignment | distance | position | gene name | gene id |
| GU305914.1 | | 170047 | 170069 | - | 0 | TCCCGCTCCCAAACCCGTCG | CGG | ||||||||[||||||||||||]PAM | NA | Intergenic | NA | NA |
| T21631 | CCGAGCTCATCATACTAAAAAGG | | 1000 | | | | | | | | | |
| Oligo fwd | CACCCCGAGCTCATCATACTAAAA | | | | | | | | | | | |
| Oligo rev | AAACTTTTAGTATGATGAGCTCGG | | | | | | | | | | | |
| Chromosome | start | | end | strand | MM | target_seq | PAM | alignment | distance | position | gene name | gene id |
| GU305914.1 | | 170021 | 170043 | - | 0 | CCGAGCTCATCATACTAAAA | AGG | ||||||||[||||||||||||]PAM | NA | Intergenic | NA | NA |
| T21632 | CTCTCCCGTCCCGCAGCCGACGG | | 1000 | | | | | | | | | |
| Oligo fwd | CACCCTCTCCCGTCCCGCAGCCGA | | | | | | | | | | | |
| Oligo rev | AAACTCGGCTGCGGGACGGGAGAG | | | | | | | | | | | |
| Chromosome | start | | end | strand | MM | target_seq | PAM | alignment | distance | position | gene name | gene id |
| GU305914.1 | | 169990 | 170012 | - | 0 | CTCTCCCGTCCCGCAGCCGA | CGG | ||||||||[||||||||||||]PAM | NA | Intergenic | NA | NA |
| T21633 | TCCCGCAGCCGACGGACGCTAGG | | 1000 | | | | | | | | | |
| Oligo fwd | CACCTCCCGCAGCCGACGGACGCT | | | | | | | | | | | |
| Oligo rev | AAACAGCGTCCGTCGGCTGCGGGA | | | | | | | | | | | |
| Chromosome | start | | end | strand | MM | target_seq | PAM | alignment | distance | position | gene name | gene id |
| GU305914.1 | | 169982 | 170004 | - | 0 | TCCCGCAGCCGACGGACGCT | AGG | ||||||||[||||||||||||]PAM | NA | Intergenic | NA | NA |
| T21634 | TAGGCCCGCGAAAGTGCCCAAGG | | 1000 | | | | | | | | | |
| Oligo fwd | CACCTAGGCCCGCGAAAGTGCCCA | | | | | | | | | | | |
| Oligo rev | AAACTGGGCACTTTCGCGGGCCTA | | | | | | | | | | | |
| Chromosome | start | | end | strand | MM | target_seq | PAM | alignment | distance | position | gene name | gene id |
| GU305914.1 | | 169963 | 169985 | - | 0 | TAGGCCCGCGAAAGTGCCCA | AGG | ||||||||[||||||||||||]PAM | NA | Intergenic | NA | NA |
| T21635 | ACCGAAACTCATCATCAAGAAGG | | 1000 | | | | | | | | | |
| Oligo fwd | CACCACCGAAACTCATCATCAAGA | | | | | | | | | | | |
| Oligo rev | AAACTCTTGATGATGAGTTTCGGT | | | | | | | | | | | |
| Chromosome | start | | end | strand | MM | target_seq | PAM | alignment | distance | position | gene name | gene id |
| GU305914.1 | | 169939 | 169961 | - | 0 | ACCGAAACTCATCATCAAGA | AGG | ||||||||[||||||||||||]PAM | NA | Intergenic | NA | NA |
| T21636 | GGTACCACGCGCACCGATCGTGG | | 1000 | | | | | | | | | |
| Oligo fwd | CACCGGTACCACGCGCACCGATCG | | | | | | | | | | | |
| Oligo rev | AAACCGATCGGTGCGCGTGGTACC | | | | | | | | | | | |
| Chromosome | start | | end | strand | MM | target_seq | PAM | alignment | distance | position | gene name | gene id |
| GU305914.1 | | 169918 | 169940 | - | 0 | GGTACCACGCGCACCGATCG | TGG | ||||||||[||||||||||||]PAM | NA | Intergenic | NA | NA |
| T21637 | CACGCGCACCGATCGTGGTGAGG | | 1000 | | | | | | | | | |
| Oligo fwd | CACCCACGCGCACCGATCGTGGTG | | | | | | | | | | | |
| Oligo rev | AAACCACCACGATCGGTGCGCGTG | | | | | | | | | | | |
| Chromosome | start | | end | strand | MM | target_seq | PAM | alignment | distance | position | gene name | gene id |
| GU305914.1 | | 169913 | 169935 | - | 0 | CACGCGCACCGATCGTGGTG | AGG | ||||||||[||||||||||||]PAM | NA | Intergenic | NA | NA |
| T21638 | CGCACCGATCGTGGTGAGGCCGG | | 1000 | | | | | | | | | |
| Oligo fwd | CACCCGCACCGATCGTGGTGAGGC | | | | | | | | | | | |
| Oligo rev | AAACGCCTCACCACGATCGGTGCG | | | | | | | | | | | |
| Chromosome | start | | end | strand | MM | target_seq | PAM | alignment | distance | position | gene name | gene id |
| GU305914.1 | | 169909 | 169931 | - | 0 | CGCACCGATCGTGGTGAGGC | CGG | ||||||||[||||||||||||]PAM | NA | Intergenic | NA | NA |
| T21639 | TGGTGAGGCCGGTTCCGACGAGG | | 1000 | | | | | | | | | |
| Oligo fwd | CACCTGGTGAGGCCGGTTCCGACG | | | | | | | | | | | |
| Oligo rev | AAACCGTCGGAACCGGCCTCACCA | | | | | | | | | | | |
| Chromosome | start | | end | strand | MM | target_seq | PAM | alignment | distance | position | gene name | gene id |
| GU305914.1 | | 169898 | 169920 | - | 0 | TGGTGAGGCCGGTTCCGACG | AGG | ||||||||[||||||||||||]PAM | NA | Intergenic | NA | NA |
| T21640 | GACGAGGTTAAGCGACGTGACGG | | 1000 | | | | | | | | | |
| Oligo fwd | CACCGACGAGGTTAAGCGACGTGA | | | | | | | | | | | |
| Oligo rev | AAACTCACGTCGCTTAACCTCGTC | | | | | | | | | | | |
| Chromosome | start | | end | strand | MM | target_seq | PAM | alignment | distance | position | gene name | gene id |
| GU305914.1 | | 169882 | 169904 | - | 0 | GACGAGGTTAAGCGACGTGA | CGG | ||||||||[||||||||||||]PAM | NA | Intergenic | NA | NA |
| T21641 | ACGTGACGGCCCCCGCGAACCGG | | 1000 | | | | | | | | | |
| Oligo fwd | CACCACGTGACGGCCCCCGCGAAC | | | | | | | | | | | |
| Oligo rev | AAACGTTCGCGGGGGCCGTCACGT | | | | | | | | | | | |
| Chromosome | start | | end | strand | MM | target_seq | PAM | alignment | distance | position | gene name | gene id |
| GU305914.1 | | 169868 | 169890 | - | 0 | ACGTGACGGCCCCCGCGAAC | CGG | ||||||||[||||||||||||]PAM | NA | Intergenic | NA | NA |
| T21642 | GACGGCCCCCGCGAACCGGCCGG | | 1000 | | | | | | | | | |
| Oligo fwd | CACCGACGGCCCCCGCGAACCGGC | | | | | | | | | | | |
| Oligo rev | AAACGCCGGTTCGCGGGGGCCGTC | | | | | | | | | | | |
| Chromosome | start | | end | strand | MM | target_seq | PAM | alignment | distance | position | gene name | gene id |
| GU305914.1 | | 169864 | 169886 | - | 0 | GACGGCCCCCGCGAACCGGC | CGG | ||||||||[||||||||||||]PAM | NA | Intergenic | NA | NA |
| T21643 | ACCGGCCGGAGCCGCGACAGCGG | | 1000 | | | | | | | | | |

Oligo fwd CACCACCGGCCGGAGCCGCGACAG
Oligo rev AAACCTGTCGCGGCTCCGGCCGGT
Chromosome start end strand MM target_seq PAM alignment distance position gene name gene id
GU305914.1 169850 169872 - 0 ACCGGCCGGAGCCGCGACAG CGG ||||||||[||||||||||||]PAM NA Intergenic NA NA

T21644 ACAGCGCGCAGAAACCGAGACGG 1000
Oligo fwd CACCACAGCGCGCAGAAACCGAGA
Oligo rev AAACTCTCGGTTTCTGCGCGCTGT
Chromosome start end strand MM target_seq PAM alignment distance position gene name gene id
GU305914.1 169825 169847 - 0 ACAGCGCGCAGAAACCGAGA CGG ||||||||[||||||||||||]PAM NA Intergenic NA NA

T21645 GCGCGCAGAAACCGAGACGGCGG 1000
Oligo fwd CACCGCGCGCAGAAACCGAGACGG
Oligo rev AAACCCGTCTCGGTTTCTGCGCGC
Chromosome start end strand MM target_seq PAM alignment distance position gene name gene id
GU305914.1 169822 169844 - 0 GCGCGCAGAAACCGAGACGG CGG ||||||||[||||||||||||]PAM NA Intergenic NA NA

T21646 CGCAGAAACCGAGACGGCGGCGG 1000
Oligo fwd CACCCGCAGAAACCGAGACGGCGG
Oligo rev AAACCCGCCGTCTCGGTTTCTGCG
Chromosome start end strand MM target_seq PAM alignment distance position gene name gene id
GU305914.1 169819 169841 - 0 CGCAGAAACCGAGACGGCGG CGG ||||||||[||||||||||||]PAM NA Intergenic NA NA

T21647 GGCGGCGTCTTCTTGCACGACGG 1000
Oligo fwd CACCGGCGGCGTCTTCTTGCACGA
Oligo rev AAACTCGTGCAAGAAGACGCCGCC
Chromosome start end strand MM target_seq PAM alignment distance position gene name gene id
GU305914.1 169801 169823 - 0 GGCGGCGTCTTCTTGCACGA CGG ||||||||[||||||||||||]PAM NA Intergenic NA NA

T21648 CAAGAAGATACCGAAGATCGTGG 1000
Oligo fwd CACCCAAGAAGATACCGAAGATCG
Oligo rev AAACCGATCTTCGGTATCTTCTTG
Chromosome start end strand MM target_seq PAM alignment distance position gene name gene id
GU305914.1 169765 169787 - 0 CAAGAAGATACCGAAGATCG TGG ||||||||[||||||||||||]PAM NA Intergenic NA NA

T21649 CGTGGTACCCGCCACGATCACGG 1000
Oligo fwd CACCCGTGGTACCCGCCACGATCA
Oligo rev AAACTGATCGTGGCGGGTACCACG
Chromosome start end strand MM target_seq PAM alignment distance position gene name gene id
GU305914.1 169747 169769 - 0 CGTGGTACCCGCCACGATCA CGG ||||||||[||||||||||||]PAM NA Intergenic NA NA

T21650 CACGGCGAAGAGAAAGACCGCGG 1000
Oligo fwd CACCCACGGCGAAGAGAAAGACCG
Oligo rev AAACCGGTCTTTCTCTTCGCCGTG
Chromosome start end strand MM target_seq PAM alignment distance position gene name gene id
GU305914.1 169729 169751 - 0 CACGGCGAAGAGAAAGACCG CGG ||||||||[||||||||||||]PAM NA Intergenic NA NA

T21651 GCGAAGAGAAAGACCGCGGAAGG 1000
Oligo fwd CACCGCGAAGAGAAAGACCGCGGA
Oligo rev AAACTCCGCGGTCTTTCTCTTCGC
Chromosome start end strand MM target_seq PAM alignment distance position gene name gene id
GU305914.1 169725 169747 - 0 GCGAAGAGAAAGACCGCGGA AGG ||||||||[||||||||||||]PAM NA Intergenic NA NA

T21652 AAAGACCGCGGAAGGCGCCCCGG 1000
Oligo fwd CACCAAAGACCGCGGAAGGCGCCC
Oligo rev AAACGGGCGCCTTCCGCGGTCTTT
Chromosome start end strand MM target_seq PAM alignment distance position gene name gene id
GU305914.1 169717 169739 - 0 AAAGACCGCGGAAGGCGCCC CGG ||||||||[||||||||||||]PAM NA Intergenic NA NA

T21653 GGCGCCCCGGACGAAGAGTTCGG 1000
Oligo fwd CACCGGCGCCCCGGACGAAGAGTT
Oligo rev AAACAACTCTTCGTCCGGGGCGCC
Chromosome start end strand MM target_seq PAM alignment distance position gene name gene id
GU305914.1 169704 169726 - 0 GGCGCCCCGGACGAAGAGTT CGG ||||||||[||||||||||||]PAM NA Intergenic NA NA

T21654 TCGGAAACGATTTATTGAAGAGG 1000
Oligo fwd CACCTCGGAAACGATTTATTGAAG
Oligo rev AAACCTTCAATAAATCGTTTCCGA
Chromosome start end strand MM target_seq PAM alignment distance position gene name gene id
GU305914.1 169685 169707 - 0 TCGGAAACGATTTATTGAAG AGG ||||||||[||||||||||||]PAM NA Intergenic NA NA

T21655 CGATTTATTGAAGAGGCTCGCGG 1000
Oligo fwd CACCCGATTTATTGAAGAGGCTCG
Oligo rev AAACCGAGCCTCTTCAATAAATCG
Chromosome start end strand MM target_seq PAM alignment distance position gene name gene id
GU305914.1 169678 169700 - 0 CGATTTATTGAAGAGGCTCG CGG ||||||||[||||||||||||]PAM NA Intergenic NA NA

T21656 TCGCGGACCTGTGCAGCGACAGG 1000
Oligo fwd CACCTCGCGGACCTGTGCAGCGAC
Oligo rev AAACGTCGCTGCACAGGTCCGCGA
Chromosome start end strand MM target_seq PAM alignment distance position gene name gene id
GU305914.1 169661 169683 - 0 TCGCGGACCTGTGCAGCGAC AGG ||||||||[||||||||||||]PAM NA Intergenic NA NA

T21657 CGACAGGCATACCGACGTCTCGG 1000
Oligo fwd CACCCGACAGGCATACCGACGTCT
Oligo rev AAACAGACGTCGGTATGCCTGTCG
Chromosome start end strand MM target_seq PAM alignment distance position gene name gene id
GU305914.1 169645 169667 - 0 CGACAGGCATACCGACGTCT CGG ||||||||[||||||||||||]PAM NA Intergenic NA NA

T21658 ATACCGACGTCTCGGACATACGG 1000
Oligo fwd CACCATACCGACGTCTCGGACATA
Oligo rev AAACTATGTCCGAGACGTCGGTAT
Chromosome start end strand MM target_seq PAM alignment distance position gene name gene id
GU305914.1 169637 169659 - 0 ATACCGACGTCTCGGACATA CGG ||||||||[||||||||||||]PAM NA Intergenic NA NA

T21659 CTCGTGTCAGAAGTTTCGAACGG 1000
Oligo fwd CACCCTCGTGTCAGAAGTTTCGAA
Oligo rev AAACTTCGAAACTTCTGACACGAG
Chromosome start end strand MM target_seq PAM alignment distance position gene name gene id
GU305914.1 169606 169628 - 0 CTCGTGTCAGAAGTTTCGAA CGG ||||||||[||||||||||||]PAM NA Intergenic NA NA

T21660 CAGAAGTTTCGAACGGCGCGCGG 1000
Oligo fwd CACCCAGAAGTTTCGAACGGCGCG
Oligo rev AAACCGCGCCGTTCGAAACTTCTG
Chromosome start end strand MM target_seq PAM alignment distance position gene name gene id
GU305914.1 169599 169621 - 0 CAGAAGTTTCGAACGGCGCG CGG ||||||||[||||||||||||]PAM NA Intergenic NA NA

T21661 AGAAGTTTCGAACGGCGCGCGGG 1000
Oligo fwd CACCAGAAGTTTCGAACGGCGCGC
Oligo rev AAACGCGCGCCGTTCGAAACTTCT
Chromosome start end strand MM target_seq PAM alignment distance position gene name gene id
GU305914.1 169598 169620 - 0 AGAAGTTTCGAACGGCGCGC GGG ||||||||[||||||||||||]PAM NA Intergenic NA NA
GU305914.1 203057 203079 - 4 AGGAGGTTCGGACGGCGCTC GGG ||-||-||[||-|||||||-|]PAM NA Intergenic NA NA

T21662 GCGCGGGACTTACAACGCGCTGG 1000
Oligo fwd CACCGCGCGGGACTTACAACGCGC
Oligo rev AAACGCGCGTTGTAAGTCCCGCGC
Chromosome start end strand MM target_seq PAM alignment distance position gene name gene id
GU305914.1 169582 169604 - 0 GCGCGGGACTTACAACGCGC TGG ||||||||[||||||||||||]PAM NA Intergenic NA NA

T21663 CCCCAACCTGAAGATCGTGATGG 1000
Oligo fwd CACCCCCCAACCTGAAGATCGTGA
Oligo rev AAACTCACGATCTTCAGGTTGGGG
Chromosome start end strand MM target_seq PAM alignment distance position gene name gene id
GU305914.1 169552 169574 - 0 CCCCAACCTGAAGATCGTGA TGG ||||||||[||||||||||||]PAM NA Intergenic NA NA

T21664 CCCAACCTGAAGATCGTGATGGG 1000
Oligo fwd CACCCCCAACCTGAAGATCGTGAT
Oligo rev AAACATCACGATCTTCAGGTTGGG
Chromosome start end strand MM target_seq PAM alignment distance position gene name gene id
GU305914.1 169551 169573 - 0 CCCAACCTGAAGATCGTGAT GGG ||||||||[||||||||||||]PAM NA Intergenic NA NA

T21665 GATGGGCGCGCAGATGCTGACGG 1000
Oligo fwd CACCGATGGGCGCGCAGATGCTGA
Oligo rev AAACTCAGCATCTGCGCGCCCATC
Chromosome start end strand MM target_seq PAM alignment distance position gene name gene id
GU305914.1 169534 169556 - 0 GATGGGCGCGCAGATGCTGA CGG ||||||||[||||||||||||]PAM NA Intergenic NA NA

T21666 ACGAGATCAACTACCAACCGAGG 1000
Oligo fwd CACCACGAGATCAACTACCAACCG
Oligo rev AAACCGGTTGGTAGTTGATCTCGT
Chromosome start end strand MM target_seq PAM alignment distance position gene name gene id
GU305914.1 169478 169500 - 0 ACGAGATCAACTACCAACCG AGG ||||||||[||||||||||||]PAM NA Intergenic NA NA

T21667 GATCAACTACCAACCGAGGACGG 1000
Oligo fwd CACCGATCAACTACCAACCGAGGA
Oligo rev AAACTCCTCGGTTGGTAGTTGATC
Chromosome start end strand MM target_seq PAM alignment distance position gene name gene id
GU305914.1 169474 169496 - 0 GATCAACTACCAACCGAGGA CGG ||||||||[||||||||||||]PAM NA Intergenic NA NA

T21668 CAACTACCAACCGAGGACGGCGG 1000
Oligo fwd CACCCAACTACCAACCGAGGACGG
Oligo rev AAACCCGTCCTCGGTTGGTAGTTG
Chromosome start end strand MM target_seq PAM alignment distance position gene name gene id
GU305914.1 169471 169493 - 0 CAACTACCAACCGAGGACGG CGG ||||||||[||||||||||||]PAM NA Intergenic NA NA

T21669 GCGAACACTCACATCGAGACCGG 1000
Oligo fwd CACCGCGAACACTCACATCGAGAC
Oligo rev AAACGTCTCGATGTGAGTGTTCGC
Chromosome start end strand MM target_seq PAM alignment distance position gene name gene id
GU305914.1 169449 169471 - 0 GCGAACACTCACATCGAGAC CGG ||||||||[||||||||||||]PAM NA Intergenic NA NA

T21670 TCGTGCGCTCGCGAGCGTCTCGG 1000
Oligo fwd CACCTCGTGCGCTCGCGAGCGTCT
Oligo rev AAACAGACGCTCGCGAGCGCACGA
Chromosome start end strand MM target_seq PAM alignment distance position gene name gene id
GU305914.1 169412 169434 - 0 TCGTGCGCTCGCGAGCGTCT CGG ||||||||[||||||||||||]PAM NA Intergenic NA NA

T21671 CGTGCGCTCGCGAGCGTCTCGGG 1000
Oligo fwd CACCCGTGCGCTCGCGAGCGTCTC
Oligo rev AAACGAGACGCTCGCGAGCGCACG
Chromosome start end strand MM target_seq PAM alignment distance position gene name gene id
GU305914.1 169411 169433 - 0 CGTGCGCTCGCGAGCGTCTC GGG ||||||||[||||||||||||]PAM NA Intergenic NA NA

T21672 GCGAGCGTCTCGGGAACTGCTGG 1000
Oligo fwd CACCGCGAGCGTCTCGGGAACTGC
Oligo rev AAACGCAGTTCCCGAGACGCTCGC
Chromosome start end strand MM target_seq PAM alignment distance position gene name gene id

GU305914.1 169402 169424 - 0 GCGAGCGTCTCGGGAACTGC TGG ||||||||[||||||||||||]PAM NA Intergenic NA NA

T21673 GTCTCGGGAACTGCTGGCCGCGG 1000
Oligo fwd CACCGTCTCGGGAACTGCTGGCCG
Oligo rev AAACCGGCCAGCAGTTCCCGAGAC
Chromosome start end strand MM target_seq PAM alignment distance position gene name gene id
GU305914.1 169396 169418 - 0 GTCTCGGGAACTGCTGGCCG CGG ||||||||[||||||||||||]PAM NA Intergenic NA NA

T21674 GGCCGCGGCCCGCGTCTGTCAGG 1000
Oligo fwd CACCGGCCGCGGCCCGCGTCTGTC
Oligo rev AAACGACAGACGCGGGCCGCGGCC
Chromosome start end strand MM target_seq PAM alignment distance position gene name gene id
GU305914.1 169381 169403 - 0 GGCCGCGGCCCGCGTCTGTC AGG ||||||||[||||||||||||]PAM NA Intergenic NA NA

T21675 GGCCCGCGTCTGTCAGGACGTGG 1000
Oligo fwd CACCGGCCCGCGTCTGTCAGGACG
Oligo rev AAACCGTCCTGACAGACGCGGGCC
Chromosome start end strand MM target_seq PAM alignment distance position gene name gene id
GU305914.1 169375 169397 - 0 GGCCCGCGTCTGTCAGGACG TGG ||||||||[||||||||||||]PAM NA Intergenic NA NA
GU305914.1 19075 19097 + 4 GGCCCACATCTGTGACGACG AGG |||||-|-[|||||-|-||||]PAM NA Intergenic NA NA

T21676 TCAGGACGTGGCCAAGAAGACGG 1000
Oligo fwd CACCTCAGGACGTGGCCAAGAAGA
Oligo rev AAACTCTTCTTGGCCACGTCCTGA
Chromosome start end strand MM target_seq PAM alignment distance position gene name gene id
GU305914.1 169363 169385 - 0 TCAGGACGTGGCCAAGAAGA CGG ||||||||[||||||||||||]PAM NA Intergenic NA NA

T21677 CAGGACGTGGCCAAGAAGACGGG 1000
Oligo fwd CACCCAGGACGTGGCCAAGAAGAC
Oligo rev AAACGTCTTCTTGGCCACGTCCTG
Chromosome start end strand MM target_seq PAM alignment distance position gene name gene id
GU305914.1 169362 169384 - 0 CAGGACGTGGCCAAGAAGAC GGG ||||||||[||||||||||||]PAM NA Intergenic NA NA

T21678 AATTCAGTTTCTCTACGATCAGG 1000
Oligo fwd CACCAATTCAGTTTCTCTACGATC
Oligo rev AAACGATCGTAGAGAAACTGAATT
Chromosome start end strand MM target_seq PAM alignment distance position gene name gene id
GU305914.1 169328 169350 - 0 AATTCAGTTTCTCTACGATC AGG ||||||||[||||||||||||]PAM NA Intergenic NA NA

T21679 CCATCAACTATTTAACCTCTCGG 1000
Oligo fwd CACCCCATCAACTATTTAACCTCT
Oligo rev AAACAGAGGTTAAATAGTTGATGG
Chromosome start end strand MM target_seq PAM alignment distance position gene name gene id
GU305914.1 169270 169292 - 0 CCATCAACTATTTAACCTCT CGG ||||||||[||||||||||||]PAM NA Intergenic NA NA

T21680 TCTCGGACACGCTCGCCGTGAGG 1000
Oligo fwd CACCTCTCGGACACGCTCGCCGTG
Oligo rev AAACCACGGCGAGCGTGTCCGAGA
Chromosome start end strand MM target_seq PAM alignment distance position gene name gene id
GU305914.1 169253 169275 - 0 TCTCGGACACGCTCGCCGTG AGG ||||||||[||||||||||||]PAM NA Intergenic NA NA

T21681 CTCGGACACGCTCGCCGTGAGGG 1000
Oligo fwd CACCCTCGGACACGCTCGCCGTGA
Oligo rev AAACTCACGGCGAGCGTGTCCGAG
Chromosome start end strand MM target_seq PAM alignment distance position gene name gene id
GU305914.1 169252 169274 - 0 CTCGGACACGCTCGCCGTGA GGG ||||||||[||||||||||||]PAM NA Intergenic NA NA
GU305914.1 188902 188924 - 4 CGGCGACACGGTCGCCGTGA CGG |---||||[||-|||||||||]PAM NA Intergenic NA NA

T21682 CACGACATCATGTTTCGTGTCGG 1000
Oligo fwd CACCCACGACATCATGTTTCGTGT
Oligo rev AAACACACGAAACATGATGTCGTG
Chromosome start end strand MM target_seq PAM alignment distance position gene name gene id
GU305914.1 169229 169251 - 0 CACGACATCATGTTTCGTGT CGG ||||||||[||||||||||||]PAM NA Intergenic NA NA

T21683 CTGCGTCGCGTACGCCGTGACGG 1000
Oligo fwd CACCCTGCGTCGCGTACGCCGTGA
Oligo rev AAACTCACGGCGTACGCGACGCAG
Chromosome start end strand MM target_seq PAM alignment distance position gene name gene id
GU305914.1 169197 169219 - 0 CTGCGTCGCGTACGCCGTGA CGG ||||||||[||||||||||||]PAM NA Intergenic NA NA

T21684 GACGGCCGTATACATAACACCGG 1000
Oligo fwd CACCGACGGCCGTATACATAACAC
Oligo rev AAACGTGTTATGTATACGGCCGTC
Chromosome start end strand MM target_seq PAM alignment distance position gene name gene id
GU305914.1 169179 169201 - 0 GACGGCCGTATACATAACAC CGG ||||||||[||||||||||||]PAM NA Intergenic NA NA

T21685 CACCTCCAGACTCATTCGCCTGG 1000
Oligo fwd CACCCACCTCCAGACTCATTCGCC
Oligo rev AAACGGCGAATGAGTCTGGAGGTG
Chromosome start end strand MM target_seq PAM alignment distance position gene name gene id
GU305914.1 169148 169170 - 0 CACCTCCAGACTCATTCGCC TGG ||||||||[||||||||||||]PAM NA Intergenic NA NA

T21686 CTCCAGACTCATTCGCCTGGTGG 1000
Oligo fwd CACCCTCCAGACTCATTCGCCTGG
Oligo rev AAACCCAGGCGAATGAGTCTGGAG
Chromosome start end strand MM target_seq PAM alignment distance position gene name gene id
GU305914.1 169145 169167 - 0 CTCCAGACTCATTCGCCTGG TGG ||||||||[||||||||||||]PAM NA Intergenic NA NA

T21687 CAGACTCATTCGCCTGGTGGAGG 1000

| | | | | | | | | | | | |
|---|---|---|---|---|---|---|---|---|---|---|---|
| Oligo fwd | CACCCAGACTCATTCGCCTGGTGG | | | | | | | | | | |
| Oligo rev | AAACCCACCAGGCGAATGAGTCTG | | | | | | | | | | |
| Chromosome | start | end | strand | MM | target_seq | PAM | alignment | distance | position | gene name | gene id |
| GU305914.1 | 169142 | 169164 | - | 0 | CAGACTCATTCGCCTGGTGG | AGG | \|\|\|\|\|\|\|\|[\|\|\|\|\|\|\|\|\|\|\|\|]PAM | NA | Intergenic | NA | NA |
| T21688 | AGACTCATTCGCCTGGTGGAGGG | 1000 | | | | | | | | | |
| Oligo fwd | CACCAGACTCATTCGCCTGGTGGA | | | | | | | | | | |
| Oligo rev | AAACTCCACCAGGCGAATGAGTCT | | | | | | | | | | |
| Chromosome | start | end | strand | MM | target_seq | PAM | alignment | distance | position | gene name | gene id |
| GU305914.1 | 169141 | 169163 | - | 0 | AGACTCATTCGCCTGGTGGA | GGG | \|\|\|\|\|\|\|\|[\|\|\|\|\|\|\|\|\|\|\|\|]PAM | NA | Intergenic | NA | NA |
| T21689 | CATACCCAAACAGCCCAAGCTGG | 1000 | | | | | | | | | |
| Oligo fwd | CACCCATACCCAAACAGCCCAAGC | | | | | | | | | | |
| Oligo rev | AAACGCTTGGGCTGTTTGGGTATG | | | | | | | | | | |
| Chromosome | start | end | strand | MM | target_seq | PAM | alignment | distance | position | gene name | gene id |
| GU305914.1 | 169083 | 169105 | - | 0 | CATACCCAAACAGCCCAAGC | TGG | \|\|\|\|\|\|\|\|[\|\|\|\|\|\|\|\|\|\|\|\|]PAM | NA | Intergenic | NA | NA |
| T21690 | AAACAGCCCAAGCTGGATAGAGG | 1000 | | | | | | | | | |
| Oligo fwd | CACCAAACAGCCCAAGCTGGATAG | | | | | | | | | | |
| Oligo rev | AAACCTATCCAGCTTGGGCTGTTT | | | | | | | | | | |
| Chromosome | start | end | strand | MM | target_seq | PAM | alignment | distance | position | gene name | gene id |
| GU305914.1 | 169076 | 169098 | - | 0 | AAACAGCCCAAGCTGGATAG | AGG | \|\|\|\|\|\|\|\|[\|\|\|\|\|\|\|\|\|\|\|\|]PAM | NA | Intergenic | NA | NA |
| T21691 | AGGACTGAGAGAAGTTCAATCGG | 1000 | | | | | | | | | |
| Oligo fwd | CACCAGGACTGAGAGAAGTTCAAT | | | | | | | | | | |
| Oligo rev | AAACATTGAACTTCTCTCAGTCCT | | | | | | | | | | |
| Chromosome | start | end | strand | MM | target_seq | PAM | alignment | distance | position | gene name | gene id |
| GU305914.1 | 169056 | 169078 | - | 0 | AGGACTGAGAGAAGTTCAAT | CGG | \|\|\|\|\|\|\|\|[\|\|\|\|\|\|\|\|\|\|\|\|]PAM | NA | Intergenic | NA | NA |
| T21692 | AGAGAAGTTCAATCGGATCCAGG | 1000 | | | | | | | | | |
| Oligo fwd | CACCAGAGAAGTTCAATCGGATCC | | | | | | | | | | |
| Oligo rev | AAACGGATCCGATTGAACTTCTCT | | | | | | | | | | |
| Chromosome | start | end | strand | MM | target_seq | PAM | alignment | distance | position | gene name | gene id |
| GU305914.1 | 169049 | 169071 | - | 0 | AGAGAAGTTCAATCGGATCC | AGG | \|\|\|\|\|\|\|\|[\|\|\|\|\|\|\|\|\|\|\|\|]PAM | NA | Intergenic | NA | NA |
| T21693 | GGATCCAGGCACCTCTCAACCGG | 1000 | | | | | | | | | |
| Oligo fwd | CACCGGATCCAGGCACCTCTCAAC | | | | | | | | | | |
| Oligo rev | AAACGTTGAGAGGTGCCTGGATCC | | | | | | | | | | |
| Chromosome | start | end | strand | MM | target_seq | PAM | alignment | distance | position | gene name | gene id |
| GU305914.1 | 169035 | 169057 | - | 0 | GGATCCAGGCACCTCTCAAC | CGG | \|\|\|\|\|\|\|\|[\|\|\|\|\|\|\|\|\|\|\|\|]PAM | NA | Intergenic | NA | NA |
| T21694 | CCCTGCACAAACCGCACAAGCGG | 1000 | | | | | | | | | |
| Oligo fwd | CACCCCCTGCACAAACCGCACAAG | | | | | | | | | | |
| Oligo rev | AAACCTTGTGCGGTTTGTGCAGGG | | | | | | | | | | |
| Chromosome | start | end | strand | MM | target_seq | PAM | alignment | distance | position | gene name | gene id |
| GU305914.1 | 168954 | 168976 | - | 0 | CCCTGCACAAACCGCACAAG | CGG | \|\|\|\|\|\|\|\|[\|\|\|\|\|\|\|\|\|\|\|\|]PAM | NA | Intergenic | NA | NA |
| T21695 | CCTGCACAAACCGCACAAGCGGG | 1000 | | | | | | | | | |
| Oligo fwd | CACCCCTGCACAAACCGCACAAGC | | | | | | | | | | |
| Oligo rev | AAACGCTTGTGCGGTTTGTGCAGG | | | | | | | | | | |
| Chromosome | start | end | strand | MM | target_seq | PAM | alignment | distance | position | gene name | gene id |
| GU305914.1 | 168953 | 168975 | - | 0 | CCTGCACAAACCGCACAAGC | GGG | \|\|\|\|\|\|\|\|[\|\|\|\|\|\|\|\|\|\|\|\|]PAM | NA | Intergenic | NA | NA |
| T21696 | ACCGCACAAGCGGGCAAGAACGG | 1000 | | | | | | | | | |
| Oligo fwd | CACCACCGCACAAGCGGGCAAGAA | | | | | | | | | | |
| Oligo rev | AAACTTCTTGCCCGCTTGTGCGGT | | | | | | | | | | |
| Chromosome | start | end | strand | MM | target_seq | PAM | alignment | distance | position | gene name | gene id |
| GU305914.1 | 168944 | 168966 | - | 0 | ACCGCACAAGCGGGCAAGAA | CGG | \|\|\|\|\|\|\|\|[\|\|\|\|\|\|\|\|\|\|\|\|]PAM | NA | Intergenic | NA | NA |
| T21697 | CAAGAACGGCAACGAGACGACGG | 1000 | | | | | | | | | |
| Oligo fwd | CACCCAAGAACGGCAACGAGACGA | | | | | | | | | | |
| Oligo rev | AAACTCGTCTCGTTGCCGTTCTTG | | | | | | | | | | |
| Chromosome | start | end | strand | MM | target_seq | PAM | alignment | distance | position | gene name | gene id |
| GU305914.1 | 168930 | 168952 | - | 0 | CAAGAACGGCAACGAGACGA | CGG | \|\|\|\|\|\|\|\|[\|\|\|\|\|\|\|\|\|\|\|\|]PAM | NA | Intergenic | NA | NA |
| T21698 | CGAGACGACGGTCGCCGCGCCGG | 1000 | | | | | | | | | |
| Oligo fwd | CACCCGAGACGACGGTCGCCGCGC | | | | | | | | | | |
| Oligo rev | AAACGCGCGGCGACCGTCGTCTCG | | | | | | | | | | |
| Chromosome | start | end | strand | MM | target_seq | PAM | alignment | distance | position | gene name | gene id |
| GU305914.1 | 168918 | 168940 | - | 0 | CGAGACGACGGTCGCCGCGC | CGG | \|\|\|\|\|\|\|\|[\|\|\|\|\|\|\|\|\|\|\|\|]PAM | NA | Intergenic | NA | NA |
| GU305914.1 | 47435 | 47457 | - | 4 | CGTGCCGACGGCGGCCGCGC | TGG | \|\|-\|-\|\|\|[\|\|\|--\|\|\|\|\|\|\|]PAM | NA | Intergenic | NA | NA |
| T21699 | CGCCGCGCCGGCTCCGCCCGCGG | 1000 | | | | | | | | | |
| Oligo fwd | CACCCGCCGCGCCGGCTCCGCCCG | | | | | | | | | | |
| Oligo rev | AAACCGGGCGGAGCCGGCGCGGCG | | | | | | | | | | |
| Chromosome | start | end | strand | MM | target_seq | PAM | alignment | distance | position | gene name | gene id |
| GU305914.1 | 168906 | 168928 | - | 0 | CGCCGCGCCGGCTCCGCCCG | CGG | \|\|\|\|\|\|\|\|[\|\|\|\|\|\|\|\|\|\|\|\|]PAM | NA | Intergenic | NA | NA |
| T21700 | CTGCCTCCGCAGCTAGCACTCGG | 1000 | | | | | | | | | |
| Oligo fwd | CACCCTGCCTCCGCAGCTAGCACT | | | | | | | | | | |
| Oligo rev | AAACAGTGCTAGCTGCGGAGGCAG | | | | | | | | | | |
| Chromosome | start | end | strand | MM | target_seq | PAM | alignment | distance | position | gene name | gene id |
| GU305914.1 | 168862 | 168884 | - | 0 | CTGCCTCCGCAGCTAGCACT | CGG | \|\|\|\|\|\|\|\|[\|\|\|\|\|\|\|\|\|\|\|\|]PAM | NA | Intergenic | NA | NA |
| T21701 | TGCCTCCGCAGCTAGCACTCGGG | 1000 | | | | | | | | | |
| Oligo fwd | CACCTGCCTCCGCAGCTAGCACTC | | | | | | | | | | |
| Oligo rev | AAACGAGTGCTAGCTGCGGAGGCA | | | | | | | | | | |
| Chromosome | start | end | strand | MM | target_seq | PAM | alignment | distance | position | gene name | gene id |
| GU305914.1 | 168861 | 168883 | - | 0 | TGCCTCCGCAGCTAGCACTC | GGG | \|\|\|\|\|\|\|\|[\|\|\|\|\|\|\|\|\|\|\|\|]PAM | NA | Intergenic | NA | NA |

| | | | | | | | | | | | |
|---|---|---|---|---|---|---|---|---|---|---|---|
| T21702 | CCACCACCGCCGCTAAAAAACGG | 1000 | | | | | | | | | |
| Oligo fwd | CACCCCACCACCGCCGCTAAAAAA | | | | | | | | | | |
| Oligo rev | AAACTTTTTTAGCGGCGGTGGTGG | | | | | | | | | | |
| Chromosome | start | end | strand | MM | target_seq | PAM | alignment | distance | position | gene name | gene id |
| GU305914.1 | 168805 | 168827 | - | 0 | CCACCACCGCCGCTAAAAAA | CGG | \|\|\|\|\|\|\|\|[\|\|\|\|\|\|\|\|\|\|\|\|]PAM | NA | Intergenic | NA | NA |

| | | | | | | | | | | | |
|---|---|---|---|---|---|---|---|---|---|---|---|
| T21703 | CACCACCGCCGCTAAAAAACGGG | 1000 | | | | | | | | | |
| Oligo fwd | CACCCACCACCGCCGCTAAAAAAC | | | | | | | | | | |
| Oligo rev | AAACGTTTTTTAGCGGCGGTGGTG | | | | | | | | | | |
| Chromosome | start | end | strand | MM | target_seq | PAM | alignment | distance | position | gene name | gene id |
| GU305914.1 | 168804 | 168826 | - | 0 | CACCACCGCCGCTAAAAAAC | GGG | \|\|\|\|\|\|\|\|[\|\|\|\|\|\|\|\|\|\|\|\|]PAM | NA | Intergenic | NA | NA |

| | | | | | | | | | | | |
|---|---|---|---|---|---|---|---|---|---|---|---|
| T21704 | ATACTATCTCGAGCGAGCGACGG | 1000 | | | | | | | | | |
| Oligo fwd | CACCATACTATCTCGAGCGAGCGA | | | | | | | | | | |
| Oligo rev | AAACTCGCTCGCTCGAGATAGTAT | | | | | | | | | | |
| Chromosome | start | end | strand | MM | target_seq | PAM | alignment | distance | position | gene name | gene id |
| GU305914.1 | 168773 | 168795 | - | 0 | ATACTATCTCGAGCGAGCGA | CGG | \|\|\|\|\|\|\|\|[\|\|\|\|\|\|\|\|\|\|\|\|]PAM | NA | Intergenic | NA | NA |

| | | | | | | | | | | | |
|---|---|---|---|---|---|---|---|---|---|---|---|
| T21705 | TACTATCTCGAGCGAGCGACGGG | 1000 | | | | | | | | | |
| Oligo fwd | CACCTACTATCTCGAGCGAGCGAC | | | | | | | | | | |
| Oligo rev | AAACGTCGCTCGCTCGAGATAGTA | | | | | | | | | | |
| Chromosome | start | end | strand | MM | target_seq | PAM | alignment | distance | position | gene name | gene id |
| GU305914.1 | 168772 | 168794 | - | 0 | TACTATCTCGAGCGAGCGAC | GGG | \|\|\|\|\|\|\|\|[\|\|\|\|\|\|\|\|\|\|\|\|]PAM | NA | Intergenic | NA | NA |

| | | | | | | | | | | | |
|---|---|---|---|---|---|---|---|---|---|---|---|
| T21706 | AAGATCTTAACCCCCACCGACGG | 1000 | | | | | | | | | |
| Oligo fwd | CACCAAGATCTTAACCCCCACCGA | | | | | | | | | | |
| Oligo rev | AAACTCGGTGGGGGTTAAGATCTT | | | | | | | | | | |
| Chromosome | start | end | strand | MM | target_seq | PAM | alignment | distance | position | gene name | gene id |
| GU305914.1 | 168746 | 168768 | - | 0 | AAGATCTTAACCCCCACCGA | CGG | \|\|\|\|\|\|\|\|[\|\|\|\|\|\|\|\|\|\|\|\|]PAM | NA | Intergenic | NA | NA |

| | | | | | | | | | | | |
|---|---|---|---|---|---|---|---|---|---|---|---|
| T21707 | AGATCTTAACCCCCACCGACGGG | 1000 | | | | | | | | | |
| Oligo fwd | CACCAGATCTTAACCCCCACCGAC | | | | | | | | | | |
| Oligo rev | AAACGTCGGTGGGGGTTAAGATCT | | | | | | | | | | |
| Chromosome | start | end | strand | MM | target_seq | PAM | alignment | distance | position | gene name | gene id |
| GU305914.1 | 168745 | 168767 | - | 0 | AGATCTTAACCCCCACCGAC | GGG | \|\|\|\|\|\|\|\|[\|\|\|\|\|\|\|\|\|\|\|\|]PAM | NA | Intergenic | NA | NA |

| | | | | | | | | | | | |
|---|---|---|---|---|---|---|---|---|---|---|---|
| T21708 | AATCTACGCCTCTTCGTCATCGG | 1000 | | | | | | | | | |
| Oligo fwd | CACCAATCTACGCCTCTTCGTCAT | | | | | | | | | | |
| Oligo rev | AAACATGACGAAGAGGCGTAGATT | | | | | | | | | | |
| Chromosome | start | end | strand | MM | target_seq | PAM | alignment | distance | position | gene name | gene id |
| GU305914.1 | 168711 | 168733 | - | 0 | AATCTACGCCTCTTCGTCAT | CGG | \|\|\|\|\|\|\|\|[\|\|\|\|\|\|\|\|\|\|\|\|]PAM | NA | Intergenic | NA | NA |

| | | | | | | | | | | | |
|---|---|---|---|---|---|---|---|---|---|---|---|
| T21709 | CTCTTCGTCATCGGACTCTTCGG | 1000 | | | | | | | | | |
| Oligo fwd | CACCCTCTTCGTCATCGGACTCTT | | | | | | | | | | |
| Oligo rev | AAACAAGAGTCCGATGACGAAGAG | | | | | | | | | | |
| Chromosome | start | end | strand | MM | target_seq | PAM | alignment | distance | position | gene name | gene id |
| GU305914.1 | 168702 | 168724 | - | 0 | CTCTTCGTCATCGGACTCTT | CGG | \|\|\|\|\|\|\|\|[\|\|\|\|\|\|\|\|\|\|\|\|]PAM | NA | Intergenic | NA | NA |

| | | | | | | | | | | | |
|---|---|---|---|---|---|---|---|---|---|---|---|
| T21710 | GGATATCTCCTCAGACGACTCGG | 1000 | | | | | | | | | |
| Oligo fwd | CACCGGATATCTCCTCAGACGACT | | | | | | | | | | |
| Oligo rev | AAACAGTCGTCTGAGGAGATATCC | | | | | | | | | | |
| Chromosome | start | end | strand | MM | target_seq | PAM | alignment | distance | position | gene name | gene id |
| GU305914.1 | 168681 | 168703 | - | 0 | GGATATCTCCTCAGACGACT | CGG | \|\|\|\|\|\|\|\|[\|\|\|\|\|\|\|\|\|\|\|\|]PAM | NA | Intergenic | NA | NA |

| | | | | | | | | | | | |
|---|---|---|---|---|---|---|---|---|---|---|---|
| T21711 | GTCTAACTCGTCCTTACCACCGG | 1000 | | | | | | | | | |
| Oligo fwd | CACCGTCTAACTCGTCCTTACCAC | | | | | | | | | | |
| Oligo rev | AAACGTGGTAAGGACGAGTTAGAC | | | | | | | | | | |
| Chromosome | start | end | strand | MM | target_seq | PAM | alignment | distance | position | gene name | gene id |
| GU305914.1 | 168627 | 168649 | - | 0 | GTCTAACTCGTCCTTACCAC | CGG | \|\|\|\|\|\|\|\|[\|\|\|\|\|\|\|\|\|\|\|\|]PAM | NA | Intergenic | NA | NA |

| | | | | | | | | | | | |
|---|---|---|---|---|---|---|---|---|---|---|---|
| T21712 | TCTAACTCGTCCTTACCACCGGG | 1000 | | | | | | | | | |
| Oligo fwd | CACCTCTAACTCGTCCTTACCACC | | | | | | | | | | |
| Oligo rev | AAACGGTGGTAAGGACGAGTTAGA | | | | | | | | | | |
| Chromosome | start | end | strand | MM | target_seq | PAM | alignment | distance | position | gene name | gene id |
| GU305914.1 | 168626 | 168648 | - | 0 | TCTAACTCGTCCTTACCACC | GGG | \|\|\|\|\|\|\|\|[\|\|\|\|\|\|\|\|\|\|\|\|]PAM | NA | Intergenic | NA | NA |

| | | | | | | | | | | | |
|---|---|---|---|---|---|---|---|---|---|---|---|
| T21713 | CTCGTCCTTACCACCGGGATCGG | 1000 | | | | | | | | | |
| Oligo fwd | CACCCTCGTCCTTACCACCGGGAT | | | | | | | | | | |
| Oligo rev | AAACATCCCGGTGGTAAGGACGAG | | | | | | | | | | |
| Chromosome | start | end | strand | MM | target_seq | PAM | alignment | distance | position | gene name | gene id |
| GU305914.1 | 168621 | 168643 | - | 0 | CTCGTCCTTACCACCGGGAT | CGG | \|\|\|\|\|\|\|\|[\|\|\|\|\|\|\|\|\|\|\|\|]PAM | NA | Intergenic | NA | NA |

| | | | | | | | | | | | |
|---|---|---|---|---|---|---|---|---|---|---|---|
| T21714 | TCCATCCTCCTCCGACTCAGAGG | 1000 | | | | | | | | | |
| Oligo fwd | CACCTCCATCCTCCTCCGACTCAG | | | | | | | | | | |
| Oligo rev | AAACCTGAGTCGGAGGAGGATGGA | | | | | | | | | | |
| Chromosome | start | end | strand | MM | target_seq | PAM | alignment | distance | position | gene name | gene id |
| GU305914.1 | 168543 | 168565 | - | 0 | TCCATCCTCCTCCGACTCAG | AGG | \|\|\|\|\|\|\|\|[\|\|\|\|\|\|\|\|\|\|\|\|]PAM | NA | Intergenic | NA | NA |

| | | | | | | | | | | | |
|---|---|---|---|---|---|---|---|---|---|---|---|
| T21715 | ATCATCGTCATCTACCAGCTCGG | 1000 | | | | | | | | | |
| Oligo fwd | CACCATCATCGTCATCTACCAGCT | | | | | | | | | | |
| Oligo rev | AAACAGCTGGTAGATGACGATGAT | | | | | | | | | | |
| Chromosome | start | end | strand | MM | target_seq | PAM | alignment | distance | position | gene name | gene id |
| GU305914.1 | 168501 | 168523 | - | 0 | ATCATCGTCATCTACCAGCT | CGG | \|\|\|\|\|\|\|\|[\|\|\|\|\|\|\|\|\|\|\|\|]PAM | NA | Intergenic | NA | NA |

| | | | | | | | | | | | |
|---|---|---|---|---|---|---|---|---|---|---|---|
| T21716 | TCGTCATCTACCAGCTCGGACGG | 1000 | | | | | | | | | |
| Oligo fwd | CACCTCGTCATCTACCAGCTCGGA | | | | | | | | | | |
| Oligo rev | AAACTCCGAGCTGGTAGATGACGA | | | | | | | | | | |
| Chromosome | start | end | strand | MM | target_seq | PAM | alignment | distance | position | gene name | gene id |

GU305914.1 168497 168519 - 0 TCGTCATCTACCAGCTCGGA CGG ||||||||[||||||||||||]PAM NA Intergenic NA NA

T21717 CAGCTCGGACGGCGCTTCATCGG 1000
Oligo fwd CACCCAGCTCGGACGGCGCTTCAT
Oligo rev AAACATGAAGCGCCGTCCGAGCTG
Chromosome start end strand MM target_seq PAM alignment distance position gene name gene id
GU305914.1 168486 168508 - 0 CAGCTCGGACGGCGCTTCAT CGG ||||||||[||||||||||||]PAM NA Intergenic NA NA

T21718 GGACGGCGCTTCATCGGATTCGG 1000
Oligo fwd CACCGGACGGCGCTTCATCGGATT
Oligo rev AAACAATCCGATGAAGCGCCGTCC
Chromosome start end strand MM target_seq PAM alignment distance position gene name gene id
GU305914.1 168480 168502 - 0 GGACGGCGCTTCATCGGATT CGG ||||||||[||||||||||||]PAM NA Intergenic NA NA

T21719 CGGCGCTTCATCGGATTCGGAGG 1000
Oligo fwd CACCCGGCGCTTCATCGGATTCGG
Oligo rev AAACCCGAATCCGATGAAGCGCCG
Chromosome start end strand MM target_seq PAM alignment distance position gene name gene id
GU305914.1 168477 168499 - 0 CGGCGCTTCATCGGATTCGG AGG ||||||||[||||||||||||]PAM NA Intergenic NA NA

T21720 GGATTCGGAGGCGCCATCTTCGG 1000
Oligo fwd CACCGGATTCGGAGGCGCCATCTT
Oligo rev AAACAAGATGGCGCCTCCGAATCC
Chromosome start end strand MM target_seq PAM alignment distance position gene name gene id
GU305914.1 168465 168487 - 0 GGATTCGGAGGCGCCATCTT CGG ||||||||[||||||||||||]PAM NA Intergenic NA NA

T21721 GAGGCGCCATCTTCGGATTCTGG 1000
Oligo fwd CACCGAGGCGCCATCTTCGGATTC
Oligo rev AAACGAATCCGAAGATGGCGCCTC
Chromosome start end strand MM target_seq PAM alignment distance position gene name gene id
GU305914.1 168458 168480 - 0 GAGGCGCCATCTTCGGATTC TGG ||||||||[||||||||||||]PAM NA Intergenic NA NA

T21722 GATTCTGGATCCGAATCCATAGG 1000
Oligo fwd CACCGATTCTGGATCCGAATCCAT
Oligo rev AAACATGGATTCGGATCCAGAATC
Chromosome start end strand MM target_seq PAM alignment distance position gene name gene id
GU305914.1 168443 168465 - 0 GATTCTGGATCCGAATCCAT AGG ||||||||[||||||||||||]PAM NA Intergenic NA NA

T21723 AGGCTCGACGTCGCCCAAATCGG 1000
Oligo fwd CACCAGGCTCGACGTCGCCCAAAT
Oligo rev AAACATTTGGGCGACGTCGAGCCT
Chromosome start end strand MM target_seq PAM alignment distance position gene name gene id
GU305914.1 168423 168445 - 0 AGGCTCGACGTCGCCCAAAT CGG ||||||||[||||||||||||]PAM NA Intergenic NA NA

T21724 GACGTCGCCCAAATCGGAATCGG 1000
Oligo fwd CACCGACGTCGCCCAAATCGGAAT
Oligo rev AAACATTCCGATTTGGGCGACGTC
Chromosome start end strand MM target_seq PAM alignment distance position gene name gene id
GU305914.1 168417 168439 - 0 GACGTCGCCCAAATCGGAAT CGG ||||||||[||||||||||||]PAM NA Intergenic NA NA

T21725 CGCGCGAGTTCGCCGCAACCAGG 1000
Oligo fwd CACCCGCGCGAGTTCGCCGCAACC
Oligo rev AAACGGTTGCGGCGAACTCGCGCG
Chromosome start end strand MM target_seq PAM alignment distance position gene name gene id
GU305914.1 168332 168354 - 0 CGCGCGAGTTCGCCGCAACC AGG ||||||||[||||||||||||]PAM NA Intergenic NA NA

T21726 AGCCCCCGCGAACACAGAACCGG 1000
Oligo fwd CACCAGCCCCCGCGAACACAGAAC
Oligo rev AAACGTTCTGTGTTCGCGGGGGCT
Chromosome start end strand MM target_seq PAM alignment distance position gene name gene id
GU305914.1 168297 168319 - 0 AGCCCCCGCGAACACAGAAC CGG ||||||||[||||||||||||]PAM NA Intergenic NA NA

T21727 ACCGGCCCGAGACGCGTCGACGG 1000
Oligo fwd CACCACCGGCCCGAGACGCGTCGA
Oligo rev AAACTCGACGCGTCTCGGGCCGGT
Chromosome start end strand MM target_seq PAM alignment distance position gene name gene id
GU305914.1 168279 168301 - 0 ACCGGCCCGAGACGCGTCGA CGG ||||||||[||||||||||||]PAM NA Intergenic NA NA

T21728 GATGTTATCCGCCTCGATCGTGG 1000
Oligo fwd CACCGATGTTATCCGCCTCGATCG
Oligo rev AAACCGATCGAGGCGGATAACATC
Chromosome start end strand MM target_seq PAM alignment distance position gene name gene id
GU305914.1 168243 168265 - 0 GATGTTATCCGCCTCGATCG TGG ||||||||[||||||||||||]PAM NA Intergenic NA NA

T21729 TGCCAAGACTACTCCTCAGCAGG 1000
Oligo fwd CACCTGCCAAGACTACTCCTCAGC
Oligo rev AAACGCTGAGGAGTAGTCTTGGCA
Chromosome start end strand MM target_seq PAM alignment distance position gene name gene id
GU305914.1 168195 168217 - 0 TGCCAAGACTACTCCTCAGC AGG ||||||||[||||||||||||]PAM NA Intergenic NA NA

T21730 CAGCAGGTAACTAACGCCACTGG 1000
Oligo fwd CACCCAGCAGGTAACTAACGCCAC
Oligo rev AAACGTGGCGTTAGTTACCTGCTG
Chromosome start end strand MM target_seq PAM alignment distance position gene name gene id
GU305914.1 168179 168201 - 0 CAGCAGGTAACTAACGCCAC TGG ||||||||[||||||||||||]PAM NA Intergenic NA NA

T21731 CAGGTAACTAACGCCACTGGCGG 1000
Oligo fwd CACCCAGGTAACTAACGCCACTGG
Oligo rev AAACCCAGTGGCGTTAGTTACCTG

<table>
<tr><td>Chromosome</td><td>start</td><td>end</td><td>strand</td><td>MM</td><td>target_seq</td><td>PAM</td><td>alignment</td><td>distance</td><td>position</td><td>gene name</td><td>gene id</td></tr>
<tr><td>GU305914.1</td><td>168176</td><td>168198</td><td>-</td><td>0</td><td>CAGGTAACTAACGCCACTGG</td><td>CGG</td><td>||||||||[||||||||||||]PAM</td><td>NA</td><td>Intergenic</td><td>NA</td><td>NA</td></tr>
<tr><td></td><td></td><td></td><td></td><td></td><td></td><td></td><td></td><td></td><td></td><td></td><td></td></tr>
<tr><td>T21732</td><td>CACTGGCGGCGATGATGCGTTGG</td><td>1000</td><td></td><td></td><td></td><td></td><td></td><td></td><td></td><td></td><td></td></tr>
<tr><td>Oligo fwd</td><td>CACCCACTGGCGGCGATGATGCGT</td><td></td><td></td><td></td><td></td><td></td><td></td><td></td><td></td><td></td><td></td></tr>
<tr><td>Oligo rev</td><td>AAACACGCATCATCGCCGCCAGTG</td><td></td><td></td><td></td><td></td><td></td><td></td><td></td><td></td><td></td><td></td></tr>
<tr><td>Chromosome</td><td>start</td><td>end</td><td>strand</td><td>MM</td><td>target_seq</td><td>PAM</td><td>alignment</td><td>distance</td><td>position</td><td>gene name</td><td>gene id</td></tr>
<tr><td>GU305914.1</td><td>168162</td><td>168184</td><td>-</td><td>0</td><td>CACTGGCGGCGATGATGCGT</td><td>TGG</td><td>||||||||[||||||||||||]PAM</td><td>NA</td><td>Intergenic</td><td>NA</td><td>NA</td></tr>
<tr><td></td><td></td><td></td><td></td><td></td><td></td><td></td><td></td><td></td><td></td><td></td><td></td></tr>
<tr><td>T21733</td><td>GCGTTGGTAGTATTGACTGTCGG</td><td>1000</td><td></td><td></td><td></td><td></td><td></td><td></td><td></td><td></td><td></td></tr>
<tr><td>Oligo fwd</td><td>CACCGCGTTGGTAGTATTGACTGT</td><td></td><td></td><td></td><td></td><td></td><td></td><td></td><td></td><td></td><td></td></tr>
<tr><td>Oligo rev</td><td>AAACACAGTCAATACTACCAACGC</td><td></td><td></td><td></td><td></td><td></td><td></td><td></td><td></td><td></td><td></td></tr>
<tr><td>Chromosome</td><td>start</td><td>end</td><td>strand</td><td>MM</td><td>target_seq</td><td>PAM</td><td>alignment</td><td>distance</td><td>position</td><td>gene name</td><td>gene id</td></tr>
<tr><td>GU305914.1</td><td>168146</td><td>168168</td><td>-</td><td>0</td><td>GCGTTGGTAGTATTGACTGT</td><td>CGG</td><td>||||||||[||||||||||||]PAM</td><td>NA</td><td>Intergenic</td><td>NA</td><td>NA</td></tr>
<tr><td></td><td></td><td></td><td></td><td></td><td></td><td></td><td></td><td></td><td></td><td></td><td></td></tr>
<tr><td>T21734</td><td>ACTCTCCGCTCTCCGATAACAGG</td><td>1000</td><td></td><td></td><td></td><td></td><td></td><td></td><td></td><td></td><td></td></tr>
<tr><td>Oligo fwd</td><td>CACCACTCTCCGCTCTCCGATAAC</td><td></td><td></td><td></td><td></td><td></td><td></td><td></td><td></td><td></td><td></td></tr>
<tr><td>Oligo rev</td><td>AAACGTTATCGGAGAGCGGAGAGT</td><td></td><td></td><td></td><td></td><td></td><td></td><td></td><td></td><td></td><td></td></tr>
<tr><td>Chromosome</td><td>start</td><td>end</td><td>strand</td><td>MM</td><td>target_seq</td><td>PAM</td><td>alignment</td><td>distance</td><td>position</td><td>gene name</td><td>gene id</td></tr>
<tr><td>GU305914.1</td><td>168112</td><td>168134</td><td>-</td><td>0</td><td>ACTCTCCGCTCTCCGATAAC</td><td>AGG</td><td>||||||||[||||||||||||]PAM</td><td>NA</td><td>Intergenic</td><td>NA</td><td>NA</td></tr>
<tr><td></td><td></td><td></td><td></td><td></td><td></td><td></td><td></td><td></td><td></td><td></td><td></td></tr>
<tr><td>T21735</td><td>GTCAACACAGAGTGCGACAACGG</td><td>1000</td><td></td><td></td><td></td><td></td><td></td><td></td><td></td><td></td><td></td></tr>
<tr><td>Oligo fwd</td><td>CACCGTCAACACAGAGTGCGACAA</td><td></td><td></td><td></td><td></td><td></td><td></td><td></td><td></td><td></td><td></td></tr>
<tr><td>Oligo rev</td><td>AAACTTGTCGCACTCTGTGTTGAC</td><td></td><td></td><td></td><td></td><td></td><td></td><td></td><td></td><td></td><td></td></tr>
<tr><td>Chromosome</td><td>start</td><td>end</td><td>strand</td><td>MM</td><td>target_seq</td><td>PAM</td><td>alignment</td><td>distance</td><td>position</td><td>gene name</td><td>gene id</td></tr>
<tr><td>GU305914.1</td><td>168090</td><td>168112</td><td>-</td><td>0</td><td>GTCAACACAGAGTGCGACAA</td><td>CGG</td><td>||||||||[||||||||||||]PAM</td><td>NA</td><td>Intergenic</td><td>NA</td><td>NA</td></tr>
<tr><td></td><td></td><td></td><td></td><td></td><td></td><td></td><td></td><td></td><td></td><td></td><td></td></tr>
<tr><td>T21736</td><td>GAGTGCGACAACGGTCCAGACGG</td><td>1000</td><td></td><td></td><td></td><td></td><td></td><td></td><td></td><td></td><td></td></tr>
<tr><td>Oligo fwd</td><td>CACCGAGTGCGACAACGGTCCAGA</td><td></td><td></td><td></td><td></td><td></td><td></td><td></td><td></td><td></td><td></td></tr>
<tr><td>Oligo rev</td><td>AAACTCTGGACCGTTGTCGCACTC</td><td></td><td></td><td></td><td></td><td></td><td></td><td></td><td></td><td></td><td></td></tr>
<tr><td>Chromosome</td><td>start</td><td>end</td><td>strand</td><td>MM</td><td>target_seq</td><td>PAM</td><td>alignment</td><td>distance</td><td>position</td><td>gene name</td><td>gene id</td></tr>
<tr><td>GU305914.1</td><td>168081</td><td>168103</td><td>-</td><td>0</td><td>GAGTGCGACAACGGTCCAGA</td><td>CGG</td><td>||||||||[||||||||||||]PAM</td><td>NA</td><td>Intergenic</td><td>NA</td><td>NA</td></tr>
<tr><td></td><td></td><td></td><td></td><td></td><td></td><td></td><td></td><td></td><td></td><td></td><td></td></tr>
<tr><td>T21737</td><td>ACAACGGTCCAGACGGAAAGCGG</td><td>1000</td><td></td><td></td><td></td><td></td><td></td><td></td><td></td><td></td><td></td></tr>
<tr><td>Oligo fwd</td><td>CACCACAACGGTCCAGACGGAAAG</td><td></td><td></td><td></td><td></td><td></td><td></td><td></td><td></td><td></td><td></td></tr>
<tr><td>Oligo rev</td><td>AAACCTTTCCGTCTGGACCGTTGT</td><td></td><td></td><td></td><td></td><td></td><td></td><td></td><td></td><td></td><td></td></tr>
<tr><td>Chromosome</td><td>start</td><td>end</td><td>strand</td><td>MM</td><td>target_seq</td><td>PAM</td><td>alignment</td><td>distance</td><td>position</td><td>gene name</td><td>gene id</td></tr>
<tr><td>GU305914.1</td><td>168074</td><td>168096</td><td>-</td><td>0</td><td>ACAACGGTCCAGACGGAAAG</td><td>CGG</td><td>||||||||[||||||||||||]PAM</td><td>NA</td><td>Intergenic</td><td>NA</td><td>NA</td></tr>
<tr><td></td><td></td><td></td><td></td><td></td><td></td><td></td><td></td><td></td><td></td><td></td><td></td></tr>
<tr><td>T21738</td><td>CAGACGGAAAGCGGTCACGCCGG</td><td>1000</td><td></td><td></td><td></td><td></td><td></td><td></td><td></td><td></td><td></td></tr>
<tr><td>Oligo fwd</td><td>CACCCAGACGGAAAGCGGTCACGC</td><td></td><td></td><td></td><td></td><td></td><td></td><td></td><td></td><td></td><td></td></tr>
<tr><td>Oligo rev</td><td>AAACGCGTGACCGCTTTCCGTCTG</td><td></td><td></td><td></td><td></td><td></td><td></td><td></td><td></td><td></td><td></td></tr>
<tr><td>Chromosome</td><td>start</td><td>end</td><td>strand</td><td>MM</td><td>target_seq</td><td>PAM</td><td>alignment</td><td>distance</td><td>position</td><td>gene name</td><td>gene id</td></tr>
<tr><td>GU305914.1</td><td>168065</td><td>168087</td><td>-</td><td>0</td><td>CAGACGGAAAGCGGTCACGC</td><td>CGG</td><td>||||||||[||||||||||||]PAM</td><td>NA</td><td>Intergenic</td><td>NA</td><td>NA</td></tr>
<tr><td></td><td></td><td></td><td></td><td></td><td></td><td></td><td></td><td></td><td></td><td></td><td></td></tr>
<tr><td>T21739</td><td>CCTATCGACCTTGAAGAGAGCGG</td><td>1000</td><td></td><td></td><td></td><td></td><td></td><td></td><td></td><td></td><td></td></tr>
<tr><td>Oligo fwd</td><td>CACCCCTATCGACCTTGAAGAGAG</td><td></td><td></td><td></td><td></td><td></td><td></td><td></td><td></td><td></td><td></td></tr>
<tr><td>Oligo rev</td><td>AAACCTCTCTTCAAGGTCGATAGG</td><td></td><td></td><td></td><td></td><td></td><td></td><td></td><td></td><td></td><td></td></tr>
<tr><td>Chromosome</td><td>start</td><td>end</td><td>strand</td><td>MM</td><td>target_seq</td><td>PAM</td><td>alignment</td><td>distance</td><td>position</td><td>gene name</td><td>gene id</td></tr>
<tr><td>GU305914.1</td><td>168009</td><td>168031</td><td>-</td><td>0</td><td>CCTATCGACCTTGAAGAGAG</td><td>CGG</td><td>||||||||[||||||||||||]PAM</td><td>NA</td><td>Intergenic</td><td>NA</td><td>NA</td></tr>
<tr><td></td><td></td><td></td><td></td><td></td><td></td><td></td><td></td><td></td><td></td><td></td><td></td></tr>
<tr><td>T21740</td><td>CTATCGACCTTGAAGAGAGCGGG</td><td>1000</td><td></td><td></td><td></td><td></td><td></td><td></td><td></td><td></td><td></td></tr>
<tr><td>Oligo fwd</td><td>CACCCTATCGACCTTGAAGAGAGC</td><td></td><td></td><td></td><td></td><td></td><td></td><td></td><td></td><td></td><td></td></tr>
<tr><td>Oligo rev</td><td>AAACGCTCTCTTCAAGGTCGATAG</td><td></td><td></td><td></td><td></td><td></td><td></td><td></td><td></td><td></td><td></td></tr>
<tr><td>Chromosome</td><td>start</td><td>end</td><td>strand</td><td>MM</td><td>target_seq</td><td>PAM</td><td>alignment</td><td>distance</td><td>position</td><td>gene name</td><td>gene id</td></tr>
<tr><td>GU305914.1</td><td>168008</td><td>168030</td><td>-</td><td>0</td><td>CTATCGACCTTGAAGAGAGC</td><td>GGG</td><td>||||||||[||||||||||||]PAM</td><td>NA</td><td>Intergenic</td><td>NA</td><td>NA</td></tr>
<tr><td></td><td></td><td></td><td></td><td></td><td></td><td></td><td></td><td></td><td></td><td></td><td></td></tr>
<tr><td>T21741</td><td>CAGAGAACTGCAGCAGAGCTCGG</td><td>1000</td><td></td><td></td><td></td><td></td><td></td><td></td><td></td><td></td><td></td></tr>
<tr><td>Oligo fwd</td><td>CACCCAGAGAACTGCAGCAGAGCT</td><td></td><td></td><td></td><td></td><td></td><td></td><td></td><td></td><td></td><td></td></tr>
<tr><td>Oligo rev</td><td>AAACAGCTCTGCTGCAGTTCTCTG</td><td></td><td></td><td></td><td></td><td></td><td></td><td></td><td></td><td></td><td></td></tr>
<tr><td>Chromosome</td><td>start</td><td>end</td><td>strand</td><td>MM</td><td>target_seq</td><td>PAM</td><td>alignment</td><td>distance</td><td>position</td><td>gene name</td><td>gene id</td></tr>
<tr><td>GU305914.1</td><td>167976</td><td>167998</td><td>-</td><td>0</td><td>CAGAGAACTGCAGCAGAGCT</td><td>CGG</td><td>||||||||[||||||||||||]PAM</td><td>NA</td><td>Intergenic</td><td>NA</td><td>NA</td></tr>
<tr><td></td><td></td><td></td><td></td><td></td><td></td><td></td><td></td><td></td><td></td><td></td><td></td></tr>
<tr><td>T21742</td><td>ACTGCAGCAGAGCTCGGAGACGG</td><td>1000</td><td></td><td></td><td></td><td></td><td></td><td></td><td></td><td></td><td></td></tr>
<tr><td>Oligo fwd</td><td>CACCACTGCAGCAGAGCTCGGAGA</td><td></td><td></td><td></td><td></td><td></td><td></td><td></td><td></td><td></td><td></td></tr>
<tr><td>Oligo rev</td><td>AAACTCTCCGAGCTCTGCTGCAGT</td><td></td><td></td><td></td><td></td><td></td><td></td><td></td><td></td><td></td><td></td></tr>
<tr><td>Chromosome</td><td>start</td><td>end</td><td>strand</td><td>MM</td><td>target_seq</td><td>PAM</td><td>alignment</td><td>distance</td><td>position</td><td>gene name</td><td>gene id</td></tr>
<tr><td>GU305914.1</td><td>167970</td><td>167992</td><td>-</td><td>0</td><td>ACTGCAGCAGAGCTCGGAGA</td><td>CGG</td><td>||||||||[||||||||||||]PAM</td><td>NA</td><td>Intergenic</td><td>NA</td><td>NA</td></tr>
<tr><td>GU305914.1</td><td>126347</td><td>126369</td><td>-</td><td>4</td><td>CATCCAGCAGAGCTCGCAGA</td><td>AGG</td><td>--|-||||[||||||||-|||]PAM</td><td>NA</td><td>Intergenic</td><td>NA</td><td>NA</td></tr>
<tr><td></td><td></td><td></td><td></td><td></td><td></td><td></td><td></td><td></td><td></td><td></td><td></td></tr>
<tr><td>T21743</td><td>CGACGATCACTGACGCGTCGCGG</td><td>1000</td><td></td><td></td><td></td><td></td><td></td><td></td><td></td><td></td><td></td></tr>
<tr><td>Oligo fwd</td><td>CACCCGACGATCACTGACGCGTCG</td><td></td><td></td><td></td><td></td><td></td><td></td><td></td><td></td><td></td><td></td></tr>
<tr><td>Oligo rev</td><td>AAACCGACGCGTCAGTGATCGTCG</td><td></td><td></td><td></td><td></td><td></td><td></td><td></td><td></td><td></td><td></td></tr>
<tr><td>Chromosome</td><td>start</td><td>end</td><td>strand</td><td>MM</td><td>target_seq</td><td>PAM</td><td>alignment</td><td>distance</td><td>position</td><td>gene name</td><td>gene id</td></tr>
<tr><td>GU305914.1</td><td>167947</td><td>167969</td><td>-</td><td>0</td><td>CGACGATCACTGACGCGTCG</td><td>CGG</td><td>||||||||[||||||||||||]PAM</td><td>NA</td><td>Intergenic</td><td>NA</td><td>NA</td></tr>
<tr><td></td><td></td><td></td><td></td><td></td><td></td><td></td><td></td><td></td><td></td><td></td><td></td></tr>
<tr><td>T21744</td><td>GACGATCACTGACGCGTCGCGGG</td><td>1000</td><td></td><td></td><td></td><td></td><td></td><td></td><td></td><td></td><td></td></tr>
<tr><td>Oligo fwd</td><td>CACCGACGATCACTGACGCGTCGC</td><td></td><td></td><td></td><td></td><td></td><td></td><td></td><td></td><td></td><td></td></tr>
<tr><td>Oligo rev</td><td>AAACGCGACGCGTCAGTGATCGTC</td><td></td><td></td><td></td><td></td><td></td><td></td><td></td><td></td><td></td><td></td></tr>
<tr><td>Chromosome</td><td>start</td><td>end</td><td>strand</td><td>MM</td><td>target_seq</td><td>PAM</td><td>alignment</td><td>distance</td><td>position</td><td>gene name</td><td>gene id</td></tr>
<tr><td>GU305914.1</td><td>167946</td><td>167968</td><td>-</td><td>0</td><td>GACGATCACTGACGCGTCGC</td><td>GGG</td><td>||||||||[||||||||||||]PAM</td><td>NA</td><td>Intergenic</td><td>NA</td><td>NA</td></tr>
<tr><td></td><td></td><td></td><td></td><td></td><td></td><td></td><td></td><td></td><td></td><td></td><td></td></tr>
<tr><td>T21745</td><td>CGCGTCGCGGGTTCCGAGCACGG</td><td>1000</td><td></td><td></td><td></td><td></td><td></td><td></td><td></td><td></td><td></td></tr>
<tr><td>Oligo fwd</td><td>CACCCGCGTCGCGGGTTCCGAGCA</td><td></td><td></td><td></td><td></td><td></td><td></td><td></td><td></td><td></td><td></td></tr>
<tr><td>Oligo rev</td><td>AAACTGCTCGGAACCCGCGACGCG</td><td></td><td></td><td></td><td></td><td></td><td></td><td></td><td></td><td></td><td></td></tr>
<tr><td>Chromosome</td><td>start</td><td>end</td><td>strand</td><td>MM</td><td>target_seq</td><td>PAM</td><td>alignment</td><td>distance</td><td>position</td><td>gene name</td><td>gene id</td></tr>
<tr><td>GU305914.1</td><td>167934</td><td>167956</td><td>-</td><td>0</td><td>CGCGTCGCGGGTTCCGAGCA</td><td>CGG</td><td>||||||||[||||||||||||]PAM</td><td>NA</td><td>Intergenic</td><td>NA</td><td>NA</td></tr>
<tr><td></td><td></td><td></td><td></td><td></td><td></td><td></td><td></td><td></td><td></td><td></td><td></td></tr>
<tr><td>T21746</td><td>GCGGGTTCCGAGCACGGTCACGG</td><td>1000</td><td></td><td></td><td></td><td></td><td></td><td></td><td></td><td></td><td></td></tr>
</table>

Oligo fwd CACCGCGGGTTCCGAGCACGGTCA
Oligo rev AAACTGACCGTGCTCGGAACCCGC
Chromosome start end strand MM target_seq PAM alignment distance position gene name gene id
GU305914.1 167928 167950 - 0 GCGGGTTCCGAGCACGGTCA CGG ||||||||[||||||||||||]PAM NA Intergenic NA NA

T21747 CATCACCGCCATCACCACCAAGG 1000
Oligo fwd CACCCATCACCGCCATCACCACCA
Oligo rev AAACTGGTGGTGATGGCGGTGATG
Chromosome start end strand MM target_seq PAM alignment distance position gene name gene id
GU305914.1 167814 167836 - 0 CATCACCGCCATCACCACCA AGG ||||||||[||||||||||||]PAM NA Intergenic NA NA

T21748 CTACTAATAAAGATAATGAATGG 1000
Oligo fwd CACCCTACTAATAAAGATAATGAA
Oligo rev AAACTTCATTATCTTTATTAGTAG
Chromosome start end strand MM target_seq PAM alignment distance position gene name gene id
GU305914.1 167659 167681 - 0 CTACTAATAAAGATAATGAA TGG ||||||||[||||||||||||]PAM NA Intergenic NA NA

T21749 TAGCGCCGCGTTCCGCACGCAGG 1000
Oligo fwd CACCTAGCGCCGCGTTCCGCACGC
Oligo rev AAACGCGTGCGGAACGCGGCGCTA
Chromosome start end strand MM target_seq PAM alignment distance position gene name gene id
GU305914.1 167607 167629 - 0 TAGCGCCGCGTTCCGCACGC AGG ||||||||[||||||||||||]PAM NA Intergenic NA NA

T21750 CCCACCATCACCACATTCCCCGG 1000
Oligo fwd CACCCCCACCATCACCACATTCCC
Oligo rev AAACGGGAATGTGGTGATGGTGGG
Chromosome start end strand MM target_seq PAM alignment distance position gene name gene id
GU305914.1 167558 167580 - 0 CCCACCATCACCACATTCCC CGG ||||||||[||||||||||||]PAM NA Intergenic NA NA

T21751 CCGGAGACCTATGCACTCGCAGG 1000
Oligo fwd CACCCCGGAGACCTATGCACTCGC
Oligo rev AAACGCGAGTGCATAGGTCTCCGG
Chromosome start end strand MM target_seq PAM alignment distance position gene name gene id
GU305914.1 167539 167561 - 0 CCGGAGACCTATGCACTCGC AGG ||||||||[||||||||||||]PAM NA Intergenic NA NA

T21752 AGGTACTGCCCCGCGAAATACGG 1000
Oligo fwd CACCAGGTACTGCCCCGCGAAATA
Oligo rev AAACTATTTCGCGGGGCAGTACCT
Chromosome start end strand MM target_seq PAM alignment distance position gene name gene id
GU305914.1 167519 167541 - 0 AGGTACTGCCCCGCGAAATA CGG ||||||||[||||||||||||]PAM NA Intergenic NA NA

T21753 GGTACTGCCCCGCGAAATACGGG 1000
Oligo fwd CACCGGTACTGCCCCGCGAAATAC
Oligo rev AAACGTATTTCGCGGGGCAGTACC
Chromosome start end strand MM target_seq PAM alignment distance position gene name gene id
GU305914.1 167518 167540 - 0 GGTACTGCCCCGCGAAATAC GGG ||||||||[||||||||||||]PAM NA Intergenic NA NA

T21754 CCGAGTGACTTTCAGTCGCGCGG 1000
Oligo fwd CACCCCGAGTGACTTTCAGTCGCG
Oligo rev AAACCGCGACTGAAAGTCACTCGG
Chromosome start end strand MM target_seq PAM alignment distance position gene name gene id
GU305914.1 167472 167494 - 0 CCGAGTGACTTTCAGTCGCG CGG ||||||||[||||||||||||]PAM NA Intergenic NA NA

T21755 CGACTCGCTCATTATCGATCCGG 1000
Oligo fwd CACCCGACTCGCTCATTATCGATC
Oligo rev AAACGATCGATAATGAGCGAGTCG
Chromosome start end strand MM target_seq PAM alignment distance position gene name gene id
GU305914.1 167421 167443 - 0 CGACTCGCTCATTATCGATC CGG ||||||||[||||||||||||]PAM NA Intergenic NA NA

T21756 GACTCGCTCATTATCGATCCGGG 1000
Oligo fwd CACCGACTCGCTCATTATCGATCC
Oligo rev AAACGGATCGATAATGAGCGAGTC
Chromosome start end strand MM target_seq PAM alignment distance position gene name gene id
GU305914.1 167420 167442 - 0 GACTCGCTCATTATCGATCC GGG ||||||||[||||||||||||]PAM NA Intergenic NA NA

T21757 GCTCATTATCGATCCGGGCCTGG 1000
Oligo fwd CACCGCTCATTATCGATCCGGGCC
Oligo rev AAACGGCCCGGATCGATAATGAGC
Chromosome start end strand MM target_seq PAM alignment distance position gene name gene id
GU305914.1 167415 167437 - 0 GCTCATTATCGATCCGGGCC TGG ||||||||[||||||||||||]PAM NA Intergenic NA NA

T21758 CTCATTATCGATCCGGGCCTGGG 1000
Oligo fwd CACCCTCATTATCGATCCGGGCCT
Oligo rev AAACAGGCCCGGATCGATAATGAG
Chromosome start end strand MM target_seq PAM alignment distance position gene name gene id
GU305914.1 167414 167436 - 0 CTCATTATCGATCCGGGCCT GGG ||||||||[||||||||||||]PAM NA Intergenic NA NA

T21759 CGATCCGGGCCTGGGCGCTTCGG 1000
Oligo fwd CACCCGATCCGGGCCTGGGCGCTT
Oligo rev AAACAAGCGCCCAGGCCCGGATCG
Chromosome start end strand MM target_seq PAM alignment distance position gene name gene id
GU305914.1 167406 167428 - 0 CGATCCGGGCCTGGGCGCTT CGG ||||||||[||||||||||||]PAM NA Intergenic NA NA

T21760 AAAGACTCAGTCTCTCAAACAGG 1000
Oligo fwd CACCAAAGACTCAGTCTCTCAAAC
Oligo rev AAACGTTTGAGAGACTGAGTCTTT
Chromosome start end strand MM target_seq PAM alignment distance position gene name gene id
GU305914.1 167365 167387 - 0 AAAGACTCAGTCTCTCAAAC AGG ||||||||[||||||||||||]PAM NA Intergenic NA NA

<table>
<tr><td>T21761</td><td>GTCTCTCAAACAGGATGATACGG</td><td>1000</td><td></td><td></td><td></td><td></td><td></td><td></td><td></td><td></td><td></td></tr>
<tr><td>Oligo fwd</td><td>CACCGTCTCTCAAACAGGATGATA</td><td></td><td></td><td></td><td></td><td></td><td></td><td></td><td></td><td></td><td></td></tr>
<tr><td>Oligo rev</td><td>AAACTATCATCCTGTTTGAGAGAC</td><td></td><td></td><td></td><td></td><td></td><td></td><td></td><td></td><td></td><td></td></tr>
<tr><td>Chromosome</td><td>start</td><td>end</td><td>strand</td><td>MM</td><td>target_seq</td><td>PAM</td><td>alignment</td><td>distance</td><td>position</td><td>gene name</td><td>gene id</td></tr>
<tr><td>GU305914.1</td><td>167356</td><td>167378</td><td>-</td><td>0</td><td>GTCTCTCAAACAGGATGATA</td><td>CGG</td><td>||||||||[||||||||||||]PAM</td><td>NA</td><td>Intergenic</td><td>NA</td><td>NA</td></tr>
<tr><td>T21762</td><td>GCTGTCCACACGTAACCTGCTGG</td><td>1000</td><td></td><td></td><td></td><td></td><td></td><td></td><td></td><td></td><td></td></tr>
<tr><td>Oligo fwd</td><td>CACCGCTGTCCACACGTAACCTGC</td><td></td><td></td><td></td><td></td><td></td><td></td><td></td><td></td><td></td><td></td></tr>
<tr><td>Oligo rev</td><td>AAACGCAGGTTACGTGTGGACAGC</td><td></td><td></td><td></td><td></td><td></td><td></td><td></td><td></td><td></td><td></td></tr>
<tr><td>Chromosome</td><td>start</td><td>end</td><td>strand</td><td>MM</td><td>target_seq</td><td>PAM</td><td>alignment</td><td>distance</td><td>position</td><td>gene name</td><td>gene id</td></tr>
<tr><td>GU305914.1</td><td>167260</td><td>167282</td><td>-</td><td>0</td><td>GCTGTCCACACGTAACCTGC</td><td>TGG</td><td>||||||||[||||||||||||]PAM</td><td>NA</td><td>Intergenic</td><td>NA</td><td>NA</td></tr>
<tr><td>T21763</td><td>CTGTCCACACGTAACCTGCTGGG</td><td>1000</td><td></td><td></td><td></td><td></td><td></td><td></td><td></td><td></td><td></td></tr>
<tr><td>Oligo fwd</td><td>CACCCTGTCCACACGTAACCTGCT</td><td></td><td></td><td></td><td></td><td></td><td></td><td></td><td></td><td></td><td></td></tr>
<tr><td>Oligo rev</td><td>AAACAGCAGGTTACGTGTGGACAG</td><td></td><td></td><td></td><td></td><td></td><td></td><td></td><td></td><td></td><td></td></tr>
<tr><td>Chromosome</td><td>start</td><td>end</td><td>strand</td><td>MM</td><td>target_seq</td><td>PAM</td><td>alignment</td><td>distance</td><td>position</td><td>gene name</td><td>gene id</td></tr>
<tr><td>GU305914.1</td><td>167259</td><td>167281</td><td>-</td><td>0</td><td>CTGTCCACACGTAACCTGCT</td><td>GGG</td><td>||||||||[||||||||||||]PAM</td><td>NA</td><td>Intergenic</td><td>NA</td><td>NA</td></tr>
<tr><td>T21764</td><td>TGTCCACACGTAACCTGCTGGGG</td><td>1000</td><td></td><td></td><td></td><td></td><td></td><td></td><td></td><td></td><td></td></tr>
<tr><td>Oligo fwd</td><td>CACCTGTCCACACGTAACCTGCTG</td><td></td><td></td><td></td><td></td><td></td><td></td><td></td><td></td><td></td><td></td></tr>
<tr><td>Oligo rev</td><td>AAACCAGCAGGTTACGTGTGGACA</td><td></td><td></td><td></td><td></td><td></td><td></td><td></td><td></td><td></td><td></td></tr>
<tr><td>Chromosome</td><td>start</td><td>end</td><td>strand</td><td>MM</td><td>target_seq</td><td>PAM</td><td>alignment</td><td>distance</td><td>position</td><td>gene name</td><td>gene id</td></tr>
<tr><td>GU305914.1</td><td>167258</td><td>167280</td><td>-</td><td>0</td><td>TGTCCACACGTAACCTGCTG</td><td>GGG</td><td>||||||||[||||||||||||]PAM</td><td>NA</td><td>Intergenic</td><td>NA</td><td>NA</td></tr>
<tr><td>T21765</td><td>GTCCACACGTAACCTGCTGGGGG</td><td>1000</td><td></td><td></td><td></td><td></td><td></td><td></td><td></td><td></td><td></td></tr>
<tr><td>Oligo fwd</td><td>CACCGTCCACACGTAACCTGCTGG</td><td></td><td></td><td></td><td></td><td></td><td></td><td></td><td></td><td></td><td></td></tr>
<tr><td>Oligo rev</td><td>AAACCCAGCAGGTTACGTGTGGAC</td><td></td><td></td><td></td><td></td><td></td><td></td><td></td><td></td><td></td><td></td></tr>
<tr><td>Chromosome</td><td>start</td><td>end</td><td>strand</td><td>MM</td><td>target_seq</td><td>PAM</td><td>alignment</td><td>distance</td><td>position</td><td>gene name</td><td>gene id</td></tr>
<tr><td>GU305914.1</td><td>167257</td><td>167279</td><td>-</td><td>0</td><td>GTCCACACGTAACCTGCTGG</td><td>GGG</td><td>||||||||[||||||||||||]PAM</td><td>NA</td><td>Intergenic</td><td>NA</td><td>NA</td></tr>
<tr><td>T21766</td><td>CACACGTAACCTGCTGGGGGCGG</td><td>1000</td><td></td><td></td><td></td><td></td><td></td><td></td><td></td><td></td><td></td></tr>
<tr><td>Oligo fwd</td><td>CACCCACACGTAACCTGCTGGGGG</td><td></td><td></td><td></td><td></td><td></td><td></td><td></td><td></td><td></td><td></td></tr>
<tr><td>Oligo rev</td><td>AAACCCCCCAGCAGGTTACGTGTG</td><td></td><td></td><td></td><td></td><td></td><td></td><td></td><td></td><td></td><td></td></tr>
<tr><td>Chromosome</td><td>start</td><td>end</td><td>strand</td><td>MM</td><td>target_seq</td><td>PAM</td><td>alignment</td><td>distance</td><td>position</td><td>gene name</td><td>gene id</td></tr>
<tr><td>GU305914.1</td><td>167254</td><td>167276</td><td>-</td><td>0</td><td>CACACGTAACCTGCTGGGGG</td><td>CGG</td><td>||||||||[||||||||||||]PAM</td><td>NA</td><td>Intergenic</td><td>NA</td><td>NA</td></tr>
<tr><td>T21767</td><td>GCTGGGGGCGGCGCAGTCGCAGG</td><td>1000</td><td></td><td></td><td></td><td></td><td></td><td></td><td></td><td></td><td></td></tr>
<tr><td>Oligo fwd</td><td>CACCGCTGGGGGCGGCGCAGTCGC</td><td></td><td></td><td></td><td></td><td></td><td></td><td></td><td></td><td></td><td></td></tr>
<tr><td>Oligo rev</td><td>AAACGCGACTGCGCCGCCCCCAGC</td><td></td><td></td><td></td><td></td><td></td><td></td><td></td><td></td><td></td><td></td></tr>
<tr><td>Chromosome</td><td>start</td><td>end</td><td>strand</td><td>MM</td><td>target_seq</td><td>PAM</td><td>alignment</td><td>distance</td><td>position</td><td>gene name</td><td>gene id</td></tr>
<tr><td>GU305914.1</td><td>167242</td><td>167264</td><td>-</td><td>0</td><td>GCTGGGGGCGGCGCAGTCGC</td><td>AGG</td><td>||||||||[||||||||||||]PAM</td><td>NA</td><td>Intergenic</td><td>NA</td><td>NA</td></tr>
<tr><td>T21768</td><td>GGCGCAGTCGCAGGAGTCGCCGG</td><td>1000</td><td></td><td></td><td></td><td></td><td></td><td></td><td></td><td></td><td></td></tr>
<tr><td>Oligo fwd</td><td>CACCGGCGCAGTCGCAGGAGTCGC</td><td></td><td></td><td></td><td></td><td></td><td></td><td></td><td></td><td></td><td></td></tr>
<tr><td>Oligo rev</td><td>AAACGCGACTCCTGCGACTGCGCC</td><td></td><td></td><td></td><td></td><td></td><td></td><td></td><td></td><td></td><td></td></tr>
<tr><td>Chromosome</td><td>start</td><td>end</td><td>strand</td><td>MM</td><td>target_seq</td><td>PAM</td><td>alignment</td><td>distance</td><td>position</td><td>gene name</td><td>gene id</td></tr>
<tr><td>GU305914.1</td><td>167233</td><td>167255</td><td>-</td><td>0</td><td>GGCGCAGTCGCAGGAGTCGC</td><td>CGG</td><td>||||||||[||||||||||||]PAM</td><td>NA</td><td>Intergenic</td><td>NA</td><td>NA</td></tr>
<tr><td>T21769</td><td>TCGCAGGAGTCGCCGGTCGCCGG</td><td>1000</td><td></td><td></td><td></td><td></td><td></td><td></td><td></td><td></td><td></td></tr>
<tr><td>Oligo fwd</td><td>CACCTCGCAGGAGTCGCCGGTCGC</td><td></td><td></td><td></td><td></td><td></td><td></td><td></td><td></td><td></td><td></td></tr>
<tr><td>Oligo rev</td><td>AAACGCGACCGGCGACTCCTGCGA</td><td></td><td></td><td></td><td></td><td></td><td></td><td></td><td></td><td></td><td></td></tr>
<tr><td>Chromosome</td><td>start</td><td>end</td><td>strand</td><td>MM</td><td>target_seq</td><td>PAM</td><td>alignment</td><td>distance</td><td>position</td><td>gene name</td><td>gene id</td></tr>
<tr><td>GU305914.1</td><td>167226</td><td>167248</td><td>-</td><td>0</td><td>TCGCAGGAGTCGCCGGTCGC</td><td>CGG</td><td>||||||||[||||||||||||]PAM</td><td>NA</td><td>Intergenic</td><td>NA</td><td>NA</td></tr>
<tr><td>T21770</td><td>CGCCGGTCGCCGGCGAGCGACGG</td><td>1000</td><td></td><td></td><td></td><td></td><td></td><td></td><td></td><td></td><td></td></tr>
<tr><td>Oligo fwd</td><td>CACCCGCCGGTCGCCGGCGAGCGA</td><td></td><td></td><td></td><td></td><td></td><td></td><td></td><td></td><td></td><td></td></tr>
<tr><td>Oligo rev</td><td>AAACTCGCTCGCCGGCGACCGGCG</td><td></td><td></td><td></td><td></td><td></td><td></td><td></td><td></td><td></td><td></td></tr>
<tr><td>Chromosome</td><td>start</td><td>end</td><td>strand</td><td>MM</td><td>target_seq</td><td>PAM</td><td>alignment</td><td>distance</td><td>position</td><td>gene name</td><td>gene id</td></tr>
<tr><td>GU305914.1</td><td>167216</td><td>167238</td><td>-</td><td>0</td><td>CGCCGGTCGCCGGCGAGCGA</td><td>CGG</td><td>||||||||[||||||||||||]PAM</td><td>NA</td><td>Intergenic</td><td>NA</td><td>NA</td></tr>
<tr><td>GU305914.1</td><td>100051</td><td>100073</td><td>+</td><td>4</td><td>CCCCGATCGCCGCCGTGCGA</td><td>GGG</td><td>|-|||-||[||||-||-||||]PAM</td><td>NA</td><td>Intergenic</td><td>NA</td><td>NA</td></tr>
<tr><td>T21771</td><td>GCCGGTCGCCGGCGAGCGACGGG</td><td>1000</td><td></td><td></td><td></td><td></td><td></td><td></td><td></td><td></td><td></td></tr>
<tr><td>Oligo fwd</td><td>CACCGCCGGTCGCCGGCGAGCGAC</td><td></td><td></td><td></td><td></td><td></td><td></td><td></td><td></td><td></td><td></td></tr>
<tr><td>Oligo rev</td><td>AAACGTCGCTCGCCGGCGACCGGC</td><td></td><td></td><td></td><td></td><td></td><td></td><td></td><td></td><td></td><td></td></tr>
<tr><td>Chromosome</td><td>start</td><td>end</td><td>strand</td><td>MM</td><td>target_seq</td><td>PAM</td><td>alignment</td><td>distance</td><td>position</td><td>gene name</td><td>gene id</td></tr>
<tr><td>GU305914.1</td><td>167215</td><td>167237</td><td>-</td><td>0</td><td>GCCGGTCGCCGGCGAGCGAC</td><td>GGG</td><td>||||||||[||||||||||||]PAM</td><td>NA</td><td>Intergenic</td><td>NA</td><td>NA</td></tr>
<tr><td>T21772</td><td>CGCCGGCGAGCGACGGGCGTTGG</td><td>1000</td><td></td><td></td><td></td><td></td><td></td><td></td><td></td><td></td><td></td></tr>
<tr><td>Oligo fwd</td><td>CACCCGCCGGCGAGCGACGGGCGT</td><td></td><td></td><td></td><td></td><td></td><td></td><td></td><td></td><td></td><td></td></tr>
<tr><td>Oligo rev</td><td>AAACACGCCCGTCGCTCGCCGGCG</td><td></td><td></td><td></td><td></td><td></td><td></td><td></td><td></td><td></td><td></td></tr>
<tr><td>Chromosome</td><td>start</td><td>end</td><td>strand</td><td>MM</td><td>target_seq</td><td>PAM</td><td>alignment</td><td>distance</td><td>position</td><td>gene name</td><td>gene id</td></tr>
<tr><td>GU305914.1</td><td>167209</td><td>167231</td><td>-</td><td>0</td><td>CGCCGGCGAGCGACGGGCGT</td><td>TGG</td><td>||||||||[||||||||||||]PAM</td><td>NA</td><td>Intergenic</td><td>NA</td><td>NA</td></tr>
<tr><td>T21773</td><td>CGTCGAACACAAGTGCCGCGAGG</td><td>1000</td><td></td><td></td><td></td><td></td><td></td><td></td><td></td><td></td><td></td></tr>
<tr><td>Oligo fwd</td><td>CACCCGTCGAACACAAGTGCCGCG</td><td></td><td></td><td></td><td></td><td></td><td></td><td></td><td></td><td></td><td></td></tr>
<tr><td>Oligo rev</td><td>AAACCGCGGCACTTGTGTTCGACG</td><td></td><td></td><td></td><td></td><td></td><td></td><td></td><td></td><td></td><td></td></tr>
<tr><td>Chromosome</td><td>start</td><td>end</td><td>strand</td><td>MM</td><td>target_seq</td><td>PAM</td><td>alignment</td><td>distance</td><td>position</td><td>gene name</td><td>gene id</td></tr>
<tr><td>GU305914.1</td><td>167137</td><td>167159</td><td>-</td><td>0</td><td>CGTCGAACACAAGTGCCGCG</td><td>AGG</td><td>||||||||[||||||||||||]PAM</td><td>NA</td><td>Intergenic</td><td>NA</td><td>NA</td></tr>
<tr><td>T21774</td><td>ACACAAGTGCCGCGAGGCCCTGG</td><td>1000</td><td></td><td></td><td></td><td></td><td></td><td></td><td></td><td></td><td></td></tr>
<tr><td>Oligo fwd</td><td>CACCACACAAGTGCCGCGAGGCCC</td><td></td><td></td><td></td><td></td><td></td><td></td><td></td><td></td><td></td><td></td></tr>
<tr><td>Oligo rev</td><td>AAACGGGCCTCGCGGCACTTGTGT</td><td></td><td></td><td></td><td></td><td></td><td></td><td></td><td></td><td></td><td></td></tr>
<tr><td>Chromosome</td><td>start</td><td>end</td><td>strand</td><td>MM</td><td>target_seq</td><td>PAM</td><td>alignment</td><td>distance</td><td>position</td><td>gene name</td><td>gene id</td></tr>
<tr><td>GU305914.1</td><td>167131</td><td>167153</td><td>-</td><td>0</td><td>ACACAAGTGCCGCGAGGCCC</td><td>TGG</td><td>||||||||[||||||||||||]PAM</td><td>NA</td><td>Intergenic</td><td>NA</td><td>NA</td></tr>
<tr><td>T21775</td><td>GCCCTGGCGAGCTGCTACAACGG</td><td>1000</td><td></td><td></td><td></td><td></td><td></td><td></td><td></td><td></td><td></td></tr>
<tr><td>Oligo fwd</td><td>CACCGCCCTGGCGAGCTGCTACAA</td><td></td><td></td><td></td><td></td><td></td><td></td><td></td><td></td><td></td><td></td></tr>
<tr><td>Oligo rev</td><td>AAACTTGTAGCAGCTCGCCAGGGC</td><td></td><td></td><td></td><td></td><td></td><td></td><td></td><td></td><td></td><td></td></tr>
<tr><td>Chromosome</td><td>start</td><td>end</td><td>strand</td><td>MM</td><td>target_seq</td><td>PAM</td><td>alignment</td><td>distance</td><td>position</td><td>gene name</td><td>gene id</td></tr>
</table>

| | | | | | | | | | | | |
|---|---|---|---|---|---|---|---|---|---|---|---|
| GU305914.1 | 167115 | 167137 | - | 0 | GCCCTGGCGAGCTGCTACAA | CGG | ||||||||[||||||||||||]PAM | NA | Intergenic | NA | NA |
| T21776 | CCCTGGCGAGCTGCTACAACGGG | 1000 | | | | | | | | | |
| Oligo fwd | CACCCCCTGGCGAGCTGCTACAAC | | | | | | | | | | |
| Oligo rev | AAACGTTGTAGCAGCTCGCCAGGG | | | | | | | | | | |
| Chromosome | start | end | strand | MM | target_seq | PAM | alignment | distance | position | gene name | gene id |
| GU305914.1 | 167114 | 167136 | - | 0 | CCCTGGCGAGCTGCTACAAC | GGG | ||||||||[||||||||||||]PAM | NA | Intergenic | NA | NA |
| T21777 | TTCCAACCGCTGCACGATGACGG | 1000 | | | | | | | | | |
| Oligo fwd | CACCTTCCAACCGCTGCACGATGA | | | | | | | | | | |
| Oligo rev | AAACTCATCGTGCAGCGGTTGGAA | | | | | | | | | | |
| Chromosome | start | end | strand | MM | target_seq | PAM | alignment | distance | position | gene name | gene id |
| GU305914.1 | 167085 | 167107 | - | 0 | TTCCAACCGCTGCACGATGA | CGG | ||||||||[||||||||||||]PAM | NA | Intergenic | NA | NA |
| T21778 | CGATGACGGCCCCATCCGTCCGG | 1000 | | | | | | | | | |
| Oligo fwd | CACCCGATGACGGCCCCATCCGTC | | | | | | | | | | |
| Oligo rev | AAACGACGGATGGGGCCGTCATCG | | | | | | | | | | |
| Chromosome | start | end | strand | MM | target_seq | PAM | alignment | distance | position | gene name | gene id |
| GU305914.1 | 167071 | 167093 | - | 0 | CGATGACGGCCCCATCCGTC | CGG | ||||||||[||||||||||||]PAM | NA | Intergenic | NA | NA |
| T21779 | CCATCCGTCCGGACCCGTATAGG | 1000 | | | | | | | | | |
| Oligo fwd | CACCCCATCCGTCCGGACCCGTAT | | | | | | | | | | |
| Oligo rev | AAACATACGGGTCCGGACGGATGG | | | | | | | | | | |
| Chromosome | start | end | strand | MM | target_seq | PAM | alignment | distance | position | gene name | gene id |
| GU305914.1 | 167060 | 167082 | - | 0 | CCATCCGTCCGGACCCGTAT | AGG | ||||||||[||||||||||||]PAM | NA | Intergenic | NA | NA |
| T21780 | ATAGGTTCTCGACGATGATCAGG | 1000 | | | | | | | | | |
| Oligo fwd | CACCATAGGTTCTCGACGATGATC | | | | | | | | | | |
| Oligo rev | AAACGATCATCGTCGAGAACCTAT | | | | | | | | | | |
| Chromosome | start | end | strand | MM | target_seq | PAM | alignment | distance | position | gene name | gene id |
| GU305914.1 | 167042 | 167064 | - | 0 | ATAGGTTCTCGACGATGATC | AGG | ||||||||[||||||||||||]PAM | NA | Intergenic | NA | NA |
| T21781 | CGACGATGATCAGGTTCAAGAGG | 1000 | | | | | | | | | |
| Oligo fwd | CACCCGACGATGATCAGGTTCAAG | | | | | | | | | | |
| Oligo rev | AAACCTTGAACCTGATCATCGTCG | | | | | | | | | | |
| Chromosome | start | end | strand | MM | target_seq | PAM | alignment | distance | position | gene name | gene id |
| GU305914.1 | 167033 | 167055 | - | 0 | CGACGATGATCAGGTTCAAG | AGG | ||||||||[||||||||||||]PAM | NA | Intergenic | NA | NA |
| T21782 | ATCAGGTTCAAGAGGAGCTACGG | 1000 | | | | | | | | | |
| Oligo fwd | CACCATCAGGTTCAAGAGGAGCTA | | | | | | | | | | |
| Oligo rev | AAACTAGCTCCTCTTGAACCTGAT | | | | | | | | | | |
| Chromosome | start | end | strand | MM | target_seq | PAM | alignment | distance | position | gene name | gene id |
| GU305914.1 | 167025 | 167047 | - | 0 | ATCAGGTTCAAGAGGAGCTA | CGG | ||||||||[||||||||||||]PAM | NA | Intergenic | NA | NA |
| T21783 | CGAGCTCAACGACGAGTTCCTGG | 1000 | | | | | | | | | |
| Oligo fwd | CACCCGAGCTCAACGACGAGTTCC | | | | | | | | | | |
| Oligo rev | AAACGGAACTCGTCGTTGAGCTCG | | | | | | | | | | |
| Chromosome | start | end | strand | MM | target_seq | PAM | alignment | distance | position | gene name | gene id |
| GU305914.1 | 166984 | 167006 | - | 0 | CGAGCTCAACGACGAGTTCC | TGG | ||||||||[||||||||||||]PAM | NA | Intergenic | NA | NA |
| T21784 | GTCCCTGCTCCACAACAACACGG | 1000 | | | | | | | | | |
| Oligo fwd | CACCGTCCCTGCTCCACAACAACA | | | | | | | | | | |
| Oligo rev | AAACTGTTGTTGTGGAGCAGGGAC | | | | | | | | | | |
| Chromosome | start | end | strand | MM | target_seq | PAM | alignment | distance | position | gene name | gene id |
| GU305914.1 | 166954 | 166976 | - | 0 | GTCCCTGCTCCACAACAACA | CGG | ||||||||[||||||||||||]PAM | NA | Intergenic | NA | NA |
| T21785 | GAGTTCTGCTGACGCTGATGCGG | 1000 | | | | | | | | | |
| Oligo fwd | CACCGAGTTCTGCTGACGCTGATG | | | | | | | | | | |
| Oligo rev | AAACCATCAGCGTCAGCAGAACTC | | | | | | | | | | |
| Chromosome | start | end | strand | MM | target_seq | PAM | alignment | distance | position | gene name | gene id |
| GU305914.1 | 166922 | 166944 | - | 0 | GAGTTCTGCTGACGCTGATG | CGG | ||||||||[||||||||||||]PAM | NA | Intergenic | NA | NA |
| T21786 | GGACCAGCCGCGCCAGCGACTGG | 1000 | | | | | | | | | |
| Oligo fwd | CACCGGACCAGCCGCGCCAGCGAC | | | | | | | | | | |
| Oligo rev | AAACGTCGCTGGCGCGGCTGGTCC | | | | | | | | | | |
| Chromosome | start | end | strand | MM | target_seq | PAM | alignment | distance | position | gene name | gene id |
| GU305914.1 | 166901 | 166923 | - | 0 | GGACCAGCCGCGCCAGCGAC | TGG | ||||||||[||||||||||||]PAM | NA | Intergenic | NA | NA |
| T21787 | CAGCGACTGGATGAGCTTCTTGG | 1000 | | | | | | | | | |
| Oligo fwd | CACCCAGCGACTGGATGAGCTTCT | | | | | | | | | | |
| Oligo rev | AAACAGAAGCTCATCCAGTCGCTG | | | | | | | | | | |
| Chromosome | start | end | strand | MM | target_seq | PAM | alignment | distance | position | gene name | gene id |
| GU305914.1 | 166888 | 166910 | - | 0 | CAGCGACTGGATGAGCTTCT | TGG | ||||||||[||||||||||||]PAM | NA | Intergenic | NA | NA |
| T21788 | AGCGACTGGATGAGCTTCTTGGG | 1000 | | | | | | | | | |
| Oligo fwd | CACCAGCGACTGGATGAGCTTCTT | | | | | | | | | | |
| Oligo rev | AAACAAGAAGCTCATCCAGTCGCT | | | | | | | | | | |
| Chromosome | start | end | strand | MM | target_seq | PAM | alignment | distance | position | gene name | gene id |
| GU305914.1 | 166887 | 166909 | - | 0 | AGCGACTGGATGAGCTTCTT | GGG | ||||||||[||||||||||||]PAM | NA | Intergenic | NA | NA |
| GU305914.1 | 181082 | 181104 | - | 4 | AGAGTCTGGATGAGCTACGT | AGG | ||-|-|||[||||||||-|-|]PAM | NA | Intergenic | NA | NA |
| T21789 | GCGACTGGATGAGCTTCTTGGGG | 1000 | | | | | | | | | |
| Oligo fwd | CACCGCGACTGGATGAGCTTCTTG | | | | | | | | | | |
| Oligo rev | AAACCAAGAAGCTCATCCAGTCGC | | | | | | | | | | |
| Chromosome | start | end | strand | MM | target_seq | PAM | alignment | distance | position | gene name | gene id |
| GU305914.1 | 166886 | 166908 | - | 0 | GCGACTGGATGAGCTTCTTG | GGG | ||||||||[||||||||||||]PAM | NA | Intergenic | NA | NA |
| T21790 | CGACTGGATGAGCTTCTTGGGGG | 1000 | | | | | | | | | |
| Oligo fwd | CACCCGACTGGATGAGCTTCTTGG | | | | | | | | | | |

Oligo rev AAACCCAAGAAGCTCATCCAGTCG
Chromosome start end strand MM target_seq PAM alignment distance position gene name gene id
GU305914.1 166885 166907 - 0 CGACTGGATGAGCTTCTTGG GGG ||||||||[||||||||||||]PAM NA Intergenic NA NA

T21791 GACTGGATGAGCTTCTTGGGGGG 1000
Oligo fwd CACCGACTGGATGAGCTTCTTGGG
Oligo rev AAACCCCAAGAAGCTCATCCAGTC
Chromosome start end strand MM target_seq PAM alignment distance position gene name gene id
GU305914.1 166884 166906 - 0 GACTGGATGAGCTTCTTGGG GGG ||||||||[||||||||||||]PAM NA Intergenic NA NA

T21792 CCAGTGCGACGCGCCGAAGTCGG 1000
Oligo fwd CACCCCAGTGCGACGCGCCGAAGT
Oligo rev AAACACTTCGGCGCGTCGCACTGG
Chromosome start end strand MM target_seq PAM alignment distance position gene name gene id
GU305914.1 166855 166877 - 0 CCAGTGCGACGCGCCGAAGT CGG ||||||||[||||||||||||]PAM NA Intergenic NA NA

T21793 GTGCGACGCGCCGAAGTCGGTGG 1000
Oligo fwd CACCGTGCGACGCGCCGAAGTCGG
Oligo rev AAACCCGACTTCGGCGCGTCGCAC
Chromosome start end strand MM target_seq PAM alignment distance position gene name gene id
GU305914.1 166852 166874 - 0 GTGCGACGCGCCGAAGTCGG TGG ||||||||[||||||||||||]PAM NA Intergenic NA NA

T21794 GTCGGTGGTGTTCACCTGCGTGG 1000
Oligo fwd CACCGTCGGTGGTGTTCACCTGCG
Oligo rev AAACCGCAGGTGAACACCACCGAC
Chromosome start end strand MM target_seq PAM alignment distance position gene name gene id
GU305914.1 166837 166859 - 0 GTCGGTGGTGTTCACCTGCG TGG ||||||||[||||||||||||]PAM NA Intergenic NA NA

T21795 GCTACGAGCACGACCTGATGCGG 1000
Oligo fwd CACCGCTACGAGCACGACCTGATG
Oligo rev AAACCATCAGGTCGTGCTCGTAGC
Chromosome start end strand MM target_seq PAM alignment distance position gene name gene id
GU305914.1 166805 166827 - 0 GCTACGAGCACGACCTGATG CGG ||||||||[||||||||||||]PAM NA Intergenic NA NA

T21796 CTACACGACCGACCTGTTCACGG 1000
Oligo fwd CACCCTACACGACCGACCTGTTCA
Oligo rev AAACTGAACAGGTCGGTCGTGTAG
Chromosome start end strand MM target_seq PAM alignment distance position gene name gene id
GU305914.1 166777 166799 - 0 CTACACGACCGACCTGTTCA CGG ||||||||[||||||||||||]PAM NA Intergenic NA NA

T21797 CCTGTTCACGGAGAACGTGTTGG 1000
Oligo fwd CACCCCTGTTCACGGAGAACGTGT
Oligo rev AAACACACGTTCTCCGTGAACAGG
Chromosome start end strand MM target_seq PAM alignment distance position gene name gene id
GU305914.1 166765 166787 - 0 CCTGTTCACGGAGAACGTGT TGG ||||||||[||||||||||||]PAM NA Intergenic NA NA

T21798 CTGTTCACGGAGAACGTGTTGGG 1000
Oligo fwd CACCCTGTTCACGGAGAACGTGTT
Oligo rev AAACAACACGTTCTCCGTGAACAG
Chromosome start end strand MM target_seq PAM alignment distance position gene name gene id
GU305914.1 166764 166786 - 0 CTGTTCACGGAGAACGTGTT GGG ||||||||[||||||||||||]PAM NA Intergenic NA NA

T21799 GCTACGTAACAACCACACCAAGG 1000
Oligo fwd CACCGCTACGTAACAACCACACCA
Oligo rev AAACTGGTGTGGTTGTTACGTAGC
Chromosome start end strand MM target_seq PAM alignment distance position gene name gene id
GU305914.1 166702 166724 - 0 GCTACGTAACAACCACACCA AGG ||||||||[||||||||||||]PAM NA Intergenic NA NA

T21800 ACGTAACAACCACACCAAGGCGG 1000
Oligo fwd CACCACGTAACAACCACACCAAGG
Oligo rev AAACCCTTGGTGTGGTTGTTACGT
Chromosome start end strand MM target_seq PAM alignment distance position gene name gene id
GU305914.1 166699 166721 - 0 ACGTAACAACCACACCAAGG CGG ||||||||[||||||||||||]PAM NA Intergenic NA NA

T21801 CAACCACACCAAGGCGGAGTCGG 1000
Oligo fwd CACCCAACCACACCAAGGCGGAGT
Oligo rev AAACACTCCGCCTTGGTGTGGTTG
Chromosome start end strand MM target_seq PAM alignment distance position gene name gene id
GU305914.1 166693 166715 - 0 CAACCACACCAAGGCGGAGT CGG ||||||||[||||||||||||]PAM NA Intergenic NA NA

T21802 CCACACCAAGGCGGAGTCGGTGG 1000
Oligo fwd CACCCCACACCAAGGCGGAGTCGG
Oligo rev AAACCCGACTCCGCCTTGGTGTGG
Chromosome start end strand MM target_seq PAM alignment distance position gene name gene id
GU305914.1 166690 166712 - 0 CCACACCAAGGCGGAGTCGG TGG ||||||||[||||||||||||]PAM NA Intergenic NA NA

T21803 AGCAGCATGTCCCTGCTCGACGG 1000
Oligo fwd CACCAGCAGCATGTCCCTGCTCGA
Oligo rev AAACTCGAGCAGGGACATGCTGCT
Chromosome start end strand MM target_seq PAM alignment distance position gene name gene id
GU305914.1 166653 166675 - 0 AGCAGCATGTCCCTGCTCGA CGG ||||||||[||||||||||||]PAM NA Intergenic NA NA

T21804 AACCTCCTCCGTACCATCTTAGG 1000
Oligo fwd CACCAACCTCCTCCGTACCATCTT
Oligo rev AAACAAGATGGTACGGAGGAGGTT
Chromosome start end strand MM target_seq PAM alignment distance position gene name gene id
GU305914.1 166623 166645 - 0 AACCTCCTCCGTACCATCTT AGG ||||||||[||||||||||||]PAM NA Intergenic NA NA

T21805 AGGCCATATGTCCCTAGACACGG 1000

<table>
<tr><td>Oligo fwd</td><td>CACCAGGCCATATGTCCCTAGACA</td><td></td><td></td><td></td><td></td><td></td><td></td><td></td><td></td><td></td><td></td></tr>
<tr><td>Oligo rev</td><td>AAACTGTCTAGGGACATATGGCCT</td><td></td><td></td><td></td><td></td><td></td><td></td><td></td><td></td><td></td><td></td></tr>
<tr><td>Chromosome</td><td>start</td><td>end</td><td>strand</td><td>MM</td><td>target_seq</td><td>PAM</td><td>alignment</td><td>distance</td><td>position</td><td>gene name</td><td>gene id</td></tr>
<tr><td>GU305914.1</td><td>166603</td><td>166625</td><td>-</td><td>0</td><td>AGGCCATATGTCCCTAGACA</td><td>CGG</td><td>||||||||[||||||||||||]PAM</td><td>NA</td><td>Intergenic</td><td>NA</td><td>NA</td></tr>
<tr><td></td><td></td><td></td><td></td><td></td><td></td><td></td><td></td><td></td><td></td><td></td><td></td></tr>
<tr><td>T21806</td><td>TCCCTAGACACGGACCTGATAGG</td><td>1000</td><td></td><td></td><td></td><td></td><td></td><td></td><td></td><td></td><td></td></tr>
<tr><td>Oligo fwd</td><td>CACCTCCCTAGACACGGACCTGAT</td><td></td><td></td><td></td><td></td><td></td><td></td><td></td><td></td><td></td><td></td></tr>
<tr><td>Oligo rev</td><td>AAACATCAGGTCCGTGTCTAGGGA</td><td></td><td></td><td></td><td></td><td></td><td></td><td></td><td></td><td></td><td></td></tr>
<tr><td>Chromosome</td><td>start</td><td>end</td><td>strand</td><td>MM</td><td>target_seq</td><td>PAM</td><td>alignment</td><td>distance</td><td>position</td><td>gene name</td><td>gene id</td></tr>
<tr><td>GU305914.1</td><td>166593</td><td>166615</td><td>-</td><td>0</td><td>TCCCTAGACACGGACCTGAT</td><td>AGG</td><td>||||||||[||||||||||||]PAM</td><td>NA</td><td>Intergenic</td><td>NA</td><td>NA</td></tr>
<tr><td></td><td></td><td></td><td></td><td></td><td></td><td></td><td></td><td></td><td></td><td></td><td></td></tr>
<tr><td>T21807</td><td>TAGGAGTCCTCCGCAGTTACAGG</td><td>1000</td><td></td><td></td><td></td><td></td><td></td><td></td><td></td><td></td><td></td></tr>
<tr><td>Oligo fwd</td><td>CACCTAGGAGTCCTCCGCAGTTAC</td><td></td><td></td><td></td><td></td><td></td><td></td><td></td><td></td><td></td><td></td></tr>
<tr><td>Oligo rev</td><td>AAACGTAACTGCGGAGGACTCCTA</td><td></td><td></td><td></td><td></td><td></td><td></td><td></td><td></td><td></td><td></td></tr>
<tr><td>Chromosome</td><td>start</td><td>end</td><td>strand</td><td>MM</td><td>target_seq</td><td>PAM</td><td>alignment</td><td>distance</td><td>position</td><td>gene name</td><td>gene id</td></tr>
<tr><td>GU305914.1</td><td>166574</td><td>166596</td><td>-</td><td>0</td><td>TAGGAGTCCTCCGCAGTTAC</td><td>AGG</td><td>||||||||[||||||||||||]PAM</td><td>NA</td><td>Intergenic</td><td>NA</td><td>NA</td></tr>
<tr><td></td><td></td><td></td><td></td><td></td><td></td><td></td><td></td><td></td><td></td><td></td><td></td></tr>
<tr><td>T21808</td><td>AGGAGTCCTCCGCAGTTACAGGG</td><td>1000</td><td></td><td></td><td></td><td></td><td></td><td></td><td></td><td></td><td></td></tr>
<tr><td>Oligo fwd</td><td>CACCAGGAGTCCTCCGCAGTTACA</td><td></td><td></td><td></td><td></td><td></td><td></td><td></td><td></td><td></td><td></td></tr>
<tr><td>Oligo rev</td><td>AAACTGTAACTGCGGAGGACTCCT</td><td></td><td></td><td></td><td></td><td></td><td></td><td></td><td></td><td></td><td></td></tr>
<tr><td>Chromosome</td><td>start</td><td>end</td><td>strand</td><td>MM</td><td>target_seq</td><td>PAM</td><td>alignment</td><td>distance</td><td>position</td><td>gene name</td><td>gene id</td></tr>
<tr><td>GU305914.1</td><td>166573</td><td>166595</td><td>-</td><td>0</td><td>AGGAGTCCTCCGCAGTTACA</td><td>GGG</td><td>||||||||[||||||||||||]PAM</td><td>NA</td><td>Intergenic</td><td>NA</td><td>NA</td></tr>
<tr><td></td><td></td><td></td><td></td><td></td><td></td><td></td><td></td><td></td><td></td><td></td><td></td></tr>
<tr><td>T21809</td><td>ATCAACGAAAGAGACCGTGATGG</td><td>1000</td><td></td><td></td><td></td><td></td><td></td><td></td><td></td><td></td><td></td></tr>
<tr><td>Oligo fwd</td><td>CACCATCAACGAAAGAGACCGTGA</td><td></td><td></td><td></td><td></td><td></td><td></td><td></td><td></td><td></td><td></td></tr>
<tr><td>Oligo rev</td><td>AAACTCACGGTCTCTTTCGTTGAT</td><td></td><td></td><td></td><td></td><td></td><td></td><td></td><td></td><td></td><td></td></tr>
<tr><td>Chromosome</td><td>start</td><td>end</td><td>strand</td><td>MM</td><td>target_seq</td><td>PAM</td><td>alignment</td><td>distance</td><td>position</td><td>gene name</td><td>gene id</td></tr>
<tr><td>GU305914.1</td><td>166481</td><td>166503</td><td>-</td><td>0</td><td>ATCAACGAAAGAGACCGTGA</td><td>TGG</td><td>||||||||[||||||||||||]PAM</td><td>NA</td><td>Intergenic</td><td>NA</td><td>NA</td></tr>
<tr><td></td><td></td><td></td><td></td><td></td><td></td><td></td><td></td><td></td><td></td><td></td><td></td></tr>
<tr><td>T21810</td><td>AAGAGACCGTGATGGCTCTCCGG</td><td>1000</td><td></td><td></td><td></td><td></td><td></td><td></td><td></td><td></td><td></td></tr>
<tr><td>Oligo fwd</td><td>CACCAAGAGACCGTGATGGCTCTC</td><td></td><td></td><td></td><td></td><td></td><td></td><td></td><td></td><td></td><td></td></tr>
<tr><td>Oligo rev</td><td>AAACGAGAGCCATCACGGTCTCTT</td><td></td><td></td><td></td><td></td><td></td><td></td><td></td><td></td><td></td><td></td></tr>
<tr><td>Chromosome</td><td>start</td><td>end</td><td>strand</td><td>MM</td><td>target_seq</td><td>PAM</td><td>alignment</td><td>distance</td><td>position</td><td>gene name</td><td>gene id</td></tr>
<tr><td>GU305914.1</td><td>166473</td><td>166495</td><td>-</td><td>0</td><td>AAGAGACCGTGATGGCTCTC</td><td>CGG</td><td>||||||||[||||||||||||]PAM</td><td>NA</td><td>Intergenic</td><td>NA</td><td>NA</td></tr>
<tr><td></td><td></td><td></td><td></td><td></td><td></td><td></td><td></td><td></td><td></td><td></td><td></td></tr>
<tr><td>T21811</td><td>CCGTGATGGCTCTCCGGCAGTGG</td><td>1000</td><td></td><td></td><td></td><td></td><td></td><td></td><td></td><td></td><td></td></tr>
<tr><td>Oligo fwd</td><td>CACCCCGTGATGGCTCTCCGGCAG</td><td></td><td></td><td></td><td></td><td></td><td></td><td></td><td></td><td></td><td></td></tr>
<tr><td>Oligo rev</td><td>AAACCTGCCGGAGAGCCATCACGG</td><td></td><td></td><td></td><td></td><td></td><td></td><td></td><td></td><td></td><td></td></tr>
<tr><td>Chromosome</td><td>start</td><td>end</td><td>strand</td><td>MM</td><td>target_seq</td><td>PAM</td><td>alignment</td><td>distance</td><td>position</td><td>gene name</td><td>gene id</td></tr>
<tr><td>GU305914.1</td><td>166467</td><td>166489</td><td>-</td><td>0</td><td>CCGTGATGGCTCTCCGGCAG</td><td>TGG</td><td>||||||||[||||||||||||]PAM</td><td>NA</td><td>Intergenic</td><td>NA</td><td>NA</td></tr>
<tr><td></td><td></td><td></td><td></td><td></td><td></td><td></td><td></td><td></td><td></td><td></td><td></td></tr>
<tr><td>T21812</td><td>GTGGATGCTCAGACACATCGCGG</td><td>1000</td><td></td><td></td><td></td><td></td><td></td><td></td><td></td><td></td><td></td></tr>
<tr><td>Oligo fwd</td><td>CACCGTGGATGCTCAGACACATCG</td><td></td><td></td><td></td><td></td><td></td><td></td><td></td><td></td><td></td><td></td></tr>
<tr><td>Oligo rev</td><td>AAACCGATGTGTCTGAGCATCCAC</td><td></td><td></td><td></td><td></td><td></td><td></td><td></td><td></td><td></td><td></td></tr>
<tr><td>Chromosome</td><td>start</td><td>end</td><td>strand</td><td>MM</td><td>target_seq</td><td>PAM</td><td>alignment</td><td>distance</td><td>position</td><td>gene name</td><td>gene id</td></tr>
<tr><td>GU305914.1</td><td>166448</td><td>166470</td><td>-</td><td>0</td><td>GTGGATGCTCAGACACATCG</td><td>CGG</td><td>||||||||[||||||||||||]PAM</td><td>NA</td><td>Intergenic</td><td>NA</td><td>NA</td></tr>
<tr><td></td><td></td><td></td><td></td><td></td><td></td><td></td><td></td><td></td><td></td><td></td><td></td></tr>
<tr><td>T21813</td><td>CACATCGCGGTGCACGACGTCGG</td><td>1000</td><td></td><td></td><td></td><td></td><td></td><td></td><td></td><td></td><td></td></tr>
<tr><td>Oligo fwd</td><td>CACCCACATCGCGGTGCACGACGT</td><td></td><td></td><td></td><td></td><td></td><td></td><td></td><td></td><td></td><td></td></tr>
<tr><td>Oligo rev</td><td>AAACACGTCGTGCACCGCGATGTG</td><td></td><td></td><td></td><td></td><td></td><td></td><td></td><td></td><td></td><td></td></tr>
<tr><td>Chromosome</td><td>start</td><td>end</td><td>strand</td><td>MM</td><td>target_seq</td><td>PAM</td><td>alignment</td><td>distance</td><td>position</td><td>gene name</td><td>gene id</td></tr>
<tr><td>GU305914.1</td><td>166435</td><td>166457</td><td>-</td><td>0</td><td>CACATCGCGGTGCACGACGT</td><td>CGG</td><td>||||||||[||||||||||||]PAM</td><td>NA</td><td>Intergenic</td><td>NA</td><td>NA</td></tr>
<tr><td></td><td></td><td></td><td></td><td></td><td></td><td></td><td></td><td></td><td></td><td></td><td></td></tr>
<tr><td>T21814</td><td>GACGTCGGCGCCGCTGCCGCCGG</td><td>1000</td><td></td><td></td><td></td><td></td><td></td><td></td><td></td><td></td><td></td></tr>
<tr><td>Oligo fwd</td><td>CACCGACGTCGGCGCCGCTGCCGC</td><td></td><td></td><td></td><td></td><td></td><td></td><td></td><td></td><td></td><td></td></tr>
<tr><td>Oligo rev</td><td>AAACGCGGCAGCGGCGCCGACGTC</td><td></td><td></td><td></td><td></td><td></td><td></td><td></td><td></td><td></td><td></td></tr>
<tr><td>Chromosome</td><td>start</td><td>end</td><td>strand</td><td>MM</td><td>target_seq</td><td>PAM</td><td>alignment</td><td>distance</td><td>position</td><td>gene name</td><td>gene id</td></tr>
<tr><td>GU305914.1</td><td>166420</td><td>166442</td><td>-</td><td>0</td><td>GACGTCGGCGCCGCTGCCGC</td><td>CGG</td><td>||||||||[||||||||||||]PAM</td><td>NA</td><td>Intergenic</td><td>NA</td><td>NA</td></tr>
<tr><td></td><td></td><td></td><td></td><td></td><td></td><td></td><td></td><td></td><td></td><td></td><td></td></tr>
<tr><td>T21815</td><td>TGCCGACGTGATTCATCAACAGG</td><td>1000</td><td></td><td></td><td></td><td></td><td></td><td></td><td></td><td></td><td></td></tr>
<tr><td>Oligo fwd</td><td>CACCTGCCGACGTGATTCATCAAC</td><td></td><td></td><td></td><td></td><td></td><td></td><td></td><td></td><td></td><td></td></tr>
<tr><td>Oligo rev</td><td>AAACGTTGATGAATCACGTCGGCA</td><td></td><td></td><td></td><td></td><td></td><td></td><td></td><td></td><td></td><td></td></tr>
<tr><td>Chromosome</td><td>start</td><td>end</td><td>strand</td><td>MM</td><td>target_seq</td><td>PAM</td><td>alignment</td><td>distance</td><td>position</td><td>gene name</td><td>gene id</td></tr>
<tr><td>GU305914.1</td><td>166385</td><td>166407</td><td>-</td><td>0</td><td>TGCCGACGTGATTCATCAAC</td><td>AGG</td><td>||||||||[||||||||||||]PAM</td><td>NA</td><td>Intergenic</td><td>NA</td><td>NA</td></tr>
<tr><td></td><td></td><td></td><td></td><td></td><td></td><td></td><td></td><td></td><td></td><td></td><td></td></tr>
<tr><td>T21816</td><td>CGACGTGATTCATCAACAGGCGG</td><td>1000</td><td></td><td></td><td></td><td></td><td></td><td></td><td></td><td></td><td></td></tr>
<tr><td>Oligo fwd</td><td>CACCCGACGTGATTCATCAACAGG</td><td></td><td></td><td></td><td></td><td></td><td></td><td></td><td></td><td></td><td></td></tr>
<tr><td>Oligo rev</td><td>AAACCCTGTTGATGAATCACGTCG</td><td></td><td></td><td></td><td></td><td></td><td></td><td></td><td></td><td></td><td></td></tr>
<tr><td>Chromosome</td><td>start</td><td>end</td><td>strand</td><td>MM</td><td>target_seq</td><td>PAM</td><td>alignment</td><td>distance</td><td>position</td><td>gene name</td><td>gene id</td></tr>
<tr><td>GU305914.1</td><td>166382</td><td>166404</td><td>-</td><td>0</td><td>CGACGTGATTCATCAACAGG</td><td>CGG</td><td>||||||||[||||||||||||]PAM</td><td>NA</td><td>Intergenic</td><td>NA</td><td>NA</td></tr>
<tr><td></td><td></td><td></td><td></td><td></td><td></td><td></td><td></td><td></td><td></td><td></td><td></td></tr>
<tr><td>T21817</td><td>CGTGATTCATCAACAGGCGGAGG</td><td>1000</td><td></td><td></td><td></td><td></td><td></td><td></td><td></td><td></td><td></td></tr>
<tr><td>Oligo fwd</td><td>CACCCGTGATTCATCAACAGGCGG</td><td></td><td></td><td></td><td></td><td></td><td></td><td></td><td></td><td></td><td></td></tr>
<tr><td>Oligo rev</td><td>AAACCCGCCTGTTGATGAATCACG</td><td></td><td></td><td></td><td></td><td></td><td></td><td></td><td></td><td></td><td></td></tr>
<tr><td>Chromosome</td><td>start</td><td>end</td><td>strand</td><td>MM</td><td>target_seq</td><td>PAM</td><td>alignment</td><td>distance</td><td>position</td><td>gene name</td><td>gene id</td></tr>
<tr><td>GU305914.1</td><td>166379</td><td>166401</td><td>-</td><td>0</td><td>CGTGATTCATCAACAGGCGG</td><td>AGG</td><td>||||||||[||||||||||||]PAM</td><td>NA</td><td>Intergenic</td><td>NA</td><td>NA</td></tr>
<tr><td></td><td></td><td></td><td></td><td></td><td></td><td></td><td></td><td></td><td></td><td></td><td></td></tr>
<tr><td>T21818</td><td>TCATCAACAGGCGGAGGCGCTGG</td><td>1000</td><td></td><td></td><td></td><td></td><td></td><td></td><td></td><td></td><td></td></tr>
<tr><td>Oligo fwd</td><td>CACCTCATCAACAGGCGGAGGCGC</td><td></td><td></td><td></td><td></td><td></td><td></td><td></td><td></td><td></td><td></td></tr>
<tr><td>Oligo rev</td><td>AAACGCGCCTCCGCCTGTTGATGA</td><td></td><td></td><td></td><td></td><td></td><td></td><td></td><td></td><td></td><td></td></tr>
<tr><td>Chromosome</td><td>start</td><td>end</td><td>strand</td><td>MM</td><td>target_seq</td><td>PAM</td><td>alignment</td><td>distance</td><td>position</td><td>gene name</td><td>gene id</td></tr>
<tr><td>GU305914.1</td><td>166373</td><td>166395</td><td>-</td><td>0</td><td>TCATCAACAGGCGGAGGCGC</td><td>TGG</td><td>||||||||[||||||||||||]PAM</td><td>NA</td><td>Intergenic</td><td>NA</td><td>NA</td></tr>
<tr><td></td><td></td><td></td><td></td><td></td><td></td><td></td><td></td><td></td><td></td><td></td><td></td></tr>
<tr><td>T21819</td><td>CATCAACAGGCGGAGGCGCTGGG</td><td>1000</td><td></td><td></td><td></td><td></td><td></td><td></td><td></td><td></td><td></td></tr>
<tr><td>Oligo fwd</td><td>CACCCATCAACAGGCGGAGGCGCT</td><td></td><td></td><td></td><td></td><td></td><td></td><td></td><td></td><td></td><td></td></tr>
<tr><td>Oligo rev</td><td>AAACAGCGCCTCCGCCTGTTGATG</td><td></td><td></td><td></td><td></td><td></td><td></td><td></td><td></td><td></td><td></td></tr>
<tr><td>Chromosome</td><td>start</td><td>end</td><td>strand</td><td>MM</td><td>target_seq</td><td>PAM</td><td>alignment</td><td>distance</td><td>position</td><td>gene name</td><td>gene id</td></tr>
<tr><td>GU305914.1</td><td>166372</td><td>166394</td><td>-</td><td>0</td><td>CATCAACAGGCGGAGGCGCT</td><td>GGG</td><td>||||||||[||||||||||||]PAM</td><td>NA</td><td>Intergenic</td><td>NA</td><td>NA</td></tr>
</table>

<table>
<tr><td>T21820</td><td>CGCTGGGCATCCACGAAGCCTGG</td><td>1000</td><td></td><td></td><td></td><td></td><td></td><td></td><td></td><td></td><td></td></tr>
<tr><td>Oligo fwd</td><td>CACCCGCTGGGCATCCACGAAGCC</td><td></td><td></td><td></td><td></td><td></td><td></td><td></td><td></td><td></td><td></td></tr>
<tr><td>Oligo rev</td><td>AAACGGCTTCGTGGATGCCCAGCG</td><td></td><td></td><td></td><td></td><td></td><td></td><td></td><td></td><td></td><td></td></tr>
<tr><td>Chromosome</td><td>start</td><td>end</td><td>strand</td><td>MM</td><td>target_seq</td><td>PAM</td><td>alignment</td><td>distance</td><td>position</td><td>gene name</td><td>gene id</td></tr>
<tr><td>GU305914.1</td><td>166356</td><td>166378</td><td>-</td><td>0</td><td>CGCTGGGCATCCACGAAGCC</td><td>TGG</td><td>||||||||[||||||||||||]PAM</td><td>NA</td><td>Intergenic</td><td>NA</td><td>NA</td></tr>
<tr><td>T21821</td><td>TCTGAAACTATCCGCCACGCAGG</td><td>1000</td><td></td><td></td><td></td><td></td><td></td><td></td><td></td><td></td><td></td></tr>
<tr><td>Oligo fwd</td><td>CACCTCTGAAACTATCCGCCACGC</td><td></td><td></td><td></td><td></td><td></td><td></td><td></td><td></td><td></td><td></td></tr>
<tr><td>Oligo rev</td><td>AAACGCGTGGCGGATAGTTTCAGA</td><td></td><td></td><td></td><td></td><td></td><td></td><td></td><td></td><td></td><td></td></tr>
<tr><td>Chromosome</td><td>start</td><td>end</td><td>strand</td><td>MM</td><td>target_seq</td><td>PAM</td><td>alignment</td><td>distance</td><td>position</td><td>gene name</td><td>gene id</td></tr>
<tr><td>GU305914.1</td><td>166325</td><td>166347</td><td>-</td><td>0</td><td>TCTGAAACTATCCGCCACGC</td><td>AGG</td><td>||||||||[||||||||||||]PAM</td><td>NA</td><td>Intergenic</td><td>NA</td><td>NA</td></tr>
<tr><td>T21822</td><td>CGCCACGCAGGCGTCACAGCTGG</td><td>1000</td><td></td><td></td><td></td><td></td><td></td><td></td><td></td><td></td><td></td></tr>
<tr><td>Oligo fwd</td><td>CACCCGCCACGCAGGCGTCACAGC</td><td></td><td></td><td></td><td></td><td></td><td></td><td></td><td></td><td></td><td></td></tr>
<tr><td>Oligo rev</td><td>AAACGCTGTGACGCCTGCGTGGCG</td><td></td><td></td><td></td><td></td><td></td><td></td><td></td><td></td><td></td><td></td></tr>
<tr><td>Chromosome</td><td>start</td><td>end</td><td>strand</td><td>MM</td><td>target_seq</td><td>PAM</td><td>alignment</td><td>distance</td><td>position</td><td>gene name</td><td>gene id</td></tr>
<tr><td>GU305914.1</td><td>166313</td><td>166335</td><td>-</td><td>0</td><td>CGCCACGCAGGCGTCACAGC</td><td>TGG</td><td>||||||||[||||||||||||]PAM</td><td>NA</td><td>Intergenic</td><td>NA</td><td>NA</td></tr>
<tr><td>T21823</td><td>AGCTGGTCCGCATCACCGACAGG</td><td>1000</td><td></td><td></td><td></td><td></td><td></td><td></td><td></td><td></td><td></td></tr>
<tr><td>Oligo fwd</td><td>CACCAGCTGGTCCGCATCACCGAC</td><td></td><td></td><td></td><td></td><td></td><td></td><td></td><td></td><td></td><td></td></tr>
<tr><td>Oligo rev</td><td>AAACGTCGGTGATGCGGACCAGCT</td><td></td><td></td><td></td><td></td><td></td><td></td><td></td><td></td><td></td><td></td></tr>
<tr><td>Chromosome</td><td>start</td><td>end</td><td>strand</td><td>MM</td><td>target_seq</td><td>PAM</td><td>alignment</td><td>distance</td><td>position</td><td>gene name</td><td>gene id</td></tr>
<tr><td>GU305914.1</td><td>166296</td><td>166318</td><td>-</td><td>0</td><td>AGCTGGTCCGCATCACCGAC</td><td>AGG</td><td>||||||||[||||||||||||]PAM</td><td>NA</td><td>Intergenic</td><td>NA</td><td>NA</td></tr>
<tr><td>T21824</td><td>GCTGGTCCGCATCACCGACAGGG</td><td>1000</td><td></td><td></td><td></td><td></td><td></td><td></td><td></td><td></td><td></td></tr>
<tr><td>Oligo fwd</td><td>CACCGCTGGTCCGCATCACCGACA</td><td></td><td></td><td></td><td></td><td></td><td></td><td></td><td></td><td></td><td></td></tr>
<tr><td>Oligo rev</td><td>AAACTGTCGGTGATGCGGACCAGC</td><td></td><td></td><td></td><td></td><td></td><td></td><td></td><td></td><td></td><td></td></tr>
<tr><td>Chromosome</td><td>start</td><td>end</td><td>strand</td><td>MM</td><td>target_seq</td><td>PAM</td><td>alignment</td><td>distance</td><td>position</td><td>gene name</td><td>gene id</td></tr>
<tr><td>GU305914.1</td><td>166295</td><td>166317</td><td>-</td><td>0</td><td>GCTGGTCCGCATCACCGACA</td><td>GGG</td><td>||||||||[||||||||||||]PAM</td><td>NA</td><td>Intergenic</td><td>NA</td><td>NA</td></tr>
<tr><td>T21825</td><td>CATCACCGACAGGGTAGATCAGG</td><td>1000</td><td></td><td></td><td></td><td></td><td></td><td></td><td></td><td></td><td></td></tr>
<tr><td>Oligo fwd</td><td>CACCCATCACCGACAGGGTAGATC</td><td></td><td></td><td></td><td></td><td></td><td></td><td></td><td></td><td></td><td></td></tr>
<tr><td>Oligo rev</td><td>AAACGATCTACCCTGTCGGTGATG</td><td></td><td></td><td></td><td></td><td></td><td></td><td></td><td></td><td></td><td></td></tr>
<tr><td>Chromosome</td><td>start</td><td>end</td><td>strand</td><td>MM</td><td>target_seq</td><td>PAM</td><td>alignment</td><td>distance</td><td>position</td><td>gene name</td><td>gene id</td></tr>
<tr><td>GU305914.1</td><td>166286</td><td>166308</td><td>-</td><td>0</td><td>CATCACCGACAGGGTAGATC</td><td>AGG</td><td>||||||||[||||||||||||]PAM</td><td>NA</td><td>Intergenic</td><td>NA</td><td>NA</td></tr>
<tr><td>T21826</td><td>AGAAGAGCGACGTCCACCGTTGG</td><td>1000</td><td></td><td></td><td></td><td></td><td></td><td></td><td></td><td></td><td></td></tr>
<tr><td>Oligo fwd</td><td>CACCAGAAGAGCGACGTCCACCGT</td><td></td><td></td><td></td><td></td><td></td><td></td><td></td><td></td><td></td><td></td></tr>
<tr><td>Oligo rev</td><td>AAACACGGTGGACGTCGCTCTTCT</td><td></td><td></td><td></td><td></td><td></td><td></td><td></td><td></td><td></td><td></td></tr>
<tr><td>Chromosome</td><td>start</td><td>end</td><td>strand</td><td>MM</td><td>target_seq</td><td>PAM</td><td>alignment</td><td>distance</td><td>position</td><td>gene name</td><td>gene id</td></tr>
<tr><td>GU305914.1</td><td>166236</td><td>166258</td><td>-</td><td>0</td><td>AGAAGAGCGACGTCCACCGT</td><td>TGG</td><td>||||||||[||||||||||||]PAM</td><td>NA</td><td>Intergenic</td><td>NA</td><td>NA</td></tr>
<tr><td>T21827</td><td>TTTCCCGTACGACGTGCGAGTGG</td><td>1000</td><td></td><td></td><td></td><td></td><td></td><td></td><td></td><td></td><td></td></tr>
<tr><td>Oligo fwd</td><td>CACCTTTCCCGTACGACGTGCGAG</td><td></td><td></td><td></td><td></td><td></td><td></td><td></td><td></td><td></td><td></td></tr>
<tr><td>Oligo rev</td><td>AAACCTCGCACGTCGTACGGGAAA</td><td></td><td></td><td></td><td></td><td></td><td></td><td></td><td></td><td></td><td></td></tr>
<tr><td>Chromosome</td><td>start</td><td>end</td><td>strand</td><td>MM</td><td>target_seq</td><td>PAM</td><td>alignment</td><td>distance</td><td>position</td><td>gene name</td><td>gene id</td></tr>
<tr><td>GU305914.1</td><td>166202</td><td>166224</td><td>-</td><td>0</td><td>TTTCCCGTACGACGTGCGAG</td><td>TGG</td><td>||||||||[||||||||||||]PAM</td><td>NA</td><td>Intergenic</td><td>NA</td><td>NA</td></tr>
<tr><td>T21828</td><td>GACGTGCGAGTGGTGATCCTAGG</td><td>1000</td><td></td><td></td><td></td><td></td><td></td><td></td><td></td><td></td><td></td></tr>
<tr><td>Oligo fwd</td><td>CACCGACGTGCGAGTGGTGATCCT</td><td></td><td></td><td></td><td></td><td></td><td></td><td></td><td></td><td></td><td></td></tr>
<tr><td>Oligo rev</td><td>AAACAGGATCACCACTCGCACGTC</td><td></td><td></td><td></td><td></td><td></td><td></td><td></td><td></td><td></td><td></td></tr>
<tr><td>Chromosome</td><td>start</td><td>end</td><td>strand</td><td>MM</td><td>target_seq</td><td>PAM</td><td>alignment</td><td>distance</td><td>position</td><td>gene name</td><td>gene id</td></tr>
<tr><td>GU305914.1</td><td>166192</td><td>166214</td><td>-</td><td>0</td><td>GACGTGCGAGTGGTGATCCT</td><td>AGG</td><td>||||||||[||||||||||||]PAM</td><td>NA</td><td>Intergenic</td><td>NA</td><td>NA</td></tr>
<tr><td>T21829</td><td>GGACAAGATCCGTACCACGACGG</td><td>1000</td><td></td><td></td><td></td><td></td><td></td><td></td><td></td><td></td><td></td></tr>
<tr><td>Oligo fwd</td><td>CACCGGACAAGATCCGTACCACGA</td><td></td><td></td><td></td><td></td><td></td><td></td><td></td><td></td><td></td><td></td></tr>
<tr><td>Oligo rev</td><td>AAACTCGTGGTACGGATCTTGTCC</td><td></td><td></td><td></td><td></td><td></td><td></td><td></td><td></td><td></td><td></td></tr>
<tr><td>Chromosome</td><td>start</td><td>end</td><td>strand</td><td>MM</td><td>target_seq</td><td>PAM</td><td>alignment</td><td>distance</td><td>position</td><td>gene name</td><td>gene id</td></tr>
<tr><td>GU305914.1</td><td>166171</td><td>166193</td><td>-</td><td>0</td><td>GGACAAGATCCGTACCACGA</td><td>CGG</td><td>||||||||[||||||||||||]PAM</td><td>NA</td><td>Intergenic</td><td>NA</td><td>NA</td></tr>
<tr><td>T21830</td><td>TACCACGACGGCAGCGCGTGCGG</td><td>1000</td><td></td><td></td><td></td><td></td><td></td><td></td><td></td><td></td><td></td></tr>
<tr><td>Oligo fwd</td><td>CACCTACCACGACGGCAGCGCGTG</td><td></td><td></td><td></td><td></td><td></td><td></td><td></td><td></td><td></td><td></td></tr>
<tr><td>Oligo rev</td><td>AAACCACGCGCTGCCGTCGTGGTA</td><td></td><td></td><td></td><td></td><td></td><td></td><td></td><td></td><td></td><td></td></tr>
<tr><td>Chromosome</td><td>start</td><td>end</td><td>strand</td><td>MM</td><td>target_seq</td><td>PAM</td><td>alignment</td><td>distance</td><td>position</td><td>gene name</td><td>gene id</td></tr>
<tr><td>GU305914.1</td><td>166159</td><td>166181</td><td>-</td><td>0</td><td>TACCACGACGGCAGCGCGTG</td><td>CGG</td><td>||||||||[||||||||||||]PAM</td><td>NA</td><td>Intergenic</td><td>NA</td><td>NA</td></tr>
<tr><td>T21831</td><td>AGCGCGTGCGGCCTCGCCTTCGG</td><td>1000</td><td></td><td></td><td></td><td></td><td></td><td></td><td></td><td></td><td></td></tr>
<tr><td>Oligo fwd</td><td>CACCAGCGCGTGCGGCCTCGCCTT</td><td></td><td></td><td></td><td></td><td></td><td></td><td></td><td></td><td></td><td></td></tr>
<tr><td>Oligo rev</td><td>AAACAAGGCGAGGCCGCACGCGCT</td><td></td><td></td><td></td><td></td><td></td><td></td><td></td><td></td><td></td><td></td></tr>
<tr><td>Chromosome</td><td>start</td><td>end</td><td>strand</td><td>MM</td><td>target_seq</td><td>PAM</td><td>alignment</td><td>distance</td><td>position</td><td>gene name</td><td>gene id</td></tr>
<tr><td>GU305914.1</td><td>166147</td><td>166169</td><td>-</td><td>0</td><td>AGCGCGTGCGGCCTCGCCTT</td><td>CGG</td><td>||||||||[||||||||||||]PAM</td><td>NA</td><td>Intergenic</td><td>NA</td><td>NA</td></tr>
<tr><td>T21832</td><td>GCGCGTGCGGCCTCGCCTTCGGG</td><td>1000</td><td></td><td></td><td></td><td></td><td></td><td></td><td></td><td></td><td></td></tr>
<tr><td>Oligo fwd</td><td>CACCGCGCGTGCGGCCTCGCCTTC</td><td></td><td></td><td></td><td></td><td></td><td></td><td></td><td></td><td></td><td></td></tr>
<tr><td>Oligo rev</td><td>AAACGAAGGCGAGGCCGCACGCGC</td><td></td><td></td><td></td><td></td><td></td><td></td><td></td><td></td><td></td><td></td></tr>
<tr><td>Chromosome</td><td>start</td><td>end</td><td>strand</td><td>MM</td><td>target_seq</td><td>PAM</td><td>alignment</td><td>distance</td><td>position</td><td>gene name</td><td>gene id</td></tr>
<tr><td>GU305914.1</td><td>166146</td><td>166168</td><td>-</td><td>0</td><td>GCGCGTGCGGCCTCGCCTTC</td><td>GGG</td><td>||||||||[||||||||||||]PAM</td><td>NA</td><td>Intergenic</td><td>NA</td><td>NA</td></tr>
<tr><td>T21833</td><td>CGGGACCGTGAGAGACCGTCCGG</td><td>1000</td><td></td><td></td><td></td><td></td><td></td><td></td><td></td><td></td><td></td></tr>
<tr><td>Oligo fwd</td><td>CACCCGGGACCGTGAGAGACCGTC</td><td></td><td></td><td></td><td></td><td></td><td></td><td></td><td></td><td></td><td></td></tr>
<tr><td>Oligo rev</td><td>AAACGACGGTCTCTCACGGTCCCG</td><td></td><td></td><td></td><td></td><td></td><td></td><td></td><td></td><td></td><td></td></tr>
<tr><td>Chromosome</td><td>start</td><td>end</td><td>strand</td><td>MM</td><td>target_seq</td><td>PAM</td><td>alignment</td><td>distance</td><td>position</td><td>gene name</td><td>gene id</td></tr>
<tr><td>GU305914.1</td><td>166127</td><td>166149</td><td>-</td><td>0</td><td>CGGGACCGTGAGAGACCGTC</td><td>CGG</td><td>||||||||[||||||||||||]PAM</td><td>NA</td><td>Intergenic</td><td>NA</td><td>NA</td></tr>
<tr><td>T21834</td><td>GGCCCCGCCGTCCCTAGTGACGG</td><td>1000</td><td></td><td></td><td></td><td></td><td></td><td></td><td></td><td></td><td></td></tr>
<tr><td>Oligo fwd</td><td>CACCGGCCCCGCCGTCCCTAGTGA</td><td></td><td></td><td></td><td></td><td></td><td></td><td></td><td></td><td></td><td></td></tr>
<tr><td>Oligo rev</td><td>AAACTCACTAGGGACGGCGGGGCC</td><td></td><td></td><td></td><td></td><td></td><td></td><td></td><td></td><td></td><td></td></tr>
<tr><td>Chromosome</td><td>start</td><td>end</td><td>strand</td><td>MM</td><td>target_seq</td><td>PAM</td><td>alignment</td><td>distance</td><td>position</td><td>gene name</td><td>gene id</td></tr>
<tr><td>GU305914.1</td><td>166106</td><td>166128</td><td>-</td><td>0</td><td>GGCCCCGCCGTCCCTAGTGA</td><td>CGG</td><td>||||||||[||||||||||||]PAM</td><td>NA</td><td>Intergenic</td><td>NA</td><td>NA</td></tr>
</table>

<table>
<tr><td>T21835</td><td>GTCCCTAGTGACGGTCTTCAAGG</td><td>1000</td><td></td><td></td><td></td><td></td><td></td><td></td><td></td><td></td><td></td></tr>
<tr><td>Oligo fwd</td><td>CACCGTCCCTAGTGACGGTCTTCA</td><td></td><td></td><td></td><td></td><td></td><td></td><td></td><td></td><td></td><td></td></tr>
<tr><td>Oligo rev</td><td>AAACTGAAGACCGTCACTAGGGAC</td><td></td><td></td><td></td><td></td><td></td><td></td><td></td><td></td><td></td><td></td></tr>
<tr><td>Chromosome</td><td>start</td><td>end</td><td>strand</td><td>MM</td><td>target_seq</td><td>PAM</td><td>alignment</td><td>distance</td><td>position</td><td>gene name</td><td>gene id</td></tr>
<tr><td>GU305914.1</td><td>166097</td><td>166119</td><td>-</td><td>0</td><td>GTCCCTAGTGACGGTCTTCA</td><td>AGG</td><td>||||||||[||||||||||||]PAM</td><td>NA</td><td>Intergenic</td><td>NA</td><td>NA</td></tr>
<tr><td>T21836</td><td>TGACGGTCTTCAAGGAACTCCGG</td><td>1000</td><td></td><td></td><td></td><td></td><td></td><td></td><td></td><td></td><td></td></tr>
<tr><td>Oligo fwd</td><td>CACCTGACGGTCTTCAAGGAACTC</td><td></td><td></td><td></td><td></td><td></td><td></td><td></td><td></td><td></td><td></td></tr>
<tr><td>Oligo rev</td><td>AAACGAGTTCCTTGAAGACCGTCA</td><td></td><td></td><td></td><td></td><td></td><td></td><td></td><td></td><td></td><td></td></tr>
<tr><td>Chromosome</td><td>start</td><td>end</td><td>strand</td><td>MM</td><td>target_seq</td><td>PAM</td><td>alignment</td><td>distance</td><td>position</td><td>gene name</td><td>gene id</td></tr>
<tr><td>GU305914.1</td><td>166089</td><td>166111</td><td>-</td><td>0</td><td>TGACGGTCTTCAAGGAACTC</td><td>CGG</td><td>||||||||[||||||||||||]PAM</td><td>NA</td><td>Intergenic</td><td>NA</td><td>NA</td></tr>
<tr><td>T21837</td><td>CGGTCTTCAAGGAACTCCGGCGG</td><td>1000</td><td></td><td></td><td></td><td></td><td></td><td></td><td></td><td></td><td></td></tr>
<tr><td>Oligo fwd</td><td>CACCCGGTCTTCAAGGAACTCCGG</td><td></td><td></td><td></td><td></td><td></td><td></td><td></td><td></td><td></td><td></td></tr>
<tr><td>Oligo rev</td><td>AAACCCGGAGTTCCTTGAAGACCG</td><td></td><td></td><td></td><td></td><td></td><td></td><td></td><td></td><td></td><td></td></tr>
<tr><td>Chromosome</td><td>start</td><td>end</td><td>strand</td><td>MM</td><td>target_seq</td><td>PAM</td><td>alignment</td><td>distance</td><td>position</td><td>gene name</td><td>gene id</td></tr>
<tr><td>GU305914.1</td><td>166086</td><td>166108</td><td>-</td><td>0</td><td>CGGTCTTCAAGGAACTCCGG</td><td>CGG</td><td>||||||||[||||||||||||]PAM</td><td>NA</td><td>Intergenic</td><td>NA</td><td>NA</td></tr>
<tr><td>T21838</td><td>GGAACTCCGGCGGAGCATCCCGG</td><td>1000</td><td></td><td></td><td></td><td></td><td></td><td></td><td></td><td></td><td></td></tr>
<tr><td>Oligo fwd</td><td>CACCGGAACTCCGGCGGAGCATCC</td><td></td><td></td><td></td><td></td><td></td><td></td><td></td><td></td><td></td><td></td></tr>
<tr><td>Oligo rev</td><td>AAACGGATGCTCCGCCGGAGTTCC</td><td></td><td></td><td></td><td></td><td></td><td></td><td></td><td></td><td></td><td></td></tr>
<tr><td>Chromosome</td><td>start</td><td>end</td><td>strand</td><td>MM</td><td>target_seq</td><td>PAM</td><td>alignment</td><td>distance</td><td>position</td><td>gene name</td><td>gene id</td></tr>
<tr><td>GU305914.1</td><td>166076</td><td>166098</td><td>-</td><td>0</td><td>GGAACTCCGGCGGAGCATCC</td><td>CGG</td><td>||||||||[||||||||||||]PAM</td><td>NA</td><td>Intergenic</td><td>NA</td><td>NA</td></tr>
<tr><td>T21839</td><td>GAGTTTTCGATGCCGAAGTGCGG</td><td>1000</td><td></td><td></td><td></td><td></td><td></td><td></td><td></td><td></td><td></td></tr>
<tr><td>Oligo fwd</td><td>CACCGAGTTTTCGATGCCGAAGTG</td><td></td><td></td><td></td><td></td><td></td><td></td><td></td><td></td><td></td><td></td></tr>
<tr><td>Oligo rev</td><td>AAACCACTTCGGCATCGAAAACTC</td><td></td><td></td><td></td><td></td><td></td><td></td><td></td><td></td><td></td><td></td></tr>
<tr><td>Chromosome</td><td>start</td><td>end</td><td>strand</td><td>MM</td><td>target_seq</td><td>PAM</td><td>alignment</td><td>distance</td><td>position</td><td>gene name</td><td>gene id</td></tr>
<tr><td>GU305914.1</td><td>166054</td><td>166076</td><td>-</td><td>0</td><td>GAGTTTTCGATGCCGAAGTG</td><td>CGG</td><td>||||||||[||||||||||||]PAM</td><td>NA</td><td>Intergenic</td><td>NA</td><td>NA</td></tr>
<tr><td>GU305914.1</td><td>191747</td><td>191769</td><td>+</td><td>4</td><td>GAGTCCTCGACGCCGAGGTG</td><td>CGG</td><td>||||--||[||-|||||-|||]PAM</td><td>NA</td><td>Intergenic</td><td>NA</td><td>NA</td></tr>
<tr><td>T21840</td><td>AGTTTTCGATGCCGAAGTGCGGG</td><td>1000</td><td></td><td></td><td></td><td></td><td></td><td></td><td></td><td></td><td></td></tr>
<tr><td>Oligo fwd</td><td>CACCAGTTTTCGATGCCGAAGTGC</td><td></td><td></td><td></td><td></td><td></td><td></td><td></td><td></td><td></td><td></td></tr>
<tr><td>Oligo rev</td><td>AAACGCACTTCGGCATCGAAAACT</td><td></td><td></td><td></td><td></td><td></td><td></td><td></td><td></td><td></td><td></td></tr>
<tr><td>Chromosome</td><td>start</td><td>end</td><td>strand</td><td>MM</td><td>target_seq</td><td>PAM</td><td>alignment</td><td>distance</td><td>position</td><td>gene name</td><td>gene id</td></tr>
<tr><td>GU305914.1</td><td>166053</td><td>166075</td><td>-</td><td>0</td><td>AGTTTTCGATGCCGAAGTGC</td><td>GGG</td><td>||||||||[||||||||||||]PAM</td><td>NA</td><td>Intergenic</td><td>NA</td><td>NA</td></tr>
<tr><td>T21841</td><td>GATGCCGAAGTGCGGGTGCCTGG</td><td>1000</td><td></td><td></td><td></td><td></td><td></td><td></td><td></td><td></td><td></td></tr>
<tr><td>Oligo fwd</td><td>CACCGATGCCGAAGTGCGGGTGCC</td><td></td><td></td><td></td><td></td><td></td><td></td><td></td><td></td><td></td><td></td></tr>
<tr><td>Oligo rev</td><td>AAACGGCACCCGCACTTCGGCATC</td><td></td><td></td><td></td><td></td><td></td><td></td><td></td><td></td><td></td><td></td></tr>
<tr><td>Chromosome</td><td>start</td><td>end</td><td>strand</td><td>MM</td><td>target_seq</td><td>PAM</td><td>alignment</td><td>distance</td><td>position</td><td>gene name</td><td>gene id</td></tr>
<tr><td>GU305914.1</td><td>166046</td><td>166068</td><td>-</td><td>0</td><td>GATGCCGAAGTGCGGGTGCC</td><td>TGG</td><td>||||||||[||||||||||||]PAM</td><td>NA</td><td>Intergenic</td><td>NA</td><td>NA</td></tr>
<tr><td>T21842</td><td>AGTGCGGGTGCCTGGACGCGTGG</td><td>1000</td><td></td><td></td><td></td><td></td><td></td><td></td><td></td><td></td><td></td></tr>
<tr><td>Oligo fwd</td><td>CACCAGTGCGGGTGCCTGGACGCG</td><td></td><td></td><td></td><td></td><td></td><td></td><td></td><td></td><td></td><td></td></tr>
<tr><td>Oligo rev</td><td>AAACCGCGTCCAGGCACCCGCACT</td><td></td><td></td><td></td><td></td><td></td><td></td><td></td><td></td><td></td><td></td></tr>
<tr><td>Chromosome</td><td>start</td><td>end</td><td>strand</td><td>MM</td><td>target_seq</td><td>PAM</td><td>alignment</td><td>distance</td><td>position</td><td>gene name</td><td>gene id</td></tr>
<tr><td>GU305914.1</td><td>166038</td><td>166060</td><td>-</td><td>0</td><td>AGTGCGGGTGCCTGGACGCG</td><td>TGG</td><td>||||||||[||||||||||||]PAM</td><td>NA</td><td>Intergenic</td><td>NA</td><td>NA</td></tr>
<tr><td>T21843</td><td>CCTGGACGCGTGGTGCCGCGAGG</td><td>1000</td><td></td><td></td><td></td><td></td><td></td><td></td><td></td><td></td><td></td></tr>
<tr><td>Oligo fwd</td><td>CACCCCTGGACGCGTGGTGCCGCG</td><td></td><td></td><td></td><td></td><td></td><td></td><td></td><td></td><td></td><td></td></tr>
<tr><td>Oligo rev</td><td>AAACCGCGGCACCACGCGTCCAGG</td><td></td><td></td><td></td><td></td><td></td><td></td><td></td><td></td><td></td><td></td></tr>
<tr><td>Chromosome</td><td>start</td><td>end</td><td>strand</td><td>MM</td><td>target_seq</td><td>PAM</td><td>alignment</td><td>distance</td><td>position</td><td>gene name</td><td>gene id</td></tr>
<tr><td>GU305914.1</td><td>166028</td><td>166050</td><td>-</td><td>0</td><td>CCTGGACGCGTGGTGCCGCG</td><td>AGG</td><td>||||||||[||||||||||||]PAM</td><td>NA</td><td>Intergenic</td><td>NA</td><td>NA</td></tr>
<tr><td>T21844</td><td>CTGGACGCGTGGTGCCGCGAGGG</td><td>1000</td><td></td><td></td><td></td><td></td><td></td><td></td><td></td><td></td><td></td></tr>
<tr><td>Oligo fwd</td><td>CACCCTGGACGCGTGGTGCCGCGA</td><td></td><td></td><td></td><td></td><td></td><td></td><td></td><td></td><td></td><td></td></tr>
<tr><td>Oligo rev</td><td>AAACTCGCGGCACCACGCGTCCAG</td><td></td><td></td><td></td><td></td><td></td><td></td><td></td><td></td><td></td><td></td></tr>
<tr><td>Chromosome</td><td>start</td><td>end</td><td>strand</td><td>MM</td><td>target_seq</td><td>PAM</td><td>alignment</td><td>distance</td><td>position</td><td>gene name</td><td>gene id</td></tr>
<tr><td>GU305914.1</td><td>166027</td><td>166049</td><td>-</td><td>0</td><td>CTGGACGCGTGGTGCCGCGA</td><td>GGG</td><td>||||||||[||||||||||||]PAM</td><td>NA</td><td>Intergenic</td><td>NA</td><td>NA</td></tr>
<tr><td>T21845</td><td>GGGTGTGCTGCTAATCAATACGG</td><td>1000</td><td></td><td></td><td></td><td></td><td></td><td></td><td></td><td></td><td></td></tr>
<tr><td>Oligo fwd</td><td>CACCGGGTGTGCTGCTAATCAATA</td><td></td><td></td><td></td><td></td><td></td><td></td><td></td><td></td><td></td><td></td></tr>
<tr><td>Oligo rev</td><td>AAACTATTGATTAGCAGCACACCC</td><td></td><td></td><td></td><td></td><td></td><td></td><td></td><td></td><td></td><td></td></tr>
<tr><td>Chromosome</td><td>start</td><td>end</td><td>strand</td><td>MM</td><td>target_seq</td><td>PAM</td><td>alignment</td><td>distance</td><td>position</td><td>gene name</td><td>gene id</td></tr>
<tr><td>GU305914.1</td><td>166007</td><td>166029</td><td>-</td><td>0</td><td>GGGTGTGCTGCTAATCAATA</td><td>CGG</td><td>||||||||[||||||||||||]PAM</td><td>NA</td><td>Intergenic</td><td>NA</td><td>NA</td></tr>
<tr><td>T21846</td><td>AATCAATACGGTGTTCACCGTGG</td><td>1000</td><td></td><td></td><td></td><td></td><td></td><td></td><td></td><td></td><td></td></tr>
<tr><td>Oligo fwd</td><td>CACCAATCAATACGGTGTTCACCG</td><td></td><td></td><td></td><td></td><td></td><td></td><td></td><td></td><td></td><td></td></tr>
<tr><td>Oligo rev</td><td>AAACCGGTGAACACCGTATTGATT</td><td></td><td></td><td></td><td></td><td></td><td></td><td></td><td></td><td></td><td></td></tr>
<tr><td>Chromosome</td><td>start</td><td>end</td><td>strand</td><td>MM</td><td>target_seq</td><td>PAM</td><td>alignment</td><td>distance</td><td>position</td><td>gene name</td><td>gene id</td></tr>
<tr><td>GU305914.1</td><td>165995</td><td>166017</td><td>-</td><td>0</td><td>AATCAATACGGTGTTCACCG</td><td>TGG</td><td>||||||||[||||||||||||]PAM</td><td>NA</td><td>Intergenic</td><td>NA</td><td>NA</td></tr>
<tr><td>T21847</td><td>TACGGTGTTCACCGTGGTCAAGG</td><td>1000</td><td></td><td></td><td></td><td></td><td></td><td></td><td></td><td></td><td></td></tr>
<tr><td>Oligo fwd</td><td>CACCTACGGTGTTCACCGTGGTCA</td><td></td><td></td><td></td><td></td><td></td><td></td><td></td><td></td><td></td><td></td></tr>
<tr><td>Oligo rev</td><td>AAACTGACCACGGTGAACACCGTA</td><td></td><td></td><td></td><td></td><td></td><td></td><td></td><td></td><td></td><td></td></tr>
<tr><td>Chromosome</td><td>start</td><td>end</td><td>strand</td><td>MM</td><td>target_seq</td><td>PAM</td><td>alignment</td><td>distance</td><td>position</td><td>gene name</td><td>gene id</td></tr>
<tr><td>GU305914.1</td><td>165989</td><td>166011</td><td>-</td><td>0</td><td>TACGGTGTTCACCGTGGTCA</td><td>AGG</td><td>||||||||[||||||||||||]PAM</td><td>NA</td><td>Intergenic</td><td>NA</td><td>NA</td></tr>
<tr><td>T21848</td><td>ACGGTGTTCACCGTGGTCAAGGG</td><td>1000</td><td></td><td></td><td></td><td></td><td></td><td></td><td></td><td></td><td></td></tr>
<tr><td>Oligo fwd</td><td>CACCACGGTGTTCACCGTGGTCAA</td><td></td><td></td><td></td><td></td><td></td><td></td><td></td><td></td><td></td><td></td></tr>
<tr><td>Oligo rev</td><td>AAACTTGACCACGGTGAACACCGT</td><td></td><td></td><td></td><td></td><td></td><td></td><td></td><td></td><td></td><td></td></tr>
<tr><td>Chromosome</td><td>start</td><td>end</td><td>strand</td><td>MM</td><td>target_seq</td><td>PAM</td><td>alignment</td><td>distance</td><td>position</td><td>gene name</td><td>gene id</td></tr>
<tr><td>GU305914.1</td><td>165988</td><td>166010</td><td>-</td><td>0</td><td>ACGGTGTTCACCGTGGTCAA</td><td>GGG</td><td>||||||||[||||||||||||]PAM</td><td>NA</td><td>Intergenic</td><td>NA</td><td>NA</td></tr>
<tr><td>T21849</td><td>CACCGTGGTCAAGGGACAGCCGG</td><td>1000</td><td></td><td></td><td></td><td></td><td></td><td></td><td></td><td></td><td></td></tr>
<tr><td>Oligo fwd</td><td>CACCCACCGTGGTCAAGGGACAGC</td><td></td><td></td><td></td><td></td><td></td><td></td><td></td><td></td><td></td><td></td></tr>
<tr><td>Oligo rev</td><td>AAACGCTGTCCCTTGACCACGGTG</td><td></td><td></td><td></td><td></td><td></td><td></td><td></td><td></td><td></td><td></td></tr>
</table>

<table>
<tr><td>Chromosome</td><td>start</td><td>end</td><td>strand</td><td>MM</td><td>target_seq</td><td>PAM</td><td>alignment</td><td>distance</td><td>position</td><td>gene name</td><td>gene id</td></tr>
<tr><td>GU305914.1</td><td>165980</td><td>166002</td><td>-</td><td>0</td><td>CACCGTGGTCAAGGGACAGC</td><td>CGG</td><td>||||||||[||||||||||||]PAM</td><td>NA</td><td>Intergenic</td><td>NA</td><td>NA</td></tr>
<tr><td>T21850</td><td>ACCGTGGTCAAGGGACAGCCGGG</td><td>1000</td><td></td><td></td><td></td><td></td><td></td><td></td><td></td><td></td><td></td></tr>
<tr><td>Oligo fwd</td><td>CACCACCGTGGTCAAGGGACAGCC</td><td></td><td></td><td></td><td></td><td></td><td></td><td></td><td></td><td></td><td></td></tr>
<tr><td>Oligo rev</td><td>AAACGGCTGTCCCTTGACCACGGT</td><td></td><td></td><td></td><td></td><td></td><td></td><td></td><td></td><td></td><td></td></tr>
<tr><td>Chromosome</td><td>start</td><td>end</td><td>strand</td><td>MM</td><td>target_seq</td><td>PAM</td><td>alignment</td><td>distance</td><td>position</td><td>gene name</td><td>gene id</td></tr>
<tr><td>GU305914.1</td><td>165979</td><td>166001</td><td>-</td><td>0</td><td>ACCGTGGTCAAGGGACAGCC</td><td>GGG</td><td>||||||||[||||||||||||]PAM</td><td>NA</td><td>Intergenic</td><td>NA</td><td>NA</td></tr>
<tr><td>T21851</td><td>GGGACAGCCGGGCTCACACGAGG</td><td>1000</td><td></td><td></td><td></td><td></td><td></td><td></td><td></td><td></td><td></td></tr>
<tr><td>Oligo fwd</td><td>CACCGGGACAGCCGGGCTCACACG</td><td></td><td></td><td></td><td></td><td></td><td></td><td></td><td></td><td></td><td></td></tr>
<tr><td>Oligo rev</td><td>AAACCGTGTGAGCCCGGCTGTCCC</td><td></td><td></td><td></td><td></td><td></td><td></td><td></td><td></td><td></td><td></td></tr>
<tr><td>Chromosome</td><td>start</td><td>end</td><td>strand</td><td>MM</td><td>target_seq</td><td>PAM</td><td>alignment</td><td>distance</td><td>position</td><td>gene name</td><td>gene id</td></tr>
<tr><td>GU305914.1</td><td>165968</td><td>165990</td><td>-</td><td>0</td><td>GGGACAGCCGGGCTCACACG</td><td>AGG</td><td>||||||||[||||||||||||]PAM</td><td>NA</td><td>Intergenic</td><td>NA</td><td>NA</td></tr>
<tr><td>T21852</td><td>GCCGGGCTCACACGAGGCCCTGG</td><td>1000</td><td></td><td></td><td></td><td></td><td></td><td></td><td></td><td></td><td></td></tr>
<tr><td>Oligo fwd</td><td>CACCGCCGGGCTCACACGAGGCCC</td><td></td><td></td><td></td><td></td><td></td><td></td><td></td><td></td><td></td><td></td></tr>
<tr><td>Oligo rev</td><td>AAACGGGCCTCGTGTGAGCCCGGC</td><td></td><td></td><td></td><td></td><td></td><td></td><td></td><td></td><td></td><td></td></tr>
<tr><td>Chromosome</td><td>start</td><td>end</td><td>strand</td><td>MM</td><td>target_seq</td><td>PAM</td><td>alignment</td><td>distance</td><td>position</td><td>gene name</td><td>gene id</td></tr>
<tr><td>GU305914.1</td><td>165962</td><td>165984</td><td>-</td><td>0</td><td>GCCGGGCTCACACGAGGCCC</td><td>TGG</td><td>||||||||[||||||||||||]PAM</td><td>NA</td><td>Intergenic</td><td>NA</td><td>NA</td></tr>
<tr><td>T21853</td><td>CCGGGCTCACACGAGGCCCTGGG</td><td>1000</td><td></td><td></td><td></td><td></td><td></td><td></td><td></td><td></td><td></td></tr>
<tr><td>Oligo fwd</td><td>CACCCCGGGCTCACACGAGGCCCT</td><td></td><td></td><td></td><td></td><td></td><td></td><td></td><td></td><td></td><td></td></tr>
<tr><td>Oligo rev</td><td>AAACAGGGCCTCGTGTGAGCCCGG</td><td></td><td></td><td></td><td></td><td></td><td></td><td></td><td></td><td></td><td></td></tr>
<tr><td>Chromosome</td><td>start</td><td>end</td><td>strand</td><td>MM</td><td>target_seq</td><td>PAM</td><td>alignment</td><td>distance</td><td>position</td><td>gene name</td><td>gene id</td></tr>
<tr><td>GU305914.1</td><td>165961</td><td>165983</td><td>-</td><td>0</td><td>CCGGGCTCACACGAGGCCCT</td><td>GGG</td><td>||||||||[||||||||||||]PAM</td><td>NA</td><td>Intergenic</td><td>NA</td><td>NA</td></tr>
<tr><td>T21854</td><td>GCTCACACGAGGCCCTGGGATGG</td><td>1000</td><td></td><td></td><td></td><td></td><td></td><td></td><td></td><td></td><td></td></tr>
<tr><td>Oligo fwd</td><td>CACCGCTCACACGAGGCCCTGGGA</td><td></td><td></td><td></td><td></td><td></td><td></td><td></td><td></td><td></td><td></td></tr>
<tr><td>Oligo rev</td><td>AAACTCCCAGGGCCTCGTGTGAGC</td><td></td><td></td><td></td><td></td><td></td><td></td><td></td><td></td><td></td><td></td></tr>
<tr><td>Chromosome</td><td>start</td><td>end</td><td>strand</td><td>MM</td><td>target_seq</td><td>PAM</td><td>alignment</td><td>distance</td><td>position</td><td>gene name</td><td>gene id</td></tr>
<tr><td>GU305914.1</td><td>165957</td><td>165979</td><td>-</td><td>0</td><td>GCTCACACGAGGCCCTGGGA</td><td>TGG</td><td>||||||||[||||||||||||]PAM</td><td>NA</td><td>Intergenic</td><td>NA</td><td>NA</td></tr>
<tr><td>T21855</td><td>CCTGGGATGGCAGATCCTCTCGG</td><td>1000</td><td></td><td></td><td></td><td></td><td></td><td></td><td></td><td></td><td></td></tr>
<tr><td>Oligo fwd</td><td>CACCCCTGGGATGGCAGATCCTCT</td><td></td><td></td><td></td><td></td><td></td><td></td><td></td><td></td><td></td><td></td></tr>
<tr><td>Oligo rev</td><td>AAACAGAGGATCTGCCATCCCAGG</td><td></td><td></td><td></td><td></td><td></td><td></td><td></td><td></td><td></td><td></td></tr>
<tr><td>Chromosome</td><td>start</td><td>end</td><td>strand</td><td>MM</td><td>target_seq</td><td>PAM</td><td>alignment</td><td>distance</td><td>position</td><td>gene name</td><td>gene id</td></tr>
<tr><td>GU305914.1</td><td>165944</td><td>165966</td><td>-</td><td>0</td><td>CCTGGGATGGCAGATCCTCT</td><td>CGG</td><td>||||||||[||||||||||||]PAM</td><td>NA</td><td>Intergenic</td><td>NA</td><td>NA</td></tr>
<tr><td>T21856</td><td>CCTCTCGGATCGCGTCCTGCAGG</td><td>1000</td><td></td><td></td><td></td><td></td><td></td><td></td><td></td><td></td><td></td></tr>
<tr><td>Oligo fwd</td><td>CACCCCTCTCGGATCGCGTCCTGC</td><td></td><td></td><td></td><td></td><td></td><td></td><td></td><td></td><td></td><td></td></tr>
<tr><td>Oligo rev</td><td>AAACGCAGGACGCGATCCGAGAGG</td><td></td><td></td><td></td><td></td><td></td><td></td><td></td><td></td><td></td><td></td></tr>
<tr><td>Chromosome</td><td>start</td><td>end</td><td>strand</td><td>MM</td><td>target_seq</td><td>PAM</td><td>alignment</td><td>distance</td><td>position</td><td>gene name</td><td>gene id</td></tr>
<tr><td>GU305914.1</td><td>165929</td><td>165951</td><td>-</td><td>0</td><td>CCTCTCGGATCGCGTCCTGC</td><td>AGG</td><td>||||||||[||||||||||||]PAM</td><td>NA</td><td>Intergenic</td><td>NA</td><td>NA</td></tr>
<tr><td>GU305914.1</td><td>208141</td><td>208163</td><td>-</td><td>4</td><td>TCTCTCTGAACGCGTCCCGC</td><td>CGG</td><td>-|||||-|[|-|||||||-||]PAM</td><td>NA</td><td>Intergenic</td><td>NA</td><td>NA</td></tr>
<tr><td>T21857</td><td>GCTCTGTCCGAGCAGAGAGAAGG</td><td>1000</td><td></td><td></td><td></td><td></td><td></td><td></td><td></td><td></td><td></td></tr>
<tr><td>Oligo fwd</td><td>CACCGCTCTGTCCGAGCAGAGAGA</td><td></td><td></td><td></td><td></td><td></td><td></td><td></td><td></td><td></td><td></td></tr>
<tr><td>Oligo rev</td><td>AAACTCTCTCTGCTCGGACAGAGC</td><td></td><td></td><td></td><td></td><td></td><td></td><td></td><td></td><td></td><td></td></tr>
<tr><td>Chromosome</td><td>start</td><td>end</td><td>strand</td><td>MM</td><td>target_seq</td><td>PAM</td><td>alignment</td><td>distance</td><td>position</td><td>gene name</td><td>gene id</td></tr>
<tr><td>GU305914.1</td><td>165907</td><td>165929</td><td>-</td><td>0</td><td>GCTCTGTCCGAGCAGAGAGA</td><td>AGG</td><td>||||||||[||||||||||||]PAM</td><td>NA</td><td>Intergenic</td><td>NA</td><td>NA</td></tr>
<tr><td>T21858</td><td>GTCCGAGCAGAGAGAAGGACTGG</td><td>1000</td><td></td><td></td><td></td><td></td><td></td><td></td><td></td><td></td><td></td></tr>
<tr><td>Oligo fwd</td><td>CACCGTCCGAGCAGAGAGAAGGAC</td><td></td><td></td><td></td><td></td><td></td><td></td><td></td><td></td><td></td><td></td></tr>
<tr><td>Oligo rev</td><td>AAACGTCCTTCTCTCTGCTCGGAC</td><td></td><td></td><td></td><td></td><td></td><td></td><td></td><td></td><td></td><td></td></tr>
<tr><td>Chromosome</td><td>start</td><td>end</td><td>strand</td><td>MM</td><td>target_seq</td><td>PAM</td><td>alignment</td><td>distance</td><td>position</td><td>gene name</td><td>gene id</td></tr>
<tr><td>GU305914.1</td><td>165902</td><td>165924</td><td>-</td><td>0</td><td>GTCCGAGCAGAGAGAAGGAC</td><td>TGG</td><td>||||||||[||||||||||||]PAM</td><td>NA</td><td>Intergenic</td><td>NA</td><td>NA</td></tr>
<tr><td>T21859</td><td>AAGGACTGGTTTTCTTGTTGTGG</td><td>1000</td><td></td><td></td><td></td><td></td><td></td><td></td><td></td><td></td><td></td></tr>
<tr><td>Oligo fwd</td><td>CACCAAGGACTGGTTTTCTTGTTG</td><td></td><td></td><td></td><td></td><td></td><td></td><td></td><td></td><td></td><td></td></tr>
<tr><td>Oligo rev</td><td>AAACCAACAAGAAAACCAGTCCTT</td><td></td><td></td><td></td><td></td><td></td><td></td><td></td><td></td><td></td><td></td></tr>
<tr><td>Chromosome</td><td>start</td><td>end</td><td>strand</td><td>MM</td><td>target_seq</td><td>PAM</td><td>alignment</td><td>distance</td><td>position</td><td>gene name</td><td>gene id</td></tr>
<tr><td>GU305914.1</td><td>165888</td><td>165910</td><td>-</td><td>0</td><td>AAGGACTGGTTTTCTTGTTG</td><td>TGG</td><td>||||||||[||||||||||||]PAM</td><td>NA</td><td>Intergenic</td><td>NA</td><td>NA</td></tr>
<tr><td>T21860</td><td>AGGACTGGTTTTCTTGTTGTGGG</td><td>1000</td><td></td><td></td><td></td><td></td><td></td><td></td><td></td><td></td><td></td></tr>
<tr><td>Oligo fwd</td><td>CACCAGGACTGGTTTTCTTGTTGT</td><td></td><td></td><td></td><td></td><td></td><td></td><td></td><td></td><td></td><td></td></tr>
<tr><td>Oligo rev</td><td>AAACACAACAAGAAAACCAGTCCT</td><td></td><td></td><td></td><td></td><td></td><td></td><td></td><td></td><td></td><td></td></tr>
<tr><td>Chromosome</td><td>start</td><td>end</td><td>strand</td><td>MM</td><td>target_seq</td><td>PAM</td><td>alignment</td><td>distance</td><td>position</td><td>gene name</td><td>gene id</td></tr>
<tr><td>GU305914.1</td><td>165887</td><td>165909</td><td>-</td><td>0</td><td>AGGACTGGTTTTCTTGTTGT</td><td>GGG</td><td>||||||||[||||||||||||]PAM</td><td>NA</td><td>Intergenic</td><td>NA</td><td>NA</td></tr>
<tr><td>T21861</td><td>GGACTGGTTTTCTTGTTGTGGGG</td><td>1000</td><td></td><td></td><td></td><td></td><td></td><td></td><td></td><td></td><td></td></tr>
<tr><td>Oligo fwd</td><td>CACCGGACTGGTTTTCTTGTTGTG</td><td></td><td></td><td></td><td></td><td></td><td></td><td></td><td></td><td></td><td></td></tr>
<tr><td>Oligo rev</td><td>AAACCACAACAAGAAAACCAGTCC</td><td></td><td></td><td></td><td></td><td></td><td></td><td></td><td></td><td></td><td></td></tr>
<tr><td>Chromosome</td><td>start</td><td>end</td><td>strand</td><td>MM</td><td>target_seq</td><td>PAM</td><td>alignment</td><td>distance</td><td>position</td><td>gene name</td><td>gene id</td></tr>
<tr><td>GU305914.1</td><td>165886</td><td>165908</td><td>-</td><td>0</td><td>GGACTGGTTTTCTTGTTGTG</td><td>GGG</td><td>||||||||[||||||||||||]PAM</td><td>NA</td><td>Intergenic</td><td>NA</td><td>NA</td></tr>
<tr><td>T21862</td><td>GACTGGTTTTCTTGTTGTGGGGG</td><td>1000</td><td></td><td></td><td></td><td></td><td></td><td></td><td></td><td></td><td></td></tr>
<tr><td>Oligo fwd</td><td>CACCGACTGGTTTTCTTGTTGTGG</td><td></td><td></td><td></td><td></td><td></td><td></td><td></td><td></td><td></td><td></td></tr>
<tr><td>Oligo rev</td><td>AAACCCACAACAAGAAAACCAGTC</td><td></td><td></td><td></td><td></td><td></td><td></td><td></td><td></td><td></td><td></td></tr>
<tr><td>Chromosome</td><td>start</td><td>end</td><td>strand</td><td>MM</td><td>target_seq</td><td>PAM</td><td>alignment</td><td>distance</td><td>position</td><td>gene name</td><td>gene id</td></tr>
<tr><td>GU305914.1</td><td>165885</td><td>165907</td><td>-</td><td>0</td><td>GACTGGTTTTCTTGTTGTGG</td><td>GGG</td><td>||||||||[||||||||||||]PAM</td><td>NA</td><td>Intergenic</td><td>NA</td><td>NA</td></tr>
<tr><td>T21863</td><td>TTTCTTGTTGTGGGGGCTGCAGG</td><td>1000</td><td></td><td></td><td></td><td></td><td></td><td></td><td></td><td></td><td></td></tr>
<tr><td>Oligo fwd</td><td>CACCTTTCTTGTTGTGGGGGCTGC</td><td></td><td></td><td></td><td></td><td></td><td></td><td></td><td></td><td></td><td></td></tr>
<tr><td>Oligo rev</td><td>AAACGCAGCCCCCACAACAAGAAA</td><td></td><td></td><td></td><td></td><td></td><td></td><td></td><td></td><td></td><td></td></tr>
<tr><td>Chromosome</td><td>start</td><td>end</td><td>strand</td><td>MM</td><td>target_seq</td><td>PAM</td><td>alignment</td><td>distance</td><td>position</td><td>gene name</td><td>gene id</td></tr>
<tr><td>GU305914.1</td><td>165878</td><td>165900</td><td>-</td><td>0</td><td>TTTCTTGTTGTGGGGGCTGC</td><td>AGG</td><td>||||||||[||||||||||||]PAM</td><td>NA</td><td>Intergenic</td><td>NA</td><td>NA</td></tr>
<tr><td>T21864</td><td>GGGGCTGCAGGCTCAGAAGAAGG</td><td>1000</td><td></td><td></td><td></td><td></td><td></td><td></td><td></td><td></td><td></td></tr>
</table>

| | | | | | | | | | | | |
|---|---|---|---|---|---|---|---|---|---|---|---|
| Oligo fwd | CACCGGGGCTGCAGGCTCAGAAGA | | | | | | | | | | |
| Oligo rev | AAACTCTTCTGAGCCTGCAGCCCC | | | | | | | | | | |
| Chromosome | start | end | strand | MM | target_seq | PAM | alignment | distance | position | gene name | gene id |
| GU305914.1 | 165866 | 165888 | - | 0 | GGGGCTGCAGGCTCAGAAGA | AGG | \|\|\|\|\|\|\|\|[\|\|\|\|\|\|\|\|\|\|\|\|]PAM | NA | Intergenic | NA | NA |
| T21865 | AGGAATACCTGATCGACCCGAGG | 1000 | | | | | | | | | |
| Oligo fwd | CACCAGGAATACCTGATCGACCCG | | | | | | | | | | |
| Oligo rev | AAACCGGGTCGATCAGGTATTCCT | | | | | | | | | | |
| Chromosome | start | end | strand | MM | target_seq | PAM | alignment | distance | position | gene name | gene id |
| GU305914.1 | 165846 | 165868 | - | 0 | AGGAATACCTGATCGACCCG | AGG | \|\|\|\|\|\|\|\|[\|\|\|\|\|\|\|\|\|\|\|\|]PAM | NA | Intergenic | NA | NA |
| T21866 | CACCCGTCCCCACGCGCTCAAGG | 1000 | | | | | | | | | |
| Oligo fwd | CACCCACCCGTCCCCACGCGCTCA | | | | | | | | | | |
| Oligo rev | AAACTGAGCGCGTGGGGACGGGTG | | | | | | | | | | |
| Chromosome | start | end | strand | MM | target_seq | PAM | alignment | distance | position | gene name | gene id |
| GU305914.1 | 165799 | 165821 | - | 0 | CACCCGTCCCCACGCGCTCA | AGG | \|\|\|\|\|\|\|\|[\|\|\|\|\|\|\|\|\|\|\|\|]PAM | NA | Intergenic | NA | NA |
| T21867 | AGGAGCACGCAACCCCTTCGTGG | 1000 | | | | | | | | | |
| Oligo fwd | CACCAGGAGCACGCAACCCCTTCG | | | | | | | | | | |
| Oligo rev | AAACCGAAGGGGTTGCGTGCTCCT | | | | | | | | | | |
| Chromosome | start | end | strand | MM | target_seq | PAM | alignment | distance | position | gene name | gene id |
| GU305914.1 | 165779 | 165801 | - | 0 | AGGAGCACGCAACCCCTTCG | TGG | \|\|\|\|\|\|\|\|[\|\|\|\|\|\|\|\|\|\|\|\|]PAM | NA | Intergenic | NA | NA |
| T21868 | GGAGCACGCAACCCCTTCGTGGG | 1000 | | | | | | | | | |
| Oligo fwd | CACCGGAGCACGCAACCCCTTCGT | | | | | | | | | | |
| Oligo rev | AAACACGAAGGGGTTGCGTGCTCC | | | | | | | | | | |
| Chromosome | start | end | strand | MM | target_seq | PAM | alignment | distance | position | gene name | gene id |
| GU305914.1 | 165778 | 165800 | - | 0 | GGAGCACGCAACCCCTTCGT | GGG | \|\|\|\|\|\|\|\|[\|\|\|\|\|\|\|\|\|\|\|\|]PAM | NA | Intergenic | NA | NA |
| T21869 | GAGCACGCAACCCCTTCGTGGGG | 1000 | | | | | | | | | |
| Oligo fwd | CACCGAGCACGCAACCCCTTCGTG | | | | | | | | | | |
| Oligo rev | AAACCACGAAGGGGTTGCGTGCTC | | | | | | | | | | |
| Chromosome | start | end | strand | MM | target_seq | PAM | alignment | distance | position | gene name | gene id |
| GU305914.1 | 165777 | 165799 | - | 0 | GAGCACGCAACCCCTTCGTG | GGG | \|\|\|\|\|\|\|\|[\|\|\|\|\|\|\|\|\|\|\|\|]PAM | NA | Intergenic | NA | NA |
| T21870 | GGGGAACAACCACTTCGTCCTGG | 1000 | | | | | | | | | |
| Oligo fwd | CACCGGGGAACAACCACTTCGTCC | | | | | | | | | | |
| Oligo rev | AAACGGACGAAGTGGTTGTTCCCC | | | | | | | | | | |
| Chromosome | start | end | strand | MM | target_seq | PAM | alignment | distance | position | gene name | gene id |
| GU305914.1 | 165758 | 165780 | - | 0 | GGGGAACAACCACTTCGTCC | TGG | \|\|\|\|\|\|\|\|[\|\|\|\|\|\|\|\|\|\|\|\|]PAM | NA | Intergenic | NA | NA |
| T21871 | AACGAGTATCTGAGCCGCCGCGG | 1000 | | | | | | | | | |
| Oligo fwd | CACCAACGAGTATCTGAGCCGCCG | | | | | | | | | | |
| Oligo rev | AAACCGGCGGCTCAGATACTCGTT | | | | | | | | | | |
| Chromosome | start | end | strand | MM | target_seq | PAM | alignment | distance | position | gene name | gene id |
| GU305914.1 | 165733 | 165755 | - | 0 | AACGAGTATCTGAGCCGCCG | CGG | \|\|\|\|\|\|\|\|[\|\|\|\|\|\|\|\|\|\|\|\|]PAM | NA | Intergenic | NA | NA |
| T21872 | ACGAGTATCTGAGCCGCCGCGGG | 1000 | | | | | | | | | |
| Oligo fwd | CACCACGAGTATCTGAGCCGCCGC | | | | | | | | | | |
| Oligo rev | AAACGCGGCGGCTCAGATACTCGT | | | | | | | | | | |
| Chromosome | start | end | strand | MM | target_seq | PAM | alignment | distance | position | gene name | gene id |
| GU305914.1 | 165732 | 165754 | - | 0 | ACGAGTATCTGAGCCGCCGC | GGG | \|\|\|\|\|\|\|\|[\|\|\|\|\|\|\|\|\|\|\|\|]PAM | NA | Intergenic | NA | NA |
| T21873 | CGAGTATCTGAGCCGCCGCGGGG | 1000 | | | | | | | | | |
| Oligo fwd | CACCCGAGTATCTGAGCCGCCGCG | | | | | | | | | | |
| Oligo rev | AAACCGCGGCGGCTCAGATACTCG | | | | | | | | | | |
| Chromosome | start | end | strand | MM | target_seq | PAM | alignment | distance | position | gene name | gene id |
| GU305914.1 | 165731 | 165753 | - | 0 | CGAGTATCTGAGCCGCCGCG | GGG | \|\|\|\|\|\|\|\|[\|\|\|\|\|\|\|\|\|\|\|\|]PAM | NA | Intergenic | NA | NA |
| T21874 | GCCGCGGGGAGCGCGTCGACTGG | 1000 | | | | | | | | | |
| Oligo fwd | CACCGCCGCGGGGAGCGCGTCGAC | | | | | | | | | | |
| Oligo rev | AAACGTCGACGCGCTCCCCGCGGC | | | | | | | | | | |
| Chromosome | start | end | strand | MM | target_seq | PAM | alignment | distance | position | gene name | gene id |
| GU305914.1 | 165717 | 165739 | - | 0 | GCCGCGGGGAGCGCGTCGAC | TGG | \|\|\|\|\|\|\|\|[\|\|\|\|\|\|\|\|\|\|\|\|]PAM | NA | Intergenic | NA | NA |
| GU305914.1 | 33429 | 33451 | - | 4 | GCCGGGGCGAGCGCCTCGAG | AGG | \|\|\|\|-\|\|-[\|\|\|\|\|\|-\|\|\|\|-]PAM | NA | Intergenic | NA | NA |
| T21875 | CCACCAAACACAAAAAATAAAGG | 1000 | | | | | | | | | |
| Oligo fwd | CACCCCACCAAACACAAAAAATAA | | | | | | | | | | |
| Oligo rev | AAACTTATTTTTTGTGTTTGGTGG | | | | | | | | | | |
| Chromosome | start | end | strand | MM | target_seq | PAM | alignment | distance | position | gene name | gene id |
| GU305914.1 | 165622 | 165644 | - | 0 | CCACCAAACACAAAAAATAA | AGG | \|\|\|\|\|\|\|\|[\|\|\|\|\|\|\|\|\|\|\|\|]PAM | NA | Intergenic | NA | NA |
| T21876 | TGTTTATTGTAACAAATGTGTGG | 1000 | | | | | | | | | |
| Oligo fwd | CACCTGTTTATTGTAACAAATGTG | | | | | | | | | | |
| Oligo rev | AAACCACATTTGTTACAATAAACA | | | | | | | | | | |
| Chromosome | start | end | strand | MM | target_seq | PAM | alignment | distance | position | gene name | gene id |
| GU305914.1 | 165590 | 165612 | - | 0 | TGTTTATTGTAACAAATGTG | TGG | \|\|\|\|\|\|\|\|[\|\|\|\|\|\|\|\|\|\|\|\|]PAM | NA | Intergenic | NA | NA |
| T21877 | AAATATTAATTTTATTTACATGG | 1000 | | | | | | | | | |
| Oligo fwd | CACCAAATATTAATTTTATTTACA | | | | | | | | | | |
| Oligo rev | AAACTGTAAATAAAATTAATATTT | | | | | | | | | | |
| Chromosome | start | end | strand | MM | target_seq | PAM | alignment | distance | position | gene name | gene id |
| GU305914.1 | 165546 | 165568 | - | 0 | AAATATTAATTTTATTTACA | TGG | \|\|\|\|\|\|\|\|[\|\|\|\|\|\|\|\|\|\|\|\|]PAM | NA | Intergenic | NA | NA |
| T21878 | TATGTTGTTAGCACAGACAGAGG | 1000 | | | | | | | | | |
| Oligo fwd | CACCTATGTTGTTAGCACAGACAG | | | | | | | | | | |
| Oligo rev | AAACCTGTCTGTGCTAACAACATA | | | | | | | | | | |
| Chromosome | start | end | strand | MM | target_seq | PAM | alignment | distance | position | gene name | gene id |
| GU305914.1 | 165519 | 165541 | - | 0 | TATGTTGTTAGCACAGACAG | AGG | \|\|\|\|\|\|\|\|[\|\|\|\|\|\|\|\|\|\|\|\|]PAM | NA | Intergenic | NA | NA |

T21879 ATGTTGTTAGCACAGACAGAGGG 1000
Oligo fwd CACCATGTTGTTAGCACAGACAGA
Oligo rev AAACTCTGTCTGTGCTAACAACAT
Chromosome start end strand MM target_seq PAM alignment distance position gene name gene id
GU305914.1 165518 165540 - 0 ATGTTGTTAGCACAGACAGA GGG ||||||||[||||||||||||]PAM NA Intergenic NA NA

T21880 TGTTGTTAGCACAGACAGAGGGG 1000
Oligo fwd CACCTGTTGTTAGCACAGACAGAG
Oligo rev AAACCTCTGTCTGTGCTAACAACA
Chromosome start end strand MM target_seq PAM alignment distance position gene name gene id
GU305914.1 165517 165539 - 0 TGTTGTTAGCACAGACAGAG GGG ||||||||[||||||||||||]PAM NA Intergenic NA NA

T21881 GTTGTTAGCACAGACAGAGGGGG 1000
Oligo fwd CACCGTTGTTAGCACAGACAGAGG
Oligo rev AAACCCTCTGTCTGTGCTAACAAC
Chromosome start end strand MM target_seq PAM alignment distance position gene name gene id
GU305914.1 165516 165538 - 0 GTTGTTAGCACAGACAGAGG GGG ||||||||[||||||||||||]PAM NA Intergenic NA NA

T21882 AGCACAGACAGAGGGGGACCAGG 1000
Oligo fwd CACCAGCACAGACAGAGGGGGACC
Oligo rev AAACGGTCCCCCTCTGTCTGTGCT
Chromosome start end strand MM target_seq PAM alignment distance position gene name gene id
GU305914.1 165510 165532 - 0 AGCACAGACAGAGGGGGACC AGG ||||||||[||||||||||||]PAM NA Intergenic NA NA

T21883 GCACAGACAGAGGGGGACCAGGG 1000
Oligo fwd CACCGCACAGACAGAGGGGGACCA
Oligo rev AAACTGGTCCCCCTCTGTCTGTGC
Chromosome start end strand MM target_seq PAM alignment distance position gene name gene id
GU305914.1 165509 165531 - 0 GCACAGACAGAGGGGGACCA GGG ||||||||[||||||||||||]PAM NA Intergenic NA NA

T21884 ACCAGGGATAATATCTCCATCGG 1000
Oligo fwd CACCACCAGGGATAATATCTCCAT
Oligo rev AAACATGGAGATATTATCCCTGGT
Chromosome start end strand MM target_seq PAM alignment distance position gene name gene id
GU305914.1 165493 165515 - 0 ACCAGGGATAATATCTCCAT CGG ||||||||[||||||||||||]PAM NA Intergenic NA NA

T21885 GGATAATATCTCCATCGGCCCGG 1000
Oligo fwd CACCGGATAATATCTCCATCGGCC
Oligo rev AAACGGCCGATGGAGATATTATCC
Chromosome start end strand MM target_seq PAM alignment distance position gene name gene id
GU305914.1 165488 165510 - 0 GGATAATATCTCCATCGGCC CGG ||||||||[||||||||||||]PAM NA Intergenic NA NA

T21886 AGTATACTTATCATCAATAACGG 1000
Oligo fwd CACCAGTATACTTATCATCAATAA
Oligo rev AAACTTATTGATGATAAGTATACT
Chromosome start end strand MM target_seq PAM alignment distance position gene name gene id
GU305914.1 165439 165461 - 0 AGTATACTTATCATCAATAA CGG ||||||||[||||||||||||]PAM NA Intergenic NA NA

T21887 ATACTTATCATCAATAACGGTGG 1000
Oligo fwd CACCATACTTATCATCAATAACGG
Oligo rev AAACCCGTTATTGATGATAAGTAT
Chromosome start end strand MM target_seq PAM alignment distance position gene name gene id
GU305914.1 165436 165458 - 0 ATACTTATCATCAATAACGG TGG ||||||||[||||||||||||]PAM NA Intergenic NA NA

T21888 GACATACACAATTTTTTCAGTGG 1000
Oligo fwd CACCGACATACACAATTTTTTCAG
Oligo rev AAACCTGAAAAAATTGTGTATGTC
Chromosome start end strand MM target_seq PAM alignment distance position gene name gene id
GU305914.1 165398 165420 - 0 GACATACACAATTTTTTCAG TGG ||||||||[||||||||||||]PAM NA Intergenic NA NA

T21889 CATGAATACAAGTTTTATATAGG 1000
Oligo fwd CACCCATGAATACAAGTTTTATAT
Oligo rev AAACATATAAAACTTGTATTCATG
Chromosome start end strand MM target_seq PAM alignment distance position gene name gene id
GU305914.1 165368 165390 - 0 CATGAATACAAGTTTTATAT AGG ||||||||[||||||||||||]PAM NA Intergenic NA NA

T21890 CTTGCTTGCGTGCTTACAGACGG 1000
Oligo fwd CACCCTTGCTTGCGTGCTTACAGA
Oligo rev AAACTCTGTAAGCACGCAAGCAAG
Chromosome start end strand MM target_seq PAM alignment distance position gene name gene id
GU305914.1 165276 165298 - 0 CTTGCTTGCGTGCTTACAGA CGG ||||||||[||||||||||||]PAM NA Intergenic NA NA

T21891 GTGCTTACAGACGGCAGCATCGG 1000
Oligo fwd CACCGTGCTTACAGACGGCAGCAT
Oligo rev AAACATGCTGCCGTCTGTAAGCAC
Chromosome start end strand MM target_seq PAM alignment distance position gene name gene id
GU305914.1 165267 165289 - 0 GTGCTTACAGACGGCAGCAT CGG ||||||||[||||||||||||]PAM NA Intergenic NA NA

T21892 CAGAAGACCCGAGCGAGACGAGG 1000
Oligo fwd CACCCAGAAGACCCGAGCGAGACG
Oligo rev AAACCGTCTCGCTCGGGTCTTCTG
Chromosome start end strand MM target_seq PAM alignment distance position gene name gene id
GU305914.1 165244 165266 - 0 CAGAAGACCCGAGCGAGACG AGG ||||||||[||||||||||||]PAM NA Intergenic NA NA

T21893 GACCATATCCTCTAACATAGCGG 1000
Oligo fwd CACCGACCATATCCTCTAACATAG
Oligo rev AAACCTATGTTAGAGGATATGGTC
Chromosome start end strand MM target_seq PAM alignment distance position gene name gene id

GU305914.1 165124 165146 - 0 GACCATATCCTCTAACATAG CGG ||||||||[||||||||||||]PAM NA Intergenic NA NA

T21894 CATATCCTCTAACATAGCGGCGG 1000
Oligo fwd CACCCATATCCTCTAACATAGCGG
Oligo rev AAACCCGCTATGTTAGAGGATATG
Chromosome start end strand MM target_seq PAM alignment distance position gene name gene id
GU305914.1 165121 165143 - 0 CATATCCTCTAACATAGCGG CGG ||||||||[||||||||||||]PAM NA Intergenic NA NA

T21895 TAACATAGCGGCGGCCATCGCGG 1000
Oligo fwd CACCTAACATAGCGGCGGCCATCG
Oligo rev AAACCGATGGCCGCCGCTATGTTA
Chromosome start end strand MM target_seq PAM alignment distance position gene name gene id
GU305914.1 165112 165134 - 0 TAACATAGCGGCGGCCATCG CGG ||||||||[||||||||||||]PAM NA Intergenic NA NA

T21896 GGCCATCGCGGTTTCCTCCTCGG 1000
Oligo fwd CACCGGCCATCGCGGTTTCCTCCT
Oligo rev AAACAGGAGGAAACCGCGATGGCC
Chromosome start end strand MM target_seq PAM alignment distance position gene name gene id
GU305914.1 165100 165122 - 0 GGCCATCGCGGTTTCCTCCT CGG ||||||||[||||||||||||]PAM NA Intergenic NA NA
GU305914.1 201600 201622 + 3 CGCCACCGCGGTCTCCTCCT CGG -||||-||[||||-|||||||]PAM NA Intergenic NA NA

T21897 CGCGGTTTCCTCCTCGGAAACGG 1000
Oligo fwd CACCCGCGGTTTCCTCCTCGGAAA
Oligo rev AAACTTTCCGAGGAGGAAACCGCG
Chromosome start end strand MM target_seq PAM alignment distance position gene name gene id
GU305914.1 165094 165116 - 0 CGCGGTTTCCTCCTCGGAAA CGG ||||||||[||||||||||||]PAM NA Intergenic NA NA

T21898 CTCGGAAACGGCCAGAGTGCCGG 1000
Oligo fwd CACCCTCGGAAACGGCCAGAGTGC
Oligo rev AAACGCACTCTGGCCGTTTCCGAG
Chromosome start end strand MM target_seq PAM alignment distance position gene name gene id
GU305914.1 165082 165104 - 0 CTCGGAAACGGCCAGAGTGC CGG ||||||||[||||||||||||]PAM NA Intergenic NA NA

T21899 GGAAACGGCCAGAGTGCCGGCGG 1000
Oligo fwd CACCGGAAACGGCCAGAGTGCCGG
Oligo rev AAACCCGGCACTCTGGCCGTTTCC
Chromosome start end strand MM target_seq PAM alignment distance position gene name gene id
GU305914.1 165079 165101 - 0 GGAAACGGCCAGAGTGCCGG CGG ||||||||[||||||||||||]PAM NA Intergenic NA NA

T21900 AACGGCCAGAGTGCCGGCGGTGG 1000
Oligo fwd CACCAACGGCCAGAGTGCCGGCGG
Oligo rev AAACCCGCCGGCACTCTGGCCGTT
Chromosome start end strand MM target_seq PAM alignment distance position gene name gene id
GU305914.1 165076 165098 - 0 AACGGCCAGAGTGCCGGCGG TGG ||||||||[||||||||||||]PAM NA Intergenic NA NA

T21901 TCTCCAACCCGTCCTCCGTGAGG 1000
Oligo fwd CACCTCTCCAACCCGTCCTCCGTG
Oligo rev AAACCACGGAGGACGGGTTGGAGA
Chromosome start end strand MM target_seq PAM alignment distance position gene name gene id
GU305914.1 165038 165060 - 0 TCTCCAACCCGTCCTCCGTG AGG ||||||||[||||||||||||]PAM NA Intergenic NA NA

T21902 AACCCGTCCTCCGTGAGGTCTGG 1000
Oligo fwd CACCAACCCGTCCTCCGTGAGGTC
Oligo rev AAACGACCTCACGGAGGACGGGTT
Chromosome start end strand MM target_seq PAM alignment distance position gene name gene id
GU305914.1 165033 165055 - 0 AACCCGTCCTCCGTGAGGTC TGG ||||||||[||||||||||||]PAM NA Intergenic NA NA

T21903 ACCCGTCCTCCGTGAGGTCTGGG 1000
Oligo fwd CACCACCCGTCCTCCGTGAGGTCT
Oligo rev AAACAGACCTCACGGAGGACGGGT
Chromosome start end strand MM target_seq PAM alignment distance position gene name gene id
GU305914.1 165032 165054 - 0 ACCCGTCCTCCGTGAGGTCT GGG ||||||||[||||||||||||]PAM NA Intergenic NA NA

T21904 ACTCCTGCTCCTTCCCCGCGTGG 1000
Oligo fwd CACCACTCCTGCTCCTTCCCCGCG
Oligo rev AAACCGCGGGGAAGGAGCAGGAGT
Chromosome start end strand MM target_seq PAM alignment distance position gene name gene id
GU305914.1 164992 165014 - 0 ACTCCTGCTCCTTCCCCGCG TGG ||||||||[||||||||||||]PAM NA Intergenic NA NA

T21905 CCTGCTCCTTCCCCGCGTGGTGG 1000
Oligo fwd CACCCCTGCTCCTTCCCCGCGTGG
Oligo rev AAACCCACGCGGGGAAGGAGCAGG
Chromosome start end strand MM target_seq PAM alignment distance position gene name gene id
GU305914.1 164989 165011 - 0 CCTGCTCCTTCCCCGCGTGG TGG ||||||||[||||||||||||]PAM NA Intergenic NA NA

T21906 CTGCTCCTTCCCCGCGTGGTGGG 1000
Oligo fwd CACCCTGCTCCTTCCCCGCGTGGT
Oligo rev AAACACCACGCGGGGAAGGAGCAG
Chromosome start end strand MM target_seq PAM alignment distance position gene name gene id
GU305914.1 164988 165010 - 0 CTGCTCCTTCCCCGCGTGGT GGG ||||||||[||||||||||||]PAM NA Intergenic NA NA

T21907 CCTTCCCCGCGTGGTGGGACCGG 1000
Oligo fwd CACCCCTTCCCCGCGTGGTGGGAC
Oligo rev AAACGTCCCACCACGCGGGGAAGG
Chromosome start end strand MM target_seq PAM alignment distance position gene name gene id
GU305914.1 164983 165005 - 0 CCTTCCCCGCGTGGTGGGAC CGG ||||||||[||||||||||||]PAM NA Intergenic NA NA

T21908 TCCCCGCGTGGTGGGACCGGCGG 1000
Oligo fwd CACCTCCCCGCGTGGTGGGACCGG

<table>
<tr><td>Oligo rev</td><td>AAACCCGGTCCCACCACGCGGGGA</td><td></td><td></td><td></td><td></td><td></td><td></td><td></td><td></td><td></td><td></td></tr>
<tr><td>Chromosome</td><td>start</td><td>end</td><td>strand</td><td>MM</td><td>target_seq</td><td>PAM</td><td>alignment</td><td>distance</td><td>position</td><td>gene name</td><td>gene id</td></tr>
<tr><td>GU305914.1</td><td>164980</td><td>165002</td><td>-</td><td>0</td><td>TCCCCGCGTGGTGGGACCGG</td><td>CGG</td><td>||||||||[||||||||||||]PAM</td><td>NA</td><td>Intergenic</td><td>NA</td><td>NA</td></tr>
<tr><td></td><td></td><td></td><td></td><td></td><td></td><td></td><td></td><td></td><td></td><td></td><td></td></tr>
<tr><td>T21909</td><td>CCGCGTGGTGGGACCGGCGGCGG</td><td>1000</td><td></td><td></td><td></td><td></td><td></td><td></td><td></td><td></td><td></td></tr>
<tr><td>Oligo fwd</td><td>CACCCCGCGTGGTGGGACCGGCGG</td><td></td><td></td><td></td><td></td><td></td><td></td><td></td><td></td><td></td><td></td></tr>
<tr><td>Oligo rev</td><td>AAACCCGCCGGTCCCACCACGCGG</td><td></td><td></td><td></td><td></td><td></td><td></td><td></td><td></td><td></td><td></td></tr>
<tr><td>Chromosome</td><td>start</td><td>end</td><td>strand</td><td>MM</td><td>target_seq</td><td>PAM</td><td>alignment</td><td>distance</td><td>position</td><td>gene name</td><td>gene id</td></tr>
<tr><td>GU305914.1</td><td>164977</td><td>164999</td><td>-</td><td>0</td><td>CCGCGTGGTGGGACCGGCGG</td><td>CGG</td><td>||||||||[||||||||||||]PAM</td><td>NA</td><td>Intergenic</td><td>NA</td><td>NA</td></tr>
<tr><td></td><td></td><td></td><td></td><td></td><td></td><td></td><td></td><td></td><td></td><td></td><td></td></tr>
<tr><td>T21910</td><td>CGTGGTGGGACCGGCGGCGGCGG</td><td>1000</td><td></td><td></td><td></td><td></td><td></td><td></td><td></td><td></td><td></td></tr>
<tr><td>Oligo fwd</td><td>CACCCGTGGTGGGACCGGCGGCGG</td><td></td><td></td><td></td><td></td><td></td><td></td><td></td><td></td><td></td><td></td></tr>
<tr><td>Oligo rev</td><td>AAACCCGCCGCCGGTCCCACCACG</td><td></td><td></td><td></td><td></td><td></td><td></td><td></td><td></td><td></td><td></td></tr>
<tr><td>Chromosome</td><td>start</td><td>end</td><td>strand</td><td>MM</td><td>target_seq</td><td>PAM</td><td>alignment</td><td>distance</td><td>position</td><td>gene name</td><td>gene id</td></tr>
<tr><td>GU305914.1</td><td>164974</td><td>164996</td><td>-</td><td>0</td><td>CGTGGTGGGACCGGCGGCGG</td><td>CGG</td><td>||||||||[||||||||||||]PAM</td><td>NA</td><td>Intergenic</td><td>NA</td><td>NA</td></tr>
<tr><td>GU305914.1</td><td>168820</td><td>168842</td><td>+</td><td>3</td><td>GGTGGTGGCAGCGGCGGCGG</td><td>CGG</td><td>-|||||||[-|-|||||||||]PAM</td><td>NA</td><td>Intergenic</td><td>NA</td><td>NA</td></tr>
<tr><td></td><td></td><td></td><td></td><td></td><td></td><td></td><td></td><td></td><td></td><td></td><td></td></tr>
<tr><td>T21911</td><td>GGTGGGACCGGCGGCGGCGGCGG</td><td>1000</td><td></td><td></td><td></td><td></td><td></td><td></td><td></td><td></td><td></td></tr>
<tr><td>Oligo fwd</td><td>CACCGGTGGGACCGGCGGCGGCGG</td><td></td><td></td><td></td><td></td><td></td><td></td><td></td><td></td><td></td><td></td></tr>
<tr><td>Oligo rev</td><td>AAACCCGCCGCCGCCGGTCCCACC</td><td></td><td></td><td></td><td></td><td></td><td></td><td></td><td></td><td></td><td></td></tr>
<tr><td>Chromosome</td><td>start</td><td>end</td><td>strand</td><td>MM</td><td>target_seq</td><td>PAM</td><td>alignment</td><td>distance</td><td>position</td><td>gene name</td><td>gene id</td></tr>
<tr><td>GU305914.1</td><td>164971</td><td>164993</td><td>-</td><td>0</td><td>GGTGGGACCGGCGGCGGCGG</td><td>CGG</td><td>||||||||[||||||||||||]PAM</td><td>NA</td><td>Intergenic</td><td>NA</td><td>NA</td></tr>
<tr><td>GU305914.1</td><td>67739</td><td>67761</td><td>+</td><td>4</td><td>TCGGGGAGCGGCGGCGGCGG</td><td>CGG</td><td>---||||-[||||||||||||]PAM</td><td>NA</td><td>Intergenic</td><td>NA</td><td>NA</td></tr>
<tr><td>GU305914.1</td><td>163896</td><td>163918</td><td>+</td><td>4</td><td>GGATCGAGCGGCGGCGGCGG</td><td>CGG</td><td>||---||-[||||||||||||]PAM</td><td>NA</td><td>Intergenic</td><td>NA</td><td>NA</td></tr>
<tr><td>GU305914.1</td><td>154407</td><td>154429</td><td>+</td><td>3</td><td>GGTGGCGGCGGCGGCGGCGG</td><td>CGG</td><td>|||||---[||||||||||||]PAM</td><td>NA</td><td>Intergenic</td><td>NA</td><td>NA</td></tr>
<tr><td>GU305914.1</td><td>58123</td><td>58145</td><td>-</td><td>3</td><td>GGTGGCGGCGGCGGCGGCGG</td><td>AGG</td><td>|||||---[||||||||||||]PAM</td><td>NA</td><td>Intergenic</td><td>NA</td><td>NA</td></tr>
<tr><td>GU305914.1</td><td>154404</td><td>154426</td><td>+</td><td>4</td><td>GGCGGTGGCGGCGGCGGCGG</td><td>CGG</td><td>||-||---[||||||||||||]PAM</td><td>NA</td><td>Intergenic</td><td>NA</td><td>NA</td></tr>
<tr><td>GU305914.1</td><td>58126</td><td>58148</td><td>-</td><td>4</td><td>GGCGGTGGCGGCGGCGGCGG</td><td>CGG</td><td>||-||---[||||||||||||]PAM</td><td>NA</td><td>Intergenic</td><td>NA</td><td>NA</td></tr>
<tr><td>GU305914.1</td><td>30000</td><td>30022</td><td>+</td><td>4</td><td>GGTGCTGGCGGCGGCGGCGG</td><td>CGG</td><td>||||----[||||||||||||]PAM</td><td>NA</td><td>Intergenic</td><td>NA</td><td>NA</td></tr>
<tr><td>GU305914.1</td><td>168820</td><td>168842</td><td>+</td><td>4</td><td>GGTGGTGGCAGCGGCGGCGG</td><td>CGG</td><td>|||||---[|-||||||||||]PAM</td><td>NA</td><td>Intergenic</td><td>NA</td><td>NA</td></tr>
<tr><td></td><td></td><td></td><td></td><td></td><td></td><td></td><td></td><td></td><td></td><td></td><td></td></tr>
<tr><td>T21912</td><td>GGCGGCGGCGGCAGCAGCTGCGG</td><td>1000</td><td></td><td></td><td></td><td></td><td></td><td></td><td></td><td></td><td></td></tr>
<tr><td>Oligo fwd</td><td>CACCGGCGGCGGCGGCAGCAGCTG</td><td></td><td></td><td></td><td></td><td></td><td></td><td></td><td></td><td></td><td></td></tr>
<tr><td>Oligo rev</td><td>AAACCAGCTGCTGCCGCCGCCGCC</td><td></td><td></td><td></td><td></td><td></td><td></td><td></td><td></td><td></td><td></td></tr>
<tr><td>Chromosome</td><td>start</td><td>end</td><td>strand</td><td>MM</td><td>target_seq</td><td>PAM</td><td>alignment</td><td>distance</td><td>position</td><td>gene name</td><td>gene id</td></tr>
<tr><td>GU305914.1</td><td>164959</td><td>164981</td><td>-</td><td>0</td><td>GGCGGCGGCGGCAGCAGCTG</td><td>CGG</td><td>||||||||[||||||||||||]PAM</td><td>NA</td><td>Intergenic</td><td>NA</td><td>NA</td></tr>
<tr><td>GU305914.1</td><td>128727</td><td>128749</td><td>-</td><td>3</td><td>GGTGGAGGCGGCAGCAGCAG</td><td>CGG</td><td>||-||-||[||||||||||-|]PAM</td><td>NA</td><td>Intergenic</td><td>NA</td><td>NA</td></tr>
<tr><td>GU305914.1</td><td>62680</td><td>62702</td><td>+</td><td>4</td><td>CCCGCCGGCGGCAGCAGCCG</td><td>CGG</td><td>--||-|||[||||||||||-|]PAM</td><td>NA</td><td>Intergenic</td><td>NA</td><td>NA</td></tr>
<tr><td>GU305914.1</td><td>27408</td><td>27430</td><td>+</td><td>4</td><td>GGCAGCAGCAGCAGCAGCAG</td><td>CGG</td><td>|||-||-|[|-||||||||-|]PAM</td><td>NA</td><td>Intergenic</td><td>NA</td><td>NA</td></tr>
<tr><td>GU305914.1</td><td>163908</td><td>163930</td><td>+</td><td>2</td><td>GGCGGCGGCGGCAGCGGCAG</td><td>CGG</td><td>||||||||[|||||||-||-|]PAM</td><td>NA</td><td>Intergenic</td><td>NA</td><td>NA</td></tr>
<tr><td>GU305914.1</td><td>151362</td><td>151384</td><td>+</td><td>4</td><td>GGCGACGACGGCAGCAGCGA</td><td>AGG</td><td>||||-||-[||||||||||--]PAM</td><td>NA</td><td>Intergenic</td><td>NA</td><td>NA</td></tr>
<tr><td></td><td></td><td></td><td></td><td></td><td></td><td></td><td></td><td></td><td></td><td></td><td></td></tr>
<tr><td>T21913</td><td>GCGGCGGCGGCAGCAGCTGCGGG</td><td>1000</td><td></td><td></td><td></td><td></td><td></td><td></td><td></td><td></td><td></td></tr>
<tr><td>Oligo fwd</td><td>CACCGCGGCGGCGGCAGCAGCTGC</td><td></td><td></td><td></td><td></td><td></td><td></td><td></td><td></td><td></td><td></td></tr>
<tr><td>Oligo rev</td><td>AAACGCAGCTGCTGCCGCCGCCGC</td><td></td><td></td><td></td><td></td><td></td><td></td><td></td><td></td><td></td><td></td></tr>
<tr><td>Chromosome</td><td>start</td><td>end</td><td>strand</td><td>MM</td><td>target_seq</td><td>PAM</td><td>alignment</td><td>distance</td><td>position</td><td>gene name</td><td>gene id</td></tr>
<tr><td>GU305914.1</td><td>164958</td><td>164980</td><td>-</td><td>0</td><td>GCGGCGGCGGCAGCAGCTGC</td><td>GGG</td><td>||||||||[||||||||||||]PAM</td><td>NA</td><td>Intergenic</td><td>NA</td><td>NA</td></tr>
<tr><td>GU305914.1</td><td>65720</td><td>65742</td><td>-</td><td>4</td><td>CCTGCGGCGGCAGCGGCAGC</td><td>GGG</td><td>-|-|||||[||||||-||-||]PAM</td><td>NA</td><td>Intergenic</td><td>NA</td><td>NA</td></tr>
<tr><td></td><td></td><td></td><td></td><td></td><td></td><td></td><td></td><td></td><td></td><td></td><td></td></tr>
<tr><td>T21914</td><td>GCGGCGGCAGCAGCTGCGGGTGG</td><td>1000</td><td></td><td></td><td></td><td></td><td></td><td></td><td></td><td></td><td></td></tr>
<tr><td>Oligo fwd</td><td>CACCGCGGCGGCAGCAGCTGCGGG</td><td></td><td></td><td></td><td></td><td></td><td></td><td></td><td></td><td></td><td></td></tr>
<tr><td>Oligo rev</td><td>AAACCCCGCAGCTGCTGCCGCCGC</td><td></td><td></td><td></td><td></td><td></td><td></td><td></td><td></td><td></td><td></td></tr>
<tr><td>Chromosome</td><td>start</td><td>end</td><td>strand</td><td>MM</td><td>target_seq</td><td>PAM</td><td>alignment</td><td>distance</td><td>position</td><td>gene name</td><td>gene id</td></tr>
<tr><td>GU305914.1</td><td>164955</td><td>164977</td><td>-</td><td>0</td><td>GCGGCGGCAGCAGCTGCGGG</td><td>TGG</td><td>||||||||[||||||||||||]PAM</td><td>NA</td><td>Intergenic</td><td>NA</td><td>NA</td></tr>
<tr><td>GU305914.1</td><td>68416</td><td>68438</td><td>+</td><td>4</td><td>GCTGCTGCAGCAGCTGCCCG</td><td>TGG</td><td>||-||-||[|||||||||--|]PAM</td><td>NA</td><td>Intergenic</td><td>NA</td><td>NA</td></tr>
<tr><td></td><td></td><td></td><td></td><td></td><td></td><td></td><td></td><td></td><td></td><td></td><td></td></tr>
<tr><td>T21915</td><td>CGGCGGCAGCAGCTGCGGGTGGG</td><td>1000</td><td></td><td></td><td></td><td></td><td></td><td></td><td></td><td></td><td></td></tr>
<tr><td>Oligo fwd</td><td>CACCCGGCGGCAGCAGCTGCGGGT</td><td></td><td></td><td></td><td></td><td></td><td></td><td></td><td></td><td></td><td></td></tr>
<tr><td>Oligo rev</td><td>AAACACCCGCAGCTGCTGCCGCCG</td><td></td><td></td><td></td><td></td><td></td><td></td><td></td><td></td><td></td><td></td></tr>
<tr><td>Chromosome</td><td>start</td><td>end</td><td>strand</td><td>MM</td><td>target_seq</td><td>PAM</td><td>alignment</td><td>distance</td><td>position</td><td>gene name</td><td>gene id</td></tr>
<tr><td>GU305914.1</td><td>164954</td><td>164976</td><td>-</td><td>0</td><td>CGGCGGCAGCAGCTGCGGGT</td><td>GGG</td><td>||||||||[||||||||||||]PAM</td><td>NA</td><td>Intergenic</td><td>NA</td><td>NA</td></tr>
<tr><td></td><td></td><td></td><td></td><td></td><td></td><td></td><td></td><td></td><td></td><td></td><td></td></tr>
<tr><td>T21916</td><td>GGCAGCAGCTGCGGGTGGGTCGG</td><td>1000</td><td></td><td></td><td></td><td></td><td></td><td></td><td></td><td></td><td></td></tr>
<tr><td>Oligo fwd</td><td>CACCGGCAGCAGCTGCGGGTGGGT</td><td></td><td></td><td></td><td></td><td></td><td></td><td></td><td></td><td></td><td></td></tr>
<tr><td>Oligo rev</td><td>AAACACCCACCCGCAGCTGCTGCC</td><td></td><td></td><td></td><td></td><td></td><td></td><td></td><td></td><td></td><td></td></tr>
<tr><td>Chromosome</td><td>start</td><td>end</td><td>strand</td><td>MM</td><td>target_seq</td><td>PAM</td><td>alignment</td><td>distance</td><td>position</td><td>gene name</td><td>gene id</td></tr>
<tr><td>GU305914.1</td><td>164950</td><td>164972</td><td>-</td><td>0</td><td>GGCAGCAGCTGCGGGTGGGT</td><td>CGG</td><td>||||||||[||||||||||||]PAM</td><td>NA</td><td>Intergenic</td><td>NA</td><td>NA</td></tr>
<tr><td></td><td></td><td></td><td></td><td></td><td></td><td></td><td></td><td></td><td></td><td></td><td></td></tr>
<tr><td>T21917</td><td>GCAGCTGCGGGTGGGTCGGTCGG</td><td>1000</td><td></td><td></td><td></td><td></td><td></td><td></td><td></td><td></td><td></td></tr>
<tr><td>Oligo fwd</td><td>CACCGCAGCTGCGGGTGGGTCGGT</td><td></td><td></td><td></td><td></td><td></td><td></td><td></td><td></td><td></td><td></td></tr>
<tr><td>Oligo rev</td><td>AAACACCGACCCACCCGCAGCTGC</td><td></td><td></td><td></td><td></td><td></td><td></td><td></td><td></td><td></td><td></td></tr>
<tr><td>Chromosome</td><td>start</td><td>end</td><td>strand</td><td>MM</td><td>target_seq</td><td>PAM</td><td>alignment</td><td>distance</td><td>position</td><td>gene name</td><td>gene id</td></tr>
<tr><td>GU305914.1</td><td>164946</td><td>164968</td><td>-</td><td>0</td><td>GCAGCTGCGGGTGGGTCGGT</td><td>CGG</td><td>||||||||[||||||||||||]PAM</td><td>NA</td><td>Intergenic</td><td>NA</td><td>NA</td></tr>
<tr><td></td><td></td><td></td><td></td><td></td><td></td><td></td><td></td><td></td><td></td><td></td><td></td></tr>
<tr><td>T21918</td><td>GTGGGTCGGTCGGATCGACCTGG</td><td>1000</td><td></td><td></td><td></td><td></td><td></td><td></td><td></td><td></td><td></td></tr>
<tr><td>Oligo fwd</td><td>CACCGTGGGTCGGTCGGATCGACC</td><td></td><td></td><td></td><td></td><td></td><td></td><td></td><td></td><td></td><td></td></tr>
<tr><td>Oligo rev</td><td>AAACGGTCGATCCGACCGACCCAC</td><td></td><td></td><td></td><td></td><td></td><td></td><td></td><td></td><td></td><td></td></tr>
<tr><td>Chromosome</td><td>start</td><td>end</td><td>strand</td><td>MM</td><td>target_seq</td><td>PAM</td><td>alignment</td><td>distance</td><td>position</td><td>gene name</td><td>gene id</td></tr>
<tr><td>GU305914.1</td><td>164936</td><td>164958</td><td>-</td><td>0</td><td>GTGGGTCGGTCGGATCGACC</td><td>TGG</td><td>||||||||[||||||||||||]PAM</td><td>NA</td><td>Intergenic</td><td>NA</td><td>NA</td></tr>
<tr><td></td><td></td><td></td><td></td><td></td><td></td><td></td><td></td><td></td><td></td><td></td><td></td></tr>
<tr><td>T21919</td><td>TGGGTCGGTCGGATCGACCTGGG</td><td>1000</td><td></td><td></td><td></td><td></td><td></td><td></td><td></td><td></td><td></td></tr>
<tr><td>Oligo fwd</td><td>CACCTGGGTCGGTCGGATCGACCT</td><td></td><td></td><td></td><td></td><td></td><td></td><td></td><td></td><td></td><td></td></tr>
<tr><td>Oligo rev</td><td>AAACAGGTCGATCCGACCGACCCA</td><td></td><td></td><td></td><td></td><td></td><td></td><td></td><td></td><td></td><td></td></tr>
<tr><td>Chromosome</td><td>start</td><td>end</td><td>strand</td><td>MM</td><td>target_seq</td><td>PAM</td><td>alignment</td><td>distance</td><td>position</td><td>gene name</td><td>gene id</td></tr>
<tr><td>GU305914.1</td><td>164935</td><td>164957</td><td>-</td><td>0</td><td>TGGGTCGGTCGGATCGACCT</td><td>GGG</td><td>||||||||[||||||||||||]PAM</td><td>NA</td><td>Intergenic</td><td>NA</td><td>NA</td></tr>
<tr><td></td><td></td><td></td><td></td><td></td><td></td><td></td><td></td><td></td><td></td><td></td><td></td></tr>
<tr><td>T21920</td><td>GCGACCCGACTGTTCCTCCGAGG</td><td>1000</td><td></td><td></td><td></td><td></td><td></td><td></td><td></td><td></td><td></td></tr>
<tr><td>Oligo fwd</td><td>CACCGCGACCCGACTGTTCCTCCG</td><td></td><td></td><td></td><td></td><td></td><td></td><td></td><td></td><td></td><td></td></tr>
<tr><td>Oligo rev</td><td>AAACCGGAGGAACAGTCGGGTCGC</td><td></td><td></td><td></td><td></td><td></td><td></td><td></td><td></td><td></td><td></td></tr>
</table>

Chromosome start end strand MM target_seq PAM alignment distance position gene name gene id
GU305914.1 164901 164923 - 0 GCGACCCGACTGTTCCTCCG AGG ||||||||[||||||||||||]PAM NA Intergenic NA NA

T21921 GCCTCTTCCCCGCCTGCGAGCGG 1000
Oligo fwd CACCGCCTCTTCCCCGCCTGCGAG
Oligo rev AAACCTCGCAGGCGGGGAAGAGGC
Chromosome start end strand MM target_seq PAM alignment distance position gene name gene id
GU305914.1 164872 164894 - 0 GCCTCTTCCCCGCCTGCGAG CGG ||||||||[||||||||||||]PAM NA Intergenic NA NA

T21922 TCTTCCCCGCCTGCGAGCGGCGG 1000
Oligo fwd CACCTCTTCCCCGCCTGCGAGCGG
Oligo rev AAACCCGCTCGCAGGCGGGGAAGA
Chromosome start end strand MM target_seq PAM alignment distance position gene name gene id
GU305914.1 164869 164891 - 0 TCTTCCCCGCCTGCGAGCGG CGG ||||||||[||||||||||||]PAM NA Intergenic NA NA

T21923 CCGCCTGCGAGCGGCGGCCGTGG 1000
Oligo fwd CACCCCGCCTGCGAGCGGCGGCCG
Oligo rev AAACCGGCCGCCGCTCGCAGGCGG
Chromosome start end strand MM target_seq PAM alignment distance position gene name gene id
GU305914.1 164863 164885 - 0 CCGCCTGCGAGCGGCGGCCG TGG ||||||||[||||||||||||]PAM NA Intergenic NA NA

T21924 CCTGCGAGCGGCGGCCGTGGTGG 1000
Oligo fwd CACCCCTGCGAGCGGCGGCCGTGG
Oligo rev AAACCCACGGCCGCCGCTCGCAGG
Chromosome start end strand MM target_seq PAM alignment distance position gene name gene id
GU305914.1 164860 164882 - 0 CCTGCGAGCGGCGGCCGTGG TGG ||||||||[||||||||||||]PAM NA Intergenic NA NA

T21925 AGCGGCGGCCGTGGTGGCTCTGG 1000
Oligo fwd CACCAGCGGCGGCCGTGGTGGCTC
Oligo rev AAACGAGCCACCACGGCCGCCGCT
Chromosome start end strand MM target_seq PAM alignment distance position gene name gene id
GU305914.1 164854 164876 - 0 AGCGGCGGCCGTGGTGGCTC TGG ||||||||[||||||||||||]PAM NA Intergenic NA NA
GU305914.1 185192 185214 + 4 AGCGGGCGCCCGGGTGGCTC GGG |||||--|[||--||||||||]PAM NA Intergenic NA NA
GU305914.1 221603 221625 - 4 ACCGGAGGCCGTGGTGGCGG CGG |-|||-||[||||||||||--]PAM NA Intergenic NA NA

T21926 GGTGGCTCTGGAGCGCGAAGCGG 1000
Oligo fwd CACCGGTGGCTCTGGAGCGCGAAG
Oligo rev AAACCTTCGCGCTCCAGAGCCACC
Chromosome start end strand MM target_seq PAM alignment distance position gene name gene id
GU305914.1 164842 164864 - 0 GGTGGCTCTGGAGCGCGAAG CGG ||||||||[||||||||||||]PAM NA Intergenic NA NA

T21927 GCTCTGGAGCGCGAAGCGGTCGG 1000
Oligo fwd CACCGCTCTGGAGCGCGAAGCGGT
Oligo rev AAACACCGCTTCGCGCTCCAGAGC
Chromosome start end strand MM target_seq PAM alignment distance position gene name gene id
GU305914.1 164838 164860 - 0 GCTCTGGAGCGCGAAGCGGT CGG ||||||||[||||||||||||]PAM NA Intergenic NA NA

T21928 GAAGCGGTCGGCGCCGCCGCCGG 1000
Oligo fwd CACCGAAGCGGTCGGCGCCGCCGC
Oligo rev AAACGCGGCGGCGCCGACCGCTTC
Chromosome start end strand MM target_seq PAM alignment distance position gene name gene id
GU305914.1 164826 164848 - 0 GAAGCGGTCGGCGCCGCCGC CGG ||||||||[||||||||||||]PAM NA Intergenic NA NA
GU305914.1 86843 86865 + 4 GAAGCTGCCGCCCCCGCCGC GGG |||||-|-[||-|-|||||||]PAM NA Intergenic NA NA

T21929 ACAGCGAGCTATGTCGATGAAGG 1000
Oligo fwd CACCACAGCGAGCTATGTCGATGA
Oligo rev AAACTCATCGACATAGCTCGCTGT
Chromosome start end strand MM target_seq PAM alignment distance position gene name gene id
GU305914.1 164797 164819 - 0 ACAGCGAGCTATGTCGATGA AGG ||||||||[||||||||||||]PAM NA Intergenic NA NA

T21930 CAGCGAGCTATGTCGATGAAGGG 1000
Oligo fwd CACCCAGCGAGCTATGTCGATGAA
Oligo rev AAACTTCATCGACATAGCTCGCTG
Chromosome start end strand MM target_seq PAM alignment distance position gene name gene id
GU305914.1 164796 164818 - 0 CAGCGAGCTATGTCGATGAA GGG ||||||||[||||||||||||]PAM NA Intergenic NA NA

T21931 AAGGGCCGTATCTCACATACAGG 1000
Oligo fwd CACCAAGGGCCGTATCTCACATAC
Oligo rev AAACGTATGTGAGATACGGCCCTT
Chromosome start end strand MM target_seq PAM alignment distance position gene name gene id
GU305914.1 164778 164800 - 0 AAGGGCCGTATCTCACATAC AGG ||||||||[||||||||||||]PAM NA Intergenic NA NA

T21932 ACATACAGGTTTCTGCTGAGCGG 1000
Oligo fwd CACCACATACAGGTTTCTGCTGAG
Oligo rev AAACCTCAGCAGAAACCTGTATGT
Chromosome start end strand MM target_seq PAM alignment distance position gene name gene id
GU305914.1 164764 164786 - 0 ACATACAGGTTTCTGCTGAG CGG ||||||||[||||||||||||]PAM NA Intergenic NA NA

T21933 TACAGGTTTCTGCTGAGCGGCGG 1000
Oligo fwd CACCTACAGGTTTCTGCTGAGCGG
Oligo rev AAACCCGCTCAGCAGAAACCTGTA
Chromosome start end strand MM target_seq PAM alignment distance position gene name gene id
GU305914.1 164761 164783 - 0 TACAGGTTTCTGCTGAGCGG CGG ||||||||[||||||||||||]PAM NA Intergenic NA NA

T21934 CCCTGTCGAAGATACCCCTGTGG 1000
Oligo fwd CACCCCCTGTCGAAGATACCCCTG
Oligo rev AAACCAGGGGTATCTTCGACAGGG
Chromosome start end strand MM target_seq PAM alignment distance position gene name gene id
GU305914.1 164732 164754 - 0 CCCTGTCGAAGATACCCCTG TGG ||||||||[||||||||||||]PAM NA Intergenic NA NA

T21935 TCGAAGATACCCCTGTGGTGAGG 1000
Oligo fwd CACCTCGAAGATACCCCTGTGGTG
Oligo rev AAACCACCACAGGGGTATCTTCGA
Chromosome start end strand MM target_seq PAM alignment distance position gene name gene id
GU305914.1 164727 164749 - 0 TCGAAGATACCCCTGTGGTG AGG ||||||||[||||||||||||]PAM NA Intergenic NA NA

T21936 AGGCATCCCCTCTGTCTGCGTGG 1000
Oligo fwd CACCAGGCATCCCCTCTGTCTGCG
Oligo rev AAACCGCAGACAGAGGGGATGCCT
Chromosome start end strand MM target_seq PAM alignment distance position gene name gene id
GU305914.1 164707 164729 - 0 AGGCATCCCCTCTGTCTGCG TGG ||||||||[||||||||||||]PAM NA Intergenic NA NA

T21937 CTGCGTGGTGCCTGTACAACTGG 1000
Oligo fwd CACCCTGCGTGGTGCCTGTACAAC
Oligo rev AAACGTTGTACAGGCACCACGCAG
Chromosome start end strand MM target_seq PAM alignment distance position gene name gene id
GU305914.1 164692 164714 - 0 CTGCGTGGTGCCTGTACAAC TGG ||||||||[||||||||||||]PAM NA Intergenic NA NA

T21938 GTGCCTGTACAACTGGATCCCGG 1000
Oligo fwd CACCGTGCCTGTACAACTGGATCC
Oligo rev AAACGGATCCAGTTGTACAGGCAC
Chromosome start end strand MM target_seq PAM alignment distance position gene name gene id
GU305914.1 164685 164707 - 0 GTGCCTGTACAACTGGATCC CGG ||||||||[||||||||||||]PAM NA Intergenic NA NA

T21939 TGTACAACTGGATCCCGGTACGG 1000
Oligo fwd CACCTGTACAACTGGATCCCGGTA
Oligo rev AAACTACCGGGATCCAGTTGTACA
Chromosome start end strand MM target_seq PAM alignment distance position gene name gene id
GU305914.1 164680 164702 - 0 TGTACAACTGGATCCCGGTA CGG ||||||||[||||||||||||]PAM NA Intergenic NA NA

T21940 ACAACTGGATCCCGGTACGGCGG 1000
Oligo fwd CACCACAACTGGATCCCGGTACGG
Oligo rev AAACCCGTACCGGGATCCAGTTGT
Chromosome start end strand MM target_seq PAM alignment distance position gene name gene id
GU305914.1 164677 164699 - 0 ACAACTGGATCCCGGTACGG CGG ||||||||[||||||||||||]PAM NA Intergenic NA NA

T21941 ACTGGATCCCGGTACGGCGGCGG 1000
Oligo fwd CACCACTGGATCCCGGTACGGCGG
Oligo rev AAACCCGCCGTACCGGGATCCAGT
Chromosome start end strand MM target_seq PAM alignment distance position gene name gene id
GU305914.1 164674 164696 - 0 ACTGGATCCCGGTACGGCGG CGG ||||||||[||||||||||||]PAM NA Intergenic NA NA

T21942 TACGGCGGCGGAACTCGAGCTGG 1000
Oligo fwd CACCTACGGCGGCGGAACTCGAGC
Oligo rev AAACGCTCGAGTTCCGCCGCCGTA
Chromosome start end strand MM target_seq PAM alignment distance position gene name gene id
GU305914.1 164662 164684 - 0 TACGGCGGCGGAACTCGAGC TGG ||||||||[||||||||||||]PAM NA Intergenic NA NA
GU305914.1 20641 20663 - 4 TACGGTCGCGGAACTCGTCC TGG |||||--|[|||||||||--|]PAM NA Intergenic NA NA

T21943 ACTCGTCTAAGAGTCTCATCAGG 1000
Oligo fwd CACCACTCGTCTAAGAGTCTCATC
Oligo rev AAACGATGAGACTCTTAGACGAGT
Chromosome start end strand MM target_seq PAM alignment distance position gene name gene id
GU305914.1 164624 164646 - 0 ACTCGTCTAAGAGTCTCATC AGG ||||||||[||||||||||||]PAM NA Intergenic NA NA

T21944 CTCGTCTAAGAGTCTCATCAGGG 1000
Oligo fwd CACCCTCGTCTAAGAGTCTCATCA
Oligo rev AAACTGATGAGACTCTTAGACGAG
Chromosome start end strand MM target_seq PAM alignment distance position gene name gene id
GU305914.1 164623 164645 - 0 CTCGTCTAAGAGTCTCATCA GGG ||||||||[||||||||||||]PAM NA Intergenic NA NA

T21945 TTAAAGACGCCTTCAAAACGCGG 1000
Oligo fwd CACCTTAAAGACGCCTTCAAAACG
Oligo rev AAACCGTTTTGAAGGCGTCTTTAA
Chromosome start end strand MM target_seq PAM alignment distance position gene name gene id
GU305914.1 164598 164620 - 0 TTAAAGACGCCTTCAAAACG CGG ||||||||[||||||||||||]PAM NA Intergenic NA NA

T21946 TAAAGACGCCTTCAAAACGCGGG 1000
Oligo fwd CACCTAAAGACGCCTTCAAAACGC
Oligo rev AAACGCGTTTTGAAGGCGTCTTTA
Chromosome start end strand MM target_seq PAM alignment distance position gene name gene id
GU305914.1 164597 164619 - 0 TAAAGACGCCTTCAAAACGC GGG ||||||||[||||||||||||]PAM NA Intergenic NA NA

T21947 TTGTTGTCCTCCTGTGAATTCGG 1000
Oligo fwd CACCTTGTTGTCCTCCTGTGAATT
Oligo rev AAACAATTCACAGGAGGACAACAA
Chromosome start end strand MM target_seq PAM alignment distance position gene name gene id
GU305914.1 164574 164596 - 0 TTGTTGTCCTCCTGTGAATT CGG ||||||||[||||||||||||]PAM NA Intergenic NA NA

T21948 TTGTCCTCCTGTGAATTCGGAGG 1000
Oligo fwd CACCTTGTCCTCCTGTGAATTCGG
Oligo rev AAACCCGAATTCACAGGAGGACAA
Chromosome start end strand MM target_seq PAM alignment distance position gene name gene id
GU305914.1 164571 164593 - 0 TTGTCCTCCTGTGAATTCGG AGG ||||||||[||||||||||||]PAM NA Intergenic NA NA

T21949 TTAGAAAGCCTCTGTTCCCCAGG 1000
Oligo fwd CACCTTAGAAAGCCTCTGTTCCCC
Oligo rev AAACGGGGAACAGAGGCTTTCTAA

<table>
<tr><td>Chromosome</td><td>start</td><td>end</td><td>strand</td><td>MM</td><td>target_seq</td><td>PAM</td><td>alignment</td><td>distance</td><td>position</td><td>gene name</td><td>gene id</td></tr>
<tr><td>GU305914.1</td><td>164538</td><td>164560</td><td>-</td><td>0</td><td>TTAGAAAGCCTCTGTTCCCC</td><td>AGG</td><td>||||||||[||||||||||||]PAM</td><td>NA</td><td>Intergenic</td><td>NA</td><td>NA</td></tr>
<tr><td>T21950</td><td>AAGCCTCTGTTCCCCAGGACAGG</td><td>1000</td><td></td><td></td><td></td><td></td><td></td><td></td><td></td><td></td><td></td></tr>
<tr><td>Oligo fwd</td><td>CACCAAGCCTCTGTTCCCCAGGAC</td><td></td><td></td><td></td><td></td><td></td><td></td><td></td><td></td><td></td><td></td></tr>
<tr><td>Oligo rev</td><td>AAACGTCCTGGGGAACAGAGGCTT</td><td></td><td></td><td></td><td></td><td></td><td></td><td></td><td></td><td></td><td></td></tr>
<tr><td>Chromosome</td><td>start</td><td>end</td><td>strand</td><td>MM</td><td>target_seq</td><td>PAM</td><td>alignment</td><td>distance</td><td>position</td><td>gene name</td><td>gene id</td></tr>
<tr><td>GU305914.1</td><td>164533</td><td>164555</td><td>-</td><td>0</td><td>AAGCCTCTGTTCCCCAGGAC</td><td>AGG</td><td>||||||||[||||||||||||]PAM</td><td>NA</td><td>Intergenic</td><td>NA</td><td>NA</td></tr>
<tr><td>T21951</td><td>CCGACTTCACCGCCGTCATGCGG</td><td>1000</td><td></td><td></td><td></td><td></td><td></td><td></td><td></td><td></td><td></td></tr>
<tr><td>Oligo fwd</td><td>CACCCCGACTTCACCGCCGTCATG</td><td></td><td></td><td></td><td></td><td></td><td></td><td></td><td></td><td></td><td></td></tr>
<tr><td>Oligo rev</td><td>AAACCATGACGGCGGTGAAGTCGG</td><td></td><td></td><td></td><td></td><td></td><td></td><td></td><td></td><td></td><td></td></tr>
<tr><td>Chromosome</td><td>start</td><td>end</td><td>strand</td><td>MM</td><td>target_seq</td><td>PAM</td><td>alignment</td><td>distance</td><td>position</td><td>gene name</td><td>gene id</td></tr>
<tr><td>GU305914.1</td><td>164489</td><td>164511</td><td>-</td><td>0</td><td>CCGACTTCACCGCCGTCATG</td><td>CGG</td><td>||||||||[||||||||||||]PAM</td><td>NA</td><td>Intergenic</td><td>NA</td><td>NA</td></tr>
<tr><td>T21952</td><td>CGCCGTCATGCGGCTCGTAAAGG</td><td>1000</td><td></td><td></td><td></td><td></td><td></td><td></td><td></td><td></td><td></td></tr>
<tr><td>Oligo fwd</td><td>CACCCGCCGTCATGCGGCTCGTAA</td><td></td><td></td><td></td><td></td><td></td><td></td><td></td><td></td><td></td><td></td></tr>
<tr><td>Oligo rev</td><td>AAACTTACGAGCCGCATGACGGCG</td><td></td><td></td><td></td><td></td><td></td><td></td><td></td><td></td><td></td><td></td></tr>
<tr><td>Chromosome</td><td>start</td><td>end</td><td>strand</td><td>MM</td><td>target_seq</td><td>PAM</td><td>alignment</td><td>distance</td><td>position</td><td>gene name</td><td>gene id</td></tr>
<tr><td>GU305914.1</td><td>164479</td><td>164501</td><td>-</td><td>0</td><td>CGCCGTCATGCGGCTCGTAA</td><td>AGG</td><td>||||||||[||||||||||||]PAM</td><td>NA</td><td>Intergenic</td><td>NA</td><td>NA</td></tr>
<tr><td>T21953</td><td>TCGTAAAGGCGAGATCCATGAGG</td><td>1000</td><td></td><td></td><td></td><td></td><td></td><td></td><td></td><td></td><td></td></tr>
<tr><td>Oligo fwd</td><td>CACCTCGTAAAGGCGAGATCCATG</td><td></td><td></td><td></td><td></td><td></td><td></td><td></td><td></td><td></td><td></td></tr>
<tr><td>Oligo rev</td><td>AAACCATGGATCTCGCCTTTACGA</td><td></td><td></td><td></td><td></td><td></td><td></td><td></td><td></td><td></td><td></td></tr>
<tr><td>Chromosome</td><td>start</td><td>end</td><td>strand</td><td>MM</td><td>target_seq</td><td>PAM</td><td>alignment</td><td>distance</td><td>position</td><td>gene name</td><td>gene id</td></tr>
<tr><td>GU305914.1</td><td>164465</td><td>164487</td><td>-</td><td>0</td><td>TCGTAAAGGCGAGATCCATG</td><td>AGG</td><td>||||||||[||||||||||||]PAM</td><td>NA</td><td>Intergenic</td><td>NA</td><td>NA</td></tr>
<tr><td>T21954</td><td>CGTAAAGGCGAGATCCATGAGGG</td><td>1000</td><td></td><td></td><td></td><td></td><td></td><td></td><td></td><td></td><td></td></tr>
<tr><td>Oligo fwd</td><td>CACCCGTAAAGGCGAGATCCATGA</td><td></td><td></td><td></td><td></td><td></td><td></td><td></td><td></td><td></td><td></td></tr>
<tr><td>Oligo rev</td><td>AAACTCATGGATCTCGCCTTTACG</td><td></td><td></td><td></td><td></td><td></td><td></td><td></td><td></td><td></td><td></td></tr>
<tr><td>Chromosome</td><td>start</td><td>end</td><td>strand</td><td>MM</td><td>target_seq</td><td>PAM</td><td>alignment</td><td>distance</td><td>position</td><td>gene name</td><td>gene id</td></tr>
<tr><td>GU305914.1</td><td>164464</td><td>164486</td><td>-</td><td>0</td><td>CGTAAAGGCGAGATCCATGA</td><td>GGG</td><td>||||||||[||||||||||||]PAM</td><td>NA</td><td>Intergenic</td><td>NA</td><td>NA</td></tr>
<tr><td>T21955</td><td>GAGGGCCCGCGAGTCAATTAAGG</td><td>1000</td><td></td><td></td><td></td><td></td><td></td><td></td><td></td><td></td><td></td></tr>
<tr><td>Oligo fwd</td><td>CACCGAGGGCCCGCGAGTCAATTA</td><td></td><td></td><td></td><td></td><td></td><td></td><td></td><td></td><td></td><td></td></tr>
<tr><td>Oligo rev</td><td>AAACTAATTGACTCGCGGGCCCTC</td><td></td><td></td><td></td><td></td><td></td><td></td><td></td><td></td><td></td><td></td></tr>
<tr><td>Chromosome</td><td>start</td><td>end</td><td>strand</td><td>MM</td><td>target_seq</td><td>PAM</td><td>alignment</td><td>distance</td><td>position</td><td>gene name</td><td>gene id</td></tr>
<tr><td>GU305914.1</td><td>164446</td><td>164468</td><td>-</td><td>0</td><td>GAGGGCCCGCGAGTCAATTA</td><td>AGG</td><td>||||||||[||||||||||||]PAM</td><td>NA</td><td>Intergenic</td><td>NA</td><td>NA</td></tr>
<tr><td>T21956</td><td>CATGCATGCTAGCCCACCGTCGG</td><td>1000</td><td></td><td></td><td></td><td></td><td></td><td></td><td></td><td></td><td></td></tr>
<tr><td>Oligo fwd</td><td>CACCCATGCATGCTAGCCCACCGT</td><td></td><td></td><td></td><td></td><td></td><td></td><td></td><td></td><td></td><td></td></tr>
<tr><td>Oligo rev</td><td>AAACACGGTGGGCTAGCATGCATG</td><td></td><td></td><td></td><td></td><td></td><td></td><td></td><td></td><td></td><td></td></tr>
<tr><td>Chromosome</td><td>start</td><td>end</td><td>strand</td><td>MM</td><td>target_seq</td><td>PAM</td><td>alignment</td><td>distance</td><td>position</td><td>gene name</td><td>gene id</td></tr>
<tr><td>GU305914.1</td><td>164419</td><td>164441</td><td>-</td><td>0</td><td>CATGCATGCTAGCCCACCGT</td><td>CGG</td><td>||||||||[||||||||||||]PAM</td><td>NA</td><td>Intergenic</td><td>NA</td><td>NA</td></tr>
<tr><td>T21957</td><td>GGTGAACGTTTGCTCGATCTCGG</td><td>1000</td><td></td><td></td><td></td><td></td><td></td><td></td><td></td><td></td><td></td></tr>
<tr><td>Oligo fwd</td><td>CACCGGTGAACGTTTGCTCGATCT</td><td></td><td></td><td></td><td></td><td></td><td></td><td></td><td></td><td></td><td></td></tr>
<tr><td>Oligo rev</td><td>AAACAGATCGAGCAAACGTTCACC</td><td></td><td></td><td></td><td></td><td></td><td></td><td></td><td></td><td></td><td></td></tr>
<tr><td>Chromosome</td><td>start</td><td>end</td><td>strand</td><td>MM</td><td>target_seq</td><td>PAM</td><td>alignment</td><td>distance</td><td>position</td><td>gene name</td><td>gene id</td></tr>
<tr><td>GU305914.1</td><td>164398</td><td>164420</td><td>-</td><td>0</td><td>GGTGAACGTTTGCTCGATCT</td><td>CGG</td><td>||||||||[||||||||||||]PAM</td><td>NA</td><td>Intergenic</td><td>NA</td><td>NA</td></tr>
<tr><td>T21958</td><td>CTCGATCTCGGCAGACGCCCCGG</td><td>1000</td><td></td><td></td><td></td><td></td><td></td><td></td><td></td><td></td><td></td></tr>
<tr><td>Oligo fwd</td><td>CACCCTCGATCTCGGCAGACGCCC</td><td></td><td></td><td></td><td></td><td></td><td></td><td></td><td></td><td></td><td></td></tr>
<tr><td>Oligo rev</td><td>AAACGGGCGTCTGCCGAGATCGAG</td><td></td><td></td><td></td><td></td><td></td><td></td><td></td><td></td><td></td><td></td></tr>
<tr><td>Chromosome</td><td>start</td><td>end</td><td>strand</td><td>MM</td><td>target_seq</td><td>PAM</td><td>alignment</td><td>distance</td><td>position</td><td>gene name</td><td>gene id</td></tr>
<tr><td>GU305914.1</td><td>164386</td><td>164408</td><td>-</td><td>0</td><td>CTCGATCTCGGCAGACGCCC</td><td>CGG</td><td>||||||||[||||||||||||]PAM</td><td>NA</td><td>Intergenic</td><td>NA</td><td>NA</td></tr>
<tr><td>T21959</td><td>CTCGGCAGACGCCCCGGTCTCGG</td><td>1000</td><td></td><td></td><td></td><td></td><td></td><td></td><td></td><td></td><td></td></tr>
<tr><td>Oligo fwd</td><td>CACCCTCGGCAGACGCCCCGGTCT</td><td></td><td></td><td></td><td></td><td></td><td></td><td></td><td></td><td></td><td></td></tr>
<tr><td>Oligo rev</td><td>AAACAGACCGGGGCGTCTGCCGAG</td><td></td><td></td><td></td><td></td><td></td><td></td><td></td><td></td><td></td><td></td></tr>
<tr><td>Chromosome</td><td>start</td><td>end</td><td>strand</td><td>MM</td><td>target_seq</td><td>PAM</td><td>alignment</td><td>distance</td><td>position</td><td>gene name</td><td>gene id</td></tr>
<tr><td>GU305914.1</td><td>164380</td><td>164402</td><td>-</td><td>0</td><td>CTCGGCAGACGCCCCGGTCT</td><td>CGG</td><td>||||||||[||||||||||||]PAM</td><td>NA</td><td>Intergenic</td><td>NA</td><td>NA</td></tr>
<tr><td>T21960</td><td>GGTTACCGTCGCCGTACCATCGG</td><td>1000</td><td></td><td></td><td></td><td></td><td></td><td></td><td></td><td></td><td></td></tr>
<tr><td>Oligo fwd</td><td>CACCGGTTACCGTCGCCGTACCAT</td><td></td><td></td><td></td><td></td><td></td><td></td><td></td><td></td><td></td><td></td></tr>
<tr><td>Oligo rev</td><td>AAACATGGTACGGCGACGGTAACC</td><td></td><td></td><td></td><td></td><td></td><td></td><td></td><td></td><td></td><td></td></tr>
<tr><td>Chromosome</td><td>start</td><td>end</td><td>strand</td><td>MM</td><td>target_seq</td><td>PAM</td><td>alignment</td><td>distance</td><td>position</td><td>gene name</td><td>gene id</td></tr>
<tr><td>GU305914.1</td><td>164359</td><td>164381</td><td>-</td><td>0</td><td>GGTTACCGTCGCCGTACCAT</td><td>CGG</td><td>||||||||[||||||||||||]PAM</td><td>NA</td><td>Intergenic</td><td>NA</td><td>NA</td></tr>
<tr><td>T21961</td><td>AGTTCCATCGTCTTCCTCCTCGG</td><td>1000</td><td></td><td></td><td></td><td></td><td></td><td></td><td></td><td></td><td></td></tr>
<tr><td>Oligo fwd</td><td>CACCAGTTCCATCGTCTTCCTCCT</td><td></td><td></td><td></td><td></td><td></td><td></td><td></td><td></td><td></td><td></td></tr>
<tr><td>Oligo rev</td><td>AAACAGGAGGAAGACGATGGAACT</td><td></td><td></td><td></td><td></td><td></td><td></td><td></td><td></td><td></td><td></td></tr>
<tr><td>Chromosome</td><td>start</td><td>end</td><td>strand</td><td>MM</td><td>target_seq</td><td>PAM</td><td>alignment</td><td>distance</td><td>position</td><td>gene name</td><td>gene id</td></tr>
<tr><td>GU305914.1</td><td>164323</td><td>164345</td><td>-</td><td>0</td><td>AGTTCCATCGTCTTCCTCCT</td><td>CGG</td><td>||||||||[||||||||||||]PAM</td><td>NA</td><td>Intergenic</td><td>NA</td><td>NA</td></tr>
<tr><td>T21962</td><td>TCTATCAAGTCATTAAACCCTGG</td><td>1000</td><td></td><td></td><td></td><td></td><td></td><td></td><td></td><td></td><td></td></tr>
<tr><td>Oligo fwd</td><td>CACCTCTATCAAGTCATTAAACCC</td><td></td><td></td><td></td><td></td><td></td><td></td><td></td><td></td><td></td><td></td></tr>
<tr><td>Oligo rev</td><td>AAACGGGTTTAATGACTTGATAGA</td><td></td><td></td><td></td><td></td><td></td><td></td><td></td><td></td><td></td><td></td></tr>
<tr><td>Chromosome</td><td>start</td><td>end</td><td>strand</td><td>MM</td><td>target_seq</td><td>PAM</td><td>alignment</td><td>distance</td><td>position</td><td>gene name</td><td>gene id</td></tr>
<tr><td>GU305914.1</td><td>164298</td><td>164320</td><td>-</td><td>0</td><td>TCTATCAAGTCATTAAACCC</td><td>TGG</td><td>||||||||[||||||||||||]PAM</td><td>NA</td><td>Intergenic</td><td>NA</td><td>NA</td></tr>
<tr><td>T21963</td><td>CCTGGTGTCTCTATAGTCACCGG</td><td>1000</td><td></td><td></td><td></td><td></td><td></td><td></td><td></td><td></td><td></td></tr>
<tr><td>Oligo fwd</td><td>CACCCCTGGTGTCTCTATAGTCAC</td><td></td><td></td><td></td><td></td><td></td><td></td><td></td><td></td><td></td><td></td></tr>
<tr><td>Oligo rev</td><td>AAACGTGACTATAGAGACACCAGG</td><td></td><td></td><td></td><td></td><td></td><td></td><td></td><td></td><td></td><td></td></tr>
<tr><td>Chromosome</td><td>start</td><td>end</td><td>strand</td><td>MM</td><td>target_seq</td><td>PAM</td><td>alignment</td><td>distance</td><td>position</td><td>gene name</td><td>gene id</td></tr>
<tr><td>GU305914.1</td><td>164280</td><td>164302</td><td>-</td><td>0</td><td>CCTGGTGTCTCTATAGTCAC</td><td>CGG</td><td>||||||||[||||||||||||]PAM</td><td>NA</td><td>Intergenic</td><td>NA</td><td>NA</td></tr>
<tr><td>T21964</td><td>TGTCTCTATAGTCACCGGAGAGG</td><td>1000</td><td></td><td></td><td></td><td></td><td></td><td></td><td></td><td></td><td></td></tr>
<tr><td>Oligo fwd</td><td>CACCTGTCTCTATAGTCACCGGAG</td><td></td><td></td><td></td><td></td><td></td><td></td><td></td><td></td><td></td><td></td></tr>
</table>

<table>
<tr><td>Oligo rev</td><td>AAACCTCCGGTGACTATAGAGACA</td><td></td><td></td><td></td><td></td><td></td><td></td><td></td><td></td><td></td><td></td></tr>
<tr><td>Chromosome</td><td>start</td><td>end</td><td>strand</td><td>MM</td><td>target_seq</td><td>PAM</td><td>alignment</td><td>distance</td><td>position</td><td>gene name</td><td>gene id</td></tr>
<tr><td>GU305914.1</td><td>164275</td><td>164297</td><td>-</td><td>0</td><td>TGTCTCTATAGTCACCGGAG</td><td>AGG</td><td>||||||||[||||||||||||]PAM</td><td>NA</td><td>Intergenic</td><td>NA</td><td>NA</td></tr>
<tr><td>T21965</td><td>CTCTATAGTCACCGGAGAGGAGG</td><td>1000</td><td></td><td></td><td></td><td></td><td></td><td></td><td></td><td></td><td></td></tr>
<tr><td>Oligo fwd</td><td>CACCCTCTATAGTCACCGGAGAGG</td><td></td><td></td><td></td><td></td><td></td><td></td><td></td><td></td><td></td><td></td></tr>
<tr><td>Oligo rev</td><td>AAACCCTCTCCGGTGACTATAGAG</td><td></td><td></td><td></td><td></td><td></td><td></td><td></td><td></td><td></td><td></td></tr>
<tr><td>Chromosome</td><td>start</td><td>end</td><td>strand</td><td>MM</td><td>target_seq</td><td>PAM</td><td>alignment</td><td>distance</td><td>position</td><td>gene name</td><td>gene id</td></tr>
<tr><td>GU305914.1</td><td>164272</td><td>164294</td><td>-</td><td>0</td><td>CTCTATAGTCACCGGAGAGG</td><td>AGG</td><td>||||||||[||||||||||||]PAM</td><td>NA</td><td>Intergenic</td><td>NA</td><td>NA</td></tr>
<tr><td>T21966</td><td>TCTATAGTCACCGGAGAGGAGGG</td><td>1000</td><td></td><td></td><td></td><td></td><td></td><td></td><td></td><td></td><td></td></tr>
<tr><td>Oligo fwd</td><td>CACCTCTATAGTCACCGGAGAGGA</td><td></td><td></td><td></td><td></td><td></td><td></td><td></td><td></td><td></td><td></td></tr>
<tr><td>Oligo rev</td><td>AAACTCCTCTCCGGTGACTATAGA</td><td></td><td></td><td></td><td></td><td></td><td></td><td></td><td></td><td></td><td></td></tr>
<tr><td>Chromosome</td><td>start</td><td>end</td><td>strand</td><td>MM</td><td>target_seq</td><td>PAM</td><td>alignment</td><td>distance</td><td>position</td><td>gene name</td><td>gene id</td></tr>
<tr><td>GU305914.1</td><td>164271</td><td>164293</td><td>-</td><td>0</td><td>TCTATAGTCACCGGAGAGGA</td><td>GGG</td><td>||||||||[||||||||||||]PAM</td><td>NA</td><td>Intergenic</td><td>NA</td><td>NA</td></tr>
<tr><td>T21967</td><td>ATAGTCACCGGAGAGGAGGGTGG</td><td>1000</td><td></td><td></td><td></td><td></td><td></td><td></td><td></td><td></td><td></td></tr>
<tr><td>Oligo fwd</td><td>CACCATAGTCACCGGAGAGGAGGG</td><td></td><td></td><td></td><td></td><td></td><td></td><td></td><td></td><td></td><td></td></tr>
<tr><td>Oligo rev</td><td>AAACCCCTCCTCTCCGGTGACTAT</td><td></td><td></td><td></td><td></td><td></td><td></td><td></td><td></td><td></td><td></td></tr>
<tr><td>Chromosome</td><td>start</td><td>end</td><td>strand</td><td>MM</td><td>target_seq</td><td>PAM</td><td>alignment</td><td>distance</td><td>position</td><td>gene name</td><td>gene id</td></tr>
<tr><td>GU305914.1</td><td>164268</td><td>164290</td><td>-</td><td>0</td><td>ATAGTCACCGGAGAGGAGGG</td><td>TGG</td><td>||||||||[||||||||||||]PAM</td><td>NA</td><td>Intergenic</td><td>NA</td><td>NA</td></tr>
<tr><td>T21968</td><td>TGGTAACGCACCATCGCTGACGG</td><td>1000</td><td></td><td></td><td></td><td></td><td></td><td></td><td></td><td></td><td></td></tr>
<tr><td>Oligo fwd</td><td>CACCTGGTAACGCACCATCGCTGA</td><td></td><td></td><td></td><td></td><td></td><td></td><td></td><td></td><td></td><td></td></tr>
<tr><td>Oligo rev</td><td>AAACTCAGCGATGGTGCGTTACCA</td><td></td><td></td><td></td><td></td><td></td><td></td><td></td><td></td><td></td><td></td></tr>
<tr><td>Chromosome</td><td>start</td><td>end</td><td>strand</td><td>MM</td><td>target_seq</td><td>PAM</td><td>alignment</td><td>distance</td><td>position</td><td>gene name</td><td>gene id</td></tr>
<tr><td>GU305914.1</td><td>164248</td><td>164270</td><td>-</td><td>0</td><td>TGGTAACGCACCATCGCTGA</td><td>CGG</td><td>||||||||[||||||||||||]PAM</td><td>NA</td><td>Intergenic</td><td>NA</td><td>NA</td></tr>
<tr><td>T21969</td><td>TAACGCACCATCGCTGACGGCGG</td><td>1000</td><td></td><td></td><td></td><td></td><td></td><td></td><td></td><td></td><td></td></tr>
<tr><td>Oligo fwd</td><td>CACCTAACGCACCATCGCTGACGG</td><td></td><td></td><td></td><td></td><td></td><td></td><td></td><td></td><td></td><td></td></tr>
<tr><td>Oligo rev</td><td>AAACCCGTCAGCGATGGTGCGTTA</td><td></td><td></td><td></td><td></td><td></td><td></td><td></td><td></td><td></td><td></td></tr>
<tr><td>Chromosome</td><td>start</td><td>end</td><td>strand</td><td>MM</td><td>target_seq</td><td>PAM</td><td>alignment</td><td>distance</td><td>position</td><td>gene name</td><td>gene id</td></tr>
<tr><td>GU305914.1</td><td>164245</td><td>164267</td><td>-</td><td>0</td><td>TAACGCACCATCGCTGACGG</td><td>CGG</td><td>||||||||[||||||||||||]PAM</td><td>NA</td><td>Intergenic</td><td>NA</td><td>NA</td></tr>
<tr><td>T21970</td><td>AGCAGCATAACTGTTCAAAAAGG</td><td>1000</td><td></td><td></td><td></td><td></td><td></td><td></td><td></td><td></td><td></td></tr>
<tr><td>Oligo fwd</td><td>CACCAGCAGCATAACTGTTCAAAA</td><td></td><td></td><td></td><td></td><td></td><td></td><td></td><td></td><td></td><td></td></tr>
<tr><td>Oligo rev</td><td>AAACTTTTGAACAGTTATGCTGCT</td><td></td><td></td><td></td><td></td><td></td><td></td><td></td><td></td><td></td><td></td></tr>
<tr><td>Chromosome</td><td>start</td><td>end</td><td>strand</td><td>MM</td><td>target_seq</td><td>PAM</td><td>alignment</td><td>distance</td><td>position</td><td>gene name</td><td>gene id</td></tr>
<tr><td>GU305914.1</td><td>164212</td><td>164234</td><td>-</td><td>0</td><td>AGCAGCATAACTGTTCAAAA</td><td>AGG</td><td>||||||||[||||||||||||]PAM</td><td>NA</td><td>Intergenic</td><td>NA</td><td>NA</td></tr>
<tr><td>T21971</td><td>GTCTGCATTACCATCGACAGTGG</td><td>1000</td><td></td><td></td><td></td><td></td><td></td><td></td><td></td><td></td><td></td></tr>
<tr><td>Oligo fwd</td><td>CACCGTCTGCATTACCATCGACAG</td><td></td><td></td><td></td><td></td><td></td><td></td><td></td><td></td><td></td><td></td></tr>
<tr><td>Oligo rev</td><td>AAACCTGTCGATGGTAATGCAGAC</td><td></td><td></td><td></td><td></td><td></td><td></td><td></td><td></td><td></td><td></td></tr>
<tr><td>Chromosome</td><td>start</td><td>end</td><td>strand</td><td>MM</td><td>target_seq</td><td>PAM</td><td>alignment</td><td>distance</td><td>position</td><td>gene name</td><td>gene id</td></tr>
<tr><td>GU305914.1</td><td>164173</td><td>164195</td><td>-</td><td>0</td><td>GTCTGCATTACCATCGACAG</td><td>TGG</td><td>||||||||[||||||||||||]PAM</td><td>NA</td><td>Intergenic</td><td>NA</td><td>NA</td></tr>
<tr><td>T21972</td><td>TCGACAGTGGTACTGCTCCTAGG</td><td>1000</td><td></td><td></td><td></td><td></td><td></td><td></td><td></td><td></td><td></td></tr>
<tr><td>Oligo fwd</td><td>CACCTCGACAGTGGTACTGCTCCT</td><td></td><td></td><td></td><td></td><td></td><td></td><td></td><td></td><td></td><td></td></tr>
<tr><td>Oligo rev</td><td>AAACAGGAGCAGTACCACTGTCGA</td><td></td><td></td><td></td><td></td><td></td><td></td><td></td><td></td><td></td><td></td></tr>
<tr><td>Chromosome</td><td>start</td><td>end</td><td>strand</td><td>MM</td><td>target_seq</td><td>PAM</td><td>alignment</td><td>distance</td><td>position</td><td>gene name</td><td>gene id</td></tr>
<tr><td>GU305914.1</td><td>164160</td><td>164182</td><td>-</td><td>0</td><td>TCGACAGTGGTACTGCTCCT</td><td>AGG</td><td>||||||||[||||||||||||]PAM</td><td>NA</td><td>Intergenic</td><td>NA</td><td>NA</td></tr>
<tr><td>T21973</td><td>ACAGTGGTACTGCTCCTAGGCGG</td><td>1000</td><td></td><td></td><td></td><td></td><td></td><td></td><td></td><td></td><td></td></tr>
<tr><td>Oligo fwd</td><td>CACCACAGTGGTACTGCTCCTAGG</td><td></td><td></td><td></td><td></td><td></td><td></td><td></td><td></td><td></td><td></td></tr>
<tr><td>Oligo rev</td><td>AAACCCTAGGAGCAGTACCACTGT</td><td></td><td></td><td></td><td></td><td></td><td></td><td></td><td></td><td></td><td></td></tr>
<tr><td>Chromosome</td><td>start</td><td>end</td><td>strand</td><td>MM</td><td>target_seq</td><td>PAM</td><td>alignment</td><td>distance</td><td>position</td><td>gene name</td><td>gene id</td></tr>
<tr><td>GU305914.1</td><td>164157</td><td>164179</td><td>-</td><td>0</td><td>ACAGTGGTACTGCTCCTAGG</td><td>CGG</td><td>||||||||[||||||||||||]PAM</td><td>NA</td><td>Intergenic</td><td>NA</td><td>NA</td></tr>
<tr><td>T21974</td><td>GGCGGTTCCAGATCCCTCTCCGG</td><td>1000</td><td></td><td></td><td></td><td></td><td></td><td></td><td></td><td></td><td></td></tr>
<tr><td>Oligo fwd</td><td>CACCGGCGGTTCCAGATCCCTCTC</td><td></td><td></td><td></td><td></td><td></td><td></td><td></td><td></td><td></td><td></td></tr>
<tr><td>Oligo rev</td><td>AAACGAGAGGGATCTGGAACCGCC</td><td></td><td></td><td></td><td></td><td></td><td></td><td></td><td></td><td></td><td></td></tr>
<tr><td>Chromosome</td><td>start</td><td>end</td><td>strand</td><td>MM</td><td>target_seq</td><td>PAM</td><td>alignment</td><td>distance</td><td>position</td><td>gene name</td><td>gene id</td></tr>
<tr><td>GU305914.1</td><td>164139</td><td>164161</td><td>-</td><td>0</td><td>GGCGGTTCCAGATCCCTCTC</td><td>CGG</td><td>||||||||[||||||||||||]PAM</td><td>NA</td><td>Intergenic</td><td>NA</td><td>NA</td></tr>
<tr><td>T21975</td><td>TCCAGATCCCTCTCCGGTCTCGG</td><td>1000</td><td></td><td></td><td></td><td></td><td></td><td></td><td></td><td></td><td></td></tr>
<tr><td>Oligo fwd</td><td>CACCTCCAGATCCCTCTCCGGTCT</td><td></td><td></td><td></td><td></td><td></td><td></td><td></td><td></td><td></td><td></td></tr>
<tr><td>Oligo rev</td><td>AAACAGACCGGAGAGGGATCTGGA</td><td></td><td></td><td></td><td></td><td></td><td></td><td></td><td></td><td></td><td></td></tr>
<tr><td>Chromosome</td><td>start</td><td>end</td><td>strand</td><td>MM</td><td>target_seq</td><td>PAM</td><td>alignment</td><td>distance</td><td>position</td><td>gene name</td><td>gene id</td></tr>
<tr><td>GU305914.1</td><td>164133</td><td>164155</td><td>-</td><td>0</td><td>TCCAGATCCCTCTCCGGTCT</td><td>CGG</td><td>||||||||[||||||||||||]PAM</td><td>NA</td><td>Intergenic</td><td>NA</td><td>NA</td></tr>
<tr><td>T21976</td><td>CCCTCTCCGGTCTCGGATCGTGG</td><td>1000</td><td></td><td></td><td></td><td></td><td></td><td></td><td></td><td></td><td></td></tr>
<tr><td>Oligo fwd</td><td>CACCCCCTCTCCGGTCTCGGATCG</td><td></td><td></td><td></td><td></td><td></td><td></td><td></td><td></td><td></td><td></td></tr>
<tr><td>Oligo rev</td><td>AAACCGATCCGAGACCGGAGAGGG</td><td></td><td></td><td></td><td></td><td></td><td></td><td></td><td></td><td></td><td></td></tr>
<tr><td>Chromosome</td><td>start</td><td>end</td><td>strand</td><td>MM</td><td>target_seq</td><td>PAM</td><td>alignment</td><td>distance</td><td>position</td><td>gene name</td><td>gene id</td></tr>
<tr><td>GU305914.1</td><td>164126</td><td>164148</td><td>-</td><td>0</td><td>CCCTCTCCGGTCTCGGATCG</td><td>TGG</td><td>||||||||[||||||||||||]PAM</td><td>NA</td><td>Intergenic</td><td>NA</td><td>NA</td></tr>
<tr><td>T21977</td><td>TCTCCGGTCTCGGATCGTGGTGG</td><td>1000</td><td></td><td></td><td></td><td></td><td></td><td></td><td></td><td></td><td></td></tr>
<tr><td>Oligo fwd</td><td>CACCTCTCCGGTCTCGGATCGTGG</td><td></td><td></td><td></td><td></td><td></td><td></td><td></td><td></td><td></td><td></td></tr>
<tr><td>Oligo rev</td><td>AAACCCACGATCCGAGACCGGAGA</td><td></td><td></td><td></td><td></td><td></td><td></td><td></td><td></td><td></td><td></td></tr>
<tr><td>Chromosome</td><td>start</td><td>end</td><td>strand</td><td>MM</td><td>target_seq</td><td>PAM</td><td>alignment</td><td>distance</td><td>position</td><td>gene name</td><td>gene id</td></tr>
<tr><td>GU305914.1</td><td>164123</td><td>164145</td><td>-</td><td>0</td><td>TCTCCGGTCTCGGATCGTGG</td><td>TGG</td><td>||||||||[||||||||||||]PAM</td><td>NA</td><td>Intergenic</td><td>NA</td><td>NA</td></tr>
<tr><td>T21978</td><td>CTCCGGTCTCGGATCGTGGTGGG</td><td>1000</td><td></td><td></td><td></td><td></td><td></td><td></td><td></td><td></td><td></td></tr>
<tr><td>Oligo fwd</td><td>CACCCTCCGGTCTCGGATCGTGGT</td><td></td><td></td><td></td><td></td><td></td><td></td><td></td><td></td><td></td><td></td></tr>
<tr><td>Oligo rev</td><td>AAACACCACGATCCGAGACCGGAG</td><td></td><td></td><td></td><td></td><td></td><td></td><td></td><td></td><td></td><td></td></tr>
<tr><td>Chromosome</td><td>start</td><td>end</td><td>strand</td><td>MM</td><td>target_seq</td><td>PAM</td><td>alignment</td><td>distance</td><td>position</td><td>gene name</td><td>gene id</td></tr>
<tr><td>GU305914.1</td><td>164122</td><td>164144</td><td>-</td><td>0</td><td>CTCCGGTCTCGGATCGTGGT</td><td>GGG</td><td>||||||||[||||||||||||]PAM</td><td>NA</td><td>Intergenic</td><td>NA</td><td>NA</td></tr>
<tr><td>T21979</td><td>TCGGATCGTGGTGGGTCTTCTGG</td><td>1000</td><td></td><td></td><td></td><td></td><td></td><td></td><td></td><td></td><td></td></tr>
</table>

<table>
<tr><td>Oligo fwd</td><td>CACCTCGGATCGTGGTGGGTCTTC</td><td></td><td></td><td></td><td></td><td></td><td></td><td></td><td></td><td></td><td></td></tr>
<tr><td>Oligo rev</td><td>AAACGAAGACCCACCACGATCCGA</td><td></td><td></td><td></td><td></td><td></td><td></td><td></td><td></td><td></td><td></td></tr>
<tr><td>Chromosome</td><td>start</td><td>end</td><td>strand</td><td>MM</td><td>target_seq</td><td>PAM</td><td>alignment</td><td>distance</td><td>position</td><td>gene name</td><td>gene id</td></tr>
<tr><td>GU305914.1</td><td>164114</td><td>164136</td><td>-</td><td>0</td><td>TCGGATCGTGGTGGGTCTTC</td><td>TGG</td><td>||||||||[||||||||||||]PAM</td><td>NA</td><td>Intergenic</td><td>NA</td><td>NA</td></tr>
<tr><td>T21980</td><td>GTGGTGGGTCTTCTGGCGCTTGG</td><td>1000</td><td></td><td></td><td></td><td></td><td></td><td></td><td></td><td></td><td></td></tr>
<tr><td>Oligo fwd</td><td>CACCGTGGTGGGTCTTCTGGCGCT</td><td></td><td></td><td></td><td></td><td></td><td></td><td></td><td></td><td></td><td></td></tr>
<tr><td>Oligo rev</td><td>AAACAGCGCCAGAAGACCCACCAC</td><td></td><td></td><td></td><td></td><td></td><td></td><td></td><td></td><td></td><td></td></tr>
<tr><td>Chromosome</td><td>start</td><td>end</td><td>strand</td><td>MM</td><td>target_seq</td><td>PAM</td><td>alignment</td><td>distance</td><td>position</td><td>gene name</td><td>gene id</td></tr>
<tr><td>GU305914.1</td><td>164107</td><td>164129</td><td>-</td><td>0</td><td>GTGGTGGGTCTTCTGGCGCT</td><td>TGG</td><td>||||||||[||||||||||||]PAM</td><td>NA</td><td>Intergenic</td><td>NA</td><td>NA</td></tr>
<tr><td>T21981</td><td>TGGTGGGTCTTCTGGCGCTTGGG</td><td>1000</td><td></td><td></td><td></td><td></td><td></td><td></td><td></td><td></td><td></td></tr>
<tr><td>Oligo fwd</td><td>CACCTGGTGGGTCTTCTGGCGCTT</td><td></td><td></td><td></td><td></td><td></td><td></td><td></td><td></td><td></td><td></td></tr>
<tr><td>Oligo rev</td><td>AAACAAGCGCCAGAAGACCCACCA</td><td></td><td></td><td></td><td></td><td></td><td></td><td></td><td></td><td></td><td></td></tr>
<tr><td>Chromosome</td><td>start</td><td>end</td><td>strand</td><td>MM</td><td>target_seq</td><td>PAM</td><td>alignment</td><td>distance</td><td>position</td><td>gene name</td><td>gene id</td></tr>
<tr><td>GU305914.1</td><td>164106</td><td>164128</td><td>-</td><td>0</td><td>TGGTGGGTCTTCTGGCGCTT</td><td>GGG</td><td>||||||||[||||||||||||]PAM</td><td>NA</td><td>Intergenic</td><td>NA</td><td>NA</td></tr>
<tr><td>T21982</td><td>TTGGGCTCCCCCCCGCTGCGTGG</td><td>1000</td><td></td><td></td><td></td><td></td><td></td><td></td><td></td><td></td><td></td></tr>
<tr><td>Oligo fwd</td><td>CACCTTGGGCTCCCCCCCGCTGCG</td><td></td><td></td><td></td><td></td><td></td><td></td><td></td><td></td><td></td><td></td></tr>
<tr><td>Oligo rev</td><td>AAACCGCAGCGGGGGGGAGCCCAA</td><td></td><td></td><td></td><td></td><td></td><td></td><td></td><td></td><td></td><td></td></tr>
<tr><td>Chromosome</td><td>start</td><td>end</td><td>strand</td><td>MM</td><td>target_seq</td><td>PAM</td><td>alignment</td><td>distance</td><td>position</td><td>gene name</td><td>gene id</td></tr>
<tr><td>GU305914.1</td><td>164088</td><td>164110</td><td>-</td><td>0</td><td>TTGGGCTCCCCCCCGCTGCG</td><td>TGG</td><td>||||||||[||||||||||||]PAM</td><td>NA</td><td>Intergenic</td><td>NA</td><td>NA</td></tr>
<tr><td>T21983</td><td>TAACATCATTAGTCCCGTGACGG</td><td>1000</td><td></td><td></td><td></td><td></td><td></td><td></td><td></td><td></td><td></td></tr>
<tr><td>Oligo fwd</td><td>CACCTAACATCATTAGTCCCGTGA</td><td></td><td></td><td></td><td></td><td></td><td></td><td></td><td></td><td></td><td></td></tr>
<tr><td>Oligo rev</td><td>AAACTCACGGGACTAATGATGTTA</td><td></td><td></td><td></td><td></td><td></td><td></td><td></td><td></td><td></td><td></td></tr>
<tr><td>Chromosome</td><td>start</td><td>end</td><td>strand</td><td>MM</td><td>target_seq</td><td>PAM</td><td>alignment</td><td>distance</td><td>position</td><td>gene name</td><td>gene id</td></tr>
<tr><td>GU305914.1</td><td>164046</td><td>164068</td><td>-</td><td>0</td><td>TAACATCATTAGTCCCGTGA</td><td>CGG</td><td>||||||||[||||||||||||]PAM</td><td>NA</td><td>Intergenic</td><td>NA</td><td>NA</td></tr>
<tr><td>T21984</td><td>GTCCCGTGACGGTTCACCGAAGG</td><td>1000</td><td></td><td></td><td></td><td></td><td></td><td></td><td></td><td></td><td></td></tr>
<tr><td>Oligo fwd</td><td>CACCGTCCCGTGACGGTTCACCGA</td><td></td><td></td><td></td><td></td><td></td><td></td><td></td><td></td><td></td><td></td></tr>
<tr><td>Oligo rev</td><td>AAACTCGGTGAACCGTCACGGGAC</td><td></td><td></td><td></td><td></td><td></td><td></td><td></td><td></td><td></td><td></td></tr>
<tr><td>Chromosome</td><td>start</td><td>end</td><td>strand</td><td>MM</td><td>target_seq</td><td>PAM</td><td>alignment</td><td>distance</td><td>position</td><td>gene name</td><td>gene id</td></tr>
<tr><td>GU305914.1</td><td>164035</td><td>164057</td><td>-</td><td>0</td><td>GTCCCGTGACGGTTCACCGA</td><td>AGG</td><td>||||||||[||||||||||||]PAM</td><td>NA</td><td>Intergenic</td><td>NA</td><td>NA</td></tr>
<tr><td>T21985</td><td>TGACGGTTCACCGAAGGTCGAGG</td><td>1000</td><td></td><td></td><td></td><td></td><td></td><td></td><td></td><td></td><td></td></tr>
<tr><td>Oligo fwd</td><td>CACCTGACGGTTCACCGAAGGTCG</td><td></td><td></td><td></td><td></td><td></td><td></td><td></td><td></td><td></td><td></td></tr>
<tr><td>Oligo rev</td><td>AAACCGACCTTCGGTGAACCGTCA</td><td></td><td></td><td></td><td></td><td></td><td></td><td></td><td></td><td></td><td></td></tr>
<tr><td>Chromosome</td><td>start</td><td>end</td><td>strand</td><td>MM</td><td>target_seq</td><td>PAM</td><td>alignment</td><td>distance</td><td>position</td><td>gene name</td><td>gene id</td></tr>
<tr><td>GU305914.1</td><td>164029</td><td>164051</td><td>-</td><td>0</td><td>TGACGGTTCACCGAAGGTCG</td><td>AGG</td><td>||||||||[||||||||||||]PAM</td><td>NA</td><td>Intergenic</td><td>NA</td><td>NA</td></tr>
<tr><td>T21986</td><td>GACGGTTCACCGAAGGTCGAGGG</td><td>1000</td><td></td><td></td><td></td><td></td><td></td><td></td><td></td><td></td><td></td></tr>
<tr><td>Oligo fwd</td><td>CACCGACGGTTCACCGAAGGTCGA</td><td></td><td></td><td></td><td></td><td></td><td></td><td></td><td></td><td></td><td></td></tr>
<tr><td>Oligo rev</td><td>AAACTCGACCTTCGGTGAACCGTC</td><td></td><td></td><td></td><td></td><td></td><td></td><td></td><td></td><td></td><td></td></tr>
<tr><td>Chromosome</td><td>start</td><td>end</td><td>strand</td><td>MM</td><td>target_seq</td><td>PAM</td><td>alignment</td><td>distance</td><td>position</td><td>gene name</td><td>gene id</td></tr>
<tr><td>GU305914.1</td><td>164028</td><td>164050</td><td>-</td><td>0</td><td>GACGGTTCACCGAAGGTCGA</td><td>GGG</td><td>||||||||[||||||||||||]PAM</td><td>NA</td><td>Intergenic</td><td>NA</td><td>NA</td></tr>
<tr><td>T21987</td><td>ACGGTTCACCGAAGGTCGAGGGG</td><td>1000</td><td></td><td></td><td></td><td></td><td></td><td></td><td></td><td></td><td></td></tr>
<tr><td>Oligo fwd</td><td>CACCACGGTTCACCGAAGGTCGAG</td><td></td><td></td><td></td><td></td><td></td><td></td><td></td><td></td><td></td><td></td></tr>
<tr><td>Oligo rev</td><td>AAACCTCGACCTTCGGTGAACCGT</td><td></td><td></td><td></td><td></td><td></td><td></td><td></td><td></td><td></td><td></td></tr>
<tr><td>Chromosome</td><td>start</td><td>end</td><td>strand</td><td>MM</td><td>target_seq</td><td>PAM</td><td>alignment</td><td>distance</td><td>position</td><td>gene name</td><td>gene id</td></tr>
<tr><td>GU305914.1</td><td>164027</td><td>164049</td><td>-</td><td>0</td><td>ACGGTTCACCGAAGGTCGAG</td><td>GGG</td><td>||||||||[||||||||||||]PAM</td><td>NA</td><td>Intergenic</td><td>NA</td><td>NA</td></tr>
<tr><td>GU305914.1</td><td>93038</td><td>93060</td><td>+</td><td>4</td><td>CCGGTTGACCGAAGGTCAAC</td><td>CGG</td><td>-|||||-|[|||||||||-|-]PAM</td><td>NA</td><td>Intergenic</td><td>NA</td><td>NA</td></tr>
<tr><td>T21988</td><td>CGGTTCACCGAAGGTCGAGGGGG</td><td>1000</td><td></td><td></td><td></td><td></td><td></td><td></td><td></td><td></td><td></td></tr>
<tr><td>Oligo fwd</td><td>CACCCGGTTCACCGAAGGTCGAGG</td><td></td><td></td><td></td><td></td><td></td><td></td><td></td><td></td><td></td><td></td></tr>
<tr><td>Oligo rev</td><td>AAACCCTCGACCTTCGGTGAACCG</td><td></td><td></td><td></td><td></td><td></td><td></td><td></td><td></td><td></td><td></td></tr>
<tr><td>Chromosome</td><td>start</td><td>end</td><td>strand</td><td>MM</td><td>target_seq</td><td>PAM</td><td>alignment</td><td>distance</td><td>position</td><td>gene name</td><td>gene id</td></tr>
<tr><td>GU305914.1</td><td>164026</td><td>164048</td><td>-</td><td>0</td><td>CGGTTCACCGAAGGTCGAGG</td><td>GGG</td><td>||||||||[||||||||||||]PAM</td><td>NA</td><td>Intergenic</td><td>NA</td><td>NA</td></tr>
<tr><td>T21989</td><td>GCTCCTTCTGCTTTGTACTGAGG</td><td>1000</td><td></td><td></td><td></td><td></td><td></td><td></td><td></td><td></td><td></td></tr>
<tr><td>Oligo fwd</td><td>CACCGCTCCTTCTGCTTTGTACTG</td><td></td><td></td><td></td><td></td><td></td><td></td><td></td><td></td><td></td><td></td></tr>
<tr><td>Oligo rev</td><td>AAACCAGTACAAAGCAGAAGGAGC</td><td></td><td></td><td></td><td></td><td></td><td></td><td></td><td></td><td></td><td></td></tr>
<tr><td>Chromosome</td><td>start</td><td>end</td><td>strand</td><td>MM</td><td>target_seq</td><td>PAM</td><td>alignment</td><td>distance</td><td>position</td><td>gene name</td><td>gene id</td></tr>
<tr><td>GU305914.1</td><td>163943</td><td>163965</td><td>-</td><td>0</td><td>GCTCCTTCTGCTTTGTACTG</td><td>AGG</td><td>||||||||[||||||||||||]PAM</td><td>NA</td><td>Intergenic</td><td>NA</td><td>NA</td></tr>
<tr><td>T21990</td><td>GCCGCCGCCGCCGCTCGATCCGG</td><td>1000</td><td></td><td></td><td></td><td></td><td></td><td></td><td></td><td></td><td></td></tr>
<tr><td>Oligo fwd</td><td>CACCGCCGCCGCCGCCGCTCGATC</td><td></td><td></td><td></td><td></td><td></td><td></td><td></td><td></td><td></td><td></td></tr>
<tr><td>Oligo rev</td><td>AAACGATCGAGCGGCGGCGGCGGC</td><td></td><td></td><td></td><td></td><td></td><td></td><td></td><td></td><td></td><td></td></tr>
<tr><td>Chromosome</td><td>start</td><td>end</td><td>strand</td><td>MM</td><td>target_seq</td><td>PAM</td><td>alignment</td><td>distance</td><td>position</td><td>gene name</td><td>gene id</td></tr>
<tr><td>GU305914.1</td><td>163894</td><td>163916</td><td>-</td><td>0</td><td>GCCGCCGCCGCCGCTCGATC</td><td>CGG</td><td>||||||||[||||||||||||]PAM</td><td>NA</td><td>Intergenic</td><td>NA</td><td>NA</td></tr>
<tr><td>T21991</td><td>GCTCGATCCGGAGCTACCCGAGG</td><td>1000</td><td></td><td></td><td></td><td></td><td></td><td></td><td></td><td></td><td></td></tr>
<tr><td>Oligo fwd</td><td>CACCGCTCGATCCGGAGCTACCCG</td><td></td><td></td><td></td><td></td><td></td><td></td><td></td><td></td><td></td><td></td></tr>
<tr><td>Oligo rev</td><td>AAACCGGGTAGCTCCGGATCGAGC</td><td></td><td></td><td></td><td></td><td></td><td></td><td></td><td></td><td></td><td></td></tr>
<tr><td>Chromosome</td><td>start</td><td>end</td><td>strand</td><td>MM</td><td>target_seq</td><td>PAM</td><td>alignment</td><td>distance</td><td>position</td><td>gene name</td><td>gene id</td></tr>
<tr><td>GU305914.1</td><td>163882</td><td>163904</td><td>-</td><td>0</td><td>GCTCGATCCGGAGCTACCCG</td><td>AGG</td><td>||||||||[||||||||||||]PAM</td><td>NA</td><td>Intergenic</td><td>NA</td><td>NA</td></tr>
<tr><td>T21992</td><td>CCGAGGTCCGCTCCCTATGAAGG</td><td>1000</td><td></td><td></td><td></td><td></td><td></td><td></td><td></td><td></td><td></td></tr>
<tr><td>Oligo fwd</td><td>CACCCCGAGGTCCGCTCCCTATGA</td><td></td><td></td><td></td><td></td><td></td><td></td><td></td><td></td><td></td><td></td></tr>
<tr><td>Oligo rev</td><td>AAACTCATAGGGAGCGGACCTCGG</td><td></td><td></td><td></td><td></td><td></td><td></td><td></td><td></td><td></td><td></td></tr>
<tr><td>Chromosome</td><td>start</td><td>end</td><td>strand</td><td>MM</td><td>target_seq</td><td>PAM</td><td>alignment</td><td>distance</td><td>position</td><td>gene name</td><td>gene id</td></tr>
<tr><td>GU305914.1</td><td>163865</td><td>163887</td><td>-</td><td>0</td><td>CCGAGGTCCGCTCCCTATGA</td><td>AGG</td><td>||||||||[||||||||||||]PAM</td><td>NA</td><td>Intergenic</td><td>NA</td><td>NA</td></tr>
<tr><td>T21993</td><td>CGAGGTCCGCTCCCTATGAAGGG</td><td>1000</td><td></td><td></td><td></td><td></td><td></td><td></td><td></td><td></td><td></td></tr>
<tr><td>Oligo fwd</td><td>CACCCGAGGTCCGCTCCCTATGAA</td><td></td><td></td><td></td><td></td><td></td><td></td><td></td><td></td><td></td><td></td></tr>
<tr><td>Oligo rev</td><td>AAACTTCATAGGGAGCGGACCTCG</td><td></td><td></td><td></td><td></td><td></td><td></td><td></td><td></td><td></td><td></td></tr>
<tr><td>Chromosome</td><td>start</td><td>end</td><td>strand</td><td>MM</td><td>target_seq</td><td>PAM</td><td>alignment</td><td>distance</td><td>position</td><td>gene name</td><td>gene id</td></tr>
<tr><td>GU305914.1</td><td>163864</td><td>163886</td><td>-</td><td>0</td><td>CGAGGTCCGCTCCCTATGAA</td><td>GGG</td><td>||||||||[||||||||||||]PAM</td><td>NA</td><td>Intergenic</td><td>NA</td><td>NA</td></tr>
</table>

<table>
<tr><td>T21994</td><td>CTCCCTATGAAGGGTACCGTTGG</td><td>1000</td><td></td><td></td><td></td><td></td><td></td><td></td><td></td><td></td><td></td></tr>
<tr><td>Oligo fwd</td><td>CACCCTCCCTATGAAGGGTACCGT</td><td></td><td></td><td></td><td></td><td></td><td></td><td></td><td></td><td></td><td></td></tr>
<tr><td>Oligo rev</td><td>AAACACGGTACCCTTCATAGGGAG</td><td></td><td></td><td></td><td></td><td></td><td></td><td></td><td></td><td></td><td></td></tr>
<tr><td>Chromosome</td><td>start</td><td>end</td><td>strand</td><td>MM</td><td>target_seq</td><td>PAM</td><td>alignment</td><td>distance</td><td>position</td><td>gene name</td><td>gene id</td></tr>
<tr><td>GU305914.1</td><td>163855</td><td>163877</td><td>-</td><td>0</td><td>CTCCCTATGAAGGGTACCGT</td><td>TGG</td><td>||||||||[||||||||||||]PAM</td><td>NA</td><td>Intergenic</td><td>NA</td><td>NA</td></tr>
<tr><td>T21995</td><td>AGAGTTCGAATCTCTTACGTCGG</td><td>1000</td><td></td><td></td><td></td><td></td><td></td><td></td><td></td><td></td><td></td></tr>
<tr><td>Oligo fwd</td><td>CACCAGAGTTCGAATCTCTTACGT</td><td></td><td></td><td></td><td></td><td></td><td></td><td></td><td></td><td></td><td></td></tr>
<tr><td>Oligo rev</td><td>AAACACGTAAGAGATTCGAACTCT</td><td></td><td></td><td></td><td></td><td></td><td></td><td></td><td></td><td></td><td></td></tr>
<tr><td>Chromosome</td><td>start</td><td>end</td><td>strand</td><td>MM</td><td>target_seq</td><td>PAM</td><td>alignment</td><td>distance</td><td>position</td><td>gene name</td><td>gene id</td></tr>
<tr><td>GU305914.1</td><td>163832</td><td>163854</td><td>-</td><td>0</td><td>AGAGTTCGAATCTCTTACGT</td><td>CGG</td><td>||||||||[||||||||||||]PAM</td><td>NA</td><td>Intergenic</td><td>NA</td><td>NA</td></tr>
<tr><td>T21996</td><td>TCGGTCGTCATCGTGAGATTTGG</td><td>1000</td><td></td><td></td><td></td><td></td><td></td><td></td><td></td><td></td><td></td></tr>
<tr><td>Oligo fwd</td><td>CACCTCGGTCGTCATCGTGAGATT</td><td></td><td></td><td></td><td></td><td></td><td></td><td></td><td></td><td></td><td></td></tr>
<tr><td>Oligo rev</td><td>AAACAATCTCACGATGACGACCGA</td><td></td><td></td><td></td><td></td><td></td><td></td><td></td><td></td><td></td><td></td></tr>
<tr><td>Chromosome</td><td>start</td><td>end</td><td>strand</td><td>MM</td><td>target_seq</td><td>PAM</td><td>alignment</td><td>distance</td><td>position</td><td>gene name</td><td>gene id</td></tr>
<tr><td>GU305914.1</td><td>163813</td><td>163835</td><td>-</td><td>0</td><td>TCGGTCGTCATCGTGAGATT</td><td>TGG</td><td>||||||||[||||||||||||]PAM</td><td>NA</td><td>Intergenic</td><td>NA</td><td>NA</td></tr>
<tr><td>T21997</td><td>ATTTGGACGAGTCTTTATCGCGG</td><td>1000</td><td></td><td></td><td></td><td></td><td></td><td></td><td></td><td></td><td></td></tr>
<tr><td>Oligo fwd</td><td>CACCATTTGGACGAGTCTTTATCG</td><td></td><td></td><td></td><td></td><td></td><td></td><td></td><td></td><td></td><td></td></tr>
<tr><td>Oligo rev</td><td>AAACCGATAAAGACTCGTCCAAAT</td><td></td><td></td><td></td><td></td><td></td><td></td><td></td><td></td><td></td><td></td></tr>
<tr><td>Chromosome</td><td>start</td><td>end</td><td>strand</td><td>MM</td><td>target_seq</td><td>PAM</td><td>alignment</td><td>distance</td><td>position</td><td>gene name</td><td>gene id</td></tr>
<tr><td>GU305914.1</td><td>163796</td><td>163818</td><td>-</td><td>0</td><td>ATTTGGACGAGTCTTTATCG</td><td>CGG</td><td>||||||||[||||||||||||]PAM</td><td>NA</td><td>Intergenic</td><td>NA</td><td>NA</td></tr>
<tr><td>T21998</td><td>AGAATTGAAGCCGCCGCCACCGG</td><td>1000</td><td></td><td></td><td></td><td></td><td></td><td></td><td></td><td></td><td></td></tr>
<tr><td>Oligo fwd</td><td>CACCAGAATTGAAGCCGCCGCCAC</td><td></td><td></td><td></td><td></td><td></td><td></td><td></td><td></td><td></td><td></td></tr>
<tr><td>Oligo rev</td><td>AAACGTGGCGGCGGCTTCAATTCT</td><td></td><td></td><td></td><td></td><td></td><td></td><td></td><td></td><td></td><td></td></tr>
<tr><td>Chromosome</td><td>start</td><td>end</td><td>strand</td><td>MM</td><td>target_seq</td><td>PAM</td><td>alignment</td><td>distance</td><td>position</td><td>gene name</td><td>gene id</td></tr>
<tr><td>GU305914.1</td><td>163759</td><td>163781</td><td>-</td><td>0</td><td>AGAATTGAAGCCGCCGCCAC</td><td>CGG</td><td>||||||||[||||||||||||]PAM</td><td>NA</td><td>Intergenic</td><td>NA</td><td>NA</td></tr>
<tr><td>T21999</td><td>GCCGCCACCGGCGCCGCCGACGG</td><td>1000</td><td></td><td></td><td></td><td></td><td></td><td></td><td></td><td></td><td></td></tr>
<tr><td>Oligo fwd</td><td>CACCGCCGCCACCGGCGCCGCCGA</td><td></td><td></td><td></td><td></td><td></td><td></td><td></td><td></td><td></td><td></td></tr>
<tr><td>Oligo rev</td><td>AAACTCGGCGGCGCCGGTGGCGGC</td><td></td><td></td><td></td><td></td><td></td><td></td><td></td><td></td><td></td><td></td></tr>
<tr><td>Chromosome</td><td>start</td><td>end</td><td>strand</td><td>MM</td><td>target_seq</td><td>PAM</td><td>alignment</td><td>distance</td><td>position</td><td>gene name</td><td>gene id</td></tr>
<tr><td>GU305914.1</td><td>163747</td><td>163769</td><td>-</td><td>0</td><td>GCCGCCACCGGCGCCGCCGA</td><td>CGG</td><td>||||||||[||||||||||||]PAM</td><td>NA</td><td>Intergenic</td><td>NA</td><td>NA</td></tr>
<tr><td>GU305914.1</td><td>152551</td><td>152573</td><td>-</td><td>3</td><td>GCCGGCACCGCCGCCGCCGC</td><td>AGG</td><td>||||-|||[||-||||||||-]PAM</td><td>NA</td><td>Intergenic</td><td>NA</td><td>NA</td></tr>
<tr><td>T22000</td><td>GCTCGAGCTACCCATCGAAGTGG</td><td>1000</td><td></td><td></td><td></td><td></td><td></td><td></td><td></td><td></td><td></td></tr>
<tr><td>Oligo fwd</td><td>CACCGCTCGAGCTACCCATCGAAG</td><td></td><td></td><td></td><td></td><td></td><td></td><td></td><td></td><td></td><td></td></tr>
<tr><td>Oligo rev</td><td>AAACCTTCGATGGGTAGCTCGAGC</td><td></td><td></td><td></td><td></td><td></td><td></td><td></td><td></td><td></td><td></td></tr>
<tr><td>Chromosome</td><td>start</td><td>end</td><td>strand</td><td>MM</td><td>target_seq</td><td>PAM</td><td>alignment</td><td>distance</td><td>position</td><td>gene name</td><td>gene id</td></tr>
<tr><td>GU305914.1</td><td>163723</td><td>163745</td><td>-</td><td>0</td><td>GCTCGAGCTACCCATCGAAG</td><td>TGG</td><td>||||||||[||||||||||||]PAM</td><td>NA</td><td>Intergenic</td><td>NA</td><td>NA</td></tr>
<tr><td>T22001</td><td>CATCGAAGTGGCCGACTGCACGG</td><td>1000</td><td></td><td></td><td></td><td></td><td></td><td></td><td></td><td></td><td></td></tr>
<tr><td>Oligo fwd</td><td>CACCCATCGAAGTGGCCGACTGCA</td><td></td><td></td><td></td><td></td><td></td><td></td><td></td><td></td><td></td><td></td></tr>
<tr><td>Oligo rev</td><td>AAACTGCAGTCGGCCACTTCGATG</td><td></td><td></td><td></td><td></td><td></td><td></td><td></td><td></td><td></td><td></td></tr>
<tr><td>Chromosome</td><td>start</td><td>end</td><td>strand</td><td>MM</td><td>target_seq</td><td>PAM</td><td>alignment</td><td>distance</td><td>position</td><td>gene name</td><td>gene id</td></tr>
<tr><td>GU305914.1</td><td>163711</td><td>163733</td><td>-</td><td>0</td><td>CATCGAAGTGGCCGACTGCA</td><td>CGG</td><td>||||||||[||||||||||||]PAM</td><td>NA</td><td>Intergenic</td><td>NA</td><td>NA</td></tr>
<tr><td>T22002</td><td>CGAAGTGGCCGACTGCACGGCGG</td><td>1000</td><td></td><td></td><td></td><td></td><td></td><td></td><td></td><td></td><td></td></tr>
<tr><td>Oligo fwd</td><td>CACCCGAAGTGGCCGACTGCACGG</td><td></td><td></td><td></td><td></td><td></td><td></td><td></td><td></td><td></td><td></td></tr>
<tr><td>Oligo rev</td><td>AAACCCGTGCAGTCGGCCACTTCG</td><td></td><td></td><td></td><td></td><td></td><td></td><td></td><td></td><td></td><td></td></tr>
<tr><td>Chromosome</td><td>start</td><td>end</td><td>strand</td><td>MM</td><td>target_seq</td><td>PAM</td><td>alignment</td><td>distance</td><td>position</td><td>gene name</td><td>gene id</td></tr>
<tr><td>GU305914.1</td><td>163708</td><td>163730</td><td>-</td><td>0</td><td>CGAAGTGGCCGACTGCACGG</td><td>CGG</td><td>||||||||[||||||||||||]PAM</td><td>NA</td><td>Intergenic</td><td>NA</td><td>NA</td></tr>
<tr><td>T22003</td><td>AGTGGCCGACTGCACGGCGGCGG</td><td>1000</td><td></td><td></td><td></td><td></td><td></td><td></td><td></td><td></td><td></td></tr>
<tr><td>Oligo fwd</td><td>CACCAGTGGCCGACTGCACGGCGG</td><td></td><td></td><td></td><td></td><td></td><td></td><td></td><td></td><td></td><td></td></tr>
<tr><td>Oligo rev</td><td>AAACCCGCCGTGCAGTCGGCCACT</td><td></td><td></td><td></td><td></td><td></td><td></td><td></td><td></td><td></td><td></td></tr>
<tr><td>Chromosome</td><td>start</td><td>end</td><td>strand</td><td>MM</td><td>target_seq</td><td>PAM</td><td>alignment</td><td>distance</td><td>position</td><td>gene name</td><td>gene id</td></tr>
<tr><td>GU305914.1</td><td>163705</td><td>163727</td><td>-</td><td>0</td><td>AGTGGCCGACTGCACGGCGG</td><td>CGG</td><td>||||||||[||||||||||||]PAM</td><td>NA</td><td>Intergenic</td><td>NA</td><td>NA</td></tr>
<tr><td>T22004</td><td>CTGCACGGCGGCGGCCACCGCGG</td><td>1000</td><td></td><td></td><td></td><td></td><td></td><td></td><td></td><td></td><td></td></tr>
<tr><td>Oligo fwd</td><td>CACCCTGCACGGCGGCGGCCACCG</td><td></td><td></td><td></td><td></td><td></td><td></td><td></td><td></td><td></td><td></td></tr>
<tr><td>Oligo rev</td><td>AAACCGGTGGCCGCCGCCGTGCAG</td><td></td><td></td><td></td><td></td><td></td><td></td><td></td><td></td><td></td><td></td></tr>
<tr><td>Chromosome</td><td>start</td><td>end</td><td>strand</td><td>MM</td><td>target_seq</td><td>PAM</td><td>alignment</td><td>distance</td><td>position</td><td>gene name</td><td>gene id</td></tr>
<tr><td>GU305914.1</td><td>163696</td><td>163718</td><td>-</td><td>0</td><td>CTGCACGGCGGCGGCCACCG</td><td>CGG</td><td>||||||||[||||||||||||]PAM</td><td>NA</td><td>Intergenic</td><td>NA</td><td>NA</td></tr>
<tr><td>GU305914.1</td><td>200735</td><td>200757</td><td>+</td><td>4</td><td>CTGCAGGACCGCGGTCACCG</td><td>CGG</td><td>|||||-|-[|-||||-|||||]PAM</td><td>NA</td><td>Intergenic</td><td>NA</td><td>NA</td></tr>
<tr><td>T22005</td><td>CGCGGCGTTGCCGACGCTCTCGG</td><td>1000</td><td></td><td></td><td></td><td></td><td></td><td></td><td></td><td></td><td></td></tr>
<tr><td>Oligo fwd</td><td>CACCCGCGGCGTTGCCGACGCTCT</td><td></td><td></td><td></td><td></td><td></td><td></td><td></td><td></td><td></td><td></td></tr>
<tr><td>Oligo rev</td><td>AAACAGAGCGTCGGCAACGCCGCG</td><td></td><td></td><td></td><td></td><td></td><td></td><td></td><td></td><td></td><td></td></tr>
<tr><td>Chromosome</td><td>start</td><td>end</td><td>strand</td><td>MM</td><td>target_seq</td><td>PAM</td><td>alignment</td><td>distance</td><td>position</td><td>gene name</td><td>gene id</td></tr>
<tr><td>GU305914.1</td><td>163678</td><td>163700</td><td>-</td><td>0</td><td>CGCGGCGTTGCCGACGCTCT</td><td>CGG</td><td>||||||||[||||||||||||]PAM</td><td>NA</td><td>Intergenic</td><td>NA</td><td>NA</td></tr>
<tr><td>T22006</td><td>GACGCTCTCGGCGACATCGTCGG</td><td>1000</td><td></td><td></td><td></td><td></td><td></td><td></td><td></td><td></td><td></td></tr>
<tr><td>Oligo fwd</td><td>CACCGACGCTCTCGGCGACATCGT</td><td></td><td></td><td></td><td></td><td></td><td></td><td></td><td></td><td></td><td></td></tr>
<tr><td>Oligo rev</td><td>AAACACGATGTCGCCGAGAGCGTC</td><td></td><td></td><td></td><td></td><td></td><td></td><td></td><td></td><td></td><td></td></tr>
<tr><td>Chromosome</td><td>start</td><td>end</td><td>strand</td><td>MM</td><td>target_seq</td><td>PAM</td><td>alignment</td><td>distance</td><td>position</td><td>gene name</td><td>gene id</td></tr>
<tr><td>GU305914.1</td><td>163666</td><td>163688</td><td>-</td><td>0</td><td>GACGCTCTCGGCGACATCGT</td><td>CGG</td><td>||||||||[||||||||||||]PAM</td><td>NA</td><td>Intergenic</td><td>NA</td><td>NA</td></tr>
<tr><td>T22007</td><td>CTCGGCGACATCGTCGGCTTCGG</td><td>1000</td><td></td><td></td><td></td><td></td><td></td><td></td><td></td><td></td><td></td></tr>
<tr><td>Oligo fwd</td><td>CACCCTCGGCGACATCGTCGGCTT</td><td></td><td></td><td></td><td></td><td></td><td></td><td></td><td></td><td></td><td></td></tr>
<tr><td>Oligo rev</td><td>AAACAAGCCGACGATGTCGCCGAG</td><td></td><td></td><td></td><td></td><td></td><td></td><td></td><td></td><td></td><td></td></tr>
<tr><td>Chromosome</td><td>start</td><td>end</td><td>strand</td><td>MM</td><td>target_seq</td><td>PAM</td><td>alignment</td><td>distance</td><td>position</td><td>gene name</td><td>gene id</td></tr>
<tr><td>GU305914.1</td><td>163660</td><td>163682</td><td>-</td><td>0</td><td>CTCGGCGACATCGTCGGCTT</td><td>CGG</td><td>||||||||[||||||||||||]PAM</td><td>NA</td><td>Intergenic</td><td>NA</td><td>NA</td></tr>
<tr><td>T22008</td><td>CGACATCGTCGGCTTCGGCGCGG</td><td>1000</td><td></td><td></td><td></td><td></td><td></td><td></td><td></td><td></td><td></td></tr>
<tr><td>Oligo fwd</td><td>CACCCGACATCGTCGGCTTCGGCG</td><td></td><td></td><td></td><td></td><td></td><td></td><td></td><td></td><td></td><td></td></tr>
</table>

| | | | | | | | | | | | |
|---|---|---|---|---|---|---|---|---|---|---|---|
| Oligo rev | AAACCGCCGAAGCCGACGATGTCG | | | | | | | | | | |
| Chromosome | start | end | strand | MM | target_seq | PAM | alignment | distance | position | gene name | gene id |
| GU305914.1 | 163655 | 163677 | - | 0 | CGACATCGTCGGCTTCGGCG | CGG | ||||||||[||||||||||||]PAM | NA | Intergenic | NA | NA |
| GU305914.1 | 30873 | 30895 | - | 4 | GGTCGTCGTCGGCTTCGTCG | AGG | -|-|-|||[|||||||||-||]PAM | NA | Intergenic | NA | NA |
| T22009 | CATCGTCGGCTTCGGCGCGGTGG | 1000 | | | | | | | | | |
| Oligo fwd | CACCCATCGTCGGCTTCGGCGCGG | | | | | | | | | | |
| Oligo rev | AAACCCGCGCCGAAGCCGACGATG | | | | | | | | | | |
| Chromosome | start | end | strand | MM | target_seq | PAM | alignment | distance | position | gene name | gene id |
| GU305914.1 | 163652 | 163674 | - | 0 | CATCGTCGGCTTCGGCGCGG | TGG | ||||||||[||||||||||||]PAM | NA | Intergenic | NA | NA |
| T22010 | ATCGTCGGCTTCGGCGCGGTGGG | 1000 | | | | | | | | | |
| Oligo fwd | CACCATCGTCGGCTTCGGCGCGGT | | | | | | | | | | |
| Oligo rev | AAACACCGCGCCGAAGCCGACGAT | | | | | | | | | | |
| Chromosome | start | end | strand | MM | target_seq | PAM | alignment | distance | position | gene name | gene id |
| GU305914.1 | 163651 | 163673 | - | 0 | ATCGTCGGCTTCGGCGCGGT | GGG | ||||||||[||||||||||||]PAM | NA | Intergenic | NA | NA |
| T22011 | GCTTCGGCGCGGTGGGCGCCCGG | 1000 | | | | | | | | | |
| Oligo fwd | CACCGCTTCGGCGCGGTGGGCGCC | | | | | | | | | | |
| Oligo rev | AAACGGCGCCCACCGCGCCGAAGC | | | | | | | | | | |
| Chromosome | start | end | strand | MM | target_seq | PAM | alignment | distance | position | gene name | gene id |
| GU305914.1 | 163644 | 163666 | - | 0 | GCTTCGGCGCGGTGGGCGCC | CGG | ||||||||[||||||||||||]PAM | NA | Intergenic | NA | NA |
| T22012 | CGCCCGGCGAGTGTTCGACCAGG | 1000 | | | | | | | | | |
| Oligo fwd | CACCCGCCCGGCGAGTGTTCGACC | | | | | | | | | | |
| Oligo rev | AAACGGTCGAACACTCGCCGGGCG | | | | | | | | | | |
| Chromosome | start | end | strand | MM | target_seq | PAM | alignment | distance | position | gene name | gene id |
| GU305914.1 | 163628 | 163650 | - | 0 | CGCCCGGCGAGTGTTCGACC | AGG | ||||||||[||||||||||||]PAM | NA | Intergenic | NA | NA |
| T22013 | CGTCCGCCACCAGAGTCAAGAGG | 1000 | | | | | | | | | |
| Oligo fwd | CACCCGTCCGCCACCAGAGTCAAG | | | | | | | | | | |
| Oligo rev | AAACCTTGACTCTGGTGGCGGACG | | | | | | | | | | |
| Chromosome | start | end | strand | MM | target_seq | PAM | alignment | distance | position | gene name | gene id |
| GU305914.1 | 163586 | 163608 | - | 0 | CGTCCGCCACCAGAGTCAAG | AGG | ||||||||[||||||||||||]PAM | NA | Intergenic | NA | NA |
| T22014 | CGCCACCAGAGTCAAGAGGCTGG | 1000 | | | | | | | | | |
| Oligo fwd | CACCCGCCACCAGAGTCAAGAGGC | | | | | | | | | | |
| Oligo rev | AAACGCCTCTTGACTCTGGTGGCG | | | | | | | | | | |
| Chromosome | start | end | strand | MM | target_seq | PAM | alignment | distance | position | gene name | gene id |
| GU305914.1 | 163582 | 163604 | - | 0 | CGCCACCAGAGTCAAGAGGC | TGG | ||||||||[||||||||||||]PAM | NA | Intergenic | NA | NA |
| T22015 | GAGGCTGGCGCCCGTGCCGTCGG | 1000 | | | | | | | | | |
| Oligo fwd | CACCGAGGCTGGCGCCCGTGCCGT | | | | | | | | | | |
| Oligo rev | AAACACGGCACGGGCGCCAGCCTC | | | | | | | | | | |
| Chromosome | start | end | strand | MM | target_seq | PAM | alignment | distance | position | gene name | gene id |
| GU305914.1 | 163567 | 163589 | - | 0 | GAGGCTGGCGCCCGTGCCGT | CGG | ||||||||[||||||||||||]PAM | NA | Intergenic | NA | NA |
| GU305914.1 | 121592 | 121614 | - | 4 | GCCGCTGGCGCCCGTGCGGA | TGG | |--|||||[|||||||||-|-]PAM | NA | Intergenic | NA | NA |
| T22016 | GCGCCCGTGCCGTCGGACCCCGG | 1000 | | | | | | | | | |
| Oligo fwd | CACCGCGCCCGTGCCGTCGGACCC | | | | | | | | | | |
| Oligo rev | AAACGGGTCCGACGGCACGGGCGC | | | | | | | | | | |
| Chromosome | start | end | strand | MM | target_seq | PAM | alignment | distance | position | gene name | gene id |
| GU305914.1 | 163560 | 163582 | - | 0 | GCGCCCGTGCCGTCGGACCC | CGG | ||||||||[||||||||||||]PAM | NA | Intergenic | NA | NA |
| T22017 | CCCGTGCCGTCGGACCCCGGCGG | 1000 | | | | | | | | | |
| Oligo fwd | CACCCCCGTGCCGTCGGACCCCGG | | | | | | | | | | |
| Oligo rev | AAACCCGGGGTCCGACGGCACGGG | | | | | | | | | | |
| Chromosome | start | end | strand | MM | target_seq | PAM | alignment | distance | position | gene name | gene id |
| GU305914.1 | 163557 | 163579 | - | 0 | CCCGTGCCGTCGGACCCCGG | CGG | ||||||||[||||||||||||]PAM | NA | Intergenic | NA | NA |
| T22018 | CGGCGTCGCGACCGAAGAGACGG | 1000 | | | | | | | | | |
| Oligo fwd | CACCCGGCGTCGCGACCGAAGAGA | | | | | | | | | | |
| Oligo rev | AAACTCTCTTCGGTCGCGACGCCG | | | | | | | | | | |
| Chromosome | start | end | strand | MM | target_seq | PAM | alignment | distance | position | gene name | gene id |
| GU305914.1 | 163537 | 163559 | - | 0 | CGGCGTCGCGACCGAAGAGA | CGG | ||||||||[||||||||||||]PAM | NA | Intergenic | NA | NA |
| T22019 | TGTTACAGACGATGCTCCCGCGG | 1000 | | | | | | | | | |
| Oligo fwd | CACCTGTTACAGACGATGCTCCCG | | | | | | | | | | |
| Oligo rev | AAACCGGGAGCATCGTCTGTAACA | | | | | | | | | | |
| Chromosome | start | end | strand | MM | target_seq | PAM | alignment | distance | position | gene name | gene id |
| GU305914.1 | 163511 | 163533 | - | 0 | TGTTACAGACGATGCTCCCG | CGG | ||||||||[||||||||||||]PAM | NA | Intergenic | NA | NA |
| T22020 | TACAGACGATGCTCCCGCGGTGG | 1000 | | | | | | | | | |
| Oligo fwd | CACCTACAGACGATGCTCCCGCGG | | | | | | | | | | |
| Oligo rev | AAACCCGCGGGAGCATCGTCTGTA | | | | | | | | | | |
| Chromosome | start | end | strand | MM | target_seq | PAM | alignment | distance | position | gene name | gene id |
| GU305914.1 | 163508 | 163530 | - | 0 | TACAGACGATGCTCCCGCGG | TGG | ||||||||[||||||||||||]PAM | NA | Intergenic | NA | NA |
| T22021 | AACATGTTCATGACCTTCTCCGG | 1000 | | | | | | | | | |
| Oligo fwd | CACCAACATGTTCATGACCTTCTC | | | | | | | | | | |
| Oligo rev | AAACGAGAAGGTCATGAACATGTT | | | | | | | | | | |
| Chromosome | start | end | strand | MM | target_seq | PAM | alignment | distance | position | gene name | gene id |
| GU305914.1 | 163485 | 163507 | - | 0 | AACATGTTCATGACCTTCTC | CGG | ||||||||[||||||||||||]PAM | NA | Intergenic | NA | NA |
| T22022 | CTTCTCCGGTCCGTCGTCCGAGG | 1000 | | | | | | | | | |
| Oligo fwd | CACCCTTCTCCGGTCCGTCGTCCG | | | | | | | | | | |
| Oligo rev | AAACCGGACGACGGACCGGAGAAG | | | | | | | | | | |
| Chromosome | start | end | strand | MM | target_seq | PAM | alignment | distance | position | gene name | gene id |
| GU305914.1 | 163471 | 163493 | - | 0 | CTTCTCCGGTCCGTCGTCCG | AGG | ||||||||[||||||||||||]PAM | NA | Intergenic | NA | NA |

<table>
<tr><td>T22023</td><td>TGTCCAGAGACATGCATCCCTGG</td><td>1000</td><td></td><td></td><td></td><td></td><td></td><td></td><td></td><td></td><td></td></tr>
<tr><td>Oligo fwd</td><td>CACCTGTCCAGAGACATGCATCCC</td><td></td><td></td><td></td><td></td><td></td><td></td><td></td><td></td><td></td><td></td></tr>
<tr><td>Oligo rev</td><td>AAACGGGATGCATGTCTCTGGACA</td><td></td><td></td><td></td><td></td><td></td><td></td><td></td><td></td><td></td><td></td></tr>
<tr><td>Chromosome</td><td>start</td><td>end</td><td>strand</td><td>MM</td><td>target_seq</td><td>PAM</td><td>alignment</td><td>distance</td><td>position</td><td>gene name</td><td>gene id</td></tr>
<tr><td>GU305914.1</td><td>163445</td><td>163467</td><td>-</td><td>0</td><td>TGTCCAGAGACATGCATCCC</td><td>TGG</td><td>||||||||[||||||||||||]PAM</td><td>NA</td><td>Intergenic</td><td>NA</td><td>NA</td></tr>
<tr><td>T22024</td><td>GTCCAGAGACATGCATCCCTGGG</td><td>1000</td><td></td><td></td><td></td><td></td><td></td><td></td><td></td><td></td><td></td></tr>
<tr><td>Oligo fwd</td><td>CACCGTCCAGAGACATGCATCCCT</td><td></td><td></td><td></td><td></td><td></td><td></td><td></td><td></td><td></td><td></td></tr>
<tr><td>Oligo rev</td><td>AAACAGGGATGCATGTCTCTGGAC</td><td></td><td></td><td></td><td></td><td></td><td></td><td></td><td></td><td></td><td></td></tr>
<tr><td>Chromosome</td><td>start</td><td>end</td><td>strand</td><td>MM</td><td>target_seq</td><td>PAM</td><td>alignment</td><td>distance</td><td>position</td><td>gene name</td><td>gene id</td></tr>
<tr><td>GU305914.1</td><td>163444</td><td>163466</td><td>-</td><td>0</td><td>GTCCAGAGACATGCATCCCT</td><td>GGG</td><td>||||||||[||||||||||||]PAM</td><td>NA</td><td>Intergenic</td><td>NA</td><td>NA</td></tr>
<tr><td>T22025</td><td>GCATCCCTGGGCCGCCGATCCGG</td><td>1000</td><td></td><td></td><td></td><td></td><td></td><td></td><td></td><td></td><td></td></tr>
<tr><td>Oligo fwd</td><td>CACCGCATCCCTGGGCCGCCGATC</td><td></td><td></td><td></td><td></td><td></td><td></td><td></td><td></td><td></td><td></td></tr>
<tr><td>Oligo rev</td><td>AAACGATCGGCGGCCCAGGGATGC</td><td></td><td></td><td></td><td></td><td></td><td></td><td></td><td></td><td></td><td></td></tr>
<tr><td>Chromosome</td><td>start</td><td>end</td><td>strand</td><td>MM</td><td>target_seq</td><td>PAM</td><td>alignment</td><td>distance</td><td>position</td><td>gene name</td><td>gene id</td></tr>
<tr><td>GU305914.1</td><td>163432</td><td>163454</td><td>-</td><td>0</td><td>GCATCCCTGGGCCGCCGATC</td><td>CGG</td><td>||||||||[||||||||||||]PAM</td><td>NA</td><td>Intergenic</td><td>NA</td><td>NA</td></tr>
<tr><td>T22026</td><td>CCGGACGACGAGCCGCTGTCCGG</td><td>1000</td><td></td><td></td><td></td><td></td><td></td><td></td><td></td><td></td><td></td></tr>
<tr><td>Oligo fwd</td><td>CACCCCGGACGACGAGCCGCTGTC</td><td></td><td></td><td></td><td></td><td></td><td></td><td></td><td></td><td></td><td></td></tr>
<tr><td>Oligo rev</td><td>AAACGACAGCGGCTCGTCGTCCGG</td><td></td><td></td><td></td><td></td><td></td><td></td><td></td><td></td><td></td><td></td></tr>
<tr><td>Chromosome</td><td>start</td><td>end</td><td>strand</td><td>MM</td><td>target_seq</td><td>PAM</td><td>alignment</td><td>distance</td><td>position</td><td>gene name</td><td>gene id</td></tr>
<tr><td>GU305914.1</td><td>163413</td><td>163435</td><td>-</td><td>0</td><td>CCGGACGACGAGCCGCTGTC</td><td>CGG</td><td>||||||||[||||||||||||]PAM</td><td>NA</td><td>Intergenic</td><td>NA</td><td>NA</td></tr>
<tr><td>T22027</td><td>GCTGTCCGGAGACTGCATGATGG</td><td>1000</td><td></td><td></td><td></td><td></td><td></td><td></td><td></td><td></td><td></td></tr>
<tr><td>Oligo fwd</td><td>CACCGCTGTCCGGAGACTGCATGA</td><td></td><td></td><td></td><td></td><td></td><td></td><td></td><td></td><td></td><td></td></tr>
<tr><td>Oligo rev</td><td>AAACTCATGCAGTCTCCGGACAGC</td><td></td><td></td><td></td><td></td><td></td><td></td><td></td><td></td><td></td><td></td></tr>
<tr><td>Chromosome</td><td>start</td><td>end</td><td>strand</td><td>MM</td><td>target_seq</td><td>PAM</td><td>alignment</td><td>distance</td><td>position</td><td>gene name</td><td>gene id</td></tr>
<tr><td>GU305914.1</td><td>163399</td><td>163421</td><td>-</td><td>0</td><td>GCTGTCCGGAGACTGCATGA</td><td>TGG</td><td>||||||||[||||||||||||]PAM</td><td>NA</td><td>Intergenic</td><td>NA</td><td>NA</td></tr>
<tr><td>T22028</td><td>GATGGTCGCGCACAGCCAGCCGG</td><td>1000</td><td></td><td></td><td></td><td></td><td></td><td></td><td></td><td></td><td></td></tr>
<tr><td>Oligo fwd</td><td>CACCGATGGTCGCGCACAGCCAGC</td><td></td><td></td><td></td><td></td><td></td><td></td><td></td><td></td><td></td><td></td></tr>
<tr><td>Oligo rev</td><td>AAACGCTGGCTGTGCGCGACCATC</td><td></td><td></td><td></td><td></td><td></td><td></td><td></td><td></td><td></td><td></td></tr>
<tr><td>Chromosome</td><td>start</td><td>end</td><td>strand</td><td>MM</td><td>target_seq</td><td>PAM</td><td>alignment</td><td>distance</td><td>position</td><td>gene name</td><td>gene id</td></tr>
<tr><td>GU305914.1</td><td>163381</td><td>163403</td><td>-</td><td>0</td><td>GATGGTCGCGCACAGCCAGC</td><td>CGG</td><td>||||||||[||||||||||||]PAM</td><td>NA</td><td>Intergenic</td><td>NA</td><td>NA</td></tr>
<tr><td>T22029</td><td>ATGGTCGCGCACAGCCAGCCGGG</td><td>1000</td><td></td><td></td><td></td><td></td><td></td><td></td><td></td><td></td><td></td></tr>
<tr><td>Oligo fwd</td><td>CACCATGGTCGCGCACAGCCAGCC</td><td></td><td></td><td></td><td></td><td></td><td></td><td></td><td></td><td></td><td></td></tr>
<tr><td>Oligo rev</td><td>AAACGGCTGGCTGTGCGCGACCAT</td><td></td><td></td><td></td><td></td><td></td><td></td><td></td><td></td><td></td><td></td></tr>
<tr><td>Chromosome</td><td>start</td><td>end</td><td>strand</td><td>MM</td><td>target_seq</td><td>PAM</td><td>alignment</td><td>distance</td><td>position</td><td>gene name</td><td>gene id</td></tr>
<tr><td>GU305914.1</td><td>163380</td><td>163402</td><td>-</td><td>0</td><td>ATGGTCGCGCACAGCCAGCC</td><td>GGG</td><td>||||||||[||||||||||||]PAM</td><td>NA</td><td>Intergenic</td><td>NA</td><td>NA</td></tr>
<tr><td>T22030</td><td>CCAGCCGGGCCCCCAGTTCTCGG</td><td>1000</td><td></td><td></td><td></td><td></td><td></td><td></td><td></td><td></td><td></td></tr>
<tr><td>Oligo fwd</td><td>CACCCCAGCCGGGCCCCCAGTTCT</td><td></td><td></td><td></td><td></td><td></td><td></td><td></td><td></td><td></td><td></td></tr>
<tr><td>Oligo rev</td><td>AAACAGAACTGGGGGCCCGGCTGG</td><td></td><td></td><td></td><td></td><td></td><td></td><td></td><td></td><td></td><td></td></tr>
<tr><td>Chromosome</td><td>start</td><td>end</td><td>strand</td><td>MM</td><td>target_seq</td><td>PAM</td><td>alignment</td><td>distance</td><td>position</td><td>gene name</td><td>gene id</td></tr>
<tr><td>GU305914.1</td><td>163366</td><td>163388</td><td>-</td><td>0</td><td>CCAGCCGGGCCCCCAGTTCT</td><td>CGG</td><td>||||||||[||||||||||||]PAM</td><td>NA</td><td>Intergenic</td><td>NA</td><td>NA</td></tr>
<tr><td>T22031</td><td>CGGGCCCCCAGTTCTCGGTGCGG</td><td>1000</td><td></td><td></td><td></td><td></td><td></td><td></td><td></td><td></td><td></td></tr>
<tr><td>Oligo fwd</td><td>CACCCGGGCCCCCAGTTCTCGGTG</td><td></td><td></td><td></td><td></td><td></td><td></td><td></td><td></td><td></td><td></td></tr>
<tr><td>Oligo rev</td><td>AAACCACCGAGAACTGGGGGCCCG</td><td></td><td></td><td></td><td></td><td></td><td></td><td></td><td></td><td></td><td></td></tr>
<tr><td>Chromosome</td><td>start</td><td>end</td><td>strand</td><td>MM</td><td>target_seq</td><td>PAM</td><td>alignment</td><td>distance</td><td>position</td><td>gene name</td><td>gene id</td></tr>
<tr><td>GU305914.1</td><td>163361</td><td>163383</td><td>-</td><td>0</td><td>CGGGCCCCCAGTTCTCGGTG</td><td>CGG</td><td>||||||||[||||||||||||]PAM</td><td>NA</td><td>Intergenic</td><td>NA</td><td>NA</td></tr>
<tr><td>T22032</td><td>CGTTCGTGCAGTCCAAAACGCGG</td><td>1000</td><td></td><td></td><td></td><td></td><td></td><td></td><td></td><td></td><td></td></tr>
<tr><td>Oligo fwd</td><td>CACCCGTTCGTGCAGTCCAAAACG</td><td></td><td></td><td></td><td></td><td></td><td></td><td></td><td></td><td></td><td></td></tr>
<tr><td>Oligo rev</td><td>AAACCGTTTTGGACTGCACGAACG</td><td></td><td></td><td></td><td></td><td></td><td></td><td></td><td></td><td></td><td></td></tr>
<tr><td>Chromosome</td><td>start</td><td>end</td><td>strand</td><td>MM</td><td>target_seq</td><td>PAM</td><td>alignment</td><td>distance</td><td>position</td><td>gene name</td><td>gene id</td></tr>
<tr><td>GU305914.1</td><td>163331</td><td>163353</td><td>-</td><td>0</td><td>CGTTCGTGCAGTCCAAAACG</td><td>CGG</td><td>||||||||[||||||||||||]PAM</td><td>NA</td><td>Intergenic</td><td>NA</td><td>NA</td></tr>
<tr><td>T22033</td><td>GTTCGTGCAGTCCAAAACGCGGG</td><td>1000</td><td></td><td></td><td></td><td></td><td></td><td></td><td></td><td></td><td></td></tr>
<tr><td>Oligo fwd</td><td>CACCGTTCGTGCAGTCCAAAACGC</td><td></td><td></td><td></td><td></td><td></td><td></td><td></td><td></td><td></td><td></td></tr>
<tr><td>Oligo rev</td><td>AAACGCGTTTTGGACTGCACGAAC</td><td></td><td></td><td></td><td></td><td></td><td></td><td></td><td></td><td></td><td></td></tr>
<tr><td>Chromosome</td><td>start</td><td>end</td><td>strand</td><td>MM</td><td>target_seq</td><td>PAM</td><td>alignment</td><td>distance</td><td>position</td><td>gene name</td><td>gene id</td></tr>
<tr><td>GU305914.1</td><td>163330</td><td>163352</td><td>-</td><td>0</td><td>GTTCGTGCAGTCCAAAACGC</td><td>GGG</td><td>||||||||[||||||||||||]PAM</td><td>NA</td><td>Intergenic</td><td>NA</td><td>NA</td></tr>
<tr><td>T22034</td><td>CAAAACGCGGGCGCTCTCGTAGG</td><td>1000</td><td></td><td></td><td></td><td></td><td></td><td></td><td></td><td></td><td></td></tr>
<tr><td>Oligo fwd</td><td>CACCCAAAACGCGGGCGCTCTCGT</td><td></td><td></td><td></td><td></td><td></td><td></td><td></td><td></td><td></td><td></td></tr>
<tr><td>Oligo rev</td><td>AAACACGAGAGCGCCCGCGTTTTG</td><td></td><td></td><td></td><td></td><td></td><td></td><td></td><td></td><td></td><td></td></tr>
<tr><td>Chromosome</td><td>start</td><td>end</td><td>strand</td><td>MM</td><td>target_seq</td><td>PAM</td><td>alignment</td><td>distance</td><td>position</td><td>gene name</td><td>gene id</td></tr>
<tr><td>GU305914.1</td><td>163318</td><td>163340</td><td>-</td><td>0</td><td>CAAAACGCGGGCGCTCTCGT</td><td>AGG</td><td>||||||||[||||||||||||]PAM</td><td>NA</td><td>Intergenic</td><td>NA</td><td>NA</td></tr>
<tr><td>T22035</td><td>CGCTCTCGTAGGTCCACTGACGG</td><td>1000</td><td></td><td></td><td></td><td></td><td></td><td></td><td></td><td></td><td></td></tr>
<tr><td>Oligo fwd</td><td>CACCCGCTCTCGTAGGTCCACTGA</td><td></td><td></td><td></td><td></td><td></td><td></td><td></td><td></td><td></td><td></td></tr>
<tr><td>Oligo rev</td><td>AAACTCAGTGGACCTACGAGAGCG</td><td></td><td></td><td></td><td></td><td></td><td></td><td></td><td></td><td></td><td></td></tr>
<tr><td>Chromosome</td><td>start</td><td>end</td><td>strand</td><td>MM</td><td>target_seq</td><td>PAM</td><td>alignment</td><td>distance</td><td>position</td><td>gene name</td><td>gene id</td></tr>
<tr><td>GU305914.1</td><td>163307</td><td>163329</td><td>-</td><td>0</td><td>CGCTCTCGTAGGTCCACTGA</td><td>CGG</td><td>||||||||[||||||||||||]PAM</td><td>NA</td><td>Intergenic</td><td>NA</td><td>NA</td></tr>
<tr><td>T22036</td><td>TCTCGTAGGTCCACTGACGGAGG</td><td>1000</td><td></td><td></td><td></td><td></td><td></td><td></td><td></td><td></td><td></td></tr>
<tr><td>Oligo fwd</td><td>CACCTCTCGTAGGTCCACTGACGG</td><td></td><td></td><td></td><td></td><td></td><td></td><td></td><td></td><td></td><td></td></tr>
<tr><td>Oligo rev</td><td>AAACCCGTCAGTGGACCTACGAGA</td><td></td><td></td><td></td><td></td><td></td><td></td><td></td><td></td><td></td><td></td></tr>
<tr><td>Chromosome</td><td>start</td><td>end</td><td>strand</td><td>MM</td><td>target_seq</td><td>PAM</td><td>alignment</td><td>distance</td><td>position</td><td>gene name</td><td>gene id</td></tr>
<tr><td>GU305914.1</td><td>163304</td><td>163326</td><td>-</td><td>0</td><td>TCTCGTAGGTCCACTGACGG</td><td>AGG</td><td>||||||||[||||||||||||]PAM</td><td>NA</td><td>Intergenic</td><td>NA</td><td>NA</td></tr>
<tr><td>T22037</td><td>ACTGACGGAGGTCGAACATGCGG</td><td>1000</td><td></td><td></td><td></td><td></td><td></td><td></td><td></td><td></td><td></td></tr>
<tr><td>Oligo fwd</td><td>CACCACTGACGGAGGTCGAACATG</td><td></td><td></td><td></td><td></td><td></td><td></td><td></td><td></td><td></td><td></td></tr>
<tr><td>Oligo rev</td><td>AAACCATGTTCGACCTCCGTCAGT</td><td></td><td></td><td></td><td></td><td></td><td></td><td></td><td></td><td></td><td></td></tr>
<tr><td>Chromosome</td><td>start</td><td>end</td><td>strand</td><td>MM</td><td>target_seq</td><td>PAM</td><td>alignment</td><td>distance</td><td>position</td><td>gene name</td><td>gene id</td></tr>
</table>

<table>
<tr><td>GU305914.1</td><td>163292</td><td>163314</td><td>-</td><td>0</td><td>ACTGACGGAGGTCGAACATG</td><td>CGG</td><td>||||||||[||||||||||||]PAM</td><td>NA</td><td>Intergenic</td><td>NA</td><td>NA</td></tr>
<tr><td>T22038</td><td>CTGACGGAGGTCGAACATGCGGG</td><td>1000</td><td></td><td></td><td></td><td></td><td></td><td></td><td></td><td></td><td></td></tr>
<tr><td>Oligo fwd</td><td>CACCCTGACGGAGGTCGAACATGC</td><td></td><td></td><td></td><td></td><td></td><td></td><td></td><td></td><td></td><td></td></tr>
<tr><td>Oligo rev</td><td>AAACGCATGTTCGACCTCCGTCAG</td><td></td><td></td><td></td><td></td><td></td><td></td><td></td><td></td><td></td><td></td></tr>
<tr><td>Chromosome</td><td>start</td><td>end</td><td>strand</td><td>MM</td><td>target_seq</td><td>PAM</td><td>alignment</td><td>distance</td><td>position</td><td>gene name</td><td>gene id</td></tr>
<tr><td>GU305914.1</td><td>163291</td><td>163313</td><td>-</td><td>0</td><td>CTGACGGAGGTCGAACATGC</td><td>GGG</td><td>||||||||[||||||||||||]PAM</td><td>NA</td><td>Intergenic</td><td>NA</td><td>NA</td></tr>
<tr><td>T22039</td><td>GCAGCGTCCGACTCGAGTTCCGG</td><td>1000</td><td></td><td></td><td></td><td></td><td></td><td></td><td></td><td></td><td></td></tr>
<tr><td>Oligo fwd</td><td>CACCGCAGCGTCCGACTCGAGTTC</td><td></td><td></td><td></td><td></td><td></td><td></td><td></td><td></td><td></td><td></td></tr>
<tr><td>Oligo rev</td><td>AAACGAACTCGAGTCGGACGCTGC</td><td></td><td></td><td></td><td></td><td></td><td></td><td></td><td></td><td></td><td></td></tr>
<tr><td>Chromosome</td><td>start</td><td>end</td><td>strand</td><td>MM</td><td>target_seq</td><td>PAM</td><td>alignment</td><td>distance</td><td>position</td><td>gene name</td><td>gene id</td></tr>
<tr><td>GU305914.1</td><td>163256</td><td>163278</td><td>-</td><td>0</td><td>GCAGCGTCCGACTCGAGTTC</td><td>CGG</td><td>||||||||[||||||||||||]PAM</td><td>NA</td><td>Intergenic</td><td>NA</td><td>NA</td></tr>
<tr><td>T22040</td><td>CGTCCGACTCGAGTTCCGGAAGG</td><td>1000</td><td></td><td></td><td></td><td></td><td></td><td></td><td></td><td></td><td></td></tr>
<tr><td>Oligo fwd</td><td>CACCCGTCCGACTCGAGTTCCGGA</td><td></td><td></td><td></td><td></td><td></td><td></td><td></td><td></td><td></td><td></td></tr>
<tr><td>Oligo rev</td><td>AAACTCCGGAACTCGAGTCGGACG</td><td></td><td></td><td></td><td></td><td></td><td></td><td></td><td></td><td></td><td></td></tr>
<tr><td>Chromosome</td><td>start</td><td>end</td><td>strand</td><td>MM</td><td>target_seq</td><td>PAM</td><td>alignment</td><td>distance</td><td>position</td><td>gene name</td><td>gene id</td></tr>
<tr><td>GU305914.1</td><td>163252</td><td>163274</td><td>-</td><td>0</td><td>CGTCCGACTCGAGTTCCGGA</td><td>AGG</td><td>||||||||[||||||||||||]PAM</td><td>NA</td><td>Intergenic</td><td>NA</td><td>NA</td></tr>
<tr><td>T22041</td><td>GTAACGTCGTCTCGAAACGATGG</td><td>1000</td><td></td><td></td><td></td><td></td><td></td><td></td><td></td><td></td><td></td></tr>
<tr><td>Oligo fwd</td><td>CACCGTAACGTCGTCTCGAAACGA</td><td></td><td></td><td></td><td></td><td></td><td></td><td></td><td></td><td></td><td></td></tr>
<tr><td>Oligo rev</td><td>AAACTCGTTTCGAGACGACGTTAC</td><td></td><td></td><td></td><td></td><td></td><td></td><td></td><td></td><td></td><td></td></tr>
<tr><td>Chromosome</td><td>start</td><td>end</td><td>strand</td><td>MM</td><td>target_seq</td><td>PAM</td><td>alignment</td><td>distance</td><td>position</td><td>gene name</td><td>gene id</td></tr>
<tr><td>GU305914.1</td><td>163225</td><td>163247</td><td>-</td><td>0</td><td>GTAACGTCGTCTCGAAACGA</td><td>TGG</td><td>||||||||[||||||||||||]PAM</td><td>NA</td><td>Intergenic</td><td>NA</td><td>NA</td></tr>
<tr><td>GU305914.1</td><td>54006</td><td>54028</td><td>+</td><td>4</td><td>GAAACGTTGTCTCCTAACGA</td><td>CGG</td><td>|-|||||-[|||||--|||||]PAM</td><td>NA</td><td>Intergenic</td><td>NA</td><td>NA</td></tr>
<tr><td>T22042</td><td>TAACGTCGTCTCGAAACGATGGG</td><td>1000</td><td></td><td></td><td></td><td></td><td></td><td></td><td></td><td></td><td></td></tr>
<tr><td>Oligo fwd</td><td>CACCTAACGTCGTCTCGAAACGAT</td><td></td><td></td><td></td><td></td><td></td><td></td><td></td><td></td><td></td><td></td></tr>
<tr><td>Oligo rev</td><td>AAACATCGTTTCGAGACGACGTTA</td><td></td><td></td><td></td><td></td><td></td><td></td><td></td><td></td><td></td><td></td></tr>
<tr><td>Chromosome</td><td>start</td><td>end</td><td>strand</td><td>MM</td><td>target_seq</td><td>PAM</td><td>alignment</td><td>distance</td><td>position</td><td>gene name</td><td>gene id</td></tr>
<tr><td>GU305914.1</td><td>163224</td><td>163246</td><td>-</td><td>0</td><td>TAACGTCGTCTCGAAACGAT</td><td>GGG</td><td>||||||||[||||||||||||]PAM</td><td>NA</td><td>Intergenic</td><td>NA</td><td>NA</td></tr>
<tr><td>T22043</td><td>ATGGGCGATCCGCGTCGATCTGG</td><td>1000</td><td></td><td></td><td></td><td></td><td></td><td></td><td></td><td></td><td></td></tr>
<tr><td>Oligo fwd</td><td>CACCATGGGCGATCCGCGTCGATC</td><td></td><td></td><td></td><td></td><td></td><td></td><td></td><td></td><td></td><td></td></tr>
<tr><td>Oligo rev</td><td>AAACGATCGACGCGGATCGCCCAT</td><td></td><td></td><td></td><td></td><td></td><td></td><td></td><td></td><td></td><td></td></tr>
<tr><td>Chromosome</td><td>start</td><td>end</td><td>strand</td><td>MM</td><td>target_seq</td><td>PAM</td><td>alignment</td><td>distance</td><td>position</td><td>gene name</td><td>gene id</td></tr>
<tr><td>GU305914.1</td><td>163206</td><td>163228</td><td>-</td><td>0</td><td>ATGGGCGATCCGCGTCGATC</td><td>TGG</td><td>||||||||[||||||||||||]PAM</td><td>NA</td><td>Intergenic</td><td>NA</td><td>NA</td></tr>
<tr><td>GU305914.1</td><td>55927</td><td>55949</td><td>+</td><td>3</td><td>ATAGTCGATCGGCGTCGATC</td><td>CGG</td><td>||-|-|||[||-|||||||||]PAM</td><td>NA</td><td>Intergenic</td><td>NA</td><td>NA</td></tr>
<tr><td>T22044</td><td>CGATCCGCGTCGATCTGGCGCGG</td><td>1000</td><td></td><td></td><td></td><td></td><td></td><td></td><td></td><td></td><td></td></tr>
<tr><td>Oligo fwd</td><td>CACCCGATCCGCGTCGATCTGGCG</td><td></td><td></td><td></td><td></td><td></td><td></td><td></td><td></td><td></td><td></td></tr>
<tr><td>Oligo rev</td><td>AAACCGCCAGATCGACGCGGATCG</td><td></td><td></td><td></td><td></td><td></td><td></td><td></td><td></td><td></td><td></td></tr>
<tr><td>Chromosome</td><td>start</td><td>end</td><td>strand</td><td>MM</td><td>target_seq</td><td>PAM</td><td>alignment</td><td>distance</td><td>position</td><td>gene name</td><td>gene id</td></tr>
<tr><td>GU305914.1</td><td>163201</td><td>163223</td><td>-</td><td>0</td><td>CGATCCGCGTCGATCTGGCG</td><td>CGG</td><td>||||||||[||||||||||||]PAM</td><td>NA</td><td>Intergenic</td><td>NA</td><td>NA</td></tr>
<tr><td>T22045</td><td>GCGTCGATCTGGCGCGGCCATGG</td><td>1000</td><td></td><td></td><td></td><td></td><td></td><td></td><td></td><td></td><td></td></tr>
<tr><td>Oligo fwd</td><td>CACCGCGTCGATCTGGCGCGGCCA</td><td></td><td></td><td></td><td></td><td></td><td></td><td></td><td></td><td></td><td></td></tr>
<tr><td>Oligo rev</td><td>AAACTGGCCGCGCCAGATCGACGC</td><td></td><td></td><td></td><td></td><td></td><td></td><td></td><td></td><td></td><td></td></tr>
<tr><td>Chromosome</td><td>start</td><td>end</td><td>strand</td><td>MM</td><td>target_seq</td><td>PAM</td><td>alignment</td><td>distance</td><td>position</td><td>gene name</td><td>gene id</td></tr>
<tr><td>GU305914.1</td><td>163195</td><td>163217</td><td>-</td><td>0</td><td>GCGTCGATCTGGCGCGGCCA</td><td>TGG</td><td>||||||||[||||||||||||]PAM</td><td>NA</td><td>Intergenic</td><td>NA</td><td>NA</td></tr>
<tr><td>T22046</td><td>CGCGGCCATGGTCCGAACGAAGG</td><td>1000</td><td></td><td></td><td></td><td></td><td></td><td></td><td></td><td></td><td></td></tr>
<tr><td>Oligo fwd</td><td>CACCCGCGGCCATGGTCCGAACGA</td><td></td><td></td><td></td><td></td><td></td><td></td><td></td><td></td><td></td><td></td></tr>
<tr><td>Oligo rev</td><td>AAACTCGTTCGGACCATGGCCGCG</td><td></td><td></td><td></td><td></td><td></td><td></td><td></td><td></td><td></td><td></td></tr>
<tr><td>Chromosome</td><td>start</td><td>end</td><td>strand</td><td>MM</td><td>target_seq</td><td>PAM</td><td>alignment</td><td>distance</td><td>position</td><td>gene name</td><td>gene id</td></tr>
<tr><td>GU305914.1</td><td>163183</td><td>163205</td><td>-</td><td>0</td><td>CGCGGCCATGGTCCGAACGA</td><td>AGG</td><td>||||||||[||||||||||||]PAM</td><td>NA</td><td>Intergenic</td><td>NA</td><td>NA</td></tr>
<tr><td>T22047</td><td>CCGAACGAAGGTCTTTTCACCGG</td><td>1000</td><td></td><td></td><td></td><td></td><td></td><td></td><td></td><td></td><td></td></tr>
<tr><td>Oligo fwd</td><td>CACCCCGAACGAAGGTCTTTTCAC</td><td></td><td></td><td></td><td></td><td></td><td></td><td></td><td></td><td></td><td></td></tr>
<tr><td>Oligo rev</td><td>AAACGTGAAAAGACCTTCGTTCGG</td><td></td><td></td><td></td><td></td><td></td><td></td><td></td><td></td><td></td><td></td></tr>
<tr><td>Chromosome</td><td>start</td><td>end</td><td>strand</td><td>MM</td><td>target_seq</td><td>PAM</td><td>alignment</td><td>distance</td><td>position</td><td>gene name</td><td>gene id</td></tr>
<tr><td>GU305914.1</td><td>163171</td><td>163193</td><td>-</td><td>0</td><td>CCGAACGAAGGTCTTTTCAC</td><td>CGG</td><td>||||||||[||||||||||||]PAM</td><td>NA</td><td>Intergenic</td><td>NA</td><td>NA</td></tr>
<tr><td>T22048</td><td>ACCGGTCGCGACGAGAAAAGTGG</td><td>1000</td><td></td><td></td><td></td><td></td><td></td><td></td><td></td><td></td><td></td></tr>
<tr><td>Oligo fwd</td><td>CACCACCGGTCGCGACGAGAAAAG</td><td></td><td></td><td></td><td></td><td></td><td></td><td></td><td></td><td></td><td></td></tr>
<tr><td>Oligo rev</td><td>AAACCTTTTCTCGTCGCGACCGGT</td><td></td><td></td><td></td><td></td><td></td><td></td><td></td><td></td><td></td><td></td></tr>
<tr><td>Chromosome</td><td>start</td><td>end</td><td>strand</td><td>MM</td><td>target_seq</td><td>PAM</td><td>alignment</td><td>distance</td><td>position</td><td>gene name</td><td>gene id</td></tr>
<tr><td>GU305914.1</td><td>163153</td><td>163175</td><td>-</td><td>0</td><td>ACCGGTCGCGACGAGAAAAG</td><td>TGG</td><td>||||||||[||||||||||||]PAM</td><td>NA</td><td>Intergenic</td><td>NA</td><td>NA</td></tr>
<tr><td>T22049</td><td>CCGGTCGCGACGAGAAAAGTGGG</td><td>1000</td><td></td><td></td><td></td><td></td><td></td><td></td><td></td><td></td><td></td></tr>
<tr><td>Oligo fwd</td><td>CACCCCGGTCGCGACGAGAAAAGT</td><td></td><td></td><td></td><td></td><td></td><td></td><td></td><td></td><td></td><td></td></tr>
<tr><td>Oligo rev</td><td>AAACACTTTTCTCGTCGCGACCGG</td><td></td><td></td><td></td><td></td><td></td><td></td><td></td><td></td><td></td><td></td></tr>
<tr><td>Chromosome</td><td>start</td><td>end</td><td>strand</td><td>MM</td><td>target_seq</td><td>PAM</td><td>alignment</td><td>distance</td><td>position</td><td>gene name</td><td>gene id</td></tr>
<tr><td>GU305914.1</td><td>163152</td><td>163174</td><td>-</td><td>0</td><td>CCGGTCGCGACGAGAAAAGT</td><td>GGG</td><td>||||||||[||||||||||||]PAM</td><td>NA</td><td>Intergenic</td><td>NA</td><td>NA</td></tr>
<tr><td>T22050</td><td>CACGTAATCTACCGCGACGCAGG</td><td>1000</td><td></td><td></td><td></td><td></td><td></td><td></td><td></td><td></td><td></td></tr>
<tr><td>Oligo fwd</td><td>CACCCACGTAATCTACCGCGACGC</td><td></td><td></td><td></td><td></td><td></td><td></td><td></td><td></td><td></td><td></td></tr>
<tr><td>Oligo rev</td><td>AAACGCGTCGCGGTAGATTACGTG</td><td></td><td></td><td></td><td></td><td></td><td></td><td></td><td></td><td></td><td></td></tr>
<tr><td>Chromosome</td><td>start</td><td>end</td><td>strand</td><td>MM</td><td>target_seq</td><td>PAM</td><td>alignment</td><td>distance</td><td>position</td><td>gene name</td><td>gene id</td></tr>
<tr><td>GU305914.1</td><td>163129</td><td>163151</td><td>-</td><td>0</td><td>CACGTAATCTACCGCGACGC</td><td>AGG</td><td>||||||||[||||||||||||]PAM</td><td>NA</td><td>Intergenic</td><td>NA</td><td>NA</td></tr>
<tr><td>GU305914.1</td><td>129693</td><td>129715</td><td>-</td><td>4</td><td>CACGTTCTCGAGCGCGACGC</td><td>CGG</td><td>|||||--|[|-|-||||||||]PAM</td><td>NA</td><td>Intergenic</td><td>NA</td><td>NA</td></tr>
<tr><td>T22051</td><td>ACGCAGGAACTATCAGCATTTGG</td><td>1000</td><td></td><td></td><td></td><td></td><td></td><td></td><td></td><td></td><td></td></tr>
<tr><td>Oligo fwd</td><td>CACCACGCAGGAACTATCAGCATT</td><td></td><td></td><td></td><td></td><td></td><td></td><td></td><td></td><td></td><td></td></tr>
<tr><td>Oligo rev</td><td>AAACAATGCTGATAGTTCCTGCGT</td><td></td><td></td><td></td><td></td><td></td><td></td><td></td><td></td><td></td><td></td></tr>
<tr><td>Chromosome</td><td>start</td><td>end</td><td>strand</td><td>MM</td><td>target_seq</td><td>PAM</td><td>alignment</td><td>distance</td><td>position</td><td>gene name</td><td>gene id</td></tr>
<tr><td>GU305914.1</td><td>163113</td><td>163135</td><td>-</td><td>0</td><td>ACGCAGGAACTATCAGCATT</td><td>TGG</td><td>||||||||[||||||||||||]PAM</td><td>NA</td><td>Intergenic</td><td>NA</td><td>NA</td></tr>
</table>

T22052 ATCGCAGAAGAAATGAGCCTAGG 1000
Oligo fwd CACCATCGCAGAAGAAATGAGCCT
Oligo rev AAACAGGCTCATTTCTTCTGCGAT
Chromosome start end strand MM target_seq PAM alignment distance position gene name gene id
GU305914.1 163074 163096 - 0 ATCGCAGAAGAAATGAGCCT AGG ||||||||[||||||||||||]PAM NA Intergenic NA NA

T22053 AATGAGCCTAGGATCCGCTCCGG 1000
Oligo fwd CACCAATGAGCCTAGGATCCGCTC
Oligo rev AAACGAGCGGATCCTAGGCTCATT
Chromosome start end strand MM target_seq PAM alignment distance position gene name gene id
GU305914.1 163063 163085 - 0 AATGAGCCTAGGATCCGCTC CGG ||||||||[||||||||||||]PAM NA Intergenic NA NA

T22054 CATGACGTTGTTATAACCTGCGG 1000
Oligo fwd CACCCATGACGTTGTTATAACCTG
Oligo rev AAACCAGGTTATAACAACGTCATG
Chromosome start end strand MM target_seq PAM alignment distance position gene name gene id
GU305914.1 162996 163018 - 0 CATGACGTTGTTATAACCTG CGG ||||||||[||||||||||||]PAM NA Intergenic NA NA

T22055 TTGTTATAACCTGCGGTCTCCGG 1000
Oligo fwd CACCTTGTTATAACCTGCGGTCTC
Oligo rev AAACGAGACCGCAGGTTATAACAA
Chromosome start end strand MM target_seq PAM alignment distance position gene name gene id
GU305914.1 162989 163011 - 0 TTGTTATAACCTGCGGTCTC CGG ||||||||[||||||||||||]PAM NA Intergenic NA NA

T22056 GCGTTCTTCTCGCTACCCGCCGG 1000
Oligo fwd CACCGCGTTCTTCTCGCTACCCGC
Oligo rev AAACGCGGGTAGCGAGAAGAACGC
Chromosome start end strand MM target_seq PAM alignment distance position gene name gene id
GU305914.1 162967 162989 - 0 GCGTTCTTCTCGCTACCCGC CGG ||||||||[||||||||||||]PAM NA Intergenic NA NA

T22057 CTCGCTACCCGCCGGTCGAATGG 1000
Oligo fwd CACCCTCGCTACCCGCCGGTCGAA
Oligo rev AAACTTCGACCGGCGGGTAGCGAG
Chromosome start end strand MM target_seq PAM alignment distance position gene name gene id
GU305914.1 162959 162981 - 0 CTCGCTACCCGCCGGTCGAA TGG ||||||||[||||||||||||]PAM NA Intergenic NA NA

T22058 CCGCCGGTCGAATGGCGCTCCGG 1000
Oligo fwd CACCCCGCCGGTCGAATGGCGCTC
Oligo rev AAACGAGCGCCATTCGACCGGCGG
Chromosome start end strand MM target_seq PAM alignment distance position gene name gene id
GU305914.1 162951 162973 - 0 CCGCCGGTCGAATGGCGCTC CGG ||||||||[||||||||||||]PAM NA Intergenic NA NA

T22059 CGAATGGCGCTCCGGACGCGCGG 1000
Oligo fwd CACCCGAATGGCGCTCCGGACGCG
Oligo rev AAACCGCGTCCGGAGCGCCATTCG
Chromosome start end strand MM target_seq PAM alignment distance position gene name gene id
GU305914.1 162943 162965 - 0 CGAATGGCGCTCCGGACGCG CGG ||||||||[||||||||||||]PAM NA Intergenic NA NA

T22060 GTCGAGCTCTAGACACGCCTAGG 1000
Oligo fwd CACCGTCGAGCTCTAGACACGCCT
Oligo rev AAACAGGCGTGTCTAGAGCTCGAC
Chromosome start end strand MM target_seq PAM alignment distance position gene name gene id
GU305914.1 162921 162943 - 0 GTCGAGCTCTAGACACGCCT AGG ||||||||[||||||||||||]PAM NA Intergenic NA NA

T22061 AGCTCTAGACACGCCTAGGCTGG 1000
Oligo fwd CACCAGCTCTAGACACGCCTAGGC
Oligo rev AAACGCCTAGGCGTGTCTAGAGCT
Chromosome start end strand MM target_seq PAM alignment distance position gene name gene id
GU305914.1 162917 162939 - 0 AGCTCTAGACACGCCTAGGC TGG ||||||||[||||||||||||]PAM NA Intergenic NA NA

T22062 GCTCTAGACACGCCTAGGCTGGG 1000
Oligo fwd CACCGCTCTAGACACGCCTAGGCT
Oligo rev AAACAGCCTAGGCGTGTCTAGAGC
Chromosome start end strand MM target_seq PAM alignment distance position gene name gene id
GU305914.1 162916 162938 - 0 GCTCTAGACACGCCTAGGCT GGG ||||||||[||||||||||||]PAM NA Intergenic NA NA

T22063 CACGCCTAGGCTGGGCGCCGAGG 1000
Oligo fwd CACCCACGCCTAGGCTGGGCGCCG
Oligo rev AAACCGGCGCCCAGCCTAGGCGTG
Chromosome start end strand MM target_seq PAM alignment distance position gene name gene id
GU305914.1 162908 162930 - 0 CACGCCTAGGCTGGGCGCCG AGG ||||||||[||||||||||||]PAM NA Intergenic NA NA

T22064 CAGTAATGTGCCTGTCTCCCCGG 1000
Oligo fwd CACCCAGTAATGTGCCTGTCTCCC
Oligo rev AAACGGGAGACAGGCACATTACTG
Chromosome start end strand MM target_seq PAM alignment distance position gene name gene id
GU305914.1 162884 162906 - 0 CAGTAATGTGCCTGTCTCCC CGG ||||||||[||||||||||||]PAM NA Intergenic NA NA

T22065 CCCCGGACCCGCCTAGCAGATGG 1000
Oligo fwd CACCCCCCGGACCCGCCTAGCAGA
Oligo rev AAACTCTGCTAGGCGGGTCCGGGG
Chromosome start end strand MM target_seq PAM alignment distance position gene name gene id
GU305914.1 162867 162889 - 0 CCCCGGACCCGCCTAGCAGA TGG ||||||||[||||||||||||]PAM NA Intergenic NA NA

T22066 GATGGTCTATAACCTCACCGCGG 1000
Oligo fwd CACCGATGGTCTATAACCTCACCG
Oligo rev AAACCGGTGAGGTTATAGACCATC
Chromosome start end strand MM target_seq PAM alignment distance position gene name gene id
GU305914.1 162849 162871 - 0 GATGGTCTATAACCTCACCG CGG ||||||||[||||||||||||]PAM NA Intergenic NA NA

T22067 TCGAGAGCCCGTCCGAGACCTGG 1000
Oligo fwd CACCTCGAGAGCCCGTCCGAGACC
Oligo rev AAACGGTCTCGGACGGGCTCTCGA
Chromosome start end strand MM target_seq PAM alignment distance position gene name gene id
GU305914.1 162800 162822 - 0 TCGAGAGCCCGTCCGAGACC TGG ||||||||[||||||||||||]PAM NA Intergenic NA NA

T22068 AGCCCGTCCGAGACCTGGCCCGG 1000
Oligo fwd CACCAGCCCGTCCGAGACCTGGCC
Oligo rev AAACGGCCAGGTCTCGGACGGGCT
Chromosome start end strand MM target_seq PAM alignment distance position gene name gene id
GU305914.1 162795 162817 - 0 AGCCCGTCCGAGACCTGGCC CGG ||||||||[||||||||||||]PAM NA Intergenic NA NA

T22069 CCGTCCGAGACCTGGCCCGGCGG 1000
Oligo fwd CACCCCGTCCGAGACCTGGCCCGG
Oligo rev AAACCCGGGCCAGGTCTCGGACGG
Chromosome start end strand MM target_seq PAM alignment distance position gene name gene id
GU305914.1 162792 162814 - 0 CCGTCCGAGACCTGGCCCGG CGG ||||||||[||||||||||||]PAM NA Intergenic NA NA
GU305914.1 195554 195576 + 4 GCGGCCGCGATCTGGCCCGG CGG -||-|||-[||-|||||||||]PAM NA Intergenic NA NA

T22070 ACCTGGCCCGGCGGCTCGTGCGG 1000
Oligo fwd CACCACCTGGCCCGGCGGCTCGTG
Oligo rev AAACCACGAGCCGCCGGGCCAGGT
Chromosome start end strand MM target_seq PAM alignment distance position gene name gene id
GU305914.1 162783 162805 - 0 ACCTGGCCCGGCGGCTCGTG CGG ||||||||[||||||||||||]PAM NA Intergenic NA NA

T22071 CGTGCGGATGTTCAACCTAGTGG 1000
Oligo fwd CACCCGTGCGGATGTTCAACCTAG
Oligo rev AAACCTAGGTTGAACATCCGCACG
Chromosome start end strand MM target_seq PAM alignment distance position gene name gene id
GU305914.1 162767 162789 - 0 CGTGCGGATGTTCAACCTAG TGG ||||||||[||||||||||||]PAM NA Intergenic NA NA

T22072 CCTAGTGGCGCTCCTCGTCCTGG 1000
Oligo fwd CACCCCTAGTGGCGCTCCTCGTCC
Oligo rev AAACGGACGAGGAGCGCCACTAGG
Chromosome start end strand MM target_seq PAM alignment distance position gene name gene id
GU305914.1 162752 162774 - 0 CCTAGTGGCGCTCCTCGTCC TGG ||||||||[||||||||||||]PAM NA Intergenic NA NA

T22073 GTGGCGCTCCTCGTCCTGGCCGG 1000
Oligo fwd CACCGTGGCGCTCCTCGTCCTGGC
Oligo rev AAACGCCAGGACGAGGAGCGCCAC
Chromosome start end strand MM target_seq PAM alignment distance position gene name gene id
GU305914.1 162748 162770 - 0 GTGGCGCTCCTCGTCCTGGC CGG ||||||||[||||||||||||]PAM NA Intergenic NA NA

T22074 TCGTCCTGGCCGGCCGCGCGAGG 1000
Oligo fwd CACCTCGTCCTGGCCGGCCGCGCG
Oligo rev AAACCGCGCGGCCGGCCAGGACGA
Chromosome start end strand MM target_seq PAM alignment distance position gene name gene id
GU305914.1 162738 162760 - 0 TCGTCCTGGCCGGCCGCGCG AGG ||||||||[||||||||||||]PAM NA Intergenic NA NA

T22075 CGAGGCCTTCGCCGAAGCGCAGG 1000
Oligo fwd CACCCGAGGCCTTCGCCGAAGCGC
Oligo rev AAACGCGCTTCGGCGAAGGCCTCG
Chromosome start end strand MM target_seq PAM alignment distance position gene name gene id
GU305914.1 162720 162742 - 0 CGAGGCCTTCGCCGAAGCGC AGG ||||||||[||||||||||||]PAM NA Intergenic NA NA

T22076 GAGGCCTTCGCCGAAGCGCAGGG 1000
Oligo fwd CACCGAGGCCTTCGCCGAAGCGCA
Oligo rev AAACTGCGCTTCGGCGAAGGCCTC
Chromosome start end strand MM target_seq PAM alignment distance position gene name gene id
GU305914.1 162719 162741 - 0 GAGGCCTTCGCCGAAGCGCA GGG ||||||||[||||||||||||]PAM NA Intergenic NA NA

T22077 AGGCCTTCGCCGAAGCGCAGGGG 1000
Oligo fwd CACCAGGCCTTCGCCGAAGCGCAG
Oligo rev AAACCTGCGCTTCGGCGAAGGCCT
Chromosome start end strand MM target_seq PAM alignment distance position gene name gene id
GU305914.1 162718 162740 - 0 AGGCCTTCGCCGAAGCGCAG GGG ||||||||[||||||||||||]PAM NA Intergenic NA NA

T22078 TCGCCGAAGCGCAGGGGCCTCGG 1000
Oligo fwd CACCTCGCCGAAGCGCAGGGGCCT
Oligo rev AAACAGGCCCCTGCGCTTCGGCGA
Chromosome start end strand MM target_seq PAM alignment distance position gene name gene id
GU305914.1 162712 162734 - 0 TCGCCGAAGCGCAGGGGCCT CGG ||||||||[||||||||||||]PAM NA Intergenic NA NA

T22079 GGCCTCGGCTCGAGATAGCGCGG 1000
Oligo fwd CACCGGCCTCGGCTCGAGATAGCG
Oligo rev AAACCGCTATCTCGAGCCGAGGCC
Chromosome start end strand MM target_seq PAM alignment distance position gene name gene id
GU305914.1 162697 162719 - 0 GGCCTCGGCTCGAGATAGCG CGG ||||||||[||||||||||||]PAM NA Intergenic NA NA

T22080 GCGGCGTCCGTCGCACCGTACGG 1000
Oligo fwd CACCGCGGCGTCCGTCGCACCGTA
Oligo rev AAACTACGGTGCGACGGACGCCGC
Chromosome start end strand MM target_seq PAM alignment distance position gene name gene id
GU305914.1 162678 162700 - 0 GCGGCGTCCGTCGCACCGTA CGG ||||||||[||||||||||||]PAM NA Intergenic NA NA

T22081 CCGTACGGCGTTCGAGATGCCGG 1000
Oligo fwd CACCCCGTACGGCGTTCGAGATGC
Oligo rev AAACGCATCTCGAACGCCGTACGG

<table>
<tr><td>Chromosome</td><td>start</td><td>end</td><td>strand</td><td>MM</td><td>target_seq</td><td>PAM</td><td>alignment</td><td>distance</td><td>position</td><td>gene name</td><td>gene id</td></tr>
<tr><td>GU305914.1</td><td>162663</td><td>162685</td><td>-</td><td>0</td><td>CCGTACGGCGTTCGAGATGC</td><td>CGG</td><td>||||||||[||||||||||||]PAM</td><td>NA</td><td>Intergenic</td><td>NA</td><td>NA</td></tr>
<tr><td></td><td></td><td></td><td></td><td></td><td></td><td></td><td></td><td></td><td></td><td></td><td></td></tr>
<tr><td>T22082</td><td>TACGGCGTTCGAGATGCCGGTGG</td><td>1000</td><td></td><td></td><td></td><td></td><td></td><td></td><td></td><td></td><td></td></tr>
<tr><td>Oligo fwd</td><td>CACCTACGGCGTTCGAGATGCCGG</td><td></td><td></td><td></td><td></td><td></td><td></td><td></td><td></td><td></td><td></td></tr>
<tr><td>Oligo rev</td><td>AAACCCGGCATCTCGAACGCCGTA</td><td></td><td></td><td></td><td></td><td></td><td></td><td></td><td></td><td></td><td></td></tr>
<tr><td>Chromosome</td><td>start</td><td>end</td><td>strand</td><td>MM</td><td>target_seq</td><td>PAM</td><td>alignment</td><td>distance</td><td>position</td><td>gene name</td><td>gene id</td></tr>
<tr><td>GU305914.1</td><td>162660</td><td>162682</td><td>-</td><td>0</td><td>TACGGCGTTCGAGATGCCGG</td><td>TGG</td><td>||||||||[||||||||||||]PAM</td><td>NA</td><td>Intergenic</td><td>NA</td><td>NA</td></tr>
<tr><td></td><td></td><td></td><td></td><td></td><td></td><td></td><td></td><td></td><td></td><td></td><td></td></tr>
<tr><td>T22083</td><td>GAGATGCCGGTGGCGTCGAGCGG</td><td>1000</td><td></td><td></td><td></td><td></td><td></td><td></td><td></td><td></td><td></td></tr>
<tr><td>Oligo fwd</td><td>CACCGAGATGCCGGTGGCGTCGAG</td><td></td><td></td><td></td><td></td><td></td><td></td><td></td><td></td><td></td><td></td></tr>
<tr><td>Oligo rev</td><td>AAACCTCGACGCCACCGGCATCTC</td><td></td><td></td><td></td><td></td><td></td><td></td><td></td><td></td><td></td><td></td></tr>
<tr><td>Chromosome</td><td>start</td><td>end</td><td>strand</td><td>MM</td><td>target_seq</td><td>PAM</td><td>alignment</td><td>distance</td><td>position</td><td>gene name</td><td>gene id</td></tr>
<tr><td>GU305914.1</td><td>162650</td><td>162672</td><td>-</td><td>0</td><td>GAGATGCCGGTGGCGTCGAG</td><td>CGG</td><td>||||||||[||||||||||||]PAM</td><td>NA</td><td>Intergenic</td><td>NA</td><td>NA</td></tr>
<tr><td>GU305914.1</td><td>117697</td><td>117719</td><td>-</td><td>4</td><td>GAAACACCGGTGGCGGCGAG</td><td>CGG</td><td>||-|--||[|||||||-||||]PAM</td><td>NA</td><td>Intergenic</td><td>NA</td><td>NA</td></tr>
<tr><td></td><td></td><td></td><td></td><td></td><td></td><td></td><td></td><td></td><td></td><td></td><td></td></tr>
<tr><td>T22084</td><td>AGATGCCGGTGGCGTCGAGCGGG</td><td>1000</td><td></td><td></td><td></td><td></td><td></td><td></td><td></td><td></td><td></td></tr>
<tr><td>Oligo fwd</td><td>CACCAGATGCCGGTGGCGTCGAGC</td><td></td><td></td><td></td><td></td><td></td><td></td><td></td><td></td><td></td><td></td></tr>
<tr><td>Oligo rev</td><td>AAACGCTCGACGCCACCGGCATCT</td><td></td><td></td><td></td><td></td><td></td><td></td><td></td><td></td><td></td><td></td></tr>
<tr><td>Chromosome</td><td>start</td><td>end</td><td>strand</td><td>MM</td><td>target_seq</td><td>PAM</td><td>alignment</td><td>distance</td><td>position</td><td>gene name</td><td>gene id</td></tr>
<tr><td>GU305914.1</td><td>162649</td><td>162671</td><td>-</td><td>0</td><td>AGATGCCGGTGGCGTCGAGC</td><td>GGG</td><td>||||||||[||||||||||||]PAM</td><td>NA</td><td>Intergenic</td><td>NA</td><td>NA</td></tr>
<tr><td></td><td></td><td></td><td></td><td></td><td></td><td></td><td></td><td></td><td></td><td></td><td></td></tr>
<tr><td>T22085</td><td>CGTCGAGCGGGTGTCCGAAGAGG</td><td>1000</td><td></td><td></td><td></td><td></td><td></td><td></td><td></td><td></td><td></td></tr>
<tr><td>Oligo fwd</td><td>CACCCGTCGAGCGGGTGTCCGAAG</td><td></td><td></td><td></td><td></td><td></td><td></td><td></td><td></td><td></td><td></td></tr>
<tr><td>Oligo rev</td><td>AAACCTTCGGACACCCGCTCGACG</td><td></td><td></td><td></td><td></td><td></td><td></td><td></td><td></td><td></td><td></td></tr>
<tr><td>Chromosome</td><td>start</td><td>end</td><td>strand</td><td>MM</td><td>target_seq</td><td>PAM</td><td>alignment</td><td>distance</td><td>position</td><td>gene name</td><td>gene id</td></tr>
<tr><td>GU305914.1</td><td>162637</td><td>162659</td><td>-</td><td>0</td><td>CGTCGAGCGGGTGTCCGAAG</td><td>AGG</td><td>||||||||[||||||||||||]PAM</td><td>NA</td><td>Intergenic</td><td>NA</td><td>NA</td></tr>
<tr><td></td><td></td><td></td><td></td><td></td><td></td><td></td><td></td><td></td><td></td><td></td><td></td></tr>
<tr><td>T22086</td><td>GTCGAGCGGGTGTCCGAAGAGGG</td><td>1000</td><td></td><td></td><td></td><td></td><td></td><td></td><td></td><td></td><td></td></tr>
<tr><td>Oligo fwd</td><td>CACCGTCGAGCGGGTGTCCGAAGA</td><td></td><td></td><td></td><td></td><td></td><td></td><td></td><td></td><td></td><td></td></tr>
<tr><td>Oligo rev</td><td>AAACTCTTCGGACACCCGCTCGAC</td><td></td><td></td><td></td><td></td><td></td><td></td><td></td><td></td><td></td><td></td></tr>
<tr><td>Chromosome</td><td>start</td><td>end</td><td>strand</td><td>MM</td><td>target_seq</td><td>PAM</td><td>alignment</td><td>distance</td><td>position</td><td>gene name</td><td>gene id</td></tr>
<tr><td>GU305914.1</td><td>162636</td><td>162658</td><td>-</td><td>0</td><td>GTCGAGCGGGTGTCCGAAGA</td><td>GGG</td><td>||||||||[||||||||||||]PAM</td><td>NA</td><td>Intergenic</td><td>NA</td><td>NA</td></tr>
<tr><td></td><td></td><td></td><td></td><td></td><td></td><td></td><td></td><td></td><td></td><td></td><td></td></tr>
<tr><td>T22087</td><td>TCGAGCGGGTGTCCGAAGAGGGG</td><td>1000</td><td></td><td></td><td></td><td></td><td></td><td></td><td></td><td></td><td></td></tr>
<tr><td>Oligo fwd</td><td>CACCTCGAGCGGGTGTCCGAAGAG</td><td></td><td></td><td></td><td></td><td></td><td></td><td></td><td></td><td></td><td></td></tr>
<tr><td>Oligo rev</td><td>AAACCTCTTCGGACACCCGCTCGA</td><td></td><td></td><td></td><td></td><td></td><td></td><td></td><td></td><td></td><td></td></tr>
<tr><td>Chromosome</td><td>start</td><td>end</td><td>strand</td><td>MM</td><td>target_seq</td><td>PAM</td><td>alignment</td><td>distance</td><td>position</td><td>gene name</td><td>gene id</td></tr>
<tr><td>GU305914.1</td><td>162635</td><td>162657</td><td>-</td><td>0</td><td>TCGAGCGGGTGTCCGAAGAG</td><td>GGG</td><td>||||||||[||||||||||||]PAM</td><td>NA</td><td>Intergenic</td><td>NA</td><td>NA</td></tr>
<tr><td></td><td></td><td></td><td></td><td></td><td></td><td></td><td></td><td></td><td></td><td></td><td></td></tr>
<tr><td>T22088</td><td>TGTCCGAAGAGGGGTTTTCCCGG</td><td>1000</td><td></td><td></td><td></td><td></td><td></td><td></td><td></td><td></td><td></td></tr>
<tr><td>Oligo fwd</td><td>CACCTGTCCGAAGAGGGGTTTTCC</td><td></td><td></td><td></td><td></td><td></td><td></td><td></td><td></td><td></td><td></td></tr>
<tr><td>Oligo rev</td><td>AAACGGAAAACCCCTCTTCGGACA</td><td></td><td></td><td></td><td></td><td></td><td></td><td></td><td></td><td></td><td></td></tr>
<tr><td>Chromosome</td><td>start</td><td>end</td><td>strand</td><td>MM</td><td>target_seq</td><td>PAM</td><td>alignment</td><td>distance</td><td>position</td><td>gene name</td><td>gene id</td></tr>
<tr><td>GU305914.1</td><td>162626</td><td>162648</td><td>-</td><td>0</td><td>TGTCCGAAGAGGGGTTTTCC</td><td>CGG</td><td>||||||||[||||||||||||]PAM</td><td>NA</td><td>Intergenic</td><td>NA</td><td>NA</td></tr>
<tr><td></td><td></td><td></td><td></td><td></td><td></td><td></td><td></td><td></td><td></td><td></td><td></td></tr>
<tr><td>T22089</td><td>AGGGGTTTTCCCGGTCCGATCGG</td><td>1000</td><td></td><td></td><td></td><td></td><td></td><td></td><td></td><td></td><td></td></tr>
<tr><td>Oligo fwd</td><td>CACCAGGGGTTTTCCCGGTCCGAT</td><td></td><td></td><td></td><td></td><td></td><td></td><td></td><td></td><td></td><td></td></tr>
<tr><td>Oligo rev</td><td>AAACATCGGACCGGGAAAACCCCT</td><td></td><td></td><td></td><td></td><td></td><td></td><td></td><td></td><td></td><td></td></tr>
<tr><td>Chromosome</td><td>start</td><td>end</td><td>strand</td><td>MM</td><td>target_seq</td><td>PAM</td><td>alignment</td><td>distance</td><td>position</td><td>gene name</td><td>gene id</td></tr>
<tr><td>GU305914.1</td><td>162617</td><td>162639</td><td>-</td><td>0</td><td>AGGGGTTTTCCCGGTCCGAT</td><td>CGG</td><td>||||||||[||||||||||||]PAM</td><td>NA</td><td>Intergenic</td><td>NA</td><td>NA</td></tr>
<tr><td></td><td></td><td></td><td></td><td></td><td></td><td></td><td></td><td></td><td></td><td></td><td></td></tr>
<tr><td>T22090</td><td>CCGGTCCGATCGGCTCGATGAGG</td><td>1000</td><td></td><td></td><td></td><td></td><td></td><td></td><td></td><td></td><td></td></tr>
<tr><td>Oligo fwd</td><td>CACCCCGGTCCGATCGGCTCGATG</td><td></td><td></td><td></td><td></td><td></td><td></td><td></td><td></td><td></td><td></td></tr>
<tr><td>Oligo rev</td><td>AAACCATCGAGCCGATCGGACCGG</td><td></td><td></td><td></td><td></td><td></td><td></td><td></td><td></td><td></td><td></td></tr>
<tr><td>Chromosome</td><td>start</td><td>end</td><td>strand</td><td>MM</td><td>target_seq</td><td>PAM</td><td>alignment</td><td>distance</td><td>position</td><td>gene name</td><td>gene id</td></tr>
<tr><td>GU305914.1</td><td>162607</td><td>162629</td><td>-</td><td>0</td><td>CCGGTCCGATCGGCTCGATG</td><td>AGG</td><td>||||||||[||||||||||||]PAM</td><td>NA</td><td>Intergenic</td><td>NA</td><td>NA</td></tr>
<tr><td></td><td></td><td></td><td></td><td></td><td></td><td></td><td></td><td></td><td></td><td></td><td></td></tr>
<tr><td>T22091</td><td>CGGTCCGATCGGCTCGATGAGGG</td><td>1000</td><td></td><td></td><td></td><td></td><td></td><td></td><td></td><td></td><td></td></tr>
<tr><td>Oligo fwd</td><td>CACCCGGTCCGATCGGCTCGATGA</td><td></td><td></td><td></td><td></td><td></td><td></td><td></td><td></td><td></td><td></td></tr>
<tr><td>Oligo rev</td><td>AAACTCATCGAGCCGATCGGACCG</td><td></td><td></td><td></td><td></td><td></td><td></td><td></td><td></td><td></td><td></td></tr>
<tr><td>Chromosome</td><td>start</td><td>end</td><td>strand</td><td>MM</td><td>target_seq</td><td>PAM</td><td>alignment</td><td>distance</td><td>position</td><td>gene name</td><td>gene id</td></tr>
<tr><td>GU305914.1</td><td>162606</td><td>162628</td><td>-</td><td>0</td><td>CGGTCCGATCGGCTCGATGA</td><td>GGG</td><td>||||||||[||||||||||||]PAM</td><td>NA</td><td>Intergenic</td><td>NA</td><td>NA</td></tr>
<tr><td></td><td></td><td></td><td></td><td></td><td></td><td></td><td></td><td></td><td></td><td></td><td></td></tr>
<tr><td>T22092</td><td>GGTCCGATCGGCTCGATGAGGGG</td><td>1000</td><td></td><td></td><td></td><td></td><td></td><td></td><td></td><td></td><td></td></tr>
<tr><td>Oligo fwd</td><td>CACCGGTCCGATCGGCTCGATGAG</td><td></td><td></td><td></td><td></td><td></td><td></td><td></td><td></td><td></td><td></td></tr>
<tr><td>Oligo rev</td><td>AAACCTCATCGAGCCGATCGGACC</td><td></td><td></td><td></td><td></td><td></td><td></td><td></td><td></td><td></td><td></td></tr>
<tr><td>Chromosome</td><td>start</td><td>end</td><td>strand</td><td>MM</td><td>target_seq</td><td>PAM</td><td>alignment</td><td>distance</td><td>position</td><td>gene name</td><td>gene id</td></tr>
<tr><td>GU305914.1</td><td>162605</td><td>162627</td><td>-</td><td>0</td><td>GGTCCGATCGGCTCGATGAG</td><td>GGG</td><td>||||||||[||||||||||||]PAM</td><td>NA</td><td>Intergenic</td><td>NA</td><td>NA</td></tr>
<tr><td></td><td></td><td></td><td></td><td></td><td></td><td></td><td></td><td></td><td></td><td></td><td></td></tr>
<tr><td>T22093</td><td>GTCCGATCGGCTCGATGAGGGGG</td><td>1000</td><td></td><td></td><td></td><td></td><td></td><td></td><td></td><td></td><td></td></tr>
<tr><td>Oligo fwd</td><td>CACCGTCCGATCGGCTCGATGAGG</td><td></td><td></td><td></td><td></td><td></td><td></td><td></td><td></td><td></td><td></td></tr>
<tr><td>Oligo rev</td><td>AAACCCTCATCGAGCCGATCGGAC</td><td></td><td></td><td></td><td></td><td></td><td></td><td></td><td></td><td></td><td></td></tr>
<tr><td>Chromosome</td><td>start</td><td>end</td><td>strand</td><td>MM</td><td>target_seq</td><td>PAM</td><td>alignment</td><td>distance</td><td>position</td><td>gene name</td><td>gene id</td></tr>
<tr><td>GU305914.1</td><td>162604</td><td>162626</td><td>-</td><td>0</td><td>GTCCGATCGGCTCGATGAGG</td><td>GGG</td><td>||||||||[||||||||||||]PAM</td><td>NA</td><td>Intergenic</td><td>NA</td><td>NA</td></tr>
<tr><td></td><td></td><td></td><td></td><td></td><td></td><td></td><td></td><td></td><td></td><td></td><td></td></tr>
<tr><td>T22094</td><td>ATGAGGGGGTCGAGCACCGTCGG</td><td>1000</td><td></td><td></td><td></td><td></td><td></td><td></td><td></td><td></td><td></td></tr>
<tr><td>Oligo fwd</td><td>CACCATGAGGGGGTCGAGCACCGT</td><td></td><td></td><td></td><td></td><td></td><td></td><td></td><td></td><td></td><td></td></tr>
<tr><td>Oligo rev</td><td>AAACACGGTGCTCGACCCCCTCAT</td><td></td><td></td><td></td><td></td><td></td><td></td><td></td><td></td><td></td><td></td></tr>
<tr><td>Chromosome</td><td>start</td><td>end</td><td>strand</td><td>MM</td><td>target_seq</td><td>PAM</td><td>alignment</td><td>distance</td><td>position</td><td>gene name</td><td>gene id</td></tr>
<tr><td>GU305914.1</td><td>162590</td><td>162612</td><td>-</td><td>0</td><td>ATGAGGGGGTCGAGCACCGT</td><td>CGG</td><td>||||||||[||||||||||||]PAM</td><td>NA</td><td>Intergenic</td><td>NA</td><td>NA</td></tr>
<tr><td></td><td></td><td></td><td></td><td></td><td></td><td></td><td></td><td></td><td></td><td></td><td></td></tr>
<tr><td>T22095</td><td>ACGAGCAACCGCCCGAAATGTGG</td><td>1000</td><td></td><td></td><td></td><td></td><td></td><td></td><td></td><td></td><td></td></tr>
<tr><td>Oligo fwd</td><td>CACCACGAGCAACCGCCCGAAATG</td><td></td><td></td><td></td><td></td><td></td><td></td><td></td><td></td><td></td><td></td></tr>
<tr><td>Oligo rev</td><td>AAACCATTTCGGGCGGTTGCTCGT</td><td></td><td></td><td></td><td></td><td></td><td></td><td></td><td></td><td></td><td></td></tr>
<tr><td>Chromosome</td><td>start</td><td>end</td><td>strand</td><td>MM</td><td>target_seq</td><td>PAM</td><td>alignment</td><td>distance</td><td>position</td><td>gene name</td><td>gene id</td></tr>
<tr><td>GU305914.1</td><td>162539</td><td>162561</td><td>-</td><td>0</td><td>ACGAGCAACCGCCCGAAATG</td><td>TGG</td><td>||||||||[||||||||||||]PAM</td><td>NA</td><td>Intergenic</td><td>NA</td><td>NA</td></tr>
<tr><td></td><td></td><td></td><td></td><td></td><td></td><td></td><td></td><td></td><td></td><td></td><td></td></tr>
<tr><td>T22096</td><td>CGAGCAACCGCCCGAAATGTGGG</td><td>1000</td><td></td><td></td><td></td><td></td><td></td><td></td><td></td><td></td><td></td></tr>
</table>

| | | | | | | | | | | | |
|---|---|---|---|---|---|---|---|---|---|---|---|
| Oligo fwd | CACCCGAGCAACCGCCCGAAATGT | | | | | | | | | | |
| Oligo rev | AAACACATTTCGGGCGGTTGCTCG | | | | | | | | | | |
| Chromosome | start | end | strand | MM | target_seq | PAM | alignment | distance | position | gene name | gene id |
| GU305914.1 | 162538 | 162560 | - | 0 | CGAGCAACCGCCCGAAATGT | GGG | \|\|\|\|\|\|\|\|[\|\|\|\|\|\|\|\|\|\|\|\|]PAM | NA | Intergenic | NA | NA |
| T22097 | GTTACGCACGTGTAAGTATACGG | 1000 | | | | | | | | | |
| Oligo fwd | CACCGTTACGCACGTGTAAGTATA | | | | | | | | | | |
| Oligo rev | AAACTATACTTACACGTGCGTAAC | | | | | | | | | | |
| Chromosome | start | end | strand | MM | target_seq | PAM | alignment | distance | position | gene name | gene id |
| GU305914.1 | 162504 | 162526 | - | 0 | GTTACGCACGTGTAAGTATA | CGG | \|\|\|\|\|\|\|\|[\|\|\|\|\|\|\|\|\|\|\|\|]PAM | NA | Intergenic | NA | NA |
| T22098 | ACGTGTAAGTATACGGCGTACGG | 1000 | | | | | | | | | |
| Oligo fwd | CACCACGTGTAAGTATACGGCGTA | | | | | | | | | | |
| Oligo rev | AAACTACGCCGTATACTTACACGT | | | | | | | | | | |
| Chromosome | start | end | strand | MM | target_seq | PAM | alignment | distance | position | gene name | gene id |
| GU305914.1 | 162497 | 162519 | - | 0 | ACGTGTAAGTATACGGCGTA | CGG | \|\|\|\|\|\|\|\|[\|\|\|\|\|\|\|\|\|\|\|\|]PAM | NA | Intergenic | NA | NA |
| T22099 | GGAGCTCCGTTCGTTTCTGACGG | 1000 | | | | | | | | | |
| Oligo fwd | CACCGGAGCTCCGTTCGTTTCTGA | | | | | | | | | | |
| Oligo rev | AAACTCAGAAACGAACGGAGCTCC | | | | | | | | | | |
| Chromosome | start | end | strand | MM | target_seq | PAM | alignment | distance | position | gene name | gene id |
| GU305914.1 | 162476 | 162498 | - | 0 | GGAGCTCCGTTCGTTTCTGA | CGG | \|\|\|\|\|\|\|\|[\|\|\|\|\|\|\|\|\|\|\|\|]PAM | NA | Intergenic | NA | NA |
| T22100 | GCTCCGTTCGTTTCTGACGGTGG | 1000 | | | | | | | | | |
| Oligo fwd | CACCGCTCCGTTCGTTTCTGACGG | | | | | | | | | | |
| Oligo rev | AAACCCGTCAGAAACGAACGGAGC | | | | | | | | | | |
| Chromosome | start | end | strand | MM | target_seq | PAM | alignment | distance | position | gene name | gene id |
| GU305914.1 | 162473 | 162495 | - | 0 | GCTCCGTTCGTTTCTGACGG | TGG | \|\|\|\|\|\|\|\|[\|\|\|\|\|\|\|\|\|\|\|\|]PAM | NA | Intergenic | NA | NA |
| T22101 | TGACGGTGGCTCGTGTCGAGCGG | 1000 | | | | | | | | | |
| Oligo fwd | CACCTGACGGTGGCTCGTGTCGAG | | | | | | | | | | |
| Oligo rev | AAACCTCGACACGAGCCACCGTCA | | | | | | | | | | |
| Chromosome | start | end | strand | MM | target_seq | PAM | alignment | distance | position | gene name | gene id |
| GU305914.1 | 162459 | 162481 | - | 0 | TGACGGTGGCTCGTGTCGAG | CGG | \|\|\|\|\|\|\|\|[\|\|\|\|\|\|\|\|\|\|\|\|]PAM | NA | Intergenic | NA | NA |
| T22102 | TCCGATCCTCCGATCCTCCGAGG | 1000 | | | | | | | | | |
| Oligo fwd | CACCTCCGATCCTCCGATCCTCCG | | | | | | | | | | |
| Oligo rev | AAACCGGAGGATCGGAGGATCGGA | | | | | | | | | | |
| Chromosome | start | end | strand | MM | target_seq | PAM | alignment | distance | position | gene name | gene id |
| GU305914.1 | 162433 | 162455 | - | 0 | TCCGATCCTCCGATCCTCCG | AGG | \|\|\|\|\|\|\|\|[\|\|\|\|\|\|\|\|\|\|\|\|]PAM | NA | Intergenic | NA | NA |
| T22103 | CCGATCCTCCGATCCTCCGAGGG | 1000 | | | | | | | | | |
| Oligo fwd | CACCCCGATCCTCCGATCCTCCGA | | | | | | | | | | |
| Oligo rev | AAACTCGGAGGATCGGAGGATCGG | | | | | | | | | | |
| Chromosome | start | end | strand | MM | target_seq | PAM | alignment | distance | position | gene name | gene id |
| GU305914.1 | 162432 | 162454 | - | 0 | CCGATCCTCCGATCCTCCGA | GGG | \|\|\|\|\|\|\|\|[\|\|\|\|\|\|\|\|\|\|\|\|]PAM | NA | Intergenic | NA | NA |
| T22104 | TCCGATCCTCCGAGGGCTGACGG | 1000 | | | | | | | | | |
| Oligo fwd | CACCTCCGATCCTCCGAGGGCTGA | | | | | | | | | | |
| Oligo rev | AAACTCAGCCCTCGGAGGATCGGA | | | | | | | | | | |
| Chromosome | start | end | strand | MM | target_seq | PAM | alignment | distance | position | gene name | gene id |
| GU305914.1 | 162425 | 162447 | - | 0 | TCCGATCCTCCGAGGGCTGA | CGG | \|\|\|\|\|\|\|\|[\|\|\|\|\|\|\|\|\|\|\|\|]PAM | NA | Intergenic | NA | NA |
| T22105 | GACGGACCTTCCCCGTTTTCAGG | 1000 | | | | | | | | | |
| Oligo fwd | CACCGACGGACCTTCCCCGTTTTC | | | | | | | | | | |
| Oligo rev | AAACGAAAACGGGGAAGGTCCGTC | | | | | | | | | | |
| Chromosome | start | end | strand | MM | target_seq | PAM | alignment | distance | position | gene name | gene id |
| GU305914.1 | 162407 | 162429 | - | 0 | GACGGACCTTCCCCGTTTTC | AGG | \|\|\|\|\|\|\|\|[\|\|\|\|\|\|\|\|\|\|\|\|]PAM | NA | Intergenic | NA | NA |
| T22106 | CCCCGTTTTCAGGTCCGCTTCGG | 1000 | | | | | | | | | |
| Oligo fwd | CACCCCCCGTTTTCAGGTCCGCTT | | | | | | | | | | |
| Oligo rev | AAACAAGCGGACCTGAAAACGGGG | | | | | | | | | | |
| Chromosome | start | end | strand | MM | target_seq | PAM | alignment | distance | position | gene name | gene id |
| GU305914.1 | 162397 | 162419 | - | 0 | CCCCGTTTTCAGGTCCGCTT | CGG | \|\|\|\|\|\|\|\|[\|\|\|\|\|\|\|\|\|\|\|\|]PAM | NA | Intergenic | NA | NA |
| T22107 | CTTCGGTTACTTGATCGCATCGG | 1000 | | | | | | | | | |
| Oligo fwd | CACCCTTCGGTTACTTGATCGCAT | | | | | | | | | | |
| Oligo rev | AAACATGCGATCAAGTAACCGAAG | | | | | | | | | | |
| Chromosome | start | end | strand | MM | target_seq | PAM | alignment | distance | position | gene name | gene id |
| GU305914.1 | 162380 | 162402 | - | 0 | CTTCGGTTACTTGATCGCAT | CGG | \|\|\|\|\|\|\|\|[\|\|\|\|\|\|\|\|\|\|\|\|]PAM | NA | Intergenic | NA | NA |
| T22108 | ACTTGATCGCATCGGACACGCGG | 1000 | | | | | | | | | |
| Oligo fwd | CACCACTTGATCGCATCGGACACG | | | | | | | | | | |
| Oligo rev | AAACCGTGTCCGATGCGATCAAGT | | | | | | | | | | |
| Chromosome | start | end | strand | MM | target_seq | PAM | alignment | distance | position | gene name | gene id |
| GU305914.1 | 162372 | 162394 | - | 0 | ACTTGATCGCATCGGACACG | CGG | \|\|\|\|\|\|\|\|[\|\|\|\|\|\|\|\|\|\|\|\|]PAM | NA | Intergenic | NA | NA |
| T22109 | TCGCATCGGACACGCGGACGTGG | 1000 | | | | | | | | | |
| Oligo fwd | CACCTCGCATCGGACACGCGGACG | | | | | | | | | | |
| Oligo rev | AAACCGTCCGCGTGTCCGATGCGA | | | | | | | | | | |
| Chromosome | start | end | strand | MM | target_seq | PAM | alignment | distance | position | gene name | gene id |
| GU305914.1 | 162366 | 162388 | - | 0 | TCGCATCGGACACGCGGACG | TGG | \|\|\|\|\|\|\|\|[\|\|\|\|\|\|\|\|\|\|\|\|]PAM | NA | Intergenic | NA | NA |
| T22110 | ACCCGTTCGCTTCGATCGAGCGG | 1000 | | | | | | | | | |
| Oligo fwd | CACCACCCGTTCGCTTCGATCGAG | | | | | | | | | | |
| Oligo rev | AAACCTCGATCGAAGCGAACGGGT | | | | | | | | | | |
| Chromosome | start | end | strand | MM | target_seq | PAM | alignment | distance | position | gene name | gene id |
| GU305914.1 | 162336 | 162358 | - | 0 | ACCCGTTCGCTTCGATCGAG | CGG | \|\|\|\|\|\|\|\|[\|\|\|\|\|\|\|\|\|\|\|\|]PAM | NA | Intergenic | NA | NA |

<table>
<tr><td>T22111</td><td>TCGCTTCGATCGAGCGGTCGTGG</td><td>1000</td><td></td><td></td><td></td><td></td><td></td><td></td><td></td><td></td><td></td></tr>
<tr><td>Oligo fwd</td><td>CACCTCGCTTCGATCGAGCGGTCG</td><td></td><td></td><td></td><td></td><td></td><td></td><td></td><td></td><td></td><td></td></tr>
<tr><td>Oligo rev</td><td>AAACCGACCGCTCGATCGAAGCGA</td><td></td><td></td><td></td><td></td><td></td><td></td><td></td><td></td><td></td><td></td></tr>
<tr><td>Chromosome</td><td>start</td><td>end</td><td>strand</td><td>MM</td><td>target_seq</td><td>PAM</td><td>alignment</td><td>distance</td><td>position</td><td>gene name</td><td>gene id</td></tr>
<tr><td>GU305914.1</td><td>162330</td><td>162352</td><td>-</td><td>0</td><td>TCGCTTCGATCGAGCGGTCG</td><td>TGG</td><td>||||||||[||||||||||||]PAM</td><td>NA</td><td>Intergenic</td><td>NA</td><td>NA</td></tr>
<tr><td>T22112</td><td>TCGAGCGGTCGTGGCGTTCGCGG</td><td>1000</td><td></td><td></td><td></td><td></td><td></td><td></td><td></td><td></td><td></td></tr>
<tr><td>Oligo fwd</td><td>CACCTCGAGCGGTCGTGGCGTTCG</td><td></td><td></td><td></td><td></td><td></td><td></td><td></td><td></td><td></td><td></td></tr>
<tr><td>Oligo rev</td><td>AAACCGAACGCCACGACCGCTCGA</td><td></td><td></td><td></td><td></td><td></td><td></td><td></td><td></td><td></td><td></td></tr>
<tr><td>Chromosome</td><td>start</td><td>end</td><td>strand</td><td>MM</td><td>target_seq</td><td>PAM</td><td>alignment</td><td>distance</td><td>position</td><td>gene name</td><td>gene id</td></tr>
<tr><td>GU305914.1</td><td>162321</td><td>162343</td><td>-</td><td>0</td><td>TCGAGCGGTCGTGGCGTTCG</td><td>CGG</td><td>||||||||[||||||||||||]PAM</td><td>NA</td><td>Intergenic</td><td>NA</td><td>NA</td></tr>
<tr><td>T22113</td><td>TTCGCGGCAATCAAATAACACGG</td><td>1000</td><td></td><td></td><td></td><td></td><td></td><td></td><td></td><td></td><td></td></tr>
<tr><td>Oligo fwd</td><td>CACCTTCGCGGCAATCAAATAACA</td><td></td><td></td><td></td><td></td><td></td><td></td><td></td><td></td><td></td><td></td></tr>
<tr><td>Oligo rev</td><td>AAACTGTTATTTGATTGCCGCGAA</td><td></td><td></td><td></td><td></td><td></td><td></td><td></td><td></td><td></td><td></td></tr>
<tr><td>Chromosome</td><td>start</td><td>end</td><td>strand</td><td>MM</td><td>target_seq</td><td>PAM</td><td>alignment</td><td>distance</td><td>position</td><td>gene name</td><td>gene id</td></tr>
<tr><td>GU305914.1</td><td>162305</td><td>162327</td><td>-</td><td>0</td><td>TTCGCGGCAATCAAATAACA</td><td>CGG</td><td>||||||||[||||||||||||]PAM</td><td>NA</td><td>Intergenic</td><td>NA</td><td>NA</td></tr>
<tr><td>T22114</td><td>AATCAAATAACACGGACGAGCGG</td><td>1000</td><td></td><td></td><td></td><td></td><td></td><td></td><td></td><td></td><td></td></tr>
<tr><td>Oligo fwd</td><td>CACCAATCAAATAACACGGACGAG</td><td></td><td></td><td></td><td></td><td></td><td></td><td></td><td></td><td></td><td></td></tr>
<tr><td>Oligo rev</td><td>AAACCTCGTCCGTGTTATTTGATT</td><td></td><td></td><td></td><td></td><td></td><td></td><td></td><td></td><td></td><td></td></tr>
<tr><td>Chromosome</td><td>start</td><td>end</td><td>strand</td><td>MM</td><td>target_seq</td><td>PAM</td><td>alignment</td><td>distance</td><td>position</td><td>gene name</td><td>gene id</td></tr>
<tr><td>GU305914.1</td><td>162297</td><td>162319</td><td>-</td><td>0</td><td>AATCAAATAACACGGACGAG</td><td>CGG</td><td>||||||||[||||||||||||]PAM</td><td>NA</td><td>Intergenic</td><td>NA</td><td>NA</td></tr>
<tr><td>T22115</td><td>ATCAAATAACACGGACGAGCGGG</td><td>1000</td><td></td><td></td><td></td><td></td><td></td><td></td><td></td><td></td><td></td></tr>
<tr><td>Oligo fwd</td><td>CACCATCAAATAACACGGACGAGC</td><td></td><td></td><td></td><td></td><td></td><td></td><td></td><td></td><td></td><td></td></tr>
<tr><td>Oligo rev</td><td>AAACGCTCGTCCGTGTTATTTGAT</td><td></td><td></td><td></td><td></td><td></td><td></td><td></td><td></td><td></td><td></td></tr>
<tr><td>Chromosome</td><td>start</td><td>end</td><td>strand</td><td>MM</td><td>target_seq</td><td>PAM</td><td>alignment</td><td>distance</td><td>position</td><td>gene name</td><td>gene id</td></tr>
<tr><td>GU305914.1</td><td>162296</td><td>162318</td><td>-</td><td>0</td><td>ATCAAATAACACGGACGAGC</td><td>GGG</td><td>||||||||[||||||||||||]PAM</td><td>NA</td><td>Intergenic</td><td>NA</td><td>NA</td></tr>
<tr><td>T22116</td><td>GAGCGGGTAATCGAAGCGAGCGG</td><td>1000</td><td></td><td></td><td></td><td></td><td></td><td></td><td></td><td></td><td></td></tr>
<tr><td>Oligo fwd</td><td>CACCGAGCGGGTAATCGAAGCGAG</td><td></td><td></td><td></td><td></td><td></td><td></td><td></td><td></td><td></td><td></td></tr>
<tr><td>Oligo rev</td><td>AAACCTCGCTTCGATTACCCGCTC</td><td></td><td></td><td></td><td></td><td></td><td></td><td></td><td></td><td></td><td></td></tr>
<tr><td>Chromosome</td><td>start</td><td>end</td><td>strand</td><td>MM</td><td>target_seq</td><td>PAM</td><td>alignment</td><td>distance</td><td>position</td><td>gene name</td><td>gene id</td></tr>
<tr><td>GU305914.1</td><td>162280</td><td>162302</td><td>-</td><td>0</td><td>GAGCGGGTAATCGAAGCGAG</td><td>CGG</td><td>||||||||[||||||||||||]PAM</td><td>NA</td><td>Intergenic</td><td>NA</td><td>NA</td></tr>
<tr><td>T22117</td><td>GGGTAATCGAAGCGAGCGGTCGG</td><td>1000</td><td></td><td></td><td></td><td></td><td></td><td></td><td></td><td></td><td></td></tr>
<tr><td>Oligo fwd</td><td>CACCGGGTAATCGAAGCGAGCGGT</td><td></td><td></td><td></td><td></td><td></td><td></td><td></td><td></td><td></td><td></td></tr>
<tr><td>Oligo rev</td><td>AAACACCGCTCGCTTCGATTACCC</td><td></td><td></td><td></td><td></td><td></td><td></td><td></td><td></td><td></td><td></td></tr>
<tr><td>Chromosome</td><td>start</td><td>end</td><td>strand</td><td>MM</td><td>target_seq</td><td>PAM</td><td>alignment</td><td>distance</td><td>position</td><td>gene name</td><td>gene id</td></tr>
<tr><td>GU305914.1</td><td>162276</td><td>162298</td><td>-</td><td>0</td><td>GGGTAATCGAAGCGAGCGGT</td><td>CGG</td><td>||||||||[||||||||||||]PAM</td><td>NA</td><td>Intergenic</td><td>NA</td><td>NA</td></tr>
<tr><td>T22118</td><td>GCGAGCGGTCGGTCGCTACGAGG</td><td>1000</td><td></td><td></td><td></td><td></td><td></td><td></td><td></td><td></td><td></td></tr>
<tr><td>Oligo fwd</td><td>CACCGCGAGCGGTCGGTCGCTACG</td><td></td><td></td><td></td><td></td><td></td><td></td><td></td><td></td><td></td><td></td></tr>
<tr><td>Oligo rev</td><td>AAACCGTAGCGACCGACCGCTCGC</td><td></td><td></td><td></td><td></td><td></td><td></td><td></td><td></td><td></td><td></td></tr>
<tr><td>Chromosome</td><td>start</td><td>end</td><td>strand</td><td>MM</td><td>target_seq</td><td>PAM</td><td>alignment</td><td>distance</td><td>position</td><td>gene name</td><td>gene id</td></tr>
<tr><td>GU305914.1</td><td>162265</td><td>162287</td><td>-</td><td>0</td><td>GCGAGCGGTCGGTCGCTACG</td><td>AGG</td><td>||||||||[||||||||||||]PAM</td><td>NA</td><td>Intergenic</td><td>NA</td><td>NA</td></tr>
<tr><td>T22119</td><td>CGAGCGGTCGGTCGCTACGAGGG</td><td>1000</td><td></td><td></td><td></td><td></td><td></td><td></td><td></td><td></td><td></td></tr>
<tr><td>Oligo fwd</td><td>CACCCGAGCGGTCGGTCGCTACGA</td><td></td><td></td><td></td><td></td><td></td><td></td><td></td><td></td><td></td><td></td></tr>
<tr><td>Oligo rev</td><td>AAACTCGTAGCGACCGACCGCTCG</td><td></td><td></td><td></td><td></td><td></td><td></td><td></td><td></td><td></td><td></td></tr>
<tr><td>Chromosome</td><td>start</td><td>end</td><td>strand</td><td>MM</td><td>target_seq</td><td>PAM</td><td>alignment</td><td>distance</td><td>position</td><td>gene name</td><td>gene id</td></tr>
<tr><td>GU305914.1</td><td>162264</td><td>162286</td><td>-</td><td>0</td><td>CGAGCGGTCGGTCGCTACGA</td><td>GGG</td><td>||||||||[||||||||||||]PAM</td><td>NA</td><td>Intergenic</td><td>NA</td><td>NA</td></tr>
<tr><td>T22120</td><td>GAGCGGTCGGTCGCTACGAGGGG</td><td>1000</td><td></td><td></td><td></td><td></td><td></td><td></td><td></td><td></td><td></td></tr>
<tr><td>Oligo fwd</td><td>CACCGAGCGGTCGGTCGCTACGAG</td><td></td><td></td><td></td><td></td><td></td><td></td><td></td><td></td><td></td><td></td></tr>
<tr><td>Oligo rev</td><td>AAACCTCGTAGCGACCGACCGCTC</td><td></td><td></td><td></td><td></td><td></td><td></td><td></td><td></td><td></td><td></td></tr>
<tr><td>Chromosome</td><td>start</td><td>end</td><td>strand</td><td>MM</td><td>target_seq</td><td>PAM</td><td>alignment</td><td>distance</td><td>position</td><td>gene name</td><td>gene id</td></tr>
<tr><td>GU305914.1</td><td>162263</td><td>162285</td><td>-</td><td>0</td><td>GAGCGGTCGGTCGCTACGAG</td><td>GGG</td><td>||||||||[||||||||||||]PAM</td><td>NA</td><td>Intergenic</td><td>NA</td><td>NA</td></tr>
<tr><td>GU305914.1</td><td>229696</td><td>229718</td><td>+</td><td>4</td><td>GAGCGGGTGCTCGCTACGCG</td><td>CGG</td><td>||||||--[|-||||||||-|]PAM</td><td>NA</td><td>Intergenic</td><td>NA</td><td>NA</td></tr>
<tr><td>T22121</td><td>GGTCGGTCGCTACGAGGGGTCGG</td><td>1000</td><td></td><td></td><td></td><td></td><td></td><td></td><td></td><td></td><td></td></tr>
<tr><td>Oligo fwd</td><td>CACCGGTCGGTCGCTACGAGGGGT</td><td></td><td></td><td></td><td></td><td></td><td></td><td></td><td></td><td></td><td></td></tr>
<tr><td>Oligo rev</td><td>AAACACCCCTCGTAGCGACCGACC</td><td></td><td></td><td></td><td></td><td></td><td></td><td></td><td></td><td></td><td></td></tr>
<tr><td>Chromosome</td><td>start</td><td>end</td><td>strand</td><td>MM</td><td>target_seq</td><td>PAM</td><td>alignment</td><td>distance</td><td>position</td><td>gene name</td><td>gene id</td></tr>
<tr><td>GU305914.1</td><td>162259</td><td>162281</td><td>-</td><td>0</td><td>GGTCGGTCGCTACGAGGGGT</td><td>CGG</td><td>||||||||[||||||||||||]PAM</td><td>NA</td><td>Intergenic</td><td>NA</td><td>NA</td></tr>
<tr><td>T22122</td><td>GAGAGATCACTCGTCGCGAGCGG</td><td>1000</td><td></td><td></td><td></td><td></td><td></td><td></td><td></td><td></td><td></td></tr>
<tr><td>Oligo fwd</td><td>CACCGAGAGATCACTCGTCGCGAG</td><td></td><td></td><td></td><td></td><td></td><td></td><td></td><td></td><td></td><td></td></tr>
<tr><td>Oligo rev</td><td>AAACCTCGCGACGAGTGATCTCTC</td><td></td><td></td><td></td><td></td><td></td><td></td><td></td><td></td><td></td><td></td></tr>
<tr><td>Chromosome</td><td>start</td><td>end</td><td>strand</td><td>MM</td><td>target_seq</td><td>PAM</td><td>alignment</td><td>distance</td><td>position</td><td>gene name</td><td>gene id</td></tr>
<tr><td>GU305914.1</td><td>162229</td><td>162251</td><td>-</td><td>0</td><td>GAGAGATCACTCGTCGCGAG</td><td>CGG</td><td>||||||||[||||||||||||]PAM</td><td>NA</td><td>Intergenic</td><td>NA</td><td>NA</td></tr>
<tr><td>T22123</td><td>ACGCGAGCACCGAGTCGCTACGG</td><td>1000</td><td></td><td></td><td></td><td></td><td></td><td></td><td></td><td></td><td></td></tr>
<tr><td>Oligo fwd</td><td>CACCACGCGAGCACCGAGTCGCTA</td><td></td><td></td><td></td><td></td><td></td><td></td><td></td><td></td><td></td><td></td></tr>
<tr><td>Oligo rev</td><td>AAACTAGCGACTCGGTGCTCGCGT</td><td></td><td></td><td></td><td></td><td></td><td></td><td></td><td></td><td></td><td></td></tr>
<tr><td>Chromosome</td><td>start</td><td>end</td><td>strand</td><td>MM</td><td>target_seq</td><td>PAM</td><td>alignment</td><td>distance</td><td>position</td><td>gene name</td><td>gene id</td></tr>
<tr><td>GU305914.1</td><td>162199</td><td>162221</td><td>-</td><td>0</td><td>ACGCGAGCACCGAGTCGCTA</td><td>CGG</td><td>||||||||[||||||||||||]PAM</td><td>NA</td><td>Intergenic</td><td>NA</td><td>NA</td></tr>
<tr><td>T22124</td><td>GAGCACCGAGTCGCTACGGACGG</td><td>1000</td><td></td><td></td><td></td><td></td><td></td><td></td><td></td><td></td><td></td></tr>
<tr><td>Oligo fwd</td><td>CACCGAGCACCGAGTCGCTACGGA</td><td></td><td></td><td></td><td></td><td></td><td></td><td></td><td></td><td></td><td></td></tr>
<tr><td>Oligo rev</td><td>AAACTCCGTAGCGACTCGGTGCTC</td><td></td><td></td><td></td><td></td><td></td><td></td><td></td><td></td><td></td><td></td></tr>
<tr><td>Chromosome</td><td>start</td><td>end</td><td>strand</td><td>MM</td><td>target_seq</td><td>PAM</td><td>alignment</td><td>distance</td><td>position</td><td>gene name</td><td>gene id</td></tr>
<tr><td>GU305914.1</td><td>162195</td><td>162217</td><td>-</td><td>0</td><td>GAGCACCGAGTCGCTACGGA</td><td>CGG</td><td>||||||||[||||||||||||]PAM</td><td>NA</td><td>Intergenic</td><td>NA</td><td>NA</td></tr>
<tr><td>T22125</td><td>GGTGACTCGAGACGAGTGACCGG</td><td>1000</td><td></td><td></td><td></td><td></td><td></td><td></td><td></td><td></td><td></td></tr>
<tr><td>Oligo fwd</td><td>CACCGGTGACTCGAGACGAGTGAC</td><td></td><td></td><td></td><td></td><td></td><td></td><td></td><td></td><td></td><td></td></tr>
<tr><td>Oligo rev</td><td>AAACGTCACTCGTCTCGAGTCACC</td><td></td><td></td><td></td><td></td><td></td><td></td><td></td><td></td><td></td><td></td></tr>
<tr><td>Chromosome</td><td>start</td><td>end</td><td>strand</td><td>MM</td><td>target_seq</td><td>PAM</td><td>alignment</td><td>distance</td><td>position</td><td>gene name</td><td>gene id</td></tr>
</table>

GU305914.1 162174 162196 - 0 GGTGACTCGAGACGAGTGAC CGG ||||||||[||||||||||||]PAM NA Intergenic NA NA

T22126 AGTGACCGGTCGACGAGCGTCGG 1000
Oligo fwd CACCAGTGACCGGTCGACGAGCGT
Oligo rev AAACACGCTCGTCGACCGGTCACT
Chromosome start end strand MM target_seq PAM alignment distance position gene name gene id
GU305914.1 162160 162182 - 0 AGTGACCGGTCGACGAGCGT CGG ||||||||[||||||||||||]PAM NA Intergenic NA NA

T22127 CCGGTCGACGAGCGTCGGTCAGG 1000
Oligo fwd CACCCCGGTCGACGAGCGTCGGTC
Oligo rev AAACGACCGACGCTCGTCGACCGG
Chromosome start end strand MM target_seq PAM alignment distance position gene name gene id
GU305914.1 162155 162177 - 0 CCGGTCGACGAGCGTCGGTC AGG ||||||||[||||||||||||]PAM NA Intergenic NA NA

T22128 AGCGTCGGTCAGGCCGCCGATGG 1000
Oligo fwd CACCAGCGTCGGTCAGGCCGCCGA
Oligo rev AAACTCGGCGGCCTGACCGACGCT
Chromosome start end strand MM target_seq PAM alignment distance position gene name gene id
GU305914.1 162145 162167 - 0 AGCGTCGGTCAGGCCGCCGA TGG ||||||||[||||||||||||]PAM NA Intergenic NA NA

T22129 CCATGCGCCCATCCAGCCCCCGG 1000
Oligo fwd CACCCCATGCGCCCATCCAGCCCC
Oligo rev AAACGGGGCTGGATGGGCGCATGG
Chromosome start end strand MM target_seq PAM alignment distance position gene name gene id
GU305914.1 162064 162086 - 0 CCATGCGCCCATCCAGCCCC CGG ||||||||[||||||||||||]PAM NA Intergenic NA NA

T22130 CATGCGCCCATCCAGCCCCCGGG 1000
Oligo fwd CACCCATGCGCCCATCCAGCCCCC
Oligo rev AAACGGGGGCTGGATGGGCGCATG
Chromosome start end strand MM target_seq PAM alignment distance position gene name gene id
GU305914.1 162063 162085 - 0 CATGCGCCCATCCAGCCCCC GGG ||||||||[||||||||||||]PAM NA Intergenic NA NA

T22131 TTAAAAAGAAAGTCCGTGACCGG 1000
Oligo fwd CACCTTAAAAAGAAAGTCCGTGAC
Oligo rev AAACGTCACGGACTTTCTTTTTAA
Chromosome start end strand MM target_seq PAM alignment distance position gene name gene id
GU305914.1 162040 162062 - 0 TTAAAAAGAAAGTCCGTGAC CGG ||||||||[||||||||||||]PAM NA Intergenic NA NA

T22132 TAAAAAGAAAGTCCGTGACCGGG 1000
Oligo fwd CACCTAAAAAGAAAGTCCGTGACC
Oligo rev AAACGGTCACGGACTTTCTTTTTA
Chromosome start end strand MM target_seq PAM alignment distance position gene name gene id
GU305914.1 162039 162061 - 0 TAAAAAGAAAGTCCGTGACC GGG ||||||||[||||||||||||]PAM NA Intergenic NA NA

T22133 AAGAAAGTCCGTGACCGGGTCGG 1000
Oligo fwd CACCAAGAAAGTCCGTGACCGGGT
Oligo rev AAACACCCGGTCACGGACTTTCTT
Chromosome start end strand MM target_seq PAM alignment distance position gene name gene id
GU305914.1 162035 162057 - 0 AAGAAAGTCCGTGACCGGGT CGG ||||||||[||||||||||||]PAM NA Intergenic NA NA

T22134 TCCGTGACCGGGTCGGCCGAAGG 1000
Oligo fwd CACCTCCGTGACCGGGTCGGCCGA
Oligo rev AAACTCGGCCGACCCGGTCACGGA
Chromosome start end strand MM target_seq PAM alignment distance position gene name gene id
GU305914.1 162028 162050 - 0 TCCGTGACCGGGTCGGCCGA AGG ||||||||[||||||||||||]PAM NA Intergenic NA NA

T22135 GTCGGCCGAAGGAACCCCACCGG 1000
Oligo fwd CACCGTCGGCCGAAGGAACCCCAC
Oligo rev AAACGTGGGGTTCCTTCGGCCGAC
Chromosome start end strand MM target_seq PAM alignment distance position gene name gene id
GU305914.1 162017 162039 - 0 GTCGGCCGAAGGAACCCCAC CGG ||||||||[||||||||||||]PAM NA Intergenic NA NA

T22136 CCGAAGGAACCCCACCGGCACGG 1000
Oligo fwd CACCCCGAAGGAACCCCACCGGCA
Oligo rev AAACTGCCGGTGGGGTTCCTTCGG
Chromosome start end strand MM target_seq PAM alignment distance position gene name gene id
GU305914.1 162012 162034 - 0 CCGAAGGAACCCCACCGGCA CGG ||||||||[||||||||||||]PAM NA Intergenic NA NA

T22137 CGAAGGAACCCCACCGGCACGGG 1000
Oligo fwd CACCCGAAGGAACCCCACCGGCAC
Oligo rev AAACGTGCCGGTGGGGTTCCTTCG
Chromosome start end strand MM target_seq PAM alignment distance position gene name gene id
GU305914.1 162011 162033 - 0 CGAAGGAACCCCACCGGCAC GGG ||||||||[||||||||||||]PAM NA Intergenic NA NA

T22138 GAAGGAACCCCACCGGCACGGGG 1000
Oligo fwd CACCGAAGGAACCCCACCGGCACG
Oligo rev AAACCGTGCCGGTGGGGTTCCTTC
Chromosome start end strand MM target_seq PAM alignment distance position gene name gene id
GU305914.1 162010 162032 - 0 GAAGGAACCCCACCGGCACG GGG ||||||||[||||||||||||]PAM NA Intergenic NA NA

T22139 TACTCCGCTATTATACTTTGCGG 1000
Oligo fwd CACCTACTCCGCTATTATACTTTG
Oligo rev AAACCAAAGTATAATAGCGGAGTA
Chromosome start end strand MM target_seq PAM alignment distance position gene name gene id
GU305914.1 161952 161974 - 0 TACTCCGCTATTATACTTTG CGG ||||||||[||||||||||||]PAM NA Intergenic NA NA

T22140 GCTATTATACTTTGCGGCGTCGG 1000
Oligo fwd CACCGCTATTATACTTTGCGGCGT
Oligo rev AAACACGCCGCAAAGTATAATAGC

<table>
<tr><td>Chromosome</td><td>start</td><td>end</td><td>strand</td><td>MM</td><td>target_seq</td><td>PAM</td><td>alignment</td><td>distance</td><td>position</td><td>gene name</td><td>gene id</td></tr>
<tr><td>GU305914.1</td><td>161946</td><td>161968</td><td>-</td><td>0</td><td>GCTATTATACTTTGCGGCGT</td><td>CGG</td><td>||||||||[||||||||||||]PAM</td><td>NA</td><td>Intergenic</td><td>NA</td><td>NA</td></tr>
<tr><td>T22141</td><td>GCGTCGGACGTGACCGCGCTAGG</td><td>1000</td><td></td><td></td><td></td><td></td><td></td><td></td><td></td><td></td><td></td></tr>
<tr><td>Oligo fwd</td><td>CACCGCGTCGGACGTGACCGCGCT</td><td></td><td></td><td></td><td></td><td></td><td></td><td></td><td></td><td></td><td></td></tr>
<tr><td>Oligo rev</td><td>AAACAGCGCGGTCACGTCCGACGC</td><td></td><td></td><td></td><td></td><td></td><td></td><td></td><td></td><td></td><td></td></tr>
<tr><td>Chromosome</td><td>start</td><td>end</td><td>strand</td><td>MM</td><td>target_seq</td><td>PAM</td><td>alignment</td><td>distance</td><td>position</td><td>gene name</td><td>gene id</td></tr>
<tr><td>GU305914.1</td><td>161930</td><td>161952</td><td>-</td><td>0</td><td>GCGTCGGACGTGACCGCGCT</td><td>AGG</td><td>||||||||[||||||||||||]PAM</td><td>NA</td><td>Intergenic</td><td>NA</td><td>NA</td></tr>
<tr><td>GU305914.1</td><td>167921</td><td>167943</td><td>+</td><td>4</td><td>GGGACGGCCGTGACCGTGCT</td><td>CGG</td><td>|-|-|||-[||||||||-|||]PAM</td><td>NA</td><td>Intergenic</td><td>NA</td><td>NA</td></tr>
<tr><td>T22142</td><td>CGGACGTGACCGCGCTAGGTAGG</td><td>1000</td><td></td><td></td><td></td><td></td><td></td><td></td><td></td><td></td><td></td></tr>
<tr><td>Oligo fwd</td><td>CACCCGGACGTGACCGCGCTAGGT</td><td></td><td></td><td></td><td></td><td></td><td></td><td></td><td></td><td></td><td></td></tr>
<tr><td>Oligo rev</td><td>AAACACCTAGCGCGGTCACGTCCG</td><td></td><td></td><td></td><td></td><td></td><td></td><td></td><td></td><td></td><td></td></tr>
<tr><td>Chromosome</td><td>start</td><td>end</td><td>strand</td><td>MM</td><td>target_seq</td><td>PAM</td><td>alignment</td><td>distance</td><td>position</td><td>gene name</td><td>gene id</td></tr>
<tr><td>GU305914.1</td><td>161926</td><td>161948</td><td>-</td><td>0</td><td>CGGACGTGACCGCGCTAGGT</td><td>AGG</td><td>||||||||[||||||||||||]PAM</td><td>NA</td><td>Intergenic</td><td>NA</td><td>NA</td></tr>
<tr><td>T22143</td><td>ACCGCGCTAGGTAGGATACCTGG</td><td>1000</td><td></td><td></td><td></td><td></td><td></td><td></td><td></td><td></td><td></td></tr>
<tr><td>Oligo fwd</td><td>CACCACCGCGCTAGGTAGGATACC</td><td></td><td></td><td></td><td></td><td></td><td></td><td></td><td></td><td></td><td></td></tr>
<tr><td>Oligo rev</td><td>AAACGGTATCCTACCTAGCGCGGT</td><td></td><td></td><td></td><td></td><td></td><td></td><td></td><td></td><td></td><td></td></tr>
<tr><td>Chromosome</td><td>start</td><td>end</td><td>strand</td><td>MM</td><td>target_seq</td><td>PAM</td><td>alignment</td><td>distance</td><td>position</td><td>gene name</td><td>gene id</td></tr>
<tr><td>GU305914.1</td><td>161918</td><td>161940</td><td>-</td><td>0</td><td>ACCGCGCTAGGTAGGATACC</td><td>TGG</td><td>||||||||[||||||||||||]PAM</td><td>NA</td><td>Intergenic</td><td>NA</td><td>NA</td></tr>
<tr><td>T22144</td><td>AGGTAGGATACCTGGAGTGTAGG</td><td>1000</td><td></td><td></td><td></td><td></td><td></td><td></td><td></td><td></td><td></td></tr>
<tr><td>Oligo fwd</td><td>CACCAGGTAGGATACCTGGAGTGT</td><td></td><td></td><td></td><td></td><td></td><td></td><td></td><td></td><td></td><td></td></tr>
<tr><td>Oligo rev</td><td>AAACACACTCCAGGTATCCTACCT</td><td></td><td></td><td></td><td></td><td></td><td></td><td></td><td></td><td></td><td></td></tr>
<tr><td>Chromosome</td><td>start</td><td>end</td><td>strand</td><td>MM</td><td>target_seq</td><td>PAM</td><td>alignment</td><td>distance</td><td>position</td><td>gene name</td><td>gene id</td></tr>
<tr><td>GU305914.1</td><td>161910</td><td>161932</td><td>-</td><td>0</td><td>AGGTAGGATACCTGGAGTGT</td><td>AGG</td><td>||||||||[||||||||||||]PAM</td><td>NA</td><td>Intergenic</td><td>NA</td><td>NA</td></tr>
<tr><td>T22145</td><td>AGACGCAACCTGACGCATCCCGG</td><td>1000</td><td></td><td></td><td></td><td></td><td></td><td></td><td></td><td></td><td></td></tr>
<tr><td>Oligo fwd</td><td>CACCAGACGCAACCTGACGCATCC</td><td></td><td></td><td></td><td></td><td></td><td></td><td></td><td></td><td></td><td></td></tr>
<tr><td>Oligo rev</td><td>AAACGGATGCGTCAGGTTGCGTCT</td><td></td><td></td><td></td><td></td><td></td><td></td><td></td><td></td><td></td><td></td></tr>
<tr><td>Chromosome</td><td>start</td><td>end</td><td>strand</td><td>MM</td><td>target_seq</td><td>PAM</td><td>alignment</td><td>distance</td><td>position</td><td>gene name</td><td>gene id</td></tr>
<tr><td>GU305914.1</td><td>161876</td><td>161898</td><td>-</td><td>0</td><td>AGACGCAACCTGACGCATCC</td><td>CGG</td><td>||||||||[||||||||||||]PAM</td><td>NA</td><td>Intergenic</td><td>NA</td><td>NA</td></tr>
<tr><td>T22146</td><td>GGCTAGAATATATAAACGTACGG</td><td>1000</td><td></td><td></td><td></td><td></td><td></td><td></td><td></td><td></td><td></td></tr>
<tr><td>Oligo fwd</td><td>CACCGGCTAGAATATATAAACGTA</td><td></td><td></td><td></td><td></td><td></td><td></td><td></td><td></td><td></td><td></td></tr>
<tr><td>Oligo rev</td><td>AAACTACGTTTATATATTCTAGCC</td><td></td><td></td><td></td><td></td><td></td><td></td><td></td><td></td><td></td><td></td></tr>
<tr><td>Chromosome</td><td>start</td><td>end</td><td>strand</td><td>MM</td><td>target_seq</td><td>PAM</td><td>alignment</td><td>distance</td><td>position</td><td>gene name</td><td>gene id</td></tr>
<tr><td>GU305914.1</td><td>161855</td><td>161877</td><td>-</td><td>0</td><td>GGCTAGAATATATAAACGTA</td><td>CGG</td><td>||||||||[||||||||||||]PAM</td><td>NA</td><td>Intergenic</td><td>NA</td><td>NA</td></tr>
<tr><td>T22147</td><td>AAACGTACGGTGTTACTGAAAGG</td><td>1000</td><td></td><td></td><td></td><td></td><td></td><td></td><td></td><td></td><td></td></tr>
<tr><td>Oligo fwd</td><td>CACCAAACGTACGGTGTTACTGAA</td><td></td><td></td><td></td><td></td><td></td><td></td><td></td><td></td><td></td><td></td></tr>
<tr><td>Oligo rev</td><td>AAACTTCAGTAACACCGTACGTTT</td><td></td><td></td><td></td><td></td><td></td><td></td><td></td><td></td><td></td><td></td></tr>
<tr><td>Chromosome</td><td>start</td><td>end</td><td>strand</td><td>MM</td><td>target_seq</td><td>PAM</td><td>alignment</td><td>distance</td><td>position</td><td>gene name</td><td>gene id</td></tr>
<tr><td>GU305914.1</td><td>161842</td><td>161864</td><td>-</td><td>0</td><td>AAACGTACGGTGTTACTGAA</td><td>AGG</td><td>||||||||[||||||||||||]PAM</td><td>NA</td><td>Intergenic</td><td>NA</td><td>NA</td></tr>
<tr><td>T22148</td><td>CATTATCACACCTGAGCGAACGG</td><td>1000</td><td></td><td></td><td></td><td></td><td></td><td></td><td></td><td></td><td></td></tr>
<tr><td>Oligo fwd</td><td>CACCCATTATCACACCTGAGCGAA</td><td></td><td></td><td></td><td></td><td></td><td></td><td></td><td></td><td></td><td></td></tr>
<tr><td>Oligo rev</td><td>AAACTTCGCTCAGGTGTGATAATG</td><td></td><td></td><td></td><td></td><td></td><td></td><td></td><td></td><td></td><td></td></tr>
<tr><td>Chromosome</td><td>start</td><td>end</td><td>strand</td><td>MM</td><td>target_seq</td><td>PAM</td><td>alignment</td><td>distance</td><td>position</td><td>gene name</td><td>gene id</td></tr>
<tr><td>GU305914.1</td><td>161817</td><td>161839</td><td>-</td><td>0</td><td>CATTATCACACCTGAGCGAA</td><td>CGG</td><td>||||||||[||||||||||||]PAM</td><td>NA</td><td>Intergenic</td><td>NA</td><td>NA</td></tr>
<tr><td>T22149</td><td>ACCTGAGCGAACGGCCGAGCAGG</td><td>1000</td><td></td><td></td><td></td><td></td><td></td><td></td><td></td><td></td><td></td></tr>
<tr><td>Oligo fwd</td><td>CACCACCTGAGCGAACGGCCGAGC</td><td></td><td></td><td></td><td></td><td></td><td></td><td></td><td></td><td></td><td></td></tr>
<tr><td>Oligo rev</td><td>AAACGCTCGGCCGTTCGCTCAGGT</td><td></td><td></td><td></td><td></td><td></td><td></td><td></td><td></td><td></td><td></td></tr>
<tr><td>Chromosome</td><td>start</td><td>end</td><td>strand</td><td>MM</td><td>target_seq</td><td>PAM</td><td>alignment</td><td>distance</td><td>position</td><td>gene name</td><td>gene id</td></tr>
<tr><td>GU305914.1</td><td>161808</td><td>161830</td><td>-</td><td>0</td><td>ACCTGAGCGAACGGCCGAGC</td><td>AGG</td><td>||||||||[||||||||||||]PAM</td><td>NA</td><td>Intergenic</td><td>NA</td><td>NA</td></tr>
<tr><td>T22150</td><td>GCGAACGGCCGAGCAGGCTCAGG</td><td>1000</td><td></td><td></td><td></td><td></td><td></td><td></td><td></td><td></td><td></td></tr>
<tr><td>Oligo fwd</td><td>CACCGCGAACGGCCGAGCAGGCTC</td><td></td><td></td><td></td><td></td><td></td><td></td><td></td><td></td><td></td><td></td></tr>
<tr><td>Oligo rev</td><td>AAACGAGCCTGCTCGGCCGTTCGC</td><td></td><td></td><td></td><td></td><td></td><td></td><td></td><td></td><td></td><td></td></tr>
<tr><td>Chromosome</td><td>start</td><td>end</td><td>strand</td><td>MM</td><td>target_seq</td><td>PAM</td><td>alignment</td><td>distance</td><td>position</td><td>gene name</td><td>gene id</td></tr>
<tr><td>GU305914.1</td><td>161802</td><td>161824</td><td>-</td><td>0</td><td>GCGAACGGCCGAGCAGGCTC</td><td>AGG</td><td>||||||||[||||||||||||]PAM</td><td>NA</td><td>Intergenic</td><td>NA</td><td>NA</td></tr>
<tr><td>T22151</td><td>CGGCCGAGCAGGCTCAGGTGAGG</td><td>1000</td><td></td><td></td><td></td><td></td><td></td><td></td><td></td><td></td><td></td></tr>
<tr><td>Oligo fwd</td><td>CACCCGGCCGAGCAGGCTCAGGTG</td><td></td><td></td><td></td><td></td><td></td><td></td><td></td><td></td><td></td><td></td></tr>
<tr><td>Oligo rev</td><td>AAACCACCTGAGCCTGCTCGGCCG</td><td></td><td></td><td></td><td></td><td></td><td></td><td></td><td></td><td></td><td></td></tr>
<tr><td>Chromosome</td><td>start</td><td>end</td><td>strand</td><td>MM</td><td>target_seq</td><td>PAM</td><td>alignment</td><td>distance</td><td>position</td><td>gene name</td><td>gene id</td></tr>
<tr><td>GU305914.1</td><td>161797</td><td>161819</td><td>-</td><td>0</td><td>CGGCCGAGCAGGCTCAGGTG</td><td>AGG</td><td>||||||||[||||||||||||]PAM</td><td>NA</td><td>Intergenic</td><td>NA</td><td>NA</td></tr>
<tr><td>T22152</td><td>CCGAGCAGGCTCAGGTGAGGCGG</td><td>1000</td><td></td><td></td><td></td><td></td><td></td><td></td><td></td><td></td><td></td></tr>
<tr><td>Oligo fwd</td><td>CACCCCGAGCAGGCTCAGGTGAGG</td><td></td><td></td><td></td><td></td><td></td><td></td><td></td><td></td><td></td><td></td></tr>
<tr><td>Oligo rev</td><td>AAACCCTCACCTGAGCCTGCTCGG</td><td></td><td></td><td></td><td></td><td></td><td></td><td></td><td></td><td></td><td></td></tr>
<tr><td>Chromosome</td><td>start</td><td>end</td><td>strand</td><td>MM</td><td>target_seq</td><td>PAM</td><td>alignment</td><td>distance</td><td>position</td><td>gene name</td><td>gene id</td></tr>
<tr><td>GU305914.1</td><td>161794</td><td>161816</td><td>-</td><td>0</td><td>CCGAGCAGGCTCAGGTGAGG</td><td>CGG</td><td>||||||||[||||||||||||]PAM</td><td>NA</td><td>Intergenic</td><td>NA</td><td>NA</td></tr>
<tr><td>T22153</td><td>AGCAGGCTCAGGTGAGGCGGCGG</td><td>1000</td><td></td><td></td><td></td><td></td><td></td><td></td><td></td><td></td><td></td></tr>
<tr><td>Oligo fwd</td><td>CACCAGCAGGCTCAGGTGAGGCGG</td><td></td><td></td><td></td><td></td><td></td><td></td><td></td><td></td><td></td><td></td></tr>
<tr><td>Oligo rev</td><td>AAACCCGCCTCACCTGAGCCTGCT</td><td></td><td></td><td></td><td></td><td></td><td></td><td></td><td></td><td></td><td></td></tr>
<tr><td>Chromosome</td><td>start</td><td>end</td><td>strand</td><td>MM</td><td>target_seq</td><td>PAM</td><td>alignment</td><td>distance</td><td>position</td><td>gene name</td><td>gene id</td></tr>
<tr><td>GU305914.1</td><td>161791</td><td>161813</td><td>-</td><td>0</td><td>AGCAGGCTCAGGTGAGGCGG</td><td>CGG</td><td>||||||||[||||||||||||]PAM</td><td>NA</td><td>Intergenic</td><td>NA</td><td>NA</td></tr>
<tr><td>T22154</td><td>GAGGCGGCGGACGCATCTCCCGG</td><td>1000</td><td></td><td></td><td></td><td></td><td></td><td></td><td></td><td></td><td></td></tr>
<tr><td>Oligo fwd</td><td>CACCGAGGCGGCGGACGCATCTCC</td><td></td><td></td><td></td><td></td><td></td><td></td><td></td><td></td><td></td><td></td></tr>
<tr><td>Oligo rev</td><td>AAACGGAGATGCGTCCGCCGCCTC</td><td></td><td></td><td></td><td></td><td></td><td></td><td></td><td></td><td></td><td></td></tr>
<tr><td>Chromosome</td><td>start</td><td>end</td><td>strand</td><td>MM</td><td>target_seq</td><td>PAM</td><td>alignment</td><td>distance</td><td>position</td><td>gene name</td><td>gene id</td></tr>
<tr><td>GU305914.1</td><td>161778</td><td>161800</td><td>-</td><td>0</td><td>GAGGCGGCGGACGCATCTCC</td><td>CGG</td><td>||||||||[||||||||||||]PAM</td><td>NA</td><td>Intergenic</td><td>NA</td><td>NA</td></tr>
<tr><td>T22155</td><td>GCGGCGGACGCATCTCCCGGAGG</td><td>1000</td><td></td><td></td><td></td><td></td><td></td><td></td><td></td><td></td><td></td></tr>
</table>

<table>
<tr><td>Oligo fwd</td><td>CACCGCGGCGGACGCATCTCCCGG</td><td></td><td></td><td></td><td></td><td></td><td></td><td></td><td></td><td></td><td></td></tr>
<tr><td>Oligo rev</td><td>AAACCCGGGAGATGCGTCCGCCGC</td><td></td><td></td><td></td><td></td><td></td><td></td><td></td><td></td><td></td><td></td></tr>
<tr><td>Chromosome</td><td>start</td><td>end</td><td>strand</td><td>MM</td><td>target_seq</td><td>PAM</td><td>alignment</td><td>distance</td><td>position</td><td>gene name</td><td>gene id</td></tr>
<tr><td>GU305914.1</td><td>161775</td><td>161797</td><td>-</td><td>0</td><td>GCGGCGGACGCATCTCCCGG</td><td>AGG</td><td>||||||||[||||||||||||]PAM</td><td>NA</td><td>Intergenic</td><td>NA</td><td>NA</td></tr>
<tr><td></td><td></td><td></td><td></td><td></td><td></td><td></td><td></td><td></td><td></td><td></td><td></td></tr>
<tr><td>T22156</td><td>GACGCATCTCCCGGAGGCCGAGG</td><td>1000</td><td></td><td></td><td></td><td></td><td></td><td></td><td></td><td></td><td></td></tr>
<tr><td>Oligo fwd</td><td>CACCGACGCATCTCCCGGAGGCCG</td><td></td><td></td><td></td><td></td><td></td><td></td><td></td><td></td><td></td><td></td></tr>
<tr><td>Oligo rev</td><td>AAACCGGCCTCCGGGAGATGCGTC</td><td></td><td></td><td></td><td></td><td></td><td></td><td></td><td></td><td></td><td></td></tr>
<tr><td>Chromosome</td><td>start</td><td>end</td><td>strand</td><td>MM</td><td>target_seq</td><td>PAM</td><td>alignment</td><td>distance</td><td>position</td><td>gene name</td><td>gene id</td></tr>
<tr><td>GU305914.1</td><td>161769</td><td>161791</td><td>-</td><td>0</td><td>GACGCATCTCCCGGAGGCCG</td><td>AGG</td><td>||||||||[||||||||||||]PAM</td><td>NA</td><td>Intergenic</td><td>NA</td><td>NA</td></tr>
<tr><td></td><td></td><td></td><td></td><td></td><td></td><td></td><td></td><td></td><td></td><td></td><td></td></tr>
<tr><td>T22157</td><td>GCATCTCCCGGAGGCCGAGGAGG</td><td>1000</td><td></td><td></td><td></td><td></td><td></td><td></td><td></td><td></td><td></td></tr>
<tr><td>Oligo fwd</td><td>CACCGCATCTCCCGGAGGCCGAGG</td><td></td><td></td><td></td><td></td><td></td><td></td><td></td><td></td><td></td><td></td></tr>
<tr><td>Oligo rev</td><td>AAACCCTCGGCCTCCGGGAGATGC</td><td></td><td></td><td></td><td></td><td></td><td></td><td></td><td></td><td></td><td></td></tr>
<tr><td>Chromosome</td><td>start</td><td>end</td><td>strand</td><td>MM</td><td>target_seq</td><td>PAM</td><td>alignment</td><td>distance</td><td>position</td><td>gene name</td><td>gene id</td></tr>
<tr><td>GU305914.1</td><td>161766</td><td>161788</td><td>-</td><td>0</td><td>GCATCTCCCGGAGGCCGAGG</td><td>AGG</td><td>||||||||[||||||||||||]PAM</td><td>NA</td><td>Intergenic</td><td>NA</td><td>NA</td></tr>
<tr><td></td><td></td><td></td><td></td><td></td><td></td><td></td><td></td><td></td><td></td><td></td><td></td></tr>
<tr><td>T22158</td><td>TCTCCCGGAGGCCGAGGAGGCGG</td><td>1000</td><td></td><td></td><td></td><td></td><td></td><td></td><td></td><td></td><td></td></tr>
<tr><td>Oligo fwd</td><td>CACCTCTCCCGGAGGCCGAGGAGG</td><td></td><td></td><td></td><td></td><td></td><td></td><td></td><td></td><td></td><td></td></tr>
<tr><td>Oligo rev</td><td>AAACCCTCCTCGGCCTCCGGGAGA</td><td></td><td></td><td></td><td></td><td></td><td></td><td></td><td></td><td></td><td></td></tr>
<tr><td>Chromosome</td><td>start</td><td>end</td><td>strand</td><td>MM</td><td>target_seq</td><td>PAM</td><td>alignment</td><td>distance</td><td>position</td><td>gene name</td><td>gene id</td></tr>
<tr><td>GU305914.1</td><td>161763</td><td>161785</td><td>-</td><td>0</td><td>TCTCCCGGAGGCCGAGGAGG</td><td>CGG</td><td>||||||||[||||||||||||]PAM</td><td>NA</td><td>Intergenic</td><td>NA</td><td>NA</td></tr>
<tr><td></td><td></td><td></td><td></td><td></td><td></td><td></td><td></td><td></td><td></td><td></td><td></td></tr>
<tr><td>T22159</td><td>CCCGGAGGCCGAGGAGGCGGCGG</td><td>1000</td><td></td><td></td><td></td><td></td><td></td><td></td><td></td><td></td><td></td></tr>
<tr><td>Oligo fwd</td><td>CACCCCCGGAGGCCGAGGAGGCGG</td><td></td><td></td><td></td><td></td><td></td><td></td><td></td><td></td><td></td><td></td></tr>
<tr><td>Oligo rev</td><td>AAACCCGCCTCCTCGGCCTCCGGG</td><td></td><td></td><td></td><td></td><td></td><td></td><td></td><td></td><td></td><td></td></tr>
<tr><td>Chromosome</td><td>start</td><td>end</td><td>strand</td><td>MM</td><td>target_seq</td><td>PAM</td><td>alignment</td><td>distance</td><td>position</td><td>gene name</td><td>gene id</td></tr>
<tr><td>GU305914.1</td><td>161760</td><td>161782</td><td>-</td><td>0</td><td>CCCGGAGGCCGAGGAGGCGG</td><td>CGG</td><td>||||||||[||||||||||||]PAM</td><td>NA</td><td>Intergenic</td><td>NA</td><td>NA</td></tr>
<tr><td>GU305914.1</td><td>99534</td><td>99556</td><td>-</td><td>4</td><td>CTCGTCGGGCGAGGAGGCGG</td><td>AGG</td><td>|-||--||[-|||||||||||]PAM</td><td>NA</td><td>Intergenic</td><td>NA</td><td>NA</td></tr>
<tr><td>GU305914.1</td><td>221603</td><td>221625</td><td>-</td><td>3</td><td>ACCGGAGGCCGTGGTGGCGG</td><td>CGG</td><td>-|||||||[|||-||-|||||]PAM</td><td>NA</td><td>Intergenic</td><td>NA</td><td>NA</td></tr>
<tr><td></td><td></td><td></td><td></td><td></td><td></td><td></td><td></td><td></td><td></td><td></td><td></td></tr>
<tr><td>T22160</td><td>GGAGGCCGAGGAGGCGGCGGCGG</td><td>1000</td><td></td><td></td><td></td><td></td><td></td><td></td><td></td><td></td><td></td></tr>
<tr><td>Oligo fwd</td><td>CACCGGAGGCCGAGGAGGCGGCGG</td><td></td><td></td><td></td><td></td><td></td><td></td><td></td><td></td><td></td><td></td></tr>
<tr><td>Oligo rev</td><td>AAACCCGCCGCCTCCTCGGCCTCC</td><td></td><td></td><td></td><td></td><td></td><td></td><td></td><td></td><td></td><td></td></tr>
<tr><td>Chromosome</td><td>start</td><td>end</td><td>strand</td><td>MM</td><td>target_seq</td><td>PAM</td><td>alignment</td><td>distance</td><td>position</td><td>gene name</td><td>gene id</td></tr>
<tr><td>GU305914.1</td><td>161757</td><td>161779</td><td>-</td><td>0</td><td>GGAGGCCGAGGAGGCGGCGG</td><td>CGG</td><td>||||||||[||||||||||||]PAM</td><td>NA</td><td>Intergenic</td><td>NA</td><td>NA</td></tr>
<tr><td>GU305914.1</td><td>81249</td><td>81271</td><td>-</td><td>4</td><td>CGCGATCGAGGAGGCGGCGG</td><td>CGG</td><td>-|-|--||[||||||||||||]PAM</td><td>NA</td><td>Intergenic</td><td>NA</td><td>NA</td></tr>
<tr><td>GU305914.1</td><td>154407</td><td>154429</td><td>+</td><td>4</td><td>GGTGGCGGCGGCGGCGGCGG</td><td>CGG</td><td>||-|||-|[-||-||||||||]PAM</td><td>NA</td><td>Intergenic</td><td>NA</td><td>NA</td></tr>
<tr><td>GU305914.1</td><td>48156</td><td>48178</td><td>+</td><td>4</td><td>GGCGGCGGCGGTGGCGGCGG</td><td>CGG</td><td>||-|||-|[-||-||||||||]PAM</td><td>NA</td><td>Intergenic</td><td>NA</td><td>NA</td></tr>
<tr><td>GU305914.1</td><td>48159</td><td>48181</td><td>+</td><td>4</td><td>GGCGGCGGTGGCGGCGGCGG</td><td>TGG</td><td>||-|||-|[-||-||||||||]PAM</td><td>NA</td><td>Intergenic</td><td>NA</td><td>NA</td></tr>
<tr><td>GU305914.1</td><td>58123</td><td>58145</td><td>-</td><td>4</td><td>GGTGGCGGCGGCGGCGGCGG</td><td>AGG</td><td>||-|||-|[-||-||||||||]PAM</td><td>NA</td><td>Intergenic</td><td>NA</td><td>NA</td></tr>
<tr><td>GU305914.1</td><td>140771</td><td>140793</td><td>+</td><td>4</td><td>GGATGAGGAGGAGGCGGCGT</td><td>CGG</td><td>|||-|--|[|||||||||||-]PAM</td><td>NA</td><td>Intergenic</td><td>NA</td><td>NA</td></tr>
<tr><td></td><td></td><td></td><td></td><td></td><td></td><td></td><td></td><td></td><td></td><td></td><td></td></tr>
<tr><td>T22161</td><td>GGCCGAGGAGGCGGCGGCGGAGG</td><td>1000</td><td></td><td></td><td></td><td></td><td></td><td></td><td></td><td></td><td></td></tr>
<tr><td>Oligo fwd</td><td>CACCGGCCGAGGAGGCGGCGGCGG</td><td></td><td></td><td></td><td></td><td></td><td></td><td></td><td></td><td></td><td></td></tr>
<tr><td>Oligo rev</td><td>AAACCCGCCGCCGCCTCCTCGGCC</td><td></td><td></td><td></td><td></td><td></td><td></td><td></td><td></td><td></td><td></td></tr>
<tr><td>Chromosome</td><td>start</td><td>end</td><td>strand</td><td>MM</td><td>target_seq</td><td>PAM</td><td>alignment</td><td>distance</td><td>position</td><td>gene name</td><td>gene id</td></tr>
<tr><td>GU305914.1</td><td>161754</td><td>161776</td><td>-</td><td>0</td><td>GGCCGAGGAGGCGGCGGCGG</td><td>AGG</td><td>||||||||[||||||||||||]PAM</td><td>NA</td><td>Intergenic</td><td>NA</td><td>NA</td></tr>
<tr><td>GU305914.1</td><td>81246</td><td>81268</td><td>-</td><td>2</td><td>GATCGAGGAGGCGGCGGCGG</td><td>GGG</td><td>|--|||||[||||||||||||]PAM</td><td>NA</td><td>Intergenic</td><td>NA</td><td>NA</td></tr>
<tr><td>GU305914.1</td><td>154404</td><td>154426</td><td>+</td><td>3</td><td>GGCGGTGGCGGCGGCGGCGG</td><td>CGG</td><td>|||-|-||[-|||||||||||]PAM</td><td>NA</td><td>Intergenic</td><td>NA</td><td>NA</td></tr>
<tr><td>GU305914.1</td><td>48159</td><td>48181</td><td>+</td><td>3</td><td>GGCGGCGGTGGCGGCGGCGG</td><td>TGG</td><td>|||-|-||[-|||||||||||]PAM</td><td>NA</td><td>Intergenic</td><td>NA</td><td>NA</td></tr>
<tr><td>GU305914.1</td><td>58126</td><td>58148</td><td>-</td><td>3</td><td>GGCGGTGGCGGCGGCGGCGG</td><td>CGG</td><td>|||-|-||[-|||||||||||]PAM</td><td>NA</td><td>Intergenic</td><td>NA</td><td>NA</td></tr>
<tr><td>GU305914.1</td><td>154407</td><td>154429</td><td>+</td><td>4</td><td>GGTGGCGGCGGCGGCGGCGG</td><td>CGG</td><td>||--|-||[-|||||||||||]PAM</td><td>NA</td><td>Intergenic</td><td>NA</td><td>NA</td></tr>
<tr><td>GU305914.1</td><td>67742</td><td>67764</td><td>+</td><td>4</td><td>GGGAGCGGCGGCGGCGGCGG</td><td>CGG</td><td>||--|-||[-|||||||||||]PAM</td><td>NA</td><td>Intergenic</td><td>NA</td><td>NA</td></tr>
<tr><td>GU305914.1</td><td>58123</td><td>58145</td><td>-</td><td>4</td><td>GGTGGCGGCGGCGGCGGCGG</td><td>AGG</td><td>||--|-||[-|||||||||||]PAM</td><td>NA</td><td>Intergenic</td><td>NA</td><td>NA</td></tr>
<tr><td>GU305914.1</td><td>149482</td><td>149504</td><td>-</td><td>4</td><td>GGCGACGGCGGCGGCGGCGG</td><td>CGG</td><td>|||---||[-|||||||||||]PAM</td><td>NA</td><td>Intergenic</td><td>NA</td><td>NA</td></tr>
<tr><td>GU305914.1</td><td>56773</td><td>56795</td><td>+</td><td>4</td><td>CGCCGCAGCGGCGGCGGCGG</td><td>CGG</td><td>-||||--|[-|||||||||||]PAM</td><td>NA</td><td>Intergenic</td><td>NA</td><td>NA</td></tr>
<tr><td>GU305914.1</td><td>48156</td><td>48178</td><td>+</td><td>4</td><td>GGCGGCGGCGGTGGCGGCGG</td><td>CGG</td><td>|||-|-||[-||-||||||||]PAM</td><td>NA</td><td>Intergenic</td><td>NA</td><td>NA</td></tr>
<tr><td>GU305914.1</td><td>62446</td><td>62468</td><td>+</td><td>4</td><td>GGCGGGGGTGGTGGCGGCGG</td><td>GGG</td><td>|||-|-||[-||-||||||||]PAM</td><td>NA</td><td>Intergenic</td><td>NA</td><td>NA</td></tr>
<tr><td>GU305914.1</td><td>114658</td><td>114680</td><td>-</td><td>4</td><td>GGCGGTGGTGGTGGCGGCGG</td><td>CGG</td><td>|||-|-||[-||-||||||||]PAM</td><td>NA</td><td>Intergenic</td><td>NA</td><td>NA</td></tr>
<tr><td>GU305914.1</td><td>168817</td><td>168839</td><td>+</td><td>4</td><td>GGCGGTGGTGGCAGCGGCGG</td><td>CGG</td><td>|||-|-||[-|||-|||||||]PAM</td><td>NA</td><td>Intergenic</td><td>NA</td><td>NA</td></tr>
<tr><td>GU305914.1</td><td>18653</td><td>18675</td><td>-</td><td>3</td><td>GGCAGAGGTGGCGGAGGCGG</td><td>GGG</td><td>|||-||||[-|||||-|||||]PAM</td><td>NA</td><td>Intergenic</td><td>NA</td><td>NA</td></tr>
<tr><td>GU305914.1</td><td>98352</td><td>98374</td><td>-</td><td>4</td><td>AGCCTAGGCGGCGGTGGCGG</td><td>CGG</td><td>-|||-|||[-|||||-|||||]PAM</td><td>NA</td><td>Intergenic</td><td>NA</td><td>NA</td></tr>
<tr><td>GU305914.1</td><td>86784</td><td>86806</td><td>-</td><td>4</td><td>GGAAGAGGGGGCGGGGGCGG</td><td>CGG</td><td>||--||||[-|||||-|||||]PAM</td><td>NA</td><td>Intergenic</td><td>NA</td><td>NA</td></tr>
<tr><td>GU305914.1</td><td>30006</td><td>30028</td><td>+</td><td>4</td><td>GGCGGCGGCGGCGGCGGCAG</td><td>TGG</td><td>|||-|-||[-|||||||||-|]PAM</td><td>NA</td><td>Intergenic</td><td>NA</td><td>NA</td></tr>
<tr><td>GU305914.1</td><td>149476</td><td>149498</td><td>-</td><td>4</td><td>GGCGGCGGCGGCGGCGGCAG</td><td>CGG</td><td>|||-|-||[-|||||||||-|]PAM</td><td>NA</td><td>Intergenic</td><td>NA</td><td>NA</td></tr>
<tr><td>GU305914.1</td><td>140771</td><td>140793</td><td>+</td><td>4</td><td>GGATGAGGAGGAGGCGGCGT</td><td>CGG</td><td>||--||||[|||-|||||||-]PAM</td><td>NA</td><td>Intergenic</td><td>NA</td><td>NA</td></tr>
<tr><td></td><td></td><td></td><td></td><td></td><td></td><td></td><td></td><td></td><td></td><td></td><td></td></tr>
<tr><td>T22162</td><td>GGAGGCGGCGGCGGAGGTGACGG</td><td>1000</td><td></td><td></td><td></td><td></td><td></td><td></td><td></td><td></td><td></td></tr>
<tr><td>Oligo fwd</td><td>CACCGGAGGCGGCGGCGGAGGTGA</td><td></td><td></td><td></td><td></td><td></td><td></td><td></td><td></td><td></td><td></td></tr>
<tr><td>Oligo rev</td><td>AAACTCACCTCCGCCGCCGCCTCC</td><td></td><td></td><td></td><td></td><td></td><td></td><td></td><td></td><td></td><td></td></tr>
<tr><td>Chromosome</td><td>start</td><td>end</td><td>strand</td><td>MM</td><td>target_seq</td><td>PAM</td><td>alignment</td><td>distance</td><td>position</td><td>gene name</td><td>gene id</td></tr>
<tr><td>GU305914.1</td><td>161748</td><td>161770</td><td>-</td><td>0</td><td>GGAGGCGGCGGCGGAGGTGA</td><td>CGG</td><td>||||||||[||||||||||||]PAM</td><td>NA</td><td>Intergenic</td><td>NA</td><td>NA</td></tr>
<tr><td>GU305914.1</td><td>19436</td><td>19458</td><td>+</td><td>4</td><td>GCCGGCGTCGGCGGAGGTGC</td><td>CGG</td><td>|--||||-[|||||||||||-]PAM</td><td>NA</td><td>Intergenic</td><td>NA</td><td>NA</td></tr>
<tr><td>GU305914.1</td><td>62452</td><td>62474</td><td>+</td><td>4</td><td>GGTGGTGGCGGCGGGGGTGG</td><td>CGG</td><td>||-||-||[||||||-||||-]PAM</td><td>NA</td><td>Intergenic</td><td>NA</td><td>NA</td></tr>
<tr><td>GU305914.1</td><td>62437</td><td>62459</td><td>+</td><td>4</td><td>GGCGGTGGCGGCGGGGGTGG</td><td>TGG</td><td>||-||-||[||||||-||||-]PAM</td><td>NA</td><td>Intergenic</td><td>NA</td><td>NA</td></tr>
<tr><td></td><td></td><td></td><td></td><td></td><td></td><td></td><td></td><td></td><td></td><td></td><td></td></tr>
<tr><td>T22163</td><td>CGGAGGTGACGGCGTCCCGAAGG</td><td>1000</td><td></td><td></td><td></td><td></td><td></td><td></td><td></td><td></td><td></td></tr>
<tr><td>Oligo fwd</td><td>CACCCGGAGGTGACGGCGTCCCGA</td><td></td><td></td><td></td><td></td><td></td><td></td><td></td><td></td><td></td><td></td></tr>
<tr><td>Oligo rev</td><td>AAACTCGGGACGCCGTCACCTCCG</td><td></td><td></td><td></td><td></td><td></td><td></td><td></td><td></td><td></td><td></td></tr>
<tr><td>Chromosome</td><td>start</td><td>end</td><td>strand</td><td>MM</td><td>target_seq</td><td>PAM</td><td>alignment</td><td>distance</td><td>position</td><td>gene name</td><td>gene id</td></tr>
<tr><td>GU305914.1</td><td>161737</td><td>161759</td><td>-</td><td>0</td><td>CGGAGGTGACGGCGTCCCGA</td><td>AGG</td><td>||||||||[||||||||||||]PAM</td><td>NA</td><td>Intergenic</td><td>NA</td><td>NA</td></tr>
<tr><td></td><td></td><td></td><td></td><td></td><td></td><td></td><td></td><td></td><td></td><td></td><td></td></tr>
<tr><td>T22164</td><td>GTGACGGCGTCCCGAAGGCCCGG</td><td>1000</td><td></td><td></td><td></td><td></td><td></td><td></td><td></td><td></td><td></td></tr>
<tr><td>Oligo fwd</td><td>CACCGTGACGGCGTCCCGAAGGCC</td><td></td><td></td><td></td><td></td><td></td><td></td><td></td><td></td><td></td><td></td></tr>
<tr><td>Oligo rev</td><td>AAACGGCCTTCGGGACGCCGTCAC</td><td></td><td></td><td></td><td></td><td></td><td></td><td></td><td></td><td></td><td></td></tr>
<tr><td>Chromosome</td><td>start</td><td>end</td><td>strand</td><td>MM</td><td>target_seq</td><td>PAM</td><td>alignment</td><td>distance</td><td>position</td><td>gene name</td><td>gene id</td></tr>
<tr><td>GU305914.1</td><td>161732</td><td>161754</td><td>-</td><td>0</td><td>GTGACGGCGTCCCGAAGGCC</td><td>CGG</td><td>||||||||[||||||||||||]PAM</td><td>NA</td><td>Intergenic</td><td>NA</td><td>NA</td></tr>
</table>

| | | | | | | | | | | | |
|---|---|---|---|---|---|---|---|---|---|---|---|
| T22165 | CCCGAAGGCCCGGAGCTCTCAGG | | 1000 | | | | | | | | |
| Oligo fwd | CACCCCCGAAGGCCCGGAGCTCTC | | | | | | | | | | |
| Oligo rev | AAACGAGAGCTCCGGGCCTTCGGG | | | | | | | | | | |
| Chromosome | start | | end | strand | MM | target_seq | PAM | alignment | distance | position | gene name | gene id |
| GU305914.1 | | 161722 | 161744 | - | 0 | CCCGAAGGCCCGGAGCTCTC | AGG | \|\|\|\|\|\|\|\|[\|\|\|\|\|\|\|\|\|\|\|\|]PAM | NA | Intergenic | NA | NA |

| | | | | | | | | | | | | |
|---|---|---|---|---|---|---|---|---|---|---|---|---|
| T22166 | AAGGCCCGGAGCTCTCAGGTAGG | | 1000 | | | | | | | | | |
| Oligo fwd | CACCAAGGCCCGGAGCTCTCAGGT | | | | | | | | | | | |
| Oligo rev | AAACACCTGAGAGCTCCGGGCCTT | | | | | | | | | | | |
| Chromosome | start | | end | strand | MM | target_seq | PAM | alignment | distance | position | gene name | gene id |
| GU305914.1 | | 161718 | 161740 | - | 0 | AAGGCCCGGAGCTCTCAGGT | AGG | \|\|\|\|\|\|\|\|[\|\|\|\|\|\|\|\|\|\|\|\|]PAM | NA | Intergenic | NA | NA |

| | | | | | | | | | | | | |
|---|---|---|---|---|---|---|---|---|---|---|---|---|
| T22167 | TCTTTTATCGTTCCTTTAGATGG | | 1000 | | | | | | | | | |
| Oligo fwd | CACCTCTTTTATCGTTCCTTTAGA | | | | | | | | | | | |
| Oligo rev | AAACTCTAAAGGAACGATAAAAGA | | | | | | | | | | | |
| Chromosome | start | | end | strand | MM | target_seq | PAM | alignment | distance | position | gene name | gene id |
| GU305914.1 | | 161655 | 161677 | - | 0 | TCTTTTATCGTTCCTTTAGA | TGG | \|\|\|\|\|\|\|\|[\|\|\|\|\|\|\|\|\|\|\|\|]PAM | NA | Intergenic | NA | NA |

| | | | | | | | | | | | | |
|---|---|---|---|---|---|---|---|---|---|---|---|---|
| T22168 | TACTTTTTTTTCTATGTTAGAGG | | 1000 | | | | | | | | | |
| Oligo fwd | CACCTACTTTTTTTTCTATGTTAG | | | | | | | | | | | |
| Oligo rev | AAACCTAACATAGAAAAAAAAGTA | | | | | | | | | | | |
| Chromosome | start | | end | strand | MM | target_seq | PAM | alignment | distance | position | gene name | gene id |
| GU305914.1 | | 161605 | 161627 | - | 0 | TACTTTTTTTTCTATGTTAG | AGG | \|\|\|\|\|\|\|\|[\|\|\|\|\|\|\|\|\|\|\|\|]PAM | NA | Intergenic | NA | NA |

| | | | | | | | | | | | | |
|---|---|---|---|---|---|---|---|---|---|---|---|---|
| T22169 | AAAAGACTTTTTTTTTCACTCGG | | 1000 | | | | | | | | | |
| Oligo fwd | CACCAAAAGACTTTTTTTTTCACT | | | | | | | | | | | |
| Oligo rev | AAACAGTGAAAAAAAAAGTCTTTT | | | | | | | | | | | |
| Chromosome | start | | end | strand | MM | target_seq | PAM | alignment | distance | position | gene name | gene id |
| GU305914.1 | | 161558 | 161580 | - | 0 | AAAAGACTTTTTTTTTCACT | CGG | \|\|\|\|\|\|\|\|[\|\|\|\|\|\|\|\|\|\|\|\|]PAM | NA | Intergenic | NA | NA |

| | | | | | | | | | | | | |
|---|---|---|---|---|---|---|---|---|---|---|---|---|
| T22170 | CTTTTTTTTTCACTCGGCCTCGG | | 1000 | | | | | | | | | |
| Oligo fwd | CACCCTTTTTTTTTCACTCGGCCT | | | | | | | | | | | |
| Oligo rev | AAACAGGCCGAGTGAAAAAAAAAG | | | | | | | | | | | |
| Chromosome | start | | end | strand | MM | target_seq | PAM | alignment | distance | position | gene name | gene id |
| GU305914.1 | | 161552 | 161574 | - | 0 | CTTTTTTTTTCACTCGGCCT | CGG | \|\|\|\|\|\|\|\|[\|\|\|\|\|\|\|\|\|\|\|\|]PAM | NA | Intergenic | NA | NA |
| GU305914.1 | | 161514 | 161536 | - | 4 | ATTTTTTCTCCAATCGGCCT | AGG | -\|\|\|\|\|\|-[\|-\|\|-\|\|\|\|\|\|\|]PAM | NA | Intergenic | NA | NA |

| | | | | | | | | | | | | |
|---|---|---|---|---|---|---|---|---|---|---|---|---|
| T22171 | GATCTCATTTTTTCTCCAATCGG | | 1000 | | | | | | | | | |
| Oligo fwd | CACCGATCTCATTTTTTCTCCAAT | | | | | | | | | | | |
| Oligo rev | AAACATTGGAGAAAAAATGAGATC | | | | | | | | | | | |
| Chromosome | start | | end | strand | MM | target_seq | PAM | alignment | distance | position | gene name | gene id |
| GU305914.1 | | 161520 | 161542 | - | 0 | GATCTCATTTTTTCTCCAAT | CGG | \|\|\|\|\|\|\|\|[\|\|\|\|\|\|\|\|\|\|\|\|]PAM | NA | Intergenic | NA | NA |

| | | | | | | | | | | | | |
|---|---|---|---|---|---|---|---|---|---|---|---|---|
| T22172 | ATTTTTTCTCCAATCGGCCTAGG | | 1000 | | | | | | | | | |
| Oligo fwd | CACCATTTTTTCTCCAATCGGCCT | | | | | | | | | | | |
| Oligo rev | AAACAGGCCGATTGGAGAAAAAAT | | | | | | | | | | | |
| Chromosome | start | | end | strand | MM | target_seq | PAM | alignment | distance | position | gene name | gene id |
| GU305914.1 | | 161514 | 161536 | - | 0 | ATTTTTTCTCCAATCGGCCT | AGG | \|\|\|\|\|\|\|\|[\|\|\|\|\|\|\|\|\|\|\|\|]PAM | NA | Intergenic | NA | NA |
| GU305914.1 | | 161552 | 161574 | - | 4 | CTTTTTTTTTCACTCGGCCT | CGG | -\|\|\|\|\|\|-[\|-\|\|-\|\|\|\|\|\|\|]PAM | NA | Intergenic | NA | NA |

| | | | | | | | | | | | | |
|---|---|---|---|---|---|---|---|---|---|---|---|---|
| T22173 | TCCAATCGGCCTAGGAATCCTGG | | 1000 | | | | | | | | | |
| Oligo fwd | CACCTCCAATCGGCCTAGGAATCC | | | | | | | | | | | |
| Oligo rev | AAACGGATTCCTAGGCCGATTGGA | | | | | | | | | | | |
| Chromosome | start | | end | strand | MM | target_seq | PAM | alignment | distance | position | gene name | gene id |
| GU305914.1 | | 161506 | 161528 | - | 0 | TCCAATCGGCCTAGGAATCC | TGG | \|\|\|\|\|\|\|\|[\|\|\|\|\|\|\|\|\|\|\|\|]PAM | NA | Intergenic | NA | NA |

| | | | | | | | | | | | | |
|---|---|---|---|---|---|---|---|---|---|---|---|---|
| T22174 | TCGGCCTAGGAATCCTGGCTAGG | | 1000 | | | | | | | | | |
| Oligo fwd | CACCTCGGCCTAGGAATCCTGGCT | | | | | | | | | | | |
| Oligo rev | AAACAGCCAGGATTCCTAGGCCGA | | | | | | | | | | | |
| Chromosome | start | | end | strand | MM | target_seq | PAM | alignment | distance | position | gene name | gene id |
| GU305914.1 | | 161501 | 161523 | - | 0 | TCGGCCTAGGAATCCTGGCT | AGG | \|\|\|\|\|\|\|\|[\|\|\|\|\|\|\|\|\|\|\|\|]PAM | NA | Intergenic | NA | NA |

| | | | | | | | | | | | | |
|---|---|---|---|---|---|---|---|---|---|---|---|---|
| T22175 | CAAACATGAAACTAGATACTAGG | | 1000 | | | | | | | | | |
| Oligo fwd | CACCCAAACATGAAACTAGATACT | | | | | | | | | | | |
| Oligo rev | AAACAGTATCTAGTTTCATGTTTG | | | | | | | | | | | |
| Chromosome | start | | end | strand | MM | target_seq | PAM | alignment | distance | position | gene name | gene id |
| GU305914.1 | | 161366 | 161388 | - | 0 | CAAACATGAAACTAGATACT | AGG | \|\|\|\|\|\|\|\|[\|\|\|\|\|\|\|\|\|\|\|\|]PAM | NA | Intergenic | NA | NA |

| | | | | | | | | | | | | |
|---|---|---|---|---|---|---|---|---|---|---|---|---|
| T22176 | TGCCGATAGATAGCGTCGACCGG | | 1000 | | | | | | | | | |
| Oligo fwd | CACCTGCCGATAGATAGCGTCGAC | | | | | | | | | | | |
| Oligo rev | AAACGTCGACGCTATCTATCGGCA | | | | | | | | | | | |
| Chromosome | start | | end | strand | MM | target_seq | PAM | alignment | distance | position | gene name | gene id |
| GU305914.1 | | 161307 | 161329 | - | 0 | TGCCGATAGATAGCGTCGAC | CGG | \|\|\|\|\|\|\|\|[\|\|\|\|\|\|\|\|\|\|\|\|]PAM | NA | Intergenic | NA | NA |

| | | | | | | | | | | | | |
|---|---|---|---|---|---|---|---|---|---|---|---|---|
| T22177 | GATCAAACAAAAACACATCACGG | | 1000 | | | | | | | | | |
| Oligo fwd | CACCGATCAAACAAAAACACATCA | | | | | | | | | | | |
| Oligo rev | AAACTGATGTGTTTTTGTTTGATC | | | | | | | | | | | |
| Chromosome | start | | end | strand | MM | target_seq | PAM | alignment | distance | position | gene name | gene id |
| GU305914.1 | | 161230 | 161252 | - | 0 | GATCAAACAAAAACACATCA | CGG | \|\|\|\|\|\|\|\|[\|\|\|\|\|\|\|\|\|\|\|\|]PAM | NA | Intergenic | NA | NA |

| | | | | | | | | | | | | |
|---|---|---|---|---|---|---|---|---|---|---|---|---|
| T22178 | AAACAAAAACACATCACGGCAGG | | 1000 | | | | | | | | | |
| Oligo fwd | CACCAAACAAAAACACATCACGGC | | | | | | | | | | | |
| Oligo rev | AAACGCCGTGATGTGTTTTTGTTT | | | | | | | | | | | |
| Chromosome | start | | end | strand | MM | target_seq | PAM | alignment | distance | position | gene name | gene id |
| GU305914.1 | | 161226 | 161248 | - | 0 | AAACAAAAACACATCACGGC | AGG | \|\|\|\|\|\|\|\|[\|\|\|\|\|\|\|\|\|\|\|\|]PAM | NA | Intergenic | NA | NA |

| | | | |
|---|---|---|---|
| T22179 | AACAAAAACACATCACGGCAGGG | | 1000 |
| Oligo fwd | CACCAACAAAAACACATCACGGCA | | |
| Oligo rev | AAACTGCCGTGATGTGTTTTTGTT | | |

<table>
<tr><td>Chromosome</td><td>start</td><td>end</td><td>strand</td><td>MM</td><td>target_seq</td><td>PAM</td><td>alignment</td><td>distance</td><td>position</td><td>gene name</td><td>gene id</td></tr>
<tr><td>GU305914.1</td><td>161225</td><td>161247</td><td>-</td><td>0</td><td>AACAAAAACACATCACGGCA</td><td>GGG</td><td>||||||||[||||||||||||]PAM</td><td>NA</td><td>Intergenic</td><td>NA</td><td>NA</td></tr>
<tr><td>T22180</td><td>CTCACTATCAACACACATTTGG</td><td>1000</td><td></td><td></td><td></td><td></td><td></td><td></td><td></td><td></td><td></td></tr>
<tr><td>Oligo fwd</td><td>CACCCTCACTATCAACACACATTT</td><td></td><td></td><td></td><td></td><td></td><td></td><td></td><td></td><td></td><td></td></tr>
<tr><td>Oligo rev</td><td>AAACAAATGTGTGTTGATAGTGAG</td><td></td><td></td><td></td><td></td><td></td><td></td><td></td><td></td><td></td><td></td></tr>
<tr><td>Chromosome</td><td>start</td><td>end</td><td>strand</td><td>MM</td><td>target_seq</td><td>PAM</td><td>alignment</td><td>distance</td><td>position</td><td>gene name</td><td>gene id</td></tr>
<tr><td>GU305914.1</td><td>161165</td><td>161187</td><td>-</td><td>0</td><td>CTCACTATCAACACACATTT</td><td>TGG</td><td>||||||||[||||||||||||]PAM</td><td>NA</td><td>Intergenic</td><td>NA</td><td>NA</td></tr>
<tr><td>T22181</td><td>CACTAAACTGTATGTAAACTCGG</td><td>1000</td><td></td><td></td><td></td><td></td><td></td><td></td><td></td><td></td><td></td></tr>
<tr><td>Oligo fwd</td><td>CACCCACTAAACTGTATGTAAACT</td><td></td><td></td><td></td><td></td><td></td><td></td><td></td><td></td><td></td><td></td></tr>
<tr><td>Oligo rev</td><td>AAACAGTTTACATACAGTTTAGTG</td><td></td><td></td><td></td><td></td><td></td><td></td><td></td><td></td><td></td><td></td></tr>
<tr><td>Chromosome</td><td>start</td><td>end</td><td>strand</td><td>MM</td><td>target_seq</td><td>PAM</td><td>alignment</td><td>distance</td><td>position</td><td>gene name</td><td>gene id</td></tr>
<tr><td>GU305914.1</td><td>161119</td><td>161141</td><td>-</td><td>0</td><td>CACTAAACTGTATGTAAACT</td><td>CGG</td><td>||||||||[||||||||||||]PAM</td><td>NA</td><td>Intergenic</td><td>NA</td><td>NA</td></tr>
<tr><td>T22182</td><td>ACTAAACTGTATGTAAACTCGGG</td><td>1000</td><td></td><td></td><td></td><td></td><td></td><td></td><td></td><td></td><td></td></tr>
<tr><td>Oligo fwd</td><td>CACCACTAAACTGTATGTAAACTC</td><td></td><td></td><td></td><td></td><td></td><td></td><td></td><td></td><td></td><td></td></tr>
<tr><td>Oligo rev</td><td>AAACGAGTTTACATACAGTTTAGT</td><td></td><td></td><td></td><td></td><td></td><td></td><td></td><td></td><td></td><td></td></tr>
<tr><td>Chromosome</td><td>start</td><td>end</td><td>strand</td><td>MM</td><td>target_seq</td><td>PAM</td><td>alignment</td><td>distance</td><td>position</td><td>gene name</td><td>gene id</td></tr>
<tr><td>GU305914.1</td><td>161118</td><td>161140</td><td>-</td><td>0</td><td>ACTAAACTGTATGTAAACTC</td><td>GGG</td><td>||||||||[||||||||||||]PAM</td><td>NA</td><td>Intergenic</td><td>NA</td><td>NA</td></tr>
<tr><td>T22183</td><td>CACAAAACAACAAACACATCCGG</td><td>1000</td><td></td><td></td><td></td><td></td><td></td><td></td><td></td><td></td><td></td></tr>
<tr><td>Oligo fwd</td><td>CACCCACAAAACAACAAACACATC</td><td></td><td></td><td></td><td></td><td></td><td></td><td></td><td></td><td></td><td></td></tr>
<tr><td>Oligo rev</td><td>AAACGATGTGTTTGTTGTTTTGTG</td><td></td><td></td><td></td><td></td><td></td><td></td><td></td><td></td><td></td><td></td></tr>
<tr><td>Chromosome</td><td>start</td><td>end</td><td>strand</td><td>MM</td><td>target_seq</td><td>PAM</td><td>alignment</td><td>distance</td><td>position</td><td>gene name</td><td>gene id</td></tr>
<tr><td>GU305914.1</td><td>161051</td><td>161073</td><td>-</td><td>0</td><td>CACAAAACAACAAACACATC</td><td>CGG</td><td>||||||||[||||||||||||]PAM</td><td>NA</td><td>Intergenic</td><td>NA</td><td>NA</td></tr>
<tr><td>T22184</td><td>ACAAAACAACAAACACATCCGGG</td><td>1000</td><td></td><td></td><td></td><td></td><td></td><td></td><td></td><td></td><td></td></tr>
<tr><td>Oligo fwd</td><td>CACCACAAAACAACAAACACATCC</td><td></td><td></td><td></td><td></td><td></td><td></td><td></td><td></td><td></td><td></td></tr>
<tr><td>Oligo rev</td><td>AAACGGATGTGTTTGTTGTTTTGT</td><td></td><td></td><td></td><td></td><td></td><td></td><td></td><td></td><td></td><td></td></tr>
<tr><td>Chromosome</td><td>start</td><td>end</td><td>strand</td><td>MM</td><td>target_seq</td><td>PAM</td><td>alignment</td><td>distance</td><td>position</td><td>gene name</td><td>gene id</td></tr>
<tr><td>GU305914.1</td><td>161050</td><td>161072</td><td>-</td><td>0</td><td>ACAAAACAACAAACACATCC</td><td>GGG</td><td>||||||||[||||||||||||]PAM</td><td>NA</td><td>Intergenic</td><td>NA</td><td>NA</td></tr>
<tr><td>T22185</td><td>CAAAACAACAAACACATCCGGGG</td><td>1000</td><td></td><td></td><td></td><td></td><td></td><td></td><td></td><td></td><td></td></tr>
<tr><td>Oligo fwd</td><td>CACCCAAAACAACAAACACATCCG</td><td></td><td></td><td></td><td></td><td></td><td></td><td></td><td></td><td></td><td></td></tr>
<tr><td>Oligo rev</td><td>AAACCGGATGTGTTTGTTGTTTTG</td><td></td><td></td><td></td><td></td><td></td><td></td><td></td><td></td><td></td><td></td></tr>
<tr><td>Chromosome</td><td>start</td><td>end</td><td>strand</td><td>MM</td><td>target_seq</td><td>PAM</td><td>alignment</td><td>distance</td><td>position</td><td>gene name</td><td>gene id</td></tr>
<tr><td>GU305914.1</td><td>161049</td><td>161071</td><td>-</td><td>0</td><td>CAAAACAACAAACACATCCG</td><td>GGG</td><td>||||||||[||||||||||||]PAM</td><td>NA</td><td>Intergenic</td><td>NA</td><td>NA</td></tr>
<tr><td>T22186</td><td>ACACAGAGCATCGTGTTCGTCGG</td><td>1000</td><td></td><td></td><td></td><td></td><td></td><td></td><td></td><td></td><td></td></tr>
<tr><td>Oligo fwd</td><td>CACCACACAGAGCATCGTGTTCGT</td><td></td><td></td><td></td><td></td><td></td><td></td><td></td><td></td><td></td><td></td></tr>
<tr><td>Oligo rev</td><td>AAACACGAACACGATGCTCTGTGT</td><td></td><td></td><td></td><td></td><td></td><td></td><td></td><td></td><td></td><td></td></tr>
<tr><td>Chromosome</td><td>start</td><td>end</td><td>strand</td><td>MM</td><td>target_seq</td><td>PAM</td><td>alignment</td><td>distance</td><td>position</td><td>gene name</td><td>gene id</td></tr>
<tr><td>GU305914.1</td><td>160967</td><td>160989</td><td>-</td><td>0</td><td>ACACAGAGCATCGTGTTCGT</td><td>CGG</td><td>||||||||[||||||||||||]PAM</td><td>NA</td><td>Intergenic</td><td>NA</td><td>NA</td></tr>
<tr><td>GU305914.1</td><td>196567</td><td>196589</td><td>+</td><td>4</td><td>ACACAGCCACTCGTGTTCGT</td><td>TGG</td><td>||||||--[--||||||||||]PAM</td><td>NA</td><td>Intergenic</td><td>NA</td><td>NA</td></tr>
<tr><td>T22187</td><td>CAGAGCATCGTGTTCGTCGGCGG</td><td>1000</td><td></td><td></td><td></td><td></td><td></td><td></td><td></td><td></td><td></td></tr>
<tr><td>Oligo fwd</td><td>CACCCAGAGCATCGTGTTCGTCGG</td><td></td><td></td><td></td><td></td><td></td><td></td><td></td><td></td><td></td><td></td></tr>
<tr><td>Oligo rev</td><td>AAACCCGACGAACACGATGCTCTG</td><td></td><td></td><td></td><td></td><td></td><td></td><td></td><td></td><td></td><td></td></tr>
<tr><td>Chromosome</td><td>start</td><td>end</td><td>strand</td><td>MM</td><td>target_seq</td><td>PAM</td><td>alignment</td><td>distance</td><td>position</td><td>gene name</td><td>gene id</td></tr>
<tr><td>GU305914.1</td><td>160964</td><td>160986</td><td>-</td><td>0</td><td>CAGAGCATCGTGTTCGTCGG</td><td>CGG</td><td>||||||||[||||||||||||]PAM</td><td>NA</td><td>Intergenic</td><td>NA</td><td>NA</td></tr>
<tr><td>T22188</td><td>AGAGCATCGTGTTCGTCGGCGGG</td><td>1000</td><td></td><td></td><td></td><td></td><td></td><td></td><td></td><td></td><td></td></tr>
<tr><td>Oligo fwd</td><td>CACCAGAGCATCGTGTTCGTCGGC</td><td></td><td></td><td></td><td></td><td></td><td></td><td></td><td></td><td></td><td></td></tr>
<tr><td>Oligo rev</td><td>AAACGCCGACGAACACGATGCTCT</td><td></td><td></td><td></td><td></td><td></td><td></td><td></td><td></td><td></td><td></td></tr>
<tr><td>Chromosome</td><td>start</td><td>end</td><td>strand</td><td>MM</td><td>target_seq</td><td>PAM</td><td>alignment</td><td>distance</td><td>position</td><td>gene name</td><td>gene id</td></tr>
<tr><td>GU305914.1</td><td>160963</td><td>160985</td><td>-</td><td>0</td><td>AGAGCATCGTGTTCGTCGGC</td><td>GGG</td><td>||||||||[||||||||||||]PAM</td><td>NA</td><td>Intergenic</td><td>NA</td><td>NA</td></tr>
<tr><td>T22189</td><td>CGTCGGCGGGAAACCCGTCTTGG</td><td>1000</td><td></td><td></td><td></td><td></td><td></td><td></td><td></td><td></td><td></td></tr>
<tr><td>Oligo fwd</td><td>CACCCGTCGGCGGGAAACCCGTCT</td><td></td><td></td><td></td><td></td><td></td><td></td><td></td><td></td><td></td><td></td></tr>
<tr><td>Oligo rev</td><td>AAACAGACGGGTTTCCCGCCGACG</td><td></td><td></td><td></td><td></td><td></td><td></td><td></td><td></td><td></td><td></td></tr>
<tr><td>Chromosome</td><td>start</td><td>end</td><td>strand</td><td>MM</td><td>target_seq</td><td>PAM</td><td>alignment</td><td>distance</td><td>position</td><td>gene name</td><td>gene id</td></tr>
<tr><td>GU305914.1</td><td>160950</td><td>160972</td><td>-</td><td>0</td><td>CGTCGGCGGGAAACCCGTCT</td><td>TGG</td><td>||||||||[||||||||||||]PAM</td><td>NA</td><td>Intergenic</td><td>NA</td><td>NA</td></tr>
<tr><td>T22190</td><td>GTCGGCGGGAAACCCGTCTTGGG</td><td>1000</td><td></td><td></td><td></td><td></td><td></td><td></td><td></td><td></td><td></td></tr>
<tr><td>Oligo fwd</td><td>CACCGTCGGCGGGAAACCCGTCTT</td><td></td><td></td><td></td><td></td><td></td><td></td><td></td><td></td><td></td><td></td></tr>
<tr><td>Oligo rev</td><td>AAACAAGACGGGTTTCCCGCCGAC</td><td></td><td></td><td></td><td></td><td></td><td></td><td></td><td></td><td></td><td></td></tr>
<tr><td>Chromosome</td><td>start</td><td>end</td><td>strand</td><td>MM</td><td>target_seq</td><td>PAM</td><td>alignment</td><td>distance</td><td>position</td><td>gene name</td><td>gene id</td></tr>
<tr><td>GU305914.1</td><td>160949</td><td>160971</td><td>-</td><td>0</td><td>GTCGGCGGGAAACCCGTCTT</td><td>GGG</td><td>||||||||[||||||||||||]PAM</td><td>NA</td><td>Intergenic</td><td>NA</td><td>NA</td></tr>
<tr><td>T22191</td><td>AGTTGTTACCCACCATTCCCAGG</td><td>1000</td><td></td><td></td><td></td><td></td><td></td><td></td><td></td><td></td><td></td></tr>
<tr><td>Oligo fwd</td><td>CACCAGTTGTTACCCACCATTCCC</td><td></td><td></td><td></td><td></td><td></td><td></td><td></td><td></td><td></td><td></td></tr>
<tr><td>Oligo rev</td><td>AAACGGGAATGGTGGGTAACAACT</td><td></td><td></td><td></td><td></td><td></td><td></td><td></td><td></td><td></td><td></td></tr>
<tr><td>Chromosome</td><td>start</td><td>end</td><td>strand</td><td>MM</td><td>target_seq</td><td>PAM</td><td>alignment</td><td>distance</td><td>position</td><td>gene name</td><td>gene id</td></tr>
<tr><td>GU305914.1</td><td>160836</td><td>160858</td><td>-</td><td>0</td><td>AGTTGTTACCCACCATTCCC</td><td>AGG</td><td>||||||||[||||||||||||]PAM</td><td>NA</td><td>Intergenic</td><td>NA</td><td>NA</td></tr>
<tr><td>T22192</td><td>GTTGTTACCCACCATTCCCAGGG</td><td>1000</td><td></td><td></td><td></td><td></td><td></td><td></td><td></td><td></td><td></td></tr>
<tr><td>Oligo fwd</td><td>CACCGTTGTTACCCACCATTCCCA</td><td></td><td></td><td></td><td></td><td></td><td></td><td></td><td></td><td></td><td></td></tr>
<tr><td>Oligo rev</td><td>AAACTGGGAATGGTGGGTAACAAC</td><td></td><td></td><td></td><td></td><td></td><td></td><td></td><td></td><td></td><td></td></tr>
<tr><td>Chromosome</td><td>start</td><td>end</td><td>strand</td><td>MM</td><td>target_seq</td><td>PAM</td><td>alignment</td><td>distance</td><td>position</td><td>gene name</td><td>gene id</td></tr>
<tr><td>GU305914.1</td><td>160835</td><td>160857</td><td>-</td><td>0</td><td>GTTGTTACCCACCATTCCCA</td><td>GGG</td><td>||||||||[||||||||||||]PAM</td><td>NA</td><td>Intergenic</td><td>NA</td><td>NA</td></tr>
<tr><td>T22193</td><td>CCACCATTCCCAGGGTGATCCGG</td><td>1000</td><td></td><td></td><td></td><td></td><td></td><td></td><td></td><td></td><td></td></tr>
<tr><td>Oligo fwd</td><td>CACCCCACCATTCCCAGGGTGATC</td><td></td><td></td><td></td><td></td><td></td><td></td><td></td><td></td><td></td><td></td></tr>
<tr><td>Oligo rev</td><td>AAACGATCACCCTGGGAATGGTGG</td><td></td><td></td><td></td><td></td><td></td><td></td><td></td><td></td><td></td><td></td></tr>
<tr><td>Chromosome</td><td>start</td><td>end</td><td>strand</td><td>MM</td><td>target_seq</td><td>PAM</td><td>alignment</td><td>distance</td><td>position</td><td>gene name</td><td>gene id</td></tr>
<tr><td>GU305914.1</td><td>160827</td><td>160849</td><td>-</td><td>0</td><td>CCACCATTCCCAGGGTGATC</td><td>CGG</td><td>||||||||[||||||||||||]PAM</td><td>NA</td><td>Intergenic</td><td>NA</td><td>NA</td></tr>
<tr><td>T22194</td><td>AAAAAAATAAAAAAAAAGAAAGG</td><td>1000</td><td></td><td></td><td></td><td></td><td></td><td></td><td></td><td></td><td></td></tr>
</table>

<table>
<tr><td>Oligo fwd</td><td>CACCAAAAAAATAAAAAAAAAGAA</td><td></td><td></td><td></td><td></td><td></td><td></td><td></td><td></td><td></td><td></td></tr>
<tr><td>Oligo rev</td><td>AAACTTCTTTTTTTTTATTTTTTT</td><td></td><td></td><td></td><td></td><td></td><td></td><td></td><td></td><td></td><td></td></tr>
<tr><td>Chromosome</td><td>start</td><td>end</td><td>strand</td><td>MM</td><td>target_seq</td><td>PAM</td><td>alignment</td><td>distance</td><td>position</td><td>gene name</td><td>gene id</td></tr>
<tr><td>GU305914.1</td><td>160765</td><td>160787</td><td>-</td><td>0</td><td>AAAAAAATAAAAAAAAAGAA</td><td>AGG</td><td>||||||||[||||||||||||]PAM</td><td>NA</td><td>Intergenic</td><td>NA</td><td>NA</td></tr>
<tr><td>T22195</td><td>CCTCCTCTCCTCCTCTTCGTTGG</td><td>1000</td><td></td><td></td><td></td><td></td><td></td><td></td><td></td><td></td><td></td></tr>
<tr><td>Oligo fwd</td><td>CACCCCTCCTCTCCTCCTCTTCGT</td><td></td><td></td><td></td><td></td><td></td><td></td><td></td><td></td><td></td><td></td></tr>
<tr><td>Oligo rev</td><td>AAACACGAAGAGGAGGAGAGGAGG</td><td></td><td></td><td></td><td></td><td></td><td></td><td></td><td></td><td></td><td></td></tr>
<tr><td>Chromosome</td><td>start</td><td>end</td><td>strand</td><td>MM</td><td>target_seq</td><td>PAM</td><td>alignment</td><td>distance</td><td>position</td><td>gene name</td><td>gene id</td></tr>
<tr><td>GU305914.1</td><td>160684</td><td>160706</td><td>-</td><td>0</td><td>CCTCCTCTCCTCCTCTTCGT</td><td>TGG</td><td>||||||||[||||||||||||]PAM</td><td>NA</td><td>Intergenic</td><td>NA</td><td>NA</td></tr>
<tr><td>T22196</td><td>TGTACGACTAACATCTGTATAGG</td><td>1000</td><td></td><td></td><td></td><td></td><td></td><td></td><td></td><td></td><td></td></tr>
<tr><td>Oligo fwd</td><td>CACCTGTACGACTAACATCTGTAT</td><td></td><td></td><td></td><td></td><td></td><td></td><td></td><td></td><td></td><td></td></tr>
<tr><td>Oligo rev</td><td>AAACATACAGATGTTAGTCGTACA</td><td></td><td></td><td></td><td></td><td></td><td></td><td></td><td></td><td></td><td></td></tr>
<tr><td>Chromosome</td><td>start</td><td>end</td><td>strand</td><td>MM</td><td>target_seq</td><td>PAM</td><td>alignment</td><td>distance</td><td>position</td><td>gene name</td><td>gene id</td></tr>
<tr><td>GU305914.1</td><td>160581</td><td>160603</td><td>-</td><td>0</td><td>TGTACGACTAACATCTGTAT</td><td>AGG</td><td>||||||||[||||||||||||]PAM</td><td>NA</td><td>Intergenic</td><td>NA</td><td>NA</td></tr>
<tr><td>T22197</td><td>GTGAAGCTCTCTGTTGCAGTTGG</td><td>1000</td><td></td><td></td><td></td><td></td><td></td><td></td><td></td><td></td><td></td></tr>
<tr><td>Oligo fwd</td><td>CACCGTGAAGCTCTCTGTTGCAGT</td><td></td><td></td><td></td><td></td><td></td><td></td><td></td><td></td><td></td><td></td></tr>
<tr><td>Oligo rev</td><td>AAACACTGCAACAGAGAGCTTCAC</td><td></td><td></td><td></td><td></td><td></td><td></td><td></td><td></td><td></td><td></td></tr>
<tr><td>Chromosome</td><td>start</td><td>end</td><td>strand</td><td>MM</td><td>target_seq</td><td>PAM</td><td>alignment</td><td>distance</td><td>position</td><td>gene name</td><td>gene id</td></tr>
<tr><td>GU305914.1</td><td>160517</td><td>160539</td><td>-</td><td>0</td><td>GTGAAGCTCTCTGTTGCAGT</td><td>TGG</td><td>||||||||[||||||||||||]PAM</td><td>NA</td><td>Intergenic</td><td>NA</td><td>NA</td></tr>
<tr><td>T22198</td><td>ACTAAGCACGCATAGAGCACTGG</td><td>1000</td><td></td><td></td><td></td><td></td><td></td><td></td><td></td><td></td><td></td></tr>
<tr><td>Oligo fwd</td><td>CACCACTAAGCACGCATAGAGCAC</td><td></td><td></td><td></td><td></td><td></td><td></td><td></td><td></td><td></td><td></td></tr>
<tr><td>Oligo rev</td><td>AAACGTGCTCTATGCGTGCTTAGT</td><td></td><td></td><td></td><td></td><td></td><td></td><td></td><td></td><td></td><td></td></tr>
<tr><td>Chromosome</td><td>start</td><td>end</td><td>strand</td><td>MM</td><td>target_seq</td><td>PAM</td><td>alignment</td><td>distance</td><td>position</td><td>gene name</td><td>gene id</td></tr>
<tr><td>GU305914.1</td><td>160494</td><td>160516</td><td>-</td><td>0</td><td>ACTAAGCACGCATAGAGCAC</td><td>TGG</td><td>||||||||[||||||||||||]PAM</td><td>NA</td><td>Intergenic</td><td>NA</td><td>NA</td></tr>
<tr><td>T22199</td><td>AATTAGCAGAAAAACTACAAAGG</td><td>1000</td><td></td><td></td><td></td><td></td><td></td><td></td><td></td><td></td><td></td></tr>
<tr><td>Oligo fwd</td><td>CACCAATTAGCAGAAAAACTACAA</td><td></td><td></td><td></td><td></td><td></td><td></td><td></td><td></td><td></td><td></td></tr>
<tr><td>Oligo rev</td><td>AAACTTGTAGTTTTTCTGCTAATT</td><td></td><td></td><td></td><td></td><td></td><td></td><td></td><td></td><td></td><td></td></tr>
<tr><td>Chromosome</td><td>start</td><td>end</td><td>strand</td><td>MM</td><td>target_seq</td><td>PAM</td><td>alignment</td><td>distance</td><td>position</td><td>gene name</td><td>gene id</td></tr>
<tr><td>GU305914.1</td><td>160390</td><td>160412</td><td>-</td><td>0</td><td>AATTAGCAGAAAAACTACAA</td><td>AGG</td><td>||||||||[||||||||||||]PAM</td><td>NA</td><td>Intergenic</td><td>NA</td><td>NA</td></tr>
<tr><td>T22200</td><td>TTAAAAAACAAATCGCAAAAAGG</td><td>1000</td><td></td><td></td><td></td><td></td><td></td><td></td><td></td><td></td><td></td></tr>
<tr><td>Oligo fwd</td><td>CACCTTAAAAAACAAATCGCAAAA</td><td></td><td></td><td></td><td></td><td></td><td></td><td></td><td></td><td></td><td></td></tr>
<tr><td>Oligo rev</td><td>AAACTTTTGCGATTTGTTTTTTAA</td><td></td><td></td><td></td><td></td><td></td><td></td><td></td><td></td><td></td><td></td></tr>
<tr><td>Chromosome</td><td>start</td><td>end</td><td>strand</td><td>MM</td><td>target_seq</td><td>PAM</td><td>alignment</td><td>distance</td><td>position</td><td>gene name</td><td>gene id</td></tr>
<tr><td>GU305914.1</td><td>160355</td><td>160377</td><td>-</td><td>0</td><td>TTAAAAAACAAATCGCAAAA</td><td>AGG</td><td>||||||||[||||||||||||]PAM</td><td>NA</td><td>Intergenic</td><td>NA</td><td>NA</td></tr>
<tr><td>T22201</td><td>AAAAACAAATCGCAAAAAGGAGG</td><td>1000</td><td></td><td></td><td></td><td></td><td></td><td></td><td></td><td></td><td></td></tr>
<tr><td>Oligo fwd</td><td>CACCAAAAACAAATCGCAAAAAGG</td><td></td><td></td><td></td><td></td><td></td><td></td><td></td><td></td><td></td><td></td></tr>
<tr><td>Oligo rev</td><td>AAACCCTTTTTGCGATTTGTTTTT</td><td></td><td></td><td></td><td></td><td></td><td></td><td></td><td></td><td></td><td></td></tr>
<tr><td>Chromosome</td><td>start</td><td>end</td><td>strand</td><td>MM</td><td>target_seq</td><td>PAM</td><td>alignment</td><td>distance</td><td>position</td><td>gene name</td><td>gene id</td></tr>
<tr><td>GU305914.1</td><td>160352</td><td>160374</td><td>-</td><td>0</td><td>AAAAACAAATCGCAAAAAGG</td><td>AGG</td><td>||||||||[||||||||||||]PAM</td><td>NA</td><td>Intergenic</td><td>NA</td><td>NA</td></tr>
<tr><td>T22202</td><td>AATCGCAAAAAGGAGGAATGAGG</td><td>1000</td><td></td><td></td><td></td><td></td><td></td><td></td><td></td><td></td><td></td></tr>
<tr><td>Oligo fwd</td><td>CACCAATCGCAAAAAGGAGGAATG</td><td></td><td></td><td></td><td></td><td></td><td></td><td></td><td></td><td></td><td></td></tr>
<tr><td>Oligo rev</td><td>AAACCATTCCTCCTTTTTGCGATT</td><td></td><td></td><td></td><td></td><td></td><td></td><td></td><td></td><td></td><td></td></tr>
<tr><td>Chromosome</td><td>start</td><td>end</td><td>strand</td><td>MM</td><td>target_seq</td><td>PAM</td><td>alignment</td><td>distance</td><td>position</td><td>gene name</td><td>gene id</td></tr>
<tr><td>GU305914.1</td><td>160345</td><td>160367</td><td>-</td><td>0</td><td>AATCGCAAAAAGGAGGAATG</td><td>AGG</td><td>||||||||[||||||||||||]PAM</td><td>NA</td><td>Intergenic</td><td>NA</td><td>NA</td></tr>
<tr><td>T22203</td><td>ATCGCAAAAAGGAGGAATGAGGG</td><td>1000</td><td></td><td></td><td></td><td></td><td></td><td></td><td></td><td></td><td></td></tr>
<tr><td>Oligo fwd</td><td>CACCATCGCAAAAAGGAGGAATGA</td><td></td><td></td><td></td><td></td><td></td><td></td><td></td><td></td><td></td><td></td></tr>
<tr><td>Oligo rev</td><td>AAACTCATTCCTCCTTTTTGCGAT</td><td></td><td></td><td></td><td></td><td></td><td></td><td></td><td></td><td></td><td></td></tr>
<tr><td>Chromosome</td><td>start</td><td>end</td><td>strand</td><td>MM</td><td>target_seq</td><td>PAM</td><td>alignment</td><td>distance</td><td>position</td><td>gene name</td><td>gene id</td></tr>
<tr><td>GU305914.1</td><td>160344</td><td>160366</td><td>-</td><td>0</td><td>ATCGCAAAAAGGAGGAATGA</td><td>GGG</td><td>||||||||[||||||||||||]PAM</td><td>NA</td><td>Intergenic</td><td>NA</td><td>NA</td></tr>
<tr><td>T22204</td><td>AGGAGGAATGAGGGCAATTACGG</td><td>1000</td><td></td><td></td><td></td><td></td><td></td><td></td><td></td><td></td><td></td></tr>
<tr><td>Oligo fwd</td><td>CACCAGGAGGAATGAGGGCAATTA</td><td></td><td></td><td></td><td></td><td></td><td></td><td></td><td></td><td></td><td></td></tr>
<tr><td>Oligo rev</td><td>AAACTAATTGCCCTCATTCCTCCT</td><td></td><td></td><td></td><td></td><td></td><td></td><td></td><td></td><td></td><td></td></tr>
<tr><td>Chromosome</td><td>start</td><td>end</td><td>strand</td><td>MM</td><td>target_seq</td><td>PAM</td><td>alignment</td><td>distance</td><td>position</td><td>gene name</td><td>gene id</td></tr>
<tr><td>GU305914.1</td><td>160335</td><td>160357</td><td>-</td><td>0</td><td>AGGAGGAATGAGGGCAATTA</td><td>CGG</td><td>||||||||[||||||||||||]PAM</td><td>NA</td><td>Intergenic</td><td>NA</td><td>NA</td></tr>
<tr><td>T22205</td><td>ACCCTCTGTGACTGTTCGCACGG</td><td>1000</td><td></td><td></td><td></td><td></td><td></td><td></td><td></td><td></td><td></td></tr>
<tr><td>Oligo fwd</td><td>CACCACCCTCTGTGACTGTTCGCA</td><td></td><td></td><td></td><td></td><td></td><td></td><td></td><td></td><td></td><td></td></tr>
<tr><td>Oligo rev</td><td>AAACTGCGAACAGTCACAGAGGGT</td><td></td><td></td><td></td><td></td><td></td><td></td><td></td><td></td><td></td><td></td></tr>
<tr><td>Chromosome</td><td>start</td><td>end</td><td>strand</td><td>MM</td><td>target_seq</td><td>PAM</td><td>alignment</td><td>distance</td><td>position</td><td>gene name</td><td>gene id</td></tr>
<tr><td>GU305914.1</td><td>160254</td><td>160276</td><td>-</td><td>0</td><td>ACCCTCTGTGACTGTTCGCA</td><td>CGG</td><td>||||||||[||||||||||||]PAM</td><td>NA</td><td>Intergenic</td><td>NA</td><td>NA</td></tr>
<tr><td>T22206</td><td>ACGACGAGAGAGATCTATAGAGG</td><td>1000</td><td></td><td></td><td></td><td></td><td></td><td></td><td></td><td></td><td></td></tr>
<tr><td>Oligo fwd</td><td>CACCACGACGAGAGAGATCTATAG</td><td></td><td></td><td></td><td></td><td></td><td></td><td></td><td></td><td></td><td></td></tr>
<tr><td>Oligo rev</td><td>AAACCTATAGATCTCTCTCGTCGT</td><td></td><td></td><td></td><td></td><td></td><td></td><td></td><td></td><td></td><td></td></tr>
<tr><td>Chromosome</td><td>start</td><td>end</td><td>strand</td><td>MM</td><td>target_seq</td><td>PAM</td><td>alignment</td><td>distance</td><td>position</td><td>gene name</td><td>gene id</td></tr>
<tr><td>GU305914.1</td><td>160111</td><td>160133</td><td>-</td><td>0</td><td>ACGACGAGAGAGATCTATAG</td><td>AGG</td><td>||||||||[||||||||||||]PAM</td><td>NA</td><td>Intergenic</td><td>NA</td><td>NA</td></tr>
<tr><td>T22207</td><td>CGACGAGAGAGATCTATAGAGGG</td><td>1000</td><td></td><td></td><td></td><td></td><td></td><td></td><td></td><td></td><td></td></tr>
<tr><td>Oligo fwd</td><td>CACCCGACGAGAGAGATCTATAGA</td><td></td><td></td><td></td><td></td><td></td><td></td><td></td><td></td><td></td><td></td></tr>
<tr><td>Oligo rev</td><td>AAACTCTATAGATCTCTCTCGTCG</td><td></td><td></td><td></td><td></td><td></td><td></td><td></td><td></td><td></td><td></td></tr>
<tr><td>Chromosome</td><td>start</td><td>end</td><td>strand</td><td>MM</td><td>target_seq</td><td>PAM</td><td>alignment</td><td>distance</td><td>position</td><td>gene name</td><td>gene id</td></tr>
<tr><td>GU305914.1</td><td>160110</td><td>160132</td><td>-</td><td>0</td><td>CGACGAGAGAGATCTATAGA</td><td>GGG</td><td>||||||||[||||||||||||]PAM</td><td>NA</td><td>Intergenic</td><td>NA</td><td>NA</td></tr>
<tr><td>T22208</td><td>CTCTAACGTGACGAACCGTTCGG</td><td>1000</td><td></td><td></td><td></td><td></td><td></td><td></td><td></td><td></td><td></td></tr>
<tr><td>Oligo fwd</td><td>CACCCTCTAACGTGACGAACCGTT</td><td></td><td></td><td></td><td></td><td></td><td></td><td></td><td></td><td></td><td></td></tr>
<tr><td>Oligo rev</td><td>AAACAACGGTTCGTCACGTTAGAG</td><td></td><td></td><td></td><td></td><td></td><td></td><td></td><td></td><td></td><td></td></tr>
<tr><td>Chromosome</td><td>start</td><td>end</td><td>strand</td><td>MM</td><td>target_seq</td><td>PAM</td><td>alignment</td><td>distance</td><td>position</td><td>gene name</td><td>gene id</td></tr>
<tr><td>GU305914.1</td><td>160067</td><td>160089</td><td>-</td><td>0</td><td>CTCTAACGTGACGAACCGTT</td><td>CGG</td><td>||||||||[||||||||||||]PAM</td><td>NA</td><td>Intergenic</td><td>NA</td><td>NA</td></tr>
</table>

<table>
<tr><td>T22209</td><td>AACACATTGTGCCAGCTGACTGG</td><td>1000</td><td></td><td></td><td></td><td></td><td></td><td></td><td></td><td></td><td></td></tr>
<tr><td>Oligo fwd</td><td>CACCAACACATTGTGCCAGCTGAC</td><td></td><td></td><td></td><td></td><td></td><td></td><td></td><td></td><td></td><td></td></tr>
<tr><td>Oligo rev</td><td>AAACGTCAGCTGGCACAATGTGTT</td><td></td><td></td><td></td><td></td><td></td><td></td><td></td><td></td><td></td><td></td></tr>
<tr><td>Chromosome</td><td>start</td><td>end</td><td>strand</td><td>MM</td><td>target_seq</td><td>PAM</td><td>alignment</td><td>distance</td><td>position</td><td>gene name</td><td>gene id</td></tr>
<tr><td>GU305914.1</td><td>160007</td><td>160029</td><td>-</td><td>0</td><td>AACACATTGTGCCAGCTGAC</td><td>TGG</td><td>||||||||[||||||||||||]PAM</td><td>NA</td><td>Intergenic</td><td>NA</td><td>NA</td></tr>
<tr><td></td><td></td><td></td><td></td><td></td><td></td><td></td><td></td><td></td><td></td><td></td><td></td></tr>
<tr><td>T22210</td><td>TGTGCCAGCTGACTGGCCCCTGG</td><td>1000</td><td></td><td></td><td></td><td></td><td></td><td></td><td></td><td></td><td></td></tr>
<tr><td>Oligo fwd</td><td>CACCTGTGCCAGCTGACTGGCCCC</td><td></td><td></td><td></td><td></td><td></td><td></td><td></td><td></td><td></td><td></td></tr>
<tr><td>Oligo rev</td><td>AAACGGGGCCAGTCAGCTGGCACA</td><td></td><td></td><td></td><td></td><td></td><td></td><td></td><td></td><td></td><td></td></tr>
<tr><td>Chromosome</td><td>start</td><td>end</td><td>strand</td><td>MM</td><td>target_seq</td><td>PAM</td><td>alignment</td><td>distance</td><td>position</td><td>gene name</td><td>gene id</td></tr>
<tr><td>GU305914.1</td><td>160000</td><td>160022</td><td>-</td><td>0</td><td>TGTGCCAGCTGACTGGCCCC</td><td>TGG</td><td>||||||||[||||||||||||]PAM</td><td>NA</td><td>Intergenic</td><td>NA</td><td>NA</td></tr>
<tr><td></td><td></td><td></td><td></td><td></td><td></td><td></td><td></td><td></td><td></td><td></td><td></td></tr>
<tr><td>T22211</td><td>GGCCCCTGGTATCATCTCCGTGG</td><td>1000</td><td></td><td></td><td></td><td></td><td></td><td></td><td></td><td></td><td></td></tr>
<tr><td>Oligo fwd</td><td>CACCGGCCCCTGGTATCATCTCCG</td><td></td><td></td><td></td><td></td><td></td><td></td><td></td><td></td><td></td><td></td></tr>
<tr><td>Oligo rev</td><td>AAACCGGAGATGATACCAGGGGCC</td><td></td><td></td><td></td><td></td><td></td><td></td><td></td><td></td><td></td><td></td></tr>
<tr><td>Chromosome</td><td>start</td><td>end</td><td>strand</td><td>MM</td><td>target_seq</td><td>PAM</td><td>alignment</td><td>distance</td><td>position</td><td>gene name</td><td>gene id</td></tr>
<tr><td>GU305914.1</td><td>159986</td><td>160008</td><td>-</td><td>0</td><td>GGCCCCTGGTATCATCTCCG</td><td>TGG</td><td>||||||||[||||||||||||]PAM</td><td>NA</td><td>Intergenic</td><td>NA</td><td>NA</td></tr>
<tr><td></td><td></td><td></td><td></td><td></td><td></td><td></td><td></td><td></td><td></td><td></td><td></td></tr>
<tr><td>T22212</td><td>CTCCGTGGTCGTCGTTCTATCGG</td><td>1000</td><td></td><td></td><td></td><td></td><td></td><td></td><td></td><td></td><td></td></tr>
<tr><td>Oligo fwd</td><td>CACCCTCCGTGGTCGTCGTTCTAT</td><td></td><td></td><td></td><td></td><td></td><td></td><td></td><td></td><td></td><td></td></tr>
<tr><td>Oligo rev</td><td>AAACATAGAACGACGACCACGGAG</td><td></td><td></td><td></td><td></td><td></td><td></td><td></td><td></td><td></td><td></td></tr>
<tr><td>Chromosome</td><td>start</td><td>end</td><td>strand</td><td>MM</td><td>target_seq</td><td>PAM</td><td>alignment</td><td>distance</td><td>position</td><td>gene name</td><td>gene id</td></tr>
<tr><td>GU305914.1</td><td>159971</td><td>159993</td><td>-</td><td>0</td><td>CTCCGTGGTCGTCGTTCTAT</td><td>CGG</td><td>||||||||[||||||||||||]PAM</td><td>NA</td><td>Intergenic</td><td>NA</td><td>NA</td></tr>
<tr><td></td><td></td><td></td><td></td><td></td><td></td><td></td><td></td><td></td><td></td><td></td><td></td></tr>
<tr><td>T22213</td><td>CGTGGTCGTCGTTCTATCGGCGG</td><td>1000</td><td></td><td></td><td></td><td></td><td></td><td></td><td></td><td></td><td></td></tr>
<tr><td>Oligo fwd</td><td>CACCCGTGGTCGTCGTTCTATCGG</td><td></td><td></td><td></td><td></td><td></td><td></td><td></td><td></td><td></td><td></td></tr>
<tr><td>Oligo rev</td><td>AAACCCGATAGAACGACGACCACG</td><td></td><td></td><td></td><td></td><td></td><td></td><td></td><td></td><td></td><td></td></tr>
<tr><td>Chromosome</td><td>start</td><td>end</td><td>strand</td><td>MM</td><td>target_seq</td><td>PAM</td><td>alignment</td><td>distance</td><td>position</td><td>gene name</td><td>gene id</td></tr>
<tr><td>GU305914.1</td><td>159968</td><td>159990</td><td>-</td><td>0</td><td>CGTGGTCGTCGTTCTATCGG</td><td>CGG</td><td>||||||||[||||||||||||]PAM</td><td>NA</td><td>Intergenic</td><td>NA</td><td>NA</td></tr>
<tr><td></td><td></td><td></td><td></td><td></td><td></td><td></td><td></td><td></td><td></td><td></td><td></td></tr>
<tr><td>T22214</td><td>CGTCGTTCTATCGGCGGCGACGG</td><td>1000</td><td></td><td></td><td></td><td></td><td></td><td></td><td></td><td></td><td></td></tr>
<tr><td>Oligo fwd</td><td>CACCCGTCGTTCTATCGGCGGCGA</td><td></td><td></td><td></td><td></td><td></td><td></td><td></td><td></td><td></td><td></td></tr>
<tr><td>Oligo rev</td><td>AAACTCGCCGCCGATAGAACGACG</td><td></td><td></td><td></td><td></td><td></td><td></td><td></td><td></td><td></td><td></td></tr>
<tr><td>Chromosome</td><td>start</td><td>end</td><td>strand</td><td>MM</td><td>target_seq</td><td>PAM</td><td>alignment</td><td>distance</td><td>position</td><td>gene name</td><td>gene id</td></tr>
<tr><td>GU305914.1</td><td>159962</td><td>159984</td><td>-</td><td>0</td><td>CGTCGTTCTATCGGCGGCGA</td><td>CGG</td><td>||||||||[||||||||||||]PAM</td><td>NA</td><td>Intergenic</td><td>NA</td><td>NA</td></tr>
<tr><td></td><td></td><td></td><td></td><td></td><td></td><td></td><td></td><td></td><td></td><td></td><td></td></tr>
<tr><td>T22215</td><td>CGTTCTATCGGCGGCGACGGCGG</td><td>1000</td><td></td><td></td><td></td><td></td><td></td><td></td><td></td><td></td><td></td></tr>
<tr><td>Oligo fwd</td><td>CACCCGTTCTATCGGCGGCGACGG</td><td></td><td></td><td></td><td></td><td></td><td></td><td></td><td></td><td></td><td></td></tr>
<tr><td>Oligo rev</td><td>AAACCCGTCGCCGCCGATAGAACG</td><td></td><td></td><td></td><td></td><td></td><td></td><td></td><td></td><td></td><td></td></tr>
<tr><td>Chromosome</td><td>start</td><td>end</td><td>strand</td><td>MM</td><td>target_seq</td><td>PAM</td><td>alignment</td><td>distance</td><td>position</td><td>gene name</td><td>gene id</td></tr>
<tr><td>GU305914.1</td><td>159959</td><td>159981</td><td>-</td><td>0</td><td>CGTTCTATCGGCGGCGACGG</td><td>CGG</td><td>||||||||[||||||||||||]PAM</td><td>NA</td><td>Intergenic</td><td>NA</td><td>NA</td></tr>
<tr><td></td><td></td><td></td><td></td><td></td><td></td><td></td><td></td><td></td><td></td><td></td><td></td></tr>
<tr><td>T22216</td><td>TATCGGCGGCGACGGCGGCGAGG</td><td>1000</td><td></td><td></td><td></td><td></td><td></td><td></td><td></td><td></td><td></td></tr>
<tr><td>Oligo fwd</td><td>CACCTATCGGCGGCGACGGCGGCG</td><td></td><td></td><td></td><td></td><td></td><td></td><td></td><td></td><td></td><td></td></tr>
<tr><td>Oligo rev</td><td>AAACCGCCGCCGTCGCCGCCGATA</td><td></td><td></td><td></td><td></td><td></td><td></td><td></td><td></td><td></td><td></td></tr>
<tr><td>Chromosome</td><td>start</td><td>end</td><td>strand</td><td>MM</td><td>target_seq</td><td>PAM</td><td>alignment</td><td>distance</td><td>position</td><td>gene name</td><td>gene id</td></tr>
<tr><td>GU305914.1</td><td>159954</td><td>159976</td><td>-</td><td>0</td><td>TATCGGCGGCGACGGCGGCG</td><td>AGG</td><td>||||||||[||||||||||||]PAM</td><td>NA</td><td>Intergenic</td><td>NA</td><td>NA</td></tr>
<tr><td>GU305914.1</td><td>35655</td><td>35677</td><td>+</td><td>4</td><td>CCACGGCGGCGGCGGCGGCG</td><td>GGG</td><td>---|||||[|||-||||||||]PAM</td><td>NA</td><td>Intergenic</td><td>NA</td><td>NA</td></tr>
<tr><td>GU305914.1</td><td>62430</td><td>62452</td><td>+</td><td>4</td><td>TTTCGACGGCGGTGGCGGCG</td><td>GGG</td><td>|-|||-||[|||--|||||||]PAM</td><td>NA</td><td>Intergenic</td><td>NA</td><td>NA</td></tr>
<tr><td>GU305914.1</td><td>201156</td><td>201178</td><td>+</td><td>4</td><td>GTTCGGCGTCGACGGGGGCG</td><td>CGG</td><td>--||||||[-||||||-||||]PAM</td><td>NA</td><td>Intergenic</td><td>NA</td><td>NA</td></tr>
<tr><td>GU305914.1</td><td>58121</td><td>58143</td><td>-</td><td>4</td><td>TGGCGGCGGCGGCGGCGGAG</td><td>GGG</td><td>|--|||||[|||-||||||-|]PAM</td><td>NA</td><td>Intergenic</td><td>NA</td><td>NA</td></tr>
<tr><td></td><td></td><td></td><td></td><td></td><td></td><td></td><td></td><td></td><td></td><td></td><td></td></tr>
<tr><td>T22217</td><td>GACGGCGGCGAGGAGCGCCTCGG</td><td>1000</td><td></td><td></td><td></td><td></td><td></td><td></td><td></td><td></td><td></td></tr>
<tr><td>Oligo fwd</td><td>CACCGACGGCGGCGAGGAGCGCCT</td><td></td><td></td><td></td><td></td><td></td><td></td><td></td><td></td><td></td><td></td></tr>
<tr><td>Oligo rev</td><td>AAACAGGCGCTCCTCGCCGCCGTC</td><td></td><td></td><td></td><td></td><td></td><td></td><td></td><td></td><td></td><td></td></tr>
<tr><td>Chromosome</td><td>start</td><td>end</td><td>strand</td><td>MM</td><td>target_seq</td><td>PAM</td><td>alignment</td><td>distance</td><td>position</td><td>gene name</td><td>gene id</td></tr>
<tr><td>GU305914.1</td><td>159944</td><td>159966</td><td>-</td><td>0</td><td>GACGGCGGCGAGGAGCGCCT</td><td>CGG</td><td>||||||||[||||||||||||]PAM</td><td>NA</td><td>Intergenic</td><td>NA</td><td>NA</td></tr>
<tr><td></td><td></td><td></td><td></td><td></td><td></td><td></td><td></td><td></td><td></td><td></td><td></td></tr>
<tr><td>T22218</td><td>ACGGCGGCGAGGAGCGCCTCGGG</td><td>1000</td><td></td><td></td><td></td><td></td><td></td><td></td><td></td><td></td><td></td></tr>
<tr><td>Oligo fwd</td><td>CACCACGGCGGCGAGGAGCGCCTC</td><td></td><td></td><td></td><td></td><td></td><td></td><td></td><td></td><td></td><td></td></tr>
<tr><td>Oligo rev</td><td>AAACGAGGCGCTCCTCGCCGCCGT</td><td></td><td></td><td></td><td></td><td></td><td></td><td></td><td></td><td></td><td></td></tr>
<tr><td>Chromosome</td><td>start</td><td>end</td><td>strand</td><td>MM</td><td>target_seq</td><td>PAM</td><td>alignment</td><td>distance</td><td>position</td><td>gene name</td><td>gene id</td></tr>
<tr><td>GU305914.1</td><td>159943</td><td>159965</td><td>-</td><td>0</td><td>ACGGCGGCGAGGAGCGCCTC</td><td>GGG</td><td>||||||||[||||||||||||]PAM</td><td>NA</td><td>Intergenic</td><td>NA</td><td>NA</td></tr>
<tr><td></td><td></td><td></td><td></td><td></td><td></td><td></td><td></td><td></td><td></td><td></td><td></td></tr>
<tr><td>T22219</td><td>CCTGTGTCATTAATTTTCTCCGG</td><td>1000</td><td></td><td></td><td></td><td></td><td></td><td></td><td></td><td></td><td></td></tr>
<tr><td>Oligo fwd</td><td>CACCCCTGTGTCATTAATTTTCTC</td><td></td><td></td><td></td><td></td><td></td><td></td><td></td><td></td><td></td><td></td></tr>
<tr><td>Oligo rev</td><td>AAACGAGAAAATTAATGACACAGG</td><td></td><td></td><td></td><td></td><td></td><td></td><td></td><td></td><td></td><td></td></tr>
<tr><td>Chromosome</td><td>start</td><td>end</td><td>strand</td><td>MM</td><td>target_seq</td><td>PAM</td><td>alignment</td><td>distance</td><td>position</td><td>gene name</td><td>gene id</td></tr>
<tr><td>GU305914.1</td><td>159895</td><td>159917</td><td>-</td><td>0</td><td>CCTGTGTCATTAATTTTCTC</td><td>CGG</td><td>||||||||[||||||||||||]PAM</td><td>NA</td><td>Intergenic</td><td>NA</td><td>NA</td></tr>
<tr><td></td><td></td><td></td><td></td><td></td><td></td><td></td><td></td><td></td><td></td><td></td><td></td></tr>
<tr><td>T22220</td><td>ACCCAAAAAAAAAGCAGTCTCCGG</td><td>1000</td><td></td><td></td><td></td><td></td><td></td><td></td><td></td><td></td><td></td></tr>
<tr><td>Oligo fwd</td><td>CACCACCCAAAAAAAAAGCAGTCTC</td><td></td><td></td><td></td><td></td><td></td><td></td><td></td><td></td><td></td><td></td></tr>
<tr><td>Oligo rev</td><td>AAACGAGACTGCTTTTTTTTTGGGT</td><td></td><td></td><td></td><td></td><td></td><td></td><td></td><td></td><td></td><td></td></tr>
<tr><td>Chromosome</td><td>start</td><td>end</td><td>strand</td><td>MM</td><td>target_seq</td><td>PAM</td><td>alignment</td><td>distance</td><td>position</td><td>gene name</td><td>gene id</td></tr>
<tr><td>GU305914.1</td><td>159841</td><td>159863</td><td>-</td><td>0</td><td>ACCCAAAAAAAAAGCAGTCTC</td><td>CGG</td><td>||||||||[||||||||||||]PAM</td><td>NA</td><td>Intergenic</td><td>NA</td><td>NA</td></tr>
<tr><td></td><td></td><td></td><td></td><td></td><td></td><td></td><td></td><td></td><td></td><td></td><td></td></tr>
<tr><td>T22221</td><td>CAAACAACGACGACGTCGATCGG</td><td>1000</td><td></td><td></td><td></td><td></td><td></td><td></td><td></td><td></td><td></td></tr>
<tr><td>Oligo fwd</td><td>CACCCAAACAACGACGACGTCGAT</td><td></td><td></td><td></td><td></td><td></td><td></td><td></td><td></td><td></td><td></td></tr>
<tr><td>Oligo rev</td><td>AAACATCGACGTCGTCGTTGTTTG</td><td></td><td></td><td></td><td></td><td></td><td></td><td></td><td></td><td></td><td></td></tr>
<tr><td>Chromosome</td><td>start</td><td>end</td><td>strand</td><td>MM</td><td>target_seq</td><td>PAM</td><td>alignment</td><td>distance</td><td>position</td><td>gene name</td><td>gene id</td></tr>
<tr><td>GU305914.1</td><td>159704</td><td>159726</td><td>-</td><td>0</td><td>CAAACAACGACGACGTCGAT</td><td>CGG</td><td>||||||||[||||||||||||]PAM</td><td>NA</td><td>Intergenic</td><td>NA</td><td>NA</td></tr>
<tr><td></td><td></td><td></td><td></td><td></td><td></td><td></td><td></td><td></td><td></td><td></td><td></td></tr>
<tr><td>T22222</td><td>ACAACGACGACGTCGATCGGAGG</td><td>1000</td><td></td><td></td><td></td><td></td><td></td><td></td><td></td><td></td><td></td></tr>
<tr><td>Oligo fwd</td><td>CACCACAACGACGACGTCGATCGG</td><td></td><td></td><td></td><td></td><td></td><td></td><td></td><td></td><td></td><td></td></tr>
<tr><td>Oligo rev</td><td>AAACCCGATCGACGTCGTCGTTGT</td><td></td><td></td><td></td><td></td><td></td><td></td><td></td><td></td><td></td><td></td></tr>
<tr><td>Chromosome</td><td>start</td><td>end</td><td>strand</td><td>MM</td><td>target_seq</td><td>PAM</td><td>alignment</td><td>distance</td><td>position</td><td>gene name</td><td>gene id</td></tr>
<tr><td>GU305914.1</td><td>159701</td><td>159723</td><td>-</td><td>0</td><td>ACAACGACGACGTCGATCGG</td><td>AGG</td><td>||||||||[||||||||||||]PAM</td><td>NA</td><td>Intergenic</td><td>NA</td><td>NA</td></tr>
<tr><td></td><td></td><td></td><td></td><td></td><td></td><td></td><td></td><td></td><td></td><td></td><td></td></tr>
<tr><td>T22223</td><td>TCGATCGGAGGTGTGACCGACGG</td><td>1000</td><td></td><td></td><td></td><td></td><td></td><td></td><td></td><td></td><td></td></tr>
</table>

<table>
<tr><td>Oligo fwd</td><td>CACCTCGATCGGAGGTGTGACCGA</td><td></td><td></td><td></td><td></td><td></td><td></td><td></td><td></td><td></td><td></td></tr>
<tr><td>Oligo rev</td><td>AAACTCGGTCACACCTCCGATCGA</td><td></td><td></td><td></td><td></td><td></td><td></td><td></td><td></td><td></td><td></td></tr>
<tr><td>Chromosome</td><td>start</td><td>end</td><td>strand</td><td>MM</td><td>target_seq</td><td>PAM</td><td>alignment</td><td>distance</td><td>position</td><td>gene name</td><td>gene id</td></tr>
<tr><td>GU305914.1</td><td>159689</td><td>159711</td><td>-</td><td>0</td><td>TCGATCGGAGGTGTGACCGA</td><td>CGG</td><td>||||||||[||||||||||||]PAM</td><td>NA</td><td>Intergenic</td><td>NA</td><td>NA</td></tr>
<tr><td></td><td></td><td></td><td></td><td></td><td></td><td></td><td></td><td></td><td></td><td></td><td></td></tr>
<tr><td>T22224</td><td>AACACATAATCAACTTTACATGG</td><td>1000</td><td></td><td></td><td></td><td></td><td></td><td></td><td></td><td></td><td></td></tr>
<tr><td>Oligo fwd</td><td>CACCAACACATAATCAACTTTACA</td><td></td><td></td><td></td><td></td><td></td><td></td><td></td><td></td><td></td><td></td></tr>
<tr><td>Oligo rev</td><td>AAACTGTAAAGTTGATTATGTGTT</td><td></td><td></td><td></td><td></td><td></td><td></td><td></td><td></td><td></td><td></td></tr>
<tr><td>Chromosome</td><td>start</td><td>end</td><td>strand</td><td>MM</td><td>target_seq</td><td>PAM</td><td>alignment</td><td>distance</td><td>position</td><td>gene name</td><td>gene id</td></tr>
<tr><td>GU305914.1</td><td>159593</td><td>159615</td><td>-</td><td>0</td><td>AACACATAATCAACTTTACA</td><td>TGG</td><td>||||||||[||||||||||||]PAM</td><td>NA</td><td>Intergenic</td><td>NA</td><td>NA</td></tr>
<tr><td></td><td></td><td></td><td></td><td></td><td></td><td></td><td></td><td></td><td></td><td></td><td></td></tr>
<tr><td>T22225</td><td>TATATATCATCCTGCCGCAACGG</td><td>1000</td><td></td><td></td><td></td><td></td><td></td><td></td><td></td><td></td><td></td></tr>
<tr><td>Oligo fwd</td><td>CACCTATATATCATCCTGCCGCAA</td><td></td><td></td><td></td><td></td><td></td><td></td><td></td><td></td><td></td><td></td></tr>
<tr><td>Oligo rev</td><td>AAACTTGCGGCAGGATGATATATA</td><td></td><td></td><td></td><td></td><td></td><td></td><td></td><td></td><td></td><td></td></tr>
<tr><td>Chromosome</td><td>start</td><td>end</td><td>strand</td><td>MM</td><td>target_seq</td><td>PAM</td><td>alignment</td><td>distance</td><td>position</td><td>gene name</td><td>gene id</td></tr>
<tr><td>GU305914.1</td><td>159454</td><td>159476</td><td>-</td><td>0</td><td>TATATATCATCCTGCCGCAA</td><td>CGG</td><td>||||||||[||||||||||||]PAM</td><td>NA</td><td>Intergenic</td><td>NA</td><td>NA</td></tr>
<tr><td></td><td></td><td></td><td></td><td></td><td></td><td></td><td></td><td></td><td></td><td></td><td></td></tr>
<tr><td>T22226</td><td>ATATATCATCCTGCCGCAACGGG</td><td>1000</td><td></td><td></td><td></td><td></td><td></td><td></td><td></td><td></td><td></td></tr>
<tr><td>Oligo fwd</td><td>CACCATATATCATCCTGCCGCAAC</td><td></td><td></td><td></td><td></td><td></td><td></td><td></td><td></td><td></td><td></td></tr>
<tr><td>Oligo rev</td><td>AAACGTTGCGGCAGGATGATATAT</td><td></td><td></td><td></td><td></td><td></td><td></td><td></td><td></td><td></td><td></td></tr>
<tr><td>Chromosome</td><td>start</td><td>end</td><td>strand</td><td>MM</td><td>target_seq</td><td>PAM</td><td>alignment</td><td>distance</td><td>position</td><td>gene name</td><td>gene id</td></tr>
<tr><td>GU305914.1</td><td>159453</td><td>159475</td><td>-</td><td>0</td><td>ATATATCATCCTGCCGCAAC</td><td>GGG</td><td>||||||||[||||||||||||]PAM</td><td>NA</td><td>Intergenic</td><td>NA</td><td>NA</td></tr>
<tr><td></td><td></td><td></td><td></td><td></td><td></td><td></td><td></td><td></td><td></td><td></td><td></td></tr>
<tr><td>T22227</td><td>CTCTGAGTCAATGTTCATCCAGG</td><td>1000</td><td></td><td></td><td></td><td></td><td></td><td></td><td></td><td></td><td></td></tr>
<tr><td>Oligo fwd</td><td>CACCCTCTGAGTCAATGTTCATCC</td><td></td><td></td><td></td><td></td><td></td><td></td><td></td><td></td><td></td><td></td></tr>
<tr><td>Oligo rev</td><td>AAACGGATGAACATTGACTCAGAG</td><td></td><td></td><td></td><td></td><td></td><td></td><td></td><td></td><td></td><td></td></tr>
<tr><td>Chromosome</td><td>start</td><td>end</td><td>strand</td><td>MM</td><td>target_seq</td><td>PAM</td><td>alignment</td><td>distance</td><td>position</td><td>gene name</td><td>gene id</td></tr>
<tr><td>GU305914.1</td><td>159400</td><td>159422</td><td>-</td><td>0</td><td>CTCTGAGTCAATGTTCATCC</td><td>AGG</td><td>||||||||[||||||||||||]PAM</td><td>NA</td><td>Intergenic</td><td>NA</td><td>NA</td></tr>
<tr><td></td><td></td><td></td><td></td><td></td><td></td><td></td><td></td><td></td><td></td><td></td><td></td></tr>
<tr><td>T22228</td><td>TTCTTCAACGCTTAACGATTCGG</td><td>1000</td><td></td><td></td><td></td><td></td><td></td><td></td><td></td><td></td><td></td></tr>
<tr><td>Oligo fwd</td><td>CACCTTCTTCAACGCTTAACGATT</td><td></td><td></td><td></td><td></td><td></td><td></td><td></td><td></td><td></td><td></td></tr>
<tr><td>Oligo rev</td><td>AAACAATCGTTAAGCGTTGAAGAA</td><td></td><td></td><td></td><td></td><td></td><td></td><td></td><td></td><td></td><td></td></tr>
<tr><td>Chromosome</td><td>start</td><td>end</td><td>strand</td><td>MM</td><td>target_seq</td><td>PAM</td><td>alignment</td><td>distance</td><td>position</td><td>gene name</td><td>gene id</td></tr>
<tr><td>GU305914.1</td><td>159372</td><td>159394</td><td>-</td><td>0</td><td>TTCTTCAACGCTTAACGATT</td><td>CGG</td><td>||||||||[||||||||||||]PAM</td><td>NA</td><td>Intergenic</td><td>NA</td><td>NA</td></tr>
<tr><td></td><td></td><td></td><td></td><td></td><td></td><td></td><td></td><td></td><td></td><td></td><td></td></tr>
<tr><td>T22229</td><td>CGGCTATTTGTAATACACGCCGG</td><td>1000</td><td></td><td></td><td></td><td></td><td></td><td></td><td></td><td></td><td></td></tr>
<tr><td>Oligo fwd</td><td>CACCCGGCTATTTGTAATACACGC</td><td></td><td></td><td></td><td></td><td></td><td></td><td></td><td></td><td></td><td></td></tr>
<tr><td>Oligo rev</td><td>AAACGCGTGTATTACAAATAGCCG</td><td></td><td></td><td></td><td></td><td></td><td></td><td></td><td></td><td></td><td></td></tr>
<tr><td>Chromosome</td><td>start</td><td>end</td><td>strand</td><td>MM</td><td>target_seq</td><td>PAM</td><td>alignment</td><td>distance</td><td>position</td><td>gene name</td><td>gene id</td></tr>
<tr><td>GU305914.1</td><td>159352</td><td>159374</td><td>-</td><td>0</td><td>CGGCTATTTGTAATACACGC</td><td>CGG</td><td>||||||||[||||||||||||]PAM</td><td>NA</td><td>Intergenic</td><td>NA</td><td>NA</td></tr>
<tr><td></td><td></td><td></td><td></td><td></td><td></td><td></td><td></td><td></td><td></td><td></td><td></td></tr>
<tr><td>T22230</td><td>GGCTATTTGTAATACACGCCGGG</td><td>1000</td><td></td><td></td><td></td><td></td><td></td><td></td><td></td><td></td><td></td></tr>
<tr><td>Oligo fwd</td><td>CACCGGCTATTTGTAATACACGCC</td><td></td><td></td><td></td><td></td><td></td><td></td><td></td><td></td><td></td><td></td></tr>
<tr><td>Oligo rev</td><td>AAACGGCGTGTATTACAAATAGCC</td><td></td><td></td><td></td><td></td><td></td><td></td><td></td><td></td><td></td><td></td></tr>
<tr><td>Chromosome</td><td>start</td><td>end</td><td>strand</td><td>MM</td><td>target_seq</td><td>PAM</td><td>alignment</td><td>distance</td><td>position</td><td>gene name</td><td>gene id</td></tr>
<tr><td>GU305914.1</td><td>159351</td><td>159373</td><td>-</td><td>0</td><td>GGCTATTTGTAATACACGCC</td><td>GGG</td><td>||||||||[||||||||||||]PAM</td><td>NA</td><td>Intergenic</td><td>NA</td><td>NA</td></tr>
<tr><td></td><td></td><td></td><td></td><td></td><td></td><td></td><td></td><td></td><td></td><td></td><td></td></tr>
<tr><td>T22231</td><td>TTGTAATACACGCCGGGTCGCGG</td><td>1000</td><td></td><td></td><td></td><td></td><td></td><td></td><td></td><td></td><td></td></tr>
<tr><td>Oligo fwd</td><td>CACCTTGTAATACACGCCGGGTCG</td><td></td><td></td><td></td><td></td><td></td><td></td><td></td><td></td><td></td><td></td></tr>
<tr><td>Oligo rev</td><td>AAACCGACCCGGCGTGTATTACAA</td><td></td><td></td><td></td><td></td><td></td><td></td><td></td><td></td><td></td><td></td></tr>
<tr><td>Chromosome</td><td>start</td><td>end</td><td>strand</td><td>MM</td><td>target_seq</td><td>PAM</td><td>alignment</td><td>distance</td><td>position</td><td>gene name</td><td>gene id</td></tr>
<tr><td>GU305914.1</td><td>159345</td><td>159367</td><td>-</td><td>0</td><td>TTGTAATACACGCCGGGTCG</td><td>CGG</td><td>||||||||[||||||||||||]PAM</td><td>NA</td><td>Intergenic</td><td>NA</td><td>NA</td></tr>
<tr><td></td><td></td><td></td><td></td><td></td><td></td><td></td><td></td><td></td><td></td><td></td><td></td></tr>
<tr><td>T22232</td><td>TAATACACGCCGGGTCGCGGCGG</td><td>1000</td><td></td><td></td><td></td><td></td><td></td><td></td><td></td><td></td><td></td></tr>
<tr><td>Oligo fwd</td><td>CACCTAATACACGCCGGGTCGCGG</td><td></td><td></td><td></td><td></td><td></td><td></td><td></td><td></td><td></td><td></td></tr>
<tr><td>Oligo rev</td><td>AAACCCGCGACCCGGCGTGTATTA</td><td></td><td></td><td></td><td></td><td></td><td></td><td></td><td></td><td></td><td></td></tr>
<tr><td>Chromosome</td><td>start</td><td>end</td><td>strand</td><td>MM</td><td>target_seq</td><td>PAM</td><td>alignment</td><td>distance</td><td>position</td><td>gene name</td><td>gene id</td></tr>
<tr><td>GU305914.1</td><td>159342</td><td>159364</td><td>-</td><td>0</td><td>TAATACACGCCGGGTCGCGG</td><td>CGG</td><td>||||||||[||||||||||||]PAM</td><td>NA</td><td>Intergenic</td><td>NA</td><td>NA</td></tr>
<tr><td></td><td></td><td></td><td></td><td></td><td></td><td></td><td></td><td></td><td></td><td></td><td></td></tr>
<tr><td>T22233</td><td>ACACGCCGGGTCGCGGCGGTCGG</td><td>1000</td><td></td><td></td><td></td><td></td><td></td><td></td><td></td><td></td><td></td></tr>
<tr><td>Oligo fwd</td><td>CACCACACGCCGGGTCGCGGCGGT</td><td></td><td></td><td></td><td></td><td></td><td></td><td></td><td></td><td></td><td></td></tr>
<tr><td>Oligo rev</td><td>AAACACCGCCGCGACCCGGCGTGT</td><td></td><td></td><td></td><td></td><td></td><td></td><td></td><td></td><td></td><td></td></tr>
<tr><td>Chromosome</td><td>start</td><td>end</td><td>strand</td><td>MM</td><td>target_seq</td><td>PAM</td><td>alignment</td><td>distance</td><td>position</td><td>gene name</td><td>gene id</td></tr>
<tr><td>GU305914.1</td><td>159338</td><td>159360</td><td>-</td><td>0</td><td>ACACGCCGGGTCGCGGCGGT</td><td>CGG</td><td>||||||||[||||||||||||]PAM</td><td>NA</td><td>Intergenic</td><td>NA</td><td>NA</td></tr>
<tr><td></td><td></td><td></td><td></td><td></td><td></td><td></td><td></td><td></td><td></td><td></td><td></td></tr>
<tr><td>T22234</td><td>AGATCGCGATTCTGACCACATGG</td><td>1000</td><td></td><td></td><td></td><td></td><td></td><td></td><td></td><td></td><td></td></tr>
<tr><td>Oligo fwd</td><td>CACCAGATCGCGATTCTGACCACA</td><td></td><td></td><td></td><td></td><td></td><td></td><td></td><td></td><td></td><td></td></tr>
<tr><td>Oligo rev</td><td>AAACTGTGGTCAGAATCGCGATCT</td><td></td><td></td><td></td><td></td><td></td><td></td><td></td><td></td><td></td><td></td></tr>
<tr><td>Chromosome</td><td>start</td><td>end</td><td>strand</td><td>MM</td><td>target_seq</td><td>PAM</td><td>alignment</td><td>distance</td><td>position</td><td>gene name</td><td>gene id</td></tr>
<tr><td>GU305914.1</td><td>159312</td><td>159334</td><td>-</td><td>0</td><td>AGATCGCGATTCTGACCACA</td><td>TGG</td><td>||||||||[||||||||||||]PAM</td><td>NA</td><td>Intergenic</td><td>NA</td><td>NA</td></tr>
<tr><td></td><td></td><td></td><td></td><td></td><td></td><td></td><td></td><td></td><td></td><td></td><td></td></tr>
<tr><td>T22235</td><td>GATCGCGATTCTGACCACATGGG</td><td>1000</td><td></td><td></td><td></td><td></td><td></td><td></td><td></td><td></td><td></td></tr>
<tr><td>Oligo fwd</td><td>CACCGATCGCGATTCTGACCACAT</td><td></td><td></td><td></td><td></td><td></td><td></td><td></td><td></td><td></td><td></td></tr>
<tr><td>Oligo rev</td><td>AAACATGTGGTCAGAATCGCGATC</td><td></td><td></td><td></td><td></td><td></td><td></td><td></td><td></td><td></td><td></td></tr>
<tr><td>Chromosome</td><td>start</td><td>end</td><td>strand</td><td>MM</td><td>target_seq</td><td>PAM</td><td>alignment</td><td>distance</td><td>position</td><td>gene name</td><td>gene id</td></tr>
<tr><td>GU305914.1</td><td>159311</td><td>159333</td><td>-</td><td>0</td><td>GATCGCGATTCTGACCACAT</td><td>GGG</td><td>||||||||[||||||||||||]PAM</td><td>NA</td><td>Intergenic</td><td>NA</td><td>NA</td></tr>
<tr><td></td><td></td><td></td><td></td><td></td><td></td><td></td><td></td><td></td><td></td><td></td><td></td></tr>
<tr><td>T22236</td><td>ACATGGGTCGCCGCGACCCGAGG</td><td>1000</td><td></td><td></td><td></td><td></td><td></td><td></td><td></td><td></td><td></td></tr>
<tr><td>Oligo fwd</td><td>CACCACATGGGTCGCCGCGACCCG</td><td></td><td></td><td></td><td></td><td></td><td></td><td></td><td></td><td></td><td></td></tr>
<tr><td>Oligo rev</td><td>AAACCGGGTCGCGGCGACCCATGT</td><td></td><td></td><td></td><td></td><td></td><td></td><td></td><td></td><td></td><td></td></tr>
<tr><td>Chromosome</td><td>start</td><td>end</td><td>strand</td><td>MM</td><td>target_seq</td><td>PAM</td><td>alignment</td><td>distance</td><td>position</td><td>gene name</td><td>gene id</td></tr>
<tr><td>GU305914.1</td><td>159295</td><td>159317</td><td>-</td><td>0</td><td>ACATGGGTCGCCGCGACCCG</td><td>AGG</td><td>||||||||[||||||||||||]PAM</td><td>NA</td><td>Intergenic</td><td>NA</td><td>NA</td></tr>
<tr><td></td><td></td><td></td><td></td><td></td><td></td><td></td><td></td><td></td><td></td><td></td><td></td></tr>
<tr><td>T22237</td><td>TCGCCGCGACCCGAGGTCTCTGG</td><td>1000</td><td></td><td></td><td></td><td></td><td></td><td></td><td></td><td></td><td></td></tr>
<tr><td>Oligo fwd</td><td>CACCTCGCCGCGACCCGAGGTCTC</td><td></td><td></td><td></td><td></td><td></td><td></td><td></td><td></td><td></td><td></td></tr>
<tr><td>Oligo rev</td><td>AAACGAGACCTCGGGTCGCGGCGA</td><td></td><td></td><td></td><td></td><td></td><td></td><td></td><td></td><td></td><td></td></tr>
<tr><td>Chromosome</td><td>start</td><td>end</td><td>strand</td><td>MM</td><td>target_seq</td><td>PAM</td><td>alignment</td><td>distance</td><td>position</td><td>gene name</td><td>gene id</td></tr>
<tr><td>GU305914.1</td><td>159288</td><td>159310</td><td>-</td><td>0</td><td>TCGCCGCGACCCGAGGTCTC</td><td>TGG</td><td>||||||||[||||||||||||]PAM</td><td>NA</td><td>Intergenic</td><td>NA</td><td>NA</td></tr>
</table>

T22238 CCGCGACCCGAGGTCTCTGGAGG 1000
Oligo fwd CACCCCGCGACCCGAGGTCTCTGG
Oligo rev AAACCCAGAGACCTCGGGTCGCGG
Chromosome start end strand MM target_seq PAM alignment distance position gene name gene id
GU305914.1 159285 159307 - 0 CCGCGACCCGAGGTCTCTGG AGG ||||||||[||||||||||||]PAM NA Intergenic NA NA

T22239 CGCGACCCGAGGTCTCTGGAGGG 1000
Oligo fwd CACCCGCGACCCGAGGTCTCTGGA
Oligo rev AAACTCCAGAGACCTCGGGTCGCG
Chromosome start end strand MM target_seq PAM alignment distance position gene name gene id
GU305914.1 159284 159306 - 0 CGCGACCCGAGGTCTCTGGA GGG ||||||||[||||||||||||]PAM NA Intergenic NA NA

T22240 ACCCGAGGTCTCTGGAGGGATGG 1000
Oligo fwd CACCACCCGAGGTCTCTGGAGGGA
Oligo rev AAACTCCCTCCAGAGACCTCGGGT
Chromosome start end strand MM target_seq PAM alignment distance position gene name gene id
GU305914.1 159280 159302 - 0 ACCCGAGGTCTCTGGAGGGA TGG ||||||||[||||||||||||]PAM NA Intergenic NA NA

T22241 AGGGATGGTAATTCCCCTCACGG 1000
Oligo fwd CACCAGGGATGGTAATTCCCCTCA
Oligo rev AAACTGAGGGGAATTACCATCCCT
Chromosome start end strand MM target_seq PAM alignment distance position gene name gene id
GU305914.1 159265 159287 - 0 AGGGATGGTAATTCCCCTCA CGG ||||||||[||||||||||||]PAM NA Intergenic NA NA

T22242 GGGATGGTAATTCCCCTCACGGG 1000
Oligo fwd CACCGGGATGGTAATTCCCCTCAC
Oligo rev AAACGTGAGGGGAATTACCATCCC
Chromosome start end strand MM target_seq PAM alignment distance position gene name gene id
GU305914.1 159264 159286 - 0 GGGATGGTAATTCCCCTCAC GGG ||||||||[||||||||||||]PAM NA Intergenic NA NA

T22243 GTAATTCCCCTCACGGGACAAGG 1000
Oligo fwd CACCGTAATTCCCCTCACGGGACA
Oligo rev AAACTGTCCCGTGAGGGGAATTAC
Chromosome start end strand MM target_seq PAM alignment distance position gene name gene id
GU305914.1 159258 159280 - 0 GTAATTCCCCTCACGGGACA AGG ||||||||[||||||||||||]PAM NA Intergenic NA NA

T22244 AGGAAGAAGCCTCTCGCCGACGG 1000
Oligo fwd CACCAGGAAGAAGCCTCTCGCCGA
Oligo rev AAACTCGGCGAGAGGCTTCTTCCT
Chromosome start end strand MM target_seq PAM alignment distance position gene name gene id
GU305914.1 159238 159260 - 0 AGGAAGAAGCCTCTCGCCGA CGG ||||||||[||||||||||||]PAM NA Intergenic NA NA

T22245 CAGCAATCAGACCCTTATTTCGG 1000
Oligo fwd CACCCAGCAATCAGACCCTTATTT
Oligo rev AAACAAATAAGGGTCTGATTGCTG
Chromosome start end strand MM target_seq PAM alignment distance position gene name gene id
GU305914.1 159137 159159 - 0 CAGCAATCAGACCCTTATTT CGG ||||||||[||||||||||||]PAM NA Intergenic NA NA

T22246 TTTCGGTAACGTGACCGCGTCGG 1000
Oligo fwd CACCTTTCGGTAACGTGACCGCGT
Oligo rev AAACACGCGGTCACGTTACCGAAA
Chromosome start end strand MM target_seq PAM alignment distance position gene name gene id
GU305914.1 159120 159142 - 0 TTTCGGTAACGTGACCGCGT CGG ||||||||[||||||||||||]PAM NA Intergenic NA NA

T22247 GCGATTTTCGTTTCGAAATAAGG 1000
Oligo fwd CACCGCGATTTTCGTTTCGAAATA
Oligo rev AAACTATTTCGAAACGAAAATCGC
Chromosome start end strand MM target_seq PAM alignment distance position gene name gene id
GU305914.1 159083 159105 - 0 GCGATTTTCGTTTCGAAATA AGG ||||||||[||||||||||||]PAM NA Intergenic NA NA

T22248 TGACCCCCTTCCTCGCAGCAAGG 1000
Oligo fwd CACCTGACCCCCTTCCTCGCAGCA
Oligo rev AAACTGCTGCGAGGAAGGGGGTCA
Chromosome start end strand MM target_seq PAM alignment distance position gene name gene id
GU305914.1 159041 159063 - 0 TGACCCCCTTCCTCGCAGCA AGG ||||||||[||||||||||||]PAM NA Intergenic NA NA

T22249 GACCCCCTTCCTCGCAGCAAGGG 1000
Oligo fwd CACCGACCCCCTTCCTCGCAGCAA
Oligo rev AAACTTGCTGCGAGGAAGGGGGTC
Chromosome start end strand MM target_seq PAM alignment distance position gene name gene id
GU305914.1 159040 159062 - 0 GACCCCCTTCCTCGCAGCAA GGG ||||||||[||||||||||||]PAM NA Intergenic NA NA

T22250 CCCTTCCTCGCAGCAAGGGAAGG 1000
Oligo fwd CACCCCCTTCCTCGCAGCAAGGGA
Oligo rev AAACTCCCTTGCTGCGAGGAAGGG
Chromosome start end strand MM target_seq PAM alignment distance position gene name gene id
GU305914.1 159036 159058 - 0 CCCTTCCTCGCAGCAAGGGA AGG ||||||||[||||||||||||]PAM NA Intergenic NA NA

T22251 GCAGCAAGGGAAGGTGTGTAAGG 1000
Oligo fwd CACCGCAGCAAGGGAAGGTGTGTA
Oligo rev AAACTACACACCTTCCCTTGCTGC
Chromosome start end strand MM target_seq PAM alignment distance position gene name gene id
GU305914.1 159027 159049 - 0 GCAGCAAGGGAAGGTGTGTA AGG ||||||||[||||||||||||]PAM NA Intergenic NA NA

T22252 GGACCGTGCCGTGCTCCCGATGG 1000
Oligo fwd CACCGGACCGTGCCGTGCTCCCGA
Oligo rev AAACTCGGGAGCACGGCACGGTCC
Chromosome start end strand MM target_seq PAM alignment distance position gene name gene id
GU305914.1 159006 159028 - 0 GGACCGTGCCGTGCTCCCGA TGG ||||||||[||||||||||||]PAM NA Intergenic NA NA

T22253 CGTGCTCCCGATGGCCGCGCTGG 1000
Oligo fwd CACCCGTGCTCCCGATGGCCGCGC
Oligo rev AAACGCGCGGCCATCGGGAGCACG
Chromosome start end strand MM target_seq PAM alignment distance position gene name gene id
GU305914.1 158997 159019 - 0 CGTGCTCCCGATGGCCGCGC TGG ||||||||[||||||||||||]PAM NA Intergenic NA NA

T22254 TGGCCGCGCTGGTCGCACGTCGG 1000
Oligo fwd CACCTGGCCGCGCTGGTCGCACGT
Oligo rev AAACACGTGCGACCAGCGCGGCCA
Chromosome start end strand MM target_seq PAM alignment distance position gene name gene id
GU305914.1 158986 159008 - 0 TGGCCGCGCTGGTCGCACGT CGG ||||||||[||||||||||||]PAM NA Intergenic NA NA

T22255 GGCCGCGCTGGTCGCACGTCGGG 1000
Oligo fwd CACCGGCCGCGCTGGTCGCACGTC
Oligo rev AAACGACGTGCGACCAGCGCGGCC
Chromosome start end strand MM target_seq PAM alignment distance position gene name gene id
GU305914.1 158985 159007 - 0 GGCCGCGCTGGTCGCACGTC GGG ||||||||[||||||||||||]PAM NA Intergenic NA NA

T22256 TTTGTCTAAATCCTTTTCAGTGG 1000
Oligo fwd CACCTTTGTCTAAATCCTTTTCAG
Oligo rev AAACCTGAAAAGGATTTAGACAAA
Chromosome start end strand MM target_seq PAM alignment distance position gene name gene id
GU305914.1 158931 158953 - 0 TTTGTCTAAATCCTTTTCAG TGG ||||||||[||||||||||||]PAM NA Intergenic NA NA

T22257 AGATTCTGTCCCCAGACACACGG 1000
Oligo fwd CACCAGATTCTGTCCCCAGACACA
Oligo rev AAACTGTGTCTGGGGACAGAATCT
Chromosome start end strand MM target_seq PAM alignment distance position gene name gene id
GU305914.1 158868 158890 - 0 AGATTCTGTCCCCAGACACA CGG ||||||||[||||||||||||]PAM NA Intergenic NA NA

T22258 GATTCTGTCCCCAGACACACGGG 1000
Oligo fwd CACCGATTCTGTCCCCAGACACAC
Oligo rev AAACGTGTGTCTGGGGACAGAATC
Chromosome start end strand MM target_seq PAM alignment distance position gene name gene id
GU305914.1 158867 158889 - 0 GATTCTGTCCCCAGACACAC GGG ||||||||[||||||||||||]PAM NA Intergenic NA NA

T22259 CCCAGACACACGGGTTCCCATGG 1000
Oligo fwd CACCCCCAGACACACGGGTTCCCA
Oligo rev AAACTGGGAACCCGTGTGTCTGGG
Chromosome start end strand MM target_seq PAM alignment distance position gene name gene id
GU305914.1 158858 158880 - 0 CCCAGACACACGGGTTCCCA TGG ||||||||[||||||||||||]PAM NA Intergenic NA NA

T22260 TGAGCGTGCGAGAGATGCGCCGG 1000
Oligo fwd CACCTGAGCGTGCGAGAGATGCGC
Oligo rev AAACGCGCATCTCTCGCACGCTCA
Chromosome start end strand MM target_seq PAM alignment distance position gene name gene id
GU305914.1 158800 158822 - 0 TGAGCGTGCGAGAGATGCGC CGG ||||||||[||||||||||||]PAM NA Intergenic NA NA
GU305914.1 112511 112533 + 4 TGCGCGCGCGAGAGACGCGA GGG ||-|||-|[|||||||-|||-]PAM NA Intergenic NA NA

T22261 AATCATTTATAAACAATTCACGG 1000
Oligo fwd CACCAATCATTTATAAACAATTCA
Oligo rev AAACTGAATTGTTTATAAATGATT
Chromosome start end strand MM target_seq PAM alignment distance position gene name gene id
GU305914.1 158680 158702 - 0 AATCATTTATAAACAATTCA CGG ||||||||[||||||||||||]PAM NA Intergenic NA NA

T22262 CACGGTAGCAAAAGTAGTCATGG 1000
Oligo fwd CACCCACGGTAGCAAAAGTAGTCA
Oligo rev AAACTGACTACTTTTGCTACCGTG
Chromosome start end strand MM target_seq PAM alignment distance position gene name gene id
GU305914.1 158662 158684 - 0 CACGGTAGCAAAAGTAGTCA TGG ||||||||[||||||||||||]PAM NA Intergenic NA NA

T22263 AATCAATCAGTAGTTAACCAAGG 1000
Oligo fwd CACCAATCAATCAGTAGTTAACCA
Oligo rev AAACTGGTTAACTACTGATTGATT
Chromosome start end strand MM target_seq PAM alignment distance position gene name gene id
GU305914.1 158565 158587 - 0 AATCAATCAGTAGTTAACCA AGG ||||||||[||||||||||||]PAM NA Intergenic NA NA

T22264 AATCAGTAGTTAACCAAGGTAGG 1000
Oligo fwd CACCAATCAGTAGTTAACCAAGGT
Oligo rev AAACACCTTGGTTAACTACTGATT
Chromosome start end strand MM target_seq PAM alignment distance position gene name gene id
GU305914.1 158561 158583 - 0 AATCAGTAGTTAACCAAGGT AGG ||||||||[||||||||||||]PAM NA Intergenic NA NA

T22265 TTAGATAACAAATAGTGATAAGG 1000
Oligo fwd CACCTTAGATAACAAATAGTGATA
Oligo rev AAACTATCACTATTTGTTATCTAA
Chromosome start end strand MM target_seq PAM alignment distance position gene name gene id
GU305914.1 158530 158552 - 0 TTAGATAACAAATAGTGATA AGG ||||||||[||||||||||||]PAM NA Intergenic NA NA

T22266 ATATTCCATATAGCACATAAAGG 1000
Oligo fwd CACCATATTCCATATAGCACATAA
Oligo rev AAACTTATGTGCTATATGGAATAT
Chromosome start end strand MM target_seq PAM alignment distance position gene name gene id
GU305914.1 158487 158509 - 0 ATATTCCATATAGCACATAA AGG ||||||||[||||||||||||]PAM NA Intergenic NA NA

T22267 TATTTATTTTACTCCAACTTAGG 1000
Oligo fwd CACCTATTTATTTTACTCCAACTT
Oligo rev AAACAAGTTGGAGTAAAATAAATA

Chromosome start end strand MM target_seq PAM alignment distance position gene name gene id
GU305914.1 158454 158476 - 0 TATTTATTTTACTCCAACTT AGG ||||||||[||||||||||||]PAM NA Intergenic NA NA

T22268 ATTTATTTTACTCCAACTTAGGG 1000
Oligo fwd CACCATTTATTTTACTCCAACTTA
Oligo rev AAACTAAGTTGGAGTAAAATAAAT
Chromosome start end strand MM target_seq PAM alignment distance position gene name gene id
GU305914.1 158453 158475 - 0 ATTTATTTTACTCCAACTTA GGG ||||||||[||||||||||||]PAM NA Intergenic NA NA

T22269 TATAATTGCTCAACTTTTATTGG 1000
Oligo fwd CACCTATAATTGCTCAACTTTTAT
Oligo rev AAACATAAAAGTTGAGCAATTATA
Chromosome start end strand MM target_seq PAM alignment distance position gene name gene id
GU305914.1 158358 158380 - 0 TATAATTGCTCAACTTTTAT TGG ||||||||[||||||||||||]PAM NA Intergenic NA NA

T22270 TCAACTTTTATTGGTTTTGTAGG 1000
Oligo fwd CACCTCAACTTTTATTGGTTTTGT
Oligo rev AAACACAAAACCAATAAAAGTTGA
Chromosome start end strand MM target_seq PAM alignment distance position gene name gene id
GU305914.1 158349 158371 - 0 TCAACTTTTATTGGTTTTGT AGG ||||||||[||||||||||||]PAM NA Intergenic NA NA

T22271 CTAGATTCCCACTCAATTCGTGG 1000
Oligo fwd CACCCTAGATTCCCACTCAATTCG
Oligo rev AAACCGAATTGAGTGGGAATCTAG
Chromosome start end strand MM target_seq PAM alignment distance position gene name gene id
GU305914.1 158323 158345 - 0 CTAGATTCCCACTCAATTCG TGG ||||||||[||||||||||||]PAM NA Intergenic NA NA

T22272 ATACGATTTTCCATAATTTTAGG 1000
Oligo fwd CACCATACGATTTTCCATAATTTT
Oligo rev AAACAAAATTATGGAAAATCGTAT
Chromosome start end strand MM target_seq PAM alignment distance position gene name gene id
GU305914.1 158249 158271 - 0 ATACGATTTTCCATAATTTT AGG ||||||||[||||||||||||]PAM NA Intergenic NA NA

T22273 TTTCCATAATTTTAGGCAATAGG 1000
Oligo fwd CACCTTTCCATAATTTTAGGCAAT
Oligo rev AAACATTGCCTAAAATTATGGAAA
Chromosome start end strand MM target_seq PAM alignment distance position gene name gene id
GU305914.1 158242 158264 - 0 TTTCCATAATTTTAGGCAAT AGG ||||||||[||||||||||||]PAM NA Intergenic NA NA

T22274 TTCCATAATTTTAGGCAATAGGG 1000
Oligo fwd CACCTTCCATAATTTTAGGCAATA
Oligo rev AAACTATTGCCTAAAATTATGGAA
Chromosome start end strand MM target_seq PAM alignment distance position gene name gene id
GU305914.1 158241 158263 - 0 TTCCATAATTTTAGGCAATA GGG ||||||||[||||||||||||]PAM NA Intergenic NA NA

T22275 TCCATAATTTTAGGCAATAGGGG 1000
Oligo fwd CACCTCCATAATTTTAGGCAATAG
Oligo rev AAACCTATTGCCTAAAATTATGGA
Chromosome start end strand MM target_seq PAM alignment distance position gene name gene id
GU305914.1 158240 158262 - 0 TCCATAATTTTAGGCAATAG GGG ||||||||[||||||||||||]PAM NA Intergenic NA NA

T22276 ACAGATCTAATGTTTCAGAACGG 1000
Oligo fwd CACCACAGATCTAATGTTTCAGAA
Oligo rev AAACTTCTGAAACATTAGATCTGT
Chromosome start end strand MM target_seq PAM alignment distance position gene name gene id
GU305914.1 158034 158056 - 0 ACAGATCTAATGTTTCAGAA CGG ||||||||[||||||||||||]PAM NA Intergenic NA NA

T22277 CGTTGACATCGCCGCGACCGCGG 1000
Oligo fwd CACCCGTTGACATCGCCGCGACCG
Oligo rev AAACCGGTCGCGGCGATGTCAACG
Chromosome start end strand MM target_seq PAM alignment distance position gene name gene id
GU305914.1 157982 158004 - 0 CGTTGACATCGCCGCGACCG CGG ||||||||[||||||||||||]PAM NA Intergenic NA NA

T22278 CGCCGCGACCGCGGTCCCGACGG 1000
Oligo fwd CACCCGCCGCGACCGCGGTCCCGA
Oligo rev AAACTCGGGACCGCGGTCGCGGCG
Chromosome start end strand MM target_seq PAM alignment distance position gene name gene id
GU305914.1 157973 157995 - 0 CGCCGCGACCGCGGTCCCGA CGG ||||||||[||||||||||||]PAM NA Intergenic NA NA
GU305914.1 200734 200756 - 4 CGCGGTGACCGCGGTCCTGC AGG |||-|-||[|||||||||-|-]PAM NA Intergenic NA NA

T22279 GTCCCGACGGTGATCACCTGCGG 1000
Oligo fwd CACCGTCCCGACGGTGATCACCTG
Oligo rev AAACCAGGTGATCACCGTCGGGAC
Chromosome start end strand MM target_seq PAM alignment distance position gene name gene id
GU305914.1 157960 157982 - 0 GTCCCGACGGTGATCACCTG CGG ||||||||[||||||||||||]PAM NA Intergenic NA NA

T22280 TATCCTTCGTATGCACCATACGG 1000
Oligo fwd CACCTATCCTTCGTATGCACCATA
Oligo rev AAACTATGGTGCATACGAAGGATA
Chromosome start end strand MM target_seq PAM alignment distance position gene name gene id
GU305914.1 157904 157926 - 0 TATCCTTCGTATGCACCATA CGG ||||||||[||||||||||||]PAM NA Intergenic NA NA

T22281 ATCCTTCGTATGCACCATACGGG 1000
Oligo fwd CACCATCCTTCGTATGCACCATAC
Oligo rev AAACGTATGGTGCATACGAAGGAT
Chromosome start end strand MM target_seq PAM alignment distance position gene name gene id
GU305914.1 157903 157925 - 0 ATCCTTCGTATGCACCATAC GGG ||||||||[||||||||||||]PAM NA Intergenic NA NA

T22282 TACAAAGTCACAAAGTTCGCTGG 1000

<table>
<tr><td>Oligo fwd</td><td>CACCTACAAAGTCACAAAGTTCGC</td><td></td><td></td><td></td><td></td><td></td><td></td><td></td><td></td><td></td><td></td></tr>
<tr><td>Oligo rev</td><td>AAACGCGAACTTTGTGACTTTGTA</td><td></td><td></td><td></td><td></td><td></td><td></td><td></td><td></td><td></td><td></td></tr>
<tr><td>Chromosome</td><td>start</td><td>end</td><td>strand</td><td>MM</td><td>target_seq</td><td>PAM</td><td>alignment</td><td>distance</td><td>position</td><td>gene name</td><td>gene id</td></tr>
<tr><td>GU305914.1</td><td>157815</td><td>157837</td><td>-</td><td>0</td><td>TACAAAGTCACAAAGTTCGC</td><td>TGG</td><td>||||||||[||||||||||||]PAM</td><td>NA</td><td>Intergenic</td><td>NA</td><td>NA</td></tr>
<tr><td></td><td></td><td></td><td></td><td></td><td></td><td></td><td></td><td></td><td></td><td></td><td></td></tr>
<tr><td>T22283</td><td>GTTCGTACGACGACGTACGTAGG</td><td>1000</td><td></td><td></td><td></td><td></td><td></td><td></td><td></td><td></td><td></td></tr>
<tr><td>Oligo fwd</td><td>CACCGTTCGTACGACGACGTACGT</td><td></td><td></td><td></td><td></td><td></td><td></td><td></td><td></td><td></td><td></td></tr>
<tr><td>Oligo rev</td><td>AAACACGTACGTCGTCGTACGAAC</td><td></td><td></td><td></td><td></td><td></td><td></td><td></td><td></td><td></td><td></td></tr>
<tr><td>Chromosome</td><td>start</td><td>end</td><td>strand</td><td>MM</td><td>target_seq</td><td>PAM</td><td>alignment</td><td>distance</td><td>position</td><td>gene name</td><td>gene id</td></tr>
<tr><td>GU305914.1</td><td>157723</td><td>157745</td><td>-</td><td>0</td><td>GTTCGTACGACGACGTACGT</td><td>AGG</td><td>||||||||[||||||||||||]PAM</td><td>NA</td><td>Intergenic</td><td>NA</td><td>NA</td></tr>
<tr><td></td><td></td><td></td><td></td><td></td><td></td><td></td><td></td><td></td><td></td><td></td><td></td></tr>
<tr><td>T22284</td><td>CTGTGTATCAATTAAATCAATGG</td><td>1000</td><td></td><td></td><td></td><td></td><td></td><td></td><td></td><td></td><td></td></tr>
<tr><td>Oligo fwd</td><td>CACCCTGTGTATCAATTAAATCAA</td><td></td><td></td><td></td><td></td><td></td><td></td><td></td><td></td><td></td><td></td></tr>
<tr><td>Oligo rev</td><td>AAACTTGATTTAATTGATACACAG</td><td></td><td></td><td></td><td></td><td></td><td></td><td></td><td></td><td></td><td></td></tr>
<tr><td>Chromosome</td><td>start</td><td>end</td><td>strand</td><td>MM</td><td>target_seq</td><td>PAM</td><td>alignment</td><td>distance</td><td>position</td><td>gene name</td><td>gene id</td></tr>
<tr><td>GU305914.1</td><td>157681</td><td>157703</td><td>-</td><td>0</td><td>CTGTGTATCAATTAAATCAA</td><td>TGG</td><td>||||||||[||||||||||||]PAM</td><td>NA</td><td>Intergenic</td><td>NA</td><td>NA</td></tr>
<tr><td></td><td></td><td></td><td></td><td></td><td></td><td></td><td></td><td></td><td></td><td></td><td></td></tr>
<tr><td>T22285</td><td>TACTTGTTTTAAAGTCCTCTTGG</td><td>1000</td><td></td><td></td><td></td><td></td><td></td><td></td><td></td><td></td><td></td></tr>
<tr><td>Oligo fwd</td><td>CACCTACTTGTTTTAAAGTCCTCT</td><td></td><td></td><td></td><td></td><td></td><td></td><td></td><td></td><td></td><td></td></tr>
<tr><td>Oligo rev</td><td>AAACAGAGGACTTTAAAACAAGTA</td><td></td><td></td><td></td><td></td><td></td><td></td><td></td><td></td><td></td><td></td></tr>
<tr><td>Chromosome</td><td>start</td><td>end</td><td>strand</td><td>MM</td><td>target_seq</td><td>PAM</td><td>alignment</td><td>distance</td><td>position</td><td>gene name</td><td>gene id</td></tr>
<tr><td>GU305914.1</td><td>157642</td><td>157664</td><td>-</td><td>0</td><td>TACTTGTTTTAAAGTCCTCT</td><td>TGG</td><td>||||||||[||||||||||||]PAM</td><td>NA</td><td>Intergenic</td><td>NA</td><td>NA</td></tr>
<tr><td></td><td></td><td></td><td></td><td></td><td></td><td></td><td></td><td></td><td></td><td></td><td></td></tr>
<tr><td>T22286</td><td>TTTAAAGTCCTCTTGGAATACGG</td><td>1000</td><td></td><td></td><td></td><td></td><td></td><td></td><td></td><td></td><td></td></tr>
<tr><td>Oligo fwd</td><td>CACCTTTAAAGTCCTCTTGGAATA</td><td></td><td></td><td></td><td></td><td></td><td></td><td></td><td></td><td></td><td></td></tr>
<tr><td>Oligo rev</td><td>AAACTATTCCAAGAGGACTTTAAA</td><td></td><td></td><td></td><td></td><td></td><td></td><td></td><td></td><td></td><td></td></tr>
<tr><td>Chromosome</td><td>start</td><td>end</td><td>strand</td><td>MM</td><td>target_seq</td><td>PAM</td><td>alignment</td><td>distance</td><td>position</td><td>gene name</td><td>gene id</td></tr>
<tr><td>GU305914.1</td><td>157635</td><td>157657</td><td>-</td><td>0</td><td>TTTAAAGTCCTCTTGGAATA</td><td>CGG</td><td>||||||||[||||||||||||]PAM</td><td>NA</td><td>Intergenic</td><td>NA</td><td>NA</td></tr>
<tr><td></td><td></td><td></td><td></td><td></td><td></td><td></td><td></td><td></td><td></td><td></td><td></td></tr>
<tr><td>T22287</td><td>GAAAAATACATAAAAATAGACGG</td><td>1000</td><td></td><td></td><td></td><td></td><td></td><td></td><td></td><td></td><td></td></tr>
<tr><td>Oligo fwd</td><td>CACCGAAAAATACATAAAAATAGA</td><td></td><td></td><td></td><td></td><td></td><td></td><td></td><td></td><td></td><td></td></tr>
<tr><td>Oligo rev</td><td>AAACTCTATTTTTATGTATTTTTC</td><td></td><td></td><td></td><td></td><td></td><td></td><td></td><td></td><td></td><td></td></tr>
<tr><td>Chromosome</td><td>start</td><td>end</td><td>strand</td><td>MM</td><td>target_seq</td><td>PAM</td><td>alignment</td><td>distance</td><td>position</td><td>gene name</td><td>gene id</td></tr>
<tr><td>GU305914.1</td><td>157601</td><td>157623</td><td>-</td><td>0</td><td>GAAAAATACATAAAAATAGA</td><td>CGG</td><td>||||||||[||||||||||||]PAM</td><td>NA</td><td>Intergenic</td><td>NA</td><td>NA</td></tr>
<tr><td></td><td></td><td></td><td></td><td></td><td></td><td></td><td></td><td></td><td></td><td></td><td></td></tr>
<tr><td>T22288</td><td>TGTTATACCTAGTTTTGTCAAGG</td><td>1000</td><td></td><td></td><td></td><td></td><td></td><td></td><td></td><td></td><td></td></tr>
<tr><td>Oligo fwd</td><td>CACCTGTTATACCTAGTTTTGTCA</td><td></td><td></td><td></td><td></td><td></td><td></td><td></td><td></td><td></td><td></td></tr>
<tr><td>Oligo rev</td><td>AAACTGACAAAACTAGGTATAACA</td><td></td><td></td><td></td><td></td><td></td><td></td><td></td><td></td><td></td><td></td></tr>
<tr><td>Chromosome</td><td>start</td><td>end</td><td>strand</td><td>MM</td><td>target_seq</td><td>PAM</td><td>alignment</td><td>distance</td><td>position</td><td>gene name</td><td>gene id</td></tr>
<tr><td>GU305914.1</td><td>157528</td><td>157550</td><td>-</td><td>0</td><td>TGTTATACCTAGTTTTGTCA</td><td>AGG</td><td>||||||||[||||||||||||]PAM</td><td>NA</td><td>Intergenic</td><td>NA</td><td>NA</td></tr>
<tr><td></td><td></td><td></td><td></td><td></td><td></td><td></td><td></td><td></td><td></td><td></td><td></td></tr>
<tr><td>T22289</td><td>CATAGTCACCCCCACCGAAACGG</td><td>1000</td><td></td><td></td><td></td><td></td><td></td><td></td><td></td><td></td><td></td></tr>
<tr><td>Oligo fwd</td><td>CACCCATAGTCACCCCCACCGAAA</td><td></td><td></td><td></td><td></td><td></td><td></td><td></td><td></td><td></td><td></td></tr>
<tr><td>Oligo rev</td><td>AAACTTTCGGTGGGGGTGACTATG</td><td></td><td></td><td></td><td></td><td></td><td></td><td></td><td></td><td></td><td></td></tr>
<tr><td>Chromosome</td><td>start</td><td>end</td><td>strand</td><td>MM</td><td>target_seq</td><td>PAM</td><td>alignment</td><td>distance</td><td>position</td><td>gene name</td><td>gene id</td></tr>
<tr><td>GU305914.1</td><td>157481</td><td>157503</td><td>-</td><td>0</td><td>CATAGTCACCCCCACCGAAA</td><td>CGG</td><td>||||||||[||||||||||||]PAM</td><td>NA</td><td>Intergenic</td><td>NA</td><td>NA</td></tr>
<tr><td></td><td></td><td></td><td></td><td></td><td></td><td></td><td></td><td></td><td></td><td></td><td></td></tr>
<tr><td>T22290</td><td>GTTCGCGAGTCGCGCCTCCGCGG</td><td>1000</td><td></td><td></td><td></td><td></td><td></td><td></td><td></td><td></td><td></td></tr>
<tr><td>Oligo fwd</td><td>CACCGTTCGCGAGTCGCGCCTCCG</td><td></td><td></td><td></td><td></td><td></td><td></td><td></td><td></td><td></td><td></td></tr>
<tr><td>Oligo rev</td><td>AAACCGGAGGCGCGACTCGCGAAC</td><td></td><td></td><td></td><td></td><td></td><td></td><td></td><td></td><td></td><td></td></tr>
<tr><td>Chromosome</td><td>start</td><td>end</td><td>strand</td><td>MM</td><td>target_seq</td><td>PAM</td><td>alignment</td><td>distance</td><td>position</td><td>gene name</td><td>gene id</td></tr>
<tr><td>GU305914.1</td><td>157366</td><td>157388</td><td>-</td><td>0</td><td>GTTCGCGAGTCGCGCCTCCG</td><td>CGG</td><td>||||||||[||||||||||||]PAM</td><td>NA</td><td>Intergenic</td><td>NA</td><td>NA</td></tr>
<tr><td></td><td></td><td></td><td></td><td></td><td></td><td></td><td></td><td></td><td></td><td></td><td></td></tr>
<tr><td>T22291</td><td>GCGAGTCGCGCCTCCGCGGCCGG</td><td>1000</td><td></td><td></td><td></td><td></td><td></td><td></td><td></td><td></td><td></td></tr>
<tr><td>Oligo fwd</td><td>CACCGCGAGTCGCGCCTCCGCGGC</td><td></td><td></td><td></td><td></td><td></td><td></td><td></td><td></td><td></td><td></td></tr>
<tr><td>Oligo rev</td><td>AAACGCCGCGGAGGCGCGACTCGC</td><td></td><td></td><td></td><td></td><td></td><td></td><td></td><td></td><td></td><td></td></tr>
<tr><td>Chromosome</td><td>start</td><td>end</td><td>strand</td><td>MM</td><td>target_seq</td><td>PAM</td><td>alignment</td><td>distance</td><td>position</td><td>gene name</td><td>gene id</td></tr>
<tr><td>GU305914.1</td><td>157362</td><td>157384</td><td>-</td><td>0</td><td>GCGAGTCGCGCCTCCGCGGC</td><td>CGG</td><td>||||||||[||||||||||||]PAM</td><td>NA</td><td>Intergenic</td><td>NA</td><td>NA</td></tr>
<tr><td></td><td></td><td></td><td></td><td></td><td></td><td></td><td></td><td></td><td></td><td></td><td></td></tr>
<tr><td>T22292</td><td>TCAGTTCTAACCCATCACGTCGG</td><td>1000</td><td></td><td></td><td></td><td></td><td></td><td></td><td></td><td></td><td></td></tr>
<tr><td>Oligo fwd</td><td>CACCTCAGTTCTAACCCATCACGT</td><td></td><td></td><td></td><td></td><td></td><td></td><td></td><td></td><td></td><td></td></tr>
<tr><td>Oligo rev</td><td>AAACACGTGATGGGTTAGAACTGA</td><td></td><td></td><td></td><td></td><td></td><td></td><td></td><td></td><td></td><td></td></tr>
<tr><td>Chromosome</td><td>start</td><td>end</td><td>strand</td><td>MM</td><td>target_seq</td><td>PAM</td><td>alignment</td><td>distance</td><td>position</td><td>gene name</td><td>gene id</td></tr>
<tr><td>GU305914.1</td><td>157288</td><td>157310</td><td>-</td><td>0</td><td>TCAGTTCTAACCCATCACGT</td><td>CGG</td><td>||||||||[||||||||||||]PAM</td><td>NA</td><td>Intergenic</td><td>NA</td><td>NA</td></tr>
<tr><td></td><td></td><td></td><td></td><td></td><td></td><td></td><td></td><td></td><td></td><td></td><td></td></tr>
<tr><td>T22293</td><td>CAGTTCTAACCCATCACGTCGGG</td><td>1000</td><td></td><td></td><td></td><td></td><td></td><td></td><td></td><td></td><td></td></tr>
<tr><td>Oligo fwd</td><td>CACCCAGTTCTAACCCATCACGTC</td><td></td><td></td><td></td><td></td><td></td><td></td><td></td><td></td><td></td><td></td></tr>
<tr><td>Oligo rev</td><td>AAACGACGTGATGGGTTAGAACTG</td><td></td><td></td><td></td><td></td><td></td><td></td><td></td><td></td><td></td><td></td></tr>
<tr><td>Chromosome</td><td>start</td><td>end</td><td>strand</td><td>MM</td><td>target_seq</td><td>PAM</td><td>alignment</td><td>distance</td><td>position</td><td>gene name</td><td>gene id</td></tr>
<tr><td>GU305914.1</td><td>157287</td><td>157309</td><td>-</td><td>0</td><td>CAGTTCTAACCCATCACGTC</td><td>GGG</td><td>||||||||[||||||||||||]PAM</td><td>NA</td><td>Intergenic</td><td>NA</td><td>NA</td></tr>
<tr><td></td><td></td><td></td><td></td><td></td><td></td><td></td><td></td><td></td><td></td><td></td><td></td></tr>
<tr><td>T22294</td><td>TGTTGTTCGAATTTACCACACGG</td><td>1000</td><td></td><td></td><td></td><td></td><td></td><td></td><td></td><td></td><td></td></tr>
<tr><td>Oligo fwd</td><td>CACCTGTTGTTCGAATTTACCACA</td><td></td><td></td><td></td><td></td><td></td><td></td><td></td><td></td><td></td><td></td></tr>
<tr><td>Oligo rev</td><td>AAACTGTGGTAAATTCGAACAACA</td><td></td><td></td><td></td><td></td><td></td><td></td><td></td><td></td><td></td><td></td></tr>
<tr><td>Chromosome</td><td>start</td><td>end</td><td>strand</td><td>MM</td><td>target_seq</td><td>PAM</td><td>alignment</td><td>distance</td><td>position</td><td>gene name</td><td>gene id</td></tr>
<tr><td>GU305914.1</td><td>157079</td><td>157101</td><td>-</td><td>0</td><td>TGTTGTTCGAATTTACCACA</td><td>CGG</td><td>||||||||[||||||||||||]PAM</td><td>NA</td><td>Intergenic</td><td>NA</td><td>NA</td></tr>
<tr><td></td><td></td><td></td><td></td><td></td><td></td><td></td><td></td><td></td><td></td><td></td><td></td></tr>
<tr><td>T22295</td><td>AAAATTTAGTTAGCTACGCAAGG</td><td>1000</td><td></td><td></td><td></td><td></td><td></td><td></td><td></td><td></td><td></td></tr>
<tr><td>Oligo fwd</td><td>CACCAAAATTTAGTTAGCTACGCA</td><td></td><td></td><td></td><td></td><td></td><td></td><td></td><td></td><td></td><td></td></tr>
<tr><td>Oligo rev</td><td>AAACTGCGTAGCTAACTAAATTTT</td><td></td><td></td><td></td><td></td><td></td><td></td><td></td><td></td><td></td><td></td></tr>
<tr><td>Chromosome</td><td>start</td><td>end</td><td>strand</td><td>MM</td><td>target_seq</td><td>PAM</td><td>alignment</td><td>distance</td><td>position</td><td>gene name</td><td>gene id</td></tr>
<tr><td>GU305914.1</td><td>156965</td><td>156987</td><td>-</td><td>0</td><td>AAAATTTAGTTAGCTACGCA</td><td>AGG</td><td>||||||||[||||||||||||]PAM</td><td>NA</td><td>Intergenic</td><td>NA</td><td>NA</td></tr>
<tr><td></td><td></td><td></td><td></td><td></td><td></td><td></td><td></td><td></td><td></td><td></td><td></td></tr>
<tr><td>T22296</td><td>TAAACCCTCTTCTAACTACTAGG</td><td>1000</td><td></td><td></td><td></td><td></td><td></td><td></td><td></td><td></td><td></td></tr>
<tr><td>Oligo fwd</td><td>CACCTAAACCCTCTTCTAACTACT</td><td></td><td></td><td></td><td></td><td></td><td></td><td></td><td></td><td></td><td></td></tr>
<tr><td>Oligo rev</td><td>AAACAGTAGTTAGAAGAGGGTTTA</td><td></td><td></td><td></td><td></td><td></td><td></td><td></td><td></td><td></td><td></td></tr>
<tr><td>Chromosome</td><td>start</td><td>end</td><td>strand</td><td>MM</td><td>target_seq</td><td>PAM</td><td>alignment</td><td>distance</td><td>position</td><td>gene name</td><td>gene id</td></tr>
<tr><td>GU305914.1</td><td>156856</td><td>156878</td><td>-</td><td>0</td><td>TAAACCCTCTTCTAACTACT</td><td>AGG</td><td>||||||||[||||||||||||]PAM</td><td>NA</td><td>Intergenic</td><td>NA</td><td>NA</td></tr>
</table>

<table>
<tr><td>T22297</td><td>CTAAGCAAAAGAAAGACGACAGG</td><td>1000</td><td></td><td></td><td></td><td></td><td></td><td></td><td></td><td></td><td></td></tr>
<tr><td>Oligo fwd</td><td>CACCCTAAGCAAAAGAAAGACGAC</td><td></td><td></td><td></td><td></td><td></td><td></td><td></td><td></td><td></td><td></td></tr>
<tr><td>Oligo rev</td><td>AAACGTCGTCTTTCTTTTGCTTAG</td><td></td><td></td><td></td><td></td><td></td><td></td><td></td><td></td><td></td><td></td></tr>
<tr><td>Chromosome</td><td>start</td><td>end</td><td>strand</td><td>MM</td><td>target_seq</td><td>PAM</td><td>alignment</td><td>distance</td><td>position</td><td>gene name</td><td>gene id</td></tr>
<tr><td>GU305914.1</td><td>156713</td><td>156735</td><td>-</td><td>0</td><td>CTAAGCAAAAGAAAGACGAC</td><td>AGG</td><td>||||||||[||||||||||||]PAM</td><td>NA</td><td>Intergenic</td><td>NA</td><td>NA</td></tr>
<tr><td>T22298</td><td>AGATAACATCTAAGAGTAGATGG</td><td>1000</td><td></td><td></td><td></td><td></td><td></td><td></td><td></td><td></td><td></td></tr>
<tr><td>Oligo fwd</td><td>CACCAGATAACATCTAAGAGTAGA</td><td></td><td></td><td></td><td></td><td></td><td></td><td></td><td></td><td></td><td></td></tr>
<tr><td>Oligo rev</td><td>AAACTCTACTCTTAGATGTTATCT</td><td></td><td></td><td></td><td></td><td></td><td></td><td></td><td></td><td></td><td></td></tr>
<tr><td>Chromosome</td><td>start</td><td>end</td><td>strand</td><td>MM</td><td>target_seq</td><td>PAM</td><td>alignment</td><td>distance</td><td>position</td><td>gene name</td><td>gene id</td></tr>
<tr><td>GU305914.1</td><td>156681</td><td>156703</td><td>-</td><td>0</td><td>AGATAACATCTAAGAGTAGA</td><td>TGG</td><td>||||||||[||||||||||||]PAM</td><td>NA</td><td>Intergenic</td><td>NA</td><td>NA</td></tr>
<tr><td>T22299</td><td>AAATCAAAGCATATTCAAAAAGG</td><td>1000</td><td></td><td></td><td></td><td></td><td></td><td></td><td></td><td></td><td></td></tr>
<tr><td>Oligo fwd</td><td>CACCAAATCAAAGCATATTCAAAA</td><td></td><td></td><td></td><td></td><td></td><td></td><td></td><td></td><td></td><td></td></tr>
<tr><td>Oligo rev</td><td>AAACTTTTGAATATGCTTTGATTT</td><td></td><td></td><td></td><td></td><td></td><td></td><td></td><td></td><td></td><td></td></tr>
<tr><td>Chromosome</td><td>start</td><td>end</td><td>strand</td><td>MM</td><td>target_seq</td><td>PAM</td><td>alignment</td><td>distance</td><td>position</td><td>gene name</td><td>gene id</td></tr>
<tr><td>GU305914.1</td><td>156619</td><td>156641</td><td>-</td><td>0</td><td>AAATCAAAGCATATTCAAAA</td><td>AGG</td><td>||||||||[||||||||||||]PAM</td><td>NA</td><td>Intergenic</td><td>NA</td><td>NA</td></tr>
<tr><td>T22300</td><td>GAACCGTAACAAAGCGTGTGAGG</td><td>1000</td><td></td><td></td><td></td><td></td><td></td><td></td><td></td><td></td><td></td></tr>
<tr><td>Oligo fwd</td><td>CACCGAACCGTAACAAAGCGTGTG</td><td></td><td></td><td></td><td></td><td></td><td></td><td></td><td></td><td></td><td></td></tr>
<tr><td>Oligo rev</td><td>AAACCACACGCTTTGTTACGGTTC</td><td></td><td></td><td></td><td></td><td></td><td></td><td></td><td></td><td></td><td></td></tr>
<tr><td>Chromosome</td><td>start</td><td>end</td><td>strand</td><td>MM</td><td>target_seq</td><td>PAM</td><td>alignment</td><td>distance</td><td>position</td><td>gene name</td><td>gene id</td></tr>
<tr><td>GU305914.1</td><td>156477</td><td>156499</td><td>-</td><td>0</td><td>GAACCGTAACAAAGCGTGTG</td><td>AGG</td><td>||||||||[||||||||||||]PAM</td><td>NA</td><td>Intergenic</td><td>NA</td><td>NA</td></tr>
<tr><td>T22301</td><td>AACCGTAACAAAGCGTGTGAGGG</td><td>1000</td><td></td><td></td><td></td><td></td><td></td><td></td><td></td><td></td><td></td></tr>
<tr><td>Oligo fwd</td><td>CACCAACCGTAACAAAGCGTGTGA</td><td></td><td></td><td></td><td></td><td></td><td></td><td></td><td></td><td></td><td></td></tr>
<tr><td>Oligo rev</td><td>AAACTCACACGCTTTGTTACGGTT</td><td></td><td></td><td></td><td></td><td></td><td></td><td></td><td></td><td></td><td></td></tr>
<tr><td>Chromosome</td><td>start</td><td>end</td><td>strand</td><td>MM</td><td>target_seq</td><td>PAM</td><td>alignment</td><td>distance</td><td>position</td><td>gene name</td><td>gene id</td></tr>
<tr><td>GU305914.1</td><td>156476</td><td>156498</td><td>-</td><td>0</td><td>AACCGTAACAAAGCGTGTGA</td><td>GGG</td><td>||||||||[||||||||||||]PAM</td><td>NA</td><td>Intergenic</td><td>NA</td><td>NA</td></tr>
<tr><td>T22302</td><td>ACCGTAACAAAGCGTGTGAGGGG</td><td>1000</td><td></td><td></td><td></td><td></td><td></td><td></td><td></td><td></td><td></td></tr>
<tr><td>Oligo fwd</td><td>CACCACCGTAACAAAGCGTGTGAG</td><td></td><td></td><td></td><td></td><td></td><td></td><td></td><td></td><td></td><td></td></tr>
<tr><td>Oligo rev</td><td>AAACCTCACACGCTTTGTTACGGT</td><td></td><td></td><td></td><td></td><td></td><td></td><td></td><td></td><td></td><td></td></tr>
<tr><td>Chromosome</td><td>start</td><td>end</td><td>strand</td><td>MM</td><td>target_seq</td><td>PAM</td><td>alignment</td><td>distance</td><td>position</td><td>gene name</td><td>gene id</td></tr>
<tr><td>GU305914.1</td><td>156475</td><td>156497</td><td>-</td><td>0</td><td>ACCGTAACAAAGCGTGTGAG</td><td>GGG</td><td>||||||||[||||||||||||]PAM</td><td>NA</td><td>Intergenic</td><td>NA</td><td>NA</td></tr>
<tr><td>T22303</td><td>TGAGGGGACAACCACAGATTCGG</td><td>1000</td><td></td><td></td><td></td><td></td><td></td><td></td><td></td><td></td><td></td></tr>
<tr><td>Oligo fwd</td><td>CACCTGAGGGGACAACCACAGATT</td><td></td><td></td><td></td><td></td><td></td><td></td><td></td><td></td><td></td><td></td></tr>
<tr><td>Oligo rev</td><td>AAACAATCTGTGGTTGTCCCCTCA</td><td></td><td></td><td></td><td></td><td></td><td></td><td></td><td></td><td></td><td></td></tr>
<tr><td>Chromosome</td><td>start</td><td>end</td><td>strand</td><td>MM</td><td>target_seq</td><td>PAM</td><td>alignment</td><td>distance</td><td>position</td><td>gene name</td><td>gene id</td></tr>
<tr><td>GU305914.1</td><td>156459</td><td>156481</td><td>-</td><td>0</td><td>TGAGGGGACAACCACAGATT</td><td>CGG</td><td>||||||||[||||||||||||]PAM</td><td>NA</td><td>Intergenic</td><td>NA</td><td>NA</td></tr>
<tr><td>T22304</td><td>AAAACATTTATTAAGATCTAAGG</td><td>1000</td><td></td><td></td><td></td><td></td><td></td><td></td><td></td><td></td><td></td></tr>
<tr><td>Oligo fwd</td><td>CACCAAAACATTTATTAAGATCTA</td><td></td><td></td><td></td><td></td><td></td><td></td><td></td><td></td><td></td><td></td></tr>
<tr><td>Oligo rev</td><td>AAACTAGATCTTAATAAATGTTTT</td><td></td><td></td><td></td><td></td><td></td><td></td><td></td><td></td><td></td><td></td></tr>
<tr><td>Chromosome</td><td>start</td><td>end</td><td>strand</td><td>MM</td><td>target_seq</td><td>PAM</td><td>alignment</td><td>distance</td><td>position</td><td>gene name</td><td>gene id</td></tr>
<tr><td>GU305914.1</td><td>156416</td><td>156438</td><td>-</td><td>0</td><td>AAAACATTTATTAAGATCTA</td><td>AGG</td><td>||||||||[||||||||||||]PAM</td><td>NA</td><td>Intergenic</td><td>NA</td><td>NA</td></tr>
<tr><td>T22305</td><td>TTAAGATCTAAGGAAAAACATGG</td><td>1000</td><td></td><td></td><td></td><td></td><td></td><td></td><td></td><td></td><td></td></tr>
<tr><td>Oligo fwd</td><td>CACCTTAAGATCTAAGGAAAAACA</td><td></td><td></td><td></td><td></td><td></td><td></td><td></td><td></td><td></td><td></td></tr>
<tr><td>Oligo rev</td><td>AAACTGTTTTTCCTTAGATCTTAA</td><td></td><td></td><td></td><td></td><td></td><td></td><td></td><td></td><td></td><td></td></tr>
<tr><td>Chromosome</td><td>start</td><td>end</td><td>strand</td><td>MM</td><td>target_seq</td><td>PAM</td><td>alignment</td><td>distance</td><td>position</td><td>gene name</td><td>gene id</td></tr>
<tr><td>GU305914.1</td><td>156406</td><td>156428</td><td>-</td><td>0</td><td>TTAAGATCTAAGGAAAAACA</td><td>TGG</td><td>||||||||[||||||||||||]PAM</td><td>NA</td><td>Intergenic</td><td>NA</td><td>NA</td></tr>
<tr><td>T22306</td><td>ATATCCATGCATGTATAGAGAGG</td><td>1000</td><td></td><td></td><td></td><td></td><td></td><td></td><td></td><td></td><td></td></tr>
<tr><td>Oligo fwd</td><td>CACCATATCCATGCATGTATAGAG</td><td></td><td></td><td></td><td></td><td></td><td></td><td></td><td></td><td></td><td></td></tr>
<tr><td>Oligo rev</td><td>AAACCTCTATACATGCATGGATAT</td><td></td><td></td><td></td><td></td><td></td><td></td><td></td><td></td><td></td><td></td></tr>
<tr><td>Chromosome</td><td>start</td><td>end</td><td>strand</td><td>MM</td><td>target_seq</td><td>PAM</td><td>alignment</td><td>distance</td><td>position</td><td>gene name</td><td>gene id</td></tr>
<tr><td>GU305914.1</td><td>156211</td><td>156233</td><td>-</td><td>0</td><td>ATATCCATGCATGTATAGAG</td><td>AGG</td><td>||||||||[||||||||||||]PAM</td><td>NA</td><td>Intergenic</td><td>NA</td><td>NA</td></tr>
<tr><td>T22307</td><td>AGGTTACTCTCAAATCGAAATGG</td><td>1000</td><td></td><td></td><td></td><td></td><td></td><td></td><td></td><td></td><td></td></tr>
<tr><td>Oligo fwd</td><td>CACCAGGTTACTCTCAAATCGAAA</td><td></td><td></td><td></td><td></td><td></td><td></td><td></td><td></td><td></td><td></td></tr>
<tr><td>Oligo rev</td><td>AAACTTTCGATTTGAGAGTAACCT</td><td></td><td></td><td></td><td></td><td></td><td></td><td></td><td></td><td></td><td></td></tr>
<tr><td>Chromosome</td><td>start</td><td>end</td><td>strand</td><td>MM</td><td>target_seq</td><td>PAM</td><td>alignment</td><td>distance</td><td>position</td><td>gene name</td><td>gene id</td></tr>
<tr><td>GU305914.1</td><td>156191</td><td>156213</td><td>-</td><td>0</td><td>AGGTTACTCTCAAATCGAAA</td><td>TGG</td><td>||||||||[||||||||||||]PAM</td><td>NA</td><td>Intergenic</td><td>NA</td><td>NA</td></tr>
<tr><td>T22308</td><td>GGTTACTCTCAAATCGAAATGGG</td><td>1000</td><td></td><td></td><td></td><td></td><td></td><td></td><td></td><td></td><td></td></tr>
<tr><td>Oligo fwd</td><td>CACCGGTTACTCTCAAATCGAAAT</td><td></td><td></td><td></td><td></td><td></td><td></td><td></td><td></td><td></td><td></td></tr>
<tr><td>Oligo rev</td><td>AAACATTTCGATTTGAGAGTAACC</td><td></td><td></td><td></td><td></td><td></td><td></td><td></td><td></td><td></td><td></td></tr>
<tr><td>Chromosome</td><td>start</td><td>end</td><td>strand</td><td>MM</td><td>target_seq</td><td>PAM</td><td>alignment</td><td>distance</td><td>position</td><td>gene name</td><td>gene id</td></tr>
<tr><td>GU305914.1</td><td>156190</td><td>156212</td><td>-</td><td>0</td><td>GGTTACTCTCAAATCGAAAT</td><td>GGG</td><td>||||||||[||||||||||||]PAM</td><td>NA</td><td>Intergenic</td><td>NA</td><td>NA</td></tr>
<tr><td>T22309</td><td>TCCTTCTATGTAAACTACTATGG</td><td>1000</td><td></td><td></td><td></td><td></td><td></td><td></td><td></td><td></td><td></td></tr>
<tr><td>Oligo fwd</td><td>CACCTCCTTCTATGTAAACTACTA</td><td></td><td></td><td></td><td></td><td></td><td></td><td></td><td></td><td></td><td></td></tr>
<tr><td>Oligo rev</td><td>AAACTAGTAGTTTACATAGAAGGA</td><td></td><td></td><td></td><td></td><td></td><td></td><td></td><td></td><td></td><td></td></tr>
<tr><td>Chromosome</td><td>start</td><td>end</td><td>strand</td><td>MM</td><td>target_seq</td><td>PAM</td><td>alignment</td><td>distance</td><td>position</td><td>gene name</td><td>gene id</td></tr>
<tr><td>GU305914.1</td><td>156083</td><td>156105</td><td>-</td><td>0</td><td>TCCTTCTATGTAAACTACTA</td><td>TGG</td><td>||||||||[||||||||||||]PAM</td><td>NA</td><td>Intergenic</td><td>NA</td><td>NA</td></tr>
<tr><td>T22310</td><td>CCTTCTATGTAAACTACTATGGG</td><td>1000</td><td></td><td></td><td></td><td></td><td></td><td></td><td></td><td></td><td></td></tr>
<tr><td>Oligo fwd</td><td>CACCCCTTCTATGTAAACTACTAT</td><td></td><td></td><td></td><td></td><td></td><td></td><td></td><td></td><td></td><td></td></tr>
<tr><td>Oligo rev</td><td>AAACATAGTAGTTTACATAGAAGG</td><td></td><td></td><td></td><td></td><td></td><td></td><td></td><td></td><td></td><td></td></tr>
<tr><td>Chromosome</td><td>start</td><td>end</td><td>strand</td><td>MM</td><td>target_seq</td><td>PAM</td><td>alignment</td><td>distance</td><td>position</td><td>gene name</td><td>gene id</td></tr>
<tr><td>GU305914.1</td><td>156082</td><td>156104</td><td>-</td><td>0</td><td>CCTTCTATGTAAACTACTAT</td><td>GGG</td><td>||||||||[||||||||||||]PAM</td><td>NA</td><td>Intergenic</td><td>NA</td><td>NA</td></tr>
<tr><td>T22311</td><td>TGTAAACTACTATGGGTTCTAGG</td><td>1000</td><td></td><td></td><td></td><td></td><td></td><td></td><td></td><td></td><td></td></tr>
<tr><td>Oligo fwd</td><td>CACCTGTAAACTACTATGGGTTCT</td><td></td><td></td><td></td><td></td><td></td><td></td><td></td><td></td><td></td><td></td></tr>
<tr><td>Oligo rev</td><td>AAACAGAACCCATAGTAGTTTACA</td><td></td><td></td><td></td><td></td><td></td><td></td><td></td><td></td><td></td><td></td></tr>
<tr><td>Chromosome</td><td>start</td><td>end</td><td>strand</td><td>MM</td><td>target_seq</td><td>PAM</td><td>alignment</td><td>distance</td><td>position</td><td>gene name</td><td>gene id</td></tr>
<tr><td>GU305914.1</td><td>156075</td><td>156097</td><td>-</td><td>0</td><td>TGTAAACTACTATGGGTTCT</td><td>AGG</td><td>||||||||[||||||||||||]PAM</td><td>NA</td><td>Intergenic</td><td>NA</td><td>NA</td></tr>
</table>

<table>
<tr><td>T22312</td><td>CAATAAGAAAAAAAAAAGACTGG</td><td>1000</td></tr>
<tr><td>Oligo fwd</td><td>CACCCAATAAGAAAAAAAAAAGAC</td></tr>
<tr><td>Oligo rev</td><td>AAACGTCTTTTTTTTTTCTTATTG</td></tr>
<tr><td>Chromosome</td><td>start</td><td>end</td><td>strand</td><td>MM</td><td>target_seq</td><td>PAM</td><td>alignment</td><td>distance</td><td>position</td><td>gene name</td><td>gene id</td></tr>
<tr><td>GU305914.1</td><td>155974</td><td>155996</td><td>-</td><td>0</td><td>CAATAAGAAAAAAAAAAGAC</td><td>TGG</td><td>||||||||[||||||||||||]PAM</td><td>NA</td><td>Intergenic</td><td>NA</td><td>NA</td></tr>
<tr><td>T22313</td><td>ACCAAACCCGCTCGTCAGTTTGG</td><td>1000</td></tr>
<tr><td>Oligo fwd</td><td>CACCACCAAACCCGCTCGTCAGTT</td></tr>
<tr><td>Oligo rev</td><td>AAACAACTGACGAGCGGGTTTGGT</td></tr>
<tr><td>Chromosome</td><td>start</td><td>end</td><td>strand</td><td>MM</td><td>target_seq</td><td>PAM</td><td>alignment</td><td>distance</td><td>position</td><td>gene name</td><td>gene id</td></tr>
<tr><td>GU305914.1</td><td>155883</td><td>155905</td><td>-</td><td>0</td><td>ACCAAACCCGCTCGTCAGTT</td><td>TGG</td><td>||||||||[||||||||||||]PAM</td><td>NA</td><td>Intergenic</td><td>NA</td><td>NA</td></tr>
<tr><td>T22314</td><td>CCAAACCCGCTCGTCAGTTTGGG</td><td>1000</td></tr>
<tr><td>Oligo fwd</td><td>CACCCCAAACCCGCTCGTCAGTTT</td></tr>
<tr><td>Oligo rev</td><td>AAACAAACTGACGAGCGGGTTTGG</td></tr>
<tr><td>Chromosome</td><td>start</td><td>end</td><td>strand</td><td>MM</td><td>target_seq</td><td>PAM</td><td>alignment</td><td>distance</td><td>position</td><td>gene name</td><td>gene id</td></tr>
<tr><td>GU305914.1</td><td>155882</td><td>155904</td><td>-</td><td>0</td><td>CCAAACCCGCTCGTCAGTTT</td><td>GGG</td><td>||||||||[||||||||||||]PAM</td><td>NA</td><td>Intergenic</td><td>NA</td><td>NA</td></tr>
<tr><td>T22315</td><td>CGTCAGTTTGGGTTTGTAAGAGG</td><td>1000</td></tr>
<tr><td>Oligo fwd</td><td>CACCCGTCAGTTTGGGTTTGTAAG</td></tr>
<tr><td>Oligo rev</td><td>AAACCTTACAAACCCAAACTGACG</td></tr>
<tr><td>Chromosome</td><td>start</td><td>end</td><td>strand</td><td>MM</td><td>target_seq</td><td>PAM</td><td>alignment</td><td>distance</td><td>position</td><td>gene name</td><td>gene id</td></tr>
<tr><td>GU305914.1</td><td>155871</td><td>155893</td><td>-</td><td>0</td><td>CGTCAGTTTGGGTTTGTAAG</td><td>AGG</td><td>||||||||[||||||||||||]PAM</td><td>NA</td><td>Intergenic</td><td>NA</td><td>NA</td></tr>
<tr><td>T22316</td><td>TTCATTTTTTTTGTCACATAAGG</td><td>1000</td></tr>
<tr><td>Oligo fwd</td><td>CACCTTCATTTTTTTTGTCACATA</td></tr>
<tr><td>Oligo rev</td><td>AAACTATGTGACAAAAAAAATGAA</td></tr>
<tr><td>Chromosome</td><td>start</td><td>end</td><td>strand</td><td>MM</td><td>target_seq</td><td>PAM</td><td>alignment</td><td>distance</td><td>position</td><td>gene name</td><td>gene id</td></tr>
<tr><td>GU305914.1</td><td>155823</td><td>155845</td><td>-</td><td>0</td><td>TTCATTTTTTTTGTCACATA</td><td>AGG</td><td>||||||||[||||||||||||]PAM</td><td>NA</td><td>Intergenic</td><td>NA</td><td>NA</td></tr>
<tr><td>T22317</td><td>GACAATGTCTTATAAGTAGTAGG</td><td>1000</td></tr>
<tr><td>Oligo fwd</td><td>CACCGACAATGTCTTATAAGTAGT</td></tr>
<tr><td>Oligo rev</td><td>AAACACTACTTATAAGACATTGTC</td></tr>
<tr><td>Chromosome</td><td>start</td><td>end</td><td>strand</td><td>MM</td><td>target_seq</td><td>PAM</td><td>alignment</td><td>distance</td><td>position</td><td>gene name</td><td>gene id</td></tr>
<tr><td>GU305914.1</td><td>155785</td><td>155807</td><td>-</td><td>0</td><td>GACAATGTCTTATAAGTAGT</td><td>AGG</td><td>||||||||[||||||||||||]PAM</td><td>NA</td><td>Intergenic</td><td>NA</td><td>NA</td></tr>
<tr><td>T22318</td><td>AGTAGTAGGTTTATCAAAATAGG</td><td>1000</td></tr>
<tr><td>Oligo fwd</td><td>CACCAGTAGTAGGTTTATCAAAAT</td></tr>
<tr><td>Oligo rev</td><td>AAACATTTTGATAAACCTACTACT</td></tr>
<tr><td>Chromosome</td><td>start</td><td>end</td><td>strand</td><td>MM</td><td>target_seq</td><td>PAM</td><td>alignment</td><td>distance</td><td>position</td><td>gene name</td><td>gene id</td></tr>
<tr><td>GU305914.1</td><td>155771</td><td>155793</td><td>-</td><td>0</td><td>AGTAGTAGGTTTATCAAAAT</td><td>AGG</td><td>||||||||[||||||||||||]PAM</td><td>NA</td><td>Intergenic</td><td>NA</td><td>NA</td></tr>
<tr><td>T22319</td><td>AGGTTTATCAAAATAGGAGTAGG</td><td>1000</td></tr>
<tr><td>Oligo fwd</td><td>CACCAGGTTTATCAAAATAGGAGT</td></tr>
<tr><td>Oligo rev</td><td>AAACACTCCTATTTTGATAAACCT</td></tr>
<tr><td>Chromosome</td><td>start</td><td>end</td><td>strand</td><td>MM</td><td>target_seq</td><td>PAM</td><td>alignment</td><td>distance</td><td>position</td><td>gene name</td><td>gene id</td></tr>
<tr><td>GU305914.1</td><td>155765</td><td>155787</td><td>-</td><td>0</td><td>AGGTTTATCAAAATAGGAGT</td><td>AGG</td><td>||||||||[||||||||||||]PAM</td><td>NA</td><td>Intergenic</td><td>NA</td><td>NA</td></tr>
<tr><td>T22320</td><td>CAGTAGTCATAGAGATGATAAGG</td><td>1000</td></tr>
<tr><td>Oligo fwd</td><td>CACCCAGTAGTCATAGAGATGATA</td></tr>
<tr><td>Oligo rev</td><td>AAACTATCATCTCTATGACTACTG</td></tr>
<tr><td>Chromosome</td><td>start</td><td>end</td><td>strand</td><td>MM</td><td>target_seq</td><td>PAM</td><td>alignment</td><td>distance</td><td>position</td><td>gene name</td><td>gene id</td></tr>
<tr><td>GU305914.1</td><td>155688</td><td>155710</td><td>-</td><td>0</td><td>CAGTAGTCATAGAGATGATA</td><td>AGG</td><td>||||||||[||||||||||||]PAM</td><td>NA</td><td>Intergenic</td><td>NA</td><td>NA</td></tr>
<tr><td>T22321</td><td>TCCGAATCACAATCTCCTTTAGG</td><td>1000</td></tr>
<tr><td>Oligo fwd</td><td>CACCTCCGAATCACAATCTCCTTT</td></tr>
<tr><td>Oligo rev</td><td>AAACAAAGGAGATTGTGATTCGGA</td></tr>
<tr><td>Chromosome</td><td>start</td><td>end</td><td>strand</td><td>MM</td><td>target_seq</td><td>PAM</td><td>alignment</td><td>distance</td><td>position</td><td>gene name</td><td>gene id</td></tr>
<tr><td>GU305914.1</td><td>155562</td><td>155584</td><td>-</td><td>0</td><td>TCCGAATCACAATCTCCTTT</td><td>AGG</td><td>||||||||[||||||||||||]PAM</td><td>NA</td><td>Intergenic</td><td>NA</td><td>NA</td></tr>
<tr><td>T22322</td><td>TCTCCTTTAGGAATACAGATAGG</td><td>1000</td></tr>
<tr><td>Oligo fwd</td><td>CACCTCTCCTTTAGGAATACAGAT</td></tr>
<tr><td>Oligo rev</td><td>AAACATCTGTATTCCTAAAGGAGA</td></tr>
<tr><td>Chromosome</td><td>start</td><td>end</td><td>strand</td><td>MM</td><td>target_seq</td><td>PAM</td><td>alignment</td><td>distance</td><td>position</td><td>gene name</td><td>gene id</td></tr>
<tr><td>GU305914.1</td><td>155550</td><td>155572</td><td>-</td><td>0</td><td>TCTCCTTTAGGAATACAGAT</td><td>AGG</td><td>||||||||[||||||||||||]PAM</td><td>NA</td><td>Intergenic</td><td>NA</td><td>NA</td></tr>
<tr><td>T22323</td><td>GGAATTAAATGCAAAATACCTGG</td><td>1000</td></tr>
<tr><td>Oligo fwd</td><td>CACCGGAATTAAATGCAAAATACC</td></tr>
<tr><td>Oligo rev</td><td>AAACGGTATTTTGCATTTAATTCC</td></tr>
<tr><td>Chromosome</td><td>start</td><td>end</td><td>strand</td><td>MM</td><td>target_seq</td><td>PAM</td><td>alignment</td><td>distance</td><td>position</td><td>gene name</td><td>gene id</td></tr>
<tr><td>GU305914.1</td><td>155529</td><td>155551</td><td>-</td><td>0</td><td>GGAATTAAATGCAAAATACC</td><td>TGG</td><td>||||||||[||||||||||||]PAM</td><td>NA</td><td>Intergenic</td><td>NA</td><td>NA</td></tr>
<tr><td>T22324</td><td>ATCACACACTTAGATCAGATCGG</td><td>1000</td></tr>
<tr><td>Oligo fwd</td><td>CACCATCACACACTTAGATCAGAT</td></tr>
<tr><td>Oligo rev</td><td>AAACATCTGATCTAAGTGTGTGAT</td></tr>
<tr><td>Chromosome</td><td>start</td><td>end</td><td>strand</td><td>MM</td><td>target_seq</td><td>PAM</td><td>alignment</td><td>distance</td><td>position</td><td>gene name</td><td>gene id</td></tr>
<tr><td>GU305914.1</td><td>155450</td><td>155472</td><td>-</td><td>0</td><td>ATCACACACTTAGATCAGAT</td><td>CGG</td><td>||||||||[||||||||||||]PAM</td><td>NA</td><td>Intergenic</td><td>NA</td><td>NA</td></tr>
<tr><td>T22325</td><td>TGTGAGAACAGATGTATAAAAGG</td><td>1000</td></tr>
<tr><td>Oligo fwd</td><td>CACCTGTGAGAACAGATGTATAAA</td></tr>
<tr><td>Oligo rev</td><td>AAACTTTATACATCTGTTCTCACA</td></tr>
<tr><td>Chromosome</td><td>start</td><td>end</td><td>strand</td><td>MM</td><td>target_seq</td><td>PAM</td><td>alignment</td><td>distance</td><td>position</td><td>gene name</td><td>gene id</td></tr>
<tr><td>GU305914.1</td><td>155392</td><td>155414</td><td>-</td><td>0</td><td>TGTGAGAACAGATGTATAAA</td><td>AGG</td><td>||||||||[||||||||||||]PAM</td><td>NA</td><td>Intergenic</td><td>NA</td><td>NA</td></tr>
<tr><td>T22326</td><td>GTCTGCGATTATTCCATCACAGG</td><td>1000</td></tr>
<tr><td>Oligo fwd</td><td>CACCGTCTGCGATTATTCCATCAC</td></tr>
<tr><td>Oligo rev</td><td>AAACGTGATGGAATAATCGCAGAC</td></tr>
<tr><td>Chromosome</td><td>start</td><td>end</td><td>strand</td><td>MM</td><td>target_seq</td><td>PAM</td><td>alignment</td><td>distance</td><td>position</td><td>gene name</td><td>gene id</td></tr>
</table>

<table>
<tr><td>GU305914.1</td><td>155216</td><td>155238</td><td>-</td><td>0</td><td>GTCTGCGATTATTCCATCAC</td><td>AGG</td><td>||||||||[||||||||||||]PAM</td><td>NA</td><td>Intergenic</td><td>NA</td><td>NA</td></tr>
<tr><td>T22327</td><td>GATTATTCCATCACAGGCACCGG</td><td>1000</td><td></td><td></td><td></td><td></td><td></td><td></td><td></td><td></td><td></td></tr>
<tr><td>Oligo fwd</td><td>CACCGATTATTCCATCACAGGCAC</td><td></td><td></td><td></td><td></td><td></td><td></td><td></td><td></td><td></td><td></td></tr>
<tr><td>Oligo rev</td><td>AAACGTGCCTGTGATGGAATAATC</td><td></td><td></td><td></td><td></td><td></td><td></td><td></td><td></td><td></td><td></td></tr>
<tr><td>Chromosome</td><td>start</td><td>end</td><td>strand</td><td>MM</td><td>target_seq</td><td>PAM</td><td>alignment</td><td>distance</td><td>position</td><td>gene name</td><td>gene id</td></tr>
<tr><td>GU305914.1</td><td>155210</td><td>155232</td><td>-</td><td>0</td><td>GATTATTCCATCACAGGCAC</td><td>CGG</td><td>||||||||[||||||||||||]PAM</td><td>NA</td><td>Intergenic</td><td>NA</td><td>NA</td></tr>
<tr><td>T22328</td><td>TTTTCATCATATTCACGATCCGG</td><td>1000</td><td></td><td></td><td></td><td></td><td></td><td></td><td></td><td></td><td></td></tr>
<tr><td>Oligo fwd</td><td>CACCTTTTCATCATATTCACGATC</td><td></td><td></td><td></td><td></td><td></td><td></td><td></td><td></td><td></td><td></td></tr>
<tr><td>Oligo rev</td><td>AAACGATCGTGAATATGATGAAAA</td><td></td><td></td><td></td><td></td><td></td><td></td><td></td><td></td><td></td><td></td></tr>
<tr><td>Chromosome</td><td>start</td><td>end</td><td>strand</td><td>MM</td><td>target_seq</td><td>PAM</td><td>alignment</td><td>distance</td><td>position</td><td>gene name</td><td>gene id</td></tr>
<tr><td>GU305914.1</td><td>155174</td><td>155196</td><td>-</td><td>0</td><td>TTTTCATCATATTCACGATC</td><td>CGG</td><td>||||||||[||||||||||||]PAM</td><td>NA</td><td>Intergenic</td><td>NA</td><td>NA</td></tr>
<tr><td>T22329</td><td>TCATCATATTCACGATCCGGCGG</td><td>1000</td><td></td><td></td><td></td><td></td><td></td><td></td><td></td><td></td><td></td></tr>
<tr><td>Oligo fwd</td><td>CACCTCATCATATTCACGATCCGG</td><td></td><td></td><td></td><td></td><td></td><td></td><td></td><td></td><td></td><td></td></tr>
<tr><td>Oligo rev</td><td>AAACCCGGATCGTGAATATGATGA</td><td></td><td></td><td></td><td></td><td></td><td></td><td></td><td></td><td></td><td></td></tr>
<tr><td>Chromosome</td><td>start</td><td>end</td><td>strand</td><td>MM</td><td>target_seq</td><td>PAM</td><td>alignment</td><td>distance</td><td>position</td><td>gene name</td><td>gene id</td></tr>
<tr><td>GU305914.1</td><td>155171</td><td>155193</td><td>-</td><td>0</td><td>TCATCATATTCACGATCCGG</td><td>CGG</td><td>||||||||[||||||||||||]PAM</td><td>NA</td><td>Intergenic</td><td>NA</td><td>NA</td></tr>
<tr><td>T22330</td><td>TATCCACTCCATAATTTTTTAGG</td><td>1000</td><td></td><td></td><td></td><td></td><td></td><td></td><td></td><td></td><td></td></tr>
<tr><td>Oligo fwd</td><td>CACCTATCCACTCCATAATTTTTT</td><td></td><td></td><td></td><td></td><td></td><td></td><td></td><td></td><td></td><td></td></tr>
<tr><td>Oligo rev</td><td>AAACAAAAAATTATGGAGTGGATA</td><td></td><td></td><td></td><td></td><td></td><td></td><td></td><td></td><td></td><td></td></tr>
<tr><td>Chromosome</td><td>start</td><td>end</td><td>strand</td><td>MM</td><td>target_seq</td><td>PAM</td><td>alignment</td><td>distance</td><td>position</td><td>gene name</td><td>gene id</td></tr>
<tr><td>GU305914.1</td><td>155010</td><td>155032</td><td>-</td><td>0</td><td>TATCCACTCCATAATTTTTT</td><td>AGG</td><td>||||||||[||||||||||||]PAM</td><td>NA</td><td>Intergenic</td><td>NA</td><td>NA</td></tr>
<tr><td>T22331</td><td>TAGAAGCCATCAACACGAAGCGG</td><td>1000</td><td></td><td></td><td></td><td></td><td></td><td></td><td></td><td></td><td></td></tr>
<tr><td>Oligo fwd</td><td>CACCTAGAAGCCATCAACACGAAG</td><td></td><td></td><td></td><td></td><td></td><td></td><td></td><td></td><td></td><td></td></tr>
<tr><td>Oligo rev</td><td>AAACCTTCGTGTTGATGGCTTCTA</td><td></td><td></td><td></td><td></td><td></td><td></td><td></td><td></td><td></td><td></td></tr>
<tr><td>Chromosome</td><td>start</td><td>end</td><td>strand</td><td>MM</td><td>target_seq</td><td>PAM</td><td>alignment</td><td>distance</td><td>position</td><td>gene name</td><td>gene id</td></tr>
<tr><td>GU305914.1</td><td>154816</td><td>154838</td><td>-</td><td>0</td><td>TAGAAGCCATCAACACGAAG</td><td>CGG</td><td>||||||||[||||||||||||]PAM</td><td>NA</td><td>Intergenic</td><td>NA</td><td>NA</td></tr>
<tr><td>T22332</td><td>GCCATCAACACGAAGCGGCTCGG</td><td>1000</td><td></td><td></td><td></td><td></td><td></td><td></td><td></td><td></td><td></td></tr>
<tr><td>Oligo fwd</td><td>CACCGCCATCAACACGAAGCGGCT</td><td></td><td></td><td></td><td></td><td></td><td></td><td></td><td></td><td></td><td></td></tr>
<tr><td>Oligo rev</td><td>AAACAGCCGCTTCGTGTTGATGGC</td><td></td><td></td><td></td><td></td><td></td><td></td><td></td><td></td><td></td><td></td></tr>
<tr><td>Chromosome</td><td>start</td><td>end</td><td>strand</td><td>MM</td><td>target_seq</td><td>PAM</td><td>alignment</td><td>distance</td><td>position</td><td>gene name</td><td>gene id</td></tr>
<tr><td>GU305914.1</td><td>154811</td><td>154833</td><td>-</td><td>0</td><td>GCCATCAACACGAAGCGGCT</td><td>CGG</td><td>||||||||[||||||||||||]PAM</td><td>NA</td><td>Intergenic</td><td>NA</td><td>NA</td></tr>
<tr><td>T22333</td><td>CCATCAACACGAAGCGGCTCGGG</td><td>1000</td><td></td><td></td><td></td><td></td><td></td><td></td><td></td><td></td><td></td></tr>
<tr><td>Oligo fwd</td><td>CACCCCATCAACACGAAGCGGCTC</td><td></td><td></td><td></td><td></td><td></td><td></td><td></td><td></td><td></td><td></td></tr>
<tr><td>Oligo rev</td><td>AAACGAGCCGCTTCGTGTTGATGG</td><td></td><td></td><td></td><td></td><td></td><td></td><td></td><td></td><td></td><td></td></tr>
<tr><td>Chromosome</td><td>start</td><td>end</td><td>strand</td><td>MM</td><td>target_seq</td><td>PAM</td><td>alignment</td><td>distance</td><td>position</td><td>gene name</td><td>gene id</td></tr>
<tr><td>GU305914.1</td><td>154810</td><td>154832</td><td>-</td><td>0</td><td>CCATCAACACGAAGCGGCTC</td><td>GGG</td><td>||||||||[||||||||||||]PAM</td><td>NA</td><td>Intergenic</td><td>NA</td><td>NA</td></tr>
<tr><td>T22334</td><td>TCAACACGAAGCGGCTCGGGAGG</td><td>1000</td><td></td><td></td><td></td><td></td><td></td><td></td><td></td><td></td><td></td></tr>
<tr><td>Oligo fwd</td><td>CACCTCAACACGAAGCGGCTCGGG</td><td></td><td></td><td></td><td></td><td></td><td></td><td></td><td></td><td></td><td></td></tr>
<tr><td>Oligo rev</td><td>AAACCCCGAGCCGCTTCGTGTTGA</td><td></td><td></td><td></td><td></td><td></td><td></td><td></td><td></td><td></td><td></td></tr>
<tr><td>Chromosome</td><td>start</td><td>end</td><td>strand</td><td>MM</td><td>target_seq</td><td>PAM</td><td>alignment</td><td>distance</td><td>position</td><td>gene name</td><td>gene id</td></tr>
<tr><td>GU305914.1</td><td>154807</td><td>154829</td><td>-</td><td>0</td><td>TCAACACGAAGCGGCTCGGG</td><td>AGG</td><td>||||||||[||||||||||||]PAM</td><td>NA</td><td>Intergenic</td><td>NA</td><td>NA</td></tr>
<tr><td>T22335</td><td>CCATACCCCCCATTCGCCTCTGG</td><td>1000</td><td></td><td></td><td></td><td></td><td></td><td></td><td></td><td></td><td></td></tr>
<tr><td>Oligo fwd</td><td>CACCCCATACCCCCCATTCGCCTC</td><td></td><td></td><td></td><td></td><td></td><td></td><td></td><td></td><td></td><td></td></tr>
<tr><td>Oligo rev</td><td>AAACGAGGCGAATGGGGGGTATGG</td><td></td><td></td><td></td><td></td><td></td><td></td><td></td><td></td><td></td><td></td></tr>
<tr><td>Chromosome</td><td>start</td><td>end</td><td>strand</td><td>MM</td><td>target_seq</td><td>PAM</td><td>alignment</td><td>distance</td><td>position</td><td>gene name</td><td>gene id</td></tr>
<tr><td>GU305914.1</td><td>154734</td><td>154756</td><td>-</td><td>0</td><td>CCATACCCCCCATTCGCCTC</td><td>TGG</td><td>||||||||[||||||||||||]PAM</td><td>NA</td><td>Intergenic</td><td>NA</td><td>NA</td></tr>
<tr><td>T22336</td><td>CATACCCCCCATTCGCCTCTGGG</td><td>1000</td><td></td><td></td><td></td><td></td><td></td><td></td><td></td><td></td><td></td></tr>
<tr><td>Oligo fwd</td><td>CACCCATACCCCCCATTCGCCTCT</td><td></td><td></td><td></td><td></td><td></td><td></td><td></td><td></td><td></td><td></td></tr>
<tr><td>Oligo rev</td><td>AAACAGAGGCGAATGGGGGGTATG</td><td></td><td></td><td></td><td></td><td></td><td></td><td></td><td></td><td></td><td></td></tr>
<tr><td>Chromosome</td><td>start</td><td>end</td><td>strand</td><td>MM</td><td>target_seq</td><td>PAM</td><td>alignment</td><td>distance</td><td>position</td><td>gene name</td><td>gene id</td></tr>
<tr><td>GU305914.1</td><td>154733</td><td>154755</td><td>-</td><td>0</td><td>CATACCCCCCATTCGCCTCT</td><td>GGG</td><td>||||||||[||||||||||||]PAM</td><td>NA</td><td>Intergenic</td><td>NA</td><td>NA</td></tr>
<tr><td>T22337</td><td>CCCCCCATTCGCCTCTGGGTCGG</td><td>1000</td><td></td><td></td><td></td><td></td><td></td><td></td><td></td><td></td><td></td></tr>
<tr><td>Oligo fwd</td><td>CACCCCCCCCATTCGCCTCTGGGT</td><td></td><td></td><td></td><td></td><td></td><td></td><td></td><td></td><td></td><td></td></tr>
<tr><td>Oligo rev</td><td>AAACACCCAGAGGCGAATGGGGGG</td><td></td><td></td><td></td><td></td><td></td><td></td><td></td><td></td><td></td><td></td></tr>
<tr><td>Chromosome</td><td>start</td><td>end</td><td>strand</td><td>MM</td><td>target_seq</td><td>PAM</td><td>alignment</td><td>distance</td><td>position</td><td>gene name</td><td>gene id</td></tr>
<tr><td>GU305914.1</td><td>154729</td><td>154751</td><td>-</td><td>0</td><td>CCCCCCATTCGCCTCTGGGT</td><td>CGG</td><td>||||||||[||||||||||||]PAM</td><td>NA</td><td>Intergenic</td><td>NA</td><td>NA</td></tr>
<tr><td>T22338</td><td>CCTATTTTTCTGCAAAAATAAGG</td><td>1000</td><td></td><td></td><td></td><td></td><td></td><td></td><td></td><td></td><td></td></tr>
<tr><td>Oligo fwd</td><td>CACCCCTATTTTTCTGCAAAAATA</td><td></td><td></td><td></td><td></td><td></td><td></td><td></td><td></td><td></td><td></td></tr>
<tr><td>Oligo rev</td><td>AAACTATTTTTGCAGAAAAATAGG</td><td></td><td></td><td></td><td></td><td></td><td></td><td></td><td></td><td></td><td></td></tr>
<tr><td>Chromosome</td><td>start</td><td>end</td><td>strand</td><td>MM</td><td>target_seq</td><td>PAM</td><td>alignment</td><td>distance</td><td>position</td><td>gene name</td><td>gene id</td></tr>
<tr><td>GU305914.1</td><td>154569</td><td>154591</td><td>-</td><td>0</td><td>CCTATTTTTCTGCAAAAATA</td><td>AGG</td><td>||||||||[||||||||||||]PAM</td><td>NA</td><td>Intergenic</td><td>NA</td><td>NA</td></tr>
<tr><td>T22339</td><td>CGCATCTGTACGCAATTTTCTGG</td><td>1000</td><td></td><td></td><td></td><td></td><td></td><td></td><td></td><td></td><td></td></tr>
<tr><td>Oligo fwd</td><td>CACCCGCATCTGTACGCAATTTTC</td><td></td><td></td><td></td><td></td><td></td><td></td><td></td><td></td><td></td><td></td></tr>
<tr><td>Oligo rev</td><td>AAACGAAAATTGCGTACAGATGCG</td><td></td><td></td><td></td><td></td><td></td><td></td><td></td><td></td><td></td><td></td></tr>
<tr><td>Chromosome</td><td>start</td><td>end</td><td>strand</td><td>MM</td><td>target_seq</td><td>PAM</td><td>alignment</td><td>distance</td><td>position</td><td>gene name</td><td>gene id</td></tr>
<tr><td>GU305914.1</td><td>154526</td><td>154548</td><td>-</td><td>0</td><td>CGCATCTGTACGCAATTTTC</td><td>TGG</td><td>||||||||[||||||||||||]PAM</td><td>NA</td><td>Intergenic</td><td>NA</td><td>NA</td></tr>
<tr><td>T22340</td><td>TACGCAATTTTCTGGTCTCACGG</td><td>1000</td><td></td><td></td><td></td><td></td><td></td><td></td><td></td><td></td><td></td></tr>
<tr><td>Oligo fwd</td><td>CACCTACGCAATTTTCTGGTCTCA</td><td></td><td></td><td></td><td></td><td></td><td></td><td></td><td></td><td></td><td></td></tr>
<tr><td>Oligo rev</td><td>AAACTGAGACCAGAAAATTGCGTA</td><td></td><td></td><td></td><td></td><td></td><td></td><td></td><td></td><td></td><td></td></tr>
<tr><td>Chromosome</td><td>start</td><td>end</td><td>strand</td><td>MM</td><td>target_seq</td><td>PAM</td><td>alignment</td><td>distance</td><td>position</td><td>gene name</td><td>gene id</td></tr>
<tr><td>GU305914.1</td><td>154518</td><td>154540</td><td>-</td><td>0</td><td>TACGCAATTTTCTGGTCTCA</td><td>CGG</td><td>||||||||[||||||||||||]PAM</td><td>NA</td><td>Intergenic</td><td>NA</td><td>NA</td></tr>
<tr><td>T22341</td><td>ACGCAATTTTCTGGTCTCACGGG</td><td>1000</td><td></td><td></td><td></td><td></td><td></td><td></td><td></td><td></td><td></td></tr>
<tr><td>Oligo fwd</td><td>CACCACGCAATTTTCTGGTCTCAC</td><td></td><td></td><td></td><td></td><td></td><td></td><td></td><td></td><td></td><td></td></tr>
<tr><td>Oligo rev</td><td>AAACGTGAGACCAGAAAATTGCGT</td><td></td><td></td><td></td><td></td><td></td><td></td><td></td><td></td><td></td><td></td></tr>
</table>

| | | | | | | | | | | | |
|---|---|---|---|---|---|---|---|---|---|---|---|
| Chromosome | start | end | strand | MM | target_seq | PAM | alignment | distance | position | gene name | gene id |
| GU305914.1 | 154517 | 154539 | - | 0 | ACGCAATTTTCTGGTCTCAC | GGG | \|\|\|\|\|\|\|\|[\|\|\|\|\|\|\|\|\|\|\|\|]PAM | NA | Intergenic | NA | NA |
| T22342 | TTCTGGTCTCACGGGAGTCGAGG | 1000 | | | | | | | | | |
| Oligo fwd | CACCTTCTGGTCTCACGGGAGTCG | | | | | | | | | | |
| Oligo rev | AAACCGACTCCCGTGAGACCAGAA | | | | | | | | | | |
| Chromosome | start | end | strand | MM | target_seq | PAM | alignment | distance | position | gene name | gene id |
| GU305914.1 | 154509 | 154531 | - | 0 | TTCTGGTCTCACGGGAGTCG | AGG | \|\|\|\|\|\|\|\|[\|\|\|\|\|\|\|\|\|\|\|\|]PAM | NA | Intergenic | NA | NA |
| T22343 | TCACGGGAGTCGAGGTCTCACGG | 1000 | | | | | | | | | |
| Oligo fwd | CACCTCACGGGAGTCGAGGTCTCA | | | | | | | | | | |
| Oligo rev | AAACTGAGACCTCGACTCCCGTGA | | | | | | | | | | |
| Chromosome | start | end | strand | MM | target_seq | PAM | alignment | distance | position | gene name | gene id |
| GU305914.1 | 154501 | 154523 | - | 0 | TCACGGGAGTCGAGGTCTCA | CGG | \|\|\|\|\|\|\|\|[\|\|\|\|\|\|\|\|\|\|\|\|]PAM | NA | Intergenic | NA | NA |
| T22344 | CGGGAGTCGAGGTCTCACGGCGG | 1000 | | | | | | | | | |
| Oligo fwd | CACCCGGGAGTCGAGGTCTCACGG | | | | | | | | | | |
| Oligo rev | AAACCCGTGAGACCTCGACTCCCG | | | | | | | | | | |
| Chromosome | start | end | strand | MM | target_seq | PAM | alignment | distance | position | gene name | gene id |
| GU305914.1 | 154498 | 154520 | - | 0 | CGGGAGTCGAGGTCTCACGG | CGG | \|\|\|\|\|\|\|\|[\|\|\|\|\|\|\|\|\|\|\|\|]PAM | NA | Intergenic | NA | NA |
| T22345 | TCGAGGTCTCACGGCGGCCTCGG | 1000 | | | | | | | | | |
| Oligo fwd | CACCTCGAGGTCTCACGGCGGCCT | | | | | | | | | | |
| Oligo rev | AAACAGGCCGCCGTGAGACCTCGA | | | | | | | | | | |
| Chromosome | start | end | strand | MM | target_seq | PAM | alignment | distance | position | gene name | gene id |
| GU305914.1 | 154492 | 154514 | - | 0 | TCGAGGTCTCACGGCGGCCT | CGG | \|\|\|\|\|\|\|\|[\|\|\|\|\|\|\|\|\|\|\|\|]PAM | NA | Intergenic | NA | NA |
| T22346 | CGTTCTCTCCTTTCTTTTTCAGG | 1000 | | | | | | | | | |
| Oligo fwd | CACCCGTTCTCTCCTTTCTTTTTC | | | | | | | | | | |
| Oligo rev | AAACGAAAAAGAAAGGAGAGAACG | | | | | | | | | | |
| Chromosome | start | end | strand | MM | target_seq | PAM | alignment | distance | position | gene name | gene id |
| GU305914.1 | 154461 | 154483 | - | 0 | CGTTCTCTCCTTTCTTTTTC | AGG | \|\|\|\|\|\|\|\|[\|\|\|\|\|\|\|\|\|\|\|\|]PAM | NA | Intergenic | NA | NA |
| T22347 | TTCTTTTTCAGGATGCGAGATGG | 1000 | | | | | | | | | |
| Oligo fwd | CACCTTCTTTTTCAGGATGCGAGA | | | | | | | | | | |
| Oligo rev | AAACTCTCGCATCCTGAAAAAGAA | | | | | | | | | | |
| Chromosome | start | end | strand | MM | target_seq | PAM | alignment | distance | position | gene name | gene id |
| GU305914.1 | 154450 | 154472 | - | 0 | TTCTTTTTCAGGATGCGAGA | TGG | \|\|\|\|\|\|\|\|[\|\|\|\|\|\|\|\|\|\|\|\|]PAM | NA | Intergenic | NA | NA |
| T22348 | TTCAGGATGCGAGATGGCGACGG | 1000 | | | | | | | | | |
| Oligo fwd | CACCTTCAGGATGCGAGATGGCGA | | | | | | | | | | |
| Oligo rev | AAACTCGCCATCTCGCATCCTGAA | | | | | | | | | | |
| Chromosome | start | end | strand | MM | target_seq | PAM | alignment | distance | position | gene name | gene id |
| GU305914.1 | 154444 | 154466 | - | 0 | TTCAGGATGCGAGATGGCGA | CGG | \|\|\|\|\|\|\|\|[\|\|\|\|\|\|\|\|\|\|\|\|]PAM | NA | Intergenic | NA | NA |
| T22349 | CCGCCGCCGCCGCCACCGCCCGG | 1000 | | | | | | | | | |
| Oligo fwd | CACCCCGCCGCCGCCGCCACCGCC | | | | | | | | | | |
| Oligo rev | AAACGGCGGTGGCGGCGGCGGCGG | | | | | | | | | | |
| Chromosome | start | end | strand | MM | target_seq | PAM | alignment | distance | position | gene name | gene id |
| GU305914.1 | 154401 | 154423 | - | 0 | CCGCCGCCGCCGCCACCGCC | CGG | \|\|\|\|\|\|\|\|[\|\|\|\|\|\|\|\|\|\|\|\|]PAM | NA | Intergenic | NA | NA |
| GU305914.1 | 97884 | 97906 | + | 4 | CGGTCGCTGCCGCCGCCGCC | CGG | \|-\|-\|\|\|-[\|\|\|\|\|\|-\|\|\|\|\|]PAM | NA | Intergenic | NA | NA |
| T22350 | CGCCGCCGCCGCCACCGCCCGGG | 1000 | | | | | | | | | |
| Oligo fwd | CACCCGCCGCCGCCGCCACCGCCC | | | | | | | | | | |
| Oligo rev | AAACGGGCGGTGGCGGCGGCGGCG | | | | | | | | | | |
| Chromosome | start | end | strand | MM | target_seq | PAM | alignment | distance | position | gene name | gene id |
| GU305914.1 | 154400 | 154422 | - | 0 | CGCCGCCGCCGCCACCGCCC | GGG | \|\|\|\|\|\|\|\|[\|\|\|\|\|\|\|\|\|\|\|\|]PAM | NA | Intergenic | NA | NA |
| GU305914.1 | 97885 | 97907 | + | 4 | GGTCGCTGCCGCCGCCGCCC | GGG | -\|-\|\|\|-\|[\|\|\|\|\|-\|\|\|\|\|\|]PAM | NA | Intergenic | NA | NA |
| GU305914.1 | 35655 | 35677 | - | 3 | CCCCGCCGCCGCCGCCGCCG | TGG | \|-\|\|\|\|\|\|[\|\|\|\|\|-\|\|\|\|\|-]PAM | NA | Intergenic | NA | NA |
| T22351 | GCCGCCGCCGCCACCGCCCGGGG | 1000 | | | | | | | | | |
| Oligo fwd | CACCGCCGCCGCCGCCACCGCCCG | | | | | | | | | | |
| Oligo rev | AAACCGGGCGGTGGCGGCGGCGGC | | | | | | | | | | |
| Chromosome | start | end | strand | MM | target_seq | PAM | alignment | distance | position | gene name | gene id |
| GU305914.1 | 154399 | 154421 | - | 0 | GCCGCCGCCGCCACCGCCCG | GGG | \|\|\|\|\|\|\|\|[\|\|\|\|\|\|\|\|\|\|\|\|]PAM | NA | Intergenic | NA | NA |
| GU305914.1 | 52720 | 52742 | - | 3 | TCCGCCACCGCCACCGCCAG | CGG | -\|\|\|\|\|-\|[\|\|\|\|\|\|\|\|\|\|-\|]PAM | NA | Intergenic | NA | NA |
| GU305914.1 | 62668 | 62690 | + | 4 | AGCGCCGCCGCCCCCGCCGG | CGG | --\|\|\|\|\|\|[\|\|\|\|-\|\|\|\|\|-\|]PAM | NA | Intergenic | NA | NA |
| T22352 | CCCGGGGCGATCGTCCCATGTGG | 1000 | | | | | | | | | |
| Oligo fwd | CACCCCCGGGGCGATCGTCCCATG | | | | | | | | | | |
| Oligo rev | AAACCATGGGACGATCGCCCCGGG | | | | | | | | | | |
| Chromosome | start | end | strand | MM | target_seq | PAM | alignment | distance | position | gene name | gene id |
| GU305914.1 | 154383 | 154405 | - | 0 | CCCGGGGCGATCGTCCCATG | TGG | \|\|\|\|\|\|\|\|[\|\|\|\|\|\|\|\|\|\|\|\|]PAM | NA | Intergenic | NA | NA |
| T22353 | CACACGATCCCCATGCTGCGAGG | 1000 | | | | | | | | | |
| Oligo fwd | CACCCACACGATCCCCATGCTGCG | | | | | | | | | | |
| Oligo rev | AAACCGCAGCATGGGGATCGTGTG | | | | | | | | | | |
| Chromosome | start | end | strand | MM | target_seq | PAM | alignment | distance | position | gene name | gene id |
| GU305914.1 | 154359 | 154381 | - | 0 | CACACGATCCCCATGCTGCG | AGG | \|\|\|\|\|\|\|\|[\|\|\|\|\|\|\|\|\|\|\|\|]PAM | NA | Intergenic | NA | NA |
| T22354 | ACGATCCCCATGCTGCGAGGAGG | 1000 | | | | | | | | | |
| Oligo fwd | CACCACGATCCCCATGCTGCGAGG | | | | | | | | | | |
| Oligo rev | AAACCCTCGCAGCATGGGGATCGT | | | | | | | | | | |
| Chromosome | start | end | strand | MM | target_seq | PAM | alignment | distance | position | gene name | gene id |
| GU305914.1 | 154356 | 154378 | - | 0 | ACGATCCCCATGCTGCGAGG | AGG | \|\|\|\|\|\|\|\|[\|\|\|\|\|\|\|\|\|\|\|\|]PAM | NA | Intergenic | NA | NA |
| T22355 | ATCCCCATGCTGCGAGGAGGCGG | 1000 | | | | | | | | | |
| Oligo fwd | CACCATCCCCATGCTGCGAGGAGG | | | | | | | | | | |
| Oligo rev | AAACCCTCCTCGCAGCATGGGGAT | | | | | | | | | | |

| | | | | | | | | | | | |
|---|---|---|---|---|---|---|---|---|---|---|---|
| Chromosome | start | end | strand | MM | target_seq | PAM | alignment | distance | position | gene name | gene id |
| GU305914.1 | 154353 | 154375 | - | 0 | ATCCCCATGCTGCGAGGAGG | CGG | ||||||||[||||||||||||]PAM | NA | Intergenic | NA | NA |
| GU305914.1 | 159037 | 159059 | + | 4 | CTTCCCTTGCTGCGAGGAAG | GGG | -|-|||-|[||||||||||-|]PAM | NA | Intergenic | NA | NA |
| T22356 | CATCGAACTCTGCGCGATCCAGG | 1000 | | | | | | | | | |
| Oligo fwd | CACCCATCGAACTCTGCGCGATCC | | | | | | | | | | |
| Oligo rev | AAACGGATCGCGCAGAGTTCGATG | | | | | | | | | | |
| Chromosome | start | end | strand | MM | target_seq | PAM | alignment | distance | position | gene name | gene id |
| GU305914.1 | 154327 | 154349 | - | 0 | CATCGAACTCTGCGCGATCC | AGG | ||||||||[||||||||||||]PAM | NA | Intergenic | NA | NA |
| T22357 | ATCGAACTCTGCGCGATCCAGGG | 1000 | | | | | | | | | |
| Oligo fwd | CACCATCGAACTCTGCGCGATCCA | | | | | | | | | | |
| Oligo rev | AAACTGGATCGCGCAGAGTTCGAT | | | | | | | | | | |
| Chromosome | start | end | strand | MM | target_seq | PAM | alignment | distance | position | gene name | gene id |
| GU305914.1 | 154326 | 154348 | - | 0 | ATCGAACTCTGCGCGATCCA | GGG | ||||||||[||||||||||||]PAM | NA | Intergenic | NA | NA |
| T22358 | CTCTGCGCGATCCAGGGACACGG | 1000 | | | | | | | | | |
| Oligo fwd | CACCCTCTGCGCGATCCAGGGACA | | | | | | | | | | |
| Oligo rev | AAACTGTCCCTGGATCGCGCAGAG | | | | | | | | | | |
| Chromosome | start | end | strand | MM | target_seq | PAM | alignment | distance | position | gene name | gene id |
| GU305914.1 | 154320 | 154342 | - | 0 | CTCTGCGCGATCCAGGGACA | CGG | ||||||||[||||||||||||]PAM | NA | Intergenic | NA | NA |
| T22359 | AACGATCACGACGACCGCGACGG | 1000 | | | | | | | | | |
| Oligo fwd | CACCAACGATCACGACGACCGCGA | | | | | | | | | | |
| Oligo rev | AAACTCGCGGTCGTCGTGATCGTT | | | | | | | | | | |
| Chromosome | start | end | strand | MM | target_seq | PAM | alignment | distance | position | gene name | gene id |
| GU305914.1 | 154228 | 154250 | - | 0 | AACGATCACGACGACCGCGA | CGG | ||||||||[||||||||||||]PAM | NA | Intergenic | NA | NA |
| T22360 | CGACGGCGCAGACAAGTCGCAGG | 1000 | | | | | | | | | |
| Oligo fwd | CACCCGACGGCGCAGACAAGTCGC | | | | | | | | | | |
| Oligo rev | AAACGCGACTTGTCTGCGCCGTCG | | | | | | | | | | |
| Chromosome | start | end | strand | MM | target_seq | PAM | alignment | distance | position | gene name | gene id |
| GU305914.1 | 154211 | 154233 | - | 0 | CGACGGCGCAGACAAGTCGC | AGG | ||||||||[||||||||||||]PAM | NA | Intergenic | NA | NA |
| T22361 | CAGACAAGTCGCAGGCTCCGCGG | 1000 | | | | | | | | | |
| Oligo fwd | CACCCAGACAAGTCGCAGGCTCCG | | | | | | | | | | |
| Oligo rev | AAACCGGAGCCTGCGACTTGTCTG | | | | | | | | | | |
| Chromosome | start | end | strand | MM | target_seq | PAM | alignment | distance | position | gene name | gene id |
| GU305914.1 | 154203 | 154225 | - | 0 | CAGACAAGTCGCAGGCTCCG | CGG | ||||||||[||||||||||||]PAM | NA | Intergenic | NA | NA |
| T22362 | TCCTATACCCAGACCGCCAGCGG | 1000 | | | | | | | | | |
| Oligo fwd | CACCTCCTATACCCAGACCGCCAG | | | | | | | | | | |
| Oligo rev | AAACCTGGCGGTCTGGGTATAGGA | | | | | | | | | | |
| Chromosome | start | end | strand | MM | target_seq | PAM | alignment | distance | position | gene name | gene id |
| GU305914.1 | 154153 | 154175 | - | 0 | TCCTATACCCAGACCGCCAG | CGG | ||||||||[||||||||||||]PAM | NA | Intergenic | NA | NA |
| T22363 | CCTATACCCAGACCGCCAGCGGG | 1000 | | | | | | | | | |
| Oligo fwd | CACCCCTATACCCAGACCGCCAGC | | | | | | | | | | |
| Oligo rev | AAACGCTGGCGGTCTGGGTATAGG | | | | | | | | | | |
| Chromosome | start | end | strand | MM | target_seq | PAM | alignment | distance | position | gene name | gene id |
| GU305914.1 | 154152 | 154174 | - | 0 | CCTATACCCAGACCGCCAGC | GGG | ||||||||[||||||||||||]PAM | NA | Intergenic | NA | NA |
| T22364 | CCCCGAGATACCCAGAATCGTGG | 1000 | | | | | | | | | |
| Oligo fwd | CACCCCCCGAGATACCCAGAATCG | | | | | | | | | | |
| Oligo rev | AAACCGATTCTGGGTATCTCGGGG | | | | | | | | | | |
| Chromosome | start | end | strand | MM | target_seq | PAM | alignment | distance | position | gene name | gene id |
| GU305914.1 | 154027 | 154049 | - | 0 | CCCCGAGATACCCAGAATCG | TGG | ||||||||[||||||||||||]PAM | NA | Intergenic | NA | NA |
| T22365 | GATACCCAGAATCGTGGTCGAGG | 1000 | | | | | | | | | |
| Oligo fwd | CACCGATACCCAGAATCGTGGTCG | | | | | | | | | | |
| Oligo rev | AAACCGACCACGATTCTGGGTATC | | | | | | | | | | |
| Chromosome | start | end | strand | MM | target_seq | PAM | alignment | distance | position | gene name | gene id |
| GU305914.1 | 154021 | 154043 | - | 0 | GATACCCAGAATCGTGGTCG | AGG | ||||||||[||||||||||||]PAM | NA | Intergenic | NA | NA |
| T22366 | ACCCAGAATCGTGGTCGAGGTGG | 1000 | | | | | | | | | |
| Oligo fwd | CACCACCCAGAATCGTGGTCGAGG | | | | | | | | | | |
| Oligo rev | AAACCCTCGACCACGATTCTGGGT | | | | | | | | | | |
| Chromosome | start | end | strand | MM | target_seq | PAM | alignment | distance | position | gene name | gene id |
| GU305914.1 | 154018 | 154040 | - | 0 | ACCCAGAATCGTGGTCGAGG | TGG | ||||||||[||||||||||||]PAM | NA | Intergenic | NA | NA |
| T22367 | CTCATGTCAGAAAACCAGAGTGG | 1000 | | | | | | | | | |
| Oligo fwd | CACCCTCATGTCAGAAAACCAGAG | | | | | | | | | | |
| Oligo rev | AAACCTCTGGTTTTCTGACATGAG | | | | | | | | | | |
| Chromosome | start | end | strand | MM | target_seq | PAM | alignment | distance | position | gene name | gene id |
| GU305914.1 | 153955 | 153977 | - | 0 | CTCATGTCAGAAAACCAGAG | TGG | ||||||||[||||||||||||]PAM | NA | Intergenic | NA | NA |
| T22368 | CTGTAACGCGCGTCGTATAAAGG | 1000 | | | | | | | | | |
| Oligo fwd | CACCCTGTAACGCGCGTCGTATAA | | | | | | | | | | |
| Oligo rev | AAACTTATACGACGCGCGTTACAG | | | | | | | | | | |
| Chromosome | start | end | strand | MM | target_seq | PAM | alignment | distance | position | gene name | gene id |
| GU305914.1 | 153922 | 153944 | - | 0 | CTGTAACGCGCGTCGTATAA | AGG | ||||||||[||||||||||||]PAM | NA | Intergenic | NA | NA |
| T22369 | TGTAACGCGCGTCGTATAAAGGG | 1000 | | | | | | | | | |
| Oligo fwd | CACCTGTAACGCGCGTCGTATAAA | | | | | | | | | | |
| Oligo rev | AAACTTTATACGACGCGCGTTACA | | | | | | | | | | |
| Chromosome | start | end | strand | MM | target_seq | PAM | alignment | distance | position | gene name | gene id |
| GU305914.1 | 153921 | 153943 | - | 0 | TGTAACGCGCGTCGTATAAA | GGG | ||||||||[||||||||||||]PAM | NA | Intergenic | NA | NA |
| T22370 | GTAACGCGCGTCGTATAAAGGGG | 1000 | | | | | | | | | |

<table>
<tr><td>Oligo fwd</td><td>CACCGTAACGCGCGTCGTATAAAG</td><td></td><td></td><td></td><td></td><td></td><td></td><td></td><td></td><td></td><td></td></tr>
<tr><td>Oligo rev</td><td>AAACCTTTATACGACGCGCGTTAC</td><td></td><td></td><td></td><td></td><td></td><td></td><td></td><td></td><td></td><td></td></tr>
<tr><td>Chromosome</td><td>start</td><td>end</td><td>strand</td><td>MM</td><td>target_seq</td><td>PAM</td><td>alignment</td><td>distance</td><td>position</td><td>gene name</td><td>gene id</td></tr>
<tr><td>GU305914.1</td><td>153920</td><td>153942</td><td>-</td><td>0</td><td>GTAACGCGCGTCGTATAAAG</td><td>GGG</td><td>||||||||[||||||||||||]PAM</td><td>NA</td><td>Intergenic</td><td>NA</td><td>NA</td></tr>
<tr><td></td><td></td><td></td><td></td><td></td><td></td><td></td><td></td><td></td><td></td><td></td><td></td></tr>
<tr><td>T22371</td><td>TAACGCGCGTCGTATAAAGGGGG</td><td>1000</td><td></td><td></td><td></td><td></td><td></td><td></td><td></td><td></td><td></td></tr>
<tr><td>Oligo fwd</td><td>CACCTAACGCGCGTCGTATAAAGG</td><td></td><td></td><td></td><td></td><td></td><td></td><td></td><td></td><td></td><td></td></tr>
<tr><td>Oligo rev</td><td>AAACCCTTTATACGACGCGCGTTA</td><td></td><td></td><td></td><td></td><td></td><td></td><td></td><td></td><td></td><td></td></tr>
<tr><td>Chromosome</td><td>start</td><td>end</td><td>strand</td><td>MM</td><td>target_seq</td><td>PAM</td><td>alignment</td><td>distance</td><td>position</td><td>gene name</td><td>gene id</td></tr>
<tr><td>GU305914.1</td><td>153919</td><td>153941</td><td>-</td><td>0</td><td>TAACGCGCGTCGTATAAAGG</td><td>GGG</td><td>||||||||[||||||||||||]PAM</td><td>NA</td><td>Intergenic</td><td>NA</td><td>NA</td></tr>
<tr><td></td><td></td><td></td><td></td><td></td><td></td><td></td><td></td><td></td><td></td><td></td><td></td></tr>
<tr><td>T22372</td><td>TCGTATAAAGGGGGTGATGAAGG</td><td>1000</td><td></td><td></td><td></td><td></td><td></td><td></td><td></td><td></td><td></td></tr>
<tr><td>Oligo fwd</td><td>CACCTCGTATAAAGGGGGTGATGA</td><td></td><td></td><td></td><td></td><td></td><td></td><td></td><td></td><td></td><td></td></tr>
<tr><td>Oligo rev</td><td>AAACTCATCACCCCCTTTATACGA</td><td></td><td></td><td></td><td></td><td></td><td></td><td></td><td></td><td></td><td></td></tr>
<tr><td>Chromosome</td><td>start</td><td>end</td><td>strand</td><td>MM</td><td>target_seq</td><td>PAM</td><td>alignment</td><td>distance</td><td>position</td><td>gene name</td><td>gene id</td></tr>
<tr><td>GU305914.1</td><td>153910</td><td>153932</td><td>-</td><td>0</td><td>TCGTATAAAGGGGGTGATGA</td><td>AGG</td><td>||||||||[||||||||||||]PAM</td><td>NA</td><td>Intergenic</td><td>NA</td><td>NA</td></tr>
<tr><td></td><td></td><td></td><td></td><td></td><td></td><td></td><td></td><td></td><td></td><td></td><td></td></tr>
<tr><td>T22373</td><td>GTTCTCGTACGTGTTGCGTACGG</td><td>1000</td><td></td><td></td><td></td><td></td><td></td><td></td><td></td><td></td><td></td></tr>
<tr><td>Oligo fwd</td><td>CACCGTTCTCGTACGTGTTGCGTA</td><td></td><td></td><td></td><td></td><td></td><td></td><td></td><td></td><td></td><td></td></tr>
<tr><td>Oligo rev</td><td>AAACTACGCAACACGTACGAGAAC</td><td></td><td></td><td></td><td></td><td></td><td></td><td></td><td></td><td></td><td></td></tr>
<tr><td>Chromosome</td><td>start</td><td>end</td><td>strand</td><td>MM</td><td>target_seq</td><td>PAM</td><td>alignment</td><td>distance</td><td>position</td><td>gene name</td><td>gene id</td></tr>
<tr><td>GU305914.1</td><td>153883</td><td>153905</td><td>-</td><td>0</td><td>GTTCTCGTACGTGTTGCGTA</td><td>CGG</td><td>||||||||[||||||||||||]PAM</td><td>NA</td><td>Intergenic</td><td>NA</td><td>NA</td></tr>
<tr><td></td><td></td><td></td><td></td><td></td><td></td><td></td><td></td><td></td><td></td><td></td><td></td></tr>
<tr><td>T22374</td><td>TTCTCGTACGTGTTGCGTACGGG</td><td>1000</td><td></td><td></td><td></td><td></td><td></td><td></td><td></td><td></td><td></td></tr>
<tr><td>Oligo fwd</td><td>CACCTTCTCGTACGTGTTGCGTAC</td><td></td><td></td><td></td><td></td><td></td><td></td><td></td><td></td><td></td><td></td></tr>
<tr><td>Oligo rev</td><td>AAACGTACGCAACACGTACGAGAA</td><td></td><td></td><td></td><td></td><td></td><td></td><td></td><td></td><td></td><td></td></tr>
<tr><td>Chromosome</td><td>start</td><td>end</td><td>strand</td><td>MM</td><td>target_seq</td><td>PAM</td><td>alignment</td><td>distance</td><td>position</td><td>gene name</td><td>gene id</td></tr>
<tr><td>GU305914.1</td><td>153882</td><td>153904</td><td>-</td><td>0</td><td>TTCTCGTACGTGTTGCGTAC</td><td>GGG</td><td>||||||||[||||||||||||]PAM</td><td>NA</td><td>Intergenic</td><td>NA</td><td>NA</td></tr>
<tr><td></td><td></td><td></td><td></td><td></td><td></td><td></td><td></td><td></td><td></td><td></td><td></td></tr>
<tr><td>T22375</td><td>TCTCGTACGTGTTGCGTACGGGG</td><td>1000</td><td></td><td></td><td></td><td></td><td></td><td></td><td></td><td></td><td></td></tr>
<tr><td>Oligo fwd</td><td>CACCTCTCGTACGTGTTGCGTACG</td><td></td><td></td><td></td><td></td><td></td><td></td><td></td><td></td><td></td><td></td></tr>
<tr><td>Oligo rev</td><td>AAACCGTACGCAACACGTACGAGA</td><td></td><td></td><td></td><td></td><td></td><td></td><td></td><td></td><td></td><td></td></tr>
<tr><td>Chromosome</td><td>start</td><td>end</td><td>strand</td><td>MM</td><td>target_seq</td><td>PAM</td><td>alignment</td><td>distance</td><td>position</td><td>gene name</td><td>gene id</td></tr>
<tr><td>GU305914.1</td><td>153881</td><td>153903</td><td>-</td><td>0</td><td>TCTCGTACGTGTTGCGTACG</td><td>GGG</td><td>||||||||[||||||||||||]PAM</td><td>NA</td><td>Intergenic</td><td>NA</td><td>NA</td></tr>
<tr><td></td><td></td><td></td><td></td><td></td><td></td><td></td><td></td><td></td><td></td><td></td><td></td></tr>
<tr><td>T22376</td><td>CGGGGTTAGTATTGAGAACTAGG</td><td>1000</td><td></td><td></td><td></td><td></td><td></td><td></td><td></td><td></td><td></td></tr>
<tr><td>Oligo fwd</td><td>CACCCGGGGTTAGTATTGAGAACT</td><td></td><td></td><td></td><td></td><td></td><td></td><td></td><td></td><td></td><td></td></tr>
<tr><td>Oligo rev</td><td>AAACAGTTCTCAATACTAACCCCG</td><td></td><td></td><td></td><td></td><td></td><td></td><td></td><td></td><td></td><td></td></tr>
<tr><td>Chromosome</td><td>start</td><td>end</td><td>strand</td><td>MM</td><td>target_seq</td><td>PAM</td><td>alignment</td><td>distance</td><td>position</td><td>gene name</td><td>gene id</td></tr>
<tr><td>GU305914.1</td><td>153863</td><td>153885</td><td>-</td><td>0</td><td>CGGGGTTAGTATTGAGAACT</td><td>AGG</td><td>||||||||[||||||||||||]PAM</td><td>NA</td><td>Intergenic</td><td>NA</td><td>NA</td></tr>
<tr><td></td><td></td><td></td><td></td><td></td><td></td><td></td><td></td><td></td><td></td><td></td><td></td></tr>
<tr><td>T22377</td><td>GGTTAGTATTGAGAACTAGGCGG</td><td>1000</td><td></td><td></td><td></td><td></td><td></td><td></td><td></td><td></td><td></td></tr>
<tr><td>Oligo fwd</td><td>CACCGGTTAGTATTGAGAACTAGG</td><td></td><td></td><td></td><td></td><td></td><td></td><td></td><td></td><td></td><td></td></tr>
<tr><td>Oligo rev</td><td>AAACCCTAGTTCTCAATACTAACC</td><td></td><td></td><td></td><td></td><td></td><td></td><td></td><td></td><td></td><td></td></tr>
<tr><td>Chromosome</td><td>start</td><td>end</td><td>strand</td><td>MM</td><td>target_seq</td><td>PAM</td><td>alignment</td><td>distance</td><td>position</td><td>gene name</td><td>gene id</td></tr>
<tr><td>GU305914.1</td><td>153860</td><td>153882</td><td>-</td><td>0</td><td>GGTTAGTATTGAGAACTAGG</td><td>CGG</td><td>||||||||[||||||||||||]PAM</td><td>NA</td><td>Intergenic</td><td>NA</td><td>NA</td></tr>
<tr><td></td><td></td><td></td><td></td><td></td><td></td><td></td><td></td><td></td><td></td><td></td><td></td></tr>
<tr><td>T22378</td><td>AGTATTGAGAACTAGGCGGTCGG</td><td>1000</td><td></td><td></td><td></td><td></td><td></td><td></td><td></td><td></td><td></td></tr>
<tr><td>Oligo fwd</td><td>CACCAGTATTGAGAACTAGGCGGT</td><td></td><td></td><td></td><td></td><td></td><td></td><td></td><td></td><td></td><td></td></tr>
<tr><td>Oligo rev</td><td>AAACACCGCCTAGTTCTCAATACT</td><td></td><td></td><td></td><td></td><td></td><td></td><td></td><td></td><td></td><td></td></tr>
<tr><td>Chromosome</td><td>start</td><td>end</td><td>strand</td><td>MM</td><td>target_seq</td><td>PAM</td><td>alignment</td><td>distance</td><td>position</td><td>gene name</td><td>gene id</td></tr>
<tr><td>GU305914.1</td><td>153856</td><td>153878</td><td>-</td><td>0</td><td>AGTATTGAGAACTAGGCGGT</td><td>CGG</td><td>||||||||[||||||||||||]PAM</td><td>NA</td><td>Intergenic</td><td>NA</td><td>NA</td></tr>
<tr><td></td><td></td><td></td><td></td><td></td><td></td><td></td><td></td><td></td><td></td><td></td><td></td></tr>
<tr><td>T22379</td><td>GTATTGAGAACTAGGCGGTCGGG</td><td>1000</td><td></td><td></td><td></td><td></td><td></td><td></td><td></td><td></td><td></td></tr>
<tr><td>Oligo fwd</td><td>CACCGTATTGAGAACTAGGCGGTC</td><td></td><td></td><td></td><td></td><td></td><td></td><td></td><td></td><td></td><td></td></tr>
<tr><td>Oligo rev</td><td>AAACGACCGCCTAGTTCTCAATAC</td><td></td><td></td><td></td><td></td><td></td><td></td><td></td><td></td><td></td><td></td></tr>
<tr><td>Chromosome</td><td>start</td><td>end</td><td>strand</td><td>MM</td><td>target_seq</td><td>PAM</td><td>alignment</td><td>distance</td><td>position</td><td>gene name</td><td>gene id</td></tr>
<tr><td>GU305914.1</td><td>153855</td><td>153877</td><td>-</td><td>0</td><td>GTATTGAGAACTAGGCGGTC</td><td>GGG</td><td>||||||||[||||||||||||]PAM</td><td>NA</td><td>Intergenic</td><td>NA</td><td>NA</td></tr>
<tr><td></td><td></td><td></td><td></td><td></td><td></td><td></td><td></td><td></td><td></td><td></td><td></td></tr>
<tr><td>T22380</td><td>TATTGAGAACTAGGCGGTCGGGG</td><td>1000</td><td></td><td></td><td></td><td></td><td></td><td></td><td></td><td></td><td></td></tr>
<tr><td>Oligo fwd</td><td>CACCTATTGAGAACTAGGCGGTCG</td><td></td><td></td><td></td><td></td><td></td><td></td><td></td><td></td><td></td><td></td></tr>
<tr><td>Oligo rev</td><td>AAACCGACCGCCTAGTTCTCAATA</td><td></td><td></td><td></td><td></td><td></td><td></td><td></td><td></td><td></td><td></td></tr>
<tr><td>Chromosome</td><td>start</td><td>end</td><td>strand</td><td>MM</td><td>target_seq</td><td>PAM</td><td>alignment</td><td>distance</td><td>position</td><td>gene name</td><td>gene id</td></tr>
<tr><td>GU305914.1</td><td>153854</td><td>153876</td><td>-</td><td>0</td><td>TATTGAGAACTAGGCGGTCG</td><td>GGG</td><td>||||||||[||||||||||||]PAM</td><td>NA</td><td>Intergenic</td><td>NA</td><td>NA</td></tr>
<tr><td></td><td></td><td></td><td></td><td></td><td></td><td></td><td></td><td></td><td></td><td></td><td></td></tr>
<tr><td>T22381</td><td>GGCGGTCGGGGTCCACCACCCGG</td><td>1000</td><td></td><td></td><td></td><td></td><td></td><td></td><td></td><td></td><td></td></tr>
<tr><td>Oligo fwd</td><td>CACCGGCGGTCGGGGTCCACCACC</td><td></td><td></td><td></td><td></td><td></td><td></td><td></td><td></td><td></td><td></td></tr>
<tr><td>Oligo rev</td><td>AAACGGTGGTGGACCCCGACCGCC</td><td></td><td></td><td></td><td></td><td></td><td></td><td></td><td></td><td></td><td></td></tr>
<tr><td>Chromosome</td><td>start</td><td>end</td><td>strand</td><td>MM</td><td>target_seq</td><td>PAM</td><td>alignment</td><td>distance</td><td>position</td><td>gene name</td><td>gene id</td></tr>
<tr><td>GU305914.1</td><td>153842</td><td>153864</td><td>-</td><td>0</td><td>GGCGGTCGGGGTCCACCACC</td><td>CGG</td><td>||||||||[||||||||||||]PAM</td><td>NA</td><td>Intergenic</td><td>NA</td><td>NA</td></tr>
<tr><td></td><td></td><td></td><td></td><td></td><td></td><td></td><td></td><td></td><td></td><td></td><td></td></tr>
<tr><td>T22382</td><td>GCGGTCGGGGTCCACCACCCGGG</td><td>1000</td><td></td><td></td><td></td><td></td><td></td><td></td><td></td><td></td><td></td></tr>
<tr><td>Oligo fwd</td><td>CACCGCGGTCGGGGTCCACCACCC</td><td></td><td></td><td></td><td></td><td></td><td></td><td></td><td></td><td></td><td></td></tr>
<tr><td>Oligo rev</td><td>AAACGGGTGGTGGACCCCGACCGC</td><td></td><td></td><td></td><td></td><td></td><td></td><td></td><td></td><td></td><td></td></tr>
<tr><td>Chromosome</td><td>start</td><td>end</td><td>strand</td><td>MM</td><td>target_seq</td><td>PAM</td><td>alignment</td><td>distance</td><td>position</td><td>gene name</td><td>gene id</td></tr>
<tr><td>GU305914.1</td><td>153841</td><td>153863</td><td>-</td><td>0</td><td>GCGGTCGGGGTCCACCACCC</td><td>GGG</td><td>||||||||[||||||||||||]PAM</td><td>NA</td><td>Intergenic</td><td>NA</td><td>NA</td></tr>
<tr><td></td><td></td><td></td><td></td><td></td><td></td><td></td><td></td><td></td><td></td><td></td><td></td></tr>
<tr><td>T22383</td><td>GGGAGATACCTACGTAGATCTGG</td><td>1000</td><td></td><td></td><td></td><td></td><td></td><td></td><td></td><td></td><td></td></tr>
<tr><td>Oligo fwd</td><td>CACCGGGAGATACCTACGTAGATC</td><td></td><td></td><td></td><td></td><td></td><td></td><td></td><td></td><td></td><td></td></tr>
<tr><td>Oligo rev</td><td>AAACGATCTACGTAGGTATCTCCC</td><td></td><td></td><td></td><td></td><td></td><td></td><td></td><td></td><td></td><td></td></tr>
<tr><td>Chromosome</td><td>start</td><td>end</td><td>strand</td><td>MM</td><td>target_seq</td><td>PAM</td><td>alignment</td><td>distance</td><td>position</td><td>gene name</td><td>gene id</td></tr>
<tr><td>GU305914.1</td><td>153821</td><td>153843</td><td>-</td><td>0</td><td>GGGAGATACCTACGTAGATC</td><td>TGG</td><td>||||||||[||||||||||||]PAM</td><td>NA</td><td>Intergenic</td><td>NA</td><td>NA</td></tr>
<tr><td></td><td></td><td></td><td></td><td></td><td></td><td></td><td></td><td></td><td></td><td></td><td></td></tr>
<tr><td>T22384</td><td>CGTAGATCTGGCTCATCTTCAGG</td><td>1000</td><td></td><td></td><td></td><td></td><td></td><td></td><td></td><td></td><td></td></tr>
<tr><td>Oligo fwd</td><td>CACCCGTAGATCTGGCTCATCTTC</td><td></td><td></td><td></td><td></td><td></td><td></td><td></td><td></td><td></td><td></td></tr>
<tr><td>Oligo rev</td><td>AAACGAAGATGAGCCAGATCTACG</td><td></td><td></td><td></td><td></td><td></td><td></td><td></td><td></td><td></td><td></td></tr>
<tr><td>Chromosome</td><td>start</td><td>end</td><td>strand</td><td>MM</td><td>target_seq</td><td>PAM</td><td>alignment</td><td>distance</td><td>position</td><td>gene name</td><td>gene id</td></tr>
<tr><td>GU305914.1</td><td>153809</td><td>153831</td><td>-</td><td>0</td><td>CGTAGATCTGGCTCATCTTC</td><td>AGG</td><td>||||||||[||||||||||||]PAM</td><td>NA</td><td>Intergenic</td><td>NA</td><td>NA</td></tr>
</table>

T22385 TGGCTCATCTTCAGGTTTTTAGG 1000
Oligo fwd CACCTGGCTCATCTTCAGGTTTTT
Oligo rev AAACAAAAACCTGAAGATGAGCCA
Chromosome start end strand MM target_seq PAM alignment distance position gene name gene id
GU305914.1 153801 153823 - 0 TGGCTCATCTTCAGGTTTTT AGG ||||||||[||||||||||||]PAM NA Intergenic NA NA

T22386 GGCTCATCTTCAGGTTTTTAGGG 1000
Oligo fwd CACCGGCTCATCTTCAGGTTTTTA
Oligo rev AAACTAAAAACCTGAAGATGAGCC
Chromosome start end strand MM target_seq PAM alignment distance position gene name gene id
GU305914.1 153800 153822 - 0 GGCTCATCTTCAGGTTTTTA GGG ||||||||[||||||||||||]PAM NA Intergenic NA NA

T22387 AGGGTTGTCTCCGAAATCGATGG 1000
Oligo fwd CACCAGGGTTGTCTCCGAAATCGA
Oligo rev AAACTCGATTTCGGAGACAACCCT
Chromosome start end strand MM target_seq PAM alignment distance position gene name gene id
GU305914.1 153781 153803 - 0 AGGGTTGTCTCCGAAATCGA TGG ||||||||[||||||||||||]PAM NA Intergenic NA NA

T22388 CAGACCGATGCCCTGACTCTTGG 1000
Oligo fwd CACCCAGACCGATGCCCTGACTCT
Oligo rev AAACAGAGTCAGGGCATCGGTCTG
Chromosome start end strand MM target_seq PAM alignment distance position gene name gene id
GU305914.1 153748 153770 - 0 CAGACCGATGCCCTGACTCT TGG ||||||||[||||||||||||]PAM NA Intergenic NA NA

T22389 GATGCCCTGACTCTTGGCGATGG 1000
Oligo fwd CACCGATGCCCTGACTCTTGGCGA
Oligo rev AAACTCGCCAAGAGTCAGGGCATC
Chromosome start end strand MM target_seq PAM alignment distance position gene name gene id
GU305914.1 153742 153764 - 0 GATGCCCTGACTCTTGGCGA TGG ||||||||[||||||||||||]PAM NA Intergenic NA NA

T22390 CTGACTCTTGGCGATGGTCATGG 1000
Oligo fwd CACCCTGACTCTTGGCGATGGTCA
Oligo rev AAACTGACCATCGCCAAGAGTCAG
Chromosome start end strand MM target_seq PAM alignment distance position gene name gene id
GU305914.1 153736 153758 - 0 CTGACTCTTGGCGATGGTCA TGG ||||||||[||||||||||||]PAM NA Intergenic NA NA

T22391 GATGGTCATGGCCATGCGCGAGG 1000
Oligo fwd CACCGATGGTCATGGCCATGCGCG
Oligo rev AAACCGCGCATGGCCATGACCATC
Chromosome start end strand MM target_seq PAM alignment distance position gene name gene id
GU305914.1 153724 153746 - 0 GATGGTCATGGCCATGCGCG AGG ||||||||[||||||||||||]PAM NA Intergenic NA NA

T22392 GGTCACCCCGTAATCCACCATGG 1000
Oligo fwd CACCGGTCACCCCGTAATCCACCA
Oligo rev AAACTGGTGGATTACGGGGTGACC
Chromosome start end strand MM target_seq PAM alignment distance position gene name gene id
GU305914.1 153703 153725 - 0 GGTCACCCCGTAATCCACCA TGG ||||||||[||||||||||||]PAM NA Intergenic NA NA

T22393 GCTCTTACCGTCTATCACGTCGG 1000
Oligo fwd CACCGCTCTTACCGTCTATCACGT
Oligo rev AAACACGTGATAGACGGTAAGAGC
Chromosome start end strand MM target_seq PAM alignment distance position gene name gene id
GU305914.1 153667 153689 - 0 GCTCTTACCGTCTATCACGT CGG ||||||||[||||||||||||]PAM NA Intergenic NA NA

T22394 GTCTATCACGTCGGAGAACTTGG 1000
Oligo fwd CACCGTCTATCACGTCGGAGAACT
Oligo rev AAACAGTTCTCCGACGTGATAGAC
Chromosome start end strand MM target_seq PAM alignment distance position gene name gene id
GU305914.1 153658 153680 - 0 GTCTATCACGTCGGAGAACT TGG ||||||||[||||||||||||]PAM NA Intergenic NA NA

T22395 ACTTGGACACGTTGTAGTCCAGG 1000
Oligo fwd CACCACTTGGACACGTTGTAGTCC
Oligo rev AAACGGACTACAACGTGTCCAAGT
Chromosome start end strand MM target_seq PAM alignment distance position gene name gene id
GU305914.1 153641 153663 - 0 ACTTGGACACGTTGTAGTCC AGG ||||||||[||||||||||||]PAM NA Intergenic NA NA

T22396 CGTTGTAGTCCAGGATGAGCAGG 1000
Oligo fwd CACCCGTTGTAGTCCAGGATGAGC
Oligo rev AAACGCTCATCCTGGACTACAACG
Chromosome start end strand MM target_seq PAM alignment distance position gene name gene id
GU305914.1 153632 153654 - 0 CGTTGTAGTCCAGGATGAGC AGG ||||||||[||||||||||||]PAM NA Intergenic NA NA

T22397 CAGGATGAGCAGGAAGCCGCAGG 1000
Oligo fwd CACCCAGGATGAGCAGGAAGCCGC
Oligo rev AAACGCGGCTTCCTGCTCATCCTG
Chromosome start end strand MM target_seq PAM alignment distance position gene name gene id
GU305914.1 153622 153644 - 0 CAGGATGAGCAGGAAGCCGC AGG ||||||||[||||||||||||]PAM NA Intergenic NA NA

T22398 GGCGTCCCTGACGATCAGCCTGG 1000
Oligo fwd CACCGGCGTCCCTGACGATCAGCC
Oligo rev AAACGGCTGATCGTCAGGGACGCC
Chromosome start end strand MM target_seq PAM alignment distance position gene name gene id
GU305914.1 153601 153623 - 0 GGCGTCCCTGACGATCAGCC TGG ||||||||[||||||||||||]PAM NA Intergenic NA NA

T22399 GCGTCCCTGACGATCAGCCTGGG 1000
Oligo fwd CACCGCGTCCCTGACGATCAGCCT
Oligo rev AAACAGGCTGATCGTCAGGGACGC
Chromosome start end strand MM target_seq PAM alignment distance position gene name gene id
GU305914.1 153600 153622 - 0 GCGTCCCTGACGATCAGCCT GGG ||||||||[||||||||||||]PAM NA Intergenic NA NA

T22400 CGTCCCTGACGATCAGCCTGGGG 1000
Oligo fwd CACCCGTCCCTGACGATCAGCCTG
Oligo rev AAACCAGGCTGATCGTCAGGGACG
Chromosome start end strand MM target_seq PAM alignment distance position gene name gene id
GU305914.1 153599 153621 - 0 CGTCCCTGACGATCAGCCTG GGG ||||||||[||||||||||||]PAM NA Intergenic NA NA

T22401 CCCTGACGATCAGCCTGGGGAGG 1000
Oligo fwd CACCCCCTGACGATCAGCCTGGGG
Oligo rev AAACCCCCAGGCTGATCGTCAGGG
Chromosome start end strand MM target_seq PAM alignment distance position gene name gene id
GU305914.1 153596 153618 - 0 CCCTGACGATCAGCCTGGGG AGG ||||||||[||||||||||||]PAM NA Intergenic NA NA

T22402 GCCTGGGGAGGCCCTTCTCCAGG 1000
Oligo fwd CACCGCCTGGGGAGGCCCTTCTCC
Oligo rev AAACGGAGAAGGGCCTCCCCAGGC
Chromosome start end strand MM target_seq PAM alignment distance position gene name gene id
GU305914.1 153584 153606 - 0 GCCTGGGGAGGCCCTTCTCC AGG ||||||||[||||||||||||]PAM NA Intergenic NA NA

T22403 CCAGGATCCGCCGATGTATACGG 1000
Oligo fwd CACCCCAGGATCCGCCGATGTATA
Oligo rev AAACTATACATCGGCGGATCCTGG
Chromosome start end strand MM target_seq PAM alignment distance position gene name gene id
GU305914.1 153566 153588 - 0 CCAGGATCCGCCGATGTATA CGG ||||||||[||||||||||||]PAM NA Intergenic NA NA

T22404 CCGATGTATACGGTTCCGTTCGG 1000
Oligo fwd CACCCCGATGTATACGGTTCCGTT
Oligo rev AAACAACGGAACCGTATACATCGG
Chromosome start end strand MM target_seq PAM alignment distance position gene name gene id
GU305914.1 153556 153578 - 0 CCGATGTATACGGTTCCGTT CGG ||||||||[||||||||||||]PAM NA Intergenic NA NA

T22405 CGTCCATGATGAAGACGTTGTGG 1000
Oligo fwd CACCCGTCCATGATGAAGACGTTG
Oligo rev AAACCAACGTCTTCATCATGGACG
Chromosome start end strand MM target_seq PAM alignment distance position gene name gene id
GU305914.1 153533 153555 - 0 CGTCCATGATGAAGACGTTG TGG ||||||||[||||||||||||]PAM NA Intergenic NA NA

T22406 CATGATGAAGACGTTGTGGTTGG 1000
Oligo fwd CACCCATGATGAAGACGTTGTGGT
Oligo rev AAACACCACAACGTCTTCATCATG
Chromosome start end strand MM target_seq PAM alignment distance position gene name gene id
GU305914.1 153529 153551 - 0 CATGATGAAGACGTTGTGGT TGG ||||||||[||||||||||||]PAM NA Intergenic NA NA

T22407 CTCCAGCACGTAGTTGTTGCTGG 1000
Oligo fwd CACCCTCCAGCACGTAGTTGTTGC
Oligo rev AAACGCAACAACTACGTGCTGGAG
Chromosome start end strand MM target_seq PAM alignment distance position gene name gene id
GU305914.1 153499 153521 - 0 CTCCAGCACGTAGTTGTTGC TGG ||||||||[||||||||||||]PAM NA Intergenic NA NA

T22408 TCCAGCACGTAGTTGTTGCTGGG 1000
Oligo fwd CACCTCCAGCACGTAGTTGTTGCT
Oligo rev AAACAGCAACAACTACGTGCTGGA
Chromosome start end strand MM target_seq PAM alignment distance position gene name gene id
GU305914.1 153498 153520 - 0 TCCAGCACGTAGTTGTTGCT GGG ||||||||[||||||||||||]PAM NA Intergenic NA NA

T22409 CCAGCACGTAGTTGTTGCTGGGG 1000
Oligo fwd CACCCCAGCACGTAGTTGTTGCTG
Oligo rev AAACCAGCAACAACTACGTGCTGG
Chromosome start end strand MM target_seq PAM alignment distance position gene name gene id
GU305914.1 153497 153519 - 0 CCAGCACGTAGTTGTTGCTG GGG ||||||||[||||||||||||]PAM NA Intergenic NA NA

T22410 CAGCACGTAGTTGTTGCTGGGGG 1000
Oligo fwd CACCCAGCACGTAGTTGTTGCTGG
Oligo rev AAACCCAGCAACAACTACGTGCTG
Chromosome start end strand MM target_seq PAM alignment distance position gene name gene id
GU305914.1 153496 153518 - 0 CAGCACGTAGTTGTTGCTGG GGG ||||||||[||||||||||||]PAM NA Intergenic NA NA

T22411 GTTGTTGCTGGGGGAGACGAAGG 1000
Oligo fwd CACCGTTGTTGCTGGGGGAGACGA
Oligo rev AAACTCGTCTCCCCCAGCAACAAC
Chromosome start end strand MM target_seq PAM alignment distance position gene name gene id
GU305914.1 153487 153509 - 0 GTTGTTGCTGGGGGAGACGA AGG ||||||||[||||||||||||]PAM NA Intergenic NA NA

T22412 GAAGGTCAGCATCCCCACGAAGG 1000
Oligo fwd CACCGAAGGTCAGCATCCCCACGA
Oligo rev AAACTCGTGGGGATGCTGACCTTC
Chromosome start end strand MM target_seq PAM alignment distance position gene name gene id
GU305914.1 153469 153491 - 0 GAAGGTCAGCATCCCCACGA AGG ||||||||[||||||||||||]PAM NA Intergenic NA NA

T22413 TCCCCACGAAGGATCCCGTCTGG 1000
Oligo fwd CACCTCCCCACGAAGGATCCCGTC
Oligo rev AAACGACGGGATCCTTCGTGGGGA
Chromosome start end strand MM target_seq PAM alignment distance position gene name gene id
GU305914.1 153458 153480 - 0 TCCCCACGAAGGATCCCGTC TGG ||||||||[||||||||||||]PAM NA Intergenic NA NA

T22414 CCCCACGAAGGATCCCGTCTGGG 1000
Oligo fwd CACCCCCCACGAAGGATCCCGTCT
Oligo rev AAACAGACGGGATCCTTCGTGGGG
Chromosome start end strand MM target_seq PAM alignment distance position gene name gene id

| | | | | | | | | | | | |
|---|---|---|---|---|---|---|---|---|---|---|---|
| GU305914.1 | 153457 | 153479 | - | 0 | CCCCACGAAGGATCCCGTCT | GGG | \|\|\|\|\|\|\|\|[\|\|\|\|\|\|\|\|\|\|\|\|]PAM | NA | Intergenic | NA | NA |
| T22415 | CCACACGTTCCTCCTGTTGTAGG | 1000 | | | | | | | | | |
| Oligo fwd | CACCCCACACGTTCCTCCTGTTGT | | | | | | | | | | |
| Oligo rev | AAACACAACAGGAGGAACGTGTGG | | | | | | | | | | |
| Chromosome | start | end | strand | MM | target_seq | PAM | alignment | distance | position | gene name | gene id |
| GU305914.1 | 153409 | 153431 | - | 0 | CCACACGTTCCTCCTGTTGT | AGG | \|\|\|\|\|\|\|\|[\|\|\|\|\|\|\|\|\|\|\|\|]PAM | NA | Intergenic | NA | NA |
| T22416 | TCCTGTTGTAGGTGACGAGCCGG | 1000 | | | | | | | | | |
| Oligo fwd | CACCTCCTGTTGTAGGTGACGAGC | | | | | | | | | | |
| Oligo rev | AAACGCTCGTCACCTACAACAGGA | | | | | | | | | | |
| Chromosome | start | end | strand | MM | target_seq | PAM | alignment | distance | position | gene name | gene id |
| GU305914.1 | 153398 | 153420 | - | 0 | TCCTGTTGTAGGTGACGAGC | CGG | \|\|\|\|\|\|\|\|[\|\|\|\|\|\|\|\|\|\|\|\|]PAM | NA | Intergenic | NA | NA |
| T22417 | CCTGTTGTAGGTGACGAGCCGGG | 1000 | | | | | | | | | |
| Oligo fwd | CACCCCTGTTGTAGGTGACGAGCC | | | | | | | | | | |
| Oligo rev | AAACGGCTCGTCACCTACAACAGG | | | | | | | | | | |
| Chromosome | start | end | strand | MM | target_seq | PAM | alignment | distance | position | gene name | gene id |
| GU305914.1 | 153397 | 153419 | - | 0 | CCTGTTGTAGGTGACGAGCC | GGG | \|\|\|\|\|\|\|\|[\|\|\|\|\|\|\|\|\|\|\|\|]PAM | NA | Intergenic | NA | NA |
| T22418 | GACGAGCCGGGTCTTGCCGAAGG | 1000 | | | | | | | | | |
| Oligo fwd | CACCGACGAGCCGGGTCTTGCCGA | | | | | | | | | | |
| Oligo rev | AAACTCGGCAAGACCCGGCTCGTC | | | | | | | | | | |
| Chromosome | start | end | strand | MM | target_seq | PAM | alignment | distance | position | gene name | gene id |
| GU305914.1 | 153385 | 153407 | - | 0 | GACGAGCCGGGTCTTGCCGA | AGG | \|\|\|\|\|\|\|\|[\|\|\|\|\|\|\|\|\|\|\|\|]PAM | NA | Intergenic | NA | NA |
| T22419 | GGTCTTGCCGAAGGCGCCCTTGG | 1000 | | | | | | | | | |
| Oligo fwd | CACCGGTCTTGCCGAAGGCGCCCT | | | | | | | | | | |
| Oligo rev | AAACAGGGCGCCTTCGGCAAGACC | | | | | | | | | | |
| Chromosome | start | end | strand | MM | target_seq | PAM | alignment | distance | position | gene name | gene id |
| GU305914.1 | 153376 | 153398 | - | 0 | GGTCTTGCCGAAGGCGCCCT | TGG | \|\|\|\|\|\|\|\|[\|\|\|\|\|\|\|\|\|\|\|\|]PAM | NA | Intergenic | NA | NA |
| T22420 | CTTCTGCATGATCCGATACCTGG | 1000 | | | | | | | | | |
| Oligo fwd | CACCCTTCTGCATGATCCGATACC | | | | | | | | | | |
| Oligo rev | AAACGGTATCGGATCATGCAGAAG | | | | | | | | | | |
| Chromosome | start | end | strand | MM | target_seq | PAM | alignment | distance | position | gene name | gene id |
| GU305914.1 | 153349 | 153371 | - | 0 | CTTCTGCATGATCCGATACC | TGG | \|\|\|\|\|\|\|\|[\|\|\|\|\|\|\|\|\|\|\|\|]PAM | NA | Intergenic | NA | NA |
| T22421 | ACCTGGCGAGAAAGATGTCTCGG | 1000 | | | | | | | | | |
| Oligo fwd | CACCACCTGGCGAGAAAGATGTCT | | | | | | | | | | |
| Oligo rev | AAACAGACATCTTTCTCGCCAGGT | | | | | | | | | | |
| Chromosome | start | end | strand | MM | target_seq | PAM | alignment | distance | position | gene name | gene id |
| GU305914.1 | 153332 | 153354 | - | 0 | ACCTGGCGAGAAAGATGTCT | CGG | \|\|\|\|\|\|\|\|[\|\|\|\|\|\|\|\|\|\|\|\|]PAM | NA | Intergenic | NA | NA |
| T22422 | AACGATCTCCTCGAAACTGATGG | 1000 | | | | | | | | | |
| Oligo fwd | CACCAACGATCTCCTCGAAACTGA | | | | | | | | | | |
| Oligo rev | AAACTCAGTTTCGAGGAGATCGTT | | | | | | | | | | |
| Chromosome | start | end | strand | MM | target_seq | PAM | alignment | distance | position | gene name | gene id |
| GU305914.1 | 153292 | 153314 | - | 0 | AACGATCTCCTCGAAACTGA | TGG | \|\|\|\|\|\|\|\|[\|\|\|\|\|\|\|\|\|\|\|\|]PAM | NA | Intergenic | NA | NA |
| T22423 | CTCGAAACTGATGGAGCTAACGG | 1000 | | | | | | | | | |
| Oligo fwd | CACCCTCGAAACTGATGGAGCTAA | | | | | | | | | | |
| Oligo rev | AAACTTAGCTCCATCAGTTTCGAG | | | | | | | | | | |
| Chromosome | start | end | strand | MM | target_seq | PAM | alignment | distance | position | gene name | gene id |
| GU305914.1 | 153283 | 153305 | - | 0 | CTCGAAACTGATGGAGCTAA | CGG | \|\|\|\|\|\|\|\|[\|\|\|\|\|\|\|\|\|\|\|\|]PAM | NA | Intergenic | NA | NA |
| T22424 | TCGAAACTGATGGAGCTAACGGG | 1000 | | | | | | | | | |
| Oligo fwd | CACCTCGAAACTGATGGAGCTAAC | | | | | | | | | | |
| Oligo rev | AAACGTTAGCTCCATCAGTTTCGA | | | | | | | | | | |
| Chromosome | start | end | strand | MM | target_seq | PAM | alignment | distance | position | gene name | gene id |
| GU305914.1 | 153282 | 153304 | - | 0 | TCGAAACTGATGGAGCTAAC | GGG | \|\|\|\|\|\|\|\|[\|\|\|\|\|\|\|\|\|\|\|\|]PAM | NA | Intergenic | NA | NA |
| T22425 | AAACTGATGGAGCTAACGGGCGG | 1000 | | | | | | | | | |
| Oligo fwd | CACCAAACTGATGGAGCTAACGGG | | | | | | | | | | |
| Oligo rev | AAACCCCGTTAGCTCCATCAGTTT | | | | | | | | | | |
| Chromosome | start | end | strand | MM | target_seq | PAM | alignment | distance | position | gene name | gene id |
| GU305914.1 | 153279 | 153301 | - | 0 | AAACTGATGGAGCTAACGGG | CGG | \|\|\|\|\|\|\|\|[\|\|\|\|\|\|\|\|\|\|\|\|]PAM | NA | Intergenic | NA | NA |
| T22426 | CTGATGGAGCTAACGGGCGGAGG | 1000 | | | | | | | | | |
| Oligo fwd | CACCCTGATGGAGCTAACGGGCGG | | | | | | | | | | |
| Oligo rev | AAACCCGCCCGTTAGCTCCATCAG | | | | | | | | | | |
| Chromosome | start | end | strand | MM | target_seq | PAM | alignment | distance | position | gene name | gene id |
| GU305914.1 | 153276 | 153298 | - | 0 | CTGATGGAGCTAACGGGCGG | AGG | \|\|\|\|\|\|\|\|[\|\|\|\|\|\|\|\|\|\|\|\|]PAM | NA | Intergenic | NA | NA |
| T22427 | GAACTTATCTGTGTAGATGTCGG | 1000 | | | | | | | | | |
| Oligo fwd | CACCGAACTTATCTGTGTAGATGT | | | | | | | | | | |
| Oligo rev | AAACACATCTACACAGATAAGTTC | | | | | | | | | | |
| Chromosome | start | end | strand | MM | target_seq | PAM | alignment | distance | position | gene name | gene id |
| GU305914.1 | 153235 | 153257 | - | 0 | GAACTTATCTGTGTAGATGT | CGG | \|\|\|\|\|\|\|\|[\|\|\|\|\|\|\|\|\|\|\|\|]PAM | NA | Intergenic | NA | NA |
| T22428 | ATAGCAGAGCATCTCCACGTCGG | 1000 | | | | | | | | | |
| Oligo fwd | CACCATAGCAGAGCATCTCCACGT | | | | | | | | | | |
| Oligo rev | AAACACGTGGAGATGCTCTGCTAT | | | | | | | | | | |
| Chromosome | start | end | strand | MM | target_seq | PAM | alignment | distance | position | gene name | gene id |
| GU305914.1 | 153211 | 153233 | - | 0 | ATAGCAGAGCATCTCCACGT | CGG | \|\|\|\|\|\|\|\|[\|\|\|\|\|\|\|\|\|\|\|\|]PAM | NA | Intergenic | NA | NA |
| T22429 | ATCTCCTCGTCGATTTGCCGCGG | 1000 | | | | | | | | | |
| Oligo fwd | CACCATCTCCTCGTCGATTTGCCG | | | | | | | | | | |
| Oligo rev | AAACCGGCAAATCGACGAGGAGAT | | | | | | | | | | |

| | | | | | | | | | | | |
|---|---|---|---|---|---|---|---|---|---|---|---|
| Chromosome | start | end | strand | MM | target_seq | PAM | alignment | distance | position | gene name | gene id |
| GU305914.1 | 153186 | 153208 | - | 0 | ATCTCCTCGTCGATTTGCCG | CGG | \|\|\|\|\|\|\|\|[\|\|\|\|\|\|\|\|\|\|\|\|]PAM | NA | Intergenic | NA | NA |
| T22430 | TCCTCGTCGATTTGCCGCGGCGG | 1000 | | | | | | | | | |
| Oligo fwd | CACCTCCTCGTCGATTTGCCGCGG | | | | | | | | | | |
| Oligo rev | AAACCCGCGGCAAATCGACGAGGA | | | | | | | | | | |
| Chromosome | start | end | strand | MM | target_seq | PAM | alignment | distance | position | gene name | gene id |
| GU305914.1 | 153183 | 153205 | - | 0 | TCCTCGTCGATTTGCCGCGG | CGG | \|\|\|\|\|\|\|\|[\|\|\|\|\|\|\|\|\|\|\|\|]PAM | NA | Intergenic | NA | NA |
| T22431 | GCGCCGCCGACATCGTCAGACGG | 1000 | | | | | | | | | |
| Oligo fwd | CACCGCGCCGCCGACATCGTCAGA | | | | | | | | | | |
| Oligo rev | AAACTCTGACGATGTCGGCGGCGC | | | | | | | | | | |
| Chromosome | start | end | strand | MM | target_seq | PAM | alignment | distance | position | gene name | gene id |
| GU305914.1 | 153158 | 153180 | - | 0 | GCGCCGCCGACATCGTCAGA | CGG | \|\|\|\|\|\|\|\|[\|\|\|\|\|\|\|\|\|\|\|\|]PAM | NA | Intergenic | NA | NA |
| T22432 | CGCCGCCGACATCGTCAGACGGG | 1000 | | | | | | | | | |
| Oligo fwd | CACCCGCCGCCGACATCGTCAGAC | | | | | | | | | | |
| Oligo rev | AAACGTCTGACGATGTCGGCGGCG | | | | | | | | | | |
| Chromosome | start | end | strand | MM | target_seq | PAM | alignment | distance | position | gene name | gene id |
| GU305914.1 | 153157 | 153179 | - | 0 | CGCCGCCGACATCGTCAGAC | GGG | \|\|\|\|\|\|\|\|[\|\|\|\|\|\|\|\|\|\|\|\|]PAM | NA | Intergenic | NA | NA |
| T22433 | GCCGCCGACATCGTCAGACGGGG | 1000 | | | | | | | | | |
| Oligo fwd | CACCGCCGCCGACATCGTCAGACG | | | | | | | | | | |
| Oligo rev | AAACCGTCTGACGATGTCGGCGGC | | | | | | | | | | |
| Chromosome | start | end | strand | MM | target_seq | PAM | alignment | distance | position | gene name | gene id |
| GU305914.1 | 153156 | 153178 | - | 0 | GCCGCCGACATCGTCAGACG | GGG | \|\|\|\|\|\|\|\|[\|\|\|\|\|\|\|\|\|\|\|\|]PAM | NA | Intergenic | NA | NA |
| T22434 | GACATCGTCAGACGGGGCTGCGG | 1000 | | | | | | | | | |
| Oligo fwd | CACCGACATCGTCAGACGGGGCTG | | | | | | | | | | |
| Oligo rev | AAACCAGCCCCGTCTGACGATGTC | | | | | | | | | | |
| Chromosome | start | end | strand | MM | target_seq | PAM | alignment | distance | position | gene name | gene id |
| GU305914.1 | 153150 | 153172 | - | 0 | GACATCGTCAGACGGGGCTG | CGG | \|\|\|\|\|\|\|\|[\|\|\|\|\|\|\|\|\|\|\|\|]PAM | NA | Intergenic | NA | NA |
| T22435 | TCGTCAGACGGGGCTGCGGTTGG | 1000 | | | | | | | | | |
| Oligo fwd | CACCTCGTCAGACGGGGCTGCGGT | | | | | | | | | | |
| Oligo rev | AAACACCGCAGCCCCGTCTGACGA | | | | | | | | | | |
| Chromosome | start | end | strand | MM | target_seq | PAM | alignment | distance | position | gene name | gene id |
| GU305914.1 | 153146 | 153168 | - | 0 | TCGTCAGACGGGGCTGCGGT | TGG | \|\|\|\|\|\|\|\|[\|\|\|\|\|\|\|\|\|\|\|\|]PAM | NA | Intergenic | NA | NA |
| T22436 | CGTCAGACGGGGCTGCGGTTGGG | 1000 | | | | | | | | | |
| Oligo fwd | CACCCGTCAGACGGGGCTGCGGTT | | | | | | | | | | |
| Oligo rev | AAACAACCGCAGCCCCGTCTGACG | | | | | | | | | | |
| Chromosome | start | end | strand | MM | target_seq | PAM | alignment | distance | position | gene name | gene id |
| GU305914.1 | 153145 | 153167 | - | 0 | CGTCAGACGGGGCTGCGGTT | GGG | \|\|\|\|\|\|\|\|[\|\|\|\|\|\|\|\|\|\|\|\|]PAM | NA | Intergenic | NA | NA |
| T22437 | GTCAGACGGGGCTGCGGTTGGGG | 1000 | | | | | | | | | |
| Oligo fwd | CACCGTCAGACGGGGCTGCGGTTG | | | | | | | | | | |
| Oligo rev | AAACCAACCGCAGCCCCGTCTGAC | | | | | | | | | | |
| Chromosome | start | end | strand | MM | target_seq | PAM | alignment | distance | position | gene name | gene id |
| GU305914.1 | 153144 | 153166 | - | 0 | GTCAGACGGGGCTGCGGTTG | GGG | \|\|\|\|\|\|\|\|[\|\|\|\|\|\|\|\|\|\|\|\|]PAM | NA | Intergenic | NA | NA |
| T22438 | GACGGGGCTGCGGTTGGGGTTGG | 1000 | | | | | | | | | |
| Oligo fwd | CACCGACGGGGCTGCGGTTGGGGT | | | | | | | | | | |
| Oligo rev | AAACACCCCAACCGCAGCCCCGTC | | | | | | | | | | |
| Chromosome | start | end | strand | MM | target_seq | PAM | alignment | distance | position | gene name | gene id |
| GU305914.1 | 153140 | 153162 | - | 0 | GACGGGGCTGCGGTTGGGGT | TGG | \|\|\|\|\|\|\|\|[\|\|\|\|\|\|\|\|\|\|\|\|]PAM | NA | Intergenic | NA | NA |
| T22439 | ACGGGGCTGCGGTTGGGGTTGGG | 1000 | | | | | | | | | |
| Oligo fwd | CACCACGGGGCTGCGGTTGGGGTT | | | | | | | | | | |
| Oligo rev | AAACAACCCCAACCGCAGCCCCGT | | | | | | | | | | |
| Chromosome | start | end | strand | MM | target_seq | PAM | alignment | distance | position | gene name | gene id |
| GU305914.1 | 153139 | 153161 | - | 0 | ACGGGGCTGCGGTTGGGGTT | GGG | \|\|\|\|\|\|\|\|[\|\|\|\|\|\|\|\|\|\|\|\|]PAM | NA | Intergenic | NA | NA |
| T22440 | CGGGGCTGCGGTTGGGGTTGGGG | 1000 | | | | | | | | | |
| Oligo fwd | CACCCGGGGCTGCGGTTGGGGTTG | | | | | | | | | | |
| Oligo rev | AAACCAACCCCAACCGCAGCCCCG | | | | | | | | | | |
| Chromosome | start | end | strand | MM | target_seq | PAM | alignment | distance | position | gene name | gene id |
| GU305914.1 | 153138 | 153160 | - | 0 | CGGGGCTGCGGTTGGGGTTG | GGG | \|\|\|\|\|\|\|\|[\|\|\|\|\|\|\|\|\|\|\|\|]PAM | NA | Intergenic | NA | NA |
| T22441 | GGCTGCGGTTGGGGTTGGGGTGG | 1000 | | | | | | | | | |
| Oligo fwd | CACCGGCTGCGGTTGGGGTTGGGG | | | | | | | | | | |
| Oligo rev | AAACCCCCAACCCCAACCGCAGCC | | | | | | | | | | |
| Chromosome | start | end | strand | MM | target_seq | PAM | alignment | distance | position | gene name | gene id |
| GU305914.1 | 153135 | 153157 | - | 0 | GGCTGCGGTTGGGGTTGGGG | TGG | \|\|\|\|\|\|\|\|[\|\|\|\|\|\|\|\|\|\|\|\|]PAM | NA | Intergenic | NA | NA |
| T22442 | TGCGGTTGGGGTTGGGGTGGAGG | 1000 | | | | | | | | | |
| Oligo fwd | CACCTGCGGTTGGGGTTGGGGTGG | | | | | | | | | | |
| Oligo rev | AAACCCACCCCAACCCCAACCGCA | | | | | | | | | | |
| Chromosome | start | end | strand | MM | target_seq | PAM | alignment | distance | position | gene name | gene id |
| GU305914.1 | 153132 | 153154 | - | 0 | TGCGGTTGGGGTTGGGGTGG | AGG | \|\|\|\|\|\|\|\|[\|\|\|\|\|\|\|\|\|\|\|\|]PAM | NA | Intergenic | NA | NA |
| T22443 | TGGGGTGGAGGTTGAAGCTGCGG | 1000 | | | | | | | | | |
| Oligo fwd | CACCTGGGGTGGAGGTTGAAGCTG | | | | | | | | | | |
| Oligo rev | AAACCAGCTTCAACCTCCACCCCA | | | | | | | | | | |
| Chromosome | start | end | strand | MM | target_seq | PAM | alignment | distance | position | gene name | gene id |
| GU305914.1 | 153120 | 153142 | - | 0 | TGGGGTGGAGGTTGAAGCTG | CGG | \|\|\|\|\|\|\|\|[\|\|\|\|\|\|\|\|\|\|\|\|]PAM | NA | Intergenic | NA | NA |
| T22444 | GGTGGAGGTTGAAGCTGCGGTGG | 1000 | | | | | | | | | |
| Oligo fwd | CACCGGTGGAGGTTGAAGCTGCGG | | | | | | | | | | |

<table>
<tr><td>Oligo rev</td><td>AAACCCGCAGCTTCAACCTCCACC</td><td></td><td></td><td></td><td></td><td></td><td></td><td></td><td></td><td></td><td></td></tr>
<tr><td>Chromosome</td><td>start</td><td>end</td><td>strand</td><td>MM</td><td>target_seq</td><td>PAM</td><td>alignment</td><td>distance</td><td>position</td><td>gene name</td><td>gene id</td></tr>
<tr><td>GU305914.1</td><td>153117</td><td>153139</td><td>-</td><td>0</td><td>GGTGGAGGTTGAAGCTGCGG</td><td>TGG</td><td>||||||||[||||||||||||]PAM</td><td>NA</td><td>Intergenic</td><td>NA</td><td>NA</td></tr>
<tr><td>GU305914.1</td><td>74930</td><td>74952</td><td>+</td><td>4</td><td>GGCGCAGGTTGAGGCGGCGG</td><td>TGG</td><td>||-|-|||[||||-||-||||]PAM</td><td>NA</td><td>Intergenic</td><td>NA</td><td>NA</td></tr>
<tr><td>T22445</td><td>GGTTGAAGCTGCGGTGGCTCTGG</td><td>1000</td><td></td><td></td><td></td><td></td><td></td><td></td><td></td><td></td><td></td></tr>
<tr><td>Oligo fwd</td><td>CACCGGTTGAAGCTGCGGTGGCTC</td><td></td><td></td><td></td><td></td><td></td><td></td><td></td><td></td><td></td><td></td></tr>
<tr><td>Oligo rev</td><td>AAACGAGCCACCGCAGCTTCAACC</td><td></td><td></td><td></td><td></td><td></td><td></td><td></td><td></td><td></td><td></td></tr>
<tr><td>Chromosome</td><td>start</td><td>end</td><td>strand</td><td>MM</td><td>target_seq</td><td>PAM</td><td>alignment</td><td>distance</td><td>position</td><td>gene name</td><td>gene id</td></tr>
<tr><td>GU305914.1</td><td>153111</td><td>153133</td><td>-</td><td>0</td><td>GGTTGAAGCTGCGGTGGCTC</td><td>TGG</td><td>||||||||[||||||||||||]PAM</td><td>NA</td><td>Intergenic</td><td>NA</td><td>NA</td></tr>
<tr><td>T22446</td><td>CTGCCGACCTCGATCGACGTAGG</td><td>1000</td><td></td><td></td><td></td><td></td><td></td><td></td><td></td><td></td><td></td></tr>
<tr><td>Oligo fwd</td><td>CACCCTGCCGACCTCGATCGACGT</td><td></td><td></td><td></td><td></td><td></td><td></td><td></td><td></td><td></td><td></td></tr>
<tr><td>Oligo rev</td><td>AAACACGTCGATCGAGGTCGGCAG</td><td></td><td></td><td></td><td></td><td></td><td></td><td></td><td></td><td></td><td></td></tr>
<tr><td>Chromosome</td><td>start</td><td>end</td><td>strand</td><td>MM</td><td>target_seq</td><td>PAM</td><td>alignment</td><td>distance</td><td>position</td><td>gene name</td><td>gene id</td></tr>
<tr><td>GU305914.1</td><td>153088</td><td>153110</td><td>-</td><td>0</td><td>CTGCCGACCTCGATCGACGT</td><td>AGG</td><td>||||||||[||||||||||||]PAM</td><td>NA</td><td>Intergenic</td><td>NA</td><td>NA</td></tr>
<tr><td>GU305914.1</td><td>89383</td><td>89405</td><td>-</td><td>4</td><td>CTGCCGCGCTCGGTCAACGT</td><td>GGG</td><td>||||||--[||||-||-||||]PAM</td><td>NA</td><td>Intergenic</td><td>NA</td><td>NA</td></tr>
<tr><td>T22447</td><td>GACCTCGATCGACGTAGGACAGG</td><td>1000</td><td></td><td></td><td></td><td></td><td></td><td></td><td></td><td></td><td></td></tr>
<tr><td>Oligo fwd</td><td>CACCGACCTCGATCGACGTAGGAC</td><td></td><td></td><td></td><td></td><td></td><td></td><td></td><td></td><td></td><td></td></tr>
<tr><td>Oligo rev</td><td>AAACGTCCTACGTCGATCGAGGTC</td><td></td><td></td><td></td><td></td><td></td><td></td><td></td><td></td><td></td><td></td></tr>
<tr><td>Chromosome</td><td>start</td><td>end</td><td>strand</td><td>MM</td><td>target_seq</td><td>PAM</td><td>alignment</td><td>distance</td><td>position</td><td>gene name</td><td>gene id</td></tr>
<tr><td>GU305914.1</td><td>153083</td><td>153105</td><td>-</td><td>0</td><td>GACCTCGATCGACGTAGGAC</td><td>AGG</td><td>||||||||[||||||||||||]PAM</td><td>NA</td><td>Intergenic</td><td>NA</td><td>NA</td></tr>
<tr><td>T22448</td><td>GTAGAAATCGTCCTCGTCGTCGG</td><td>1000</td><td></td><td></td><td></td><td></td><td></td><td></td><td></td><td></td><td></td></tr>
<tr><td>Oligo fwd</td><td>CACCGTAGAAATCGTCCTCGTCGT</td><td></td><td></td><td></td><td></td><td></td><td></td><td></td><td></td><td></td><td></td></tr>
<tr><td>Oligo rev</td><td>AAACACGACGAGGACGATTTCTAC</td><td></td><td></td><td></td><td></td><td></td><td></td><td></td><td></td><td></td><td></td></tr>
<tr><td>Chromosome</td><td>start</td><td>end</td><td>strand</td><td>MM</td><td>target_seq</td><td>PAM</td><td>alignment</td><td>distance</td><td>position</td><td>gene name</td><td>gene id</td></tr>
<tr><td>GU305914.1</td><td>153058</td><td>153080</td><td>-</td><td>0</td><td>GTAGAAATCGTCCTCGTCGT</td><td>CGG</td><td>||||||||[||||||||||||]PAM</td><td>NA</td><td>Intergenic</td><td>NA</td><td>NA</td></tr>
<tr><td>T22449</td><td>GAAATCGTCCTCGTCGTCGGAGG</td><td>1000</td><td></td><td></td><td></td><td></td><td></td><td></td><td></td><td></td><td></td></tr>
<tr><td>Oligo fwd</td><td>CACCGAAATCGTCCTCGTCGTCGG</td><td></td><td></td><td></td><td></td><td></td><td></td><td></td><td></td><td></td><td></td></tr>
<tr><td>Oligo rev</td><td>AAACCCGACGACGAGGACGATTTC</td><td></td><td></td><td></td><td></td><td></td><td></td><td></td><td></td><td></td><td></td></tr>
<tr><td>Chromosome</td><td>start</td><td>end</td><td>strand</td><td>MM</td><td>target_seq</td><td>PAM</td><td>alignment</td><td>distance</td><td>position</td><td>gene name</td><td>gene id</td></tr>
<tr><td>GU305914.1</td><td>153055</td><td>153077</td><td>-</td><td>0</td><td>GAAATCGTCCTCGTCGTCGG</td><td>AGG</td><td>||||||||[||||||||||||]PAM</td><td>NA</td><td>Intergenic</td><td>NA</td><td>NA</td></tr>
<tr><td>T22450</td><td>CCTCGCGCCGTCGCCGCCGTTGG</td><td>1000</td><td></td><td></td><td></td><td></td><td></td><td></td><td></td><td></td><td></td></tr>
<tr><td>Oligo fwd</td><td>CACCCCTCGCGCCGTCGCCGCCGT</td><td></td><td></td><td></td><td></td><td></td><td></td><td></td><td></td><td></td><td></td></tr>
<tr><td>Oligo rev</td><td>AAACACGGCGGCGACGGCGCGAGG</td><td></td><td></td><td></td><td></td><td></td><td></td><td></td><td></td><td></td><td></td></tr>
<tr><td>Chromosome</td><td>start</td><td>end</td><td>strand</td><td>MM</td><td>target_seq</td><td>PAM</td><td>alignment</td><td>distance</td><td>position</td><td>gene name</td><td>gene id</td></tr>
<tr><td>GU305914.1</td><td>152917</td><td>152939</td><td>-</td><td>0</td><td>CCTCGCGCCGTCGCCGCCGT</td><td>TGG</td><td>||||||||[||||||||||||]PAM</td><td>NA</td><td>Intergenic</td><td>NA</td><td>NA</td></tr>
<tr><td>T22451</td><td>TTGGCTCCGATGACGAGCGTCGG</td><td>1000</td><td></td><td></td><td></td><td></td><td></td><td></td><td></td><td></td><td></td></tr>
<tr><td>Oligo fwd</td><td>CACCTTGGCTCCGATGACGAGCGT</td><td></td><td></td><td></td><td></td><td></td><td></td><td></td><td></td><td></td><td></td></tr>
<tr><td>Oligo rev</td><td>AAACACGCTCGTCATCGGAGCCAA</td><td></td><td></td><td></td><td></td><td></td><td></td><td></td><td></td><td></td><td></td></tr>
<tr><td>Chromosome</td><td>start</td><td>end</td><td>strand</td><td>MM</td><td>target_seq</td><td>PAM</td><td>alignment</td><td>distance</td><td>position</td><td>gene name</td><td>gene id</td></tr>
<tr><td>GU305914.1</td><td>152898</td><td>152920</td><td>-</td><td>0</td><td>TTGGCTCCGATGACGAGCGT</td><td>CGG</td><td>||||||||[||||||||||||]PAM</td><td>NA</td><td>Intergenic</td><td>NA</td><td>NA</td></tr>
<tr><td>T22452</td><td>TGGCTCCGATGACGAGCGTCGGG</td><td>1000</td><td></td><td></td><td></td><td></td><td></td><td></td><td></td><td></td><td></td></tr>
<tr><td>Oligo fwd</td><td>CACCTGGCTCCGATGACGAGCGTC</td><td></td><td></td><td></td><td></td><td></td><td></td><td></td><td></td><td></td><td></td></tr>
<tr><td>Oligo rev</td><td>AAACGACGCTCGTCATCGGAGCCA</td><td></td><td></td><td></td><td></td><td></td><td></td><td></td><td></td><td></td><td></td></tr>
<tr><td>Chromosome</td><td>start</td><td>end</td><td>strand</td><td>MM</td><td>target_seq</td><td>PAM</td><td>alignment</td><td>distance</td><td>position</td><td>gene name</td><td>gene id</td></tr>
<tr><td>GU305914.1</td><td>152897</td><td>152919</td><td>-</td><td>0</td><td>TGGCTCCGATGACGAGCGTC</td><td>GGG</td><td>||||||||[||||||||||||]PAM</td><td>NA</td><td>Intergenic</td><td>NA</td><td>NA</td></tr>
<tr><td>T22453</td><td>GATGACGAGCGTCGGGATGTCGG</td><td>1000</td><td></td><td></td><td></td><td></td><td></td><td></td><td></td><td></td><td></td></tr>
<tr><td>Oligo fwd</td><td>CACCGATGACGAGCGTCGGGATGT</td><td></td><td></td><td></td><td></td><td></td><td></td><td></td><td></td><td></td><td></td></tr>
<tr><td>Oligo rev</td><td>AAACACATCCCGACGCTCGTCATC</td><td></td><td></td><td></td><td></td><td></td><td></td><td></td><td></td><td></td><td></td></tr>
<tr><td>Chromosome</td><td>start</td><td>end</td><td>strand</td><td>MM</td><td>target_seq</td><td>PAM</td><td>alignment</td><td>distance</td><td>position</td><td>gene name</td><td>gene id</td></tr>
<tr><td>GU305914.1</td><td>152890</td><td>152912</td><td>-</td><td>0</td><td>GATGACGAGCGTCGGGATGT</td><td>CGG</td><td>||||||||[||||||||||||]PAM</td><td>NA</td><td>Intergenic</td><td>NA</td><td>NA</td></tr>
<tr><td>T22454</td><td>ATGACGAGCGTCGGGATGTCGGG</td><td>1000</td><td></td><td></td><td></td><td></td><td></td><td></td><td></td><td></td><td></td></tr>
<tr><td>Oligo fwd</td><td>CACCATGACGAGCGTCGGGATGTC</td><td></td><td></td><td></td><td></td><td></td><td></td><td></td><td></td><td></td><td></td></tr>
<tr><td>Oligo rev</td><td>AAACGACATCCCGACGCTCGTCAT</td><td></td><td></td><td></td><td></td><td></td><td></td><td></td><td></td><td></td><td></td></tr>
<tr><td>Chromosome</td><td>start</td><td>end</td><td>strand</td><td>MM</td><td>target_seq</td><td>PAM</td><td>alignment</td><td>distance</td><td>position</td><td>gene name</td><td>gene id</td></tr>
<tr><td>GU305914.1</td><td>152889</td><td>152911</td><td>-</td><td>0</td><td>ATGACGAGCGTCGGGATGTC</td><td>GGG</td><td>||||||||[||||||||||||]PAM</td><td>NA</td><td>Intergenic</td><td>NA</td><td>NA</td></tr>
<tr><td>T22455</td><td>GAGCGTCGGGATGTCGGGCAAGG</td><td>1000</td><td></td><td></td><td></td><td></td><td></td><td></td><td></td><td></td><td></td></tr>
<tr><td>Oligo fwd</td><td>CACCGAGCGTCGGGATGTCGGGCA</td><td></td><td></td><td></td><td></td><td></td><td></td><td></td><td></td><td></td><td></td></tr>
<tr><td>Oligo rev</td><td>AAACTGCCCGACATCCCGACGCTC</td><td></td><td></td><td></td><td></td><td></td><td></td><td></td><td></td><td></td><td></td></tr>
<tr><td>Chromosome</td><td>start</td><td>end</td><td>strand</td><td>MM</td><td>target_seq</td><td>PAM</td><td>alignment</td><td>distance</td><td>position</td><td>gene name</td><td>gene id</td></tr>
<tr><td>GU305914.1</td><td>152884</td><td>152906</td><td>-</td><td>0</td><td>GAGCGTCGGGATGTCGGGCA</td><td>AGG</td><td>||||||||[||||||||||||]PAM</td><td>NA</td><td>Intergenic</td><td>NA</td><td>NA</td></tr>
<tr><td>T22456</td><td>AGCGTCGGGATGTCGGGCAAGGG</td><td>1000</td><td></td><td></td><td></td><td></td><td></td><td></td><td></td><td></td><td></td></tr>
<tr><td>Oligo fwd</td><td>CACCAGCGTCGGGATGTCGGGCAA</td><td></td><td></td><td></td><td></td><td></td><td></td><td></td><td></td><td></td><td></td></tr>
<tr><td>Oligo rev</td><td>AAACTTGCCCGACATCCCGACGCT</td><td></td><td></td><td></td><td></td><td></td><td></td><td></td><td></td><td></td><td></td></tr>
<tr><td>Chromosome</td><td>start</td><td>end</td><td>strand</td><td>MM</td><td>target_seq</td><td>PAM</td><td>alignment</td><td>distance</td><td>position</td><td>gene name</td><td>gene id</td></tr>
<tr><td>GU305914.1</td><td>152883</td><td>152905</td><td>-</td><td>0</td><td>AGCGTCGGGATGTCGGGCAA</td><td>GGG</td><td>||||||||[||||||||||||]PAM</td><td>NA</td><td>Intergenic</td><td>NA</td><td>NA</td></tr>
<tr><td>T22457</td><td>GCGTCGGGATGTCGGGCAAGGGG</td><td>1000</td><td></td><td></td><td></td><td></td><td></td><td></td><td></td><td></td><td></td></tr>
<tr><td>Oligo fwd</td><td>CACCGCGTCGGGATGTCGGGCAAG</td><td></td><td></td><td></td><td></td><td></td><td></td><td></td><td></td><td></td><td></td></tr>
<tr><td>Oligo rev</td><td>AAACCTTGCCCGACATCCCGACGC</td><td></td><td></td><td></td><td></td><td></td><td></td><td></td><td></td><td></td><td></td></tr>
<tr><td>Chromosome</td><td>start</td><td>end</td><td>strand</td><td>MM</td><td>target_seq</td><td>PAM</td><td>alignment</td><td>distance</td><td>position</td><td>gene name</td><td>gene id</td></tr>
<tr><td>GU305914.1</td><td>152882</td><td>152904</td><td>-</td><td>0</td><td>GCGTCGGGATGTCGGGCAAG</td><td>GGG</td><td>||||||||[||||||||||||]PAM</td><td>NA</td><td>Intergenic</td><td>NA</td><td>NA</td></tr>
<tr><td>T22458</td><td>GATGTCGGGCAAGGGGATCTCGG</td><td>1000</td><td></td><td></td><td></td><td></td><td></td><td></td><td></td><td></td><td></td></tr>
<tr><td>Oligo fwd</td><td>CACCGATGTCGGGCAAGGGGATCT</td><td></td><td></td><td></td><td></td><td></td><td></td><td></td><td></td><td></td><td></td></tr>
<tr><td>Oligo rev</td><td>AAACAGATCCCCTTGCCCGACATC</td><td></td><td></td><td></td><td></td><td></td><td></td><td></td><td></td><td></td><td></td></tr>
<tr><td>Chromosome</td><td>start</td><td>end</td><td>strand</td><td>MM</td><td>target_seq</td><td>PAM</td><td>alignment</td><td>distance</td><td>position</td><td>gene name</td><td>gene id</td></tr>
<tr><td>GU305914.1</td><td>152875</td><td>152897</td><td>-</td><td>0</td><td>GATGTCGGGCAAGGGGATCT</td><td>CGG</td><td>||||||||[||||||||||||]PAM</td><td>NA</td><td>Intergenic</td><td>NA</td><td>NA</td></tr>
</table>

<table>
<tr><td>T22459</td><td>CGGGCAAGGGGATCTCGGAGAGG</td><td>1000</td><td></td><td></td><td></td><td></td><td></td><td></td><td></td><td></td><td></td></tr>
<tr><td>Oligo fwd</td><td>CACCCGGGCAAGGGGATCTCGGAG</td><td></td><td></td><td></td><td></td><td></td><td></td><td></td><td></td><td></td><td></td></tr>
<tr><td>Oligo rev</td><td>AAACCTCCGAGATCCCCTTGCCCG</td><td></td><td></td><td></td><td></td><td></td><td></td><td></td><td></td><td></td><td></td></tr>
<tr><td>Chromosome</td><td>start</td><td>end</td><td>strand</td><td>MM</td><td>target_seq</td><td>PAM</td><td>alignment</td><td>distance</td><td>position</td><td>gene name</td><td>gene id</td></tr>
<tr><td>GU305914.1</td><td>152870</td><td>152892</td><td>-</td><td>0</td><td>CGGGCAAGGGGATCTCGGAG</td><td>AGG</td><td>||||||||[||||||||||||]PAM</td><td>NA</td><td>Intergenic</td><td>NA</td><td>NA</td></tr>
<tr><td>T22460</td><td>GGATCTCGGAGAGGTCGCGCAGG</td><td>1000</td><td></td><td></td><td></td><td></td><td></td><td></td><td></td><td></td><td></td></tr>
<tr><td>Oligo fwd</td><td>CACCGGATCTCGGAGAGGTCGCGC</td><td></td><td></td><td></td><td></td><td></td><td></td><td></td><td></td><td></td><td></td></tr>
<tr><td>Oligo rev</td><td>AAACGCGCGACCTCTCCGAGATCC</td><td></td><td></td><td></td><td></td><td></td><td></td><td></td><td></td><td></td><td></td></tr>
<tr><td>Chromosome</td><td>start</td><td>end</td><td>strand</td><td>MM</td><td>target_seq</td><td>PAM</td><td>alignment</td><td>distance</td><td>position</td><td>gene name</td><td>gene id</td></tr>
<tr><td>GU305914.1</td><td>152861</td><td>152883</td><td>-</td><td>0</td><td>GGATCTCGGAGAGGTCGCGC</td><td>AGG</td><td>||||||||[||||||||||||]PAM</td><td>NA</td><td>Intergenic</td><td>NA</td><td>NA</td></tr>
<tr><td>T22461</td><td>GAGAGGTCGCGCAGGATCTCCGG</td><td>1000</td><td></td><td></td><td></td><td></td><td></td><td></td><td></td><td></td><td></td></tr>
<tr><td>Oligo fwd</td><td>CACCGAGAGGTCGCGCAGGATCTC</td><td></td><td></td><td></td><td></td><td></td><td></td><td></td><td></td><td></td><td></td></tr>
<tr><td>Oligo rev</td><td>AAACGAGATCCTGCGCGACCTCTC</td><td></td><td></td><td></td><td></td><td></td><td></td><td></td><td></td><td></td><td></td></tr>
<tr><td>Chromosome</td><td>start</td><td>end</td><td>strand</td><td>MM</td><td>target_seq</td><td>PAM</td><td>alignment</td><td>distance</td><td>position</td><td>gene name</td><td>gene id</td></tr>
<tr><td>GU305914.1</td><td>152853</td><td>152875</td><td>-</td><td>0</td><td>GAGAGGTCGCGCAGGATCTC</td><td>CGG</td><td>||||||||[||||||||||||]PAM</td><td>NA</td><td>Intergenic</td><td>NA</td><td>NA</td></tr>
<tr><td>T22462</td><td>AGAGGTCGCGCAGGATCTCCGGG</td><td>1000</td><td></td><td></td><td></td><td></td><td></td><td></td><td></td><td></td><td></td></tr>
<tr><td>Oligo fwd</td><td>CACCAGAGGTCGCGCAGGATCTCC</td><td></td><td></td><td></td><td></td><td></td><td></td><td></td><td></td><td></td><td></td></tr>
<tr><td>Oligo rev</td><td>AAACGGAGATCCTGCGCGACCTCT</td><td></td><td></td><td></td><td></td><td></td><td></td><td></td><td></td><td></td><td></td></tr>
<tr><td>Chromosome</td><td>start</td><td>end</td><td>strand</td><td>MM</td><td>target_seq</td><td>PAM</td><td>alignment</td><td>distance</td><td>position</td><td>gene name</td><td>gene id</td></tr>
<tr><td>GU305914.1</td><td>152852</td><td>152874</td><td>-</td><td>0</td><td>AGAGGTCGCGCAGGATCTCC</td><td>GGG</td><td>||||||||[||||||||||||]PAM</td><td>NA</td><td>Intergenic</td><td>NA</td><td>NA</td></tr>
<tr><td>T22463</td><td>GAGGTCGCGCAGGATCTCCGGGG</td><td>1000</td><td></td><td></td><td></td><td></td><td></td><td></td><td></td><td></td><td></td></tr>
<tr><td>Oligo fwd</td><td>CACCGAGGTCGCGCAGGATCTCCG</td><td></td><td></td><td></td><td></td><td></td><td></td><td></td><td></td><td></td><td></td></tr>
<tr><td>Oligo rev</td><td>AAACCGGAGATCCTGCGCGACCTC</td><td></td><td></td><td></td><td></td><td></td><td></td><td></td><td></td><td></td><td></td></tr>
<tr><td>Chromosome</td><td>start</td><td>end</td><td>strand</td><td>MM</td><td>target_seq</td><td>PAM</td><td>alignment</td><td>distance</td><td>position</td><td>gene name</td><td>gene id</td></tr>
<tr><td>GU305914.1</td><td>152851</td><td>152873</td><td>-</td><td>0</td><td>GAGGTCGCGCAGGATCTCCG</td><td>GGG</td><td>||||||||[||||||||||||]PAM</td><td>NA</td><td>Intergenic</td><td>NA</td><td>NA</td></tr>
<tr><td>T22464</td><td>GTCGCGCAGGATCTCCGGGGTGG</td><td>1000</td><td></td><td></td><td></td><td></td><td></td><td></td><td></td><td></td><td></td></tr>
<tr><td>Oligo fwd</td><td>CACCGTCGCGCAGGATCTCCGGGG</td><td></td><td></td><td></td><td></td><td></td><td></td><td></td><td></td><td></td><td></td></tr>
<tr><td>Oligo rev</td><td>AAACCCCCGGAGATCCTGCGCGAC</td><td></td><td></td><td></td><td></td><td></td><td></td><td></td><td></td><td></td><td></td></tr>
<tr><td>Chromosome</td><td>start</td><td>end</td><td>strand</td><td>MM</td><td>target_seq</td><td>PAM</td><td>alignment</td><td>distance</td><td>position</td><td>gene name</td><td>gene id</td></tr>
<tr><td>GU305914.1</td><td>152848</td><td>152870</td><td>-</td><td>0</td><td>GTCGCGCAGGATCTCCGGGG</td><td>TGG</td><td>||||||||[||||||||||||]PAM</td><td>NA</td><td>Intergenic</td><td>NA</td><td>NA</td></tr>
<tr><td>T22465</td><td>GCAGGATCTCCGGGGTGGTGAGG</td><td>1000</td><td></td><td></td><td></td><td></td><td></td><td></td><td></td><td></td><td></td></tr>
<tr><td>Oligo fwd</td><td>CACCGCAGGATCTCCGGGGTGGTG</td><td></td><td></td><td></td><td></td><td></td><td></td><td></td><td></td><td></td><td></td></tr>
<tr><td>Oligo rev</td><td>AAACCACCACCCCGGAGATCCTGC</td><td></td><td></td><td></td><td></td><td></td><td></td><td></td><td></td><td></td><td></td></tr>
<tr><td>Chromosome</td><td>start</td><td>end</td><td>strand</td><td>MM</td><td>target_seq</td><td>PAM</td><td>alignment</td><td>distance</td><td>position</td><td>gene name</td><td>gene id</td></tr>
<tr><td>GU305914.1</td><td>152843</td><td>152865</td><td>-</td><td>0</td><td>GCAGGATCTCCGGGGTGGTG</td><td>AGG</td><td>||||||||[||||||||||||]PAM</td><td>NA</td><td>Intergenic</td><td>NA</td><td>NA</td></tr>
<tr><td>T22466</td><td>CAGGATCTCCGGGGTGGTGAGGG</td><td>1000</td><td></td><td></td><td></td><td></td><td></td><td></td><td></td><td></td><td></td></tr>
<tr><td>Oligo fwd</td><td>CACCCAGGATCTCCGGGGTGGTGA</td><td></td><td></td><td></td><td></td><td></td><td></td><td></td><td></td><td></td><td></td></tr>
<tr><td>Oligo rev</td><td>AAACTCACCACCCCGGAGATCCTG</td><td></td><td></td><td></td><td></td><td></td><td></td><td></td><td></td><td></td><td></td></tr>
<tr><td>Chromosome</td><td>start</td><td>end</td><td>strand</td><td>MM</td><td>target_seq</td><td>PAM</td><td>alignment</td><td>distance</td><td>position</td><td>gene name</td><td>gene id</td></tr>
<tr><td>GU305914.1</td><td>152842</td><td>152864</td><td>-</td><td>0</td><td>CAGGATCTCCGGGGTGGTGA</td><td>GGG</td><td>||||||||[||||||||||||]PAM</td><td>NA</td><td>Intergenic</td><td>NA</td><td>NA</td></tr>
<tr><td>T22467</td><td>AGGATCTCCGGGGTGGTGAGGGG</td><td>1000</td><td></td><td></td><td></td><td></td><td></td><td></td><td></td><td></td><td></td></tr>
<tr><td>Oligo fwd</td><td>CACCAGGATCTCCGGGGTGGTGAG</td><td></td><td></td><td></td><td></td><td></td><td></td><td></td><td></td><td></td><td></td></tr>
<tr><td>Oligo rev</td><td>AAACCTCACCACCCCGGAGATCCT</td><td></td><td></td><td></td><td></td><td></td><td></td><td></td><td></td><td></td><td></td></tr>
<tr><td>Chromosome</td><td>start</td><td>end</td><td>strand</td><td>MM</td><td>target_seq</td><td>PAM</td><td>alignment</td><td>distance</td><td>position</td><td>gene name</td><td>gene id</td></tr>
<tr><td>GU305914.1</td><td>152841</td><td>152863</td><td>-</td><td>0</td><td>AGGATCTCCGGGGTGGTGAG</td><td>GGG</td><td>||||||||[||||||||||||]PAM</td><td>NA</td><td>Intergenic</td><td>NA</td><td>NA</td></tr>
<tr><td>T22468</td><td>GGATCTCCGGGGTGGTGAGGGGG</td><td>1000</td><td></td><td></td><td></td><td></td><td></td><td></td><td></td><td></td><td></td></tr>
<tr><td>Oligo fwd</td><td>CACCGGATCTCCGGGGTGGTGAGG</td><td></td><td></td><td></td><td></td><td></td><td></td><td></td><td></td><td></td><td></td></tr>
<tr><td>Oligo rev</td><td>AAACCCTCACCACCCCGGAGATCC</td><td></td><td></td><td></td><td></td><td></td><td></td><td></td><td></td><td></td><td></td></tr>
<tr><td>Chromosome</td><td>start</td><td>end</td><td>strand</td><td>MM</td><td>target_seq</td><td>PAM</td><td>alignment</td><td>distance</td><td>position</td><td>gene name</td><td>gene id</td></tr>
<tr><td>GU305914.1</td><td>152840</td><td>152862</td><td>-</td><td>0</td><td>GGATCTCCGGGGTGGTGAGG</td><td>GGG</td><td>||||||||[||||||||||||]PAM</td><td>NA</td><td>Intergenic</td><td>NA</td><td>NA</td></tr>
<tr><td>T22469</td><td>GATCTCCGGGGTGGTGAGGGGGG</td><td>1000</td><td></td><td></td><td></td><td></td><td></td><td></td><td></td><td></td><td></td></tr>
<tr><td>Oligo fwd</td><td>CACCGATCTCCGGGGTGGTGAGGG</td><td></td><td></td><td></td><td></td><td></td><td></td><td></td><td></td><td></td><td></td></tr>
<tr><td>Oligo rev</td><td>AAACCCCTCACCACCCCGGAGATC</td><td></td><td></td><td></td><td></td><td></td><td></td><td></td><td></td><td></td><td></td></tr>
<tr><td>Chromosome</td><td>start</td><td>end</td><td>strand</td><td>MM</td><td>target_seq</td><td>PAM</td><td>alignment</td><td>distance</td><td>position</td><td>gene name</td><td>gene id</td></tr>
<tr><td>GU305914.1</td><td>152839</td><td>152861</td><td>-</td><td>0</td><td>GATCTCCGGGGTGGTGAGGG</td><td>GGG</td><td>||||||||[||||||||||||]PAM</td><td>NA</td><td>Intergenic</td><td>NA</td><td>NA</td></tr>
<tr><td>T22470</td><td>GGAGCTGTAAAAGAGATACATGG</td><td>1000</td><td></td><td></td><td></td><td></td><td></td><td></td><td></td><td></td><td></td></tr>
<tr><td>Oligo fwd</td><td>CACCGGAGCTGTAAAAGAGATACA</td><td></td><td></td><td></td><td></td><td></td><td></td><td></td><td></td><td></td><td></td></tr>
<tr><td>Oligo rev</td><td>AAACTGTATCTCTTTTACAGCTCC</td><td></td><td></td><td></td><td></td><td></td><td></td><td></td><td></td><td></td><td></td></tr>
<tr><td>Chromosome</td><td>start</td><td>end</td><td>strand</td><td>MM</td><td>target_seq</td><td>PAM</td><td>alignment</td><td>distance</td><td>position</td><td>gene name</td><td>gene id</td></tr>
<tr><td>GU305914.1</td><td>152818</td><td>152840</td><td>-</td><td>0</td><td>GGAGCTGTAAAAGAGATACA</td><td>TGG</td><td>||||||||[||||||||||||]PAM</td><td>NA</td><td>Intergenic</td><td>NA</td><td>NA</td></tr>
<tr><td>T22471</td><td>AAAAGAGCAGTCCCGAGATGAGG</td><td>1000</td><td></td><td></td><td></td><td></td><td></td><td></td><td></td><td></td><td></td></tr>
<tr><td>Oligo fwd</td><td>CACCAAAAGAGCAGTCCCGAGATG</td><td></td><td></td><td></td><td></td><td></td><td></td><td></td><td></td><td></td><td></td></tr>
<tr><td>Oligo rev</td><td>AAACCATCTCGGGACTGCTCTTTT</td><td></td><td></td><td></td><td></td><td></td><td></td><td></td><td></td><td></td><td></td></tr>
<tr><td>Chromosome</td><td>start</td><td>end</td><td>strand</td><td>MM</td><td>target_seq</td><td>PAM</td><td>alignment</td><td>distance</td><td>position</td><td>gene name</td><td>gene id</td></tr>
<tr><td>GU305914.1</td><td>152792</td><td>152814</td><td>-</td><td>0</td><td>AAAAGAGCAGTCCCGAGATG</td><td>AGG</td><td>||||||||[||||||||||||]PAM</td><td>NA</td><td>Intergenic</td><td>NA</td><td>NA</td></tr>
<tr><td>T22472</td><td>AAAGAGCAGTCCCGAGATGAGGG</td><td>1000</td><td></td><td></td><td></td><td></td><td></td><td></td><td></td><td></td><td></td></tr>
<tr><td>Oligo fwd</td><td>CACCAAAGAGCAGTCCCGAGATGA</td><td></td><td></td><td></td><td></td><td></td><td></td><td></td><td></td><td></td><td></td></tr>
<tr><td>Oligo rev</td><td>AAACTCATCTCGGGACTGCTCTTT</td><td></td><td></td><td></td><td></td><td></td><td></td><td></td><td></td><td></td><td></td></tr>
<tr><td>Chromosome</td><td>start</td><td>end</td><td>strand</td><td>MM</td><td>target_seq</td><td>PAM</td><td>alignment</td><td>distance</td><td>position</td><td>gene name</td><td>gene id</td></tr>
<tr><td>GU305914.1</td><td>152791</td><td>152813</td><td>-</td><td>0</td><td>AAAGAGCAGTCCCGAGATGA</td><td>GGG</td><td>||||||||[||||||||||||]PAM</td><td>NA</td><td>Intergenic</td><td>NA</td><td>NA</td></tr>
<tr><td>T22473</td><td>CAGTCCCGAGATGAGGGAATAGG</td><td>1000</td><td></td><td></td><td></td><td></td><td></td><td></td><td></td><td></td><td></td></tr>
<tr><td>Oligo fwd</td><td>CACCCAGTCCCGAGATGAGGGAAT</td><td></td><td></td><td></td><td></td><td></td><td></td><td></td><td></td><td></td><td></td></tr>
<tr><td>Oligo rev</td><td>AAACATTCCCTCATCTCGGGACTG</td><td></td><td></td><td></td><td></td><td></td><td></td><td></td><td></td><td></td><td></td></tr>
<tr><td>Chromosome</td><td>start</td><td>end</td><td>strand</td><td>MM</td><td>target_seq</td><td>PAM</td><td>alignment</td><td>distance</td><td>position</td><td>gene name</td><td>gene id</td></tr>
</table>

GU305914.1 152785 152807 - 0 CAGTCCCGAGATGAGGGAAT AGG ||||||||[||||||||||||]PAM NA Intergenic NA NA

T22474 GATGAGGGAATAGGCGTAGACGG 1000
Oligo fwd CACCGATGAGGGAATAGGCGTAGA
Oligo rev AAACTCTACGCCTATTCCCTCATC
Chromosome start end strand MM target_seq PAM alignment distance position gene name gene id
GU305914.1 152776 152798 - 0 GATGAGGGAATAGGCGTAGA CGG ||||||||[||||||||||||]PAM NA Intergenic NA NA

T22475 GGCGTAGACGGCCTGCTCGCAGG 1000
Oligo fwd CACCGGCGTAGACGGCCTGCTCGC
Oligo rev AAACGCGAGCAGGCCGTCTACGCC
Chromosome start end strand MM target_seq PAM alignment distance position gene name gene id
GU305914.1 152764 152786 - 0 GGCGTAGACGGCCTGCTCGC AGG ||||||||[||||||||||||]PAM NA Intergenic NA NA

T22476 GCGTAGACGGCCTGCTCGCAGGG 1000
Oligo fwd CACCGCGTAGACGGCCTGCTCGCA
Oligo rev AAACTGCGAGCAGGCCGTCTACGC
Chromosome start end strand MM target_seq PAM alignment distance position gene name gene id
GU305914.1 152763 152785 - 0 GCGTAGACGGCCTGCTCGCA GGG ||||||||[||||||||||||]PAM NA Intergenic NA NA

T22477 CGTAGACGGCCTGCTCGCAGGGG 1000
Oligo fwd CACCCGTAGACGGCCTGCTCGCAG
Oligo rev AAACCTGCGAGCAGGCCGTCTACG
Chromosome start end strand MM target_seq PAM alignment distance position gene name gene id
GU305914.1 152762 152784 - 0 CGTAGACGGCCTGCTCGCAG GGG ||||||||[||||||||||||]PAM NA Intergenic NA NA

T22478 GTAGACGGCCTGCTCGCAGGGGG 1000
Oligo fwd CACCGTAGACGGCCTGCTCGCAGG
Oligo rev AAACCCTGCGAGCAGGCCGTCTAC
Chromosome start end strand MM target_seq PAM alignment distance position gene name gene id
GU305914.1 152761 152783 - 0 GTAGACGGCCTGCTCGCAGG GGG ||||||||[||||||||||||]PAM NA Intergenic NA NA

T22479 AGGGGGTCCTCTCGATGAAGAGG 1000
Oligo fwd CACCAGGGGGTCCTCTCGATGAAG
Oligo rev AAACCTTCATCGAGAGGACCCCCT
Chromosome start end strand MM target_seq PAM alignment distance position gene name gene id
GU305914.1 152744 152766 - 0 AGGGGGTCCTCTCGATGAAG AGG ||||||||[||||||||||||]PAM NA Intergenic NA NA

T22480 CGATGAAGAGGTCCTTTTGCAGG 1000
Oligo fwd CACCCGATGAAGAGGTCCTTTTGC
Oligo rev AAACGCAAAAGGACCTCTTCATCG
Chromosome start end strand MM target_seq PAM alignment distance position gene name gene id
GU305914.1 152732 152754 - 0 CGATGAAGAGGTCCTTTTGC AGG ||||||||[||||||||||||]PAM NA Intergenic NA NA

T22481 GAGGTCCTTTTGCAGGATCTCGG 1000
Oligo fwd CACCGAGGTCCTTTTGCAGGATCT
Oligo rev AAACAGATCCTGCAAAAGGACCTC
Chromosome start end strand MM target_seq PAM alignment distance position gene name gene id
GU305914.1 152725 152747 - 0 GAGGTCCTTTTGCAGGATCT CGG ||||||||[||||||||||||]PAM NA Intergenic NA NA

T22482 GGCGAAATCGTCGAAAGTGCCGG 1000
Oligo fwd CACCGGCGAAATCGTCGAAAGTGC
Oligo rev AAACGCACTTTCGACGATTTCGCC
Chromosome start end strand MM target_seq PAM alignment distance position gene name gene id
GU305914.1 152704 152726 - 0 GGCGAAATCGTCGAAAGTGC CGG ||||||||[||||||||||||]PAM NA Intergenic NA NA

T22483 GCCGGTGTACCCGACGACGCAGG 1000
Oligo fwd CACCGCCGGTGTACCCGACGACGC
Oligo rev AAACGCGTCGTCGGGTACACCGGC
Chromosome start end strand MM target_seq PAM alignment distance position gene name gene id
GU305914.1 152686 152708 - 0 GCCGGTGTACCCGACGACGC AGG ||||||||[||||||||||||]PAM NA Intergenic NA NA

T22484 GTACCCGACGACGCAGGCCTTGG 1000
Oligo fwd CACCGTACCCGACGACGCAGGCCT
Oligo rev AAACAGGCCTGCGTCGTCGGGTAC
Chromosome start end strand MM target_seq PAM alignment distance position gene name gene id
GU305914.1 152680 152702 - 0 GTACCCGACGACGCAGGCCT TGG ||||||||[||||||||||||]PAM NA Intergenic NA NA

T22485 GACGACGCAGGCCTTGGTTTTGG 1000
Oligo fwd CACCGACGACGCAGGCCTTGGTTT
Oligo rev AAACAAACCAAGGCCTGCGTCGTC
Chromosome start end strand MM target_seq PAM alignment distance position gene name gene id
GU305914.1 152674 152696 - 0 GACGACGCAGGCCTTGGTTT TGG ||||||||[||||||||||||]PAM NA Intergenic NA NA

T22486 GGTTTTGGCCGTGACGCCGACGG 1000
Oligo fwd CACCGGTTTTGGCCGTGACGCCGA
Oligo rev AAACTCGGCGTCACGGCCAAAACC
Chromosome start end strand MM target_seq PAM alignment distance position gene name gene id
GU305914.1 152659 152681 - 0 GGTTTTGGCCGTGACGCCGA CGG ||||||||[||||||||||||]PAM NA Intergenic NA NA

T22487 GACGGCGCTGTCCCTGATGAAGG 1000
Oligo fwd CACCGACGGCGCTGTCCCTGATGA
Oligo rev AAACTCATCAGGGACAGCGCCGTC
Chromosome start end strand MM target_seq PAM alignment distance position gene name gene id
GU305914.1 152641 152663 - 0 GACGGCGCTGTCCCTGATGA AGG ||||||||[||||||||||||]PAM NA Intergenic NA NA

T22488 TGTCCCTGATGAAGGTGATGCGG 1000
Oligo fwd CACCTGTCCCTGATGAAGGTGATG
Oligo rev AAACCATCACCTTCATCAGGGACA

Chromosome start end strand MM target_seq PAM alignment distance position gene name gene id
GU305914.1 152633 152655 - 0 TGTCCCTGATGAAGGTGATG CGG ||||||||[||||||||||||]PAM NA Intergenic NA NA

T22489 CCTGATGAAGGTGATGCGGCAGG 1000
Oligo fwd CACCCCTGATGAAGGTGATGCGGC
Oligo rev AAACGCCGCATCACCTTCATCAGG
Chromosome start end strand MM target_seq PAM alignment distance position gene name gene id
GU305914.1 152629 152651 - 0 CCTGATGAAGGTGATGCGGC AGG ||||||||[||||||||||||]PAM NA Intergenic NA NA

T22490 AGGTGATGCGGCAGGTGAGCAGG 1000
Oligo fwd CACCAGGTGATGCGGCAGGTGAGC
Oligo rev AAACGCTCACCTGCCGCATCACCT
Chromosome start end strand MM target_seq PAM alignment distance position gene name gene id
GU305914.1 152621 152643 - 0 AGGTGATGCGGCAGGTGAGC AGG ||||||||[||||||||||||]PAM NA Intergenic NA NA

T22491 GGTGATGCGGCAGGTGAGCAGGG 1000
Oligo fwd CACCGGTGATGCGGCAGGTGAGCA
Oligo rev AAACTGCTCACCTGCCGCATCACC
Chromosome start end strand MM target_seq PAM alignment distance position gene name gene id
GU305914.1 152620 152642 - 0 GGTGATGCGGCAGGTGAGCA GGG ||||||||[||||||||||||]PAM NA Intergenic NA NA

T22492 CGTTGATCATCTCCTCCTCGAGG 1000
Oligo fwd CACCCGTTGATCATCTCCTCCTCG
Oligo rev AAACCGAGGAGGAGATGATCAACG
Chromosome start end strand MM target_seq PAM alignment distance position gene name gene id
GU305914.1 152594 152616 - 0 CGTTGATCATCTCCTCCTCG AGG ||||||||[||||||||||||]PAM NA Intergenic NA NA

T22493 GAAGCCGTCATCGTCCCCGCCGG 1000
Oligo fwd CACCGAAGCCGTCATCGTCCCCGC
Oligo rev AAACGCGGGGACGATGACGGCTTC
Chromosome start end strand MM target_seq PAM alignment distance position gene name gene id
GU305914.1 152569 152591 - 0 GAAGCCGTCATCGTCCCCGC CGG ||||||||[||||||||||||]PAM NA Intergenic NA NA

T22494 GCCGGCACCGCCGCCGCCGCAGG 1000
Oligo fwd CACCGCCGGCACCGCCGCCGCCGC
Oligo rev AAACGCGGCGGCGGCGGTGCCGGC
Chromosome start end strand MM target_seq PAM alignment distance position gene name gene id
GU305914.1 152551 152573 - 0 GCCGGCACCGCCGCCGCCGC AGG ||||||||[||||||||||||]PAM NA Intergenic NA NA
GU305914.1 164964 164986 + 4 GCTGCTGCCGCCGCCGCCGC CGG ||-|---|[||||||||||||]PAM NA Intergenic NA NA
GU305914.1 24620 24642 + 4 GTCGCCGCCACCGCCGCCGC CGG |-||-|-|[|-||||||||||]PAM NA Intergenic NA NA
GU305914.1 19437 19459 - 3 TCCGGCACCTCCGCCGACGC CGG -|||||||[|-||||||-|||]PAM NA Intergenic NA NA
GU305914.1 48151 48173 - 2 GCCGCCACCGCCGCCGCCAC AGG ||||-|||[||||||||||-|]PAM NA Intergenic NA NA
GU305914.1 163747 163769 - 3 GCCGCCACCGGCGCCGCCGA CGG ||||-|||[||-||||||||-]PAM NA Intergenic NA NA

T22495 CACCGCCGCCGCCGCAGGACTGG 1000
Oligo fwd CACCCACCGCCGCCGCCGCAGGAC
Oligo rev AAACGTCCTGCGGCGGCGGCGGTG
Chromosome start end strand MM target_seq PAM alignment distance position gene name gene id
GU305914.1 152546 152568 - 0 CACCGCCGCCGCCGCAGGAC TGG ||||||||[||||||||||||]PAM NA Intergenic NA NA

T22496 ACCGCCGCCGCCGCAGGACTGGG 1000
Oligo fwd CACCACCGCCGCCGCCGCAGGACT
Oligo rev AAACAGTCCTGCGGCGGCGGCGGT
Chromosome start end strand MM target_seq PAM alignment distance position gene name gene id
GU305914.1 152545 152567 - 0 ACCGCCGCCGCCGCAGGACT GGG ||||||||[||||||||||||]PAM NA Intergenic NA NA
GU305914.1 56560 56582 - 4 TCCGTCGCCGCAGCAGTACT AGG -|||-|||[|||-||||-|||]PAM NA Intergenic NA NA

T22497 ACTGGGCGAGCTCCTCGACCTGG 1000
Oligo fwd CACCACTGGGCGAGCTCCTCGACC
Oligo rev AAACGGTCGAGGAGCTCGCCCAGT
Chromosome start end strand MM target_seq PAM alignment distance position gene name gene id
GU305914.1 152528 152550 - 0 ACTGGGCGAGCTCCTCGACC TGG ||||||||[||||||||||||]PAM NA Intergenic NA NA

T22498 CGACCTGGATGTCCTCCGACAGG 1000
Oligo fwd CACCCGACCTGGATGTCCTCCGAC
Oligo rev AAACGTCGGAGGACATCCAGGTCG
Chromosome start end strand MM target_seq PAM alignment distance position gene name gene id
GU305914.1 152513 152535 - 0 CGACCTGGATGTCCTCCGAC AGG ||||||||[||||||||||||]PAM NA Intergenic NA NA

T22499 GGATGTCCTCCGACAGGTTATGG 1000
Oligo fwd CACCGGATGTCCTCCGACAGGTTA
Oligo rev AAACTAACCTGTCGGAGGACATCC
Chromosome start end strand MM target_seq PAM alignment distance position gene name gene id
GU305914.1 152507 152529 - 0 GGATGTCCTCCGACAGGTTA TGG ||||||||[||||||||||||]PAM NA Intergenic NA NA

T22500 GTCCTCCGACAGGTTATGGTCGG 1000
Oligo fwd CACCGTCCTCCGACAGGTTATGGT
Oligo rev AAACACCATAACCTGTCGGAGGAC
Chromosome start end strand MM target_seq PAM alignment distance position gene name gene id
GU305914.1 152503 152525 - 0 GTCCTCCGACAGGTTATGGT CGG ||||||||[||||||||||||]PAM NA Intergenic NA NA

T22501 ACTGCGAGTAGTTGCTGATGCGG 1000
Oligo fwd CACCACTGCGAGTAGTTGCTGATG
Oligo rev AAACCATCAGCAACTACTCGCAGT
Chromosome start end strand MM target_seq PAM alignment distance position gene name gene id
GU305914.1 152477 152499 - 0 ACTGCGAGTAGTTGCTGATG CGG ||||||||[||||||||||||]PAM NA Intergenic NA NA

T22502 CTGCGAGTAGTTGCTGATGCGGG 1000
Oligo fwd CACCCTGCGAGTAGTTGCTGATGC

Oligo rev AAACGCATCAGCAACTACTCGCAG
Chromosome start end strand MM target_seq PAM alignment distance position gene name gene id
GU305914.1 152476 152498 - 0 CTGCGAGTAGTTGCTGATGC GGG ||||||||[||||||||||||]PAM NA Intergenic NA NA

T22503 AGTAGTTGCTGATGCGGGCCAGG 1000
Oligo fwd CACCAGTAGTTGCTGATGCGGGCC
Oligo rev AAACGGCCCGCATCAGCAACTACT
Chromosome start end strand MM target_seq PAM alignment distance position gene name gene id
GU305914.1 152471 152493 - 0 AGTAGTTGCTGATGCGGGCC AGG ||||||||[||||||||||||]PAM NA Intergenic NA NA

T22504 TGATGCGGGCCAGGTTCTTGCGG 1000
Oligo fwd CACCTGATGCGGGCCAGGTTCTTG
Oligo rev AAACCAAGAACCTGGCCCGCATCA
Chromosome start end strand MM target_seq PAM alignment distance position gene name gene id
GU305914.1 152462 152484 - 0 TGATGCGGGCCAGGTTCTTG CGG ||||||||[||||||||||||]PAM NA Intergenic NA NA

T22505 GGCCAGGTTCTTGCGGAACCAGG 1000
Oligo fwd CACCGGCCAGGTTCTTGCGGAACC
Oligo rev AAACGGTTCCGCAAGAACCTGGCC
Chromosome start end strand MM target_seq PAM alignment distance position gene name gene id
GU305914.1 152455 152477 - 0 GGCCAGGTTCTTGCGGAACC AGG ||||||||[||||||||||||]PAM NA Intergenic NA NA

T22506 TTCTTGCGGAACCAGGTCTCCGG 1000
Oligo fwd CACCTTCTTGCGGAACCAGGTCTC
Oligo rev AAACGAGACCTGGTTCCGCAAGAA
Chromosome start end strand MM target_seq PAM alignment distance position gene name gene id
GU305914.1 152448 152470 - 0 TTCTTGCGGAACCAGGTCTC CGG ||||||||[||||||||||||]PAM NA Intergenic NA NA

T22507 CGGAACCAGGTCTCCGGACGCGG 1000
Oligo fwd CACCCGGAACCAGGTCTCCGGACG
Oligo rev AAACCGTCCGGAGACCTGGTTCCG
Chromosome start end strand MM target_seq PAM alignment distance position gene name gene id
GU305914.1 152442 152464 - 0 CGGAACCAGGTCTCCGGACG CGG ||||||||[||||||||||||]PAM NA Intergenic NA NA

T22508 CCAGGTCTCCGGACGCGGCGTGG 1000
Oligo fwd CACCCCAGGTCTCCGGACGCGGCG
Oligo rev AAACCGCCGCGTCCGGAGACCTGG
Chromosome start end strand MM target_seq PAM alignment distance position gene name gene id
GU305914.1 152437 152459 - 0 CCAGGTCTCCGGACGCGGCG TGG ||||||||[||||||||||||]PAM NA Intergenic NA NA

T22509 CAGGTCTCCGGACGCGGCGTGGG 1000
Oligo fwd CACCCAGGTCTCCGGACGCGGCGT
Oligo rev AAACACGCCGCGTCCGGAGACCTG
Chromosome start end strand MM target_seq PAM alignment distance position gene name gene id
GU305914.1 152436 152458 - 0 CAGGTCTCCGGACGCGGCGT GGG ||||||||[||||||||||||]PAM NA Intergenic NA NA

T22510 CGGACGCGGCGTGGGCTCCCCGG 1000
Oligo fwd CACCCGGACGCGGCGTGGGCTCCC
Oligo rev AAACGGGAGCCCACGCCGCGTCCG
Chromosome start end strand MM target_seq PAM alignment distance position gene name gene id
GU305914.1 152428 152450 - 0 CGGACGCGGCGTGGGCTCCC CGG ||||||||[||||||||||||]PAM NA Intergenic NA NA

T22511 GGGCTCCCCGGTGCACATGCTGG 1000
Oligo fwd CACCGGGCTCCCCGGTGCACATGC
Oligo rev AAACGCATGTGCACCGGGGAGCCC
Chromosome start end strand MM target_seq PAM alignment distance position gene name gene id
GU305914.1 152416 152438 - 0 GGGCTCCCCGGTGCACATGC TGG ||||||||[||||||||||||]PAM NA Intergenic NA NA

T22512 ACATGCTGGCACAGTACTCGTGG 1000
Oligo fwd CACCACATGCTGGCACAGTACTCG
Oligo rev AAACCGAGTACTGTGCCAGCATGT
Chromosome start end strand MM target_seq PAM alignment distance position gene name gene id
GU305914.1 152402 152424 - 0 ACATGCTGGCACAGTACTCG TGG ||||||||[||||||||||||]PAM NA Intergenic NA NA

T22513 GCTGGCACAGTACTCGTGGAAGG 1000
Oligo fwd CACCGCTGGCACAGTACTCGTGGA
Oligo rev AAACTCCACGAGTACTGTGCCAGC
Chromosome start end strand MM target_seq PAM alignment distance position gene name gene id
GU305914.1 152398 152420 - 0 GCTGGCACAGTACTCGTGGA AGG ||||||||[||||||||||||]PAM NA Intergenic NA NA

T22514 CGTGGAAGGCCGAGTTGTTGAGG 1000
Oligo fwd CACCCGTGGAAGGCCGAGTTGTTG
Oligo rev AAACCAACAACTCGGCCTTCCACG
Chromosome start end strand MM target_seq PAM alignment distance position gene name gene id
GU305914.1 152384 152406 - 0 CGTGGAAGGCCGAGTTGTTG AGG ||||||||[||||||||||||]PAM NA Intergenic NA NA

T22515 TTGTTGAGGACGCAGTAGACCGG 1000
Oligo fwd CACCTTGTTGAGGACGCAGTAGAC
Oligo rev AAACGTCTACTGCGTCCTCAACAA
Chromosome start end strand MM target_seq PAM alignment distance position gene name gene id
GU305914.1 152370 152392 - 0 TTGTTGAGGACGCAGTAGAC CGG ||||||||[||||||||||||]PAM NA Intergenic NA NA

T22516 AGACCGGCACCTCGAAGATGAGG 1000
Oligo fwd CACCAGACCGGCACCTCGAAGATG
Oligo rev AAACCATCTTCGAGGTGCCGGTCT
Chromosome start end strand MM target_seq PAM alignment distance position gene name gene id
GU305914.1 152354 152376 - 0 AGACCGGCACCTCGAAGATG AGG ||||||||[||||||||||||]PAM NA Intergenic NA NA

T22517 GCACCTCGAAGATGAGGTGCTGG 1000

| | | | | | | | | | | | | |
|---|---|---|---|---|---|---|---|---|---|---|---|---|
| Oligo fwd | CACCGCACCTCGAAGATGAGGTGC | | | | | | | | | | | |
| Oligo rev | AAACGCACCTCATCTTCGAGGTGC | | | | | | | | | | | |
| Chromosome | start | | end | strand | MM | target_seq | PAM | alignment | distance | position | gene name | gene id |
| GU305914.1 | | 152348 | 152370 | - | 0 | GCACCTCGAAGATGAGGTGC | TGG | \|\|\|\|\|\|\|\|[\|\|\|\|\|\|\|\|\|\|\|\|]PAM | NA | Intergenic | NA | NA |
| T22518 | TGAGGTGCTGGCTGTAGATGCGG | | 1000 | | | | | | | | | |
| Oligo fwd | CACCTGAGGTGCTGGCTGTAGATG | | | | | | | | | | | |
| Oligo rev | AAACCATCTACAGCCAGCACCTCA | | | | | | | | | | | |
| Chromosome | start | | end | strand | MM | target_seq | PAM | alignment | distance | position | gene name | gene id |
| GU305914.1 | | 152336 | 152358 | - | 0 | TGAGGTGCTGGCTGTAGATG | CGG | \|\|\|\|\|\|\|\|[\|\|\|\|\|\|\|\|\|\|\|\|]PAM | NA | Intergenic | NA | NA |
| T22519 | GCTGGCTGTAGATGCGGACGCGG | | 1000 | | | | | | | | | |
| Oligo fwd | CACCGCTGGCTGTAGATGCGGACG | | | | | | | | | | | |
| Oligo rev | AAACCGTCCGCATCTACAGCCAGC | | | | | | | | | | | |
| Chromosome | start | | end | strand | MM | target_seq | PAM | alignment | distance | position | gene name | gene id |
| GU305914.1 | | 152330 | 152352 | - | 0 | GCTGGCTGTAGATGCGGACG | CGG | \|\|\|\|\|\|\|\|[\|\|\|\|\|\|\|\|\|\|\|\|]PAM | NA | Intergenic | NA | NA |
| T22520 | TGTAGATGCGGACGCGGTCGTGG | | 1000 | | | | | | | | | |
| Oligo fwd | CACCTGTAGATGCGGACGCGGTCG | | | | | | | | | | | |
| Oligo rev | AAACCGACCGCGTCCGCATCTACA | | | | | | | | | | | |
| Chromosome | start | | end | strand | MM | target_seq | PAM | alignment | distance | position | gene name | gene id |
| GU305914.1 | | 152324 | 152346 | - | 0 | TGTAGATGCGGACGCGGTCG | TGG | \|\|\|\|\|\|\|\|[\|\|\|\|\|\|\|\|\|\|\|\|]PAM | NA | Intergenic | NA | NA |
| T22521 | CGTGAAGTACCTCTTGACCTCGG | | 1000 | | | | | | | | | |
| Oligo fwd | CACCCGTGAAGTACCTCTTGACCT | | | | | | | | | | | |
| Oligo rev | AAACAGGTCAAGAGGTACTTCACG | | | | | | | | | | | |
| Chromosome | start | | end | strand | MM | target_seq | PAM | alignment | distance | position | gene name | gene id |
| GU305914.1 | | 152296 | 152318 | - | 0 | CGTGAAGTACCTCTTGACCT | CGG | \|\|\|\|\|\|\|\|[\|\|\|\|\|\|\|\|\|\|\|\|]PAM | NA | Intergenic | NA | NA |
| T22522 | CCTCGGCGTGCGAGATGAAGAGG | | 1000 | | | | | | | | | |
| Oligo fwd | CACCCCTCGGCGTGCGAGATGAAG | | | | | | | | | | | |
| Oligo rev | AAACCTTCATCTCGCACGCCGAGG | | | | | | | | | | | |
| Chromosome | start | | end | strand | MM | target_seq | PAM | alignment | distance | position | gene name | gene id |
| GU305914.1 | | 152279 | 152301 | - | 0 | CCTCGGCGTGCGAGATGAAG | AGG | \|\|\|\|\|\|\|\|[\|\|\|\|\|\|\|\|\|\|\|\|]PAM | NA | Intergenic | NA | NA |
| T22523 | CGGCGTGCGAGATGAAGAGGCGG | | 1000 | | | | | | | | | |
| Oligo fwd | CACCCGGCGTGCGAGATGAAGAGG | | | | | | | | | | | |
| Oligo rev | AAACCCTCTTCATCTCGCACGCCG | | | | | | | | | | | |
| Chromosome | start | | end | strand | MM | target_seq | PAM | alignment | distance | position | gene name | gene id |
| GU305914.1 | | 152276 | 152298 | - | 0 | CGGCGTGCGAGATGAAGAGG | CGG | \|\|\|\|\|\|\|\|[\|\|\|\|\|\|\|\|\|\|\|\|]PAM | NA | Intergenic | NA | NA |
| T22524 | GGCGTGCGAGATGAAGAGGCGGG | | 1000 | | | | | | | | | |
| Oligo fwd | CACCGGCGTGCGAGATGAAGAGGC | | | | | | | | | | | |
| Oligo rev | AAACGCCTCTTCATCTCGCACGCC | | | | | | | | | | | |
| Chromosome | start | | end | strand | MM | target_seq | PAM | alignment | distance | position | gene name | gene id |
| GU305914.1 | | 152275 | 152297 | - | 0 | GGCGTGCGAGATGAAGAGGC | GGG | \|\|\|\|\|\|\|\|[\|\|\|\|\|\|\|\|\|\|\|\|]PAM | NA | Intergenic | NA | NA |
| GU305914.1 | | 199319 | 199341 | - | 4 | GGGTGGCGAGATGGAGAGGC | CGG | \|\|---\|\|\|[\|\|\|\|\|-\|\|\|\|\|\|]PAM | NA | Intergenic | NA | NA |
| T22525 | GAAGAGGCGGGTGACATCGATGG | | 1000 | | | | | | | | | |
| Oligo fwd | CACCGAAGAGGCGGGTGACATCGA | | | | | | | | | | | |
| Oligo rev | AAACTCGATGTCACCCGCCTCTTC | | | | | | | | | | | |
| Chromosome | start | | end | strand | MM | target_seq | PAM | alignment | distance | position | gene name | gene id |
| GU305914.1 | | 152263 | 152285 | - | 0 | GAAGAGGCGGGTGACATCGA | TGG | \|\|\|\|\|\|\|\|[\|\|\|\|\|\|\|\|\|\|\|\|]PAM | NA | Intergenic | NA | NA |
| T22526 | GTGACATCGATGGCGTGCGCCGG | | 1000 | | | | | | | | | |
| Oligo fwd | CACCGTGACATCGATGGCGTGCGC | | | | | | | | | | | |
| Oligo rev | AAACGCGCACGCCATCGATGTCAC | | | | | | | | | | | |
| Chromosome | start | | end | strand | MM | target_seq | PAM | alignment | distance | position | gene name | gene id |
| GU305914.1 | | 152253 | 152275 | - | 0 | GTGACATCGATGGCGTGCGC | CGG | \|\|\|\|\|\|\|\|[\|\|\|\|\|\|\|\|\|\|\|\|]PAM | NA | Intergenic | NA | NA |
| T22527 | TGACATCGATGGCGTGCGCCGGG | | 1000 | | | | | | | | | |
| Oligo fwd | CACCTGACATCGATGGCGTGCGCC | | | | | | | | | | | |
| Oligo rev | AAACGGCGCACGCCATCGATGTCA | | | | | | | | | | | |
| Chromosome | start | | end | strand | MM | target_seq | PAM | alignment | distance | position | gene name | gene id |
| GU305914.1 | | 152252 | 152274 | - | 0 | TGACATCGATGGCGTGCGCC | GGG | \|\|\|\|\|\|\|\|[\|\|\|\|\|\|\|\|\|\|\|\|]PAM | NA | Intergenic | NA | NA |
| T22528 | CGTGCGCCGGGTCCCTGATGAGG | | 1000 | | | | | | | | | |
| Oligo fwd | CACCCGTGCGCCGGGTCCCTGATG | | | | | | | | | | | |
| Oligo rev | AAACCATCAGGGACCCGGCGCACG | | | | | | | | | | | |
| Chromosome | start | | end | strand | MM | target_seq | PAM | alignment | distance | position | gene name | gene id |
| GU305914.1 | | 152240 | 152262 | - | 0 | CGTGCGCCGGGTCCCTGATG | AGG | \|\|\|\|\|\|\|\|[\|\|\|\|\|\|\|\|\|\|\|\|]PAM | NA | Intergenic | NA | NA |
| T22529 | CGCCGGGTCCCTGATGAGGCCGG | | 1000 | | | | | | | | | |
| Oligo fwd | CACCCGCCGGGTCCCTGATGAGGC | | | | | | | | | | | |
| Oligo rev | AAACGCCTCATCAGGGACCCGGCG | | | | | | | | | | | |
| Chromosome | start | | end | strand | MM | target_seq | PAM | alignment | distance | position | gene name | gene id |
| GU305914.1 | | 152236 | 152258 | - | 0 | CGCCGGGTCCCTGATGAGGC | CGG | \|\|\|\|\|\|\|\|[\|\|\|\|\|\|\|\|\|\|\|\|]PAM | NA | Intergenic | NA | NA |
| T22530 | CGGGTCCCTGATGAGGCCGGCGG | | 1000 | | | | | | | | | |
| Oligo fwd | CACCCGGGTCCCTGATGAGGCCGG | | | | | | | | | | | |
| Oligo rev | AAACCCGGCCTCATCAGGGACCCG | | | | | | | | | | | |
| Chromosome | start | | end | strand | MM | target_seq | PAM | alignment | distance | position | gene name | gene id |
| GU305914.1 | | 152233 | 152255 | - | 0 | CGGGTCCCTGATGAGGCCGG | CGG | \|\|\|\|\|\|\|\|[\|\|\|\|\|\|\|\|\|\|\|\|]PAM | NA | Intergenic | NA | NA |
| T22531 | GGGTCCCTGATGAGGCCGGCGGG | | 1000 | | | | | | | | | |
| Oligo fwd | CACCGGGTCCCTGATGAGGCCGGC | | | | | | | | | | | |
| Oligo rev | AAACGCCGGCCTCATCAGGGACCC | | | | | | | | | | | |
| Chromosome | start | | end | strand | MM | target_seq | PAM | alignment | distance | position | gene name | gene id |
| GU305914.1 | | 152232 | 152254 | - | 0 | GGGTCCCTGATGAGGCCGGC | GGG | \|\|\|\|\|\|\|\|[\|\|\|\|\|\|\|\|\|\|\|\|]PAM | NA | Intergenic | NA | NA |

| | | | | | | | | | | | |
|---|---|---|---|---|---|---|---|---|---|---|---|
| T22532 | GGTCCCTGATGAGGCCGGCGGGG | 1000 | | | | | | | | | |
| Oligo fwd | CACCGGTCCCTGATGAGGCCGGCG | | | | | | | | | | |
| Oligo rev | AAACCGCCGGCCTCATCAGGGACC | | | | | | | | | | |
| Chromosome | start | end | strand | MM | target_seq | PAM | alignment | distance | position | gene name | gene id |
| GU305914.1 | 152231 | 152253 | - | 0 | GGTCCCTGATGAGGCCGGCG | GGG | \|\|\|\|\|\|\|\|[\|\|\|\|\|\|\|\|\|\|\|\|]PAM | NA | Intergenic | NA | NA |

| | | | | | | | | | | | |
|---|---|---|---|---|---|---|---|---|---|---|---|
| T22533 | CGGGGCGCACGAACCTGTCGAGG | 1000 | | | | | | | | | |
| Oligo fwd | CACCCGGGGCGCACGAACCTGTCG | | | | | | | | | | |
| Oligo rev | AAACCGACAGGTTCGTGCGCCCCG | | | | | | | | | | |
| Chromosome | start | end | strand | MM | target_seq | PAM | alignment | distance | position | gene name | gene id |
| GU305914.1 | 152213 | 152235 | - | 0 | CGGGGCGCACGAACCTGTCG | AGG | \|\|\|\|\|\|\|\|[\|\|\|\|\|\|\|\|\|\|\|\|]PAM | NA | Intergenic | NA | NA |

| | | | | | | | | | | | |
|---|---|---|---|---|---|---|---|---|---|---|---|
| T22534 | GTCGAGGTACTCGACGTGCTCGG | 1000 | | | | | | | | | |
| Oligo fwd | CACCGTCGAGGTACTCGACGTGCT | | | | | | | | | | |
| Oligo rev | AAACAGCACGTCGAGTACCTCGAC | | | | | | | | | | |
| Chromosome | start | end | strand | MM | target_seq | PAM | alignment | distance | position | gene name | gene id |
| GU305914.1 | 152197 | 152219 | - | 0 | GTCGAGGTACTCGACGTGCT | CGG | \|\|\|\|\|\|\|\|[\|\|\|\|\|\|\|\|\|\|\|\|]PAM | NA | Intergenic | NA | NA |

| | | | | | | | | | | | |
|---|---|---|---|---|---|---|---|---|---|---|---|
| T22535 | ACTCGACGTGCTCGGACTTGAGG | 1000 | | | | | | | | | |
| Oligo fwd | CACCACTCGACGTGCTCGGACTTG | | | | | | | | | | |
| Oligo rev | AAACCAAGTCCGAGCACGTCGAGT | | | | | | | | | | |
| Chromosome | start | end | strand | MM | target_seq | PAM | alignment | distance | position | gene name | gene id |
| GU305914.1 | 152189 | 152211 | - | 0 | ACTCGACGTGCTCGGACTTG | AGG | \|\|\|\|\|\|\|\|[\|\|\|\|\|\|\|\|\|\|\|\|]PAM | NA | Intergenic | NA | NA |

| | | | | | | | | | | | |
|---|---|---|---|---|---|---|---|---|---|---|---|
| T22536 | CTCGACGTGCTCGGACTTGAGGG | 1000 | | | | | | | | | |
| Oligo fwd | CACCCTCGACGTGCTCGGACTTGA | | | | | | | | | | |
| Oligo rev | AAACTCAAGTCCGAGCACGTCGAG | | | | | | | | | | |
| Chromosome | start | end | strand | MM | target_seq | PAM | alignment | distance | position | gene name | gene id |
| GU305914.1 | 152188 | 152210 | - | 0 | CTCGACGTGCTCGGACTTGA | GGG | \|\|\|\|\|\|\|\|[\|\|\|\|\|\|\|\|\|\|\|\|]PAM | NA | Intergenic | NA | NA |

| | | | | | | | | | | | |
|---|---|---|---|---|---|---|---|---|---|---|---|
| T22537 | TCGACGTGCTCGGACTTGAGGGG | 1000 | | | | | | | | | |
| Oligo fwd | CACCTCGACGTGCTCGGACTTGAG | | | | | | | | | | |
| Oligo rev | AAACCTCAAGTCCGAGCACGTCGA | | | | | | | | | | |
| Chromosome | start | end | strand | MM | target_seq | PAM | alignment | distance | position | gene name | gene id |
| GU305914.1 | 152187 | 152209 | - | 0 | TCGACGTGCTCGGACTTGAG | GGG | \|\|\|\|\|\|\|\|[\|\|\|\|\|\|\|\|\|\|\|\|]PAM | NA | Intergenic | NA | NA |

| | | | | | | | | | | | |
|---|---|---|---|---|---|---|---|---|---|---|---|
| T22538 | CGAACTCGATGTGCTTGAGCAGG | 1000 | | | | | | | | | |
| Oligo fwd | CACCCGAACTCGATGTGCTTGAGC | | | | | | | | | | |
| Oligo rev | AAACGCTCAAGCACATCGAGTTCG | | | | | | | | | | |
| Chromosome | start | end | strand | MM | target_seq | PAM | alignment | distance | position | gene name | gene id |
| GU305914.1 | 152159 | 152181 | - | 0 | CGAACTCGATGTGCTTGAGC | AGG | \|\|\|\|\|\|\|\|[\|\|\|\|\|\|\|\|\|\|\|\|]PAM | NA | Intergenic | NA | NA |

| | | | | | | | | | | | |
|---|---|---|---|---|---|---|---|---|---|---|---|
| T22539 | TGAGCAGGTCGCCGAACTCGAGG | 1000 | | | | | | | | | |
| Oligo fwd | CACCTGAGCAGGTCGCCGAACTCG | | | | | | | | | | |
| Oligo rev | AAACCGAGTTCGGCGACCTGCTCA | | | | | | | | | | |
| Chromosome | start | end | strand | MM | target_seq | PAM | alignment | distance | position | gene name | gene id |
| GU305914.1 | 152144 | 152166 | - | 0 | TGAGCAGGTCGCCGAACTCG | AGG | \|\|\|\|\|\|\|\|[\|\|\|\|\|\|\|\|\|\|\|\|]PAM | NA | Intergenic | NA | NA |

| | | | | | | | | | | | |
|---|---|---|---|---|---|---|---|---|---|---|---|
| T22540 | GGTCGCCGAACTCGAGGTCGAGG | 1000 | | | | | | | | | |
| Oligo fwd | CACCGGTCGCCGAACTCGAGGTCG | | | | | | | | | | |
| Oligo rev | AAACCGACCTCGAGTTCGGCGACC | | | | | | | | | | |
| Chromosome | start | end | strand | MM | target_seq | PAM | alignment | distance | position | gene name | gene id |
| GU305914.1 | 152138 | 152160 | - | 0 | GGTCGCCGAACTCGAGGTCG | AGG | \|\|\|\|\|\|\|\|[\|\|\|\|\|\|\|\|\|\|\|\|]PAM | NA | Intergenic | NA | NA |

| | | | | | | | | | | | |
|---|---|---|---|---|---|---|---|---|---|---|---|
| T22541 | TGATGAACATGACCCAGTTGTGG | 1000 | | | | | | | | | |
| Oligo fwd | CACCTGATGAACATGACCCAGTTG | | | | | | | | | | |
| Oligo rev | AAACCAACTGGGTCATGTTCATCA | | | | | | | | | | |
| Chromosome | start | end | strand | MM | target_seq | PAM | alignment | distance | position | gene name | gene id |
| GU305914.1 | 152102 | 152124 | - | 0 | TGATGAACATGACCCAGTTG | TGG | \|\|\|\|\|\|\|\|[\|\|\|\|\|\|\|\|\|\|\|\|]PAM | NA | Intergenic | NA | NA |

| | | | | | | | | | | | |
|---|---|---|---|---|---|---|---|---|---|---|---|
| T22542 | GATGAACATGACCCAGTTGTGGG | 1000 | | | | | | | | | |
| Oligo fwd | CACCGATGAACATGACCCAGTTGT | | | | | | | | | | |
| Oligo rev | AAACACAACTGGGTCATGTTCATC | | | | | | | | | | |
| Chromosome | start | end | strand | MM | target_seq | PAM | alignment | distance | position | gene name | gene id |
| GU305914.1 | 152101 | 152123 | - | 0 | GATGAACATGACCCAGTTGT | GGG | \|\|\|\|\|\|\|\|[\|\|\|\|\|\|\|\|\|\|\|\|]PAM | NA | Intergenic | NA | NA |

| | | | | | | | | | | | |
|---|---|---|---|---|---|---|---|---|---|---|---|
| T22543 | GTGGGCGACGTCGCAGTACTCGG | 1000 | | | | | | | | | |
| Oligo fwd | CACCGTGGGCGACGTCGCAGTACT | | | | | | | | | | |
| Oligo rev | AAACAGTACTGCGACGTCGCCCAC | | | | | | | | | | |
| Chromosome | start | end | strand | MM | target_seq | PAM | alignment | distance | position | gene name | gene id |
| GU305914.1 | 152083 | 152105 | - | 0 | GTGGGCGACGTCGCAGTACT | CGG | \|\|\|\|\|\|\|\|[\|\|\|\|\|\|\|\|\|\|\|\|]PAM | NA | Intergenic | NA | NA |

| | | | | | | | | | | | |
|---|---|---|---|---|---|---|---|---|---|---|---|
| T22544 | CGACGTCGCAGTACTCGGAGAGG | 1000 | | | | | | | | | |
| Oligo fwd | CACCCGACGTCGCAGTACTCGGAG | | | | | | | | | | |
| Oligo rev | AAACCTCCGAGTACTGCGACGTCG | | | | | | | | | | |
| Chromosome | start | end | strand | MM | target_seq | PAM | alignment | distance | position | gene name | gene id |
| GU305914.1 | 152078 | 152100 | - | 0 | CGACGTCGCAGTACTCGGAG | AGG | \|\|\|\|\|\|\|\|[\|\|\|\|\|\|\|\|\|\|\|\|]PAM | NA | Intergenic | NA | NA |

| | | | | | | | | | | | |
|---|---|---|---|---|---|---|---|---|---|---|---|
| T22545 | TCGCAGTACTCGGAGAGGACCGG | 1000 | | | | | | | | | |
| Oligo fwd | CACCTCGCAGTACTCGGAGAGGAC | | | | | | | | | | |
| Oligo rev | AAACGTCCTCTCCGAGTACTGCGA | | | | | | | | | | |
| Chromosome | start | end | strand | MM | target_seq | PAM | alignment | distance | position | gene name | gene id |
| GU305914.1 | 152073 | 152095 | - | 0 | TCGCAGTACTCGGAGAGGAC | CGG | \|\|\|\|\|\|\|\|[\|\|\|\|\|\|\|\|\|\|\|\|]PAM | NA | Intergenic | NA | NA |

| | | | | | | | | | | | |
|---|---|---|---|---|---|---|---|---|---|---|---|
| T22546 | CGCAGTACTCGGAGAGGACCGGG | 1000 | | | | | | | | | |
| Oligo fwd | CACCCGCAGTACTCGGAGAGGACC | | | | | | | | | | |
| Oligo rev | AAACGGTCCTCTCCGAGTACTGCG | | | | | | | | | | |
| Chromosome | start | end | strand | MM | target_seq | PAM | alignment | distance | position | gene name | gene id |

<table>
<tr><td>GU305914.1</td><td></td><td>152072</td><td>152094</td><td>-</td><td>0</td><td>CGCAGTACTCGGAGAGGACC</td><td>GGG</td><td>||||||||[||||||||||||]PAM</td><td>NA</td><td>Intergenic</td><td>NA</td><td>NA</td></tr>
<tr><td>T22547</td><td>GACCGGGTCGCTGATGAGCGCGG</td><td></td><td>1000</td><td></td><td></td><td></td><td></td><td></td><td></td><td></td><td></td><td></td></tr>
<tr><td>Oligo fwd</td><td>CACCGACCGGGTCGCTGATGAGCG</td><td></td><td></td><td></td><td></td><td></td><td></td><td></td><td></td><td></td><td></td><td></td></tr>
<tr><td>Oligo rev</td><td>AAACCGCTCATCAGCGACCCGGTC</td><td></td><td></td><td></td><td></td><td></td><td></td><td></td><td></td><td></td><td></td><td></td></tr>
<tr><td>Chromosome</td><td>start</td><td></td><td>end</td><td>strand</td><td>MM</td><td>target_seq</td><td>PAM</td><td>alignment</td><td>distance</td><td>position</td><td>gene name</td><td>gene id</td></tr>
<tr><td>GU305914.1</td><td></td><td>152056</td><td>152078</td><td>-</td><td>0</td><td>GACCGGGTCGCTGATGAGCG</td><td>CGG</td><td>||||||||[||||||||||||]PAM</td><td>NA</td><td>Intergenic</td><td>NA</td><td>NA</td></tr>
<tr><td>T22548</td><td>GGTCGCTGATGAGCGCGGAGAGG</td><td></td><td>1000</td><td></td><td></td><td></td><td></td><td></td><td></td><td></td><td></td><td></td></tr>
<tr><td>Oligo fwd</td><td>CACCGGTCGCTGATGAGCGCGGAG</td><td></td><td></td><td></td><td></td><td></td><td></td><td></td><td></td><td></td><td></td><td></td></tr>
<tr><td>Oligo rev</td><td>AAACCTCCGCGCTCATCAGCGACC</td><td></td><td></td><td></td><td></td><td></td><td></td><td></td><td></td><td></td><td></td><td></td></tr>
<tr><td>Chromosome</td><td>start</td><td></td><td>end</td><td>strand</td><td>MM</td><td>target_seq</td><td>PAM</td><td>alignment</td><td>distance</td><td>position</td><td>gene name</td><td>gene id</td></tr>
<tr><td>GU305914.1</td><td></td><td>152051</td><td>152073</td><td>-</td><td>0</td><td>GGTCGCTGATGAGCGCGGAG</td><td>AGG</td><td>||||||||[||||||||||||]PAM</td><td>NA</td><td>Intergenic</td><td>NA</td><td>NA</td></tr>
<tr><td>T22549</td><td>GCACGTCCCGATTCTGCGTGCGG</td><td></td><td>1000</td><td></td><td></td><td></td><td></td><td></td><td></td><td></td><td></td><td></td></tr>
<tr><td>Oligo fwd</td><td>CACCGCACGTCCCGATTCTGCGTG</td><td></td><td></td><td></td><td></td><td></td><td></td><td></td><td></td><td></td><td></td><td></td></tr>
<tr><td>Oligo rev</td><td>AAACCACGCAGAATCGGGACGTGC</td><td></td><td></td><td></td><td></td><td></td><td></td><td></td><td></td><td></td><td></td><td></td></tr>
<tr><td>Chromosome</td><td>start</td><td></td><td>end</td><td>strand</td><td>MM</td><td>target_seq</td><td>PAM</td><td>alignment</td><td>distance</td><td>position</td><td>gene name</td><td>gene id</td></tr>
<tr><td>GU305914.1</td><td></td><td>152012</td><td>152034</td><td>-</td><td>0</td><td>GCACGTCCCGATTCTGCGTG</td><td>CGG</td><td>||||||||[||||||||||||]PAM</td><td>NA</td><td>Intergenic</td><td>NA</td><td>NA</td></tr>
<tr><td>GU305914.1</td><td></td><td>63764</td><td>63786</td><td>-</td><td>4</td><td>TCACGGCCCGCTTCTTCGTG</td><td>CGG</td><td>-||||-||[||-||||-||||]PAM</td><td>NA</td><td>Intergenic</td><td>NA</td><td>NA</td></tr>
<tr><td>T22550</td><td>CGTCCCGATTCTGCGTGCGGTGG</td><td></td><td>1000</td><td></td><td></td><td></td><td></td><td></td><td></td><td></td><td></td><td></td></tr>
<tr><td>Oligo fwd</td><td>CACCCGTCCCGATTCTGCGTGCGG</td><td></td><td></td><td></td><td></td><td></td><td></td><td></td><td></td><td></td><td></td><td></td></tr>
<tr><td>Oligo rev</td><td>AAACCCGCACGCAGAATCGGGACG</td><td></td><td></td><td></td><td></td><td></td><td></td><td></td><td></td><td></td><td></td><td></td></tr>
<tr><td>Chromosome</td><td>start</td><td></td><td>end</td><td>strand</td><td>MM</td><td>target_seq</td><td>PAM</td><td>alignment</td><td>distance</td><td>position</td><td>gene name</td><td>gene id</td></tr>
<tr><td>GU305914.1</td><td></td><td>152009</td><td>152031</td><td>-</td><td>0</td><td>CGTCCCGATTCTGCGTGCGG</td><td>TGG</td><td>||||||||[||||||||||||]PAM</td><td>NA</td><td>Intergenic</td><td>NA</td><td>NA</td></tr>
<tr><td>T22551</td><td>TCTGCGTGCGGTGGTCGTAGCGG</td><td></td><td>1000</td><td></td><td></td><td></td><td></td><td></td><td></td><td></td><td></td><td></td></tr>
<tr><td>Oligo fwd</td><td>CACCTCTGCGTGCGGTGGTCGTAG</td><td></td><td></td><td></td><td></td><td></td><td></td><td></td><td></td><td></td><td></td><td></td></tr>
<tr><td>Oligo rev</td><td>AAACCTACGACCACCGCACGCAGA</td><td></td><td></td><td></td><td></td><td></td><td></td><td></td><td></td><td></td><td></td><td></td></tr>
<tr><td>Chromosome</td><td>start</td><td></td><td>end</td><td>strand</td><td>MM</td><td>target_seq</td><td>PAM</td><td>alignment</td><td>distance</td><td>position</td><td>gene name</td><td>gene id</td></tr>
<tr><td>GU305914.1</td><td></td><td>152000</td><td>152022</td><td>-</td><td>0</td><td>TCTGCGTGCGGTGGTCGTAG</td><td>CGG</td><td>||||||||[||||||||||||]PAM</td><td>NA</td><td>Intergenic</td><td>NA</td><td>NA</td></tr>
<tr><td>T22552</td><td>GGTGGTCGTAGCGGCTCTCGAGG</td><td></td><td>1000</td><td></td><td></td><td></td><td></td><td></td><td></td><td></td><td></td><td></td></tr>
<tr><td>Oligo fwd</td><td>CACCGGTGGTCGTAGCGGCTCTCG</td><td></td><td></td><td></td><td></td><td></td><td></td><td></td><td></td><td></td><td></td><td></td></tr>
<tr><td>Oligo rev</td><td>AAACCGAGAGCCGCTACGACCACC</td><td></td><td></td><td></td><td></td><td></td><td></td><td></td><td></td><td></td><td></td><td></td></tr>
<tr><td>Chromosome</td><td>start</td><td></td><td>end</td><td>strand</td><td>MM</td><td>target_seq</td><td>PAM</td><td>alignment</td><td>distance</td><td>position</td><td>gene name</td><td>gene id</td></tr>
<tr><td>GU305914.1</td><td></td><td>151991</td><td>152013</td><td>-</td><td>0</td><td>GGTGGTCGTAGCGGCTCTCG</td><td>AGG</td><td>||||||||[||||||||||||]PAM</td><td>NA</td><td>Intergenic</td><td>NA</td><td>NA</td></tr>
<tr><td>GU305914.1</td><td></td><td>197382</td><td>197404</td><td>+</td><td>4</td><td>GTTCCTCGTAGCGGCTCGCG</td><td>GGG</td><td>|-|--|||[|||||||||-||]PAM</td><td>NA</td><td>Intergenic</td><td>NA</td><td>NA</td></tr>
<tr><td>T22553</td><td>GTGGTCGTAGCGGCTCTCGAGGG</td><td></td><td>1000</td><td></td><td></td><td></td><td></td><td></td><td></td><td></td><td></td><td></td></tr>
<tr><td>Oligo fwd</td><td>CACCGTGGTCGTAGCGGCTCTCGA</td><td></td><td></td><td></td><td></td><td></td><td></td><td></td><td></td><td></td><td></td><td></td></tr>
<tr><td>Oligo rev</td><td>AAACTCGAGAGCCGCTACGACCAC</td><td></td><td></td><td></td><td></td><td></td><td></td><td></td><td></td><td></td><td></td><td></td></tr>
<tr><td>Chromosome</td><td>start</td><td></td><td>end</td><td>strand</td><td>MM</td><td>target_seq</td><td>PAM</td><td>alignment</td><td>distance</td><td>position</td><td>gene name</td><td>gene id</td></tr>
<tr><td>GU305914.1</td><td></td><td>151990</td><td>152012</td><td>-</td><td>0</td><td>GTGGTCGTAGCGGCTCTCGA</td><td>GGG</td><td>||||||||[||||||||||||]PAM</td><td>NA</td><td>Intergenic</td><td>NA</td><td>NA</td></tr>
<tr><td>T22554</td><td>GTAGCGGCTCTCGAGGGCCTCGG</td><td></td><td>1000</td><td></td><td></td><td></td><td></td><td></td><td></td><td></td><td></td><td></td></tr>
<tr><td>Oligo fwd</td><td>CACCGTAGCGGCTCTCGAGGGCCT</td><td></td><td></td><td></td><td></td><td></td><td></td><td></td><td></td><td></td><td></td><td></td></tr>
<tr><td>Oligo rev</td><td>AAACAGGCCCTCGAGAGCCGCTAC</td><td></td><td></td><td></td><td></td><td></td><td></td><td></td><td></td><td></td><td></td><td></td></tr>
<tr><td>Chromosome</td><td>start</td><td></td><td>end</td><td>strand</td><td>MM</td><td>target_seq</td><td>PAM</td><td>alignment</td><td>distance</td><td>position</td><td>gene name</td><td>gene id</td></tr>
<tr><td>GU305914.1</td><td></td><td>151984</td><td>152006</td><td>-</td><td>0</td><td>GTAGCGGCTCTCGAGGGCCT</td><td>CGG</td><td>||||||||[||||||||||||]PAM</td><td>NA</td><td>Intergenic</td><td>NA</td><td>NA</td></tr>
<tr><td>T22555</td><td>CTCGAGGGCCTCGGACTGCGTGG</td><td></td><td>1000</td><td></td><td></td><td></td><td></td><td></td><td></td><td></td><td></td><td></td></tr>
<tr><td>Oligo fwd</td><td>CACCCTCGAGGGCCTCGGACTGCG</td><td></td><td></td><td></td><td></td><td></td><td></td><td></td><td></td><td></td><td></td><td></td></tr>
<tr><td>Oligo rev</td><td>AAACCGCAGTCCGAGGCCCTCGAG</td><td></td><td></td><td></td><td></td><td></td><td></td><td></td><td></td><td></td><td></td><td></td></tr>
<tr><td>Chromosome</td><td>start</td><td></td><td>end</td><td>strand</td><td>MM</td><td>target_seq</td><td>PAM</td><td>alignment</td><td>distance</td><td>position</td><td>gene name</td><td>gene id</td></tr>
<tr><td>GU305914.1</td><td></td><td>151975</td><td>151997</td><td>-</td><td>0</td><td>CTCGAGGGCCTCGGACTGCG</td><td>TGG</td><td>||||||||[||||||||||||]PAM</td><td>NA</td><td>Intergenic</td><td>NA</td><td>NA</td></tr>
<tr><td>T22556</td><td>TCGAGGGCCTCGGACTGCGTGGG</td><td></td><td>1000</td><td></td><td></td><td></td><td></td><td></td><td></td><td></td><td></td><td></td></tr>
<tr><td>Oligo fwd</td><td>CACCTCGAGGGCCTCGGACTGCGT</td><td></td><td></td><td></td><td></td><td></td><td></td><td></td><td></td><td></td><td></td><td></td></tr>
<tr><td>Oligo rev</td><td>AAACACGCAGTCCGAGGCCCTCGA</td><td></td><td></td><td></td><td></td><td></td><td></td><td></td><td></td><td></td><td></td><td></td></tr>
<tr><td>Chromosome</td><td>start</td><td></td><td>end</td><td>strand</td><td>MM</td><td>target_seq</td><td>PAM</td><td>alignment</td><td>distance</td><td>position</td><td>gene name</td><td>gene id</td></tr>
<tr><td>GU305914.1</td><td></td><td>151974</td><td>151996</td><td>-</td><td>0</td><td>TCGAGGGCCTCGGACTGCGT</td><td>GGG</td><td>||||||||[||||||||||||]PAM</td><td>NA</td><td>Intergenic</td><td>NA</td><td>NA</td></tr>
<tr><td>T22557</td><td>CGACCCGACGCAGACGATGCAGG</td><td></td><td>1000</td><td></td><td></td><td></td><td></td><td></td><td></td><td></td><td></td><td></td></tr>
<tr><td>Oligo fwd</td><td>CACCCGACCCGACGCAGACGATGC</td><td></td><td></td><td></td><td></td><td></td><td></td><td></td><td></td><td></td><td></td><td></td></tr>
<tr><td>Oligo rev</td><td>AAACGCATCGTCTGCGTCGGGTCG</td><td></td><td></td><td></td><td></td><td></td><td></td><td></td><td></td><td></td><td></td><td></td></tr>
<tr><td>Chromosome</td><td>start</td><td></td><td>end</td><td>strand</td><td>MM</td><td>target_seq</td><td>PAM</td><td>alignment</td><td>distance</td><td>position</td><td>gene name</td><td>gene id</td></tr>
<tr><td>GU305914.1</td><td></td><td>151951</td><td>151973</td><td>-</td><td>0</td><td>CGACCCGACGCAGACGATGC</td><td>AGG</td><td>||||||||[||||||||||||]PAM</td><td>NA</td><td>Intergenic</td><td>NA</td><td>NA</td></tr>
<tr><td>T22558</td><td>GACCCGACGCAGACGATGCAGGG</td><td></td><td>1000</td><td></td><td></td><td></td><td></td><td></td><td></td><td></td><td></td><td></td></tr>
<tr><td>Oligo fwd</td><td>CACCGACCCGACGCAGACGATGCA</td><td></td><td></td><td></td><td></td><td></td><td></td><td></td><td></td><td></td><td></td><td></td></tr>
<tr><td>Oligo rev</td><td>AAACTGCATCGTCTGCGTCGGGTC</td><td></td><td></td><td></td><td></td><td></td><td></td><td></td><td></td><td></td><td></td><td></td></tr>
<tr><td>Chromosome</td><td>start</td><td></td><td>end</td><td>strand</td><td>MM</td><td>target_seq</td><td>PAM</td><td>alignment</td><td>distance</td><td>position</td><td>gene name</td><td>gene id</td></tr>
<tr><td>GU305914.1</td><td></td><td>151950</td><td>151972</td><td>-</td><td>0</td><td>GACCCGACGCAGACGATGCA</td><td>GGG</td><td>||||||||[||||||||||||]PAM</td><td>NA</td><td>Intergenic</td><td>NA</td><td>NA</td></tr>
<tr><td>T22559</td><td>CGATGCAGGGCGCCGCGCGCTGG</td><td></td><td>1000</td><td></td><td></td><td></td><td></td><td></td><td></td><td></td><td></td><td></td></tr>
<tr><td>Oligo fwd</td><td>CACCCGATGCAGGGCGCCGCGCGC</td><td></td><td></td><td></td><td></td><td></td><td></td><td></td><td></td><td></td><td></td><td></td></tr>
<tr><td>Oligo rev</td><td>AAACGCGCGCGGCGCCCTGCATCG</td><td></td><td></td><td></td><td></td><td></td><td></td><td></td><td></td><td></td><td></td><td></td></tr>
<tr><td>Chromosome</td><td>start</td><td></td><td>end</td><td>strand</td><td>MM</td><td>target_seq</td><td>PAM</td><td>alignment</td><td>distance</td><td>position</td><td>gene name</td><td>gene id</td></tr>
<tr><td>GU305914.1</td><td></td><td>151937</td><td>151959</td><td>-</td><td>0</td><td>CGATGCAGGGCGCCGCGCGC</td><td>TGG</td><td>||||||||[||||||||||||]PAM</td><td>NA</td><td>Intergenic</td><td>NA</td><td>NA</td></tr>
<tr><td>T22560</td><td>TGCAGGGCGCCGCGCGCTGGCGG</td><td></td><td>1000</td><td></td><td></td><td></td><td></td><td></td><td></td><td></td><td></td><td></td></tr>
<tr><td>Oligo fwd</td><td>CACCTGCAGGGCGCCGCGCGCTGG</td><td></td><td></td><td></td><td></td><td></td><td></td><td></td><td></td><td></td><td></td><td></td></tr>
<tr><td>Oligo rev</td><td>AAACCCAGCGCGCGGCGCCCTGCA</td><td></td><td></td><td></td><td></td><td></td><td></td><td></td><td></td><td></td><td></td><td></td></tr>
<tr><td>Chromosome</td><td>start</td><td></td><td>end</td><td>strand</td><td>MM</td><td>target_seq</td><td>PAM</td><td>alignment</td><td>distance</td><td>position</td><td>gene name</td><td>gene id</td></tr>
<tr><td>GU305914.1</td><td></td><td>151934</td><td>151956</td><td>-</td><td>0</td><td>TGCAGGGCGCCGCGCGCTGG</td><td>CGG</td><td>||||||||[||||||||||||]PAM</td><td>NA</td><td>Intergenic</td><td>NA</td><td>NA</td></tr>
<tr><td>T22561</td><td>GGTAGAGCTCGCTGTCGTTGAGG</td><td></td><td>1000</td><td></td><td></td><td></td><td></td><td></td><td></td><td></td><td></td><td></td></tr>
</table>

Oligo fwd CACCGGTAGAGCTCGCTGTCGTTG
Oligo rev AAACCAACGACAGCGAGCTCTACC
Chromosome start end strand MM target_seq PAM alignment distance position gene name gene id
GU305914.1 151913 151935 - 0 GGTAGAGCTCGCTGTCGTTG AGG ||||||||[||||||||||||]PAM NA Intergenic NA NA

T22562 GTAGAGCTCGCTGTCGTTGAGGG 1000
Oligo fwd CACCGTAGAGCTCGCTGTCGTTGA
Oligo rev AAACTCAACGACAGCGAGCTCTAC
Chromosome start end strand MM target_seq PAM alignment distance position gene name gene id
GU305914.1 151912 151934 - 0 GTAGAGCTCGCTGTCGTTGA GGG ||||||||[||||||||||||]PAM NA Intergenic NA NA

T22563 CGACGACGTGCAGCATGTGTCGG 1000
Oligo fwd CACCCGACGACGTGCAGCATGTGT
Oligo rev AAACACACATGCTGCACGTCGTCG
Chromosome start end strand MM target_seq PAM alignment distance position gene name gene id
GU305914.1 151865 151887 - 0 CGACGACGTGCAGCATGTGT CGG ||||||||[||||||||||||]PAM NA Intergenic NA NA

T22564 CGACGTGCAGCATGTGTCGGAGG 1000
Oligo fwd CACCCGACGTGCAGCATGTGTCGG
Oligo rev AAACCCGACACATGCTGCACGTCG
Chromosome start end strand MM target_seq PAM alignment distance position gene name gene id
GU305914.1 151862 151884 - 0 CGACGTGCAGCATGTGTCGG AGG ||||||||[||||||||||||]PAM NA Intergenic NA NA

T22565 CGATGACGATGATGTTGCTGCGG 1000
Oligo fwd CACCCGATGACGATGATGTTGCTG
Oligo rev AAACCAGCAACATCATCGTCATCG
Chromosome start end strand MM target_seq PAM alignment distance position gene name gene id
GU305914.1 151823 151845 - 0 CGATGACGATGATGTTGCTG CGG ||||||||[||||||||||||]PAM NA Intergenic NA NA

T22566 CGATGATGTTGCTGCGGCAGAGG 1000
Oligo fwd CACCCGATGATGTTGCTGCGGCAG
Oligo rev AAACCTGCCGCAGCAACATCATCG
Chromosome start end strand MM target_seq PAM alignment distance position gene name gene id
GU305914.1 151817 151839 - 0 CGATGATGTTGCTGCGGCAG AGG ||||||||[||||||||||||]PAM NA Intergenic NA NA

T22567 GTTGCTGCGGCAGAGGTCGTCGG 1000
Oligo fwd CACCGTTGCTGCGGCAGAGGTCGT
Oligo rev AAACACGACCTCTGCCGCAGCAAC
Chromosome start end strand MM target_seq PAM alignment distance position gene name gene id
GU305914.1 151810 151832 - 0 GTTGCTGCGGCAGAGGTCGT CGG ||||||||[||||||||||||]PAM NA Intergenic NA NA
GU305914.1 24053 24075 + 4 GTAGACGCGACAGAGGTCGT CGG ||-|--||[|-||||||||||]PAM NA Intergenic NA NA

T22568 AGAGGTCGTCGGTGTCCTTGCGG 1000
Oligo fwd CACCAGAGGTCGTCGGTGTCCTTG
Oligo rev AAACCAAGGACACCGACGACCTCT
Chromosome start end strand MM target_seq PAM alignment distance position gene name gene id
GU305914.1 151799 151821 - 0 AGAGGTCGTCGGTGTCCTTG CGG ||||||||[||||||||||||]PAM NA Intergenic NA NA

T22569 TGCGGCCCTTGCCGCCGTCCAGG 1000
Oligo fwd CACCTGCGGCCCTTGCCGCCGTCC
Oligo rev AAACGGACGGCGGCAAGGGCCGCA
Chromosome start end strand MM target_seq PAM alignment distance position gene name gene id
GU305914.1 151781 151803 - 0 TGCGGCCCTTGCCGCCGTCC AGG ||||||||[||||||||||||]PAM NA Intergenic NA NA

T22570 GCGGCCCTTGCCGCCGTCCAGGG 1000
Oligo fwd CACCGCGGCCCTTGCCGCCGTCCA
Oligo rev AAACTGGACGGCGGCAAGGGCCGC
Chromosome start end strand MM target_seq PAM alignment distance position gene name gene id
GU305914.1 151780 151802 - 0 GCGGCCCTTGCCGCCGTCCA GGG ||||||||[||||||||||||]PAM NA Intergenic NA NA

T22571 CCTTGCCGCCGTCCAGGGCGCGG 1000
Oligo fwd CACCCCTTGCCGCCGTCCAGGGCG
Oligo rev AAACCGCCCTGGACGGCGGCAAGG
Chromosome start end strand MM target_seq PAM alignment distance position gene name gene id
GU305914.1 151775 151797 - 0 CCTTGCCGCCGTCCAGGGCG CGG ||||||||[||||||||||||]PAM NA Intergenic NA NA

T22572 GCCGCCGTCCAGGGCGCGGATGG 1000
Oligo fwd CACCGCCGCCGTCCAGGGCGCGGA
Oligo rev AAACTCCGCGCCCTGGACGGCGGC
Chromosome start end strand MM target_seq PAM alignment distance position gene name gene id
GU305914.1 151771 151793 - 0 GCCGCCGTCCAGGGCGCGGA TGG ||||||||[||||||||||||]PAM NA Intergenic NA NA

T22573 CAGGGCGCGGATGGCGATGTCGG 1000
Oligo fwd CACCCAGGGCGCGGATGGCGATGT
Oligo rev AAACACATCGCCATCCGCGCCCTG
Chromosome start end strand MM target_seq PAM alignment distance position gene name gene id
GU305914.1 151762 151784 - 0 CAGGGCGCGGATGGCGATGT CGG ||||||||[||||||||||||]PAM NA Intergenic NA NA

T22574 ATGGCGATGTCGGAGACGACCGG 1000
Oligo fwd CACCATGGCGATGTCGGAGACGAC
Oligo rev AAACGTCGTCTCCGACATCGCCAT
Chromosome start end strand MM target_seq PAM alignment distance position gene name gene id
GU305914.1 151752 151774 - 0 ATGGCGATGTCGGAGACGAC CGG ||||||||[||||||||||||]PAM NA Intergenic NA NA

T22575 GTCGGAGACGACCGGCCAGTAGG 1000
Oligo fwd CACCGTCGGAGACGACCGGCCAGT
Oligo rev AAACACTGGCCGGTCGTCTCCGAC
Chromosome start end strand MM target_seq PAM alignment distance position gene name gene id
GU305914.1 151744 151766 - 0 GTCGGAGACGACCGGCCAGT AGG ||||||||[||||||||||||]PAM NA Intergenic NA NA

<table>
<tr><td>T22576</td><td>AGAGATCGCGCCACTGCTGCTGG</td><td>1000</td><td></td><td></td><td></td><td></td><td></td><td></td><td></td><td></td><td></td></tr>
<tr><td>Oligo fwd</td><td>CACCAGAGATCGCGCCACTGCTGC</td><td></td><td></td><td></td><td></td><td></td><td></td><td></td><td></td><td></td><td></td></tr>
<tr><td>Oligo rev</td><td>AAACGCAGCAGTGGCGCGATCTCT</td><td></td><td></td><td></td><td></td><td></td><td></td><td></td><td></td><td></td><td></td></tr>
<tr><td>Chromosome</td><td>start</td><td>end</td><td>strand</td><td>MM</td><td>target_seq</td><td>PAM</td><td>alignment</td><td>distance</td><td>position</td><td>gene name</td><td>gene id</td></tr>
<tr><td>GU305914.1</td><td>151718</td><td>151740</td><td>-</td><td>0</td><td>AGAGATCGCGCCACTGCTGC</td><td>TGG</td><td>||||||||[||||||||||||]PAM</td><td>NA</td><td>Intergenic</td><td>NA</td><td>NA</td></tr>
<tr><td>T22577</td><td>GCGCCACTGCTGCTGGCAGATGG</td><td>1000</td><td></td><td></td><td></td><td></td><td></td><td></td><td></td><td></td><td></td></tr>
<tr><td>Oligo fwd</td><td>CACCGCGCCACTGCTGCTGGCAGA</td><td></td><td></td><td></td><td></td><td></td><td></td><td></td><td></td><td></td><td></td></tr>
<tr><td>Oligo rev</td><td>AAACTCTGCCAGCAGCAGTGGCGC</td><td></td><td></td><td></td><td></td><td></td><td></td><td></td><td></td><td></td><td></td></tr>
<tr><td>Chromosome</td><td>start</td><td>end</td><td>strand</td><td>MM</td><td>target_seq</td><td>PAM</td><td>alignment</td><td>distance</td><td>position</td><td>gene name</td><td>gene id</td></tr>
<tr><td>GU305914.1</td><td>151711</td><td>151733</td><td>-</td><td>0</td><td>GCGCCACTGCTGCTGGCAGA</td><td>TGG</td><td>||||||||[||||||||||||]PAM</td><td>NA</td><td>Intergenic</td><td>NA</td><td>NA</td></tr>
<tr><td>T22578</td><td>CGCCACTGCTGCTGGCAGATGGG</td><td>1000</td><td></td><td></td><td></td><td></td><td></td><td></td><td></td><td></td><td></td></tr>
<tr><td>Oligo fwd</td><td>CACCCGCCACTGCTGCTGGCAGAT</td><td></td><td></td><td></td><td></td><td></td><td></td><td></td><td></td><td></td><td></td></tr>
<tr><td>Oligo rev</td><td>AAACATCTGCCAGCAGCAGTGGCG</td><td></td><td></td><td></td><td></td><td></td><td></td><td></td><td></td><td></td><td></td></tr>
<tr><td>Chromosome</td><td>start</td><td>end</td><td>strand</td><td>MM</td><td>target_seq</td><td>PAM</td><td>alignment</td><td>distance</td><td>position</td><td>gene name</td><td>gene id</td></tr>
<tr><td>GU305914.1</td><td>151710</td><td>151732</td><td>-</td><td>0</td><td>CGCCACTGCTGCTGGCAGAT</td><td>GGG</td><td>||||||||[||||||||||||]PAM</td><td>NA</td><td>Intergenic</td><td>NA</td><td>NA</td></tr>
<tr><td>T22579</td><td>GCCACTGCTGCTGGCAGATGGGG</td><td>1000</td><td></td><td></td><td></td><td></td><td></td><td></td><td></td><td></td><td></td></tr>
<tr><td>Oligo fwd</td><td>CACCGCCACTGCTGCTGGCAGATG</td><td></td><td></td><td></td><td></td><td></td><td></td><td></td><td></td><td></td><td></td></tr>
<tr><td>Oligo rev</td><td>AAACCATCTGCCAGCAGCAGTGGC</td><td></td><td></td><td></td><td></td><td></td><td></td><td></td><td></td><td></td><td></td></tr>
<tr><td>Chromosome</td><td>start</td><td>end</td><td>strand</td><td>MM</td><td>target_seq</td><td>PAM</td><td>alignment</td><td>distance</td><td>position</td><td>gene name</td><td>gene id</td></tr>
<tr><td>GU305914.1</td><td>151709</td><td>151731</td><td>-</td><td>0</td><td>GCCACTGCTGCTGGCAGATG</td><td>GGG</td><td>||||||||[||||||||||||]PAM</td><td>NA</td><td>Intergenic</td><td>NA</td><td>NA</td></tr>
<tr><td>T22580</td><td>CGACGTCGTCGCGCCGCCTGAGG</td><td>1000</td><td></td><td></td><td></td><td></td><td></td><td></td><td></td><td></td><td></td></tr>
<tr><td>Oligo fwd</td><td>CACCCGACGTCGTCGCGCCGCCTG</td><td></td><td></td><td></td><td></td><td></td><td></td><td></td><td></td><td></td><td></td></tr>
<tr><td>Oligo rev</td><td>AAACCAGGCGGCGCGACGACGTCG</td><td></td><td></td><td></td><td></td><td></td><td></td><td></td><td></td><td></td><td></td></tr>
<tr><td>Chromosome</td><td>start</td><td>end</td><td>strand</td><td>MM</td><td>target_seq</td><td>PAM</td><td>alignment</td><td>distance</td><td>position</td><td>gene name</td><td>gene id</td></tr>
<tr><td>GU305914.1</td><td>151667</td><td>151689</td><td>-</td><td>0</td><td>CGACGTCGTCGCGCCGCCTG</td><td>AGG</td><td>||||||||[||||||||||||]PAM</td><td>NA</td><td>Intergenic</td><td>NA</td><td>NA</td></tr>
<tr><td>T22581</td><td>CGCGCCGCCTGAGGTGAACGCGG</td><td>1000</td><td></td><td></td><td></td><td></td><td></td><td></td><td></td><td></td><td></td></tr>
<tr><td>Oligo fwd</td><td>CACCCGCGCCGCCTGAGGTGAACG</td><td></td><td></td><td></td><td></td><td></td><td></td><td></td><td></td><td></td><td></td></tr>
<tr><td>Oligo rev</td><td>AAACCGTTCACCTCAGGCGGCGCG</td><td></td><td></td><td></td><td></td><td></td><td></td><td></td><td></td><td></td><td></td></tr>
<tr><td>Chromosome</td><td>start</td><td>end</td><td>strand</td><td>MM</td><td>target_seq</td><td>PAM</td><td>alignment</td><td>distance</td><td>position</td><td>gene name</td><td>gene id</td></tr>
<tr><td>GU305914.1</td><td>151658</td><td>151680</td><td>-</td><td>0</td><td>CGCGCCGCCTGAGGTGAACG</td><td>CGG</td><td>||||||||[||||||||||||]PAM</td><td>NA</td><td>Intergenic</td><td>NA</td><td>NA</td></tr>
<tr><td>T22582</td><td>CCGCCTGAGGTGAACGCGGTCGG</td><td>1000</td><td></td><td></td><td></td><td></td><td></td><td></td><td></td><td></td><td></td></tr>
<tr><td>Oligo fwd</td><td>CACCCCGCCTGAGGTGAACGCGGT</td><td></td><td></td><td></td><td></td><td></td><td></td><td></td><td></td><td></td><td></td></tr>
<tr><td>Oligo rev</td><td>AAACACCGCGTTCACCTCAGGCGG</td><td></td><td></td><td></td><td></td><td></td><td></td><td></td><td></td><td></td><td></td></tr>
<tr><td>Chromosome</td><td>start</td><td>end</td><td>strand</td><td>MM</td><td>target_seq</td><td>PAM</td><td>alignment</td><td>distance</td><td>position</td><td>gene name</td><td>gene id</td></tr>
<tr><td>GU305914.1</td><td>151654</td><td>151676</td><td>-</td><td>0</td><td>CCGCCTGAGGTGAACGCGGT</td><td>CGG</td><td>||||||||[||||||||||||]PAM</td><td>NA</td><td>Intergenic</td><td>NA</td><td>NA</td></tr>
<tr><td>T22583</td><td>TGAGGTGAACGCGGTCGGCGAGG</td><td>1000</td><td></td><td></td><td></td><td></td><td></td><td></td><td></td><td></td><td></td></tr>
<tr><td>Oligo fwd</td><td>CACCTGAGGTGAACGCGGTCGGCG</td><td></td><td></td><td></td><td></td><td></td><td></td><td></td><td></td><td></td><td></td></tr>
<tr><td>Oligo rev</td><td>AAACCGCCGACCGCGTTCACCTCA</td><td></td><td></td><td></td><td></td><td></td><td></td><td></td><td></td><td></td><td></td></tr>
<tr><td>Chromosome</td><td>start</td><td>end</td><td>strand</td><td>MM</td><td>target_seq</td><td>PAM</td><td>alignment</td><td>distance</td><td>position</td><td>gene name</td><td>gene id</td></tr>
<tr><td>GU305914.1</td><td>151649</td><td>151671</td><td>-</td><td>0</td><td>TGAGGTGAACGCGGTCGGCG</td><td>AGG</td><td>||||||||[||||||||||||]PAM</td><td>NA</td><td>Intergenic</td><td>NA</td><td>NA</td></tr>
<tr><td>GU305914.1</td><td>114495</td><td>114517</td><td>-</td><td>4</td><td>TGAGGTAGACGCAGTCGGCC</td><td>GGG</td><td>||||||--[||||-||||||-]PAM</td><td>NA</td><td>Intergenic</td><td>NA</td><td>NA</td></tr>
<tr><td>T22584</td><td>GAGGTGAACGCGGTCGGCGAGGG</td><td>1000</td><td></td><td></td><td></td><td></td><td></td><td></td><td></td><td></td><td></td></tr>
<tr><td>Oligo fwd</td><td>CACCGAGGTGAACGCGGTCGGCGA</td><td></td><td></td><td></td><td></td><td></td><td></td><td></td><td></td><td></td><td></td></tr>
<tr><td>Oligo rev</td><td>AAACTCGCCGACCGCGTTCACCTC</td><td></td><td></td><td></td><td></td><td></td><td></td><td></td><td></td><td></td><td></td></tr>
<tr><td>Chromosome</td><td>start</td><td>end</td><td>strand</td><td>MM</td><td>target_seq</td><td>PAM</td><td>alignment</td><td>distance</td><td>position</td><td>gene name</td><td>gene id</td></tr>
<tr><td>GU305914.1</td><td>151648</td><td>151670</td><td>-</td><td>0</td><td>GAGGTGAACGCGGTCGGCGA</td><td>GGG</td><td>||||||||[||||||||||||]PAM</td><td>NA</td><td>Intergenic</td><td>NA</td><td>NA</td></tr>
<tr><td>T22585</td><td>AAACCCAAAAGTCCTGAAGATGG</td><td>1000</td><td></td><td></td><td></td><td></td><td></td><td></td><td></td><td></td><td></td></tr>
<tr><td>Oligo fwd</td><td>CACCAAACCCAAAAGTCCTGAAGA</td><td></td><td></td><td></td><td></td><td></td><td></td><td></td><td></td><td></td><td></td></tr>
<tr><td>Oligo rev</td><td>AAACTCTTCAGGACTTTTGGGTTT</td><td></td><td></td><td></td><td></td><td></td><td></td><td></td><td></td><td></td><td></td></tr>
<tr><td>Chromosome</td><td>start</td><td>end</td><td>strand</td><td>MM</td><td>target_seq</td><td>PAM</td><td>alignment</td><td>distance</td><td>position</td><td>gene name</td><td>gene id</td></tr>
<tr><td>GU305914.1</td><td>151609</td><td>151631</td><td>-</td><td>0</td><td>AAACCCAAAAGTCCTGAAGA</td><td>TGG</td><td>||||||||[||||||||||||]PAM</td><td>NA</td><td>Intergenic</td><td>NA</td><td>NA</td></tr>
<tr><td>T22586</td><td>GAAGATGGTCTTGATCTGAGCGG</td><td>1000</td><td></td><td></td><td></td><td></td><td></td><td></td><td></td><td></td><td></td></tr>
<tr><td>Oligo fwd</td><td>CACCGAAGATGGTCTTGATCTGAG</td><td></td><td></td><td></td><td></td><td></td><td></td><td></td><td></td><td></td><td></td></tr>
<tr><td>Oligo rev</td><td>AAACCTCAGATCAAGACCATCTTC</td><td></td><td></td><td></td><td></td><td></td><td></td><td></td><td></td><td></td><td></td></tr>
<tr><td>Chromosome</td><td>start</td><td>end</td><td>strand</td><td>MM</td><td>target_seq</td><td>PAM</td><td>alignment</td><td>distance</td><td>position</td><td>gene name</td><td>gene id</td></tr>
<tr><td>GU305914.1</td><td>151594</td><td>151616</td><td>-</td><td>0</td><td>GAAGATGGTCTTGATCTGAG</td><td>CGG</td><td>||||||||[||||||||||||]PAM</td><td>NA</td><td>Intergenic</td><td>NA</td><td>NA</td></tr>
<tr><td>T22587</td><td>TGATCTGAGCGGATCTGCTGCGG</td><td>1000</td><td></td><td></td><td></td><td></td><td></td><td></td><td></td><td></td><td></td></tr>
<tr><td>Oligo fwd</td><td>CACCTGATCTGAGCGGATCTGCTG</td><td></td><td></td><td></td><td></td><td></td><td></td><td></td><td></td><td></td><td></td></tr>
<tr><td>Oligo rev</td><td>AAACCAGCAGATCCGCTCAGATCA</td><td></td><td></td><td></td><td></td><td></td><td></td><td></td><td></td><td></td><td></td></tr>
<tr><td>Chromosome</td><td>start</td><td>end</td><td>strand</td><td>MM</td><td>target_seq</td><td>PAM</td><td>alignment</td><td>distance</td><td>position</td><td>gene name</td><td>gene id</td></tr>
<tr><td>GU305914.1</td><td>151583</td><td>151605</td><td>-</td><td>0</td><td>TGATCTGAGCGGATCTGCTG</td><td>CGG</td><td>||||||||[||||||||||||]PAM</td><td>NA</td><td>Intergenic</td><td>NA</td><td>NA</td></tr>
<tr><td>T22588</td><td>GGATCTGCTGCGGTTGAGCGCGG</td><td>1000</td><td></td><td></td><td></td><td></td><td></td><td></td><td></td><td></td><td></td></tr>
<tr><td>Oligo fwd</td><td>CACCGGATCTGCTGCGGTTGAGCG</td><td></td><td></td><td></td><td></td><td></td><td></td><td></td><td></td><td></td><td></td></tr>
<tr><td>Oligo rev</td><td>AAACCGCTCAACCGCAGCAGATCC</td><td></td><td></td><td></td><td></td><td></td><td></td><td></td><td></td><td></td><td></td></tr>
<tr><td>Chromosome</td><td>start</td><td>end</td><td>strand</td><td>MM</td><td>target_seq</td><td>PAM</td><td>alignment</td><td>distance</td><td>position</td><td>gene name</td><td>gene id</td></tr>
<tr><td>GU305914.1</td><td>151573</td><td>151595</td><td>-</td><td>0</td><td>GGATCTGCTGCGGTTGAGCG</td><td>CGG</td><td>||||||||[||||||||||||]PAM</td><td>NA</td><td>Intergenic</td><td>NA</td><td>NA</td></tr>
<tr><td>T22589</td><td>TGAGCGCGGAACTGAGCGCCTGG</td><td>1000</td><td></td><td></td><td></td><td></td><td></td><td></td><td></td><td></td><td></td></tr>
<tr><td>Oligo fwd</td><td>CACCTGAGCGCGGAACTGAGCGCC</td><td></td><td></td><td></td><td></td><td></td><td></td><td></td><td></td><td></td><td></td></tr>
<tr><td>Oligo rev</td><td>AAACGGCGCTCAGTTCCGCGCTCA</td><td></td><td></td><td></td><td></td><td></td><td></td><td></td><td></td><td></td><td></td></tr>
<tr><td>Chromosome</td><td>start</td><td>end</td><td>strand</td><td>MM</td><td>target_seq</td><td>PAM</td><td>alignment</td><td>distance</td><td>position</td><td>gene name</td><td>gene id</td></tr>
<tr><td>GU305914.1</td><td>151559</td><td>151581</td><td>-</td><td>0</td><td>TGAGCGCGGAACTGAGCGCC</td><td>TGG</td><td>||||||||[||||||||||||]PAM</td><td>NA</td><td>Intergenic</td><td>NA</td><td>NA</td></tr>
<tr><td>T22590</td><td>GAGCGCGGAACTGAGCGCCTGGG</td><td>1000</td><td></td><td></td><td></td><td></td><td></td><td></td><td></td><td></td><td></td></tr>
<tr><td>Oligo fwd</td><td>CACCGAGCGCGGAACTGAGCGCCT</td><td></td><td></td><td></td><td></td><td></td><td></td><td></td><td></td><td></td><td></td></tr>
<tr><td>Oligo rev</td><td>AAACAGGCGCTCAGTTCCGCGCTC</td><td></td><td></td><td></td><td></td><td></td><td></td><td></td><td></td><td></td><td></td></tr>
</table>

<table>
<tr><td>Chromosome</td><td>start</td><td>end</td><td>strand</td><td>MM</td><td>target_seq</td><td>PAM</td><td>alignment</td><td>distance</td><td>position</td><td>gene name</td><td>gene id</td></tr>
<tr><td>GU305914.1</td><td>151558</td><td>151580</td><td>-</td><td>0</td><td>GAGCGCGGAACTGAGCGCCT</td><td>GGG</td><td>||||||||[||||||||||||]PAM</td><td>NA</td><td>Intergenic</td><td>NA</td><td>NA</td></tr>
<tr><td></td><td></td><td></td><td></td><td></td><td></td><td></td><td></td><td></td><td></td><td></td><td></td></tr>
<tr><td>T22591</td><td>CGCGGAACTGAGCGCCTGGGAGG</td><td>1000</td><td></td><td></td><td></td><td></td><td></td><td></td><td></td><td></td><td></td></tr>
<tr><td>Oligo fwd</td><td>CACCCGCGGAACTGAGCGCCTGGG</td><td></td><td></td><td></td><td></td><td></td><td></td><td></td><td></td><td></td><td></td></tr>
<tr><td>Oligo rev</td><td>AAACCCCAGGCGCTCAGTTCCGCG</td><td></td><td></td><td></td><td></td><td></td><td></td><td></td><td></td><td></td><td></td></tr>
<tr><td>Chromosome</td><td>start</td><td>end</td><td>strand</td><td>MM</td><td>target_seq</td><td>PAM</td><td>alignment</td><td>distance</td><td>position</td><td>gene name</td><td>gene id</td></tr>
<tr><td>GU305914.1</td><td>151555</td><td>151577</td><td>-</td><td>0</td><td>CGCGGAACTGAGCGCCTGGG</td><td>AGG</td><td>||||||||[||||||||||||]PAM</td><td>NA</td><td>Intergenic</td><td>NA</td><td>NA</td></tr>
<tr><td></td><td></td><td></td><td></td><td></td><td></td><td></td><td></td><td></td><td></td><td></td><td></td></tr>
<tr><td>T22592</td><td>GAGCGCCTGGGAGGAGATCACGG</td><td>1000</td><td></td><td></td><td></td><td></td><td></td><td></td><td></td><td></td><td></td></tr>
<tr><td>Oligo fwd</td><td>CACCGAGCGCCTGGGAGGAGATCA</td><td></td><td></td><td></td><td></td><td></td><td></td><td></td><td></td><td></td><td></td></tr>
<tr><td>Oligo rev</td><td>AAACTGATCTCCTCCCAGGCGCTC</td><td></td><td></td><td></td><td></td><td></td><td></td><td></td><td></td><td></td><td></td></tr>
<tr><td>Chromosome</td><td>start</td><td>end</td><td>strand</td><td>MM</td><td>target_seq</td><td>PAM</td><td>alignment</td><td>distance</td><td>position</td><td>gene name</td><td>gene id</td></tr>
<tr><td>GU305914.1</td><td>151546</td><td>151568</td><td>-</td><td>0</td><td>GAGCGCCTGGGAGGAGATCA</td><td>CGG</td><td>||||||||[||||||||||||]PAM</td><td>NA</td><td>Intergenic</td><td>NA</td><td>NA</td></tr>
<tr><td></td><td></td><td></td><td></td><td></td><td></td><td></td><td></td><td></td><td></td><td></td><td></td></tr>
<tr><td>T22593</td><td>TGCCCGTGATCACGCAGTCGAGG</td><td>1000</td><td></td><td></td><td></td><td></td><td></td><td></td><td></td><td></td><td></td></tr>
<tr><td>Oligo fwd</td><td>CACCTGCCCGTGATCACGCAGTCG</td><td></td><td></td><td></td><td></td><td></td><td></td><td></td><td></td><td></td><td></td></tr>
<tr><td>Oligo rev</td><td>AAACCGACTGCGTGATCACGGGCA</td><td></td><td></td><td></td><td></td><td></td><td></td><td></td><td></td><td></td><td></td></tr>
<tr><td>Chromosome</td><td>start</td><td>end</td><td>strand</td><td>MM</td><td>target_seq</td><td>PAM</td><td>alignment</td><td>distance</td><td>position</td><td>gene name</td><td>gene id</td></tr>
<tr><td>GU305914.1</td><td>151520</td><td>151542</td><td>-</td><td>0</td><td>TGCCCGTGATCACGCAGTCG</td><td>AGG</td><td>||||||||[||||||||||||]PAM</td><td>NA</td><td>Intergenic</td><td>NA</td><td>NA</td></tr>
<tr><td></td><td></td><td></td><td></td><td></td><td></td><td></td><td></td><td></td><td></td><td></td><td></td></tr>
<tr><td>T22594</td><td>GATCACGCAGTCGAGGTTAGCGG</td><td>1000</td><td></td><td></td><td></td><td></td><td></td><td></td><td></td><td></td><td></td></tr>
<tr><td>Oligo fwd</td><td>CACCGATCACGCAGTCGAGGTTAG</td><td></td><td></td><td></td><td></td><td></td><td></td><td></td><td></td><td></td><td></td></tr>
<tr><td>Oligo rev</td><td>AAACCTAACCTCGACTGCGTGATC</td><td></td><td></td><td></td><td></td><td></td><td></td><td></td><td></td><td></td><td></td></tr>
<tr><td>Chromosome</td><td>start</td><td>end</td><td>strand</td><td>MM</td><td>target_seq</td><td>PAM</td><td>alignment</td><td>distance</td><td>position</td><td>gene name</td><td>gene id</td></tr>
<tr><td>GU305914.1</td><td>151513</td><td>151535</td><td>-</td><td>0</td><td>GATCACGCAGTCGAGGTTAG</td><td>CGG</td><td>||||||||[||||||||||||]PAM</td><td>NA</td><td>Intergenic</td><td>NA</td><td>NA</td></tr>
<tr><td></td><td></td><td></td><td></td><td></td><td></td><td></td><td></td><td></td><td></td><td></td><td></td></tr>
<tr><td>T22595</td><td>CGCAGTCGAGGTTAGCGGCCAGG</td><td>1000</td><td></td><td></td><td></td><td></td><td></td><td></td><td></td><td></td><td></td></tr>
<tr><td>Oligo fwd</td><td>CACCCGCAGTCGAGGTTAGCGGCC</td><td></td><td></td><td></td><td></td><td></td><td></td><td></td><td></td><td></td><td></td></tr>
<tr><td>Oligo rev</td><td>AAACGGCCGCTAACCTCGACTGCG</td><td></td><td></td><td></td><td></td><td></td><td></td><td></td><td></td><td></td><td></td></tr>
<tr><td>Chromosome</td><td>start</td><td>end</td><td>strand</td><td>MM</td><td>target_seq</td><td>PAM</td><td>alignment</td><td>distance</td><td>position</td><td>gene name</td><td>gene id</td></tr>
<tr><td>GU305914.1</td><td>151508</td><td>151530</td><td>-</td><td>0</td><td>CGCAGTCGAGGTTAGCGGCC</td><td>AGG</td><td>||||||||[||||||||||||]PAM</td><td>NA</td><td>Intergenic</td><td>NA</td><td>NA</td></tr>
<tr><td></td><td></td><td></td><td></td><td></td><td></td><td></td><td></td><td></td><td></td><td></td><td></td></tr>
<tr><td>T22596</td><td>CGAGGTTAGCGGCCAGGACCTGG</td><td>1000</td><td></td><td></td><td></td><td></td><td></td><td></td><td></td><td></td><td></td></tr>
<tr><td>Oligo fwd</td><td>CACCCGAGGTTAGCGGCCAGGACC</td><td></td><td></td><td></td><td></td><td></td><td></td><td></td><td></td><td></td><td></td></tr>
<tr><td>Oligo rev</td><td>AAACGGTCCTGGCCGCTAACCTCG</td><td></td><td></td><td></td><td></td><td></td><td></td><td></td><td></td><td></td><td></td></tr>
<tr><td>Chromosome</td><td>start</td><td>end</td><td>strand</td><td>MM</td><td>target_seq</td><td>PAM</td><td>alignment</td><td>distance</td><td>position</td><td>gene name</td><td>gene id</td></tr>
<tr><td>GU305914.1</td><td>151502</td><td>151524</td><td>-</td><td>0</td><td>CGAGGTTAGCGGCCAGGACC</td><td>TGG</td><td>||||||||[||||||||||||]PAM</td><td>NA</td><td>Intergenic</td><td>NA</td><td>NA</td></tr>
<tr><td></td><td></td><td></td><td></td><td></td><td></td><td></td><td></td><td></td><td></td><td></td><td></td></tr>
<tr><td>T22597</td><td>GGCCAGGACCTGGACGCTCGAGG</td><td>1000</td><td></td><td></td><td></td><td></td><td></td><td></td><td></td><td></td><td></td></tr>
<tr><td>Oligo fwd</td><td>CACCGGCCAGGACCTGGACGCTCG</td><td></td><td></td><td></td><td></td><td></td><td></td><td></td><td></td><td></td><td></td></tr>
<tr><td>Oligo rev</td><td>AAACCGAGCGTCCAGGTCCTGGCC</td><td></td><td></td><td></td><td></td><td></td><td></td><td></td><td></td><td></td><td></td></tr>
<tr><td>Chromosome</td><td>start</td><td>end</td><td>strand</td><td>MM</td><td>target_seq</td><td>PAM</td><td>alignment</td><td>distance</td><td>position</td><td>gene name</td><td>gene id</td></tr>
<tr><td>GU305914.1</td><td>151492</td><td>151514</td><td>-</td><td>0</td><td>GGCCAGGACCTGGACGCTCG</td><td>AGG</td><td>||||||||[||||||||||||]PAM</td><td>NA</td><td>Intergenic</td><td>NA</td><td>NA</td></tr>
<tr><td></td><td></td><td></td><td></td><td></td><td></td><td></td><td></td><td></td><td></td><td></td><td></td></tr>
<tr><td>T22598</td><td>GCTCGAGGTCTTCCCCGCGCCGG</td><td>1000</td><td></td><td></td><td></td><td></td><td></td><td></td><td></td><td></td><td></td></tr>
<tr><td>Oligo fwd</td><td>CACCGCTCGAGGTCTTCCCCGCGC</td><td></td><td></td><td></td><td></td><td></td><td></td><td></td><td></td><td></td><td></td></tr>
<tr><td>Oligo rev</td><td>AAACGCGCGGGGAAGACCTCGAGC</td><td></td><td></td><td></td><td></td><td></td><td></td><td></td><td></td><td></td><td></td></tr>
<tr><td>Chromosome</td><td>start</td><td>end</td><td>strand</td><td>MM</td><td>target_seq</td><td>PAM</td><td>alignment</td><td>distance</td><td>position</td><td>gene name</td><td>gene id</td></tr>
<tr><td>GU305914.1</td><td>151477</td><td>151499</td><td>-</td><td>0</td><td>GCTCGAGGTCTTCCCCGCGC</td><td>CGG</td><td>||||||||[||||||||||||]PAM</td><td>NA</td><td>Intergenic</td><td>NA</td><td>NA</td></tr>
<tr><td></td><td></td><td></td><td></td><td></td><td></td><td></td><td></td><td></td><td></td><td></td><td></td></tr>
<tr><td>T22599</td><td>CGAGGTCTTCCCCGCGCCGGCGG</td><td>1000</td><td></td><td></td><td></td><td></td><td></td><td></td><td></td><td></td><td></td></tr>
<tr><td>Oligo fwd</td><td>CACCCGAGGTCTTCCCCGCGCCGG</td><td></td><td></td><td></td><td></td><td></td><td></td><td></td><td></td><td></td><td></td></tr>
<tr><td>Oligo rev</td><td>AAACCCGGCGCGGGGAAGACCTCG</td><td></td><td></td><td></td><td></td><td></td><td></td><td></td><td></td><td></td><td></td></tr>
<tr><td>Chromosome</td><td>start</td><td>end</td><td>strand</td><td>MM</td><td>target_seq</td><td>PAM</td><td>alignment</td><td>distance</td><td>position</td><td>gene name</td><td>gene id</td></tr>
<tr><td>GU305914.1</td><td>151474</td><td>151496</td><td>-</td><td>0</td><td>CGAGGTCTTCCCCGCGCCGG</td><td>CGG</td><td>||||||||[||||||||||||]PAM</td><td>NA</td><td>Intergenic</td><td>NA</td><td>NA</td></tr>
<tr><td></td><td></td><td></td><td></td><td></td><td></td><td></td><td></td><td></td><td></td><td></td><td></td></tr>
<tr><td>T22600</td><td>CTTCCCCGCGCCGGCGGTACCGG</td><td>1000</td><td></td><td></td><td></td><td></td><td></td><td></td><td></td><td></td><td></td></tr>
<tr><td>Oligo fwd</td><td>CACCCTTCCCCGCGCCGGCGGTAC</td><td></td><td></td><td></td><td></td><td></td><td></td><td></td><td></td><td></td><td></td></tr>
<tr><td>Oligo rev</td><td>AAACGTACCGCCGGCGCGGGGAAG</td><td></td><td></td><td></td><td></td><td></td><td></td><td></td><td></td><td></td><td></td></tr>
<tr><td>Chromosome</td><td>start</td><td>end</td><td>strand</td><td>MM</td><td>target_seq</td><td>PAM</td><td>alignment</td><td>distance</td><td>position</td><td>gene name</td><td>gene id</td></tr>
<tr><td>GU305914.1</td><td>151468</td><td>151490</td><td>-</td><td>0</td><td>CTTCCCCGCGCCGGCGGTAC</td><td>CGG</td><td>||||||||[||||||||||||]PAM</td><td>NA</td><td>Intergenic</td><td>NA</td><td>NA</td></tr>
<tr><td></td><td></td><td></td><td></td><td></td><td></td><td></td><td></td><td></td><td></td><td></td><td></td></tr>
<tr><td>T22601</td><td>CGCCGGCGGTACCGGTGATCAGG</td><td>1000</td><td></td><td></td><td></td><td></td><td></td><td></td><td></td><td></td><td></td></tr>
<tr><td>Oligo fwd</td><td>CACCCGCCGGCGGTACCGGTGATC</td><td></td><td></td><td></td><td></td><td></td><td></td><td></td><td></td><td></td><td></td></tr>
<tr><td>Oligo rev</td><td>AAACGATCACCGGTACCGCCGGCG</td><td></td><td></td><td></td><td></td><td></td><td></td><td></td><td></td><td></td><td></td></tr>
<tr><td>Chromosome</td><td>start</td><td>end</td><td>strand</td><td>MM</td><td>target_seq</td><td>PAM</td><td>alignment</td><td>distance</td><td>position</td><td>gene name</td><td>gene id</td></tr>
<tr><td>GU305914.1</td><td>151460</td><td>151482</td><td>-</td><td>0</td><td>CGCCGGCGGTACCGGTGATC</td><td>AGG</td><td>||||||||[||||||||||||]PAM</td><td>NA</td><td>Intergenic</td><td>NA</td><td>NA</td></tr>
<tr><td></td><td></td><td></td><td></td><td></td><td></td><td></td><td></td><td></td><td></td><td></td><td></td></tr>
<tr><td>T22602</td><td>CGGCGGTACCGGTGATCAGGAGG</td><td>1000</td><td></td><td></td><td></td><td></td><td></td><td></td><td></td><td></td><td></td></tr>
<tr><td>Oligo fwd</td><td>CACCCGGCGGTACCGGTGATCAGG</td><td></td><td></td><td></td><td></td><td></td><td></td><td></td><td></td><td></td><td></td></tr>
<tr><td>Oligo rev</td><td>AAACCCTGATCACCGGTACCGCCG</td><td></td><td></td><td></td><td></td><td></td><td></td><td></td><td></td><td></td><td></td></tr>
<tr><td>Chromosome</td><td>start</td><td>end</td><td>strand</td><td>MM</td><td>target_seq</td><td>PAM</td><td>alignment</td><td>distance</td><td>position</td><td>gene name</td><td>gene id</td></tr>
<tr><td>GU305914.1</td><td>151457</td><td>151479</td><td>-</td><td>0</td><td>CGGCGGTACCGGTGATCAGG</td><td>AGG</td><td>||||||||[||||||||||||]PAM</td><td>NA</td><td>Intergenic</td><td>NA</td><td>NA</td></tr>
<tr><td></td><td></td><td></td><td></td><td></td><td></td><td></td><td></td><td></td><td></td><td></td><td></td></tr>
<tr><td>T22603</td><td>GGTACCGGTGATCAGGAGGCAGG</td><td>1000</td><td></td><td></td><td></td><td></td><td></td><td></td><td></td><td></td><td></td></tr>
<tr><td>Oligo fwd</td><td>CACCGGTACCGGTGATCAGGAGGC</td><td></td><td></td><td></td><td></td><td></td><td></td><td></td><td></td><td></td><td></td></tr>
<tr><td>Oligo rev</td><td>AAACGCCTCCTGATCACCGGTACC</td><td></td><td></td><td></td><td></td><td></td><td></td><td></td><td></td><td></td><td></td></tr>
<tr><td>Chromosome</td><td>start</td><td>end</td><td>strand</td><td>MM</td><td>target_seq</td><td>PAM</td><td>alignment</td><td>distance</td><td>position</td><td>gene name</td><td>gene id</td></tr>
<tr><td>GU305914.1</td><td>151453</td><td>151475</td><td>-</td><td>0</td><td>GGTACCGGTGATCAGGAGGC</td><td>AGG</td><td>||||||||[||||||||||||]PAM</td><td>NA</td><td>Intergenic</td><td>NA</td><td>NA</td></tr>
<tr><td></td><td></td><td></td><td></td><td></td><td></td><td></td><td></td><td></td><td></td><td></td><td></td></tr>
<tr><td>T22604</td><td>GGTGATCAGGAGGCAGGAGAAGG</td><td>1000</td><td></td><td></td><td></td><td></td><td></td><td></td><td></td><td></td><td></td></tr>
<tr><td>Oligo fwd</td><td>CACCGGTGATCAGGAGGCAGGAGA</td><td></td><td></td><td></td><td></td><td></td><td></td><td></td><td></td><td></td><td></td></tr>
<tr><td>Oligo rev</td><td>AAACTCTCCTGCCTCCTGATCACC</td><td></td><td></td><td></td><td></td><td></td><td></td><td></td><td></td><td></td><td></td></tr>
<tr><td>Chromosome</td><td>start</td><td>end</td><td>strand</td><td>MM</td><td>target_seq</td><td>PAM</td><td>alignment</td><td>distance</td><td>position</td><td>gene name</td><td>gene id</td></tr>
<tr><td>GU305914.1</td><td>151447</td><td>151469</td><td>-</td><td>0</td><td>GGTGATCAGGAGGCAGGAGA</td><td>AGG</td><td>||||||||[||||||||||||]PAM</td><td>NA</td><td>Intergenic</td><td>NA</td><td>NA</td></tr>
<tr><td></td><td></td><td></td><td></td><td></td><td></td><td></td><td></td><td></td><td></td><td></td><td></td></tr>
<tr><td>T22605</td><td>GTGATCAGGAGGCAGGAGAAGGG</td><td>1000</td><td></td><td></td><td></td><td></td><td></td><td></td><td></td><td></td><td></td></tr>
<tr><td>Oligo fwd</td><td>CACCGTGATCAGGAGGCAGGAGAA</td><td></td><td></td><td></td><td></td><td></td><td></td><td></td><td></td><td></td><td></td></tr>
</table>

Oligo rev AAACTTCTCCTGCCTCCTGATCAC
Chromosome start end strand MM target_seq PAM alignment distance position gene name gene id
GU305914.1 151446 151468 - 0 GTGATCAGGAGGCAGGAGAA GGG ||||||||[||||||||||||]PAM NA Intergenic NA NA

T22606 TCAGGAGGCAGGAGAAGGGTAGG 1000
Oligo fwd CACCTCAGGAGGCAGGAGAAGGGT
Oligo rev AAACACCCTTCTCCTGCCTCCTGA
Chromosome start end strand MM target_seq PAM alignment distance position gene name gene id
GU305914.1 151442 151464 - 0 TCAGGAGGCAGGAGAAGGGT AGG ||||||||[||||||||||||]PAM NA Intergenic NA NA

T22607 GAGGCAGGAGAAGGGTAGGAAGG 1000
Oligo fwd CACCGAGGCAGGAGAAGGGTAGGA
Oligo rev AAACTCCTACCCTTCTCCTGCCTC
Chromosome start end strand MM target_seq PAM alignment distance position gene name gene id
GU305914.1 151438 151460 - 0 GAGGCAGGAGAAGGGTAGGA AGG ||||||||[||||||||||||]PAM NA Intergenic NA NA
GU305914.1 82326 82348 - 4 GATGCTGGAGGAGGGGAGGA AGG ||-||-||[||-||||-||||]PAM NA Intergenic NA NA

T22608 AGGCAGGAGAAGGGTAGGAAGGG 1000
Oligo fwd CACCAGGCAGGAGAAGGGTAGGAA
Oligo rev AAACTTCCTACCCTTCTCCTGCCT
Chromosome start end strand MM target_seq PAM alignment distance position gene name gene id
GU305914.1 151437 151459 - 0 AGGCAGGAGAAGGGTAGGAA GGG ||||||||[||||||||||||]PAM NA Intergenic NA NA

T22609 CAGGAGAAGGGTAGGAAGGGCGG 1000
Oligo fwd CACCCAGGAGAAGGGTAGGAAGGG
Oligo rev AAACCCCTTCCTACCCTTCTCCTG
Chromosome start end strand MM target_seq PAM alignment distance position gene name gene id
GU305914.1 151434 151456 - 0 CAGGAGAAGGGTAGGAAGGG CGG ||||||||[||||||||||||]PAM NA Intergenic NA NA

T22610 AGAAGGGTAGGAAGGGCGGCCGG 1000
Oligo fwd CACCAGAAGGGTAGGAAGGGCGGC
Oligo rev AAACGCCGCCCTTCCTACCCTTCT
Chromosome start end strand MM target_seq PAM alignment distance position gene name gene id
GU305914.1 151430 151452 - 0 AGAAGGGTAGGAAGGGCGGC CGG ||||||||[||||||||||||]PAM NA Intergenic NA NA

T22611 GCGGCCGGCGCGCCAGCTCGCGG 1000
Oligo fwd CACCGCGGCCGGCGCGCCAGCTCG
Oligo rev AAACCGAGCTGGCGCGCCGGCCGC
Chromosome start end strand MM target_seq PAM alignment distance position gene name gene id
GU305914.1 151415 151437 - 0 GCGGCCGGCGCGCCAGCTCG CGG ||||||||[||||||||||||]PAM NA Intergenic NA NA

T22612 GCGGTCGCCGACGATCATCGCGG 1000
Oligo fwd CACCGCGGTCGCCGACGATCATCG
Oligo rev AAACCGATGATCGTCGGCGACCGC
Chromosome start end strand MM target_seq PAM alignment distance position gene name gene id
GU305914.1 151396 151418 - 0 GCGGTCGCCGACGATCATCG CGG ||||||||[||||||||||||]PAM NA Intergenic NA NA

T22613 GCTGCCGTCGTCGCCGAGCTCGG 1000
Oligo fwd CACCGCTGCCGTCGTCGCCGAGCT
Oligo rev AAACAGCTCGGCGACGACGGCAGC
Chromosome start end strand MM target_seq PAM alignment distance position gene name gene id
GU305914.1 151354 151376 - 0 GCTGCCGTCGTCGCCGAGCT CGG ||||||||[||||||||||||]PAM NA Intergenic NA NA

T22614 CTGCCGTCGTCGCCGAGCTCGGG 1000
Oligo fwd CACCCTGCCGTCGTCGCCGAGCTC
Oligo rev AAACGAGCTCGGCGACGACGGCAG
Chromosome start end strand MM target_seq PAM alignment distance position gene name gene id
GU305914.1 151353 151375 - 0 CTGCCGTCGTCGCCGAGCTC GGG ||||||||[||||||||||||]PAM NA Intergenic NA NA

T22615 CGCCGAGCTCGGGCGCCACCAGG 1000
Oligo fwd CACCCGCCGAGCTCGGGCGCCACC
Oligo rev AAACGGTGGCGCCCGAGCTCGGCG
Chromosome start end strand MM target_seq PAM alignment distance position gene name gene id
GU305914.1 151343 151365 - 0 CGCCGAGCTCGGGCGCCACC AGG ||||||||[||||||||||||]PAM NA Intergenic NA NA

T22616 GGGCGCCACCAGGCCCGCGCAGG 1000
Oligo fwd CACCGGGCGCCACCAGGCCCGCGC
Oligo rev AAACGCGCGGGCCTGGTGGCGCCC
Chromosome start end strand MM target_seq PAM alignment distance position gene name gene id
GU305914.1 151333 151355 - 0 GGGCGCCACCAGGCCCGCGC AGG ||||||||[||||||||||||]PAM NA Intergenic NA NA

T22617 GGCGCCACCAGGCCCGCGCAGGG 1000
Oligo fwd CACCGGCGCCACCAGGCCCGCGCA
Oligo rev AAACTGCGCGGGCCTGGTGGCGCC
Chromosome start end strand MM target_seq PAM alignment distance position gene name gene id
GU305914.1 151332 151354 - 0 GGCGCCACCAGGCCCGCGCA GGG ||||||||[||||||||||||]PAM NA Intergenic NA NA

T22618 CCGCGCAGGGATCCAACATGTGG 1000
Oligo fwd CACCCCGCGCAGGGATCCAACATG
Oligo rev AAACCATGTTGGATCCCTGCGCGG
Chromosome start end strand MM target_seq PAM alignment distance position gene name gene id
GU305914.1 151319 151341 - 0 CCGCGCAGGGATCCAACATG TGG ||||||||[||||||||||||]PAM NA Intergenic NA NA

T22619 CGCAGGGATCCAACATGTGGCGG 1000
Oligo fwd CACCCGCAGGGATCCAACATGTGG
Oligo rev AAACCCACATGTTGGATCCCTGCG
Chromosome start end strand MM target_seq PAM alignment distance position gene name gene id
GU305914.1 151316 151338 - 0 CGCAGGGATCCAACATGTGG CGG ||||||||[||||||||||||]PAM NA Intergenic NA NA

| | | | | | | | | | | | |
|---|---|---|---|---|---|---|---|---|---|---|---|
| T22620 | AGTCCTTGTACAGAGATTCGCGG | 1000 | | | | | | | | | |
| Oligo fwd | CACCAGTCCTTGTACAGAGATTCG | | | | | | | | | | |
| Oligo rev | AAACCGAATCTCTGTACAAGGACT | | | | | | | | | | |
| Chromosome | start | end | strand | MM | target_seq | PAM | alignment | distance | position | gene name | gene id |
| GU305914.1 | 151289 | 151311 | - | 0 | AGTCCTTGTACAGAGATTCG | CGG | \|\|\|\|\|\|\|\|[\|\|\|\|\|\|\|\|\|\|\|\|]PAM | NA | Intergenic | NA | NA |
| T22621 | GTCCTTGTACAGAGATTCGCGGG | 1000 | | | | | | | | | |
| Oligo fwd | CACCGTCCTTGTACAGAGATTCGC | | | | | | | | | | |
| Oligo rev | AAACGCGAATCTCTGTACAAGGAC | | | | | | | | | | |
| Chromosome | start | end | strand | MM | target_seq | PAM | alignment | distance | position | gene name | gene id |
| GU305914.1 | 151288 | 151310 | - | 0 | GTCCTTGTACAGAGATTCGC | GGG | \|\|\|\|\|\|\|\|[\|\|\|\|\|\|\|\|\|\|\|\|]PAM | NA | Intergenic | NA | NA |
| T22622 | TTCGCGGGAGAACCGCTTCAAGG | 1000 | | | | | | | | | |
| Oligo fwd | CACCTTCGCGGGAGAACCGCTTCA | | | | | | | | | | |
| Oligo rev | AAACTGAAGCGGTTCTCCCGCGAA | | | | | | | | | | |
| Chromosome | start | end | strand | MM | target_seq | PAM | alignment | distance | position | gene name | gene id |
| GU305914.1 | 151273 | 151295 | - | 0 | TTCGCGGGAGAACCGCTTCA | AGG | \|\|\|\|\|\|\|\|[\|\|\|\|\|\|\|\|\|\|\|\|]PAM | NA | Intergenic | NA | NA |
| T22623 | TCAAGGCCAGCGACTTGACGCGG | 1000 | | | | | | | | | |
| Oligo fwd | CACCTCAAGGCCAGCGACTTGACG | | | | | | | | | | |
| Oligo rev | AAACCGTCAAGTCGCTGGCCTTGA | | | | | | | | | | |
| Chromosome | start | end | strand | MM | target_seq | PAM | alignment | distance | position | gene name | gene id |
| GU305914.1 | 151256 | 151278 | - | 0 | TCAAGGCCAGCGACTTGACG | CGG | \|\|\|\|\|\|\|\|[\|\|\|\|\|\|\|\|\|\|\|\|]PAM | NA | Intergenic | NA | NA |
| T22624 | AACGATCCGCTCGATCTTCGAGG | 1000 | | | | | | | | | |
| Oligo fwd | CACCAACGATCCGCTCGATCTTCG | | | | | | | | | | |
| Oligo rev | AAACCGAAGATCGAGCGGATCGTT | | | | | | | | | | |
| Chromosome | start | end | strand | MM | target_seq | PAM | alignment | distance | position | gene name | gene id |
| GU305914.1 | 151231 | 151253 | - | 0 | AACGATCCGCTCGATCTTCG | AGG | \|\|\|\|\|\|\|\|[\|\|\|\|\|\|\|\|\|\|\|\|]PAM | NA | Intergenic | NA | NA |
| T22625 | GATCCGCTCGATCTTCGAGGCGG | 1000 | | | | | | | | | |
| Oligo fwd | CACCGATCCGCTCGATCTTCGAGG | | | | | | | | | | |
| Oligo rev | AAACCCTCGAAGATCGAGCGGATC | | | | | | | | | | |
| Chromosome | start | end | strand | MM | target_seq | PAM | alignment | distance | position | gene name | gene id |
| GU305914.1 | 151228 | 151250 | - | 0 | GATCCGCTCGATCTTCGAGG | CGG | \|\|\|\|\|\|\|\|[\|\|\|\|\|\|\|\|\|\|\|\|]PAM | NA | Intergenic | NA | NA |
| T22626 | TCGAGGCGGACGACATATTCAGG | 1000 | | | | | | | | | |
| Oligo fwd | CACCTCGAGGCGGACGACATATTC | | | | | | | | | | |
| Oligo rev | AAACGAATATGTCGTCCGCCTCGA | | | | | | | | | | |
| Chromosome | start | end | strand | MM | target_seq | PAM | alignment | distance | position | gene name | gene id |
| GU305914.1 | 151214 | 151236 | - | 0 | TCGAGGCGGACGACATATTC | AGG | \|\|\|\|\|\|\|\|[\|\|\|\|\|\|\|\|\|\|\|\|]PAM | NA | Intergenic | NA | NA |
| T22627 | GTCGTACCTCGACAACCCGCCGG | 1000 | | | | | | | | | |
| Oligo fwd | CACCGTCGTACCTCGACAACCCGC | | | | | | | | | | |
| Oligo rev | AAACGCGGGTTGTCGAGGTACGAC | | | | | | | | | | |
| Chromosome | start | end | strand | MM | target_seq | PAM | alignment | distance | position | gene name | gene id |
| GU305914.1 | 151180 | 151202 | - | 0 | GTCGTACCTCGACAACCCGC | CGG | \|\|\|\|\|\|\|\|[\|\|\|\|\|\|\|\|\|\|\|\|]PAM | NA | Intergenic | NA | NA |
| T22628 | CAACCCGCCGGAGCCGCCGTCGG | 1000 | | | | | | | | | |
| Oligo fwd | CACCCAACCCGCCGGAGCCGCCGT | | | | | | | | | | |
| Oligo rev | AAACACGGCGGCTCCGGCGGGTTG | | | | | | | | | | |
| Chromosome | start | end | strand | MM | target_seq | PAM | alignment | distance | position | gene name | gene id |
| GU305914.1 | 151168 | 151190 | - | 0 | CAACCCGCCGGAGCCGCCGT | CGG | \|\|\|\|\|\|\|\|[\|\|\|\|\|\|\|\|\|\|\|\|]PAM | NA | Intergenic | NA | NA |
| T22629 | GTCGGACCCCTTGTTCCCGACGG | 1000 | | | | | | | | | |
| Oligo fwd | CACCGTCGGACCCCTTGTTCCCGA | | | | | | | | | | |
| Oligo rev | AAACTCGGGAACAAGGGGTCCGAC | | | | | | | | | | |
| Chromosome | start | end | strand | MM | target_seq | PAM | alignment | distance | position | gene name | gene id |
| GU305914.1 | 151150 | 151172 | - | 0 | GTCGGACCCCTTGTTCCCGA | CGG | \|\|\|\|\|\|\|\|[\|\|\|\|\|\|\|\|\|\|\|\|]PAM | NA | Intergenic | NA | NA |
| T22630 | GTTCCCGACGGACTCGTCGCTGG | 1000 | | | | | | | | | |
| Oligo fwd | CACCGTTCCCGACGGACTCGTCGC | | | | | | | | | | |
| Oligo rev | AAACGCGACGAGTCCGTCGGGAAC | | | | | | | | | | |
| Chromosome | start | end | strand | MM | target_seq | PAM | alignment | distance | position | gene name | gene id |
| GU305914.1 | 151138 | 151160 | - | 0 | GTTCCCGACGGACTCGTCGC | TGG | \|\|\|\|\|\|\|\|[\|\|\|\|\|\|\|\|\|\|\|\|]PAM | NA | Intergenic | NA | NA |
| T22631 | GACCTCTTCTCCATGATCAACGG | 1000 | | | | | | | | | |
| Oligo fwd | CACCGACCTCTTCTCCATGATCAA | | | | | | | | | | |
| Oligo rev | AAACTTGATCATGGAGAAGAGGTC | | | | | | | | | | |
| Chromosome | start | end | strand | MM | target_seq | PAM | alignment | distance | position | gene name | gene id |
| GU305914.1 | 151116 | 151138 | - | 0 | GACCTCTTCTCCATGATCAA | CGG | \|\|\|\|\|\|\|\|[\|\|\|\|\|\|\|\|\|\|\|\|]PAM | NA | Intergenic | NA | NA |
| T22632 | CTTCTCCATGATCAACGGCACGG | 1000 | | | | | | | | | |
| Oligo fwd | CACCCTTCTCCATGATCAACGGCA | | | | | | | | | | |
| Oligo rev | AAACTGCCGTTGATCATGGAGAAG | | | | | | | | | | |
| Chromosome | start | end | strand | MM | target_seq | PAM | alignment | distance | position | gene name | gene id |
| GU305914.1 | 151111 | 151133 | - | 0 | CTTCTCCATGATCAACGGCA | CGG | \|\|\|\|\|\|\|\|[\|\|\|\|\|\|\|\|\|\|\|\|]PAM | NA | Intergenic | NA | NA |
| T22633 | CTCCATGATCAACGGCACGGAGG | 1000 | | | | | | | | | |
| Oligo fwd | CACCCTCCATGATCAACGGCACGG | | | | | | | | | | |
| Oligo rev | AAACCCGTGCCGTTGATCATGGAG | | | | | | | | | | |
| Chromosome | start | end | strand | MM | target_seq | PAM | alignment | distance | position | gene name | gene id |
| GU305914.1 | 151108 | 151130 | - | 0 | CTCCATGATCAACGGCACGG | AGG | \|\|\|\|\|\|\|\|[\|\|\|\|\|\|\|\|\|\|\|\|]PAM | NA | Intergenic | NA | NA |
| T22634 | TCCATGATCAACGGCACGGAGGG | 1000 | | | | | | | | | |
| Oligo fwd | CACCTCCATGATCAACGGCACGGA | | | | | | | | | | |
| Oligo rev | AAACTCCGTGCCGTTGATCATGGA | | | | | | | | | | |
| Chromosome | start | end | strand | MM | target_seq | PAM | alignment | distance | position | gene name | gene id |
| GU305914.1 | 151107 | 151129 | - | 0 | TCCATGATCAACGGCACGGA | GGG | \|\|\|\|\|\|\|\|[\|\|\|\|\|\|\|\|\|\|\|\|]PAM | NA | Intergenic | NA | NA |

<table>
<tr><td>T22635</td><td>GGCACGGAGGGCGCGTGTATCGG</td><td>1000</td><td></td><td></td><td></td><td></td><td></td><td></td><td></td><td></td><td></td></tr>
<tr><td>Oligo fwd</td><td>CACCGGCACGGAGGGCGCGTGTAT</td><td></td><td></td><td></td><td></td><td></td><td></td><td></td><td></td><td></td><td></td></tr>
<tr><td>Oligo rev</td><td>AAACATACACGCGCCCTCCGTGCC</td><td></td><td></td><td></td><td></td><td></td><td></td><td></td><td></td><td></td><td></td></tr>
<tr><td>Chromosome</td><td>start</td><td>end</td><td>strand</td><td>MM</td><td>target_seq</td><td>PAM</td><td>alignment</td><td>distance</td><td>position</td><td>gene name</td><td>gene id</td></tr>
<tr><td>GU305914.1</td><td>151095</td><td>151117</td><td>-</td><td>0</td><td>GGCACGGAGGGCGCGTGTAT</td><td>CGG</td><td>||||||||[||||||||||||]PAM</td><td>NA</td><td>Intergenic</td><td>NA</td><td>NA</td></tr>
<tr><td>T22636</td><td>GCACGGAGGGCGCGTGTATCGGG</td><td>1000</td><td></td><td></td><td></td><td></td><td></td><td></td><td></td><td></td><td></td></tr>
<tr><td>Oligo fwd</td><td>CACCGCACGGAGGGCGCGTGTATC</td><td></td><td></td><td></td><td></td><td></td><td></td><td></td><td></td><td></td><td></td></tr>
<tr><td>Oligo rev</td><td>AAACGATACACGCGCCCTCCGTGC</td><td></td><td></td><td></td><td></td><td></td><td></td><td></td><td></td><td></td><td></td></tr>
<tr><td>Chromosome</td><td>start</td><td>end</td><td>strand</td><td>MM</td><td>target_seq</td><td>PAM</td><td>alignment</td><td>distance</td><td>position</td><td>gene name</td><td>gene id</td></tr>
<tr><td>GU305914.1</td><td>151094</td><td>151116</td><td>-</td><td>0</td><td>GCACGGAGGGCGCGTGTATC</td><td>GGG</td><td>||||||||[||||||||||||]PAM</td><td>NA</td><td>Intergenic</td><td>NA</td><td>NA</td></tr>
<tr><td>T22637</td><td>AGATCTTCTACGCGATGATGCGG</td><td>1000</td><td></td><td></td><td></td><td></td><td></td><td></td><td></td><td></td><td></td></tr>
<tr><td>Oligo fwd</td><td>CACCAGATCTTCTACGCGATGATG</td><td></td><td></td><td></td><td></td><td></td><td></td><td></td><td></td><td></td><td></td></tr>
<tr><td>Oligo rev</td><td>AAACCATCATCGCGTAGAAGATCT</td><td></td><td></td><td></td><td></td><td></td><td></td><td></td><td></td><td></td><td></td></tr>
<tr><td>Chromosome</td><td>start</td><td>end</td><td>strand</td><td>MM</td><td>target_seq</td><td>PAM</td><td>alignment</td><td>distance</td><td>position</td><td>gene name</td><td>gene id</td></tr>
<tr><td>GU305914.1</td><td>151025</td><td>151047</td><td>-</td><td>0</td><td>AGATCTTCTACGCGATGATG</td><td>CGG</td><td>||||||||[||||||||||||]PAM</td><td>NA</td><td>Intergenic</td><td>NA</td><td>NA</td></tr>
<tr><td>T22638</td><td>ATGATGCGGTTCCTGCTGAACGG</td><td>1000</td><td></td><td></td><td></td><td></td><td></td><td></td><td></td><td></td><td></td></tr>
<tr><td>Oligo fwd</td><td>CACCATGATGCGGTTCCTGCTGAA</td><td></td><td></td><td></td><td></td><td></td><td></td><td></td><td></td><td></td><td></td></tr>
<tr><td>Oligo rev</td><td>AAACTTCAGCAGGAACCGCATCAT</td><td></td><td></td><td></td><td></td><td></td><td></td><td></td><td></td><td></td><td></td></tr>
<tr><td>Chromosome</td><td>start</td><td>end</td><td>strand</td><td>MM</td><td>target_seq</td><td>PAM</td><td>alignment</td><td>distance</td><td>position</td><td>gene name</td><td>gene id</td></tr>
<tr><td>GU305914.1</td><td>151011</td><td>151033</td><td>-</td><td>0</td><td>ATGATGCGGTTCCTGCTGAA</td><td>CGG</td><td>||||||||[||||||||||||]PAM</td><td>NA</td><td>Intergenic</td><td>NA</td><td>NA</td></tr>
<tr><td>T22639</td><td>CCTGCTGAACGGCATCAGCGTGG</td><td>1000</td><td></td><td></td><td></td><td></td><td></td><td></td><td></td><td></td><td></td></tr>
<tr><td>Oligo fwd</td><td>CACCCCTGCTGAACGGCATCAGCG</td><td></td><td></td><td></td><td></td><td></td><td></td><td></td><td></td><td></td><td></td></tr>
<tr><td>Oligo rev</td><td>AAACCGCTGATGCCGTTCAGCAGG</td><td></td><td></td><td></td><td></td><td></td><td></td><td></td><td></td><td></td><td></td></tr>
<tr><td>Chromosome</td><td>start</td><td>end</td><td>strand</td><td>MM</td><td>target_seq</td><td>PAM</td><td>alignment</td><td>distance</td><td>position</td><td>gene name</td><td>gene id</td></tr>
<tr><td>GU305914.1</td><td>151000</td><td>151022</td><td>-</td><td>0</td><td>CCTGCTGAACGGCATCAGCG</td><td>TGG</td><td>||||||||[||||||||||||]PAM</td><td>NA</td><td>Intergenic</td><td>NA</td><td>NA</td></tr>
<tr><td>T22640</td><td>CTGCTGAACGGCATCAGCGTGGG</td><td>1000</td><td></td><td></td><td></td><td></td><td></td><td></td><td></td><td></td><td></td></tr>
<tr><td>Oligo fwd</td><td>CACCCTGCTGAACGGCATCAGCGT</td><td></td><td></td><td></td><td></td><td></td><td></td><td></td><td></td><td></td><td></td></tr>
<tr><td>Oligo rev</td><td>AAACACGCTGATGCCGTTCAGCAG</td><td></td><td></td><td></td><td></td><td></td><td></td><td></td><td></td><td></td><td></td></tr>
<tr><td>Chromosome</td><td>start</td><td>end</td><td>strand</td><td>MM</td><td>target_seq</td><td>PAM</td><td>alignment</td><td>distance</td><td>position</td><td>gene name</td><td>gene id</td></tr>
<tr><td>GU305914.1</td><td>150999</td><td>151021</td><td>-</td><td>0</td><td>CTGCTGAACGGCATCAGCGT</td><td>GGG</td><td>||||||||[||||||||||||]PAM</td><td>NA</td><td>Intergenic</td><td>NA</td><td>NA</td></tr>
<tr><td>T22641</td><td>CAGCGTGGGCGAGCTGTCGACGG</td><td>1000</td><td></td><td></td><td></td><td></td><td></td><td></td><td></td><td></td><td></td></tr>
<tr><td>Oligo fwd</td><td>CACCCAGCGTGGGCGAGCTGTCGA</td><td></td><td></td><td></td><td></td><td></td><td></td><td></td><td></td><td></td><td></td></tr>
<tr><td>Oligo rev</td><td>AAACTCGACAGCTCGCCCACGCTG</td><td></td><td></td><td></td><td></td><td></td><td></td><td></td><td></td><td></td><td></td></tr>
<tr><td>Chromosome</td><td>start</td><td>end</td><td>strand</td><td>MM</td><td>target_seq</td><td>PAM</td><td>alignment</td><td>distance</td><td>position</td><td>gene name</td><td>gene id</td></tr>
<tr><td>GU305914.1</td><td>150985</td><td>151007</td><td>-</td><td>0</td><td>CAGCGTGGGCGAGCTGTCGA</td><td>CGG</td><td>||||||||[||||||||||||]PAM</td><td>NA</td><td>Intergenic</td><td>NA</td><td>NA</td></tr>
<tr><td>T22642</td><td>TGGGCGAGCTGTCGACGGCCTGG</td><td>1000</td><td></td><td></td><td></td><td></td><td></td><td></td><td></td><td></td><td></td></tr>
<tr><td>Oligo fwd</td><td>CACCTGGGCGAGCTGTCGACGGCC</td><td></td><td></td><td></td><td></td><td></td><td></td><td></td><td></td><td></td><td></td></tr>
<tr><td>Oligo rev</td><td>AAACGGCCGTCGACAGCTCGCCCA</td><td></td><td></td><td></td><td></td><td></td><td></td><td></td><td></td><td></td><td></td></tr>
<tr><td>Chromosome</td><td>start</td><td>end</td><td>strand</td><td>MM</td><td>target_seq</td><td>PAM</td><td>alignment</td><td>distance</td><td>position</td><td>gene name</td><td>gene id</td></tr>
<tr><td>GU305914.1</td><td>150980</td><td>151002</td><td>-</td><td>0</td><td>TGGGCGAGCTGTCGACGGCC</td><td>TGG</td><td>||||||||[||||||||||||]PAM</td><td>NA</td><td>Intergenic</td><td>NA</td><td>NA</td></tr>
<tr><td>T22643</td><td>GGGCGAGCTGTCGACGGCCTGGG</td><td>1000</td><td></td><td></td><td></td><td></td><td></td><td></td><td></td><td></td><td></td></tr>
<tr><td>Oligo fwd</td><td>CACCGGGCGAGCTGTCGACGGCCT</td><td></td><td></td><td></td><td></td><td></td><td></td><td></td><td></td><td></td><td></td></tr>
<tr><td>Oligo rev</td><td>AAACAGGCCGTCGACAGCTCGCCC</td><td></td><td></td><td></td><td></td><td></td><td></td><td></td><td></td><td></td><td></td></tr>
<tr><td>Chromosome</td><td>start</td><td>end</td><td>strand</td><td>MM</td><td>target_seq</td><td>PAM</td><td>alignment</td><td>distance</td><td>position</td><td>gene name</td><td>gene id</td></tr>
<tr><td>GU305914.1</td><td>150979</td><td>151001</td><td>-</td><td>0</td><td>GGGCGAGCTGTCGACGGCCT</td><td>GGG</td><td>||||||||[||||||||||||]PAM</td><td>NA</td><td>Intergenic</td><td>NA</td><td>NA</td></tr>
<tr><td>T22644</td><td>CGACGGCCTGGGCCTCCCACCGG</td><td>1000</td><td></td><td></td><td></td><td></td><td></td><td></td><td></td><td></td><td></td></tr>
<tr><td>Oligo fwd</td><td>CACCCGACGGCCTGGGCCTCCCAC</td><td></td><td></td><td></td><td></td><td></td><td></td><td></td><td></td><td></td><td></td></tr>
<tr><td>Oligo rev</td><td>AAACGTGGGAGGCCCAGGCCGTCG</td><td></td><td></td><td></td><td></td><td></td><td></td><td></td><td></td><td></td><td></td></tr>
<tr><td>Chromosome</td><td>start</td><td>end</td><td>strand</td><td>MM</td><td>target_seq</td><td>PAM</td><td>alignment</td><td>distance</td><td>position</td><td>gene name</td><td>gene id</td></tr>
<tr><td>GU305914.1</td><td>150968</td><td>150990</td><td>-</td><td>0</td><td>CGACGGCCTGGGCCTCCCAC</td><td>CGG</td><td>||||||||[||||||||||||]PAM</td><td>NA</td><td>Intergenic</td><td>NA</td><td>NA</td></tr>
<tr><td>T22645</td><td>CGGCCTGGGCCTCCCACCGGCGG</td><td>1000</td><td></td><td></td><td></td><td></td><td></td><td></td><td></td><td></td><td></td></tr>
<tr><td>Oligo fwd</td><td>CACCCGGCCTGGGCCTCCCACCGG</td><td></td><td></td><td></td><td></td><td></td><td></td><td></td><td></td><td></td><td></td></tr>
<tr><td>Oligo rev</td><td>AAACCCGGTGGGAGGCCCAGGCCG</td><td></td><td></td><td></td><td></td><td></td><td></td><td></td><td></td><td></td><td></td></tr>
<tr><td>Chromosome</td><td>start</td><td>end</td><td>strand</td><td>MM</td><td>target_seq</td><td>PAM</td><td>alignment</td><td>distance</td><td>position</td><td>gene name</td><td>gene id</td></tr>
<tr><td>GU305914.1</td><td>150965</td><td>150987</td><td>-</td><td>0</td><td>CGGCCTGGGCCTCCCACCGG</td><td>CGG</td><td>||||||||[||||||||||||]PAM</td><td>NA</td><td>Intergenic</td><td>NA</td><td>NA</td></tr>
<tr><td>T22646</td><td>CCTGGGCCTCCCACCGGCGGCGG</td><td>1000</td><td></td><td></td><td></td><td></td><td></td><td></td><td></td><td></td><td></td></tr>
<tr><td>Oligo fwd</td><td>CACCCCTGGGCCTCCCACCGGCGG</td><td></td><td></td><td></td><td></td><td></td><td></td><td></td><td></td><td></td><td></td></tr>
<tr><td>Oligo rev</td><td>AAACCCGCCGGTGGGAGGCCCAGG</td><td></td><td></td><td></td><td></td><td></td><td></td><td></td><td></td><td></td><td></td></tr>
<tr><td>Chromosome</td><td>start</td><td>end</td><td>strand</td><td>MM</td><td>target_seq</td><td>PAM</td><td>alignment</td><td>distance</td><td>position</td><td>gene name</td><td>gene id</td></tr>
<tr><td>GU305914.1</td><td>150962</td><td>150984</td><td>-</td><td>0</td><td>CCTGGGCCTCCCACCGGCGG</td><td>CGG</td><td>||||||||[||||||||||||]PAM</td><td>NA</td><td>Intergenic</td><td>NA</td><td>NA</td></tr>
<tr><td>T22647</td><td>CCGGCGGCGGTTCGTGCGCGCGG</td><td>1000</td><td></td><td></td><td></td><td></td><td></td><td></td><td></td><td></td><td></td></tr>
<tr><td>Oligo fwd</td><td>CACCCCGGCGGCGGTTCGTGCGCG</td><td></td><td></td><td></td><td></td><td></td><td></td><td></td><td></td><td></td><td></td></tr>
<tr><td>Oligo rev</td><td>AAACCGCGCACGAACCGCCGCCGG</td><td></td><td></td><td></td><td></td><td></td><td></td><td></td><td></td><td></td><td></td></tr>
<tr><td>Chromosome</td><td>start</td><td>end</td><td>strand</td><td>MM</td><td>target_seq</td><td>PAM</td><td>alignment</td><td>distance</td><td>position</td><td>gene name</td><td>gene id</td></tr>
<tr><td>GU305914.1</td><td>150949</td><td>150971</td><td>-</td><td>0</td><td>CCGGCGGCGGTTCGTGCGCG</td><td>CGG</td><td>||||||||[||||||||||||]PAM</td><td>NA</td><td>Intergenic</td><td>NA</td><td>NA</td></tr>
<tr><td>T22648</td><td>GCGGCGGTTCGTGCGCGCGGAGG</td><td>1000</td><td></td><td></td><td></td><td></td><td></td><td></td><td></td><td></td><td></td></tr>
<tr><td>Oligo fwd</td><td>CACCGCGGCGGTTCGTGCGCGCGG</td><td></td><td></td><td></td><td></td><td></td><td></td><td></td><td></td><td></td><td></td></tr>
<tr><td>Oligo rev</td><td>AAACCCGCGCGCACGAACCGCCGC</td><td></td><td></td><td></td><td></td><td></td><td></td><td></td><td></td><td></td><td></td></tr>
<tr><td>Chromosome</td><td>start</td><td>end</td><td>strand</td><td>MM</td><td>target_seq</td><td>PAM</td><td>alignment</td><td>distance</td><td>position</td><td>gene name</td><td>gene id</td></tr>
<tr><td>GU305914.1</td><td>150946</td><td>150968</td><td>-</td><td>0</td><td>GCGGCGGTTCGTGCGCGCGG</td><td>AGG</td><td>||||||||[||||||||||||]PAM</td><td>NA</td><td>Intergenic</td><td>NA</td><td>NA</td></tr>
<tr><td>T22649</td><td>GTTCGTGCGCGCGGAGGACGAGG</td><td>1000</td><td></td><td></td><td></td><td></td><td></td><td></td><td></td><td></td><td></td></tr>
<tr><td>Oligo fwd</td><td>CACCGTTCGTGCGCGCGGAGGACG</td><td></td><td></td><td></td><td></td><td></td><td></td><td></td><td></td><td></td><td></td></tr>
<tr><td>Oligo rev</td><td>AAACCGTCCTCCGCGCGCACGAAC</td><td></td><td></td><td></td><td></td><td></td><td></td><td></td><td></td><td></td><td></td></tr>
<tr><td>Chromosome</td><td>start</td><td>end</td><td>strand</td><td>MM</td><td>target_seq</td><td>PAM</td><td>alignment</td><td>distance</td><td>position</td><td>gene name</td><td>gene id</td></tr>
</table>

GU305914.1 150940 150962 - 0 GTTCGTGCGCGCGGAGGACG AGG ||||||||[||||||||||||]PAM NA Intergenic NA NA

T22650 TTCGTGCGCGCGGAGGACGAGGG 1000
Oligo fwd CACCTTCGTGCGCGCGGAGGACGA
Oligo rev AAACTCGTCCTCCGCGCGCACGAA
Chromosome start end strand MM target_seq PAM alignment distance position gene name gene id
GU305914.1 150939 150961 - 0 TTCGTGCGCGCGGAGGACGA GGG ||||||||[||||||||||||]PAM NA Intergenic NA NA

T22651 GTGCGCGCGGAGGACGAGGGCGG 1000
Oligo fwd CACCGTGCGCGCGGAGGACGAGGG
Oligo rev AAACCCCTCGTCCTCCGCGCGCAC
Chromosome start end strand MM target_seq PAM alignment distance position gene name gene id
GU305914.1 150936 150958 - 0 GTGCGCGCGGAGGACGAGGG CGG ||||||||[||||||||||||]PAM NA Intergenic NA NA

T22652 CGCGGAGGACGAGGGCGGCGCGG 1000
Oligo fwd CACCCGCGGAGGACGAGGGCGGCG
Oligo rev AAACCGCCGCCCTCGTCCTCCGCG
Chromosome start end strand MM target_seq PAM alignment distance position gene name gene id
GU305914.1 150931 150953 - 0 CGCGGAGGACGAGGGCGGCG CGG ||||||||[||||||||||||]PAM NA Intergenic NA NA
GU305914.1 65123 65145 + 3 CGACGAGGAGGAGGGCGGCG AGG ||--||||[|-||||||||||]PAM NA Intergenic NA NA

T22653 GGACGAGGGCGGCGCGGCGCTGG 1000
Oligo fwd CACCGGACGAGGGCGGCGCGGCGC
Oligo rev AAACGCGCCGCGCCGCCCTCGTCC
Chromosome start end strand MM target_seq PAM alignment distance position gene name gene id
GU305914.1 150925 150947 - 0 GGACGAGGGCGGCGCGGCGC TGG ||||||||[||||||||||||]PAM NA Intergenic NA NA
GU305914.1 201100 201122 + 4 GGGCGACGGCGTCTCGGCGC GGG ||-|||-|[|||-|-||||||]PAM NA Intergenic NA NA

T22654 CGCTGGAACAGTTCGAGATCTGG 1000
Oligo fwd CACCCGCTGGAACAGTTCGAGATC
Oligo rev AAACGATCTCGAACTGTTCCAGCG
Chromosome start end strand MM target_seq PAM alignment distance position gene name gene id
GU305914.1 150908 150930 - 0 CGCTGGAACAGTTCGAGATC TGG ||||||||[||||||||||||]PAM NA Intergenic NA NA

T22655 GCTGGAACAGTTCGAGATCTGGG 1000
Oligo fwd CACCGCTGGAACAGTTCGAGATCT
Oligo rev AAACAGATCTCGAACTGTTCCAGC
Chromosome start end strand MM target_seq PAM alignment distance position gene name gene id
GU305914.1 150907 150929 - 0 GCTGGAACAGTTCGAGATCT GGG ||||||||[||||||||||||]PAM NA Intergenic NA NA

T22656 GGAACAGTTCGAGATCTGGGCGG 1000
Oligo fwd CACCGGAACAGTTCGAGATCTGGG
Oligo rev AAACCCCAGATCTCGAACTGTTCC
Chromosome start end strand MM target_seq PAM alignment distance position gene name gene id
GU305914.1 150904 150926 - 0 GGAACAGTTCGAGATCTGGG CGG ||||||||[||||||||||||]PAM NA Intergenic NA NA

T22657 CGCGCTGAAACACACGATCGTGG 1000
Oligo fwd CACCCGCGCTGAAACACACGATCG
Oligo rev AAACCGATCGTGTGTTTCAGCGCG
Chromosome start end strand MM target_seq PAM alignment distance position gene name gene id
GU305914.1 150880 150902 - 0 CGCGCTGAAACACACGATCG TGG ||||||||[||||||||||||]PAM NA Intergenic NA NA

T22658 GGACAGCATAGCCGCGCTGCTGG 1000
Oligo fwd CACCGGACAGCATAGCCGCGCTGC
Oligo rev AAACGCAGCGCGGCTATGCTGTCC
Chromosome start end strand MM target_seq PAM alignment distance position gene name gene id
GU305914.1 150859 150881 - 0 GGACAGCATAGCCGCGCTGC TGG ||||||||[||||||||||||]PAM NA Intergenic NA NA

T22659 GAAGCTTATATACACGTACGCGG 1000
Oligo fwd CACCGAAGCTTATATACACGTACG
Oligo rev AAACCGTACGTGTATATAAGCTTC
Chromosome start end strand MM target_seq PAM alignment distance position gene name gene id
GU305914.1 150835 150857 - 0 GAAGCTTATATACACGTACG CGG ||||||||[||||||||||||]PAM NA Intergenic NA NA

T22660 GCTTATATACACGTACGCGGCGG 1000
Oligo fwd CACCGCTTATATACACGTACGCGG
Oligo rev AAACCCGCGTACGTGTATATAAGC
Chromosome start end strand MM target_seq PAM alignment distance position gene name gene id
GU305914.1 150832 150854 - 0 GCTTATATACACGTACGCGG CGG ||||||||[||||||||||||]PAM NA Intergenic NA NA

T22661 ACACGTACGCGGCGGACGACAGG 1000
Oligo fwd CACCACACGTACGCGGCGGACGAC
Oligo rev AAACGTCGTCCGCCGCGTACGTGT
Chromosome start end strand MM target_seq PAM alignment distance position gene name gene id
GU305914.1 150824 150846 - 0 ACACGTACGCGGCGGACGAC AGG ||||||||[||||||||||||]PAM NA Intergenic NA NA

T22662 GGTACTGTCGCTACGTAGACTGG 1000
Oligo fwd CACCGGTACTGTCGCTACGTAGAC
Oligo rev AAACGTCTACGTAGCGACAGTACC
Chromosome start end strand MM target_seq PAM alignment distance position gene name gene id
GU305914.1 150803 150825 - 0 GGTACTGTCGCTACGTAGAC TGG ||||||||[||||||||||||]PAM NA Intergenic NA NA

T22663 CTACGTAGACTGGATCGTGTCGG 1000
Oligo fwd CACCCTACGTAGACTGGATCGTGT
Oligo rev AAACACACGATCCAGTCTACGTAG
Chromosome start end strand MM target_seq PAM alignment distance position gene name gene id
GU305914.1 150793 150815 - 0 CTACGTAGACTGGATCGTGT CGG ||||||||[||||||||||||]PAM NA Intergenic NA NA

T22664 CGTAGACTGGATCGTGTCGGTGG 1000

Oligo fwd CACCCGTAGACTGGATCGTGTCGG
Oligo rev AAACCCGACACGATCCAGTCTACG
Chromosome start end strand MM target_seq PAM alignment distance position gene name gene id
GU305914.1 150790 150812 - 0 CGTAGACTGGATCGTGTCGG TGG ||||||||[||||||||||||]PAM NA Intergenic NA NA

T22665 GTAGACTGGATCGTGTCGGTGGG 1000
Oligo fwd CACCGTAGACTGGATCGTGTCGGT
Oligo rev AAACACCGACACGATCCAGTCTAC
Chromosome start end strand MM target_seq PAM alignment distance position gene name gene id
GU305914.1 150789 150811 - 0 GTAGACTGGATCGTGTCGGT GGG ||||||||[||||||||||||]PAM NA Intergenic NA NA

T22666 TAGACTGGATCGTGTCGGTGGGG 1000
Oligo fwd CACCTAGACTGGATCGTGTCGGTG
Oligo rev AAACCACCGACACGATCCAGTCTA
Chromosome start end strand MM target_seq PAM alignment distance position gene name gene id
GU305914.1 150788 150810 - 0 TAGACTGGATCGTGTCGGTG GGG ||||||||[||||||||||||]PAM NA Intergenic NA NA

T22667 AGACTGGATCGTGTCGGTGGGGG 1000
Oligo fwd CACCAGACTGGATCGTGTCGGTGG
Oligo rev AAACCCACCGACACGATCCAGTCT
Chromosome start end strand MM target_seq PAM alignment distance position gene name gene id
GU305914.1 150787 150809 - 0 AGACTGGATCGTGTCGGTGG GGG ||||||||[||||||||||||]PAM NA Intergenic NA NA

T22668 CTGGATCGTGTCGGTGGGGGTGG 1000
Oligo fwd CACCCTGGATCGTGTCGGTGGGGG
Oligo rev AAACCCCCCACCGACACGATCCAG
Chromosome start end strand MM target_seq PAM alignment distance position gene name gene id
GU305914.1 150784 150806 - 0 CTGGATCGTGTCGGTGGGGG TGG ||||||||[||||||||||||]PAM NA Intergenic NA NA

T22669 GGGGGTGGTTCCCATCGCCGAGG 1000
Oligo fwd CACCGGGGGTGGTTCCCATCGCCG
Oligo rev AAACCGGCGATGGGAACCACCCCC
Chromosome start end strand MM target_seq PAM alignment distance position gene name gene id
GU305914.1 150769 150791 - 0 GGGGGTGGTTCCCATCGCCG AGG ||||||||[||||||||||||]PAM NA Intergenic NA NA

T22670 TCCCATCGCCGAGGTACGCACGG 1000
Oligo fwd CACCTCCCATCGCCGAGGTACGCA
Oligo rev AAACTGCGTACCTCGGCGATGGGA
Chromosome start end strand MM target_seq PAM alignment distance position gene name gene id
GU305914.1 150760 150782 - 0 TCCCATCGCCGAGGTACGCA CGG ||||||||[||||||||||||]PAM NA Intergenic NA NA

T22671 CCGAGGTACGCACGGCCGAGCGG 1000
Oligo fwd CACCCCGAGGTACGCACGGCCGAG
Oligo rev AAACCTCGGCCGTGCGTACCTCGG
Chromosome start end strand MM target_seq PAM alignment distance position gene name gene id
GU305914.1 150752 150774 - 0 CCGAGGTACGCACGGCCGAG CGG ||||||||[||||||||||||]PAM NA Intergenic NA NA

T22672 CGAGGTACGCACGGCCGAGCGGG 1000
Oligo fwd CACCCGAGGTACGCACGGCCGAGC
Oligo rev AAACGCTCGGCCGTGCGTACCTCG
Chromosome start end strand MM target_seq PAM alignment distance position gene name gene id
GU305914.1 150751 150773 - 0 CGAGGTACGCACGGCCGAGC GGG ||||||||[||||||||||||]PAM NA Intergenic NA NA

T22673 ACGCACGGCCGAGCGGGAGAAGG 1000
Oligo fwd CACCACGCACGGCCGAGCGGGAGA
Oligo rev AAACTCTCCCGCTCGGCCGTGCGT
Chromosome start end strand MM target_seq PAM alignment distance position gene name gene id
GU305914.1 150745 150767 - 0 ACGCACGGCCGAGCGGGAGA AGG ||||||||[||||||||||||]PAM NA Intergenic NA NA

T22674 CACGGCCGAGCGGGAGAAGGCGG 1000
Oligo fwd CACCCACGGCCGAGCGGGAGAAGG
Oligo rev AAACCCTTCTCCCGCTCGGCCGTG
Chromosome start end strand MM target_seq PAM alignment distance position gene name gene id
GU305914.1 150742 150764 - 0 CACGGCCGAGCGGGAGAAGG CGG ||||||||[||||||||||||]PAM NA Intergenic NA NA

T22675 GCGGGAGAAGGCGGTCGATGCGG 1000
Oligo fwd CACCGCGGGAGAAGGCGGTCGATG
Oligo rev AAACCATCGACCGCCTTCTCCCGC
Chromosome start end strand MM target_seq PAM alignment distance position gene name gene id
GU305914.1 150733 150755 - 0 GCGGGAGAAGGCGGTCGATG CGG ||||||||[||||||||||||]PAM NA Intergenic NA NA

T22676 AGGCGGTCGATGCGGCGCAGCGG 1000
Oligo fwd CACCAGGCGGTCGATGCGGCGCAG
Oligo rev AAACCTGCGCCGCATCGACCGCCT
Chromosome start end strand MM target_seq PAM alignment distance position gene name gene id
GU305914.1 150725 150747 - 0 AGGCGGTCGATGCGGCGCAG CGG ||||||||[||||||||||||]PAM NA Intergenic NA NA

T22677 TGCGGCGCAGCGGCGCTTCCTGG 1000
Oligo fwd CACCTGCGGCGCAGCGGCGCTTCC
Oligo rev AAACGGAAGCGCCGCTGCGCCGCA
Chromosome start end strand MM target_seq PAM alignment distance position gene name gene id
GU305914.1 150715 150737 - 0 TGCGGCGCAGCGGCGCTTCC TGG ||||||||[||||||||||||]PAM NA Intergenic NA NA

T22678 GCAGCGGCGCTTCCTGGCCGAGG 1000
Oligo fwd CACCGCAGCGGCGCTTCCTGGCCG
Oligo rev AAACCGGCCAGGAAGCGCCGCTGC
Chromosome start end strand MM target_seq PAM alignment distance position gene name gene id
GU305914.1 150709 150731 - 0 GCAGCGGCGCTTCCTGGCCG AGG ||||||||[||||||||||||]PAM NA Intergenic NA NA

<table>
<tr><td>T22679</td><td>GCGGCGCTTCCTGGCCGAGGTGG</td><td>1000</td><td></td><td></td><td></td><td></td><td></td><td></td><td></td><td></td><td></td></tr>
<tr><td>Oligo fwd</td><td>CACCGCGGCGCTTCCTGGCCGAGG</td><td></td><td></td><td></td><td></td><td></td><td></td><td></td><td></td><td></td><td></td></tr>
<tr><td>Oligo rev</td><td>AAACCCTCGGCCAGGAAGCGCCGC</td><td></td><td></td><td></td><td></td><td></td><td></td><td></td><td></td><td></td><td></td></tr>
<tr><td>Chromosome</td><td>start</td><td>end</td><td>strand</td><td>MM</td><td>target_seq</td><td>PAM</td><td>alignment</td><td>distance</td><td>position</td><td>gene name</td><td>gene id</td></tr>
<tr><td>GU305914.1</td><td>150706</td><td>150728</td><td>-</td><td>0</td><td>GCGGCGCTTCCTGGCCGAGG</td><td>TGG</td><td>||||||||[||||||||||||]PAM</td><td>NA</td><td>Intergenic</td><td>NA</td><td>NA</td></tr>
<tr><td>T22680</td><td>GGTGGCCGAGTGTCACCTGTTGG</td><td>1000</td><td></td><td></td><td></td><td></td><td></td><td></td><td></td><td></td><td></td></tr>
<tr><td>Oligo fwd</td><td>CACCGGTGGCCGAGTGTCACCTGT</td><td></td><td></td><td></td><td></td><td></td><td></td><td></td><td></td><td></td><td></td></tr>
<tr><td>Oligo rev</td><td>AAACACAGGTGACACTCGGCCACC</td><td></td><td></td><td></td><td></td><td></td><td></td><td></td><td></td><td></td><td></td></tr>
<tr><td>Chromosome</td><td>start</td><td>end</td><td>strand</td><td>MM</td><td>target_seq</td><td>PAM</td><td>alignment</td><td>distance</td><td>position</td><td>gene name</td><td>gene id</td></tr>
<tr><td>GU305914.1</td><td>150688</td><td>150710</td><td>-</td><td>0</td><td>GGTGGCCGAGTGTCACCTGT</td><td>TGG</td><td>||||||||[||||||||||||]PAM</td><td>NA</td><td>Intergenic</td><td>NA</td><td>NA</td></tr>
<tr><td>T22681</td><td>CCGAGTGTCACCTGTTGGACAGG</td><td>1000</td><td></td><td></td><td></td><td></td><td></td><td></td><td></td><td></td><td></td></tr>
<tr><td>Oligo fwd</td><td>CACCCCGAGTGTCACCTGTTGGAC</td><td></td><td></td><td></td><td></td><td></td><td></td><td></td><td></td><td></td><td></td></tr>
<tr><td>Oligo rev</td><td>AAACGTCCAACAGGTGACACTCGG</td><td></td><td></td><td></td><td></td><td></td><td></td><td></td><td></td><td></td><td></td></tr>
<tr><td>Chromosome</td><td>start</td><td>end</td><td>strand</td><td>MM</td><td>target_seq</td><td>PAM</td><td>alignment</td><td>distance</td><td>position</td><td>gene name</td><td>gene id</td></tr>
<tr><td>GU305914.1</td><td>150683</td><td>150705</td><td>-</td><td>0</td><td>CCGAGTGTCACCTGTTGGAC</td><td>AGG</td><td>||||||||[||||||||||||]PAM</td><td>NA</td><td>Intergenic</td><td>NA</td><td>NA</td></tr>
<tr><td>T22682</td><td>GTGTCACCTGTTGGACAGGCCGG</td><td>1000</td><td></td><td></td><td></td><td></td><td></td><td></td><td></td><td></td><td></td></tr>
<tr><td>Oligo fwd</td><td>CACCGTGTCACCTGTTGGACAGGC</td><td></td><td></td><td></td><td></td><td></td><td></td><td></td><td></td><td></td><td></td></tr>
<tr><td>Oligo rev</td><td>AAACGCCTGTCCAACAGGTGACAC</td><td></td><td></td><td></td><td></td><td></td><td></td><td></td><td></td><td></td><td></td></tr>
<tr><td>Chromosome</td><td>start</td><td>end</td><td>strand</td><td>MM</td><td>target_seq</td><td>PAM</td><td>alignment</td><td>distance</td><td>position</td><td>gene name</td><td>gene id</td></tr>
<tr><td>GU305914.1</td><td>150679</td><td>150701</td><td>-</td><td>0</td><td>GTGTCACCTGTTGGACAGGC</td><td>CGG</td><td>||||||||[||||||||||||]PAM</td><td>NA</td><td>Intergenic</td><td>NA</td><td>NA</td></tr>
<tr><td>T22683</td><td>GGCCGGACCCGCTGCGCGCGCGG</td><td>1000</td><td></td><td></td><td></td><td></td><td></td><td></td><td></td><td></td><td></td></tr>
<tr><td>Oligo fwd</td><td>CACCGGCCGGACCCGCTGCGCGCG</td><td></td><td></td><td></td><td></td><td></td><td></td><td></td><td></td><td></td><td></td></tr>
<tr><td>Oligo rev</td><td>AAACCGCGCGCAGCGGGTCCGGCC</td><td></td><td></td><td></td><td></td><td></td><td></td><td></td><td></td><td></td><td></td></tr>
<tr><td>Chromosome</td><td>start</td><td>end</td><td>strand</td><td>MM</td><td>target_seq</td><td>PAM</td><td>alignment</td><td>distance</td><td>position</td><td>gene name</td><td>gene id</td></tr>
<tr><td>GU305914.1</td><td>150662</td><td>150684</td><td>-</td><td>0</td><td>GGCCGGACCCGCTGCGCGCG</td><td>CGG</td><td>||||||||[||||||||||||]PAM</td><td>NA</td><td>Intergenic</td><td>NA</td><td>NA</td></tr>
<tr><td>T22684</td><td>CGCTGCGCGCGCGGACGCTGCGG</td><td>1000</td><td></td><td></td><td></td><td></td><td></td><td></td><td></td><td></td><td></td></tr>
<tr><td>Oligo fwd</td><td>CACCCGCTGCGCGCGCGGACGCTG</td><td></td><td></td><td></td><td></td><td></td><td></td><td></td><td></td><td></td><td></td></tr>
<tr><td>Oligo rev</td><td>AAACCAGCGTCCGCGCGCGCAGCG</td><td></td><td></td><td></td><td></td><td></td><td></td><td></td><td></td><td></td><td></td></tr>
<tr><td>Chromosome</td><td>start</td><td>end</td><td>strand</td><td>MM</td><td>target_seq</td><td>PAM</td><td>alignment</td><td>distance</td><td>position</td><td>gene name</td><td>gene id</td></tr>
<tr><td>GU305914.1</td><td>150653</td><td>150675</td><td>-</td><td>0</td><td>CGCTGCGCGCGCGGACGCTG</td><td>CGG</td><td>||||||||[||||||||||||]PAM</td><td>NA</td><td>Intergenic</td><td>NA</td><td>NA</td></tr>
<tr><td>T22685</td><td>GCTGCGCGCGCGGACGCTGCGGG</td><td>1000</td><td></td><td></td><td></td><td></td><td></td><td></td><td></td><td></td><td></td></tr>
<tr><td>Oligo fwd</td><td>CACCGCTGCGCGCGCGGACGCTGC</td><td></td><td></td><td></td><td></td><td></td><td></td><td></td><td></td><td></td><td></td></tr>
<tr><td>Oligo rev</td><td>AAACGCAGCGTCCGCGCGCGCAGC</td><td></td><td></td><td></td><td></td><td></td><td></td><td></td><td></td><td></td><td></td></tr>
<tr><td>Chromosome</td><td>start</td><td>end</td><td>strand</td><td>MM</td><td>target_seq</td><td>PAM</td><td>alignment</td><td>distance</td><td>position</td><td>gene name</td><td>gene id</td></tr>
<tr><td>GU305914.1</td><td>150652</td><td>150674</td><td>-</td><td>0</td><td>GCTGCGCGCGCGGACGCTGC</td><td>GGG</td><td>||||||||[||||||||||||]PAM</td><td>NA</td><td>Intergenic</td><td>NA</td><td>NA</td></tr>
<tr><td>T22686</td><td>GACGCTGCGGGCGTGCGTGTCGG</td><td>1000</td><td></td><td></td><td></td><td></td><td></td><td></td><td></td><td></td><td></td></tr>
<tr><td>Oligo fwd</td><td>CACCGACGCTGCGGGCGTGCGTGT</td><td></td><td></td><td></td><td></td><td></td><td></td><td></td><td></td><td></td><td></td></tr>
<tr><td>Oligo rev</td><td>AAACACACGCACGCCCGCAGCGTC</td><td></td><td></td><td></td><td></td><td></td><td></td><td></td><td></td><td></td><td></td></tr>
<tr><td>Chromosome</td><td>start</td><td>end</td><td>strand</td><td>MM</td><td>target_seq</td><td>PAM</td><td>alignment</td><td>distance</td><td>position</td><td>gene name</td><td>gene id</td></tr>
<tr><td>GU305914.1</td><td>150640</td><td>150662</td><td>-</td><td>0</td><td>GACGCTGCGGGCGTGCGTGT</td><td>CGG</td><td>||||||||[||||||||||||]PAM</td><td>NA</td><td>Intergenic</td><td>NA</td><td>NA</td></tr>
<tr><td>T22687</td><td>GCGTGTCGGCGCTGATGACCCGG</td><td>1000</td><td></td><td></td><td></td><td></td><td></td><td></td><td></td><td></td><td></td></tr>
<tr><td>Oligo fwd</td><td>CACCGCGTGTCGGCGCTGATGACC</td><td></td><td></td><td></td><td></td><td></td><td></td><td></td><td></td><td></td><td></td></tr>
<tr><td>Oligo rev</td><td>AAACGGTCATCAGCGCCGACACGC</td><td></td><td></td><td></td><td></td><td></td><td></td><td></td><td></td><td></td><td></td></tr>
<tr><td>Chromosome</td><td>start</td><td>end</td><td>strand</td><td>MM</td><td>target_seq</td><td>PAM</td><td>alignment</td><td>distance</td><td>position</td><td>gene name</td><td>gene id</td></tr>
<tr><td>GU305914.1</td><td>150626</td><td>150648</td><td>-</td><td>0</td><td>GCGTGTCGGCGCTGATGACC</td><td>CGG</td><td>||||||||[||||||||||||]PAM</td><td>NA</td><td>Intergenic</td><td>NA</td><td>NA</td></tr>
<tr><td>T22688</td><td>CGTGTCGGCGCTGATGACCCGGG</td><td>1000</td><td></td><td></td><td></td><td></td><td></td><td></td><td></td><td></td><td></td></tr>
<tr><td>Oligo fwd</td><td>CACCCGTGTCGGCGCTGATGACCC</td><td></td><td></td><td></td><td></td><td></td><td></td><td></td><td></td><td></td><td></td></tr>
<tr><td>Oligo rev</td><td>AAACGGGTCATCAGCGCCGACACG</td><td></td><td></td><td></td><td></td><td></td><td></td><td></td><td></td><td></td><td></td></tr>
<tr><td>Chromosome</td><td>start</td><td>end</td><td>strand</td><td>MM</td><td>target_seq</td><td>PAM</td><td>alignment</td><td>distance</td><td>position</td><td>gene name</td><td>gene id</td></tr>
<tr><td>GU305914.1</td><td>150625</td><td>150647</td><td>-</td><td>0</td><td>CGTGTCGGCGCTGATGACCC</td><td>GGG</td><td>||||||||[||||||||||||]PAM</td><td>NA</td><td>Intergenic</td><td>NA</td><td>NA</td></tr>
<tr><td>GU305914.1</td><td>90510</td><td>90532</td><td>+</td><td>3</td><td>CGTGTCGCCGCTCGTGACCC</td><td>CGG</td><td>|||||||-[||||--||||||]PAM</td><td>NA</td><td>Intergenic</td><td>NA</td><td>NA</td></tr>
<tr><td>T22689</td><td>GTCGGCGCTGATGACCCGGGAGG</td><td>1000</td><td></td><td></td><td></td><td></td><td></td><td></td><td></td><td></td><td></td></tr>
<tr><td>Oligo fwd</td><td>CACCGTCGGCGCTGATGACCCGGG</td><td></td><td></td><td></td><td></td><td></td><td></td><td></td><td></td><td></td><td></td></tr>
<tr><td>Oligo rev</td><td>AAACCCCGGGTCATCAGCGCCGAC</td><td></td><td></td><td></td><td></td><td></td><td></td><td></td><td></td><td></td><td></td></tr>
<tr><td>Chromosome</td><td>start</td><td>end</td><td>strand</td><td>MM</td><td>target_seq</td><td>PAM</td><td>alignment</td><td>distance</td><td>position</td><td>gene name</td><td>gene id</td></tr>
<tr><td>GU305914.1</td><td>150622</td><td>150644</td><td>-</td><td>0</td><td>GTCGGCGCTGATGACCCGGG</td><td>AGG</td><td>||||||||[||||||||||||]PAM</td><td>NA</td><td>Intergenic</td><td>NA</td><td>NA</td></tr>
<tr><td>T22690</td><td>TGCCCGACATCCGCATCCACCGG</td><td>1000</td><td></td><td></td><td></td><td></td><td></td><td></td><td></td><td></td><td></td></tr>
<tr><td>Oligo fwd</td><td>CACCTGCCCGACATCCGCATCCAC</td><td></td><td></td><td></td><td></td><td></td><td></td><td></td><td></td><td></td><td></td></tr>
<tr><td>Oligo rev</td><td>AAACGTGGATGCGGATGTCGGGCA</td><td></td><td></td><td></td><td></td><td></td><td></td><td></td><td></td><td></td><td></td></tr>
<tr><td>Chromosome</td><td>start</td><td>end</td><td>strand</td><td>MM</td><td>target_seq</td><td>PAM</td><td>alignment</td><td>distance</td><td>position</td><td>gene name</td><td>gene id</td></tr>
<tr><td>GU305914.1</td><td>150590</td><td>150612</td><td>-</td><td>0</td><td>TGCCCGACATCCGCATCCAC</td><td>CGG</td><td>||||||||[||||||||||||]PAM</td><td>NA</td><td>Intergenic</td><td>NA</td><td>NA</td></tr>
<tr><td>T22691</td><td>ATCCACCGGCTGAAGAGCAACGG</td><td>1000</td><td></td><td></td><td></td><td></td><td></td><td></td><td></td><td></td><td></td></tr>
<tr><td>Oligo fwd</td><td>CACCATCCACCGGCTGAAGAGCAA</td><td></td><td></td><td></td><td></td><td></td><td></td><td></td><td></td><td></td><td></td></tr>
<tr><td>Oligo rev</td><td>AAACTTGCTCTTCAGCCGGTGGAT</td><td></td><td></td><td></td><td></td><td></td><td></td><td></td><td></td><td></td><td></td></tr>
<tr><td>Chromosome</td><td>start</td><td>end</td><td>strand</td><td>MM</td><td>target_seq</td><td>PAM</td><td>alignment</td><td>distance</td><td>position</td><td>gene name</td><td>gene id</td></tr>
<tr><td>GU305914.1</td><td>150576</td><td>150598</td><td>-</td><td>0</td><td>ATCCACCGGCTGAAGAGCAA</td><td>CGG</td><td>||||||||[||||||||||||]PAM</td><td>NA</td><td>Intergenic</td><td>NA</td><td>NA</td></tr>
<tr><td>T22692</td><td>TCCACCGGCTGAAGAGCAACGGG</td><td>1000</td><td></td><td></td><td></td><td></td><td></td><td></td><td></td><td></td><td></td></tr>
<tr><td>Oligo fwd</td><td>CACCTCCACCGGCTGAAGAGCAAC</td><td></td><td></td><td></td><td></td><td></td><td></td><td></td><td></td><td></td><td></td></tr>
<tr><td>Oligo rev</td><td>AAACGTTGCTCTTCAGCCGGTGGA</td><td></td><td></td><td></td><td></td><td></td><td></td><td></td><td></td><td></td><td></td></tr>
<tr><td>Chromosome</td><td>start</td><td>end</td><td>strand</td><td>MM</td><td>target_seq</td><td>PAM</td><td>alignment</td><td>distance</td><td>position</td><td>gene name</td><td>gene id</td></tr>
<tr><td>GU305914.1</td><td>150575</td><td>150597</td><td>-</td><td>0</td><td>TCCACCGGCTGAAGAGCAAC</td><td>GGG</td><td>||||||||[||||||||||||]PAM</td><td>NA</td><td>Intergenic</td><td>NA</td><td>NA</td></tr>
<tr><td>T22693</td><td>CGGGCACATCGAGTGCTTCTCGG</td><td>1000</td><td></td><td></td><td></td><td></td><td></td><td></td><td></td><td></td><td></td></tr>
<tr><td>Oligo fwd</td><td>CACCCGGGCACATCGAGTGCTTCT</td><td></td><td></td><td></td><td></td><td></td><td></td><td></td><td></td><td></td><td></td></tr>
<tr><td>Oligo rev</td><td>AAACAGAAGCACTCGATGTGCCCG</td><td></td><td></td><td></td><td></td><td></td><td></td><td></td><td></td><td></td><td></td></tr>
<tr><td>Chromosome</td><td>start</td><td>end</td><td>strand</td><td>MM</td><td>target_seq</td><td>PAM</td><td>alignment</td><td>distance</td><td>position</td><td>gene name</td><td>gene id</td></tr>
</table>

GU305914.1 150556 150578 - 0 CGGGCACATCGAGTGCTTCT CGG ||||||||[||||||||||||]PAM NA Intergenic NA NA

T22694 GGGCACATCGAGTGCTTCTCGGG 1000
Oligo fwd CACCGGGCACATCGAGTGCTTCTC
Oligo rev AAACGAGAAGCACTCGATGTGCCC
Chromosome start end strand MM target_seq PAM alignment distance position gene name gene id
GU305914.1 150555 150577 - 0 GGGCACATCGAGTGCTTCTC GGG ||||||||[||||||||||||]PAM NA Intergenic NA NA

T22695 TCGAGTGCTTCTCGGGCAAGCGG 1000
Oligo fwd CACCTCGAGTGCTTCTCGGGCAAG
Oligo rev AAACCTTGCCCGAGAAGCACTCGA
Chromosome start end strand MM target_seq PAM alignment distance position gene name gene id
GU305914.1 150548 150570 - 0 TCGAGTGCTTCTCGGGCAAG CGG ||||||||[||||||||||||]PAM NA Intergenic NA NA

T22696 AGTGCTTCTCGGGCAAGCGGCGG 1000
Oligo fwd CACCAGTGCTTCTCGGGCAAGCGG
Oligo rev AAACCCGCTTGCCCGAGAAGCACT
Chromosome start end strand MM target_seq PAM alignment distance position gene name gene id
GU305914.1 150545 150567 - 0 AGTGCTTCTCGGGCAAGCGG CGG ||||||||[||||||||||||]PAM NA Intergenic NA NA

T22697 GCTGAAGCGTTTCATCTACGCGG 1000
Oligo fwd CACCGCTGAAGCGTTTCATCTACG
Oligo rev AAACCGTAGATGAAACGCTTCAGC
Chromosome start end strand MM target_seq PAM alignment distance position gene name gene id
GU305914.1 150523 150545 - 0 GCTGAAGCGTTTCATCTACG CGG ||||||||[||||||||||||]PAM NA Intergenic NA NA

T22698 CGCGGAGCCCACGATCCTCGAGG 1000
Oligo fwd CACCCGCGGAGCCCACGATCCTCG
Oligo rev AAACCGAGGATCGTGGGCTCCGCG
Chromosome start end strand MM target_seq PAM alignment distance position gene name gene id
GU305914.1 150505 150527 - 0 CGCGGAGCCCACGATCCTCG AGG ||||||||[||||||||||||]PAM NA Intergenic NA NA

T22699 GGAGCCCACGATCCTCGAGGAGG 1000
Oligo fwd CACCGGAGCCCACGATCCTCGAGG
Oligo rev AAACCCTCGAGGATCGTGGGCTCC
Chromosome start end strand MM target_seq PAM alignment distance position gene name gene id
GU305914.1 150502 150524 - 0 GGAGCCCACGATCCTCGAGG AGG ||||||||[||||||||||||]PAM NA Intergenic NA NA

T22700 CCACGATCCTCGAGGAGGAGCGG 1000
Oligo fwd CACCCCACGATCCTCGAGGAGGAG
Oligo rev AAACCTCCTCCTCGAGGATCGTGG
Chromosome start end strand MM target_seq PAM alignment distance position gene name gene id
GU305914.1 150497 150519 - 0 CCACGATCCTCGAGGAGGAG CGG ||||||||[||||||||||||]PAM NA Intergenic NA NA

T22701 GCTGATCCTGACGACGCCGCTGG 1000
Oligo fwd CACCGCTGATCCTGACGACGCCGC
Oligo rev AAACGCGGCGTCGTCAGGATCAGC
Chromosome start end strand MM target_seq PAM alignment distance position gene name gene id
GU305914.1 150475 150497 - 0 GCTGATCCTGACGACGCCGC TGG ||||||||[||||||||||||]PAM NA Intergenic NA NA

T22702 TCCTGACGACGCCGCTGGCCCGG 1000
Oligo fwd CACCTCCTGACGACGCCGCTGGCC
Oligo rev AAACGGCCAGCGGCGTCGTCAGGA
Chromosome start end strand MM target_seq PAM alignment distance position gene name gene id
GU305914.1 150470 150492 - 0 TCCTGACGACGCCGCTGGCC CGG ||||||||[||||||||||||]PAM NA Intergenic NA NA

T22703 GCTGAACACCAACCCGATCAAGG 1000
Oligo fwd CACCGCTGAACACCAACCCGATCA
Oligo rev AAACTGATCGGGTTGGTGTTCAGC
Chromosome start end strand MM target_seq PAM alignment distance position gene name gene id
GU305914.1 150388 150410 - 0 GCTGAACACCAACCCGATCA AGG ||||||||[||||||||||||]PAM NA Intergenic NA NA

T22704 GAACACCAACCCGATCAAGGTGG 1000
Oligo fwd CACCGAACACCAACCCGATCAAGG
Oligo rev AAACCCTTGATCGGGTTGGTGTTC
Chromosome start end strand MM target_seq PAM alignment distance position gene name gene id
GU305914.1 150385 150407 - 0 GAACACCAACCCGATCAAGG TGG ||||||||[||||||||||||]PAM NA Intergenic NA NA

T22705 TCAAGGTGGTGACCACGAGCCGG 1000
Oligo fwd CACCTCAAGGTGGTGACCACGAGC
Oligo rev AAACGCTCGTGGTCACCACCTTGA
Chromosome start end strand MM target_seq PAM alignment distance position gene name gene id
GU305914.1 150371 150393 - 0 TCAAGGTGGTGACCACGAGC CGG ||||||||[||||||||||||]PAM NA Intergenic NA NA

T22706 GATGAACACGAAGCGCATCGTGG 1000
Oligo fwd CACCGATGAACACGAAGCGCATCG
Oligo rev AAACCGATGCGCTTCGTGTTCATC
Chromosome start end strand MM target_seq PAM alignment distance position gene name gene id
GU305914.1 150343 150365 - 0 GATGAACACGAAGCGCATCG TGG ||||||||[||||||||||||]PAM NA Intergenic NA NA

T22707 GAAGCGCATCGTGGAGCTGATGG 1000
Oligo fwd CACCGAAGCGCATCGTGGAGCTGA
Oligo rev AAACTCAGCTCCACGATGCGCTTC
Chromosome start end strand MM target_seq PAM alignment distance position gene name gene id
GU305914.1 150334 150356 - 0 GAAGCGCATCGTGGAGCTGA TGG ||||||||[||||||||||||]PAM NA Intergenic NA NA

T22708 TCGTGGAGCTGATGGAGAAGCGG 1000
Oligo fwd CACCTCGTGGAGCTGATGGAGAAG
Oligo rev AAACCTTCTCCATCAGCTCCACGA

<table>
<tr><td>Chromosome</td><td>start</td><td>end</td><td>strand</td><td>MM</td><td>target_seq</td><td>PAM</td><td>alignment</td><td>distance</td><td>position</td><td>gene name</td><td>gene id</td></tr>
<tr><td>GU305914.1</td><td>150326</td><td>150348</td><td>-</td><td>0</td><td>TCGTGGAGCTGATGGAGAAG</td><td>CGG</td><td>||||||||[||||||||||||]PAM</td><td>NA</td><td>Intergenic</td><td>NA</td><td>NA</td></tr>
<tr><td></td><td></td><td></td><td></td><td></td><td></td><td></td><td></td><td></td><td></td><td></td><td></td></tr>
<tr><td>T22709</td><td>CGTGGAGCTGATGGAGAAGCGGG</td><td>1000</td><td></td><td></td><td></td><td></td><td></td><td></td><td></td><td></td><td></td></tr>
<tr><td>Oligo fwd</td><td>CACCCGTGGAGCTGATGGAGAAGC</td><td></td><td></td><td></td><td></td><td></td><td></td><td></td><td></td><td></td><td></td></tr>
<tr><td>Oligo rev</td><td>AAACGCTTCTCCATCAGCTCCACG</td><td></td><td></td><td></td><td></td><td></td><td></td><td></td><td></td><td></td><td></td></tr>
<tr><td>Chromosome</td><td>start</td><td>end</td><td>strand</td><td>MM</td><td>target_seq</td><td>PAM</td><td>alignment</td><td>distance</td><td>position</td><td>gene name</td><td>gene id</td></tr>
<tr><td>GU305914.1</td><td>150325</td><td>150347</td><td>-</td><td>0</td><td>CGTGGAGCTGATGGAGAAGC</td><td>GGG</td><td>||||||||[||||||||||||]PAM</td><td>NA</td><td>Intergenic</td><td>NA</td><td>NA</td></tr>
<tr><td></td><td></td><td></td><td></td><td></td><td></td><td></td><td></td><td></td><td></td><td></td><td></td></tr>
<tr><td>T22710</td><td>AGCTGATGGAGAAGCGGGACCGG</td><td>1000</td><td></td><td></td><td></td><td></td><td></td><td></td><td></td><td></td><td></td></tr>
<tr><td>Oligo fwd</td><td>CACCAGCTGATGGAGAAGCGGGAC</td><td></td><td></td><td></td><td></td><td></td><td></td><td></td><td></td><td></td><td></td></tr>
<tr><td>Oligo rev</td><td>AAACGTCCCGCTTCTCCATCAGCT</td><td></td><td></td><td></td><td></td><td></td><td></td><td></td><td></td><td></td><td></td></tr>
<tr><td>Chromosome</td><td>start</td><td>end</td><td>strand</td><td>MM</td><td>target_seq</td><td>PAM</td><td>alignment</td><td>distance</td><td>position</td><td>gene name</td><td>gene id</td></tr>
<tr><td>GU305914.1</td><td>150320</td><td>150342</td><td>-</td><td>0</td><td>AGCTGATGGAGAAGCGGGAC</td><td>CGG</td><td>||||||||[||||||||||||]PAM</td><td>NA</td><td>Intergenic</td><td>NA</td><td>NA</td></tr>
<tr><td></td><td></td><td></td><td></td><td></td><td></td><td></td><td></td><td></td><td></td><td></td><td></td></tr>
<tr><td>T22711</td><td>GATGGAGAAGCGGGACCGGCAGG</td><td>1000</td><td></td><td></td><td></td><td></td><td></td><td></td><td></td><td></td><td></td></tr>
<tr><td>Oligo fwd</td><td>CACCGATGGAGAAGCGGGACCGGC</td><td></td><td></td><td></td><td></td><td></td><td></td><td></td><td></td><td></td><td></td></tr>
<tr><td>Oligo rev</td><td>AAACGCCGGTCCCGCTTCTCCATC</td><td></td><td></td><td></td><td></td><td></td><td></td><td></td><td></td><td></td><td></td></tr>
<tr><td>Chromosome</td><td>start</td><td>end</td><td>strand</td><td>MM</td><td>target_seq</td><td>PAM</td><td>alignment</td><td>distance</td><td>position</td><td>gene name</td><td>gene id</td></tr>
<tr><td>GU305914.1</td><td>150316</td><td>150338</td><td>-</td><td>0</td><td>GATGGAGAAGCGGGACCGGC</td><td>AGG</td><td>||||||||[||||||||||||]PAM</td><td>NA</td><td>Intergenic</td><td>NA</td><td>NA</td></tr>
<tr><td></td><td></td><td></td><td></td><td></td><td></td><td></td><td></td><td></td><td></td><td></td><td></td></tr>
<tr><td>T22712</td><td>GGAGAAGCGGGACCGGCAGGTGG</td><td>1000</td><td></td><td></td><td></td><td></td><td></td><td></td><td></td><td></td><td></td></tr>
<tr><td>Oligo fwd</td><td>CACCGGAGAAGCGGGACCGGCAGG</td><td></td><td></td><td></td><td></td><td></td><td></td><td></td><td></td><td></td><td></td></tr>
<tr><td>Oligo rev</td><td>AAACCCTGCCGGTCCCGCTTCTCC</td><td></td><td></td><td></td><td></td><td></td><td></td><td></td><td></td><td></td><td></td></tr>
<tr><td>Chromosome</td><td>start</td><td>end</td><td>strand</td><td>MM</td><td>target_seq</td><td>PAM</td><td>alignment</td><td>distance</td><td>position</td><td>gene name</td><td>gene id</td></tr>
<tr><td>GU305914.1</td><td>150313</td><td>150335</td><td>-</td><td>0</td><td>GGAGAAGCGGGACCGGCAGG</td><td>TGG</td><td>||||||||[||||||||||||]PAM</td><td>NA</td><td>Intergenic</td><td>NA</td><td>NA</td></tr>
<tr><td>GU305914.1</td><td>72580</td><td>72602</td><td>+</td><td>3</td><td>GGAGAAGGTGGAACGGCAGG</td><td>TGG</td><td>|||||||-[-|||-|||||||]PAM</td><td>NA</td><td>Intergenic</td><td>NA</td><td>NA</td></tr>
<tr><td></td><td></td><td></td><td></td><td></td><td></td><td></td><td></td><td></td><td></td><td></td><td></td></tr>
<tr><td>T22713</td><td>TCCGACAGCAAGTCGAAGATCGG</td><td>1000</td><td></td><td></td><td></td><td></td><td></td><td></td><td></td><td></td><td></td></tr>
<tr><td>Oligo fwd</td><td>CACCTCCGACAGCAAGTCGAAGAT</td><td></td><td></td><td></td><td></td><td></td><td></td><td></td><td></td><td></td><td></td></tr>
<tr><td>Oligo rev</td><td>AAACATCTTCGACTTGCTGTCGGA</td><td></td><td></td><td></td><td></td><td></td><td></td><td></td><td></td><td></td><td></td></tr>
<tr><td>Chromosome</td><td>start</td><td>end</td><td>strand</td><td>MM</td><td>target_seq</td><td>PAM</td><td>alignment</td><td>distance</td><td>position</td><td>gene name</td><td>gene id</td></tr>
<tr><td>GU305914.1</td><td>150252</td><td>150274</td><td>-</td><td>0</td><td>TCCGACAGCAAGTCGAAGAT</td><td>CGG</td><td>||||||||[||||||||||||]PAM</td><td>NA</td><td>Intergenic</td><td>NA</td><td>NA</td></tr>
<tr><td></td><td></td><td></td><td></td><td></td><td></td><td></td><td></td><td></td><td></td><td></td><td></td></tr>
<tr><td>T22714</td><td>CAGCAAGTCGAAGATCGGACTGG</td><td>1000</td><td></td><td></td><td></td><td></td><td></td><td></td><td></td><td></td><td></td></tr>
<tr><td>Oligo fwd</td><td>CACCCAGCAAGTCGAAGATCGGAC</td><td></td><td></td><td></td><td></td><td></td><td></td><td></td><td></td><td></td><td></td></tr>
<tr><td>Oligo rev</td><td>AAACGTCCGATCTTCGACTTGCTG</td><td></td><td></td><td></td><td></td><td></td><td></td><td></td><td></td><td></td><td></td></tr>
<tr><td>Chromosome</td><td>start</td><td>end</td><td>strand</td><td>MM</td><td>target_seq</td><td>PAM</td><td>alignment</td><td>distance</td><td>position</td><td>gene name</td><td>gene id</td></tr>
<tr><td>GU305914.1</td><td>150247</td><td>150269</td><td>-</td><td>0</td><td>CAGCAAGTCGAAGATCGGAC</td><td>TGG</td><td>||||||||[||||||||||||]PAM</td><td>NA</td><td>Intergenic</td><td>NA</td><td>NA</td></tr>
<tr><td></td><td></td><td></td><td></td><td></td><td></td><td></td><td></td><td></td><td></td><td></td><td></td></tr>
<tr><td>T22715</td><td>CAAGTCGAAGATCGGACTGGAGG</td><td>1000</td><td></td><td></td><td></td><td></td><td></td><td></td><td></td><td></td><td></td></tr>
<tr><td>Oligo fwd</td><td>CACCCAAGTCGAAGATCGGACTGG</td><td></td><td></td><td></td><td></td><td></td><td></td><td></td><td></td><td></td><td></td></tr>
<tr><td>Oligo rev</td><td>AAACCCAGTCCGATCTTCGACTTG</td><td></td><td></td><td></td><td></td><td></td><td></td><td></td><td></td><td></td><td></td></tr>
<tr><td>Chromosome</td><td>start</td><td>end</td><td>strand</td><td>MM</td><td>target_seq</td><td>PAM</td><td>alignment</td><td>distance</td><td>position</td><td>gene name</td><td>gene id</td></tr>
<tr><td>GU305914.1</td><td>150244</td><td>150266</td><td>-</td><td>0</td><td>CAAGTCGAAGATCGGACTGG</td><td>AGG</td><td>||||||||[||||||||||||]PAM</td><td>NA</td><td>Intergenic</td><td>NA</td><td>NA</td></tr>
<tr><td></td><td></td><td></td><td></td><td></td><td></td><td></td><td></td><td></td><td></td><td></td><td></td></tr>
<tr><td>T22716</td><td>GAAGATCGGACTGGAGGACTCGG</td><td>1000</td><td></td><td></td><td></td><td></td><td></td><td></td><td></td><td></td><td></td></tr>
<tr><td>Oligo fwd</td><td>CACCGAAGATCGGACTGGAGGACT</td><td></td><td></td><td></td><td></td><td></td><td></td><td></td><td></td><td></td><td></td></tr>
<tr><td>Oligo rev</td><td>AAACAGTCCTCCAGTCCGATCTTC</td><td></td><td></td><td></td><td></td><td></td><td></td><td></td><td></td><td></td><td></td></tr>
<tr><td>Chromosome</td><td>start</td><td>end</td><td>strand</td><td>MM</td><td>target_seq</td><td>PAM</td><td>alignment</td><td>distance</td><td>position</td><td>gene name</td><td>gene id</td></tr>
<tr><td>GU305914.1</td><td>150238</td><td>150260</td><td>-</td><td>0</td><td>GAAGATCGGACTGGAGGACT</td><td>CGG</td><td>||||||||[||||||||||||]PAM</td><td>NA</td><td>Intergenic</td><td>NA</td><td>NA</td></tr>
<tr><td></td><td></td><td></td><td></td><td></td><td></td><td></td><td></td><td></td><td></td><td></td><td></td></tr>
<tr><td>T22717</td><td>GATCGGACTGGAGGACTCGGTGG</td><td>1000</td><td></td><td></td><td></td><td></td><td></td><td></td><td></td><td></td><td></td></tr>
<tr><td>Oligo fwd</td><td>CACCGATCGGACTGGAGGACTCGG</td><td></td><td></td><td></td><td></td><td></td><td></td><td></td><td></td><td></td><td></td></tr>
<tr><td>Oligo rev</td><td>AAACCCGAGTCCTCCAGTCCGATC</td><td></td><td></td><td></td><td></td><td></td><td></td><td></td><td></td><td></td><td></td></tr>
<tr><td>Chromosome</td><td>start</td><td>end</td><td>strand</td><td>MM</td><td>target_seq</td><td>PAM</td><td>alignment</td><td>distance</td><td>position</td><td>gene name</td><td>gene id</td></tr>
<tr><td>GU305914.1</td><td>150235</td><td>150257</td><td>-</td><td>0</td><td>GATCGGACTGGAGGACTCGG</td><td>TGG</td><td>||||||||[||||||||||||]PAM</td><td>NA</td><td>Intergenic</td><td>NA</td><td>NA</td></tr>
<tr><td></td><td></td><td></td><td></td><td></td><td></td><td></td><td></td><td></td><td></td><td></td><td></td></tr>
<tr><td>T22718</td><td>CTCGGTGGAGTCTTTCCTGCAGG</td><td>1000</td><td></td><td></td><td></td><td></td><td></td><td></td><td></td><td></td><td></td></tr>
<tr><td>Oligo fwd</td><td>CACCCTCGGTGGAGTCTTTCCTGC</td><td></td><td></td><td></td><td></td><td></td><td></td><td></td><td></td><td></td><td></td></tr>
<tr><td>Oligo rev</td><td>AAACGCAGGAAAGACTCCACCGAG</td><td></td><td></td><td></td><td></td><td></td><td></td><td></td><td></td><td></td><td></td></tr>
<tr><td>Chromosome</td><td>start</td><td>end</td><td>strand</td><td>MM</td><td>target_seq</td><td>PAM</td><td>alignment</td><td>distance</td><td>position</td><td>gene name</td><td>gene id</td></tr>
<tr><td>GU305914.1</td><td>150220</td><td>150242</td><td>-</td><td>0</td><td>CTCGGTGGAGTCTTTCCTGC</td><td>AGG</td><td>||||||||[||||||||||||]PAM</td><td>NA</td><td>Intergenic</td><td>NA</td><td>NA</td></tr>
<tr><td></td><td></td><td></td><td></td><td></td><td></td><td></td><td></td><td></td><td></td><td></td><td></td></tr>
<tr><td>T22719</td><td>GCAGGACCTGACGCCGTCTGTGG</td><td>1000</td><td></td><td></td><td></td><td></td><td></td><td></td><td></td><td></td><td></td></tr>
<tr><td>Oligo fwd</td><td>CACCGCAGGACCTGACGCCGTCTG</td><td></td><td></td><td></td><td></td><td></td><td></td><td></td><td></td><td></td><td></td></tr>
<tr><td>Oligo rev</td><td>AAACCAGACGGCGTCAGGTCCTGC</td><td></td><td></td><td></td><td></td><td></td><td></td><td></td><td></td><td></td><td></td></tr>
<tr><td>Chromosome</td><td>start</td><td>end</td><td>strand</td><td>MM</td><td>target_seq</td><td>PAM</td><td>alignment</td><td>distance</td><td>position</td><td>gene name</td><td>gene id</td></tr>
<tr><td>GU305914.1</td><td>150202</td><td>150224</td><td>-</td><td>0</td><td>GCAGGACCTGACGCCGTCTG</td><td>TGG</td><td>||||||||[||||||||||||]PAM</td><td>NA</td><td>Intergenic</td><td>NA</td><td>NA</td></tr>
<tr><td></td><td></td><td></td><td></td><td></td><td></td><td></td><td></td><td></td><td></td><td></td><td></td></tr>
<tr><td>T22720</td><td>CCGCGCGCCCCTCATCCAACCGG</td><td>1000</td><td></td><td></td><td></td><td></td><td></td><td></td><td></td><td></td><td></td></tr>
<tr><td>Oligo fwd</td><td>CACCCCGCGCGCCCCTCATCCAAC</td><td></td><td></td><td></td><td></td><td></td><td></td><td></td><td></td><td></td><td></td></tr>
<tr><td>Oligo rev</td><td>AAACGTTGGATGAGGGGCGCGCGG</td><td></td><td></td><td></td><td></td><td></td><td></td><td></td><td></td><td></td><td></td></tr>
<tr><td>Chromosome</td><td>start</td><td>end</td><td>strand</td><td>MM</td><td>target_seq</td><td>PAM</td><td>alignment</td><td>distance</td><td>position</td><td>gene name</td><td>gene id</td></tr>
<tr><td>GU305914.1</td><td>150157</td><td>150179</td><td>-</td><td>0</td><td>CCGCGCGCCCCTCATCCAAC</td><td>CGG</td><td>||||||||[||||||||||||]PAM</td><td>NA</td><td>Intergenic</td><td>NA</td><td>NA</td></tr>
<tr><td></td><td></td><td></td><td></td><td></td><td></td><td></td><td></td><td></td><td></td><td></td><td></td></tr>
<tr><td>T22721</td><td>CCTCATCCAACCGGCACCCTCGG</td><td>1000</td><td></td><td></td><td></td><td></td><td></td><td></td><td></td><td></td><td></td></tr>
<tr><td>Oligo fwd</td><td>CACCCCTCATCCAACCGGCACCCT</td><td></td><td></td><td></td><td></td><td></td><td></td><td></td><td></td><td></td><td></td></tr>
<tr><td>Oligo rev</td><td>AAACAGGGTGCCGGTTGGATGAGG</td><td></td><td></td><td></td><td></td><td></td><td></td><td></td><td></td><td></td><td></td></tr>
<tr><td>Chromosome</td><td>start</td><td>end</td><td>strand</td><td>MM</td><td>target_seq</td><td>PAM</td><td>alignment</td><td>distance</td><td>position</td><td>gene name</td><td>gene id</td></tr>
<tr><td>GU305914.1</td><td>150148</td><td>150170</td><td>-</td><td>0</td><td>CCTCATCCAACCGGCACCCT</td><td>CGG</td><td>||||||||[||||||||||||]PAM</td><td>NA</td><td>Intergenic</td><td>NA</td><td>NA</td></tr>
<tr><td></td><td></td><td></td><td></td><td></td><td></td><td></td><td></td><td></td><td></td><td></td><td></td></tr>
<tr><td>T22722</td><td>CTCATCCAACCGGCACCCTCGGG</td><td>1000</td><td></td><td></td><td></td><td></td><td></td><td></td><td></td><td></td><td></td></tr>
<tr><td>Oligo fwd</td><td>CACCCTCATCCAACCGGCACCCTC</td><td></td><td></td><td></td><td></td><td></td><td></td><td></td><td></td><td></td><td></td></tr>
<tr><td>Oligo rev</td><td>AAACGAGGGTGCCGGTTGGATGAG</td><td></td><td></td><td></td><td></td><td></td><td></td><td></td><td></td><td></td><td></td></tr>
<tr><td>Chromosome</td><td>start</td><td>end</td><td>strand</td><td>MM</td><td>target_seq</td><td>PAM</td><td>alignment</td><td>distance</td><td>position</td><td>gene name</td><td>gene id</td></tr>
<tr><td>GU305914.1</td><td>150147</td><td>150169</td><td>-</td><td>0</td><td>CTCATCCAACCGGCACCCTC</td><td>GGG</td><td>||||||||[||||||||||||]PAM</td><td>NA</td><td>Intergenic</td><td>NA</td><td>NA</td></tr>
<tr><td></td><td></td><td></td><td></td><td></td><td></td><td></td><td></td><td></td><td></td><td></td><td></td></tr>
<tr><td>T22723</td><td>CCAACCGGCACCCTCGGGTTCGG</td><td>1000</td><td></td><td></td><td></td><td></td><td></td><td></td><td></td><td></td><td></td></tr>
</table>

<table>
<tr><td>Oligo fwd</td><td>CACCCCAACCGGCACCCTCGGGTT</td><td></td><td></td><td></td><td></td><td></td><td></td><td></td><td></td><td></td><td></td></tr>
<tr><td>Oligo rev</td><td>AAACAACCCGAGGGTGCCGGTTGG</td><td></td><td></td><td></td><td></td><td></td><td></td><td></td><td></td><td></td><td></td></tr>
<tr><td>Chromosome</td><td>start</td><td>end</td><td>strand</td><td>MM</td><td>target_seq</td><td>PAM</td><td>alignment</td><td>distance</td><td>position</td><td>gene name</td><td>gene id</td></tr>
<tr><td>GU305914.1</td><td>150142</td><td>150164</td><td>-</td><td>0</td><td>CCAACCGGCACCCTCGGGTT</td><td>CGG</td><td>||||||||[||||||||||||]PAM</td><td>NA</td><td>Intergenic</td><td>NA</td><td>NA</td></tr>
<tr><td>T22724</td><td>CAACCGGCACCCTCGGGTTCGGG</td><td>1000</td><td></td><td></td><td></td><td></td><td></td><td></td><td></td><td></td><td></td></tr>
<tr><td>Oligo fwd</td><td>CACCCAACCGGCACCCTCGGGTTC</td><td></td><td></td><td></td><td></td><td></td><td></td><td></td><td></td><td></td><td></td></tr>
<tr><td>Oligo rev</td><td>AAACGAACCCGAGGGTGCCGGTTG</td><td></td><td></td><td></td><td></td><td></td><td></td><td></td><td></td><td></td><td></td></tr>
<tr><td>Chromosome</td><td>start</td><td>end</td><td>strand</td><td>MM</td><td>target_seq</td><td>PAM</td><td>alignment</td><td>distance</td><td>position</td><td>gene name</td><td>gene id</td></tr>
<tr><td>GU305914.1</td><td>150141</td><td>150163</td><td>-</td><td>0</td><td>CAACCGGCACCCTCGGGTTC</td><td>GGG</td><td>||||||||[||||||||||||]PAM</td><td>NA</td><td>Intergenic</td><td>NA</td><td>NA</td></tr>
<tr><td>T22725</td><td>ACCCTCGGGTTCGGGCGCGCAGG</td><td>1000</td><td></td><td></td><td></td><td></td><td></td><td></td><td></td><td></td><td></td></tr>
<tr><td>Oligo fwd</td><td>CACCACCCTCGGGTTCGGGCGCGC</td><td></td><td></td><td></td><td></td><td></td><td></td><td></td><td></td><td></td><td></td></tr>
<tr><td>Oligo rev</td><td>AAACGCGCGCCCGAACCCGAGGGT</td><td></td><td></td><td></td><td></td><td></td><td></td><td></td><td></td><td></td><td></td></tr>
<tr><td>Chromosome</td><td>start</td><td>end</td><td>strand</td><td>MM</td><td>target_seq</td><td>PAM</td><td>alignment</td><td>distance</td><td>position</td><td>gene name</td><td>gene id</td></tr>
<tr><td>GU305914.1</td><td>150133</td><td>150155</td><td>-</td><td>0</td><td>ACCCTCGGGTTCGGGCGCGC</td><td>AGG</td><td>||||||||[||||||||||||]PAM</td><td>NA</td><td>Intergenic</td><td>NA</td><td>NA</td></tr>
<tr><td>T22726</td><td>AGGACATCCGAGAGCTCTTTAGG</td><td>1000</td><td></td><td></td><td></td><td></td><td></td><td></td><td></td><td></td><td></td></tr>
<tr><td>Oligo fwd</td><td>CACCAGGACATCCGAGAGCTCTTT</td><td></td><td></td><td></td><td></td><td></td><td></td><td></td><td></td><td></td><td></td></tr>
<tr><td>Oligo rev</td><td>AAACAAAGAGCTCTCGGATGTCCT</td><td></td><td></td><td></td><td></td><td></td><td></td><td></td><td></td><td></td><td></td></tr>
<tr><td>Chromosome</td><td>start</td><td>end</td><td>strand</td><td>MM</td><td>target_seq</td><td>PAM</td><td>alignment</td><td>distance</td><td>position</td><td>gene name</td><td>gene id</td></tr>
<tr><td>GU305914.1</td><td>150113</td><td>150135</td><td>-</td><td>0</td><td>AGGACATCCGAGAGCTCTTT</td><td>AGG</td><td>||||||||[||||||||||||]PAM</td><td>NA</td><td>Intergenic</td><td>NA</td><td>NA</td></tr>
<tr><td>GU305914.1</td><td>75495</td><td>75517</td><td>-</td><td>4</td><td>GGGTCATCCGTGAGCTCTTC</td><td>GGG</td><td>-||-||||[||-||||||||-]PAM</td><td>NA</td><td>Intergenic</td><td>NA</td><td>NA</td></tr>
<tr><td>T22727</td><td>CCGAGAGCTCTTTAGGCGACAGG</td><td>1000</td><td></td><td></td><td></td><td></td><td></td><td></td><td></td><td></td><td></td></tr>
<tr><td>Oligo fwd</td><td>CACCCCGAGAGCTCTTTAGGCGAC</td><td></td><td></td><td></td><td></td><td></td><td></td><td></td><td></td><td></td><td></td></tr>
<tr><td>Oligo rev</td><td>AAACGTCGCCTAAAGAGCTCTCGG</td><td></td><td></td><td></td><td></td><td></td><td></td><td></td><td></td><td></td><td></td></tr>
<tr><td>Chromosome</td><td>start</td><td>end</td><td>strand</td><td>MM</td><td>target_seq</td><td>PAM</td><td>alignment</td><td>distance</td><td>position</td><td>gene name</td><td>gene id</td></tr>
<tr><td>GU305914.1</td><td>150106</td><td>150128</td><td>-</td><td>0</td><td>CCGAGAGCTCTTTAGGCGAC</td><td>AGG</td><td>||||||||[||||||||||||]PAM</td><td>NA</td><td>Intergenic</td><td>NA</td><td>NA</td></tr>
<tr><td>T22728</td><td>TCTTTAGGCGACAGGTCATCAGG</td><td>1000</td><td></td><td></td><td></td><td></td><td></td><td></td><td></td><td></td><td></td></tr>
<tr><td>Oligo fwd</td><td>CACCTCTTTAGGCGACAGGTCATC</td><td></td><td></td><td></td><td></td><td></td><td></td><td></td><td></td><td></td><td></td></tr>
<tr><td>Oligo rev</td><td>AAACGATGACCTGTCGCCTAAAGA</td><td></td><td></td><td></td><td></td><td></td><td></td><td></td><td></td><td></td><td></td></tr>
<tr><td>Chromosome</td><td>start</td><td>end</td><td>strand</td><td>MM</td><td>target_seq</td><td>PAM</td><td>alignment</td><td>distance</td><td>position</td><td>gene name</td><td>gene id</td></tr>
<tr><td>GU305914.1</td><td>150098</td><td>150120</td><td>-</td><td>0</td><td>TCTTTAGGCGACAGGTCATC</td><td>AGG</td><td>||||||||[||||||||||||]PAM</td><td>NA</td><td>Intergenic</td><td>NA</td><td>NA</td></tr>
<tr><td>T22729</td><td>GCGACAGGTCATCAGGTGCCTGG</td><td>1000</td><td></td><td></td><td></td><td></td><td></td><td></td><td></td><td></td><td></td></tr>
<tr><td>Oligo fwd</td><td>CACCGCGACAGGTCATCAGGTGCC</td><td></td><td></td><td></td><td></td><td></td><td></td><td></td><td></td><td></td><td></td></tr>
<tr><td>Oligo rev</td><td>AAACGGCACCTGATGACCTGTCGC</td><td></td><td></td><td></td><td></td><td></td><td></td><td></td><td></td><td></td><td></td></tr>
<tr><td>Chromosome</td><td>start</td><td>end</td><td>strand</td><td>MM</td><td>target_seq</td><td>PAM</td><td>alignment</td><td>distance</td><td>position</td><td>gene name</td><td>gene id</td></tr>
<tr><td>GU305914.1</td><td>150091</td><td>150113</td><td>-</td><td>0</td><td>GCGACAGGTCATCAGGTGCC</td><td>TGG</td><td>||||||||[||||||||||||]PAM</td><td>NA</td><td>Intergenic</td><td>NA</td><td>NA</td></tr>
<tr><td>T22730</td><td>ACAGGTCATCAGGTGCCTGGAGG</td><td>1000</td><td></td><td></td><td></td><td></td><td></td><td></td><td></td><td></td><td></td></tr>
<tr><td>Oligo fwd</td><td>CACCACAGGTCATCAGGTGCCTGG</td><td></td><td></td><td></td><td></td><td></td><td></td><td></td><td></td><td></td><td></td></tr>
<tr><td>Oligo rev</td><td>AAACCCAGGCACCTGATGACCTGT</td><td></td><td></td><td></td><td></td><td></td><td></td><td></td><td></td><td></td><td></td></tr>
<tr><td>Chromosome</td><td>start</td><td>end</td><td>strand</td><td>MM</td><td>target_seq</td><td>PAM</td><td>alignment</td><td>distance</td><td>position</td><td>gene name</td><td>gene id</td></tr>
<tr><td>GU305914.1</td><td>150088</td><td>150110</td><td>-</td><td>0</td><td>ACAGGTCATCAGGTGCCTGG</td><td>AGG</td><td>||||||||[||||||||||||]PAM</td><td>NA</td><td>Intergenic</td><td>NA</td><td>NA</td></tr>
<tr><td>T22731</td><td>CCAGATCCAAGACCATGTCGAGG</td><td>1000</td><td></td><td></td><td></td><td></td><td></td><td></td><td></td><td></td><td></td></tr>
<tr><td>Oligo fwd</td><td>CACCCCAGATCCAAGACCATGTCG</td><td></td><td></td><td></td><td></td><td></td><td></td><td></td><td></td><td></td><td></td></tr>
<tr><td>Oligo rev</td><td>AAACCGACATGGTCTTGGATCTGG</td><td></td><td></td><td></td><td></td><td></td><td></td><td></td><td></td><td></td><td></td></tr>
<tr><td>Chromosome</td><td>start</td><td>end</td><td>strand</td><td>MM</td><td>target_seq</td><td>PAM</td><td>alignment</td><td>distance</td><td>position</td><td>gene name</td><td>gene id</td></tr>
<tr><td>GU305914.1</td><td>150064</td><td>150086</td><td>-</td><td>0</td><td>CCAGATCCAAGACCATGTCG</td><td>AGG</td><td>||||||||[||||||||||||]PAM</td><td>NA</td><td>Intergenic</td><td>NA</td><td>NA</td></tr>
<tr><td>T22732</td><td>TGAAGCTGCTGAACAAGACGTGG</td><td>1000</td><td></td><td></td><td></td><td></td><td></td><td></td><td></td><td></td><td></td></tr>
<tr><td>Oligo fwd</td><td>CACCTGAAGCTGCTGAACAAGACG</td><td></td><td></td><td></td><td></td><td></td><td></td><td></td><td></td><td></td><td></td></tr>
<tr><td>Oligo rev</td><td>AAACCGTCTTGTTCAGCAGCTTCA</td><td></td><td></td><td></td><td></td><td></td><td></td><td></td><td></td><td></td><td></td></tr>
<tr><td>Chromosome</td><td>start</td><td>end</td><td>strand</td><td>MM</td><td>target_seq</td><td>PAM</td><td>alignment</td><td>distance</td><td>position</td><td>gene name</td><td>gene id</td></tr>
<tr><td>GU305914.1</td><td>150029</td><td>150051</td><td>-</td><td>0</td><td>TGAAGCTGCTGAACAAGACG</td><td>TGG</td><td>||||||||[||||||||||||]PAM</td><td>NA</td><td>Intergenic</td><td>NA</td><td>NA</td></tr>
<tr><td>T22733</td><td>GAAGCTGCTGAACAAGACGTGGG</td><td>1000</td><td></td><td></td><td></td><td></td><td></td><td></td><td></td><td></td><td></td></tr>
<tr><td>Oligo fwd</td><td>CACCGAAGCTGCTGAACAAGACGT</td><td></td><td></td><td></td><td></td><td></td><td></td><td></td><td></td><td></td><td></td></tr>
<tr><td>Oligo rev</td><td>AAACACGTCTTGTTCAGCAGCTTC</td><td></td><td></td><td></td><td></td><td></td><td></td><td></td><td></td><td></td><td></td></tr>
<tr><td>Chromosome</td><td>start</td><td>end</td><td>strand</td><td>MM</td><td>target_seq</td><td>PAM</td><td>alignment</td><td>distance</td><td>position</td><td>gene name</td><td>gene id</td></tr>
<tr><td>GU305914.1</td><td>150028</td><td>150050</td><td>-</td><td>0</td><td>GAAGCTGCTGAACAAGACGT</td><td>GGG</td><td>||||||||[||||||||||||]PAM</td><td>NA</td><td>Intergenic</td><td>NA</td><td>NA</td></tr>
<tr><td>T22734</td><td>GCGCGAGCTGCGCGACGCGCTGG</td><td>1000</td><td></td><td></td><td></td><td></td><td></td><td></td><td></td><td></td><td></td></tr>
<tr><td>Oligo fwd</td><td>CACCGCGCGAGCTGCGCGACGCGC</td><td></td><td></td><td></td><td></td><td></td><td></td><td></td><td></td><td></td><td></td></tr>
<tr><td>Oligo rev</td><td>AAACGCGCGTCGCGCAGCTCGCGC</td><td></td><td></td><td></td><td></td><td></td><td></td><td></td><td></td><td></td><td></td></tr>
<tr><td>Chromosome</td><td>start</td><td>end</td><td>strand</td><td>MM</td><td>target_seq</td><td>PAM</td><td>alignment</td><td>distance</td><td>position</td><td>gene name</td><td>gene id</td></tr>
<tr><td>GU305914.1</td><td>149995</td><td>150017</td><td>-</td><td>0</td><td>GCGCGAGCTGCGCGACGCGC</td><td>TGG</td><td>||||||||[||||||||||||]PAM</td><td>NA</td><td>Intergenic</td><td>NA</td><td>NA</td></tr>
<tr><td>T22735</td><td>GCTGGACCGCTATGAGTCCGAGG</td><td>1000</td><td></td><td></td><td></td><td></td><td></td><td></td><td></td><td></td><td></td></tr>
<tr><td>Oligo fwd</td><td>CACCGCTGGACCGCTATGAGTCCG</td><td></td><td></td><td></td><td></td><td></td><td></td><td></td><td></td><td></td><td></td></tr>
<tr><td>Oligo rev</td><td>AAACCGGACTCATAGCGGTCCAGC</td><td></td><td></td><td></td><td></td><td></td><td></td><td></td><td></td><td></td><td></td></tr>
<tr><td>Chromosome</td><td>start</td><td>end</td><td>strand</td><td>MM</td><td>target_seq</td><td>PAM</td><td>alignment</td><td>distance</td><td>position</td><td>gene name</td><td>gene id</td></tr>
<tr><td>GU305914.1</td><td>149977</td><td>149999</td><td>-</td><td>0</td><td>GCTGGACCGCTATGAGTCCG</td><td>AGG</td><td>||||||||[||||||||||||]PAM</td><td>NA</td><td>Intergenic</td><td>NA</td><td>NA</td></tr>
<tr><td>T22736</td><td>CTGGACCGCTATGAGTCCGAGGG</td><td>1000</td><td></td><td></td><td></td><td></td><td></td><td></td><td></td><td></td><td></td></tr>
<tr><td>Oligo fwd</td><td>CACCCTGGACCGCTATGAGTCCGA</td><td></td><td></td><td></td><td></td><td></td><td></td><td></td><td></td><td></td><td></td></tr>
<tr><td>Oligo rev</td><td>AAACTCGGACTCATAGCGGTCCAG</td><td></td><td></td><td></td><td></td><td></td><td></td><td></td><td></td><td></td><td></td></tr>
<tr><td>Chromosome</td><td>start</td><td>end</td><td>strand</td><td>MM</td><td>target_seq</td><td>PAM</td><td>alignment</td><td>distance</td><td>position</td><td>gene name</td><td>gene id</td></tr>
<tr><td>GU305914.1</td><td>149976</td><td>149998</td><td>-</td><td>0</td><td>CTGGACCGCTATGAGTCCGA</td><td>GGG</td><td>||||||||[||||||||||||]PAM</td><td>NA</td><td>Intergenic</td><td>NA</td><td>NA</td></tr>
<tr><td>T22737</td><td>ACCGCTATGAGTCCGAGGGCCGG</td><td>1000</td><td></td><td></td><td></td><td></td><td></td><td></td><td></td><td></td><td></td></tr>
<tr><td>Oligo fwd</td><td>CACCACCGCTATGAGTCCGAGGGC</td><td></td><td></td><td></td><td></td><td></td><td></td><td></td><td></td><td></td><td></td></tr>
<tr><td>Oligo rev</td><td>AAACGCCCTCGGACTCATAGCGGT</td><td></td><td></td><td></td><td></td><td></td><td></td><td></td><td></td><td></td><td></td></tr>
<tr><td>Chromosome</td><td>start</td><td>end</td><td>strand</td><td>MM</td><td>target_seq</td><td>PAM</td><td>alignment</td><td>distance</td><td>position</td><td>gene name</td><td>gene id</td></tr>
<tr><td>GU305914.1</td><td>149972</td><td>149994</td><td>-</td><td>0</td><td>ACCGCTATGAGTCCGAGGGC</td><td>CGG</td><td>||||||||[||||||||||||]PAM</td><td>NA</td><td>Intergenic</td><td>NA</td><td>NA</td></tr>
</table>

| | | | | | | | | | | | |
|---|---|---|---|---|---|---|---|---|---|---|---|
| T22738 | TATGAGTCCGAGGGCCGGCGCGG | 1000 | | | | | | | | | |
| Oligo fwd | CACCTATGAGTCCGAGGGCCGGCG | | | | | | | | | | |
| Oligo rev | AAACCGCCGGCCCTCGGACTCATA | | | | | | | | | | |
| Chromosome | start | end | strand | MM | target_seq | PAM | alignment | distance | position | gene name | gene id |
| GU305914.1 | 149967 | 149989 | - | 0 | TATGAGTCCGAGGGCCGGCG | CGG | \|\|\|\|\|\|\|\|[\|\|\|\|\|\|\|\|\|\|\|\|]PAM | NA | Intergenic | NA | NA |
| T22739 | TCCGAGGGCCGGCGCGGCCGCGG | 1000 | | | | | | | | | |
| Oligo fwd | CACCTCCGAGGGCCGGCGCGGCCG | | | | | | | | | | |
| Oligo rev | AAACCGGCCGCGCCGGCCCTCGGA | | | | | | | | | | |
| Chromosome | start | end | strand | MM | target_seq | PAM | alignment | distance | position | gene name | gene id |
| GU305914.1 | 149961 | 149983 | - | 0 | TCCGAGGGCCGGCGCGGCCG | CGG | \|\|\|\|\|\|\|\|[\|\|\|\|\|\|\|\|\|\|\|\|]PAM | NA | Intergenic | NA | NA |
| T22740 | CCGGCGCGGCCGCGGCCCCCGG | 1000 | | | | | | | | | |
| Oligo fwd | CACCCCGGCGCGGCCGCGGCCCCC | | | | | | | | | | |
| Oligo rev | AAACGGGGGCCGCGGCCGCGCCGG | | | | | | | | | | |
| Chromosome | start | end | strand | MM | target_seq | PAM | alignment | distance | position | gene name | gene id |
| GU305914.1 | 149953 | 149975 | - | 0 | CCGGCGCGGCCGCGGCCCCC | CGG | \|\|\|\|\|\|\|\|[\|\|\|\|\|\|\|\|\|\|\|\|]PAM | NA | Intergenic | NA | NA |
| GU305914.1 | 55167 | 55189 | - | 4 | TCGGCGCCGCCGCGGGCGCC | AGG | -\|\|\|\|\|\|-[\|\|\|\|\|\|\|-\|-\|\|]PAM | NA | Intergenic | NA | NA |
| T22741 | GGCGTTCGACCTGCAGACCCTGG | 1000 | | | | | | | | | |
| Oligo fwd | CACCGGCGTTCGACCTGCAGACCC | | | | | | | | | | |
| Oligo rev | AAACGGGTCTGCAGGTCGAACGCC | | | | | | | | | | |
| Chromosome | start | end | strand | MM | target_seq | PAM | alignment | distance | position | gene name | gene id |
| GU305914.1 | 149932 | 149954 | - | 0 | GGCGTTCGACCTGCAGACCC | TGG | \|\|\|\|\|\|\|\|[\|\|\|\|\|\|\|\|\|\|\|\|]PAM | NA | Intergenic | NA | NA |
| T22742 | CGACCTGCAGACCCTGGACACGG | 1000 | | | | | | | | | |
| Oligo fwd | CACCCGACCTGCAGACCCTGGACA | | | | | | | | | | |
| Oligo rev | AAACTGTCCAGGGTCTGCAGGTCG | | | | | | | | | | |
| Chromosome | start | end | strand | MM | target_seq | PAM | alignment | distance | position | gene name | gene id |
| GU305914.1 | 149926 | 149948 | - | 0 | CGACCTGCAGACCCTGGACA | CGG | \|\|\|\|\|\|\|\|[\|\|\|\|\|\|\|\|\|\|\|\|]PAM | NA | Intergenic | NA | NA |
| T22743 | TGGACACGGTCAACGCGCTGCGG | 1000 | | | | | | | | | |
| Oligo fwd | CACCTGGACACGGTCAACGCGCTG | | | | | | | | | | |
| Oligo rev | AAACCAGCGCGTTGACCGTGTCCA | | | | | | | | | | |
| Chromosome | start | end | strand | MM | target_seq | PAM | alignment | distance | position | gene name | gene id |
| GU305914.1 | 149912 | 149934 | - | 0 | TGGACACGGTCAACGCGCTG | CGG | \|\|\|\|\|\|\|\|[\|\|\|\|\|\|\|\|\|\|\|\|]PAM | NA | Intergenic | NA | NA |
| T22744 | CAACGCGCTGCGGCGCGTGCAGG | 1000 | | | | | | | | | |
| Oligo fwd | CACCCAACGCGCTGCGGCGCGTGC | | | | | | | | | | |
| Oligo rev | AAACGCACGCGCCGCAGCGCGTTG | | | | | | | | | | |
| Chromosome | start | end | strand | MM | target_seq | PAM | alignment | distance | position | gene name | gene id |
| GU305914.1 | 149902 | 149924 | - | 0 | CAACGCGCTGCGGCGCGTGC | AGG | \|\|\|\|\|\|\|\|[\|\|\|\|\|\|\|\|\|\|\|\|]PAM | NA | Intergenic | NA | NA |
| T22745 | AACGCGCTGCGGCGCGTGCAGGG | 1000 | | | | | | | | | |
| Oligo fwd | CACCAACGCGCTGCGGCGCGTGCA | | | | | | | | | | |
| Oligo rev | AAACTGCACGCGCCGCAGCGCGTT | | | | | | | | | | |
| Chromosome | start | end | strand | MM | target_seq | PAM | alignment | distance | position | gene name | gene id |
| GU305914.1 | 149901 | 149923 | - | 0 | AACGCGCTGCGGCGCGTGCA | GGG | \|\|\|\|\|\|\|\|[\|\|\|\|\|\|\|\|\|\|\|\|]PAM | NA | Intergenic | NA | NA |
| T22746 | GCGCGTGCAGGGCCTGCCGACGG | 1000 | | | | | | | | | |
| Oligo fwd | CACCGCGCGTGCAGGGCCTGCCGA | | | | | | | | | | |
| Oligo rev | AAACTCGGCAGGCCCTGCACGCGC | | | | | | | | | | |
| Chromosome | start | end | strand | MM | target_seq | PAM | alignment | distance | position | gene name | gene id |
| GU305914.1 | 149890 | 149912 | - | 0 | GCGCGTGCAGGGCCTGCCGA | CGG | \|\|\|\|\|\|\|\|[\|\|\|\|\|\|\|\|\|\|\|\|]PAM | NA | Intergenic | NA | NA |
| T22747 | GCAGGGCCTGCCGACGGCGCCGG | 1000 | | | | | | | | | |
| Oligo fwd | CACCGCAGGGCCTGCCGACGGCGC | | | | | | | | | | |
| Oligo rev | AAACGCGCCGTCGGCAGGCCCTGC | | | | | | | | | | |
| Chromosome | start | end | strand | MM | target_seq | PAM | alignment | distance | position | gene name | gene id |
| GU305914.1 | 149884 | 149906 | - | 0 | GCAGGGCCTGCCGACGGCGC | CGG | \|\|\|\|\|\|\|\|[\|\|\|\|\|\|\|\|\|\|\|\|]PAM | NA | Intergenic | NA | NA |
| T22748 | CCTGCCGACGGCGCCGGTGACGG | 1000 | | | | | | | | | |
| Oligo fwd | CACCCCTGCCGACGGCGCCGGTGA | | | | | | | | | | |
| Oligo rev | AAACTCACCGGCGCCGTCGGCAGG | | | | | | | | | | |
| Chromosome | start | end | strand | MM | target_seq | PAM | alignment | distance | position | gene name | gene id |
| GU305914.1 | 149878 | 149900 | - | 0 | CCTGCCGACGGCGCCGGTGA | CGG | \|\|\|\|\|\|\|\|[\|\|\|\|\|\|\|\|\|\|\|\|]PAM | NA | Intergenic | NA | NA |
| T22749 | GCCGACGGCGCCGGTGACGGTGG | 1000 | | | | | | | | | |
| Oligo fwd | CACCGCCGACGGCGCCGGTGACGG | | | | | | | | | | |
| Oligo rev | AAACCCGTCACCGGCGCCGTCGGC | | | | | | | | | | |
| Chromosome | start | end | strand | MM | target_seq | PAM | alignment | distance | position | gene name | gene id |
| GU305914.1 | 149875 | 149897 | - | 0 | GCCGACGGCGCCGGTGACGG | TGG | \|\|\|\|\|\|\|\|[\|\|\|\|\|\|\|\|\|\|\|\|]PAM | NA | Intergenic | NA | NA |
| GU305914.1 | 31155 | 31177 | - | 4 | GGAGACGGAGACGGTGACGG | TGG | \|--\|\|\|\|\|[-\|-\|\|\|\|\|\|\|\|\|]PAM | NA | Intergenic | NA | NA |
| GU305914.1 | 163749 | 163771 | + | 3 | GTCGGCGGCGCCGGTGGCGG | CGG | \|-\|\|-\|\|\|[\|\|\|\|\|\|\|\|-\|\|\|]PAM | NA | Intergenic | NA | NA |
| T22750 | CGGTGACGGTGGACGACAACCGG | 1000 | | | | | | | | | |
| Oligo fwd | CACCCGGTGACGGTGGACGACAAC | | | | | | | | | | |
| Oligo rev | AAACGTTGTCGTCCACCGTCACCG | | | | | | | | | | |
| Chromosome | start | end | strand | MM | target_seq | PAM | alignment | distance | position | gene name | gene id |
| GU305914.1 | 149864 | 149886 | - | 0 | CGGTGACGGTGGACGACAAC | CGG | \|\|\|\|\|\|\|\|[\|\|\|\|\|\|\|\|\|\|\|\|]PAM | NA | Intergenic | NA | NA |
| T22751 | GGTGACGGTGGACGACAACCGGG | 1000 | | | | | | | | | |
| Oligo fwd | CACCGGTGACGGTGGACGACAACC | | | | | | | | | | |
| Oligo rev | AAACGGTTGTCGTCCACCGTCACC | | | | | | | | | | |
| Chromosome | start | end | strand | MM | target_seq | PAM | alignment | distance | position | gene name | gene id |
| GU305914.1 | 149863 | 149885 | - | 0 | GGTGACGGTGGACGACAACC | GGG | \|\|\|\|\|\|\|\|[\|\|\|\|\|\|\|\|\|\|\|\|]PAM | NA | Intergenic | NA | NA |
| T22752 | CGTACCCGACGAGCGCGAGTCGG | 1000 | | | | | | | | | |

Oligo fwd CACCCGTACCCGACGAGCGCGAGT
Oligo rev AAACACTCGCGCTCGTCGGGTACG
Chromosome start end strand MM target_seq PAM alignment distance position gene name gene id
GU305914.1 149812 149834 - 0 CGTACCCGACGAGCGCGAGT CGG ||||||||[||||||||||||]PAM NA Intergenic NA NA

T22753 ACGAGCGCCTGTCGCGTCTCTGG 1000
Oligo fwd CACCACGAGCGCCTGTCGCGTCTC
Oligo rev AAACGAGACGCGACAGGCGCTCGT
Chromosome start end strand MM target_seq PAM alignment distance position gene name gene id
GU305914.1 149789 149811 - 0 ACGAGCGCCTGTCGCGTCTC TGG ||||||||[||||||||||||]PAM NA Intergenic NA NA

T22754 CGAGCGCCTGTCGCGTCTCTGGG 1000
Oligo fwd CACCCGAGCGCCTGTCGCGTCTCT
Oligo rev AAACAGAGACGCGACAGGCGCTCG
Chromosome start end strand MM target_seq PAM alignment distance position gene name gene id
GU305914.1 149788 149810 - 0 CGAGCGCCTGTCGCGTCTCT GGG ||||||||[||||||||||||]PAM NA Intergenic NA NA

T22755 TTCGATGTTTCAAGTTCCGACGG 1000
Oligo fwd CACCTTCGATGTTTCAAGTTCCGA
Oligo rev AAACTCGGAACTTGAAACATCGAA
Chromosome start end strand MM target_seq PAM alignment distance position gene name gene id
GU305914.1 149753 149775 - 0 TTCGATGTTTCAAGTTCCGA CGG ||||||||[||||||||||||]PAM NA Intergenic NA NA

T22756 CCGACGGAACGTGACGAACCAGG 1000
Oligo fwd CACCCCGACGGAACGTGACGAACC
Oligo rev AAACGGTTCGTCACGTTCCGTCGG
Chromosome start end strand MM target_seq PAM alignment distance position gene name gene id
GU305914.1 149737 149759 - 0 CCGACGGAACGTGACGAACC AGG ||||||||[||||||||||||]PAM NA Intergenic NA NA

T22757 CGACGGAACGTGACGAACCAGGG 1000
Oligo fwd CACCCGACGGAACGTGACGAACCA
Oligo rev AAACTGGTTCGTCACGTTCCGTCG
Chromosome start end strand MM target_seq PAM alignment distance position gene name gene id
GU305914.1 149736 149758 - 0 CGACGGAACGTGACGAACCA GGG ||||||||[||||||||||||]PAM NA Intergenic NA NA

T22758 CGTGACGAACCAGGGCGCCGAGG 1000
Oligo fwd CACCCGTGACGAACCAGGGCGCCG
Oligo rev AAACCGGCGCCCTGGTTCGTCACG
Chromosome start end strand MM target_seq PAM alignment distance position gene name gene id
GU305914.1 149728 149750 - 0 CGTGACGAACCAGGGCGCCG AGG ||||||||[||||||||||||]PAM NA Intergenic NA NA

T22759 CGTGCTGCTCCCGTTCCTCACGG 1000
Oligo fwd CACCCGTGCTGCTCCCGTTCCTCA
Oligo rev AAACTGAGGAACGGGAGCAGCACG
Chromosome start end strand MM target_seq PAM alignment distance position gene name gene id
GU305914.1 149671 149693 - 0 CGTGCTGCTCCCGTTCCTCA CGG ||||||||[||||||||||||]PAM NA Intergenic NA NA

T22760 GCTGCTCCCGTTCCTCACGGCGG 1000
Oligo fwd CACCGCTGCTCCCGTTCCTCACGG
Oligo rev AAACCCGTGAGGAACGGGAGCAGC
Chromosome start end strand MM target_seq PAM alignment distance position gene name gene id
GU305914.1 149668 149690 - 0 GCTGCTCCCGTTCCTCACGG CGG ||||||||[||||||||||||]PAM NA Intergenic NA NA

T22761 GGTCATCGAGTTCCCCATGTTGG 1000
Oligo fwd CACCGGTCATCGAGTTCCCCATGT
Oligo rev AAACACATGGGGAACTCGATGACC
Chromosome start end strand MM target_seq PAM alignment distance position gene name gene id
GU305914.1 149647 149669 - 0 GGTCATCGAGTTCCCCATGT TGG ||||||||[||||||||||||]PAM NA Intergenic NA NA

T22762 ATTTCTGTCGCTCAGCGAGTTGG 1000
Oligo fwd CACCATTTCTGTCGCTCAGCGAGT
Oligo rev AAACACTCGCTGAGCGACAGAAAT
Chromosome start end strand MM target_seq PAM alignment distance position gene name gene id
GU305914.1 149602 149624 - 0 ATTTCTGTCGCTCAGCGAGT TGG ||||||||[||||||||||||]PAM NA Intergenic NA NA

T22763 GCGTCCAAGCCTTTTTAGAGAGG 1000
Oligo fwd CACCGCGTCCAAGCCTTTTTAGAG
Oligo rev AAACCTCTAAAAAGGCTTGGACGC
Chromosome start end strand MM target_seq PAM alignment distance position gene name gene id
GU305914.1 149510 149532 - 0 GCGTCCAAGCCTTTTTAGAG AGG ||||||||[||||||||||||]PAM NA Intergenic NA NA

T22764 CGTCCAAGCCTTTTTAGAGAGGG 1000
Oligo fwd CACCCGTCCAAGCCTTTTTAGAGA
Oligo rev AAACTCTCTAAAAAGGCTTGGACG
Chromosome start end strand MM target_seq PAM alignment distance position gene name gene id
GU305914.1 149509 149531 - 0 CGTCCAAGCCTTTTTAGAGA GGG ||||||||[||||||||||||]PAM NA Intergenic NA NA

T22765 AGCCTTTTTAGAGAGGGAGCAGG 1000
Oligo fwd CACCAGCCTTTTTAGAGAGGGAGC
Oligo rev AAACGCTCCCTCTCTAAAAAGGCT
Chromosome start end strand MM target_seq PAM alignment distance position gene name gene id
GU305914.1 149503 149525 - 0 AGCCTTTTTAGAGAGGGAGC AGG ||||||||[||||||||||||]PAM NA Intergenic NA NA

T22766 TTTAGAGAGGGAGCAGGCGACGG 1000
Oligo fwd CACCTTTAGAGAGGGAGCAGGCGA
Oligo rev AAACTCGCCTGCTCCCTCTCTAAA
Chromosome start end strand MM target_seq PAM alignment distance position gene name gene id
GU305914.1 149497 149519 - 0 TTTAGAGAGGGAGCAGGCGA CGG ||||||||[||||||||||||]PAM NA Intergenic NA NA

T22767 AGAGAGGGAGCAGGCGACGGCGG 1000
Oligo fwd CACCAGAGAGGGAGCAGGCGACGG
Oligo rev AAACCCGTCGCCTGCTCCCTCTCT

| Chromosome | start | end | strand | MM | target_seq | PAM | alignment | distance | position | gene name | gene id |
|---|---|---|---|---|---|---|---|---|---|---|---|
| GU305914.1 | 149494 | 149516 | - | 0 | AGAGAGGGAGCAGGCGACGG | CGG | ||||||||[||||||||||||]PAM | NA | Intergenic | NA | NA |

T22768 GAGGGAGCAGGCGACGGCGGCGG 1000
Oligo fwd CACCGAGGGAGCAGGCGACGGCGG
Oligo rev AAACCCGCCGTCGCCTGCTCCCTC

| Chromosome | start | end | strand | MM | target_seq | PAM | alignment | distance | position | gene name | gene id |
|---|---|---|---|---|---|---|---|---|---|---|---|
| GU305914.1 | 149491 | 149513 | - | 0 | GAGGGAGCAGGCGACGGCGG | CGG | ||||||||[||||||||||||]PAM | NA | Intergenic | NA | NA |
| GU305914.1 | 81246 | 81268 | - | 4 | GATCGAGGAGGCGGCGGCGG | GGG | ||--|||-[|||||-||||||]PAM | NA | Intergenic | NA | NA |
| GU305914.1 | 149289 | 149311 | + | 4 | TCGGGAGCGGGCGACGACGG | CGG | --||||||[-|||||||-|||]PAM | NA | Intergenic | NA | NA |

T22769 GGAGCAGGCGACGGCGGCGGCGG 1000
Oligo fwd CACCGGAGCAGGCGACGGCGGCGG
Oligo rev AAACCCGCCGCCGTCGCCTGCTCC

| Chromosome | start | end | strand | MM | target_seq | PAM | alignment | distance | position | gene name | gene id |
|---|---|---|---|---|---|---|---|---|---|---|---|
| GU305914.1 | 149488 | 149510 | - | 0 | GGAGCAGGCGACGGCGGCGG | CGG | ||||||||[||||||||||||]PAM | NA | Intergenic | NA | NA |
| GU305914.1 | 176454 | 176476 | + | 4 | GACGATGGCGACGGCGGCGG | AGG | |--|--||[||||||||||||]PAM | NA | Intergenic | NA | NA |
| GU305914.1 | 30000 | 30022 | + | 3 | GGTGCTGGCGGCGGCGGCGG | CGG | ||-||-||[||-|||||||||]PAM | NA | Intergenic | NA | NA |
| GU305914.1 | 154404 | 154426 | + | 4 | GGCGGTGGCGGCGGCGGCGG | CGG | ||-|--||[||-|||||||||]PAM | NA | Intergenic | NA | NA |
| GU305914.1 | 154407 | 154429 | + | 4 | GGTGGCGGCGGCGGCGGCGG | CGG | ||-|--||[||-|||||||||]PAM | NA | Intergenic | NA | NA |
| GU305914.1 | 58123 | 58145 | - | 4 | GGTGGCGGCGGCGGCGGCGG | AGG | ||-|--||[||-|||||||||]PAM | NA | Intergenic | NA | NA |
| GU305914.1 | 149482 | 149504 | - | 4 | GGCGACGGCGGCGGCGGCGG | CGG | ||-|--||[||-|||||||||]PAM | NA | Intergenic | NA | NA |
| GU305914.1 | 58126 | 58148 | - | 4 | GGCGGTGGCGGCGGCGGCGG | CGG | ||-|--||[||-|||||||||]PAM | NA | Intergenic | NA | NA |
| GU305914.1 | 149292 | 149314 | + | 2 | GGAGCGGGCGACGACGGCGG | CGG | |||||-||[|||||-||||||]PAM | NA | Intergenic | NA | NA |
| GU305914.1 | 163896 | 163918 | + | 4 | GGATCGAGCGGCGGCGGCGG | CGG | |||-|--|[||-|||||||||]PAM | NA | Intergenic | NA | NA |
| GU305914.1 | 201344 | 201366 | - | 4 | GGATCAGTCTCCGGCGGCGG | TGG | |||-|||-[|--|||||||||]PAM | NA | Intergenic | NA | NA |
| GU305914.1 | 90262 | 90284 | - | 4 | GAGGCGGGCGACGGCGGAGG | CGG | |--||-||[|||||||||-||]PAM | NA | Intergenic | NA | NA |

T22770 GCAGGCGACGGCGGCGGCGGCGG 1000
Oligo fwd CACCGCAGGCGACGGCGGCGGCGG
Oligo rev AAACCCGCCGCCGCCGTCGCCTGC

| Chromosome | start | end | strand | MM | target_seq | PAM | alignment | distance | position | gene name | gene id |
|---|---|---|---|---|---|---|---|---|---|---|---|
| GU305914.1 | 149485 | 149507 | - | 0 | GCAGGCGACGGCGGCGGCGG | CGG | ||||||||[||||||||||||]PAM | NA | Intergenic | NA | NA |
| GU305914.1 | 154407 | 154429 | + | 3 | GGTGGCGGCGGCGGCGGCGG | CGG | |--||||-[||||||||||||]PAM | NA | Intergenic | NA | NA |
| GU305914.1 | 58123 | 58145 | - | 3 | GGTGGCGGCGGCGGCGGCGG | AGG | |--||||-[||||||||||||]PAM | NA | Intergenic | NA | NA |
| GU305914.1 | 67742 | 67764 | + | 4 | GGGAGCGGCGGCGGCGGCGG | CGG | |---|||-[||||||||||||]PAM | NA | Intergenic | NA | NA |
| GU305914.1 | 149482 | 149504 | - | 4 | GGCGACGGCGGCGGCGGCGG | CGG | |--|-||-[||||||||||||]PAM | NA | Intergenic | NA | NA |
| GU305914.1 | 154404 | 154426 | + | 4 | GGCGGTGGCGGCGGCGGCGG | CGG | |--||-|-[||||||||||||]PAM | NA | Intergenic | NA | NA |
| GU305914.1 | 58126 | 58148 | - | 4 | GGCGGTGGCGGCGGCGGCGG | CGG | |--||-|-[||||||||||||]PAM | NA | Intergenic | NA | NA |
| GU305914.1 | 58129 | 58151 | - | 3 | GCGGGCGGTGGCGGCGGCGG | CGG | ||-||||-[-|||||||||||]PAM | NA | Intergenic | NA | NA |
| GU305914.1 | 154401 | 154423 | + | 4 | CCGGGCGGTGGCGGCGGCGG | CGG | -|-||||-[-|||||||||||]PAM | NA | Intergenic | NA | NA |
| GU305914.1 | 48159 | 48181 | + | 4 | GGCGGCGGTGGCGGCGGCGG | TGG | |--||||-[-|||||||||||]PAM | NA | Intergenic | NA | NA |
| GU305914.1 | 48156 | 48178 | + | 4 | GGCGGCGGCGGTGGCGGCGG | CGG | |--||||-[|||-||||||||]PAM | NA | Intergenic | NA | NA |
| GU305914.1 | 90259 | 90281 | - | 2 | GCGGGCGACGGCGGAGGCGG | CGG | ||-|||||[||||||-|||||]PAM | NA | Intergenic | NA | NA |
| GU305914.1 | 110125 | 110147 | - | 3 | GGACGCGACGGCGGCCGCGG | CGG | |-|-||||[|||||||-||||]PAM | NA | Intergenic | NA | NA |
| GU305914.1 | 62672 | 62694 | - | 4 | GCTGCCGCCGGCGGGGGCGG | CGG | ||-|-||-[||||||-|||||]PAM | NA | Intergenic | NA | NA |
| GU305914.1 | 30006 | 30028 | + | 4 | GGCGGCGGCGGCGGCGGCAG | TGG | |--||||-[||||||||||-|]PAM | NA | Intergenic | NA | NA |
| GU305914.1 | 149476 | 149498 | - | 4 | GGCGGCGGCGGCGGCGGCAG | CGG | |--||||-[||||||||||-|]PAM | NA | Intergenic | NA | NA |
| GU305914.1 | 113924 | 113946 | - | 4 | GTCGGCGTCGGCGGCGGCAG | CGG | |--||||-[||||||||||-|]PAM | NA | Intergenic | NA | NA |
| GU305914.1 | 128981 | 129003 | + | 4 | GCAGCCGTCGGCAGCGGCAG | CGG | ||||-||-[||||-|||||-|]PAM | NA | Intergenic | NA | NA |

T22771 GGCGACGGCGGCGGCGGCGGCGG 1000
Oligo fwd CACCGGCGACGGCGGCGGCGGCGG
Oligo rev AAACCCGCCGCCGCCGCCGTCGCC

| Chromosome | start | end | strand | MM | target_seq | PAM | alignment | distance | position | gene name | gene id |
|---|---|---|---|---|---|---|---|---|---|---|---|
| GU305914.1 | 149482 | 149504 | - | 0 | GGCGACGGCGGCGGCGGCGG | CGG | ||||||||[||||||||||||]PAM | NA | Intergenic | NA | NA |
| GU305914.1 | 154407 | 154429 | + | 2 | GGTGGCGGCGGCGGCGGCGG | CGG | ||-|-|||[||||||||||||]PAM | NA | Intergenic | NA | NA |
| GU305914.1 | 58123 | 58145 | - | 2 | GGTGGCGGCGGCGGCGGCGG | AGG | ||-|-|||[||||||||||||]PAM | NA | Intergenic | NA | NA |
| GU305914.1 | 35653 | 35675 | + | 3 | ATCCACGGCGGCGGCGGCGG | CGG | --|-||||[||||||||||||]PAM | NA | Intergenic | NA | NA |
| GU305914.1 | 154404 | 154426 | + | 2 | GGCGGTGGCGGCGGCGGCGG | CGG | ||||--||[||||||||||||]PAM | NA | Intergenic | NA | NA |
| GU305914.1 | 58126 | 58148 | - | 2 | GGCGGTGGCGGCGGCGGCGG | CGG | ||||--||[||||||||||||]PAM | NA | Intergenic | NA | NA |
| GU305914.1 | 67742 | 67764 | + | 3 | GGGAGCGGCGGCGGCGGCGG | CGG | ||---|||[||||||||||||]PAM | NA | Intergenic | NA | NA |
| GU305914.1 | 30000 | 30022 | + | 3 | GGTGCTGGCGGCGGCGGCGG | CGG | ||-|--||[||||||||||||]PAM | NA | Intergenic | NA | NA |
| GU305914.1 | 48159 | 48181 | + | 2 | GGCGGCGGTGGCGGCGGCGG | TGG | ||||-|||[-|||||||||||]PAM | NA | Intergenic | NA | NA |
| GU305914.1 | 56773 | 56795 | + | 4 | CGCCGCAGCGGCGGCGGCGG | CGG | -||--|-|[||||||||||||]PAM | NA | Intergenic | NA | NA |
| GU305914.1 | 149485 | 149507 | - | 4 | GCAGGCGACGGCGGCGGCGG | CGG | |--|-||-[||||||||||||]PAM | NA | Intergenic | NA | NA |
| GU305914.1 | 58129 | 58151 | - | 4 | GCGGGCGGTGGCGGCGGCGG | CGG | |--|-|||[-|||||||||||]PAM | NA | Intergenic | NA | NA |
| GU305914.1 | 24627 | 24649 | - | 4 | CGCAACACCGGCGGCGGCGG | TGG | -||-||--[||||||||||||]PAM | NA | Intergenic | NA | NA |
| GU305914.1 | 48156 | 48178 | + | 2 | GGCGGCGGCGGTGGCGGCGG | CGG | ||||-|||[|||-||||||||]PAM | NA | Intergenic | NA | NA |
| GU305914.1 | 62431 | 62453 | + | 3 | TTCGACGGCGGTGGCGGCGG | GGG | --||||||[|||-||||||||]PAM | NA | Intergenic | NA | NA |
| GU305914.1 | 176454 | 176476 | + | 3 | GACGATGGCGACGGCGGCGG | AGG | |-|||-||[||-|||||||||]PAM | NA | Intergenic | NA | NA |
| GU305914.1 | 73792 | 73814 | - | 3 | GGCGCCAGCAGCGGCGGCGG | AGG | ||||-|-|[|-||||||||||]PAM | NA | Intergenic | NA | NA |
| GU305914.1 | 161754 | 161776 | - | 4 | GGCCGAGGAGGCGGCGGCGG | AGG | |||---||[-|||||||||||]PAM | NA | Intergenic | NA | NA |
| GU305914.1 | 168820 | 168842 | + | 4 | GGTGGTGGCAGCGGCGGCGG | CGG | ||-|--||[|-||||||||||]PAM | NA | Intergenic | NA | NA |
| GU305914.1 | 149488 | 149510 | - | 4 | GGAGCAGGCGACGGCGGCGG | CGG | ||-|--||[||-|||||||||]PAM | NA | Intergenic | NA | NA |
| GU305914.1 | 62446 | 62468 | + | 4 | GGCGGGGGTGGTGGCGGCGG | GGG | ||||--||[-||-||||||||]PAM | NA | Intergenic | NA | NA |
| GU305914.1 | 114658 | 114680 | - | 4 | GGCGGTGGTGGTGGCGGCGG | CGG | ||||--||[-||-||||||||]PAM | NA | Intergenic | NA | NA |
| GU305914.1 | 48153 | 48175 | + | 4 | TGTGGCGGCGGCGGTGGCGG | CGG | -|-|-|||[||||||-|||||]PAM | NA | Intergenic | NA | NA |
| GU305914.1 | 168817 | 168839 | + | 4 | GGCGGTGGTGGCAGCGGCGG | CGG | ||||--||[-|||-|||||||]PAM | NA | Intergenic | NA | NA |
| GU305914.1 | 62155 | 62177 | + | 2 | GGCGACGGCAGCGGCAGCGG | CGG | ||||||||[|-|||||-||||]PAM | NA | Intergenic | NA | NA |
| GU305914.1 | 163749 | 163771 | + | 4 | GTCGGCGGCGCCGGTGGCGG | CGG | |-||-|||[||-|||-|||||]PAM | NA | Intergenic | NA | NA |
| GU305914.1 | 163911 | 163933 | + | 3 | GGCGGCGGCAGCGGCAGCGG | AGG | ||||-|||[|-|||||-||||]PAM | NA | Intergenic | NA | NA |
| GU305914.1 | 45255 | 45277 | - | 4 | GGTGGCGGTGGCGGCAGCGG | CGG | ||-|-|||[-||||||-||||]PAM | NA | Intergenic | NA | NA |
| GU305914.1 | 30006 | 30028 | + | 2 | GGCGGCGGCGGCGGCGGCAG | TGG | ||||-|||[||||||||||-|]PAM | NA | Intergenic | NA | NA |
| GU305914.1 | 149476 | 149498 | - | 2 | GGCGGCGGCGGCGGCGGCAG | CGG | ||||-|||[||||||||||-|]PAM | NA | Intergenic | NA | NA |
| GU305914.1 | 163902 | 163924 | + | 3 | AGCGGCGGCGGCGGCGGCAG | CGG | -|||-|||[||||||||||-|]PAM | NA | Intergenic | NA | NA |

<table>
<tr><td>GU305914.1</td><td>62434</td><td>62456</td><td>+</td><td>4</td><td>GACGGCGGTGGCGGCGGGGG</td><td>TGG</td><td>|-||-|||[-||||||||-||]PAM</td><td>NA</td><td>Intergenic</td><td>NA</td><td>NA</td></tr>
<tr><td>GU305914.1</td><td>113924</td><td>113946</td><td>-</td><td>4</td><td>GTCGGCGTCGGCGGCGGCAG</td><td>CGG</td><td>|-||-||-[||||||||||-|]PAM</td><td>NA</td><td>Intergenic</td><td>NA</td><td>NA</td></tr>
<tr><td>GU305914.1</td><td>152553</td><td>152575</td><td>+</td><td>4</td><td>TGCGGCGGCGGCGGTGCCGG</td><td>CGG</td><td>-|||-|||[||||||-|-|||]PAM</td><td>NA</td><td>Intergenic</td><td>NA</td><td>NA</td></tr>
<tr><td>GU305914.1</td><td>90250</td><td>90272</td><td>-</td><td>4</td><td>GGCGGAGGCGGCGGAGCCGG</td><td>CGG</td><td>||||--||[||||||-|-|||]PAM</td><td>NA</td><td>Intergenic</td><td>NA</td><td>NA</td></tr>
<tr><td>GU305914.1</td><td>166410</td><td>166432</td><td>+</td><td>4</td><td>GACGTCGGCGCCGGCGGCAG</td><td>CGG</td><td>|-||-|||[||-|||||||-|]PAM</td><td>NA</td><td>Intergenic</td><td>NA</td><td>NA</td></tr>
<tr><td>GU305914.1</td><td>163908</td><td>163930</td><td>+</td><td>3</td><td>GGCGGCGGCGGCAGCGGCAG</td><td>CGG</td><td>||||-|||[||||-|||||-|]PAM</td><td>NA</td><td>Intergenic</td><td>NA</td><td>NA</td></tr>
<tr><td>GU305914.1</td><td>225498</td><td>225520</td><td>+</td><td>4</td><td>GCCGAGGGCGGCGGTGGTGG</td><td>TGG</td><td>|-|||-||[||||||-||-||]PAM</td><td>NA</td><td>Intergenic</td><td>NA</td><td>NA</td></tr>
<tr><td>GU305914.1</td><td>62437</td><td>62459</td><td>+</td><td>4</td><td>GGCGGTGGCGGCGGGGGTGG</td><td>TGG</td><td>||||--||[||||||-||-||]PAM</td><td>NA</td><td>Intergenic</td><td>NA</td><td>NA</td></tr>
<tr><td>GU305914.1</td><td>113918</td><td>113940</td><td>-</td><td>4</td><td>GTCGGCGGCGGCAGCGGCGA</td><td>CGG</td><td>|-||-|||[||||-||||||-]PAM</td><td>NA</td><td>Intergenic</td><td>NA</td><td>NA</td></tr>
<tr><td>GU305914.1</td><td>113912</td><td>113934</td><td>-</td><td>4</td><td>GGCGGCAGCGGCGACGGCGT</td><td>CGG</td><td>||||-|-|[|||||-|||||-]PAM</td><td>NA</td><td>Intergenic</td><td>NA</td><td>NA</td></tr>
<tr><td>GU305914.1</td><td>168829</td><td>168851</td><td>+</td><td>4</td><td>AGCGGCGGCGGCGGCGACAG</td><td>CGG</td><td>-|||-|||[||||||||-|-|]PAM</td><td>NA</td><td>Intergenic</td><td>NA</td><td>NA</td></tr>
<tr><td></td></tr>
<tr><td>T22772</td><td>GGCGGCGGCGGCGGCGGCAGCGG</td><td>1000</td></tr>
<tr><td>Oligo fwd</td><td>CACCGGCGGCGGCGGCGGCGGCAG</td></tr>
<tr><td>Oligo rev</td><td>AAACCTGCCGCCGCCGCCGCCGCC</td></tr>
<tr><td>Chromosome</td><td>start</td><td>end</td><td>strand</td><td>MM</td><td>target_seq</td><td>PAM</td><td>alignment</td><td>distance</td><td>position</td><td>gene name</td><td>gene id</td></tr>
<tr><td>GU305914.1</td><td>30006</td><td>30028</td><td>+</td><td>0</td><td>GGCGGCGGCGGCGGCGGCAG</td><td>TGG</td><td>||||||||[||||||||||||]PAM</td><td>NA</td><td>Intergenic</td><td>NA</td><td>NA</td></tr>
<tr><td>GU305914.1</td><td>149476</td><td>149498</td><td>-</td><td>0</td><td>GGCGGCGGCGGCGGCGGCAG</td><td>CGG</td><td>||||||||[||||||||||||]PAM</td><td>NA</td><td>Intergenic</td><td>NA</td><td>NA</td></tr>
<tr><td>GU305914.1</td><td>163902</td><td>163924</td><td>+</td><td>1</td><td>AGCGGCGGCGGCGGCGGCAG</td><td>CGG</td><td>-|||||||[||||||||||||]PAM</td><td>NA</td><td>Intergenic</td><td>NA</td><td>NA</td></tr>
<tr><td>GU305914.1</td><td>113924</td><td>113946</td><td>-</td><td>2</td><td>GTCGGCGTCGGCGGCGGCAG</td><td>CGG</td><td>|-|||||-[||||||||||||]PAM</td><td>NA</td><td>Intergenic</td><td>NA</td><td>NA</td></tr>
<tr><td>GU305914.1</td><td>163908</td><td>163930</td><td>+</td><td>1</td><td>GGCGGCGGCGGCAGCGGCAG</td><td>CGG</td><td>||||||||[||||-|||||||]PAM</td><td>NA</td><td>Intergenic</td><td>NA</td><td>NA</td></tr>
<tr><td>GU305914.1</td><td>166410</td><td>166432</td><td>+</td><td>3</td><td>GACGTCGGCGCCGGCGGCAG</td><td>CGG</td><td>|-||-|||[||-|||||||||]PAM</td><td>NA</td><td>Intergenic</td><td>NA</td><td>NA</td></tr>
<tr><td>GU305914.1</td><td>45258</td><td>45280</td><td>-</td><td>4</td><td>AACGGTGGCGGTGGCGGCAG</td><td>CGG</td><td>--|||-||[|||-||||||||]PAM</td><td>NA</td><td>Intergenic</td><td>NA</td><td>NA</td></tr>
<tr><td>GU305914.1</td><td>65721</td><td>65743</td><td>-</td><td>4</td><td>TCCTGCGGCGGCAGCGGCAG</td><td>CGG</td><td>--|-||||[||||-|||||||]PAM</td><td>NA</td><td>Intergenic</td><td>NA</td><td>NA</td></tr>
<tr><td>GU305914.1</td><td>62152</td><td>62174</td><td>+</td><td>4</td><td>AGGGGCGACGGCAGCGGCAG</td><td>CGG</td><td>-|-||||-[||||-|||||||]PAM</td><td>NA</td><td>Intergenic</td><td>NA</td><td>NA</td></tr>
<tr><td>GU305914.1</td><td>168829</td><td>168851</td><td>+</td><td>2</td><td>AGCGGCGGCGGCGGCGACAG</td><td>CGG</td><td>-|||||||[||||||||-|||]PAM</td><td>NA</td><td>Intergenic</td><td>NA</td><td>NA</td></tr>
<tr><td>GU305914.1</td><td>24557</td><td>24579</td><td>+</td><td>3</td><td>GGCGGCAGCGACGACGGCAG</td><td>CGG</td><td>||||||-|[||-||-||||||]PAM</td><td>NA</td><td>Intergenic</td><td>NA</td><td>NA</td></tr>
<tr><td>GU305914.1</td><td>163920</td><td>163942</td><td>+</td><td>4</td><td>AGCGGCAGCGGAGGCAGCAG</td><td>CGG</td><td>-|||||-|[|||-|||-||||]PAM</td><td>NA</td><td>Intergenic</td><td>NA</td><td>NA</td></tr>
<tr><td>GU305914.1</td><td>154407</td><td>154429</td><td>+</td><td>2</td><td>GGTGGCGGCGGCGGCGGCGG</td><td>CGG</td><td>||-|||||[||||||||||-|]PAM</td><td>NA</td><td>Intergenic</td><td>NA</td><td>NA</td></tr>
<tr><td>GU305914.1</td><td>58123</td><td>58145</td><td>-</td><td>2</td><td>GGTGGCGGCGGCGGCGGCGG</td><td>AGG</td><td>||-|||||[||||||||||-|]PAM</td><td>NA</td><td>Intergenic</td><td>NA</td><td>NA</td></tr>
<tr><td>GU305914.1</td><td>128727</td><td>128749</td><td>-</td><td>4</td><td>GGTGGAGGCGGCAGCAGCAG</td><td>CGG</td><td>||-||-||[||||-||-||||]PAM</td><td>NA</td><td>Intergenic</td><td>NA</td><td>NA</td></tr>
<tr><td>GU305914.1</td><td>149482</td><td>149504</td><td>-</td><td>2</td><td>GGCGACGGCGGCGGCGGCGG</td><td>CGG</td><td>||||-|||[||||||||||-|]PAM</td><td>NA</td><td>Intergenic</td><td>NA</td><td>NA</td></tr>
<tr><td>GU305914.1</td><td>154404</td><td>154426</td><td>+</td><td>2</td><td>GGCGGTGGCGGCGGCGGCGG</td><td>CGG</td><td>|||||-||[||||||||||-|]PAM</td><td>NA</td><td>Intergenic</td><td>NA</td><td>NA</td></tr>
<tr><td>GU305914.1</td><td>58126</td><td>58148</td><td>-</td><td>2</td><td>GGCGGTGGCGGCGGCGGCGG</td><td>CGG</td><td>|||||-||[||||||||||-|]PAM</td><td>NA</td><td>Intergenic</td><td>NA</td><td>NA</td></tr>
<tr><td>GU305914.1</td><td>67742</td><td>67764</td><td>+</td><td>3</td><td>GGGAGCGGCGGCGGCGGCGG</td><td>CGG</td><td>||--||||[||||||||||-|]PAM</td><td>NA</td><td>Intergenic</td><td>NA</td><td>NA</td></tr>
<tr><td>GU305914.1</td><td>48159</td><td>48181</td><td>+</td><td>2</td><td>GGCGGCGGTGGCGGCGGCGG</td><td>TGG</td><td>||||||||[-|||||||||-|]PAM</td><td>NA</td><td>Intergenic</td><td>NA</td><td>NA</td></tr>
<tr><td>GU305914.1</td><td>56773</td><td>56795</td><td>+</td><td>4</td><td>CGCCGCAGCGGCGGCGGCGG</td><td>CGG</td><td>-||-||-|[||||||||||-|]PAM</td><td>NA</td><td>Intergenic</td><td>NA</td><td>NA</td></tr>
<tr><td>GU305914.1</td><td>30000</td><td>30022</td><td>+</td><td>4</td><td>GGTGCTGGCGGCGGCGGCGG</td><td>CGG</td><td>||-|--||[||||||||||-|]PAM</td><td>NA</td><td>Intergenic</td><td>NA</td><td>NA</td></tr>
<tr><td>GU305914.1</td><td>149485</td><td>149507</td><td>-</td><td>4</td><td>GCAGGCGACGGCGGCGGCGG</td><td>CGG</td><td>|--||||-[||||||||||-|]PAM</td><td>NA</td><td>Intergenic</td><td>NA</td><td>NA</td></tr>
<tr><td>GU305914.1</td><td>58129</td><td>58151</td><td>-</td><td>4</td><td>GCGGGCGGTGGCGGCGGCGG</td><td>CGG</td><td>|--|||||[-|||||||||-|]PAM</td><td>NA</td><td>Intergenic</td><td>NA</td><td>NA</td></tr>
<tr><td>GU305914.1</td><td>48156</td><td>48178</td><td>+</td><td>2</td><td>GGCGGCGGCGGTGGCGGCGG</td><td>CGG</td><td>||||||||[|||-||||||-|]PAM</td><td>NA</td><td>Intergenic</td><td>NA</td><td>NA</td></tr>
<tr><td>GU305914.1</td><td>161754</td><td>161776</td><td>-</td><td>4</td><td>GGCCGAGGAGGCGGCGGCGG</td><td>AGG</td><td>|||-|-||[-|||||||||-|]PAM</td><td>NA</td><td>Intergenic</td><td>NA</td><td>NA</td></tr>
<tr><td>GU305914.1</td><td>168820</td><td>168842</td><td>+</td><td>4</td><td>GGTGGTGGCAGCGGCGGCGG</td><td>CGG</td><td>||-||-||[|-||||||||-|]PAM</td><td>NA</td><td>Intergenic</td><td>NA</td><td>NA</td></tr>
<tr><td>GU305914.1</td><td>73792</td><td>73814</td><td>-</td><td>4</td><td>GGCGCCAGCAGCGGCGGCGG</td><td>AGG</td><td>||||-|-|[|-||||||||-|]PAM</td><td>NA</td><td>Intergenic</td><td>NA</td><td>NA</td></tr>
<tr><td>GU305914.1</td><td>71753</td><td>71775</td><td>+</td><td>4</td><td>AGCAGCAGCGGCGGCGGCAA</td><td>CGG</td><td>-||-||-|[|||||||||||-]PAM</td><td>NA</td><td>Intergenic</td><td>NA</td><td>NA</td></tr>
<tr><td>GU305914.1</td><td>48153</td><td>48175</td><td>+</td><td>4</td><td>TGTGGCGGCGGCGGTGGCGG</td><td>CGG</td><td>-|-|||||[||||||-|||-|]PAM</td><td>NA</td><td>Intergenic</td><td>NA</td><td>NA</td></tr>
<tr><td></td></tr>
<tr><td>T22773</td><td>GCGGCGGCGGCGGCGGCAGCGGG</td><td>1000</td></tr>
<tr><td>Oligo fwd</td><td>CACCGCGGCGGCGGCGGCGGCAGC</td></tr>
<tr><td>Oligo rev</td><td>AAACGCTGCCGCCGCCGCCGCCGC</td></tr>
<tr><td>Chromosome</td><td>start</td><td>end</td><td>strand</td><td>MM</td><td>target_seq</td><td>PAM</td><td>alignment</td><td>distance</td><td>position</td><td>gene name</td><td>gene id</td></tr>
<tr><td>GU305914.1</td><td>149475</td><td>149497</td><td>-</td><td>0</td><td>GCGGCGGCGGCGGCGGCAGC</td><td>GGG</td><td>||||||||[||||||||||||]PAM</td><td>NA</td><td>Intergenic</td><td>NA</td><td>NA</td></tr>
<tr><td>GU305914.1</td><td>65720</td><td>65742</td><td>-</td><td>3</td><td>CCTGCGGCGGCAGCGGCAGC</td><td>GGG</td><td>-|-|||||[|||-||||||||]PAM</td><td>NA</td><td>Intergenic</td><td>NA</td><td>NA</td></tr>
<tr><td>GU305914.1</td><td>168830</td><td>168852</td><td>+</td><td>1</td><td>GCGGCGGCGGCGGCGACAGC</td><td>GGG</td><td>||||||||[|||||||-||||]PAM</td><td>NA</td><td>Intergenic</td><td>NA</td><td>NA</td></tr>
<tr><td>GU305914.1</td><td>225977</td><td>225999</td><td>-</td><td>2</td><td>GCGACGGCGGCGGCGGGAGC</td><td>CGG</td><td>|||-||||[||||||||-|||]PAM</td><td>NA</td><td>Intergenic</td><td>NA</td><td>NA</td></tr>
<tr><td>GU305914.1</td><td>67743</td><td>67765</td><td>+</td><td>3</td><td>GGAGCGGCGGCGGCGGCGGC</td><td>GGG</td><td>|--|||||[|||||||||-||]PAM</td><td>NA</td><td>Intergenic</td><td>NA</td><td>NA</td></tr>
<tr><td>GU305914.1</td><td>35654</td><td>35676</td><td>+</td><td>4</td><td>TCCACGGCGGCGGCGGCGGC</td><td>GGG</td><td>-|--||||[|||||||||-||]PAM</td><td>NA</td><td>Intergenic</td><td>NA</td><td>NA</td></tr>
<tr><td>GU305914.1</td><td>71754</td><td>71776</td><td>+</td><td>3</td><td>GCAGCAGCGGCGGCGGCAAC</td><td>GGG</td><td>||-||-||[||||||||||-|]PAM</td><td>NA</td><td>Intergenic</td><td>NA</td><td>NA</td></tr>
<tr><td>GU305914.1</td><td>86780</td><td>86802</td><td>-</td><td>4</td><td>GAGGGGGCGGGGGCGGCGGC</td><td>AGG</td><td>|-||-|||[||-||||||-||]PAM</td><td>NA</td><td>Intergenic</td><td>NA</td><td>NA</td></tr>
<tr><td>GU305914.1</td><td>114657</td><td>114679</td><td>-</td><td>4</td><td>GCGGTGGTGGTGGCGGCGGC</td><td>GGG</td><td>||||-||-[||-||||||-||]PAM</td><td>NA</td><td>Intergenic</td><td>NA</td><td>NA</td></tr>
<tr><td>GU305914.1</td><td>58122</td><td>58144</td><td>-</td><td>3</td><td>GTGGCGGCGGCGGCGGCGGA</td><td>GGG</td><td>|-||||||[|||||||||-|-]PAM</td><td>NA</td><td>Intergenic</td><td>NA</td><td>NA</td></tr>
<tr><td>GU305914.1</td><td>48160</td><td>48182</td><td>+</td><td>3</td><td>GCGGCGGTGGCGGCGGCGGT</td><td>GGG</td><td>|||||||-[|||||||||-|-]PAM</td><td>NA</td><td>Intergenic</td><td>NA</td><td>NA</td></tr>
<tr><td></td></tr>
<tr><td>T22774</td><td>CAGCAGCAGCGCCGAGCGAGCGG</td><td>1000</td></tr>
<tr><td>Oligo fwd</td><td>CACCCAGCAGCAGCGCCGAGCGAG</td></tr>
<tr><td>Oligo rev</td><td>AAACCTCGCTCGGCGCTGCTGCTG</td></tr>
<tr><td>Chromosome</td><td>start</td><td>end</td><td>strand</td><td>MM</td><td>target_seq</td><td>PAM</td><td>alignment</td><td>distance</td><td>position</td><td>gene name</td><td>gene id</td></tr>
<tr><td>GU305914.1</td><td>149444</td><td>149466</td><td>-</td><td>0</td><td>CAGCAGCAGCGCCGAGCGAG</td><td>CGG</td><td>||||||||[||||||||||||]PAM</td><td>NA</td><td>Intergenic</td><td>NA</td><td>NA</td></tr>
<tr><td></td></tr>
<tr><td>T22775</td><td>GCAGCGCCGAGCGAGCGGATCGG</td><td>1000</td></tr>
<tr><td>Oligo fwd</td><td>CACCGCAGCGCCGAGCGAGCGGAT</td></tr>
<tr><td>Oligo rev</td><td>AAACATCCGCTCGCTCGGCGCTGC</td></tr>
<tr><td>Chromosome</td><td>start</td><td>end</td><td>strand</td><td>MM</td><td>target_seq</td><td>PAM</td><td>alignment</td><td>distance</td><td>position</td><td>gene name</td><td>gene id</td></tr>
<tr><td>GU305914.1</td><td>149439</td><td>149461</td><td>-</td><td>0</td><td>GCAGCGCCGAGCGAGCGGAT</td><td>CGG</td><td>||||||||[||||||||||||]PAM</td><td>NA</td><td>Intergenic</td><td>NA</td><td>NA</td></tr>
<tr><td></td></tr>
<tr><td>T22776</td><td>GAGCGGATCGGACGAGCGCCAGG</td><td>1000</td></tr>
<tr><td>Oligo fwd</td><td>CACCGAGCGGATCGGACGAGCGCC</td></tr>
<tr><td>Oligo rev</td><td>AAACGGCGCTCGTCCGATCCGCTC</td></tr>
<tr><td>Chromosome</td><td>start</td><td>end</td><td>strand</td><td>MM</td><td>target_seq</td><td>PAM</td><td>alignment</td><td>distance</td><td>position</td><td>gene name</td><td>gene id</td></tr>
<tr><td>GU305914.1</td><td>149427</td><td>149449</td><td>-</td><td>0</td><td>GAGCGGATCGGACGAGCGCC</td><td>AGG</td><td>||||||||[||||||||||||]PAM</td><td>NA</td><td>Intergenic</td><td>NA</td><td>NA</td></tr>
<tr><td></td></tr>
<tr><td>T22777</td><td>GATCGGACGAGCGCCAGGCCAGG</td><td>1000</td></tr>
<tr><td>Oligo fwd</td><td>CACCGATCGGACGAGCGCCAGGCC</td></tr>
<tr><td>Oligo rev</td><td>AAACGGCCTGGCGCTCGTCCGATC</td></tr>
<tr><td>Chromosome</td><td>start</td><td>end</td><td>strand</td><td>MM</td><td>target_seq</td><td>PAM</td><td>alignment</td><td>distance</td><td>position</td><td>gene name</td><td>gene id</td></tr>
<tr><td>GU305914.1</td><td>149422</td><td>149444</td><td>-</td><td>0</td><td>GATCGGACGAGCGCCAGGCC</td><td>AGG</td><td>||||||||[||||||||||||]PAM</td><td>NA</td><td>Intergenic</td><td>NA</td><td>NA</td></tr>
<tr><td></td></tr>
<tr><td>T22778</td><td>CGAGCGCCAGGCCAGGTCTCCGG</td><td>1000</td></tr>
<tr><td>Oligo fwd</td><td>CACCCGAGCGCCAGGCCAGGTCTC</td></tr>
</table>

<table>
<tr><td>Oligo rev</td><td>AAACGAGACCTGGCCTGGCGCTCG</td><td></td><td></td><td></td><td></td><td></td><td></td><td></td><td></td><td></td><td></td></tr>
<tr><td>Chromosome</td><td>start</td><td>end</td><td>strand</td><td>MM</td><td>target_seq</td><td>PAM</td><td>alignment</td><td>distance</td><td>position</td><td>gene name</td><td>gene id</td></tr>
<tr><td>GU305914.1</td><td>149415</td><td>149437</td><td>-</td><td>0</td><td>CGAGCGCCAGGCCAGGTCTC</td><td>CGG</td><td>||||||||[||||||||||||]PAM</td><td>NA</td><td>Intergenic</td><td>NA</td><td>NA</td></tr>
<tr><td>T22779</td><td>CCGGCCCCCCGACGAAGATCAGG</td><td>1000</td><td></td><td></td><td></td><td></td><td></td><td></td><td></td><td></td><td></td></tr>
<tr><td>Oligo fwd</td><td>CACCCCGGCCCCCCGACGAAGATC</td><td></td><td></td><td></td><td></td><td></td><td></td><td></td><td></td><td></td><td></td></tr>
<tr><td>Oligo rev</td><td>AAACGATCTTCGTCGGGGGGCCGG</td><td></td><td></td><td></td><td></td><td></td><td></td><td></td><td></td><td></td><td></td></tr>
<tr><td>Chromosome</td><td>start</td><td>end</td><td>strand</td><td>MM</td><td>target_seq</td><td>PAM</td><td>alignment</td><td>distance</td><td>position</td><td>gene name</td><td>gene id</td></tr>
<tr><td>GU305914.1</td><td>149396</td><td>149418</td><td>-</td><td>0</td><td>CCGGCCCCCCGACGAAGATC</td><td>AGG</td><td>||||||||[||||||||||||]PAM</td><td>NA</td><td>Intergenic</td><td>NA</td><td>NA</td></tr>
<tr><td>T22780</td><td>CGACTCGACGAAACGACACCAGG</td><td>1000</td><td></td><td></td><td></td><td></td><td></td><td></td><td></td><td></td><td></td></tr>
<tr><td>Oligo fwd</td><td>CACCCGACTCGACGAAACGACACC</td><td></td><td></td><td></td><td></td><td></td><td></td><td></td><td></td><td></td><td></td></tr>
<tr><td>Oligo rev</td><td>AAACGGTGTCGTTTCGTCGAGTCG</td><td></td><td></td><td></td><td></td><td></td><td></td><td></td><td></td><td></td><td></td></tr>
<tr><td>Chromosome</td><td>start</td><td>end</td><td>strand</td><td>MM</td><td>target_seq</td><td>PAM</td><td>alignment</td><td>distance</td><td>position</td><td>gene name</td><td>gene id</td></tr>
<tr><td>GU305914.1</td><td>149373</td><td>149395</td><td>-</td><td>0</td><td>CGACTCGACGAAACGACACC</td><td>AGG</td><td>||||||||[||||||||||||]PAM</td><td>NA</td><td>Intergenic</td><td>NA</td><td>NA</td></tr>
<tr><td>T22781</td><td>CGACGAAACGACACCAGGCACGG</td><td>1000</td><td></td><td></td><td></td><td></td><td></td><td></td><td></td><td></td><td></td></tr>
<tr><td>Oligo fwd</td><td>CACCCGACGAAACGACACCAGGCA</td><td></td><td></td><td></td><td></td><td></td><td></td><td></td><td></td><td></td><td></td></tr>
<tr><td>Oligo rev</td><td>AAACTGCCTGGTGTCGTTTCGTCG</td><td></td><td></td><td></td><td></td><td></td><td></td><td></td><td></td><td></td><td></td></tr>
<tr><td>Chromosome</td><td>start</td><td>end</td><td>strand</td><td>MM</td><td>target_seq</td><td>PAM</td><td>alignment</td><td>distance</td><td>position</td><td>gene name</td><td>gene id</td></tr>
<tr><td>GU305914.1</td><td>149368</td><td>149390</td><td>-</td><td>0</td><td>CGACGAAACGACACCAGGCA</td><td>CGG</td><td>||||||||[||||||||||||]PAM</td><td>NA</td><td>Intergenic</td><td>NA</td><td>NA</td></tr>
<tr><td>T22782</td><td>ATCGTGACGACGACCCCGATCGG</td><td>1000</td><td></td><td></td><td></td><td></td><td></td><td></td><td></td><td></td><td></td></tr>
<tr><td>Oligo fwd</td><td>CACCATCGTGACGACGACCCCGAT</td><td></td><td></td><td></td><td></td><td></td><td></td><td></td><td></td><td></td><td></td></tr>
<tr><td>Oligo rev</td><td>AAACATCGGGGTCGTCGTCACGAT</td><td></td><td></td><td></td><td></td><td></td><td></td><td></td><td></td><td></td><td></td></tr>
<tr><td>Chromosome</td><td>start</td><td>end</td><td>strand</td><td>MM</td><td>target_seq</td><td>PAM</td><td>alignment</td><td>distance</td><td>position</td><td>gene name</td><td>gene id</td></tr>
<tr><td>GU305914.1</td><td>149319</td><td>149341</td><td>-</td><td>0</td><td>ATCGTGACGACGACCCCGAT</td><td>CGG</td><td>||||||||[||||||||||||]PAM</td><td>NA</td><td>Intergenic</td><td>NA</td><td>NA</td></tr>
<tr><td>T22783</td><td>GCCGTCGTCGCCCGCTCCCGAGG</td><td>1000</td><td></td><td></td><td></td><td></td><td></td><td></td><td></td><td></td><td></td></tr>
<tr><td>Oligo fwd</td><td>CACCGCCGTCGTCGCCCGCTCCCG</td><td></td><td></td><td></td><td></td><td></td><td></td><td></td><td></td><td></td><td></td></tr>
<tr><td>Oligo rev</td><td>AAACCGGGAGCGGGCGACGACGGC</td><td></td><td></td><td></td><td></td><td></td><td></td><td></td><td></td><td></td><td></td></tr>
<tr><td>Chromosome</td><td>start</td><td>end</td><td>strand</td><td>MM</td><td>target_seq</td><td>PAM</td><td>alignment</td><td>distance</td><td>position</td><td>gene name</td><td>gene id</td></tr>
<tr><td>GU305914.1</td><td>149287</td><td>149309</td><td>-</td><td>0</td><td>GCCGTCGTCGCCCGCTCCCG</td><td>AGG</td><td>||||||||[||||||||||||]PAM</td><td>NA</td><td>Intergenic</td><td>NA</td><td>NA</td></tr>
<tr><td>T22784</td><td>CCGCTCCCGAGGTCAGCCCGAGG</td><td>1000</td><td></td><td></td><td></td><td></td><td></td><td></td><td></td><td></td><td></td></tr>
<tr><td>Oligo fwd</td><td>CACCCCGCTCCCGAGGTCAGCCCG</td><td></td><td></td><td></td><td></td><td></td><td></td><td></td><td></td><td></td><td></td></tr>
<tr><td>Oligo rev</td><td>AAACCGGGCTGACCTCGGGAGCGG</td><td></td><td></td><td></td><td></td><td></td><td></td><td></td><td></td><td></td><td></td></tr>
<tr><td>Chromosome</td><td>start</td><td>end</td><td>strand</td><td>MM</td><td>target_seq</td><td>PAM</td><td>alignment</td><td>distance</td><td>position</td><td>gene name</td><td>gene id</td></tr>
<tr><td>GU305914.1</td><td>149276</td><td>149298</td><td>-</td><td>0</td><td>CCGCTCCCGAGGTCAGCCCG</td><td>AGG</td><td>||||||||[||||||||||||]PAM</td><td>NA</td><td>Intergenic</td><td>NA</td><td>NA</td></tr>
<tr><td>T22785</td><td>CGCAATGTCCAGCACCTTAATGG</td><td>1000</td><td></td><td></td><td></td><td></td><td></td><td></td><td></td><td></td><td></td></tr>
<tr><td>Oligo fwd</td><td>CACCCGCAATGTCCAGCACCTTAA</td><td></td><td></td><td></td><td></td><td></td><td></td><td></td><td></td><td></td><td></td></tr>
<tr><td>Oligo rev</td><td>AAACTTAAGGTGCTGGACATTGCG</td><td></td><td></td><td></td><td></td><td></td><td></td><td></td><td></td><td></td><td></td></tr>
<tr><td>Chromosome</td><td>start</td><td>end</td><td>strand</td><td>MM</td><td>target_seq</td><td>PAM</td><td>alignment</td><td>distance</td><td>position</td><td>gene name</td><td>gene id</td></tr>
<tr><td>GU305914.1</td><td>149214</td><td>149236</td><td>-</td><td>0</td><td>CGCAATGTCCAGCACCTTAA</td><td>TGG</td><td>||||||||[||||||||||||]PAM</td><td>NA</td><td>Intergenic</td><td>NA</td><td>NA</td></tr>
<tr><td>T22786</td><td>TGTCCAGCACCTTAATGGTTAGG</td><td>1000</td><td></td><td></td><td></td><td></td><td></td><td></td><td></td><td></td><td></td></tr>
<tr><td>Oligo fwd</td><td>CACCTGTCCAGCACCTTAATGGTT</td><td></td><td></td><td></td><td></td><td></td><td></td><td></td><td></td><td></td><td></td></tr>
<tr><td>Oligo rev</td><td>AAACAACCATTAAGGTGCTGGACA</td><td></td><td></td><td></td><td></td><td></td><td></td><td></td><td></td><td></td><td></td></tr>
<tr><td>Chromosome</td><td>start</td><td>end</td><td>strand</td><td>MM</td><td>target_seq</td><td>PAM</td><td>alignment</td><td>distance</td><td>position</td><td>gene name</td><td>gene id</td></tr>
<tr><td>GU305914.1</td><td>149209</td><td>149231</td><td>-</td><td>0</td><td>TGTCCAGCACCTTAATGGTT</td><td>AGG</td><td>||||||||[||||||||||||]PAM</td><td>NA</td><td>Intergenic</td><td>NA</td><td>NA</td></tr>
<tr><td>T22787</td><td>GTCCAGCACCTTAATGGTTAGGG</td><td>1000</td><td></td><td></td><td></td><td></td><td></td><td></td><td></td><td></td><td></td></tr>
<tr><td>Oligo fwd</td><td>CACCGTCCAGCACCTTAATGGTTA</td><td></td><td></td><td></td><td></td><td></td><td></td><td></td><td></td><td></td><td></td></tr>
<tr><td>Oligo rev</td><td>AAACTAACCATTAAGGTGCTGGAC</td><td></td><td></td><td></td><td></td><td></td><td></td><td></td><td></td><td></td><td></td></tr>
<tr><td>Chromosome</td><td>start</td><td>end</td><td>strand</td><td>MM</td><td>target_seq</td><td>PAM</td><td>alignment</td><td>distance</td><td>position</td><td>gene name</td><td>gene id</td></tr>
<tr><td>GU305914.1</td><td>149208</td><td>149230</td><td>-</td><td>0</td><td>GTCCAGCACCTTAATGGTTA</td><td>GGG</td><td>||||||||[||||||||||||]PAM</td><td>NA</td><td>Intergenic</td><td>NA</td><td>NA</td></tr>
<tr><td>T22788</td><td>TCCAGCACCTTAATGGTTAGGGG</td><td>1000</td><td></td><td></td><td></td><td></td><td></td><td></td><td></td><td></td><td></td></tr>
<tr><td>Oligo fwd</td><td>CACCTCCAGCACCTTAATGGTTAG</td><td></td><td></td><td></td><td></td><td></td><td></td><td></td><td></td><td></td><td></td></tr>
<tr><td>Oligo rev</td><td>AAACCTAACCATTAAGGTGCTGGA</td><td></td><td></td><td></td><td></td><td></td><td></td><td></td><td></td><td></td><td></td></tr>
<tr><td>Chromosome</td><td>start</td><td>end</td><td>strand</td><td>MM</td><td>target_seq</td><td>PAM</td><td>alignment</td><td>distance</td><td>position</td><td>gene name</td><td>gene id</td></tr>
<tr><td>GU305914.1</td><td>149207</td><td>149229</td><td>-</td><td>0</td><td>TCCAGCACCTTAATGGTTAG</td><td>GGG</td><td>||||||||[||||||||||||]PAM</td><td>NA</td><td>Intergenic</td><td>NA</td><td>NA</td></tr>
<tr><td>T22789</td><td>CTTAATGGTTAGGGGCGCCATGG</td><td>1000</td><td></td><td></td><td></td><td></td><td></td><td></td><td></td><td></td><td></td></tr>
<tr><td>Oligo fwd</td><td>CACCCTTAATGGTTAGGGGCGCCA</td><td></td><td></td><td></td><td></td><td></td><td></td><td></td><td></td><td></td><td></td></tr>
<tr><td>Oligo rev</td><td>AAACTGGCGCCCCTAACCATTAAG</td><td></td><td></td><td></td><td></td><td></td><td></td><td></td><td></td><td></td><td></td></tr>
<tr><td>Chromosome</td><td>start</td><td>end</td><td>strand</td><td>MM</td><td>target_seq</td><td>PAM</td><td>alignment</td><td>distance</td><td>position</td><td>gene name</td><td>gene id</td></tr>
<tr><td>GU305914.1</td><td>149199</td><td>149221</td><td>-</td><td>0</td><td>CTTAATGGTTAGGGGCGCCA</td><td>TGG</td><td>||||||||[||||||||||||]PAM</td><td>NA</td><td>Intergenic</td><td>NA</td><td>NA</td></tr>
<tr><td>T22790</td><td>AATGGTTAGGGGCGCCATGGAGG</td><td>1000</td><td></td><td></td><td></td><td></td><td></td><td></td><td></td><td></td><td></td></tr>
<tr><td>Oligo fwd</td><td>CACCAATGGTTAGGGGCGCCATGG</td><td></td><td></td><td></td><td></td><td></td><td></td><td></td><td></td><td></td><td></td></tr>
<tr><td>Oligo rev</td><td>AAACCCATGGCGCCCCTAACCATT</td><td></td><td></td><td></td><td></td><td></td><td></td><td></td><td></td><td></td><td></td></tr>
<tr><td>Chromosome</td><td>start</td><td>end</td><td>strand</td><td>MM</td><td>target_seq</td><td>PAM</td><td>alignment</td><td>distance</td><td>position</td><td>gene name</td><td>gene id</td></tr>
<tr><td>GU305914.1</td><td>149196</td><td>149218</td><td>-</td><td>0</td><td>AATGGTTAGGGGCGCCATGG</td><td>AGG</td><td>||||||||[||||||||||||]PAM</td><td>NA</td><td>Intergenic</td><td>NA</td><td>NA</td></tr>
<tr><td>T22791</td><td>CCATGGAGGTGCACAACGACCGG</td><td>1000</td><td></td><td></td><td></td><td></td><td></td><td></td><td></td><td></td><td></td></tr>
<tr><td>Oligo fwd</td><td>CACCCCATGGAGGTGCACAACGAC</td><td></td><td></td><td></td><td></td><td></td><td></td><td></td><td></td><td></td><td></td></tr>
<tr><td>Oligo rev</td><td>AAACGTCGTTGTGCACCTCCATGG</td><td></td><td></td><td></td><td></td><td></td><td></td><td></td><td></td><td></td><td></td></tr>
<tr><td>Chromosome</td><td>start</td><td>end</td><td>strand</td><td>MM</td><td>target_seq</td><td>PAM</td><td>alignment</td><td>distance</td><td>position</td><td>gene name</td><td>gene id</td></tr>
<tr><td>GU305914.1</td><td>149182</td><td>149204</td><td>-</td><td>0</td><td>CCATGGAGGTGCACAACGAC</td><td>CGG</td><td>||||||||[||||||||||||]PAM</td><td>NA</td><td>Intergenic</td><td>NA</td><td>NA</td></tr>
<tr><td>T22792</td><td>CATGGAGGTGCACAACGACCGGG</td><td>1000</td><td></td><td></td><td></td><td></td><td></td><td></td><td></td><td></td><td></td></tr>
<tr><td>Oligo fwd</td><td>CACCCATGGAGGTGCACAACGACC</td><td></td><td></td><td></td><td></td><td></td><td></td><td></td><td></td><td></td><td></td></tr>
<tr><td>Oligo rev</td><td>AAACGGTCGTTGTGCACCTCCATG</td><td></td><td></td><td></td><td></td><td></td><td></td><td></td><td></td><td></td><td></td></tr>
<tr><td>Chromosome</td><td>start</td><td>end</td><td>strand</td><td>MM</td><td>target_seq</td><td>PAM</td><td>alignment</td><td>distance</td><td>position</td><td>gene name</td><td>gene id</td></tr>
<tr><td>GU305914.1</td><td>149181</td><td>149203</td><td>-</td><td>0</td><td>CATGGAGGTGCACAACGACC</td><td>GGG</td><td>||||||||[||||||||||||]PAM</td><td>NA</td><td>Intergenic</td><td>NA</td><td>NA</td></tr>
<tr><td>T22793</td><td>CCGGGACTACGTCCGACTGATGG</td><td>1000</td><td></td><td></td><td></td><td></td><td></td><td></td><td></td><td></td><td></td></tr>
</table>

<table>
<tr><td>Oligo fwd</td><td>CACCCCGGGACTACGTCCGACTGA</td><td></td><td></td><td></td><td></td><td></td><td></td><td></td><td></td><td></td><td></td></tr>
<tr><td>Oligo rev</td><td>AAACTCAGTCGGACGTAGTCCCGG</td><td></td><td></td><td></td><td></td><td></td><td></td><td></td><td></td><td></td><td></td></tr>
<tr><td>Chromosome</td><td>start</td><td>end</td><td>strand</td><td>MM</td><td>target_seq</td><td>PAM</td><td>alignment</td><td>distance</td><td>position</td><td>gene name</td><td>gene id</td></tr>
<tr><td>GU305914.1</td><td>149163</td><td>149185</td><td>-</td><td>0</td><td>CCGGGACTACGTCCGACTGA</td><td>TGG</td><td>||||||||[||||||||||||]PAM</td><td>NA</td><td>Intergenic</td><td>NA</td><td>NA</td></tr>
<tr><td></td><td></td><td></td><td></td><td></td><td></td><td></td><td></td><td></td><td></td><td></td><td></td></tr>
<tr><td>T22794</td><td>CTACGTCCGACTGATGGCGCCGG</td><td>1000</td><td></td><td></td><td></td><td></td><td></td><td></td><td></td><td></td><td></td></tr>
<tr><td>Oligo fwd</td><td>CACCCTACGTCCGACTGATGGCGC</td><td></td><td></td><td></td><td></td><td></td><td></td><td></td><td></td><td></td><td></td></tr>
<tr><td>Oligo rev</td><td>AAACGCGCCATCAGTCGGACGTAG</td><td></td><td></td><td></td><td></td><td></td><td></td><td></td><td></td><td></td><td></td></tr>
<tr><td>Chromosome</td><td>start</td><td>end</td><td>strand</td><td>MM</td><td>target_seq</td><td>PAM</td><td>alignment</td><td>distance</td><td>position</td><td>gene name</td><td>gene id</td></tr>
<tr><td>GU305914.1</td><td>149157</td><td>149179</td><td>-</td><td>0</td><td>CTACGTCCGACTGATGGCGC</td><td>CGG</td><td>||||||||[||||||||||||]PAM</td><td>NA</td><td>Intergenic</td><td>NA</td><td>NA</td></tr>
<tr><td></td><td></td><td></td><td></td><td></td><td></td><td></td><td></td><td></td><td></td><td></td><td></td></tr>
<tr><td>T22795</td><td>GCCGGAGATCGTGAGCATCACGG</td><td>1000</td><td></td><td></td><td></td><td></td><td></td><td></td><td></td><td></td><td></td></tr>
<tr><td>Oligo fwd</td><td>CACCGCCGGAGATCGTGAGCATCA</td><td></td><td></td><td></td><td></td><td></td><td></td><td></td><td></td><td></td><td></td></tr>
<tr><td>Oligo rev</td><td>AAACTGATGCTCACGATCTCCGGC</td><td></td><td></td><td></td><td></td><td></td><td></td><td></td><td></td><td></td><td></td></tr>
<tr><td>Chromosome</td><td>start</td><td>end</td><td>strand</td><td>MM</td><td>target_seq</td><td>PAM</td><td>alignment</td><td>distance</td><td>position</td><td>gene name</td><td>gene id</td></tr>
<tr><td>GU305914.1</td><td>149139</td><td>149161</td><td>-</td><td>0</td><td>GCCGGAGATCGTGAGCATCA</td><td>CGG</td><td>||||||||[||||||||||||]PAM</td><td>NA</td><td>Intergenic</td><td>NA</td><td>NA</td></tr>
<tr><td></td><td></td><td></td><td></td><td></td><td></td><td></td><td></td><td></td><td></td><td></td><td></td></tr>
<tr><td>T22796</td><td>GATCGTGAGCATCACGGTGTCGG</td><td>1000</td><td></td><td></td><td></td><td></td><td></td><td></td><td></td><td></td><td></td></tr>
<tr><td>Oligo fwd</td><td>CACCGATCGTGAGCATCACGGTGT</td><td></td><td></td><td></td><td></td><td></td><td></td><td></td><td></td><td></td><td></td></tr>
<tr><td>Oligo rev</td><td>AAACACACCGTGATGCTCACGATC</td><td></td><td></td><td></td><td></td><td></td><td></td><td></td><td></td><td></td><td></td></tr>
<tr><td>Chromosome</td><td>start</td><td>end</td><td>strand</td><td>MM</td><td>target_seq</td><td>PAM</td><td>alignment</td><td>distance</td><td>position</td><td>gene name</td><td>gene id</td></tr>
<tr><td>GU305914.1</td><td>149133</td><td>149155</td><td>-</td><td>0</td><td>GATCGTGAGCATCACGGTGT</td><td>CGG</td><td>||||||||[||||||||||||]PAM</td><td>NA</td><td>Intergenic</td><td>NA</td><td>NA</td></tr>
<tr><td>GU305914.1</td><td>58451</td><td>58473</td><td>-</td><td>4</td><td>GAACCTGCGCATCACGGTGC</td><td>AGG</td><td>||-|-||-[|||||||||||-]PAM</td><td>NA</td><td>Intergenic</td><td>NA</td><td>NA</td></tr>
<tr><td></td><td></td><td></td><td></td><td></td><td></td><td></td><td></td><td></td><td></td><td></td><td></td></tr>
<tr><td>T22797</td><td>TGAGCATCACGGTGTCGGACAGG</td><td>1000</td><td></td><td></td><td></td><td></td><td></td><td></td><td></td><td></td><td></td></tr>
<tr><td>Oligo fwd</td><td>CACCTGAGCATCACGGTGTCGGAC</td><td></td><td></td><td></td><td></td><td></td><td></td><td></td><td></td><td></td><td></td></tr>
<tr><td>Oligo rev</td><td>AAACGTCCGACACCGTGATGCTCA</td><td></td><td></td><td></td><td></td><td></td><td></td><td></td><td></td><td></td><td></td></tr>
<tr><td>Chromosome</td><td>start</td><td>end</td><td>strand</td><td>MM</td><td>target_seq</td><td>PAM</td><td>alignment</td><td>distance</td><td>position</td><td>gene name</td><td>gene id</td></tr>
<tr><td>GU305914.1</td><td>149128</td><td>149150</td><td>-</td><td>0</td><td>TGAGCATCACGGTGTCGGAC</td><td>AGG</td><td>||||||||[||||||||||||]PAM</td><td>NA</td><td>Intergenic</td><td>NA</td><td>NA</td></tr>
<tr><td></td><td></td><td></td><td></td><td></td><td></td><td></td><td></td><td></td><td></td><td></td><td></td></tr>
<tr><td>T22798</td><td>CGGTGTCGGACAGGCAGATCTGG</td><td>1000</td><td></td><td></td><td></td><td></td><td></td><td></td><td></td><td></td><td></td></tr>
<tr><td>Oligo fwd</td><td>CACCCGGTGTCGGACAGGCAGATC</td><td></td><td></td><td></td><td></td><td></td><td></td><td></td><td></td><td></td><td></td></tr>
<tr><td>Oligo rev</td><td>AAACGATCTGCCTGTCCGACACCG</td><td></td><td></td><td></td><td></td><td></td><td></td><td></td><td></td><td></td><td></td></tr>
<tr><td>Chromosome</td><td>start</td><td>end</td><td>strand</td><td>MM</td><td>target_seq</td><td>PAM</td><td>alignment</td><td>distance</td><td>position</td><td>gene name</td><td>gene id</td></tr>
<tr><td>GU305914.1</td><td>149119</td><td>149141</td><td>-</td><td>0</td><td>CGGTGTCGGACAGGCAGATC</td><td>TGG</td><td>||||||||[||||||||||||]PAM</td><td>NA</td><td>Intergenic</td><td>NA</td><td>NA</td></tr>
<tr><td></td><td></td><td></td><td></td><td></td><td></td><td></td><td></td><td></td><td></td><td></td><td></td></tr>
<tr><td>T22799</td><td>GATCTGGTTCCACACCGAGTCGG</td><td>1000</td><td></td><td></td><td></td><td></td><td></td><td></td><td></td><td></td><td></td></tr>
<tr><td>Oligo fwd</td><td>CACCGATCTGGTTCCACACCGAGT</td><td></td><td></td><td></td><td></td><td></td><td></td><td></td><td></td><td></td><td></td></tr>
<tr><td>Oligo rev</td><td>AAACACTCGGTGTGGAACCAGATC</td><td></td><td></td><td></td><td></td><td></td><td></td><td></td><td></td><td></td><td></td></tr>
<tr><td>Chromosome</td><td>start</td><td>end</td><td>strand</td><td>MM</td><td>target_seq</td><td>PAM</td><td>alignment</td><td>distance</td><td>position</td><td>gene name</td><td>gene id</td></tr>
<tr><td>GU305914.1</td><td>149103</td><td>149125</td><td>-</td><td>0</td><td>GATCTGGTTCCACACCGAGT</td><td>CGG</td><td>||||||||[||||||||||||]PAM</td><td>NA</td><td>Intergenic</td><td>NA</td><td>NA</td></tr>
<tr><td></td><td></td><td></td><td></td><td></td><td></td><td></td><td></td><td></td><td></td><td></td><td></td></tr>
<tr><td>T22800</td><td>ATCTGGTTCCACACCGAGTCGGG</td><td>1000</td><td></td><td></td><td></td><td></td><td></td><td></td><td></td><td></td><td></td></tr>
<tr><td>Oligo fwd</td><td>CACCATCTGGTTCCACACCGAGTC</td><td></td><td></td><td></td><td></td><td></td><td></td><td></td><td></td><td></td><td></td></tr>
<tr><td>Oligo rev</td><td>AAACGACTCGGTGTGGAACCAGAT</td><td></td><td></td><td></td><td></td><td></td><td></td><td></td><td></td><td></td><td></td></tr>
<tr><td>Chromosome</td><td>start</td><td>end</td><td>strand</td><td>MM</td><td>target_seq</td><td>PAM</td><td>alignment</td><td>distance</td><td>position</td><td>gene name</td><td>gene id</td></tr>
<tr><td>GU305914.1</td><td>149102</td><td>149124</td><td>-</td><td>0</td><td>ATCTGGTTCCACACCGAGTC</td><td>GGG</td><td>||||||||[||||||||||||]PAM</td><td>NA</td><td>Intergenic</td><td>NA</td><td>NA</td></tr>
<tr><td></td><td></td><td></td><td></td><td></td><td></td><td></td><td></td><td></td><td></td><td></td><td></td></tr>
<tr><td>T22801</td><td>CCCCAACGCGCAGTACCGCGCGG</td><td>1000</td><td></td><td></td><td></td><td></td><td></td><td></td><td></td><td></td><td></td></tr>
<tr><td>Oligo fwd</td><td>CACCCCCCAACGCGCAGTACCGCG</td><td></td><td></td><td></td><td></td><td></td><td></td><td></td><td></td><td></td><td></td></tr>
<tr><td>Oligo rev</td><td>AAACCGCGGTACTGCGCGTTGGGG</td><td></td><td></td><td></td><td></td><td></td><td></td><td></td><td></td><td></td><td></td></tr>
<tr><td>Chromosome</td><td>start</td><td>end</td><td>strand</td><td>MM</td><td>target_seq</td><td>PAM</td><td>alignment</td><td>distance</td><td>position</td><td>gene name</td><td>gene id</td></tr>
<tr><td>GU305914.1</td><td>149070</td><td>149092</td><td>-</td><td>0</td><td>CCCCAACGCGCAGTACCGCG</td><td>CGG</td><td>||||||||[||||||||||||]PAM</td><td>NA</td><td>Intergenic</td><td>NA</td><td>NA</td></tr>
<tr><td></td><td></td><td></td><td></td><td></td><td></td><td></td><td></td><td></td><td></td><td></td><td></td></tr>
<tr><td>T22802</td><td>CGCGCAGTACCGCGCGGACTCGG</td><td>1000</td><td></td><td></td><td></td><td></td><td></td><td></td><td></td><td></td><td></td></tr>
<tr><td>Oligo fwd</td><td>CACCCGCGCAGTACCGCGCGGACT</td><td></td><td></td><td></td><td></td><td></td><td></td><td></td><td></td><td></td><td></td></tr>
<tr><td>Oligo rev</td><td>AAACAGTCCGCGCGGTACTGCGCG</td><td></td><td></td><td></td><td></td><td></td><td></td><td></td><td></td><td></td><td></td></tr>
<tr><td>Chromosome</td><td>start</td><td>end</td><td>strand</td><td>MM</td><td>target_seq</td><td>PAM</td><td>alignment</td><td>distance</td><td>position</td><td>gene name</td><td>gene id</td></tr>
<tr><td>GU305914.1</td><td>149064</td><td>149086</td><td>-</td><td>0</td><td>CGCGCAGTACCGCGCGGACT</td><td>CGG</td><td>||||||||[||||||||||||]PAM</td><td>NA</td><td>Intergenic</td><td>NA</td><td>NA</td></tr>
<tr><td></td><td></td><td></td><td></td><td></td><td></td><td></td><td></td><td></td><td></td><td></td><td></td></tr>
<tr><td>T22803</td><td>ACCGCGCGGACTCGGACTGCCGG</td><td>1000</td><td></td><td></td><td></td><td></td><td></td><td></td><td></td><td></td><td></td></tr>
<tr><td>Oligo fwd</td><td>CACCACCGCGCGGACTCGGACTGC</td><td></td><td></td><td></td><td></td><td></td><td></td><td></td><td></td><td></td><td></td></tr>
<tr><td>Oligo rev</td><td>AAACGCAGTCCGAGTCCGCGCGGT</td><td></td><td></td><td></td><td></td><td></td><td></td><td></td><td></td><td></td><td></td></tr>
<tr><td>Chromosome</td><td>start</td><td>end</td><td>strand</td><td>MM</td><td>target_seq</td><td>PAM</td><td>alignment</td><td>distance</td><td>position</td><td>gene name</td><td>gene id</td></tr>
<tr><td>GU305914.1</td><td>149056</td><td>149078</td><td>-</td><td>0</td><td>ACCGCGCGGACTCGGACTGC</td><td>CGG</td><td>||||||||[||||||||||||]PAM</td><td>NA</td><td>Intergenic</td><td>NA</td><td>NA</td></tr>
<tr><td></td><td></td><td></td><td></td><td></td><td></td><td></td><td></td><td></td><td></td><td></td><td></td></tr>
<tr><td>T22804</td><td>GGACTGCCGGTCGTCGTTCCTGG</td><td>1000</td><td></td><td></td><td></td><td></td><td></td><td></td><td></td><td></td><td></td></tr>
<tr><td>Oligo fwd</td><td>CACCGGACTGCCGGTCGTCGTTCC</td><td></td><td></td><td></td><td></td><td></td><td></td><td></td><td></td><td></td><td></td></tr>
<tr><td>Oligo rev</td><td>AAACGGAACGACGACCGGCAGTCC</td><td></td><td></td><td></td><td></td><td></td><td></td><td></td><td></td><td></td><td></td></tr>
<tr><td>Chromosome</td><td>start</td><td>end</td><td>strand</td><td>MM</td><td>target_seq</td><td>PAM</td><td>alignment</td><td>distance</td><td>position</td><td>gene name</td><td>gene id</td></tr>
<tr><td>GU305914.1</td><td>149043</td><td>149065</td><td>-</td><td>0</td><td>GGACTGCCGGTCGTCGTTCC</td><td>TGG</td><td>||||||||[||||||||||||]PAM</td><td>NA</td><td>Intergenic</td><td>NA</td><td>NA</td></tr>
<tr><td></td><td></td><td></td><td></td><td></td><td></td><td></td><td></td><td></td><td></td><td></td><td></td></tr>
<tr><td>T22805</td><td>GACTGCCGGTCGTCGTTCCTGGG</td><td>1000</td><td></td><td></td><td></td><td></td><td></td><td></td><td></td><td></td><td></td></tr>
<tr><td>Oligo fwd</td><td>CACCGACTGCCGGTCGTCGTTCCT</td><td></td><td></td><td></td><td></td><td></td><td></td><td></td><td></td><td></td><td></td></tr>
<tr><td>Oligo rev</td><td>AAACAGGAACGACGACCGGCAGTC</td><td></td><td></td><td></td><td></td><td></td><td></td><td></td><td></td><td></td><td></td></tr>
<tr><td>Chromosome</td><td>start</td><td>end</td><td>strand</td><td>MM</td><td>target_seq</td><td>PAM</td><td>alignment</td><td>distance</td><td>position</td><td>gene name</td><td>gene id</td></tr>
<tr><td>GU305914.1</td><td>149042</td><td>149064</td><td>-</td><td>0</td><td>GACTGCCGGTCGTCGTTCCT</td><td>GGG</td><td>||||||||[||||||||||||]PAM</td><td>NA</td><td>Intergenic</td><td>NA</td><td>NA</td></tr>
<tr><td></td><td></td><td></td><td></td><td></td><td></td><td></td><td></td><td></td><td></td><td></td><td></td></tr>
<tr><td>T22806</td><td>ATTCTGCCTGTTCTTCATCCTGG</td><td>1000</td><td></td><td></td><td></td><td></td><td></td><td></td><td></td><td></td><td></td></tr>
<tr><td>Oligo fwd</td><td>CACCATTCTGCCTGTTCTTCATCC</td><td></td><td></td><td></td><td></td><td></td><td></td><td></td><td></td><td></td><td></td></tr>
<tr><td>Oligo rev</td><td>AAACGGATGAAGAACAGGCAGAAT</td><td></td><td></td><td></td><td></td><td></td><td></td><td></td><td></td><td></td><td></td></tr>
<tr><td>Chromosome</td><td>start</td><td>end</td><td>strand</td><td>MM</td><td>target_seq</td><td>PAM</td><td>alignment</td><td>distance</td><td>position</td><td>gene name</td><td>gene id</td></tr>
<tr><td>GU305914.1</td><td>149019</td><td>149041</td><td>-</td><td>0</td><td>ATTCTGCCTGTTCTTCATCC</td><td>TGG</td><td>||||||||[||||||||||||]PAM</td><td>NA</td><td>Intergenic</td><td>NA</td><td>NA</td></tr>
<tr><td></td><td></td><td></td><td></td><td></td><td></td><td></td><td></td><td></td><td></td><td></td><td></td></tr>
<tr><td>T22807</td><td>GTTCTTCATCCTGGACGCCGAGG</td><td>1000</td><td></td><td></td><td></td><td></td><td></td><td></td><td></td><td></td><td></td></tr>
<tr><td>Oligo fwd</td><td>CACCGTTCTTCATCCTGGACGCCG</td><td></td><td></td><td></td><td></td><td></td><td></td><td></td><td></td><td></td><td></td></tr>
<tr><td>Oligo rev</td><td>AAACCGGCGTCCAGGATGAAGAAC</td><td></td><td></td><td></td><td></td><td></td><td></td><td></td><td></td><td></td><td></td></tr>
<tr><td>Chromosome</td><td>start</td><td>end</td><td>strand</td><td>MM</td><td>target_seq</td><td>PAM</td><td>alignment</td><td>distance</td><td>position</td><td>gene name</td><td>gene id</td></tr>
<tr><td>GU305914.1</td><td>149010</td><td>149032</td><td>-</td><td>0</td><td>GTTCTTCATCCTGGACGCCG</td><td>AGG</td><td>||||||||[||||||||||||]PAM</td><td>NA</td><td>Intergenic</td><td>NA</td><td>NA</td></tr>
</table>

| | | | | | | | | | | | |
|---|---|---|---|---|---|---|---|---|---|---|---|
| GU305914.1 | 187153 | 187175 | + | 4 | GTTTATCATGGTGGACGCCG | GGG | \|\|\|--\|\|\|[\|--\|\|\|\|\|\|\|\|\|]PAM | NA | Intergenic | NA | NA |
| T22808 | GGACGCCGAGGACGCCCTGTCGG | 1000 | | | | | | | | | |
| Oligo fwd | CACCGGACGCCGAGGACGCCCTGT | | | | | | | | | | |
| Oligo rev | AAACACAGGGCGTCCTCGGCGTCC | | | | | | | | | | |
| Chromosome | start | end | strand | MM | target_seq | PAM | alignment | distance | position | gene name | gene id |
| GU305914.1 | 148998 | 149020 | - | 0 | GGACGCCGAGGACGCCCTGT | CGG | \|\|\|\|\|\|\|\|[\|\|\|\|\|\|\|\|\|\|\|\|]PAM | NA | Intergenic | NA | NA |
| T22809 | AGGACGCCCTGTCGGAGCTGCGG | 1000 | | | | | | | | | |
| Oligo fwd | CACCAGGACGCCCTGTCGGAGCTG | | | | | | | | | | |
| Oligo rev | AAACCAGCTCCGACAGGGCGTCCT | | | | | | | | | | |
| Chromosome | start | end | strand | MM | target_seq | PAM | alignment | distance | position | gene name | gene id |
| GU305914.1 | 148990 | 149012 | - | 0 | AGGACGCCCTGTCGGAGCTG | CGG | \|\|\|\|\|\|\|\|[\|\|\|\|\|\|\|\|\|\|\|\|]PAM | NA | Intergenic | NA | NA |
| T22810 | AGCTGCGGCTGTCCTCGATCCGG | 1000 | | | | | | | | | |
| Oligo fwd | CACCAGCTGCGGCTGTCCTCGATC | | | | | | | | | | |
| Oligo rev | AAACGATCGAGGACAGCCGCAGCT | | | | | | | | | | |
| Chromosome | start | end | strand | MM | target_seq | PAM | alignment | distance | position | gene name | gene id |
| GU305914.1 | 148975 | 148997 | - | 0 | AGCTGCGGCTGTCCTCGATC | CGG | \|\|\|\|\|\|\|\|[\|\|\|\|\|\|\|\|\|\|\|\|]PAM | NA | Intergenic | NA | NA |
| T22811 | CCTCGATCCGGACCAAGCACCGG | 1000 | | | | | | | | | |
| Oligo fwd | CACCCCTCGATCCGGACCAAGCAC | | | | | | | | | | |
| Oligo rev | AAACGTGCTTGGTCCGGATCGAGG | | | | | | | | | | |
| Chromosome | start | end | strand | MM | target_seq | PAM | alignment | distance | position | gene name | gene id |
| GU305914.1 | 148963 | 148985 | - | 0 | CCTCGATCCGGACCAAGCAC | CGG | \|\|\|\|\|\|\|\|[\|\|\|\|\|\|\|\|\|\|\|\|]PAM | NA | Intergenic | NA | NA |
| T22812 | GTTCCGCCCGAAGACCAACACGG | 1000 | | | | | | | | | |
| Oligo fwd | CACCGTTCCGCCCGAAGACCAACA | | | | | | | | | | |
| Oligo rev | AAACTGTTGGTCTTCGGGCGGAAC | | | | | | | | | | |
| Chromosome | start | end | strand | MM | target_seq | PAM | alignment | distance | position | gene name | gene id |
| GU305914.1 | 148932 | 148954 | - | 0 | GTTCCGCCCGAAGACCAACA | CGG | \|\|\|\|\|\|\|\|[\|\|\|\|\|\|\|\|\|\|\|\|]PAM | NA | Intergenic | NA | NA |
| T22813 | GCCGCTGTCCAGACAGACGCTGG | 1000 | | | | | | | | | |
| Oligo fwd | CACCGCCGCTGTCCAGACAGACGC | | | | | | | | | | |
| Oligo rev | AAACGCGTCTGTCTGGACAGCGGC | | | | | | | | | | |
| Chromosome | start | end | strand | MM | target_seq | PAM | alignment | distance | position | gene name | gene id |
| GU305914.1 | 148869 | 148891 | - | 0 | GCCGCTGTCCAGACAGACGC | TGG | \|\|\|\|\|\|\|\|[\|\|\|\|\|\|\|\|\|\|\|\|]PAM | NA | Intergenic | NA | NA |
| T22814 | CAGACAGACGCTGGAGCACCTGG | 1000 | | | | | | | | | |
| Oligo fwd | CACCCAGACAGACGCTGGAGCACC | | | | | | | | | | |
| Oligo rev | AAACGGTGCTCCAGCGTCTGTCTG | | | | | | | | | | |
| Chromosome | start | end | strand | MM | target_seq | PAM | alignment | distance | position | gene name | gene id |
| GU305914.1 | 148860 | 148882 | - | 0 | CAGACAGACGCTGGAGCACC | TGG | \|\|\|\|\|\|\|\|[\|\|\|\|\|\|\|\|\|\|\|\|]PAM | NA | Intergenic | NA | NA |
| T22815 | CACCTGGTGACCTTCCTGACCGG | 1000 | | | | | | | | | |
| Oligo fwd | CACCCACCTGGTGACCTTCCTGAC | | | | | | | | | | |
| Oligo rev | AAACGTCAGGAAGGTCACCAGGTG | | | | | | | | | | |
| Chromosome | start | end | strand | MM | target_seq | PAM | alignment | distance | position | gene name | gene id |
| GU305914.1 | 148844 | 148866 | - | 0 | CACCTGGTGACCTTCCTGAC | CGG | \|\|\|\|\|\|\|\|[\|\|\|\|\|\|\|\|\|\|\|\|]PAM | NA | Intergenic | NA | NA |
| T22816 | TGACCTTCCTGACCGGCACCAGG | 1000 | | | | | | | | | |
| Oligo fwd | CACCTGACCTTCCTGACCGGCACC | | | | | | | | | | |
| Oligo rev | AAACGGTGCCGGTCAGGAAGGTCA | | | | | | | | | | |
| Chromosome | start | end | strand | MM | target_seq | PAM | alignment | distance | position | gene name | gene id |
| GU305914.1 | 148837 | 148859 | - | 0 | TGACCTTCCTGACCGGCACC | AGG | \|\|\|\|\|\|\|\|[\|\|\|\|\|\|\|\|\|\|\|\|]PAM | NA | Intergenic | NA | NA |
| T22817 | GACCGGCACCAGGCCCCGCACGG | 1000 | | | | | | | | | |
| Oligo fwd | CACCGACCGGCACCAGGCCCCGCA | | | | | | | | | | |
| Oligo rev | AAACTGCGGGGCCTGGTGCCGGTC | | | | | | | | | | |
| Chromosome | start | end | strand | MM | target_seq | PAM | alignment | distance | position | gene name | gene id |
| GU305914.1 | 148827 | 148849 | - | 0 | GACCGGCACCAGGCCCCGCA | CGG | \|\|\|\|\|\|\|\|[\|\|\|\|\|\|\|\|\|\|\|\|]PAM | NA | Intergenic | NA | NA |
| T22818 | ACCGGCACCAGGCCCCGCACGGG | 1000 | | | | | | | | | |
| Oligo fwd | CACCACCGGCACCAGGCCCCGCAC | | | | | | | | | | |
| Oligo rev | AAACGTGCGGGGCCTGGTGCCGGT | | | | | | | | | | |
| Chromosome | start | end | strand | MM | target_seq | PAM | alignment | distance | position | gene name | gene id |
| GU305914.1 | 148826 | 148848 | - | 0 | ACCGGCACCAGGCCCCGCAC | GGG | \|\|\|\|\|\|\|\|[\|\|\|\|\|\|\|\|\|\|\|\|]PAM | NA | Intergenic | NA | NA |
| T22819 | CCGGCACCAGGCCCCGCACGGGG | 1000 | | | | | | | | | |
| Oligo fwd | CACCCCGGCACCAGGCCCCGCACG | | | | | | | | | | |
| Oligo rev | AAACCGTGCGGGGCCTGGTGCCGG | | | | | | | | | | |
| Chromosome | start | end | strand | MM | target_seq | PAM | alignment | distance | position | gene name | gene id |
| GU305914.1 | 148825 | 148847 | - | 0 | CCGGCACCAGGCCCCGCACG | GGG | \|\|\|\|\|\|\|\|[\|\|\|\|\|\|\|\|\|\|\|\|]PAM | NA | Intergenic | NA | NA |
| T22820 | GGCCCCGCACGGGGCTGACCCGG | 1000 | | | | | | | | | |
| Oligo fwd | CACCGGCCCCGCACGGGGCTGACC | | | | | | | | | | |
| Oligo rev | AAACGGTCAGCCCCGTGCGGGGCC | | | | | | | | | | |
| Chromosome | start | end | strand | MM | target_seq | PAM | alignment | distance | position | gene name | gene id |
| GU305914.1 | 148816 | 148838 | - | 0 | GGCCCCGCACGGGGCTGACC | CGG | \|\|\|\|\|\|\|\|[\|\|\|\|\|\|\|\|\|\|\|\|]PAM | NA | Intergenic | NA | NA |
| T22821 | CTGCAAGAGCTGCATCAAGCTGG | 1000 | | | | | | | | | |
| Oligo fwd | CACCCTGCAAGAGCTGCATCAAGC | | | | | | | | | | |
| Oligo rev | AAACGCTTGATGCAGCTCTTGCAG | | | | | | | | | | |
| Chromosome | start | end | strand | MM | target_seq | PAM | alignment | distance | position | gene name | gene id |
| GU305914.1 | 148788 | 148810 | - | 0 | CTGCAAGAGCTGCATCAAGC | TGG | \|\|\|\|\|\|\|\|[\|\|\|\|\|\|\|\|\|\|\|\|]PAM | NA | Intergenic | NA | NA |
| T22822 | GTACCTGTTCTTCGACGAGACGG | 1000 | | | | | | | | | |
| Oligo fwd | CACCGTACCTGTTCTTCGACGAGA | | | | | | | | | | |
| Oligo rev | AAACTCTCGTCGAAGAACAGGTAC | | | | | | | | | | |

Chromosome start end strand MM target_seq PAM alignment distance position gene name gene id
GU305914.1 148752 148774 - 0 GTACCTGTTCTTCGACGAGA CGG ||||||||[||||||||||||]PAM NA Intergenic NA NA

T22823 TCTTCGACGAGACGGACCCGAGG 1000
Oligo fwd CACCTCTTCGACGAGACGGACCCG
Oligo rev AAACCGGGTCCGTCTCGTCGAAGA
Chromosome start end strand MM target_seq PAM alignment distance position gene name gene id
GU305914.1 148744 148766 - 0 TCTTCGACGAGACGGACCCG AGG ||||||||[||||||||||||]PAM NA Intergenic NA NA

T22824 CGAGATCTGCATGCTGTACAAGG 1000
Oligo fwd CACCCGAGATCTGCATGCTGTACA
Oligo rev AAACTGTACAGCATGCAGATCTCG
Chromosome start end strand MM target_seq PAM alignment distance position gene name gene id
GU305914.1 148704 148726 - 0 CGAGATCTGCATGCTGTACA AGG ||||||||[||||||||||||]PAM NA Intergenic NA NA

T22825 CAAGGAGACCCAACGCGCCCAGG 1000
Oligo fwd CACCCAAGGAGACCCAACGCGCCC
Oligo rev AAACGGGCGCGTTGGGTCTCCTTG
Chromosome start end strand MM target_seq PAM alignment distance position gene name gene id
GU305914.1 148686 148708 - 0 CAAGGAGACCCAACGCGCCC AGG ||||||||[||||||||||||]PAM NA Intergenic NA NA

T22826 ACGCGCCCAGGCGACCATGCTGG 1000
Oligo fwd CACCACGCGCCCAGGCGACCATGC
Oligo rev AAACGCATGGTCGCCTGGGCGCGT
Chromosome start end strand MM target_seq PAM alignment distance position gene name gene id
GU305914.1 148674 148696 - 0 ACGCGCCCAGGCGACCATGC TGG ||||||||[||||||||||||]PAM NA Intergenic NA NA

T22827 ATGCTGGCCGAGACGTACTTCGG 1000
Oligo fwd CACCATGCTGGCCGAGACGTACTT
Oligo rev AAACAAGTACGTCTCGGCCAGCAT
Chromosome start end strand MM target_seq PAM alignment distance position gene name gene id
GU305914.1 148658 148680 - 0 ATGCTGGCCGAGACGTACTT CGG ||||||||[||||||||||||]PAM NA Intergenic NA NA

T22828 CGAGACGTACTTCGGCGTCCAGG 1000
Oligo fwd CACCCGAGACGTACTTCGGCGTCC
Oligo rev AAACGGACGCCGAAGTACGTCTCG
Chromosome start end strand MM target_seq PAM alignment distance position gene name gene id
GU305914.1 148650 148672 - 0 CGAGACGTACTTCGGCGTCC AGG ||||||||[||||||||||||]PAM NA Intergenic NA NA

T22829 GGACATCAGCAGCATGTCTCTGG 1000
Oligo fwd CACCGGACATCAGCAGCATGTCTC
Oligo rev AAACGAGACATGCTGCTGATGTCC
Chromosome start end strand MM target_seq PAM alignment distance position gene name gene id
GU305914.1 148629 148651 - 0 GGACATCAGCAGCATGTCTC TGG ||||||||[||||||||||||]PAM NA Intergenic NA NA

T22830 TCTGGTGTCGCTGACGCTGACGG 1000
Oligo fwd CACCTCTGGTGTCGCTGACGCTGA
Oligo rev AAACTCAGCGTCAGCGACACCAGA
Chromosome start end strand MM target_seq PAM alignment distance position gene name gene id
GU305914.1 148611 148633 - 0 TCTGGTGTCGCTGACGCTGA CGG ||||||||[||||||||||||]PAM NA Intergenic NA NA

T22831 TGTCGCTGACGCTGACGGACCGG 1000
Oligo fwd CACCTGTCGCTGACGCTGACGGAC
Oligo rev AAACGTCCGTCAGCGTCAGCGACA
Chromosome start end strand MM target_seq PAM alignment distance position gene name gene id
GU305914.1 148606 148628 - 0 TGTCGCTGACGCTGACGGAC CGG ||||||||[||||||||||||]PAM NA Intergenic NA NA

T22832 GTCGCTGACGCTGACGGACCGGG 1000
Oligo fwd CACCGTCGCTGACGCTGACGGACC
Oligo rev AAACGGTCCGTCAGCGTCAGCGAC
Chromosome start end strand MM target_seq PAM alignment distance position gene name gene id
GU305914.1 148605 148627 - 0 GTCGCTGACGCTGACGGACC GGG ||||||||[||||||||||||]PAM NA Intergenic NA NA

T22833 GCTGACGGACCGGGTGACCAAGG 1000
Oligo fwd CACCGCTGACGGACCGGGTGACCA
Oligo rev AAACTGGTCACCCGGTCCGTCAGC
Chromosome start end strand MM target_seq PAM alignment distance position gene name gene id
GU305914.1 148596 148618 - 0 GCTGACGGACCGGGTGACCA AGG ||||||||[||||||||||||]PAM NA Intergenic NA NA

T22834 ACGGACCGGGTGACCAAGGACGG 1000
Oligo fwd CACCACGGACCGGGTGACCAAGGA
Oligo rev AAACTCCTTGGTCACCCGGTCCGT
Chromosome start end strand MM target_seq PAM alignment distance position gene name gene id
GU305914.1 148592 148614 - 0 ACGGACCGGGTGACCAAGGA CGG ||||||||[||||||||||||]PAM NA Intergenic NA NA

T22835 GACCAAGGACGGCGACACCGTGG 1000
Oligo fwd CACCGACCAAGGACGGCGACACCG
Oligo rev AAACCGGTGTCGCCGTCCTTGGTC
Chromosome start end strand MM target_seq PAM alignment distance position gene name gene id
GU305914.1 148581 148603 - 0 GACCAAGGACGGCGACACCG TGG ||||||||[||||||||||||]PAM NA Intergenic NA NA

T22836 ACCAAGGACGGCGACACCGTGGG 1000
Oligo fwd CACCACCAAGGACGGCGACACCGT
Oligo rev AAACACGGTGTCGCCGTCCTTGGT
Chromosome start end strand MM target_seq PAM alignment distance position gene name gene id
GU305914.1 148580 148602 - 0 ACCAAGGACGGCGACACCGT GGG ||||||||[||||||||||||]PAM NA Intergenic NA NA

T22837 CGGCGACACCGTGGGCGACCTGG 1000
Oligo fwd CACCCGGCGACACCGTGGGCGACC

<table>
<tr><td>Oligo rev</td><td>AAACGGTCGCCCACGGTGTCGCCG</td><td></td><td></td><td></td><td></td><td></td><td></td><td></td><td></td><td></td><td></td></tr>
<tr><td>Chromosome</td><td>start</td><td>end</td><td>strand</td><td>MM</td><td>target_seq</td><td>PAM</td><td>alignment</td><td>distance</td><td>position</td><td>gene name</td><td>gene id</td></tr>
<tr><td>GU305914.1</td><td>148572</td><td>148594</td><td>-</td><td>0</td><td>CGGCGACACCGTGGGCGACC</td><td>TGG</td><td>||||||||[||||||||||||]PAM</td><td>NA</td><td>Intergenic</td><td>NA</td><td>NA</td></tr>
<tr><td>GU305914.1</td><td>46331</td><td>46353</td><td>+</td><td>4</td><td>GGTCGAGGCCGTGGGCGACC</td><td>TGG</td><td>-|-|||--[||||||||||||]PAM</td><td>NA</td><td>Intergenic</td><td>NA</td><td>NA</td></tr>
<tr><td></td><td></td><td></td><td></td><td></td><td></td><td></td><td></td><td></td><td></td><td></td><td></td></tr>
<tr><td>T22838</td><td>CGACACCGTGGGCGACCTGGCGG</td><td>1000</td><td></td><td></td><td></td><td></td><td></td><td></td><td></td><td></td><td></td></tr>
<tr><td>Oligo fwd</td><td>CACCCGACACCGTGGGCGACCTGG</td><td></td><td></td><td></td><td></td><td></td><td></td><td></td><td></td><td></td><td></td></tr>
<tr><td>Oligo rev</td><td>AAACCCAGGTCGCCCACGGTGTCG</td><td></td><td></td><td></td><td></td><td></td><td></td><td></td><td></td><td></td><td></td></tr>
<tr><td>Chromosome</td><td>start</td><td>end</td><td>strand</td><td>MM</td><td>target_seq</td><td>PAM</td><td>alignment</td><td>distance</td><td>position</td><td>gene name</td><td>gene id</td></tr>
<tr><td>GU305914.1</td><td>148569</td><td>148591</td><td>-</td><td>0</td><td>CGACACCGTGGGCGACCTGG</td><td>CGG</td><td>||||||||[||||||||||||]PAM</td><td>NA</td><td>Intergenic</td><td>NA</td><td>NA</td></tr>
<tr><td></td><td></td><td></td><td></td><td></td><td></td><td></td><td></td><td></td><td></td><td></td><td></td></tr>
<tr><td>T22839</td><td>CGTGGGCGACCTGGCGGCCGAGG</td><td>1000</td><td></td><td></td><td></td><td></td><td></td><td></td><td></td><td></td><td></td></tr>
<tr><td>Oligo fwd</td><td>CACCCGTGGGCGACCTGGCGGCCG</td><td></td><td></td><td></td><td></td><td></td><td></td><td></td><td></td><td></td><td></td></tr>
<tr><td>Oligo rev</td><td>AAACCGGCCGCCAGGTCGCCCACG</td><td></td><td></td><td></td><td></td><td></td><td></td><td></td><td></td><td></td><td></td></tr>
<tr><td>Chromosome</td><td>start</td><td>end</td><td>strand</td><td>MM</td><td>target_seq</td><td>PAM</td><td>alignment</td><td>distance</td><td>position</td><td>gene name</td><td>gene id</td></tr>
<tr><td>GU305914.1</td><td>148563</td><td>148585</td><td>-</td><td>0</td><td>CGTGGGCGACCTGGCGGCCG</td><td>AGG</td><td>||||||||[||||||||||||]PAM</td><td>NA</td><td>Intergenic</td><td>NA</td><td>NA</td></tr>
<tr><td></td><td></td><td></td><td></td><td></td><td></td><td></td><td></td><td></td><td></td><td></td><td></td></tr>
<tr><td>T22840</td><td>TAACGTGTTTTACGTGCCACTGG</td><td>1000</td><td></td><td></td><td></td><td></td><td></td><td></td><td></td><td></td><td></td></tr>
<tr><td>Oligo fwd</td><td>CACCTAACGTGTTTTACGTGCCAC</td><td></td><td></td><td></td><td></td><td></td><td></td><td></td><td></td><td></td><td></td></tr>
<tr><td>Oligo rev</td><td>AAACGTGGCACGTAAAACACGTTA</td><td></td><td></td><td></td><td></td><td></td><td></td><td></td><td></td><td></td><td></td></tr>
<tr><td>Chromosome</td><td>start</td><td>end</td><td>strand</td><td>MM</td><td>target_seq</td><td>PAM</td><td>alignment</td><td>distance</td><td>position</td><td>gene name</td><td>gene id</td></tr>
<tr><td>GU305914.1</td><td>148524</td><td>148546</td><td>-</td><td>0</td><td>TAACGTGTTTTACGTGCCAC</td><td>TGG</td><td>||||||||[||||||||||||]PAM</td><td>NA</td><td>Intergenic</td><td>NA</td><td>NA</td></tr>
<tr><td></td><td></td><td></td><td></td><td></td><td></td><td></td><td></td><td></td><td></td><td></td><td></td></tr>
<tr><td>T22841</td><td>AACGTGTTTTACGTGCCACTGGG</td><td>1000</td><td></td><td></td><td></td><td></td><td></td><td></td><td></td><td></td><td></td></tr>
<tr><td>Oligo fwd</td><td>CACCAACGTGTTTTACGTGCCACT</td><td></td><td></td><td></td><td></td><td></td><td></td><td></td><td></td><td></td><td></td></tr>
<tr><td>Oligo rev</td><td>AAACAGTGGCACGTAAAACACGTT</td><td></td><td></td><td></td><td></td><td></td><td></td><td></td><td></td><td></td><td></td></tr>
<tr><td>Chromosome</td><td>start</td><td>end</td><td>strand</td><td>MM</td><td>target_seq</td><td>PAM</td><td>alignment</td><td>distance</td><td>position</td><td>gene name</td><td>gene id</td></tr>
<tr><td>GU305914.1</td><td>148523</td><td>148545</td><td>-</td><td>0</td><td>AACGTGTTTTACGTGCCACT</td><td>GGG</td><td>||||||||[||||||||||||]PAM</td><td>NA</td><td>Intergenic</td><td>NA</td><td>NA</td></tr>
<tr><td></td><td></td><td></td><td></td><td></td><td></td><td></td><td></td><td></td><td></td><td></td><td></td></tr>
<tr><td>T22842</td><td>ACGTGTTTTACGTGCCACTGGGG</td><td>1000</td><td></td><td></td><td></td><td></td><td></td><td></td><td></td><td></td><td></td></tr>
<tr><td>Oligo fwd</td><td>CACCACGTGTTTTACGTGCCACTG</td><td></td><td></td><td></td><td></td><td></td><td></td><td></td><td></td><td></td><td></td></tr>
<tr><td>Oligo rev</td><td>AAACCAGTGGCACGTAAAACACGT</td><td></td><td></td><td></td><td></td><td></td><td></td><td></td><td></td><td></td><td></td></tr>
<tr><td>Chromosome</td><td>start</td><td>end</td><td>strand</td><td>MM</td><td>target_seq</td><td>PAM</td><td>alignment</td><td>distance</td><td>position</td><td>gene name</td><td>gene id</td></tr>
<tr><td>GU305914.1</td><td>148522</td><td>148544</td><td>-</td><td>0</td><td>ACGTGTTTTACGTGCCACTG</td><td>GGG</td><td>||||||||[||||||||||||]PAM</td><td>NA</td><td>Intergenic</td><td>NA</td><td>NA</td></tr>
<tr><td></td><td></td><td></td><td></td><td></td><td></td><td></td><td></td><td></td><td></td><td></td><td></td></tr>
<tr><td>T22843</td><td>TACGTGCCACTGGGGACCAACGG</td><td>1000</td><td></td><td></td><td></td><td></td><td></td><td></td><td></td><td></td><td></td></tr>
<tr><td>Oligo fwd</td><td>CACCTACGTGCCACTGGGGACCAA</td><td></td><td></td><td></td><td></td><td></td><td></td><td></td><td></td><td></td><td></td></tr>
<tr><td>Oligo rev</td><td>AAACTTGGTCCCCAGTGGCACGTA</td><td></td><td></td><td></td><td></td><td></td><td></td><td></td><td></td><td></td><td></td></tr>
<tr><td>Chromosome</td><td>start</td><td>end</td><td>strand</td><td>MM</td><td>target_seq</td><td>PAM</td><td>alignment</td><td>distance</td><td>position</td><td>gene name</td><td>gene id</td></tr>
<tr><td>GU305914.1</td><td>148514</td><td>148536</td><td>-</td><td>0</td><td>TACGTGCCACTGGGGACCAA</td><td>CGG</td><td>||||||||[||||||||||||]PAM</td><td>NA</td><td>Intergenic</td><td>NA</td><td>NA</td></tr>
<tr><td></td><td></td><td></td><td></td><td></td><td></td><td></td><td></td><td></td><td></td><td></td><td></td></tr>
<tr><td>T22844</td><td>CCACTGGGGACCAACGGTTCCGG</td><td>1000</td><td></td><td></td><td></td><td></td><td></td><td></td><td></td><td></td><td></td></tr>
<tr><td>Oligo fwd</td><td>CACCCCACTGGGGACCAACGGTTC</td><td></td><td></td><td></td><td></td><td></td><td></td><td></td><td></td><td></td><td></td></tr>
<tr><td>Oligo rev</td><td>AAACGAACCGTTGGTCCCCAGTGG</td><td></td><td></td><td></td><td></td><td></td><td></td><td></td><td></td><td></td><td></td></tr>
<tr><td>Chromosome</td><td>start</td><td>end</td><td>strand</td><td>MM</td><td>target_seq</td><td>PAM</td><td>alignment</td><td>distance</td><td>position</td><td>gene name</td><td>gene id</td></tr>
<tr><td>GU305914.1</td><td>148508</td><td>148530</td><td>-</td><td>0</td><td>CCACTGGGGACCAACGGTTC</td><td>CGG</td><td>||||||||[||||||||||||]PAM</td><td>NA</td><td>Intergenic</td><td>NA</td><td>NA</td></tr>
<tr><td></td><td></td><td></td><td></td><td></td><td></td><td></td><td></td><td></td><td></td><td></td><td></td></tr>
<tr><td>T22845</td><td>GTTCCGGTGTCGTGACCCGCTGG</td><td>1000</td><td></td><td></td><td></td><td></td><td></td><td></td><td></td><td></td><td></td></tr>
<tr><td>Oligo fwd</td><td>CACCGTTCCGGTGTCGTGACCCGC</td><td></td><td></td><td></td><td></td><td></td><td></td><td></td><td></td><td></td><td></td></tr>
<tr><td>Oligo rev</td><td>AAACGCGGGTCACGACACCGGAAC</td><td></td><td></td><td></td><td></td><td></td><td></td><td></td><td></td><td></td><td></td></tr>
<tr><td>Chromosome</td><td>start</td><td>end</td><td>strand</td><td>MM</td><td>target_seq</td><td>PAM</td><td>alignment</td><td>distance</td><td>position</td><td>gene name</td><td>gene id</td></tr>
<tr><td>GU305914.1</td><td>148492</td><td>148514</td><td>-</td><td>0</td><td>GTTCCGGTGTCGTGACCCGC</td><td>TGG</td><td>||||||||[||||||||||||]PAM</td><td>NA</td><td>Intergenic</td><td>NA</td><td>NA</td></tr>
<tr><td></td><td></td><td></td><td></td><td></td><td></td><td></td><td></td><td></td><td></td><td></td><td></td></tr>
<tr><td>T22846</td><td>AATACCACCACCACTACTCCCGG</td><td>1000</td><td></td><td></td><td></td><td></td><td></td><td></td><td></td><td></td><td></td></tr>
<tr><td>Oligo fwd</td><td>CACCAATACCACCACCACTACTCC</td><td></td><td></td><td></td><td></td><td></td><td></td><td></td><td></td><td></td><td></td></tr>
<tr><td>Oligo rev</td><td>AAACGGAGTAGTGGTGGTGGTATT</td><td></td><td></td><td></td><td></td><td></td><td></td><td></td><td></td><td></td><td></td></tr>
<tr><td>Chromosome</td><td>start</td><td>end</td><td>strand</td><td>MM</td><td>target_seq</td><td>PAM</td><td>alignment</td><td>distance</td><td>position</td><td>gene name</td><td>gene id</td></tr>
<tr><td>GU305914.1</td><td>148319</td><td>148341</td><td>-</td><td>0</td><td>AATACCACCACCACTACTCC</td><td>CGG</td><td>||||||||[||||||||||||]PAM</td><td>NA</td><td>Intergenic</td><td>NA</td><td>NA</td></tr>
<tr><td></td><td></td><td></td><td></td><td></td><td></td><td></td><td></td><td></td><td></td><td></td><td></td></tr>
<tr><td>T22847</td><td>TGAAAATGAAGCATATACAGTGG</td><td>1000</td><td></td><td></td><td></td><td></td><td></td><td></td><td></td><td></td><td></td></tr>
<tr><td>Oligo fwd</td><td>CACCTGAAAATGAAGCATATACAG</td><td></td><td></td><td></td><td></td><td></td><td></td><td></td><td></td><td></td><td></td></tr>
<tr><td>Oligo rev</td><td>AAACCTGTATATGCTTCATTTTCA</td><td></td><td></td><td></td><td></td><td></td><td></td><td></td><td></td><td></td><td></td></tr>
<tr><td>Chromosome</td><td>start</td><td>end</td><td>strand</td><td>MM</td><td>target_seq</td><td>PAM</td><td>alignment</td><td>distance</td><td>position</td><td>gene name</td><td>gene id</td></tr>
<tr><td>GU305914.1</td><td>148208</td><td>148230</td><td>-</td><td>0</td><td>TGAAAATGAAGCATATACAG</td><td>TGG</td><td>||||||||[||||||||||||]PAM</td><td>NA</td><td>Intergenic</td><td>NA</td><td>NA</td></tr>
<tr><td></td><td></td><td></td><td></td><td></td><td></td><td></td><td></td><td></td><td></td><td></td><td></td></tr>
<tr><td>T22848</td><td>TAGTTAGTTATAGTTAGTTATGG</td><td>1000</td><td></td><td></td><td></td><td></td><td></td><td></td><td></td><td></td><td></td></tr>
<tr><td>Oligo fwd</td><td>CACCTAGTTAGTTATAGTTAGTTA</td><td></td><td></td><td></td><td></td><td></td><td></td><td></td><td></td><td></td><td></td></tr>
<tr><td>Oligo rev</td><td>AAACTAACTAACTATAACTAACTA</td><td></td><td></td><td></td><td></td><td></td><td></td><td></td><td></td><td></td><td></td></tr>
<tr><td>Chromosome</td><td>start</td><td>end</td><td>strand</td><td>MM</td><td>target_seq</td><td>PAM</td><td>alignment</td><td>distance</td><td>position</td><td>gene name</td><td>gene id</td></tr>
<tr><td>GU305914.1</td><td>148185</td><td>148207</td><td>-</td><td>0</td><td>TAGTTAGTTATAGTTAGTTA</td><td>TGG</td><td>||||||||[||||||||||||]PAM</td><td>NA</td><td>Intergenic</td><td>NA</td><td>NA</td></tr>
<tr><td></td><td></td><td></td><td></td><td></td><td></td><td></td><td></td><td></td><td></td><td></td><td></td></tr>
<tr><td>T22849</td><td>CAGTCAGTTCATAGAAAAGAAGG</td><td>1000</td><td></td><td></td><td></td><td></td><td></td><td></td><td></td><td></td><td></td></tr>
<tr><td>Oligo fwd</td><td>CACCCAGTCAGTTCATAGAAAAGA</td><td></td><td></td><td></td><td></td><td></td><td></td><td></td><td></td><td></td><td></td></tr>
<tr><td>Oligo rev</td><td>AAACTCTTTTCTATGAACTGACTG</td><td></td><td></td><td></td><td></td><td></td><td></td><td></td><td></td><td></td><td></td></tr>
<tr><td>Chromosome</td><td>start</td><td>end</td><td>strand</td><td>MM</td><td>target_seq</td><td>PAM</td><td>alignment</td><td>distance</td><td>position</td><td>gene name</td><td>gene id</td></tr>
<tr><td>GU305914.1</td><td>148112</td><td>148134</td><td>-</td><td>0</td><td>CAGTCAGTTCATAGAAAAGA</td><td>AGG</td><td>||||||||[||||||||||||]PAM</td><td>NA</td><td>Intergenic</td><td>NA</td><td>NA</td></tr>
<tr><td></td><td></td><td></td><td></td><td></td><td></td><td></td><td></td><td></td><td></td><td></td><td></td></tr>
<tr><td>T22850</td><td>CATCGTTAAACTCTTCGATCAGG</td><td>1000</td><td></td><td></td><td></td><td></td><td></td><td></td><td></td><td></td><td></td></tr>
<tr><td>Oligo fwd</td><td>CACCCATCGTTAAACTCTTCGATC</td><td></td><td></td><td></td><td></td><td></td><td></td><td></td><td></td><td></td><td></td></tr>
<tr><td>Oligo rev</td><td>AAACGATCGAAGAGTTTAACGATG</td><td></td><td></td><td></td><td></td><td></td><td></td><td></td><td></td><td></td><td></td></tr>
<tr><td>Chromosome</td><td>start</td><td>end</td><td>strand</td><td>MM</td><td>target_seq</td><td>PAM</td><td>alignment</td><td>distance</td><td>position</td><td>gene name</td><td>gene id</td></tr>
<tr><td>GU305914.1</td><td>148047</td><td>148069</td><td>-</td><td>0</td><td>CATCGTTAAACTCTTCGATC</td><td>AGG</td><td>||||||||[||||||||||||]PAM</td><td>NA</td><td>Intergenic</td><td>NA</td><td>NA</td></tr>
<tr><td></td><td></td><td></td><td></td><td></td><td></td><td></td><td></td><td></td><td></td><td></td><td></td></tr>
<tr><td>T22851</td><td>TTCGATCAGGTTGTGCGTGTCGG</td><td>1000</td><td></td><td></td><td></td><td></td><td></td><td></td><td></td><td></td><td></td></tr>
<tr><td>Oligo fwd</td><td>CACCTTCGATCAGGTTGTGCGTGT</td><td></td><td></td><td></td><td></td><td></td><td></td><td></td><td></td><td></td><td></td></tr>
<tr><td>Oligo rev</td><td>AAACACACGCACAACCTGATCGAA</td><td></td><td></td><td></td><td></td><td></td><td></td><td></td><td></td><td></td><td></td></tr>
<tr><td>Chromosome</td><td>start</td><td>end</td><td>strand</td><td>MM</td><td>target_seq</td><td>PAM</td><td>alignment</td><td>distance</td><td>position</td><td>gene name</td><td>gene id</td></tr>
<tr><td>GU305914.1</td><td>148034</td><td>148056</td><td>-</td><td>0</td><td>TTCGATCAGGTTGTGCGTGT</td><td>CGG</td><td>||||||||[||||||||||||]PAM</td><td>NA</td><td>Intergenic</td><td>NA</td><td>NA</td></tr>
</table>

<table>
<tr><td>T22852</td><td>GCGTTCCACCACCATGCCGACGG</td><td>1000</td><td></td><td></td><td></td><td></td><td></td><td></td><td></td><td></td><td></td></tr>
<tr><td>Oligo fwd</td><td>CACCGCGTTCCACCACCATGCCGA</td><td></td><td></td><td></td><td></td><td></td><td></td><td></td><td></td><td></td><td></td></tr>
<tr><td>Oligo rev</td><td>AAACTCGGCATGGTGGTGGAACGC</td><td></td><td></td><td></td><td></td><td></td><td></td><td></td><td></td><td></td><td></td></tr>
<tr><td>Chromosome</td><td>start</td><td>end</td><td>strand</td><td>MM</td><td>target_seq</td><td>PAM</td><td>alignment</td><td>distance</td><td>position</td><td>gene name</td><td>gene id</td></tr>
<tr><td>GU305914.1</td><td>148010</td><td>148032</td><td>-</td><td>0</td><td>GCGTTCCACCACCATGCCGA</td><td>CGG</td><td>||||||||[||||||||||||]PAM</td><td>NA</td><td>Intergenic</td><td>NA</td><td>NA</td></tr>
<tr><td>GU305914.1</td><td>61549</td><td>61571</td><td>+</td><td>4</td><td>GGCGTCCACCACCATGCCGT</td><td>AGG</td><td>|---||||[|||||||||||-]PAM</td><td>NA</td><td>Intergenic</td><td>NA</td><td>NA</td></tr>
<tr><td>T22853</td><td>CACCACCATGCCGACGGTGTTGG</td><td>1000</td><td></td><td></td><td></td><td></td><td></td><td></td><td></td><td></td><td></td></tr>
<tr><td>Oligo fwd</td><td>CACCCACCACCATGCCGACGGTGT</td><td></td><td></td><td></td><td></td><td></td><td></td><td></td><td></td><td></td><td></td></tr>
<tr><td>Oligo rev</td><td>AAACACACCGTCGGCATGGTGGTG</td><td></td><td></td><td></td><td></td><td></td><td></td><td></td><td></td><td></td><td></td></tr>
<tr><td>Chromosome</td><td>start</td><td>end</td><td>strand</td><td>MM</td><td>target_seq</td><td>PAM</td><td>alignment</td><td>distance</td><td>position</td><td>gene name</td><td>gene id</td></tr>
<tr><td>GU305914.1</td><td>148004</td><td>148026</td><td>-</td><td>0</td><td>CACCACCATGCCGACGGTGT</td><td>TGG</td><td>||||||||[||||||||||||]PAM</td><td>NA</td><td>Intergenic</td><td>NA</td><td>NA</td></tr>
<tr><td>T22854</td><td>CATGCCGACGGTGTTGGCCGTGG</td><td>1000</td><td></td><td></td><td></td><td></td><td></td><td></td><td></td><td></td><td></td></tr>
<tr><td>Oligo fwd</td><td>CACCCATGCCGACGGTGTTGGCCG</td><td></td><td></td><td></td><td></td><td></td><td></td><td></td><td></td><td></td><td></td></tr>
<tr><td>Oligo rev</td><td>AAACCGGCCAACACCGTCGGCATG</td><td></td><td></td><td></td><td></td><td></td><td></td><td></td><td></td><td></td><td></td></tr>
<tr><td>Chromosome</td><td>start</td><td>end</td><td>strand</td><td>MM</td><td>target_seq</td><td>PAM</td><td>alignment</td><td>distance</td><td>position</td><td>gene name</td><td>gene id</td></tr>
<tr><td>GU305914.1</td><td>147998</td><td>148020</td><td>-</td><td>0</td><td>CATGCCGACGGTGTTGGCCG</td><td>TGG</td><td>||||||||[||||||||||||]PAM</td><td>NA</td><td>Intergenic</td><td>NA</td><td>NA</td></tr>
<tr><td>T22855</td><td>TGTTGGCCGTGGTCTCCCTGAGG</td><td>1000</td><td></td><td></td><td></td><td></td><td></td><td></td><td></td><td></td><td></td></tr>
<tr><td>Oligo fwd</td><td>CACCTGTTGGCCGTGGTCTCCCTG</td><td></td><td></td><td></td><td></td><td></td><td></td><td></td><td></td><td></td><td></td></tr>
<tr><td>Oligo rev</td><td>AAACCAGGGAGACCACGGCCAACA</td><td></td><td></td><td></td><td></td><td></td><td></td><td></td><td></td><td></td><td></td></tr>
<tr><td>Chromosome</td><td>start</td><td>end</td><td>strand</td><td>MM</td><td>target_seq</td><td>PAM</td><td>alignment</td><td>distance</td><td>position</td><td>gene name</td><td>gene id</td></tr>
<tr><td>GU305914.1</td><td>147987</td><td>148009</td><td>-</td><td>0</td><td>TGTTGGCCGTGGTCTCCCTG</td><td>AGG</td><td>||||||||[||||||||||||]PAM</td><td>NA</td><td>Intergenic</td><td>NA</td><td>NA</td></tr>
<tr><td>T22856</td><td>GTTGGCCGTGGTCTCCCTGAGGG</td><td>1000</td><td></td><td></td><td></td><td></td><td></td><td></td><td></td><td></td><td></td></tr>
<tr><td>Oligo fwd</td><td>CACCGTTGGCCGTGGTCTCCCTGA</td><td></td><td></td><td></td><td></td><td></td><td></td><td></td><td></td><td></td><td></td></tr>
<tr><td>Oligo rev</td><td>AAACTCAGGGAGACCACGGCCAAC</td><td></td><td></td><td></td><td></td><td></td><td></td><td></td><td></td><td></td><td></td></tr>
<tr><td>Chromosome</td><td>start</td><td>end</td><td>strand</td><td>MM</td><td>target_seq</td><td>PAM</td><td>alignment</td><td>distance</td><td>position</td><td>gene name</td><td>gene id</td></tr>
<tr><td>GU305914.1</td><td>147986</td><td>148008</td><td>-</td><td>0</td><td>GTTGGCCGTGGTCTCCCTGA</td><td>GGG</td><td>||||||||[||||||||||||]PAM</td><td>NA</td><td>Intergenic</td><td>NA</td><td>NA</td></tr>
<tr><td>T22857</td><td>CTCCCTGAGGGCCCGTAGCGCGG</td><td>1000</td><td></td><td></td><td></td><td></td><td></td><td></td><td></td><td></td><td></td></tr>
<tr><td>Oligo fwd</td><td>CACCCTCCCTGAGGGCCCGTAGCG</td><td></td><td></td><td></td><td></td><td></td><td></td><td></td><td></td><td></td><td></td></tr>
<tr><td>Oligo rev</td><td>AAACCGCTACGGGCCCTCAGGGAG</td><td></td><td></td><td></td><td></td><td></td><td></td><td></td><td></td><td></td><td></td></tr>
<tr><td>Chromosome</td><td>start</td><td>end</td><td>strand</td><td>MM</td><td>target_seq</td><td>PAM</td><td>alignment</td><td>distance</td><td>position</td><td>gene name</td><td>gene id</td></tr>
<tr><td>GU305914.1</td><td>147974</td><td>147996</td><td>-</td><td>0</td><td>CTCCCTGAGGGCCCGTAGCG</td><td>CGG</td><td>||||||||[||||||||||||]PAM</td><td>NA</td><td>Intergenic</td><td>NA</td><td>NA</td></tr>
<tr><td>T22858</td><td>GGCGTCCAGCTCATCCGAGCCGG</td><td>1000</td><td></td><td></td><td></td><td></td><td></td><td></td><td></td><td></td><td></td></tr>
<tr><td>Oligo fwd</td><td>CACCGGCGTCCAGCTCATCCGAGC</td><td></td><td></td><td></td><td></td><td></td><td></td><td></td><td></td><td></td><td></td></tr>
<tr><td>Oligo rev</td><td>AAACGCTCGGATGAGCTGGACGCC</td><td></td><td></td><td></td><td></td><td></td><td></td><td></td><td></td><td></td><td></td></tr>
<tr><td>Chromosome</td><td>start</td><td>end</td><td>strand</td><td>MM</td><td>target_seq</td><td>PAM</td><td>alignment</td><td>distance</td><td>position</td><td>gene name</td><td>gene id</td></tr>
<tr><td>GU305914.1</td><td>147953</td><td>147975</td><td>-</td><td>0</td><td>GGCGTCCAGCTCATCCGAGC</td><td>CGG</td><td>||||||||[||||||||||||]PAM</td><td>NA</td><td>Intergenic</td><td>NA</td><td>NA</td></tr>
<tr><td>GU305914.1</td><td>76466</td><td>76488</td><td>-</td><td>4</td><td>GGCGGCGAGCTCATCCGCGG</td><td>CGG</td><td>||||-|-|[|||||||||-|-]PAM</td><td>NA</td><td>Intergenic</td><td>NA</td><td>NA</td></tr>
<tr><td>T22859</td><td>GTCCAGCTCATCCGAGCCGGTGG</td><td>1000</td><td></td><td></td><td></td><td></td><td></td><td></td><td></td><td></td><td></td></tr>
<tr><td>Oligo fwd</td><td>CACCGTCCAGCTCATCCGAGCCGG</td><td></td><td></td><td></td><td></td><td></td><td></td><td></td><td></td><td></td><td></td></tr>
<tr><td>Oligo rev</td><td>AAACCCGGCTCGGATGAGCTGGAC</td><td></td><td></td><td></td><td></td><td></td><td></td><td></td><td></td><td></td><td></td></tr>
<tr><td>Chromosome</td><td>start</td><td>end</td><td>strand</td><td>MM</td><td>target_seq</td><td>PAM</td><td>alignment</td><td>distance</td><td>position</td><td>gene name</td><td>gene id</td></tr>
<tr><td>GU305914.1</td><td>147950</td><td>147972</td><td>-</td><td>0</td><td>GTCCAGCTCATCCGAGCCGG</td><td>TGG</td><td>||||||||[||||||||||||]PAM</td><td>NA</td><td>Intergenic</td><td>NA</td><td>NA</td></tr>
<tr><td>T22860</td><td>GTCAGTGACCTCCTGCTTGATGG</td><td>1000</td><td></td><td></td><td></td><td></td><td></td><td></td><td></td><td></td><td></td></tr>
<tr><td>Oligo fwd</td><td>CACCGTCAGTGACCTCCTGCTTGA</td><td></td><td></td><td></td><td></td><td></td><td></td><td></td><td></td><td></td><td></td></tr>
<tr><td>Oligo rev</td><td>AAACTCAAGCAGGAGGTCACTGAC</td><td></td><td></td><td></td><td></td><td></td><td></td><td></td><td></td><td></td><td></td></tr>
<tr><td>Chromosome</td><td>start</td><td>end</td><td>strand</td><td>MM</td><td>target_seq</td><td>PAM</td><td>alignment</td><td>distance</td><td>position</td><td>gene name</td><td>gene id</td></tr>
<tr><td>GU305914.1</td><td>147908</td><td>147930</td><td>-</td><td>0</td><td>GTCAGTGACCTCCTGCTTGA</td><td>TGG</td><td>||||||||[||||||||||||]PAM</td><td>NA</td><td>Intergenic</td><td>NA</td><td>NA</td></tr>
<tr><td>T22861</td><td>CGTGCTGTTGTTGACGACCGCGG</td><td>1000</td><td></td><td></td><td></td><td></td><td></td><td></td><td></td><td></td><td></td></tr>
<tr><td>Oligo fwd</td><td>CACCCGTGCTGTTGTTGACGACCG</td><td></td><td></td><td></td><td></td><td></td><td></td><td></td><td></td><td></td><td></td></tr>
<tr><td>Oligo rev</td><td>AAACCGGTCGTCAACAACAGCACG</td><td></td><td></td><td></td><td></td><td></td><td></td><td></td><td></td><td></td><td></td></tr>
<tr><td>Chromosome</td><td>start</td><td>end</td><td>strand</td><td>MM</td><td>target_seq</td><td>PAM</td><td>alignment</td><td>distance</td><td>position</td><td>gene name</td><td>gene id</td></tr>
<tr><td>GU305914.1</td><td>147866</td><td>147888</td><td>-</td><td>0</td><td>CGTGCTGTTGTTGACGACCG</td><td>CGG</td><td>||||||||[||||||||||||]PAM</td><td>NA</td><td>Intergenic</td><td>NA</td><td>NA</td></tr>
<tr><td>T22862</td><td>GTTGTTGACGACCGCGGTGTTGG</td><td>1000</td><td></td><td></td><td></td><td></td><td></td><td></td><td></td><td></td><td></td></tr>
<tr><td>Oligo fwd</td><td>CACCGTTGTTGACGACCGCGGTGT</td><td></td><td></td><td></td><td></td><td></td><td></td><td></td><td></td><td></td><td></td></tr>
<tr><td>Oligo rev</td><td>AAACACACCGCGGTCGTCAACAAC</td><td></td><td></td><td></td><td></td><td></td><td></td><td></td><td></td><td></td><td></td></tr>
<tr><td>Chromosome</td><td>start</td><td>end</td><td>strand</td><td>MM</td><td>target_seq</td><td>PAM</td><td>alignment</td><td>distance</td><td>position</td><td>gene name</td><td>gene id</td></tr>
<tr><td>GU305914.1</td><td>147860</td><td>147882</td><td>-</td><td>0</td><td>GTTGTTGACGACCGCGGTGT</td><td>TGG</td><td>||||||||[||||||||||||]PAM</td><td>NA</td><td>Intergenic</td><td>NA</td><td>NA</td></tr>
<tr><td>T22863</td><td>TGACGACCGCGGTGTTGGTCTGG</td><td>1000</td><td></td><td></td><td></td><td></td><td></td><td></td><td></td><td></td><td></td></tr>
<tr><td>Oligo fwd</td><td>CACCTGACGACCGCGGTGTTGGTC</td><td></td><td></td><td></td><td></td><td></td><td></td><td></td><td></td><td></td><td></td></tr>
<tr><td>Oligo rev</td><td>AAACGACCAACACCGCGGTCGTCA</td><td></td><td></td><td></td><td></td><td></td><td></td><td></td><td></td><td></td><td></td></tr>
<tr><td>Chromosome</td><td>start</td><td>end</td><td>strand</td><td>MM</td><td>target_seq</td><td>PAM</td><td>alignment</td><td>distance</td><td>position</td><td>gene name</td><td>gene id</td></tr>
<tr><td>GU305914.1</td><td>147855</td><td>147877</td><td>-</td><td>0</td><td>TGACGACCGCGGTGTTGGTC</td><td>TGG</td><td>||||||||[||||||||||||]PAM</td><td>NA</td><td>Intergenic</td><td>NA</td><td>NA</td></tr>
<tr><td>T22864</td><td>TGTTGGTCTGGTCCTGCGAGAGG</td><td>1000</td><td></td><td></td><td></td><td></td><td></td><td></td><td></td><td></td><td></td></tr>
<tr><td>Oligo fwd</td><td>CACCTGTTGGTCTGGTCCTGCGAG</td><td></td><td></td><td></td><td></td><td></td><td></td><td></td><td></td><td></td><td></td></tr>
<tr><td>Oligo rev</td><td>AAACCTCGCAGGACCAGACCAACA</td><td></td><td></td><td></td><td></td><td></td><td></td><td></td><td></td><td></td><td></td></tr>
<tr><td>Chromosome</td><td>start</td><td>end</td><td>strand</td><td>MM</td><td>target_seq</td><td>PAM</td><td>alignment</td><td>distance</td><td>position</td><td>gene name</td><td>gene id</td></tr>
<tr><td>GU305914.1</td><td>147843</td><td>147865</td><td>-</td><td>0</td><td>TGTTGGTCTGGTCCTGCGAG</td><td>AGG</td><td>||||||||[||||||||||||]PAM</td><td>NA</td><td>Intergenic</td><td>NA</td><td>NA</td></tr>
<tr><td>T22865</td><td>GGACGATGATGACCTTCTCGAGG</td><td>1000</td><td></td><td></td><td></td><td></td><td></td><td></td><td></td><td></td><td></td></tr>
<tr><td>Oligo fwd</td><td>CACCGGACGATGATGACCTTCTCG</td><td></td><td></td><td></td><td></td><td></td><td></td><td></td><td></td><td></td><td></td></tr>
<tr><td>Oligo rev</td><td>AAACCGAGAAGGTCATCATCGTCC</td><td></td><td></td><td></td><td></td><td></td><td></td><td></td><td></td><td></td><td></td></tr>
<tr><td>Chromosome</td><td>start</td><td>end</td><td>strand</td><td>MM</td><td>target_seq</td><td>PAM</td><td>alignment</td><td>distance</td><td>position</td><td>gene name</td><td>gene id</td></tr>
<tr><td>GU305914.1</td><td>147822</td><td>147844</td><td>-</td><td>0</td><td>GGACGATGATGACCTTCTCG</td><td>AGG</td><td>||||||||[||||||||||||]PAM</td><td>NA</td><td>Intergenic</td><td>NA</td><td>NA</td></tr>
<tr><td>T22866</td><td>GACGATGATGACCTTCTCGAGGG</td><td>1000</td><td></td><td></td><td></td><td></td><td></td><td></td><td></td><td></td><td></td></tr>
<tr><td>Oligo fwd</td><td>CACCGACGATGATGACCTTCTCGA</td><td></td><td></td><td></td><td></td><td></td><td></td><td></td><td></td><td></td><td></td></tr>
<tr><td>Oligo rev</td><td>AAACTCGAGAAGGTCATCATCGTC</td><td></td><td></td><td></td><td></td><td></td><td></td><td></td><td></td><td></td><td></td></tr>
</table>

| | | | | | | | | | | | |
|---|---|---|---|---|---|---|---|---|---|---|---|
| Chromosome | start | end | strand | MM | target_seq | PAM | alignment | distance | position | gene name | gene id |
| GU305914.1 | 147821 | 147843 | - | 0 | GACGATGATGACCTTCTCGA | GGG | \|\|\|\|\|\|\|\|[\|\|\|\|\|\|\|\|\|\|\|\|]PAM | NA | Intergenic | NA | NA |
| GU305914.1 | 81370 | 81392 | + | 4 | GTCGAAGATGACCTTCCTGA | GGG | \|-\|\|\|-\|\|[\|\|\|\|\|\|\|\|--\|\|]PAM | NA | Intergenic | NA | NA |
| T22867 | GGTCACCCTGAGCACCTCGAAGG | 1000 | | | | | | | | | |
| Oligo fwd | CACCGGTCACCCTGAGCACCTCGA | | | | | | | | | | |
| Oligo rev | AAACTCGAGGTGCTCAGGGTGACC | | | | | | | | | | |
| Chromosome | start | end | strand | MM | target_seq | PAM | alignment | distance | position | gene name | gene id |
| GU305914.1 | 147800 | 147822 | - | 0 | GGTCACCCTGAGCACCTCGA | AGG | \|\|\|\|\|\|\|\|[\|\|\|\|\|\|\|\|\|\|\|\|]PAM | NA | Intergenic | NA | NA |
| GU305914.1 | 186642 | 186664 | - | 4 | GATCACGCCGAGAACCTCGA | GGG | \|-\|\|\|\|-\|[-\|\|\|-\|\|\|\|\|\|\|]PAM | NA | Intergenic | NA | NA |
| T22868 | TGTAGCAGATGTTGTCGTGCAGG | 1000 | | | | | | | | | |
| Oligo fwd | CACCTGTAGCAGATGTTGTCGTGC | | | | | | | | | | |
| Oligo rev | AAACGCACGACAACATCTGCTACA | | | | | | | | | | |
| Chromosome | start | end | strand | MM | target_seq | PAM | alignment | distance | position | gene name | gene id |
| GU305914.1 | 147759 | 147781 | - | 0 | TGTAGCAGATGTTGTCGTGC | AGG | \|\|\|\|\|\|\|\|[\|\|\|\|\|\|\|\|\|\|\|\|]PAM | NA | Intergenic | NA | NA |
| T22869 | GTAGCAGATGTTGTCGTGCAGGG | 1000 | | | | | | | | | |
| Oligo fwd | CACCGTAGCAGATGTTGTCGTGCA | | | | | | | | | | |
| Oligo rev | AAACTGCACGACAACATCTGCTAC | | | | | | | | | | |
| Chromosome | start | end | strand | MM | target_seq | PAM | alignment | distance | position | gene name | gene id |
| GU305914.1 | 147758 | 147780 | - | 0 | GTAGCAGATGTTGTCGTGCA | GGG | \|\|\|\|\|\|\|\|[\|\|\|\|\|\|\|\|\|\|\|\|]PAM | NA | Intergenic | NA | NA |
| T22870 | TGTCGTGCAGGGCCCCGTCCAGG | 1000 | | | | | | | | | |
| Oligo fwd | CACCTGTCGTGCAGGGCCCCGTCC | | | | | | | | | | |
| Oligo rev | AAACGGACGGGGCCCTGCACGACA | | | | | | | | | | |
| Chromosome | start | end | strand | MM | target_seq | PAM | alignment | distance | position | gene name | gene id |
| GU305914.1 | 147747 | 147769 | - | 0 | TGTCGTGCAGGGCCCCGTCC | AGG | \|\|\|\|\|\|\|\|[\|\|\|\|\|\|\|\|\|\|\|\|]PAM | NA | Intergenic | NA | NA |
| T22871 | GTCGTGCAGGGCCCCGTCCAGGG | 1000 | | | | | | | | | |
| Oligo fwd | CACCGTCGTGCAGGGCCCCGTCCA | | | | | | | | | | |
| Oligo rev | AAACTGGACGGGGCCCTGCACGAC | | | | | | | | | | |
| Chromosome | start | end | strand | MM | target_seq | PAM | alignment | distance | position | gene name | gene id |
| GU305914.1 | 147746 | 147768 | - | 0 | GTCGTGCAGGGCCCCGTCCA | GGG | \|\|\|\|\|\|\|\|[\|\|\|\|\|\|\|\|\|\|\|\|]PAM | NA | Intergenic | NA | NA |
| T22872 | TCGTGCAGGGCCCCGTCCAGGGG | 1000 | | | | | | | | | |
| Oligo fwd | CACCTCGTGCAGGGCCCCGTCCAG | | | | | | | | | | |
| Oligo rev | AAACCTGGACGGGGCCCTGCACGA | | | | | | | | | | |
| Chromosome | start | end | strand | MM | target_seq | PAM | alignment | distance | position | gene name | gene id |
| GU305914.1 | 147745 | 147767 | - | 0 | TCGTGCAGGGCCCCGTCCAG | GGG | \|\|\|\|\|\|\|\|[\|\|\|\|\|\|\|\|\|\|\|\|]PAM | NA | Intergenic | NA | NA |
| T22873 | GCAGGGCCCCGTCCAGGGGCTGG | 1000 | | | | | | | | | |
| Oligo fwd | CACCGCAGGGCCCCGTCCAGGGGC | | | | | | | | | | |
| Oligo rev | AAACGCCCCTGGACGGGGCCCTGC | | | | | | | | | | |
| Chromosome | start | end | strand | MM | target_seq | PAM | alignment | distance | position | gene name | gene id |
| GU305914.1 | 147741 | 147763 | - | 0 | GCAGGGCCCCGTCCAGGGGC | TGG | \|\|\|\|\|\|\|\|[\|\|\|\|\|\|\|\|\|\|\|\|]PAM | NA | Intergenic | NA | NA |
| T22874 | GGCCCCGTCCAGGGGCTGGACGG | 1000 | | | | | | | | | |
| Oligo fwd | CACCGGCCCCGTCCAGGGGCTGGA | | | | | | | | | | |
| Oligo rev | AAACTCCAGCCCCTGGACGGGGCC | | | | | | | | | | |
| Chromosome | start | end | strand | MM | target_seq | PAM | alignment | distance | position | gene name | gene id |
| GU305914.1 | 147737 | 147759 | - | 0 | GGCCCCGTCCAGGGGCTGGA | CGG | \|\|\|\|\|\|\|\|[\|\|\|\|\|\|\|\|\|\|\|\|]PAM | NA | Intergenic | NA | NA |
| T22875 | GGGGCTGGACGGCCACCATGCGG | 1000 | | | | | | | | | |
| Oligo fwd | CACCGGGGCTGGACGGCCACCATG | | | | | | | | | | |
| Oligo rev | AAACCATGGTGGCCGTCCAGCCCC | | | | | | | | | | |
| Chromosome | start | end | strand | MM | target_seq | PAM | alignment | distance | position | gene name | gene id |
| GU305914.1 | 147726 | 147748 | - | 0 | GGGGCTGGACGGCCACCATG | CGG | \|\|\|\|\|\|\|\|[\|\|\|\|\|\|\|\|\|\|\|\|]PAM | NA | Intergenic | NA | NA |
| T22876 | CCACCATGCGGCCGCAGACGTGG | 1000 | | | | | | | | | |
| Oligo fwd | CACCCCACCATGCGGCCGCAGACG | | | | | | | | | | |
| Oligo rev | AAACCGTCTGCGGCCGCATGGTGG | | | | | | | | | | |
| Chromosome | start | end | strand | MM | target_seq | PAM | alignment | distance | position | gene name | gene id |
| GU305914.1 | 147714 | 147736 | - | 0 | CCACCATGCGGCCGCAGACG | TGG | \|\|\|\|\|\|\|\|[\|\|\|\|\|\|\|\|\|\|\|\|]PAM | NA | Intergenic | NA | NA |
| T22877 | GTGGATCCCGCAGAACTCGTTGG | 1000 | | | | | | | | | |
| Oligo fwd | CACCGTGGATCCCGCAGAACTCGT | | | | | | | | | | |
| Oligo rev | AAACACGAGTTCTGCGGGATCCAC | | | | | | | | | | |
| Chromosome | start | end | strand | MM | target_seq | PAM | alignment | distance | position | gene name | gene id |
| GU305914.1 | 147695 | 147717 | - | 0 | GTGGATCCCGCAGAACTCGT | TGG | \|\|\|\|\|\|\|\|[\|\|\|\|\|\|\|\|\|\|\|\|]PAM | NA | Intergenic | NA | NA |
| T22878 | CCATGACGCCGTCGTGCTGCTGG | 1000 | | | | | | | | | |
| Oligo fwd | CACCCCATGACGCCGTCGTGCTGC | | | | | | | | | | |
| Oligo rev | AAACGCAGCACGACGGCGTCATGG | | | | | | | | | | |
| Chromosome | start | end | strand | MM | target_seq | PAM | alignment | distance | position | gene name | gene id |
| GU305914.1 | 147669 | 147691 | - | 0 | CCATGACGCCGTCGTGCTGC | TGG | \|\|\|\|\|\|\|\|[\|\|\|\|\|\|\|\|\|\|\|\|]PAM | NA | Intergenic | NA | NA |
| T22879 | CGCCGTCGTGCTGCTGGACGCGG | 1000 | | | | | | | | | |
| Oligo fwd | CACCCGCCGTCGTGCTGCTGGACG | | | | | | | | | | |
| Oligo rev | AAACCGTCCAGCAGCACGACGGCG | | | | | | | | | | |
| Chromosome | start | end | strand | MM | target_seq | PAM | alignment | distance | position | gene name | gene id |
| GU305914.1 | 147663 | 147685 | - | 0 | CGCCGTCGTGCTGCTGGACG | CGG | \|\|\|\|\|\|\|\|[\|\|\|\|\|\|\|\|\|\|\|\|]PAM | NA | Intergenic | NA | NA |
| GU305914.1 | 125138 | 125160 | - | 3 | CGACGACGTGCCGCTGGACG | CGG | \|\|-\|\|-\|\|[\|\|\|-\|\|\|\|\|\|\|\|]PAM | NA | Intergenic | NA | NA |
| T22880 | CTGCTGGACGCGGTTGCGTCCGG | 1000 | | | | | | | | | |
| Oligo fwd | CACCCTGCTGGACGCGGTTGCGTC | | | | | | | | | | |
| Oligo rev | AAACGACGCAACCGCGTCCAGCAG | | | | | | | | | | |
| Chromosome | start | end | strand | MM | target_seq | PAM | alignment | distance | position | gene name | gene id |
| GU305914.1 | 147653 | 147675 | - | 0 | CTGCTGGACGCGGTTGCGTC | CGG | \|\|\|\|\|\|\|\|[\|\|\|\|\|\|\|\|\|\|\|\|]PAM | NA | Intergenic | NA | NA |

<table>
<tr><td>T22881</td><td>TGCTGGACGCGGTTGCGTCCGGG</td><td>1000</td></tr>
<tr><td>Oligo fwd</td><td>CACCTGCTGGACGCGGTTGCGTCC</td></tr>
<tr><td>Oligo rev</td><td>AAACGGACGCAACCGCGTCCAGCA</td></tr>
<tr><td>Chromosome</td><td>start</td><td>end</td><td>strand</td><td>MM</td><td>target_seq</td><td>PAM</td><td>alignment</td><td>distance</td><td>position</td><td>gene name</td><td>gene id</td></tr>
<tr><td>GU305914.1</td><td>147652</td><td>147674</td><td>-</td><td>0</td><td>TGCTGGACGCGGTTGCGTCC</td><td>GGG</td><td>||||||||[||||||||||||]PAM</td><td>NA</td><td>Intergenic</td><td>NA</td><td>NA</td></tr>
<tr><td>T22882</td><td>GCTGGACGCGGTTGCGTCCGGGG</td><td>1000</td></tr>
<tr><td>Oligo fwd</td><td>CACCGCTGGACGCGGTTGCGTCCG</td></tr>
<tr><td>Oligo rev</td><td>AAACCGGACGCAACCGCGTCCAGC</td></tr>
<tr><td>Chromosome</td><td>start</td><td>end</td><td>strand</td><td>MM</td><td>target_seq</td><td>PAM</td><td>alignment</td><td>distance</td><td>position</td><td>gene name</td><td>gene id</td></tr>
<tr><td>GU305914.1</td><td>147651</td><td>147673</td><td>-</td><td>0</td><td>GCTGGACGCGGTTGCGTCCG</td><td>GGG</td><td>||||||||[||||||||||||]PAM</td><td>NA</td><td>Intergenic</td><td>NA</td><td>NA</td></tr>
<tr><td>T22883</td><td>CTGGACGCGGTTGCGTCCGGGGG</td><td>1000</td></tr>
<tr><td>Oligo fwd</td><td>CACCCTGGACGCGGTTGCGTCCGG</td></tr>
<tr><td>Oligo rev</td><td>AAACCCGGACGCAACCGCGTCCAG</td></tr>
<tr><td>Chromosome</td><td>start</td><td>end</td><td>strand</td><td>MM</td><td>target_seq</td><td>PAM</td><td>alignment</td><td>distance</td><td>position</td><td>gene name</td><td>gene id</td></tr>
<tr><td>GU305914.1</td><td>147650</td><td>147672</td><td>-</td><td>0</td><td>CTGGACGCGGTTGCGTCCGG</td><td>GGG</td><td>||||||||[||||||||||||]PAM</td><td>NA</td><td>Intergenic</td><td>NA</td><td>NA</td></tr>
<tr><td>T22884</td><td>CGAACCTCTGCGCCCAGAAGCGG</td><td>1000</td></tr>
<tr><td>Oligo fwd</td><td>CACCCGAACCTCTGCGCCCAGAAG</td></tr>
<tr><td>Oligo rev</td><td>AAACCTTCTGGGCGCAGAGGTTCG</td></tr>
<tr><td>Chromosome</td><td>start</td><td>end</td><td>strand</td><td>MM</td><td>target_seq</td><td>PAM</td><td>alignment</td><td>distance</td><td>position</td><td>gene name</td><td>gene id</td></tr>
<tr><td>GU305914.1</td><td>147627</td><td>147649</td><td>-</td><td>0</td><td>CGAACCTCTGCGCCCAGAAG</td><td>CGG</td><td>||||||||[||||||||||||]PAM</td><td>NA</td><td>Intergenic</td><td>NA</td><td>NA</td></tr>
<tr><td>T22885</td><td>GAACCTCTGCGCCCAGAAGCGGG</td><td>1000</td></tr>
<tr><td>Oligo fwd</td><td>CACCGAACCTCTGCGCCCAGAAGC</td></tr>
<tr><td>Oligo rev</td><td>AAACGCTTCTGGGCGCAGAGGTTC</td></tr>
<tr><td>Chromosome</td><td>start</td><td>end</td><td>strand</td><td>MM</td><td>target_seq</td><td>PAM</td><td>alignment</td><td>distance</td><td>position</td><td>gene name</td><td>gene id</td></tr>
<tr><td>GU305914.1</td><td>147626</td><td>147648</td><td>-</td><td>0</td><td>GAACCTCTGCGCCCAGAAGC</td><td>GGG</td><td>||||||||[||||||||||||]PAM</td><td>NA</td><td>Intergenic</td><td>NA</td><td>NA</td></tr>
<tr><td>T22886</td><td>GAAGCGGGTCTCGTGCCTCCTGG</td><td>1000</td></tr>
<tr><td>Oligo fwd</td><td>CACCGAAGCGGGTCTCGTGCCTCC</td></tr>
<tr><td>Oligo rev</td><td>AAACGGAGGCACGAGACCCGCTTC</td></tr>
<tr><td>Chromosome</td><td>start</td><td>end</td><td>strand</td><td>MM</td><td>target_seq</td><td>PAM</td><td>alignment</td><td>distance</td><td>position</td><td>gene name</td><td>gene id</td></tr>
<tr><td>GU305914.1</td><td>147611</td><td>147633</td><td>-</td><td>0</td><td>GAAGCGGGTCTCGTGCCTCC</td><td>TGG</td><td>||||||||[||||||||||||]PAM</td><td>NA</td><td>Intergenic</td><td>NA</td><td>NA</td></tr>
<tr><td>T22887</td><td>TCTCGTGCCTCCTGGCCGAGAGG</td><td>1000</td></tr>
<tr><td>Oligo fwd</td><td>CACCTCTCGTGCCTCCTGGCCGAG</td></tr>
<tr><td>Oligo rev</td><td>AAACCTCGGCCAGGAGGCACGAGA</td></tr>
<tr><td>Chromosome</td><td>start</td><td>end</td><td>strand</td><td>MM</td><td>target_seq</td><td>PAM</td><td>alignment</td><td>distance</td><td>position</td><td>gene name</td><td>gene id</td></tr>
<tr><td>GU305914.1</td><td>147603</td><td>147625</td><td>-</td><td>0</td><td>TCTCGTGCCTCCTGGCCGAG</td><td>AGG</td><td>||||||||[||||||||||||]PAM</td><td>NA</td><td>Intergenic</td><td>NA</td><td>NA</td></tr>
<tr><td>T22888</td><td>CGAGAGGCGCAGCATGTGCGTGG</td><td>1000</td></tr>
<tr><td>Oligo fwd</td><td>CACCCGAGAGGCGCAGCATGTGCG</td></tr>
<tr><td>Oligo rev</td><td>AAACCGCACATGCTGCGCCTCTCG</td></tr>
<tr><td>Chromosome</td><td>start</td><td>end</td><td>strand</td><td>MM</td><td>target_seq</td><td>PAM</td><td>alignment</td><td>distance</td><td>position</td><td>gene name</td><td>gene id</td></tr>
<tr><td>GU305914.1</td><td>147587</td><td>147609</td><td>-</td><td>0</td><td>CGAGAGGCGCAGCATGTGCG</td><td>TGG</td><td>||||||||[||||||||||||]PAM</td><td>NA</td><td>Intergenic</td><td>NA</td><td>NA</td></tr>
<tr><td>T22889</td><td>GCATGTGCGTGGCCATGAAGCGG</td><td>1000</td></tr>
<tr><td>Oligo fwd</td><td>CACCGCATGTGCGTGGCCATGAAG</td></tr>
<tr><td>Oligo rev</td><td>AAACCTTCATGGCCACGCACATGC</td></tr>
<tr><td>Chromosome</td><td>start</td><td>end</td><td>strand</td><td>MM</td><td>target_seq</td><td>PAM</td><td>alignment</td><td>distance</td><td>position</td><td>gene name</td><td>gene id</td></tr>
<tr><td>GU305914.1</td><td>147576</td><td>147598</td><td>-</td><td>0</td><td>GCATGTGCGTGGCCATGAAG</td><td>CGG</td><td>||||||||[||||||||||||]PAM</td><td>NA</td><td>Intergenic</td><td>NA</td><td>NA</td></tr>
<tr><td>T22890</td><td>GAAGCGGATGATCTTGTCTTTGG</td><td>1000</td></tr>
<tr><td>Oligo fwd</td><td>CACCGAAGCGGATGATCTTGTCTT</td></tr>
<tr><td>Oligo rev</td><td>AAACAAGACAAGATCATCCGCTTC</td></tr>
<tr><td>Chromosome</td><td>start</td><td>end</td><td>strand</td><td>MM</td><td>target_seq</td><td>PAM</td><td>alignment</td><td>distance</td><td>position</td><td>gene name</td><td>gene id</td></tr>
<tr><td>GU305914.1</td><td>147560</td><td>147582</td><td>-</td><td>0</td><td>GAAGCGGATGATCTTGTCTT</td><td>TGG</td><td>||||||||[||||||||||||]PAM</td><td>NA</td><td>Intergenic</td><td>NA</td><td>NA</td></tr>
<tr><td>T22891</td><td>CTTGTCTTTGGCGCCCCAATCGG</td><td>1000</td></tr>
<tr><td>Oligo fwd</td><td>CACCCTTGTCTTTGGCGCCCCAAT</td></tr>
<tr><td>Oligo rev</td><td>AAACATTGGGGCGCCAAAGACAAG</td></tr>
<tr><td>Chromosome</td><td>start</td><td>end</td><td>strand</td><td>MM</td><td>target_seq</td><td>PAM</td><td>alignment</td><td>distance</td><td>position</td><td>gene name</td><td>gene id</td></tr>
<tr><td>GU305914.1</td><td>147548</td><td>147570</td><td>-</td><td>0</td><td>CTTGTCTTTGGCGCCCCAAT</td><td>CGG</td><td>||||||||[||||||||||||]PAM</td><td>NA</td><td>Intergenic</td><td>NA</td><td>NA</td></tr>
<tr><td>T22892</td><td>TTGTCTTTGGCGCCCCAATCGGG</td><td>1000</td></tr>
<tr><td>Oligo fwd</td><td>CACCTTGTCTTTGGCGCCCCAATC</td></tr>
<tr><td>Oligo rev</td><td>AAACGATTGGGGCGCCAAAGACAA</td></tr>
<tr><td>Chromosome</td><td>start</td><td>end</td><td>strand</td><td>MM</td><td>target_seq</td><td>PAM</td><td>alignment</td><td>distance</td><td>position</td><td>gene name</td><td>gene id</td></tr>
<tr><td>GU305914.1</td><td>147547</td><td>147569</td><td>-</td><td>0</td><td>TTGTCTTTGGCGCCCCAATC</td><td>GGG</td><td>||||||||[||||||||||||]PAM</td><td>NA</td><td>Intergenic</td><td>NA</td><td>NA</td></tr>
<tr><td>T22893</td><td>TGTCTTTGGCGCCCCAATCGGGG</td><td>1000</td></tr>
<tr><td>Oligo fwd</td><td>CACCTGTCTTTGGCGCCCCAATCG</td></tr>
<tr><td>Oligo rev</td><td>AAACCGATTGGGGCGCCAAAGACA</td></tr>
<tr><td>Chromosome</td><td>start</td><td>end</td><td>strand</td><td>MM</td><td>target_seq</td><td>PAM</td><td>alignment</td><td>distance</td><td>position</td><td>gene name</td><td>gene id</td></tr>
<tr><td>GU305914.1</td><td>147546</td><td>147568</td><td>-</td><td>0</td><td>TGTCTTTGGCGCCCCAATCG</td><td>GGG</td><td>||||||||[||||||||||||]PAM</td><td>NA</td><td>Intergenic</td><td>NA</td><td>NA</td></tr>
<tr><td>T22894</td><td>GTCTTTGGCGCCCCAATCGGGGG</td><td>1000</td></tr>
<tr><td>Oligo fwd</td><td>CACCGTCTTTGGCGCCCCAATCGG</td></tr>
<tr><td>Oligo rev</td><td>AAACCCGATTGGGGCGCCAAAGAC</td></tr>
<tr><td>Chromosome</td><td>start</td><td>end</td><td>strand</td><td>MM</td><td>target_seq</td><td>PAM</td><td>alignment</td><td>distance</td><td>position</td><td>gene name</td><td>gene id</td></tr>
<tr><td>GU305914.1</td><td>147545</td><td>147567</td><td>-</td><td>0</td><td>GTCTTTGGCGCCCCAATCGG</td><td>GGG</td><td>||||||||[||||||||||||]PAM</td><td>NA</td><td>Intergenic</td><td>NA</td><td>NA</td></tr>
<tr><td>T22895</td><td>TGGCGCCCCAATCGGGGGACTGG</td><td>1000</td></tr>
<tr><td>Oligo fwd</td><td>CACCTGGCGCCCCAATCGGGGGAC</td></tr>
<tr><td>Oligo rev</td><td>AAACGTCCCCCGATTGGGGCGCCA</td></tr>
<tr><td>Chromosome</td><td>start</td><td>end</td><td>strand</td><td>MM</td><td>target_seq</td><td>PAM</td><td>alignment</td><td>distance</td><td>position</td><td>gene name</td><td>gene id</td></tr>
</table>

<table>
<tr><td>GU305914.1</td><td></td><td>147540</td><td>147562</td><td>-</td><td>0</td><td>TGGCGCCCCAATCGGGGGAC</td><td>TGG</td><td>||||||||[||||||||||||]PAM</td><td>NA</td><td>Intergenic</td><td>NA</td><td>NA</td></tr>
<tr><td>T22896</td><td>GCCCCAATCGGGGGACTGGAAGG</td><td></td><td>1000</td><td></td><td></td><td></td><td></td><td></td><td></td><td></td><td></td><td></td></tr>
<tr><td>Oligo fwd</td><td>CACCGCCCCAATCGGGGGACTGGA</td><td></td><td></td><td></td><td></td><td></td><td></td><td></td><td></td><td></td><td></td><td></td></tr>
<tr><td>Oligo rev</td><td>AAACTCCAGTCCCCCGATTGGGGC</td><td></td><td></td><td></td><td></td><td></td><td></td><td></td><td></td><td></td><td></td><td></td></tr>
<tr><td>Chromosome</td><td>start</td><td></td><td>end</td><td>strand</td><td>MM</td><td>target_seq</td><td>PAM</td><td>alignment</td><td>distance</td><td>position</td><td>gene name</td><td>gene id</td></tr>
<tr><td>GU305914.1</td><td></td><td>147536</td><td>147558</td><td>-</td><td>0</td><td>GCCCCAATCGGGGGACTGGA</td><td>AGG</td><td>||||||||[||||||||||||]PAM</td><td>NA</td><td>Intergenic</td><td>NA</td><td>NA</td></tr>
<tr><td>T22897</td><td>GGACTGGAAGGCGCAGAAGACGG</td><td></td><td>1000</td><td></td><td></td><td></td><td></td><td></td><td></td><td></td><td></td><td></td></tr>
<tr><td>Oligo fwd</td><td>CACCGGACTGGAAGGCGCAGAAGA</td><td></td><td></td><td></td><td></td><td></td><td></td><td></td><td></td><td></td><td></td><td></td></tr>
<tr><td>Oligo rev</td><td>AAACTCTTCTGCGCCTTCCAGTCC</td><td></td><td></td><td></td><td></td><td></td><td></td><td></td><td></td><td></td><td></td><td></td></tr>
<tr><td>Chromosome</td><td>start</td><td></td><td>end</td><td>strand</td><td>MM</td><td>target_seq</td><td>PAM</td><td>alignment</td><td>distance</td><td>position</td><td>gene name</td><td>gene id</td></tr>
<tr><td>GU305914.1</td><td></td><td>147524</td><td>147546</td><td>-</td><td>0</td><td>GGACTGGAAGGCGCAGAAGA</td><td>CGG</td><td>||||||||[||||||||||||]PAM</td><td>NA</td><td>Intergenic</td><td>NA</td><td>NA</td></tr>
<tr><td>GU305914.1</td><td></td><td>128343</td><td>128365</td><td>-</td><td>4</td><td>GTGGTGGAAGGCGGAGAAGA</td><td>CGG</td><td>|---||||[|||||-||||||]PAM</td><td>NA</td><td>Intergenic</td><td>NA</td><td>NA</td></tr>
<tr><td>T22898</td><td>GAAGGCGCAGAAGACGGCCTCGG</td><td></td><td>1000</td><td></td><td></td><td></td><td></td><td></td><td></td><td></td><td></td><td></td></tr>
<tr><td>Oligo fwd</td><td>CACCGAAGGCGCAGAAGACGGCCT</td><td></td><td></td><td></td><td></td><td></td><td></td><td></td><td></td><td></td><td></td><td></td></tr>
<tr><td>Oligo rev</td><td>AAACAGGCCGTCTTCTGCGCCTTC</td><td></td><td></td><td></td><td></td><td></td><td></td><td></td><td></td><td></td><td></td><td></td></tr>
<tr><td>Chromosome</td><td>start</td><td></td><td>end</td><td>strand</td><td>MM</td><td>target_seq</td><td>PAM</td><td>alignment</td><td>distance</td><td>position</td><td>gene name</td><td>gene id</td></tr>
<tr><td>GU305914.1</td><td></td><td>147518</td><td>147540</td><td>-</td><td>0</td><td>GAAGGCGCAGAAGACGGCCT</td><td>CGG</td><td>||||||||[||||||||||||]PAM</td><td>NA</td><td>Intergenic</td><td>NA</td><td>NA</td></tr>
<tr><td>GU305914.1</td><td></td><td>52925</td><td>52947</td><td>+</td><td>4</td><td>AAAGGCGAAGAAGACGCCCG</td><td>CGG</td><td>-||||||-[||||||||-||-]PAM</td><td>NA</td><td>Intergenic</td><td>NA</td><td>NA</td></tr>
<tr><td>T22899</td><td>AGAAGACGGCCTCGGAGAGCAGG</td><td></td><td>1000</td><td></td><td></td><td></td><td></td><td></td><td></td><td></td><td></td><td></td></tr>
<tr><td>Oligo fwd</td><td>CACCAGAAGACGGCCTCGGAGAGC</td><td></td><td></td><td></td><td></td><td></td><td></td><td></td><td></td><td></td><td></td><td></td></tr>
<tr><td>Oligo rev</td><td>AAACGCTCTCCGAGGCCGTCTTCT</td><td></td><td></td><td></td><td></td><td></td><td></td><td></td><td></td><td></td><td></td><td></td></tr>
<tr><td>Chromosome</td><td>start</td><td></td><td>end</td><td>strand</td><td>MM</td><td>target_seq</td><td>PAM</td><td>alignment</td><td>distance</td><td>position</td><td>gene name</td><td>gene id</td></tr>
<tr><td>GU305914.1</td><td></td><td>147510</td><td>147532</td><td>-</td><td>0</td><td>AGAAGACGGCCTCGGAGAGC</td><td>AGG</td><td>||||||||[||||||||||||]PAM</td><td>NA</td><td>Intergenic</td><td>NA</td><td>NA</td></tr>
<tr><td>T22900</td><td>GGCCTCGGAGAGCAGGCGTCCGG</td><td></td><td>1000</td><td></td><td></td><td></td><td></td><td></td><td></td><td></td><td></td><td></td></tr>
<tr><td>Oligo fwd</td><td>CACCGGCCTCGGAGAGCAGGCGTC</td><td></td><td></td><td></td><td></td><td></td><td></td><td></td><td></td><td></td><td></td><td></td></tr>
<tr><td>Oligo rev</td><td>AAACGACGCCTGCTCTCCGAGGCC</td><td></td><td></td><td></td><td></td><td></td><td></td><td></td><td></td><td></td><td></td><td></td></tr>
<tr><td>Chromosome</td><td>start</td><td></td><td>end</td><td>strand</td><td>MM</td><td>target_seq</td><td>PAM</td><td>alignment</td><td>distance</td><td>position</td><td>gene name</td><td>gene id</td></tr>
<tr><td>GU305914.1</td><td></td><td>147503</td><td>147525</td><td>-</td><td>0</td><td>GGCCTCGGAGAGCAGGCGTC</td><td>CGG</td><td>||||||||[||||||||||||]PAM</td><td>NA</td><td>Intergenic</td><td>NA</td><td>NA</td></tr>
<tr><td>T22901</td><td>CTCGGAGAGCAGGCGTCCGGTGG</td><td></td><td>1000</td><td></td><td></td><td></td><td></td><td></td><td></td><td></td><td></td><td></td></tr>
<tr><td>Oligo fwd</td><td>CACCCTCGGAGAGCAGGCGTCCGG</td><td></td><td></td><td></td><td></td><td></td><td></td><td></td><td></td><td></td><td></td><td></td></tr>
<tr><td>Oligo rev</td><td>AAACCCGGACGCCTGCTCTCCGAG</td><td></td><td></td><td></td><td></td><td></td><td></td><td></td><td></td><td></td><td></td><td></td></tr>
<tr><td>Chromosome</td><td>start</td><td></td><td>end</td><td>strand</td><td>MM</td><td>target_seq</td><td>PAM</td><td>alignment</td><td>distance</td><td>position</td><td>gene name</td><td>gene id</td></tr>
<tr><td>GU305914.1</td><td></td><td>147500</td><td>147522</td><td>-</td><td>0</td><td>CTCGGAGAGCAGGCGTCCGG</td><td>TGG</td><td>||||||||[||||||||||||]PAM</td><td>NA</td><td>Intergenic</td><td>NA</td><td>NA</td></tr>
<tr><td>T22902</td><td>TCCGGTGGCGCCGACCCATCCGG</td><td></td><td>1000</td><td></td><td></td><td></td><td></td><td></td><td></td><td></td><td></td><td></td></tr>
<tr><td>Oligo fwd</td><td>CACCTCCGGTGGCGCCGACCCATC</td><td></td><td></td><td></td><td></td><td></td><td></td><td></td><td></td><td></td><td></td><td></td></tr>
<tr><td>Oligo rev</td><td>AAACGATGGGTCGGCGCCACCGGA</td><td></td><td></td><td></td><td></td><td></td><td></td><td></td><td></td><td></td><td></td><td></td></tr>
<tr><td>Chromosome</td><td>start</td><td></td><td>end</td><td>strand</td><td>MM</td><td>target_seq</td><td>PAM</td><td>alignment</td><td>distance</td><td>position</td><td>gene name</td><td>gene id</td></tr>
<tr><td>GU305914.1</td><td></td><td>147485</td><td>147507</td><td>-</td><td>0</td><td>TCCGGTGGCGCCGACCCATC</td><td>CGG</td><td>||||||||[||||||||||||]PAM</td><td>NA</td><td>Intergenic</td><td>NA</td><td>NA</td></tr>
<tr><td>GU305914.1</td><td></td><td>159297</td><td>159319</td><td>+</td><td>4</td><td>TCGGGTCGCGGCGACCCATG</td><td>TGG</td><td>||-|||-|[||-||||||||-]PAM</td><td>NA</td><td>Intergenic</td><td>NA</td><td>NA</td></tr>
<tr><td>T22903</td><td>CCGGTGGCGCCGACCCATCCGGG</td><td></td><td>1000</td><td></td><td></td><td></td><td></td><td></td><td></td><td></td><td></td><td></td></tr>
<tr><td>Oligo fwd</td><td>CACCCCGGTGGCGCCGACCCATCC</td><td></td><td></td><td></td><td></td><td></td><td></td><td></td><td></td><td></td><td></td><td></td></tr>
<tr><td>Oligo rev</td><td>AAACGGATGGGTCGGCGCCACCGG</td><td></td><td></td><td></td><td></td><td></td><td></td><td></td><td></td><td></td><td></td><td></td></tr>
<tr><td>Chromosome</td><td>start</td><td></td><td>end</td><td>strand</td><td>MM</td><td>target_seq</td><td>PAM</td><td>alignment</td><td>distance</td><td>position</td><td>gene name</td><td>gene id</td></tr>
<tr><td>GU305914.1</td><td></td><td>147484</td><td>147506</td><td>-</td><td>0</td><td>CCGGTGGCGCCGACCCATCC</td><td>GGG</td><td>||||||||[||||||||||||]PAM</td><td>NA</td><td>Intergenic</td><td>NA</td><td>NA</td></tr>
<tr><td>T22904</td><td>CGGTGGCGCCGACCCATCCGGGG</td><td></td><td>1000</td><td></td><td></td><td></td><td></td><td></td><td></td><td></td><td></td><td></td></tr>
<tr><td>Oligo fwd</td><td>CACCCGGTGGCGCCGACCCATCCG</td><td></td><td></td><td></td><td></td><td></td><td></td><td></td><td></td><td></td><td></td><td></td></tr>
<tr><td>Oligo rev</td><td>AAACCGGATGGGTCGGCGCCACCG</td><td></td><td></td><td></td><td></td><td></td><td></td><td></td><td></td><td></td><td></td><td></td></tr>
<tr><td>Chromosome</td><td>start</td><td></td><td>end</td><td>strand</td><td>MM</td><td>target_seq</td><td>PAM</td><td>alignment</td><td>distance</td><td>position</td><td>gene name</td><td>gene id</td></tr>
<tr><td>GU305914.1</td><td></td><td>147483</td><td>147505</td><td>-</td><td>0</td><td>CGGTGGCGCCGACCCATCCG</td><td>GGG</td><td>||||||||[||||||||||||]PAM</td><td>NA</td><td>Intergenic</td><td>NA</td><td>NA</td></tr>
<tr><td>T22905</td><td>GGTGGCGCCGACCCATCCGGGGG</td><td></td><td>1000</td><td></td><td></td><td></td><td></td><td></td><td></td><td></td><td></td><td></td></tr>
<tr><td>Oligo fwd</td><td>CACCGGTGGCGCCGACCCATCCGG</td><td></td><td></td><td></td><td></td><td></td><td></td><td></td><td></td><td></td><td></td><td></td></tr>
<tr><td>Oligo rev</td><td>AAACCCGGATGGGTCGGCGCCACC</td><td></td><td></td><td></td><td></td><td></td><td></td><td></td><td></td><td></td><td></td><td></td></tr>
<tr><td>Chromosome</td><td>start</td><td></td><td>end</td><td>strand</td><td>MM</td><td>target_seq</td><td>PAM</td><td>alignment</td><td>distance</td><td>position</td><td>gene name</td><td>gene id</td></tr>
<tr><td>GU305914.1</td><td></td><td>147482</td><td>147504</td><td>-</td><td>0</td><td>GGTGGCGCCGACCCATCCGG</td><td>GGG</td><td>||||||||[||||||||||||]PAM</td><td>NA</td><td>Intergenic</td><td>NA</td><td>NA</td></tr>
<tr><td>GU305914.1</td><td></td><td>176217</td><td>176239</td><td>+</td><td>4</td><td>GGTTGCGACGACCCGTACGG</td><td>AGG</td><td>|||-|||-[||||||-|-|||]PAM</td><td>NA</td><td>Intergenic</td><td>NA</td><td>NA</td></tr>
<tr><td>T22906</td><td>GGCGCCGACCCATCCGGGGGTGG</td><td></td><td>1000</td><td></td><td></td><td></td><td></td><td></td><td></td><td></td><td></td><td></td></tr>
<tr><td>Oligo fwd</td><td>CACCGGCGCCGACCCATCCGGGGG</td><td></td><td></td><td></td><td></td><td></td><td></td><td></td><td></td><td></td><td></td><td></td></tr>
<tr><td>Oligo rev</td><td>AAACCCCCCGGATGGGTCGGCGCC</td><td></td><td></td><td></td><td></td><td></td><td></td><td></td><td></td><td></td><td></td><td></td></tr>
<tr><td>Chromosome</td><td>start</td><td></td><td>end</td><td>strand</td><td>MM</td><td>target_seq</td><td>PAM</td><td>alignment</td><td>distance</td><td>position</td><td>gene name</td><td>gene id</td></tr>
<tr><td>GU305914.1</td><td></td><td>147479</td><td>147501</td><td>-</td><td>0</td><td>GGCGCCGACCCATCCGGGGG</td><td>TGG</td><td>||||||||[||||||||||||]PAM</td><td>NA</td><td>Intergenic</td><td>NA</td><td>NA</td></tr>
<tr><td>T22907</td><td>GCGCCGACCCATCCGGGGGTGGG</td><td></td><td>1000</td><td></td><td></td><td></td><td></td><td></td><td></td><td></td><td></td><td></td></tr>
<tr><td>Oligo fwd</td><td>CACCGCGCCGACCCATCCGGGGGT</td><td></td><td></td><td></td><td></td><td></td><td></td><td></td><td></td><td></td><td></td><td></td></tr>
<tr><td>Oligo rev</td><td>AAACACCCCCGGATGGGTCGGCGC</td><td></td><td></td><td></td><td></td><td></td><td></td><td></td><td></td><td></td><td></td><td></td></tr>
<tr><td>Chromosome</td><td>start</td><td></td><td>end</td><td>strand</td><td>MM</td><td>target_seq</td><td>PAM</td><td>alignment</td><td>distance</td><td>position</td><td>gene name</td><td>gene id</td></tr>
<tr><td>GU305914.1</td><td></td><td>147478</td><td>147500</td><td>-</td><td>0</td><td>GCGCCGACCCATCCGGGGGT</td><td>GGG</td><td>||||||||[||||||||||||]PAM</td><td>NA</td><td>Intergenic</td><td>NA</td><td>NA</td></tr>
<tr><td>T22908</td><td>CGCCGACCCATCCGGGGGTGGGG</td><td></td><td>1000</td><td></td><td></td><td></td><td></td><td></td><td></td><td></td><td></td><td></td></tr>
<tr><td>Oligo fwd</td><td>CACCCGCCGACCCATCCGGGGGTG</td><td></td><td></td><td></td><td></td><td></td><td></td><td></td><td></td><td></td><td></td><td></td></tr>
<tr><td>Oligo rev</td><td>AAACCACCCCCGGATGGGTCGGCG</td><td></td><td></td><td></td><td></td><td></td><td></td><td></td><td></td><td></td><td></td><td></td></tr>
<tr><td>Chromosome</td><td>start</td><td></td><td>end</td><td>strand</td><td>MM</td><td>target_seq</td><td>PAM</td><td>alignment</td><td>distance</td><td>position</td><td>gene name</td><td>gene id</td></tr>
<tr><td>GU305914.1</td><td></td><td>147477</td><td>147499</td><td>-</td><td>0</td><td>CGCCGACCCATCCGGGGGTG</td><td>GGG</td><td>||||||||[||||||||||||]PAM</td><td>NA</td><td>Intergenic</td><td>NA</td><td>NA</td></tr>
<tr><td>T22909</td><td>GGGGAAGTACTTCTGCTCGTCGG</td><td></td><td>1000</td><td></td><td></td><td></td><td></td><td></td><td></td><td></td><td></td><td></td></tr>
<tr><td>Oligo fwd</td><td>CACCGGGGAAGTACTTCTGCTCGT</td><td></td><td></td><td></td><td></td><td></td><td></td><td></td><td></td><td></td><td></td><td></td></tr>
<tr><td>Oligo rev</td><td>AAACACGAGCAGAAGTACTTCCCC</td><td></td><td></td><td></td><td></td><td></td><td></td><td></td><td></td><td></td><td></td><td></td></tr>
<tr><td>Chromosome</td><td>start</td><td></td><td>end</td><td>strand</td><td>MM</td><td>target_seq</td><td>PAM</td><td>alignment</td><td>distance</td><td>position</td><td>gene name</td><td>gene id</td></tr>
<tr><td>GU305914.1</td><td></td><td>147458</td><td>147480</td><td>-</td><td>0</td><td>GGGGAAGTACTTCTGCTCGT</td><td>CGG</td><td>||||||||[||||||||||||]PAM</td><td>NA</td><td>Intergenic</td><td>NA</td><td>NA</td></tr>
</table>

<table>
<tr><td>T22910</td><td>GGGAAGTACTTCTGCTCGTCGGG</td><td>1000</td><td></td><td></td><td></td><td></td><td></td><td></td><td></td><td></td><td></td></tr>
<tr><td>Oligo fwd</td><td>CACCGGGAAGTACTTCTGCTCGTC</td><td></td><td></td><td></td><td></td><td></td><td></td><td></td><td></td><td></td><td></td></tr>
<tr><td>Oligo rev</td><td>AAACGACGAGCAGAAGTACTTCCC</td><td></td><td></td><td></td><td></td><td></td><td></td><td></td><td></td><td></td><td></td></tr>
<tr><td>Chromosome</td><td>start</td><td>end</td><td>strand</td><td>MM</td><td>target_seq</td><td>PAM</td><td>alignment</td><td>distance</td><td>position</td><td>gene name</td><td>gene id</td></tr>
<tr><td>GU305914.1</td><td>147457</td><td>147479</td><td>-</td><td>0</td><td>GGGAAGTACTTCTGCTCGTC</td><td>GGG</td><td>||||||||[||||||||||||]PAM</td><td>NA</td><td>Intergenic</td><td>NA</td><td>NA</td></tr>
<tr><td>T22911</td><td>GGAAGTACTTCTGCTCGTCGGGG</td><td>1000</td><td></td><td></td><td></td><td></td><td></td><td></td><td></td><td></td><td></td></tr>
<tr><td>Oligo fwd</td><td>CACCGGAAGTACTTCTGCTCGTCG</td><td></td><td></td><td></td><td></td><td></td><td></td><td></td><td></td><td></td><td></td></tr>
<tr><td>Oligo rev</td><td>AAACCGACGAGCAGAAGTACTTCC</td><td></td><td></td><td></td><td></td><td></td><td></td><td></td><td></td><td></td><td></td></tr>
<tr><td>Chromosome</td><td>start</td><td>end</td><td>strand</td><td>MM</td><td>target_seq</td><td>PAM</td><td>alignment</td><td>distance</td><td>position</td><td>gene name</td><td>gene id</td></tr>
<tr><td>GU305914.1</td><td>147456</td><td>147478</td><td>-</td><td>0</td><td>GGAAGTACTTCTGCTCGTCG</td><td>GGG</td><td>||||||||[||||||||||||]PAM</td><td>NA</td><td>Intergenic</td><td>NA</td><td>NA</td></tr>
<tr><td>T22912</td><td>CTTCTGCTCGTCGGGGAAGCCGG</td><td>1000</td><td></td><td></td><td></td><td></td><td></td><td></td><td></td><td></td><td></td></tr>
<tr><td>Oligo fwd</td><td>CACCCTTCTGCTCGTCGGGGAAGC</td><td></td><td></td><td></td><td></td><td></td><td></td><td></td><td></td><td></td><td></td></tr>
<tr><td>Oligo rev</td><td>AAACGCTTCCCCGACGAGCAGAAG</td><td></td><td></td><td></td><td></td><td></td><td></td><td></td><td></td><td></td><td></td></tr>
<tr><td>Chromosome</td><td>start</td><td>end</td><td>strand</td><td>MM</td><td>target_seq</td><td>PAM</td><td>alignment</td><td>distance</td><td>position</td><td>gene name</td><td>gene id</td></tr>
<tr><td>GU305914.1</td><td>147449</td><td>147471</td><td>-</td><td>0</td><td>CTTCTGCTCGTCGGGGAAGC</td><td>CGG</td><td>||||||||[||||||||||||]PAM</td><td>NA</td><td>Intergenic</td><td>NA</td><td>NA</td></tr>
<tr><td>T22913</td><td>CGTCGGGGAAGCCGGAGAGCAGG</td><td>1000</td><td></td><td></td><td></td><td></td><td></td><td></td><td></td><td></td><td></td></tr>
<tr><td>Oligo fwd</td><td>CACCCGTCGGGGAAGCCGGAGAGC</td><td></td><td></td><td></td><td></td><td></td><td></td><td></td><td></td><td></td><td></td></tr>
<tr><td>Oligo rev</td><td>AAACGCTCTCCGGCTTCCCCGACG</td><td></td><td></td><td></td><td></td><td></td><td></td><td></td><td></td><td></td><td></td></tr>
<tr><td>Chromosome</td><td>start</td><td>end</td><td>strand</td><td>MM</td><td>target_seq</td><td>PAM</td><td>alignment</td><td>distance</td><td>position</td><td>gene name</td><td>gene id</td></tr>
<tr><td>GU305914.1</td><td>147441</td><td>147463</td><td>-</td><td>0</td><td>CGTCGGGGAAGCCGGAGAGC</td><td>AGG</td><td>||||||||[||||||||||||]PAM</td><td>NA</td><td>Intergenic</td><td>NA</td><td>NA</td></tr>
<tr><td>T22914</td><td>GCCGGAGAGCAGGTACTTGCCGG</td><td>1000</td><td></td><td></td><td></td><td></td><td></td><td></td><td></td><td></td><td></td></tr>
<tr><td>Oligo fwd</td><td>CACCGCCGGAGAGCAGGTACTTGC</td><td></td><td></td><td></td><td></td><td></td><td></td><td></td><td></td><td></td><td></td></tr>
<tr><td>Oligo rev</td><td>AAACGCAAGTACCTGCTCTCCGGC</td><td></td><td></td><td></td><td></td><td></td><td></td><td></td><td></td><td></td><td></td></tr>
<tr><td>Chromosome</td><td>start</td><td>end</td><td>strand</td><td>MM</td><td>target_seq</td><td>PAM</td><td>alignment</td><td>distance</td><td>position</td><td>gene name</td><td>gene id</td></tr>
<tr><td>GU305914.1</td><td>147431</td><td>147453</td><td>-</td><td>0</td><td>GCCGGAGAGCAGGTACTTGC</td><td>CGG</td><td>||||||||[||||||||||||]PAM</td><td>NA</td><td>Intergenic</td><td>NA</td><td>NA</td></tr>
<tr><td>T22915</td><td>GGAGAGCAGGTACTTGCCGGTGG</td><td>1000</td><td></td><td></td><td></td><td></td><td></td><td></td><td></td><td></td><td></td></tr>
<tr><td>Oligo fwd</td><td>CACCGGAGAGCAGGTACTTGCCGG</td><td></td><td></td><td></td><td></td><td></td><td></td><td></td><td></td><td></td><td></td></tr>
<tr><td>Oligo rev</td><td>AAACCCGGCAAGTACCTGCTCTCC</td><td></td><td></td><td></td><td></td><td></td><td></td><td></td><td></td><td></td><td></td></tr>
<tr><td>Chromosome</td><td>start</td><td>end</td><td>strand</td><td>MM</td><td>target_seq</td><td>PAM</td><td>alignment</td><td>distance</td><td>position</td><td>gene name</td><td>gene id</td></tr>
<tr><td>GU305914.1</td><td>147428</td><td>147450</td><td>-</td><td>0</td><td>GGAGAGCAGGTACTTGCCGG</td><td>TGG</td><td>||||||||[||||||||||||]PAM</td><td>NA</td><td>Intergenic</td><td>NA</td><td>NA</td></tr>
<tr><td>T22916</td><td>GGCGAAGAGCACGCCGCTCATGG</td><td>1000</td><td></td><td></td><td></td><td></td><td></td><td></td><td></td><td></td><td></td></tr>
<tr><td>Oligo fwd</td><td>CACCGGCGAAGAGCACGCCGCTCA</td><td></td><td></td><td></td><td></td><td></td><td></td><td></td><td></td><td></td><td></td></tr>
<tr><td>Oligo rev</td><td>AAACTGAGCGGCGTGCTCTTCGCC</td><td></td><td></td><td></td><td></td><td></td><td></td><td></td><td></td><td></td><td></td></tr>
<tr><td>Chromosome</td><td>start</td><td>end</td><td>strand</td><td>MM</td><td>target_seq</td><td>PAM</td><td>alignment</td><td>distance</td><td>position</td><td>gene name</td><td>gene id</td></tr>
<tr><td>GU305914.1</td><td>147407</td><td>147429</td><td>-</td><td>0</td><td>GGCGAAGAGCACGCCGCTCA</td><td>TGG</td><td>||||||||[||||||||||||]PAM</td><td>NA</td><td>Intergenic</td><td>NA</td><td>NA</td></tr>
<tr><td>T22917</td><td>GAGCACGCCGCTCATGGCCACGG</td><td>1000</td><td></td><td></td><td></td><td></td><td></td><td></td><td></td><td></td><td></td></tr>
<tr><td>Oligo fwd</td><td>CACCGAGCACGCCGCTCATGGCCA</td><td></td><td></td><td></td><td></td><td></td><td></td><td></td><td></td><td></td><td></td></tr>
<tr><td>Oligo rev</td><td>AAACTGGCCATGAGCGGCGTGCTC</td><td></td><td></td><td></td><td></td><td></td><td></td><td></td><td></td><td></td><td></td></tr>
<tr><td>Chromosome</td><td>start</td><td>end</td><td>strand</td><td>MM</td><td>target_seq</td><td>PAM</td><td>alignment</td><td>distance</td><td>position</td><td>gene name</td><td>gene id</td></tr>
<tr><td>GU305914.1</td><td>147401</td><td>147423</td><td>-</td><td>0</td><td>GAGCACGCCGCTCATGGCCA</td><td>CGG</td><td>||||||||[||||||||||||]PAM</td><td>NA</td><td>Intergenic</td><td>NA</td><td>NA</td></tr>
<tr><td>T22918</td><td>CGCTCATGGCCACGGTGATGCGG</td><td>1000</td><td></td><td></td><td></td><td></td><td></td><td></td><td></td><td></td><td></td></tr>
<tr><td>Oligo fwd</td><td>CACCCGCTCATGGCCACGGTGATG</td><td></td><td></td><td></td><td></td><td></td><td></td><td></td><td></td><td></td><td></td></tr>
<tr><td>Oligo rev</td><td>AAACCATCACCGTGGCCATGAGCG</td><td></td><td></td><td></td><td></td><td></td><td></td><td></td><td></td><td></td><td></td></tr>
<tr><td>Chromosome</td><td>start</td><td>end</td><td>strand</td><td>MM</td><td>target_seq</td><td>PAM</td><td>alignment</td><td>distance</td><td>position</td><td>gene name</td><td>gene id</td></tr>
<tr><td>GU305914.1</td><td>147393</td><td>147415</td><td>-</td><td>0</td><td>CGCTCATGGCCACGGTGATG</td><td>CGG</td><td>||||||||[||||||||||||]PAM</td><td>NA</td><td>Intergenic</td><td>NA</td><td>NA</td></tr>
<tr><td>T22919</td><td>GGCCACGGTGATGCGGCAGACGG</td><td>1000</td><td></td><td></td><td></td><td></td><td></td><td></td><td></td><td></td><td></td></tr>
<tr><td>Oligo fwd</td><td>CACCGGCCACGGTGATGCGGCAGA</td><td></td><td></td><td></td><td></td><td></td><td></td><td></td><td></td><td></td><td></td></tr>
<tr><td>Oligo rev</td><td>AAACTCTGCCGCATCACCGTGGCC</td><td></td><td></td><td></td><td></td><td></td><td></td><td></td><td></td><td></td><td></td></tr>
<tr><td>Chromosome</td><td>start</td><td>end</td><td>strand</td><td>MM</td><td>target_seq</td><td>PAM</td><td>alignment</td><td>distance</td><td>position</td><td>gene name</td><td>gene id</td></tr>
<tr><td>GU305914.1</td><td>147386</td><td>147408</td><td>-</td><td>0</td><td>GGCCACGGTGATGCGGCAGA</td><td>CGG</td><td>||||||||[||||||||||||]PAM</td><td>NA</td><td>Intergenic</td><td>NA</td><td>NA</td></tr>
<tr><td>T22920</td><td>GCCACGGTGATGCGGCAGACGGG</td><td>1000</td><td></td><td></td><td></td><td></td><td></td><td></td><td></td><td></td><td></td></tr>
<tr><td>Oligo fwd</td><td>CACCGCCACGGTGATGCGGCAGAC</td><td></td><td></td><td></td><td></td><td></td><td></td><td></td><td></td><td></td><td></td></tr>
<tr><td>Oligo rev</td><td>AAACGTCTGCCGCATCACCGTGGC</td><td></td><td></td><td></td><td></td><td></td><td></td><td></td><td></td><td></td><td></td></tr>
<tr><td>Chromosome</td><td>start</td><td>end</td><td>strand</td><td>MM</td><td>target_seq</td><td>PAM</td><td>alignment</td><td>distance</td><td>position</td><td>gene name</td><td>gene id</td></tr>
<tr><td>GU305914.1</td><td>147385</td><td>147407</td><td>-</td><td>0</td><td>GCCACGGTGATGCGGCAGAC</td><td>GGG</td><td>||||||||[||||||||||||]PAM</td><td>NA</td><td>Intergenic</td><td>NA</td><td>NA</td></tr>
<tr><td>T22921</td><td>CCACGGTGATGCGGCAGACGGGG</td><td>1000</td><td></td><td></td><td></td><td></td><td></td><td></td><td></td><td></td><td></td></tr>
<tr><td>Oligo fwd</td><td>CACCCCACGGTGATGCGGCAGACG</td><td></td><td></td><td></td><td></td><td></td><td></td><td></td><td></td><td></td><td></td></tr>
<tr><td>Oligo rev</td><td>AAACCGTCTGCCGCATCACCGTGG</td><td></td><td></td><td></td><td></td><td></td><td></td><td></td><td></td><td></td><td></td></tr>
<tr><td>Chromosome</td><td>start</td><td>end</td><td>strand</td><td>MM</td><td>target_seq</td><td>PAM</td><td>alignment</td><td>distance</td><td>position</td><td>gene name</td><td>gene id</td></tr>
<tr><td>GU305914.1</td><td>147384</td><td>147406</td><td>-</td><td>0</td><td>CCACGGTGATGCGGCAGACG</td><td>GGG</td><td>||||||||[||||||||||||]PAM</td><td>NA</td><td>Intergenic</td><td>NA</td><td>NA</td></tr>
<tr><td>T22922</td><td>TGATGCGGCAGACGGGGCAGAGG</td><td>1000</td><td></td><td></td><td></td><td></td><td></td><td></td><td></td><td></td><td></td></tr>
<tr><td>Oligo fwd</td><td>CACCTGATGCGGCAGACGGGGCAG</td><td></td><td></td><td></td><td></td><td></td><td></td><td></td><td></td><td></td><td></td></tr>
<tr><td>Oligo rev</td><td>AAACCTGCCCCGTCTGCCGCATCA</td><td></td><td></td><td></td><td></td><td></td><td></td><td></td><td></td><td></td><td></td></tr>
<tr><td>Chromosome</td><td>start</td><td>end</td><td>strand</td><td>MM</td><td>target_seq</td><td>PAM</td><td>alignment</td><td>distance</td><td>position</td><td>gene name</td><td>gene id</td></tr>
<tr><td>GU305914.1</td><td>147378</td><td>147400</td><td>-</td><td>0</td><td>TGATGCGGCAGACGGGGCAG</td><td>AGG</td><td>||||||||[||||||||||||]PAM</td><td>NA</td><td>Intergenic</td><td>NA</td><td>NA</td></tr>
<tr><td>T22923</td><td>GGCAGACGGGGCAGAGGTACTGG</td><td>1000</td><td></td><td></td><td></td><td></td><td></td><td></td><td></td><td></td><td></td></tr>
<tr><td>Oligo fwd</td><td>CACCGGCAGACGGGGCAGAGGTAC</td><td></td><td></td><td></td><td></td><td></td><td></td><td></td><td></td><td></td><td></td></tr>
<tr><td>Oligo rev</td><td>AAACGTACCTCTGCCCCGTCTGCC</td><td></td><td></td><td></td><td></td><td></td><td></td><td></td><td></td><td></td><td></td></tr>
<tr><td>Chromosome</td><td>start</td><td>end</td><td>strand</td><td>MM</td><td>target_seq</td><td>PAM</td><td>alignment</td><td>distance</td><td>position</td><td>gene name</td><td>gene id</td></tr>
<tr><td>GU305914.1</td><td>147372</td><td>147394</td><td>-</td><td>0</td><td>GGCAGACGGGGCAGAGGTAC</td><td>TGG</td><td>||||||||[||||||||||||]PAM</td><td>NA</td><td>Intergenic</td><td>NA</td><td>NA</td></tr>
<tr><td>T22924</td><td>AGACGGGGCAGAGGTACTGGAGG</td><td>1000</td><td></td><td></td><td></td><td></td><td></td><td></td><td></td><td></td><td></td></tr>
<tr><td>Oligo fwd</td><td>CACCAGACGGGGCAGAGGTACTGG</td><td></td><td></td><td></td><td></td><td></td><td></td><td></td><td></td><td></td><td></td></tr>
<tr><td>Oligo rev</td><td>AAACCCAGTACCTCTGCCCCGTCT</td><td></td><td></td><td></td><td></td><td></td><td></td><td></td><td></td><td></td><td></td></tr>
<tr><td>Chromosome</td><td>start</td><td>end</td><td>strand</td><td>MM</td><td>target_seq</td><td>PAM</td><td>alignment</td><td>distance</td><td>position</td><td>gene name</td><td>gene id</td></tr>
</table>

<table>
<tr><td>GU305914.1</td><td>147369</td><td>147391</td><td>-</td><td>0</td><td>AGACGGGGCAGAGGTACTGG</td><td>AGG</td><td>||||||||[||||||||||||]PAM</td><td>NA</td><td>Intergenic</td><td>NA</td><td>NA</td></tr>
<tr><td>T22925</td><td>GTACTGGAGGCCGAAGAGCTCGG</td><td>1000</td><td></td><td></td><td></td><td></td><td></td><td></td><td></td><td></td><td></td></tr>
<tr><td>Oligo fwd</td><td>CACCGTACTGGAGGCCGAAGAGCT</td><td></td><td></td><td></td><td></td><td></td><td></td><td></td><td></td><td></td><td></td></tr>
<tr><td>Oligo rev</td><td>AAACAGCTCTTCGGCCTCCAGTAC</td><td></td><td></td><td></td><td></td><td></td><td></td><td></td><td></td><td></td><td></td></tr>
<tr><td>Chromosome</td><td>start</td><td>end</td><td>strand</td><td>MM</td><td>target_seq</td><td>PAM</td><td>alignment</td><td>distance</td><td>position</td><td>gene name</td><td>gene id</td></tr>
<tr><td>GU305914.1</td><td>147356</td><td>147378</td><td>-</td><td>0</td><td>GTACTGGAGGCCGAAGAGCT</td><td>CGG</td><td>||||||||[||||||||||||]PAM</td><td>NA</td><td>Intergenic</td><td>NA</td><td>NA</td></tr>
<tr><td>T22926</td><td>GAAGAGCTCGGCCCAGACCCTGG</td><td>1000</td><td></td><td></td><td></td><td></td><td></td><td></td><td></td><td></td><td></td></tr>
<tr><td>Oligo fwd</td><td>CACCGAAGAGCTCGGCCCAGACCC</td><td></td><td></td><td></td><td></td><td></td><td></td><td></td><td></td><td></td><td></td></tr>
<tr><td>Oligo rev</td><td>AAACGGGTCTGGGCCGAGCTCTTC</td><td></td><td></td><td></td><td></td><td></td><td></td><td></td><td></td><td></td><td></td></tr>
<tr><td>Chromosome</td><td>start</td><td>end</td><td>strand</td><td>MM</td><td>target_seq</td><td>PAM</td><td>alignment</td><td>distance</td><td>position</td><td>gene name</td><td>gene id</td></tr>
<tr><td>GU305914.1</td><td>147344</td><td>147366</td><td>-</td><td>0</td><td>GAAGAGCTCGGCCCAGACCC</td><td>TGG</td><td>||||||||[||||||||||||]PAM</td><td>NA</td><td>Intergenic</td><td>NA</td><td>NA</td></tr>
<tr><td>T22927</td><td>GCTCGGCCCAGACCCTGGCCAGG</td><td>1000</td><td></td><td></td><td></td><td></td><td></td><td></td><td></td><td></td><td></td></tr>
<tr><td>Oligo fwd</td><td>CACCGCTCGGCCCAGACCCTGGCC</td><td></td><td></td><td></td><td></td><td></td><td></td><td></td><td></td><td></td><td></td></tr>
<tr><td>Oligo rev</td><td>AAACGGCCAGGGTCTGGGCCGAGC</td><td></td><td></td><td></td><td></td><td></td><td></td><td></td><td></td><td></td><td></td></tr>
<tr><td>Chromosome</td><td>start</td><td>end</td><td>strand</td><td>MM</td><td>target_seq</td><td>PAM</td><td>alignment</td><td>distance</td><td>position</td><td>gene name</td><td>gene id</td></tr>
<tr><td>GU305914.1</td><td>147339</td><td>147361</td><td>-</td><td>0</td><td>GCTCGGCCCAGACCCTGGCC</td><td>AGG</td><td>||||||||[||||||||||||]PAM</td><td>NA</td><td>Intergenic</td><td>NA</td><td>NA</td></tr>
<tr><td>T22928</td><td>GGCCCAGACCCTGGCCAGGTTGG</td><td>1000</td><td></td><td></td><td></td><td></td><td></td><td></td><td></td><td></td><td></td></tr>
<tr><td>Oligo fwd</td><td>CACCGGCCCAGACCCTGGCCAGGT</td><td></td><td></td><td></td><td></td><td></td><td></td><td></td><td></td><td></td><td></td></tr>
<tr><td>Oligo rev</td><td>AAACACCTGGCCAGGGTCTGGGCC</td><td></td><td></td><td></td><td></td><td></td><td></td><td></td><td></td><td></td><td></td></tr>
<tr><td>Chromosome</td><td>start</td><td>end</td><td>strand</td><td>MM</td><td>target_seq</td><td>PAM</td><td>alignment</td><td>distance</td><td>position</td><td>gene name</td><td>gene id</td></tr>
<tr><td>GU305914.1</td><td>147335</td><td>147357</td><td>-</td><td>0</td><td>GGCCCAGACCCTGGCCAGGT</td><td>TGG</td><td>||||||||[||||||||||||]PAM</td><td>NA</td><td>Intergenic</td><td>NA</td><td>NA</td></tr>
<tr><td>T22929</td><td>GGCCAGGTTGGCCCTGACGATGG</td><td>1000</td><td></td><td></td><td></td><td></td><td></td><td></td><td></td><td></td><td></td></tr>
<tr><td>Oligo fwd</td><td>CACCGGCCAGGTTGGCCCTGACGA</td><td></td><td></td><td></td><td></td><td></td><td></td><td></td><td></td><td></td><td></td></tr>
<tr><td>Oligo rev</td><td>AAACTCGTCAGGGCCAACCTGGCC</td><td></td><td></td><td></td><td></td><td></td><td></td><td></td><td></td><td></td><td></td></tr>
<tr><td>Chromosome</td><td>start</td><td>end</td><td>strand</td><td>MM</td><td>target_seq</td><td>PAM</td><td>alignment</td><td>distance</td><td>position</td><td>gene name</td><td>gene id</td></tr>
<tr><td>GU305914.1</td><td>147323</td><td>147345</td><td>-</td><td>0</td><td>GGCCAGGTTGGCCCTGACGA</td><td>TGG</td><td>||||||||[||||||||||||]PAM</td><td>NA</td><td>Intergenic</td><td>NA</td><td>NA</td></tr>
<tr><td>T22930</td><td>CGATGGTGTCGAAGTTGACCTGG</td><td>1000</td><td></td><td></td><td></td><td></td><td></td><td></td><td></td><td></td><td></td></tr>
<tr><td>Oligo fwd</td><td>CACCCGATGGTGTCGAAGTTGACC</td><td></td><td></td><td></td><td></td><td></td><td></td><td></td><td></td><td></td><td></td></tr>
<tr><td>Oligo rev</td><td>AAACGGTCAACTTCGACACCATCG</td><td></td><td></td><td></td><td></td><td></td><td></td><td></td><td></td><td></td><td></td></tr>
<tr><td>Chromosome</td><td>start</td><td>end</td><td>strand</td><td>MM</td><td>target_seq</td><td>PAM</td><td>alignment</td><td>distance</td><td>position</td><td>gene name</td><td>gene id</td></tr>
<tr><td>GU305914.1</td><td>147306</td><td>147328</td><td>-</td><td>0</td><td>CGATGGTGTCGAAGTTGACC</td><td>TGG</td><td>||||||||[||||||||||||]PAM</td><td>NA</td><td>Intergenic</td><td>NA</td><td>NA</td></tr>
<tr><td>GU305914.1</td><td>121721</td><td>121743</td><td>+</td><td>4</td><td>TGAAGGTGGCGAAGTTCACC</td><td>AGG</td><td>-||-||||[-|||||||-|||]PAM</td><td>NA</td><td>Intergenic</td><td>NA</td><td>NA</td></tr>
<tr><td>T22931</td><td>GTTGACCTGGTCGCAGTCGACGG</td><td>1000</td><td></td><td></td><td></td><td></td><td></td><td></td><td></td><td></td><td></td></tr>
<tr><td>Oligo fwd</td><td>CACCGTTGACCTGGTCGCAGTCGA</td><td></td><td></td><td></td><td></td><td></td><td></td><td></td><td></td><td></td><td></td></tr>
<tr><td>Oligo rev</td><td>AAACTCGACTGCGACCAGGTCAAC</td><td></td><td></td><td></td><td></td><td></td><td></td><td></td><td></td><td></td><td></td></tr>
<tr><td>Chromosome</td><td>start</td><td>end</td><td>strand</td><td>MM</td><td>target_seq</td><td>PAM</td><td>alignment</td><td>distance</td><td>position</td><td>gene name</td><td>gene id</td></tr>
<tr><td>GU305914.1</td><td>147293</td><td>147315</td><td>-</td><td>0</td><td>GTTGACCTGGTCGCAGTCGA</td><td>CGG</td><td>||||||||[||||||||||||]PAM</td><td>NA</td><td>Intergenic</td><td>NA</td><td>NA</td></tr>
<tr><td>T22932</td><td>TTGACCTGGTCGCAGTCGACGGG</td><td>1000</td><td></td><td></td><td></td><td></td><td></td><td></td><td></td><td></td><td></td></tr>
<tr><td>Oligo fwd</td><td>CACCTTGACCTGGTCGCAGTCGAC</td><td></td><td></td><td></td><td></td><td></td><td></td><td></td><td></td><td></td><td></td></tr>
<tr><td>Oligo rev</td><td>AAACGTCGACTGCGACCAGGTCAA</td><td></td><td></td><td></td><td></td><td></td><td></td><td></td><td></td><td></td><td></td></tr>
<tr><td>Chromosome</td><td>start</td><td>end</td><td>strand</td><td>MM</td><td>target_seq</td><td>PAM</td><td>alignment</td><td>distance</td><td>position</td><td>gene name</td><td>gene id</td></tr>
<tr><td>GU305914.1</td><td>147292</td><td>147314</td><td>-</td><td>0</td><td>TTGACCTGGTCGCAGTCGAC</td><td>GGG</td><td>||||||||[||||||||||||]PAM</td><td>NA</td><td>Intergenic</td><td>NA</td><td>NA</td></tr>
<tr><td>T22933</td><td>TGACCTGGTCGCAGTCGACGGGG</td><td>1000</td><td></td><td></td><td></td><td></td><td></td><td></td><td></td><td></td><td></td></tr>
<tr><td>Oligo fwd</td><td>CACCTGACCTGGTCGCAGTCGACG</td><td></td><td></td><td></td><td></td><td></td><td></td><td></td><td></td><td></td><td></td></tr>
<tr><td>Oligo rev</td><td>AAACCGTCGACTGCGACCAGGTCA</td><td></td><td></td><td></td><td></td><td></td><td></td><td></td><td></td><td></td><td></td></tr>
<tr><td>Chromosome</td><td>start</td><td>end</td><td>strand</td><td>MM</td><td>target_seq</td><td>PAM</td><td>alignment</td><td>distance</td><td>position</td><td>gene name</td><td>gene id</td></tr>
<tr><td>GU305914.1</td><td>147291</td><td>147313</td><td>-</td><td>0</td><td>TGACCTGGTCGCAGTCGACG</td><td>GGG</td><td>||||||||[||||||||||||]PAM</td><td>NA</td><td>Intergenic</td><td>NA</td><td>NA</td></tr>
<tr><td>T22934</td><td>CGAAGATCTCGCAGTCGTCCCGG</td><td>1000</td><td></td><td></td><td></td><td></td><td></td><td></td><td></td><td></td><td></td></tr>
<tr><td>Oligo fwd</td><td>CACCCGAAGATCTCGCAGTCGTCC</td><td></td><td></td><td></td><td></td><td></td><td></td><td></td><td></td><td></td><td></td></tr>
<tr><td>Oligo rev</td><td>AAACGGACGACTGCGAGATCTTCG</td><td></td><td></td><td></td><td></td><td></td><td></td><td></td><td></td><td></td><td></td></tr>
<tr><td>Chromosome</td><td>start</td><td>end</td><td>strand</td><td>MM</td><td>target_seq</td><td>PAM</td><td>alignment</td><td>distance</td><td>position</td><td>gene name</td><td>gene id</td></tr>
<tr><td>GU305914.1</td><td>147261</td><td>147283</td><td>-</td><td>0</td><td>CGAAGATCTCGCAGTCGTCC</td><td>CGG</td><td>||||||||[||||||||||||]PAM</td><td>NA</td><td>Intergenic</td><td>NA</td><td>NA</td></tr>
<tr><td>T22935</td><td>TCTCGCAGTCGTCCCGGACCAGG</td><td>1000</td><td></td><td></td><td></td><td></td><td></td><td></td><td></td><td></td><td></td></tr>
<tr><td>Oligo fwd</td><td>CACCTCTCGCAGTCGTCCCGGACC</td><td></td><td></td><td></td><td></td><td></td><td></td><td></td><td></td><td></td><td></td></tr>
<tr><td>Oligo rev</td><td>AAACGGTCCGGGACGACTGCGAGA</td><td></td><td></td><td></td><td></td><td></td><td></td><td></td><td></td><td></td><td></td></tr>
<tr><td>Chromosome</td><td>start</td><td>end</td><td>strand</td><td>MM</td><td>target_seq</td><td>PAM</td><td>alignment</td><td>distance</td><td>position</td><td>gene name</td><td>gene id</td></tr>
<tr><td>GU305914.1</td><td>147255</td><td>147277</td><td>-</td><td>0</td><td>TCTCGCAGTCGTCCCGGACC</td><td>AGG</td><td>||||||||[||||||||||||]PAM</td><td>NA</td><td>Intergenic</td><td>NA</td><td>NA</td></tr>
<tr><td>T22936</td><td>GTCGTCCCGGACCAGGATCCAGG</td><td>1000</td><td></td><td></td><td></td><td></td><td></td><td></td><td></td><td></td><td></td></tr>
<tr><td>Oligo fwd</td><td>CACCGTCGTCCCGGACCAGGATCC</td><td></td><td></td><td></td><td></td><td></td><td></td><td></td><td></td><td></td><td></td></tr>
<tr><td>Oligo rev</td><td>AAACGGATCCTGGTCCGGGACGAC</td><td></td><td></td><td></td><td></td><td></td><td></td><td></td><td></td><td></td><td></td></tr>
<tr><td>Chromosome</td><td>start</td><td>end</td><td>strand</td><td>MM</td><td>target_seq</td><td>PAM</td><td>alignment</td><td>distance</td><td>position</td><td>gene name</td><td>gene id</td></tr>
<tr><td>GU305914.1</td><td>147248</td><td>147270</td><td>-</td><td>0</td><td>GTCGTCCCGGACCAGGATCC</td><td>AGG</td><td>||||||||[||||||||||||]PAM</td><td>NA</td><td>Intergenic</td><td>NA</td><td>NA</td></tr>
<tr><td>T22937</td><td>TCGTCCCGGACCAGGATCCAGGG</td><td>1000</td><td></td><td></td><td></td><td></td><td></td><td></td><td></td><td></td><td></td></tr>
<tr><td>Oligo fwd</td><td>CACCTCGTCCCGGACCAGGATCCA</td><td></td><td></td><td></td><td></td><td></td><td></td><td></td><td></td><td></td><td></td></tr>
<tr><td>Oligo rev</td><td>AAACTGGATCCTGGTCCGGGACGA</td><td></td><td></td><td></td><td></td><td></td><td></td><td></td><td></td><td></td><td></td></tr>
<tr><td>Chromosome</td><td>start</td><td>end</td><td>strand</td><td>MM</td><td>target_seq</td><td>PAM</td><td>alignment</td><td>distance</td><td>position</td><td>gene name</td><td>gene id</td></tr>
<tr><td>GU305914.1</td><td>147247</td><td>147269</td><td>-</td><td>0</td><td>TCGTCCCGGACCAGGATCCA</td><td>GGG</td><td>||||||||[||||||||||||]PAM</td><td>NA</td><td>Intergenic</td><td>NA</td><td>NA</td></tr>
<tr><td>T22938</td><td>CGTCCCGGACCAGGATCCAGGGG</td><td>1000</td><td></td><td></td><td></td><td></td><td></td><td></td><td></td><td></td><td></td></tr>
<tr><td>Oligo fwd</td><td>CACCCGTCCCGGACCAGGATCCAG</td><td></td><td></td><td></td><td></td><td></td><td></td><td></td><td></td><td></td><td></td></tr>
<tr><td>Oligo rev</td><td>AAACCTGGATCCTGGTCCGGGACG</td><td></td><td></td><td></td><td></td><td></td><td></td><td></td><td></td><td></td><td></td></tr>
<tr><td>Chromosome</td><td>start</td><td>end</td><td>strand</td><td>MM</td><td>target_seq</td><td>PAM</td><td>alignment</td><td>distance</td><td>position</td><td>gene name</td><td>gene id</td></tr>
<tr><td>GU305914.1</td><td>147246</td><td>147268</td><td>-</td><td>0</td><td>CGTCCCGGACCAGGATCCAG</td><td>GGG</td><td>||||||||[||||||||||||]PAM</td><td>NA</td><td>Intergenic</td><td>NA</td><td>NA</td></tr>
<tr><td>T22939</td><td>GGGGTAACCCTCGTCTTCGCAGG</td><td>1000</td><td></td><td></td><td></td><td></td><td></td><td></td><td></td><td></td><td></td></tr>
<tr><td>Oligo fwd</td><td>CACCGGGGTAACCCTCGTCTTCGC</td><td></td><td></td><td></td><td></td><td></td><td></td><td></td><td></td><td></td><td></td></tr>
</table>

<table>
<tr><td>Oligo rev</td><td>AAACGCGAAGACGAGGGTTACCCC</td><td></td><td></td><td></td><td></td><td></td><td></td><td></td><td></td><td></td><td></td></tr>
<tr><td>Chromosome</td><td>start</td><td>end</td><td>strand</td><td>MM</td><td>target_seq</td><td>PAM</td><td>alignment</td><td>distance</td><td>position</td><td>gene name</td><td>gene id</td></tr>
<tr><td>GU305914.1</td><td>147227</td><td>147249</td><td>-</td><td>0</td><td>GGGGTAACCCTCGTCTTCGC</td><td>AGG</td><td>||||||||[||||||||||||]PAM</td><td>NA</td><td>Intergenic</td><td>NA</td><td>NA</td></tr>
<tr><td></td><td></td><td></td><td></td><td></td><td></td><td></td><td></td><td></td><td></td><td></td><td></td></tr>
<tr><td>T22940</td><td>AGTTTTCGAGAGAGCGCACGCGG</td><td>1000</td><td></td><td></td><td></td><td></td><td></td><td></td><td></td><td></td><td></td></tr>
<tr><td>Oligo fwd</td><td>CACCAGTTTTCGAGAGAGCGCACG</td><td></td><td></td><td></td><td></td><td></td><td></td><td></td><td></td><td></td><td></td></tr>
<tr><td>Oligo rev</td><td>AAACCGTGCGCTCTCTCGAAAACT</td><td></td><td></td><td></td><td></td><td></td><td></td><td></td><td></td><td></td><td></td></tr>
<tr><td>Chromosome</td><td>start</td><td>end</td><td>strand</td><td>MM</td><td>target_seq</td><td>PAM</td><td>alignment</td><td>distance</td><td>position</td><td>gene name</td><td>gene id</td></tr>
<tr><td>GU305914.1</td><td>147180</td><td>147202</td><td>-</td><td>0</td><td>AGTTTTCGAGAGAGCGCACG</td><td>CGG</td><td>||||||||[||||||||||||]PAM</td><td>NA</td><td>Intergenic</td><td>NA</td><td>NA</td></tr>
<tr><td></td><td></td><td></td><td></td><td></td><td></td><td></td><td></td><td></td><td></td><td></td><td></td></tr>
<tr><td>T22941</td><td>TTTCGAGAGAGCGCACGCGGCGG</td><td>1000</td><td></td><td></td><td></td><td></td><td></td><td></td><td></td><td></td><td></td></tr>
<tr><td>Oligo fwd</td><td>CACCTTTCGAGAGAGCGCACGCGG</td><td></td><td></td><td></td><td></td><td></td><td></td><td></td><td></td><td></td><td></td></tr>
<tr><td>Oligo rev</td><td>AAACCCGCGTGCGCTCTCTCGAAA</td><td></td><td></td><td></td><td></td><td></td><td></td><td></td><td></td><td></td><td></td></tr>
<tr><td>Chromosome</td><td>start</td><td>end</td><td>strand</td><td>MM</td><td>target_seq</td><td>PAM</td><td>alignment</td><td>distance</td><td>position</td><td>gene name</td><td>gene id</td></tr>
<tr><td>GU305914.1</td><td>147177</td><td>147199</td><td>-</td><td>0</td><td>TTTCGAGAGAGCGCACGCGG</td><td>CGG</td><td>||||||||[||||||||||||]PAM</td><td>NA</td><td>Intergenic</td><td>NA</td><td>NA</td></tr>
<tr><td></td><td></td><td></td><td></td><td></td><td></td><td></td><td></td><td></td><td></td><td></td><td></td></tr>
<tr><td>T22942</td><td>CGCACGCGGCGGAAGATGCGCGG</td><td>1000</td><td></td><td></td><td></td><td></td><td></td><td></td><td></td><td></td><td></td></tr>
<tr><td>Oligo fwd</td><td>CACCCGCACGCGGCGGAAGATGCG</td><td></td><td></td><td></td><td></td><td></td><td></td><td></td><td></td><td></td><td></td></tr>
<tr><td>Oligo rev</td><td>AAACCGCATCTTCCGCCGCGTGCG</td><td></td><td></td><td></td><td></td><td></td><td></td><td></td><td></td><td></td><td></td></tr>
<tr><td>Chromosome</td><td>start</td><td>end</td><td>strand</td><td>MM</td><td>target_seq</td><td>PAM</td><td>alignment</td><td>distance</td><td>position</td><td>gene name</td><td>gene id</td></tr>
<tr><td>GU305914.1</td><td>147166</td><td>147188</td><td>-</td><td>0</td><td>CGCACGCGGCGGAAGATGCG</td><td>CGG</td><td>||||||||[||||||||||||]PAM</td><td>NA</td><td>Intergenic</td><td>NA</td><td>NA</td></tr>
<tr><td></td><td></td><td></td><td></td><td></td><td></td><td></td><td></td><td></td><td></td><td></td><td></td></tr>
<tr><td>T22943</td><td>GCACGCGGCGGAAGATGCGCGGG</td><td>1000</td><td></td><td></td><td></td><td></td><td></td><td></td><td></td><td></td><td></td></tr>
<tr><td>Oligo fwd</td><td>CACCGCACGCGGCGGAAGATGCGC</td><td></td><td></td><td></td><td></td><td></td><td></td><td></td><td></td><td></td><td></td></tr>
<tr><td>Oligo rev</td><td>AAACGCGCATCTTCCGCCGCGTGC</td><td></td><td></td><td></td><td></td><td></td><td></td><td></td><td></td><td></td><td></td></tr>
<tr><td>Chromosome</td><td>start</td><td>end</td><td>strand</td><td>MM</td><td>target_seq</td><td>PAM</td><td>alignment</td><td>distance</td><td>position</td><td>gene name</td><td>gene id</td></tr>
<tr><td>GU305914.1</td><td>147165</td><td>147187</td><td>-</td><td>0</td><td>GCACGCGGCGGAAGATGCGC</td><td>GGG</td><td>||||||||[||||||||||||]PAM</td><td>NA</td><td>Intergenic</td><td>NA</td><td>NA</td></tr>
<tr><td></td><td></td><td></td><td></td><td></td><td></td><td></td><td></td><td></td><td></td><td></td><td></td></tr>
<tr><td>T22944</td><td>CACGCGGCGGAAGATGCGCGGGG</td><td>1000</td><td></td><td></td><td></td><td></td><td></td><td></td><td></td><td></td><td></td></tr>
<tr><td>Oligo fwd</td><td>CACCCACGCGGCGGAAGATGCGCG</td><td></td><td></td><td></td><td></td><td></td><td></td><td></td><td></td><td></td><td></td></tr>
<tr><td>Oligo rev</td><td>AAACCGCGCATCTTCCGCCGCGTG</td><td></td><td></td><td></td><td></td><td></td><td></td><td></td><td></td><td></td><td></td></tr>
<tr><td>Chromosome</td><td>start</td><td>end</td><td>strand</td><td>MM</td><td>target_seq</td><td>PAM</td><td>alignment</td><td>distance</td><td>position</td><td>gene name</td><td>gene id</td></tr>
<tr><td>GU305914.1</td><td>147164</td><td>147186</td><td>-</td><td>0</td><td>CACGCGGCGGAAGATGCGCG</td><td>GGG</td><td>||||||||[||||||||||||]PAM</td><td>NA</td><td>Intergenic</td><td>NA</td><td>NA</td></tr>
<tr><td>GU305914.1</td><td>3812</td><td>3834</td><td>+</td><td>4</td><td>GATGCTGCGGAAGATGAGCG</td><td>AGG</td><td>-|-||-||[||||||||-|||]PAM</td><td>NA</td><td>Intergenic</td><td>NA</td><td>NA</td></tr>
<tr><td></td><td></td><td></td><td></td><td></td><td></td><td></td><td></td><td></td><td></td><td></td><td></td></tr>
<tr><td>T22945</td><td>GGAAGATGCGCGGGGAGCTGTGG</td><td>1000</td><td></td><td></td><td></td><td></td><td></td><td></td><td></td><td></td><td></td></tr>
<tr><td>Oligo fwd</td><td>CACCGGAAGATGCGCGGGGAGCTG</td><td></td><td></td><td></td><td></td><td></td><td></td><td></td><td></td><td></td><td></td></tr>
<tr><td>Oligo rev</td><td>AAACCAGCTCCCCGCGCATCTTCC</td><td></td><td></td><td></td><td></td><td></td><td></td><td></td><td></td><td></td><td></td></tr>
<tr><td>Chromosome</td><td>start</td><td>end</td><td>strand</td><td>MM</td><td>target_seq</td><td>PAM</td><td>alignment</td><td>distance</td><td>position</td><td>gene name</td><td>gene id</td></tr>
<tr><td>GU305914.1</td><td>147156</td><td>147178</td><td>-</td><td>0</td><td>GGAAGATGCGCGGGGAGCTG</td><td>TGG</td><td>||||||||[||||||||||||]PAM</td><td>NA</td><td>Intergenic</td><td>NA</td><td>NA</td></tr>
<tr><td></td><td></td><td></td><td></td><td></td><td></td><td></td><td></td><td></td><td></td><td></td><td></td></tr>
<tr><td>T22946</td><td>GAAGATGCGCGGGGAGCTGTGGG</td><td>1000</td><td></td><td></td><td></td><td></td><td></td><td></td><td></td><td></td><td></td></tr>
<tr><td>Oligo fwd</td><td>CACCGAAGATGCGCGGGGAGCTGT</td><td></td><td></td><td></td><td></td><td></td><td></td><td></td><td></td><td></td><td></td></tr>
<tr><td>Oligo rev</td><td>AAACACAGCTCCCCGCGCATCTTC</td><td></td><td></td><td></td><td></td><td></td><td></td><td></td><td></td><td></td><td></td></tr>
<tr><td>Chromosome</td><td>start</td><td>end</td><td>strand</td><td>MM</td><td>target_seq</td><td>PAM</td><td>alignment</td><td>distance</td><td>position</td><td>gene name</td><td>gene id</td></tr>
<tr><td>GU305914.1</td><td>147155</td><td>147177</td><td>-</td><td>0</td><td>GAAGATGCGCGGGGAGCTGT</td><td>GGG</td><td>||||||||[||||||||||||]PAM</td><td>NA</td><td>Intergenic</td><td>NA</td><td>NA</td></tr>
<tr><td></td><td></td><td></td><td></td><td></td><td></td><td></td><td></td><td></td><td></td><td></td><td></td></tr>
<tr><td>T22947</td><td>TGCGCGGGGAGCTGTGGGCGAGG</td><td>1000</td><td></td><td></td><td></td><td></td><td></td><td></td><td></td><td></td><td></td></tr>
<tr><td>Oligo fwd</td><td>CACCTGCGCGGGGAGCTGTGGGCG</td><td></td><td></td><td></td><td></td><td></td><td></td><td></td><td></td><td></td><td></td></tr>
<tr><td>Oligo rev</td><td>AAACCGCCCACAGCTCCCCGCGCA</td><td></td><td></td><td></td><td></td><td></td><td></td><td></td><td></td><td></td><td></td></tr>
<tr><td>Chromosome</td><td>start</td><td>end</td><td>strand</td><td>MM</td><td>target_seq</td><td>PAM</td><td>alignment</td><td>distance</td><td>position</td><td>gene name</td><td>gene id</td></tr>
<tr><td>GU305914.1</td><td>147150</td><td>147172</td><td>-</td><td>0</td><td>TGCGCGGGGAGCTGTGGGCG</td><td>AGG</td><td>||||||||[||||||||||||]PAM</td><td>NA</td><td>Intergenic</td><td>NA</td><td>NA</td></tr>
<tr><td></td><td></td><td></td><td></td><td></td><td></td><td></td><td></td><td></td><td></td><td></td><td></td></tr>
<tr><td>T22948</td><td>GGGAGCTGTGGGCGAGGCGTTGG</td><td>1000</td><td></td><td></td><td></td><td></td><td></td><td></td><td></td><td></td><td></td></tr>
<tr><td>Oligo fwd</td><td>CACCGGGAGCTGTGGGCGAGGCGT</td><td></td><td></td><td></td><td></td><td></td><td></td><td></td><td></td><td></td><td></td></tr>
<tr><td>Oligo rev</td><td>AAACACGCCTCGCCCACAGCTCCC</td><td></td><td></td><td></td><td></td><td></td><td></td><td></td><td></td><td></td><td></td></tr>
<tr><td>Chromosome</td><td>start</td><td>end</td><td>strand</td><td>MM</td><td>target_seq</td><td>PAM</td><td>alignment</td><td>distance</td><td>position</td><td>gene name</td><td>gene id</td></tr>
<tr><td>GU305914.1</td><td>147144</td><td>147166</td><td>-</td><td>0</td><td>GGGAGCTGTGGGCGAGGCGT</td><td>TGG</td><td>||||||||[||||||||||||]PAM</td><td>NA</td><td>Intergenic</td><td>NA</td><td>NA</td></tr>
<tr><td></td><td></td><td></td><td></td><td></td><td></td><td></td><td></td><td></td><td></td><td></td><td></td></tr>
<tr><td>T22949</td><td>AGCTGTGGGCGAGGCGTTGGAGG</td><td>1000</td><td></td><td></td><td></td><td></td><td></td><td></td><td></td><td></td><td></td></tr>
<tr><td>Oligo fwd</td><td>CACCAGCTGTGGGCGAGGCGTTGG</td><td></td><td></td><td></td><td></td><td></td><td></td><td></td><td></td><td></td><td></td></tr>
<tr><td>Oligo rev</td><td>AAACCCAACGCCTCGCCCACAGCT</td><td></td><td></td><td></td><td></td><td></td><td></td><td></td><td></td><td></td><td></td></tr>
<tr><td>Chromosome</td><td>start</td><td>end</td><td>strand</td><td>MM</td><td>target_seq</td><td>PAM</td><td>alignment</td><td>distance</td><td>position</td><td>gene name</td><td>gene id</td></tr>
<tr><td>GU305914.1</td><td>147141</td><td>147163</td><td>-</td><td>0</td><td>AGCTGTGGGCGAGGCGTTGG</td><td>AGG</td><td>||||||||[||||||||||||]PAM</td><td>NA</td><td>Intergenic</td><td>NA</td><td>NA</td></tr>
<tr><td></td><td></td><td></td><td></td><td></td><td></td><td></td><td></td><td></td><td></td><td></td><td></td></tr>
<tr><td>T22950</td><td>GGAAAGCGTCGATCATCATGCGG</td><td>1000</td><td></td><td></td><td></td><td></td><td></td><td></td><td></td><td></td><td></td></tr>
<tr><td>Oligo fwd</td><td>CACCGGAAAGCGTCGATCATCATG</td><td></td><td></td><td></td><td></td><td></td><td></td><td></td><td></td><td></td><td></td></tr>
<tr><td>Oligo rev</td><td>AAACCATGATGATCGACGCTTTCC</td><td></td><td></td><td></td><td></td><td></td><td></td><td></td><td></td><td></td><td></td></tr>
<tr><td>Chromosome</td><td>start</td><td>end</td><td>strand</td><td>MM</td><td>target_seq</td><td>PAM</td><td>alignment</td><td>distance</td><td>position</td><td>gene name</td><td>gene id</td></tr>
<tr><td>GU305914.1</td><td>147120</td><td>147142</td><td>-</td><td>0</td><td>GGAAAGCGTCGATCATCATG</td><td>CGG</td><td>||||||||[||||||||||||]PAM</td><td>NA</td><td>Intergenic</td><td>NA</td><td>NA</td></tr>
<tr><td></td><td></td><td></td><td></td><td></td><td></td><td></td><td></td><td></td><td></td><td></td><td></td></tr>
<tr><td>T22951</td><td>CGATCATCATGCGGTCGACACGG</td><td>1000</td><td></td><td></td><td></td><td></td><td></td><td></td><td></td><td></td><td></td></tr>
<tr><td>Oligo fwd</td><td>CACCCGATCATCATGCGGTCGACA</td><td></td><td></td><td></td><td></td><td></td><td></td><td></td><td></td><td></td><td></td></tr>
<tr><td>Oligo rev</td><td>AAACTGTCGACCGCATGATGATCG</td><td></td><td></td><td></td><td></td><td></td><td></td><td></td><td></td><td></td><td></td></tr>
<tr><td>Chromosome</td><td>start</td><td>end</td><td>strand</td><td>MM</td><td>target_seq</td><td>PAM</td><td>alignment</td><td>distance</td><td>position</td><td>gene name</td><td>gene id</td></tr>
<tr><td>GU305914.1</td><td>147111</td><td>147133</td><td>-</td><td>0</td><td>CGATCATCATGCGGTCGACA</td><td>CGG</td><td>||||||||[||||||||||||]PAM</td><td>NA</td><td>Intergenic</td><td>NA</td><td>NA</td></tr>
<tr><td></td><td></td><td></td><td></td><td></td><td></td><td></td><td></td><td></td><td></td><td></td><td></td></tr>
<tr><td>T22952</td><td>CGCGACGTCGCTGAGCACGAAGG</td><td>1000</td><td></td><td></td><td></td><td></td><td></td><td></td><td></td><td></td><td></td></tr>
<tr><td>Oligo fwd</td><td>CACCCGCGACGTCGCTGAGCACGA</td><td></td><td></td><td></td><td></td><td></td><td></td><td></td><td></td><td></td><td></td></tr>
<tr><td>Oligo rev</td><td>AAACTCGTGCTCAGCGACGTCGCG</td><td></td><td></td><td></td><td></td><td></td><td></td><td></td><td></td><td></td><td></td></tr>
<tr><td>Chromosome</td><td>start</td><td>end</td><td>strand</td><td>MM</td><td>target_seq</td><td>PAM</td><td>alignment</td><td>distance</td><td>position</td><td>gene name</td><td>gene id</td></tr>
<tr><td>GU305914.1</td><td>147086</td><td>147108</td><td>-</td><td>0</td><td>CGCGACGTCGCTGAGCACGA</td><td>AGG</td><td>||||||||[||||||||||||]PAM</td><td>NA</td><td>Intergenic</td><td>NA</td><td>NA</td></tr>
<tr><td></td><td></td><td></td><td></td><td></td><td></td><td></td><td></td><td></td><td></td><td></td><td></td></tr>
<tr><td>T22953</td><td>GCGACGTCGCTGAGCACGAAGGG</td><td>1000</td><td></td><td></td><td></td><td></td><td></td><td></td><td></td><td></td><td></td></tr>
<tr><td>Oligo fwd</td><td>CACCGCGACGTCGCTGAGCACGAA</td><td></td><td></td><td></td><td></td><td></td><td></td><td></td><td></td><td></td><td></td></tr>
<tr><td>Oligo rev</td><td>AAACTTCGTGCTCAGCGACGTCGC</td><td></td><td></td><td></td><td></td><td></td><td></td><td></td><td></td><td></td><td></td></tr>
<tr><td>Chromosome</td><td>start</td><td>end</td><td>strand</td><td>MM</td><td>target_seq</td><td>PAM</td><td>alignment</td><td>distance</td><td>position</td><td>gene name</td><td>gene id</td></tr>
<tr><td>GU305914.1</td><td>147085</td><td>147107</td><td>-</td><td>0</td><td>GCGACGTCGCTGAGCACGAA</td><td>GGG</td><td>||||||||[||||||||||||]PAM</td><td>NA</td><td>Intergenic</td><td>NA</td><td>NA</td></tr>
<tr><td>GU305914.1</td><td>58285</td><td>58307</td><td>+</td><td>4</td><td>CCCACGTCGCCGCGCACGAA</td><td>GGG</td><td>-|-|||||[||-|-|||||||]PAM</td><td>NA</td><td>Intergenic</td><td>NA</td><td>NA</td></tr>
</table>

T22954 GCTGAGCACGAAGGGAAATTCGG 1000
Oligo fwd CACCGCTGAGCACGAAGGGAAATT
Oligo rev AAACAATTTCCCTTCGTGCTCAGC
Chromosome start end strand MM target_seq PAM alignment distance position gene name gene id
GU305914.1 147077 147099 - 0 GCTGAGCACGAAGGGAAATT CGG ||||||||[||||||||||||]PAM NA Intergenic NA NA

T22955 CTGAGCACGAAGGGAAATTCGGG 1000
Oligo fwd CACCCTGAGCACGAAGGGAAATTC
Oligo rev AAACGAATTTCCCTTCGTGCTCAG
Chromosome start end strand MM target_seq PAM alignment distance position gene name gene id
GU305914.1 147076 147098 - 0 CTGAGCACGAAGGGAAATTC GGG ||||||||[||||||||||||]PAM NA Intergenic NA NA

T22956 CACGAAGGGAAATTCGGGCCCGG 1000
Oligo fwd CACCCACGAAGGGAAATTCGGGCC
Oligo rev AAACGGCCCGAATTTCCCTTCGTG
Chromosome start end strand MM target_seq PAM alignment distance position gene name gene id
GU305914.1 147071 147093 - 0 CACGAAGGGAAATTCGGGCC CGG ||||||||[||||||||||||]PAM NA Intergenic NA NA

T22957 ACGAAGGGAAATTCGGGCCCGGG 1000
Oligo fwd CACCACGAAGGGAAATTCGGGCCC
Oligo rev AAACGGGCCCGAATTTCCCTTCGT
Chromosome start end strand MM target_seq PAM alignment distance position gene name gene id
GU305914.1 147070 147092 - 0 ACGAAGGGAAATTCGGGCCC GGG ||||||||[||||||||||||]PAM NA Intergenic NA NA

T22958 AGGGAAATTCGGGCCCGGGCAGG 1000
Oligo fwd CACCAGGGAAATTCGGGCCCGGGC
Oligo rev AAACGCCCGGGCCCGAATTTCCCT
Chromosome start end strand MM target_seq PAM alignment distance position gene name gene id
GU305914.1 147066 147088 - 0 AGGGAAATTCGGGCCCGGGC AGG ||||||||[||||||||||||]PAM NA Intergenic NA NA

T22959 GCCCGGGCAGGCACATCCGCAGG 1000
Oligo fwd CACCGCCCGGGCAGGCACATCCGC
Oligo rev AAACGCGGATGTGCCTGCCCGGGC
Chromosome start end strand MM target_seq PAM alignment distance position gene name gene id
GU305914.1 147054 147076 - 0 GCCCGGGCAGGCACATCCGC AGG ||||||||[||||||||||||]PAM NA Intergenic NA NA

T22960 GGGCAGGCACATCCGCAGGACGG 1000
Oligo fwd CACCGGGCAGGCACATCCGCAGGA
Oligo rev AAACTCCTGCGGATGTGCCTGCCC
Chromosome start end strand MM target_seq PAM alignment distance position gene name gene id
GU305914.1 147050 147072 - 0 GGGCAGGCACATCCGCAGGA CGG ||||||||[||||||||||||]PAM NA Intergenic NA NA

T22961 GGCGAACCTGAGATCGACGATGG 1000
Oligo fwd CACCGGCGAACCTGAGATCGACGA
Oligo rev AAACTCGTCGATCTCAGGTTCGCC
Chromosome start end strand MM target_seq PAM alignment distance position gene name gene id
GU305914.1 147029 147051 - 0 GGCGAACCTGAGATCGACGA TGG ||||||||[||||||||||||]PAM NA Intergenic NA NA

T22962 TGAGATCGACGATGGAGATGCGG 1000
Oligo fwd CACCTGAGATCGACGATGGAGATG
Oligo rev AAACCATCTCCATCGTCGATCTCA
Chromosome start end strand MM target_seq PAM alignment distance position gene name gene id
GU305914.1 147021 147043 - 0 TGAGATCGACGATGGAGATG CGG ||||||||[||||||||||||]PAM NA Intergenic NA NA

T22963 CGCCGACGCCCACATAGATGTGG 1000
Oligo fwd CACCCGCCGACGCCCACATAGATG
Oligo rev AAACCATCTATGTGGGCGTCGGCG
Chromosome start end strand MM target_seq PAM alignment distance position gene name gene id
GU305914.1 146991 147013 - 0 CGCCGACGCCCACATAGATG TGG ||||||||[||||||||||||]PAM NA Intergenic NA NA
GU305914.1 146982 147004 + 4 CTCCGTCGCCCACATCTATG TGG |-|||-||[|||||||--|||]PAM NA Intergenic NA NA

T22964 GCCGACGCCCACATAGATGTGGG 1000
Oligo fwd CACCGCCGACGCCCACATAGATGT
Oligo rev AAACACATCTATGTGGGCGTCGGC
Chromosome start end strand MM target_seq PAM alignment distance position gene name gene id
GU305914.1 146990 147012 - 0 GCCGACGCCCACATAGATGT GGG ||||||||[||||||||||||]PAM NA Intergenic NA NA
GU305914.1 146983 147005 + 4 TCCGTCGCCCACATCTATGT GGG -|||-|||[||||||--||||]PAM NA Intergenic NA NA

T22965 GCCCACATAGATGTGGGCGACGG 1000
Oligo fwd CACCGCCCACATAGATGTGGGCGA
Oligo rev AAACTCGCCCACATCTATGTGGGC
Chromosome start end strand MM target_seq PAM alignment distance position gene name gene id
GU305914.1 146984 147006 - 0 GCCCACATAGATGTGGGCGA CGG ||||||||[||||||||||||]PAM NA Intergenic NA NA

T22966 GGAGCTCTCGTCGTCGAGCGCGG 1000
Oligo fwd CACCGGAGCTCTCGTCGTCGAGCG
Oligo rev AAACCGCTCGACGACGAGAGCTCC
Chromosome start end strand MM target_seq PAM alignment distance position gene name gene id
GU305914.1 146963 146985 - 0 GGAGCTCTCGTCGTCGAGCG CGG ||||||||[||||||||||||]PAM NA Intergenic NA NA

T22967 GCGCGGCGTCGCCCATGTCGTGG 1000
Oligo fwd CACCGCGCGGCGTCGCCCATGTCG
Oligo rev AAACCGACATGGGCGACGCCGCGC
Chromosome start end strand MM target_seq PAM alignment distance position gene name gene id
GU305914.1 146946 146968 - 0 GCGCGGCGTCGCCCATGTCG TGG ||||||||[||||||||||||]PAM NA Intergenic NA NA

T22968 CGCGGCGTCGCCCATGTCGTGGG 1000
Oligo fwd CACCCGCGGCGTCGCCCATGTCGT

Oligo rev AAACACGACATGGGCGACGCCGCG
Chromosome start end strand MM target_seq PAM alignment distance position gene name gene id
GU305914.1 146945 146967 - 0 CGCGGCGTCGCCCATGTCGT GGG ||||||||[||||||||||||]PAM NA Intergenic NA NA

T22969 GTCGTGGGTAACGCAGAGCCTGG 1000
Oligo fwd CACCGTCGTGGGTAACGCAGAGCC
Oligo rev AAACGGCTCTGCGTTACCCACGAC
Chromosome start end strand MM target_seq PAM alignment distance position gene name gene id
GU305914.1 146930 146952 - 0 GTCGTGGGTAACGCAGAGCC TGG ||||||||[||||||||||||]PAM NA Intergenic NA NA

T22970 GATGCGAAAGTGCCTCATGTCGG 1000
Oligo fwd CACCGATGCGAAAGTGCCTCATGT
Oligo rev AAACACATGAGGCACTTTCGCATC
Chromosome start end strand MM target_seq PAM alignment distance position gene name gene id
GU305914.1 146903 146925 - 0 GATGCGAAAGTGCCTCATGT CGG ||||||||[||||||||||||]PAM NA Intergenic NA NA

T22971 GCCTCATGTCGGCGTAGTGCAGG 1000
Oligo fwd CACCGCCTCATGTCGGCGTAGTGC
Oligo rev AAACGCACTACGCCGACATGAGGC
Chromosome start end strand MM target_seq PAM alignment distance position gene name gene id
GU305914.1 146892 146914 - 0 GCCTCATGTCGGCGTAGTGC AGG ||||||||[||||||||||||]PAM NA Intergenic NA NA

T22972 GTCGATATCGTACATCCTGACGG 1000
Oligo fwd CACCGTCGATATCGTACATCCTGA
Oligo rev AAACTCAGGATGTACGATATCGAC
Chromosome start end strand MM target_seq PAM alignment distance position gene name gene id
GU305914.1 146870 146892 - 0 GTCGATATCGTACATCCTGA CGG ||||||||[||||||||||||]PAM NA Intergenic NA NA

T22973 CGTACATCCTGACGGCGTCCAGG 1000
Oligo fwd CACCCGTACATCCTGACGGCGTCC
Oligo rev AAACGGACGCCGTCAGGATGTACG
Chromosome start end strand MM target_seq PAM alignment distance position gene name gene id
GU305914.1 146862 146884 - 0 CGTACATCCTGACGGCGTCC AGG ||||||||[||||||||||||]PAM NA Intergenic NA NA

T22974 GTACATCCTGACGGCGTCCAGGG 1000
Oligo fwd CACCGTACATCCTGACGGCGTCCA
Oligo rev AAACTGGACGCCGTCAGGATGTAC
Chromosome start end strand MM target_seq PAM alignment distance position gene name gene id
GU305914.1 146861 146883 - 0 GTACATCCTGACGGCGTCCA GGG ||||||||[||||||||||||]PAM NA Intergenic NA NA

T22975 GACGGCGTCCAGGGTGTCGACGG 1000
Oligo fwd CACCGACGGCGTCCAGGGTGTCGA
Oligo rev AAACTCGACACCCTGGACGCCGTC
Chromosome start end strand MM target_seq PAM alignment distance position gene name gene id
GU305914.1 146852 146874 - 0 GACGGCGTCCAGGGTGTCGA CGG ||||||||[||||||||||||]PAM NA Intergenic NA NA
GU305914.1 30886 30908 + 4 GACGACGACCAGCCTGTCGA CGG ||||-||-[||||--||||||]PAM NA Intergenic NA NA
GU305914.1 25777 25799 + 4 GACGAGGACCAGGGTGTCGT CGG ||||--|-[|||||||||||-]PAM NA Intergenic NA NA

T22976 CAGGGTGTCGACGGCGTTGCCGG 1000
Oligo fwd CACCCAGGGTGTCGACGGCGTTGC
Oligo rev AAACGCAACGCCGTCGACACCCTG
Chromosome start end strand MM target_seq PAM alignment distance position gene name gene id
GU305914.1 146843 146865 - 0 CAGGGTGTCGACGGCGTTGC CGG ||||||||[||||||||||||]PAM NA Intergenic NA NA

T22977 AGGGTGTCGACGGCGTTGCCGGG 1000
Oligo fwd CACCAGGGTGTCGACGGCGTTGCC
Oligo rev AAACGGCAACGCCGTCGACACCCT
Chromosome start end strand MM target_seq PAM alignment distance position gene name gene id
GU305914.1 146842 146864 - 0 AGGGTGTCGACGGCGTTGCC GGG ||||||||[||||||||||||]PAM NA Intergenic NA NA

T22978 TCGACGGCGTTGCCGGGCTCCGG 1000
Oligo fwd CACCTCGACGGCGTTGCCGGGCTC
Oligo rev AAACGAGCCCGGCAACGCCGTCGA
Chromosome start end strand MM target_seq PAM alignment distance position gene name gene id
GU305914.1 146836 146858 - 0 TCGACGGCGTTGCCGGGCTC CGG ||||||||[||||||||||||]PAM NA Intergenic NA NA

T22979 TTGCCGGGCTCCGGCACGAGCGG 1000
Oligo fwd CACCTTGCCGGGCTCCGGCACGAG
Oligo rev AAACCTCGTGCCGGAGCCCGGCAA
Chromosome start end strand MM target_seq PAM alignment distance position gene name gene id
GU305914.1 146827 146849 - 0 TTGCCGGGCTCCGGCACGAG CGG ||||||||[||||||||||||]PAM NA Intergenic NA NA

T22980 ACCCGCGACGCCGACGATGTCGG 1000
Oligo fwd CACCACCCGCGACGCCGACGATGT
Oligo rev AAACACATCGTCGGCGTCGCGGGT
Chromosome start end strand MM target_seq PAM alignment distance position gene name gene id
GU305914.1 146798 146820 - 0 ACCCGCGACGCCGACGATGT CGG ||||||||[||||||||||||]PAM NA Intergenic NA NA
GU305914.1 61965 61987 - 4 CCCCCCGACCCCGACGATGA CGG -|||-|||[|-|||||||||-]PAM NA Intergenic NA NA

T22981 CCCGCGACGCCGACGATGTCGGG 1000
Oligo fwd CACCCCCGCGACGCCGACGATGTC
Oligo rev AAACGACATCGTCGGCGTCGCGGG
Chromosome start end strand MM target_seq PAM alignment distance position gene name gene id
GU305914.1 146797 146819 - 0 CCCGCGACGCCGACGATGTC GGG ||||||||[||||||||||||]PAM NA Intergenic NA NA

T22982 CGACGCCGACGATGTCGGGCAGG 1000
Oligo fwd CACCCGACGCCGACGATGTCGGGC
Oligo rev AAACGCCCGACATCGTCGGCGTCG
Chromosome start end strand MM target_seq PAM alignment distance position gene name gene id

| | | | | | | | | | | | |
|---|---|---|---|---|---|---|---|---|---|---|---|
| GU305914.1 | | 146793 | 146815 | - | 0 | CGACGCCGACGATGTCGGGC | AGG | ||||||||[||||||||||||]PAM | NA | Intergenic | NA | NA |
| T22983 | GATGTCGGGCAGGCCGACCAAGG | 1000 | | | | | | | | | |
| Oligo fwd | CACCGATGTCGGGCAGGCCGACCA | | | | | | | | | | |
| Oligo rev | AAACTGGTCGGCCTGCCCGACATC | | | | | | | | | | |
| Chromosome | start | end | strand | MM | target_seq | PAM | alignment | distance | position | gene name | gene id |
| GU305914.1 | 146783 | 146805 | - | 0 | GATGTCGGGCAGGCCGACCA | AGG | ||||||||[||||||||||||]PAM | NA | Intergenic | NA | NA |
| T22984 | CAAGGACCTGATCTCGTTGATGG | 1000 | | | | | | | | | |
| Oligo fwd | CACCCAAGGACCTGATCTCGTTGA | | | | | | | | | | |
| Oligo rev | AAACTCAACGAGATCAGGTCCTTG | | | | | | | | | | |
| Chromosome | start | end | strand | MM | target_seq | PAM | alignment | distance | position | gene name | gene id |
| GU305914.1 | 146765 | 146787 | - | 0 | CAAGGACCTGATCTCGTTGA | TGG | ||||||||[||||||||||||]PAM | NA | Intergenic | NA | NA |
| T22985 | CCTGATCTCGTTGATGGCGTCGG | 1000 | | | | | | | | | |
| Oligo fwd | CACCCCTGATCTCGTTGATGGCGT | | | | | | | | | | |
| Oligo rev | AAACACGCCATCAACGAGATCAGG | | | | | | | | | | |
| Chromosome | start | end | strand | MM | target_seq | PAM | alignment | distance | position | gene name | gene id |
| GU305914.1 | 146759 | 146781 | - | 0 | CCTGATCTCGTTGATGGCGT | CGG | ||||||||[||||||||||||]PAM | NA | Intergenic | NA | NA |
| T22986 | CGTATTCATGTCGTAGCGCCAGG | 1000 | | | | | | | | | |
| Oligo fwd | CACCCGTATTCATGTCGTAGCGCC | | | | | | | | | | |
| Oligo rev | AAACGGCGCTACGACATGAATACG | | | | | | | | | | |
| Chromosome | start | end | strand | MM | target_seq | PAM | alignment | distance | position | gene name | gene id |
| GU305914.1 | 146726 | 146748 | - | 0 | CGTATTCATGTCGTAGCGCC | AGG | ||||||||[||||||||||||]PAM | NA | Intergenic | NA | NA |
| T22987 | GTATTCATGTCGTAGCGCCAGGG | 1000 | | | | | | | | | |
| Oligo fwd | CACCGTATTCATGTCGTAGCGCCA | | | | | | | | | | |
| Oligo rev | AAACTGGCGCTACGACATGAATAC | | | | | | | | | | |
| Chromosome | start | end | strand | MM | target_seq | PAM | alignment | distance | position | gene name | gene id |
| GU305914.1 | 146725 | 146747 | - | 0 | GTATTCATGTCGTAGCGCCA | GGG | ||||||||[||||||||||||]PAM | NA | Intergenic | NA | NA |
| T22988 | CATGTCGTAGCGCCAGGGCACGG | 1000 | | | | | | | | | |
| Oligo fwd | CACCCATGTCGTAGCGCCAGGGCA | | | | | | | | | | |
| Oligo rev | AAACTGCCCTGGCGCTACGACATG | | | | | | | | | | |
| Chromosome | start | end | strand | MM | target_seq | PAM | alignment | distance | position | gene name | gene id |
| GU305914.1 | 146720 | 146742 | - | 0 | CATGTCGTAGCGCCAGGGCA | CGG | ||||||||[||||||||||||]PAM | NA | Intergenic | NA | NA |
| T22989 | GCGCCAGGGCACGGAGTAGACGG | 1000 | | | | | | | | | |
| Oligo fwd | CACCGCGCCAGGGCACGGAGTAGA | | | | | | | | | | |
| Oligo rev | AAACTCTACTCCGTGCCCTGGCGC | | | | | | | | | | |
| Chromosome | start | end | strand | MM | target_seq | PAM | alignment | distance | position | gene name | gene id |
| GU305914.1 | 146711 | 146733 | - | 0 | GCGCCAGGGCACGGAGTAGA | CGG | ||||||||[||||||||||||]PAM | NA | Intergenic | NA | NA |
| GU305914.1 | 29450 | 29472 | + | 3 | GGGCCAGGGCAGGGAGAAGA | CGG | |-||||||[|||-||||-|||]PAM | NA | Intergenic | NA | NA |
| T22990 | GGGCACGGAGTAGACGGCGACGG | 1000 | | | | | | | | | |
| Oligo fwd | CACCGGGCACGGAGTAGACGGCGA | | | | | | | | | | |
| Oligo rev | AAACTCGCCGTCTACTCCGTGCCC | | | | | | | | | | |
| Chromosome | start | end | strand | MM | target_seq | PAM | alignment | distance | position | gene name | gene id |
| GU305914.1 | 146705 | 146727 | - | 0 | GGGCACGGAGTAGACGGCGA | CGG | ||||||||[||||||||||||]PAM | NA | Intergenic | NA | NA |
| T22991 | CACGGAGTAGACGGCGACGGCGG | 1000 | | | | | | | | | |
| Oligo fwd | CACCCACGGAGTAGACGGCGACGG | | | | | | | | | | |
| Oligo rev | AAACCCGTCGCCGTCTACTCCGTG | | | | | | | | | | |
| Chromosome | start | end | strand | MM | target_seq | PAM | alignment | distance | position | gene name | gene id |
| GU305914.1 | 146702 | 146724 | - | 0 | CACGGAGTAGACGGCGACGG | CGG | ||||||||[||||||||||||]PAM | NA | Intergenic | NA | NA |
| T22992 | ACGGAGTAGACGGCGACGGCGGG | 1000 | | | | | | | | | |
| Oligo fwd | CACCACGGAGTAGACGGCGACGGC | | | | | | | | | | |
| Oligo rev | AAACGCCGTCGCCGTCTACTCCGT | | | | | | | | | | |
| Chromosome | start | end | strand | MM | target_seq | PAM | alignment | distance | position | gene name | gene id |
| GU305914.1 | 146701 | 146723 | - | 0 | ACGGAGTAGACGGCGACGGC | GGG | ||||||||[||||||||||||]PAM | NA | Intergenic | NA | NA |
| GU305914.1 | 109329 | 109351 | + | 4 | ACGGAGGTGGCGGGGACGGC | GGG | ||||||--[|-|||-||||||]PAM | NA | Intergenic | NA | NA |
| T22993 | CGGAGTAGACGGCGACGGCGGGG | 1000 | | | | | | | | | |
| Oligo fwd | CACCCGGAGTAGACGGCGACGGCG | | | | | | | | | | |
| Oligo rev | AAACCGCCGTCGCCGTCTACTCCG | | | | | | | | | | |
| Chromosome | start | end | strand | MM | target_seq | PAM | alignment | distance | position | gene name | gene id |
| GU305914.1 | 146700 | 146722 | - | 0 | CGGAGTAGACGGCGACGGCG | GGG | ||||||||[||||||||||||]PAM | NA | Intergenic | NA | NA |
| T22994 | GACGGCGACGGCGGGGAAGCCGG | 1000 | | | | | | | | | |
| Oligo fwd | CACCGACGGCGACGGCGGGGAAGC | | | | | | | | | | |
| Oligo rev | AAACGCTTCCCCGCCGTCGCCGTC | | | | | | | | | | |
| Chromosome | start | end | strand | MM | target_seq | PAM | alignment | distance | position | gene name | gene id |
| GU305914.1 | 146693 | 146715 | - | 0 | GACGGCGACGGCGGGGAAGC | CGG | ||||||||[||||||||||||]PAM | NA | Intergenic | NA | NA |
| T22995 | ACGGCGACGGCGGGGAAGCCGGG | 1000 | | | | | | | | | |
| Oligo fwd | CACCACGGCGACGGCGGGGAAGCC | | | | | | | | | | |
| Oligo rev | AAACGGCTTCCCCGCCGTCGCCGT | | | | | | | | | | |
| Chromosome | start | end | strand | MM | target_seq | PAM | alignment | distance | position | gene name | gene id |
| GU305914.1 | 146692 | 146714 | - | 0 | ACGGCGACGGCGGGGAAGCC | GGG | ||||||||[||||||||||||]PAM | NA | Intergenic | NA | NA |
| T22996 | CGGCGACGGCGGGGAAGCCGGGG | 1000 | | | | | | | | | |
| Oligo fwd | CACCCGGCGACGGCGGGGAAGCCG | | | | | | | | | | |
| Oligo rev | AAACCGGCTTCCCCGCCGTCGCCG | | | | | | | | | | |
| Chromosome | start | end | strand | MM | target_seq | PAM | alignment | distance | position | gene name | gene id |
| GU305914.1 | 146691 | 146713 | - | 0 | CGGCGACGGCGGGGAAGCCG | GGG | ||||||||[||||||||||||]PAM | NA | Intergenic | NA | NA |
| T22997 | GGGGAAGCCGGGGAAGAAGACGG | 1000 | | | | | | | | | |

Oligo fwd CACCGGGGAAGCCGGGGAAGAAGA
Oligo rev AAACTCTTCTTCCCCGGCTTCCCC
Chromosome start end strand MM target_seq PAM alignment distance position gene name gene id
GU305914.1 146681 146703 - 0 GGGGAAGCCGGGGAAGAAGA CGG ||||||||[||||||||||||]PAM NA Intergenic NA NA

T22998 CCGGGGAAGAAGACGGAGCGCGG 1000
Oligo fwd CACCCCGGGGAAGAAGACGGAGCG
Oligo rev AAACCGCTCCGTCTTCTTCCCCGG
Chromosome start end strand MM target_seq PAM alignment distance position gene name gene id
GU305914.1 146674 146696 - 0 CCGGGGAAGAAGACGGAGCG CGG ||||||||[||||||||||||]PAM NA Intergenic NA NA

T22999 CGGGGAAGAAGACGGAGCGCGGG 1000
Oligo fwd CACCCGGGGAAGAAGACGGAGCGC
Oligo rev AAACGCGCTCCGTCTTCTTCCCCG
Chromosome start end strand MM target_seq PAM alignment distance position gene name gene id
GU305914.1 146673 146695 - 0 CGGGGAAGAAGACGGAGCGC GGG ||||||||[||||||||||||]PAM NA Intergenic NA NA

T23000 GAAGACGGAGCGCGGGCCCCCGG 1000
Oligo fwd CACCGAAGACGGAGCGCGGGCCCC
Oligo rev AAACGGGGCCCGCGCTCCGTCTTC
Chromosome start end strand MM target_seq PAM alignment distance position gene name gene id
GU305914.1 146666 146688 - 0 GAAGACGGAGCGCGGGCCCC CGG ||||||||[||||||||||||]PAM NA Intergenic NA NA

T23001 GCCCCCGGAGCGCAGCATATTGG 1000
Oligo fwd CACCGCCCCCGGAGCGCAGCATAT
Oligo rev AAACATATGCTGCGCTCCGGGGGC
Chromosome start end strand MM target_seq PAM alignment distance position gene name gene id
GU305914.1 146651 146673 - 0 GCCCCCGGAGCGCAGCATAT TGG ||||||||[||||||||||||]PAM NA Intergenic NA NA

T23002 CCCCCGGAGCGCAGCATATTGGG 1000
Oligo fwd CACCCCCCCGGAGCGCAGCATATT
Oligo rev AAACAATATGCTGCGCTCCGGGGG
Chromosome start end strand MM target_seq PAM alignment distance position gene name gene id
GU305914.1 146650 146672 - 0 CCCCCGGAGCGCAGCATATT GGG ||||||||[||||||||||||]PAM NA Intergenic NA NA

T23003 GGGCCCGAAGAAGTCGAAGACGG 1000
Oligo fwd CACCGGGCCCGAAGAAGTCGAAGA
Oligo rev AAACTCTTCGACTTCTTCGGGCCC
Chromosome start end strand MM target_seq PAM alignment distance position gene name gene id
GU305914.1 146630 146652 - 0 GGGCCCGAAGAAGTCGAAGA CGG ||||||||[||||||||||||]PAM NA Intergenic NA NA
GU305914.1 77403 77425 - 4 GTGCGCGTGGAAGTCGAAGA TGG |-||-||-[-|||||||||||]PAM NA Intergenic NA NA

T23004 GGCCCGAAGAAGTCGAAGACGGG 1000
Oligo fwd CACCGGCCCGAAGAAGTCGAAGAC
Oligo rev AAACGTCTTCGACTTCTTCGGGCC
Chromosome start end strand MM target_seq PAM alignment distance position gene name gene id
GU305914.1 146629 146651 - 0 GGCCCGAAGAAGTCGAAGAC GGG ||||||||[||||||||||||]PAM NA Intergenic NA NA

T23005 GCCCGAAGAAGTCGAAGACGGGG 1000
Oligo fwd CACCGCCCGAAGAAGTCGAAGACG
Oligo rev AAACCGTCTTCGACTTCTTCGGGC
Chromosome start end strand MM target_seq PAM alignment distance position gene name gene id
GU305914.1 146628 146650 - 0 GCCCGAAGAAGTCGAAGACG GGG ||||||||[||||||||||||]PAM NA Intergenic NA NA

T23006 AGAAGTCGAAGACGGGGTAGAGG 1000
Oligo fwd CACCAGAAGTCGAAGACGGGGTAG
Oligo rev AAACCTACCCCGTCTTCGACTTCT
Chromosome start end strand MM target_seq PAM alignment distance position gene name gene id
GU305914.1 146622 146644 - 0 AGAAGTCGAAGACGGGGTAG AGG ||||||||[||||||||||||]PAM NA Intergenic NA NA

T23007 GTCGAAGACGGGGTAGAGGCCGG 1000
Oligo fwd CACCGTCGAAGACGGGGTAGAGGC
Oligo rev AAACGCCTCTACCCCGTCTTCGAC
Chromosome start end strand MM target_seq PAM alignment distance position gene name gene id
GU305914.1 146618 146640 - 0 GTCGAAGACGGGGTAGAGGC CGG ||||||||[||||||||||||]PAM NA Intergenic NA NA

T23008 CGCCGTAGACCTCCTTGAAGAGG 1000
Oligo fwd CACCCGCCGTAGACCTCCTTGAAG
Oligo rev AAACCTTCAAGGAGGTCTACGGCG
Chromosome start end strand MM target_seq PAM alignment distance position gene name gene id
GU305914.1 146589 146611 - 0 CGCCGTAGACCTCCTTGAAG AGG ||||||||[||||||||||||]PAM NA Intergenic NA NA
GU305914.1 213358 213380 + 4 CCCGGTAGACCTCCTCGAAA CGG |-|-||||[||||||||-|||-]PAM NA Intergenic NA NA

T23009 CTCCTTGAAGAGGCGCTGCATGG 1000
Oligo fwd CACCCTCCTTGAAGAGGCGCTGCA
Oligo rev AAACTGCAGCGCCTCTTCAAGGAG
Chromosome start end strand MM target_seq PAM alignment distance position gene name gene id
GU305914.1 146579 146601 - 0 CTCCTTGAAGAGGCGCTGCA TGG ||||||||[||||||||||||]PAM NA Intergenic NA NA

T23010 TGGCATCGCGCAGTAAGACGAGG 1000
Oligo fwd CACCTGGCATCGCGCAGTAAGACG
Oligo rev AAACCGTCTTACTGCGCGATGCCA
Chromosome start end strand MM target_seq PAM alignment distance position gene name gene id
GU305914.1 146559 146581 - 0 TGGCATCGCGCAGTAAGACG AGG ||||||||[||||||||||||]PAM NA Intergenic NA NA

T23011 CGCGCAGTAAGACGAGGCACTGG 1000
Oligo fwd CACCCGCGCAGTAAGACGAGGCAC
Oligo rev AAACGTGCCTCGTCTTACTGCGCG
Chromosome start end strand MM target_seq PAM alignment distance position gene name gene id

<table>
<tr><td>GU305914.1</td><td>146553</td><td>146575</td><td>-</td><td>0</td><td>CGCGCAGTAAGACGAGGCAC</td><td>TGG</td><td>||||||||[||||||||||||]PAM</td><td>NA</td><td>Intergenic</td><td>NA</td><td>NA</td></tr>
<tr><td>T23012</td><td>CTGGTCGTCCGAGAAGACGAAGG</td><td>1000</td><td></td><td></td><td></td><td></td><td></td><td></td><td></td><td></td><td></td></tr>
<tr><td>Oligo fwd</td><td>CACCCTGGTCGTCCGAGAAGACGA</td><td></td><td></td><td></td><td></td><td></td><td></td><td></td><td></td><td></td><td></td></tr>
<tr><td>Oligo rev</td><td>AAACTCGTCTTCTCGGACGACCAG</td><td></td><td></td><td></td><td></td><td></td><td></td><td></td><td></td><td></td><td></td></tr>
<tr><td>Chromosome</td><td>start</td><td>end</td><td>strand</td><td>MM</td><td>target_seq</td><td>PAM</td><td>alignment</td><td>distance</td><td>position</td><td>gene name</td><td>gene id</td></tr>
<tr><td>GU305914.1</td><td>146534</td><td>146556</td><td>-</td><td>0</td><td>CTGGTCGTCCGAGAAGACGA</td><td>AGG</td><td>||||||||[||||||||||||]PAM</td><td>NA</td><td>Intergenic</td><td>NA</td><td>NA</td></tr>
<tr><td>T23013</td><td>CGTCCGAGAAGACGAAGGCCAGG</td><td>1000</td><td></td><td></td><td></td><td></td><td></td><td></td><td></td><td></td><td></td></tr>
<tr><td>Oligo fwd</td><td>CACCCGTCCGAGAAGACGAAGGCC</td><td></td><td></td><td></td><td></td><td></td><td></td><td></td><td></td><td></td><td></td></tr>
<tr><td>Oligo rev</td><td>AAACGGCCTTCGTCTTCTCGGACG</td><td></td><td></td><td></td><td></td><td></td><td></td><td></td><td></td><td></td><td></td></tr>
<tr><td>Chromosome</td><td>start</td><td>end</td><td>strand</td><td>MM</td><td>target_seq</td><td>PAM</td><td>alignment</td><td>distance</td><td>position</td><td>gene name</td><td>gene id</td></tr>
<tr><td>GU305914.1</td><td>146529</td><td>146551</td><td>-</td><td>0</td><td>CGTCCGAGAAGACGAAGGCC</td><td>AGG</td><td>||||||||[||||||||||||]PAM</td><td>NA</td><td>Intergenic</td><td>NA</td><td>NA</td></tr>
<tr><td>T23014</td><td>ACGAAGGCCAGGAACTCGCGCGG</td><td>1000</td><td></td><td></td><td></td><td></td><td></td><td></td><td></td><td></td><td></td></tr>
<tr><td>Oligo fwd</td><td>CACCACGAAGGCCAGGAACTCGCG</td><td></td><td></td><td></td><td></td><td></td><td></td><td></td><td></td><td></td><td></td></tr>
<tr><td>Oligo rev</td><td>AAACCGCGAGTTCCTGGCCTTCGT</td><td></td><td></td><td></td><td></td><td></td><td></td><td></td><td></td><td></td><td></td></tr>
<tr><td>Chromosome</td><td>start</td><td>end</td><td>strand</td><td>MM</td><td>target_seq</td><td>PAM</td><td>alignment</td><td>distance</td><td>position</td><td>gene name</td><td>gene id</td></tr>
<tr><td>GU305914.1</td><td>146518</td><td>146540</td><td>-</td><td>0</td><td>ACGAAGGCCAGGAACTCGCG</td><td>CGG</td><td>||||||||[||||||||||||]PAM</td><td>NA</td><td>Intergenic</td><td>NA</td><td>NA</td></tr>
<tr><td>T23015</td><td>GGAACTCGCGCGGAAAGCACAGG</td><td>1000</td><td></td><td></td><td></td><td></td><td></td><td></td><td></td><td></td><td></td></tr>
<tr><td>Oligo fwd</td><td>CACCGGAACTCGCGCGGAAAGCAC</td><td></td><td></td><td></td><td></td><td></td><td></td><td></td><td></td><td></td><td></td></tr>
<tr><td>Oligo rev</td><td>AAACGTGCTTTCCGCGCGAGTTCC</td><td></td><td></td><td></td><td></td><td></td><td></td><td></td><td></td><td></td><td></td></tr>
<tr><td>Chromosome</td><td>start</td><td>end</td><td>strand</td><td>MM</td><td>target_seq</td><td>PAM</td><td>alignment</td><td>distance</td><td>position</td><td>gene name</td><td>gene id</td></tr>
<tr><td>GU305914.1</td><td>146508</td><td>146530</td><td>-</td><td>0</td><td>GGAACTCGCGCGGAAAGCAC</td><td>AGG</td><td>||||||||[||||||||||||]PAM</td><td>NA</td><td>Intergenic</td><td>NA</td><td>NA</td></tr>
<tr><td>T23016</td><td>TGCCGCGCCACTTGACATCCTGG</td><td>1000</td><td></td><td></td><td></td><td></td><td></td><td></td><td></td><td></td><td></td></tr>
<tr><td>Oligo fwd</td><td>CACCTGCCGCGCCACTTGACATCC</td><td></td><td></td><td></td><td></td><td></td><td></td><td></td><td></td><td></td><td></td></tr>
<tr><td>Oligo rev</td><td>AAACGGATGTCAAGTGGCGCGGCA</td><td></td><td></td><td></td><td></td><td></td><td></td><td></td><td></td><td></td><td></td></tr>
<tr><td>Chromosome</td><td>start</td><td>end</td><td>strand</td><td>MM</td><td>target_seq</td><td>PAM</td><td>alignment</td><td>distance</td><td>position</td><td>gene name</td><td>gene id</td></tr>
<tr><td>GU305914.1</td><td>146475</td><td>146497</td><td>-</td><td>0</td><td>TGCCGCGCCACTTGACATCC</td><td>TGG</td><td>||||||||[||||||||||||]PAM</td><td>NA</td><td>Intergenic</td><td>NA</td><td>NA</td></tr>
<tr><td>T23017</td><td>ATCCTGGATCACGAAGACGTCGG</td><td>1000</td><td></td><td></td><td></td><td></td><td></td><td></td><td></td><td></td><td></td></tr>
<tr><td>Oligo fwd</td><td>CACCATCCTGGATCACGAAGACGT</td><td></td><td></td><td></td><td></td><td></td><td></td><td></td><td></td><td></td><td></td></tr>
<tr><td>Oligo rev</td><td>AAACACGTCTTCGTGATCCAGGAT</td><td></td><td></td><td></td><td></td><td></td><td></td><td></td><td></td><td></td><td></td></tr>
<tr><td>Chromosome</td><td>start</td><td>end</td><td>strand</td><td>MM</td><td>target_seq</td><td>PAM</td><td>alignment</td><td>distance</td><td>position</td><td>gene name</td><td>gene id</td></tr>
<tr><td>GU305914.1</td><td>146459</td><td>146481</td><td>-</td><td>0</td><td>ATCCTGGATCACGAAGACGT</td><td>CGG</td><td>||||||||[||||||||||||]PAM</td><td>NA</td><td>Intergenic</td><td>NA</td><td>NA</td></tr>
<tr><td>T23018</td><td>GATCACGAAGACGTCGGAGACGG</td><td>1000</td><td></td><td></td><td></td><td></td><td></td><td></td><td></td><td></td><td></td></tr>
<tr><td>Oligo fwd</td><td>CACCGATCACGAAGACGTCGGAGA</td><td></td><td></td><td></td><td></td><td></td><td></td><td></td><td></td><td></td><td></td></tr>
<tr><td>Oligo rev</td><td>AAACTCTCCGACGTCTTCGTGATC</td><td></td><td></td><td></td><td></td><td></td><td></td><td></td><td></td><td></td><td></td></tr>
<tr><td>Chromosome</td><td>start</td><td>end</td><td>strand</td><td>MM</td><td>target_seq</td><td>PAM</td><td>alignment</td><td>distance</td><td>position</td><td>gene name</td><td>gene id</td></tr>
<tr><td>GU305914.1</td><td>146453</td><td>146475</td><td>-</td><td>0</td><td>GATCACGAAGACGTCGGAGA</td><td>CGG</td><td>||||||||[||||||||||||]PAM</td><td>NA</td><td>Intergenic</td><td>NA</td><td>NA</td></tr>
<tr><td>T23019</td><td>GACGTCGGAGACGGAGAACTTGG</td><td>1000</td><td></td><td></td><td></td><td></td><td></td><td></td><td></td><td></td><td></td></tr>
<tr><td>Oligo fwd</td><td>CACCGACGTCGGAGACGGAGAACT</td><td></td><td></td><td></td><td></td><td></td><td></td><td></td><td></td><td></td><td></td></tr>
<tr><td>Oligo rev</td><td>AAACAGTTCTCCGTCTCCGACGTC</td><td></td><td></td><td></td><td></td><td></td><td></td><td></td><td></td><td></td><td></td></tr>
<tr><td>Chromosome</td><td>start</td><td>end</td><td>strand</td><td>MM</td><td>target_seq</td><td>PAM</td><td>alignment</td><td>distance</td><td>position</td><td>gene name</td><td>gene id</td></tr>
<tr><td>GU305914.1</td><td>146444</td><td>146466</td><td>-</td><td>0</td><td>GACGTCGGAGACGGAGAACT</td><td>TGG</td><td>||||||||[||||||||||||]PAM</td><td>NA</td><td>Intergenic</td><td>NA</td><td>NA</td></tr>
<tr><td>T23020</td><td>GACGGAGAACTTGGCGCCCACGG</td><td>1000</td><td></td><td></td><td></td><td></td><td></td><td></td><td></td><td></td><td></td></tr>
<tr><td>Oligo fwd</td><td>CACCGACGGAGAACTTGGCGCCCA</td><td></td><td></td><td></td><td></td><td></td><td></td><td></td><td></td><td></td><td></td></tr>
<tr><td>Oligo rev</td><td>AAACTGGGCGCCAAGTTCTCCGTC</td><td></td><td></td><td></td><td></td><td></td><td></td><td></td><td></td><td></td><td></td></tr>
<tr><td>Chromosome</td><td>start</td><td>end</td><td>strand</td><td>MM</td><td>target_seq</td><td>PAM</td><td>alignment</td><td>distance</td><td>position</td><td>gene name</td><td>gene id</td></tr>
<tr><td>GU305914.1</td><td>146435</td><td>146457</td><td>-</td><td>0</td><td>GACGGAGAACTTGGCGCCCA</td><td>CGG</td><td>||||||||[||||||||||||]PAM</td><td>NA</td><td>Intergenic</td><td>NA</td><td>NA</td></tr>
<tr><td>T23021</td><td>GGAGAACTTGGCGCCCACGGCGG</td><td>1000</td><td></td><td></td><td></td><td></td><td></td><td></td><td></td><td></td><td></td></tr>
<tr><td>Oligo fwd</td><td>CACCGGAGAACTTGGCGCCCACGG</td><td></td><td></td><td></td><td></td><td></td><td></td><td></td><td></td><td></td><td></td></tr>
<tr><td>Oligo rev</td><td>AAACCCGTGGGCGCCAAGTTCTCC</td><td></td><td></td><td></td><td></td><td></td><td></td><td></td><td></td><td></td><td></td></tr>
<tr><td>Chromosome</td><td>start</td><td>end</td><td>strand</td><td>MM</td><td>target_seq</td><td>PAM</td><td>alignment</td><td>distance</td><td>position</td><td>gene name</td><td>gene id</td></tr>
<tr><td>GU305914.1</td><td>146432</td><td>146454</td><td>-</td><td>0</td><td>GGAGAACTTGGCGCCCACGG</td><td>CGG</td><td>||||||||[||||||||||||]PAM</td><td>NA</td><td>Intergenic</td><td>NA</td><td>NA</td></tr>
<tr><td>T23022</td><td>GAACTTGGCGCCCACGGCGGAGG</td><td>1000</td><td></td><td></td><td></td><td></td><td></td><td></td><td></td><td></td><td></td></tr>
<tr><td>Oligo fwd</td><td>CACCGAACTTGGCGCCCACGGCGG</td><td></td><td></td><td></td><td></td><td></td><td></td><td></td><td></td><td></td><td></td></tr>
<tr><td>Oligo rev</td><td>AAACCCGCCGTGGGCGCCAAGTTC</td><td></td><td></td><td></td><td></td><td></td><td></td><td></td><td></td><td></td><td></td></tr>
<tr><td>Chromosome</td><td>start</td><td>end</td><td>strand</td><td>MM</td><td>target_seq</td><td>PAM</td><td>alignment</td><td>distance</td><td>position</td><td>gene name</td><td>gene id</td></tr>
<tr><td>GU305914.1</td><td>146429</td><td>146451</td><td>-</td><td>0</td><td>GAACTTGGCGCCCACGGCGG</td><td>AGG</td><td>||||||||[||||||||||||]PAM</td><td>NA</td><td>Intergenic</td><td>NA</td><td>NA</td></tr>
<tr><td>T23023</td><td>AACTTGGCGCCCACGGCGGAGGG</td><td>1000</td><td></td><td></td><td></td><td></td><td></td><td></td><td></td><td></td><td></td></tr>
<tr><td>Oligo fwd</td><td>CACCAACTTGGCGCCCACGGCGGA</td><td></td><td></td><td></td><td></td><td></td><td></td><td></td><td></td><td></td><td></td></tr>
<tr><td>Oligo rev</td><td>AAACTCCGCCGTGGGCGCCAAGTT</td><td></td><td></td><td></td><td></td><td></td><td></td><td></td><td></td><td></td><td></td></tr>
<tr><td>Chromosome</td><td>start</td><td>end</td><td>strand</td><td>MM</td><td>target_seq</td><td>PAM</td><td>alignment</td><td>distance</td><td>position</td><td>gene name</td><td>gene id</td></tr>
<tr><td>GU305914.1</td><td>146428</td><td>146450</td><td>-</td><td>0</td><td>AACTTGGCGCCCACGGCGGA</td><td>GGG</td><td>||||||||[||||||||||||]PAM</td><td>NA</td><td>Intergenic</td><td>NA</td><td>NA</td></tr>
<tr><td>T23024</td><td>TGAACTTTTTGTTACCGATGCGG</td><td>1000</td><td></td><td></td><td></td><td></td><td></td><td></td><td></td><td></td><td></td></tr>
<tr><td>Oligo fwd</td><td>CACCTGAACTTTTTGTTACCGATG</td><td></td><td></td><td></td><td></td><td></td><td></td><td></td><td></td><td></td><td></td></tr>
<tr><td>Oligo rev</td><td>AAACCATCGGTAACAAAAAGTTCA</td><td></td><td></td><td></td><td></td><td></td><td></td><td></td><td></td><td></td><td></td></tr>
<tr><td>Chromosome</td><td>start</td><td>end</td><td>strand</td><td>MM</td><td>target_seq</td><td>PAM</td><td>alignment</td><td>distance</td><td>position</td><td>gene name</td><td>gene id</td></tr>
<tr><td>GU305914.1</td><td>146394</td><td>146416</td><td>-</td><td>0</td><td>TGAACTTTTTGTTACCGATG</td><td>CGG</td><td>||||||||[||||||||||||]PAM</td><td>NA</td><td>Intergenic</td><td>NA</td><td>NA</td></tr>
<tr><td>T23025</td><td>CTTTTTGTTACCGATGCGGATGG</td><td>1000</td><td></td><td></td><td></td><td></td><td></td><td></td><td></td><td></td><td></td></tr>
<tr><td>Oligo fwd</td><td>CACCCTTTTTGTTACCGATGCGGA</td><td></td><td></td><td></td><td></td><td></td><td></td><td></td><td></td><td></td><td></td></tr>
<tr><td>Oligo rev</td><td>AAACTCCGCATCGGTAACAAAAAG</td><td></td><td></td><td></td><td></td><td></td><td></td><td></td><td></td><td></td><td></td></tr>
<tr><td>Chromosome</td><td>start</td><td>end</td><td>strand</td><td>MM</td><td>target_seq</td><td>PAM</td><td>alignment</td><td>distance</td><td>position</td><td>gene name</td><td>gene id</td></tr>
<tr><td>GU305914.1</td><td>146390</td><td>146412</td><td>-</td><td>0</td><td>CTTTTTGTTACCGATGCGGA</td><td>TGG</td><td>||||||||[||||||||||||]PAM</td><td>NA</td><td>Intergenic</td><td>NA</td><td>NA</td></tr>
<tr><td>T23026</td><td>GTTACCGATGCGGATGGAGAAGG</td><td>1000</td><td></td><td></td><td></td><td></td><td></td><td></td><td></td><td></td><td></td></tr>
<tr><td>Oligo fwd</td><td>CACCGTTACCGATGCGGATGGAGA</td><td></td><td></td><td></td><td></td><td></td><td></td><td></td><td></td><td></td><td></td></tr>
<tr><td>Oligo rev</td><td>AAACTCTCCATCCGCATCGGTAAC</td><td></td><td></td><td></td><td></td><td></td><td></td><td></td><td></td><td></td><td></td></tr>
</table>

<table>
<tr><td>Chromosome</td><td>start</td><td>end</td><td>strand</td><td>MM</td><td>target_seq</td><td>PAM</td><td>alignment</td><td>distance</td><td>position</td><td>gene name</td><td>gene id</td></tr>
<tr><td>GU305914.1</td><td>146384</td><td>146406</td><td>-</td><td>0</td><td>GTTACCGATGCGGATGGAGA</td><td>AGG</td><td>||||||||[||||||||||||]PAM</td><td>NA</td><td>Intergenic</td><td>NA</td><td>NA</td></tr>
<tr><td></td><td></td><td></td><td></td><td></td><td></td><td></td><td></td><td></td><td></td><td></td><td></td></tr>
<tr><td>T23027</td><td>TGTCGCGTGAGACGAAGCCGCGG</td><td>1000</td><td></td><td></td><td></td><td></td><td></td><td></td><td></td><td></td><td></td></tr>
<tr><td>Oligo fwd</td><td>CACCTGTCGCGTGAGACGAAGCCG</td><td></td><td></td><td></td><td></td><td></td><td></td><td></td><td></td><td></td><td></td></tr>
<tr><td>Oligo rev</td><td>AAACCGGCTTCGTCTCACGCGACA</td><td></td><td></td><td></td><td></td><td></td><td></td><td></td><td></td><td></td><td></td></tr>
<tr><td>Chromosome</td><td>start</td><td>end</td><td>strand</td><td>MM</td><td>target_seq</td><td>PAM</td><td>alignment</td><td>distance</td><td>position</td><td>gene name</td><td>gene id</td></tr>
<tr><td>GU305914.1</td><td>146358</td><td>146380</td><td>-</td><td>0</td><td>TGTCGCGTGAGACGAAGCCG</td><td>CGG</td><td>||||||||[||||||||||||]PAM</td><td>NA</td><td>Intergenic</td><td>NA</td><td>NA</td></tr>
<tr><td></td><td></td><td></td><td></td><td></td><td></td><td></td><td></td><td></td><td></td><td></td><td></td></tr>
<tr><td>T23028</td><td>CGCGTGAGACGAAGCCGCGGCGG</td><td>1000</td><td></td><td></td><td></td><td></td><td></td><td></td><td></td><td></td><td></td></tr>
<tr><td>Oligo fwd</td><td>CACCCGCGTGAGACGAAGCCGCGG</td><td></td><td></td><td></td><td></td><td></td><td></td><td></td><td></td><td></td><td></td></tr>
<tr><td>Oligo rev</td><td>AAACCCGCGGCTTCGTCTCACGCG</td><td></td><td></td><td></td><td></td><td></td><td></td><td></td><td></td><td></td><td></td></tr>
<tr><td>Chromosome</td><td>start</td><td>end</td><td>strand</td><td>MM</td><td>target_seq</td><td>PAM</td><td>alignment</td><td>distance</td><td>position</td><td>gene name</td><td>gene id</td></tr>
<tr><td>GU305914.1</td><td>146355</td><td>146377</td><td>-</td><td>0</td><td>CGCGTGAGACGAAGCCGCGG</td><td>CGG</td><td>||||||||[||||||||||||]PAM</td><td>NA</td><td>Intergenic</td><td>NA</td><td>NA</td></tr>
<tr><td></td><td></td><td></td><td></td><td></td><td></td><td></td><td></td><td></td><td></td><td></td><td></td></tr>
<tr><td>T23029</td><td>GCCGCGGCGGAGCTTCCTGTAGG</td><td>1000</td><td></td><td></td><td></td><td></td><td></td><td></td><td></td><td></td><td></td></tr>
<tr><td>Oligo fwd</td><td>CACCGCCGCGGCGGAGCTTCCTGT</td><td></td><td></td><td></td><td></td><td></td><td></td><td></td><td></td><td></td><td></td></tr>
<tr><td>Oligo rev</td><td>AAACACAGGAAGCTCCGCCGCGGC</td><td></td><td></td><td></td><td></td><td></td><td></td><td></td><td></td><td></td><td></td></tr>
<tr><td>Chromosome</td><td>start</td><td>end</td><td>strand</td><td>MM</td><td>target_seq</td><td>PAM</td><td>alignment</td><td>distance</td><td>position</td><td>gene name</td><td>gene id</td></tr>
<tr><td>GU305914.1</td><td>146342</td><td>146364</td><td>-</td><td>0</td><td>GCCGCGGCGGAGCTTCCTGT</td><td>AGG</td><td>||||||||[||||||||||||]PAM</td><td>NA</td><td>Intergenic</td><td>NA</td><td>NA</td></tr>
<tr><td></td><td></td><td></td><td></td><td></td><td></td><td></td><td></td><td></td><td></td><td></td><td></td></tr>
<tr><td>T23030</td><td>TCCTGTAGGCGTGCGTGTTGCGG</td><td>1000</td><td></td><td></td><td></td><td></td><td></td><td></td><td></td><td></td><td></td></tr>
<tr><td>Oligo fwd</td><td>CACCTCCTGTAGGCGTGCGTGTTG</td><td></td><td></td><td></td><td></td><td></td><td></td><td></td><td></td><td></td><td></td></tr>
<tr><td>Oligo rev</td><td>AAACCAACACGCACGCCTACAGGA</td><td></td><td></td><td></td><td></td><td></td><td></td><td></td><td></td><td></td><td></td></tr>
<tr><td>Chromosome</td><td>start</td><td>end</td><td>strand</td><td>MM</td><td>target_seq</td><td>PAM</td><td>alignment</td><td>distance</td><td>position</td><td>gene name</td><td>gene id</td></tr>
<tr><td>GU305914.1</td><td>146328</td><td>146350</td><td>-</td><td>0</td><td>TCCTGTAGGCGTGCGTGTTG</td><td>CGG</td><td>||||||||[||||||||||||]PAM</td><td>NA</td><td>Intergenic</td><td>NA</td><td>NA</td></tr>
<tr><td></td><td></td><td></td><td></td><td></td><td></td><td></td><td></td><td></td><td></td><td></td><td></td></tr>
<tr><td>T23031</td><td>AGGCGTGCGTGTTGCGGTAGCGG</td><td>1000</td><td></td><td></td><td></td><td></td><td></td><td></td><td></td><td></td><td></td></tr>
<tr><td>Oligo fwd</td><td>CACCAGGCGTGCGTGTTGCGGTAG</td><td></td><td></td><td></td><td></td><td></td><td></td><td></td><td></td><td></td><td></td></tr>
<tr><td>Oligo rev</td><td>AAACCTACCGCAACACGCACGCCT</td><td></td><td></td><td></td><td></td><td></td><td></td><td></td><td></td><td></td><td></td></tr>
<tr><td>Chromosome</td><td>start</td><td>end</td><td>strand</td><td>MM</td><td>target_seq</td><td>PAM</td><td>alignment</td><td>distance</td><td>position</td><td>gene name</td><td>gene id</td></tr>
<tr><td>GU305914.1</td><td>146322</td><td>146344</td><td>-</td><td>0</td><td>AGGCGTGCGTGTTGCGGTAG</td><td>CGG</td><td>||||||||[||||||||||||]PAM</td><td>NA</td><td>Intergenic</td><td>NA</td><td>NA</td></tr>
<tr><td></td><td></td><td></td><td></td><td></td><td></td><td></td><td></td><td></td><td></td><td></td><td></td></tr>
<tr><td>T23032</td><td>GCGGTAGCGGCACTCGTCCTCGG</td><td>1000</td><td></td><td></td><td></td><td></td><td></td><td></td><td></td><td></td><td></td></tr>
<tr><td>Oligo fwd</td><td>CACCGCGGTAGCGGCACTCGTCCT</td><td></td><td></td><td></td><td></td><td></td><td></td><td></td><td></td><td></td><td></td></tr>
<tr><td>Oligo rev</td><td>AAACAGGACGAGTGCCGCTACCGC</td><td></td><td></td><td></td><td></td><td></td><td></td><td></td><td></td><td></td><td></td></tr>
<tr><td>Chromosome</td><td>start</td><td>end</td><td>strand</td><td>MM</td><td>target_seq</td><td>PAM</td><td>alignment</td><td>distance</td><td>position</td><td>gene name</td><td>gene id</td></tr>
<tr><td>GU305914.1</td><td>146309</td><td>146331</td><td>-</td><td>0</td><td>GCGGTAGCGGCACTCGTCCT</td><td>CGG</td><td>||||||||[||||||||||||]PAM</td><td>NA</td><td>Intergenic</td><td>NA</td><td>NA</td></tr>
<tr><td></td><td></td><td></td><td></td><td></td><td></td><td></td><td></td><td></td><td></td><td></td><td></td></tr>
<tr><td>T23033</td><td>GCACTCGTCCTCGGCATCCTCGG</td><td>1000</td><td></td><td></td><td></td><td></td><td></td><td></td><td></td><td></td><td></td></tr>
<tr><td>Oligo fwd</td><td>CACCGCACTCGTCCTCGGCATCCT</td><td></td><td></td><td></td><td></td><td></td><td></td><td></td><td></td><td></td><td></td></tr>
<tr><td>Oligo rev</td><td>AAACAGGATGCCGAGGACGAGTGC</td><td></td><td></td><td></td><td></td><td></td><td></td><td></td><td></td><td></td><td></td></tr>
<tr><td>Chromosome</td><td>start</td><td>end</td><td>strand</td><td>MM</td><td>target_seq</td><td>PAM</td><td>alignment</td><td>distance</td><td>position</td><td>gene name</td><td>gene id</td></tr>
<tr><td>GU305914.1</td><td>146300</td><td>146322</td><td>-</td><td>0</td><td>GCACTCGTCCTCGGCATCCT</td><td>CGG</td><td>||||||||[||||||||||||]PAM</td><td>NA</td><td>Intergenic</td><td>NA</td><td>NA</td></tr>
<tr><td></td><td></td><td></td><td></td><td></td><td></td><td></td><td></td><td></td><td></td><td></td><td></td></tr>
<tr><td>T23034</td><td>GTCCTCGGCATCCTCGGTCGTGG</td><td>1000</td><td></td><td></td><td></td><td></td><td></td><td></td><td></td><td></td><td></td></tr>
<tr><td>Oligo fwd</td><td>CACCGTCCTCGGCATCCTCGGTCG</td><td></td><td></td><td></td><td></td><td></td><td></td><td></td><td></td><td></td><td></td></tr>
<tr><td>Oligo rev</td><td>AAACCGACCGAGGATGCCGAGGAC</td><td></td><td></td><td></td><td></td><td></td><td></td><td></td><td></td><td></td><td></td></tr>
<tr><td>Chromosome</td><td>start</td><td>end</td><td>strand</td><td>MM</td><td>target_seq</td><td>PAM</td><td>alignment</td><td>distance</td><td>position</td><td>gene name</td><td>gene id</td></tr>
<tr><td>GU305914.1</td><td>146294</td><td>146316</td><td>-</td><td>0</td><td>GTCCTCGGCATCCTCGGTCG</td><td>TGG</td><td>||||||||[||||||||||||]PAM</td><td>NA</td><td>Intergenic</td><td>NA</td><td>NA</td></tr>
<tr><td></td><td></td><td></td><td></td><td></td><td></td><td></td><td></td><td></td><td></td><td></td><td></td></tr>
<tr><td>T23035</td><td>ATCCTCGGTCGTGGCGCCTTCGG</td><td>1000</td><td></td><td></td><td></td><td></td><td></td><td></td><td></td><td></td><td></td></tr>
<tr><td>Oligo fwd</td><td>CACCATCCTCGGTCGTGGCGCCTT</td><td></td><td></td><td></td><td></td><td></td><td></td><td></td><td></td><td></td><td></td></tr>
<tr><td>Oligo rev</td><td>AAACAAGGCGCCACGACCGAGGAT</td><td></td><td></td><td></td><td></td><td></td><td></td><td></td><td></td><td></td><td></td></tr>
<tr><td>Chromosome</td><td>start</td><td>end</td><td>strand</td><td>MM</td><td>target_seq</td><td>PAM</td><td>alignment</td><td>distance</td><td>position</td><td>gene name</td><td>gene id</td></tr>
<tr><td>GU305914.1</td><td>146285</td><td>146307</td><td>-</td><td>0</td><td>ATCCTCGGTCGTGGCGCCTT</td><td>CGG</td><td>||||||||[||||||||||||]PAM</td><td>NA</td><td>Intergenic</td><td>NA</td><td>NA</td></tr>
<tr><td></td><td></td><td></td><td></td><td></td><td></td><td></td><td></td><td></td><td></td><td></td><td></td></tr>
<tr><td>T23036</td><td>CGTGGCGCCTTCGGCGATGATGG</td><td>1000</td><td></td><td></td><td></td><td></td><td></td><td></td><td></td><td></td><td></td></tr>
<tr><td>Oligo fwd</td><td>CACCCGTGGCGCCTTCGGCGATGA</td><td></td><td></td><td></td><td></td><td></td><td></td><td></td><td></td><td></td><td></td></tr>
<tr><td>Oligo rev</td><td>AAACTCATCGCCGAAGGCGCCACG</td><td></td><td></td><td></td><td></td><td></td><td></td><td></td><td></td><td></td><td></td></tr>
<tr><td>Chromosome</td><td>start</td><td>end</td><td>strand</td><td>MM</td><td>target_seq</td><td>PAM</td><td>alignment</td><td>distance</td><td>position</td><td>gene name</td><td>gene id</td></tr>
<tr><td>GU305914.1</td><td>146276</td><td>146298</td><td>-</td><td>0</td><td>CGTGGCGCCTTCGGCGATGA</td><td>TGG</td><td>||||||||[||||||||||||]PAM</td><td>NA</td><td>Intergenic</td><td>NA</td><td>NA</td></tr>
<tr><td></td><td></td><td></td><td></td><td></td><td></td><td></td><td></td><td></td><td></td><td></td><td></td></tr>
<tr><td>T23037</td><td>CGCCTTCGGCGATGATGGACTGG</td><td>1000</td><td></td><td></td><td></td><td></td><td></td><td></td><td></td><td></td><td></td></tr>
<tr><td>Oligo fwd</td><td>CACCCGCCTTCGGCGATGATGGAC</td><td></td><td></td><td></td><td></td><td></td><td></td><td></td><td></td><td></td><td></td></tr>
<tr><td>Oligo rev</td><td>AAACGTCCATCATCGCCGAAGGCG</td><td></td><td></td><td></td><td></td><td></td><td></td><td></td><td></td><td></td><td></td></tr>
<tr><td>Chromosome</td><td>start</td><td>end</td><td>strand</td><td>MM</td><td>target_seq</td><td>PAM</td><td>alignment</td><td>distance</td><td>position</td><td>gene name</td><td>gene id</td></tr>
<tr><td>GU305914.1</td><td>146271</td><td>146293</td><td>-</td><td>0</td><td>CGCCTTCGGCGATGATGGAC</td><td>TGG</td><td>||||||||[||||||||||||]PAM</td><td>NA</td><td>Intergenic</td><td>NA</td><td>NA</td></tr>
<tr><td></td><td></td><td></td><td></td><td></td><td></td><td></td><td></td><td></td><td></td><td></td><td></td></tr>
<tr><td>T23038</td><td>GATGGACTGGATCGCCGTACTGG</td><td>1000</td><td></td><td></td><td></td><td></td><td></td><td></td><td></td><td></td><td></td></tr>
<tr><td>Oligo fwd</td><td>CACCGATGGACTGGATCGCCGTAC</td><td></td><td></td><td></td><td></td><td></td><td></td><td></td><td></td><td></td><td></td></tr>
<tr><td>Oligo rev</td><td>AAACGTACGGCGATCCAGTCCATC</td><td></td><td></td><td></td><td></td><td></td><td></td><td></td><td></td><td></td><td></td></tr>
<tr><td>Chromosome</td><td>start</td><td>end</td><td>strand</td><td>MM</td><td>target_seq</td><td>PAM</td><td>alignment</td><td>distance</td><td>position</td><td>gene name</td><td>gene id</td></tr>
<tr><td>GU305914.1</td><td>146258</td><td>146280</td><td>-</td><td>0</td><td>GATGGACTGGATCGCCGTAC</td><td>TGG</td><td>||||||||[||||||||||||]PAM</td><td>NA</td><td>Intergenic</td><td>NA</td><td>NA</td></tr>
<tr><td></td><td></td><td></td><td></td><td></td><td></td><td></td><td></td><td></td><td></td><td></td><td></td></tr>
<tr><td>T23039</td><td>CGTACTGGTCTCGATCGAGACGG</td><td>1000</td><td></td><td></td><td></td><td></td><td></td><td></td><td></td><td></td><td></td></tr>
<tr><td>Oligo fwd</td><td>CACCCGTACTGGTCTCGATCGAGA</td><td></td><td></td><td></td><td></td><td></td><td></td><td></td><td></td><td></td><td></td></tr>
<tr><td>Oligo rev</td><td>AAACTCTCGATCGAGACCAGTACG</td><td></td><td></td><td></td><td></td><td></td><td></td><td></td><td></td><td></td><td></td></tr>
<tr><td>Chromosome</td><td>start</td><td>end</td><td>strand</td><td>MM</td><td>target_seq</td><td>PAM</td><td>alignment</td><td>distance</td><td>position</td><td>gene name</td><td>gene id</td></tr>
<tr><td>GU305914.1</td><td>146243</td><td>146265</td><td>-</td><td>0</td><td>CGTACTGGTCTCGATCGAGA</td><td>CGG</td><td>||||||||[||||||||||||]PAM</td><td>NA</td><td>Intergenic</td><td>NA</td><td>NA</td></tr>
<tr><td></td><td></td><td></td><td></td><td></td><td></td><td></td><td></td><td></td><td></td><td></td><td></td></tr>
<tr><td>T23040</td><td>TCGAGACGGCGTCCGCCATCTGG</td><td>1000</td><td></td><td></td><td></td><td></td><td></td><td></td><td></td><td></td><td></td></tr>
<tr><td>Oligo fwd</td><td>CACCTCGAGACGGCGTCCGCCATC</td><td></td><td></td><td></td><td></td><td></td><td></td><td></td><td></td><td></td><td></td></tr>
<tr><td>Oligo rev</td><td>AAACGATGGCGGACGCCGTCTCGA</td><td></td><td></td><td></td><td></td><td></td><td></td><td></td><td></td><td></td><td></td></tr>
<tr><td>Chromosome</td><td>start</td><td>end</td><td>strand</td><td>MM</td><td>target_seq</td><td>PAM</td><td>alignment</td><td>distance</td><td>position</td><td>gene name</td><td>gene id</td></tr>
<tr><td>GU305914.1</td><td>146229</td><td>146251</td><td>-</td><td>0</td><td>TCGAGACGGCGTCCGCCATC</td><td>TGG</td><td>||||||||[||||||||||||]PAM</td><td>NA</td><td>Intergenic</td><td>NA</td><td>NA</td></tr>
<tr><td></td><td></td><td></td><td></td><td></td><td></td><td></td><td></td><td></td><td></td><td></td><td></td></tr>
<tr><td>T23041</td><td>GGCGTCCGCCATCTGGTGATAGG</td><td>1000</td><td></td><td></td><td></td><td></td><td></td><td></td><td></td><td></td><td></td></tr>
<tr><td>Oligo fwd</td><td>CACCGGCGTCCGCCATCTGGTGAT</td><td></td><td></td><td></td><td></td><td></td><td></td><td></td><td></td><td></td><td></td></tr>
</table>

Oligo rev AAACATCACCAGATGGCGGACGCC
Chromosome start end strand MM target_seq PAM alignment distance position gene name gene id
GU305914.1 146222 146244 - 0 GGCGTCCGCCATCTGGTGAT AGG ||||||||[||||||||||||]PAM NA Intergenic NA NA

T23042 CATCTGGTGATAGGAGCCGATGG 1000
Oligo fwd CACCCATCTGGTGATAGGAGCCGA
Oligo rev AAACTCGGCTCCTATCACCAGATG
Chromosome start end strand MM target_seq PAM alignment distance position gene name gene id
GU305914.1 146213 146235 - 0 CATCTGGTGATAGGAGCCGA TGG ||||||||[||||||||||||]PAM NA Intergenic NA NA

T23043 GGTGATAGGAGCCGATGGCTCGG 1000
Oligo fwd CACCGGTGATAGGAGCCGATGGCT
Oligo rev AAACAGCCATCGGCTCCTATCACC
Chromosome start end strand MM target_seq PAM alignment distance position gene name gene id
GU305914.1 146208 146230 - 0 GGTGATAGGAGCCGATGGCT CGG ||||||||[||||||||||||]PAM NA Intergenic NA NA

T23044 CTCGGATCCTGCCGACCGTCAGG 1000
Oligo fwd CACCCTCGGATCCTGCCGACCGTC
Oligo rev AAACGACGGTCGGCAGGATCCGAG
Chromosome start end strand MM target_seq PAM alignment distance position gene name gene id
GU305914.1 146190 146212 - 0 CTCGGATCCTGCCGACCGTC AGG ||||||||[||||||||||||]PAM NA Intergenic NA NA

T23045 TCGGATCCTGCCGACCGTCAGGG 1000
Oligo fwd CACCTCGGATCCTGCCGACCGTCA
Oligo rev AAACTGACGGTCGGCAGGATCCGA
Chromosome start end strand MM target_seq PAM alignment distance position gene name gene id
GU305914.1 146189 146211 - 0 TCGGATCCTGCCGACCGTCA GGG ||||||||[||||||||||||]PAM NA Intergenic NA NA

T23046 GATCCTGCCGACCGTCAGGGCGG 1000
Oligo fwd CACCGATCCTGCCGACCGTCAGGG
Oligo rev AAACCCCTGACGGTCGGCAGGATC
Chromosome start end strand MM target_seq PAM alignment distance position gene name gene id
GU305914.1 146186 146208 - 0 GATCCTGCCGACCGTCAGGG CGG ||||||||[||||||||||||]PAM NA Intergenic NA NA

T23047 TGCCGACCGTCAGGGCGGCCAGG 1000
Oligo fwd CACCTGCCGACCGTCAGGGCGGCC
Oligo rev AAACGGCCGCCCTGACGGTCGGCA
Chromosome start end strand MM target_seq PAM alignment distance position gene name gene id
GU305914.1 146181 146203 - 0 TGCCGACCGTCAGGGCGGCC AGG ||||||||[||||||||||||]PAM NA Intergenic NA NA

T23048 GCCGACCGTCAGGGCGGCCAGGG 1000
Oligo fwd CACCGCCGACCGTCAGGGCGGCCA
Oligo rev AAACTGGCCGCCCTGACGGTCGGC
Chromosome start end strand MM target_seq PAM alignment distance position gene name gene id
GU305914.1 146180 146202 - 0 GCCGACCGTCAGGGCGGCCA GGG ||||||||[||||||||||||]PAM NA Intergenic NA NA

T23049 TCAGGGCGGCCAGGGTCTGCAGG 1000
Oligo fwd CACCTCAGGGCGGCCAGGGTCTGC
Oligo rev AAACGCAGACCCTGGCCGCCCTGA
Chromosome start end strand MM target_seq PAM alignment distance position gene name gene id
GU305914.1 146172 146194 - 0 TCAGGGCGGCCAGGGTCTGC AGG ||||||||[||||||||||||]PAM NA Intergenic NA NA

T23050 CGCGCACATTCTCCCATCTCCGG 1000
Oligo fwd CACCCGCGCACATTCTCCCATCTC
Oligo rev AAACGAGATGGGAGAATGTGCGCG
Chromosome start end strand MM target_seq PAM alignment distance position gene name gene id
GU305914.1 146133 146155 - 0 CGCGCACATTCTCCCATCTC CGG ||||||||[||||||||||||]PAM NA Intergenic NA NA

T23051 GCGCACATTCTCCCATCTCCGGG 1000
Oligo fwd CACCGCGCACATTCTCCCATCTCC
Oligo rev AAACGGAGATGGGAGAATGTGCGC
Chromosome start end strand MM target_seq PAM alignment distance position gene name gene id
GU305914.1 146132 146154 - 0 GCGCACATTCTCCCATCTCC GGG ||||||||[||||||||||||]PAM NA Intergenic NA NA

T23052 TCTCCCATCTCCGGGAGCCAAGG 1000
Oligo fwd CACCTCTCCCATCTCCGGGAGCCA
Oligo rev AAACTGGCTCCCGGAGATGGGAGA
Chromosome start end strand MM target_seq PAM alignment distance position gene name gene id
GU305914.1 146124 146146 - 0 TCTCCCATCTCCGGGAGCCA AGG ||||||||[||||||||||||]PAM NA Intergenic NA NA

T23053 GCCAAGGATCTCGACGCCGTCGG 1000
Oligo fwd CACCGCCAAGGATCTCGACGCCGT
Oligo rev AAACACGGCGTCGAGATCCTTGGC
Chromosome start end strand MM target_seq PAM alignment distance position gene name gene id
GU305914.1 146108 146130 - 0 GCCAAGGATCTCGACGCCGT CGG ||||||||[||||||||||||]PAM NA Intergenic NA NA

T23054 GCCGTCGGAGAAGATCTCGACGG 1000
Oligo fwd CACCGCCGTCGGAGAAGATCTCGA
Oligo rev AAACTCGAGATCTTCTCCGACGGC
Chromosome start end strand MM target_seq PAM alignment distance position gene name gene id
GU305914.1 146093 146115 - 0 GCCGTCGGAGAAGATCTCGA CGG ||||||||[||||||||||||]PAM NA Intergenic NA NA

T23055 CCGTCGGAGAAGATCTCGACGGG 1000
Oligo fwd CACCCCGTCGGAGAAGATCTCGAC
Oligo rev AAACGTCGAGATCTTCTCCGACGG
Chromosome start end strand MM target_seq PAM alignment distance position gene name gene id
GU305914.1 146092 146114 - 0 CCGTCGGAGAAGATCTCGAC GGG ||||||||[||||||||||||]PAM NA Intergenic NA NA

T23056 CGGGCGTGAAAAACCTAGTCAGG 1000

<table>
<tr><td>Oligo fwd</td><td>CACCCGGGCGTGAAAAACCTAGTC</td><td></td><td></td><td></td><td></td><td></td><td></td><td></td><td></td><td></td><td></td></tr>
<tr><td>Oligo rev</td><td>AAACGACTAGGTTTTTCACGCCCG</td><td></td><td></td><td></td><td></td><td></td><td></td><td></td><td></td><td></td><td></td></tr>
<tr><td>Chromosome</td><td>start</td><td>end</td><td>strand</td><td>MM</td><td>target_seq</td><td>PAM</td><td>alignment</td><td>distance</td><td>position</td><td>gene name</td><td>gene id</td></tr>
<tr><td>GU305914.1</td><td>146073</td><td>146095</td><td>-</td><td>0</td><td>CGGGCGTGAAAAACCTAGTC</td><td>AGG</td><td>||||||||[||||||||||||]PAM</td><td>NA</td><td>Intergenic</td><td>NA</td><td>NA</td></tr>
<tr><td>T23057</td><td>GGGCGTGAAAAACCTAGTCAGGG</td><td>1000</td><td></td><td></td><td></td><td></td><td></td><td></td><td></td><td></td><td></td></tr>
<tr><td>Oligo fwd</td><td>CACCGGGCGTGAAAAACCTAGTCA</td><td></td><td></td><td></td><td></td><td></td><td></td><td></td><td></td><td></td><td></td></tr>
<tr><td>Oligo rev</td><td>AAACTGACTAGGTTTTTCACGCCC</td><td></td><td></td><td></td><td></td><td></td><td></td><td></td><td></td><td></td><td></td></tr>
<tr><td>Chromosome</td><td>start</td><td>end</td><td>strand</td><td>MM</td><td>target_seq</td><td>PAM</td><td>alignment</td><td>distance</td><td>position</td><td>gene name</td><td>gene id</td></tr>
<tr><td>GU305914.1</td><td>146072</td><td>146094</td><td>-</td><td>0</td><td>GGGCGTGAAAAACCTAGTCA</td><td>GGG</td><td>||||||||[||||||||||||]PAM</td><td>NA</td><td>Intergenic</td><td>NA</td><td>NA</td></tr>
<tr><td>T23058</td><td>GGCGTGAAAAACCTAGTCAGGGG</td><td>1000</td><td></td><td></td><td></td><td></td><td></td><td></td><td></td><td></td><td></td></tr>
<tr><td>Oligo fwd</td><td>CACCGGCGTGAAAAACCTAGTCAG</td><td></td><td></td><td></td><td></td><td></td><td></td><td></td><td></td><td></td><td></td></tr>
<tr><td>Oligo rev</td><td>AAACCTGACTAGGTTTTTCACGCC</td><td></td><td></td><td></td><td></td><td></td><td></td><td></td><td></td><td></td><td></td></tr>
<tr><td>Chromosome</td><td>start</td><td>end</td><td>strand</td><td>MM</td><td>target_seq</td><td>PAM</td><td>alignment</td><td>distance</td><td>position</td><td>gene name</td><td>gene id</td></tr>
<tr><td>GU305914.1</td><td>146071</td><td>146093</td><td>-</td><td>0</td><td>GGCGTGAAAAACCTAGTCAG</td><td>GGG</td><td>||||||||[||||||||||||]PAM</td><td>NA</td><td>Intergenic</td><td>NA</td><td>NA</td></tr>
<tr><td>T23059</td><td>CTAGTCAGGGGCTCGAAGTCCGG</td><td>1000</td><td></td><td></td><td></td><td></td><td></td><td></td><td></td><td></td><td></td></tr>
<tr><td>Oligo fwd</td><td>CACCCTAGTCAGGGGCTCGAAGTC</td><td></td><td></td><td></td><td></td><td></td><td></td><td></td><td></td><td></td><td></td></tr>
<tr><td>Oligo rev</td><td>AAACGACTTCGAGCCCCTGACTAG</td><td></td><td></td><td></td><td></td><td></td><td></td><td></td><td></td><td></td><td></td></tr>
<tr><td>Chromosome</td><td>start</td><td>end</td><td>strand</td><td>MM</td><td>target_seq</td><td>PAM</td><td>alignment</td><td>distance</td><td>position</td><td>gene name</td><td>gene id</td></tr>
<tr><td>GU305914.1</td><td>146059</td><td>146081</td><td>-</td><td>0</td><td>CTAGTCAGGGGCTCGAAGTC</td><td>CGG</td><td>||||||||[||||||||||||]PAM</td><td>NA</td><td>Intergenic</td><td>NA</td><td>NA</td></tr>
<tr><td>T23060</td><td>AGTCCGGCCCGACGTGCATGAGG</td><td>1000</td><td></td><td></td><td></td><td></td><td></td><td></td><td></td><td></td><td></td></tr>
<tr><td>Oligo fwd</td><td>CACCAGTCCGGCCCGACGTGCATG</td><td></td><td></td><td></td><td></td><td></td><td></td><td></td><td></td><td></td><td></td></tr>
<tr><td>Oligo rev</td><td>AAACCATGCACGTCGGGCCGGACT</td><td></td><td></td><td></td><td></td><td></td><td></td><td></td><td></td><td></td><td></td></tr>
<tr><td>Chromosome</td><td>start</td><td>end</td><td>strand</td><td>MM</td><td>target_seq</td><td>PAM</td><td>alignment</td><td>distance</td><td>position</td><td>gene name</td><td>gene id</td></tr>
<tr><td>GU305914.1</td><td>146043</td><td>146065</td><td>-</td><td>0</td><td>AGTCCGGCCCGACGTGCATG</td><td>AGG</td><td>||||||||[||||||||||||]PAM</td><td>NA</td><td>Intergenic</td><td>NA</td><td>NA</td></tr>
<tr><td>T23061</td><td>TCTGTGCGAGCGCCCGCGACCGG</td><td>1000</td><td></td><td></td><td></td><td></td><td></td><td></td><td></td><td></td><td></td></tr>
<tr><td>Oligo fwd</td><td>CACCTCTGTGCGAGCGCCCGCGAC</td><td></td><td></td><td></td><td></td><td></td><td></td><td></td><td></td><td></td><td></td></tr>
<tr><td>Oligo rev</td><td>AAACGTCGCGGGCGCTCGCACAGA</td><td></td><td></td><td></td><td></td><td></td><td></td><td></td><td></td><td></td><td></td></tr>
<tr><td>Chromosome</td><td>start</td><td>end</td><td>strand</td><td>MM</td><td>target_seq</td><td>PAM</td><td>alignment</td><td>distance</td><td>position</td><td>gene name</td><td>gene id</td></tr>
<tr><td>GU305914.1</td><td>146019</td><td>146041</td><td>-</td><td>0</td><td>TCTGTGCGAGCGCCCGCGAC</td><td>CGG</td><td>||||||||[||||||||||||]PAM</td><td>NA</td><td>Intergenic</td><td>NA</td><td>NA</td></tr>
<tr><td>T23062</td><td>CGACCGGCGCGCGACGCGATCGG</td><td>1000</td><td></td><td></td><td></td><td></td><td></td><td></td><td></td><td></td><td></td></tr>
<tr><td>Oligo fwd</td><td>CACCCGACCGGCGCGCGACGCGAT</td><td></td><td></td><td></td><td></td><td></td><td></td><td></td><td></td><td></td><td></td></tr>
<tr><td>Oligo rev</td><td>AAACATCGCGTCGCGCGCCGGTCG</td><td></td><td></td><td></td><td></td><td></td><td></td><td></td><td></td><td></td><td></td></tr>
<tr><td>Chromosome</td><td>start</td><td>end</td><td>strand</td><td>MM</td><td>target_seq</td><td>PAM</td><td>alignment</td><td>distance</td><td>position</td><td>gene name</td><td>gene id</td></tr>
<tr><td>GU305914.1</td><td>146003</td><td>146025</td><td>-</td><td>0</td><td>CGACCGGCGCGCGACGCGAT</td><td>CGG</td><td>||||||||[||||||||||||]PAM</td><td>NA</td><td>Intergenic</td><td>NA</td><td>NA</td></tr>
<tr><td>T23063</td><td>TCCTACCCCCGACCCGTTAGCGG</td><td>1000</td><td></td><td></td><td></td><td></td><td></td><td></td><td></td><td></td><td></td></tr>
<tr><td>Oligo fwd</td><td>CACCTCCTACCCCCGACCCGTTAG</td><td></td><td></td><td></td><td></td><td></td><td></td><td></td><td></td><td></td><td></td></tr>
<tr><td>Oligo rev</td><td>AAACCTAACGGGTCGGGGGTAGGA</td><td></td><td></td><td></td><td></td><td></td><td></td><td></td><td></td><td></td><td></td></tr>
<tr><td>Chromosome</td><td>start</td><td>end</td><td>strand</td><td>MM</td><td>target_seq</td><td>PAM</td><td>alignment</td><td>distance</td><td>position</td><td>gene name</td><td>gene id</td></tr>
<tr><td>GU305914.1</td><td>145940</td><td>145962</td><td>-</td><td>0</td><td>TCCTACCCCCGACCCGTTAG</td><td>CGG</td><td>||||||||[||||||||||||]PAM</td><td>NA</td><td>Intergenic</td><td>NA</td><td>NA</td></tr>
<tr><td>T23064</td><td>TACCCCCGACCCGTTAGCGGCGG</td><td>1000</td><td></td><td></td><td></td><td></td><td></td><td></td><td></td><td></td><td></td></tr>
<tr><td>Oligo fwd</td><td>CACCTACCCCCGACCCGTTAGCGG</td><td></td><td></td><td></td><td></td><td></td><td></td><td></td><td></td><td></td><td></td></tr>
<tr><td>Oligo rev</td><td>AAACCCGCTAACGGGTCGGGGGTA</td><td></td><td></td><td></td><td></td><td></td><td></td><td></td><td></td><td></td><td></td></tr>
<tr><td>Chromosome</td><td>start</td><td>end</td><td>strand</td><td>MM</td><td>target_seq</td><td>PAM</td><td>alignment</td><td>distance</td><td>position</td><td>gene name</td><td>gene id</td></tr>
<tr><td>GU305914.1</td><td>145937</td><td>145959</td><td>-</td><td>0</td><td>TACCCCCGACCCGTTAGCGG</td><td>CGG</td><td>||||||||[||||||||||||]PAM</td><td>NA</td><td>Intergenic</td><td>NA</td><td>NA</td></tr>
<tr><td>T23065</td><td>TTAGCGGCGGTCGTCGAATCCGG</td><td>1000</td><td></td><td></td><td></td><td></td><td></td><td></td><td></td><td></td><td></td></tr>
<tr><td>Oligo fwd</td><td>CACCTTAGCGGCGGTCGTCGAATC</td><td></td><td></td><td></td><td></td><td></td><td></td><td></td><td></td><td></td><td></td></tr>
<tr><td>Oligo rev</td><td>AAACGATTCGACGACCGCCGCTAA</td><td></td><td></td><td></td><td></td><td></td><td></td><td></td><td></td><td></td><td></td></tr>
<tr><td>Chromosome</td><td>start</td><td>end</td><td>strand</td><td>MM</td><td>target_seq</td><td>PAM</td><td>alignment</td><td>distance</td><td>position</td><td>gene name</td><td>gene id</td></tr>
<tr><td>GU305914.1</td><td>145924</td><td>145946</td><td>-</td><td>0</td><td>TTAGCGGCGGTCGTCGAATC</td><td>CGG</td><td>||||||||[||||||||||||]PAM</td><td>NA</td><td>Intergenic</td><td>NA</td><td>NA</td></tr>
<tr><td>T23066</td><td>TAGCGGCGGTCGTCGAATCCGGG</td><td>1000</td><td></td><td></td><td></td><td></td><td></td><td></td><td></td><td></td><td></td></tr>
<tr><td>Oligo fwd</td><td>CACCTAGCGGCGGTCGTCGAATCC</td><td></td><td></td><td></td><td></td><td></td><td></td><td></td><td></td><td></td><td></td></tr>
<tr><td>Oligo rev</td><td>AAACGGATTCGACGACCGCCGCTA</td><td></td><td></td><td></td><td></td><td></td><td></td><td></td><td></td><td></td><td></td></tr>
<tr><td>Chromosome</td><td>start</td><td>end</td><td>strand</td><td>MM</td><td>target_seq</td><td>PAM</td><td>alignment</td><td>distance</td><td>position</td><td>gene name</td><td>gene id</td></tr>
<tr><td>GU305914.1</td><td>145923</td><td>145945</td><td>-</td><td>0</td><td>TAGCGGCGGTCGTCGAATCC</td><td>GGG</td><td>||||||||[||||||||||||]PAM</td><td>NA</td><td>Intergenic</td><td>NA</td><td>NA</td></tr>
<tr><td>T23067</td><td>TCGAATCCGGGATCGTGTCGAGG</td><td>1000</td><td></td><td></td><td></td><td></td><td></td><td></td><td></td><td></td><td></td></tr>
<tr><td>Oligo fwd</td><td>CACCTCGAATCCGGGATCGTGTCG</td><td></td><td></td><td></td><td></td><td></td><td></td><td></td><td></td><td></td><td></td></tr>
<tr><td>Oligo rev</td><td>AAACCGACACGATCCCGGATTCGA</td><td></td><td></td><td></td><td></td><td></td><td></td><td></td><td></td><td></td><td></td></tr>
<tr><td>Chromosome</td><td>start</td><td>end</td><td>strand</td><td>MM</td><td>target_seq</td><td>PAM</td><td>alignment</td><td>distance</td><td>position</td><td>gene name</td><td>gene id</td></tr>
<tr><td>GU305914.1</td><td>145911</td><td>145933</td><td>-</td><td>0</td><td>TCGAATCCGGGATCGTGTCG</td><td>AGG</td><td>||||||||[||||||||||||]PAM</td><td>NA</td><td>Intergenic</td><td>NA</td><td>NA</td></tr>
<tr><td>T23068</td><td>CGAATCCGGGATCGTGTCGAGGG</td><td>1000</td><td></td><td></td><td></td><td></td><td></td><td></td><td></td><td></td><td></td></tr>
<tr><td>Oligo fwd</td><td>CACCCGAATCCGGGATCGTGTCGA</td><td></td><td></td><td></td><td></td><td></td><td></td><td></td><td></td><td></td><td></td></tr>
<tr><td>Oligo rev</td><td>AAACTCGACACGATCCCGGATTCG</td><td></td><td></td><td></td><td></td><td></td><td></td><td></td><td></td><td></td><td></td></tr>
<tr><td>Chromosome</td><td>start</td><td>end</td><td>strand</td><td>MM</td><td>target_seq</td><td>PAM</td><td>alignment</td><td>distance</td><td>position</td><td>gene name</td><td>gene id</td></tr>
<tr><td>GU305914.1</td><td>145910</td><td>145932</td><td>-</td><td>0</td><td>CGAATCCGGGATCGTGTCGA</td><td>GGG</td><td>||||||||[||||||||||||]PAM</td><td>NA</td><td>Intergenic</td><td>NA</td><td>NA</td></tr>
<tr><td>T23069</td><td>CCGACGTGTAATTAGCGAGCAGG</td><td>1000</td><td></td><td></td><td></td><td></td><td></td><td></td><td></td><td></td><td></td></tr>
<tr><td>Oligo fwd</td><td>CACCCCGACGTGTAATTAGCGAGC</td><td></td><td></td><td></td><td></td><td></td><td></td><td></td><td></td><td></td><td></td></tr>
<tr><td>Oligo rev</td><td>AAACGCTCGCTAATTACACGTCGG</td><td></td><td></td><td></td><td></td><td></td><td></td><td></td><td></td><td></td><td></td></tr>
<tr><td>Chromosome</td><td>start</td><td>end</td><td>strand</td><td>MM</td><td>target_seq</td><td>PAM</td><td>alignment</td><td>distance</td><td>position</td><td>gene name</td><td>gene id</td></tr>
<tr><td>GU305914.1</td><td>145872</td><td>145894</td><td>-</td><td>0</td><td>CCGACGTGTAATTAGCGAGC</td><td>AGG</td><td>||||||||[||||||||||||]PAM</td><td>NA</td><td>Intergenic</td><td>NA</td><td>NA</td></tr>
<tr><td>T23070</td><td>CGTGTAATTAGCGAGCAGGTAGG</td><td>1000</td><td></td><td></td><td></td><td></td><td></td><td></td><td></td><td></td><td></td></tr>
<tr><td>Oligo fwd</td><td>CACCCGTGTAATTAGCGAGCAGGT</td><td></td><td></td><td></td><td></td><td></td><td></td><td></td><td></td><td></td><td></td></tr>
<tr><td>Oligo rev</td><td>AAACACCTGCTCGCTAATTACACG</td><td></td><td></td><td></td><td></td><td></td><td></td><td></td><td></td><td></td><td></td></tr>
<tr><td>Chromosome</td><td>start</td><td>end</td><td>strand</td><td>MM</td><td>target_seq</td><td>PAM</td><td>alignment</td><td>distance</td><td>position</td><td>gene name</td><td>gene id</td></tr>
<tr><td>GU305914.1</td><td>145868</td><td>145890</td><td>-</td><td>0</td><td>CGTGTAATTAGCGAGCAGGT</td><td>AGG</td><td>||||||||[||||||||||||]PAM</td><td>NA</td><td>Intergenic</td><td>NA</td><td>NA</td></tr>
</table>

<table>
<tr><td>T23071</td><td>GTAGGTGCGAAGCTCCCCTTCGG</td><td>1000</td></tr>
<tr><td>Oligo fwd</td><td>CACCGTAGGTGCGAAGCTCCCCTT</td></tr>
<tr><td>Oligo rev</td><td>AAACAAGGGGAGCTTCGCACCTAC</td></tr>
<tr><td>Chromosome</td><td>start</td><td>end</td><td>strand</td><td>MM</td><td>target_seq</td><td>PAM</td><td>alignment</td><td>distance</td><td>position</td><td>gene name</td><td>gene id</td></tr>
<tr><td>GU305914.1</td><td>145850</td><td>145872</td><td>-</td><td>0</td><td>GTAGGTGCGAAGCTCCCCTT</td><td>CGG</td><td>||||||||[||||||||||||]PAM</td><td>NA</td><td>Intergenic</td><td>NA</td><td>NA</td></tr>
<tr><td>T23072</td><td>GGTGCGAAGCTCCCCTTCGGCGG</td><td>1000</td></tr>
<tr><td>Oligo fwd</td><td>CACCGGTGCGAAGCTCCCCTTCGG</td></tr>
<tr><td>Oligo rev</td><td>AAACCCGAAGGGGAGCTTCGCACC</td></tr>
<tr><td>Chromosome</td><td>start</td><td>end</td><td>strand</td><td>MM</td><td>target_seq</td><td>PAM</td><td>alignment</td><td>distance</td><td>position</td><td>gene name</td><td>gene id</td></tr>
<tr><td>GU305914.1</td><td>145847</td><td>145869</td><td>-</td><td>0</td><td>GGTGCGAAGCTCCCCTTCGG</td><td>CGG</td><td>||||||||[||||||||||||]PAM</td><td>NA</td><td>Intergenic</td><td>NA</td><td>NA</td></tr>
<tr><td>T23073</td><td>GTGCGAAGCTCCCCTTCGGCGGG</td><td>1000</td></tr>
<tr><td>Oligo fwd</td><td>CACCGTGCGAAGCTCCCCTTCGGC</td></tr>
<tr><td>Oligo rev</td><td>AAACGCCGAAGGGGAGCTTCGCAC</td></tr>
<tr><td>Chromosome</td><td>start</td><td>end</td><td>strand</td><td>MM</td><td>target_seq</td><td>PAM</td><td>alignment</td><td>distance</td><td>position</td><td>gene name</td><td>gene id</td></tr>
<tr><td>GU305914.1</td><td>145846</td><td>145868</td><td>-</td><td>0</td><td>GTGCGAAGCTCCCCTTCGGC</td><td>GGG</td><td>||||||||[||||||||||||]PAM</td><td>NA</td><td>Intergenic</td><td>NA</td><td>NA</td></tr>
<tr><td>T23074</td><td>TGCGAAGCTCCCCTTCGGCGGGG</td><td>1000</td></tr>
<tr><td>Oligo fwd</td><td>CACCTGCGAAGCTCCCCTTCGGCG</td></tr>
<tr><td>Oligo rev</td><td>AAACCGCCGAAGGGGAGCTTCGCA</td></tr>
<tr><td>Chromosome</td><td>start</td><td>end</td><td>strand</td><td>MM</td><td>target_seq</td><td>PAM</td><td>alignment</td><td>distance</td><td>position</td><td>gene name</td><td>gene id</td></tr>
<tr><td>GU305914.1</td><td>145845</td><td>145867</td><td>-</td><td>0</td><td>TGCGAAGCTCCCCTTCGGCG</td><td>GGG</td><td>||||||||[||||||||||||]PAM</td><td>NA</td><td>Intergenic</td><td>NA</td><td>NA</td></tr>
<tr><td>T23075</td><td>CCTTCGGCGGGGATATTGACCGG</td><td>1000</td></tr>
<tr><td>Oligo fwd</td><td>CACCCCTTCGGCGGGGATATTGAC</td></tr>
<tr><td>Oligo rev</td><td>AAACGTCAATATCCCCGCCGAAGG</td></tr>
<tr><td>Chromosome</td><td>start</td><td>end</td><td>strand</td><td>MM</td><td>target_seq</td><td>PAM</td><td>alignment</td><td>distance</td><td>position</td><td>gene name</td><td>gene id</td></tr>
<tr><td>GU305914.1</td><td>145834</td><td>145856</td><td>-</td><td>0</td><td>CCTTCGGCGGGGATATTGAC</td><td>CGG</td><td>||||||||[||||||||||||]PAM</td><td>NA</td><td>Intergenic</td><td>NA</td><td>NA</td></tr>
<tr><td>T23076</td><td>CTTCGGCGGGGATATTGACCGGG</td><td>1000</td></tr>
<tr><td>Oligo fwd</td><td>CACCCTTCGGCGGGGATATTGACC</td></tr>
<tr><td>Oligo rev</td><td>AAACGGTCAATATCCCCGCCGAAG</td></tr>
<tr><td>Chromosome</td><td>start</td><td>end</td><td>strand</td><td>MM</td><td>target_seq</td><td>PAM</td><td>alignment</td><td>distance</td><td>position</td><td>gene name</td><td>gene id</td></tr>
<tr><td>GU305914.1</td><td>145833</td><td>145855</td><td>-</td><td>0</td><td>CTTCGGCGGGGATATTGACC</td><td>GGG</td><td>||||||||[||||||||||||]PAM</td><td>NA</td><td>Intergenic</td><td>NA</td><td>NA</td></tr>
<tr><td>T23077</td><td>GACCGGGAAAACGAAGAGTCTGG</td><td>1000</td></tr>
<tr><td>Oligo fwd</td><td>CACCGACCGGGAAAACGAAGAGTC</td></tr>
<tr><td>Oligo rev</td><td>AAACGACTCTTCGTTTTCCCGGTC</td></tr>
<tr><td>Chromosome</td><td>start</td><td>end</td><td>strand</td><td>MM</td><td>target_seq</td><td>PAM</td><td>alignment</td><td>distance</td><td>position</td><td>gene name</td><td>gene id</td></tr>
<tr><td>GU305914.1</td><td>145817</td><td>145839</td><td>-</td><td>0</td><td>GACCGGGAAAACGAAGAGTC</td><td>TGG</td><td>||||||||[||||||||||||]PAM</td><td>NA</td><td>Intergenic</td><td>NA</td><td>NA</td></tr>
<tr><td>T23078</td><td>GGTCACGACGCGATCTCCGTCGG</td><td>1000</td></tr>
<tr><td>Oligo fwd</td><td>CACCGGTCACGACGCGATCTCCGT</td></tr>
<tr><td>Oligo rev</td><td>AAACACGGAGATCGCGTCGTGACC</td></tr>
<tr><td>Chromosome</td><td>start</td><td>end</td><td>strand</td><td>MM</td><td>target_seq</td><td>PAM</td><td>alignment</td><td>distance</td><td>position</td><td>gene name</td><td>gene id</td></tr>
<tr><td>GU305914.1</td><td>145796</td><td>145818</td><td>-</td><td>0</td><td>GGTCACGACGCGATCTCCGT</td><td>CGG</td><td>||||||||[||||||||||||]PAM</td><td>NA</td><td>Intergenic</td><td>NA</td><td>NA</td></tr>
<tr><td>T23079</td><td>CTTCGCAGAAGAAACACTGCAGG</td><td>1000</td></tr>
<tr><td>Oligo fwd</td><td>CACCCTTCGCAGAAGAAACACTGC</td></tr>
<tr><td>Oligo rev</td><td>AAACGCAGTGTTTCTTCTGCGAAG</td></tr>
<tr><td>Chromosome</td><td>start</td><td>end</td><td>strand</td><td>MM</td><td>target_seq</td><td>PAM</td><td>alignment</td><td>distance</td><td>position</td><td>gene name</td><td>gene id</td></tr>
<tr><td>GU305914.1</td><td>145770</td><td>145792</td><td>-</td><td>0</td><td>CTTCGCAGAAGAAACACTGC</td><td>AGG</td><td>||||||||[||||||||||||]PAM</td><td>NA</td><td>Intergenic</td><td>NA</td><td>NA</td></tr>
<tr><td>T23080</td><td>GCAGAAGAAACACTGCAGGATGG</td><td>1000</td></tr>
<tr><td>Oligo fwd</td><td>CACCGCAGAAGAAACACTGCAGGA</td></tr>
<tr><td>Oligo rev</td><td>AAACTCCTGCAGTGTTTCTTCTGC</td></tr>
<tr><td>Chromosome</td><td>start</td><td>end</td><td>strand</td><td>MM</td><td>target_seq</td><td>PAM</td><td>alignment</td><td>distance</td><td>position</td><td>gene name</td><td>gene id</td></tr>
<tr><td>GU305914.1</td><td>145766</td><td>145788</td><td>-</td><td>0</td><td>GCAGAAGAAACACTGCAGGA</td><td>TGG</td><td>||||||||[||||||||||||]PAM</td><td>NA</td><td>Intergenic</td><td>NA</td><td>NA</td></tr>
<tr><td>T23081</td><td>AGAAACACTGCAGGATGGCCTGG</td><td>1000</td></tr>
<tr><td>Oligo fwd</td><td>CACCAGAAACACTGCAGGATGGCC</td></tr>
<tr><td>Oligo rev</td><td>AAACGGCCATCCTGCAGTGTTTCT</td></tr>
<tr><td>Chromosome</td><td>start</td><td>end</td><td>strand</td><td>MM</td><td>target_seq</td><td>PAM</td><td>alignment</td><td>distance</td><td>position</td><td>gene name</td><td>gene id</td></tr>
<tr><td>GU305914.1</td><td>145761</td><td>145783</td><td>-</td><td>0</td><td>AGAAACACTGCAGGATGGCC</td><td>TGG</td><td>||||||||[||||||||||||]PAM</td><td>NA</td><td>Intergenic</td><td>NA</td><td>NA</td></tr>
<tr><td>T23082</td><td>CAGGATGGCCTGGTTCTCCTCGG</td><td>1000</td></tr>
<tr><td>Oligo fwd</td><td>CACCCAGGATGGCCTGGTTCTCCT</td></tr>
<tr><td>Oligo rev</td><td>AAACAGGAGAACCAGGCCATCCTG</td></tr>
<tr><td>Chromosome</td><td>start</td><td>end</td><td>strand</td><td>MM</td><td>target_seq</td><td>PAM</td><td>alignment</td><td>distance</td><td>position</td><td>gene name</td><td>gene id</td></tr>
<tr><td>GU305914.1</td><td>145751</td><td>145773</td><td>-</td><td>0</td><td>CAGGATGGCCTGGTTCTCCT</td><td>CGG</td><td>||||||||[||||||||||||]PAM</td><td>NA</td><td>Intergenic</td><td>NA</td><td>NA</td></tr>
<tr><td>T23083</td><td>AGGATGGCCTGGTTCTCCTCGGG</td><td>1000</td></tr>
<tr><td>Oligo fwd</td><td>CACCAGGATGGCCTGGTTCTCCTC</td></tr>
<tr><td>Oligo rev</td><td>AAACGAGGAGAACCAGGCCATCCT</td></tr>
<tr><td>Chromosome</td><td>start</td><td>end</td><td>strand</td><td>MM</td><td>target_seq</td><td>PAM</td><td>alignment</td><td>distance</td><td>position</td><td>gene name</td><td>gene id</td></tr>
<tr><td>GU305914.1</td><td>145750</td><td>145772</td><td>-</td><td>0</td><td>AGGATGGCCTGGTTCTCCTC</td><td>GGG</td><td>||||||||[||||||||||||]PAM</td><td>NA</td><td>Intergenic</td><td>NA</td><td>NA</td></tr>
<tr><td>T23084</td><td>TGGCCTGGTTCTCCTCGGGCAGG</td><td>1000</td></tr>
<tr><td>Oligo fwd</td><td>CACCTGGCCTGGTTCTCCTCGGGC</td></tr>
<tr><td>Oligo rev</td><td>AAACGCCCGAGGAGAACCAGGCCA</td></tr>
<tr><td>Chromosome</td><td>start</td><td>end</td><td>strand</td><td>MM</td><td>target_seq</td><td>PAM</td><td>alignment</td><td>distance</td><td>position</td><td>gene name</td><td>gene id</td></tr>
<tr><td>GU305914.1</td><td>145746</td><td>145768</td><td>-</td><td>0</td><td>TGGCCTGGTTCTCCTCGGGC</td><td>AGG</td><td>||||||||[||||||||||||]PAM</td><td>NA</td><td>Intergenic</td><td>NA</td><td>NA</td></tr>
<tr><td>T23085</td><td>CTCGGGCAGGAAAGCATAGATGG</td><td>1000</td></tr>
<tr><td>Oligo fwd</td><td>CACCCTCGGGCAGGAAAGCATAGA</td></tr>
<tr><td>Oligo rev</td><td>AAACTCTATGCTTTCCTGCCCGAG</td></tr>
<tr><td>Chromosome</td><td>start</td><td>end</td><td>strand</td><td>MM</td><td>target_seq</td><td>PAM</td><td>alignment</td><td>distance</td><td>position</td><td>gene name</td><td>gene id</td></tr>
<tr><td>GU305914.1</td><td>145733</td><td>145755</td><td>-</td><td>0</td><td>CTCGGGCAGGAAAGCATAGA</td><td>TGG</td><td>||||||||[||||||||||||]PAM</td><td>NA</td><td>Intergenic</td><td>NA</td><td>NA</td></tr>
</table>

<table>
<tr><td>T23086</td><td>GGAAAGCATAGATGGACAGATGG</td><td>1000</td><td></td><td></td><td></td><td></td><td></td><td></td><td></td><td></td><td></td></tr>
<tr><td>Oligo fwd</td><td>CACCGGAAAGCATAGATGGACAGA</td><td></td><td></td><td></td><td></td><td></td><td></td><td></td><td></td><td></td><td></td></tr>
<tr><td>Oligo rev</td><td>AAACTCTGTCCATCTATGCTTTCC</td><td></td><td></td><td></td><td></td><td></td><td></td><td></td><td></td><td></td><td></td></tr>
<tr><td>Chromosome</td><td>start</td><td>end</td><td>strand</td><td>MM</td><td>target_seq</td><td>PAM</td><td>alignment</td><td>distance</td><td>position</td><td>gene name</td><td>gene id</td></tr>
<tr><td>GU305914.1</td><td>145725</td><td>145747</td><td>-</td><td>0</td><td>GGAAAGCATAGATGGACAGA</td><td>TGG</td><td>||||||||[||||||||||||]PAM</td><td>NA</td><td>Intergenic</td><td>NA</td><td>NA</td></tr>
<tr><td>T23087</td><td>GCATAGATGGACAGATGGACCGG</td><td>1000</td><td></td><td></td><td></td><td></td><td></td><td></td><td></td><td></td><td></td></tr>
<tr><td>Oligo fwd</td><td>CACCGCATAGATGGACAGATGGAC</td><td></td><td></td><td></td><td></td><td></td><td></td><td></td><td></td><td></td><td></td></tr>
<tr><td>Oligo rev</td><td>AAACGTCCATCTGTCCATCTATGC</td><td></td><td></td><td></td><td></td><td></td><td></td><td></td><td></td><td></td><td></td></tr>
<tr><td>Chromosome</td><td>start</td><td>end</td><td>strand</td><td>MM</td><td>target_seq</td><td>PAM</td><td>alignment</td><td>distance</td><td>position</td><td>gene name</td><td>gene id</td></tr>
<tr><td>GU305914.1</td><td>145720</td><td>145742</td><td>-</td><td>0</td><td>GCATAGATGGACAGATGGAC</td><td>CGG</td><td>||||||||[||||||||||||]PAM</td><td>NA</td><td>Intergenic</td><td>NA</td><td>NA</td></tr>
<tr><td>T23088</td><td>CGAGAGTCTGAGACAATGTGTGG</td><td>1000</td><td></td><td></td><td></td><td></td><td></td><td></td><td></td><td></td><td></td></tr>
<tr><td>Oligo fwd</td><td>CACCCGAGAGTCTGAGACAATGTG</td><td></td><td></td><td></td><td></td><td></td><td></td><td></td><td></td><td></td><td></td></tr>
<tr><td>Oligo rev</td><td>AAACCACATTGTCTCAGACTCTCG</td><td></td><td></td><td></td><td></td><td></td><td></td><td></td><td></td><td></td><td></td></tr>
<tr><td>Chromosome</td><td>start</td><td>end</td><td>strand</td><td>MM</td><td>target_seq</td><td>PAM</td><td>alignment</td><td>distance</td><td>position</td><td>gene name</td><td>gene id</td></tr>
<tr><td>GU305914.1</td><td>145647</td><td>145669</td><td>-</td><td>0</td><td>CGAGAGTCTGAGACAATGTG</td><td>TGG</td><td>||||||||[||||||||||||]PAM</td><td>NA</td><td>Intergenic</td><td>NA</td><td>NA</td></tr>
<tr><td>T23089</td><td>ATGTGTGGTGTGATCCCAGCAGG</td><td>1000</td><td></td><td></td><td></td><td></td><td></td><td></td><td></td><td></td><td></td></tr>
<tr><td>Oligo fwd</td><td>CACCATGTGTGGTGTGATCCCAGC</td><td></td><td></td><td></td><td></td><td></td><td></td><td></td><td></td><td></td><td></td></tr>
<tr><td>Oligo rev</td><td>AAACGCTGGGATCACACCACACAT</td><td></td><td></td><td></td><td></td><td></td><td></td><td></td><td></td><td></td><td></td></tr>
<tr><td>Chromosome</td><td>start</td><td>end</td><td>strand</td><td>MM</td><td>target_seq</td><td>PAM</td><td>alignment</td><td>distance</td><td>position</td><td>gene name</td><td>gene id</td></tr>
<tr><td>GU305914.1</td><td>145632</td><td>145654</td><td>-</td><td>0</td><td>ATGTGTGGTGTGATCCCAGC</td><td>AGG</td><td>||||||||[||||||||||||]PAM</td><td>NA</td><td>Intergenic</td><td>NA</td><td>NA</td></tr>
<tr><td>T23090</td><td>TGTGATCCCAGCAGGAGCCCTGG</td><td>1000</td><td></td><td></td><td></td><td></td><td></td><td></td><td></td><td></td><td></td></tr>
<tr><td>Oligo fwd</td><td>CACCTGTGATCCCAGCAGGAGCCC</td><td></td><td></td><td></td><td></td><td></td><td></td><td></td><td></td><td></td><td></td></tr>
<tr><td>Oligo rev</td><td>AAACGGGCTCCTGCTGGGATCACA</td><td></td><td></td><td></td><td></td><td></td><td></td><td></td><td></td><td></td><td></td></tr>
<tr><td>Chromosome</td><td>start</td><td>end</td><td>strand</td><td>MM</td><td>target_seq</td><td>PAM</td><td>alignment</td><td>distance</td><td>position</td><td>gene name</td><td>gene id</td></tr>
<tr><td>GU305914.1</td><td>145624</td><td>145646</td><td>-</td><td>0</td><td>TGTGATCCCAGCAGGAGCCC</td><td>TGG</td><td>||||||||[||||||||||||]PAM</td><td>NA</td><td>Intergenic</td><td>NA</td><td>NA</td></tr>
<tr><td>T23091</td><td>GTGATCCCAGCAGGAGCCCTGGG</td><td>1000</td><td></td><td></td><td></td><td></td><td></td><td></td><td></td><td></td><td></td></tr>
<tr><td>Oligo fwd</td><td>CACCGTGATCCCAGCAGGAGCCCT</td><td></td><td></td><td></td><td></td><td></td><td></td><td></td><td></td><td></td><td></td></tr>
<tr><td>Oligo rev</td><td>AAACAGGGCTCCTGCTGGGATCAC</td><td></td><td></td><td></td><td></td><td></td><td></td><td></td><td></td><td></td><td></td></tr>
<tr><td>Chromosome</td><td>start</td><td>end</td><td>strand</td><td>MM</td><td>target_seq</td><td>PAM</td><td>alignment</td><td>distance</td><td>position</td><td>gene name</td><td>gene id</td></tr>
<tr><td>GU305914.1</td><td>145623</td><td>145645</td><td>-</td><td>0</td><td>GTGATCCCAGCAGGAGCCCT</td><td>GGG</td><td>||||||||[||||||||||||]PAM</td><td>NA</td><td>Intergenic</td><td>NA</td><td>NA</td></tr>
<tr><td>T23092</td><td>AGCCCTGGGATTTATAGTACCGG</td><td>1000</td><td></td><td></td><td></td><td></td><td></td><td></td><td></td><td></td><td></td></tr>
<tr><td>Oligo fwd</td><td>CACCAGCCCTGGGATTTATAGTAC</td><td></td><td></td><td></td><td></td><td></td><td></td><td></td><td></td><td></td><td></td></tr>
<tr><td>Oligo rev</td><td>AAACGTACTATAAATCCCAGGGCT</td><td></td><td></td><td></td><td></td><td></td><td></td><td></td><td></td><td></td><td></td></tr>
<tr><td>Chromosome</td><td>start</td><td>end</td><td>strand</td><td>MM</td><td>target_seq</td><td>PAM</td><td>alignment</td><td>distance</td><td>position</td><td>gene name</td><td>gene id</td></tr>
<tr><td>GU305914.1</td><td>145609</td><td>145631</td><td>-</td><td>0</td><td>AGCCCTGGGATTTATAGTAC</td><td>CGG</td><td>||||||||[||||||||||||]PAM</td><td>NA</td><td>Intergenic</td><td>NA</td><td>NA</td></tr>
<tr><td>T23093</td><td>GCCCTGGGATTTATAGTACCGGG</td><td>1000</td><td></td><td></td><td></td><td></td><td></td><td></td><td></td><td></td><td></td></tr>
<tr><td>Oligo fwd</td><td>CACCGCCCTGGGATTTATAGTACC</td><td></td><td></td><td></td><td></td><td></td><td></td><td></td><td></td><td></td><td></td></tr>
<tr><td>Oligo rev</td><td>AAACGGTACTATAAATCCCAGGGC</td><td></td><td></td><td></td><td></td><td></td><td></td><td></td><td></td><td></td><td></td></tr>
<tr><td>Chromosome</td><td>start</td><td>end</td><td>strand</td><td>MM</td><td>target_seq</td><td>PAM</td><td>alignment</td><td>distance</td><td>position</td><td>gene name</td><td>gene id</td></tr>
<tr><td>GU305914.1</td><td>145608</td><td>145630</td><td>-</td><td>0</td><td>GCCCTGGGATTTATAGTACC</td><td>GGG</td><td>||||||||[||||||||||||]PAM</td><td>NA</td><td>Intergenic</td><td>NA</td><td>NA</td></tr>
<tr><td>T23094</td><td>CCCTGGGATTTATAGTACCGGGG</td><td>1000</td><td></td><td></td><td></td><td></td><td></td><td></td><td></td><td></td><td></td></tr>
<tr><td>Oligo fwd</td><td>CACCCCCTGGGATTTATAGTACCG</td><td></td><td></td><td></td><td></td><td></td><td></td><td></td><td></td><td></td><td></td></tr>
<tr><td>Oligo rev</td><td>AAACCGGTACTATAAATCCCAGGG</td><td></td><td></td><td></td><td></td><td></td><td></td><td></td><td></td><td></td><td></td></tr>
<tr><td>Chromosome</td><td>start</td><td>end</td><td>strand</td><td>MM</td><td>target_seq</td><td>PAM</td><td>alignment</td><td>distance</td><td>position</td><td>gene name</td><td>gene id</td></tr>
<tr><td>GU305914.1</td><td>145607</td><td>145629</td><td>-</td><td>0</td><td>CCCTGGGATTTATAGTACCG</td><td>GGG</td><td>||||||||[||||||||||||]PAM</td><td>NA</td><td>Intergenic</td><td>NA</td><td>NA</td></tr>
<tr><td>T23095</td><td>CCTGGGATTTATAGTACCGGGGG</td><td>1000</td><td></td><td></td><td></td><td></td><td></td><td></td><td></td><td></td><td></td></tr>
<tr><td>Oligo fwd</td><td>CACCCCTGGGATTTATAGTACCGG</td><td></td><td></td><td></td><td></td><td></td><td></td><td></td><td></td><td></td><td></td></tr>
<tr><td>Oligo rev</td><td>AAACCCGGTACTATAAATCCCAGG</td><td></td><td></td><td></td><td></td><td></td><td></td><td></td><td></td><td></td><td></td></tr>
<tr><td>Chromosome</td><td>start</td><td>end</td><td>strand</td><td>MM</td><td>target_seq</td><td>PAM</td><td>alignment</td><td>distance</td><td>position</td><td>gene name</td><td>gene id</td></tr>
<tr><td>GU305914.1</td><td>145606</td><td>145628</td><td>-</td><td>0</td><td>CCTGGGATTTATAGTACCGG</td><td>GGG</td><td>||||||||[||||||||||||]PAM</td><td>NA</td><td>Intergenic</td><td>NA</td><td>NA</td></tr>
<tr><td>T23096</td><td>TACCGGGGGCAGCCGAGATGAGG</td><td>1000</td><td></td><td></td><td></td><td></td><td></td><td></td><td></td><td></td><td></td></tr>
<tr><td>Oligo fwd</td><td>CACCTACCGGGGGCAGCCGAGATG</td><td></td><td></td><td></td><td></td><td></td><td></td><td></td><td></td><td></td><td></td></tr>
<tr><td>Oligo rev</td><td>AAACCATCTCGGCTGCCCCCGGTA</td><td></td><td></td><td></td><td></td><td></td><td></td><td></td><td></td><td></td><td></td></tr>
<tr><td>Chromosome</td><td>start</td><td>end</td><td>strand</td><td>MM</td><td>target_seq</td><td>PAM</td><td>alignment</td><td>distance</td><td>position</td><td>gene name</td><td>gene id</td></tr>
<tr><td>GU305914.1</td><td>145592</td><td>145614</td><td>-</td><td>0</td><td>TACCGGGGGCAGCCGAGATG</td><td>AGG</td><td>||||||||[||||||||||||]PAM</td><td>NA</td><td>Intergenic</td><td>NA</td><td>NA</td></tr>
<tr><td>T23097</td><td>GCCCTGTGTCGAGTGTCGCTCGG</td><td>1000</td><td></td><td></td><td></td><td></td><td></td><td></td><td></td><td></td><td></td></tr>
<tr><td>Oligo fwd</td><td>CACCGCCCTGTGTCGAGTGTCGCT</td><td></td><td></td><td></td><td></td><td></td><td></td><td></td><td></td><td></td><td></td></tr>
<tr><td>Oligo rev</td><td>AAACAGCGACACTCGACACAGGGC</td><td></td><td></td><td></td><td></td><td></td><td></td><td></td><td></td><td></td><td></td></tr>
<tr><td>Chromosome</td><td>start</td><td>end</td><td>strand</td><td>MM</td><td>target_seq</td><td>PAM</td><td>alignment</td><td>distance</td><td>position</td><td>gene name</td><td>gene id</td></tr>
<tr><td>GU305914.1</td><td>145568</td><td>145590</td><td>-</td><td>0</td><td>GCCCTGTGTCGAGTGTCGCT</td><td>CGG</td><td>||||||||[||||||||||||]PAM</td><td>NA</td><td>Intergenic</td><td>NA</td><td>NA</td></tr>
<tr><td>GU305914.1</td><td>51611</td><td>51633</td><td>+</td><td>4</td><td>GACCTGTCTCGAGGGTCGAT</td><td>TGG</td><td>|-|||||-[|||||-||||-|]PAM</td><td>NA</td><td>Intergenic</td><td>NA</td><td>NA</td></tr>
<tr><td>T23098</td><td>GTGTCGCTCGGCACCCGAAGCGG</td><td>1000</td><td></td><td></td><td></td><td></td><td></td><td></td><td></td><td></td><td></td></tr>
<tr><td>Oligo fwd</td><td>CACCGTGTCGCTCGGCACCCGAAG</td><td></td><td></td><td></td><td></td><td></td><td></td><td></td><td></td><td></td><td></td></tr>
<tr><td>Oligo rev</td><td>AAACCTTCGGGTGCCGAGCGACAC</td><td></td><td></td><td></td><td></td><td></td><td></td><td></td><td></td><td></td><td></td></tr>
<tr><td>Chromosome</td><td>start</td><td>end</td><td>strand</td><td>MM</td><td>target_seq</td><td>PAM</td><td>alignment</td><td>distance</td><td>position</td><td>gene name</td><td>gene id</td></tr>
<tr><td>GU305914.1</td><td>145556</td><td>145578</td><td>-</td><td>0</td><td>GTGTCGCTCGGCACCCGAAG</td><td>CGG</td><td>||||||||[||||||||||||]PAM</td><td>NA</td><td>Intergenic</td><td>NA</td><td>NA</td></tr>
<tr><td>T23099</td><td>GCGACGTTACCCCGTCGTCTCGG</td><td>1000</td><td></td><td></td><td></td><td></td><td></td><td></td><td></td><td></td><td></td></tr>
<tr><td>Oligo fwd</td><td>CACCGCGACGTTACCCCGTCGTCT</td><td></td><td></td><td></td><td></td><td></td><td></td><td></td><td></td><td></td><td></td></tr>
<tr><td>Oligo rev</td><td>AAACAGACGACGGGGTAACGTCGC</td><td></td><td></td><td></td><td></td><td></td><td></td><td></td><td></td><td></td><td></td></tr>
<tr><td>Chromosome</td><td>start</td><td>end</td><td>strand</td><td>MM</td><td>target_seq</td><td>PAM</td><td>alignment</td><td>distance</td><td>position</td><td>gene name</td><td>gene id</td></tr>
<tr><td>GU305914.1</td><td>145534</td><td>145556</td><td>-</td><td>0</td><td>GCGACGTTACCCCGTCGTCT</td><td>CGG</td><td>||||||||[||||||||||||]PAM</td><td>NA</td><td>Intergenic</td><td>NA</td><td>NA</td></tr>
<tr><td>T23100</td><td>GTTACCCCGTCGTCTCGGCGAGG</td><td>1000</td><td></td><td></td><td></td><td></td><td></td><td></td><td></td><td></td><td></td></tr>
<tr><td>Oligo fwd</td><td>CACCGTTACCCCGTCGTCTCGGCG</td><td></td><td></td><td></td><td></td><td></td><td></td><td></td><td></td><td></td><td></td></tr>
<tr><td>Oligo rev</td><td>AAACCGCCGAGACGACGGGGTAAC</td><td></td><td></td><td></td><td></td><td></td><td></td><td></td><td></td><td></td><td></td></tr>
</table>

<table>
<tr><td>Chromosome</td><td>start</td><td>end</td><td>strand</td><td>MM</td><td>target_seq</td><td>PAM</td><td>alignment</td><td>distance</td><td>position</td><td>gene name</td><td>gene id</td></tr>
<tr><td>GU305914.1</td><td>145529</td><td>145551</td><td>-</td><td>0</td><td>GTTACCCCGTCGTCTCGGCG</td><td>AGG</td><td>||||||||[||||||||||||]PAM</td><td>NA</td><td>Intergenic</td><td>NA</td><td>NA</td></tr>
<tr><td>T23101</td><td>CCCGTCGTCTCGGCGAGGCTCGG</td><td>1000</td><td></td><td></td><td></td><td></td><td></td><td></td><td></td><td></td><td></td></tr>
<tr><td>Oligo fwd</td><td>CACCCCCGTCGTCTCGGCGAGGCT</td><td></td><td></td><td></td><td></td><td></td><td></td><td></td><td></td><td></td><td></td></tr>
<tr><td>Oligo rev</td><td>AAACAGCCTCGCCGAGACGACGGG</td><td></td><td></td><td></td><td></td><td></td><td></td><td></td><td></td><td></td><td></td></tr>
<tr><td>Chromosome</td><td>start</td><td>end</td><td>strand</td><td>MM</td><td>target_seq</td><td>PAM</td><td>alignment</td><td>distance</td><td>position</td><td>gene name</td><td>gene id</td></tr>
<tr><td>GU305914.1</td><td>145524</td><td>145546</td><td>-</td><td>0</td><td>CCCGTCGTCTCGGCGAGGCT</td><td>CGG</td><td>||||||||[||||||||||||]PAM</td><td>NA</td><td>Intergenic</td><td>NA</td><td>NA</td></tr>
<tr><td>T23102</td><td>TCGTCTCGGCGAGGCTCGGACGG</td><td>1000</td><td></td><td></td><td></td><td></td><td></td><td></td><td></td><td></td><td></td></tr>
<tr><td>Oligo fwd</td><td>CACCTCGTCTCGGCGAGGCTCGGA</td><td></td><td></td><td></td><td></td><td></td><td></td><td></td><td></td><td></td><td></td></tr>
<tr><td>Oligo rev</td><td>AAACTCCGAGCCTCGCCGAGACGA</td><td></td><td></td><td></td><td></td><td></td><td></td><td></td><td></td><td></td><td></td></tr>
<tr><td>Chromosome</td><td>start</td><td>end</td><td>strand</td><td>MM</td><td>target_seq</td><td>PAM</td><td>alignment</td><td>distance</td><td>position</td><td>gene name</td><td>gene id</td></tr>
<tr><td>GU305914.1</td><td>145520</td><td>145542</td><td>-</td><td>0</td><td>TCGTCTCGGCGAGGCTCGGA</td><td>CGG</td><td>||||||||[||||||||||||]PAM</td><td>NA</td><td>Intergenic</td><td>NA</td><td>NA</td></tr>
<tr><td>T23103</td><td>AGGCTCGGACGGTCTCTCTAAGG</td><td>1000</td><td></td><td></td><td></td><td></td><td></td><td></td><td></td><td></td><td></td></tr>
<tr><td>Oligo fwd</td><td>CACCAGGCTCGGACGGTCTCTCTA</td><td></td><td></td><td></td><td></td><td></td><td></td><td></td><td></td><td></td><td></td></tr>
<tr><td>Oligo rev</td><td>AAACTAGAGAGACCGTCCGAGCCT</td><td></td><td></td><td></td><td></td><td></td><td></td><td></td><td></td><td></td><td></td></tr>
<tr><td>Chromosome</td><td>start</td><td>end</td><td>strand</td><td>MM</td><td>target_seq</td><td>PAM</td><td>alignment</td><td>distance</td><td>position</td><td>gene name</td><td>gene id</td></tr>
<tr><td>GU305914.1</td><td>145509</td><td>145531</td><td>-</td><td>0</td><td>AGGCTCGGACGGTCTCTCTA</td><td>AGG</td><td>||||||||[||||||||||||]PAM</td><td>NA</td><td>Intergenic</td><td>NA</td><td>NA</td></tr>
<tr><td>T23104</td><td>TTGACCCGCCCCGTCGACCGCGG</td><td>1000</td><td></td><td></td><td></td><td></td><td></td><td></td><td></td><td></td><td></td></tr>
<tr><td>Oligo fwd</td><td>CACCTTGACCCGCCCCGTCGACCG</td><td></td><td></td><td></td><td></td><td></td><td></td><td></td><td></td><td></td><td></td></tr>
<tr><td>Oligo rev</td><td>AAACCGGTCGACGGGGCGGGTCAA</td><td></td><td></td><td></td><td></td><td></td><td></td><td></td><td></td><td></td><td></td></tr>
<tr><td>Chromosome</td><td>start</td><td>end</td><td>strand</td><td>MM</td><td>target_seq</td><td>PAM</td><td>alignment</td><td>distance</td><td>position</td><td>gene name</td><td>gene id</td></tr>
<tr><td>GU305914.1</td><td>145473</td><td>145495</td><td>-</td><td>0</td><td>TTGACCCGCCCCGTCGACCG</td><td>CGG</td><td>||||||||[||||||||||||]PAM</td><td>NA</td><td>Intergenic</td><td>NA</td><td>NA</td></tr>
<tr><td>T23105</td><td>GACACGAGACACCTGCACGATGG</td><td>1000</td><td></td><td></td><td></td><td></td><td></td><td></td><td></td><td></td><td></td></tr>
<tr><td>Oligo fwd</td><td>CACCGACACGAGACACCTGCACGA</td><td></td><td></td><td></td><td></td><td></td><td></td><td></td><td></td><td></td><td></td></tr>
<tr><td>Oligo rev</td><td>AAACTCGTGCAGGTGTCTCGTGTC</td><td></td><td></td><td></td><td></td><td></td><td></td><td></td><td></td><td></td><td></td></tr>
<tr><td>Chromosome</td><td>start</td><td>end</td><td>strand</td><td>MM</td><td>target_seq</td><td>PAM</td><td>alignment</td><td>distance</td><td>position</td><td>gene name</td><td>gene id</td></tr>
<tr><td>GU305914.1</td><td>145451</td><td>145473</td><td>-</td><td>0</td><td>GACACGAGACACCTGCACGA</td><td>TGG</td><td>||||||||[||||||||||||]PAM</td><td>NA</td><td>Intergenic</td><td>NA</td><td>NA</td></tr>
<tr><td>T23106</td><td>CACCTGCACGATGGCGAAAGCGG</td><td>1000</td><td></td><td></td><td></td><td></td><td></td><td></td><td></td><td></td><td></td></tr>
<tr><td>Oligo fwd</td><td>CACCCACCTGCACGATGGCGAAAG</td><td></td><td></td><td></td><td></td><td></td><td></td><td></td><td></td><td></td><td></td></tr>
<tr><td>Oligo rev</td><td>AAACCTTTCGCCATCGTGCAGGTG</td><td></td><td></td><td></td><td></td><td></td><td></td><td></td><td></td><td></td><td></td></tr>
<tr><td>Chromosome</td><td>start</td><td>end</td><td>strand</td><td>MM</td><td>target_seq</td><td>PAM</td><td>alignment</td><td>distance</td><td>position</td><td>gene name</td><td>gene id</td></tr>
<tr><td>GU305914.1</td><td>145442</td><td>145464</td><td>-</td><td>0</td><td>CACCTGCACGATGGCGAAAG</td><td>CGG</td><td>||||||||[||||||||||||]PAM</td><td>NA</td><td>Intergenic</td><td>NA</td><td>NA</td></tr>
<tr><td>T23107</td><td>ACCTGCACGATGGCGAAAGCGGG</td><td>1000</td><td></td><td></td><td></td><td></td><td></td><td></td><td></td><td></td><td></td></tr>
<tr><td>Oligo fwd</td><td>CACCACCTGCACGATGGCGAAAGC</td><td></td><td></td><td></td><td></td><td></td><td></td><td></td><td></td><td></td><td></td></tr>
<tr><td>Oligo rev</td><td>AAACGCTTTCGCCATCGTGCAGGT</td><td></td><td></td><td></td><td></td><td></td><td></td><td></td><td></td><td></td><td></td></tr>
<tr><td>Chromosome</td><td>start</td><td>end</td><td>strand</td><td>MM</td><td>target_seq</td><td>PAM</td><td>alignment</td><td>distance</td><td>position</td><td>gene name</td><td>gene id</td></tr>
<tr><td>GU305914.1</td><td>145441</td><td>145463</td><td>-</td><td>0</td><td>ACCTGCACGATGGCGAAAGC</td><td>GGG</td><td>||||||||[||||||||||||]PAM</td><td>NA</td><td>Intergenic</td><td>NA</td><td>NA</td></tr>
<tr><td>T23108</td><td>CGTCATGACGCTCTCCCACGTGG</td><td>1000</td><td></td><td></td><td></td><td></td><td></td><td></td><td></td><td></td><td></td></tr>
<tr><td>Oligo fwd</td><td>CACCCGTCATGACGCTCTCCCACG</td><td></td><td></td><td></td><td></td><td></td><td></td><td></td><td></td><td></td><td></td></tr>
<tr><td>Oligo rev</td><td>AAACCGTGGGAGAGCGTCATGACG</td><td></td><td></td><td></td><td></td><td></td><td></td><td></td><td></td><td></td><td></td></tr>
<tr><td>Chromosome</td><td>start</td><td>end</td><td>strand</td><td>MM</td><td>target_seq</td><td>PAM</td><td>alignment</td><td>distance</td><td>position</td><td>gene name</td><td>gene id</td></tr>
<tr><td>GU305914.1</td><td>145418</td><td>145440</td><td>-</td><td>0</td><td>CGTCATGACGCTCTCCCACG</td><td>TGG</td><td>||||||||[||||||||||||]PAM</td><td>NA</td><td>Intergenic</td><td>NA</td><td>NA</td></tr>
<tr><td>T23109</td><td>ACGTGGACCGCATGAACCTGCGG</td><td>1000</td><td></td><td></td><td></td><td></td><td></td><td></td><td></td><td></td><td></td></tr>
<tr><td>Oligo fwd</td><td>CACCACGTGGACCGCATGAACCTG</td><td></td><td></td><td></td><td></td><td></td><td></td><td></td><td></td><td></td><td></td></tr>
<tr><td>Oligo rev</td><td>AAACCAGGTTCATGCGGTCCACGT</td><td></td><td></td><td></td><td></td><td></td><td></td><td></td><td></td><td></td><td></td></tr>
<tr><td>Chromosome</td><td>start</td><td>end</td><td>strand</td><td>MM</td><td>target_seq</td><td>PAM</td><td>alignment</td><td>distance</td><td>position</td><td>gene name</td><td>gene id</td></tr>
<tr><td>GU305914.1</td><td>145401</td><td>145423</td><td>-</td><td>0</td><td>ACGTGGACCGCATGAACCTG</td><td>CGG</td><td>||||||||[||||||||||||]PAM</td><td>NA</td><td>Intergenic</td><td>NA</td><td>NA</td></tr>
<tr><td>T23110</td><td>ACCGCATGAACCTGCGGACGTGG</td><td>1000</td><td></td><td></td><td></td><td></td><td></td><td></td><td></td><td></td><td></td></tr>
<tr><td>Oligo fwd</td><td>CACCACCGCATGAACCTGCGGACG</td><td></td><td></td><td></td><td></td><td></td><td></td><td></td><td></td><td></td><td></td></tr>
<tr><td>Oligo rev</td><td>AAACCGTCCGCAGGTTCATGCGGT</td><td></td><td></td><td></td><td></td><td></td><td></td><td></td><td></td><td></td><td></td></tr>
<tr><td>Chromosome</td><td>start</td><td>end</td><td>strand</td><td>MM</td><td>target_seq</td><td>PAM</td><td>alignment</td><td>distance</td><td>position</td><td>gene name</td><td>gene id</td></tr>
<tr><td>GU305914.1</td><td>145395</td><td>145417</td><td>-</td><td>0</td><td>ACCGCATGAACCTGCGGACG</td><td>TGG</td><td>||||||||[||||||||||||]PAM</td><td>NA</td><td>Intergenic</td><td>NA</td><td>NA</td></tr>
<tr><td>T23111</td><td>GAACCTGCGGACGTGGACGATGG</td><td>1000</td><td></td><td></td><td></td><td></td><td></td><td></td><td></td><td></td><td></td></tr>
<tr><td>Oligo fwd</td><td>CACCGAACCTGCGGACGTGGACGA</td><td></td><td></td><td></td><td></td><td></td><td></td><td></td><td></td><td></td><td></td></tr>
<tr><td>Oligo rev</td><td>AAACTCGTCCACGTCCGCAGGTTC</td><td></td><td></td><td></td><td></td><td></td><td></td><td></td><td></td><td></td><td></td></tr>
<tr><td>Chromosome</td><td>start</td><td>end</td><td>strand</td><td>MM</td><td>target_seq</td><td>PAM</td><td>alignment</td><td>distance</td><td>position</td><td>gene name</td><td>gene id</td></tr>
<tr><td>GU305914.1</td><td>145388</td><td>145410</td><td>-</td><td>0</td><td>GAACCTGCGGACGTGGACGA</td><td>TGG</td><td>||||||||[||||||||||||]PAM</td><td>NA</td><td>Intergenic</td><td>NA</td><td>NA</td></tr>
<tr><td>T23112</td><td>GCTGTCGTTCGTGAACATCGTGG</td><td>1000</td><td></td><td></td><td></td><td></td><td></td><td></td><td></td><td></td><td></td></tr>
<tr><td>Oligo fwd</td><td>CACCGCTGTCGTTCGTGAACATCG</td><td></td><td></td><td></td><td></td><td></td><td></td><td></td><td></td><td></td><td></td></tr>
<tr><td>Oligo rev</td><td>AAACCGATGTTCACGAACGACAGC</td><td></td><td></td><td></td><td></td><td></td><td></td><td></td><td></td><td></td><td></td></tr>
<tr><td>Chromosome</td><td>start</td><td>end</td><td>strand</td><td>MM</td><td>target_seq</td><td>PAM</td><td>alignment</td><td>distance</td><td>position</td><td>gene name</td><td>gene id</td></tr>
<tr><td>GU305914.1</td><td>145349</td><td>145371</td><td>-</td><td>0</td><td>GCTGTCGTTCGTGAACATCG</td><td>TGG</td><td>||||||||[||||||||||||]PAM</td><td>NA</td><td>Intergenic</td><td>NA</td><td>NA</td></tr>
<tr><td>T23113</td><td>CGTGAACATCGTGGTGTTCTCGG</td><td>1000</td><td></td><td></td><td></td><td></td><td></td><td></td><td></td><td></td><td></td></tr>
<tr><td>Oligo fwd</td><td>CACCCGTGAACATCGTGGTGTTCT</td><td></td><td></td><td></td><td></td><td></td><td></td><td></td><td></td><td></td><td></td></tr>
<tr><td>Oligo rev</td><td>AAACAGAACACCACGATGTTCACG</td><td></td><td></td><td></td><td></td><td></td><td></td><td></td><td></td><td></td><td></td></tr>
<tr><td>Chromosome</td><td>start</td><td>end</td><td>strand</td><td>MM</td><td>target_seq</td><td>PAM</td><td>alignment</td><td>distance</td><td>position</td><td>gene name</td><td>gene id</td></tr>
<tr><td>GU305914.1</td><td>145340</td><td>145362</td><td>-</td><td>0</td><td>CGTGAACATCGTGGTGTTCT</td><td>CGG</td><td>||||||||[||||||||||||]PAM</td><td>NA</td><td>Intergenic</td><td>NA</td><td>NA</td></tr>
<tr><td>T23114</td><td>GAACATCGTGGTGTTCTCGGTGG</td><td>1000</td><td></td><td></td><td></td><td></td><td></td><td></td><td></td><td></td><td></td></tr>
<tr><td>Oligo fwd</td><td>CACCGAACATCGTGGTGTTCTCGG</td><td></td><td></td><td></td><td></td><td></td><td></td><td></td><td></td><td></td><td></td></tr>
<tr><td>Oligo rev</td><td>AAACCCGAGAACACCACGATGTTC</td><td></td><td></td><td></td><td></td><td></td><td></td><td></td><td></td><td></td><td></td></tr>
<tr><td>Chromosome</td><td>start</td><td>end</td><td>strand</td><td>MM</td><td>target_seq</td><td>PAM</td><td>alignment</td><td>distance</td><td>position</td><td>gene name</td><td>gene id</td></tr>
<tr><td>GU305914.1</td><td>145337</td><td>145359</td><td>-</td><td>0</td><td>GAACATCGTGGTGTTCTCGG</td><td>TGG</td><td>||||||||[||||||||||||]PAM</td><td>NA</td><td>Intergenic</td><td>NA</td><td>NA</td></tr>
<tr><td>T23115</td><td>CATCGTGGTGTTCTCGGTGGCGG</td><td>1000</td><td></td><td></td><td></td><td></td><td></td><td></td><td></td><td></td><td></td></tr>
<tr><td>Oligo fwd</td><td>CACCCATCGTGGTGTTCTCGGTGG</td><td></td><td></td><td></td><td></td><td></td><td></td><td></td><td></td><td></td><td></td></tr>
</table>

<table>
<tr><td>Oligo rev</td><td>AAACCCACCGAGAACACCACGATG</td><td></td><td></td><td></td><td></td><td></td><td></td><td></td><td></td><td></td><td></td></tr>
<tr><td>Chromosome</td><td>start</td><td>end</td><td>strand</td><td>MM</td><td>target_seq</td><td>PAM</td><td>alignment</td><td>distance</td><td>position</td><td>gene name</td><td>gene id</td></tr>
<tr><td>GU305914.1</td><td>145334</td><td>145356</td><td>-</td><td>0</td><td>CATCGTGGTGTTCTCGGTGG</td><td>CGG</td><td>||||||||[||||||||||||]PAM</td><td>NA</td><td>Intergenic</td><td>NA</td><td>NA</td></tr>
<tr><td>T23116</td><td>TCGGTGGCGGCGCACTTCCCCGG</td><td>1000</td><td></td><td></td><td></td><td></td><td></td><td></td><td></td><td></td><td></td></tr>
<tr><td>Oligo fwd</td><td>CACCTCGGTGGCGGCGCACTTCCC</td><td></td><td></td><td></td><td></td><td></td><td></td><td></td><td></td><td></td><td></td></tr>
<tr><td>Oligo rev</td><td>AAACGGGAAGTGCGCCGCCACCGA</td><td></td><td></td><td></td><td></td><td></td><td></td><td></td><td></td><td></td><td></td></tr>
<tr><td>Chromosome</td><td>start</td><td>end</td><td>strand</td><td>MM</td><td>target_seq</td><td>PAM</td><td>alignment</td><td>distance</td><td>position</td><td>gene name</td><td>gene id</td></tr>
<tr><td>GU305914.1</td><td>145321</td><td>145343</td><td>-</td><td>0</td><td>TCGGTGGCGGCGCACTTCCC</td><td>CGG</td><td>||||||||[||||||||||||]PAM</td><td>NA</td><td>Intergenic</td><td>NA</td><td>NA</td></tr>
<tr><td>T23117</td><td>GCGGCGCACTTCCCCGGCATCGG</td><td>1000</td><td></td><td></td><td></td><td></td><td></td><td></td><td></td><td></td><td></td></tr>
<tr><td>Oligo fwd</td><td>CACCGCGGCGCACTTCCCCGGCAT</td><td></td><td></td><td></td><td></td><td></td><td></td><td></td><td></td><td></td><td></td></tr>
<tr><td>Oligo rev</td><td>AAACATGCCGGGGAAGTGCGCCGC</td><td></td><td></td><td></td><td></td><td></td><td></td><td></td><td></td><td></td><td></td></tr>
<tr><td>Chromosome</td><td>start</td><td>end</td><td>strand</td><td>MM</td><td>target_seq</td><td>PAM</td><td>alignment</td><td>distance</td><td>position</td><td>gene name</td><td>gene id</td></tr>
<tr><td>GU305914.1</td><td>145315</td><td>145337</td><td>-</td><td>0</td><td>GCGGCGCACTTCCCCGGCAT</td><td>CGG</td><td>||||||||[||||||||||||]PAM</td><td>NA</td><td>Intergenic</td><td>NA</td><td>NA</td></tr>
<tr><td>T23118</td><td>TCTGACGCCGCAGCTGTTCCTGG</td><td>1000</td><td></td><td></td><td></td><td></td><td></td><td></td><td></td><td></td><td></td></tr>
<tr><td>Oligo fwd</td><td>CACCTCTGACGCCGCAGCTGTTCC</td><td></td><td></td><td></td><td></td><td></td><td></td><td></td><td></td><td></td><td></td></tr>
<tr><td>Oligo rev</td><td>AAACGGAACAGCTGCGGCGTCAGA</td><td></td><td></td><td></td><td></td><td></td><td></td><td></td><td></td><td></td><td></td></tr>
<tr><td>Chromosome</td><td>start</td><td>end</td><td>strand</td><td>MM</td><td>target_seq</td><td>PAM</td><td>alignment</td><td>distance</td><td>position</td><td>gene name</td><td>gene id</td></tr>
<tr><td>GU305914.1</td><td>145220</td><td>145242</td><td>-</td><td>0</td><td>TCTGACGCCGCAGCTGTTCC</td><td>TGG</td><td>||||||||[||||||||||||]PAM</td><td>NA</td><td>Intergenic</td><td>NA</td><td>NA</td></tr>
<tr><td>T23119</td><td>CATCGTGTACGTGATCTTCACGG</td><td>1000</td><td></td><td></td><td></td><td></td><td></td><td></td><td></td><td></td><td></td></tr>
<tr><td>Oligo fwd</td><td>CACCCATCGTGTACGTGATCTTCA</td><td></td><td></td><td></td><td></td><td></td><td></td><td></td><td></td><td></td><td></td></tr>
<tr><td>Oligo rev</td><td>AAACTGAAGATCACGTACACGATG</td><td></td><td></td><td></td><td></td><td></td><td></td><td></td><td></td><td></td><td></td></tr>
<tr><td>Chromosome</td><td>start</td><td>end</td><td>strand</td><td>MM</td><td>target_seq</td><td>PAM</td><td>alignment</td><td>distance</td><td>position</td><td>gene name</td><td>gene id</td></tr>
<tr><td>GU305914.1</td><td>145184</td><td>145206</td><td>-</td><td>0</td><td>CATCGTGTACGTGATCTTCA</td><td>CGG</td><td>||||||||[||||||||||||]PAM</td><td>NA</td><td>Intergenic</td><td>NA</td><td>NA</td></tr>
<tr><td>T23120</td><td>TGAGCTACTACATCGTGTGCTGG</td><td>1000</td><td></td><td></td><td></td><td></td><td></td><td></td><td></td><td></td><td></td></tr>
<tr><td>Oligo fwd</td><td>CACCTGAGCTACTACATCGTGTGC</td><td></td><td></td><td></td><td></td><td></td><td></td><td></td><td></td><td></td><td></td></tr>
<tr><td>Oligo rev</td><td>AAACGCACACGATGTAGTAGCTCA</td><td></td><td></td><td></td><td></td><td></td><td></td><td></td><td></td><td></td><td></td></tr>
<tr><td>Chromosome</td><td>start</td><td>end</td><td>strand</td><td>MM</td><td>target_seq</td><td>PAM</td><td>alignment</td><td>distance</td><td>position</td><td>gene name</td><td>gene id</td></tr>
<tr><td>GU305914.1</td><td>145140</td><td>145162</td><td>-</td><td>0</td><td>TGAGCTACTACATCGTGTGC</td><td>TGG</td><td>||||||||[||||||||||||]PAM</td><td>NA</td><td>Intergenic</td><td>NA</td><td>NA</td></tr>
<tr><td>T23121</td><td>GAGCTACTACATCGTGTGCTGGG</td><td>1000</td><td></td><td></td><td></td><td></td><td></td><td></td><td></td><td></td><td></td></tr>
<tr><td>Oligo fwd</td><td>CACCGAGCTACTACATCGTGTGCT</td><td></td><td></td><td></td><td></td><td></td><td></td><td></td><td></td><td></td><td></td></tr>
<tr><td>Oligo rev</td><td>AAACAGCACACGATGTAGTAGCTC</td><td></td><td></td><td></td><td></td><td></td><td></td><td></td><td></td><td></td><td></td></tr>
<tr><td>Chromosome</td><td>start</td><td>end</td><td>strand</td><td>MM</td><td>target_seq</td><td>PAM</td><td>alignment</td><td>distance</td><td>position</td><td>gene name</td><td>gene id</td></tr>
<tr><td>GU305914.1</td><td>145139</td><td>145161</td><td>-</td><td>0</td><td>GAGCTACTACATCGTGTGCT</td><td>GGG</td><td>||||||||[||||||||||||]PAM</td><td>NA</td><td>Intergenic</td><td>NA</td><td>NA</td></tr>
<tr><td>T23122</td><td>ATCTACTTCCGCAGCGAGCACGG</td><td>1000</td><td></td><td></td><td></td><td></td><td></td><td></td><td></td><td></td><td></td></tr>
<tr><td>Oligo fwd</td><td>CACCATCTACTTCCGCAGCGAGCA</td><td></td><td></td><td></td><td></td><td></td><td></td><td></td><td></td><td></td><td></td></tr>
<tr><td>Oligo rev</td><td>AAACTGCTCGCTGCGGAAGTAGAT</td><td></td><td></td><td></td><td></td><td></td><td></td><td></td><td></td><td></td><td></td></tr>
<tr><td>Chromosome</td><td>start</td><td>end</td><td>strand</td><td>MM</td><td>target_seq</td><td>PAM</td><td>alignment</td><td>distance</td><td>position</td><td>gene name</td><td>gene id</td></tr>
<tr><td>GU305914.1</td><td>145111</td><td>145133</td><td>-</td><td>0</td><td>ATCTACTTCCGCAGCGAGCA</td><td>CGG</td><td>||||||||[||||||||||||]PAM</td><td>NA</td><td>Intergenic</td><td>NA</td><td>NA</td></tr>
<tr><td>T23123</td><td>CCGCAGCGAGCACGGCACGCAGG</td><td>1000</td><td></td><td></td><td></td><td></td><td></td><td></td><td></td><td></td><td></td></tr>
<tr><td>Oligo fwd</td><td>CACCCCGCAGCGAGCACGGCACGC</td><td></td><td></td><td></td><td></td><td></td><td></td><td></td><td></td><td></td><td></td></tr>
<tr><td>Oligo rev</td><td>AAACGCGTGCCGTGCTCGCTGCGG</td><td></td><td></td><td></td><td></td><td></td><td></td><td></td><td></td><td></td><td></td></tr>
<tr><td>Chromosome</td><td>start</td><td>end</td><td>strand</td><td>MM</td><td>target_seq</td><td>PAM</td><td>alignment</td><td>distance</td><td>position</td><td>gene name</td><td>gene id</td></tr>
<tr><td>GU305914.1</td><td>145103</td><td>145125</td><td>-</td><td>0</td><td>CCGCAGCGAGCACGGCACGC</td><td>AGG</td><td>||||||||[||||||||||||]PAM</td><td>NA</td><td>Intergenic</td><td>NA</td><td>NA</td></tr>
<tr><td>T23124</td><td>GACCCGCGACATCAACTTCATGG</td><td>1000</td><td></td><td></td><td></td><td></td><td></td><td></td><td></td><td></td><td></td></tr>
<tr><td>Oligo fwd</td><td>CACCGACCCGCGACATCAACTTCA</td><td></td><td></td><td></td><td></td><td></td><td></td><td></td><td></td><td></td><td></td></tr>
<tr><td>Oligo rev</td><td>AAACTGAAGTTGATGTCGCGGGTC</td><td></td><td></td><td></td><td></td><td></td><td></td><td></td><td></td><td></td><td></td></tr>
<tr><td>Chromosome</td><td>start</td><td>end</td><td>strand</td><td>MM</td><td>target_seq</td><td>PAM</td><td>alignment</td><td>distance</td><td>position</td><td>gene name</td><td>gene id</td></tr>
<tr><td>GU305914.1</td><td>145070</td><td>145092</td><td>-</td><td>0</td><td>GACCCGCGACATCAACTTCA</td><td>TGG</td><td>||||||||[||||||||||||]PAM</td><td>NA</td><td>Intergenic</td><td>NA</td><td>NA</td></tr>
<tr><td>T23125</td><td>ACCCGCGACATCAACTTCATGGG</td><td>1000</td><td></td><td></td><td></td><td></td><td></td><td></td><td></td><td></td><td></td></tr>
<tr><td>Oligo fwd</td><td>CACCACCCGCGACATCAACTTCAT</td><td></td><td></td><td></td><td></td><td></td><td></td><td></td><td></td><td></td><td></td></tr>
<tr><td>Oligo rev</td><td>AAACATGAAGTTGATGTCGCGGGT</td><td></td><td></td><td></td><td></td><td></td><td></td><td></td><td></td><td></td><td></td></tr>
<tr><td>Chromosome</td><td>start</td><td>end</td><td>strand</td><td>MM</td><td>target_seq</td><td>PAM</td><td>alignment</td><td>distance</td><td>position</td><td>gene name</td><td>gene id</td></tr>
<tr><td>GU305914.1</td><td>145069</td><td>145091</td><td>-</td><td>0</td><td>ACCCGCGACATCAACTTCAT</td><td>GGG</td><td>||||||||[||||||||||||]PAM</td><td>NA</td><td>Intergenic</td><td>NA</td><td>NA</td></tr>
<tr><td>T23126</td><td>CATCAACTTCATGGGCGACTCGG</td><td>1000</td><td></td><td></td><td></td><td></td><td></td><td></td><td></td><td></td><td></td></tr>
<tr><td>Oligo fwd</td><td>CACCCATCAACTTCATGGGCGACT</td><td></td><td></td><td></td><td></td><td></td><td></td><td></td><td></td><td></td><td></td></tr>
<tr><td>Oligo rev</td><td>AAACAGTCGCCCATGAAGTTGATG</td><td></td><td></td><td></td><td></td><td></td><td></td><td></td><td></td><td></td><td></td></tr>
<tr><td>Chromosome</td><td>start</td><td>end</td><td>strand</td><td>MM</td><td>target_seq</td><td>PAM</td><td>alignment</td><td>distance</td><td>position</td><td>gene name</td><td>gene id</td></tr>
<tr><td>GU305914.1</td><td>145061</td><td>145083</td><td>-</td><td>0</td><td>CATCAACTTCATGGGCGACT</td><td>CGG</td><td>||||||||[||||||||||||]PAM</td><td>NA</td><td>Intergenic</td><td>NA</td><td>NA</td></tr>
<tr><td>GU305914.1</td><td>100411</td><td>100433</td><td>-</td><td>4</td><td>GTTCAACTTCGTGGGCGACG</td><td>TGG</td><td>--||||||[||-||||||||-]PAM</td><td>NA</td><td>Intergenic</td><td>NA</td><td>NA</td></tr>
<tr><td>T23127</td><td>CTTCACCTTCGTGCTGACGATGG</td><td>1000</td><td></td><td></td><td></td><td></td><td></td><td></td><td></td><td></td><td></td></tr>
<tr><td>Oligo fwd</td><td>CACCCTTCACCTTCGTGCTGACGA</td><td></td><td></td><td></td><td></td><td></td><td></td><td></td><td></td><td></td><td></td></tr>
<tr><td>Oligo rev</td><td>AAACTCGTCAGCACGAAGGTGAAG</td><td></td><td></td><td></td><td></td><td></td><td></td><td></td><td></td><td></td><td></td></tr>
<tr><td>Chromosome</td><td>start</td><td>end</td><td>strand</td><td>MM</td><td>target_seq</td><td>PAM</td><td>alignment</td><td>distance</td><td>position</td><td>gene name</td><td>gene id</td></tr>
<tr><td>GU305914.1</td><td>145031</td><td>145053</td><td>-</td><td>0</td><td>CTTCACCTTCGTGCTGACGA</td><td>TGG</td><td>||||||||[||||||||||||]PAM</td><td>NA</td><td>Intergenic</td><td>NA</td><td>NA</td></tr>
<tr><td>GU305914.1</td><td>21988</td><td>22010</td><td>-</td><td>4</td><td>CCTCACCGTCCTGCCGACGA</td><td>CGG</td><td>|-|||||-[||-|||-|||||]PAM</td><td>NA</td><td>Intergenic</td><td>NA</td><td>NA</td></tr>
<tr><td>T23128</td><td>GTCGTTCCGCCTACCGTCCATGG</td><td>1000</td><td></td><td></td><td></td><td></td><td></td><td></td><td></td><td></td><td></td></tr>
<tr><td>Oligo fwd</td><td>CACCGTCGTTCCGCCTACCGTCCA</td><td></td><td></td><td></td><td></td><td></td><td></td><td></td><td></td><td></td><td></td></tr>
<tr><td>Oligo rev</td><td>AAACTGGACGGTAGGCGGAACGAC</td><td></td><td></td><td></td><td></td><td></td><td></td><td></td><td></td><td></td><td></td></tr>
<tr><td>Chromosome</td><td>start</td><td>end</td><td>strand</td><td>MM</td><td>target_seq</td><td>PAM</td><td>alignment</td><td>distance</td><td>position</td><td>gene name</td><td>gene id</td></tr>
<tr><td>GU305914.1</td><td>144980</td><td>145002</td><td>-</td><td>0</td><td>GTCGTTCCGCCTACCGTCCA</td><td>TGG</td><td>||||||||[||||||||||||]PAM</td><td>NA</td><td>Intergenic</td><td>NA</td><td>NA</td></tr>
<tr><td>T23129</td><td>CATGTACTTTATGTGCCTGACGG</td><td>1000</td><td></td><td></td><td></td><td></td><td></td><td></td><td></td><td></td><td></td></tr>
<tr><td>Oligo fwd</td><td>CACCCATGTACTTTATGTGCCTGA</td><td></td><td></td><td></td><td></td><td></td><td></td><td></td><td></td><td></td><td></td></tr>
<tr><td>Oligo rev</td><td>AAACTCAGGCACATAAAGTACATG</td><td></td><td></td><td></td><td></td><td></td><td></td><td></td><td></td><td></td><td></td></tr>
<tr><td>Chromosome</td><td>start</td><td>end</td><td>strand</td><td>MM</td><td>target_seq</td><td>PAM</td><td>alignment</td><td>distance</td><td>position</td><td>gene name</td><td>gene id</td></tr>
<tr><td>GU305914.1</td><td>144941</td><td>144963</td><td>-</td><td>0</td><td>CATGTACTTTATGTGCCTGA</td><td>CGG</td><td>||||||||[||||||||||||]PAM</td><td>NA</td><td>Intergenic</td><td>NA</td><td>NA</td></tr>
</table>

| | | | | | | | | | | | |
|---|---|---|---|---|---|---|---|---|---|---|---|
| T23130 | TATGTGCCTGACGGCCTTCGTGG | 1000 | | | | | | | | | |
| Oligo fwd | CACCTATGTGCCTGACGGCCTTCG | | | | | | | | | | |
| Oligo rev | AAACCGAAGGCCGTCAGGCACATA | | | | | | | | | | |
| Chromosome | start | end | strand | MM | target_seq | PAM | alignment | distance | position | gene name | gene id |
| GU305914.1 | 144932 | 144954 | - | 0 | TATGTGCCTGACGGCCTTCG | TGG | \|\|\|\|\|\|\|\|[\|\|\|\|\|\|\|\|\|\|\|\|]PAM | NA | Intergenic | NA | NA |
| T23131 | GAAGATCCACCCGAAGCTGCAGG | 1000 | | | | | | | | | |
| Oligo fwd | CACCGAAGATCCACCCGAAGCTGC | | | | | | | | | | |
| Oligo rev | AAACGCAGCTTCGGGTGGATCTTC | | | | | | | | | | |
| Chromosome | start | end | strand | MM | target_seq | PAM | alignment | distance | position | gene name | gene id |
| GU305914.1 | 144857 | 144879 | - | 0 | GAAGATCCACCCGAAGCTGC | AGG | \|\|\|\|\|\|\|\|[\|\|\|\|\|\|\|\|\|\|\|\|]PAM | NA | Intergenic | NA | NA |
| T23132 | AAGATCCACCCGAAGCTGCAGGG | 1000 | | | | | | | | | |
| Oligo fwd | CACCAAGATCCACCCGAAGCTGCA | | | | | | | | | | |
| Oligo rev | AAACTGCAGCTTCGGGTGGATCTT | | | | | | | | | | |
| Chromosome | start | end | strand | MM | target_seq | PAM | alignment | distance | position | gene name | gene id |
| GU305914.1 | 144856 | 144878 | - | 0 | AAGATCCACCCGAAGCTGCA | GGG | \|\|\|\|\|\|\|\|[\|\|\|\|\|\|\|\|\|\|\|\|]PAM | NA | Intergenic | NA | NA |
| T23133 | AGATCCACCCGAAGCTGCAGGGG | 1000 | | | | | | | | | |
| Oligo fwd | CACCAGATCCACCCGAAGCTGCAG | | | | | | | | | | |
| Oligo rev | AAACCTGCAGCTTCGGGTGGATCT | | | | | | | | | | |
| Chromosome | start | end | strand | MM | target_seq | PAM | alignment | distance | position | gene name | gene id |
| GU305914.1 | 144855 | 144877 | - | 0 | AGATCCACCCGAAGCTGCAG | GGG | \|\|\|\|\|\|\|\|[\|\|\|\|\|\|\|\|\|\|\|\|]PAM | NA | Intergenic | NA | NA |
| T23134 | TGCAGGGGACCATCCGCTATCGG | 1000 | | | | | | | | | |
| Oligo fwd | CACCTGCAGGGGACCATCCGCTAT | | | | | | | | | | |
| Oligo rev | AAACATAGCGGATGGTCCCCTGCA | | | | | | | | | | |
| Chromosome | start | end | strand | MM | target_seq | PAM | alignment | distance | position | gene name | gene id |
| GU305914.1 | 144840 | 144862 | - | 0 | TGCAGGGGACCATCCGCTAT | CGG | \|\|\|\|\|\|\|\|[\|\|\|\|\|\|\|\|\|\|\|\|]PAM | NA | Intergenic | NA | NA |
| T23135 | GGGGACCATCCGCTATCGGACGG | 1000 | | | | | | | | | |
| Oligo fwd | CACCGGGGACCATCCGCTATCGGA | | | | | | | | | | |
| Oligo rev | AAACTCCGATAGCGGATGGTCCCC | | | | | | | | | | |
| Chromosome | start | end | strand | MM | target_seq | PAM | alignment | distance | position | gene name | gene id |
| GU305914.1 | 144836 | 144858 | - | 0 | GGGGACCATCCGCTATCGGA | CGG | \|\|\|\|\|\|\|\|[\|\|\|\|\|\|\|\|\|\|\|\|]PAM | NA | Intergenic | NA | NA |
| T23136 | GACCATCCGCTATCGGACGGCGG | 1000 | | | | | | | | | |
| Oligo fwd | CACCGACCATCCGCTATCGGACGG | | | | | | | | | | |
| Oligo rev | AAACCCGTCCGATAGCGGATGGTC | | | | | | | | | | |
| Chromosome | start | end | strand | MM | target_seq | PAM | alignment | distance | position | gene name | gene id |
| GU305914.1 | 144833 | 144855 | - | 0 | GACCATCCGCTATCGGACGG | CGG | \|\|\|\|\|\|\|\|[\|\|\|\|\|\|\|\|\|\|\|\|]PAM | NA | Intergenic | NA | NA |
| T23137 | CATCCGCTATCGGACGGCGGTGG | 1000 | | | | | | | | | |
| Oligo fwd | CACCCATCCGCTATCGGACGGCGG | | | | | | | | | | |
| Oligo rev | AAACCCGCCGTCCGATAGCGGATG | | | | | | | | | | |
| Chromosome | start | end | strand | MM | target_seq | PAM | alignment | distance | position | gene name | gene id |
| GU305914.1 | 144830 | 144852 | - | 0 | CATCCGCTATCGGACGGCGG | TGG | \|\|\|\|\|\|\|\|[\|\|\|\|\|\|\|\|\|\|\|\|]PAM | NA | Intergenic | NA | NA |
| T23138 | GAACCTGACCCAGCTGATCTTGG | 1000 | | | | | | | | | |
| Oligo fwd | CACCGAACCTGACCCAGCTGATCT | | | | | | | | | | |
| Oligo rev | AAACAGATCAGCTGGGTCAGGTTC | | | | | | | | | | |
| Chromosome | start | end | strand | MM | target_seq | PAM | alignment | distance | position | gene name | gene id |
| GU305914.1 | 144806 | 144828 | - | 0 | GAACCTGACCCAGCTGATCT | TGG | \|\|\|\|\|\|\|\|[\|\|\|\|\|\|\|\|\|\|\|\|]PAM | NA | Intergenic | NA | NA |
| GU305914.1 | 31606 | 31628 | - | 4 | CAACCTCACGCAGCTCATCT | CGG | -\|\|\|\|\|-\|[\|-\|\|\|\|\|-\|\|\|\|]PAM | NA | Intergenic | NA | NA |
| T23139 | AACCTGACCCAGCTGATCTTGGG | 1000 | | | | | | | | | |
| Oligo fwd | CACCAACCTGACCCAGCTGATCTT | | | | | | | | | | |
| Oligo rev | AAACAAGATCAGCTGGGTCAGGTT | | | | | | | | | | |
| Chromosome | start | end | strand | MM | target_seq | PAM | alignment | distance | position | gene name | gene id |
| GU305914.1 | 144805 | 144827 | - | 0 | AACCTGACCCAGCTGATCTT | GGG | \|\|\|\|\|\|\|\|[\|\|\|\|\|\|\|\|\|\|\|\|]PAM | NA | Intergenic | NA | NA |
| T23140 | GATCTTGGGCTTCGCGACCATGG | 1000 | | | | | | | | | |
| Oligo fwd | CACCGATCTTGGGCTTCGCGACCA | | | | | | | | | | |
| Oligo rev | AAACTGGTCGCGAAGCCCAAGATC | | | | | | | | | | |
| Chromosome | start | end | strand | MM | target_seq | PAM | alignment | distance | position | gene name | gene id |
| GU305914.1 | 144791 | 144813 | - | 0 | GATCTTGGGCTTCGCGACCA | TGG | \|\|\|\|\|\|\|\|[\|\|\|\|\|\|\|\|\|\|\|\|]PAM | NA | Intergenic | NA | NA |
| T23141 | GGGCTTCGCGACCATGGTGCTGG | 1000 | | | | | | | | | |
| Oligo fwd | CACCGGGCTTCGCGACCATGGTGC | | | | | | | | | | |
| Oligo rev | AAACGCACCATGGTCGCGAAGCCC | | | | | | | | | | |
| Chromosome | start | end | strand | MM | target_seq | PAM | alignment | distance | position | gene name | gene id |
| GU305914.1 | 144785 | 144807 | - | 0 | GGGCTTCGCGACCATGGTGC | TGG | \|\|\|\|\|\|\|\|[\|\|\|\|\|\|\|\|\|\|\|\|]PAM | NA | Intergenic | NA | NA |
| T23142 | CATGGTGCTGGCCATGTCGCTGG | 1000 | | | | | | | | | |
| Oligo fwd | CACCCATGGTGCTGGCCATGTCGC | | | | | | | | | | |
| Oligo rev | AAACGCGACATGGCCAGCACCATG | | | | | | | | | | |
| Chromosome | start | end | strand | MM | target_seq | PAM | alignment | distance | position | gene name | gene id |
| GU305914.1 | 144773 | 144795 | - | 0 | CATGGTGCTGGCCATGTCGC | TGG | \|\|\|\|\|\|\|\|[\|\|\|\|\|\|\|\|\|\|\|\|]PAM | NA | Intergenic | NA | NA |
| GU305914.1 | 197602 | 197624 | + | 4 | CAGTGTGCTGTCCATCTCGC | CGG | \|\|--\|\|\|\|[\|\|-\|\|\|\|-\|\|\|\|]PAM | NA | Intergenic | NA | NA |
| T23143 | CTGGCCATGTCGCTGGCGCTCGG | 1000 | | | | | | | | | |
| Oligo fwd | CACCCTGGCCATGTCGCTGGCGCT | | | | | | | | | | |
| Oligo rev | AAACAGCGCCAGCGACATGGCCAG | | | | | | | | | | |
| Chromosome | start | end | strand | MM | target_seq | PAM | alignment | distance | position | gene name | gene id |
| GU305914.1 | 144766 | 144788 | - | 0 | CTGGCCATGTCGCTGGCGCT | CGG | \|\|\|\|\|\|\|\|[\|\|\|\|\|\|\|\|\|\|\|\|]PAM | NA | Intergenic | NA | NA |
| T23144 | ATGTCGCTGGCGCTCGGCTTCGG | 1000 | | | | | | | | | |
| Oligo fwd | CACCATGTCGCTGGCGCTCGGCTT | | | | | | | | | | |

<table>
<tr><td>Oligo rev</td><td>AAACAAGCCGAGCGCCAGCGACAT</td><td></td><td></td><td></td><td></td><td></td><td></td><td></td><td></td><td></td><td></td></tr>
<tr><td>Chromosome</td><td>start</td><td>end</td><td>strand</td><td>MM</td><td>target_seq</td><td>PAM</td><td>alignment</td><td>distance</td><td>position</td><td>gene name</td><td>gene id</td></tr>
<tr><td>GU305914.1</td><td>144760</td><td>144782</td><td>-</td><td>0</td><td>ATGTCGCTGGCGCTCGGCTT</td><td>CGG</td><td>||||||||[||||||||||||]PAM</td><td>NA</td><td>Intergenic</td><td>NA</td><td>NA</td></tr>
<tr><td>T23145</td><td>CAACAGCTTTTTCGTGAAGACGG</td><td>1000</td><td></td><td></td><td></td><td></td><td></td><td></td><td></td><td></td><td></td></tr>
<tr><td>Oligo fwd</td><td>CACCCAACAGCTTTTTCGTGAAGA</td><td></td><td></td><td></td><td></td><td></td><td></td><td></td><td></td><td></td><td></td></tr>
<tr><td>Oligo rev</td><td>AAACTCTTCACGAAAAAGCTGTTG</td><td></td><td></td><td></td><td></td><td></td><td></td><td></td><td></td><td></td><td></td></tr>
<tr><td>Chromosome</td><td>start</td><td>end</td><td>strand</td><td>MM</td><td>target_seq</td><td>PAM</td><td>alignment</td><td>distance</td><td>position</td><td>gene name</td><td>gene id</td></tr>
<tr><td>GU305914.1</td><td>144737</td><td>144759</td><td>-</td><td>0</td><td>CAACAGCTTTTTCGTGAAGA</td><td>CGG</td><td>||||||||[||||||||||||]PAM</td><td>NA</td><td>Intergenic</td><td>NA</td><td>NA</td></tr>
<tr><td>T23146</td><td>AAGACGGCGCACGTCGTGTTCGG</td><td>1000</td><td></td><td></td><td></td><td></td><td></td><td></td><td></td><td></td><td></td></tr>
<tr><td>Oligo fwd</td><td>CACCAAGACGGCGCACGTCGTGTT</td><td></td><td></td><td></td><td></td><td></td><td></td><td></td><td></td><td></td><td></td></tr>
<tr><td>Oligo rev</td><td>AAACAACACGACGTGCGCCGTCTT</td><td></td><td></td><td></td><td></td><td></td><td></td><td></td><td></td><td></td><td></td></tr>
<tr><td>Chromosome</td><td>start</td><td>end</td><td>strand</td><td>MM</td><td>target_seq</td><td>PAM</td><td>alignment</td><td>distance</td><td>position</td><td>gene name</td><td>gene id</td></tr>
<tr><td>GU305914.1</td><td>144721</td><td>144743</td><td>-</td><td>0</td><td>AAGACGGCGCACGTCGTGTT</td><td>CGG</td><td>||||||||[||||||||||||]PAM</td><td>NA</td><td>Intergenic</td><td>NA</td><td>NA</td></tr>
<tr><td>GU305914.1</td><td>91438</td><td>91460</td><td>-</td><td>4</td><td>TACACGGGGCACGTCGTGTG</td><td>CGG</td><td>-|-||||-[|||||||||||-]PAM</td><td>NA</td><td>Intergenic</td><td>NA</td><td>NA</td></tr>
<tr><td>T23147</td><td>GCACGTCGTGTTCGGCGCCATGG</td><td>1000</td><td></td><td></td><td></td><td></td><td></td><td></td><td></td><td></td><td></td></tr>
<tr><td>Oligo fwd</td><td>CACCGCACGTCGTGTTCGGCGCCA</td><td></td><td></td><td></td><td></td><td></td><td></td><td></td><td></td><td></td><td></td></tr>
<tr><td>Oligo rev</td><td>AAACTGGCGCCGAACACGACGTGC</td><td></td><td></td><td></td><td></td><td></td><td></td><td></td><td></td><td></td><td></td></tr>
<tr><td>Chromosome</td><td>start</td><td>end</td><td>strand</td><td>MM</td><td>target_seq</td><td>PAM</td><td>alignment</td><td>distance</td><td>position</td><td>gene name</td><td>gene id</td></tr>
<tr><td>GU305914.1</td><td>144713</td><td>144735</td><td>-</td><td>0</td><td>GCACGTCGTGTTCGGCGCCA</td><td>TGG</td><td>||||||||[||||||||||||]PAM</td><td>NA</td><td>Intergenic</td><td>NA</td><td>NA</td></tr>
<tr><td>T23148</td><td>CATGGTCGCCTTCGCGATCGTGG</td><td>1000</td><td></td><td></td><td></td><td></td><td></td><td></td><td></td><td></td><td></td></tr>
<tr><td>Oligo fwd</td><td>CACCCATGGTCGCCTTCGCGATCG</td><td></td><td></td><td></td><td></td><td></td><td></td><td></td><td></td><td></td><td></td></tr>
<tr><td>Oligo rev</td><td>AAACCGATCGCGAAGGCGACCATG</td><td></td><td></td><td></td><td></td><td></td><td></td><td></td><td></td><td></td><td></td></tr>
<tr><td>Chromosome</td><td>start</td><td>end</td><td>strand</td><td>MM</td><td>target_seq</td><td>PAM</td><td>alignment</td><td>distance</td><td>position</td><td>gene name</td><td>gene id</td></tr>
<tr><td>GU305914.1</td><td>144695</td><td>144717</td><td>-</td><td>0</td><td>CATGGTCGCCTTCGCGATCG</td><td>TGG</td><td>||||||||[||||||||||||]PAM</td><td>NA</td><td>Intergenic</td><td>NA</td><td>NA</td></tr>
<tr><td>T23149</td><td>GTACTTCTCGATCATCGAGTCGG</td><td>1000</td><td></td><td></td><td></td><td></td><td></td><td></td><td></td><td></td><td></td></tr>
<tr><td>Oligo fwd</td><td>CACCGTACTTCTCGATCATCGAGT</td><td></td><td></td><td></td><td></td><td></td><td></td><td></td><td></td><td></td><td></td></tr>
<tr><td>Oligo rev</td><td>AAACACTCGATGATCGAGAAGTAC</td><td></td><td></td><td></td><td></td><td></td><td></td><td></td><td></td><td></td><td></td></tr>
<tr><td>Chromosome</td><td>start</td><td>end</td><td>strand</td><td>MM</td><td>target_seq</td><td>PAM</td><td>alignment</td><td>distance</td><td>position</td><td>gene name</td><td>gene id</td></tr>
<tr><td>GU305914.1</td><td>144665</td><td>144687</td><td>-</td><td>0</td><td>GTACTTCTCGATCATCGAGT</td><td>CGG</td><td>||||||||[||||||||||||]PAM</td><td>NA</td><td>Intergenic</td><td>NA</td><td>NA</td></tr>
<tr><td>T23150</td><td>GGTGCTGTCGCGCTACATGAAGG</td><td>1000</td><td></td><td></td><td></td><td></td><td></td><td></td><td></td><td></td><td></td></tr>
<tr><td>Oligo fwd</td><td>CACCGGTGCTGTCGCGCTACATGA</td><td></td><td></td><td></td><td></td><td></td><td></td><td></td><td></td><td></td><td></td></tr>
<tr><td>Oligo rev</td><td>AAACTCATGTAGCGCGACAGCACC</td><td></td><td></td><td></td><td></td><td></td><td></td><td></td><td></td><td></td><td></td></tr>
<tr><td>Chromosome</td><td>start</td><td>end</td><td>strand</td><td>MM</td><td>target_seq</td><td>PAM</td><td>alignment</td><td>distance</td><td>position</td><td>gene name</td><td>gene id</td></tr>
<tr><td>GU305914.1</td><td>144644</td><td>144666</td><td>-</td><td>0</td><td>GGTGCTGTCGCGCTACATGA</td><td>AGG</td><td>||||||||[||||||||||||]PAM</td><td>NA</td><td>Intergenic</td><td>NA</td><td>NA</td></tr>
<tr><td>T23151</td><td>CGCTACATGAAGGTGCAGTTCGG</td><td>1000</td><td></td><td></td><td></td><td></td><td></td><td></td><td></td><td></td><td></td></tr>
<tr><td>Oligo fwd</td><td>CACCCGCTACATGAAGGTGCAGTT</td><td></td><td></td><td></td><td></td><td></td><td></td><td></td><td></td><td></td><td></td></tr>
<tr><td>Oligo rev</td><td>AAACAACTGCACCTTCATGTAGCG</td><td></td><td></td><td></td><td></td><td></td><td></td><td></td><td></td><td></td><td></td></tr>
<tr><td>Chromosome</td><td>start</td><td>end</td><td>strand</td><td>MM</td><td>target_seq</td><td>PAM</td><td>alignment</td><td>distance</td><td>position</td><td>gene name</td><td>gene id</td></tr>
<tr><td>GU305914.1</td><td>144634</td><td>144656</td><td>-</td><td>0</td><td>CGCTACATGAAGGTGCAGTT</td><td>CGG</td><td>||||||||[||||||||||||]PAM</td><td>NA</td><td>Intergenic</td><td>NA</td><td>NA</td></tr>
<tr><td>T23152</td><td>GTGCAGTTCGGCTACCACATCGG</td><td>1000</td><td></td><td></td><td></td><td></td><td></td><td></td><td></td><td></td><td></td></tr>
<tr><td>Oligo fwd</td><td>CACCGTGCAGTTCGGCTACCACAT</td><td></td><td></td><td></td><td></td><td></td><td></td><td></td><td></td><td></td><td></td></tr>
<tr><td>Oligo rev</td><td>AAACATGTGGTAGCCGAACTGCAC</td><td></td><td></td><td></td><td></td><td></td><td></td><td></td><td></td><td></td><td></td></tr>
<tr><td>Chromosome</td><td>start</td><td>end</td><td>strand</td><td>MM</td><td>target_seq</td><td>PAM</td><td>alignment</td><td>distance</td><td>position</td><td>gene name</td><td>gene id</td></tr>
<tr><td>GU305914.1</td><td>144622</td><td>144644</td><td>-</td><td>0</td><td>GTGCAGTTCGGCTACCACAT</td><td>CGG</td><td>||||||||[||||||||||||]PAM</td><td>NA</td><td>Intergenic</td><td>NA</td><td>NA</td></tr>
<tr><td>T23153</td><td>TGCAGTTCGGCTACCACATCGGG</td><td>1000</td><td></td><td></td><td></td><td></td><td></td><td></td><td></td><td></td><td></td></tr>
<tr><td>Oligo fwd</td><td>CACCTGCAGTTCGGCTACCACATC</td><td></td><td></td><td></td><td></td><td></td><td></td><td></td><td></td><td></td><td></td></tr>
<tr><td>Oligo rev</td><td>AAACGATGTGGTAGCCGAACTGCA</td><td></td><td></td><td></td><td></td><td></td><td></td><td></td><td></td><td></td><td></td></tr>
<tr><td>Chromosome</td><td>start</td><td>end</td><td>strand</td><td>MM</td><td>target_seq</td><td>PAM</td><td>alignment</td><td>distance</td><td>position</td><td>gene name</td><td>gene id</td></tr>
<tr><td>GU305914.1</td><td>144621</td><td>144643</td><td>-</td><td>0</td><td>TGCAGTTCGGCTACCACATC</td><td>GGG</td><td>||||||||[||||||||||||]PAM</td><td>NA</td><td>Intergenic</td><td>NA</td><td>NA</td></tr>
<tr><td>T23154</td><td>CTACCACATCGGGACGATACTGG</td><td>1000</td><td></td><td></td><td></td><td></td><td></td><td></td><td></td><td></td><td></td></tr>
<tr><td>Oligo fwd</td><td>CACCCTACCACATCGGGACGATAC</td><td></td><td></td><td></td><td></td><td></td><td></td><td></td><td></td><td></td><td></td></tr>
<tr><td>Oligo rev</td><td>AAACGTATCGTCCCGATGTGGTAG</td><td></td><td></td><td></td><td></td><td></td><td></td><td></td><td></td><td></td><td></td></tr>
<tr><td>Chromosome</td><td>start</td><td>end</td><td>strand</td><td>MM</td><td>target_seq</td><td>PAM</td><td>alignment</td><td>distance</td><td>position</td><td>gene name</td><td>gene id</td></tr>
<tr><td>GU305914.1</td><td>144611</td><td>144633</td><td>-</td><td>0</td><td>CTACCACATCGGGACGATAC</td><td>TGG</td><td>||||||||[||||||||||||]PAM</td><td>NA</td><td>Intergenic</td><td>NA</td><td>NA</td></tr>
<tr><td>T23155</td><td>TACCACATCGGGACGATACTGGG</td><td>1000</td><td></td><td></td><td></td><td></td><td></td><td></td><td></td><td></td><td></td></tr>
<tr><td>Oligo fwd</td><td>CACCTACCACATCGGGACGATACT</td><td></td><td></td><td></td><td></td><td></td><td></td><td></td><td></td><td></td><td></td></tr>
<tr><td>Oligo rev</td><td>AAACAGTATCGTCCCGATGTGGTA</td><td></td><td></td><td></td><td></td><td></td><td></td><td></td><td></td><td></td><td></td></tr>
<tr><td>Chromosome</td><td>start</td><td>end</td><td>strand</td><td>MM</td><td>target_seq</td><td>PAM</td><td>alignment</td><td>distance</td><td>position</td><td>gene name</td><td>gene id</td></tr>
<tr><td>GU305914.1</td><td>144610</td><td>144632</td><td>-</td><td>0</td><td>TACCACATCGGGACGATACT</td><td>GGG</td><td>||||||||[||||||||||||]PAM</td><td>NA</td><td>Intergenic</td><td>NA</td><td>NA</td></tr>
<tr><td>T23156</td><td>GGGACGATACTGGGCGTCTGCGG</td><td>1000</td><td></td><td></td><td></td><td></td><td></td><td></td><td></td><td></td><td></td></tr>
<tr><td>Oligo fwd</td><td>CACCGGGACGATACTGGGCGTCTG</td><td></td><td></td><td></td><td></td><td></td><td></td><td></td><td></td><td></td><td></td></tr>
<tr><td>Oligo rev</td><td>AAACCAGACGCCCAGTATCGTCCC</td><td></td><td></td><td></td><td></td><td></td><td></td><td></td><td></td><td></td><td></td></tr>
<tr><td>Chromosome</td><td>start</td><td>end</td><td>strand</td><td>MM</td><td>target_seq</td><td>PAM</td><td>alignment</td><td>distance</td><td>position</td><td>gene name</td><td>gene id</td></tr>
<tr><td>GU305914.1</td><td>144601</td><td>144623</td><td>-</td><td>0</td><td>GGGACGATACTGGGCGTCTG</td><td>CGG</td><td>||||||||[||||||||||||]PAM</td><td>NA</td><td>Intergenic</td><td>NA</td><td>NA</td></tr>
<tr><td>T23157</td><td>GTACCCCATCATCCGCTACGAGG</td><td>1000</td><td></td><td></td><td></td><td></td><td></td><td></td><td></td><td></td><td></td></tr>
<tr><td>Oligo fwd</td><td>CACCGTACCCCATCATCCGCTACG</td><td></td><td></td><td></td><td></td><td></td><td></td><td></td><td></td><td></td><td></td></tr>
<tr><td>Oligo rev</td><td>AAACCGTAGCGGATGATGGGGTAC</td><td></td><td></td><td></td><td></td><td></td><td></td><td></td><td></td><td></td><td></td></tr>
<tr><td>Chromosome</td><td>start</td><td>end</td><td>strand</td><td>MM</td><td>target_seq</td><td>PAM</td><td>alignment</td><td>distance</td><td>position</td><td>gene name</td><td>gene id</td></tr>
<tr><td>GU305914.1</td><td>144572</td><td>144594</td><td>-</td><td>0</td><td>GTACCCCATCATCCGCTACG</td><td>AGG</td><td>||||||||[||||||||||||]PAM</td><td>NA</td><td>Intergenic</td><td>NA</td><td>NA</td></tr>
<tr><td>T23158</td><td>TGAACGCCTCGAGCTACGCGAGG</td><td>1000</td><td></td><td></td><td></td><td></td><td></td><td></td><td></td><td></td><td></td></tr>
<tr><td>Oligo fwd</td><td>CACCTGAACGCCTCGAGCTACGCG</td><td></td><td></td><td></td><td></td><td></td><td></td><td></td><td></td><td></td><td></td></tr>
<tr><td>Oligo rev</td><td>AAACCGCGTAGCTCGAGGCGTTCA</td><td></td><td></td><td></td><td></td><td></td><td></td><td></td><td></td><td></td><td></td></tr>
<tr><td>Chromosome</td><td>start</td><td>end</td><td>strand</td><td>MM</td><td>target_seq</td><td>PAM</td><td>alignment</td><td>distance</td><td>position</td><td>gene name</td><td>gene id</td></tr>
<tr><td>GU305914.1</td><td>144546</td><td>144568</td><td>-</td><td>0</td><td>TGAACGCCTCGAGCTACGCG</td><td>AGG</td><td>||||||||[||||||||||||]PAM</td><td>NA</td><td>Intergenic</td><td>NA</td><td>NA</td></tr>
</table>

T23159 GAACGCCTCGAGCTACGCGAGGG 1000
Oligo fwd CACCGAACGCCTCGAGCTACGCGA
Oligo rev AAACTCGCGTAGCTCGAGGCGTTC
Chromosome start end strand MM target_seq PAM alignment distance position gene name gene id
GU305914.1 144545 144567 - 0 GAACGCCTCGAGCTACGCGA GGG ||||||||[||||||||||||]PAM NA Intergenic NA NA

T23160 TACGCGAGGGACATCAACATCGG 1000
Oligo fwd CACCTACGCGAGGGACATCAACAT
Oligo rev AAACATGTTGATGTCCCTCGCGTA
Chromosome start end strand MM target_seq PAM alignment distance position gene name gene id
GU305914.1 144532 144554 - 0 TACGCGAGGGACATCAACAT CGG ||||||||[||||||||||||]PAM NA Intergenic NA NA

T23161 GGACATCAACATCGGCATCACGG 1000
Oligo fwd CACCGGACATCAACATCGGCATCA
Oligo rev AAACTGATGCCGATGTTGATGTCC
Chromosome start end strand MM target_seq PAM alignment distance position gene name gene id
GU305914.1 144524 144546 - 0 GGACATCAACATCGGCATCA CGG ||||||||[||||||||||||]PAM NA Intergenic NA NA

T23162 GGTGCTGCTACTGCTCTGCGTGG 1000
Oligo fwd CACCGGTGCTGCTACTGCTCTGCG
Oligo rev AAACCGCAGAGCAGTAGCAGCACC
Chromosome start end strand MM target_seq PAM alignment distance position gene name gene id
GU305914.1 144503 144525 - 0 GGTGCTGCTACTGCTCTGCG TGG ||||||||[||||||||||||]PAM NA Intergenic NA NA

T23163 ACTGCTCTGCGTGGCGTTCTCGG 1000
Oligo fwd CACCACTGCTCTGCGTGGCGTTCT
Oligo rev AAACAGAACGCCACGCAGAGCAGT
Chromosome start end strand MM target_seq PAM alignment distance position gene name gene id
GU305914.1 144494 144516 - 0 ACTGCTCTGCGTGGCGTTCT CGG ||||||||[||||||||||||]PAM NA Intergenic NA NA

T23164 GGCGTTCTCGGTCATCCGTACGG 1000
Oligo fwd CACCGGCGTTCTCGGTCATCCGTA
Oligo rev AAACTACGGATGACCGAGAACGCC
Chromosome start end strand MM target_seq PAM alignment distance position gene name gene id
GU305914.1 144482 144504 - 0 GGCGTTCTCGGTCATCCGTA CGG ||||||||[||||||||||||]PAM NA Intergenic NA NA

T23165 TCTCGGTCATCCGTACGGTGAGG 1000
Oligo fwd CACCTCTCGGTCATCCGTACGGTG
Oligo rev AAACCACCGTACGGATGACCGAGA
Chromosome start end strand MM target_seq PAM alignment distance position gene name gene id
GU305914.1 144477 144499 - 0 TCTCGGTCATCCGTACGGTG AGG ||||||||[||||||||||||]PAM NA Intergenic NA NA

T23166 CAACAAGCGATATCGAGCCCTGG 1000
Oligo fwd CACCCAACAAGCGATATCGAGCCC
Oligo rev AAACGGGCTCGATATCGCTTGTTG
Chromosome start end strand MM target_seq PAM alignment distance position gene name gene id
GU305914.1 144440 144462 - 0 CAACAAGCGATATCGAGCCC TGG ||||||||[||||||||||||]PAM NA Intergenic NA NA

T23167 GCGATATCGAGCCCTGGCCCTGG 1000
Oligo fwd CACCGCGATATCGAGCCCTGGCCC
Oligo rev AAACGGGCCAGGGCTCGATATCGC
Chromosome start end strand MM target_seq PAM alignment distance position gene name gene id
GU305914.1 144434 144456 - 0 GCGATATCGAGCCCTGGCCC TGG ||||||||[||||||||||||]PAM NA Intergenic NA NA

T23168 CTCCCTTTCTCCCCCCCTCACGG 1000
Oligo fwd CACCCTCCCTTTCTCCCCCCCTCA
Oligo rev AAACTGAGGGGGGGAGAAAGGGAG
Chromosome start end strand MM target_seq PAM alignment distance position gene name gene id
GU305914.1 144177 144199 - 0 CTCCCTTTCTCCCCCCCTCA CGG ||||||||[||||||||||||]PAM NA Intergenic NA NA

T23169 ATCGATAGATAGATATCACAAGG 1000
Oligo fwd CACCATCGATAGATAGATATCACA
Oligo rev AAACTGTGATATCTATCTATCGAT
Chromosome start end strand MM target_seq PAM alignment distance position gene name gene id
GU305914.1 144151 144173 - 0 ATCGATAGATAGATATCACA AGG ||||||||[||||||||||||]PAM NA Intergenic NA NA

T23170 CCTGACTTTTTTCTTCACGCCGG 1000
Oligo fwd CACCCCTGACTTTTTTCTTCACGC
Oligo rev AAACGCGTGAAGAAAAAAGTCAGG
Chromosome start end strand MM target_seq PAM alignment distance position gene name gene id
GU305914.1 144127 144149 - 0 CCTGACTTTTTTCTTCACGC CGG ||||||||[||||||||||||]PAM NA Intergenic NA NA

T23171 CTTCACGCCGGCGCCGTTCTTGG 1000
Oligo fwd CACCCTTCACGCCGGCGCCGTTCT
Oligo rev AAACAGAACGGCGCCGGCGTGAAG
Chromosome start end strand MM target_seq PAM alignment distance position gene name gene id
GU305914.1 144115 144137 - 0 CTTCACGCCGGCGCCGTTCT TGG ||||||||[||||||||||||]PAM NA Intergenic NA NA

T23172 CACGCCGGCGCCGTTCTTGGCGG 1000
Oligo fwd CACCCACGCCGGCGCCGTTCTTGG
Oligo rev AAACCCAAGAACGGCGCCGGCGTG
Chromosome start end strand MM target_seq PAM alignment distance position gene name gene id
GU305914.1 144112 144134 - 0 CACGCCGGCGCCGTTCTTGG CGG ||||||||[||||||||||||]PAM NA Intergenic NA NA

T23173 TTGTTCGCCCCGAGCTGTATCGG 1000
Oligo fwd CACCTTGTTCGCCCCGAGCTGTAT
Oligo rev AAACATACAGCTCGGGGCGAACAA
Chromosome start end strand MM target_seq PAM alignment distance position gene name gene id
GU305914.1 144057 144079 - 0 TTGTTCGCCCCGAGCTGTAT CGG ||||||||[||||||||||||]PAM NA Intergenic NA NA

| | | | | | | | | | | | |
|---|---|---|---|---|---|---|---|---|---|---|---|
| T23174 | GAGCTGTATCGGAGTGACCACGG | 1000 | | | | | | | | | |
| Oligo fwd | CACCGAGCTGTATCGGAGTGACCA | | | | | | | | | | |
| Oligo rev | AAACTGGTCACTCCGATACAGCTC | | | | | | | | | | |
| Chromosome | start | end | strand | MM | target_seq | PAM | alignment | distance | position | gene name | gene id |
| GU305914.1 | 144046 | 144068 | - | 0 | GAGCTGTATCGGAGTGACCA | CGG | \|\|\|\|\|\|\|\|[\|\|\|\|\|\|\|\|\|\|\|\|]PAM | NA | Intergenic | NA | NA |
| T23175 | GTATCGGAGTGACCACGGTCAGG | 1000 | | | | | | | | | |
| Oligo fwd | CACCGTATCGGAGTGACCACGGTC | | | | | | | | | | |
| Oligo rev | AAACGACCGTGGTCACTCCGATAC | | | | | | | | | | |
| Chromosome | start | end | strand | MM | target_seq | PAM | alignment | distance | position | gene name | gene id |
| GU305914.1 | 144041 | 144063 | - | 0 | GTATCGGAGTGACCACGGTC | AGG | \|\|\|\|\|\|\|\|[\|\|\|\|\|\|\|\|\|\|\|\|]PAM | NA | Intergenic | NA | NA |
| T23176 | TATCGGAGTGACCACGGTCAGGG | 1000 | | | | | | | | | |
| Oligo fwd | CACCTATCGGAGTGACCACGGTCA | | | | | | | | | | |
| Oligo rev | AAACTGACCGTGGTCACTCCGATA | | | | | | | | | | |
| Chromosome | start | end | strand | MM | target_seq | PAM | alignment | distance | position | gene name | gene id |
| GU305914.1 | 144040 | 144062 | - | 0 | TATCGGAGTGACCACGGTCA | GGG | \|\|\|\|\|\|\|\|[\|\|\|\|\|\|\|\|\|\|\|\|]PAM | NA | Intergenic | NA | NA |
| T23177 | ATCGGAGTGACCACGGTCAGGGG | 1000 | | | | | | | | | |
| Oligo fwd | CACCATCGGAGTGACCACGGTCAG | | | | | | | | | | |
| Oligo rev | AAACCTGACCGTGGTCACTCCGAT | | | | | | | | | | |
| Chromosome | start | end | strand | MM | target_seq | PAM | alignment | distance | position | gene name | gene id |
| GU305914.1 | 144039 | 144061 | - | 0 | ATCGGAGTGACCACGGTCAG | GGG | \|\|\|\|\|\|\|\|[\|\|\|\|\|\|\|\|\|\|\|\|]PAM | NA | Intergenic | NA | NA |
| T23178 | GGGTATAGCGCTATTCTTGTCGG | 1000 | | | | | | | | | |
| Oligo fwd | CACCGGGTATAGCGCTATTCTTGT | | | | | | | | | | |
| Oligo rev | AAACACAAGAATAGCGCTATACCC | | | | | | | | | | |
| Chromosome | start | end | strand | MM | target_seq | PAM | alignment | distance | position | gene name | gene id |
| GU305914.1 | 144019 | 144041 | - | 0 | GGGTATAGCGCTATTCTTGT | CGG | \|\|\|\|\|\|\|\|[\|\|\|\|\|\|\|\|\|\|\|\|]PAM | NA | Intergenic | NA | NA |
| T23179 | GCTATTCTTGTCGGAAAGTAAGG | 1000 | | | | | | | | | |
| Oligo fwd | CACCGCTATTCTTGTCGGAAAGTA | | | | | | | | | | |
| Oligo rev | AAACTACTTTCCGACAAGAATAGC | | | | | | | | | | |
| Chromosome | start | end | strand | MM | target_seq | PAM | alignment | distance | position | gene name | gene id |
| GU305914.1 | 144010 | 144032 | - | 0 | GCTATTCTTGTCGGAAAGTA | AGG | \|\|\|\|\|\|\|\|[\|\|\|\|\|\|\|\|\|\|\|\|]PAM | NA | Intergenic | NA | NA |
| T23180 | CTATTCTTGTCGGAAAGTAAGGG | 1000 | | | | | | | | | |
| Oligo fwd | CACCCTATTCTTGTCGGAAAGTAA | | | | | | | | | | |
| Oligo rev | AAACTTACTTTCCGACAAGAATAG | | | | | | | | | | |
| Chromosome | start | end | strand | MM | target_seq | PAM | alignment | distance | position | gene name | gene id |
| GU305914.1 | 144009 | 144031 | - | 0 | CTATTCTTGTCGGAAAGTAA | GGG | \|\|\|\|\|\|\|\|[\|\|\|\|\|\|\|\|\|\|\|\|]PAM | NA | Intergenic | NA | NA |
| T23181 | TATTCTTGTCGGAAAGTAAGGGG | 1000 | | | | | | | | | |
| Oligo fwd | CACCTATTCTTGTCGGAAAGTAAG | | | | | | | | | | |
| Oligo rev | AAACCTTACTTTCCGACAAGAATA | | | | | | | | | | |
| Chromosome | start | end | strand | MM | target_seq | PAM | alignment | distance | position | gene name | gene id |
| GU305914.1 | 144008 | 144030 | - | 0 | TATTCTTGTCGGAAAGTAAG | GGG | \|\|\|\|\|\|\|\|[\|\|\|\|\|\|\|\|\|\|\|\|]PAM | NA | Intergenic | NA | NA |
| T23182 | ATTCTTGTCGGAAAGTAAGGGGG | 1000 | | | | | | | | | |
| Oligo fwd | CACCATTCTTGTCGGAAAGTAAGG | | | | | | | | | | |
| Oligo rev | AAACCCTTACTTTCCGACAAGAAT | | | | | | | | | | |
| Chromosome | start | end | strand | MM | target_seq | PAM | alignment | distance | position | gene name | gene id |
| GU305914.1 | 144007 | 144029 | - | 0 | ATTCTTGTCGGAAAGTAAGG | GGG | \|\|\|\|\|\|\|\|[\|\|\|\|\|\|\|\|\|\|\|\|]PAM | NA | Intergenic | NA | NA |
| T23183 | CACGAAGTCGTCTCGCTCGCCGG | 1000 | | | | | | | | | |
| Oligo fwd | CACCCACGAAGTCGTCTCGCTCGC | | | | | | | | | | |
| Oligo rev | AAACGCGAGCGAGACGACTTCGTG | | | | | | | | | | |
| Chromosome | start | end | strand | MM | target_seq | PAM | alignment | distance | position | gene name | gene id |
| GU305914.1 | 143968 | 143990 | - | 0 | CACGAAGTCGTCTCGCTCGC | CGG | \|\|\|\|\|\|\|\|[\|\|\|\|\|\|\|\|\|\|\|\|]PAM | NA | Intergenic | NA | NA |
| T23184 | CTCGCTCGCCGGCGCCGACCTGG | 1000 | | | | | | | | | |
| Oligo fwd | CACCCTCGCTCGCCGGCGCCGACC | | | | | | | | | | |
| Oligo rev | AAACGGTCGGCGCCGGCGAGCGAG | | | | | | | | | | |
| Chromosome | start | end | strand | MM | target_seq | PAM | alignment | distance | position | gene name | gene id |
| GU305914.1 | 143957 | 143979 | - | 0 | CTCGCTCGCCGGCGCCGACC | TGG | \|\|\|\|\|\|\|\|[\|\|\|\|\|\|\|\|\|\|\|\|]PAM | NA | Intergenic | NA | NA |
| T23185 | ATCCGTCAGCACAGAGAATTCGG | 1000 | | | | | | | | | |
| Oligo fwd | CACCATCCGTCAGCACAGAGAATT | | | | | | | | | | |
| Oligo rev | AAACAATTCTCTGTGCTGACGGAT | | | | | | | | | | |
| Chromosome | start | end | strand | MM | target_seq | PAM | alignment | distance | position | gene name | gene id |
| GU305914.1 | 143917 | 143939 | - | 0 | ATCCGTCAGCACAGAGAATT | CGG | \|\|\|\|\|\|\|\|[\|\|\|\|\|\|\|\|\|\|\|\|]PAM | NA | Intergenic | NA | NA |
| T23186 | GCACAGAGAATTCGGTCCCGAGG | 1000 | | | | | | | | | |
| Oligo fwd | CACCGCACAGAGAATTCGGTCCCG | | | | | | | | | | |
| Oligo rev | AAACCGGGACCGAATTCTCTGTGC | | | | | | | | | | |
| Chromosome | start | end | strand | MM | target_seq | PAM | alignment | distance | position | gene name | gene id |
| GU305914.1 | 143909 | 143931 | - | 0 | GCACAGAGAATTCGGTCCCG | AGG | \|\|\|\|\|\|\|\|[\|\|\|\|\|\|\|\|\|\|\|\|]PAM | NA | Intergenic | NA | NA |
| T23187 | GAGGTCCACGCGTCGCCCCGAGG | 1000 | | | | | | | | | |
| Oligo fwd | CACCGAGGTCCACGCGTCGCCCCG | | | | | | | | | | |
| Oligo rev | AAACCGGGGCGACGCGTGGACCTC | | | | | | | | | | |
| Chromosome | start | end | strand | MM | target_seq | PAM | alignment | distance | position | gene name | gene id |
| GU305914.1 | 143890 | 143912 | - | 0 | GAGGTCCACGCGTCGCCCCG | AGG | \|\|\|\|\|\|\|\|[\|\|\|\|\|\|\|\|\|\|\|\|]PAM | NA | Intergenic | NA | NA |
| T23188 | GTCGCCCCGAGGAATCCTTCAGG | 1000 | | | | | | | | | |
| Oligo fwd | CACCGTCGCCCCGAGGAATCCTTC | | | | | | | | | | |
| Oligo rev | AAACGAAGGATTCCTCGGGGCGAC | | | | | | | | | | |
| Chromosome | start | end | strand | MM | target_seq | PAM | alignment | distance | position | gene name | gene id |

<table>
<tr><td>GU305914.1</td><td>143879</td><td>143901</td><td>-</td><td>0</td><td>GTCGCCCCGAGGAATCCTTC</td><td>AGG</td><td>||||||||[||||||||||||]PAM</td><td>NA</td><td>Intergenic</td><td>NA</td><td>NA</td></tr>
<tr><td>T23189</td><td>TCGCCCCGAGGAATCCTTCAGGG</td><td>1000</td></tr>
<tr><td>Oligo fwd</td><td>CACCTCGCCCCGAGGAATCCTTCA</td></tr>
<tr><td>Oligo rev</td><td>AAACTGAAGGATTCCTCGGGGCGA</td></tr>
<tr><td>Chromosome</td><td>start</td><td>end</td><td>strand</td><td>MM</td><td>target_seq</td><td>PAM</td><td>alignment</td><td>distance</td><td>position</td><td>gene name</td><td>gene id</td></tr>
<tr><td>GU305914.1</td><td>143878</td><td>143900</td><td>-</td><td>0</td><td>TCGCCCCGAGGAATCCTTCA</td><td>GGG</td><td>||||||||[||||||||||||]PAM</td><td>NA</td><td>Intergenic</td><td>NA</td><td>NA</td></tr>
<tr><td>T23190</td><td>GGAATCCTTCAGGGAGTCCGCGG</td><td>1000</td></tr>
<tr><td>Oligo fwd</td><td>CACCGGAATCCTTCAGGGAGTCCG</td></tr>
<tr><td>Oligo rev</td><td>AAACCGGACTCCCTGAAGGATTCC</td></tr>
<tr><td>Chromosome</td><td>start</td><td>end</td><td>strand</td><td>MM</td><td>target_seq</td><td>PAM</td><td>alignment</td><td>distance</td><td>position</td><td>gene name</td><td>gene id</td></tr>
<tr><td>GU305914.1</td><td>143869</td><td>143891</td><td>-</td><td>0</td><td>GGAATCCTTCAGGGAGTCCG</td><td>CGG</td><td>||||||||[||||||||||||]PAM</td><td>NA</td><td>Intergenic</td><td>NA</td><td>NA</td></tr>
<tr><td>T23191</td><td>CTTCAGGGAGTCCGCGGCGTAGG</td><td>1000</td></tr>
<tr><td>Oligo fwd</td><td>CACCCTTCAGGGAGTCCGCGGCGT</td></tr>
<tr><td>Oligo rev</td><td>AAACACGCCGCGGACTCCCTGAAG</td></tr>
<tr><td>Chromosome</td><td>start</td><td>end</td><td>strand</td><td>MM</td><td>target_seq</td><td>PAM</td><td>alignment</td><td>distance</td><td>position</td><td>gene name</td><td>gene id</td></tr>
<tr><td>GU305914.1</td><td>143863</td><td>143885</td><td>-</td><td>0</td><td>CTTCAGGGAGTCCGCGGCGT</td><td>AGG</td><td>||||||||[||||||||||||]PAM</td><td>NA</td><td>Intergenic</td><td>NA</td><td>NA</td></tr>
<tr><td>T23192</td><td>TTCAGGGAGTCCGCGGCGTAGGG</td><td>1000</td></tr>
<tr><td>Oligo fwd</td><td>CACCTTCAGGGAGTCCGCGGCGTA</td></tr>
<tr><td>Oligo rev</td><td>AAACTACGCCGCGGACTCCCTGAA</td></tr>
<tr><td>Chromosome</td><td>start</td><td>end</td><td>strand</td><td>MM</td><td>target_seq</td><td>PAM</td><td>alignment</td><td>distance</td><td>position</td><td>gene name</td><td>gene id</td></tr>
<tr><td>GU305914.1</td><td>143862</td><td>143884</td><td>-</td><td>0</td><td>TTCAGGGAGTCCGCGGCGTA</td><td>GGG</td><td>||||||||[||||||||||||]PAM</td><td>NA</td><td>Intergenic</td><td>NA</td><td>NA</td></tr>
<tr><td>T23193</td><td>AGTCCGCGGCGTAGGGATGCAGG</td><td>1000</td></tr>
<tr><td>Oligo fwd</td><td>CACCAGTCCGCGGCGTAGGGATGC</td></tr>
<tr><td>Oligo rev</td><td>AAACGCATCCCTACGCCGCGGACT</td></tr>
<tr><td>Chromosome</td><td>start</td><td>end</td><td>strand</td><td>MM</td><td>target_seq</td><td>PAM</td><td>alignment</td><td>distance</td><td>position</td><td>gene name</td><td>gene id</td></tr>
<tr><td>GU305914.1</td><td>143855</td><td>143877</td><td>-</td><td>0</td><td>AGTCCGCGGCGTAGGGATGC</td><td>AGG</td><td>||||||||[||||||||||||]PAM</td><td>NA</td><td>Intergenic</td><td>NA</td><td>NA</td></tr>
<tr><td>T23194</td><td>TGTCCCCCTCCCAGACGTCGAGG</td><td>1000</td></tr>
<tr><td>Oligo fwd</td><td>CACCTGTCCCCCTCCCAGACGTCG</td></tr>
<tr><td>Oligo rev</td><td>AAACCGACGTCTGGGAGGGGGACA</td></tr>
<tr><td>Chromosome</td><td>start</td><td>end</td><td>strand</td><td>MM</td><td>target_seq</td><td>PAM</td><td>alignment</td><td>distance</td><td>position</td><td>gene name</td><td>gene id</td></tr>
<tr><td>GU305914.1</td><td>143772</td><td>143794</td><td>-</td><td>0</td><td>TGTCCCCCTCCCAGACGTCG</td><td>AGG</td><td>||||||||[||||||||||||]PAM</td><td>NA</td><td>Intergenic</td><td>NA</td><td>NA</td></tr>
<tr><td>T23195</td><td>GGACGCTGCTCACAGCATCCCGG</td><td>1000</td></tr>
<tr><td>Oligo fwd</td><td>CACCGGACGCTGCTCACAGCATCC</td></tr>
<tr><td>Oligo rev</td><td>AAACGGATGCTGTGAGCAGCGTCC</td></tr>
<tr><td>Chromosome</td><td>start</td><td>end</td><td>strand</td><td>MM</td><td>target_seq</td><td>PAM</td><td>alignment</td><td>distance</td><td>position</td><td>gene name</td><td>gene id</td></tr>
<tr><td>GU305914.1</td><td>143751</td><td>143773</td><td>-</td><td>0</td><td>GGACGCTGCTCACAGCATCC</td><td>CGG</td><td>||||||||[||||||||||||]PAM</td><td>NA</td><td>Intergenic</td><td>NA</td><td>NA</td></tr>
<tr><td>T23196</td><td>GACGCTGCTCACAGCATCCCGGG</td><td>1000</td></tr>
<tr><td>Oligo fwd</td><td>CACCGACGCTGCTCACAGCATCCC</td></tr>
<tr><td>Oligo rev</td><td>AAACGGGATGCTGTGAGCAGCGTC</td></tr>
<tr><td>Chromosome</td><td>start</td><td>end</td><td>strand</td><td>MM</td><td>target_seq</td><td>PAM</td><td>alignment</td><td>distance</td><td>position</td><td>gene name</td><td>gene id</td></tr>
<tr><td>GU305914.1</td><td>143750</td><td>143772</td><td>-</td><td>0</td><td>GACGCTGCTCACAGCATCCC</td><td>GGG</td><td>||||||||[||||||||||||]PAM</td><td>NA</td><td>Intergenic</td><td>NA</td><td>NA</td></tr>
<tr><td>T23197</td><td>TCACAGCATCCCGGGCGAAGTGG</td><td>1000</td></tr>
<tr><td>Oligo fwd</td><td>CACCTCACAGCATCCCGGGCGAAG</td></tr>
<tr><td>Oligo rev</td><td>AAACCTTCGCCCGGGATGCTGTGA</td></tr>
<tr><td>Chromosome</td><td>start</td><td>end</td><td>strand</td><td>MM</td><td>target_seq</td><td>PAM</td><td>alignment</td><td>distance</td><td>position</td><td>gene name</td><td>gene id</td></tr>
<tr><td>GU305914.1</td><td>143742</td><td>143764</td><td>-</td><td>0</td><td>TCACAGCATCCCGGGCGAAG</td><td>TGG</td><td>||||||||[||||||||||||]PAM</td><td>NA</td><td>Intergenic</td><td>NA</td><td>NA</td></tr>
<tr><td>T23198</td><td>CACAGCATCCCGGGCGAAGTGGG</td><td>1000</td></tr>
<tr><td>Oligo fwd</td><td>CACCCACAGCATCCCGGGCGAAGT</td></tr>
<tr><td>Oligo rev</td><td>AAACACTTCGCCCGGGATGCTGTG</td></tr>
<tr><td>Chromosome</td><td>start</td><td>end</td><td>strand</td><td>MM</td><td>target_seq</td><td>PAM</td><td>alignment</td><td>distance</td><td>position</td><td>gene name</td><td>gene id</td></tr>
<tr><td>GU305914.1</td><td>143741</td><td>143763</td><td>-</td><td>0</td><td>CACAGCATCCCGGGCGAAGT</td><td>GGG</td><td>||||||||[||||||||||||]PAM</td><td>NA</td><td>Intergenic</td><td>NA</td><td>NA</td></tr>
<tr><td>T23199</td><td>ACAGCATCCCGGGCGAAGTGGGG</td><td>1000</td></tr>
<tr><td>Oligo fwd</td><td>CACCACAGCATCCCGGGCGAAGTG</td></tr>
<tr><td>Oligo rev</td><td>AAACCACTTCGCCCGGGATGCTGT</td></tr>
<tr><td>Chromosome</td><td>start</td><td>end</td><td>strand</td><td>MM</td><td>target_seq</td><td>PAM</td><td>alignment</td><td>distance</td><td>position</td><td>gene name</td><td>gene id</td></tr>
<tr><td>GU305914.1</td><td>143740</td><td>143762</td><td>-</td><td>0</td><td>ACAGCATCCCGGGCGAAGTG</td><td>GGG</td><td>||||||||[||||||||||||]PAM</td><td>NA</td><td>Intergenic</td><td>NA</td><td>NA</td></tr>
<tr><td>T23200</td><td>GAAGTGGGGATCCAGACGCACGG</td><td>1000</td></tr>
<tr><td>Oligo fwd</td><td>CACCGAAGTGGGGATCCAGACGCA</td></tr>
<tr><td>Oligo rev</td><td>AAACTGCGTCTGGATCCCCACTTC</td></tr>
<tr><td>Chromosome</td><td>start</td><td>end</td><td>strand</td><td>MM</td><td>target_seq</td><td>PAM</td><td>alignment</td><td>distance</td><td>position</td><td>gene name</td><td>gene id</td></tr>
<tr><td>GU305914.1</td><td>143726</td><td>143748</td><td>-</td><td>0</td><td>GAAGTGGGGATCCAGACGCA</td><td>CGG</td><td>||||||||[||||||||||||]PAM</td><td>NA</td><td>Intergenic</td><td>NA</td><td>NA</td></tr>
<tr><td>T23201</td><td>AAGTGGGGATCCAGACGCACGGG</td><td>1000</td></tr>
<tr><td>Oligo fwd</td><td>CACCAAGTGGGGATCCAGACGCAC</td></tr>
<tr><td>Oligo rev</td><td>AAACGTGCGTCTGGATCCCCACTT</td></tr>
<tr><td>Chromosome</td><td>start</td><td>end</td><td>strand</td><td>MM</td><td>target_seq</td><td>PAM</td><td>alignment</td><td>distance</td><td>position</td><td>gene name</td><td>gene id</td></tr>
<tr><td>GU305914.1</td><td>143725</td><td>143747</td><td>-</td><td>0</td><td>AAGTGGGGATCCAGACGCAC</td><td>GGG</td><td>||||||||[||||||||||||]PAM</td><td>NA</td><td>Intergenic</td><td>NA</td><td>NA</td></tr>
<tr><td>T23202</td><td>GGCGTCACGACGACGCAGAGCGG</td><td>1000</td></tr>
<tr><td>Oligo fwd</td><td>CACCGGCGTCACGACGACGCAGAG</td></tr>
<tr><td>Oligo rev</td><td>AAACCTCTGCGTCGTCGTGACGCC</td></tr>
<tr><td>Chromosome</td><td>start</td><td>end</td><td>strand</td><td>MM</td><td>target_seq</td><td>PAM</td><td>alignment</td><td>distance</td><td>position</td><td>gene name</td><td>gene id</td></tr>
<tr><td>GU305914.1</td><td>143704</td><td>143726</td><td>-</td><td>0</td><td>GGCGTCACGACGACGCAGAG</td><td>CGG</td><td>||||||||[||||||||||||]PAM</td><td>NA</td><td>Intergenic</td><td>NA</td><td>NA</td></tr>
<tr><td>T23203</td><td>GAGCGGCACCTCGTCGCAGTCGG</td><td>1000</td></tr>
<tr><td>Oligo fwd</td><td>CACCGAGCGGCACCTCGTCGCAGT</td></tr>
<tr><td>Oligo rev</td><td>AAACACTGCGACGAGGTGCCGCTC</td></tr>
</table>

<table>
<tr><td>Chromosome</td><td>start</td><td>end</td><td>strand</td><td>MM</td><td>target_seq</td><td>PAM</td><td>alignment</td><td>distance</td><td>position</td><td>gene name</td><td>gene id</td></tr>
<tr><td>GU305914.1</td><td>143687</td><td>143709</td><td>-</td><td>0</td><td>GAGCGGCACCTCGTCGCAGT</td><td>CGG</td><td>||||||||[||||||||||||]PAM</td><td>NA</td><td>Intergenic</td><td>NA</td><td>NA</td></tr>
<tr><td>T23204</td><td>GCACCTCGTCGCAGTCGGAACGG</td><td>1000</td><td></td><td></td><td></td><td></td><td></td><td></td><td></td><td></td><td></td></tr>
<tr><td>Oligo fwd</td><td>CACCGCACCTCGTCGCAGTCGGAA</td><td></td><td></td><td></td><td></td><td></td><td></td><td></td><td></td><td></td><td></td></tr>
<tr><td>Oligo rev</td><td>AAACTTCCGACTGCGACGAGGTGC</td><td></td><td></td><td></td><td></td><td></td><td></td><td></td><td></td><td></td><td></td></tr>
<tr><td>Chromosome</td><td>start</td><td>end</td><td>strand</td><td>MM</td><td>target_seq</td><td>PAM</td><td>alignment</td><td>distance</td><td>position</td><td>gene name</td><td>gene id</td></tr>
<tr><td>GU305914.1</td><td>143682</td><td>143704</td><td>-</td><td>0</td><td>GCACCTCGTCGCAGTCGGAA</td><td>CGG</td><td>||||||||[||||||||||||]PAM</td><td>NA</td><td>Intergenic</td><td>NA</td><td>NA</td></tr>
<tr><td>T23205</td><td>CTCGTCCCGTTTTCTCAGTATGG</td><td>1000</td><td></td><td></td><td></td><td></td><td></td><td></td><td></td><td></td><td></td></tr>
<tr><td>Oligo fwd</td><td>CACCCTCGTCCCGTTTTCTCAGTA</td><td></td><td></td><td></td><td></td><td></td><td></td><td></td><td></td><td></td><td></td></tr>
<tr><td>Oligo rev</td><td>AAACTACTGAGAAAACGGGACGAG</td><td></td><td></td><td></td><td></td><td></td><td></td><td></td><td></td><td></td><td></td></tr>
<tr><td>Chromosome</td><td>start</td><td>end</td><td>strand</td><td>MM</td><td>target_seq</td><td>PAM</td><td>alignment</td><td>distance</td><td>position</td><td>gene name</td><td>gene id</td></tr>
<tr><td>GU305914.1</td><td>143624</td><td>143646</td><td>-</td><td>0</td><td>CTCGTCCCGTTTTCTCAGTA</td><td>TGG</td><td>||||||||[||||||||||||]PAM</td><td>NA</td><td>Intergenic</td><td>NA</td><td>NA</td></tr>
<tr><td>T23206</td><td>TTCTCAGTATGGCCGAGACCAGG</td><td>1000</td><td></td><td></td><td></td><td></td><td></td><td></td><td></td><td></td><td></td></tr>
<tr><td>Oligo fwd</td><td>CACCTTCTCAGTATGGCCGAGACC</td><td></td><td></td><td></td><td></td><td></td><td></td><td></td><td></td><td></td><td></td></tr>
<tr><td>Oligo rev</td><td>AAACGGTCTCGGCCATACTGAGAA</td><td></td><td></td><td></td><td></td><td></td><td></td><td></td><td></td><td></td><td></td></tr>
<tr><td>Chromosome</td><td>start</td><td>end</td><td>strand</td><td>MM</td><td>target_seq</td><td>PAM</td><td>alignment</td><td>distance</td><td>position</td><td>gene name</td><td>gene id</td></tr>
<tr><td>GU305914.1</td><td>143613</td><td>143635</td><td>-</td><td>0</td><td>TTCTCAGTATGGCCGAGACC</td><td>AGG</td><td>||||||||[||||||||||||]PAM</td><td>NA</td><td>Intergenic</td><td>NA</td><td>NA</td></tr>
<tr><td>T23207</td><td>TATGGCCGAGACCAGGTACACGG</td><td>1000</td><td></td><td></td><td></td><td></td><td></td><td></td><td></td><td></td><td></td></tr>
<tr><td>Oligo fwd</td><td>CACCTATGGCCGAGACCAGGTACA</td><td></td><td></td><td></td><td></td><td></td><td></td><td></td><td></td><td></td><td></td></tr>
<tr><td>Oligo rev</td><td>AAACTGTACCTGGTCTCGGCCATA</td><td></td><td></td><td></td><td></td><td></td><td></td><td></td><td></td><td></td><td></td></tr>
<tr><td>Chromosome</td><td>start</td><td>end</td><td>strand</td><td>MM</td><td>target_seq</td><td>PAM</td><td>alignment</td><td>distance</td><td>position</td><td>gene name</td><td>gene id</td></tr>
<tr><td>GU305914.1</td><td>143606</td><td>143628</td><td>-</td><td>0</td><td>TATGGCCGAGACCAGGTACA</td><td>CGG</td><td>||||||||[||||||||||||]PAM</td><td>NA</td><td>Intergenic</td><td>NA</td><td>NA</td></tr>
<tr><td>T23208</td><td>GGCCGAGACCAGGTACACGGAGG</td><td>1000</td><td></td><td></td><td></td><td></td><td></td><td></td><td></td><td></td><td></td></tr>
<tr><td>Oligo fwd</td><td>CACCGGCCGAGACCAGGTACACGG</td><td></td><td></td><td></td><td></td><td></td><td></td><td></td><td></td><td></td><td></td></tr>
<tr><td>Oligo rev</td><td>AAACCCGTGTACCTGGTCTCGGCC</td><td></td><td></td><td></td><td></td><td></td><td></td><td></td><td></td><td></td><td></td></tr>
<tr><td>Chromosome</td><td>start</td><td>end</td><td>strand</td><td>MM</td><td>target_seq</td><td>PAM</td><td>alignment</td><td>distance</td><td>position</td><td>gene name</td><td>gene id</td></tr>
<tr><td>GU305914.1</td><td>143603</td><td>143625</td><td>-</td><td>0</td><td>GGCCGAGACCAGGTACACGG</td><td>AGG</td><td>||||||||[||||||||||||]PAM</td><td>NA</td><td>Intergenic</td><td>NA</td><td>NA</td></tr>
<tr><td>T23209</td><td>GCCGAGACCAGGTACACGGAGGG</td><td>1000</td><td></td><td></td><td></td><td></td><td></td><td></td><td></td><td></td><td></td></tr>
<tr><td>Oligo fwd</td><td>CACCGCCGAGACCAGGTACACGGA</td><td></td><td></td><td></td><td></td><td></td><td></td><td></td><td></td><td></td><td></td></tr>
<tr><td>Oligo rev</td><td>AAACTCCGTGTACCTGGTCTCGGC</td><td></td><td></td><td></td><td></td><td></td><td></td><td></td><td></td><td></td><td></td></tr>
<tr><td>Chromosome</td><td>start</td><td>end</td><td>strand</td><td>MM</td><td>target_seq</td><td>PAM</td><td>alignment</td><td>distance</td><td>position</td><td>gene name</td><td>gene id</td></tr>
<tr><td>GU305914.1</td><td>143602</td><td>143624</td><td>-</td><td>0</td><td>GCCGAGACCAGGTACACGGA</td><td>GGG</td><td>||||||||[||||||||||||]PAM</td><td>NA</td><td>Intergenic</td><td>NA</td><td>NA</td></tr>
<tr><td>T23210</td><td>GTACACGGAGGGTAGTTCGTCGG</td><td>1000</td><td></td><td></td><td></td><td></td><td></td><td></td><td></td><td></td><td></td></tr>
<tr><td>Oligo fwd</td><td>CACCGTACACGGAGGGTAGTTCGT</td><td></td><td></td><td></td><td></td><td></td><td></td><td></td><td></td><td></td><td></td></tr>
<tr><td>Oligo rev</td><td>AAACACGAACTACCCTCCGTGTAC</td><td></td><td></td><td></td><td></td><td></td><td></td><td></td><td></td><td></td><td></td></tr>
<tr><td>Chromosome</td><td>start</td><td>end</td><td>strand</td><td>MM</td><td>target_seq</td><td>PAM</td><td>alignment</td><td>distance</td><td>position</td><td>gene name</td><td>gene id</td></tr>
<tr><td>GU305914.1</td><td>143591</td><td>143613</td><td>-</td><td>0</td><td>GTACACGGAGGGTAGTTCGT</td><td>CGG</td><td>||||||||[||||||||||||]PAM</td><td>NA</td><td>Intergenic</td><td>NA</td><td>NA</td></tr>
<tr><td>T23211</td><td>TACACGGAGGGTAGTTCGTCGGG</td><td>1000</td><td></td><td></td><td></td><td></td><td></td><td></td><td></td><td></td><td></td></tr>
<tr><td>Oligo fwd</td><td>CACCTACACGGAGGGTAGTTCGTC</td><td></td><td></td><td></td><td></td><td></td><td></td><td></td><td></td><td></td><td></td></tr>
<tr><td>Oligo rev</td><td>AAACGACGAACTACCCTCCGTGTA</td><td></td><td></td><td></td><td></td><td></td><td></td><td></td><td></td><td></td><td></td></tr>
<tr><td>Chromosome</td><td>start</td><td>end</td><td>strand</td><td>MM</td><td>target_seq</td><td>PAM</td><td>alignment</td><td>distance</td><td>position</td><td>gene name</td><td>gene id</td></tr>
<tr><td>GU305914.1</td><td>143590</td><td>143612</td><td>-</td><td>0</td><td>TACACGGAGGGTAGTTCGTC</td><td>GGG</td><td>||||||||[||||||||||||]PAM</td><td>NA</td><td>Intergenic</td><td>NA</td><td>NA</td></tr>
<tr><td>T23212</td><td>TTCGTCGGGAGACATACGTTCGG</td><td>1000</td><td></td><td></td><td></td><td></td><td></td><td></td><td></td><td></td><td></td></tr>
<tr><td>Oligo fwd</td><td>CACCTTCGTCGGGAGACATACGTT</td><td></td><td></td><td></td><td></td><td></td><td></td><td></td><td></td><td></td><td></td></tr>
<tr><td>Oligo rev</td><td>AAACAACGTATGTCTCCCGACGAA</td><td></td><td></td><td></td><td></td><td></td><td></td><td></td><td></td><td></td><td></td></tr>
<tr><td>Chromosome</td><td>start</td><td>end</td><td>strand</td><td>MM</td><td>target_seq</td><td>PAM</td><td>alignment</td><td>distance</td><td>position</td><td>gene name</td><td>gene id</td></tr>
<tr><td>GU305914.1</td><td>143576</td><td>143598</td><td>-</td><td>0</td><td>TTCGTCGGGAGACATACGTT</td><td>CGG</td><td>||||||||[||||||||||||]PAM</td><td>NA</td><td>Intergenic</td><td>NA</td><td>NA</td></tr>
<tr><td>T23213</td><td>TCGTCGGGAGACATACGTTCGGG</td><td>1000</td><td></td><td></td><td></td><td></td><td></td><td></td><td></td><td></td><td></td></tr>
<tr><td>Oligo fwd</td><td>CACCTCGTCGGGAGACATACGTTC</td><td></td><td></td><td></td><td></td><td></td><td></td><td></td><td></td><td></td><td></td></tr>
<tr><td>Oligo rev</td><td>AAACGAACGTATGTCTCCCGACGA</td><td></td><td></td><td></td><td></td><td></td><td></td><td></td><td></td><td></td><td></td></tr>
<tr><td>Chromosome</td><td>start</td><td>end</td><td>strand</td><td>MM</td><td>target_seq</td><td>PAM</td><td>alignment</td><td>distance</td><td>position</td><td>gene name</td><td>gene id</td></tr>
<tr><td>GU305914.1</td><td>143575</td><td>143597</td><td>-</td><td>0</td><td>TCGTCGGGAGACATACGTTC</td><td>GGG</td><td>||||||||[||||||||||||]PAM</td><td>NA</td><td>Intergenic</td><td>NA</td><td>NA</td></tr>
<tr><td>T23214</td><td>ACTGACTCAACACATAGTGCTGG</td><td>1000</td><td></td><td></td><td></td><td></td><td></td><td></td><td></td><td></td><td></td></tr>
<tr><td>Oligo fwd</td><td>CACCACTGACTCAACACATAGTGC</td><td></td><td></td><td></td><td></td><td></td><td></td><td></td><td></td><td></td><td></td></tr>
<tr><td>Oligo rev</td><td>AAACGCACTATGTGTTGAGTCAGT</td><td></td><td></td><td></td><td></td><td></td><td></td><td></td><td></td><td></td><td></td></tr>
<tr><td>Chromosome</td><td>start</td><td>end</td><td>strand</td><td>MM</td><td>target_seq</td><td>PAM</td><td>alignment</td><td>distance</td><td>position</td><td>gene name</td><td>gene id</td></tr>
<tr><td>GU305914.1</td><td>143529</td><td>143551</td><td>-</td><td>0</td><td>ACTGACTCAACACATAGTGC</td><td>TGG</td><td>||||||||[||||||||||||]PAM</td><td>NA</td><td>Intergenic</td><td>NA</td><td>NA</td></tr>
<tr><td>T23215</td><td>TCAACACATAGTGCTGGACCAGG</td><td>1000</td><td></td><td></td><td></td><td></td><td></td><td></td><td></td><td></td><td></td></tr>
<tr><td>Oligo fwd</td><td>CACCTCAACACATAGTGCTGGACC</td><td></td><td></td><td></td><td></td><td></td><td></td><td></td><td></td><td></td><td></td></tr>
<tr><td>Oligo rev</td><td>AAACGGTCCAGCACTATGTGTTGA</td><td></td><td></td><td></td><td></td><td></td><td></td><td></td><td></td><td></td><td></td></tr>
<tr><td>Chromosome</td><td>start</td><td>end</td><td>strand</td><td>MM</td><td>target_seq</td><td>PAM</td><td>alignment</td><td>distance</td><td>position</td><td>gene name</td><td>gene id</td></tr>
<tr><td>GU305914.1</td><td>143523</td><td>143545</td><td>-</td><td>0</td><td>TCAACACATAGTGCTGGACC</td><td>AGG</td><td>||||||||[||||||||||||]PAM</td><td>NA</td><td>Intergenic</td><td>NA</td><td>NA</td></tr>
<tr><td>T23216</td><td>CAACACATAGTGCTGGACCAGGG</td><td>1000</td><td></td><td></td><td></td><td></td><td></td><td></td><td></td><td></td><td></td></tr>
<tr><td>Oligo fwd</td><td>CACCCAACACATAGTGCTGGACCA</td><td></td><td></td><td></td><td></td><td></td><td></td><td></td><td></td><td></td><td></td></tr>
<tr><td>Oligo rev</td><td>AAACTGGTCCAGCACTATGTGTTG</td><td></td><td></td><td></td><td></td><td></td><td></td><td></td><td></td><td></td><td></td></tr>
<tr><td>Chromosome</td><td>start</td><td>end</td><td>strand</td><td>MM</td><td>target_seq</td><td>PAM</td><td>alignment</td><td>distance</td><td>position</td><td>gene name</td><td>gene id</td></tr>
<tr><td>GU305914.1</td><td>143522</td><td>143544</td><td>-</td><td>0</td><td>CAACACATAGTGCTGGACCA</td><td>GGG</td><td>||||||||[||||||||||||]PAM</td><td>NA</td><td>Intergenic</td><td>NA</td><td>NA</td></tr>
<tr><td>T23217</td><td>ACCAGGGTCTGACAGTAGTACGG</td><td>1000</td><td></td><td></td><td></td><td></td><td></td><td></td><td></td><td></td><td></td></tr>
<tr><td>Oligo fwd</td><td>CACCACCAGGGTCTGACAGTAGTA</td><td></td><td></td><td></td><td></td><td></td><td></td><td></td><td></td><td></td><td></td></tr>
<tr><td>Oligo rev</td><td>AAACTACTACTGTCAGACCCTGGT</td><td></td><td></td><td></td><td></td><td></td><td></td><td></td><td></td><td></td><td></td></tr>
<tr><td>Chromosome</td><td>start</td><td>end</td><td>strand</td><td>MM</td><td>target_seq</td><td>PAM</td><td>alignment</td><td>distance</td><td>position</td><td>gene name</td><td>gene id</td></tr>
<tr><td>GU305914.1</td><td>143506</td><td>143528</td><td>-</td><td>0</td><td>ACCAGGGTCTGACAGTAGTA</td><td>CGG</td><td>||||||||[||||||||||||]PAM</td><td>NA</td><td>Intergenic</td><td>NA</td><td>NA</td></tr>
<tr><td>T23218</td><td>CCAGGGTCTGACAGTAGTACGGG</td><td>1000</td><td></td><td></td><td></td><td></td><td></td><td></td><td></td><td></td><td></td></tr>
<tr><td>Oligo fwd</td><td>CACCCCAGGGTCTGACAGTAGTAC</td><td></td><td></td><td></td><td></td><td></td><td></td><td></td><td></td><td></td><td></td></tr>
</table>

Oligo rev AAACGTACTACTGTCAGACCCTGG
Chromosome start end strand MM target_seq PAM alignment distance position gene name gene id
GU305914.1 143505 143527 - 0 CCAGGGTCTGACAGTAGTAC GGG ||||||||[||||||||||||]PAM NA Intergenic NA NA

T23219 TGACAGTAGTACGGGTGACGCGG 1000
Oligo fwd CACCTGACAGTAGTACGGGTGACG
Oligo rev AAACCGTCACCCGTACTACTGTCA
Chromosome start end strand MM target_seq PAM alignment distance position gene name gene id
GU305914.1 143497 143519 - 0 TGACAGTAGTACGGGTGACG CGG ||||||||[||||||||||||]PAM NA Intergenic NA NA

T23220 GACAGTAGTACGGGTGACGCGGG 1000
Oligo fwd CACCGACAGTAGTACGGGTGACGC
Oligo rev AAACGCGTCACCCGTACTACTGTC
Chromosome start end strand MM target_seq PAM alignment distance position gene name gene id
GU305914.1 143496 143518 - 0 GACAGTAGTACGGGTGACGC GGG ||||||||[||||||||||||]PAM NA Intergenic NA NA

T23221 GTGACGCGGGTTAGCGAAGACGG 1000
Oligo fwd CACCGTGACGCGGGTTAGCGAAGA
Oligo rev AAACTCTTCGCTAACCCGCGTCAC
Chromosome start end strand MM target_seq PAM alignment distance position gene name gene id
GU305914.1 143483 143505 - 0 GTGACGCGGGTTAGCGAAGA CGG ||||||||[||||||||||||]PAM NA Intergenic NA NA

T23222 TGACGCGGGTTAGCGAAGACGGG 1000
Oligo fwd CACCTGACGCGGGTTAGCGAAGAC
Oligo rev AAACGTCTTCGCTAACCCGCGTCA
Chromosome start end strand MM target_seq PAM alignment distance position gene name gene id
GU305914.1 143482 143504 - 0 TGACGCGGGTTAGCGAAGAC GGG ||||||||[||||||||||||]PAM NA Intergenic NA NA

T23223 CGGGTTAGCGAAGACGGGCGCGG 1000
Oligo fwd CACCCGGGTTAGCGAAGACGGGCG
Oligo rev AAACCGCCCGTCTTCGCTAACCCG
Chromosome start end strand MM target_seq PAM alignment distance position gene name gene id
GU305914.1 143477 143499 - 0 CGGGTTAGCGAAGACGGGCG CGG ||||||||[||||||||||||]PAM NA Intergenic NA NA

T23224 GAAGACGGGCGCGGCGAACGCGG 1000
Oligo fwd CACCGAAGACGGGCGCGGCGAACG
Oligo rev AAACCGTTCGCCGCGCCCGTCTTC
Chromosome start end strand MM target_seq PAM alignment distance position gene name gene id
GU305914.1 143468 143490 - 0 GAAGACGGGCGCGGCGAACG CGG ||||||||[||||||||||||]PAM NA Intergenic NA NA

T23225 GCGCGGCGAACGCGGCCTTGAGG 1000
Oligo fwd CACCGCGCGGCGAACGCGGCCTTG
Oligo rev AAACCAAGGCCGCGTTCGCCGCGC
Chromosome start end strand MM target_seq PAM alignment distance position gene name gene id
GU305914.1 143460 143482 - 0 GCGCGGCGAACGCGGCCTTG AGG ||||||||[||||||||||||]PAM NA Intergenic NA NA

T23226 CGGCCTTGAGGAACAGATCTAGG 1000
Oligo fwd CACCCGGCCTTGAGGAACAGATCT
Oligo rev AAACAGATCTGTTCCTCAAGGCCG
Chromosome start end strand MM target_seq PAM alignment distance position gene name gene id
GU305914.1 143448 143470 - 0 CGGCCTTGAGGAACAGATCT AGG ||||||||[||||||||||||]PAM NA Intergenic NA NA

T23227 AGGAACAGATCTAGGTAGAGCGG 1000
Oligo fwd CACCAGGAACAGATCTAGGTAGAG
Oligo rev AAACCTCTACCTAGATCTGTTCCT
Chromosome start end strand MM target_seq PAM alignment distance position gene name gene id
GU305914.1 143440 143462 - 0 AGGAACAGATCTAGGTAGAG CGG ||||||||[||||||||||||]PAM NA Intergenic NA NA

T23228 GGAACAGATCTAGGTAGAGCGGG 1000
Oligo fwd CACCGGAACAGATCTAGGTAGAGC
Oligo rev AAACGCTCTACCTAGATCTGTTCC
Chromosome start end strand MM target_seq PAM alignment distance position gene name gene id
GU305914.1 143439 143461 - 0 GGAACAGATCTAGGTAGAGC GGG ||||||||[||||||||||||]PAM NA Intergenic NA NA

T23229 GAACAGATCTAGGTAGAGCGGGG 1000
Oligo fwd CACCGAACAGATCTAGGTAGAGCG
Oligo rev AAACCGCTCTACCTAGATCTGTTC
Chromosome start end strand MM target_seq PAM alignment distance position gene name gene id
GU305914.1 143438 143460 - 0 GAACAGATCTAGGTAGAGCG GGG ||||||||[||||||||||||]PAM NA Intergenic NA NA

T23230 AACAGATCTAGGTAGAGCGGGGG 1000
Oligo fwd CACCAACAGATCTAGGTAGAGCGG
Oligo rev AAACCCGCTCTACCTAGATCTGTT
Chromosome start end strand MM target_seq PAM alignment distance position gene name gene id
GU305914.1 143437 143459 - 0 AACAGATCTAGGTAGAGCGG GGG ||||||||[||||||||||||]PAM NA Intergenic NA NA

T23231 TAGGTAGAGCGGGGGCCGCTCGG 1000
Oligo fwd CACCTAGGTAGAGCGGGGGCCGCT
Oligo rev AAACAGCGGCCCCCGCTCTACCTA
Chromosome start end strand MM target_seq PAM alignment distance position gene name gene id
GU305914.1 143429 143451 - 0 TAGGTAGAGCGGGGGCCGCT CGG ||||||||[||||||||||||]PAM NA Intergenic NA NA

T23232 GTAGAGCGGGGGCCGCTCGGCGG 1000
Oligo fwd CACCGTAGAGCGGGGGCCGCTCGG
Oligo rev AAACCCGAGCGGCCCCCGCTCTAC
Chromosome start end strand MM target_seq PAM alignment distance position gene name gene id
GU305914.1 143426 143448 - 0 GTAGAGCGGGGGCCGCTCGG CGG ||||||||[||||||||||||]PAM NA Intergenic NA NA

T23233 GCGGGGGCCGCTCGGCGGACAGG 1000

<table>
<tr><td>Oligo fwd</td><td>CACCGCGGGGGCCGCTCGGCGGAC</td><td></td><td></td><td></td><td></td><td></td><td></td><td></td><td></td><td></td><td></td></tr>
<tr><td>Oligo rev</td><td>AAACGTCCGCCGAGCGGCCCCCGC</td><td></td><td></td><td></td><td></td><td></td><td></td><td></td><td></td><td></td><td></td></tr>
<tr><td>Chromosome</td><td>start</td><td>end</td><td>strand</td><td>MM</td><td>target_seq</td><td>PAM</td><td>alignment</td><td>distance</td><td>position</td><td>gene name</td><td>gene id</td></tr>
<tr><td>GU305914.1</td><td>143421</td><td>143443</td><td>-</td><td>0</td><td>GCGGGGGCCGCTCGGCGGAC</td><td>AGG</td><td>||||||||[||||||||||||]PAM</td><td>NA</td><td>Intergenic</td><td>NA</td><td>NA</td></tr>
<tr><td></td><td></td><td></td><td></td><td></td><td></td><td></td><td></td><td></td><td></td><td></td><td></td></tr>
<tr><td>T23234</td><td>CTCGTCGTCATCACCGACGCAGG</td><td>1000</td><td></td><td></td><td></td><td></td><td></td><td></td><td></td><td></td><td></td></tr>
<tr><td>Oligo fwd</td><td>CACCCTCGTCGTCATCACCGACGC</td><td></td><td></td><td></td><td></td><td></td><td></td><td></td><td></td><td></td><td></td></tr>
<tr><td>Oligo rev</td><td>AAACGCGTCGGTGATGACGACGAG</td><td></td><td></td><td></td><td></td><td></td><td></td><td></td><td></td><td></td><td></td></tr>
<tr><td>Chromosome</td><td>start</td><td>end</td><td>strand</td><td>MM</td><td>target_seq</td><td>PAM</td><td>alignment</td><td>distance</td><td>position</td><td>gene name</td><td>gene id</td></tr>
<tr><td>GU305914.1</td><td>143390</td><td>143412</td><td>-</td><td>0</td><td>CTCGTCGTCATCACCGACGC</td><td>AGG</td><td>||||||||[||||||||||||]PAM</td><td>NA</td><td>Intergenic</td><td>NA</td><td>NA</td></tr>
<tr><td>GU305914.1</td><td>173742</td><td>173764</td><td>+</td><td>4</td><td>CTCGACGCCAGCACCGAAGC</td><td>GGG</td><td>||||-||-[||-||||||-||]PAM</td><td>NA</td><td>Intergenic</td><td>NA</td><td>NA</td></tr>
<tr><td></td><td></td><td></td><td></td><td></td><td></td><td></td><td></td><td></td><td></td><td></td><td></td></tr>
<tr><td>T23235</td><td>GTCATCACCGACGCAGGTCGCGG</td><td>1000</td><td></td><td></td><td></td><td></td><td></td><td></td><td></td><td></td><td></td></tr>
<tr><td>Oligo fwd</td><td>CACCGTCATCACCGACGCAGGTCG</td><td></td><td></td><td></td><td></td><td></td><td></td><td></td><td></td><td></td><td></td></tr>
<tr><td>Oligo rev</td><td>AAACCGACCTGCGTCGGTGATGAC</td><td></td><td></td><td></td><td></td><td></td><td></td><td></td><td></td><td></td><td></td></tr>
<tr><td>Chromosome</td><td>start</td><td>end</td><td>strand</td><td>MM</td><td>target_seq</td><td>PAM</td><td>alignment</td><td>distance</td><td>position</td><td>gene name</td><td>gene id</td></tr>
<tr><td>GU305914.1</td><td>143384</td><td>143406</td><td>-</td><td>0</td><td>GTCATCACCGACGCAGGTCG</td><td>CGG</td><td>||||||||[||||||||||||]PAM</td><td>NA</td><td>Intergenic</td><td>NA</td><td>NA</td></tr>
<tr><td></td><td></td><td></td><td></td><td></td><td></td><td></td><td></td><td></td><td></td><td></td><td></td></tr>
<tr><td>T23236</td><td>ACCGACGCAGGTCGCGGCGCCGG</td><td>1000</td><td></td><td></td><td></td><td></td><td></td><td></td><td></td><td></td><td></td></tr>
<tr><td>Oligo fwd</td><td>CACCACCGACGCAGGTCGCGGCGC</td><td></td><td></td><td></td><td></td><td></td><td></td><td></td><td></td><td></td><td></td></tr>
<tr><td>Oligo rev</td><td>AAACGCGCCGCGACCTGCGTCGGT</td><td></td><td></td><td></td><td></td><td></td><td></td><td></td><td></td><td></td><td></td></tr>
<tr><td>Chromosome</td><td>start</td><td>end</td><td>strand</td><td>MM</td><td>target_seq</td><td>PAM</td><td>alignment</td><td>distance</td><td>position</td><td>gene name</td><td>gene id</td></tr>
<tr><td>GU305914.1</td><td>143378</td><td>143400</td><td>-</td><td>0</td><td>ACCGACGCAGGTCGCGGCGC</td><td>CGG</td><td>||||||||[||||||||||||]PAM</td><td>NA</td><td>Intergenic</td><td>NA</td><td>NA</td></tr>
<tr><td></td><td></td><td></td><td></td><td></td><td></td><td></td><td></td><td></td><td></td><td></td><td></td></tr>
<tr><td>T23237</td><td>CGCGGCGCCGGTCTCGCCCTCGG</td><td>1000</td><td></td><td></td><td></td><td></td><td></td><td></td><td></td><td></td><td></td></tr>
<tr><td>Oligo fwd</td><td>CACCCGCGGCGCCGGTCTCGCCCT</td><td></td><td></td><td></td><td></td><td></td><td></td><td></td><td></td><td></td><td></td></tr>
<tr><td>Oligo rev</td><td>AAACAGGGCGAGACCGGCGCCGCG</td><td></td><td></td><td></td><td></td><td></td><td></td><td></td><td></td><td></td><td></td></tr>
<tr><td>Chromosome</td><td>start</td><td>end</td><td>strand</td><td>MM</td><td>target_seq</td><td>PAM</td><td>alignment</td><td>distance</td><td>position</td><td>gene name</td><td>gene id</td></tr>
<tr><td>GU305914.1</td><td>143366</td><td>143388</td><td>-</td><td>0</td><td>CGCGGCGCCGGTCTCGCCCT</td><td>CGG</td><td>||||||||[||||||||||||]PAM</td><td>NA</td><td>Intergenic</td><td>NA</td><td>NA</td></tr>
<tr><td></td><td></td><td></td><td></td><td></td><td></td><td></td><td></td><td></td><td></td><td></td><td></td></tr>
<tr><td>T23238</td><td>GGCGCCGGTCTCGCCCTCGGCGG</td><td>1000</td><td></td><td></td><td></td><td></td><td></td><td></td><td></td><td></td><td></td></tr>
<tr><td>Oligo fwd</td><td>CACCGGCGCCGGTCTCGCCCTCGG</td><td></td><td></td><td></td><td></td><td></td><td></td><td></td><td></td><td></td><td></td></tr>
<tr><td>Oligo rev</td><td>AAACCCGAGGGCGAGACCGGCGCC</td><td></td><td></td><td></td><td></td><td></td><td></td><td></td><td></td><td></td><td></td></tr>
<tr><td>Chromosome</td><td>start</td><td>end</td><td>strand</td><td>MM</td><td>target_seq</td><td>PAM</td><td>alignment</td><td>distance</td><td>position</td><td>gene name</td><td>gene id</td></tr>
<tr><td>GU305914.1</td><td>143363</td><td>143385</td><td>-</td><td>0</td><td>GGCGCCGGTCTCGCCCTCGG</td><td>CGG</td><td>||||||||[||||||||||||]PAM</td><td>NA</td><td>Intergenic</td><td>NA</td><td>NA</td></tr>
<tr><td></td><td></td><td></td><td></td><td></td><td></td><td></td><td></td><td></td><td></td><td></td><td></td></tr>
<tr><td>T23239</td><td>GCGCGCGTCGAAAACGAAGACGG</td><td>1000</td><td></td><td></td><td></td><td></td><td></td><td></td><td></td><td></td><td></td></tr>
<tr><td>Oligo fwd</td><td>CACCGCGCGCGTCGAAAACGAAGA</td><td></td><td></td><td></td><td></td><td></td><td></td><td></td><td></td><td></td><td></td></tr>
<tr><td>Oligo rev</td><td>AAACTCTTCGTTTTCGACGCGCGC</td><td></td><td></td><td></td><td></td><td></td><td></td><td></td><td></td><td></td><td></td></tr>
<tr><td>Chromosome</td><td>start</td><td>end</td><td>strand</td><td>MM</td><td>target_seq</td><td>PAM</td><td>alignment</td><td>distance</td><td>position</td><td>gene name</td><td>gene id</td></tr>
<tr><td>GU305914.1</td><td>143339</td><td>143361</td><td>-</td><td>0</td><td>GCGCGCGTCGAAAACGAAGA</td><td>CGG</td><td>||||||||[||||||||||||]PAM</td><td>NA</td><td>Intergenic</td><td>NA</td><td>NA</td></tr>
<tr><td></td><td></td><td></td><td></td><td></td><td></td><td></td><td></td><td></td><td></td><td></td><td></td></tr>
<tr><td>T23240</td><td>CGCGTCGAAAACGAAGACGGTGG</td><td>1000</td><td></td><td></td><td></td><td></td><td></td><td></td><td></td><td></td><td></td></tr>
<tr><td>Oligo fwd</td><td>CACCCGCGTCGAAAACGAAGACGG</td><td></td><td></td><td></td><td></td><td></td><td></td><td></td><td></td><td></td><td></td></tr>
<tr><td>Oligo rev</td><td>AAACCCGTCTTCGTTTTCGACGCG</td><td></td><td></td><td></td><td></td><td></td><td></td><td></td><td></td><td></td><td></td></tr>
<tr><td>Chromosome</td><td>start</td><td>end</td><td>strand</td><td>MM</td><td>target_seq</td><td>PAM</td><td>alignment</td><td>distance</td><td>position</td><td>gene name</td><td>gene id</td></tr>
<tr><td>GU305914.1</td><td>143336</td><td>143358</td><td>-</td><td>0</td><td>CGCGTCGAAAACGAAGACGG</td><td>TGG</td><td>||||||||[||||||||||||]PAM</td><td>NA</td><td>Intergenic</td><td>NA</td><td>NA</td></tr>
<tr><td></td><td></td><td></td><td></td><td></td><td></td><td></td><td></td><td></td><td></td><td></td><td></td></tr>
<tr><td>T23241</td><td>CGAAAACGAAGACGGTGGAGAGG</td><td>1000</td><td></td><td></td><td></td><td></td><td></td><td></td><td></td><td></td><td></td></tr>
<tr><td>Oligo fwd</td><td>CACCCGAAAACGAAGACGGTGGAG</td><td></td><td></td><td></td><td></td><td></td><td></td><td></td><td></td><td></td><td></td></tr>
<tr><td>Oligo rev</td><td>AAACCTCCACCGTCTTCGTTTTCG</td><td></td><td></td><td></td><td></td><td></td><td></td><td></td><td></td><td></td><td></td></tr>
<tr><td>Chromosome</td><td>start</td><td>end</td><td>strand</td><td>MM</td><td>target_seq</td><td>PAM</td><td>alignment</td><td>distance</td><td>position</td><td>gene name</td><td>gene id</td></tr>
<tr><td>GU305914.1</td><td>143331</td><td>143353</td><td>-</td><td>0</td><td>CGAAAACGAAGACGGTGGAG</td><td>AGG</td><td>||||||||[||||||||||||]PAM</td><td>NA</td><td>Intergenic</td><td>NA</td><td>NA</td></tr>
<tr><td></td><td></td><td></td><td></td><td></td><td></td><td></td><td></td><td></td><td></td><td></td><td></td></tr>
<tr><td>T23242</td><td>CCTCCGAGTTCTCCTTGATGAGG</td><td>1000</td><td></td><td></td><td></td><td></td><td></td><td></td><td></td><td></td><td></td></tr>
<tr><td>Oligo fwd</td><td>CACCCCTCCGAGTTCTCCTTGATG</td><td></td><td></td><td></td><td></td><td></td><td></td><td></td><td></td><td></td><td></td></tr>
<tr><td>Oligo rev</td><td>AAACCATCAAGGAGAACTCGGAGG</td><td></td><td></td><td></td><td></td><td></td><td></td><td></td><td></td><td></td><td></td></tr>
<tr><td>Chromosome</td><td>start</td><td>end</td><td>strand</td><td>MM</td><td>target_seq</td><td>PAM</td><td>alignment</td><td>distance</td><td>position</td><td>gene name</td><td>gene id</td></tr>
<tr><td>GU305914.1</td><td>143307</td><td>143329</td><td>-</td><td>0</td><td>CCTCCGAGTTCTCCTTGATG</td><td>AGG</td><td>||||||||[||||||||||||]PAM</td><td>NA</td><td>Intergenic</td><td>NA</td><td>NA</td></tr>
<tr><td></td><td></td><td></td><td></td><td></td><td></td><td></td><td></td><td></td><td></td><td></td><td></td></tr>
<tr><td>T23243</td><td>TGATGAGGCCGTCGATCCAGAGG</td><td>1000</td><td></td><td></td><td></td><td></td><td></td><td></td><td></td><td></td><td></td></tr>
<tr><td>Oligo fwd</td><td>CACCTGATGAGGCCGTCGATCCAG</td><td></td><td></td><td></td><td></td><td></td><td></td><td></td><td></td><td></td><td></td></tr>
<tr><td>Oligo rev</td><td>AAACCTGGATCGACGGCCTCATCA</td><td></td><td></td><td></td><td></td><td></td><td></td><td></td><td></td><td></td><td></td></tr>
<tr><td>Chromosome</td><td>start</td><td>end</td><td>strand</td><td>MM</td><td>target_seq</td><td>PAM</td><td>alignment</td><td>distance</td><td>position</td><td>gene name</td><td>gene id</td></tr>
<tr><td>GU305914.1</td><td>143292</td><td>143314</td><td>-</td><td>0</td><td>TGATGAGGCCGTCGATCCAG</td><td>AGG</td><td>||||||||[||||||||||||]PAM</td><td>NA</td><td>Intergenic</td><td>NA</td><td>NA</td></tr>
<tr><td></td><td></td><td></td><td></td><td></td><td></td><td></td><td></td><td></td><td></td><td></td><td></td></tr>
<tr><td>T23244</td><td>GGCCGTCGATCCAGAGGCGCAGG</td><td>1000</td><td></td><td></td><td></td><td></td><td></td><td></td><td></td><td></td><td></td></tr>
<tr><td>Oligo fwd</td><td>CACCGGCCGTCGATCCAGAGGCGC</td><td></td><td></td><td></td><td></td><td></td><td></td><td></td><td></td><td></td><td></td></tr>
<tr><td>Oligo rev</td><td>AAACGCGCCTCTGGATCGACGGCC</td><td></td><td></td><td></td><td></td><td></td><td></td><td></td><td></td><td></td><td></td></tr>
<tr><td>Chromosome</td><td>start</td><td>end</td><td>strand</td><td>MM</td><td>target_seq</td><td>PAM</td><td>alignment</td><td>distance</td><td>position</td><td>gene name</td><td>gene id</td></tr>
<tr><td>GU305914.1</td><td>143286</td><td>143308</td><td>-</td><td>0</td><td>GGCCGTCGATCCAGAGGCGC</td><td>AGG</td><td>||||||||[||||||||||||]PAM</td><td>NA</td><td>Intergenic</td><td>NA</td><td>NA</td></tr>
<tr><td></td><td></td><td></td><td></td><td></td><td></td><td></td><td></td><td></td><td></td><td></td><td></td></tr>
<tr><td>T23245</td><td>GCCGTCGATCCAGAGGCGCAGGG</td><td>1000</td><td></td><td></td><td></td><td></td><td></td><td></td><td></td><td></td><td></td></tr>
<tr><td>Oligo fwd</td><td>CACCGCCGTCGATCCAGAGGCGCA</td><td></td><td></td><td></td><td></td><td></td><td></td><td></td><td></td><td></td><td></td></tr>
<tr><td>Oligo rev</td><td>AAACTGCGCCTCTGGATCGACGGC</td><td></td><td></td><td></td><td></td><td></td><td></td><td></td><td></td><td></td><td></td></tr>
<tr><td>Chromosome</td><td>start</td><td>end</td><td>strand</td><td>MM</td><td>target_seq</td><td>PAM</td><td>alignment</td><td>distance</td><td>position</td><td>gene name</td><td>gene id</td></tr>
<tr><td>GU305914.1</td><td>143285</td><td>143307</td><td>-</td><td>0</td><td>GCCGTCGATCCAGAGGCGCA</td><td>GGG</td><td>||||||||[||||||||||||]PAM</td><td>NA</td><td>Intergenic</td><td>NA</td><td>NA</td></tr>
<tr><td></td><td></td><td></td><td></td><td></td><td></td><td></td><td></td><td></td><td></td><td></td><td></td></tr>
<tr><td>T23246</td><td>CAGGGACTCCGAAACGACGCCGG</td><td>1000</td><td></td><td></td><td></td><td></td><td></td><td></td><td></td><td></td><td></td></tr>
<tr><td>Oligo fwd</td><td>CACCCAGGGACTCCGAAACGACGC</td><td></td><td></td><td></td><td></td><td></td><td></td><td></td><td></td><td></td><td></td></tr>
<tr><td>Oligo rev</td><td>AAACGCGTCGTTTCGGAGTCCCTG</td><td></td><td></td><td></td><td></td><td></td><td></td><td></td><td></td><td></td><td></td></tr>
<tr><td>Chromosome</td><td>start</td><td>end</td><td>strand</td><td>MM</td><td>target_seq</td><td>PAM</td><td>alignment</td><td>distance</td><td>position</td><td>gene name</td><td>gene id</td></tr>
<tr><td>GU305914.1</td><td>143267</td><td>143289</td><td>-</td><td>0</td><td>CAGGGACTCCGAAACGACGC</td><td>CGG</td><td>||||||||[||||||||||||]PAM</td><td>NA</td><td>Intergenic</td><td>NA</td><td>NA</td></tr>
<tr><td></td><td></td><td></td><td></td><td></td><td></td><td></td><td></td><td></td><td></td><td></td><td></td></tr>
<tr><td>T23247</td><td>TGCACGACTGCCGAAAGACGCGG</td><td>1000</td><td></td><td></td><td></td><td></td><td></td><td></td><td></td><td></td><td></td></tr>
<tr><td>Oligo fwd</td><td>CACCTGCACGACTGCCGAAAGACG</td><td></td><td></td><td></td><td></td><td></td><td></td><td></td><td></td><td></td><td></td></tr>
<tr><td>Oligo rev</td><td>AAACCGTCTTTCGGCAGTCGTGCA</td><td></td><td></td><td></td><td></td><td></td><td></td><td></td><td></td><td></td><td></td></tr>
<tr><td>Chromosome</td><td>start</td><td>end</td><td>strand</td><td>MM</td><td>target_seq</td><td>PAM</td><td>alignment</td><td>distance</td><td>position</td><td>gene name</td><td>gene id</td></tr>
<tr><td>GU305914.1</td><td>143232</td><td>143254</td><td>-</td><td>0</td><td>TGCACGACTGCCGAAAGACG</td><td>CGG</td><td>||||||||[||||||||||||]PAM</td><td>NA</td><td>Intergenic</td><td>NA</td><td>NA</td></tr>
</table>

T23248 CTGCCGAAAGACGCGGTCGTAGG 1000
Oligo fwd CACCCTGCCGAAAGACGCGGTCGT
Oligo rev AAACACGACCGCGTCTTTCGGCAG
Chromosome start end strand MM target_seq PAM alignment distance position gene name gene id
GU305914.1 143225 143247 - 0 CTGCCGAAAGACGCGGTCGT AGG ||||||||[||||||||||||]PAM NA Intergenic NA NA

T23249 GGACACCAGACACTCGCGCGAGG 1000
Oligo fwd CACCGGACACCAGACACTCGCGCG
Oligo rev AAACCGCGCGAGTGTCTGGTGTCC
Chromosome start end strand MM target_seq PAM alignment distance position gene name gene id
GU305914.1 143204 143226 - 0 GGACACCAGACACTCGCGCG AGG ||||||||[||||||||||||]PAM NA Intergenic NA NA

T23250 CACCAGACACTCGCGCGAGGTGG 1000
Oligo fwd CACCCACCAGACACTCGCGCGAGG
Oligo rev AAACCCTCGCGCGAGTGTCTGGTG
Chromosome start end strand MM target_seq PAM alignment distance position gene name gene id
GU305914.1 143201 143223 - 0 CACCAGACACTCGCGCGAGG TGG ||||||||[||||||||||||]PAM NA Intergenic NA NA

T23251 ACCAGACACTCGCGCGAGGTGGG 1000
Oligo fwd CACCACCAGACACTCGCGCGAGGT
Oligo rev AAACACCTCGCGCGAGTGTCTGGT
Chromosome start end strand MM target_seq PAM alignment distance position gene name gene id
GU305914.1 143200 143222 - 0 ACCAGACACTCGCGCGAGGT GGG ||||||||[||||||||||||]PAM NA Intergenic NA NA

T23252 ACTCGCGCGAGGTGGGCAGCTGG 1000
Oligo fwd CACCACTCGCGCGAGGTGGGCAGC
Oligo rev AAACGCTGCCCACCTCGCGCGAGT
Chromosome start end strand MM target_seq PAM alignment distance position gene name gene id
GU305914.1 143193 143215 - 0 ACTCGCGCGAGGTGGGCAGC TGG ||||||||[||||||||||||]PAM NA Intergenic NA NA

T23253 GCGAGGTGGGCAGCTGGCCCTGG 1000
Oligo fwd CACCGCGAGGTGGGCAGCTGGCCC
Oligo rev AAACGGGCCAGCTGCCCACCTCGC
Chromosome start end strand MM target_seq PAM alignment distance position gene name gene id
GU305914.1 143187 143209 - 0 GCGAGGTGGGCAGCTGGCCC TGG ||||||||[||||||||||||]PAM NA Intergenic NA NA

T23254 TGGGCAGCTGGCCCTGGTGCCGG 1000
Oligo fwd CACCTGGGCAGCTGGCCCTGGTGC
Oligo rev AAACGCACCAGGGCCAGCTGCCCA
Chromosome start end strand MM target_seq PAM alignment distance position gene name gene id
GU305914.1 143181 143203 - 0 TGGGCAGCTGGCCCTGGTGC CGG ||||||||[||||||||||||]PAM NA Intergenic NA NA

T23255 GCCCTGGTGCCGGAACTCGACGG 1000
Oligo fwd CACCGCCCTGGTGCCGGAACTCGA
Oligo rev AAACTCGAGTTCCGGCACCAGGGC
Chromosome start end strand MM target_seq PAM alignment distance position gene name gene id
GU305914.1 143171 143193 - 0 GCCCTGGTGCCGGAACTCGA CGG ||||||||[||||||||||||]PAM NA Intergenic NA NA

T23256 TGGTGCCGGAACTCGACGGCCGG 1000
Oligo fwd CACCTGGTGCCGGAACTCGACGGC
Oligo rev AAACGCCGTCGAGTTCCGGCACCA
Chromosome start end strand MM target_seq PAM alignment distance position gene name gene id
GU305914.1 143167 143189 - 0 TGGTGCCGGAACTCGACGGC CGG ||||||||[||||||||||||]PAM NA Intergenic NA NA

T23257 GGAACTCGACGGCCGGCGTGTGG 1000
Oligo fwd CACCGGAACTCGACGGCCGGCGTG
Oligo rev AAACCACGCCGGCCGTCGAGTTCC
Chromosome start end strand MM target_seq PAM alignment distance position gene name gene id
GU305914.1 143160 143182 - 0 GGAACTCGACGGCCGGCGTG TGG ||||||||[||||||||||||]PAM NA Intergenic NA NA

T23258 ACTCGACGGCCGGCGTGTGGTGG 1000
Oligo fwd CACCACTCGACGGCCGGCGTGTGG
Oligo rev AAACCCACACGCCGGCCGTCGAGT
Chromosome start end strand MM target_seq PAM alignment distance position gene name gene id
GU305914.1 143157 143179 - 0 ACTCGACGGCCGGCGTGTGG TGG ||||||||[||||||||||||]PAM NA Intergenic NA NA

T23259 CGGCCGGCGTGTGGTGGTCCAGG 1000
Oligo fwd CACCCGGCCGGCGTGTGGTGGTCC
Oligo rev AAACGGACCACCACACGCCGGCCG
Chromosome start end strand MM target_seq PAM alignment distance position gene name gene id
GU305914.1 143151 143173 - 0 CGGCCGGCGTGTGGTGGTCC AGG ||||||||[||||||||||||]PAM NA Intergenic NA NA

T23260 GTGGTGGTCCAGGATGAAATCGG 1000
Oligo fwd CACCGTGGTGGTCCAGGATGAAAT
Oligo rev AAACATTTCATCCTGGACCACCAC
Chromosome start end strand MM target_seq PAM alignment distance position gene name gene id
GU305914.1 143141 143163 - 0 GTGGTGGTCCAGGATGAAAT CGG ||||||||[||||||||||||]PAM NA Intergenic NA NA

T23261 GGATGAAATCGGCGAAGCTCTGG 1000
Oligo fwd CACCGGATGAAATCGGCGAAGCTC
Oligo rev AAACGAGCTTCGCCGATTTCATCC
Chromosome start end strand MM target_seq PAM alignment distance position gene name gene id
GU305914.1 143130 143152 - 0 GGATGAAATCGGCGAAGCTC TGG ||||||||[||||||||||||]PAM NA Intergenic NA NA

T23262 ATCGGCGAAGCTCTGGAGACCGG 1000
Oligo fwd CACCATCGGCGAAGCTCTGGAGAC
Oligo rev AAACGTCTCCAGAGCTTCGCCGAT
Chromosome start end strand MM target_seq PAM alignment distance position gene name gene id

GU305914.1 143123 143145 - 0 ATCGGCGAAGCTCTGGAGAC CGG ||||||||[||||||||||||]PAM NA Intergenic NA NA

T23263 TCGGCGAAGCTCTGGAGACCGGG 1000
Oligo fwd CACCTCGGCGAAGCTCTGGAGACC
Oligo rev AAACGGTCTCCAGAGCTTCGCCGA
Chromosome start end strand MM target_seq PAM alignment distance position gene name gene id
GU305914.1 143122 143144 - 0 TCGGCGAAGCTCTGGAGACC GGG ||||||||[||||||||||||]PAM NA Intergenic NA NA

T23264 CGGCGAAGCTCTGGAGACCGGGG 1000
Oligo fwd CACCCGGCGAAGCTCTGGAGACCG
Oligo rev AAACCGGTCTCCAGAGCTTCGCCG
Chromosome start end strand MM target_seq PAM alignment distance position gene name gene id
GU305914.1 143121 143143 - 0 CGGCGAAGCTCTGGAGACCG GGG ||||||||[||||||||||||]PAM NA Intergenic NA NA

T23265 TCTGGAGACCGGGGTCGTCCAGG 1000
Oligo fwd CACCTCTGGAGACCGGGGTCGTCC
Oligo rev AAACGGACGACCCCGGTCTCCAGA
Chromosome start end strand MM target_seq PAM alignment distance position gene name gene id
GU305914.1 143112 143134 - 0 TCTGGAGACCGGGGTCGTCC AGG ||||||||[||||||||||||]PAM NA Intergenic NA NA

T23266 CCAGGAGCCCCTGCACCGCGCGG 1000
Oligo fwd CACCCCAGGAGCCCCTGCACCGCG
Oligo rev AAACCGCGGTGCAGGGGCTCCTGG
Chromosome start end strand MM target_seq PAM alignment distance position gene name gene id
GU305914.1 143094 143116 - 0 CCAGGAGCCCCTGCACCGCG CGG ||||||||[||||||||||||]PAM NA Intergenic NA NA

T23267 CGCGGTCGTCGCGCGACCGCAGG 1000
Oligo fwd CACCCGCGGTCGTCGCGCGACCGC
Oligo rev AAACGCGGTCGCGCGACGACCGCG
Chromosome start end strand MM target_seq PAM alignment distance position gene name gene id
GU305914.1 143076 143098 - 0 CGCGGTCGTCGCGCGACCGC AGG ||||||||[||||||||||||]PAM NA Intergenic NA NA

T23268 GCGACCGCAGGTACTTGAAGCGG 1000
Oligo fwd CACCGCGACCGCAGGTACTTGAAG
Oligo rev AAACCTTCAAGTACCTGCGGTCGC
Chromosome start end strand MM target_seq PAM alignment distance position gene name gene id
GU305914.1 143064 143086 - 0 GCGACCGCAGGTACTTGAAG CGG ||||||||[||||||||||||]PAM NA Intergenic NA NA

T23269 GCGGCATTTGATCTCGAAGATGG 1000
Oligo fwd CACCGCGGCATTTGATCTCGAAGA
Oligo rev AAACTCTTCGAGATCAAATGCCGC
Chromosome start end strand MM target_seq PAM alignment distance position gene name gene id
GU305914.1 143045 143067 - 0 GCGGCATTTGATCTCGAAGA TGG ||||||||[||||||||||||]PAM NA Intergenic NA NA

T23270 ATCTCGAAGATGGCCGCCCCCGG 1000
Oligo fwd CACCATCTCGAAGATGGCCGCCCC
Oligo rev AAACGGGGCGGCCATCTTCGAGAT
Chromosome start end strand MM target_seq PAM alignment distance position gene name gene id
GU305914.1 143035 143057 - 0 ATCTCGAAGATGGCCGCCCC CGG ||||||||[||||||||||||]PAM NA Intergenic NA NA

T23271 CCCCCGGCGCCACCACCAGCAGG 1000
Oligo fwd CACCCCCCCGGCGCCACCACCAGC
Oligo rev AAACGCTGGTGGTGGCGCCGGGGG
Chromosome start end strand MM target_seq PAM alignment distance position gene name gene id
GU305914.1 143019 143041 - 0 CCCCCGGCGCCACCACCAGC AGG ||||||||[||||||||||||]PAM NA Intergenic NA NA

T23272 CCAGCAGGCCGTCGTCGTCGCGG 1000
Oligo fwd CACCCCAGCAGGCCGTCGTCGTCG
Oligo rev AAACCGACGACGACGGCCTGCTGG
Chromosome start end strand MM target_seq PAM alignment distance position gene name gene id
GU305914.1 143004 143026 - 0 CCAGCAGGCCGTCGTCGTCG CGG ||||||||[||||||||||||]PAM NA Intergenic NA NA

T23273 CAGCAGGCCGTCGTCGTCGCGGG 1000
Oligo fwd CACCCAGCAGGCCGTCGTCGTCGC
Oligo rev AAACGCGACGACGACGGCCTGCTG
Chromosome start end strand MM target_seq PAM alignment distance position gene name gene id
GU305914.1 143003 143025 - 0 CAGCAGGCCGTCGTCGTCGC GGG ||||||||[||||||||||||]PAM NA Intergenic NA NA

T23274 ACCGCTACAGAAGTCGAGCGAGG 1000
Oligo fwd CACCACCGCTACAGAAGTCGAGCG
Oligo rev AAACCGCTCGACTTCTGTAGCGGT
Chromosome start end strand MM target_seq PAM alignment distance position gene name gene id
GU305914.1 142976 142998 - 0 ACCGCTACAGAAGTCGAGCG AGG ||||||||[||||||||||||]PAM NA Intergenic NA NA

T23275 AGAAGTCGAGCGAGGCGCCCAGG 1000
Oligo fwd CACCAGAAGTCGAGCGAGGCGCCC
Oligo rev AAACGGGCGCCTCGCTCGACTTCT
Chromosome start end strand MM target_seq PAM alignment distance position gene name gene id
GU305914.1 142968 142990 - 0 AGAAGTCGAGCGAGGCGCCC AGG ||||||||[||||||||||||]PAM NA Intergenic NA NA

T23276 GCGCCCAGGACCCCCGACGCCGG 1000
Oligo fwd CACCGCGCCCAGGACCCCCGACGC
Oligo rev AAACGCGTCGGGGGTCCTGGGCGC
Chromosome start end strand MM target_seq PAM alignment distance position gene name gene id
GU305914.1 142954 142976 - 0 GCGCCCAGGACCCCCGACGC CGG ||||||||[||||||||||||]PAM NA Intergenic NA NA

T23277 CGCCCAGGACCCCCGACGCCGGG 1000
Oligo fwd CACCCGCCCAGGACCCCCGACGCC
Oligo rev AAACGGCGTCGGGGGTCCTGGGCG

| | | | | | | | | | | | |
|---|---|---|---|---|---|---|---|---|---|---|---|
| Chromosome | start | end | strand | MM | target_seq | PAM | alignment | distance | position | gene name | gene id |
| GU305914.1 | 142953 | 142975 | - | 0 | CGCCCAGGACCCCCGACGCC | GGG | \|\|\|\|\|\|\|\|[\|\|\|\|\|\|\|\|\|\|\|\|]PAM | NA | Intergenic | NA | NA |
| T23278 | GGACCCCCGACGCCGGGTCCAGG | 1000 | | | | | | | | | |
| Oligo fwd | CACCGGACCCCCGACGCCGGGTCC | | | | | | | | | | |
| Oligo rev | AAACGGACCCGGCGTCGGGGGTCC | | | | | | | | | | |
| Chromosome | start | end | strand | MM | target_seq | PAM | alignment | distance | position | gene name | gene id |
| GU305914.1 | 142947 | 142969 | - | 0 | GGACCCCCGACGCCGGGTCC | AGG | \|\|\|\|\|\|\|\|[\|\|\|\|\|\|\|\|\|\|\|\|]PAM | NA | Intergenic | NA | NA |
| T23279 | GCTCACTCCCCCTCGCCCCTCGG | 1000 | | | | | | | | | |
| Oligo fwd | CACCGCTCACTCCCCCTCGCCCCT | | | | | | | | | | |
| Oligo rev | AAACAGGGGCGAGGGGGAGTGAGC | | | | | | | | | | |
| Chromosome | start | end | strand | MM | target_seq | PAM | alignment | distance | position | gene name | gene id |
| GU305914.1 | 142910 | 142932 | - | 0 | GCTCACTCCCCCTCGCCCCT | CGG | \|\|\|\|\|\|\|\|[\|\|\|\|\|\|\|\|\|\|\|\|]PAM | NA | Intergenic | NA | NA |
| T23280 | CCTGAGCACCTGCGCGATCGCGG | 1000 | | | | | | | | | |
| Oligo fwd | CACCCCTGAGCACCTGCGCGATCG | | | | | | | | | | |
| Oligo rev | AAACCGATCGCGCAGGTGCTCAGG | | | | | | | | | | |
| Chromosome | start | end | strand | MM | target_seq | PAM | alignment | distance | position | gene name | gene id |
| GU305914.1 | 142877 | 142899 | - | 0 | CCTGAGCACCTGCGCGATCG | CGG | \|\|\|\|\|\|\|\|[\|\|\|\|\|\|\|\|\|\|\|\|]PAM | NA | Intergenic | NA | NA |
| T23281 | AGCGCACCGCTTCGCTGCTGTGG | 1000 | | | | | | | | | |
| Oligo fwd | CACCAGCGCACCGCTTCGCTGCTG | | | | | | | | | | |
| Oligo rev | AAACCAGCAGCGAAGCGGTGCGCT | | | | | | | | | | |
| Chromosome | start | end | strand | MM | target_seq | PAM | alignment | distance | position | gene name | gene id |
| GU305914.1 | 142836 | 142858 | - | 0 | AGCGCACCGCTTCGCTGCTG | TGG | \|\|\|\|\|\|\|\|[\|\|\|\|\|\|\|\|\|\|\|\|]PAM | NA | Intergenic | NA | NA |
| T23282 | AACATCCAGATCCTTCCTGAAGG | 1000 | | | | | | | | | |
| Oligo fwd | CACCAACATCCAGATCCTTCCTGA | | | | | | | | | | |
| Oligo rev | AAACTCAGGAAGGATCTGGATGTT | | | | | | | | | | |
| Chromosome | start | end | strand | MM | target_seq | PAM | alignment | distance | position | gene name | gene id |
| GU305914.1 | 142811 | 142833 | - | 0 | AACATCCAGATCCTTCCTGA | AGG | \|\|\|\|\|\|\|\|[\|\|\|\|\|\|\|\|\|\|\|\|]PAM | NA | Intergenic | NA | NA |
| T23283 | AGGCGATCTTACGCGTCGCGAGG | 1000 | | | | | | | | | |
| Oligo fwd | CACCAGGCGATCTTACGCGTCGCG | | | | | | | | | | |
| Oligo rev | AAACCGCGACGCGTAAGATCGCCT | | | | | | | | | | |
| Chromosome | start | end | strand | MM | target_seq | PAM | alignment | distance | position | gene name | gene id |
| GU305914.1 | 142791 | 142813 | - | 0 | AGGCGATCTTACGCGTCGCG | AGG | \|\|\|\|\|\|\|\|[\|\|\|\|\|\|\|\|\|\|\|\|]PAM | NA | Intergenic | NA | NA |
| T23284 | GTGAAACCGCGTCGCTGACACGG | 1000 | | | | | | | | | |
| Oligo fwd | CACCGTGAAACCGCGTCGCTGACA | | | | | | | | | | |
| Oligo rev | AAACTGTCAGCGACGCGGTTTCAC | | | | | | | | | | |
| Chromosome | start | end | strand | MM | target_seq | PAM | alignment | distance | position | gene name | gene id |
| GU305914.1 | 142763 | 142785 | - | 0 | GTGAAACCGCGTCGCTGACA | CGG | \|\|\|\|\|\|\|\|[\|\|\|\|\|\|\|\|\|\|\|\|]PAM | NA | Intergenic | NA | NA |
| T23285 | GACGCGCAGCGCGTGCCAGACGG | 1000 | | | | | | | | | |
| Oligo fwd | CACCGACGCGCAGCGCGTGCCAGA | | | | | | | | | | |
| Oligo rev | AAACTCTGGCACGCGCTGCGCGTC | | | | | | | | | | |
| Chromosome | start | end | strand | MM | target_seq | PAM | alignment | distance | position | gene name | gene id |
| GU305914.1 | 142736 | 142758 | - | 0 | GACGCGCAGCGCGTGCCAGA | CGG | \|\|\|\|\|\|\|\|[\|\|\|\|\|\|\|\|\|\|\|\|]PAM | NA | Intergenic | NA | NA |
| T23286 | ACGCGCAGCGCGTGCCAGACGGG | 1000 | | | | | | | | | |
| Oligo fwd | CACCACGCGCAGCGCGTGCCAGAC | | | | | | | | | | |
| Oligo rev | AAACGTCTGGCACGCGCTGCGCGT | | | | | | | | | | |
| Chromosome | start | end | strand | MM | target_seq | PAM | alignment | distance | position | gene name | gene id |
| GU305914.1 | 142735 | 142757 | - | 0 | ACGCGCAGCGCGTGCCAGAC | GGG | \|\|\|\|\|\|\|\|[\|\|\|\|\|\|\|\|\|\|\|\|]PAM | NA | Intergenic | NA | NA |
| T23287 | CGCGCAGCGCGTGCCAGACGGGG | 1000 | | | | | | | | | |
| Oligo fwd | CACCCGCGCAGCGCGTGCCAGACG | | | | | | | | | | |
| Oligo rev | AAACCGTCTGGCACGCGCTGCGCG | | | | | | | | | | |
| Chromosome | start | end | strand | MM | target_seq | PAM | alignment | distance | position | gene name | gene id |
| GU305914.1 | 142734 | 142756 | - | 0 | CGCGCAGCGCGTGCCAGACG | GGG | \|\|\|\|\|\|\|\|[\|\|\|\|\|\|\|\|\|\|\|\|]PAM | NA | Intergenic | NA | NA |
| T23288 | CGATCGCCTTCATGACCCTGAGG | 1000 | | | | | | | | | |
| Oligo fwd | CACCCGATCGCCTTCATGACCCTG | | | | | | | | | | |
| Oligo rev | AAACCAGGGTCATGAAGGCGATCG | | | | | | | | | | |
| Chromosome | start | end | strand | MM | target_seq | PAM | alignment | distance | position | gene name | gene id |
| GU305914.1 | 142683 | 142705 | - | 0 | CGATCGCCTTCATGACCCTG | AGG | \|\|\|\|\|\|\|\|[\|\|\|\|\|\|\|\|\|\|\|\|]PAM | NA | Intergenic | NA | NA |
| T23289 | CCTTCATGACCCTGAGGAGCCGG | 1000 | | | | | | | | | |
| Oligo fwd | CACCCCTTCATGACCCTGAGGAGC | | | | | | | | | | |
| Oligo rev | AAACGCTCCTCAGGGTCATGAAGG | | | | | | | | | | |
| Chromosome | start | end | strand | MM | target_seq | PAM | alignment | distance | position | gene name | gene id |
| GU305914.1 | 142677 | 142699 | - | 0 | CCTTCATGACCCTGAGGAGC | CGG | \|\|\|\|\|\|\|\|[\|\|\|\|\|\|\|\|\|\|\|\|]PAM | NA | Intergenic | NA | NA |
| T23290 | ACCCTGAGGAGCCGGCGCGCCGG | 1000 | | | | | | | | | |
| Oligo fwd | CACCACCCTGAGGAGCCGGCGCGC | | | | | | | | | | |
| Oligo rev | AAACGCGCGCCGGCTCCTCAGGGT | | | | | | | | | | |
| Chromosome | start | end | strand | MM | target_seq | PAM | alignment | distance | position | gene name | gene id |
| GU305914.1 | 142669 | 142691 | - | 0 | ACCCTGAGGAGCCGGCGCGC | CGG | \|\|\|\|\|\|\|\|[\|\|\|\|\|\|\|\|\|\|\|\|]PAM | NA | Intergenic | NA | NA |
| T23291 | TGAGGAGCCGGCGCGCCGGCAGG | 1000 | | | | | | | | | |
| Oligo fwd | CACCTGAGGAGCCGGCGCGCCGGC | | | | | | | | | | |
| Oligo rev | AAACGCCGGCGCGCCGGCTCCTCA | | | | | | | | | | |
| Chromosome | start | end | strand | MM | target_seq | PAM | alignment | distance | position | gene name | gene id |
| GU305914.1 | 142665 | 142687 | - | 0 | TGAGGAGCCGGCGCGCCGGC | AGG | \|\|\|\|\|\|\|\|[\|\|\|\|\|\|\|\|\|\|\|\|]PAM | NA | Intergenic | NA | NA |
| T23292 | GCGCGCCGGCAGGCGAAAGATGG | 1000 | | | | | | | | | |
| Oligo fwd | CACCGCGCGCCGGCAGGCGAAAGA | | | | | | | | | | |

Oligo rev AAACTCTTTCGCCTGCCGGCGCGC
Chromosome start end strand MM target_seq PAM alignment distance position gene name gene id
GU305914.1 142655 142677 - 0 GCGCGCCGGCAGGCGAAAGA TGG ||||||||[||||||||||||]PAM NA Intergenic NA NA

T23293 ACCCCCGACCGCATCTCTGACGG 1000
Oligo fwd CACCACCCCCGACCGCATCTCTGA
Oligo rev AAACTCAGAGATGCGGTCGGGGGT
Chromosome start end strand MM target_seq PAM alignment distance position gene name gene id
GU305914.1 142601 142623 - 0 ACCCCCGACCGCATCTCTGA CGG ||||||||[||||||||||||]PAM NA Intergenic NA NA

T23294 TCTGACGGTGCTCTCGCCGCCGG 1000
Oligo fwd CACCTCTGACGGTGCTCTCGCCGC
Oligo rev AAACGCGGCGAGAGCACCGTCAGA
Chromosome start end strand MM target_seq PAM alignment distance position gene name gene id
GU305914.1 142586 142608 - 0 TCTGACGGTGCTCTCGCCGC CGG ||||||||[||||||||||||]PAM NA Intergenic NA NA

T23295 CGCCGGCGTTCGAGACCTCGCGG 1000
Oligo fwd CACCCGCCGGCGTTCGAGACCTCG
Oligo rev AAACCGAGGTCTCGAACGCCGGCG
Chromosome start end strand MM target_seq PAM alignment distance position gene name gene id
GU305914.1 142569 142591 - 0 CGCCGGCGTTCGAGACCTCG CGG ||||||||[||||||||||||]PAM NA Intergenic NA NA

T23296 CAAGAAATTAGTAGCCAACCCGG 1000
Oligo fwd CACCCAAGAAATTAGTAGCCAACC
Oligo rev AAACGGTTGGCTACTAATTTCTTG
Chromosome start end strand MM target_seq PAM alignment distance position gene name gene id
GU305914.1 142535 142557 - 0 CAAGAAATTAGTAGCCAACC CGG ||||||||[||||||||||||]PAM NA Intergenic NA NA

T23297 AAGAAATTAGTAGCCAACCCGGG 1000
Oligo fwd CACCAAGAAATTAGTAGCCAACCC
Oligo rev AAACGGGTTGGCTACTAATTTCTT
Chromosome start end strand MM target_seq PAM alignment distance position gene name gene id
GU305914.1 142534 142556 - 0 AAGAAATTAGTAGCCAACCC GGG ||||||||[||||||||||||]PAM NA Intergenic NA NA

T23298 ACCCGGGCCCATCGTGTGTCAGG 1000
Oligo fwd CACCACCCGGGCCCATCGTGTGTC
Oligo rev AAACGACACACGATGGGCCCGGGT
Chromosome start end strand MM target_seq PAM alignment distance position gene name gene id
GU305914.1 142518 142540 - 0 ACCCGGGCCCATCGTGTGTC AGG ||||||||[||||||||||||]PAM NA Intergenic NA NA

T23299 ATACCGACCGAAAATACGCCAGG 1000
Oligo fwd CACCATACCGACCGAAAATACGCC
Oligo rev AAACGGCGTATTTTCGGTCGGTAT
Chromosome start end strand MM target_seq PAM alignment distance position gene name gene id
GU305914.1 142482 142504 - 0 ATACCGACCGAAAATACGCC AGG ||||||||[||||||||||||]PAM NA Intergenic NA NA

T23300 CGCCAGGCGCAACGCGCACACGG 1000
Oligo fwd CACCCGCCAGGCGCAACGCGCACA
Oligo rev AAACTGTGCGCGTTGCGCCTGGCG
Chromosome start end strand MM target_seq PAM alignment distance position gene name gene id
GU305914.1 142466 142488 - 0 CGCCAGGCGCAACGCGCACA CGG ||||||||[||||||||||||]PAM NA Intergenic NA NA

T23301 GCCAGGCGCAACGCGCACACGGG 1000
Oligo fwd CACCGCCAGGCGCAACGCGCACAC
Oligo rev AAACGTGTGCGCGTTGCGCCTGGC
Chromosome start end strand MM target_seq PAM alignment distance position gene name gene id
GU305914.1 142465 142487 - 0 GCCAGGCGCAACGCGCACAC GGG ||||||||[||||||||||||]PAM NA Intergenic NA NA

T23302 CCAGGCGCAACGCGCACACGGGG 1000
Oligo fwd CACCCCAGGCGCAACGCGCACACG
Oligo rev AAACCGTGTGCGCGTTGCGCCTGG
Chromosome start end strand MM target_seq PAM alignment distance position gene name gene id
GU305914.1 142464 142486 - 0 CCAGGCGCAACGCGCACACG GGG ||||||||[||||||||||||]PAM NA Intergenic NA NA

T23303 CAACGCGCACACGGGGACGCTGG 1000
Oligo fwd CACCCAACGCGCACACGGGGACGC
Oligo rev AAACGCGTCCCCGTGTGCGCGTTG
Chromosome start end strand MM target_seq PAM alignment distance position gene name gene id
GU305914.1 142457 142479 - 0 CAACGCGCACACGGGGACGC TGG ||||||||[||||||||||||]PAM NA Intergenic NA NA

T23304 AAACACGCTCCTTGATATCTAGG 1000
Oligo fwd CACCAAACACGCTCCTTGATATCT
Oligo rev AAACAGATATCAAGGAGCGTGTTT
Chromosome start end strand MM target_seq PAM alignment distance position gene name gene id
GU305914.1 142434 142456 - 0 AAACACGCTCCTTGATATCT AGG ||||||||[||||||||||||]PAM NA Intergenic NA NA

T23305 CACGCTCCTTGATATCTAGGAGG 1000
Oligo fwd CACCCACGCTCCTTGATATCTAGG
Oligo rev AAACCCTAGATATCAAGGAGCGTG
Chromosome start end strand MM target_seq PAM alignment distance position gene name gene id
GU305914.1 142431 142453 - 0 CACGCTCCTTGATATCTAGG AGG ||||||||[||||||||||||]PAM NA Intergenic NA NA

T23306 TGATATCTAGGAGGTTATACAGG 1000
Oligo fwd CACCTGATATCTAGGAGGTTATAC
Oligo rev AAACGTATAACCTCCTAGATATCA
Chromosome start end strand MM target_seq PAM alignment distance position gene name gene id
GU305914.1 142422 142444 - 0 TGATATCTAGGAGGTTATAC AGG ||||||||[||||||||||||]PAM NA Intergenic NA NA

T23307 GTAAATATCAAAAGATTCGCTGG 1000

<table>
<tr><td>Oligo fwd</td><td>CACCGTAAATATCAAAAGATTCGC</td><td></td><td></td><td></td><td></td><td></td><td></td><td></td><td></td><td></td><td></td></tr>
<tr><td>Oligo rev</td><td>AAACGCGAATCTTTTGATATTTAC</td><td></td><td></td><td></td><td></td><td></td><td></td><td></td><td></td><td></td><td></td></tr>
<tr><td>Chromosome</td><td>start</td><td>end</td><td>strand</td><td>MM</td><td>target_seq</td><td>PAM</td><td>alignment</td><td>distance</td><td>position</td><td>gene name</td><td>gene id</td></tr>
<tr><td>GU305914.1</td><td>142397</td><td>142419</td><td>-</td><td>0</td><td>GTAAATATCAAAAGATTCGC</td><td>TGG</td><td>||||||||[||||||||||||]PAM</td><td>NA</td><td>Intergenic</td><td>NA</td><td>NA</td></tr>
<tr><td></td><td></td><td></td><td></td><td></td><td></td><td></td><td></td><td></td><td></td><td></td><td></td></tr>
<tr><td>T23308</td><td>GCTCGCTCTGATCGAAGACGAGG</td><td>1000</td><td></td><td></td><td></td><td></td><td></td><td></td><td></td><td></td><td></td></tr>
<tr><td>Oligo fwd</td><td>CACCGCTCGCTCTGATCGAAGACG</td><td></td><td></td><td></td><td></td><td></td><td></td><td></td><td></td><td></td><td></td></tr>
<tr><td>Oligo rev</td><td>AAACCGTCTTCGATCAGAGCGAGC</td><td></td><td></td><td></td><td></td><td></td><td></td><td></td><td></td><td></td><td></td></tr>
<tr><td>Chromosome</td><td>start</td><td>end</td><td>strand</td><td>MM</td><td>target_seq</td><td>PAM</td><td>alignment</td><td>distance</td><td>position</td><td>gene name</td><td>gene id</td></tr>
<tr><td>GU305914.1</td><td>142344</td><td>142366</td><td>-</td><td>0</td><td>GCTCGCTCTGATCGAAGACG</td><td>AGG</td><td>||||||||[||||||||||||]PAM</td><td>NA</td><td>Intergenic</td><td>NA</td><td>NA</td></tr>
<tr><td></td><td></td><td></td><td></td><td></td><td></td><td></td><td></td><td></td><td></td><td></td><td></td></tr>
<tr><td>T23309</td><td>GACGAGGCACTCGTTCCGTTCGG</td><td>1000</td><td></td><td></td><td></td><td></td><td></td><td></td><td></td><td></td><td></td></tr>
<tr><td>Oligo fwd</td><td>CACCGACGAGGCACTCGTTCCGTT</td><td></td><td></td><td></td><td></td><td></td><td></td><td></td><td></td><td></td><td></td></tr>
<tr><td>Oligo rev</td><td>AAACAACGGAACGAGTGCCTCGTC</td><td></td><td></td><td></td><td></td><td></td><td></td><td></td><td></td><td></td><td></td></tr>
<tr><td>Chromosome</td><td>start</td><td>end</td><td>strand</td><td>MM</td><td>target_seq</td><td>PAM</td><td>alignment</td><td>distance</td><td>position</td><td>gene name</td><td>gene id</td></tr>
<tr><td>GU305914.1</td><td>142328</td><td>142350</td><td>-</td><td>0</td><td>GACGAGGCACTCGTTCCGTT</td><td>CGG</td><td>||||||||[||||||||||||]PAM</td><td>NA</td><td>Intergenic</td><td>NA</td><td>NA</td></tr>
<tr><td></td><td></td><td></td><td></td><td></td><td></td><td></td><td></td><td></td><td></td><td></td><td></td></tr>
<tr><td>T23310</td><td>CGAACGCGCTGTCGATATTATGG</td><td>1000</td><td></td><td></td><td></td><td></td><td></td><td></td><td></td><td></td><td></td></tr>
<tr><td>Oligo fwd</td><td>CACCCGAACGCGCTGTCGATATTA</td><td></td><td></td><td></td><td></td><td></td><td></td><td></td><td></td><td></td><td></td></tr>
<tr><td>Oligo rev</td><td>AAACTAATATCGACAGCGCGTTCG</td><td></td><td></td><td></td><td></td><td></td><td></td><td></td><td></td><td></td><td></td></tr>
<tr><td>Chromosome</td><td>start</td><td>end</td><td>strand</td><td>MM</td><td>target_seq</td><td>PAM</td><td>alignment</td><td>distance</td><td>position</td><td>gene name</td><td>gene id</td></tr>
<tr><td>GU305914.1</td><td>142299</td><td>142321</td><td>-</td><td>0</td><td>CGAACGCGCTGTCGATATTA</td><td>TGG</td><td>||||||||[||||||||||||]PAM</td><td>NA</td><td>Intergenic</td><td>NA</td><td>NA</td></tr>
<tr><td></td><td></td><td></td><td></td><td></td><td></td><td></td><td></td><td></td><td></td><td></td><td></td></tr>
<tr><td>T23311</td><td>GAACGCGCTGTCGATATTATGGG</td><td>1000</td><td></td><td></td><td></td><td></td><td></td><td></td><td></td><td></td><td></td></tr>
<tr><td>Oligo fwd</td><td>CACCGAACGCGCTGTCGATATTAT</td><td></td><td></td><td></td><td></td><td></td><td></td><td></td><td></td><td></td><td></td></tr>
<tr><td>Oligo rev</td><td>AAACATAATATCGACAGCGCGTTC</td><td></td><td></td><td></td><td></td><td></td><td></td><td></td><td></td><td></td><td></td></tr>
<tr><td>Chromosome</td><td>start</td><td>end</td><td>strand</td><td>MM</td><td>target_seq</td><td>PAM</td><td>alignment</td><td>distance</td><td>position</td><td>gene name</td><td>gene id</td></tr>
<tr><td>GU305914.1</td><td>142298</td><td>142320</td><td>-</td><td>0</td><td>GAACGCGCTGTCGATATTAT</td><td>GGG</td><td>||||||||[||||||||||||]PAM</td><td>NA</td><td>Intergenic</td><td>NA</td><td>NA</td></tr>
<tr><td></td><td></td><td></td><td></td><td></td><td></td><td></td><td></td><td></td><td></td><td></td><td></td></tr>
<tr><td>T23312</td><td>ACGTCGTCGACGATCCGCCGCGG</td><td>1000</td><td></td><td></td><td></td><td></td><td></td><td></td><td></td><td></td><td></td></tr>
<tr><td>Oligo fwd</td><td>CACCACGTCGTCGACGATCCGCCG</td><td></td><td></td><td></td><td></td><td></td><td></td><td></td><td></td><td></td><td></td></tr>
<tr><td>Oligo rev</td><td>AAACCGGCGGATCGTCGACGACGT</td><td></td><td></td><td></td><td></td><td></td><td></td><td></td><td></td><td></td><td></td></tr>
<tr><td>Chromosome</td><td>start</td><td>end</td><td>strand</td><td>MM</td><td>target_seq</td><td>PAM</td><td>alignment</td><td>distance</td><td>position</td><td>gene name</td><td>gene id</td></tr>
<tr><td>GU305914.1</td><td>142246</td><td>142268</td><td>-</td><td>0</td><td>ACGTCGTCGACGATCCGCCG</td><td>CGG</td><td>||||||||[||||||||||||]PAM</td><td>NA</td><td>Intergenic</td><td>NA</td><td>NA</td></tr>
<tr><td></td><td></td><td></td><td></td><td></td><td></td><td></td><td></td><td></td><td></td><td></td><td></td></tr>
<tr><td>T23313</td><td>GTCGTCATCGCCGTCGCCTCCGG</td><td>1000</td><td></td><td></td><td></td><td></td><td></td><td></td><td></td><td></td><td></td></tr>
<tr><td>Oligo fwd</td><td>CACCGTCGTCATCGCCGTCGCCTC</td><td></td><td></td><td></td><td></td><td></td><td></td><td></td><td></td><td></td><td></td></tr>
<tr><td>Oligo rev</td><td>AAACGAGGCGACGGCGATGACGAC</td><td></td><td></td><td></td><td></td><td></td><td></td><td></td><td></td><td></td><td></td></tr>
<tr><td>Chromosome</td><td>start</td><td>end</td><td>strand</td><td>MM</td><td>target_seq</td><td>PAM</td><td>alignment</td><td>distance</td><td>position</td><td>gene name</td><td>gene id</td></tr>
<tr><td>GU305914.1</td><td>142219</td><td>142241</td><td>-</td><td>0</td><td>GTCGTCATCGCCGTCGCCTC</td><td>CGG</td><td>||||||||[||||||||||||]PAM</td><td>NA</td><td>Intergenic</td><td>NA</td><td>NA</td></tr>
<tr><td></td><td></td><td></td><td></td><td></td><td></td><td></td><td></td><td></td><td></td><td></td><td></td></tr>
<tr><td>T23314</td><td>GTCATCGCCGTCGCCTCCGGAGG</td><td>1000</td><td></td><td></td><td></td><td></td><td></td><td></td><td></td><td></td><td></td></tr>
<tr><td>Oligo fwd</td><td>CACCGTCATCGCCGTCGCCTCCGG</td><td></td><td></td><td></td><td></td><td></td><td></td><td></td><td></td><td></td><td></td></tr>
<tr><td>Oligo rev</td><td>AAACCCGGAGGCGACGGCGATGAC</td><td></td><td></td><td></td><td></td><td></td><td></td><td></td><td></td><td></td><td></td></tr>
<tr><td>Chromosome</td><td>start</td><td>end</td><td>strand</td><td>MM</td><td>target_seq</td><td>PAM</td><td>alignment</td><td>distance</td><td>position</td><td>gene name</td><td>gene id</td></tr>
<tr><td>GU305914.1</td><td>142216</td><td>142238</td><td>-</td><td>0</td><td>GTCATCGCCGTCGCCTCCGG</td><td>AGG</td><td>||||||||[||||||||||||]PAM</td><td>NA</td><td>Intergenic</td><td>NA</td><td>NA</td></tr>
<tr><td>GU305914.1</td><td>60614</td><td>60636</td><td>-</td><td>4</td><td>GGCGTCTCCGTCGTCTCCGG</td><td>CGG</td><td>|-|-||-|[|||||-||||||]PAM</td><td>NA</td><td>Intergenic</td><td>NA</td><td>NA</td></tr>
<tr><td></td><td></td><td></td><td></td><td></td><td></td><td></td><td></td><td></td><td></td><td></td><td></td></tr>
<tr><td>T23315</td><td>TCATCGCCGTCGCCTCCGGAGGG</td><td>1000</td><td></td><td></td><td></td><td></td><td></td><td></td><td></td><td></td><td></td></tr>
<tr><td>Oligo fwd</td><td>CACCTCATCGCCGTCGCCTCCGGA</td><td></td><td></td><td></td><td></td><td></td><td></td><td></td><td></td><td></td><td></td></tr>
<tr><td>Oligo rev</td><td>AAACTCCGGAGGCGACGGCGATGA</td><td></td><td></td><td></td><td></td><td></td><td></td><td></td><td></td><td></td><td></td></tr>
<tr><td>Chromosome</td><td>start</td><td>end</td><td>strand</td><td>MM</td><td>target_seq</td><td>PAM</td><td>alignment</td><td>distance</td><td>position</td><td>gene name</td><td>gene id</td></tr>
<tr><td>GU305914.1</td><td>142215</td><td>142237</td><td>-</td><td>0</td><td>TCATCGCCGTCGCCTCCGGA</td><td>GGG</td><td>||||||||[||||||||||||]PAM</td><td>NA</td><td>Intergenic</td><td>NA</td><td>NA</td></tr>
<tr><td></td><td></td><td></td><td></td><td></td><td></td><td></td><td></td><td></td><td></td><td></td><td></td></tr>
<tr><td>T23316</td><td>TCGCCGTCGCCTCCGGAGGGAGG</td><td>1000</td><td></td><td></td><td></td><td></td><td></td><td></td><td></td><td></td><td></td></tr>
<tr><td>Oligo fwd</td><td>CACCTCGCCGTCGCCTCCGGAGGG</td><td></td><td></td><td></td><td></td><td></td><td></td><td></td><td></td><td></td><td></td></tr>
<tr><td>Oligo rev</td><td>AAACCCCTCCGGAGGCGACGGCGA</td><td></td><td></td><td></td><td></td><td></td><td></td><td></td><td></td><td></td><td></td></tr>
<tr><td>Chromosome</td><td>start</td><td>end</td><td>strand</td><td>MM</td><td>target_seq</td><td>PAM</td><td>alignment</td><td>distance</td><td>position</td><td>gene name</td><td>gene id</td></tr>
<tr><td>GU305914.1</td><td>142212</td><td>142234</td><td>-</td><td>0</td><td>TCGCCGTCGCCTCCGGAGGG</td><td>AGG</td><td>||||||||[||||||||||||]PAM</td><td>NA</td><td>Intergenic</td><td>NA</td><td>NA</td></tr>
<tr><td></td><td></td><td></td><td></td><td></td><td></td><td></td><td></td><td></td><td></td><td></td><td></td></tr>
<tr><td>T23317</td><td>AGGACCGTCGTCGCCATATCTGG</td><td>1000</td><td></td><td></td><td></td><td></td><td></td><td></td><td></td><td></td><td></td></tr>
<tr><td>Oligo fwd</td><td>CACCAGGACCGTCGTCGCCATATC</td><td></td><td></td><td></td><td></td><td></td><td></td><td></td><td></td><td></td><td></td></tr>
<tr><td>Oligo rev</td><td>AAACGATATGGCGACGACGGTCCT</td><td></td><td></td><td></td><td></td><td></td><td></td><td></td><td></td><td></td><td></td></tr>
<tr><td>Chromosome</td><td>start</td><td>end</td><td>strand</td><td>MM</td><td>target_seq</td><td>PAM</td><td>alignment</td><td>distance</td><td>position</td><td>gene name</td><td>gene id</td></tr>
<tr><td>GU305914.1</td><td>142192</td><td>142214</td><td>-</td><td>0</td><td>AGGACCGTCGTCGCCATATC</td><td>TGG</td><td>||||||||[||||||||||||]PAM</td><td>NA</td><td>Intergenic</td><td>NA</td><td>NA</td></tr>
<tr><td></td><td></td><td></td><td></td><td></td><td></td><td></td><td></td><td></td><td></td><td></td><td></td></tr>
<tr><td>T23318</td><td>ATATCTGGATTTCGCCGACCCGG</td><td>1000</td><td></td><td></td><td></td><td></td><td></td><td></td><td></td><td></td><td></td></tr>
<tr><td>Oligo fwd</td><td>CACCATATCTGGATTTCGCCGACC</td><td></td><td></td><td></td><td></td><td></td><td></td><td></td><td></td><td></td><td></td></tr>
<tr><td>Oligo rev</td><td>AAACGGTCGGCGAAATCCAGATAT</td><td></td><td></td><td></td><td></td><td></td><td></td><td></td><td></td><td></td><td></td></tr>
<tr><td>Chromosome</td><td>start</td><td>end</td><td>strand</td><td>MM</td><td>target_seq</td><td>PAM</td><td>alignment</td><td>distance</td><td>position</td><td>gene name</td><td>gene id</td></tr>
<tr><td>GU305914.1</td><td>142177</td><td>142199</td><td>-</td><td>0</td><td>ATATCTGGATTTCGCCGACC</td><td>CGG</td><td>||||||||[||||||||||||]PAM</td><td>NA</td><td>Intergenic</td><td>NA</td><td>NA</td></tr>
<tr><td></td><td></td><td></td><td></td><td></td><td></td><td></td><td></td><td></td><td></td><td></td><td></td></tr>
<tr><td>T23319</td><td>TCTGGATTTCGCCGACCCGGAGG</td><td>1000</td><td></td><td></td><td></td><td></td><td></td><td></td><td></td><td></td><td></td></tr>
<tr><td>Oligo fwd</td><td>CACCTCTGGATTTCGCCGACCCGG</td><td></td><td></td><td></td><td></td><td></td><td></td><td></td><td></td><td></td><td></td></tr>
<tr><td>Oligo rev</td><td>AAACCCGGGTCGGCGAAATCCAGA</td><td></td><td></td><td></td><td></td><td></td><td></td><td></td><td></td><td></td><td></td></tr>
<tr><td>Chromosome</td><td>start</td><td>end</td><td>strand</td><td>MM</td><td>target_seq</td><td>PAM</td><td>alignment</td><td>distance</td><td>position</td><td>gene name</td><td>gene id</td></tr>
<tr><td>GU305914.1</td><td>142174</td><td>142196</td><td>-</td><td>0</td><td>TCTGGATTTCGCCGACCCGG</td><td>AGG</td><td>||||||||[||||||||||||]PAM</td><td>NA</td><td>Intergenic</td><td>NA</td><td>NA</td></tr>
<tr><td></td><td></td><td></td><td></td><td></td><td></td><td></td><td></td><td></td><td></td><td></td><td></td></tr>
<tr><td>T23320</td><td>CGCCGACCCGGAGGACGATCAGG</td><td>1000</td><td></td><td></td><td></td><td></td><td></td><td></td><td></td><td></td><td></td></tr>
<tr><td>Oligo fwd</td><td>CACCCGCCGACCCGGAGGACGATC</td><td></td><td></td><td></td><td></td><td></td><td></td><td></td><td></td><td></td><td></td></tr>
<tr><td>Oligo rev</td><td>AAACGATCGTCCTCCGGGTCGGCG</td><td></td><td></td><td></td><td></td><td></td><td></td><td></td><td></td><td></td><td></td></tr>
<tr><td>Chromosome</td><td>start</td><td>end</td><td>strand</td><td>MM</td><td>target_seq</td><td>PAM</td><td>alignment</td><td>distance</td><td>position</td><td>gene name</td><td>gene id</td></tr>
<tr><td>GU305914.1</td><td>142165</td><td>142187</td><td>-</td><td>0</td><td>CGCCGACCCGGAGGACGATC</td><td>AGG</td><td>||||||||[||||||||||||]PAM</td><td>NA</td><td>Intergenic</td><td>NA</td><td>NA</td></tr>
<tr><td></td><td></td><td></td><td></td><td></td><td></td><td></td><td></td><td></td><td></td><td></td><td></td></tr>
<tr><td>T23321</td><td>GCCGACCCGGAGGACGATCAGGG</td><td>1000</td><td></td><td></td><td></td><td></td><td></td><td></td><td></td><td></td><td></td></tr>
<tr><td>Oligo fwd</td><td>CACCGCCGACCCGGAGGACGATCA</td><td></td><td></td><td></td><td></td><td></td><td></td><td></td><td></td><td></td><td></td></tr>
<tr><td>Oligo rev</td><td>AAACTGATCGTCCTCCGGGTCGGC</td><td></td><td></td><td></td><td></td><td></td><td></td><td></td><td></td><td></td><td></td></tr>
<tr><td>Chromosome</td><td>start</td><td>end</td><td>strand</td><td>MM</td><td>target_seq</td><td>PAM</td><td>alignment</td><td>distance</td><td>position</td><td>gene name</td><td>gene id</td></tr>
<tr><td>GU305914.1</td><td>142164</td><td>142186</td><td>-</td><td>0</td><td>GCCGACCCGGAGGACGATCA</td><td>GGG</td><td>||||||||[||||||||||||]PAM</td><td>NA</td><td>Intergenic</td><td>NA</td><td>NA</td></tr>
</table>

<table>
<tr><td>GU305914.1</td><td>118304</td><td>118326</td><td>-</td><td>4</td><td>GCCGTCGCGGTGGACGATTA</td><td>CGG</td><td>||||-|-|[||-|||||||-|]PAM</td><td>NA</td><td>Intergenic</td><td>NA</td><td>NA</td></tr>
<tr><td>T23322</td><td>GATCAGGGAAAGATCTTGTACGG</td><td>1000</td><td></td><td></td><td></td><td></td><td></td><td></td><td></td><td></td><td></td></tr>
<tr><td>Oligo fwd</td><td>CACCGATCAGGGAAAGATCTTGTA</td><td></td><td></td><td></td><td></td><td></td><td></td><td></td><td></td><td></td><td></td></tr>
<tr><td>Oligo rev</td><td>AAACTACAAGATCTTTCCCTGATC</td><td></td><td></td><td></td><td></td><td></td><td></td><td></td><td></td><td></td><td></td></tr>
<tr><td>Chromosome</td><td>start</td><td>end</td><td>strand</td><td>MM</td><td>target_seq</td><td>PAM</td><td>alignment</td><td>distance</td><td>position</td><td>gene name</td><td>gene id</td></tr>
<tr><td>GU305914.1</td><td>142149</td><td>142171</td><td>-</td><td>0</td><td>GATCAGGGAAAGATCTTGTA</td><td>CGG</td><td>||||||||[||||||||||||]PAM</td><td>NA</td><td>Intergenic</td><td>NA</td><td>NA</td></tr>
<tr><td>T23323</td><td>TCGCACGTATGTTATCGAGAAGG</td><td>1000</td><td></td><td></td><td></td><td></td><td></td><td></td><td></td><td></td><td></td></tr>
<tr><td>Oligo fwd</td><td>CACCTCGCACGTATGTTATCGAGA</td><td></td><td></td><td></td><td></td><td></td><td></td><td></td><td></td><td></td><td></td></tr>
<tr><td>Oligo rev</td><td>AAACTCTCGATAACATACGTGCGA</td><td></td><td></td><td></td><td></td><td></td><td></td><td></td><td></td><td></td><td></td></tr>
<tr><td>Chromosome</td><td>start</td><td>end</td><td>strand</td><td>MM</td><td>target_seq</td><td>PAM</td><td>alignment</td><td>distance</td><td>position</td><td>gene name</td><td>gene id</td></tr>
<tr><td>GU305914.1</td><td>142088</td><td>142110</td><td>-</td><td>0</td><td>TCGCACGTATGTTATCGAGA</td><td>AGG</td><td>||||||||[||||||||||||]PAM</td><td>NA</td><td>Intergenic</td><td>NA</td><td>NA</td></tr>
<tr><td>T23324</td><td>ACCGTCCTCCGTCTCCGACGCGG</td><td>1000</td><td></td><td></td><td></td><td></td><td></td><td></td><td></td><td></td><td></td></tr>
<tr><td>Oligo fwd</td><td>CACCACCGTCCTCCGTCTCCGACG</td><td></td><td></td><td></td><td></td><td></td><td></td><td></td><td></td><td></td><td></td></tr>
<tr><td>Oligo rev</td><td>AAACCGTCGGAGACGGAGGACGGT</td><td></td><td></td><td></td><td></td><td></td><td></td><td></td><td></td><td></td><td></td></tr>
<tr><td>Chromosome</td><td>start</td><td>end</td><td>strand</td><td>MM</td><td>target_seq</td><td>PAM</td><td>alignment</td><td>distance</td><td>position</td><td>gene name</td><td>gene id</td></tr>
<tr><td>GU305914.1</td><td>142060</td><td>142082</td><td>-</td><td>0</td><td>ACCGTCCTCCGTCTCCGACG</td><td>CGG</td><td>||||||||[||||||||||||]PAM</td><td>NA</td><td>Intergenic</td><td>NA</td><td>NA</td></tr>
<tr><td>T23325</td><td>CCGTCTCCGACGCGGCTAACAGG</td><td>1000</td><td></td><td></td><td></td><td></td><td></td><td></td><td></td><td></td><td></td></tr>
<tr><td>Oligo fwd</td><td>CACCCCGTCTCCGACGCGGCTAAC</td><td></td><td></td><td></td><td></td><td></td><td></td><td></td><td></td><td></td><td></td></tr>
<tr><td>Oligo rev</td><td>AAACGTTAGCCGCGTCGGAGACGG</td><td></td><td></td><td></td><td></td><td></td><td></td><td></td><td></td><td></td><td></td></tr>
<tr><td>Chromosome</td><td>start</td><td>end</td><td>strand</td><td>MM</td><td>target_seq</td><td>PAM</td><td>alignment</td><td>distance</td><td>position</td><td>gene name</td><td>gene id</td></tr>
<tr><td>GU305914.1</td><td>142052</td><td>142074</td><td>-</td><td>0</td><td>CCGTCTCCGACGCGGCTAAC</td><td>AGG</td><td>||||||||[||||||||||||]PAM</td><td>NA</td><td>Intergenic</td><td>NA</td><td>NA</td></tr>
<tr><td>T23326</td><td>CCGACGCGGCTAACAGGAGCCGG</td><td>1000</td><td></td><td></td><td></td><td></td><td></td><td></td><td></td><td></td><td></td></tr>
<tr><td>Oligo fwd</td><td>CACCCCGACGCGGCTAACAGGAGC</td><td></td><td></td><td></td><td></td><td></td><td></td><td></td><td></td><td></td><td></td></tr>
<tr><td>Oligo rev</td><td>AAACGCTCCTGTTAGCCGCGTCGG</td><td></td><td></td><td></td><td></td><td></td><td></td><td></td><td></td><td></td><td></td></tr>
<tr><td>Chromosome</td><td>start</td><td>end</td><td>strand</td><td>MM</td><td>target_seq</td><td>PAM</td><td>alignment</td><td>distance</td><td>position</td><td>gene name</td><td>gene id</td></tr>
<tr><td>GU305914.1</td><td>142046</td><td>142068</td><td>-</td><td>0</td><td>CCGACGCGGCTAACAGGAGC</td><td>CGG</td><td>||||||||[||||||||||||]PAM</td><td>NA</td><td>Intergenic</td><td>NA</td><td>NA</td></tr>
<tr><td>T23327</td><td>TTTTATAGACGCTCCGAAAACGG</td><td>1000</td><td></td><td></td><td></td><td></td><td></td><td></td><td></td><td></td><td></td></tr>
<tr><td>Oligo fwd</td><td>CACCTTTTATAGACGCTCCGAAAA</td><td></td><td></td><td></td><td></td><td></td><td></td><td></td><td></td><td></td><td></td></tr>
<tr><td>Oligo rev</td><td>AAACTTTTCGGAGCGTCTATAAAA</td><td></td><td></td><td></td><td></td><td></td><td></td><td></td><td></td><td></td><td></td></tr>
<tr><td>Chromosome</td><td>start</td><td>end</td><td>strand</td><td>MM</td><td>target_seq</td><td>PAM</td><td>alignment</td><td>distance</td><td>position</td><td>gene name</td><td>gene id</td></tr>
<tr><td>GU305914.1</td><td>141989</td><td>142011</td><td>-</td><td>0</td><td>TTTTATAGACGCTCCGAAAA</td><td>CGG</td><td>||||||||[||||||||||||]PAM</td><td>NA</td><td>Intergenic</td><td>NA</td><td>NA</td></tr>
<tr><td>T23328</td><td>ATAGACGCTCCGAAAACGGTCGG</td><td>1000</td><td></td><td></td><td></td><td></td><td></td><td></td><td></td><td></td><td></td></tr>
<tr><td>Oligo fwd</td><td>CACCATAGACGCTCCGAAAACGGT</td><td></td><td></td><td></td><td></td><td></td><td></td><td></td><td></td><td></td><td></td></tr>
<tr><td>Oligo rev</td><td>AAACACCGTTTTCGGAGCGTCTAT</td><td></td><td></td><td></td><td></td><td></td><td></td><td></td><td></td><td></td><td></td></tr>
<tr><td>Chromosome</td><td>start</td><td>end</td><td>strand</td><td>MM</td><td>target_seq</td><td>PAM</td><td>alignment</td><td>distance</td><td>position</td><td>gene name</td><td>gene id</td></tr>
<tr><td>GU305914.1</td><td>141985</td><td>142007</td><td>-</td><td>0</td><td>ATAGACGCTCCGAAAACGGT</td><td>CGG</td><td>||||||||[||||||||||||]PAM</td><td>NA</td><td>Intergenic</td><td>NA</td><td>NA</td></tr>
<tr><td>T23329</td><td>TAGACGCTCCGAAAACGGTCGGG</td><td>1000</td><td></td><td></td><td></td><td></td><td></td><td></td><td></td><td></td><td></td></tr>
<tr><td>Oligo fwd</td><td>CACCTAGACGCTCCGAAAACGGTC</td><td></td><td></td><td></td><td></td><td></td><td></td><td></td><td></td><td></td><td></td></tr>
<tr><td>Oligo rev</td><td>AAACGACCGTTTTCGGAGCGTCTA</td><td></td><td></td><td></td><td></td><td></td><td></td><td></td><td></td><td></td><td></td></tr>
<tr><td>Chromosome</td><td>start</td><td>end</td><td>strand</td><td>MM</td><td>target_seq</td><td>PAM</td><td>alignment</td><td>distance</td><td>position</td><td>gene name</td><td>gene id</td></tr>
<tr><td>GU305914.1</td><td>141984</td><td>142006</td><td>-</td><td>0</td><td>TAGACGCTCCGAAAACGGTC</td><td>GGG</td><td>||||||||[||||||||||||]PAM</td><td>NA</td><td>Intergenic</td><td>NA</td><td>NA</td></tr>
<tr><td>T23330</td><td>TAGTCACACAACCGCGCCATAGG</td><td>1000</td><td></td><td></td><td></td><td></td><td></td><td></td><td></td><td></td><td></td></tr>
<tr><td>Oligo fwd</td><td>CACCTAGTCACACAACCGCGCCAT</td><td></td><td></td><td></td><td></td><td></td><td></td><td></td><td></td><td></td><td></td></tr>
<tr><td>Oligo rev</td><td>AAACATGGCGCGGTTGTGTGACTA</td><td></td><td></td><td></td><td></td><td></td><td></td><td></td><td></td><td></td><td></td></tr>
<tr><td>Chromosome</td><td>start</td><td>end</td><td>strand</td><td>MM</td><td>target_seq</td><td>PAM</td><td>alignment</td><td>distance</td><td>position</td><td>gene name</td><td>gene id</td></tr>
<tr><td>GU305914.1</td><td>141933</td><td>141955</td><td>-</td><td>0</td><td>TAGTCACACAACCGCGCCAT</td><td>AGG</td><td>||||||||[||||||||||||]PAM</td><td>NA</td><td>Intergenic</td><td>NA</td><td>NA</td></tr>
<tr><td>T23331</td><td>CATGTCGTCGCTGCCCAGCAAGG</td><td>1000</td><td></td><td></td><td></td><td></td><td></td><td></td><td></td><td></td><td></td></tr>
<tr><td>Oligo fwd</td><td>CACCCATGTCGTCGCTGCCCAGCA</td><td></td><td></td><td></td><td></td><td></td><td></td><td></td><td></td><td></td><td></td></tr>
<tr><td>Oligo rev</td><td>AAACTGCTGGGCAGCGACGACATG</td><td></td><td></td><td></td><td></td><td></td><td></td><td></td><td></td><td></td><td></td></tr>
<tr><td>Chromosome</td><td>start</td><td>end</td><td>strand</td><td>MM</td><td>target_seq</td><td>PAM</td><td>alignment</td><td>distance</td><td>position</td><td>gene name</td><td>gene id</td></tr>
<tr><td>GU305914.1</td><td>141904</td><td>141926</td><td>-</td><td>0</td><td>CATGTCGTCGCTGCCCAGCA</td><td>AGG</td><td>||||||||[||||||||||||]PAM</td><td>NA</td><td>Intergenic</td><td>NA</td><td>NA</td></tr>
<tr><td>GU305914.1</td><td>60241</td><td>60263</td><td>+</td><td>4</td><td>CAGGTCGACGCTGGCCCGCA</td><td>CGG</td><td>||-||||-[|||||-||-|||]PAM</td><td>NA</td><td>Intergenic</td><td>NA</td><td>NA</td></tr>
<tr><td>T23332</td><td>GTCGTCGCTGCCCAGCAAGGTGG</td><td>1000</td><td></td><td></td><td></td><td></td><td></td><td></td><td></td><td></td><td></td></tr>
<tr><td>Oligo fwd</td><td>CACCGTCGTCGCTGCCCAGCAAGG</td><td></td><td></td><td></td><td></td><td></td><td></td><td></td><td></td><td></td><td></td></tr>
<tr><td>Oligo rev</td><td>AAACCCTTGCTGGGCAGCGACGAC</td><td></td><td></td><td></td><td></td><td></td><td></td><td></td><td></td><td></td><td></td></tr>
<tr><td>Chromosome</td><td>start</td><td>end</td><td>strand</td><td>MM</td><td>target_seq</td><td>PAM</td><td>alignment</td><td>distance</td><td>position</td><td>gene name</td><td>gene id</td></tr>
<tr><td>GU305914.1</td><td>141901</td><td>141923</td><td>-</td><td>0</td><td>GTCGTCGCTGCCCAGCAAGG</td><td>TGG</td><td>||||||||[||||||||||||]PAM</td><td>NA</td><td>Intergenic</td><td>NA</td><td>NA</td></tr>
<tr><td>T23333</td><td>GTCGCTGCCCAGCAAGGTGGCGG</td><td>1000</td><td></td><td></td><td></td><td></td><td></td><td></td><td></td><td></td><td></td></tr>
<tr><td>Oligo fwd</td><td>CACCGTCGCTGCCCAGCAAGGTGG</td><td></td><td></td><td></td><td></td><td></td><td></td><td></td><td></td><td></td><td></td></tr>
<tr><td>Oligo rev</td><td>AAACCCACCTTGCTGGGCAGCGAC</td><td></td><td></td><td></td><td></td><td></td><td></td><td></td><td></td><td></td><td></td></tr>
<tr><td>Chromosome</td><td>start</td><td>end</td><td>strand</td><td>MM</td><td>target_seq</td><td>PAM</td><td>alignment</td><td>distance</td><td>position</td><td>gene name</td><td>gene id</td></tr>
<tr><td>GU305914.1</td><td>141898</td><td>141920</td><td>-</td><td>0</td><td>GTCGCTGCCCAGCAAGGTGG</td><td>CGG</td><td>||||||||[||||||||||||]PAM</td><td>NA</td><td>Intergenic</td><td>NA</td><td>NA</td></tr>
<tr><td>T23334</td><td>GCCCAGCAAGGTGGCGGTGTAGG</td><td>1000</td><td></td><td></td><td></td><td></td><td></td><td></td><td></td><td></td><td></td></tr>
<tr><td>Oligo fwd</td><td>CACCGCCCAGCAAGGTGGCGGTGT</td><td></td><td></td><td></td><td></td><td></td><td></td><td></td><td></td><td></td><td></td></tr>
<tr><td>Oligo rev</td><td>AAACACACCGCCACCTTGCTGGGC</td><td></td><td></td><td></td><td></td><td></td><td></td><td></td><td></td><td></td><td></td></tr>
<tr><td>Chromosome</td><td>start</td><td>end</td><td>strand</td><td>MM</td><td>target_seq</td><td>PAM</td><td>alignment</td><td>distance</td><td>position</td><td>gene name</td><td>gene id</td></tr>
<tr><td>GU305914.1</td><td>141892</td><td>141914</td><td>-</td><td>0</td><td>GCCCAGCAAGGTGGCGGTGT</td><td>AGG</td><td>||||||||[||||||||||||]PAM</td><td>NA</td><td>Intergenic</td><td>NA</td><td>NA</td></tr>
<tr><td>T23335</td><td>AGGTGGCGGTGTAGGCCAGCTGG</td><td>1000</td><td></td><td></td><td></td><td></td><td></td><td></td><td></td><td></td><td></td></tr>
<tr><td>Oligo fwd</td><td>CACCAGGTGGCGGTGTAGGCCAGC</td><td></td><td></td><td></td><td></td><td></td><td></td><td></td><td></td><td></td><td></td></tr>
<tr><td>Oligo rev</td><td>AAACGCTGGCCTACACCGCCACCT</td><td></td><td></td><td></td><td></td><td></td><td></td><td></td><td></td><td></td><td></td></tr>
<tr><td>Chromosome</td><td>start</td><td>end</td><td>strand</td><td>MM</td><td>target_seq</td><td>PAM</td><td>alignment</td><td>distance</td><td>position</td><td>gene name</td><td>gene id</td></tr>
<tr><td>GU305914.1</td><td>141884</td><td>141906</td><td>-</td><td>0</td><td>AGGTGGCGGTGTAGGCCAGC</td><td>TGG</td><td>||||||||[||||||||||||]PAM</td><td>NA</td><td>Intergenic</td><td>NA</td><td>NA</td></tr>
<tr><td>T23336</td><td>CCACCTCGTCCACCGAGTGCCGG</td><td>1000</td><td></td><td></td><td></td><td></td><td></td><td></td><td></td><td></td><td></td></tr>
<tr><td>Oligo fwd</td><td>CACCCCACCTCGTCCACCGAGTGC</td><td></td><td></td><td></td><td></td><td></td><td></td><td></td><td></td><td></td><td></td></tr>
</table>

Oligo rev AAACGCACTCGGTGGACGAGGTGG
Chromosome start end strand MM target_seq PAM alignment distance position gene name gene id
GU305914.1 141833 141855 - 0 CCACCTCGTCCACCGAGTGC CGG ||||||||[||||||||||||]PAM NA Intergenic NA NA

T23337 CACCTCGTCCACCGAGTGCCGGG 1000
Oligo fwd CACCCACCTCGTCCACCGAGTGCC
Oligo rev AAACGGCACTCGGTGGACGAGGTG
Chromosome start end strand MM target_seq PAM alignment distance position gene name gene id
GU305914.1 141832 141854 - 0 CACCTCGTCCACCGAGTGCC GGG ||||||||[||||||||||||]PAM NA Intergenic NA NA

T23338 CGTCCACCGAGTGCCGGGCGAGG 1000
Oligo fwd CACCCGTCCACCGAGTGCCGGGCG
Oligo rev AAACCGCCCGGCACTCGGTGGACG
Chromosome start end strand MM target_seq PAM alignment distance position gene name gene id
GU305914.1 141827 141849 - 0 CGTCCACCGAGTGCCGGGCG AGG ||||||||[||||||||||||]PAM NA Intergenic NA NA

T23339 GTCCACCGAGTGCCGGGCGAGGG 1000
Oligo fwd CACCGTCCACCGAGTGCCGGGCGA
Oligo rev AAACTCGCCCGGCACTCGGTGGAC
Chromosome start end strand MM target_seq PAM alignment distance position gene name gene id
GU305914.1 141826 141848 - 0 GTCCACCGAGTGCCGGGCGA GGG ||||||||[||||||||||||]PAM NA Intergenic NA NA

T23340 GTGCCGGGCGAGGGCGTCGCAGG 1000
Oligo fwd CACCGTGCCGGGCGAGGGCGTCGC
Oligo rev AAACGCGACGCCCTCGCCCGGCAC
Chromosome start end strand MM target_seq PAM alignment distance position gene name gene id
GU305914.1 141817 141839 - 0 GTGCCGGGCGAGGGCGTCGC AGG ||||||||[||||||||||||]PAM NA Intergenic NA NA

T23341 GACGCCGAAGAGCGCGTCCTCGG 1000
Oligo fwd CACCGACGCCGAAGAGCGCGTCCT
Oligo rev AAACAGGACGCGCTCTTCGGCGTC
Chromosome start end strand MM target_seq PAM alignment distance position gene name gene id
GU305914.1 141784 141806 - 0 GACGCCGAAGAGCGCGTCCT CGG ||||||||[||||||||||||]PAM NA Intergenic NA NA

T23342 GCCGAAGAGCGCGTCCTCGGAGG 1000
Oligo fwd CACCGCCGAAGAGCGCGTCCTCGG
Oligo rev AAACCCGAGGACGCGCTCTTCGGC
Chromosome start end strand MM target_seq PAM alignment distance position gene name gene id
GU305914.1 141781 141803 - 0 GCCGAAGAGCGCGTCCTCGG AGG ||||||||[||||||||||||]PAM NA Intergenic NA NA

T23343 TGAGCTGTCCGCGCTCGTCCAGG 1000
Oligo fwd CACCTGAGCTGTCCGCGCTCGTCC
Oligo rev AAACGGACGAGCGCGGACAGCTCA
Chromosome start end strand MM target_seq PAM alignment distance position gene name gene id
GU305914.1 141746 141768 - 0 TGAGCTGTCCGCGCTCGTCC AGG ||||||||[||||||||||||]PAM NA Intergenic NA NA

T23344 GCTCGTCCAGGACCCTGACCAGG 1000
Oligo fwd CACCGCTCGTCCAGGACCCTGACC
Oligo rev AAACGGTCAGGGTCCTGGACGAGC
Chromosome start end strand MM target_seq PAM alignment distance position gene name gene id
GU305914.1 141734 141756 - 0 GCTCGTCCAGGACCCTGACC AGG ||||||||[||||||||||||]PAM NA Intergenic NA NA

T23345 GACCCTGACCAGGCAGAAAGCGG 1000
Oligo fwd CACCGACCCTGACCAGGCAGAAAG
Oligo rev AAACCTTTCTGCCTGGTCAGGGTC
Chromosome start end strand MM target_seq PAM alignment distance position gene name gene id
GU305914.1 141724 141746 - 0 GACCCTGACCAGGCAGAAAG CGG ||||||||[||||||||||||]PAM NA Intergenic NA NA

T23346 AGAAAGCGGCCACGTGTCCCAGG 1000
Oligo fwd CACCAGAAAGCGGCCACGTGTCCC
Oligo rev AAACGGGACACGTGGCCGCTTTCT
Chromosome start end strand MM target_seq PAM alignment distance position gene name gene id
GU305914.1 141710 141732 - 0 AGAAAGCGGCCACGTGTCCC AGG ||||||||[||||||||||||]PAM NA Intergenic NA NA

T23347 GAAAGCGGCCACGTGTCCCAGGG 1000
Oligo fwd CACCGAAAGCGGCCACGTGTCCCA
Oligo rev AAACTGGGACACGTGGCCGCTTTC
Chromosome start end strand MM target_seq PAM alignment distance position gene name gene id
GU305914.1 141709 141731 - 0 GAAAGCGGCCACGTGTCCCA GGG ||||||||[||||||||||||]PAM NA Intergenic NA NA

T23348 CCACGTGTCCCAGGGCGCAGAGG 1000
Oligo fwd CACCCCACGTGTCCCAGGGCGCAG
Oligo rev AAACCTGCGCCCTGGGACACGTGG
Chromosome start end strand MM target_seq PAM alignment distance position gene name gene id
GU305914.1 141701 141723 - 0 CCACGTGTCCCAGGGCGCAG AGG ||||||||[||||||||||||]PAM NA Intergenic NA NA

T23349 AGAGGTCCGAGACGCAGAAGCGG 1000
Oligo fwd CACCAGAGGTCCGAGACGCAGAAG
Oligo rev AAACCTTCTGCGTCTCGGACCTCT
Chromosome start end strand MM target_seq PAM alignment distance position gene name gene id
GU305914.1 141683 141705 - 0 AGAGGTCCGAGACGCAGAAG CGG ||||||||[||||||||||||]PAM NA Intergenic NA NA

T23350 GACGCAGAAGCGGTTCGCCGAGG 1000
Oligo fwd CACCGACGCAGAAGCGGTTCGCCG
Oligo rev AAACCGGCGAACCGCTTCTGCGTC
Chromosome start end strand MM target_seq PAM alignment distance position gene name gene id
GU305914.1 141673 141695 - 0 GACGCAGAAGCGGTTCGCCG AGG ||||||||[||||||||||||]PAM NA Intergenic NA NA

T23351 ACGCAGAAGCGGTTCGCCGAGGG 1000

Oligo fwd CACCACGCAGAAGCGGTTCGCCGA
Oligo rev AAACTCGGCGAACCGCTTCTGCGT
Chromosome start end strand MM target_seq PAM alignment distance position gene name gene id
GU305914.1 141672 141694 - 0 ACGCAGAAGCGGTTCGCCGA GGG ||||||||[||||||||||||]PAM NA Intergenic NA NA

T23352 GAAGCGGTTCGCCGAGGGCTTGG 1000
Oligo fwd CACCGAAGCGGTTCGCCGAGGGCT
Oligo rev AAACAGCCCTCGGCGAACCGCTTC
Chromosome start end strand MM target_seq PAM alignment distance position gene name gene id
GU305914.1 141667 141689 - 0 GAAGCGGTTCGCCGAGGGCT TGG ||||||||[||||||||||||]PAM NA Intergenic NA NA

T23353 AAGCGGTTCGCCGAGGGCTTGGG 1000
Oligo fwd CACCAAGCGGTTCGCCGAGGGCTT
Oligo rev AAACAAGCCCTCGGCGAACCGCTT
Chromosome start end strand MM target_seq PAM alignment distance position gene name gene id
GU305914.1 141666 141688 - 0 AAGCGGTTCGCCGAGGGCTT GGG ||||||||[||||||||||||]PAM NA Intergenic NA NA

T23354 AGCGGTTCGCCGAGGGCTTGGGG 1000
Oligo fwd CACCAGCGGTTCGCCGAGGGCTTG
Oligo rev AAACCAAGCCCTCGGCGAACCGCT
Chromosome start end strand MM target_seq PAM alignment distance position gene name gene id
GU305914.1 141665 141687 - 0 AGCGGTTCGCCGAGGGCTTG GGG ||||||||[||||||||||||]PAM NA Intergenic NA NA

T23355 GGCTTGGGGAACTGCTTGCGCGG 1000
Oligo fwd CACCGGCTTGGGGAACTGCTTGCG
Oligo rev AAACCGCAAGCAGTTCCCCAAGCC
Chromosome start end strand MM target_seq PAM alignment distance position gene name gene id
GU305914.1 141651 141673 - 0 GGCTTGGGGAACTGCTTGCG CGG ||||||||[||||||||||||]PAM NA Intergenic NA NA

T23356 GAACTGCTTGCGCGGCTCGATGG 1000
Oligo fwd CACCGAACTGCTTGCGCGGCTCGA
Oligo rev AAACTCGAGCCGCGCAAGCAGTTC
Chromosome start end strand MM target_seq PAM alignment distance position gene name gene id
GU305914.1 141643 141665 - 0 GAACTGCTTGCGCGGCTCGA TGG ||||||||[||||||||||||]PAM NA Intergenic NA NA
GU305914.1 133107 133129 + 4 GATCGGCTTGCGCGGCCAGA GGG ||-|-|||[||||||||--||]PAM NA Intergenic NA NA

T23357 CAGAGTTTCTGCAGCATGAGCGG 1000
Oligo fwd CACCCAGAGTTTCTGCAGCATGAG
Oligo rev AAACCTCATGCTGCAGAAACTCTG
Chromosome start end strand MM target_seq PAM alignment distance position gene name gene id
GU305914.1 141618 141640 - 0 CAGAGTTTCTGCAGCATGAG CGG ||||||||[||||||||||||]PAM NA Intergenic NA NA

T23358 CTGCAGCATGAGCGGCTTGAAGG 1000
Oligo fwd CACCCTGCAGCATGAGCGGCTTGA
Oligo rev AAACTCAAGCCGCTCATGCTGCAG
Chromosome start end strand MM target_seq PAM alignment distance position gene name gene id
GU305914.1 141610 141632 - 0 CTGCAGCATGAGCGGCTTGA AGG ||||||||[||||||||||||]PAM NA Intergenic NA NA

T23359 CAGCATGAGCGGCTTGAAGGCGG 1000
Oligo fwd CACCCAGCATGAGCGGCTTGAAGG
Oligo rev AAACCCTTCAAGCCGCTCATGCTG
Chromosome start end strand MM target_seq PAM alignment distance position gene name gene id
GU305914.1 141607 141629 - 0 CAGCATGAGCGGCTTGAAGG CGG ||||||||[||||||||||||]PAM NA Intergenic NA NA

T23360 AGCATGAGCGGCTTGAAGGCGGG 1000
Oligo fwd CACCAGCATGAGCGGCTTGAAGGC
Oligo rev AAACGCCTTCAAGCCGCTCATGCT
Chromosome start end strand MM target_seq PAM alignment distance position gene name gene id
GU305914.1 141606 141628 - 0 AGCATGAGCGGCTTGAAGGC GGG ||||||||[||||||||||||]PAM NA Intergenic NA NA

T23361 TGAGCGGCTTGAAGGCGGGATGG 1000
Oligo fwd CACCTGAGCGGCTTGAAGGCGGGA
Oligo rev AAACTCCCGCCTTCAAGCCGCTCA
Chromosome start end strand MM target_seq PAM alignment distance position gene name gene id
GU305914.1 141602 141624 - 0 TGAGCGGCTTGAAGGCGGGA TGG ||||||||[||||||||||||]PAM NA Intergenic NA NA

T23362 CGGGATGGTAGATGTCCGTGAGG 1000
Oligo fwd CACCCGGGATGGTAGATGTCCGTG
Oligo rev AAACCACGGACATCTACCATCCCG
Chromosome start end strand MM target_seq PAM alignment distance position gene name gene id
GU305914.1 141587 141609 - 0 CGGGATGGTAGATGTCCGTG AGG ||||||||[||||||||||||]PAM NA Intergenic NA NA

T23363 GTAGATGTCCGTGAGGTAGTAGG 1000
Oligo fwd CACCGTAGATGTCCGTGAGGTAGT
Oligo rev AAACACTACCTCACGGACATCTAC
Chromosome start end strand MM target_seq PAM alignment distance position gene name gene id
GU305914.1 141580 141602 - 0 GTAGATGTCCGTGAGGTAGT AGG ||||||||[||||||||||||]PAM NA Intergenic NA NA

T23364 CGTGAGGTAGTAGGCGCTCTTGG 1000
Oligo fwd CACCCGTGAGGTAGTAGGCGCTCT
Oligo rev AAACAGAGCGCCTACTACCTCACG
Chromosome start end strand MM target_seq PAM alignment distance position gene name gene id
GU305914.1 141571 141593 - 0 CGTGAGGTAGTAGGCGCTCT TGG ||||||||[||||||||||||]PAM NA Intergenic NA NA

T23365 GTGAGGTAGTAGGCGCTCTTGGG 1000
Oligo fwd CACCGTGAGGTAGTAGGCGCTCTT
Oligo rev AAACAAGAGCGCCTACTACCTCAC
Chromosome start end strand MM target_seq PAM alignment distance position gene name gene id
GU305914.1 141570 141592 - 0 GTGAGGTAGTAGGCGCTCTT GGG ||||||||[||||||||||||]PAM NA Intergenic NA NA

T23366 GCTCTTGGGCAGCGCCCTGACGG 1000
Oligo fwd CACCGCTCTTGGGCAGCGCCCTGA
Oligo rev AAACTCAGGGCGCTGCCCAAGAGC
Chromosome start end strand MM target_seq PAM alignment distance position gene name gene id
GU305914.1 141556 141578 - 0 GCTCTTGGGCAGCGCCCTGA CGG ||||||||[||||||||||||]PAM NA Intergenic NA NA

T23367 GGGCAGCGCCCTGACGGTCTTGG 1000
Oligo fwd CACCGGGCAGCGCCCTGACGGTCT
Oligo rev AAACAGACCGTCAGGGCGCTGCCC
Chromosome start end strand MM target_seq PAM alignment distance position gene name gene id
GU305914.1 141550 141572 - 0 GGGCAGCGCCCTGACGGTCT TGG ||||||||[||||||||||||]PAM NA Intergenic NA NA

T23368 CCCTGACGGTCTTGGTCTGCTGG 1000
Oligo fwd CACCCCCTGACGGTCTTGGTCTGC
Oligo rev AAACGCAGACCAAGACCGTCAGGG
Chromosome start end strand MM target_seq PAM alignment distance position gene name gene id
GU305914.1 141542 141564 - 0 CCCTGACGGTCTTGGTCTGC TGG ||||||||[||||||||||||]PAM NA Intergenic NA NA

T23369 GAAGACCACGACGCAGTGTCCGG 1000
Oligo fwd CACCGAAGACCACGACGCAGTGTC
Oligo rev AAACGACACTGCGTCGTGGTCTTC
Chromosome start end strand MM target_seq PAM alignment distance position gene name gene id
GU305914.1 141520 141542 - 0 GAAGACCACGACGCAGTGTC CGG ||||||||[||||||||||||]PAM NA Intergenic NA NA
GU305914.1 69574 69596 + 4 GAAGAACTCGACGCAGTCGC TGG |||||-|-[|||||||||--|]PAM NA Intergenic NA NA

T23370 GCAGTGTCCGGTAGTACCCTCGG 1000
Oligo fwd CACCGCAGTGTCCGGTAGTACCCT
Oligo rev AAACAGGGTACTACCGGACACTGC
Chromosome start end strand MM target_seq PAM alignment distance position gene name gene id
GU305914.1 141508 141530 - 0 GCAGTGTCCGGTAGTACCCT CGG ||||||||[||||||||||||]PAM NA Intergenic NA NA

T23371 CCTCGGTGTGACACTGCGACAGG 1000
Oligo fwd CACCCCTCGGTGTGACACTGCGAC
Oligo rev AAACGTCGCAGTGTCACACCGAGG
Chromosome start end strand MM target_seq PAM alignment distance position gene name gene id
GU305914.1 141491 141513 - 0 CCTCGGTGTGACACTGCGAC AGG ||||||||[||||||||||||]PAM NA Intergenic NA NA

T23372 ACGCGCGATCTGCCTGTCGTCGG 1000
Oligo fwd CACCACGCGCGATCTGCCTGTCGT
Oligo rev AAACACGACAGGCAGATCGCGCGT
Chromosome start end strand MM target_seq PAM alignment distance position gene name gene id
GU305914.1 141448 141470 - 0 ACGCGCGATCTGCCTGTCGT CGG ||||||||[||||||||||||]PAM NA Intergenic NA NA

T23373 CGATCTGCCTGTCGTCGGCGCGG 1000
Oligo fwd CACCCGATCTGCCTGTCGTCGGCG
Oligo rev AAACCGCCGACGACAGGCAGATCG
Chromosome start end strand MM target_seq PAM alignment distance position gene name gene id
GU305914.1 141443 141465 - 0 CGATCTGCCTGTCGTCGGCG CGG ||||||||[||||||||||||]PAM NA Intergenic NA NA

T23374 TAGTTGATGAGCACGTTCATAGG 1000
Oligo fwd CACCTAGTTGATGAGCACGTTCAT
Oligo rev AAACATGAACGTGCTCATCAACTA
Chromosome start end strand MM target_seq PAM alignment distance position gene name gene id
GU305914.1 141417 141439 - 0 TAGTTGATGAGCACGTTCAT AGG ||||||||[||||||||||||]PAM NA Intergenic NA NA

T23375 AGTGACCGACGCCGCAACGCAGG 1000
Oligo fwd CACCAGTGACCGACGCCGCAACGC
Oligo rev AAACGCGTTGCGGCGTCGGTCACT
Chromosome start end strand MM target_seq PAM alignment distance position gene name gene id
GU305914.1 141383 141405 - 0 AGTGACCGACGCCGCAACGC AGG ||||||||[||||||||||||]PAM NA Intergenic NA NA

T23376 CGACGCCGCAACGCAGGTTGAGG 1000
Oligo fwd CACCCGACGCCGCAACGCAGGTTG
Oligo rev AAACCAACCTGCGTTGCGGCGTCG
Chromosome start end strand MM target_seq PAM alignment distance position gene name gene id
GU305914.1 141377 141399 - 0 CGACGCCGCAACGCAGGTTG AGG ||||||||[||||||||||||]PAM NA Intergenic NA NA

T23377 GTTGAGGAACCTGAGCGCGTCGG 1000
Oligo fwd CACCGTTGAGGAACCTGAGCGCGT
Oligo rev AAACACGCGCTCAGGTTCCTCAAC
Chromosome start end strand MM target_seq PAM alignment distance position gene name gene id
GU305914.1 141361 141383 - 0 GTTGAGGAACCTGAGCGCGT CGG ||||||||[||||||||||||]PAM NA Intergenic NA NA

T23378 CGCGTCGGCGATGCCGCTGAAGG 1000
Oligo fwd CACCCGCGTCGGCGATGCCGCTGA
Oligo rev AAACTCAGCGGCATCGCCGACGCG
Chromosome start end strand MM target_seq PAM alignment distance position gene name gene id
GU305914.1 141346 141368 - 0 CGCGTCGGCGATGCCGCTGA AGG ||||||||[||||||||||||]PAM NA Intergenic NA NA

T23379 CGGCGATGCCGCTGAAGGCGCGG 1000
Oligo fwd CACCCGGCGATGCCGCTGAAGGCG
Oligo rev AAACCGCCTTCAGCGGCATCGCCG
Chromosome start end strand MM target_seq PAM alignment distance position gene name gene id
GU305914.1 141341 141363 - 0 CGGCGATGCCGCTGAAGGCG CGG ||||||||[||||||||||||]PAM NA Intergenic NA NA

T23380 CGATGCCGCTGAAGGCGCGGCGG 1000
Oligo fwd CACCCGATGCCGCTGAAGGCGCGG
Oligo rev AAACCCGCGCCTTCAGCGGCATCG

<table>
<tr><td>Chromosome</td><td>start</td><td>end</td><td>strand</td><td>MM</td><td>target_seq</td><td>PAM</td><td>alignment</td><td>distance</td><td>position</td><td>gene name</td><td>gene id</td></tr>
<tr><td>GU305914.1</td><td>141338</td><td>141360</td><td>-</td><td>0</td><td>CGATGCCGCTGAAGGCGCGG</td><td>CGG</td><td>||||||||[||||||||||||]PAM</td><td>NA</td><td>Intergenic</td><td>NA</td><td>NA</td></tr>
<tr><td>T23381</td><td>GCTGAAGGCGCGGCGGTAGCTGG</td><td>1000</td><td></td><td></td><td></td><td></td><td></td><td></td><td></td><td></td><td></td></tr>
<tr><td>Oligo fwd</td><td>CACCGCTGAAGGCGCGGCGGTAGC</td><td></td><td></td><td></td><td></td><td></td><td></td><td></td><td></td><td></td><td></td></tr>
<tr><td>Oligo rev</td><td>AAACGCTACCGCCGCGCCTTCAGC</td><td></td><td></td><td></td><td></td><td></td><td></td><td></td><td></td><td></td><td></td></tr>
<tr><td>Chromosome</td><td>start</td><td>end</td><td>strand</td><td>MM</td><td>target_seq</td><td>PAM</td><td>alignment</td><td>distance</td><td>position</td><td>gene name</td><td>gene id</td></tr>
<tr><td>GU305914.1</td><td>141331</td><td>141353</td><td>-</td><td>0</td><td>GCTGAAGGCGCGGCGGTAGC</td><td>TGG</td><td>||||||||[||||||||||||]PAM</td><td>NA</td><td>Intergenic</td><td>NA</td><td>NA</td></tr>
<tr><td>T23382</td><td>AGGCGCGGCGGTAGCTGGCCAGG</td><td>1000</td><td></td><td></td><td></td><td></td><td></td><td></td><td></td><td></td><td></td></tr>
<tr><td>Oligo fwd</td><td>CACCAGGCGCGGCGGTAGCTGGCC</td><td></td><td></td><td></td><td></td><td></td><td></td><td></td><td></td><td></td><td></td></tr>
<tr><td>Oligo rev</td><td>AAACGGCCAGCTACCGCCGCGCCT</td><td></td><td></td><td></td><td></td><td></td><td></td><td></td><td></td><td></td><td></td></tr>
<tr><td>Chromosome</td><td>start</td><td>end</td><td>strand</td><td>MM</td><td>target_seq</td><td>PAM</td><td>alignment</td><td>distance</td><td>position</td><td>gene name</td><td>gene id</td></tr>
<tr><td>GU305914.1</td><td>141326</td><td>141348</td><td>-</td><td>0</td><td>AGGCGCGGCGGTAGCTGGCC</td><td>AGG</td><td>||||||||[||||||||||||]PAM</td><td>NA</td><td>Intergenic</td><td>NA</td><td>NA</td></tr>
<tr><td>T23383</td><td>GGCCCGCGTAGTGCCAGCCCCGG</td><td>1000</td><td></td><td></td><td></td><td></td><td></td><td></td><td></td><td></td><td></td></tr>
<tr><td>Oligo fwd</td><td>CACCGGCCCGCGTAGTGCCAGCCC</td><td></td><td></td><td></td><td></td><td></td><td></td><td></td><td></td><td></td><td></td></tr>
<tr><td>Oligo rev</td><td>AAACGGGCTGGCACTACGCGGGCC</td><td></td><td></td><td></td><td></td><td></td><td></td><td></td><td></td><td></td><td></td></tr>
<tr><td>Chromosome</td><td>start</td><td>end</td><td>strand</td><td>MM</td><td>target_seq</td><td>PAM</td><td>alignment</td><td>distance</td><td>position</td><td>gene name</td><td>gene id</td></tr>
<tr><td>GU305914.1</td><td>141305</td><td>141327</td><td>-</td><td>0</td><td>GGCCCGCGTAGTGCCAGCCC</td><td>CGG</td><td>||||||||[||||||||||||]PAM</td><td>NA</td><td>Intergenic</td><td>NA</td><td>NA</td></tr>
<tr><td>T23384</td><td>GCCAGCCCCGGTAGTTGTAGAGG</td><td>1000</td><td></td><td></td><td></td><td></td><td></td><td></td><td></td><td></td><td></td></tr>
<tr><td>Oligo fwd</td><td>CACCGCCAGCCCCGGTAGTTGTAG</td><td></td><td></td><td></td><td></td><td></td><td></td><td></td><td></td><td></td><td></td></tr>
<tr><td>Oligo rev</td><td>AAACCTACAACTACCGGGGCTGGC</td><td></td><td></td><td></td><td></td><td></td><td></td><td></td><td></td><td></td><td></td></tr>
<tr><td>Chromosome</td><td>start</td><td>end</td><td>strand</td><td>MM</td><td>target_seq</td><td>PAM</td><td>alignment</td><td>distance</td><td>position</td><td>gene name</td><td>gene id</td></tr>
<tr><td>GU305914.1</td><td>141293</td><td>141315</td><td>-</td><td>0</td><td>GCCAGCCCCGGTAGTTGTAG</td><td>AGG</td><td>||||||||[||||||||||||]PAM</td><td>NA</td><td>Intergenic</td><td>NA</td><td>NA</td></tr>
<tr><td>T23385</td><td>GTAGAGGTCCCGATCGAAACTGG</td><td>1000</td><td></td><td></td><td></td><td></td><td></td><td></td><td></td><td></td><td></td></tr>
<tr><td>Oligo fwd</td><td>CACCGTAGAGGTCCCGATCGAAAC</td><td></td><td></td><td></td><td></td><td></td><td></td><td></td><td></td><td></td><td></td></tr>
<tr><td>Oligo rev</td><td>AAACGTTTCGATCGGGACCTCTAC</td><td></td><td></td><td></td><td></td><td></td><td></td><td></td><td></td><td></td><td></td></tr>
<tr><td>Chromosome</td><td>start</td><td>end</td><td>strand</td><td>MM</td><td>target_seq</td><td>PAM</td><td>alignment</td><td>distance</td><td>position</td><td>gene name</td><td>gene id</td></tr>
<tr><td>GU305914.1</td><td>141277</td><td>141299</td><td>-</td><td>0</td><td>GTAGAGGTCCCGATCGAAAC</td><td>TGG</td><td>||||||||[||||||||||||]PAM</td><td>NA</td><td>Intergenic</td><td>NA</td><td>NA</td></tr>
<tr><td>T23386</td><td>CGAAACTGGCGAACGACACGAGG</td><td>1000</td><td></td><td></td><td></td><td></td><td></td><td></td><td></td><td></td><td></td></tr>
<tr><td>Oligo fwd</td><td>CACCCGAAACTGGCGAACGACACG</td><td></td><td></td><td></td><td></td><td></td><td></td><td></td><td></td><td></td><td></td></tr>
<tr><td>Oligo rev</td><td>AAACCGTGTCGTTCGCCAGTTTCG</td><td></td><td></td><td></td><td></td><td></td><td></td><td></td><td></td><td></td><td></td></tr>
<tr><td>Chromosome</td><td>start</td><td>end</td><td>strand</td><td>MM</td><td>target_seq</td><td>PAM</td><td>alignment</td><td>distance</td><td>position</td><td>gene name</td><td>gene id</td></tr>
<tr><td>GU305914.1</td><td>141263</td><td>141285</td><td>-</td><td>0</td><td>CGAAACTGGCGAACGACACG</td><td>AGG</td><td>||||||||[||||||||||||]PAM</td><td>NA</td><td>Intergenic</td><td>NA</td><td>NA</td></tr>
<tr><td>T23387</td><td>ACGACACGAGGTTGTGTCTCAGG</td><td>1000</td><td></td><td></td><td></td><td></td><td></td><td></td><td></td><td></td><td></td></tr>
<tr><td>Oligo fwd</td><td>CACCACGACACGAGGTTGTGTCTC</td><td></td><td></td><td></td><td></td><td></td><td></td><td></td><td></td><td></td><td></td></tr>
<tr><td>Oligo rev</td><td>AAACGAGACACAACCTCGTGTCGT</td><td></td><td></td><td></td><td></td><td></td><td></td><td></td><td></td><td></td><td></td></tr>
<tr><td>Chromosome</td><td>start</td><td>end</td><td>strand</td><td>MM</td><td>target_seq</td><td>PAM</td><td>alignment</td><td>distance</td><td>position</td><td>gene name</td><td>gene id</td></tr>
<tr><td>GU305914.1</td><td>141251</td><td>141273</td><td>-</td><td>0</td><td>ACGACACGAGGTTGTGTCTC</td><td>AGG</td><td>||||||||[||||||||||||]PAM</td><td>NA</td><td>Intergenic</td><td>NA</td><td>NA</td></tr>
<tr><td>T23388</td><td>GTTGTGTCTCAGGCAGCTGCCGG</td><td>1000</td><td></td><td></td><td></td><td></td><td></td><td></td><td></td><td></td><td></td></tr>
<tr><td>Oligo fwd</td><td>CACCGTTGTGTCTCAGGCAGCTGC</td><td></td><td></td><td></td><td></td><td></td><td></td><td></td><td></td><td></td><td></td></tr>
<tr><td>Oligo rev</td><td>AAACGCAGCTGCCTGAGACACAAC</td><td></td><td></td><td></td><td></td><td></td><td></td><td></td><td></td><td></td><td></td></tr>
<tr><td>Chromosome</td><td>start</td><td>end</td><td>strand</td><td>MM</td><td>target_seq</td><td>PAM</td><td>alignment</td><td>distance</td><td>position</td><td>gene name</td><td>gene id</td></tr>
<tr><td>GU305914.1</td><td>141241</td><td>141263</td><td>-</td><td>0</td><td>GTTGTGTCTCAGGCAGCTGC</td><td>CGG</td><td>||||||||[||||||||||||]PAM</td><td>NA</td><td>Intergenic</td><td>NA</td><td>NA</td></tr>
<tr><td>T23389</td><td>GTGTCTCAGGCAGCTGCCGGTGG</td><td>1000</td><td></td><td></td><td></td><td></td><td></td><td></td><td></td><td></td><td></td></tr>
<tr><td>Oligo fwd</td><td>CACCGTGTCTCAGGCAGCTGCCGG</td><td></td><td></td><td></td><td></td><td></td><td></td><td></td><td></td><td></td><td></td></tr>
<tr><td>Oligo rev</td><td>AAACCCGGCAGCTGCCTGAGACAC</td><td></td><td></td><td></td><td></td><td></td><td></td><td></td><td></td><td></td><td></td></tr>
<tr><td>Chromosome</td><td>start</td><td>end</td><td>strand</td><td>MM</td><td>target_seq</td><td>PAM</td><td>alignment</td><td>distance</td><td>position</td><td>gene name</td><td>gene id</td></tr>
<tr><td>GU305914.1</td><td>141238</td><td>141260</td><td>-</td><td>0</td><td>GTGTCTCAGGCAGCTGCCGG</td><td>TGG</td><td>||||||||[||||||||||||]PAM</td><td>NA</td><td>Intergenic</td><td>NA</td><td>NA</td></tr>
<tr><td>T23390</td><td>GGCAGCTGCCGGTGGCCACCAGG</td><td>1000</td><td></td><td></td><td></td><td></td><td></td><td></td><td></td><td></td><td></td></tr>
<tr><td>Oligo fwd</td><td>CACCGGCAGCTGCCGGTGGCCACC</td><td></td><td></td><td></td><td></td><td></td><td></td><td></td><td></td><td></td><td></td></tr>
<tr><td>Oligo rev</td><td>AAACGGTGGCCACCGGCAGCTGCC</td><td></td><td></td><td></td><td></td><td></td><td></td><td></td><td></td><td></td><td></td></tr>
<tr><td>Chromosome</td><td>start</td><td>end</td><td>strand</td><td>MM</td><td>target_seq</td><td>PAM</td><td>alignment</td><td>distance</td><td>position</td><td>gene name</td><td>gene id</td></tr>
<tr><td>GU305914.1</td><td>141230</td><td>141252</td><td>-</td><td>0</td><td>GGCAGCTGCCGGTGGCCACC</td><td>AGG</td><td>||||||||[||||||||||||]PAM</td><td>NA</td><td>Intergenic</td><td>NA</td><td>NA</td></tr>
<tr><td>T23391</td><td>CGCTATCGTCCAGTTCTCCCCGG</td><td>1000</td><td></td><td></td><td></td><td></td><td></td><td></td><td></td><td></td><td></td></tr>
<tr><td>Oligo fwd</td><td>CACCCGCTATCGTCCAGTTCTCCC</td><td></td><td></td><td></td><td></td><td></td><td></td><td></td><td></td><td></td><td></td></tr>
<tr><td>Oligo rev</td><td>AAACGGGAGAACTGGACGATAGCG</td><td></td><td></td><td></td><td></td><td></td><td></td><td></td><td></td><td></td><td></td></tr>
<tr><td>Chromosome</td><td>start</td><td>end</td><td>strand</td><td>MM</td><td>target_seq</td><td>PAM</td><td>alignment</td><td>distance</td><td>position</td><td>gene name</td><td>gene id</td></tr>
<tr><td>GU305914.1</td><td>141194</td><td>141216</td><td>-</td><td>0</td><td>CGCTATCGTCCAGTTCTCCC</td><td>CGG</td><td>||||||||[||||||||||||]PAM</td><td>NA</td><td>Intergenic</td><td>NA</td><td>NA</td></tr>
<tr><td>T23392</td><td>GTCCAGTTCTCCCCGGTAGCCGG</td><td>1000</td><td></td><td></td><td></td><td></td><td></td><td></td><td></td><td></td><td></td></tr>
<tr><td>Oligo fwd</td><td>CACCGTCCAGTTCTCCCCGGTAGC</td><td></td><td></td><td></td><td></td><td></td><td></td><td></td><td></td><td></td><td></td></tr>
<tr><td>Oligo rev</td><td>AAACGCTACCGGGGAGAACTGGAC</td><td></td><td></td><td></td><td></td><td></td><td></td><td></td><td></td><td></td><td></td></tr>
<tr><td>Chromosome</td><td>start</td><td>end</td><td>strand</td><td>MM</td><td>target_seq</td><td>PAM</td><td>alignment</td><td>distance</td><td>position</td><td>gene name</td><td>gene id</td></tr>
<tr><td>GU305914.1</td><td>141187</td><td>141209</td><td>-</td><td>0</td><td>GTCCAGTTCTCCCCGGTAGC</td><td>CGG</td><td>||||||||[||||||||||||]PAM</td><td>NA</td><td>Intergenic</td><td>NA</td><td>NA</td></tr>
<tr><td>T23393</td><td>CTCCCCGGTAGCCGGCGTCCTGG</td><td>1000</td><td></td><td></td><td></td><td></td><td></td><td></td><td></td><td></td><td></td></tr>
<tr><td>Oligo fwd</td><td>CACCCTCCCCGGTAGCCGGCGTCC</td><td></td><td></td><td></td><td></td><td></td><td></td><td></td><td></td><td></td><td></td></tr>
<tr><td>Oligo rev</td><td>AAACGGACGCCGGCTACCGGGGAG</td><td></td><td></td><td></td><td></td><td></td><td></td><td></td><td></td><td></td><td></td></tr>
<tr><td>Chromosome</td><td>start</td><td>end</td><td>strand</td><td>MM</td><td>target_seq</td><td>PAM</td><td>alignment</td><td>distance</td><td>position</td><td>gene name</td><td>gene id</td></tr>
<tr><td>GU305914.1</td><td>141179</td><td>141201</td><td>-</td><td>0</td><td>CTCCCCGGTAGCCGGCGTCC</td><td>TGG</td><td>||||||||[||||||||||||]PAM</td><td>NA</td><td>Intergenic</td><td>NA</td><td>NA</td></tr>
<tr><td>T23394</td><td>TCCCCGGTAGCCGGCGTCCTGGG</td><td>1000</td><td></td><td></td><td></td><td></td><td></td><td></td><td></td><td></td><td></td></tr>
<tr><td>Oligo fwd</td><td>CACCTCCCCGGTAGCCGGCGTCCT</td><td></td><td></td><td></td><td></td><td></td><td></td><td></td><td></td><td></td><td></td></tr>
<tr><td>Oligo rev</td><td>AAACAGGACGCCGGCTACCGGGGA</td><td></td><td></td><td></td><td></td><td></td><td></td><td></td><td></td><td></td><td></td></tr>
<tr><td>Chromosome</td><td>start</td><td>end</td><td>strand</td><td>MM</td><td>target_seq</td><td>PAM</td><td>alignment</td><td>distance</td><td>position</td><td>gene name</td><td>gene id</td></tr>
<tr><td>GU305914.1</td><td>141178</td><td>141200</td><td>-</td><td>0</td><td>TCCCCGGTAGCCGGCGTCCT</td><td>GGG</td><td>||||||||[||||||||||||]PAM</td><td>NA</td><td>Intergenic</td><td>NA</td><td>NA</td></tr>
<tr><td>T23395</td><td>GGTAGCCGGCGTCCTGGGCTCGG</td><td>1000</td><td></td><td></td><td></td><td></td><td></td><td></td><td></td><td></td><td></td></tr>
<tr><td>Oligo fwd</td><td>CACCGGTAGCCGGCGTCCTGGGCT</td><td></td><td></td><td></td><td></td><td></td><td></td><td></td><td></td><td></td><td></td></tr>
</table>

Oligo rev AAACAGCCCAGGACGCCGGCTACC
Chromosome start end strand MM target_seq PAM alignment distance position gene name gene id
GU305914.1 141173 141195 - 0 GGTAGCCGGCGTCCTGGGCT CGG ||||||||[||||||||||||]PAM NA Intergenic NA NA

T23396 GTAGCCGGCGTCCTGGGCTCGGG 1000
Oligo fwd CACCGTAGCCGGCGTCCTGGGCTC
Oligo rev AAACGAGCCCAGGACGCCGGCTAC
Chromosome start end strand MM target_seq PAM alignment distance position gene name gene id
GU305914.1 141172 141194 - 0 GTAGCCGGCGTCCTGGGCTC GGG ||||||||[||||||||||||]PAM NA Intergenic NA NA

T23397 TACTACGCCCGAGATCCAGACGG 1000
Oligo fwd CACCTACTACGCCCGAGATCCAGA
Oligo rev AAACTCTGGATCTCGGGCGTAGTA
Chromosome start end strand MM target_seq PAM alignment distance position gene name gene id
GU305914.1 141145 141167 - 0 TACTACGCCCGAGATCCAGA CGG ||||||||[||||||||||||]PAM NA Intergenic NA NA

T23398 CCGAGATCCAGACGGTCAACAGG 1000
Oligo fwd CACCCCGAGATCCAGACGGTCAAC
Oligo rev AAACGTTGACCGTCTGGATCTCGG
Chromosome start end strand MM target_seq PAM alignment distance position gene name gene id
GU305914.1 141137 141159 - 0 CCGAGATCCAGACGGTCAAC AGG ||||||||[||||||||||||]PAM NA Intergenic NA NA

T23399 CGAGATCCAGACGGTCAACAGGG 1000
Oligo fwd CACCCGAGATCCAGACGGTCAACA
Oligo rev AAACTGTTGACCGTCTGGATCTCG
Chromosome start end strand MM target_seq PAM alignment distance position gene name gene id
GU305914.1 141136 141158 - 0 CGAGATCCAGACGGTCAACA GGG ||||||||[||||||||||||]PAM NA Intergenic NA NA

T23400 GGTCAACAGGGTCTCGTTGATGG 1000
Oligo fwd CACCGGTCAACAGGGTCTCGTTGA
Oligo rev AAACTCAACGAGACCCTGTTGACC
Chromosome start end strand MM target_seq PAM alignment distance position gene name gene id
GU305914.1 141124 141146 - 0 GGTCAACAGGGTCTCGTTGA TGG ||||||||[||||||||||||]PAM NA Intergenic NA NA

T23401 GGTCTCGTTGATGGACTCGCTGG 1000
Oligo fwd CACCGGTCTCGTTGATGGACTCGC
Oligo rev AAACGCGAGTCCATCAACGAGACC
Chromosome start end strand MM target_seq PAM alignment distance position gene name gene id
GU305914.1 141115 141137 - 0 GGTCTCGTTGATGGACTCGC TGG ||||||||[||||||||||||]PAM NA Intergenic NA NA

T23402 CTCGTTGATGGACTCGCTGGCGG 1000
Oligo fwd CACCCTCGTTGATGGACTCGCTGG
Oligo rev AAACCCAGCGAGTCCATCAACGAG
Chromosome start end strand MM target_seq PAM alignment distance position gene name gene id
GU305914.1 141112 141134 - 0 CTCGTTGATGGACTCGCTGG CGG ||||||||[||||||||||||]PAM NA Intergenic NA NA

T23403 GTTGATGGACTCGCTGGCGGCGG 1000
Oligo fwd CACCGTTGATGGACTCGCTGGCGG
Oligo rev AAACCCGCCAGCGAGTCCATCAAC
Chromosome start end strand MM target_seq PAM alignment distance position gene name gene id
GU305914.1 141109 141131 - 0 GTTGATGGACTCGCTGGCGG CGG ||||||||[||||||||||||]PAM NA Intergenic NA NA

T23404 ACTCGCTGGCGGCGGCCTTCAGG 1000
Oligo fwd CACCACTCGCTGGCGGCGGCCTTC
Oligo rev AAACGAAGGCCGCCGCCAGCGAGT
Chromosome start end strand MM target_seq PAM alignment distance position gene name gene id
GU305914.1 141101 141123 - 0 ACTCGCTGGCGGCGGCCTTC AGG ||||||||[||||||||||||]PAM NA Intergenic NA NA

T23405 CTCGCTGGCGGCGGCCTTCAGGG 1000
Oligo fwd CACCCTCGCTGGCGGCGGCCTTCA
Oligo rev AAACTGAAGGCCGCCGCCAGCGAG
Chromosome start end strand MM target_seq PAM alignment distance position gene name gene id
GU305914.1 141100 141122 - 0 CTCGCTGGCGGCGGCCTTCA GGG ||||||||[||||||||||||]PAM NA Intergenic NA NA

T23406 CTTCAGGGCCGTCTTCTTGTCGG 1000
Oligo fwd CACCCTTCAGGGCCGTCTTCTTGT
Oligo rev AAACACAAGAAGACGGCCCTGAAG
Chromosome start end strand MM target_seq PAM alignment distance position gene name gene id
GU305914.1 141085 141107 - 0 CTTCAGGGCCGTCTTCTTGT CGG ||||||||[||||||||||||]PAM NA Intergenic NA NA

T23407 CGACAGAAACTCTCGTCATACGG 1000
Oligo fwd CACCCGACAGAAACTCTCGTCATA
Oligo rev AAACTATGACGAGAGTTTCTGTCG
Chromosome start end strand MM target_seq PAM alignment distance position gene name gene id
GU305914.1 140970 140992 - 0 CGACAGAAACTCTCGTCATA CGG ||||||||[||||||||||||]PAM NA Intergenic NA NA

T23408 CTCGTCATACGGCACGTACACGG 1000
Oligo fwd CACCCTCGTCATACGGCACGTACA
Oligo rev AAACTGTACGTGCCGTATGACGAG
Chromosome start end strand MM target_seq PAM alignment distance position gene name gene id
GU305914.1 140959 140981 - 0 CTCGTCATACGGCACGTACA CGG ||||||||[||||||||||||]PAM NA Intergenic NA NA

T23409 TCGTCATACGGCACGTACACGGG 1000
Oligo fwd CACCTCGTCATACGGCACGTACAC
Oligo rev AAACGTGTACGTGCCGTATGACGA
Chromosome start end strand MM target_seq PAM alignment distance position gene name gene id
GU305914.1 140958 140980 - 0 TCGTCATACGGCACGTACAC GGG ||||||||[||||||||||||]PAM NA Intergenic NA NA

T23410 CGTCATACGGCACGTACACGGGG 1000

<table>
<tr><td>Oligo fwd</td><td>CACCCGTCATACGGCACGTACACG</td></tr>
<tr><td>Oligo rev</td><td>AAACCGTGTACGTGCCGTATGACG</td></tr>
<tr><td>Chromosome</td><td>start</td><td></td><td>end</td><td>strand</td><td>MM</td><td>target_seq</td><td>PAM</td><td>alignment</td><td>distance</td><td>position</td><td>gene name</td><td>gene id</td></tr>
<tr><td>GU305914.1</td><td></td><td>140957</td><td>140979</td><td>-</td><td>0</td><td>CGTCATACGGCACGTACACG</td><td>GGG</td><td>||||||||[||||||||||||]PAM</td><td>NA</td><td>Intergenic</td><td>NA</td><td>NA</td></tr>
<tr><td></td></tr>
<tr><td>T23411</td><td>CATACGGCACGTACACGGGGAGG</td><td></td><td>1000</td></tr>
<tr><td>Oligo fwd</td><td>CACCCATACGGCACGTACACGGGG</td></tr>
<tr><td>Oligo rev</td><td>AAACCCCCGTGTACGTGCCGTATG</td></tr>
<tr><td>Chromosome</td><td>start</td><td></td><td>end</td><td>strand</td><td>MM</td><td>target_seq</td><td>PAM</td><td>alignment</td><td>distance</td><td>position</td><td>gene name</td><td>gene id</td></tr>
<tr><td>GU305914.1</td><td></td><td>140954</td><td>140976</td><td>-</td><td>0</td><td>CATACGGCACGTACACGGGG</td><td>AGG</td><td>||||||||[||||||||||||]PAM</td><td>NA</td><td>Intergenic</td><td>NA</td><td>NA</td></tr>
<tr><td></td></tr>
<tr><td>T23412</td><td>ACTACCCGTCTCGAGCAGCACGG</td><td></td><td>1000</td></tr>
<tr><td>Oligo fwd</td><td>CACCACTACCCGTCTCGAGCAGCA</td></tr>
<tr><td>Oligo rev</td><td>AAACTGCTGCTCGAGACGGGTAGT</td></tr>
<tr><td>Chromosome</td><td>start</td><td></td><td>end</td><td>strand</td><td>MM</td><td>target_seq</td><td>PAM</td><td>alignment</td><td>distance</td><td>position</td><td>gene name</td><td>gene id</td></tr>
<tr><td>GU305914.1</td><td></td><td>140926</td><td>140948</td><td>-</td><td>0</td><td>ACTACCCGTCTCGAGCAGCA</td><td>CGG</td><td>||||||||[||||||||||||]PAM</td><td>NA</td><td>Intergenic</td><td>NA</td><td>NA</td></tr>
<tr><td></td></tr>
<tr><td>T23413</td><td>CTACCCGTCTCGAGCAGCACGGG</td><td></td><td>1000</td></tr>
<tr><td>Oligo fwd</td><td>CACCCTACCCGTCTCGAGCAGCAC</td></tr>
<tr><td>Oligo rev</td><td>AAACGTGCTGCTCGAGACGGGTAG</td></tr>
<tr><td>Chromosome</td><td>start</td><td></td><td>end</td><td>strand</td><td>MM</td><td>target_seq</td><td>PAM</td><td>alignment</td><td>distance</td><td>position</td><td>gene name</td><td>gene id</td></tr>
<tr><td>GU305914.1</td><td></td><td>140925</td><td>140947</td><td>-</td><td>0</td><td>CTACCCGTCTCGAGCAGCAC</td><td>GGG</td><td>||||||||[||||||||||||]PAM</td><td>NA</td><td>Intergenic</td><td>NA</td><td>NA</td></tr>
<tr><td></td></tr>
<tr><td>T23414</td><td>TCTCGAGCAGCACGGGCTGTCGG</td><td></td><td>1000</td></tr>
<tr><td>Oligo fwd</td><td>CACCTCTCGAGCAGCACGGGCTGT</td></tr>
<tr><td>Oligo rev</td><td>AAACACAGCCCGTGCTGCTCGAGA</td></tr>
<tr><td>Chromosome</td><td>start</td><td></td><td>end</td><td>strand</td><td>MM</td><td>target_seq</td><td>PAM</td><td>alignment</td><td>distance</td><td>position</td><td>gene name</td><td>gene id</td></tr>
<tr><td>GU305914.1</td><td></td><td>140918</td><td>140940</td><td>-</td><td>0</td><td>TCTCGAGCAGCACGGGCTGT</td><td>CGG</td><td>||||||||[||||||||||||]PAM</td><td>NA</td><td>Intergenic</td><td>NA</td><td>NA</td></tr>
<tr><td></td></tr>
<tr><td>T23415</td><td>CGTGACATAACATTTCAACCCGG</td><td></td><td>1000</td></tr>
<tr><td>Oligo fwd</td><td>CACCCGTGACATAACATTTCAACC</td></tr>
<tr><td>Oligo rev</td><td>AAACGGTTGAAATGTTATGTCACG</td></tr>
<tr><td>Chromosome</td><td>start</td><td></td><td>end</td><td>strand</td><td>MM</td><td>target_seq</td><td>PAM</td><td>alignment</td><td>distance</td><td>position</td><td>gene name</td><td>gene id</td></tr>
<tr><td>GU305914.1</td><td></td><td>140881</td><td>140903</td><td>-</td><td>0</td><td>CGTGACATAACATTTCAACC</td><td>CGG</td><td>||||||||[||||||||||||]PAM</td><td>NA</td><td>Intergenic</td><td>NA</td><td>NA</td></tr>
<tr><td></td></tr>
<tr><td>T23416</td><td>GTGACATAACATTTCAACCCGGG</td><td></td><td>1000</td></tr>
<tr><td>Oligo fwd</td><td>CACCGTGACATAACATTTCAACCC</td></tr>
<tr><td>Oligo rev</td><td>AAACGGGTTGAAATGTTATGTCAC</td></tr>
<tr><td>Chromosome</td><td>start</td><td></td><td>end</td><td>strand</td><td>MM</td><td>target_seq</td><td>PAM</td><td>alignment</td><td>distance</td><td>position</td><td>gene name</td><td>gene id</td></tr>
<tr><td>GU305914.1</td><td></td><td>140880</td><td>140902</td><td>-</td><td>0</td><td>GTGACATAACATTTCAACCC</td><td>GGG</td><td>||||||||[||||||||||||]PAM</td><td>NA</td><td>Intergenic</td><td>NA</td><td>NA</td></tr>
<tr><td></td></tr>
<tr><td>T23417</td><td>TAACATTTCAACCCGGGAAACGG</td><td></td><td>1000</td></tr>
<tr><td>Oligo fwd</td><td>CACCTAACATTTCAACCCGGGAAA</td></tr>
<tr><td>Oligo rev</td><td>AAACTTTCCCGGGTTGAAATGTTA</td></tr>
<tr><td>Chromosome</td><td>start</td><td></td><td>end</td><td>strand</td><td>MM</td><td>target_seq</td><td>PAM</td><td>alignment</td><td>distance</td><td>position</td><td>gene name</td><td>gene id</td></tr>
<tr><td>GU305914.1</td><td></td><td>140874</td><td>140896</td><td>-</td><td>0</td><td>TAACATTTCAACCCGGGAAA</td><td>CGG</td><td>||||||||[||||||||||||]PAM</td><td>NA</td><td>Intergenic</td><td>NA</td><td>NA</td></tr>
<tr><td></td></tr>
<tr><td>T23418</td><td>AACATTTCAACCCGGGAAACGGG</td><td></td><td>1000</td></tr>
<tr><td>Oligo fwd</td><td>CACCAACATTTCAACCCGGGAAAC</td></tr>
<tr><td>Oligo rev</td><td>AAACGTTTCCCGGGTTGAAATGTT</td></tr>
<tr><td>Chromosome</td><td>start</td><td></td><td>end</td><td>strand</td><td>MM</td><td>target_seq</td><td>PAM</td><td>alignment</td><td>distance</td><td>position</td><td>gene name</td><td>gene id</td></tr>
<tr><td>GU305914.1</td><td></td><td>140873</td><td>140895</td><td>-</td><td>0</td><td>AACATTTCAACCCGGGAAAC</td><td>GGG</td><td>||||||||[||||||||||||]PAM</td><td>NA</td><td>Intergenic</td><td>NA</td><td>NA</td></tr>
<tr><td></td></tr>
<tr><td>T23419</td><td>ACATTTCAACCCGGGAAACGGGG</td><td></td><td>1000</td></tr>
<tr><td>Oligo fwd</td><td>CACCACATTTCAACCCGGGAAACG</td></tr>
<tr><td>Oligo rev</td><td>AAACCGTTTCCCGGGTTGAAATGT</td></tr>
<tr><td>Chromosome</td><td>start</td><td></td><td>end</td><td>strand</td><td>MM</td><td>target_seq</td><td>PAM</td><td>alignment</td><td>distance</td><td>position</td><td>gene name</td><td>gene id</td></tr>
<tr><td>GU305914.1</td><td></td><td>140872</td><td>140894</td><td>-</td><td>0</td><td>ACATTTCAACCCGGGAAACG</td><td>GGG</td><td>||||||||[||||||||||||]PAM</td><td>NA</td><td>Intergenic</td><td>NA</td><td>NA</td></tr>
<tr><td></td></tr>
<tr><td>T23420</td><td>CCACTCCAGCCCGCCGCTAGAGG</td><td></td><td>1000</td></tr>
<tr><td>Oligo fwd</td><td>CACCCCACTCCAGCCCGCCGCTAG</td></tr>
<tr><td>Oligo rev</td><td>AAACCTAGCGGCGGGCTGGAGTGG</td></tr>
<tr><td>Chromosome</td><td>start</td><td></td><td>end</td><td>strand</td><td>MM</td><td>target_seq</td><td>PAM</td><td>alignment</td><td>distance</td><td>position</td><td>gene name</td><td>gene id</td></tr>
<tr><td>GU305914.1</td><td></td><td>140797</td><td>140819</td><td>-</td><td>0</td><td>CCACTCCAGCCCGCCGCTAG</td><td>AGG</td><td>||||||||[||||||||||||]PAM</td><td>NA</td><td>Intergenic</td><td>NA</td><td>NA</td></tr>
<tr><td></td></tr>
<tr><td>T23421</td><td>CTCCCCCCGTCACCTTACTTCGG</td><td></td><td>1000</td></tr>
<tr><td>Oligo fwd</td><td>CACCCTCCCCCCGTCACCTTACTT</td></tr>
<tr><td>Oligo rev</td><td>AAACAAGTAAGGTGACGGGGGGAG</td></tr>
<tr><td>Chromosome</td><td>start</td><td></td><td>end</td><td>strand</td><td>MM</td><td>target_seq</td><td>PAM</td><td>alignment</td><td>distance</td><td>position</td><td>gene name</td><td>gene id</td></tr>
<tr><td>GU305914.1</td><td></td><td>140696</td><td>140718</td><td>-</td><td>0</td><td>CTCCCCCCGTCACCTTACTT</td><td>CGG</td><td>||||||||[||||||||||||]PAM</td><td>NA</td><td>Intergenic</td><td>NA</td><td>NA</td></tr>
<tr><td></td></tr>
<tr><td>T23422</td><td>CCGTCACCTTACTTCGGCCGCGG</td><td></td><td>1000</td></tr>
<tr><td>Oligo fwd</td><td>CACCCCGTCACCTTACTTCGGCCG</td></tr>
<tr><td>Oligo rev</td><td>AAACCGGCCGAAGTAAGGTGACGG</td></tr>
<tr><td>Chromosome</td><td>start</td><td></td><td>end</td><td>strand</td><td>MM</td><td>target_seq</td><td>PAM</td><td>alignment</td><td>distance</td><td>position</td><td>gene name</td><td>gene id</td></tr>
<tr><td>GU305914.1</td><td></td><td>140690</td><td>140712</td><td>-</td><td>0</td><td>CCGTCACCTTACTTCGGCCG</td><td>CGG</td><td>||||||||[||||||||||||]PAM</td><td>NA</td><td>Intergenic</td><td>NA</td><td>NA</td></tr>
<tr><td></td></tr>
<tr><td>T23423</td><td>TTCGGCCGCGGCAACAGCCGCGG</td><td></td><td>1000</td></tr>
<tr><td>Oligo fwd</td><td>CACCTTCGGCCGCGGCAACAGCCG</td></tr>
<tr><td>Oligo rev</td><td>AAACCGGCTGTTGCCGCGGCCGAA</td></tr>
<tr><td>Chromosome</td><td>start</td><td></td><td>end</td><td>strand</td><td>MM</td><td>target_seq</td><td>PAM</td><td>alignment</td><td>distance</td><td>position</td><td>gene name</td><td>gene id</td></tr>
<tr><td>GU305914.1</td><td></td><td>140678</td><td>140700</td><td>-</td><td>0</td><td>TTCGGCCGCGGCAACAGCCG</td><td>CGG</td><td>||||||||[||||||||||||]PAM</td><td>NA</td><td>Intergenic</td><td>NA</td><td>NA</td></tr>
<tr><td></td></tr>
<tr><td>T23424</td><td>TGCTAATTCCTAAAGAACACAGG</td><td></td><td>1000</td></tr>
<tr><td>Oligo fwd</td><td>CACCTGCTAATTCCTAAAGAACAC</td></tr>
<tr><td>Oligo rev</td><td>AAACGTGTTCTTTAGGAATTAGCA</td></tr>
<tr><td>Chromosome</td><td>start</td><td></td><td>end</td><td>strand</td><td>MM</td><td>target_seq</td><td>PAM</td><td>alignment</td><td>distance</td><td>position</td><td>gene name</td><td>gene id</td></tr>
<tr><td>GU305914.1</td><td></td><td>140642</td><td>140664</td><td>-</td><td>0</td><td>TGCTAATTCCTAAAGAACAC</td><td>AGG</td><td>||||||||[||||||||||||]PAM</td><td>NA</td><td>Intergenic</td><td>NA</td><td>NA</td></tr>
</table>

T23425 GCTAATTCCTAAAGAACACAGGG 1000
Oligo fwd CACCGCTAATTCCTAAAGAACACA
Oligo rev AAACTGTGTTCTTTAGGAATTAGC
Chromosome start end strand MM target_seq PAM alignment distance position gene name gene id
GU305914.1 140641 140663 - 0 GCTAATTCCTAAAGAACACA GGG ||||||||[||||||||||||]PAM NA Intergenic NA NA

T23426 CTAATTCCTAAAGAACACAGGGG 1000
Oligo fwd CACCCTAATTCCTAAAGAACACAG
Oligo rev AAACCTGTGTTCTTTAGGAATTAG
Chromosome start end strand MM target_seq PAM alignment distance position gene name gene id
GU305914.1 140640 140662 - 0 CTAATTCCTAAAGAACACAG GGG ||||||||[||||||||||||]PAM NA Intergenic NA NA

T23427 CTAAAGAACACAGGGGACACCGG 1000
Oligo fwd CACCCTAAAGAACACAGGGGACAC
Oligo rev AAACGTGTCCCCTGTGTTCTTTAG
Chromosome start end strand MM target_seq PAM alignment distance position gene name gene id
GU305914.1 140633 140655 - 0 CTAAAGAACACAGGGGACAC CGG ||||||||[||||||||||||]PAM NA Intergenic NA NA

T23428 TAAAGAACACAGGGGACACCGGG 1000
Oligo fwd CACCTAAAGAACACAGGGGACACC
Oligo rev AAACGGTGTCCCCTGTGTTCTTTA
Chromosome start end strand MM target_seq PAM alignment distance position gene name gene id
GU305914.1 140632 140654 - 0 TAAAGAACACAGGGGACACC GGG ||||||||[||||||||||||]PAM NA Intergenic NA NA

T23429 GGGACACCGGGCGACCGACACGG 1000
Oligo fwd CACCGGGACACCGGGCGACCGACA
Oligo rev AAACTGTCGGTCGCCCGGTGTCCC
Chromosome start end strand MM target_seq PAM alignment distance position gene name gene id
GU305914.1 140620 140642 - 0 GGGACACCGGGCGACCGACA CGG ||||||||[||||||||||||]PAM NA Intergenic NA NA

T23430 GACCGACACGGAATCCCCAATGG 1000
Oligo fwd CACCGACCGACACGGAATCCCCAA
Oligo rev AAACTTGGGGATTCCGTGTCGGTC
Chromosome start end strand MM target_seq PAM alignment distance position gene name gene id
GU305914.1 140608 140630 - 0 GACCGACACGGAATCCCCAA TGG ||||||||[||||||||||||]PAM NA Intergenic NA NA

T23431 GCAGAGCAAGTGTCTCTCACCGG 1000
Oligo fwd CACCGCAGAGCAAGTGTCTCTCAC
Oligo rev AAACGTGAGAGACACTTGCTCTGC
Chromosome start end strand MM target_seq PAM alignment distance position gene name gene id
GU305914.1 140572 140594 - 0 GCAGAGCAAGTGTCTCTCAC CGG ||||||||[||||||||||||]PAM NA Intergenic NA NA

T23432 CAGAGCAAGTGTCTCTCACCGGG 1000
Oligo fwd CACCCAGAGCAAGTGTCTCTCACC
Oligo rev AAACGGTGAGAGACACTTGCTCTG
Chromosome start end strand MM target_seq PAM alignment distance position gene name gene id
GU305914.1 140571 140593 - 0 CAGAGCAAGTGTCTCTCACC GGG ||||||||[||||||||||||]PAM NA Intergenic NA NA

T23433 AGAGCAAGTGTCTCTCACCGGGG 1000
Oligo fwd CACCAGAGCAAGTGTCTCTCACCG
Oligo rev AAACCGGTGAGAGACACTTGCTCT
Chromosome start end strand MM target_seq PAM alignment distance position gene name gene id
GU305914.1 140570 140592 - 0 AGAGCAAGTGTCTCTCACCG GGG ||||||||[||||||||||||]PAM NA Intergenic NA NA

T23434 GAGCAAGTGTCTCTCACCGGGGG 1000
Oligo fwd CACCGAGCAAGTGTCTCTCACCGG
Oligo rev AAACCCGGTGAGAGACACTTGCTC
Chromosome start end strand MM target_seq PAM alignment distance position gene name gene id
GU305914.1 140569 140591 - 0 GAGCAAGTGTCTCTCACCGG GGG ||||||||[||||||||||||]PAM NA Intergenic NA NA

T23435 GTGTCTCTCACCGGGGGTACAGG 1000
Oligo fwd CACCGTGTCTCTCACCGGGGGTAC
Oligo rev AAACGTACCCCCGGTGAGAGACAC
Chromosome start end strand MM target_seq PAM alignment distance position gene name gene id
GU305914.1 140563 140585 - 0 GTGTCTCTCACCGGGGGTAC AGG ||||||||[||||||||||||]PAM NA Intergenic NA NA

T23436 CTCACCGGGGGTACAGGCGCAGG 1000
Oligo fwd CACCCTCACCGGGGGTACAGGCGC
Oligo rev AAACGCGCCTGTACCCCCGGTGAG
Chromosome start end strand MM target_seq PAM alignment distance position gene name gene id
GU305914.1 140557 140579 - 0 CTCACCGGGGGTACAGGCGC AGG ||||||||[||||||||||||]PAM NA Intergenic NA NA

T23437 CGGGGGTACAGGCGCAGGAATGG 1000
Oligo fwd CACCCGGGGGTACAGGCGCAGGAA
Oligo rev AAACTTCCTGCGCCTGTACCCCCG
Chromosome start end strand MM target_seq PAM alignment distance position gene name gene id
GU305914.1 140552 140574 - 0 CGGGGGTACAGGCGCAGGAA TGG ||||||||[||||||||||||]PAM NA Intergenic NA NA

T23438 TGTGTATCCATCCTCGCCCGCGG 1000
Oligo fwd CACCTGTGTATCCATCCTCGCCCG
Oligo rev AAACCGGGCGAGGATGGATACACA
Chromosome start end strand MM target_seq PAM alignment distance position gene name gene id
GU305914.1 140521 140543 - 0 TGTGTATCCATCCTCGCCCG CGG ||||||||[||||||||||||]PAM NA Intergenic NA NA

T23439 GTATCCATCCTCGCCCGCGGTGG 1000
Oligo fwd CACCGTATCCATCCTCGCCCGCGG
Oligo rev AAACCCGCGGGCGAGGATGGATAC
Chromosome start end strand MM target_seq PAM alignment distance position gene name gene id
GU305914.1 140518 140540 - 0 GTATCCATCCTCGCCCGCGG TGG ||||||||[||||||||||||]PAM NA Intergenic NA NA

T23440 GGAAGAATCATCAGTGCGTTTGG 1000
Oligo fwd CACCGGAAGAATCATCAGTGCGTT
Oligo rev AAACAACGCACTGATGATTCTTCC
Chromosome start end strand MM target_seq PAM alignment distance position gene name gene id
GU305914.1 140497 140519 - 0 GGAAGAATCATCAGTGCGTT TGG ||||||||[||||||||||||]PAM NA Intergenic NA NA

T23441 GAAGAATCATCAGTGCGTTTGGG 1000
Oligo fwd CACCGAAGAATCATCAGTGCGTTT
Oligo rev AAACAAACGCACTGATGATTCTTC
Chromosome start end strand MM target_seq PAM alignment distance position gene name gene id
GU305914.1 140496 140518 - 0 GAAGAATCATCAGTGCGTTT GGG ||||||||[||||||||||||]PAM NA Intergenic NA NA

T23442 ATCAGTGCGTTTGGGACTATCGG 1000
Oligo fwd CACCATCAGTGCGTTTGGGACTAT
Oligo rev AAACATAGTCCCAAACGCACTGAT
Chromosome start end strand MM target_seq PAM alignment distance position gene name gene id
GU305914.1 140488 140510 - 0 ATCAGTGCGTTTGGGACTAT CGG ||||||||[||||||||||||]PAM NA Intergenic NA NA

T23443 GGGACTATCGGATACGACATCGG 1000
Oligo fwd CACCGGGACTATCGGATACGACAT
Oligo rev AAACATGTCGTATCCGATAGTCCC
Chromosome start end strand MM target_seq PAM alignment distance position gene name gene id
GU305914.1 140476 140498 - 0 GGGACTATCGGATACGACAT CGG ||||||||[||||||||||||]PAM NA Intergenic NA NA

T23444 ACTATCGGATACGACATCGGCGG 1000
Oligo fwd CACCACTATCGGATACGACATCGG
Oligo rev AAACCCGATGTCGTATCCGATAGT
Chromosome start end strand MM target_seq PAM alignment distance position gene name gene id
GU305914.1 140473 140495 - 0 ACTATCGGATACGACATCGG CGG ||||||||[||||||||||||]PAM NA Intergenic NA NA

T23445 CGTCTTGCGCCGCGACCTGCTGG 1000
Oligo fwd CACCCGTCTTGCGCCGCGACCTGC
Oligo rev AAACGCAGGTCGCGGCGCAAGACG
Chromosome start end strand MM target_seq PAM alignment distance position gene name gene id
GU305914.1 140449 140471 - 0 CGTCTTGCGCCGCGACCTGC TGG ||||||||[||||||||||||]PAM NA Intergenic NA NA

T23446 CTTGCGCCGCGACCTGCTGGTGG 1000
Oligo fwd CACCCTTGCGCCGCGACCTGCTGG
Oligo rev AAACCCAGCAGGTCGCGGCGCAAG
Chromosome start end strand MM target_seq PAM alignment distance position gene name gene id
GU305914.1 140446 140468 - 0 CTTGCGCCGCGACCTGCTGG TGG ||||||||[||||||||||||]PAM NA Intergenic NA NA

T23447 ACTGCCGCTGTTGTTGTTACTGG 1000
Oligo fwd CACCACTGCCGCTGTTGTTGTTAC
Oligo rev AAACGTAACAACAACAGCGGCAGT
Chromosome start end strand MM target_seq PAM alignment distance position gene name gene id
GU305914.1 140416 140438 - 0 ACTGCCGCTGTTGTTGTTAC TGG ||||||||[||||||||||||]PAM NA Intergenic NA NA

T23448 GTTACTGGTATCGTTGTCGACGG 1000
Oligo fwd CACCGTTACTGGTATCGTTGTCGA
Oligo rev AAACTCGACAACGATACCAGTAAC
Chromosome start end strand MM target_seq PAM alignment distance position gene name gene id
GU305914.1 140401 140423 - 0 GTTACTGGTATCGTTGTCGA CGG ||||||||[||||||||||||]PAM NA Intergenic NA NA

T23449 GTTGTCGACGGCGCCATCGTCGG 1000
Oligo fwd CACCGTTGTCGACGGCGCCATCGT
Oligo rev AAACACGATGGCGCCGTCGACAAC
Chromosome start end strand MM target_seq PAM alignment distance position gene name gene id
GU305914.1 140389 140411 - 0 GTTGTCGACGGCGCCATCGT CGG ||||||||[||||||||||||]PAM NA Intergenic NA NA

T23450 CAGTTTTTCCAGATCGCTCACGG 1000
Oligo fwd CACCCAGTTTTTCCAGATCGCTCA
Oligo rev AAACTGAGCGATCTGGAAAAACTG
Chromosome start end strand MM target_seq PAM alignment distance position gene name gene id
GU305914.1 140335 140357 - 0 CAGTTTTTCCAGATCGCTCA CGG ||||||||[||||||||||||]PAM NA Intergenic NA NA

T23451 CTCACGGCGCGTCGCCTCAACGG 1000
Oligo fwd CACCCTCACGGCGCGTCGCCTCAA
Oligo rev AAACTTGAGGCGACGCGCCGTGAG
Chromosome start end strand MM target_seq PAM alignment distance position gene name gene id
GU305914.1 140319 140341 - 0 CTCACGGCGCGTCGCCTCAA CGG ||||||||[||||||||||||]PAM NA Intergenic NA NA

T23452 GCGTCGCCTCAACGGCTTTCTGG 1000
Oligo fwd CACCGCGTCGCCTCAACGGCTTTC
Oligo rev AAACGAAAGCCGTTGAGGCGACGC
Chromosome start end strand MM target_seq PAM alignment distance position gene name gene id
GU305914.1 140311 140333 - 0 GCGTCGCCTCAACGGCTTTC TGG ||||||||[||||||||||||]PAM NA Intergenic NA NA
GU305914.1 93643 93665 + 4 GCCTCGGCTCCCCGGCTTTC CGG ||-|||-|[||--||||||||]PAM NA Intergenic NA NA

T23453 CGTCGCCTCAACGGCTTTCTGGG 1000
Oligo fwd CACCCGTCGCCTCAACGGCTTTCT
Oligo rev AAACAGAAAGCCGTTGAGGCGACG
Chromosome start end strand MM target_seq PAM alignment distance position gene name gene id
GU305914.1 140310 140332 - 0 CGTCGCCTCAACGGCTTTCT GGG ||||||||[||||||||||||]PAM NA Intergenic NA NA

T23454 CTCAACGGCTTTCTGGGAGCCGG 1000
Oligo fwd CACCCTCAACGGCTTTCTGGGAGC
Oligo rev AAACGCTCCCAGAAAGCCGTTGAG

| | | | | | | | | | | | |
|---|---|---|---|---|---|---|---|---|---|---|---|
| Chromosome | start | end | strand | MM | target_seq | PAM | alignment | distance | position | gene name | gene id |
| GU305914.1 | 140304 | 140326 | - | 0 | CTCAACGGCTTTCTGGGAGC | CGG | ||||||||[||||||||||||]PAM | NA | Intergenic | NA | NA |
| T23455 | AACGGCTTTCTGGGAGCCGGAGG | 1000 | | | | | | | | | |
| Oligo fwd | CACCAACGGCTTTCTGGGAGCCGG | | | | | | | | | | |
| Oligo rev | AAACCCGGCTCCCAGAAAGCCGTT | | | | | | | | | | |
| Chromosome | start | end | strand | MM | target_seq | PAM | alignment | distance | position | gene name | gene id |
| GU305914.1 | 140301 | 140323 | - | 0 | AACGGCTTTCTGGGAGCCGG | AGG | ||||||||[||||||||||||]PAM | NA | Intergenic | NA | NA |
| T23456 | ACGGCTTTCTGGGAGCCGGAGGG | 1000 | | | | | | | | | |
| Oligo fwd | CACCACGGCTTTCTGGGAGCCGGA | | | | | | | | | | |
| Oligo rev | AAACTCCGGCTCCCAGAAAGCCGT | | | | | | | | | | |
| Chromosome | start | end | strand | MM | target_seq | PAM | alignment | distance | position | gene name | gene id |
| GU305914.1 | 140300 | 140322 | - | 0 | ACGGCTTTCTGGGAGCCGGA | GGG | ||||||||[||||||||||||]PAM | NA | Intergenic | NA | NA |
| T23457 | GGGAGCCGGAGGGACGACGACGG | 1000 | | | | | | | | | |
| Oligo fwd | CACCGGGAGCCGGAGGGACGACGA | | | | | | | | | | |
| Oligo rev | AAACTCGTCGTCCCTCCGGCTCCC | | | | | | | | | | |
| Chromosome | start | end | strand | MM | target_seq | PAM | alignment | distance | position | gene name | gene id |
| GU305914.1 | 140290 | 140312 | - | 0 | GGGAGCCGGAGGGACGACGA | CGG | ||||||||[||||||||||||]PAM | NA | Intergenic | NA | NA |
| T23458 | GGAGCCGGAGGGACGACGACGGG | 1000 | | | | | | | | | |
| Oligo fwd | CACCGGAGCCGGAGGGACGACGAC | | | | | | | | | | |
| Oligo rev | AAACGTCGTCGTCCCTCCGGCTCC | | | | | | | | | | |
| Chromosome | start | end | strand | MM | target_seq | PAM | alignment | distance | position | gene name | gene id |
| GU305914.1 | 140289 | 140311 | - | 0 | GGAGCCGGAGGGACGACGAC | GGG | ||||||||[||||||||||||]PAM | NA | Intergenic | NA | NA |
| T23459 | GACGACGGGAGCGAACCCGACGG | 1000 | | | | | | | | | |
| Oligo fwd | CACCGACGACGGGAGCGAACCCGA | | | | | | | | | | |
| Oligo rev | AAACTCGGGTTCGCTCCCGTCGTC | | | | | | | | | | |
| Chromosome | start | end | strand | MM | target_seq | PAM | alignment | distance | position | gene name | gene id |
| GU305914.1 | 140275 | 140297 | - | 0 | GACGACGGGAGCGAACCCGA | CGG | ||||||||[||||||||||||]PAM | NA | Intergenic | NA | NA |
| T23460 | CCGACGGAGCCGTCGCTTCTCGG | 1000 | | | | | | | | | |
| Oligo fwd | CACCCCGACGGAGCCGTCGCTTCT | | | | | | | | | | |
| Oligo rev | AAACAGAAGCGACGGCTCCGTCGG | | | | | | | | | | |
| Chromosome | start | end | strand | MM | target_seq | PAM | alignment | distance | position | gene name | gene id |
| GU305914.1 | 140259 | 140281 | - | 0 | CCGACGGAGCCGTCGCTTCT | CGG | ||||||||[||||||||||||]PAM | NA | Intergenic | NA | NA |
| T23461 | ACGGAGCCGTCGCTTCTCGGCGG | 1000 | | | | | | | | | |
| Oligo fwd | CACCACGGAGCCGTCGCTTCTCGG | | | | | | | | | | |
| Oligo rev | AAACCCGAGAAGCGACGGCTCCGT | | | | | | | | | | |
| Chromosome | start | end | strand | MM | target_seq | PAM | alignment | distance | position | gene name | gene id |
| GU305914.1 | 140256 | 140278 | - | 0 | ACGGAGCCGTCGCTTCTCGG | CGG | ||||||||[||||||||||||]PAM | NA | Intergenic | NA | NA |
| T23462 | CGGCGTCAGCTCCACGCTCATGG | 1000 | | | | | | | | | |
| Oligo fwd | CACCCGGCGTCAGCTCCACGCTCA | | | | | | | | | | |
| Oligo rev | AAACTGAGCGTGGAGCTGACGCCG | | | | | | | | | | |
| Chromosome | start | end | strand | MM | target_seq | PAM | alignment | distance | position | gene name | gene id |
| GU305914.1 | 140236 | 140258 | - | 0 | CGGCGTCAGCTCCACGCTCA | TGG | ||||||||[||||||||||||]PAM | NA | Intergenic | NA | NA |
| GU305914.1 | 38128 | 38150 | - | 4 | CGCCGACAGCTCCAGCCTCA | CGG | ||-||-||[||||||--||||]PAM | NA | Intergenic | NA | NA |
| T23463 | CAGCTCCACGCTCATGGCGACGG | 1000 | | | | | | | | | |
| Oligo fwd | CACCCAGCTCCACGCTCATGGCGA | | | | | | | | | | |
| Oligo rev | AAACTCGCCATGAGCGTGGAGCTG | | | | | | | | | | |
| Chromosome | start | end | strand | MM | target_seq | PAM | alignment | distance | position | gene name | gene id |
| GU305914.1 | 140230 | 140252 | - | 0 | CAGCTCCACGCTCATGGCGA | CGG | ||||||||[||||||||||||]PAM | NA | Intergenic | NA | NA |
| T23464 | CTCCACGCTCATGGCGACGGCGG | 1000 | | | | | | | | | |
| Oligo fwd | CACCCTCCACGCTCATGGCGACGG | | | | | | | | | | |
| Oligo rev | AAACCCGTCGCCATGAGCGTGGAG | | | | | | | | | | |
| Chromosome | start | end | strand | MM | target_seq | PAM | alignment | distance | position | gene name | gene id |
| GU305914.1 | 140227 | 140249 | - | 0 | CTCCACGCTCATGGCGACGG | CGG | ||||||||[||||||||||||]PAM | NA | Intergenic | NA | NA |
| T23465 | TCCACGCTCATGGCGACGGCGGG | 1000 | | | | | | | | | |
| Oligo fwd | CACCTCCACGCTCATGGCGACGGC | | | | | | | | | | |
| Oligo rev | AAACGCCGTCGCCATGAGCGTGGA | | | | | | | | | | |
| Chromosome | start | end | strand | MM | target_seq | PAM | alignment | distance | position | gene name | gene id |
| GU305914.1 | 140226 | 140248 | - | 0 | TCCACGCTCATGGCGACGGC | GGG | ||||||||[||||||||||||]PAM | NA | Intergenic | NA | NA |
| T23466 | CCACGCTCATGGCGACGGCGGGG | 1000 | | | | | | | | | |
| Oligo fwd | CACCCCACGCTCATGGCGACGGCG | | | | | | | | | | |
| Oligo rev | AAACCGCCGTCGCCATGAGCGTGG | | | | | | | | | | |
| Chromosome | start | end | strand | MM | target_seq | PAM | alignment | distance | position | gene name | gene id |
| GU305914.1 | 140225 | 140247 | - | 0 | CCACGCTCATGGCGACGGCG | GGG | ||||||||[||||||||||||]PAM | NA | Intergenic | NA | NA |
| GU305914.1 | 173163 | 173185 | - | 4 | CAACGATCACGGCGACGGCA | CGG | |-|||-||[|-|||||||||-]PAM | NA | Intergenic | NA | NA |
| T23467 | ACGGCGGGGTTTTTTTCTCGCGG | 1000 | | | | | | | | | |
| Oligo fwd | CACCACGGCGGGGTTTTTTTCTCG | | | | | | | | | | |
| Oligo rev | AAACCGAGAAAAAAACCCCGCCGT | | | | | | | | | | |
| Chromosome | start | end | strand | MM | target_seq | PAM | alignment | distance | position | gene name | gene id |
| GU305914.1 | 140211 | 140233 | - | 0 | ACGGCGGGGTTTTTTTCTCG | CGG | ||||||||[||||||||||||]PAM | NA | Intergenic | NA | NA |
| T23468 | CCGTTCGTCGACCGACCCCTCGG | 1000 | | | | | | | | | |
| Oligo fwd | CACCCCGTTCGTCGACCGACCCCT | | | | | | | | | | |
| Oligo rev | AAACAGGGGTCGGTCGACGAACGG | | | | | | | | | | |
| Chromosome | start | end | strand | MM | target_seq | PAM | alignment | distance | position | gene name | gene id |
| GU305914.1 | 140155 | 140177 | - | 0 | CCGTTCGTCGACCGACCCCT | CGG | ||||||||[||||||||||||]PAM | NA | Intergenic | NA | NA |

<table>
<tr><td>T23469</td><td>CGTTCGTCGACCGACCCCTCGGG</td><td>1000</td><td></td><td></td><td></td><td></td><td></td><td></td><td></td><td></td><td></td></tr>
<tr><td>Oligo fwd</td><td>CACCCGTTCGTCGACCGACCCCTC</td><td></td><td></td><td></td><td></td><td></td><td></td><td></td><td></td><td></td><td></td></tr>
<tr><td>Oligo rev</td><td>AAACGAGGGGTCGGTCGACGAACG</td><td></td><td></td><td></td><td></td><td></td><td></td><td></td><td></td><td></td><td></td></tr>
<tr><td>Chromosome</td><td>start</td><td>end</td><td>strand</td><td>MM</td><td>target_seq</td><td>PAM</td><td>alignment</td><td>distance</td><td>position</td><td>gene name</td><td>gene id</td></tr>
<tr><td>GU305914.1</td><td>140154</td><td>140176</td><td>-</td><td>0</td><td>CGTTCGTCGACCGACCCCTC</td><td>GGG</td><td>||||||||[||||||||||||]PAM</td><td>NA</td><td>Intergenic</td><td>NA</td><td>NA</td></tr>
<tr><td>T23470</td><td>GTTCGTCGACCGACCCCTCGGGG</td><td>1000</td><td></td><td></td><td></td><td></td><td></td><td></td><td></td><td></td><td></td></tr>
<tr><td>Oligo fwd</td><td>CACCGTTCGTCGACCGACCCCTCG</td><td></td><td></td><td></td><td></td><td></td><td></td><td></td><td></td><td></td><td></td></tr>
<tr><td>Oligo rev</td><td>AAACCGAGGGGTCGGTCGACGAAC</td><td></td><td></td><td></td><td></td><td></td><td></td><td></td><td></td><td></td><td></td></tr>
<tr><td>Chromosome</td><td>start</td><td>end</td><td>strand</td><td>MM</td><td>target_seq</td><td>PAM</td><td>alignment</td><td>distance</td><td>position</td><td>gene name</td><td>gene id</td></tr>
<tr><td>GU305914.1</td><td>140153</td><td>140175</td><td>-</td><td>0</td><td>GTTCGTCGACCGACCCCTCG</td><td>GGG</td><td>||||||||[||||||||||||]PAM</td><td>NA</td><td>Intergenic</td><td>NA</td><td>NA</td></tr>
<tr><td>T23471</td><td>TTCGTCGACCGACCCCTCGGGGG</td><td>1000</td><td></td><td></td><td></td><td></td><td></td><td></td><td></td><td></td><td></td></tr>
<tr><td>Oligo fwd</td><td>CACCTTCGTCGACCGACCCCTCGG</td><td></td><td></td><td></td><td></td><td></td><td></td><td></td><td></td><td></td><td></td></tr>
<tr><td>Oligo rev</td><td>AAACCCGAGGGGTCGGTCGACGAA</td><td></td><td></td><td></td><td></td><td></td><td></td><td></td><td></td><td></td><td></td></tr>
<tr><td>Chromosome</td><td>start</td><td>end</td><td>strand</td><td>MM</td><td>target_seq</td><td>PAM</td><td>alignment</td><td>distance</td><td>position</td><td>gene name</td><td>gene id</td></tr>
<tr><td>GU305914.1</td><td>140152</td><td>140174</td><td>-</td><td>0</td><td>TTCGTCGACCGACCCCTCGG</td><td>GGG</td><td>||||||||[||||||||||||]PAM</td><td>NA</td><td>Intergenic</td><td>NA</td><td>NA</td></tr>
<tr><td>T23472</td><td>GGGATGCGATCACCCTCACCCGG</td><td>1000</td><td></td><td></td><td></td><td></td><td></td><td></td><td></td><td></td><td></td></tr>
<tr><td>Oligo fwd</td><td>CACCGGGATGCGATCACCCTCACC</td><td></td><td></td><td></td><td></td><td></td><td></td><td></td><td></td><td></td><td></td></tr>
<tr><td>Oligo rev</td><td>AAACGGTGAGGGTGATCGCATCCC</td><td></td><td></td><td></td><td></td><td></td><td></td><td></td><td></td><td></td><td></td></tr>
<tr><td>Chromosome</td><td>start</td><td>end</td><td>strand</td><td>MM</td><td>target_seq</td><td>PAM</td><td>alignment</td><td>distance</td><td>position</td><td>gene name</td><td>gene id</td></tr>
<tr><td>GU305914.1</td><td>140132</td><td>140154</td><td>-</td><td>0</td><td>GGGATGCGATCACCCTCACC</td><td>CGG</td><td>||||||||[||||||||||||]PAM</td><td>NA</td><td>Intergenic</td><td>NA</td><td>NA</td></tr>
<tr><td>T23473</td><td>CCTCACCCGGACCACCGTCACGG</td><td>1000</td><td></td><td></td><td></td><td></td><td></td><td></td><td></td><td></td><td></td></tr>
<tr><td>Oligo fwd</td><td>CACCCCTCACCCGGACCACCGTCA</td><td></td><td></td><td></td><td></td><td></td><td></td><td></td><td></td><td></td><td></td></tr>
<tr><td>Oligo rev</td><td>AAACTGACGGTGGTCCGGGTGAGG</td><td></td><td></td><td></td><td></td><td></td><td></td><td></td><td></td><td></td><td></td></tr>
<tr><td>Chromosome</td><td>start</td><td>end</td><td>strand</td><td>MM</td><td>target_seq</td><td>PAM</td><td>alignment</td><td>distance</td><td>position</td><td>gene name</td><td>gene id</td></tr>
<tr><td>GU305914.1</td><td>140119</td><td>140141</td><td>-</td><td>0</td><td>CCTCACCCGGACCACCGTCA</td><td>CGG</td><td>||||||||[||||||||||||]PAM</td><td>NA</td><td>Intergenic</td><td>NA</td><td>NA</td></tr>
<tr><td>T23474</td><td>GATCCGAGCGCGCAACACAGCGG</td><td>1000</td><td></td><td></td><td></td><td></td><td></td><td></td><td></td><td></td><td></td></tr>
<tr><td>Oligo fwd</td><td>CACCGATCCGAGCGCGCAACACAG</td><td></td><td></td><td></td><td></td><td></td><td></td><td></td><td></td><td></td><td></td></tr>
<tr><td>Oligo rev</td><td>AAACCTGTGTTGCGCGCTCGGATC</td><td></td><td></td><td></td><td></td><td></td><td></td><td></td><td></td><td></td><td></td></tr>
<tr><td>Chromosome</td><td>start</td><td>end</td><td>strand</td><td>MM</td><td>target_seq</td><td>PAM</td><td>alignment</td><td>distance</td><td>position</td><td>gene name</td><td>gene id</td></tr>
<tr><td>GU305914.1</td><td>140057</td><td>140079</td><td>-</td><td>0</td><td>GATCCGAGCGCGCAACACAG</td><td>CGG</td><td>||||||||[||||||||||||]PAM</td><td>NA</td><td>Intergenic</td><td>NA</td><td>NA</td></tr>
<tr><td>T23475</td><td>GAACACACTACGAAGTTCCTCGG</td><td>1000</td><td></td><td></td><td></td><td></td><td></td><td></td><td></td><td></td><td></td></tr>
<tr><td>Oligo fwd</td><td>CACCGAACACACTACGAAGTTCCT</td><td></td><td></td><td></td><td></td><td></td><td></td><td></td><td></td><td></td><td></td></tr>
<tr><td>Oligo rev</td><td>AAACAGGAACTTCGTAGTGTGTTC</td><td></td><td></td><td></td><td></td><td></td><td></td><td></td><td></td><td></td><td></td></tr>
<tr><td>Chromosome</td><td>start</td><td>end</td><td>strand</td><td>MM</td><td>target_seq</td><td>PAM</td><td>alignment</td><td>distance</td><td>position</td><td>gene name</td><td>gene id</td></tr>
<tr><td>GU305914.1</td><td>139972</td><td>139994</td><td>-</td><td>0</td><td>GAACACACTACGAAGTTCCT</td><td>CGG</td><td>||||||||[||||||||||||]PAM</td><td>NA</td><td>Intergenic</td><td>NA</td><td>NA</td></tr>
<tr><td>T23476</td><td>CTCGGCCACGTCCCTCATCTCGG</td><td>1000</td><td></td><td></td><td></td><td></td><td></td><td></td><td></td><td></td><td></td></tr>
<tr><td>Oligo fwd</td><td>CACCCTCGGCCACGTCCCTCATCT</td><td></td><td></td><td></td><td></td><td></td><td></td><td></td><td></td><td></td><td></td></tr>
<tr><td>Oligo rev</td><td>AAACAGATGAGGGACGTGGCCGAG</td><td></td><td></td><td></td><td></td><td></td><td></td><td></td><td></td><td></td><td></td></tr>
<tr><td>Chromosome</td><td>start</td><td>end</td><td>strand</td><td>MM</td><td>target_seq</td><td>PAM</td><td>alignment</td><td>distance</td><td>position</td><td>gene name</td><td>gene id</td></tr>
<tr><td>GU305914.1</td><td>139954</td><td>139976</td><td>-</td><td>0</td><td>CTCGGCCACGTCCCTCATCT</td><td>CGG</td><td>||||||||[||||||||||||]PAM</td><td>NA</td><td>Intergenic</td><td>NA</td><td>NA</td></tr>
<tr><td>T23477</td><td>CCGCATCTGCGCCTTGCCAACGG</td><td>1000</td><td></td><td></td><td></td><td></td><td></td><td></td><td></td><td></td><td></td></tr>
<tr><td>Oligo fwd</td><td>CACCCCGCATCTGCGCCTTGCCAA</td><td></td><td></td><td></td><td></td><td></td><td></td><td></td><td></td><td></td><td></td></tr>
<tr><td>Oligo rev</td><td>AAACTTGGCAAGGCGCAGATGCGG</td><td></td><td></td><td></td><td></td><td></td><td></td><td></td><td></td><td></td><td></td></tr>
<tr><td>Chromosome</td><td>start</td><td>end</td><td>strand</td><td>MM</td><td>target_seq</td><td>PAM</td><td>alignment</td><td>distance</td><td>position</td><td>gene name</td><td>gene id</td></tr>
<tr><td>GU305914.1</td><td>139846</td><td>139868</td><td>-</td><td>0</td><td>CCGCATCTGCGCCTTGCCAA</td><td>CGG</td><td>||||||||[||||||||||||]PAM</td><td>NA</td><td>Intergenic</td><td>NA</td><td>NA</td></tr>
<tr><td>T23478</td><td>CTTGCCAACGGCGATCGCCGCGG</td><td>1000</td><td></td><td></td><td></td><td></td><td></td><td></td><td></td><td></td><td></td></tr>
<tr><td>Oligo fwd</td><td>CACCCTTGCCAACGGCGATCGCCG</td><td></td><td></td><td></td><td></td><td></td><td></td><td></td><td></td><td></td><td></td></tr>
<tr><td>Oligo rev</td><td>AAACCGGCGATCGCCGTTGGCAAG</td><td></td><td></td><td></td><td></td><td></td><td></td><td></td><td></td><td></td><td></td></tr>
<tr><td>Chromosome</td><td>start</td><td>end</td><td>strand</td><td>MM</td><td>target_seq</td><td>PAM</td><td>alignment</td><td>distance</td><td>position</td><td>gene name</td><td>gene id</td></tr>
<tr><td>GU305914.1</td><td>139834</td><td>139856</td><td>-</td><td>0</td><td>CTTGCCAACGGCGATCGCCG</td><td>CGG</td><td>||||||||[||||||||||||]PAM</td><td>NA</td><td>Intergenic</td><td>NA</td><td>NA</td></tr>
<tr><td>T23479</td><td>GAGTGCTCTGCGCGCATTCTCGG</td><td>1000</td><td></td><td></td><td></td><td></td><td></td><td></td><td></td><td></td><td></td></tr>
<tr><td>Oligo fwd</td><td>CACCGAGTGCTCTGCGCGCATTCT</td><td></td><td></td><td></td><td></td><td></td><td></td><td></td><td></td><td></td><td></td></tr>
<tr><td>Oligo rev</td><td>AAACAGAATGCGCGCAGAGCACTC</td><td></td><td></td><td></td><td></td><td></td><td></td><td></td><td></td><td></td><td></td></tr>
<tr><td>Chromosome</td><td>start</td><td>end</td><td>strand</td><td>MM</td><td>target_seq</td><td>PAM</td><td>alignment</td><td>distance</td><td>position</td><td>gene name</td><td>gene id</td></tr>
<tr><td>GU305914.1</td><td>139802</td><td>139824</td><td>-</td><td>0</td><td>GAGTGCTCTGCGCGCATTCT</td><td>CGG</td><td>||||||||[||||||||||||]PAM</td><td>NA</td><td>Intergenic</td><td>NA</td><td>NA</td></tr>
<tr><td>T23480</td><td>TCGGATACTCGCTCGCGTCTTGG</td><td>1000</td><td></td><td></td><td></td><td></td><td></td><td></td><td></td><td></td><td></td></tr>
<tr><td>Oligo fwd</td><td>CACCTCGGATACTCGCTCGCGTCT</td><td></td><td></td><td></td><td></td><td></td><td></td><td></td><td></td><td></td><td></td></tr>
<tr><td>Oligo rev</td><td>AAACAGACGCGAGCGAGTATCCGA</td><td></td><td></td><td></td><td></td><td></td><td></td><td></td><td></td><td></td><td></td></tr>
<tr><td>Chromosome</td><td>start</td><td>end</td><td>strand</td><td>MM</td><td>target_seq</td><td>PAM</td><td>alignment</td><td>distance</td><td>position</td><td>gene name</td><td>gene id</td></tr>
<tr><td>GU305914.1</td><td>139783</td><td>139805</td><td>-</td><td>0</td><td>TCGGATACTCGCTCGCGTCT</td><td>TGG</td><td>||||||||[||||||||||||]PAM</td><td>NA</td><td>Intergenic</td><td>NA</td><td>NA</td></tr>
<tr><td>T23481</td><td>TGATCCCCTGAATACGCACCAGG</td><td>1000</td><td></td><td></td><td></td><td></td><td></td><td></td><td></td><td></td><td></td></tr>
<tr><td>Oligo fwd</td><td>CACCTGATCCCCTGAATACGCACC</td><td></td><td></td><td></td><td></td><td></td><td></td><td></td><td></td><td></td><td></td></tr>
<tr><td>Oligo rev</td><td>AAACGGTGCGTATTCAGGGGATCA</td><td></td><td></td><td></td><td></td><td></td><td></td><td></td><td></td><td></td><td></td></tr>
<tr><td>Chromosome</td><td>start</td><td>end</td><td>strand</td><td>MM</td><td>target_seq</td><td>PAM</td><td>alignment</td><td>distance</td><td>position</td><td>gene name</td><td>gene id</td></tr>
<tr><td>GU305914.1</td><td>139748</td><td>139770</td><td>-</td><td>0</td><td>TGATCCCCTGAATACGCACC</td><td>AGG</td><td>||||||||[||||||||||||]PAM</td><td>NA</td><td>Intergenic</td><td>NA</td><td>NA</td></tr>
<tr><td>T23482</td><td>GATCCCCTGAATACGCACCAGGG</td><td>1000</td><td></td><td></td><td></td><td></td><td></td><td></td><td></td><td></td><td></td></tr>
<tr><td>Oligo fwd</td><td>CACCGATCCCCTGAATACGCACCA</td><td></td><td></td><td></td><td></td><td></td><td></td><td></td><td></td><td></td><td></td></tr>
<tr><td>Oligo rev</td><td>AAACTGGTGCGTATTCAGGGGATC</td><td></td><td></td><td></td><td></td><td></td><td></td><td></td><td></td><td></td><td></td></tr>
<tr><td>Chromosome</td><td>start</td><td>end</td><td>strand</td><td>MM</td><td>target_seq</td><td>PAM</td><td>alignment</td><td>distance</td><td>position</td><td>gene name</td><td>gene id</td></tr>
<tr><td>GU305914.1</td><td>139747</td><td>139769</td><td>-</td><td>0</td><td>GATCCCCTGAATACGCACCA</td><td>GGG</td><td>||||||||[||||||||||||]PAM</td><td>NA</td><td>Intergenic</td><td>NA</td><td>NA</td></tr>
<tr><td>T23483</td><td>ATCCCCTGAATACGCACCAGGGG</td><td>1000</td><td></td><td></td><td></td><td></td><td></td><td></td><td></td><td></td><td></td></tr>
<tr><td>Oligo fwd</td><td>CACCATCCCCTGAATACGCACCAG</td><td></td><td></td><td></td><td></td><td></td><td></td><td></td><td></td><td></td><td></td></tr>
<tr><td>Oligo rev</td><td>AAACCTGGTGCGTATTCAGGGGAT</td><td></td><td></td><td></td><td></td><td></td><td></td><td></td><td></td><td></td><td></td></tr>
<tr><td>Chromosome</td><td>start</td><td>end</td><td>strand</td><td>MM</td><td>target_seq</td><td>PAM</td><td>alignment</td><td>distance</td><td>position</td><td>gene name</td><td>gene id</td></tr>
<tr><td>GU305914.1</td><td>139746</td><td>139768</td><td>-</td><td>0</td><td>ATCCCCTGAATACGCACCAG</td><td>GGG</td><td>||||||||[||||||||||||]PAM</td><td>NA</td><td>Intergenic</td><td>NA</td><td>NA</td></tr>
</table>

<table>
<tr><td>T23484</td><td>ATACGCACCAGGGGATGATCAGG</td><td>1000</td><td></td><td></td><td></td><td></td><td></td><td></td><td></td><td></td><td></td></tr>
<tr><td>Oligo fwd</td><td>CACCATACGCACCAGGGGATGATC</td><td></td><td></td><td></td><td></td><td></td><td></td><td></td><td></td><td></td><td></td></tr>
<tr><td>Oligo rev</td><td>AAACGATCATCCCCTGGTGCGTAT</td><td></td><td></td><td></td><td></td><td></td><td></td><td></td><td></td><td></td><td></td></tr>
<tr><td>Chromosome</td><td>start</td><td>end</td><td>strand</td><td>MM</td><td>target_seq</td><td>PAM</td><td>alignment</td><td>distance</td><td>position</td><td>gene name</td><td>gene id</td></tr>
<tr><td>GU305914.1</td><td>139737</td><td>139759</td><td>-</td><td>0</td><td>ATACGCACCAGGGGATGATC</td><td>AGG</td><td>||||||||[||||||||||||]PAM</td><td>NA</td><td>Intergenic</td><td>NA</td><td>NA</td></tr>
<tr><td>T23485</td><td>ATCGCTTCTGCCGTTTCTCCAGG</td><td>1000</td><td></td><td></td><td></td><td></td><td></td><td></td><td></td><td></td><td></td></tr>
<tr><td>Oligo fwd</td><td>CACCATCGCTTCTGCCGTTTCTCC</td><td></td><td></td><td></td><td></td><td></td><td></td><td></td><td></td><td></td><td></td></tr>
<tr><td>Oligo rev</td><td>AAACGGAGAAACGGCAGAAGCGAT</td><td></td><td></td><td></td><td></td><td></td><td></td><td></td><td></td><td></td><td></td></tr>
<tr><td>Chromosome</td><td>start</td><td>end</td><td>strand</td><td>MM</td><td>target_seq</td><td>PAM</td><td>alignment</td><td>distance</td><td>position</td><td>gene name</td><td>gene id</td></tr>
<tr><td>GU305914.1</td><td>139694</td><td>139716</td><td>-</td><td>0</td><td>ATCGCTTCTGCCGTTTCTCC</td><td>AGG</td><td>||||||||[||||||||||||]PAM</td><td>NA</td><td>Intergenic</td><td>NA</td><td>NA</td></tr>
<tr><td>T23486</td><td>CAGCGCACTCTCGCTGCGCGCGG</td><td>1000</td><td></td><td></td><td></td><td></td><td></td><td></td><td></td><td></td><td></td></tr>
<tr><td>Oligo fwd</td><td>CACCCAGCGCACTCTCGCTGCGCG</td><td></td><td></td><td></td><td></td><td></td><td></td><td></td><td></td><td></td><td></td></tr>
<tr><td>Oligo rev</td><td>AAACCGCGCAGCGAGAGTGCGCTG</td><td></td><td></td><td></td><td></td><td></td><td></td><td></td><td></td><td></td><td></td></tr>
<tr><td>Chromosome</td><td>start</td><td>end</td><td>strand</td><td>MM</td><td>target_seq</td><td>PAM</td><td>alignment</td><td>distance</td><td>position</td><td>gene name</td><td>gene id</td></tr>
<tr><td>GU305914.1</td><td>139648</td><td>139670</td><td>-</td><td>0</td><td>CAGCGCACTCTCGCTGCGCG</td><td>CGG</td><td>||||||||[||||||||||||]PAM</td><td>NA</td><td>Intergenic</td><td>NA</td><td>NA</td></tr>
<tr><td>T23487</td><td>AGCGCACTCTCGCTGCGCGCGGG</td><td>1000</td><td></td><td></td><td></td><td></td><td></td><td></td><td></td><td></td><td></td></tr>
<tr><td>Oligo fwd</td><td>CACCAGCGCACTCTCGCTGCGCGC</td><td></td><td></td><td></td><td></td><td></td><td></td><td></td><td></td><td></td><td></td></tr>
<tr><td>Oligo rev</td><td>AAACGCGCGCAGCGAGAGTGCGCT</td><td></td><td></td><td></td><td></td><td></td><td></td><td></td><td></td><td></td><td></td></tr>
<tr><td>Chromosome</td><td>start</td><td>end</td><td>strand</td><td>MM</td><td>target_seq</td><td>PAM</td><td>alignment</td><td>distance</td><td>position</td><td>gene name</td><td>gene id</td></tr>
<tr><td>GU305914.1</td><td>139647</td><td>139669</td><td>-</td><td>0</td><td>AGCGCACTCTCGCTGCGCGC</td><td>GGG</td><td>||||||||[||||||||||||]PAM</td><td>NA</td><td>Intergenic</td><td>NA</td><td>NA</td></tr>
<tr><td>T23488</td><td>ACCCCATCAGACCGCGTCGTCGG</td><td>1000</td><td></td><td></td><td></td><td></td><td></td><td></td><td></td><td></td><td></td></tr>
<tr><td>Oligo fwd</td><td>CACCACCCCATCAGACCGCGTCGT</td><td></td><td></td><td></td><td></td><td></td><td></td><td></td><td></td><td></td><td></td></tr>
<tr><td>Oligo rev</td><td>AAACACGACGCGGTCTGATGGGGT</td><td></td><td></td><td></td><td></td><td></td><td></td><td></td><td></td><td></td><td></td></tr>
<tr><td>Chromosome</td><td>start</td><td>end</td><td>strand</td><td>MM</td><td>target_seq</td><td>PAM</td><td>alignment</td><td>distance</td><td>position</td><td>gene name</td><td>gene id</td></tr>
<tr><td>GU305914.1</td><td>139616</td><td>139638</td><td>-</td><td>0</td><td>ACCCCATCAGACCGCGTCGT</td><td>CGG</td><td>||||||||[||||||||||||]PAM</td><td>NA</td><td>Intergenic</td><td>NA</td><td>NA</td></tr>
<tr><td>T23489</td><td>CCATCAGACCGCGTCGTCGGTGG</td><td>1000</td><td></td><td></td><td></td><td></td><td></td><td></td><td></td><td></td><td></td></tr>
<tr><td>Oligo fwd</td><td>CACCCCATCAGACCGCGTCGTCGG</td><td></td><td></td><td></td><td></td><td></td><td></td><td></td><td></td><td></td><td></td></tr>
<tr><td>Oligo rev</td><td>AAACCCGACGACGCGGTCTGATGG</td><td></td><td></td><td></td><td></td><td></td><td></td><td></td><td></td><td></td><td></td></tr>
<tr><td>Chromosome</td><td>start</td><td>end</td><td>strand</td><td>MM</td><td>target_seq</td><td>PAM</td><td>alignment</td><td>distance</td><td>position</td><td>gene name</td><td>gene id</td></tr>
<tr><td>GU305914.1</td><td>139613</td><td>139635</td><td>-</td><td>0</td><td>CCATCAGACCGCGTCGTCGG</td><td>TGG</td><td>||||||||[||||||||||||]PAM</td><td>NA</td><td>Intergenic</td><td>NA</td><td>NA</td></tr>
<tr><td>T23490</td><td>TCAGACCGCGTCGTCGGTGGTGG</td><td>1000</td><td></td><td></td><td></td><td></td><td></td><td></td><td></td><td></td><td></td></tr>
<tr><td>Oligo fwd</td><td>CACCTCAGACCGCGTCGTCGGTGG</td><td></td><td></td><td></td><td></td><td></td><td></td><td></td><td></td><td></td><td></td></tr>
<tr><td>Oligo rev</td><td>AAACCCACCGACGACGCGGTCTGA</td><td></td><td></td><td></td><td></td><td></td><td></td><td></td><td></td><td></td><td></td></tr>
<tr><td>Chromosome</td><td>start</td><td>end</td><td>strand</td><td>MM</td><td>target_seq</td><td>PAM</td><td>alignment</td><td>distance</td><td>position</td><td>gene name</td><td>gene id</td></tr>
<tr><td>GU305914.1</td><td>139610</td><td>139632</td><td>-</td><td>0</td><td>TCAGACCGCGTCGTCGGTGG</td><td>TGG</td><td>||||||||[||||||||||||]PAM</td><td>NA</td><td>Intergenic</td><td>NA</td><td>NA</td></tr>
<tr><td>T23491</td><td>GGTGGAGAGCGCCGAATCGATGG</td><td>1000</td><td></td><td></td><td></td><td></td><td></td><td></td><td></td><td></td><td></td></tr>
<tr><td>Oligo fwd</td><td>CACCGGTGGAGAGCGCCGAATCGA</td><td></td><td></td><td></td><td></td><td></td><td></td><td></td><td></td><td></td><td></td></tr>
<tr><td>Oligo rev</td><td>AAACTCGATTCGGCGCTCTCCACC</td><td></td><td></td><td></td><td></td><td></td><td></td><td></td><td></td><td></td><td></td></tr>
<tr><td>Chromosome</td><td>start</td><td>end</td><td>strand</td><td>MM</td><td>target_seq</td><td>PAM</td><td>alignment</td><td>distance</td><td>position</td><td>gene name</td><td>gene id</td></tr>
<tr><td>GU305914.1</td><td>139592</td><td>139614</td><td>-</td><td>0</td><td>GGTGGAGAGCGCCGAATCGA</td><td>TGG</td><td>||||||||[||||||||||||]PAM</td><td>NA</td><td>Intergenic</td><td>NA</td><td>NA</td></tr>
<tr><td>T23492</td><td>CCTGCGCTCCTCGCCGTCGTTGG</td><td>1000</td><td></td><td></td><td></td><td></td><td></td><td></td><td></td><td></td><td></td></tr>
<tr><td>Oligo fwd</td><td>CACCCCTGCGCTCCTCGCCGTCGT</td><td></td><td></td><td></td><td></td><td></td><td></td><td></td><td></td><td></td><td></td></tr>
<tr><td>Oligo rev</td><td>AAACACGACGGCGAGGAGCGCAGG</td><td></td><td></td><td></td><td></td><td></td><td></td><td></td><td></td><td></td><td></td></tr>
<tr><td>Chromosome</td><td>start</td><td>end</td><td>strand</td><td>MM</td><td>target_seq</td><td>PAM</td><td>alignment</td><td>distance</td><td>position</td><td>gene name</td><td>gene id</td></tr>
<tr><td>GU305914.1</td><td>139553</td><td>139575</td><td>-</td><td>0</td><td>CCTGCGCTCCTCGCCGTCGT</td><td>TGG</td><td>||||||||[||||||||||||]PAM</td><td>NA</td><td>Intergenic</td><td>NA</td><td>NA</td></tr>
<tr><td>T23493</td><td>CGTTGGAGTCGCAGAGCGCTCGG</td><td>1000</td><td></td><td></td><td></td><td></td><td></td><td></td><td></td><td></td><td></td></tr>
<tr><td>Oligo fwd</td><td>CACCCGTTGGAGTCGCAGAGCGCT</td><td></td><td></td><td></td><td></td><td></td><td></td><td></td><td></td><td></td><td></td></tr>
<tr><td>Oligo rev</td><td>AAACAGCGCTCTGCGACTCCAACG</td><td></td><td></td><td></td><td></td><td></td><td></td><td></td><td></td><td></td><td></td></tr>
<tr><td>Chromosome</td><td>start</td><td>end</td><td>strand</td><td>MM</td><td>target_seq</td><td>PAM</td><td>alignment</td><td>distance</td><td>position</td><td>gene name</td><td>gene id</td></tr>
<tr><td>GU305914.1</td><td>139536</td><td>139558</td><td>-</td><td>0</td><td>CGTTGGAGTCGCAGAGCGCT</td><td>CGG</td><td>||||||||[||||||||||||]PAM</td><td>NA</td><td>Intergenic</td><td>NA</td><td>NA</td></tr>
<tr><td>T23494</td><td>CAGCGTCCCGATGTCGTAGACGG</td><td>1000</td><td></td><td></td><td></td><td></td><td></td><td></td><td></td><td></td><td></td></tr>
<tr><td>Oligo fwd</td><td>CACCCAGCGTCCCGATGTCGTAGA</td><td></td><td></td><td></td><td></td><td></td><td></td><td></td><td></td><td></td><td></td></tr>
<tr><td>Oligo rev</td><td>AAACTCTACGACATCGGGACGCTG</td><td></td><td></td><td></td><td></td><td></td><td></td><td></td><td></td><td></td><td></td></tr>
<tr><td>Chromosome</td><td>start</td><td>end</td><td>strand</td><td>MM</td><td>target_seq</td><td>PAM</td><td>alignment</td><td>distance</td><td>position</td><td>gene name</td><td>gene id</td></tr>
<tr><td>GU305914.1</td><td>139511</td><td>139533</td><td>-</td><td>0</td><td>CAGCGTCCCGATGTCGTAGA</td><td>CGG</td><td>||||||||[||||||||||||]PAM</td><td>NA</td><td>Intergenic</td><td>NA</td><td>NA</td></tr>
<tr><td>T23495</td><td>AGCGTCCCGATGTCGTAGACGGG</td><td>1000</td><td></td><td></td><td></td><td></td><td></td><td></td><td></td><td></td><td></td></tr>
<tr><td>Oligo fwd</td><td>CACCAGCGTCCCGATGTCGTAGAC</td><td></td><td></td><td></td><td></td><td></td><td></td><td></td><td></td><td></td><td></td></tr>
<tr><td>Oligo rev</td><td>AAACGTCTACGACATCGGGACGCT</td><td></td><td></td><td></td><td></td><td></td><td></td><td></td><td></td><td></td><td></td></tr>
<tr><td>Chromosome</td><td>start</td><td>end</td><td>strand</td><td>MM</td><td>target_seq</td><td>PAM</td><td>alignment</td><td>distance</td><td>position</td><td>gene name</td><td>gene id</td></tr>
<tr><td>GU305914.1</td><td>139510</td><td>139532</td><td>-</td><td>0</td><td>AGCGTCCCGATGTCGTAGAC</td><td>GGG</td><td>||||||||[||||||||||||]PAM</td><td>NA</td><td>Intergenic</td><td>NA</td><td>NA</td></tr>
<tr><td>T23496</td><td>CGATGTCGTAGACGGGAAACAGG</td><td>1000</td><td></td><td></td><td></td><td></td><td></td><td></td><td></td><td></td><td></td></tr>
<tr><td>Oligo fwd</td><td>CACCCGATGTCGTAGACGGGAAAC</td><td></td><td></td><td></td><td></td><td></td><td></td><td></td><td></td><td></td><td></td></tr>
<tr><td>Oligo rev</td><td>AAACGTTTCCCGTCTACGACATCG</td><td></td><td></td><td></td><td></td><td></td><td></td><td></td><td></td><td></td><td></td></tr>
<tr><td>Chromosome</td><td>start</td><td>end</td><td>strand</td><td>MM</td><td>target_seq</td><td>PAM</td><td>alignment</td><td>distance</td><td>position</td><td>gene name</td><td>gene id</td></tr>
<tr><td>GU305914.1</td><td>139503</td><td>139525</td><td>-</td><td>0</td><td>CGATGTCGTAGACGGGAAAC</td><td>AGG</td><td>||||||||[||||||||||||]PAM</td><td>NA</td><td>Intergenic</td><td>NA</td><td>NA</td></tr>
<tr><td>T23497</td><td>GATGTCGTAGACGGGAAACAGGG</td><td>1000</td><td></td><td></td><td></td><td></td><td></td><td></td><td></td><td></td><td></td></tr>
<tr><td>Oligo fwd</td><td>CACCGATGTCGTAGACGGGAAACA</td><td></td><td></td><td></td><td></td><td></td><td></td><td></td><td></td><td></td><td></td></tr>
<tr><td>Oligo rev</td><td>AAACTGTTTCCCGTCTACGACATC</td><td></td><td></td><td></td><td></td><td></td><td></td><td></td><td></td><td></td><td></td></tr>
<tr><td>Chromosome</td><td>start</td><td>end</td><td>strand</td><td>MM</td><td>target_seq</td><td>PAM</td><td>alignment</td><td>distance</td><td>position</td><td>gene name</td><td>gene id</td></tr>
<tr><td>GU305914.1</td><td>139502</td><td>139524</td><td>-</td><td>0</td><td>GATGTCGTAGACGGGAAACA</td><td>GGG</td><td>||||||||[||||||||||||]PAM</td><td>NA</td><td>Intergenic</td><td>NA</td><td>NA</td></tr>
<tr><td>T23498</td><td>GTCGTAGACGGGAAACAGGGAGG</td><td>1000</td><td></td><td></td><td></td><td></td><td></td><td></td><td></td><td></td><td></td></tr>
<tr><td>Oligo fwd</td><td>CACCGTCGTAGACGGGAAACAGGG</td><td></td><td></td><td></td><td></td><td></td><td></td><td></td><td></td><td></td><td></td></tr>
<tr><td>Oligo rev</td><td>AAACCCCTGTTTCCCGTCTACGAC</td><td></td><td></td><td></td><td></td><td></td><td></td><td></td><td></td><td></td><td></td></tr>
<tr><td>Chromosome</td><td>start</td><td>end</td><td>strand</td><td>MM</td><td>target_seq</td><td>PAM</td><td>alignment</td><td>distance</td><td>position</td><td>gene name</td><td>gene id</td></tr>
</table>

<table>
<tr><td>GU305914.1</td><td>139499</td><td>139521</td><td>-</td><td>0</td><td>GTCGTAGACGGGAAACAGGG</td><td>AGG</td><td>||||||||[||||||||||||]PAM</td><td>NA</td><td>Intergenic</td><td>NA</td><td>NA</td></tr>
<tr><td>T23499</td><td>AGGGAGGCGACGCGCTGTCCCGG</td><td>1000</td><td></td><td></td><td></td><td></td><td></td><td></td><td></td><td></td><td></td></tr>
<tr><td>Oligo fwd</td><td>CACCAGGGAGGCGACGCGCTGTCC</td><td></td><td></td><td></td><td></td><td></td><td></td><td></td><td></td><td></td><td></td></tr>
<tr><td>Oligo rev</td><td>AAACGGACAGCGCGTCGCCTCCCT</td><td></td><td></td><td></td><td></td><td></td><td></td><td></td><td></td><td></td><td></td></tr>
<tr><td>Chromosome</td><td>start</td><td>end</td><td>strand</td><td>MM</td><td>target_seq</td><td>PAM</td><td>alignment</td><td>distance</td><td>position</td><td>gene name</td><td>gene id</td></tr>
<tr><td>GU305914.1</td><td>139483</td><td>139505</td><td>-</td><td>0</td><td>AGGGAGGCGACGCGCTGTCC</td><td>CGG</td><td>||||||||[||||||||||||]PAM</td><td>NA</td><td>Intergenic</td><td>NA</td><td>NA</td></tr>
<tr><td>T23500</td><td>AGGCGACGCGCTGTCCCGGCAGG</td><td>1000</td><td></td><td></td><td></td><td></td><td></td><td></td><td></td><td></td><td></td></tr>
<tr><td>Oligo fwd</td><td>CACCAGGCGACGCGCTGTCCCGGC</td><td></td><td></td><td></td><td></td><td></td><td></td><td></td><td></td><td></td><td></td></tr>
<tr><td>Oligo rev</td><td>AAACGCCGGGACAGCGCGTCGCCT</td><td></td><td></td><td></td><td></td><td></td><td></td><td></td><td></td><td></td><td></td></tr>
<tr><td>Chromosome</td><td>start</td><td>end</td><td>strand</td><td>MM</td><td>target_seq</td><td>PAM</td><td>alignment</td><td>distance</td><td>position</td><td>gene name</td><td>gene id</td></tr>
<tr><td>GU305914.1</td><td>139479</td><td>139501</td><td>-</td><td>0</td><td>AGGCGACGCGCTGTCCCGGC</td><td>AGG</td><td>||||||||[||||||||||||]PAM</td><td>NA</td><td>Intergenic</td><td>NA</td><td>NA</td></tr>
<tr><td>GU305914.1</td><td>97037</td><td>97059</td><td>+</td><td>4</td><td>AGACGGCGCGCTGTCGCCGC</td><td>GGG</td><td>||-||-||[|||||||-|-||]PAM</td><td>NA</td><td>Intergenic</td><td>NA</td><td>NA</td></tr>
<tr><td>T23501</td><td>GTCCCGGCAGGACGTGACTGAGG</td><td>1000</td><td></td><td></td><td></td><td></td><td></td><td></td><td></td><td></td><td></td></tr>
<tr><td>Oligo fwd</td><td>CACCGTCCCGGCAGGACGTGACTG</td><td></td><td></td><td></td><td></td><td></td><td></td><td></td><td></td><td></td><td></td></tr>
<tr><td>Oligo rev</td><td>AAACCAGTCACGTCCTGCCGGGAC</td><td></td><td></td><td></td><td></td><td></td><td></td><td></td><td></td><td></td><td></td></tr>
<tr><td>Chromosome</td><td>start</td><td>end</td><td>strand</td><td>MM</td><td>target_seq</td><td>PAM</td><td>alignment</td><td>distance</td><td>position</td><td>gene name</td><td>gene id</td></tr>
<tr><td>GU305914.1</td><td>139467</td><td>139489</td><td>-</td><td>0</td><td>GTCCCGGCAGGACGTGACTG</td><td>AGG</td><td>||||||||[||||||||||||]PAM</td><td>NA</td><td>Intergenic</td><td>NA</td><td>NA</td></tr>
<tr><td>T23502</td><td>TCCCGGCAGGACGTGACTGAGGG</td><td>1000</td><td></td><td></td><td></td><td></td><td></td><td></td><td></td><td></td><td></td></tr>
<tr><td>Oligo fwd</td><td>CACCTCCCGGCAGGACGTGACTGA</td><td></td><td></td><td></td><td></td><td></td><td></td><td></td><td></td><td></td><td></td></tr>
<tr><td>Oligo rev</td><td>AAACTCAGTCACGTCCTGCCGGGA</td><td></td><td></td><td></td><td></td><td></td><td></td><td></td><td></td><td></td><td></td></tr>
<tr><td>Chromosome</td><td>start</td><td>end</td><td>strand</td><td>MM</td><td>target_seq</td><td>PAM</td><td>alignment</td><td>distance</td><td>position</td><td>gene name</td><td>gene id</td></tr>
<tr><td>GU305914.1</td><td>139466</td><td>139488</td><td>-</td><td>0</td><td>TCCCGGCAGGACGTGACTGA</td><td>GGG</td><td>||||||||[||||||||||||]PAM</td><td>NA</td><td>Intergenic</td><td>NA</td><td>NA</td></tr>
<tr><td>T23503</td><td>CGACAGTCCTTGATGATGATCGG</td><td>1000</td><td></td><td></td><td></td><td></td><td></td><td></td><td></td><td></td><td></td></tr>
<tr><td>Oligo fwd</td><td>CACCCGACAGTCCTTGATGATGAT</td><td></td><td></td><td></td><td></td><td></td><td></td><td></td><td></td><td></td><td></td></tr>
<tr><td>Oligo rev</td><td>AAACATCATCATCAAGGACTGTCG</td><td></td><td></td><td></td><td></td><td></td><td></td><td></td><td></td><td></td><td></td></tr>
<tr><td>Chromosome</td><td>start</td><td>end</td><td>strand</td><td>MM</td><td>target_seq</td><td>PAM</td><td>alignment</td><td>distance</td><td>position</td><td>gene name</td><td>gene id</td></tr>
<tr><td>GU305914.1</td><td>139423</td><td>139445</td><td>-</td><td>0</td><td>CGACAGTCCTTGATGATGAT</td><td>CGG</td><td>||||||||[||||||||||||]PAM</td><td>NA</td><td>Intergenic</td><td>NA</td><td>NA</td></tr>
<tr><td>T23504</td><td>GACAGTCCTTGATGATGATCGGG</td><td>1000</td><td></td><td></td><td></td><td></td><td></td><td></td><td></td><td></td><td></td></tr>
<tr><td>Oligo fwd</td><td>CACCGACAGTCCTTGATGATGATC</td><td></td><td></td><td></td><td></td><td></td><td></td><td></td><td></td><td></td><td></td></tr>
<tr><td>Oligo rev</td><td>AAACGATCATCATCAAGGACTGTC</td><td></td><td></td><td></td><td></td><td></td><td></td><td></td><td></td><td></td><td></td></tr>
<tr><td>Chromosome</td><td>start</td><td>end</td><td>strand</td><td>MM</td><td>target_seq</td><td>PAM</td><td>alignment</td><td>distance</td><td>position</td><td>gene name</td><td>gene id</td></tr>
<tr><td>GU305914.1</td><td>139422</td><td>139444</td><td>-</td><td>0</td><td>GACAGTCCTTGATGATGATC</td><td>GGG</td><td>||||||||[||||||||||||]PAM</td><td>NA</td><td>Intergenic</td><td>NA</td><td>NA</td></tr>
<tr><td>T23505</td><td>CTTGATGATGATCGGGCTGTAGG</td><td>1000</td><td></td><td></td><td></td><td></td><td></td><td></td><td></td><td></td><td></td></tr>
<tr><td>Oligo fwd</td><td>CACCCTTGATGATGATCGGGCTGT</td><td></td><td></td><td></td><td></td><td></td><td></td><td></td><td></td><td></td><td></td></tr>
<tr><td>Oligo rev</td><td>AAACACAGCCCGATCATCATCAAG</td><td></td><td></td><td></td><td></td><td></td><td></td><td></td><td></td><td></td><td></td></tr>
<tr><td>Chromosome</td><td>start</td><td>end</td><td>strand</td><td>MM</td><td>target_seq</td><td>PAM</td><td>alignment</td><td>distance</td><td>position</td><td>gene name</td><td>gene id</td></tr>
<tr><td>GU305914.1</td><td>139415</td><td>139437</td><td>-</td><td>0</td><td>CTTGATGATGATCGGGCTGT</td><td>AGG</td><td>||||||||[||||||||||||]PAM</td><td>NA</td><td>Intergenic</td><td>NA</td><td>NA</td></tr>
<tr><td>T23506</td><td>CGTCGACATCGCTCTTAGCGAGG</td><td>1000</td><td></td><td></td><td></td><td></td><td></td><td></td><td></td><td></td><td></td></tr>
<tr><td>Oligo fwd</td><td>CACCCGTCGACATCGCTCTTAGCG</td><td></td><td></td><td></td><td></td><td></td><td></td><td></td><td></td><td></td><td></td></tr>
<tr><td>Oligo rev</td><td>AAACCGCTAAGAGCGATGTCGACG</td><td></td><td></td><td></td><td></td><td></td><td></td><td></td><td></td><td></td><td></td></tr>
<tr><td>Chromosome</td><td>start</td><td>end</td><td>strand</td><td>MM</td><td>target_seq</td><td>PAM</td><td>alignment</td><td>distance</td><td>position</td><td>gene name</td><td>gene id</td></tr>
<tr><td>GU305914.1</td><td>139386</td><td>139408</td><td>-</td><td>0</td><td>CGTCGACATCGCTCTTAGCG</td><td>AGG</td><td>||||||||[||||||||||||]PAM</td><td>NA</td><td>Intergenic</td><td>NA</td><td>NA</td></tr>
<tr><td>T23507</td><td>TAGCGAGGTCCAGACACACACGG</td><td>1000</td><td></td><td></td><td></td><td></td><td></td><td></td><td></td><td></td><td></td></tr>
<tr><td>Oligo fwd</td><td>CACCTAGCGAGGTCCAGACACACA</td><td></td><td></td><td></td><td></td><td></td><td></td><td></td><td></td><td></td><td></td></tr>
<tr><td>Oligo rev</td><td>AAACTGTGTGTCTGGACCTCGCTA</td><td></td><td></td><td></td><td></td><td></td><td></td><td></td><td></td><td></td><td></td></tr>
<tr><td>Chromosome</td><td>start</td><td>end</td><td>strand</td><td>MM</td><td>target_seq</td><td>PAM</td><td>alignment</td><td>distance</td><td>position</td><td>gene name</td><td>gene id</td></tr>
<tr><td>GU305914.1</td><td>139371</td><td>139393</td><td>-</td><td>0</td><td>TAGCGAGGTCCAGACACACA</td><td>CGG</td><td>||||||||[||||||||||||]PAM</td><td>NA</td><td>Intergenic</td><td>NA</td><td>NA</td></tr>
<tr><td>T23508</td><td>ATATAAAGCACAAAACACGCAGG</td><td>1000</td><td></td><td></td><td></td><td></td><td></td><td></td><td></td><td></td><td></td></tr>
<tr><td>Oligo fwd</td><td>CACCATATAAAGCACAAAACACGC</td><td></td><td></td><td></td><td></td><td></td><td></td><td></td><td></td><td></td><td></td></tr>
<tr><td>Oligo rev</td><td>AAACGCGTGTTTTGTGCTTTATAT</td><td></td><td></td><td></td><td></td><td></td><td></td><td></td><td></td><td></td><td></td></tr>
<tr><td>Chromosome</td><td>start</td><td>end</td><td>strand</td><td>MM</td><td>target_seq</td><td>PAM</td><td>alignment</td><td>distance</td><td>position</td><td>gene name</td><td>gene id</td></tr>
<tr><td>GU305914.1</td><td>139340</td><td>139362</td><td>-</td><td>0</td><td>ATATAAAGCACAAAACACGC</td><td>AGG</td><td>||||||||[||||||||||||]PAM</td><td>NA</td><td>Intergenic</td><td>NA</td><td>NA</td></tr>
<tr><td>T23509</td><td>CACGCAGGTGCGAGACGAGACGG</td><td>1000</td><td></td><td></td><td></td><td></td><td></td><td></td><td></td><td></td><td></td></tr>
<tr><td>Oligo fwd</td><td>CACCCACGCAGGTGCGAGACGAGA</td><td></td><td></td><td></td><td></td><td></td><td></td><td></td><td></td><td></td><td></td></tr>
<tr><td>Oligo rev</td><td>AAACTCTCGTCTCGCACCTGCGTG</td><td></td><td></td><td></td><td></td><td></td><td></td><td></td><td></td><td></td><td></td></tr>
<tr><td>Chromosome</td><td>start</td><td>end</td><td>strand</td><td>MM</td><td>target_seq</td><td>PAM</td><td>alignment</td><td>distance</td><td>position</td><td>gene name</td><td>gene id</td></tr>
<tr><td>GU305914.1</td><td>139325</td><td>139347</td><td>-</td><td>0</td><td>CACGCAGGTGCGAGACGAGA</td><td>CGG</td><td>||||||||[||||||||||||]PAM</td><td>NA</td><td>Intergenic</td><td>NA</td><td>NA</td></tr>
<tr><td>T23510</td><td>ACGCAGGTGCGAGACGAGACGGG</td><td>1000</td><td></td><td></td><td></td><td></td><td></td><td></td><td></td><td></td><td></td></tr>
<tr><td>Oligo fwd</td><td>CACCACGCAGGTGCGAGACGAGAC</td><td></td><td></td><td></td><td></td><td></td><td></td><td></td><td></td><td></td><td></td></tr>
<tr><td>Oligo rev</td><td>AAACGTCTCGTCTCGCACCTGCGT</td><td></td><td></td><td></td><td></td><td></td><td></td><td></td><td></td><td></td><td></td></tr>
<tr><td>Chromosome</td><td>start</td><td>end</td><td>strand</td><td>MM</td><td>target_seq</td><td>PAM</td><td>alignment</td><td>distance</td><td>position</td><td>gene name</td><td>gene id</td></tr>
<tr><td>GU305914.1</td><td>139324</td><td>139346</td><td>-</td><td>0</td><td>ACGCAGGTGCGAGACGAGAC</td><td>GGG</td><td>||||||||[||||||||||||]PAM</td><td>NA</td><td>Intergenic</td><td>NA</td><td>NA</td></tr>
<tr><td>T23511</td><td>GCGAGACGAGACGGGCCAGACGG</td><td>1000</td><td></td><td></td><td></td><td></td><td></td><td></td><td></td><td></td><td></td></tr>
<tr><td>Oligo fwd</td><td>CACCGCGAGACGAGACGGGCCAGA</td><td></td><td></td><td></td><td></td><td></td><td></td><td></td><td></td><td></td><td></td></tr>
<tr><td>Oligo rev</td><td>AAACTCTGGCCCGTCTCGTCTCGC</td><td></td><td></td><td></td><td></td><td></td><td></td><td></td><td></td><td></td><td></td></tr>
<tr><td>Chromosome</td><td>start</td><td>end</td><td>strand</td><td>MM</td><td>target_seq</td><td>PAM</td><td>alignment</td><td>distance</td><td>position</td><td>gene name</td><td>gene id</td></tr>
<tr><td>GU305914.1</td><td>139316</td><td>139338</td><td>-</td><td>0</td><td>GCGAGACGAGACGGGCCAGA</td><td>CGG</td><td>||||||||[||||||||||||]PAM</td><td>NA</td><td>Intergenic</td><td>NA</td><td>NA</td></tr>
<tr><td>T23512</td><td>CGAGACGAGACGGGCCAGACGGG</td><td>1000</td><td></td><td></td><td></td><td></td><td></td><td></td><td></td><td></td><td></td></tr>
<tr><td>Oligo fwd</td><td>CACCCGAGACGAGACGGGCCAGAC</td><td></td><td></td><td></td><td></td><td></td><td></td><td></td><td></td><td></td><td></td></tr>
<tr><td>Oligo rev</td><td>AAACGTCTGGCCCGTCTCGTCTCG</td><td></td><td></td><td></td><td></td><td></td><td></td><td></td><td></td><td></td><td></td></tr>
<tr><td>Chromosome</td><td>start</td><td>end</td><td>strand</td><td>MM</td><td>target_seq</td><td>PAM</td><td>alignment</td><td>distance</td><td>position</td><td>gene name</td><td>gene id</td></tr>
<tr><td>GU305914.1</td><td>139315</td><td>139337</td><td>-</td><td>0</td><td>CGAGACGAGACGGGCCAGAC</td><td>GGG</td><td>||||||||[||||||||||||]PAM</td><td>NA</td><td>Intergenic</td><td>NA</td><td>NA</td></tr>
<tr><td>T23513</td><td>CGGGCCAGACGGGCACGCTGCGG</td><td>1000</td><td></td><td></td><td></td><td></td><td></td><td></td><td></td><td></td><td></td></tr>
<tr><td>Oligo fwd</td><td>CACCCGGGCCAGACGGGCACGCTG</td><td></td><td></td><td></td><td></td><td></td><td></td><td></td><td></td><td></td><td></td></tr>
</table>

| | | | | | | | | | | | | |
|---|---|---|---|---|---|---|---|---|---|---|---|---|
| Oligo rev | AAACCAGCGTGCCCGTCTGGCCCG | | | | | | | | | | | |
| Chromosome | | start | end | strand | MM | target_seq | PAM | alignment | distance | position | gene name | gene id |
| GU305914.1 | | 139305 | 139327 | - | 0 | CGGGCCAGACGGGCACGCTG | CGG | \|\|\|\|\|\|\|\|[\|\|\|\|\|\|\|\|\|\|\|\|]PAM | NA | Intergenic | NA | NA |
| T23514 | CCAGACGGGCACGCTGCGGTCGG | | 1000 | | | | | | | | | |
| Oligo fwd | CACCCCAGACGGGCACGCTGCGGT | | | | | | | | | | | |
| Oligo rev | AAACACCGCAGCGTGCCCGTCTGG | | | | | | | | | | | |
| Chromosome | | start | end | strand | MM | target_seq | PAM | alignment | distance | position | gene name | gene id |
| GU305914.1 | | 139301 | 139323 | - | 0 | CCAGACGGGCACGCTGCGGT | CGG | \|\|\|\|\|\|\|\|[\|\|\|\|\|\|\|\|\|\|\|\|]PAM | NA | Intergenic | NA | NA |
| T23515 | CGCTGCGGTCGGAAAACCTCTGG | | 1000 | | | | | | | | | |
| Oligo fwd | CACCCGCTGCGGTCGGAAAACCTC | | | | | | | | | | | |
| Oligo rev | AAACGAGGTTTTCCGACCGCAGCG | | | | | | | | | | | |
| Chromosome | | start | end | strand | MM | target_seq | PAM | alignment | distance | position | gene name | gene id |
| GU305914.1 | | 139290 | 139312 | - | 0 | CGCTGCGGTCGGAAAACCTC | TGG | \|\|\|\|\|\|\|\|[\|\|\|\|\|\|\|\|\|\|\|\|]PAM | NA | Intergenic | NA | NA |
| T23516 | TGCGGTCGGAAAACCTCTGGCGG | | 1000 | | | | | | | | | |
| Oligo fwd | CACCTGCGGTCGGAAAACCTCTGG | | | | | | | | | | | |
| Oligo rev | AAACCCAGAGGTTTTCCGACCGCA | | | | | | | | | | | |
| Chromosome | | start | end | strand | MM | target_seq | PAM | alignment | distance | position | gene name | gene id |
| GU305914.1 | | 139287 | 139309 | - | 0 | TGCGGTCGGAAAACCTCTGG | CGG | \|\|\|\|\|\|\|\|[\|\|\|\|\|\|\|\|\|\|\|\|]PAM | NA | Intergenic | NA | NA |
| T23517 | GTCGGAAAACCTCTGGCGGACGG | | 1000 | | | | | | | | | |
| Oligo fwd | CACCGTCGGAAAACCTCTGGCGGA | | | | | | | | | | | |
| Oligo rev | AAACTCCGCCAGAGGTTTTCCGAC | | | | | | | | | | | |
| Chromosome | | start | end | strand | MM | target_seq | PAM | alignment | distance | position | gene name | gene id |
| GU305914.1 | | 139283 | 139305 | - | 0 | GTCGGAAAACCTCTGGCGGA | CGG | \|\|\|\|\|\|\|\|[\|\|\|\|\|\|\|\|\|\|\|\|]PAM | NA | Intergenic | NA | NA |
| T23518 | ACTGCAGCGTCTTCTTGACGCGG | | 1000 | | | | | | | | | |
| Oligo fwd | CACCACTGCAGCGTCTTCTTGACG | | | | | | | | | | | |
| Oligo rev | AAACCGTCAAGAAGACGCTGCAGT | | | | | | | | | | | |
| Chromosome | | start | end | strand | MM | target_seq | PAM | alignment | distance | position | gene name | gene id |
| GU305914.1 | | 139260 | 139282 | - | 0 | ACTGCAGCGTCTTCTTGACG | CGG | \|\|\|\|\|\|\|\|[\|\|\|\|\|\|\|\|\|\|\|\|]PAM | NA | Intergenic | NA | NA |
| T23519 | CGTCTTCTTGACGCGGTCCATGG | | 1000 | | | | | | | | | |
| Oligo fwd | CACCCGTCTTCTTGACGCGGTCCA | | | | | | | | | | | |
| Oligo rev | AAACTGGACCGCGTCAAGAAGACG | | | | | | | | | | | |
| Chromosome | | start | end | strand | MM | target_seq | PAM | alignment | distance | position | gene name | gene id |
| GU305914.1 | | 139253 | 139275 | - | 0 | CGTCTTCTTGACGCGGTCCA | TGG | \|\|\|\|\|\|\|\|[\|\|\|\|\|\|\|\|\|\|\|\|]PAM | NA | Intergenic | NA | NA |
| T23520 | CCCGAGCTCGTAGCACCGCGTGG | | 1000 | | | | | | | | | |
| Oligo fwd | CACCCCCGAGCTCGTAGCACCGCG | | | | | | | | | | | |
| Oligo rev | AAACCGCGGTGCTACGAGCTCGGG | | | | | | | | | | | |
| Chromosome | | start | end | strand | MM | target_seq | PAM | alignment | distance | position | gene name | gene id |
| GU305914.1 | | 139223 | 139245 | - | 0 | CCCGAGCTCGTAGCACCGCG | TGG | \|\|\|\|\|\|\|\|[\|\|\|\|\|\|\|\|\|\|\|\|]PAM | NA | Intergenic | NA | NA |
| T23521 | CCGAGCTCGTAGCACCGCGTGGG | | 1000 | | | | | | | | | |
| Oligo fwd | CACCCCGAGCTCGTAGCACCGCGT | | | | | | | | | | | |
| Oligo rev | AAACACGCGGTGCTACGAGCTCGG | | | | | | | | | | | |
| Chromosome | | start | end | strand | MM | target_seq | PAM | alignment | distance | position | gene name | gene id |
| GU305914.1 | | 139222 | 139244 | - | 0 | CCGAGCTCGTAGCACCGCGT | GGG | \|\|\|\|\|\|\|\|[\|\|\|\|\|\|\|\|\|\|\|\|]PAM | NA | Intergenic | NA | NA |
| T23522 | CTCGTAGCACCGCGTGGGCTCGG | | 1000 | | | | | | | | | |
| Oligo fwd | CACCCTCGTAGCACCGCGTGGGCT | | | | | | | | | | | |
| Oligo rev | AAACAGCCCACGCGGTGCTACGAG | | | | | | | | | | | |
| Chromosome | | start | end | strand | MM | target_seq | PAM | alignment | distance | position | gene name | gene id |
| GU305914.1 | | 139217 | 139239 | - | 0 | CTCGTAGCACCGCGTGGGCT | CGG | \|\|\|\|\|\|\|\|[\|\|\|\|\|\|\|\|\|\|\|\|]PAM | NA | Intergenic | NA | NA |
| T23523 | CCACACGTACACGTAGAGCTCGG | | 1000 | | | | | | | | | |
| Oligo fwd | CACCCCACACGTACACGTAGAGCT | | | | | | | | | | | |
| Oligo rev | AAACAGCTCTACGTGTACGTGTGG | | | | | | | | | | | |
| Chromosome | | start | end | strand | MM | target_seq | PAM | alignment | distance | position | gene name | gene id |
| GU305914.1 | | 139172 | 139194 | - | 0 | CCACACGTACACGTAGAGCT | CGG | \|\|\|\|\|\|\|\|[\|\|\|\|\|\|\|\|\|\|\|\|]PAM | NA | Intergenic | NA | NA |
| T23524 | CGTACACGTAGAGCTCGGCCAGG | | 1000 | | | | | | | | | |
| Oligo fwd | CACCCGTACACGTAGAGCTCGGCC | | | | | | | | | | | |
| Oligo rev | AAACGGCCGAGCTCTACGTGTACG | | | | | | | | | | | |
| Chromosome | | start | end | strand | MM | target_seq | PAM | alignment | distance | position | gene name | gene id |
| GU305914.1 | | 139167 | 139189 | - | 0 | CGTACACGTAGAGCTCGGCC | AGG | \|\|\|\|\|\|\|\|[\|\|\|\|\|\|\|\|\|\|\|\|]PAM | NA | Intergenic | NA | NA |
| T23525 | GAGCTCGGCCAGGTTGATGTAGG | | 1000 | | | | | | | | | |
| Oligo fwd | CACCGAGCTCGGCCAGGTTGATGT | | | | | | | | | | | |
| Oligo rev | AAACACATCAACCTGGCCGAGCTC | | | | | | | | | | | |
| Chromosome | | start | end | strand | MM | target_seq | PAM | alignment | distance | position | gene name | gene id |
| GU305914.1 | | 139157 | 139179 | - | 0 | GAGCTCGGCCAGGTTGATGT | AGG | \|\|\|\|\|\|\|\|[\|\|\|\|\|\|\|\|\|\|\|\|]PAM | NA | Intergenic | NA | NA |
| T23526 | GTTGATGTAGGCCGAGTAGTCGG | | 1000 | | | | | | | | | |
| Oligo fwd | CACCGTTGATGTAGGCCGAGTAGT | | | | | | | | | | | |
| Oligo rev | AAACACTACTCGGCCTACATCAAC | | | | | | | | | | | |
| Chromosome | | start | end | strand | MM | target_seq | PAM | alignment | distance | position | gene name | gene id |
| GU305914.1 | | 139145 | 139167 | - | 0 | GTTGATGTAGGCCGAGTAGT | CGG | \|\|\|\|\|\|\|\|[\|\|\|\|\|\|\|\|\|\|\|\|]PAM | NA | Intergenic | NA | NA |
| T23527 | GTAGGCCGAGTAGTCGGTGATGG | | 1000 | | | | | | | | | |
| Oligo fwd | CACCGTAGGCCGAGTAGTCGGTGA | | | | | | | | | | | |
| Oligo rev | AAACTCACCGACTACTCGGCCTAC | | | | | | | | | | | |
| Chromosome | | start | end | strand | MM | target_seq | PAM | alignment | distance | position | gene name | gene id |
| GU305914.1 | | 139139 | 139161 | - | 0 | GTAGGCCGAGTAGTCGGTGA | TGG | \|\|\|\|\|\|\|\|[\|\|\|\|\|\|\|\|\|\|\|\|]PAM | NA | Intergenic | NA | NA |
| T23528 | TAGGCCGAGTAGTCGGTGATGGG | | 1000 | | | | | | | | | |

<table>
<tr><td>Oligo fwd</td><td>CACCTAGGCCGAGTAGTCGGTGAT</td><td></td><td></td><td></td><td></td><td></td><td></td><td></td><td></td><td></td><td></td></tr>
<tr><td>Oligo rev</td><td>AAACATCACCGACTACTCGGCCTA</td><td></td><td></td><td></td><td></td><td></td><td></td><td></td><td></td><td></td><td></td></tr>
<tr><td>Chromosome</td><td>start</td><td>end</td><td>strand</td><td>MM</td><td>target_seq</td><td>PAM</td><td>alignment</td><td>distance</td><td>position</td><td>gene name</td><td>gene id</td></tr>
<tr><td>GU305914.1</td><td>139138</td><td>139160</td><td>-</td><td>0</td><td>TAGGCCGAGTAGTCGGTGAT</td><td>GGG</td><td>||||||||[||||||||||||]PAM</td><td>NA</td><td>Intergenic</td><td>NA</td><td>NA</td></tr>
<tr><td></td><td></td><td></td><td></td><td></td><td></td><td></td><td></td><td></td><td></td><td></td><td></td></tr>
<tr><td>T23529</td><td>AGGCCGAGTAGTCGGTGATGGGG</td><td>1000</td><td></td><td></td><td></td><td></td><td></td><td></td><td></td><td></td><td></td></tr>
<tr><td>Oligo fwd</td><td>CACCAGGCCGAGTAGTCGGTGATG</td><td></td><td></td><td></td><td></td><td></td><td></td><td></td><td></td><td></td><td></td></tr>
<tr><td>Oligo rev</td><td>AAACCATCACCGACTACTCGGCCT</td><td></td><td></td><td></td><td></td><td></td><td></td><td></td><td></td><td></td><td></td></tr>
<tr><td>Chromosome</td><td>start</td><td>end</td><td>strand</td><td>MM</td><td>target_seq</td><td>PAM</td><td>alignment</td><td>distance</td><td>position</td><td>gene name</td><td>gene id</td></tr>
<tr><td>GU305914.1</td><td>139137</td><td>139159</td><td>-</td><td>0</td><td>AGGCCGAGTAGTCGGTGATG</td><td>GGG</td><td>||||||||[||||||||||||]PAM</td><td>NA</td><td>Intergenic</td><td>NA</td><td>NA</td></tr>
<tr><td></td><td></td><td></td><td></td><td></td><td></td><td></td><td></td><td></td><td></td><td></td><td></td></tr>
<tr><td>T23530</td><td>GGTGATGGGGTCGTAGAAGACGG</td><td>1000</td><td></td><td></td><td></td><td></td><td></td><td></td><td></td><td></td><td></td></tr>
<tr><td>Oligo fwd</td><td>CACCGGTGATGGGGTCGTAGAAGA</td><td></td><td></td><td></td><td></td><td></td><td></td><td></td><td></td><td></td><td></td></tr>
<tr><td>Oligo rev</td><td>AAACTCTTCTACGACCCCATCACC</td><td></td><td></td><td></td><td></td><td></td><td></td><td></td><td></td><td></td><td></td></tr>
<tr><td>Chromosome</td><td>start</td><td>end</td><td>strand</td><td>MM</td><td>target_seq</td><td>PAM</td><td>alignment</td><td>distance</td><td>position</td><td>gene name</td><td>gene id</td></tr>
<tr><td>GU305914.1</td><td>139124</td><td>139146</td><td>-</td><td>0</td><td>GGTGATGGGGTCGTAGAAGA</td><td>CGG</td><td>||||||||[||||||||||||]PAM</td><td>NA</td><td>Intergenic</td><td>NA</td><td>NA</td></tr>
<tr><td></td><td></td><td></td><td></td><td></td><td></td><td></td><td></td><td></td><td></td><td></td><td></td></tr>
<tr><td>T23531</td><td>GTGATGGGGTCGTAGAAGACGGG</td><td>1000</td><td></td><td></td><td></td><td></td><td></td><td></td><td></td><td></td><td></td></tr>
<tr><td>Oligo fwd</td><td>CACCGTGATGGGGTCGTAGAAGAC</td><td></td><td></td><td></td><td></td><td></td><td></td><td></td><td></td><td></td><td></td></tr>
<tr><td>Oligo rev</td><td>AAACGTCTTCTACGACCCCATCAC</td><td></td><td></td><td></td><td></td><td></td><td></td><td></td><td></td><td></td><td></td></tr>
<tr><td>Chromosome</td><td>start</td><td>end</td><td>strand</td><td>MM</td><td>target_seq</td><td>PAM</td><td>alignment</td><td>distance</td><td>position</td><td>gene name</td><td>gene id</td></tr>
<tr><td>GU305914.1</td><td>139123</td><td>139145</td><td>-</td><td>0</td><td>GTGATGGGGTCGTAGAAGAC</td><td>GGG</td><td>||||||||[||||||||||||]PAM</td><td>NA</td><td>Intergenic</td><td>NA</td><td>NA</td></tr>
<tr><td></td><td></td><td></td><td></td><td></td><td></td><td></td><td></td><td></td><td></td><td></td><td></td></tr>
<tr><td>T23532</td><td>TGATGGGGTCGTAGAAGACGGGG</td><td>1000</td><td></td><td></td><td></td><td></td><td></td><td></td><td></td><td></td><td></td></tr>
<tr><td>Oligo fwd</td><td>CACCTGATGGGGTCGTAGAAGACG</td><td></td><td></td><td></td><td></td><td></td><td></td><td></td><td></td><td></td><td></td></tr>
<tr><td>Oligo rev</td><td>AAACCGTCTTCTACGACCCCATCA</td><td></td><td></td><td></td><td></td><td></td><td></td><td></td><td></td><td></td><td></td></tr>
<tr><td>Chromosome</td><td>start</td><td>end</td><td>strand</td><td>MM</td><td>target_seq</td><td>PAM</td><td>alignment</td><td>distance</td><td>position</td><td>gene name</td><td>gene id</td></tr>
<tr><td>GU305914.1</td><td>139122</td><td>139144</td><td>-</td><td>0</td><td>TGATGGGGTCGTAGAAGACG</td><td>GGG</td><td>||||||||[||||||||||||]PAM</td><td>NA</td><td>Intergenic</td><td>NA</td><td>NA</td></tr>
<tr><td></td><td></td><td></td><td></td><td></td><td></td><td></td><td></td><td></td><td></td><td></td><td></td></tr>
<tr><td>T23533</td><td>GATGGGGTCGTAGAAGACGGGGG</td><td>1000</td><td></td><td></td><td></td><td></td><td></td><td></td><td></td><td></td><td></td></tr>
<tr><td>Oligo fwd</td><td>CACCGATGGGGTCGTAGAAGACGG</td><td></td><td></td><td></td><td></td><td></td><td></td><td></td><td></td><td></td><td></td></tr>
<tr><td>Oligo rev</td><td>AAACCCGTCTTCTACGACCCCATC</td><td></td><td></td><td></td><td></td><td></td><td></td><td></td><td></td><td></td><td></td></tr>
<tr><td>Chromosome</td><td>start</td><td>end</td><td>strand</td><td>MM</td><td>target_seq</td><td>PAM</td><td>alignment</td><td>distance</td><td>position</td><td>gene name</td><td>gene id</td></tr>
<tr><td>GU305914.1</td><td>139121</td><td>139143</td><td>-</td><td>0</td><td>GATGGGGTCGTAGAAGACGG</td><td>GGG</td><td>||||||||[||||||||||||]PAM</td><td>NA</td><td>Intergenic</td><td>NA</td><td>NA</td></tr>
<tr><td></td><td></td><td></td><td></td><td></td><td></td><td></td><td></td><td></td><td></td><td></td><td></td></tr>
<tr><td>T23534</td><td>ATGGGGTCGTAGAAGACGGGGGG</td><td>1000</td><td></td><td></td><td></td><td></td><td></td><td></td><td></td><td></td><td></td></tr>
<tr><td>Oligo fwd</td><td>CACCATGGGGTCGTAGAAGACGGG</td><td></td><td></td><td></td><td></td><td></td><td></td><td></td><td></td><td></td><td></td></tr>
<tr><td>Oligo rev</td><td>AAACCCCGTCTTCTACGACCCCAT</td><td></td><td></td><td></td><td></td><td></td><td></td><td></td><td></td><td></td><td></td></tr>
<tr><td>Chromosome</td><td>start</td><td>end</td><td>strand</td><td>MM</td><td>target_seq</td><td>PAM</td><td>alignment</td><td>distance</td><td>position</td><td>gene name</td><td>gene id</td></tr>
<tr><td>GU305914.1</td><td>139120</td><td>139142</td><td>-</td><td>0</td><td>ATGGGGTCGTAGAAGACGGG</td><td>GGG</td><td>||||||||[||||||||||||]PAM</td><td>NA</td><td>Intergenic</td><td>NA</td><td>NA</td></tr>
<tr><td></td><td></td><td></td><td></td><td></td><td></td><td></td><td></td><td></td><td></td><td></td><td></td></tr>
<tr><td>T23535</td><td>GGTCGTAGAAGACGGGGGGCAGG</td><td>1000</td><td></td><td></td><td></td><td></td><td></td><td></td><td></td><td></td><td></td></tr>
<tr><td>Oligo fwd</td><td>CACCGGTCGTAGAAGACGGGGGGC</td><td></td><td></td><td></td><td></td><td></td><td></td><td></td><td></td><td></td><td></td></tr>
<tr><td>Oligo rev</td><td>AAACGCCCCCCGTCTTCTACGACC</td><td></td><td></td><td></td><td></td><td></td><td></td><td></td><td></td><td></td><td></td></tr>
<tr><td>Chromosome</td><td>start</td><td>end</td><td>strand</td><td>MM</td><td>target_seq</td><td>PAM</td><td>alignment</td><td>distance</td><td>position</td><td>gene name</td><td>gene id</td></tr>
<tr><td>GU305914.1</td><td>139116</td><td>139138</td><td>-</td><td>0</td><td>GGTCGTAGAAGACGGGGGGC</td><td>AGG</td><td>||||||||[||||||||||||]PAM</td><td>NA</td><td>Intergenic</td><td>NA</td><td>NA</td></tr>
<tr><td></td><td></td><td></td><td></td><td></td><td></td><td></td><td></td><td></td><td></td><td></td><td></td></tr>
<tr><td>T23536</td><td>GAAGACGGGGGGCAGGAGCACGG</td><td>1000</td><td></td><td></td><td></td><td></td><td></td><td></td><td></td><td></td><td></td></tr>
<tr><td>Oligo fwd</td><td>CACCGAAGACGGGGGGCAGGAGCA</td><td></td><td></td><td></td><td></td><td></td><td></td><td></td><td></td><td></td><td></td></tr>
<tr><td>Oligo rev</td><td>AAACTGCTCCTGCCCCCCGTCTTC</td><td></td><td></td><td></td><td></td><td></td><td></td><td></td><td></td><td></td><td></td></tr>
<tr><td>Chromosome</td><td>start</td><td>end</td><td>strand</td><td>MM</td><td>target_seq</td><td>PAM</td><td>alignment</td><td>distance</td><td>position</td><td>gene name</td><td>gene id</td></tr>
<tr><td>GU305914.1</td><td>139109</td><td>139131</td><td>-</td><td>0</td><td>GAAGACGGGGGGCAGGAGCA</td><td>CGG</td><td>||||||||[||||||||||||]PAM</td><td>NA</td><td>Intergenic</td><td>NA</td><td>NA</td></tr>
<tr><td></td><td></td><td></td><td></td><td></td><td></td><td></td><td></td><td></td><td></td><td></td><td></td></tr>
<tr><td>T23537</td><td>AAGACGGGGGGCAGGAGCACGGG</td><td>1000</td><td></td><td></td><td></td><td></td><td></td><td></td><td></td><td></td><td></td></tr>
<tr><td>Oligo fwd</td><td>CACCAAGACGGGGGGCAGGAGCAC</td><td></td><td></td><td></td><td></td><td></td><td></td><td></td><td></td><td></td><td></td></tr>
<tr><td>Oligo rev</td><td>AAACGTGCTCCTGCCCCCCGTCTT</td><td></td><td></td><td></td><td></td><td></td><td></td><td></td><td></td><td></td><td></td></tr>
<tr><td>Chromosome</td><td>start</td><td>end</td><td>strand</td><td>MM</td><td>target_seq</td><td>PAM</td><td>alignment</td><td>distance</td><td>position</td><td>gene name</td><td>gene id</td></tr>
<tr><td>GU305914.1</td><td>139108</td><td>139130</td><td>-</td><td>0</td><td>AAGACGGGGGGCAGGAGCAC</td><td>GGG</td><td>||||||||[||||||||||||]PAM</td><td>NA</td><td>Intergenic</td><td>NA</td><td>NA</td></tr>
<tr><td></td><td></td><td></td><td></td><td></td><td></td><td></td><td></td><td></td><td></td><td></td><td></td></tr>
<tr><td>T23538</td><td>AGACGGGGGGCAGGAGCACGGGG</td><td>1000</td><td></td><td></td><td></td><td></td><td></td><td></td><td></td><td></td><td></td></tr>
<tr><td>Oligo fwd</td><td>CACCAGACGGGGGGCAGGAGCACG</td><td></td><td></td><td></td><td></td><td></td><td></td><td></td><td></td><td></td><td></td></tr>
<tr><td>Oligo rev</td><td>AAACCGTGCTCCTGCCCCCCGTCT</td><td></td><td></td><td></td><td></td><td></td><td></td><td></td><td></td><td></td><td></td></tr>
<tr><td>Chromosome</td><td>start</td><td>end</td><td>strand</td><td>MM</td><td>target_seq</td><td>PAM</td><td>alignment</td><td>distance</td><td>position</td><td>gene name</td><td>gene id</td></tr>
<tr><td>GU305914.1</td><td>139107</td><td>139129</td><td>-</td><td>0</td><td>AGACGGGGGGCAGGAGCACG</td><td>GGG</td><td>||||||||[||||||||||||]PAM</td><td>NA</td><td>Intergenic</td><td>NA</td><td>NA</td></tr>
<tr><td>GU305914.1</td><td>115019</td><td>115041</td><td>+</td><td>4</td><td>GTACGGGGGGCAAGAGAACG</td><td>CGG</td><td>--||||||[||||-|||-|||]PAM</td><td>NA</td><td>Intergenic</td><td>NA</td><td>NA</td></tr>
<tr><td></td><td></td><td></td><td></td><td></td><td></td><td></td><td></td><td></td><td></td><td></td><td></td></tr>
<tr><td>T23539</td><td>GGTAAGACACCTCACACTCTCGG</td><td>1000</td><td></td><td></td><td></td><td></td><td></td><td></td><td></td><td></td><td></td></tr>
<tr><td>Oligo fwd</td><td>CACCGGTAAGACACCTCACACTCT</td><td></td><td></td><td></td><td></td><td></td><td></td><td></td><td></td><td></td><td></td></tr>
<tr><td>Oligo rev</td><td>AAACAGAGTGTGAGGTGTCTTACC</td><td></td><td></td><td></td><td></td><td></td><td></td><td></td><td></td><td></td><td></td></tr>
<tr><td>Chromosome</td><td>start</td><td>end</td><td>strand</td><td>MM</td><td>target_seq</td><td>PAM</td><td>alignment</td><td>distance</td><td>position</td><td>gene name</td><td>gene id</td></tr>
<tr><td>GU305914.1</td><td>139086</td><td>139108</td><td>-</td><td>0</td><td>GGTAAGACACCTCACACTCT</td><td>CGG</td><td>||||||||[||||||||||||]PAM</td><td>NA</td><td>Intergenic</td><td>NA</td><td>NA</td></tr>
<tr><td></td><td></td><td></td><td></td><td></td><td></td><td></td><td></td><td></td><td></td><td></td><td></td></tr>
<tr><td>T23540</td><td>CTCTCGGTCGCGCCCGAAACAGG</td><td>1000</td><td></td><td></td><td></td><td></td><td></td><td></td><td></td><td></td><td></td></tr>
<tr><td>Oligo fwd</td><td>CACCCTCTCGGTCGCGCCCGAAAC</td><td></td><td></td><td></td><td></td><td></td><td></td><td></td><td></td><td></td><td></td></tr>
<tr><td>Oligo rev</td><td>AAACGTTTCGGGCGCGACCGAGAG</td><td></td><td></td><td></td><td></td><td></td><td></td><td></td><td></td><td></td><td></td></tr>
<tr><td>Chromosome</td><td>start</td><td>end</td><td>strand</td><td>MM</td><td>target_seq</td><td>PAM</td><td>alignment</td><td>distance</td><td>position</td><td>gene name</td><td>gene id</td></tr>
<tr><td>GU305914.1</td><td>139070</td><td>139092</td><td>-</td><td>0</td><td>CTCTCGGTCGCGCCCGAAAC</td><td>AGG</td><td>||||||||[||||||||||||]PAM</td><td>NA</td><td>Intergenic</td><td>NA</td><td>NA</td></tr>
<tr><td></td><td></td><td></td><td></td><td></td><td></td><td></td><td></td><td></td><td></td><td></td><td></td></tr>
<tr><td>T23541</td><td>CGGTCGCGCCCGAAACAGGTAGG</td><td>1000</td><td></td><td></td><td></td><td></td><td></td><td></td><td></td><td></td><td></td></tr>
<tr><td>Oligo fwd</td><td>CACCCGGTCGCGCCCGAAACAGGT</td><td></td><td></td><td></td><td></td><td></td><td></td><td></td><td></td><td></td><td></td></tr>
<tr><td>Oligo rev</td><td>AAACACCTGTTTCGGGCGCGACCG</td><td></td><td></td><td></td><td></td><td></td><td></td><td></td><td></td><td></td><td></td></tr>
<tr><td>Chromosome</td><td>start</td><td>end</td><td>strand</td><td>MM</td><td>target_seq</td><td>PAM</td><td>alignment</td><td>distance</td><td>position</td><td>gene name</td><td>gene id</td></tr>
<tr><td>GU305914.1</td><td>139066</td><td>139088</td><td>-</td><td>0</td><td>CGGTCGCGCCCGAAACAGGT</td><td>AGG</td><td>||||||||[||||||||||||]PAM</td><td>NA</td><td>Intergenic</td><td>NA</td><td>NA</td></tr>
<tr><td></td><td></td><td></td><td></td><td></td><td></td><td></td><td></td><td></td><td></td><td></td><td></td></tr>
<tr><td>T23542</td><td>GGTCGCGCCCGAAACAGGTAGGG</td><td>1000</td><td></td><td></td><td></td><td></td><td></td><td></td><td></td><td></td><td></td></tr>
<tr><td>Oligo fwd</td><td>CACCGGTCGCGCCCGAAACAGGTA</td><td></td><td></td><td></td><td></td><td></td><td></td><td></td><td></td><td></td><td></td></tr>
<tr><td>Oligo rev</td><td>AAACTACCTGTTTCGGGCGCGACC</td><td></td><td></td><td></td><td></td><td></td><td></td><td></td><td></td><td></td><td></td></tr>
<tr><td>Chromosome</td><td>start</td><td>end</td><td>strand</td><td>MM</td><td>target_seq</td><td>PAM</td><td>alignment</td><td>distance</td><td>position</td><td>gene name</td><td>gene id</td></tr>
<tr><td>GU305914.1</td><td>139065</td><td>139087</td><td>-</td><td>0</td><td>GGTCGCGCCCGAAACAGGTA</td><td>GGG</td><td>||||||||[||||||||||||]PAM</td><td>NA</td><td>Intergenic</td><td>NA</td><td>NA</td></tr>
</table>

| | | | | | | | | | | | |
|---|---|---|---|---|---|---|---|---|---|---|---|
| T23543 | GGGTTCTGACACACGCTCTGCGG | 1000 | | | | | | | | | |
| Oligo fwd | CACCGGGTTCTGACACACGCTCTG | | | | | | | | | | |
| Oligo rev | AAACCAGAGCGTGTGTCAGAACCC | | | | | | | | | | |
| Chromosome | start | end | strand | MM | target_seq | PAM | alignment | distance | position | gene name | gene id |
| GU305914.1 | 139045 | 139067 | - | 0 | GGGTTCTGACACACGCTCTG | CGG | \|\|\|\|\|\|\|\|[\|\|\|\|\|\|\|\|\|\|\|\|]PAM | NA | Intergenic | NA | NA |

| | | | | | | | | | | | |
|---|---|---|---|---|---|---|---|---|---|---|---|
| T23544 | GCAGAACAGCAGCCAGAGCACGG | 1000 | | | | | | | | | |
| Oligo fwd | CACCGCAGAACAGCAGCCAGAGCA | | | | | | | | | | |
| Oligo rev | AAACTGCTCTGGCTGCTGTTCTGC | | | | | | | | | | |
| Chromosome | start | end | strand | MM | target_seq | PAM | alignment | distance | position | gene name | gene id |
| GU305914.1 | 139019 | 139041 | - | 0 | GCAGAACAGCAGCCAGAGCA | CGG | \|\|\|\|\|\|\|\|[\|\|\|\|\|\|\|\|\|\|\|\|]PAM | NA | Intergenic | NA | NA |
| GU305914.1 | 110290 | 110312 | - | 4 | GAGGAACAGCAGGCAGAGGA | CGG | \|--\|\|\|\|\|[\|\|\|\|-\|\|\|\|\|-\|]PAM | NA | Intergenic | NA | NA |

| | | | | | | | | | | | |
|---|---|---|---|---|---|---|---|---|---|---|---|
| T23545 | CGACATCCTGCACGTTATTGTGG | 1000 | | | | | | | | | |
| Oligo fwd | CACCCGACATCCTGCACGTTATTG | | | | | | | | | | |
| Oligo rev | AAACCAATAACGTGCAGGATGTCG | | | | | | | | | | |
| Chromosome | start | end | strand | MM | target_seq | PAM | alignment | distance | position | gene name | gene id |
| GU305914.1 | 138993 | 139015 | - | 0 | CGACATCCTGCACGTTATTG | TGG | \|\|\|\|\|\|\|\|[\|\|\|\|\|\|\|\|\|\|\|\|]PAM | NA | Intergenic | NA | NA |

| | | | | | | | | | | | |
|---|---|---|---|---|---|---|---|---|---|---|---|
| T23546 | TTATTGTGGTCTAAGATCCCCGG | 1000 | | | | | | | | | |
| Oligo fwd | CACCTTATTGTGGTCTAAGATCCC | | | | | | | | | | |
| Oligo rev | AAACGGGATCTTAGACCACAATAA | | | | | | | | | | |
| Chromosome | start | end | strand | MM | target_seq | PAM | alignment | distance | position | gene name | gene id |
| GU305914.1 | 138979 | 139001 | - | 0 | TTATTGTGGTCTAAGATCCC | CGG | \|\|\|\|\|\|\|\|[\|\|\|\|\|\|\|\|\|\|\|\|]PAM | NA | Intergenic | NA | NA |

| | | | | | | | | | | | |
|---|---|---|---|---|---|---|---|---|---|---|---|
| T23547 | CCGGCGTGTCCACGAACCTGAGG | 1000 | | | | | | | | | |
| Oligo fwd | CACCCCGGCGTGTCCACGAACCTG | | | | | | | | | | |
| Oligo rev | AAACCAGGTTCGTGGACACGCCGG | | | | | | | | | | |
| Chromosome | start | end | strand | MM | target_seq | PAM | alignment | distance | position | gene name | gene id |
| GU305914.1 | 138960 | 138982 | - | 0 | CCGGCGTGTCCACGAACCTG | AGG | \|\|\|\|\|\|\|\|[\|\|\|\|\|\|\|\|\|\|\|\|]PAM | NA | Intergenic | NA | NA |

| | | | | | | | | | | | |
|---|---|---|---|---|---|---|---|---|---|---|---|
| T23548 | CTCCACGATCGCCCGCTTATCGG | 1000 | | | | | | | | | |
| Oligo fwd | CACCCTCCACGATCGCCCGCTTAT | | | | | | | | | | |
| Oligo rev | AAACATAAGCGGGCGATCGTGGAG | | | | | | | | | | |
| Chromosome | start | end | strand | MM | target_seq | PAM | alignment | distance | position | gene name | gene id |
| GU305914.1 | 138932 | 138954 | - | 0 | CTCCACGATCGCCCGCTTAT | CGG | \|\|\|\|\|\|\|\|[\|\|\|\|\|\|\|\|\|\|\|\|]PAM | NA | Intergenic | NA | NA |

| | | | | | | | | | | | |
|---|---|---|---|---|---|---|---|---|---|---|---|
| T23549 | TCCACGATCGCCCGCTTATCGGG | 1000 | | | | | | | | | |
| Oligo fwd | CACCTCCACGATCGCCCGCTTATC | | | | | | | | | | |
| Oligo rev | AAACGATAAGCGGGCGATCGTGGA | | | | | | | | | | |
| Chromosome | start | end | strand | MM | target_seq | PAM | alignment | distance | position | gene name | gene id |
| GU305914.1 | 138931 | 138953 | - | 0 | TCCACGATCGCCCGCTTATC | GGG | \|\|\|\|\|\|\|\|[\|\|\|\|\|\|\|\|\|\|\|\|]PAM | NA | Intergenic | NA | NA |

| | | | | | | | | | | | |
|---|---|---|---|---|---|---|---|---|---|---|---|
| T23550 | TTCCTTAGTGTAAAGCACGTAGG | 1000 | | | | | | | | | |
| Oligo fwd | CACCTTCCTTAGTGTAAAGCACGT | | | | | | | | | | |
| Oligo rev | AAACACGTGCTTTACACTAAGGAA | | | | | | | | | | |
| Chromosome | start | end | strand | MM | target_seq | PAM | alignment | distance | position | gene name | gene id |
| GU305914.1 | 138887 | 138909 | - | 0 | TTCCTTAGTGTAAAGCACGT | AGG | \|\|\|\|\|\|\|\|[\|\|\|\|\|\|\|\|\|\|\|\|]PAM | NA | Intergenic | NA | NA |

| | | | | | | | | | | | |
|---|---|---|---|---|---|---|---|---|---|---|---|
| T23551 | ACCGATTTCGCTACATATCCCGG | 1000 | | | | | | | | | |
| Oligo fwd | CACCACCGATTTCGCTACATATCC | | | | | | | | | | |
| Oligo rev | AAACGGATATGTAGCGAAATCGGT | | | | | | | | | | |
| Chromosome | start | end | strand | MM | target_seq | PAM | alignment | distance | position | gene name | gene id |
| GU305914.1 | 138826 | 138848 | - | 0 | ACCGATTTCGCTACATATCC | CGG | \|\|\|\|\|\|\|\|[\|\|\|\|\|\|\|\|\|\|\|\|]PAM | NA | Intergenic | NA | NA |

| | | | | | | | | | | | |
|---|---|---|---|---|---|---|---|---|---|---|---|
| T23552 | TTTCGCTACATATCCCGGAGCGG | 1000 | | | | | | | | | |
| Oligo fwd | CACCTTTCGCTACATATCCCGGAG | | | | | | | | | | |
| Oligo rev | AAACCTCCGGGATATGTAGCGAAA | | | | | | | | | | |
| Chromosome | start | end | strand | MM | target_seq | PAM | alignment | distance | position | gene name | gene id |
| GU305914.1 | 138821 | 138843 | - | 0 | TTTCGCTACATATCCCGGAG | CGG | \|\|\|\|\|\|\|\|[\|\|\|\|\|\|\|\|\|\|\|\|]PAM | NA | Intergenic | NA | NA |

| | | | | | | | | | | | |
|---|---|---|---|---|---|---|---|---|---|---|---|
| T23553 | GCTACATATCCCGGAGCGGACGG | 1000 | | | | | | | | | |
| Oligo fwd | CACCGCTACATATCCCGGAGCGGA | | | | | | | | | | |
| Oligo rev | AAACTCCGCTCCGGGATATGTAGC | | | | | | | | | | |
| Chromosome | start | end | strand | MM | target_seq | PAM | alignment | distance | position | gene name | gene id |
| GU305914.1 | 138817 | 138839 | - | 0 | GCTACATATCCCGGAGCGGA | CGG | \|\|\|\|\|\|\|\|[\|\|\|\|\|\|\|\|\|\|\|\|]PAM | NA | Intergenic | NA | NA |

| | | | | | | | | | | | |
|---|---|---|---|---|---|---|---|---|---|---|---|
| T23554 | GAACGCGTTACCGCTACCTGCGG | 1000 | | | | | | | | | |
| Oligo fwd | CACCGAACGCGTTACCGCTACCTG | | | | | | | | | | |
| Oligo rev | AAACCAGGTAGCGGTAACGCGTTC | | | | | | | | | | |
| Chromosome | start | end | strand | MM | target_seq | PAM | alignment | distance | position | gene name | gene id |
| GU305914.1 | 138737 | 138759 | - | 0 | GAACGCGTTACCGCTACCTG | CGG | \|\|\|\|\|\|\|\|[\|\|\|\|\|\|\|\|\|\|\|\|]PAM | NA | Intergenic | NA | NA |

| | | | | | | | | | | | |
|---|---|---|---|---|---|---|---|---|---|---|---|
| T23555 | GCGTTACCGCTACCTGCGGACGG | 1000 | | | | | | | | | |
| Oligo fwd | CACCGCGTTACCGCTACCTGCGGA | | | | | | | | | | |
| Oligo rev | AAACTCCGCAGGTAGCGGTAACGC | | | | | | | | | | |
| Chromosome | start | end | strand | MM | target_seq | PAM | alignment | distance | position | gene name | gene id |
| GU305914.1 | 138733 | 138755 | - | 0 | GCGTTACCGCTACCTGCGGA | CGG | \|\|\|\|\|\|\|\|[\|\|\|\|\|\|\|\|\|\|\|\|]PAM | NA | Intergenic | NA | NA |

| | | | | | | | | | | | |
|---|---|---|---|---|---|---|---|---|---|---|---|
| T23556 | TCGCCCGCCCCGACCCCTCGAGG | 1000 | | | | | | | | | |
| Oligo fwd | CACCTCGCCCGCCCCGACCCCTCG | | | | | | | | | | |
| Oligo rev | AAACCGAGGGGTCGGGGCGGGCGA | | | | | | | | | | |
| Chromosome | start | end | strand | MM | target_seq | PAM | alignment | distance | position | gene name | gene id |
| GU305914.1 | 138678 | 138700 | - | 0 | TCGCCCGCCCCGACCCCTCG | AGG | \|\|\|\|\|\|\|\|[\|\|\|\|\|\|\|\|\|\|\|\|]PAM | NA | Intergenic | NA | NA |

| | | |
|---|---|---|
| T23557 | CCCGACCCCTCGAGGACGCACGG | 1000 |
| Oligo fwd | CACCCCCGACCCCTCGAGGACGCA | |
| Oligo rev | AAACTGCGTCCTCGAGGGGTCGGG | |

<table>
<tr><td>Chromosome</td><td>start</td><td>end</td><td>strand</td><td>MM</td><td>target_seq</td><td>PAM</td><td>alignment</td><td>distance</td><td>position</td><td>gene name</td><td>gene id</td></tr>
<tr><td>GU305914.1</td><td>138670</td><td>138692</td><td>-</td><td>0</td><td>CCCGACCCCTCGAGGACGCA</td><td>CGG</td><td>||||||||[||||||||||||]PAM</td><td>NA</td><td>Intergenic</td><td>NA</td><td>NA</td></tr>
<tr><td></td><td></td><td></td><td></td><td></td><td></td><td></td><td></td><td></td><td></td><td></td><td></td></tr>
<tr><td>T23558</td><td>GACCCCTCGAGGACGCACGGCGG</td><td>1000</td><td></td><td></td><td></td><td></td><td></td><td></td><td></td><td></td><td></td></tr>
<tr><td>Oligo fwd</td><td>CACCGACCCCTCGAGGACGCACGG</td><td></td><td></td><td></td><td></td><td></td><td></td><td></td><td></td><td></td><td></td></tr>
<tr><td>Oligo rev</td><td>AAACCCGTGCGTCCTCGAGGGGTC</td><td></td><td></td><td></td><td></td><td></td><td></td><td></td><td></td><td></td><td></td></tr>
<tr><td>Chromosome</td><td>start</td><td>end</td><td>strand</td><td>MM</td><td>target_seq</td><td>PAM</td><td>alignment</td><td>distance</td><td>position</td><td>gene name</td><td>gene id</td></tr>
<tr><td>GU305914.1</td><td>138667</td><td>138689</td><td>-</td><td>0</td><td>GACCCCTCGAGGACGCACGG</td><td>CGG</td><td>||||||||[||||||||||||]PAM</td><td>NA</td><td>Intergenic</td><td>NA</td><td>NA</td></tr>
<tr><td>GU305914.1</td><td>41415</td><td>41437</td><td>-</td><td>4</td><td>GTCCGCTCGAGGGCGCGCGG</td><td>CGG</td><td>|-||-|||[||||-|||-|||]PAM</td><td>NA</td><td>Intergenic</td><td>NA</td><td>NA</td></tr>
<tr><td></td><td></td><td></td><td></td><td></td><td></td><td></td><td></td><td></td><td></td><td></td><td></td></tr>
<tr><td>T23559</td><td>ACGGCGGTTGTTGTTCCGTCCGG</td><td>1000</td><td></td><td></td><td></td><td></td><td></td><td></td><td></td><td></td><td></td></tr>
<tr><td>Oligo fwd</td><td>CACCACGGCGGTTGTTGTTCCGTC</td><td></td><td></td><td></td><td></td><td></td><td></td><td></td><td></td><td></td><td></td></tr>
<tr><td>Oligo rev</td><td>AAACGACGGAACAACAACCGCCGT</td><td></td><td></td><td></td><td></td><td></td><td></td><td></td><td></td><td></td><td></td></tr>
<tr><td>Chromosome</td><td>start</td><td>end</td><td>strand</td><td>MM</td><td>target_seq</td><td>PAM</td><td>alignment</td><td>distance</td><td>position</td><td>gene name</td><td>gene id</td></tr>
<tr><td>GU305914.1</td><td>138651</td><td>138673</td><td>-</td><td>0</td><td>ACGGCGGTTGTTGTTCCGTC</td><td>CGG</td><td>||||||||[||||||||||||]PAM</td><td>NA</td><td>Intergenic</td><td>NA</td><td>NA</td></tr>
<tr><td></td><td></td><td></td><td></td><td></td><td></td><td></td><td></td><td></td><td></td><td></td><td></td></tr>
<tr><td>T23560</td><td>TTGTTGTTCCGTCCGGTCCTTGG</td><td>1000</td><td></td><td></td><td></td><td></td><td></td><td></td><td></td><td></td><td></td></tr>
<tr><td>Oligo fwd</td><td>CACCTTGTTGTTCCGTCCGGTCCT</td><td></td><td></td><td></td><td></td><td></td><td></td><td></td><td></td><td></td><td></td></tr>
<tr><td>Oligo rev</td><td>AAACAGGACCGGACGGAACAACAA</td><td></td><td></td><td></td><td></td><td></td><td></td><td></td><td></td><td></td><td></td></tr>
<tr><td>Chromosome</td><td>start</td><td>end</td><td>strand</td><td>MM</td><td>target_seq</td><td>PAM</td><td>alignment</td><td>distance</td><td>position</td><td>gene name</td><td>gene id</td></tr>
<tr><td>GU305914.1</td><td>138644</td><td>138666</td><td>-</td><td>0</td><td>TTGTTGTTCCGTCCGGTCCT</td><td>TGG</td><td>||||||||[||||||||||||]PAM</td><td>NA</td><td>Intergenic</td><td>NA</td><td>NA</td></tr>
<tr><td></td><td></td><td></td><td></td><td></td><td></td><td></td><td></td><td></td><td></td><td></td><td></td></tr>
<tr><td>T23561</td><td>TGTTGTTCCGTCCGGTCCTTGGG</td><td>1000</td><td></td><td></td><td></td><td></td><td></td><td></td><td></td><td></td><td></td></tr>
<tr><td>Oligo fwd</td><td>CACCTGTTGTTCCGTCCGGTCCTT</td><td></td><td></td><td></td><td></td><td></td><td></td><td></td><td></td><td></td><td></td></tr>
<tr><td>Oligo rev</td><td>AAACAAGGACCGGACGGAACAACA</td><td></td><td></td><td></td><td></td><td></td><td></td><td></td><td></td><td></td><td></td></tr>
<tr><td>Chromosome</td><td>start</td><td>end</td><td>strand</td><td>MM</td><td>target_seq</td><td>PAM</td><td>alignment</td><td>distance</td><td>position</td><td>gene name</td><td>gene id</td></tr>
<tr><td>GU305914.1</td><td>138643</td><td>138665</td><td>-</td><td>0</td><td>TGTTGTTCCGTCCGGTCCTT</td><td>GGG</td><td>||||||||[||||||||||||]PAM</td><td>NA</td><td>Intergenic</td><td>NA</td><td>NA</td></tr>
<tr><td></td><td></td><td></td><td></td><td></td><td></td><td></td><td></td><td></td><td></td><td></td><td></td></tr>
<tr><td>T23562</td><td>CCGGTCCTTGGGCAGCGCGTTGG</td><td>1000</td><td></td><td></td><td></td><td></td><td></td><td></td><td></td><td></td><td></td></tr>
<tr><td>Oligo fwd</td><td>CACCCCGGTCCTTGGGCAGCGCGT</td><td></td><td></td><td></td><td></td><td></td><td></td><td></td><td></td><td></td><td></td></tr>
<tr><td>Oligo rev</td><td>AAACACGCGCTGCCCAAGGACCGG</td><td></td><td></td><td></td><td></td><td></td><td></td><td></td><td></td><td></td><td></td></tr>
<tr><td>Chromosome</td><td>start</td><td>end</td><td>strand</td><td>MM</td><td>target_seq</td><td>PAM</td><td>alignment</td><td>distance</td><td>position</td><td>gene name</td><td>gene id</td></tr>
<tr><td>GU305914.1</td><td>138632</td><td>138654</td><td>-</td><td>0</td><td>CCGGTCCTTGGGCAGCGCGT</td><td>TGG</td><td>||||||||[||||||||||||]PAM</td><td>NA</td><td>Intergenic</td><td>NA</td><td>NA</td></tr>
<tr><td>GU305914.1</td><td>110791</td><td>110813</td><td>-</td><td>4</td><td>CCTGTGCTTGGGCAGCGTGA</td><td>TGG</td><td>||-||-||[|||||||||-|-]PAM</td><td>NA</td><td>Intergenic</td><td>NA</td><td>NA</td></tr>
<tr><td></td><td></td><td></td><td></td><td></td><td></td><td></td><td></td><td></td><td></td><td></td><td></td></tr>
<tr><td>T23563</td><td>GACGAGCAAGAAGTTGCCCCCGG</td><td>1000</td><td></td><td></td><td></td><td></td><td></td><td></td><td></td><td></td><td></td></tr>
<tr><td>Oligo fwd</td><td>CACCGACGAGCAAGAAGTTGCCCC</td><td></td><td></td><td></td><td></td><td></td><td></td><td></td><td></td><td></td><td></td></tr>
<tr><td>Oligo rev</td><td>AAACGGGGCAACTTCTTGCTCGTC</td><td></td><td></td><td></td><td></td><td></td><td></td><td></td><td></td><td></td><td></td></tr>
<tr><td>Chromosome</td><td>start</td><td>end</td><td>strand</td><td>MM</td><td>target_seq</td><td>PAM</td><td>alignment</td><td>distance</td><td>position</td><td>gene name</td><td>gene id</td></tr>
<tr><td>GU305914.1</td><td>138608</td><td>138630</td><td>-</td><td>0</td><td>GACGAGCAAGAAGTTGCCCC</td><td>CGG</td><td>||||||||[||||||||||||]PAM</td><td>NA</td><td>Intergenic</td><td>NA</td><td>NA</td></tr>
<tr><td></td><td></td><td></td><td></td><td></td><td></td><td></td><td></td><td></td><td></td><td></td><td></td></tr>
<tr><td>T23564</td><td>TGCCCCCGGCCGTCTCGATCAGG</td><td>1000</td><td></td><td></td><td></td><td></td><td></td><td></td><td></td><td></td><td></td></tr>
<tr><td>Oligo fwd</td><td>CACCTGCCCCCGGCCGTCTCGATC</td><td></td><td></td><td></td><td></td><td></td><td></td><td></td><td></td><td></td><td></td></tr>
<tr><td>Oligo rev</td><td>AAACGATCGAGACGGCCGGGGGCA</td><td></td><td></td><td></td><td></td><td></td><td></td><td></td><td></td><td></td><td></td></tr>
<tr><td>Chromosome</td><td>start</td><td>end</td><td>strand</td><td>MM</td><td>target_seq</td><td>PAM</td><td>alignment</td><td>distance</td><td>position</td><td>gene name</td><td>gene id</td></tr>
<tr><td>GU305914.1</td><td>138594</td><td>138616</td><td>-</td><td>0</td><td>TGCCCCCGGCCGTCTCGATC</td><td>AGG</td><td>||||||||[||||||||||||]PAM</td><td>NA</td><td>Intergenic</td><td>NA</td><td>NA</td></tr>
<tr><td></td><td></td><td></td><td></td><td></td><td></td><td></td><td></td><td></td><td></td><td></td><td></td></tr>
<tr><td>T23565</td><td>GATCAGGCGAAACCTGTCCCTGG</td><td>1000</td><td></td><td></td><td></td><td></td><td></td><td></td><td></td><td></td><td></td></tr>
<tr><td>Oligo fwd</td><td>CACCGATCAGGCGAAACCTGTCCC</td><td></td><td></td><td></td><td></td><td></td><td></td><td></td><td></td><td></td><td></td></tr>
<tr><td>Oligo rev</td><td>AAACGGGACAGGTTTCGCCTGATC</td><td></td><td></td><td></td><td></td><td></td><td></td><td></td><td></td><td></td><td></td></tr>
<tr><td>Chromosome</td><td>start</td><td>end</td><td>strand</td><td>MM</td><td>target_seq</td><td>PAM</td><td>alignment</td><td>distance</td><td>position</td><td>gene name</td><td>gene id</td></tr>
<tr><td>GU305914.1</td><td>138578</td><td>138600</td><td>-</td><td>0</td><td>GATCAGGCGAAACCTGTCCC</td><td>TGG</td><td>||||||||[||||||||||||]PAM</td><td>NA</td><td>Intergenic</td><td>NA</td><td>NA</td></tr>
<tr><td></td><td></td><td></td><td></td><td></td><td></td><td></td><td></td><td></td><td></td><td></td><td></td></tr>
<tr><td>T23566</td><td>ATCAGGCGAAACCTGTCCCTGGG</td><td>1000</td><td></td><td></td><td></td><td></td><td></td><td></td><td></td><td></td><td></td></tr>
<tr><td>Oligo fwd</td><td>CACCATCAGGCGAAACCTGTCCCT</td><td></td><td></td><td></td><td></td><td></td><td></td><td></td><td></td><td></td><td></td></tr>
<tr><td>Oligo rev</td><td>AAACAGGGACAGGTTTCGCCTGAT</td><td></td><td></td><td></td><td></td><td></td><td></td><td></td><td></td><td></td><td></td></tr>
<tr><td>Chromosome</td><td>start</td><td>end</td><td>strand</td><td>MM</td><td>target_seq</td><td>PAM</td><td>alignment</td><td>distance</td><td>position</td><td>gene name</td><td>gene id</td></tr>
<tr><td>GU305914.1</td><td>138577</td><td>138599</td><td>-</td><td>0</td><td>ATCAGGCGAAACCTGTCCCT</td><td>GGG</td><td>||||||||[||||||||||||]PAM</td><td>NA</td><td>Intergenic</td><td>NA</td><td>NA</td></tr>
<tr><td></td><td></td><td></td><td></td><td></td><td></td><td></td><td></td><td></td><td></td><td></td><td></td></tr>
<tr><td>T23567</td><td>GCGAAACCTGTCCCTGGGTGAGG</td><td>1000</td><td></td><td></td><td></td><td></td><td></td><td></td><td></td><td></td><td></td></tr>
<tr><td>Oligo fwd</td><td>CACCGCGAAACCTGTCCCTGGGTG</td><td></td><td></td><td></td><td></td><td></td><td></td><td></td><td></td><td></td><td></td></tr>
<tr><td>Oligo rev</td><td>AAACCACCCAGGGACAGGTTTCGC</td><td></td><td></td><td></td><td></td><td></td><td></td><td></td><td></td><td></td><td></td></tr>
<tr><td>Chromosome</td><td>start</td><td>end</td><td>strand</td><td>MM</td><td>target_seq</td><td>PAM</td><td>alignment</td><td>distance</td><td>position</td><td>gene name</td><td>gene id</td></tr>
<tr><td>GU305914.1</td><td>138572</td><td>138594</td><td>-</td><td>0</td><td>GCGAAACCTGTCCCTGGGTG</td><td>AGG</td><td>||||||||[||||||||||||]PAM</td><td>NA</td><td>Intergenic</td><td>NA</td><td>NA</td></tr>
<tr><td></td><td></td><td></td><td></td><td></td><td></td><td></td><td></td><td></td><td></td><td></td><td></td></tr>
<tr><td>T23568</td><td>AAACCTGTCCCTGGGTGAGGAGG</td><td>1000</td><td></td><td></td><td></td><td></td><td></td><td></td><td></td><td></td><td></td></tr>
<tr><td>Oligo fwd</td><td>CACCAAACCTGTCCCTGGGTGAGG</td><td></td><td></td><td></td><td></td><td></td><td></td><td></td><td></td><td></td><td></td></tr>
<tr><td>Oligo rev</td><td>AAACCCTCACCCAGGGACAGGTTT</td><td></td><td></td><td></td><td></td><td></td><td></td><td></td><td></td><td></td><td></td></tr>
<tr><td>Chromosome</td><td>start</td><td>end</td><td>strand</td><td>MM</td><td>target_seq</td><td>PAM</td><td>alignment</td><td>distance</td><td>position</td><td>gene name</td><td>gene id</td></tr>
<tr><td>GU305914.1</td><td>138569</td><td>138591</td><td>-</td><td>0</td><td>AAACCTGTCCCTGGGTGAGG</td><td>AGG</td><td>||||||||[||||||||||||]PAM</td><td>NA</td><td>Intergenic</td><td>NA</td><td>NA</td></tr>
<tr><td></td><td></td><td></td><td></td><td></td><td></td><td></td><td></td><td></td><td></td><td></td><td></td></tr>
<tr><td>T23569</td><td>AGGAGGCCTCGCACGCGATCAGG</td><td>1000</td><td></td><td></td><td></td><td></td><td></td><td></td><td></td><td></td><td></td></tr>
<tr><td>Oligo fwd</td><td>CACCAGGAGGCCTCGCACGCGATC</td><td></td><td></td><td></td><td></td><td></td><td></td><td></td><td></td><td></td><td></td></tr>
<tr><td>Oligo rev</td><td>AAACGATCGCGTGCGAGGCCTCCT</td><td></td><td></td><td></td><td></td><td></td><td></td><td></td><td></td><td></td><td></td></tr>
<tr><td>Chromosome</td><td>start</td><td>end</td><td>strand</td><td>MM</td><td>target_seq</td><td>PAM</td><td>alignment</td><td>distance</td><td>position</td><td>gene name</td><td>gene id</td></tr>
<tr><td>GU305914.1</td><td>138552</td><td>138574</td><td>-</td><td>0</td><td>AGGAGGCCTCGCACGCGATC</td><td>AGG</td><td>||||||||[||||||||||||]PAM</td><td>NA</td><td>Intergenic</td><td>NA</td><td>NA</td></tr>
<tr><td></td><td></td><td></td><td></td><td></td><td></td><td></td><td></td><td></td><td></td><td></td><td></td></tr>
<tr><td>T23570</td><td>GGAGGCCTCGCACGCGATCAGGG</td><td>1000</td><td></td><td></td><td></td><td></td><td></td><td></td><td></td><td></td><td></td></tr>
<tr><td>Oligo fwd</td><td>CACCGGAGGCCTCGCACGCGATCA</td><td></td><td></td><td></td><td></td><td></td><td></td><td></td><td></td><td></td><td></td></tr>
<tr><td>Oligo rev</td><td>AAACTGATCGCGTGCGAGGCCTCC</td><td></td><td></td><td></td><td></td><td></td><td></td><td></td><td></td><td></td><td></td></tr>
<tr><td>Chromosome</td><td>start</td><td>end</td><td>strand</td><td>MM</td><td>target_seq</td><td>PAM</td><td>alignment</td><td>distance</td><td>position</td><td>gene name</td><td>gene id</td></tr>
<tr><td>GU305914.1</td><td>138551</td><td>138573</td><td>-</td><td>0</td><td>GGAGGCCTCGCACGCGATCA</td><td>GGG</td><td>||||||||[||||||||||||]PAM</td><td>NA</td><td>Intergenic</td><td>NA</td><td>NA</td></tr>
<tr><td></td><td></td><td></td><td></td><td></td><td></td><td></td><td></td><td></td><td></td><td></td><td></td></tr>
<tr><td>T23571</td><td>CGCGATCAGGGCCGCGTCCACGG</td><td>1000</td><td></td><td></td><td></td><td></td><td></td><td></td><td></td><td></td><td></td></tr>
<tr><td>Oligo fwd</td><td>CACCCGCGATCAGGGCCGCGTCCA</td><td></td><td></td><td></td><td></td><td></td><td></td><td></td><td></td><td></td><td></td></tr>
<tr><td>Oligo rev</td><td>AAACTGGACGCGGCCCTGATCGCG</td><td></td><td></td><td></td><td></td><td></td><td></td><td></td><td></td><td></td><td></td></tr>
<tr><td>Chromosome</td><td>start</td><td>end</td><td>strand</td><td>MM</td><td>target_seq</td><td>PAM</td><td>alignment</td><td>distance</td><td>position</td><td>gene name</td><td>gene id</td></tr>
<tr><td>GU305914.1</td><td>138539</td><td>138561</td><td>-</td><td>0</td><td>CGCGATCAGGGCCGCGTCCA</td><td>CGG</td><td>||||||||[||||||||||||]PAM</td><td>NA</td><td>Intergenic</td><td>NA</td><td>NA</td></tr>
</table>

<table>
<tr><td>T23572</td><td>GTACCTGTTGACGTCCGCCTCGG</td><td></td><td>1000</td><td></td><td></td><td></td><td></td><td></td><td></td><td></td><td></td><td></td></tr>
<tr><td>Oligo fwd</td><td>CACCGTACCTGTTGACGTCCGCCT</td><td></td><td></td><td></td><td></td><td></td><td></td><td></td><td></td><td></td><td></td><td></td></tr>
<tr><td>Oligo rev</td><td>AAACAGGCGGACGTCAACAGGTAC</td><td></td><td></td><td></td><td></td><td></td><td></td><td></td><td></td><td></td><td></td><td></td></tr>
<tr><td>Chromosome</td><td>start</td><td></td><td>end</td><td>strand</td><td>MM</td><td>target_seq</td><td>PAM</td><td>alignment</td><td>distance</td><td>position</td><td>gene name</td><td>gene id</td></tr>
<tr><td>GU305914.1</td><td></td><td>138509</td><td>138531</td><td>-</td><td>0</td><td>GTACCTGTTGACGTCCGCCT</td><td>CGG</td><td>||||||||[||||||||||||]PAM</td><td>NA</td><td>Intergenic</td><td>NA</td><td>NA</td></tr>
<tr><td>T23573</td><td>GGACACCCGACCCTCCTCGACGG</td><td></td><td>1000</td><td></td><td></td><td></td><td></td><td></td><td></td><td></td><td></td><td></td></tr>
<tr><td>Oligo fwd</td><td>CACCGGACACCCGACCCTCCTCGA</td><td></td><td></td><td></td><td></td><td></td><td></td><td></td><td></td><td></td><td></td><td></td></tr>
<tr><td>Oligo rev</td><td>AAACTCGAGGAGGGTCGGGTGTCC</td><td></td><td></td><td></td><td></td><td></td><td></td><td></td><td></td><td></td><td></td><td></td></tr>
<tr><td>Chromosome</td><td>start</td><td></td><td>end</td><td>strand</td><td>MM</td><td>target_seq</td><td>PAM</td><td>alignment</td><td>distance</td><td>position</td><td>gene name</td><td>gene id</td></tr>
<tr><td>GU305914.1</td><td></td><td>138488</td><td>138510</td><td>-</td><td>0</td><td>GGACACCCGACCCTCCTCGA</td><td>CGG</td><td>||||||||[||||||||||||]PAM</td><td>NA</td><td>Intergenic</td><td>NA</td><td>NA</td></tr>
<tr><td>T23574</td><td>CACCCGACCCTCCTCGACGGCGG</td><td></td><td>1000</td><td></td><td></td><td></td><td></td><td></td><td></td><td></td><td></td><td></td></tr>
<tr><td>Oligo fwd</td><td>CACCCACCCGACCCTCCTCGACGG</td><td></td><td></td><td></td><td></td><td></td><td></td><td></td><td></td><td></td><td></td><td></td></tr>
<tr><td>Oligo rev</td><td>AAACCCGTCGAGGAGGGTCGGGTG</td><td></td><td></td><td></td><td></td><td></td><td></td><td></td><td></td><td></td><td></td><td></td></tr>
<tr><td>Chromosome</td><td>start</td><td></td><td>end</td><td>strand</td><td>MM</td><td>target_seq</td><td>PAM</td><td>alignment</td><td>distance</td><td>position</td><td>gene name</td><td>gene id</td></tr>
<tr><td>GU305914.1</td><td></td><td>138485</td><td>138507</td><td>-</td><td>0</td><td>CACCCGACCCTCCTCGACGG</td><td>CGG</td><td>||||||||[||||||||||||]PAM</td><td>NA</td><td>Intergenic</td><td>NA</td><td>NA</td></tr>
<tr><td>T23575</td><td>CGCCTCGACATCGCAGAGCGCGG</td><td></td><td>1000</td><td></td><td></td><td></td><td></td><td></td><td></td><td></td><td></td><td></td></tr>
<tr><td>Oligo fwd</td><td>CACCCGCCTCGACATCGCAGAGCG</td><td></td><td></td><td></td><td></td><td></td><td></td><td></td><td></td><td></td><td></td><td></td></tr>
<tr><td>Oligo rev</td><td>AAACCGCTCTGCGATGTCGAGGCG</td><td></td><td></td><td></td><td></td><td></td><td></td><td></td><td></td><td></td><td></td><td></td></tr>
<tr><td>Chromosome</td><td>start</td><td></td><td>end</td><td>strand</td><td>MM</td><td>target_seq</td><td>PAM</td><td>alignment</td><td>distance</td><td>position</td><td>gene name</td><td>gene id</td></tr>
<tr><td>GU305914.1</td><td></td><td>138458</td><td>138480</td><td>-</td><td>0</td><td>CGCCTCGACATCGCAGAGCG</td><td>CGG</td><td>||||||||[||||||||||||]PAM</td><td>NA</td><td>Intergenic</td><td>NA</td><td>NA</td></tr>
<tr><td>T23576</td><td>GCAGAGCGCGGTCGAGACGTCGG</td><td></td><td>1000</td><td></td><td></td><td></td><td></td><td></td><td></td><td></td><td></td><td></td></tr>
<tr><td>Oligo fwd</td><td>CACCGCAGAGCGCGGTCGAGACGT</td><td></td><td></td><td></td><td></td><td></td><td></td><td></td><td></td><td></td><td></td><td></td></tr>
<tr><td>Oligo rev</td><td>AAACACGTCTCGACCGCGCTCTGC</td><td></td><td></td><td></td><td></td><td></td><td></td><td></td><td></td><td></td><td></td><td></td></tr>
<tr><td>Chromosome</td><td>start</td><td></td><td>end</td><td>strand</td><td>MM</td><td>target_seq</td><td>PAM</td><td>alignment</td><td>distance</td><td>position</td><td>gene name</td><td>gene id</td></tr>
<tr><td>GU305914.1</td><td></td><td>138446</td><td>138468</td><td>-</td><td>0</td><td>GCAGAGCGCGGTCGAGACGT</td><td>CGG</td><td>||||||||[||||||||||||]PAM</td><td>NA</td><td>Intergenic</td><td>NA</td><td>NA</td></tr>
<tr><td>T23577</td><td>GTCGAGACGTCGGTCGCCCGTGG</td><td></td><td>1000</td><td></td><td></td><td></td><td></td><td></td><td></td><td></td><td></td><td></td></tr>
<tr><td>Oligo fwd</td><td>CACCGTCGAGACGTCGGTCGCCCG</td><td></td><td></td><td></td><td></td><td></td><td></td><td></td><td></td><td></td><td></td><td></td></tr>
<tr><td>Oligo rev</td><td>AAACCGGGCGACCGACGTCTCGAC</td><td></td><td></td><td></td><td></td><td></td><td></td><td></td><td></td><td></td><td></td><td></td></tr>
<tr><td>Chromosome</td><td>start</td><td></td><td>end</td><td>strand</td><td>MM</td><td>target_seq</td><td>PAM</td><td>alignment</td><td>distance</td><td>position</td><td>gene name</td><td>gene id</td></tr>
<tr><td>GU305914.1</td><td></td><td>138436</td><td>138458</td><td>-</td><td>0</td><td>GTCGAGACGTCGGTCGCCCG</td><td>TGG</td><td>||||||||[||||||||||||]PAM</td><td>NA</td><td>Intergenic</td><td>NA</td><td>NA</td></tr>
<tr><td>T23578</td><td>GCCACCAGCGTAAACACCGCCGG</td><td></td><td>1000</td><td></td><td></td><td></td><td></td><td></td><td></td><td></td><td></td><td></td></tr>
<tr><td>Oligo fwd</td><td>CACCGCCACCAGCGTAAACACCGC</td><td></td><td></td><td></td><td></td><td></td><td></td><td></td><td></td><td></td><td></td><td></td></tr>
<tr><td>Oligo rev</td><td>AAACGCGGTGTTTACGCTGGTGGC</td><td></td><td></td><td></td><td></td><td></td><td></td><td></td><td></td><td></td><td></td><td></td></tr>
<tr><td>Chromosome</td><td>start</td><td></td><td>end</td><td>strand</td><td>MM</td><td>target_seq</td><td>PAM</td><td>alignment</td><td>distance</td><td>position</td><td>gene name</td><td>gene id</td></tr>
<tr><td>GU305914.1</td><td></td><td>138392</td><td>138414</td><td>-</td><td>0</td><td>GCCACCAGCGTAAACACCGC</td><td>CGG</td><td>||||||||[||||||||||||]PAM</td><td>NA</td><td>Intergenic</td><td>NA</td><td>NA</td></tr>
<tr><td>T23579</td><td>CACCGACGCGATCGCCGTCGCGG</td><td></td><td>1000</td><td></td><td></td><td></td><td></td><td></td><td></td><td></td><td></td><td></td></tr>
<tr><td>Oligo fwd</td><td>CACCCACCGACGCGATCGCCGTCG</td><td></td><td></td><td></td><td></td><td></td><td></td><td></td><td></td><td></td><td></td><td></td></tr>
<tr><td>Oligo rev</td><td>AAACCGACGGCGATCGCGTCGGTG</td><td></td><td></td><td></td><td></td><td></td><td></td><td></td><td></td><td></td><td></td><td></td></tr>
<tr><td>Chromosome</td><td>start</td><td></td><td>end</td><td>strand</td><td>MM</td><td>target_seq</td><td>PAM</td><td>alignment</td><td>distance</td><td>position</td><td>gene name</td><td>gene id</td></tr>
<tr><td>GU305914.1</td><td></td><td>138353</td><td>138375</td><td>-</td><td>0</td><td>CACCGACGCGATCGCCGTCG</td><td>CGG</td><td>||||||||[||||||||||||]PAM</td><td>NA</td><td>Intergenic</td><td>NA</td><td>NA</td></tr>
<tr><td>GU305914.1</td><td></td><td>90107</td><td>90129</td><td>-</td><td>4</td><td>GATCGACGCGATCACCGGCG</td><td>AGG</td><td>-|-|||||[|||||-|||-||]PAM</td><td>NA</td><td>Intergenic</td><td>NA</td><td>NA</td></tr>
<tr><td>T23580</td><td>GCCCACGACCGTGATCACCACGG</td><td></td><td>1000</td><td></td><td></td><td></td><td></td><td></td><td></td><td></td><td></td><td></td></tr>
<tr><td>Oligo fwd</td><td>CACCGCCCACGACCGTGATCACCA</td><td></td><td></td><td></td><td></td><td></td><td></td><td></td><td></td><td></td><td></td><td></td></tr>
<tr><td>Oligo rev</td><td>AAACTGGTGATCACGGTCGTGGGC</td><td></td><td></td><td></td><td></td><td></td><td></td><td></td><td></td><td></td><td></td><td></td></tr>
<tr><td>Chromosome</td><td>start</td><td></td><td>end</td><td>strand</td><td>MM</td><td>target_seq</td><td>PAM</td><td>alignment</td><td>distance</td><td>position</td><td>gene name</td><td>gene id</td></tr>
<tr><td>GU305914.1</td><td></td><td>138290</td><td>138312</td><td>-</td><td>0</td><td>GCCCACGACCGTGATCACCA</td><td>CGG</td><td>||||||||[||||||||||||]PAM</td><td>NA</td><td>Intergenic</td><td>NA</td><td>NA</td></tr>
<tr><td>T23581</td><td>CACCACGGAGTTTCCCGACGTGG</td><td></td><td>1000</td><td></td><td></td><td></td><td></td><td></td><td></td><td></td><td></td><td></td></tr>
<tr><td>Oligo fwd</td><td>CACCCACCACGGAGTTTCCCGACG</td><td></td><td></td><td></td><td></td><td></td><td></td><td></td><td></td><td></td><td></td><td></td></tr>
<tr><td>Oligo rev</td><td>AAACCGTCGGGAAACTCCGTGGTG</td><td></td><td></td><td></td><td></td><td></td><td></td><td></td><td></td><td></td><td></td><td></td></tr>
<tr><td>Chromosome</td><td>start</td><td></td><td>end</td><td>strand</td><td>MM</td><td>target_seq</td><td>PAM</td><td>alignment</td><td>distance</td><td>position</td><td>gene name</td><td>gene id</td></tr>
<tr><td>GU305914.1</td><td></td><td>138275</td><td>138297</td><td>-</td><td>0</td><td>CACCACGGAGTTTCCCGACG</td><td>TGG</td><td>||||||||[||||||||||||]PAM</td><td>NA</td><td>Intergenic</td><td>NA</td><td>NA</td></tr>
<tr><td>T23582</td><td>ACCACGGAGTTTCCCGACGTGGG</td><td></td><td>1000</td><td></td><td></td><td></td><td></td><td></td><td></td><td></td><td></td><td></td></tr>
<tr><td>Oligo fwd</td><td>CACCACCACGGAGTTTCCCGACGT</td><td></td><td></td><td></td><td></td><td></td><td></td><td></td><td></td><td></td><td></td><td></td></tr>
<tr><td>Oligo rev</td><td>AAACACGTCGGGAAACTCCGTGGT</td><td></td><td></td><td></td><td></td><td></td><td></td><td></td><td></td><td></td><td></td><td></td></tr>
<tr><td>Chromosome</td><td>start</td><td></td><td>end</td><td>strand</td><td>MM</td><td>target_seq</td><td>PAM</td><td>alignment</td><td>distance</td><td>position</td><td>gene name</td><td>gene id</td></tr>
<tr><td>GU305914.1</td><td></td><td>138274</td><td>138296</td><td>-</td><td>0</td><td>ACCACGGAGTTTCCCGACGT</td><td>GGG</td><td>||||||||[||||||||||||]PAM</td><td>NA</td><td>Intergenic</td><td>NA</td><td>NA</td></tr>
<tr><td>GU305914.1</td><td></td><td>99336</td><td>99358</td><td>-</td><td>4</td><td>GCCACGGTGTTTACCGCCGT</td><td>CGG</td><td>-||||||-[||||-|||-|||]PAM</td><td>NA</td><td>Intergenic</td><td>NA</td><td>NA</td></tr>
<tr><td>T23583</td><td>GCGATATCAGACCGTACACCCGG</td><td></td><td>1000</td><td></td><td></td><td></td><td></td><td></td><td></td><td></td><td></td><td></td></tr>
<tr><td>Oligo fwd</td><td>CACCGCGATATCAGACCGTACACC</td><td></td><td></td><td></td><td></td><td></td><td></td><td></td><td></td><td></td><td></td><td></td></tr>
<tr><td>Oligo rev</td><td>AAACGGTGTACGGTCTGATATCGC</td><td></td><td></td><td></td><td></td><td></td><td></td><td></td><td></td><td></td><td></td><td></td></tr>
<tr><td>Chromosome</td><td>start</td><td></td><td>end</td><td>strand</td><td>MM</td><td>target_seq</td><td>PAM</td><td>alignment</td><td>distance</td><td>position</td><td>gene name</td><td>gene id</td></tr>
<tr><td>GU305914.1</td><td></td><td>138236</td><td>138258</td><td>-</td><td>0</td><td>GCGATATCAGACCGTACACC</td><td>CGG</td><td>||||||||[||||||||||||]PAM</td><td>NA</td><td>Intergenic</td><td>NA</td><td>NA</td></tr>
<tr><td>T23584</td><td>TCAGACCGTACACCCGGAGCTGG</td><td></td><td>1000</td><td></td><td></td><td></td><td></td><td></td><td></td><td></td><td></td><td></td></tr>
<tr><td>Oligo fwd</td><td>CACCTCAGACCGTACACCCGGAGC</td><td></td><td></td><td></td><td></td><td></td><td></td><td></td><td></td><td></td><td></td><td></td></tr>
<tr><td>Oligo rev</td><td>AAACGCTCCGGGTGTACGGTCTGA</td><td></td><td></td><td></td><td></td><td></td><td></td><td></td><td></td><td></td><td></td><td></td></tr>
<tr><td>Chromosome</td><td>start</td><td></td><td>end</td><td>strand</td><td>MM</td><td>target_seq</td><td>PAM</td><td>alignment</td><td>distance</td><td>position</td><td>gene name</td><td>gene id</td></tr>
<tr><td>GU305914.1</td><td></td><td>138230</td><td>138252</td><td>-</td><td>0</td><td>TCAGACCGTACACCCGGAGC</td><td>TGG</td><td>||||||||[||||||||||||]PAM</td><td>NA</td><td>Intergenic</td><td>NA</td><td>NA</td></tr>
<tr><td>T23585</td><td>CAGACCGTACACCCGGAGCTGGG</td><td></td><td>1000</td><td></td><td></td><td></td><td></td><td></td><td></td><td></td><td></td><td></td></tr>
<tr><td>Oligo fwd</td><td>CACCCAGACCGTACACCCGGAGCT</td><td></td><td></td><td></td><td></td><td></td><td></td><td></td><td></td><td></td><td></td><td></td></tr>
<tr><td>Oligo rev</td><td>AAACAGCTCCGGGTGTACGGTCTG</td><td></td><td></td><td></td><td></td><td></td><td></td><td></td><td></td><td></td><td></td><td></td></tr>
<tr><td>Chromosome</td><td>start</td><td></td><td>end</td><td>strand</td><td>MM</td><td>target_seq</td><td>PAM</td><td>alignment</td><td>distance</td><td>position</td><td>gene name</td><td>gene id</td></tr>
<tr><td>GU305914.1</td><td></td><td>138229</td><td>138251</td><td>-</td><td>0</td><td>CAGACCGTACACCCGGAGCT</td><td>GGG</td><td>||||||||[||||||||||||]PAM</td><td>NA</td><td>Intergenic</td><td>NA</td><td>NA</td></tr>
<tr><td>T23586</td><td>CCAAGTTCTGCGTGCAATACCGG</td><td></td><td>1000</td><td></td><td></td><td></td><td></td><td></td><td></td><td></td><td></td><td></td></tr>
<tr><td>Oligo fwd</td><td>CACCCCAAGTTCTGCGTGCAATAC</td><td></td><td></td><td></td><td></td><td></td><td></td><td></td><td></td><td></td><td></td><td></td></tr>
<tr><td>Oligo rev</td><td>AAACGTATTGCACGCAGAACTTGG</td><td></td><td></td><td></td><td></td><td></td><td></td><td></td><td></td><td></td><td></td><td></td></tr>
</table>

Chromosome start end strand MM target_seq PAM alignment distance position gene name gene id
GU305914.1 138168 138190 - 0 CCAAGTTCTGCGTGCAATAC CGG ||||||||[||||||||||||]PAM NA Intergenic NA NA

T23587 CAAGTTCTGCGTGCAATACCGGG 1000
Oligo fwd CACCCAAGTTCTGCGTGCAATACC
Oligo rev AAACGGTATTGCACGCAGAACTTG
Chromosome start end strand MM target_seq PAM alignment distance position gene name gene id
GU305914.1 138167 138189 - 0 CAAGTTCTGCGTGCAATACC GGG ||||||||[||||||||||||]PAM NA Intergenic NA NA

T23588 GAAGATCACCCCGACGCTGTCGG 1000
Oligo fwd CACCGAAGATCACCCCGACGCTGT
Oligo rev AAACACAGCGTCGGGGTGATCTTC
Chromosome start end strand MM target_seq PAM alignment distance position gene name gene id
GU305914.1 138110 138132 - 0 GAAGATCACCCCGACGCTGT CGG ||||||||[||||||||||||]PAM NA Intergenic NA NA

T23589 GATCACCCCGACGCTGTCGGCGG 1000
Oligo fwd CACCGATCACCCCGACGCTGTCGG
Oligo rev AAACCCGACAGCGTCGGGGTGATC
Chromosome start end strand MM target_seq PAM alignment distance position gene name gene id
GU305914.1 138107 138129 - 0 GATCACCCCGACGCTGTCGG CGG ||||||||[||||||||||||]PAM NA Intergenic NA NA

T23590 GCTGTCGGCGGCGATCGACGCGG 1000
Oligo fwd CACCGCTGTCGGCGGCGATCGACG
Oligo rev AAACCGTCGATCGCCGCCGACAGC
Chromosome start end strand MM target_seq PAM alignment distance position gene name gene id
GU305914.1 138095 138117 - 0 GCTGTCGGCGGCGATCGACG CGG ||||||||[||||||||||||]PAM NA Intergenic NA NA
GU305914.1 218360 218382 - 4 GCTGGTGGCCGCGATCGTCG CGG ||||--||[|-|||||||-||]PAM NA Intergenic NA NA

T23591 CAAGTTTCAGCCGTGTACCCCGG 1000
Oligo fwd CACCCAAGTTTCAGCCGTGTACCC
Oligo rev AAACGGGTACACGGCTGAAACTTG
Chromosome start end strand MM target_seq PAM alignment distance position gene name gene id
GU305914.1 138071 138093 - 0 CAAGTTTCAGCCGTGTACCC CGG ||||||||[||||||||||||]PAM NA Intergenic NA NA

T23592 GCCGTGTACCCCGGAACAGCTGG 1000
Oligo fwd CACCGCCGTGTACCCCGGAACAGC
Oligo rev AAACGCTGTTCCGGGGTACACGGC
Chromosome start end strand MM target_seq PAM alignment distance position gene name gene id
GU305914.1 138062 138084 - 0 GCCGTGTACCCCGGAACAGC TGG ||||||||[||||||||||||]PAM NA Intergenic NA NA

T23593 CTCCGACTTATATAGAGATCCGG 1000
Oligo fwd CACCCTCCGACTTATATAGAGATC
Oligo rev AAACGATCTCTATATAAGTCGGAG
Chromosome start end strand MM target_seq PAM alignment distance position gene name gene id
GU305914.1 138014 138036 - 0 CTCCGACTTATATAGAGATC CGG ||||||||[||||||||||||]PAM NA Intergenic NA NA

T23594 GCAGATAGTGAACTTCCTACAGG 1000
Oligo fwd CACCGCAGATAGTGAACTTCCTAC
Oligo rev AAACGTAGGAAGTTCACTATCTGC
Chromosome start end strand MM target_seq PAM alignment distance position gene name gene id
GU305914.1 137984 138006 - 0 GCAGATAGTGAACTTCCTAC AGG ||||||||[||||||||||||]PAM NA Intergenic NA NA

T23595 TCCTACAGGACTTCGAGAACTGG 1000
Oligo fwd CACCTCCTACAGGACTTCGAGAAC
Oligo rev AAACGTTCTCGAAGTCCTGTAGGA
Chromosome start end strand MM target_seq PAM alignment distance position gene name gene id
GU305914.1 137970 137992 - 0 TCCTACAGGACTTCGAGAAC TGG ||||||||[||||||||||||]PAM NA Intergenic NA NA

T23596 TTCGAGAACTGGATCGCAACCGG 1000
Oligo fwd CACCTTCGAGAACTGGATCGCAAC
Oligo rev AAACGTTGCGATCCAGTTCTCGAA
Chromosome start end strand MM target_seq PAM alignment distance position gene name gene id
GU305914.1 137959 137981 - 0 TTCGAGAACTGGATCGCAAC CGG ||||||||[||||||||||||]PAM NA Intergenic NA NA

T23597 AGAACTGCCCGAGCGAACGACGG 1000
Oligo fwd CACCAGAACTGCCCGAGCGAACGA
Oligo rev AAACTCGTTCGCTCGGGCAGTTCT
Chromosome start end strand MM target_seq PAM alignment distance position gene name gene id
GU305914.1 137928 137950 - 0 AGAACTGCCCGAGCGAACGA CGG ||||||||[||||||||||||]PAM NA Intergenic NA NA

T23598 CACTACCAAGAACCCCACGCTGG 1000
Oligo fwd CACCCACTACCAAGAACCCCACGC
Oligo rev AAACGCGTGGGGTTCTTGGTAGTG
Chromosome start end strand MM target_seq PAM alignment distance position gene name gene id
GU305914.1 137843 137865 - 0 CACTACCAAGAACCCCACGC TGG ||||||||[||||||||||||]PAM NA Intergenic NA NA

T23599 CACGCTGGCCAACCGCGTGACGG 1000
Oligo fwd CACCCACGCTGGCCAACCGCGTGA
Oligo rev AAACTCACGCGGTTGGCCAGCGTG
Chromosome start end strand MM target_seq PAM alignment distance position gene name gene id
GU305914.1 137828 137850 - 0 CACGCTGGCCAACCGCGTGA CGG ||||||||[||||||||||||]PAM NA Intergenic NA NA

T23600 GATAGACCTGTTCAAGCAGAAGG 1000
Oligo fwd CACCGATAGACCTGTTCAAGCAGA
Oligo rev AAACTCTGCTTGAACAGGTCTATC
Chromosome start end strand MM target_seq PAM alignment distance position gene name gene id
GU305914.1 137759 137781 - 0 GATAGACCTGTTCAAGCAGA AGG ||||||||[||||||||||||]PAM NA Intergenic NA NA

T23601 CCTGTTCAAGCAGAAGGCCACGG 1000

<table>
<tr><td>Oligo fwd</td><td>CACCCCTGTTCAAGCAGAAGGCCA</td></tr>
<tr><td>Oligo rev</td><td>AAACTGGCCTTCTGCTTGAACAGG</td></tr>
<tr><td>Chromosome</td><td>start</td><td>end</td><td>strand</td><td>MM</td><td>target_seq</td><td>PAM</td><td>alignment</td><td>distance</td><td>position</td><td>gene name</td><td>gene id</td></tr>
<tr><td>GU305914.1</td><td>137753</td><td>137775</td><td>-</td><td>0</td><td>CCTGTTCAAGCAGAAGGCCA</td><td>CGG</td><td>||||||||[||||||||||||]PAM</td><td>NA</td><td>Intergenic</td><td>NA</td><td>NA</td></tr>
<tr><td></td></tr>
<tr><td>T23602</td><td>CCACGGTGTTCCTCGTGCCCCGG</td><td>1000</td></tr>
<tr><td>Oligo fwd</td><td>CACCCCACGGTGTTCCTCGTGCCC</td></tr>
<tr><td>Oligo rev</td><td>AAACGGGCACGAGGAACACCGTGG</td></tr>
<tr><td>Chromosome</td><td>start</td><td>end</td><td>strand</td><td>MM</td><td>target_seq</td><td>PAM</td><td>alignment</td><td>distance</td><td>position</td><td>gene name</td><td>gene id</td></tr>
<tr><td>GU305914.1</td><td>137736</td><td>137758</td><td>-</td><td>0</td><td>CCACGGTGTTCCTCGTGCCC</td><td>CGG</td><td>||||||||[||||||||||||]PAM</td><td>NA</td><td>Intergenic</td><td>NA</td><td>NA</td></tr>
<tr><td></td></tr>
<tr><td>T23603</td><td>CGGTGTTCCTCGTGCCCCGGCGG</td><td>1000</td></tr>
<tr><td>Oligo fwd</td><td>CACCCGGTGTTCCTCGTGCCCCGG</td></tr>
<tr><td>Oligo rev</td><td>AAACCCGGGGCACGAGGAACACCG</td></tr>
<tr><td>Chromosome</td><td>start</td><td>end</td><td>strand</td><td>MM</td><td>target_seq</td><td>PAM</td><td>alignment</td><td>distance</td><td>position</td><td>gene name</td><td>gene id</td></tr>
<tr><td>GU305914.1</td><td>137733</td><td>137755</td><td>-</td><td>0</td><td>CGGTGTTCCTCGTGCCCCGG</td><td>CGG</td><td>||||||||[||||||||||||]PAM</td><td>NA</td><td>Intergenic</td><td>NA</td><td>NA</td></tr>
<tr><td></td></tr>
<tr><td>T23604</td><td>TTCCTCGTGCCCCGGCGGCACGG</td><td>1000</td></tr>
<tr><td>Oligo fwd</td><td>CACCTTCCTCGTGCCCCGGCGGCA</td></tr>
<tr><td>Oligo rev</td><td>AAACTGCCGCCGGGGCACGAGGAA</td></tr>
<tr><td>Chromosome</td><td>start</td><td>end</td><td>strand</td><td>MM</td><td>target_seq</td><td>PAM</td><td>alignment</td><td>distance</td><td>position</td><td>gene name</td><td>gene id</td></tr>
<tr><td>GU305914.1</td><td>137728</td><td>137750</td><td>-</td><td>0</td><td>TTCCTCGTGCCCCGGCGGCA</td><td>CGG</td><td>||||||||[||||||||||||]PAM</td><td>NA</td><td>Intergenic</td><td>NA</td><td>NA</td></tr>
<tr><td></td></tr>
<tr><td>T23605</td><td>TCCTCGTGCCCCGGCGGCACGGG</td><td>1000</td></tr>
<tr><td>Oligo fwd</td><td>CACCTCCTCGTGCCCCGGCGGCAC</td></tr>
<tr><td>Oligo rev</td><td>AAACGTGCCGCCGGGGCACGAGGA</td></tr>
<tr><td>Chromosome</td><td>start</td><td>end</td><td>strand</td><td>MM</td><td>target_seq</td><td>PAM</td><td>alignment</td><td>distance</td><td>position</td><td>gene name</td><td>gene id</td></tr>
<tr><td>GU305914.1</td><td>137727</td><td>137749</td><td>-</td><td>0</td><td>TCCTCGTGCCCCGGCGGCAC</td><td>GGG</td><td>||||||||[||||||||||||]PAM</td><td>NA</td><td>Intergenic</td><td>NA</td><td>NA</td></tr>
<tr><td></td></tr>
<tr><td>T23606</td><td>CCCGGCGGCACGGGAAGACCTGG</td><td>1000</td></tr>
<tr><td>Oligo fwd</td><td>CACCCCCGGCGGCACGGGAAGACC</td></tr>
<tr><td>Oligo rev</td><td>AAACGGTCTTCCCGTGCCGCCGGG</td></tr>
<tr><td>Chromosome</td><td>start</td><td>end</td><td>strand</td><td>MM</td><td>target_seq</td><td>PAM</td><td>alignment</td><td>distance</td><td>position</td><td>gene name</td><td>gene id</td></tr>
<tr><td>GU305914.1</td><td>137718</td><td>137740</td><td>-</td><td>0</td><td>CCCGGCGGCACGGGAAGACC</td><td>TGG</td><td>||||||||[||||||||||||]PAM</td><td>NA</td><td>Intergenic</td><td>NA</td><td>NA</td></tr>
<tr><td></td></tr>
<tr><td>T23607</td><td>TTCCTACTGAAGAACATCATCGG</td><td>1000</td></tr>
<tr><td>Oligo fwd</td><td>CACCTTCCTACTGAAGAACATCAT</td></tr>
<tr><td>Oligo rev</td><td>AAACATGATGTTCTTCAGTAGGAA</td></tr>
<tr><td>Chromosome</td><td>start</td><td>end</td><td>strand</td><td>MM</td><td>target_seq</td><td>PAM</td><td>alignment</td><td>distance</td><td>position</td><td>gene name</td><td>gene id</td></tr>
<tr><td>GU305914.1</td><td>137674</td><td>137696</td><td>-</td><td>0</td><td>TTCCTACTGAAGAACATCAT</td><td>CGG</td><td>||||||||[||||||||||||]PAM</td><td>NA</td><td>Intergenic</td><td>NA</td><td>NA</td></tr>
<tr><td></td></tr>
<tr><td>T23608</td><td>TCCTACTGAAGAACATCATCGGG</td><td>1000</td></tr>
<tr><td>Oligo fwd</td><td>CACCTCCTACTGAAGAACATCATC</td></tr>
<tr><td>Oligo rev</td><td>AAACGATGATGTTCTTCAGTAGGA</td></tr>
<tr><td>Chromosome</td><td>start</td><td>end</td><td>strand</td><td>MM</td><td>target_seq</td><td>PAM</td><td>alignment</td><td>distance</td><td>position</td><td>gene name</td><td>gene id</td></tr>
<tr><td>GU305914.1</td><td>137673</td><td>137695</td><td>-</td><td>0</td><td>TCCTACTGAAGAACATCATC</td><td>GGG</td><td>||||||||[||||||||||||]PAM</td><td>NA</td><td>Intergenic</td><td>NA</td><td>NA</td></tr>
<tr><td></td></tr>
<tr><td>T23609</td><td>AACATCATCGGGATCAGCATCGG</td><td>1000</td></tr>
<tr><td>Oligo fwd</td><td>CACCAACATCATCGGGATCAGCAT</td></tr>
<tr><td>Oligo rev</td><td>AAACATGCTGATCCCGATGATGTT</td></tr>
<tr><td>Chromosome</td><td>start</td><td>end</td><td>strand</td><td>MM</td><td>target_seq</td><td>PAM</td><td>alignment</td><td>distance</td><td>position</td><td>gene name</td><td>gene id</td></tr>
<tr><td>GU305914.1</td><td>137662</td><td>137684</td><td>-</td><td>0</td><td>AACATCATCGGGATCAGCAT</td><td>CGG</td><td>||||||||[||||||||||||]PAM</td><td>NA</td><td>Intergenic</td><td>NA</td><td>NA</td></tr>
<tr><td></td></tr>
<tr><td>T23610</td><td>CGGGATCAGCATCGGCTACGTGG</td><td>1000</td></tr>
<tr><td>Oligo fwd</td><td>CACCCGGGATCAGCATCGGCTACG</td></tr>
<tr><td>Oligo rev</td><td>AAACCGTAGCCGATGCTGATCCCG</td></tr>
<tr><td>Chromosome</td><td>start</td><td>end</td><td>strand</td><td>MM</td><td>target_seq</td><td>PAM</td><td>alignment</td><td>distance</td><td>position</td><td>gene name</td><td>gene id</td></tr>
<tr><td>GU305914.1</td><td>137654</td><td>137676</td><td>-</td><td>0</td><td>CGGGATCAGCATCGGCTACG</td><td>TGG</td><td>||||||||[||||||||||||]PAM</td><td>NA</td><td>Intergenic</td><td>NA</td><td>NA</td></tr>
<tr><td></td></tr>
<tr><td>T23611</td><td>CTCCCAGTACGTGCTCAAAGAGG</td><td>1000</td></tr>
<tr><td>Oligo fwd</td><td>CACCCTCCCAGTACGTGCTCAAAG</td></tr>
<tr><td>Oligo rev</td><td>AAACCTTTGAGCACGTACTGGGAG</td></tr>
<tr><td>Chromosome</td><td>start</td><td>end</td><td>strand</td><td>MM</td><td>target_seq</td><td>PAM</td><td>alignment</td><td>distance</td><td>position</td><td>gene name</td><td>gene id</td></tr>
<tr><td>GU305914.1</td><td>137615</td><td>137637</td><td>-</td><td>0</td><td>CTCCCAGTACGTGCTCAAAG</td><td>AGG</td><td>||||||||[||||||||||||]PAM</td><td>NA</td><td>Intergenic</td><td>NA</td><td>NA</td></tr>
<tr><td></td></tr>
<tr><td>T23612</td><td>AGGTCGAGTTTCGCTGCAAGCGG</td><td>1000</td></tr>
<tr><td>Oligo fwd</td><td>CACCAGGTCGAGTTTCGCTGCAAG</td></tr>
<tr><td>Oligo rev</td><td>AAACCTTGCAGCGAAACTCGACCT</td></tr>
<tr><td>Chromosome</td><td>start</td><td>end</td><td>strand</td><td>MM</td><td>target_seq</td><td>PAM</td><td>alignment</td><td>distance</td><td>position</td><td>gene name</td><td>gene id</td></tr>
<tr><td>GU305914.1</td><td>137595</td><td>137617</td><td>-</td><td>0</td><td>AGGTCGAGTTTCGCTGCAAG</td><td>CGG</td><td>||||||||[||||||||||||]PAM</td><td>NA</td><td>Intergenic</td><td>NA</td><td>NA</td></tr>
<tr><td></td></tr>
<tr><td>T23613</td><td>TCGCTGCAAGCGGATGTTCTCGG</td><td>1000</td></tr>
<tr><td>Oligo fwd</td><td>CACCTCGCTGCAAGCGGATGTTCT</td></tr>
<tr><td>Oligo rev</td><td>AAACAGAACATCCGCTTGCAGCGA</td></tr>
<tr><td>Chromosome</td><td>start</td><td>end</td><td>strand</td><td>MM</td><td>target_seq</td><td>PAM</td><td>alignment</td><td>distance</td><td>position</td><td>gene name</td><td>gene id</td></tr>
<tr><td>GU305914.1</td><td>137585</td><td>137607</td><td>-</td><td>0</td><td>TCGCTGCAAGCGGATGTTCT</td><td>CGG</td><td>||||||||[||||||||||||]PAM</td><td>NA</td><td>Intergenic</td><td>NA</td><td>NA</td></tr>
<tr><td></td></tr>
<tr><td>T23614</td><td>CGCTGCAAGCGGATGTTCTCGGG</td><td>1000</td></tr>
<tr><td>Oligo fwd</td><td>CACCCGCTGCAAGCGGATGTTCTC</td></tr>
<tr><td>Oligo rev</td><td>AAACGAGAACATCCGCTTGCAGCG</td></tr>
<tr><td>Chromosome</td><td>start</td><td>end</td><td>strand</td><td>MM</td><td>target_seq</td><td>PAM</td><td>alignment</td><td>distance</td><td>position</td><td>gene name</td><td>gene id</td></tr>
<tr><td>GU305914.1</td><td>137584</td><td>137606</td><td>-</td><td>0</td><td>CGCTGCAAGCGGATGTTCTC</td><td>GGG</td><td>||||||||[||||||||||||]PAM</td><td>NA</td><td>Intergenic</td><td>NA</td><td>NA</td></tr>
<tr><td></td></tr>
<tr><td>T23615</td><td>GCAAGCGGATGTTCTCGGGCCGG</td><td>1000</td></tr>
<tr><td>Oligo fwd</td><td>CACCGCAAGCGGATGTTCTCGGGC</td></tr>
<tr><td>Oligo rev</td><td>AAACGCCCGAGAACATCCGCTTGC</td></tr>
<tr><td>Chromosome</td><td>start</td><td>end</td><td>strand</td><td>MM</td><td>target_seq</td><td>PAM</td><td>alignment</td><td>distance</td><td>position</td><td>gene name</td><td>gene id</td></tr>
<tr><td>GU305914.1</td><td>137580</td><td>137602</td><td>-</td><td>0</td><td>GCAAGCGGATGTTCTCGGGC</td><td>CGG</td><td>||||||||[||||||||||||]PAM</td><td>NA</td><td>Intergenic</td><td>NA</td><td>NA</td></tr>
</table>

<table>
<tr><td>T23616</td><td>GATGTTCTCGGGCCGGTTCATGG</td><td>1000</td><td></td><td></td><td></td><td></td><td></td><td></td><td></td><td></td><td></td></tr>
<tr><td>Oligo fwd</td><td>CACCGATGTTCTCGGGCCGGTTCA</td><td></td><td></td><td></td><td></td><td></td><td></td><td></td><td></td><td></td><td></td></tr>
<tr><td>Oligo rev</td><td>AAACTGAACCGGCCCGAGAACATC</td><td></td><td></td><td></td><td></td><td></td><td></td><td></td><td></td><td></td><td></td></tr>
<tr><td>Chromosome</td><td>start</td><td>end</td><td>strand</td><td>MM</td><td>target_seq</td><td>PAM</td><td>alignment</td><td>distance</td><td>position</td><td>gene name</td><td>gene id</td></tr>
<tr><td>GU305914.1</td><td>137573</td><td>137595</td><td>-</td><td>0</td><td>GATGTTCTCGGGCCGGTTCA</td><td>TGG</td><td>||||||||[||||||||||||]PAM</td><td>NA</td><td>Intergenic</td><td>NA</td><td>NA</td></tr>
<tr><td>T23617</td><td>CCGGTTCATGGTCGAGAACAAGG</td><td>1000</td><td></td><td></td><td></td><td></td><td></td><td></td><td></td><td></td><td></td></tr>
<tr><td>Oligo fwd</td><td>CACCCCGGTTCATGGTCGAGAACA</td><td></td><td></td><td></td><td></td><td></td><td></td><td></td><td></td><td></td><td></td></tr>
<tr><td>Oligo rev</td><td>AAACTGTTCTCGACCATGAACCGG</td><td></td><td></td><td></td><td></td><td></td><td></td><td></td><td></td><td></td><td></td></tr>
<tr><td>Chromosome</td><td>start</td><td>end</td><td>strand</td><td>MM</td><td>target_seq</td><td>PAM</td><td>alignment</td><td>distance</td><td>position</td><td>gene name</td><td>gene id</td></tr>
<tr><td>GU305914.1</td><td>137561</td><td>137583</td><td>-</td><td>0</td><td>CCGGTTCATGGTCGAGAACA</td><td>AGG</td><td>||||||||[||||||||||||]PAM</td><td>NA</td><td>Intergenic</td><td>NA</td><td>NA</td></tr>
<tr><td>GU305914.1</td><td>132507</td><td>132529</td><td>-</td><td>4</td><td>CTCGTTCATGGACGAGATCA</td><td>TGG</td><td>|--|||||[|||-|||||-||]PAM</td><td>NA</td><td>Intergenic</td><td>NA</td><td>NA</td></tr>
<tr><td>T23618</td><td>CAAGGATAATGTCATCAGCGTGG</td><td>1000</td><td></td><td></td><td></td><td></td><td></td><td></td><td></td><td></td><td></td></tr>
<tr><td>Oligo fwd</td><td>CACCCAAGGATAATGTCATCAGCG</td><td></td><td></td><td></td><td></td><td></td><td></td><td></td><td></td><td></td><td></td></tr>
<tr><td>Oligo rev</td><td>AAACCGCTGATGACATTATCCTTG</td><td></td><td></td><td></td><td></td><td></td><td></td><td></td><td></td><td></td><td></td></tr>
<tr><td>Chromosome</td><td>start</td><td>end</td><td>strand</td><td>MM</td><td>target_seq</td><td>PAM</td><td>alignment</td><td>distance</td><td>position</td><td>gene name</td><td>gene id</td></tr>
<tr><td>GU305914.1</td><td>137543</td><td>137565</td><td>-</td><td>0</td><td>CAAGGATAATGTCATCAGCG</td><td>TGG</td><td>||||||||[||||||||||||]PAM</td><td>NA</td><td>Intergenic</td><td>NA</td><td>NA</td></tr>
<tr><td>T23619</td><td>CCACAAGATCGCCAAGAGCACGG</td><td>1000</td><td></td><td></td><td></td><td></td><td></td><td></td><td></td><td></td><td></td></tr>
<tr><td>Oligo fwd</td><td>CACCCCACAAGATCGCCAAGAGCA</td><td></td><td></td><td></td><td></td><td></td><td></td><td></td><td></td><td></td><td></td></tr>
<tr><td>Oligo rev</td><td>AAACTGCTCTTGGCGATCTTGTGG</td><td></td><td></td><td></td><td></td><td></td><td></td><td></td><td></td><td></td><td></td></tr>
<tr><td>Chromosome</td><td>start</td><td>end</td><td>strand</td><td>MM</td><td>target_seq</td><td>PAM</td><td>alignment</td><td>distance</td><td>position</td><td>gene name</td><td>gene id</td></tr>
<tr><td>GU305914.1</td><td>137519</td><td>137541</td><td>-</td><td>0</td><td>CCACAAGATCGCCAAGAGCA</td><td>CGG</td><td>||||||||[||||||||||||]PAM</td><td>NA</td><td>Intergenic</td><td>NA</td><td>NA</td></tr>
<tr><td>T23620</td><td>TAACGTGAGTATATCTCAGACGG</td><td>1000</td><td></td><td></td><td></td><td></td><td></td><td></td><td></td><td></td><td></td></tr>
<tr><td>Oligo fwd</td><td>CACCTAACGTGAGTATATCTCAGA</td><td></td><td></td><td></td><td></td><td></td><td></td><td></td><td></td><td></td><td></td></tr>
<tr><td>Oligo rev</td><td>AAACTCTGAGATATACTCACGTTA</td><td></td><td></td><td></td><td></td><td></td><td></td><td></td><td></td><td></td><td></td></tr>
<tr><td>Chromosome</td><td>start</td><td>end</td><td>strand</td><td>MM</td><td>target_seq</td><td>PAM</td><td>alignment</td><td>distance</td><td>position</td><td>gene name</td><td>gene id</td></tr>
<tr><td>GU305914.1</td><td>137471</td><td>137493</td><td>-</td><td>0</td><td>TAACGTGAGTATATCTCAGA</td><td>CGG</td><td>||||||||[||||||||||||]PAM</td><td>NA</td><td>Intergenic</td><td>NA</td><td>NA</td></tr>
<tr><td>T23621</td><td>AAAAAACCAGACAAGATATGCGG</td><td>1000</td><td></td><td></td><td></td><td></td><td></td><td></td><td></td><td></td><td></td></tr>
<tr><td>Oligo fwd</td><td>CACCAAAAAACCAGACAAGATATG</td><td></td><td></td><td></td><td></td><td></td><td></td><td></td><td></td><td></td><td></td></tr>
<tr><td>Oligo rev</td><td>AAACCATATCTTGTCTGGTTTTTT</td><td></td><td></td><td></td><td></td><td></td><td></td><td></td><td></td><td></td><td></td></tr>
<tr><td>Chromosome</td><td>start</td><td>end</td><td>strand</td><td>MM</td><td>target_seq</td><td>PAM</td><td>alignment</td><td>distance</td><td>position</td><td>gene name</td><td>gene id</td></tr>
<tr><td>GU305914.1</td><td>137446</td><td>137468</td><td>-</td><td>0</td><td>AAAAAACCAGACAAGATATG</td><td>CGG</td><td>||||||||[||||||||||||]PAM</td><td>NA</td><td>Intergenic</td><td>NA</td><td>NA</td></tr>
<tr><td>T23622</td><td>AAAAACCAGACAAGATATGCGGG</td><td>1000</td><td></td><td></td><td></td><td></td><td></td><td></td><td></td><td></td><td></td></tr>
<tr><td>Oligo fwd</td><td>CACCAAAAACCAGACAAGATATGC</td><td></td><td></td><td></td><td></td><td></td><td></td><td></td><td></td><td></td><td></td></tr>
<tr><td>Oligo rev</td><td>AAACGCATATCTTGTCTGGTTTTT</td><td></td><td></td><td></td><td></td><td></td><td></td><td></td><td></td><td></td><td></td></tr>
<tr><td>Chromosome</td><td>start</td><td>end</td><td>strand</td><td>MM</td><td>target_seq</td><td>PAM</td><td>alignment</td><td>distance</td><td>position</td><td>gene name</td><td>gene id</td></tr>
<tr><td>GU305914.1</td><td>137445</td><td>137467</td><td>-</td><td>0</td><td>AAAAACCAGACAAGATATGC</td><td>GGG</td><td>||||||||[||||||||||||]PAM</td><td>NA</td><td>Intergenic</td><td>NA</td><td>NA</td></tr>
<tr><td>T23623</td><td>AACCAGACAAGATATGCGGGAGG</td><td>1000</td><td></td><td></td><td></td><td></td><td></td><td></td><td></td><td></td><td></td></tr>
<tr><td>Oligo fwd</td><td>CACCAACCAGACAAGATATGCGGG</td><td></td><td></td><td></td><td></td><td></td><td></td><td></td><td></td><td></td><td></td></tr>
<tr><td>Oligo rev</td><td>AAACCCCGCATATCTTGTCTGGTT</td><td></td><td></td><td></td><td></td><td></td><td></td><td></td><td></td><td></td><td></td></tr>
<tr><td>Chromosome</td><td>start</td><td>end</td><td>strand</td><td>MM</td><td>target_seq</td><td>PAM</td><td>alignment</td><td>distance</td><td>position</td><td>gene name</td><td>gene id</td></tr>
<tr><td>GU305914.1</td><td>137442</td><td>137464</td><td>-</td><td>0</td><td>AACCAGACAAGATATGCGGG</td><td>AGG</td><td>||||||||[||||||||||||]PAM</td><td>NA</td><td>Intergenic</td><td>NA</td><td>NA</td></tr>
<tr><td>T23624</td><td>ACCAGACAAGATATGCGGGAGGG</td><td>1000</td><td></td><td></td><td></td><td></td><td></td><td></td><td></td><td></td><td></td></tr>
<tr><td>Oligo fwd</td><td>CACCACCAGACAAGATATGCGGGA</td><td></td><td></td><td></td><td></td><td></td><td></td><td></td><td></td><td></td><td></td></tr>
<tr><td>Oligo rev</td><td>AAACTCCCGCATATCTTGTCTGGT</td><td></td><td></td><td></td><td></td><td></td><td></td><td></td><td></td><td></td><td></td></tr>
<tr><td>Chromosome</td><td>start</td><td>end</td><td>strand</td><td>MM</td><td>target_seq</td><td>PAM</td><td>alignment</td><td>distance</td><td>position</td><td>gene name</td><td>gene id</td></tr>
<tr><td>GU305914.1</td><td>137441</td><td>137463</td><td>-</td><td>0</td><td>ACCAGACAAGATATGCGGGA</td><td>GGG</td><td>||||||||[||||||||||||]PAM</td><td>NA</td><td>Intergenic</td><td>NA</td><td>NA</td></tr>
<tr><td>T23625</td><td>AGACAAGATATGCGGGAGGGAGG</td><td>1000</td><td></td><td></td><td></td><td></td><td></td><td></td><td></td><td></td><td></td></tr>
<tr><td>Oligo fwd</td><td>CACCAGACAAGATATGCGGGAGGG</td><td></td><td></td><td></td><td></td><td></td><td></td><td></td><td></td><td></td><td></td></tr>
<tr><td>Oligo rev</td><td>AAACCCCTCCCGCATATCTTGTCT</td><td></td><td></td><td></td><td></td><td></td><td></td><td></td><td></td><td></td><td></td></tr>
<tr><td>Chromosome</td><td>start</td><td>end</td><td>strand</td><td>MM</td><td>target_seq</td><td>PAM</td><td>alignment</td><td>distance</td><td>position</td><td>gene name</td><td>gene id</td></tr>
<tr><td>GU305914.1</td><td>137438</td><td>137460</td><td>-</td><td>0</td><td>AGACAAGATATGCGGGAGGG</td><td>AGG</td><td>||||||||[||||||||||||]PAM</td><td>NA</td><td>Intergenic</td><td>NA</td><td>NA</td></tr>
<tr><td>T23626</td><td>TTTTATTATTCATTGTGCCATGG</td><td>1000</td><td></td><td></td><td></td><td></td><td></td><td></td><td></td><td></td><td></td></tr>
<tr><td>Oligo fwd</td><td>CACCTTTTATTATTCATTGTGCCA</td><td></td><td></td><td></td><td></td><td></td><td></td><td></td><td></td><td></td><td></td></tr>
<tr><td>Oligo rev</td><td>AAACTGGCACAATGAATAATAAAA</td><td></td><td></td><td></td><td></td><td></td><td></td><td></td><td></td><td></td><td></td></tr>
<tr><td>Chromosome</td><td>start</td><td>end</td><td>strand</td><td>MM</td><td>target_seq</td><td>PAM</td><td>alignment</td><td>distance</td><td>position</td><td>gene name</td><td>gene id</td></tr>
<tr><td>GU305914.1</td><td>137405</td><td>137427</td><td>-</td><td>0</td><td>TTTTATTATTCATTGTGCCA</td><td>TGG</td><td>||||||||[||||||||||||]PAM</td><td>NA</td><td>Intergenic</td><td>NA</td><td>NA</td></tr>
<tr><td>T23627</td><td>ATTATTCATTGTGCCATGGATGG</td><td>1000</td><td></td><td></td><td></td><td></td><td></td><td></td><td></td><td></td><td></td></tr>
<tr><td>Oligo fwd</td><td>CACCATTATTCATTGTGCCATGGA</td><td></td><td></td><td></td><td></td><td></td><td></td><td></td><td></td><td></td><td></td></tr>
<tr><td>Oligo rev</td><td>AAACTCCATGGCACAATGAATAAT</td><td></td><td></td><td></td><td></td><td></td><td></td><td></td><td></td><td></td><td></td></tr>
<tr><td>Chromosome</td><td>start</td><td>end</td><td>strand</td><td>MM</td><td>target_seq</td><td>PAM</td><td>alignment</td><td>distance</td><td>position</td><td>gene name</td><td>gene id</td></tr>
<tr><td>GU305914.1</td><td>137401</td><td>137423</td><td>-</td><td>0</td><td>ATTATTCATTGTGCCATGGA</td><td>TGG</td><td>||||||||[||||||||||||]PAM</td><td>NA</td><td>Intergenic</td><td>NA</td><td>NA</td></tr>
<tr><td>T23628</td><td>TTATTCATTGTGCCATGGATGGG</td><td>1000</td><td></td><td></td><td></td><td></td><td></td><td></td><td></td><td></td><td></td></tr>
<tr><td>Oligo fwd</td><td>CACCTTATTCATTGTGCCATGGAT</td><td></td><td></td><td></td><td></td><td></td><td></td><td></td><td></td><td></td><td></td></tr>
<tr><td>Oligo rev</td><td>AAACATCCATGGCACAATGAATAA</td><td></td><td></td><td></td><td></td><td></td><td></td><td></td><td></td><td></td><td></td></tr>
<tr><td>Chromosome</td><td>start</td><td>end</td><td>strand</td><td>MM</td><td>target_seq</td><td>PAM</td><td>alignment</td><td>distance</td><td>position</td><td>gene name</td><td>gene id</td></tr>
<tr><td>GU305914.1</td><td>137400</td><td>137422</td><td>-</td><td>0</td><td>TTATTCATTGTGCCATGGAT</td><td>GGG</td><td>||||||||[||||||||||||]PAM</td><td>NA</td><td>Intergenic</td><td>NA</td><td>NA</td></tr>
<tr><td>T23629</td><td>TCATTGTGCCATGGATGGGTTGG</td><td>1000</td><td></td><td></td><td></td><td></td><td></td><td></td><td></td><td></td><td></td></tr>
<tr><td>Oligo fwd</td><td>CACCTCATTGTGCCATGGATGGGT</td><td></td><td></td><td></td><td></td><td></td><td></td><td></td><td></td><td></td><td></td></tr>
<tr><td>Oligo rev</td><td>AAACACCCATCCATGGCACAATGA</td><td></td><td></td><td></td><td></td><td></td><td></td><td></td><td></td><td></td><td></td></tr>
<tr><td>Chromosome</td><td>start</td><td>end</td><td>strand</td><td>MM</td><td>target_seq</td><td>PAM</td><td>alignment</td><td>distance</td><td>position</td><td>gene name</td><td>gene id</td></tr>
<tr><td>GU305914.1</td><td>137396</td><td>137418</td><td>-</td><td>0</td><td>TCATTGTGCCATGGATGGGT</td><td>TGG</td><td>||||||||[||||||||||||]PAM</td><td>NA</td><td>Intergenic</td><td>NA</td><td>NA</td></tr>
<tr><td>T23630</td><td>TGGTTGATTTGTATGTATGTCGG</td><td>1000</td><td></td><td></td><td></td><td></td><td></td><td></td><td></td><td></td><td></td></tr>
<tr><td>Oligo fwd</td><td>CACCTGGTTGATTTGTATGTATGT</td><td></td><td></td><td></td><td></td><td></td><td></td><td></td><td></td><td></td><td></td></tr>
<tr><td>Oligo rev</td><td>AAACACATACATACAAATCAACCA</td><td></td><td></td><td></td><td></td><td></td><td></td><td></td><td></td><td></td><td></td></tr>
<tr><td>Chromosome</td><td>start</td><td>end</td><td>strand</td><td>MM</td><td>target_seq</td><td>PAM</td><td>alignment</td><td>distance</td><td>position</td><td>gene name</td><td>gene id</td></tr>
</table>

<table>
<tr><td>GU305914.1</td><td>137376</td><td>137398</td><td>-</td><td>0</td><td>TGGTTGATTTGTATGTATGT</td><td>CGG</td><td>||||||||[||||||||||||]PAM</td><td>NA</td><td>Intergenic</td><td>NA</td><td>NA</td></tr>
<tr><td>T23631</td><td>GAGAACGATCTTTTTGAGTAAGG</td><td>1000</td><td colspan="9"></td></tr>
<tr><td>Oligo fwd</td><td>CACCGAGAACGATCTTTTTGAGTA</td><td colspan="10"></td></tr>
<tr><td>Oligo rev</td><td>AAACTACTCAAAAAGATCGTTCTC</td><td colspan="10"></td></tr>
<tr><td>Chromosome</td><td>start</td><td>end</td><td>strand</td><td>MM</td><td>target_seq</td><td>PAM</td><td>alignment</td><td>distance</td><td>position</td><td>gene name</td><td>gene id</td></tr>
<tr><td>GU305914.1</td><td>137337</td><td>137359</td><td>-</td><td>0</td><td>GAGAACGATCTTTTTGAGTA</td><td>AGG</td><td>||||||||[||||||||||||]PAM</td><td>NA</td><td>Intergenic</td><td>NA</td><td>NA</td></tr>
<tr><td>T23632</td><td>AGAACGATCTTTTTGAGTAAGGG</td><td>1000</td><td colspan="9"></td></tr>
<tr><td>Oligo fwd</td><td>CACCAGAACGATCTTTTTGAGTAA</td><td colspan="10"></td></tr>
<tr><td>Oligo rev</td><td>AAACTTACTCAAAAAGATCGTTCT</td><td colspan="10"></td></tr>
<tr><td>Chromosome</td><td>start</td><td>end</td><td>strand</td><td>MM</td><td>target_seq</td><td>PAM</td><td>alignment</td><td>distance</td><td>position</td><td>gene name</td><td>gene id</td></tr>
<tr><td>GU305914.1</td><td>137336</td><td>137358</td><td>-</td><td>0</td><td>AGAACGATCTTTTTGAGTAA</td><td>GGG</td><td>||||||||[||||||||||||]PAM</td><td>NA</td><td>Intergenic</td><td>NA</td><td>NA</td></tr>
<tr><td>T23633</td><td>GATCTTTTTGAGTAAGGGACAGG</td><td>1000</td><td colspan="9"></td></tr>
<tr><td>Oligo fwd</td><td>CACCGATCTTTTTGAGTAAGGGAC</td><td colspan="10"></td></tr>
<tr><td>Oligo rev</td><td>AAACGTCCCTTACTCAAAAAGATC</td><td colspan="10"></td></tr>
<tr><td>Chromosome</td><td>start</td><td>end</td><td>strand</td><td>MM</td><td>target_seq</td><td>PAM</td><td>alignment</td><td>distance</td><td>position</td><td>gene name</td><td>gene id</td></tr>
<tr><td>GU305914.1</td><td>137331</td><td>137353</td><td>-</td><td>0</td><td>GATCTTTTTGAGTAAGGGAC</td><td>AGG</td><td>||||||||[||||||||||||]PAM</td><td>NA</td><td>Intergenic</td><td>NA</td><td>NA</td></tr>
<tr><td>T23634</td><td>GGGACAGGCCATGATGATCATGG</td><td>1000</td><td colspan="9"></td></tr>
<tr><td>Oligo fwd</td><td>CACCGGGACAGGCCATGATGATCA</td><td colspan="10"></td></tr>
<tr><td>Oligo rev</td><td>AAACTGATCATCATGGCCTGTCCC</td><td colspan="10"></td></tr>
<tr><td>Chromosome</td><td>start</td><td>end</td><td>strand</td><td>MM</td><td>target_seq</td><td>PAM</td><td>alignment</td><td>distance</td><td>position</td><td>gene name</td><td>gene id</td></tr>
<tr><td>GU305914.1</td><td>137316</td><td>137338</td><td>-</td><td>0</td><td>GGGACAGGCCATGATGATCA</td><td>TGG</td><td>||||||||[||||||||||||]PAM</td><td>NA</td><td>Intergenic</td><td>NA</td><td>NA</td></tr>
<tr><td>T23635</td><td>ACAGGCCATGATGATCATGGCGG</td><td>1000</td><td colspan="9"></td></tr>
<tr><td>Oligo fwd</td><td>CACCACAGGCCATGATGATCATGG</td><td colspan="10"></td></tr>
<tr><td>Oligo rev</td><td>AAACCCATGATCATCATGGCCTGT</td><td colspan="10"></td></tr>
<tr><td>Chromosome</td><td>start</td><td>end</td><td>strand</td><td>MM</td><td>target_seq</td><td>PAM</td><td>alignment</td><td>distance</td><td>position</td><td>gene name</td><td>gene id</td></tr>
<tr><td>GU305914.1</td><td>137313</td><td>137335</td><td>-</td><td>0</td><td>ACAGGCCATGATGATCATGG</td><td>CGG</td><td>||||||||[||||||||||||]PAM</td><td>NA</td><td>Intergenic</td><td>NA</td><td>NA</td></tr>
<tr><td>T23636</td><td>ATGATCATGGCGGACAGCGTCGG</td><td>1000</td><td colspan="9"></td></tr>
<tr><td>Oligo fwd</td><td>CACCATGATCATGGCGGACAGCGT</td><td colspan="10"></td></tr>
<tr><td>Oligo rev</td><td>AAACACGCTGTCCGCCATGATCAT</td><td colspan="10"></td></tr>
<tr><td>Chromosome</td><td>start</td><td>end</td><td>strand</td><td>MM</td><td>target_seq</td><td>PAM</td><td>alignment</td><td>distance</td><td>position</td><td>gene name</td><td>gene id</td></tr>
<tr><td>GU305914.1</td><td>137303</td><td>137325</td><td>-</td><td>0</td><td>ATGATCATGGCGGACAGCGT</td><td>CGG</td><td>||||||||[||||||||||||]PAM</td><td>NA</td><td>Intergenic</td><td>NA</td><td>NA</td></tr>
<tr><td>T23637</td><td>TTCTATCTTCAAGAGCTGCCAGG</td><td>1000</td><td colspan="9"></td></tr>
<tr><td>Oligo fwd</td><td>CACCTTCTATCTTCAAGAGCTGCC</td><td colspan="10"></td></tr>
<tr><td>Oligo rev</td><td>AAACGGCAGCTCTTGAAGATAGAA</td><td colspan="10"></td></tr>
<tr><td>Chromosome</td><td>start</td><td>end</td><td>strand</td><td>MM</td><td>target_seq</td><td>PAM</td><td>alignment</td><td>distance</td><td>position</td><td>gene name</td><td>gene id</td></tr>
<tr><td>GU305914.1</td><td>137280</td><td>137302</td><td>-</td><td>0</td><td>TTCTATCTTCAAGAGCTGCC</td><td>AGG</td><td>||||||||[||||||||||||]PAM</td><td>NA</td><td>Intergenic</td><td>NA</td><td>NA</td></tr>
<tr><td>T23638</td><td>TCTATCTTCAAGAGCTGCCAGGG</td><td>1000</td><td colspan="9"></td></tr>
<tr><td>Oligo fwd</td><td>CACCTCTATCTTCAAGAGCTGCCA</td><td colspan="10"></td></tr>
<tr><td>Oligo rev</td><td>AAACTGGCAGCTCTTGAAGATAGA</td><td colspan="10"></td></tr>
<tr><td>Chromosome</td><td>start</td><td>end</td><td>strand</td><td>MM</td><td>target_seq</td><td>PAM</td><td>alignment</td><td>distance</td><td>position</td><td>gene name</td><td>gene id</td></tr>
<tr><td>GU305914.1</td><td>137279</td><td>137301</td><td>-</td><td>0</td><td>TCTATCTTCAAGAGCTGCCA</td><td>GGG</td><td>||||||||[||||||||||||]PAM</td><td>NA</td><td>Intergenic</td><td>NA</td><td>NA</td></tr>
<tr><td>T23639</td><td>CTTCAAGAGCTGCCAGGGCTCGG</td><td>1000</td><td colspan="9"></td></tr>
<tr><td>Oligo fwd</td><td>CACCCTTCAAGAGCTGCCAGGGCT</td><td colspan="10"></td></tr>
<tr><td>Oligo rev</td><td>AAACAGCCCTGGCAGCTCTTGAAG</td><td colspan="10"></td></tr>
<tr><td>Chromosome</td><td>start</td><td>end</td><td>strand</td><td>MM</td><td>target_seq</td><td>PAM</td><td>alignment</td><td>distance</td><td>position</td><td>gene name</td><td>gene id</td></tr>
<tr><td>GU305914.1</td><td>137274</td><td>137296</td><td>-</td><td>0</td><td>CTTCAAGAGCTGCCAGGGCT</td><td>CGG</td><td>||||||||[||||||||||||]PAM</td><td>NA</td><td>Intergenic</td><td>NA</td><td>NA</td></tr>
<tr><td>T23640</td><td>TGCCAGGGCTCGGTCCTCAGCGG</td><td>1000</td><td colspan="9"></td></tr>
<tr><td>Oligo fwd</td><td>CACCTGCCAGGGCTCGGTCCTCAG</td><td colspan="10"></td></tr>
<tr><td>Oligo rev</td><td>AAACCTGAGGACCGAGCCCTGGCA</td><td colspan="10"></td></tr>
<tr><td>Chromosome</td><td>start</td><td>end</td><td>strand</td><td>MM</td><td>target_seq</td><td>PAM</td><td>alignment</td><td>distance</td><td>position</td><td>gene name</td><td>gene id</td></tr>
<tr><td>GU305914.1</td><td>137264</td><td>137286</td><td>-</td><td>0</td><td>TGCCAGGGCTCGGTCCTCAG</td><td>CGG</td><td>||||||||[||||||||||||]PAM</td><td>NA</td><td>Intergenic</td><td>NA</td><td>NA</td></tr>
<tr><td>T23641</td><td>GCCAGGGCTCGGTCCTCAGCGGG</td><td>1000</td><td colspan="9"></td></tr>
<tr><td>Oligo fwd</td><td>CACCGCCAGGGCTCGGTCCTCAGC</td><td colspan="10"></td></tr>
<tr><td>Oligo rev</td><td>AAACGCTGAGGACCGAGCCCTGGC</td><td colspan="10"></td></tr>
<tr><td>Chromosome</td><td>start</td><td>end</td><td>strand</td><td>MM</td><td>target_seq</td><td>PAM</td><td>alignment</td><td>distance</td><td>position</td><td>gene name</td><td>gene id</td></tr>
<tr><td>GU305914.1</td><td>137263</td><td>137285</td><td>-</td><td>0</td><td>GCCAGGGCTCGGTCCTCAGC</td><td>GGG</td><td>||||||||[||||||||||||]PAM</td><td>NA</td><td>Intergenic</td><td>NA</td><td>NA</td></tr>
<tr><td>T23642</td><td>CTCCTCGCCCTCGCTGTTCCTGG</td><td>1000</td><td colspan="9"></td></tr>
<tr><td>Oligo fwd</td><td>CACCCTCCTCGCCCTCGCTGTTCC</td><td colspan="10"></td></tr>
<tr><td>Oligo rev</td><td>AAACGGAACAGCGAGGGCGAGGAG</td><td colspan="10"></td></tr>
<tr><td>Chromosome</td><td>start</td><td>end</td><td>strand</td><td>MM</td><td>target_seq</td><td>PAM</td><td>alignment</td><td>distance</td><td>position</td><td>gene name</td><td>gene id</td></tr>
<tr><td>GU305914.1</td><td>137238</td><td>137260</td><td>-</td><td>0</td><td>CTCCTCGCCCTCGCTGTTCC</td><td>TGG</td><td>||||||||[||||||||||||]PAM</td><td>NA</td><td>Intergenic</td><td>NA</td><td>NA</td></tr>
<tr><td>T23643</td><td>CCTCGCTGTTCCTGGCGCCGAGG</td><td>1000</td><td colspan="9"></td></tr>
<tr><td>Oligo fwd</td><td>CACCCCTCGCTGTTCCTGGCGCCG</td><td colspan="10"></td></tr>
<tr><td>Oligo rev</td><td>AAACCGGCGCCAGGAACAGCGAGG</td><td colspan="10"></td></tr>
<tr><td>Chromosome</td><td>start</td><td>end</td><td>strand</td><td>MM</td><td>target_seq</td><td>PAM</td><td>alignment</td><td>distance</td><td>position</td><td>gene name</td><td>gene id</td></tr>
<tr><td>GU305914.1</td><td>137230</td><td>137252</td><td>-</td><td>0</td><td>CCTCGCTGTTCCTGGCGCCG</td><td>AGG</td><td>||||||||[||||||||||||]PAM</td><td>NA</td><td>Intergenic</td><td>NA</td><td>NA</td></tr>
<tr><td>T23644</td><td>GCCGAGGTAGTCAGATATCGTGG</td><td>1000</td><td colspan="9"></td></tr>
<tr><td>Oligo fwd</td><td>CACCGCCGAGGTAGTCAGATATCG</td><td colspan="10"></td></tr>
<tr><td>Oligo rev</td><td>AAACCGATATCTGACTACCTCGGC</td><td colspan="10"></td></tr>
<tr><td>Chromosome</td><td>start</td><td>end</td><td>strand</td><td>MM</td><td>target_seq</td><td>PAM</td><td>alignment</td><td>distance</td><td>position</td><td>gene name</td><td>gene id</td></tr>
<tr><td>GU305914.1</td><td>137214</td><td>137236</td><td>-</td><td>0</td><td>GCCGAGGTAGTCAGATATCG</td><td>TGG</td><td>||||||||[||||||||||||]PAM</td><td>NA</td><td>Intergenic</td><td>NA</td><td>NA</td></tr>
<tr><td>T23645</td><td>AGGTAGTCAGATATCGTGGCCGG</td><td>1000</td><td colspan="9"></td></tr>
<tr><td>Oligo fwd</td><td>CACCAGGTAGTCAGATATCGTGGC</td><td colspan="10"></td></tr>
<tr><td>Oligo rev</td><td>AAACGCCACGATATCTGACTACCT</td><td colspan="10"></td></tr>
</table>

Chromosome start end strand MM target_seq PAM alignment distance position gene name gene id
GU305914.1 137210 137232 - 0 AGGTAGTCAGATATCGTGGC CGG ||||||||[||||||||||||]PAM NA Intergenic NA NA

T23646 CGTCGTAGCGCCGCACGAAGAGG 1000
Oligo fwd CACCCGTCGTAGCGCCGCACGAAG
Oligo rev AAACCTTCGTGCGGCGCTACGACG
Chromosome start end strand MM target_seq PAM alignment distance position gene name gene id
GU305914.1 137173 137195 - 0 CGTCGTAGCGCCGCACGAAG AGG ||||||||[||||||||||||]PAM NA Intergenic NA NA

T23647 TCGCACCGTCTTCCAGTTCCAGG 1000
Oligo fwd CACCTCGCACCGTCTTCCAGTTCC
Oligo rev AAACGGAACTGGAAGACGGTGCGA
Chromosome start end strand MM target_seq PAM alignment distance position gene name gene id
GU305914.1 137140 137162 - 0 TCGCACCGTCTTCCAGTTCC AGG ||||||||[||||||||||||]PAM NA Intergenic NA NA

T23648 CCGTCTTCCAGTTCCAGGTAAGG 1000
Oligo fwd CACCCCGTCTTCCAGTTCCAGGTA
Oligo rev AAACTACCTGGAACTGGAAGACGG
Chromosome start end strand MM target_seq PAM alignment distance position gene name gene id
GU305914.1 137135 137157 - 0 CCGTCTTCCAGTTCCAGGTA AGG ||||||||[||||||||||||]PAM NA Intergenic NA NA

T23649 CCAGCTTCTGACGCTGACCACGG 1000
Oligo fwd CACCCCAGCTTCTGACGCTGACCA
Oligo rev AAACTGGTCAGCGTCAGAAGCTGG
Chromosome start end strand MM target_seq PAM alignment distance position gene name gene id
GU305914.1 137109 137131 - 0 CCAGCTTCTGACGCTGACCA CGG ||||||||[||||||||||||]PAM NA Intergenic NA NA
GU305914.1 52475 52497 - 4 ACAACTTCTGACCCTGACCG TGG -||-||||[||||-|||||||-]PAM NA Intergenic NA NA

T23650 ACGGCATCCGTCCCGCGACACGG 1000
Oligo fwd CACCACGGCATCCGTCCCGCGACA
Oligo rev AAACTGTCGCGGGACGGATGCCGT
Chromosome start end strand MM target_seq PAM alignment distance position gene name gene id
GU305914.1 137090 137112 - 0 ACGGCATCCGTCCCGCGACA CGG ||||||||[||||||||||||]PAM NA Intergenic NA NA

T23651 ACAGAGACACACGCCACTGTCGG 1000
Oligo fwd CACCACAGAGACACACGCCACTGT
Oligo rev AAACACAGTGGCGTGTGTCTCTGT
Chromosome start end strand MM target_seq PAM alignment distance position gene name gene id
GU305914.1 137061 137083 - 0 ACAGAGACACACGCCACTGT CGG ||||||||[||||||||||||]PAM NA Intergenic NA NA

T23652 CAGAGACACACGCCACTGTCGGG 1000
Oligo fwd CACCCAGAGACACACGCCACTGTC
Oligo rev AAACGACAGTGGCGTGTGTCTCTG
Chromosome start end strand MM target_seq PAM alignment distance position gene name gene id
GU305914.1 137060 137082 - 0 CAGAGACACACGCCACTGTC GGG ||||||||[||||||||||||]PAM NA Intergenic NA NA

T23653 AGAGACACACGCCACTGTCGGGG 1000
Oligo fwd CACCAGAGACACACGCCACTGTCG
Oligo rev AAACCGACAGTGGCGTGTGTCTCT
Chromosome start end strand MM target_seq PAM alignment distance position gene name gene id
GU305914.1 137059 137081 - 0 AGAGACACACGCCACTGTCG GGG ||||||||[||||||||||||]PAM NA Intergenic NA NA

T23654 CGGGGACGCAGCCGACGAAGCGG 1000
Oligo fwd CACCCGGGGACGCAGCCGACGAAG
Oligo rev AAACCTTCGTCGGCTGCGTCCCCG
Chromosome start end strand MM target_seq PAM alignment distance position gene name gene id
GU305914.1 137041 137063 - 0 CGGGGACGCAGCCGACGAAG CGG ||||||||[||||||||||||]PAM NA Intergenic NA NA

T23655 GACGCAGCCGACGAAGCGGCCGG 1000
Oligo fwd CACCGACGCAGCCGACGAAGCGGC
Oligo rev AAACGCCGCTTCGTCGGCTGCGTC
Chromosome start end strand MM target_seq PAM alignment distance position gene name gene id
GU305914.1 137037 137059 - 0 GACGCAGCCGACGAAGCGGC CGG ||||||||[||||||||||||]PAM NA Intergenic NA NA
GU305914.1 60598 60620 + 4 GGTACAGCCGACGAAGCCGC CGG |---||||[|||||||||-||]PAM NA Intergenic NA NA

T23656 GCAGCCGACGAAGCGGCCGGTGG 1000
Oligo fwd CACCGCAGCCGACGAAGCGGCCGG
Oligo rev AAACCCGGCCGCTTCGTCGGCTGC
Chromosome start end strand MM target_seq PAM alignment distance position gene name gene id
GU305914.1 137034 137056 - 0 GCAGCCGACGAAGCGGCCGG TGG ||||||||[||||||||||||]PAM NA Intergenic NA NA

T23657 CAGCCGACGAAGCGGCCGGTGGG 1000
Oligo fwd CACCCAGCCGACGAAGCGGCCGGT
Oligo rev AAACACCGGCCGCTTCGTCGGCTG
Chromosome start end strand MM target_seq PAM alignment distance position gene name gene id
GU305914.1 137033 137055 - 0 CAGCCGACGAAGCGGCCGGT GGG ||||||||[||||||||||||]PAM NA Intergenic NA NA

T23658 AGCCGACGAAGCGGCCGGTGGGG 1000
Oligo fwd CACCAGCCGACGAAGCGGCCGGTG
Oligo rev AAACCACCGGCCGCTTCGTCGGCT
Chromosome start end strand MM target_seq PAM alignment distance position gene name gene id
GU305914.1 137032 137054 - 0 AGCCGACGAAGCGGCCGGTG GGG ||||||||[||||||||||||]PAM NA Intergenic NA NA

T23659 GGTGGGGTCGACGTGCCTGCCGG 1000
Oligo fwd CACCGGTGGGGTCGACGTGCCTGC
Oligo rev AAACGCAGGCACGTCGACCCCACC
Chromosome start end strand MM target_seq PAM alignment distance position gene name gene id
GU305914.1 137016 137038 - 0 GGTGGGGTCGACGTGCCTGC CGG ||||||||[||||||||||||]PAM NA Intergenic NA NA

| | | | | | | | | | | | |
|---|---|---|---|---|---|---|---|---|---|---|---|
| T23660 | TCGACAGAATCCGCACTTGTCGG | 1000 | | | | | | | | | |
| Oligo fwd | CACCTCGACAGAATCCGCACTTGT | | | | | | | | | | |
| Oligo rev | AAACACAAGTGCGGATTCTGTCGA | | | | | | | | | | |
| Chromosome | start | end | strand | MM | target_seq | PAM | alignment | distance | position | gene name | gene id |
| GU305914.1 | 136989 | 137011 | - | 0 | TCGACAGAATCCGCACTTGT | CGG | \|\|\|\|\|\|\|\|[\|\|\|\|\|\|\|\|\|\|\|\|]PAM | NA | Intergenic | NA | NA |

| | | | | | | | | | | | |
|---|---|---|---|---|---|---|---|---|---|---|---|
| T23661 | TCCGCACTTGTCGGCCTCGCAGG | 1000 | | | | | | | | | |
| Oligo fwd | CACCTCCGCACTTGTCGGCCTCGC | | | | | | | | | | |
| Oligo rev | AAACGCGAGGCCGACAAGTGCGGA | | | | | | | | | | |
| Chromosome | start | end | strand | MM | target_seq | PAM | alignment | distance | position | gene name | gene id |
| GU305914.1 | 136980 | 137002 | - | 0 | TCCGCACTTGTCGGCCTCGC | AGG | \|\|\|\|\|\|\|\|[\|\|\|\|\|\|\|\|\|\|\|\|]PAM | NA | Intergenic | NA | NA |

| | | | | | | | | | | | |
|---|---|---|---|---|---|---|---|---|---|---|---|
| T23662 | CACAGTTTCGCTGTAGATCTTGG | 1000 | | | | | | | | | |
| Oligo fwd | CACCCACAGTTTCGCTGTAGATCT | | | | | | | | | | |
| Oligo rev | AAACAGATCTACAGCGAAACTGTG | | | | | | | | | | |
| Chromosome | start | end | strand | MM | target_seq | PAM | alignment | distance | position | gene name | gene id |
| GU305914.1 | 136956 | 136978 | - | 0 | CACAGTTTCGCTGTAGATCT | TGG | \|\|\|\|\|\|\|\|[\|\|\|\|\|\|\|\|\|\|\|\|]PAM | NA | Intergenic | NA | NA |

| | | | | | | | | | | | |
|---|---|---|---|---|---|---|---|---|---|---|---|
| T23663 | TTTCGCTGTAGATCTTGGCCAGG | 1000 | | | | | | | | | |
| Oligo fwd | CACCTTTCGCTGTAGATCTTGGCC | | | | | | | | | | |
| Oligo rev | AAACGGCCAAGATCTACAGCGAAA | | | | | | | | | | |
| Chromosome | start | end | strand | MM | target_seq | PAM | alignment | distance | position | gene name | gene id |
| GU305914.1 | 136951 | 136973 | - | 0 | TTTCGCTGTAGATCTTGGCC | AGG | \|\|\|\|\|\|\|\|[\|\|\|\|\|\|\|\|\|\|\|\|]PAM | NA | Intergenic | NA | NA |

| | | | | | | | | | | | |
|---|---|---|---|---|---|---|---|---|---|---|---|
| T23664 | TTCGCTGTAGATCTTGGCCAGGG | 1000 | | | | | | | | | |
| Oligo fwd | CACCTTCGCTGTAGATCTTGGCCA | | | | | | | | | | |
| Oligo rev | AAACTGGCCAAGATCTACAGCGAA | | | | | | | | | | |
| Chromosome | start | end | strand | MM | target_seq | PAM | alignment | distance | position | gene name | gene id |
| GU305914.1 | 136950 | 136972 | - | 0 | TTCGCTGTAGATCTTGGCCA | GGG | \|\|\|\|\|\|\|\|[\|\|\|\|\|\|\|\|\|\|\|\|]PAM | NA | Intergenic | NA | NA |

| | | | | | | | | | | | |
|---|---|---|---|---|---|---|---|---|---|---|---|
| T23665 | CTGTAGATCTTGGCCAGGGACGG | 1000 | | | | | | | | | |
| Oligo fwd | CACCCTGTAGATCTTGGCCAGGGA | | | | | | | | | | |
| Oligo rev | AAACTCCCTGGCCAAGATCTACAG | | | | | | | | | | |
| Chromosome | start | end | strand | MM | target_seq | PAM | alignment | distance | position | gene name | gene id |
| GU305914.1 | 136946 | 136968 | - | 0 | CTGTAGATCTTGGCCAGGGA | CGG | \|\|\|\|\|\|\|\|[\|\|\|\|\|\|\|\|\|\|\|\|]PAM | NA | Intergenic | NA | NA |

| | | | | | | | | | | | |
|---|---|---|---|---|---|---|---|---|---|---|---|
| T23666 | TGTAGATCTTGGCCAGGGACGGG | 1000 | | | | | | | | | |
| Oligo fwd | CACCTGTAGATCTTGGCCAGGGAC | | | | | | | | | | |
| Oligo rev | AAACGTCCCTGGCCAAGATCTACA | | | | | | | | | | |
| Chromosome | start | end | strand | MM | target_seq | PAM | alignment | distance | position | gene name | gene id |
| GU305914.1 | 136945 | 136967 | - | 0 | TGTAGATCTTGGCCAGGGAC | GGG | \|\|\|\|\|\|\|\|[\|\|\|\|\|\|\|\|\|\|\|\|]PAM | NA | Intergenic | NA | NA |

| | | | | | | | | | | | |
|---|---|---|---|---|---|---|---|---|---|---|---|
| T23667 | GATCTTGGCCAGGGACGGGAAGG | 1000 | | | | | | | | | |
| Oligo fwd | CACCGATCTTGGCCAGGGACGGGA | | | | | | | | | | |
| Oligo rev | AAACTCCCGTCCCTGGCCAAGATC | | | | | | | | | | |
| Chromosome | start | end | strand | MM | target_seq | PAM | alignment | distance | position | gene name | gene id |
| GU305914.1 | 136941 | 136963 | - | 0 | GATCTTGGCCAGGGACGGGA | AGG | \|\|\|\|\|\|\|\|[\|\|\|\|\|\|\|\|\|\|\|\|]PAM | NA | Intergenic | NA | NA |

| | | | | | | | | | | | |
|---|---|---|---|---|---|---|---|---|---|---|---|
| T23668 | GAAGGCGAGCCGATCGCAGCAGG | 1000 | | | | | | | | | |
| Oligo fwd | CACCGAAGGCGAGCCGATCGCAGC | | | | | | | | | | |
| Oligo rev | AAACGCTGCGATCGGCTCGCCTTC | | | | | | | | | | |
| Chromosome | start | end | strand | MM | target_seq | PAM | alignment | distance | position | gene name | gene id |
| GU305914.1 | 136923 | 136945 | - | 0 | GAAGGCGAGCCGATCGCAGC | AGG | \|\|\|\|\|\|\|\|[\|\|\|\|\|\|\|\|\|\|\|\|]PAM | NA | Intergenic | NA | NA |

| | | | | | | | | | | | |
|---|---|---|---|---|---|---|---|---|---|---|---|
| T23669 | GGCGAGCCGATCGCAGCAGGCGG | 1000 | | | | | | | | | |
| Oligo fwd | CACCGGCGAGCCGATCGCAGCAGG | | | | | | | | | | |
| Oligo rev | AAACCCTGCTGCGATCGGCTCGCC | | | | | | | | | | |
| Chromosome | start | end | strand | MM | target_seq | PAM | alignment | distance | position | gene name | gene id |
| GU305914.1 | 136920 | 136942 | - | 0 | GGCGAGCCGATCGCAGCAGG | CGG | \|\|\|\|\|\|\|\|[\|\|\|\|\|\|\|\|\|\|\|\|]PAM | NA | Intergenic | NA | NA |

| | | | | | | | | | | | |
|---|---|---|---|---|---|---|---|---|---|---|---|
| T23670 | CGGTAAAGACGTCGAAGCAGAGG | 1000 | | | | | | | | | |
| Oligo fwd | CACCCGGTAAAGACGTCGAAGCAG | | | | | | | | | | |
| Oligo rev | AAACCTGCTTCGACGTCTTTACCG | | | | | | | | | | |
| Chromosome | start | end | strand | MM | target_seq | PAM | alignment | distance | position | gene name | gene id |
| GU305914.1 | 136900 | 136922 | - | 0 | CGGTAAAGACGTCGAAGCAG | AGG | \|\|\|\|\|\|\|\|[\|\|\|\|\|\|\|\|\|\|\|\|]PAM | NA | Intergenic | NA | NA |

| | | | | | | | | | | | |
|---|---|---|---|---|---|---|---|---|---|---|---|
| T23671 | ACCATCCTTGATCTGCACCCAGG | 1000 | | | | | | | | | |
| Oligo fwd | CACCACCATCCTTGATCTGCACCC | | | | | | | | | | |
| Oligo rev | AAACGGGTGCAGATCAAGGATGGT | | | | | | | | | | |
| Chromosome | start | end | strand | MM | target_seq | PAM | alignment | distance | position | gene name | gene id |
| GU305914.1 | 136851 | 136873 | - | 0 | ACCATCCTTGATCTGCACCC | AGG | \|\|\|\|\|\|\|\|[\|\|\|\|\|\|\|\|\|\|\|\|]PAM | NA | Intergenic | NA | NA |

| | | | | | | | | | | | |
|---|---|---|---|---|---|---|---|---|---|---|---|
| T23672 | GCCAACGCCACTGAGTCCCACGG | 1000 | | | | | | | | | |
| Oligo fwd | CACCGCCAACGCCACTGAGTCCCA | | | | | | | | | | |
| Oligo rev | AAACTGGGACTCAGTGGCGTTGGC | | | | | | | | | | |
| Chromosome | start | end | strand | MM | target_seq | PAM | alignment | distance | position | gene name | gene id |
| GU305914.1 | 136827 | 136849 | - | 0 | GCCAACGCCACTGAGTCCCA | CGG | \|\|\|\|\|\|\|\|[\|\|\|\|\|\|\|\|\|\|\|\|]PAM | NA | Intergenic | NA | NA |

| | | | | | | | | | | | |
|---|---|---|---|---|---|---|---|---|---|---|---|
| T23673 | GAGTCCCACGGTCACGATGCCGG | 1000 | | | | | | | | | |
| Oligo fwd | CACCGAGTCCCACGGTCACGATGC | | | | | | | | | | |
| Oligo rev | AAACGCATCGTGACCGTGGGACTC | | | | | | | | | | |
| Chromosome | start | end | strand | MM | target_seq | PAM | alignment | distance | position | gene name | gene id |
| GU305914.1 | 136815 | 136837 | - | 0 | GAGTCCCACGGTCACGATGC | CGG | \|\|\|\|\|\|\|\|[\|\|\|\|\|\|\|\|\|\|\|\|]PAM | NA | Intergenic | NA | NA |

| | | | | | | | | | | | |
|---|---|---|---|---|---|---|---|---|---|---|---|
| T23674 | GCCGGAACCCTGCATAGTGACGG | 1000 | | | | | | | | | |
| Oligo fwd | CACCGCCGGAACCCTGCATAGTGA | | | | | | | | | | |
| Oligo rev | AAACTCACTATGCAGGGTTCCGGC | | | | | | | | | | |
| Chromosome | start | end | strand | MM | target_seq | PAM | alignment | distance | position | gene name | gene id |
| GU305914.1 | 136797 | 136819 | - | 0 | GCCGGAACCCTGCATAGTGA | CGG | \|\|\|\|\|\|\|\|[\|\|\|\|\|\|\|\|\|\|\|\|]PAM | NA | Intergenic | NA | NA |

| | | | | | | | | | | | |
|---|---|---|---|---|---|---|---|---|---|---|---|
| T23675 | GGAACCCTGCATAGTGACGGTGG | 1000 | | | | | | | | | |
| Oligo fwd | CACCGGAACCCTGCATAGTGACGG | | | | | | | | | | |
| Oligo rev | AAACCCGTCACTATGCAGGGTTCC | | | | | | | | | | |
| Chromosome | start | end | strand | MM | target_seq | PAM | alignment | distance | position | gene name | gene id |
| GU305914.1 | 136794 | 136816 | - | 0 | GGAACCCTGCATAGTGACGG | TGG | \|\|\|\|\|\|\|\|[\|\|\|\|\|\|\|\|\|\|\|\|]PAM | NA | Intergenic | NA | NA |
| T23676 | ACCCTGCATAGTGACGGTGGCGG | 1000 | | | | | | | | | |
| Oligo fwd | CACCACCCTGCATAGTGACGGTGG | | | | | | | | | | |
| Oligo rev | AAACCCACCGTCACTATGCAGGGT | | | | | | | | | | |
| Chromosome | start | end | strand | MM | target_seq | PAM | alignment | distance | position | gene name | gene id |
| GU305914.1 | 136791 | 136813 | - | 0 | ACCCTGCATAGTGACGGTGG | CGG | \|\|\|\|\|\|\|\|[\|\|\|\|\|\|\|\|\|\|\|\|]PAM | NA | Intergenic | NA | NA |
| T23677 | AGTGACGGTGGCGGCCTCTTCGG | 1000 | | | | | | | | | |
| Oligo fwd | CACCAGTGACGGTGGCGGCCTCTT | | | | | | | | | | |
| Oligo rev | AAACAAGAGGCCGCCACCGTCACT | | | | | | | | | | |
| Chromosome | start | end | strand | MM | target_seq | PAM | alignment | distance | position | gene name | gene id |
| GU305914.1 | 136782 | 136804 | - | 0 | AGTGACGGTGGCGGCCTCTT | CGG | \|\|\|\|\|\|\|\|[\|\|\|\|\|\|\|\|\|\|\|\|]PAM | NA | Intergenic | NA | NA |
| T23678 | GTGACGGTGGCGGCCTCTTCGGG | 1000 | | | | | | | | | |
| Oligo fwd | CACCGTGACGGTGGCGGCCTCTTC | | | | | | | | | | |
| Oligo rev | AAACGAAGAGGCCGCCACCGTCAC | | | | | | | | | | |
| Chromosome | start | end | strand | MM | target_seq | PAM | alignment | distance | position | gene name | gene id |
| GU305914.1 | 136781 | 136803 | - | 0 | GTGACGGTGGCGGCCTCTTC | GGG | \|\|\|\|\|\|\|\|[\|\|\|\|\|\|\|\|\|\|\|\|]PAM | NA | Intergenic | NA | NA |
| T23679 | GCGTGACCACACTCGCGTTCTGG | 1000 | | | | | | | | | |
| Oligo fwd | CACCGCGTGACCACACTCGCGTTC | | | | | | | | | | |
| Oligo rev | AAACGAACGCGAGTGTGGTCACGC | | | | | | | | | | |
| Chromosome | start | end | strand | MM | target_seq | PAM | alignment | distance | position | gene name | gene id |
| GU305914.1 | 136759 | 136781 | - | 0 | GCGTGACCACACTCGCGTTC | TGG | \|\|\|\|\|\|\|\|[\|\|\|\|\|\|\|\|\|\|\|\|]PAM | NA | Intergenic | NA | NA |
| T23680 | CGCGTTCTGGTAGATGACGCTGG | 1000 | | | | | | | | | |
| Oligo fwd | CACCCGCGTTCTGGTAGATGACGC | | | | | | | | | | |
| Oligo rev | AAACGCGTCATCTACCAGAACGCG | | | | | | | | | | |
| Chromosome | start | end | strand | MM | target_seq | PAM | alignment | distance | position | gene name | gene id |
| GU305914.1 | 136746 | 136768 | - | 0 | CGCGTTCTGGTAGATGACGC | TGG | \|\|\|\|\|\|\|\|[\|\|\|\|\|\|\|\|\|\|\|\|]PAM | NA | Intergenic | NA | NA |
| T23681 | GCGTTCTGGTAGATGACGCTGGG | 1000 | | | | | | | | | |
| Oligo fwd | CACCGCGTTCTGGTAGATGACGCT | | | | | | | | | | |
| Oligo rev | AAACAGCGTCATCTACCAGAACGC | | | | | | | | | | |
| Chromosome | start | end | strand | MM | target_seq | PAM | alignment | distance | position | gene name | gene id |
| GU305914.1 | 136745 | 136767 | - | 0 | GCGTTCTGGTAGATGACGCT | GGG | \|\|\|\|\|\|\|\|[\|\|\|\|\|\|\|\|\|\|\|\|]PAM | NA | Intergenic | NA | NA |
| T23682 | CGTTCTGGTAGATGACGCTGGGG | 1000 | | | | | | | | | |
| Oligo fwd | CACCCGTTCTGGTAGATGACGCTG | | | | | | | | | | |
| Oligo rev | AAACCAGCGTCATCTACCAGAACG | | | | | | | | | | |
| Chromosome | start | end | strand | MM | target_seq | PAM | alignment | distance | position | gene name | gene id |
| GU305914.1 | 136744 | 136766 | - | 0 | CGTTCTGGTAGATGACGCTG | GGG | \|\|\|\|\|\|\|\|[\|\|\|\|\|\|\|\|\|\|\|\|]PAM | NA | Intergenic | NA | NA |
| GU305914.1 | 177596 | 177618 | + | 4 | CGTTCTCATAGTTGACGATG | AGG | \|\|\|\|\|\|--[\|\|\|-\|\|\|\|\|-\|\|]PAM | NA | Intergenic | NA | NA |
| T23683 | GACGCTGGGGTCCAGACCGCTGG | 1000 | | | | | | | | | |
| Oligo fwd | CACCGACGCTGGGGTCCAGACCGC | | | | | | | | | | |
| Oligo rev | AAACGCGGTCTGGACCCCAGCGTC | | | | | | | | | | |
| Chromosome | start | end | strand | MM | target_seq | PAM | alignment | distance | position | gene name | gene id |
| GU305914.1 | 136731 | 136753 | - | 0 | GACGCTGGGGTCCAGACCGC | TGG | \|\|\|\|\|\|\|\|[\|\|\|\|\|\|\|\|\|\|\|\|]PAM | NA | Intergenic | NA | NA |
| T23684 | CTGGAGCCGCCGATGCTCTTAGG | 1000 | | | | | | | | | |
| Oligo fwd | CACCCTGGAGCCGCCGATGCTCTT | | | | | | | | | | |
| Oligo rev | AAACAAGAGCATCGGCGGCTCCAG | | | | | | | | | | |
| Chromosome | start | end | strand | MM | target_seq | PAM | alignment | distance | position | gene name | gene id |
| GU305914.1 | 136712 | 136734 | - | 0 | CTGGAGCCGCCGATGCTCTT | AGG | \|\|\|\|\|\|\|\|[\|\|\|\|\|\|\|\|\|\|\|\|]PAM | NA | Intergenic | NA | NA |
| T23685 | GAGCCGCCGATGCTCTTAGGCGG | 1000 | | | | | | | | | |
| Oligo fwd | CACCGAGCCGCCGATGCTCTTAGG | | | | | | | | | | |
| Oligo rev | AAACCCTAAGAGCATCGGCGGCTC | | | | | | | | | | |
| Chromosome | start | end | strand | MM | target_seq | PAM | alignment | distance | position | gene name | gene id |
| GU305914.1 | 136709 | 136731 | - | 0 | GAGCCGCCGATGCTCTTAGG | CGG | \|\|\|\|\|\|\|\|[\|\|\|\|\|\|\|\|\|\|\|\|]PAM | NA | Intergenic | NA | NA |
| T23686 | GCTCTTAGGCGGCAACATGTAGG | 1000 | | | | | | | | | |
| Oligo fwd | CACCGCTCTTAGGCGGCAACATGT | | | | | | | | | | |
| Oligo rev | AAACACATGTTGCCGCCTAAGAGC | | | | | | | | | | |
| Chromosome | start | end | strand | MM | target_seq | PAM | alignment | distance | position | gene name | gene id |
| GU305914.1 | 136698 | 136720 | - | 0 | GCTCTTAGGCGGCAACATGT | AGG | \|\|\|\|\|\|\|\|[\|\|\|\|\|\|\|\|\|\|\|\|]PAM | NA | Intergenic | NA | NA |
| T23687 | CTCTTAGGCGGCAACATGTAGGG | 1000 | | | | | | | | | |
| Oligo fwd | CACCCTCTTAGGCGGCAACATGTA | | | | | | | | | | |
| Oligo rev | AAACTACATGTTGCCGCCTAAGAG | | | | | | | | | | |
| Chromosome | start | end | strand | MM | target_seq | PAM | alignment | distance | position | gene name | gene id |
| GU305914.1 | 136697 | 136719 | - | 0 | CTCTTAGGCGGCAACATGTA | GGG | \|\|\|\|\|\|\|\|[\|\|\|\|\|\|\|\|\|\|\|\|]PAM | NA | Intergenic | NA | NA |
| T23688 | GGGCGAGAATCCTAGCTTGACGG | 1000 | | | | | | | | | |
| Oligo fwd | CACCGGGCGAGAATCCTAGCTTGA | | | | | | | | | | |
| Oligo rev | AAACTCAAGCTAGGATTCTCGCCC | | | | | | | | | | |
| Chromosome | start | end | strand | MM | target_seq | PAM | alignment | distance | position | gene name | gene id |
| GU305914.1 | 136677 | 136699 | - | 0 | GGGCGAGAATCCTAGCTTGA | CGG | \|\|\|\|\|\|\|\|[\|\|\|\|\|\|\|\|\|\|\|\|]PAM | NA | Intergenic | NA | NA |
| T23689 | GGCGAGAATCCTAGCTTGACGGG | 1000 | | | | | | | | | |
| Oligo fwd | CACCGGCGAGAATCCTAGCTTGAC | | | | | | | | | | |
| Oligo rev | AAACGTCAAGCTAGGATTCTCGCC | | | | | | | | | | |

Chromosome start end strand MM target_seq PAM alignment distance position gene name gene id
GU305914.1 136676 136698 - 0 GGCGAGAATCCTAGCTTGAC GGG ||||||||[||||||||||||]PAM NA Intergenic NA NA

T23690 GCGAGAATCCTAGCTTGACGGGG 1000
Oligo fwd CACCGCGAGAATCCTAGCTTGACG
Oligo rev AAACCGTCAAGCTAGGATTCTCGC
Chromosome start end strand MM target_seq PAM alignment distance position gene name gene id
GU305914.1 136675 136697 - 0 GCGAGAATCCTAGCTTGACG GGG ||||||||[||||||||||||]PAM NA Intergenic NA NA

T23691 TCCTAGCTTGACGGGGATGATGG 1000
Oligo fwd CACCTCCTAGCTTGACGGGGATGA
Oligo rev AAACTCATCCCCGTCAAGCTAGGA
Chromosome start end strand MM target_seq PAM alignment distance position gene name gene id
GU305914.1 136668 136690 - 0 TCCTAGCTTGACGGGGATGA TGG ||||||||[||||||||||||]PAM NA Intergenic NA NA

T23692 CCTAGCTTGACGGGGATGATGGG 1000
Oligo fwd CACCCCTAGCTTGACGGGGATGAT
Oligo rev AAACATCATCCCCGTCAAGCTAGG
Chromosome start end strand MM target_seq PAM alignment distance position gene name gene id
GU305914.1 136667 136689 - 0 CCTAGCTTGACGGGGATGAT GGG ||||||||[||||||||||||]PAM NA Intergenic NA NA

T23693 CTAGCTTGACGGGGATGATGGGG 1000
Oligo fwd CACCCTAGCTTGACGGGGATGATG
Oligo rev AAACCATCATCCCCGTCAAGCTAG
Chromosome start end strand MM target_seq PAM alignment distance position gene name gene id
GU305914.1 136666 136688 - 0 CTAGCTTGACGGGGATGATG GGG ||||||||[||||||||||||]PAM NA Intergenic NA NA

T23694 TAGCTTGACGGGGATGATGGGGG 1000
Oligo fwd CACCTAGCTTGACGGGGATGATGG
Oligo rev AAACCCATCATCCCCGTCAAGCTA
Chromosome start end strand MM target_seq PAM alignment distance position gene name gene id
GU305914.1 136665 136687 - 0 TAGCTTGACGGGGATGATGG GGG ||||||||[||||||||||||]PAM NA Intergenic NA NA

T23695 TGACGGGGATGATGGGGGCCAGG 1000
Oligo fwd CACCTGACGGGGATGATGGGGGCC
Oligo rev AAACGGCCCCCATCATCCCCGTCA
Chromosome start end strand MM target_seq PAM alignment distance position gene name gene id
GU305914.1 136660 136682 - 0 TGACGGGGATGATGGGGGCC AGG ||||||||[||||||||||||]PAM NA Intergenic NA NA

T23696 CAGGAGACGCATCACGTAGACGG 1000
Oligo fwd CACCCAGGAGACGCATCACGTAGA
Oligo rev AAACTCTACGTGATGCGTCTCCTG
Chromosome start end strand MM target_seq PAM alignment distance position gene name gene id
GU305914.1 136641 136663 - 0 CAGGAGACGCATCACGTAGA CGG ||||||||[||||||||||||]PAM NA Intergenic NA NA

T23697 AGGAGACGCATCACGTAGACGGG 1000
Oligo fwd CACCAGGAGACGCATCACGTAGAC
Oligo rev AAACGTCTACGTGATGCGTCTCCT
Chromosome start end strand MM target_seq PAM alignment distance position gene name gene id
GU305914.1 136640 136662 - 0 AGGAGACGCATCACGTAGAC GGG ||||||||[||||||||||||]PAM NA Intergenic NA NA

T23698 GCATCACGTAGACGGGCTCCTGG 1000
Oligo fwd CACCGCATCACGTAGACGGGCTCC
Oligo rev AAACGGAGCCCGTCTACGTGATGC
Chromosome start end strand MM target_seq PAM alignment distance position gene name gene id
GU305914.1 136633 136655 - 0 GCATCACGTAGACGGGCTCC TGG ||||||||[||||||||||||]PAM NA Intergenic NA NA

T23699 CTGCAGCCGAACTGACCCACCGG 1000
Oligo fwd CACCCTGCAGCCGAACTGACCCAC
Oligo rev AAACGTGGGTCAGTTCGGCTGCAG
Chromosome start end strand MM target_seq PAM alignment distance position gene name gene id
GU305914.1 136574 136596 - 0 CTGCAGCCGAACTGACCCAC CGG ||||||||[||||||||||||]PAM NA Intergenic NA NA

T23700 GTCACGTCTGAAAAGCATCATGG 1000
Oligo fwd CACCGTCACGTCTGAAAAGCATCA
Oligo rev AAACTGATGCTTTTCAGACGTGAC
Chromosome start end strand MM target_seq PAM alignment distance position gene name gene id
GU305914.1 136515 136537 - 0 GTCACGTCTGAAAAGCATCA TGG ||||||||[||||||||||||]PAM NA Intergenic NA NA

T23701 ACGTCTGAAAAGCATCATGGAGG 1000
Oligo fwd CACCACGTCTGAAAAGCATCATGG
Oligo rev AAACCCATGATGCTTTTCAGACGT
Chromosome start end strand MM target_seq PAM alignment distance position gene name gene id
GU305914.1 136512 136534 - 0 ACGTCTGAAAAGCATCATGG AGG ||||||||[||||||||||||]PAM NA Intergenic NA NA

T23702 TCTGAAAAGCATCATGGAGGCGG 1000
Oligo fwd CACCTCTGAAAAGCATCATGGAGG
Oligo rev AAACCCTCCATGATGCTTTTCAGA
Chromosome start end strand MM target_seq PAM alignment distance position gene name gene id
GU305914.1 136509 136531 - 0 TCTGAAAAGCATCATGGAGG CGG ||||||||[||||||||||||]PAM NA Intergenic NA NA

T23703 GGCGAGACATCTCCCGTCGTCGG 1000
Oligo fwd CACCGGCGAGACATCTCCCGTCGT
Oligo rev AAACACGACGGGAGATGTCTCGCC
Chromosome start end strand MM target_seq PAM alignment distance position gene name gene id
GU305914.1 136488 136510 - 0 GGCGAGACATCTCCCGTCGT CGG ||||||||[||||||||||||]PAM NA Intergenic NA NA

T23704 GAGACATCTCCCGTCGTCGGTGG 1000
Oligo fwd CACCGAGACATCTCCCGTCGTCGG

<table>
<tr><td>Oligo rev</td><td>AAACCCGACGACGGGAGATGTCTC</td><td></td><td></td><td></td><td></td><td></td><td></td><td></td><td></td><td></td><td></td></tr>
<tr><td>Chromosome</td><td>start</td><td>end</td><td>strand</td><td>MM</td><td>target_seq</td><td>PAM</td><td>alignment</td><td>distance</td><td>position</td><td>gene name</td><td>gene id</td></tr>
<tr><td>GU305914.1</td><td>136485</td><td>136507</td><td>-</td><td>0</td><td>GAGACATCTCCCGTCGTCGG</td><td>TGG</td><td>||||||||[||||||||||||]PAM</td><td>NA</td><td>Intergenic</td><td>NA</td><td>NA</td></tr>
<tr><td></td><td></td><td></td><td></td><td></td><td></td><td></td><td></td><td></td><td></td><td></td><td></td></tr>
<tr><td>T23705</td><td>ACATCTCCCGTCGTCGGTGGAGG</td><td>1000</td><td></td><td></td><td></td><td></td><td></td><td></td><td></td><td></td><td></td></tr>
<tr><td>Oligo fwd</td><td>CACCACATCTCCCGTCGTCGGTGG</td><td></td><td></td><td></td><td></td><td></td><td></td><td></td><td></td><td></td><td></td></tr>
<tr><td>Oligo rev</td><td>AAACCCACCGACGACGGGAGATGT</td><td></td><td></td><td></td><td></td><td></td><td></td><td></td><td></td><td></td><td></td></tr>
<tr><td>Chromosome</td><td>start</td><td>end</td><td>strand</td><td>MM</td><td>target_seq</td><td>PAM</td><td>alignment</td><td>distance</td><td>position</td><td>gene name</td><td>gene id</td></tr>
<tr><td>GU305914.1</td><td>136482</td><td>136504</td><td>-</td><td>0</td><td>ACATCTCCCGTCGTCGGTGG</td><td>AGG</td><td>||||||||[||||||||||||]PAM</td><td>NA</td><td>Intergenic</td><td>NA</td><td>NA</td></tr>
<tr><td></td><td></td><td></td><td></td><td></td><td></td><td></td><td></td><td></td><td></td><td></td><td></td></tr>
<tr><td>T23706</td><td>GAGGAGACCTTCCCCAGCCCCGG</td><td>1000</td><td></td><td></td><td></td><td></td><td></td><td></td><td></td><td></td><td></td></tr>
<tr><td>Oligo fwd</td><td>CACCGAGGAGACCTTCCCCAGCCC</td><td></td><td></td><td></td><td></td><td></td><td></td><td></td><td></td><td></td><td></td></tr>
<tr><td>Oligo rev</td><td>AAACGGGCTGGGGAAGGTCTCCTC</td><td></td><td></td><td></td><td></td><td></td><td></td><td></td><td></td><td></td><td></td></tr>
<tr><td>Chromosome</td><td>start</td><td>end</td><td>strand</td><td>MM</td><td>target_seq</td><td>PAM</td><td>alignment</td><td>distance</td><td>position</td><td>gene name</td><td>gene id</td></tr>
<tr><td>GU305914.1</td><td>136463</td><td>136485</td><td>-</td><td>0</td><td>GAGGAGACCTTCCCCAGCCC</td><td>CGG</td><td>||||||||[||||||||||||]PAM</td><td>NA</td><td>Intergenic</td><td>NA</td><td>NA</td></tr>
<tr><td></td><td></td><td></td><td></td><td></td><td></td><td></td><td></td><td></td><td></td><td></td><td></td></tr>
<tr><td>T23707</td><td>CCTTCCCCAGCCCCGGCGACTGG</td><td>1000</td><td></td><td></td><td></td><td></td><td></td><td></td><td></td><td></td><td></td></tr>
<tr><td>Oligo fwd</td><td>CACCCCTTCCCCAGCCCCGGCGAC</td><td></td><td></td><td></td><td></td><td></td><td></td><td></td><td></td><td></td><td></td></tr>
<tr><td>Oligo rev</td><td>AAACGTCGCCGGGGCTGGGGAAGG</td><td></td><td></td><td></td><td></td><td></td><td></td><td></td><td></td><td></td><td></td></tr>
<tr><td>Chromosome</td><td>start</td><td>end</td><td>strand</td><td>MM</td><td>target_seq</td><td>PAM</td><td>alignment</td><td>distance</td><td>position</td><td>gene name</td><td>gene id</td></tr>
<tr><td>GU305914.1</td><td>136456</td><td>136478</td><td>-</td><td>0</td><td>CCTTCCCCAGCCCCGGCGAC</td><td>TGG</td><td>||||||||[||||||||||||]PAM</td><td>NA</td><td>Intergenic</td><td>NA</td><td>NA</td></tr>
<tr><td></td><td></td><td></td><td></td><td></td><td></td><td></td><td></td><td></td><td></td><td></td><td></td></tr>
<tr><td>T23708</td><td>CCCCAGCCCCGGCGACTGGCTGG</td><td>1000</td><td></td><td></td><td></td><td></td><td></td><td></td><td></td><td></td><td></td></tr>
<tr><td>Oligo fwd</td><td>CACCCCCCAGCCCCGGCGACTGGC</td><td></td><td></td><td></td><td></td><td></td><td></td><td></td><td></td><td></td><td></td></tr>
<tr><td>Oligo rev</td><td>AAACGCCAGTCGCCGGGGCTGGGG</td><td></td><td></td><td></td><td></td><td></td><td></td><td></td><td></td><td></td><td></td></tr>
<tr><td>Chromosome</td><td>start</td><td>end</td><td>strand</td><td>MM</td><td>target_seq</td><td>PAM</td><td>alignment</td><td>distance</td><td>position</td><td>gene name</td><td>gene id</td></tr>
<tr><td>GU305914.1</td><td>136452</td><td>136474</td><td>-</td><td>0</td><td>CCCCAGCCCCGGCGACTGGC</td><td>TGG</td><td>||||||||[||||||||||||]PAM</td><td>NA</td><td>Intergenic</td><td>NA</td><td>NA</td></tr>
<tr><td></td><td></td><td></td><td></td><td></td><td></td><td></td><td></td><td></td><td></td><td></td><td></td></tr>
<tr><td>T23709</td><td>CCCCGGCGACTGGCTGGTGACGG</td><td>1000</td><td></td><td></td><td></td><td></td><td></td><td></td><td></td><td></td><td></td></tr>
<tr><td>Oligo fwd</td><td>CACCCCCCGGCGACTGGCTGGTGA</td><td></td><td></td><td></td><td></td><td></td><td></td><td></td><td></td><td></td><td></td></tr>
<tr><td>Oligo rev</td><td>AAACTCACCAGCCAGTCGCCGGGG</td><td></td><td></td><td></td><td></td><td></td><td></td><td></td><td></td><td></td><td></td></tr>
<tr><td>Chromosome</td><td>start</td><td>end</td><td>strand</td><td>MM</td><td>target_seq</td><td>PAM</td><td>alignment</td><td>distance</td><td>position</td><td>gene name</td><td>gene id</td></tr>
<tr><td>GU305914.1</td><td>136446</td><td>136468</td><td>-</td><td>0</td><td>CCCCGGCGACTGGCTGGTGA</td><td>CGG</td><td>||||||||[||||||||||||]PAM</td><td>NA</td><td>Intergenic</td><td>NA</td><td>NA</td></tr>
<tr><td></td><td></td><td></td><td></td><td></td><td></td><td></td><td></td><td></td><td></td><td></td><td></td></tr>
<tr><td>T23710</td><td>GCGACTGGCTGGTGACGGCGCGG</td><td>1000</td><td></td><td></td><td></td><td></td><td></td><td></td><td></td><td></td><td></td></tr>
<tr><td>Oligo fwd</td><td>CACCGCGACTGGCTGGTGACGGCG</td><td></td><td></td><td></td><td></td><td></td><td></td><td></td><td></td><td></td><td></td></tr>
<tr><td>Oligo rev</td><td>AAACCGCCGTCACCAGCCAGTCGC</td><td></td><td></td><td></td><td></td><td></td><td></td><td></td><td></td><td></td><td></td></tr>
<tr><td>Chromosome</td><td>start</td><td>end</td><td>strand</td><td>MM</td><td>target_seq</td><td>PAM</td><td>alignment</td><td>distance</td><td>position</td><td>gene name</td><td>gene id</td></tr>
<tr><td>GU305914.1</td><td>136441</td><td>136463</td><td>-</td><td>0</td><td>GCGACTGGCTGGTGACGGCG</td><td>CGG</td><td>||||||||[||||||||||||]PAM</td><td>NA</td><td>Intergenic</td><td>NA</td><td>NA</td></tr>
<tr><td>GU305914.1</td><td>117425</td><td>117447</td><td>+</td><td>4</td><td>GCGCCAGGCAGGTGATGGCG</td><td>GGG</td><td>|||-|-||[|-|||||-||||]PAM</td><td>NA</td><td>Intergenic</td><td>NA</td><td>NA</td></tr>
<tr><td></td><td></td><td></td><td></td><td></td><td></td><td></td><td></td><td></td><td></td><td></td><td></td></tr>
<tr><td>T23711</td><td>TGGTGACGGCGCGGAACTCTCGG</td><td>1000</td><td></td><td></td><td></td><td></td><td></td><td></td><td></td><td></td><td></td></tr>
<tr><td>Oligo fwd</td><td>CACCTGGTGACGGCGCGGAACTCT</td><td></td><td></td><td></td><td></td><td></td><td></td><td></td><td></td><td></td><td></td></tr>
<tr><td>Oligo rev</td><td>AAACAGAGTTCCGCGCCGTCACCA</td><td></td><td></td><td></td><td></td><td></td><td></td><td></td><td></td><td></td><td></td></tr>
<tr><td>Chromosome</td><td>start</td><td>end</td><td>strand</td><td>MM</td><td>target_seq</td><td>PAM</td><td>alignment</td><td>distance</td><td>position</td><td>gene name</td><td>gene id</td></tr>
<tr><td>GU305914.1</td><td>136432</td><td>136454</td><td>-</td><td>0</td><td>TGGTGACGGCGCGGAACTCT</td><td>CGG</td><td>||||||||[||||||||||||]PAM</td><td>NA</td><td>Intergenic</td><td>NA</td><td>NA</td></tr>
<tr><td></td><td></td><td></td><td></td><td></td><td></td><td></td><td></td><td></td><td></td><td></td><td></td></tr>
<tr><td>T23712</td><td>GCATCCAACAACACTCCCACTGG</td><td>1000</td><td></td><td></td><td></td><td></td><td></td><td></td><td></td><td></td><td></td></tr>
<tr><td>Oligo fwd</td><td>CACCGCATCCAACAACACTCCCAC</td><td></td><td></td><td></td><td></td><td></td><td></td><td></td><td></td><td></td><td></td></tr>
<tr><td>Oligo rev</td><td>AAACGTGGGAGTGTTGTTGGATGC</td><td></td><td></td><td></td><td></td><td></td><td></td><td></td><td></td><td></td><td></td></tr>
<tr><td>Chromosome</td><td>start</td><td>end</td><td>strand</td><td>MM</td><td>target_seq</td><td>PAM</td><td>alignment</td><td>distance</td><td>position</td><td>gene name</td><td>gene id</td></tr>
<tr><td>GU305914.1</td><td>136387</td><td>136409</td><td>-</td><td>0</td><td>GCATCCAACAACACTCCCAC</td><td>TGG</td><td>||||||||[||||||||||||]PAM</td><td>NA</td><td>Intergenic</td><td>NA</td><td>NA</td></tr>
<tr><td></td><td></td><td></td><td></td><td></td><td></td><td></td><td></td><td></td><td></td><td></td><td></td></tr>
<tr><td>T23713</td><td>GATGCTTCTCAAACTTTTCACGG</td><td>1000</td><td></td><td></td><td></td><td></td><td></td><td></td><td></td><td></td><td></td></tr>
<tr><td>Oligo fwd</td><td>CACCGATGCTTCTCAAACTTTTCA</td><td></td><td></td><td></td><td></td><td></td><td></td><td></td><td></td><td></td><td></td></tr>
<tr><td>Oligo rev</td><td>AAACTGAAAAGTTTGAGAAGCATC</td><td></td><td></td><td></td><td></td><td></td><td></td><td></td><td></td><td></td><td></td></tr>
<tr><td>Chromosome</td><td>start</td><td>end</td><td>strand</td><td>MM</td><td>target_seq</td><td>PAM</td><td>alignment</td><td>distance</td><td>position</td><td>gene name</td><td>gene id</td></tr>
<tr><td>GU305914.1</td><td>136353</td><td>136375</td><td>-</td><td>0</td><td>GATGCTTCTCAAACTTTTCA</td><td>CGG</td><td>||||||||[||||||||||||]PAM</td><td>NA</td><td>Intergenic</td><td>NA</td><td>NA</td></tr>
<tr><td></td><td></td><td></td><td></td><td></td><td></td><td></td><td></td><td></td><td></td><td></td><td></td></tr>
<tr><td>T23714</td><td>CAAACTTTTCACGGATAGTCTGG</td><td>1000</td><td></td><td></td><td></td><td></td><td></td><td></td><td></td><td></td><td></td></tr>
<tr><td>Oligo fwd</td><td>CACCCAAACTTTTCACGGATAGTC</td><td></td><td></td><td></td><td></td><td></td><td></td><td></td><td></td><td></td><td></td></tr>
<tr><td>Oligo rev</td><td>AAACGACTATCCGTGAAAAGTTTG</td><td></td><td></td><td></td><td></td><td></td><td></td><td></td><td></td><td></td><td></td></tr>
<tr><td>Chromosome</td><td>start</td><td>end</td><td>strand</td><td>MM</td><td>target_seq</td><td>PAM</td><td>alignment</td><td>distance</td><td>position</td><td>gene name</td><td>gene id</td></tr>
<tr><td>GU305914.1</td><td>136344</td><td>136366</td><td>-</td><td>0</td><td>CAAACTTTTCACGGATAGTC</td><td>TGG</td><td>||||||||[||||||||||||]PAM</td><td>NA</td><td>Intergenic</td><td>NA</td><td>NA</td></tr>
<tr><td></td><td></td><td></td><td></td><td></td><td></td><td></td><td></td><td></td><td></td><td></td><td></td></tr>
<tr><td>T23715</td><td>GGATAGTCTGGACGTCGCCATGG</td><td>1000</td><td></td><td></td><td></td><td></td><td></td><td></td><td></td><td></td><td></td></tr>
<tr><td>Oligo fwd</td><td>CACCGGATAGTCTGGACGTCGCCA</td><td></td><td></td><td></td><td></td><td></td><td></td><td></td><td></td><td></td><td></td></tr>
<tr><td>Oligo rev</td><td>AAACTGGCGACGTCCAGACTATCC</td><td></td><td></td><td></td><td></td><td></td><td></td><td></td><td></td><td></td><td></td></tr>
<tr><td>Chromosome</td><td>start</td><td>end</td><td>strand</td><td>MM</td><td>target_seq</td><td>PAM</td><td>alignment</td><td>distance</td><td>position</td><td>gene name</td><td>gene id</td></tr>
<tr><td>GU305914.1</td><td>136332</td><td>136354</td><td>-</td><td>0</td><td>GGATAGTCTGGACGTCGCCA</td><td>TGG</td><td>||||||||[||||||||||||]PAM</td><td>NA</td><td>Intergenic</td><td>NA</td><td>NA</td></tr>
<tr><td></td><td></td><td></td><td></td><td></td><td></td><td></td><td></td><td></td><td></td><td></td><td></td></tr>
<tr><td>T23716</td><td>CGCCATGGTCTACCTGCAGCTGG</td><td>1000</td><td></td><td></td><td></td><td></td><td></td><td></td><td></td><td></td><td></td></tr>
<tr><td>Oligo fwd</td><td>CACCCGCCATGGTCTACCTGCAGC</td><td></td><td></td><td></td><td></td><td></td><td></td><td></td><td></td><td></td><td></td></tr>
<tr><td>Oligo rev</td><td>AAACGCTGCAGGTAGACCATGGCG</td><td></td><td></td><td></td><td></td><td></td><td></td><td></td><td></td><td></td><td></td></tr>
<tr><td>Chromosome</td><td>start</td><td>end</td><td>strand</td><td>MM</td><td>target_seq</td><td>PAM</td><td>alignment</td><td>distance</td><td>position</td><td>gene name</td><td>gene id</td></tr>
<tr><td>GU305914.1</td><td>136317</td><td>136339</td><td>-</td><td>0</td><td>CGCCATGGTCTACCTGCAGC</td><td>TGG</td><td>||||||||[||||||||||||]PAM</td><td>NA</td><td>Intergenic</td><td>NA</td><td>NA</td></tr>
<tr><td></td><td></td><td></td><td></td><td></td><td></td><td></td><td></td><td></td><td></td><td></td><td></td></tr>
<tr><td>T23717</td><td>ATGGTCTACCTGCAGCTGGAAGG</td><td>1000</td><td></td><td></td><td></td><td></td><td></td><td></td><td></td><td></td><td></td></tr>
<tr><td>Oligo fwd</td><td>CACCATGGTCTACCTGCAGCTGGA</td><td></td><td></td><td></td><td></td><td></td><td></td><td></td><td></td><td></td><td></td></tr>
<tr><td>Oligo rev</td><td>AAACTCCAGCTGCAGGTAGACCAT</td><td></td><td></td><td></td><td></td><td></td><td></td><td></td><td></td><td></td><td></td></tr>
<tr><td>Chromosome</td><td>start</td><td>end</td><td>strand</td><td>MM</td><td>target_seq</td><td>PAM</td><td>alignment</td><td>distance</td><td>position</td><td>gene name</td><td>gene id</td></tr>
<tr><td>GU305914.1</td><td>136313</td><td>136335</td><td>-</td><td>0</td><td>ATGGTCTACCTGCAGCTGGA</td><td>AGG</td><td>||||||||[||||||||||||]PAM</td><td>NA</td><td>Intergenic</td><td>NA</td><td>NA</td></tr>
<tr><td></td><td></td><td></td><td></td><td></td><td></td><td></td><td></td><td></td><td></td><td></td><td></td></tr>
<tr><td>T23718</td><td>GAAGGTCGCGCAGAAGCCGCCGG</td><td>1000</td><td></td><td></td><td></td><td></td><td></td><td></td><td></td><td></td><td></td></tr>
<tr><td>Oligo fwd</td><td>CACCGAAGGTCGCGCAGAAGCCGC</td><td></td><td></td><td></td><td></td><td></td><td></td><td></td><td></td><td></td><td></td></tr>
<tr><td>Oligo rev</td><td>AAACGCGGCTTCTGCGCGACCTTC</td><td></td><td></td><td></td><td></td><td></td><td></td><td></td><td></td><td></td><td></td></tr>
<tr><td>Chromosome</td><td>start</td><td>end</td><td>strand</td><td>MM</td><td>target_seq</td><td>PAM</td><td>alignment</td><td>distance</td><td>position</td><td>gene name</td><td>gene id</td></tr>
<tr><td>GU305914.1</td><td>136295</td><td>136317</td><td>-</td><td>0</td><td>GAAGGTCGCGCAGAAGCCGC</td><td>CGG</td><td>||||||||[||||||||||||]PAM</td><td>NA</td><td>Intergenic</td><td>NA</td><td>NA</td></tr>
<tr><td>GU305914.1</td><td>128325</td><td>128347</td><td>-</td><td>4</td><td>GACGGTCTCGAAGAAGTCGC</td><td>GGG</td><td>||-||||-[||-|||||-|||]PAM</td><td>NA</td><td>Intergenic</td><td>NA</td><td>NA</td></tr>
</table>

T23719 AAGGTCGCGCAGAAGCCGCCGGG 1000
Oligo fwd CACCAAGGTCGCGCAGAAGCCGCC
Oligo rev AAACGGCGGCTTCTGCGCGACCTT
Chromosome start end strand MM target_seq PAM alignment distance position gene name gene id
GU305914.1 136294 136316 - 0 AAGGTCGCGCAGAAGCCGCC GGG ||||||||[||||||||||||]PAM NA Intergenic NA NA

T23720 CGCCGGGAAGCACCCAGAAGAGG 1000
Oligo fwd CACCCGCCGGGAAGCACCCAGAAG
Oligo rev AAACCTTCTGGGTGCTTCCCGGCG
Chromosome start end strand MM target_seq PAM alignment distance position gene name gene id
GU305914.1 136278 136300 - 0 CGCCGGGAAGCACCCAGAAG AGG ||||||||[||||||||||||]PAM NA Intergenic NA NA

T23721 GAAGCACCCAGAAGAGGCTGCGG 1000
Oligo fwd CACCGAAGCACCCAGAAGAGGCTG
Oligo rev AAACCAGCCTCTTCTGGGTGCTTC
Chromosome start end strand MM target_seq PAM alignment distance position gene name gene id
GU305914.1 136272 136294 - 0 GAAGCACCCAGAAGAGGCTG CGG ||||||||[||||||||||||]PAM NA Intergenic NA NA

T23722 GAAGAGGCTGCGGTCCTCGAAGG 1000
Oligo fwd CACCGAAGAGGCTGCGGTCCTCGA
Oligo rev AAACTCGAGGACCGCAGCCTCTTC
Chromosome start end strand MM target_seq PAM alignment distance position gene name gene id
GU305914.1 136262 136284 - 0 GAAGAGGCTGCGGTCCTCGA AGG ||||||||[||||||||||||]PAM NA Intergenic NA NA

T23723 TGCGGTCCTCGAAGGCGTAGAGG 1000
Oligo fwd CACCTGCGGTCCTCGAAGGCGTAG
Oligo rev AAACCTACGCCTTCGAGGACCGCA
Chromosome start end strand MM target_seq PAM alignment distance position gene name gene id
GU305914.1 136254 136276 - 0 TGCGGTCCTCGAAGGCGTAG AGG ||||||||[||||||||||||]PAM NA Intergenic NA NA

T23724 GGCGTAGAGGACACAGACGTCGG 1000
Oligo fwd CACCGGCGTAGAGGACACAGACGT
Oligo rev AAACACGTCTGTGTCCTCTACGCC
Chromosome start end strand MM target_seq PAM alignment distance position gene name gene id
GU305914.1 136241 136263 - 0 GGCGTAGAGGACACAGACGT CGG ||||||||[||||||||||||]PAM NA Intergenic NA NA

T23725 TGTCGAGAAAAGATCGCCAGAGG 1000
Oligo fwd CACCTGTCGAGAAAAGATCGCCAG
Oligo rev AAACCTGGCGATCTTTTCTCGACA
Chromosome start end strand MM target_seq PAM alignment distance position gene name gene id
GU305914.1 136188 136210 - 0 TGTCGAGAAAAGATCGCCAG AGG ||||||||[||||||||||||]PAM NA Intergenic NA NA

T23726 ACCGACCCTCATGAGCTTTATGG 1000
Oligo fwd CACCACCGACCCTCATGAGCTTTA
Oligo rev AAACTAAAGCTCATGAGGGTCGGT
Chromosome start end strand MM target_seq PAM alignment distance position gene name gene id
GU305914.1 136157 136179 - 0 ACCGACCCTCATGAGCTTTA TGG ||||||||[||||||||||||]PAM NA Intergenic NA NA

T23727 CCGACCCTCATGAGCTTTATGGG 1000
Oligo fwd CACCCCGACCCTCATGAGCTTTAT
Oligo rev AAACATAAAGCTCATGAGGGTCGG
Chromosome start end strand MM target_seq PAM alignment distance position gene name gene id
GU305914.1 136156 136178 - 0 CCGACCCTCATGAGCTTTAT GGG ||||||||[||||||||||||]PAM NA Intergenic NA NA

T23728 CTGCGCGTACCAGATCACATCGG 1000
Oligo fwd CACCCTGCGCGTACCAGATCACAT
Oligo rev AAACATGTGATCTGGTACGCGCAG
Chromosome start end strand MM target_seq PAM alignment distance position gene name gene id
GU305914.1 136112 136134 - 0 CTGCGCGTACCAGATCACAT CGG ||||||||[||||||||||||]PAM NA Intergenic NA NA

T23729 CGACACGACACCCATCGACGAGG 1000
Oligo fwd CACCCGACACGACACCCATCGACG
Oligo rev AAACCGTCGATGGGTGTCGTGTCG
Chromosome start end strand MM target_seq PAM alignment distance position gene name gene id
GU305914.1 136083 136105 - 0 CGACACGACACCCATCGACG AGG ||||||||[||||||||||||]PAM NA Intergenic NA NA

T23730 TCGCCGACGCCGATGAAGATCGG 1000
Oligo fwd CACCTCGCCGACGCCGATGAAGAT
Oligo rev AAACATCTTCATCGGCGTCGGCGA
Chromosome start end strand MM target_seq PAM alignment distance position gene name gene id
GU305914.1 136052 136074 - 0 TCGCCGACGCCGATGAAGAT CGG ||||||||[||||||||||||]PAM NA Intergenic NA NA

T23731 CCGACGCCGATGAAGATCGGAGG 1000
Oligo fwd CACCCCGACGCCGATGAAGATCGG
Oligo rev AAACCCGATCTTCATCGGCGTCGG
Chromosome start end strand MM target_seq PAM alignment distance position gene name gene id
GU305914.1 136049 136071 - 0 CCGACGCCGATGAAGATCGG AGG ||||||||[||||||||||||]PAM NA Intergenic NA NA

T23732 CTTAATAATGCAGCCCCCTCCGG 1000
Oligo fwd CACCCTTAATAATGCAGCCCCCTC
Oligo rev AAACGAGGGGGCTGCATTATTAAG
Chromosome start end strand MM target_seq PAM alignment distance position gene name gene id
GU305914.1 136019 136041 - 0 CTTAATAATGCAGCCCCCTC CGG ||||||||[||||||||||||]PAM NA Intergenic NA NA

T23733 TCCGGTGCCGTTCACGAGCCAGG 1000
Oligo fwd CACCTCCGGTGCCGTTCACGAGCC
Oligo rev AAACGGCTCGTGAACGGCACCGGA
Chromosome start end strand MM target_seq PAM alignment distance position gene name gene id

| | | | | | | | | | | | | |
|---|---|---|---|---|---|---|---|---|---|---|---|---|
| GU305914.1 | | 136001 | 136023 | - | 0 | TCCGGTGCCGTTCACGAGCC | AGG | \|\|\|\|\|\|\|\|[\|\|\|\|\|\|\|\|\|\|\|\|]PAM | NA | Intergenic | NA | NA |
| T23734 | CAGATCCCCCGCCCGCAGCCCGG | | 1000 | | | | | | | | | |
| Oligo fwd | CACCCAGATCCCCCGCCCGCAGCC | | | | | | | | | | | |
| Oligo rev | AAACGGCTGCGGGCGGGGGATCTG | | | | | | | | | | | |
| Chromosome | start | | end | strand | MM | target_seq | PAM | alignment | distance | position | gene name | gene id |
| GU305914.1 | | 135950 | 135972 | - | 0 | CAGATCCCCCGCCCGCAGCC | CGG | \|\|\|\|\|\|\|\|[\|\|\|\|\|\|\|\|\|\|\|\|]PAM | NA | Intergenic | NA | NA |
| T23735 | GAGCCCCCCGACGCGCGTGACGG | | 1000 | | | | | | | | | |
| Oligo fwd | CACCGAGCCCCCCGACGCGCGTGA | | | | | | | | | | | |
| Oligo rev | AAACTCACGCGCGTCGGGGGGCTC | | | | | | | | | | | |
| Chromosome | start | | end | strand | MM | target_seq | PAM | alignment | distance | position | gene name | gene id |
| GU305914.1 | | 135899 | 135921 | - | 0 | GAGCCCCCCGACGCGCGTGA | CGG | \|\|\|\|\|\|\|\|[\|\|\|\|\|\|\|\|\|\|\|\|]PAM | NA | Intergenic | NA | NA |
| T23736 | CCCGACGCGCGTGACGGCCGCGG | | 1000 | | | | | | | | | |
| Oligo fwd | CACCCCCGACGCGCGTGACGGCCG | | | | | | | | | | | |
| Oligo rev | AAACCGGCCGTCACGCGCGTCGGG | | | | | | | | | | | |
| Chromosome | start | | end | strand | MM | target_seq | PAM | alignment | distance | position | gene name | gene id |
| GU305914.1 | | 135893 | 135915 | - | 0 | CCCGACGCGCGTGACGGCCG | CGG | \|\|\|\|\|\|\|\|[\|\|\|\|\|\|\|\|\|\|\|\|]PAM | NA | Intergenic | NA | NA |
| T23737 | TGACGGCCGCGGCGAGCTGTCGG | | 1000 | | | | | | | | | |
| Oligo fwd | CACCTGACGGCCGCGGCGAGCTGT | | | | | | | | | | | |
| Oligo rev | AAACACAGCTCGCCGCGGCCGTCA | | | | | | | | | | | |
| Chromosome | start | | end | strand | MM | target_seq | PAM | alignment | distance | position | gene name | gene id |
| GU305914.1 | | 135882 | 135904 | - | 0 | TGACGGCCGCGGCGAGCTGT | CGG | \|\|\|\|\|\|\|\|[\|\|\|\|\|\|\|\|\|\|\|\|]PAM | NA | Intergenic | NA | NA |
| T23738 | GGCCGCGGCGAGCTGTCGGCTGG | | 1000 | | | | | | | | | |
| Oligo fwd | CACCGGCCGCGGCGAGCTGTCGGC | | | | | | | | | | | |
| Oligo rev | AAACGCCGACAGCTCGCCGCGGCC | | | | | | | | | | | |
| Chromosome | start | | end | strand | MM | target_seq | PAM | alignment | distance | position | gene name | gene id |
| GU305914.1 | | 135878 | 135900 | - | 0 | GGCCGCGGCGAGCTGTCGGC | TGG | \|\|\|\|\|\|\|\|[\|\|\|\|\|\|\|\|\|\|\|\|]PAM | NA | Intergenic | NA | NA |
| T23739 | CGCGGCGAGCTGTCGGCTGGTGG | | 1000 | | | | | | | | | |
| Oligo fwd | CACCCGCGGCGAGCTGTCGGCTGG | | | | | | | | | | | |
| Oligo rev | AAACCCAGCCGACAGCTCGCCGCG | | | | | | | | | | | |
| Chromosome | start | | end | strand | MM | target_seq | PAM | alignment | distance | position | gene name | gene id |
| GU305914.1 | | 135875 | 135897 | - | 0 | CGCGGCGAGCTGTCGGCTGG | TGG | \|\|\|\|\|\|\|\|[\|\|\|\|\|\|\|\|\|\|\|\|]PAM | NA | Intergenic | NA | NA |
| T23740 | GGCGCCGTAGACCCGCGTGACGG | | 1000 | | | | | | | | | |
| Oligo fwd | CACCGGCGCCGTAGACCCGCGTGA | | | | | | | | | | | |
| Oligo rev | AAACTCACGCGGGTCTACGGCGCC | | | | | | | | | | | |
| Chromosome | start | | end | strand | MM | target_seq | PAM | alignment | distance | position | gene name | gene id |
| GU305914.1 | | 135854 | 135876 | - | 0 | GGCGCCGTAGACCCGCGTGA | CGG | \|\|\|\|\|\|\|\|[\|\|\|\|\|\|\|\|\|\|\|\|]PAM | NA | Intergenic | NA | NA |
| T23741 | GCGTGACGGCCGTCTCCCAGCGG | | 1000 | | | | | | | | | |
| Oligo fwd | CACCGCGTGACGGCCGTCTCCCAG | | | | | | | | | | | |
| Oligo rev | AAACCTGGGAGACGGCCGTCACGC | | | | | | | | | | | |
| Chromosome | start | | end | strand | MM | target_seq | PAM | alignment | distance | position | gene name | gene id |
| GU305914.1 | | 135840 | 135862 | - | 0 | GCGTGACGGCCGTCTCCCAG | CGG | \|\|\|\|\|\|\|\|[\|\|\|\|\|\|\|\|\|\|\|\|]PAM | NA | Intergenic | NA | NA |
| T23742 | CCAGCGGCGCGTGTGTTCGAAGG | | 1000 | | | | | | | | | |
| Oligo fwd | CACCCCAGCGGCGCGTGTGTTCGA | | | | | | | | | | | |
| Oligo rev | AAACTCGAACACACGCGCCGCTGG | | | | | | | | | | | |
| Chromosome | start | | end | strand | MM | target_seq | PAM | alignment | distance | position | gene name | gene id |
| GU305914.1 | | 135824 | 135846 | - | 0 | CCAGCGGCGCGTGTGTTCGA | AGG | \|\|\|\|\|\|\|\|[\|\|\|\|\|\|\|\|\|\|\|\|]PAM | NA | Intergenic | NA | NA |
| GU305914.1 | | 167138 | 167160 | + | 4 | CTCGCGGCACTTGTGTTCGA | CGG | \|--\|\|\|\|\|[-\|-\|\|\|\|\|\|\|\|\|]PAM | NA | Intergenic | NA | NA |
| T23743 | GCGGCGCGTGTGTTCGAAGGAGG | | 1000 | | | | | | | | | |
| Oligo fwd | CACCGCGGCGCGTGTGTTCGAAGG | | | | | | | | | | | |
| Oligo rev | AAACCCTTCGAACACACGCGCCGC | | | | | | | | | | | |
| Chromosome | start | | end | strand | MM | target_seq | PAM | alignment | distance | position | gene name | gene id |
| GU305914.1 | | 135821 | 135843 | - | 0 | GCGGCGCGTGTGTTCGAAGG | AGG | \|\|\|\|\|\|\|\|[\|\|\|\|\|\|\|\|\|\|\|\|]PAM | NA | Intergenic | NA | NA |
| GU305914.1 | | 167141 | 167163 | + | 3 | GCGGCACTTGTGTTCGACGG | TGG | \|\|\|\|\|-\|-[\|\|\|\|\|\|\|\|\|-\|\|]PAM | NA | Intergenic | NA | NA |
| T23744 | CGTGTGTTCGAAGGAGGTGACGG | | 1000 | | | | | | | | | |
| Oligo fwd | CACCCGTGTGTTCGAAGGAGGTGA | | | | | | | | | | | |
| Oligo rev | AAACTCACCTCCTTCGAACACACG | | | | | | | | | | | |
| Chromosome | start | | end | strand | MM | target_seq | PAM | alignment | distance | position | gene name | gene id |
| GU305914.1 | | 135815 | 135837 | - | 0 | CGTGTGTTCGAAGGAGGTGA | CGG | \|\|\|\|\|\|\|\|[\|\|\|\|\|\|\|\|\|\|\|\|]PAM | NA | Intergenic | NA | NA |
| T23745 | CCGTCCGACGTCGACGTCGACGG | | 1000 | | | | | | | | | |
| Oligo fwd | CACCCCGTCCGACGTCGACGTCGA | | | | | | | | | | | |
| Oligo rev | AAACTCGACGTCGACGTCGGACGG | | | | | | | | | | | |
| Chromosome | start | | end | strand | MM | target_seq | PAM | alignment | distance | position | gene name | gene id |
| GU305914.1 | | 135773 | 135795 | - | 0 | CCGTCCGACGTCGACGTCGA | CGG | \|\|\|\|\|\|\|\|[\|\|\|\|\|\|\|\|\|\|\|\|]PAM | NA | Intergenic | NA | NA |
| T23746 | GACGTCGACGGTCGATCCGATGG | | 1000 | | | | | | | | | |
| Oligo fwd | CACCGACGTCGACGGTCGATCCGA | | | | | | | | | | | |
| Oligo rev | AAACTCGGATCGACCGTCGACGTC | | | | | | | | | | | |
| Chromosome | start | | end | strand | MM | target_seq | PAM | alignment | distance | position | gene name | gene id |
| GU305914.1 | | 135761 | 135783 | - | 0 | GACGTCGACGGTCGATCCGA | TGG | \|\|\|\|\|\|\|\|[\|\|\|\|\|\|\|\|\|\|\|\|]PAM | NA | Intergenic | NA | NA |
| T23747 | GACGGTCGATCCGATGGACTCGG | | 1000 | | | | | | | | | |
| Oligo fwd | CACCGACGGTCGATCCGATGGACT | | | | | | | | | | | |
| Oligo rev | AAACAGTCCATCGGATCGACCGTC | | | | | | | | | | | |
| Chromosome | start | | end | strand | MM | target_seq | PAM | alignment | distance | position | gene name | gene id |
| GU305914.1 | | 135755 | 135777 | - | 0 | GACGGTCGATCCGATGGACT | CGG | \|\|\|\|\|\|\|\|[\|\|\|\|\|\|\|\|\|\|\|\|]PAM | NA | Intergenic | NA | NA |
| T23748 | GGACTCGGAGTACCAGATCACGG | | 1000 | | | | | | | | | |

<table>
<tr><td>Oligo fwd</td><td>CACCGGACTCGGAGTACCAGATCA</td><td></td><td></td><td></td><td></td><td></td><td></td><td></td><td></td><td></td><td></td><td></td></tr>
<tr><td>Oligo rev</td><td>AAACTGATCTGGTACTCCGAGTCC</td><td></td><td></td><td></td><td></td><td></td><td></td><td></td><td></td><td></td><td></td><td></td></tr>
<tr><td>Chromosome</td><td>start</td><td></td><td>end</td><td>strand</td><td>MM</td><td>target_seq</td><td>PAM</td><td>alignment</td><td>distance</td><td>position</td><td>gene name</td><td>gene id</td></tr>
<tr><td>GU305914.1</td><td></td><td>135740</td><td>135762</td><td>-</td><td>0</td><td>GGACTCGGAGTACCAGATCA</td><td>CGG</td><td>||||||||[||||||||||||]PAM</td><td>NA</td><td>Intergenic</td><td>NA</td><td>NA</td></tr>
<tr><td>T23749</td><td>CACGGCGAACCTGCGCGAGCCGG</td><td></td><td>1000</td><td></td><td></td><td></td><td></td><td></td><td></td><td></td><td></td><td></td></tr>
<tr><td>Oligo fwd</td><td>CACCCACGGCGAACCTGCGCGAGC</td><td></td><td></td><td></td><td></td><td></td><td></td><td></td><td></td><td></td><td></td><td></td></tr>
<tr><td>Oligo rev</td><td>AAACGCTCGCGCAGGTTCGCCGTG</td><td></td><td></td><td></td><td></td><td></td><td></td><td></td><td></td><td></td><td></td><td></td></tr>
<tr><td>Chromosome</td><td>start</td><td></td><td>end</td><td>strand</td><td>MM</td><td>target_seq</td><td>PAM</td><td>alignment</td><td>distance</td><td>position</td><td>gene name</td><td>gene id</td></tr>
<tr><td>GU305914.1</td><td></td><td>135722</td><td>135744</td><td>-</td><td>0</td><td>CACGGCGAACCTGCGCGAGC</td><td>CGG</td><td>||||||||[||||||||||||]PAM</td><td>NA</td><td>Intergenic</td><td>NA</td><td>NA</td></tr>
<tr><td>T23750</td><td>TGCGCGAGCCGGTGACCTCCTGG</td><td></td><td>1000</td><td></td><td></td><td></td><td></td><td></td><td></td><td></td><td></td><td></td></tr>
<tr><td>Oligo fwd</td><td>CACCTGCGCGAGCCGGTGACCTCC</td><td></td><td></td><td></td><td></td><td></td><td></td><td></td><td></td><td></td><td></td><td></td></tr>
<tr><td>Oligo rev</td><td>AAACGGAGGTCACCGGCTCGCGCA</td><td></td><td></td><td></td><td></td><td></td><td></td><td></td><td></td><td></td><td></td><td></td></tr>
<tr><td>Chromosome</td><td>start</td><td></td><td>end</td><td>strand</td><td>MM</td><td>target_seq</td><td>PAM</td><td>alignment</td><td>distance</td><td>position</td><td>gene name</td><td>gene id</td></tr>
<tr><td>GU305914.1</td><td></td><td>135711</td><td>135733</td><td>-</td><td>0</td><td>TGCGCGAGCCGGTGACCTCC</td><td>TGG</td><td>||||||||[||||||||||||]PAM</td><td>NA</td><td>Intergenic</td><td>NA</td><td>NA</td></tr>
<tr><td>T23751</td><td>GCGCGAGCCGGTGACCTCCTGGG</td><td></td><td>1000</td><td></td><td></td><td></td><td></td><td></td><td></td><td></td><td></td><td></td></tr>
<tr><td>Oligo fwd</td><td>CACCGCGCGAGCCGGTGACCTCCT</td><td></td><td></td><td></td><td></td><td></td><td></td><td></td><td></td><td></td><td></td><td></td></tr>
<tr><td>Oligo rev</td><td>AAACAGGAGGTCACCGGCTCGCGC</td><td></td><td></td><td></td><td></td><td></td><td></td><td></td><td></td><td></td><td></td><td></td></tr>
<tr><td>Chromosome</td><td>start</td><td></td><td>end</td><td>strand</td><td>MM</td><td>target_seq</td><td>PAM</td><td>alignment</td><td>distance</td><td>position</td><td>gene name</td><td>gene id</td></tr>
<tr><td>GU305914.1</td><td></td><td>135710</td><td>135732</td><td>-</td><td>0</td><td>GCGCGAGCCGGTGACCTCCT</td><td>GGG</td><td>||||||||[||||||||||||]PAM</td><td>NA</td><td>Intergenic</td><td>NA</td><td>NA</td></tr>
<tr><td>T23752</td><td>CTGGGCGTACGAGATGCGAATGG</td><td></td><td>1000</td><td></td><td></td><td></td><td></td><td></td><td></td><td></td><td></td><td></td></tr>
<tr><td>Oligo fwd</td><td>CACCCTGGGCGTACGAGATGCGAA</td><td></td><td></td><td></td><td></td><td></td><td></td><td></td><td></td><td></td><td></td><td></td></tr>
<tr><td>Oligo rev</td><td>AAACTTCGCATCTCGTACGCCCAG</td><td></td><td></td><td></td><td></td><td></td><td></td><td></td><td></td><td></td><td></td><td></td></tr>
<tr><td>Chromosome</td><td>start</td><td></td><td>end</td><td>strand</td><td>MM</td><td>target_seq</td><td>PAM</td><td>alignment</td><td>distance</td><td>position</td><td>gene name</td><td>gene id</td></tr>
<tr><td>GU305914.1</td><td></td><td>135692</td><td>135714</td><td>-</td><td>0</td><td>CTGGGCGTACGAGATGCGAA</td><td>TGG</td><td>||||||||[||||||||||||]PAM</td><td>NA</td><td>Intergenic</td><td>NA</td><td>NA</td></tr>
<tr><td>T23753</td><td>CGTTGTCGCAGCCGAACCAGCGG</td><td></td><td>1000</td><td></td><td></td><td></td><td></td><td></td><td></td><td></td><td></td><td></td></tr>
<tr><td>Oligo fwd</td><td>CACCCGTTGTCGCAGCCGAACCAG</td><td></td><td></td><td></td><td></td><td></td><td></td><td></td><td></td><td></td><td></td><td></td></tr>
<tr><td>Oligo rev</td><td>AAACCTGGTTCGGCTGCGACAACG</td><td></td><td></td><td></td><td></td><td></td><td></td><td></td><td></td><td></td><td></td><td></td></tr>
<tr><td>Chromosome</td><td>start</td><td></td><td>end</td><td>strand</td><td>MM</td><td>target_seq</td><td>PAM</td><td>alignment</td><td>distance</td><td>position</td><td>gene name</td><td>gene id</td></tr>
<tr><td>GU305914.1</td><td></td><td>135669</td><td>135691</td><td>-</td><td>0</td><td>CGTTGTCGCAGCCGAACCAG</td><td>CGG</td><td>||||||||[||||||||||||]PAM</td><td>NA</td><td>Intergenic</td><td>NA</td><td>NA</td></tr>
<tr><td>T23754</td><td>AGCCGAACCAGCGGTCCGATCGG</td><td></td><td>1000</td><td></td><td></td><td></td><td></td><td></td><td></td><td></td><td></td><td></td></tr>
<tr><td>Oligo fwd</td><td>CACCAGCCGAACCAGCGGTCCGAT</td><td></td><td></td><td></td><td></td><td></td><td></td><td></td><td></td><td></td><td></td><td></td></tr>
<tr><td>Oligo rev</td><td>AAACATCGGACCGCTGGTTCGGCT</td><td></td><td></td><td></td><td></td><td></td><td></td><td></td><td></td><td></td><td></td><td></td></tr>
<tr><td>Chromosome</td><td>start</td><td></td><td>end</td><td>strand</td><td>MM</td><td>target_seq</td><td>PAM</td><td>alignment</td><td>distance</td><td>position</td><td>gene name</td><td>gene id</td></tr>
<tr><td>GU305914.1</td><td></td><td>135660</td><td>135682</td><td>-</td><td>0</td><td>AGCCGAACCAGCGGTCCGAT</td><td>CGG</td><td>||||||||[||||||||||||]PAM</td><td>NA</td><td>Intergenic</td><td>NA</td><td>NA</td></tr>
<tr><td>T23755</td><td>GCGGTCCGATCGGCTCCTGACGG</td><td></td><td>1000</td><td></td><td></td><td></td><td></td><td></td><td></td><td></td><td></td><td></td></tr>
<tr><td>Oligo fwd</td><td>CACCGCGGTCCGATCGGCTCCTGA</td><td></td><td></td><td></td><td></td><td></td><td></td><td></td><td></td><td></td><td></td><td></td></tr>
<tr><td>Oligo rev</td><td>AAACTCAGGAGCCGATCGGACCGC</td><td></td><td></td><td></td><td></td><td></td><td></td><td></td><td></td><td></td><td></td><td></td></tr>
<tr><td>Chromosome</td><td>start</td><td></td><td>end</td><td>strand</td><td>MM</td><td>target_seq</td><td>PAM</td><td>alignment</td><td>distance</td><td>position</td><td>gene name</td><td>gene id</td></tr>
<tr><td>GU305914.1</td><td></td><td>135650</td><td>135672</td><td>-</td><td>0</td><td>GCGGTCCGATCGGCTCCTGA</td><td>CGG</td><td>||||||||[||||||||||||]PAM</td><td>NA</td><td>Intergenic</td><td>NA</td><td>NA</td></tr>
<tr><td>T23756</td><td>CGGTCCGATCGGCTCCTGACGGG</td><td></td><td>1000</td><td></td><td></td><td></td><td></td><td></td><td></td><td></td><td></td><td></td></tr>
<tr><td>Oligo fwd</td><td>CACCCGGTCCGATCGGCTCCTGAC</td><td></td><td></td><td></td><td></td><td></td><td></td><td></td><td></td><td></td><td></td><td></td></tr>
<tr><td>Oligo rev</td><td>AAACGTCAGGAGCCGATCGGACCG</td><td></td><td></td><td></td><td></td><td></td><td></td><td></td><td></td><td></td><td></td><td></td></tr>
<tr><td>Chromosome</td><td>start</td><td></td><td>end</td><td>strand</td><td>MM</td><td>target_seq</td><td>PAM</td><td>alignment</td><td>distance</td><td>position</td><td>gene name</td><td>gene id</td></tr>
<tr><td>GU305914.1</td><td></td><td>135649</td><td>135671</td><td>-</td><td>0</td><td>CGGTCCGATCGGCTCCTGAC</td><td>GGG</td><td>||||||||[||||||||||||]PAM</td><td>NA</td><td>Intergenic</td><td>NA</td><td>NA</td></tr>
<tr><td>T23757</td><td>GGTCCGATCGGCTCCTGACGGGG</td><td></td><td>1000</td><td></td><td></td><td></td><td></td><td></td><td></td><td></td><td></td><td></td></tr>
<tr><td>Oligo fwd</td><td>CACCGGTCCGATCGGCTCCTGACG</td><td></td><td></td><td></td><td></td><td></td><td></td><td></td><td></td><td></td><td></td><td></td></tr>
<tr><td>Oligo rev</td><td>AAACCGTCAGGAGCCGATCGGACC</td><td></td><td></td><td></td><td></td><td></td><td></td><td></td><td></td><td></td><td></td><td></td></tr>
<tr><td>Chromosome</td><td>start</td><td></td><td>end</td><td>strand</td><td>MM</td><td>target_seq</td><td>PAM</td><td>alignment</td><td>distance</td><td>position</td><td>gene name</td><td>gene id</td></tr>
<tr><td>GU305914.1</td><td></td><td>135648</td><td>135670</td><td>-</td><td>0</td><td>GGTCCGATCGGCTCCTGACG</td><td>GGG</td><td>||||||||[||||||||||||]PAM</td><td>NA</td><td>Intergenic</td><td>NA</td><td>NA</td></tr>
<tr><td>T23758</td><td>CGATCGGCTCCTGACGGGGTCGG</td><td></td><td>1000</td><td></td><td></td><td></td><td></td><td></td><td></td><td></td><td></td><td></td></tr>
<tr><td>Oligo fwd</td><td>CACCCGATCGGCTCCTGACGGGGT</td><td></td><td></td><td></td><td></td><td></td><td></td><td></td><td></td><td></td><td></td><td></td></tr>
<tr><td>Oligo rev</td><td>AAACACCCCGTCAGGAGCCGATCG</td><td></td><td></td><td></td><td></td><td></td><td></td><td></td><td></td><td></td><td></td><td></td></tr>
<tr><td>Chromosome</td><td>start</td><td></td><td>end</td><td>strand</td><td>MM</td><td>target_seq</td><td>PAM</td><td>alignment</td><td>distance</td><td>position</td><td>gene name</td><td>gene id</td></tr>
<tr><td>GU305914.1</td><td></td><td>135644</td><td>135666</td><td>-</td><td>0</td><td>CGATCGGCTCCTGACGGGGT</td><td>CGG</td><td>||||||||[||||||||||||]PAM</td><td>NA</td><td>Intergenic</td><td>NA</td><td>NA</td></tr>
<tr><td>T23759</td><td>GATCGGCTCCTGACGGGGTCGGG</td><td></td><td>1000</td><td></td><td></td><td></td><td></td><td></td><td></td><td></td><td></td><td></td></tr>
<tr><td>Oligo fwd</td><td>CACCGATCGGCTCCTGACGGGGTC</td><td></td><td></td><td></td><td></td><td></td><td></td><td></td><td></td><td></td><td></td><td></td></tr>
<tr><td>Oligo rev</td><td>AAACGACCCCGTCAGGAGCCGATC</td><td></td><td></td><td></td><td></td><td></td><td></td><td></td><td></td><td></td><td></td><td></td></tr>
<tr><td>Chromosome</td><td>start</td><td></td><td>end</td><td>strand</td><td>MM</td><td>target_seq</td><td>PAM</td><td>alignment</td><td>distance</td><td>position</td><td>gene name</td><td>gene id</td></tr>
<tr><td>GU305914.1</td><td></td><td>135643</td><td>135665</td><td>-</td><td>0</td><td>GATCGGCTCCTGACGGGGTC</td><td>GGG</td><td>||||||||[||||||||||||]PAM</td><td>NA</td><td>Intergenic</td><td>NA</td><td>NA</td></tr>
<tr><td>GU305914.1</td><td></td><td>35112</td><td>35134</td><td>-</td><td>4</td><td>TATCGTCTCCGAACGGGGTC</td><td>TGG</td><td>-||||-||[||--||||||||]PAM</td><td>NA</td><td>Intergenic</td><td>NA</td><td>NA</td></tr>
<tr><td>T23760</td><td>GCTCCTGACGGGGTCGGGCGCGG</td><td></td><td>1000</td><td></td><td></td><td></td><td></td><td></td><td></td><td></td><td></td><td></td></tr>
<tr><td>Oligo fwd</td><td>CACCGCTCCTGACGGGGTCGGGCG</td><td></td><td></td><td></td><td></td><td></td><td></td><td></td><td></td><td></td><td></td><td></td></tr>
<tr><td>Oligo rev</td><td>AAACCGCCCGACCCCGTCAGGAGC</td><td></td><td></td><td></td><td></td><td></td><td></td><td></td><td></td><td></td><td></td><td></td></tr>
<tr><td>Chromosome</td><td>start</td><td></td><td>end</td><td>strand</td><td>MM</td><td>target_seq</td><td>PAM</td><td>alignment</td><td>distance</td><td>position</td><td>gene name</td><td>gene id</td></tr>
<tr><td>GU305914.1</td><td></td><td>135638</td><td>135660</td><td>-</td><td>0</td><td>GCTCCTGACGGGGTCGGGCG</td><td>CGG</td><td>||||||||[||||||||||||]PAM</td><td>NA</td><td>Intergenic</td><td>NA</td><td>NA</td></tr>
<tr><td>T23761</td><td>CTGACGGGGTCGGGCGCGGCCGG</td><td></td><td>1000</td><td></td><td></td><td></td><td></td><td></td><td></td><td></td><td></td><td></td></tr>
<tr><td>Oligo fwd</td><td>CACCCTGACGGGGTCGGGCGCGGC</td><td></td><td></td><td></td><td></td><td></td><td></td><td></td><td></td><td></td><td></td><td></td></tr>
<tr><td>Oligo rev</td><td>AAACGCCGCGCCCGACCCCGTCAG</td><td></td><td></td><td></td><td></td><td></td><td></td><td></td><td></td><td></td><td></td><td></td></tr>
<tr><td>Chromosome</td><td>start</td><td></td><td>end</td><td>strand</td><td>MM</td><td>target_seq</td><td>PAM</td><td>alignment</td><td>distance</td><td>position</td><td>gene name</td><td>gene id</td></tr>
<tr><td>GU305914.1</td><td></td><td>135634</td><td>135656</td><td>-</td><td>0</td><td>CTGACGGGGTCGGGCGCGGC</td><td>CGG</td><td>||||||||[||||||||||||]PAM</td><td>NA</td><td>Intergenic</td><td>NA</td><td>NA</td></tr>
<tr><td>GU305914.1</td><td></td><td>102744</td><td>102766</td><td>-</td><td>4</td><td>GGGAGGTGGTCGGGCGCGGC</td><td>GGG</td><td>--||-|-|[||||||||||||]PAM</td><td>NA</td><td>Intergenic</td><td>NA</td><td>NA</td></tr>
<tr><td>T23762</td><td>CCTGAAGAACTCGCCGCGTCTGG</td><td></td><td>1000</td><td></td><td></td><td></td><td></td><td></td><td></td><td></td><td></td><td></td></tr>
<tr><td>Oligo fwd</td><td>CACCCCTGAAGAACTCGCCGCGTC</td><td></td><td></td><td></td><td></td><td></td><td></td><td></td><td></td><td></td><td></td><td></td></tr>
<tr><td>Oligo rev</td><td>AAACGACGCGGCGAGTTCTTCAGG</td><td></td><td></td><td></td><td></td><td></td><td></td><td></td><td></td><td></td><td></td><td></td></tr>
<tr><td>Chromosome</td><td>start</td><td></td><td>end</td><td>strand</td><td>MM</td><td>target_seq</td><td>PAM</td><td>alignment</td><td>distance</td><td>position</td><td>gene name</td><td>gene id</td></tr>
</table>

| | | | | | | | | | | | | |
|---|---|---|---|---|---|---|---|---|---|---|---|---|
| GU305914.1 | | 135599 | 135621 | - | 0 | CCTGAAGAACTCGCCGCGTC | TGG | \|\|\|\|\|\|\|\|[\|\|\|\|\|\|\|\|\|\|\|\|]PAM | NA | Intergenic | NA | NA |
| T23763 | CGCGTCTGGCCTTGATCTCGCGG | | 1000 | | | | | | | | | |
| Oligo fwd | CACCCGCGTCTGGCCTTGATCTCG | | | | | | | | | | | |
| Oligo rev | AAACCGAGATCAAGGCCAGACGCG | | | | | | | | | | | |
| Chromosome | start | | end | strand | MM | target_seq | PAM | alignment | distance | position | gene name | gene id |
| GU305914.1 | | 135585 | 135607 | - | 0 | CGCGTCTGGCCTTGATCTCG | CGG | \|\|\|\|\|\|\|\|[\|\|\|\|\|\|\|\|\|\|\|\|]PAM | NA | Intergenic | NA | NA |
| GU305914.1 | | 47776 | 47798 | + | 4 | CGAGCGTGGCCTTGATCACG | GGG | \|\|-\|--\|\|[\|\|\|\|\|\|\|\|\|-\|\|]PAM | NA | Intergenic | NA | NA |
| T23764 | GATCTCGCGGCCCCTGACCTTGG | | 1000 | | | | | | | | | |
| Oligo fwd | CACCGATCTCGCGGCCCCTGACCT | | | | | | | | | | | |
| Oligo rev | AAACAGGTCAGGGGCCGCGAGATC | | | | | | | | | | | |
| Chromosome | start | | end | strand | MM | target_seq | PAM | alignment | distance | position | gene name | gene id |
| GU305914.1 | | 135572 | 135594 | - | 0 | GATCTCGCGGCCCCTGACCT | TGG | \|\|\|\|\|\|\|\|[\|\|\|\|\|\|\|\|\|\|\|\|]PAM | NA | Intergenic | NA | NA |
| T23765 | CTCGCGGCCCCTGACCTTGGAGG | | 1000 | | | | | | | | | |
| Oligo fwd | CACCCTCGCGGCCCCTGACCTTGG | | | | | | | | | | | |
| Oligo rev | AAACCCAAGGTCAGGGGCCGCGAG | | | | | | | | | | | |
| Chromosome | start | | end | strand | MM | target_seq | PAM | alignment | distance | position | gene name | gene id |
| GU305914.1 | | 135569 | 135591 | - | 0 | CTCGCGGCCCCTGACCTTGG | AGG | \|\|\|\|\|\|\|\|[\|\|\|\|\|\|\|\|\|\|\|\|]PAM | NA | Intergenic | NA | NA |
| T23766 | TCGCGGCCCCTGACCTTGGAGGG | | 1000 | | | | | | | | | |
| Oligo fwd | CACCTCGCGGCCCCTGACCTTGGA | | | | | | | | | | | |
| Oligo rev | AAACTCCAAGGTCAGGGGCCGCGA | | | | | | | | | | | |
| Chromosome | start | | end | strand | MM | target_seq | PAM | alignment | distance | position | gene name | gene id |
| GU305914.1 | | 135568 | 135590 | - | 0 | TCGCGGCCCCTGACCTTGGA | GGG | \|\|\|\|\|\|\|\|[\|\|\|\|\|\|\|\|\|\|\|\|]PAM | NA | Intergenic | NA | NA |
| T23767 | CGGCCCCTGACCTTGGAGGGCGG | | 1000 | | | | | | | | | |
| Oligo fwd | CACCCGGCCCCTGACCTTGGAGGG | | | | | | | | | | | |
| Oligo rev | AAACCCCTCCAAGGTCAGGGGCCG | | | | | | | | | | | |
| Chromosome | start | | end | strand | MM | target_seq | PAM | alignment | distance | position | gene name | gene id |
| GU305914.1 | | 135565 | 135587 | - | 0 | CGGCCCCTGACCTTGGAGGG | CGG | \|\|\|\|\|\|\|\|[\|\|\|\|\|\|\|\|\|\|\|\|]PAM | NA | Intergenic | NA | NA |
| T23768 | ACCTTGGAGGGCGGCTCGAGCGG | | 1000 | | | | | | | | | |
| Oligo fwd | CACCACCTTGGAGGGCGGCTCGAG | | | | | | | | | | | |
| Oligo rev | AAACCTCGAGCCGCCCTCCAAGGT | | | | | | | | | | | |
| Chromosome | start | | end | strand | MM | target_seq | PAM | alignment | distance | position | gene name | gene id |
| GU305914.1 | | 135556 | 135578 | - | 0 | ACCTTGGAGGGCGGCTCGAG | CGG | \|\|\|\|\|\|\|\|[\|\|\|\|\|\|\|\|\|\|\|\|]PAM | NA | Intergenic | NA | NA |
| T23769 | CCTTGGAGGGCGGCTCGAGCGGG | | 1000 | | | | | | | | | |
| Oligo fwd | CACCCCTTGGAGGGCGGCTCGAGC | | | | | | | | | | | |
| Oligo rev | AAACGCTCGAGCCGCCCTCCAAGG | | | | | | | | | | | |
| Chromosome | start | | end | strand | MM | target_seq | PAM | alignment | distance | position | gene name | gene id |
| GU305914.1 | | 135555 | 135577 | - | 0 | CCTTGGAGGGCGGCTCGAGC | GGG | \|\|\|\|\|\|\|\|[\|\|\|\|\|\|\|\|\|\|\|\|]PAM | NA | Intergenic | NA | NA |
| T23770 | CGGCTCGAGCGGGCCGAAGTCGG | | 1000 | | | | | | | | | |
| Oligo fwd | CACCCGGCTCGAGCGGGCCGAAGT | | | | | | | | | | | |
| Oligo rev | AAACACTTCGGCCCGCTCGAGCCG | | | | | | | | | | | |
| Chromosome | start | | end | strand | MM | target_seq | PAM | alignment | distance | position | gene name | gene id |
| GU305914.1 | | 135545 | 135567 | - | 0 | CGGCTCGAGCGGGCCGAAGT | CGG | \|\|\|\|\|\|\|\|[\|\|\|\|\|\|\|\|\|\|\|\|]PAM | NA | Intergenic | NA | NA |
| T23771 | CTCGAGCGGGCCGAAGTCGGCGG | | 1000 | | | | | | | | | |
| Oligo fwd | CACCCTCGAGCGGGCCGAAGTCGG | | | | | | | | | | | |
| Oligo rev | AAACCCGACTTCGGCCCGCTCGAG | | | | | | | | | | | |
| Chromosome | start | | end | strand | MM | target_seq | PAM | alignment | distance | position | gene name | gene id |
| GU305914.1 | | 135542 | 135564 | - | 0 | CTCGAGCGGGCCGAAGTCGG | CGG | \|\|\|\|\|\|\|\|[\|\|\|\|\|\|\|\|\|\|\|\|]PAM | NA | Intergenic | NA | NA |
| T23772 | GAGCGGGCCGAAGTCGGCGGCGG | | 1000 | | | | | | | | | |
| Oligo fwd | CACCGAGCGGGCCGAAGTCGGCGG | | | | | | | | | | | |
| Oligo rev | AAACCCGCCGACTTCGGCCCGCTC | | | | | | | | | | | |
| Chromosome | start | | end | strand | MM | target_seq | PAM | alignment | distance | position | gene name | gene id |
| GU305914.1 | | 135539 | 135561 | - | 0 | GAGCGGGCCGAAGTCGGCGG | CGG | \|\|\|\|\|\|\|\|[\|\|\|\|\|\|\|\|\|\|\|\|]PAM | NA | Intergenic | NA | NA |
| GU305914.1 | | 225492 | 225514 | + | 4 | GACCGAGCCGAGGGCGGCGG | TGG | \|\|-\|\|-\|\|[\|\|\|-\|-\|\|\|\|\|\|]PAM | NA | Intergenic | NA | NA |
| T23773 | GACGCGTTTCAGCGCGAACCTGG | | 1000 | | | | | | | | | |
| Oligo fwd | CACCGACGCGTTTCAGCGCGAACC | | | | | | | | | | | |
| Oligo rev | AAACGGTTCGCGCTGAAACGCGTC | | | | | | | | | | | |
| Chromosome | start | | end | strand | MM | target_seq | PAM | alignment | distance | position | gene name | gene id |
| GU305914.1 | | 135515 | 135537 | - | 0 | GACGCGTTTCAGCGCGAACC | TGG | \|\|\|\|\|\|\|\|[\|\|\|\|\|\|\|\|\|\|\|\|]PAM | NA | Intergenic | NA | NA |
| T23774 | TCAGCGCGAACCTGGCCAGCCGG | | 1000 | | | | | | | | | |
| Oligo fwd | CACCTCAGCGCGAACCTGGCCAGC | | | | | | | | | | | |
| Oligo rev | AAACGCTGGCCAGGTTCGCGCTGA | | | | | | | | | | | |
| Chromosome | start | | end | strand | MM | target_seq | PAM | alignment | distance | position | gene name | gene id |
| GU305914.1 | | 135507 | 135529 | - | 0 | TCAGCGCGAACCTGGCCAGC | CGG | \|\|\|\|\|\|\|\|[\|\|\|\|\|\|\|\|\|\|\|\|]PAM | NA | Intergenic | NA | NA |
| T23775 | CAGCCGGTCCTCCATCTCCTCGG | | 1000 | | | | | | | | | |
| Oligo fwd | CACCCAGCCGGTCCTCCATCTCCT | | | | | | | | | | | |
| Oligo rev | AAACAGGAGATGGAGGACCGGCTG | | | | | | | | | | | |
| Chromosome | start | | end | strand | MM | target_seq | PAM | alignment | distance | position | gene name | gene id |
| GU305914.1 | | 135491 | 135513 | - | 0 | CAGCCGGTCCTCCATCTCCT | CGG | \|\|\|\|\|\|\|\|[\|\|\|\|\|\|\|\|\|\|\|\|]PAM | NA | Intergenic | NA | NA |
| T23776 | TCGCCGAGATCGAAGAAACCCGG | | 1000 | | | | | | | | | |
| Oligo fwd | CACCTCGCCGAGATCGAAGAAACC | | | | | | | | | | | |
| Oligo rev | AAACGGTTTCTTCGATCTCGGCGA | | | | | | | | | | | |
| Chromosome | start | | end | strand | MM | target_seq | PAM | alignment | distance | position | gene name | gene id |
| GU305914.1 | | 135454 | 135476 | - | 0 | TCGCCGAGATCGAAGAAACC | CGG | \|\|\|\|\|\|\|\|[\|\|\|\|\|\|\|\|\|\|\|\|]PAM | NA | Intergenic | NA | NA |
| T23777 | CGCCGAGATCGAAGAAACCCGGG | | 1000 | | | | | | | | | |

<table>
<tr><td>Oligo fwd</td><td>CACCCGCCGAGATCGAAGAAACCC</td><td></td><td></td><td></td><td></td><td></td><td></td><td></td><td></td><td></td><td></td></tr>
<tr><td>Oligo rev</td><td>AAACGGGTTTCTTCGATCTCGGCG</td><td></td><td></td><td></td><td></td><td></td><td></td><td></td><td></td><td></td><td></td></tr>
<tr><td>Chromosome</td><td>start</td><td>end</td><td>strand</td><td>MM</td><td>target_seq</td><td>PAM</td><td>alignment</td><td>distance</td><td>position</td><td>gene name</td><td>gene id</td></tr>
<tr><td>GU305914.1</td><td>135453</td><td>135475</td><td>-</td><td>0</td><td>CGCCGAGATCGAAGAAACCC</td><td>GGG</td><td>||||||||[||||||||||||]PAM</td><td>NA</td><td>Intergenic</td><td>NA</td><td>NA</td></tr>
<tr><td></td><td></td><td></td><td></td><td></td><td></td><td></td><td></td><td></td><td></td><td></td><td></td></tr>
<tr><td>T23778</td><td>ATCGAAGAAACCCGGGCCGTCGG</td><td>1000</td><td></td><td></td><td></td><td></td><td></td><td></td><td></td><td></td><td></td></tr>
<tr><td>Oligo fwd</td><td>CACCATCGAAGAAACCCGGGCCGT</td><td></td><td></td><td></td><td></td><td></td><td></td><td></td><td></td><td></td><td></td></tr>
<tr><td>Oligo rev</td><td>AAACACGGCCCGGGTTTCTTCGAT</td><td></td><td></td><td></td><td></td><td></td><td></td><td></td><td></td><td></td><td></td></tr>
<tr><td>Chromosome</td><td>start</td><td>end</td><td>strand</td><td>MM</td><td>target_seq</td><td>PAM</td><td>alignment</td><td>distance</td><td>position</td><td>gene name</td><td>gene id</td></tr>
<tr><td>GU305914.1</td><td>135446</td><td>135468</td><td>-</td><td>0</td><td>ATCGAAGAAACCCGGGCCGT</td><td>CGG</td><td>||||||||[||||||||||||]PAM</td><td>NA</td><td>Intergenic</td><td>NA</td><td>NA</td></tr>
<tr><td></td><td></td><td></td><td></td><td></td><td></td><td></td><td></td><td></td><td></td><td></td><td></td></tr>
<tr><td>T23779</td><td>GAAGAAACCCGGGCCGTCGGCGG</td><td>1000</td><td></td><td></td><td></td><td></td><td></td><td></td><td></td><td></td><td></td></tr>
<tr><td>Oligo fwd</td><td>CACCGAAGAAACCCGGGCCGTCGG</td><td></td><td></td><td></td><td></td><td></td><td></td><td></td><td></td><td></td><td></td></tr>
<tr><td>Oligo rev</td><td>AAACCCGACGGCCCGGGTTTCTTC</td><td></td><td></td><td></td><td></td><td></td><td></td><td></td><td></td><td></td><td></td></tr>
<tr><td>Chromosome</td><td>start</td><td>end</td><td>strand</td><td>MM</td><td>target_seq</td><td>PAM</td><td>alignment</td><td>distance</td><td>position</td><td>gene name</td><td>gene id</td></tr>
<tr><td>GU305914.1</td><td>135443</td><td>135465</td><td>-</td><td>0</td><td>GAAGAAACCCGGGCCGTCGG</td><td>CGG</td><td>||||||||[||||||||||||]PAM</td><td>NA</td><td>Intergenic</td><td>NA</td><td>NA</td></tr>
<tr><td>GU305914.1</td><td>184984</td><td>185006</td><td>-</td><td>4</td><td>GTAAAAACCCGGGCGGTCCG</td><td>CGG</td><td>|-|-||||[||||||-|||-|]PAM</td><td>NA</td><td>Intergenic</td><td>NA</td><td>NA</td></tr>
<tr><td></td><td></td><td></td><td></td><td></td><td></td><td></td><td></td><td></td><td></td><td></td><td></td></tr>
<tr><td>T23780</td><td>GAAACCCGGGCCGTCGGCGGCGG</td><td>1000</td><td></td><td></td><td></td><td></td><td></td><td></td><td></td><td></td><td></td></tr>
<tr><td>Oligo fwd</td><td>CACCGAAACCCGGGCCGTCGGCGG</td><td></td><td></td><td></td><td></td><td></td><td></td><td></td><td></td><td></td><td></td></tr>
<tr><td>Oligo rev</td><td>AAACCCGCCGACGGCCCGGGTTTC</td><td></td><td></td><td></td><td></td><td></td><td></td><td></td><td></td><td></td><td></td></tr>
<tr><td>Chromosome</td><td>start</td><td>end</td><td>strand</td><td>MM</td><td>target_seq</td><td>PAM</td><td>alignment</td><td>distance</td><td>position</td><td>gene name</td><td>gene id</td></tr>
<tr><td>GU305914.1</td><td>135440</td><td>135462</td><td>-</td><td>0</td><td>GAAACCCGGGCCGTCGGCGG</td><td>CGG</td><td>||||||||[||||||||||||]PAM</td><td>NA</td><td>Intergenic</td><td>NA</td><td>NA</td></tr>
<tr><td></td><td></td><td></td><td></td><td></td><td></td><td></td><td></td><td></td><td></td><td></td><td></td></tr>
<tr><td>T23781</td><td>GGCGGCGCCCAACAGTCCCTCGG</td><td>1000</td><td></td><td></td><td></td><td></td><td></td><td></td><td></td><td></td><td></td></tr>
<tr><td>Oligo fwd</td><td>CACCGGCGGCGCCCAACAGTCCCT</td><td></td><td></td><td></td><td></td><td></td><td></td><td></td><td></td><td></td><td></td></tr>
<tr><td>Oligo rev</td><td>AAACAGGGACTGTTGGGCGCCGCC</td><td></td><td></td><td></td><td></td><td></td><td></td><td></td><td></td><td></td><td></td></tr>
<tr><td>Chromosome</td><td>start</td><td>end</td><td>strand</td><td>MM</td><td>target_seq</td><td>PAM</td><td>alignment</td><td>distance</td><td>position</td><td>gene name</td><td>gene id</td></tr>
<tr><td>GU305914.1</td><td>135422</td><td>135444</td><td>-</td><td>0</td><td>GGCGGCGCCCAACAGTCCCT</td><td>CGG</td><td>||||||||[||||||||||||]PAM</td><td>NA</td><td>Intergenic</td><td>NA</td><td>NA</td></tr>
<tr><td></td><td></td><td></td><td></td><td></td><td></td><td></td><td></td><td></td><td></td><td></td><td></td></tr>
<tr><td>T23782</td><td>GCGGCGCCCAACAGTCCCTCGGG</td><td>1000</td><td></td><td></td><td></td><td></td><td></td><td></td><td></td><td></td><td></td></tr>
<tr><td>Oligo fwd</td><td>CACCGCGGCGCCCAACAGTCCCTC</td><td></td><td></td><td></td><td></td><td></td><td></td><td></td><td></td><td></td><td></td></tr>
<tr><td>Oligo rev</td><td>AAACGAGGGACTGTTGGGCGCCGC</td><td></td><td></td><td></td><td></td><td></td><td></td><td></td><td></td><td></td><td></td></tr>
<tr><td>Chromosome</td><td>start</td><td>end</td><td>strand</td><td>MM</td><td>target_seq</td><td>PAM</td><td>alignment</td><td>distance</td><td>position</td><td>gene name</td><td>gene id</td></tr>
<tr><td>GU305914.1</td><td>135421</td><td>135443</td><td>-</td><td>0</td><td>GCGGCGCCCAACAGTCCCTC</td><td>GGG</td><td>||||||||[||||||||||||]PAM</td><td>NA</td><td>Intergenic</td><td>NA</td><td>NA</td></tr>
<tr><td></td><td></td><td></td><td></td><td></td><td></td><td></td><td></td><td></td><td></td><td></td><td></td></tr>
<tr><td>T23783</td><td>CATCACCCGCCGCGCCCGCCCGG</td><td>1000</td><td></td><td></td><td></td><td></td><td></td><td></td><td></td><td></td><td></td></tr>
<tr><td>Oligo fwd</td><td>CACCCATCACCCGCCGCGCCCGCC</td><td></td><td></td><td></td><td></td><td></td><td></td><td></td><td></td><td></td><td></td></tr>
<tr><td>Oligo rev</td><td>AAACGGCGGGCGCGGCGGGTGATG</td><td></td><td></td><td></td><td></td><td></td><td></td><td></td><td></td><td></td><td></td></tr>
<tr><td>Chromosome</td><td>start</td><td>end</td><td>strand</td><td>MM</td><td>target_seq</td><td>PAM</td><td>alignment</td><td>distance</td><td>position</td><td>gene name</td><td>gene id</td></tr>
<tr><td>GU305914.1</td><td>135366</td><td>135388</td><td>-</td><td>0</td><td>CATCACCCGCCGCGCCCGCC</td><td>CGG</td><td>||||||||[||||||||||||]PAM</td><td>NA</td><td>Intergenic</td><td>NA</td><td>NA</td></tr>
<tr><td></td><td></td><td></td><td></td><td></td><td></td><td></td><td></td><td></td><td></td><td></td><td></td></tr>
<tr><td>T23784</td><td>CCGCCGCGCCCGCCCGGTGATGG</td><td>1000</td><td></td><td></td><td></td><td></td><td></td><td></td><td></td><td></td><td></td></tr>
<tr><td>Oligo fwd</td><td>CACCCCGCCGCGCCCGCCCGGTGA</td><td></td><td></td><td></td><td></td><td></td><td></td><td></td><td></td><td></td><td></td></tr>
<tr><td>Oligo rev</td><td>AAACTCACCGGGCGGGCGCGGCGG</td><td></td><td></td><td></td><td></td><td></td><td></td><td></td><td></td><td></td><td></td></tr>
<tr><td>Chromosome</td><td>start</td><td>end</td><td>strand</td><td>MM</td><td>target_seq</td><td>PAM</td><td>alignment</td><td>distance</td><td>position</td><td>gene name</td><td>gene id</td></tr>
<tr><td>GU305914.1</td><td>135360</td><td>135382</td><td>-</td><td>0</td><td>CCGCCGCGCCCGCCCGGTGA</td><td>TGG</td><td>||||||||[||||||||||||]PAM</td><td>NA</td><td>Intergenic</td><td>NA</td><td>NA</td></tr>
<tr><td></td><td></td><td></td><td></td><td></td><td></td><td></td><td></td><td></td><td></td><td></td><td></td></tr>
<tr><td>T23785</td><td>GCGCCCGCCCGGTGATGGTGAGG</td><td>1000</td><td></td><td></td><td></td><td></td><td></td><td></td><td></td><td></td><td></td></tr>
<tr><td>Oligo fwd</td><td>CACCGCGCCCGCCCGGTGATGGTG</td><td></td><td></td><td></td><td></td><td></td><td></td><td></td><td></td><td></td><td></td></tr>
<tr><td>Oligo rev</td><td>AAACCACCATCACCGGGCGGGCGC</td><td></td><td></td><td></td><td></td><td></td><td></td><td></td><td></td><td></td><td></td></tr>
<tr><td>Chromosome</td><td>start</td><td>end</td><td>strand</td><td>MM</td><td>target_seq</td><td>PAM</td><td>alignment</td><td>distance</td><td>position</td><td>gene name</td><td>gene id</td></tr>
<tr><td>GU305914.1</td><td>135355</td><td>135377</td><td>-</td><td>0</td><td>GCGCCCGCCCGGTGATGGTG</td><td>AGG</td><td>||||||||[||||||||||||]PAM</td><td>NA</td><td>Intergenic</td><td>NA</td><td>NA</td></tr>
<tr><td>GU305914.1</td><td>117425</td><td>117447</td><td>+</td><td>4</td><td>GCGCCAGGCAGGTGATGGCG</td><td>GGG</td><td>|||||-|-[|-||||||||-|]PAM</td><td>NA</td><td>Intergenic</td><td>NA</td><td>NA</td></tr>
<tr><td></td><td></td><td></td><td></td><td></td><td></td><td></td><td></td><td></td><td></td><td></td><td></td></tr>
<tr><td>T23786</td><td>ATGATGCAGCAGCCGACTACTGG</td><td>1000</td><td></td><td></td><td></td><td></td><td></td><td></td><td></td><td></td><td></td></tr>
<tr><td>Oligo fwd</td><td>CACCATGATGCAGCAGCCGACTAC</td><td></td><td></td><td></td><td></td><td></td><td></td><td></td><td></td><td></td><td></td></tr>
<tr><td>Oligo rev</td><td>AAACGTAGTCGGCTGCTGCATCAT</td><td></td><td></td><td></td><td></td><td></td><td></td><td></td><td></td><td></td><td></td></tr>
<tr><td>Chromosome</td><td>start</td><td>end</td><td>strand</td><td>MM</td><td>target_seq</td><td>PAM</td><td>alignment</td><td>distance</td><td>position</td><td>gene name</td><td>gene id</td></tr>
<tr><td>GU305914.1</td><td>135329</td><td>135351</td><td>-</td><td>0</td><td>ATGATGCAGCAGCCGACTAC</td><td>TGG</td><td>||||||||[||||||||||||]PAM</td><td>NA</td><td>Intergenic</td><td>NA</td><td>NA</td></tr>
<tr><td></td><td></td><td></td><td></td><td></td><td></td><td></td><td></td><td></td><td></td><td></td><td></td></tr>
<tr><td>T23787</td><td>GCAGCCGACTACTGGCGCTGCGG</td><td>1000</td><td></td><td></td><td></td><td></td><td></td><td></td><td></td><td></td><td></td></tr>
<tr><td>Oligo fwd</td><td>CACCGCAGCCGACTACTGGCGCTG</td><td></td><td></td><td></td><td></td><td></td><td></td><td></td><td></td><td></td><td></td></tr>
<tr><td>Oligo rev</td><td>AAACCAGCGCCAGTAGTCGGCTGC</td><td></td><td></td><td></td><td></td><td></td><td></td><td></td><td></td><td></td><td></td></tr>
<tr><td>Chromosome</td><td>start</td><td>end</td><td>strand</td><td>MM</td><td>target_seq</td><td>PAM</td><td>alignment</td><td>distance</td><td>position</td><td>gene name</td><td>gene id</td></tr>
<tr><td>GU305914.1</td><td>135321</td><td>135343</td><td>-</td><td>0</td><td>GCAGCCGACTACTGGCGCTG</td><td>CGG</td><td>||||||||[||||||||||||]PAM</td><td>NA</td><td>Intergenic</td><td>NA</td><td>NA</td></tr>
<tr><td></td><td></td><td></td><td></td><td></td><td></td><td></td><td></td><td></td><td></td><td></td><td></td></tr>
<tr><td>T23788</td><td>CAGCCGACTACTGGCGCTGCGGG</td><td>1000</td><td></td><td></td><td></td><td></td><td></td><td></td><td></td><td></td><td></td></tr>
<tr><td>Oligo fwd</td><td>CACCCAGCCGACTACTGGCGCTGC</td><td></td><td></td><td></td><td></td><td></td><td></td><td></td><td></td><td></td><td></td></tr>
<tr><td>Oligo rev</td><td>AAACGCAGCGCCAGTAGTCGGCTG</td><td></td><td></td><td></td><td></td><td></td><td></td><td></td><td></td><td></td><td></td></tr>
<tr><td>Chromosome</td><td>start</td><td>end</td><td>strand</td><td>MM</td><td>target_seq</td><td>PAM</td><td>alignment</td><td>distance</td><td>position</td><td>gene name</td><td>gene id</td></tr>
<tr><td>GU305914.1</td><td>135320</td><td>135342</td><td>-</td><td>0</td><td>CAGCCGACTACTGGCGCTGC</td><td>GGG</td><td>||||||||[||||||||||||]PAM</td><td>NA</td><td>Intergenic</td><td>NA</td><td>NA</td></tr>
<tr><td></td><td></td><td></td><td></td><td></td><td></td><td></td><td></td><td></td><td></td><td></td><td></td></tr>
<tr><td>T23789</td><td>AGCCGACTACTGGCGCTGCGGGG</td><td>1000</td><td></td><td></td><td></td><td></td><td></td><td></td><td></td><td></td><td></td></tr>
<tr><td>Oligo fwd</td><td>CACCAGCCGACTACTGGCGCTGCG</td><td></td><td></td><td></td><td></td><td></td><td></td><td></td><td></td><td></td><td></td></tr>
<tr><td>Oligo rev</td><td>AAACCGCAGCGCCAGTAGTCGGCT</td><td></td><td></td><td></td><td></td><td></td><td></td><td></td><td></td><td></td><td></td></tr>
<tr><td>Chromosome</td><td>start</td><td>end</td><td>strand</td><td>MM</td><td>target_seq</td><td>PAM</td><td>alignment</td><td>distance</td><td>position</td><td>gene name</td><td>gene id</td></tr>
<tr><td>GU305914.1</td><td>135319</td><td>135341</td><td>-</td><td>0</td><td>AGCCGACTACTGGCGCTGCG</td><td>GGG</td><td>||||||||[||||||||||||]PAM</td><td>NA</td><td>Intergenic</td><td>NA</td><td>NA</td></tr>
<tr><td></td><td></td><td></td><td></td><td></td><td></td><td></td><td></td><td></td><td></td><td></td><td></td></tr>
<tr><td>T23790</td><td>GCGCTGCGGGGCCCGCTCCGCGG</td><td>1000</td><td></td><td></td><td></td><td></td><td></td><td></td><td></td><td></td><td></td></tr>
<tr><td>Oligo fwd</td><td>CACCGCGCTGCGGGGCCCGCTCCG</td><td></td><td></td><td></td><td></td><td></td><td></td><td></td><td></td><td></td><td></td></tr>
<tr><td>Oligo rev</td><td>AAACCGGAGCGGGCCCCGCAGCGC</td><td></td><td></td><td></td><td></td><td></td><td></td><td></td><td></td><td></td><td></td></tr>
<tr><td>Chromosome</td><td>start</td><td>end</td><td>strand</td><td>MM</td><td>target_seq</td><td>PAM</td><td>alignment</td><td>distance</td><td>position</td><td>gene name</td><td>gene id</td></tr>
<tr><td>GU305914.1</td><td>135307</td><td>135329</td><td>-</td><td>0</td><td>GCGCTGCGGGGCCCGCTCCG</td><td>CGG</td><td>||||||||[||||||||||||]PAM</td><td>NA</td><td>Intergenic</td><td>NA</td><td>NA</td></tr>
<tr><td>GU305914.1</td><td>191574</td><td>191596</td><td>+</td><td>4</td><td>GCACTTCGGGGCCCTCTCTG</td><td>AGG</td><td>||-||-||[||||||-|||-|]PAM</td><td>NA</td><td>Intergenic</td><td>NA</td><td>NA</td></tr>
<tr><td></td><td></td><td></td><td></td><td></td><td></td><td></td><td></td><td></td><td></td><td></td><td></td></tr>
<tr><td>T23791</td><td>GCCTGACCTCTCCGAAAAAGCGG</td><td>1000</td><td></td><td></td><td></td><td></td><td></td><td></td><td></td><td></td><td></td></tr>
<tr><td>Oligo fwd</td><td>CACCGCCTGACCTCTCCGAAAAAG</td><td></td><td></td><td></td><td></td><td></td><td></td><td></td><td></td><td></td><td></td></tr>
<tr><td>Oligo rev</td><td>AAACCTTTTTCGGAGAGGTCAGGC</td><td></td><td></td><td></td><td></td><td></td><td></td><td></td><td></td><td></td><td></td></tr>
</table>

| | | | | | | | | | | | |
|---|---|---|---|---|---|---|---|---|---|---|---|
| Chromosome | start | end | strand | MM | target_seq | PAM | alignment | distance | position | gene name | gene id |
| GU305914.1 | 135273 | 135295 | - | 0 | GCCTGACCTCTCCGAAAAAG | CGG | \|\|\|\|\|\|\|\|[\|\|\|\|\|\|\|\|\|\|\|\|]PAM | NA | Intergenic | NA | NA |
| T23792 | TTCGCGCAGAAGTCTGACGAAGG | 1000 | | | | | | | | | |
| Oligo fwd | CACCTTCGCGCAGAAGTCTGACGA | | | | | | | | | | |
| Oligo rev | AAACTCGTCAGACTTCTGCGCGAA | | | | | | | | | | |
| Chromosome | start | end | strand | MM | target_seq | PAM | alignment | distance | position | gene name | gene id |
| GU305914.1 | 135245 | 135267 | - | 0 | TTCGCGCAGAAGTCTGACGA | AGG | \|\|\|\|\|\|\|\|[\|\|\|\|\|\|\|\|\|\|\|\|]PAM | NA | Intergenic | NA | NA |
| T23793 | CCTTCCACCACTCGACGTCGCGG | 1000 | | | | | | | | | |
| Oligo fwd | CACCCCTTCCACCACTCGACGTCG | | | | | | | | | | |
| Oligo rev | AAACCGACGTCGAGTGGTGGAAGG | | | | | | | | | | |
| Chromosome | start | end | strand | MM | target_seq | PAM | alignment | distance | position | gene name | gene id |
| GU305914.1 | 135165 | 135187 | - | 0 | CCTTCCACCACTCGACGTCG | CGG | \|\|\|\|\|\|\|\|[\|\|\|\|\|\|\|\|\|\|\|\|]PAM | NA | Intergenic | NA | NA |
| T23794 | TCCACCACTCGACGTCGCGGAGG | 1000 | | | | | | | | | |
| Oligo fwd | CACCTCCACCACTCGACGTCGCGG | | | | | | | | | | |
| Oligo rev | AAACCCGCGACGTCGAGTGGTGGA | | | | | | | | | | |
| Chromosome | start | end | strand | MM | target_seq | PAM | alignment | distance | position | gene name | gene id |
| GU305914.1 | 135162 | 135184 | - | 0 | TCCACCACTCGACGTCGCGG | AGG | \|\|\|\|\|\|\|\|[\|\|\|\|\|\|\|\|\|\|\|\|]PAM | NA | Intergenic | NA | NA |
| T23795 | GAGGTGCAGCACCACGCTGAAGG | 1000 | | | | | | | | | |
| Oligo fwd | CACCGAGGTGCAGCACCACGCTGA | | | | | | | | | | |
| Oligo rev | AAACTCAGCGTGGTGCTGCACCTC | | | | | | | | | | |
| Chromosome | start | end | strand | MM | target_seq | PAM | alignment | distance | position | gene name | gene id |
| GU305914.1 | 135143 | 135165 | - | 0 | GAGGTGCAGCACCACGCTGA | AGG | \|\|\|\|\|\|\|\|[\|\|\|\|\|\|\|\|\|\|\|\|]PAM | NA | Intergenic | NA | NA |
| GU305914.1 | 77692 | 77714 | + | 4 | GATGTTCAACACGACGCTGA | CGG | \|\|-\|\|-\|\|[-\|\|\|-\|\|\|\|\|\|\|]PAM | NA | Intergenic | NA | NA |
| T23796 | TCTCGTCGCCGCCGCCGTGCAGG | 1000 | | | | | | | | | |
| Oligo fwd | CACCTCTCGTCGCCGCCGCCGTGC | | | | | | | | | | |
| Oligo rev | AAACGCACGGCGGCGGCGACGAGA | | | | | | | | | | |
| Chromosome | start | end | strand | MM | target_seq | PAM | alignment | distance | position | gene name | gene id |
| GU305914.1 | 135102 | 135124 | - | 0 | TCTCGTCGCCGCCGCCGTGC | AGG | \|\|\|\|\|\|\|\|[\|\|\|\|\|\|\|\|\|\|\|\|]PAM | NA | Intergenic | NA | NA |
| T23797 | CTCGTCGCCGCCGCCGTGCAGGG | 1000 | | | | | | | | | |
| Oligo fwd | CACCCTCGTCGCCGCCGCCGTGCA | | | | | | | | | | |
| Oligo rev | AAACTGCACGGCGGCGGCGACGAG | | | | | | | | | | |
| Chromosome | start | end | strand | MM | target_seq | PAM | alignment | distance | position | gene name | gene id |
| GU305914.1 | 135101 | 135123 | - | 0 | CTCGTCGCCGCCGCCGTGCA | GGG | \|\|\|\|\|\|\|\|[\|\|\|\|\|\|\|\|\|\|\|\|]PAM | NA | Intergenic | NA | NA |
| T23798 | GCCGCCGCCGTGCAGGGTCCAGG | 1000 | | | | | | | | | |
| Oligo fwd | CACCGCCGCCGCCGTGCAGGGTCC | | | | | | | | | | |
| Oligo rev | AAACGGACCCTGCACGGCGGCGGC | | | | | | | | | | |
| Chromosome | start | end | strand | MM | target_seq | PAM | alignment | distance | position | gene name | gene id |
| GU305914.1 | 135095 | 135117 | - | 0 | GCCGCCGCCGTGCAGGGTCC | AGG | \|\|\|\|\|\|\|\|[\|\|\|\|\|\|\|\|\|\|\|\|]PAM | NA | Intergenic | NA | NA |
| T23799 | CCGCCGCCGTGCAGGGTCCAGGG | 1000 | | | | | | | | | |
| Oligo fwd | CACCCCGCCGCCGTGCAGGGTCCA | | | | | | | | | | |
| Oligo rev | AAACTGGACCCTGCACGGCGGCGG | | | | | | | | | | |
| Chromosome | start | end | strand | MM | target_seq | PAM | alignment | distance | position | gene name | gene id |
| GU305914.1 | 135094 | 135116 | - | 0 | CCGCCGCCGTGCAGGGTCCA | GGG | \|\|\|\|\|\|\|\|[\|\|\|\|\|\|\|\|\|\|\|\|]PAM | NA | Intergenic | NA | NA |
| T23800 | CAGGGTCCAGGGCCCGTCTCCGG | 1000 | | | | | | | | | |
| Oligo fwd | CACCCAGGGTCCAGGGCCCGTCTC | | | | | | | | | | |
| Oligo rev | AAACGAGACGGGCCCTGGACCCTG | | | | | | | | | | |
| Chromosome | start | end | strand | MM | target_seq | PAM | alignment | distance | position | gene name | gene id |
| GU305914.1 | 135083 | 135105 | - | 0 | CAGGGTCCAGGGCCCGTCTC | CGG | \|\|\|\|\|\|\|\|[\|\|\|\|\|\|\|\|\|\|\|\|]PAM | NA | Intergenic | NA | NA |
| T23801 | AGGGTCCAGGGCCCGTCTCCGGG | 1000 | | | | | | | | | |
| Oligo fwd | CACCAGGGTCCAGGGCCCGTCTCC | | | | | | | | | | |
| Oligo rev | AAACGGAGACGGGCCCTGGACCCT | | | | | | | | | | |
| Chromosome | start | end | strand | MM | target_seq | PAM | alignment | distance | position | gene name | gene id |
| GU305914.1 | 135082 | 135104 | - | 0 | AGGGTCCAGGGCCCGTCTCC | GGG | \|\|\|\|\|\|\|\|[\|\|\|\|\|\|\|\|\|\|\|\|]PAM | NA | Intergenic | NA | NA |
| T23802 | GGGTCCAGGGCCCGTCTCCGGGG | 1000 | | | | | | | | | |
| Oligo fwd | CACCGGGTCCAGGGCCCGTCTCCG | | | | | | | | | | |
| Oligo rev | AAACCGGAGACGGGCCCTGGACCC | | | | | | | | | | |
| Chromosome | start | end | strand | MM | target_seq | PAM | alignment | distance | position | gene name | gene id |
| GU305914.1 | 135081 | 135103 | - | 0 | GGGTCCAGGGCCCGTCTCCG | GGG | \|\|\|\|\|\|\|\|[\|\|\|\|\|\|\|\|\|\|\|\|]PAM | NA | Intergenic | NA | NA |
| T23803 | GGTCCAGGGCCCGTCTCCGGGGG | 1000 | | | | | | | | | |
| Oligo fwd | CACCGGTCCAGGGCCCGTCTCCGG | | | | | | | | | | |
| Oligo rev | AAACCCGGAGACGGGCCCTGGACC | | | | | | | | | | |
| Chromosome | start | end | strand | MM | target_seq | PAM | alignment | distance | position | gene name | gene id |
| GU305914.1 | 135080 | 135102 | - | 0 | GGTCCAGGGCCCGTCTCCGG | GGG | \|\|\|\|\|\|\|\|[\|\|\|\|\|\|\|\|\|\|\|\|]PAM | NA | Intergenic | NA | NA |
| T23804 | CCAGGGCCCGTCTCCGGGGGAGG | 1000 | | | | | | | | | |
| Oligo fwd | CACCCCAGGGCCCGTCTCCGGGGG | | | | | | | | | | |
| Oligo rev | AAACCCCCCGGAGACGGGCCCTGG | | | | | | | | | | |
| Chromosome | start | end | strand | MM | target_seq | PAM | alignment | distance | position | gene name | gene id |
| GU305914.1 | 135077 | 135099 | - | 0 | CCAGGGCCCGTCTCCGGGGG | AGG | \|\|\|\|\|\|\|\|[\|\|\|\|\|\|\|\|\|\|\|\|]PAM | NA | Intergenic | NA | NA |
| T23805 | CGTCTCCGGGGGAGGCCGAGAGG | 1000 | | | | | | | | | |
| Oligo fwd | CACCCGTCTCCGGGGGAGGCCGAG | | | | | | | | | | |
| Oligo rev | AAACCTCGGCCTCCCCCGGAGACG | | | | | | | | | | |
| Chromosome | start | end | strand | MM | target_seq | PAM | alignment | distance | position | gene name | gene id |
| GU305914.1 | 135069 | 135091 | - | 0 | CGTCTCCGGGGGAGGCCGAG | AGG | \|\|\|\|\|\|\|\|[\|\|\|\|\|\|\|\|\|\|\|\|]PAM | NA | Intergenic | NA | NA |
| T23806 | AGAGGTGCCTGCACTTGTCGTGG | 1000 | | | | | | | | | |

<table>
<tr><td>Oligo fwd</td><td>CACCAGAGGTGCCTGCACTTGTCG</td><td></td><td></td><td></td><td></td><td></td><td></td><td></td><td></td><td></td><td></td></tr>
<tr><td>Oligo rev</td><td>AAACCGACAAGTGCAGGCACCTCT</td><td></td><td></td><td></td><td></td><td></td><td></td><td></td><td></td><td></td><td></td></tr>
<tr><td>Chromosome</td><td>start</td><td>end</td><td>strand</td><td>MM</td><td>target_seq</td><td>PAM</td><td>alignment</td><td>distance</td><td>position</td><td>gene name</td><td>gene id</td></tr>
<tr><td>GU305914.1</td><td>135051</td><td>135073</td><td>-</td><td>0</td><td>AGAGGTGCCTGCACTTGTCG</td><td>TGG</td><td>||||||||[||||||||||||]PAM</td><td>NA</td><td>Intergenic</td><td>NA</td><td>NA</td></tr>
<tr><td></td><td></td><td></td><td></td><td></td><td></td><td></td><td></td><td></td><td></td><td></td><td></td></tr>
<tr><td>T23807</td><td>GACTCTTGAGCACGAACTCGCGG</td><td>1000</td><td></td><td></td><td></td><td></td><td></td><td></td><td></td><td></td><td></td></tr>
<tr><td>Oligo fwd</td><td>CACCGACTCTTGAGCACGAACTCG</td><td></td><td></td><td></td><td></td><td></td><td></td><td></td><td></td><td></td><td></td></tr>
<tr><td>Oligo rev</td><td>AAACCGAGTTCGTGCTCAAGAGTC</td><td></td><td></td><td></td><td></td><td></td><td></td><td></td><td></td><td></td><td></td></tr>
<tr><td>Chromosome</td><td>start</td><td>end</td><td>strand</td><td>MM</td><td>target_seq</td><td>PAM</td><td>alignment</td><td>distance</td><td>position</td><td>gene name</td><td>gene id</td></tr>
<tr><td>GU305914.1</td><td>135024</td><td>135046</td><td>-</td><td>0</td><td>GACTCTTGAGCACGAACTCG</td><td>CGG</td><td>||||||||[||||||||||||]PAM</td><td>NA</td><td>Intergenic</td><td>NA</td><td>NA</td></tr>
<tr><td></td><td></td><td></td><td></td><td></td><td></td><td></td><td></td><td></td><td></td><td></td><td></td></tr>
<tr><td>T23808</td><td>GCGGTCCAGACGCTCGAAGTAGG</td><td>1000</td><td></td><td></td><td></td><td></td><td></td><td></td><td></td><td></td><td></td></tr>
<tr><td>Oligo fwd</td><td>CACCGCGGTCCAGACGCTCGAAGT</td><td></td><td></td><td></td><td></td><td></td><td></td><td></td><td></td><td></td><td></td></tr>
<tr><td>Oligo rev</td><td>AAACACTTCGAGCGTCTGGACCGC</td><td></td><td></td><td></td><td></td><td></td><td></td><td></td><td></td><td></td><td></td></tr>
<tr><td>Chromosome</td><td>start</td><td>end</td><td>strand</td><td>MM</td><td>target_seq</td><td>PAM</td><td>alignment</td><td>distance</td><td>position</td><td>gene name</td><td>gene id</td></tr>
<tr><td>GU305914.1</td><td>135005</td><td>135027</td><td>-</td><td>0</td><td>GCGGTCCAGACGCTCGAAGT</td><td>AGG</td><td>||||||||[||||||||||||]PAM</td><td>NA</td><td>Intergenic</td><td>NA</td><td>NA</td></tr>
<tr><td></td><td></td><td></td><td></td><td></td><td></td><td></td><td></td><td></td><td></td><td></td><td></td></tr>
<tr><td>T23809</td><td>CGGTCCAGACGCTCGAAGTAGGG</td><td>1000</td><td></td><td></td><td></td><td></td><td></td><td></td><td></td><td></td><td></td></tr>
<tr><td>Oligo fwd</td><td>CACCCGGTCCAGACGCTCGAAGTA</td><td></td><td></td><td></td><td></td><td></td><td></td><td></td><td></td><td></td><td></td></tr>
<tr><td>Oligo rev</td><td>AAACTACTTCGAGCGTCTGGACCG</td><td></td><td></td><td></td><td></td><td></td><td></td><td></td><td></td><td></td><td></td></tr>
<tr><td>Chromosome</td><td>start</td><td>end</td><td>strand</td><td>MM</td><td>target_seq</td><td>PAM</td><td>alignment</td><td>distance</td><td>position</td><td>gene name</td><td>gene id</td></tr>
<tr><td>GU305914.1</td><td>135004</td><td>135026</td><td>-</td><td>0</td><td>CGGTCCAGACGCTCGAAGTA</td><td>GGG</td><td>||||||||[||||||||||||]PAM</td><td>NA</td><td>Intergenic</td><td>NA</td><td>NA</td></tr>
<tr><td></td><td></td><td></td><td></td><td></td><td></td><td></td><td></td><td></td><td></td><td></td><td></td></tr>
<tr><td>T23810</td><td>GGTCCAGACGCTCGAAGTAGGGG</td><td>1000</td><td></td><td></td><td></td><td></td><td></td><td></td><td></td><td></td><td></td></tr>
<tr><td>Oligo fwd</td><td>CACCGGTCCAGACGCTCGAAGTAG</td><td></td><td></td><td></td><td></td><td></td><td></td><td></td><td></td><td></td><td></td></tr>
<tr><td>Oligo rev</td><td>AAACCTACTTCGAGCGTCTGGACC</td><td></td><td></td><td></td><td></td><td></td><td></td><td></td><td></td><td></td><td></td></tr>
<tr><td>Chromosome</td><td>start</td><td>end</td><td>strand</td><td>MM</td><td>target_seq</td><td>PAM</td><td>alignment</td><td>distance</td><td>position</td><td>gene name</td><td>gene id</td></tr>
<tr><td>GU305914.1</td><td>135003</td><td>135025</td><td>-</td><td>0</td><td>GGTCCAGACGCTCGAAGTAG</td><td>GGG</td><td>||||||||[||||||||||||]PAM</td><td>NA</td><td>Intergenic</td><td>NA</td><td>NA</td></tr>
<tr><td></td><td></td><td></td><td></td><td></td><td></td><td></td><td></td><td></td><td></td><td></td><td></td></tr>
<tr><td>T23811</td><td>GATGTGCTCCCAGCGCGCGTCGG</td><td>1000</td><td></td><td></td><td></td><td></td><td></td><td></td><td></td><td></td><td></td></tr>
<tr><td>Oligo fwd</td><td>CACCGATGTGCTCCCAGCGCGCGT</td><td></td><td></td><td></td><td></td><td></td><td></td><td></td><td></td><td></td><td></td></tr>
<tr><td>Oligo rev</td><td>AAACACGCGCGCTGGGAGCACATC</td><td></td><td></td><td></td><td></td><td></td><td></td><td></td><td></td><td></td><td></td></tr>
<tr><td>Chromosome</td><td>start</td><td>end</td><td>strand</td><td>MM</td><td>target_seq</td><td>PAM</td><td>alignment</td><td>distance</td><td>position</td><td>gene name</td><td>gene id</td></tr>
<tr><td>GU305914.1</td><td>134981</td><td>135003</td><td>-</td><td>0</td><td>GATGTGCTCCCAGCGCGCGT</td><td>CGG</td><td>||||||||[||||||||||||]PAM</td><td>NA</td><td>Intergenic</td><td>NA</td><td>NA</td></tr>
<tr><td></td><td></td><td></td><td></td><td></td><td></td><td></td><td></td><td></td><td></td><td></td><td></td></tr>
<tr><td>T23812</td><td>ATGTGCTCCCAGCGCGCGTCGGG</td><td>1000</td><td></td><td></td><td></td><td></td><td></td><td></td><td></td><td></td><td></td></tr>
<tr><td>Oligo fwd</td><td>CACCATGTGCTCCCAGCGCGCGTC</td><td></td><td></td><td></td><td></td><td></td><td></td><td></td><td></td><td></td><td></td></tr>
<tr><td>Oligo rev</td><td>AAACGACGCGCGCTGGGAGCACAT</td><td></td><td></td><td></td><td></td><td></td><td></td><td></td><td></td><td></td><td></td></tr>
<tr><td>Chromosome</td><td>start</td><td>end</td><td>strand</td><td>MM</td><td>target_seq</td><td>PAM</td><td>alignment</td><td>distance</td><td>position</td><td>gene name</td><td>gene id</td></tr>
<tr><td>GU305914.1</td><td>134980</td><td>135002</td><td>-</td><td>0</td><td>ATGTGCTCCCAGCGCGCGTC</td><td>GGG</td><td>||||||||[||||||||||||]PAM</td><td>NA</td><td>Intergenic</td><td>NA</td><td>NA</td></tr>
<tr><td></td><td></td><td></td><td></td><td></td><td></td><td></td><td></td><td></td><td></td><td></td><td></td></tr>
<tr><td>T23813</td><td>CTCCCAGCGCGCGTCGGGTCTGG</td><td>1000</td><td></td><td></td><td></td><td></td><td></td><td></td><td></td><td></td><td></td></tr>
<tr><td>Oligo fwd</td><td>CACCCTCCCAGCGCGCGTCGGGTC</td><td></td><td></td><td></td><td></td><td></td><td></td><td></td><td></td><td></td><td></td></tr>
<tr><td>Oligo rev</td><td>AAACGACCCGACGCGCGCTGGGAG</td><td></td><td></td><td></td><td></td><td></td><td></td><td></td><td></td><td></td><td></td></tr>
<tr><td>Chromosome</td><td>start</td><td>end</td><td>strand</td><td>MM</td><td>target_seq</td><td>PAM</td><td>alignment</td><td>distance</td><td>position</td><td>gene name</td><td>gene id</td></tr>
<tr><td>GU305914.1</td><td>134975</td><td>134997</td><td>-</td><td>0</td><td>CTCCCAGCGCGCGTCGGGTC</td><td>TGG</td><td>||||||||[||||||||||||]PAM</td><td>NA</td><td>Intergenic</td><td>NA</td><td>NA</td></tr>
<tr><td></td><td></td><td></td><td></td><td></td><td></td><td></td><td></td><td></td><td></td><td></td><td></td></tr>
<tr><td>T23814</td><td>TCTTGAAGCACAGCGTGTGCAGG</td><td>1000</td><td></td><td></td><td></td><td></td><td></td><td></td><td></td><td></td><td></td></tr>
<tr><td>Oligo fwd</td><td>CACCTCTTGAAGCACAGCGTGTGC</td><td></td><td></td><td></td><td></td><td></td><td></td><td></td><td></td><td></td><td></td></tr>
<tr><td>Oligo rev</td><td>AAACGCACACGCTGTGCTTCAAGA</td><td></td><td></td><td></td><td></td><td></td><td></td><td></td><td></td><td></td><td></td></tr>
<tr><td>Chromosome</td><td>start</td><td>end</td><td>strand</td><td>MM</td><td>target_seq</td><td>PAM</td><td>alignment</td><td>distance</td><td>position</td><td>gene name</td><td>gene id</td></tr>
<tr><td>GU305914.1</td><td>134946</td><td>134968</td><td>-</td><td>0</td><td>TCTTGAAGCACAGCGTGTGC</td><td>AGG</td><td>||||||||[||||||||||||]PAM</td><td>NA</td><td>Intergenic</td><td>NA</td><td>NA</td></tr>
<tr><td></td><td></td><td></td><td></td><td></td><td></td><td></td><td></td><td></td><td></td><td></td><td></td></tr>
<tr><td>T23815</td><td>TGTGCAGGCCCTCGAACTCCCGG</td><td>1000</td><td></td><td></td><td></td><td></td><td></td><td></td><td></td><td></td><td></td></tr>
<tr><td>Oligo fwd</td><td>CACCTGTGCAGGCCCTCGAACTCC</td><td></td><td></td><td></td><td></td><td></td><td></td><td></td><td></td><td></td><td></td></tr>
<tr><td>Oligo rev</td><td>AAACGGAGTTCGAGGGCCTGCACA</td><td></td><td></td><td></td><td></td><td></td><td></td><td></td><td></td><td></td><td></td></tr>
<tr><td>Chromosome</td><td>start</td><td>end</td><td>strand</td><td>MM</td><td>target_seq</td><td>PAM</td><td>alignment</td><td>distance</td><td>position</td><td>gene name</td><td>gene id</td></tr>
<tr><td>GU305914.1</td><td>134931</td><td>134953</td><td>-</td><td>0</td><td>TGTGCAGGCCCTCGAACTCC</td><td>CGG</td><td>||||||||[||||||||||||]PAM</td><td>NA</td><td>Intergenic</td><td>NA</td><td>NA</td></tr>
<tr><td></td><td></td><td></td><td></td><td></td><td></td><td></td><td></td><td></td><td></td><td></td><td></td></tr>
<tr><td>T23816</td><td>CAGGCCCTCGAACTCCCGGTAGG</td><td>1000</td><td></td><td></td><td></td><td></td><td></td><td></td><td></td><td></td><td></td></tr>
<tr><td>Oligo fwd</td><td>CACCCAGGCCCTCGAACTCCCGGT</td><td></td><td></td><td></td><td></td><td></td><td></td><td></td><td></td><td></td><td></td></tr>
<tr><td>Oligo rev</td><td>AAACACCGGGAGTTCGAGGGCCTG</td><td></td><td></td><td></td><td></td><td></td><td></td><td></td><td></td><td></td><td></td></tr>
<tr><td>Chromosome</td><td>start</td><td>end</td><td>strand</td><td>MM</td><td>target_seq</td><td>PAM</td><td>alignment</td><td>distance</td><td>position</td><td>gene name</td><td>gene id</td></tr>
<tr><td>GU305914.1</td><td>134927</td><td>134949</td><td>-</td><td>0</td><td>CAGGCCCTCGAACTCCCGGT</td><td>AGG</td><td>||||||||[||||||||||||]PAM</td><td>NA</td><td>Intergenic</td><td>NA</td><td>NA</td></tr>
<tr><td></td><td></td><td></td><td></td><td></td><td></td><td></td><td></td><td></td><td></td><td></td><td></td></tr>
<tr><td>T23817</td><td>TCAGCGAAAACGACGTGTACCGG</td><td>1000</td><td></td><td></td><td></td><td></td><td></td><td></td><td></td><td></td><td></td></tr>
<tr><td>Oligo fwd</td><td>CACCTCAGCGAAAACGACGTGTAC</td><td></td><td></td><td></td><td></td><td></td><td></td><td></td><td></td><td></td><td></td></tr>
<tr><td>Oligo rev</td><td>AAACGTACACGTCGTTTTCGCTGA</td><td></td><td></td><td></td><td></td><td></td><td></td><td></td><td></td><td></td><td></td></tr>
<tr><td>Chromosome</td><td>start</td><td>end</td><td>strand</td><td>MM</td><td>target_seq</td><td>PAM</td><td>alignment</td><td>distance</td><td>position</td><td>gene name</td><td>gene id</td></tr>
<tr><td>GU305914.1</td><td>134893</td><td>134915</td><td>-</td><td>0</td><td>TCAGCGAAAACGACGTGTAC</td><td>CGG</td><td>||||||||[||||||||||||]PAM</td><td>NA</td><td>Intergenic</td><td>NA</td><td>NA</td></tr>
<tr><td></td><td></td><td></td><td></td><td></td><td></td><td></td><td></td><td></td><td></td><td></td><td></td></tr>
<tr><td>T23818</td><td>GTACCGGCACGAGCCTGTCGCGG</td><td>1000</td><td></td><td></td><td></td><td></td><td></td><td></td><td></td><td></td><td></td></tr>
<tr><td>Oligo fwd</td><td>CACCGTACCGGCACGAGCCTGTCG</td><td></td><td></td><td></td><td></td><td></td><td></td><td></td><td></td><td></td><td></td></tr>
<tr><td>Oligo rev</td><td>AAACCGACAGGCTCGTGCCGGTAC</td><td></td><td></td><td></td><td></td><td></td><td></td><td></td><td></td><td></td><td></td></tr>
<tr><td>Chromosome</td><td>start</td><td>end</td><td>strand</td><td>MM</td><td>target_seq</td><td>PAM</td><td>alignment</td><td>distance</td><td>position</td><td>gene name</td><td>gene id</td></tr>
<tr><td>GU305914.1</td><td>134877</td><td>134899</td><td>-</td><td>0</td><td>GTACCGGCACGAGCCTGTCG</td><td>CGG</td><td>||||||||[||||||||||||]PAM</td><td>NA</td><td>Intergenic</td><td>NA</td><td>NA</td></tr>
<tr><td></td><td></td><td></td><td></td><td></td><td></td><td></td><td></td><td></td><td></td><td></td><td></td></tr>
<tr><td>T23819</td><td>GTCGCGGTCGTCCTCGCTAGCGG</td><td>1000</td><td></td><td></td><td></td><td></td><td></td><td></td><td></td><td></td><td></td></tr>
<tr><td>Oligo fwd</td><td>CACCGTCGCGGTCGTCCTCGCTAG</td><td></td><td></td><td></td><td></td><td></td><td></td><td></td><td></td><td></td><td></td></tr>
<tr><td>Oligo rev</td><td>AAACCTAGCGAGGACGACCGCGAC</td><td></td><td></td><td></td><td></td><td></td><td></td><td></td><td></td><td></td><td></td></tr>
<tr><td>Chromosome</td><td>start</td><td>end</td><td>strand</td><td>MM</td><td>target_seq</td><td>PAM</td><td>alignment</td><td>distance</td><td>position</td><td>gene name</td><td>gene id</td></tr>
<tr><td>GU305914.1</td><td>134861</td><td>134883</td><td>-</td><td>0</td><td>GTCGCGGTCGTCCTCGCTAG</td><td>CGG</td><td>||||||||[||||||||||||]PAM</td><td>NA</td><td>Intergenic</td><td>NA</td><td>NA</td></tr>
<tr><td>GU305914.1</td><td>79716</td><td>79738</td><td>-</td><td>4</td><td>GGAGCGGTCGTCCTCGCACG</td><td>AGG</td><td>|--|||||[|||||||||--|]PAM</td><td>NA</td><td>Intergenic</td><td>NA</td><td>NA</td></tr>
<tr><td></td><td></td><td></td><td></td><td></td><td></td><td></td><td></td><td></td><td></td><td></td><td></td></tr>
<tr><td>T23820</td><td>TAGCGGCCTGTTCGAAACGAAGG</td><td>1000</td><td></td><td></td><td></td><td></td><td></td><td></td><td></td><td></td><td></td></tr>
<tr><td>Oligo fwd</td><td>CACCTAGCGGCCTGTTCGAAACGA</td><td></td><td></td><td></td><td></td><td></td><td></td><td></td><td></td><td></td><td></td></tr>
<tr><td>Oligo rev</td><td>AAACTCGTTTCGAACAGGCCGCTA</td><td></td><td></td><td></td><td></td><td></td><td></td><td></td><td></td><td></td><td></td></tr>
<tr><td>Chromosome</td><td>start</td><td>end</td><td>strand</td><td>MM</td><td>target_seq</td><td>PAM</td><td>alignment</td><td>distance</td><td>position</td><td>gene name</td><td>gene id</td></tr>
<tr><td>GU305914.1</td><td>134844</td><td>134866</td><td>-</td><td>0</td><td>TAGCGGCCTGTTCGAAACGA</td><td>AGG</td><td>||||||||[||||||||||||]PAM</td><td>NA</td><td>Intergenic</td><td>NA</td><td>NA</td></tr>
</table>

<table>
<tr><td>T23821</td><td>ATTCGAAACGCATCCGTTTCAGG</td><td>1000</td><td></td><td></td><td></td><td></td><td></td><td></td><td></td><td></td><td></td></tr>
<tr><td>Oligo fwd</td><td>CACCATTCGAAACGCATCCGTTTC</td><td></td><td></td><td></td><td></td><td></td><td></td><td></td><td></td><td></td><td></td></tr>
<tr><td>Oligo rev</td><td>AAACGAAACGGATGCGTTTCGAAT</td><td></td><td></td><td></td><td></td><td></td><td></td><td></td><td></td><td></td><td></td></tr>
<tr><td>Chromosome</td><td>start</td><td>end</td><td>strand</td><td>MM</td><td>target_seq</td><td>PAM</td><td>alignment</td><td>distance</td><td>position</td><td>gene name</td><td>gene id</td></tr>
<tr><td>GU305914.1</td><td>134810</td><td>134832</td><td>-</td><td>0</td><td>ATTCGAAACGCATCCGTTTC</td><td>AGG</td><td>||||||||[||||||||||||]PAM</td><td>NA</td><td>Intergenic</td><td>NA</td><td>NA</td></tr>
<tr><td>T23822</td><td>CCTCCGACTGACTTTGATCACGG</td><td>1000</td><td></td><td></td><td></td><td></td><td></td><td></td><td></td><td></td><td></td></tr>
<tr><td>Oligo fwd</td><td>CACCCCTCCGACTGACTTTGATCA</td><td></td><td></td><td></td><td></td><td></td><td></td><td></td><td></td><td></td><td></td></tr>
<tr><td>Oligo rev</td><td>AAACTGATCAAAGTCAGTCGGAGG</td><td></td><td></td><td></td><td></td><td></td><td></td><td></td><td></td><td></td><td></td></tr>
<tr><td>Chromosome</td><td>start</td><td>end</td><td>strand</td><td>MM</td><td>target_seq</td><td>PAM</td><td>alignment</td><td>distance</td><td>position</td><td>gene name</td><td>gene id</td></tr>
<tr><td>GU305914.1</td><td>134770</td><td>134792</td><td>-</td><td>0</td><td>CCTCCGACTGACTTTGATCA</td><td>CGG</td><td>||||||||[||||||||||||]PAM</td><td>NA</td><td>Intergenic</td><td>NA</td><td>NA</td></tr>
<tr><td>T23823</td><td>GATCACGGTCGAGTCGTATTTGG</td><td>1000</td><td></td><td></td><td></td><td></td><td></td><td></td><td></td><td></td><td></td></tr>
<tr><td>Oligo fwd</td><td>CACCGATCACGGTCGAGTCGTATT</td><td></td><td></td><td></td><td></td><td></td><td></td><td></td><td></td><td></td><td></td></tr>
<tr><td>Oligo rev</td><td>AAACAATACGACTCGACCGTGATC</td><td></td><td></td><td></td><td></td><td></td><td></td><td></td><td></td><td></td><td></td></tr>
<tr><td>Chromosome</td><td>start</td><td>end</td><td>strand</td><td>MM</td><td>target_seq</td><td>PAM</td><td>alignment</td><td>distance</td><td>position</td><td>gene name</td><td>gene id</td></tr>
<tr><td>GU305914.1</td><td>134755</td><td>134777</td><td>-</td><td>0</td><td>GATCACGGTCGAGTCGTATT</td><td>TGG</td><td>||||||||[||||||||||||]PAM</td><td>NA</td><td>Intergenic</td><td>NA</td><td>NA</td></tr>
<tr><td>T23824</td><td>CGAGTCGTATTTGGTCACCCTGG</td><td>1000</td><td></td><td></td><td></td><td></td><td></td><td></td><td></td><td></td><td></td></tr>
<tr><td>Oligo fwd</td><td>CACCCGAGTCGTATTTGGTCACCC</td><td></td><td></td><td></td><td></td><td></td><td></td><td></td><td></td><td></td><td></td></tr>
<tr><td>Oligo rev</td><td>AAACGGGTGACCAAATACGACTCG</td><td></td><td></td><td></td><td></td><td></td><td></td><td></td><td></td><td></td><td></td></tr>
<tr><td>Chromosome</td><td>start</td><td>end</td><td>strand</td><td>MM</td><td>target_seq</td><td>PAM</td><td>alignment</td><td>distance</td><td>position</td><td>gene name</td><td>gene id</td></tr>
<tr><td>GU305914.1</td><td>134746</td><td>134768</td><td>-</td><td>0</td><td>CGAGTCGTATTTGGTCACCC</td><td>TGG</td><td>||||||||[||||||||||||]PAM</td><td>NA</td><td>Intergenic</td><td>NA</td><td>NA</td></tr>
<tr><td>T23825</td><td>CAGCTGACCTATGACGCTGATGG</td><td>1000</td><td></td><td></td><td></td><td></td><td></td><td></td><td></td><td></td><td></td></tr>
<tr><td>Oligo fwd</td><td>CACCCAGCTGACCTATGACGCTGA</td><td></td><td></td><td></td><td></td><td></td><td></td><td></td><td></td><td></td><td></td></tr>
<tr><td>Oligo rev</td><td>AAACTCAGCGTCATAGGTCAGCTG</td><td></td><td></td><td></td><td></td><td></td><td></td><td></td><td></td><td></td><td></td></tr>
<tr><td>Chromosome</td><td>start</td><td>end</td><td>strand</td><td>MM</td><td>target_seq</td><td>PAM</td><td>alignment</td><td>distance</td><td>position</td><td>gene name</td><td>gene id</td></tr>
<tr><td>GU305914.1</td><td>134695</td><td>134717</td><td>-</td><td>0</td><td>CAGCTGACCTATGACGCTGA</td><td>TGG</td><td>||||||||[||||||||||||]PAM</td><td>NA</td><td>Intergenic</td><td>NA</td><td>NA</td></tr>
<tr><td>T23826</td><td>AGCTGACCTATGACGCTGATGGG</td><td>1000</td><td></td><td></td><td></td><td></td><td></td><td></td><td></td><td></td><td></td></tr>
<tr><td>Oligo fwd</td><td>CACCAGCTGACCTATGACGCTGAT</td><td></td><td></td><td></td><td></td><td></td><td></td><td></td><td></td><td></td><td></td></tr>
<tr><td>Oligo rev</td><td>AAACATCAGCGTCATAGGTCAGCT</td><td></td><td></td><td></td><td></td><td></td><td></td><td></td><td></td><td></td><td></td></tr>
<tr><td>Chromosome</td><td>start</td><td>end</td><td>strand</td><td>MM</td><td>target_seq</td><td>PAM</td><td>alignment</td><td>distance</td><td>position</td><td>gene name</td><td>gene id</td></tr>
<tr><td>GU305914.1</td><td>134694</td><td>134716</td><td>-</td><td>0</td><td>AGCTGACCTATGACGCTGAT</td><td>GGG</td><td>||||||||[||||||||||||]PAM</td><td>NA</td><td>Intergenic</td><td>NA</td><td>NA</td></tr>
<tr><td>T23827</td><td>CGTTATGCTGCTTGAAGACGCGG</td><td>1000</td><td></td><td></td><td></td><td></td><td></td><td></td><td></td><td></td><td></td></tr>
<tr><td>Oligo fwd</td><td>CACCCGTTATGCTGCTTGAAGACG</td><td></td><td></td><td></td><td></td><td></td><td></td><td></td><td></td><td></td><td></td></tr>
<tr><td>Oligo rev</td><td>AAACCGTCTTCAAGCAGCATAACG</td><td></td><td></td><td></td><td></td><td></td><td></td><td></td><td></td><td></td><td></td></tr>
<tr><td>Chromosome</td><td>start</td><td>end</td><td>strand</td><td>MM</td><td>target_seq</td><td>PAM</td><td>alignment</td><td>distance</td><td>position</td><td>gene name</td><td>gene id</td></tr>
<tr><td>GU305914.1</td><td>134657</td><td>134679</td><td>-</td><td>0</td><td>CGTTATGCTGCTTGAAGACG</td><td>CGG</td><td>||||||||[||||||||||||]PAM</td><td>NA</td><td>Intergenic</td><td>NA</td><td>NA</td></tr>
<tr><td>T23828</td><td>AAAATAGATCGCATTCTCCGTGG</td><td>1000</td><td></td><td></td><td></td><td></td><td></td><td></td><td></td><td></td><td></td></tr>
<tr><td>Oligo fwd</td><td>CACCAAAATAGATCGCATTCTCCG</td><td></td><td></td><td></td><td></td><td></td><td></td><td></td><td></td><td></td><td></td></tr>
<tr><td>Oligo rev</td><td>AAACCGGAGAATGCGATCTATTTT</td><td></td><td></td><td></td><td></td><td></td><td></td><td></td><td></td><td></td><td></td></tr>
<tr><td>Chromosome</td><td>start</td><td>end</td><td>strand</td><td>MM</td><td>target_seq</td><td>PAM</td><td>alignment</td><td>distance</td><td>position</td><td>gene name</td><td>gene id</td></tr>
<tr><td>GU305914.1</td><td>134626</td><td>134648</td><td>-</td><td>0</td><td>AAAATAGATCGCATTCTCCG</td><td>TGG</td><td>||||||||[||||||||||||]PAM</td><td>NA</td><td>Intergenic</td><td>NA</td><td>NA</td></tr>
<tr><td>T23829</td><td>AAATAGATCGCATTCTCCGTGGG</td><td>1000</td><td></td><td></td><td></td><td></td><td></td><td></td><td></td><td></td><td></td></tr>
<tr><td>Oligo fwd</td><td>CACCAAATAGATCGCATTCTCCGT</td><td></td><td></td><td></td><td></td><td></td><td></td><td></td><td></td><td></td><td></td></tr>
<tr><td>Oligo rev</td><td>AAACACGGAGAATGCGATCTATTT</td><td></td><td></td><td></td><td></td><td></td><td></td><td></td><td></td><td></td><td></td></tr>
<tr><td>Chromosome</td><td>start</td><td>end</td><td>strand</td><td>MM</td><td>target_seq</td><td>PAM</td><td>alignment</td><td>distance</td><td>position</td><td>gene name</td><td>gene id</td></tr>
<tr><td>GU305914.1</td><td>134625</td><td>134647</td><td>-</td><td>0</td><td>AAATAGATCGCATTCTCCGT</td><td>GGG</td><td>||||||||[||||||||||||]PAM</td><td>NA</td><td>Intergenic</td><td>NA</td><td>NA</td></tr>
<tr><td>T23830</td><td>TCCCTATGACATCCGCGTAGTGG</td><td>1000</td><td></td><td></td><td></td><td></td><td></td><td></td><td></td><td></td><td></td></tr>
<tr><td>Oligo fwd</td><td>CACCTCCCTATGACATCCGCGTAG</td><td></td><td></td><td></td><td></td><td></td><td></td><td></td><td></td><td></td><td></td></tr>
<tr><td>Oligo rev</td><td>AAACCTACGCGGATGTCATAGGGA</td><td></td><td></td><td></td><td></td><td></td><td></td><td></td><td></td><td></td><td></td></tr>
<tr><td>Chromosome</td><td>start</td><td>end</td><td>strand</td><td>MM</td><td>target_seq</td><td>PAM</td><td>alignment</td><td>distance</td><td>position</td><td>gene name</td><td>gene id</td></tr>
<tr><td>GU305914.1</td><td>134573</td><td>134595</td><td>-</td><td>0</td><td>TCCCTATGACATCCGCGTAG</td><td>TGG</td><td>||||||||[||||||||||||]PAM</td><td>NA</td><td>Intergenic</td><td>NA</td><td>NA</td></tr>
<tr><td>T23831</td><td>CGCGTAGTGGTCAGCGTTCCTGG</td><td>1000</td><td></td><td></td><td></td><td></td><td></td><td></td><td></td><td></td><td></td></tr>
<tr><td>Oligo fwd</td><td>CACCCGCGTAGTGGTCAGCGTTCC</td><td></td><td></td><td></td><td></td><td></td><td></td><td></td><td></td><td></td><td></td></tr>
<tr><td>Oligo rev</td><td>AAACGGAACGCTGACCACTACGCG</td><td></td><td></td><td></td><td></td><td></td><td></td><td></td><td></td><td></td><td></td></tr>
<tr><td>Chromosome</td><td>start</td><td>end</td><td>strand</td><td>MM</td><td>target_seq</td><td>PAM</td><td>alignment</td><td>distance</td><td>position</td><td>gene name</td><td>gene id</td></tr>
<tr><td>GU305914.1</td><td>134560</td><td>134582</td><td>-</td><td>0</td><td>CGCGTAGTGGTCAGCGTTCC</td><td>TGG</td><td>||||||||[||||||||||||]PAM</td><td>NA</td><td>Intergenic</td><td>NA</td><td>NA</td></tr>
<tr><td>T23832</td><td>GTCAGCGTTCCTGGTGAGATAGG</td><td>1000</td><td></td><td></td><td></td><td></td><td></td><td></td><td></td><td></td><td></td></tr>
<tr><td>Oligo fwd</td><td>CACCGTCAGCGTTCCTGGTGAGAT</td><td></td><td></td><td></td><td></td><td></td><td></td><td></td><td></td><td></td><td></td></tr>
<tr><td>Oligo rev</td><td>AAACATCTCACCAGGAACGCTGAC</td><td></td><td></td><td></td><td></td><td></td><td></td><td></td><td></td><td></td><td></td></tr>
<tr><td>Chromosome</td><td>start</td><td>end</td><td>strand</td><td>MM</td><td>target_seq</td><td>PAM</td><td>alignment</td><td>distance</td><td>position</td><td>gene name</td><td>gene id</td></tr>
<tr><td>GU305914.1</td><td>134551</td><td>134573</td><td>-</td><td>0</td><td>GTCAGCGTTCCTGGTGAGAT</td><td>AGG</td><td>||||||||[||||||||||||]PAM</td><td>NA</td><td>Intergenic</td><td>NA</td><td>NA</td></tr>
<tr><td>T23833</td><td>GGTGAGATAGGTGTAGACGTAGG</td><td>1000</td><td></td><td></td><td></td><td></td><td></td><td></td><td></td><td></td><td></td></tr>
<tr><td>Oligo fwd</td><td>CACCGGTGAGATAGGTGTAGACGT</td><td></td><td></td><td></td><td></td><td></td><td></td><td></td><td></td><td></td><td></td></tr>
<tr><td>Oligo rev</td><td>AAACACGTCTACACCTATCTCACC</td><td></td><td></td><td></td><td></td><td></td><td></td><td></td><td></td><td></td><td></td></tr>
<tr><td>Chromosome</td><td>start</td><td>end</td><td>strand</td><td>MM</td><td>target_seq</td><td>PAM</td><td>alignment</td><td>distance</td><td>position</td><td>gene name</td><td>gene id</td></tr>
<tr><td>GU305914.1</td><td>134539</td><td>134561</td><td>-</td><td>0</td><td>GGTGAGATAGGTGTAGACGT</td><td>AGG</td><td>||||||||[||||||||||||]PAM</td><td>NA</td><td>Intergenic</td><td>NA</td><td>NA</td></tr>
<tr><td>T23834</td><td>ACATTGATCTCGTCGACGTGCGG</td><td>1000</td><td></td><td></td><td></td><td></td><td></td><td></td><td></td><td></td><td></td></tr>
<tr><td>Oligo fwd</td><td>CACCACATTGATCTCGTCGACGTG</td><td></td><td></td><td></td><td></td><td></td><td></td><td></td><td></td><td></td><td></td></tr>
<tr><td>Oligo rev</td><td>AAACCACGTCGACGAGATCAATGT</td><td></td><td></td><td></td><td></td><td></td><td></td><td></td><td></td><td></td><td></td></tr>
<tr><td>Chromosome</td><td>start</td><td>end</td><td>strand</td><td>MM</td><td>target_seq</td><td>PAM</td><td>alignment</td><td>distance</td><td>position</td><td>gene name</td><td>gene id</td></tr>
<tr><td>GU305914.1</td><td>134499</td><td>134521</td><td>-</td><td>0</td><td>ACATTGATCTCGTCGACGTG</td><td>CGG</td><td>||||||||[||||||||||||]PAM</td><td>NA</td><td>Intergenic</td><td>NA</td><td>NA</td></tr>
<tr><td>T23835</td><td>GATCTCGTCGACGTGCGGCTCGG</td><td>1000</td><td></td><td></td><td></td><td></td><td></td><td></td><td></td><td></td><td></td></tr>
<tr><td>Oligo fwd</td><td>CACCGATCTCGTCGACGTGCGGCT</td><td></td><td></td><td></td><td></td><td></td><td></td><td></td><td></td><td></td><td></td></tr>
<tr><td>Oligo rev</td><td>AAACAGCCGCACGTCGACGAGATC</td><td></td><td></td><td></td><td></td><td></td><td></td><td></td><td></td><td></td><td></td></tr>
<tr><td>Chromosome</td><td>start</td><td>end</td><td>strand</td><td>MM</td><td>target_seq</td><td>PAM</td><td>alignment</td><td>distance</td><td>position</td><td>gene name</td><td>gene id</td></tr>
</table>

| | | | | | | | | | | | |
|---|---|---|---|---|---|---|---|---|---|---|---|
| GU305914.1 | | 134494 | 134516 | - | 0 | GATCTCGTCGACGTGCGGCT | CGG | \|\|\|\|\|\|\|\|[\|\|\|\|\|\|\|\|\|\|\|\|]PAM | NA | Intergenic | NA | NA |
| T23836 | GTCGACGTGCGGCTCGGTGACGG | | 1000 | | | | | | | | | |
| Oligo fwd | CACCGTCGACGTGCGGCTCGGTGA | | | | | | | | | | | |
| Oligo rev | AAACTCACCGAGCCGCACGTCGAC | | | | | | | | | | | |
| Chromosome | start | | end | strand | MM | target_seq | PAM | alignment | distance | position | gene name | gene id |
| GU305914.1 | | 134488 | 134510 | - | 0 | GTCGACGTGCGGCTCGGTGA | CGG | \|\|\|\|\|\|\|\|[\|\|\|\|\|\|\|\|\|\|\|\|]PAM | NA | Intergenic | NA | NA |
| GU305914.1 | | 126288 | 126310 | + | 4 | GTCCATCTGCGTCTCGGTGA | AGG | \|\|\|-\|--\|[\|\|\|-\|\|\|\|\|\|\|\|]PAM | NA | Intergenic | NA | NA |
| T23837 | TCGACGTGCGGCTCGGTGACGGG | | 1000 | | | | | | | | | |
| Oligo fwd | CACCTCGACGTGCGGCTCGGTGAC | | | | | | | | | | | |
| Oligo rev | AAACGTCACCGAGCCGCACGTCGA | | | | | | | | | | | |
| Chromosome | start | | end | strand | MM | target_seq | PAM | alignment | distance | position | gene name | gene id |
| GU305914.1 | | 134487 | 134509 | - | 0 | TCGACGTGCGGCTCGGTGAC | GGG | \|\|\|\|\|\|\|\|[\|\|\|\|\|\|\|\|\|\|\|\|]PAM | NA | Intergenic | NA | NA |
| T23838 | GCGGCTCGGTGACGGGTTCCAGG | | 1000 | | | | | | | | | |
| Oligo fwd | CACCGCGGCTCGGTGACGGGTTCC | | | | | | | | | | | |
| Oligo rev | AAACGGAACCCGTCACCGAGCCGC | | | | | | | | | | | |
| Chromosome | start | | end | strand | MM | target_seq | PAM | alignment | distance | position | gene name | gene id |
| GU305914.1 | | 134480 | 134502 | - | 0 | GCGGCTCGGTGACGGGTTCC | AGG | \|\|\|\|\|\|\|\|[\|\|\|\|\|\|\|\|\|\|\|\|]PAM | NA | Intergenic | NA | NA |
| T23839 | CGGCTCGGTGACGGGTTCCAGGG | | 1000 | | | | | | | | | |
| Oligo fwd | CACCCGGCTCGGTGACGGGTTCCA | | | | | | | | | | | |
| Oligo rev | AAACTGGAACCCGTCACCGAGCCG | | | | | | | | | | | |
| Chromosome | start | | end | strand | MM | target_seq | PAM | alignment | distance | position | gene name | gene id |
| GU305914.1 | | 134479 | 134501 | - | 0 | CGGCTCGGTGACGGGTTCCA | GGG | \|\|\|\|\|\|\|\|[\|\|\|\|\|\|\|\|\|\|\|\|]PAM | NA | Intergenic | NA | NA |
| T23840 | CAGGGCGCTCACGCAGCCCCCGG | | 1000 | | | | | | | | | |
| Oligo fwd | CACCCAGGGCGCTCACGCAGCCCC | | | | | | | | | | | |
| Oligo rev | AAACGGGGCTGCGTGAGCGCCCTG | | | | | | | | | | | |
| Chromosome | start | | end | strand | MM | target_seq | PAM | alignment | distance | position | gene name | gene id |
| GU305914.1 | | 134461 | 134483 | - | 0 | CAGGGCGCTCACGCAGCCCC | CGG | \|\|\|\|\|\|\|\|[\|\|\|\|\|\|\|\|\|\|\|\|]PAM | NA | Intergenic | NA | NA |
| T23841 | AGGGCGCTCACGCAGCCCCCGGG | | 1000 | | | | | | | | | |
| Oligo fwd | CACCAGGGCGCTCACGCAGCCCCC | | | | | | | | | | | |
| Oligo rev | AAACGGGGGCTGCGTGAGCGCCCT | | | | | | | | | | | |
| Chromosome | start | | end | strand | MM | target_seq | PAM | alignment | distance | position | gene name | gene id |
| GU305914.1 | | 134460 | 134482 | - | 0 | AGGGCGCTCACGCAGCCCCC | GGG | \|\|\|\|\|\|\|\|[\|\|\|\|\|\|\|\|\|\|\|\|]PAM | NA | Intergenic | NA | NA |
| T23842 | GGGCGCTCACGCAGCCCCCGGGG | | 1000 | | | | | | | | | |
| Oligo fwd | CACCGGGCGCTCACGCAGCCCCCG | | | | | | | | | | | |
| Oligo rev | AAACCGGGGGCTGCGTGAGCGCCC | | | | | | | | | | | |
| Chromosome | start | | end | strand | MM | target_seq | PAM | alignment | distance | position | gene name | gene id |
| GU305914.1 | | 134459 | 134481 | - | 0 | GGGCGCTCACGCAGCCCCCG | GGG | \|\|\|\|\|\|\|\|[\|\|\|\|\|\|\|\|\|\|\|\|]PAM | NA | Intergenic | NA | NA |
| T23843 | CACGCAGCCCCCGGGGAAAACGG | | 1000 | | | | | | | | | |
| Oligo fwd | CACCCACGCAGCCCCCGGGGAAAA | | | | | | | | | | | |
| Oligo rev | AAACTTTTCCCCGGGGGCTGCGTG | | | | | | | | | | | |
| Chromosome | start | | end | strand | MM | target_seq | PAM | alignment | distance | position | gene name | gene id |
| GU305914.1 | | 134452 | 134474 | - | 0 | CACGCAGCCCCCGGGGAAAA | CGG | \|\|\|\|\|\|\|\|[\|\|\|\|\|\|\|\|\|\|\|\|]PAM | NA | Intergenic | NA | NA |
| T23844 | CCCGGGGAAAACGGAGCCGTAGG | | 1000 | | | | | | | | | |
| Oligo fwd | CACCCCCGGGGAAAACGGAGCCGT | | | | | | | | | | | |
| Oligo rev | AAACACGGCTCCGTTTTCCCCGGG | | | | | | | | | | | |
| Chromosome | start | | end | strand | MM | target_seq | PAM | alignment | distance | position | gene name | gene id |
| GU305914.1 | | 134443 | 134465 | - | 0 | CCCGGGGAAAACGGAGCCGT | AGG | \|\|\|\|\|\|\|\|[\|\|\|\|\|\|\|\|\|\|\|\|]PAM | NA | Intergenic | NA | NA |
| T23845 | GGAAAACGGAGCCGTAGGTGAGG | | 1000 | | | | | | | | | |
| Oligo fwd | CACCGGAAAACGGAGCCGTAGGTG | | | | | | | | | | | |
| Oligo rev | AAACCACCTACGGCTCCGTTTTCC | | | | | | | | | | | |
| Chromosome | start | | end | strand | MM | target_seq | PAM | alignment | distance | position | gene name | gene id |
| GU305914.1 | | 134438 | 134460 | - | 0 | GGAAAACGGAGCCGTAGGTG | AGG | \|\|\|\|\|\|\|\|[\|\|\|\|\|\|\|\|\|\|\|\|]PAM | NA | Intergenic | NA | NA |
| T23846 | GAGGCCCGTCTTGATGCAGACGG | | 1000 | | | | | | | | | |
| Oligo fwd | CACCGAGGCCCGTCTTGATGCAGA | | | | | | | | | | | |
| Oligo rev | AAACTCTGCATCAAGACGGGCCTC | | | | | | | | | | | |
| Chromosome | start | | end | strand | MM | target_seq | PAM | alignment | distance | position | gene name | gene id |
| GU305914.1 | | 134419 | 134441 | - | 0 | GAGGCCCGTCTTGATGCAGA | CGG | \|\|\|\|\|\|\|\|[\|\|\|\|\|\|\|\|\|\|\|\|]PAM | NA | Intergenic | NA | NA |
| T23847 | AGACGGAGCCGTCCTGCGTGTGG | | 1000 | | | | | | | | | |
| Oligo fwd | CACCAGACGGAGCCGTCCTGCGTG | | | | | | | | | | | |
| Oligo rev | AAACCACGCAGGACGGCTCCGTCT | | | | | | | | | | | |
| Chromosome | start | | end | strand | MM | target_seq | PAM | alignment | distance | position | gene name | gene id |
| GU305914.1 | | 134402 | 134424 | - | 0 | AGACGGAGCCGTCCTGCGTG | TGG | \|\|\|\|\|\|\|\|[\|\|\|\|\|\|\|\|\|\|\|\|]PAM | NA | Intergenic | NA | NA |
| T23848 | AGTCGTGCCGTAGATCGCAGAGG | | 1000 | | | | | | | | | |
| Oligo fwd | CACCAGTCGTGCCGTAGATCGCAG | | | | | | | | | | | |
| Oligo rev | AAACCTGCGATCTACGGCACGACT | | | | | | | | | | | |
| Chromosome | start | | end | strand | MM | target_seq | PAM | alignment | distance | position | gene name | gene id |
| GU305914.1 | | 134369 | 134391 | - | 0 | AGTCGTGCCGTAGATCGCAG | AGG | \|\|\|\|\|\|\|\|[\|\|\|\|\|\|\|\|\|\|\|\|]PAM | NA | Intergenic | NA | NA |
| T23849 | GTAGATCGCAGAGGTGAGTGCGG | | 1000 | | | | | | | | | |
| Oligo fwd | CACCGTAGATCGCAGAGGTGAGTG | | | | | | | | | | | |
| Oligo rev | AAACCACTCACCTCTGCGATCTAC | | | | | | | | | | | |
| Chromosome | start | | end | strand | MM | target_seq | PAM | alignment | distance | position | gene name | gene id |
| GU305914.1 | | 134360 | 134382 | - | 0 | GTAGATCGCAGAGGTGAGTG | CGG | \|\|\|\|\|\|\|\|[\|\|\|\|\|\|\|\|\|\|\|\|]PAM | NA | Intergenic | NA | NA |
| T23850 | CGCACACGTACATATTCCTGAGG | | 1000 | | | | | | | | | |
| Oligo fwd | CACCCGCACACGTACATATTCCTG | | | | | | | | | | | |

<table>
<tr><td>Oligo rev</td><td>AAACCAGGAATATGTACGTGTGCG</td><td></td><td></td><td></td><td></td><td></td><td></td><td></td><td></td><td></td><td></td></tr>
<tr><td>Chromosome</td><td>start</td><td>end</td><td>strand</td><td>MM</td><td>target_seq</td><td>PAM</td><td>alignment</td><td>distance</td><td>position</td><td>gene name</td><td>gene id</td></tr>
<tr><td>GU305914.1</td><td>134327</td><td>134349</td><td>-</td><td>0</td><td>CGCACACGTACATATTCCTG</td><td>AGG</td><td>||||||||[||||||||||||]PAM</td><td>NA</td><td>Intergenic</td><td>NA</td><td>NA</td></tr>
<tr><td>T23851</td><td>AGGTTGAGCTCCTGCGTCAGCGG</td><td>1000</td><td></td><td></td><td></td><td></td><td></td><td></td><td></td><td></td><td></td></tr>
<tr><td>Oligo fwd</td><td>CACCAGGTTGAGCTCCTGCGTCAG</td><td></td><td></td><td></td><td></td><td></td><td></td><td></td><td></td><td></td><td></td></tr>
<tr><td>Oligo rev</td><td>AAACCTGACGCAGGAGCTCAACCT</td><td></td><td></td><td></td><td></td><td></td><td></td><td></td><td></td><td></td><td></td></tr>
<tr><td>Chromosome</td><td>start</td><td>end</td><td>strand</td><td>MM</td><td>target_seq</td><td>PAM</td><td>alignment</td><td>distance</td><td>position</td><td>gene name</td><td>gene id</td></tr>
<tr><td>GU305914.1</td><td>134307</td><td>134329</td><td>-</td><td>0</td><td>AGGTTGAGCTCCTGCGTCAG</td><td>CGG</td><td>||||||||[||||||||||||]PAM</td><td>NA</td><td>Intergenic</td><td>NA</td><td>NA</td></tr>
<tr><td>T23852</td><td>GGTTGAGCTCCTGCGTCAGCGGG</td><td>1000</td><td></td><td></td><td></td><td></td><td></td><td></td><td></td><td></td><td></td></tr>
<tr><td>Oligo fwd</td><td>CACCGGTTGAGCTCCTGCGTCAGC</td><td></td><td></td><td></td><td></td><td></td><td></td><td></td><td></td><td></td><td></td></tr>
<tr><td>Oligo rev</td><td>AAACGCTGACGCAGGAGCTCAACC</td><td></td><td></td><td></td><td></td><td></td><td></td><td></td><td></td><td></td><td></td></tr>
<tr><td>Chromosome</td><td>start</td><td>end</td><td>strand</td><td>MM</td><td>target_seq</td><td>PAM</td><td>alignment</td><td>distance</td><td>position</td><td>gene name</td><td>gene id</td></tr>
<tr><td>GU305914.1</td><td>134306</td><td>134328</td><td>-</td><td>0</td><td>GGTTGAGCTCCTGCGTCAGC</td><td>GGG</td><td>||||||||[||||||||||||]PAM</td><td>NA</td><td>Intergenic</td><td>NA</td><td>NA</td></tr>
<tr><td>T23853</td><td>CACTACCGCCCCCGCGTGCGTGG</td><td>1000</td><td></td><td></td><td></td><td></td><td></td><td></td><td></td><td></td><td></td></tr>
<tr><td>Oligo fwd</td><td>CACCCACTACCGCCCCCGCGTGCG</td><td></td><td></td><td></td><td></td><td></td><td></td><td></td><td></td><td></td><td></td></tr>
<tr><td>Oligo rev</td><td>AAACCGCACGCGGGGGCGGTAGTG</td><td></td><td></td><td></td><td></td><td></td><td></td><td></td><td></td><td></td><td></td></tr>
<tr><td>Chromosome</td><td>start</td><td>end</td><td>strand</td><td>MM</td><td>target_seq</td><td>PAM</td><td>alignment</td><td>distance</td><td>position</td><td>gene name</td><td>gene id</td></tr>
<tr><td>GU305914.1</td><td>134237</td><td>134259</td><td>-</td><td>0</td><td>CACTACCGCCCCCGCGTGCG</td><td>TGG</td><td>||||||||[||||||||||||]PAM</td><td>NA</td><td>Intergenic</td><td>NA</td><td>NA</td></tr>
<tr><td>GU305914.1</td><td>21335</td><td>21357</td><td>+</td><td>3</td><td>CACGACCGCCTCGGCGTGCG</td><td>CGG</td><td>|||-||||[||-|-|||||||]PAM</td><td>NA</td><td>Intergenic</td><td>NA</td><td>NA</td></tr>
<tr><td>T23854</td><td>ACTACCGCCCCCGCGTGCGTGGG</td><td>1000</td><td></td><td></td><td></td><td></td><td></td><td></td><td></td><td></td><td></td></tr>
<tr><td>Oligo fwd</td><td>CACCACTACCGCCCCCGCGTGCGT</td><td></td><td></td><td></td><td></td><td></td><td></td><td></td><td></td><td></td><td></td></tr>
<tr><td>Oligo rev</td><td>AAACACGCACGCGGGGGCGGTAGT</td><td></td><td></td><td></td><td></td><td></td><td></td><td></td><td></td><td></td><td></td></tr>
<tr><td>Chromosome</td><td>start</td><td>end</td><td>strand</td><td>MM</td><td>target_seq</td><td>PAM</td><td>alignment</td><td>distance</td><td>position</td><td>gene name</td><td>gene id</td></tr>
<tr><td>GU305914.1</td><td>134236</td><td>134258</td><td>-</td><td>0</td><td>ACTACCGCCCCCGCGTGCGT</td><td>GGG</td><td>||||||||[||||||||||||]PAM</td><td>NA</td><td>Intergenic</td><td>NA</td><td>NA</td></tr>
<tr><td>T23855</td><td>CGCCCCCGCGTGCGTGGGCACGG</td><td>1000</td><td></td><td></td><td></td><td></td><td></td><td></td><td></td><td></td><td></td></tr>
<tr><td>Oligo fwd</td><td>CACCCGCCCCCGCGTGCGTGGGCA</td><td></td><td></td><td></td><td></td><td></td><td></td><td></td><td></td><td></td><td></td></tr>
<tr><td>Oligo rev</td><td>AAACTGCCCACGCACGCGGGGGCG</td><td></td><td></td><td></td><td></td><td></td><td></td><td></td><td></td><td></td><td></td></tr>
<tr><td>Chromosome</td><td>start</td><td>end</td><td>strand</td><td>MM</td><td>target_seq</td><td>PAM</td><td>alignment</td><td>distance</td><td>position</td><td>gene name</td><td>gene id</td></tr>
<tr><td>GU305914.1</td><td>134231</td><td>134253</td><td>-</td><td>0</td><td>CGCCCCCGCGTGCGTGGGCA</td><td>CGG</td><td>||||||||[||||||||||||]PAM</td><td>NA</td><td>Intergenic</td><td>NA</td><td>NA</td></tr>
<tr><td>T23856</td><td>GCCCCCGCGTGCGTGGGCACGGG</td><td>1000</td><td></td><td></td><td></td><td></td><td></td><td></td><td></td><td></td><td></td></tr>
<tr><td>Oligo fwd</td><td>CACCGCCCCCGCGTGCGTGGGCAC</td><td></td><td></td><td></td><td></td><td></td><td></td><td></td><td></td><td></td><td></td></tr>
<tr><td>Oligo rev</td><td>AAACGTGCCCACGCACGCGGGGGC</td><td></td><td></td><td></td><td></td><td></td><td></td><td></td><td></td><td></td><td></td></tr>
<tr><td>Chromosome</td><td>start</td><td>end</td><td>strand</td><td>MM</td><td>target_seq</td><td>PAM</td><td>alignment</td><td>distance</td><td>position</td><td>gene name</td><td>gene id</td></tr>
<tr><td>GU305914.1</td><td>134230</td><td>134252</td><td>-</td><td>0</td><td>GCCCCCGCGTGCGTGGGCAC</td><td>GGG</td><td>||||||||[||||||||||||]PAM</td><td>NA</td><td>Intergenic</td><td>NA</td><td>NA</td></tr>
<tr><td>T23857</td><td>GCGTGCGTGGGCACGGGCCCCGG</td><td>1000</td><td></td><td></td><td></td><td></td><td></td><td></td><td></td><td></td><td></td></tr>
<tr><td>Oligo fwd</td><td>CACCGCGTGCGTGGGCACGGGCCC</td><td></td><td></td><td></td><td></td><td></td><td></td><td></td><td></td><td></td><td></td></tr>
<tr><td>Oligo rev</td><td>AAACGGGCCCGTGCCCACGCACGC</td><td></td><td></td><td></td><td></td><td></td><td></td><td></td><td></td><td></td><td></td></tr>
<tr><td>Chromosome</td><td>start</td><td>end</td><td>strand</td><td>MM</td><td>target_seq</td><td>PAM</td><td>alignment</td><td>distance</td><td>position</td><td>gene name</td><td>gene id</td></tr>
<tr><td>GU305914.1</td><td>134224</td><td>134246</td><td>-</td><td>0</td><td>GCGTGCGTGGGCACGGGCCC</td><td>CGG</td><td>||||||||[||||||||||||]PAM</td><td>NA</td><td>Intergenic</td><td>NA</td><td>NA</td></tr>
<tr><td>T23858</td><td>CGGGCCCCGGCCGCCGATATAGG</td><td>1000</td><td></td><td></td><td></td><td></td><td></td><td></td><td></td><td></td><td></td></tr>
<tr><td>Oligo fwd</td><td>CACCCGGGCCCCGGCCGCCGATAT</td><td></td><td></td><td></td><td></td><td></td><td></td><td></td><td></td><td></td><td></td></tr>
<tr><td>Oligo rev</td><td>AAACATATCGGCGGCCGGGGCCCG</td><td></td><td></td><td></td><td></td><td></td><td></td><td></td><td></td><td></td><td></td></tr>
<tr><td>Chromosome</td><td>start</td><td>end</td><td>strand</td><td>MM</td><td>target_seq</td><td>PAM</td><td>alignment</td><td>distance</td><td>position</td><td>gene name</td><td>gene id</td></tr>
<tr><td>GU305914.1</td><td>134211</td><td>134233</td><td>-</td><td>0</td><td>CGGGCCCCGGCCGCCGATAT</td><td>AGG</td><td>||||||||[||||||||||||]PAM</td><td>NA</td><td>Intergenic</td><td>NA</td><td>NA</td></tr>
<tr><td>T23859</td><td>GATATAGGCTTGCCCGTCCGCGG</td><td>1000</td><td></td><td></td><td></td><td></td><td></td><td></td><td></td><td></td><td></td></tr>
<tr><td>Oligo fwd</td><td>CACCGATATAGGCTTGCCCGTCCG</td><td></td><td></td><td></td><td></td><td></td><td></td><td></td><td></td><td></td><td></td></tr>
<tr><td>Oligo rev</td><td>AAACCGGACGGGCAAGCCTATATC</td><td></td><td></td><td></td><td></td><td></td><td></td><td></td><td></td><td></td><td></td></tr>
<tr><td>Chromosome</td><td>start</td><td>end</td><td>strand</td><td>MM</td><td>target_seq</td><td>PAM</td><td>alignment</td><td>distance</td><td>position</td><td>gene name</td><td>gene id</td></tr>
<tr><td>GU305914.1</td><td>134196</td><td>134218</td><td>-</td><td>0</td><td>GATATAGGCTTGCCCGTCCG</td><td>CGG</td><td>||||||||[||||||||||||]PAM</td><td>NA</td><td>Intergenic</td><td>NA</td><td>NA</td></tr>
<tr><td>T23860</td><td>ATATAGGCTTGCCCGTCCGCGGG</td><td>1000</td><td></td><td></td><td></td><td></td><td></td><td></td><td></td><td></td><td></td></tr>
<tr><td>Oligo fwd</td><td>CACCATATAGGCTTGCCCGTCCGC</td><td></td><td></td><td></td><td></td><td></td><td></td><td></td><td></td><td></td><td></td></tr>
<tr><td>Oligo rev</td><td>AAACGCGGACGGGCAAGCCTATAT</td><td></td><td></td><td></td><td></td><td></td><td></td><td></td><td></td><td></td><td></td></tr>
<tr><td>Chromosome</td><td>start</td><td>end</td><td>strand</td><td>MM</td><td>target_seq</td><td>PAM</td><td>alignment</td><td>distance</td><td>position</td><td>gene name</td><td>gene id</td></tr>
<tr><td>GU305914.1</td><td>134195</td><td>134217</td><td>-</td><td>0</td><td>ATATAGGCTTGCCCGTCCGC</td><td>GGG</td><td>||||||||[||||||||||||]PAM</td><td>NA</td><td>Intergenic</td><td>NA</td><td>NA</td></tr>
<tr><td>T23861</td><td>GCTTGCCCGTCCGCGGGTCTCGG</td><td>1000</td><td></td><td></td><td></td><td></td><td></td><td></td><td></td><td></td><td></td></tr>
<tr><td>Oligo fwd</td><td>CACCGCTTGCCCGTCCGCGGGTCT</td><td></td><td></td><td></td><td></td><td></td><td></td><td></td><td></td><td></td><td></td></tr>
<tr><td>Oligo rev</td><td>AAACAGACCCGCGGACGGGCAAGC</td><td></td><td></td><td></td><td></td><td></td><td></td><td></td><td></td><td></td><td></td></tr>
<tr><td>Chromosome</td><td>start</td><td>end</td><td>strand</td><td>MM</td><td>target_seq</td><td>PAM</td><td>alignment</td><td>distance</td><td>position</td><td>gene name</td><td>gene id</td></tr>
<tr><td>GU305914.1</td><td>134189</td><td>134211</td><td>-</td><td>0</td><td>GCTTGCCCGTCCGCGGGTCT</td><td>CGG</td><td>||||||||[||||||||||||]PAM</td><td>NA</td><td>Intergenic</td><td>NA</td><td>NA</td></tr>
<tr><td>T23862</td><td>AGAACATAGTCAGTCCTTCGAGG</td><td>1000</td><td></td><td></td><td></td><td></td><td></td><td></td><td></td><td></td><td></td></tr>
<tr><td>Oligo fwd</td><td>CACCAGAACATAGTCAGTCCTTCG</td><td></td><td></td><td></td><td></td><td></td><td></td><td></td><td></td><td></td><td></td></tr>
<tr><td>Oligo rev</td><td>AAACCGAAGGACTGACTATGTTCT</td><td></td><td></td><td></td><td></td><td></td><td></td><td></td><td></td><td></td><td></td></tr>
<tr><td>Chromosome</td><td>start</td><td>end</td><td>strand</td><td>MM</td><td>target_seq</td><td>PAM</td><td>alignment</td><td>distance</td><td>position</td><td>gene name</td><td>gene id</td></tr>
<tr><td>GU305914.1</td><td>134159</td><td>134181</td><td>-</td><td>0</td><td>AGAACATAGTCAGTCCTTCG</td><td>AGG</td><td>||||||||[||||||||||||]PAM</td><td>NA</td><td>Intergenic</td><td>NA</td><td>NA</td></tr>
<tr><td>T23863</td><td>ACATAGTCAGTCCTTCGAGGCGG</td><td>1000</td><td></td><td></td><td></td><td></td><td></td><td></td><td></td><td></td><td></td></tr>
<tr><td>Oligo fwd</td><td>CACCACATAGTCAGTCCTTCGAGG</td><td></td><td></td><td></td><td></td><td></td><td></td><td></td><td></td><td></td><td></td></tr>
<tr><td>Oligo rev</td><td>AAACCCTCGAAGGACTGACTATGT</td><td></td><td></td><td></td><td></td><td></td><td></td><td></td><td></td><td></td><td></td></tr>
<tr><td>Chromosome</td><td>start</td><td>end</td><td>strand</td><td>MM</td><td>target_seq</td><td>PAM</td><td>alignment</td><td>distance</td><td>position</td><td>gene name</td><td>gene id</td></tr>
<tr><td>GU305914.1</td><td>134156</td><td>134178</td><td>-</td><td>0</td><td>ACATAGTCAGTCCTTCGAGG</td><td>CGG</td><td>||||||||[||||||||||||]PAM</td><td>NA</td><td>Intergenic</td><td>NA</td><td>NA</td></tr>
<tr><td>T23864</td><td>TTCGAGGCGGTCGCTCCGCGAGG</td><td>1000</td><td></td><td></td><td></td><td></td><td></td><td></td><td></td><td></td><td></td></tr>
<tr><td>Oligo fwd</td><td>CACCTTCGAGGCGGTCGCTCCGCG</td><td></td><td></td><td></td><td></td><td></td><td></td><td></td><td></td><td></td><td></td></tr>
<tr><td>Oligo rev</td><td>AAACCGCGGAGCGACCGCCTCGAA</td><td></td><td></td><td></td><td></td><td></td><td></td><td></td><td></td><td></td><td></td></tr>
<tr><td>Chromosome</td><td>start</td><td>end</td><td>strand</td><td>MM</td><td>target_seq</td><td>PAM</td><td>alignment</td><td>distance</td><td>position</td><td>gene name</td><td>gene id</td></tr>
<tr><td>GU305914.1</td><td>134143</td><td>134165</td><td>-</td><td>0</td><td>TTCGAGGCGGTCGCTCCGCG</td><td>AGG</td><td>||||||||[||||||||||||]PAM</td><td>NA</td><td>Intergenic</td><td>NA</td><td>NA</td></tr>
</table>

<table>
<tr><td>T23865</td><td>TCGAGGCGGTCGCTCCGCGAGGG</td><td>1000</td><td></td><td></td><td></td><td></td><td></td><td></td><td></td><td></td><td></td></tr>
<tr><td>Oligo fwd</td><td>CACCTCGAGGCGGTCGCTCCGCGA</td><td></td><td></td><td></td><td></td><td></td><td></td><td></td><td></td><td></td><td></td></tr>
<tr><td>Oligo rev</td><td>AAACTCGCGGAGCGACCGCCTCGA</td><td></td><td></td><td></td><td></td><td></td><td></td><td></td><td></td><td></td><td></td></tr>
<tr><td>Chromosome</td><td>start</td><td>end</td><td>strand</td><td>MM</td><td>target_seq</td><td>PAM</td><td>alignment</td><td>distance</td><td>position</td><td>gene name</td><td>gene id</td></tr>
<tr><td>GU305914.1</td><td>134142</td><td>134164</td><td>-</td><td>0</td><td>TCGAGGCGGTCGCTCCGCGA</td><td>GGG</td><td>||||||||[||||||||||||]PAM</td><td>NA</td><td>Intergenic</td><td>NA</td><td>NA</td></tr>
<tr><td>GU305914.1</td><td>53912</td><td>53934</td><td>+</td><td>3</td><td>TCGCGGCGGTCGCGCCGGGA</td><td>TGG</td><td>|||-||||[|||||-|||-||]PAM</td><td>NA</td><td>Intergenic</td><td>NA</td><td>NA</td></tr>
<tr><td></td><td></td><td></td><td></td><td></td><td></td><td></td><td></td><td></td><td></td><td></td><td></td></tr>
<tr><td>T23866</td><td>AGGCGGTCGCTCCGCGAGGGTGG</td><td>1000</td><td></td><td></td><td></td><td></td><td></td><td></td><td></td><td></td><td></td></tr>
<tr><td>Oligo fwd</td><td>CACCAGGCGGTCGCTCCGCGAGGG</td><td></td><td></td><td></td><td></td><td></td><td></td><td></td><td></td><td></td><td></td></tr>
<tr><td>Oligo rev</td><td>AAACCCCTCGCGGAGCGACCGCCT</td><td></td><td></td><td></td><td></td><td></td><td></td><td></td><td></td><td></td><td></td></tr>
<tr><td>Chromosome</td><td>start</td><td>end</td><td>strand</td><td>MM</td><td>target_seq</td><td>PAM</td><td>alignment</td><td>distance</td><td>position</td><td>gene name</td><td>gene id</td></tr>
<tr><td>GU305914.1</td><td>134139</td><td>134161</td><td>-</td><td>0</td><td>AGGCGGTCGCTCCGCGAGGG</td><td>TGG</td><td>||||||||[||||||||||||]PAM</td><td>NA</td><td>Intergenic</td><td>NA</td><td>NA</td></tr>
<tr><td></td><td></td><td></td><td></td><td></td><td></td><td></td><td></td><td></td><td></td><td></td><td></td></tr>
<tr><td>T23867</td><td>GGGTGGCCCGTCGTTCCGCCCGG</td><td>1000</td><td></td><td></td><td></td><td></td><td></td><td></td><td></td><td></td><td></td></tr>
<tr><td>Oligo fwd</td><td>CACCGGGTGGCCCGTCGTTCCGCC</td><td></td><td></td><td></td><td></td><td></td><td></td><td></td><td></td><td></td><td></td></tr>
<tr><td>Oligo rev</td><td>AAACGGCGGAACGACGGGCCACCC</td><td></td><td></td><td></td><td></td><td></td><td></td><td></td><td></td><td></td><td></td></tr>
<tr><td>Chromosome</td><td>start</td><td>end</td><td>strand</td><td>MM</td><td>target_seq</td><td>PAM</td><td>alignment</td><td>distance</td><td>position</td><td>gene name</td><td>gene id</td></tr>
<tr><td>GU305914.1</td><td>134122</td><td>134144</td><td>-</td><td>0</td><td>GGGTGGCCCGTCGTTCCGCC</td><td>CGG</td><td>||||||||[||||||||||||]PAM</td><td>NA</td><td>Intergenic</td><td>NA</td><td>NA</td></tr>
<tr><td></td><td></td><td></td><td></td><td></td><td></td><td></td><td></td><td></td><td></td><td></td><td></td></tr>
<tr><td>T23868</td><td>CGCCGCAACGTGATCAAATAAGG</td><td>1000</td><td></td><td></td><td></td><td></td><td></td><td></td><td></td><td></td><td></td></tr>
<tr><td>Oligo fwd</td><td>CACCCGCCGCAACGTGATCAAATA</td><td></td><td></td><td></td><td></td><td></td><td></td><td></td><td></td><td></td><td></td></tr>
<tr><td>Oligo rev</td><td>AAACTATTTGATCACGTTGCGGCG</td><td></td><td></td><td></td><td></td><td></td><td></td><td></td><td></td><td></td><td></td></tr>
<tr><td>Chromosome</td><td>start</td><td>end</td><td>strand</td><td>MM</td><td>target_seq</td><td>PAM</td><td>alignment</td><td>distance</td><td>position</td><td>gene name</td><td>gene id</td></tr>
<tr><td>GU305914.1</td><td>134045</td><td>134067</td><td>-</td><td>0</td><td>CGCCGCAACGTGATCAAATA</td><td>AGG</td><td>||||||||[||||||||||||]PAM</td><td>NA</td><td>Intergenic</td><td>NA</td><td>NA</td></tr>
<tr><td></td><td></td><td></td><td></td><td></td><td></td><td></td><td></td><td></td><td></td><td></td><td></td></tr>
<tr><td>T23869</td><td>CGAGATCCAAGACCCGCAGTCGG</td><td>1000</td><td></td><td></td><td></td><td></td><td></td><td></td><td></td><td></td><td></td></tr>
<tr><td>Oligo fwd</td><td>CACCCGAGATCCAAGACCCGCAGT</td><td></td><td></td><td></td><td></td><td></td><td></td><td></td><td></td><td></td><td></td></tr>
<tr><td>Oligo rev</td><td>AAACACTGCGGGTCTTGGATCTCG</td><td></td><td></td><td></td><td></td><td></td><td></td><td></td><td></td><td></td><td></td></tr>
<tr><td>Chromosome</td><td>start</td><td>end</td><td>strand</td><td>MM</td><td>target_seq</td><td>PAM</td><td>alignment</td><td>distance</td><td>position</td><td>gene name</td><td>gene id</td></tr>
<tr><td>GU305914.1</td><td>134016</td><td>134038</td><td>-</td><td>0</td><td>CGAGATCCAAGACCCGCAGT</td><td>CGG</td><td>||||||||[||||||||||||]PAM</td><td>NA</td><td>Intergenic</td><td>NA</td><td>NA</td></tr>
<tr><td></td><td></td><td></td><td></td><td></td><td></td><td></td><td></td><td></td><td></td><td></td><td></td></tr>
<tr><td>T23870</td><td>GAGATCCAAGACCCGCAGTCGGG</td><td>1000</td><td></td><td></td><td></td><td></td><td></td><td></td><td></td><td></td><td></td></tr>
<tr><td>Oligo fwd</td><td>CACCGAGATCCAAGACCCGCAGTC</td><td></td><td></td><td></td><td></td><td></td><td></td><td></td><td></td><td></td><td></td></tr>
<tr><td>Oligo rev</td><td>AAACGACTGCGGGTCTTGGATCTC</td><td></td><td></td><td></td><td></td><td></td><td></td><td></td><td></td><td></td><td></td></tr>
<tr><td>Chromosome</td><td>start</td><td>end</td><td>strand</td><td>MM</td><td>target_seq</td><td>PAM</td><td>alignment</td><td>distance</td><td>position</td><td>gene name</td><td>gene id</td></tr>
<tr><td>GU305914.1</td><td>134015</td><td>134037</td><td>-</td><td>0</td><td>GAGATCCAAGACCCGCAGTC</td><td>GGG</td><td>||||||||[||||||||||||]PAM</td><td>NA</td><td>Intergenic</td><td>NA</td><td>NA</td></tr>
<tr><td></td><td></td><td></td><td></td><td></td><td></td><td></td><td></td><td></td><td></td><td></td><td></td></tr>
<tr><td>T23871</td><td>GACCCGCAGTCGGGATTCCCCGG</td><td>1000</td><td></td><td></td><td></td><td></td><td></td><td></td><td></td><td></td><td></td></tr>
<tr><td>Oligo fwd</td><td>CACCGACCCGCAGTCGGGATTCCC</td><td></td><td></td><td></td><td></td><td></td><td></td><td></td><td></td><td></td><td></td></tr>
<tr><td>Oligo rev</td><td>AAACGGGAATCCCGACTGCGGGTC</td><td></td><td></td><td></td><td></td><td></td><td></td><td></td><td></td><td></td><td></td></tr>
<tr><td>Chromosome</td><td>start</td><td>end</td><td>strand</td><td>MM</td><td>target_seq</td><td>PAM</td><td>alignment</td><td>distance</td><td>position</td><td>gene name</td><td>gene id</td></tr>
<tr><td>GU305914.1</td><td>134006</td><td>134028</td><td>-</td><td>0</td><td>GACCCGCAGTCGGGATTCCC</td><td>CGG</td><td>||||||||[||||||||||||]PAM</td><td>NA</td><td>Intergenic</td><td>NA</td><td>NA</td></tr>
<tr><td></td><td></td><td></td><td></td><td></td><td></td><td></td><td></td><td></td><td></td><td></td><td></td></tr>
<tr><td>T23872</td><td>ACCCGCAGTCGGGATTCCCCGGG</td><td>1000</td><td></td><td></td><td></td><td></td><td></td><td></td><td></td><td></td><td></td></tr>
<tr><td>Oligo fwd</td><td>CACCACCCGCAGTCGGGATTCCCC</td><td></td><td></td><td></td><td></td><td></td><td></td><td></td><td></td><td></td><td></td></tr>
<tr><td>Oligo rev</td><td>AAACGGGGAATCCCGACTGCGGGT</td><td></td><td></td><td></td><td></td><td></td><td></td><td></td><td></td><td></td><td></td></tr>
<tr><td>Chromosome</td><td>start</td><td>end</td><td>strand</td><td>MM</td><td>target_seq</td><td>PAM</td><td>alignment</td><td>distance</td><td>position</td><td>gene name</td><td>gene id</td></tr>
<tr><td>GU305914.1</td><td>134005</td><td>134027</td><td>-</td><td>0</td><td>ACCCGCAGTCGGGATTCCCC</td><td>GGG</td><td>||||||||[||||||||||||]PAM</td><td>NA</td><td>Intergenic</td><td>NA</td><td>NA</td></tr>
<tr><td></td><td></td><td></td><td></td><td></td><td></td><td></td><td></td><td></td><td></td><td></td><td></td></tr>
<tr><td>T23873</td><td>GAAGTCGCACGCCTCCCCGATGG</td><td>1000</td><td></td><td></td><td></td><td></td><td></td><td></td><td></td><td></td><td></td></tr>
<tr><td>Oligo fwd</td><td>CACCGAAGTCGCACGCCTCCCCGA</td><td></td><td></td><td></td><td></td><td></td><td></td><td></td><td></td><td></td><td></td></tr>
<tr><td>Oligo rev</td><td>AAACTCGGGGAGGCGTGCGACTTC</td><td></td><td></td><td></td><td></td><td></td><td></td><td></td><td></td><td></td><td></td></tr>
<tr><td>Chromosome</td><td>start</td><td>end</td><td>strand</td><td>MM</td><td>target_seq</td><td>PAM</td><td>alignment</td><td>distance</td><td>position</td><td>gene name</td><td>gene id</td></tr>
<tr><td>GU305914.1</td><td>133973</td><td>133995</td><td>-</td><td>0</td><td>GAAGTCGCACGCCTCCCCGA</td><td>TGG</td><td>||||||||[||||||||||||]PAM</td><td>NA</td><td>Intergenic</td><td>NA</td><td>NA</td></tr>
<tr><td></td><td></td><td></td><td></td><td></td><td></td><td></td><td></td><td></td><td></td><td></td><td></td></tr>
<tr><td>T23874</td><td>GCACGCCTCCCCGATGGACTCGG</td><td>1000</td><td></td><td></td><td></td><td></td><td></td><td></td><td></td><td></td><td></td></tr>
<tr><td>Oligo fwd</td><td>CACCGCACGCCTCCCCGATGGACT</td><td></td><td></td><td></td><td></td><td></td><td></td><td></td><td></td><td></td><td></td></tr>
<tr><td>Oligo rev</td><td>AAACAGTCCATCGGGGAGGCGTGC</td><td></td><td></td><td></td><td></td><td></td><td></td><td></td><td></td><td></td><td></td></tr>
<tr><td>Chromosome</td><td>start</td><td>end</td><td>strand</td><td>MM</td><td>target_seq</td><td>PAM</td><td>alignment</td><td>distance</td><td>position</td><td>gene name</td><td>gene id</td></tr>
<tr><td>GU305914.1</td><td>133967</td><td>133989</td><td>-</td><td>0</td><td>GCACGCCTCCCCGATGGACT</td><td>CGG</td><td>||||||||[||||||||||||]PAM</td><td>NA</td><td>Intergenic</td><td>NA</td><td>NA</td></tr>
<tr><td></td><td></td><td></td><td></td><td></td><td></td><td></td><td></td><td></td><td></td><td></td><td></td></tr>
<tr><td>T23875</td><td>CGAGACGAAACACGTCTTCCAGG</td><td>1000</td><td></td><td></td><td></td><td></td><td></td><td></td><td></td><td></td><td></td></tr>
<tr><td>Oligo fwd</td><td>CACCCGAGACGAAACACGTCTTCC</td><td></td><td></td><td></td><td></td><td></td><td></td><td></td><td></td><td></td><td></td></tr>
<tr><td>Oligo rev</td><td>AAACGGAAGACGTGTTTCGTCTCG</td><td></td><td></td><td></td><td></td><td></td><td></td><td></td><td></td><td></td><td></td></tr>
<tr><td>Chromosome</td><td>start</td><td>end</td><td>strand</td><td>MM</td><td>target_seq</td><td>PAM</td><td>alignment</td><td>distance</td><td>position</td><td>gene name</td><td>gene id</td></tr>
<tr><td>GU305914.1</td><td>133944</td><td>133966</td><td>-</td><td>0</td><td>CGAGACGAAACACGTCTTCC</td><td>AGG</td><td>||||||||[||||||||||||]PAM</td><td>NA</td><td>Intergenic</td><td>NA</td><td>NA</td></tr>
<tr><td></td><td></td><td></td><td></td><td></td><td></td><td></td><td></td><td></td><td></td><td></td><td></td></tr>
<tr><td>T23876</td><td>GAGACGAAACACGTCTTCCAGGG</td><td>1000</td><td></td><td></td><td></td><td></td><td></td><td></td><td></td><td></td><td></td></tr>
<tr><td>Oligo fwd</td><td>CACCGAGACGAAACACGTCTTCCA</td><td></td><td></td><td></td><td></td><td></td><td></td><td></td><td></td><td></td><td></td></tr>
<tr><td>Oligo rev</td><td>AAACTGGAAGACGTGTTTCGTCTC</td><td></td><td></td><td></td><td></td><td></td><td></td><td></td><td></td><td></td><td></td></tr>
<tr><td>Chromosome</td><td>start</td><td>end</td><td>strand</td><td>MM</td><td>target_seq</td><td>PAM</td><td>alignment</td><td>distance</td><td>position</td><td>gene name</td><td>gene id</td></tr>
<tr><td>GU305914.1</td><td>133943</td><td>133965</td><td>-</td><td>0</td><td>GAGACGAAACACGTCTTCCA</td><td>GGG</td><td>||||||||[||||||||||||]PAM</td><td>NA</td><td>Intergenic</td><td>NA</td><td>NA</td></tr>
<tr><td></td><td></td><td></td><td></td><td></td><td></td><td></td><td></td><td></td><td></td><td></td><td></td></tr>
<tr><td>T23877</td><td>CAGGGTCTCGTGCTCGATGCCGG</td><td>1000</td><td></td><td></td><td></td><td></td><td></td><td></td><td></td><td></td><td></td></tr>
<tr><td>Oligo fwd</td><td>CACCCAGGGTCTCGTGCTCGATGC</td><td></td><td></td><td></td><td></td><td></td><td></td><td></td><td></td><td></td><td></td></tr>
<tr><td>Oligo rev</td><td>AAACGCATCGAGCACGAGACCCTG</td><td></td><td></td><td></td><td></td><td></td><td></td><td></td><td></td><td></td><td></td></tr>
<tr><td>Chromosome</td><td>start</td><td>end</td><td>strand</td><td>MM</td><td>target_seq</td><td>PAM</td><td>alignment</td><td>distance</td><td>position</td><td>gene name</td><td>gene id</td></tr>
<tr><td>GU305914.1</td><td>133925</td><td>133947</td><td>-</td><td>0</td><td>CAGGGTCTCGTGCTCGATGC</td><td>CGG</td><td>||||||||[||||||||||||]PAM</td><td>NA</td><td>Intergenic</td><td>NA</td><td>NA</td></tr>
<tr><td></td><td></td><td></td><td></td><td></td><td></td><td></td><td></td><td></td><td></td><td></td><td></td></tr>
<tr><td>T23878</td><td>CGGCCTTCCTGAGCTCGCCGAGG</td><td>1000</td><td></td><td></td><td></td><td></td><td></td><td></td><td></td><td></td><td></td></tr>
<tr><td>Oligo fwd</td><td>CACCCGGCCTTCCTGAGCTCGCCG</td><td></td><td></td><td></td><td></td><td></td><td></td><td></td><td></td><td></td><td></td></tr>
<tr><td>Oligo rev</td><td>AAACCGGCGAGCTCAGGAAGGCCG</td><td></td><td></td><td></td><td></td><td></td><td></td><td></td><td></td><td></td><td></td></tr>
<tr><td>Chromosome</td><td>start</td><td>end</td><td>strand</td><td>MM</td><td>target_seq</td><td>PAM</td><td>alignment</td><td>distance</td><td>position</td><td>gene name</td><td>gene id</td></tr>
<tr><td>GU305914.1</td><td>133905</td><td>133927</td><td>-</td><td>0</td><td>CGGCCTTCCTGAGCTCGCCG</td><td>AGG</td><td>||||||||[||||||||||||]PAM</td><td>NA</td><td>Intergenic</td><td>NA</td><td>NA</td></tr>
<tr><td></td><td></td><td></td><td></td><td></td><td></td><td></td><td></td><td></td><td></td><td></td><td></td></tr>
<tr><td>T23879</td><td>GAGGATCACATTCATGCTGCTGG</td><td>1000</td><td></td><td></td><td></td><td></td><td></td><td></td><td></td><td></td><td></td></tr>
<tr><td>Oligo fwd</td><td>CACCGAGGATCACATTCATGCTGC</td><td></td><td></td><td></td><td></td><td></td><td></td><td></td><td></td><td></td><td></td></tr>
<tr><td>Oligo rev</td><td>AAACGCAGCATGAATGTGATCCTC</td><td></td><td></td><td></td><td></td><td></td><td></td><td></td><td></td><td></td><td></td></tr>
<tr><td>Chromosome</td><td>start</td><td>end</td><td>strand</td><td>MM</td><td>target_seq</td><td>PAM</td><td>alignment</td><td>distance</td><td>position</td><td>gene name</td><td>gene id</td></tr>
</table>

<table>
<tr><td>GU305914.1</td><td>133886</td><td>133908</td><td>-</td><td>0</td><td>GAGGATCACATTCATGCTGC</td><td>TGG</td><td>||||||||[||||||||||||]PAM</td><td>NA</td><td>Intergenic</td><td>NA</td><td>NA</td></tr>
<tr><td>T23880</td><td>AGGATCACATTCATGCTGCTGGG</td><td>1000</td><td></td><td></td><td></td><td></td><td></td><td></td><td></td><td></td><td></td></tr>
<tr><td>Oligo fwd</td><td>CACCAGGATCACATTCATGCTGCT</td><td></td><td></td><td></td><td></td><td></td><td></td><td></td><td></td><td></td><td></td></tr>
<tr><td>Oligo rev</td><td>AAACAGCAGCATGAATGTGATCCT</td><td></td><td></td><td></td><td></td><td></td><td></td><td></td><td></td><td></td><td></td></tr>
<tr><td>Chromosome</td><td>start</td><td>end</td><td>strand</td><td>MM</td><td>target_seq</td><td>PAM</td><td>alignment</td><td>distance</td><td>position</td><td>gene name</td><td>gene id</td></tr>
<tr><td>GU305914.1</td><td>133885</td><td>133907</td><td>-</td><td>0</td><td>AGGATCACATTCATGCTGCT</td><td>GGG</td><td>||||||||[||||||||||||]PAM</td><td>NA</td><td>Intergenic</td><td>NA</td><td>NA</td></tr>
<tr><td>T23881</td><td>CTGTGTTTTGCGCTCGCGAGCGG</td><td>1000</td><td></td><td></td><td></td><td></td><td></td><td></td><td></td><td></td><td></td></tr>
<tr><td>Oligo fwd</td><td>CACCCTGTGTTTTGCGCTCGCGAG</td><td></td><td></td><td></td><td></td><td></td><td></td><td></td><td></td><td></td><td></td></tr>
<tr><td>Oligo rev</td><td>AAACCTCGCGAGCGCAAAACACAG</td><td></td><td></td><td></td><td></td><td></td><td></td><td></td><td></td><td></td><td></td></tr>
<tr><td>Chromosome</td><td>start</td><td>end</td><td>strand</td><td>MM</td><td>target_seq</td><td>PAM</td><td>alignment</td><td>distance</td><td>position</td><td>gene name</td><td>gene id</td></tr>
<tr><td>GU305914.1</td><td>133855</td><td>133877</td><td>-</td><td>0</td><td>CTGTGTTTTGCGCTCGCGAG</td><td>CGG</td><td>||||||||[||||||||||||]PAM</td><td>NA</td><td>Intergenic</td><td>NA</td><td>NA</td></tr>
<tr><td>T23882</td><td>TGTTTTGCGCTCGCGAGCGGCGG</td><td>1000</td><td></td><td></td><td></td><td></td><td></td><td></td><td></td><td></td><td></td></tr>
<tr><td>Oligo fwd</td><td>CACCTGTTTTGCGCTCGCGAGCGG</td><td></td><td></td><td></td><td></td><td></td><td></td><td></td><td></td><td></td><td></td></tr>
<tr><td>Oligo rev</td><td>AAACCCGCTCGCGAGCGCAAAACA</td><td></td><td></td><td></td><td></td><td></td><td></td><td></td><td></td><td></td><td></td></tr>
<tr><td>Chromosome</td><td>start</td><td>end</td><td>strand</td><td>MM</td><td>target_seq</td><td>PAM</td><td>alignment</td><td>distance</td><td>position</td><td>gene name</td><td>gene id</td></tr>
<tr><td>GU305914.1</td><td>133852</td><td>133874</td><td>-</td><td>0</td><td>TGTTTTGCGCTCGCGAGCGG</td><td>CGG</td><td>||||||||[||||||||||||]PAM</td><td>NA</td><td>Intergenic</td><td>NA</td><td>NA</td></tr>
<tr><td>T23883</td><td>TGCGCTCGCGAGCGGCGGCACGG</td><td>1000</td><td></td><td></td><td></td><td></td><td></td><td></td><td></td><td></td><td></td></tr>
<tr><td>Oligo fwd</td><td>CACCTGCGCTCGCGAGCGGCGGCA</td><td></td><td></td><td></td><td></td><td></td><td></td><td></td><td></td><td></td><td></td></tr>
<tr><td>Oligo rev</td><td>AAACTGCCGCCGCTCGCGAGCGCA</td><td></td><td></td><td></td><td></td><td></td><td></td><td></td><td></td><td></td><td></td></tr>
<tr><td>Chromosome</td><td>start</td><td>end</td><td>strand</td><td>MM</td><td>target_seq</td><td>PAM</td><td>alignment</td><td>distance</td><td>position</td><td>gene name</td><td>gene id</td></tr>
<tr><td>GU305914.1</td><td>133847</td><td>133869</td><td>-</td><td>0</td><td>TGCGCTCGCGAGCGGCGGCA</td><td>CGG</td><td>||||||||[||||||||||||]PAM</td><td>NA</td><td>Intergenic</td><td>NA</td><td>NA</td></tr>
<tr><td>T23884</td><td>GCTCGCGAGCGGCGGCACGGCGG</td><td>1000</td><td></td><td></td><td></td><td></td><td></td><td></td><td></td><td></td><td></td></tr>
<tr><td>Oligo fwd</td><td>CACCGCTCGCGAGCGGCGGCACGG</td><td></td><td></td><td></td><td></td><td></td><td></td><td></td><td></td><td></td><td></td></tr>
<tr><td>Oligo rev</td><td>AAACCCGTGCCGCCGCTCGCGAGC</td><td></td><td></td><td></td><td></td><td></td><td></td><td></td><td></td><td></td><td></td></tr>
<tr><td>Chromosome</td><td>start</td><td>end</td><td>strand</td><td>MM</td><td>target_seq</td><td>PAM</td><td>alignment</td><td>distance</td><td>position</td><td>gene name</td><td>gene id</td></tr>
<tr><td>GU305914.1</td><td>133844</td><td>133866</td><td>-</td><td>0</td><td>GCTCGCGAGCGGCGGCACGG</td><td>CGG</td><td>||||||||[||||||||||||]PAM</td><td>NA</td><td>Intergenic</td><td>NA</td><td>NA</td></tr>
<tr><td>T23885</td><td>GCACGGCGGACTTATATACCTGG</td><td>1000</td><td></td><td></td><td></td><td></td><td></td><td></td><td></td><td></td><td></td></tr>
<tr><td>Oligo fwd</td><td>CACCGCACGGCGGACTTATATACC</td><td></td><td></td><td></td><td></td><td></td><td></td><td></td><td></td><td></td><td></td></tr>
<tr><td>Oligo rev</td><td>AAACGGTATATAAGTCCGCCGTGC</td><td></td><td></td><td></td><td></td><td></td><td></td><td></td><td></td><td></td><td></td></tr>
<tr><td>Chromosome</td><td>start</td><td>end</td><td>strand</td><td>MM</td><td>target_seq</td><td>PAM</td><td>alignment</td><td>distance</td><td>position</td><td>gene name</td><td>gene id</td></tr>
<tr><td>GU305914.1</td><td>133830</td><td>133852</td><td>-</td><td>0</td><td>GCACGGCGGACTTATATACC</td><td>TGG</td><td>||||||||[||||||||||||]PAM</td><td>NA</td><td>Intergenic</td><td>NA</td><td>NA</td></tr>
<tr><td>T23886</td><td>GGACTTATATACCTGGATACAGG</td><td>1000</td><td></td><td></td><td></td><td></td><td></td><td></td><td></td><td></td><td></td></tr>
<tr><td>Oligo fwd</td><td>CACCGGACTTATATACCTGGATAC</td><td></td><td></td><td></td><td></td><td></td><td></td><td></td><td></td><td></td><td></td></tr>
<tr><td>Oligo rev</td><td>AAACGTATCCAGGTATATAAGTCC</td><td></td><td></td><td></td><td></td><td></td><td></td><td></td><td></td><td></td><td></td></tr>
<tr><td>Chromosome</td><td>start</td><td>end</td><td>strand</td><td>MM</td><td>target_seq</td><td>PAM</td><td>alignment</td><td>distance</td><td>position</td><td>gene name</td><td>gene id</td></tr>
<tr><td>GU305914.1</td><td>133823</td><td>133845</td><td>-</td><td>0</td><td>GGACTTATATACCTGGATAC</td><td>AGG</td><td>||||||||[||||||||||||]PAM</td><td>NA</td><td>Intergenic</td><td>NA</td><td>NA</td></tr>
<tr><td>T23887</td><td>GAGCAACATCACGTCGCCCTCGG</td><td>1000</td><td></td><td></td><td></td><td></td><td></td><td></td><td></td><td></td><td></td></tr>
<tr><td>Oligo fwd</td><td>CACCGAGCAACATCACGTCGCCCT</td><td></td><td></td><td></td><td></td><td></td><td></td><td></td><td></td><td></td><td></td></tr>
<tr><td>Oligo rev</td><td>AAACAGGGCGACGTGATGTTGCTC</td><td></td><td></td><td></td><td></td><td></td><td></td><td></td><td></td><td></td><td></td></tr>
<tr><td>Chromosome</td><td>start</td><td>end</td><td>strand</td><td>MM</td><td>target_seq</td><td>PAM</td><td>alignment</td><td>distance</td><td>position</td><td>gene name</td><td>gene id</td></tr>
<tr><td>GU305914.1</td><td>133793</td><td>133815</td><td>-</td><td>0</td><td>GAGCAACATCACGTCGCCCT</td><td>CGG</td><td>||||||||[||||||||||||]PAM</td><td>NA</td><td>Intergenic</td><td>NA</td><td>NA</td></tr>
<tr><td>T23888</td><td>TACCATGAACGTCACCATCTCGG</td><td>1000</td><td></td><td></td><td></td><td></td><td></td><td></td><td></td><td></td><td></td></tr>
<tr><td>Oligo fwd</td><td>CACCTACCATGAACGTCACCATCT</td><td></td><td></td><td></td><td></td><td></td><td></td><td></td><td></td><td></td><td></td></tr>
<tr><td>Oligo rev</td><td>AAACAGATGGTGACGTTCATGGTA</td><td></td><td></td><td></td><td></td><td></td><td></td><td></td><td></td><td></td><td></td></tr>
<tr><td>Chromosome</td><td>start</td><td>end</td><td>strand</td><td>MM</td><td>target_seq</td><td>PAM</td><td>alignment</td><td>distance</td><td>position</td><td>gene name</td><td>gene id</td></tr>
<tr><td>GU305914.1</td><td>133763</td><td>133785</td><td>-</td><td>0</td><td>TACCATGAACGTCACCATCT</td><td>CGG</td><td>||||||||[||||||||||||]PAM</td><td>NA</td><td>Intergenic</td><td>NA</td><td>NA</td></tr>
<tr><td>T23889</td><td>CACCATCTCGGCGACCGACTCGG</td><td>1000</td><td></td><td></td><td></td><td></td><td></td><td></td><td></td><td></td><td></td></tr>
<tr><td>Oligo fwd</td><td>CACCCACCATCTCGGCGACCGACT</td><td></td><td></td><td></td><td></td><td></td><td></td><td></td><td></td><td></td><td></td></tr>
<tr><td>Oligo rev</td><td>AAACAGTCGGTCGCCGAGATGGTG</td><td></td><td></td><td></td><td></td><td></td><td></td><td></td><td></td><td></td><td></td></tr>
<tr><td>Chromosome</td><td>start</td><td>end</td><td>strand</td><td>MM</td><td>target_seq</td><td>PAM</td><td>alignment</td><td>distance</td><td>position</td><td>gene name</td><td>gene id</td></tr>
<tr><td>GU305914.1</td><td>133751</td><td>133773</td><td>-</td><td>0</td><td>CACCATCTCGGCGACCGACT</td><td>CGG</td><td>||||||||[||||||||||||]PAM</td><td>NA</td><td>Intergenic</td><td>NA</td><td>NA</td></tr>
<tr><td>T23890</td><td>GCGACCGACTCGGACCGTAGCGG</td><td>1000</td><td></td><td></td><td></td><td></td><td></td><td></td><td></td><td></td><td></td></tr>
<tr><td>Oligo fwd</td><td>CACCGCGACCGACTCGGACCGTAG</td><td></td><td></td><td></td><td></td><td></td><td></td><td></td><td></td><td></td><td></td></tr>
<tr><td>Oligo rev</td><td>AAACCTACGGTCCGAGTCGGTCGC</td><td></td><td></td><td></td><td></td><td></td><td></td><td></td><td></td><td></td><td></td></tr>
<tr><td>Chromosome</td><td>start</td><td>end</td><td>strand</td><td>MM</td><td>target_seq</td><td>PAM</td><td>alignment</td><td>distance</td><td>position</td><td>gene name</td><td>gene id</td></tr>
<tr><td>GU305914.1</td><td>133741</td><td>133763</td><td>-</td><td>0</td><td>GCGACCGACTCGGACCGTAG</td><td>CGG</td><td>||||||||[||||||||||||]PAM</td><td>NA</td><td>Intergenic</td><td>NA</td><td>NA</td></tr>
<tr><td>T23891</td><td>CACTTTCGCCTACGTGACCGAGG</td><td>1000</td><td></td><td></td><td></td><td></td><td></td><td></td><td></td><td></td><td></td></tr>
<tr><td>Oligo fwd</td><td>CACCCACTTTCGCCTACGTGACCG</td><td></td><td></td><td></td><td></td><td></td><td></td><td></td><td></td><td></td><td></td></tr>
<tr><td>Oligo rev</td><td>AAACCGGTCACGTAGGCGAAAGTG</td><td></td><td></td><td></td><td></td><td></td><td></td><td></td><td></td><td></td><td></td></tr>
<tr><td>Chromosome</td><td>start</td><td>end</td><td>strand</td><td>MM</td><td>target_seq</td><td>PAM</td><td>alignment</td><td>distance</td><td>position</td><td>gene name</td><td>gene id</td></tr>
<tr><td>GU305914.1</td><td>133703</td><td>133725</td><td>-</td><td>0</td><td>CACTTTCGCCTACGTGACCG</td><td>AGG</td><td>||||||||[||||||||||||]PAM</td><td>NA</td><td>Intergenic</td><td>NA</td><td>NA</td></tr>
<tr><td>T23892</td><td>TTTCGCCTACGTGACCGAGGCGG</td><td>1000</td><td></td><td></td><td></td><td></td><td></td><td></td><td></td><td></td><td></td></tr>
<tr><td>Oligo fwd</td><td>CACCTTTCGCCTACGTGACCGAGG</td><td></td><td></td><td></td><td></td><td></td><td></td><td></td><td></td><td></td><td></td></tr>
<tr><td>Oligo rev</td><td>AAACCCTCGGTCACGTAGGCGAAA</td><td></td><td></td><td></td><td></td><td></td><td></td><td></td><td></td><td></td><td></td></tr>
<tr><td>Chromosome</td><td>start</td><td>end</td><td>strand</td><td>MM</td><td>target_seq</td><td>PAM</td><td>alignment</td><td>distance</td><td>position</td><td>gene name</td><td>gene id</td></tr>
<tr><td>GU305914.1</td><td>133700</td><td>133722</td><td>-</td><td>0</td><td>TTTCGCCTACGTGACCGAGG</td><td>CGG</td><td>||||||||[||||||||||||]PAM</td><td>NA</td><td>Intergenic</td><td>NA</td><td>NA</td></tr>
<tr><td>T23893</td><td>GGCGGAACAATCGCAGAACGAGG</td><td>1000</td><td></td><td></td><td></td><td></td><td></td><td></td><td></td><td></td><td></td></tr>
<tr><td>Oligo fwd</td><td>CACCGGCGGAACAATCGCAGAACG</td><td></td><td></td><td></td><td></td><td></td><td></td><td></td><td></td><td></td><td></td></tr>
<tr><td>Oligo rev</td><td>AAACCGTTCTGCGATTGTTCCGCC</td><td></td><td></td><td></td><td></td><td></td><td></td><td></td><td></td><td></td><td></td></tr>
<tr><td>Chromosome</td><td>start</td><td>end</td><td>strand</td><td>MM</td><td>target_seq</td><td>PAM</td><td>alignment</td><td>distance</td><td>position</td><td>gene name</td><td>gene id</td></tr>
<tr><td>GU305914.1</td><td>133682</td><td>133704</td><td>-</td><td>0</td><td>GGCGGAACAATCGCAGAACG</td><td>AGG</td><td>||||||||[||||||||||||]PAM</td><td>NA</td><td>Intergenic</td><td>NA</td><td>NA</td></tr>
<tr><td>T23894</td><td>CGACGCGCCCGAATACGACACGG</td><td>1000</td><td></td><td></td><td></td><td></td><td></td><td></td><td></td><td></td><td></td></tr>
<tr><td>Oligo fwd</td><td>CACCCGACGCGCCCGAATACGACA</td><td></td><td></td><td></td><td></td><td></td><td></td><td></td><td></td><td></td><td></td></tr>
<tr><td>Oligo rev</td><td>AAACTGTCGTATTCGGGCGCGTCG</td><td></td><td></td><td></td><td></td><td></td><td></td><td></td><td></td><td></td><td></td></tr>
</table>

Chromosome start end strand MM target_seq PAM alignment distance position gene name gene id
GU305914.1 133658 133680 - 0 CGACGCGCCCGAATACGACA CGG ||||||||[||||||||||||]PAM NA Intergenic NA NA

T23895 TCGCGCCCGTACTCTTCAACTGG 1000
Oligo fwd CACCTCGCGCCCGTACTCTTCAAC
Oligo rev AAACGTTGAAGAGTACGGGCGCGA
Chromosome start end strand MM target_seq PAM alignment distance position gene name gene id
GU305914.1 133611 133633 - 0 TCGCGCCCGTACTCTTCAAC TGG ||||||||[||||||||||||]PAM NA Intergenic NA NA

T23896 TTCAACTGGTTTCGCGTCACCGG 1000
Oligo fwd CACCTTCAACTGGTTTCGCGTCAC
Oligo rev AAACGTGACGCGAAACCAGTTGAA
Chromosome start end strand MM target_seq PAM alignment distance position gene name gene id
GU305914.1 133597 133619 - 0 TTCAACTGGTTTCGCGTCAC CGG ||||||||[||||||||||||]PAM NA Intergenic NA NA

T23897 CCGGCAAGAACAACACCGCCAGG 1000
Oligo fwd CACCCCGGCAAGAACAACACCGCC
Oligo rev AAACGGCGGTGTTGTTCTTGCCGG
Chromosome start end strand MM target_seq PAM alignment distance position gene name gene id
GU305914.1 133578 133600 - 0 CCGGCAAGAACAACACCGCC AGG ||||||||[||||||||||||]PAM NA Intergenic NA NA

T23898 AGAACAACACCGCCAGGTTCAGG 1000
Oligo fwd CACCAGAACAACACCGCCAGGTTC
Oligo rev AAACGAACCTGGCGGTGTTGTTCT
Chromosome start end strand MM target_seq PAM alignment distance position gene name gene id
GU305914.1 133572 133594 - 0 AGAACAACACCGCCAGGTTC AGG ||||||||[||||||||||||]PAM NA Intergenic NA NA

T23899 CAACACCGCCAGGTTCAGGACGG 1000
Oligo fwd CACCCAACACCGCCAGGTTCAGGA
Oligo rev AAACTCCTGAACCTGGCGGTGTTG
Chromosome start end strand MM target_seq PAM alignment distance position gene name gene id
GU305914.1 133568 133590 - 0 CAACACCGCCAGGTTCAGGA CGG ||||||||[||||||||||||]PAM NA Intergenic NA NA

T23900 CATCACAAAGCGTTCTCACGTGG 1000
Oligo fwd CACCCATCACAAAGCGTTCTCACG
Oligo rev AAACCGTGAGAACGCTTTGTGATG
Chromosome start end strand MM target_seq PAM alignment distance position gene name gene id
GU305914.1 133382 133404 - 0 CATCACAAAGCGTTCTCACG TGG ||||||||[||||||||||||]PAM NA Intergenic NA NA

T23901 ATCACAAAGCGTTCTCACGTGGG 1000
Oligo fwd CACCATCACAAAGCGTTCTCACGT
Oligo rev AAACACGTGAGAACGCTTTGTGAT
Chromosome start end strand MM target_seq PAM alignment distance position gene name gene id
GU305914.1 133381 133403 - 0 ATCACAAAGCGTTCTCACGT GGG ||||||||[||||||||||||]PAM NA Intergenic NA NA

T23902 CTGCCTCACTTACCGTCGCGCGG 1000
Oligo fwd CACCCTGCCTCACTTACCGTCGCG
Oligo rev AAACCGCGACGGTAAGTGAGGCAG
Chromosome start end strand MM target_seq PAM alignment distance position gene name gene id
GU305914.1 133334 133356 - 0 CTGCCTCACTTACCGTCGCG CGG ||||||||[||||||||||||]PAM NA Intergenic NA NA

T23903 CCGTCGCGCGGCCGATCCCGAGG 1000
Oligo fwd CACCCCGTCGCGCGGCCGATCCCG
Oligo rev AAACCGGGATCGGCCGCGCGACGG
Chromosome start end strand MM target_seq PAM alignment distance position gene name gene id
GU305914.1 133322 133344 - 0 CCGTCGCGCGGCCGATCCCG AGG ||||||||[||||||||||||]PAM NA Intergenic NA NA

T23904 TCCCGAGGTCCAGCTCGCCGAGG 1000
Oligo fwd CACCTCCCGAGGTCCAGCTCGCCG
Oligo rev AAACCGGCGAGCTGGACCTCGGGA
Chromosome start end strand MM target_seq PAM alignment distance position gene name gene id
GU305914.1 133307 133329 - 0 TCCCGAGGTCCAGCTCGCCG AGG ||||||||[||||||||||||]PAM NA Intergenic NA NA

T23905 CCCGAGGTCCAGCTCGCCGAGGG 1000
Oligo fwd CACCCCCGAGGTCCAGCTCGCCGA
Oligo rev AAACTCGGCGAGCTGGACCTCGGG
Chromosome start end strand MM target_seq PAM alignment distance position gene name gene id
GU305914.1 133306 133328 - 0 CCCGAGGTCCAGCTCGCCGA GGG ||||||||[||||||||||||]PAM NA Intergenic NA NA

T23906 CCAGCTCGCCGAGGGAACCGTGG 1000
Oligo fwd CACCCCAGCTCGCCGAGGGAACCG
Oligo rev AAACCGGTTCCCTCGGCGAGCTGG
Chromosome start end strand MM target_seq PAM alignment distance position gene name gene id
GU305914.1 133298 133320 - 0 CCAGCTCGCCGAGGGAACCG TGG ||||||||[||||||||||||]PAM NA Intergenic NA NA

T23907 CAGCTCGCCGAGGGAACCGTGGG 1000
Oligo fwd CACCCAGCTCGCCGAGGGAACCGT
Oligo rev AAACACGGTTCCCTCGGCGAGCTG
Chromosome start end strand MM target_seq PAM alignment distance position gene name gene id
GU305914.1 133297 133319 - 0 CAGCTCGCCGAGGGAACCGT GGG ||||||||[||||||||||||]PAM NA Intergenic NA NA

T23908 AACCGTGGGCAGAGAACCCCAGG 1000
Oligo fwd CACCAACCGTGGGCAGAGAACCCC
Oligo rev AAACGGGGTTCTCTGCCCACGGTT
Chromosome start end strand MM target_seq PAM alignment distance position gene name gene id
GU305914.1 133283 133305 - 0 AACCGTGGGCAGAGAACCCC AGG ||||||||[||||||||||||]PAM NA Intergenic NA NA

T23909 CGTGGGCAGAGAACCCCAGGAGG 1000
Oligo fwd CACCCGTGGGCAGAGAACCCCAGG

<table>
<tr><td>Oligo rev</td><td>AAACCCTGGGGTTCTCTGCCCACG</td><td></td><td></td><td></td><td></td><td></td><td></td><td></td><td></td><td></td><td></td></tr>
<tr><td>Chromosome</td><td>start</td><td>end</td><td>strand</td><td>MM</td><td>target_seq</td><td>PAM</td><td>alignment</td><td>distance</td><td>position</td><td>gene name</td><td>gene id</td></tr>
<tr><td>GU305914.1</td><td>133280</td><td>133302</td><td>-</td><td>0</td><td>CGTGGGCAGAGAACCCCAGG</td><td>AGG</td><td>||||||||[||||||||||||]PAM</td><td>NA</td><td>Intergenic</td><td>NA</td><td>NA</td></tr>
<tr><td>T23910</td><td>ATCGCCTTCCTCTTCTGCATCGG</td><td>1000</td><td></td><td></td><td></td><td></td><td></td><td></td><td></td><td></td><td></td></tr>
<tr><td>Oligo fwd</td><td>CACCATCGCCTTCCTCTTCTGCAT</td><td></td><td></td><td></td><td></td><td></td><td></td><td></td><td></td><td></td><td></td></tr>
<tr><td>Oligo rev</td><td>AAACATGCAGAAGAGGAAGGCGAT</td><td></td><td></td><td></td><td></td><td></td><td></td><td></td><td></td><td></td><td></td></tr>
<tr><td>Chromosome</td><td>start</td><td>end</td><td>strand</td><td>MM</td><td>target_seq</td><td>PAM</td><td>alignment</td><td>distance</td><td>position</td><td>gene name</td><td>gene id</td></tr>
<tr><td>GU305914.1</td><td>133234</td><td>133256</td><td>-</td><td>0</td><td>ATCGCCTTCCTCTTCTGCAT</td><td>CGG</td><td>||||||||[||||||||||||]PAM</td><td>NA</td><td>Intergenic</td><td>NA</td><td>NA</td></tr>
<tr><td>T23911</td><td>CTTCCTCTTCTGCATCGGCACGG</td><td>1000</td><td></td><td></td><td></td><td></td><td></td><td></td><td></td><td></td><td></td></tr>
<tr><td>Oligo fwd</td><td>CACCCTTCCTCTTCTGCATCGGCA</td><td></td><td></td><td></td><td></td><td></td><td></td><td></td><td></td><td></td><td></td></tr>
<tr><td>Oligo rev</td><td>AAACTGCCGATGCAGAAGAGGAAG</td><td></td><td></td><td></td><td></td><td></td><td></td><td></td><td></td><td></td><td></td></tr>
<tr><td>Chromosome</td><td>start</td><td>end</td><td>strand</td><td>MM</td><td>target_seq</td><td>PAM</td><td>alignment</td><td>distance</td><td>position</td><td>gene name</td><td>gene id</td></tr>
<tr><td>GU305914.1</td><td>133229</td><td>133251</td><td>-</td><td>0</td><td>CTTCCTCTTCTGCATCGGCA</td><td>CGG</td><td>||||||||[||||||||||||]PAM</td><td>NA</td><td>Intergenic</td><td>NA</td><td>NA</td></tr>
<tr><td>T23912</td><td>TTCCTCTTCTGCATCGGCACGGG</td><td>1000</td><td></td><td></td><td></td><td></td><td></td><td></td><td></td><td></td><td></td></tr>
<tr><td>Oligo fwd</td><td>CACCTTCCTCTTCTGCATCGGCAC</td><td></td><td></td><td></td><td></td><td></td><td></td><td></td><td></td><td></td><td></td></tr>
<tr><td>Oligo rev</td><td>AAACGTGCCGATGCAGAAGAGGAA</td><td></td><td></td><td></td><td></td><td></td><td></td><td></td><td></td><td></td><td></td></tr>
<tr><td>Chromosome</td><td>start</td><td>end</td><td>strand</td><td>MM</td><td>target_seq</td><td>PAM</td><td>alignment</td><td>distance</td><td>position</td><td>gene name</td><td>gene id</td></tr>
<tr><td>GU305914.1</td><td>133228</td><td>133250</td><td>-</td><td>0</td><td>TTCCTCTTCTGCATCGGCAC</td><td>GGG</td><td>||||||||[||||||||||||]PAM</td><td>NA</td><td>Intergenic</td><td>NA</td><td>NA</td></tr>
<tr><td>T23913</td><td>CCCGTACTACCAGAAGAAGTCGG</td><td>1000</td><td></td><td></td><td></td><td></td><td></td><td></td><td></td><td></td><td></td></tr>
<tr><td>Oligo fwd</td><td>CACCCCCGTACTACCAGAAGAAGT</td><td></td><td></td><td></td><td></td><td></td><td></td><td></td><td></td><td></td><td></td></tr>
<tr><td>Oligo rev</td><td>AAACACTTCTTCTGGTAGTACGGG</td><td></td><td></td><td></td><td></td><td></td><td></td><td></td><td></td><td></td><td></td></tr>
<tr><td>Chromosome</td><td>start</td><td>end</td><td>strand</td><td>MM</td><td>target_seq</td><td>PAM</td><td>alignment</td><td>distance</td><td>position</td><td>gene name</td><td>gene id</td></tr>
<tr><td>GU305914.1</td><td>133175</td><td>133197</td><td>-</td><td>0</td><td>CCCGTACTACCAGAAGAAGT</td><td>CGG</td><td>||||||||[||||||||||||]PAM</td><td>NA</td><td>Intergenic</td><td>NA</td><td>NA</td></tr>
<tr><td>T23914</td><td>GAAAAAGAACAGAGACGACAAGG</td><td>1000</td><td></td><td></td><td></td><td></td><td></td><td></td><td></td><td></td><td></td></tr>
<tr><td>Oligo fwd</td><td>CACCGAAAAAGAACAGAGACGACA</td><td></td><td></td><td></td><td></td><td></td><td></td><td></td><td></td><td></td><td></td></tr>
<tr><td>Oligo rev</td><td>AAACTGTCGTCTCTGTTCTTTTTC</td><td></td><td></td><td></td><td></td><td></td><td></td><td></td><td></td><td></td><td></td></tr>
<tr><td>Chromosome</td><td>start</td><td>end</td><td>strand</td><td>MM</td><td>target_seq</td><td>PAM</td><td>alignment</td><td>distance</td><td>position</td><td>gene name</td><td>gene id</td></tr>
<tr><td>GU305914.1</td><td>133133</td><td>133155</td><td>-</td><td>0</td><td>GAAAAAGAACAGAGACGACA</td><td>AGG</td><td>||||||||[||||||||||||]PAM</td><td>NA</td><td>Intergenic</td><td>NA</td><td>NA</td></tr>
<tr><td>T23915</td><td>GAGACGACAAGGCGACCCTCTGG</td><td>1000</td><td></td><td></td><td></td><td></td><td></td><td></td><td></td><td></td><td></td></tr>
<tr><td>Oligo fwd</td><td>CACCGAGACGACAAGGCGACCCTC</td><td></td><td></td><td></td><td></td><td></td><td></td><td></td><td></td><td></td><td></td></tr>
<tr><td>Oligo rev</td><td>AAACGAGGGTCGCCTTGTCGTCTC</td><td></td><td></td><td></td><td></td><td></td><td></td><td></td><td></td><td></td><td></td></tr>
<tr><td>Chromosome</td><td>start</td><td>end</td><td>strand</td><td>MM</td><td>target_seq</td><td>PAM</td><td>alignment</td><td>distance</td><td>position</td><td>gene name</td><td>gene id</td></tr>
<tr><td>GU305914.1</td><td>133122</td><td>133144</td><td>-</td><td>0</td><td>GAGACGACAAGGCGACCCTC</td><td>TGG</td><td>||||||||[||||||||||||]PAM</td><td>NA</td><td>Intergenic</td><td>NA</td><td>NA</td></tr>
<tr><td>T23916</td><td>GCCGCGCAAGCCGATCTTTACGG</td><td>1000</td><td></td><td></td><td></td><td></td><td></td><td></td><td></td><td></td><td></td></tr>
<tr><td>Oligo fwd</td><td>CACCGCCGCGCAAGCCGATCTTTA</td><td></td><td></td><td></td><td></td><td></td><td></td><td></td><td></td><td></td><td></td></tr>
<tr><td>Oligo rev</td><td>AAACTAAAGATCGGCTTGCGCGGC</td><td></td><td></td><td></td><td></td><td></td><td></td><td></td><td></td><td></td><td></td></tr>
<tr><td>Chromosome</td><td>start</td><td>end</td><td>strand</td><td>MM</td><td>target_seq</td><td>PAM</td><td>alignment</td><td>distance</td><td>position</td><td>gene name</td><td>gene id</td></tr>
<tr><td>GU305914.1</td><td>133100</td><td>133122</td><td>-</td><td>0</td><td>GCCGCGCAAGCCGATCTTTA</td><td>CGG</td><td>||||||||[||||||||||||]PAM</td><td>NA</td><td>Intergenic</td><td>NA</td><td>NA</td></tr>
<tr><td>T23917</td><td>GAATATTACGACTCGCCGCCCGG</td><td>1000</td><td></td><td></td><td></td><td></td><td></td><td></td><td></td><td></td><td></td></tr>
<tr><td>Oligo fwd</td><td>CACCGAATATTACGACTCGCCGCC</td><td></td><td></td><td></td><td></td><td></td><td></td><td></td><td></td><td></td><td></td></tr>
<tr><td>Oligo rev</td><td>AAACGGCGGCGAGTCGTAATATTC</td><td></td><td></td><td></td><td></td><td></td><td></td><td></td><td></td><td></td><td></td></tr>
<tr><td>Chromosome</td><td>start</td><td>end</td><td>strand</td><td>MM</td><td>target_seq</td><td>PAM</td><td>alignment</td><td>distance</td><td>position</td><td>gene name</td><td>gene id</td></tr>
<tr><td>GU305914.1</td><td>133075</td><td>133097</td><td>-</td><td>0</td><td>GAATATTACGACTCGCCGCC</td><td>CGG</td><td>||||||||[||||||||||||]PAM</td><td>NA</td><td>Intergenic</td><td>NA</td><td>NA</td></tr>
<tr><td>T23918</td><td>AATATTACGACTCGCCGCCCGGG</td><td>1000</td><td></td><td></td><td></td><td></td><td></td><td></td><td></td><td></td><td></td></tr>
<tr><td>Oligo fwd</td><td>CACCAATATTACGACTCGCCGCCC</td><td></td><td></td><td></td><td></td><td></td><td></td><td></td><td></td><td></td><td></td></tr>
<tr><td>Oligo rev</td><td>AAACGGGCGGCGAGTCGTAATATT</td><td></td><td></td><td></td><td></td><td></td><td></td><td></td><td></td><td></td><td></td></tr>
<tr><td>Chromosome</td><td>start</td><td>end</td><td>strand</td><td>MM</td><td>target_seq</td><td>PAM</td><td>alignment</td><td>distance</td><td>position</td><td>gene name</td><td>gene id</td></tr>
<tr><td>GU305914.1</td><td>133074</td><td>133096</td><td>-</td><td>0</td><td>AATATTACGACTCGCCGCCC</td><td>GGG</td><td>||||||||[||||||||||||]PAM</td><td>NA</td><td>Intergenic</td><td>NA</td><td>NA</td></tr>
<tr><td>T23919</td><td>CATCAAGAAGCCCTACGCCGTGG</td><td>1000</td><td></td><td></td><td></td><td></td><td></td><td></td><td></td><td></td><td></td></tr>
<tr><td>Oligo fwd</td><td>CACCCATCAAGAAGCCCTACGCCG</td><td></td><td></td><td></td><td></td><td></td><td></td><td></td><td></td><td></td><td></td></tr>
<tr><td>Oligo rev</td><td>AAACCGGCGTAGGGCTTCTTGATG</td><td></td><td></td><td></td><td></td><td></td><td></td><td></td><td></td><td></td><td></td></tr>
<tr><td>Chromosome</td><td>start</td><td>end</td><td>strand</td><td>MM</td><td>target_seq</td><td>PAM</td><td>alignment</td><td>distance</td><td>position</td><td>gene name</td><td>gene id</td></tr>
<tr><td>GU305914.1</td><td>133049</td><td>133071</td><td>-</td><td>0</td><td>CATCAAGAAGCCCTACGCCG</td><td>TGG</td><td>||||||||[||||||||||||]PAM</td><td>NA</td><td>Intergenic</td><td>NA</td><td>NA</td></tr>
<tr><td>T23920</td><td>GCCCTACGCCGTGGTCAACCCGG</td><td>1000</td><td></td><td></td><td></td><td></td><td></td><td></td><td></td><td></td><td></td></tr>
<tr><td>Oligo fwd</td><td>CACCGCCCTACGCCGTGGTCAACC</td><td></td><td></td><td></td><td></td><td></td><td></td><td></td><td></td><td></td><td></td></tr>
<tr><td>Oligo rev</td><td>AAACGGTTGACCACGGCGTAGGGC</td><td></td><td></td><td></td><td></td><td></td><td></td><td></td><td></td><td></td><td></td></tr>
<tr><td>Chromosome</td><td>start</td><td>end</td><td>strand</td><td>MM</td><td>target_seq</td><td>PAM</td><td>alignment</td><td>distance</td><td>position</td><td>gene name</td><td>gene id</td></tr>
<tr><td>GU305914.1</td><td>133040</td><td>133062</td><td>-</td><td>0</td><td>GCCCTACGCCGTGGTCAACC</td><td>CGG</td><td>||||||||[||||||||||||]PAM</td><td>NA</td><td>Intergenic</td><td>NA</td><td>NA</td></tr>
<tr><td>T23921</td><td>CCCTACGCCGTGGTCAACCCGGG</td><td>1000</td><td></td><td></td><td></td><td></td><td></td><td></td><td></td><td></td><td></td></tr>
<tr><td>Oligo fwd</td><td>CACCCCCTACGCCGTGGTCAACCC</td><td></td><td></td><td></td><td></td><td></td><td></td><td></td><td></td><td></td><td></td></tr>
<tr><td>Oligo rev</td><td>AAACGGGTTGACCACGGCGTAGGG</td><td></td><td></td><td></td><td></td><td></td><td></td><td></td><td></td><td></td><td></td></tr>
<tr><td>Chromosome</td><td>start</td><td>end</td><td>strand</td><td>MM</td><td>target_seq</td><td>PAM</td><td>alignment</td><td>distance</td><td>position</td><td>gene name</td><td>gene id</td></tr>
<tr><td>GU305914.1</td><td>133039</td><td>133061</td><td>-</td><td>0</td><td>CCCTACGCCGTGGTCAACCC</td><td>GGG</td><td>||||||||[||||||||||||]PAM</td><td>NA</td><td>Intergenic</td><td>NA</td><td>NA</td></tr>
<tr><td>T23922</td><td>CCTACGCCGTGGTCAACCCGGGG</td><td>1000</td><td></td><td></td><td></td><td></td><td></td><td></td><td></td><td></td><td></td></tr>
<tr><td>Oligo fwd</td><td>CACCCCTACGCCGTGGTCAACCCG</td><td></td><td></td><td></td><td></td><td></td><td></td><td></td><td></td><td></td><td></td></tr>
<tr><td>Oligo rev</td><td>AAACCGGGTTGACCACGGCGTAGG</td><td></td><td></td><td></td><td></td><td></td><td></td><td></td><td></td><td></td><td></td></tr>
<tr><td>Chromosome</td><td>start</td><td>end</td><td>strand</td><td>MM</td><td>target_seq</td><td>PAM</td><td>alignment</td><td>distance</td><td>position</td><td>gene name</td><td>gene id</td></tr>
<tr><td>GU305914.1</td><td>133038</td><td>133060</td><td>-</td><td>0</td><td>CCTACGCCGTGGTCAACCCG</td><td>GGG</td><td>||||||||[||||||||||||]PAM</td><td>NA</td><td>Intergenic</td><td>NA</td><td>NA</td></tr>
<tr><td>T23923</td><td>GGGTTCATGTACAAAGACTGAGG</td><td>1000</td><td></td><td></td><td></td><td></td><td></td><td></td><td></td><td></td><td></td></tr>
<tr><td>Oligo fwd</td><td>CACCGGGTTCATGTACAAAGACTG</td><td></td><td></td><td></td><td></td><td></td><td></td><td></td><td></td><td></td><td></td></tr>
<tr><td>Oligo rev</td><td>AAACCAGTCTTTGTACATGAACCC</td><td></td><td></td><td></td><td></td><td></td><td></td><td></td><td></td><td></td><td></td></tr>
<tr><td>Chromosome</td><td>start</td><td>end</td><td>strand</td><td>MM</td><td>target_seq</td><td>PAM</td><td>alignment</td><td>distance</td><td>position</td><td>gene name</td><td>gene id</td></tr>
<tr><td>GU305914.1</td><td>133018</td><td>133040</td><td>-</td><td>0</td><td>GGGTTCATGTACAAAGACTG</td><td>AGG</td><td>||||||||[||||||||||||]PAM</td><td>NA</td><td>Intergenic</td><td>NA</td><td>NA</td></tr>
<tr><td>T23924</td><td>GGTTCATGTACAAAGACTGAGGG</td><td>1000</td><td></td><td></td><td></td><td></td><td></td><td></td><td></td><td></td><td></td></tr>
</table>

<table>
<tr><td>Oligo fwd</td><td>CACCGGTTCATGTACAAAGACTGA</td><td></td><td></td><td></td><td></td><td></td><td></td><td></td><td></td><td></td><td></td></tr>
<tr><td>Oligo rev</td><td>AAACTCAGTCTTTGTACATGAACC</td><td></td><td></td><td></td><td></td><td></td><td></td><td></td><td></td><td></td><td></td></tr>
<tr><td>Chromosome</td><td>start</td><td>end</td><td>strand</td><td>MM</td><td>target_seq</td><td>PAM</td><td>alignment</td><td>distance</td><td>position</td><td>gene name</td><td>gene id</td></tr>
<tr><td>GU305914.1</td><td>133017</td><td>133039</td><td>-</td><td>0</td><td>GGTTCATGTACAAAGACTGA</td><td>GGG</td><td>||||||||[||||||||||||]PAM</td><td>NA</td><td>Intergenic</td><td>NA</td><td>NA</td></tr>
<tr><td></td><td></td><td></td><td></td><td></td><td></td><td></td><td></td><td></td><td></td><td></td><td></td></tr>
<tr><td>T23925</td><td>GTTCATGTACAAAGACTGAGGGG</td><td>1000</td><td></td><td></td><td></td><td></td><td></td><td></td><td></td><td></td><td></td></tr>
<tr><td>Oligo fwd</td><td>CACCGTTCATGTACAAAGACTGAG</td><td></td><td></td><td></td><td></td><td></td><td></td><td></td><td></td><td></td><td></td></tr>
<tr><td>Oligo rev</td><td>AAACCTCAGTCTTTGTACATGAAC</td><td></td><td></td><td></td><td></td><td></td><td></td><td></td><td></td><td></td><td></td></tr>
<tr><td>Chromosome</td><td>start</td><td>end</td><td>strand</td><td>MM</td><td>target_seq</td><td>PAM</td><td>alignment</td><td>distance</td><td>position</td><td>gene name</td><td>gene id</td></tr>
<tr><td>GU305914.1</td><td>133016</td><td>133038</td><td>-</td><td>0</td><td>GTTCATGTACAAAGACTGAG</td><td>GGG</td><td>||||||||[||||||||||||]PAM</td><td>NA</td><td>Intergenic</td><td>NA</td><td>NA</td></tr>
<tr><td></td><td></td><td></td><td></td><td></td><td></td><td></td><td></td><td></td><td></td><td></td><td></td></tr>
<tr><td>T23926</td><td>ACCAGCCGCCCCGACCCCGTCGG</td><td>1000</td><td></td><td></td><td></td><td></td><td></td><td></td><td></td><td></td><td></td></tr>
<tr><td>Oligo fwd</td><td>CACCACCAGCCGCCCCGACCCCGT</td><td></td><td></td><td></td><td></td><td></td><td></td><td></td><td></td><td></td><td></td></tr>
<tr><td>Oligo rev</td><td>AAACACGGGGTCGGGGCGGCTGGT</td><td></td><td></td><td></td><td></td><td></td><td></td><td></td><td></td><td></td><td></td></tr>
<tr><td>Chromosome</td><td>start</td><td>end</td><td>strand</td><td>MM</td><td>target_seq</td><td>PAM</td><td>alignment</td><td>distance</td><td>position</td><td>gene name</td><td>gene id</td></tr>
<tr><td>GU305914.1</td><td>132936</td><td>132958</td><td>-</td><td>0</td><td>ACCAGCCGCCCCGACCCCGT</td><td>CGG</td><td>||||||||[||||||||||||]PAM</td><td>NA</td><td>Intergenic</td><td>NA</td><td>NA</td></tr>
<tr><td></td><td></td><td></td><td></td><td></td><td></td><td></td><td></td><td></td><td></td><td></td><td></td></tr>
<tr><td>T23927</td><td>CCAGCCGCCCCGACCCCGTCGGG</td><td>1000</td><td></td><td></td><td></td><td></td><td></td><td></td><td></td><td></td><td></td></tr>
<tr><td>Oligo fwd</td><td>CACCCCAGCCGCCCCGACCCCGTC</td><td></td><td></td><td></td><td></td><td></td><td></td><td></td><td></td><td></td><td></td></tr>
<tr><td>Oligo rev</td><td>AAACGACGGGGTCGGGGCGGCTGG</td><td></td><td></td><td></td><td></td><td></td><td></td><td></td><td></td><td></td><td></td></tr>
<tr><td>Chromosome</td><td>start</td><td>end</td><td>strand</td><td>MM</td><td>target_seq</td><td>PAM</td><td>alignment</td><td>distance</td><td>position</td><td>gene name</td><td>gene id</td></tr>
<tr><td>GU305914.1</td><td>132935</td><td>132957</td><td>-</td><td>0</td><td>CCAGCCGCCCCGACCCCGTC</td><td>GGG</td><td>||||||||[||||||||||||]PAM</td><td>NA</td><td>Intergenic</td><td>NA</td><td>NA</td></tr>
<tr><td></td><td></td><td></td><td></td><td></td><td></td><td></td><td></td><td></td><td></td><td></td><td></td></tr>
<tr><td>T23928</td><td>TTCCTCACGCAGAGCATCCGCGG</td><td>1000</td><td></td><td></td><td></td><td></td><td></td><td></td><td></td><td></td><td></td></tr>
<tr><td>Oligo fwd</td><td>CACCTTCCTCACGCAGAGCATCCG</td><td></td><td></td><td></td><td></td><td></td><td></td><td></td><td></td><td></td><td></td></tr>
<tr><td>Oligo rev</td><td>AAACCGGATGCTCTGCGTGAGGAA</td><td></td><td></td><td></td><td></td><td></td><td></td><td></td><td></td><td></td><td></td></tr>
<tr><td>Chromosome</td><td>start</td><td>end</td><td>strand</td><td>MM</td><td>target_seq</td><td>PAM</td><td>alignment</td><td>distance</td><td>position</td><td>gene name</td><td>gene id</td></tr>
<tr><td>GU305914.1</td><td>132860</td><td>132882</td><td>-</td><td>0</td><td>TTCCTCACGCAGAGCATCCG</td><td>CGG</td><td>||||||||[||||||||||||]PAM</td><td>NA</td><td>Intergenic</td><td>NA</td><td>NA</td></tr>
<tr><td></td><td></td><td></td><td></td><td></td><td></td><td></td><td></td><td></td><td></td><td></td><td></td></tr>
<tr><td>T23929</td><td>TCCTCACGCAGAGCATCCGCGGG</td><td>1000</td><td></td><td></td><td></td><td></td><td></td><td></td><td></td><td></td><td></td></tr>
<tr><td>Oligo fwd</td><td>CACCTCCTCACGCAGAGCATCCGC</td><td></td><td></td><td></td><td></td><td></td><td></td><td></td><td></td><td></td><td></td></tr>
<tr><td>Oligo rev</td><td>AAACGCGGATGCTCTGCGTGAGGA</td><td></td><td></td><td></td><td></td><td></td><td></td><td></td><td></td><td></td><td></td></tr>
<tr><td>Chromosome</td><td>start</td><td>end</td><td>strand</td><td>MM</td><td>target_seq</td><td>PAM</td><td>alignment</td><td>distance</td><td>position</td><td>gene name</td><td>gene id</td></tr>
<tr><td>GU305914.1</td><td>132859</td><td>132881</td><td>-</td><td>0</td><td>TCCTCACGCAGAGCATCCGC</td><td>GGG</td><td>||||||||[||||||||||||]PAM</td><td>NA</td><td>Intergenic</td><td>NA</td><td>NA</td></tr>
<tr><td></td><td></td><td></td><td></td><td></td><td></td><td></td><td></td><td></td><td></td><td></td><td></td></tr>
<tr><td>T23930</td><td>GAACTTCAACCTCTTGATCGTGG</td><td>1000</td><td></td><td></td><td></td><td></td><td></td><td></td><td></td><td></td><td></td></tr>
<tr><td>Oligo fwd</td><td>CACCGAACTTCAACCTCTTGATCG</td><td></td><td></td><td></td><td></td><td></td><td></td><td></td><td></td><td></td><td></td></tr>
<tr><td>Oligo rev</td><td>AAACCGATCAAGAGGTTGAAGTTC</td><td></td><td></td><td></td><td></td><td></td><td></td><td></td><td></td><td></td><td></td></tr>
<tr><td>Chromosome</td><td>start</td><td>end</td><td>strand</td><td>MM</td><td>target_seq</td><td>PAM</td><td>alignment</td><td>distance</td><td>position</td><td>gene name</td><td>gene id</td></tr>
<tr><td>GU305914.1</td><td>132834</td><td>132856</td><td>-</td><td>0</td><td>GAACTTCAACCTCTTGATCG</td><td>TGG</td><td>||||||||[||||||||||||]PAM</td><td>NA</td><td>Intergenic</td><td>NA</td><td>NA</td></tr>
<tr><td></td><td></td><td></td><td></td><td></td><td></td><td></td><td></td><td></td><td></td><td></td><td></td></tr>
<tr><td>T23931</td><td>CAACCTCTTGATCGTGGACGAGG</td><td>1000</td><td></td><td></td><td></td><td></td><td></td><td></td><td></td><td></td><td></td></tr>
<tr><td>Oligo fwd</td><td>CACCCAACCTCTTGATCGTGGACG</td><td></td><td></td><td></td><td></td><td></td><td></td><td></td><td></td><td></td><td></td></tr>
<tr><td>Oligo rev</td><td>AAACCGTCCACGATCAAGAGGTTG</td><td></td><td></td><td></td><td></td><td></td><td></td><td></td><td></td><td></td><td></td></tr>
<tr><td>Chromosome</td><td>start</td><td>end</td><td>strand</td><td>MM</td><td>target_seq</td><td>PAM</td><td>alignment</td><td>distance</td><td>position</td><td>gene name</td><td>gene id</td></tr>
<tr><td>GU305914.1</td><td>132828</td><td>132850</td><td>-</td><td>0</td><td>CAACCTCTTGATCGTGGACG</td><td>AGG</td><td>||||||||[||||||||||||]PAM</td><td>NA</td><td>Intergenic</td><td>NA</td><td>NA</td></tr>
<tr><td></td><td></td><td></td><td></td><td></td><td></td><td></td><td></td><td></td><td></td><td></td><td></td></tr>
<tr><td>T23932</td><td>CGAGGCACACTTCATCAAGAAGG</td><td>1000</td><td></td><td></td><td></td><td></td><td></td><td></td><td></td><td></td><td></td></tr>
<tr><td>Oligo fwd</td><td>CACCCGAGGCACACTTCATCAAGA</td><td></td><td></td><td></td><td></td><td></td><td></td><td></td><td></td><td></td><td></td></tr>
<tr><td>Oligo rev</td><td>AAACTCTTGATGAAGTGTGCCTCG</td><td></td><td></td><td></td><td></td><td></td><td></td><td></td><td></td><td></td><td></td></tr>
<tr><td>Chromosome</td><td>start</td><td>end</td><td>strand</td><td>MM</td><td>target_seq</td><td>PAM</td><td>alignment</td><td>distance</td><td>position</td><td>gene name</td><td>gene id</td></tr>
<tr><td>GU305914.1</td><td>132810</td><td>132832</td><td>-</td><td>0</td><td>CGAGGCACACTTCATCAAGA</td><td>AGG</td><td>||||||||[||||||||||||]PAM</td><td>NA</td><td>Intergenic</td><td>NA</td><td>NA</td></tr>
<tr><td></td><td></td><td></td><td></td><td></td><td></td><td></td><td></td><td></td><td></td><td></td><td></td></tr>
<tr><td>T23933</td><td>GGCACACTTCATCAAGAAGGAGG</td><td>1000</td><td></td><td></td><td></td><td></td><td></td><td></td><td></td><td></td><td></td></tr>
<tr><td>Oligo fwd</td><td>CACCGGCACACTTCATCAAGAAGG</td><td></td><td></td><td></td><td></td><td></td><td></td><td></td><td></td><td></td><td></td></tr>
<tr><td>Oligo rev</td><td>AAACCCTTCTTGATGAAGTGTGCC</td><td></td><td></td><td></td><td></td><td></td><td></td><td></td><td></td><td></td><td></td></tr>
<tr><td>Chromosome</td><td>start</td><td>end</td><td>strand</td><td>MM</td><td>target_seq</td><td>PAM</td><td>alignment</td><td>distance</td><td>position</td><td>gene name</td><td>gene id</td></tr>
<tr><td>GU305914.1</td><td>132807</td><td>132829</td><td>-</td><td>0</td><td>GGCACACTTCATCAAGAAGG</td><td>AGG</td><td>||||||||[||||||||||||]PAM</td><td>NA</td><td>Intergenic</td><td>NA</td><td>NA</td></tr>
<tr><td></td><td></td><td></td><td></td><td></td><td></td><td></td><td></td><td></td><td></td><td></td><td></td></tr>
<tr><td>T23934</td><td>GGAGGCCTTTAACACGATCCTGG</td><td>1000</td><td></td><td></td><td></td><td></td><td></td><td></td><td></td><td></td><td></td></tr>
<tr><td>Oligo fwd</td><td>CACCGGAGGCCTTTAACACGATCC</td><td></td><td></td><td></td><td></td><td></td><td></td><td></td><td></td><td></td><td></td></tr>
<tr><td>Oligo rev</td><td>AAACGGATCGTGTTAAAGGCCTCC</td><td></td><td></td><td></td><td></td><td></td><td></td><td></td><td></td><td></td><td></td></tr>
<tr><td>Chromosome</td><td>start</td><td>end</td><td>strand</td><td>MM</td><td>target_seq</td><td>PAM</td><td>alignment</td><td>distance</td><td>position</td><td>gene name</td><td>gene id</td></tr>
<tr><td>GU305914.1</td><td>132789</td><td>132811</td><td>-</td><td>0</td><td>GGAGGCCTTTAACACGATCC</td><td>TGG</td><td>||||||||[||||||||||||]PAM</td><td>NA</td><td>Intergenic</td><td>NA</td><td>NA</td></tr>
<tr><td></td><td></td><td></td><td></td><td></td><td></td><td></td><td></td><td></td><td></td><td></td><td></td></tr>
<tr><td>T23935</td><td>GAGGCCTTTAACACGATCCTGGG</td><td>1000</td><td></td><td></td><td></td><td></td><td></td><td></td><td></td><td></td><td></td></tr>
<tr><td>Oligo fwd</td><td>CACCGAGGCCTTTAACACGATCCT</td><td></td><td></td><td></td><td></td><td></td><td></td><td></td><td></td><td></td><td></td></tr>
<tr><td>Oligo rev</td><td>AAACAGGATCGTGTTAAAGGCCTC</td><td></td><td></td><td></td><td></td><td></td><td></td><td></td><td></td><td></td><td></td></tr>
<tr><td>Chromosome</td><td>start</td><td>end</td><td>strand</td><td>MM</td><td>target_seq</td><td>PAM</td><td>alignment</td><td>distance</td><td>position</td><td>gene name</td><td>gene id</td></tr>
<tr><td>GU305914.1</td><td>132788</td><td>132810</td><td>-</td><td>0</td><td>GAGGCCTTTAACACGATCCT</td><td>GGG</td><td>||||||||[||||||||||||]PAM</td><td>NA</td><td>Intergenic</td><td>NA</td><td>NA</td></tr>
<tr><td></td><td></td><td></td><td></td><td></td><td></td><td></td><td></td><td></td><td></td><td></td><td></td></tr>
<tr><td>T23936</td><td>TAACACGATCCTGGGCTTCCTGG</td><td>1000</td><td></td><td></td><td></td><td></td><td></td><td></td><td></td><td></td><td></td></tr>
<tr><td>Oligo fwd</td><td>CACCTAACACGATCCTGGGCTTCC</td><td></td><td></td><td></td><td></td><td></td><td></td><td></td><td></td><td></td><td></td></tr>
<tr><td>Oligo rev</td><td>AAACGGAAGCCCAGGATCGTGTTA</td><td></td><td></td><td></td><td></td><td></td><td></td><td></td><td></td><td></td><td></td></tr>
<tr><td>Chromosome</td><td>start</td><td>end</td><td>strand</td><td>MM</td><td>target_seq</td><td>PAM</td><td>alignment</td><td>distance</td><td>position</td><td>gene name</td><td>gene id</td></tr>
<tr><td>GU305914.1</td><td>132780</td><td>132802</td><td>-</td><td>0</td><td>TAACACGATCCTGGGCTTCC</td><td>TGG</td><td>||||||||[||||||||||||]PAM</td><td>NA</td><td>Intergenic</td><td>NA</td><td>NA</td></tr>
<tr><td></td><td></td><td></td><td></td><td></td><td></td><td></td><td></td><td></td><td></td><td></td><td></td></tr>
<tr><td>T23937</td><td>CTTCCTGACGAAGCTGACCTCGG</td><td>1000</td><td></td><td></td><td></td><td></td><td></td><td></td><td></td><td></td><td></td></tr>
<tr><td>Oligo fwd</td><td>CACCCTTCCTGACGAAGCTGACCT</td><td></td><td></td><td></td><td></td><td></td><td></td><td></td><td></td><td></td><td></td></tr>
<tr><td>Oligo rev</td><td>AAACAGGTCAGCTTCGTCAGGAAG</td><td></td><td></td><td></td><td></td><td></td><td></td><td></td><td></td><td></td><td></td></tr>
<tr><td>Chromosome</td><td>start</td><td>end</td><td>strand</td><td>MM</td><td>target_seq</td><td>PAM</td><td>alignment</td><td>distance</td><td>position</td><td>gene name</td><td>gene id</td></tr>
<tr><td>GU305914.1</td><td>132696</td><td>132718</td><td>-</td><td>0</td><td>CTTCCTGACGAAGCTGACCT</td><td>CGG</td><td>||||||||[||||||||||||]PAM</td><td>NA</td><td>Intergenic</td><td>NA</td><td>NA</td></tr>
<tr><td></td><td></td><td></td><td></td><td></td><td></td><td></td><td></td><td></td><td></td><td></td><td></td></tr>
<tr><td>T23938</td><td>GCCCTTCGACATGCTGAACGTGG</td><td>1000</td><td></td><td></td><td></td><td></td><td></td><td></td><td></td><td></td><td></td></tr>
<tr><td>Oligo fwd</td><td>CACCGCCCTTCGACATGCTGAACG</td><td></td><td></td><td></td><td></td><td></td><td></td><td></td><td></td><td></td><td></td></tr>
<tr><td>Oligo rev</td><td>AAACCGTTCAGCATGTCGAAGGGC</td><td></td><td></td><td></td><td></td><td></td><td></td><td></td><td></td><td></td><td></td></tr>
<tr><td>Chromosome</td><td>start</td><td>end</td><td>strand</td><td>MM</td><td>target_seq</td><td>PAM</td><td>alignment</td><td>distance</td><td>position</td><td>gene name</td><td>gene id</td></tr>
<tr><td>GU305914.1</td><td>132672</td><td>132694</td><td>-</td><td>0</td><td>GCCCTTCGACATGCTGAACG</td><td>TGG</td><td>||||||||[||||||||||||]PAM</td><td>NA</td><td>Intergenic</td><td>NA</td><td>NA</td></tr>
</table>

<table>
<tr><td>T23939</td><td>CGTGGTATCCTACGTGTGCGAGG</td><td>1000</td><td></td><td></td><td></td><td></td><td></td><td></td><td></td><td></td><td></td></tr>
<tr><td>Oligo fwd</td><td>CACCCGTGGTATCCTACGTGTGCG</td><td></td><td></td><td></td><td></td><td></td><td></td><td></td><td></td><td></td><td></td></tr>
<tr><td>Oligo rev</td><td>AAACCGCACACGTAGGATACCACG</td><td></td><td></td><td></td><td></td><td></td><td></td><td></td><td></td><td></td><td></td></tr>
<tr><td>Chromosome</td><td>start</td><td>end</td><td>strand</td><td>MM</td><td>target_seq</td><td>PAM</td><td>alignment</td><td>distance</td><td>position</td><td>gene name</td><td>gene id</td></tr>
<tr><td>GU305914.1</td><td>132654</td><td>132676</td><td>-</td><td>0</td><td>CGTGGTATCCTACGTGTGCG</td><td>AGG</td><td>||||||||[||||||||||||]PAM</td><td>NA</td><td>Intergenic</td><td>NA</td><td>NA</td></tr>
<tr><td>T23940</td><td>CGTGTGCGAGGAGCACATCCAGG</td><td>1000</td><td></td><td></td><td></td><td></td><td></td><td></td><td></td><td></td><td></td></tr>
<tr><td>Oligo fwd</td><td>CACCCGTGTGCGAGGAGCACATCC</td><td></td><td></td><td></td><td></td><td></td><td></td><td></td><td></td><td></td><td></td></tr>
<tr><td>Oligo rev</td><td>AAACGGATGTGCTCCTCGCACACG</td><td></td><td></td><td></td><td></td><td></td><td></td><td></td><td></td><td></td><td></td></tr>
<tr><td>Chromosome</td><td>start</td><td>end</td><td>strand</td><td>MM</td><td>target_seq</td><td>PAM</td><td>alignment</td><td>distance</td><td>position</td><td>gene name</td><td>gene id</td></tr>
<tr><td>GU305914.1</td><td>132642</td><td>132664</td><td>-</td><td>0</td><td>CGTGTGCGAGGAGCACATCC</td><td>AGG</td><td>||||||||[||||||||||||]PAM</td><td>NA</td><td>Intergenic</td><td>NA</td><td>NA</td></tr>
<tr><td>T23941</td><td>GGAGCACATCCAGGCGTTCTCGG</td><td>1000</td><td></td><td></td><td></td><td></td><td></td><td></td><td></td><td></td><td></td></tr>
<tr><td>Oligo fwd</td><td>CACCGGAGCACATCCAGGCGTTCT</td><td></td><td></td><td></td><td></td><td></td><td></td><td></td><td></td><td></td><td></td></tr>
<tr><td>Oligo rev</td><td>AAACAGAACGCCTGGATGTGCTCC</td><td></td><td></td><td></td><td></td><td></td><td></td><td></td><td></td><td></td><td></td></tr>
<tr><td>Chromosome</td><td>start</td><td>end</td><td>strand</td><td>MM</td><td>target_seq</td><td>PAM</td><td>alignment</td><td>distance</td><td>position</td><td>gene name</td><td>gene id</td></tr>
<tr><td>GU305914.1</td><td>132633</td><td>132655</td><td>-</td><td>0</td><td>GGAGCACATCCAGGCGTTCT</td><td>CGG</td><td>||||||||[||||||||||||]PAM</td><td>NA</td><td>Intergenic</td><td>NA</td><td>NA</td></tr>
<tr><td>T23942</td><td>CATCCAGGCGTTCTCGGAGAAGG</td><td>1000</td><td></td><td></td><td></td><td></td><td></td><td></td><td></td><td></td><td></td></tr>
<tr><td>Oligo fwd</td><td>CACCCATCCAGGCGTTCTCGGAGA</td><td></td><td></td><td></td><td></td><td></td><td></td><td></td><td></td><td></td><td></td></tr>
<tr><td>Oligo rev</td><td>AAACTCTCCGAGAACGCCTGGATG</td><td></td><td></td><td></td><td></td><td></td><td></td><td></td><td></td><td></td><td></td></tr>
<tr><td>Chromosome</td><td>start</td><td>end</td><td>strand</td><td>MM</td><td>target_seq</td><td>PAM</td><td>alignment</td><td>distance</td><td>position</td><td>gene name</td><td>gene id</td></tr>
<tr><td>GU305914.1</td><td>132627</td><td>132649</td><td>-</td><td>0</td><td>CATCCAGGCGTTCTCGGAGA</td><td>AGG</td><td>||||||||[||||||||||||]PAM</td><td>NA</td><td>Intergenic</td><td>NA</td><td>NA</td></tr>
<tr><td>T23943</td><td>ATCCAGGCGTTCTCGGAGAAGGG</td><td>1000</td><td></td><td></td><td></td><td></td><td></td><td></td><td></td><td></td><td></td></tr>
<tr><td>Oligo fwd</td><td>CACCATCCAGGCGTTCTCGGAGAA</td><td></td><td></td><td></td><td></td><td></td><td></td><td></td><td></td><td></td><td></td></tr>
<tr><td>Oligo rev</td><td>AAACTTCTCCGAGAACGCCTGGAT</td><td></td><td></td><td></td><td></td><td></td><td></td><td></td><td></td><td></td><td></td></tr>
<tr><td>Chromosome</td><td>start</td><td>end</td><td>strand</td><td>MM</td><td>target_seq</td><td>PAM</td><td>alignment</td><td>distance</td><td>position</td><td>gene name</td><td>gene id</td></tr>
<tr><td>GU305914.1</td><td>132626</td><td>132648</td><td>-</td><td>0</td><td>ATCCAGGCGTTCTCGGAGAA</td><td>GGG</td><td>||||||||[||||||||||||]PAM</td><td>NA</td><td>Intergenic</td><td>NA</td><td>NA</td></tr>
<tr><td>T23944</td><td>CTCGGAGAAGGGCGACGCGACGG</td><td>1000</td><td></td><td></td><td></td><td></td><td></td><td></td><td></td><td></td><td></td></tr>
<tr><td>Oligo fwd</td><td>CACCCTCGGAGAAGGGCGACGCGA</td><td></td><td></td><td></td><td></td><td></td><td></td><td></td><td></td><td></td><td></td></tr>
<tr><td>Oligo rev</td><td>AAACTCGCGTCGCCCTTCTCCGAG</td><td></td><td></td><td></td><td></td><td></td><td></td><td></td><td></td><td></td><td></td></tr>
<tr><td>Chromosome</td><td>start</td><td>end</td><td>strand</td><td>MM</td><td>target_seq</td><td>PAM</td><td>alignment</td><td>distance</td><td>position</td><td>gene name</td><td>gene id</td></tr>
<tr><td>GU305914.1</td><td>132615</td><td>132637</td><td>-</td><td>0</td><td>CTCGGAGAAGGGCGACGCGA</td><td>CGG</td><td>||||||||[||||||||||||]PAM</td><td>NA</td><td>Intergenic</td><td>NA</td><td>NA</td></tr>
<tr><td>T23945</td><td>GACCTTCATCACGCTGAACTCGG</td><td>1000</td><td></td><td></td><td></td><td></td><td></td><td></td><td></td><td></td><td></td></tr>
<tr><td>Oligo fwd</td><td>CACCGACCTTCATCACGCTGAACT</td><td></td><td></td><td></td><td></td><td></td><td></td><td></td><td></td><td></td><td></td></tr>
<tr><td>Oligo rev</td><td>AAACAGTTCAGCGTGATGAAGGTC</td><td></td><td></td><td></td><td></td><td></td><td></td><td></td><td></td><td></td><td></td></tr>
<tr><td>Chromosome</td><td>start</td><td>end</td><td>strand</td><td>MM</td><td>target_seq</td><td>PAM</td><td>alignment</td><td>distance</td><td>position</td><td>gene name</td><td>gene id</td></tr>
<tr><td>GU305914.1</td><td>132564</td><td>132586</td><td>-</td><td>0</td><td>GACCTTCATCACGCTGAACT</td><td>CGG</td><td>||||||||[||||||||||||]PAM</td><td>NA</td><td>Intergenic</td><td>NA</td><td>NA</td></tr>
<tr><td>T23946</td><td>TCACGCTGAACTCGGACGTGCGG</td><td>1000</td><td></td><td></td><td></td><td></td><td></td><td></td><td></td><td></td><td></td></tr>
<tr><td>Oligo fwd</td><td>CACCTCACGCTGAACTCGGACGTG</td><td></td><td></td><td></td><td></td><td></td><td></td><td></td><td></td><td></td><td></td></tr>
<tr><td>Oligo rev</td><td>AAACCACGTCCGAGTTCAGCGTGA</td><td></td><td></td><td></td><td></td><td></td><td></td><td></td><td></td><td></td><td></td></tr>
<tr><td>Chromosome</td><td>start</td><td>end</td><td>strand</td><td>MM</td><td>target_seq</td><td>PAM</td><td>alignment</td><td>distance</td><td>position</td><td>gene name</td><td>gene id</td></tr>
<tr><td>GU305914.1</td><td>132556</td><td>132578</td><td>-</td><td>0</td><td>TCACGCTGAACTCGGACGTG</td><td>CGG</td><td>||||||||[||||||||||||]PAM</td><td>NA</td><td>Intergenic</td><td>NA</td><td>NA</td></tr>
<tr><td>T23947</td><td>GAACTCGGACGTGCGGAAGACGG</td><td>1000</td><td></td><td></td><td></td><td></td><td></td><td></td><td></td><td></td><td></td></tr>
<tr><td>Oligo fwd</td><td>CACCGAACTCGGACGTGCGGAAGA</td><td></td><td></td><td></td><td></td><td></td><td></td><td></td><td></td><td></td><td></td></tr>
<tr><td>Oligo rev</td><td>AAACTCTTCCGCACGTCCGAGTTC</td><td></td><td></td><td></td><td></td><td></td><td></td><td></td><td></td><td></td><td></td></tr>
<tr><td>Chromosome</td><td>start</td><td>end</td><td>strand</td><td>MM</td><td>target_seq</td><td>PAM</td><td>alignment</td><td>distance</td><td>position</td><td>gene name</td><td>gene id</td></tr>
<tr><td>GU305914.1</td><td>132549</td><td>132571</td><td>-</td><td>0</td><td>GAACTCGGACGTGCGGAAGA</td><td>CGG</td><td>||||||||[||||||||||||]PAM</td><td>NA</td><td>Intergenic</td><td>NA</td><td>NA</td></tr>
<tr><td>GU305914.1</td><td>135784</td><td>135806</td><td>+</td><td>4</td><td>GACGTCGGACGGGCGGACGA</td><td>CGG</td><td>||--||||[|||-|||||-||]PAM</td><td>NA</td><td>Intergenic</td><td>NA</td><td>NA</td></tr>
<tr><td>T23948</td><td>ACGGCGAACATGTTTATGCCCGG</td><td>1000</td><td></td><td></td><td></td><td></td><td></td><td></td><td></td><td></td><td></td></tr>
<tr><td>Oligo fwd</td><td>CACCACGGCGAACATGTTTATGCC</td><td></td><td></td><td></td><td></td><td></td><td></td><td></td><td></td><td></td><td></td></tr>
<tr><td>Oligo rev</td><td>AAACGGCATAAACATGTTCGCCGT</td><td></td><td></td><td></td><td></td><td></td><td></td><td></td><td></td><td></td><td></td></tr>
<tr><td>Chromosome</td><td>start</td><td>end</td><td>strand</td><td>MM</td><td>target_seq</td><td>PAM</td><td>alignment</td><td>distance</td><td>position</td><td>gene name</td><td>gene id</td></tr>
<tr><td>GU305914.1</td><td>132530</td><td>132552</td><td>-</td><td>0</td><td>ACGGCGAACATGTTTATGCC</td><td>CGG</td><td>||||||||[||||||||||||]PAM</td><td>NA</td><td>Intergenic</td><td>NA</td><td>NA</td></tr>
<tr><td>T23949</td><td>GTTTATGCCCGGCTCGTTCATGG</td><td>1000</td><td></td><td></td><td></td><td></td><td></td><td></td><td></td><td></td><td></td></tr>
<tr><td>Oligo fwd</td><td>CACCGTTTATGCCCGGCTCGTTCA</td><td></td><td></td><td></td><td></td><td></td><td></td><td></td><td></td><td></td><td></td></tr>
<tr><td>Oligo rev</td><td>AAACTGAACGAGCCGGGCATAAAC</td><td></td><td></td><td></td><td></td><td></td><td></td><td></td><td></td><td></td><td></td></tr>
<tr><td>Chromosome</td><td>start</td><td>end</td><td>strand</td><td>MM</td><td>target_seq</td><td>PAM</td><td>alignment</td><td>distance</td><td>position</td><td>gene name</td><td>gene id</td></tr>
<tr><td>GU305914.1</td><td>132519</td><td>132541</td><td>-</td><td>0</td><td>GTTTATGCCCGGCTCGTTCA</td><td>TGG</td><td>||||||||[||||||||||||]PAM</td><td>NA</td><td>Intergenic</td><td>NA</td><td>NA</td></tr>
<tr><td>T23950</td><td>CTCGTTCATGGACGAGATCATGG</td><td>1000</td><td></td><td></td><td></td><td></td><td></td><td></td><td></td><td></td><td></td></tr>
<tr><td>Oligo fwd</td><td>CACCCTCGTTCATGGACGAGATCA</td><td></td><td></td><td></td><td></td><td></td><td></td><td></td><td></td><td></td><td></td></tr>
<tr><td>Oligo rev</td><td>AAACTGATCTCGTCCATGAACGAG</td><td></td><td></td><td></td><td></td><td></td><td></td><td></td><td></td><td></td><td></td></tr>
<tr><td>Chromosome</td><td>start</td><td>end</td><td>strand</td><td>MM</td><td>target_seq</td><td>PAM</td><td>alignment</td><td>distance</td><td>position</td><td>gene name</td><td>gene id</td></tr>
<tr><td>GU305914.1</td><td>132507</td><td>132529</td><td>-</td><td>0</td><td>CTCGTTCATGGACGAGATCA</td><td>TGG</td><td>||||||||[||||||||||||]PAM</td><td>NA</td><td>Intergenic</td><td>NA</td><td>NA</td></tr>
<tr><td>GU305914.1</td><td>137561</td><td>137583</td><td>-</td><td>4</td><td>CCGGTTCATGGTCGAGAACA</td><td>AGG</td><td>|--|||||[|||-|||||-||]PAM</td><td>NA</td><td>Intergenic</td><td>NA</td><td>NA</td></tr>
<tr><td>T23951</td><td>TCGTTCATGGACGAGATCATGGG</td><td>1000</td><td></td><td></td><td></td><td></td><td></td><td></td><td></td><td></td><td></td></tr>
<tr><td>Oligo fwd</td><td>CACCTCGTTCATGGACGAGATCAT</td><td></td><td></td><td></td><td></td><td></td><td></td><td></td><td></td><td></td><td></td></tr>
<tr><td>Oligo rev</td><td>AAACATGATCTCGTCCATGAACGA</td><td></td><td></td><td></td><td></td><td></td><td></td><td></td><td></td><td></td><td></td></tr>
<tr><td>Chromosome</td><td>start</td><td>end</td><td>strand</td><td>MM</td><td>target_seq</td><td>PAM</td><td>alignment</td><td>distance</td><td>position</td><td>gene name</td><td>gene id</td></tr>
<tr><td>GU305914.1</td><td>132506</td><td>132528</td><td>-</td><td>0</td><td>TCGTTCATGGACGAGATCAT</td><td>GGG</td><td>||||||||[||||||||||||]PAM</td><td>NA</td><td>Intergenic</td><td>NA</td><td>NA</td></tr>
<tr><td>T23952</td><td>TTCATGGACGAGATCATGGGCGG</td><td>1000</td><td></td><td></td><td></td><td></td><td></td><td></td><td></td><td></td><td></td></tr>
<tr><td>Oligo fwd</td><td>CACCTTCATGGACGAGATCATGGG</td><td></td><td></td><td></td><td></td><td></td><td></td><td></td><td></td><td></td><td></td></tr>
<tr><td>Oligo rev</td><td>AAACCCCATGATCTCGTCCATGAA</td><td></td><td></td><td></td><td></td><td></td><td></td><td></td><td></td><td></td><td></td></tr>
<tr><td>Chromosome</td><td>start</td><td>end</td><td>strand</td><td>MM</td><td>target_seq</td><td>PAM</td><td>alignment</td><td>distance</td><td>position</td><td>gene name</td><td>gene id</td></tr>
<tr><td>GU305914.1</td><td>132503</td><td>132525</td><td>-</td><td>0</td><td>TTCATGGACGAGATCATGGG</td><td>CGG</td><td>||||||||[||||||||||||]PAM</td><td>NA</td><td>Intergenic</td><td>NA</td><td>NA</td></tr>
<tr><td>T23953</td><td>CGGCACGAACAAGATCAACGAGG</td><td>1000</td><td></td><td></td><td></td><td></td><td></td><td></td><td></td><td></td><td></td></tr>
<tr><td>Oligo fwd</td><td>CACCCGGCACGAACAAGATCAACG</td><td></td><td></td><td></td><td></td><td></td><td></td><td></td><td></td><td></td><td></td></tr>
<tr><td>Oligo rev</td><td>AAACCGTTGATCTTGTTCGTGCCG</td><td></td><td></td><td></td><td></td><td></td><td></td><td></td><td></td><td></td><td></td></tr>
</table>

<table>
<tr><td>Chromosome</td><td>start</td><td>end</td><td>strand</td><td>MM</td><td>target_seq</td><td>PAM</td><td>alignment</td><td>distance</td><td>position</td><td>gene name</td><td>gene id</td></tr>
<tr><td>GU305914.1</td><td>132483</td><td>132505</td><td>-</td><td>0</td><td>CGGCACGAACAAGATCAACG</td><td>AGG</td><td>||||||||[||||||||||||]PAM</td><td>NA</td><td>Intergenic</td><td>NA</td><td>NA</td></tr>
<tr><td></td><td></td><td></td><td></td><td></td><td></td><td></td><td></td><td></td><td></td><td></td><td></td></tr>
<tr><td>T23954</td><td>GAACAAGATCAACGAGGAGACGG</td><td>1000</td><td></td><td></td><td></td><td></td><td></td><td></td><td></td><td></td><td></td></tr>
<tr><td>Oligo fwd</td><td>CACCGAACAAGATCAACGAGGAGA</td><td></td><td></td><td></td><td></td><td></td><td></td><td></td><td></td><td></td><td></td></tr>
<tr><td>Oligo rev</td><td>AAACTCTCCTCGTTGATCTTGTTC</td><td></td><td></td><td></td><td></td><td></td><td></td><td></td><td></td><td></td><td></td></tr>
<tr><td>Chromosome</td><td>start</td><td>end</td><td>strand</td><td>MM</td><td>target_seq</td><td>PAM</td><td>alignment</td><td>distance</td><td>position</td><td>gene name</td><td>gene id</td></tr>
<tr><td>GU305914.1</td><td>132477</td><td>132499</td><td>-</td><td>0</td><td>GAACAAGATCAACGAGGAGA</td><td>CGG</td><td>||||||||[||||||||||||]PAM</td><td>NA</td><td>Intergenic</td><td>NA</td><td>NA</td></tr>
<tr><td></td><td></td><td></td><td></td><td></td><td></td><td></td><td></td><td></td><td></td><td></td><td></td></tr>
<tr><td>T23955</td><td>GATCACCGACGAGAGCCGCGAGG</td><td>1000</td><td></td><td></td><td></td><td></td><td></td><td></td><td></td><td></td><td></td></tr>
<tr><td>Oligo fwd</td><td>CACCGATCACCGACGAGAGCCGCG</td><td></td><td></td><td></td><td></td><td></td><td></td><td></td><td></td><td></td><td></td></tr>
<tr><td>Oligo rev</td><td>AAACCGCGGCTCTCGTCGGTGATC</td><td></td><td></td><td></td><td></td><td></td><td></td><td></td><td></td><td></td><td></td></tr>
<tr><td>Chromosome</td><td>start</td><td>end</td><td>strand</td><td>MM</td><td>target_seq</td><td>PAM</td><td>alignment</td><td>distance</td><td>position</td><td>gene name</td><td>gene id</td></tr>
<tr><td>GU305914.1</td><td>132450</td><td>132472</td><td>-</td><td>0</td><td>GATCACCGACGAGAGCCGCG</td><td>AGG</td><td>||||||||[||||||||||||]PAM</td><td>NA</td><td>Intergenic</td><td>NA</td><td>NA</td></tr>
<tr><td></td><td></td><td></td><td></td><td></td><td></td><td></td><td></td><td></td><td></td><td></td><td></td></tr>
<tr><td>T23956</td><td>CCCGCAGTTCCACCCTCACCTGG</td><td>1000</td><td></td><td></td><td></td><td></td><td></td><td></td><td></td><td></td><td></td></tr>
<tr><td>Oligo fwd</td><td>CACCCCCGCAGTTCCACCCTCACC</td><td></td><td></td><td></td><td></td><td></td><td></td><td></td><td></td><td></td><td></td></tr>
<tr><td>Oligo rev</td><td>AAACGGTGAGGGTGGAACTGCGGG</td><td></td><td></td><td></td><td></td><td></td><td></td><td></td><td></td><td></td><td></td></tr>
<tr><td>Chromosome</td><td>start</td><td>end</td><td>strand</td><td>MM</td><td>target_seq</td><td>PAM</td><td>alignment</td><td>distance</td><td>position</td><td>gene name</td><td>gene id</td></tr>
<tr><td>GU305914.1</td><td>132396</td><td>132418</td><td>-</td><td>0</td><td>CCCGCAGTTCCACCCTCACC</td><td>TGG</td><td>||||||||[||||||||||||]PAM</td><td>NA</td><td>Intergenic</td><td>NA</td><td>NA</td></tr>
<tr><td></td><td></td><td></td><td></td><td></td><td></td><td></td><td></td><td></td><td></td><td></td><td></td></tr>
<tr><td>T23957</td><td>CCGCAGTTCCACCCTCACCTGGG</td><td>1000</td><td></td><td></td><td></td><td></td><td></td><td></td><td></td><td></td><td></td></tr>
<tr><td>Oligo fwd</td><td>CACCCCGCAGTTCCACCCTCACCT</td><td></td><td></td><td></td><td></td><td></td><td></td><td></td><td></td><td></td><td></td></tr>
<tr><td>Oligo rev</td><td>AAACAGGTGAGGGTGGAACTGCGG</td><td></td><td></td><td></td><td></td><td></td><td></td><td></td><td></td><td></td><td></td></tr>
<tr><td>Chromosome</td><td>start</td><td>end</td><td>strand</td><td>MM</td><td>target_seq</td><td>PAM</td><td>alignment</td><td>distance</td><td>position</td><td>gene name</td><td>gene id</td></tr>
<tr><td>GU305914.1</td><td>132395</td><td>132417</td><td>-</td><td>0</td><td>CCGCAGTTCCACCCTCACCT</td><td>GGG</td><td>||||||||[||||||||||||]PAM</td><td>NA</td><td>Intergenic</td><td>NA</td><td>NA</td></tr>
<tr><td>GU305914.1</td><td>33923</td><td>33945</td><td>-</td><td>4</td><td>CCGCGGTCTCACCCTCGCCT</td><td>GGG</td><td>||||-||-[-|||||||-|||]PAM</td><td>NA</td><td>Intergenic</td><td>NA</td><td>NA</td></tr>
<tr><td></td><td></td><td></td><td></td><td></td><td></td><td></td><td></td><td></td><td></td><td></td><td></td></tr>
<tr><td>T23958</td><td>CGCGATCCTGAGCGTCTACGTGG</td><td>1000</td><td></td><td></td><td></td><td></td><td></td><td></td><td></td><td></td><td></td></tr>
<tr><td>Oligo fwd</td><td>CACCCGCGATCCTGAGCGTCTACG</td><td></td><td></td><td></td><td></td><td></td><td></td><td></td><td></td><td></td><td></td></tr>
<tr><td>Oligo rev</td><td>AAACCGTAGACGCTCAGGATCGCG</td><td></td><td></td><td></td><td></td><td></td><td></td><td></td><td></td><td></td><td></td></tr>
<tr><td>Chromosome</td><td>start</td><td>end</td><td>strand</td><td>MM</td><td>target_seq</td><td>PAM</td><td>alignment</td><td>distance</td><td>position</td><td>gene name</td><td>gene id</td></tr>
<tr><td>GU305914.1</td><td>132372</td><td>132394</td><td>-</td><td>0</td><td>CGCGATCCTGAGCGTCTACG</td><td>TGG</td><td>||||||||[||||||||||||]PAM</td><td>NA</td><td>Intergenic</td><td>NA</td><td>NA</td></tr>
<tr><td></td><td></td><td></td><td></td><td></td><td></td><td></td><td></td><td></td><td></td><td></td><td></td></tr>
<tr><td>T23959</td><td>CCTGAGCGTCTACGTGGACCCGG</td><td>1000</td><td></td><td></td><td></td><td></td><td></td><td></td><td></td><td></td><td></td></tr>
<tr><td>Oligo fwd</td><td>CACCCCTGAGCGTCTACGTGGACC</td><td></td><td></td><td></td><td></td><td></td><td></td><td></td><td></td><td></td><td></td></tr>
<tr><td>Oligo rev</td><td>AAACGGTCCACGTAGACGCTCAGG</td><td></td><td></td><td></td><td></td><td></td><td></td><td></td><td></td><td></td><td></td></tr>
<tr><td>Chromosome</td><td>start</td><td>end</td><td>strand</td><td>MM</td><td>target_seq</td><td>PAM</td><td>alignment</td><td>distance</td><td>position</td><td>gene name</td><td>gene id</td></tr>
<tr><td>GU305914.1</td><td>132366</td><td>132388</td><td>-</td><td>0</td><td>CCTGAGCGTCTACGTGGACC</td><td>CGG</td><td>||||||||[||||||||||||]PAM</td><td>NA</td><td>Intergenic</td><td>NA</td><td>NA</td></tr>
<tr><td></td><td></td><td></td><td></td><td></td><td></td><td></td><td></td><td></td><td></td><td></td><td></td></tr>
<tr><td>T23960</td><td>ACCTCGAACCGCCGCGCGTCCGG</td><td>1000</td><td></td><td></td><td></td><td></td><td></td><td></td><td></td><td></td><td></td></tr>
<tr><td>Oligo fwd</td><td>CACCACCTCGAACCGCCGCGCGTC</td><td></td><td></td><td></td><td></td><td></td><td></td><td></td><td></td><td></td><td></td></tr>
<tr><td>Oligo rev</td><td>AAACGACGCGCGGCGGTTCGAGGT</td><td></td><td></td><td></td><td></td><td></td><td></td><td></td><td></td><td></td><td></td></tr>
<tr><td>Chromosome</td><td>start</td><td>end</td><td>strand</td><td>MM</td><td>target_seq</td><td>PAM</td><td>alignment</td><td>distance</td><td>position</td><td>gene name</td><td>gene id</td></tr>
<tr><td>GU305914.1</td><td>132338</td><td>132360</td><td>-</td><td>0</td><td>ACCTCGAACCGCCGCGCGTC</td><td>CGG</td><td>||||||||[||||||||||||]PAM</td><td>NA</td><td>Intergenic</td><td>NA</td><td>NA</td></tr>
<tr><td></td><td></td><td></td><td></td><td></td><td></td><td></td><td></td><td></td><td></td><td></td><td></td></tr>
<tr><td>T23961</td><td>AACCGCCGCGCGTCCGGCACCGG</td><td>1000</td><td></td><td></td><td></td><td></td><td></td><td></td><td></td><td></td><td></td></tr>
<tr><td>Oligo fwd</td><td>CACCAACCGCCGCGCGTCCGGCAC</td><td></td><td></td><td></td><td></td><td></td><td></td><td></td><td></td><td></td><td></td></tr>
<tr><td>Oligo rev</td><td>AAACGTGCCGGACGCGCGGCGGTT</td><td></td><td></td><td></td><td></td><td></td><td></td><td></td><td></td><td></td><td></td></tr>
<tr><td>Chromosome</td><td>start</td><td>end</td><td>strand</td><td>MM</td><td>target_seq</td><td>PAM</td><td>alignment</td><td>distance</td><td>position</td><td>gene name</td><td>gene id</td></tr>
<tr><td>GU305914.1</td><td>132332</td><td>132354</td><td>-</td><td>0</td><td>AACCGCCGCGCGTCCGGCAC</td><td>CGG</td><td>||||||||[||||||||||||]PAM</td><td>NA</td><td>Intergenic</td><td>NA</td><td>NA</td></tr>
<tr><td></td><td></td><td></td><td></td><td></td><td></td><td></td><td></td><td></td><td></td><td></td><td></td></tr>
<tr><td>T23962</td><td>CCGCGCGTCCGGCACCGGCGTGG</td><td>1000</td><td></td><td></td><td></td><td></td><td></td><td></td><td></td><td></td><td></td></tr>
<tr><td>Oligo fwd</td><td>CACCCCGCGCGTCCGGCACCGGCG</td><td></td><td></td><td></td><td></td><td></td><td></td><td></td><td></td><td></td><td></td></tr>
<tr><td>Oligo rev</td><td>AAACCGCCGGTGCCGGACGCGCGG</td><td></td><td></td><td></td><td></td><td></td><td></td><td></td><td></td><td></td><td></td></tr>
<tr><td>Chromosome</td><td>start</td><td>end</td><td>strand</td><td>MM</td><td>target_seq</td><td>PAM</td><td>alignment</td><td>distance</td><td>position</td><td>gene name</td><td>gene id</td></tr>
<tr><td>GU305914.1</td><td>132327</td><td>132349</td><td>-</td><td>0</td><td>CCGCGCGTCCGGCACCGGCG</td><td>TGG</td><td>||||||||[||||||||||||]PAM</td><td>NA</td><td>Intergenic</td><td>NA</td><td>NA</td></tr>
<tr><td>GU305914.1</td><td>30041</td><td>30063</td><td>+</td><td>3</td><td>CTGCGCGACCGTCACCGGCG</td><td>CGG</td><td>|-|||||-[|||-||||||||]PAM</td><td>NA</td><td>Intergenic</td><td>NA</td><td>NA</td></tr>
<tr><td></td><td></td><td></td><td></td><td></td><td></td><td></td><td></td><td></td><td></td><td></td><td></td></tr>
<tr><td>T23963</td><td>GTCCGGCACCGGCGTGGCAGCGG</td><td>1000</td><td></td><td></td><td></td><td></td><td></td><td></td><td></td><td></td><td></td></tr>
<tr><td>Oligo fwd</td><td>CACCGTCCGGCACCGGCGTGGCAG</td><td></td><td></td><td></td><td></td><td></td><td></td><td></td><td></td><td></td><td></td></tr>
<tr><td>Oligo rev</td><td>AAACCTGCCACGCCGGTGCCGGAC</td><td></td><td></td><td></td><td></td><td></td><td></td><td></td><td></td><td></td><td></td></tr>
<tr><td>Chromosome</td><td>start</td><td>end</td><td>strand</td><td>MM</td><td>target_seq</td><td>PAM</td><td>alignment</td><td>distance</td><td>position</td><td>gene name</td><td>gene id</td></tr>
<tr><td>GU305914.1</td><td>132321</td><td>132343</td><td>-</td><td>0</td><td>GTCCGGCACCGGCGTGGCAG</td><td>CGG</td><td>||||||||[||||||||||||]PAM</td><td>NA</td><td>Intergenic</td><td>NA</td><td>NA</td></tr>
<tr><td></td><td></td><td></td><td></td><td></td><td></td><td></td><td></td><td></td><td></td><td></td><td></td></tr>
<tr><td>T23964</td><td>CGGCACCGGCGTGGCAGCGGTGG</td><td>1000</td><td></td><td></td><td></td><td></td><td></td><td></td><td></td><td></td><td></td></tr>
<tr><td>Oligo fwd</td><td>CACCCGGCACCGGCGTGGCAGCGG</td><td></td><td></td><td></td><td></td><td></td><td></td><td></td><td></td><td></td><td></td></tr>
<tr><td>Oligo rev</td><td>AAACCCGCTGCCACGCCGGTGCCG</td><td></td><td></td><td></td><td></td><td></td><td></td><td></td><td></td><td></td><td></td></tr>
<tr><td>Chromosome</td><td>start</td><td>end</td><td>strand</td><td>MM</td><td>target_seq</td><td>PAM</td><td>alignment</td><td>distance</td><td>position</td><td>gene name</td><td>gene id</td></tr>
<tr><td>GU305914.1</td><td>132318</td><td>132340</td><td>-</td><td>0</td><td>CGGCACCGGCGTGGCAGCGG</td><td>TGG</td><td>||||||||[||||||||||||]PAM</td><td>NA</td><td>Intergenic</td><td>NA</td><td>NA</td></tr>
<tr><td></td><td></td><td></td><td></td><td></td><td></td><td></td><td></td><td></td><td></td><td></td><td></td></tr>
<tr><td>T23965</td><td>GGCACCGGCGTGGCAGCGGTGGG</td><td>1000</td><td></td><td></td><td></td><td></td><td></td><td></td><td></td><td></td><td></td></tr>
<tr><td>Oligo fwd</td><td>CACCGGCACCGGCGTGGCAGCGGT</td><td></td><td></td><td></td><td></td><td></td><td></td><td></td><td></td><td></td><td></td></tr>
<tr><td>Oligo rev</td><td>AAACACCGCTGCCACGCCGGTGCC</td><td></td><td></td><td></td><td></td><td></td><td></td><td></td><td></td><td></td><td></td></tr>
<tr><td>Chromosome</td><td>start</td><td>end</td><td>strand</td><td>MM</td><td>target_seq</td><td>PAM</td><td>alignment</td><td>distance</td><td>position</td><td>gene name</td><td>gene id</td></tr>
<tr><td>GU305914.1</td><td>132317</td><td>132339</td><td>-</td><td>0</td><td>GGCACCGGCGTGGCAGCGGT</td><td>GGG</td><td>||||||||[||||||||||||]PAM</td><td>NA</td><td>Intergenic</td><td>NA</td><td>NA</td></tr>
<tr><td></td><td></td><td></td><td></td><td></td><td></td><td></td><td></td><td></td><td></td><td></td><td></td></tr>
<tr><td>T23966</td><td>GCACCGGCGTGGCAGCGGTGGGG</td><td>1000</td><td></td><td></td><td></td><td></td><td></td><td></td><td></td><td></td><td></td></tr>
<tr><td>Oligo fwd</td><td>CACCGCACCGGCGTGGCAGCGGTG</td><td></td><td></td><td></td><td></td><td></td><td></td><td></td><td></td><td></td><td></td></tr>
<tr><td>Oligo rev</td><td>AAACCACCGCTGCCACGCCGGTGC</td><td></td><td></td><td></td><td></td><td></td><td></td><td></td><td></td><td></td><td></td></tr>
<tr><td>Chromosome</td><td>start</td><td>end</td><td>strand</td><td>MM</td><td>target_seq</td><td>PAM</td><td>alignment</td><td>distance</td><td>position</td><td>gene name</td><td>gene id</td></tr>
<tr><td>GU305914.1</td><td>132316</td><td>132338</td><td>-</td><td>0</td><td>GCACCGGCGTGGCAGCGGTG</td><td>GGG</td><td>||||||||[||||||||||||]PAM</td><td>NA</td><td>Intergenic</td><td>NA</td><td>NA</td></tr>
<tr><td></td><td></td><td></td><td></td><td></td><td></td><td></td><td></td><td></td><td></td><td></td><td></td></tr>
<tr><td>T23967</td><td>CGCGACCAGTTTATCGTGTACGG</td><td>1000</td><td></td><td></td><td></td><td></td><td></td><td></td><td></td><td></td><td></td></tr>
<tr><td>Oligo fwd</td><td>CACCCGCGACCAGTTTATCGTGTA</td><td></td><td></td><td></td><td></td><td></td><td></td><td></td><td></td><td></td><td></td></tr>
<tr><td>Oligo rev</td><td>AAACTACACGATAAACTGGTCGCG</td><td></td><td></td><td></td><td></td><td></td><td></td><td></td><td></td><td></td><td></td></tr>
<tr><td>Chromosome</td><td>start</td><td>end</td><td>strand</td><td>MM</td><td>target_seq</td><td>PAM</td><td>alignment</td><td>distance</td><td>position</td><td>gene name</td><td>gene id</td></tr>
<tr><td>GU305914.1</td><td>132287</td><td>132309</td><td>-</td><td>0</td><td>CGCGACCAGTTTATCGTGTA</td><td>CGG</td><td>||||||||[||||||||||||]PAM</td><td>NA</td><td>Intergenic</td><td>NA</td><td>NA</td></tr>
</table>

| | | | | | | | | | | | |
|---|---|---|---|---|---|---|---|---|---|---|---|
| T23968 | GCGACCAGTTTATCGTGTACGGG | 1000 | | | | | | | | | |
| Oligo fwd | CACCGCGACCAGTTTATCGTGTAC | | | | | | | | | | |
| Oligo rev | AAACGTACACGATAAACTGGTCGC | | | | | | | | | | |
| Chromosome | start | end | strand | MM | target_seq | PAM | alignment | distance | position | gene name | gene id |
| GU305914.1 | 132286 | 132308 | - | 0 | GCGACCAGTTTATCGTGTAC | GGG | ||||||||[||||||||||||]PAM | NA | Intergenic | NA | NA |

| | | | | | | | | | | | |
|---|---|---|---|---|---|---|---|---|---|---|---|
| T23969 | CCAGTTTATCGTGTACGGGTTGG | 1000 | | | | | | | | | |
| Oligo fwd | CACCCCAGTTTATCGTGTACGGGT | | | | | | | | | | |
| Oligo rev | AAACACCCGTACACGATAAACTGG | | | | | | | | | | |
| Chromosome | start | end | strand | MM | target_seq | PAM | alignment | distance | position | gene name | gene id |
| GU305914.1 | 132282 | 132304 | - | 0 | CCAGTTTATCGTGTACGGGT | TGG | ||||||||[||||||||||||]PAM | NA | Intergenic | NA | NA |

| | | | | | | | | | | | |
|---|---|---|---|---|---|---|---|---|---|---|---|
| T23970 | GTTGGAGCACTACTTCCTGAAGG | 1000 | | | | | | | | | |
| Oligo fwd | CACCGTTGGAGCACTACTTCCTGA | | | | | | | | | | |
| Oligo rev | AAACTCAGGAAGTAGTGCTCCAAC | | | | | | | | | | |
| Chromosome | start | end | strand | MM | target_seq | PAM | alignment | distance | position | gene name | gene id |
| GU305914.1 | 132264 | 132286 | - | 0 | GTTGGAGCACTACTTCCTGA | AGG | ||||||||[||||||||||||]PAM | NA | Intergenic | NA | NA |

| | | | | | | | | | | | |
|---|---|---|---|---|---|---|---|---|---|---|---|
| T23971 | CTACTTCCTGAAGGACCTGCTGG | 1000 | | | | | | | | | |
| Oligo fwd | CACCCTACTTCCTGAAGGACCTGC | | | | | | | | | | |
| Oligo rev | AAACGCAGGTCCTTCAGGAAGTAG | | | | | | | | | | |
| Chromosome | start | end | strand | MM | target_seq | PAM | alignment | distance | position | gene name | gene id |
| GU305914.1 | 132255 | 132277 | - | 0 | CTACTTCCTGAAGGACCTGC | TGG | ||||||||[||||||||||||]PAM | NA | Intergenic | NA | NA |

| | | | | | | | | | | | |
|---|---|---|---|---|---|---|---|---|---|---|---|
| T23972 | GAAGGACCTGCTGGACAGCTCGG | 1000 | | | | | | | | | |
| Oligo fwd | CACCGAAGGACCTGCTGGACAGCT | | | | | | | | | | |
| Oligo rev | AAACAGCTGTCCAGCAGGTCCTTC | | | | | | | | | | |
| Chromosome | start | end | strand | MM | target_seq | PAM | alignment | distance | position | gene name | gene id |
| GU305914.1 | 132246 | 132268 | - | 0 | GAAGGACCTGCTGGACAGCT | CGG | ||||||||[||||||||||||]PAM | NA | Intergenic | NA | NA |

| | | | | | | | | | | | |
|---|---|---|---|---|---|---|---|---|---|---|---|
| T23973 | CAGCTCGGAGACGTCGATCGCGG | 1000 | | | | | | | | | |
| Oligo fwd | CACCCAGCTCGGAGACGTCGATCG | | | | | | | | | | |
| Oligo rev | AAACCGATCGACGTCTCCGAGCTG | | | | | | | | | | |
| Chromosome | start | end | strand | MM | target_seq | PAM | alignment | distance | position | gene name | gene id |
| GU305914.1 | 132231 | 132253 | - | 0 | CAGCTCGGAGACGTCGATCG | CGG | ||||||||[||||||||||||]PAM | NA | Intergenic | NA | NA |

| | | | | | | | | | | | |
|---|---|---|---|---|---|---|---|---|---|---|---|
| T23974 | CCTGCACCCGTTCCTGTCACAGG | 1000 | | | | | | | | | |
| Oligo fwd | CACCCCTGCACCCGTTCCTGTCAC | | | | | | | | | | |
| Oligo rev | AAACGTGACAGGAACGGGTGCAGG | | | | | | | | | | |
| Chromosome | start | end | strand | MM | target_seq | PAM | alignment | distance | position | gene name | gene id |
| GU305914.1 | 132174 | 132196 | - | 0 | CCTGCACCCGTTCCTGTCAC | AGG | ||||||||[||||||||||||]PAM | NA | Intergenic | NA | NA |

| | | | | | | | | | | | |
|---|---|---|---|---|---|---|---|---|---|---|---|
| T23975 | ACCCGTTCCTGTCACAGGTGCGG | 1000 | | | | | | | | | |
| Oligo fwd | CACCACCCGTTCCTGTCACAGGTG | | | | | | | | | | |
| Oligo rev | AAACCACCTGTGACAGGAACGGGT | | | | | | | | | | |
| Chromosome | start | end | strand | MM | target_seq | PAM | alignment | distance | position | gene name | gene id |
| GU305914.1 | 132169 | 132191 | - | 0 | ACCCGTTCCTGTCACAGGTG | CGG | ||||||||[||||||||||||]PAM | NA | Intergenic | NA | NA |

| | | | | | | | | | | | |
|---|---|---|---|---|---|---|---|---|---|---|---|
| T23976 | CCCGTTCCTGTCACAGGTGCGGG | 1000 | | | | | | | | | |
| Oligo fwd | CACCCCCGTTCCTGTCACAGGTGC | | | | | | | | | | |
| Oligo rev | AAACGCACCTGTGACAGGAACGGG | | | | | | | | | | |
| Chromosome | start | end | strand | MM | target_seq | PAM | alignment | distance | position | gene name | gene id |
| GU305914.1 | 132168 | 132190 | - | 0 | CCCGTTCCTGTCACAGGTGC | GGG | ||||||||[||||||||||||]PAM | NA | Intergenic | NA | NA |
| GU305914.1 | 18756 | 18778 | - | 4 | CCCGCTACTGTCACTGCTGC | TGG | ||||-|-|[||||||-|-|||]PAM | NA | Intergenic | NA | NA |

| | | | | | | | | | | | |
|---|---|---|---|---|---|---|---|---|---|---|---|
| T23977 | CAGGTGCGGGTGACGATCGAAGG | 1000 | | | | | | | | | |
| Oligo fwd | CACCCAGGTGCGGGTGACGATCGA | | | | | | | | | | |
| Oligo rev | AAACTCGATCGTCACCCGCACCTG | | | | | | | | | | |
| Chromosome | start | end | strand | MM | target_seq | PAM | alignment | distance | position | gene name | gene id |
| GU305914.1 | 132155 | 132177 | - | 0 | CAGGTGCGGGTGACGATCGA | AGG | ||||||||[||||||||||||]PAM | NA | Intergenic | NA | NA |

| | | | | | | | | | | | |
|---|---|---|---|---|---|---|---|---|---|---|---|
| T23978 | GATCGAAGGCAACTCGAACCAGG | 1000 | | | | | | | | | |
| Oligo fwd | CACCGATCGAAGGCAACTCGAACC | | | | | | | | | | |
| Oligo rev | AAACGGTTCGAGTTGCCTTCGATC | | | | | | | | | | |
| Chromosome | start | end | strand | MM | target_seq | PAM | alignment | distance | position | gene name | gene id |
| GU305914.1 | 132141 | 132163 | - | 0 | GATCGAAGGCAACTCGAACC | AGG | ||||||||[||||||||||||]PAM | NA | Intergenic | NA | NA |

| | | | | | | | | | | | |
|---|---|---|---|---|---|---|---|---|---|---|---|
| T23979 | CGAAGGCAACTCGAACCAGGCGG | 1000 | | | | | | | | | |
| Oligo fwd | CACCCGAAGGCAACTCGAACCAGG | | | | | | | | | | |
| Oligo rev | AAACCCTGGTTCGAGTTGCCTTCG | | | | | | | | | | |
| Chromosome | start | end | strand | MM | target_seq | PAM | alignment | distance | position | gene name | gene id |
| GU305914.1 | 132138 | 132160 | - | 0 | CGAAGGCAACTCGAACCAGG | CGG | ||||||||[||||||||||||]PAM | NA | Intergenic | NA | NA |

| | | | | | | | | | | | |
|---|---|---|---|---|---|---|---|---|---|---|---|
| T23980 | AGGCAACTCGAACCAGGCGGCGG | 1000 | | | | | | | | | |
| Oligo fwd | CACCAGGCAACTCGAACCAGGCGG | | | | | | | | | | |
| Oligo rev | AAACCCGCCTGGTTCGAGTTGCCT | | | | | | | | | | |
| Chromosome | start | end | strand | MM | target_seq | PAM | alignment | distance | position | gene name | gene id |
| GU305914.1 | 132135 | 132157 | - | 0 | AGGCAACTCGAACCAGGCGG | CGG | ||||||||[||||||||||||]PAM | NA | Intergenic | NA | NA |

| | | | | | | | | | | | |
|---|---|---|---|---|---|---|---|---|---|---|---|
| T23981 | CAACTCGAACCAGGCGGCGGCGG | 1000 | | | | | | | | | |
| Oligo fwd | CACCCAACTCGAACCAGGCGGCGG | | | | | | | | | | |
| Oligo rev | AAACCCGCCGCCTGGTTCGAGTTG | | | | | | | | | | |
| Chromosome | start | end | strand | MM | target_seq | PAM | alignment | distance | position | gene name | gene id |
| GU305914.1 | 132132 | 132154 | - | 0 | CAACTCGAACCAGGCGGCGG | CGG | ||||||||[||||||||||||]PAM | NA | Intergenic | NA | NA |

| | | | | | | | | | | | |
|---|---|---|---|---|---|---|---|---|---|---|---|
| T23982 | CATCAAACACAACCTGCTGTCGG | 1000 | | | | | | | | | |
| Oligo fwd | CACCCATCAAACACAACCTGCTGT | | | | | | | | | | |
| Oligo rev | AAACACAGCAGGTTGTGTTTGATG | | | | | | | | | | |
| Chromosome | start | end | strand | MM | target_seq | PAM | alignment | distance | position | gene name | gene id |

| | | | | | | | | | | | |
|---|---|---|---|---|---|---|---|---|---|---|---|
| GU305914.1 | | 132093 | 132115 | - | 0 | CATCAAACACAACCTGCTGT | CGG | \|\|\|\|\|\|\|\|[\|\|\|\|\|\|\|\|\|\|\|\|]PAM | NA | Intergenic | NA | NA |
| T23983 | GACGCTGTTCTACCACAGCCCGG | | 1000 | | | | | | | | | |
| Oligo fwd | CACCGACGCTGTTCTACCACAGCC | | | | | | | | | | | |
| Oligo rev | AAACGGCTGTGGTAGAACAGCGTC | | | | | | | | | | | |
| Chromosome | start | | end | strand | MM | target_seq | PAM | alignment | distance | position | gene name | gene id |
| GU305914.1 | | 132060 | 132082 | - | 0 | GACGCTGTTCTACCACAGCC | CGG | \|\|\|\|\|\|\|\|[\|\|\|\|\|\|\|\|\|\|\|\|]PAM | NA | Intergenic | NA | NA |
| T23984 | GAACAGAGACAAGAGACTCGCGG | | 1000 | | | | | | | | | |
| Oligo fwd | CACCGAACAGAGACAAGAGACTCG | | | | | | | | | | | |
| Oligo rev | AAACCGAGTCTCTTGTCTCTGTTC | | | | | | | | | | | |
| Chromosome | start | | end | strand | MM | target_seq | PAM | alignment | distance | position | gene name | gene id |
| GU305914.1 | | 132003 | 132025 | - | 0 | GAACAGAGACAAGAGACTCG | CGG | \|\|\|\|\|\|\|\|[\|\|\|\|\|\|\|\|\|\|\|\|]PAM | NA | Intergenic | NA | NA |
| T23985 | AGACAAGAGACTCGCGGTCGAGG | | 1000 | | | | | | | | | |
| Oligo fwd | CACCAGACAAGAGACTCGCGGTCG | | | | | | | | | | | |
| Oligo rev | AAACCGACCGCGAGTCTCTTGTCT | | | | | | | | | | | |
| Chromosome | start | | end | strand | MM | target_seq | PAM | alignment | distance | position | gene name | gene id |
| GU305914.1 | | 131997 | 132019 | - | 0 | AGACAAGAGACTCGCGGTCG | AGG | \|\|\|\|\|\|\|\|[\|\|\|\|\|\|\|\|\|\|\|\|]PAM | NA | Intergenic | NA | NA |
| T23986 | GTTTAATTCGTCGTACATCAAGG | | 1000 | | | | | | | | | |
| Oligo fwd | CACCGTTTAATTCGTCGTACATCA | | | | | | | | | | | |
| Oligo rev | AAACTGATGTACGACGAATTAAAC | | | | | | | | | | | |
| Chromosome | start | | end | strand | MM | target_seq | PAM | alignment | distance | position | gene name | gene id |
| GU305914.1 | | 131961 | 131983 | - | 0 | GTTTAATTCGTCGTACATCA | AGG | \|\|\|\|\|\|\|\|[\|\|\|\|\|\|\|\|\|\|\|\|]PAM | NA | Intergenic | NA | NA |
| T23987 | GTCGTACATCAAGGCCTCCCAGG | | 1000 | | | | | | | | | |
| Oligo fwd | CACCGTCGTACATCAAGGCCTCCC | | | | | | | | | | | |
| Oligo rev | AAACGGGAGGCCTTGATGTACGAC | | | | | | | | | | | |
| Chromosome | start | | end | strand | MM | target_seq | PAM | alignment | distance | position | gene name | gene id |
| GU305914.1 | | 131952 | 131974 | - | 0 | GTCGTACATCAAGGCCTCCC | AGG | \|\|\|\|\|\|\|\|[\|\|\|\|\|\|\|\|\|\|\|\|]PAM | NA | Intergenic | NA | NA |
| GU305914.1 | | 196911 | 196933 | - | 4 | GTCGAACCTGAAGGCCTGCC | TGG | \|\|\|\|-\|\|-[\|-\|\|\|\|\|\|\|-\|\|]PAM | NA | Intergenic | NA | NA |
| T23988 | CATCAAGCTGAGTTACGACCCGG | | 1000 | | | | | | | | | |
| Oligo fwd | CACCCATCAAGCTGAGTTACGACC | | | | | | | | | | | |
| Oligo rev | AAACGGTCGTAACTCAGCTTGATG | | | | | | | | | | | |
| Chromosome | start | | end | strand | MM | target_seq | PAM | alignment | distance | position | gene name | gene id |
| GU305914.1 | | 131913 | 131935 | - | 0 | CATCAAGCTGAGTTACGACC | CGG | \|\|\|\|\|\|\|\|[\|\|\|\|\|\|\|\|\|\|\|\|]PAM | NA | Intergenic | NA | NA |
| T23989 | ACGTGACATACAGCGCGAAACGG | | 1000 | | | | | | | | | |
| Oligo fwd | CACCACGTGACATACAGCGCGAAA | | | | | | | | | | | |
| Oligo rev | AAACTTTCGCGCTGTATGTCACGT | | | | | | | | | | | |
| Chromosome | start | | end | strand | MM | target_seq | PAM | alignment | distance | position | gene name | gene id |
| GU305914.1 | | 131836 | 131858 | - | 0 | ACGTGACATACAGCGCGAAA | CGG | \|\|\|\|\|\|\|\|[\|\|\|\|\|\|\|\|\|\|\|\|]PAM | NA | Intergenic | NA | NA |
| T23990 | CGCGAAACGGAACAACCAGTCGG | | 1000 | | | | | | | | | |
| Oligo fwd | CACCCGCGAAACGGAACAACCAGT | | | | | | | | | | | |
| Oligo rev | AAACACTGGTTGTTCCGTTTCGCG | | | | | | | | | | | |
| Chromosome | start | | end | strand | MM | target_seq | PAM | alignment | distance | position | gene name | gene id |
| GU305914.1 | | 131823 | 131845 | - | 0 | CGCGAAACGGAACAACCAGT | CGG | \|\|\|\|\|\|\|\|[\|\|\|\|\|\|\|\|\|\|\|\|]PAM | NA | Intergenic | NA | NA |
| T23991 | GAACAACCAGTCGGACGACCTGG | | 1000 | | | | | | | | | |
| Oligo fwd | CACCGAACAACCAGTCGGACGACC | | | | | | | | | | | |
| Oligo rev | AAACGGTCGTCCGACTGGTTGTTC | | | | | | | | | | | |
| Chromosome | start | | end | strand | MM | target_seq | PAM | alignment | distance | position | gene name | gene id |
| GU305914.1 | | 131814 | 131836 | - | 0 | GAACAACCAGTCGGACGACC | TGG | \|\|\|\|\|\|\|\|[\|\|\|\|\|\|\|\|\|\|\|\|]PAM | NA | Intergenic | NA | NA |
| T23992 | CAACCAGTCGGACGACCTGGTGG | | 1000 | | | | | | | | | |
| Oligo fwd | CACCCAACCAGTCGGACGACCTGG | | | | | | | | | | | |
| Oligo rev | AAACCCAGGTCGTCCGACTGGTTG | | | | | | | | | | | |
| Chromosome | start | | end | strand | MM | target_seq | PAM | alignment | distance | position | gene name | gene id |
| GU305914.1 | | 131811 | 131833 | - | 0 | CAACCAGTCGGACGACCTGG | TGG | \|\|\|\|\|\|\|\|[\|\|\|\|\|\|\|\|\|\|\|\|]PAM | NA | Intergenic | NA | NA |
| T23993 | CCTGGTGGTCGCCCTGATCATGG | | 1000 | | | | | | | | | |
| Oligo fwd | CACCCCTGGTGGTCGCCCTGATCA | | | | | | | | | | | |
| Oligo rev | AAACTGATCAGGGCGACCACCAGG | | | | | | | | | | | |
| Chromosome | start | | end | strand | MM | target_seq | PAM | alignment | distance | position | gene name | gene id |
| GU305914.1 | | 131796 | 131818 | - | 0 | CCTGGTGGTCGCCCTGATCA | TGG | \|\|\|\|\|\|\|\|[\|\|\|\|\|\|\|\|\|\|\|\|]PAM | NA | Intergenic | NA | NA |
| T23994 | GGCCGTGTACATGTGCTCGCCGG | | 1000 | | | | | | | | | |
| Oligo fwd | CACCGGCCGTGTACATGTGCTCGC | | | | | | | | | | | |
| Oligo rev | AAACGCGAGCACATGTACACGGCC | | | | | | | | | | | |
| Chromosome | start | | end | strand | MM | target_seq | PAM | alignment | distance | position | gene name | gene id |
| GU305914.1 | | 131775 | 131797 | - | 0 | GGCCGTGTACATGTGCTCGC | CGG | \|\|\|\|\|\|\|\|[\|\|\|\|\|\|\|\|\|\|\|\|]PAM | NA | Intergenic | NA | NA |
| T23995 | TGTACATGTGCTCGCCGGAGAGG | | 1000 | | | | | | | | | |
| Oligo fwd | CACCTGTACATGTGCTCGCCGGAG | | | | | | | | | | | |
| Oligo rev | AAACCTCCGGCGAGCACATGTACA | | | | | | | | | | | |
| Chromosome | start | | end | strand | MM | target_seq | PAM | alignment | distance | position | gene name | gene id |
| GU305914.1 | | 131770 | 131792 | - | 0 | TGTACATGTGCTCGCCGGAG | AGG | \|\|\|\|\|\|\|\|[\|\|\|\|\|\|\|\|\|\|\|\|]PAM | NA | Intergenic | NA | NA |
| T23996 | TCAATCGATCGATCGATCGAAGG | | 1000 | | | | | | | | | |
| Oligo fwd | CACCTCAATCGATCGATCGATCGA | | | | | | | | | | | |
| Oligo rev | AAACTCGATCGATCGATCGATTGA | | | | | | | | | | | |
| Chromosome | start | | end | strand | MM | target_seq | PAM | alignment | distance | position | gene name | gene id |
| GU305914.1 | | 131723 | 131745 | - | 0 | TCAATCGATCGATCGATCGA | AGG | \|\|\|\|\|\|\|\|[\|\|\|\|\|\|\|\|\|\|\|\|]PAM | NA | Intergenic | NA | NA |
| T23997 | CAATCGATCGATCGATCGAAGGG | | 1000 | | | | | | | | | |
| Oligo fwd | CACCCAATCGATCGATCGATCGAA | | | | | | | | | | | |

<table>
<tr><td>Oligo rev</td><td>AAACTTCGATCGATCGATCGATTG</td><td></td><td></td><td></td><td></td><td></td><td></td><td></td><td></td><td></td><td></td></tr>
<tr><td>Chromosome</td><td>start</td><td>end</td><td>strand</td><td>MM</td><td>target_seq</td><td>PAM</td><td>alignment</td><td>distance</td><td>position</td><td>gene name</td><td>gene id</td></tr>
<tr><td>GU305914.1</td><td>131722</td><td>131744</td><td>-</td><td>0</td><td>CAATCGATCGATCGATCGAA</td><td>GGG</td><td>||||||||[||||||||||||]PAM</td><td>NA</td><td>Intergenic</td><td>NA</td><td>NA</td></tr>
<tr><td></td><td></td><td></td><td></td><td></td><td></td><td></td><td></td><td></td><td></td><td></td><td></td></tr>
<tr><td>T23998</td><td>GATCACGTCGTCCCGATCGACGG</td><td>1000</td><td></td><td></td><td></td><td></td><td></td><td></td><td></td><td></td><td></td></tr>
<tr><td>Oligo fwd</td><td>CACCGATCACGTCGTCCCGATCGA</td><td></td><td></td><td></td><td></td><td></td><td></td><td></td><td></td><td></td><td></td></tr>
<tr><td>Oligo rev</td><td>AAACTCGATCGGGACGACGTGATC</td><td></td><td></td><td></td><td></td><td></td><td></td><td></td><td></td><td></td><td></td></tr>
<tr><td>Chromosome</td><td>start</td><td>end</td><td>strand</td><td>MM</td><td>target_seq</td><td>PAM</td><td>alignment</td><td>distance</td><td>position</td><td>gene name</td><td>gene id</td></tr>
<tr><td>GU305914.1</td><td>131587</td><td>131609</td><td>-</td><td>0</td><td>GATCACGTCGTCCCGATCGA</td><td>CGG</td><td>||||||||[||||||||||||]PAM</td><td>NA</td><td>Intergenic</td><td>NA</td><td>NA</td></tr>
<tr><td></td><td></td><td></td><td></td><td></td><td></td><td></td><td></td><td></td><td></td><td></td><td></td></tr>
<tr><td>T23999</td><td>ATCACGTCGTCCCGATCGACGGG</td><td>1000</td><td></td><td></td><td></td><td></td><td></td><td></td><td></td><td></td><td></td></tr>
<tr><td>Oligo fwd</td><td>CACCATCACGTCGTCCCGATCGAC</td><td></td><td></td><td></td><td></td><td></td><td></td><td></td><td></td><td></td><td></td></tr>
<tr><td>Oligo rev</td><td>AAACGTCGATCGGGACGACGTGAT</td><td></td><td></td><td></td><td></td><td></td><td></td><td></td><td></td><td></td><td></td></tr>
<tr><td>Chromosome</td><td>start</td><td>end</td><td>strand</td><td>MM</td><td>target_seq</td><td>PAM</td><td>alignment</td><td>distance</td><td>position</td><td>gene name</td><td>gene id</td></tr>
<tr><td>GU305914.1</td><td>131586</td><td>131608</td><td>-</td><td>0</td><td>ATCACGTCGTCCCGATCGAC</td><td>GGG</td><td>||||||||[||||||||||||]PAM</td><td>NA</td><td>Intergenic</td><td>NA</td><td>NA</td></tr>
<tr><td></td><td></td><td></td><td></td><td></td><td></td><td></td><td></td><td></td><td></td><td></td><td></td></tr>
<tr><td>T24000</td><td>CCGATCGACGGGACCCGTCATGG</td><td>1000</td><td></td><td></td><td></td><td></td><td></td><td></td><td></td><td></td><td></td></tr>
<tr><td>Oligo fwd</td><td>CACCCCGATCGACGGGACCCGTCA</td><td></td><td></td><td></td><td></td><td></td><td></td><td></td><td></td><td></td><td></td></tr>
<tr><td>Oligo rev</td><td>AAACTGACGGGTCCCGTCGATCGG</td><td></td><td></td><td></td><td></td><td></td><td></td><td></td><td></td><td></td><td></td></tr>
<tr><td>Chromosome</td><td>start</td><td>end</td><td>strand</td><td>MM</td><td>target_seq</td><td>PAM</td><td>alignment</td><td>distance</td><td>position</td><td>gene name</td><td>gene id</td></tr>
<tr><td>GU305914.1</td><td>131575</td><td>131597</td><td>-</td><td>0</td><td>CCGATCGACGGGACCCGTCA</td><td>TGG</td><td>||||||||[||||||||||||]PAM</td><td>NA</td><td>Intergenic</td><td>NA</td><td>NA</td></tr>
<tr><td></td><td></td><td></td><td></td><td></td><td></td><td></td><td></td><td></td><td></td><td></td><td></td></tr>
<tr><td>T24001</td><td>CGATCGACGGGACCCGTCATGGG</td><td>1000</td><td></td><td></td><td></td><td></td><td></td><td></td><td></td><td></td><td></td></tr>
<tr><td>Oligo fwd</td><td>CACCCGATCGACGGGACCCGTCAT</td><td></td><td></td><td></td><td></td><td></td><td></td><td></td><td></td><td></td><td></td></tr>
<tr><td>Oligo rev</td><td>AAACATGACGGGTCCCGTCGATCG</td><td></td><td></td><td></td><td></td><td></td><td></td><td></td><td></td><td></td><td></td></tr>
<tr><td>Chromosome</td><td>start</td><td>end</td><td>strand</td><td>MM</td><td>target_seq</td><td>PAM</td><td>alignment</td><td>distance</td><td>position</td><td>gene name</td><td>gene id</td></tr>
<tr><td>GU305914.1</td><td>131574</td><td>131596</td><td>-</td><td>0</td><td>CGATCGACGGGACCCGTCAT</td><td>GGG</td><td>||||||||[||||||||||||]PAM</td><td>NA</td><td>Intergenic</td><td>NA</td><td>NA</td></tr>
<tr><td></td><td></td><td></td><td></td><td></td><td></td><td></td><td></td><td></td><td></td><td></td><td></td></tr>
<tr><td>T24002</td><td>GATCGACGGGACCCGTCATGGGG</td><td>1000</td><td></td><td></td><td></td><td></td><td></td><td></td><td></td><td></td><td></td></tr>
<tr><td>Oligo fwd</td><td>CACCGATCGACGGGACCCGTCATG</td><td></td><td></td><td></td><td></td><td></td><td></td><td></td><td></td><td></td><td></td></tr>
<tr><td>Oligo rev</td><td>AAACCATGACGGGTCCCGTCGATC</td><td></td><td></td><td></td><td></td><td></td><td></td><td></td><td></td><td></td><td></td></tr>
<tr><td>Chromosome</td><td>start</td><td>end</td><td>strand</td><td>MM</td><td>target_seq</td><td>PAM</td><td>alignment</td><td>distance</td><td>position</td><td>gene name</td><td>gene id</td></tr>
<tr><td>GU305914.1</td><td>131573</td><td>131595</td><td>-</td><td>0</td><td>GATCGACGGGACCCGTCATG</td><td>GGG</td><td>||||||||[||||||||||||]PAM</td><td>NA</td><td>Intergenic</td><td>NA</td><td>NA</td></tr>
<tr><td></td><td></td><td></td><td></td><td></td><td></td><td></td><td></td><td></td><td></td><td></td><td></td></tr>
<tr><td>T24003</td><td>CGCCGTCGCTCACCAACCCCCGG</td><td>1000</td><td></td><td></td><td></td><td></td><td></td><td></td><td></td><td></td><td></td></tr>
<tr><td>Oligo fwd</td><td>CACCCGCCGTCGCTCACCAACCCC</td><td></td><td></td><td></td><td></td><td></td><td></td><td></td><td></td><td></td><td></td></tr>
<tr><td>Oligo rev</td><td>AAACGGGGTTGGTGAGCGACGGCG</td><td></td><td></td><td></td><td></td><td></td><td></td><td></td><td></td><td></td><td></td></tr>
<tr><td>Chromosome</td><td>start</td><td>end</td><td>strand</td><td>MM</td><td>target_seq</td><td>PAM</td><td>alignment</td><td>distance</td><td>position</td><td>gene name</td><td>gene id</td></tr>
<tr><td>GU305914.1</td><td>131525</td><td>131547</td><td>-</td><td>0</td><td>CGCCGTCGCTCACCAACCCC</td><td>CGG</td><td>||||||||[||||||||||||]PAM</td><td>NA</td><td>Intergenic</td><td>NA</td><td>NA</td></tr>
<tr><td></td><td></td><td></td><td></td><td></td><td></td><td></td><td></td><td></td><td></td><td></td><td></td></tr>
<tr><td>T24004</td><td>CGTCGCTCACCAACCCCCGGAGG</td><td>1000</td><td></td><td></td><td></td><td></td><td></td><td></td><td></td><td></td><td></td></tr>
<tr><td>Oligo fwd</td><td>CACCCGTCGCTCACCAACCCCCGG</td><td></td><td></td><td></td><td></td><td></td><td></td><td></td><td></td><td></td><td></td></tr>
<tr><td>Oligo rev</td><td>AAACCCGGGGGTTGGTGAGCGACG</td><td></td><td></td><td></td><td></td><td></td><td></td><td></td><td></td><td></td><td></td></tr>
<tr><td>Chromosome</td><td>start</td><td>end</td><td>strand</td><td>MM</td><td>target_seq</td><td>PAM</td><td>alignment</td><td>distance</td><td>position</td><td>gene name</td><td>gene id</td></tr>
<tr><td>GU305914.1</td><td>131522</td><td>131544</td><td>-</td><td>0</td><td>CGTCGCTCACCAACCCCCGG</td><td>AGG</td><td>||||||||[||||||||||||]PAM</td><td>NA</td><td>Intergenic</td><td>NA</td><td>NA</td></tr>
<tr><td></td><td></td><td></td><td></td><td></td><td></td><td></td><td></td><td></td><td></td><td></td><td></td></tr>
<tr><td>T24005</td><td>GTCGCTCACCAACCCCCGGAGGG</td><td>1000</td><td></td><td></td><td></td><td></td><td></td><td></td><td></td><td></td><td></td></tr>
<tr><td>Oligo fwd</td><td>CACCGTCGCTCACCAACCCCCGGA</td><td></td><td></td><td></td><td></td><td></td><td></td><td></td><td></td><td></td><td></td></tr>
<tr><td>Oligo rev</td><td>AAACTCCGGGGGTTGGTGAGCGAC</td><td></td><td></td><td></td><td></td><td></td><td></td><td></td><td></td><td></td><td></td></tr>
<tr><td>Chromosome</td><td>start</td><td>end</td><td>strand</td><td>MM</td><td>target_seq</td><td>PAM</td><td>alignment</td><td>distance</td><td>position</td><td>gene name</td><td>gene id</td></tr>
<tr><td>GU305914.1</td><td>131521</td><td>131543</td><td>-</td><td>0</td><td>GTCGCTCACCAACCCCCGGA</td><td>GGG</td><td>||||||||[||||||||||||]PAM</td><td>NA</td><td>Intergenic</td><td>NA</td><td>NA</td></tr>
<tr><td></td><td></td><td></td><td></td><td></td><td></td><td></td><td></td><td></td><td></td><td></td><td></td></tr>
<tr><td>T24006</td><td>CACCAACCCCCGGAGGGAGAAGG</td><td>1000</td><td></td><td></td><td></td><td></td><td></td><td></td><td></td><td></td><td></td></tr>
<tr><td>Oligo fwd</td><td>CACCCACCAACCCCCGGAGGGAGA</td><td></td><td></td><td></td><td></td><td></td><td></td><td></td><td></td><td></td><td></td></tr>
<tr><td>Oligo rev</td><td>AAACTCTCCCTCCGGGGGTTGGTG</td><td></td><td></td><td></td><td></td><td></td><td></td><td></td><td></td><td></td><td></td></tr>
<tr><td>Chromosome</td><td>start</td><td>end</td><td>strand</td><td>MM</td><td>target_seq</td><td>PAM</td><td>alignment</td><td>distance</td><td>position</td><td>gene name</td><td>gene id</td></tr>
<tr><td>GU305914.1</td><td>131515</td><td>131537</td><td>-</td><td>0</td><td>CACCAACCCCCGGAGGGAGA</td><td>AGG</td><td>||||||||[||||||||||||]PAM</td><td>NA</td><td>Intergenic</td><td>NA</td><td>NA</td></tr>
<tr><td></td><td></td><td></td><td></td><td></td><td></td><td></td><td></td><td></td><td></td><td></td><td></td></tr>
<tr><td>T24007</td><td>GCGCAGAACGTCCGCCGCCACGG</td><td>1000</td><td></td><td></td><td></td><td></td><td></td><td></td><td></td><td></td><td></td></tr>
<tr><td>Oligo fwd</td><td>CACCGCGCAGAACGTCCGCCGCCA</td><td></td><td></td><td></td><td></td><td></td><td></td><td></td><td></td><td></td><td></td></tr>
<tr><td>Oligo rev</td><td>AAACTGGCGGCGGACGTTCTGCGC</td><td></td><td></td><td></td><td></td><td></td><td></td><td></td><td></td><td></td><td></td></tr>
<tr><td>Chromosome</td><td>start</td><td>end</td><td>strand</td><td>MM</td><td>target_seq</td><td>PAM</td><td>alignment</td><td>distance</td><td>position</td><td>gene name</td><td>gene id</td></tr>
<tr><td>GU305914.1</td><td>131491</td><td>131513</td><td>-</td><td>0</td><td>GCGCAGAACGTCCGCCGCCA</td><td>CGG</td><td>||||||||[||||||||||||]PAM</td><td>NA</td><td>Intergenic</td><td>NA</td><td>NA</td></tr>
<tr><td></td><td></td><td></td><td></td><td></td><td></td><td></td><td></td><td></td><td></td><td></td><td></td></tr>
<tr><td>T24008</td><td>GACCCCGACATCGACTTCTGCGG</td><td>1000</td><td></td><td></td><td></td><td></td><td></td><td></td><td></td><td></td><td></td></tr>
<tr><td>Oligo fwd</td><td>CACCGACCCCGACATCGACTTCTG</td><td></td><td></td><td></td><td></td><td></td><td></td><td></td><td></td><td></td><td></td></tr>
<tr><td>Oligo rev</td><td>AAACCAGAAGTCGATGTCGGGGTC</td><td></td><td></td><td></td><td></td><td></td><td></td><td></td><td></td><td></td><td></td></tr>
<tr><td>Chromosome</td><td>start</td><td>end</td><td>strand</td><td>MM</td><td>target_seq</td><td>PAM</td><td>alignment</td><td>distance</td><td>position</td><td>gene name</td><td>gene id</td></tr>
<tr><td>GU305914.1</td><td>131458</td><td>131480</td><td>-</td><td>0</td><td>GACCCCGACATCGACTTCTG</td><td>CGG</td><td>||||||||[||||||||||||]PAM</td><td>NA</td><td>Intergenic</td><td>NA</td><td>NA</td></tr>
<tr><td>GU305914.1</td><td>146626</td><td>146648</td><td>+</td><td>4</td><td>TACCCCGTCTTCGACTTCTT</td><td>CGG</td><td>-||||||-[|-|||||||||-]PAM</td><td>NA</td><td>Intergenic</td><td>NA</td><td>NA</td></tr>
<tr><td></td><td></td><td></td><td></td><td></td><td></td><td></td><td></td><td></td><td></td><td></td><td></td></tr>
<tr><td>T24009</td><td>AATCTCGCGCCGCCGATGTCAGG</td><td>1000</td><td></td><td></td><td></td><td></td><td></td><td></td><td></td><td></td><td></td></tr>
<tr><td>Oligo fwd</td><td>CACCAATCTCGCGCCGCCGATGTC</td><td></td><td></td><td></td><td></td><td></td><td></td><td></td><td></td><td></td><td></td></tr>
<tr><td>Oligo rev</td><td>AAACGACATCGGCGGCGCGAGATT</td><td></td><td></td><td></td><td></td><td></td><td></td><td></td><td></td><td></td><td></td></tr>
<tr><td>Chromosome</td><td>start</td><td>end</td><td>strand</td><td>MM</td><td>target_seq</td><td>PAM</td><td>alignment</td><td>distance</td><td>position</td><td>gene name</td><td>gene id</td></tr>
<tr><td>GU305914.1</td><td>131429</td><td>131451</td><td>-</td><td>0</td><td>AATCTCGCGCCGCCGATGTC</td><td>AGG</td><td>||||||||[||||||||||||]PAM</td><td>NA</td><td>Intergenic</td><td>NA</td><td>NA</td></tr>
<tr><td></td><td></td><td></td><td></td><td></td><td></td><td></td><td></td><td></td><td></td><td></td><td></td></tr>
<tr><td>T24010</td><td>ATCTCGCGCCGCCGATGTCAGGG</td><td>1000</td><td></td><td></td><td></td><td></td><td></td><td></td><td></td><td></td><td></td></tr>
<tr><td>Oligo fwd</td><td>CACCATCTCGCGCCGCCGATGTCA</td><td></td><td></td><td></td><td></td><td></td><td></td><td></td><td></td><td></td><td></td></tr>
<tr><td>Oligo rev</td><td>AAACTGACATCGGCGGCGCGAGAT</td><td></td><td></td><td></td><td></td><td></td><td></td><td></td><td></td><td></td><td></td></tr>
<tr><td>Chromosome</td><td>start</td><td>end</td><td>strand</td><td>MM</td><td>target_seq</td><td>PAM</td><td>alignment</td><td>distance</td><td>position</td><td>gene name</td><td>gene id</td></tr>
<tr><td>GU305914.1</td><td>131428</td><td>131450</td><td>-</td><td>0</td><td>ATCTCGCGCCGCCGATGTCA</td><td>GGG</td><td>||||||||[||||||||||||]PAM</td><td>NA</td><td>Intergenic</td><td>NA</td><td>NA</td></tr>
<tr><td></td><td></td><td></td><td></td><td></td><td></td><td></td><td></td><td></td><td></td><td></td><td></td></tr>
<tr><td>T24011</td><td>TCTCGCGCCGCCGATGTCAGGGG</td><td>1000</td><td></td><td></td><td></td><td></td><td></td><td></td><td></td><td></td><td></td></tr>
<tr><td>Oligo fwd</td><td>CACCTCTCGCGCCGCCGATGTCAG</td><td></td><td></td><td></td><td></td><td></td><td></td><td></td><td></td><td></td><td></td></tr>
<tr><td>Oligo rev</td><td>AAACCTGACATCGGCGGCGCGAGA</td><td></td><td></td><td></td><td></td><td></td><td></td><td></td><td></td><td></td><td></td></tr>
<tr><td>Chromosome</td><td>start</td><td>end</td><td>strand</td><td>MM</td><td>target_seq</td><td>PAM</td><td>alignment</td><td>distance</td><td>position</td><td>gene name</td><td>gene id</td></tr>
<tr><td>GU305914.1</td><td>131427</td><td>131449</td><td>-</td><td>0</td><td>TCTCGCGCCGCCGATGTCAG</td><td>GGG</td><td>||||||||[||||||||||||]PAM</td><td>NA</td><td>Intergenic</td><td>NA</td><td>NA</td></tr>
</table>

<table>
<tr><td>T24012</td><td>CTCGCGCCGCCGATGTCAGGGGG</td><td>1000</td><td></td><td></td><td></td><td></td><td></td><td></td><td></td><td></td><td></td></tr>
<tr><td>Oligo fwd</td><td>CACCCTCGCGCCGCCGATGTCAGG</td><td></td><td></td><td></td><td></td><td></td><td></td><td></td><td></td><td></td><td></td></tr>
<tr><td>Oligo rev</td><td>AAACCCTGACATCGGCGGCGCGAG</td><td></td><td></td><td></td><td></td><td></td><td></td><td></td><td></td><td></td><td></td></tr>
<tr><td>Chromosome</td><td>start</td><td>end</td><td>strand</td><td>MM</td><td>target_seq</td><td>PAM</td><td>alignment</td><td>distance</td><td>position</td><td>gene name</td><td>gene id</td></tr>
<tr><td>GU305914.1</td><td>131426</td><td>131448</td><td>-</td><td>0</td><td>CTCGCGCCGCCGATGTCAGG</td><td>GGG</td><td>||||||||[||||||||||||]PAM</td><td>NA</td><td>Intergenic</td><td>NA</td><td>NA</td></tr>
<tr><td>T24013</td><td>CGCCGCCGATGTCAGGGGGTCGG</td><td>1000</td><td></td><td></td><td></td><td></td><td></td><td></td><td></td><td></td><td></td></tr>
<tr><td>Oligo fwd</td><td>CACCCGCCGCCGATGTCAGGGGGT</td><td></td><td></td><td></td><td></td><td></td><td></td><td></td><td></td><td></td><td></td></tr>
<tr><td>Oligo rev</td><td>AAACACCCCCTGACATCGGCGGCG</td><td></td><td></td><td></td><td></td><td></td><td></td><td></td><td></td><td></td><td></td></tr>
<tr><td>Chromosome</td><td>start</td><td>end</td><td>strand</td><td>MM</td><td>target_seq</td><td>PAM</td><td>alignment</td><td>distance</td><td>position</td><td>gene name</td><td>gene id</td></tr>
<tr><td>GU305914.1</td><td>131422</td><td>131444</td><td>-</td><td>0</td><td>CGCCGCCGATGTCAGGGGGT</td><td>CGG</td><td>||||||||[||||||||||||]PAM</td><td>NA</td><td>Intergenic</td><td>NA</td><td>NA</td></tr>
<tr><td>T24014</td><td>CATATCCAGACGCCGCTCTACGG</td><td>1000</td><td></td><td></td><td></td><td></td><td></td><td></td><td></td><td></td><td></td></tr>
<tr><td>Oligo fwd</td><td>CACCCATATCCAGACGCCGCTCTA</td><td></td><td></td><td></td><td></td><td></td><td></td><td></td><td></td><td></td><td></td></tr>
<tr><td>Oligo rev</td><td>AAACTAGAGCGGCGTCTGGATATG</td><td></td><td></td><td></td><td></td><td></td><td></td><td></td><td></td><td></td><td></td></tr>
<tr><td>Chromosome</td><td>start</td><td>end</td><td>strand</td><td>MM</td><td>target_seq</td><td>PAM</td><td>alignment</td><td>distance</td><td>position</td><td>gene name</td><td>gene id</td></tr>
<tr><td>GU305914.1</td><td>131398</td><td>131420</td><td>-</td><td>0</td><td>CATATCCAGACGCCGCTCTA</td><td>CGG</td><td>||||||||[||||||||||||]PAM</td><td>NA</td><td>Intergenic</td><td>NA</td><td>NA</td></tr>
<tr><td>T24015</td><td>ATATCCAGACGCCGCTCTACGGG</td><td>1000</td><td></td><td></td><td></td><td></td><td></td><td></td><td></td><td></td><td></td></tr>
<tr><td>Oligo fwd</td><td>CACCATATCCAGACGCCGCTCTAC</td><td></td><td></td><td></td><td></td><td></td><td></td><td></td><td></td><td></td><td></td></tr>
<tr><td>Oligo rev</td><td>AAACGTAGAGCGGCGTCTGGATAT</td><td></td><td></td><td></td><td></td><td></td><td></td><td></td><td></td><td></td><td></td></tr>
<tr><td>Chromosome</td><td>start</td><td>end</td><td>strand</td><td>MM</td><td>target_seq</td><td>PAM</td><td>alignment</td><td>distance</td><td>position</td><td>gene name</td><td>gene id</td></tr>
<tr><td>GU305914.1</td><td>131397</td><td>131419</td><td>-</td><td>0</td><td>ATATCCAGACGCCGCTCTAC</td><td>GGG</td><td>||||||||[||||||||||||]PAM</td><td>NA</td><td>Intergenic</td><td>NA</td><td>NA</td></tr>
<tr><td>T24016</td><td>TCCAGACGCCGCTCTACGGGCGG</td><td>1000</td><td></td><td></td><td></td><td></td><td></td><td></td><td></td><td></td><td></td></tr>
<tr><td>Oligo fwd</td><td>CACCTCCAGACGCCGCTCTACGGG</td><td></td><td></td><td></td><td></td><td></td><td></td><td></td><td></td><td></td><td></td></tr>
<tr><td>Oligo rev</td><td>AAACCCCGTAGAGCGGCGTCTGGA</td><td></td><td></td><td></td><td></td><td></td><td></td><td></td><td></td><td></td><td></td></tr>
<tr><td>Chromosome</td><td>start</td><td>end</td><td>strand</td><td>MM</td><td>target_seq</td><td>PAM</td><td>alignment</td><td>distance</td><td>position</td><td>gene name</td><td>gene id</td></tr>
<tr><td>GU305914.1</td><td>131394</td><td>131416</td><td>-</td><td>0</td><td>TCCAGACGCCGCTCTACGGG</td><td>CGG</td><td>||||||||[||||||||||||]PAM</td><td>NA</td><td>Intergenic</td><td>NA</td><td>NA</td></tr>
<tr><td>T24017</td><td>GACGCCGCTCTACGGGCGGCAGG</td><td>1000</td><td></td><td></td><td></td><td></td><td></td><td></td><td></td><td></td><td></td></tr>
<tr><td>Oligo fwd</td><td>CACCGACGCCGCTCTACGGGCGGC</td><td></td><td></td><td></td><td></td><td></td><td></td><td></td><td></td><td></td><td></td></tr>
<tr><td>Oligo rev</td><td>AAACGCCGCCCGTAGAGCGGCGTC</td><td></td><td></td><td></td><td></td><td></td><td></td><td></td><td></td><td></td><td></td></tr>
<tr><td>Chromosome</td><td>start</td><td>end</td><td>strand</td><td>MM</td><td>target_seq</td><td>PAM</td><td>alignment</td><td>distance</td><td>position</td><td>gene name</td><td>gene id</td></tr>
<tr><td>GU305914.1</td><td>131390</td><td>131412</td><td>-</td><td>0</td><td>GACGCCGCTCTACGGGCGGC</td><td>AGG</td><td>||||||||[||||||||||||]PAM</td><td>NA</td><td>Intergenic</td><td>NA</td><td>NA</td></tr>
<tr><td>T24018</td><td>CCGCTCTACGGGCGGCAGGTAGG</td><td>1000</td><td></td><td></td><td></td><td></td><td></td><td></td><td></td><td></td><td></td></tr>
<tr><td>Oligo fwd</td><td>CACCCCGCTCTACGGGCGGCAGGT</td><td></td><td></td><td></td><td></td><td></td><td></td><td></td><td></td><td></td><td></td></tr>
<tr><td>Oligo rev</td><td>AAACACCTGCCGCCCGTAGAGCGG</td><td></td><td></td><td></td><td></td><td></td><td></td><td></td><td></td><td></td><td></td></tr>
<tr><td>Chromosome</td><td>start</td><td>end</td><td>strand</td><td>MM</td><td>target_seq</td><td>PAM</td><td>alignment</td><td>distance</td><td>position</td><td>gene name</td><td>gene id</td></tr>
<tr><td>GU305914.1</td><td>131386</td><td>131408</td><td>-</td><td>0</td><td>CCGCTCTACGGGCGGCAGGT</td><td>AGG</td><td>||||||||[||||||||||||]PAM</td><td>NA</td><td>Intergenic</td><td>NA</td><td>NA</td></tr>
<tr><td>T24019</td><td>AGTACTCCTGAATCCTCCTGAGG</td><td>1000</td><td></td><td></td><td></td><td></td><td></td><td></td><td></td><td></td><td></td></tr>
<tr><td>Oligo fwd</td><td>CACCAGTACTCCTGAATCCTCCTG</td><td></td><td></td><td></td><td></td><td></td><td></td><td></td><td></td><td></td><td></td></tr>
<tr><td>Oligo rev</td><td>AAACCAGGAGGATTCAGGAGTACT</td><td></td><td></td><td></td><td></td><td></td><td></td><td></td><td></td><td></td><td></td></tr>
<tr><td>Chromosome</td><td>start</td><td>end</td><td>strand</td><td>MM</td><td>target_seq</td><td>PAM</td><td>alignment</td><td>distance</td><td>position</td><td>gene name</td><td>gene id</td></tr>
<tr><td>GU305914.1</td><td>131360</td><td>131382</td><td>-</td><td>0</td><td>AGTACTCCTGAATCCTCCTG</td><td>AGG</td><td>||||||||[||||||||||||]PAM</td><td>NA</td><td>Intergenic</td><td>NA</td><td>NA</td></tr>
<tr><td>T24020</td><td>CCTGAATCCTCCTGAGGATCAGG</td><td>1000</td><td></td><td></td><td></td><td></td><td></td><td></td><td></td><td></td><td></td></tr>
<tr><td>Oligo fwd</td><td>CACCCCTGAATCCTCCTGAGGATC</td><td></td><td></td><td></td><td></td><td></td><td></td><td></td><td></td><td></td><td></td></tr>
<tr><td>Oligo rev</td><td>AAACGATCCTCAGGAGGATTCAGG</td><td></td><td></td><td></td><td></td><td></td><td></td><td></td><td></td><td></td><td></td></tr>
<tr><td>Chromosome</td><td>start</td><td>end</td><td>strand</td><td>MM</td><td>target_seq</td><td>PAM</td><td>alignment</td><td>distance</td><td>position</td><td>gene name</td><td>gene id</td></tr>
<tr><td>GU305914.1</td><td>131354</td><td>131376</td><td>-</td><td>0</td><td>CCTGAATCCTCCTGAGGATC</td><td>AGG</td><td>||||||||[||||||||||||]PAM</td><td>NA</td><td>Intergenic</td><td>NA</td><td>NA</td></tr>
<tr><td>T24021</td><td>CTGAATCCTCCTGAGGATCAGGG</td><td>1000</td><td></td><td></td><td></td><td></td><td></td><td></td><td></td><td></td><td></td></tr>
<tr><td>Oligo fwd</td><td>CACCCTGAATCCTCCTGAGGATCA</td><td></td><td></td><td></td><td></td><td></td><td></td><td></td><td></td><td></td><td></td></tr>
<tr><td>Oligo rev</td><td>AAACTGATCCTCAGGAGGATTCAG</td><td></td><td></td><td></td><td></td><td></td><td></td><td></td><td></td><td></td><td></td></tr>
<tr><td>Chromosome</td><td>start</td><td>end</td><td>strand</td><td>MM</td><td>target_seq</td><td>PAM</td><td>alignment</td><td>distance</td><td>position</td><td>gene name</td><td>gene id</td></tr>
<tr><td>GU305914.1</td><td>131353</td><td>131375</td><td>-</td><td>0</td><td>CTGAATCCTCCTGAGGATCA</td><td>GGG</td><td>||||||||[||||||||||||]PAM</td><td>NA</td><td>Intergenic</td><td>NA</td><td>NA</td></tr>
<tr><td>T24022</td><td>CAGGGCGAGCTGCGTACCGTTGG</td><td>1000</td><td></td><td></td><td></td><td></td><td></td><td></td><td></td><td></td><td></td></tr>
<tr><td>Oligo fwd</td><td>CACCCAGGGCGAGCTGCGTACCGT</td><td></td><td></td><td></td><td></td><td></td><td></td><td></td><td></td><td></td><td></td></tr>
<tr><td>Oligo rev</td><td>AAACACGGTACGCAGCTCGCCCTG</td><td></td><td></td><td></td><td></td><td></td><td></td><td></td><td></td><td></td><td></td></tr>
<tr><td>Chromosome</td><td>start</td><td>end</td><td>strand</td><td>MM</td><td>target_seq</td><td>PAM</td><td>alignment</td><td>distance</td><td>position</td><td>gene name</td><td>gene id</td></tr>
<tr><td>GU305914.1</td><td>131335</td><td>131357</td><td>-</td><td>0</td><td>CAGGGCGAGCTGCGTACCGT</td><td>TGG</td><td>||||||||[||||||||||||]PAM</td><td>NA</td><td>Intergenic</td><td>NA</td><td>NA</td></tr>
<tr><td>T24023</td><td>GAGCACCCCCACTATCTCATTGG</td><td>1000</td><td></td><td></td><td></td><td></td><td></td><td></td><td></td><td></td><td></td></tr>
<tr><td>Oligo fwd</td><td>CACCGAGCACCCCCACTATCTCAT</td><td></td><td></td><td></td><td></td><td></td><td></td><td></td><td></td><td></td><td></td></tr>
<tr><td>Oligo rev</td><td>AAACATGAGATAGTGGGGGTGCTC</td><td></td><td></td><td></td><td></td><td></td><td></td><td></td><td></td><td></td><td></td></tr>
<tr><td>Chromosome</td><td>start</td><td>end</td><td>strand</td><td>MM</td><td>target_seq</td><td>PAM</td><td>alignment</td><td>distance</td><td>position</td><td>gene name</td><td>gene id</td></tr>
<tr><td>GU305914.1</td><td>131254</td><td>131276</td><td>-</td><td>0</td><td>GAGCACCCCCACTATCTCAT</td><td>TGG</td><td>||||||||[||||||||||||]PAM</td><td>NA</td><td>Intergenic</td><td>NA</td><td>NA</td></tr>
<tr><td>T24024</td><td>TGTCCTCTCCGAGTCCACAGAGG</td><td>1000</td><td></td><td></td><td></td><td></td><td></td><td></td><td></td><td></td><td></td></tr>
<tr><td>Oligo fwd</td><td>CACCTGTCCTCTCCGAGTCCACAG</td><td></td><td></td><td></td><td></td><td></td><td></td><td></td><td></td><td></td><td></td></tr>
<tr><td>Oligo rev</td><td>AAACCTGTGGACTCGGAGAGGACA</td><td></td><td></td><td></td><td></td><td></td><td></td><td></td><td></td><td></td><td></td></tr>
<tr><td>Chromosome</td><td>start</td><td>end</td><td>strand</td><td>MM</td><td>target_seq</td><td>PAM</td><td>alignment</td><td>distance</td><td>position</td><td>gene name</td><td>gene id</td></tr>
<tr><td>GU305914.1</td><td>131225</td><td>131247</td><td>-</td><td>0</td><td>TGTCCTCTCCGAGTCCACAG</td><td>AGG</td><td>||||||||[||||||||||||]PAM</td><td>NA</td><td>Intergenic</td><td>NA</td><td>NA</td></tr>
<tr><td>T24025</td><td>TCCACAGAGGCAGACCTCCGAGG</td><td>1000</td><td></td><td></td><td></td><td></td><td></td><td></td><td></td><td></td><td></td></tr>
<tr><td>Oligo fwd</td><td>CACCTCCACAGAGGCAGACCTCCG</td><td></td><td></td><td></td><td></td><td></td><td></td><td></td><td></td><td></td><td></td></tr>
<tr><td>Oligo rev</td><td>AAACCGGAGGTCTGCCTCTGTGGA</td><td></td><td></td><td></td><td></td><td></td><td></td><td></td><td></td><td></td><td></td></tr>
<tr><td>Chromosome</td><td>start</td><td>end</td><td>strand</td><td>MM</td><td>target_seq</td><td>PAM</td><td>alignment</td><td>distance</td><td>position</td><td>gene name</td><td>gene id</td></tr>
<tr><td>GU305914.1</td><td>131212</td><td>131234</td><td>-</td><td>0</td><td>TCCACAGAGGCAGACCTCCG</td><td>AGG</td><td>||||||||[||||||||||||]PAM</td><td>NA</td><td>Intergenic</td><td>NA</td><td>NA</td></tr>
<tr><td>T24026</td><td>CGAGGCGAACATGTTGCCGATGG</td><td>1000</td><td></td><td></td><td></td><td></td><td></td><td></td><td></td><td></td><td></td></tr>
<tr><td>Oligo fwd</td><td>CACCCGAGGCGAACATGTTGCCGA</td><td></td><td></td><td></td><td></td><td></td><td></td><td></td><td></td><td></td><td></td></tr>
<tr><td>Oligo rev</td><td>AAACTCGGCAACATGTTCGCCTCG</td><td></td><td></td><td></td><td></td><td></td><td></td><td></td><td></td><td></td><td></td></tr>
<tr><td>Chromosome</td><td>start</td><td>end</td><td>strand</td><td>MM</td><td>target_seq</td><td>PAM</td><td>alignment</td><td>distance</td><td>position</td><td>gene name</td><td>gene id</td></tr>
<tr><td>GU305914.1</td><td>131194</td><td>131216</td><td>-</td><td>0</td><td>CGAGGCGAACATGTTGCCGA</td><td>TGG</td><td>||||||||[||||||||||||]PAM</td><td>NA</td><td>Intergenic</td><td>NA</td><td>NA</td></tr>
</table>

| | | | | | | | | | | | |
|---|---|---|---|---|---|---|---|---|---|---|---|
| GU305914.1 | 113052 | 113074 | + | 4 | CGAGAAGAAGATGTTGCCGT | CGG | \|\|\|\|--\|\|[\|-\|\|\|\|\|\|\|\|\|-]PAM | NA | Intergenic | NA | NA |
| T24027 | GAGGCGAACATGTTGCCGATGGG | 1000 | | | | | | | | | |
| Oligo fwd | CACCGAGGCGAACATGTTGCCGAT | | | | | | | | | | |
| Oligo rev | AAACATCGGCAACATGTTCGCCTC | | | | | | | | | | |
| Chromosome | start | end | strand | MM | target_seq | PAM | alignment | distance | position | gene name | gene id |
| GU305914.1 | 131193 | 131215 | - | 0 | GAGGCGAACATGTTGCCGAT | GGG | \|\|\|\|\|\|\|\|[\|\|\|\|\|\|\|\|\|\|\|\|]PAM | NA | Intergenic | NA | NA |
| T24028 | AGGCGAACATGTTGCCGATGGGG | 1000 | | | | | | | | | |
| Oligo fwd | CACCAGGCGAACATGTTGCCGATG | | | | | | | | | | |
| Oligo rev | AAACCATCGGCAACATGTTCGCCT | | | | | | | | | | |
| Chromosome | start | end | strand | MM | target_seq | PAM | alignment | distance | position | gene name | gene id |
| GU305914.1 | 131192 | 131214 | - | 0 | AGGCGAACATGTTGCCGATG | GGG | \|\|\|\|\|\|\|\|[\|\|\|\|\|\|\|\|\|\|\|\|]PAM | NA | Intergenic | NA | NA |
| T24029 | GGCGAACATGTTGCCGATGGGGG | 1000 | | | | | | | | | |
| Oligo fwd | CACCGGCGAACATGTTGCCGATGG | | | | | | | | | | |
| Oligo rev | AAACCCATCGGCAACATGTTCGCC | | | | | | | | | | |
| Chromosome | start | end | strand | MM | target_seq | PAM | alignment | distance | position | gene name | gene id |
| GU305914.1 | 131191 | 131213 | - | 0 | GGCGAACATGTTGCCGATGG | GGG | \|\|\|\|\|\|\|\|[\|\|\|\|\|\|\|\|\|\|\|\|]PAM | NA | Intergenic | NA | NA |
| T24030 | GACCAACAAGAGTCTCTTCCTGG | 1000 | | | | | | | | | |
| Oligo fwd | CACCGACCAACAAGAGTCTCTTCC | | | | | | | | | | |
| Oligo rev | AAACGGAAGAGACTCTTGTTGGTC | | | | | | | | | | |
| Chromosome | start | end | strand | MM | target_seq | PAM | alignment | distance | position | gene name | gene id |
| GU305914.1 | 131161 | 131183 | - | 0 | GACCAACAAGAGTCTCTTCC | TGG | \|\|\|\|\|\|\|\|[\|\|\|\|\|\|\|\|\|\|\|\|]PAM | NA | Intergenic | NA | NA |
| T24031 | CAACAAGAGTCTCTTCCTGGCGG | 1000 | | | | | | | | | |
| Oligo fwd | CACCCAACAAGAGTCTCTTCCTGG | | | | | | | | | | |
| Oligo rev | AAACCCAGGAAGAGACTCTTGTTG | | | | | | | | | | |
| Chromosome | start | end | strand | MM | target_seq | PAM | alignment | distance | position | gene name | gene id |
| GU305914.1 | 131158 | 131180 | - | 0 | CAACAAGAGTCTCTTCCTGG | CGG | \|\|\|\|\|\|\|\|[\|\|\|\|\|\|\|\|\|\|\|\|]PAM | NA | Intergenic | NA | NA |
| T24032 | CGAGCTCCACACACCTCATGAGG | 1000 | | | | | | | | | |
| Oligo fwd | CACCCGAGCTCCACACACCTCATG | | | | | | | | | | |
| Oligo rev | AAACCATGAGGTGTGTGGAGCTCG | | | | | | | | | | |
| Chromosome | start | end | strand | MM | target_seq | PAM | alignment | distance | position | gene name | gene id |
| GU305914.1 | 131135 | 131157 | - | 0 | CGAGCTCCACACACCTCATG | AGG | \|\|\|\|\|\|\|\|[\|\|\|\|\|\|\|\|\|\|\|\|]PAM | NA | Intergenic | NA | NA |
| T24033 | ACCTCATGAGGACCTCACGCCGG | 1000 | | | | | | | | | |
| Oligo fwd | CACCACCTCATGAGGACCTCACGC | | | | | | | | | | |
| Oligo rev | AAACGCGTGAGGTCCTCATGAGGT | | | | | | | | | | |
| Chromosome | start | end | strand | MM | target_seq | PAM | alignment | distance | position | gene name | gene id |
| GU305914.1 | 131123 | 131145 | - | 0 | ACCTCATGAGGACCTCACGC | CGG | \|\|\|\|\|\|\|\|[\|\|\|\|\|\|\|\|\|\|\|\|]PAM | NA | Intergenic | NA | NA |
| T24034 | GGACCTCACGCCGGCGCCCGAGG | 1000 | | | | | | | | | |
| Oligo fwd | CACCGGACCTCACGCCGGCGCCCG | | | | | | | | | | |
| Oligo rev | AAACCGGGCGCCGGCGTGAGGTCC | | | | | | | | | | |
| Chromosome | start | end | strand | MM | target_seq | PAM | alignment | distance | position | gene name | gene id |
| GU305914.1 | 131114 | 131136 | - | 0 | GGACCTCACGCCGGCGCCCG | AGG | \|\|\|\|\|\|\|\|[\|\|\|\|\|\|\|\|\|\|\|\|]PAM | NA | Intergenic | NA | NA |
| T24035 | CTCACGCCGGCGCCCGAGGCTGG | 1000 | | | | | | | | | |
| Oligo fwd | CACCCTCACGCCGGCGCCCGAGGC | | | | | | | | | | |
| Oligo rev | AAACGCCTCGGGCGCCGGCGTGAG | | | | | | | | | | |
| Chromosome | start | end | strand | MM | target_seq | PAM | alignment | distance | position | gene name | gene id |
| GU305914.1 | 131110 | 131132 | - | 0 | CTCACGCCGGCGCCCGAGGC | TGG | \|\|\|\|\|\|\|\|[\|\|\|\|\|\|\|\|\|\|\|\|]PAM | NA | Intergenic | NA | NA |
| T24036 | GGTCTCCAGTTCGCGCTCGATGG | 1000 | | | | | | | | | |
| Oligo fwd | CACCGGTCTCCAGTTCGCGCTCGA | | | | | | | | | | |
| Oligo rev | AAACTCGAGCGCGAACTGGAGACC | | | | | | | | | | |
| Chromosome | start | end | strand | MM | target_seq | PAM | alignment | distance | position | gene name | gene id |
| GU305914.1 | 131089 | 131111 | - | 0 | GGTCTCCAGTTCGCGCTCGA | TGG | \|\|\|\|\|\|\|\|[\|\|\|\|\|\|\|\|\|\|\|\|]PAM | NA | Intergenic | NA | NA |
| T24037 | GCTCGATGGCGAGCAGTCGCAGG | 1000 | | | | | | | | | |
| Oligo fwd | CACCGCTCGATGGCGAGCAGTCGC | | | | | | | | | | |
| Oligo rev | AAACGCGACTGCTCGCCATCGAGC | | | | | | | | | | |
| Chromosome | start | end | strand | MM | target_seq | PAM | alignment | distance | position | gene name | gene id |
| GU305914.1 | 131075 | 131097 | - | 0 | GCTCGATGGCGAGCAGTCGC | AGG | \|\|\|\|\|\|\|\|[\|\|\|\|\|\|\|\|\|\|\|\|]PAM | NA | Intergenic | NA | NA |
| T24038 | GGCGAGCAGTCGCAGGTAGACGG | 1000 | | | | | | | | | |
| Oligo fwd | CACCGGCGAGCAGTCGCAGGTAGA | | | | | | | | | | |
| Oligo rev | AAACTCTACCTGCGACTGCTCGCC | | | | | | | | | | |
| Chromosome | start | end | strand | MM | target_seq | PAM | alignment | distance | position | gene name | gene id |
| GU305914.1 | 131068 | 131090 | - | 0 | GGCGAGCAGTCGCAGGTAGA | CGG | \|\|\|\|\|\|\|\|[\|\|\|\|\|\|\|\|\|\|\|\|]PAM | NA | Intergenic | NA | NA |
| T24039 | CAGTCGCAGGTAGACGGCCGAGG | 1000 | | | | | | | | | |
| Oligo fwd | CACCCAGTCGCAGGTAGACGGCCG | | | | | | | | | | |
| Oligo rev | AAACCGGCCGTCTACCTGCGACTG | | | | | | | | | | |
| Chromosome | start | end | strand | MM | target_seq | PAM | alignment | distance | position | gene name | gene id |
| GU305914.1 | 131062 | 131084 | - | 0 | CAGTCGCAGGTAGACGGCCG | AGG | \|\|\|\|\|\|\|\|[\|\|\|\|\|\|\|\|\|\|\|\|]PAM | NA | Intergenic | NA | NA |
| T24040 | AGACGGCCGAGGCGCGCTCCAGG | 1000 | | | | | | | | | |
| Oligo fwd | CACCAGACGGCCGAGGCGCGCTCC | | | | | | | | | | |
| Oligo rev | AAACGGAGCGCGCCTCGGCCGTCT | | | | | | | | | | |
| Chromosome | start | end | strand | MM | target_seq | PAM | alignment | distance | position | gene name | gene id |
| GU305914.1 | 131051 | 131073 | - | 0 | AGACGGCCGAGGCGCGCTCC | AGG | \|\|\|\|\|\|\|\|[\|\|\|\|\|\|\|\|\|\|\|\|]PAM | NA | Intergenic | NA | NA |
| T24041 | CGAGGCGCGCTCCAGGAAGATGG | 1000 | | | | | | | | | |
| Oligo fwd | CACCCGAGGCGCGCTCCAGGAAGA | | | | | | | | | | |
| Oligo rev | AAACTCTTCCTGGAGCGCGCCTCG | | | | | | | | | | |

| | | | | | | | | | | | |
|---|---|---|---|---|---|---|---|---|---|---|---|
| Chromosome | start | end | strand | MM | target_seq | PAM | alignment | distance | position | gene name | gene id |
| GU305914.1 | 131044 | 131066 | - | 0 | CGAGGCGCGCTCCAGGAAGA | TGG | \|\|\|\|\|\|\|\|[\|\|\|\|\|\|\|\|\|\|\|\|]PAM | NA | Intergenic | NA | NA |
| T24042 | GTCGAGCCGTAGCCTCTCGTCGG | 1000 | | | | | | | | | |
| Oligo fwd | CACCGTCGAGCCGTAGCCTCTCGT | | | | | | | | | | |
| Oligo rev | AAACACGAGAGGCTACGGCTCGAC | | | | | | | | | | |
| Chromosome | start | end | strand | MM | target_seq | PAM | alignment | distance | position | gene name | gene id |
| GU305914.1 | 131011 | 131033 | - | 0 | GTCGAGCCGTAGCCTCTCGT | CGG | \|\|\|\|\|\|\|\|[\|\|\|\|\|\|\|\|\|\|\|\|]PAM | NA | Intergenic | NA | NA |
| T24043 | TCTCGTCGGTGTACTGCTCGCGG | 1000 | | | | | | | | | |
| Oligo fwd | CACCTCTCGTCGGTGTACTGCTCG | | | | | | | | | | |
| Oligo rev | AAACCGAGCAGTACACCGACGAGA | | | | | | | | | | |
| Chromosome | start | end | strand | MM | target_seq | PAM | alignment | distance | position | gene name | gene id |
| GU305914.1 | 130997 | 131019 | - | 0 | TCTCGTCGGTGTACTGCTCG | CGG | \|\|\|\|\|\|\|\|[\|\|\|\|\|\|\|\|\|\|\|\|]PAM | NA | Intergenic | NA | NA |
| T24044 | GCTCGCGGTCGCCCACGACGAGG | 1000 | | | | | | | | | |
| Oligo fwd | CACCGCTCGCGGTCGCCCACGACG | | | | | | | | | | |
| Oligo rev | AAACCGTCGTGGGCGACCGCGAGC | | | | | | | | | | |
| Chromosome | start | end | strand | MM | target_seq | PAM | alignment | distance | position | gene name | gene id |
| GU305914.1 | 130982 | 131004 | - | 0 | GCTCGCGGTCGCCCACGACG | AGG | \|\|\|\|\|\|\|\|[\|\|\|\|\|\|\|\|\|\|\|\|]PAM | NA | Intergenic | NA | NA |
| GU305914.1 | 223123 | 223145 | + | 4 | GGTCGATGTCGCACACGACG | AGG | \|-\|\|\|--\|[\|\|\|\|-\|\|\|\|\|\|\|]PAM | NA | Intergenic | NA | NA |
| T24045 | CTCGCGGTCGCCCACGACGAGGG | 1000 | | | | | | | | | |
| Oligo fwd | CACCCTCGCGGTCGCCCACGACGA | | | | | | | | | | |
| Oligo rev | AAACTCGTCGTGGGCGACCGCGAG | | | | | | | | | | |
| Chromosome | start | end | strand | MM | target_seq | PAM | alignment | distance | position | gene name | gene id |
| GU305914.1 | 130981 | 131003 | - | 0 | CTCGCGGTCGCCCACGACGA | GGG | \|\|\|\|\|\|\|\|[\|\|\|\|\|\|\|\|\|\|\|\|]PAM | NA | Intergenic | NA | NA |
| T24046 | TCGCGGTCGCCCACGACGAGGGG | 1000 | | | | | | | | | |
| Oligo fwd | CACCTCGCGGTCGCCCACGACGAG | | | | | | | | | | |
| Oligo rev | AAACCTCGTCGTGGGCGACCGCGA | | | | | | | | | | |
| Chromosome | start | end | strand | MM | target_seq | PAM | alignment | distance | position | gene name | gene id |
| GU305914.1 | 130980 | 131002 | - | 0 | TCGCGGTCGCCCACGACGAG | GGG | \|\|\|\|\|\|\|\|[\|\|\|\|\|\|\|\|\|\|\|\|]PAM | NA | Intergenic | NA | NA |
| T24047 | GCCCACGACGAGGGGCACATAGG | 1000 | | | | | | | | | |
| Oligo fwd | CACCGCCCACGACGAGGGGCACAT | | | | | | | | | | |
| Oligo rev | AAACATGTGCCCCTCGTCGTGGGC | | | | | | | | | | |
| Chromosome | start | end | strand | MM | target_seq | PAM | alignment | distance | position | gene name | gene id |
| GU305914.1 | 130972 | 130994 | - | 0 | GCCCACGACGAGGGGCACAT | AGG | \|\|\|\|\|\|\|\|[\|\|\|\|\|\|\|\|\|\|\|\|]PAM | NA | Intergenic | NA | NA |
| T24048 | ATAGGCCCCGATGTCGATGTTGG | 1000 | | | | | | | | | |
| Oligo fwd | CACCATAGGCCCCGATGTCGATGT | | | | | | | | | | |
| Oligo rev | AAACACATCGACATCGGGGCCTAT | | | | | | | | | | |
| Chromosome | start | end | strand | MM | target_seq | PAM | alignment | distance | position | gene name | gene id |
| GU305914.1 | 130954 | 130976 | - | 0 | ATAGGCCCCGATGTCGATGT | TGG | \|\|\|\|\|\|\|\|[\|\|\|\|\|\|\|\|\|\|\|\|]PAM | NA | Intergenic | NA | NA |
| T24049 | GGCCCCGATGTCGATGTTGGTGG | 1000 | | | | | | | | | |
| Oligo fwd | CACCGGCCCCGATGTCGATGTTGG | | | | | | | | | | |
| Oligo rev | AAACCCAACATCGACATCGGGGCC | | | | | | | | | | |
| Chromosome | start | end | strand | MM | target_seq | PAM | alignment | distance | position | gene name | gene id |
| GU305914.1 | 130951 | 130973 | - | 0 | GGCCCCGATGTCGATGTTGG | TGG | \|\|\|\|\|\|\|\|[\|\|\|\|\|\|\|\|\|\|\|\|]PAM | NA | Intergenic | NA | NA |
| T24050 | GCCCCGATGTCGATGTTGGTGGG | 1000 | | | | | | | | | |
| Oligo fwd | CACCGCCCCGATGTCGATGTTGGT | | | | | | | | | | |
| Oligo rev | AAACACCAACATCGACATCGGGGC | | | | | | | | | | |
| Chromosome | start | end | strand | MM | target_seq | PAM | alignment | distance | position | gene name | gene id |
| GU305914.1 | 130950 | 130972 | - | 0 | GCCCCGATGTCGATGTTGGT | GGG | \|\|\|\|\|\|\|\|[\|\|\|\|\|\|\|\|\|\|\|\|]PAM | NA | Intergenic | NA | NA |
| T24051 | TCGCCGAAGAGTCTGACGCAAGG | 1000 | | | | | | | | | |
| Oligo fwd | CACCTCGCCGAAGAGTCTGACGCA | | | | | | | | | | |
| Oligo rev | AAACTGCGTCAGACTCTTCGGCGA | | | | | | | | | | |
| Chromosome | start | end | strand | MM | target_seq | PAM | alignment | distance | position | gene name | gene id |
| GU305914.1 | 130926 | 130948 | - | 0 | TCGCCGAAGAGTCTGACGCA | AGG | \|\|\|\|\|\|\|\|[\|\|\|\|\|\|\|\|\|\|\|\|]PAM | NA | Intergenic | NA | NA |
| T24052 | CGCCGAAGAGTCTGACGCAAGGG | 1000 | | | | | | | | | |
| Oligo fwd | CACCCGCCGAAGAGTCTGACGCAA | | | | | | | | | | |
| Oligo rev | AAACTTGCGTCAGACTCTTCGGCG | | | | | | | | | | |
| Chromosome | start | end | strand | MM | target_seq | PAM | alignment | distance | position | gene name | gene id |
| GU305914.1 | 130925 | 130947 | - | 0 | CGCCGAAGAGTCTGACGCAA | GGG | \|\|\|\|\|\|\|\|[\|\|\|\|\|\|\|\|\|\|\|\|]PAM | NA | Intergenic | NA | NA |
| T24053 | GAAGCCGATCATGCCGACGTTGG | 1000 | | | | | | | | | |
| Oligo fwd | CACCGAAGCCGATCATGCCGACGT | | | | | | | | | | |
| Oligo rev | AAACACGTCGGCATGATCGGCTTC | | | | | | | | | | |
| Chromosome | start | end | strand | MM | target_seq | PAM | alignment | distance | position | gene name | gene id |
| GU305914.1 | 130903 | 130925 | - | 0 | GAAGCCGATCATGCCGACGT | TGG | \|\|\|\|\|\|\|\|[\|\|\|\|\|\|\|\|\|\|\|\|]PAM | NA | Intergenic | NA | NA |
| T24054 | AAGCCGATCATGCCGACGTTGGG | 1000 | | | | | | | | | |
| Oligo fwd | CACCAAGCCGATCATGCCGACGTT | | | | | | | | | | |
| Oligo rev | AAACAACGTCGGCATGATCGGCTT | | | | | | | | | | |
| Chromosome | start | end | strand | MM | target_seq | PAM | alignment | distance | position | gene name | gene id |
| GU305914.1 | 130902 | 130924 | - | 0 | AAGCCGATCATGCCGACGTT | GGG | \|\|\|\|\|\|\|\|[\|\|\|\|\|\|\|\|\|\|\|\|]PAM | NA | Intergenic | NA | NA |
| T24055 | CATGCCGACGTTGGGCATGACGG | 1000 | | | | | | | | | |
| Oligo fwd | CACCCATGCCGACGTTGGGCATGA | | | | | | | | | | |
| Oligo rev | AAACTCATGCCCAACGTCGGCATG | | | | | | | | | | |
| Chromosome | start | end | strand | MM | target_seq | PAM | alignment | distance | position | gene name | gene id |
| GU305914.1 | 130894 | 130916 | - | 0 | CATGCCGACGTTGGGCATGA | CGG | \|\|\|\|\|\|\|\|[\|\|\|\|\|\|\|\|\|\|\|\|]PAM | NA | Intergenic | NA | NA |
| T24056 | ATGCCGACGTTGGGCATGACGGG | 1000 | | | | | | | | | |

<table>
<tr><td>Oligo fwd</td><td>CACCATGCCGACGTTGGGCATGAC</td><td></td><td></td><td></td><td></td><td></td><td></td><td></td><td></td><td></td><td></td><td></td></tr>
<tr><td>Oligo rev</td><td>AAACGTCATGCCCAACGTCGGCAT</td><td></td><td></td><td></td><td></td><td></td><td></td><td></td><td></td><td></td><td></td><td></td></tr>
<tr><td>Chromosome</td><td>start</td><td></td><td>end</td><td>strand</td><td>MM</td><td>target_seq</td><td>PAM</td><td>alignment</td><td>distance</td><td>position</td><td>gene name</td><td>gene id</td></tr>
<tr><td>GU305914.1</td><td></td><td>130893</td><td>130915</td><td>-</td><td>0</td><td>ATGCCGACGTTGGGCATGAC</td><td>GGG</td><td>||||||||[||||||||||||]PAM</td><td>NA</td><td>Intergenic</td><td>NA</td><td>NA</td></tr>
<tr><td>T24057</td><td>GACGTTGGGCATGACGGGCGTGG</td><td></td><td>1000</td><td></td><td></td><td></td><td></td><td></td><td></td><td></td><td></td><td></td></tr>
<tr><td>Oligo fwd</td><td>CACCGACGTTGGGCATGACGGGCG</td><td></td><td></td><td></td><td></td><td></td><td></td><td></td><td></td><td></td><td></td><td></td></tr>
<tr><td>Oligo rev</td><td>AAACCGCCCGTCATGCCCAACGTC</td><td></td><td></td><td></td><td></td><td></td><td></td><td></td><td></td><td></td><td></td><td></td></tr>
<tr><td>Chromosome</td><td>start</td><td></td><td>end</td><td>strand</td><td>MM</td><td>target_seq</td><td>PAM</td><td>alignment</td><td>distance</td><td>position</td><td>gene name</td><td>gene id</td></tr>
<tr><td>GU305914.1</td><td></td><td>130888</td><td>130910</td><td>-</td><td>0</td><td>GACGTTGGGCATGACGGGCG</td><td>TGG</td><td>||||||||[||||||||||||]PAM</td><td>NA</td><td>Intergenic</td><td>NA</td><td>NA</td></tr>
<tr><td>T24058</td><td>ACGTTGGGCATGACGGGCGTGGG</td><td></td><td>1000</td><td></td><td></td><td></td><td></td><td></td><td></td><td></td><td></td><td></td></tr>
<tr><td>Oligo fwd</td><td>CACCACGTTGGGCATGACGGGCGT</td><td></td><td></td><td></td><td></td><td></td><td></td><td></td><td></td><td></td><td></td><td></td></tr>
<tr><td>Oligo rev</td><td>AAACACGCCCGTCATGCCCAACGT</td><td></td><td></td><td></td><td></td><td></td><td></td><td></td><td></td><td></td><td></td><td></td></tr>
<tr><td>Chromosome</td><td>start</td><td></td><td>end</td><td>strand</td><td>MM</td><td>target_seq</td><td>PAM</td><td>alignment</td><td>distance</td><td>position</td><td>gene name</td><td>gene id</td></tr>
<tr><td>GU305914.1</td><td></td><td>130887</td><td>130909</td><td>-</td><td>0</td><td>ACGTTGGGCATGACGGGCGT</td><td>GGG</td><td>||||||||[||||||||||||]PAM</td><td>NA</td><td>Intergenic</td><td>NA</td><td>NA</td></tr>
<tr><td>T24059</td><td>CATGACGGGCGTGGGCTGCCTGG</td><td></td><td>1000</td><td></td><td></td><td></td><td></td><td></td><td></td><td></td><td></td><td></td></tr>
<tr><td>Oligo fwd</td><td>CACCCATGACGGGCGTGGGCTGCC</td><td></td><td></td><td></td><td></td><td></td><td></td><td></td><td></td><td></td><td></td><td></td></tr>
<tr><td>Oligo rev</td><td>AAACGGCAGCCCACGCCCGTCATG</td><td></td><td></td><td></td><td></td><td></td><td></td><td></td><td></td><td></td><td></td><td></td></tr>
<tr><td>Chromosome</td><td>start</td><td></td><td>end</td><td>strand</td><td>MM</td><td>target_seq</td><td>PAM</td><td>alignment</td><td>distance</td><td>position</td><td>gene name</td><td>gene id</td></tr>
<tr><td>GU305914.1</td><td></td><td>130879</td><td>130901</td><td>-</td><td>0</td><td>CATGACGGGCGTGGGCTGCC</td><td>TGG</td><td>||||||||[||||||||||||]PAM</td><td>NA</td><td>Intergenic</td><td>NA</td><td>NA</td></tr>
<tr><td>T24060</td><td>GACGGGCGTGGGCTGCCTGGCGG</td><td></td><td>1000</td><td></td><td></td><td></td><td></td><td></td><td></td><td></td><td></td><td></td></tr>
<tr><td>Oligo fwd</td><td>CACCGACGGGCGTGGGCTGCCTGG</td><td></td><td></td><td></td><td></td><td></td><td></td><td></td><td></td><td></td><td></td><td></td></tr>
<tr><td>Oligo rev</td><td>AAACCCAGGCAGCCCACGCCCGTC</td><td></td><td></td><td></td><td></td><td></td><td></td><td></td><td></td><td></td><td></td><td></td></tr>
<tr><td>Chromosome</td><td>start</td><td></td><td>end</td><td>strand</td><td>MM</td><td>target_seq</td><td>PAM</td><td>alignment</td><td>distance</td><td>position</td><td>gene name</td><td>gene id</td></tr>
<tr><td>GU305914.1</td><td></td><td>130876</td><td>130898</td><td>-</td><td>0</td><td>GACGGGCGTGGGCTGCCTGG</td><td>CGG</td><td>||||||||[||||||||||||]PAM</td><td>NA</td><td>Intergenic</td><td>NA</td><td>NA</td></tr>
<tr><td>T24061</td><td>GGGCGTGGGCTGCCTGGCGGCGG</td><td></td><td>1000</td><td></td><td></td><td></td><td></td><td></td><td></td><td></td><td></td><td></td></tr>
<tr><td>Oligo fwd</td><td>CACCGGGCGTGGGCTGCCTGGCGG</td><td></td><td></td><td></td><td></td><td></td><td></td><td></td><td></td><td></td><td></td><td></td></tr>
<tr><td>Oligo rev</td><td>AAACCCGCCAGGCAGCCCACGCCC</td><td></td><td></td><td></td><td></td><td></td><td></td><td></td><td></td><td></td><td></td><td></td></tr>
<tr><td>Chromosome</td><td>start</td><td></td><td>end</td><td>strand</td><td>MM</td><td>target_seq</td><td>PAM</td><td>alignment</td><td>distance</td><td>position</td><td>gene name</td><td>gene id</td></tr>
<tr><td>GU305914.1</td><td></td><td>130873</td><td>130895</td><td>-</td><td>0</td><td>GGGCGTGGGCTGCCTGGCGG</td><td>CGG</td><td>||||||||[||||||||||||]PAM</td><td>NA</td><td>Intergenic</td><td>NA</td><td>NA</td></tr>
<tr><td>T24062</td><td>GGCGTGGGCTGCCTGGCGGCGGG</td><td></td><td>1000</td><td></td><td></td><td></td><td></td><td></td><td></td><td></td><td></td><td></td></tr>
<tr><td>Oligo fwd</td><td>CACCGGCGTGGGCTGCCTGGCGGC</td><td></td><td></td><td></td><td></td><td></td><td></td><td></td><td></td><td></td><td></td><td></td></tr>
<tr><td>Oligo rev</td><td>AAACGCCGCCAGGCAGCCCACGCC</td><td></td><td></td><td></td><td></td><td></td><td></td><td></td><td></td><td></td><td></td><td></td></tr>
<tr><td>Chromosome</td><td>start</td><td></td><td>end</td><td>strand</td><td>MM</td><td>target_seq</td><td>PAM</td><td>alignment</td><td>distance</td><td>position</td><td>gene name</td><td>gene id</td></tr>
<tr><td>GU305914.1</td><td></td><td>130872</td><td>130894</td><td>-</td><td>0</td><td>GGCGTGGGCTGCCTGGCGGC</td><td>GGG</td><td>||||||||[||||||||||||]PAM</td><td>NA</td><td>Intergenic</td><td>NA</td><td>NA</td></tr>
<tr><td>T24063</td><td>CCTGGCGGCGGGCGCGTTGCTGG</td><td></td><td>1000</td><td></td><td></td><td></td><td></td><td></td><td></td><td></td><td></td><td></td></tr>
<tr><td>Oligo fwd</td><td>CACCCCTGGCGGCGGGCGCGTTGC</td><td></td><td></td><td></td><td></td><td></td><td></td><td></td><td></td><td></td><td></td><td></td></tr>
<tr><td>Oligo rev</td><td>AAACGCAACGCGCCCGCCGCCAGG</td><td></td><td></td><td></td><td></td><td></td><td></td><td></td><td></td><td></td><td></td><td></td></tr>
<tr><td>Chromosome</td><td>start</td><td></td><td>end</td><td>strand</td><td>MM</td><td>target_seq</td><td>PAM</td><td>alignment</td><td>distance</td><td>position</td><td>gene name</td><td>gene id</td></tr>
<tr><td>GU305914.1</td><td></td><td>130861</td><td>130883</td><td>-</td><td>0</td><td>CCTGGCGGCGGGCGCGTTGC</td><td>TGG</td><td>||||||||[||||||||||||]PAM</td><td>NA</td><td>Intergenic</td><td>NA</td><td>NA</td></tr>
<tr><td>T24064</td><td>AGCTTGATGCCGACGTCCTTCGG</td><td></td><td>1000</td><td></td><td></td><td></td><td></td><td></td><td></td><td></td><td></td><td></td></tr>
<tr><td>Oligo fwd</td><td>CACCAGCTTGATGCCGACGTCCTT</td><td></td><td></td><td></td><td></td><td></td><td></td><td></td><td></td><td></td><td></td><td></td></tr>
<tr><td>Oligo rev</td><td>AAACAAGGACGTCGGCATCAAGCT</td><td></td><td></td><td></td><td></td><td></td><td></td><td></td><td></td><td></td><td></td><td></td></tr>
<tr><td>Chromosome</td><td>start</td><td></td><td>end</td><td>strand</td><td>MM</td><td>target_seq</td><td>PAM</td><td>alignment</td><td>distance</td><td>position</td><td>gene name</td><td>gene id</td></tr>
<tr><td>GU305914.1</td><td></td><td>130827</td><td>130849</td><td>-</td><td>0</td><td>AGCTTGATGCCGACGTCCTT</td><td>CGG</td><td>||||||||[||||||||||||]PAM</td><td>NA</td><td>Intergenic</td><td>NA</td><td>NA</td></tr>
<tr><td>T24065</td><td>ACGTCCTTCGGCCAGATGCCCGG</td><td></td><td>1000</td><td></td><td></td><td></td><td></td><td></td><td></td><td></td><td></td><td></td></tr>
<tr><td>Oligo fwd</td><td>CACCACGTCCTTCGGCCAGATGCC</td><td></td><td></td><td></td><td></td><td></td><td></td><td></td><td></td><td></td><td></td><td></td></tr>
<tr><td>Oligo rev</td><td>AAACGGCATCTGGCCGAAGGACGT</td><td></td><td></td><td></td><td></td><td></td><td></td><td></td><td></td><td></td><td></td><td></td></tr>
<tr><td>Chromosome</td><td>start</td><td></td><td>end</td><td>strand</td><td>MM</td><td>target_seq</td><td>PAM</td><td>alignment</td><td>distance</td><td>position</td><td>gene name</td><td>gene id</td></tr>
<tr><td>GU305914.1</td><td></td><td>130815</td><td>130837</td><td>-</td><td>0</td><td>ACGTCCTTCGGCCAGATGCC</td><td>CGG</td><td>||||||||[||||||||||||]PAM</td><td>NA</td><td>Intergenic</td><td>NA</td><td>NA</td></tr>
<tr><td>T24066</td><td>CGGCCAGATGCCCGGAGTCTTGG</td><td></td><td>1000</td><td></td><td></td><td></td><td></td><td></td><td></td><td></td><td></td><td></td></tr>
<tr><td>Oligo fwd</td><td>CACCCGGCCAGATGCCCGGAGTCT</td><td></td><td></td><td></td><td></td><td></td><td></td><td></td><td></td><td></td><td></td><td></td></tr>
<tr><td>Oligo rev</td><td>AAACAGACTCCGGGCATCTGGCCG</td><td></td><td></td><td></td><td></td><td></td><td></td><td></td><td></td><td></td><td></td><td></td></tr>
<tr><td>Chromosome</td><td>start</td><td></td><td>end</td><td>strand</td><td>MM</td><td>target_seq</td><td>PAM</td><td>alignment</td><td>distance</td><td>position</td><td>gene name</td><td>gene id</td></tr>
<tr><td>GU305914.1</td><td></td><td>130807</td><td>130829</td><td>-</td><td>0</td><td>CGGCCAGATGCCCGGAGTCT</td><td>TGG</td><td>||||||||[||||||||||||]PAM</td><td>NA</td><td>Intergenic</td><td>NA</td><td>NA</td></tr>
<tr><td>T24067</td><td>GATGCCCGGAGTCTTGGACCAGG</td><td></td><td>1000</td><td></td><td></td><td></td><td></td><td></td><td></td><td></td><td></td><td></td></tr>
<tr><td>Oligo fwd</td><td>CACCGATGCCCGGAGTCTTGGACC</td><td></td><td></td><td></td><td></td><td></td><td></td><td></td><td></td><td></td><td></td><td></td></tr>
<tr><td>Oligo rev</td><td>AAACGGTCCAAGACTCCGGGCATC</td><td></td><td></td><td></td><td></td><td></td><td></td><td></td><td></td><td></td><td></td><td></td></tr>
<tr><td>Chromosome</td><td>start</td><td></td><td>end</td><td>strand</td><td>MM</td><td>target_seq</td><td>PAM</td><td>alignment</td><td>distance</td><td>position</td><td>gene name</td><td>gene id</td></tr>
<tr><td>GU305914.1</td><td></td><td>130801</td><td>130823</td><td>-</td><td>0</td><td>GATGCCCGGAGTCTTGGACC</td><td>AGG</td><td>||||||||[||||||||||||]PAM</td><td>NA</td><td>Intergenic</td><td>NA</td><td>NA</td></tr>
<tr><td>T24068</td><td>GACGTGCGAGATGCCGCCGTTGG</td><td></td><td>1000</td><td></td><td></td><td></td><td></td><td></td><td></td><td></td><td></td><td></td></tr>
<tr><td>Oligo fwd</td><td>CACCGACGTGCGAGATGCCGCCGT</td><td></td><td></td><td></td><td></td><td></td><td></td><td></td><td></td><td></td><td></td><td></td></tr>
<tr><td>Oligo rev</td><td>AAACACGGCGGCATCTCGCACGTC</td><td></td><td></td><td></td><td></td><td></td><td></td><td></td><td></td><td></td><td></td><td></td></tr>
<tr><td>Chromosome</td><td>start</td><td></td><td>end</td><td>strand</td><td>MM</td><td>target_seq</td><td>PAM</td><td>alignment</td><td>distance</td><td>position</td><td>gene name</td><td>gene id</td></tr>
<tr><td>GU305914.1</td><td></td><td>130777</td><td>130799</td><td>-</td><td>0</td><td>GACGTGCGAGATGCCGCCGT</td><td>TGG</td><td>||||||||[||||||||||||]PAM</td><td>NA</td><td>Intergenic</td><td>NA</td><td>NA</td></tr>
<tr><td>T24069</td><td>ACGTGCGAGATGCCGCCGTTGGG</td><td></td><td>1000</td><td></td><td></td><td></td><td></td><td></td><td></td><td></td><td></td><td></td></tr>
<tr><td>Oligo fwd</td><td>CACCACGTGCGAGATGCCGCCGTT</td><td></td><td></td><td></td><td></td><td></td><td></td><td></td><td></td><td></td><td></td><td></td></tr>
<tr><td>Oligo rev</td><td>AAACAACGGCGGCATCTCGCACGT</td><td></td><td></td><td></td><td></td><td></td><td></td><td></td><td></td><td></td><td></td><td></td></tr>
<tr><td>Chromosome</td><td>start</td><td></td><td>end</td><td>strand</td><td>MM</td><td>target_seq</td><td>PAM</td><td>alignment</td><td>distance</td><td>position</td><td>gene name</td><td>gene id</td></tr>
<tr><td>GU305914.1</td><td></td><td>130776</td><td>130798</td><td>-</td><td>0</td><td>ACGTGCGAGATGCCGCCGTT</td><td>GGG</td><td>||||||||[||||||||||||]PAM</td><td>NA</td><td>Intergenic</td><td>NA</td><td>NA</td></tr>
<tr><td>T24070</td><td>CGTGCGAGATGCCGCCGTTGGGG</td><td></td><td>1000</td><td></td><td></td><td></td><td></td><td></td><td></td><td></td><td></td><td></td></tr>
<tr><td>Oligo fwd</td><td>CACCCGTGCGAGATGCCGCCGTTG</td><td></td><td></td><td></td><td></td><td></td><td></td><td></td><td></td><td></td><td></td><td></td></tr>
<tr><td>Oligo rev</td><td>AAACCAACGGCGGCATCTCGCACG</td><td></td><td></td><td></td><td></td><td></td><td></td><td></td><td></td><td></td><td></td><td></td></tr>
<tr><td>Chromosome</td><td>start</td><td></td><td>end</td><td>strand</td><td>MM</td><td>target_seq</td><td>PAM</td><td>alignment</td><td>distance</td><td>position</td><td>gene name</td><td>gene id</td></tr>
<tr><td>GU305914.1</td><td></td><td>130775</td><td>130797</td><td>-</td><td>0</td><td>CGTGCGAGATGCCGCCGTTG</td><td>GGG</td><td>||||||||[||||||||||||]PAM</td><td>NA</td><td>Intergenic</td><td>NA</td><td>NA</td></tr>
</table>

<table>
<tr><td>T24071</td><td>GATGCCGCCGTTGGGGTCCCTGG</td><td>1000</td><td></td><td></td><td></td><td></td><td></td><td></td><td></td><td></td><td></td></tr>
<tr><td>Oligo fwd</td><td>CACCGATGCCGCCGTTGGGGTCCC</td><td></td><td></td><td></td><td></td><td></td><td></td><td></td><td></td><td></td><td></td></tr>
<tr><td>Oligo rev</td><td>AAACGGGACCCCAACGGCGGCATC</td><td></td><td></td><td></td><td></td><td></td><td></td><td></td><td></td><td></td><td></td></tr>
<tr><td>Chromosome</td><td>start</td><td>end</td><td>strand</td><td>MM</td><td>target_seq</td><td>PAM</td><td>alignment</td><td>distance</td><td>position</td><td>gene name</td><td>gene id</td></tr>
<tr><td>GU305914.1</td><td>130768</td><td>130790</td><td>-</td><td>0</td><td>GATGCCGCCGTTGGGGTCCC</td><td>TGG</td><td>||||||||[||||||||||||]PAM</td><td>NA</td><td>Intergenic</td><td>NA</td><td>NA</td></tr>
<tr><td></td><td></td><td></td><td></td><td></td><td></td><td></td><td></td><td></td><td></td><td></td><td></td></tr>
<tr><td>T24072</td><td>TGGGGTCCCTGGCGCGCACGAGG</td><td>1000</td><td></td><td></td><td></td><td></td><td></td><td></td><td></td><td></td><td></td></tr>
<tr><td>Oligo fwd</td><td>CACCTGGGGTCCCTGGCGCGCACG</td><td></td><td></td><td></td><td></td><td></td><td></td><td></td><td></td><td></td><td></td></tr>
<tr><td>Oligo rev</td><td>AAACCGTGCGCGCCAGGGACCCCA</td><td></td><td></td><td></td><td></td><td></td><td></td><td></td><td></td><td></td><td></td></tr>
<tr><td>Chromosome</td><td>start</td><td>end</td><td>strand</td><td>MM</td><td>target_seq</td><td>PAM</td><td>alignment</td><td>distance</td><td>position</td><td>gene name</td><td>gene id</td></tr>
<tr><td>GU305914.1</td><td>130757</td><td>130779</td><td>-</td><td>0</td><td>TGGGGTCCCTGGCGCGCACG</td><td>AGG</td><td>||||||||[||||||||||||]PAM</td><td>NA</td><td>Intergenic</td><td>NA</td><td>NA</td></tr>
<tr><td></td><td></td><td></td><td></td><td></td><td></td><td></td><td></td><td></td><td></td><td></td><td></td></tr>
<tr><td>T24073</td><td>GGTCCCTGGCGCGCACGAGGAGG</td><td>1000</td><td></td><td></td><td></td><td></td><td></td><td></td><td></td><td></td><td></td></tr>
<tr><td>Oligo fwd</td><td>CACCGGTCCCTGGCGCGCACGAGG</td><td></td><td></td><td></td><td></td><td></td><td></td><td></td><td></td><td></td><td></td></tr>
<tr><td>Oligo rev</td><td>AAACCCTCGTGCGCGCCAGGGACC</td><td></td><td></td><td></td><td></td><td></td><td></td><td></td><td></td><td></td><td></td></tr>
<tr><td>Chromosome</td><td>start</td><td>end</td><td>strand</td><td>MM</td><td>target_seq</td><td>PAM</td><td>alignment</td><td>distance</td><td>position</td><td>gene name</td><td>gene id</td></tr>
<tr><td>GU305914.1</td><td>130754</td><td>130776</td><td>-</td><td>0</td><td>GGTCCCTGGCGCGCACGAGG</td><td>AGG</td><td>||||||||[||||||||||||]PAM</td><td>NA</td><td>Intergenic</td><td>NA</td><td>NA</td></tr>
<tr><td></td><td></td><td></td><td></td><td></td><td></td><td></td><td></td><td></td><td></td><td></td><td></td></tr>
<tr><td>T24074</td><td>GCGCGCACGAGGAGGACGCGCGG</td><td>1000</td><td></td><td></td><td></td><td></td><td></td><td></td><td></td><td></td><td></td></tr>
<tr><td>Oligo fwd</td><td>CACCGCGCGCACGAGGAGGACGCG</td><td></td><td></td><td></td><td></td><td></td><td></td><td></td><td></td><td></td><td></td></tr>
<tr><td>Oligo rev</td><td>AAACCGCGTCCTCCTCGTGCGCGC</td><td></td><td></td><td></td><td></td><td></td><td></td><td></td><td></td><td></td><td></td></tr>
<tr><td>Chromosome</td><td>start</td><td>end</td><td>strand</td><td>MM</td><td>target_seq</td><td>PAM</td><td>alignment</td><td>distance</td><td>position</td><td>gene name</td><td>gene id</td></tr>
<tr><td>GU305914.1</td><td>130746</td><td>130768</td><td>-</td><td>0</td><td>GCGCGCACGAGGAGGACGCG</td><td>CGG</td><td>||||||||[||||||||||||]PAM</td><td>NA</td><td>Intergenic</td><td>NA</td><td>NA</td></tr>
<tr><td></td><td></td><td></td><td></td><td></td><td></td><td></td><td></td><td></td><td></td><td></td><td></td></tr>
<tr><td>T24075</td><td>CGGCTCGATGAGCATCTCCATGG</td><td>1000</td><td></td><td></td><td></td><td></td><td></td><td></td><td></td><td></td><td></td></tr>
<tr><td>Oligo fwd</td><td>CACCCGGCTCGATGAGCATCTCCA</td><td></td><td></td><td></td><td></td><td></td><td></td><td></td><td></td><td></td><td></td></tr>
<tr><td>Oligo rev</td><td>AAACTGGAGATGCTCATCGAGCCG</td><td></td><td></td><td></td><td></td><td></td><td></td><td></td><td></td><td></td><td></td></tr>
<tr><td>Chromosome</td><td>start</td><td>end</td><td>strand</td><td>MM</td><td>target_seq</td><td>PAM</td><td>alignment</td><td>distance</td><td>position</td><td>gene name</td><td>gene id</td></tr>
<tr><td>GU305914.1</td><td>130726</td><td>130748</td><td>-</td><td>0</td><td>CGGCTCGATGAGCATCTCCA</td><td>TGG</td><td>||||||||[||||||||||||]PAM</td><td>NA</td><td>Intergenic</td><td>NA</td><td>NA</td></tr>
<tr><td></td><td></td><td></td><td></td><td></td><td></td><td></td><td></td><td></td><td></td><td></td><td></td></tr>
<tr><td>T24076</td><td>AGCATCTCCATGGTGCCGTCCGG</td><td>1000</td><td></td><td></td><td></td><td></td><td></td><td></td><td></td><td></td><td></td></tr>
<tr><td>Oligo fwd</td><td>CACCAGCATCTCCATGGTGCCGTC</td><td></td><td></td><td></td><td></td><td></td><td></td><td></td><td></td><td></td><td></td></tr>
<tr><td>Oligo rev</td><td>AAACGACGGCACCATGGAGATGCT</td><td></td><td></td><td></td><td></td><td></td><td></td><td></td><td></td><td></td><td></td></tr>
<tr><td>Chromosome</td><td>start</td><td>end</td><td>strand</td><td>MM</td><td>target_seq</td><td>PAM</td><td>alignment</td><td>distance</td><td>position</td><td>gene name</td><td>gene id</td></tr>
<tr><td>GU305914.1</td><td>130716</td><td>130738</td><td>-</td><td>0</td><td>AGCATCTCCATGGTGCCGTC</td><td>CGG</td><td>||||||||[||||||||||||]PAM</td><td>NA</td><td>Intergenic</td><td>NA</td><td>NA</td></tr>
<tr><td></td><td></td><td></td><td></td><td></td><td></td><td></td><td></td><td></td><td></td><td></td><td></td></tr>
<tr><td>T24077</td><td>GACGTAGATCTGCGCGCAGTAGG</td><td>1000</td><td></td><td></td><td></td><td></td><td></td><td></td><td></td><td></td><td></td></tr>
<tr><td>Oligo fwd</td><td>CACCGACGTAGATCTGCGCGCAGT</td><td></td><td></td><td></td><td></td><td></td><td></td><td></td><td></td><td></td><td></td></tr>
<tr><td>Oligo rev</td><td>AAACACTGCGCGCAGATCTACGTC</td><td></td><td></td><td></td><td></td><td></td><td></td><td></td><td></td><td></td><td></td></tr>
<tr><td>Chromosome</td><td>start</td><td>end</td><td>strand</td><td>MM</td><td>target_seq</td><td>PAM</td><td>alignment</td><td>distance</td><td>position</td><td>gene name</td><td>gene id</td></tr>
<tr><td>GU305914.1</td><td>130675</td><td>130697</td><td>-</td><td>0</td><td>GACGTAGATCTGCGCGCAGT</td><td>AGG</td><td>||||||||[||||||||||||]PAM</td><td>NA</td><td>Intergenic</td><td>NA</td><td>NA</td></tr>
<tr><td></td><td></td><td></td><td></td><td></td><td></td><td></td><td></td><td></td><td></td><td></td><td></td></tr>
<tr><td>T24078</td><td>CTGCGCGCAGTAGGTGACGCCGG</td><td>1000</td><td></td><td></td><td></td><td></td><td></td><td></td><td></td><td></td><td></td></tr>
<tr><td>Oligo fwd</td><td>CACCCTGCGCGCAGTAGGTGACGC</td><td></td><td></td><td></td><td></td><td></td><td></td><td></td><td></td><td></td><td></td></tr>
<tr><td>Oligo rev</td><td>AAACGCGTCACCTACTGCGCGCAG</td><td></td><td></td><td></td><td></td><td></td><td></td><td></td><td></td><td></td><td></td></tr>
<tr><td>Chromosome</td><td>start</td><td>end</td><td>strand</td><td>MM</td><td>target_seq</td><td>PAM</td><td>alignment</td><td>distance</td><td>position</td><td>gene name</td><td>gene id</td></tr>
<tr><td>GU305914.1</td><td>130666</td><td>130688</td><td>-</td><td>0</td><td>CTGCGCGCAGTAGGTGACGC</td><td>CGG</td><td>||||||||[||||||||||||]PAM</td><td>NA</td><td>Intergenic</td><td>NA</td><td>NA</td></tr>
<tr><td></td><td></td><td></td><td></td><td></td><td></td><td></td><td></td><td></td><td></td><td></td><td></td></tr>
<tr><td>T24079</td><td>CGCAGTAGGTGACGCCGGTGAGG</td><td>1000</td><td></td><td></td><td></td><td></td><td></td><td></td><td></td><td></td><td></td></tr>
<tr><td>Oligo fwd</td><td>CACCCGCAGTAGGTGACGCCGGTG</td><td></td><td></td><td></td><td></td><td></td><td></td><td></td><td></td><td></td><td></td></tr>
<tr><td>Oligo rev</td><td>AAACCACCGGCGTCACCTACTGCG</td><td></td><td></td><td></td><td></td><td></td><td></td><td></td><td></td><td></td><td></td></tr>
<tr><td>Chromosome</td><td>start</td><td>end</td><td>strand</td><td>MM</td><td>target_seq</td><td>PAM</td><td>alignment</td><td>distance</td><td>position</td><td>gene name</td><td>gene id</td></tr>
<tr><td>GU305914.1</td><td>130661</td><td>130683</td><td>-</td><td>0</td><td>CGCAGTAGGTGACGCCGGTG</td><td>AGG</td><td>||||||||[||||||||||||]PAM</td><td>NA</td><td>Intergenic</td><td>NA</td><td>NA</td></tr>
<tr><td></td><td></td><td></td><td></td><td></td><td></td><td></td><td></td><td></td><td></td><td></td><td></td></tr>
<tr><td>T24080</td><td>CGCCGGTGAGGACGAAGATGAGG</td><td>1000</td><td></td><td></td><td></td><td></td><td></td><td></td><td></td><td></td><td></td></tr>
<tr><td>Oligo fwd</td><td>CACCCGCCGGTGAGGACGAAGATG</td><td></td><td></td><td></td><td></td><td></td><td></td><td></td><td></td><td></td><td></td></tr>
<tr><td>Oligo rev</td><td>AAACCATCTTCGTCCTCACCGGCG</td><td></td><td></td><td></td><td></td><td></td><td></td><td></td><td></td><td></td><td></td></tr>
<tr><td>Chromosome</td><td>start</td><td>end</td><td>strand</td><td>MM</td><td>target_seq</td><td>PAM</td><td>alignment</td><td>distance</td><td>position</td><td>gene name</td><td>gene id</td></tr>
<tr><td>GU305914.1</td><td>130649</td><td>130671</td><td>-</td><td>0</td><td>CGCCGGTGAGGACGAAGATG</td><td>AGG</td><td>||||||||[||||||||||||]PAM</td><td>NA</td><td>Intergenic</td><td>NA</td><td>NA</td></tr>
<tr><td>GU305914.1</td><td>99829</td><td>99851</td><td>+</td><td>4</td><td>CGGCGGGGACGATGAAGATG</td><td>GGG</td><td>||-|||-|[|-||-|||||||]PAM</td><td>NA</td><td>Intergenic</td><td>NA</td><td>NA</td></tr>
<tr><td></td><td></td><td></td><td></td><td></td><td></td><td></td><td></td><td></td><td></td><td></td><td></td></tr>
<tr><td>T24081</td><td>GGACGAAGATGAGGTCACTGTGG</td><td>1000</td><td></td><td></td><td></td><td></td><td></td><td></td><td></td><td></td><td></td></tr>
<tr><td>Oligo fwd</td><td>CACCGGACGAAGATGAGGTCACTG</td><td></td><td></td><td></td><td></td><td></td><td></td><td></td><td></td><td></td><td></td></tr>
<tr><td>Oligo rev</td><td>AAACCAGTGACCTCATCTTCGTCC</td><td></td><td></td><td></td><td></td><td></td><td></td><td></td><td></td><td></td><td></td></tr>
<tr><td>Chromosome</td><td>start</td><td>end</td><td>strand</td><td>MM</td><td>target_seq</td><td>PAM</td><td>alignment</td><td>distance</td><td>position</td><td>gene name</td><td>gene id</td></tr>
<tr><td>GU305914.1</td><td>130640</td><td>130662</td><td>-</td><td>0</td><td>GGACGAAGATGAGGTCACTG</td><td>TGG</td><td>||||||||[||||||||||||]PAM</td><td>NA</td><td>Intergenic</td><td>NA</td><td>NA</td></tr>
<tr><td></td><td></td><td></td><td></td><td></td><td></td><td></td><td></td><td></td><td></td><td></td><td></td></tr>
<tr><td>T24082</td><td>ACTGTGGCTGTGCCAGTAGATGG</td><td>1000</td><td></td><td></td><td></td><td></td><td></td><td></td><td></td><td></td><td></td></tr>
<tr><td>Oligo fwd</td><td>CACCACTGTGGCTGTGCCAGTAGA</td><td></td><td></td><td></td><td></td><td></td><td></td><td></td><td></td><td></td><td></td></tr>
<tr><td>Oligo rev</td><td>AAACTCTACTGGCACAGCCACAGT</td><td></td><td></td><td></td><td></td><td></td><td></td><td></td><td></td><td></td><td></td></tr>
<tr><td>Chromosome</td><td>start</td><td>end</td><td>strand</td><td>MM</td><td>target_seq</td><td>PAM</td><td>alignment</td><td>distance</td><td>position</td><td>gene name</td><td>gene id</td></tr>
<tr><td>GU305914.1</td><td>130624</td><td>130646</td><td>-</td><td>0</td><td>ACTGTGGCTGTGCCAGTAGA</td><td>TGG</td><td>||||||||[||||||||||||]PAM</td><td>NA</td><td>Intergenic</td><td>NA</td><td>NA</td></tr>
<tr><td></td><td></td><td></td><td></td><td></td><td></td><td></td><td></td><td></td><td></td><td></td><td></td></tr>
<tr><td>T24083</td><td>GTGCCAGTAGATGGAGAAGCCGG</td><td>1000</td><td></td><td></td><td></td><td></td><td></td><td></td><td></td><td></td><td></td></tr>
<tr><td>Oligo fwd</td><td>CACCGTGCCAGTAGATGGAGAAGC</td><td></td><td></td><td></td><td></td><td></td><td></td><td></td><td></td><td></td><td></td></tr>
<tr><td>Oligo rev</td><td>AAACGCTTCTCCATCTACTGGCAC</td><td></td><td></td><td></td><td></td><td></td><td></td><td></td><td></td><td></td><td></td></tr>
<tr><td>Chromosome</td><td>start</td><td>end</td><td>strand</td><td>MM</td><td>target_seq</td><td>PAM</td><td>alignment</td><td>distance</td><td>position</td><td>gene name</td><td>gene id</td></tr>
<tr><td>GU305914.1</td><td>130615</td><td>130637</td><td>-</td><td>0</td><td>GTGCCAGTAGATGGAGAAGC</td><td>CGG</td><td>||||||||[||||||||||||]PAM</td><td>NA</td><td>Intergenic</td><td>NA</td><td>NA</td></tr>
<tr><td></td><td></td><td></td><td></td><td></td><td></td><td></td><td></td><td></td><td></td><td></td><td></td></tr>
<tr><td>T24084</td><td>CCAGTAGATGGAGAAGCCGGCGG</td><td>1000</td><td></td><td></td><td></td><td></td><td></td><td></td><td></td><td></td><td></td></tr>
<tr><td>Oligo fwd</td><td>CACCCCAGTAGATGGAGAAGCCGG</td><td></td><td></td><td></td><td></td><td></td><td></td><td></td><td></td><td></td><td></td></tr>
<tr><td>Oligo rev</td><td>AAACCCGGCTTCTCCATCTACTGG</td><td></td><td></td><td></td><td></td><td></td><td></td><td></td><td></td><td></td><td></td></tr>
<tr><td>Chromosome</td><td>start</td><td>end</td><td>strand</td><td>MM</td><td>target_seq</td><td>PAM</td><td>alignment</td><td>distance</td><td>position</td><td>gene name</td><td>gene id</td></tr>
<tr><td>GU305914.1</td><td>130612</td><td>130634</td><td>-</td><td>0</td><td>CCAGTAGATGGAGAAGCCGG</td><td>CGG</td><td>||||||||[||||||||||||]PAM</td><td>NA</td><td>Intergenic</td><td>NA</td><td>NA</td></tr>
<tr><td></td><td></td><td></td><td></td><td></td><td></td><td></td><td></td><td></td><td></td><td></td><td></td></tr>
<tr><td>T24085</td><td>AGATGGAGAAGCCGGCGGCGCGG</td><td>1000</td><td></td><td></td><td></td><td></td><td></td><td></td><td></td><td></td><td></td></tr>
<tr><td>Oligo fwd</td><td>CACCAGATGGAGAAGCCGGCGGCG</td><td></td><td></td><td></td><td></td><td></td><td></td><td></td><td></td><td></td><td></td></tr>
<tr><td>Oligo rev</td><td>AAACCGCCGCCGGCTTCTCCATCT</td><td></td><td></td><td></td><td></td><td></td><td></td><td></td><td></td><td></td><td></td></tr>
<tr><td>Chromosome</td><td>start</td><td>end</td><td>strand</td><td>MM</td><td>target_seq</td><td>PAM</td><td>alignment</td><td>distance</td><td>position</td><td>gene name</td><td>gene id</td></tr>
</table>

<table>
<tr><td>GU305914.1</td><td></td><td>130607</td><td>130629</td><td>-</td><td>0</td><td>AGATGGAGAAGCCGGCGGCG</td><td>CGG</td><td>||||||||[||||||||||||]PAM</td><td>NA</td><td>Intergenic</td><td>NA</td><td>NA</td></tr>
<tr><td>GU305914.1</td><td></td><td>81247</td><td>81269</td><td>-</td><td>4</td><td>CGATCGAGGAGGCGGCGGCG</td><td>GGG</td><td>-|||-|||[-||-||||||||]PAM</td><td>NA</td><td>Intergenic</td><td>NA</td><td>NA</td></tr>
<tr><td>GU305914.1</td><td></td><td>71196</td><td>71218</td><td>+</td><td>4</td><td>ATCTGGAGAAGCTGTCGGCG</td><td>CGG</td><td>|--|||||[||||-|-|||||]PAM</td><td>NA</td><td>Intergenic</td><td>NA</td><td>NA</td></tr>
<tr><td>T24086</td><td>GGCGCGGCGCCGACAGCGTCCGG</td><td></td><td>1000</td><td></td><td></td><td></td><td></td><td></td><td></td><td></td><td></td><td></td></tr>
<tr><td>Oligo fwd</td><td>CACCGGCGCGGCGCCGACAGCGTC</td><td></td><td></td><td></td><td></td><td></td><td></td><td></td><td></td><td></td><td></td><td></td></tr>
<tr><td>Oligo rev</td><td>AAACGACGCTGTCGGCGCCGCGCC</td><td></td><td></td><td></td><td></td><td></td><td></td><td></td><td></td><td></td><td></td><td></td></tr>
<tr><td>Chromosome</td><td>start</td><td></td><td>end</td><td>strand</td><td>MM</td><td>target_seq</td><td>PAM</td><td>alignment</td><td>distance</td><td>position</td><td>gene name</td><td>gene id</td></tr>
<tr><td>GU305914.1</td><td></td><td>130591</td><td>130613</td><td>-</td><td>0</td><td>GGCGCGGCGCCGACAGCGTC</td><td>CGG</td><td>||||||||[||||||||||||]PAM</td><td>NA</td><td>Intergenic</td><td>NA</td><td>NA</td></tr>
<tr><td>GU305914.1</td><td></td><td>138101</td><td>138123</td><td>+</td><td>4</td><td>CGATCGCCGCCGACAGCGTC</td><td>GGG</td><td>-|--||-|[||||||||||||]PAM</td><td>NA</td><td>Intergenic</td><td>NA</td><td>NA</td></tr>
<tr><td>T24087</td><td>TGACAAAGAGCGAGAGCTCCCGG</td><td></td><td>1000</td><td></td><td></td><td></td><td></td><td></td><td></td><td></td><td></td><td></td></tr>
<tr><td>Oligo fwd</td><td>CACCTGACAAAGAGCGAGAGCTCC</td><td></td><td></td><td></td><td></td><td></td><td></td><td></td><td></td><td></td><td></td><td></td></tr>
<tr><td>Oligo rev</td><td>AAACGGAGCTCTCGCTCTTTGTCA</td><td></td><td></td><td></td><td></td><td></td><td></td><td></td><td></td><td></td><td></td><td></td></tr>
<tr><td>Chromosome</td><td>start</td><td></td><td>end</td><td>strand</td><td>MM</td><td>target_seq</td><td>PAM</td><td>alignment</td><td>distance</td><td>position</td><td>gene name</td><td>gene id</td></tr>
<tr><td>GU305914.1</td><td></td><td>130568</td><td>130590</td><td>-</td><td>0</td><td>TGACAAAGAGCGAGAGCTCC</td><td>CGG</td><td>||||||||[||||||||||||]PAM</td><td>NA</td><td>Intergenic</td><td>NA</td><td>NA</td></tr>
<tr><td>T24088</td><td>GACAAAGAGCGAGAGCTCCCGGG</td><td></td><td>1000</td><td></td><td></td><td></td><td></td><td></td><td></td><td></td><td></td><td></td></tr>
<tr><td>Oligo fwd</td><td>CACCGACAAAGAGCGAGAGCTCCC</td><td></td><td></td><td></td><td></td><td></td><td></td><td></td><td></td><td></td><td></td><td></td></tr>
<tr><td>Oligo rev</td><td>AAACGGGAGCTCTCGCTCTTTGTC</td><td></td><td></td><td></td><td></td><td></td><td></td><td></td><td></td><td></td><td></td><td></td></tr>
<tr><td>Chromosome</td><td>start</td><td></td><td>end</td><td>strand</td><td>MM</td><td>target_seq</td><td>PAM</td><td>alignment</td><td>distance</td><td>position</td><td>gene name</td><td>gene id</td></tr>
<tr><td>GU305914.1</td><td></td><td>130567</td><td>130589</td><td>-</td><td>0</td><td>GACAAAGAGCGAGAGCTCCC</td><td>GGG</td><td>||||||||[||||||||||||]PAM</td><td>NA</td><td>Intergenic</td><td>NA</td><td>NA</td></tr>
<tr><td>T24089</td><td>GAGCTCCCGGGAGCTGACGAAGG</td><td></td><td>1000</td><td></td><td></td><td></td><td></td><td></td><td></td><td></td><td></td><td></td></tr>
<tr><td>Oligo fwd</td><td>CACCGAGCTCCCGGGAGCTGACGA</td><td></td><td></td><td></td><td></td><td></td><td></td><td></td><td></td><td></td><td></td><td></td></tr>
<tr><td>Oligo rev</td><td>AAACTCGTCAGCTCCCGGGAGCTC</td><td></td><td></td><td></td><td></td><td></td><td></td><td></td><td></td><td></td><td></td><td></td></tr>
<tr><td>Chromosome</td><td>start</td><td></td><td>end</td><td>strand</td><td>MM</td><td>target_seq</td><td>PAM</td><td>alignment</td><td>distance</td><td>position</td><td>gene name</td><td>gene id</td></tr>
<tr><td>GU305914.1</td><td></td><td>130555</td><td>130577</td><td>-</td><td>0</td><td>GAGCTCCCGGGAGCTGACGA</td><td>AGG</td><td>||||||||[||||||||||||]PAM</td><td>NA</td><td>Intergenic</td><td>NA</td><td>NA</td></tr>
<tr><td>T24090</td><td>CTCCCGGGAGCTGACGAAGGCGG</td><td></td><td>1000</td><td></td><td></td><td></td><td></td><td></td><td></td><td></td><td></td><td></td></tr>
<tr><td>Oligo fwd</td><td>CACCCTCCCGGGAGCTGACGAAGG</td><td></td><td></td><td></td><td></td><td></td><td></td><td></td><td></td><td></td><td></td><td></td></tr>
<tr><td>Oligo rev</td><td>AAACCCTTCGTCAGCTCCCGGGAG</td><td></td><td></td><td></td><td></td><td></td><td></td><td></td><td></td><td></td><td></td><td></td></tr>
<tr><td>Chromosome</td><td>start</td><td></td><td>end</td><td>strand</td><td>MM</td><td>target_seq</td><td>PAM</td><td>alignment</td><td>distance</td><td>position</td><td>gene name</td><td>gene id</td></tr>
<tr><td>GU305914.1</td><td></td><td>130552</td><td>130574</td><td>-</td><td>0</td><td>CTCCCGGGAGCTGACGAAGG</td><td>CGG</td><td>||||||||[||||||||||||]PAM</td><td>NA</td><td>Intergenic</td><td>NA</td><td>NA</td></tr>
<tr><td>T24091</td><td>TCCTTCTGATCAGAGAGACCAGG</td><td></td><td>1000</td><td></td><td></td><td></td><td></td><td></td><td></td><td></td><td></td><td></td></tr>
<tr><td>Oligo fwd</td><td>CACCTCCTTCTGATCAGAGAGACC</td><td></td><td></td><td></td><td></td><td></td><td></td><td></td><td></td><td></td><td></td><td></td></tr>
<tr><td>Oligo rev</td><td>AAACGGTCTCTCTGATCAGAAGGA</td><td></td><td></td><td></td><td></td><td></td><td></td><td></td><td></td><td></td><td></td><td></td></tr>
<tr><td>Chromosome</td><td>start</td><td></td><td>end</td><td>strand</td><td>MM</td><td>target_seq</td><td>PAM</td><td>alignment</td><td>distance</td><td>position</td><td>gene name</td><td>gene id</td></tr>
<tr><td>GU305914.1</td><td></td><td>130490</td><td>130512</td><td>-</td><td>0</td><td>TCCTTCTGATCAGAGAGACC</td><td>AGG</td><td>||||||||[||||||||||||]PAM</td><td>NA</td><td>Intergenic</td><td>NA</td><td>NA</td></tr>
<tr><td>T24092</td><td>CCTTCTGATCAGAGAGACCAGGG</td><td></td><td>1000</td><td></td><td></td><td></td><td></td><td></td><td></td><td></td><td></td><td></td></tr>
<tr><td>Oligo fwd</td><td>CACCCCTTCTGATCAGAGAGACCA</td><td></td><td></td><td></td><td></td><td></td><td></td><td></td><td></td><td></td><td></td><td></td></tr>
<tr><td>Oligo rev</td><td>AAACTGGTCTCTCTGATCAGAAGG</td><td></td><td></td><td></td><td></td><td></td><td></td><td></td><td></td><td></td><td></td><td></td></tr>
<tr><td>Chromosome</td><td>start</td><td></td><td>end</td><td>strand</td><td>MM</td><td>target_seq</td><td>PAM</td><td>alignment</td><td>distance</td><td>position</td><td>gene name</td><td>gene id</td></tr>
<tr><td>GU305914.1</td><td></td><td>130489</td><td>130511</td><td>-</td><td>0</td><td>CCTTCTGATCAGAGAGACCA</td><td>GGG</td><td>||||||||[||||||||||||]PAM</td><td>NA</td><td>Intergenic</td><td>NA</td><td>NA</td></tr>
<tr><td>T24093</td><td>CTTCTGATCAGAGAGACCAGGGG</td><td></td><td>1000</td><td></td><td></td><td></td><td></td><td></td><td></td><td></td><td></td><td></td></tr>
<tr><td>Oligo fwd</td><td>CACCCTTCTGATCAGAGAGACCAG</td><td></td><td></td><td></td><td></td><td></td><td></td><td></td><td></td><td></td><td></td><td></td></tr>
<tr><td>Oligo rev</td><td>AAACCTGGTCTCTCTGATCAGAAG</td><td></td><td></td><td></td><td></td><td></td><td></td><td></td><td></td><td></td><td></td><td></td></tr>
<tr><td>Chromosome</td><td>start</td><td></td><td>end</td><td>strand</td><td>MM</td><td>target_seq</td><td>PAM</td><td>alignment</td><td>distance</td><td>position</td><td>gene name</td><td>gene id</td></tr>
<tr><td>GU305914.1</td><td></td><td>130488</td><td>130510</td><td>-</td><td>0</td><td>CTTCTGATCAGAGAGACCAG</td><td>GGG</td><td>||||||||[||||||||||||]PAM</td><td>NA</td><td>Intergenic</td><td>NA</td><td>NA</td></tr>
<tr><td>T24094</td><td>GATCAGAGAGACCAGGGGTTTGG</td><td></td><td>1000</td><td></td><td></td><td></td><td></td><td></td><td></td><td></td><td></td><td></td></tr>
<tr><td>Oligo fwd</td><td>CACCGATCAGAGAGACCAGGGGTT</td><td></td><td></td><td></td><td></td><td></td><td></td><td></td><td></td><td></td><td></td><td></td></tr>
<tr><td>Oligo rev</td><td>AAACAACCCCTGGTCTCTCTGATC</td><td></td><td></td><td></td><td></td><td></td><td></td><td></td><td></td><td></td><td></td><td></td></tr>
<tr><td>Chromosome</td><td>start</td><td></td><td>end</td><td>strand</td><td>MM</td><td>target_seq</td><td>PAM</td><td>alignment</td><td>distance</td><td>position</td><td>gene name</td><td>gene id</td></tr>
<tr><td>GU305914.1</td><td></td><td>130483</td><td>130505</td><td>-</td><td>0</td><td>GATCAGAGAGACCAGGGGTT</td><td>TGG</td><td>||||||||[||||||||||||]PAM</td><td>NA</td><td>Intergenic</td><td>NA</td><td>NA</td></tr>
<tr><td>T24095</td><td>GTAGAACTCGTGTTCCATGATGG</td><td></td><td>1000</td><td></td><td></td><td></td><td></td><td></td><td></td><td></td><td></td><td></td></tr>
<tr><td>Oligo fwd</td><td>CACCGTAGAACTCGTGTTCCATGA</td><td></td><td></td><td></td><td></td><td></td><td></td><td></td><td></td><td></td><td></td><td></td></tr>
<tr><td>Oligo rev</td><td>AAACTCATGGAACACGAGTTCTAC</td><td></td><td></td><td></td><td></td><td></td><td></td><td></td><td></td><td></td><td></td><td></td></tr>
<tr><td>Chromosome</td><td>start</td><td></td><td>end</td><td>strand</td><td>MM</td><td>target_seq</td><td>PAM</td><td>alignment</td><td>distance</td><td>position</td><td>gene name</td><td>gene id</td></tr>
<tr><td>GU305914.1</td><td></td><td>130459</td><td>130481</td><td>-</td><td>0</td><td>GTAGAACTCGTGTTCCATGA</td><td>TGG</td><td>||||||||[||||||||||||]PAM</td><td>NA</td><td>Intergenic</td><td>NA</td><td>NA</td></tr>
<tr><td>T24096</td><td>GGTCCCGTCGCTCATGAGCACGG</td><td></td><td>1000</td><td></td><td></td><td></td><td></td><td></td><td></td><td></td><td></td><td></td></tr>
<tr><td>Oligo fwd</td><td>CACCGGTCCCGTCGCTCATGAGCA</td><td></td><td></td><td></td><td></td><td></td><td></td><td></td><td></td><td></td><td></td><td></td></tr>
<tr><td>Oligo rev</td><td>AAACTGCTCATGAGCGACGGGACC</td><td></td><td></td><td></td><td></td><td></td><td></td><td></td><td></td><td></td><td></td><td></td></tr>
<tr><td>Chromosome</td><td>start</td><td></td><td>end</td><td>strand</td><td>MM</td><td>target_seq</td><td>PAM</td><td>alignment</td><td>distance</td><td>position</td><td>gene name</td><td>gene id</td></tr>
<tr><td>GU305914.1</td><td></td><td>130438</td><td>130460</td><td>-</td><td>0</td><td>GGTCCCGTCGCTCATGAGCA</td><td>CGG</td><td>||||||||[||||||||||||]PAM</td><td>NA</td><td>Intergenic</td><td>NA</td><td>NA</td></tr>
<tr><td>T24097</td><td>CACGGCCCCGTCGCACCAGATGG</td><td></td><td>1000</td><td></td><td></td><td></td><td></td><td></td><td></td><td></td><td></td><td></td></tr>
<tr><td>Oligo fwd</td><td>CACCCACGGCCCCGTCGCACCAGA</td><td></td><td></td><td></td><td></td><td></td><td></td><td></td><td></td><td></td><td></td><td></td></tr>
<tr><td>Oligo rev</td><td>AAACTCTGGTGCGACGGGGCCGTG</td><td></td><td></td><td></td><td></td><td></td><td></td><td></td><td></td><td></td><td></td><td></td></tr>
<tr><td>Chromosome</td><td>start</td><td></td><td>end</td><td>strand</td><td>MM</td><td>target_seq</td><td>PAM</td><td>alignment</td><td>distance</td><td>position</td><td>gene name</td><td>gene id</td></tr>
<tr><td>GU305914.1</td><td></td><td>130420</td><td>130442</td><td>-</td><td>0</td><td>CACGGCCCCGTCGCACCAGA</td><td>TGG</td><td>||||||||[||||||||||||]PAM</td><td>NA</td><td>Intergenic</td><td>NA</td><td>NA</td></tr>
<tr><td>T24098</td><td>CCCGTCGCACCAGATGGCGCCGG</td><td></td><td>1000</td><td></td><td></td><td></td><td></td><td></td><td></td><td></td><td></td><td></td></tr>
<tr><td>Oligo fwd</td><td>CACCCCCGTCGCACCAGATGGCGC</td><td></td><td></td><td></td><td></td><td></td><td></td><td></td><td></td><td></td><td></td><td></td></tr>
<tr><td>Oligo rev</td><td>AAACGCGCCATCTGGTGCGACGGG</td><td></td><td></td><td></td><td></td><td></td><td></td><td></td><td></td><td></td><td></td><td></td></tr>
<tr><td>Chromosome</td><td>start</td><td></td><td>end</td><td>strand</td><td>MM</td><td>target_seq</td><td>PAM</td><td>alignment</td><td>distance</td><td>position</td><td>gene name</td><td>gene id</td></tr>
<tr><td>GU305914.1</td><td></td><td>130414</td><td>130436</td><td>-</td><td>0</td><td>CCCGTCGCACCAGATGGCGC</td><td>CGG</td><td>||||||||[||||||||||||]PAM</td><td>NA</td><td>Intergenic</td><td>NA</td><td>NA</td></tr>
<tr><td>T24099</td><td>CCGTCGCACCAGATGGCGCCGGG</td><td></td><td>1000</td><td></td><td></td><td></td><td></td><td></td><td></td><td></td><td></td><td></td></tr>
<tr><td>Oligo fwd</td><td>CACCCCGTCGCACCAGATGGCGCC</td><td></td><td></td><td></td><td></td><td></td><td></td><td></td><td></td><td></td><td></td><td></td></tr>
<tr><td>Oligo rev</td><td>AAACGGCGCCATCTGGTGCGACGG</td><td></td><td></td><td></td><td></td><td></td><td></td><td></td><td></td><td></td><td></td><td></td></tr>
<tr><td>Chromosome</td><td>start</td><td></td><td>end</td><td>strand</td><td>MM</td><td>target_seq</td><td>PAM</td><td>alignment</td><td>distance</td><td>position</td><td>gene name</td><td>gene id</td></tr>
<tr><td>GU305914.1</td><td></td><td>130413</td><td>130435</td><td>-</td><td>0</td><td>CCGTCGCACCAGATGGCGCC</td><td>GGG</td><td>||||||||[||||||||||||]PAM</td><td>NA</td><td>Intergenic</td><td>NA</td><td>NA</td></tr>
</table>

<table>
<tr><td>T24100</td><td>CGTCGCACCAGATGGCGCCGGGG</td><td>1000</td><td></td><td></td><td></td><td></td><td></td><td></td><td></td><td></td><td></td></tr>
<tr><td>Oligo fwd</td><td>CACCCGTCGCACCAGATGGCGCCG</td><td></td><td></td><td></td><td></td><td></td><td></td><td></td><td></td><td></td><td></td></tr>
<tr><td>Oligo rev</td><td>AAACCGGCGCCATCTGGTGCGACG</td><td></td><td></td><td></td><td></td><td></td><td></td><td></td><td></td><td></td><td></td></tr>
<tr><td>Chromosome</td><td>start</td><td>end</td><td>strand</td><td>MM</td><td>target_seq</td><td>PAM</td><td>alignment</td><td>distance</td><td>position</td><td>gene name</td><td>gene id</td></tr>
<tr><td>GU305914.1</td><td>130412</td><td>130434</td><td>-</td><td>0</td><td>CGTCGCACCAGATGGCGCCG</td><td>GGG</td><td>||||||||[||||||||||||]PAM</td><td>NA</td><td>Intergenic</td><td>NA</td><td>NA</td></tr>
<tr><td>T24101</td><td>ACCAGATGGCGCCGGGGTCGCGG</td><td>1000</td><td></td><td></td><td></td><td></td><td></td><td></td><td></td><td></td><td></td></tr>
<tr><td>Oligo fwd</td><td>CACCACCAGATGGCGCCGGGGTCG</td><td></td><td></td><td></td><td></td><td></td><td></td><td></td><td></td><td></td><td></td></tr>
<tr><td>Oligo rev</td><td>AAACCGACCCCGGCGCCATCTGGT</td><td></td><td></td><td></td><td></td><td></td><td></td><td></td><td></td><td></td><td></td></tr>
<tr><td>Chromosome</td><td>start</td><td>end</td><td>strand</td><td>MM</td><td>target_seq</td><td>PAM</td><td>alignment</td><td>distance</td><td>position</td><td>gene name</td><td>gene id</td></tr>
<tr><td>GU305914.1</td><td>130406</td><td>130428</td><td>-</td><td>0</td><td>ACCAGATGGCGCCGGGGTCG</td><td>CGG</td><td>||||||||[||||||||||||]PAM</td><td>NA</td><td>Intergenic</td><td>NA</td><td>NA</td></tr>
<tr><td>T24102</td><td>GCCGGGGTCGCGGATAGCGACGG</td><td>1000</td><td></td><td></td><td></td><td></td><td></td><td></td><td></td><td></td><td></td></tr>
<tr><td>Oligo fwd</td><td>CACCGCCGGGGTCGCGGATAGCGA</td><td></td><td></td><td></td><td></td><td></td><td></td><td></td><td></td><td></td><td></td></tr>
<tr><td>Oligo rev</td><td>AAACTCGCTATCCGCGACCCCGGC</td><td></td><td></td><td></td><td></td><td></td><td></td><td></td><td></td><td></td><td></td></tr>
<tr><td>Chromosome</td><td>start</td><td>end</td><td>strand</td><td>MM</td><td>target_seq</td><td>PAM</td><td>alignment</td><td>distance</td><td>position</td><td>gene name</td><td>gene id</td></tr>
<tr><td>GU305914.1</td><td>130396</td><td>130418</td><td>-</td><td>0</td><td>GCCGGGGTCGCGGATAGCGA</td><td>CGG</td><td>||||||||[||||||||||||]PAM</td><td>NA</td><td>Intergenic</td><td>NA</td><td>NA</td></tr>
<tr><td>T24103</td><td>CCTCTCGCCGTTGATGCTGCCGG</td><td>1000</td><td></td><td></td><td></td><td></td><td></td><td></td><td></td><td></td><td></td></tr>
<tr><td>Oligo fwd</td><td>CACCCCTCTCGCCGTTGATGCTGC</td><td></td><td></td><td></td><td></td><td></td><td></td><td></td><td></td><td></td><td></td></tr>
<tr><td>Oligo rev</td><td>AAACGCAGCATCAACGGCGAGAGG</td><td></td><td></td><td></td><td></td><td></td><td></td><td></td><td></td><td></td><td></td></tr>
<tr><td>Chromosome</td><td>start</td><td>end</td><td>strand</td><td>MM</td><td>target_seq</td><td>PAM</td><td>alignment</td><td>distance</td><td>position</td><td>gene name</td><td>gene id</td></tr>
<tr><td>GU305914.1</td><td>130363</td><td>130385</td><td>-</td><td>0</td><td>CCTCTCGCCGTTGATGCTGC</td><td>CGG</td><td>||||||||[||||||||||||]PAM</td><td>NA</td><td>Intergenic</td><td>NA</td><td>NA</td></tr>
<tr><td>T24104</td><td>GTCTTCTTCGTCCATGACGCCGG</td><td>1000</td><td></td><td></td><td></td><td></td><td></td><td></td><td></td><td></td><td></td></tr>
<tr><td>Oligo fwd</td><td>CACCGTCTTCTTCGTCCATGACGC</td><td></td><td></td><td></td><td></td><td></td><td></td><td></td><td></td><td></td><td></td></tr>
<tr><td>Oligo rev</td><td>AAACGCGTCATGGACGAAGAAGAC</td><td></td><td></td><td></td><td></td><td></td><td></td><td></td><td></td><td></td><td></td></tr>
<tr><td>Chromosome</td><td>start</td><td>end</td><td>strand</td><td>MM</td><td>target_seq</td><td>PAM</td><td>alignment</td><td>distance</td><td>position</td><td>gene name</td><td>gene id</td></tr>
<tr><td>GU305914.1</td><td>130339</td><td>130361</td><td>-</td><td>0</td><td>GTCTTCTTCGTCCATGACGC</td><td>CGG</td><td>||||||||[||||||||||||]PAM</td><td>NA</td><td>Intergenic</td><td>NA</td><td>NA</td></tr>
<tr><td>T24105</td><td>TCCATGACGCCGGACCGCCGCGG</td><td>1000</td><td></td><td></td><td></td><td></td><td></td><td></td><td></td><td></td><td></td></tr>
<tr><td>Oligo fwd</td><td>CACCTCCATGACGCCGGACCGCCG</td><td></td><td></td><td></td><td></td><td></td><td></td><td></td><td></td><td></td><td></td></tr>
<tr><td>Oligo rev</td><td>AAACCGGCGGTCCGGCGTCATGGA</td><td></td><td></td><td></td><td></td><td></td><td></td><td></td><td></td><td></td><td></td></tr>
<tr><td>Chromosome</td><td>start</td><td>end</td><td>strand</td><td>MM</td><td>target_seq</td><td>PAM</td><td>alignment</td><td>distance</td><td>position</td><td>gene name</td><td>gene id</td></tr>
<tr><td>GU305914.1</td><td>130329</td><td>130351</td><td>-</td><td>0</td><td>TCCATGACGCCGGACCGCCG</td><td>CGG</td><td>||||||||[||||||||||||]PAM</td><td>NA</td><td>Intergenic</td><td>NA</td><td>NA</td></tr>
<tr><td>T24106</td><td>ATGACGCCGGACCGCCGCGGCGG</td><td>1000</td><td></td><td></td><td></td><td></td><td></td><td></td><td></td><td></td><td></td></tr>
<tr><td>Oligo fwd</td><td>CACCATGACGCCGGACCGCCGCGG</td><td></td><td></td><td></td><td></td><td></td><td></td><td></td><td></td><td></td><td></td></tr>
<tr><td>Oligo rev</td><td>AAACCCGCGGCGGTCCGGCGTCAT</td><td></td><td></td><td></td><td></td><td></td><td></td><td></td><td></td><td></td><td></td></tr>
<tr><td>Chromosome</td><td>start</td><td>end</td><td>strand</td><td>MM</td><td>target_seq</td><td>PAM</td><td>alignment</td><td>distance</td><td>position</td><td>gene name</td><td>gene id</td></tr>
<tr><td>GU305914.1</td><td>130326</td><td>130348</td><td>-</td><td>0</td><td>ATGACGCCGGACCGCCGCGG</td><td>CGG</td><td>||||||||[||||||||||||]PAM</td><td>NA</td><td>Intergenic</td><td>NA</td><td>NA</td></tr>
<tr><td>T24107</td><td>ACGCCGGACCGCCGCGGCGGCGG</td><td>1000</td><td></td><td></td><td></td><td></td><td></td><td></td><td></td><td></td><td></td></tr>
<tr><td>Oligo fwd</td><td>CACCACGCCGGACCGCCGCGGCGG</td><td></td><td></td><td></td><td></td><td></td><td></td><td></td><td></td><td></td><td></td></tr>
<tr><td>Oligo rev</td><td>AAACCCGCCGCGGCGGTCCGGCGT</td><td></td><td></td><td></td><td></td><td></td><td></td><td></td><td></td><td></td><td></td></tr>
<tr><td>Chromosome</td><td>start</td><td>end</td><td>strand</td><td>MM</td><td>target_seq</td><td>PAM</td><td>alignment</td><td>distance</td><td>position</td><td>gene name</td><td>gene id</td></tr>
<tr><td>GU305914.1</td><td>130323</td><td>130345</td><td>-</td><td>0</td><td>ACGCCGGACCGCCGCGGCGG</td><td>CGG</td><td>||||||||[||||||||||||]PAM</td><td>NA</td><td>Intergenic</td><td>NA</td><td>NA</td></tr>
<tr><td>T24108</td><td>TGACAGTCTATCTCTATCGCCGG</td><td>1000</td><td></td><td></td><td></td><td></td><td></td><td></td><td></td><td></td><td></td></tr>
<tr><td>Oligo fwd</td><td>CACCTGACAGTCTATCTCTATCGC</td><td></td><td></td><td></td><td></td><td></td><td></td><td></td><td></td><td></td><td></td></tr>
<tr><td>Oligo rev</td><td>AAACGCGATAGAGATAGACTGTCA</td><td></td><td></td><td></td><td></td><td></td><td></td><td></td><td></td><td></td><td></td></tr>
<tr><td>Chromosome</td><td>start</td><td>end</td><td>strand</td><td>MM</td><td>target_seq</td><td>PAM</td><td>alignment</td><td>distance</td><td>position</td><td>gene name</td><td>gene id</td></tr>
<tr><td>GU305914.1</td><td>130258</td><td>130280</td><td>-</td><td>0</td><td>TGACAGTCTATCTCTATCGC</td><td>CGG</td><td>||||||||[||||||||||||]PAM</td><td>NA</td><td>Intergenic</td><td>NA</td><td>NA</td></tr>
<tr><td>T24109</td><td>TCGCCGGCTCGTCGCTGCCTCGG</td><td>1000</td><td></td><td></td><td></td><td></td><td></td><td></td><td></td><td></td><td></td></tr>
<tr><td>Oligo fwd</td><td>CACCTCGCCGGCTCGTCGCTGCCT</td><td></td><td></td><td></td><td></td><td></td><td></td><td></td><td></td><td></td><td></td></tr>
<tr><td>Oligo rev</td><td>AAACAGGCAGCGACGAGCCGGCGA</td><td></td><td></td><td></td><td></td><td></td><td></td><td></td><td></td><td></td><td></td></tr>
<tr><td>Chromosome</td><td>start</td><td>end</td><td>strand</td><td>MM</td><td>target_seq</td><td>PAM</td><td>alignment</td><td>distance</td><td>position</td><td>gene name</td><td>gene id</td></tr>
<tr><td>GU305914.1</td><td>130242</td><td>130264</td><td>-</td><td>0</td><td>TCGCCGGCTCGTCGCTGCCT</td><td>CGG</td><td>||||||||[||||||||||||]PAM</td><td>NA</td><td>Intergenic</td><td>NA</td><td>NA</td></tr>
<tr><td>T24110</td><td>CGCCGGCTCGTCGCTGCCTCGGG</td><td>1000</td><td></td><td></td><td></td><td></td><td></td><td></td><td></td><td></td><td></td></tr>
<tr><td>Oligo fwd</td><td>CACCCGCCGGCTCGTCGCTGCCTC</td><td></td><td></td><td></td><td></td><td></td><td></td><td></td><td></td><td></td><td></td></tr>
<tr><td>Oligo rev</td><td>AAACGAGGCAGCGACGAGCCGGCG</td><td></td><td></td><td></td><td></td><td></td><td></td><td></td><td></td><td></td><td></td></tr>
<tr><td>Chromosome</td><td>start</td><td>end</td><td>strand</td><td>MM</td><td>target_seq</td><td>PAM</td><td>alignment</td><td>distance</td><td>position</td><td>gene name</td><td>gene id</td></tr>
<tr><td>GU305914.1</td><td>130241</td><td>130263</td><td>-</td><td>0</td><td>CGCCGGCTCGTCGCTGCCTC</td><td>GGG</td><td>||||||||[||||||||||||]PAM</td><td>NA</td><td>Intergenic</td><td>NA</td><td>NA</td></tr>
<tr><td>T24111</td><td>GCCGGCTCGTCGCTGCCTCGGGG</td><td>1000</td><td></td><td></td><td></td><td></td><td></td><td></td><td></td><td></td><td></td></tr>
<tr><td>Oligo fwd</td><td>CACCGCCGGCTCGTCGCTGCCTCG</td><td></td><td></td><td></td><td></td><td></td><td></td><td></td><td></td><td></td><td></td></tr>
<tr><td>Oligo rev</td><td>AAACCGAGGCAGCGACGAGCCGGC</td><td></td><td></td><td></td><td></td><td></td><td></td><td></td><td></td><td></td><td></td></tr>
<tr><td>Chromosome</td><td>start</td><td>end</td><td>strand</td><td>MM</td><td>target_seq</td><td>PAM</td><td>alignment</td><td>distance</td><td>position</td><td>gene name</td><td>gene id</td></tr>
<tr><td>GU305914.1</td><td>130240</td><td>130262</td><td>-</td><td>0</td><td>GCCGGCTCGTCGCTGCCTCG</td><td>GGG</td><td>||||||||[||||||||||||]PAM</td><td>NA</td><td>Intergenic</td><td>NA</td><td>NA</td></tr>
<tr><td>T24112</td><td>GGCTCGTCGCTGCCTCGGGGCGG</td><td>1000</td><td></td><td></td><td></td><td></td><td></td><td></td><td></td><td></td><td></td></tr>
<tr><td>Oligo fwd</td><td>CACCGGCTCGTCGCTGCCTCGGGG</td><td></td><td></td><td></td><td></td><td></td><td></td><td></td><td></td><td></td><td></td></tr>
<tr><td>Oligo rev</td><td>AAACCCCCGAGGCAGCGACGAGCC</td><td></td><td></td><td></td><td></td><td></td><td></td><td></td><td></td><td></td><td></td></tr>
<tr><td>Chromosome</td><td>start</td><td>end</td><td>strand</td><td>MM</td><td>target_seq</td><td>PAM</td><td>alignment</td><td>distance</td><td>position</td><td>gene name</td><td>gene id</td></tr>
<tr><td>GU305914.1</td><td>130237</td><td>130259</td><td>-</td><td>0</td><td>GGCTCGTCGCTGCCTCGGGG</td><td>CGG</td><td>||||||||[||||||||||||]PAM</td><td>NA</td><td>Intergenic</td><td>NA</td><td>NA</td></tr>
<tr><td>T24113</td><td>CGCTCGCGTGAGCGCCTCTTTGG</td><td>1000</td><td></td><td></td><td></td><td></td><td></td><td></td><td></td><td></td><td></td></tr>
<tr><td>Oligo fwd</td><td>CACCCGCTCGCGTGAGCGCCTCTT</td><td></td><td></td><td></td><td></td><td></td><td></td><td></td><td></td><td></td><td></td></tr>
<tr><td>Oligo rev</td><td>AAACAAGAGGCGCTCACGCGAGCG</td><td></td><td></td><td></td><td></td><td></td><td></td><td></td><td></td><td></td><td></td></tr>
<tr><td>Chromosome</td><td>start</td><td>end</td><td>strand</td><td>MM</td><td>target_seq</td><td>PAM</td><td>alignment</td><td>distance</td><td>position</td><td>gene name</td><td>gene id</td></tr>
<tr><td>GU305914.1</td><td>130191</td><td>130213</td><td>-</td><td>0</td><td>CGCTCGCGTGAGCGCCTCTT</td><td>TGG</td><td>||||||||[||||||||||||]PAM</td><td>NA</td><td>Intergenic</td><td>NA</td><td>NA</td></tr>
<tr><td>T24114</td><td>CGCCTCTTTGGTCCTGTCGATGG</td><td>1000</td><td></td><td></td><td></td><td></td><td></td><td></td><td></td><td></td><td></td></tr>
<tr><td>Oligo fwd</td><td>CACCCGCCTCTTTGGTCCTGTCGA</td><td></td><td></td><td></td><td></td><td></td><td></td><td></td><td></td><td></td><td></td></tr>
<tr><td>Oligo rev</td><td>AAACTCGACAGGACCAAAGAGGCG</td><td></td><td></td><td></td><td></td><td></td><td></td><td></td><td></td><td></td><td></td></tr>
<tr><td>Chromosome</td><td>start</td><td>end</td><td>strand</td><td>MM</td><td>target_seq</td><td>PAM</td><td>alignment</td><td>distance</td><td>position</td><td>gene name</td><td>gene id</td></tr>
<tr><td>GU305914.1</td><td>130179</td><td>130201</td><td>-</td><td>0</td><td>CGCCTCTTTGGTCCTGTCGA</td><td>TGG</td><td>||||||||[||||||||||||]PAM</td><td>NA</td><td>Intergenic</td><td>NA</td><td>NA</td></tr>
</table>

T24115 CAGCCGATGAGCATCGACGCAGG 1000
Oligo fwd CACCCAGCCGATGAGCATCGACGC
Oligo rev AAACGCGTCGATGCTCATCGGCTG
Chromosome start end strand MM target_seq PAM alignment distance position gene name gene id
GU305914.1 130113 130135 - 0 CAGCCGATGAGCATCGACGC AGG ||||||||[||||||||||||]PAM NA Intergenic NA NA

T24116 CGACGCAGGCGTGATCAACGTGG 1000
Oligo fwd CACCCGACGCAGGCGTGATCAACG
Oligo rev AAACCGTTGATCACGCCTGCGTCG
Chromosome start end strand MM target_seq PAM alignment distance position gene name gene id
GU305914.1 130099 130121 - 0 CGACGCAGGCGTGATCAACG TGG ||||||||[||||||||||||]PAM NA Intergenic NA NA

T24117 CGTGATCAACGTGGCGACAGAGG 1000
Oligo fwd CACCCGTGATCAACGTGGCGACAG
Oligo rev AAACCTGTCGCCACGTTGATCACG
Chromosome start end strand MM target_seq PAM alignment distance position gene name gene id
GU305914.1 130090 130112 - 0 CGTGATCAACGTGGCGACAG AGG ||||||||[||||||||||||]PAM NA Intergenic NA NA

T24118 GTGATCAACGTGGCGACAGAGGG 1000
Oligo fwd CACCGTGATCAACGTGGCGACAGA
Oligo rev AAACTCTGTCGCCACGTTGATCAC
Chromosome start end strand MM target_seq PAM alignment distance position gene name gene id
GU305914.1 130089 130111 - 0 GTGATCAACGTGGCGACAGA GGG ||||||||[||||||||||||]PAM NA Intergenic NA NA

T24119 CGTGGCGACAGAGGGTCAAGAGG 1000
Oligo fwd CACCCGTGGCGACAGAGGGTCAAG
Oligo rev AAACCTTGACCCTCTGTCGCCACG
Chromosome start end strand MM target_seq PAM alignment distance position gene name gene id
GU305914.1 130081 130103 - 0 CGTGGCGACAGAGGGTCAAG AGG ||||||||[||||||||||||]PAM NA Intergenic NA NA

T24120 GCGACAGAGGGTCAAGAGGTTGG 1000
Oligo fwd CACCGCGACAGAGGGTCAAGAGGT
Oligo rev AAACACCTCTTGACCCTCTGTCGC
Chromosome start end strand MM target_seq PAM alignment distance position gene name gene id
GU305914.1 130077 130099 - 0 GCGACAGAGGGTCAAGAGGT TGG ||||||||[||||||||||||]PAM NA Intergenic NA NA
GU305914.1 55056 55078 - 4 GCTTCGGAAGGTCAAGAGGT AGG ||--|-||[-|||||||||||]PAM NA Intergenic NA NA

T24121 TGATTTTTTTGTTCTTGACGAGG 1000
Oligo fwd CACCTGATTTTTTTGTTCTTGACG
Oligo rev AAACCGTCAAGAACAAAAAAATCA
Chromosome start end strand MM target_seq PAM alignment distance position gene name gene id
GU305914.1 130030 130052 - 0 TGATTTTTTTGTTCTTGACG AGG ||||||||[||||||||||||]PAM NA Intergenic NA NA

T24122 TTGTTCTTGACGAGGACTTTTGG 1000
Oligo fwd CACCTTGTTCTTGACGAGGACTTT
Oligo rev AAACAAAGTCCTCGTCAAGAACAA
Chromosome start end strand MM target_seq PAM alignment distance position gene name gene id
GU305914.1 130022 130044 - 0 TTGTTCTTGACGAGGACTTT TGG ||||||||[||||||||||||]PAM NA Intergenic NA NA

T24123 TGTTCTTGACGAGGACTTTTGGG 1000
Oligo fwd CACCTGTTCTTGACGAGGACTTTT
Oligo rev AAACAAAAGTCCTCGTCAAGAACA
Chromosome start end strand MM target_seq PAM alignment distance position gene name gene id
GU305914.1 130021 130043 - 0 TGTTCTTGACGAGGACTTTT GGG ||||||||[||||||||||||]PAM NA Intergenic NA NA

T24124 GAGGACTTTTGGGACGTTATTGG 1000
Oligo fwd CACCGAGGACTTTTGGGACGTTAT
Oligo rev AAACATAACGTCCCAAAAGTCCTC
Chromosome start end strand MM target_seq PAM alignment distance position gene name gene id
GU305914.1 130011 130033 - 0 GAGGACTTTTGGGACGTTAT TGG ||||||||[||||||||||||]PAM NA Intergenic NA NA

T24125 AGCGCATGTGTGTTTGATCTTGG 1000
Oligo fwd CACCAGCGCATGTGTGTTTGATCT
Oligo rev AAACAGATCAAACACACATGCGCT
Chromosome start end strand MM target_seq PAM alignment distance position gene name gene id
GU305914.1 129969 129991 - 0 AGCGCATGTGTGTTTGATCT TGG ||||||||[||||||||||||]PAM NA Intergenic NA NA

T24126 TGTGATAATCGAAGACGCCCTGG 1000
Oligo fwd CACCTGTGATAATCGAAGACGCCC
Oligo rev AAACGGGCGTCTTCGATTATCACA
Chromosome start end strand MM target_seq PAM alignment distance position gene name gene id
GU305914.1 129928 129950 - 0 TGTGATAATCGAAGACGCCC TGG ||||||||[||||||||||||]PAM NA Intergenic NA NA

T24127 AGACGCCCTGGCGTATAGCGCGG 1000
Oligo fwd CACCAGACGCCCTGGCGTATAGCG
Oligo rev AAACCGCTATACGCCAGGGCGTCT
Chromosome start end strand MM target_seq PAM alignment distance position gene name gene id
GU305914.1 129916 129938 - 0 AGACGCCCTGGCGTATAGCG CGG ||||||||[||||||||||||]PAM NA Intergenic NA NA

T24128 GTATAGCGCGGAAGATGAGCCGG 1000
Oligo fwd CACCGTATAGCGCGGAAGATGAGC
Oligo rev AAACGCTCATCTTCCGCGCTATAC
Chromosome start end strand MM target_seq PAM alignment distance position gene name gene id
GU305914.1 129904 129926 - 0 GTATAGCGCGGAAGATGAGC CGG ||||||||[||||||||||||]PAM NA Intergenic NA NA

T24129 GCGGAAGATGAGCCGGTCGACGG 1000
Oligo fwd CACCGCGGAAGATGAGCCGGTCGA
Oligo rev AAACTCGACCGGCTCATCTTCCGC

Chromosome start end strand MM target_seq PAM alignment distance position gene name gene id
GU305914.1 129897 129919 - 0 GCGGAAGATGAGCCGGTCGA CGG ||||||||[||||||||||||]PAM NA Intergenic NA NA

T24130 TGTGTTCCAGACCGCAGATGCGG 1000
Oligo fwd CACCTGTGTTCCAGACCGCAGATG
Oligo rev AAACCATCTGCGGTCTGGAACACA
Chromosome start end strand MM target_seq PAM alignment distance position gene name gene id
GU305914.1 129868 129890 - 0 TGTGTTCCAGACCGCAGATG CGG ||||||||[||||||||||||]PAM NA Intergenic NA NA

T24131 ATAAACTGCTTACTCAAAAACGG 1000
Oligo fwd CACCATAAACTGCTTACTCAAAAA
Oligo rev AAACTTTTTGAGTAAGCAGTTTAT
Chromosome start end strand MM target_seq PAM alignment distance position gene name gene id
GU305914.1 129821 129843 - 0 ATAAACTGCTTACTCAAAAA CGG ||||||||[||||||||||||]PAM NA Intergenic NA NA

T24132 GCTTACTCAAAAACGGTATCTGG 1000
Oligo fwd CACCGCTTACTCAAAAACGGTATC
Oligo rev AAACGATACCGTTTTTGAGTAAGC
Chromosome start end strand MM target_seq PAM alignment distance position gene name gene id
GU305914.1 129814 129836 - 0 GCTTACTCAAAAACGGTATC TGG ||||||||[||||||||||||]PAM NA Intergenic NA NA

T24133 CGGTATCTGGTAGCAATTTCTGG 1000
Oligo fwd CACCCGGTATCTGGTAGCAATTTC
Oligo rev AAACGAAATTGCTACCAGATACCG
Chromosome start end strand MM target_seq PAM alignment distance position gene name gene id
GU305914.1 129801 129823 - 0 CGGTATCTGGTAGCAATTTC TGG ||||||||[||||||||||||]PAM NA Intergenic NA NA

T24134 GCAATTTCTGGTCTCGAGCACGG 1000
Oligo fwd CACCGCAATTTCTGGTCTCGAGCA
Oligo rev AAACTGCTCGAGACCAGAAATTGC
Chromosome start end strand MM target_seq PAM alignment distance position gene name gene id
GU305914.1 129789 129811 - 0 GCAATTTCTGGTCTCGAGCA CGG ||||||||[||||||||||||]PAM NA Intergenic NA NA

T24135 TCTCGAGCACGGTAAAGACGCGG 1000
Oligo fwd CACCTCTCGAGCACGGTAAAGACG
Oligo rev AAACCGTCTTTACCGTGCTCGAGA
Chromosome start end strand MM target_seq PAM alignment distance position gene name gene id
GU305914.1 129778 129800 - 0 TCTCGAGCACGGTAAAGACG CGG ||||||||[||||||||||||]PAM NA Intergenic NA NA

T24136 GCACGGTAAAGACGCGGTGCAGG 1000
Oligo fwd CACCGCACGGTAAAGACGCGGTGC
Oligo rev AAACGCACCGCGTCTTTACCGTGC
Chromosome start end strand MM target_seq PAM alignment distance position gene name gene id
GU305914.1 129772 129794 - 0 GCACGGTAAAGACGCGGTGC AGG ||||||||[||||||||||||]PAM NA Intergenic NA NA

T24137 CACGGTAAAGACGCGGTGCAGGG 1000
Oligo fwd CACCCACGGTAAAGACGCGGTGCA
Oligo rev AAACTGCACCGCGTCTTTACCGTG
Chromosome start end strand MM target_seq PAM alignment distance position gene name gene id
GU305914.1 129771 129793 - 0 CACGGTAAAGACGCGGTGCA GGG ||||||||[||||||||||||]PAM NA Intergenic NA NA

T24138 GGGTGTAGTCGACGCAGAGCTGG 1000
Oligo fwd CACCGGGTGTAGTCGACGCAGAGC
Oligo rev AAACGCTCTGCGTCGACTACACCC
Chromosome start end strand MM target_seq PAM alignment distance position gene name gene id
GU305914.1 129751 129773 - 0 GGGTGTAGTCGACGCAGAGC TGG ||||||||[||||||||||||]PAM NA Intergenic NA NA

T24139 GACGCAGAGCTGGTCGATGCTGG 1000
Oligo fwd CACCGACGCAGAGCTGGTCGATGC
Oligo rev AAACGCATCGACCAGCTCTGCGTC
Chromosome start end strand MM target_seq PAM alignment distance position gene name gene id
GU305914.1 129741 129763 - 0 GACGCAGAGCTGGTCGATGC TGG ||||||||[||||||||||||]PAM NA Intergenic NA NA

T24140 CTGGTCGATGCTGGTCACGTAGG 1000
Oligo fwd CACCCTGGTCGATGCTGGTCACGT
Oligo rev AAACACGTGACCAGCATCGACCAG
Chromosome start end strand MM target_seq PAM alignment distance position gene name gene id
GU305914.1 129732 129754 - 0 CTGGTCGATGCTGGTCACGT AGG ||||||||[||||||||||||]PAM NA Intergenic NA NA

T24141 CACGTTCTCGAGCGCGACGCCGG 1000
Oligo fwd CACCCACGTTCTCGAGCGCGACGC
Oligo rev AAACGCGTCGCGCTCGAGAACGTG
Chromosome start end strand MM target_seq PAM alignment distance position gene name gene id
GU305914.1 129693 129715 - 0 CACGTTCTCGAGCGCGACGC CGG ||||||||[||||||||||||]PAM NA Intergenic NA NA
GU305914.1 163129 163151 - 4 CACGTAATCTACCGCGACGC AGG |||||--|[|-|-||||||||]PAM NA Intergenic NA NA

T24142 AGCGCGACGCCGGAGACCTTAGG 1000
Oligo fwd CACCAGCGCGACGCCGGAGACCTT
Oligo rev AAACAAGGTCTCCGGCGTCGCGCT
Chromosome start end strand MM target_seq PAM alignment distance position gene name gene id
GU305914.1 129683 129705 - 0 AGCGCGACGCCGGAGACCTT AGG ||||||||[||||||||||||]PAM NA Intergenic NA NA

T24143 GCCGGAGACCTTAGGTATGATGG 1000
Oligo fwd CACCGCCGGAGACCTTAGGTATGA
Oligo rev AAACTCATACCTAAGGTCTCCGGC
Chromosome start end strand MM target_seq PAM alignment distance position gene name gene id
GU305914.1 129675 129697 - 0 GCCGGAGACCTTAGGTATGA TGG ||||||||[||||||||||||]PAM NA Intergenic NA NA

T24144 GGAGACCTTAGGTATGATGGCGG 1000

Oligo fwd CACCGGAGACCTTAGGTATGATGG
Oligo rev AAACCCATCATACCTAAGGTCTCC
Chromosome start end strand MM target_seq PAM alignment distance position gene name gene id
GU305914.1 129672 129694 - 0 GGAGACCTTAGGTATGATGG CGG ||||||||[||||||||||||]PAM NA Intergenic NA NA

T24145 CTTAGGTATGATGGCGGTCGCGG 1000
Oligo fwd CACCCTTAGGTATGATGGCGGTCG
Oligo rev AAACCGACCGCCATCATACCTAAG
Chromosome start end strand MM target_seq PAM alignment distance position gene name gene id
GU305914.1 129666 129688 - 0 CTTAGGTATGATGGCGGTCG CGG ||||||||[||||||||||||]PAM NA Intergenic NA NA

T24146 CCAGCCGACGTTAATGTTGTCGG 1000
Oligo fwd CACCCCAGCCGACGTTAATGTTGT
Oligo rev AAACACAACATTAACGTCGGCTGG
Chromosome start end strand MM target_seq PAM alignment distance position gene name gene id
GU305914.1 129636 129658 - 0 CCAGCCGACGTTAATGTTGT CGG ||||||||[||||||||||||]PAM NA Intergenic NA NA

T24147 GTCGGTGTTCGAGATCATGCAGG 1000
Oligo fwd CACCGTCGGTGTTCGAGATCATGC
Oligo rev AAACGCATGATCTCGAACACCGAC
Chromosome start end strand MM target_seq PAM alignment distance position gene name gene id
GU305914.1 129618 129640 - 0 GTCGGTGTTCGAGATCATGC AGG ||||||||[||||||||||||]PAM NA Intergenic NA NA

T24148 TCGGTGTTCGAGATCATGCAGGG 1000
Oligo fwd CACCTCGGTGTTCGAGATCATGCA
Oligo rev AAACTGCATGATCTCGAACACCGA
Chromosome start end strand MM target_seq PAM alignment distance position gene name gene id
GU305914.1 129617 129639 - 0 TCGGTGTTCGAGATCATGCA GGG ||||||||[||||||||||||]PAM NA Intergenic NA NA

T24149 CGGTGTTCGAGATCATGCAGGGG 1000
Oligo fwd CACCCGGTGTTCGAGATCATGCAG
Oligo rev AAACCTGCATGATCTCGAACACCG
Chromosome start end strand MM target_seq PAM alignment distance position gene name gene id
GU305914.1 129616 129638 - 0 CGGTGTTCGAGATCATGCAG GGG ||||||||[||||||||||||]PAM NA Intergenic NA NA

T24150 CGAGATCATGCAGGGGAAGTTGG 1000
Oligo fwd CACCCGAGATCATGCAGGGGAAGT
Oligo rev AAACACTTCCCCTGCATGATCTCG
Chromosome start end strand MM target_seq PAM alignment distance position gene name gene id
GU305914.1 129609 129631 - 0 CGAGATCATGCAGGGGAAGT TGG ||||||||[||||||||||||]PAM NA Intergenic NA NA

T24151 GATCATGCAGGGGAAGTTGGTGG 1000
Oligo fwd CACCGATCATGCAGGGGAAGTTGG
Oligo rev AAACCCAACTTCCCCTGCATGATC
Chromosome start end strand MM target_seq PAM alignment distance position gene name gene id
GU305914.1 129606 129628 - 0 GATCATGCAGGGGAAGTTGG TGG ||||||||[||||||||||||]PAM NA Intergenic NA NA

T24152 CCAGTAGACCTTATCCCGCTCGG 1000
Oligo fwd CACCCCAGTAGACCTTATCCCGCT
Oligo rev AAACAGCGGGATAAGGTCTACTGG
Chromosome start end strand MM target_seq PAM alignment distance position gene name gene id
GU305914.1 129582 129604 - 0 CCAGTAGACCTTATCCCGCT CGG ||||||||[||||||||||||]PAM NA Intergenic NA NA

T24153 AGACCTTATCCCGCTCGGATCGG 1000
Oligo fwd CACCAGACCTTATCCCGCTCGGAT
Oligo rev AAACATCCGAGCGGGATAAGGTCT
Chromosome start end strand MM target_seq PAM alignment distance position gene name gene id
GU305914.1 129577 129599 - 0 AGACCTTATCCCGCTCGGAT CGG ||||||||[||||||||||||]PAM NA Intergenic NA NA

T24154 GCTCGGATCGGATGACCCCCTGG 1000
Oligo fwd CACCGCTCGGATCGGATGACCCCC
Oligo rev AAACGGGGGTCATCCGATCCGAGC
Chromosome start end strand MM target_seq PAM alignment distance position gene name gene id
GU305914.1 129565 129587 - 0 GCTCGGATCGGATGACCCCC TGG ||||||||[||||||||||||]PAM NA Intergenic NA NA

T24155 GATCCCGTGTACGTCTAGCGAGG 1000
Oligo fwd CACCGATCCCGTGTACGTCTAGCG
Oligo rev AAACCGCTAGACGTACACGGGATC
Chromosome start end strand MM target_seq PAM alignment distance position gene name gene id
GU305914.1 129537 129559 - 0 GATCCCGTGTACGTCTAGCG AGG ||||||||[||||||||||||]PAM NA Intergenic NA NA

T24156 CGAGGAGTTACTATTGCACACGG 1000
Oligo fwd CACCCGAGGAGTTACTATTGCACA
Oligo rev AAACTGTGCAATAGTAACTCCTCG
Chromosome start end strand MM target_seq PAM alignment distance position gene name gene id
GU305914.1 129519 129541 - 0 CGAGGAGTTACTATTGCACA CGG ||||||||[||||||||||||]PAM NA Intergenic NA NA

T24157 GTTACTATTGCACACGGACGCGG 1000
Oligo fwd CACCGTTACTATTGCACACGGACG
Oligo rev AAACCGTCCGTGTGCAATAGTAAC
Chromosome start end strand MM target_seq PAM alignment distance position gene name gene id
GU305914.1 129513 129535 - 0 GTTACTATTGCACACGGACG CGG ||||||||[||||||||||||]PAM NA Intergenic NA NA

T24158 TTACTATTGCACACGGACGCGGG 1000
Oligo fwd CACCTTACTATTGCACACGGACGC
Oligo rev AAACGCGTCCGTGTGCAATAGTAA
Chromosome start end strand MM target_seq PAM alignment distance position gene name gene id
GU305914.1 129512 129534 - 0 TTACTATTGCACACGGACGC GGG ||||||||[||||||||||||]PAM NA Intergenic NA NA

<table>
<tr><td>T24159</td><td>CATATAGCTTATAGACACTGAGG</td><td>1000</td><td></td><td></td><td></td><td></td><td></td><td></td><td></td><td></td><td></td></tr>
<tr><td>Oligo fwd</td><td>CACCCATATAGCTTATAGACACTG</td><td></td><td></td><td></td><td></td><td></td><td></td><td></td><td></td><td></td><td></td></tr>
<tr><td>Oligo rev</td><td>AAACCAGTGTCTATAAGCTATATG</td><td></td><td></td><td></td><td></td><td></td><td></td><td></td><td></td><td></td><td></td></tr>
<tr><td>Chromosome</td><td>start</td><td>end</td><td>strand</td><td>MM</td><td>target_seq</td><td>PAM</td><td>alignment</td><td>distance</td><td>position</td><td>gene name</td><td>gene id</td></tr>
<tr><td>GU305914.1</td><td>129466</td><td>129488</td><td>-</td><td>0</td><td>CATATAGCTTATAGACACTG</td><td>AGG</td><td>||||||||[||||||||||||]PAM</td><td>NA</td><td>Intergenic</td><td>NA</td><td>NA</td></tr>
<tr><td>T24160</td><td>ATATAGCTTATAGACACTGAGGG</td><td>1000</td><td></td><td></td><td></td><td></td><td></td><td></td><td></td><td></td><td></td></tr>
<tr><td>Oligo fwd</td><td>CACCATATAGCTTATAGACACTGA</td><td></td><td></td><td></td><td></td><td></td><td></td><td></td><td></td><td></td><td></td></tr>
<tr><td>Oligo rev</td><td>AAACTCAGTGTCTATAAGCTATAT</td><td></td><td></td><td></td><td></td><td></td><td></td><td></td><td></td><td></td><td></td></tr>
<tr><td>Chromosome</td><td>start</td><td>end</td><td>strand</td><td>MM</td><td>target_seq</td><td>PAM</td><td>alignment</td><td>distance</td><td>position</td><td>gene name</td><td>gene id</td></tr>
<tr><td>GU305914.1</td><td>129465</td><td>129487</td><td>-</td><td>0</td><td>ATATAGCTTATAGACACTGA</td><td>GGG</td><td>||||||||[||||||||||||]PAM</td><td>NA</td><td>Intergenic</td><td>NA</td><td>NA</td></tr>
<tr><td>T24161</td><td>TATAGACACTGAGGGCGTAGAGG</td><td>1000</td><td></td><td></td><td></td><td></td><td></td><td></td><td></td><td></td><td></td></tr>
<tr><td>Oligo fwd</td><td>CACCTATAGACACTGAGGGCGTAG</td><td></td><td></td><td></td><td></td><td></td><td></td><td></td><td></td><td></td><td></td></tr>
<tr><td>Oligo rev</td><td>AAACCTACGCCCTCAGTGTCTATA</td><td></td><td></td><td></td><td></td><td></td><td></td><td></td><td></td><td></td><td></td></tr>
<tr><td>Chromosome</td><td>start</td><td>end</td><td>strand</td><td>MM</td><td>target_seq</td><td>PAM</td><td>alignment</td><td>distance</td><td>position</td><td>gene name</td><td>gene id</td></tr>
<tr><td>GU305914.1</td><td>129457</td><td>129479</td><td>-</td><td>0</td><td>TATAGACACTGAGGGCGTAG</td><td>AGG</td><td>||||||||[||||||||||||]PAM</td><td>NA</td><td>Intergenic</td><td>NA</td><td>NA</td></tr>
<tr><td>T24162</td><td>TGAGGGCGTAGAGGAACTCCTGG</td><td>1000</td><td></td><td></td><td></td><td></td><td></td><td></td><td></td><td></td><td></td></tr>
<tr><td>Oligo fwd</td><td>CACCTGAGGGCGTAGAGGAACTCC</td><td></td><td></td><td></td><td></td><td></td><td></td><td></td><td></td><td></td><td></td></tr>
<tr><td>Oligo rev</td><td>AAACGGAGTTCCTCTACGCCCTCA</td><td></td><td></td><td></td><td></td><td></td><td></td><td></td><td></td><td></td><td></td></tr>
<tr><td>Chromosome</td><td>start</td><td>end</td><td>strand</td><td>MM</td><td>target_seq</td><td>PAM</td><td>alignment</td><td>distance</td><td>position</td><td>gene name</td><td>gene id</td></tr>
<tr><td>GU305914.1</td><td>129448</td><td>129470</td><td>-</td><td>0</td><td>TGAGGGCGTAGAGGAACTCC</td><td>TGG</td><td>||||||||[||||||||||||]PAM</td><td>NA</td><td>Intergenic</td><td>NA</td><td>NA</td></tr>
<tr><td>T24163</td><td>GGCGTAGAGGAACTCCTGGTAGG</td><td>1000</td><td></td><td></td><td></td><td></td><td></td><td></td><td></td><td></td><td></td></tr>
<tr><td>Oligo fwd</td><td>CACCGGCGTAGAGGAACTCCTGGT</td><td></td><td></td><td></td><td></td><td></td><td></td><td></td><td></td><td></td><td></td></tr>
<tr><td>Oligo rev</td><td>AAACACCAGGAGTTCCTCTACGCC</td><td></td><td></td><td></td><td></td><td></td><td></td><td></td><td></td><td></td><td></td></tr>
<tr><td>Chromosome</td><td>start</td><td>end</td><td>strand</td><td>MM</td><td>target_seq</td><td>PAM</td><td>alignment</td><td>distance</td><td>position</td><td>gene name</td><td>gene id</td></tr>
<tr><td>GU305914.1</td><td>129444</td><td>129466</td><td>-</td><td>0</td><td>GGCGTAGAGGAACTCCTGGT</td><td>AGG</td><td>||||||||[||||||||||||]PAM</td><td>NA</td><td>Intergenic</td><td>NA</td><td>NA</td></tr>
<tr><td>T24164</td><td>TCACGTTTAGAGAGTTTGTGTGG</td><td>1000</td><td></td><td></td><td></td><td></td><td></td><td></td><td></td><td></td><td></td></tr>
<tr><td>Oligo fwd</td><td>CACCTCACGTTTAGAGAGTTTGTG</td><td></td><td></td><td></td><td></td><td></td><td></td><td></td><td></td><td></td><td></td></tr>
<tr><td>Oligo rev</td><td>AAACCACAAACTCTCTAAACGTGA</td><td></td><td></td><td></td><td></td><td></td><td></td><td></td><td></td><td></td><td></td></tr>
<tr><td>Chromosome</td><td>start</td><td>end</td><td>strand</td><td>MM</td><td>target_seq</td><td>PAM</td><td>alignment</td><td>distance</td><td>position</td><td>gene name</td><td>gene id</td></tr>
<tr><td>GU305914.1</td><td>129409</td><td>129431</td><td>-</td><td>0</td><td>TCACGTTTAGAGAGTTTGTG</td><td>TGG</td><td>||||||||[||||||||||||]PAM</td><td>NA</td><td>Intergenic</td><td>NA</td><td>NA</td></tr>
<tr><td>T24165</td><td>CACGTTTAGAGAGTTTGTGTGGG</td><td>1000</td><td></td><td></td><td></td><td></td><td></td><td></td><td></td><td></td><td></td></tr>
<tr><td>Oligo fwd</td><td>CACCCACGTTTAGAGAGTTTGTGT</td><td></td><td></td><td></td><td></td><td></td><td></td><td></td><td></td><td></td><td></td></tr>
<tr><td>Oligo rev</td><td>AAACACACAAACTCTCTAAACGTG</td><td></td><td></td><td></td><td></td><td></td><td></td><td></td><td></td><td></td><td></td></tr>
<tr><td>Chromosome</td><td>start</td><td>end</td><td>strand</td><td>MM</td><td>target_seq</td><td>PAM</td><td>alignment</td><td>distance</td><td>position</td><td>gene name</td><td>gene id</td></tr>
<tr><td>GU305914.1</td><td>129408</td><td>129430</td><td>-</td><td>0</td><td>CACGTTTAGAGAGTTTGTGT</td><td>GGG</td><td>||||||||[||||||||||||]PAM</td><td>NA</td><td>Intergenic</td><td>NA</td><td>NA</td></tr>
<tr><td>T24166</td><td>ACGAATGTGCTGACCGTATATGG</td><td>1000</td><td></td><td></td><td></td><td></td><td></td><td></td><td></td><td></td><td></td></tr>
<tr><td>Oligo fwd</td><td>CACCACGAATGTGCTGACCGTATA</td><td></td><td></td><td></td><td></td><td></td><td></td><td></td><td></td><td></td><td></td></tr>
<tr><td>Oligo rev</td><td>AAACTATACGGTCAGCACATTCGT</td><td></td><td></td><td></td><td></td><td></td><td></td><td></td><td></td><td></td><td></td></tr>
<tr><td>Chromosome</td><td>start</td><td>end</td><td>strand</td><td>MM</td><td>target_seq</td><td>PAM</td><td>alignment</td><td>distance</td><td>position</td><td>gene name</td><td>gene id</td></tr>
<tr><td>GU305914.1</td><td>129371</td><td>129393</td><td>-</td><td>0</td><td>ACGAATGTGCTGACCGTATA</td><td>TGG</td><td>||||||||[||||||||||||]PAM</td><td>NA</td><td>Intergenic</td><td>NA</td><td>NA</td></tr>
<tr><td>T24167</td><td>CGAATGTGCTGACCGTATATGGG</td><td>1000</td><td></td><td></td><td></td><td></td><td></td><td></td><td></td><td></td><td></td></tr>
<tr><td>Oligo fwd</td><td>CACCCGAATGTGCTGACCGTATAT</td><td></td><td></td><td></td><td></td><td></td><td></td><td></td><td></td><td></td><td></td></tr>
<tr><td>Oligo rev</td><td>AAACATATACGGTCAGCACATTCG</td><td></td><td></td><td></td><td></td><td></td><td></td><td></td><td></td><td></td><td></td></tr>
<tr><td>Chromosome</td><td>start</td><td>end</td><td>strand</td><td>MM</td><td>target_seq</td><td>PAM</td><td>alignment</td><td>distance</td><td>position</td><td>gene name</td><td>gene id</td></tr>
<tr><td>GU305914.1</td><td>129370</td><td>129392</td><td>-</td><td>0</td><td>CGAATGTGCTGACCGTATAT</td><td>GGG</td><td>||||||||[||||||||||||]PAM</td><td>NA</td><td>Intergenic</td><td>NA</td><td>NA</td></tr>
<tr><td>T24168</td><td>GAATGTGCTGACCGTATATGGGG</td><td>1000</td><td></td><td></td><td></td><td></td><td></td><td></td><td></td><td></td><td></td></tr>
<tr><td>Oligo fwd</td><td>CACCGAATGTGCTGACCGTATATG</td><td></td><td></td><td></td><td></td><td></td><td></td><td></td><td></td><td></td><td></td></tr>
<tr><td>Oligo rev</td><td>AAACCATATACGGTCAGCACATTC</td><td></td><td></td><td></td><td></td><td></td><td></td><td></td><td></td><td></td><td></td></tr>
<tr><td>Chromosome</td><td>start</td><td>end</td><td>strand</td><td>MM</td><td>target_seq</td><td>PAM</td><td>alignment</td><td>distance</td><td>position</td><td>gene name</td><td>gene id</td></tr>
<tr><td>GU305914.1</td><td>129369</td><td>129391</td><td>-</td><td>0</td><td>GAATGTGCTGACCGTATATG</td><td>GGG</td><td>||||||||[||||||||||||]PAM</td><td>NA</td><td>Intergenic</td><td>NA</td><td>NA</td></tr>
<tr><td>T24169</td><td>AATGTGCTGACCGTATATGGGGG</td><td>1000</td><td></td><td></td><td></td><td></td><td></td><td></td><td></td><td></td><td></td></tr>
<tr><td>Oligo fwd</td><td>CACCAATGTGCTGACCGTATATGG</td><td></td><td></td><td></td><td></td><td></td><td></td><td></td><td></td><td></td><td></td></tr>
<tr><td>Oligo rev</td><td>AAACCCATATACGGTCAGCACATT</td><td></td><td></td><td></td><td></td><td></td><td></td><td></td><td></td><td></td><td></td></tr>
<tr><td>Chromosome</td><td>start</td><td>end</td><td>strand</td><td>MM</td><td>target_seq</td><td>PAM</td><td>alignment</td><td>distance</td><td>position</td><td>gene name</td><td>gene id</td></tr>
<tr><td>GU305914.1</td><td>129368</td><td>129390</td><td>-</td><td>0</td><td>AATGTGCTGACCGTATATGG</td><td>GGG</td><td>||||||||[||||||||||||]PAM</td><td>NA</td><td>Intergenic</td><td>NA</td><td>NA</td></tr>
<tr><td>T24170</td><td>CGTATATGGGGGTTCTTTGAAGG</td><td>1000</td><td></td><td></td><td></td><td></td><td></td><td></td><td></td><td></td><td></td></tr>
<tr><td>Oligo fwd</td><td>CACCCGTATATGGGGGTTCTTTGA</td><td></td><td></td><td></td><td></td><td></td><td></td><td></td><td></td><td></td><td></td></tr>
<tr><td>Oligo rev</td><td>AAACTCAAAGAACCCCCATATACG</td><td></td><td></td><td></td><td></td><td></td><td></td><td></td><td></td><td></td><td></td></tr>
<tr><td>Chromosome</td><td>start</td><td>end</td><td>strand</td><td>MM</td><td>target_seq</td><td>PAM</td><td>alignment</td><td>distance</td><td>position</td><td>gene name</td><td>gene id</td></tr>
<tr><td>GU305914.1</td><td>129357</td><td>129379</td><td>-</td><td>0</td><td>CGTATATGGGGGTTCTTTGA</td><td>AGG</td><td>||||||||[||||||||||||]PAM</td><td>NA</td><td>Intergenic</td><td>NA</td><td>NA</td></tr>
<tr><td>T24171</td><td>AAGCTGTGAGTCATGTTTCGAGG</td><td>1000</td><td></td><td></td><td></td><td></td><td></td><td></td><td></td><td></td><td></td></tr>
<tr><td>Oligo fwd</td><td>CACCAAGCTGTGAGTCATGTTTCG</td><td></td><td></td><td></td><td></td><td></td><td></td><td></td><td></td><td></td><td></td></tr>
<tr><td>Oligo rev</td><td>AAACCGAAACATGACTCACAGCTT</td><td></td><td></td><td></td><td></td><td></td><td></td><td></td><td></td><td></td><td></td></tr>
<tr><td>Chromosome</td><td>start</td><td>end</td><td>strand</td><td>MM</td><td>target_seq</td><td>PAM</td><td>alignment</td><td>distance</td><td>position</td><td>gene name</td><td>gene id</td></tr>
<tr><td>GU305914.1</td><td>129317</td><td>129339</td><td>-</td><td>0</td><td>AAGCTGTGAGTCATGTTTCG</td><td>AGG</td><td>||||||||[||||||||||||]PAM</td><td>NA</td><td>Intergenic</td><td>NA</td><td>NA</td></tr>
<tr><td>T24172</td><td>AGCTGTGAGTCATGTTTCGAGGG</td><td>1000</td><td></td><td></td><td></td><td></td><td></td><td></td><td></td><td></td><td></td></tr>
<tr><td>Oligo fwd</td><td>CACCAGCTGTGAGTCATGTTTCGA</td><td></td><td></td><td></td><td></td><td></td><td></td><td></td><td></td><td></td><td></td></tr>
<tr><td>Oligo rev</td><td>AAACTCGAAACATGACTCACAGCT</td><td></td><td></td><td></td><td></td><td></td><td></td><td></td><td></td><td></td><td></td></tr>
<tr><td>Chromosome</td><td>start</td><td>end</td><td>strand</td><td>MM</td><td>target_seq</td><td>PAM</td><td>alignment</td><td>distance</td><td>position</td><td>gene name</td><td>gene id</td></tr>
<tr><td>GU305914.1</td><td>129316</td><td>129338</td><td>-</td><td>0</td><td>AGCTGTGAGTCATGTTTCGA</td><td>GGG</td><td>||||||||[||||||||||||]PAM</td><td>NA</td><td>Intergenic</td><td>NA</td><td>NA</td></tr>
<tr><td>T24173</td><td>GACATAAAACGCAGACAACCTGG</td><td>1000</td><td></td><td></td><td></td><td></td><td></td><td></td><td></td><td></td><td></td></tr>
<tr><td>Oligo fwd</td><td>CACCGACATAAAACGCAGACAACC</td><td></td><td></td><td></td><td></td><td></td><td></td><td></td><td></td><td></td><td></td></tr>
<tr><td>Oligo rev</td><td>AAACGGTTGTCTGCGTTTTATGTC</td><td></td><td></td><td></td><td></td><td></td><td></td><td></td><td></td><td></td><td></td></tr>
<tr><td>Chromosome</td><td>start</td><td>end</td><td>strand</td><td>MM</td><td>target_seq</td><td>PAM</td><td>alignment</td><td>distance</td><td>position</td><td>gene name</td><td>gene id</td></tr>
<tr><td>GU305914.1</td><td>129294</td><td>129316</td><td>-</td><td>0</td><td>GACATAAAACGCAGACAACC</td><td>TGG</td><td>||||||||[||||||||||||]PAM</td><td>NA</td><td>Intergenic</td><td>NA</td><td>NA</td></tr>
</table>

| | | | | | | | | | | | | |
|---|---|---|---|---|---|---|---|---|---|---|---|---|
| T24174 | ACATAAAACGCAGACAACCTGGG | | 1000 | | | | | | | | | |
| Oligo fwd | CACCACATAAAACGCAGACAACCT | | | | | | | | | | | |
| Oligo rev | AAACAGGTTGTCTGCGTTTTATGT | | | | | | | | | | | |
| Chromosome | start | | end | strand | MM | target_seq | PAM | alignment | distance | position | gene name | gene id |
| GU305914.1 | | 129293 | 129315 | - | 0 | ACATAAAACGCAGACAACCT | GGG | \|\|\|\|\|\|\|\|[\|\|\|\|\|\|\|\|\|\|\|\|]PAM | NA | Intergenic | NA | NA |

| | | | | | | | | | | | | |
|---|---|---|---|---|---|---|---|---|---|---|---|---|
| T24175 | CAACCTGGGCGTCTGCACGATGG | | 1000 | | | | | | | | | |
| Oligo fwd | CACCCAACCTGGGCGTCTGCACGA | | | | | | | | | | | |
| Oligo rev | AAACTCGTGCAGACGCCCAGGTTG | | | | | | | | | | | |
| Chromosome | start | | end | strand | MM | target_seq | PAM | alignment | distance | position | gene name | gene id |
| GU305914.1 | | 129279 | 129301 | - | 0 | CAACCTGGGCGTCTGCACGA | TGG | \|\|\|\|\|\|\|\|[\|\|\|\|\|\|\|\|\|\|\|\|]PAM | NA | Intergenic | NA | NA |

| | | | | | | | | | | | | |
|---|---|---|---|---|---|---|---|---|---|---|---|---|
| T24176 | TCTGCACGATGGTCGCTTTGAGG | | 1000 | | | | | | | | | |
| Oligo fwd | CACCTCTGCACGATGGTCGCTTTG | | | | | | | | | | | |
| Oligo rev | AAACCAAAGCGACCATCGTGCAGA | | | | | | | | | | | |
| Chromosome | start | | end | strand | MM | target_seq | PAM | alignment | distance | position | gene name | gene id |
| GU305914.1 | | 129268 | 129290 | - | 0 | TCTGCACGATGGTCGCTTTG | AGG | \|\|\|\|\|\|\|\|[\|\|\|\|\|\|\|\|\|\|\|\|]PAM | NA | Intergenic | NA | NA |

| | | | | | | | | | | | | |
|---|---|---|---|---|---|---|---|---|---|---|---|---|
| T24177 | TCGCTTTGAGGTTGATGACGCGG | | 1000 | | | | | | | | | |
| Oligo fwd | CACCTCGCTTTGAGGTTGATGACG | | | | | | | | | | | |
| Oligo rev | AAACCGTCATCAACCTCAAAGCGA | | | | | | | | | | | |
| Chromosome | start | | end | strand | MM | target_seq | PAM | alignment | distance | position | gene name | gene id |
| GU305914.1 | | 129256 | 129278 | - | 0 | TCGCTTTGAGGTTGATGACG | CGG | \|\|\|\|\|\|\|\|[\|\|\|\|\|\|\|\|\|\|\|\|]PAM | NA | Intergenic | NA | NA |

| | | | | | | | | | | | | |
|---|---|---|---|---|---|---|---|---|---|---|---|---|
| T24178 | TTTGAGGTTGATGACGCGGTTGG | | 1000 | | | | | | | | | |
| Oligo fwd | CACCTTTGAGGTTGATGACGCGGT | | | | | | | | | | | |
| Oligo rev | AAACACCGCGTCATCAACCTCAAA | | | | | | | | | | | |
| Chromosome | start | | end | strand | MM | target_seq | PAM | alignment | distance | position | gene name | gene id |
| GU305914.1 | | 129252 | 129274 | - | 0 | TTTGAGGTTGATGACGCGGT | TGG | \|\|\|\|\|\|\|\|[\|\|\|\|\|\|\|\|\|\|\|\|]PAM | NA | Intergenic | NA | NA |

| | | | | | | | | | | | | |
|---|---|---|---|---|---|---|---|---|---|---|---|---|
| T24179 | GATGACGCGGTTGGTGTTGAAGG | | 1000 | | | | | | | | | |
| Oligo fwd | CACCGATGACGCGGTTGGTGTTGA | | | | | | | | | | | |
| Oligo rev | AAACTCAACACCAACCGCGTCATC | | | | | | | | | | | |
| Chromosome | start | | end | strand | MM | target_seq | PAM | alignment | distance | position | gene name | gene id |
| GU305914.1 | | 129243 | 129265 | - | 0 | GATGACGCGGTTGGTGTTGA | AGG | \|\|\|\|\|\|\|\|[\|\|\|\|\|\|\|\|\|\|\|\|]PAM | NA | Intergenic | NA | NA |

| | | | | | | | | | | | | |
|---|---|---|---|---|---|---|---|---|---|---|---|---|
| T24180 | GTTGGTGTTGAAGGCATTGACGG | | 1000 | | | | | | | | | |
| Oligo fwd | CACCGTTGGTGTTGAAGGCATTGA | | | | | | | | | | | |
| Oligo rev | AAACTCAATGCCTTCAACACCAAC | | | | | | | | | | | |
| Chromosome | start | | end | strand | MM | target_seq | PAM | alignment | distance | position | gene name | gene id |
| GU305914.1 | | 129234 | 129256 | - | 0 | GTTGGTGTTGAAGGCATTGA | CGG | \|\|\|\|\|\|\|\|[\|\|\|\|\|\|\|\|\|\|\|\|]PAM | NA | Intergenic | NA | NA |

| | | | | | | | | | | | | |
|---|---|---|---|---|---|---|---|---|---|---|---|---|
| T24181 | GGCCACCCGATTGAGCGAGACGG | | 1000 | | | | | | | | | |
| Oligo fwd | CACCGGCCACCCGATTGAGCGAGA | | | | | | | | | | | |
| Oligo rev | AAACTCTCGCTCAATCGGGTGGCC | | | | | | | | | | | |
| Chromosome | start | | end | strand | MM | target_seq | PAM | alignment | distance | position | gene name | gene id |
| GU305914.1 | | 129213 | 129235 | - | 0 | GGCCACCCGATTGAGCGAGA | CGG | \|\|\|\|\|\|\|\|[\|\|\|\|\|\|\|\|\|\|\|\|]PAM | NA | Intergenic | NA | NA |

| | | | | | | | | | | | | |
|---|---|---|---|---|---|---|---|---|---|---|---|---|
| T24182 | GATCTTACGTGCGCCGTAAACGG | | 1000 | | | | | | | | | |
| Oligo fwd | CACCGATCTTACGTGCGCCGTAAA | | | | | | | | | | | |
| Oligo rev | AAACTTTACGGCGCACGTAAGATC | | | | | | | | | | | |
| Chromosome | start | | end | strand | MM | target_seq | PAM | alignment | distance | position | gene name | gene id |
| GU305914.1 | | 129159 | 129181 | - | 0 | GATCTTACGTGCGCCGTAAA | CGG | \|\|\|\|\|\|\|\|[\|\|\|\|\|\|\|\|\|\|\|\|]PAM | NA | Intergenic | NA | NA |

| | | | | | | | | | | | | |
|---|---|---|---|---|---|---|---|---|---|---|---|---|
| T24183 | GCCCTCGCGTCGCACGCCCTCGG | | 1000 | | | | | | | | | |
| Oligo fwd | CACCGCCCTCGCGTCGCACGCCCT | | | | | | | | | | | |
| Oligo rev | AAACAGGGCGTGCGACGCGAGGGC | | | | | | | | | | | |
| Chromosome | start | | end | strand | MM | target_seq | PAM | alignment | distance | position | gene name | gene id |
| GU305914.1 | | 129129 | 129151 | - | 0 | GCCCTCGCGTCGCACGCCCT | CGG | \|\|\|\|\|\|\|\|[\|\|\|\|\|\|\|\|\|\|\|\|]PAM | NA | Intergenic | NA | NA |

| | | | | | | | | | | | | |
|---|---|---|---|---|---|---|---|---|---|---|---|---|
| T24184 | CGGCCGCACGATGTCTGACGAGG | | 1000 | | | | | | | | | |
| Oligo fwd | CACCCGGCCGCACGATGTCTGACG | | | | | | | | | | | |
| Oligo rev | AAACCGTCAGACATCGTGCGGCCG | | | | | | | | | | | |
| Chromosome | start | | end | strand | MM | target_seq | PAM | alignment | distance | position | gene name | gene id |
| GU305914.1 | | 129109 | 129131 | - | 0 | CGGCCGCACGATGTCTGACG | AGG | \|\|\|\|\|\|\|\|[\|\|\|\|\|\|\|\|\|\|\|\|]PAM | NA | Intergenic | NA | NA |

| | | | | | | | | | | | | |
|---|---|---|---|---|---|---|---|---|---|---|---|---|
| T24185 | CACGATGTCTGACGAGGCATCGG | | 1000 | | | | | | | | | |
| Oligo fwd | CACCCACGATGTCTGACGAGGCAT | | | | | | | | | | | |
| Oligo rev | AAACATGCCTCGTCAGACATCGTG | | | | | | | | | | | |
| Chromosome | start | | end | strand | MM | target_seq | PAM | alignment | distance | position | gene name | gene id |
| GU305914.1 | | 129103 | 129125 | - | 0 | CACGATGTCTGACGAGGCAT | CGG | \|\|\|\|\|\|\|\|[\|\|\|\|\|\|\|\|\|\|\|\|]PAM | NA | Intergenic | NA | NA |

| | | | | | | | | | | | | |
|---|---|---|---|---|---|---|---|---|---|---|---|---|
| T24186 | TGTCTGACGAGGCATCGGTACGG | | 1000 | | | | | | | | | |
| Oligo fwd | CACCTGTCTGACGAGGCATCGGTA | | | | | | | | | | | |
| Oligo rev | AAACTACCGATGCCTCGTCAGACA | | | | | | | | | | | |
| Chromosome | start | | end | strand | MM | target_seq | PAM | alignment | distance | position | gene name | gene id |
| GU305914.1 | | 129098 | 129120 | - | 0 | TGTCTGACGAGGCATCGGTA | CGG | \|\|\|\|\|\|\|\|[\|\|\|\|\|\|\|\|\|\|\|\|]PAM | NA | Intergenic | NA | NA |

| | | | | | | | | | | | | |
|---|---|---|---|---|---|---|---|---|---|---|---|---|
| T24187 | GTCTGACGAGGCATCGGTACGGG | | 1000 | | | | | | | | | |
| Oligo fwd | CACCGTCTGACGAGGCATCGGTAC | | | | | | | | | | | |
| Oligo rev | AAACGTACCGATGCCTCGTCAGAC | | | | | | | | | | | |
| Chromosome | start | | end | strand | MM | target_seq | PAM | alignment | distance | position | gene name | gene id |
| GU305914.1 | | 129097 | 129119 | - | 0 | GTCTGACGAGGCATCGGTAC | GGG | \|\|\|\|\|\|\|\|[\|\|\|\|\|\|\|\|\|\|\|\|]PAM | NA | Intergenic | NA | NA |

| | | | | | | | | | | | | |
|---|---|---|---|---|---|---|---|---|---|---|---|---|
| T24188 | TACGGGTCGAAATCGAAACTCGG | | 1000 | | | | | | | | | |
| Oligo fwd | CACCTACGGGTCGAAATCGAAACT | | | | | | | | | | | |
| Oligo rev | AAACAGTTTCGATTTCGACCCGTA | | | | | | | | | | | |
| Chromosome | start | | end | strand | MM | target_seq | PAM | alignment | distance | position | gene name | gene id |

<table>
<tr><td>GU305914.1</td><td></td><td>129080</td><td>129102</td><td>-</td><td>0</td><td>TACGGGTCGAAATCGAAACT</td><td>CGG</td><td>||||||||[||||||||||||]PAM</td><td>NA</td><td>Intergenic</td><td>NA</td><td>NA</td></tr>
<tr><td>T24189</td><td>GTCGAAATCGAAACTCGGCACGG</td><td></td><td>1000</td><td></td><td></td><td></td><td></td><td></td><td></td><td></td><td></td><td></td></tr>
<tr><td>Oligo fwd</td><td>CACCGTCGAAATCGAAACTCGGCA</td><td></td><td></td><td></td><td></td><td></td><td></td><td></td><td></td><td></td><td></td><td></td></tr>
<tr><td>Oligo rev</td><td>AAACTGCCGAGTTTCGATTTCGAC</td><td></td><td></td><td></td><td></td><td></td><td></td><td></td><td></td><td></td><td></td><td></td></tr>
<tr><td>Chromosome</td><td>start</td><td></td><td>end</td><td>strand</td><td>MM</td><td>target_seq</td><td>PAM</td><td>alignment</td><td>distance</td><td>position</td><td>gene name</td><td>gene id</td></tr>
<tr><td>GU305914.1</td><td></td><td>129075</td><td>129097</td><td>-</td><td>0</td><td>GTCGAAATCGAAACTCGGCA</td><td>CGG</td><td>||||||||[||||||||||||]PAM</td><td>NA</td><td>Intergenic</td><td>NA</td><td>NA</td></tr>
<tr><td>T24190</td><td>TCGAAACTCGGCACGGCTACTGG</td><td></td><td>1000</td><td></td><td></td><td></td><td></td><td></td><td></td><td></td><td></td><td></td></tr>
<tr><td>Oligo fwd</td><td>CACCTCGAAACTCGGCACGGCTAC</td><td></td><td></td><td></td><td></td><td></td><td></td><td></td><td></td><td></td><td></td><td></td></tr>
<tr><td>Oligo rev</td><td>AAACGTAGCCGTGCCGAGTTTCGA</td><td></td><td></td><td></td><td></td><td></td><td></td><td></td><td></td><td></td><td></td><td></td></tr>
<tr><td>Chromosome</td><td>start</td><td></td><td>end</td><td>strand</td><td>MM</td><td>target_seq</td><td>PAM</td><td>alignment</td><td>distance</td><td>position</td><td>gene name</td><td>gene id</td></tr>
<tr><td>GU305914.1</td><td></td><td>129068</td><td>129090</td><td>-</td><td>0</td><td>TCGAAACTCGGCACGGCTAC</td><td>TGG</td><td>||||||||[||||||||||||]PAM</td><td>NA</td><td>Intergenic</td><td>NA</td><td>NA</td></tr>
<tr><td>T24191</td><td>TGGCTCGTCCGCCTTCACGTCGG</td><td></td><td>1000</td><td></td><td></td><td></td><td></td><td></td><td></td><td></td><td></td><td></td></tr>
<tr><td>Oligo fwd</td><td>CACCTGGCTCGTCCGCCTTCACGT</td><td></td><td></td><td></td><td></td><td></td><td></td><td></td><td></td><td></td><td></td><td></td></tr>
<tr><td>Oligo rev</td><td>AAACACGTGAAGGCGGACGAGCCA</td><td></td><td></td><td></td><td></td><td></td><td></td><td></td><td></td><td></td><td></td><td></td></tr>
<tr><td>Chromosome</td><td>start</td><td></td><td>end</td><td>strand</td><td>MM</td><td>target_seq</td><td>PAM</td><td>alignment</td><td>distance</td><td>position</td><td>gene name</td><td>gene id</td></tr>
<tr><td>GU305914.1</td><td></td><td>129048</td><td>129070</td><td>-</td><td>0</td><td>TGGCTCGTCCGCCTTCACGT</td><td>CGG</td><td>||||||||[||||||||||||]PAM</td><td>NA</td><td>Intergenic</td><td>NA</td><td>NA</td></tr>
<tr><td>T24192</td><td>CACGTCGGTCTGAACGCCAGAGG</td><td></td><td>1000</td><td></td><td></td><td></td><td></td><td></td><td></td><td></td><td></td><td></td></tr>
<tr><td>Oligo fwd</td><td>CACCCACGTCGGTCTGAACGCCAG</td><td></td><td></td><td></td><td></td><td></td><td></td><td></td><td></td><td></td><td></td><td></td></tr>
<tr><td>Oligo rev</td><td>AAACCTGGCGTTCAGACCGACGTG</td><td></td><td></td><td></td><td></td><td></td><td></td><td></td><td></td><td></td><td></td><td></td></tr>
<tr><td>Chromosome</td><td>start</td><td></td><td>end</td><td>strand</td><td>MM</td><td>target_seq</td><td>PAM</td><td>alignment</td><td>distance</td><td>position</td><td>gene name</td><td>gene id</td></tr>
<tr><td>GU305914.1</td><td></td><td>129033</td><td>129055</td><td>-</td><td>0</td><td>CACGTCGGTCTGAACGCCAG</td><td>AGG</td><td>||||||||[||||||||||||]PAM</td><td>NA</td><td>Intergenic</td><td>NA</td><td>NA</td></tr>
<tr><td>T24193</td><td>CTGAACGCCAGAGGTGCCGCCGG</td><td></td><td>1000</td><td></td><td></td><td></td><td></td><td></td><td></td><td></td><td></td><td></td></tr>
<tr><td>Oligo fwd</td><td>CACCCTGAACGCCAGAGGTGCCGC</td><td></td><td></td><td></td><td></td><td></td><td></td><td></td><td></td><td></td><td></td><td></td></tr>
<tr><td>Oligo rev</td><td>AAACGCGGCACCTCTGGCGTTCAG</td><td></td><td></td><td></td><td></td><td></td><td></td><td></td><td></td><td></td><td></td><td></td></tr>
<tr><td>Chromosome</td><td>start</td><td></td><td>end</td><td>strand</td><td>MM</td><td>target_seq</td><td>PAM</td><td>alignment</td><td>distance</td><td>position</td><td>gene name</td><td>gene id</td></tr>
<tr><td>GU305914.1</td><td></td><td>129024</td><td>129046</td><td>-</td><td>0</td><td>CTGAACGCCAGAGGTGCCGC</td><td>CGG</td><td>||||||||[||||||||||||]PAM</td><td>NA</td><td>Intergenic</td><td>NA</td><td>NA</td></tr>
<tr><td>T24194</td><td>GCCAGAGGTGCCGCCGGCGACGG</td><td></td><td>1000</td><td></td><td></td><td></td><td></td><td></td><td></td><td></td><td></td><td></td></tr>
<tr><td>Oligo fwd</td><td>CACCGCCAGAGGTGCCGCCGGCGA</td><td></td><td></td><td></td><td></td><td></td><td></td><td></td><td></td><td></td><td></td><td></td></tr>
<tr><td>Oligo rev</td><td>AAACTCGCCGGCGGCACCTCTGGC</td><td></td><td></td><td></td><td></td><td></td><td></td><td></td><td></td><td></td><td></td><td></td></tr>
<tr><td>Chromosome</td><td>start</td><td></td><td>end</td><td>strand</td><td>MM</td><td>target_seq</td><td>PAM</td><td>alignment</td><td>distance</td><td>position</td><td>gene name</td><td>gene id</td></tr>
<tr><td>GU305914.1</td><td></td><td>129018</td><td>129040</td><td>-</td><td>0</td><td>GCCAGAGGTGCCGCCGGCGA</td><td>CGG</td><td>||||||||[||||||||||||]PAM</td><td>NA</td><td>Intergenic</td><td>NA</td><td>NA</td></tr>
<tr><td>T24195</td><td>GCTGCCGCTGCCGCTGCCGACGG</td><td></td><td>1000</td><td></td><td></td><td></td><td></td><td></td><td></td><td></td><td></td><td></td></tr>
<tr><td>Oligo fwd</td><td>CACCGCTGCCGCTGCCGCTGCCGA</td><td></td><td></td><td></td><td></td><td></td><td></td><td></td><td></td><td></td><td></td><td></td></tr>
<tr><td>Oligo rev</td><td>AAACTCGGCAGCGGCAGCGGCAGC</td><td></td><td></td><td></td><td></td><td></td><td></td><td></td><td></td><td></td><td></td><td></td></tr>
<tr><td>Chromosome</td><td>start</td><td></td><td>end</td><td>strand</td><td>MM</td><td>target_seq</td><td>PAM</td><td>alignment</td><td>distance</td><td>position</td><td>gene name</td><td>gene id</td></tr>
<tr><td>GU305914.1</td><td></td><td>128985</td><td>129007</td><td>-</td><td>0</td><td>GCTGCCGCTGCCGCTGCCGA</td><td>CGG</td><td>||||||||[||||||||||||]PAM</td><td>NA</td><td>Intergenic</td><td>NA</td><td>NA</td></tr>
<tr><td>T24196</td><td>GCCGCTGCCGACGGCTGCCTCGG</td><td></td><td>1000</td><td></td><td></td><td></td><td></td><td></td><td></td><td></td><td></td><td></td></tr>
<tr><td>Oligo fwd</td><td>CACCGCCGCTGCCGACGGCTGCCT</td><td></td><td></td><td></td><td></td><td></td><td></td><td></td><td></td><td></td><td></td><td></td></tr>
<tr><td>Oligo rev</td><td>AAACAGGCAGCCGTCGGCAGCGGC</td><td></td><td></td><td></td><td></td><td></td><td></td><td></td><td></td><td></td><td></td><td></td></tr>
<tr><td>Chromosome</td><td>start</td><td></td><td>end</td><td>strand</td><td>MM</td><td>target_seq</td><td>PAM</td><td>alignment</td><td>distance</td><td>position</td><td>gene name</td><td>gene id</td></tr>
<tr><td>GU305914.1</td><td></td><td>128976</td><td>128998</td><td>-</td><td>0</td><td>GCCGCTGCCGACGGCTGCCT</td><td>CGG</td><td>||||||||[||||||||||||]PAM</td><td>NA</td><td>Intergenic</td><td>NA</td><td>NA</td></tr>
<tr><td>T24197</td><td>GCTGCCGACGGCTGCCTCGGCGG</td><td></td><td>1000</td><td></td><td></td><td></td><td></td><td></td><td></td><td></td><td></td><td></td></tr>
<tr><td>Oligo fwd</td><td>CACCGCTGCCGACGGCTGCCTCGG</td><td></td><td></td><td></td><td></td><td></td><td></td><td></td><td></td><td></td><td></td><td></td></tr>
<tr><td>Oligo rev</td><td>AAACCCGAGGCAGCCGTCGGCAGC</td><td></td><td></td><td></td><td></td><td></td><td></td><td></td><td></td><td></td><td></td><td></td></tr>
<tr><td>Chromosome</td><td>start</td><td></td><td>end</td><td>strand</td><td>MM</td><td>target_seq</td><td>PAM</td><td>alignment</td><td>distance</td><td>position</td><td>gene name</td><td>gene id</td></tr>
<tr><td>GU305914.1</td><td></td><td>128973</td><td>128995</td><td>-</td><td>0</td><td>GCTGCCGACGGCTGCCTCGG</td><td>CGG</td><td>||||||||[||||||||||||]PAM</td><td>NA</td><td>Intergenic</td><td>NA</td><td>NA</td></tr>
<tr><td>GU305914.1</td><td></td><td>24134</td><td>24156</td><td>+</td><td>4</td><td>GGCGCCGACGTCGGCCTCGG</td><td>CGG</td><td>|--|||||[||-|-|||||||]PAM</td><td>NA</td><td>Intergenic</td><td>NA</td><td>NA</td></tr>
<tr><td>T24198</td><td>CCAGCCGCCTCTCCCTGAAGCGG</td><td></td><td>1000</td><td></td><td></td><td></td><td></td><td></td><td></td><td></td><td></td><td></td></tr>
<tr><td>Oligo fwd</td><td>CACCCCAGCCGCCTCTCCCTGAAG</td><td></td><td></td><td></td><td></td><td></td><td></td><td></td><td></td><td></td><td></td><td></td></tr>
<tr><td>Oligo rev</td><td>AAACCTTCAGGGAGAGGCGGCTGG</td><td></td><td></td><td></td><td></td><td></td><td></td><td></td><td></td><td></td><td></td><td></td></tr>
<tr><td>Chromosome</td><td>start</td><td></td><td>end</td><td>strand</td><td>MM</td><td>target_seq</td><td>PAM</td><td>alignment</td><td>distance</td><td>position</td><td>gene name</td><td>gene id</td></tr>
<tr><td>GU305914.1</td><td></td><td>128950</td><td>128972</td><td>-</td><td>0</td><td>CCAGCCGCCTCTCCCTGAAG</td><td>CGG</td><td>||||||||[||||||||||||]PAM</td><td>NA</td><td>Intergenic</td><td>NA</td><td>NA</td></tr>
<tr><td>T24199</td><td>AGCGGAGCCGAACCAGAGCCAGG</td><td></td><td>1000</td><td></td><td></td><td></td><td></td><td></td><td></td><td></td><td></td><td></td></tr>
<tr><td>Oligo fwd</td><td>CACCAGCGGAGCCGAACCAGAGCC</td><td></td><td></td><td></td><td></td><td></td><td></td><td></td><td></td><td></td><td></td><td></td></tr>
<tr><td>Oligo rev</td><td>AAACGGCTCTGGTTCGGCTCCGCT</td><td></td><td></td><td></td><td></td><td></td><td></td><td></td><td></td><td></td><td></td><td></td></tr>
<tr><td>Chromosome</td><td>start</td><td></td><td>end</td><td>strand</td><td>MM</td><td>target_seq</td><td>PAM</td><td>alignment</td><td>distance</td><td>position</td><td>gene name</td><td>gene id</td></tr>
<tr><td>GU305914.1</td><td></td><td>128932</td><td>128954</td><td>-</td><td>0</td><td>AGCGGAGCCGAACCAGAGCC</td><td>AGG</td><td>||||||||[||||||||||||]PAM</td><td>NA</td><td>Intergenic</td><td>NA</td><td>NA</td></tr>
<tr><td>T24200</td><td>CTGAGTCGCTGTACTGCTGACGG</td><td></td><td>1000</td><td></td><td></td><td></td><td></td><td></td><td></td><td></td><td></td><td></td></tr>
<tr><td>Oligo fwd</td><td>CACCCTGAGTCGCTGTACTGCTGA</td><td></td><td></td><td></td><td></td><td></td><td></td><td></td><td></td><td></td><td></td><td></td></tr>
<tr><td>Oligo rev</td><td>AAACTCAGCAGTACAGCGACTCAG</td><td></td><td></td><td></td><td></td><td></td><td></td><td></td><td></td><td></td><td></td><td></td></tr>
<tr><td>Chromosome</td><td>start</td><td></td><td>end</td><td>strand</td><td>MM</td><td>target_seq</td><td>PAM</td><td>alignment</td><td>distance</td><td>position</td><td>gene name</td><td>gene id</td></tr>
<tr><td>GU305914.1</td><td></td><td>128886</td><td>128908</td><td>-</td><td>0</td><td>CTGAGTCGCTGTACTGCTGA</td><td>CGG</td><td>||||||||[||||||||||||]PAM</td><td>NA</td><td>Intergenic</td><td>NA</td><td>NA</td></tr>
<tr><td>T24201</td><td>AGTCGCTGTACTGCTGACGGCGG</td><td></td><td>1000</td><td></td><td></td><td></td><td></td><td></td><td></td><td></td><td></td><td></td></tr>
<tr><td>Oligo fwd</td><td>CACCAGTCGCTGTACTGCTGACGG</td><td></td><td></td><td></td><td></td><td></td><td></td><td></td><td></td><td></td><td></td><td></td></tr>
<tr><td>Oligo rev</td><td>AAACCCGTCAGCAGTACAGCGACT</td><td></td><td></td><td></td><td></td><td></td><td></td><td></td><td></td><td></td><td></td><td></td></tr>
<tr><td>Chromosome</td><td>start</td><td></td><td>end</td><td>strand</td><td>MM</td><td>target_seq</td><td>PAM</td><td>alignment</td><td>distance</td><td>position</td><td>gene name</td><td>gene id</td></tr>
<tr><td>GU305914.1</td><td></td><td>128883</td><td>128905</td><td>-</td><td>0</td><td>AGTCGCTGTACTGCTGACGG</td><td>CGG</td><td>||||||||[||||||||||||]PAM</td><td>NA</td><td>Intergenic</td><td>NA</td><td>NA</td></tr>
<tr><td>T24202</td><td>CTGACCTCCGCCAACGACCGTGG</td><td></td><td>1000</td><td></td><td></td><td></td><td></td><td></td><td></td><td></td><td></td><td></td></tr>
<tr><td>Oligo fwd</td><td>CACCCTGACCTCCGCCAACGACCG</td><td></td><td></td><td></td><td></td><td></td><td></td><td></td><td></td><td></td><td></td><td></td></tr>
<tr><td>Oligo rev</td><td>AAACCGGTCGTTGGCGGAGGTCAG</td><td></td><td></td><td></td><td></td><td></td><td></td><td></td><td></td><td></td><td></td><td></td></tr>
<tr><td>Chromosome</td><td>start</td><td></td><td>end</td><td>strand</td><td>MM</td><td>target_seq</td><td>PAM</td><td>alignment</td><td>distance</td><td>position</td><td>gene name</td><td>gene id</td></tr>
<tr><td>GU305914.1</td><td></td><td>128856</td><td>128878</td><td>-</td><td>0</td><td>CTGACCTCCGCCAACGACCG</td><td>TGG</td><td>||||||||[||||||||||||]PAM</td><td>NA</td><td>Intergenic</td><td>NA</td><td>NA</td></tr>
<tr><td>T24203</td><td>TGACCTCCGCCAACGACCGTGGG</td><td></td><td>1000</td><td></td><td></td><td></td><td></td><td></td><td></td><td></td><td></td><td></td></tr>
<tr><td>Oligo fwd</td><td>CACCTGACCTCCGCCAACGACCGT</td><td></td><td></td><td></td><td></td><td></td><td></td><td></td><td></td><td></td><td></td><td></td></tr>
</table>

<table>
<tr><td>Oligo rev</td><td>AAACACGGTCGTTGGCGGAGGTCA</td><td></td><td></td><td></td><td></td><td></td><td></td><td></td><td></td><td></td><td></td></tr>
<tr><td>Chromosome</td><td>start</td><td>end</td><td>strand</td><td>MM</td><td>target_seq</td><td>PAM</td><td>alignment</td><td>distance</td><td>position</td><td>gene name</td><td>gene id</td></tr>
<tr><td>GU305914.1</td><td>128855</td><td>128877</td><td>-</td><td>0</td><td>TGACCTCCGCCAACGACCGT</td><td>GGG</td><td>||||||||[||||||||||||]PAM</td><td>NA</td><td>Intergenic</td><td>NA</td><td>NA</td></tr>
<tr><td>T24204</td><td>GACCTCCGCCAACGACCGTGGGG</td><td>1000</td><td></td><td></td><td></td><td></td><td></td><td></td><td></td><td></td><td></td></tr>
<tr><td>Oligo fwd</td><td>CACCGACCTCCGCCAACGACCGTG</td><td></td><td></td><td></td><td></td><td></td><td></td><td></td><td></td><td></td><td></td></tr>
<tr><td>Oligo rev</td><td>AAACCACGGTCGTTGGCGGAGGTC</td><td></td><td></td><td></td><td></td><td></td><td></td><td></td><td></td><td></td><td></td></tr>
<tr><td>Chromosome</td><td>start</td><td>end</td><td>strand</td><td>MM</td><td>target_seq</td><td>PAM</td><td>alignment</td><td>distance</td><td>position</td><td>gene name</td><td>gene id</td></tr>
<tr><td>GU305914.1</td><td>128854</td><td>128876</td><td>-</td><td>0</td><td>GACCTCCGCCAACGACCGTG</td><td>GGG</td><td>||||||||[||||||||||||]PAM</td><td>NA</td><td>Intergenic</td><td>NA</td><td>NA</td></tr>
<tr><td>T24205</td><td>ACCTCCGCCAACGACCGTGGGGG</td><td>1000</td><td></td><td></td><td></td><td></td><td></td><td></td><td></td><td></td><td></td></tr>
<tr><td>Oligo fwd</td><td>CACCACCTCCGCCAACGACCGTGG</td><td></td><td></td><td></td><td></td><td></td><td></td><td></td><td></td><td></td><td></td></tr>
<tr><td>Oligo rev</td><td>AAACCCACGGTCGTTGGCGGAGGT</td><td></td><td></td><td></td><td></td><td></td><td></td><td></td><td></td><td></td><td></td></tr>
<tr><td>Chromosome</td><td>start</td><td>end</td><td>strand</td><td>MM</td><td>target_seq</td><td>PAM</td><td>alignment</td><td>distance</td><td>position</td><td>gene name</td><td>gene id</td></tr>
<tr><td>GU305914.1</td><td>128853</td><td>128875</td><td>-</td><td>0</td><td>ACCTCCGCCAACGACCGTGG</td><td>GGG</td><td>||||||||[||||||||||||]PAM</td><td>NA</td><td>Intergenic</td><td>NA</td><td>NA</td></tr>
<tr><td>T24206</td><td>TCCGCCAACGACCGTGGGGGCGG</td><td>1000</td><td></td><td></td><td></td><td></td><td></td><td></td><td></td><td></td><td></td></tr>
<tr><td>Oligo fwd</td><td>CACCTCCGCCAACGACCGTGGGGG</td><td></td><td></td><td></td><td></td><td></td><td></td><td></td><td></td><td></td><td></td></tr>
<tr><td>Oligo rev</td><td>AAACCCCCCACGGTCGTTGGCGGA</td><td></td><td></td><td></td><td></td><td></td><td></td><td></td><td></td><td></td><td></td></tr>
<tr><td>Chromosome</td><td>start</td><td>end</td><td>strand</td><td>MM</td><td>target_seq</td><td>PAM</td><td>alignment</td><td>distance</td><td>position</td><td>gene name</td><td>gene id</td></tr>
<tr><td>GU305914.1</td><td>128850</td><td>128872</td><td>-</td><td>0</td><td>TCCGCCAACGACCGTGGGGG</td><td>CGG</td><td>||||||||[||||||||||||]PAM</td><td>NA</td><td>Intergenic</td><td>NA</td><td>NA</td></tr>
<tr><td>T24207</td><td>CCAACGACCGTGGGGGCGGCTGG</td><td>1000</td><td></td><td></td><td></td><td></td><td></td><td></td><td></td><td></td><td></td></tr>
<tr><td>Oligo fwd</td><td>CACCCCAACGACCGTGGGGGCGGC</td><td></td><td></td><td></td><td></td><td></td><td></td><td></td><td></td><td></td><td></td></tr>
<tr><td>Oligo rev</td><td>AAACGCCGCCCCCACGGTCGTTGG</td><td></td><td></td><td></td><td></td><td></td><td></td><td></td><td></td><td></td><td></td></tr>
<tr><td>Chromosome</td><td>start</td><td>end</td><td>strand</td><td>MM</td><td>target_seq</td><td>PAM</td><td>alignment</td><td>distance</td><td>position</td><td>gene name</td><td>gene id</td></tr>
<tr><td>GU305914.1</td><td>128846</td><td>128868</td><td>-</td><td>0</td><td>CCAACGACCGTGGGGGCGGC</td><td>TGG</td><td>||||||||[||||||||||||]PAM</td><td>NA</td><td>Intergenic</td><td>NA</td><td>NA</td></tr>
<tr><td>T24208</td><td>GTGGGGGCGGCTGGCGCGTGCGG</td><td>1000</td><td></td><td></td><td></td><td></td><td></td><td></td><td></td><td></td><td></td></tr>
<tr><td>Oligo fwd</td><td>CACCGTGGGGGCGGCTGGCGCGTG</td><td></td><td></td><td></td><td></td><td></td><td></td><td></td><td></td><td></td><td></td></tr>
<tr><td>Oligo rev</td><td>AAACCACGCGCCAGCCGCCCCCAC</td><td></td><td></td><td></td><td></td><td></td><td></td><td></td><td></td><td></td><td></td></tr>
<tr><td>Chromosome</td><td>start</td><td>end</td><td>strand</td><td>MM</td><td>target_seq</td><td>PAM</td><td>alignment</td><td>distance</td><td>position</td><td>gene name</td><td>gene id</td></tr>
<tr><td>GU305914.1</td><td>128837</td><td>128859</td><td>-</td><td>0</td><td>GTGGGGGCGGCTGGCGCGTG</td><td>CGG</td><td>||||||||[||||||||||||]PAM</td><td>NA</td><td>Intergenic</td><td>NA</td><td>NA</td></tr>
<tr><td>T24209</td><td>CGCCGACGTCGCTGCCGCTGCGG</td><td>1000</td><td></td><td></td><td></td><td></td><td></td><td></td><td></td><td></td><td></td></tr>
<tr><td>Oligo fwd</td><td>CACCCGCCGACGTCGCTGCCGCTG</td><td></td><td></td><td></td><td></td><td></td><td></td><td></td><td></td><td></td><td></td></tr>
<tr><td>Oligo rev</td><td>AAACCAGCGGCAGCGACGTCGGCG</td><td></td><td></td><td></td><td></td><td></td><td></td><td></td><td></td><td></td><td></td></tr>
<tr><td>Chromosome</td><td>start</td><td>end</td><td>strand</td><td>MM</td><td>target_seq</td><td>PAM</td><td>alignment</td><td>distance</td><td>position</td><td>gene name</td><td>gene id</td></tr>
<tr><td>GU305914.1</td><td>128802</td><td>128824</td><td>-</td><td>0</td><td>CGCCGACGTCGCTGCCGCTG</td><td>CGG</td><td>||||||||[||||||||||||]PAM</td><td>NA</td><td>Intergenic</td><td>NA</td><td>NA</td></tr>
<tr><td>GU305914.1</td><td>56775</td><td>56797</td><td>-</td><td>4</td><td>TGCCGCCGCCGCCGCCGCTG</td><td>CGG</td><td>-||||-||[-|||-||||||||]PAM</td><td>NA</td><td>Intergenic</td><td>NA</td><td>NA</td></tr>
<tr><td>T24210</td><td>GCCGACGTCGCTGCCGCTGCGGG</td><td>1000</td><td></td><td></td><td></td><td></td><td></td><td></td><td></td><td></td><td></td></tr>
<tr><td>Oligo fwd</td><td>CACCGCCGACGTCGCTGCCGCTGC</td><td></td><td></td><td></td><td></td><td></td><td></td><td></td><td></td><td></td><td></td></tr>
<tr><td>Oligo rev</td><td>AAACGCAGCGGCAGCGACGTCGGC</td><td></td><td></td><td></td><td></td><td></td><td></td><td></td><td></td><td></td><td></td></tr>
<tr><td>Chromosome</td><td>start</td><td>end</td><td>strand</td><td>MM</td><td>target_seq</td><td>PAM</td><td>alignment</td><td>distance</td><td>position</td><td>gene name</td><td>gene id</td></tr>
<tr><td>GU305914.1</td><td>128801</td><td>128823</td><td>-</td><td>0</td><td>GCCGACGTCGCTGCCGCTGC</td><td>GGG</td><td>||||||||[||||||||||||]PAM</td><td>NA</td><td>Intergenic</td><td>NA</td><td>NA</td></tr>
<tr><td>GU305914.1</td><td>62096</td><td>62118</td><td>+</td><td>4</td><td>GTCGCCATCGCTGCTGCTGC</td><td>TGG</td><td>|-||-|-|[||||||-|||||]PAM</td><td>NA</td><td>Intergenic</td><td>NA</td><td>NA</td></tr>
<tr><td>T24211</td><td>ACCACTCTCCGTCGACGCCGCGG</td><td>1000</td><td></td><td></td><td></td><td></td><td></td><td></td><td></td><td></td><td></td></tr>
<tr><td>Oligo fwd</td><td>CACCACCACTCTCCGTCGACGCCG</td><td></td><td></td><td></td><td></td><td></td><td></td><td></td><td></td><td></td><td></td></tr>
<tr><td>Oligo rev</td><td>AAACCGGCGTCGACGGAGAGTGGT</td><td></td><td></td><td></td><td></td><td></td><td></td><td></td><td></td><td></td><td></td></tr>
<tr><td>Chromosome</td><td>start</td><td>end</td><td>strand</td><td>MM</td><td>target_seq</td><td>PAM</td><td>alignment</td><td>distance</td><td>position</td><td>gene name</td><td>gene id</td></tr>
<tr><td>GU305914.1</td><td>128763</td><td>128785</td><td>-</td><td>0</td><td>ACCACTCTCCGTCGACGCCG</td><td>CGG</td><td>||||||||[||||||||||||]PAM</td><td>NA</td><td>Intergenic</td><td>NA</td><td>NA</td></tr>
<tr><td>T24212</td><td>ACTCTCCGTCGACGCCGCGGAGG</td><td>1000</td><td></td><td></td><td></td><td></td><td></td><td></td><td></td><td></td><td></td></tr>
<tr><td>Oligo fwd</td><td>CACCACTCTCCGTCGACGCCGCGG</td><td></td><td></td><td></td><td></td><td></td><td></td><td></td><td></td><td></td><td></td></tr>
<tr><td>Oligo rev</td><td>AAACCCGCGGCGTCGACGGAGAGT</td><td></td><td></td><td></td><td></td><td></td><td></td><td></td><td></td><td></td><td></td></tr>
<tr><td>Chromosome</td><td>start</td><td>end</td><td>strand</td><td>MM</td><td>target_seq</td><td>PAM</td><td>alignment</td><td>distance</td><td>position</td><td>gene name</td><td>gene id</td></tr>
<tr><td>GU305914.1</td><td>128760</td><td>128782</td><td>-</td><td>0</td><td>ACTCTCCGTCGACGCCGCGG</td><td>AGG</td><td>||||||||[||||||||||||]PAM</td><td>NA</td><td>Intergenic</td><td>NA</td><td>NA</td></tr>
<tr><td>T24213</td><td>CGCCGCGGAGGCATCGTCGTCGG</td><td>1000</td><td></td><td></td><td></td><td></td><td></td><td></td><td></td><td></td><td></td></tr>
<tr><td>Oligo fwd</td><td>CACCCGCCGCGGAGGCATCGTCGT</td><td></td><td></td><td></td><td></td><td></td><td></td><td></td><td></td><td></td><td></td></tr>
<tr><td>Oligo rev</td><td>AAACACGACGATGCCTCCGCGGCG</td><td></td><td></td><td></td><td></td><td></td><td></td><td></td><td></td><td></td><td></td></tr>
<tr><td>Chromosome</td><td>start</td><td>end</td><td>strand</td><td>MM</td><td>target_seq</td><td>PAM</td><td>alignment</td><td>distance</td><td>position</td><td>gene name</td><td>gene id</td></tr>
<tr><td>GU305914.1</td><td>128748</td><td>128770</td><td>-</td><td>0</td><td>CGCCGCGGAGGCATCGTCGT</td><td>CGG</td><td>||||||||[||||||||||||]PAM</td><td>NA</td><td>Intergenic</td><td>NA</td><td>NA</td></tr>
<tr><td>T24214</td><td>CGCGGAGGCATCGTCGTCGGTGG</td><td>1000</td><td></td><td></td><td></td><td></td><td></td><td></td><td></td><td></td><td></td></tr>
<tr><td>Oligo fwd</td><td>CACCCGCGGAGGCATCGTCGTCGG</td><td></td><td></td><td></td><td></td><td></td><td></td><td></td><td></td><td></td><td></td></tr>
<tr><td>Oligo rev</td><td>AAACCCGACGACGATGCCTCCGCG</td><td></td><td></td><td></td><td></td><td></td><td></td><td></td><td></td><td></td><td></td></tr>
<tr><td>Chromosome</td><td>start</td><td>end</td><td>strand</td><td>MM</td><td>target_seq</td><td>PAM</td><td>alignment</td><td>distance</td><td>position</td><td>gene name</td><td>gene id</td></tr>
<tr><td>GU305914.1</td><td>128745</td><td>128767</td><td>-</td><td>0</td><td>CGCGGAGGCATCGTCGTCGG</td><td>TGG</td><td>||||||||[||||||||||||]PAM</td><td>NA</td><td>Intergenic</td><td>NA</td><td>NA</td></tr>
<tr><td>GU305914.1</td><td>128803</td><td>128825</td><td>+</td><td>4</td><td>CGCAGCGGCAGCGACGTCGG</td><td>CGG</td><td>|||-|-||[||-||-||||||]PAM</td><td>NA</td><td>Intergenic</td><td>NA</td><td>NA</td></tr>
<tr><td>T24215</td><td>GGAGGCATCGTCGTCGGTGGAGG</td><td>1000</td><td></td><td></td><td></td><td></td><td></td><td></td><td></td><td></td><td></td></tr>
<tr><td>Oligo fwd</td><td>CACCGGAGGCATCGTCGTCGGTGG</td><td></td><td></td><td></td><td></td><td></td><td></td><td></td><td></td><td></td><td></td></tr>
<tr><td>Oligo rev</td><td>AAACCCACCGACGACGATGCCTCC</td><td></td><td></td><td></td><td></td><td></td><td></td><td></td><td></td><td></td><td></td></tr>
<tr><td>Chromosome</td><td>start</td><td>end</td><td>strand</td><td>MM</td><td>target_seq</td><td>PAM</td><td>alignment</td><td>distance</td><td>position</td><td>gene name</td><td>gene id</td></tr>
<tr><td>GU305914.1</td><td>128742</td><td>128764</td><td>-</td><td>0</td><td>GGAGGCATCGTCGTCGGTGG</td><td>AGG</td><td>||||||||[||||||||||||]PAM</td><td>NA</td><td>Intergenic</td><td>NA</td><td>NA</td></tr>
<tr><td>T24216</td><td>GGCATCGTCGTCGGTGGAGGCGG</td><td>1000</td><td></td><td></td><td></td><td></td><td></td><td></td><td></td><td></td><td></td></tr>
<tr><td>Oligo fwd</td><td>CACCGGCATCGTCGTCGGTGGAGG</td><td></td><td></td><td></td><td></td><td></td><td></td><td></td><td></td><td></td><td></td></tr>
<tr><td>Oligo rev</td><td>AAACCCTCCACCGACGACGATGCC</td><td></td><td></td><td></td><td></td><td></td><td></td><td></td><td></td><td></td><td></td></tr>
<tr><td>Chromosome</td><td>start</td><td>end</td><td>strand</td><td>MM</td><td>target_seq</td><td>PAM</td><td>alignment</td><td>distance</td><td>position</td><td>gene name</td><td>gene id</td></tr>
<tr><td>GU305914.1</td><td>128739</td><td>128761</td><td>-</td><td>0</td><td>GGCATCGTCGTCGGTGGAGG</td><td>CGG</td><td>||||||||[||||||||||||]PAM</td><td>NA</td><td>Intergenic</td><td>NA</td><td>NA</td></tr>
<tr><td>T24217</td><td>GGTGGAGGCGGCAGCAGCAGCGG</td><td>1000</td><td></td><td></td><td></td><td></td><td></td><td></td><td></td><td></td><td></td></tr>
<tr><td>Oligo fwd</td><td>CACCGGTGGAGGCGGCAGCAGCAG</td><td></td><td></td><td></td><td></td><td></td><td></td><td></td><td></td><td></td><td></td></tr>
<tr><td>Oligo rev</td><td>AAACCTGCTGCTGCCGCCTCCACC</td><td></td><td></td><td></td><td></td><td></td><td></td><td></td><td></td><td></td><td></td></tr>
<tr><td>Chromosome</td><td>start</td><td>end</td><td>strand</td><td>MM</td><td>target_seq</td><td>PAM</td><td>alignment</td><td>distance</td><td>position</td><td>gene name</td><td>gene id</td></tr>
</table>

| | | | | | | | | | | | |
|---|---|---|---|---|---|---|---|---|---|---|---|
| GU305914.1 | 128727 | 128749 | - | 0 | GGTGGAGGCGGCAGCAGCAG | CGG | \|\|\|\|\|\|\|\|[\|\|\|\|\|\|\|\|\|\|\|\|]PAM | NA | Intergenic | NA | NA |
| GU305914.1 | 163908 | 163930 | + | 3 | GGCGGCGGCGGCAGCGGCAG | CGG | \|\|-\|\|-\|\|[\|\|\|\|\|\|\|-\|\|\|\|]PAM | NA | Intergenic | NA | NA |
| GU305914.1 | 30006 | 30028 | + | 4 | GGCGGCGGCGGCGGCGGCAG | TGG | \|\|-\|\|-\|\|[\|\|\|\|-\|\|-\|\|\|\|]PAM | NA | Intergenic | NA | NA |
| GU305914.1 | 149476 | 149498 | - | 4 | GGCGGCGGCGGCGGCGGCAG | CGG | \|\|-\|\|-\|\|[\|\|\|\|-\|\|-\|\|\|\|]PAM | NA | Intergenic | NA | NA |
| GU305914.1 | 164959 | 164981 | - | 3 | GGCGGCGGCGGCAGCAGCTG | CGG | \|\|-\|\|-\|\|[\|\|\|\|\|\|\|\|\|\|-\|]PAM | NA | Intergenic | NA | NA |
| T24218 | GGCGGCAGCAGCAGCGGCGCTGG | 1000 | | | | | | | | | |
| Oligo fwd | CACCGGCGGCAGCAGCAGCGGCGC | | | | | | | | | | |
| Oligo rev | AAACGCGCCGCTGCTGCTGCCGCC | | | | | | | | | | |
| Chromosome | start | end | strand | MM | target_seq | PAM | alignment | distance | position | gene name | gene id |
| GU305914.1 | 128721 | 128743 | - | 0 | GGCGGCAGCAGCAGCGGCGC | TGG | \|\|\|\|\|\|\|\|[\|\|\|\|\|\|\|\|\|\|\|\|]PAM | NA | Intergenic | NA | NA |
| GU305914.1 | 71747 | 71769 | + | 4 | CTCAGCAGCAGCAGCGGCGG | CGG | --\|-\|\|\|\|[\|\|\|\|\|\|\|\|\|\|\|-]PAM | NA | Intergenic | NA | NA |
| GU305914.1 | 113918 | 113940 | - | 4 | GTCGGCGGCGGCAGCGGCGA | CGG | \|-\|\|\|\|-\|[\|-\|\|\|\|\|\|\|\|\|-]PAM | NA | Intergenic | NA | NA |
| GU305914.1 | 73792 | 73814 | - | 3 | GGCGCCAGCAGCGGCGGCGG | AGG | \|\|\|\|-\|\|\|[\|\|\|\|-\|\|\|\|\|\|-]PAM | NA | Intergenic | NA | NA |
| GU305914.1 | 27411 | 27433 | + | 4 | AGCAGCAGCAGCAGCAGCGG | CGG | -\|\|-\|\|\|\|[\|\|\|\|\|\|\|-\|\|\|-]PAM | NA | Intergenic | NA | NA |
| T24219 | AGCAGCAGCGGCGCTGGCAGTGG | 1000 | | | | | | | | | |
| Oligo fwd | CACCAGCAGCAGCGGCGCTGGCAG | | | | | | | | | | |
| Oligo rev | AAACCTGCCAGCGCCGCTGCTGCT | | | | | | | | | | |
| Chromosome | start | end | strand | MM | target_seq | PAM | alignment | distance | position | gene name | gene id |
| GU305914.1 | 128715 | 128737 | - | 0 | AGCAGCAGCGGCGCTGGCAG | TGG | \|\|\|\|\|\|\|\|[\|\|\|\|\|\|\|\|\|\|\|\|]PAM | NA | Intergenic | NA | NA |
| GU305914.1 | 163902 | 163924 | + | 4 | AGCGGCGGCGGCGGCGGCAG | CGG | \|\|\|-\|\|-\|[\|\|\|\|\|--\|\|\|\|\|]PAM | NA | Intergenic | NA | NA |
| T24220 | TGGTAGCCTGATCGCGCCCGCGG | 1000 | | | | | | | | | |
| Oligo fwd | CACCTGGTAGCCTGATCGCGCCCG | | | | | | | | | | |
| Oligo rev | AAACCGGGCGCGATCAGGCTACCA | | | | | | | | | | |
| Chromosome | start | end | strand | MM | target_seq | PAM | alignment | distance | position | gene name | gene id |
| GU305914.1 | 128695 | 128717 | - | 0 | TGGTAGCCTGATCGCGCCCG | CGG | \|\|\|\|\|\|\|\|[\|\|\|\|\|\|\|\|\|\|\|\|]PAM | NA | Intergenic | NA | NA |
| T24221 | GCCTCCGCCGCGCTCAGACTCGG | 1000 | | | | | | | | | |
| Oligo fwd | CACCGCCTCCGCCGCGCTCAGACT | | | | | | | | | | |
| Oligo rev | AAACAGTCTGAGCGCGGCGGAGGC | | | | | | | | | | |
| Chromosome | start | end | strand | MM | target_seq | PAM | alignment | distance | position | gene name | gene id |
| GU305914.1 | 128670 | 128692 | - | 0 | GCCTCCGCCGCGCTCAGACT | CGG | \|\|\|\|\|\|\|\|[\|\|\|\|\|\|\|\|\|\|\|\|]PAM | NA | Intergenic | NA | NA |
| T24222 | TCCGCCGCGCTCAGACTCGGTGG | 1000 | | | | | | | | | |
| Oligo fwd | CACCTCCGCCGCGCTCAGACTCGG | | | | | | | | | | |
| Oligo rev | AAACCCGAGTCTGAGCGCGGCGGA | | | | | | | | | | |
| Chromosome | start | end | strand | MM | target_seq | PAM | alignment | distance | position | gene name | gene id |
| GU305914.1 | 128667 | 128689 | - | 0 | TCCGCCGCGCTCAGACTCGG | TGG | \|\|\|\|\|\|\|\|[\|\|\|\|\|\|\|\|\|\|\|\|]PAM | NA | Intergenic | NA | NA |
| T24223 | GCCGCGCTCAGACTCGGTGGCGG | 1000 | | | | | | | | | |
| Oligo fwd | CACCGCCGCGCTCAGACTCGGTGG | | | | | | | | | | |
| Oligo rev | AAACCCACCGAGTCTGAGCGCGGC | | | | | | | | | | |
| Chromosome | start | end | strand | MM | target_seq | PAM | alignment | distance | position | gene name | gene id |
| GU305914.1 | 128664 | 128686 | - | 0 | GCCGCGCTCAGACTCGGTGG | CGG | \|\|\|\|\|\|\|\|[\|\|\|\|\|\|\|\|\|\|\|\|]PAM | NA | Intergenic | NA | NA |
| T24224 | GCGCCGCCGCCGCTGCCGCTCGG | 1000 | | | | | | | | | |
| Oligo fwd | CACCGCGCCGCCGCCGCTGCCGCT | | | | | | | | | | |
| Oligo rev | AAACAGCGGCAGCGGCGGCGGCGC | | | | | | | | | | |
| Chromosome | start | end | strand | MM | target_seq | PAM | alignment | distance | position | gene name | gene id |
| GU305914.1 | 128642 | 128664 | - | 0 | GCGCCGCCGCCGCTGCCGCT | CGG | \|\|\|\|\|\|\|\|[\|\|\|\|\|\|\|\|\|\|\|\|]PAM | NA | Intergenic | NA | NA |
| T24225 | TGCCGCTCGGCCCCCACAGCCGG | 1000 | | | | | | | | | |
| Oligo fwd | CACCTGCCGCTCGGCCCCCACAGC | | | | | | | | | | |
| Oligo rev | AAACGCTGTGGGGGCCGAGCGGCA | | | | | | | | | | |
| Chromosome | start | end | strand | MM | target_seq | PAM | alignment | distance | position | gene name | gene id |
| GU305914.1 | 128629 | 128651 | - | 0 | TGCCGCTCGGCCCCCACAGC | CGG | \|\|\|\|\|\|\|\|[\|\|\|\|\|\|\|\|\|\|\|\|]PAM | NA | Intergenic | NA | NA |
| T24226 | TCTCGCCGCCGAAGCCCCGATGG | 1000 | | | | | | | | | |
| Oligo fwd | CACCTCTCGCCGCCGAAGCCCCGA | | | | | | | | | | |
| Oligo rev | AAACTCGGGGCTTCGGCGGCGAGA | | | | | | | | | | |
| Chromosome | start | end | strand | MM | target_seq | PAM | alignment | distance | position | gene name | gene id |
| GU305914.1 | 128605 | 128627 | - | 0 | TCTCGCCGCCGAAGCCCCGA | TGG | \|\|\|\|\|\|\|\|[\|\|\|\|\|\|\|\|\|\|\|\|]PAM | NA | Intergenic | NA | NA |
| T24227 | CTCGCCGCCGAAGCCCCGATGGG | 1000 | | | | | | | | | |
| Oligo fwd | CACCCTCGCCGCCGAAGCCCCGAT | | | | | | | | | | |
| Oligo rev | AAACATCGGGGCTTCGGCGGCGAG | | | | | | | | | | |
| Chromosome | start | end | strand | MM | target_seq | PAM | alignment | distance | position | gene name | gene id |
| GU305914.1 | 128604 | 128626 | - | 0 | CTCGCCGCCGAAGCCCCGAT | GGG | \|\|\|\|\|\|\|\|[\|\|\|\|\|\|\|\|\|\|\|\|]PAM | NA | Intergenic | NA | NA |
| T24228 | GAAGCCCCGATGGGCGCACCAGG | 1000 | | | | | | | | | |
| Oligo fwd | CACCGAAGCCCCGATGGGCGCACC | | | | | | | | | | |
| Oligo rev | AAACGGTGCGCCCATCGGGGCTTC | | | | | | | | | | |
| Chromosome | start | end | strand | MM | target_seq | PAM | alignment | distance | position | gene name | gene id |
| GU305914.1 | 128595 | 128617 | - | 0 | GAAGCCCCGATGGGCGCACC | AGG | \|\|\|\|\|\|\|\|[\|\|\|\|\|\|\|\|\|\|\|\|]PAM | NA | Intergenic | NA | NA |
| T24229 | ATGGGCGCACCAGGCATAGTAGG | 1000 | | | | | | | | | |
| Oligo fwd | CACCATGGGCGCACCAGGCATAGT | | | | | | | | | | |
| Oligo rev | AAACACTATGCCTGGTGCGCCCAT | | | | | | | | | | |
| Chromosome | start | end | strand | MM | target_seq | PAM | alignment | distance | position | gene name | gene id |
| GU305914.1 | 128586 | 128608 | - | 0 | ATGGGCGCACCAGGCATAGT | AGG | \|\|\|\|\|\|\|\|[\|\|\|\|\|\|\|\|\|\|\|\|]PAM | NA | Intergenic | NA | NA |
| T24230 | AGGCATAGTAGGTGTAGAACAGG | 1000 | | | | | | | | | |
| Oligo fwd | CACCAGGCATAGTAGGTGTAGAAC | | | | | | | | | | |
| Oligo rev | AAACGTTCTACACCTACTATGCCT | | | | | | | | | | |
| Chromosome | start | end | strand | MM | target_seq | PAM | alignment | distance | position | gene name | gene id |
| GU305914.1 | 128575 | 128597 | - | 0 | AGGCATAGTAGGTGTAGAAC | AGG | \|\|\|\|\|\|\|\|[\|\|\|\|\|\|\|\|\|\|\|\|]PAM | NA | Intergenic | NA | NA |

<table>
<tr><td>T24231</td><td>GGTTCCTCCAGATCACGTCGTGG</td><td>1000</td><td></td><td></td><td></td><td></td><td></td><td></td><td></td><td></td><td></td></tr>
<tr><td>Oligo fwd</td><td>CACCGGTTCCTCCAGATCACGTCG</td><td></td><td></td><td></td><td></td><td></td><td></td><td></td><td></td><td></td><td></td></tr>
<tr><td>Oligo rev</td><td>AAACCGACGTGATCTGGAGGAACC</td><td></td><td></td><td></td><td></td><td></td><td></td><td></td><td></td><td></td><td></td></tr>
<tr><td>Chromosome</td><td>start</td><td>end</td><td>strand</td><td>MM</td><td>target_seq</td><td>PAM</td><td>alignment</td><td>distance</td><td>position</td><td>gene name</td><td>gene id</td></tr>
<tr><td>GU305914.1</td><td>128554</td><td>128576</td><td>-</td><td>0</td><td>GGTTCCTCCAGATCACGTCG</td><td>TGG</td><td>||||||||[||||||||||||]PAM</td><td>NA</td><td>Intergenic</td><td>NA</td><td>NA</td></tr>
<tr><td>T24232</td><td>GTGGTCCGCGAACAGAACGCTGG</td><td>1000</td><td></td><td></td><td></td><td></td><td></td><td></td><td></td><td></td><td></td></tr>
<tr><td>Oligo fwd</td><td>CACCGTGGTCCGCGAACAGAACGC</td><td></td><td></td><td></td><td></td><td></td><td></td><td></td><td></td><td></td><td></td></tr>
<tr><td>Oligo rev</td><td>AAACGCGTTCTGTTCGCGGACCAC</td><td></td><td></td><td></td><td></td><td></td><td></td><td></td><td></td><td></td><td></td></tr>
<tr><td>Chromosome</td><td>start</td><td>end</td><td>strand</td><td>MM</td><td>target_seq</td><td>PAM</td><td>alignment</td><td>distance</td><td>position</td><td>gene name</td><td>gene id</td></tr>
<tr><td>GU305914.1</td><td>128535</td><td>128557</td><td>-</td><td>0</td><td>GTGGTCCGCGAACAGAACGC</td><td>TGG</td><td>||||||||[||||||||||||]PAM</td><td>NA</td><td>Intergenic</td><td>NA</td><td>NA</td></tr>
<tr><td>T24233</td><td>CATTTTTGAGTTCGTCCTTCCGG</td><td>1000</td><td></td><td></td><td></td><td></td><td></td><td></td><td></td><td></td><td></td></tr>
<tr><td>Oligo fwd</td><td>CACCCATTTTTGAGTTCGTCCTTC</td><td></td><td></td><td></td><td></td><td></td><td></td><td></td><td></td><td></td><td></td></tr>
<tr><td>Oligo rev</td><td>AAACGAAGGACGAACTCAAAAATG</td><td></td><td></td><td></td><td></td><td></td><td></td><td></td><td></td><td></td><td></td></tr>
<tr><td>Chromosome</td><td>start</td><td>end</td><td>strand</td><td>MM</td><td>target_seq</td><td>PAM</td><td>alignment</td><td>distance</td><td>position</td><td>gene name</td><td>gene id</td></tr>
<tr><td>GU305914.1</td><td>128494</td><td>128516</td><td>-</td><td>0</td><td>CATTTTTGAGTTCGTCCTTC</td><td>CGG</td><td>||||||||[||||||||||||]PAM</td><td>NA</td><td>Intergenic</td><td>NA</td><td>NA</td></tr>
<tr><td>T24234</td><td>CCTTCCGGCTCTTAGTACATCGG</td><td>1000</td><td></td><td></td><td></td><td></td><td></td><td></td><td></td><td></td><td></td></tr>
<tr><td>Oligo fwd</td><td>CACCCCTTCCGGCTCTTAGTACAT</td><td></td><td></td><td></td><td></td><td></td><td></td><td></td><td></td><td></td><td></td></tr>
<tr><td>Oligo rev</td><td>AAACATGTACTAAGAGCCGGAAGG</td><td></td><td></td><td></td><td></td><td></td><td></td><td></td><td></td><td></td><td></td></tr>
<tr><td>Chromosome</td><td>start</td><td>end</td><td>strand</td><td>MM</td><td>target_seq</td><td>PAM</td><td>alignment</td><td>distance</td><td>position</td><td>gene name</td><td>gene id</td></tr>
<tr><td>GU305914.1</td><td>128479</td><td>128501</td><td>-</td><td>0</td><td>CCTTCCGGCTCTTAGTACAT</td><td>CGG</td><td>||||||||[||||||||||||]PAM</td><td>NA</td><td>Intergenic</td><td>NA</td><td>NA</td></tr>
<tr><td>T24235</td><td>GGACTTCGAGCTCGAAGTAAAGG</td><td>1000</td><td></td><td></td><td></td><td></td><td></td><td></td><td></td><td></td><td></td></tr>
<tr><td>Oligo fwd</td><td>CACCGGACTTCGAGCTCGAAGTAA</td><td></td><td></td><td></td><td></td><td></td><td></td><td></td><td></td><td></td><td></td></tr>
<tr><td>Oligo rev</td><td>AAACTTACTTCGAGCTCGAAGTCC</td><td></td><td></td><td></td><td></td><td></td><td></td><td></td><td></td><td></td><td></td></tr>
<tr><td>Chromosome</td><td>start</td><td>end</td><td>strand</td><td>MM</td><td>target_seq</td><td>PAM</td><td>alignment</td><td>distance</td><td>position</td><td>gene name</td><td>gene id</td></tr>
<tr><td>GU305914.1</td><td>128458</td><td>128480</td><td>-</td><td>0</td><td>GGACTTCGAGCTCGAAGTAA</td><td>AGG</td><td>||||||||[||||||||||||]PAM</td><td>NA</td><td>Intergenic</td><td>NA</td><td>NA</td></tr>
<tr><td>T24236</td><td>CGCGACATCGATATCCATCGCGG</td><td>1000</td><td></td><td></td><td></td><td></td><td></td><td></td><td></td><td></td><td></td></tr>
<tr><td>Oligo fwd</td><td>CACCCGCGACATCGATATCCATCG</td><td></td><td></td><td></td><td></td><td></td><td></td><td></td><td></td><td></td><td></td></tr>
<tr><td>Oligo rev</td><td>AAACCGATGGATATCGATGTCGCG</td><td></td><td></td><td></td><td></td><td></td><td></td><td></td><td></td><td></td><td></td></tr>
<tr><td>Chromosome</td><td>start</td><td>end</td><td>strand</td><td>MM</td><td>target_seq</td><td>PAM</td><td>alignment</td><td>distance</td><td>position</td><td>gene name</td><td>gene id</td></tr>
<tr><td>GU305914.1</td><td>128420</td><td>128442</td><td>-</td><td>0</td><td>CGCGACATCGATATCCATCG</td><td>CGG</td><td>||||||||[||||||||||||]PAM</td><td>NA</td><td>Intergenic</td><td>NA</td><td>NA</td></tr>
<tr><td>T24237</td><td>GATATCCATCGCGGAAACAGCGG</td><td>1000</td><td></td><td></td><td></td><td></td><td></td><td></td><td></td><td></td><td></td></tr>
<tr><td>Oligo fwd</td><td>CACCGATATCCATCGCGGAAACAG</td><td></td><td></td><td></td><td></td><td></td><td></td><td></td><td></td><td></td><td></td></tr>
<tr><td>Oligo rev</td><td>AAACCTGTTTCCGCGATGGATATC</td><td></td><td></td><td></td><td></td><td></td><td></td><td></td><td></td><td></td><td></td></tr>
<tr><td>Chromosome</td><td>start</td><td>end</td><td>strand</td><td>MM</td><td>target_seq</td><td>PAM</td><td>alignment</td><td>distance</td><td>position</td><td>gene name</td><td>gene id</td></tr>
<tr><td>GU305914.1</td><td>128411</td><td>128433</td><td>-</td><td>0</td><td>GATATCCATCGCGGAAACAG</td><td>CGG</td><td>||||||||[||||||||||||]PAM</td><td>NA</td><td>Intergenic</td><td>NA</td><td>NA</td></tr>
<tr><td>T24238</td><td>TCCATCGCGGAAACAGCGGCTGG</td><td>1000</td><td></td><td></td><td></td><td></td><td></td><td></td><td></td><td></td><td></td></tr>
<tr><td>Oligo fwd</td><td>CACCTCCATCGCGGAAACAGCGGC</td><td></td><td></td><td></td><td></td><td></td><td></td><td></td><td></td><td></td><td></td></tr>
<tr><td>Oligo rev</td><td>AAACGCCGCTGTTTCCGCGATGGA</td><td></td><td></td><td></td><td></td><td></td><td></td><td></td><td></td><td></td><td></td></tr>
<tr><td>Chromosome</td><td>start</td><td>end</td><td>strand</td><td>MM</td><td>target_seq</td><td>PAM</td><td>alignment</td><td>distance</td><td>position</td><td>gene name</td><td>gene id</td></tr>
<tr><td>GU305914.1</td><td>128407</td><td>128429</td><td>-</td><td>0</td><td>TCCATCGCGGAAACAGCGGC</td><td>TGG</td><td>||||||||[||||||||||||]PAM</td><td>NA</td><td>Intergenic</td><td>NA</td><td>NA</td></tr>
<tr><td>T24239</td><td>ATCGCGGAAACAGCGGCTGGCGG</td><td>1000</td><td></td><td></td><td></td><td></td><td></td><td></td><td></td><td></td><td></td></tr>
<tr><td>Oligo fwd</td><td>CACCATCGCGGAAACAGCGGCTGG</td><td></td><td></td><td></td><td></td><td></td><td></td><td></td><td></td><td></td><td></td></tr>
<tr><td>Oligo rev</td><td>AAACCCAGCCGCTGTTTCCGCGAT</td><td></td><td></td><td></td><td></td><td></td><td></td><td></td><td></td><td></td><td></td></tr>
<tr><td>Chromosome</td><td>start</td><td>end</td><td>strand</td><td>MM</td><td>target_seq</td><td>PAM</td><td>alignment</td><td>distance</td><td>position</td><td>gene name</td><td>gene id</td></tr>
<tr><td>GU305914.1</td><td>128404</td><td>128426</td><td>-</td><td>0</td><td>ATCGCGGAAACAGCGGCTGG</td><td>CGG</td><td>||||||||[||||||||||||]PAM</td><td>NA</td><td>Intergenic</td><td>NA</td><td>NA</td></tr>
<tr><td>T24240</td><td>GGAAACAGCGGCTGGCGGCCCGG</td><td>1000</td><td></td><td></td><td></td><td></td><td></td><td></td><td></td><td></td><td></td></tr>
<tr><td>Oligo fwd</td><td>CACCGGAAACAGCGGCTGGCGGCC</td><td></td><td></td><td></td><td></td><td></td><td></td><td></td><td></td><td></td><td></td></tr>
<tr><td>Oligo rev</td><td>AAACGGCCGCCAGCCGCTGTTTCC</td><td></td><td></td><td></td><td></td><td></td><td></td><td></td><td></td><td></td><td></td></tr>
<tr><td>Chromosome</td><td>start</td><td>end</td><td>strand</td><td>MM</td><td>target_seq</td><td>PAM</td><td>alignment</td><td>distance</td><td>position</td><td>gene name</td><td>gene id</td></tr>
<tr><td>GU305914.1</td><td>128399</td><td>128421</td><td>-</td><td>0</td><td>GGAAACAGCGGCTGGCGGCC</td><td>CGG</td><td>||||||||[||||||||||||]PAM</td><td>NA</td><td>Intergenic</td><td>NA</td><td>NA</td></tr>
<tr><td>T24241</td><td>GCATGACCTTACCTGTCTCGTGG</td><td>1000</td><td></td><td></td><td></td><td></td><td></td><td></td><td></td><td></td><td></td></tr>
<tr><td>Oligo fwd</td><td>CACCGCATGACCTTACCTGTCTCG</td><td></td><td></td><td></td><td></td><td></td><td></td><td></td><td></td><td></td><td></td></tr>
<tr><td>Oligo rev</td><td>AAACCGAGACAGGTAAGGTCATGC</td><td></td><td></td><td></td><td></td><td></td><td></td><td></td><td></td><td></td><td></td></tr>
<tr><td>Chromosome</td><td>start</td><td>end</td><td>strand</td><td>MM</td><td>target_seq</td><td>PAM</td><td>alignment</td><td>distance</td><td>position</td><td>gene name</td><td>gene id</td></tr>
<tr><td>GU305914.1</td><td>128362</td><td>128384</td><td>-</td><td>0</td><td>GCATGACCTTACCTGTCTCG</td><td>TGG</td><td>||||||||[||||||||||||]PAM</td><td>NA</td><td>Intergenic</td><td>NA</td><td>NA</td></tr>
<tr><td>T24242</td><td>TGACCTTACCTGTCTCGTGGTGG</td><td>1000</td><td></td><td></td><td></td><td></td><td></td><td></td><td></td><td></td><td></td></tr>
<tr><td>Oligo fwd</td><td>CACCTGACCTTACCTGTCTCGTGG</td><td></td><td></td><td></td><td></td><td></td><td></td><td></td><td></td><td></td><td></td></tr>
<tr><td>Oligo rev</td><td>AAACCCACGAGACAGGTAAGGTCA</td><td></td><td></td><td></td><td></td><td></td><td></td><td></td><td></td><td></td><td></td></tr>
<tr><td>Chromosome</td><td>start</td><td>end</td><td>strand</td><td>MM</td><td>target_seq</td><td>PAM</td><td>alignment</td><td>distance</td><td>position</td><td>gene name</td><td>gene id</td></tr>
<tr><td>GU305914.1</td><td>128359</td><td>128381</td><td>-</td><td>0</td><td>TGACCTTACCTGTCTCGTGG</td><td>TGG</td><td>||||||||[||||||||||||]PAM</td><td>NA</td><td>Intergenic</td><td>NA</td><td>NA</td></tr>
<tr><td>T24243</td><td>CTTACCTGTCTCGTGGTGGAAGG</td><td>1000</td><td></td><td></td><td></td><td></td><td></td><td></td><td></td><td></td><td></td></tr>
<tr><td>Oligo fwd</td><td>CACCCTTACCTGTCTCGTGGTGGA</td><td></td><td></td><td></td><td></td><td></td><td></td><td></td><td></td><td></td><td></td></tr>
<tr><td>Oligo rev</td><td>AAACTCCACCACGAGACAGGTAAG</td><td></td><td></td><td></td><td></td><td></td><td></td><td></td><td></td><td></td><td></td></tr>
<tr><td>Chromosome</td><td>start</td><td>end</td><td>strand</td><td>MM</td><td>target_seq</td><td>PAM</td><td>alignment</td><td>distance</td><td>position</td><td>gene name</td><td>gene id</td></tr>
<tr><td>GU305914.1</td><td>128355</td><td>128377</td><td>-</td><td>0</td><td>CTTACCTGTCTCGTGGTGGA</td><td>AGG</td><td>||||||||[||||||||||||]PAM</td><td>NA</td><td>Intergenic</td><td>NA</td><td>NA</td></tr>
<tr><td>T24244</td><td>ACCTGTCTCGTGGTGGAAGGCGG</td><td>1000</td><td></td><td></td><td></td><td></td><td></td><td></td><td></td><td></td><td></td></tr>
<tr><td>Oligo fwd</td><td>CACCACCTGTCTCGTGGTGGAAGG</td><td></td><td></td><td></td><td></td><td></td><td></td><td></td><td></td><td></td><td></td></tr>
<tr><td>Oligo rev</td><td>AAACCCTTCCACCACGAGACAGGT</td><td></td><td></td><td></td><td></td><td></td><td></td><td></td><td></td><td></td><td></td></tr>
<tr><td>Chromosome</td><td>start</td><td>end</td><td>strand</td><td>MM</td><td>target_seq</td><td>PAM</td><td>alignment</td><td>distance</td><td>position</td><td>gene name</td><td>gene id</td></tr>
<tr><td>GU305914.1</td><td>128352</td><td>128374</td><td>-</td><td>0</td><td>ACCTGTCTCGTGGTGGAAGG</td><td>CGG</td><td>||||||||[||||||||||||]PAM</td><td>NA</td><td>Intergenic</td><td>NA</td><td>NA</td></tr>
<tr><td>T24245</td><td>GTGGTGGAAGGCGGAGAAGACGG</td><td>1000</td><td></td><td></td><td></td><td></td><td></td><td></td><td></td><td></td><td></td></tr>
<tr><td>Oligo fwd</td><td>CACCGTGGTGGAAGGCGGAGAAGA</td><td></td><td></td><td></td><td></td><td></td><td></td><td></td><td></td><td></td><td></td></tr>
<tr><td>Oligo rev</td><td>AAACTCTTCTCCGCCTTCCACCAC</td><td></td><td></td><td></td><td></td><td></td><td></td><td></td><td></td><td></td><td></td></tr>
<tr><td>Chromosome</td><td>start</td><td>end</td><td>strand</td><td>MM</td><td>target_seq</td><td>PAM</td><td>alignment</td><td>distance</td><td>position</td><td>gene name</td><td>gene id</td></tr>
<tr><td>GU305914.1</td><td>128343</td><td>128365</td><td>-</td><td>0</td><td>GTGGTGGAAGGCGGAGAAGA</td><td>CGG</td><td>||||||||[||||||||||||]PAM</td><td>NA</td><td>Intergenic</td><td>NA</td><td>NA</td></tr>
</table>

| | | | | | | | | | | | |
|---|---|---|---|---|---|---|---|---|---|---|---|
| GU305914.1 | 147524 | 147546 | - | 4 | GGACTGGAAGGCGCAGAAGA | CGG | \|---\|\|\|\|[\|\|\|\|\|-\|\|\|\|\|\|]PAM | NA | Intergenic | NA | NA |
| T24246 | AGACGGTCTCGAAGAAGTCGCGG | 1000 | | | | | | | | | |
| Oligo fwd | CACCAGACGGTCTCGAAGAAGTCG | | | | | | | | | | |
| Oligo rev | AAACCGACTTCTTCGAGACCGTCT | | | | | | | | | | |
| Chromosome | start | end | strand | MM | target_seq | PAM | alignment | distance | position | gene name | gene id |
| GU305914.1 | 128326 | 128348 | - | 0 | AGACGGTCTCGAAGAAGTCG | CGG | \|\|\|\|\|\|\|\|[\|\|\|\|\|\|\|\|\|\|\|\|]PAM | NA | Intergenic | NA | NA |
| T24247 | GACGGTCTCGAAGAAGTCGCGGG | 1000 | | | | | | | | | |
| Oligo fwd | CACCGACGGTCTCGAAGAAGTCGC | | | | | | | | | | |
| Oligo rev | AAACGCGACTTCTTCGAGACCGTC | | | | | | | | | | |
| Chromosome | start | end | strand | MM | target_seq | PAM | alignment | distance | position | gene name | gene id |
| GU305914.1 | 128325 | 128347 | - | 0 | GACGGTCTCGAAGAAGTCGC | GGG | \|\|\|\|\|\|\|\|[\|\|\|\|\|\|\|\|\|\|\|\|]PAM | NA | Intergenic | NA | NA |
| GU305914.1 | 136295 | 136317 | - | 4 | GAAGGTCGCGCAGAAGCCGC | CGG | \|\|-\|\|\|\|-[\|\|-\|\|\|\|\|-\|\|\|]PAM | NA | Intergenic | NA | NA |
| T24248 | AGTCGCGGGTGACCGCATTACGG | 1000 | | | | | | | | | |
| Oligo fwd | CACCAGTCGCGGGTGACCGCATTA | | | | | | | | | | |
| Oligo rev | AAACTAATGCGGTCACCCGCGACT | | | | | | | | | | |
| Chromosome | start | end | strand | MM | target_seq | PAM | alignment | distance | position | gene name | gene id |
| GU305914.1 | 128311 | 128333 | - | 0 | AGTCGCGGGTGACCGCATTA | CGG | \|\|\|\|\|\|\|\|[\|\|\|\|\|\|\|\|\|\|\|\|]PAM | NA | Intergenic | NA | NA |
| T24249 | GCGGGTGACCGCATTACGGTAGG | 1000 | | | | | | | | | |
| Oligo fwd | CACCGCGGGTGACCGCATTACGGT | | | | | | | | | | |
| Oligo rev | AAACACCGTAATGCGGTCACCCGC | | | | | | | | | | |
| Chromosome | start | end | strand | MM | target_seq | PAM | alignment | distance | position | gene name | gene id |
| GU305914.1 | 128307 | 128329 | - | 0 | GCGGGTGACCGCATTACGGT | AGG | \|\|\|\|\|\|\|\|[\|\|\|\|\|\|\|\|\|\|\|\|]PAM | NA | Intergenic | NA | NA |
| T24250 | AGAGCTTCTGCAGCTCGAGCAGG | 1000 | | | | | | | | | |
| Oligo fwd | CACCAGAGCTTCTGCAGCTCGAGC | | | | | | | | | | |
| Oligo rev | AAACGCTCGAGCTGCAGAAGCTCT | | | | | | | | | | |
| Chromosome | start | end | strand | MM | target_seq | PAM | alignment | distance | position | gene name | gene id |
| GU305914.1 | 128272 | 128294 | - | 0 | AGAGCTTCTGCAGCTCGAGC | AGG | \|\|\|\|\|\|\|\|[\|\|\|\|\|\|\|\|\|\|\|\|]PAM | NA | Intergenic | NA | NA |
| T24251 | TCTGCAGCTCGAGCAGGAAGAGG | 1000 | | | | | | | | | |
| Oligo fwd | CACCTCTGCAGCTCGAGCAGGAAG | | | | | | | | | | |
| Oligo rev | AAACCTTCCTGCTCGAGCTGCAGA | | | | | | | | | | |
| Chromosome | start | end | strand | MM | target_seq | PAM | alignment | distance | position | gene name | gene id |
| GU305914.1 | 128266 | 128288 | - | 0 | TCTGCAGCTCGAGCAGGAAG | AGG | \|\|\|\|\|\|\|\|[\|\|\|\|\|\|\|\|\|\|\|\|]PAM | NA | Intergenic | NA | NA |
| T24252 | GAGCAGGAAGAGGTCGTAGTTGG | 1000 | | | | | | | | | |
| Oligo fwd | CACCGAGCAGGAAGAGGTCGTAGT | | | | | | | | | | |
| Oligo rev | AAACACTACGACCTCTTCCTGCTC | | | | | | | | | | |
| Chromosome | start | end | strand | MM | target_seq | PAM | alignment | distance | position | gene name | gene id |
| GU305914.1 | 128256 | 128278 | - | 0 | GAGCAGGAAGAGGTCGTAGT | TGG | \|\|\|\|\|\|\|\|[\|\|\|\|\|\|\|\|\|\|\|\|]PAM | NA | Intergenic | NA | NA |
| GU305914.1 | 28667 | 28689 | - | 4 | TTGCAGGAACAGCTCGTAGT | TGG | --\|\|\|\|\|\|[\|-\|\|-\|\|\|\|\|\|\|]PAM | NA | Intergenic | NA | NA |
| T24253 | GGAAGAGGTCGTAGTTGGCGCGG | 1000 | | | | | | | | | |
| Oligo fwd | CACCGGAAGAGGTCGTAGTTGGCG | | | | | | | | | | |
| Oligo rev | AAACCGCCAACTACGACCTCTTCC | | | | | | | | | | |
| Chromosome | start | end | strand | MM | target_seq | PAM | alignment | distance | position | gene name | gene id |
| GU305914.1 | 128251 | 128273 | - | 0 | GGAAGAGGTCGTAGTTGGCG | CGG | \|\|\|\|\|\|\|\|[\|\|\|\|\|\|\|\|\|\|\|\|]PAM | NA | Intergenic | NA | NA |
| T24254 | AGAGGTCGTAGTTGGCGCGGCGG | 1000 | | | | | | | | | |
| Oligo fwd | CACCAGAGGTCGTAGTTGGCGCGG | | | | | | | | | | |
| Oligo rev | AAACCCGCGCCAACTACGACCTCT | | | | | | | | | | |
| Chromosome | start | end | strand | MM | target_seq | PAM | alignment | distance | position | gene name | gene id |
| GU305914.1 | 128248 | 128270 | - | 0 | AGAGGTCGTAGTTGGCGCGG | CGG | \|\|\|\|\|\|\|\|[\|\|\|\|\|\|\|\|\|\|\|\|]PAM | NA | Intergenic | NA | NA |
| T24255 | GAGGTCGTAGTTGGCGCGGCGGG | 1000 | | | | | | | | | |
| Oligo fwd | CACCGAGGTCGTAGTTGGCGCGGC | | | | | | | | | | |
| Oligo rev | AAACGCCGCGCCAACTACGACCTC | | | | | | | | | | |
| Chromosome | start | end | strand | MM | target_seq | PAM | alignment | distance | position | gene name | gene id |
| GU305914.1 | 128247 | 128269 | - | 0 | GAGGTCGTAGTTGGCGCGGC | GGG | \|\|\|\|\|\|\|\|[\|\|\|\|\|\|\|\|\|\|\|\|]PAM | NA | Intergenic | NA | NA |
| T24256 | GTTGGCGCGGCGGGCCACCTCGG | 1000 | | | | | | | | | |
| Oligo fwd | CACCGTTGGCGCGGCGGGCCACCT | | | | | | | | | | |
| Oligo rev | AAACAGGTGGCCCGCCGCGCCAAC | | | | | | | | | | |
| Chromosome | start | end | strand | MM | target_seq | PAM | alignment | distance | position | gene name | gene id |
| GU305914.1 | 128238 | 128260 | - | 0 | GTTGGCGCGGCGGGCCACCT | CGG | \|\|\|\|\|\|\|\|[\|\|\|\|\|\|\|\|\|\|\|\|]PAM | NA | Intergenic | NA | NA |
| T24257 | CGGGCCACCTCGGTGACCCGCGG | 1000 | | | | | | | | | |
| Oligo fwd | CACCCGGGCCACCTCGGTGACCCG | | | | | | | | | | |
| Oligo rev | AAACCGGGTCACCGAGGTGGCCCG | | | | | | | | | | |
| Chromosome | start | end | strand | MM | target_seq | PAM | alignment | distance | position | gene name | gene id |
| GU305914.1 | 128228 | 128250 | - | 0 | CGGGCCACCTCGGTGACCCG | CGG | \|\|\|\|\|\|\|\|[\|\|\|\|\|\|\|\|\|\|\|\|]PAM | NA | Intergenic | NA | NA |
| T24258 | GCCACCTCGGTGACCCGCGGCGG | 1000 | | | | | | | | | |
| Oligo fwd | CACCGCCACCTCGGTGACCCGCGG | | | | | | | | | | |
| Oligo rev | AAACCCGCGGGTCACCGAGGTGGC | | | | | | | | | | |
| Chromosome | start | end | strand | MM | target_seq | PAM | alignment | distance | position | gene name | gene id |
| GU305914.1 | 128225 | 128247 | - | 0 | GCCACCTCGGTGACCCGCGG | CGG | \|\|\|\|\|\|\|\|[\|\|\|\|\|\|\|\|\|\|\|\|]PAM | NA | Intergenic | NA | NA |
| T24259 | CCTCGGTGACCCGCGGCGGCCGG | 1000 | | | | | | | | | |
| Oligo fwd | CACCCCTCGGTGACCCGCGGCGGC | | | | | | | | | | |
| Oligo rev | AAACGCCGCCGCGGGTCACCGAGG | | | | | | | | | | |
| Chromosome | start | end | strand | MM | target_seq | PAM | alignment | distance | position | gene name | gene id |
| GU305914.1 | 128221 | 128243 | - | 0 | CCTCGGTGACCCGCGGCGGC | CGG | \|\|\|\|\|\|\|\|[\|\|\|\|\|\|\|\|\|\|\|\|]PAM | NA | Intergenic | NA | NA |
| T24260 | CTCGGTGACCCGCGGCGGCCGGG | 1000 | | | | | | | | | |

<table>
<tr><td>Oligo fwd</td><td>CACCCTCGGTGACCCGCGGCGGCC</td><td></td><td></td><td></td><td></td><td></td><td></td><td></td><td></td><td></td><td></td></tr>
<tr><td>Oligo rev</td><td>AAACGGCCGCCGCGGGTCACCGAG</td><td></td><td></td><td></td><td></td><td></td><td></td><td></td><td></td><td></td><td></td></tr>
<tr><td>Chromosome</td><td>start</td><td>end</td><td>strand</td><td>MM</td><td>target_seq</td><td>PAM</td><td>alignment</td><td>distance</td><td>position</td><td>gene name</td><td>gene id</td></tr>
<tr><td>GU305914.1</td><td>128220</td><td>128242</td><td>-</td><td>0</td><td>CTCGGTGACCCGCGGCGGCC</td><td>GGG</td><td>||||||||[||||||||||||]PAM</td><td>NA</td><td>Intergenic</td><td>NA</td><td>NA</td></tr>
<tr><td>T24261</td><td>TCTTGTAGTGCACGCCGACGCGG</td><td>1000</td><td></td><td></td><td></td><td></td><td></td><td></td><td></td><td></td><td></td></tr>
<tr><td>Oligo fwd</td><td>CACCTCTTGTAGTGCACGCCGACG</td><td></td><td></td><td></td><td></td><td></td><td></td><td></td><td></td><td></td><td></td></tr>
<tr><td>Oligo rev</td><td>AAACCGTCGGCGTGCACTACAAGA</td><td></td><td></td><td></td><td></td><td></td><td></td><td></td><td></td><td></td><td></td></tr>
<tr><td>Chromosome</td><td>start</td><td>end</td><td>strand</td><td>MM</td><td>target_seq</td><td>PAM</td><td>alignment</td><td>distance</td><td>position</td><td>gene name</td><td>gene id</td></tr>
<tr><td>GU305914.1</td><td>128179</td><td>128201</td><td>-</td><td>0</td><td>TCTTGTAGTGCACGCCGACG</td><td>CGG</td><td>||||||||[||||||||||||]PAM</td><td>NA</td><td>Intergenic</td><td>NA</td><td>NA</td></tr>
<tr><td>T24262</td><td>CTTGTAGTGCACGCCGACGCGGG</td><td>1000</td><td></td><td></td><td></td><td></td><td></td><td></td><td></td><td></td><td></td></tr>
<tr><td>Oligo fwd</td><td>CACCCTTGTAGTGCACGCCGACGC</td><td></td><td></td><td></td><td></td><td></td><td></td><td></td><td></td><td></td><td></td></tr>
<tr><td>Oligo rev</td><td>AAACGCGTCGGCGTGCACTACAAG</td><td></td><td></td><td></td><td></td><td></td><td></td><td></td><td></td><td></td><td></td></tr>
<tr><td>Chromosome</td><td>start</td><td>end</td><td>strand</td><td>MM</td><td>target_seq</td><td>PAM</td><td>alignment</td><td>distance</td><td>position</td><td>gene name</td><td>gene id</td></tr>
<tr><td>GU305914.1</td><td>128178</td><td>128200</td><td>-</td><td>0</td><td>CTTGTAGTGCACGCCGACGC</td><td>GGG</td><td>||||||||[||||||||||||]PAM</td><td>NA</td><td>Intergenic</td><td>NA</td><td>NA</td></tr>
<tr><td>T24263</td><td>GTAGTGCACGCCGACGCGGGCGG</td><td>1000</td><td></td><td></td><td></td><td></td><td></td><td></td><td></td><td></td><td></td></tr>
<tr><td>Oligo fwd</td><td>CACCGTAGTGCACGCCGACGCGGG</td><td></td><td></td><td></td><td></td><td></td><td></td><td></td><td></td><td></td><td></td></tr>
<tr><td>Oligo rev</td><td>AAACCCCGCGTCGGCGTGCACTAC</td><td></td><td></td><td></td><td></td><td></td><td></td><td></td><td></td><td></td><td></td></tr>
<tr><td>Chromosome</td><td>start</td><td>end</td><td>strand</td><td>MM</td><td>target_seq</td><td>PAM</td><td>alignment</td><td>distance</td><td>position</td><td>gene name</td><td>gene id</td></tr>
<tr><td>GU305914.1</td><td>128175</td><td>128197</td><td>-</td><td>0</td><td>GTAGTGCACGCCGACGCGGG</td><td>CGG</td><td>||||||||[||||||||||||]PAM</td><td>NA</td><td>Intergenic</td><td>NA</td><td>NA</td></tr>
<tr><td>T24264</td><td>CACGCCGACGCGGGCGGCGATGG</td><td>1000</td><td></td><td></td><td></td><td></td><td></td><td></td><td></td><td></td><td></td></tr>
<tr><td>Oligo fwd</td><td>CACCCACGCCGACGCGGGCGGCGA</td><td></td><td></td><td></td><td></td><td></td><td></td><td></td><td></td><td></td><td></td></tr>
<tr><td>Oligo rev</td><td>AAACTCGCCGCCCGCGTCGGCGTG</td><td></td><td></td><td></td><td></td><td></td><td></td><td></td><td></td><td></td><td></td></tr>
<tr><td>Chromosome</td><td>start</td><td>end</td><td>strand</td><td>MM</td><td>target_seq</td><td>PAM</td><td>alignment</td><td>distance</td><td>position</td><td>gene name</td><td>gene id</td></tr>
<tr><td>GU305914.1</td><td>128169</td><td>128191</td><td>-</td><td>0</td><td>CACGCCGACGCGGGCGGCGA</td><td>TGG</td><td>||||||||[||||||||||||]PAM</td><td>NA</td><td>Intergenic</td><td>NA</td><td>NA</td></tr>
<tr><td>T24265</td><td>CCGACGCGGGCGGCGATGGCCGG</td><td>1000</td><td></td><td></td><td></td><td></td><td></td><td></td><td></td><td></td><td></td></tr>
<tr><td>Oligo fwd</td><td>CACCCCGACGCGGGCGGCGATGGC</td><td></td><td></td><td></td><td></td><td></td><td></td><td></td><td></td><td></td><td></td></tr>
<tr><td>Oligo rev</td><td>AAACGCCATCGCCGCCCGCGTCGG</td><td></td><td></td><td></td><td></td><td></td><td></td><td></td><td></td><td></td><td></td></tr>
<tr><td>Chromosome</td><td>start</td><td>end</td><td>strand</td><td>MM</td><td>target_seq</td><td>PAM</td><td>alignment</td><td>distance</td><td>position</td><td>gene name</td><td>gene id</td></tr>
<tr><td>GU305914.1</td><td>128165</td><td>128187</td><td>-</td><td>0</td><td>CCGACGCGGGCGGCGATGGC</td><td>CGG</td><td>||||||||[||||||||||||]PAM</td><td>NA</td><td>Intergenic</td><td>NA</td><td>NA</td></tr>
<tr><td>T24266</td><td>GGGCGGCGATGGCCGGACCGAGG</td><td>1000</td><td></td><td></td><td></td><td></td><td></td><td></td><td></td><td></td><td></td></tr>
<tr><td>Oligo fwd</td><td>CACCGGGCGGCGATGGCCGGACCG</td><td></td><td></td><td></td><td></td><td></td><td></td><td></td><td></td><td></td><td></td></tr>
<tr><td>Oligo rev</td><td>AAACCGGTCCGGCCATCGCCGCCC</td><td></td><td></td><td></td><td></td><td></td><td></td><td></td><td></td><td></td><td></td></tr>
<tr><td>Chromosome</td><td>start</td><td>end</td><td>strand</td><td>MM</td><td>target_seq</td><td>PAM</td><td>alignment</td><td>distance</td><td>position</td><td>gene name</td><td>gene id</td></tr>
<tr><td>GU305914.1</td><td>128158</td><td>128180</td><td>-</td><td>0</td><td>GGGCGGCGATGGCCGGACCG</td><td>AGG</td><td>||||||||[||||||||||||]PAM</td><td>NA</td><td>Intergenic</td><td>NA</td><td>NA</td></tr>
<tr><td>T24267</td><td>GATGGCCGGACCGAGGTGCGCGG</td><td>1000</td><td></td><td></td><td></td><td></td><td></td><td></td><td></td><td></td><td></td></tr>
<tr><td>Oligo fwd</td><td>CACCGATGGCCGGACCGAGGTGCG</td><td></td><td></td><td></td><td></td><td></td><td></td><td></td><td></td><td></td><td></td></tr>
<tr><td>Oligo rev</td><td>AAACCGCACCTCGGTCCGGCCATC</td><td></td><td></td><td></td><td></td><td></td><td></td><td></td><td></td><td></td><td></td></tr>
<tr><td>Chromosome</td><td>start</td><td>end</td><td>strand</td><td>MM</td><td>target_seq</td><td>PAM</td><td>alignment</td><td>distance</td><td>position</td><td>gene name</td><td>gene id</td></tr>
<tr><td>GU305914.1</td><td>128151</td><td>128173</td><td>-</td><td>0</td><td>GATGGCCGGACCGAGGTGCG</td><td>CGG</td><td>||||||||[||||||||||||]PAM</td><td>NA</td><td>Intergenic</td><td>NA</td><td>NA</td></tr>
<tr><td>T24268</td><td>CGGACCGAGGTGCGCGGTCATGG</td><td>1000</td><td></td><td></td><td></td><td></td><td></td><td></td><td></td><td></td><td></td></tr>
<tr><td>Oligo fwd</td><td>CACCCGGACCGAGGTGCGCGGTCA</td><td></td><td></td><td></td><td></td><td></td><td></td><td></td><td></td><td></td><td></td></tr>
<tr><td>Oligo rev</td><td>AAACTGACCGCGCACCTCGGTCCG</td><td></td><td></td><td></td><td></td><td></td><td></td><td></td><td></td><td></td><td></td></tr>
<tr><td>Chromosome</td><td>start</td><td>end</td><td>strand</td><td>MM</td><td>target_seq</td><td>PAM</td><td>alignment</td><td>distance</td><td>position</td><td>gene name</td><td>gene id</td></tr>
<tr><td>GU305914.1</td><td>128145</td><td>128167</td><td>-</td><td>0</td><td>CGGACCGAGGTGCGCGGTCA</td><td>TGG</td><td>||||||||[||||||||||||]PAM</td><td>NA</td><td>Intergenic</td><td>NA</td><td>NA</td></tr>
<tr><td>T24269</td><td>GAGGTGCGCGGTCATGGAGTTGG</td><td>1000</td><td></td><td></td><td></td><td></td><td></td><td></td><td></td><td></td><td></td></tr>
<tr><td>Oligo fwd</td><td>CACCGAGGTGCGCGGTCATGGAGT</td><td></td><td></td><td></td><td></td><td></td><td></td><td></td><td></td><td></td><td></td></tr>
<tr><td>Oligo rev</td><td>AAACACTCCATGACCGCGCACCTC</td><td></td><td></td><td></td><td></td><td></td><td></td><td></td><td></td><td></td><td></td></tr>
<tr><td>Chromosome</td><td>start</td><td>end</td><td>strand</td><td>MM</td><td>target_seq</td><td>PAM</td><td>alignment</td><td>distance</td><td>position</td><td>gene name</td><td>gene id</td></tr>
<tr><td>GU305914.1</td><td>128139</td><td>128161</td><td>-</td><td>0</td><td>GAGGTGCGCGGTCATGGAGT</td><td>TGG</td><td>||||||||[||||||||||||]PAM</td><td>NA</td><td>Intergenic</td><td>NA</td><td>NA</td></tr>
<tr><td>T24270</td><td>GTTGGCGCAGTCGTCGATCTCGG</td><td>1000</td><td></td><td></td><td></td><td></td><td></td><td></td><td></td><td></td><td></td></tr>
<tr><td>Oligo fwd</td><td>CACCGTTGGCGCAGTCGTCGATCT</td><td></td><td></td><td></td><td></td><td></td><td></td><td></td><td></td><td></td><td></td></tr>
<tr><td>Oligo rev</td><td>AAACAGATCGACGACTGCGCCAAC</td><td></td><td></td><td></td><td></td><td></td><td></td><td></td><td></td><td></td><td></td></tr>
<tr><td>Chromosome</td><td>start</td><td>end</td><td>strand</td><td>MM</td><td>target_seq</td><td>PAM</td><td>alignment</td><td>distance</td><td>position</td><td>gene name</td><td>gene id</td></tr>
<tr><td>GU305914.1</td><td>128121</td><td>128143</td><td>-</td><td>0</td><td>GTTGGCGCAGTCGTCGATCT</td><td>CGG</td><td>||||||||[||||||||||||]PAM</td><td>NA</td><td>Intergenic</td><td>NA</td><td>NA</td></tr>
<tr><td>T24271</td><td>TTGGCGCAGTCGTCGATCTCGGG</td><td>1000</td><td></td><td></td><td></td><td></td><td></td><td></td><td></td><td></td><td></td></tr>
<tr><td>Oligo fwd</td><td>CACCTTGGCGCAGTCGTCGATCTC</td><td></td><td></td><td></td><td></td><td></td><td></td><td></td><td></td><td></td><td></td></tr>
<tr><td>Oligo rev</td><td>AAACGAGATCGACGACTGCGCCAA</td><td></td><td></td><td></td><td></td><td></td><td></td><td></td><td></td><td></td><td></td></tr>
<tr><td>Chromosome</td><td>start</td><td>end</td><td>strand</td><td>MM</td><td>target_seq</td><td>PAM</td><td>alignment</td><td>distance</td><td>position</td><td>gene name</td><td>gene id</td></tr>
<tr><td>GU305914.1</td><td>128120</td><td>128142</td><td>-</td><td>0</td><td>TTGGCGCAGTCGTCGATCTC</td><td>GGG</td><td>||||||||[||||||||||||]PAM</td><td>NA</td><td>Intergenic</td><td>NA</td><td>NA</td></tr>
<tr><td>T24272</td><td>GCAGTCGTCGATCTCGGGCACGG</td><td>1000</td><td></td><td></td><td></td><td></td><td></td><td></td><td></td><td></td><td></td></tr>
<tr><td>Oligo fwd</td><td>CACCGCAGTCGTCGATCTCGGGCA</td><td></td><td></td><td></td><td></td><td></td><td></td><td></td><td></td><td></td><td></td></tr>
<tr><td>Oligo rev</td><td>AAACTGCCCGAGATCGACGACTGC</td><td></td><td></td><td></td><td></td><td></td><td></td><td></td><td></td><td></td><td></td></tr>
<tr><td>Chromosome</td><td>start</td><td>end</td><td>strand</td><td>MM</td><td>target_seq</td><td>PAM</td><td>alignment</td><td>distance</td><td>position</td><td>gene name</td><td>gene id</td></tr>
<tr><td>GU305914.1</td><td>128115</td><td>128137</td><td>-</td><td>0</td><td>GCAGTCGTCGATCTCGGGCA</td><td>CGG</td><td>||||||||[||||||||||||]PAM</td><td>NA</td><td>Intergenic</td><td>NA</td><td>NA</td></tr>
<tr><td>T24273</td><td>CGTCGATCTCGGGCACGGTGAGG</td><td>1000</td><td></td><td></td><td></td><td></td><td></td><td></td><td></td><td></td><td></td></tr>
<tr><td>Oligo fwd</td><td>CACCCGTCGATCTCGGGCACGGTG</td><td></td><td></td><td></td><td></td><td></td><td></td><td></td><td></td><td></td><td></td></tr>
<tr><td>Oligo rev</td><td>AAACCACCGTGCCCGAGATCGACG</td><td></td><td></td><td></td><td></td><td></td><td></td><td></td><td></td><td></td><td></td></tr>
<tr><td>Chromosome</td><td>start</td><td>end</td><td>strand</td><td>MM</td><td>target_seq</td><td>PAM</td><td>alignment</td><td>distance</td><td>position</td><td>gene name</td><td>gene id</td></tr>
<tr><td>GU305914.1</td><td>128110</td><td>128132</td><td>-</td><td>0</td><td>CGTCGATCTCGGGCACGGTG</td><td>AGG</td><td>||||||||[||||||||||||]PAM</td><td>NA</td><td>Intergenic</td><td>NA</td><td>NA</td></tr>
<tr><td>T24274</td><td>CTCGGGCACGGTGAGGTACCTGG</td><td>1000</td><td></td><td></td><td></td><td></td><td></td><td></td><td></td><td></td><td></td></tr>
<tr><td>Oligo fwd</td><td>CACCCTCGGGCACGGTGAGGTACC</td><td></td><td></td><td></td><td></td><td></td><td></td><td></td><td></td><td></td><td></td></tr>
<tr><td>Oligo rev</td><td>AAACGGTACCTCACCGTGCCCGAG</td><td></td><td></td><td></td><td></td><td></td><td></td><td></td><td></td><td></td><td></td></tr>
<tr><td>Chromosome</td><td>start</td><td>end</td><td>strand</td><td>MM</td><td>target_seq</td><td>PAM</td><td>alignment</td><td>distance</td><td>position</td><td>gene name</td><td>gene id</td></tr>
<tr><td>GU305914.1</td><td>128103</td><td>128125</td><td>-</td><td>0</td><td>CTCGGGCACGGTGAGGTACC</td><td>TGG</td><td>||||||||[||||||||||||]PAM</td><td>NA</td><td>Intergenic</td><td>NA</td><td>NA</td></tr>
<tr><td>GU305914.1</td><td>81183</td><td>81205</td><td>-</td><td>3</td><td>CTCGCGGACGGTGGGGTACC</td><td>AGG</td><td>||||-|-|[|||||-||||||]PAM</td><td>NA</td><td>Intergenic</td><td>NA</td><td>NA</td></tr>
</table>

T24275 GCACGGTGAGGTACCTGGCGAGG 1000
Oligo fwd CACCGCACGGTGAGGTACCTGGCG
Oligo rev AAACCGCCAGGTACCTCACCGTGC
Chromosome start end strand MM target_seq PAM alignment distance position gene name gene id
GU305914.1 128098 128120 - 0 GCACGGTGAGGTACCTGGCG AGG ||||||||[||||||||||||]PAM NA Intergenic NA NA

T24276 GGTACCTGGCGAGGCAGACGAGG 1000
Oligo fwd CACCGGTACCTGGCGAGGCAGACG
Oligo rev AAACCGTCTGCCTCGCCAGGTACC
Chromosome start end strand MM target_seq PAM alignment distance position gene name gene id
GU305914.1 128089 128111 - 0 GGTACCTGGCGAGGCAGACG AGG ||||||||[||||||||||||]PAM NA Intergenic NA NA
GU305914.1 80220 80242 + 4 CGTCCCTGGCGCGGCAGAGG CGG -||-||||[|||-||||||-|]PAM NA Intergenic NA NA

T24277 GGCGAGGCAGACGAGGAAATAGG 1000
Oligo fwd CACCGGCGAGGCAGACGAGGAAAT
Oligo rev AAACATTTCCTCGTCTGCCTCGCC
Chromosome start end strand MM target_seq PAM alignment distance position gene name gene id
GU305914.1 128082 128104 - 0 GGCGAGGCAGACGAGGAAAT AGG ||||||||[||||||||||||]PAM NA Intergenic NA NA

T24278 GCGAGGCAGACGAGGAAATAGGG 1000
Oligo fwd CACCGCGAGGCAGACGAGGAAATA
Oligo rev AAACTATTTCCTCGTCTGCCTCGC
Chromosome start end strand MM target_seq PAM alignment distance position gene name gene id
GU305914.1 128081 128103 - 0 GCGAGGCAGACGAGGAAATA GGG ||||||||[||||||||||||]PAM NA Intergenic NA NA

T24279 CGAGGCAGACGAGGAAATAGGGG 1000
Oligo fwd CACCCGAGGCAGACGAGGAAATAG
Oligo rev AAACCTATTTCCTCGTCTGCCTCG
Chromosome start end strand MM target_seq PAM alignment distance position gene name gene id
GU305914.1 128080 128102 - 0 CGAGGCAGACGAGGAAATAG GGG ||||||||[||||||||||||]PAM NA Intergenic NA NA

T24280 GGGGTAGTACTCGATAAAGACGG 1000
Oligo fwd CACCGGGGTAGTACTCGATAAAGA
Oligo rev AAACTCTTTATCGAGTACTACCCC
Chromosome start end strand MM target_seq PAM alignment distance position gene name gene id
GU305914.1 128061 128083 - 0 GGGGTAGTACTCGATAAAGA CGG ||||||||[||||||||||||]PAM NA Intergenic NA NA

T24281 AGTACTCGATAAAGACGGACTGG 1000
Oligo fwd CACCAGTACTCGATAAAGACGGAC
Oligo rev AAACGTCCGTCTTTATCGAGTACT
Chromosome start end strand MM target_seq PAM alignment distance position gene name gene id
GU305914.1 128056 128078 - 0 AGTACTCGATAAAGACGGAC TGG ||||||||[||||||||||||]PAM NA Intergenic NA NA

T24282 GATAAAGACGGACTGGTGCTTGG 1000
Oligo fwd CACCGATAAAGACGGACTGGTGCT
Oligo rev AAACAGCACCAGTCCGTCTTTATC
Chromosome start end strand MM target_seq PAM alignment distance position gene name gene id
GU305914.1 128049 128071 - 0 GATAAAGACGGACTGGTGCT TGG ||||||||[||||||||||||]PAM NA Intergenic NA NA

T24283 ATAAAGACGGACTGGTGCTTGGG 1000
Oligo fwd CACCATAAAGACGGACTGGTGCTT
Oligo rev AAACAAGCACCAGTCCGTCTTTAT
Chromosome start end strand MM target_seq PAM alignment distance position gene name gene id
GU305914.1 128048 128070 - 0 ATAAAGACGGACTGGTGCTT GGG ||||||||[||||||||||||]PAM NA Intergenic NA NA

T24284 GGACTGGTGCTTGGGCAGCACGG 1000
Oligo fwd CACCGGACTGGTGCTTGGGCAGCA
Oligo rev AAACTGCTGCCCAAGCACCAGTCC
Chromosome start end strand MM target_seq PAM alignment distance position gene name gene id
GU305914.1 128040 128062 - 0 GGACTGGTGCTTGGGCAGCA CGG ||||||||[||||||||||||]PAM NA Intergenic NA NA

T24285 GCAGCACGGCCATGACATACCGG 1000
Oligo fwd CACCGCAGCACGGCCATGACATAC
Oligo rev AAACGTATGTCATGGCCGTGCTGC
Chromosome start end strand MM target_seq PAM alignment distance position gene name gene id
GU305914.1 128026 128048 - 0 GCAGCACGGCCATGACATAC CGG ||||||||[||||||||||||]PAM NA Intergenic NA NA

T24286 CACGGCCATGACATACCGGTCGG 1000
Oligo fwd CACCCACGGCCATGACATACCGGT
Oligo rev AAACACCGGTATGTCATGGCCGTG
Chromosome start end strand MM target_seq PAM alignment distance position gene name gene id
GU305914.1 128022 128044 - 0 CACGGCCATGACATACCGGT CGG ||||||||[||||||||||||]PAM NA Intergenic NA NA

T24287 GACATACCGGTCGGAGCGAAAGG 1000
Oligo fwd CACCGACATACCGGTCGGAGCGAA
Oligo rev AAACTTCGCTCCGACCGGTATGTC
Chromosome start end strand MM target_seq PAM alignment distance position gene name gene id
GU305914.1 128013 128035 - 0 GACATACCGGTCGGAGCGAA AGG ||||||||[||||||||||||]PAM NA Intergenic NA NA

T24288 GTCGGAGCGAAAGGCGTTACAGG 1000
Oligo fwd CACCGTCGGAGCGAAAGGCGTTAC
Oligo rev AAACGTAACGCCTTTCGCTCCGAC
Chromosome start end strand MM target_seq PAM alignment distance position gene name gene id
GU305914.1 128004 128026 - 0 GTCGGAGCGAAAGGCGTTAC AGG ||||||||[||||||||||||]PAM NA Intergenic NA NA

T24289 GTTACAGGCGCAGAAGACGTCGG 1000
Oligo fwd CACCGTTACAGGCGCAGAAGACGT
Oligo rev AAACACGTCTTCTGCGCCTGTAAC

<table>
<tr><td>Chromosome</td><td>start</td><td>end</td><td>strand</td><td>MM</td><td>target_seq</td><td>PAM</td><td>alignment</td><td>distance</td><td>position</td><td>gene name</td><td>gene id</td></tr>
<tr><td>GU305914.1</td><td>127989</td><td>128011</td><td>-</td><td>0</td><td>GTTACAGGCGCAGAAGACGT</td><td>CGG</td><td>||||||||[||||||||||||]PAM</td><td>NA</td><td>Intergenic</td><td>NA</td><td>NA</td></tr>
<tr><td></td><td></td><td></td><td></td><td></td><td></td><td></td><td></td><td></td><td></td><td></td><td></td></tr>
<tr><td>T24290</td><td>AGGCGCAGAAGACGTCGGTCAGG</td><td>1000</td><td></td><td></td><td></td><td></td><td></td><td></td><td></td><td></td><td></td></tr>
<tr><td>Oligo fwd</td><td>CACCAGGCGCAGAAGACGTCGGTC</td><td></td><td></td><td></td><td></td><td></td><td></td><td></td><td></td><td></td><td></td></tr>
<tr><td>Oligo rev</td><td>AAACGACCGACGTCTTCTGCGCCT</td><td></td><td></td><td></td><td></td><td></td><td></td><td></td><td></td><td></td><td></td></tr>
<tr><td>Chromosome</td><td>start</td><td>end</td><td>strand</td><td>MM</td><td>target_seq</td><td>PAM</td><td>alignment</td><td>distance</td><td>position</td><td>gene name</td><td>gene id</td></tr>
<tr><td>GU305914.1</td><td>127984</td><td>128006</td><td>-</td><td>0</td><td>AGGCGCAGAAGACGTCGGTC</td><td>AGG</td><td>||||||||[||||||||||||]PAM</td><td>NA</td><td>Intergenic</td><td>NA</td><td>NA</td></tr>
<tr><td></td><td></td><td></td><td></td><td></td><td></td><td></td><td></td><td></td><td></td><td></td><td></td></tr>
<tr><td>T24291</td><td>GGTCAGGCGCACGTTCCACTCGG</td><td>1000</td><td></td><td></td><td></td><td></td><td></td><td></td><td></td><td></td><td></td></tr>
<tr><td>Oligo fwd</td><td>CACCGGTCAGGCGCACGTTCCACT</td><td></td><td></td><td></td><td></td><td></td><td></td><td></td><td></td><td></td><td></td></tr>
<tr><td>Oligo rev</td><td>AAACAGTGGAACGTGCGCCTGACC</td><td></td><td></td><td></td><td></td><td></td><td></td><td></td><td></td><td></td><td></td></tr>
<tr><td>Chromosome</td><td>start</td><td>end</td><td>strand</td><td>MM</td><td>target_seq</td><td>PAM</td><td>alignment</td><td>distance</td><td>position</td><td>gene name</td><td>gene id</td></tr>
<tr><td>GU305914.1</td><td>127968</td><td>127990</td><td>-</td><td>0</td><td>GGTCAGGCGCACGTTCCACT</td><td>CGG</td><td>||||||||[||||||||||||]PAM</td><td>NA</td><td>Intergenic</td><td>NA</td><td>NA</td></tr>
<tr><td></td><td></td><td></td><td></td><td></td><td></td><td></td><td></td><td></td><td></td><td></td><td></td></tr>
<tr><td>T24292</td><td>CAGCCCCCGAAACACGTGCGTGG</td><td>1000</td><td></td><td></td><td></td><td></td><td></td><td></td><td></td><td></td><td></td></tr>
<tr><td>Oligo fwd</td><td>CACCCAGCCCCCGAAACACGTGCG</td><td></td><td></td><td></td><td></td><td></td><td></td><td></td><td></td><td></td><td></td></tr>
<tr><td>Oligo rev</td><td>AAACCGCACGTGTTTCGGGGGCTG</td><td></td><td></td><td></td><td></td><td></td><td></td><td></td><td></td><td></td><td></td></tr>
<tr><td>Chromosome</td><td>start</td><td>end</td><td>strand</td><td>MM</td><td>target_seq</td><td>PAM</td><td>alignment</td><td>distance</td><td>position</td><td>gene name</td><td>gene id</td></tr>
<tr><td>GU305914.1</td><td>127935</td><td>127957</td><td>-</td><td>0</td><td>CAGCCCCCGAAACACGTGCG</td><td>TGG</td><td>||||||||[||||||||||||]PAM</td><td>NA</td><td>Intergenic</td><td>NA</td><td>NA</td></tr>
<tr><td></td><td></td><td></td><td></td><td></td><td></td><td></td><td></td><td></td><td></td><td></td><td></td></tr>
<tr><td>T24293</td><td>GAAACACGTGCGTGGTGTAGTGG</td><td>1000</td><td></td><td></td><td></td><td></td><td></td><td></td><td></td><td></td><td></td></tr>
<tr><td>Oligo fwd</td><td>CACCGAAACACGTGCGTGGTGTAG</td><td></td><td></td><td></td><td></td><td></td><td></td><td></td><td></td><td></td><td></td></tr>
<tr><td>Oligo rev</td><td>AAACCTACACCACGCACGTGTTTC</td><td></td><td></td><td></td><td></td><td></td><td></td><td></td><td></td><td></td><td></td></tr>
<tr><td>Chromosome</td><td>start</td><td>end</td><td>strand</td><td>MM</td><td>target_seq</td><td>PAM</td><td>alignment</td><td>distance</td><td>position</td><td>gene name</td><td>gene id</td></tr>
<tr><td>GU305914.1</td><td>127927</td><td>127949</td><td>-</td><td>0</td><td>GAAACACGTGCGTGGTGTAG</td><td>TGG</td><td>||||||||[||||||||||||]PAM</td><td>NA</td><td>Intergenic</td><td>NA</td><td>NA</td></tr>
<tr><td></td><td></td><td></td><td></td><td></td><td></td><td></td><td></td><td></td><td></td><td></td><td></td></tr>
<tr><td>T24294</td><td>GTCGTGCAACAACGTGAACATGG</td><td>1000</td><td></td><td></td><td></td><td></td><td></td><td></td><td></td><td></td><td></td></tr>
<tr><td>Oligo fwd</td><td>CACCGTCGTGCAACAACGTGAACA</td><td></td><td></td><td></td><td></td><td></td><td></td><td></td><td></td><td></td><td></td></tr>
<tr><td>Oligo rev</td><td>AAACTGTTCACGTTGTTGCACGAC</td><td></td><td></td><td></td><td></td><td></td><td></td><td></td><td></td><td></td><td></td></tr>
<tr><td>Chromosome</td><td>start</td><td>end</td><td>strand</td><td>MM</td><td>target_seq</td><td>PAM</td><td>alignment</td><td>distance</td><td>position</td><td>gene name</td><td>gene id</td></tr>
<tr><td>GU305914.1</td><td>127905</td><td>127927</td><td>-</td><td>0</td><td>GTCGTGCAACAACGTGAACA</td><td>TGG</td><td>||||||||[||||||||||||]PAM</td><td>NA</td><td>Intergenic</td><td>NA</td><td>NA</td></tr>
<tr><td></td><td></td><td></td><td></td><td></td><td></td><td></td><td></td><td></td><td></td><td></td><td></td></tr>
<tr><td>T24295</td><td>GAACATGGTCCCGCCGCCCTCGG</td><td>1000</td><td></td><td></td><td></td><td></td><td></td><td></td><td></td><td></td><td></td></tr>
<tr><td>Oligo fwd</td><td>CACCGAACATGGTCCCGCCGCCCT</td><td></td><td></td><td></td><td></td><td></td><td></td><td></td><td></td><td></td><td></td></tr>
<tr><td>Oligo rev</td><td>AAACAGGGCGGCGGGACCATGTTC</td><td></td><td></td><td></td><td></td><td></td><td></td><td></td><td></td><td></td><td></td></tr>
<tr><td>Chromosome</td><td>start</td><td>end</td><td>strand</td><td>MM</td><td>target_seq</td><td>PAM</td><td>alignment</td><td>distance</td><td>position</td><td>gene name</td><td>gene id</td></tr>
<tr><td>GU305914.1</td><td>127890</td><td>127912</td><td>-</td><td>0</td><td>GAACATGGTCCCGCCGCCCT</td><td>CGG</td><td>||||||||[||||||||||||]PAM</td><td>NA</td><td>Intergenic</td><td>NA</td><td>NA</td></tr>
<tr><td></td><td></td><td></td><td></td><td></td><td></td><td></td><td></td><td></td><td></td><td></td><td></td></tr>
<tr><td>T24296</td><td>CGCCGCCCTCGGCGCCGCAGCGG</td><td>1000</td><td></td><td></td><td></td><td></td><td></td><td></td><td></td><td></td><td></td></tr>
<tr><td>Oligo fwd</td><td>CACCCGCCGCCCTCGGCGCCGCAG</td><td></td><td></td><td></td><td></td><td></td><td></td><td></td><td></td><td></td><td></td></tr>
<tr><td>Oligo rev</td><td>AAACCTGCGGCGCCGAGGGCGGCG</td><td></td><td></td><td></td><td></td><td></td><td></td><td></td><td></td><td></td><td></td></tr>
<tr><td>Chromosome</td><td>start</td><td>end</td><td>strand</td><td>MM</td><td>target_seq</td><td>PAM</td><td>alignment</td><td>distance</td><td>position</td><td>gene name</td><td>gene id</td></tr>
<tr><td>GU305914.1</td><td>127879</td><td>127901</td><td>-</td><td>0</td><td>CGCCGCCCTCGGCGCCGCAG</td><td>CGG</td><td>||||||||[||||||||||||]PAM</td><td>NA</td><td>Intergenic</td><td>NA</td><td>NA</td></tr>
<tr><td></td><td></td><td></td><td></td><td></td><td></td><td></td><td></td><td></td><td></td><td></td><td></td></tr>
<tr><td>T24297</td><td>CGCCGCAGCGGCAGCGCGAGCGG</td><td>1000</td><td></td><td></td><td></td><td></td><td></td><td></td><td></td><td></td><td></td></tr>
<tr><td>Oligo fwd</td><td>CACCCGCCGCAGCGGCAGCGCGAG</td><td></td><td></td><td></td><td></td><td></td><td></td><td></td><td></td><td></td><td></td></tr>
<tr><td>Oligo rev</td><td>AAACCTCGCGCTGCCGCTGCGGCG</td><td></td><td></td><td></td><td></td><td></td><td></td><td></td><td></td><td></td><td></td></tr>
<tr><td>Chromosome</td><td>start</td><td>end</td><td>strand</td><td>MM</td><td>target_seq</td><td>PAM</td><td>alignment</td><td>distance</td><td>position</td><td>gene name</td><td>gene id</td></tr>
<tr><td>GU305914.1</td><td>127867</td><td>127889</td><td>-</td><td>0</td><td>CGCCGCAGCGGCAGCGCGAG</td><td>CGG</td><td>||||||||[||||||||||||]PAM</td><td>NA</td><td>Intergenic</td><td>NA</td><td>NA</td></tr>
<tr><td></td><td></td><td></td><td></td><td></td><td></td><td></td><td></td><td></td><td></td><td></td><td></td></tr>
<tr><td>T24298</td><td>GAGCGGTGCACGTAGAGCGCCGG</td><td>1000</td><td></td><td></td><td></td><td></td><td></td><td></td><td></td><td></td><td></td></tr>
<tr><td>Oligo fwd</td><td>CACCGAGCGGTGCACGTAGAGCGC</td><td></td><td></td><td></td><td></td><td></td><td></td><td></td><td></td><td></td><td></td></tr>
<tr><td>Oligo rev</td><td>AAACGCGCTCTACGTGCACCGCTC</td><td></td><td></td><td></td><td></td><td></td><td></td><td></td><td></td><td></td><td></td></tr>
<tr><td>Chromosome</td><td>start</td><td>end</td><td>strand</td><td>MM</td><td>target_seq</td><td>PAM</td><td>alignment</td><td>distance</td><td>position</td><td>gene name</td><td>gene id</td></tr>
<tr><td>GU305914.1</td><td>127850</td><td>127872</td><td>-</td><td>0</td><td>GAGCGGTGCACGTAGAGCGC</td><td>CGG</td><td>||||||||[||||||||||||]PAM</td><td>NA</td><td>Intergenic</td><td>NA</td><td>NA</td></tr>
<tr><td></td><td></td><td></td><td></td><td></td><td></td><td></td><td></td><td></td><td></td><td></td><td></td></tr>
<tr><td>T24299</td><td>AGAGCGCCGGTCCCGAGCAGCGG</td><td>1000</td><td></td><td></td><td></td><td></td><td></td><td></td><td></td><td></td><td></td></tr>
<tr><td>Oligo fwd</td><td>CACCAGAGCGCCGGTCCCGAGCAG</td><td></td><td></td><td></td><td></td><td></td><td></td><td></td><td></td><td></td><td></td></tr>
<tr><td>Oligo rev</td><td>AAACCTGCTCGGGACCGGCGCTCT</td><td></td><td></td><td></td><td></td><td></td><td></td><td></td><td></td><td></td><td></td></tr>
<tr><td>Chromosome</td><td>start</td><td>end</td><td>strand</td><td>MM</td><td>target_seq</td><td>PAM</td><td>alignment</td><td>distance</td><td>position</td><td>gene name</td><td>gene id</td></tr>
<tr><td>GU305914.1</td><td>127837</td><td>127859</td><td>-</td><td>0</td><td>AGAGCGCCGGTCCCGAGCAG</td><td>CGG</td><td>||||||||[||||||||||||]PAM</td><td>NA</td><td>Intergenic</td><td>NA</td><td>NA</td></tr>
<tr><td></td><td></td><td></td><td></td><td></td><td></td><td></td><td></td><td></td><td></td><td></td><td></td></tr>
<tr><td>T24300</td><td>CGCCGGTCCCGAGCAGCGGTAGG</td><td>1000</td><td></td><td></td><td></td><td></td><td></td><td></td><td></td><td></td><td></td></tr>
<tr><td>Oligo fwd</td><td>CACCCGCCGGTCCCGAGCAGCGGT</td><td></td><td></td><td></td><td></td><td></td><td></td><td></td><td></td><td></td><td></td></tr>
<tr><td>Oligo rev</td><td>AAACACCGCTGCTCGGGACCGGCG</td><td></td><td></td><td></td><td></td><td></td><td></td><td></td><td></td><td></td><td></td></tr>
<tr><td>Chromosome</td><td>start</td><td>end</td><td>strand</td><td>MM</td><td>target_seq</td><td>PAM</td><td>alignment</td><td>distance</td><td>position</td><td>gene name</td><td>gene id</td></tr>
<tr><td>GU305914.1</td><td>127833</td><td>127855</td><td>-</td><td>0</td><td>CGCCGGTCCCGAGCAGCGGT</td><td>AGG</td><td>||||||||[||||||||||||]PAM</td><td>NA</td><td>Intergenic</td><td>NA</td><td>NA</td></tr>
<tr><td></td><td></td><td></td><td></td><td></td><td></td><td></td><td></td><td></td><td></td><td></td><td></td></tr>
<tr><td>T24301</td><td>CGAGCAGCGGTAGGCGCACTTGG</td><td>1000</td><td></td><td></td><td></td><td></td><td></td><td></td><td></td><td></td><td></td></tr>
<tr><td>Oligo fwd</td><td>CACCCGAGCAGCGGTAGGCGCACT</td><td></td><td></td><td></td><td></td><td></td><td></td><td></td><td></td><td></td><td></td></tr>
<tr><td>Oligo rev</td><td>AAACAGTGCGCCTACCGCTGCTCG</td><td></td><td></td><td></td><td></td><td></td><td></td><td></td><td></td><td></td><td></td></tr>
<tr><td>Chromosome</td><td>start</td><td>end</td><td>strand</td><td>MM</td><td>target_seq</td><td>PAM</td><td>alignment</td><td>distance</td><td>position</td><td>gene name</td><td>gene id</td></tr>
<tr><td>GU305914.1</td><td>127824</td><td>127846</td><td>-</td><td>0</td><td>CGAGCAGCGGTAGGCGCACT</td><td>TGG</td><td>||||||||[||||||||||||]PAM</td><td>NA</td><td>Intergenic</td><td>NA</td><td>NA</td></tr>
<tr><td></td><td></td><td></td><td></td><td></td><td></td><td></td><td></td><td></td><td></td><td></td><td></td></tr>
<tr><td>T24302</td><td>AGCGGTAGGCGCACTTGGCGAGG</td><td>1000</td><td></td><td></td><td></td><td></td><td></td><td></td><td></td><td></td><td></td></tr>
<tr><td>Oligo fwd</td><td>CACCAGCGGTAGGCGCACTTGGCG</td><td></td><td></td><td></td><td></td><td></td><td></td><td></td><td></td><td></td><td></td></tr>
<tr><td>Oligo rev</td><td>AAACCGCCAAGTGCGCCTACCGCT</td><td></td><td></td><td></td><td></td><td></td><td></td><td></td><td></td><td></td><td></td></tr>
<tr><td>Chromosome</td><td>start</td><td>end</td><td>strand</td><td>MM</td><td>target_seq</td><td>PAM</td><td>alignment</td><td>distance</td><td>position</td><td>gene name</td><td>gene id</td></tr>
<tr><td>GU305914.1</td><td>127819</td><td>127841</td><td>-</td><td>0</td><td>AGCGGTAGGCGCACTTGGCG</td><td>AGG</td><td>||||||||[||||||||||||]PAM</td><td>NA</td><td>Intergenic</td><td>NA</td><td>NA</td></tr>
<tr><td></td><td></td><td></td><td></td><td></td><td></td><td></td><td></td><td></td><td></td><td></td><td></td></tr>
<tr><td>T24303</td><td>GTCCCACTCGTCGCAGTAGCAGG</td><td>1000</td><td></td><td></td><td></td><td></td><td></td><td></td><td></td><td></td><td></td></tr>
<tr><td>Oligo fwd</td><td>CACCGTCCCACTCGTCGCAGTAGC</td><td></td><td></td><td></td><td></td><td></td><td></td><td></td><td></td><td></td><td></td></tr>
<tr><td>Oligo rev</td><td>AAACGCTACTGCGACGAGTGGGAC</td><td></td><td></td><td></td><td></td><td></td><td></td><td></td><td></td><td></td><td></td></tr>
<tr><td>Chromosome</td><td>start</td><td>end</td><td>strand</td><td>MM</td><td>target_seq</td><td>PAM</td><td>alignment</td><td>distance</td><td>position</td><td>gene name</td><td>gene id</td></tr>
<tr><td>GU305914.1</td><td>127785</td><td>127807</td><td>-</td><td>0</td><td>GTCCCACTCGTCGCAGTAGC</td><td>AGG</td><td>||||||||[||||||||||||]PAM</td><td>NA</td><td>Intergenic</td><td>NA</td><td>NA</td></tr>
<tr><td></td><td></td><td></td><td></td><td></td><td></td><td></td><td></td><td></td><td></td><td></td><td></td></tr>
<tr><td>T24304</td><td>TCCCACTCGTCGCAGTAGCAGGG</td><td>1000</td><td></td><td></td><td></td><td></td><td></td><td></td><td></td><td></td><td></td></tr>
<tr><td>Oligo fwd</td><td>CACCTCCCACTCGTCGCAGTAGCA</td><td></td><td></td><td></td><td></td><td></td><td></td><td></td><td></td><td></td><td></td></tr>
</table>

<table>
<tr><td>Oligo rev</td><td>AAACTGCTACTGCGACGAGTGGGA</td><td></td><td></td><td></td><td></td><td></td><td></td><td></td><td></td><td></td><td></td></tr>
<tr><td>Chromosome</td><td>start</td><td>end</td><td>strand</td><td>MM</td><td>target_seq</td><td>PAM</td><td>alignment</td><td>distance</td><td>position</td><td>gene name</td><td>gene id</td></tr>
<tr><td>GU305914.1</td><td>127784</td><td>127806</td><td>-</td><td>0</td><td>TCCCACTCGTCGCAGTAGCA</td><td>GGG</td><td>||||||||[||||||||||||]PAM</td><td>NA</td><td>Intergenic</td><td>NA</td><td>NA</td></tr>
<tr><td></td><td></td><td></td><td></td><td></td><td></td><td></td><td></td><td></td><td></td><td></td><td></td></tr>
<tr><td>T24305</td><td>CTCGTCGCAGTAGCAGGGCACGG</td><td>1000</td><td></td><td></td><td></td><td></td><td></td><td></td><td></td><td></td><td></td></tr>
<tr><td>Oligo fwd</td><td>CACCCTCGTCGCAGTAGCAGGGCA</td><td></td><td></td><td></td><td></td><td></td><td></td><td></td><td></td><td></td><td></td></tr>
<tr><td>Oligo rev</td><td>AAACTGCCCTGCTACTGCGACGAG</td><td></td><td></td><td></td><td></td><td></td><td></td><td></td><td></td><td></td><td></td></tr>
<tr><td>Chromosome</td><td>start</td><td>end</td><td>strand</td><td>MM</td><td>target_seq</td><td>PAM</td><td>alignment</td><td>distance</td><td>position</td><td>gene name</td><td>gene id</td></tr>
<tr><td>GU305914.1</td><td>127779</td><td>127801</td><td>-</td><td>0</td><td>CTCGTCGCAGTAGCAGGGCA</td><td>CGG</td><td>||||||||[||||||||||||]PAM</td><td>NA</td><td>Intergenic</td><td>NA</td><td>NA</td></tr>
<tr><td>GU305914.1</td><td>61867</td><td>61889</td><td>+</td><td>4</td><td>GCCGTGGCAGTAGCAGCGCA</td><td>CGG</td><td>--|||-||[||||||||-|||]PAM</td><td>NA</td><td>Intergenic</td><td>NA</td><td>NA</td></tr>
<tr><td></td><td></td><td></td><td></td><td></td><td></td><td></td><td></td><td></td><td></td><td></td><td></td></tr>
<tr><td>T24306</td><td>GCAGTAGCAGGGCACGGCGACGG</td><td>1000</td><td></td><td></td><td></td><td></td><td></td><td></td><td></td><td></td><td></td></tr>
<tr><td>Oligo fwd</td><td>CACCGCAGTAGCAGGGCACGGCGA</td><td></td><td></td><td></td><td></td><td></td><td></td><td></td><td></td><td></td><td></td></tr>
<tr><td>Oligo rev</td><td>AAACTCGCCGTGCCCTGCTACTGC</td><td></td><td></td><td></td><td></td><td></td><td></td><td></td><td></td><td></td><td></td></tr>
<tr><td>Chromosome</td><td>start</td><td>end</td><td>strand</td><td>MM</td><td>target_seq</td><td>PAM</td><td>alignment</td><td>distance</td><td>position</td><td>gene name</td><td>gene id</td></tr>
<tr><td>GU305914.1</td><td>127773</td><td>127795</td><td>-</td><td>0</td><td>GCAGTAGCAGGGCACGGCGA</td><td>CGG</td><td>||||||||[||||||||||||]PAM</td><td>NA</td><td>Intergenic</td><td>NA</td><td>NA</td></tr>
<tr><td></td><td></td><td></td><td></td><td></td><td></td><td></td><td></td><td></td><td></td><td></td><td></td></tr>
<tr><td>T24307</td><td>CAGTAGCAGGGCACGGCGACGGG</td><td>1000</td><td></td><td></td><td></td><td></td><td></td><td></td><td></td><td></td><td></td></tr>
<tr><td>Oligo fwd</td><td>CACCCAGTAGCAGGGCACGGCGAC</td><td></td><td></td><td></td><td></td><td></td><td></td><td></td><td></td><td></td><td></td></tr>
<tr><td>Oligo rev</td><td>AAACGTCGCCGTGCCCTGCTACTG</td><td></td><td></td><td></td><td></td><td></td><td></td><td></td><td></td><td></td><td></td></tr>
<tr><td>Chromosome</td><td>start</td><td>end</td><td>strand</td><td>MM</td><td>target_seq</td><td>PAM</td><td>alignment</td><td>distance</td><td>position</td><td>gene name</td><td>gene id</td></tr>
<tr><td>GU305914.1</td><td>127772</td><td>127794</td><td>-</td><td>0</td><td>CAGTAGCAGGGCACGGCGAC</td><td>GGG</td><td>||||||||[||||||||||||]PAM</td><td>NA</td><td>Intergenic</td><td>NA</td><td>NA</td></tr>
<tr><td></td><td></td><td></td><td></td><td></td><td></td><td></td><td></td><td></td><td></td><td></td><td></td></tr>
<tr><td>T24308</td><td>AGTAGCAGGGCACGGCGACGGGG</td><td>1000</td><td></td><td></td><td></td><td></td><td></td><td></td><td></td><td></td><td></td></tr>
<tr><td>Oligo fwd</td><td>CACCAGTAGCAGGGCACGGCGACG</td><td></td><td></td><td></td><td></td><td></td><td></td><td></td><td></td><td></td><td></td></tr>
<tr><td>Oligo rev</td><td>AAACCGTCGCCGTGCCCTGCTACT</td><td></td><td></td><td></td><td></td><td></td><td></td><td></td><td></td><td></td><td></td></tr>
<tr><td>Chromosome</td><td>start</td><td>end</td><td>strand</td><td>MM</td><td>target_seq</td><td>PAM</td><td>alignment</td><td>distance</td><td>position</td><td>gene name</td><td>gene id</td></tr>
<tr><td>GU305914.1</td><td>127771</td><td>127793</td><td>-</td><td>0</td><td>AGTAGCAGGGCACGGCGACG</td><td>GGG</td><td>||||||||[||||||||||||]PAM</td><td>NA</td><td>Intergenic</td><td>NA</td><td>NA</td></tr>
<tr><td></td><td></td><td></td><td></td><td></td><td></td><td></td><td></td><td></td><td></td><td></td><td></td></tr>
<tr><td>T24309</td><td>CGGCGACGGGGCCCATGAGCAGG</td><td>1000</td><td></td><td></td><td></td><td></td><td></td><td></td><td></td><td></td><td></td></tr>
<tr><td>Oligo fwd</td><td>CACCCGGCGACGGGGCCCATGAGC</td><td></td><td></td><td></td><td></td><td></td><td></td><td></td><td></td><td></td><td></td></tr>
<tr><td>Oligo rev</td><td>AAACGCTCATGGGCCCCGTCGCCG</td><td></td><td></td><td></td><td></td><td></td><td></td><td></td><td></td><td></td><td></td></tr>
<tr><td>Chromosome</td><td>start</td><td>end</td><td>strand</td><td>MM</td><td>target_seq</td><td>PAM</td><td>alignment</td><td>distance</td><td>position</td><td>gene name</td><td>gene id</td></tr>
<tr><td>GU305914.1</td><td>127759</td><td>127781</td><td>-</td><td>0</td><td>CGGCGACGGGGCCCATGAGC</td><td>AGG</td><td>||||||||[||||||||||||]PAM</td><td>NA</td><td>Intergenic</td><td>NA</td><td>NA</td></tr>
<tr><td></td><td></td><td></td><td></td><td></td><td></td><td></td><td></td><td></td><td></td><td></td><td></td></tr>
<tr><td>T24310</td><td>CGACGGGGCCCATGAGCAGGCGG</td><td>1000</td><td></td><td></td><td></td><td></td><td></td><td></td><td></td><td></td><td></td></tr>
<tr><td>Oligo fwd</td><td>CACCCGACGGGGCCCATGAGCAGG</td><td></td><td></td><td></td><td></td><td></td><td></td><td></td><td></td><td></td><td></td></tr>
<tr><td>Oligo rev</td><td>AAACCCTGCTCATGGGCCCCGTCG</td><td></td><td></td><td></td><td></td><td></td><td></td><td></td><td></td><td></td><td></td></tr>
<tr><td>Chromosome</td><td>start</td><td>end</td><td>strand</td><td>MM</td><td>target_seq</td><td>PAM</td><td>alignment</td><td>distance</td><td>position</td><td>gene name</td><td>gene id</td></tr>
<tr><td>GU305914.1</td><td>127756</td><td>127778</td><td>-</td><td>0</td><td>CGACGGGGCCCATGAGCAGG</td><td>CGG</td><td>||||||||[||||||||||||]PAM</td><td>NA</td><td>Intergenic</td><td>NA</td><td>NA</td></tr>
<tr><td></td><td></td><td></td><td></td><td></td><td></td><td></td><td></td><td></td><td></td><td></td><td></td></tr>
<tr><td>T24311</td><td>GGCCCATGAGCAGGCGGCAGAGG</td><td>1000</td><td></td><td></td><td></td><td></td><td></td><td></td><td></td><td></td><td></td></tr>
<tr><td>Oligo fwd</td><td>CACCGGCCCATGAGCAGGCGGCAG</td><td></td><td></td><td></td><td></td><td></td><td></td><td></td><td></td><td></td><td></td></tr>
<tr><td>Oligo rev</td><td>AAACCTGCCGCCTGCTCATGGGCC</td><td></td><td></td><td></td><td></td><td></td><td></td><td></td><td></td><td></td><td></td></tr>
<tr><td>Chromosome</td><td>start</td><td>end</td><td>strand</td><td>MM</td><td>target_seq</td><td>PAM</td><td>alignment</td><td>distance</td><td>position</td><td>gene name</td><td>gene id</td></tr>
<tr><td>GU305914.1</td><td>127750</td><td>127772</td><td>-</td><td>0</td><td>GGCCCATGAGCAGGCGGCAG</td><td>AGG</td><td>||||||||[||||||||||||]PAM</td><td>NA</td><td>Intergenic</td><td>NA</td><td>NA</td></tr>
<tr><td></td><td></td><td></td><td></td><td></td><td></td><td></td><td></td><td></td><td></td><td></td><td></td></tr>
<tr><td>T24312</td><td>GACGCGCGCCTTGTCCTCGTCGG</td><td>1000</td><td></td><td></td><td></td><td></td><td></td><td></td><td></td><td></td><td></td></tr>
<tr><td>Oligo fwd</td><td>CACCGACGCGCGCCTTGTCCTCGT</td><td></td><td></td><td></td><td></td><td></td><td></td><td></td><td></td><td></td><td></td></tr>
<tr><td>Oligo rev</td><td>AAACACGAGGACAAGGCGCGCGTC</td><td></td><td></td><td></td><td></td><td></td><td></td><td></td><td></td><td></td><td></td></tr>
<tr><td>Chromosome</td><td>start</td><td>end</td><td>strand</td><td>MM</td><td>target_seq</td><td>PAM</td><td>alignment</td><td>distance</td><td>position</td><td>gene name</td><td>gene id</td></tr>
<tr><td>GU305914.1</td><td>127725</td><td>127747</td><td>-</td><td>0</td><td>GACGCGCGCCTTGTCCTCGT</td><td>CGG</td><td>||||||||[||||||||||||]PAM</td><td>NA</td><td>Intergenic</td><td>NA</td><td>NA</td></tr>
<tr><td></td><td></td><td></td><td></td><td></td><td></td><td></td><td></td><td></td><td></td><td></td><td></td></tr>
<tr><td>T24313</td><td>GCGCGCCTTGTCCTCGTCGGTGG</td><td>1000</td><td></td><td></td><td></td><td></td><td></td><td></td><td></td><td></td><td></td></tr>
<tr><td>Oligo fwd</td><td>CACCGCGCGCCTTGTCCTCGTCGG</td><td></td><td></td><td></td><td></td><td></td><td></td><td></td><td></td><td></td><td></td></tr>
<tr><td>Oligo rev</td><td>AAACCCGACGAGGACAAGGCGCGC</td><td></td><td></td><td></td><td></td><td></td><td></td><td></td><td></td><td></td><td></td></tr>
<tr><td>Chromosome</td><td>start</td><td>end</td><td>strand</td><td>MM</td><td>target_seq</td><td>PAM</td><td>alignment</td><td>distance</td><td>position</td><td>gene name</td><td>gene id</td></tr>
<tr><td>GU305914.1</td><td>127722</td><td>127744</td><td>-</td><td>0</td><td>GCGCGCCTTGTCCTCGTCGG</td><td>TGG</td><td>||||||||[||||||||||||]PAM</td><td>NA</td><td>Intergenic</td><td>NA</td><td>NA</td></tr>
<tr><td></td><td></td><td></td><td></td><td></td><td></td><td></td><td></td><td></td><td></td><td></td><td></td></tr>
<tr><td>T24314</td><td>CTTGTCCTCGTCGGTGGCGCCGG</td><td>1000</td><td></td><td></td><td></td><td></td><td></td><td></td><td></td><td></td><td></td></tr>
<tr><td>Oligo fwd</td><td>CACCCTTGTCCTCGTCGGTGGCGC</td><td></td><td></td><td></td><td></td><td></td><td></td><td></td><td></td><td></td><td></td></tr>
<tr><td>Oligo rev</td><td>AAACGCGCCACCGACGAGGACAAG</td><td></td><td></td><td></td><td></td><td></td><td></td><td></td><td></td><td></td><td></td></tr>
<tr><td>Chromosome</td><td>start</td><td>end</td><td>strand</td><td>MM</td><td>target_seq</td><td>PAM</td><td>alignment</td><td>distance</td><td>position</td><td>gene name</td><td>gene id</td></tr>
<tr><td>GU305914.1</td><td>127716</td><td>127738</td><td>-</td><td>0</td><td>CTTGTCCTCGTCGGTGGCGC</td><td>CGG</td><td>||||||||[||||||||||||]PAM</td><td>NA</td><td>Intergenic</td><td>NA</td><td>NA</td></tr>
<tr><td>GU305914.1</td><td>73186</td><td>73208</td><td>+</td><td>4</td><td>CTCTTCCTCGTCGGTGCCGG</td><td>CGG</td><td>||--||||[||||||||-||-]PAM</td><td>NA</td><td>Intergenic</td><td>NA</td><td>NA</td></tr>
<tr><td></td><td></td><td></td><td></td><td></td><td></td><td></td><td></td><td></td><td></td><td></td><td></td></tr>
<tr><td>T24315</td><td>TTGTCCTCGTCGGTGGCGCCGGG</td><td>1000</td><td></td><td></td><td></td><td></td><td></td><td></td><td></td><td></td><td></td></tr>
<tr><td>Oligo fwd</td><td>CACCTTGTCCTCGTCGGTGGCGCC</td><td></td><td></td><td></td><td></td><td></td><td></td><td></td><td></td><td></td><td></td></tr>
<tr><td>Oligo rev</td><td>AAACGGCGCCACCGACGAGGACAA</td><td></td><td></td><td></td><td></td><td></td><td></td><td></td><td></td><td></td><td></td></tr>
<tr><td>Chromosome</td><td>start</td><td>end</td><td>strand</td><td>MM</td><td>target_seq</td><td>PAM</td><td>alignment</td><td>distance</td><td>position</td><td>gene name</td><td>gene id</td></tr>
<tr><td>GU305914.1</td><td>127715</td><td>127737</td><td>-</td><td>0</td><td>TTGTCCTCGTCGGTGGCGCC</td><td>GGG</td><td>||||||||[||||||||||||]PAM</td><td>NA</td><td>Intergenic</td><td>NA</td><td>NA</td></tr>
<tr><td></td><td></td><td></td><td></td><td></td><td></td><td></td><td></td><td></td><td></td><td></td><td></td></tr>
<tr><td>T24316</td><td>TGTCCTCGTCGGTGGCGCCGGGG</td><td>1000</td><td></td><td></td><td></td><td></td><td></td><td></td><td></td><td></td><td></td></tr>
<tr><td>Oligo fwd</td><td>CACCTGTCCTCGTCGGTGGCGCCG</td><td></td><td></td><td></td><td></td><td></td><td></td><td></td><td></td><td></td><td></td></tr>
<tr><td>Oligo rev</td><td>AAACCGGCGCCACCGACGAGGACA</td><td></td><td></td><td></td><td></td><td></td><td></td><td></td><td></td><td></td><td></td></tr>
<tr><td>Chromosome</td><td>start</td><td>end</td><td>strand</td><td>MM</td><td>target_seq</td><td>PAM</td><td>alignment</td><td>distance</td><td>position</td><td>gene name</td><td>gene id</td></tr>
<tr><td>GU305914.1</td><td>127714</td><td>127736</td><td>-</td><td>0</td><td>TGTCCTCGTCGGTGGCGCCG</td><td>GGG</td><td>||||||||[||||||||||||]PAM</td><td>NA</td><td>Intergenic</td><td>NA</td><td>NA</td></tr>
<tr><td></td><td></td><td></td><td></td><td></td><td></td><td></td><td></td><td></td><td></td><td></td><td></td></tr>
<tr><td>T24317</td><td>CTCGTCGGTGGCGCCGGGGTCGG</td><td>1000</td><td></td><td></td><td></td><td></td><td></td><td></td><td></td><td></td><td></td></tr>
<tr><td>Oligo fwd</td><td>CACCCTCGTCGGTGGCGCCGGGGT</td><td></td><td></td><td></td><td></td><td></td><td></td><td></td><td></td><td></td><td></td></tr>
<tr><td>Oligo rev</td><td>AAACACCCCGGCGCCACCGACGAG</td><td></td><td></td><td></td><td></td><td></td><td></td><td></td><td></td><td></td><td></td></tr>
<tr><td>Chromosome</td><td>start</td><td>end</td><td>strand</td><td>MM</td><td>target_seq</td><td>PAM</td><td>alignment</td><td>distance</td><td>position</td><td>gene name</td><td>gene id</td></tr>
<tr><td>GU305914.1</td><td>127710</td><td>127732</td><td>-</td><td>0</td><td>CTCGTCGGTGGCGCCGGGGT</td><td>CGG</td><td>||||||||[||||||||||||]PAM</td><td>NA</td><td>Intergenic</td><td>NA</td><td>NA</td></tr>
<tr><td></td><td></td><td></td><td></td><td></td><td></td><td></td><td></td><td></td><td></td><td></td><td></td></tr>
<tr><td>T24318</td><td>TCGTCGGTGGCGCCGGGGTCGGG</td><td>1000</td><td></td><td></td><td></td><td></td><td></td><td></td><td></td><td></td><td></td></tr>
<tr><td>Oligo fwd</td><td>CACCTCGTCGGTGGCGCCGGGGTC</td><td></td><td></td><td></td><td></td><td></td><td></td><td></td><td></td><td></td><td></td></tr>
<tr><td>Oligo rev</td><td>AAACGACCCCGGCGCCACCGACGA</td><td></td><td></td><td></td><td></td><td></td><td></td><td></td><td></td><td></td><td></td></tr>
<tr><td>Chromosome</td><td>start</td><td>end</td><td>strand</td><td>MM</td><td>target_seq</td><td>PAM</td><td>alignment</td><td>distance</td><td>position</td><td>gene name</td><td>gene id</td></tr>
<tr><td>GU305914.1</td><td>127709</td><td>127731</td><td>-</td><td>0</td><td>TCGTCGGTGGCGCCGGGGTC</td><td>GGG</td><td>||||||||[||||||||||||]PAM</td><td>NA</td><td>Intergenic</td><td>NA</td><td>NA</td></tr>
</table>

<table>
<tr><td>T24319</td><td>GGCGCCGGGGTCGGGTCCCGCGG</td><td>1000</td><td></td><td></td><td></td><td></td><td></td><td></td><td></td><td></td><td></td></tr>
<tr><td>Oligo fwd</td><td>CACCGGCGCCGGGGTCGGGTCCCG</td><td></td><td></td><td></td><td></td><td></td><td></td><td></td><td></td><td></td><td></td></tr>
<tr><td>Oligo rev</td><td>AAACCGGGACCCGACCCCGGCGCC</td><td></td><td></td><td></td><td></td><td></td><td></td><td></td><td></td><td></td><td></td></tr>
<tr><td>Chromosome</td><td>start</td><td>end</td><td>strand</td><td>MM</td><td>target_seq</td><td>PAM</td><td>alignment</td><td>distance</td><td>position</td><td>gene name</td><td>gene id</td></tr>
<tr><td>GU305914.1</td><td>127701</td><td>127723</td><td>-</td><td>0</td><td>GGCGCCGGGGTCGGGTCCCG</td><td>CGG</td><td>||||||||[||||||||||||]PAM</td><td>NA</td><td>Intergenic</td><td>NA</td><td>NA</td></tr>
<tr><td>T24320</td><td>CATCAGCGTCCTGAAGATCGAGG</td><td>1000</td><td></td><td></td><td></td><td></td><td></td><td></td><td></td><td></td><td></td></tr>
<tr><td>Oligo fwd</td><td>CACCCATCAGCGTCCTGAAGATCG</td><td></td><td></td><td></td><td></td><td></td><td></td><td></td><td></td><td></td><td></td></tr>
<tr><td>Oligo rev</td><td>AAACCGATCTTCAGGACGCTGATG</td><td></td><td></td><td></td><td></td><td></td><td></td><td></td><td></td><td></td><td></td></tr>
<tr><td>Chromosome</td><td>start</td><td>end</td><td>strand</td><td>MM</td><td>target_seq</td><td>PAM</td><td>alignment</td><td>distance</td><td>position</td><td>gene name</td><td>gene id</td></tr>
<tr><td>GU305914.1</td><td>127677</td><td>127699</td><td>-</td><td>0</td><td>CATCAGCGTCCTGAAGATCG</td><td>AGG</td><td>||||||||[||||||||||||]PAM</td><td>NA</td><td>Intergenic</td><td>NA</td><td>NA</td></tr>
<tr><td>T24321</td><td>GGAGATGTGCTCCTCGTCGACGG</td><td>1000</td><td></td><td></td><td></td><td></td><td></td><td></td><td></td><td></td><td></td></tr>
<tr><td>Oligo fwd</td><td>CACCGGAGATGTGCTCCTCGTCGA</td><td></td><td></td><td></td><td></td><td></td><td></td><td></td><td></td><td></td><td></td></tr>
<tr><td>Oligo rev</td><td>AAACTCGACGAGGAGCACATCTCC</td><td></td><td></td><td></td><td></td><td></td><td></td><td></td><td></td><td></td><td></td></tr>
<tr><td>Chromosome</td><td>start</td><td>end</td><td>strand</td><td>MM</td><td>target_seq</td><td>PAM</td><td>alignment</td><td>distance</td><td>position</td><td>gene name</td><td>gene id</td></tr>
<tr><td>GU305914.1</td><td>127656</td><td>127678</td><td>-</td><td>0</td><td>GGAGATGTGCTCCTCGTCGA</td><td>CGG</td><td>||||||||[||||||||||||]PAM</td><td>NA</td><td>Intergenic</td><td>NA</td><td>NA</td></tr>
<tr><td>T24322</td><td>GTGCTCCTCGTCGACGGTGACGG</td><td>1000</td><td></td><td></td><td></td><td></td><td></td><td></td><td></td><td></td><td></td></tr>
<tr><td>Oligo fwd</td><td>CACCGTGCTCCTCGTCGACGGTGA</td><td></td><td></td><td></td><td></td><td></td><td></td><td></td><td></td><td></td><td></td></tr>
<tr><td>Oligo rev</td><td>AAACTCACCGTCGACGAGGAGCAC</td><td></td><td></td><td></td><td></td><td></td><td></td><td></td><td></td><td></td><td></td></tr>
<tr><td>Chromosome</td><td>start</td><td>end</td><td>strand</td><td>MM</td><td>target_seq</td><td>PAM</td><td>alignment</td><td>distance</td><td>position</td><td>gene name</td><td>gene id</td></tr>
<tr><td>GU305914.1</td><td>127650</td><td>127672</td><td>-</td><td>0</td><td>GTGCTCCTCGTCGACGGTGA</td><td>CGG</td><td>||||||||[||||||||||||]PAM</td><td>NA</td><td>Intergenic</td><td>NA</td><td>NA</td></tr>
<tr><td>T24323</td><td>CCTCGTCGACGGTGACGGCCTGG</td><td>1000</td><td></td><td></td><td></td><td></td><td></td><td></td><td></td><td></td><td></td></tr>
<tr><td>Oligo fwd</td><td>CACCCCTCGTCGACGGTGACGGCC</td><td></td><td></td><td></td><td></td><td></td><td></td><td></td><td></td><td></td><td></td></tr>
<tr><td>Oligo rev</td><td>AAACGGCCGTCACCGTCGACGAGG</td><td></td><td></td><td></td><td></td><td></td><td></td><td></td><td></td><td></td><td></td></tr>
<tr><td>Chromosome</td><td>start</td><td>end</td><td>strand</td><td>MM</td><td>target_seq</td><td>PAM</td><td>alignment</td><td>distance</td><td>position</td><td>gene name</td><td>gene id</td></tr>
<tr><td>GU305914.1</td><td>127645</td><td>127667</td><td>-</td><td>0</td><td>CCTCGTCGACGGTGACGGCC</td><td>TGG</td><td>||||||||[||||||||||||]PAM</td><td>NA</td><td>Intergenic</td><td>NA</td><td>NA</td></tr>
<tr><td>T24324</td><td>CTCGTCGACGGTGACGGCCTGGG</td><td>1000</td><td></td><td></td><td></td><td></td><td></td><td></td><td></td><td></td><td></td></tr>
<tr><td>Oligo fwd</td><td>CACCCTCGTCGACGGTGACGGCCT</td><td></td><td></td><td></td><td></td><td></td><td></td><td></td><td></td><td></td><td></td></tr>
<tr><td>Oligo rev</td><td>AAACAGGCCGTCACCGTCGACGAG</td><td></td><td></td><td></td><td></td><td></td><td></td><td></td><td></td><td></td><td></td></tr>
<tr><td>Chromosome</td><td>start</td><td>end</td><td>strand</td><td>MM</td><td>target_seq</td><td>PAM</td><td>alignment</td><td>distance</td><td>position</td><td>gene name</td><td>gene id</td></tr>
<tr><td>GU305914.1</td><td>127644</td><td>127666</td><td>-</td><td>0</td><td>CTCGTCGACGGTGACGGCCT</td><td>GGG</td><td>||||||||[||||||||||||]PAM</td><td>NA</td><td>Intergenic</td><td>NA</td><td>NA</td></tr>
<tr><td>GU305914.1</td><td>170135</td><td>170157</td><td>+</td><td>4</td><td>TTCGGGGACGGTGACGGGCT</td><td>CGG</td><td>-|||--||[|||||||||-||]PAM</td><td>NA</td><td>Intergenic</td><td>NA</td><td>NA</td></tr>
<tr><td>T24325</td><td>CGACGGTGACGGCCTGGGAGAGG</td><td>1000</td><td></td><td></td><td></td><td></td><td></td><td></td><td></td><td></td><td></td></tr>
<tr><td>Oligo fwd</td><td>CACCCGACGGTGACGGCCTGGGAG</td><td></td><td></td><td></td><td></td><td></td><td></td><td></td><td></td><td></td><td></td></tr>
<tr><td>Oligo rev</td><td>AAACCTCCCAGGCCGTCACCGTCG</td><td></td><td></td><td></td><td></td><td></td><td></td><td></td><td></td><td></td><td></td></tr>
<tr><td>Chromosome</td><td>start</td><td>end</td><td>strand</td><td>MM</td><td>target_seq</td><td>PAM</td><td>alignment</td><td>distance</td><td>position</td><td>gene name</td><td>gene id</td></tr>
<tr><td>GU305914.1</td><td>127639</td><td>127661</td><td>-</td><td>0</td><td>CGACGGTGACGGCCTGGGAG</td><td>AGG</td><td>||||||||[||||||||||||]PAM</td><td>NA</td><td>Intergenic</td><td>NA</td><td>NA</td></tr>
<tr><td>T24326</td><td>CGGTGACGGCCTGGGAGAGGCGG</td><td>1000</td><td></td><td></td><td></td><td></td><td></td><td></td><td></td><td></td><td></td></tr>
<tr><td>Oligo fwd</td><td>CACCCGGTGACGGCCTGGGAGAGG</td><td></td><td></td><td></td><td></td><td></td><td></td><td></td><td></td><td></td><td></td></tr>
<tr><td>Oligo rev</td><td>AAACCCTCTCCCAGGCCGTCACCG</td><td></td><td></td><td></td><td></td><td></td><td></td><td></td><td></td><td></td><td></td></tr>
<tr><td>Chromosome</td><td>start</td><td>end</td><td>strand</td><td>MM</td><td>target_seq</td><td>PAM</td><td>alignment</td><td>distance</td><td>position</td><td>gene name</td><td>gene id</td></tr>
<tr><td>GU305914.1</td><td>127636</td><td>127658</td><td>-</td><td>0</td><td>CGGTGACGGCCTGGGAGAGG</td><td>CGG</td><td>||||||||[||||||||||||]PAM</td><td>NA</td><td>Intergenic</td><td>NA</td><td>NA</td></tr>
<tr><td>T24327</td><td>GGTGACGGCCTGGGAGAGGCGGG</td><td>1000</td><td></td><td></td><td></td><td></td><td></td><td></td><td></td><td></td><td></td></tr>
<tr><td>Oligo fwd</td><td>CACCGGTGACGGCCTGGGAGAGGC</td><td></td><td></td><td></td><td></td><td></td><td></td><td></td><td></td><td></td><td></td></tr>
<tr><td>Oligo rev</td><td>AAACGCCTCTCCCAGGCCGTCACC</td><td></td><td></td><td></td><td></td><td></td><td></td><td></td><td></td><td></td><td></td></tr>
<tr><td>Chromosome</td><td>start</td><td>end</td><td>strand</td><td>MM</td><td>target_seq</td><td>PAM</td><td>alignment</td><td>distance</td><td>position</td><td>gene name</td><td>gene id</td></tr>
<tr><td>GU305914.1</td><td>127635</td><td>127657</td><td>-</td><td>0</td><td>GGTGACGGCCTGGGAGAGGC</td><td>GGG</td><td>||||||||[||||||||||||]PAM</td><td>NA</td><td>Intergenic</td><td>NA</td><td>NA</td></tr>
<tr><td>T24328</td><td>GAGAGGCGGGAGTCCTCGAACGG</td><td>1000</td><td></td><td></td><td></td><td></td><td></td><td></td><td></td><td></td><td></td></tr>
<tr><td>Oligo fwd</td><td>CACCGAGAGGCGGGAGTCCTCGAA</td><td></td><td></td><td></td><td></td><td></td><td></td><td></td><td></td><td></td><td></td></tr>
<tr><td>Oligo rev</td><td>AAACTTCGAGGACTCCCGCCTCTC</td><td></td><td></td><td></td><td></td><td></td><td></td><td></td><td></td><td></td><td></td></tr>
<tr><td>Chromosome</td><td>start</td><td>end</td><td>strand</td><td>MM</td><td>target_seq</td><td>PAM</td><td>alignment</td><td>distance</td><td>position</td><td>gene name</td><td>gene id</td></tr>
<tr><td>GU305914.1</td><td>127622</td><td>127644</td><td>-</td><td>0</td><td>GAGAGGCGGGAGTCCTCGAA</td><td>CGG</td><td>||||||||[||||||||||||]PAM</td><td>NA</td><td>Intergenic</td><td>NA</td><td>NA</td></tr>
<tr><td>T24329</td><td>CCTCGAACGGCGAGATCTCTCGG</td><td>1000</td><td></td><td></td><td></td><td></td><td></td><td></td><td></td><td></td><td></td></tr>
<tr><td>Oligo fwd</td><td>CACCCCTCGAACGGCGAGATCTCT</td><td></td><td></td><td></td><td></td><td></td><td></td><td></td><td></td><td></td><td></td></tr>
<tr><td>Oligo rev</td><td>AAACAGAGATCTCGCCGTTCGAGG</td><td></td><td></td><td></td><td></td><td></td><td></td><td></td><td></td><td></td><td></td></tr>
<tr><td>Chromosome</td><td>start</td><td>end</td><td>strand</td><td>MM</td><td>target_seq</td><td>PAM</td><td>alignment</td><td>distance</td><td>position</td><td>gene name</td><td>gene id</td></tr>
<tr><td>GU305914.1</td><td>127609</td><td>127631</td><td>-</td><td>0</td><td>CCTCGAACGGCGAGATCTCT</td><td>CGG</td><td>||||||||[||||||||||||]PAM</td><td>NA</td><td>Intergenic</td><td>NA</td><td>NA</td></tr>
<tr><td>T24330</td><td>CTCGAACGGCGAGATCTCTCGGG</td><td>1000</td><td></td><td></td><td></td><td></td><td></td><td></td><td></td><td></td><td></td></tr>
<tr><td>Oligo fwd</td><td>CACCCTCGAACGGCGAGATCTCTC</td><td></td><td></td><td></td><td></td><td></td><td></td><td></td><td></td><td></td><td></td></tr>
<tr><td>Oligo rev</td><td>AAACGAGAGATCTCGCCGTTCGAG</td><td></td><td></td><td></td><td></td><td></td><td></td><td></td><td></td><td></td><td></td></tr>
<tr><td>Chromosome</td><td>start</td><td>end</td><td>strand</td><td>MM</td><td>target_seq</td><td>PAM</td><td>alignment</td><td>distance</td><td>position</td><td>gene name</td><td>gene id</td></tr>
<tr><td>GU305914.1</td><td>127608</td><td>127630</td><td>-</td><td>0</td><td>CTCGAACGGCGAGATCTCTC</td><td>GGG</td><td>||||||||[||||||||||||]PAM</td><td>NA</td><td>Intergenic</td><td>NA</td><td>NA</td></tr>
<tr><td>T24331</td><td>GATCTCTCGGGTGAGATTGTAGG</td><td>1000</td><td></td><td></td><td></td><td></td><td></td><td></td><td></td><td></td><td></td></tr>
<tr><td>Oligo fwd</td><td>CACCGATCTCTCGGGTGAGATTGT</td><td></td><td></td><td></td><td></td><td></td><td></td><td></td><td></td><td></td><td></td></tr>
<tr><td>Oligo rev</td><td>AAACACAATCTCACCCGAGAGATC</td><td></td><td></td><td></td><td></td><td></td><td></td><td></td><td></td><td></td><td></td></tr>
<tr><td>Chromosome</td><td>start</td><td>end</td><td>strand</td><td>MM</td><td>target_seq</td><td>PAM</td><td>alignment</td><td>distance</td><td>position</td><td>gene name</td><td>gene id</td></tr>
<tr><td>GU305914.1</td><td>127596</td><td>127618</td><td>-</td><td>0</td><td>GATCTCTCGGGTGAGATTGT</td><td>AGG</td><td>||||||||[||||||||||||]PAM</td><td>NA</td><td>Intergenic</td><td>NA</td><td>NA</td></tr>
<tr><td>T24332</td><td>TTGTAGGAGTTGACGCAGACCGG</td><td>1000</td><td></td><td></td><td></td><td></td><td></td><td></td><td></td><td></td><td></td></tr>
<tr><td>Oligo fwd</td><td>CACCTTGTAGGAGTTGACGCAGAC</td><td></td><td></td><td></td><td></td><td></td><td></td><td></td><td></td><td></td><td></td></tr>
<tr><td>Oligo rev</td><td>AAACGTCTGCGTCAACTCCTACAA</td><td></td><td></td><td></td><td></td><td></td><td></td><td></td><td></td><td></td><td></td></tr>
<tr><td>Chromosome</td><td>start</td><td>end</td><td>strand</td><td>MM</td><td>target_seq</td><td>PAM</td><td>alignment</td><td>distance</td><td>position</td><td>gene name</td><td>gene id</td></tr>
<tr><td>GU305914.1</td><td>127580</td><td>127602</td><td>-</td><td>0</td><td>TTGTAGGAGTTGACGCAGAC</td><td>CGG</td><td>||||||||[||||||||||||]PAM</td><td>NA</td><td>Intergenic</td><td>NA</td><td>NA</td></tr>
<tr><td>T24333</td><td>GCGCTCTTGACGCGCGCAGTCGG</td><td>1000</td><td></td><td></td><td></td><td></td><td></td><td></td><td></td><td></td><td></td></tr>
<tr><td>Oligo fwd</td><td>CACCGCGCTCTTGACGCGCGCAGT</td><td></td><td></td><td></td><td></td><td></td><td></td><td></td><td></td><td></td><td></td></tr>
<tr><td>Oligo rev</td><td>AAACACTGCGCGCGTCAAGAGCGC</td><td></td><td></td><td></td><td></td><td></td><td></td><td></td><td></td><td></td><td></td></tr>
</table>

<table>
<tr><td>Chromosome</td><td>start</td><td>end</td><td>strand</td><td>MM</td><td>target_seq</td><td>PAM</td><td>alignment</td><td>distance</td><td>position</td><td>gene name</td><td>gene id</td></tr>
<tr><td>GU305914.1</td><td>127556</td><td>127578</td><td>-</td><td>0</td><td>GCGCTCTTGACGCGCGCAGT</td><td>CGG</td><td>||||||||[||||||||||||]PAM</td><td>NA</td><td>Intergenic</td><td>NA</td><td>NA</td></tr>
<tr><td></td><td></td><td></td><td></td><td></td><td></td><td></td><td></td><td></td><td></td><td></td><td></td></tr>
<tr><td>T24334</td><td>GGCGACTTAACTATTAAAGCCGG</td><td>1000</td><td></td><td></td><td></td><td></td><td></td><td></td><td></td><td></td><td></td></tr>
<tr><td>Oligo fwd</td><td>CACCGGCGACTTAACTATTAAAGC</td><td></td><td></td><td></td><td></td><td></td><td></td><td></td><td></td><td></td><td></td></tr>
<tr><td>Oligo rev</td><td>AAACGCTTTAATAGTTAAGTCGCC</td><td></td><td></td><td></td><td></td><td></td><td></td><td></td><td></td><td></td><td></td></tr>
<tr><td>Chromosome</td><td>start</td><td>end</td><td>strand</td><td>MM</td><td>target_seq</td><td>PAM</td><td>alignment</td><td>distance</td><td>position</td><td>gene name</td><td>gene id</td></tr>
<tr><td>GU305914.1</td><td>127535</td><td>127557</td><td>-</td><td>0</td><td>GGCGACTTAACTATTAAAGC</td><td>CGG</td><td>||||||||[||||||||||||]PAM</td><td>NA</td><td>Intergenic</td><td>NA</td><td>NA</td></tr>
<tr><td></td><td></td><td></td><td></td><td></td><td></td><td></td><td></td><td></td><td></td><td></td><td></td></tr>
<tr><td>T24335</td><td>TATTAAAGCCGGATCGAGACAGG</td><td>1000</td><td></td><td></td><td></td><td></td><td></td><td></td><td></td><td></td><td></td></tr>
<tr><td>Oligo fwd</td><td>CACCTATTAAAGCCGGATCGAGAC</td><td></td><td></td><td></td><td></td><td></td><td></td><td></td><td></td><td></td><td></td></tr>
<tr><td>Oligo rev</td><td>AAACGTCTCGATCCGGCTTTAATA</td><td></td><td></td><td></td><td></td><td></td><td></td><td></td><td></td><td></td><td></td></tr>
<tr><td>Chromosome</td><td>start</td><td>end</td><td>strand</td><td>MM</td><td>target_seq</td><td>PAM</td><td>alignment</td><td>distance</td><td>position</td><td>gene name</td><td>gene id</td></tr>
<tr><td>GU305914.1</td><td>127524</td><td>127546</td><td>-</td><td>0</td><td>TATTAAAGCCGGATCGAGAC</td><td>AGG</td><td>||||||||[||||||||||||]PAM</td><td>NA</td><td>Intergenic</td><td>NA</td><td>NA</td></tr>
<tr><td></td><td></td><td></td><td></td><td></td><td></td><td></td><td></td><td></td><td></td><td></td><td></td></tr>
<tr><td>T24336</td><td>TCGAGACAGGACAGAGACGCCGG</td><td>1000</td><td></td><td></td><td></td><td></td><td></td><td></td><td></td><td></td><td></td></tr>
<tr><td>Oligo fwd</td><td>CACCTCGAGACAGGACAGAGACGC</td><td></td><td></td><td></td><td></td><td></td><td></td><td></td><td></td><td></td><td></td></tr>
<tr><td>Oligo rev</td><td>AAACGCGTCTCTGTCCTGTCTCGA</td><td></td><td></td><td></td><td></td><td></td><td></td><td></td><td></td><td></td><td></td></tr>
<tr><td>Chromosome</td><td>start</td><td>end</td><td>strand</td><td>MM</td><td>target_seq</td><td>PAM</td><td>alignment</td><td>distance</td><td>position</td><td>gene name</td><td>gene id</td></tr>
<tr><td>GU305914.1</td><td>127511</td><td>127533</td><td>-</td><td>0</td><td>TCGAGACAGGACAGAGACGC</td><td>CGG</td><td>||||||||[||||||||||||]PAM</td><td>NA</td><td>Intergenic</td><td>NA</td><td>NA</td></tr>
<tr><td></td><td></td><td></td><td></td><td></td><td></td><td></td><td></td><td></td><td></td><td></td><td></td></tr>
<tr><td>T24337</td><td>GCCACCGCGCCCGATCCTCGCGG</td><td>1000</td><td></td><td></td><td></td><td></td><td></td><td></td><td></td><td></td><td></td></tr>
<tr><td>Oligo fwd</td><td>CACCGCCACCGCGCCCGATCCTCG</td><td></td><td></td><td></td><td></td><td></td><td></td><td></td><td></td><td></td><td></td></tr>
<tr><td>Oligo rev</td><td>AAACCGAGGATCGGGCGCGGTGGC</td><td></td><td></td><td></td><td></td><td></td><td></td><td></td><td></td><td></td><td></td></tr>
<tr><td>Chromosome</td><td>start</td><td>end</td><td>strand</td><td>MM</td><td>target_seq</td><td>PAM</td><td>alignment</td><td>distance</td><td>position</td><td>gene name</td><td>gene id</td></tr>
<tr><td>GU305914.1</td><td>127460</td><td>127482</td><td>-</td><td>0</td><td>GCCACCGCGCCCGATCCTCG</td><td>CGG</td><td>||||||||[||||||||||||]PAM</td><td>NA</td><td>Intergenic</td><td>NA</td><td>NA</td></tr>
<tr><td></td><td></td><td></td><td></td><td></td><td></td><td></td><td></td><td></td><td></td><td></td><td></td></tr>
<tr><td>T24338</td><td>GCTCCCGAACCACGTCCCCGAGG</td><td>1000</td><td></td><td></td><td></td><td></td><td></td><td></td><td></td><td></td><td></td></tr>
<tr><td>Oligo fwd</td><td>CACCGCTCCCGAACCACGTCCCCG</td><td></td><td></td><td></td><td></td><td></td><td></td><td></td><td></td><td></td><td></td></tr>
<tr><td>Oligo rev</td><td>AAACCGGGGACGTGGTTCGGGAGC</td><td></td><td></td><td></td><td></td><td></td><td></td><td></td><td></td><td></td><td></td></tr>
<tr><td>Chromosome</td><td>start</td><td>end</td><td>strand</td><td>MM</td><td>target_seq</td><td>PAM</td><td>alignment</td><td>distance</td><td>position</td><td>gene name</td><td>gene id</td></tr>
<tr><td>GU305914.1</td><td>127432</td><td>127454</td><td>-</td><td>0</td><td>GCTCCCGAACCACGTCCCCG</td><td>AGG</td><td>||||||||[||||||||||||]PAM</td><td>NA</td><td>Intergenic</td><td>NA</td><td>NA</td></tr>
<tr><td></td><td></td><td></td><td></td><td></td><td></td><td></td><td></td><td></td><td></td><td></td><td></td></tr>
<tr><td>T24339</td><td>CTCCCGAACCACGTCCCCGAGGG</td><td>1000</td><td></td><td></td><td></td><td></td><td></td><td></td><td></td><td></td><td></td></tr>
<tr><td>Oligo fwd</td><td>CACCCTCCCGAACCACGTCCCCGA</td><td></td><td></td><td></td><td></td><td></td><td></td><td></td><td></td><td></td><td></td></tr>
<tr><td>Oligo rev</td><td>AAACTCGGGGACGTGGTTCGGGAG</td><td></td><td></td><td></td><td></td><td></td><td></td><td></td><td></td><td></td><td></td></tr>
<tr><td>Chromosome</td><td>start</td><td>end</td><td>strand</td><td>MM</td><td>target_seq</td><td>PAM</td><td>alignment</td><td>distance</td><td>position</td><td>gene name</td><td>gene id</td></tr>
<tr><td>GU305914.1</td><td>127431</td><td>127453</td><td>-</td><td>0</td><td>CTCCCGAACCACGTCCCCGA</td><td>GGG</td><td>||||||||[||||||||||||]PAM</td><td>NA</td><td>Intergenic</td><td>NA</td><td>NA</td></tr>
<tr><td></td><td></td><td></td><td></td><td></td><td></td><td></td><td></td><td></td><td></td><td></td><td></td></tr>
<tr><td>T24340</td><td>AGAGCGAGAGAGAGACCGAGAGG</td><td>1000</td><td></td><td></td><td></td><td></td><td></td><td></td><td></td><td></td><td></td></tr>
<tr><td>Oligo fwd</td><td>CACCAGAGCGAGAGAGAGACCGAG</td><td></td><td></td><td></td><td></td><td></td><td></td><td></td><td></td><td></td><td></td></tr>
<tr><td>Oligo rev</td><td>AAACCTCGGTCTCTCTCTCGCTCT</td><td></td><td></td><td></td><td></td><td></td><td></td><td></td><td></td><td></td><td></td></tr>
<tr><td>Chromosome</td><td>start</td><td>end</td><td>strand</td><td>MM</td><td>target_seq</td><td>PAM</td><td>alignment</td><td>distance</td><td>position</td><td>gene name</td><td>gene id</td></tr>
<tr><td>GU305914.1</td><td>127398</td><td>127420</td><td>-</td><td>0</td><td>AGAGCGAGAGAGAGACCGAG</td><td>AGG</td><td>||||||||[||||||||||||]PAM</td><td>NA</td><td>Intergenic</td><td>NA</td><td>NA</td></tr>
<tr><td></td><td></td><td></td><td></td><td></td><td></td><td></td><td></td><td></td><td></td><td></td><td></td></tr>
<tr><td>T24341</td><td>GAGCGAGAGAGAGACCGAGAGGG</td><td>1000</td><td></td><td></td><td></td><td></td><td></td><td></td><td></td><td></td><td></td></tr>
<tr><td>Oligo fwd</td><td>CACCGAGCGAGAGAGAGACCGAGA</td><td></td><td></td><td></td><td></td><td></td><td></td><td></td><td></td><td></td><td></td></tr>
<tr><td>Oligo rev</td><td>AAACTCTCGGTCTCTCTCTCGCTC</td><td></td><td></td><td></td><td></td><td></td><td></td><td></td><td></td><td></td><td></td></tr>
<tr><td>Chromosome</td><td>start</td><td>end</td><td>strand</td><td>MM</td><td>target_seq</td><td>PAM</td><td>alignment</td><td>distance</td><td>position</td><td>gene name</td><td>gene id</td></tr>
<tr><td>GU305914.1</td><td>127397</td><td>127419</td><td>-</td><td>0</td><td>GAGCGAGAGAGAGACCGAGA</td><td>GGG</td><td>||||||||[||||||||||||]PAM</td><td>NA</td><td>Intergenic</td><td>NA</td><td>NA</td></tr>
<tr><td></td><td></td><td></td><td></td><td></td><td></td><td></td><td></td><td></td><td></td><td></td><td></td></tr>
<tr><td>T24342</td><td>GAGAGGGCGATCGCCGCCGCCGG</td><td>1000</td><td></td><td></td><td></td><td></td><td></td><td></td><td></td><td></td><td></td></tr>
<tr><td>Oligo fwd</td><td>CACCGAGAGGGCGATCGCCGCCGC</td><td></td><td></td><td></td><td></td><td></td><td></td><td></td><td></td><td></td><td></td></tr>
<tr><td>Oligo rev</td><td>AAACGCGGCGGCGATCGCCCTCTC</td><td></td><td></td><td></td><td></td><td></td><td></td><td></td><td></td><td></td><td></td></tr>
<tr><td>Chromosome</td><td>start</td><td>end</td><td>strand</td><td>MM</td><td>target_seq</td><td>PAM</td><td>alignment</td><td>distance</td><td>position</td><td>gene name</td><td>gene id</td></tr>
<tr><td>GU305914.1</td><td>127381</td><td>127403</td><td>-</td><td>0</td><td>GAGAGGGCGATCGCCGCCGC</td><td>CGG</td><td>||||||||[||||||||||||]PAM</td><td>NA</td><td>Intergenic</td><td>NA</td><td>NA</td></tr>
<tr><td></td><td></td><td></td><td></td><td></td><td></td><td></td><td></td><td></td><td></td><td></td><td></td></tr>
<tr><td>T24343</td><td>CGGTAGCGCCGCCGCCGCGATGG</td><td>1000</td><td></td><td></td><td></td><td></td><td></td><td></td><td></td><td></td><td></td></tr>
<tr><td>Oligo fwd</td><td>CACCCGGTAGCGCCGCCGCCGCGA</td><td></td><td></td><td></td><td></td><td></td><td></td><td></td><td></td><td></td><td></td></tr>
<tr><td>Oligo rev</td><td>AAACTCGCGGCGGCGGCGCTACCG</td><td></td><td></td><td></td><td></td><td></td><td></td><td></td><td></td><td></td><td></td></tr>
<tr><td>Chromosome</td><td>start</td><td>end</td><td>strand</td><td>MM</td><td>target_seq</td><td>PAM</td><td>alignment</td><td>distance</td><td>position</td><td>gene name</td><td>gene id</td></tr>
<tr><td>GU305914.1</td><td>127361</td><td>127383</td><td>-</td><td>0</td><td>CGGTAGCGCCGCCGCCGCGA</td><td>TGG</td><td>||||||||[||||||||||||]PAM</td><td>NA</td><td>Intergenic</td><td>NA</td><td>NA</td></tr>
<tr><td>GU305914.1</td><td>114687</td><td>114709</td><td>-</td><td>4</td><td>AAGTACCGCCACCGCCGCGA</td><td>TGG</td><td>--|||-||[||-|||||||||]PAM</td><td>NA</td><td>Intergenic</td><td>NA</td><td>NA</td></tr>
<tr><td></td><td></td><td></td><td></td><td></td><td></td><td></td><td></td><td></td><td></td><td></td><td></td></tr>
<tr><td>T24344</td><td>GGTAGCGCCGCCGCCGCGATGGG</td><td>1000</td><td></td><td></td><td></td><td></td><td></td><td></td><td></td><td></td><td></td></tr>
<tr><td>Oligo fwd</td><td>CACCGGTAGCGCCGCCGCCGCGAT</td><td></td><td></td><td></td><td></td><td></td><td></td><td></td><td></td><td></td><td></td></tr>
<tr><td>Oligo rev</td><td>AAACATCGCGGCGGCGGCGCTACC</td><td></td><td></td><td></td><td></td><td></td><td></td><td></td><td></td><td></td><td></td></tr>
<tr><td>Chromosome</td><td>start</td><td>end</td><td>strand</td><td>MM</td><td>target_seq</td><td>PAM</td><td>alignment</td><td>distance</td><td>position</td><td>gene name</td><td>gene id</td></tr>
<tr><td>GU305914.1</td><td>127360</td><td>127382</td><td>-</td><td>0</td><td>GGTAGCGCCGCCGCCGCGAT</td><td>GGG</td><td>||||||||[||||||||||||]PAM</td><td>NA</td><td>Intergenic</td><td>NA</td><td>NA</td></tr>
<tr><td></td><td></td><td></td><td></td><td></td><td></td><td></td><td></td><td></td><td></td><td></td><td></td></tr>
<tr><td>T24345</td><td>GTAGCGCCGCCGCCGCGATGGGG</td><td>1000</td><td></td><td></td><td></td><td></td><td></td><td></td><td></td><td></td><td></td></tr>
<tr><td>Oligo fwd</td><td>CACCGTAGCGCCGCCGCCGCGATG</td><td></td><td></td><td></td><td></td><td></td><td></td><td></td><td></td><td></td><td></td></tr>
<tr><td>Oligo rev</td><td>AAACCATCGCGGCGGCGGCGCTAC</td><td></td><td></td><td></td><td></td><td></td><td></td><td></td><td></td><td></td><td></td></tr>
<tr><td>Chromosome</td><td>start</td><td>end</td><td>strand</td><td>MM</td><td>target_seq</td><td>PAM</td><td>alignment</td><td>distance</td><td>position</td><td>gene name</td><td>gene id</td></tr>
<tr><td>GU305914.1</td><td>127359</td><td>127381</td><td>-</td><td>0</td><td>GTAGCGCCGCCGCCGCGATG</td><td>GGG</td><td>||||||||[||||||||||||]PAM</td><td>NA</td><td>Intergenic</td><td>NA</td><td>NA</td></tr>
<tr><td></td><td></td><td></td><td></td><td></td><td></td><td></td><td></td><td></td><td></td><td></td><td></td></tr>
<tr><td>T24346</td><td>TAGCGCCGCCGCCGCGATGGGGG</td><td>1000</td><td></td><td></td><td></td><td></td><td></td><td></td><td></td><td></td><td></td></tr>
<tr><td>Oligo fwd</td><td>CACCTAGCGCCGCCGCCGCGATGG</td><td></td><td></td><td></td><td></td><td></td><td></td><td></td><td></td><td></td><td></td></tr>
<tr><td>Oligo rev</td><td>AAACCCATCGCGGCGGCGGCGCTA</td><td></td><td></td><td></td><td></td><td></td><td></td><td></td><td></td><td></td><td></td></tr>
<tr><td>Chromosome</td><td>start</td><td>end</td><td>strand</td><td>MM</td><td>target_seq</td><td>PAM</td><td>alignment</td><td>distance</td><td>position</td><td>gene name</td><td>gene id</td></tr>
<tr><td>GU305914.1</td><td>127358</td><td>127380</td><td>-</td><td>0</td><td>TAGCGCCGCCGCCGCGATGG</td><td>GGG</td><td>||||||||[||||||||||||]PAM</td><td>NA</td><td>Intergenic</td><td>NA</td><td>NA</td></tr>
<tr><td></td><td></td><td></td><td></td><td></td><td></td><td></td><td></td><td></td><td></td><td></td><td></td></tr>
<tr><td>T24347</td><td>CCGCCGCGATGGGGGAGAACTGG</td><td>1000</td><td></td><td></td><td></td><td></td><td></td><td></td><td></td><td></td><td></td></tr>
<tr><td>Oligo fwd</td><td>CACCCCGCCGCGATGGGGGAGAAC</td><td></td><td></td><td></td><td></td><td></td><td></td><td></td><td></td><td></td><td></td></tr>
<tr><td>Oligo rev</td><td>AAACGTTCTCCCCCATCGCGGCGG</td><td></td><td></td><td></td><td></td><td></td><td></td><td></td><td></td><td></td><td></td></tr>
<tr><td>Chromosome</td><td>start</td><td>end</td><td>strand</td><td>MM</td><td>target_seq</td><td>PAM</td><td>alignment</td><td>distance</td><td>position</td><td>gene name</td><td>gene id</td></tr>
<tr><td>GU305914.1</td><td>127350</td><td>127372</td><td>-</td><td>0</td><td>CCGCCGCGATGGGGGAGAAC</td><td>TGG</td><td>||||||||[||||||||||||]PAM</td><td>NA</td><td>Intergenic</td><td>NA</td><td>NA</td></tr>
<tr><td></td><td></td><td></td><td></td><td></td><td></td><td></td><td></td><td></td><td></td><td></td><td></td></tr>
<tr><td>T24348</td><td>CGCGATGGGGGAGAACTGGACGG</td><td>1000</td><td></td><td></td><td></td><td></td><td></td><td></td><td></td><td></td><td></td></tr>
</table>

| | | | | | | | | | | | |
|---|---|---|---|---|---|---|---|---|---|---|---|
| Oligo fwd | CACCCGCGATGGGGGAGAACTGGA | | | | | | | | | | |
| Oligo rev | AAACTCCAGTTCTCCCCCATCGCG | | | | | | | | | | |
| Chromosome | start | end | strand | MM | target_seq | PAM | alignment | distance | position | gene name | gene id |
| GU305914.1 | 127346 | 127368 | - | 0 | CGCGATGGGGGAGAACTGGA | CGG | \|\|\|\|\|\|\|\|[\|\|\|\|\|\|\|\|\|\|\|\|]PAM | NA | Intergenic | NA | NA |
| T24349 | GGGGGAGAACTGGACGGCGACGG | 1000 | | | | | | | | | |
| Oligo fwd | CACCGGGGGAGAACTGGACGGCGA | | | | | | | | | | |
| Oligo rev | AAACTCGCCGTCCAGTTCTCCCCC | | | | | | | | | | |
| Chromosome | start | end | strand | MM | target_seq | PAM | alignment | distance | position | gene name | gene id |
| GU305914.1 | 127340 | 127362 | - | 0 | GGGGGAGAACTGGACGGCGA | CGG | \|\|\|\|\|\|\|\|[\|\|\|\|\|\|\|\|\|\|\|\|]PAM | NA | Intergenic | NA | NA |
| T24350 | GACGCACATCAAACTGAGCGTGG | 1000 | | | | | | | | | |
| Oligo fwd | CACCGACGCACATCAAACTGAGCG | | | | | | | | | | |
| Oligo rev | AAACCGCTCAGTTTGATGTGCGTC | | | | | | | | | | |
| Chromosome | start | end | strand | MM | target_seq | PAM | alignment | distance | position | gene name | gene id |
| GU305914.1 | 127280 | 127302 | - | 0 | GACGCACATCAAACTGAGCG | TGG | \|\|\|\|\|\|\|\|[\|\|\|\|\|\|\|\|\|\|\|\|]PAM | NA | Intergenic | NA | NA |
| T24351 | ACGCACATCAAACTGAGCGTGGG | 1000 | | | | | | | | | |
| Oligo fwd | CACCACGCACATCAAACTGAGCGT | | | | | | | | | | |
| Oligo rev | AAACACGCTCAGTTTGATGTGCGT | | | | | | | | | | |
| Chromosome | start | end | strand | MM | target_seq | PAM | alignment | distance | position | gene name | gene id |
| GU305914.1 | 127279 | 127301 | - | 0 | ACGCACATCAAACTGAGCGT | GGG | \|\|\|\|\|\|\|\|[\|\|\|\|\|\|\|\|\|\|\|\|]PAM | NA | Intergenic | NA | NA |
| T24352 | CGCACATCAAACTGAGCGTGGGG | 1000 | | | | | | | | | |
| Oligo fwd | CACCCGCACATCAAACTGAGCGTG | | | | | | | | | | |
| Oligo rev | AAACCACGCTCAGTTTGATGTGCG | | | | | | | | | | |
| Chromosome | start | end | strand | MM | target_seq | PAM | alignment | distance | position | gene name | gene id |
| GU305914.1 | 127278 | 127300 | - | 0 | CGCACATCAAACTGAGCGTG | GGG | \|\|\|\|\|\|\|\|[\|\|\|\|\|\|\|\|\|\|\|\|]PAM | NA | Intergenic | NA | NA |
| T24353 | GCACATCAAACTGAGCGTGGGGG | 1000 | | | | | | | | | |
| Oligo fwd | CACCGCACATCAAACTGAGCGTGG | | | | | | | | | | |
| Oligo rev | AAACCCACGCTCAGTTTGATGTGC | | | | | | | | | | |
| Chromosome | start | end | strand | MM | target_seq | PAM | alignment | distance | position | gene name | gene id |
| GU305914.1 | 127277 | 127299 | - | 0 | GCACATCAAACTGAGCGTGG | GGG | \|\|\|\|\|\|\|\|[\|\|\|\|\|\|\|\|\|\|\|\|]PAM | NA | Intergenic | NA | NA |
| T24354 | CATCAAACTGAGCGTGGGGGAGG | 1000 | | | | | | | | | |
| Oligo fwd | CACCCATCAAACTGAGCGTGGGGG | | | | | | | | | | |
| Oligo rev | AAACCCCCCACGCTCAGTTTGATG | | | | | | | | | | |
| Chromosome | start | end | strand | MM | target_seq | PAM | alignment | distance | position | gene name | gene id |
| GU305914.1 | 127274 | 127296 | - | 0 | CATCAAACTGAGCGTGGGGG | AGG | \|\|\|\|\|\|\|\|[\|\|\|\|\|\|\|\|\|\|\|\|]PAM | NA | Intergenic | NA | NA |
| T24355 | AACAACTTTCGCCTGTACTACGG | 1000 | | | | | | | | | |
| Oligo fwd | CACCAACAACTTTCGCCTGTACTA | | | | | | | | | | |
| Oligo rev | AAACTAGTACAGGCGAAAGTTGTT | | | | | | | | | | |
| Chromosome | start | end | strand | MM | target_seq | PAM | alignment | distance | position | gene name | gene id |
| GU305914.1 | 127243 | 127265 | - | 0 | AACAACTTTCGCCTGTACTA | CGG | \|\|\|\|\|\|\|\|[\|\|\|\|\|\|\|\|\|\|\|\|]PAM | NA | Intergenic | NA | NA |
| T24356 | ACAACTTTCGCCTGTACTACGGG | 1000 | | | | | | | | | |
| Oligo fwd | CACCACAACTTTCGCCTGTACTAC | | | | | | | | | | |
| Oligo rev | AAACGTAGTACAGGCGAAAGTTGT | | | | | | | | | | |
| Chromosome | start | end | strand | MM | target_seq | PAM | alignment | distance | position | gene name | gene id |
| GU305914.1 | 127242 | 127264 | - | 0 | ACAACTTTCGCCTGTACTAC | GGG | \|\|\|\|\|\|\|\|[\|\|\|\|\|\|\|\|\|\|\|\|]PAM | NA | Intergenic | NA | NA |
| T24357 | CAACTTTCGCCTGTACTACGGGG | 1000 | | | | | | | | | |
| Oligo fwd | CACCCAACTTTCGCCTGTACTACG | | | | | | | | | | |
| Oligo rev | AAACCGTAGTACAGGCGAAAGTTG | | | | | | | | | | |
| Chromosome | start | end | strand | MM | target_seq | PAM | alignment | distance | position | gene name | gene id |
| GU305914.1 | 127241 | 127263 | - | 0 | CAACTTTCGCCTGTACTACG | GGG | \|\|\|\|\|\|\|\|[\|\|\|\|\|\|\|\|\|\|\|\|]PAM | NA | Intergenic | NA | NA |
| T24358 | ACTACGGGGACGACCCCGAACGG | 1000 | | | | | | | | | |
| Oligo fwd | CACCACTACGGGGACGACCCCGAA | | | | | | | | | | |
| Oligo rev | AAACTTCGGGGTCGTCCCCGTAGT | | | | | | | | | | |
| Chromosome | start | end | strand | MM | target_seq | PAM | alignment | distance | position | gene name | gene id |
| GU305914.1 | 127227 | 127249 | - | 0 | ACTACGGGGACGACCCCGAA | CGG | \|\|\|\|\|\|\|\|[\|\|\|\|\|\|\|\|\|\|\|\|]PAM | NA | Intergenic | NA | NA |
| T24359 | ACGGTACAACCTGAGCTTCGAGG | 1000 | | | | | | | | | |
| Oligo fwd | CACCACGGTACAACCTGAGCTTCG | | | | | | | | | | |
| Oligo rev | AAACCGAAGCTCAGGTTGTACCGT | | | | | | | | | | |
| Chromosome | start | end | strand | MM | target_seq | PAM | alignment | distance | position | gene name | gene id |
| GU305914.1 | 127208 | 127230 | - | 0 | ACGGTACAACCTGAGCTTCG | AGG | \|\|\|\|\|\|\|\|[\|\|\|\|\|\|\|\|\|\|\|\|]PAM | NA | Intergenic | NA | NA |
| T24360 | CTGAGCTTCGAGGCGATCTTCGG | 1000 | | | | | | | | | |
| Oligo fwd | CACCCTGAGCTTCGAGGCGATCTT | | | | | | | | | | |
| Oligo rev | AAACAAGATCGCCTCGAAGCTCAG | | | | | | | | | | |
| Chromosome | start | end | strand | MM | target_seq | PAM | alignment | distance | position | gene name | gene id |
| GU305914.1 | 127198 | 127220 | - | 0 | CTGAGCTTCGAGGCGATCTT | CGG | \|\|\|\|\|\|\|\|[\|\|\|\|\|\|\|\|\|\|\|\|]PAM | NA | Intergenic | NA | NA |
| T24361 | CGTACTGCAACAAGATCGAGTGG | 1000 | | | | | | | | | |
| Oligo fwd | CACCCGTACTGCAACAAGATCGAG | | | | | | | | | | |
| Oligo rev | AAACCTCGATCTTGTTGCAGTACG | | | | | | | | | | |
| Chromosome | start | end | strand | MM | target_seq | PAM | alignment | distance | position | gene name | gene id |
| GU305914.1 | 127173 | 127195 | - | 0 | CGTACTGCAACAAGATCGAG | TGG | \|\|\|\|\|\|\|\|[\|\|\|\|\|\|\|\|\|\|\|\|]PAM | NA | Intergenic | NA | NA |
| T24362 | GTACTGCAACAAGATCGAGTGGG | 1000 | | | | | | | | | |
| Oligo fwd | CACCGTACTGCAACAAGATCGAGT | | | | | | | | | | |
| Oligo rev | AAACACTCGATCTTGTTGCAGTAC | | | | | | | | | | |
| Chromosome | start | end | strand | MM | target_seq | PAM | alignment | distance | position | gene name | gene id |
| GU305914.1 | 127172 | 127194 | - | 0 | GTACTGCAACAAGATCGAGT | GGG | \|\|\|\|\|\|\|\|[\|\|\|\|\|\|\|\|\|\|\|\|]PAM | NA | Intergenic | NA | NA |

| | | | | | | | | | | | |
|---|---|---|---|---|---|---|---|---|---|---|---|
| T24363 | GATCGAGTGGGTGACCTTCCTGG | 1000 | | | | | | | | | |
| Oligo fwd | CACCGATCGAGTGGGTGACCTTCC | | | | | | | | | | |
| Oligo rev | AAACGGAAGGTCACCCACTCGATC | | | | | | | | | | |
| Chromosome | start | end | strand | MM | target_seq | PAM | alignment | distance | position | gene name | gene id |
| GU305914.1 | 127160 | 127182 | - | 0 | GATCGAGTGGGTGACCTTCC | TGG | \|\|\|\|\|\|\|\|[\|\|\|\|\|\|\|\|\|\|\|\|]PAM | NA | Intergenic | NA | NA |
| T24364 | ATCGAGTGGGTGACCTTCCTGGG | 1000 | | | | | | | | | |
| Oligo fwd | CACCATCGAGTGGGTGACCTTCCT | | | | | | | | | | |
| Oligo rev | AAACAGGAAGGTCACCCACTCGAT | | | | | | | | | | |
| Chromosome | start | end | strand | MM | target_seq | PAM | alignment | distance | position | gene name | gene id |
| GU305914.1 | 127159 | 127181 | - | 0 | ATCGAGTGGGTGACCTTCCT | GGG | \|\|\|\|\|\|\|\|[\|\|\|\|\|\|\|\|\|\|\|\|]PAM | NA | Intergenic | NA | NA |
| T24365 | GTGGGTGACCTTCCTGGGAACGG | 1000 | | | | | | | | | |
| Oligo fwd | CACCGTGGGTGACCTTCCTGGGAA | | | | | | | | | | |
| Oligo rev | AAACTTCCCAGGAAGGTCACCCAC | | | | | | | | | | |
| Chromosome | start | end | strand | MM | target_seq | PAM | alignment | distance | position | gene name | gene id |
| GU305914.1 | 127154 | 127176 | - | 0 | GTGGGTGACCTTCCTGGGAA | CGG | \|\|\|\|\|\|\|\|[\|\|\|\|\|\|\|\|\|\|\|\|]PAM | NA | Intergenic | NA | NA |
| T24366 | GACCTTCCTGGGAACGGCGTTGG | 1000 | | | | | | | | | |
| Oligo fwd | CACCGACCTTCCTGGGAACGGCGT | | | | | | | | | | |
| Oligo rev | AAACACGCCGTTCCCAGGAAGGTC | | | | | | | | | | |
| Chromosome | start | end | strand | MM | target_seq | PAM | alignment | distance | position | gene name | gene id |
| GU305914.1 | 127148 | 127170 | - | 0 | GACCTTCCTGGGAACGGCGT | TGG | \|\|\|\|\|\|\|\|[\|\|\|\|\|\|\|\|\|\|\|\|]PAM | NA | Intergenic | NA | NA |
| T24367 | CCTGGGAACGGCGTTGGCGACGG | 1000 | | | | | | | | | |
| Oligo fwd | CACCCCTGGGAACGGCGTTGGCGA | | | | | | | | | | |
| Oligo rev | AAACTCGCCAACGCCGTTCCCAGG | | | | | | | | | | |
| Chromosome | start | end | strand | MM | target_seq | PAM | alignment | distance | position | gene name | gene id |
| GU305914.1 | 127142 | 127164 | - | 0 | CCTGGGAACGGCGTTGGCGA | CGG | \|\|\|\|\|\|\|\|[\|\|\|\|\|\|\|\|\|\|\|\|]PAM | NA | Intergenic | NA | NA |
| T24368 | GGGAACGGCGTTGGCGACGGCGG | 1000 | | | | | | | | | |
| Oligo fwd | CACCGGGAACGGCGTTGGCGACGG | | | | | | | | | | |
| Oligo rev | AAACCCGTCGCCAACGCCGTTCCC | | | | | | | | | | |
| Chromosome | start | end | strand | MM | target_seq | PAM | alignment | distance | position | gene name | gene id |
| GU305914.1 | 127139 | 127161 | - | 0 | GGGAACGGCGTTGGCGACGG | CGG | \|\|\|\|\|\|\|\|[\|\|\|\|\|\|\|\|\|\|\|\|]PAM | NA | Intergenic | NA | NA |
| GU305914.1 | 26379 | 26401 | + | 4 | GTCATCGGCGCTGGCGACGG | CGG | \|--\|-\|\|\|[\|\|-\|\|\|\|\|\|\|\|\|]PAM | NA | Intergenic | NA | NA |
| GU305914.1 | 134259 | 134281 | + | 4 | GGGAAGTGCGATGGCGGCGG | AGG | \|\|\|\|\|--\|[\|\|-\|\|\|\|\|-\|\|\|]PAM | NA | Intergenic | NA | NA |
| T24369 | CGACCTGAACAAAATGACTACGG | 1000 | | | | | | | | | |
| Oligo fwd | CACCCGACCTGAACAAAATGACTA | | | | | | | | | | |
| Oligo rev | AAACTAGTCATTTTGTTCAGGTCG | | | | | | | | | | |
| Chromosome | start | end | strand | MM | target_seq | PAM | alignment | distance | position | gene name | gene id |
| GU305914.1 | 127097 | 127119 | - | 0 | CGACCTGAACAAAATGACTA | CGG | \|\|\|\|\|\|\|\|[\|\|\|\|\|\|\|\|\|\|\|\|]PAM | NA | Intergenic | NA | NA |
| T24370 | GACCTGAACAAAATGACTACGGG | 1000 | | | | | | | | | |
| Oligo fwd | CACCGACCTGAACAAAATGACTAC | | | | | | | | | | |
| Oligo rev | AAACGTAGTCATTTTGTTCAGGTC | | | | | | | | | | |
| Chromosome | start | end | strand | MM | target_seq | PAM | alignment | distance | position | gene name | gene id |
| GU305914.1 | 127096 | 127118 | - | 0 | GACCTGAACAAAATGACTAC | GGG | \|\|\|\|\|\|\|\|[\|\|\|\|\|\|\|\|\|\|\|\|]PAM | NA | Intergenic | NA | NA |
| T24371 | CAAGATGCTTTTTTATATACAGG | 1000 | | | | | | | | | |
| Oligo fwd | CACCCAAGATGCTTTTTTATATAC | | | | | | | | | | |
| Oligo rev | AAACGTATATAAAAAAGCATCTTG | | | | | | | | | | |
| Chromosome | start | end | strand | MM | target_seq | PAM | alignment | distance | position | gene name | gene id |
| GU305914.1 | 127073 | 127095 | - | 0 | CAAGATGCTTTTTTATATAC | AGG | \|\|\|\|\|\|\|\|[\|\|\|\|\|\|\|\|\|\|\|\|]PAM | NA | Intergenic | NA | NA |
| T24372 | TTATATACAGGTCCCTCGAGTGG | 1000 | | | | | | | | | |
| Oligo fwd | CACCTTATATACAGGTCCCTCGAG | | | | | | | | | | |
| Oligo rev | AAACCTCGAGGGACCTGTATATAA | | | | | | | | | | |
| Chromosome | start | end | strand | MM | target_seq | PAM | alignment | distance | position | gene name | gene id |
| GU305914.1 | 127061 | 127083 | - | 0 | TTATATACAGGTCCCTCGAG | TGG | \|\|\|\|\|\|\|\|[\|\|\|\|\|\|\|\|\|\|\|\|]PAM | NA | Intergenic | NA | NA |
| T24373 | CAGGTCCCTCGAGTGGCCACCGG | 1000 | | | | | | | | | |
| Oligo fwd | CACCCAGGTCCCTCGAGTGGCCAC | | | | | | | | | | |
| Oligo rev | AAACGTGGCCACTCGAGGGACCTG | | | | | | | | | | |
| Chromosome | start | end | strand | MM | target_seq | PAM | alignment | distance | position | gene name | gene id |
| GU305914.1 | 127054 | 127076 | - | 0 | CAGGTCCCTCGAGTGGCCAC | CGG | \|\|\|\|\|\|\|\|[\|\|\|\|\|\|\|\|\|\|\|\|]PAM | NA | Intergenic | NA | NA |
| T24374 | AGGTCCCTCGAGTGGCCACCGGG | 1000 | | | | | | | | | |
| Oligo fwd | CACCAGGTCCCTCGAGTGGCCACC | | | | | | | | | | |
| Oligo rev | AAACGGTGGCCACTCGAGGGACCT | | | | | | | | | | |
| Chromosome | start | end | strand | MM | target_seq | PAM | alignment | distance | position | gene name | gene id |
| GU305914.1 | 127053 | 127075 | - | 0 | AGGTCCCTCGAGTGGCCACC | GGG | \|\|\|\|\|\|\|\|[\|\|\|\|\|\|\|\|\|\|\|\|]PAM | NA | Intergenic | NA | NA |
| T24375 | GGTCCCTCGAGTGGCCACCGGGG | 1000 | | | | | | | | | |
| Oligo fwd | CACCGGTCCCTCGAGTGGCCACCG | | | | | | | | | | |
| Oligo rev | AAACCGGTGGCCACTCGAGGGACC | | | | | | | | | | |
| Chromosome | start | end | strand | MM | target_seq | PAM | alignment | distance | position | gene name | gene id |
| GU305914.1 | 127052 | 127074 | - | 0 | GGTCCCTCGAGTGGCCACCG | GGG | \|\|\|\|\|\|\|\|[\|\|\|\|\|\|\|\|\|\|\|\|]PAM | NA | Intergenic | NA | NA |
| T24376 | CCTCGAGTGGCCACCGGGGCCGG | 1000 | | | | | | | | | |
| Oligo fwd | CACCCCTCGAGTGGCCACCGGGGC | | | | | | | | | | |
| Oligo rev | AAACGCCCCGGTGGCCACTCGAGG | | | | | | | | | | |
| Chromosome | start | end | strand | MM | target_seq | PAM | alignment | distance | position | gene name | gene id |
| GU305914.1 | 127048 | 127070 | - | 0 | CCTCGAGTGGCCACCGGGGC | CGG | \|\|\|\|\|\|\|\|[\|\|\|\|\|\|\|\|\|\|\|\|]PAM | NA | Intergenic | NA | NA |
| T24377 | CTCGAGTGGCCACCGGGGCCGGG | 1000 | | | | | | | | | |
| Oligo fwd | CACCCTCGAGTGGCCACCGGGGCC | | | | | | | | | | |
| Oligo rev | AAACGGCCCCGGTGGCCACTCGAG | | | | | | | | | | |

<table>
<tr><td>Chromosome</td><td>start</td><td>end</td><td>strand</td><td>MM</td><td>target_seq</td><td>PAM</td><td>alignment</td><td>distance</td><td>position</td><td>gene name</td><td>gene id</td></tr>
<tr><td>GU305914.1</td><td>127047</td><td>127069</td><td>-</td><td>0</td><td>CTCGAGTGGCCACCGGGGCC</td><td>GGG</td><td>||||||||[||||||||||||]PAM</td><td>NA</td><td>Intergenic</td><td>NA</td><td>NA</td></tr>
<tr><td>T24378</td><td>CCCCACCTCGCGACAGACGACGG</td><td>1000</td><td></td><td></td><td></td><td></td><td></td><td></td><td></td><td></td><td></td></tr>
<tr><td>Oligo fwd</td><td>CACCCCCCACCTCGCGACAGACGA</td><td></td><td></td><td></td><td></td><td></td><td></td><td></td><td></td><td></td><td></td></tr>
<tr><td>Oligo rev</td><td>AAACTCGTCTGTCGCGAGGTGGGG</td><td></td><td></td><td></td><td></td><td></td><td></td><td></td><td></td><td></td><td></td></tr>
<tr><td>Chromosome</td><td>start</td><td>end</td><td>strand</td><td>MM</td><td>target_seq</td><td>PAM</td><td>alignment</td><td>distance</td><td>position</td><td>gene name</td><td>gene id</td></tr>
<tr><td>GU305914.1</td><td>127022</td><td>127044</td><td>-</td><td>0</td><td>CCCCACCTCGCGACAGACGA</td><td>CGG</td><td>||||||||[||||||||||||]PAM</td><td>NA</td><td>Intergenic</td><td>NA</td><td>NA</td></tr>
<tr><td>T24379</td><td>CTCGCGACAGACGACGGTGATGG</td><td>1000</td><td></td><td></td><td></td><td></td><td></td><td></td><td></td><td></td><td></td></tr>
<tr><td>Oligo fwd</td><td>CACCCTCGCGACAGACGACGGTGA</td><td></td><td></td><td></td><td></td><td></td><td></td><td></td><td></td><td></td><td></td></tr>
<tr><td>Oligo rev</td><td>AAACTCACCGTCGTCTGTCGCGAG</td><td></td><td></td><td></td><td></td><td></td><td></td><td></td><td></td><td></td><td></td></tr>
<tr><td>Chromosome</td><td>start</td><td>end</td><td>strand</td><td>MM</td><td>target_seq</td><td>PAM</td><td>alignment</td><td>distance</td><td>position</td><td>gene name</td><td>gene id</td></tr>
<tr><td>GU305914.1</td><td>127016</td><td>127038</td><td>-</td><td>0</td><td>CTCGCGACAGACGACGGTGA</td><td>TGG</td><td>||||||||[||||||||||||]PAM</td><td>NA</td><td>Intergenic</td><td>NA</td><td>NA</td></tr>
<tr><td>T24380</td><td>CGCCTGTCTGACCCACCTGAAGG</td><td>1000</td><td></td><td></td><td></td><td></td><td></td><td></td><td></td><td></td><td></td></tr>
<tr><td>Oligo fwd</td><td>CACCCGCCTGTCTGACCCACCTGA</td><td></td><td></td><td></td><td></td><td></td><td></td><td></td><td></td><td></td><td></td></tr>
<tr><td>Oligo rev</td><td>AAACTCAGGTGGGTCAGACAGGCG</td><td></td><td></td><td></td><td></td><td></td><td></td><td></td><td></td><td></td><td></td></tr>
<tr><td>Chromosome</td><td>start</td><td>end</td><td>strand</td><td>MM</td><td>target_seq</td><td>PAM</td><td>alignment</td><td>distance</td><td>position</td><td>gene name</td><td>gene id</td></tr>
<tr><td>GU305914.1</td><td>126941</td><td>126963</td><td>-</td><td>0</td><td>CGCCTGTCTGACCCACCTGA</td><td>AGG</td><td>||||||||[||||||||||||]PAM</td><td>NA</td><td>Intergenic</td><td>NA</td><td>NA</td></tr>
<tr><td>T24381</td><td>CCACCTGAAGGAGACCTTCGAGG</td><td>1000</td><td></td><td></td><td></td><td></td><td></td><td></td><td></td><td></td><td></td></tr>
<tr><td>Oligo fwd</td><td>CACCCCACCTGAAGGAGACCTTCG</td><td></td><td></td><td></td><td></td><td></td><td></td><td></td><td></td><td></td><td></td></tr>
<tr><td>Oligo rev</td><td>AAACCGAAGGTCTCCTTCAGGTGG</td><td></td><td></td><td></td><td></td><td></td><td></td><td></td><td></td><td></td><td></td></tr>
<tr><td>Chromosome</td><td>start</td><td>end</td><td>strand</td><td>MM</td><td>target_seq</td><td>PAM</td><td>alignment</td><td>distance</td><td>position</td><td>gene name</td><td>gene id</td></tr>
<tr><td>GU305914.1</td><td>126929</td><td>126951</td><td>-</td><td>0</td><td>CCACCTGAAGGAGACCTTCG</td><td>AGG</td><td>||||||||[||||||||||||]PAM</td><td>NA</td><td>Intergenic</td><td>NA</td><td>NA</td></tr>
<tr><td>GU305914.1</td><td>147786</td><td>147808</td><td>+</td><td>4</td><td>CTACGTCAAGAAGACCTTCG</td><td>AGG</td><td>|-||-|-|[||-|||||||||]PAM</td><td>NA</td><td>Intergenic</td><td>NA</td><td>NA</td></tr>
<tr><td>T24382</td><td>GACCTTCGAGGACACGCTGCTGG</td><td>1000</td><td></td><td></td><td></td><td></td><td></td><td></td><td></td><td></td><td></td></tr>
<tr><td>Oligo fwd</td><td>CACCGACCTTCGAGGACACGCTGC</td><td></td><td></td><td></td><td></td><td></td><td></td><td></td><td></td><td></td><td></td></tr>
<tr><td>Oligo rev</td><td>AAACGCAGCGTGTCCTCGAAGGTC</td><td></td><td></td><td></td><td></td><td></td><td></td><td></td><td></td><td></td><td></td></tr>
<tr><td>Chromosome</td><td>start</td><td>end</td><td>strand</td><td>MM</td><td>target_seq</td><td>PAM</td><td>alignment</td><td>distance</td><td>position</td><td>gene name</td><td>gene id</td></tr>
<tr><td>GU305914.1</td><td>126917</td><td>126939</td><td>-</td><td>0</td><td>GACCTTCGAGGACACGCTGC</td><td>TGG</td><td>||||||||[||||||||||||]PAM</td><td>NA</td><td>Intergenic</td><td>NA</td><td>NA</td></tr>
<tr><td>T24383</td><td>GAACGCCGACGCCGTCAACACGG</td><td>1000</td><td></td><td></td><td></td><td></td><td></td><td></td><td></td><td></td><td></td></tr>
<tr><td>Oligo fwd</td><td>CACCGAACGCCGACGCCGTCAACA</td><td></td><td></td><td></td><td></td><td></td><td></td><td></td><td></td><td></td><td></td></tr>
<tr><td>Oligo rev</td><td>AAACTGTTGACGGCGTCGGCGTTC</td><td></td><td></td><td></td><td></td><td></td><td></td><td></td><td></td><td></td><td></td></tr>
<tr><td>Chromosome</td><td>start</td><td>end</td><td>strand</td><td>MM</td><td>target_seq</td><td>PAM</td><td>alignment</td><td>distance</td><td>position</td><td>gene name</td><td>gene id</td></tr>
<tr><td>GU305914.1</td><td>126884</td><td>126906</td><td>-</td><td>0</td><td>GAACGCCGACGCCGTCAACA</td><td>CGG</td><td>||||||||[||||||||||||]PAM</td><td>NA</td><td>Intergenic</td><td>NA</td><td>NA</td></tr>
<tr><td>T24384</td><td>CGTCAACACGGTGTTGCGTGCGG</td><td>1000</td><td></td><td></td><td></td><td></td><td></td><td></td><td></td><td></td><td></td></tr>
<tr><td>Oligo fwd</td><td>CACCCGTCAACACGGTGTTGCGTG</td><td></td><td></td><td></td><td></td><td></td><td></td><td></td><td></td><td></td><td></td></tr>
<tr><td>Oligo rev</td><td>AAACCACGCAACACCGTGTTGACG</td><td></td><td></td><td></td><td></td><td></td><td></td><td></td><td></td><td></td><td></td></tr>
<tr><td>Chromosome</td><td>start</td><td>end</td><td>strand</td><td>MM</td><td>target_seq</td><td>PAM</td><td>alignment</td><td>distance</td><td>position</td><td>gene name</td><td>gene id</td></tr>
<tr><td>GU305914.1</td><td>126872</td><td>126894</td><td>-</td><td>0</td><td>CGTCAACACGGTGTTGCGTG</td><td>CGG</td><td>||||||||[||||||||||||]PAM</td><td>NA</td><td>Intergenic</td><td>NA</td><td>NA</td></tr>
<tr><td>T24385</td><td>GTTGCGTGCGGTGAAAAACACGG</td><td>1000</td><td></td><td></td><td></td><td></td><td></td><td></td><td></td><td></td><td></td></tr>
<tr><td>Oligo fwd</td><td>CACCGTTGCGTGCGGTGAAAAACA</td><td></td><td></td><td></td><td></td><td></td><td></td><td></td><td></td><td></td><td></td></tr>
<tr><td>Oligo rev</td><td>AAACTGTTTTTCACCGCACGCAAC</td><td></td><td></td><td></td><td></td><td></td><td></td><td></td><td></td><td></td><td></td></tr>
<tr><td>Chromosome</td><td>start</td><td>end</td><td>strand</td><td>MM</td><td>target_seq</td><td>PAM</td><td>alignment</td><td>distance</td><td>position</td><td>gene name</td><td>gene id</td></tr>
<tr><td>GU305914.1</td><td>126860</td><td>126882</td><td>-</td><td>0</td><td>GTTGCGTGCGGTGAAAAACA</td><td>CGG</td><td>||||||||[||||||||||||]PAM</td><td>NA</td><td>Intergenic</td><td>NA</td><td>NA</td></tr>
<tr><td>T24386</td><td>GAAAAACACGGCCGACGCGATGG</td><td>1000</td><td></td><td></td><td></td><td></td><td></td><td></td><td></td><td></td><td></td></tr>
<tr><td>Oligo fwd</td><td>CACCGAAAAACACGGCCGACGCGA</td><td></td><td></td><td></td><td></td><td></td><td></td><td></td><td></td><td></td><td></td></tr>
<tr><td>Oligo rev</td><td>AAACTCGCGTCGGCCGTGTTTTTC</td><td></td><td></td><td></td><td></td><td></td><td></td><td></td><td></td><td></td><td></td></tr>
<tr><td>Chromosome</td><td>start</td><td>end</td><td>strand</td><td>MM</td><td>target_seq</td><td>PAM</td><td>alignment</td><td>distance</td><td>position</td><td>gene name</td><td>gene id</td></tr>
<tr><td>GU305914.1</td><td>126848</td><td>126870</td><td>-</td><td>0</td><td>GAAAAACACGGCCGACGCGA</td><td>TGG</td><td>||||||||[||||||||||||]PAM</td><td>NA</td><td>Intergenic</td><td>NA</td><td>NA</td></tr>
<tr><td>T24387</td><td>ACGGCCGACGCGATGGAGCGCGG</td><td>1000</td><td></td><td></td><td></td><td></td><td></td><td></td><td></td><td></td><td></td></tr>
<tr><td>Oligo fwd</td><td>CACCACGGCCGACGCGATGGAGCG</td><td></td><td></td><td></td><td></td><td></td><td></td><td></td><td></td><td></td><td></td></tr>
<tr><td>Oligo rev</td><td>AAACCGCTCCATCGCGTCGGCCGT</td><td></td><td></td><td></td><td></td><td></td><td></td><td></td><td></td><td></td><td></td></tr>
<tr><td>Chromosome</td><td>start</td><td>end</td><td>strand</td><td>MM</td><td>target_seq</td><td>PAM</td><td>alignment</td><td>distance</td><td>position</td><td>gene name</td><td>gene id</td></tr>
<tr><td>GU305914.1</td><td>126841</td><td>126863</td><td>-</td><td>0</td><td>ACGGCCGACGCGATGGAGCG</td><td>CGG</td><td>||||||||[||||||||||||]PAM</td><td>NA</td><td>Intergenic</td><td>NA</td><td>NA</td></tr>
<tr><td>T24388</td><td>CTTCGTGTTGCGTACGCTGATGG</td><td>1000</td><td></td><td></td><td></td><td></td><td></td><td></td><td></td><td></td><td></td></tr>
<tr><td>Oligo fwd</td><td>CACCCTTCGTGTTGCGTACGCTGA</td><td></td><td></td><td></td><td></td><td></td><td></td><td></td><td></td><td></td><td></td></tr>
<tr><td>Oligo rev</td><td>AAACTCAGCGTACGCAACACGAAG</td><td></td><td></td><td></td><td></td><td></td><td></td><td></td><td></td><td></td><td></td></tr>
<tr><td>Chromosome</td><td>start</td><td>end</td><td>strand</td><td>MM</td><td>target_seq</td><td>PAM</td><td>alignment</td><td>distance</td><td>position</td><td>gene name</td><td>gene id</td></tr>
<tr><td>GU305914.1</td><td>126770</td><td>126792</td><td>-</td><td>0</td><td>CTTCGTGTTGCGTACGCTGA</td><td>TGG</td><td>||||||||[||||||||||||]PAM</td><td>NA</td><td>Intergenic</td><td>NA</td><td>NA</td></tr>
<tr><td>T24389</td><td>TTGCGTACGCTGATGGAGCACGG</td><td>1000</td><td></td><td></td><td></td><td></td><td></td><td></td><td></td><td></td><td></td></tr>
<tr><td>Oligo fwd</td><td>CACCTTGCGTACGCTGATGGAGCA</td><td></td><td></td><td></td><td></td><td></td><td></td><td></td><td></td><td></td><td></td></tr>
<tr><td>Oligo rev</td><td>AAACTGCTCCATCAGCGTACGCAA</td><td></td><td></td><td></td><td></td><td></td><td></td><td></td><td></td><td></td><td></td></tr>
<tr><td>Chromosome</td><td>start</td><td>end</td><td>strand</td><td>MM</td><td>target_seq</td><td>PAM</td><td>alignment</td><td>distance</td><td>position</td><td>gene name</td><td>gene id</td></tr>
<tr><td>GU305914.1</td><td>126763</td><td>126785</td><td>-</td><td>0</td><td>TTGCGTACGCTGATGGAGCA</td><td>CGG</td><td>||||||||[||||||||||||]PAM</td><td>NA</td><td>Intergenic</td><td>NA</td><td>NA</td></tr>
<tr><td>T24390</td><td>TGCGTACGCTGATGGAGCACGGG</td><td>1000</td><td></td><td></td><td></td><td></td><td></td><td></td><td></td><td></td><td></td></tr>
<tr><td>Oligo fwd</td><td>CACCTGCGTACGCTGATGGAGCAC</td><td></td><td></td><td></td><td></td><td></td><td></td><td></td><td></td><td></td><td></td></tr>
<tr><td>Oligo rev</td><td>AAACGTGCTCCATCAGCGTACGCA</td><td></td><td></td><td></td><td></td><td></td><td></td><td></td><td></td><td></td><td></td></tr>
<tr><td>Chromosome</td><td>start</td><td>end</td><td>strand</td><td>MM</td><td>target_seq</td><td>PAM</td><td>alignment</td><td>distance</td><td>position</td><td>gene name</td><td>gene id</td></tr>
<tr><td>GU305914.1</td><td>126762</td><td>126784</td><td>-</td><td>0</td><td>TGCGTACGCTGATGGAGCAC</td><td>GGG</td><td>||||||||[||||||||||||]PAM</td><td>NA</td><td>Intergenic</td><td>NA</td><td>NA</td></tr>
<tr><td>T24391</td><td>TGGAGCACGGGACGATCGCGCGG</td><td>1000</td><td></td><td></td><td></td><td></td><td></td><td></td><td></td><td></td><td></td></tr>
<tr><td>Oligo fwd</td><td>CACCTGGAGCACGGGACGATCGCG</td><td></td><td></td><td></td><td></td><td></td><td></td><td></td><td></td><td></td><td></td></tr>
<tr><td>Oligo rev</td><td>AAACCGCGATCGTCCCGTGCTCCA</td><td></td><td></td><td></td><td></td><td></td><td></td><td></td><td></td><td></td><td></td></tr>
<tr><td>Chromosome</td><td>start</td><td>end</td><td>strand</td><td>MM</td><td>target_seq</td><td>PAM</td><td>alignment</td><td>distance</td><td>position</td><td>gene name</td><td>gene id</td></tr>
<tr><td>GU305914.1</td><td>126750</td><td>126772</td><td>-</td><td>0</td><td>TGGAGCACGGGACGATCGCG</td><td>CGG</td><td>||||||||[||||||||||||]PAM</td><td>NA</td><td>Intergenic</td><td>NA</td><td>NA</td></tr>
<tr><td>T24392</td><td>CGGGACGATCGCGCGGCGCATGG</td><td>1000</td><td></td><td></td><td></td><td></td><td></td><td></td><td></td><td></td><td></td></tr>
</table>

| | | | | | | | | | | | |
|---|---|---|---|---|---|---|---|---|---|---|---|
| Oligo fwd | CACCCGGGACGATCGCGCGGCGCA | | | | | | | | | | |
| Oligo rev | AAACTGCGCCGCGCGATCGTCCCG | | | | | | | | | | |
| Chromosome | start | end | strand | MM | target_seq | PAM | alignment | distance | position | gene name | gene id |
| GU305914.1 | 126743 | 126765 | - | 0 | CGGGACGATCGCGCGGCGCA | TGG | \|\|\|\|\|\|\|\|[\|\|\|\|\|\|\|\|\|\|\|\|]PAM | NA | Intergenic | NA | NA |
| GU305914.1 | 143086 | 143108 | + | 3 | CGCGACGACCGCGCGGTGCA | GGG | \|\|-\|\|\|\|\|[-\|\|\|\|\|\|\|-\|\|\|]PAM | NA | Intergenic | NA | NA |
| T24393 | TCGCGCGGCGCATGGCGACGCGG | 1000 | | | | | | | | | |
| Oligo fwd | CACCTCGCGCGGCGCATGGCGACG | | | | | | | | | | |
| Oligo rev | AAACCGTCGCCATGCGCCGCGCGA | | | | | | | | | | |
| Chromosome | start | end | strand | MM | target_seq | PAM | alignment | distance | position | gene name | gene id |
| GU305914.1 | 126735 | 126757 | - | 0 | TCGCGCGGCGCATGGCGACG | CGG | \|\|\|\|\|\|\|\|[\|\|\|\|\|\|\|\|\|\|\|\|]PAM | NA | Intergenic | NA | NA |
| T24394 | CGCGCGGCGCATGGCGACGCGGG | 1000 | | | | | | | | | |
| Oligo fwd | CACCCGCGCGGCGCATGGCGACGC | | | | | | | | | | |
| Oligo rev | AAACGCGTCGCCATGCGCCGCGCG | | | | | | | | | | |
| Chromosome | start | end | strand | MM | target_seq | PAM | alignment | distance | position | gene name | gene id |
| GU305914.1 | 126734 | 126756 | - | 0 | CGCGCGGCGCATGGCGACGC | GGG | \|\|\|\|\|\|\|\|[\|\|\|\|\|\|\|\|\|\|\|\|]PAM | NA | Intergenic | NA | NA |
| GU305914.1 | 74988 | 75010 | + | 4 | GGCGCGGAGGAAGGCGACGC | AGG | -\|\|\|\|\|\|-[\|-\|-\|\|\|\|\|\|\|\|]PAM | NA | Intergenic | NA | NA |
| T24395 | GCAGCGCGCCAACATCGCGCAGG | 1000 | | | | | | | | | |
| Oligo fwd | CACCGCAGCGCGCCAACATCGCGC | | | | | | | | | | |
| Oligo rev | AAACGCGCGATGTTGGCGCGCTGC | | | | | | | | | | |
| Chromosome | start | end | strand | MM | target_seq | PAM | alignment | distance | position | gene name | gene id |
| GU305914.1 | 126710 | 126732 | - | 0 | GCAGCGCGCCAACATCGCGC | AGG | \|\|\|\|\|\|\|\|[\|\|\|\|\|\|\|\|\|\|\|\|]PAM | NA | Intergenic | NA | NA |
| T24396 | CAGCGCGCCAACATCGCGCAGGG | 1000 | | | | | | | | | |
| Oligo fwd | CACCCAGCGCGCCAACATCGCGCA | | | | | | | | | | |
| Oligo rev | AAACTGCGCGATGTTGGCGCGCTG | | | | | | | | | | |
| Chromosome | start | end | strand | MM | target_seq | PAM | alignment | distance | position | gene name | gene id |
| GU305914.1 | 126709 | 126731 | - | 0 | CAGCGCGCCAACATCGCGCA | GGG | \|\|\|\|\|\|\|\|[\|\|\|\|\|\|\|\|\|\|\|\|]PAM | NA | Intergenic | NA | NA |
| T24397 | GGGCTTCAAGAGCAAGATGCTGG | 1000 | | | | | | | | | |
| Oligo fwd | CACCGGGCTTCAAGAGCAAGATGC | | | | | | | | | | |
| Oligo rev | AAACGCATCTTGCTCTTGAAGCCC | | | | | | | | | | |
| Chromosome | start | end | strand | MM | target_seq | PAM | alignment | distance | position | gene name | gene id |
| GU305914.1 | 126689 | 126711 | - | 0 | GGGCTTCAAGAGCAAGATGC | TGG | \|\|\|\|\|\|\|\|[\|\|\|\|\|\|\|\|\|\|\|\|]PAM | NA | Intergenic | NA | NA |
| T24398 | GCTGGCCACCATCTTCCTGCTGG | 1000 | | | | | | | | | |
| Oligo fwd | CACCGCTGGCCACCATCTTCCTGC | | | | | | | | | | |
| Oligo rev | AAACGCAGGAAGATGGTGGCCAGC | | | | | | | | | | |
| Chromosome | start | end | strand | MM | target_seq | PAM | alignment | distance | position | gene name | gene id |
| GU305914.1 | 126671 | 126693 | - | 0 | GCTGGCCACCATCTTCCTGC | TGG | \|\|\|\|\|\|\|\|[\|\|\|\|\|\|\|\|\|\|\|\|]PAM | NA | Intergenic | NA | NA |
| T24399 | TCTTCCTGCTGGACCGCAGCCGG | 1000 | | | | | | | | | |
| Oligo fwd | CACCTCTTCCTGCTGGACCGCAGC | | | | | | | | | | |
| Oligo rev | AAACGCTGCGGTCCAGCAGGAAGA | | | | | | | | | | |
| Chromosome | start | end | strand | MM | target_seq | PAM | alignment | distance | position | gene name | gene id |
| GU305914.1 | 126660 | 126682 | - | 0 | TCTTCCTGCTGGACCGCAGC | CGG | \|\|\|\|\|\|\|\|[\|\|\|\|\|\|\|\|\|\|\|\|]PAM | NA | Intergenic | NA | NA |
| T24400 | CTTCCTGCTGGACCGCAGCCGGG | 1000 | | | | | | | | | |
| Oligo fwd | CACCCTTCCTGCTGGACCGCAGCC | | | | | | | | | | |
| Oligo rev | AAACGGCTGCGGTCCAGCAGGAAG | | | | | | | | | | |
| Chromosome | start | end | strand | MM | target_seq | PAM | alignment | distance | position | gene name | gene id |
| GU305914.1 | 126659 | 126681 | - | 0 | CTTCCTGCTGGACCGCAGCC | GGG | \|\|\|\|\|\|\|\|[\|\|\|\|\|\|\|\|\|\|\|\|]PAM | NA | Intergenic | NA | NA |
| T24401 | CTGGACCGCAGCCGGGACCGCGG | 1000 | | | | | | | | | |
| Oligo fwd | CACCCTGGACCGCAGCCGGGACCG | | | | | | | | | | |
| Oligo rev | AAACCGGTCCCGGCTGCGGTCCAG | | | | | | | | | | |
| Chromosome | start | end | strand | MM | target_seq | PAM | alignment | distance | position | gene name | gene id |
| GU305914.1 | 126652 | 126674 | - | 0 | CTGGACCGCAGCCGGGACCG | CGG | \|\|\|\|\|\|\|\|[\|\|\|\|\|\|\|\|\|\|\|\|]PAM | NA | Intergenic | NA | NA |
| T24402 | TGGACCGCAGCCGGGACCGCGGG | 1000 | | | | | | | | | |
| Oligo fwd | CACCTGGACCGCAGCCGGGACCGC | | | | | | | | | | |
| Oligo rev | AAACGCGGTCCCGGCTGCGGTCCA | | | | | | | | | | |
| Chromosome | start | end | strand | MM | target_seq | PAM | alignment | distance | position | gene name | gene id |
| GU305914.1 | 126651 | 126673 | - | 0 | TGGACCGCAGCCGGGACCGC | GGG | \|\|\|\|\|\|\|\|[\|\|\|\|\|\|\|\|\|\|\|\|]PAM | NA | Intergenic | NA | NA |
| T24403 | CGGGCAGCTGACGCGTTACCTGG | 1000 | | | | | | | | | |
| Oligo fwd | CACCCGGGCAGCTGACGCGTTACC | | | | | | | | | | |
| Oligo rev | AAACGGTAACGCGTCAGCTGCCCG | | | | | | | | | | |
| Chromosome | start | end | strand | MM | target_seq | PAM | alignment | distance | position | gene name | gene id |
| GU305914.1 | 126632 | 126654 | - | 0 | CGGGCAGCTGACGCGTTACC | TGG | \|\|\|\|\|\|\|\|[\|\|\|\|\|\|\|\|\|\|\|\|]PAM | NA | Intergenic | NA | NA |
| T24404 | GCGTTACCTGGACATGCTGACGG | 1000 | | | | | | | | | |
| Oligo fwd | CACCGCGTTACCTGGACATGCTGA | | | | | | | | | | |
| Oligo rev | AAACTCAGCATGTCCAGGTAACGC | | | | | | | | | | |
| Chromosome | start | end | strand | MM | target_seq | PAM | alignment | distance | position | gene name | gene id |
| GU305914.1 | 126620 | 126642 | - | 0 | GCGTTACCTGGACATGCTGA | CGG | \|\|\|\|\|\|\|\|[\|\|\|\|\|\|\|\|\|\|\|\|]PAM | NA | Intergenic | NA | NA |
| T24405 | CATGCTGACGGACTGCGTGACGG | 1000 | | | | | | | | | |
| Oligo fwd | CACCCATGCTGACGGACTGCGTGA | | | | | | | | | | |
| Oligo rev | AAACTCACGCAGTCCGTCAGCATG | | | | | | | | | | |
| Chromosome | start | end | strand | MM | target_seq | PAM | alignment | distance | position | gene name | gene id |
| GU305914.1 | 126608 | 126630 | - | 0 | CATGCTGACGGACTGCGTGA | CGG | \|\|\|\|\|\|\|\|[\|\|\|\|\|\|\|\|\|\|\|\|]PAM | NA | Intergenic | NA | NA |
| T24406 | CTGCGTGACGGAGAGCATCCTGG | 1000 | | | | | | | | | |
| Oligo fwd | CACCCTGCGTGACGGAGAGCATCC | | | | | | | | | | |
| Oligo rev | AAACGGATGCTCTCCGTCACGCAG | | | | | | | | | | |
| Chromosome | start | end | strand | MM | target_seq | PAM | alignment | distance | position | gene name | gene id |

| | | | | | | | | | | | |
|---|---|---|---|---|---|---|---|---|---|---|---|
| GU305914.1 | | 126596 | 126618 | - | 0 | CTGCGTGACGGAGAGCATCC | TGG | \|\|\|\|\|\|\|\|[\|\|\|\|\|\|\|\|\|\|\|\|]PAM | NA | Intergenic | NA | NA |
| T24407 | GGAGAGCATCCTGGACAACCCGG | 1000 | | | | | | | | | |
| Oligo fwd | CACCGGAGAGCATCCTGGACAACC | | | | | | | | | | |
| Oligo rev | AAACGGTTGTCCAGGATGCTCTCC | | | | | | | | | | |
| Chromosome | start | end | strand | MM | target_seq | PAM | alignment | distance | position | gene name | gene id |
| GU305914.1 | 126587 | 126609 | - | 0 | GGAGAGCATCCTGGACAACC | CGG | \|\|\|\|\|\|\|\|[\|\|\|\|\|\|\|\|\|\|\|\|]PAM | NA | Intergenic | NA | NA |
| T24408 | GGACAACCCGGAGACCTACACGG | 1000 | | | | | | | | | |
| Oligo fwd | CACCGGACAACCCGGAGACCTACA | | | | | | | | | | |
| Oligo rev | AAACTGTAGGTCTCCGGGTTGTCC | | | | | | | | | | |
| Chromosome | start | end | strand | MM | target_seq | PAM | alignment | distance | position | gene name | gene id |
| GU305914.1 | 126575 | 126597 | - | 0 | GGACAACCCGGAGACCTACA | CGG | \|\|\|\|\|\|\|\|[\|\|\|\|\|\|\|\|\|\|\|\|]PAM | NA | Intergenic | NA | NA |
| T24409 | CAACCCGGAGACCTACACGGTGG | 1000 | | | | | | | | | |
| Oligo fwd | CACCCAACCCGGAGACCTACACGG | | | | | | | | | | |
| Oligo rev | AAACCCGTGTAGGTCTCCGGGTTG | | | | | | | | | | |
| Chromosome | start | end | strand | MM | target_seq | PAM | alignment | distance | position | gene name | gene id |
| GU305914.1 | 126572 | 126594 | - | 0 | CAACCCGGAGACCTACACGG | TGG | \|\|\|\|\|\|\|\|[\|\|\|\|\|\|\|\|\|\|\|\|]PAM | NA | Intergenic | NA | NA |
| T24410 | AACCCGGAGACCTACACGGTGGG | 1000 | | | | | | | | | |
| Oligo fwd | CACCAACCCGGAGACCTACACGGT | | | | | | | | | | |
| Oligo rev | AAACACCGTGTAGGTCTCCGGGTT | | | | | | | | | | |
| Chromosome | start | end | strand | MM | target_seq | PAM | alignment | distance | position | gene name | gene id |
| GU305914.1 | 126571 | 126593 | - | 0 | AACCCGGAGACCTACACGGT | GGG | \|\|\|\|\|\|\|\|[\|\|\|\|\|\|\|\|\|\|\|\|]PAM | NA | Intergenic | NA | NA |
| T24411 | CCGGAGACCTACACGGTGGGCGG | 1000 | | | | | | | | | |
| Oligo fwd | CACCCCGGAGACCTACACGGTGGG | | | | | | | | | | |
| Oligo rev | AAACCCCACCGTGTAGGTCTCCGG | | | | | | | | | | |
| Chromosome | start | end | strand | MM | target_seq | PAM | alignment | distance | position | gene name | gene id |
| GU305914.1 | 126568 | 126590 | - | 0 | CCGGAGACCTACACGGTGGG | CGG | \|\|\|\|\|\|\|\|[\|\|\|\|\|\|\|\|\|\|\|\|]PAM | NA | Intergenic | NA | NA |
| T24412 | GAGACCTACACGGTGGGCGGCGG | 1000 | | | | | | | | | |
| Oligo fwd | CACCGAGACCTACACGGTGGGCGG | | | | | | | | | | |
| Oligo rev | AAACCCGCCCACCGTGTAGGTCTC | | | | | | | | | | |
| Chromosome | start | end | strand | MM | target_seq | PAM | alignment | distance | position | gene name | gene id |
| GU305914.1 | 126565 | 126587 | - | 0 | GAGACCTACACGGTGGGCGG | CGG | \|\|\|\|\|\|\|\|[\|\|\|\|\|\|\|\|\|\|\|\|]PAM | NA | Intergenic | NA | NA |
| T24413 | AGACCTACACGGTGGGCGGCGGG | 1000 | | | | | | | | | |
| Oligo fwd | CACCAGACCTACACGGTGGGCGGC | | | | | | | | | | |
| Oligo rev | AAACGCCGCCCACCGTGTAGGTCT | | | | | | | | | | |
| Chromosome | start | end | strand | MM | target_seq | PAM | alignment | distance | position | gene name | gene id |
| GU305914.1 | 126564 | 126586 | - | 0 | AGACCTACACGGTGGGCGGC | GGG | \|\|\|\|\|\|\|\|[\|\|\|\|\|\|\|\|\|\|\|\|]PAM | NA | Intergenic | NA | NA |
| T24414 | GACCTACACGGTGGGCGGCGGGG | 1000 | | | | | | | | | |
| Oligo fwd | CACCGACCTACACGGTGGGCGGCG | | | | | | | | | | |
| Oligo rev | AAACCGCCGCCCACCGTGTAGGTC | | | | | | | | | | |
| Chromosome | start | end | strand | MM | target_seq | PAM | alignment | distance | position | gene name | gene id |
| GU305914.1 | 126563 | 126585 | - | 0 | GACCTACACGGTGGGCGGCG | GGG | \|\|\|\|\|\|\|\|[\|\|\|\|\|\|\|\|\|\|\|\|]PAM | NA | Intergenic | NA | NA |
| T24415 | GGTGGGCGGCGGGGAGCGCCTGG | 1000 | | | | | | | | | |
| Oligo fwd | CACCGGTGGGCGGCGGGGAGCGCC | | | | | | | | | | |
| Oligo rev | AAACGGCGCTCCCCGCCGCCCACC | | | | | | | | | | |
| Chromosome | start | end | strand | MM | target_seq | PAM | alignment | distance | position | gene name | gene id |
| GU305914.1 | 126554 | 126576 | - | 0 | GGTGGGCGGCGGGGAGCGCC | TGG | \|\|\|\|\|\|\|\|[\|\|\|\|\|\|\|\|\|\|\|\|]PAM | NA | Intergenic | NA | NA |
| GU305914.1 | 83269 | 83291 | - | 4 | AGCGGGCCGCCGGGAGCGCC | TGG | -\|-\|\|\|\|-[\|\|-\|\|\|\|\|\|\|\|\|]PAM | NA | Intergenic | NA | NA |
| T24416 | GGGCGGCGGGGAGCGCCTGGCGG | 1000 | | | | | | | | | |
| Oligo fwd | CACCGGGCGGCGGGGAGCGCCTGG | | | | | | | | | | |
| Oligo rev | AAACCCAGGCGCTCCCCGCCGCCC | | | | | | | | | | |
| Chromosome | start | end | strand | MM | target_seq | PAM | alignment | distance | position | gene name | gene id |
| GU305914.1 | 126551 | 126573 | - | 0 | GGGCGGCGGGGAGCGCCTGG | CGG | \|\|\|\|\|\|\|\|[\|\|\|\|\|\|\|\|\|\|\|\|]PAM | NA | Intergenic | NA | NA |
| T24417 | GGCGGCGGGGAGCGCCTGGCGGG | 1000 | | | | | | | | | |
| Oligo fwd | CACCGGCGGCGGGGAGCGCCTGGC | | | | | | | | | | |
| Oligo rev | AAACGCCAGGCGCTCCCCGCCGCC | | | | | | | | | | |
| Chromosome | start | end | strand | MM | target_seq | PAM | alignment | distance | position | gene name | gene id |
| GU305914.1 | 126550 | 126572 | - | 0 | GGCGGCGGGGAGCGCCTGGC | GGG | \|\|\|\|\|\|\|\|[\|\|\|\|\|\|\|\|\|\|\|\|]PAM | NA | Intergenic | NA | NA |
| T24418 | GATCGTGAGCACGCACACCGTGG | 1000 | | | | | | | | | |
| Oligo fwd | CACCGATCGTGAGCACGCACACCG | | | | | | | | | | |
| Oligo rev | AAACCGGTGTGCGTGCTCACGATC | | | | | | | | | | |
| Chromosome | start | end | strand | MM | target_seq | PAM | alignment | distance | position | gene name | gene id |
| GU305914.1 | 126524 | 126546 | - | 0 | GATCGTGAGCACGCACACCG | TGG | \|\|\|\|\|\|\|\|[\|\|\|\|\|\|\|\|\|\|\|\|]PAM | NA | Intergenic | NA | NA |
| T24419 | GAGCACGCACACCGTGGTGCAGG | 1000 | | | | | | | | | |
| Oligo fwd | CACCGAGCACGCACACCGTGGTGC | | | | | | | | | | |
| Oligo rev | AAACGCACCACGGTGTGCGTGCTC | | | | | | | | | | |
| Chromosome | start | end | strand | MM | target_seq | PAM | alignment | distance | position | gene name | gene id |
| GU305914.1 | 126518 | 126540 | - | 0 | GAGCACGCACACCGTGGTGC | AGG | \|\|\|\|\|\|\|\|[\|\|\|\|\|\|\|\|\|\|\|\|]PAM | NA | Intergenic | NA | NA |
| GU305914.1 | 77832 | 77854 | - | 4 | CAGCAGGCACACCGTGGTCG | TGG | -\|\|\|\|-\|\|[\|\|\|\|\|\|\|\|\|\|--]PAM | NA | Intergenic | NA | NA |
| T24420 | GCAGGCGCTGCTGAACGCGCTGG | 1000 | | | | | | | | | |
| Oligo fwd | CACCGCAGGCGCTGCTGAACGCGC | | | | | | | | | | |
| Oligo rev | AAACGCGCGTTCAGCAGCGCCTGC | | | | | | | | | | |
| Chromosome | start | end | strand | MM | target_seq | PAM | alignment | distance | position | gene name | gene id |
| GU305914.1 | 126500 | 126522 | - | 0 | GCAGGCGCTGCTGAACGCGC | TGG | \|\|\|\|\|\|\|\|[\|\|\|\|\|\|\|\|\|\|\|\|]PAM | NA | Intergenic | NA | NA |
| T24421 | CAGGCGCTGCTGAACGCGCTGGG | 1000 | | | | | | | | | |

| | | | | | | | | | | | |
|---|---|---|---|---|---|---|---|---|---|---|---|
| Oligo fwd | CACCCAGGCGCTGCTGAACGCGCT | | | | | | | | | | |
| Oligo rev | AAACAGCGCGTTCAGCAGCGCCTG | | | | | | | | | | |
| Chromosome | start | end | strand | MM | target_seq | PAM | alignment | distance | position | gene name | gene id |
| GU305914.1 | 126499 | 126521 | - | 0 | CAGGCGCTGCTGAACGCGCT | GGG | \|\|\|\|\|\|\|\|[\|\|\|\|\|\|\|\|\|\|\|\|]PAM | NA | Intergenic | NA | NA |
| T24422 | GCGCTGCTGAACGCGCTGGGCGG | 1000 | | | | | | | | | |
| Oligo fwd | CACCGCGCTGCTGAACGCGCTGGG | | | | | | | | | | |
| Oligo rev | AAACCCCAGCGCGTTCAGCAGCGC | | | | | | | | | | |
| Chromosome | start | end | strand | MM | target_seq | PAM | alignment | distance | position | gene name | gene id |
| GU305914.1 | 126496 | 126518 | - | 0 | GCGCTGCTGAACGCGCTGGG | CGG | \|\|\|\|\|\|\|\|[\|\|\|\|\|\|\|\|\|\|\|\|]PAM | NA | Intergenic | NA | NA |
| GU305914.1 | 66165 | 66187 | + | 4 | TCGTTCCTGCACGCGCTGGG | GGG | -\|\|-\|-\|\|[\|-\|\|\|\|\|\|\|\|\|\|]PAM | NA | Intergenic | NA | NA |
| T24423 | GGGCGGCTCGATCCGCCGCACGG | 1000 | | | | | | | | | |
| Oligo fwd | CACCGGGCGGCTCGATCCGCCGCA | | | | | | | | | | |
| Oligo rev | AAACTGCGGCGGATCGAGCCGCCC | | | | | | | | | | |
| Chromosome | start | end | strand | MM | target_seq | PAM | alignment | distance | position | gene name | gene id |
| GU305914.1 | 126479 | 126501 | - | 0 | GGGCGGCTCGATCCGCCGCA | CGG | \|\|\|\|\|\|\|\|[\|\|\|\|\|\|\|\|\|\|\|\|]PAM | NA | Intergenic | NA | NA |
| T24424 | GGCGGCTCGATCCGCCGCACGGG | 1000 | | | | | | | | | |
| Oligo fwd | CACCGGCGGCTCGATCCGCCGCAC | | | | | | | | | | |
| Oligo rev | AAACGTGCGGCGGATCGAGCCGCC | | | | | | | | | | |
| Chromosome | start | end | strand | MM | target_seq | PAM | alignment | distance | position | gene name | gene id |
| GU305914.1 | 126478 | 126500 | - | 0 | GGCGGCTCGATCCGCCGCAC | GGG | \|\|\|\|\|\|\|\|[\|\|\|\|\|\|\|\|\|\|\|\|]PAM | NA | Intergenic | NA | NA |
| T24425 | GTGAAGACGCCCGCCTCGTACGG | 1000 | | | | | | | | | |
| Oligo fwd | CACCGTGAAGACGCCCGCCTCGTA | | | | | | | | | | |
| Oligo rev | AAACTACGAGGCGGGCGTCTTCAC | | | | | | | | | | |
| Chromosome | start | end | strand | MM | target_seq | PAM | alignment | distance | position | gene name | gene id |
| GU305914.1 | 126454 | 126476 | - | 0 | GTGAAGACGCCCGCCTCGTA | CGG | \|\|\|\|\|\|\|\|[\|\|\|\|\|\|\|\|\|\|\|\|]PAM | NA | Intergenic | NA | NA |
| T24426 | CGGCAAGTTCGTGCTGAGCAAGG | 1000 | | | | | | | | | |
| Oligo fwd | CACCCGGCAAGTTCGTGCTGAGCA | | | | | | | | | | |
| Oligo rev | AAACTGCTCAGCACGAACTTGCCG | | | | | | | | | | |
| Chromosome | start | end | strand | MM | target_seq | PAM | alignment | distance | position | gene name | gene id |
| GU305914.1 | 126434 | 126456 | - | 0 | CGGCAAGTTCGTGCTGAGCA | AGG | \|\|\|\|\|\|\|\|[\|\|\|\|\|\|\|\|\|\|\|\|]PAM | NA | Intergenic | NA | NA |
| T24427 | GAGCAAGGAGAACGCCGTGACGG | 1000 | | | | | | | | | |
| Oligo fwd | CACCGAGCAAGGAGAACGCCGTGA | | | | | | | | | | |
| Oligo rev | AAACTCACGGCGTTCTCCTTGCTC | | | | | | | | | | |
| Chromosome | start | end | strand | MM | target_seq | PAM | alignment | distance | position | gene name | gene id |
| GU305914.1 | 126419 | 126441 | - | 0 | GAGCAAGGAGAACGCCGTGA | CGG | \|\|\|\|\|\|\|\|[\|\|\|\|\|\|\|\|\|\|\|\|]PAM | NA | Intergenic | NA | NA |
| GU305914.1 | 120635 | 120657 | - | 4 | GAGCGAGAAGAACGCGGCGA | AGG | \|\|\|\|-\|\|-[\|\|\|\|\|\|\|-\|-\|\|]PAM | NA | Intergenic | NA | NA |
| T24428 | GATCGCGCACCACGCGATCATGG | 1000 | | | | | | | | | |
| Oligo fwd | CACCGATCGCGCACCACGCGATCA | | | | | | | | | | |
| Oligo rev | AAACTGATCGCGTGGTGCGCGATC | | | | | | | | | | |
| Chromosome | start | end | strand | MM | target_seq | PAM | alignment | distance | position | gene name | gene id |
| GU305914.1 | 126395 | 126417 | - | 0 | GATCGCGCACCACGCGATCA | TGG | \|\|\|\|\|\|\|\|[\|\|\|\|\|\|\|\|\|\|\|\|]PAM | NA | Intergenic | NA | NA |
| T24429 | CGCGCACCACGCGATCATGGCGG | 1000 | | | | | | | | | |
| Oligo fwd | CACCCGCGCACCACGCGATCATGG | | | | | | | | | | |
| Oligo rev | AAACCCATGATCGCGTGGTGCGCG | | | | | | | | | | |
| Chromosome | start | end | strand | MM | target_seq | PAM | alignment | distance | position | gene name | gene id |
| GU305914.1 | 126392 | 126414 | - | 0 | CGCGCACCACGCGATCATGG | CGG | \|\|\|\|\|\|\|\|[\|\|\|\|\|\|\|\|\|\|\|\|]PAM | NA | Intergenic | NA | NA |
| T24430 | CATCCAGCAGAGCTCGCAGAAGG | 1000 | | | | | | | | | |
| Oligo fwd | CACCCATCCAGCAGAGCTCGCAGA | | | | | | | | | | |
| Oligo rev | AAACTCTGCGAGCTCTGCTGGATG | | | | | | | | | | |
| Chromosome | start | end | strand | MM | target_seq | PAM | alignment | distance | position | gene name | gene id |
| GU305914.1 | 126347 | 126369 | - | 0 | CATCCAGCAGAGCTCGCAGA | AGG | \|\|\|\|\|\|\|\|[\|\|\|\|\|\|\|\|\|\|\|\|]PAM | NA | Intergenic | NA | NA |
| GU305914.1 | 102541 | 102563 | - | 4 | GATCCTGCCGACCTCGCAGA | GGG | -\|\|\|\|-\|\|[-\|\|-\|\|\|\|\|\|\|\|]PAM | NA | Intergenic | NA | NA |
| GU305914.1 | 167970 | 167992 | - | 4 | ACTGCAGCAGAGCTCGGAGA | CGG | --\|-\|\|\|\|[\|\|\|\|\|\|\|\|-\|\|\|]PAM | NA | Intergenic | NA | NA |
| T24431 | CTTACCCGAGTCGCAGTTCCTGG | 1000 | | | | | | | | | |
| Oligo fwd | CACCCTTACCCGAGTCGCAGTTCC | | | | | | | | | | |
| Oligo rev | AAACGGAACTGCGACTCGGGTAAG | | | | | | | | | | |
| Chromosome | start | end | strand | MM | target_seq | PAM | alignment | distance | position | gene name | gene id |
| GU305914.1 | 126323 | 126345 | - | 0 | CTTACCCGAGTCGCAGTTCC | TGG | \|\|\|\|\|\|\|\|[\|\|\|\|\|\|\|\|\|\|\|\|]PAM | NA | Intergenic | NA | NA |
| T24432 | AGTCGCAGTTCCTGGACCAGCGG | 1000 | | | | | | | | | |
| Oligo fwd | CACCAGTCGCAGTTCCTGGACCAG | | | | | | | | | | |
| Oligo rev | AAACCTGGTCCAGGAACTGCGACT | | | | | | | | | | |
| Chromosome | start | end | strand | MM | target_seq | PAM | alignment | distance | position | gene name | gene id |
| GU305914.1 | 126315 | 126337 | - | 0 | AGTCGCAGTTCCTGGACCAG | CGG | \|\|\|\|\|\|\|\|[\|\|\|\|\|\|\|\|\|\|\|\|]PAM | NA | Intergenic | NA | NA |
| T24433 | GACCTTCACCGAGACGCAGATGG | 1000 | | | | | | | | | |
| Oligo fwd | CACCGACCTTCACCGAGACGCAGA | | | | | | | | | | |
| Oligo rev | AAACTCTGCGTCTCGGTGAAGGTC | | | | | | | | | | |
| Chromosome | start | end | strand | MM | target_seq | PAM | alignment | distance | position | gene name | gene id |
| GU305914.1 | 126290 | 126312 | - | 0 | GACCTTCACCGAGACGCAGA | TGG | \|\|\|\|\|\|\|\|[\|\|\|\|\|\|\|\|\|\|\|\|]PAM | NA | Intergenic | NA | NA |
| T24434 | GACGCAGATGGACGTGCTGAAGG | 1000 | | | | | | | | | |
| Oligo fwd | CACCGACGCAGATGGACGTGCTGA | | | | | | | | | | |
| Oligo rev | AAACTCAGCACGTCCATCTGCGTC | | | | | | | | | | |
| Chromosome | start | end | strand | MM | target_seq | PAM | alignment | distance | position | gene name | gene id |
| GU305914.1 | 126278 | 126300 | - | 0 | GACGCAGATGGACGTGCTGA | AGG | \|\|\|\|\|\|\|\|[\|\|\|\|\|\|\|\|\|\|\|\|]PAM | NA | Intergenic | NA | NA |
| T24435 | GCAGATGGACGTGCTGAAGGTGG | 1000 | | | | | | | | | |
| Oligo fwd | CACCGCAGATGGACGTGCTGAAGG | | | | | | | | | | |

Oligo rev AAACCCTTCAGCACGTCCATCTGC
Chromosome start end strand MM target_seq PAM alignment distance position gene name gene id
GU305914.1 126275 126297 - 0 GCAGATGGACGTGCTGAAGG TGG ||||||||[||||||||||||]PAM NA Intergenic NA NA
GU305914.1 82453 82475 + 4 AGAGATGGACACGCTGAAGG GGG --||||||[||--||||||||]PAM NA Intergenic NA NA

T24436 CAGATGGACGTGCTGAAGGTGGG 1000
Oligo fwd CACCCAGATGGACGTGCTGAAGGT
Oligo rev AAACACCTTCAGCACGTCCATCTG
Chromosome start end strand MM target_seq PAM alignment distance position gene name gene id
GU305914.1 126274 126296 - 0 CAGATGGACGTGCTGAAGGT GGG ||||||||[||||||||||||]PAM NA Intergenic NA NA

T24437 ACGTGCTGAAGGTGGGCGAGCGG 1000
Oligo fwd CACCACGTGCTGAAGGTGGGCGAG
Oligo rev AAACCTCGCCCACCTTCAGCACGT
Chromosome start end strand MM target_seq PAM alignment distance position gene name gene id
GU305914.1 126267 126289 - 0 ACGTGCTGAAGGTGGGCGAG CGG ||||||||[||||||||||||]PAM NA Intergenic NA NA

T24438 GAAGGTGGGCGAGCGGCTCGTGG 1000
Oligo fwd CACCGAAGGTGGGCGAGCGGCTCG
Oligo rev AAACCGAGCCGCTCGCCCACCTTC
Chromosome start end strand MM target_seq PAM alignment distance position gene name gene id
GU305914.1 126260 126282 - 0 GAAGGTGGGCGAGCGGCTCG TGG ||||||||[||||||||||||]PAM NA Intergenic NA NA

T24439 GGGCGAGCGGCTCGTGGCGCTGG 1000
Oligo fwd CACCGGGCGAGCGGCTCGTGGCGC
Oligo rev AAACGCGCCACGAGCCGCTCGCCC
Chromosome start end strand MM target_seq PAM alignment distance position gene name gene id
GU305914.1 126254 126276 - 0 GGGCGAGCGGCTCGTGGCGC TGG ||||||||[||||||||||||]PAM NA Intergenic NA NA

T24440 GGCGCTGGAGCACCTGCGAAAGG 1000
Oligo fwd CACCGGCGCTGGAGCACCTGCGAA
Oligo rev AAACTTCGCAGGTGCTCCAGCGCC
Chromosome start end strand MM target_seq PAM alignment distance position gene name gene id
GU305914.1 126239 126261 - 0 GGCGCTGGAGCACCTGCGAA AGG ||||||||[||||||||||||]PAM NA Intergenic NA NA

T24441 GTACAAGAACACAGACGTGCAGG 1000
Oligo fwd CACCGTACAAGAACACAGACGTGC
Oligo rev AAACGCACGTCTGTGTTCTTGTAC
Chromosome start end strand MM target_seq PAM alignment distance position gene name gene id
GU305914.1 126215 126237 - 0 GTACAAGAACACAGACGTGC AGG ||||||||[||||||||||||]PAM NA Intergenic NA NA

T24442 CACAGACGTGCAGGACCCGCTGG 1000
Oligo fwd CACCCACAGACGTGCAGGACCCGC
Oligo rev AAACGCGGGTCCTGCACGTCTGTG
Chromosome start end strand MM target_seq PAM alignment distance position gene name gene id
GU305914.1 126206 126228 - 0 CACAGACGTGCAGGACCCGC TGG ||||||||[||||||||||||]PAM NA Intergenic NA NA

T24443 ACGTGCAGGACCCGCTGGAGCGG 1000
Oligo fwd CACCACGTGCAGGACCCGCTGGAG
Oligo rev AAACCTCCAGCGGGTCCTGCACGT
Chromosome start end strand MM target_seq PAM alignment distance position gene name gene id
GU305914.1 126201 126223 - 0 ACGTGCAGGACCCGCTGGAG CGG ||||||||[||||||||||||]PAM NA Intergenic NA NA

T24444 CGTGCAGGACCCGCTGGAGCGGG 1000
Oligo fwd CACCCGTGCAGGACCCGCTGGAGC
Oligo rev AAACGCTCCAGCGGGTCCTGCACG
Chromosome start end strand MM target_seq PAM alignment distance position gene name gene id
GU305914.1 126200 126222 - 0 CGTGCAGGACCCGCTGGAGC GGG ||||||||[||||||||||||]PAM NA Intergenic NA NA

T24445 CGAGCTCACCTTCTACTTTCCGG 1000
Oligo fwd CACCCGAGCTCACCTTCTACTTTC
Oligo rev AAACGAAAGTAGAAGGTGAGCTCG
Chromosome start end strand MM target_seq PAM alignment distance position gene name gene id
GU305914.1 126173 126195 - 0 CGAGCTCACCTTCTACTTTC CGG ||||||||[||||||||||||]PAM NA Intergenic NA NA

T24446 CTCACCTTCTACTTTCCGGTCGG 1000
Oligo fwd CACCCTCACCTTCTACTTTCCGGT
Oligo rev AAACACCGGAAAGTAGAAGGTGAG
Chromosome start end strand MM target_seq PAM alignment distance position gene name gene id
GU305914.1 126169 126191 - 0 CTCACCTTCTACTTTCCGGT CGG ||||||||[||||||||||||]PAM NA Intergenic NA NA

T24447 GTCGGCCTGCACGTGCCGAGCGG 1000
Oligo fwd CACCGTCGGCCTGCACGTGCCGAG
Oligo rev AAACCTCGGCACGTGCAGGCCGAC
Chromosome start end strand MM target_seq PAM alignment distance position gene name gene id
GU305914.1 126151 126173 - 0 GTCGGCCTGCACGTGCCGAG CGG ||||||||[||||||||||||]PAM NA Intergenic NA NA

T24448 GCCTGCACGTGCCGAGCGGCCGG 1000
Oligo fwd CACCGCCTGCACGTGCCGAGCGGC
Oligo rev AAACGCCGCTCGGCACGTGCAGGC
Chromosome start end strand MM target_seq PAM alignment distance position gene name gene id
GU305914.1 126147 126169 - 0 GCCTGCACGTGCCGAGCGGC CGG ||||||||[||||||||||||]PAM NA Intergenic NA NA

T24449 CCTGCACGTGCCGAGCGGCCGGG 1000
Oligo fwd CACCCCTGCACGTGCCGAGCGGCC
Oligo rev AAACGGCCGCTCGGCACGTGCAGG
Chromosome start end strand MM target_seq PAM alignment distance position gene name gene id
GU305914.1 126146 126168 - 0 CCTGCACGTGCCGAGCGGCC GGG ||||||||[||||||||||||]PAM NA Intergenic NA NA

<table>
<tr><td>T24450</td><td>GAGCGGCCGGGCCTACAGCACGG</td><td>1000</td></tr>
<tr><td>Oligo fwd</td><td>CACCGAGCGGCCGGGCCTACAGCA</td></tr>
<tr><td>Oligo rev</td><td>AAACTGCTGTAGGCCCGGCCGCTC</td></tr>
<tr><td>Chromosome</td><td>start</td><td>end</td><td>strand</td><td>MM</td><td>target_seq</td><td>PAM</td><td>alignment</td><td>distance</td><td>position</td><td>gene name</td><td>gene id</td></tr>
<tr><td>GU305914.1</td><td>126134</td><td>126156</td><td>-</td><td>0</td><td>GAGCGGCCGGGCCTACAGCA</td><td>CGG</td><td>||||||||[||||||||||||]PAM</td><td>NA</td><td>Intergenic</td><td>NA</td><td>NA</td></tr>
<tr><td>T24451</td><td>CGGCCGGGCCTACAGCACGGCGG</td><td>1000</td></tr>
<tr><td>Oligo fwd</td><td>CACCCGGCCGGGCCTACAGCACGG</td></tr>
<tr><td>Oligo rev</td><td>AAACCCGTGCTGTAGGCCCGGCCG</td></tr>
<tr><td>Chromosome</td><td>start</td><td>end</td><td>strand</td><td>MM</td><td>target_seq</td><td>PAM</td><td>alignment</td><td>distance</td><td>position</td><td>gene name</td><td>gene id</td></tr>
<tr><td>GU305914.1</td><td>126131</td><td>126153</td><td>-</td><td>0</td><td>CGGCCGGGCCTACAGCACGG</td><td>CGG</td><td>||||||||[||||||||||||]PAM</td><td>NA</td><td>Intergenic</td><td>NA</td><td>NA</td></tr>
<tr><td>T24452</td><td>GGCGGAGAACAAGATCAAACTGG</td><td>1000</td></tr>
<tr><td>Oligo fwd</td><td>CACCGGCGGAGAACAAGATCAAAC</td></tr>
<tr><td>Oligo rev</td><td>AAACGTTTGATCTTGTTCTCCGCC</td></tr>
<tr><td>Chromosome</td><td>start</td><td>end</td><td>strand</td><td>MM</td><td>target_seq</td><td>PAM</td><td>alignment</td><td>distance</td><td>position</td><td>gene name</td><td>gene id</td></tr>
<tr><td>GU305914.1</td><td>126113</td><td>126135</td><td>-</td><td>0</td><td>GGCGGAGAACAAGATCAAAC</td><td>TGG</td><td>||||||||[||||||||||||]PAM</td><td>NA</td><td>Intergenic</td><td>NA</td><td>NA</td></tr>
<tr><td>T24453</td><td>GGAGAACAAGATCAAACTGGTGG</td><td>1000</td></tr>
<tr><td>Oligo fwd</td><td>CACCGGAGAACAAGATCAAACTGG</td></tr>
<tr><td>Oligo rev</td><td>AAACCCAGTTTGATCTTGTTCTCC</td></tr>
<tr><td>Chromosome</td><td>start</td><td>end</td><td>strand</td><td>MM</td><td>target_seq</td><td>PAM</td><td>alignment</td><td>distance</td><td>position</td><td>gene name</td><td>gene id</td></tr>
<tr><td>GU305914.1</td><td>126110</td><td>126132</td><td>-</td><td>0</td><td>GGAGAACAAGATCAAACTGG</td><td>TGG</td><td>||||||||[||||||||||||]PAM</td><td>NA</td><td>Intergenic</td><td>NA</td><td>NA</td></tr>
<tr><td>T24454</td><td>CAAGATCAAACTGGTGGACACGG</td><td>1000</td></tr>
<tr><td>Oligo fwd</td><td>CACCCAAGATCAAACTGGTGGACA</td></tr>
<tr><td>Oligo rev</td><td>AAACTGTCCACCAGTTTGATCTTG</td></tr>
<tr><td>Chromosome</td><td>start</td><td>end</td><td>strand</td><td>MM</td><td>target_seq</td><td>PAM</td><td>alignment</td><td>distance</td><td>position</td><td>gene name</td><td>gene id</td></tr>
<tr><td>GU305914.1</td><td>126104</td><td>126126</td><td>-</td><td>0</td><td>CAAGATCAAACTGGTGGACA</td><td>CGG</td><td>||||||||[||||||||||||]PAM</td><td>NA</td><td>Intergenic</td><td>NA</td><td>NA</td></tr>
<tr><td>T24455</td><td>GACCGTGTACTTCTACAACAAGG</td><td>1000</td></tr>
<tr><td>Oligo fwd</td><td>CACCGACCGTGTACTTCTACAACA</td></tr>
<tr><td>Oligo rev</td><td>AAACTGTTGTAGAAGTACACGGTC</td></tr>
<tr><td>Chromosome</td><td>start</td><td>end</td><td>strand</td><td>MM</td><td>target_seq</td><td>PAM</td><td>alignment</td><td>distance</td><td>position</td><td>gene name</td><td>gene id</td></tr>
<tr><td>GU305914.1</td><td>126062</td><td>126084</td><td>-</td><td>0</td><td>GACCGTGTACTTCTACAACA</td><td>AGG</td><td>||||||||[||||||||||||]PAM</td><td>NA</td><td>Intergenic</td><td>NA</td><td>NA</td></tr>
<tr><td>T24456</td><td>TGTACTTCTACAACAAGGACAGG</td><td>1000</td></tr>
<tr><td>Oligo fwd</td><td>CACCTGTACTTCTACAACAAGGAC</td></tr>
<tr><td>Oligo rev</td><td>AAACGTCCTTGTTGTAGAAGTACA</td></tr>
<tr><td>Chromosome</td><td>start</td><td>end</td><td>strand</td><td>MM</td><td>target_seq</td><td>PAM</td><td>alignment</td><td>distance</td><td>position</td><td>gene name</td><td>gene id</td></tr>
<tr><td>GU305914.1</td><td>126057</td><td>126079</td><td>-</td><td>0</td><td>TGTACTTCTACAACAAGGAC</td><td>AGG</td><td>||||||||[||||||||||||]PAM</td><td>NA</td><td>Intergenic</td><td>NA</td><td>NA</td></tr>
<tr><td>T24457</td><td>CCAGCGCATCTCGCACGCCGAGG</td><td>1000</td></tr>
<tr><td>Oligo fwd</td><td>CACCCCAGCGCATCTCGCACGCCG</td></tr>
<tr><td>Oligo rev</td><td>AAACCGGCGTGCGAGATGCGCTGG</td></tr>
<tr><td>Chromosome</td><td>start</td><td>end</td><td>strand</td><td>MM</td><td>target_seq</td><td>PAM</td><td>alignment</td><td>distance</td><td>position</td><td>gene name</td><td>gene id</td></tr>
<tr><td>GU305914.1</td><td>126029</td><td>126051</td><td>-</td><td>0</td><td>CCAGCGCATCTCGCACGCCG</td><td>AGG</td><td>||||||||[||||||||||||]PAM</td><td>NA</td><td>Intergenic</td><td>NA</td><td>NA</td></tr>
<tr><td>GU305914.1</td><td>152279</td><td>152301</td><td>+</td><td>4</td><td>CCTCTTCATCTCGCACGCCG</td><td>AGG</td><td>||----||[||||||||||||]PAM</td><td>NA</td><td>Intergenic</td><td>NA</td><td>NA</td></tr>
<tr><td>T24458</td><td>CACCCCGCGATACACGACGCCGG</td><td>1000</td></tr>
<tr><td>Oligo fwd</td><td>CACCCACCCCGCGATACACGACGC</td></tr>
<tr><td>Oligo rev</td><td>AAACGCGTCGTGTATCGCGGGGTG</td></tr>
<tr><td>Chromosome</td><td>start</td><td>end</td><td>strand</td><td>MM</td><td>target_seq</td><td>PAM</td><td>alignment</td><td>distance</td><td>position</td><td>gene name</td><td>gene id</td></tr>
<tr><td>GU305914.1</td><td>125989</td><td>126011</td><td>-</td><td>0</td><td>CACCCCGCGATACACGACGC</td><td>CGG</td><td>||||||||[||||||||||||]PAM</td><td>NA</td><td>Intergenic</td><td>NA</td><td>NA</td></tr>
<tr><td>T24459</td><td>ACCCCGCGATACACGACGCCGGG</td><td>1000</td></tr>
<tr><td>Oligo fwd</td><td>CACCACCCCGCGATACACGACGCC</td></tr>
<tr><td>Oligo rev</td><td>AAACGGCGTCGTGTATCGCGGGGT</td></tr>
<tr><td>Chromosome</td><td>start</td><td>end</td><td>strand</td><td>MM</td><td>target_seq</td><td>PAM</td><td>alignment</td><td>distance</td><td>position</td><td>gene name</td><td>gene id</td></tr>
<tr><td>GU305914.1</td><td>125988</td><td>126010</td><td>-</td><td>0</td><td>ACCCCGCGATACACGACGCC</td><td>GGG</td><td>||||||||[||||||||||||]PAM</td><td>NA</td><td>Intergenic</td><td>NA</td><td>NA</td></tr>
<tr><td>T24460</td><td>ACACGACGCCGGGCCCTGTCTGG</td><td>1000</td></tr>
<tr><td>Oligo fwd</td><td>CACCACACGACGCCGGGCCCTGTC</td></tr>
<tr><td>Oligo rev</td><td>AAACGACAGGGCCCGGCGTCGTGT</td></tr>
<tr><td>Chromosome</td><td>start</td><td>end</td><td>strand</td><td>MM</td><td>target_seq</td><td>PAM</td><td>alignment</td><td>distance</td><td>position</td><td>gene name</td><td>gene id</td></tr>
<tr><td>GU305914.1</td><td>125978</td><td>126000</td><td>-</td><td>0</td><td>ACACGACGCCGGGCCCTGTC</td><td>TGG</td><td>||||||||[||||||||||||]PAM</td><td>NA</td><td>Intergenic</td><td>NA</td><td>NA</td></tr>
<tr><td>T24461</td><td>CGACGCCGGGCCCTGTCTGGAGG</td><td>1000</td></tr>
<tr><td>Oligo fwd</td><td>CACCCGACGCCGGGCCCTGTCTGG</td></tr>
<tr><td>Oligo rev</td><td>AAACCCAGACAGGGCCCGGCGTCG</td></tr>
<tr><td>Chromosome</td><td>start</td><td>end</td><td>strand</td><td>MM</td><td>target_seq</td><td>PAM</td><td>alignment</td><td>distance</td><td>position</td><td>gene name</td><td>gene id</td></tr>
<tr><td>GU305914.1</td><td>125975</td><td>125997</td><td>-</td><td>0</td><td>CGACGCCGGGCCCTGTCTGG</td><td>AGG</td><td>||||||||[||||||||||||]PAM</td><td>NA</td><td>Intergenic</td><td>NA</td><td>NA</td></tr>
<tr><td>T24462</td><td>CTGTCTGGAGGCGTTCGCGCAGG</td><td>1000</td></tr>
<tr><td>Oligo fwd</td><td>CACCCTGTCTGGAGGCGTTCGCGC</td></tr>
<tr><td>Oligo rev</td><td>AAACGCGCGAACGCCTCCAGACAG</td></tr>
<tr><td>Chromosome</td><td>start</td><td>end</td><td>strand</td><td>MM</td><td>target_seq</td><td>PAM</td><td>alignment</td><td>distance</td><td>position</td><td>gene name</td><td>gene id</td></tr>
<tr><td>GU305914.1</td><td>125963</td><td>125985</td><td>-</td><td>0</td><td>CTGTCTGGAGGCGTTCGCGC</td><td>AGG</td><td>||||||||[||||||||||||]PAM</td><td>NA</td><td>Intergenic</td><td>NA</td><td>NA</td></tr>
<tr><td>T24463</td><td>TCTGGAGGCGTTCGCGCAGGCGG</td><td>1000</td></tr>
<tr><td>Oligo fwd</td><td>CACCTCTGGAGGCGTTCGCGCAGG</td></tr>
<tr><td>Oligo rev</td><td>AAACCCTGCGCGAACGCCTCCAGA</td></tr>
<tr><td>Chromosome</td><td>start</td><td>end</td><td>strand</td><td>MM</td><td>target_seq</td><td>PAM</td><td>alignment</td><td>distance</td><td>position</td><td>gene name</td><td>gene id</td></tr>
<tr><td>GU305914.1</td><td>125960</td><td>125982</td><td>-</td><td>0</td><td>TCTGGAGGCGTTCGCGCAGG</td><td>CGG</td><td>||||||||[||||||||||||]PAM</td><td>NA</td><td>Intergenic</td><td>NA</td><td>NA</td></tr>
<tr><td>T24464</td><td>CTGGAGGCGTTCGCGCAGGCGGG</td><td>1000</td></tr>
<tr><td>Oligo fwd</td><td>CACCCTGGAGGCGTTCGCGCAGGC</td></tr>
<tr><td>Oligo rev</td><td>AAACGCCTGCGCGAACGCCTCCAG</td></tr>
<tr><td>Chromosome</td><td>start</td><td>end</td><td>strand</td><td>MM</td><td>target_seq</td><td>PAM</td><td>alignment</td><td>distance</td><td>position</td><td>gene name</td><td>gene id</td></tr>
</table>

GU305914.1 125959 125981 - 0 CTGGAGGCGTTCGCGCAGGC GGG ||||||||[||||||||||||]PAM NA Intergenic NA NA

T24465 CGCGCAGGCGGGCCCGCCCCAGG 1000
Oligo fwd CACCCGCGCAGGCGGGCCCGCCCC
Oligo rev AAACGGGGCGGGCCCGCCTGCGCG
Chromosome start end strand MM target_seq PAM alignment distance position gene name gene id
GU305914.1 125948 125970 - 0 CGCGCAGGCGGGCCCGCCCC AGG ||||||||[||||||||||||]PAM NA Intergenic NA NA

T24466 GCGCAGGCGGGCCCGCCCCAGGG 1000
Oligo fwd CACCGCGCAGGCGGGCCCGCCCCA
Oligo rev AAACTGGGGCGGGCCCGCCTGCGC
Chromosome start end strand MM target_seq PAM alignment distance position gene name gene id
GU305914.1 125947 125969 - 0 GCGCAGGCGGGCCCGCCCCA GGG ||||||||[||||||||||||]PAM NA Intergenic NA NA
GU305914.1 96852 96874 + 4 GCGTCGGCGGGCTCGCCCGA CGG |||--|||[||||-|||||-|]PAM NA Intergenic NA NA

T24467 CGCAGGCGGGCCCGCCCCAGGGG 1000
Oligo fwd CACCCGCAGGCGGGCCCGCCCCAG
Oligo rev AAACCTGGGGCGGGCCCGCCTGCG
Chromosome start end strand MM target_seq PAM alignment distance position gene name gene id
GU305914.1 125946 125968 - 0 CGCAGGCGGGCCCGCCCCAG GGG ||||||||[||||||||||||]PAM NA Intergenic NA NA

T24468 GCAGGCGGGCCCGCCCCAGGGGG 1000
Oligo fwd CACCGCAGGCGGGCCCGCCCCAGG
Oligo rev AAACCCTGGGGCGGGCCCGCCTGC
Chromosome start end strand MM target_seq PAM alignment distance position gene name gene id
GU305914.1 125945 125967 - 0 GCAGGCGGGCCCGCCCCAGG GGG ||||||||[||||||||||||]PAM NA Intergenic NA NA

T24469 GCCCGCCCCAGGGGGATGATCGG 1000
Oligo fwd CACCGCCCGCCCCAGGGGGATGAT
Oligo rev AAACATCATCCCCCTGGGGCGGGC
Chromosome start end strand MM target_seq PAM alignment distance position gene name gene id
GU305914.1 125937 125959 - 0 GCCCGCCCCAGGGGGATGAT CGG ||||||||[||||||||||||]PAM NA Intergenic NA NA

T24470 CCCGCCCCAGGGGGATGATCGGG 1000
Oligo fwd CACCCCCGCCCCAGGGGGATGATC
Oligo rev AAACGATCATCCCCCTGGGGCGGG
Chromosome start end strand MM target_seq PAM alignment distance position gene name gene id
GU305914.1 125936 125958 - 0 CCCGCCCCAGGGGGATGATC GGG ||||||||[||||||||||||]PAM NA Intergenic NA NA

T24471 CCCAGGGGGATGATCGGGTCAGG 1000
Oligo fwd CACCCCCAGGGGGATGATCGGGTC
Oligo rev AAACGACCCGATCATCCCCCTGGG
Chromosome start end strand MM target_seq PAM alignment distance position gene name gene id
GU305914.1 125931 125953 - 0 CCCAGGGGGATGATCGGGTC AGG ||||||||[||||||||||||]PAM NA Intergenic NA NA

T24472 CCAGGGGGATGATCGGGTCAGGG 1000
Oligo fwd CACCCCAGGGGGATGATCGGGTCA
Oligo rev AAACTGACCCGATCATCCCCCTGG
Chromosome start end strand MM target_seq PAM alignment distance position gene name gene id
GU305914.1 125930 125952 - 0 CCAGGGGGATGATCGGGTCA GGG ||||||||[||||||||||||]PAM NA Intergenic NA NA

T24473 ATCGGGTCAGGGCGCTCTGCCGG 1000
Oligo fwd CACCATCGGGTCAGGGCGCTCTGC
Oligo rev AAACGCAGAGCGCCCTGACCCGAT
Chromosome start end strand MM target_seq PAM alignment distance position gene name gene id
GU305914.1 125919 125941 - 0 ATCGGGTCAGGGCGCTCTGC CGG ||||||||[||||||||||||]PAM NA Intergenic NA NA

T24474 TCTGCCGGCGAGAGTTCGTCAGG 1000
Oligo fwd CACCTCTGCCGGCGAGAGTTCGTC
Oligo rev AAACGACGAACTCTCGCCGGCAGA
Chromosome start end strand MM target_seq PAM alignment distance position gene name gene id
GU305914.1 125904 125926 - 0 TCTGCCGGCGAGAGTTCGTC AGG ||||||||[||||||||||||]PAM NA Intergenic NA NA

T24475 CTGCCGGCGAGAGTTCGTCAGGG 1000
Oligo fwd CACCCTGCCGGCGAGAGTTCGTCA
Oligo rev AAACTGACGAACTCTCGCCGGCAG
Chromosome start end strand MM target_seq PAM alignment distance position gene name gene id
GU305914.1 125903 125925 - 0 CTGCCGGCGAGAGTTCGTCA GGG ||||||||[||||||||||||]PAM NA Intergenic NA NA

T24476 AGAGTTCGTCAGGGAGCACATGG 1000
Oligo fwd CACCAGAGTTCGTCAGGGAGCACA
Oligo rev AAACTGTGCTCCCTGACGAACTCT
Chromosome start end strand MM target_seq PAM alignment distance position gene name gene id
GU305914.1 125894 125916 - 0 AGAGTTCGTCAGGGAGCACA TGG ||||||||[||||||||||||]PAM NA Intergenic NA NA

T24477 AGCACATGGCACACGCCACGCGG 1000
Oligo fwd CACCAGCACATGGCACACGCCACG
Oligo rev AAACCGTGGCGTGTGCCATGTGCT
Chromosome start end strand MM target_seq PAM alignment distance position gene name gene id
GU305914.1 125880 125902 - 0 AGCACATGGCACACGCCACG CGG ||||||||[||||||||||||]PAM NA Intergenic NA NA

T24478 ACATGGCACACGCCACGCGGCGG 1000
Oligo fwd CACCACATGGCACACGCCACGCGG
Oligo rev AAACCCGCGTGGCGTGTGCCATGT
Chromosome start end strand MM target_seq PAM alignment distance position gene name gene id
GU305914.1 125877 125899 - 0 ACATGGCACACGCCACGCGG CGG ||||||||[||||||||||||]PAM NA Intergenic NA NA

T24479 GCGGCTCGTGCACTTCTACCAGG 1000
Oligo fwd CACCGCGGCTCGTGCACTTCTACC

<table>
<tr><td>Oligo rev</td><td>AAACGGTAGAAGTGCACGAGCCGC</td><td></td><td></td><td></td><td></td><td></td><td></td><td></td><td></td><td></td><td></td></tr>
<tr><td>Chromosome</td><td>start</td><td>end</td><td>strand</td><td>MM</td><td>target_seq</td><td>PAM</td><td>alignment</td><td>distance</td><td>position</td><td>gene name</td><td>gene id</td></tr>
<tr><td>GU305914.1</td><td>125858</td><td>125880</td><td>-</td><td>0</td><td>GCGGCTCGTGCACTTCTACC</td><td>AGG</td><td>||||||||[||||||||||||]PAM</td><td>NA</td><td>Intergenic</td><td>NA</td><td>NA</td></tr>
<tr><td></td><td></td><td></td><td></td><td></td><td></td><td></td><td></td><td></td><td></td><td></td><td></td></tr>
<tr><td>T24480</td><td>GCGCATCGACCCGCCGCGCACGG</td><td>1000</td><td></td><td></td><td></td><td></td><td></td><td></td><td></td><td></td><td></td></tr>
<tr><td>Oligo fwd</td><td>CACCGCGCATCGACCCGCCGCGCA</td><td></td><td></td><td></td><td></td><td></td><td></td><td></td><td></td><td></td><td></td></tr>
<tr><td>Oligo rev</td><td>AAACTGCGCGGCGGGTCGATGCGC</td><td></td><td></td><td></td><td></td><td></td><td></td><td></td><td></td><td></td><td></td></tr>
<tr><td>Chromosome</td><td>start</td><td>end</td><td>strand</td><td>MM</td><td>target_seq</td><td>PAM</td><td>alignment</td><td>distance</td><td>position</td><td>gene name</td><td>gene id</td></tr>
<tr><td>GU305914.1</td><td>125834</td><td>125856</td><td>-</td><td>0</td><td>GCGCATCGACCCGCCGCGCA</td><td>CGG</td><td>||||||||[||||||||||||]PAM</td><td>NA</td><td>Intergenic</td><td>NA</td><td>NA</td></tr>
<tr><td></td><td></td><td></td><td></td><td></td><td></td><td></td><td></td><td></td><td></td><td></td><td></td></tr>
<tr><td>T24481</td><td>CCCGCCGCGCACGGCCAACGAGG</td><td>1000</td><td></td><td></td><td></td><td></td><td></td><td></td><td></td><td></td><td></td></tr>
<tr><td>Oligo fwd</td><td>CACCCCCGCCGCGCACGGCCAACG</td><td></td><td></td><td></td><td></td><td></td><td></td><td></td><td></td><td></td><td></td></tr>
<tr><td>Oligo rev</td><td>AAACCGTTGGCCGTGCGCGGCGGG</td><td></td><td></td><td></td><td></td><td></td><td></td><td></td><td></td><td></td><td></td></tr>
<tr><td>Chromosome</td><td>start</td><td>end</td><td>strand</td><td>MM</td><td>target_seq</td><td>PAM</td><td>alignment</td><td>distance</td><td>position</td><td>gene name</td><td>gene id</td></tr>
<tr><td>GU305914.1</td><td>125825</td><td>125847</td><td>-</td><td>0</td><td>CCCGCCGCGCACGGCCAACG</td><td>AGG</td><td>||||||||[||||||||||||]PAM</td><td>NA</td><td>Intergenic</td><td>NA</td><td>NA</td></tr>
<tr><td>GU305914.1</td><td>89384</td><td>89406</td><td>-</td><td>4</td><td>GCTGCCGCGCTCGGTCAACG</td><td>TGG</td><td>-|-|||||[||-|||-|||||]PAM</td><td>NA</td><td>Intergenic</td><td>NA</td><td>NA</td></tr>
<tr><td></td><td></td><td></td><td></td><td></td><td></td><td></td><td></td><td></td><td></td><td></td><td></td></tr>
<tr><td>T24482</td><td>GAAGCACGATTTCAGCACCAAGG</td><td>1000</td><td></td><td></td><td></td><td></td><td></td><td></td><td></td><td></td><td></td></tr>
<tr><td>Oligo fwd</td><td>CACCGAAGCACGATTTCAGCACCA</td><td></td><td></td><td></td><td></td><td></td><td></td><td></td><td></td><td></td><td></td></tr>
<tr><td>Oligo rev</td><td>AAACTGGTGCTGAAATCGTGCTTC</td><td></td><td></td><td></td><td></td><td></td><td></td><td></td><td></td><td></td><td></td></tr>
<tr><td>Chromosome</td><td>start</td><td>end</td><td>strand</td><td>MM</td><td>target_seq</td><td>PAM</td><td>alignment</td><td>distance</td><td>position</td><td>gene name</td><td>gene id</td></tr>
<tr><td>GU305914.1</td><td>125801</td><td>125823</td><td>-</td><td>0</td><td>GAAGCACGATTTCAGCACCA</td><td>AGG</td><td>||||||||[||||||||||||]PAM</td><td>NA</td><td>Intergenic</td><td>NA</td><td>NA</td></tr>
<tr><td></td><td></td><td></td><td></td><td></td><td></td><td></td><td></td><td></td><td></td><td></td><td></td></tr>
<tr><td>T24483</td><td>CAGCACCAAGGAGTTCGCCAAGG</td><td>1000</td><td></td><td></td><td></td><td></td><td></td><td></td><td></td><td></td><td></td></tr>
<tr><td>Oligo fwd</td><td>CACCCAGCACCAAGGAGTTCGCCA</td><td></td><td></td><td></td><td></td><td></td><td></td><td></td><td></td><td></td><td></td></tr>
<tr><td>Oligo rev</td><td>AAACTGGCGAACTCCTTGGTGCTG</td><td></td><td></td><td></td><td></td><td></td><td></td><td></td><td></td><td></td><td></td></tr>
<tr><td>Chromosome</td><td>start</td><td>end</td><td>strand</td><td>MM</td><td>target_seq</td><td>PAM</td><td>alignment</td><td>distance</td><td>position</td><td>gene name</td><td>gene id</td></tr>
<tr><td>GU305914.1</td><td>125789</td><td>125811</td><td>-</td><td>0</td><td>CAGCACCAAGGAGTTCGCCA</td><td>AGG</td><td>||||||||[||||||||||||]PAM</td><td>NA</td><td>Intergenic</td><td>NA</td><td>NA</td></tr>
<tr><td></td><td></td><td></td><td></td><td></td><td></td><td></td><td></td><td></td><td></td><td></td><td></td></tr>
<tr><td>T24484</td><td>CACCAAGGAGTTCGCCAAGGTGG</td><td>1000</td><td></td><td></td><td></td><td></td><td></td><td></td><td></td><td></td><td></td></tr>
<tr><td>Oligo fwd</td><td>CACCCACCAAGGAGTTCGCCAAGG</td><td></td><td></td><td></td><td></td><td></td><td></td><td></td><td></td><td></td><td></td></tr>
<tr><td>Oligo rev</td><td>AAACCCTTGGCGAACTCCTTGGTG</td><td></td><td></td><td></td><td></td><td></td><td></td><td></td><td></td><td></td><td></td></tr>
<tr><td>Chromosome</td><td>start</td><td>end</td><td>strand</td><td>MM</td><td>target_seq</td><td>PAM</td><td>alignment</td><td>distance</td><td>position</td><td>gene name</td><td>gene id</td></tr>
<tr><td>GU305914.1</td><td>125786</td><td>125808</td><td>-</td><td>0</td><td>CACCAAGGAGTTCGCCAAGG</td><td>TGG</td><td>||||||||[||||||||||||]PAM</td><td>NA</td><td>Intergenic</td><td>NA</td><td>NA</td></tr>
<tr><td></td><td></td><td></td><td></td><td></td><td></td><td></td><td></td><td></td><td></td><td></td><td></td></tr>
<tr><td>T24485</td><td>CTTCGACTTCTGCTTCCACACGG</td><td>1000</td><td></td><td></td><td></td><td></td><td></td><td></td><td></td><td></td><td></td></tr>
<tr><td>Oligo fwd</td><td>CACCCTTCGACTTCTGCTTCCACA</td><td></td><td></td><td></td><td></td><td></td><td></td><td></td><td></td><td></td><td></td></tr>
<tr><td>Oligo rev</td><td>AAACTGTGGAAGCAGAAGTCGAAG</td><td></td><td></td><td></td><td></td><td></td><td></td><td></td><td></td><td></td><td></td></tr>
<tr><td>Chromosome</td><td>start</td><td>end</td><td>strand</td><td>MM</td><td>target_seq</td><td>PAM</td><td>alignment</td><td>distance</td><td>position</td><td>gene name</td><td>gene id</td></tr>
<tr><td>GU305914.1</td><td>125729</td><td>125751</td><td>-</td><td>0</td><td>CTTCGACTTCTGCTTCCACA</td><td>CGG</td><td>||||||||[||||||||||||]PAM</td><td>NA</td><td>Intergenic</td><td>NA</td><td>NA</td></tr>
<tr><td></td><td></td><td></td><td></td><td></td><td></td><td></td><td></td><td></td><td></td><td></td><td></td></tr>
<tr><td>T24486</td><td>TTCTGCTTCCACACGGAGAACGG</td><td>1000</td><td></td><td></td><td></td><td></td><td></td><td></td><td></td><td></td><td></td></tr>
<tr><td>Oligo fwd</td><td>CACCTTCTGCTTCCACACGGAGAA</td><td></td><td></td><td></td><td></td><td></td><td></td><td></td><td></td><td></td><td></td></tr>
<tr><td>Oligo rev</td><td>AAACTTCTCCGTGTGGAAGCAGAA</td><td></td><td></td><td></td><td></td><td></td><td></td><td></td><td></td><td></td><td></td></tr>
<tr><td>Chromosome</td><td>start</td><td>end</td><td>strand</td><td>MM</td><td>target_seq</td><td>PAM</td><td>alignment</td><td>distance</td><td>position</td><td>gene name</td><td>gene id</td></tr>
<tr><td>GU305914.1</td><td>125722</td><td>125744</td><td>-</td><td>0</td><td>TTCTGCTTCCACACGGAGAA</td><td>CGG</td><td>||||||||[||||||||||||]PAM</td><td>NA</td><td>Intergenic</td><td>NA</td><td>NA</td></tr>
<tr><td></td><td></td><td></td><td></td><td></td><td></td><td></td><td></td><td></td><td></td><td></td><td></td></tr>
<tr><td>T24487</td><td>TCTGCTTCCACACGGAGAACGGG</td><td>1000</td><td></td><td></td><td></td><td></td><td></td><td></td><td></td><td></td><td></td></tr>
<tr><td>Oligo fwd</td><td>CACCTCTGCTTCCACACGGAGAAC</td><td></td><td></td><td></td><td></td><td></td><td></td><td></td><td></td><td></td><td></td></tr>
<tr><td>Oligo rev</td><td>AAACGTTCTCCGTGTGGAAGCAGA</td><td></td><td></td><td></td><td></td><td></td><td></td><td></td><td></td><td></td><td></td></tr>
<tr><td>Chromosome</td><td>start</td><td>end</td><td>strand</td><td>MM</td><td>target_seq</td><td>PAM</td><td>alignment</td><td>distance</td><td>position</td><td>gene name</td><td>gene id</td></tr>
<tr><td>GU305914.1</td><td>125721</td><td>125743</td><td>-</td><td>0</td><td>TCTGCTTCCACACGGAGAAC</td><td>GGG</td><td>||||||||[||||||||||||]PAM</td><td>NA</td><td>Intergenic</td><td>NA</td><td>NA</td></tr>
<tr><td></td><td></td><td></td><td></td><td></td><td></td><td></td><td></td><td></td><td></td><td></td><td></td></tr>
<tr><td>T24488</td><td>CTTCCACACGGAGAACGGGCAGG</td><td>1000</td><td></td><td></td><td></td><td></td><td></td><td></td><td></td><td></td><td></td></tr>
<tr><td>Oligo fwd</td><td>CACCCTTCCACACGGAGAACGGGC</td><td></td><td></td><td></td><td></td><td></td><td></td><td></td><td></td><td></td><td></td></tr>
<tr><td>Oligo rev</td><td>AAACGCCCGTTCTCCGTGTGGAAG</td><td></td><td></td><td></td><td></td><td></td><td></td><td></td><td></td><td></td><td></td></tr>
<tr><td>Chromosome</td><td>start</td><td>end</td><td>strand</td><td>MM</td><td>target_seq</td><td>PAM</td><td>alignment</td><td>distance</td><td>position</td><td>gene name</td><td>gene id</td></tr>
<tr><td>GU305914.1</td><td>125717</td><td>125739</td><td>-</td><td>0</td><td>CTTCCACACGGAGAACGGGC</td><td>AGG</td><td>||||||||[||||||||||||]PAM</td><td>NA</td><td>Intergenic</td><td>NA</td><td>NA</td></tr>
<tr><td></td><td></td><td></td><td></td><td></td><td></td><td></td><td></td><td></td><td></td><td></td><td></td></tr>
<tr><td>T24489</td><td>ACACGGAGAACGGGCAGGTGCGG</td><td>1000</td><td></td><td></td><td></td><td></td><td></td><td></td><td></td><td></td><td></td></tr>
<tr><td>Oligo fwd</td><td>CACCACACGGAGAACGGGCAGGTG</td><td></td><td></td><td></td><td></td><td></td><td></td><td></td><td></td><td></td><td></td></tr>
<tr><td>Oligo rev</td><td>AAACCACCTGCCCGTTCTCCGTGT</td><td></td><td></td><td></td><td></td><td></td><td></td><td></td><td></td><td></td><td></td></tr>
<tr><td>Chromosome</td><td>start</td><td>end</td><td>strand</td><td>MM</td><td>target_seq</td><td>PAM</td><td>alignment</td><td>distance</td><td>position</td><td>gene name</td><td>gene id</td></tr>
<tr><td>GU305914.1</td><td>125712</td><td>125734</td><td>-</td><td>0</td><td>ACACGGAGAACGGGCAGGTG</td><td>CGG</td><td>||||||||[||||||||||||]PAM</td><td>NA</td><td>Intergenic</td><td>NA</td><td>NA</td></tr>
<tr><td></td><td></td><td></td><td></td><td></td><td></td><td></td><td></td><td></td><td></td><td></td><td></td></tr>
<tr><td>T24490</td><td>GCTGTGCACGCCGCGCATCATGG</td><td>1000</td><td></td><td></td><td></td><td></td><td></td><td></td><td></td><td></td><td></td></tr>
<tr><td>Oligo fwd</td><td>CACCGCTGTGCACGCCGCGCATCA</td><td></td><td></td><td></td><td></td><td></td><td></td><td></td><td></td><td></td><td></td></tr>
<tr><td>Oligo rev</td><td>AAACTGATGCGCGGCGTGCACAGC</td><td></td><td></td><td></td><td></td><td></td><td></td><td></td><td></td><td></td><td></td></tr>
<tr><td>Chromosome</td><td>start</td><td>end</td><td>strand</td><td>MM</td><td>target_seq</td><td>PAM</td><td>alignment</td><td>distance</td><td>position</td><td>gene name</td><td>gene id</td></tr>
<tr><td>GU305914.1</td><td>125687</td><td>125709</td><td>-</td><td>0</td><td>GCTGTGCACGCCGCGCATCA</td><td>TGG</td><td>||||||||[||||||||||||]PAM</td><td>NA</td><td>Intergenic</td><td>NA</td><td>NA</td></tr>
<tr><td></td><td></td><td></td><td></td><td></td><td></td><td></td><td></td><td></td><td></td><td></td><td></td></tr>
<tr><td>T24491</td><td>GTGCACGCCGCGCATCATGGTGG</td><td>1000</td><td></td><td></td><td></td><td></td><td></td><td></td><td></td><td></td><td></td></tr>
<tr><td>Oligo fwd</td><td>CACCGTGCACGCCGCGCATCATGG</td><td></td><td></td><td></td><td></td><td></td><td></td><td></td><td></td><td></td><td></td></tr>
<tr><td>Oligo rev</td><td>AAACCCATGATGCGCGGCGTGCAC</td><td></td><td></td><td></td><td></td><td></td><td></td><td></td><td></td><td></td><td></td></tr>
<tr><td>Chromosome</td><td>start</td><td>end</td><td>strand</td><td>MM</td><td>target_seq</td><td>PAM</td><td>alignment</td><td>distance</td><td>position</td><td>gene name</td><td>gene id</td></tr>
<tr><td>GU305914.1</td><td>125684</td><td>125706</td><td>-</td><td>0</td><td>GTGCACGCCGCGCATCATGG</td><td>TGG</td><td>||||||||[||||||||||||]PAM</td><td>NA</td><td>Intergenic</td><td>NA</td><td>NA</td></tr>
<tr><td></td><td></td><td></td><td></td><td></td><td></td><td></td><td></td><td></td><td></td><td></td><td></td></tr>
<tr><td>T24492</td><td>TGCACGCCGCGCATCATGGTGGG</td><td>1000</td><td></td><td></td><td></td><td></td><td></td><td></td><td></td><td></td><td></td></tr>
<tr><td>Oligo fwd</td><td>CACCTGCACGCCGCGCATCATGGT</td><td></td><td></td><td></td><td></td><td></td><td></td><td></td><td></td><td></td><td></td></tr>
<tr><td>Oligo rev</td><td>AAACACCATGATGCGCGGCGTGCA</td><td></td><td></td><td></td><td></td><td></td><td></td><td></td><td></td><td></td><td></td></tr>
<tr><td>Chromosome</td><td>start</td><td>end</td><td>strand</td><td>MM</td><td>target_seq</td><td>PAM</td><td>alignment</td><td>distance</td><td>position</td><td>gene name</td><td>gene id</td></tr>
<tr><td>GU305914.1</td><td>125683</td><td>125705</td><td>-</td><td>0</td><td>TGCACGCCGCGCATCATGGT</td><td>GGG</td><td>||||||||[||||||||||||]PAM</td><td>NA</td><td>Intergenic</td><td>NA</td><td>NA</td></tr>
<tr><td></td><td></td><td></td><td></td><td></td><td></td><td></td><td></td><td></td><td></td><td></td><td></td></tr>
<tr><td>T24493</td><td>CATGGTGGGTAACCTGCCCGAGG</td><td>1000</td><td></td><td></td><td></td><td></td><td></td><td></td><td></td><td></td><td></td></tr>
<tr><td>Oligo fwd</td><td>CACCCATGGTGGGTAACCTGCCCG</td><td></td><td></td><td></td><td></td><td></td><td></td><td></td><td></td><td></td><td></td></tr>
<tr><td>Oligo rev</td><td>AAACCGGGCAGGTTACCCACCATG</td><td></td><td></td><td></td><td></td><td></td><td></td><td></td><td></td><td></td><td></td></tr>
<tr><td>Chromosome</td><td>start</td><td>end</td><td>strand</td><td>MM</td><td>target_seq</td><td>PAM</td><td>alignment</td><td>distance</td><td>position</td><td>gene name</td><td>gene id</td></tr>
<tr><td>GU305914.1</td><td>125669</td><td>125691</td><td>-</td><td>0</td><td>CATGGTGGGTAACCTGCCCG</td><td>AGG</td><td>||||||||[||||||||||||]PAM</td><td>NA</td><td>Intergenic</td><td>NA</td><td>NA</td></tr>
</table>

T24494 GGGTAACCTGCCCGAGGCGCTGG 1000
Oligo fwd CACCGGGTAACCTGCCCGAGGCGC
Oligo rev AAACGCGCCTCGGGCAGGTTACCC
Chromosome start end strand MM target_seq PAM alignment distance position gene name gene id
GU305914.1 125663 125685 - 0 GGGTAACCTGCCCGAGGCGC TGG ||||||||[||||||||||||]PAM NA Intergenic NA NA

T24495 CCTGCCCGAGGCGCTGGCGCCGG 1000
Oligo fwd CACCCCTGCCCGAGGCGCTGGCGC
Oligo rev AAACGCGCCAGCGCCTCGGGCAGG
Chromosome start end strand MM target_seq PAM alignment distance position gene name gene id
GU305914.1 125657 125679 - 0 CCTGCCCGAGGCGCTGGCGC CGG ||||||||[||||||||||||]PAM NA Intergenic NA NA

T24496 CGGCCGACTTCCACGACCTGCGG 1000
Oligo fwd CACCCGGCCGACTTCCACGACCTG
Oligo rev AAACCAGGTCGTGGAAGTCGGCCG
Chromosome start end strand MM target_seq PAM alignment distance position gene name gene id
GU305914.1 125637 125659 - 0 CGGCCGACTTCCACGACCTG CGG ||||||||[||||||||||||]PAM NA Intergenic NA NA

T24497 GGCCGACTTCCACGACCTGCGGG 1000
Oligo fwd CACCGGCCGACTTCCACGACCTGC
Oligo rev AAACGCAGGTCGTGGAAGTCGGCC
Chromosome start end strand MM target_seq PAM alignment distance position gene name gene id
GU305914.1 125636 125658 - 0 GGCCGACTTCCACGACCTGC GGG ||||||||[||||||||||||]PAM NA Intergenic NA NA

T24498 CCACGACCTGCGGGCGAAGCAGG 1000
Oligo fwd CACCCCACGACCTGCGGGCGAAGC
Oligo rev AAACGCTTCGCCCGCAGGTCGTGG
Chromosome start end strand MM target_seq PAM alignment distance position gene name gene id
GU305914.1 125627 125649 - 0 CCACGACCTGCGGGCGAAGC AGG ||||||||[||||||||||||]PAM NA Intergenic NA NA

T24499 CCTGCGGGCGAAGCAGGCGCTGG 1000
Oligo fwd CACCCCTGCGGGCGAAGCAGGCGC
Oligo rev AAACGCGCCTGCTTCGCCCGCAGG
Chromosome start end strand MM target_seq PAM alignment distance position gene name gene id
GU305914.1 125621 125643 - 0 CCTGCGGGCGAAGCAGGCGC TGG ||||||||[||||||||||||]PAM NA Intergenic NA NA

T24500 GCAGGCGCTGGAGCTGACCAAGG 1000
Oligo fwd CACCGCAGGCGCTGGAGCTGACCA
Oligo rev AAACTGGTCAGCTCCAGCGCCTGC
Chromosome start end strand MM target_seq PAM alignment distance position gene name gene id
GU305914.1 125609 125631 - 0 GCAGGCGCTGGAGCTGACCA AGG ||||||||[||||||||||||]PAM NA Intergenic NA NA

T24501 GCTGACCAAGGTGCGCGCGCCGG 1000
Oligo fwd CACCGCTGACCAAGGTGCGCGCGC
Oligo rev AAACGCGCGCGCACCTTGGTCAGC
Chromosome start end strand MM target_seq PAM alignment distance position gene name gene id
GU305914.1 125597 125619 - 0 GCTGACCAAGGTGCGCGCGC CGG ||||||||[||||||||||||]PAM NA Intergenic NA NA

T24502 GACCAAGGTGCGCGCGCCGGAGG 1000
Oligo fwd CACCGACCAAGGTGCGCGCGCCGG
Oligo rev AAACCCGGCGCGCGCACCTTGGTC
Chromosome start end strand MM target_seq PAM alignment distance position gene name gene id
GU305914.1 125594 125616 - 0 GACCAAGGTGCGCGCGCCGG AGG ||||||||[||||||||||||]PAM NA Intergenic NA NA

T24503 ACCAAGGTGCGCGCGCCGGAGGG 1000
Oligo fwd CACCACCAAGGTGCGCGCGCCGGA
Oligo rev AAACTCCGGCGCGCGCACCTTGGT
Chromosome start end strand MM target_seq PAM alignment distance position gene name gene id
GU305914.1 125593 125615 - 0 ACCAAGGTGCGCGCGCCGGA GGG ||||||||[||||||||||||]PAM NA Intergenic NA NA

T24504 GCGCGCGCCGGAGGGTCACGAGG 1000
Oligo fwd CACCGCGCGCGCCGGAGGGTCACG
Oligo rev AAACCGTGACCCTCCGGCGCGCGC
Chromosome start end strand MM target_seq PAM alignment distance position gene name gene id
GU305914.1 125585 125607 - 0 GCGCGCGCCGGAGGGTCACG AGG ||||||||[||||||||||||]PAM NA Intergenic NA NA

T24505 GGGTCACGAGGCGACGCTGCAGG 1000
Oligo fwd CACCGGGTCACGAGGCGACGCTGC
Oligo rev AAACGCAGCGTCGCCTCGTGACCC
Chromosome start end strand MM target_seq PAM alignment distance position gene name gene id
GU305914.1 125573 125595 - 0 GGGTCACGAGGCGACGCTGC AGG ||||||||[||||||||||||]PAM NA Intergenic NA NA

T24506 AGGCGACGCTGCAGGTGCTGCGG 1000
Oligo fwd CACCAGGCGACGCTGCAGGTGCTG
Oligo rev AAACCAGCACCTGCAGCGTCGCCT
Chromosome start end strand MM target_seq PAM alignment distance position gene name gene id
GU305914.1 125565 125587 - 0 AGGCGACGCTGCAGGTGCTG CGG ||||||||[||||||||||||]PAM NA Intergenic NA NA

T24507 GGCGACGCTGCAGGTGCTGCGGG 1000
Oligo fwd CACCGGCGACGCTGCAGGTGCTGC
Oligo rev AAACGCAGCACCTGCAGCGTCGCC
Chromosome start end strand MM target_seq PAM alignment distance position gene name gene id
GU305914.1 125564 125586 - 0 GGCGACGCTGCAGGTGCTGC GGG ||||||||[||||||||||||]PAM NA Intergenic NA NA

T24508 GGTGCTGCGGGCCTCGCTGACGG 1000
Oligo fwd CACCGGTGCTGCGGGCCTCGCTGA
Oligo rev AAACTCAGCGAGGCCCGCAGCACC
Chromosome start end strand MM target_seq PAM alignment distance position gene name gene id
GU305914.1 125552 125574 - 0 GGTGCTGCGGGCCTCGCTGA CGG ||||||||[||||||||||||]PAM NA Intergenic NA NA

<table>
<tr><td>T24509</td><td>GCTGACGGACCACCAGTACCCGG</td><td>1000</td><td></td><td></td><td></td><td></td><td></td><td></td><td></td><td></td><td></td></tr>
<tr><td>Oligo fwd</td><td>CACCGCTGACGGACCACCAGTACC</td><td></td><td></td><td></td><td></td><td></td><td></td><td></td><td></td><td></td><td></td></tr>
<tr><td>Oligo rev</td><td>AAACGGTACTGGTGGTCCGTCAGC</td><td></td><td></td><td></td><td></td><td></td><td></td><td></td><td></td><td></td><td></td></tr>
<tr><td>Chromosome</td><td>start</td><td>end</td><td>strand</td><td>MM</td><td>target_seq</td><td>PAM</td><td>alignment</td><td>distance</td><td>position</td><td>gene name</td><td>gene id</td></tr>
<tr><td>GU305914.1</td><td>125537</td><td>125559</td><td>-</td><td>0</td><td>GCTGACGGACCACCAGTACC</td><td>CGG</td><td>||||||||[||||||||||||]PAM</td><td>NA</td><td>Intergenic</td><td>NA</td><td>NA</td></tr>
<tr><td>T24510</td><td>CTGATAGAGTCTCTGATCCACGG</td><td>1000</td><td></td><td></td><td></td><td></td><td></td><td></td><td></td><td></td><td></td></tr>
<tr><td>Oligo fwd</td><td>CACCCTGATAGAGTCTCTGATCCA</td><td></td><td></td><td></td><td></td><td></td><td></td><td></td><td></td><td></td><td></td></tr>
<tr><td>Oligo rev</td><td>AAACTGGATCAGAGACTCTATCAG</td><td></td><td></td><td></td><td></td><td></td><td></td><td></td><td></td><td></td><td></td></tr>
<tr><td>Chromosome</td><td>start</td><td>end</td><td>strand</td><td>MM</td><td>target_seq</td><td>PAM</td><td>alignment</td><td>distance</td><td>position</td><td>gene name</td><td>gene id</td></tr>
<tr><td>GU305914.1</td><td>125503</td><td>125525</td><td>-</td><td>0</td><td>CTGATAGAGTCTCTGATCCA</td><td>CGG</td><td>||||||||[||||||||||||]PAM</td><td>NA</td><td>Intergenic</td><td>NA</td><td>NA</td></tr>
<tr><td>T24511</td><td>GTCTCTGATCCACGGCGACCCGG</td><td>1000</td><td></td><td></td><td></td><td></td><td></td><td></td><td></td><td></td><td></td></tr>
<tr><td>Oligo fwd</td><td>CACCGTCTCTGATCCACGGCGACC</td><td></td><td></td><td></td><td></td><td></td><td></td><td></td><td></td><td></td><td></td></tr>
<tr><td>Oligo rev</td><td>AAACGGTCGCCGTGGATCAGAGAC</td><td></td><td></td><td></td><td></td><td></td><td></td><td></td><td></td><td></td><td></td></tr>
<tr><td>Chromosome</td><td>start</td><td>end</td><td>strand</td><td>MM</td><td>target_seq</td><td>PAM</td><td>alignment</td><td>distance</td><td>position</td><td>gene name</td><td>gene id</td></tr>
<tr><td>GU305914.1</td><td>125495</td><td>125517</td><td>-</td><td>0</td><td>GTCTCTGATCCACGGCGACC</td><td>CGG</td><td>||||||||[||||||||||||]PAM</td><td>NA</td><td>Intergenic</td><td>NA</td><td>NA</td></tr>
<tr><td>T24512</td><td>TCTGATCCACGGCGACCCGGCGG</td><td>1000</td><td></td><td></td><td></td><td></td><td></td><td></td><td></td><td></td><td></td></tr>
<tr><td>Oligo fwd</td><td>CACCTCTGATCCACGGCGACCCGG</td><td></td><td></td><td></td><td></td><td></td><td></td><td></td><td></td><td></td><td></td></tr>
<tr><td>Oligo rev</td><td>AAACCCGGGTCGCCGTGGATCAGA</td><td></td><td></td><td></td><td></td><td></td><td></td><td></td><td></td><td></td><td></td></tr>
<tr><td>Chromosome</td><td>start</td><td>end</td><td>strand</td><td>MM</td><td>target_seq</td><td>PAM</td><td>alignment</td><td>distance</td><td>position</td><td>gene name</td><td>gene id</td></tr>
<tr><td>GU305914.1</td><td>125492</td><td>125514</td><td>-</td><td>0</td><td>TCTGATCCACGGCGACCCGG</td><td>CGG</td><td>||||||||[||||||||||||]PAM</td><td>NA</td><td>Intergenic</td><td>NA</td><td>NA</td></tr>
<tr><td>T24513</td><td>CGACCCGGCGGCCTTCGAGACGG</td><td>1000</td><td></td><td></td><td></td><td></td><td></td><td></td><td></td><td></td><td></td></tr>
<tr><td>Oligo fwd</td><td>CACCCGACCCGGCGGCCTTCGAGA</td><td></td><td></td><td></td><td></td><td></td><td></td><td></td><td></td><td></td><td></td></tr>
<tr><td>Oligo rev</td><td>AAACTCTCGAAGGCCGCCGGGTCG</td><td></td><td></td><td></td><td></td><td></td><td></td><td></td><td></td><td></td><td></td></tr>
<tr><td>Chromosome</td><td>start</td><td>end</td><td>strand</td><td>MM</td><td>target_seq</td><td>PAM</td><td>alignment</td><td>distance</td><td>position</td><td>gene name</td><td>gene id</td></tr>
<tr><td>GU305914.1</td><td>125480</td><td>125502</td><td>-</td><td>0</td><td>CGACCCGGCGGCCTTCGAGA</td><td>CGG</td><td>||||||||[||||||||||||]PAM</td><td>NA</td><td>Intergenic</td><td>NA</td><td>NA</td></tr>
<tr><td>T24514</td><td>GACCCGGCGGCCTTCGAGACGGG</td><td>1000</td><td></td><td></td><td></td><td></td><td></td><td></td><td></td><td></td><td></td></tr>
<tr><td>Oligo fwd</td><td>CACCGACCCGGCGGCCTTCGAGAC</td><td></td><td></td><td></td><td></td><td></td><td></td><td></td><td></td><td></td><td></td></tr>
<tr><td>Oligo rev</td><td>AAACGTCTCGAAGGCCGCCGGGTC</td><td></td><td></td><td></td><td></td><td></td><td></td><td></td><td></td><td></td><td></td></tr>
<tr><td>Chromosome</td><td>start</td><td>end</td><td>strand</td><td>MM</td><td>target_seq</td><td>PAM</td><td>alignment</td><td>distance</td><td>position</td><td>gene name</td><td>gene id</td></tr>
<tr><td>GU305914.1</td><td>125479</td><td>125501</td><td>-</td><td>0</td><td>GACCCGGCGGCCTTCGAGAC</td><td>GGG</td><td>||||||||[||||||||||||]PAM</td><td>NA</td><td>Intergenic</td><td>NA</td><td>NA</td></tr>
<tr><td>T24515</td><td>CTTCGAGACGGGCATCGAGCTGG</td><td>1000</td><td></td><td></td><td></td><td></td><td></td><td></td><td></td><td></td><td></td></tr>
<tr><td>Oligo fwd</td><td>CACCCTTCGAGACGGGCATCGAGC</td><td></td><td></td><td></td><td></td><td></td><td></td><td></td><td></td><td></td><td></td></tr>
<tr><td>Oligo rev</td><td>AAACGCTCGATGCCCGTCTCGAAG</td><td></td><td></td><td></td><td></td><td></td><td></td><td></td><td></td><td></td><td></td></tr>
<tr><td>Chromosome</td><td>start</td><td>end</td><td>strand</td><td>MM</td><td>target_seq</td><td>PAM</td><td>alignment</td><td>distance</td><td>position</td><td>gene name</td><td>gene id</td></tr>
<tr><td>GU305914.1</td><td>125468</td><td>125490</td><td>-</td><td>0</td><td>CTTCGAGACGGGCATCGAGC</td><td>TGG</td><td>||||||||[||||||||||||]PAM</td><td>NA</td><td>Intergenic</td><td>NA</td><td>NA</td></tr>
<tr><td>T24516</td><td>CGCGCTGCGTGAACAACTACTGG</td><td>1000</td><td></td><td></td><td></td><td></td><td></td><td></td><td></td><td></td><td></td></tr>
<tr><td>Oligo fwd</td><td>CACCCGCGCTGCGTGAACAACTAC</td><td></td><td></td><td></td><td></td><td></td><td></td><td></td><td></td><td></td><td></td></tr>
<tr><td>Oligo rev</td><td>AAACGTAGTTGTTCACGCAGCGCG</td><td></td><td></td><td></td><td></td><td></td><td></td><td></td><td></td><td></td><td></td></tr>
<tr><td>Chromosome</td><td>start</td><td>end</td><td>strand</td><td>MM</td><td>target_seq</td><td>PAM</td><td>alignment</td><td>distance</td><td>position</td><td>gene name</td><td>gene id</td></tr>
<tr><td>GU305914.1</td><td>125442</td><td>125464</td><td>-</td><td>0</td><td>CGCGCTGCGTGAACAACTAC</td><td>TGG</td><td>||||||||[||||||||||||]PAM</td><td>NA</td><td>Intergenic</td><td>NA</td><td>NA</td></tr>
<tr><td>T24517</td><td>TGAACAACTACTGGCGTCAGCGG</td><td>1000</td><td></td><td></td><td></td><td></td><td></td><td></td><td></td><td></td><td></td></tr>
<tr><td>Oligo fwd</td><td>CACCTGAACAACTACTGGCGTCAG</td><td></td><td></td><td></td><td></td><td></td><td></td><td></td><td></td><td></td><td></td></tr>
<tr><td>Oligo rev</td><td>AAACCTGACGCCAGTAGTTGTTCA</td><td></td><td></td><td></td><td></td><td></td><td></td><td></td><td></td><td></td><td></td></tr>
<tr><td>Chromosome</td><td>start</td><td>end</td><td>strand</td><td>MM</td><td>target_seq</td><td>PAM</td><td>alignment</td><td>distance</td><td>position</td><td>gene name</td><td>gene id</td></tr>
<tr><td>GU305914.1</td><td>125433</td><td>125455</td><td>-</td><td>0</td><td>TGAACAACTACTGGCGTCAG</td><td>CGG</td><td>||||||||[||||||||||||]PAM</td><td>NA</td><td>Intergenic</td><td>NA</td><td>NA</td></tr>
<tr><td>T24518</td><td>GAACAACTACTGGCGTCAGCGGG</td><td>1000</td><td></td><td></td><td></td><td></td><td></td><td></td><td></td><td></td><td></td></tr>
<tr><td>Oligo fwd</td><td>CACCGAACAACTACTGGCGTCAGC</td><td></td><td></td><td></td><td></td><td></td><td></td><td></td><td></td><td></td><td></td></tr>
<tr><td>Oligo rev</td><td>AAACGCTGACGCCAGTAGTTGTTC</td><td></td><td></td><td></td><td></td><td></td><td></td><td></td><td></td><td></td><td></td></tr>
<tr><td>Chromosome</td><td>start</td><td>end</td><td>strand</td><td>MM</td><td>target_seq</td><td>PAM</td><td>alignment</td><td>distance</td><td>position</td><td>gene name</td><td>gene id</td></tr>
<tr><td>GU305914.1</td><td>125432</td><td>125454</td><td>-</td><td>0</td><td>GAACAACTACTGGCGTCAGC</td><td>GGG</td><td>||||||||[||||||||||||]PAM</td><td>NA</td><td>Intergenic</td><td>NA</td><td>NA</td></tr>
<tr><td>T24519</td><td>AACAACTACTGGCGTCAGCGGGG</td><td>1000</td><td></td><td></td><td></td><td></td><td></td><td></td><td></td><td></td><td></td></tr>
<tr><td>Oligo fwd</td><td>CACCAACAACTACTGGCGTCAGCG</td><td></td><td></td><td></td><td></td><td></td><td></td><td></td><td></td><td></td><td></td></tr>
<tr><td>Oligo rev</td><td>AAACCGCTGACGCCAGTAGTTGTT</td><td></td><td></td><td></td><td></td><td></td><td></td><td></td><td></td><td></td><td></td></tr>
<tr><td>Chromosome</td><td>start</td><td>end</td><td>strand</td><td>MM</td><td>target_seq</td><td>PAM</td><td>alignment</td><td>distance</td><td>position</td><td>gene name</td><td>gene id</td></tr>
<tr><td>GU305914.1</td><td>125431</td><td>125453</td><td>-</td><td>0</td><td>AACAACTACTGGCGTCAGCG</td><td>GGG</td><td>||||||||[||||||||||||]PAM</td><td>NA</td><td>Intergenic</td><td>NA</td><td>NA</td></tr>
<tr><td>T24520</td><td>CTGGCGTCAGCGGGGCCTGCTGG</td><td>1000</td><td></td><td></td><td></td><td></td><td></td><td></td><td></td><td></td><td></td></tr>
<tr><td>Oligo fwd</td><td>CACCCTGGCGTCAGCGGGGCCTGC</td><td></td><td></td><td></td><td></td><td></td><td></td><td></td><td></td><td></td><td></td></tr>
<tr><td>Oligo rev</td><td>AAACGCAGGCCCCGCTGACGCCAG</td><td></td><td></td><td></td><td></td><td></td><td></td><td></td><td></td><td></td><td></td></tr>
<tr><td>Chromosome</td><td>start</td><td>end</td><td>strand</td><td>MM</td><td>target_seq</td><td>PAM</td><td>alignment</td><td>distance</td><td>position</td><td>gene name</td><td>gene id</td></tr>
<tr><td>GU305914.1</td><td>125423</td><td>125445</td><td>-</td><td>0</td><td>CTGGCGTCAGCGGGGCCTGC</td><td>TGG</td><td>||||||||[||||||||||||]PAM</td><td>NA</td><td>Intergenic</td><td>NA</td><td>NA</td></tr>
<tr><td>T24521</td><td>GTTCGCCAACAGCTACGACATGG</td><td>1000</td><td></td><td></td><td></td><td></td><td></td><td></td><td></td><td></td><td></td></tr>
<tr><td>Oligo fwd</td><td>CACCGTTCGCCAACAGCTACGACA</td><td></td><td></td><td></td><td></td><td></td><td></td><td></td><td></td><td></td><td></td></tr>
<tr><td>Oligo rev</td><td>AAACTGTCGTAGCTGTTGGCGAAC</td><td></td><td></td><td></td><td></td><td></td><td></td><td></td><td></td><td></td><td></td></tr>
<tr><td>Chromosome</td><td>start</td><td>end</td><td>strand</td><td>MM</td><td>target_seq</td><td>PAM</td><td>alignment</td><td>distance</td><td>position</td><td>gene name</td><td>gene id</td></tr>
<tr><td>GU305914.1</td><td>125399</td><td>125421</td><td>-</td><td>0</td><td>GTTCGCCAACAGCTACGACA</td><td>TGG</td><td>||||||||[||||||||||||]PAM</td><td>NA</td><td>Intergenic</td><td>NA</td><td>NA</td></tr>
<tr><td>T24522</td><td>CCAACAGCTACGACATGGTGCGG</td><td>1000</td><td></td><td></td><td></td><td></td><td></td><td></td><td></td><td></td><td></td></tr>
<tr><td>Oligo fwd</td><td>CACCCCAACAGCTACGACATGGTG</td><td></td><td></td><td></td><td></td><td></td><td></td><td></td><td></td><td></td><td></td></tr>
<tr><td>Oligo rev</td><td>AAACCACCATGTCGTAGCTGTTGG</td><td></td><td></td><td></td><td></td><td></td><td></td><td></td><td></td><td></td><td></td></tr>
<tr><td>Chromosome</td><td>start</td><td>end</td><td>strand</td><td>MM</td><td>target_seq</td><td>PAM</td><td>alignment</td><td>distance</td><td>position</td><td>gene name</td><td>gene id</td></tr>
<tr><td>GU305914.1</td><td>125394</td><td>125416</td><td>-</td><td>0</td><td>CCAACAGCTACGACATGGTG</td><td>CGG</td><td>||||||||[||||||||||||]PAM</td><td>NA</td><td>Intergenic</td><td>NA</td><td>NA</td></tr>
<tr><td>T24523</td><td>TGGTGCGGCTGATCGCCACGCGG</td><td>1000</td><td></td><td></td><td></td><td></td><td></td><td></td><td></td><td></td><td></td></tr>
<tr><td>Oligo fwd</td><td>CACCTGGTGCGGCTGATCGCCACG</td><td></td><td></td><td></td><td></td><td></td><td></td><td></td><td></td><td></td><td></td></tr>
<tr><td>Oligo rev</td><td>AAACCGTGGCGATCAGCCGCACCA</td><td></td><td></td><td></td><td></td><td></td><td></td><td></td><td></td><td></td><td></td></tr>
<tr><td>Chromosome</td><td>start</td><td>end</td><td>strand</td><td>MM</td><td>target_seq</td><td>PAM</td><td>alignment</td><td>distance</td><td>position</td><td>gene name</td><td>gene id</td></tr>
</table>

| | | | | | | | | | | | |
|---|---|---|---|---|---|---|---|---|---|---|---|
| GU305914.1 | 125379 | 125401 | - | 0 | TGGTGCGGCTGATCGCCACG | CGG | \|\|\|\|\|\|\|\|[\|\|\|\|\|\|\|\|\|\|\|\|]PAM | NA | Intergenic | NA | NA |
| T24524 | GCGGCTGATCGCCACGCGGCTGG | 1000 | | | | | | | | | |
| Oligo fwd | CACCGCGGCTGATCGCCACGCGGC | | | | | | | | | | |
| Oligo rev | AAACGCCGCGTGGCGATCAGCCGC | | | | | | | | | | |
| Chromosome | start | end | strand | MM | target_seq | PAM | alignment | distance | position | gene name | gene id |
| GU305914.1 | 125375 | 125397 | - | 0 | GCGGCTGATCGCCACGCGGC | TGG | \|\|\|\|\|\|\|\|[\|\|\|\|\|\|\|\|\|\|\|\|]PAM | NA | Intergenic | NA | NA |
| T24525 | CGGCTGATCGCCACGCGGCTGGG | 1000 | | | | | | | | | |
| Oligo fwd | CACCCGGCTGATCGCCACGCGGCT | | | | | | | | | | |
| Oligo rev | AAACAGCCGCGTGGCGATCAGCCG | | | | | | | | | | |
| Chromosome | start | end | strand | MM | target_seq | PAM | alignment | distance | position | gene name | gene id |
| GU305914.1 | 125374 | 125396 | - | 0 | CGGCTGATCGCCACGCGGCT | GGG | \|\|\|\|\|\|\|\|[\|\|\|\|\|\|\|\|\|\|\|\|]PAM | NA | Intergenic | NA | NA |
| T24526 | ATCGCCACGCGGCTGGGCGACGG | 1000 | | | | | | | | | |
| Oligo fwd | CACCATCGCCACGCGGCTGGGCGA | | | | | | | | | | |
| Oligo rev | AAACTCGCCCAGCCGCGTGGCGAT | | | | | | | | | | |
| Chromosome | start | end | strand | MM | target_seq | PAM | alignment | distance | position | gene name | gene id |
| GU305914.1 | 125368 | 125390 | - | 0 | ATCGCCACGCGGCTGGGCGA | CGG | \|\|\|\|\|\|\|\|[\|\|\|\|\|\|\|\|\|\|\|\|]PAM | NA | Intergenic | NA | NA |
| T24527 | CACGCGGCTGGGCGACGGCGCGG | 1000 | | | | | | | | | |
| Oligo fwd | CACCCACGCGGCTGGGCGACGGCG | | | | | | | | | | |
| Oligo rev | AAACCGCCGTCGCCCAGCCGCGTG | | | | | | | | | | |
| Chromosome | start | end | strand | MM | target_seq | PAM | alignment | distance | position | gene name | gene id |
| GU305914.1 | 125363 | 125385 | - | 0 | CACGCGGCTGGGCGACGGCG | CGG | \|\|\|\|\|\|\|\|[\|\|\|\|\|\|\|\|\|\|\|\|]PAM | NA | Intergenic | NA | NA |
| T24528 | GCGGCTGGGCGACGGCGCGGTGG | 1000 | | | | | | | | | |
| Oligo fwd | CACCGCGGCTGGGCGACGGCGCGG | | | | | | | | | | |
| Oligo rev | AAACCCGCGCCGTCGCCCAGCCGC | | | | | | | | | | |
| Chromosome | start | end | strand | MM | target_seq | PAM | alignment | distance | position | gene name | gene id |
| GU305914.1 | 125360 | 125382 | - | 0 | GCGGCTGGGCGACGGCGCGG | TGG | \|\|\|\|\|\|\|\|[\|\|\|\|\|\|\|\|\|\|\|\|]PAM | NA | Intergenic | NA | NA |
| GU305914.1 | 103961 | 103983 | - | 4 | GCTGCTGCGCGACGGTCCGG | AGG | \|\|-\|\|\|\|-[\|\|\|\|\|\|\|--\|\|\|]PAM | NA | Intergenic | NA | NA |
| T24529 | GGGCGACGGCGCGGTGGTGCCGG | 1000 | | | | | | | | | |
| Oligo fwd | CACCGGGCGACGGCGCGGTGGTGC | | | | | | | | | | |
| Oligo rev | AAACGCACCACCGCGCCGTCGCCC | | | | | | | | | | |
| Chromosome | start | end | strand | MM | target_seq | PAM | alignment | distance | position | gene name | gene id |
| GU305914.1 | 125354 | 125376 | - | 0 | GGGCGACGGCGCGGTGGTGC | CGG | \|\|\|\|\|\|\|\|[\|\|\|\|\|\|\|\|\|\|\|\|]PAM | NA | Intergenic | NA | NA |
| GU305914.1 | 60453 | 60475 | - | 4 | GGTCCGCGACGCGGTGGTGC | TGG | \|\|-\|--\|\|[-\|\|\|\|\|\|\|\|\|\|\|]PAM | NA | Intergenic | NA | NA |
| GU305914.1 | 57528 | 57550 | + | 4 | TGGCGATGGCGCGGCGGTGG | TGG | -\|\|\|\|\|-\|[\|\|\|\|\|\|-\|\|\|\|-]PAM | NA | Intergenic | NA | NA |
| T24530 | CGACGGCGCGGTGGTGCCGGCGG | 1000 | | | | | | | | | |
| Oligo fwd | CACCCGACGGCGCGGTGGTGCCGG | | | | | | | | | | |
| Oligo rev | AAACCCGGCACCACCGCGCCGTCG | | | | | | | | | | |
| Chromosome | start | end | strand | MM | target_seq | PAM | alignment | distance | position | gene name | gene id |
| GU305914.1 | 125351 | 125373 | - | 0 | CGACGGCGCGGTGGTGCCGG | CGG | \|\|\|\|\|\|\|\|[\|\|\|\|\|\|\|\|\|\|\|\|]PAM | NA | Intergenic | NA | NA |
| T24531 | GCAACCTGCTGTCCATCACACGG | 1000 | | | | | | | | | |
| Oligo fwd | CACCGCAACCTGCTGTCCATCACA | | | | | | | | | | |
| Oligo rev | AAACTGTGATGGACAGCAGGTTGC | | | | | | | | | | |
| Chromosome | start | end | strand | MM | target_seq | PAM | alignment | distance | position | gene name | gene id |
| GU305914.1 | 125313 | 125335 | - | 0 | GCAACCTGCTGTCCATCACA | CGG | \|\|\|\|\|\|\|\|[\|\|\|\|\|\|\|\|\|\|\|\|]PAM | NA | Intergenic | NA | NA |
| T24532 | GCTGTCCATCACACGGTTCGTGG | 1000 | | | | | | | | | |
| Oligo fwd | CACCGCTGTCCATCACACGGTTCG | | | | | | | | | | |
| Oligo rev | AAACCGAACCGTGTGATGGACAGC | | | | | | | | | | |
| Chromosome | start | end | strand | MM | target_seq | PAM | alignment | distance | position | gene name | gene id |
| GU305914.1 | 125306 | 125328 | - | 0 | GCTGTCCATCACACGGTTCG | TGG | \|\|\|\|\|\|\|\|[\|\|\|\|\|\|\|\|\|\|\|\|]PAM | NA | Intergenic | NA | NA |
| T24533 | CCATCACACGGTTCGTGGCGCGG | 1000 | | | | | | | | | |
| Oligo fwd | CACCCCATCACACGGTTCGTGGCG | | | | | | | | | | |
| Oligo rev | AAACCGCCACGAACCGTGTGATGG | | | | | | | | | | |
| Chromosome | start | end | strand | MM | target_seq | PAM | alignment | distance | position | gene name | gene id |
| GU305914.1 | 125301 | 125323 | - | 0 | CCATCACACGGTTCGTGGCG | CGG | \|\|\|\|\|\|\|\|[\|\|\|\|\|\|\|\|\|\|\|\|]PAM | NA | Intergenic | NA | NA |
| T24534 | GGCGCGGACCTGCGAACTGACGG | 1000 | | | | | | | | | |
| Oligo fwd | CACCGGCGCGGACCTGCGAACTGA | | | | | | | | | | |
| Oligo rev | AAACTCAGTTCGCAGGTCCGCGCC | | | | | | | | | | |
| Chromosome | start | end | strand | MM | target_seq | PAM | alignment | distance | position | gene name | gene id |
| GU305914.1 | 125285 | 125307 | - | 0 | GGCGCGGACCTGCGAACTGA | CGG | \|\|\|\|\|\|\|\|[\|\|\|\|\|\|\|\|\|\|\|\|]PAM | NA | Intergenic | NA | NA |
| T24535 | GCGCGGACCTGCGAACTGACGGG | 1000 | | | | | | | | | |
| Oligo fwd | CACCGCGCGGACCTGCGAACTGAC | | | | | | | | | | |
| Oligo rev | AAACGTCAGTTCGCAGGTCCGCGC | | | | | | | | | | |
| Chromosome | start | end | strand | MM | target_seq | PAM | alignment | distance | position | gene name | gene id |
| GU305914.1 | 125284 | 125306 | - | 0 | GCGCGGACCTGCGAACTGAC | GGG | \|\|\|\|\|\|\|\|[\|\|\|\|\|\|\|\|\|\|\|\|]PAM | NA | Intergenic | NA | NA |
| T24536 | TGCGAACTGACGGGCCTGAACGG | 1000 | | | | | | | | | |
| Oligo fwd | CACCTGCGAACTGACGGGCCTGAA | | | | | | | | | | |
| Oligo rev | AAACTTCAGGCCCGTCAGTTCGCA | | | | | | | | | | |
| Chromosome | start | end | strand | MM | target_seq | PAM | alignment | distance | position | gene name | gene id |
| GU305914.1 | 125275 | 125297 | - | 0 | TGCGAACTGACGGGCCTGAA | CGG | \|\|\|\|\|\|\|\|[\|\|\|\|\|\|\|\|\|\|\|\|]PAM | NA | Intergenic | NA | NA |
| T24537 | GCGAACTGACGGGCCTGAACGGG | 1000 | | | | | | | | | |
| Oligo fwd | CACCGCGAACTGACGGGCCTGAAC | | | | | | | | | | |
| Oligo rev | AAACGTTCAGGCCCGTCAGTTCGC | | | | | | | | | | |
| Chromosome | start | end | strand | MM | target_seq | PAM | alignment | distance | position | gene name | gene id |
| GU305914.1 | 125274 | 125296 | - | 0 | GCGAACTGACGGGCCTGAAC | GGG | \|\|\|\|\|\|\|\|[\|\|\|\|\|\|\|\|\|\|\|\|]PAM | NA | Intergenic | NA | NA |

T24538 AACTGACGGGCCTGAACGGGCGG 1000
Oligo fwd CACCAACTGACGGGCCTGAACGGG
Oligo rev AAACCCCGTTCAGGCCCGTCAGTT
Chromosome start end strand MM target_seq PAM alignment distance position gene name gene id
GU305914.1 125271 125293 - 0 AACTGACGGGCCTGAACGGG CGG ||||||||[||||||||||||]PAM NA Intergenic NA NA

T24539 GCTGTGTGACGAGCCGCTGCTGG 1000
Oligo fwd CACCGCTGTGTGACGAGCCGCTGC
Oligo rev AAACGCAGCGGCTCGTCACACAGC
Chromosome start end strand MM target_seq PAM alignment distance position gene name gene id
GU305914.1 125249 125271 - 0 GCTGTGTGACGAGCCGCTGC TGG ||||||||[||||||||||||]PAM NA Intergenic NA NA

T24540 CGCTGCACGACCCGCGCCTCTGG 1000
Oligo fwd CACCCGCTGCACGACCCGCGCCTC
Oligo rev AAACGAGGCGCGGGTCGTGCAGCG
Chromosome start end strand MM target_seq PAM alignment distance position gene name gene id
GU305914.1 125214 125236 - 0 CGCTGCACGACCCGCGCCTC TGG ||||||||[||||||||||||]PAM NA Intergenic NA NA

T24541 CCTCTGGCCGCCCTTCGTGCAGG 1000
Oligo fwd CACCCCTCTGGCCGCCCTTCGTGC
Oligo rev AAACGCACGAAGGGCGGCCAGAGG
Chromosome start end strand MM target_seq PAM alignment distance position gene name gene id
GU305914.1 125198 125220 - 0 CCTCTGGCCGCCCTTCGTGC AGG ||||||||[||||||||||||]PAM NA Intergenic NA NA

T24542 GCTGCCGCGCAACGCGAACCTGG 1000
Oligo fwd CACCGCTGCCGCGCAACGCGAACC
Oligo rev AAACGGTTCGCGTTGCGCGGCAGC
Chromosome start end strand MM target_seq PAM alignment distance position gene name gene id
GU305914.1 125174 125196 - 0 GCTGCCGCGCAACGCGAACC TGG ||||||||[||||||||||||]PAM NA Intergenic NA NA

T24543 CAACGCGAACCTGGTGCGCGTGG 1000
Oligo fwd CACCCAACGCGAACCTGGTGCGCG
Oligo rev AAACCGCGCACCAGGTTCGCGTTG
Chromosome start end strand MM target_seq PAM alignment distance position gene name gene id
GU305914.1 125165 125187 - 0 CAACGCGAACCTGGTGCGCG TGG ||||||||[||||||||||||]PAM NA Intergenic NA NA

T24544 CGCGAACCTGGTGCGCGTGGTGG 1000
Oligo fwd CACCCGCGAACCTGGTGCGCGTGG
Oligo rev AAACCCACGCGCACCAGGTTCGCG
Chromosome start end strand MM target_seq PAM alignment distance position gene name gene id
GU305914.1 125162 125184 - 0 CGCGAACCTGGTGCGCGTGG TGG ||||||||[||||||||||||]PAM NA Intergenic NA NA

T24545 GGTGGCCGACGACGTGCCGCTGG 1000
Oligo fwd CACCGGTGGCCGACGACGTGCCGC
Oligo rev AAACGCGGCACGTCGTCGGCCACC
Chromosome start end strand MM target_seq PAM alignment distance position gene name gene id
GU305914.1 125144 125166 - 0 GGTGGCCGACGACGTGCCGC TGG ||||||||[||||||||||||]PAM NA Intergenic NA NA
GU305914.1 116174 116196 - 4 GGACGCCGACAACGTGCCGT CGG ||--||||[||-||||||||-]PAM NA Intergenic NA NA

T24546 CGACGACGTGCCGCTGGACGCGG 1000
Oligo fwd CACCCGACGACGTGCCGCTGGACG
Oligo rev AAACCGTCCAGCGGCACGTCGTCG
Chromosome start end strand MM target_seq PAM alignment distance position gene name gene id
GU305914.1 125138 125160 - 0 CGACGACGTGCCGCTGGACG CGG ||||||||[||||||||||||]PAM NA Intergenic NA NA
GU305914.1 147663 147685 - 3 CGCCGTCGTGCTGCTGGACG CGG ||-||-||[|||-||||||||]PAM NA Intergenic NA NA

T24547 GCTGGACGCGGCGCACATCGAGG 1000
Oligo fwd CACCGCTGGACGCGGCGCACATCG
Oligo rev AAACCGATGTGCGCCGCGTCCAGC
Chromosome start end strand MM target_seq PAM alignment distance position gene name gene id
GU305914.1 125126 125148 - 0 GCTGGACGCGGCGCACATCG AGG ||||||||[||||||||||||]PAM NA Intergenic NA NA

T24548 GCACATCGAGGAGCGCAACCCGG 1000
Oligo fwd CACCGCACATCGAGGAGCGCAACC
Oligo rev AAACGGTTGCGCTCCTCGATGTGC
Chromosome start end strand MM target_seq PAM alignment distance position gene name gene id
GU305914.1 125114 125136 - 0 GCACATCGAGGAGCGCAACC CGG ||||||||[||||||||||||]PAM NA Intergenic NA NA
GU305914.1 198935 198957 - 4 GCACCTGGTGGCGCGCAACC TGG ||||-|-|[-||-||||||||]PAM NA Intergenic NA NA

T24549 CACATCGAGGAGCGCAACCCGGG 1000
Oligo fwd CACCCACATCGAGGAGCGCAACCC
Oligo rev AAACGGGTTGCGCTCCTCGATGTG
Chromosome start end strand MM target_seq PAM alignment distance position gene name gene id
GU305914.1 125113 125135 - 0 CACATCGAGGAGCGCAACCC GGG ||||||||[||||||||||||]PAM NA Intergenic NA NA

T24550 GGAGCGCAACCCGGGCACCTCGG 1000
Oligo fwd CACCGGAGCGCAACCCGGGCACCT
Oligo rev AAACAGGTGCCCGGGTTGCGCTCC
Chromosome start end strand MM target_seq PAM alignment distance position gene name gene id
GU305914.1 125105 125127 - 0 GGAGCGCAACCCGGGCACCT CGG ||||||||[||||||||||||]PAM NA Intergenic NA NA

T24551 CAACCCGGGCACCTCGGACGTGG 1000
Oligo fwd CACCCAACCCGGGCACCTCGGACG
Oligo rev AAACCGTCCGAGGTGCCCGGGTTG
Chromosome start end strand MM target_seq PAM alignment distance position gene name gene id
GU305914.1 125099 125121 - 0 CAACCCGGGCACCTCGGACG TGG ||||||||[||||||||||||]PAM NA Intergenic NA NA

T24552 CGGGCACCTCGGACGTGGCGCGG 1000
Oligo fwd CACCCGGGCACCTCGGACGTGGCG

| | | | | | | | | | | | |
|---|---|---|---|---|---|---|---|---|---|---|---|
| Oligo rev | AAACCGCCACGTCCGAGGTGCCCG | | | | | | | | | | |
| Chromosome | start | end | strand | MM | target_seq | PAM | alignment | distance | position | gene name | gene id |
| GU305914.1 | 125094 | 125116 | - | 0 | CGGGCACCTCGGACGTGGCG | CGG | ||||||||[||||||||||||]PAM | NA | Intergenic | NA | NA |
| T24553 | CGTGGCGCGGATGATCGCGATGG | 1000 | | | | | | | | | |
| Oligo fwd | CACCCGTGGCGCGGATGATCGCGA | | | | | | | | | | |
| Oligo rev | AAACTCGCGATCATCCGCGCCACG | | | | | | | | | | |
| Chromosome | start | end | strand | MM | target_seq | PAM | alignment | distance | position | gene name | gene id |
| GU305914.1 | 125081 | 125103 | - | 0 | CGTGGCGCGGATGATCGCGA | TGG | ||||||||[||||||||||||]PAM | NA | Intergenic | NA | NA |
| T24554 | GCGGATGATCGCGATGGACCAGG | 1000 | | | | | | | | | |
| Oligo fwd | CACCGCGGATGATCGCGATGGACC | | | | | | | | | | |
| Oligo rev | AAACGGTCCATCGCGATCATCCGC | | | | | | | | | | |
| Chromosome | start | end | strand | MM | target_seq | PAM | alignment | distance | position | gene name | gene id |
| GU305914.1 | 125075 | 125097 | - | 0 | GCGGATGATCGCGATGGACC | AGG | ||||||||[||||||||||||]PAM | NA | Intergenic | NA | NA |
| T24555 | GATGATCGCGATGGACCAGGCGG | 1000 | | | | | | | | | |
| Oligo fwd | CACCGATGATCGCGATGGACCAGG | | | | | | | | | | |
| Oligo rev | AAACCCTGGTCCATCGCGATCATC | | | | | | | | | | |
| Chromosome | start | end | strand | MM | target_seq | PAM | alignment | distance | position | gene name | gene id |
| GU305914.1 | 125072 | 125094 | - | 0 | GATGATCGCGATGGACCAGG | CGG | ||||||||[||||||||||||]PAM | NA | Intergenic | NA | NA |
| T24556 | CCAGGCGGAGCCGCTGTTCGTGG | 1000 | | | | | | | | | |
| Oligo fwd | CACCCCAGGCGGAGCCGCTGTTCG | | | | | | | | | | |
| Oligo rev | AAACCGAACAGCGGCTCCGCCTGG | | | | | | | | | | |
| Chromosome | start | end | strand | MM | target_seq | PAM | alignment | distance | position | gene name | gene id |
| GU305914.1 | 125057 | 125079 | - | 0 | CCAGGCGGAGCCGCTGTTCG | TGG | ||||||||[||||||||||||]PAM | NA | Intergenic | NA | NA |
| T24557 | AGCCGCTGTTCGTGGACGCCCGG | 1000 | | | | | | | | | |
| Oligo fwd | CACCAGCCGCTGTTCGTGGACGCC | | | | | | | | | | |
| Oligo rev | AAACGGCGTCCACGAACAGCGGCT | | | | | | | | | | |
| Chromosome | start | end | strand | MM | target_seq | PAM | alignment | distance | position | gene name | gene id |
| GU305914.1 | 125049 | 125071 | - | 0 | AGCCGCTGTTCGTGGACGCC | CGG | ||||||||[||||||||||||]PAM | NA | Intergenic | NA | NA |
| T24558 | CGCTGTTCGTGGACGCCCGGCGG | 1000 | | | | | | | | | |
| Oligo fwd | CACCCGCTGTTCGTGGACGCCCGG | | | | | | | | | | |
| Oligo rev | AAACCCGGGCGTCCACGAACAGCG | | | | | | | | | | |
| Chromosome | start | end | strand | MM | target_seq | PAM | alignment | distance | position | gene name | gene id |
| GU305914.1 | 125046 | 125068 | - | 0 | CGCTGTTCGTGGACGCCCGG | CGG | ||||||||[||||||||||||]PAM | NA | Intergenic | NA | NA |
| T24559 | CGTGGACGCCCGGCGGACGTCGG | 1000 | | | | | | | | | |
| Oligo fwd | CACCCGTGGACGCCCGGCGGACGT | | | | | | | | | | |
| Oligo rev | AAACACGTCCGCCGGGCGTCCACG | | | | | | | | | | |
| Chromosome | start | end | strand | MM | target_seq | PAM | alignment | distance | position | gene name | gene id |
| GU305914.1 | 125039 | 125061 | - | 0 | CGTGGACGCCCGGCGGACGT | CGG | ||||||||[||||||||||||]PAM | NA | Intergenic | NA | NA |
| T24560 | CGCCCGGCGGACGTCGGACGAGG | 1000 | | | | | | | | | |
| Oligo fwd | CACCCGCCCGGCGGACGTCGGACG | | | | | | | | | | |
| Oligo rev | AAACCGTCCGACGTCCGCCGGGCG | | | | | | | | | | |
| Chromosome | start | end | strand | MM | target_seq | PAM | alignment | distance | position | gene name | gene id |
| GU305914.1 | 125033 | 125055 | - | 0 | CGCCCGGCGGACGTCGGACG | AGG | ||||||||[||||||||||||]PAM | NA | Intergenic | NA | NA |
| T24561 | GCGGACGTCGGACGAGGAGATGG | 1000 | | | | | | | | | |
| Oligo fwd | CACCGCGGACGTCGGACGAGGAGA | | | | | | | | | | |
| Oligo rev | AAACTCTCCTCGTCCGACGTCCGC | | | | | | | | | | |
| Chromosome | start | end | strand | MM | target_seq | PAM | alignment | distance | position | gene name | gene id |
| GU305914.1 | 125027 | 125049 | - | 0 | GCGGACGTCGGACGAGGAGA | TGG | ||||||||[||||||||||||]PAM | NA | Intergenic | NA | NA |
| T24562 | GACGTCGGACGAGGAGATGGTGG | 1000 | | | | | | | | | |
| Oligo fwd | CACCGACGTCGGACGAGGAGATGG | | | | | | | | | | |
| Oligo rev | AAACCCATCTCCTCGTCCGACGTC | | | | | | | | | | |
| Chromosome | start | end | strand | MM | target_seq | PAM | alignment | distance | position | gene name | gene id |
| GU305914.1 | 125024 | 125046 | - | 0 | GACGTCGGACGAGGAGATGG | TGG | ||||||||[||||||||||||]PAM | NA | Intergenic | NA | NA |
| GU305914.1 | 67767 | 67789 | - | 4 | GGCGGCGGAAGAAGAGATGG | AGG | |-||-|||[|-||-|||||||]PAM | NA | Intergenic | NA | NA |
| T24563 | CGAGGAGATGGTGGCGCAGAAGG | 1000 | | | | | | | | | |
| Oligo fwd | CACCCGAGGAGATGGTGGCGCAGA | | | | | | | | | | |
| Oligo rev | AAACTCTGCGCCACCATCTCCTCG | | | | | | | | | | |
| Chromosome | start | end | strand | MM | target_seq | PAM | alignment | distance | position | gene name | gene id |
| GU305914.1 | 125015 | 125037 | - | 0 | CGAGGAGATGGTGGCGCAGA | AGG | ||||||||[||||||||||||]PAM | NA | Intergenic | NA | NA |
| T24564 | GAAGGTATACTACCTGTGCCTGG | 1000 | | | | | | | | | |
| Oligo fwd | CACCGAAGGTATACTACCTGTGCC | | | | | | | | | | |
| Oligo rev | AAACGGCACAGGTAGTATACCTTC | | | | | | | | | | |
| Chromosome | start | end | strand | MM | target_seq | PAM | alignment | distance | position | gene name | gene id |
| GU305914.1 | 124997 | 125019 | - | 0 | GAAGGTATACTACCTGTGCC | TGG | ||||||||[||||||||||||]PAM | NA | Intergenic | NA | NA |
| T24565 | GTGCTGAACAACCACGCGTGCGG | 1000 | | | | | | | | | |
| Oligo fwd | CACCGTGCTGAACAACCACGCGTG | | | | | | | | | | |
| Oligo rev | AAACCACGCGTGGTTGTTCAGCAC | | | | | | | | | | |
| Chromosome | start | end | strand | MM | target_seq | PAM | alignment | distance | position | gene name | gene id |
| GU305914.1 | 124966 | 124988 | - | 0 | GTGCTGAACAACCACGCGTG | CGG | ||||||||[||||||||||||]PAM | NA | Intergenic | NA | NA |
| T24566 | GAACAACCACGCGTGCGGCGCGG | 1000 | | | | | | | | | |
| Oligo fwd | CACCGAACAACCACGCGTGCGGCG | | | | | | | | | | |
| Oligo rev | AAACCGCCGCACGCGTGGTTGTTC | | | | | | | | | | |
| Chromosome | start | end | strand | MM | target_seq | PAM | alignment | distance | position | gene name | gene id |
| GU305914.1 | 124961 | 124983 | - | 0 | GAACAACCACGCGTGCGGCG | CGG | ||||||||[||||||||||||]PAM | NA | Intergenic | NA | NA |

T24567 AACAACCACGCGTGCGGCGCGGG 1000
Oligo fwd CACCAACAACCACGCGTGCGGCGC
Oligo rev AAACGCGCCGCACGCGTGGTTGTT
Chromosome start end strand MM target_seq PAM alignment distance position gene name gene id
GU305914.1 124960 124982 - 0 AACAACCACGCGTGCGGCGC GGG ||||||||[||||||||||||]PAM NA Intergenic NA NA

T24568 CCTGAACCTGAAGCACCTGCTGG 1000
Oligo fwd CACCCCTGAACCTGAAGCACCTGC
Oligo rev AAACGCAGGTGCTTCAGGTTCAGG
Chromosome start end strand MM target_seq PAM alignment distance position gene name gene id
GU305914.1 124937 124959 - 0 CCTGAACCTGAAGCACCTGC TGG ||||||||[||||||||||||]PAM NA Intergenic NA NA

T24569 CTACACCAAGTTCTTCCTGACGG 1000
Oligo fwd CACCCTACACCAAGTTCTTCCTGA
Oligo rev AAACTCAGGAAGAACTTGGTGTAG
Chromosome start end strand MM target_seq PAM alignment distance position gene name gene id
GU305914.1 124904 124926 - 0 CTACACCAAGTTCTTCCTGA CGG ||||||||[||||||||||||]PAM NA Intergenic NA NA

T24570 CACCAAGTTCTTCCTGACGGCGG 1000
Oligo fwd CACCCACCAAGTTCTTCCTGACGG
Oligo rev AAACCCGTCAGGAAGAACTTGGTG
Chromosome start end strand MM target_seq PAM alignment distance position gene name gene id
GU305914.1 124901 124923 - 0 CACCAAGTTCTTCCTGACGG CGG ||||||||[||||||||||||]PAM NA Intergenic NA NA

T24571 GTTCTTCCTGACGGCGGACCCGG 1000
Oligo fwd CACCGTTCTTCCTGACGGCGGACC
Oligo rev AAACGGTCCGCCGTCAGGAAGAAC
Chromosome start end strand MM target_seq PAM alignment distance position gene name gene id
GU305914.1 124895 124917 - 0 GTTCTTCCTGACGGCGGACC CGG ||||||||[||||||||||||]PAM NA Intergenic NA NA

T24572 GGCGGACCCGGACTCGCTGACGG 1000
Oligo fwd CACCGGCGGACCCGGACTCGCTGA
Oligo rev AAACTCAGCGAGTCCGGGTCCGCC
Chromosome start end strand MM target_seq PAM alignment distance position gene name gene id
GU305914.1 124883 124905 - 0 GGCGGACCCGGACTCGCTGA CGG ||||||||[||||||||||||]PAM NA Intergenic NA NA

T24573 GACCCGGACTCGCTGACGGCCGG 1000
Oligo fwd CACCGACCCGGACTCGCTGACGGC
Oligo rev AAACGCCGTCAGCGAGTCCGGGTC
Chromosome start end strand MM target_seq PAM alignment distance position gene name gene id
GU305914.1 124879 124901 - 0 GACCCGGACTCGCTGACGGC CGG ||||||||[||||||||||||]PAM NA Intergenic NA NA

T24574 GGACTCGCTGACGGCCGGCGAGG 1000
Oligo fwd CACCGGACTCGCTGACGGCCGGCG
Oligo rev AAACCGCCGGCCGTCAGCGAGTCC
Chromosome start end strand MM target_seq PAM alignment distance position gene name gene id
GU305914.1 124874 124896 - 0 GGACTCGCTGACGGCCGGCG AGG ||||||||[||||||||||||]PAM NA Intergenic NA NA

T24575 CTCGCTGACGGCCGGCGAGGAGG 1000
Oligo fwd CACCCTCGCTGACGGCCGGCGAGG
Oligo rev AAACCCTCGCCGGCCGTCAGCGAG
Chromosome start end strand MM target_seq PAM alignment distance position gene name gene id
GU305914.1 124871 124893 - 0 CTCGCTGACGGCCGGCGAGG AGG ||||||||[||||||||||||]PAM NA Intergenic NA NA

T24576 GCTGACCAACAACCCGCTGCTGG 1000
Oligo fwd CACCGCTGACCAACAACCCGCTGC
Oligo rev AAACGCAGCGGGTTGTTGGTCAGC
Chromosome start end strand MM target_seq PAM alignment distance position gene name gene id
GU305914.1 124847 124869 - 0 GCTGACCAACAACCCGCTGC TGG ||||||||[||||||||||||]PAM NA Intergenic NA NA

T24577 CAACCCGCTGCTGGCCGCGCTGG 1000
Oligo fwd CACCCAACCCGCTGCTGGCCGCGC
Oligo rev AAACGCGCGGCCAGCAGCGGGTTG
Chromosome start end strand MM target_seq PAM alignment distance position gene name gene id
GU305914.1 124838 124860 - 0 CAACCCGCTGCTGGCCGCGC TGG ||||||||[||||||||||||]PAM NA Intergenic NA NA

T24578 CGCTGCTGGCCGCGCTGGTGCGG 1000
Oligo fwd CACCCGCTGCTGGCCGCGCTGGTG
Oligo rev AAACCACCAGCGCGGCCAGCAGCG
Chromosome start end strand MM target_seq PAM alignment distance position gene name gene id
GU305914.1 124833 124855 - 0 CGCTGCTGGCCGCGCTGGTG CGG ||||||||[||||||||||||]PAM NA Intergenic NA NA

T24579 GCTGCTGGCCGCGCTGGTGCGGG 1000
Oligo fwd CACCGCTGCTGGCCGCGCTGGTGC
Oligo rev AAACGCACCAGCGCGGCCAGCAGC
Chromosome start end strand MM target_seq PAM alignment distance position gene name gene id
GU305914.1 124832 124854 - 0 GCTGCTGGCCGCGCTGGTGC GGG ||||||||[||||||||||||]PAM NA Intergenic NA NA

T24580 GGCCGCGCTGGTGCGGGACGTGG 1000
Oligo fwd CACCGGCCGCGCTGGTGCGGGACG
Oligo rev AAACCGTCCCGCACCAGCGCGGCC
Chromosome start end strand MM target_seq PAM alignment distance position gene name gene id
GU305914.1 124826 124848 - 0 GGCCGCGCTGGTGCGGGACG TGG ||||||||[||||||||||||]PAM NA Intergenic NA NA
GU305914.1 201464 201486 - 4 GTCGGCGCTGCTGCGTGACG AGG |-|-||||[||-||||-||||]PAM NA Intergenic NA NA

T24581 GCTGGTGCGGGACGTGGCGACGG 1000
Oligo fwd CACCGCTGGTGCGGGACGTGGCGA
Oligo rev AAACTCGCCACGTCCCGCACCAGC
Chromosome start end strand MM target_seq PAM alignment distance position gene name gene id

<table>
<tr><td>GU305914.1</td><td>124820</td><td>124842</td><td>-</td><td>0</td><td>GCTGGTGCGGGACGTGGCGA</td><td>CGG</td><td>||||||||[||||||||||||]PAM</td><td>NA</td><td>Intergenic</td><td>NA</td><td>NA</td></tr>
<tr><td>T24582</td><td>GGCGACGGACGAGAACGTGACGG</td><td>1000</td><td></td><td></td><td></td><td></td><td></td><td></td><td></td><td></td><td></td></tr>
<tr><td>Oligo fwd</td><td>CACCGGCGACGGACGAGAACGTGA</td><td></td><td></td><td></td><td></td><td></td><td></td><td></td><td></td><td></td><td></td></tr>
<tr><td>Oligo rev</td><td>AAACTCACGTTCTCGTCCGTCGCC</td><td></td><td></td><td></td><td></td><td></td><td></td><td></td><td></td><td></td><td></td></tr>
<tr><td>Chromosome</td><td>start</td><td>end</td><td>strand</td><td>MM</td><td>target_seq</td><td>PAM</td><td>alignment</td><td>distance</td><td>position</td><td>gene name</td><td>gene id</td></tr>
<tr><td>GU305914.1</td><td>124805</td><td>124827</td><td>-</td><td>0</td><td>GGCGACGGACGAGAACGTGA</td><td>CGG</td><td>||||||||[||||||||||||]PAM</td><td>NA</td><td>Intergenic</td><td>NA</td><td>NA</td></tr>
<tr><td>T24583</td><td>CGAGAACGTGACGGCGAACCAGG</td><td>1000</td><td></td><td></td><td></td><td></td><td></td><td></td><td></td><td></td><td></td></tr>
<tr><td>Oligo fwd</td><td>CACCCGAGAACGTGACGGCGAACC</td><td></td><td></td><td></td><td></td><td></td><td></td><td></td><td></td><td></td><td></td></tr>
<tr><td>Oligo rev</td><td>AAACGGTTCGCCGTCACGTTCTCG</td><td></td><td></td><td></td><td></td><td></td><td></td><td></td><td></td><td></td><td></td></tr>
<tr><td>Chromosome</td><td>start</td><td>end</td><td>strand</td><td>MM</td><td>target_seq</td><td>PAM</td><td>alignment</td><td>distance</td><td>position</td><td>gene name</td><td>gene id</td></tr>
<tr><td>GU305914.1</td><td>124796</td><td>124818</td><td>-</td><td>0</td><td>CGAGAACGTGACGGCGAACC</td><td>AGG</td><td>||||||||[||||||||||||]PAM</td><td>NA</td><td>Intergenic</td><td>NA</td><td>NA</td></tr>
<tr><td>T24584</td><td>GAACGTGACGGCGAACCAGGCGG</td><td>1000</td><td></td><td></td><td></td><td></td><td></td><td></td><td></td><td></td><td></td></tr>
<tr><td>Oligo fwd</td><td>CACCGAACGTGACGGCGAACCAGG</td><td></td><td></td><td></td><td></td><td></td><td></td><td></td><td></td><td></td><td></td></tr>
<tr><td>Oligo rev</td><td>AAACCCTGGTTCGCCGTCACGTTC</td><td></td><td></td><td></td><td></td><td></td><td></td><td></td><td></td><td></td><td></td></tr>
<tr><td>Chromosome</td><td>start</td><td>end</td><td>strand</td><td>MM</td><td>target_seq</td><td>PAM</td><td>alignment</td><td>distance</td><td>position</td><td>gene name</td><td>gene id</td></tr>
<tr><td>GU305914.1</td><td>124793</td><td>124815</td><td>-</td><td>0</td><td>GAACGTGACGGCGAACCAGG</td><td>CGG</td><td>||||||||[||||||||||||]PAM</td><td>NA</td><td>Intergenic</td><td>NA</td><td>NA</td></tr>
<tr><td>T24585</td><td>GACGGCGAACCAGGCGGCCGAGG</td><td>1000</td><td></td><td></td><td></td><td></td><td></td><td></td><td></td><td></td><td></td></tr>
<tr><td>Oligo fwd</td><td>CACCGACGGCGAACCAGGCGGCCG</td><td></td><td></td><td></td><td></td><td></td><td></td><td></td><td></td><td></td><td></td></tr>
<tr><td>Oligo rev</td><td>AAACCGGCCGCCTGGTTCGCCGTC</td><td></td><td></td><td></td><td></td><td></td><td></td><td></td><td></td><td></td><td></td></tr>
<tr><td>Chromosome</td><td>start</td><td>end</td><td>strand</td><td>MM</td><td>target_seq</td><td>PAM</td><td>alignment</td><td>distance</td><td>position</td><td>gene name</td><td>gene id</td></tr>
<tr><td>GU305914.1</td><td>124787</td><td>124809</td><td>-</td><td>0</td><td>GACGGCGAACCAGGCGGCCG</td><td>AGG</td><td>||||||||[||||||||||||]PAM</td><td>NA</td><td>Intergenic</td><td>NA</td><td>NA</td></tr>
<tr><td>T24586</td><td>GGCCGAGGAGCTGTTCCACCTGG</td><td>1000</td><td></td><td></td><td></td><td></td><td></td><td></td><td></td><td></td><td></td></tr>
<tr><td>Oligo fwd</td><td>CACCGGCCGAGGAGCTGTTCCACC</td><td></td><td></td><td></td><td></td><td></td><td></td><td></td><td></td><td></td><td></td></tr>
<tr><td>Oligo rev</td><td>AAACGGTGGAACAGCTCCTCGGCC</td><td></td><td></td><td></td><td></td><td></td><td></td><td></td><td></td><td></td><td></td></tr>
<tr><td>Chromosome</td><td>start</td><td>end</td><td>strand</td><td>MM</td><td>target_seq</td><td>PAM</td><td>alignment</td><td>distance</td><td>position</td><td>gene name</td><td>gene id</td></tr>
<tr><td>GU305914.1</td><td>124772</td><td>124794</td><td>-</td><td>0</td><td>GGCCGAGGAGCTGTTCCACC</td><td>TGG</td><td>||||||||[||||||||||||]PAM</td><td>NA</td><td>Intergenic</td><td>NA</td><td>NA</td></tr>
<tr><td>T24587</td><td>CGAGGAGCTGTTCCACCTGGTGG</td><td>1000</td><td></td><td></td><td></td><td></td><td></td><td></td><td></td><td></td><td></td></tr>
<tr><td>Oligo fwd</td><td>CACCCGAGGAGCTGTTCCACCTGG</td><td></td><td></td><td></td><td></td><td></td><td></td><td></td><td></td><td></td><td></td></tr>
<tr><td>Oligo rev</td><td>AAACCCAGGTGGAACAGCTCCTCG</td><td></td><td></td><td></td><td></td><td></td><td></td><td></td><td></td><td></td><td></td></tr>
<tr><td>Chromosome</td><td>start</td><td>end</td><td>strand</td><td>MM</td><td>target_seq</td><td>PAM</td><td>alignment</td><td>distance</td><td>position</td><td>gene name</td><td>gene id</td></tr>
<tr><td>GU305914.1</td><td>124769</td><td>124791</td><td>-</td><td>0</td><td>CGAGGAGCTGTTCCACCTGG</td><td>TGG</td><td>||||||||[||||||||||||]PAM</td><td>NA</td><td>Intergenic</td><td>NA</td><td>NA</td></tr>
<tr><td>T24588</td><td>GCCCGAGAACGCGCAGATGCTGG</td><td>1000</td><td></td><td></td><td></td><td></td><td></td><td></td><td></td><td></td><td></td></tr>
<tr><td>Oligo fwd</td><td>CACCGCCCGAGAACGCGCAGATGC</td><td></td><td></td><td></td><td></td><td></td><td></td><td></td><td></td><td></td><td></td></tr>
<tr><td>Oligo rev</td><td>AAACGCATCTGCGCGTTCTCGGGC</td><td></td><td></td><td></td><td></td><td></td><td></td><td></td><td></td><td></td><td></td></tr>
<tr><td>Chromosome</td><td>start</td><td>end</td><td>strand</td><td>MM</td><td>target_seq</td><td>PAM</td><td>alignment</td><td>distance</td><td>position</td><td>gene name</td><td>gene id</td></tr>
<tr><td>GU305914.1</td><td>124739</td><td>124761</td><td>-</td><td>0</td><td>GCCCGAGAACGCGCAGATGC</td><td>TGG</td><td>||||||||[||||||||||||]PAM</td><td>NA</td><td>Intergenic</td><td>NA</td><td>NA</td></tr>
<tr><td>T24589</td><td>GCAGATGCTGGAGATCCGCGCGG</td><td>1000</td><td></td><td></td><td></td><td></td><td></td><td></td><td></td><td></td><td></td></tr>
<tr><td>Oligo fwd</td><td>CACCGCAGATGCTGGAGATCCGCG</td><td></td><td></td><td></td><td></td><td></td><td></td><td></td><td></td><td></td><td></td></tr>
<tr><td>Oligo rev</td><td>AAACCGCGGATCTCCAGCATCTGC</td><td></td><td></td><td></td><td></td><td></td><td></td><td></td><td></td><td></td><td></td></tr>
<tr><td>Chromosome</td><td>start</td><td>end</td><td>strand</td><td>MM</td><td>target_seq</td><td>PAM</td><td>alignment</td><td>distance</td><td>position</td><td>gene name</td><td>gene id</td></tr>
<tr><td>GU305914.1</td><td>124727</td><td>124749</td><td>-</td><td>0</td><td>GCAGATGCTGGAGATCCGCG</td><td>CGG</td><td>||||||||[||||||||||||]PAM</td><td>NA</td><td>Intergenic</td><td>NA</td><td>NA</td></tr>
<tr><td>T24590</td><td>GCTGGAGATCCGCGCGGCCCTGG</td><td>1000</td><td></td><td></td><td></td><td></td><td></td><td></td><td></td><td></td><td></td></tr>
<tr><td>Oligo fwd</td><td>CACCGCTGGAGATCCGCGCGGCCC</td><td></td><td></td><td></td><td></td><td></td><td></td><td></td><td></td><td></td><td></td></tr>
<tr><td>Oligo rev</td><td>AAACGGGCCGCGCGGATCTCCAGC</td><td></td><td></td><td></td><td></td><td></td><td></td><td></td><td></td><td></td><td></td></tr>
<tr><td>Chromosome</td><td>start</td><td>end</td><td>strand</td><td>MM</td><td>target_seq</td><td>PAM</td><td>alignment</td><td>distance</td><td>position</td><td>gene name</td><td>gene id</td></tr>
<tr><td>GU305914.1</td><td>124721</td><td>124743</td><td>-</td><td>0</td><td>GCTGGAGATCCGCGCGGCCC</td><td>TGG</td><td>||||||||[||||||||||||]PAM</td><td>NA</td><td>Intergenic</td><td>NA</td><td>NA</td></tr>
<tr><td>T24591</td><td>GATCCGCGCGGCCCTGGACCCGG</td><td>1000</td><td></td><td></td><td></td><td></td><td></td><td></td><td></td><td></td><td></td></tr>
<tr><td>Oligo fwd</td><td>CACCGATCCGCGCGGCCCTGGACC</td><td></td><td></td><td></td><td></td><td></td><td></td><td></td><td></td><td></td><td></td></tr>
<tr><td>Oligo rev</td><td>AAACGGTCCAGGGCCGCGCGGATC</td><td></td><td></td><td></td><td></td><td></td><td></td><td></td><td></td><td></td><td></td></tr>
<tr><td>Chromosome</td><td>start</td><td>end</td><td>strand</td><td>MM</td><td>target_seq</td><td>PAM</td><td>alignment</td><td>distance</td><td>position</td><td>gene name</td><td>gene id</td></tr>
<tr><td>GU305914.1</td><td>124715</td><td>124737</td><td>-</td><td>0</td><td>GATCCGCGCGGCCCTGGACC</td><td>CGG</td><td>||||||||[||||||||||||]PAM</td><td>NA</td><td>Intergenic</td><td>NA</td><td>NA</td></tr>
<tr><td>GU305914.1</td><td>197196</td><td>197218</td><td>-</td><td>4</td><td>GATCCTGGTGGCCCCGGACC</td><td>TGG</td><td>|||||--|[-|||||-|||||]PAM</td><td>NA</td><td>Intergenic</td><td>NA</td><td>NA</td></tr>
<tr><td>T24592</td><td>CGGCCCTGGACCCGGCGCAGCGG</td><td>1000</td><td></td><td></td><td></td><td></td><td></td><td></td><td></td><td></td><td></td></tr>
<tr><td>Oligo fwd</td><td>CACCCGGCCCTGGACCCGGCGCAG</td><td></td><td></td><td></td><td></td><td></td><td></td><td></td><td></td><td></td><td></td></tr>
<tr><td>Oligo rev</td><td>AAACCTGCGCCGGGTCCAGGGCCG</td><td></td><td></td><td></td><td></td><td></td><td></td><td></td><td></td><td></td><td></td></tr>
<tr><td>Chromosome</td><td>start</td><td>end</td><td>strand</td><td>MM</td><td>target_seq</td><td>PAM</td><td>alignment</td><td>distance</td><td>position</td><td>gene name</td><td>gene id</td></tr>
<tr><td>GU305914.1</td><td>124707</td><td>124729</td><td>-</td><td>0</td><td>CGGCCCTGGACCCGGCGCAG</td><td>CGG</td><td>||||||||[||||||||||||]PAM</td><td>NA</td><td>Intergenic</td><td>NA</td><td>NA</td></tr>
<tr><td>T24593</td><td>CTGGACCCGGCGCAGCGGCACGG</td><td>1000</td><td></td><td></td><td></td><td></td><td></td><td></td><td></td><td></td><td></td></tr>
<tr><td>Oligo fwd</td><td>CACCCTGGACCCGGCGCAGCGGCA</td><td></td><td></td><td></td><td></td><td></td><td></td><td></td><td></td><td></td><td></td></tr>
<tr><td>Oligo rev</td><td>AAACTGCCGCTGCGCCGGGTCCAG</td><td></td><td></td><td></td><td></td><td></td><td></td><td></td><td></td><td></td><td></td></tr>
<tr><td>Chromosome</td><td>start</td><td>end</td><td>strand</td><td>MM</td><td>target_seq</td><td>PAM</td><td>alignment</td><td>distance</td><td>position</td><td>gene name</td><td>gene id</td></tr>
<tr><td>GU305914.1</td><td>124702</td><td>124724</td><td>-</td><td>0</td><td>CTGGACCCGGCGCAGCGGCA</td><td>CGG</td><td>||||||||[||||||||||||]PAM</td><td>NA</td><td>Intergenic</td><td>NA</td><td>NA</td></tr>
<tr><td>T24594</td><td>GCAGCGGCACGGCGCGCCGTCGG</td><td>1000</td><td></td><td></td><td></td><td></td><td></td><td></td><td></td><td></td><td></td></tr>
<tr><td>Oligo fwd</td><td>CACCGCAGCGGCACGGCGCGCCGT</td><td></td><td></td><td></td><td></td><td></td><td></td><td></td><td></td><td></td><td></td></tr>
<tr><td>Oligo rev</td><td>AAACACGGCGCGCCGTGCCGCTGC</td><td></td><td></td><td></td><td></td><td></td><td></td><td></td><td></td><td></td><td></td></tr>
<tr><td>Chromosome</td><td>start</td><td>end</td><td>strand</td><td>MM</td><td>target_seq</td><td>PAM</td><td>alignment</td><td>distance</td><td>position</td><td>gene name</td><td>gene id</td></tr>
<tr><td>GU305914.1</td><td>124691</td><td>124713</td><td>-</td><td>0</td><td>GCAGCGGCACGGCGCGCCGT</td><td>CGG</td><td>||||||||[||||||||||||]PAM</td><td>NA</td><td>Intergenic</td><td>NA</td><td>NA</td></tr>
<tr><td>T24595</td><td>GCGGCACGGCGCGCCGTCGGCGG</td><td>1000</td><td></td><td></td><td></td><td></td><td></td><td></td><td></td><td></td><td></td></tr>
<tr><td>Oligo fwd</td><td>CACCGCGGCACGGCGCGCCGTCGG</td><td></td><td></td><td></td><td></td><td></td><td></td><td></td><td></td><td></td><td></td></tr>
<tr><td>Oligo rev</td><td>AAACCCGACGGCGCGCCGTGCCGC</td><td></td><td></td><td></td><td></td><td></td><td></td><td></td><td></td><td></td><td></td></tr>
<tr><td>Chromosome</td><td>start</td><td>end</td><td>strand</td><td>MM</td><td>target_seq</td><td>PAM</td><td>alignment</td><td>distance</td><td>position</td><td>gene name</td><td>gene id</td></tr>
<tr><td>GU305914.1</td><td>124688</td><td>124710</td><td>-</td><td>0</td><td>GCGGCACGGCGCGCCGTCGG</td><td>CGG</td><td>||||||||[||||||||||||]PAM</td><td>NA</td><td>Intergenic</td><td>NA</td><td>NA</td></tr>
<tr><td>T24596</td><td>CGGCACGGCGCGCCGTCGGCGGG</td><td>1000</td><td></td><td></td><td></td><td></td><td></td><td></td><td></td><td></td><td></td></tr>
<tr><td>Oligo fwd</td><td>CACCCGGCACGGCGCGCCGTCGGC</td><td></td><td></td><td></td><td></td><td></td><td></td><td></td><td></td><td></td><td></td></tr>
</table>

<table>
<tr><td>Oligo rev</td><td>AAACGCCGACGGCGCGCCGTGCCG</td><td></td><td></td><td></td><td></td><td></td><td></td><td></td><td></td><td></td><td></td></tr>
<tr><td>Chromosome</td><td>start</td><td>end</td><td>strand</td><td>MM</td><td>target_seq</td><td>PAM</td><td>alignment</td><td>distance</td><td>position</td><td>gene name</td><td>gene id</td></tr>
<tr><td>GU305914.1</td><td>124687</td><td>124709</td><td>-</td><td>0</td><td>CGGCACGGCGCGCCGTCGGC</td><td>GGG</td><td>||||||||[||||||||||||]PAM</td><td>NA</td><td>Intergenic</td><td>NA</td><td>NA</td></tr>
<tr><td></td><td></td><td></td><td></td><td></td><td></td><td></td><td></td><td></td><td></td><td></td><td></td></tr>
<tr><td>T24597</td><td>CTGCAGCACGTCCTGTACAACGG</td><td>1000</td><td></td><td></td><td></td><td></td><td></td><td></td><td></td><td></td><td></td></tr>
<tr><td>Oligo fwd</td><td>CACCCTGCAGCACGTCCTGTACAA</td><td></td><td></td><td></td><td></td><td></td><td></td><td></td><td></td><td></td><td></td></tr>
<tr><td>Oligo rev</td><td>AAACTTGTACAGGACGTGCTGCAG</td><td></td><td></td><td></td><td></td><td></td><td></td><td></td><td></td><td></td><td></td></tr>
<tr><td>Chromosome</td><td>start</td><td>end</td><td>strand</td><td>MM</td><td>target_seq</td><td>PAM</td><td>alignment</td><td>distance</td><td>position</td><td>gene name</td><td>gene id</td></tr>
<tr><td>GU305914.1</td><td>124654</td><td>124676</td><td>-</td><td>0</td><td>CTGCAGCACGTCCTGTACAA</td><td>CGG</td><td>||||||||[||||||||||||]PAM</td><td>NA</td><td>Intergenic</td><td>NA</td><td>NA</td></tr>
<tr><td></td><td></td><td></td><td></td><td></td><td></td><td></td><td></td><td></td><td></td><td></td><td></td></tr>
<tr><td>T24598</td><td>TGCAGCACGTCCTGTACAACGGG</td><td>1000</td><td></td><td></td><td></td><td></td><td></td><td></td><td></td><td></td><td></td></tr>
<tr><td>Oligo fwd</td><td>CACCTGCAGCACGTCCTGTACAAC</td><td></td><td></td><td></td><td></td><td></td><td></td><td></td><td></td><td></td><td></td></tr>
<tr><td>Oligo rev</td><td>AAACGTTGTACAGGACGTGCTGCA</td><td></td><td></td><td></td><td></td><td></td><td></td><td></td><td></td><td></td><td></td></tr>
<tr><td>Chromosome</td><td>start</td><td>end</td><td>strand</td><td>MM</td><td>target_seq</td><td>PAM</td><td>alignment</td><td>distance</td><td>position</td><td>gene name</td><td>gene id</td></tr>
<tr><td>GU305914.1</td><td>124653</td><td>124675</td><td>-</td><td>0</td><td>TGCAGCACGTCCTGTACAAC</td><td>GGG</td><td>||||||||[||||||||||||]PAM</td><td>NA</td><td>Intergenic</td><td>NA</td><td>NA</td></tr>
<tr><td></td><td></td><td></td><td></td><td></td><td></td><td></td><td></td><td></td><td></td><td></td><td></td></tr>
<tr><td>T24599</td><td>GACCGTGCCGAAGCTGCTGCAGG</td><td>1000</td><td></td><td></td><td></td><td></td><td></td><td></td><td></td><td></td><td></td></tr>
<tr><td>Oligo fwd</td><td>CACCGACCGTGCCGAAGCTGCTGC</td><td></td><td></td><td></td><td></td><td></td><td></td><td></td><td></td><td></td><td></td></tr>
<tr><td>Oligo rev</td><td>AAACGCAGCAGCTTCGGCACGGTC</td><td></td><td></td><td></td><td></td><td></td><td></td><td></td><td></td><td></td><td></td></tr>
<tr><td>Chromosome</td><td>start</td><td>end</td><td>strand</td><td>MM</td><td>target_seq</td><td>PAM</td><td>alignment</td><td>distance</td><td>position</td><td>gene name</td><td>gene id</td></tr>
<tr><td>GU305914.1</td><td>124619</td><td>124641</td><td>-</td><td>0</td><td>GACCGTGCCGAAGCTGCTGC</td><td>AGG</td><td>||||||||[||||||||||||]PAM</td><td>NA</td><td>Intergenic</td><td>NA</td><td>NA</td></tr>
<tr><td></td><td></td><td></td><td></td><td></td><td></td><td></td><td></td><td></td><td></td><td></td><td></td></tr>
<tr><td>T24600</td><td>GCTGCTGCAGGAGTACCTGACGG</td><td>1000</td><td></td><td></td><td></td><td></td><td></td><td></td><td></td><td></td><td></td></tr>
<tr><td>Oligo fwd</td><td>CACCGCTGCTGCAGGAGTACCTGA</td><td></td><td></td><td></td><td></td><td></td><td></td><td></td><td></td><td></td><td></td></tr>
<tr><td>Oligo rev</td><td>AAACTCAGGTACTCCTGCAGCAGC</td><td></td><td></td><td></td><td></td><td></td><td></td><td></td><td></td><td></td><td></td></tr>
<tr><td>Chromosome</td><td>start</td><td>end</td><td>strand</td><td>MM</td><td>target_seq</td><td>PAM</td><td>alignment</td><td>distance</td><td>position</td><td>gene name</td><td>gene id</td></tr>
<tr><td>GU305914.1</td><td>124607</td><td>124629</td><td>-</td><td>0</td><td>GCTGCTGCAGGAGTACCTGA</td><td>CGG</td><td>||||||||[||||||||||||]PAM</td><td>NA</td><td>Intergenic</td><td>NA</td><td>NA</td></tr>
<tr><td>GU305914.1</td><td>65696</td><td>65718</td><td>+</td><td>4</td><td>GCTGGACCAGGAGGACCTGA</td><td>CGG</td><td>||||---|[|||||-||||||]PAM</td><td>NA</td><td>Intergenic</td><td>NA</td><td>NA</td></tr>
<tr><td></td><td></td><td></td><td></td><td></td><td></td><td></td><td></td><td></td><td></td><td></td><td></td></tr>
<tr><td>T24601</td><td>TGACGGTGATCCCCTTCCACCGG</td><td>1000</td><td></td><td></td><td></td><td></td><td></td><td></td><td></td><td></td><td></td></tr>
<tr><td>Oligo fwd</td><td>CACCTGACGGTGATCCCCTTCCAC</td><td></td><td></td><td></td><td></td><td></td><td></td><td></td><td></td><td></td><td></td></tr>
<tr><td>Oligo rev</td><td>AAACGTGGAAGGGGATCACCGTCA</td><td></td><td></td><td></td><td></td><td></td><td></td><td></td><td></td><td></td><td></td></tr>
<tr><td>Chromosome</td><td>start</td><td>end</td><td>strand</td><td>MM</td><td>target_seq</td><td>PAM</td><td>alignment</td><td>distance</td><td>position</td><td>gene name</td><td>gene id</td></tr>
<tr><td>GU305914.1</td><td>124590</td><td>124612</td><td>-</td><td>0</td><td>TGACGGTGATCCCCTTCCAC</td><td>CGG</td><td>||||||||[||||||||||||]PAM</td><td>NA</td><td>Intergenic</td><td>NA</td><td>NA</td></tr>
<tr><td></td><td></td><td></td><td></td><td></td><td></td><td></td><td></td><td></td><td></td><td></td><td></td></tr>
<tr><td>T24602</td><td>CCCCTTCCACCGGTTCTACTCGG</td><td>1000</td><td></td><td></td><td></td><td></td><td></td><td></td><td></td><td></td><td></td></tr>
<tr><td>Oligo fwd</td><td>CACCCCCCTTCCACCGGTTCTACT</td><td></td><td></td><td></td><td></td><td></td><td></td><td></td><td></td><td></td><td></td></tr>
<tr><td>Oligo rev</td><td>AAACAGTAGAACCGGTGGAAGGGG</td><td></td><td></td><td></td><td></td><td></td><td></td><td></td><td></td><td></td><td></td></tr>
<tr><td>Chromosome</td><td>start</td><td>end</td><td>strand</td><td>MM</td><td>target_seq</td><td>PAM</td><td>alignment</td><td>distance</td><td>position</td><td>gene name</td><td>gene id</td></tr>
<tr><td>GU305914.1</td><td>124580</td><td>124602</td><td>-</td><td>0</td><td>CCCCTTCCACCGGTTCTACT</td><td>CGG</td><td>||||||||[||||||||||||]PAM</td><td>NA</td><td>Intergenic</td><td>NA</td><td>NA</td></tr>
<tr><td></td><td></td><td></td><td></td><td></td><td></td><td></td><td></td><td></td><td></td><td></td><td></td></tr>
<tr><td>T24603</td><td>CCACCGGTTCTACTCGGACCCGG</td><td>1000</td><td></td><td></td><td></td><td></td><td></td><td></td><td></td><td></td><td></td></tr>
<tr><td>Oligo fwd</td><td>CACCCCACCGGTTCTACTCGGACC</td><td></td><td></td><td></td><td></td><td></td><td></td><td></td><td></td><td></td><td></td></tr>
<tr><td>Oligo rev</td><td>AAACGGTCCGAGTAGAACCGGTGG</td><td></td><td></td><td></td><td></td><td></td><td></td><td></td><td></td><td></td><td></td></tr>
<tr><td>Chromosome</td><td>start</td><td>end</td><td>strand</td><td>MM</td><td>target_seq</td><td>PAM</td><td>alignment</td><td>distance</td><td>position</td><td>gene name</td><td>gene id</td></tr>
<tr><td>GU305914.1</td><td>124574</td><td>124596</td><td>-</td><td>0</td><td>CCACCGGTTCTACTCGGACC</td><td>CGG</td><td>||||||||[||||||||||||]PAM</td><td>NA</td><td>Intergenic</td><td>NA</td><td>NA</td></tr>
<tr><td></td><td></td><td></td><td></td><td></td><td></td><td></td><td></td><td></td><td></td><td></td><td></td></tr>
<tr><td>T24604</td><td>CACCGGTTCTACTCGGACCCGGG</td><td>1000</td><td></td><td></td><td></td><td></td><td></td><td></td><td></td><td></td><td></td></tr>
<tr><td>Oligo fwd</td><td>CACCCACCGGTTCTACTCGGACCC</td><td></td><td></td><td></td><td></td><td></td><td></td><td></td><td></td><td></td><td></td></tr>
<tr><td>Oligo rev</td><td>AAACGGGTCCGAGTAGAACCGGTG</td><td></td><td></td><td></td><td></td><td></td><td></td><td></td><td></td><td></td><td></td></tr>
<tr><td>Chromosome</td><td>start</td><td>end</td><td>strand</td><td>MM</td><td>target_seq</td><td>PAM</td><td>alignment</td><td>distance</td><td>position</td><td>gene name</td><td>gene id</td></tr>
<tr><td>GU305914.1</td><td>124573</td><td>124595</td><td>-</td><td>0</td><td>CACCGGTTCTACTCGGACCC</td><td>GGG</td><td>||||||||[||||||||||||]PAM</td><td>NA</td><td>Intergenic</td><td>NA</td><td>NA</td></tr>
<tr><td></td><td></td><td></td><td></td><td></td><td></td><td></td><td></td><td></td><td></td><td></td><td></td></tr>
<tr><td>T24605</td><td>GTTCTACTCGGACCCGGGACTGG</td><td>1000</td><td></td><td></td><td></td><td></td><td></td><td></td><td></td><td></td><td></td></tr>
<tr><td>Oligo fwd</td><td>CACCGTTCTACTCGGACCCGGGAC</td><td></td><td></td><td></td><td></td><td></td><td></td><td></td><td></td><td></td><td></td></tr>
<tr><td>Oligo rev</td><td>AAACGTCCCGGGTCCGAGTAGAAC</td><td></td><td></td><td></td><td></td><td></td><td></td><td></td><td></td><td></td><td></td></tr>
<tr><td>Chromosome</td><td>start</td><td>end</td><td>strand</td><td>MM</td><td>target_seq</td><td>PAM</td><td>alignment</td><td>distance</td><td>position</td><td>gene name</td><td>gene id</td></tr>
<tr><td>GU305914.1</td><td>124568</td><td>124590</td><td>-</td><td>0</td><td>GTTCTACTCGGACCCGGGAC</td><td>TGG</td><td>||||||||[||||||||||||]PAM</td><td>NA</td><td>Intergenic</td><td>NA</td><td>NA</td></tr>
<tr><td></td><td></td><td></td><td></td><td></td><td></td><td></td><td></td><td></td><td></td><td></td><td></td></tr>
<tr><td>T24606</td><td>CTACTCGGACCCGGGACTGGCGG</td><td>1000</td><td></td><td></td><td></td><td></td><td></td><td></td><td></td><td></td><td></td></tr>
<tr><td>Oligo fwd</td><td>CACCCTACTCGGACCCGGGACTGG</td><td></td><td></td><td></td><td></td><td></td><td></td><td></td><td></td><td></td><td></td></tr>
<tr><td>Oligo rev</td><td>AAACCCAGTCCCGGGTCCGAGTAG</td><td></td><td></td><td></td><td></td><td></td><td></td><td></td><td></td><td></td><td></td></tr>
<tr><td>Chromosome</td><td>start</td><td>end</td><td>strand</td><td>MM</td><td>target_seq</td><td>PAM</td><td>alignment</td><td>distance</td><td>position</td><td>gene name</td><td>gene id</td></tr>
<tr><td>GU305914.1</td><td>124565</td><td>124587</td><td>-</td><td>0</td><td>CTACTCGGACCCGGGACTGG</td><td>CGG</td><td>||||||||[||||||||||||]PAM</td><td>NA</td><td>Intergenic</td><td>NA</td><td>NA</td></tr>
<tr><td></td><td></td><td></td><td></td><td></td><td></td><td></td><td></td><td></td><td></td><td></td><td></td></tr>
<tr><td>T24607</td><td>GGACCCGGGACTGGCGGCGACGG</td><td>1000</td><td></td><td></td><td></td><td></td><td></td><td></td><td></td><td></td><td></td></tr>
<tr><td>Oligo fwd</td><td>CACCGGACCCGGGACTGGCGGCGA</td><td></td><td></td><td></td><td></td><td></td><td></td><td></td><td></td><td></td><td></td></tr>
<tr><td>Oligo rev</td><td>AAACTCGCCGCCAGTCCCGGGTCC</td><td></td><td></td><td></td><td></td><td></td><td></td><td></td><td></td><td></td><td></td></tr>
<tr><td>Chromosome</td><td>start</td><td>end</td><td>strand</td><td>MM</td><td>target_seq</td><td>PAM</td><td>alignment</td><td>distance</td><td>position</td><td>gene name</td><td>gene id</td></tr>
<tr><td>GU305914.1</td><td>124559</td><td>124581</td><td>-</td><td>0</td><td>GGACCCGGGACTGGCGGCGA</td><td>CGG</td><td>||||||||[||||||||||||]PAM</td><td>NA</td><td>Intergenic</td><td>NA</td><td>NA</td></tr>
<tr><td></td><td></td><td></td><td></td><td></td><td></td><td></td><td></td><td></td><td></td><td></td><td></td></tr>
<tr><td>T24608</td><td>ACGACTTCCCGCAGTACCAGCGG</td><td>1000</td><td></td><td></td><td></td><td></td><td></td><td></td><td></td><td></td><td></td></tr>
<tr><td>Oligo fwd</td><td>CACCACGACTTCCCGCAGTACCAG</td><td></td><td></td><td></td><td></td><td></td><td></td><td></td><td></td><td></td><td></td></tr>
<tr><td>Oligo rev</td><td>AAACCTGGTACTGCGGGAAGTCGT</td><td></td><td></td><td></td><td></td><td></td><td></td><td></td><td></td><td></td><td></td></tr>
<tr><td>Chromosome</td><td>start</td><td>end</td><td>strand</td><td>MM</td><td>target_seq</td><td>PAM</td><td>alignment</td><td>distance</td><td>position</td><td>gene name</td><td>gene id</td></tr>
<tr><td>GU305914.1</td><td>124509</td><td>124531</td><td>-</td><td>0</td><td>ACGACTTCCCGCAGTACCAG</td><td>CGG</td><td>||||||||[||||||||||||]PAM</td><td>NA</td><td>Intergenic</td><td>NA</td><td>NA</td></tr>
<tr><td></td><td></td><td></td><td></td><td></td><td></td><td></td><td></td><td></td><td></td><td></td><td></td></tr>
<tr><td>T24609</td><td>CCGCAGTACCAGCGGTGCGACGG</td><td>1000</td><td></td><td></td><td></td><td></td><td></td><td></td><td></td><td></td><td></td></tr>
<tr><td>Oligo fwd</td><td>CACCCCGCAGTACCAGCGGTGCGA</td><td></td><td></td><td></td><td></td><td></td><td></td><td></td><td></td><td></td><td></td></tr>
<tr><td>Oligo rev</td><td>AAACTCGCACCGCTGGTACTGCGG</td><td></td><td></td><td></td><td></td><td></td><td></td><td></td><td></td><td></td><td></td></tr>
<tr><td>Chromosome</td><td>start</td><td>end</td><td>strand</td><td>MM</td><td>target_seq</td><td>PAM</td><td>alignment</td><td>distance</td><td>position</td><td>gene name</td><td>gene id</td></tr>
<tr><td>GU305914.1</td><td>124501</td><td>124523</td><td>-</td><td>0</td><td>CCGCAGTACCAGCGGTGCGA</td><td>CGG</td><td>||||||||[||||||||||||]PAM</td><td>NA</td><td>Intergenic</td><td>NA</td><td>NA</td></tr>
<tr><td></td><td></td><td></td><td></td><td></td><td></td><td></td><td></td><td></td><td></td><td></td><td></td></tr>
<tr><td>T24610</td><td>CAGTACCAGCGGTGCGACGGCGG</td><td>1000</td><td></td><td></td><td></td><td></td><td></td><td></td><td></td><td></td><td></td></tr>
<tr><td>Oligo fwd</td><td>CACCCAGTACCAGCGGTGCGACGG</td><td></td><td></td><td></td><td></td><td></td><td></td><td></td><td></td><td></td><td></td></tr>
<tr><td>Oligo rev</td><td>AAACCCGTCGCACCGCTGGTACTG</td><td></td><td></td><td></td><td></td><td></td><td></td><td></td><td></td><td></td><td></td></tr>
<tr><td>Chromosome</td><td>start</td><td>end</td><td>strand</td><td>MM</td><td>target_seq</td><td>PAM</td><td>alignment</td><td>distance</td><td>position</td><td>gene name</td><td>gene id</td></tr>
<tr><td>GU305914.1</td><td>124498</td><td>124520</td><td>-</td><td>0</td><td>CAGTACCAGCGGTGCGACGG</td><td>CGG</td><td>||||||||[||||||||||||]PAM</td><td>NA</td><td>Intergenic</td><td>NA</td><td>NA</td></tr>
</table>

| | | | | | | | | | | | |
|---|---|---|---|---|---|---|---|---|---|---|---|
| T24611 | TCGCGCACGAGTACCACCACTGG | 1000 | | | | | | | | | |
| Oligo fwd | CACCTCGCGCACGAGTACCACCAC | | | | | | | | | | |
| Oligo rev | AAACGTGGTGGTACTCGTGCGCGA | | | | | | | | | | |
| Chromosome | start | end | strand | MM | target_seq | PAM | alignment | distance | position | gene name | gene id |
| GU305914.1 | 124455 | 124477 | - | 0 | TCGCGCACGAGTACCACCAC | TGG | \|\|\|\|\|\|\|\|[\|\|\|\|\|\|\|\|\|\|\|\|]PAM | NA | Intergenic | NA | NA |

| | | | | | | | | | | | |
|---|---|---|---|---|---|---|---|---|---|---|---|
| T24612 | CACGCCGTTCTCGTGCTACTCGG | 1000 | | | | | | | | | |
| Oligo fwd | CACCCACGCCGTTCTCGTGCTACT | | | | | | | | | | |
| Oligo rev | AAACAGTAGCACGAGAACGGCGTG | | | | | | | | | | |
| Chromosome | start | end | strand | MM | target_seq | PAM | alignment | distance | position | gene name | gene id |
| GU305914.1 | 124427 | 124449 | - | 0 | CACGCCGTTCTCGTGCTACT | CGG | \|\|\|\|\|\|\|\|[\|\|\|\|\|\|\|\|\|\|\|\|]PAM | NA | Intergenic | NA | NA |

| | | | | | | | | | | | |
|---|---|---|---|---|---|---|---|---|---|---|---|
| T24613 | GCCGTTCTCGTGCTACTCGGCGG | 1000 | | | | | | | | | |
| Oligo fwd | CACCGCCGTTCTCGTGCTACTCGG | | | | | | | | | | |
| Oligo rev | AAACCCGAGTAGCACGAGAACGGC | | | | | | | | | | |
| Chromosome | start | end | strand | MM | target_seq | PAM | alignment | distance | position | gene name | gene id |
| GU305914.1 | 124424 | 124446 | - | 0 | GCCGTTCTCGTGCTACTCGG | CGG | \|\|\|\|\|\|\|\|[\|\|\|\|\|\|\|\|\|\|\|\|]PAM | NA | Intergenic | NA | NA |

| | | | | | | | | | | | |
|---|---|---|---|---|---|---|---|---|---|---|---|
| T24614 | GGCGGCCTGCGCGCACACCCTGG | 1000 | | | | | | | | | |
| Oligo fwd | CACCGGCGGCCTGCGCGCACACCC | | | | | | | | | | |
| Oligo rev | AAACGGGTGTGCGCGCAGGCCGCC | | | | | | | | | | |
| Chromosome | start | end | strand | MM | target_seq | PAM | alignment | distance | position | gene name | gene id |
| GU305914.1 | 124406 | 124428 | - | 0 | GGCGGCCTGCGCGCACACCC | TGG | \|\|\|\|\|\|\|\|[\|\|\|\|\|\|\|\|\|\|\|\|]PAM | NA | Intergenic | NA | NA |

| | | | | | | | | | | | |
|---|---|---|---|---|---|---|---|---|---|---|---|
| T24615 | CTGCGCGCACACCCTGGAGTCGG | 1000 | | | | | | | | | |
| Oligo fwd | CACCCTGCGCGCACACCCTGGAGT | | | | | | | | | | |
| Oligo rev | AAACACTCCAGGGTGTGCGCGCAG | | | | | | | | | | |
| Chromosome | start | end | strand | MM | target_seq | PAM | alignment | distance | position | gene name | gene id |
| GU305914.1 | 124400 | 124422 | - | 0 | CTGCGCGCACACCCTGGAGT | CGG | \|\|\|\|\|\|\|\|[\|\|\|\|\|\|\|\|\|\|\|\|]PAM | NA | Intergenic | NA | NA |

| | | | | | | | | | | | |
|---|---|---|---|---|---|---|---|---|---|---|---|
| T24616 | CCTGGAGTCGGTGTTGACGCTGG | 1000 | | | | | | | | | |
| Oligo fwd | CACCCCTGGAGTCGGTGTTGACGC | | | | | | | | | | |
| Oligo rev | AAACGCGTCAACACCGACTCCAGG | | | | | | | | | | |
| Chromosome | start | end | strand | MM | target_seq | PAM | alignment | distance | position | gene name | gene id |
| GU305914.1 | 124388 | 124410 | - | 0 | CCTGGAGTCGGTGTTGACGC | TGG | \|\|\|\|\|\|\|\|[\|\|\|\|\|\|\|\|\|\|\|\|]PAM | NA | Intergenic | NA | NA |

| | | | | | | | | | | | |
|---|---|---|---|---|---|---|---|---|---|---|---|
| T24617 | CATGCATCACAAGATGTCCCCGG | 1000 | | | | | | | | | |
| Oligo fwd | CACCCATGCATCACAAGATGTCCC | | | | | | | | | | |
| Oligo rev | AAACGGGACATCTTGTGATGCATG | | | | | | | | | | |
| Chromosome | start | end | strand | MM | target_seq | PAM | alignment | distance | position | gene name | gene id |
| GU305914.1 | 124361 | 124383 | - | 0 | CATGCATCACAAGATGTCCC | CGG | \|\|\|\|\|\|\|\|[\|\|\|\|\|\|\|\|\|\|\|\|]PAM | NA | Intergenic | NA | NA |

| | | | | | | | | | | | |
|---|---|---|---|---|---|---|---|---|---|---|---|
| T24618 | GATGTCCCCGGTGTCGATCGCGG | 1000 | | | | | | | | | |
| Oligo fwd | CACCGATGTCCCCGGTGTCGATCG | | | | | | | | | | |
| Oligo rev | AAACCGATCGACACCGGGGACATC | | | | | | | | | | |
| Chromosome | start | end | strand | MM | target_seq | PAM | alignment | distance | position | gene name | gene id |
| GU305914.1 | 124349 | 124371 | - | 0 | GATGTCCCCGGTGTCGATCG | CGG | \|\|\|\|\|\|\|\|[\|\|\|\|\|\|\|\|\|\|\|\|]PAM | NA | Intergenic | NA | NA |
| GU305914.1 | 191725 | 191747 | + | 4 | GATGTGCTCGGTGACGACCG | AGG | \|\|\|\|\|-\|-[\|\|\|\|\|-\|\|\|-\|\|]PAM | NA | Intergenic | NA | NA |

| | | | | | | | | | | | |
|---|---|---|---|---|---|---|---|---|---|---|---|
| T24619 | TGTCGATCGCGGCGCTGAGCCGG | 1000 | | | | | | | | | |
| Oligo fwd | CACCTGTCGATCGCGGCGCTGAGC | | | | | | | | | | |
| Oligo rev | AAACGCTCAGCGCCGCGATCGACA | | | | | | | | | | |
| Chromosome | start | end | strand | MM | target_seq | PAM | alignment | distance | position | gene name | gene id |
| GU305914.1 | 124338 | 124360 | - | 0 | TGTCGATCGCGGCGCTGAGC | CGG | \|\|\|\|\|\|\|\|[\|\|\|\|\|\|\|\|\|\|\|\|]PAM | NA | Intergenic | NA | NA |
| GU305914.1 | 55833 | 55855 | + | 4 | TGGCGGTCTCGGCGCCGAGC | AGG | \|\|-\|\|-\|\|[-\|\|\|\|\|\|-\|\|\|\|]PAM | NA | Intergenic | NA | NA |

| | | | | | | | | | | | |
|---|---|---|---|---|---|---|---|---|---|---|---|
| T24620 | GATCGCGGCGCTGAGCCGGATGG | 1000 | | | | | | | | | |
| Oligo fwd | CACCGATCGCGGCGCTGAGCCGGA | | | | | | | | | | |
| Oligo rev | AAACTCCGGCTCAGCGCCGCGATC | | | | | | | | | | |
| Chromosome | start | end | strand | MM | target_seq | PAM | alignment | distance | position | gene name | gene id |
| GU305914.1 | 124334 | 124356 | - | 0 | GATCGCGGCGCTGAGCCGGA | TGG | \|\|\|\|\|\|\|\|[\|\|\|\|\|\|\|\|\|\|\|\|]PAM | NA | Intergenic | NA | NA |

| | | | | | | | | | | | |
|---|---|---|---|---|---|---|---|---|---|---|---|
| T24621 | ATCGCGGCGCTGAGCCGGATGGG | 1000 | | | | | | | | | |
| Oligo fwd | CACCATCGCGGCGCTGAGCCGGAT | | | | | | | | | | |
| Oligo rev | AAACATCCGGCTCAGCGCCGCGAT | | | | | | | | | | |
| Chromosome | start | end | strand | MM | target_seq | PAM | alignment | distance | position | gene name | gene id |
| GU305914.1 | 124333 | 124355 | - | 0 | ATCGCGGCGCTGAGCCGGAT | GGG | \|\|\|\|\|\|\|\|[\|\|\|\|\|\|\|\|\|\|\|\|]PAM | NA | Intergenic | NA | NA |

| | | | | | | | | | | | |
|---|---|---|---|---|---|---|---|---|---|---|---|
| T24622 | GAGCCGGATGGGCCTGCACCCGG | 1000 | | | | | | | | | |
| Oligo fwd | CACCGAGCCGGATGGGCCTGCACC | | | | | | | | | | |
| Oligo rev | AAACGGTGCAGGCCCATCCGGCTC | | | | | | | | | | |
| Chromosome | start | end | strand | MM | target_seq | PAM | alignment | distance | position | gene name | gene id |
| GU305914.1 | 124322 | 124344 | - | 0 | GAGCCGGATGGGCCTGCACC | CGG | \|\|\|\|\|\|\|\|[\|\|\|\|\|\|\|\|\|\|\|\|]PAM | NA | Intergenic | NA | NA |

| | | | | | | | | | | | |
|---|---|---|---|---|---|---|---|---|---|---|---|
| T24623 | AGCCGGATGGGCCTGCACCCGGG | 1000 | | | | | | | | | |
| Oligo fwd | CACCAGCCGGATGGGCCTGCACCC | | | | | | | | | | |
| Oligo rev | AAACGGGTGCAGGCCCATCCGGCT | | | | | | | | | | |
| Chromosome | start | end | strand | MM | target_seq | PAM | alignment | distance | position | gene name | gene id |
| GU305914.1 | 124321 | 124343 | - | 0 | AGCCGGATGGGCCTGCACCC | GGG | \|\|\|\|\|\|\|\|[\|\|\|\|\|\|\|\|\|\|\|\|]PAM | NA | Intergenic | NA | NA |

| | | | | | | | | | | | |
|---|---|---|---|---|---|---|---|---|---|---|---|
| T24624 | CCCGGGCTTCGCGCTGACCGTGG | 1000 | | | | | | | | | |
| Oligo fwd | CACCCCCGGGCTTCGCGCTGACCG | | | | | | | | | | |
| Oligo rev | AAACCGGTCAGCGCGAAGCCCGGG | | | | | | | | | | |
| Chromosome | start | end | strand | MM | target_seq | PAM | alignment | distance | position | gene name | gene id |
| GU305914.1 | 124304 | 124326 | - | 0 | CCCGGGCTTCGCGCTGACCG | TGG | \|\|\|\|\|\|\|\|[\|\|\|\|\|\|\|\|\|\|\|\|]PAM | NA | Intergenic | NA | NA |

| | | |
|---|---|---|
| T24625 | GCTTCGCGCTGACCGTGGTGCGG | 1000 |
| Oligo fwd | CACCGCTTCGCGCTGACCGTGGTG | |
| Oligo rev | AAACCACCACGGTCAGCGCGAAGC | |

<table>
<tr><td>Chromosome</td><td>start</td><td>end</td><td>strand</td><td>MM</td><td>target_seq</td><td>PAM</td><td>alignment</td><td>distance</td><td>position</td><td>gene name</td><td>gene id</td></tr>
<tr><td>GU305914.1</td><td>124299</td><td>124321</td><td>-</td><td>0</td><td>GCTTCGCGCTGACCGTGGTG</td><td>CGG</td><td>||||||||[||||||||||||]PAM</td><td>NA</td><td>Intergenic</td><td>NA</td><td>NA</td></tr>
<tr><td></td><td></td><td></td><td></td><td></td><td></td><td></td><td></td><td></td><td></td><td></td><td></td></tr>
<tr><td>T24626</td><td>CGCGCTGACCGTGGTGCGGACGG</td><td>1000</td><td></td><td></td><td></td><td></td><td></td><td></td><td></td><td></td><td></td></tr>
<tr><td>Oligo fwd</td><td>CACCCGCGCTGACCGTGGTGCGGA</td><td></td><td></td><td></td><td></td><td></td><td></td><td></td><td></td><td></td><td></td></tr>
<tr><td>Oligo rev</td><td>AAACTCCGCACCACGGTCAGCGCG</td><td></td><td></td><td></td><td></td><td></td><td></td><td></td><td></td><td></td><td></td></tr>
<tr><td>Chromosome</td><td>start</td><td>end</td><td>strand</td><td>MM</td><td>target_seq</td><td>PAM</td><td>alignment</td><td>distance</td><td>position</td><td>gene name</td><td>gene id</td></tr>
<tr><td>GU305914.1</td><td>124295</td><td>124317</td><td>-</td><td>0</td><td>CGCGCTGACCGTGGTGCGGA</td><td>CGG</td><td>||||||||[||||||||||||]PAM</td><td>NA</td><td>Intergenic</td><td>NA</td><td>NA</td></tr>
<tr><td>GU305914.1</td><td>19474</td><td>19496</td><td>+</td><td>4</td><td>GGCGATGAGCGTGGTGGGGA</td><td>CGG</td><td>-|||-|||[-|||||||-|||]PAM</td><td>NA</td><td>Intergenic</td><td>NA</td><td>NA</td></tr>
<tr><td></td><td></td><td></td><td></td><td></td><td></td><td></td><td></td><td></td><td></td><td></td><td></td></tr>
<tr><td>T24627</td><td>GCGGACGGACACGTTCGAGACGG</td><td>1000</td><td></td><td></td><td></td><td></td><td></td><td></td><td></td><td></td><td></td></tr>
<tr><td>Oligo fwd</td><td>CACCGCGGACGGACACGTTCGAGA</td><td></td><td></td><td></td><td></td><td></td><td></td><td></td><td></td><td></td><td></td></tr>
<tr><td>Oligo rev</td><td>AAACTCTCGAACGTGTCCGTCCGC</td><td></td><td></td><td></td><td></td><td></td><td></td><td></td><td></td><td></td><td></td></tr>
<tr><td>Chromosome</td><td>start</td><td>end</td><td>strand</td><td>MM</td><td>target_seq</td><td>PAM</td><td>alignment</td><td>distance</td><td>position</td><td>gene name</td><td>gene id</td></tr>
<tr><td>GU305914.1</td><td>124280</td><td>124302</td><td>-</td><td>0</td><td>GCGGACGGACACGTTCGAGA</td><td>CGG</td><td>||||||||[||||||||||||]PAM</td><td>NA</td><td>Intergenic</td><td>NA</td><td>NA</td></tr>
<tr><td></td><td></td><td></td><td></td><td></td><td></td><td></td><td></td><td></td><td></td><td></td><td></td></tr>
<tr><td>T24628</td><td>CACGCTGCTGTACAGCACCAAGG</td><td>1000</td><td></td><td></td><td></td><td></td><td></td><td></td><td></td><td></td><td></td></tr>
<tr><td>Oligo fwd</td><td>CACCCACGCTGCTGTACAGCACCA</td><td></td><td></td><td></td><td></td><td></td><td></td><td></td><td></td><td></td><td></td></tr>
<tr><td>Oligo rev</td><td>AAACTGGTGCTGTACAGCAGCGTG</td><td></td><td></td><td></td><td></td><td></td><td></td><td></td><td></td><td></td><td></td></tr>
<tr><td>Chromosome</td><td>start</td><td>end</td><td>strand</td><td>MM</td><td>target_seq</td><td>PAM</td><td>alignment</td><td>distance</td><td>position</td><td>gene name</td><td>gene id</td></tr>
<tr><td>GU305914.1</td><td>124256</td><td>124278</td><td>-</td><td>0</td><td>CACGCTGCTGTACAGCACCA</td><td>AGG</td><td>||||||||[||||||||||||]PAM</td><td>NA</td><td>Intergenic</td><td>NA</td><td>NA</td></tr>
<tr><td></td><td></td><td></td><td></td><td></td><td></td><td></td><td></td><td></td><td></td><td></td><td></td></tr>
<tr><td>T24629</td><td>GTACAGCACCAAGGCGTCGACGG</td><td>1000</td><td></td><td></td><td></td><td></td><td></td><td></td><td></td><td></td><td></td></tr>
<tr><td>Oligo fwd</td><td>CACCGTACAGCACCAAGGCGTCGA</td><td></td><td></td><td></td><td></td><td></td><td></td><td></td><td></td><td></td><td></td></tr>
<tr><td>Oligo rev</td><td>AAACTCGACGCCTTGGTGCTGTAC</td><td></td><td></td><td></td><td></td><td></td><td></td><td></td><td></td><td></td><td></td></tr>
<tr><td>Chromosome</td><td>start</td><td>end</td><td>strand</td><td>MM</td><td>target_seq</td><td>PAM</td><td>alignment</td><td>distance</td><td>position</td><td>gene name</td><td>gene id</td></tr>
<tr><td>GU305914.1</td><td>124247</td><td>124269</td><td>-</td><td>0</td><td>GTACAGCACCAAGGCGTCGA</td><td>CGG</td><td>||||||||[||||||||||||]PAM</td><td>NA</td><td>Intergenic</td><td>NA</td><td>NA</td></tr>
<tr><td></td><td></td><td></td><td></td><td></td><td></td><td></td><td></td><td></td><td></td><td></td><td></td></tr>
<tr><td>T24630</td><td>CAGCACCAAGGCGTCGACGGCGG</td><td>1000</td><td></td><td></td><td></td><td></td><td></td><td></td><td></td><td></td><td></td></tr>
<tr><td>Oligo fwd</td><td>CACCCAGCACCAAGGCGTCGACGG</td><td></td><td></td><td></td><td></td><td></td><td></td><td></td><td></td><td></td><td></td></tr>
<tr><td>Oligo rev</td><td>AAACCCGTCGACGCCTTGGTGCTG</td><td></td><td></td><td></td><td></td><td></td><td></td><td></td><td></td><td></td><td></td></tr>
<tr><td>Chromosome</td><td>start</td><td>end</td><td>strand</td><td>MM</td><td>target_seq</td><td>PAM</td><td>alignment</td><td>distance</td><td>position</td><td>gene name</td><td>gene id</td></tr>
<tr><td>GU305914.1</td><td>124244</td><td>124266</td><td>-</td><td>0</td><td>CAGCACCAAGGCGTCGACGG</td><td>CGG</td><td>||||||||[||||||||||||]PAM</td><td>NA</td><td>Intergenic</td><td>NA</td><td>NA</td></tr>
<tr><td></td><td></td><td></td><td></td><td></td><td></td><td></td><td></td><td></td><td></td><td></td><td></td></tr>
<tr><td>T24631</td><td>CAACACGCCGATCGTGACCAAGG</td><td>1000</td><td></td><td></td><td></td><td></td><td></td><td></td><td></td><td></td><td></td></tr>
<tr><td>Oligo fwd</td><td>CACCCAACACGCCGATCGTGACCA</td><td></td><td></td><td></td><td></td><td></td><td></td><td></td><td></td><td></td><td></td></tr>
<tr><td>Oligo rev</td><td>AAACTGGTCACGATCGGCGTGTTG</td><td></td><td></td><td></td><td></td><td></td><td></td><td></td><td></td><td></td><td></td></tr>
<tr><td>Chromosome</td><td>start</td><td>end</td><td>strand</td><td>MM</td><td>target_seq</td><td>PAM</td><td>alignment</td><td>distance</td><td>position</td><td>gene name</td><td>gene id</td></tr>
<tr><td>GU305914.1</td><td>124214</td><td>124236</td><td>-</td><td>0</td><td>CAACACGCCGATCGTGACCA</td><td>AGG</td><td>||||||||[||||||||||||]PAM</td><td>NA</td><td>Intergenic</td><td>NA</td><td>NA</td></tr>
<tr><td>GU305914.1</td><td>222623</td><td>222645</td><td>+</td><td>4</td><td>GTACACGACGATCGTGACGA</td><td>TGG</td><td>--|||||-[||||||||||-|]PAM</td><td>NA</td><td>Intergenic</td><td>NA</td><td>NA</td></tr>
<tr><td></td><td></td><td></td><td></td><td></td><td></td><td></td><td></td><td></td><td></td><td></td><td></td></tr>
<tr><td>T24632</td><td>CACGCCGATCGTGACCAAGGAGG</td><td>1000</td><td></td><td></td><td></td><td></td><td></td><td></td><td></td><td></td><td></td></tr>
<tr><td>Oligo fwd</td><td>CACCCACGCCGATCGTGACCAAGG</td><td></td><td></td><td></td><td></td><td></td><td></td><td></td><td></td><td></td><td></td></tr>
<tr><td>Oligo rev</td><td>AAACCCTTGGTCACGATCGGCGTG</td><td></td><td></td><td></td><td></td><td></td><td></td><td></td><td></td><td></td><td></td></tr>
<tr><td>Chromosome</td><td>start</td><td>end</td><td>strand</td><td>MM</td><td>target_seq</td><td>PAM</td><td>alignment</td><td>distance</td><td>position</td><td>gene name</td><td>gene id</td></tr>
<tr><td>GU305914.1</td><td>124211</td><td>124233</td><td>-</td><td>0</td><td>CACGCCGATCGTGACCAAGG</td><td>AGG</td><td>||||||||[||||||||||||]PAM</td><td>NA</td><td>Intergenic</td><td>NA</td><td>NA</td></tr>
<tr><td></td><td></td><td></td><td></td><td></td><td></td><td></td><td></td><td></td><td></td><td></td><td></td></tr>
<tr><td>T24633</td><td>CGATCGTGACCAAGGAGGACCGG</td><td>1000</td><td></td><td></td><td></td><td></td><td></td><td></td><td></td><td></td><td></td></tr>
<tr><td>Oligo fwd</td><td>CACCCGATCGTGACCAAGGAGGAC</td><td></td><td></td><td></td><td></td><td></td><td></td><td></td><td></td><td></td><td></td></tr>
<tr><td>Oligo rev</td><td>AAACGTCCTCCTTGGTCACGATCG</td><td></td><td></td><td></td><td></td><td></td><td></td><td></td><td></td><td></td><td></td></tr>
<tr><td>Chromosome</td><td>start</td><td>end</td><td>strand</td><td>MM</td><td>target_seq</td><td>PAM</td><td>alignment</td><td>distance</td><td>position</td><td>gene name</td><td>gene id</td></tr>
<tr><td>GU305914.1</td><td>124206</td><td>124228</td><td>-</td><td>0</td><td>CGATCGTGACCAAGGAGGAC</td><td>CGG</td><td>||||||||[||||||||||||]PAM</td><td>NA</td><td>Intergenic</td><td>NA</td><td>NA</td></tr>
<tr><td>GU305914.1</td><td>70032</td><td>70054</td><td>+</td><td>4</td><td>TGAGGCTGACCAAGGAGGAC</td><td>AGG</td><td>-||---||[||||||||||||]PAM</td><td>NA</td><td>Intergenic</td><td>NA</td><td>NA</td></tr>
<tr><td></td><td></td><td></td><td></td><td></td><td></td><td></td><td></td><td></td><td></td><td></td><td></td></tr>
<tr><td>T24634</td><td>GATCGTGACCAAGGAGGACCGGG</td><td>1000</td><td></td><td></td><td></td><td></td><td></td><td></td><td></td><td></td><td></td></tr>
<tr><td>Oligo fwd</td><td>CACCGATCGTGACCAAGGAGGACC</td><td></td><td></td><td></td><td></td><td></td><td></td><td></td><td></td><td></td><td></td></tr>
<tr><td>Oligo rev</td><td>AAACGGTCCTCCTTGGTCACGATC</td><td></td><td></td><td></td><td></td><td></td><td></td><td></td><td></td><td></td><td></td></tr>
<tr><td>Chromosome</td><td>start</td><td>end</td><td>strand</td><td>MM</td><td>target_seq</td><td>PAM</td><td>alignment</td><td>distance</td><td>position</td><td>gene name</td><td>gene id</td></tr>
<tr><td>GU305914.1</td><td>124205</td><td>124227</td><td>-</td><td>0</td><td>GATCGTGACCAAGGAGGACC</td><td>GGG</td><td>||||||||[||||||||||||]PAM</td><td>NA</td><td>Intergenic</td><td>NA</td><td>NA</td></tr>
<tr><td></td><td></td><td></td><td></td><td></td><td></td><td></td><td></td><td></td><td></td><td></td><td></td></tr>
<tr><td>T24635</td><td>GGAGGACCGGGACATCAACACGG</td><td>1000</td><td></td><td></td><td></td><td></td><td></td><td></td><td></td><td></td><td></td></tr>
<tr><td>Oligo fwd</td><td>CACCGGAGGACCGGGACATCAACA</td><td></td><td></td><td></td><td></td><td></td><td></td><td></td><td></td><td></td><td></td></tr>
<tr><td>Oligo rev</td><td>AAACTGTTGATGTCCCGGTCCTCC</td><td></td><td></td><td></td><td></td><td></td><td></td><td></td><td></td><td></td><td></td></tr>
<tr><td>Chromosome</td><td>start</td><td>end</td><td>strand</td><td>MM</td><td>target_seq</td><td>PAM</td><td>alignment</td><td>distance</td><td>position</td><td>gene name</td><td>gene id</td></tr>
<tr><td>GU305914.1</td><td>124193</td><td>124215</td><td>-</td><td>0</td><td>GGAGGACCGGGACATCAACA</td><td>CGG</td><td>||||||||[||||||||||||]PAM</td><td>NA</td><td>Intergenic</td><td>NA</td><td>NA</td></tr>
<tr><td></td><td></td><td></td><td></td><td></td><td></td><td></td><td></td><td></td><td></td><td></td><td></td></tr>
<tr><td>T24636</td><td>CGTGAGCCAGAACATCAACACGG</td><td>1000</td><td></td><td></td><td></td><td></td><td></td><td></td><td></td><td></td><td></td></tr>
<tr><td>Oligo fwd</td><td>CACCCGTGAGCCAGAACATCAACA</td><td></td><td></td><td></td><td></td><td></td><td></td><td></td><td></td><td></td><td></td></tr>
<tr><td>Oligo rev</td><td>AAACTGTTGATGTTCTGGCTCACG</td><td></td><td></td><td></td><td></td><td></td><td></td><td></td><td></td><td></td><td></td></tr>
<tr><td>Chromosome</td><td>start</td><td>end</td><td>strand</td><td>MM</td><td>target_seq</td><td>PAM</td><td>alignment</td><td>distance</td><td>position</td><td>gene name</td><td>gene id</td></tr>
<tr><td>GU305914.1</td><td>124163</td><td>124185</td><td>-</td><td>0</td><td>CGTGAGCCAGAACATCAACA</td><td>CGG</td><td>||||||||[||||||||||||]PAM</td><td>NA</td><td>Intergenic</td><td>NA</td><td>NA</td></tr>
<tr><td></td><td></td><td></td><td></td><td></td><td></td><td></td><td></td><td></td><td></td><td></td><td></td></tr>
<tr><td>T24637</td><td>GAGCCAGAACATCAACACGGTGG</td><td>1000</td><td></td><td></td><td></td><td></td><td></td><td></td><td></td><td></td><td></td></tr>
<tr><td>Oligo fwd</td><td>CACCGAGCCAGAACATCAACACGG</td><td></td><td></td><td></td><td></td><td></td><td></td><td></td><td></td><td></td><td></td></tr>
<tr><td>Oligo rev</td><td>AAACCCGTGTTGATGTTCTGGCTC</td><td></td><td></td><td></td><td></td><td></td><td></td><td></td><td></td><td></td><td></td></tr>
<tr><td>Chromosome</td><td>start</td><td>end</td><td>strand</td><td>MM</td><td>target_seq</td><td>PAM</td><td>alignment</td><td>distance</td><td>position</td><td>gene name</td><td>gene id</td></tr>
<tr><td>GU305914.1</td><td>124160</td><td>124182</td><td>-</td><td>0</td><td>GAGCCAGAACATCAACACGG</td><td>TGG</td><td>||||||||[||||||||||||]PAM</td><td>NA</td><td>Intergenic</td><td>NA</td><td>NA</td></tr>
<tr><td>GU305914.1</td><td>91827</td><td>91849</td><td>+</td><td>4</td><td>GAGGAAGAACATCCAGACGG</td><td>CGG</td><td>|||--|||[|||||-|-||||]PAM</td><td>NA</td><td>Intergenic</td><td>NA</td><td>NA</td></tr>
<tr><td></td><td></td><td></td><td></td><td></td><td></td><td></td><td></td><td></td><td></td><td></td><td></td></tr>
<tr><td>T24638</td><td>GAACATCAACACGGTGGAGATGG</td><td>1000</td><td></td><td></td><td></td><td></td><td></td><td></td><td></td><td></td><td></td></tr>
<tr><td>Oligo fwd</td><td>CACCGAACATCAACACGGTGGAGA</td><td></td><td></td><td></td><td></td><td></td><td></td><td></td><td></td><td></td><td></td></tr>
<tr><td>Oligo rev</td><td>AAACTCTCCACCGTGTTGATGTTC</td><td></td><td></td><td></td><td></td><td></td><td></td><td></td><td></td><td></td><td></td></tr>
<tr><td>Chromosome</td><td>start</td><td>end</td><td>strand</td><td>MM</td><td>target_seq</td><td>PAM</td><td>alignment</td><td>distance</td><td>position</td><td>gene name</td><td>gene id</td></tr>
<tr><td>GU305914.1</td><td>124154</td><td>124176</td><td>-</td><td>0</td><td>GAACATCAACACGGTGGAGA</td><td>TGG</td><td>||||||||[||||||||||||]PAM</td><td>NA</td><td>Intergenic</td><td>NA</td><td>NA</td></tr>
<tr><td></td><td></td><td></td><td></td><td></td><td></td><td></td><td></td><td></td><td></td><td></td><td></td></tr>
<tr><td>T24639</td><td>AACATCAACACGGTGGAGATGGG</td><td>1000</td><td></td><td></td><td></td><td></td><td></td><td></td><td></td><td></td><td></td></tr>
<tr><td>Oligo fwd</td><td>CACCAACATCAACACGGTGGAGAT</td><td></td><td></td><td></td><td></td><td></td><td></td><td></td><td></td><td></td><td></td></tr>
<tr><td>Oligo rev</td><td>AAACATCTCCACCGTGTTGATGTT</td><td></td><td></td><td></td><td></td><td></td><td></td><td></td><td></td><td></td><td></td></tr>
<tr><td>Chromosome</td><td>start</td><td>end</td><td>strand</td><td>MM</td><td>target_seq</td><td>PAM</td><td>alignment</td><td>distance</td><td>position</td><td>gene name</td><td>gene id</td></tr>
</table>

<table>
<tr><td>GU305914.1</td><td>124153</td><td>124175</td><td>-</td><td>0</td><td>AACATCAACACGGTGGAGAT</td><td>GGG</td><td>||||||||[||||||||||||]PAM</td><td>NA</td><td>Intergenic</td><td>NA</td><td>NA</td></tr>
<tr><td></td><td></td><td></td><td></td><td></td><td></td><td></td><td></td><td></td><td></td><td></td><td></td></tr>
<tr><td>T24640</td><td>CAACACGGTGGAGATGGGCCTGG</td><td>1000</td><td></td><td></td><td></td><td></td><td></td><td></td><td></td><td></td><td></td></tr>
<tr><td>Oligo fwd</td><td>CACCCAACACGGTGGAGATGGGCC</td><td></td><td></td><td></td><td></td><td></td><td></td><td></td><td></td><td></td><td></td></tr>
<tr><td>Oligo rev</td><td>AAACGGCCCATCTCCACCGTGTTG</td><td></td><td></td><td></td><td></td><td></td><td></td><td></td><td></td><td></td><td></td></tr>
<tr><td>Chromosome</td><td>start</td><td>end</td><td>strand</td><td>MM</td><td>target_seq</td><td>PAM</td><td>alignment</td><td>distance</td><td>position</td><td>gene name</td><td>gene id</td></tr>
<tr><td>GU305914.1</td><td>124148</td><td>124170</td><td>-</td><td>0</td><td>CAACACGGTGGAGATGGGCC</td><td>TGG</td><td>||||||||[||||||||||||]PAM</td><td>NA</td><td>Intergenic</td><td>NA</td><td>NA</td></tr>
<tr><td></td><td></td><td></td><td></td><td></td><td></td><td></td><td></td><td></td><td></td><td></td><td></td></tr>
<tr><td>T24641</td><td>AACACGGTGGAGATGGGCCTGGG</td><td>1000</td><td></td><td></td><td></td><td></td><td></td><td></td><td></td><td></td><td></td></tr>
<tr><td>Oligo fwd</td><td>CACCAACACGGTGGAGATGGGCCT</td><td></td><td></td><td></td><td></td><td></td><td></td><td></td><td></td><td></td><td></td></tr>
<tr><td>Oligo rev</td><td>AAACAGGCCCATCTCCACCGTGTT</td><td></td><td></td><td></td><td></td><td></td><td></td><td></td><td></td><td></td><td></td></tr>
<tr><td>Chromosome</td><td>start</td><td>end</td><td>strand</td><td>MM</td><td>target_seq</td><td>PAM</td><td>alignment</td><td>distance</td><td>position</td><td>gene name</td><td>gene id</td></tr>
<tr><td>GU305914.1</td><td>124147</td><td>124169</td><td>-</td><td>0</td><td>AACACGGTGGAGATGGGCCT</td><td>GGG</td><td>||||||||[||||||||||||]PAM</td><td>NA</td><td>Intergenic</td><td>NA</td><td>NA</td></tr>
<tr><td></td><td></td><td></td><td></td><td></td><td></td><td></td><td></td><td></td><td></td><td></td><td></td></tr>
<tr><td>T24642</td><td>ACACGGTGGAGATGGGCCTGGGG</td><td>1000</td><td></td><td></td><td></td><td></td><td></td><td></td><td></td><td></td><td></td></tr>
<tr><td>Oligo fwd</td><td>CACCACACGGTGGAGATGGGCCTG</td><td></td><td></td><td></td><td></td><td></td><td></td><td></td><td></td><td></td><td></td></tr>
<tr><td>Oligo rev</td><td>AAACCAGGCCCATCTCCACCGTGT</td><td></td><td></td><td></td><td></td><td></td><td></td><td></td><td></td><td></td><td></td></tr>
<tr><td>Chromosome</td><td>start</td><td>end</td><td>strand</td><td>MM</td><td>target_seq</td><td>PAM</td><td>alignment</td><td>distance</td><td>position</td><td>gene name</td><td>gene id</td></tr>
<tr><td>GU305914.1</td><td>124146</td><td>124168</td><td>-</td><td>0</td><td>ACACGGTGGAGATGGGCCTG</td><td>GGG</td><td>||||||||[||||||||||||]PAM</td><td>NA</td><td>Intergenic</td><td>NA</td><td>NA</td></tr>
<tr><td></td><td></td><td></td><td></td><td></td><td></td><td></td><td></td><td></td><td></td><td></td><td></td></tr>
<tr><td>T24643</td><td>GTGGAGATGGGCCTGGGGTACGG</td><td>1000</td><td></td><td></td><td></td><td></td><td></td><td></td><td></td><td></td><td></td></tr>
<tr><td>Oligo fwd</td><td>CACCGTGGAGATGGGCCTGGGGTA</td><td></td><td></td><td></td><td></td><td></td><td></td><td></td><td></td><td></td><td></td></tr>
<tr><td>Oligo rev</td><td>AAACTACCCCAGGCCCATCTCCAC</td><td></td><td></td><td></td><td></td><td></td><td></td><td></td><td></td><td></td><td></td></tr>
<tr><td>Chromosome</td><td>start</td><td>end</td><td>strand</td><td>MM</td><td>target_seq</td><td>PAM</td><td>alignment</td><td>distance</td><td>position</td><td>gene name</td><td>gene id</td></tr>
<tr><td>GU305914.1</td><td>124141</td><td>124163</td><td>-</td><td>0</td><td>GTGGAGATGGGCCTGGGGTA</td><td>CGG</td><td>||||||||[||||||||||||]PAM</td><td>NA</td><td>Intergenic</td><td>NA</td><td>NA</td></tr>
<tr><td></td><td></td><td></td><td></td><td></td><td></td><td></td><td></td><td></td><td></td><td></td><td></td></tr>
<tr><td>T24644</td><td>TGGAGATGGGCCTGGGGTACGGG</td><td>1000</td><td></td><td></td><td></td><td></td><td></td><td></td><td></td><td></td><td></td></tr>
<tr><td>Oligo fwd</td><td>CACCTGGAGATGGGCCTGGGGTAC</td><td></td><td></td><td></td><td></td><td></td><td></td><td></td><td></td><td></td><td></td></tr>
<tr><td>Oligo rev</td><td>AAACGTACCCCAGGCCCATCTCCA</td><td></td><td></td><td></td><td></td><td></td><td></td><td></td><td></td><td></td><td></td></tr>
<tr><td>Chromosome</td><td>start</td><td>end</td><td>strand</td><td>MM</td><td>target_seq</td><td>PAM</td><td>alignment</td><td>distance</td><td>position</td><td>gene name</td><td>gene id</td></tr>
<tr><td>GU305914.1</td><td>124140</td><td>124162</td><td>-</td><td>0</td><td>TGGAGATGGGCCTGGGGTAC</td><td>GGG</td><td>||||||||[||||||||||||]PAM</td><td>NA</td><td>Intergenic</td><td>NA</td><td>NA</td></tr>
<tr><td></td><td></td><td></td><td></td><td></td><td></td><td></td><td></td><td></td><td></td><td></td><td></td></tr>
<tr><td>T24645</td><td>GGAGATGGGCCTGGGGTACGGGG</td><td>1000</td><td></td><td></td><td></td><td></td><td></td><td></td><td></td><td></td><td></td></tr>
<tr><td>Oligo fwd</td><td>CACCGGAGATGGGCCTGGGGTACG</td><td></td><td></td><td></td><td></td><td></td><td></td><td></td><td></td><td></td><td></td></tr>
<tr><td>Oligo rev</td><td>AAACCGTACCCCAGGCCCATCTCC</td><td></td><td></td><td></td><td></td><td></td><td></td><td></td><td></td><td></td><td></td></tr>
<tr><td>Chromosome</td><td>start</td><td>end</td><td>strand</td><td>MM</td><td>target_seq</td><td>PAM</td><td>alignment</td><td>distance</td><td>position</td><td>gene name</td><td>gene id</td></tr>
<tr><td>GU305914.1</td><td>124139</td><td>124161</td><td>-</td><td>0</td><td>GGAGATGGGCCTGGGGTACG</td><td>GGG</td><td>||||||||[||||||||||||]PAM</td><td>NA</td><td>Intergenic</td><td>NA</td><td>NA</td></tr>
<tr><td>GU305914.1</td><td>93579</td><td>93601</td><td>-</td><td>4</td><td>GGTGCTGGTCCTGGGGTCCG</td><td>GGG</td><td>||-|-|||[-||||||||-||]PAM</td><td>NA</td><td>Intergenic</td><td>NA</td><td>NA</td></tr>
<tr><td></td><td></td><td></td><td></td><td></td><td></td><td></td><td></td><td></td><td></td><td></td><td></td></tr>
<tr><td>T24646</td><td>GTACGGGGCGACGACGTGCACGG</td><td>1000</td><td></td><td></td><td></td><td></td><td></td><td></td><td></td><td></td><td></td></tr>
<tr><td>Oligo fwd</td><td>CACCGTACGGGGCGACGACGTGCA</td><td></td><td></td><td></td><td></td><td></td><td></td><td></td><td></td><td></td><td></td></tr>
<tr><td>Oligo rev</td><td>AAACTGCACGTCGTCGCCCCGTAC</td><td></td><td></td><td></td><td></td><td></td><td></td><td></td><td></td><td></td><td></td></tr>
<tr><td>Chromosome</td><td>start</td><td>end</td><td>strand</td><td>MM</td><td>target_seq</td><td>PAM</td><td>alignment</td><td>distance</td><td>position</td><td>gene name</td><td>gene id</td></tr>
<tr><td>GU305914.1</td><td>124124</td><td>124146</td><td>-</td><td>0</td><td>GTACGGGGCGACGACGTGCA</td><td>CGG</td><td>||||||||[||||||||||||]PAM</td><td>NA</td><td>Intergenic</td><td>NA</td><td>NA</td></tr>
<tr><td></td><td></td><td></td><td></td><td></td><td></td><td></td><td></td><td></td><td></td><td></td><td></td></tr>
<tr><td>T24647</td><td>CGACGTGCACGGCGCACCTGCGG</td><td>1000</td><td></td><td></td><td></td><td></td><td></td><td></td><td></td><td></td><td></td></tr>
<tr><td>Oligo fwd</td><td>CACCCGACGTGCACGGCGCACCTG</td><td></td><td></td><td></td><td></td><td></td><td></td><td></td><td></td><td></td><td></td></tr>
<tr><td>Oligo rev</td><td>AAACCAGGTGCGCCGTGCACGTCG</td><td></td><td></td><td></td><td></td><td></td><td></td><td></td><td></td><td></td><td></td></tr>
<tr><td>Chromosome</td><td>start</td><td>end</td><td>strand</td><td>MM</td><td>target_seq</td><td>PAM</td><td>alignment</td><td>distance</td><td>position</td><td>gene name</td><td>gene id</td></tr>
<tr><td>GU305914.1</td><td>124113</td><td>124135</td><td>-</td><td>0</td><td>CGACGTGCACGGCGCACCTG</td><td>CGG</td><td>||||||||[||||||||||||]PAM</td><td>NA</td><td>Intergenic</td><td>NA</td><td>NA</td></tr>
<tr><td></td><td></td><td></td><td></td><td></td><td></td><td></td><td></td><td></td><td></td><td></td><td></td></tr>
<tr><td>T24648</td><td>CGGCGCACCTGCGGCGCGTGCGG</td><td>1000</td><td></td><td></td><td></td><td></td><td></td><td></td><td></td><td></td><td></td></tr>
<tr><td>Oligo fwd</td><td>CACCCGGCGCACCTGCGGCGCGTG</td><td></td><td></td><td></td><td></td><td></td><td></td><td></td><td></td><td></td><td></td></tr>
<tr><td>Oligo rev</td><td>AAACCACGCGCCGCAGGTGCGCCG</td><td></td><td></td><td></td><td></td><td></td><td></td><td></td><td></td><td></td><td></td></tr>
<tr><td>Chromosome</td><td>start</td><td>end</td><td>strand</td><td>MM</td><td>target_seq</td><td>PAM</td><td>alignment</td><td>distance</td><td>position</td><td>gene name</td><td>gene id</td></tr>
<tr><td>GU305914.1</td><td>124104</td><td>124126</td><td>-</td><td>0</td><td>CGGCGCACCTGCGGCGCGTG</td><td>CGG</td><td>||||||||[||||||||||||]PAM</td><td>NA</td><td>Intergenic</td><td>NA</td><td>NA</td></tr>
<tr><td></td><td></td><td></td><td></td><td></td><td></td><td></td><td></td><td></td><td></td><td></td><td></td></tr>
<tr><td>T24649</td><td>GCGGCGCGTGCGGAGCGACATGG</td><td>1000</td><td></td><td></td><td></td><td></td><td></td><td></td><td></td><td></td><td></td></tr>
<tr><td>Oligo fwd</td><td>CACCGCGGCGCGTGCGGAGCGACA</td><td></td><td></td><td></td><td></td><td></td><td></td><td></td><td></td><td></td><td></td></tr>
<tr><td>Oligo rev</td><td>AAACTGTCGCTCCGCACGCGCCGC</td><td></td><td></td><td></td><td></td><td></td><td></td><td></td><td></td><td></td><td></td></tr>
<tr><td>Chromosome</td><td>start</td><td>end</td><td>strand</td><td>MM</td><td>target_seq</td><td>PAM</td><td>alignment</td><td>distance</td><td>position</td><td>gene name</td><td>gene id</td></tr>
<tr><td>GU305914.1</td><td>124094</td><td>124116</td><td>-</td><td>0</td><td>GCGGCGCGTGCGGAGCGACA</td><td>TGG</td><td>||||||||[||||||||||||]PAM</td><td>NA</td><td>Intergenic</td><td>NA</td><td>NA</td></tr>
<tr><td></td><td></td><td></td><td></td><td></td><td></td><td></td><td></td><td></td><td></td><td></td><td></td></tr>
<tr><td>T24650</td><td>CGGCGCGTGCGGAGCGACATGGG</td><td>1000</td><td></td><td></td><td></td><td></td><td></td><td></td><td></td><td></td><td></td></tr>
<tr><td>Oligo fwd</td><td>CACCCGGCGCGTGCGGAGCGACAT</td><td></td><td></td><td></td><td></td><td></td><td></td><td></td><td></td><td></td><td></td></tr>
<tr><td>Oligo rev</td><td>AAACATGTCGCTCCGCACGCGCCG</td><td></td><td></td><td></td><td></td><td></td><td></td><td></td><td></td><td></td><td></td></tr>
<tr><td>Chromosome</td><td>start</td><td>end</td><td>strand</td><td>MM</td><td>target_seq</td><td>PAM</td><td>alignment</td><td>distance</td><td>position</td><td>gene name</td><td>gene id</td></tr>
<tr><td>GU305914.1</td><td>124093</td><td>124115</td><td>-</td><td>0</td><td>CGGCGCGTGCGGAGCGACAT</td><td>GGG</td><td>||||||||[||||||||||||]PAM</td><td>NA</td><td>Intergenic</td><td>NA</td><td>NA</td></tr>
<tr><td></td><td></td><td></td><td></td><td></td><td></td><td></td><td></td><td></td><td></td><td></td><td></td></tr>
<tr><td>T24651</td><td>TGCGGAGCGACATGGGCTCGCGG</td><td>1000</td><td></td><td></td><td></td><td></td><td></td><td></td><td></td><td></td><td></td></tr>
<tr><td>Oligo fwd</td><td>CACCTGCGGAGCGACATGGGCTCG</td><td></td><td></td><td></td><td></td><td></td><td></td><td></td><td></td><td></td><td></td></tr>
<tr><td>Oligo rev</td><td>AAACCGAGCCCATGTCGCTCCGCA</td><td></td><td></td><td></td><td></td><td></td><td></td><td></td><td></td><td></td><td></td></tr>
<tr><td>Chromosome</td><td>start</td><td>end</td><td>strand</td><td>MM</td><td>target_seq</td><td>PAM</td><td>alignment</td><td>distance</td><td>position</td><td>gene name</td><td>gene id</td></tr>
<tr><td>GU305914.1</td><td>124086</td><td>124108</td><td>-</td><td>0</td><td>TGCGGAGCGACATGGGCTCG</td><td>CGG</td><td>||||||||[||||||||||||]PAM</td><td>NA</td><td>Intergenic</td><td>NA</td><td>NA</td></tr>
<tr><td></td><td></td><td></td><td></td><td></td><td></td><td></td><td></td><td></td><td></td><td></td><td></td></tr>
<tr><td>T24652</td><td>CGACATGGGCTCGCGGATGCAGG</td><td>1000</td><td></td><td></td><td></td><td></td><td></td><td></td><td></td><td></td><td></td></tr>
<tr><td>Oligo fwd</td><td>CACCCGACATGGGCTCGCGGATGC</td><td></td><td></td><td></td><td></td><td></td><td></td><td></td><td></td><td></td><td></td></tr>
<tr><td>Oligo rev</td><td>AAACGCATCCGCGAGCCCATGTCG</td><td></td><td></td><td></td><td></td><td></td><td></td><td></td><td></td><td></td><td></td></tr>
<tr><td>Chromosome</td><td>start</td><td>end</td><td>strand</td><td>MM</td><td>target_seq</td><td>PAM</td><td>alignment</td><td>distance</td><td>position</td><td>gene name</td><td>gene id</td></tr>
<tr><td>GU305914.1</td><td>124079</td><td>124101</td><td>-</td><td>0</td><td>CGACATGGGCTCGCGGATGC</td><td>AGG</td><td>||||||||[||||||||||||]PAM</td><td>NA</td><td>Intergenic</td><td>NA</td><td>NA</td></tr>
<tr><td></td><td></td><td></td><td></td><td></td><td></td><td></td><td></td><td></td><td></td><td></td><td></td></tr>
<tr><td>T24653</td><td>GCGGATGCAGGACCTGTTCCAGG</td><td>1000</td><td></td><td></td><td></td><td></td><td></td><td></td><td></td><td></td><td></td></tr>
<tr><td>Oligo fwd</td><td>CACCGCGGATGCAGGACCTGTTCC</td><td></td><td></td><td></td><td></td><td></td><td></td><td></td><td></td><td></td><td></td></tr>
<tr><td>Oligo rev</td><td>AAACGGAACAGGTCCTGCATCCGC</td><td></td><td></td><td></td><td></td><td></td><td></td><td></td><td></td><td></td><td></td></tr>
<tr><td>Chromosome</td><td>start</td><td>end</td><td>strand</td><td>MM</td><td>target_seq</td><td>PAM</td><td>alignment</td><td>distance</td><td>position</td><td>gene name</td><td>gene id</td></tr>
<tr><td>GU305914.1</td><td>124067</td><td>124089</td><td>-</td><td>0</td><td>GCGGATGCAGGACCTGTTCC</td><td>AGG</td><td>||||||||[||||||||||||]PAM</td><td>NA</td><td>Intergenic</td><td>NA</td><td>NA</td></tr>
<tr><td></td><td></td><td></td><td></td><td></td><td></td><td></td><td></td><td></td><td></td><td></td><td></td></tr>
<tr><td>T24654</td><td>GATGCACGTGTATCGCAACGAGG</td><td>1000</td><td></td><td></td><td></td><td></td><td></td><td></td><td></td><td></td><td></td></tr>
<tr><td>Oligo fwd</td><td>CACCGATGCACGTGTATCGCAACG</td><td></td><td></td><td></td><td></td><td></td><td></td><td></td><td></td><td></td><td></td></tr>
</table>

<table>
<tr><td>Oligo rev</td><td>AAACCGTTGCGATACACGTGCATC</td></tr>
<tr><td>Chromosome</td><td>start</td><td>end</td><td>strand</td><td>MM</td><td>target_seq</td><td>PAM</td><td>alignment</td><td>distance</td><td>position</td><td>gene name</td><td>gene id</td></tr>
<tr><td>GU305914.1</td><td>124037</td><td>124059</td><td>-</td><td>0</td><td>GATGCACGTGTATCGCAACG</td><td>AGG</td><td>||||||||[||||||||||||]PAM</td><td>NA</td><td>Intergenic</td><td>NA</td><td>NA</td></tr>
<tr><td>T24655</td><td>CGTGTATCGCAACGAGGACGTGG</td><td>1000</td></tr>
<tr><td>Oligo fwd</td><td>CACCCGTGTATCGCAACGAGGACG</td></tr>
<tr><td>Oligo rev</td><td>AAACCGTCCTCGTTGCGATACACG</td></tr>
<tr><td>Chromosome</td><td>start</td><td>end</td><td>strand</td><td>MM</td><td>target_seq</td><td>PAM</td><td>alignment</td><td>distance</td><td>position</td><td>gene name</td><td>gene id</td></tr>
<tr><td>GU305914.1</td><td>124031</td><td>124053</td><td>-</td><td>0</td><td>CGTGTATCGCAACGAGGACG</td><td>TGG</td><td>||||||||[||||||||||||]PAM</td><td>NA</td><td>Intergenic</td><td>NA</td><td>NA</td></tr>
<tr><td>T24656</td><td>GCAACGAGGACGTGGACGCCTGG</td><td>1000</td></tr>
<tr><td>Oligo fwd</td><td>CACCGCAACGAGGACGTGGACGCC</td></tr>
<tr><td>Oligo rev</td><td>AAACGGCGTCCACGTCCTCGTTGC</td></tr>
<tr><td>Chromosome</td><td>start</td><td>end</td><td>strand</td><td>MM</td><td>target_seq</td><td>PAM</td><td>alignment</td><td>distance</td><td>position</td><td>gene name</td><td>gene id</td></tr>
<tr><td>GU305914.1</td><td>124023</td><td>124045</td><td>-</td><td>0</td><td>GCAACGAGGACGTGGACGCC</td><td>TGG</td><td>||||||||[||||||||||||]PAM</td><td>NA</td><td>Intergenic</td><td>NA</td><td>NA</td></tr>
<tr><td>T24657</td><td>CAACGAGGACGTGGACGCCTGGG</td><td>1000</td></tr>
<tr><td>Oligo fwd</td><td>CACCCAACGAGGACGTGGACGCCT</td></tr>
<tr><td>Oligo rev</td><td>AAACAGGCGTCCACGTCCTCGTTG</td></tr>
<tr><td>Chromosome</td><td>start</td><td>end</td><td>strand</td><td>MM</td><td>target_seq</td><td>PAM</td><td>alignment</td><td>distance</td><td>position</td><td>gene name</td><td>gene id</td></tr>
<tr><td>GU305914.1</td><td>124022</td><td>124044</td><td>-</td><td>0</td><td>CAACGAGGACGTGGACGCCT</td><td>GGG</td><td>||||||||[||||||||||||]PAM</td><td>NA</td><td>Intergenic</td><td>NA</td><td>NA</td></tr>
<tr><td>T24658</td><td>CGTGGACGCCTGGGTCCGACAGG</td><td>1000</td></tr>
<tr><td>Oligo fwd</td><td>CACCCGTGGACGCCTGGGTCCGAC</td></tr>
<tr><td>Oligo rev</td><td>AAACGTCGGACCCAGGCGTCCACG</td></tr>
<tr><td>Chromosome</td><td>start</td><td>end</td><td>strand</td><td>MM</td><td>target_seq</td><td>PAM</td><td>alignment</td><td>distance</td><td>position</td><td>gene name</td><td>gene id</td></tr>
<tr><td>GU305914.1</td><td>124013</td><td>124035</td><td>-</td><td>0</td><td>CGTGGACGCCTGGGTCCGAC</td><td>AGG</td><td>||||||||[||||||||||||]PAM</td><td>NA</td><td>Intergenic</td><td>NA</td><td>NA</td></tr>
<tr><td>T24659</td><td>GCCTGGGTCCGACAGGCGACCGG</td><td>1000</td></tr>
<tr><td>Oligo fwd</td><td>CACCGCCTGGGTCCGACAGGCGAC</td></tr>
<tr><td>Oligo rev</td><td>AAACGTCGCCTGTCGGACCCAGGC</td></tr>
<tr><td>Chromosome</td><td>start</td><td>end</td><td>strand</td><td>MM</td><td>target_seq</td><td>PAM</td><td>alignment</td><td>distance</td><td>position</td><td>gene name</td><td>gene id</td></tr>
<tr><td>GU305914.1</td><td>124006</td><td>124028</td><td>-</td><td>0</td><td>GCCTGGGTCCGACAGGCGAC</td><td>CGG</td><td>||||||||[||||||||||||]PAM</td><td>NA</td><td>Intergenic</td><td>NA</td><td>NA</td></tr>
<tr><td>T24660</td><td>TCCGACAGGCGACCGGCGCGCGG</td><td>1000</td></tr>
<tr><td>Oligo fwd</td><td>CACCTCCGACAGGCGACCGGCGCG</td></tr>
<tr><td>Oligo rev</td><td>AAACCGCGCCGGTCGCCTGTCGGA</td></tr>
<tr><td>Chromosome</td><td>start</td><td>end</td><td>strand</td><td>MM</td><td>target_seq</td><td>PAM</td><td>alignment</td><td>distance</td><td>position</td><td>gene name</td><td>gene id</td></tr>
<tr><td>GU305914.1</td><td>123999</td><td>124021</td><td>-</td><td>0</td><td>TCCGACAGGCGACCGGCGCG</td><td>CGG</td><td>||||||||[||||||||||||]PAM</td><td>NA</td><td>Intergenic</td><td>NA</td><td>NA</td></tr>
<tr><td>T24661</td><td>GACCGGCGCGCGGCGCACCGAGG</td><td>1000</td></tr>
<tr><td>Oligo fwd</td><td>CACCGACCGGCGCGCGGCGCACCG</td></tr>
<tr><td>Oligo rev</td><td>AAACCGGTGCGCCGCGCGCCGGTC</td></tr>
<tr><td>Chromosome</td><td>start</td><td>end</td><td>strand</td><td>MM</td><td>target_seq</td><td>PAM</td><td>alignment</td><td>distance</td><td>position</td><td>gene name</td><td>gene id</td></tr>
<tr><td>GU305914.1</td><td>123989</td><td>124011</td><td>-</td><td>0</td><td>GACCGGCGCGCGGCGCACCG</td><td>AGG</td><td>||||||||[||||||||||||]PAM</td><td>NA</td><td>Intergenic</td><td>NA</td><td>NA</td></tr>
<tr><td>T24662</td><td>CGCGCGGCGCACCGAGGTGCTGG</td><td>1000</td></tr>
<tr><td>Oligo fwd</td><td>CACCCGCGCGGCGCACCGAGGTGC</td></tr>
<tr><td>Oligo rev</td><td>AAACGCACCTCGGTGCGCCGCGCG</td></tr>
<tr><td>Chromosome</td><td>start</td><td>end</td><td>strand</td><td>MM</td><td>target_seq</td><td>PAM</td><td>alignment</td><td>distance</td><td>position</td><td>gene name</td><td>gene id</td></tr>
<tr><td>GU305914.1</td><td>123983</td><td>124005</td><td>-</td><td>0</td><td>CGCGCGGCGCACCGAGGTGC</td><td>TGG</td><td>||||||||[||||||||||||]PAM</td><td>NA</td><td>Intergenic</td><td>NA</td><td>NA</td></tr>
<tr><td>T24663</td><td>GCGCACCGAGGTGCTGGACTCGG</td><td>1000</td></tr>
<tr><td>Oligo fwd</td><td>CACCGCGCACCGAGGTGCTGGACT</td></tr>
<tr><td>Oligo rev</td><td>AAACAGTCCAGCACCTCGGTGCGC</td></tr>
<tr><td>Chromosome</td><td>start</td><td>end</td><td>strand</td><td>MM</td><td>target_seq</td><td>PAM</td><td>alignment</td><td>distance</td><td>position</td><td>gene name</td><td>gene id</td></tr>
<tr><td>GU305914.1</td><td>123977</td><td>123999</td><td>-</td><td>0</td><td>GCGCACCGAGGTGCTGGACT</td><td>CGG</td><td>||||||||[||||||||||||]PAM</td><td>NA</td><td>Intergenic</td><td>NA</td><td>NA</td></tr>
<tr><td>T24664</td><td>CACCGAGGTGCTGGACTCGGAGG</td><td>1000</td></tr>
<tr><td>Oligo fwd</td><td>CACCCACCGAGGTGCTGGACTCGG</td></tr>
<tr><td>Oligo rev</td><td>AAACCCGAGTCCAGCACCTCGGTG</td></tr>
<tr><td>Chromosome</td><td>start</td><td>end</td><td>strand</td><td>MM</td><td>target_seq</td><td>PAM</td><td>alignment</td><td>distance</td><td>position</td><td>gene name</td><td>gene id</td></tr>
<tr><td>GU305914.1</td><td>123974</td><td>123996</td><td>-</td><td>0</td><td>CACCGAGGTGCTGGACTCGG</td><td>AGG</td><td>||||||||[||||||||||||]PAM</td><td>NA</td><td>Intergenic</td><td>NA</td><td>NA</td></tr>
<tr><td>GU305914.1</td><td>69505</td><td>69527</td><td>+</td><td>4</td><td>CGCCGACCTGCTGGACTCGA</td><td>CGG</td><td>|-||||--[|||||||||||-]PAM</td><td>NA</td><td>Intergenic</td><td>NA</td><td>NA</td></tr>
<tr><td>T24665</td><td>GCCATCTCGATCCTGACCTTCGG</td><td>1000</td></tr>
<tr><td>Oligo fwd</td><td>CACCGCCATCTCGATCCTGACCTT</td></tr>
<tr><td>Oligo rev</td><td>AAACAAGGTCAGGATCGAGATGGC</td></tr>
<tr><td>Chromosome</td><td>start</td><td>end</td><td>strand</td><td>MM</td><td>target_seq</td><td>PAM</td><td>alignment</td><td>distance</td><td>position</td><td>gene name</td><td>gene id</td></tr>
<tr><td>GU305914.1</td><td>123952</td><td>123974</td><td>-</td><td>0</td><td>GCCATCTCGATCCTGACCTT</td><td>CGG</td><td>||||||||[||||||||||||]PAM</td><td>NA</td><td>Intergenic</td><td>NA</td><td>NA</td></tr>
<tr><td>T24666</td><td>CCATCTCGATCCTGACCTTCGGG</td><td>1000</td></tr>
<tr><td>Oligo fwd</td><td>CACCCCATCTCGATCCTGACCTTC</td></tr>
<tr><td>Oligo rev</td><td>AAACGAAGGTCAGGATCGAGATGG</td></tr>
<tr><td>Chromosome</td><td>start</td><td>end</td><td>strand</td><td>MM</td><td>target_seq</td><td>PAM</td><td>alignment</td><td>distance</td><td>position</td><td>gene name</td><td>gene id</td></tr>
<tr><td>GU305914.1</td><td>123951</td><td>123973</td><td>-</td><td>0</td><td>CCATCTCGATCCTGACCTTC</td><td>GGG</td><td>||||||||[||||||||||||]PAM</td><td>NA</td><td>Intergenic</td><td>NA</td><td>NA</td></tr>
<tr><td>T24667</td><td>CCTGACCTTCGGGCGCAAGACGG</td><td>1000</td></tr>
<tr><td>Oligo fwd</td><td>CACCCCTGACCTTCGGGCGCAAGA</td></tr>
<tr><td>Oligo rev</td><td>AAACTCTTGCGCCCGAAGGTCAGG</td></tr>
<tr><td>Chromosome</td><td>start</td><td>end</td><td>strand</td><td>MM</td><td>target_seq</td><td>PAM</td><td>alignment</td><td>distance</td><td>position</td><td>gene name</td><td>gene id</td></tr>
<tr><td>GU305914.1</td><td>123941</td><td>123963</td><td>-</td><td>0</td><td>CCTGACCTTCGGGCGCAAGA</td><td>CGG</td><td>||||||||[||||||||||||]PAM</td><td>NA</td><td>Intergenic</td><td>NA</td><td>NA</td></tr>
<tr><td>T24668</td><td>CTTCGGGCGCAAGACGGACAAGG</td><td>1000</td></tr>
<tr><td>Oligo fwd</td><td>CACCCTTCGGGCGCAAGACGGACA</td></tr>
<tr><td>Oligo rev</td><td>AAACTGTCCGTCTTGCGCCCGAAG</td></tr>
<tr><td>Chromosome</td><td>start</td><td>end</td><td>strand</td><td>MM</td><td>target_seq</td><td>PAM</td><td>alignment</td><td>distance</td><td>position</td><td>gene name</td><td>gene id</td></tr>
<tr><td>GU305914.1</td><td>123935</td><td>123957</td><td>-</td><td>0</td><td>CTTCGGGCGCAAGACGGACA</td><td>AGG</td><td>||||||||[||||||||||||]PAM</td><td>NA</td><td>Intergenic</td><td>NA</td><td>NA</td></tr>
</table>

| | | | | | | | | | | | |
|---|---|---|---|---|---|---|---|---|---|---|---|
| T24669 | TTCGGGCGCAAGACGGACAAGGG | 1000 | | | | | | | | | |
| Oligo fwd | CACCTTCGGGCGCAAGACGGACAA | | | | | | | | | | |
| Oligo rev | AAACTTGTCCGTCTTGCGCCCGAA | | | | | | | | | | |
| Chromosome | start | end | strand | MM | target_seq | PAM | alignment | distance | position | gene name | gene id |
| GU305914.1 | 123934 | 123956 | - | 0 | TTCGGGCGCAAGACGGACAA | GGG | ||||||||[||||||||||||]PAM | NA | Intergenic | NA | NA |
| T24670 | GGGCGCAAGACGGACAAGGGCGG | 1000 | | | | | | | | | |
| Oligo fwd | CACCGGGCGCAAGACGGACAAGGG | | | | | | | | | | |
| Oligo rev | AAACCCCTTGTCCGTCTTGCGCCC | | | | | | | | | | |
| Chromosome | start | end | strand | MM | target_seq | PAM | alignment | distance | position | gene name | gene id |
| GU305914.1 | 123931 | 123953 | - | 0 | GGGCGCAAGACGGACAAGGG | CGG | ||||||||[||||||||||||]PAM | NA | Intergenic | NA | NA |
| T24671 | CAAGACGGACAAGGGCGGCCCGG | 1000 | | | | | | | | | |
| Oligo fwd | CACCCAAGACGGACAAGGGCGGCC | | | | | | | | | | |
| Oligo rev | AAACGGCCGCCCTTGTCCGTCTTG | | | | | | | | | | |
| Chromosome | start | end | strand | MM | target_seq | PAM | alignment | distance | position | gene name | gene id |
| GU305914.1 | 123926 | 123948 | - | 0 | CAAGACGGACAAGGGCGGCC | CGG | ||||||||[||||||||||||]PAM | NA | Intergenic | NA | NA |
| T24672 | GGCGGCCCGGCGCTGCTGCACGG | 1000 | | | | | | | | | |
| Oligo fwd | CACCGGCGGCCCGGCGCTGCTGCA | | | | | | | | | | |
| Oligo rev | AAACTGCAGCAGCGCCGGGCCGCC | | | | | | | | | | |
| Chromosome | start | end | strand | MM | target_seq | PAM | alignment | distance | position | gene name | gene id |
| GU305914.1 | 123913 | 123935 | - | 0 | GGCGGCCCGGCGCTGCTGCA | CGG | ||||||||[||||||||||||]PAM | NA | Intergenic | NA | NA |
| T24673 | GCGGCCCGGCGCTGCTGCACGGG | 1000 | | | | | | | | | |
| Oligo fwd | CACCGCGGCCCGGCGCTGCTGCAC | | | | | | | | | | |
| Oligo rev | AAACGTGCAGCAGCGCCGGGCCGC | | | | | | | | | | |
| Chromosome | start | end | strand | MM | target_seq | PAM | alignment | distance | position | gene name | gene id |
| GU305914.1 | 123912 | 123934 | - | 0 | GCGGCCCGGCGCTGCTGCAC | GGG | ||||||||[||||||||||||]PAM | NA | Intergenic | NA | NA |
| T24674 | CGGGCAGCGCGCCACGTGCGAGG | 1000 | | | | | | | | | |
| Oligo fwd | CACCCGGGCAGCGCGCCACGTGCG | | | | | | | | | | |
| Oligo rev | AAACCGCACGTGGCGCGCTGCCCG | | | | | | | | | | |
| Chromosome | start | end | strand | MM | target_seq | PAM | alignment | distance | position | gene name | gene id |
| GU305914.1 | 123893 | 123915 | - | 0 | CGGGCAGCGCGCCACGTGCG | AGG | ||||||||[||||||||||||]PAM | NA | Intergenic | NA | NA |
| GU305914.1 | 521 | 543 | + | 3 | CGGGCCGCGCGCCGCGTCCG | CGG | |||||-||[|||||-|||-||]PAM | NA | Intergenic | NA | NA |
| T24675 | GTGCGAGGTGATCCTGACGCCGG | 1000 | | | | | | | | | |
| Oligo fwd | CACCGTGCGAGGTGATCCTGACGC | | | | | | | | | | |
| Oligo rev | AAACGCGTCAGGATCACCTCGCAC | | | | | | | | | | |
| Chromosome | start | end | strand | MM | target_seq | PAM | alignment | distance | position | gene name | gene id |
| GU305914.1 | 123878 | 123900 | - | 0 | GTGCGAGGTGATCCTGACGC | CGG | ||||||||[||||||||||||]PAM | NA | Intergenic | NA | NA |
| T24676 | GGTGATCCTGACGCCGGTGTCGG | 1000 | | | | | | | | | |
| Oligo fwd | CACCGGTGATCCTGACGCCGGTGT | | | | | | | | | | |
| Oligo rev | AAACACACCGGCGTCAGGATCACC | | | | | | | | | | |
| Chromosome | start | end | strand | MM | target_seq | PAM | alignment | distance | position | gene name | gene id |
| GU305914.1 | 123872 | 123894 | - | 0 | GGTGATCCTGACGCCGGTGT | CGG | ||||||||[||||||||||||]PAM | NA | Intergenic | NA | NA |
| T24677 | GACGCCGGTGTCGGCCGACCTGG | 1000 | | | | | | | | | |
| Oligo fwd | CACCGACGCCGGTGTCGGCCGACC | | | | | | | | | | |
| Oligo rev | AAACGGTCGGCCGACACCGGCGTC | | | | | | | | | | |
| Chromosome | start | end | strand | MM | target_seq | PAM | alignment | distance | position | gene name | gene id |
| GU305914.1 | 123863 | 123885 | - | 0 | GACGCCGGTGTCGGCCGACC | TGG | ||||||||[||||||||||||]PAM | NA | Intergenic | NA | NA |
| T24678 | CGCTACCCGAACAACCCGCGCGG | 1000 | | | | | | | | | |
| Oligo fwd | CACCCGCTACCCGAACAACCCGCG | | | | | | | | | | |
| Oligo rev | AAACCGCGGGTTGTTCGGGTAGCG | | | | | | | | | | |
| Chromosome | start | end | strand | MM | target_seq | PAM | alignment | distance | position | gene name | gene id |
| GU305914.1 | 123832 | 123854 | - | 0 | CGCTACCCGAACAACCCGCG | CGG | ||||||||[||||||||||||]PAM | NA | Intergenic | NA | NA |
| T24679 | GCTACCCGAACAACCCGCGCGGG | 1000 | | | | | | | | | |
| Oligo fwd | CACCGCTACCCGAACAACCCGCGC | | | | | | | | | | |
| Oligo rev | AAACGCGCGGGTTGTTCGGGTAGC | | | | | | | | | | |
| Chromosome | start | end | strand | MM | target_seq | PAM | alignment | distance | position | gene name | gene id |
| GU305914.1 | 123831 | 123853 | - | 0 | GCTACCCGAACAACCCGCGC | GGG | ||||||||[||||||||||||]PAM | NA | Intergenic | NA | NA |
| T24680 | CGGGCGCTCGTCGTCGATGCTGG | 1000 | | | | | | | | | |
| Oligo fwd | CACCCGGGCGCTCGTCGTCGATGC | | | | | | | | | | |
| Oligo rev | AAACGCATCGACGACGAGCGCCCG | | | | | | | | | | |
| Chromosome | start | end | strand | MM | target_seq | PAM | alignment | distance | position | gene name | gene id |
| GU305914.1 | 123812 | 123834 | - | 0 | CGGGCGCTCGTCGTCGATGC | TGG | ||||||||[||||||||||||]PAM | NA | Intergenic | NA | NA |
| T24681 | GGGCGCTCGTCGTCGATGCTGGG | 1000 | | | | | | | | | |
| Oligo fwd | CACCGGGCGCTCGTCGTCGATGCT | | | | | | | | | | |
| Oligo rev | AAACAGCATCGACGACGAGCGCCC | | | | | | | | | | |
| Chromosome | start | end | strand | MM | target_seq | PAM | alignment | distance | position | gene name | gene id |
| GU305914.1 | 123811 | 123833 | - | 0 | GGGCGCTCGTCGTCGATGCT | GGG | ||||||||[||||||||||||]PAM | NA | Intergenic | NA | NA |
| T24682 | GGGCGTCGACCCGTACGACGAGG | 1000 | | | | | | | | | |
| Oligo fwd | CACCGGGCGTCGACCCGTACGACG | | | | | | | | | | |
| Oligo rev | AAACCGTCGTACGGGTCGACGCCC | | | | | | | | | | |
| Chromosome | start | end | strand | MM | target_seq | PAM | alignment | distance | position | gene name | gene id |
| GU305914.1 | 123791 | 123813 | - | 0 | GGGCGTCGACCCGTACGACG | AGG | ||||||||[||||||||||||]PAM | NA | Intergenic | NA | NA |
| T24683 | CGACCCGTACGACGAGGACGCGG | 1000 | | | | | | | | | |
| Oligo fwd | CACCCGACCCGTACGACGAGGACG | | | | | | | | | | |
| Oligo rev | AAACCGTCCTCGTCGTACGGGTCG | | | | | | | | | | |
| Chromosome | start | end | strand | MM | target_seq | PAM | alignment | distance | position | gene name | gene id |

<table>
<tr><td>GU305914.1</td><td>123785</td><td>123807</td><td>-</td><td>0</td><td>CGACCCGTACGACGAGGACG</td><td>CGG</td><td>||||||||[||||||||||||]PAM</td><td>NA</td><td>Intergenic</td><td>NA</td><td>NA</td></tr>
<tr><td>GU305914.1</td><td>59739</td><td>59761</td><td>-</td><td>2</td><td>CGACTCGGACGACGAGGACG</td><td>AGG</td><td>||||-||-[||||||||||||]PAM</td><td>NA</td><td>Intergenic</td><td>NA</td><td>NA</td></tr>
<tr><td>GU305914.1</td><td>116060</td><td>116082</td><td>-</td><td>3</td><td>CGACTCGTACGAGGAGGAGG</td><td>CGG</td><td>||||-|||[||||-|||||-|]PAM</td><td>NA</td><td>Intergenic</td><td>NA</td><td>NA</td></tr>
<tr><td></td></tr>
<tr><td>T24684</td><td>GTACGACGAGGACGCGGCGCTGG</td><td>1000</td></tr>
<tr><td>Oligo fwd</td><td>CACCGTACGACGAGGACGCGGCGC</td></tr>
<tr><td>Oligo rev</td><td>AAACGCGCCGCGTCCTCGTCGTAC</td></tr>
<tr><td>Chromosome</td><td>start</td><td>end</td><td>strand</td><td>MM</td><td>target_seq</td><td>PAM</td><td>alignment</td><td>distance</td><td>position</td><td>gene name</td><td>gene id</td></tr>
<tr><td>GU305914.1</td><td>123779</td><td>123801</td><td>-</td><td>0</td><td>GTACGACGAGGACGCGGCGC</td><td>TGG</td><td>||||||||[||||||||||||]PAM</td><td>NA</td><td>Intergenic</td><td>NA</td><td>NA</td></tr>
<tr><td></td></tr>
<tr><td>T24685</td><td>CCTGTACGACCACACCAGCCCGG</td><td>1000</td></tr>
<tr><td>Oligo fwd</td><td>CACCCCTGTACGACCACACCAGCC</td></tr>
<tr><td>Oligo rev</td><td>AAACGGCTGGTGTGGTCGTACAGG</td></tr>
<tr><td>Chromosome</td><td>start</td><td>end</td><td>strand</td><td>MM</td><td>target_seq</td><td>PAM</td><td>alignment</td><td>distance</td><td>position</td><td>gene name</td><td>gene id</td></tr>
<tr><td>GU305914.1</td><td>123752</td><td>123774</td><td>-</td><td>0</td><td>CCTGTACGACCACACCAGCC</td><td>CGG</td><td>||||||||[||||||||||||]PAM</td><td>NA</td><td>Intergenic</td><td>NA</td><td>NA</td></tr>
<tr><td></td></tr>
<tr><td>T24686</td><td>TCGTGTCGACCAACAACCCGTGG</td><td>1000</td></tr>
<tr><td>Oligo fwd</td><td>CACCTCGTGTCGACCAACAACCCG</td></tr>
<tr><td>Oligo rev</td><td>AAACCGGGTTGTTGGTCGACACGA</td></tr>
<tr><td>Chromosome</td><td>start</td><td>end</td><td>strand</td><td>MM</td><td>target_seq</td><td>PAM</td><td>alignment</td><td>distance</td><td>position</td><td>gene name</td><td>gene id</td></tr>
<tr><td>GU305914.1</td><td>123717</td><td>123739</td><td>-</td><td>0</td><td>TCGTGTCGACCAACAACCCG</td><td>TGG</td><td>||||||||[||||||||||||]PAM</td><td>NA</td><td>Intergenic</td><td>NA</td><td>NA</td></tr>
<tr><td></td></tr>
<tr><td>T24687</td><td>CGTGTCGACCAACAACCCGTGGG</td><td>1000</td></tr>
<tr><td>Oligo fwd</td><td>CACCCGTGTCGACCAACAACCCGT</td></tr>
<tr><td>Oligo rev</td><td>AAACACGGGTTGTTGGTCGACACG</td></tr>
<tr><td>Chromosome</td><td>start</td><td>end</td><td>strand</td><td>MM</td><td>target_seq</td><td>PAM</td><td>alignment</td><td>distance</td><td>position</td><td>gene name</td><td>gene id</td></tr>
<tr><td>GU305914.1</td><td>123716</td><td>123738</td><td>-</td><td>0</td><td>CGTGTCGACCAACAACCCGT</td><td>GGG</td><td>||||||||[||||||||||||]PAM</td><td>NA</td><td>Intergenic</td><td>NA</td><td>NA</td></tr>
<tr><td></td></tr>
<tr><td>T24688</td><td>GTGTCGACCAACAACCCGTGGGG</td><td>1000</td></tr>
<tr><td>Oligo fwd</td><td>CACCGTGTCGACCAACAACCCGTG</td></tr>
<tr><td>Oligo rev</td><td>AAACCACGGGTTGTTGGTCGACAC</td></tr>
<tr><td>Chromosome</td><td>start</td><td>end</td><td>strand</td><td>MM</td><td>target_seq</td><td>PAM</td><td>alignment</td><td>distance</td><td>position</td><td>gene name</td><td>gene id</td></tr>
<tr><td>GU305914.1</td><td>123715</td><td>123737</td><td>-</td><td>0</td><td>GTGTCGACCAACAACCCGTG</td><td>GGG</td><td>||||||||[||||||||||||]PAM</td><td>NA</td><td>Intergenic</td><td>NA</td><td>NA</td></tr>
<tr><td></td></tr>
<tr><td>T24689</td><td>AACCCGTGGGGCTCGCAGCGCGG</td><td>1000</td></tr>
<tr><td>Oligo fwd</td><td>CACCAACCCGTGGGGCTCGCAGCG</td></tr>
<tr><td>Oligo rev</td><td>AAACCGCTGCGAGCCCCACGGGTT</td></tr>
<tr><td>Chromosome</td><td>start</td><td>end</td><td>strand</td><td>MM</td><td>target_seq</td><td>PAM</td><td>alignment</td><td>distance</td><td>position</td><td>gene name</td><td>gene id</td></tr>
<tr><td>GU305914.1</td><td>123703</td><td>123725</td><td>-</td><td>0</td><td>AACCCGTGGGGCTCGCAGCG</td><td>CGG</td><td>||||||||[||||||||||||]PAM</td><td>NA</td><td>Intergenic</td><td>NA</td><td>NA</td></tr>
<tr><td></td></tr>
<tr><td>T24690</td><td>ACCCGTGGGGCTCGCAGCGCGGG</td><td>1000</td></tr>
<tr><td>Oligo fwd</td><td>CACCACCCGTGGGGCTCGCAGCGC</td></tr>
<tr><td>Oligo rev</td><td>AAACGCGCTGCGAGCCCCACGGGT</td></tr>
<tr><td>Chromosome</td><td>start</td><td>end</td><td>strand</td><td>MM</td><td>target_seq</td><td>PAM</td><td>alignment</td><td>distance</td><td>position</td><td>gene name</td><td>gene id</td></tr>
<tr><td>GU305914.1</td><td>123702</td><td>123724</td><td>-</td><td>0</td><td>ACCCGTGGGGCTCGCAGCGC</td><td>GGG</td><td>||||||||[||||||||||||]PAM</td><td>NA</td><td>Intergenic</td><td>NA</td><td>NA</td></tr>
<tr><td></td></tr>
<tr><td>T24691</td><td>GGGCTCGCAGCGCGGGTCGCTGG</td><td>1000</td></tr>
<tr><td>Oligo fwd</td><td>CACCGGGCTCGCAGCGCGGGTCGC</td></tr>
<tr><td>Oligo rev</td><td>AAACGCGACCCGCGCTGCGAGCCC</td></tr>
<tr><td>Chromosome</td><td>start</td><td>end</td><td>strand</td><td>MM</td><td>target_seq</td><td>PAM</td><td>alignment</td><td>distance</td><td>position</td><td>gene name</td><td>gene id</td></tr>
<tr><td>GU305914.1</td><td>123695</td><td>123717</td><td>-</td><td>0</td><td>GGGCTCGCAGCGCGGGTCGC</td><td>TGG</td><td>||||||||[||||||||||||]PAM</td><td>NA</td><td>Intergenic</td><td>NA</td><td>NA</td></tr>
<tr><td></td></tr>
<tr><td>T24692</td><td>GGCTCGCAGCGCGGGTCGCTGGG</td><td>1000</td></tr>
<tr><td>Oligo fwd</td><td>CACCGGCTCGCAGCGCGGGTCGCT</td></tr>
<tr><td>Oligo rev</td><td>AAACAGCGACCCGCGCTGCGAGCC</td></tr>
<tr><td>Chromosome</td><td>start</td><td>end</td><td>strand</td><td>MM</td><td>target_seq</td><td>PAM</td><td>alignment</td><td>distance</td><td>position</td><td>gene name</td><td>gene id</td></tr>
<tr><td>GU305914.1</td><td>123694</td><td>123716</td><td>-</td><td>0</td><td>GGCTCGCAGCGCGGGTCGCT</td><td>GGG</td><td>||||||||[||||||||||||]PAM</td><td>NA</td><td>Intergenic</td><td>NA</td><td>NA</td></tr>
<tr><td></td></tr>
<tr><td>T24693</td><td>CACCCGCAACCGCGAGAAGCTGG</td><td>1000</td></tr>
<tr><td>Oligo fwd</td><td>CACCCACCCGCAACCGCGAGAAGC</td></tr>
<tr><td>Oligo rev</td><td>AAACGCTTCTCGCGGTTGCGGGTG</td></tr>
<tr><td>Chromosome</td><td>start</td><td>end</td><td>strand</td><td>MM</td><td>target_seq</td><td>PAM</td><td>alignment</td><td>distance</td><td>position</td><td>gene name</td><td>gene id</td></tr>
<tr><td>GU305914.1</td><td>123656</td><td>123678</td><td>-</td><td>0</td><td>CACCCGCAACCGCGAGAAGC</td><td>TGG</td><td>||||||||[||||||||||||]PAM</td><td>NA</td><td>Intergenic</td><td>NA</td><td>NA</td></tr>
<tr><td></td></tr>
<tr><td>T24694</td><td>ACCCGCAACCGCGAGAAGCTGGG</td><td>1000</td></tr>
<tr><td>Oligo fwd</td><td>CACCACCCGCAACCGCGAGAAGCT</td></tr>
<tr><td>Oligo rev</td><td>AAACAGCTTCTCGCGGTTGCGGGT</td></tr>
<tr><td>Chromosome</td><td>start</td><td>end</td><td>strand</td><td>MM</td><td>target_seq</td><td>PAM</td><td>alignment</td><td>distance</td><td>position</td><td>gene name</td><td>gene id</td></tr>
<tr><td>GU305914.1</td><td>123655</td><td>123677</td><td>-</td><td>0</td><td>ACCCGCAACCGCGAGAAGCT</td><td>GGG</td><td>||||||||[||||||||||||]PAM</td><td>NA</td><td>Intergenic</td><td>NA</td><td>NA</td></tr>
<tr><td></td></tr>
<tr><td>T24695</td><td>GTGCGCGCAGTTCTTCACGACGG</td><td>1000</td></tr>
<tr><td>Oligo fwd</td><td>CACCGTGCGCGCAGTTCTTCACGA</td></tr>
<tr><td>Oligo rev</td><td>AAACTCGTGAAGAACTGCGCGCAC</td></tr>
<tr><td>Chromosome</td><td>start</td><td>end</td><td>strand</td><td>MM</td><td>target_seq</td><td>PAM</td><td>alignment</td><td>distance</td><td>position</td><td>gene name</td><td>gene id</td></tr>
<tr><td>GU305914.1</td><td>123608</td><td>123630</td><td>-</td><td>0</td><td>GTGCGCGCAGTTCTTCACGA</td><td>CGG</td><td>||||||||[||||||||||||]PAM</td><td>NA</td><td>Intergenic</td><td>NA</td><td>NA</td></tr>
<tr><td></td></tr>
<tr><td>T24696</td><td>GAACAAGACCCTGTTCAAGACGG</td><td>1000</td></tr>
<tr><td>Oligo fwd</td><td>CACCGAACAAGACCCTGTTCAAGA</td></tr>
<tr><td>Oligo rev</td><td>AAACTCTTGAACAGGGTCTTGTTC</td></tr>
<tr><td>Chromosome</td><td>start</td><td>end</td><td>strand</td><td>MM</td><td>target_seq</td><td>PAM</td><td>alignment</td><td>distance</td><td>position</td><td>gene name</td><td>gene id</td></tr>
<tr><td>GU305914.1</td><td>123569</td><td>123591</td><td>-</td><td>0</td><td>GAACAAGACCCTGTTCAAGA</td><td>CGG</td><td>||||||||[||||||||||||]PAM</td><td>NA</td><td>Intergenic</td><td>NA</td><td>NA</td></tr>
<tr><td></td></tr>
<tr><td>T24697</td><td>CAAGACCCTGTTCAAGACGGTGG</td><td>1000</td></tr>
<tr><td>Oligo fwd</td><td>CACCCAAGACCCTGTTCAAGACGG</td></tr>
<tr><td>Oligo rev</td><td>AAACCCGTCTTGAACAGGGTCTTG</td></tr>
<tr><td>Chromosome</td><td>start</td><td>end</td><td>strand</td><td>MM</td><td>target_seq</td><td>PAM</td><td>alignment</td><td>distance</td><td>position</td><td>gene name</td><td>gene id</td></tr>
<tr><td>GU305914.1</td><td>123566</td><td>123588</td><td>-</td><td>0</td><td>CAAGACCCTGTTCAAGACGG</td><td>TGG</td><td>||||||||[||||||||||||]PAM</td><td>NA</td><td>Intergenic</td><td>NA</td><td>NA</td></tr>
<tr><td></td></tr>
<tr><td>T24698</td><td>GACCCTGTTCAAGACGGTGGAGG</td><td>1000</td></tr>
</table>

<table>
<tr><td>Oligo fwd</td><td>CACCGACCCTGTTCAAGACGGTGG</td><td></td><td></td><td></td><td></td><td></td><td></td><td></td><td></td><td></td><td></td></tr>
<tr><td>Oligo rev</td><td>AAACCCACCGTCTTGAACAGGGTC</td><td></td><td></td><td></td><td></td><td></td><td></td><td></td><td></td><td></td><td></td></tr>
<tr><td>Chromosome</td><td>start</td><td>end</td><td>strand</td><td>MM</td><td>target_seq</td><td>PAM</td><td>alignment</td><td>distance</td><td>position</td><td>gene name</td><td>gene id</td></tr>
<tr><td>GU305914.1</td><td>123563</td><td>123585</td><td>-</td><td>0</td><td>GACCCTGTTCAAGACGGTGG</td><td>AGG</td><td>||||||||[||||||||||||]PAM</td><td>NA</td><td>Intergenic</td><td>NA</td><td>NA</td></tr>
<tr><td></td><td></td><td></td><td></td><td></td><td></td><td></td><td></td><td></td><td></td><td></td><td></td></tr>
<tr><td>T24699</td><td>GGAGTACCTGAACCGCTCTCAGG</td><td>1000</td><td></td><td></td><td></td><td></td><td></td><td></td><td></td><td></td><td></td></tr>
<tr><td>Oligo fwd</td><td>CACCGGAGTACCTGAACCGCTCTC</td><td></td><td></td><td></td><td></td><td></td><td></td><td></td><td></td><td></td><td></td></tr>
<tr><td>Oligo rev</td><td>AAACGAGAGCGGTTCAGGTACTCC</td><td></td><td></td><td></td><td></td><td></td><td></td><td></td><td></td><td></td><td></td></tr>
<tr><td>Chromosome</td><td>start</td><td>end</td><td>strand</td><td>MM</td><td>target_seq</td><td>PAM</td><td>alignment</td><td>distance</td><td>position</td><td>gene name</td><td>gene id</td></tr>
<tr><td>GU305914.1</td><td>123542</td><td>123564</td><td>-</td><td>0</td><td>GGAGTACCTGAACCGCTCTC</td><td>AGG</td><td>||||||||[||||||||||||]PAM</td><td>NA</td><td>Intergenic</td><td>NA</td><td>NA</td></tr>
<tr><td>GU305914.1</td><td>57664</td><td>57686</td><td>-</td><td>4</td><td>GGAGGATCTGAACCCCCCTC</td><td>AGG</td><td>||||-|-|[||||||-|-|||]PAM</td><td>NA</td><td>Intergenic</td><td>NA</td><td>NA</td></tr>
<tr><td></td><td></td><td></td><td></td><td></td><td></td><td></td><td></td><td></td><td></td><td></td><td></td></tr>
<tr><td>T24700</td><td>CGCTCTCAGGACTGCATCCACGG</td><td>1000</td><td></td><td></td><td></td><td></td><td></td><td></td><td></td><td></td><td></td></tr>
<tr><td>Oligo fwd</td><td>CACCCGCTCTCAGGACTGCATCCA</td><td></td><td></td><td></td><td></td><td></td><td></td><td></td><td></td><td></td><td></td></tr>
<tr><td>Oligo rev</td><td>AAACTGGATGCAGTCCTGAGAGCG</td><td></td><td></td><td></td><td></td><td></td><td></td><td></td><td></td><td></td><td></td></tr>
<tr><td>Chromosome</td><td>start</td><td>end</td><td>strand</td><td>MM</td><td>target_seq</td><td>PAM</td><td>alignment</td><td>distance</td><td>position</td><td>gene name</td><td>gene id</td></tr>
<tr><td>GU305914.1</td><td>123529</td><td>123551</td><td>-</td><td>0</td><td>CGCTCTCAGGACTGCATCCA</td><td>CGG</td><td>||||||||[||||||||||||]PAM</td><td>NA</td><td>Intergenic</td><td>NA</td><td>NA</td></tr>
<tr><td></td><td></td><td></td><td></td><td></td><td></td><td></td><td></td><td></td><td></td><td></td><td></td></tr>
<tr><td>T24701</td><td>GGACTGCATCCACGGCGAGACGG</td><td>1000</td><td></td><td></td><td></td><td></td><td></td><td></td><td></td><td></td><td></td></tr>
<tr><td>Oligo fwd</td><td>CACCGGACTGCATCCACGGCGAGA</td><td></td><td></td><td></td><td></td><td></td><td></td><td></td><td></td><td></td><td></td></tr>
<tr><td>Oligo rev</td><td>AAACTCTCGCCGTGGATGCAGTCC</td><td></td><td></td><td></td><td></td><td></td><td></td><td></td><td></td><td></td><td></td></tr>
<tr><td>Chromosome</td><td>start</td><td>end</td><td>strand</td><td>MM</td><td>target_seq</td><td>PAM</td><td>alignment</td><td>distance</td><td>position</td><td>gene name</td><td>gene id</td></tr>
<tr><td>GU305914.1</td><td>123521</td><td>123543</td><td>-</td><td>0</td><td>GGACTGCATCCACGGCGAGA</td><td>CGG</td><td>||||||||[||||||||||||]PAM</td><td>NA</td><td>Intergenic</td><td>NA</td><td>NA</td></tr>
<tr><td></td><td></td><td></td><td></td><td></td><td></td><td></td><td></td><td></td><td></td><td></td><td></td></tr>
<tr><td>T24702</td><td>GGACCTGCAGTACATCTGCGTGG</td><td>1000</td><td></td><td></td><td></td><td></td><td></td><td></td><td></td><td></td><td></td></tr>
<tr><td>Oligo fwd</td><td>CACCGGACCTGCAGTACATCTGCG</td><td></td><td></td><td></td><td></td><td></td><td></td><td></td><td></td><td></td><td></td></tr>
<tr><td>Oligo rev</td><td>AAACCGCAGATGTACTGCAGGTCC</td><td></td><td></td><td></td><td></td><td></td><td></td><td></td><td></td><td></td><td></td></tr>
<tr><td>Chromosome</td><td>start</td><td>end</td><td>strand</td><td>MM</td><td>target_seq</td><td>PAM</td><td>alignment</td><td>distance</td><td>position</td><td>gene name</td><td>gene id</td></tr>
<tr><td>GU305914.1</td><td>123500</td><td>123522</td><td>-</td><td>0</td><td>GGACCTGCAGTACATCTGCG</td><td>TGG</td><td>||||||||[||||||||||||]PAM</td><td>NA</td><td>Intergenic</td><td>NA</td><td>NA</td></tr>
<tr><td></td><td></td><td></td><td></td><td></td><td></td><td></td><td></td><td></td><td></td><td></td><td></td></tr>
<tr><td>T24703</td><td>CCTGCAGTACATCTGCGTGGAGG</td><td>1000</td><td></td><td></td><td></td><td></td><td></td><td></td><td></td><td></td><td></td></tr>
<tr><td>Oligo fwd</td><td>CACCCCTGCAGTACATCTGCGTGG</td><td></td><td></td><td></td><td></td><td></td><td></td><td></td><td></td><td></td><td></td></tr>
<tr><td>Oligo rev</td><td>AAACCCACGCAGATGTACTGCAGG</td><td></td><td></td><td></td><td></td><td></td><td></td><td></td><td></td><td></td><td></td></tr>
<tr><td>Chromosome</td><td>start</td><td>end</td><td>strand</td><td>MM</td><td>target_seq</td><td>PAM</td><td>alignment</td><td>distance</td><td>position</td><td>gene name</td><td>gene id</td></tr>
<tr><td>GU305914.1</td><td>123497</td><td>123519</td><td>-</td><td>0</td><td>CCTGCAGTACATCTGCGTGG</td><td>AGG</td><td>||||||||[||||||||||||]PAM</td><td>NA</td><td>Intergenic</td><td>NA</td><td>NA</td></tr>
<tr><td></td><td></td><td></td><td></td><td></td><td></td><td></td><td></td><td></td><td></td><td></td><td></td></tr>
<tr><td>T24704</td><td>CTGCAGTACATCTGCGTGGAGGG</td><td>1000</td><td></td><td></td><td></td><td></td><td></td><td></td><td></td><td></td><td></td></tr>
<tr><td>Oligo fwd</td><td>CACCCTGCAGTACATCTGCGTGGA</td><td></td><td></td><td></td><td></td><td></td><td></td><td></td><td></td><td></td><td></td></tr>
<tr><td>Oligo rev</td><td>AAACTCCACGCAGATGTACTGCAG</td><td></td><td></td><td></td><td></td><td></td><td></td><td></td><td></td><td></td><td></td></tr>
<tr><td>Chromosome</td><td>start</td><td>end</td><td>strand</td><td>MM</td><td>target_seq</td><td>PAM</td><td>alignment</td><td>distance</td><td>position</td><td>gene name</td><td>gene id</td></tr>
<tr><td>GU305914.1</td><td>123496</td><td>123518</td><td>-</td><td>0</td><td>CTGCAGTACATCTGCGTGGA</td><td>GGG</td><td>||||||||[||||||||||||]PAM</td><td>NA</td><td>Intergenic</td><td>NA</td><td>NA</td></tr>
<tr><td></td><td></td><td></td><td></td><td></td><td></td><td></td><td></td><td></td><td></td><td></td><td></td></tr>
<tr><td>T24705</td><td>GGAGGGCACCAACTCGATCGTGG</td><td>1000</td><td></td><td></td><td></td><td></td><td></td><td></td><td></td><td></td><td></td></tr>
<tr><td>Oligo fwd</td><td>CACCGGAGGGCACCAACTCGATCG</td><td></td><td></td><td></td><td></td><td></td><td></td><td></td><td></td><td></td><td></td></tr>
<tr><td>Oligo rev</td><td>AAACCGATCGAGTTGGTGCCCTCC</td><td></td><td></td><td></td><td></td><td></td><td></td><td></td><td></td><td></td><td></td></tr>
<tr><td>Chromosome</td><td>start</td><td>end</td><td>strand</td><td>MM</td><td>target_seq</td><td>PAM</td><td>alignment</td><td>distance</td><td>position</td><td>gene name</td><td>gene id</td></tr>
<tr><td>GU305914.1</td><td>123479</td><td>123501</td><td>-</td><td>0</td><td>GGAGGGCACCAACTCGATCG</td><td>TGG</td><td>||||||||[||||||||||||]PAM</td><td>NA</td><td>Intergenic</td><td>NA</td><td>NA</td></tr>
<tr><td></td><td></td><td></td><td></td><td></td><td></td><td></td><td></td><td></td><td></td><td></td><td></td></tr>
<tr><td>T24706</td><td>GAAGCCCTGTCGCTTCCTCCAGG</td><td>1000</td><td></td><td></td><td></td><td></td><td></td><td></td><td></td><td></td><td></td></tr>
<tr><td>Oligo fwd</td><td>CACCGAAGCCCTGTCGCTTCCTCC</td><td></td><td></td><td></td><td></td><td></td><td></td><td></td><td></td><td></td><td></td></tr>
<tr><td>Oligo rev</td><td>AAACGGAGGAAGCGACAGGGCTTC</td><td></td><td></td><td></td><td></td><td></td><td></td><td></td><td></td><td></td><td></td></tr>
<tr><td>Chromosome</td><td>start</td><td>end</td><td>strand</td><td>MM</td><td>target_seq</td><td>PAM</td><td>alignment</td><td>distance</td><td>position</td><td>gene name</td><td>gene id</td></tr>
<tr><td>GU305914.1</td><td>123455</td><td>123477</td><td>-</td><td>0</td><td>GAAGCCCTGTCGCTTCCTCC</td><td>AGG</td><td>||||||||[||||||||||||]PAM</td><td>NA</td><td>Intergenic</td><td>NA</td><td>NA</td></tr>
<tr><td></td><td></td><td></td><td></td><td></td><td></td><td></td><td></td><td></td><td></td><td></td><td></td></tr>
<tr><td>T24707</td><td>GGAAGCCCTCACGCAACACACGG</td><td>1000</td><td></td><td></td><td></td><td></td><td></td><td></td><td></td><td></td><td></td></tr>
<tr><td>Oligo fwd</td><td>CACCGGAAGCCCTCACGCAACACA</td><td></td><td></td><td></td><td></td><td></td><td></td><td></td><td></td><td></td><td></td></tr>
<tr><td>Oligo rev</td><td>AAACTGTGTTGCGTGAGGGCTTCC</td><td></td><td></td><td></td><td></td><td></td><td></td><td></td><td></td><td></td><td></td></tr>
<tr><td>Chromosome</td><td>start</td><td>end</td><td>strand</td><td>MM</td><td>target_seq</td><td>PAM</td><td>alignment</td><td>distance</td><td>position</td><td>gene name</td><td>gene id</td></tr>
<tr><td>GU305914.1</td><td>123434</td><td>123456</td><td>-</td><td>0</td><td>GGAAGCCCTCACGCAACACA</td><td>CGG</td><td>||||||||[||||||||||||]PAM</td><td>NA</td><td>Intergenic</td><td>NA</td><td>NA</td></tr>
<tr><td></td><td></td><td></td><td></td><td></td><td></td><td></td><td></td><td></td><td></td><td></td><td></td></tr>
<tr><td>T24708</td><td>GAAGCCCTCACGCAACACACGGG</td><td>1000</td><td></td><td></td><td></td><td></td><td></td><td></td><td></td><td></td><td></td></tr>
<tr><td>Oligo fwd</td><td>CACCGAAGCCCTCACGCAACACAC</td><td></td><td></td><td></td><td></td><td></td><td></td><td></td><td></td><td></td><td></td></tr>
<tr><td>Oligo rev</td><td>AAACGTGTGTTGCGTGAGGGCTTC</td><td></td><td></td><td></td><td></td><td></td><td></td><td></td><td></td><td></td><td></td></tr>
<tr><td>Chromosome</td><td>start</td><td>end</td><td>strand</td><td>MM</td><td>target_seq</td><td>PAM</td><td>alignment</td><td>distance</td><td>position</td><td>gene name</td><td>gene id</td></tr>
<tr><td>GU305914.1</td><td>123433</td><td>123455</td><td>-</td><td>0</td><td>GAAGCCCTCACGCAACACAC</td><td>GGG</td><td>||||||||[||||||||||||]PAM</td><td>NA</td><td>Intergenic</td><td>NA</td><td>NA</td></tr>
<tr><td></td><td></td><td></td><td></td><td></td><td></td><td></td><td></td><td></td><td></td><td></td><td></td></tr>
<tr><td>T24709</td><td>GCAACACACGGGCACGACGCAGG</td><td>1000</td><td></td><td></td><td></td><td></td><td></td><td></td><td></td><td></td><td></td></tr>
<tr><td>Oligo fwd</td><td>CACCGCAACACACGGGCACGACGC</td><td></td><td></td><td></td><td></td><td></td><td></td><td></td><td></td><td></td><td></td></tr>
<tr><td>Oligo rev</td><td>AAACGCGTCGTGCCCGTGTGTTGC</td><td></td><td></td><td></td><td></td><td></td><td></td><td></td><td></td><td></td><td></td></tr>
<tr><td>Chromosome</td><td>start</td><td>end</td><td>strand</td><td>MM</td><td>target_seq</td><td>PAM</td><td>alignment</td><td>distance</td><td>position</td><td>gene name</td><td>gene id</td></tr>
<tr><td>GU305914.1</td><td>123422</td><td>123444</td><td>-</td><td>0</td><td>GCAACACACGGGCACGACGC</td><td>AGG</td><td>||||||||[||||||||||||]PAM</td><td>NA</td><td>Intergenic</td><td>NA</td><td>NA</td></tr>
<tr><td></td><td></td><td></td><td></td><td></td><td></td><td></td><td></td><td></td><td></td><td></td><td></td></tr>
<tr><td>T24710</td><td>GGGCACGACGCAGGCCCTGATGG</td><td>1000</td><td></td><td></td><td></td><td></td><td></td><td></td><td></td><td></td><td></td></tr>
<tr><td>Oligo fwd</td><td>CACCGGGCACGACGCAGGCCCTGA</td><td></td><td></td><td></td><td></td><td></td><td></td><td></td><td></td><td></td><td></td></tr>
<tr><td>Oligo rev</td><td>AAACTCAGGGCCTGCGTCGTGCCC</td><td></td><td></td><td></td><td></td><td></td><td></td><td></td><td></td><td></td><td></td></tr>
<tr><td>Chromosome</td><td>start</td><td>end</td><td>strand</td><td>MM</td><td>target_seq</td><td>PAM</td><td>alignment</td><td>distance</td><td>position</td><td>gene name</td><td>gene id</td></tr>
<tr><td>GU305914.1</td><td>123413</td><td>123435</td><td>-</td><td>0</td><td>GGGCACGACGCAGGCCCTGA</td><td>TGG</td><td>||||||||[||||||||||||]PAM</td><td>NA</td><td>Intergenic</td><td>NA</td><td>NA</td></tr>
<tr><td></td><td></td><td></td><td></td><td></td><td></td><td></td><td></td><td></td><td></td><td></td><td></td></tr>
<tr><td>T24711</td><td>CCTGATGGAGAGCCAGCTGAAGG</td><td>1000</td><td></td><td></td><td></td><td></td><td></td><td></td><td></td><td></td><td></td></tr>
<tr><td>Oligo fwd</td><td>CACCCCTGATGGAGAGCCAGCTGA</td><td></td><td></td><td></td><td></td><td></td><td></td><td></td><td></td><td></td><td></td></tr>
<tr><td>Oligo rev</td><td>AAACTCAGCTGGCTCTCCATCAGG</td><td></td><td></td><td></td><td></td><td></td><td></td><td></td><td></td><td></td><td></td></tr>
<tr><td>Chromosome</td><td>start</td><td>end</td><td>strand</td><td>MM</td><td>target_seq</td><td>PAM</td><td>alignment</td><td>distance</td><td>position</td><td>gene name</td><td>gene id</td></tr>
<tr><td>GU305914.1</td><td>123398</td><td>123420</td><td>-</td><td>0</td><td>CCTGATGGAGAGCCAGCTGA</td><td>AGG</td><td>||||||||[||||||||||||]PAM</td><td>NA</td><td>Intergenic</td><td>NA</td><td>NA</td></tr>
<tr><td></td><td></td><td></td><td></td><td></td><td></td><td></td><td></td><td></td><td></td><td></td><td></td></tr>
<tr><td>T24712</td><td>CTGATGGAGAGCCAGCTGAAGGG</td><td>1000</td><td></td><td></td><td></td><td></td><td></td><td></td><td></td><td></td><td></td></tr>
<tr><td>Oligo fwd</td><td>CACCCTGATGGAGAGCCAGCTGAA</td><td></td><td></td><td></td><td></td><td></td><td></td><td></td><td></td><td></td><td></td></tr>
<tr><td>Oligo rev</td><td>AAACTTCAGCTGGCTCTCCATCAG</td><td></td><td></td><td></td><td></td><td></td><td></td><td></td><td></td><td></td><td></td></tr>
<tr><td>Chromosome</td><td>start</td><td>end</td><td>strand</td><td>MM</td><td>target_seq</td><td>PAM</td><td>alignment</td><td>distance</td><td>position</td><td>gene name</td><td>gene id</td></tr>
<tr><td>GU305914.1</td><td>123397</td><td>123419</td><td>-</td><td>0</td><td>CTGATGGAGAGCCAGCTGAA</td><td>GGG</td><td>||||||||[||||||||||||]PAM</td><td>NA</td><td>Intergenic</td><td>NA</td><td>NA</td></tr>
</table>

<table>
<tr><td>T24713</td><td>GCTGAAGGGCACGAGCAAACTGG</td><td>1000</td><td></td><td></td><td></td><td></td><td></td><td></td><td></td><td></td><td></td></tr>
<tr><td>Oligo fwd</td><td>CACCGCTGAAGGGCACGAGCAAAC</td><td></td><td></td><td></td><td></td><td></td><td></td><td></td><td></td><td></td><td></td></tr>
<tr><td>Oligo rev</td><td>AAACGTTTGCTCGTGCCCTTCAGC</td><td></td><td></td><td></td><td></td><td></td><td></td><td></td><td></td><td></td><td></td></tr>
<tr><td>Chromosome</td><td>start</td><td>end</td><td>strand</td><td>MM</td><td>target_seq</td><td>PAM</td><td>alignment</td><td>distance</td><td>position</td><td>gene name</td><td>gene id</td></tr>
<tr><td>GU305914.1</td><td>123383</td><td>123405</td><td>-</td><td>0</td><td>GCTGAAGGGCACGAGCAAAC</td><td>TGG</td><td>||||||||[||||||||||||]PAM</td><td>NA</td><td>Intergenic</td><td>NA</td><td>NA</td></tr>
<tr><td>T24714</td><td>CTGAAGGGCACGAGCAAACTGGG</td><td>1000</td><td></td><td></td><td></td><td></td><td></td><td></td><td></td><td></td><td></td></tr>
<tr><td>Oligo fwd</td><td>CACCCTGAAGGGCACGAGCAAACT</td><td></td><td></td><td></td><td></td><td></td><td></td><td></td><td></td><td></td><td></td></tr>
<tr><td>Oligo rev</td><td>AAACAGTTTGCTCGTGCCCTTCAG</td><td></td><td></td><td></td><td></td><td></td><td></td><td></td><td></td><td></td><td></td></tr>
<tr><td>Chromosome</td><td>start</td><td>end</td><td>strand</td><td>MM</td><td>target_seq</td><td>PAM</td><td>alignment</td><td>distance</td><td>position</td><td>gene name</td><td>gene id</td></tr>
<tr><td>GU305914.1</td><td>123382</td><td>123404</td><td>-</td><td>0</td><td>CTGAAGGGCACGAGCAAACT</td><td>GGG</td><td>||||||||[||||||||||||]PAM</td><td>NA</td><td>Intergenic</td><td>NA</td><td>NA</td></tr>
<tr><td>T24715</td><td>GGGCACGAGCAAACTGGGACTGG</td><td>1000</td><td></td><td></td><td></td><td></td><td></td><td></td><td></td><td></td><td></td></tr>
<tr><td>Oligo fwd</td><td>CACCGGGCACGAGCAAACTGGGAC</td><td></td><td></td><td></td><td></td><td></td><td></td><td></td><td></td><td></td><td></td></tr>
<tr><td>Oligo rev</td><td>AAACGTCCCAGTTTGCTCGTGCCC</td><td></td><td></td><td></td><td></td><td></td><td></td><td></td><td></td><td></td><td></td></tr>
<tr><td>Chromosome</td><td>start</td><td>end</td><td>strand</td><td>MM</td><td>target_seq</td><td>PAM</td><td>alignment</td><td>distance</td><td>position</td><td>gene name</td><td>gene id</td></tr>
<tr><td>GU305914.1</td><td>123377</td><td>123399</td><td>-</td><td>0</td><td>GGGCACGAGCAAACTGGGAC</td><td>TGG</td><td>||||||||[||||||||||||]PAM</td><td>NA</td><td>Intergenic</td><td>NA</td><td>NA</td></tr>
<tr><td>T24716</td><td>GGACTGGACGAAACCCACTACGG</td><td>1000</td><td></td><td></td><td></td><td></td><td></td><td></td><td></td><td></td><td></td></tr>
<tr><td>Oligo fwd</td><td>CACCGGACTGGACGAAACCCACTA</td><td></td><td></td><td></td><td></td><td></td><td></td><td></td><td></td><td></td><td></td></tr>
<tr><td>Oligo rev</td><td>AAACTAGTGGGTTTCGTCCAGTCC</td><td></td><td></td><td></td><td></td><td></td><td></td><td></td><td></td><td></td><td></td></tr>
<tr><td>Chromosome</td><td>start</td><td>end</td><td>strand</td><td>MM</td><td>target_seq</td><td>PAM</td><td>alignment</td><td>distance</td><td>position</td><td>gene name</td><td>gene id</td></tr>
<tr><td>GU305914.1</td><td>123361</td><td>123383</td><td>-</td><td>0</td><td>GGACTGGACGAAACCCACTA</td><td>CGG</td><td>||||||||[||||||||||||]PAM</td><td>NA</td><td>Intergenic</td><td>NA</td><td>NA</td></tr>
<tr><td>T24717</td><td>CACTACGGCAACTACAGCATCGG</td><td>1000</td><td></td><td></td><td></td><td></td><td></td><td></td><td></td><td></td><td></td></tr>
<tr><td>Oligo fwd</td><td>CACCCACTACGGCAACTACAGCAT</td><td></td><td></td><td></td><td></td><td></td><td></td><td></td><td></td><td></td><td></td></tr>
<tr><td>Oligo rev</td><td>AAACATGCTGTAGTTGCCGTAGTG</td><td></td><td></td><td></td><td></td><td></td><td></td><td></td><td></td><td></td><td></td></tr>
<tr><td>Chromosome</td><td>start</td><td>end</td><td>strand</td><td>MM</td><td>target_seq</td><td>PAM</td><td>alignment</td><td>distance</td><td>position</td><td>gene name</td><td>gene id</td></tr>
<tr><td>GU305914.1</td><td>123346</td><td>123368</td><td>-</td><td>0</td><td>CACTACGGCAACTACAGCAT</td><td>CGG</td><td>||||||||[||||||||||||]PAM</td><td>NA</td><td>Intergenic</td><td>NA</td><td>NA</td></tr>
<tr><td>T24718</td><td>TTATTTAACTCGTGATCGAGCGG</td><td>1000</td><td></td><td></td><td></td><td></td><td></td><td></td><td></td><td></td><td></td></tr>
<tr><td>Oligo fwd</td><td>CACCTTATTTAACTCGTGATCGAG</td><td></td><td></td><td></td><td></td><td></td><td></td><td></td><td></td><td></td><td></td></tr>
<tr><td>Oligo rev</td><td>AAACCTCGATCACGAGTTAAATAA</td><td></td><td></td><td></td><td></td><td></td><td></td><td></td><td></td><td></td><td></td></tr>
<tr><td>Chromosome</td><td>start</td><td>end</td><td>strand</td><td>MM</td><td>target_seq</td><td>PAM</td><td>alignment</td><td>distance</td><td>position</td><td>gene name</td><td>gene id</td></tr>
<tr><td>GU305914.1</td><td>123295</td><td>123317</td><td>-</td><td>0</td><td>TTATTTAACTCGTGATCGAG</td><td>CGG</td><td>||||||||[||||||||||||]PAM</td><td>NA</td><td>Intergenic</td><td>NA</td><td>NA</td></tr>
<tr><td>T24719</td><td>TATTTAACTCGTGATCGAGCGGG</td><td>1000</td><td></td><td></td><td></td><td></td><td></td><td></td><td></td><td></td><td></td></tr>
<tr><td>Oligo fwd</td><td>CACCTATTTAACTCGTGATCGAGC</td><td></td><td></td><td></td><td></td><td></td><td></td><td></td><td></td><td></td><td></td></tr>
<tr><td>Oligo rev</td><td>AAACGCTCGATCACGAGTTAAATA</td><td></td><td></td><td></td><td></td><td></td><td></td><td></td><td></td><td></td><td></td></tr>
<tr><td>Chromosome</td><td>start</td><td>end</td><td>strand</td><td>MM</td><td>target_seq</td><td>PAM</td><td>alignment</td><td>distance</td><td>position</td><td>gene name</td><td>gene id</td></tr>
<tr><td>GU305914.1</td><td>123294</td><td>123316</td><td>-</td><td>0</td><td>TATTTAACTCGTGATCGAGC</td><td>GGG</td><td>||||||||[||||||||||||]PAM</td><td>NA</td><td>Intergenic</td><td>NA</td><td>NA</td></tr>
<tr><td>T24720</td><td>ATCGAGCGGGTCCTCCCGAGCGG</td><td>1000</td><td></td><td></td><td></td><td></td><td></td><td></td><td></td><td></td><td></td></tr>
<tr><td>Oligo fwd</td><td>CACCATCGAGCGGGTCCTCCCGAG</td><td></td><td></td><td></td><td></td><td></td><td></td><td></td><td></td><td></td><td></td></tr>
<tr><td>Oligo rev</td><td>AAACCTCGGGAGGACCCGCTCGAT</td><td></td><td></td><td></td><td></td><td></td><td></td><td></td><td></td><td></td><td></td></tr>
<tr><td>Chromosome</td><td>start</td><td>end</td><td>strand</td><td>MM</td><td>target_seq</td><td>PAM</td><td>alignment</td><td>distance</td><td>position</td><td>gene name</td><td>gene id</td></tr>
<tr><td>GU305914.1</td><td>123281</td><td>123303</td><td>-</td><td>0</td><td>ATCGAGCGGGTCCTCCCGAG</td><td>CGG</td><td>||||||||[||||||||||||]PAM</td><td>NA</td><td>Intergenic</td><td>NA</td><td>NA</td></tr>
<tr><td>T24721</td><td>AGCGGGTCCTCCCGAGCGGAAGG</td><td>1000</td><td></td><td></td><td></td><td></td><td></td><td></td><td></td><td></td><td></td></tr>
<tr><td>Oligo fwd</td><td>CACCAGCGGGTCCTCCCGAGCGGA</td><td></td><td></td><td></td><td></td><td></td><td></td><td></td><td></td><td></td><td></td></tr>
<tr><td>Oligo rev</td><td>AAACTCCGCTCGGGAGGACCCGCT</td><td></td><td></td><td></td><td></td><td></td><td></td><td></td><td></td><td></td><td></td></tr>
<tr><td>Chromosome</td><td>start</td><td>end</td><td>strand</td><td>MM</td><td>target_seq</td><td>PAM</td><td>alignment</td><td>distance</td><td>position</td><td>gene name</td><td>gene id</td></tr>
<tr><td>GU305914.1</td><td>123277</td><td>123299</td><td>-</td><td>0</td><td>AGCGGGTCCTCCCGAGCGGA</td><td>AGG</td><td>||||||||[||||||||||||]PAM</td><td>NA</td><td>Intergenic</td><td>NA</td><td>NA</td></tr>
<tr><td>T24722</td><td>AGCGGAAGGCGCAGCGCCGTCGG</td><td>1000</td><td></td><td></td><td></td><td></td><td></td><td></td><td></td><td></td><td></td></tr>
<tr><td>Oligo fwd</td><td>CACCAGCGGAAGGCGCAGCGCCGT</td><td></td><td></td><td></td><td></td><td></td><td></td><td></td><td></td><td></td><td></td></tr>
<tr><td>Oligo rev</td><td>AAACACGGCGCTGCGCCTTCCGCT</td><td></td><td></td><td></td><td></td><td></td><td></td><td></td><td></td><td></td><td></td></tr>
<tr><td>Chromosome</td><td>start</td><td>end</td><td>strand</td><td>MM</td><td>target_seq</td><td>PAM</td><td>alignment</td><td>distance</td><td>position</td><td>gene name</td><td>gene id</td></tr>
<tr><td>GU305914.1</td><td>123263</td><td>123285</td><td>-</td><td>0</td><td>AGCGGAAGGCGCAGCGCCGT</td><td>CGG</td><td>||||||||[||||||||||||]PAM</td><td>NA</td><td>Intergenic</td><td>NA</td><td>NA</td></tr>
<tr><td>T24723</td><td>CCGTCGGACGTCACGCTCGCCGG</td><td>1000</td><td></td><td></td><td></td><td></td><td></td><td></td><td></td><td></td><td></td></tr>
<tr><td>Oligo fwd</td><td>CACCCCGTCGGACGTCACGCTCGC</td><td></td><td></td><td></td><td></td><td></td><td></td><td></td><td></td><td></td><td></td></tr>
<tr><td>Oligo rev</td><td>AAACGCGAGCGTGACGTCCGACGG</td><td></td><td></td><td></td><td></td><td></td><td></td><td></td><td></td><td></td><td></td></tr>
<tr><td>Chromosome</td><td>start</td><td>end</td><td>strand</td><td>MM</td><td>target_seq</td><td>PAM</td><td>alignment</td><td>distance</td><td>position</td><td>gene name</td><td>gene id</td></tr>
<tr><td>GU305914.1</td><td>123247</td><td>123269</td><td>-</td><td>0</td><td>CCGTCGGACGTCACGCTCGC</td><td>CGG</td><td>||||||||[||||||||||||]PAM</td><td>NA</td><td>Intergenic</td><td>NA</td><td>NA</td></tr>
<tr><td>GU305914.1</td><td>62171</td><td>62193</td><td>+</td><td>4</td><td>GCGGCGGTCGTTACGCTCGC</td><td>GGG</td><td>-||-|||-[|||-||||||||]PAM</td><td>NA</td><td>Intergenic</td><td>NA</td><td>NA</td></tr>
<tr><td>T24724</td><td>GAGACCGACACCCACGACCATGG</td><td>1000</td><td></td><td></td><td></td><td></td><td></td><td></td><td></td><td></td><td></td></tr>
<tr><td>Oligo fwd</td><td>CACCGAGACCGACACCCACGACCA</td><td></td><td></td><td></td><td></td><td></td><td></td><td></td><td></td><td></td><td></td></tr>
<tr><td>Oligo rev</td><td>AAACTGGTCGTGGGTGTCGGTCTC</td><td></td><td></td><td></td><td></td><td></td><td></td><td></td><td></td><td></td><td></td></tr>
<tr><td>Chromosome</td><td>start</td><td>end</td><td>strand</td><td>MM</td><td>target_seq</td><td>PAM</td><td>alignment</td><td>distance</td><td>position</td><td>gene name</td><td>gene id</td></tr>
<tr><td>GU305914.1</td><td>123225</td><td>123247</td><td>-</td><td>0</td><td>GAGACCGACACCCACGACCA</td><td>TGG</td><td>||||||||[||||||||||||]PAM</td><td>NA</td><td>Intergenic</td><td>NA</td><td>NA</td></tr>
<tr><td>T24725</td><td>ACCCACGACCATGGAAACGACGG</td><td>1000</td><td></td><td></td><td></td><td></td><td></td><td></td><td></td><td></td><td></td></tr>
<tr><td>Oligo fwd</td><td>CACCACCCACGACCATGGAAACGA</td><td></td><td></td><td></td><td></td><td></td><td></td><td></td><td></td><td></td><td></td></tr>
<tr><td>Oligo rev</td><td>AAACTCGTTTCCATGGTCGTGGGT</td><td></td><td></td><td></td><td></td><td></td><td></td><td></td><td></td><td></td><td></td></tr>
<tr><td>Chromosome</td><td>start</td><td>end</td><td>strand</td><td>MM</td><td>target_seq</td><td>PAM</td><td>alignment</td><td>distance</td><td>position</td><td>gene name</td><td>gene id</td></tr>
<tr><td>GU305914.1</td><td>123216</td><td>123238</td><td>-</td><td>0</td><td>ACCCACGACCATGGAAACGA</td><td>CGG</td><td>||||||||[||||||||||||]PAM</td><td>NA</td><td>Intergenic</td><td>NA</td><td>NA</td></tr>
<tr><td>T24726</td><td>GACCATGGAAACGACGGTTTTGG</td><td>1000</td><td></td><td></td><td></td><td></td><td></td><td></td><td></td><td></td><td></td></tr>
<tr><td>Oligo fwd</td><td>CACCGACCATGGAAACGACGGTTT</td><td></td><td></td><td></td><td></td><td></td><td></td><td></td><td></td><td></td><td></td></tr>
<tr><td>Oligo rev</td><td>AAACAAACCGTCGTTTCCATGGTC</td><td></td><td></td><td></td><td></td><td></td><td></td><td></td><td></td><td></td><td></td></tr>
<tr><td>Chromosome</td><td>start</td><td>end</td><td>strand</td><td>MM</td><td>target_seq</td><td>PAM</td><td>alignment</td><td>distance</td><td>position</td><td>gene name</td><td>gene id</td></tr>
<tr><td>GU305914.1</td><td>123210</td><td>123232</td><td>-</td><td>0</td><td>GACCATGGAAACGACGGTTT</td><td>TGG</td><td>||||||||[||||||||||||]PAM</td><td>NA</td><td>Intergenic</td><td>NA</td><td>NA</td></tr>
<tr><td>T24727</td><td>TTCGAGCAGCGCCTCACGACCGG</td><td>1000</td><td></td><td></td><td></td><td></td><td></td><td></td><td></td><td></td><td></td></tr>
<tr><td>Oligo fwd</td><td>CACCTTCGAGCAGCGCCTCACGAC</td><td></td><td></td><td></td><td></td><td></td><td></td><td></td><td></td><td></td><td></td></tr>
<tr><td>Oligo rev</td><td>AAACGTCGTGAGGCGCTGCTCGAA</td><td></td><td></td><td></td><td></td><td></td><td></td><td></td><td></td><td></td><td></td></tr>
</table>

| Chromosome | start | end | strand | MM | target_seq | PAM | alignment | distance | position | gene name | gene id |
|---|---|---|---|---|---|---|---|---|---|---|---|
| GU305914.1 | 123182 | 123204 | - | 0 | TTCGAGCAGCGCCTCACGAC | CGG | \|\|\|\|\|\|\|\|[\|\|\|\|\|\|\|\|\|\|\|\|]PAM | NA | Intergenic | NA | NA |
| T24728 | TCGAGCAGCGCCTCACGACCGGG | 1000 | | | | | | | | | |
| Oligo fwd | CACCTCGAGCAGCGCCTCACGACC | | | | | | | | | | |
| Oligo rev | AAACGGTCGTGAGGCGCTGCTCGA | | | | | | | | | | |
| Chromosome | start | end | strand | MM | target_seq | PAM | alignment | distance | position | gene name | gene id |
| GU305914.1 | 123181 | 123203 | - | 0 | TCGAGCAGCGCCTCACGACC | GGG | \|\|\|\|\|\|\|\|[\|\|\|\|\|\|\|\|\|\|\|\|]PAM | NA | Intergenic | NA | NA |
| T24729 | CGAGCAGCGCCTCACGACCGGGG | 1000 | | | | | | | | | |
| Oligo fwd | CACCCGAGCAGCGCCTCACGACCG | | | | | | | | | | |
| Oligo rev | AAACCGGTCGTGAGGCGCTGCTCG | | | | | | | | | | |
| Chromosome | start | end | strand | MM | target_seq | PAM | alignment | distance | position | gene name | gene id |
| GU305914.1 | 123180 | 123202 | - | 0 | CGAGCAGCGCCTCACGACCG | GGG | \|\|\|\|\|\|\|\|[\|\|\|\|\|\|\|\|\|\|\|\|]PAM | NA | Intergenic | NA | NA |
| T24730 | GCGCCTCACGACCGGGGACGTGG | 1000 | | | | | | | | | |
| Oligo fwd | CACCGCGCCTCACGACCGGGGACG | | | | | | | | | | |
| Oligo rev | AAACCGTCCCCGGTCGTGAGGCGC | | | | | | | | | | |
| Chromosome | start | end | strand | MM | target_seq | PAM | alignment | distance | position | gene name | gene id |
| GU305914.1 | 123174 | 123196 | - | 0 | GCGCCTCACGACCGGGGACG | TGG | \|\|\|\|\|\|\|\|[\|\|\|\|\|\|\|\|\|\|\|\|]PAM | NA | Intergenic | NA | NA |
| T24731 | CGCCTCACGACCGGGGACGTGGG | 1000 | | | | | | | | | |
| Oligo fwd | CACCCGCCTCACGACCGGGGACGT | | | | | | | | | | |
| Oligo rev | AAACACGTCCCCGGTCGTGAGGCG | | | | | | | | | | |
| Chromosome | start | end | strand | MM | target_seq | PAM | alignment | distance | position | gene name | gene id |
| GU305914.1 | 123173 | 123195 | - | 0 | CGCCTCACGACCGGGGACGT | GGG | \|\|\|\|\|\|\|\|[\|\|\|\|\|\|\|\|\|\|\|\|]PAM | NA | Intergenic | NA | NA |
| T24732 | GCCTCACGACCGGGGACGTGGGG | 1000 | | | | | | | | | |
| Oligo fwd | CACCGCCTCACGACCGGGGACGTG | | | | | | | | | | |
| Oligo rev | AAACCACGTCCCCGGTCGTGAGGC | | | | | | | | | | |
| Chromosome | start | end | strand | MM | target_seq | PAM | alignment | distance | position | gene name | gene id |
| GU305914.1 | 123172 | 123194 | - | 0 | GCCTCACGACCGGGGACGTG | GGG | \|\|\|\|\|\|\|\|[\|\|\|\|\|\|\|\|\|\|\|\|]PAM | NA | Intergenic | NA | NA |
| T24733 | GGGACGTGGGGAAGCTGTCGCGG | 1000 | | | | | | | | | |
| Oligo fwd | CACCGGGACGTGGGGAAGCTGTCG | | | | | | | | | | |
| Oligo rev | AAACCGACAGCTTCCCCACGTCCC | | | | | | | | | | |
| Chromosome | start | end | strand | MM | target_seq | PAM | alignment | distance | position | gene name | gene id |
| GU305914.1 | 123160 | 123182 | - | 0 | GGGACGTGGGGAAGCTGTCG | CGG | \|\|\|\|\|\|\|\|[\|\|\|\|\|\|\|\|\|\|\|\|]PAM | NA | Intergenic | NA | NA |
| T24734 | GGGAAGCTGTCGCGGCTGATCGG | 1000 | | | | | | | | | |
| Oligo fwd | CACCGGGAAGCTGTCGCGGCTGAT | | | | | | | | | | |
| Oligo rev | AAACATCAGCCGCGACAGCTTCCC | | | | | | | | | | |
| Chromosome | start | end | strand | MM | target_seq | PAM | alignment | distance | position | gene name | gene id |
| GU305914.1 | 123152 | 123174 | - | 0 | GGGAAGCTGTCGCGGCTGAT | CGG | \|\|\|\|\|\|\|\|[\|\|\|\|\|\|\|\|\|\|\|\|]PAM | NA | Intergenic | NA | NA |
| T24735 | GCTGTCGCGGCTGATCGGCGCGG | 1000 | | | | | | | | | |
| Oligo fwd | CACCGCTGTCGCGGCTGATCGGCG | | | | | | | | | | |
| Oligo rev | AAACCGCCGATCAGCCGCGACAGC | | | | | | | | | | |
| Chromosome | start | end | strand | MM | target_seq | PAM | alignment | distance | position | gene name | gene id |
| GU305914.1 | 123147 | 123169 | - | 0 | GCTGTCGCGGCTGATCGGCG | CGG | \|\|\|\|\|\|\|\|[\|\|\|\|\|\|\|\|\|\|\|\|]PAM | NA | Intergenic | NA | NA |
| T24736 | CCCCTACCGCCACCATCTGCTGG | 1000 | | | | | | | | | |
| Oligo fwd | CACCCCCCTACCGCCACCATCTGC | | | | | | | | | | |
| Oligo rev | AAACGCAGATGGTGGCGGTAGGGG | | | | | | | | | | |
| Chromosome | start | end | strand | MM | target_seq | PAM | alignment | distance | position | gene name | gene id |
| GU305914.1 | 123114 | 123136 | - | 0 | CCCCTACCGCCACCATCTGC | TGG | \|\|\|\|\|\|\|\|[\|\|\|\|\|\|\|\|\|\|\|\|]PAM | NA | Intergenic | NA | NA |
| T24737 | CCCTACCGCCACCATCTGCTGGG | 1000 | | | | | | | | | |
| Oligo fwd | CACCCCCTACCGCCACCATCTGCT | | | | | | | | | | |
| Oligo rev | AAACAGCAGATGGTGGCGGTAGGG | | | | | | | | | | |
| Chromosome | start | end | strand | MM | target_seq | PAM | alignment | distance | position | gene name | gene id |
| GU305914.1 | 123113 | 123135 | - | 0 | CCCTACCGCCACCATCTGCT | GGG | \|\|\|\|\|\|\|\|[\|\|\|\|\|\|\|\|\|\|\|\|]PAM | NA | Intergenic | NA | NA |
| T24738 | CCATCTGCTGGGCAGCTCGCAGG | 1000 | | | | | | | | | |
| Oligo fwd | CACCCCATCTGCTGGGCAGCTCGC | | | | | | | | | | |
| Oligo rev | AAACGCGAGCTGCCCAGCAGATGG | | | | | | | | | | |
| Chromosome | start | end | strand | MM | target_seq | PAM | alignment | distance | position | gene name | gene id |
| GU305914.1 | 123102 | 123124 | - | 0 | CCATCTGCTGGGCAGCTCGC | AGG | \|\|\|\|\|\|\|\|[\|\|\|\|\|\|\|\|\|\|\|\|]PAM | NA | Intergenic | NA | NA |
| T24739 | TCTGCTGGGCAGCTCGCAGGTGG | 1000 | | | | | | | | | |
| Oligo fwd | CACCTCTGCTGGGCAGCTCGCAGG | | | | | | | | | | |
| Oligo rev | AAACCCTGCGAGCTGCCCAGCAGA | | | | | | | | | | |
| Chromosome | start | end | strand | MM | target_seq | PAM | alignment | distance | position | gene name | gene id |
| GU305914.1 | 123099 | 123121 | - | 0 | TCTGCTGGGCAGCTCGCAGG | TGG | \|\|\|\|\|\|\|\|[\|\|\|\|\|\|\|\|\|\|\|\|]PAM | NA | Intergenic | NA | NA |
| T24740 | CTGCTGGGCAGCTCGCAGGTGGG | 1000 | | | | | | | | | |
| Oligo fwd | CACCCTGCTGGGCAGCTCGCAGGT | | | | | | | | | | |
| Oligo rev | AAACACCTGCGAGCTGCCCAGCAG | | | | | | | | | | |
| Chromosome | start | end | strand | MM | target_seq | PAM | alignment | distance | position | gene name | gene id |
| GU305914.1 | 123098 | 123120 | - | 0 | CTGCTGGGCAGCTCGCAGGT | GGG | \|\|\|\|\|\|\|\|[\|\|\|\|\|\|\|\|\|\|\|\|]PAM | NA | Intergenic | NA | NA |
| T24741 | TGCTGGGCAGCTCGCAGGTGGGG | 1000 | | | | | | | | | |
| Oligo fwd | CACCTGCTGGGCAGCTCGCAGGTG | | | | | | | | | | |
| Oligo rev | AAACCACCTGCGAGCTGCCCAGCA | | | | | | | | | | |
| Chromosome | start | end | strand | MM | target_seq | PAM | alignment | distance | position | gene name | gene id |
| GU305914.1 | 123097 | 123119 | - | 0 | TGCTGGGCAGCTCGCAGGTG | GGG | \|\|\|\|\|\|\|\|[\|\|\|\|\|\|\|\|\|\|\|\|]PAM | NA | Intergenic | NA | NA |
| T24742 | GGGCAGCTCGCAGGTGGGGCTGG | 1000 | | | | | | | | | |
| Oligo fwd | CACCGGGCAGCTCGCAGGTGGGGC | | | | | | | | | | |

| | | | | | | | | | | | |
|---|---|---|---|---|---|---|---|---|---|---|---|
| Oligo rev | AAACGCCCCACCTGCGAGCTGCCC | | | | | | | | | | |
| Chromosome | start | end | strand | MM | target_seq | PAM | alignment | distance | position | gene name | gene id |
| GU305914.1 | 123093 | 123115 | - | 0 | GGGCAGCTCGCAGGTGGGGC | TGG | \|\|\|\|\|\|\|\|[\|\|\|\|\|\|\|\|\|\|\|\|]PAM | NA | Intergenic | NA | NA |
| T24743 | CTCGCAGGTGGGGCTGGACGCGG | 1000 | | | | | | | | | |
| Oligo fwd | CACCCTCGCAGGTGGGGCTGGACG | | | | | | | | | | |
| Oligo rev | AAACCGTCCAGCCCCACCTGCGAG | | | | | | | | | | |
| Chromosome | start | end | strand | MM | target_seq | PAM | alignment | distance | position | gene name | gene id |
| GU305914.1 | 123087 | 123109 | - | 0 | CTCGCAGGTGGGGCTGGACG | CGG | \|\|\|\|\|\|\|\|[\|\|\|\|\|\|\|\|\|\|\|\|]PAM | NA | Intergenic | NA | NA |
| T24744 | GGGGCTGGACGCGGTCGTCAAGG | 1000 | | | | | | | | | |
| Oligo fwd | CACCGGGGCTGGACGCGGTCGTCA | | | | | | | | | | |
| Oligo rev | AAACTGACGACCGCGTCCAGCCCC | | | | | | | | | | |
| Chromosome | start | end | strand | MM | target_seq | PAM | alignment | distance | position | gene name | gene id |
| GU305914.1 | 123078 | 123100 | - | 0 | GGGGCTGGACGCGGTCGTCA | AGG | \|\|\|\|\|\|\|\|[\|\|\|\|\|\|\|\|\|\|\|\|]PAM | NA | Intergenic | NA | NA |
| T24745 | ACTACTCGCGCATGCGAGCGCGG | 1000 | | | | | | | | | |
| Oligo fwd | CACCACTACTCGCGCATGCGAGCG | | | | | | | | | | |
| Oligo rev | AAACCGCTCGCATGCGCGAGTAGT | | | | | | | | | | |
| Chromosome | start | end | strand | MM | target_seq | PAM | alignment | distance | position | gene name | gene id |
| GU305914.1 | 123043 | 123065 | - | 0 | ACTACTCGCGCATGCGAGCG | CGG | \|\|\|\|\|\|\|\|[\|\|\|\|\|\|\|\|\|\|\|\|]PAM | NA | Intergenic | NA | NA |
| T24746 | AGATGACTCTCACCATCATGCGG | 1000 | | | | | | | | | |
| Oligo fwd | CACCAGATGACTCTCACCATCATG | | | | | | | | | | |
| Oligo rev | AAACCATGATGGTGAGAGTCATCT | | | | | | | | | | |
| Chromosome | start | end | strand | MM | target_seq | PAM | alignment | distance | position | gene name | gene id |
| GU305914.1 | 123013 | 123035 | - | 0 | AGATGACTCTCACCATCATG | CGG | \|\|\|\|\|\|\|\|[\|\|\|\|\|\|\|\|\|\|\|\|]PAM | NA | Intergenic | NA | NA |
| T24747 | CACCATCATGCGGCGCGTCGAGG | 1000 | | | | | | | | | |
| Oligo fwd | CACCCACCATCATGCGGCGCGTCG | | | | | | | | | | |
| Oligo rev | AAACCGACGCGCCGCATGATGGTG | | | | | | | | | | |
| Chromosome | start | end | strand | MM | target_seq | PAM | alignment | distance | position | gene name | gene id |
| GU305914.1 | 123003 | 123025 | - | 0 | CACCATCATGCGGCGCGTCG | AGG | \|\|\|\|\|\|\|\|[\|\|\|\|\|\|\|\|\|\|\|\|]PAM | NA | Intergenic | NA | NA |
| GU305914.1 | 37409 | 37431 | - | 4 | CACGGTAATGCGTCGCGTCG | GGG | \|\|\|--\|-\|[\|\|\|\|-\|\|\|\|\|\|\|]PAM | NA | Intergenic | NA | NA |
| T24748 | ACCATCATGCGGCGCGTCGAGGG | 1000 | | | | | | | | | |
| Oligo fwd | CACCACCATCATGCGGCGCGTCGA | | | | | | | | | | |
| Oligo rev | AAACTCGACGCGCCGCATGATGGT | | | | | | | | | | |
| Chromosome | start | end | strand | MM | target_seq | PAM | alignment | distance | position | gene name | gene id |
| GU305914.1 | 123002 | 123024 | - | 0 | ACCATCATGCGGCGCGTCGA | GGG | \|\|\|\|\|\|\|\|[\|\|\|\|\|\|\|\|\|\|\|\|]PAM | NA | Intergenic | NA | NA |
| T24749 | CGAGGGTAACCAGATGATCCTGG | 1000 | | | | | | | | | |
| Oligo fwd | CACCCGAGGGTAACCAGATGATCC | | | | | | | | | | |
| Oligo rev | AAACGGATCATCTGGTTACCCTCG | | | | | | | | | | |
| Chromosome | start | end | strand | MM | target_seq | PAM | alignment | distance | position | gene name | gene id |
| GU305914.1 | 122985 | 123007 | - | 0 | CGAGGGTAACCAGATGATCC | TGG | \|\|\|\|\|\|\|\|[\|\|\|\|\|\|\|\|\|\|\|\|]PAM | NA | Intergenic | NA | NA |
| T24750 | GAGGGTAACCAGATGATCCTGGG | 1000 | | | | | | | | | |
| Oligo fwd | CACCGAGGGTAACCAGATGATCCT | | | | | | | | | | |
| Oligo rev | AAACAGGATCATCTGGTTACCCTC | | | | | | | | | | |
| Chromosome | start | end | strand | MM | target_seq | PAM | alignment | distance | position | gene name | gene id |
| GU305914.1 | 122984 | 123006 | - | 0 | GAGGGTAACCAGATGATCCT | GGG | \|\|\|\|\|\|\|\|[\|\|\|\|\|\|\|\|\|\|\|\|]PAM | NA | Intergenic | NA | NA |
| T24751 | ATCCTGGGCGTCCCGACGCACGG | 1000 | | | | | | | | | |
| Oligo fwd | CACCATCCTGGGCGTCCCGACGCA | | | | | | | | | | |
| Oligo rev | AAACTGCGTCGGGACGCCCAGGAT | | | | | | | | | | |
| Chromosome | start | end | strand | MM | target_seq | PAM | alignment | distance | position | gene name | gene id |
| GU305914.1 | 122969 | 122991 | - | 0 | ATCCTGGGCGTCCCGACGCA | CGG | \|\|\|\|\|\|\|\|[\|\|\|\|\|\|\|\|\|\|\|\|]PAM | NA | Intergenic | NA | NA |
| T24752 | GTGCTACACCATCCGCAACACGG | 1000 | | | | | | | | | |
| Oligo fwd | CACCGTGCTACACCATCCGCAACA | | | | | | | | | | |
| Oligo rev | AAACTGTTGCGGATGGTGTAGCAC | | | | | | | | | | |
| Chromosome | start | end | strand | MM | target_seq | PAM | alignment | distance | position | gene name | gene id |
| GU305914.1 | 122943 | 122965 | - | 0 | GTGCTACACCATCCGCAACA | CGG | \|\|\|\|\|\|\|\|[\|\|\|\|\|\|\|\|\|\|\|\|]PAM | NA | Intergenic | NA | NA |
| T24753 | TGCTACACCATCCGCAACACGGG | 1000 | | | | | | | | | |
| Oligo fwd | CACCTGCTACACCATCCGCAACAC | | | | | | | | | | |
| Oligo rev | AAACGTGTTGCGGATGGTGTAGCA | | | | | | | | | | |
| Chromosome | start | end | strand | MM | target_seq | PAM | alignment | distance | position | gene name | gene id |
| GU305914.1 | 122942 | 122964 | - | 0 | TGCTACACCATCCGCAACAC | GGG | \|\|\|\|\|\|\|\|[\|\|\|\|\|\|\|\|\|\|\|\|]PAM | NA | Intergenic | NA | NA |
| T24754 | GCAACACGGGTCCCGTGAGCTGG | 1000 | | | | | | | | | |
| Oligo fwd | CACCGCAACACGGGTCCCGTGAGC | | | | | | | | | | |
| Oligo rev | AAACGCTCACGGGACCCGTGTTGC | | | | | | | | | | |
| Chromosome | start | end | strand | MM | target_seq | PAM | alignment | distance | position | gene name | gene id |
| GU305914.1 | 122929 | 122951 | - | 0 | GCAACACGGGTCCCGTGAGC | TGG | \|\|\|\|\|\|\|\|[\|\|\|\|\|\|\|\|\|\|\|\|]PAM | NA | Intergenic | NA | NA |
| T24755 | CAACACGGGTCCCGTGAGCTGGG | 1000 | | | | | | | | | |
| Oligo fwd | CACCCAACACGGGTCCCGTGAGCT | | | | | | | | | | |
| Oligo rev | AAACAGCTCACGGGACCCGTGTTG | | | | | | | | | | |
| Chromosome | start | end | strand | MM | target_seq | PAM | alignment | distance | position | gene name | gene id |
| GU305914.1 | 122928 | 122950 | - | 0 | CAACACGGGTCCCGTGAGCT | GGG | \|\|\|\|\|\|\|\|[\|\|\|\|\|\|\|\|\|\|\|\|]PAM | NA | Intergenic | NA | NA |
| T24756 | GGGTCCCGTGAGCTGGGAGAAGG | 1000 | | | | | | | | | |
| Oligo fwd | CACCGGGTCCCGTGAGCTGGGAGA | | | | | | | | | | |
| Oligo rev | AAACTCTCCCAGCTCACGGGACCC | | | | | | | | | | |
| Chromosome | start | end | strand | MM | target_seq | PAM | alignment | distance | position | gene name | gene id |
| GU305914.1 | 122922 | 122944 | - | 0 | GGGTCCCGTGAGCTGGGAGA | AGG | \|\|\|\|\|\|\|\|[\|\|\|\|\|\|\|\|\|\|\|\|]PAM | NA | Intergenic | NA | NA |

| | | | | | | | | | | | |
|---|---|---|---|---|---|---|---|---|---|---|---|
| T24757 | GGTCCCGTGAGCTGGGAGAAGGG | 1000 | | | | | | | | | |
| Oligo fwd | CACCGGTCCCGTGAGCTGGGAGAA | | | | | | | | | | |
| Oligo rev | AAACTTCTCCCAGCTCACGGGACC | | | | | | | | | | |
| Chromosome | start | end | strand | MM | target_seq | PAM | alignment | distance | position | gene name | gene id |
| GU305914.1 | 122921 | 122943 | - | 0 | GGTCCCGTGAGCTGGGAGAA | GGG | \|\|\|\|\|\|\|\|[\|\|\|\|\|\|\|\|\|\|\|\|]PAM | NA | Intergenic | NA | NA |
| T24758 | CACCCTGCCGCCCGTCTTCTCGG | 1000 | | | | | | | | | |
| Oligo fwd | CACCCACCCTGCCGCCCGTCTTCT | | | | | | | | | | |
| Oligo rev | AAACAGAAGACGGGCGGCAGGGTG | | | | | | | | | | |
| Chromosome | start | end | strand | MM | target_seq | PAM | alignment | distance | position | gene name | gene id |
| GU305914.1 | 122886 | 122908 | - | 0 | CACCCTGCCGCCCGTCTTCT | CGG | \|\|\|\|\|\|\|\|[\|\|\|\|\|\|\|\|\|\|\|\|]PAM | NA | Intergenic | NA | NA |
| T24759 | ACCCTGCCGCCCGTCTTCTCGGG | 1000 | | | | | | | | | |
| Oligo fwd | CACCACCCTGCCGCCCGTCTTCTC | | | | | | | | | | |
| Oligo rev | AAACGAGAAGACGGGCGGCAGGGT | | | | | | | | | | |
| Chromosome | start | end | strand | MM | target_seq | PAM | alignment | distance | position | gene name | gene id |
| GU305914.1 | 122885 | 122907 | - | 0 | ACCCTGCCGCCCGTCTTCTC | GGG | \|\|\|\|\|\|\|\|[\|\|\|\|\|\|\|\|\|\|\|\|]PAM | NA | Intergenic | NA | NA |
| T24760 | GCCGCCCGTCTTCTCGGGAGAGG | 1000 | | | | | | | | | |
| Oligo fwd | CACCGCCGCCCGTCTTCTCGGGAG | | | | | | | | | | |
| Oligo rev | AAACCTCCCGAGAAGACGGGCGGC | | | | | | | | | | |
| Chromosome | start | end | strand | MM | target_seq | PAM | alignment | distance | position | gene name | gene id |
| GU305914.1 | 122880 | 122902 | - | 0 | GCCGCCCGTCTTCTCGGGAG | AGG | \|\|\|\|\|\|\|\|[\|\|\|\|\|\|\|\|\|\|\|\|]PAM | NA | Intergenic | NA | NA |
| T24761 | CGTCTTCTCGGGAGAGGTGACGG | 1000 | | | | | | | | | |
| Oligo fwd | CACCCGTCTTCTCGGGAGAGGTGA | | | | | | | | | | |
| Oligo rev | AAACTCACCTCTCCCGAGAAGACG | | | | | | | | | | |
| Chromosome | start | end | strand | MM | target_seq | PAM | alignment | distance | position | gene name | gene id |
| GU305914.1 | 122874 | 122896 | - | 0 | CGTCTTCTCGGGAGAGGTGA | CGG | \|\|\|\|\|\|\|\|[\|\|\|\|\|\|\|\|\|\|\|\|]PAM | NA | Intergenic | NA | NA |
| T24762 | GTCTTCTCGGGAGAGGTGACGGG | 1000 | | | | | | | | | |
| Oligo fwd | CACCGTCTTCTCGGGAGAGGTGAC | | | | | | | | | | |
| Oligo rev | AAACGTCACCTCTCCCGAGAAGAC | | | | | | | | | | |
| Chromosome | start | end | strand | MM | target_seq | PAM | alignment | distance | position | gene name | gene id |
| GU305914.1 | 122873 | 122895 | - | 0 | GTCTTCTCGGGAGAGGTGAC | GGG | \|\|\|\|\|\|\|\|[\|\|\|\|\|\|\|\|\|\|\|\|]PAM | NA | Intergenic | NA | NA |
| T24763 | GACGGGCCTCGTCTCCGTCTCGG | 1000 | | | | | | | | | |
| Oligo fwd | CACCGACGGGCCTCGTCTCCGTCT | | | | | | | | | | |
| Oligo rev | AAACAGACGGAGACGAGGCCCGTC | | | | | | | | | | |
| Chromosome | start | end | strand | MM | target_seq | PAM | alignment | distance | position | gene name | gene id |
| GU305914.1 | 122856 | 122878 | - | 0 | GACGGGCCTCGTCTCCGTCT | CGG | \|\|\|\|\|\|\|\|[\|\|\|\|\|\|\|\|\|\|\|\|]PAM | NA | Intergenic | NA | NA |
| T24764 | GCCTCGTCTCCGTCTCGGACTGG | 1000 | | | | | | | | | |
| Oligo fwd | CACCGCCTCGTCTCCGTCTCGGAC | | | | | | | | | | |
| Oligo rev | AAACGTCCGAGACGGAGACGAGGC | | | | | | | | | | |
| Chromosome | start | end | strand | MM | target_seq | PAM | alignment | distance | position | gene name | gene id |
| GU305914.1 | 122851 | 122873 | - | 0 | GCCTCGTCTCCGTCTCGGAC | TGG | \|\|\|\|\|\|\|\|[\|\|\|\|\|\|\|\|\|\|\|\|]PAM | NA | Intergenic | NA | NA |
| T24765 | CCTCGTCTCCGTCTCGGACTGGG | 1000 | | | | | | | | | |
| Oligo fwd | CACCCCTCGTCTCCGTCTCGGACT | | | | | | | | | | |
| Oligo rev | AAACAGTCCGAGACGGAGACGAGG | | | | | | | | | | |
| Chromosome | start | end | strand | MM | target_seq | PAM | alignment | distance | position | gene name | gene id |
| GU305914.1 | 122850 | 122872 | - | 0 | CCTCGTCTCCGTCTCGGACT | GGG | \|\|\|\|\|\|\|\|[\|\|\|\|\|\|\|\|\|\|\|\|]PAM | NA | Intergenic | NA | NA |
| GU305914.1 | 31162 | 31184 | + | 3 | CACCGTCTCCGTCTCCGACT | CGG | \|--\|\|\|\|\|[\|\|\|\|\|\|\|-\|\|\|\|]PAM | NA | Intergenic | NA | NA |
| T24766 | ACTGGGACCTCGTGCTGCCCTGG | 1000 | | | | | | | | | |
| Oligo fwd | CACCACTGGGACCTCGTGCTGCCC | | | | | | | | | | |
| Oligo rev | AAACGGGCAGCACGAGGTCCCAGT | | | | | | | | | | |
| Chromosome | start | end | strand | MM | target_seq | PAM | alignment | distance | position | gene name | gene id |
| GU305914.1 | 122833 | 122855 | - | 0 | ACTGGGACCTCGTGCTGCCC | TGG | \|\|\|\|\|\|\|\|[\|\|\|\|\|\|\|\|\|\|\|\|]PAM | NA | Intergenic | NA | NA |
| T24767 | GCTGCCCTGGATCGTGCCCATGG | 1000 | | | | | | | | | |
| Oligo fwd | CACCGCTGCCCTGGATCGTGCCCA | | | | | | | | | | |
| Oligo rev | AAACTGGGCACGATCCAGGGCAGC | | | | | | | | | | |
| Chromosome | start | end | strand | MM | target_seq | PAM | alignment | distance | position | gene name | gene id |
| GU305914.1 | 122820 | 122842 | - | 0 | GCTGCCCTGGATCGTGCCCA | TGG | \|\|\|\|\|\|\|\|[\|\|\|\|\|\|\|\|\|\|\|\|]PAM | NA | Intergenic | NA | NA |
| T24768 | CTGGATCGTGCCCATGGCGCTGG | 1000 | | | | | | | | | |
| Oligo fwd | CACCCTGGATCGTGCCCATGGCGC | | | | | | | | | | |
| Oligo rev | AAACGCGCCATGGGCACGATCCAG | | | | | | | | | | |
| Chromosome | start | end | strand | MM | target_seq | PAM | alignment | distance | position | gene name | gene id |
| GU305914.1 | 122814 | 122836 | - | 0 | CTGGATCGTGCCCATGGCGC | TGG | \|\|\|\|\|\|\|\|[\|\|\|\|\|\|\|\|\|\|\|\|]PAM | NA | Intergenic | NA | NA |
| T24769 | CAACCAGCGCATGATGATGCTGG | 1000 | | | | | | | | | |
| Oligo fwd | CACCCAACCAGCGCATGATGATGC | | | | | | | | | | |
| Oligo rev | AAACGCATCATCATGCGCTGGTTG | | | | | | | | | | |
| Chromosome | start | end | strand | MM | target_seq | PAM | alignment | distance | position | gene name | gene id |
| GU305914.1 | 122781 | 122803 | - | 0 | CAACCAGCGCATGATGATGC | TGG | \|\|\|\|\|\|\|\|[\|\|\|\|\|\|\|\|\|\|\|\|]PAM | NA | Intergenic | NA | NA |
| T24770 | GTCGCTCGACCGCTCGCACGAGG | 1000 | | | | | | | | | |
| Oligo fwd | CACCGTCGCTCGACCGCTCGCACG | | | | | | | | | | |
| Oligo rev | AAACCGTGCGAGCGGTCGAGCGAC | | | | | | | | | | |
| Chromosome | start | end | strand | MM | target_seq | PAM | alignment | distance | position | gene name | gene id |
| GU305914.1 | 122751 | 122773 | - | 0 | GTCGCTCGACCGCTCGCACG | AGG | \|\|\|\|\|\|\|\|[\|\|\|\|\|\|\|\|\|\|\|\|]PAM | NA | Intergenic | NA | NA |
| T24771 | GCTCGACCGCTCGCACGAGGAGG | 1000 | | | | | | | | | |
| Oligo fwd | CACCGCTCGACCGCTCGCACGAGG | | | | | | | | | | |
| Oligo rev | AAACCCTCGTGCGAGCGGTCGAGC | | | | | | | | | | |
| Chromosome | start | end | strand | MM | target_seq | PAM | alignment | distance | position | gene name | gene id |

| | | | | | | | | | | | |
|---|---|---|---|---|---|---|---|---|---|---|---|
| GU305914.1 | 122748 | 122770 | - | 0 | GCTCGACCGCTCGCACGAGG | AGG | \|\|\|\|\|\|\|\|[\|\|\|\|\|\|\|\|\|\|\|\|]PAM | NA | Intergenic | NA | NA |
| T24772 | CTCGCACGAGGAGGTGCGCGCGG | 1000 | | | | | | | | | |
| Oligo fwd | CACCCTCGCACGAGGAGGTGCGCG | | | | | | | | | | |
| Oligo rev | AAACCGCGCACCTCCTCGTGCGAG | | | | | | | | | | |
| Chromosome | start | end | strand | MM | target_seq | PAM | alignment | distance | position | gene name | gene id |
| GU305914.1 | 122739 | 122761 | - | 0 | CTCGCACGAGGAGGTGCGCG | CGG | \|\|\|\|\|\|\|\|[\|\|\|\|\|\|\|\|\|\|\|\|]PAM | NA | Intergenic | NA | NA |
| T24773 | GCGCGGCGACCGCCCAGCTGCGG | 1000 | | | | | | | | | |
| Oligo fwd | CACCGCGCGGCGACCGCCCAGCTG | | | | | | | | | | |
| Oligo rev | AAACCAGCTGGGCGGTCGCCGCGC | | | | | | | | | | |
| Chromosome | start | end | strand | MM | target_seq | PAM | alignment | distance | position | gene name | gene id |
| GU305914.1 | 122722 | 122744 | - | 0 | GCGCGGCGACCGCCCAGCTG | CGG | \|\|\|\|\|\|\|\|[\|\|\|\|\|\|\|\|\|\|\|\|]PAM | NA | Intergenic | NA | NA |
| T24774 | CGCGGCGACCGCCCAGCTGCGGG | 1000 | | | | | | | | | |
| Oligo fwd | CACCCGCGGCGACCGCCCAGCTGC | | | | | | | | | | |
| Oligo rev | AAACGCAGCTGGGCGGTCGCCGCG | | | | | | | | | | |
| Chromosome | start | end | strand | MM | target_seq | PAM | alignment | distance | position | gene name | gene id |
| GU305914.1 | 122721 | 122743 | - | 0 | CGCGGCGACCGCCCAGCTGC | GGG | \|\|\|\|\|\|\|\|[\|\|\|\|\|\|\|\|\|\|\|\|]PAM | NA | Intergenic | NA | NA |
| T24775 | AGCTGCGGGTCGTGCGCTACCGG | 1000 | | | | | | | | | |
| Oligo fwd | CACCAGCTGCGGGTCGTGCGCTAC | | | | | | | | | | |
| Oligo rev | AAACGTAGCGCACGACCCGCAGCT | | | | | | | | | | |
| Chromosome | start | end | strand | MM | target_seq | PAM | alignment | distance | position | gene name | gene id |
| GU305914.1 | 122707 | 122729 | - | 0 | AGCTGCGGGTCGTGCGCTAC | CGG | \|\|\|\|\|\|\|\|[\|\|\|\|\|\|\|\|\|\|\|\|]PAM | NA | Intergenic | NA | NA |
| T24776 | GCTGCGGGTCGTGCGCTACCGGG | 1000 | | | | | | | | | |
| Oligo fwd | CACCGCTGCGGGTCGTGCGCTACC | | | | | | | | | | |
| Oligo rev | AAACGGTAGCGCACGACCCGCAGC | | | | | | | | | | |
| Chromosome | start | end | strand | MM | target_seq | PAM | alignment | distance | position | gene name | gene id |
| GU305914.1 | 122706 | 122728 | - | 0 | GCTGCGGGTCGTGCGCTACC | GGG | \|\|\|\|\|\|\|\|[\|\|\|\|\|\|\|\|\|\|\|\|]PAM | NA | Intergenic | NA | NA |
| T24777 | CGAGATCACGATCGACGACACGG | 1000 | | | | | | | | | |
| Oligo fwd | CACCCGAGATCACGATCGACGACA | | | | | | | | | | |
| Oligo rev | AAACTGTCGTCGATCGTGATCTCG | | | | | | | | | | |
| Chromosome | start | end | strand | MM | target_seq | PAM | alignment | distance | position | gene name | gene id |
| GU305914.1 | 122664 | 122686 | - | 0 | CGAGATCACGATCGACGACA | CGG | \|\|\|\|\|\|\|\|[\|\|\|\|\|\|\|\|\|\|\|\|]PAM | NA | Intergenic | NA | NA |
| GU305914.1 | 90509 | 90531 | - | 4 | CGGGGTCACGAGCGGCGACA | CGG | \|\|-\|-\|\|\|[\|\|\|-\|\|-\|\|\|\|\|]PAM | NA | Intergenic | NA | NA |
| T24778 | GATCGCCAACCTATCCTCCGAGG | 1000 | | | | | | | | | |
| Oligo fwd | CACCGATCGCCAACCTATCCTCCG | | | | | | | | | | |
| Oligo rev | AAACCGGAGGATAGGTTGGCGATC | | | | | | | | | | |
| Chromosome | start | end | strand | MM | target_seq | PAM | alignment | distance | position | gene name | gene id |
| GU305914.1 | 122601 | 122623 | - | 0 | GATCGCCAACCTATCCTCCG | AGG | \|\|\|\|\|\|\|\|[\|\|\|\|\|\|\|\|\|\|\|\|]PAM | NA | Intergenic | NA | NA |
| T24779 | CCTATCCTCCGAGGTGACGCTGG | 1000 | | | | | | | | | |
| Oligo fwd | CACCCCTATCCTCCGAGGTGACGC | | | | | | | | | | |
| Oligo rev | AAACGCGTCACCTCGGAGGATAGG | | | | | | | | | | |
| Chromosome | start | end | strand | MM | target_seq | PAM | alignment | distance | position | gene name | gene id |
| GU305914.1 | 122592 | 122614 | - | 0 | CCTATCCTCCGAGGTGACGC | TGG | \|\|\|\|\|\|\|\|[\|\|\|\|\|\|\|\|\|\|\|\|]PAM | NA | Intergenic | NA | NA |
| T24780 | AGGTGACGCTGGCCTACGTGCGG | 1000 | | | | | | | | | |
| Oligo fwd | CACCAGGTGACGCTGGCCTACGTG | | | | | | | | | | |
| Oligo rev | AAACCACGTAGGCCAGCGTCACCT | | | | | | | | | | |
| Chromosome | start | end | strand | MM | target_seq | PAM | alignment | distance | position | gene name | gene id |
| GU305914.1 | 122581 | 122603 | - | 0 | AGGTGACGCTGGCCTACGTG | CGG | \|\|\|\|\|\|\|\|[\|\|\|\|\|\|\|\|\|\|\|\|]PAM | NA | Intergenic | NA | NA |
| T24781 | GCTGGCCTACGTGCGGAAGCTGG | 1000 | | | | | | | | | |
| Oligo fwd | CACCGCTGGCCTACGTGCGGAAGC | | | | | | | | | | |
| Oligo rev | AAACGCTTCCGCACGTAGGCCAGC | | | | | | | | | | |
| Chromosome | start | end | strand | MM | target_seq | PAM | alignment | distance | position | gene name | gene id |
| GU305914.1 | 122574 | 122596 | - | 0 | GCTGGCCTACGTGCGGAAGC | TGG | \|\|\|\|\|\|\|\|[\|\|\|\|\|\|\|\|\|\|\|\|]PAM | NA | Intergenic | NA | NA |
| T24782 | CTACGTGCGGAAGCTGGCCCTGG | 1000 | | | | | | | | | |
| Oligo fwd | CACCCTACGTGCGGAAGCTGGCCC | | | | | | | | | | |
| Oligo rev | AAACGGGCCAGCTTCCGCACGTAG | | | | | | | | | | |
| Chromosome | start | end | strand | MM | target_seq | PAM | alignment | distance | position | gene name | gene id |
| GU305914.1 | 122568 | 122590 | - | 0 | CTACGTGCGGAAGCTGGCCC | TGG | \|\|\|\|\|\|\|\|[\|\|\|\|\|\|\|\|\|\|\|\|]PAM | NA | Intergenic | NA | NA |
| T24783 | CGTGCGGAAGCTGGCCCTGGAGG | 1000 | | | | | | | | | |
| Oligo fwd | CACCCGTGCGGAAGCTGGCCCTGG | | | | | | | | | | |
| Oligo rev | AAACCCAGGGCCAGCTTCCGCACG | | | | | | | | | | |
| Chromosome | start | end | strand | MM | target_seq | PAM | alignment | distance | position | gene name | gene id |
| GU305914.1 | 122565 | 122587 | - | 0 | CGTGCGGAAGCTGGCCCTGG | AGG | \|\|\|\|\|\|\|\|[\|\|\|\|\|\|\|\|\|\|\|\|]PAM | NA | Intergenic | NA | NA |
| T24784 | CATGCTCCTCATGAAATGTCAGG | 1000 | | | | | | | | | |
| Oligo fwd | CACCCATGCTCCTCATGAAATGTC | | | | | | | | | | |
| Oligo rev | AAACGACATTTCATGAGGAGCATG | | | | | | | | | | |
| Chromosome | start | end | strand | MM | target_seq | PAM | alignment | distance | position | gene name | gene id |
| GU305914.1 | 122535 | 122557 | - | 0 | CATGCTCCTCATGAAATGTC | AGG | \|\|\|\|\|\|\|\|[\|\|\|\|\|\|\|\|\|\|\|\|]PAM | NA | Intergenic | NA | NA |
| T24785 | ATGAAATGTCAGGAGATCCTCGG | 1000 | | | | | | | | | |
| Oligo fwd | CACCATGAAATGTCAGGAGATCCT | | | | | | | | | | |
| Oligo rev | AAACAGGATCTCCTGACATTTCAT | | | | | | | | | | |
| Chromosome | start | end | strand | MM | target_seq | PAM | alignment | distance | position | gene name | gene id |
| GU305914.1 | 122525 | 122547 | - | 0 | ATGAAATGTCAGGAGATCCT | CGG | \|\|\|\|\|\|\|\|[\|\|\|\|\|\|\|\|\|\|\|\|]PAM | NA | Intergenic | NA | NA |
| T24786 | AATGTCAGGAGATCCTCGGCCGG | 1000 | | | | | | | | | |
| Oligo fwd | CACCAATGTCAGGAGATCCTCGGC | | | | | | | | | | |

<table>
<tr><td>Oligo rev</td><td>AAACGCCGAGGATCTCCTGACATT</td><td></td><td></td><td></td><td></td><td></td><td></td><td></td><td></td><td></td><td></td></tr>
<tr><td>Chromosome</td><td>start</td><td>end</td><td>strand</td><td>MM</td><td>target_seq</td><td>PAM</td><td>alignment</td><td>distance</td><td>position</td><td>gene name</td><td>gene id</td></tr>
<tr><td>GU305914.1</td><td>122521</td><td>122543</td><td>-</td><td>0</td><td>AATGTCAGGAGATCCTCGGC</td><td>CGG</td><td>||||||||[||||||||||||]PAM</td><td>NA</td><td>Intergenic</td><td>NA</td><td>NA</td></tr>
<tr><td>GU305914.1</td><td>223050</td><td>223072</td><td>+</td><td>4</td><td>ATCGTCAGGAGCTCCTCGCC</td><td>CGG</td><td>|--|||||[|||-||||||-|]PAM</td><td>NA</td><td>Intergenic</td><td>NA</td><td>NA</td></tr>
<tr><td></td><td></td><td></td><td></td><td></td><td></td><td></td><td></td><td></td><td></td><td></td><td></td></tr>
<tr><td>T24787</td><td>GTCAGGAGATCCTCGGCCGGCGG</td><td>1000</td><td></td><td></td><td></td><td></td><td></td><td></td><td></td><td></td><td></td></tr>
<tr><td>Oligo fwd</td><td>CACCGTCAGGAGATCCTCGGCCGG</td><td></td><td></td><td></td><td></td><td></td><td></td><td></td><td></td><td></td><td></td></tr>
<tr><td>Oligo rev</td><td>AAACCCGGCCGAGGATCTCCTGAC</td><td></td><td></td><td></td><td></td><td></td><td></td><td></td><td></td><td></td><td></td></tr>
<tr><td>Chromosome</td><td>start</td><td>end</td><td>strand</td><td>MM</td><td>target_seq</td><td>PAM</td><td>alignment</td><td>distance</td><td>position</td><td>gene name</td><td>gene id</td></tr>
<tr><td>GU305914.1</td><td>122518</td><td>122540</td><td>-</td><td>0</td><td>GTCAGGAGATCCTCGGCCGG</td><td>CGG</td><td>||||||||[||||||||||||]PAM</td><td>NA</td><td>Intergenic</td><td>NA</td><td>NA</td></tr>
<tr><td></td><td></td><td></td><td></td><td></td><td></td><td></td><td></td><td></td><td></td><td></td><td></td></tr>
<tr><td>T24788</td><td>CCTCGGCCGGCGGATGCCCCAGG</td><td>1000</td><td></td><td></td><td></td><td></td><td></td><td></td><td></td><td></td><td></td></tr>
<tr><td>Oligo fwd</td><td>CACCCCTCGGCCGGCGGATGCCCC</td><td></td><td></td><td></td><td></td><td></td><td></td><td></td><td></td><td></td><td></td></tr>
<tr><td>Oligo rev</td><td>AAACGGGGCATCCGCCGGCCGAGG</td><td></td><td></td><td></td><td></td><td></td><td></td><td></td><td></td><td></td><td></td></tr>
<tr><td>Chromosome</td><td>start</td><td>end</td><td>strand</td><td>MM</td><td>target_seq</td><td>PAM</td><td>alignment</td><td>distance</td><td>position</td><td>gene name</td><td>gene id</td></tr>
<tr><td>GU305914.1</td><td>122508</td><td>122530</td><td>-</td><td>0</td><td>CCTCGGCCGGCGGATGCCCC</td><td>AGG</td><td>||||||||[||||||||||||]PAM</td><td>NA</td><td>Intergenic</td><td>NA</td><td>NA</td></tr>
<tr><td></td><td></td><td></td><td></td><td></td><td></td><td></td><td></td><td></td><td></td><td></td><td></td></tr>
<tr><td>T24789</td><td>GGCCGGCGGATGCCCCAGGTCGG</td><td>1000</td><td></td><td></td><td></td><td></td><td></td><td></td><td></td><td></td><td></td></tr>
<tr><td>Oligo fwd</td><td>CACCGGCCGGCGGATGCCCCAGGT</td><td></td><td></td><td></td><td></td><td></td><td></td><td></td><td></td><td></td><td></td></tr>
<tr><td>Oligo rev</td><td>AAACACCTGGGGCATCCGCCGGCC</td><td></td><td></td><td></td><td></td><td></td><td></td><td></td><td></td><td></td><td></td></tr>
<tr><td>Chromosome</td><td>start</td><td>end</td><td>strand</td><td>MM</td><td>target_seq</td><td>PAM</td><td>alignment</td><td>distance</td><td>position</td><td>gene name</td><td>gene id</td></tr>
<tr><td>GU305914.1</td><td>122504</td><td>122526</td><td>-</td><td>0</td><td>GGCCGGCGGATGCCCCAGGT</td><td>CGG</td><td>||||||||[||||||||||||]PAM</td><td>NA</td><td>Intergenic</td><td>NA</td><td>NA</td></tr>
<tr><td></td><td></td><td></td><td></td><td></td><td></td><td></td><td></td><td></td><td></td><td></td><td></td></tr>
<tr><td>T24790</td><td>CGGATGCCCCAGGTCGGTGTCGG</td><td>1000</td><td></td><td></td><td></td><td></td><td></td><td></td><td></td><td></td><td></td></tr>
<tr><td>Oligo fwd</td><td>CACCCGGATGCCCCAGGTCGGTGT</td><td></td><td></td><td></td><td></td><td></td><td></td><td></td><td></td><td></td><td></td></tr>
<tr><td>Oligo rev</td><td>AAACACACCGACCTGGGGCATCCG</td><td></td><td></td><td></td><td></td><td></td><td></td><td></td><td></td><td></td><td></td></tr>
<tr><td>Chromosome</td><td>start</td><td>end</td><td>strand</td><td>MM</td><td>target_seq</td><td>PAM</td><td>alignment</td><td>distance</td><td>position</td><td>gene name</td><td>gene id</td></tr>
<tr><td>GU305914.1</td><td>122498</td><td>122520</td><td>-</td><td>0</td><td>CGGATGCCCCAGGTCGGTGT</td><td>CGG</td><td>||||||||[||||||||||||]PAM</td><td>NA</td><td>Intergenic</td><td>NA</td><td>NA</td></tr>
<tr><td></td><td></td><td></td><td></td><td></td><td></td><td></td><td></td><td></td><td></td><td></td><td></td></tr>
<tr><td>T24791</td><td>GCCCCAGGTCGGTGTCGGCGCGG</td><td>1000</td><td></td><td></td><td></td><td></td><td></td><td></td><td></td><td></td><td></td></tr>
<tr><td>Oligo fwd</td><td>CACCGCCCCAGGTCGGTGTCGGCG</td><td></td><td></td><td></td><td></td><td></td><td></td><td></td><td></td><td></td><td></td></tr>
<tr><td>Oligo rev</td><td>AAACCGCCGACACCGACCTGGGGC</td><td></td><td></td><td></td><td></td><td></td><td></td><td></td><td></td><td></td><td></td></tr>
<tr><td>Chromosome</td><td>start</td><td>end</td><td>strand</td><td>MM</td><td>target_seq</td><td>PAM</td><td>alignment</td><td>distance</td><td>position</td><td>gene name</td><td>gene id</td></tr>
<tr><td>GU305914.1</td><td>122493</td><td>122515</td><td>-</td><td>0</td><td>GCCCCAGGTCGGTGTCGGCG</td><td>CGG</td><td>||||||||[||||||||||||]PAM</td><td>NA</td><td>Intergenic</td><td>NA</td><td>NA</td></tr>
<tr><td></td><td></td><td></td><td></td><td></td><td></td><td></td><td></td><td></td><td></td><td></td><td></td></tr>
<tr><td>T24792</td><td>CCCCAGGTCGGTGTCGGCGCGGG</td><td>1000</td><td></td><td></td><td></td><td></td><td></td><td></td><td></td><td></td><td></td></tr>
<tr><td>Oligo fwd</td><td>CACCCCCCAGGTCGGTGTCGGCGC</td><td></td><td></td><td></td><td></td><td></td><td></td><td></td><td></td><td></td><td></td></tr>
<tr><td>Oligo rev</td><td>AAACGCGCCGACACCGACCTGGGG</td><td></td><td></td><td></td><td></td><td></td><td></td><td></td><td></td><td></td><td></td></tr>
<tr><td>Chromosome</td><td>start</td><td>end</td><td>strand</td><td>MM</td><td>target_seq</td><td>PAM</td><td>alignment</td><td>distance</td><td>position</td><td>gene name</td><td>gene id</td></tr>
<tr><td>GU305914.1</td><td>122492</td><td>122514</td><td>-</td><td>0</td><td>CCCCAGGTCGGTGTCGGCGC</td><td>GGG</td><td>||||||||[||||||||||||]PAM</td><td>NA</td><td>Intergenic</td><td>NA</td><td>NA</td></tr>
<tr><td></td><td></td><td></td><td></td><td></td><td></td><td></td><td></td><td></td><td></td><td></td><td></td></tr>
<tr><td>T24793</td><td>GGTGTCGGCGCGGGCAGCAGCGG</td><td>1000</td><td></td><td></td><td></td><td></td><td></td><td></td><td></td><td></td><td></td></tr>
<tr><td>Oligo fwd</td><td>CACCGGTGTCGGCGCGGGCAGCAG</td><td></td><td></td><td></td><td></td><td></td><td></td><td></td><td></td><td></td><td></td></tr>
<tr><td>Oligo rev</td><td>AAACCTGCTGCCCGCGCCGACACC</td><td></td><td></td><td></td><td></td><td></td><td></td><td></td><td></td><td></td><td></td></tr>
<tr><td>Chromosome</td><td>start</td><td>end</td><td>strand</td><td>MM</td><td>target_seq</td><td>PAM</td><td>alignment</td><td>distance</td><td>position</td><td>gene name</td><td>gene id</td></tr>
<tr><td>GU305914.1</td><td>122483</td><td>122505</td><td>-</td><td>0</td><td>GGTGTCGGCGCGGGCAGCAG</td><td>CGG</td><td>||||||||[||||||||||||]PAM</td><td>NA</td><td>Intergenic</td><td>NA</td><td>NA</td></tr>
<tr><td>GU305914.1</td><td>166410</td><td>166432</td><td>+</td><td>4</td><td>GACGTCGGCGCCGGCGGCAG</td><td>CGG</td><td>|--|||||[|||-|||-||||]PAM</td><td>NA</td><td>Intergenic</td><td>NA</td><td>NA</td></tr>
<tr><td></td><td></td><td></td><td></td><td></td><td></td><td></td><td></td><td></td><td></td><td></td><td></td></tr>
<tr><td>T24794</td><td>GCGCGGGCAGCAGCGGCGACAGG</td><td>1000</td><td></td><td></td><td></td><td></td><td></td><td></td><td></td><td></td><td></td></tr>
<tr><td>Oligo fwd</td><td>CACCGCGCGGGCAGCAGCGGCGAC</td><td></td><td></td><td></td><td></td><td></td><td></td><td></td><td></td><td></td><td></td></tr>
<tr><td>Oligo rev</td><td>AAACGTCGCCGCTGCTGCCCGCGC</td><td></td><td></td><td></td><td></td><td></td><td></td><td></td><td></td><td></td><td></td></tr>
<tr><td>Chromosome</td><td>start</td><td>end</td><td>strand</td><td>MM</td><td>target_seq</td><td>PAM</td><td>alignment</td><td>distance</td><td>position</td><td>gene name</td><td>gene id</td></tr>
<tr><td>GU305914.1</td><td>122476</td><td>122498</td><td>-</td><td>0</td><td>GCGCGGGCAGCAGCGGCGAC</td><td>AGG</td><td>||||||||[||||||||||||]PAM</td><td>NA</td><td>Intergenic</td><td>NA</td><td>NA</td></tr>
<tr><td></td><td></td><td></td><td></td><td></td><td></td><td></td><td></td><td></td><td></td><td></td><td></td></tr>
<tr><td>T24795</td><td>CGGCGACAGGAACGACCCGCCGG</td><td>1000</td><td></td><td></td><td></td><td></td><td></td><td></td><td></td><td></td><td></td></tr>
<tr><td>Oligo fwd</td><td>CACCCGGCGACAGGAACGACCCGC</td><td></td><td></td><td></td><td></td><td></td><td></td><td></td><td></td><td></td><td></td></tr>
<tr><td>Oligo rev</td><td>AAACGCGGGTCGTTCCTGTCGCCG</td><td></td><td></td><td></td><td></td><td></td><td></td><td></td><td></td><td></td><td></td></tr>
<tr><td>Chromosome</td><td>start</td><td>end</td><td>strand</td><td>MM</td><td>target_seq</td><td>PAM</td><td>alignment</td><td>distance</td><td>position</td><td>gene name</td><td>gene id</td></tr>
<tr><td>GU305914.1</td><td>122463</td><td>122485</td><td>-</td><td>0</td><td>CGGCGACAGGAACGACCCGC</td><td>CGG</td><td>||||||||[||||||||||||]PAM</td><td>NA</td><td>Intergenic</td><td>NA</td><td>NA</td></tr>
<tr><td>GU305914.1</td><td>201132</td><td>201154</td><td>+</td><td>4</td><td>GGGCGACGGGGACGACCCGG</td><td>TGG</td><td>-||||||-[||-||||||||-]PAM</td><td>NA</td><td>Intergenic</td><td>NA</td><td>NA</td></tr>
<tr><td></td><td></td><td></td><td></td><td></td><td></td><td></td><td></td><td></td><td></td><td></td><td></td></tr>
<tr><td>T24796</td><td>ACGACCCGCCGGCCCGCTCGAGG</td><td>1000</td><td></td><td></td><td></td><td></td><td></td><td></td><td></td><td></td><td></td></tr>
<tr><td>Oligo fwd</td><td>CACCACGACCCGCCGGCCCGCTCG</td><td></td><td></td><td></td><td></td><td></td><td></td><td></td><td></td><td></td><td></td></tr>
<tr><td>Oligo rev</td><td>AAACCGAGCGGGCCGGCGGGTCGT</td><td></td><td></td><td></td><td></td><td></td><td></td><td></td><td></td><td></td><td></td></tr>
<tr><td>Chromosome</td><td>start</td><td>end</td><td>strand</td><td>MM</td><td>target_seq</td><td>PAM</td><td>alignment</td><td>distance</td><td>position</td><td>gene name</td><td>gene id</td></tr>
<tr><td>GU305914.1</td><td>122452</td><td>122474</td><td>-</td><td>0</td><td>ACGACCCGCCGGCCCGCTCG</td><td>AGG</td><td>||||||||[||||||||||||]PAM</td><td>NA</td><td>Intergenic</td><td>NA</td><td>NA</td></tr>
<tr><td></td><td></td><td></td><td></td><td></td><td></td><td></td><td></td><td></td><td></td><td></td><td></td></tr>
<tr><td>T24797</td><td>CTACAACATCACCCCCACCGAGG</td><td>1000</td><td></td><td></td><td></td><td></td><td></td><td></td><td></td><td></td><td></td></tr>
<tr><td>Oligo fwd</td><td>CACCCTACAACATCACCCCCACCG</td><td></td><td></td><td></td><td></td><td></td><td></td><td></td><td></td><td></td><td></td></tr>
<tr><td>Oligo rev</td><td>AAACCGGTGGGGGTGATGTTGTAG</td><td></td><td></td><td></td><td></td><td></td><td></td><td></td><td></td><td></td><td></td></tr>
<tr><td>Chromosome</td><td>start</td><td>end</td><td>strand</td><td>MM</td><td>target_seq</td><td>PAM</td><td>alignment</td><td>distance</td><td>position</td><td>gene name</td><td>gene id</td></tr>
<tr><td>GU305914.1</td><td>122424</td><td>122446</td><td>-</td><td>0</td><td>CTACAACATCACCCCCACCG</td><td>AGG</td><td>||||||||[||||||||||||]PAM</td><td>NA</td><td>Intergenic</td><td>NA</td><td>NA</td></tr>
<tr><td></td><td></td><td></td><td></td><td></td><td></td><td></td><td></td><td></td><td></td><td></td><td></td></tr>
<tr><td>T24798</td><td>CCGCCCTGTTCGTCATGATCCGG</td><td>1000</td><td></td><td></td><td></td><td></td><td></td><td></td><td></td><td></td><td></td></tr>
<tr><td>Oligo fwd</td><td>CACCCCGCCCTGTTCGTCATGATC</td><td></td><td></td><td></td><td></td><td></td><td></td><td></td><td></td><td></td><td></td></tr>
<tr><td>Oligo rev</td><td>AAACGATCATGACGAACAGGGCGG</td><td></td><td></td><td></td><td></td><td></td><td></td><td></td><td></td><td></td><td></td></tr>
<tr><td>Chromosome</td><td>start</td><td>end</td><td>strand</td><td>MM</td><td>target_seq</td><td>PAM</td><td>alignment</td><td>distance</td><td>position</td><td>gene name</td><td>gene id</td></tr>
<tr><td>GU305914.1</td><td>122386</td><td>122408</td><td>-</td><td>0</td><td>CCGCCCTGTTCGTCATGATC</td><td>CGG</td><td>||||||||[||||||||||||]PAM</td><td>NA</td><td>Intergenic</td><td>NA</td><td>NA</td></tr>
<tr><td></td><td></td><td></td><td></td><td></td><td></td><td></td><td></td><td></td><td></td><td></td><td></td></tr>
<tr><td>T24799</td><td>CGTCATGATCCGGCAGATCACGG</td><td>1000</td><td></td><td></td><td></td><td></td><td></td><td></td><td></td><td></td><td></td></tr>
<tr><td>Oligo fwd</td><td>CACCCGTCATGATCCGGCAGATCA</td><td></td><td></td><td></td><td></td><td></td><td></td><td></td><td></td><td></td><td></td></tr>
<tr><td>Oligo rev</td><td>AAACTGATCTGCCGGATCATGACG</td><td></td><td></td><td></td><td></td><td></td><td></td><td></td><td></td><td></td><td></td></tr>
<tr><td>Chromosome</td><td>start</td><td>end</td><td>strand</td><td>MM</td><td>target_seq</td><td>PAM</td><td>alignment</td><td>distance</td><td>position</td><td>gene name</td><td>gene id</td></tr>
<tr><td>GU305914.1</td><td>122376</td><td>122398</td><td>-</td><td>0</td><td>CGTCATGATCCGGCAGATCA</td><td>CGG</td><td>||||||||[||||||||||||]PAM</td><td>NA</td><td>Intergenic</td><td>NA</td><td>NA</td></tr>
<tr><td></td><td></td><td></td><td></td><td></td><td></td><td></td><td></td><td></td><td></td><td></td><td></td></tr>
<tr><td>T24800</td><td>CTCCGAGCAGCCCGCGTTCCTGG</td><td>1000</td><td></td><td></td><td></td><td></td><td></td><td></td><td></td><td></td><td></td></tr>
<tr><td>Oligo fwd</td><td>CACCCTCCGAGCAGCCCGCGTTCC</td><td></td><td></td><td></td><td></td><td></td><td></td><td></td><td></td><td></td><td></td></tr>
<tr><td>Oligo rev</td><td>AAACGGAACGCGGGCTGCTCGGAG</td><td></td><td></td><td></td><td></td><td></td><td></td><td></td><td></td><td></td><td></td></tr>
<tr><td>Chromosome</td><td>start</td><td>end</td><td>strand</td><td>MM</td><td>target_seq</td><td>PAM</td><td>alignment</td><td>distance</td><td>position</td><td>gene name</td><td>gene id</td></tr>
</table>

GU305914.1 122346 122368 - 0 CTCCGAGCAGCCCGCGTTCC TGG ||||||||[||||||||||||]PAM NA Intergenic NA NA

T24801 CGTGTCCCCCGACGACAAATCGG 1000
Oligo fwd CACCCGTGTCCCCCGACGACAAAT
Oligo rev AAACATTTGTCGTCGGGGGACACG
Chromosome start end strand MM target_seq PAM alignment distance position gene name gene id
GU305914.1 122316 122338 - 0 CGTGTCCCCCGACGACAAAT CGG ||||||||[||||||||||||]PAM NA Intergenic NA NA

T24802 ATCGGCCCTGTGCATCTACAAGG 1000
Oligo fwd CACCATCGGCCCTGTGCATCTACA
Oligo rev AAACTGTAGATGCACAGGGCCGAT
Chromosome start end strand MM target_seq PAM alignment distance position gene name gene id
GU305914.1 122298 122320 - 0 ATCGGCCCTGTGCATCTACA AGG ||||||||[||||||||||||]PAM NA Intergenic NA NA

T24803 TCGGCCCTGTGCATCTACAAGGG 1000
Oligo fwd CACCTCGGCCCTGTGCATCTACAA
Oligo rev AAACTTGTAGATGCACAGGGCCGA
Chromosome start end strand MM target_seq PAM alignment distance position gene name gene id
GU305914.1 122297 122319 - 0 TCGGCCCTGTGCATCTACAA GGG ||||||||[||||||||||||]PAM NA Intergenic NA NA

T24804 GCATCTACAAGGGTTGAGACCGG 1000
Oligo fwd CACCGCATCTACAAGGGTTGAGAC
Oligo rev AAACGTCTCAACCCTTGTAGATGC
Chromosome start end strand MM target_seq PAM alignment distance position gene name gene id
GU305914.1 122287 122309 - 0 GCATCTACAAGGGTTGAGAC CGG ||||||||[||||||||||||]PAM NA Intergenic NA NA

T24805 ACAAGGGTTGAGACCGGCAGCGG 1000
Oligo fwd CACCACAAGGGTTGAGACCGGCAG
Oligo rev AAACCTGCCGGTCTCAACCCTTGT
Chromosome start end strand MM target_seq PAM alignment distance position gene name gene id
GU305914.1 122281 122303 - 0 ACAAGGGTTGAGACCGGCAG CGG ||||||||[||||||||||||]PAM NA Intergenic NA NA

T24806 GGGTTGAGACCGGCAGCGGACGG 1000
Oligo fwd CACCGGGTTGAGACCGGCAGCGGA
Oligo rev AAACTCCGCTGCCGGTCTCAACCC
Chromosome start end strand MM target_seq PAM alignment distance position gene name gene id
GU305914.1 122277 122299 - 0 GGGTTGAGACCGGCAGCGGA CGG ||||||||[||||||||||||]PAM NA Intergenic NA NA

T24807 CGACCTTCGACAACACCGCGAGG 1000
Oligo fwd CACCCGACCTTCGACAACACCGCG
Oligo rev AAACCGCGGTGTTGTCGAAGGTCG
Chromosome start end strand MM target_seq PAM alignment distance position gene name gene id
GU305914.1 122221 122243 - 0 CGACCTTCGACAACACCGCG AGG ||||||||[||||||||||||]PAM NA Intergenic NA NA

T24808 GTCGTATCCCAACCCCCGAGCGG 1000
Oligo fwd CACCGTCGTATCCCAACCCCCGAG
Oligo rev AAACCTCGGGGGTTGGGATACGAC
Chromosome start end strand MM target_seq PAM alignment distance position gene name gene id
GU305914.1 122189 122211 - 0 GTCGTATCCCAACCCCCGAG CGG ||||||||[||||||||||||]PAM NA Intergenic NA NA

T24809 CACTTATAAGAGAAAGTGCGCGG 1000
Oligo fwd CACCCACTTATAAGAGAAAGTGCG
Oligo rev AAACCGCACTTTCTCTTATAAGTG
Chromosome start end strand MM target_seq PAM alignment distance position gene name gene id
GU305914.1 122092 122114 - 0 CACTTATAAGAGAAAGTGCG CGG ||||||||[||||||||||||]PAM NA Intergenic NA NA

T24810 CCGCCACCTCCGTCCGCAGAAGG 1000
Oligo fwd CACCCCGCCACCTCCGTCCGCAGA
Oligo rev AAACTCTGCGGACGGAGGTGGCGG
Chromosome start end strand MM target_seq PAM alignment distance position gene name gene id
GU305914.1 122046 122068 - 0 CCGCCACCTCCGTCCGCAGA AGG ||||||||[||||||||||||]PAM NA Intergenic NA NA

T24811 CTCCGTCCGCAGAAGGAGAGAGG 1000
Oligo fwd CACCCTCCGTCCGCAGAAGGAGAG
Oligo rev AAACCTCTCCTTCTGCGGACGGAG
Chromosome start end strand MM target_seq PAM alignment distance position gene name gene id
GU305914.1 122039 122061 - 0 CTCCGTCCGCAGAAGGAGAG AGG ||||||||[||||||||||||]PAM NA Intergenic NA NA

T24812 TCCGTCCGCAGAAGGAGAGAGGG 1000
Oligo fwd CACCTCCGTCCGCAGAAGGAGAGA
Oligo rev AAACTCTCTCCTTCTGCGGACGGA
Chromosome start end strand MM target_seq PAM alignment distance position gene name gene id
GU305914.1 122038 122060 - 0 TCCGTCCGCAGAAGGAGAGA GGG ||||||||[||||||||||||]PAM NA Intergenic NA NA
GU305914.1 85622 85644 - 4 CGCGTCCGCAGAAACAGAGA CGG --||||||[|||||--|||||]PAM NA Intergenic NA NA

T24813 GCAGAAGGAGAGAGGGATAGAGG 1000
Oligo fwd CACCGCAGAAGGAGAGAGGGATAG
Oligo rev AAACCTATCCCTCTCTCCTTCTGC
Chromosome start end strand MM target_seq PAM alignment distance position gene name gene id
GU305914.1 122031 122053 - 0 GCAGAAGGAGAGAGGGATAG AGG ||||||||[||||||||||||]PAM NA Intergenic NA NA

T24814 CAGAAGGAGAGAGGGATAGAGGG 1000
Oligo fwd CACCCAGAAGGAGAGAGGGATAGA
Oligo rev AAACTCTATCCCTCTCTCCTTCTG
Chromosome start end strand MM target_seq PAM alignment distance position gene name gene id
GU305914.1 122030 122052 - 0 CAGAAGGAGAGAGGGATAGA GGG ||||||||[||||||||||||]PAM NA Intergenic NA NA

T24815 TTTTTTTTGACACACTCTCACGG 1000
Oligo fwd CACCTTTTTTTTGACACACTCTCA

<table>
<tr><td>Oligo rev</td><td>AAACTGAGAGTGTGTCAAAAAAAA</td><td></td><td></td><td></td><td></td><td></td><td></td><td></td><td></td><td></td><td></td></tr>
<tr><td>Chromosome</td><td>start</td><td>end</td><td>strand</td><td>MM</td><td>target_seq</td><td>PAM</td><td>alignment</td><td>distance</td><td>position</td><td>gene name</td><td>gene id</td></tr>
<tr><td>GU305914.1</td><td>121983</td><td>122005</td><td>-</td><td>0</td><td>TTTTTTTGACACACTCTCA</td><td>CGG</td><td>||||||||[||||||||||||]PAM</td><td>NA</td><td>Intergenic</td><td>NA</td><td>NA</td></tr>
<tr><td>T24816</td><td>TCTTCCTTCGTTCGACATGTCGG</td><td>1000</td><td></td><td></td><td></td><td></td><td></td><td></td><td></td><td></td><td></td></tr>
<tr><td>Oligo fwd</td><td>CACCTCTTCCTTCGTTCGACATGT</td><td></td><td></td><td></td><td></td><td></td><td></td><td></td><td></td><td></td><td></td></tr>
<tr><td>Oligo rev</td><td>AAACACATGTCGAACGAAGGAAGA</td><td></td><td></td><td></td><td></td><td></td><td></td><td></td><td></td><td></td><td></td></tr>
<tr><td>Chromosome</td><td>start</td><td>end</td><td>strand</td><td>MM</td><td>target_seq</td><td>PAM</td><td>alignment</td><td>distance</td><td>position</td><td>gene name</td><td>gene id</td></tr>
<tr><td>GU305914.1</td><td>121943</td><td>121965</td><td>-</td><td>0</td><td>TCTTCCTTCGTTCGACATGT</td><td>CGG</td><td>||||||||[||||||||||||]PAM</td><td>NA</td><td>Intergenic</td><td>NA</td><td>NA</td></tr>
<tr><td>T24817</td><td>GTCAACGTTTACTTGCCGAGCGG</td><td>1000</td><td></td><td></td><td></td><td></td><td></td><td></td><td></td><td></td><td></td></tr>
<tr><td>Oligo fwd</td><td>CACCGTCAACGTTTACTTGCCGAG</td><td></td><td></td><td></td><td></td><td></td><td></td><td></td><td></td><td></td><td></td></tr>
<tr><td>Oligo rev</td><td>AAACCTCGGCAAGTAAACGTTGAC</td><td></td><td></td><td></td><td></td><td></td><td></td><td></td><td></td><td></td><td></td></tr>
<tr><td>Chromosome</td><td>start</td><td>end</td><td>strand</td><td>MM</td><td>target_seq</td><td>PAM</td><td>alignment</td><td>distance</td><td>position</td><td>gene name</td><td>gene id</td></tr>
<tr><td>GU305914.1</td><td>121921</td><td>121943</td><td>-</td><td>0</td><td>GTCAACGTTTACTTGCCGAG</td><td>CGG</td><td>||||||||[||||||||||||]PAM</td><td>NA</td><td>Intergenic</td><td>NA</td><td>NA</td></tr>
<tr><td>T24818</td><td>CGGCACGAGCTCCTTGTTCGCGG</td><td>1000</td><td></td><td></td><td></td><td></td><td></td><td></td><td></td><td></td><td></td></tr>
<tr><td>Oligo fwd</td><td>CACCCGGCACGAGCTCCTTGTTCG</td><td></td><td></td><td></td><td></td><td></td><td></td><td></td><td></td><td></td><td></td></tr>
<tr><td>Oligo rev</td><td>AAACCGAACAAGGAGCTCGTGCCG</td><td></td><td></td><td></td><td></td><td></td><td></td><td></td><td></td><td></td><td></td></tr>
<tr><td>Chromosome</td><td>start</td><td>end</td><td>strand</td><td>MM</td><td>target_seq</td><td>PAM</td><td>alignment</td><td>distance</td><td>position</td><td>gene name</td><td>gene id</td></tr>
<tr><td>GU305914.1</td><td>121901</td><td>121923</td><td>-</td><td>0</td><td>CGGCACGAGCTCCTTGTTCG</td><td>CGG</td><td>||||||||[||||||||||||]PAM</td><td>NA</td><td>Intergenic</td><td>NA</td><td>NA</td></tr>
<tr><td>T24819</td><td>GGCACGAGCTCCTTGTTCGCGGG</td><td>1000</td><td></td><td></td><td></td><td></td><td></td><td></td><td></td><td></td><td></td></tr>
<tr><td>Oligo fwd</td><td>CACCGGCACGAGCTCCTTGTTCGC</td><td></td><td></td><td></td><td></td><td></td><td></td><td></td><td></td><td></td><td></td></tr>
<tr><td>Oligo rev</td><td>AAACGCGAACAAGGAGCTCGTGCC</td><td></td><td></td><td></td><td></td><td></td><td></td><td></td><td></td><td></td><td></td></tr>
<tr><td>Chromosome</td><td>start</td><td>end</td><td>strand</td><td>MM</td><td>target_seq</td><td>PAM</td><td>alignment</td><td>distance</td><td>position</td><td>gene name</td><td>gene id</td></tr>
<tr><td>GU305914.1</td><td>121900</td><td>121922</td><td>-</td><td>0</td><td>GGCACGAGCTCCTTGTTCGC</td><td>GGG</td><td>||||||||[||||||||||||]PAM</td><td>NA</td><td>Intergenic</td><td>NA</td><td>NA</td></tr>
<tr><td>T24820</td><td>GCACGAGCTCCTTGTTCGCGGGG</td><td>1000</td><td></td><td></td><td></td><td></td><td></td><td></td><td></td><td></td><td></td></tr>
<tr><td>Oligo fwd</td><td>CACCGCACGAGCTCCTTGTTCGCG</td><td></td><td></td><td></td><td></td><td></td><td></td><td></td><td></td><td></td><td></td></tr>
<tr><td>Oligo rev</td><td>AAACCGCGAACAAGGAGCTCGTGC</td><td></td><td></td><td></td><td></td><td></td><td></td><td></td><td></td><td></td><td></td></tr>
<tr><td>Chromosome</td><td>start</td><td>end</td><td>strand</td><td>MM</td><td>target_seq</td><td>PAM</td><td>alignment</td><td>distance</td><td>position</td><td>gene name</td><td>gene id</td></tr>
<tr><td>GU305914.1</td><td>121899</td><td>121921</td><td>-</td><td>0</td><td>GCACGAGCTCCTTGTTCGCG</td><td>GGG</td><td>||||||||[||||||||||||]PAM</td><td>NA</td><td>Intergenic</td><td>NA</td><td>NA</td></tr>
<tr><td>T24821</td><td>GGGGAGTCTGTCTTACCTGCTGG</td><td>1000</td><td></td><td></td><td></td><td></td><td></td><td></td><td></td><td></td><td></td></tr>
<tr><td>Oligo fwd</td><td>CACCGGGGAGTCTGTCTTACCTGC</td><td></td><td></td><td></td><td></td><td></td><td></td><td></td><td></td><td></td><td></td></tr>
<tr><td>Oligo rev</td><td>AAACGCAGGTAAGACAGACTCCCC</td><td></td><td></td><td></td><td></td><td></td><td></td><td></td><td></td><td></td><td></td></tr>
<tr><td>Chromosome</td><td>start</td><td>end</td><td>strand</td><td>MM</td><td>target_seq</td><td>PAM</td><td>alignment</td><td>distance</td><td>position</td><td>gene name</td><td>gene id</td></tr>
<tr><td>GU305914.1</td><td>121880</td><td>121902</td><td>-</td><td>0</td><td>GGGGAGTCTGTCTTACCTGC</td><td>TGG</td><td>||||||||[||||||||||||]PAM</td><td>NA</td><td>Intergenic</td><td>NA</td><td>NA</td></tr>
<tr><td>T24822</td><td>GTCCCCGTGACTGTAGATTTCGG</td><td>1000</td><td></td><td></td><td></td><td></td><td></td><td></td><td></td><td></td><td></td></tr>
<tr><td>Oligo fwd</td><td>CACCGTCCCCGTGACTGTAGATTT</td><td></td><td></td><td></td><td></td><td></td><td></td><td></td><td></td><td></td><td></td></tr>
<tr><td>Oligo rev</td><td>AAACAAATCTACAGTCACGGGGAC</td><td></td><td></td><td></td><td></td><td></td><td></td><td></td><td></td><td></td><td></td></tr>
<tr><td>Chromosome</td><td>start</td><td>end</td><td>strand</td><td>MM</td><td>target_seq</td><td>PAM</td><td>alignment</td><td>distance</td><td>position</td><td>gene name</td><td>gene id</td></tr>
<tr><td>GU305914.1</td><td>121849</td><td>121871</td><td>-</td><td>0</td><td>GTCCCCGTGACTGTAGATTT</td><td>CGG</td><td>||||||||[||||||||||||]PAM</td><td>NA</td><td>Intergenic</td><td>NA</td><td>NA</td></tr>
<tr><td>T24823</td><td>GAGACCAAATTCCTCCAGACCGG</td><td>1000</td><td></td><td></td><td></td><td></td><td></td><td></td><td></td><td></td><td></td></tr>
<tr><td>Oligo fwd</td><td>CACCGAGACCAAATTCCTCCAGAC</td><td></td><td></td><td></td><td></td><td></td><td></td><td></td><td></td><td></td><td></td></tr>
<tr><td>Oligo rev</td><td>AAACGTCTGGAGGAATTTGGTCTC</td><td></td><td></td><td></td><td></td><td></td><td></td><td></td><td></td><td></td><td></td></tr>
<tr><td>Chromosome</td><td>start</td><td>end</td><td>strand</td><td>MM</td><td>target_seq</td><td>PAM</td><td>alignment</td><td>distance</td><td>position</td><td>gene name</td><td>gene id</td></tr>
<tr><td>GU305914.1</td><td>121819</td><td>121841</td><td>-</td><td>0</td><td>GAGACCAAATTCCTCCAGAC</td><td>CGG</td><td>||||||||[||||||||||||]PAM</td><td>NA</td><td>Intergenic</td><td>NA</td><td>NA</td></tr>
<tr><td>T24824</td><td>AGACCAAATTCCTCCAGACCGGG</td><td>1000</td><td></td><td></td><td></td><td></td><td></td><td></td><td></td><td></td><td></td></tr>
<tr><td>Oligo fwd</td><td>CACCAGACCAAATTCCTCCAGACC</td><td></td><td></td><td></td><td></td><td></td><td></td><td></td><td></td><td></td><td></td></tr>
<tr><td>Oligo rev</td><td>AAACGGTCTGGAGGAATTTGGTCT</td><td></td><td></td><td></td><td></td><td></td><td></td><td></td><td></td><td></td><td></td></tr>
<tr><td>Chromosome</td><td>start</td><td>end</td><td>strand</td><td>MM</td><td>target_seq</td><td>PAM</td><td>alignment</td><td>distance</td><td>position</td><td>gene name</td><td>gene id</td></tr>
<tr><td>GU305914.1</td><td>121818</td><td>121840</td><td>-</td><td>0</td><td>AGACCAAATTCCTCCAGACC</td><td>GGG</td><td>||||||||[||||||||||||]PAM</td><td>NA</td><td>Intergenic</td><td>NA</td><td>NA</td></tr>
<tr><td>T24825</td><td>AATTCCTCCAGACCGGGATCCGG</td><td>1000</td><td></td><td></td><td></td><td></td><td></td><td></td><td></td><td></td><td></td></tr>
<tr><td>Oligo fwd</td><td>CACCAATTCCTCCAGACCGGGATC</td><td></td><td></td><td></td><td></td><td></td><td></td><td></td><td></td><td></td><td></td></tr>
<tr><td>Oligo rev</td><td>AAACGATCCCGGTCTGGAGGAATT</td><td></td><td></td><td></td><td></td><td></td><td></td><td></td><td></td><td></td><td></td></tr>
<tr><td>Chromosome</td><td>start</td><td>end</td><td>strand</td><td>MM</td><td>target_seq</td><td>PAM</td><td>alignment</td><td>distance</td><td>position</td><td>gene name</td><td>gene id</td></tr>
<tr><td>GU305914.1</td><td>121812</td><td>121834</td><td>-</td><td>0</td><td>AATTCCTCCAGACCGGGATC</td><td>CGG</td><td>||||||||[||||||||||||]PAM</td><td>NA</td><td>Intergenic</td><td>NA</td><td>NA</td></tr>
<tr><td>T24826</td><td>CGGGATCCGGCTGAGTCCGCAGG</td><td>1000</td><td></td><td></td><td></td><td></td><td></td><td></td><td></td><td></td><td></td></tr>
<tr><td>Oligo fwd</td><td>CACCCGGGATCCGGCTGAGTCCGC</td><td></td><td></td><td></td><td></td><td></td><td></td><td></td><td></td><td></td><td></td></tr>
<tr><td>Oligo rev</td><td>AAACGCGGACTCAGCCGGATCCCG</td><td></td><td></td><td></td><td></td><td></td><td></td><td></td><td></td><td></td><td></td></tr>
<tr><td>Chromosome</td><td>start</td><td>end</td><td>strand</td><td>MM</td><td>target_seq</td><td>PAM</td><td>alignment</td><td>distance</td><td>position</td><td>gene name</td><td>gene id</td></tr>
<tr><td>GU305914.1</td><td>121799</td><td>121821</td><td>-</td><td>0</td><td>CGGGATCCGGCTGAGTCCGC</td><td>AGG</td><td>||||||||[||||||||||||]PAM</td><td>NA</td><td>Intergenic</td><td>NA</td><td>NA</td></tr>
<tr><td>T24827</td><td>GGCGCCCTGCGTCCTGTGTGTGG</td><td>1000</td><td></td><td></td><td></td><td></td><td></td><td></td><td></td><td></td><td></td></tr>
<tr><td>Oligo fwd</td><td>CACCGGCGCCCTGCGTCCTGTGTG</td><td></td><td></td><td></td><td></td><td></td><td></td><td></td><td></td><td></td><td></td></tr>
<tr><td>Oligo rev</td><td>AAACCACACAGGACGCAGGGCGCC</td><td></td><td></td><td></td><td></td><td></td><td></td><td></td><td></td><td></td><td></td></tr>
<tr><td>Chromosome</td><td>start</td><td>end</td><td>strand</td><td>MM</td><td>target_seq</td><td>PAM</td><td>alignment</td><td>distance</td><td>position</td><td>gene name</td><td>gene id</td></tr>
<tr><td>GU305914.1</td><td>121778</td><td>121800</td><td>-</td><td>0</td><td>GGCGCCCTGCGTCCTGTGTG</td><td>TGG</td><td>||||||||[||||||||||||]PAM</td><td>NA</td><td>Intergenic</td><td>NA</td><td>NA</td></tr>
<tr><td>T24828</td><td>GCGCCCTGCGTCCTGTGTGTGGG</td><td>1000</td><td></td><td></td><td></td><td></td><td></td><td></td><td></td><td></td><td></td></tr>
<tr><td>Oligo fwd</td><td>CACCGCGCCCTGCGTCCTGTGTGT</td><td></td><td></td><td></td><td></td><td></td><td></td><td></td><td></td><td></td><td></td></tr>
<tr><td>Oligo rev</td><td>AAACACACACAGGACGCAGGGCGC</td><td></td><td></td><td></td><td></td><td></td><td></td><td></td><td></td><td></td><td></td></tr>
<tr><td>Chromosome</td><td>start</td><td>end</td><td>strand</td><td>MM</td><td>target_seq</td><td>PAM</td><td>alignment</td><td>distance</td><td>position</td><td>gene name</td><td>gene id</td></tr>
<tr><td>GU305914.1</td><td>121777</td><td>121799</td><td>-</td><td>0</td><td>GCGCCCTGCGTCCTGTGTGT</td><td>GGG</td><td>||||||||[||||||||||||]PAM</td><td>NA</td><td>Intergenic</td><td>NA</td><td>NA</td></tr>
<tr><td>T24829</td><td>CGCCCTGCGTCCTGTGTGTGGGG</td><td>1000</td><td></td><td></td><td></td><td></td><td></td><td></td><td></td><td></td><td></td></tr>
<tr><td>Oligo fwd</td><td>CACCCGCCCTGCGTCCTGTGTGTG</td><td></td><td></td><td></td><td></td><td></td><td></td><td></td><td></td><td></td><td></td></tr>
<tr><td>Oligo rev</td><td>AAACCACACACAGGACGCAGGGCG</td><td></td><td></td><td></td><td></td><td></td><td></td><td></td><td></td><td></td><td></td></tr>
<tr><td>Chromosome</td><td>start</td><td>end</td><td>strand</td><td>MM</td><td>target_seq</td><td>PAM</td><td>alignment</td><td>distance</td><td>position</td><td>gene name</td><td>gene id</td></tr>
<tr><td>GU305914.1</td><td>121776</td><td>121798</td><td>-</td><td>0</td><td>CGCCCTGCGTCCTGTGTGTG</td><td>GGG</td><td>||||||||[||||||||||||]PAM</td><td>NA</td><td>Intergenic</td><td>NA</td><td>NA</td></tr>
<tr><td>T24830</td><td>TGTGTGTGGGGAAGATGTACCGG</td><td>1000</td><td></td><td></td><td></td><td></td><td></td><td></td><td></td><td></td><td></td></tr>
</table>

<table>
<tr><td>Oligo fwd</td><td>CACCTGTGTGTGGGGAAGATGTAC</td><td></td><td></td><td></td><td></td><td></td><td></td><td></td><td></td><td></td><td></td></tr>
<tr><td>Oligo rev</td><td>AAACGTACATCTTCCCCACACACA</td><td></td><td></td><td></td><td></td><td></td><td></td><td></td><td></td><td></td><td></td></tr>
<tr><td>Chromosome</td><td>start</td><td>end</td><td>strand</td><td>MM</td><td>target_seq</td><td>PAM</td><td>alignment</td><td>distance</td><td>position</td><td>gene name</td><td>gene id</td></tr>
<tr><td>GU305914.1</td><td>121764</td><td>121786</td><td>-</td><td>0</td><td>TGTGTGTGGGGAAGATGTAC</td><td>CGG</td><td>||||||||[||||||||||||]PAM</td><td>NA</td><td>Intergenic</td><td>NA</td><td>NA</td></tr>
<tr><td>T24831</td><td>GGGGAAGATGTACCGGCACCCGG</td><td>1000</td><td></td><td></td><td></td><td></td><td></td><td></td><td></td><td></td><td></td></tr>
<tr><td>Oligo fwd</td><td>CACCGGGGAAGATGTACCGGCACC</td><td></td><td></td><td></td><td></td><td></td><td></td><td></td><td></td><td></td><td></td></tr>
<tr><td>Oligo rev</td><td>AAACGGTGCCGGTACATCTTCCCC</td><td></td><td></td><td></td><td></td><td></td><td></td><td></td><td></td><td></td><td></td></tr>
<tr><td>Chromosome</td><td>start</td><td>end</td><td>strand</td><td>MM</td><td>target_seq</td><td>PAM</td><td>alignment</td><td>distance</td><td>position</td><td>gene name</td><td>gene id</td></tr>
<tr><td>GU305914.1</td><td>121757</td><td>121779</td><td>-</td><td>0</td><td>GGGGAAGATGTACCGGCACC</td><td>CGG</td><td>||||||||[||||||||||||]PAM</td><td>NA</td><td>Intergenic</td><td>NA</td><td>NA</td></tr>
<tr><td>T24832</td><td>CCCGGACAACCACCCACACCTGG</td><td>1000</td><td></td><td></td><td></td><td></td><td></td><td></td><td></td><td></td><td></td></tr>
<tr><td>Oligo fwd</td><td>CACCCCCGGACAACCACCCACACC</td><td></td><td></td><td></td><td></td><td></td><td></td><td></td><td></td><td></td><td></td></tr>
<tr><td>Oligo rev</td><td>AAACGGTGTGGGTGGTTGTCCGGG</td><td></td><td></td><td></td><td></td><td></td><td></td><td></td><td></td><td></td><td></td></tr>
<tr><td>Chromosome</td><td>start</td><td>end</td><td>strand</td><td>MM</td><td>target_seq</td><td>PAM</td><td>alignment</td><td>distance</td><td>position</td><td>gene name</td><td>gene id</td></tr>
<tr><td>GU305914.1</td><td>121739</td><td>121761</td><td>-</td><td>0</td><td>CCCGGACAACCACCCACACC</td><td>TGG</td><td>||||||||[||||||||||||]PAM</td><td>NA</td><td>Intergenic</td><td>NA</td><td>NA</td></tr>
<tr><td>T24833</td><td>CTTCGCCACCTTCACGCCCGCGG</td><td>1000</td><td></td><td></td><td></td><td></td><td></td><td></td><td></td><td></td><td></td></tr>
<tr><td>Oligo fwd</td><td>CACCCTTCGCCACCTTCACGCCCG</td><td></td><td></td><td></td><td></td><td></td><td></td><td></td><td></td><td></td><td></td></tr>
<tr><td>Oligo rev</td><td>AAACCGGGCGTGAAGGTGGCGAAG</td><td></td><td></td><td></td><td></td><td></td><td></td><td></td><td></td><td></td><td></td></tr>
<tr><td>Chromosome</td><td>start</td><td>end</td><td>strand</td><td>MM</td><td>target_seq</td><td>PAM</td><td>alignment</td><td>distance</td><td>position</td><td>gene name</td><td>gene id</td></tr>
<tr><td>GU305914.1</td><td>121712</td><td>121734</td><td>-</td><td>0</td><td>CTTCGCCACCTTCACGCCCG</td><td>CGG</td><td>||||||||[||||||||||||]PAM</td><td>NA</td><td>Intergenic</td><td>NA</td><td>NA</td></tr>
<tr><td>T24834</td><td>CGCGGACGCGCCGACCGACCTGG</td><td>1000</td><td></td><td></td><td></td><td></td><td></td><td></td><td></td><td></td><td></td></tr>
<tr><td>Oligo fwd</td><td>CACCCGCGGACGCGCCGACCGACC</td><td></td><td></td><td></td><td></td><td></td><td></td><td></td><td></td><td></td><td></td></tr>
<tr><td>Oligo rev</td><td>AAACGGTCGGTCGGCGCGTCCGCG</td><td></td><td></td><td></td><td></td><td></td><td></td><td></td><td></td><td></td><td></td></tr>
<tr><td>Chromosome</td><td>start</td><td>end</td><td>strand</td><td>MM</td><td>target_seq</td><td>PAM</td><td>alignment</td><td>distance</td><td>position</td><td>gene name</td><td>gene id</td></tr>
<tr><td>GU305914.1</td><td>121694</td><td>121716</td><td>-</td><td>0</td><td>CGCGGACGCGCCGACCGACC</td><td>TGG</td><td>||||||||[||||||||||||]PAM</td><td>NA</td><td>Intergenic</td><td>NA</td><td>NA</td></tr>
<tr><td>T24835</td><td>CATCCGCTGAAGCCGCCGTCCGG</td><td>1000</td><td></td><td></td><td></td><td></td><td></td><td></td><td></td><td></td><td></td></tr>
<tr><td>Oligo fwd</td><td>CACCCATCCGCTGAAGCCGCCGTC</td><td></td><td></td><td></td><td></td><td></td><td></td><td></td><td></td><td></td><td></td></tr>
<tr><td>Oligo rev</td><td>AAACGACGGCGGCTTCAGCGGATG</td><td></td><td></td><td></td><td></td><td></td><td></td><td></td><td></td><td></td><td></td></tr>
<tr><td>Chromosome</td><td>start</td><td>end</td><td>strand</td><td>MM</td><td>target_seq</td><td>PAM</td><td>alignment</td><td>distance</td><td>position</td><td>gene name</td><td>gene id</td></tr>
<tr><td>GU305914.1</td><td>121645</td><td>121667</td><td>-</td><td>0</td><td>CATCCGCTGAAGCCGCCGTC</td><td>CGG</td><td>||||||||[||||||||||||]PAM</td><td>NA</td><td>Intergenic</td><td>NA</td><td>NA</td></tr>
<tr><td>T24836</td><td>CCGCTGAAGCCGCCGTCCGGCGG</td><td>1000</td><td></td><td></td><td></td><td></td><td></td><td></td><td></td><td></td><td></td></tr>
<tr><td>Oligo fwd</td><td>CACCCCGCTGAAGCCGCCGTCCGG</td><td></td><td></td><td></td><td></td><td></td><td></td><td></td><td></td><td></td><td></td></tr>
<tr><td>Oligo rev</td><td>AAACCCGGACGGCGGCTTCAGCGG</td><td></td><td></td><td></td><td></td><td></td><td></td><td></td><td></td><td></td><td></td></tr>
<tr><td>Chromosome</td><td>start</td><td>end</td><td>strand</td><td>MM</td><td>target_seq</td><td>PAM</td><td>alignment</td><td>distance</td><td>position</td><td>gene name</td><td>gene id</td></tr>
<tr><td>GU305914.1</td><td>121642</td><td>121664</td><td>-</td><td>0</td><td>CCGCTGAAGCCGCCGTCCGG</td><td>CGG</td><td>||||||||[||||||||||||]PAM</td><td>NA</td><td>Intergenic</td><td>NA</td><td>NA</td></tr>
<tr><td>GU305914.1</td><td>90202</td><td>90224</td><td>+</td><td>3</td><td>CCGCTGGCGCCGCCGGCCGG</td><td>AGG</td><td>||||||--[|||||||-||||]PAM</td><td>NA</td><td>Intergenic</td><td>NA</td><td>NA</td></tr>
<tr><td>T24837</td><td>CGCTGAAGCCGCCGTCCGGCGGG</td><td>1000</td><td></td><td></td><td></td><td></td><td></td><td></td><td></td><td></td><td></td></tr>
<tr><td>Oligo fwd</td><td>CACCCGCTGAAGCCGCCGTCCGGC</td><td></td><td></td><td></td><td></td><td></td><td></td><td></td><td></td><td></td><td></td></tr>
<tr><td>Oligo rev</td><td>AAACGCCGGACGGCGGCTTCAGCG</td><td></td><td></td><td></td><td></td><td></td><td></td><td></td><td></td><td></td><td></td></tr>
<tr><td>Chromosome</td><td>start</td><td>end</td><td>strand</td><td>MM</td><td>target_seq</td><td>PAM</td><td>alignment</td><td>distance</td><td>position</td><td>gene name</td><td>gene id</td></tr>
<tr><td>GU305914.1</td><td>121641</td><td>121663</td><td>-</td><td>0</td><td>CGCTGAAGCCGCCGTCCGGC</td><td>GGG</td><td>||||||||[||||||||||||]PAM</td><td>NA</td><td>Intergenic</td><td>NA</td><td>NA</td></tr>
<tr><td>T24838</td><td>GCCGCCGTCCGGCGGGCAGCTGG</td><td>1000</td><td></td><td></td><td></td><td></td><td></td><td></td><td></td><td></td><td></td></tr>
<tr><td>Oligo fwd</td><td>CACCGCCGCCGTCCGGCGGGCAGC</td><td></td><td></td><td></td><td></td><td></td><td></td><td></td><td></td><td></td><td></td></tr>
<tr><td>Oligo rev</td><td>AAACGCTGCCCGCCGGACGGCGGC</td><td></td><td></td><td></td><td></td><td></td><td></td><td></td><td></td><td></td><td></td></tr>
<tr><td>Chromosome</td><td>start</td><td>end</td><td>strand</td><td>MM</td><td>target_seq</td><td>PAM</td><td>alignment</td><td>distance</td><td>position</td><td>gene name</td><td>gene id</td></tr>
<tr><td>GU305914.1</td><td>121634</td><td>121656</td><td>-</td><td>0</td><td>GCCGCCGTCCGGCGGGCAGC</td><td>TGG</td><td>||||||||[||||||||||||]PAM</td><td>NA</td><td>Intergenic</td><td>NA</td><td>NA</td></tr>
<tr><td>T24839</td><td>GATCGTGTTCTGCGTGCCGCTGG</td><td>1000</td><td></td><td></td><td></td><td></td><td></td><td></td><td></td><td></td><td></td></tr>
<tr><td>Oligo fwd</td><td>CACCGATCGTGTTCTGCGTGCCGC</td><td></td><td></td><td></td><td></td><td></td><td></td><td></td><td></td><td></td><td></td></tr>
<tr><td>Oligo rev</td><td>AAACGCGGCACGCAGAACACGATC</td><td></td><td></td><td></td><td></td><td></td><td></td><td></td><td></td><td></td><td></td></tr>
<tr><td>Chromosome</td><td>start</td><td>end</td><td>strand</td><td>MM</td><td>target_seq</td><td>PAM</td><td>alignment</td><td>distance</td><td>position</td><td>gene name</td><td>gene id</td></tr>
<tr><td>GU305914.1</td><td>121607</td><td>121629</td><td>-</td><td>0</td><td>GATCGTGTTCTGCGTGCCGC</td><td>TGG</td><td>||||||||[||||||||||||]PAM</td><td>NA</td><td>Intergenic</td><td>NA</td><td>NA</td></tr>
<tr><td>T24840</td><td>GCGTGCCGCTGGCGCCCGTGCGG</td><td>1000</td><td></td><td></td><td></td><td></td><td></td><td></td><td></td><td></td><td></td></tr>
<tr><td>Oligo fwd</td><td>CACCGCGTGCCGCTGGCGCCCGTG</td><td></td><td></td><td></td><td></td><td></td><td></td><td></td><td></td><td></td><td></td></tr>
<tr><td>Oligo rev</td><td>AAACCACGGGCGCCAGCGGCACGC</td><td></td><td></td><td></td><td></td><td></td><td></td><td></td><td></td><td></td><td></td></tr>
<tr><td>Chromosome</td><td>start</td><td>end</td><td>strand</td><td>MM</td><td>target_seq</td><td>PAM</td><td>alignment</td><td>distance</td><td>position</td><td>gene name</td><td>gene id</td></tr>
<tr><td>GU305914.1</td><td>121596</td><td>121618</td><td>-</td><td>0</td><td>GCGTGCCGCTGGCGCCCGTG</td><td>CGG</td><td>||||||||[||||||||||||]PAM</td><td>NA</td><td>Intergenic</td><td>NA</td><td>NA</td></tr>
<tr><td>T24841</td><td>GCCGCTGGCGCCCGTGCGGATGG</td><td>1000</td><td></td><td></td><td></td><td></td><td></td><td></td><td></td><td></td><td></td></tr>
<tr><td>Oligo fwd</td><td>CACCGCCGCTGGCGCCCGTGCGGA</td><td></td><td></td><td></td><td></td><td></td><td></td><td></td><td></td><td></td><td></td></tr>
<tr><td>Oligo rev</td><td>AAACTCCGCACGGGCGCCAGCGGC</td><td></td><td></td><td></td><td></td><td></td><td></td><td></td><td></td><td></td><td></td></tr>
<tr><td>Chromosome</td><td>start</td><td>end</td><td>strand</td><td>MM</td><td>target_seq</td><td>PAM</td><td>alignment</td><td>distance</td><td>position</td><td>gene name</td><td>gene id</td></tr>
<tr><td>GU305914.1</td><td>121592</td><td>121614</td><td>-</td><td>0</td><td>GCCGCTGGCGCCCGTGCGGA</td><td>TGG</td><td>||||||||[||||||||||||]PAM</td><td>NA</td><td>Intergenic</td><td>NA</td><td>NA</td></tr>
<tr><td>GU305914.1</td><td>163567</td><td>163589</td><td>-</td><td>4</td><td>GAGGCTGGCGCCCGTGCCGT</td><td>CGG</td><td>|--|||||[|||||||||-|-]PAM</td><td>NA</td><td>Intergenic</td><td>NA</td><td>NA</td></tr>
<tr><td>T24842</td><td>TGGACCTGCTGTGCGTGCAGAGG</td><td>1000</td><td></td><td></td><td></td><td></td><td></td><td></td><td></td><td></td><td></td></tr>
<tr><td>Oligo fwd</td><td>CACCTGGACCTGCTGTGCGTGCAG</td><td></td><td></td><td></td><td></td><td></td><td></td><td></td><td></td><td></td><td></td></tr>
<tr><td>Oligo rev</td><td>AAACCTGCACGCACAGCAGGTCCA</td><td></td><td></td><td></td><td></td><td></td><td></td><td></td><td></td><td></td><td></td></tr>
<tr><td>Chromosome</td><td>start</td><td>end</td><td>strand</td><td>MM</td><td>target_seq</td><td>PAM</td><td>alignment</td><td>distance</td><td>position</td><td>gene name</td><td>gene id</td></tr>
<tr><td>GU305914.1</td><td>121572</td><td>121594</td><td>-</td><td>0</td><td>TGGACCTGCTGTGCGTGCAG</td><td>AGG</td><td>||||||||[||||||||||||]PAM</td><td>NA</td><td>Intergenic</td><td>NA</td><td>NA</td></tr>
<tr><td>T24843</td><td>GACTAGACGAGCCCCGAGTGCGG</td><td>1000</td><td></td><td></td><td></td><td></td><td></td><td></td><td></td><td></td><td></td></tr>
<tr><td>Oligo fwd</td><td>CACCGACTAGACGAGCCCCGAGTG</td><td></td><td></td><td></td><td></td><td></td><td></td><td></td><td></td><td></td><td></td></tr>
<tr><td>Oligo rev</td><td>AAACCACTCGGGGCTCGTCTAGTC</td><td></td><td></td><td></td><td></td><td></td><td></td><td></td><td></td><td></td><td></td></tr>
<tr><td>Chromosome</td><td>start</td><td>end</td><td>strand</td><td>MM</td><td>target_seq</td><td>PAM</td><td>alignment</td><td>distance</td><td>position</td><td>gene name</td><td>gene id</td></tr>
<tr><td>GU305914.1</td><td>121548</td><td>121570</td><td>-</td><td>0</td><td>GACTAGACGAGCCCCGAGTG</td><td>CGG</td><td>||||||||[||||||||||||]PAM</td><td>NA</td><td>Intergenic</td><td>NA</td><td>NA</td></tr>
<tr><td>T24844</td><td>ACTAGACGAGCCCCGAGTGCGGG</td><td>1000</td><td></td><td></td><td></td><td></td><td></td><td></td><td></td><td></td><td></td></tr>
<tr><td>Oligo fwd</td><td>CACCACTAGACGAGCCCCGAGTGC</td><td></td><td></td><td></td><td></td><td></td><td></td><td></td><td></td><td></td><td></td></tr>
<tr><td>Oligo rev</td><td>AAACGCACTCGGGGCTCGTCTAGT</td><td></td><td></td><td></td><td></td><td></td><td></td><td></td><td></td><td></td><td></td></tr>
<tr><td>Chromosome</td><td>start</td><td>end</td><td>strand</td><td>MM</td><td>target_seq</td><td>PAM</td><td>alignment</td><td>distance</td><td>position</td><td>gene name</td><td>gene id</td></tr>
</table>

<table>
<tr><td>GU305914.1</td><td>121547</td><td>121569</td><td>-</td><td>0</td><td>ACTAGACGAGCCCCGAGTGC</td><td>GGG</td><td>||||||||[||||||||||||]PAM</td><td>NA</td><td>Intergenic</td><td>NA</td><td>NA</td></tr>
<tr><td>T24845</td><td>CTAGACGAGCCCCGAGTGCGGGG</td><td>1000</td><td></td><td></td><td></td><td></td><td></td><td></td><td></td><td></td><td></td></tr>
<tr><td>Oligo fwd</td><td>CACCCTAGACGAGCCCCGAGTGCG</td><td></td><td></td><td></td><td></td><td></td><td></td><td></td><td></td><td></td><td></td></tr>
<tr><td>Oligo rev</td><td>AAACCGCACTCGGGGCTCGTCTAG</td><td></td><td></td><td></td><td></td><td></td><td></td><td></td><td></td><td></td><td></td></tr>
<tr><td>Chromosome</td><td>start</td><td>end</td><td>strand</td><td>MM</td><td>target_seq</td><td>PAM</td><td>alignment</td><td>distance</td><td>position</td><td>gene name</td><td>gene id</td></tr>
<tr><td>GU305914.1</td><td>121546</td><td>121568</td><td>-</td><td>0</td><td>CTAGACGAGCCCCGAGTGCG</td><td>GGG</td><td>||||||||[||||||||||||]PAM</td><td>NA</td><td>Intergenic</td><td>NA</td><td>NA</td></tr>
<tr><td>T24846</td><td>CCGAGTGCGGGGCCGCTGCCCGG</td><td>1000</td><td></td><td></td><td></td><td></td><td></td><td></td><td></td><td></td><td></td></tr>
<tr><td>Oligo fwd</td><td>CACCCCGAGTGCGGGGCCGCTGCC</td><td></td><td></td><td></td><td></td><td></td><td></td><td></td><td></td><td></td><td></td></tr>
<tr><td>Oligo rev</td><td>AAACGGCAGCGGCCCCGCACTCGG</td><td></td><td></td><td></td><td></td><td></td><td></td><td></td><td></td><td></td><td></td></tr>
<tr><td>Chromosome</td><td>start</td><td>end</td><td>strand</td><td>MM</td><td>target_seq</td><td>PAM</td><td>alignment</td><td>distance</td><td>position</td><td>gene name</td><td>gene id</td></tr>
<tr><td>GU305914.1</td><td>121535</td><td>121557</td><td>-</td><td>0</td><td>CCGAGTGCGGGGCCGCTGCC</td><td>CGG</td><td>||||||||[||||||||||||]PAM</td><td>NA</td><td>Intergenic</td><td>NA</td><td>NA</td></tr>
<tr><td>T24847</td><td>CCGCTGCCCGGTGAGCATCCTGG</td><td>1000</td><td></td><td></td><td></td><td></td><td></td><td></td><td></td><td></td><td></td></tr>
<tr><td>Oligo fwd</td><td>CACCCCGCTGCCCGGTGAGCATCC</td><td></td><td></td><td></td><td></td><td></td><td></td><td></td><td></td><td></td><td></td></tr>
<tr><td>Oligo rev</td><td>AAACGGATGCTCACCGGGCAGCGG</td><td></td><td></td><td></td><td></td><td></td><td></td><td></td><td></td><td></td><td></td></tr>
<tr><td>Chromosome</td><td>start</td><td>end</td><td>strand</td><td>MM</td><td>target_seq</td><td>PAM</td><td>alignment</td><td>distance</td><td>position</td><td>gene name</td><td>gene id</td></tr>
<tr><td>GU305914.1</td><td>121523</td><td>121545</td><td>-</td><td>0</td><td>CCGCTGCCCGGTGAGCATCC</td><td>TGG</td><td>||||||||[||||||||||||]PAM</td><td>NA</td><td>Intergenic</td><td>NA</td><td>NA</td></tr>
<tr><td>T24848</td><td>CGGTGAGCATCCTGGCGATGCGG</td><td>1000</td><td></td><td></td><td></td><td></td><td></td><td></td><td></td><td></td><td></td></tr>
<tr><td>Oligo fwd</td><td>CACCCGGTGAGCATCCTGGCGATG</td><td></td><td></td><td></td><td></td><td></td><td></td><td></td><td></td><td></td><td></td></tr>
<tr><td>Oligo rev</td><td>AAACCATCGCCAGGATGCTCACCG</td><td></td><td></td><td></td><td></td><td></td><td></td><td></td><td></td><td></td><td></td></tr>
<tr><td>Chromosome</td><td>start</td><td>end</td><td>strand</td><td>MM</td><td>target_seq</td><td>PAM</td><td>alignment</td><td>distance</td><td>position</td><td>gene name</td><td>gene id</td></tr>
<tr><td>GU305914.1</td><td>121515</td><td>121537</td><td>-</td><td>0</td><td>CGGTGAGCATCCTGGCGATG</td><td>CGG</td><td>||||||||[||||||||||||]PAM</td><td>NA</td><td>Intergenic</td><td>NA</td><td>NA</td></tr>
<tr><td>T24849</td><td>GCGGACGAGCAAGAGCACGCTGG</td><td>1000</td><td></td><td></td><td></td><td></td><td></td><td></td><td></td><td></td><td></td></tr>
<tr><td>Oligo fwd</td><td>CACCGCGGACGAGCAAGAGCACGC</td><td></td><td></td><td></td><td></td><td></td><td></td><td></td><td></td><td></td><td></td></tr>
<tr><td>Oligo rev</td><td>AAACGCGTGCTCTTGCTCGTCCGC</td><td></td><td></td><td></td><td></td><td></td><td></td><td></td><td></td><td></td><td></td></tr>
<tr><td>Chromosome</td><td>start</td><td>end</td><td>strand</td><td>MM</td><td>target_seq</td><td>PAM</td><td>alignment</td><td>distance</td><td>position</td><td>gene name</td><td>gene id</td></tr>
<tr><td>GU305914.1</td><td>121496</td><td>121518</td><td>-</td><td>0</td><td>GCGGACGAGCAAGAGCACGC</td><td>TGG</td><td>||||||||[||||||||||||]PAM</td><td>NA</td><td>Intergenic</td><td>NA</td><td>NA</td></tr>
<tr><td>T24850</td><td>CAAGAGCACGCTGGACCTAGAGG</td><td>1000</td><td></td><td></td><td></td><td></td><td></td><td></td><td></td><td></td><td></td></tr>
<tr><td>Oligo fwd</td><td>CACCCAAGAGCACGCTGGACCTAG</td><td></td><td></td><td></td><td></td><td></td><td></td><td></td><td></td><td></td><td></td></tr>
<tr><td>Oligo rev</td><td>AAACCTAGGTCCAGCGTGCTCTTG</td><td></td><td></td><td></td><td></td><td></td><td></td><td></td><td></td><td></td><td></td></tr>
<tr><td>Chromosome</td><td>start</td><td>end</td><td>strand</td><td>MM</td><td>target_seq</td><td>PAM</td><td>alignment</td><td>distance</td><td>position</td><td>gene name</td><td>gene id</td></tr>
<tr><td>GU305914.1</td><td>121487</td><td>121509</td><td>-</td><td>0</td><td>CAAGAGCACGCTGGACCTAG</td><td>AGG</td><td>||||||||[||||||||||||]PAM</td><td>NA</td><td>Intergenic</td><td>NA</td><td>NA</td></tr>
<tr><td>T24851</td><td>GAGCACGCTGGACCTAGAGGTGG</td><td>1000</td><td></td><td></td><td></td><td></td><td></td><td></td><td></td><td></td><td></td></tr>
<tr><td>Oligo fwd</td><td>CACCGAGCACGCTGGACCTAGAGG</td><td></td><td></td><td></td><td></td><td></td><td></td><td></td><td></td><td></td><td></td></tr>
<tr><td>Oligo rev</td><td>AAACCCTCTAGGTCCAGCGTGCTC</td><td></td><td></td><td></td><td></td><td></td><td></td><td></td><td></td><td></td><td></td></tr>
<tr><td>Chromosome</td><td>start</td><td>end</td><td>strand</td><td>MM</td><td>target_seq</td><td>PAM</td><td>alignment</td><td>distance</td><td>position</td><td>gene name</td><td>gene id</td></tr>
<tr><td>GU305914.1</td><td>121484</td><td>121506</td><td>-</td><td>0</td><td>GAGCACGCTGGACCTAGAGG</td><td>TGG</td><td>||||||||[||||||||||||]PAM</td><td>NA</td><td>Intergenic</td><td>NA</td><td>NA</td></tr>
<tr><td>T24852</td><td>CACGCTGGACCTAGAGGTGGCGG</td><td>1000</td><td></td><td></td><td></td><td></td><td></td><td></td><td></td><td></td><td></td></tr>
<tr><td>Oligo fwd</td><td>CACCCACGCTGGACCTAGAGGTGG</td><td></td><td></td><td></td><td></td><td></td><td></td><td></td><td></td><td></td><td></td></tr>
<tr><td>Oligo rev</td><td>AAACCCACCTCTAGGTCCAGCGTG</td><td></td><td></td><td></td><td></td><td></td><td></td><td></td><td></td><td></td><td></td></tr>
<tr><td>Chromosome</td><td>start</td><td>end</td><td>strand</td><td>MM</td><td>target_seq</td><td>PAM</td><td>alignment</td><td>distance</td><td>position</td><td>gene name</td><td>gene id</td></tr>
<tr><td>GU305914.1</td><td>121481</td><td>121503</td><td>-</td><td>0</td><td>CACGCTGGACCTAGAGGTGG</td><td>CGG</td><td>||||||||[||||||||||||]PAM</td><td>NA</td><td>Intergenic</td><td>NA</td><td>NA</td></tr>
<tr><td>T24853</td><td>AGAGGTGGCGGCGTCGAACCTGG</td><td>1000</td><td></td><td></td><td></td><td></td><td></td><td></td><td></td><td></td><td></td></tr>
<tr><td>Oligo fwd</td><td>CACCAGAGGTGGCGGCGTCGAACC</td><td></td><td></td><td></td><td></td><td></td><td></td><td></td><td></td><td></td><td></td></tr>
<tr><td>Oligo rev</td><td>AAACGGTTCGACGCCGCCACCTCT</td><td></td><td></td><td></td><td></td><td></td><td></td><td></td><td></td><td></td><td></td></tr>
<tr><td>Chromosome</td><td>start</td><td>end</td><td>strand</td><td>MM</td><td>target_seq</td><td>PAM</td><td>alignment</td><td>distance</td><td>position</td><td>gene name</td><td>gene id</td></tr>
<tr><td>GU305914.1</td><td>121469</td><td>121491</td><td>-</td><td>0</td><td>AGAGGTGGCGGCGTCGAACC</td><td>TGG</td><td>||||||||[||||||||||||]PAM</td><td>NA</td><td>Intergenic</td><td>NA</td><td>NA</td></tr>
<tr><td>T24854</td><td>TGGCGGCGTCGAACCTGGCGTGG</td><td>1000</td><td></td><td></td><td></td><td></td><td></td><td></td><td></td><td></td><td></td></tr>
<tr><td>Oligo fwd</td><td>CACCTGGCGGCGTCGAACCTGGCG</td><td></td><td></td><td></td><td></td><td></td><td></td><td></td><td></td><td></td><td></td></tr>
<tr><td>Oligo rev</td><td>AAACCGCCAGGTTCGACGCCGCCA</td><td></td><td></td><td></td><td></td><td></td><td></td><td></td><td></td><td></td><td></td></tr>
<tr><td>Chromosome</td><td>start</td><td>end</td><td>strand</td><td>MM</td><td>target_seq</td><td>PAM</td><td>alignment</td><td>distance</td><td>position</td><td>gene name</td><td>gene id</td></tr>
<tr><td>GU305914.1</td><td>121464</td><td>121486</td><td>-</td><td>0</td><td>TGGCGGCGTCGAACCTGGCG</td><td>TGG</td><td>||||||||[||||||||||||]PAM</td><td>NA</td><td>Intergenic</td><td>NA</td><td>NA</td></tr>
<tr><td>T24855</td><td>CGGCGTCGAACCTGGCGTGGCGG</td><td>1000</td><td></td><td></td><td></td><td></td><td></td><td></td><td></td><td></td><td></td></tr>
<tr><td>Oligo fwd</td><td>CACCCGGCGTCGAACCTGGCGTGG</td><td></td><td></td><td></td><td></td><td></td><td></td><td></td><td></td><td></td><td></td></tr>
<tr><td>Oligo rev</td><td>AAACCCACGCCAGGTTCGACGCCG</td><td></td><td></td><td></td><td></td><td></td><td></td><td></td><td></td><td></td><td></td></tr>
<tr><td>Chromosome</td><td>start</td><td>end</td><td>strand</td><td>MM</td><td>target_seq</td><td>PAM</td><td>alignment</td><td>distance</td><td>position</td><td>gene name</td><td>gene id</td></tr>
<tr><td>GU305914.1</td><td>121461</td><td>121483</td><td>-</td><td>0</td><td>CGGCGTCGAACCTGGCGTGG</td><td>CGG</td><td>||||||||[||||||||||||]PAM</td><td>NA</td><td>Intergenic</td><td>NA</td><td>NA</td></tr>
<tr><td>T24856</td><td>GCGTGGCGGCCGTTCAGCCGCGG</td><td>1000</td><td></td><td></td><td></td><td></td><td></td><td></td><td></td><td></td><td></td></tr>
<tr><td>Oligo fwd</td><td>CACCGCGTGGCGGCCGTTCAGCCG</td><td></td><td></td><td></td><td></td><td></td><td></td><td></td><td></td><td></td><td></td></tr>
<tr><td>Oligo rev</td><td>AAACCGGCTGAACGGCCGCCACGC</td><td></td><td></td><td></td><td></td><td></td><td></td><td></td><td></td><td></td><td></td></tr>
<tr><td>Chromosome</td><td>start</td><td>end</td><td>strand</td><td>MM</td><td>target_seq</td><td>PAM</td><td>alignment</td><td>distance</td><td>position</td><td>gene name</td><td>gene id</td></tr>
<tr><td>GU305914.1</td><td>121447</td><td>121469</td><td>-</td><td>0</td><td>GCGTGGCGGCCGTTCAGCCG</td><td>CGG</td><td>||||||||[||||||||||||]PAM</td><td>NA</td><td>Intergenic</td><td>NA</td><td>NA</td></tr>
<tr><td>T24857</td><td>GGCCGTTCAGCCGCGGCAACAGG</td><td>1000</td><td></td><td></td><td></td><td></td><td></td><td></td><td></td><td></td><td></td></tr>
<tr><td>Oligo fwd</td><td>CACCGGCCGTTCAGCCGCGGCAAC</td><td></td><td></td><td></td><td></td><td></td><td></td><td></td><td></td><td></td><td></td></tr>
<tr><td>Oligo rev</td><td>AAACGTTGCCGCGGCTGAACGGCC</td><td></td><td></td><td></td><td></td><td></td><td></td><td></td><td></td><td></td><td></td></tr>
<tr><td>Chromosome</td><td>start</td><td>end</td><td>strand</td><td>MM</td><td>target_seq</td><td>PAM</td><td>alignment</td><td>distance</td><td>position</td><td>gene name</td><td>gene id</td></tr>
<tr><td>GU305914.1</td><td>121440</td><td>121462</td><td>-</td><td>0</td><td>GGCCGTTCAGCCGCGGCAAC</td><td>AGG</td><td>||||||||[||||||||||||]PAM</td><td>NA</td><td>Intergenic</td><td>NA</td><td>NA</td></tr>
<tr><td>T24858</td><td>CAGCCGCGGCAACAGGCTGACGG</td><td>1000</td><td></td><td></td><td></td><td></td><td></td><td></td><td></td><td></td><td></td></tr>
<tr><td>Oligo fwd</td><td>CACCCAGCCGCGGCAACAGGCTGA</td><td></td><td></td><td></td><td></td><td></td><td></td><td></td><td></td><td></td><td></td></tr>
<tr><td>Oligo rev</td><td>AAACTCAGCCTGTTGCCGCGGCTG</td><td></td><td></td><td></td><td></td><td></td><td></td><td></td><td></td><td></td><td></td></tr>
<tr><td>Chromosome</td><td>start</td><td>end</td><td>strand</td><td>MM</td><td>target_seq</td><td>PAM</td><td>alignment</td><td>distance</td><td>position</td><td>gene name</td><td>gene id</td></tr>
<tr><td>GU305914.1</td><td>121433</td><td>121455</td><td>-</td><td>0</td><td>CAGCCGCGGCAACAGGCTGA</td><td>CGG</td><td>||||||||[||||||||||||]PAM</td><td>NA</td><td>Intergenic</td><td>NA</td><td>NA</td></tr>
<tr><td>T24859</td><td>CGGCAACAGGCTGACGGCCTCGG</td><td>1000</td><td></td><td></td><td></td><td></td><td></td><td></td><td></td><td></td><td></td></tr>
<tr><td>Oligo fwd</td><td>CACCCGGCAACAGGCTGACGGCCT</td><td></td><td></td><td></td><td></td><td></td><td></td><td></td><td></td><td></td><td></td></tr>
<tr><td>Oligo rev</td><td>AAACAGGCCGTCAGCCTGTTGCCG</td><td></td><td></td><td></td><td></td><td></td><td></td><td></td><td></td><td></td><td></td></tr>
</table>

<table>
<tr><td>Chromosome</td><td>start</td><td>end</td><td>strand</td><td>MM</td><td>target_seq</td><td>PAM</td><td>alignment</td><td>distance</td><td>position</td><td>gene name</td><td>gene id</td></tr>
<tr><td>GU305914.1</td><td>121427</td><td>121449</td><td>-</td><td>0</td><td>CGGCAACAGGCTGACGGCCT</td><td>CGG</td><td>||||||||[||||||||||||]PAM</td><td>NA</td><td>Intergenic</td><td>NA</td><td>NA</td></tr>
<tr><td>T24860</td><td>GCTGACGGCCTCGGTGAGCCTGG</td><td>1000</td></tr>
<tr><td>Oligo fwd</td><td>CACCGCTGACGGCCTCGGTGAGCC</td></tr>
<tr><td>Oligo rev</td><td>AAACGGCTCACCGAGGCCGTCAGC</td></tr>
<tr><td>Chromosome</td><td>start</td><td>end</td><td>strand</td><td>MM</td><td>target_seq</td><td>PAM</td><td>alignment</td><td>distance</td><td>position</td><td>gene name</td><td>gene id</td></tr>
<tr><td>GU305914.1</td><td>121418</td><td>121440</td><td>-</td><td>0</td><td>GCTGACGGCCTCGGTGAGCC</td><td>TGG</td><td>||||||||[||||||||||||]PAM</td><td>NA</td><td>Intergenic</td><td>NA</td><td>NA</td></tr>
<tr><td>T24861</td><td>ACGTGTCGTCGACGAACATGCGG</td><td>1000</td></tr>
<tr><td>Oligo fwd</td><td>CACCACGTGTCGTCGACGAACATG</td></tr>
<tr><td>Oligo rev</td><td>AAACCATGTTCGTCGACGACACGT</td></tr>
<tr><td>Chromosome</td><td>start</td><td>end</td><td>strand</td><td>MM</td><td>target_seq</td><td>PAM</td><td>alignment</td><td>distance</td><td>position</td><td>gene name</td><td>gene id</td></tr>
<tr><td>GU305914.1</td><td>121395</td><td>121417</td><td>-</td><td>0</td><td>ACGTGTCGTCGACGAACATG</td><td>CGG</td><td>||||||||[||||||||||||]PAM</td><td>NA</td><td>Intergenic</td><td>NA</td><td>NA</td></tr>
<tr><td>T24862</td><td>CGTCCTCTCCTGCTCGCACTCGG</td><td>1000</td></tr>
<tr><td>Oligo fwd</td><td>CACCCGTCCTCTCCTGCTCGCACT</td></tr>
<tr><td>Oligo rev</td><td>AAACAGTGCGAGCAGGAGAGGACG</td></tr>
<tr><td>Chromosome</td><td>start</td><td>end</td><td>strand</td><td>MM</td><td>target_seq</td><td>PAM</td><td>alignment</td><td>distance</td><td>position</td><td>gene name</td><td>gene id</td></tr>
<tr><td>GU305914.1</td><td>121355</td><td>121377</td><td>-</td><td>0</td><td>CGTCCTCTCCTGCTCGCACT</td><td>CGG</td><td>||||||||[||||||||||||]PAM</td><td>NA</td><td>Intergenic</td><td>NA</td><td>NA</td></tr>
<tr><td>T24863</td><td>CTCGCACTCGGACATCTACGTGG</td><td>1000</td></tr>
<tr><td>Oligo fwd</td><td>CACCCTCGCACTCGGACATCTACG</td></tr>
<tr><td>Oligo rev</td><td>AAACCGTAGATGTCCGAGTGCGAG</td></tr>
<tr><td>Chromosome</td><td>start</td><td>end</td><td>strand</td><td>MM</td><td>target_seq</td><td>PAM</td><td>alignment</td><td>distance</td><td>position</td><td>gene name</td><td>gene id</td></tr>
<tr><td>GU305914.1</td><td>121343</td><td>121365</td><td>-</td><td>0</td><td>CTCGCACTCGGACATCTACG</td><td>TGG</td><td>||||||||[||||||||||||]PAM</td><td>NA</td><td>Intergenic</td><td>NA</td><td>NA</td></tr>
<tr><td>T24864</td><td>CTCGGACATCTACGTGGACCAGG</td><td>1000</td></tr>
<tr><td>Oligo fwd</td><td>CACCCTCGGACATCTACGTGGACC</td></tr>
<tr><td>Oligo rev</td><td>AAACGGTCCACGTAGATGTCCGAG</td></tr>
<tr><td>Chromosome</td><td>start</td><td>end</td><td>strand</td><td>MM</td><td>target_seq</td><td>PAM</td><td>alignment</td><td>distance</td><td>position</td><td>gene name</td><td>gene id</td></tr>
<tr><td>GU305914.1</td><td>121337</td><td>121359</td><td>-</td><td>0</td><td>CTCGGACATCTACGTGGACC</td><td>AGG</td><td>||||||||[||||||||||||]PAM</td><td>NA</td><td>Intergenic</td><td>NA</td><td>NA</td></tr>
<tr><td>GU305914.1</td><td>46988</td><td>47010</td><td>+</td><td>4</td><td>CCTGGACATCGAGGTGGACC</td><td>CGG</td><td>|--|||||[||-|-|||||||]PAM</td><td>NA</td><td>Intergenic</td><td>NA</td><td>NA</td></tr>
<tr><td>T24865</td><td>CATCTACGTGGACCAGGTGACGG</td><td>1000</td></tr>
<tr><td>Oligo fwd</td><td>CACCCATCTACGTGGACCAGGTGA</td></tr>
<tr><td>Oligo rev</td><td>AAACTCACCTGGTCCACGTAGATG</td></tr>
<tr><td>Chromosome</td><td>start</td><td>end</td><td>strand</td><td>MM</td><td>target_seq</td><td>PAM</td><td>alignment</td><td>distance</td><td>position</td><td>gene name</td><td>gene id</td></tr>
<tr><td>GU305914.1</td><td>121331</td><td>121353</td><td>-</td><td>0</td><td>CATCTACGTGGACCAGGTGA</td><td>CGG</td><td>||||||||[||||||||||||]PAM</td><td>NA</td><td>Intergenic</td><td>NA</td><td>NA</td></tr>
<tr><td>T24866</td><td>CTACGTGGACCAGGTGACGGTGG</td><td>1000</td></tr>
<tr><td>Oligo fwd</td><td>CACCCTACGTGGACCAGGTGACGG</td></tr>
<tr><td>Oligo rev</td><td>AAACCCGTCACCTGGTCCACGTAG</td></tr>
<tr><td>Chromosome</td><td>start</td><td>end</td><td>strand</td><td>MM</td><td>target_seq</td><td>PAM</td><td>alignment</td><td>distance</td><td>position</td><td>gene name</td><td>gene id</td></tr>
<tr><td>GU305914.1</td><td>121328</td><td>121350</td><td>-</td><td>0</td><td>CTACGTGGACCAGGTGACGG</td><td>TGG</td><td>||||||||[||||||||||||]PAM</td><td>NA</td><td>Intergenic</td><td>NA</td><td>NA</td></tr>
<tr><td>T24867</td><td>GGTGACGGTGGTCCAGAGCTCGG</td><td>1000</td></tr>
<tr><td>Oligo fwd</td><td>CACCGGTGACGGTGGTCCAGAGCT</td></tr>
<tr><td>Oligo rev</td><td>AAACAGCTCTGGACCACCGTCACC</td></tr>
<tr><td>Chromosome</td><td>start</td><td>end</td><td>strand</td><td>MM</td><td>target_seq</td><td>PAM</td><td>alignment</td><td>distance</td><td>position</td><td>gene name</td><td>gene id</td></tr>
<tr><td>GU305914.1</td><td>121316</td><td>121338</td><td>-</td><td>0</td><td>GGTGACGGTGGTCCAGAGCT</td><td>CGG</td><td>||||||||[||||||||||||]PAM</td><td>NA</td><td>Intergenic</td><td>NA</td><td>NA</td></tr>
<tr><td>GU305914.1</td><td>64738</td><td>64760</td><td>-</td><td>4</td><td>GATGTCGGTGGTGCAGATCT</td><td>CGG</td><td>|-||-|||[||||-||||-||]PAM</td><td>NA</td><td>Intergenic</td><td>NA</td><td>NA</td></tr>
<tr><td>T24868</td><td>GTGACGGTGGTCCAGAGCTCGGG</td><td>1000</td></tr>
<tr><td>Oligo fwd</td><td>CACCGTGACGGTGGTCCAGAGCTC</td></tr>
<tr><td>Oligo rev</td><td>AAACGAGCTCTGGACCACCGTCAC</td></tr>
<tr><td>Chromosome</td><td>start</td><td>end</td><td>strand</td><td>MM</td><td>target_seq</td><td>PAM</td><td>alignment</td><td>distance</td><td>position</td><td>gene name</td><td>gene id</td></tr>
<tr><td>GU305914.1</td><td>121315</td><td>121337</td><td>-</td><td>0</td><td>GTGACGGTGGTCCAGAGCTC</td><td>GGG</td><td>||||||||[||||||||||||]PAM</td><td>NA</td><td>Intergenic</td><td>NA</td><td>NA</td></tr>
<tr><td>T24869</td><td>TGACGGTGGTCCAGAGCTCGGGG</td><td>1000</td></tr>
<tr><td>Oligo fwd</td><td>CACCTGACGGTGGTCCAGAGCTCG</td></tr>
<tr><td>Oligo rev</td><td>AAACCGAGCTCTGGACCACCGTCA</td></tr>
<tr><td>Chromosome</td><td>start</td><td>end</td><td>strand</td><td>MM</td><td>target_seq</td><td>PAM</td><td>alignment</td><td>distance</td><td>position</td><td>gene name</td><td>gene id</td></tr>
<tr><td>GU305914.1</td><td>121314</td><td>121336</td><td>-</td><td>0</td><td>TGACGGTGGTCCAGAGCTCG</td><td>GGG</td><td>||||||||[||||||||||||]PAM</td><td>NA</td><td>Intergenic</td><td>NA</td><td>NA</td></tr>
<tr><td>T24870</td><td>GACGGTGGTCCAGAGCTCGGGGG</td><td>1000</td></tr>
<tr><td>Oligo fwd</td><td>CACCGACGGTGGTCCAGAGCTCGG</td></tr>
<tr><td>Oligo rev</td><td>AAACCCGAGCTCTGGACCACCGTC</td></tr>
<tr><td>Chromosome</td><td>start</td><td>end</td><td>strand</td><td>MM</td><td>target_seq</td><td>PAM</td><td>alignment</td><td>distance</td><td>position</td><td>gene name</td><td>gene id</td></tr>
<tr><td>GU305914.1</td><td>121313</td><td>121335</td><td>-</td><td>0</td><td>GACGGTGGTCCAGAGCTCGG</td><td>GGG</td><td>||||||||[||||||||||||]PAM</td><td>NA</td><td>Intergenic</td><td>NA</td><td>NA</td></tr>
<tr><td>T24871</td><td>GGTGGTCCAGAGCTCGGGGGTGG</td><td>1000</td></tr>
<tr><td>Oligo fwd</td><td>CACCGGTGGTCCAGAGCTCGGGGG</td></tr>
<tr><td>Oligo rev</td><td>AAACCCCCCGAGCTCTGGACCACC</td></tr>
<tr><td>Chromosome</td><td>start</td><td>end</td><td>strand</td><td>MM</td><td>target_seq</td><td>PAM</td><td>alignment</td><td>distance</td><td>position</td><td>gene name</td><td>gene id</td></tr>
<tr><td>GU305914.1</td><td>121310</td><td>121332</td><td>-</td><td>0</td><td>GGTGGTCCAGAGCTCGGGGG</td><td>TGG</td><td>||||||||[||||||||||||]PAM</td><td>NA</td><td>Intergenic</td><td>NA</td><td>NA</td></tr>
<tr><td>T24872</td><td>GACCAAGTACGCGACGATCGCGG</td><td>1000</td></tr>
<tr><td>Oligo fwd</td><td>CACCGACCAAGTACGCGACGATCG</td></tr>
<tr><td>Oligo rev</td><td>AAACCGATCGTCGCGTACTTGGTC</td></tr>
<tr><td>Chromosome</td><td>start</td><td>end</td><td>strand</td><td>MM</td><td>target_seq</td><td>PAM</td><td>alignment</td><td>distance</td><td>position</td><td>gene name</td><td>gene id</td></tr>
<tr><td>GU305914.1</td><td>121271</td><td>121293</td><td>-</td><td>0</td><td>GACCAAGTACGCGACGATCG</td><td>CGG</td><td>||||||||[||||||||||||]PAM</td><td>NA</td><td>Intergenic</td><td>NA</td><td>NA</td></tr>
<tr><td>GU305914.1</td><td>218352</td><td>218374</td><td>+</td><td>4</td><td>CACTAAGACCGCGACGATCG</td><td>CGG</td><td>-||-|||-[-|||||||||||]PAM</td><td>NA</td><td>Intergenic</td><td>NA</td><td>NA</td></tr>
<tr><td>T24873</td><td>CTTCAAGATCCTGTTCCGCGAGG</td><td>1000</td></tr>
<tr><td>Oligo fwd</td><td>CACCCTTCAAGATCCTGTTCCGCG</td></tr>
<tr><td>Oligo rev</td><td>AAACCGCGGAACAGGATCTTGAAG</td></tr>
<tr><td>Chromosome</td><td>start</td><td>end</td><td>strand</td><td>MM</td><td>target_seq</td><td>PAM</td><td>alignment</td><td>distance</td><td>position</td><td>gene name</td><td>gene id</td></tr>
<tr><td>GU305914.1</td><td>121232</td><td>121254</td><td>-</td><td>0</td><td>CTTCAAGATCCTGTTCCGCG</td><td>AGG</td><td>||||||||[||||||||||||]PAM</td><td>NA</td><td>Intergenic</td><td>NA</td><td>NA</td></tr>
</table>

| | | | | | | | | | | | |
|---|---|---|---|---|---|---|---|---|---|---|---|
| T24874 | CAAGATCCTGTTCCGCGAGGAGG | 1000 | | | | | | | | | |
| Oligo fwd | CACCCAAGATCCTGTTCCGCGAGG | | | | | | | | | | |
| Oligo rev | AAACCCTCGCGGAACAGGATCTTG | | | | | | | | | | |
| Chromosome | start | end | strand | MM | target_seq | PAM | alignment | distance | position | gene name | gene id |
| GU305914.1 | 121229 | 121251 | - | 0 | CAAGATCCTGTTCCGCGAGG | AGG | \|\|\|\|\|\|\|\|[\|\|\|\|\|\|\|\|\|\|\|\|]PAM | NA | Intergenic | NA | NA |
| T24875 | CGAGGAGGACGACGCGCAGCTGG | 1000 | | | | | | | | | |
| Oligo fwd | CACCCGAGGAGGACGACGCGCAGC | | | | | | | | | | |
| Oligo rev | AAACGCTGCGCGTCGTCCTCCTCG | | | | | | | | | | |
| Chromosome | start | end | strand | MM | target_seq | PAM | alignment | distance | position | gene name | gene id |
| GU305914.1 | 121214 | 121236 | - | 0 | CGAGGAGGACGACGCGCAGC | TGG | \|\|\|\|\|\|\|\|[\|\|\|\|\|\|\|\|\|\|\|\|]PAM | NA | Intergenic | NA | NA |
| GU305914.1 | 66392 | 66414 | + | 4 | CGCGGCGGACAACGCGGAGC | TGG | \|\|-\|\|-\|\|[\|\|-\|\|\|\|\|-\|\|\|]PAM | NA | Intergenic | NA | NA |
| T24876 | ACGCGCAGCTGGAAGACAAGCGG | 1000 | | | | | | | | | |
| Oligo fwd | CACCACGCGCAGCTGGAAGACAAG | | | | | | | | | | |
| Oligo rev | AAACCTTGTCTTCCAGCTGCGCGT | | | | | | | | | | |
| Chromosome | start | end | strand | MM | target_seq | PAM | alignment | distance | position | gene name | gene id |
| GU305914.1 | 121203 | 121225 | - | 0 | ACGCGCAGCTGGAAGACAAG | CGG | \|\|\|\|\|\|\|\|[\|\|\|\|\|\|\|\|\|\|\|\|]PAM | NA | Intergenic | NA | NA |
| T24877 | GATCACGTTCCCCCGCTATCCGG | 1000 | | | | | | | | | |
| Oligo fwd | CACCGATCACGTTCCCCCGCTATC | | | | | | | | | | |
| Oligo rev | AAACGATAGCGGGGGAACGTGATC | | | | | | | | | | |
| Chromosome | start | end | strand | MM | target_seq | PAM | alignment | distance | position | gene name | gene id |
| GU305914.1 | 121181 | 121203 | - | 0 | GATCACGTTCCCCCGCTATC | CGG | \|\|\|\|\|\|\|\|[\|\|\|\|\|\|\|\|\|\|\|\|]PAM | NA | Intergenic | NA | NA |
| T24878 | GTGTGCACGCCGCACAAGTACGG | 1000 | | | | | | | | | |
| Oligo fwd | CACCGTGTGCACGCCGCACAAGTA | | | | | | | | | | |
| Oligo rev | AAACTACTTGTGCGGCGTGCACAC | | | | | | | | | | |
| Chromosome | start | end | strand | MM | target_seq | PAM | alignment | distance | position | gene name | gene id |
| GU305914.1 | 121150 | 121172 | - | 0 | GTGTGCACGCCGCACAAGTA | CGG | \|\|\|\|\|\|\|\|[\|\|\|\|\|\|\|\|\|\|\|\|]PAM | NA | Intergenic | NA | NA |
| T24879 | CGCCGCACAAGTACGGACTGCGG | 1000 | | | | | | | | | |
| Oligo fwd | CACCCGCCGCACAAGTACGGACTG | | | | | | | | | | |
| Oligo rev | AAACCAGTCCGTACTTGTGCGGCG | | | | | | | | | | |
| Chromosome | start | end | strand | MM | target_seq | PAM | alignment | distance | position | gene name | gene id |
| GU305914.1 | 121143 | 121165 | - | 0 | CGCCGCACAAGTACGGACTG | CGG | \|\|\|\|\|\|\|\|[\|\|\|\|\|\|\|\|\|\|\|\|]PAM | NA | Intergenic | NA | NA |
| T24880 | GCCGCACAAGTACGGACTGCGGG | 1000 | | | | | | | | | |
| Oligo fwd | CACCGCCGCACAAGTACGGACTGC | | | | | | | | | | |
| Oligo rev | AAACGCAGTCCGTACTTGTGCGGC | | | | | | | | | | |
| Chromosome | start | end | strand | MM | target_seq | PAM | alignment | distance | position | gene name | gene id |
| GU305914.1 | 121142 | 121164 | - | 0 | GCCGCACAAGTACGGACTGC | GGG | \|\|\|\|\|\|\|\|[\|\|\|\|\|\|\|\|\|\|\|\|]PAM | NA | Intergenic | NA | NA |
| T24881 | GCGAGACGCCGCCCTGACCCCGG | 1000 | | | | | | | | | |
| Oligo fwd | CACCGCGAGACGCCGCCCTGACCC | | | | | | | | | | |
| Oligo rev | AAACGGGTCAGGGCGGCGTCTCGC | | | | | | | | | | |
| Chromosome | start | end | strand | MM | target_seq | PAM | alignment | distance | position | gene name | gene id |
| GU305914.1 | 121109 | 121131 | - | 0 | GCGAGACGCCGCCCTGACCC | CGG | \|\|\|\|\|\|\|\|[\|\|\|\|\|\|\|\|\|\|\|\|]PAM | NA | Intergenic | NA | NA |
| T24882 | CGAGACGCCGCCCTGACCCCGGG | 1000 | | | | | | | | | |
| Oligo fwd | CACCCGAGACGCCGCCCTGACCCC | | | | | | | | | | |
| Oligo rev | AAACGGGGTCAGGGCGGCGTCTCG | | | | | | | | | | |
| Chromosome | start | end | strand | MM | target_seq | PAM | alignment | distance | position | gene name | gene id |
| GU305914.1 | 121108 | 121130 | - | 0 | CGAGACGCCGCCCTGACCCC | GGG | \|\|\|\|\|\|\|\|[\|\|\|\|\|\|\|\|\|\|\|\|]PAM | NA | Intergenic | NA | NA |
| T24883 | CGCCCTGACCCCGGGCAAGCCGG | 1000 | | | | | | | | | |
| Oligo fwd | CACCCGCCCTGACCCCGGGCAAGC | | | | | | | | | | |
| Oligo rev | AAACGCTTGCCCGGGGTCAGGGCG | | | | | | | | | | |
| Chromosome | start | end | strand | MM | target_seq | PAM | alignment | distance | position | gene name | gene id |
| GU305914.1 | 121100 | 121122 | - | 0 | CGCCCTGACCCCGGGCAAGC | CGG | \|\|\|\|\|\|\|\|[\|\|\|\|\|\|\|\|\|\|\|\|]PAM | NA | Intergenic | NA | NA |
| T24884 | TGTCGATCAGCTACTTCTCGAGG | 1000 | | | | | | | | | |
| Oligo fwd | CACCTGTCGATCAGCTACTTCTCG | | | | | | | | | | |
| Oligo rev | AAACCGAGAAGTAGCTGATCGACA | | | | | | | | | | |
| Chromosome | start | end | strand | MM | target_seq | PAM | alignment | distance | position | gene name | gene id |
| GU305914.1 | 121065 | 121087 | - | 0 | TGTCGATCAGCTACTTCTCG | AGG | \|\|\|\|\|\|\|\|[\|\|\|\|\|\|\|\|\|\|\|\|]PAM | NA | Intergenic | NA | NA |
| T24885 | GTCGATCAGCTACTTCTCGAGGG | 1000 | | | | | | | | | |
| Oligo fwd | CACCGTCGATCAGCTACTTCTCGA | | | | | | | | | | |
| Oligo rev | AAACTCGAGAAGTAGCTGATCGAC | | | | | | | | | | |
| Chromosome | start | end | strand | MM | target_seq | PAM | alignment | distance | position | gene name | gene id |
| GU305914.1 | 121064 | 121086 | - | 0 | GTCGATCAGCTACTTCTCGA | GGG | \|\|\|\|\|\|\|\|[\|\|\|\|\|\|\|\|\|\|\|\|]PAM | NA | Intergenic | NA | NA |
| T24886 | CTTCTCGAGGGCCGCGTACGCGG | 1000 | | | | | | | | | |
| Oligo fwd | CACCCTTCTCGAGGGCCGCGTACG | | | | | | | | | | |
| Oligo rev | AAACCGTACGCGGCCCTCGAGAAG | | | | | | | | | | |
| Chromosome | start | end | strand | MM | target_seq | PAM | alignment | distance | position | gene name | gene id |
| GU305914.1 | 121052 | 121074 | - | 0 | CTTCTCGAGGGCCGCGTACG | CGG | \|\|\|\|\|\|\|\|[\|\|\|\|\|\|\|\|\|\|\|\|]PAM | NA | Intergenic | NA | NA |
| T24887 | TTCTCGAGGGCCGCGTACGCGGG | 1000 | | | | | | | | | |
| Oligo fwd | CACCTTCTCGAGGGCCGCGTACGC | | | | | | | | | | |
| Oligo rev | AAACGCGTACGCGGCCCTCGAGAA | | | | | | | | | | |
| Chromosome | start | end | strand | MM | target_seq | PAM | alignment | distance | position | gene name | gene id |
| GU305914.1 | 121051 | 121073 | - | 0 | TTCTCGAGGGCCGCGTACGC | GGG | \|\|\|\|\|\|\|\|[\|\|\|\|\|\|\|\|\|\|\|\|]PAM | NA | Intergenic | NA | NA |
| T24888 | GTTCACGCCGCTGTACCGTCCGG | 1000 | | | | | | | | | |
| Oligo fwd | CACCGTTCACGCCGCTGTACCGTC | | | | | | | | | | |
| Oligo rev | AAACGACGGTACAGCGGCGTGAAC | | | | | | | | | | |

<table>
<tr><td>Chromosome</td><td>start</td><td>end</td><td>strand</td><td>MM</td><td>target_seq</td><td>PAM</td><td>alignment</td><td>distance</td><td>position</td><td>gene name</td><td>gene id</td></tr>
<tr><td>GU305914.1</td><td>121025</td><td>121047</td><td>-</td><td>0</td><td>GTTCACGCCGCTGTACCGTC</td><td>CGG</td><td>||||||||[||||||||||||]PAM</td><td>NA</td><td>Intergenic</td><td>NA</td><td>NA</td></tr>
<tr><td></td><td></td><td></td><td></td><td></td><td></td><td></td><td></td><td></td><td></td><td></td><td></td></tr>
<tr><td>T24889</td><td>TTCACGCCGCTGTACCGTCCGGG</td><td>1000</td><td></td><td></td><td></td><td></td><td></td><td></td><td></td><td></td><td></td></tr>
<tr><td>Oligo fwd</td><td>CACCTTCACGCCGCTGTACCGTCC</td><td></td><td></td><td></td><td></td><td></td><td></td><td></td><td></td><td></td><td></td></tr>
<tr><td>Oligo rev</td><td>AAACGGACGGTACAGCGGCGTGAA</td><td></td><td></td><td></td><td></td><td></td><td></td><td></td><td></td><td></td><td></td></tr>
<tr><td>Chromosome</td><td>start</td><td>end</td><td>strand</td><td>MM</td><td>target_seq</td><td>PAM</td><td>alignment</td><td>distance</td><td>position</td><td>gene name</td><td>gene id</td></tr>
<tr><td>GU305914.1</td><td>121024</td><td>121046</td><td>-</td><td>0</td><td>TTCACGCCGCTGTACCGTCC</td><td>GGG</td><td>||||||||[||||||||||||]PAM</td><td>NA</td><td>Intergenic</td><td>NA</td><td>NA</td></tr>
<tr><td></td><td></td><td></td><td></td><td></td><td></td><td></td><td></td><td></td><td></td><td></td><td></td></tr>
<tr><td>T24890</td><td>TCACGCCGCTGTACCGTCCGGGG</td><td>1000</td><td></td><td></td><td></td><td></td><td></td><td></td><td></td><td></td><td></td></tr>
<tr><td>Oligo fwd</td><td>CACCTCACGCCGCTGTACCGTCCG</td><td></td><td></td><td></td><td></td><td></td><td></td><td></td><td></td><td></td><td></td></tr>
<tr><td>Oligo rev</td><td>AAACCGGACGGTACAGCGGCGTGA</td><td></td><td></td><td></td><td></td><td></td><td></td><td></td><td></td><td></td><td></td></tr>
<tr><td>Chromosome</td><td>start</td><td>end</td><td>strand</td><td>MM</td><td>target_seq</td><td>PAM</td><td>alignment</td><td>distance</td><td>position</td><td>gene name</td><td>gene id</td></tr>
<tr><td>GU305914.1</td><td>121023</td><td>121045</td><td>-</td><td>0</td><td>TCACGCCGCTGTACCGTCCG</td><td>GGG</td><td>||||||||[||||||||||||]PAM</td><td>NA</td><td>Intergenic</td><td>NA</td><td>NA</td></tr>
<tr><td></td><td></td><td></td><td></td><td></td><td></td><td></td><td></td><td></td><td></td><td></td><td></td></tr>
<tr><td>T24891</td><td>TGAGCTGCCAGATCGTACCGTGG</td><td>1000</td><td></td><td></td><td></td><td></td><td></td><td></td><td></td><td></td><td></td></tr>
<tr><td>Oligo fwd</td><td>CACCTGAGCTGCCAGATCGTACCG</td><td></td><td></td><td></td><td></td><td></td><td></td><td></td><td></td><td></td><td></td></tr>
<tr><td>Oligo rev</td><td>AAACCGGTACGATCTGGCAGCTCA</td><td></td><td></td><td></td><td></td><td></td><td></td><td></td><td></td><td></td><td></td></tr>
<tr><td>Chromosome</td><td>start</td><td>end</td><td>strand</td><td>MM</td><td>target_seq</td><td>PAM</td><td>alignment</td><td>distance</td><td>position</td><td>gene name</td><td>gene id</td></tr>
<tr><td>GU305914.1</td><td>120999</td><td>121021</td><td>-</td><td>0</td><td>TGAGCTGCCAGATCGTACCG</td><td>TGG</td><td>||||||||[||||||||||||]PAM</td><td>NA</td><td>Intergenic</td><td>NA</td><td>NA</td></tr>
<tr><td></td><td></td><td></td><td></td><td></td><td></td><td></td><td></td><td></td><td></td><td></td><td></td></tr>
<tr><td>T24892</td><td>CTGCCAGATCGTACCGTGGAAGG</td><td>1000</td><td></td><td></td><td></td><td></td><td></td><td></td><td></td><td></td><td></td></tr>
<tr><td>Oligo fwd</td><td>CACCCTGCCAGATCGTACCGTGGA</td><td></td><td></td><td></td><td></td><td></td><td></td><td></td><td></td><td></td><td></td></tr>
<tr><td>Oligo rev</td><td>AAACTCCACGGTACGATCTGGCAG</td><td></td><td></td><td></td><td></td><td></td><td></td><td></td><td></td><td></td><td></td></tr>
<tr><td>Chromosome</td><td>start</td><td>end</td><td>strand</td><td>MM</td><td>target_seq</td><td>PAM</td><td>alignment</td><td>distance</td><td>position</td><td>gene name</td><td>gene id</td></tr>
<tr><td>GU305914.1</td><td>120995</td><td>121017</td><td>-</td><td>0</td><td>CTGCCAGATCGTACCGTGGA</td><td>AGG</td><td>||||||||[||||||||||||]PAM</td><td>NA</td><td>Intergenic</td><td>NA</td><td>NA</td></tr>
<tr><td></td><td></td><td></td><td></td><td></td><td></td><td></td><td></td><td></td><td></td><td></td><td></td></tr>
<tr><td>T24893</td><td>CGTACCGTGGAAGGAGCGTCAGG</td><td>1000</td><td></td><td></td><td></td><td></td><td></td><td></td><td></td><td></td><td></td></tr>
<tr><td>Oligo fwd</td><td>CACCCGTACCGTGGAAGGAGCGTC</td><td></td><td></td><td></td><td></td><td></td><td></td><td></td><td></td><td></td><td></td></tr>
<tr><td>Oligo rev</td><td>AAACGACGCTCCTTCCACGGTACG</td><td></td><td></td><td></td><td></td><td></td><td></td><td></td><td></td><td></td><td></td></tr>
<tr><td>Chromosome</td><td>start</td><td>end</td><td>strand</td><td>MM</td><td>target_seq</td><td>PAM</td><td>alignment</td><td>distance</td><td>position</td><td>gene name</td><td>gene id</td></tr>
<tr><td>GU305914.1</td><td>120986</td><td>121008</td><td>-</td><td>0</td><td>CGTACCGTGGAAGGAGCGTC</td><td>AGG</td><td>||||||||[||||||||||||]PAM</td><td>NA</td><td>Intergenic</td><td>NA</td><td>NA</td></tr>
<tr><td></td><td></td><td></td><td></td><td></td><td></td><td></td><td></td><td></td><td></td><td></td><td></td></tr>
<tr><td>T24894</td><td>GCGTCAGGTGCTGAGCATGACGG</td><td>1000</td><td></td><td></td><td></td><td></td><td></td><td></td><td></td><td></td><td></td></tr>
<tr><td>Oligo fwd</td><td>CACCGCGTCAGGTGCTGAGCATGA</td><td></td><td></td><td></td><td></td><td></td><td></td><td></td><td></td><td></td><td></td></tr>
<tr><td>Oligo rev</td><td>AAACTCATGCTCAGCACCTGACGC</td><td></td><td></td><td></td><td></td><td></td><td></td><td></td><td></td><td></td><td></td></tr>
<tr><td>Chromosome</td><td>start</td><td>end</td><td>strand</td><td>MM</td><td>target_seq</td><td>PAM</td><td>alignment</td><td>distance</td><td>position</td><td>gene name</td><td>gene id</td></tr>
<tr><td>GU305914.1</td><td>120971</td><td>120993</td><td>-</td><td>0</td><td>GCGTCAGGTGCTGAGCATGA</td><td>CGG</td><td>||||||||[||||||||||||]PAM</td><td>NA</td><td>Intergenic</td><td>NA</td><td>NA</td></tr>
<tr><td></td><td></td><td></td><td></td><td></td><td></td><td></td><td></td><td></td><td></td><td></td><td></td></tr>
<tr><td>T24895</td><td>AGGTGCTGAGCATGACGGTGCGG</td><td>1000</td><td></td><td></td><td></td><td></td><td></td><td></td><td></td><td></td><td></td></tr>
<tr><td>Oligo fwd</td><td>CACCAGGTGCTGAGCATGACGGTG</td><td></td><td></td><td></td><td></td><td></td><td></td><td></td><td></td><td></td><td></td></tr>
<tr><td>Oligo rev</td><td>AAACCACCGTCATGCTCAGCACCT</td><td></td><td></td><td></td><td></td><td></td><td></td><td></td><td></td><td></td><td></td></tr>
<tr><td>Chromosome</td><td>start</td><td>end</td><td>strand</td><td>MM</td><td>target_seq</td><td>PAM</td><td>alignment</td><td>distance</td><td>position</td><td>gene name</td><td>gene id</td></tr>
<tr><td>GU305914.1</td><td>120966</td><td>120988</td><td>-</td><td>0</td><td>AGGTGCTGAGCATGACGGTG</td><td>CGG</td><td>||||||||[||||||||||||]PAM</td><td>NA</td><td>Intergenic</td><td>NA</td><td>NA</td></tr>
<tr><td></td><td></td><td></td><td></td><td></td><td></td><td></td><td></td><td></td><td></td><td></td><td></td></tr>
<tr><td>T24896</td><td>GGTGCGGTCGTCGTTTACGACGG</td><td>1000</td><td></td><td></td><td></td><td></td><td></td><td></td><td></td><td></td><td></td></tr>
<tr><td>Oligo fwd</td><td>CACCGGTGCGGTCGTCGTTTACGA</td><td></td><td></td><td></td><td></td><td></td><td></td><td></td><td></td><td></td><td></td></tr>
<tr><td>Oligo rev</td><td>AAACTCGTAAACGACGACCGCACC</td><td></td><td></td><td></td><td></td><td></td><td></td><td></td><td></td><td></td><td></td></tr>
<tr><td>Chromosome</td><td>start</td><td>end</td><td>strand</td><td>MM</td><td>target_seq</td><td>PAM</td><td>alignment</td><td>distance</td><td>position</td><td>gene name</td><td>gene id</td></tr>
<tr><td>GU305914.1</td><td>120950</td><td>120972</td><td>-</td><td>0</td><td>GGTGCGGTCGTCGTTTACGA</td><td>CGG</td><td>||||||||[||||||||||||]PAM</td><td>NA</td><td>Intergenic</td><td>NA</td><td>NA</td></tr>
<tr><td></td><td></td><td></td><td></td><td></td><td></td><td></td><td></td><td></td><td></td><td></td><td></td></tr>
<tr><td>T24897</td><td>GTTTACGACGGAGCTGAAAGCGG</td><td>1000</td><td></td><td></td><td></td><td></td><td></td><td></td><td></td><td></td><td></td></tr>
<tr><td>Oligo fwd</td><td>CACCGTTTACGACGGAGCTGAAAG</td><td></td><td></td><td></td><td></td><td></td><td></td><td></td><td></td><td></td><td></td></tr>
<tr><td>Oligo rev</td><td>AAACCTTTCAGCTCCGTCGTAAAC</td><td></td><td></td><td></td><td></td><td></td><td></td><td></td><td></td><td></td><td></td></tr>
<tr><td>Chromosome</td><td>start</td><td>end</td><td>strand</td><td>MM</td><td>target_seq</td><td>PAM</td><td>alignment</td><td>distance</td><td>position</td><td>gene name</td><td>gene id</td></tr>
<tr><td>GU305914.1</td><td>120938</td><td>120960</td><td>-</td><td>0</td><td>GTTTACGACGGAGCTGAAAG</td><td>CGG</td><td>||||||||[||||||||||||]PAM</td><td>NA</td><td>Intergenic</td><td>NA</td><td>NA</td></tr>
<tr><td></td><td></td><td></td><td></td><td></td><td></td><td></td><td></td><td></td><td></td><td></td><td></td></tr>
<tr><td>T24898</td><td>TTTACGACGGAGCTGAAAGCGGG</td><td>1000</td><td></td><td></td><td></td><td></td><td></td><td></td><td></td><td></td><td></td></tr>
<tr><td>Oligo fwd</td><td>CACCTTTACGACGGAGCTGAAAGC</td><td></td><td></td><td></td><td></td><td></td><td></td><td></td><td></td><td></td><td></td></tr>
<tr><td>Oligo rev</td><td>AAACGCTTTCAGCTCCGTCGTAAA</td><td></td><td></td><td></td><td></td><td></td><td></td><td></td><td></td><td></td><td></td></tr>
<tr><td>Chromosome</td><td>start</td><td>end</td><td>strand</td><td>MM</td><td>target_seq</td><td>PAM</td><td>alignment</td><td>distance</td><td>position</td><td>gene name</td><td>gene id</td></tr>
<tr><td>GU305914.1</td><td>120937</td><td>120959</td><td>-</td><td>0</td><td>TTTACGACGGAGCTGAAAGC</td><td>GGG</td><td>||||||||[||||||||||||]PAM</td><td>NA</td><td>Intergenic</td><td>NA</td><td>NA</td></tr>
<tr><td></td><td></td><td></td><td></td><td></td><td></td><td></td><td></td><td></td><td></td><td></td><td></td></tr>
<tr><td>T24899</td><td>GCTGAAAGCGGGCGAGCCGCTGG</td><td>1000</td><td></td><td></td><td></td><td></td><td></td><td></td><td></td><td></td><td></td></tr>
<tr><td>Oligo fwd</td><td>CACCGCTGAAAGCGGGCGAGCCGC</td><td></td><td></td><td></td><td></td><td></td><td></td><td></td><td></td><td></td><td></td></tr>
<tr><td>Oligo rev</td><td>AAACGCGGCTCGCCCGCTTTCAGC</td><td></td><td></td><td></td><td></td><td></td><td></td><td></td><td></td><td></td><td></td></tr>
<tr><td>Chromosome</td><td>start</td><td>end</td><td>strand</td><td>MM</td><td>target_seq</td><td>PAM</td><td>alignment</td><td>distance</td><td>position</td><td>gene name</td><td>gene id</td></tr>
<tr><td>GU305914.1</td><td>120926</td><td>120948</td><td>-</td><td>0</td><td>GCTGAAAGCGGGCGAGCCGC</td><td>TGG</td><td>||||||||[||||||||||||]PAM</td><td>NA</td><td>Intergenic</td><td>NA</td><td>NA</td></tr>
<tr><td></td><td></td><td></td><td></td><td></td><td></td><td></td><td></td><td></td><td></td><td></td><td></td></tr>
<tr><td>T24900</td><td>CTGAAAGCGGGCGAGCCGCTGGG</td><td>1000</td><td></td><td></td><td></td><td></td><td></td><td></td><td></td><td></td><td></td></tr>
<tr><td>Oligo fwd</td><td>CACCCTGAAAGCGGGCGAGCCGCT</td><td></td><td></td><td></td><td></td><td></td><td></td><td></td><td></td><td></td><td></td></tr>
<tr><td>Oligo rev</td><td>AAACAGCGGCTCGCCCGCTTTCAG</td><td></td><td></td><td></td><td></td><td></td><td></td><td></td><td></td><td></td><td></td></tr>
<tr><td>Chromosome</td><td>start</td><td>end</td><td>strand</td><td>MM</td><td>target_seq</td><td>PAM</td><td>alignment</td><td>distance</td><td>position</td><td>gene name</td><td>gene id</td></tr>
<tr><td>GU305914.1</td><td>120925</td><td>120947</td><td>-</td><td>0</td><td>CTGAAAGCGGGCGAGCCGCT</td><td>GGG</td><td>||||||||[||||||||||||]PAM</td><td>NA</td><td>Intergenic</td><td>NA</td><td>NA</td></tr>
<tr><td></td><td></td><td></td><td></td><td></td><td></td><td></td><td></td><td></td><td></td><td></td><td></td></tr>
<tr><td>T24901</td><td>CGTGAAGAAGACAAGCCCGTCGG</td><td>1000</td><td></td><td></td><td></td><td></td><td></td><td></td><td></td><td></td><td></td></tr>
<tr><td>Oligo fwd</td><td>CACCCGTGAAGAAGACAAGCCCGT</td><td></td><td></td><td></td><td></td><td></td><td></td><td></td><td></td><td></td><td></td></tr>
<tr><td>Oligo rev</td><td>AAACACGGGCTTGTCTTCTTCACG</td><td></td><td></td><td></td><td></td><td></td><td></td><td></td><td></td><td></td><td></td></tr>
<tr><td>Chromosome</td><td>start</td><td>end</td><td>strand</td><td>MM</td><td>target_seq</td><td>PAM</td><td>alignment</td><td>distance</td><td>position</td><td>gene name</td><td>gene id</td></tr>
<tr><td>GU305914.1</td><td>120869</td><td>120891</td><td>-</td><td>0</td><td>CGTGAAGAAGACAAGCCCGT</td><td>CGG</td><td>||||||||[||||||||||||]PAM</td><td>NA</td><td>Intergenic</td><td>NA</td><td>NA</td></tr>
<tr><td></td><td></td><td></td><td></td><td></td><td></td><td></td><td></td><td></td><td></td><td></td><td></td></tr>
<tr><td>T24902</td><td>GAAGACAAGCCCGTCGGCCGCGG</td><td>1000</td><td></td><td></td><td></td><td></td><td></td><td></td><td></td><td></td><td></td></tr>
<tr><td>Oligo fwd</td><td>CACCGAAGACAAGCCCGTCGGCCG</td><td></td><td></td><td></td><td></td><td></td><td></td><td></td><td></td><td></td><td></td></tr>
<tr><td>Oligo rev</td><td>AAACCGGCCGACGGGCTTGTCTTC</td><td></td><td></td><td></td><td></td><td></td><td></td><td></td><td></td><td></td><td></td></tr>
<tr><td>Chromosome</td><td>start</td><td>end</td><td>strand</td><td>MM</td><td>target_seq</td><td>PAM</td><td>alignment</td><td>distance</td><td>position</td><td>gene name</td><td>gene id</td></tr>
<tr><td>GU305914.1</td><td>120863</td><td>120885</td><td>-</td><td>0</td><td>GAAGACAAGCCCGTCGGCCG</td><td>CGG</td><td>||||||||[||||||||||||]PAM</td><td>NA</td><td>Intergenic</td><td>NA</td><td>NA</td></tr>
<tr><td></td><td></td><td></td><td></td><td></td><td></td><td></td><td></td><td></td><td></td><td></td><td></td></tr>
<tr><td>T24903</td><td>GACAAGCCCGTCGGCCGCGGCGG</td><td>1000</td><td></td><td></td><td></td><td></td><td></td><td></td><td></td><td></td><td></td></tr>
<tr><td>Oligo fwd</td><td>CACCGACAAGCCCGTCGGCCGCGG</td><td></td><td></td><td></td><td></td><td></td><td></td><td></td><td></td><td></td><td></td></tr>
</table>

Oligo rev AAACCCGCGGCCGACGGGCTTGTC
Chromosome start end strand MM target_seq PAM alignment distance position gene name gene id
GU305914.1 120860 120882 - 0 GACAAGCCCGTCGGCCGCGG CGG ||||||||[||||||||||||]PAM NA Intergenic NA NA

T24904 CTACTCGATCCTCGTGCACCCGG 1000
Oligo fwd CACCCTACTCGATCCTCGTGCACC
Oligo rev AAACGGTGCACGAGGATCGAGTAG
Chromosome start end strand MM target_seq PAM alignment distance position gene name gene id
GU305914.1 120827 120849 - 0 CTACTCGATCCTCGTGCACC CGG ||||||||[||||||||||||]PAM NA Intergenic NA NA

T24905 CTCGTGCACCCGGCCGAGACCGG 1000
Oligo fwd CACCCTCGTGCACCCGGCCGAGAC
Oligo rev AAACGTCTCGGCCGGGTGCACGAG
Chromosome start end strand MM target_seq PAM alignment distance position gene name gene id
GU305914.1 120817 120839 - 0 CTCGTGCACCCGGCCGAGAC CGG ||||||||[||||||||||||]PAM NA Intergenic NA NA

T24906 CACCCGGCCGAGACCGGCGCCGG 1000
Oligo fwd CACCCACCCGGCCGAGACCGGCGC
Oligo rev AAACGCGCCGGTCTCGGCCGGGTG
Chromosome start end strand MM target_seq PAM alignment distance position gene name gene id
GU305914.1 120811 120833 - 0 CACCCGGCCGAGACCGGCGC CGG ||||||||[||||||||||||]PAM NA Intergenic NA NA

T24907 CCGGCCGAGACCGGCGCCGGAGG 1000
Oligo fwd CACCCCGGCCGAGACCGGCGCCGG
Oligo rev AAACCCGGCGCCGGTCTCGGCCGG
Chromosome start end strand MM target_seq PAM alignment distance position gene name gene id
GU305914.1 120808 120830 - 0 CCGGCCGAGACCGGCGCCGG AGG ||||||||[||||||||||||]PAM NA Intergenic NA NA

T24908 GCCGACCACCGCCGACACACCGG 1000
Oligo fwd CACCGCCGACCACCGCCGACACAC
Oligo rev AAACGTGTGTCGGCGGTGGTCGGC
Chromosome start end strand MM target_seq PAM alignment distance position gene name gene id
GU305914.1 120771 120793 - 0 GCCGACCACCGCCGACACAC CGG ||||||||[||||||||||||]PAM NA Intergenic NA NA

T24909 AGACCCTCGACATCGTCGAGCGG 1000
Oligo fwd CACCAGACCCTCGACATCGTCGAG
Oligo rev AAACCTCGACGATGTCGAGGGTCT
Chromosome start end strand MM target_seq PAM alignment distance position gene name gene id
GU305914.1 120748 120770 - 0 AGACCCTCGACATCGTCGAG CGG ||||||||[||||||||||||]PAM NA Intergenic NA NA

T24910 CATCGTCGAGCGGCGCCAAGAGG 1000
Oligo fwd CACCCATCGTCGAGCGGCGCCAAG
Oligo rev AAACCTTGGCGCCGCTCGACGATG
Chromosome start end strand MM target_seq PAM alignment distance position gene name gene id
GU305914.1 120738 120760 - 0 CATCGTCGAGCGGCGCCAAG AGG ||||||||[||||||||||||]PAM NA Intergenic NA NA

T24911 ATCGTCGAGCGGCGCCAAGAGGG 1000
Oligo fwd CACCATCGTCGAGCGGCGCCAAGA
Oligo rev AAACTCTTGGCGCCGCTCGACGAT
Chromosome start end strand MM target_seq PAM alignment distance position gene name gene id
GU305914.1 120737 120759 - 0 ATCGTCGAGCGGCGCCAAGA GGG ||||||||[||||||||||||]PAM NA Intergenic NA NA

T24912 TCGTCGAGCGGCGCCAAGAGGGG 1000
Oligo fwd CACCTCGTCGAGCGGCGCCAAGAG
Oligo rev AAACCTCTTGGCGCCGCTCGACGA
Chromosome start end strand MM target_seq PAM alignment distance position gene name gene id
GU305914.1 120736 120758 - 0 TCGTCGAGCGGCGCCAAGAG GGG ||||||||[||||||||||||]PAM NA Intergenic NA NA

T24913 CGTCGAGCGGCGCCAAGAGGGGG 1000
Oligo fwd CACCCGTCGAGCGGCGCCAAGAGG
Oligo rev AAACCCTCTTGGCGCCGCTCGACG
Chromosome start end strand MM target_seq PAM alignment distance position gene name gene id
GU305914.1 120735 120757 - 0 CGTCGAGCGGCGCCAAGAGG GGG ||||||||[||||||||||||]PAM NA Intergenic NA NA

T24914 AAGAGGGGGTCGATGAACCCCGG 1000
Oligo fwd CACCAAGAGGGGGTCGATGAACCC
Oligo rev AAACGGGTTCATCGACCCCCTCTT
Chromosome start end strand MM target_seq PAM alignment distance position gene name gene id
GU305914.1 120721 120743 - 0 AAGAGGGGGTCGATGAACCC CGG ||||||||[||||||||||||]PAM NA Intergenic NA NA

T24915 ACCCCGGATTTAGCGAGTTCTGG 1000
Oligo fwd CACCACCCCGGATTTAGCGAGTTC
Oligo rev AAACGAACTCGCTAAATCCGGGGT
Chromosome start end strand MM target_seq PAM alignment distance position gene name gene id
GU305914.1 120705 120727 - 0 ACCCCGGATTTAGCGAGTTC TGG ||||||||[||||||||||||]PAM NA Intergenic NA NA

T24916 TCCCGCCGCCCGACAGCAAGAGG 1000
Oligo fwd CACCTCCCGCCGCCCGACAGCAAG
Oligo rev AAACCTTGCTGTCGGGCGGCGGGA
Chromosome start end strand MM target_seq PAM alignment distance position gene name gene id
GU305914.1 120675 120697 - 0 TCCCGCCGCCCGACAGCAAG AGG ||||||||[||||||||||||]PAM NA Intergenic NA NA

T24917 CCCGCCGCCCGACAGCAAGAGGG 1000
Oligo fwd CACCCCCGCCGCCCGACAGCAAGA
Oligo rev AAACTCTTGCTGTCGGGCGGCGGG
Chromosome start end strand MM target_seq PAM alignment distance position gene name gene id
GU305914.1 120674 120696 - 0 CCCGCCGCCCGACAGCAAGA GGG ||||||||[||||||||||||]PAM NA Intergenic NA NA

T24918 CGCCCGACAGCAAGAGGGCCCGG 1000

<table>
<tr><td>Oligo fwd</td><td>CACCCGCCCGACAGCAAGAGGGCC</td><td></td><td></td><td></td><td></td><td></td><td></td><td></td><td></td><td></td><td></td></tr>
<tr><td>Oligo rev</td><td>AAACGGCCCTCTTGCTGTCGGGCG</td><td></td><td></td><td></td><td></td><td></td><td></td><td></td><td></td><td></td><td></td></tr>
<tr><td>Chromosome</td><td>start</td><td>end</td><td>strand</td><td>MM</td><td>target_seq</td><td>PAM</td><td>alignment</td><td>distance</td><td>position</td><td>gene name</td><td>gene id</td></tr>
<tr><td>GU305914.1</td><td>120669</td><td>120691</td><td>-</td><td>0</td><td>CGCCCGACAGCAAGAGGGCC</td><td>CGG</td><td>||||||||[||||||||||||]PAM</td><td>NA</td><td>Intergenic</td><td>NA</td><td>NA</td></tr>
<tr><td></td><td></td><td></td><td></td><td></td><td></td><td></td><td></td><td></td><td></td><td></td><td></td></tr>
<tr><td>T24919</td><td>ACAGCAAGAGGGCCCGGACGAGG</td><td>1000</td><td></td><td></td><td></td><td></td><td></td><td></td><td></td><td></td><td></td></tr>
<tr><td>Oligo fwd</td><td>CACCACAGCAAGAGGGCCCGGACG</td><td></td><td></td><td></td><td></td><td></td><td></td><td></td><td></td><td></td><td></td></tr>
<tr><td>Oligo rev</td><td>AAACCGTCCGGGCCCTCTTGCTGT</td><td></td><td></td><td></td><td></td><td></td><td></td><td></td><td></td><td></td><td></td></tr>
<tr><td>Chromosome</td><td>start</td><td>end</td><td>strand</td><td>MM</td><td>target_seq</td><td>PAM</td><td>alignment</td><td>distance</td><td>position</td><td>gene name</td><td>gene id</td></tr>
<tr><td>GU305914.1</td><td>120663</td><td>120685</td><td>-</td><td>0</td><td>ACAGCAAGAGGGCCCGGACG</td><td>AGG</td><td>||||||||[||||||||||||]PAM</td><td>NA</td><td>Intergenic</td><td>NA</td><td>NA</td></tr>
<tr><td></td><td></td><td></td><td></td><td></td><td></td><td></td><td></td><td></td><td></td><td></td><td></td></tr>
<tr><td>T24920</td><td>CAAGAGGGCCCGGACGAGGTCGG</td><td>1000</td><td></td><td></td><td></td><td></td><td></td><td></td><td></td><td></td><td></td></tr>
<tr><td>Oligo fwd</td><td>CACCCAAGAGGGCCCGGACGAGGT</td><td></td><td></td><td></td><td></td><td></td><td></td><td></td><td></td><td></td><td></td></tr>
<tr><td>Oligo rev</td><td>AAACACCTCGTCCGGGCCCTCTTG</td><td></td><td></td><td></td><td></td><td></td><td></td><td></td><td></td><td></td><td></td></tr>
<tr><td>Chromosome</td><td>start</td><td>end</td><td>strand</td><td>MM</td><td>target_seq</td><td>PAM</td><td>alignment</td><td>distance</td><td>position</td><td>gene name</td><td>gene id</td></tr>
<tr><td>GU305914.1</td><td>120659</td><td>120681</td><td>-</td><td>0</td><td>CAAGAGGGCCCGGACGAGGT</td><td>CGG</td><td>||||||||[||||||||||||]PAM</td><td>NA</td><td>Intergenic</td><td>NA</td><td>NA</td></tr>
<tr><td></td><td></td><td></td><td></td><td></td><td></td><td></td><td></td><td></td><td></td><td></td><td></td></tr>
<tr><td>T24921</td><td>GTCGGAGAGCGAGAAGAACGCGG</td><td>1000</td><td></td><td></td><td></td><td></td><td></td><td></td><td></td><td></td><td></td></tr>
<tr><td>Oligo fwd</td><td>CACCGTCGGAGAGCGAGAAGAACG</td><td></td><td></td><td></td><td></td><td></td><td></td><td></td><td></td><td></td><td></td></tr>
<tr><td>Oligo rev</td><td>AAACCGTTCTTCTCGCTCTCCGAC</td><td></td><td></td><td></td><td></td><td></td><td></td><td></td><td></td><td></td><td></td></tr>
<tr><td>Chromosome</td><td>start</td><td>end</td><td>strand</td><td>MM</td><td>target_seq</td><td>PAM</td><td>alignment</td><td>distance</td><td>position</td><td>gene name</td><td>gene id</td></tr>
<tr><td>GU305914.1</td><td>120641</td><td>120663</td><td>-</td><td>0</td><td>GTCGGAGAGCGAGAAGAACG</td><td>CGG</td><td>||||||||[||||||||||||]PAM</td><td>NA</td><td>Intergenic</td><td>NA</td><td>NA</td></tr>
<tr><td></td><td></td><td></td><td></td><td></td><td></td><td></td><td></td><td></td><td></td><td></td><td></td></tr>
<tr><td>T24922</td><td>GAGCGAGAAGAACGCGGCGAAGG</td><td>1000</td><td></td><td></td><td></td><td></td><td></td><td></td><td></td><td></td><td></td></tr>
<tr><td>Oligo fwd</td><td>CACCGAGCGAGAAGAACGCGGCGA</td><td></td><td></td><td></td><td></td><td></td><td></td><td></td><td></td><td></td><td></td></tr>
<tr><td>Oligo rev</td><td>AAACTCGCCGCGTTCTTCTCGCTC</td><td></td><td></td><td></td><td></td><td></td><td></td><td></td><td></td><td></td><td></td></tr>
<tr><td>Chromosome</td><td>start</td><td>end</td><td>strand</td><td>MM</td><td>target_seq</td><td>PAM</td><td>alignment</td><td>distance</td><td>position</td><td>gene name</td><td>gene id</td></tr>
<tr><td>GU305914.1</td><td>120635</td><td>120657</td><td>-</td><td>0</td><td>GAGCGAGAAGAACGCGGCGA</td><td>AGG</td><td>||||||||[||||||||||||]PAM</td><td>NA</td><td>Intergenic</td><td>NA</td><td>NA</td></tr>
<tr><td>GU305914.1</td><td>126419</td><td>126441</td><td>-</td><td>4</td><td>GAGCAAGGAGAACGCCGTGA</td><td>CGG</td><td>||||-||-[|||||||-|-||]PAM</td><td>NA</td><td>Intergenic</td><td>NA</td><td>NA</td></tr>
<tr><td></td><td></td><td></td><td></td><td></td><td></td><td></td><td></td><td></td><td></td><td></td><td></td></tr>
<tr><td>T24923</td><td>GCAGCGTCCACCCGATCCGCCGG</td><td>1000</td><td></td><td></td><td></td><td></td><td></td><td></td><td></td><td></td><td></td></tr>
<tr><td>Oligo fwd</td><td>CACCGCAGCGTCCACCCGATCCGC</td><td></td><td></td><td></td><td></td><td></td><td></td><td></td><td></td><td></td><td></td></tr>
<tr><td>Oligo rev</td><td>AAACGCGGATCGGGTGGACGCTGC</td><td></td><td></td><td></td><td></td><td></td><td></td><td></td><td></td><td></td><td></td></tr>
<tr><td>Chromosome</td><td>start</td><td>end</td><td>strand</td><td>MM</td><td>target_seq</td><td>PAM</td><td>alignment</td><td>distance</td><td>position</td><td>gene name</td><td>gene id</td></tr>
<tr><td>GU305914.1</td><td>120611</td><td>120633</td><td>-</td><td>0</td><td>GCAGCGTCCACCCGATCCGC</td><td>CGG</td><td>||||||||[||||||||||||]PAM</td><td>NA</td><td>Intergenic</td><td>NA</td><td>NA</td></tr>
<tr><td></td><td></td><td></td><td></td><td></td><td></td><td></td><td></td><td></td><td></td><td></td><td></td></tr>
<tr><td>T24924</td><td>TCCGCCGGAGAGCGAACGTCAGG</td><td>1000</td><td></td><td></td><td></td><td></td><td></td><td></td><td></td><td></td><td></td></tr>
<tr><td>Oligo fwd</td><td>CACCTCCGCCGGAGAGCGAACGTC</td><td></td><td></td><td></td><td></td><td></td><td></td><td></td><td></td><td></td><td></td></tr>
<tr><td>Oligo rev</td><td>AAACGACGTTCGCTCTCCGGCGGA</td><td></td><td></td><td></td><td></td><td></td><td></td><td></td><td></td><td></td><td></td></tr>
<tr><td>Chromosome</td><td>start</td><td>end</td><td>strand</td><td>MM</td><td>target_seq</td><td>PAM</td><td>alignment</td><td>distance</td><td>position</td><td>gene name</td><td>gene id</td></tr>
<tr><td>GU305914.1</td><td>120596</td><td>120618</td><td>-</td><td>0</td><td>TCCGCCGGAGAGCGAACGTC</td><td>AGG</td><td>||||||||[||||||||||||]PAM</td><td>NA</td><td>Intergenic</td><td>NA</td><td>NA</td></tr>
<tr><td></td><td></td><td></td><td></td><td></td><td></td><td></td><td></td><td></td><td></td><td></td><td></td></tr>
<tr><td>T24925</td><td>TCAGGCGCCGACGTACCCCGAGG</td><td>1000</td><td></td><td></td><td></td><td></td><td></td><td></td><td></td><td></td><td></td></tr>
<tr><td>Oligo fwd</td><td>CACCTCAGGCGCCGACGTACCCCG</td><td></td><td></td><td></td><td></td><td></td><td></td><td></td><td></td><td></td><td></td></tr>
<tr><td>Oligo rev</td><td>AAACCGGGGTACGTCGGCGCCTGA</td><td></td><td></td><td></td><td></td><td></td><td></td><td></td><td></td><td></td><td></td></tr>
<tr><td>Chromosome</td><td>start</td><td>end</td><td>strand</td><td>MM</td><td>target_seq</td><td>PAM</td><td>alignment</td><td>distance</td><td>position</td><td>gene name</td><td>gene id</td></tr>
<tr><td>GU305914.1</td><td>120578</td><td>120600</td><td>-</td><td>0</td><td>TCAGGCGCCGACGTACCCCG</td><td>AGG</td><td>||||||||[||||||||||||]PAM</td><td>NA</td><td>Intergenic</td><td>NA</td><td>NA</td></tr>
<tr><td></td><td></td><td></td><td></td><td></td><td></td><td></td><td></td><td></td><td></td><td></td><td></td></tr>
<tr><td>T24926</td><td>GGCGCCGACGTACCCCGAGGTGG</td><td>1000</td><td></td><td></td><td></td><td></td><td></td><td></td><td></td><td></td><td></td></tr>
<tr><td>Oligo fwd</td><td>CACCGGCGCCGACGTACCCCGAGG</td><td></td><td></td><td></td><td></td><td></td><td></td><td></td><td></td><td></td><td></td></tr>
<tr><td>Oligo rev</td><td>AAACCCTCGGGGTACGTCGGCGCC</td><td></td><td></td><td></td><td></td><td></td><td></td><td></td><td></td><td></td><td></td></tr>
<tr><td>Chromosome</td><td>start</td><td>end</td><td>strand</td><td>MM</td><td>target_seq</td><td>PAM</td><td>alignment</td><td>distance</td><td>position</td><td>gene name</td><td>gene id</td></tr>
<tr><td>GU305914.1</td><td>120575</td><td>120597</td><td>-</td><td>0</td><td>GGCGCCGACGTACCCCGAGG</td><td>TGG</td><td>||||||||[||||||||||||]PAM</td><td>NA</td><td>Intergenic</td><td>NA</td><td>NA</td></tr>
<tr><td></td><td></td><td></td><td></td><td></td><td></td><td></td><td></td><td></td><td></td><td></td><td></td></tr>
<tr><td>T24927</td><td>CCCCGAGGTGGAGAGATTCTCGG</td><td>1000</td><td></td><td></td><td></td><td></td><td></td><td></td><td></td><td></td><td></td></tr>
<tr><td>Oligo fwd</td><td>CACCCCCCGAGGTGGAGAGATTCT</td><td></td><td></td><td></td><td></td><td></td><td></td><td></td><td></td><td></td><td></td></tr>
<tr><td>Oligo rev</td><td>AAACAGAATCTCTCCACCTCGGGG</td><td></td><td></td><td></td><td></td><td></td><td></td><td></td><td></td><td></td><td></td></tr>
<tr><td>Chromosome</td><td>start</td><td>end</td><td>strand</td><td>MM</td><td>target_seq</td><td>PAM</td><td>alignment</td><td>distance</td><td>position</td><td>gene name</td><td>gene id</td></tr>
<tr><td>GU305914.1</td><td>120563</td><td>120585</td><td>-</td><td>0</td><td>CCCCGAGGTGGAGAGATTCT</td><td>CGG</td><td>||||||||[||||||||||||]PAM</td><td>NA</td><td>Intergenic</td><td>NA</td><td>NA</td></tr>
<tr><td></td><td></td><td></td><td></td><td></td><td></td><td></td><td></td><td></td><td></td><td></td><td></td></tr>
<tr><td>T24928</td><td>GAGATTCTCGGAGCACGACGAGG</td><td>1000</td><td></td><td></td><td></td><td></td><td></td><td></td><td></td><td></td><td></td></tr>
<tr><td>Oligo fwd</td><td>CACCGAGATTCTCGGAGCACGACG</td><td></td><td></td><td></td><td></td><td></td><td></td><td></td><td></td><td></td><td></td></tr>
<tr><td>Oligo rev</td><td>AAACCGTCGTGCTCCGAGAATCTC</td><td></td><td></td><td></td><td></td><td></td><td></td><td></td><td></td><td></td><td></td></tr>
<tr><td>Chromosome</td><td>start</td><td>end</td><td>strand</td><td>MM</td><td>target_seq</td><td>PAM</td><td>alignment</td><td>distance</td><td>position</td><td>gene name</td><td>gene id</td></tr>
<tr><td>GU305914.1</td><td>120551</td><td>120573</td><td>-</td><td>0</td><td>GAGATTCTCGGAGCACGACG</td><td>AGG</td><td>||||||||[||||||||||||]PAM</td><td>NA</td><td>Intergenic</td><td>NA</td><td>NA</td></tr>
<tr><td></td><td></td><td></td><td></td><td></td><td></td><td></td><td></td><td></td><td></td><td></td><td></td></tr>
<tr><td>T24929</td><td>GGCCCTAGAGATCGTGCCGACGG</td><td>1000</td><td></td><td></td><td></td><td></td><td></td><td></td><td></td><td></td><td></td></tr>
<tr><td>Oligo fwd</td><td>CACCGGCCCTAGAGATCGTGCCGA</td><td></td><td></td><td></td><td></td><td></td><td></td><td></td><td></td><td></td><td></td></tr>
<tr><td>Oligo rev</td><td>AAACTCGGCACGATCTCTAGGGCC</td><td></td><td></td><td></td><td></td><td></td><td></td><td></td><td></td><td></td><td></td></tr>
<tr><td>Chromosome</td><td>start</td><td>end</td><td>strand</td><td>MM</td><td>target_seq</td><td>PAM</td><td>alignment</td><td>distance</td><td>position</td><td>gene name</td><td>gene id</td></tr>
<tr><td>GU305914.1</td><td>120530</td><td>120552</td><td>-</td><td>0</td><td>GGCCCTAGAGATCGTGCCGA</td><td>CGG</td><td>||||||||[||||||||||||]PAM</td><td>NA</td><td>Intergenic</td><td>NA</td><td>NA</td></tr>
<tr><td></td><td></td><td></td><td></td><td></td><td></td><td></td><td></td><td></td><td></td><td></td><td></td></tr>
<tr><td>T24930</td><td>GCCCTAGAGATCGTGCCGACGGG</td><td>1000</td><td></td><td></td><td></td><td></td><td></td><td></td><td></td><td></td><td></td></tr>
<tr><td>Oligo fwd</td><td>CACCGCCCTAGAGATCGTGCCGAC</td><td></td><td></td><td></td><td></td><td></td><td></td><td></td><td></td><td></td><td></td></tr>
<tr><td>Oligo rev</td><td>AAACGTCGGCACGATCTCTAGGGC</td><td></td><td></td><td></td><td></td><td></td><td></td><td></td><td></td><td></td><td></td></tr>
<tr><td>Chromosome</td><td>start</td><td>end</td><td>strand</td><td>MM</td><td>target_seq</td><td>PAM</td><td>alignment</td><td>distance</td><td>position</td><td>gene name</td><td>gene id</td></tr>
<tr><td>GU305914.1</td><td>120529</td><td>120551</td><td>-</td><td>0</td><td>GCCCTAGAGATCGTGCCGAC</td><td>GGG</td><td>||||||||[||||||||||||]PAM</td><td>NA</td><td>Intergenic</td><td>NA</td><td>NA</td></tr>
<tr><td>GU305914.1</td><td>224626</td><td>224648</td><td>-</td><td>4</td><td>GCCCTGGGGATCGTGCTGAT</td><td>CGG</td><td>|||||-|-[||||||||-||-]PAM</td><td>NA</td><td>Intergenic</td><td>NA</td><td>NA</td></tr>
<tr><td></td><td></td><td></td><td></td><td></td><td></td><td></td><td></td><td></td><td></td><td></td><td></td></tr>
<tr><td>T24931</td><td>CGAGACTCGCCGATCACGCGCGG</td><td>1000</td><td></td><td></td><td></td><td></td><td></td><td></td><td></td><td></td><td></td></tr>
<tr><td>Oligo fwd</td><td>CACCCGAGACTCGCCGATCACGCG</td><td></td><td></td><td></td><td></td><td></td><td></td><td></td><td></td><td></td><td></td></tr>
<tr><td>Oligo rev</td><td>AAACCGCGTGATCGGCGAGTCTCG</td><td></td><td></td><td></td><td></td><td></td><td></td><td></td><td></td><td></td><td></td></tr>
<tr><td>Chromosome</td><td>start</td><td>end</td><td>strand</td><td>MM</td><td>target_seq</td><td>PAM</td><td>alignment</td><td>distance</td><td>position</td><td>gene name</td><td>gene id</td></tr>
<tr><td>GU305914.1</td><td>120454</td><td>120476</td><td>-</td><td>0</td><td>CGAGACTCGCCGATCACGCG</td><td>CGG</td><td>||||||||[||||||||||||]PAM</td><td>NA</td><td>Intergenic</td><td>NA</td><td>NA</td></tr>
<tr><td></td><td></td><td></td><td></td><td></td><td></td><td></td><td></td><td></td><td></td><td></td><td></td></tr>
<tr><td>T24932</td><td>CGCGGTCCGCCCAAGAAGATCGG</td><td>1000</td><td></td><td></td><td></td><td></td><td></td><td></td><td></td><td></td><td></td></tr>
<tr><td>Oligo fwd</td><td>CACCCGCGGTCCGCCCAAGAAGAT</td><td></td><td></td><td></td><td></td><td></td><td></td><td></td><td></td><td></td><td></td></tr>
<tr><td>Oligo rev</td><td>AAACATCTTCTTGGGCGGACCGCG</td><td></td><td></td><td></td><td></td><td></td><td></td><td></td><td></td><td></td><td></td></tr>
<tr><td>Chromosome</td><td>start</td><td>end</td><td>strand</td><td>MM</td><td>target_seq</td><td>PAM</td><td>alignment</td><td>distance</td><td>position</td><td>gene name</td><td>gene id</td></tr>
</table>

GU305914.1 120436 120458 - 0 CGCGGTCCGCCCAAGAAGAT CGG ||||||||[||||||||||||]PAM NA Intergenic NA NA

T24933 GTCCGCCCAAGAAGATCGGCAGG 1000
Oligo fwd CACCGTCCGCCCAAGAAGATCGGC
Oligo rev AAACGCCGATCTTCTTGGGCGGAC
Chromosome start end strand MM target_seq PAM alignment distance position gene name gene id
GU305914.1 120432 120454 - 0 GTCCGCCCAAGAAGATCGGC AGG ||||||||[||||||||||||]PAM NA Intergenic NA NA

T24934 TCCGCCCAAGAAGATCGGCAGGG 1000
Oligo fwd CACCTCCGCCCAAGAAGATCGGCA
Oligo rev AAACTGCCGATCTTCTTGGGCGGA
Chromosome start end strand MM target_seq PAM alignment distance position gene name gene id
GU305914.1 120431 120453 - 0 TCCGCCCAAGAAGATCGGCA GGG ||||||||[||||||||||||]PAM NA Intergenic NA NA

T24935 GAAGATCGGCAGGGCGAGCGCGG 1000
Oligo fwd CACCGAAGATCGGCAGGGCGAGCG
Oligo rev AAACCGCTCGCCCTGCCGATCTTC
Chromosome start end strand MM target_seq PAM alignment distance position gene name gene id
GU305914.1 120422 120444 - 0 GAAGATCGGCAGGGCGAGCG CGG ||||||||[||||||||||||]PAM NA Intergenic NA NA

T24936 CGGCGTCGCTCCCCTCGCCCAGG 1000
Oligo fwd CACCCGGCGTCGCTCCCCTCGCCC
Oligo rev AAACGGGCGAGGGGAGCGACGCCG
Chromosome start end strand MM target_seq PAM alignment distance position gene name gene id
GU305914.1 120402 120424 - 0 CGGCGTCGCTCCCCTCGCCC AGG ||||||||[||||||||||||]PAM NA Intergenic NA NA

T24937 CGTCGCTCCCCTCGCCCAGGTGG 1000
Oligo fwd CACCCGTCGCTCCCCTCGCCCAGG
Oligo rev AAACCCTGGGCGAGGGGAGCGACG
Chromosome start end strand MM target_seq PAM alignment distance position gene name gene id
GU305914.1 120399 120421 - 0 CGTCGCTCCCCTCGCCCAGG TGG ||||||||[||||||||||||]PAM NA Intergenic NA NA

T24938 CCCCTCGCCCAGGTGGCACTCGG 1000
Oligo fwd CACCCCCCTCGCCCAGGTGGCACT
Oligo rev AAACAGTGCCACCTGGGCGAGGGG
Chromosome start end strand MM target_seq PAM alignment distance position gene name gene id
GU305914.1 120392 120414 - 0 CCCCTCGCCCAGGTGGCACT CGG ||||||||[||||||||||||]PAM NA Intergenic NA NA

T24939 TGGCACTCGGAATTGTGCGTCGG 1000
Oligo fwd CACCTGGCACTCGGAATTGTGCGT
Oligo rev AAACACGCACAATTCCGAGTGCCA
Chromosome start end strand MM target_seq PAM alignment distance position gene name gene id
GU305914.1 120379 120401 - 0 TGGCACTCGGAATTGTGCGT CGG ||||||||[||||||||||||]PAM NA Intergenic NA NA

T24940 GGAATTGTGCGTCGGAACGCTGG 1000
Oligo fwd CACCGGAATTGTGCGTCGGAACGC
Oligo rev AAACGCGTTCCGACGCACAATTCC
Chromosome start end strand MM target_seq PAM alignment distance position gene name gene id
GU305914.1 120371 120393 - 0 GGAATTGTGCGTCGGAACGC TGG ||||||||[||||||||||||]PAM NA Intergenic NA NA

T24941 TTGTGCGTCGGAACGCTGGTAGG 1000
Oligo fwd CACCTTGTGCGTCGGAACGCTGGT
Oligo rev AAACACCAGCGTTCCGACGCACAA
Chromosome start end strand MM target_seq PAM alignment distance position gene name gene id
GU305914.1 120367 120389 - 0 TTGTGCGTCGGAACGCTGGT AGG ||||||||[||||||||||||]PAM NA Intergenic NA NA

T24942 TGTGCGTCGGAACGCTGGTAGGG 1000
Oligo fwd CACCTGTGCGTCGGAACGCTGGTA
Oligo rev AAACTACCAGCGTTCCGACGCACA
Chromosome start end strand MM target_seq PAM alignment distance position gene name gene id
GU305914.1 120366 120388 - 0 TGTGCGTCGGAACGCTGGTA GGG ||||||||[||||||||||||]PAM NA Intergenic NA NA

T24943 CGTCGGAACGCTGGTAGGGACGG 1000
Oligo fwd CACCCGTCGGAACGCTGGTAGGGA
Oligo rev AAACTCCCTACCAGCGTTCCGACG
Chromosome start end strand MM target_seq PAM alignment distance position gene name gene id
GU305914.1 120362 120384 - 0 CGTCGGAACGCTGGTAGGGA CGG ||||||||[||||||||||||]PAM NA Intergenic NA NA

T24944 AACGCTGGTAGGGACGGCGCCGG 1000
Oligo fwd CACCAACGCTGGTAGGGACGGCGC
Oligo rev AAACGCGCCGTCCCTACCAGCGTT
Chromosome start end strand MM target_seq PAM alignment distance position gene name gene id
GU305914.1 120356 120378 - 0 AACGCTGGTAGGGACGGCGC CGG ||||||||[||||||||||||]PAM NA Intergenic NA NA

T24945 CCGGCGAGTCAAGCGCCGTACGG 1000
Oligo fwd CACCCCGGCGAGTCAAGCGCCGTA
Oligo rev AAACTACGGCGCTTGACTCGCCGG
Chromosome start end strand MM target_seq PAM alignment distance position gene name gene id
GU305914.1 120337 120359 - 0 CCGGCGAGTCAAGCGCCGTA CGG ||||||||[||||||||||||]PAM NA Intergenic NA NA

T24946 GCCTCCGTTCTCCAACATCCTGG 1000
Oligo fwd CACCGCCTCCGTTCTCCAACATCC
Oligo rev AAACGGATGTTGGAGAACGGAGGC
Chromosome start end strand MM target_seq PAM alignment distance position gene name gene id
GU305914.1 120308 120330 - 0 GCCTCCGTTCTCCAACATCC TGG ||||||||[||||||||||||]PAM NA Intergenic NA NA

T24947 CTCCAACATCCTGGCGTTGCAGG 1000
Oligo fwd CACCCTCCAACATCCTGGCGTTGC
Oligo rev AAACGCAACGCCAGGATGTTGGAG

<table>
<tr><td>Chromosome</td><td>start</td><td>end</td><td>strand</td><td>MM</td><td>target_seq</td><td>PAM</td><td>alignment</td><td>distance</td><td>position</td><td>gene name</td><td>gene id</td></tr>
<tr><td>GU305914.1</td><td>120299</td><td>120321</td><td>-</td><td>0</td><td>CTCCAACATCCTGGCGTTGC</td><td>AGG</td><td>||||||||[||||||||||||]PAM</td><td>NA</td><td>Intergenic</td><td>NA</td><td>NA</td></tr>
<tr><td>T24948</td><td>TCCTCAGCTCCAGCTTCAGCCGG</td><td>1000</td><td></td><td></td><td></td><td></td><td></td><td></td><td></td><td></td><td></td></tr>
<tr><td>Oligo fwd</td><td>CACCTCCTCAGCTCCAGCTTCAGC</td><td></td><td></td><td></td><td></td><td></td><td></td><td></td><td></td><td></td><td></td></tr>
<tr><td>Oligo rev</td><td>AAACGCTGAAGCTGGAGCTGAGGA</td><td></td><td></td><td></td><td></td><td></td><td></td><td></td><td></td><td></td><td></td></tr>
<tr><td>Chromosome</td><td>start</td><td>end</td><td>strand</td><td>MM</td><td>target_seq</td><td>PAM</td><td>alignment</td><td>distance</td><td>position</td><td>gene name</td><td>gene id</td></tr>
<tr><td>GU305914.1</td><td>120248</td><td>120270</td><td>-</td><td>0</td><td>TCCTCAGCTCCAGCTTCAGC</td><td>CGG</td><td>||||||||[||||||||||||]PAM</td><td>NA</td><td>Intergenic</td><td>NA</td><td>NA</td></tr>
<tr><td>T24949</td><td>GCCGTGTTCTTCCACCCGCTCGG</td><td>1000</td><td></td><td></td><td></td><td></td><td></td><td></td><td></td><td></td><td></td></tr>
<tr><td>Oligo fwd</td><td>CACCGCCGTGTTCTTCCACCCGCT</td><td></td><td></td><td></td><td></td><td></td><td></td><td></td><td></td><td></td><td></td></tr>
<tr><td>Oligo rev</td><td>AAACAGCGGGTGGAAGAACACGGC</td><td></td><td></td><td></td><td></td><td></td><td></td><td></td><td></td><td></td><td></td></tr>
<tr><td>Chromosome</td><td>start</td><td>end</td><td>strand</td><td>MM</td><td>target_seq</td><td>PAM</td><td>alignment</td><td>distance</td><td>position</td><td>gene name</td><td>gene id</td></tr>
<tr><td>GU305914.1</td><td>120217</td><td>120239</td><td>-</td><td>0</td><td>GCCGTGTTCTTCCACCCGCT</td><td>CGG</td><td>||||||||[||||||||||||]PAM</td><td>NA</td><td>Intergenic</td><td>NA</td><td>NA</td></tr>
<tr><td>T24950</td><td>ACCCGCTCGGCGCGCTCCAGAGG</td><td>1000</td><td></td><td></td><td></td><td></td><td></td><td></td><td></td><td></td><td></td></tr>
<tr><td>Oligo fwd</td><td>CACCACCCGCTCGGCGCGCTCCAG</td><td></td><td></td><td></td><td></td><td></td><td></td><td></td><td></td><td></td><td></td></tr>
<tr><td>Oligo rev</td><td>AAACCTGGAGCGCGCCGAGCGGGT</td><td></td><td></td><td></td><td></td><td></td><td></td><td></td><td></td><td></td><td></td></tr>
<tr><td>Chromosome</td><td>start</td><td>end</td><td>strand</td><td>MM</td><td>target_seq</td><td>PAM</td><td>alignment</td><td>distance</td><td>position</td><td>gene name</td><td>gene id</td></tr>
<tr><td>GU305914.1</td><td>120204</td><td>120226</td><td>-</td><td>0</td><td>ACCCGCTCGGCGCGCTCCAG</td><td>AGG</td><td>||||||||[||||||||||||]PAM</td><td>NA</td><td>Intergenic</td><td>NA</td><td>NA</td></tr>
<tr><td>T24951</td><td>AGGCTGCAGCAGAACATCTGAGG</td><td>1000</td><td></td><td></td><td></td><td></td><td></td><td></td><td></td><td></td><td></td></tr>
<tr><td>Oligo fwd</td><td>CACCAGGCTGCAGCAGAACATCTG</td><td></td><td></td><td></td><td></td><td></td><td></td><td></td><td></td><td></td><td></td></tr>
<tr><td>Oligo rev</td><td>AAACCAGATGTTCTGCTGCAGCCT</td><td></td><td></td><td></td><td></td><td></td><td></td><td></td><td></td><td></td><td></td></tr>
<tr><td>Chromosome</td><td>start</td><td>end</td><td>strand</td><td>MM</td><td>target_seq</td><td>PAM</td><td>alignment</td><td>distance</td><td>position</td><td>gene name</td><td>gene id</td></tr>
<tr><td>GU305914.1</td><td>120184</td><td>120206</td><td>-</td><td>0</td><td>AGGCTGCAGCAGAACATCTG</td><td>AGG</td><td>||||||||[||||||||||||]PAM</td><td>NA</td><td>Intergenic</td><td>NA</td><td>NA</td></tr>
<tr><td>T24952</td><td>ACAAACAGACAGAGCCCGATCGG</td><td>1000</td><td></td><td></td><td></td><td></td><td></td><td></td><td></td><td></td><td></td></tr>
<tr><td>Oligo fwd</td><td>CACCACAAACAGACAGAGCCCGAT</td><td></td><td></td><td></td><td></td><td></td><td></td><td></td><td></td><td></td><td></td></tr>
<tr><td>Oligo rev</td><td>AAACATCGGGCTCTGTCTGTTTGT</td><td></td><td></td><td></td><td></td><td></td><td></td><td></td><td></td><td></td><td></td></tr>
<tr><td>Chromosome</td><td>start</td><td>end</td><td>strand</td><td>MM</td><td>target_seq</td><td>PAM</td><td>alignment</td><td>distance</td><td>position</td><td>gene name</td><td>gene id</td></tr>
<tr><td>GU305914.1</td><td>120155</td><td>120177</td><td>-</td><td>0</td><td>ACAAACAGACAGAGCCCGAT</td><td>CGG</td><td>||||||||[||||||||||||]PAM</td><td>NA</td><td>Intergenic</td><td>NA</td><td>NA</td></tr>
<tr><td>T24953</td><td>AGATGAACGCCATGAACAACAGG</td><td>1000</td><td></td><td></td><td></td><td></td><td></td><td></td><td></td><td></td><td></td></tr>
<tr><td>Oligo fwd</td><td>CACCAGATGAACGCCATGAACAAC</td><td></td><td></td><td></td><td></td><td></td><td></td><td></td><td></td><td></td><td></td></tr>
<tr><td>Oligo rev</td><td>AAACGTTGTTCATGGCGTTCATCT</td><td></td><td></td><td></td><td></td><td></td><td></td><td></td><td></td><td></td><td></td></tr>
<tr><td>Chromosome</td><td>start</td><td>end</td><td>strand</td><td>MM</td><td>target_seq</td><td>PAM</td><td>alignment</td><td>distance</td><td>position</td><td>gene name</td><td>gene id</td></tr>
<tr><td>GU305914.1</td><td>120127</td><td>120149</td><td>-</td><td>0</td><td>AGATGAACGCCATGAACAAC</td><td>AGG</td><td>||||||||[||||||||||||]PAM</td><td>NA</td><td>Intergenic</td><td>NA</td><td>NA</td></tr>
<tr><td>T24954</td><td>GACCGAAGTGATCAATCCCTTGG</td><td>1000</td><td></td><td></td><td></td><td></td><td></td><td></td><td></td><td></td><td></td></tr>
<tr><td>Oligo fwd</td><td>CACCGACCGAAGTGATCAATCCCT</td><td></td><td></td><td></td><td></td><td></td><td></td><td></td><td></td><td></td><td></td></tr>
<tr><td>Oligo rev</td><td>AAACAGGGATTGATCACTTCGGTC</td><td></td><td></td><td></td><td></td><td></td><td></td><td></td><td></td><td></td><td></td></tr>
<tr><td>Chromosome</td><td>start</td><td>end</td><td>strand</td><td>MM</td><td>target_seq</td><td>PAM</td><td>alignment</td><td>distance</td><td>position</td><td>gene name</td><td>gene id</td></tr>
<tr><td>GU305914.1</td><td>120105</td><td>120127</td><td>-</td><td>0</td><td>GACCGAAGTGATCAATCCCT</td><td>TGG</td><td>||||||||[||||||||||||]PAM</td><td>NA</td><td>Intergenic</td><td>NA</td><td>NA</td></tr>
<tr><td>T24955</td><td>GGACTCCCGCACCTCGTCGATGG</td><td>1000</td><td></td><td></td><td></td><td></td><td></td><td></td><td></td><td></td><td></td></tr>
<tr><td>Oligo fwd</td><td>CACCGGACTCCCGCACCTCGTCGA</td><td></td><td></td><td></td><td></td><td></td><td></td><td></td><td></td><td></td><td></td></tr>
<tr><td>Oligo rev</td><td>AAACTCGACGAGGTGCGGGAGTCC</td><td></td><td></td><td></td><td></td><td></td><td></td><td></td><td></td><td></td><td></td></tr>
<tr><td>Chromosome</td><td>start</td><td>end</td><td>strand</td><td>MM</td><td>target_seq</td><td>PAM</td><td>alignment</td><td>distance</td><td>position</td><td>gene name</td><td>gene id</td></tr>
<tr><td>GU305914.1</td><td>120084</td><td>120106</td><td>-</td><td>0</td><td>GGACTCCCGCACCTCGTCGA</td><td>TGG</td><td>||||||||[||||||||||||]PAM</td><td>NA</td><td>Intergenic</td><td>NA</td><td>NA</td></tr>
<tr><td>T24956</td><td>CTCCCGCACCTCGTCGATGGTGG</td><td>1000</td><td></td><td></td><td></td><td></td><td></td><td></td><td></td><td></td><td></td></tr>
<tr><td>Oligo fwd</td><td>CACCCTCCCGCACCTCGTCGATGG</td><td></td><td></td><td></td><td></td><td></td><td></td><td></td><td></td><td></td><td></td></tr>
<tr><td>Oligo rev</td><td>AAACCCATCGACGAGGTGCGGGAG</td><td></td><td></td><td></td><td></td><td></td><td></td><td></td><td></td><td></td><td></td></tr>
<tr><td>Chromosome</td><td>start</td><td>end</td><td>strand</td><td>MM</td><td>target_seq</td><td>PAM</td><td>alignment</td><td>distance</td><td>position</td><td>gene name</td><td>gene id</td></tr>
<tr><td>GU305914.1</td><td>120081</td><td>120103</td><td>-</td><td>0</td><td>CTCCCGCACCTCGTCGATGG</td><td>TGG</td><td>||||||||[||||||||||||]PAM</td><td>NA</td><td>Intergenic</td><td>NA</td><td>NA</td></tr>
<tr><td>T24957</td><td>TCCCGCACCTCGTCGATGGTGGG</td><td>1000</td><td></td><td></td><td></td><td></td><td></td><td></td><td></td><td></td><td></td></tr>
<tr><td>Oligo fwd</td><td>CACCTCCCGCACCTCGTCGATGGT</td><td></td><td></td><td></td><td></td><td></td><td></td><td></td><td></td><td></td><td></td></tr>
<tr><td>Oligo rev</td><td>AAACACCATCGACGAGGTGCGGGA</td><td></td><td></td><td></td><td></td><td></td><td></td><td></td><td></td><td></td><td></td></tr>
<tr><td>Chromosome</td><td>start</td><td>end</td><td>strand</td><td>MM</td><td>target_seq</td><td>PAM</td><td>alignment</td><td>distance</td><td>position</td><td>gene name</td><td>gene id</td></tr>
<tr><td>GU305914.1</td><td>120080</td><td>120102</td><td>-</td><td>0</td><td>TCCCGCACCTCGTCGATGGT</td><td>GGG</td><td>||||||||[||||||||||||]PAM</td><td>NA</td><td>Intergenic</td><td>NA</td><td>NA</td></tr>
<tr><td>T24958</td><td>GGGCTCGACGAAGATCCTGCTGG</td><td>1000</td><td></td><td></td><td></td><td></td><td></td><td></td><td></td><td></td><td></td></tr>
<tr><td>Oligo fwd</td><td>CACCGGGCTCGACGAAGATCCTGC</td><td></td><td></td><td></td><td></td><td></td><td></td><td></td><td></td><td></td><td></td></tr>
<tr><td>Oligo rev</td><td>AAACGCAGGATCTTCGTCGAGCCC</td><td></td><td></td><td></td><td></td><td></td><td></td><td></td><td></td><td></td><td></td></tr>
<tr><td>Chromosome</td><td>start</td><td>end</td><td>strand</td><td>MM</td><td>target_seq</td><td>PAM</td><td>alignment</td><td>distance</td><td>position</td><td>gene name</td><td>gene id</td></tr>
<tr><td>GU305914.1</td><td>120060</td><td>120082</td><td>-</td><td>0</td><td>GGGCTCGACGAAGATCCTGC</td><td>TGG</td><td>||||||||[||||||||||||]PAM</td><td>NA</td><td>Intergenic</td><td>NA</td><td>NA</td></tr>
<tr><td>T24959</td><td>TGCTGGCGAGTTTCCTCCAGCGG</td><td>1000</td><td></td><td></td><td></td><td></td><td></td><td></td><td></td><td></td><td></td></tr>
<tr><td>Oligo fwd</td><td>CACCTGCTGGCGAGTTTCCTCCAG</td><td></td><td></td><td></td><td></td><td></td><td></td><td></td><td></td><td></td><td></td></tr>
<tr><td>Oligo rev</td><td>AAACCTGGAGGAAACTCGCCAGCA</td><td></td><td></td><td></td><td></td><td></td><td></td><td></td><td></td><td></td><td></td></tr>
<tr><td>Chromosome</td><td>start</td><td>end</td><td>strand</td><td>MM</td><td>target_seq</td><td>PAM</td><td>alignment</td><td>distance</td><td>position</td><td>gene name</td><td>gene id</td></tr>
<tr><td>GU305914.1</td><td>120043</td><td>120065</td><td>-</td><td>0</td><td>TGCTGGCGAGTTTCCTCCAG</td><td>CGG</td><td>||||||||[||||||||||||]PAM</td><td>NA</td><td>Intergenic</td><td>NA</td><td>NA</td></tr>
<tr><td>T24960</td><td>GCTGGCGAGTTTCCTCCAGCGGG</td><td>1000</td><td></td><td></td><td></td><td></td><td></td><td></td><td></td><td></td><td></td></tr>
<tr><td>Oligo fwd</td><td>CACCGCTGGCGAGTTTCCTCCAGC</td><td></td><td></td><td></td><td></td><td></td><td></td><td></td><td></td><td></td><td></td></tr>
<tr><td>Oligo rev</td><td>AAACGCTGGAGGAAACTCGCCAGC</td><td></td><td></td><td></td><td></td><td></td><td></td><td></td><td></td><td></td><td></td></tr>
<tr><td>Chromosome</td><td>start</td><td>end</td><td>strand</td><td>MM</td><td>target_seq</td><td>PAM</td><td>alignment</td><td>distance</td><td>position</td><td>gene name</td><td>gene id</td></tr>
<tr><td>GU305914.1</td><td>120042</td><td>120064</td><td>-</td><td>0</td><td>GCTGGCGAGTTTCCTCCAGC</td><td>GGG</td><td>||||||||[||||||||||||]PAM</td><td>NA</td><td>Intergenic</td><td>NA</td><td>NA</td></tr>
<tr><td>T24961</td><td>GCGGGTCTTCTTCGAGCCCGAGG</td><td>1000</td><td></td><td></td><td></td><td></td><td></td><td></td><td></td><td></td><td></td></tr>
<tr><td>Oligo fwd</td><td>CACCGCGGGTCTTCTTCGAGCCCG</td><td></td><td></td><td></td><td></td><td></td><td></td><td></td><td></td><td></td><td></td></tr>
<tr><td>Oligo rev</td><td>AAACCGGGCTCGAAGAAGACCCGC</td><td></td><td></td><td></td><td></td><td></td><td></td><td></td><td></td><td></td><td></td></tr>
<tr><td>Chromosome</td><td>start</td><td>end</td><td>strand</td><td>MM</td><td>target_seq</td><td>PAM</td><td>alignment</td><td>distance</td><td>position</td><td>gene name</td><td>gene id</td></tr>
<tr><td>GU305914.1</td><td>120024</td><td>120046</td><td>-</td><td>0</td><td>GCGGGTCTTCTTCGAGCCCG</td><td>AGG</td><td>||||||||[||||||||||||]PAM</td><td>NA</td><td>Intergenic</td><td>NA</td><td>NA</td></tr>
<tr><td>T24962</td><td>GGAGCTGAAAGTGTTCTCGACGG</td><td>1000</td><td></td><td></td><td></td><td></td><td></td><td></td><td></td><td></td><td></td></tr>
<tr><td>Oligo fwd</td><td>CACCGGAGCTGAAAGTGTTCTCGA</td><td></td><td></td><td></td><td></td><td></td><td></td><td></td><td></td><td></td><td></td></tr>
</table>

| | | | | | | | | | | | |
|---|---|---|---|---|---|---|---|---|---|---|---|
| Oligo rev | AAACTCGAGAACACTTTCAGCTCC | | | | | | | | | | |
| Chromosome | start | end | strand | MM | target_seq | PAM | alignment | distance | position | gene name | gene id |
| GU305914.1 | 120003 | 120025 | - | 0 | GGAGCTGAAAGTGTTCTCGA | CGG | \|\|\|\|\|\|\|\|[\|\|\|\|\|\|\|\|\|\|\|\|]PAM | NA | Intergenic | NA | NA |
| T24963 | GAGCTGAAAGTGTTCTCGACGGG | 1000 | | | | | | | | | |
| Oligo fwd | CACCGAGCTGAAAGTGTTCTCGAC | | | | | | | | | | |
| Oligo rev | AAACGTCGAGAACACTTTCAGCTC | | | | | | | | | | |
| Chromosome | start | end | strand | MM | target_seq | PAM | alignment | distance | position | gene name | gene id |
| GU305914.1 | 120002 | 120024 | - | 0 | GAGCTGAAAGTGTTCTCGAC | GGG | \|\|\|\|\|\|\|\|[\|\|\|\|\|\|\|\|\|\|\|\|]PAM | NA | Intergenic | NA | NA |
| T24964 | CCTGTGCGTGTGTCGCGACGAGG | 1000 | | | | | | | | | |
| Oligo fwd | CACCCCTGTGCGTGTGTCGCGACG | | | | | | | | | | |
| Oligo rev | AAACCGTCGCGACACACGCACAGG | | | | | | | | | | |
| Chromosome | start | end | strand | MM | target_seq | PAM | alignment | distance | position | gene name | gene id |
| GU305914.1 | 119949 | 119971 | - | 0 | CCTGTGCGTGTGTCGCGACG | AGG | \|\|\|\|\|\|\|\|[\|\|\|\|\|\|\|\|\|\|\|\|]PAM | NA | Intergenic | NA | NA |
| T24965 | CTGTGCGTGTGTCGCGACGAGGG | 1000 | | | | | | | | | |
| Oligo fwd | CACCCTGTGCGTGTGTCGCGACGA | | | | | | | | | | |
| Oligo rev | AAACTCGTCGCGACACACGCACAG | | | | | | | | | | |
| Chromosome | start | end | strand | MM | target_seq | PAM | alignment | distance | position | gene name | gene id |
| GU305914.1 | 119948 | 119970 | - | 0 | CTGTGCGTGTGTCGCGACGA | GGG | \|\|\|\|\|\|\|\|[\|\|\|\|\|\|\|\|\|\|\|\|]PAM | NA | Intergenic | NA | NA |
| T24966 | GTGTCGCGACGAGGGCATGATGG | 1000 | | | | | | | | | |
| Oligo fwd | CACCGTGTCGCGACGAGGGCATGA | | | | | | | | | | |
| Oligo rev | AAACTCATGCCCTCGTCGCGACAC | | | | | | | | | | |
| Chromosome | start | end | strand | MM | target_seq | PAM | alignment | distance | position | gene name | gene id |
| GU305914.1 | 119940 | 119962 | - | 0 | GTGTCGCGACGAGGGCATGA | TGG | \|\|\|\|\|\|\|\|[\|\|\|\|\|\|\|\|\|\|\|\|]PAM | NA | Intergenic | NA | NA |
| T24967 | GGGCATGATGGTACCCTTCCAGG | 1000 | | | | | | | | | |
| Oligo fwd | CACCGGGCATGATGGTACCCTTCC | | | | | | | | | | |
| Oligo rev | AAACGGAAGGGTACCATCATGCCC | | | | | | | | | | |
| Chromosome | start | end | strand | MM | target_seq | PAM | alignment | distance | position | gene name | gene id |
| GU305914.1 | 119928 | 119950 | - | 0 | GGGCATGATGGTACCCTTCC | AGG | \|\|\|\|\|\|\|\|[\|\|\|\|\|\|\|\|\|\|\|\|]PAM | NA | Intergenic | NA | NA |
| T24968 | TACGCCGTGAACACCCAGTTCGG | 1000 | | | | | | | | | |
| Oligo fwd | CACCTACGCCGTGAACACCCAGTT | | | | | | | | | | |
| Oligo rev | AAACAACTGGGTGTTCACGGCGTA | | | | | | | | | | |
| Chromosome | start | end | strand | MM | target_seq | PAM | alignment | distance | position | gene name | gene id |
| GU305914.1 | 119873 | 119895 | - | 0 | TACGCCGTGAACACCCAGTT | CGG | \|\|\|\|\|\|\|\|[\|\|\|\|\|\|\|\|\|\|\|\|]PAM | NA | Intergenic | NA | NA |
| T24969 | GTGAACACCCAGTTCGGTATCGG | 1000 | | | | | | | | | |
| Oligo fwd | CACCGTGAACACCCAGTTCGGTAT | | | | | | | | | | |
| Oligo rev | AAACATACCGAACTGGGTGTTCAC | | | | | | | | | | |
| Chromosome | start | end | strand | MM | target_seq | PAM | alignment | distance | position | gene name | gene id |
| GU305914.1 | 119867 | 119889 | - | 0 | GTGAACACCCAGTTCGGTAT | CGG | \|\|\|\|\|\|\|\|[\|\|\|\|\|\|\|\|\|\|\|\|]PAM | NA | Intergenic | NA | NA |
| T24970 | TATCGGCTCGCTCCCCGTCCCGG | 1000 | | | | | | | | | |
| Oligo fwd | CACCTATCGGCTCGCTCCCCGTCC | | | | | | | | | | |
| Oligo rev | AAACGGACGGGGAGCGAGCCGATA | | | | | | | | | | |
| Chromosome | start | end | strand | MM | target_seq | PAM | alignment | distance | position | gene name | gene id |
| GU305914.1 | 119850 | 119872 | - | 0 | TATCGGCTCGCTCCCCGTCC | CGG | \|\|\|\|\|\|\|\|[\|\|\|\|\|\|\|\|\|\|\|\|]PAM | NA | Intergenic | NA | NA |
| T24971 | CGGCACAGAACTCCCAGCCGCGG | 1000 | | | | | | | | | |
| Oligo fwd | CACCCGGCACAGAACTCCCAGCCG | | | | | | | | | | |
| Oligo rev | AAACCGGCTGGGAGTTCTGTGCCG | | | | | | | | | | |
| Chromosome | start | end | strand | MM | target_seq | PAM | alignment | distance | position | gene name | gene id |
| GU305914.1 | 119830 | 119852 | - | 0 | CGGCACAGAACTCCCAGCCG | CGG | \|\|\|\|\|\|\|\|[\|\|\|\|\|\|\|\|\|\|\|\|]PAM | NA | Intergenic | NA | NA |
| T24972 | CCCAGCCGCGGAAACAGACTCGG | 1000 | | | | | | | | | |
| Oligo fwd | CACCCCCAGCCGCGGAAACAGACT | | | | | | | | | | |
| Oligo rev | AAACAGTCTGTTTCCGCGGCTGGG | | | | | | | | | | |
| Chromosome | start | end | strand | MM | target_seq | PAM | alignment | distance | position | gene name | gene id |
| GU305914.1 | 119818 | 119840 | - | 0 | CCCAGCCGCGGAAACAGACT | CGG | \|\|\|\|\|\|\|\|[\|\|\|\|\|\|\|\|\|\|\|\|]PAM | NA | Intergenic | NA | NA |
| GU305914.1 | 174975 | 174997 | - | 4 | CCGAGCGGCGGAAACGGACA | CGG | \|\|-\|\|\|-\|[\|\|\|\|\|\|\|-\|\|\|-]PAM | NA | Intergenic | NA | NA |
| T24973 | GCTGAGATCCAACAAGTACCTGG | 1000 | | | | | | | | | |
| Oligo fwd | CACCGCTGAGATCCAACAAGTACC | | | | | | | | | | |
| Oligo rev | AAACGGTACTTGTTGGATCTCAGC | | | | | | | | | | |
| Chromosome | start | end | strand | MM | target_seq | PAM | alignment | distance | position | gene name | gene id |
| GU305914.1 | 119793 | 119815 | - | 0 | GCTGAGATCCAACAAGTACC | TGG | \|\|\|\|\|\|\|\|[\|\|\|\|\|\|\|\|\|\|\|\|]PAM | NA | Intergenic | NA | NA |
| T24974 | CTGAGATCCAACAAGTACCTGGG | 1000 | | | | | | | | | |
| Oligo fwd | CACCCTGAGATCCAACAAGTACCT | | | | | | | | | | |
| Oligo rev | AAACAGGTACTTGTTGGATCTCAG | | | | | | | | | | |
| Chromosome | start | end | strand | MM | target_seq | PAM | alignment | distance | position | gene name | gene id |
| GU305914.1 | 119792 | 119814 | - | 0 | CTGAGATCCAACAAGTACCT | GGG | \|\|\|\|\|\|\|\|[\|\|\|\|\|\|\|\|\|\|\|\|]PAM | NA | Intergenic | NA | NA |
| T24975 | GGGTTTCTGCGTGTTCGCCCTGG | 1000 | | | | | | | | | |
| Oligo fwd | CACCGGGTTTCTGCGTGTTCGCCC | | | | | | | | | | |
| Oligo rev | AAACGGGCGAACACGCAGAAACCC | | | | | | | | | | |
| Chromosome | start | end | strand | MM | target_seq | PAM | alignment | distance | position | gene name | gene id |
| GU305914.1 | 119772 | 119794 | - | 0 | GGGTTTCTGCGTGTTCGCCC | TGG | \|\|\|\|\|\|\|\|[\|\|\|\|\|\|\|\|\|\|\|\|]PAM | NA | Intergenic | NA | NA |
| T24976 | CGCCGCCCATGCTGATCGCCAGG | 1000 | | | | | | | | | |
| Oligo fwd | CACCCGCCGCCCATGCTGATCGCC | | | | | | | | | | |
| Oligo rev | AAACGGCGATCAGCATGGGCGGCG | | | | | | | | | | |
| Chromosome | start | end | strand | MM | target_seq | PAM | alignment | distance | position | gene name | gene id |
| GU305914.1 | 119731 | 119753 | - | 0 | CGCCGCCCATGCTGATCGCC | AGG | \|\|\|\|\|\|\|\|[\|\|\|\|\|\|\|\|\|\|\|\|]PAM | NA | Intergenic | NA | NA |

<table>
<tr><td>T24977</td><td>GCCGCCCATGCTGATCGCCAGGG</td><td></td><td>1000</td><td></td><td></td><td></td><td></td><td></td><td></td><td></td><td></td></tr>
<tr><td>Oligo fwd</td><td>CACCGCCGCCCATGCTGATCGCCA</td><td></td><td></td><td></td><td></td><td></td><td></td><td></td><td></td><td></td><td></td></tr>
<tr><td>Oligo rev</td><td>AAACTGGCGATCAGCATGGGCGGC</td><td></td><td></td><td></td><td></td><td></td><td></td><td></td><td></td><td></td><td></td></tr>
<tr><td>Chromosome</td><td>start</td><td>end</td><td>strand</td><td>MM</td><td>target_seq</td><td>PAM</td><td>alignment</td><td>distance</td><td>position</td><td>gene name</td><td>gene id</td></tr>
<tr><td>GU305914.1</td><td>119730</td><td>119752</td><td>-</td><td>0</td><td>GCCGCCCATGCTGATCGCCA</td><td>GGG</td><td>||||||||[||||||||||||]PAM</td><td>NA</td><td>Intergenic</td><td>NA</td><td>NA</td></tr>
<tr><td>T24978</td><td>GCTGATCGCCAGGGTGAACTCGG</td><td></td><td>1000</td><td></td><td></td><td></td><td></td><td></td><td></td><td></td><td></td></tr>
<tr><td>Oligo fwd</td><td>CACCGCTGATCGCCAGGGTGAACT</td><td></td><td></td><td></td><td></td><td></td><td></td><td></td><td></td><td></td><td></td></tr>
<tr><td>Oligo rev</td><td>AAACAGTTCACCCTGGCGATCAGC</td><td></td><td></td><td></td><td></td><td></td><td></td><td></td><td></td><td></td><td></td></tr>
<tr><td>Chromosome</td><td>start</td><td>end</td><td>strand</td><td>MM</td><td>target_seq</td><td>PAM</td><td>alignment</td><td>distance</td><td>position</td><td>gene name</td><td>gene id</td></tr>
<tr><td>GU305914.1</td><td>119721</td><td>119743</td><td>-</td><td>0</td><td>GCTGATCGCCAGGGTGAACT</td><td>CGG</td><td>||||||||[||||||||||||]PAM</td><td>NA</td><td>Intergenic</td><td>NA</td><td>NA</td></tr>
<tr><td>T24979</td><td>CGCCAGGGTGAACTCGGTGAAGG</td><td></td><td>1000</td><td></td><td></td><td></td><td></td><td></td><td></td><td></td><td></td></tr>
<tr><td>Oligo fwd</td><td>CACCCGCCAGGGTGAACTCGGTGA</td><td></td><td></td><td></td><td></td><td></td><td></td><td></td><td></td><td></td><td></td></tr>
<tr><td>Oligo rev</td><td>AAACTCACCGAGTTCACCCTGGCG</td><td></td><td></td><td></td><td></td><td></td><td></td><td></td><td></td><td></td><td></td></tr>
<tr><td>Chromosome</td><td>start</td><td>end</td><td>strand</td><td>MM</td><td>target_seq</td><td>PAM</td><td>alignment</td><td>distance</td><td>position</td><td>gene name</td><td>gene id</td></tr>
<tr><td>GU305914.1</td><td>119715</td><td>119737</td><td>-</td><td>0</td><td>CGCCAGGGTGAACTCGGTGA</td><td>AGG</td><td>||||||||[||||||||||||]PAM</td><td>NA</td><td>Intergenic</td><td>NA</td><td>NA</td></tr>
<tr><td>T24980</td><td>CAGGGTGAACTCGGTGAAGGTGG</td><td></td><td>1000</td><td></td><td></td><td></td><td></td><td></td><td></td><td></td><td></td></tr>
<tr><td>Oligo fwd</td><td>CACCCAGGGTGAACTCGGTGAAGG</td><td></td><td></td><td></td><td></td><td></td><td></td><td></td><td></td><td></td><td></td></tr>
<tr><td>Oligo rev</td><td>AAACCCTTCACCGAGTTCACCCTG</td><td></td><td></td><td></td><td></td><td></td><td></td><td></td><td></td><td></td><td></td></tr>
<tr><td>Chromosome</td><td>start</td><td>end</td><td>strand</td><td>MM</td><td>target_seq</td><td>PAM</td><td>alignment</td><td>distance</td><td>position</td><td>gene name</td><td>gene id</td></tr>
<tr><td>GU305914.1</td><td>119712</td><td>119734</td><td>-</td><td>0</td><td>CAGGGTGAACTCGGTGAAGG</td><td>TGG</td><td>||||||||[||||||||||||]PAM</td><td>NA</td><td>Intergenic</td><td>NA</td><td>NA</td></tr>
<tr><td>T24981</td><td>CTCGGTGAAGGTGGACGCTTCGG</td><td></td><td>1000</td><td></td><td></td><td></td><td></td><td></td><td></td><td></td><td></td></tr>
<tr><td>Oligo fwd</td><td>CACCCTCGGTGAAGGTGGACGCTT</td><td></td><td></td><td></td><td></td><td></td><td></td><td></td><td></td><td></td><td></td></tr>
<tr><td>Oligo rev</td><td>AAACAAGCGTCCACCTTCACCGAG</td><td></td><td></td><td></td><td></td><td></td><td></td><td></td><td></td><td></td><td></td></tr>
<tr><td>Chromosome</td><td>start</td><td>end</td><td>strand</td><td>MM</td><td>target_seq</td><td>PAM</td><td>alignment</td><td>distance</td><td>position</td><td>gene name</td><td>gene id</td></tr>
<tr><td>GU305914.1</td><td>119703</td><td>119725</td><td>-</td><td>0</td><td>CTCGGTGAAGGTGGACGCTT</td><td>CGG</td><td>||||||||[||||||||||||]PAM</td><td>NA</td><td>Intergenic</td><td>NA</td><td>NA</td></tr>
<tr><td>T24982</td><td>GGTGAAGGTGGACGCTTCGGCGG</td><td></td><td>1000</td><td></td><td></td><td></td><td></td><td></td><td></td><td></td><td></td></tr>
<tr><td>Oligo fwd</td><td>CACCGGTGAAGGTGGACGCTTCGG</td><td></td><td></td><td></td><td></td><td></td><td></td><td></td><td></td><td></td><td></td></tr>
<tr><td>Oligo rev</td><td>AAACCCGAAGCGTCCACCTTCACC</td><td></td><td></td><td></td><td></td><td></td><td></td><td></td><td></td><td></td><td></td></tr>
<tr><td>Chromosome</td><td>start</td><td>end</td><td>strand</td><td>MM</td><td>target_seq</td><td>PAM</td><td>alignment</td><td>distance</td><td>position</td><td>gene name</td><td>gene id</td></tr>
<tr><td>GU305914.1</td><td>119700</td><td>119722</td><td>-</td><td>0</td><td>GGTGAAGGTGGACGCTTCGG</td><td>CGG</td><td>||||||||[||||||||||||]PAM</td><td>NA</td><td>Intergenic</td><td>NA</td><td>NA</td></tr>
<tr><td>T24983</td><td>GGACGCTTCGGCGGTCCGCCAGG</td><td></td><td>1000</td><td></td><td></td><td></td><td></td><td></td><td></td><td></td><td></td></tr>
<tr><td>Oligo fwd</td><td>CACCGGACGCTTCGGCGGTCCGCC</td><td></td><td></td><td></td><td></td><td></td><td></td><td></td><td></td><td></td><td></td></tr>
<tr><td>Oligo rev</td><td>AAACGGCGGACCGCCGAAGCGTCC</td><td></td><td></td><td></td><td></td><td></td><td></td><td></td><td></td><td></td><td></td></tr>
<tr><td>Chromosome</td><td>start</td><td>end</td><td>strand</td><td>MM</td><td>target_seq</td><td>PAM</td><td>alignment</td><td>distance</td><td>position</td><td>gene name</td><td>gene id</td></tr>
<tr><td>GU305914.1</td><td>119691</td><td>119713</td><td>-</td><td>0</td><td>GGACGCTTCGGCGGTCCGCC</td><td>AGG</td><td>||||||||[||||||||||||]PAM</td><td>NA</td><td>Intergenic</td><td>NA</td><td>NA</td></tr>
<tr><td>T24984</td><td>TTCGGCGGTCCGCCAGGCGTCGG</td><td></td><td>1000</td><td></td><td></td><td></td><td></td><td></td><td></td><td></td><td></td></tr>
<tr><td>Oligo fwd</td><td>CACCTTCGGCGGTCCGCCAGGCGT</td><td></td><td></td><td></td><td></td><td></td><td></td><td></td><td></td><td></td><td></td></tr>
<tr><td>Oligo rev</td><td>AAACACGCCTGGCGGACCGCCGAA</td><td></td><td></td><td></td><td></td><td></td><td></td><td></td><td></td><td></td><td></td></tr>
<tr><td>Chromosome</td><td>start</td><td>end</td><td>strand</td><td>MM</td><td>target_seq</td><td>PAM</td><td>alignment</td><td>distance</td><td>position</td><td>gene name</td><td>gene id</td></tr>
<tr><td>GU305914.1</td><td>119685</td><td>119707</td><td>-</td><td>0</td><td>TTCGGCGGTCCGCCAGGCGT</td><td>CGG</td><td>||||||||[||||||||||||]PAM</td><td>NA</td><td>Intergenic</td><td>NA</td><td>NA</td></tr>
<tr><td>T24985</td><td>GGTCCGCCAGGCGTCGGTCTCGG</td><td></td><td>1000</td><td></td><td></td><td></td><td></td><td></td><td></td><td></td><td></td></tr>
<tr><td>Oligo fwd</td><td>CACCGGTCCGCCAGGCGTCGGTCT</td><td></td><td></td><td></td><td></td><td></td><td></td><td></td><td></td><td></td><td></td></tr>
<tr><td>Oligo rev</td><td>AAACAGACCGACGCCTGGCGGACC</td><td></td><td></td><td></td><td></td><td></td><td></td><td></td><td></td><td></td><td></td></tr>
<tr><td>Chromosome</td><td>start</td><td>end</td><td>strand</td><td>MM</td><td>target_seq</td><td>PAM</td><td>alignment</td><td>distance</td><td>position</td><td>gene name</td><td>gene id</td></tr>
<tr><td>GU305914.1</td><td>119679</td><td>119701</td><td>-</td><td>0</td><td>GGTCCGCCAGGCGTCGGTCT</td><td>CGG</td><td>||||||||[||||||||||||]PAM</td><td>NA</td><td>Intergenic</td><td>NA</td><td>NA</td></tr>
<tr><td>T24986</td><td>CTCGGCGACCACCACGCGCACGG</td><td></td><td>1000</td><td></td><td></td><td></td><td></td><td></td><td></td><td></td><td></td></tr>
<tr><td>Oligo fwd</td><td>CACCCTCGGCGACCACCACGCGCA</td><td></td><td></td><td></td><td></td><td></td><td></td><td></td><td></td><td></td><td></td></tr>
<tr><td>Oligo rev</td><td>AAACTGCGCGTGGTGGTCGCCGAG</td><td></td><td></td><td></td><td></td><td></td><td></td><td></td><td></td><td></td><td></td></tr>
<tr><td>Chromosome</td><td>start</td><td>end</td><td>strand</td><td>MM</td><td>target_seq</td><td>PAM</td><td>alignment</td><td>distance</td><td>position</td><td>gene name</td><td>gene id</td></tr>
<tr><td>GU305914.1</td><td>119661</td><td>119683</td><td>-</td><td>0</td><td>CTCGGCGACCACCACGCGCA</td><td>CGG</td><td>||||||||[||||||||||||]PAM</td><td>NA</td><td>Intergenic</td><td>NA</td><td>NA</td></tr>
<tr><td>T24987</td><td>TCGGCGACCACCACGCGCACGGG</td><td></td><td>1000</td><td></td><td></td><td></td><td></td><td></td><td></td><td></td><td></td></tr>
<tr><td>Oligo fwd</td><td>CACCTCGGCGACCACCACGCGCAC</td><td></td><td></td><td></td><td></td><td></td><td></td><td></td><td></td><td></td><td></td></tr>
<tr><td>Oligo rev</td><td>AAACGTGCGCGTGGTGGTCGCCGA</td><td></td><td></td><td></td><td></td><td></td><td></td><td></td><td></td><td></td><td></td></tr>
<tr><td>Chromosome</td><td>start</td><td>end</td><td>strand</td><td>MM</td><td>target_seq</td><td>PAM</td><td>alignment</td><td>distance</td><td>position</td><td>gene name</td><td>gene id</td></tr>
<tr><td>GU305914.1</td><td>119660</td><td>119682</td><td>-</td><td>0</td><td>TCGGCGACCACCACGCGCAC</td><td>GGG</td><td>||||||||[||||||||||||]PAM</td><td>NA</td><td>Intergenic</td><td>NA</td><td>NA</td></tr>
<tr><td>T24988</td><td>CGGCGACCACCACGCGCACGGGG</td><td></td><td>1000</td><td></td><td></td><td></td><td></td><td></td><td></td><td></td><td></td></tr>
<tr><td>Oligo fwd</td><td>CACCCGGCGACCACCACGCGCACG</td><td></td><td></td><td></td><td></td><td></td><td></td><td></td><td></td><td></td><td></td></tr>
<tr><td>Oligo rev</td><td>AAACCGTGCGCGTGGTGGTCGCCG</td><td></td><td></td><td></td><td></td><td></td><td></td><td></td><td></td><td></td><td></td></tr>
<tr><td>Chromosome</td><td>start</td><td>end</td><td>strand</td><td>MM</td><td>target_seq</td><td>PAM</td><td>alignment</td><td>distance</td><td>position</td><td>gene name</td><td>gene id</td></tr>
<tr><td>GU305914.1</td><td>119659</td><td>119681</td><td>-</td><td>0</td><td>CGGCGACCACCACGCGCACG</td><td>GGG</td><td>||||||||[||||||||||||]PAM</td><td>NA</td><td>Intergenic</td><td>NA</td><td>NA</td></tr>
<tr><td>GU305914.1</td><td>106135</td><td>106157</td><td>-</td><td>4</td><td>CGTCGAGCACCCCCCGCACG</td><td>CGG</td><td>||-|||-|[|||-|-||||||]PAM</td><td>NA</td><td>Intergenic</td><td>NA</td><td>NA</td></tr>
<tr><td>GU305914.1</td><td>125156</td><td>125178</td><td>+</td><td>3</td><td>CGTCGGCCACCACGCGCACC</td><td>AGG</td><td>||-||-||[|||||||||||-]PAM</td><td>NA</td><td>Intergenic</td><td>NA</td><td>NA</td></tr>
<tr><td>T24989</td><td>GCACGGGGATGAAGACCGTCAGG</td><td></td><td>1000</td><td></td><td></td><td></td><td></td><td></td><td></td><td></td><td></td></tr>
<tr><td>Oligo fwd</td><td>CACCGCACGGGGATGAAGACCGTC</td><td></td><td></td><td></td><td></td><td></td><td></td><td></td><td></td><td></td><td></td></tr>
<tr><td>Oligo rev</td><td>AAACGACGGTCTTCATCCCCGTGC</td><td></td><td></td><td></td><td></td><td></td><td></td><td></td><td></td><td></td><td></td></tr>
<tr><td>Chromosome</td><td>start</td><td>end</td><td>strand</td><td>MM</td><td>target_seq</td><td>PAM</td><td>alignment</td><td>distance</td><td>position</td><td>gene name</td><td>gene id</td></tr>
<tr><td>GU305914.1</td><td>119644</td><td>119666</td><td>-</td><td>0</td><td>GCACGGGGATGAAGACCGTC</td><td>AGG</td><td>||||||||[||||||||||||]PAM</td><td>NA</td><td>Intergenic</td><td>NA</td><td>NA</td></tr>
<tr><td>T24990</td><td>GATGAAGACCGTCAGGATGACGG</td><td></td><td>1000</td><td></td><td></td><td></td><td></td><td></td><td></td><td></td><td></td></tr>
<tr><td>Oligo fwd</td><td>CACCGATGAAGACCGTCAGGATGA</td><td></td><td></td><td></td><td></td><td></td><td></td><td></td><td></td><td></td><td></td></tr>
<tr><td>Oligo rev</td><td>AAACTCATCCTGACGGTCTTCATC</td><td></td><td></td><td></td><td></td><td></td><td></td><td></td><td></td><td></td><td></td></tr>
<tr><td>Chromosome</td><td>start</td><td>end</td><td>strand</td><td>MM</td><td>target_seq</td><td>PAM</td><td>alignment</td><td>distance</td><td>position</td><td>gene name</td><td>gene id</td></tr>
<tr><td>GU305914.1</td><td>119637</td><td>119659</td><td>-</td><td>0</td><td>GATGAAGACCGTCAGGATGA</td><td>CGG</td><td>||||||||[||||||||||||]PAM</td><td>NA</td><td>Intergenic</td><td>NA</td><td>NA</td></tr>
<tr><td>T24991</td><td>CGTCAGGATGACGGTCCACAAGG</td><td></td><td>1000</td><td></td><td></td><td></td><td></td><td></td><td></td><td></td><td></td></tr>
<tr><td>Oligo fwd</td><td>CACCCGTCAGGATGACGGTCCACA</td><td></td><td></td><td></td><td></td><td></td><td></td><td></td><td></td><td></td><td></td></tr>
<tr><td>Oligo rev</td><td>AAACTGTGGACCGTCATCCTGACG</td><td></td><td></td><td></td><td></td><td></td><td></td><td></td><td></td><td></td><td></td></tr>
</table>

Chromosome start end strand MM target_seq PAM alignment distance position gene name gene id
GU305914.1 119628 119650 - 0 CGTCAGGATGACGGTCCACA AGG ||||||||[||||||||||||]PAM NA Intergenic NA NA

T24992 TGACGGTCCACAAGGTGCGCTGG 1000
Oligo fwd CACCTGACGGTCCACAAGGTGCGC
Oligo rev AAACGCGCACCTTGTGGACCGTCA
Chromosome start end strand MM target_seq PAM alignment distance position gene name gene id
GU305914.1 119620 119642 - 0 TGACGGTCCACAAGGTGCGC TGG ||||||||[||||||||||||]PAM NA Intergenic NA NA

T24993 GACGGTCCACAAGGTGCGCTGGG 1000
Oligo fwd CACCGACGGTCCACAAGGTGCGCT
Oligo rev AAACAGCGCACCTTGTGGACCGTC
Chromosome start end strand MM target_seq PAM alignment distance position gene name gene id
GU305914.1 119619 119641 - 0 GACGGTCCACAAGGTGCGCT GGG ||||||||[||||||||||||]PAM NA Intergenic NA NA

T24994 GGTGCGCTGGGCCAACGATCCGG 1000
Oligo fwd CACCGGTGCGCTGGGCCAACGATC
Oligo rev AAACGATCGTTGGCCCAGCGCACC
Chromosome start end strand MM target_seq PAM alignment distance position gene name gene id
GU305914.1 119607 119629 - 0 GGTGCGCTGGGCCAACGATC CGG ||||||||[||||||||||||]PAM NA Intergenic NA NA

T24995 GGACTCCAAACGCTTCAAGAAGG 1000
Oligo fwd CACCGGACTCCAAACGCTTCAAGA
Oligo rev AAACTCTTGAAGCGTTTGGAGTCC
Chromosome start end strand MM target_seq PAM alignment distance position gene name gene id
GU305914.1 119586 119608 - 0 GGACTCCAAACGCTTCAAGA AGG ||||||||[||||||||||||]PAM NA Intergenic NA NA

T24996 CAAGAAGGCCATCGTGTCTCTGG 1000
Oligo fwd CACCCAAGAAGGCCATCGTGTCTC
Oligo rev AAACGAGACACGATGGCCTTCTTG
Chromosome start end strand MM target_seq PAM alignment distance position gene name gene id
GU305914.1 119571 119593 - 0 CAAGAAGGCCATCGTGTCTC TGG ||||||||[||||||||||||]PAM NA Intergenic NA NA

T24997 TCGTGTCTCTGGAGCTGCCCAGG 1000
Oligo fwd CACCTCGTGTCTCTGGAGCTGCCC
Oligo rev AAACGGGCAGCTCCAGAGACACGA
Chromosome start end strand MM target_seq PAM alignment distance position gene name gene id
GU305914.1 119560 119582 - 0 TCGTGTCTCTGGAGCTGCCC AGG ||||||||[||||||||||||]PAM NA Intergenic NA NA

T24998 CGTGTCTCTGGAGCTGCCCAGGG 1000
Oligo fwd CACCCGTGTCTCTGGAGCTGCCCA
Oligo rev AAACTGGGCAGCTCCAGAGACACG
Chromosome start end strand MM target_seq PAM alignment distance position gene name gene id
GU305914.1 119559 119581 - 0 CGTGTCTCTGGAGCTGCCCA GGG ||||||||[||||||||||||]PAM NA Intergenic NA NA

T24999 CCTCGACCCGAAGACCTGCTCGG 1000
Oligo fwd CACCCCTCGACCCGAAGACCTGCT
Oligo rev AAACAGCAGGTCTTCGGGTCGAGG
Chromosome start end strand MM target_seq PAM alignment distance position gene name gene id
GU305914.1 119523 119545 - 0 CCTCGACCCGAAGACCTGCT CGG ||||||||[||||||||||||]PAM NA Intergenic NA NA

T25000 CGTCAATTTCTACCAGATGACGG 1000
Oligo fwd CACCCGTCAATTTCTACCAGATGA
Oligo rev AAACTCATCTGGTAGAAATTGACG
Chromosome start end strand MM target_seq PAM alignment distance position gene name gene id
GU305914.1 119493 119515 - 0 CGTCAATTTCTACCAGATGA CGG ||||||||[||||||||||||]PAM NA Intergenic NA NA

T25001 CGACGATCTACCTGACGCTGCGG 1000
Oligo fwd CACCCGACGATCTACCTGACGCTG
Oligo rev AAACCAGCGTCAGGTAGATCGTCG
Chromosome start end strand MM target_seq PAM alignment distance position gene name gene id
GU305914.1 119455 119477 - 0 CGACGATCTACCTGACGCTG CGG ||||||||[||||||||||||]PAM NA Intergenic NA NA

T25002 CCTGACGCTGCGGTACGCGAAGG 1000
Oligo fwd CACCCCTGACGCTGCGGTACGCGA
Oligo rev AAACTCGCGTACCGCAGCGTCAGG
Chromosome start end strand MM target_seq PAM alignment distance position gene name gene id
GU305914.1 119445 119467 - 0 CCTGACGCTGCGGTACGCGA AGG ||||||||[||||||||||||]PAM NA Intergenic NA NA

T25003 CTGCGGTACGCGAAGGCCAACGG 1000
Oligo fwd CACCCTGCGGTACGCGAAGGCCAA
Oligo rev AAACTTGGCCTTCGCGTACCGCAG
Chromosome start end strand MM target_seq PAM alignment distance position gene name gene id
GU305914.1 119438 119460 - 0 CTGCGGTACGCGAAGGCCAA CGG ||||||||[||||||||||||]PAM NA Intergenic NA NA

T25004 CACGCCTCCAGACTCGCTCACGG 1000
Oligo fwd CACCCACGCCTCCAGACTCGCTCA
Oligo rev AAACTGAGCGAGTCTGGAGGCGTG
Chromosome start end strand MM target_seq PAM alignment distance position gene name gene id
GU305914.1 119415 119437 - 0 CACGCCTCCAGACTCGCTCA CGG ||||||||[||||||||||||]PAM NA Intergenic NA NA

T25005 AGACTCGCTCACGGCCATCCTGG 1000
Oligo fwd CACCAGACTCGCTCACGGCCATCC
Oligo rev AAACGGATGGCCGTGAGCGAGTCT
Chromosome start end strand MM target_seq PAM alignment distance position gene name gene id
GU305914.1 119406 119428 - 0 AGACTCGCTCACGGCCATCC TGG ||||||||[||||||||||||]PAM NA Intergenic NA NA

T25006 CTCGCTCACGGCCATCCTGGCGG 1000
Oligo fwd CACCCTCGCTCACGGCCATCCTGG

<table>
<tr><td>Oligo rev</td><td>AAACCCAGGATGGCCGTGAGCGAG</td><td></td><td></td><td></td><td></td><td></td><td></td><td></td><td></td><td></td><td></td></tr>
<tr><td>Chromosome</td><td>start</td><td>end</td><td>strand</td><td>MM</td><td>target_seq</td><td>PAM</td><td>alignment</td><td>distance</td><td>position</td><td>gene name</td><td>gene id</td></tr>
<tr><td>GU305914.1</td><td>119403</td><td>119425</td><td>-</td><td>0</td><td>CTCGCTCACGGCCATCCTGG</td><td>CGG</td><td>||||||||[||||||||||||]PAM</td><td>NA</td><td>Intergenic</td><td>NA</td><td>NA</td></tr>
<tr><td>T25007</td><td>GCTCACGGCCATCCTGGCGGTGG</td><td>1000</td><td></td><td></td><td></td><td></td><td></td><td></td><td></td><td></td><td></td></tr>
<tr><td>Oligo fwd</td><td>CACCGCTCACGGCCATCCTGGCGG</td><td></td><td></td><td></td><td></td><td></td><td></td><td></td><td></td><td></td><td></td></tr>
<tr><td>Oligo rev</td><td>AAACCCGCCAGGATGGCCGTGAGC</td><td></td><td></td><td></td><td></td><td></td><td></td><td></td><td></td><td></td><td></td></tr>
<tr><td>Chromosome</td><td>start</td><td>end</td><td>strand</td><td>MM</td><td>target_seq</td><td>PAM</td><td>alignment</td><td>distance</td><td>position</td><td>gene name</td><td>gene id</td></tr>
<tr><td>GU305914.1</td><td>119400</td><td>119422</td><td>-</td><td>0</td><td>GCTCACGGCCATCCTGGCGG</td><td>TGG</td><td>||||||||[||||||||||||]PAM</td><td>NA</td><td>Intergenic</td><td>NA</td><td>NA</td></tr>
<tr><td>T25008</td><td>CTCACGGCCATCCTGGCGGTGGG</td><td>1000</td><td></td><td></td><td></td><td></td><td></td><td></td><td></td><td></td><td></td></tr>
<tr><td>Oligo fwd</td><td>CACCCTCACGGCCATCCTGGCGGT</td><td></td><td></td><td></td><td></td><td></td><td></td><td></td><td></td><td></td><td></td></tr>
<tr><td>Oligo rev</td><td>AAACACCGCCAGGATGGCCGTGAG</td><td></td><td></td><td></td><td></td><td></td><td></td><td></td><td></td><td></td><td></td></tr>
<tr><td>Chromosome</td><td>start</td><td>end</td><td>strand</td><td>MM</td><td>target_seq</td><td>PAM</td><td>alignment</td><td>distance</td><td>position</td><td>gene name</td><td>gene id</td></tr>
<tr><td>GU305914.1</td><td>119399</td><td>119421</td><td>-</td><td>0</td><td>CTCACGGCCATCCTGGCGGT</td><td>GGG</td><td>||||||||[||||||||||||]PAM</td><td>NA</td><td>Intergenic</td><td>NA</td><td>NA</td></tr>
<tr><td>T25009</td><td>GGTGGGTAACCGCGCGAAGAAGG</td><td>1000</td><td></td><td></td><td></td><td></td><td></td><td></td><td></td><td></td><td></td></tr>
<tr><td>Oligo fwd</td><td>CACCGGTGGGTAACCGCGCGAAGA</td><td></td><td></td><td></td><td></td><td></td><td></td><td></td><td></td><td></td><td></td></tr>
<tr><td>Oligo rev</td><td>AAACTCTTCGCGCGGTTACCCACC</td><td></td><td></td><td></td><td></td><td></td><td></td><td></td><td></td><td></td><td></td></tr>
<tr><td>Chromosome</td><td>start</td><td>end</td><td>strand</td><td>MM</td><td>target_seq</td><td>PAM</td><td>alignment</td><td>distance</td><td>position</td><td>gene name</td><td>gene id</td></tr>
<tr><td>GU305914.1</td><td>119382</td><td>119404</td><td>-</td><td>0</td><td>GGTGGGTAACCGCGCGAAGA</td><td>AGG</td><td>||||||||[||||||||||||]PAM</td><td>NA</td><td>Intergenic</td><td>NA</td><td>NA</td></tr>
<tr><td>T25010</td><td>GGTGAACCTGAAGCGTAGCCTGG</td><td>1000</td><td></td><td></td><td></td><td></td><td></td><td></td><td></td><td></td><td></td></tr>
<tr><td>Oligo fwd</td><td>CACCGGTGAACCTGAAGCGTAGCC</td><td></td><td></td><td></td><td></td><td></td><td></td><td></td><td></td><td></td><td></td></tr>
<tr><td>Oligo rev</td><td>AAACGGCTACGCTTCAGGTTCACC</td><td></td><td></td><td></td><td></td><td></td><td></td><td></td><td></td><td></td><td></td></tr>
<tr><td>Chromosome</td><td>start</td><td>end</td><td>strand</td><td>MM</td><td>target_seq</td><td>PAM</td><td>alignment</td><td>distance</td><td>position</td><td>gene name</td><td>gene id</td></tr>
<tr><td>GU305914.1</td><td>119361</td><td>119383</td><td>-</td><td>0</td><td>GGTGAACCTGAAGCGTAGCC</td><td>TGG</td><td>||||||||[||||||||||||]PAM</td><td>NA</td><td>Intergenic</td><td>NA</td><td>NA</td></tr>
<tr><td>T25011</td><td>CCCTTTTCTGAAACCGCTCCCGG</td><td>1000</td><td></td><td></td><td></td><td></td><td></td><td></td><td></td><td></td><td></td></tr>
<tr><td>Oligo fwd</td><td>CACCCCCTTTTCTGAAACCGCTCC</td><td></td><td></td><td></td><td></td><td></td><td></td><td></td><td></td><td></td><td></td></tr>
<tr><td>Oligo rev</td><td>AAACGGAGCGGTTTCAGAAAAGGG</td><td></td><td></td><td></td><td></td><td></td><td></td><td></td><td></td><td></td><td></td></tr>
<tr><td>Chromosome</td><td>start</td><td>end</td><td>strand</td><td>MM</td><td>target_seq</td><td>PAM</td><td>alignment</td><td>distance</td><td>position</td><td>gene name</td><td>gene id</td></tr>
<tr><td>GU305914.1</td><td>119337</td><td>119359</td><td>-</td><td>0</td><td>CCCTTTTCTGAAACCGCTCC</td><td>CGG</td><td>||||||||[||||||||||||]PAM</td><td>NA</td><td>Intergenic</td><td>NA</td><td>NA</td></tr>
<tr><td>T25012</td><td>CTGAAACCGCTCCCGGTGAACGG</td><td>1000</td><td></td><td></td><td></td><td></td><td></td><td></td><td></td><td></td><td></td></tr>
<tr><td>Oligo fwd</td><td>CACCCTGAAACCGCTCCCGGTGAA</td><td></td><td></td><td></td><td></td><td></td><td></td><td></td><td></td><td></td><td></td></tr>
<tr><td>Oligo rev</td><td>AAACTTCACCGGGAGCGGTTTCAG</td><td></td><td></td><td></td><td></td><td></td><td></td><td></td><td></td><td></td><td></td></tr>
<tr><td>Chromosome</td><td>start</td><td>end</td><td>strand</td><td>MM</td><td>target_seq</td><td>PAM</td><td>alignment</td><td>distance</td><td>position</td><td>gene name</td><td>gene id</td></tr>
<tr><td>GU305914.1</td><td>119330</td><td>119352</td><td>-</td><td>0</td><td>CTGAAACCGCTCCCGGTGAA</td><td>CGG</td><td>||||||||[||||||||||||]PAM</td><td>NA</td><td>Intergenic</td><td>NA</td><td>NA</td></tr>
<tr><td>T25013</td><td>TGAAACCGCTCCCGGTGAACGGG</td><td>1000</td><td></td><td></td><td></td><td></td><td></td><td></td><td></td><td></td><td></td></tr>
<tr><td>Oligo fwd</td><td>CACCTGAAACCGCTCCCGGTGAAC</td><td></td><td></td><td></td><td></td><td></td><td></td><td></td><td></td><td></td><td></td></tr>
<tr><td>Oligo rev</td><td>AAACGTTCACCGGGAGCGGTTTCA</td><td></td><td></td><td></td><td></td><td></td><td></td><td></td><td></td><td></td><td></td></tr>
<tr><td>Chromosome</td><td>start</td><td>end</td><td>strand</td><td>MM</td><td>target_seq</td><td>PAM</td><td>alignment</td><td>distance</td><td>position</td><td>gene name</td><td>gene id</td></tr>
<tr><td>GU305914.1</td><td>119329</td><td>119351</td><td>-</td><td>0</td><td>TGAAACCGCTCCCGGTGAAC</td><td>GGG</td><td>||||||||[||||||||||||]PAM</td><td>NA</td><td>Intergenic</td><td>NA</td><td>NA</td></tr>
<tr><td>T25014</td><td>CCCGGTGAACGGGTTCCGCGTGG</td><td>1000</td><td></td><td></td><td></td><td></td><td></td><td></td><td></td><td></td><td></td></tr>
<tr><td>Oligo fwd</td><td>CACCCCCGGTGAACGGGTTCCGCG</td><td></td><td></td><td></td><td></td><td></td><td></td><td></td><td></td><td></td><td></td></tr>
<tr><td>Oligo rev</td><td>AAACCGCGGAACCCGTTCACCGGG</td><td></td><td></td><td></td><td></td><td></td><td></td><td></td><td></td><td></td><td></td></tr>
<tr><td>Chromosome</td><td>start</td><td>end</td><td>strand</td><td>MM</td><td>target_seq</td><td>PAM</td><td>alignment</td><td>distance</td><td>position</td><td>gene name</td><td>gene id</td></tr>
<tr><td>GU305914.1</td><td>119319</td><td>119341</td><td>-</td><td>0</td><td>CCCGGTGAACGGGTTCCGCG</td><td>TGG</td><td>||||||||[||||||||||||]PAM</td><td>NA</td><td>Intergenic</td><td>NA</td><td>NA</td></tr>
<tr><td>T25015</td><td>TGCCGCGCACCGTGCAGCTGAGG</td><td>1000</td><td></td><td></td><td></td><td></td><td></td><td></td><td></td><td></td><td></td></tr>
<tr><td>Oligo fwd</td><td>CACCTGCCGCGCACCGTGCAGCTG</td><td></td><td></td><td></td><td></td><td></td><td></td><td></td><td></td><td></td><td></td></tr>
<tr><td>Oligo rev</td><td>AAACCAGCTGCACGGTGCGCGGCA</td><td></td><td></td><td></td><td></td><td></td><td></td><td></td><td></td><td></td><td></td></tr>
<tr><td>Chromosome</td><td>start</td><td>end</td><td>strand</td><td>MM</td><td>target_seq</td><td>PAM</td><td>alignment</td><td>distance</td><td>position</td><td>gene name</td><td>gene id</td></tr>
<tr><td>GU305914.1</td><td>119293</td><td>119315</td><td>-</td><td>0</td><td>TGCCGCGCACCGTGCAGCTG</td><td>AGG</td><td>||||||||[||||||||||||]PAM</td><td>NA</td><td>Intergenic</td><td>NA</td><td>NA</td></tr>
<tr><td>T25016</td><td>GCGCACCGTGCAGCTGAGGACGG</td><td>1000</td><td></td><td></td><td></td><td></td><td></td><td></td><td></td><td></td><td></td></tr>
<tr><td>Oligo fwd</td><td>CACCGCGCACCGTGCAGCTGAGGA</td><td></td><td></td><td></td><td></td><td></td><td></td><td></td><td></td><td></td><td></td></tr>
<tr><td>Oligo rev</td><td>AAACTCCTCAGCTGCACGGTGCGC</td><td></td><td></td><td></td><td></td><td></td><td></td><td></td><td></td><td></td><td></td></tr>
<tr><td>Chromosome</td><td>start</td><td>end</td><td>strand</td><td>MM</td><td>target_seq</td><td>PAM</td><td>alignment</td><td>distance</td><td>position</td><td>gene name</td><td>gene id</td></tr>
<tr><td>GU305914.1</td><td>119289</td><td>119311</td><td>-</td><td>0</td><td>GCGCACCGTGCAGCTGAGGA</td><td>CGG</td><td>||||||||[||||||||||||]PAM</td><td>NA</td><td>Intergenic</td><td>NA</td><td>NA</td></tr>
<tr><td>T25017</td><td>CGCACCGTGCAGCTGAGGACGGG</td><td>1000</td><td></td><td></td><td></td><td></td><td></td><td></td><td></td><td></td><td></td></tr>
<tr><td>Oligo fwd</td><td>CACCCGCACCGTGCAGCTGAGGAC</td><td></td><td></td><td></td><td></td><td></td><td></td><td></td><td></td><td></td><td></td></tr>
<tr><td>Oligo rev</td><td>AAACGTCCTCAGCTGCACGGTGCG</td><td></td><td></td><td></td><td></td><td></td><td></td><td></td><td></td><td></td><td></td></tr>
<tr><td>Chromosome</td><td>start</td><td>end</td><td>strand</td><td>MM</td><td>target_seq</td><td>PAM</td><td>alignment</td><td>distance</td><td>position</td><td>gene name</td><td>gene id</td></tr>
<tr><td>GU305914.1</td><td>119288</td><td>119310</td><td>-</td><td>0</td><td>CGCACCGTGCAGCTGAGGAC</td><td>GGG</td><td>||||||||[||||||||||||]PAM</td><td>NA</td><td>Intergenic</td><td>NA</td><td>NA</td></tr>
<tr><td>T25018</td><td>GTACCACGCGTCCAACCACGCGG</td><td>1000</td><td></td><td></td><td></td><td></td><td></td><td></td><td></td><td></td><td></td></tr>
<tr><td>Oligo fwd</td><td>CACCGTACCACGCGTCCAACCACG</td><td></td><td></td><td></td><td></td><td></td><td></td><td></td><td></td><td></td><td></td></tr>
<tr><td>Oligo rev</td><td>AAACCGTGGTTGGACGCGTGGTAC</td><td></td><td></td><td></td><td></td><td></td><td></td><td></td><td></td><td></td><td></td></tr>
<tr><td>Chromosome</td><td>start</td><td>end</td><td>strand</td><td>MM</td><td>target_seq</td><td>PAM</td><td>alignment</td><td>distance</td><td>position</td><td>gene name</td><td>gene id</td></tr>
<tr><td>GU305914.1</td><td>119235</td><td>119257</td><td>-</td><td>0</td><td>GTACCACGCGTCCAACCACG</td><td>CGG</td><td>||||||||[||||||||||||]PAM</td><td>NA</td><td>Intergenic</td><td>NA</td><td>NA</td></tr>
<tr><td>T25019</td><td>CTTCATCCCTTACCTGATCCCGG</td><td>1000</td><td></td><td></td><td></td><td></td><td></td><td></td><td></td><td></td><td></td></tr>
<tr><td>Oligo fwd</td><td>CACCCTTCATCCCTTACCTGATCC</td><td></td><td></td><td></td><td></td><td></td><td></td><td></td><td></td><td></td><td></td></tr>
<tr><td>Oligo rev</td><td>AAACGGATCAGGTAAGGGATGAAG</td><td></td><td></td><td></td><td></td><td></td><td></td><td></td><td></td><td></td><td></td></tr>
<tr><td>Chromosome</td><td>start</td><td>end</td><td>strand</td><td>MM</td><td>target_seq</td><td>PAM</td><td>alignment</td><td>distance</td><td>position</td><td>gene name</td><td>gene id</td></tr>
<tr><td>GU305914.1</td><td>119208</td><td>119230</td><td>-</td><td>0</td><td>CTTCATCCCTTACCTGATCC</td><td>CGG</td><td>||||||||[||||||||||||]PAM</td><td>NA</td><td>Intergenic</td><td>NA</td><td>NA</td></tr>
<tr><td>T25020</td><td>TTCATCCCTTACCTGATCCCGGG</td><td>1000</td><td></td><td></td><td></td><td></td><td></td><td></td><td></td><td></td><td></td></tr>
<tr><td>Oligo fwd</td><td>CACCTTCATCCCTTACCTGATCCC</td><td></td><td></td><td></td><td></td><td></td><td></td><td></td><td></td><td></td><td></td></tr>
<tr><td>Oligo rev</td><td>AAACGGGATCAGGTAAGGGATGAA</td><td></td><td></td><td></td><td></td><td></td><td></td><td></td><td></td><td></td><td></td></tr>
<tr><td>Chromosome</td><td>start</td><td>end</td><td>strand</td><td>MM</td><td>target_seq</td><td>PAM</td><td>alignment</td><td>distance</td><td>position</td><td>gene name</td><td>gene id</td></tr>
<tr><td>GU305914.1</td><td>119207</td><td>119229</td><td>-</td><td>0</td><td>TTCATCCCTTACCTGATCCC</td><td>GGG</td><td>||||||||[||||||||||||]PAM</td><td>NA</td><td>Intergenic</td><td>NA</td><td>NA</td></tr>
<tr><td>T25021</td><td>CCCTTACCTGATCCCGGGTCTGG</td><td>1000</td><td></td><td></td><td></td><td></td><td></td><td></td><td></td><td></td><td></td></tr>
</table>

<table>
<tr><td>Oligo fwd</td><td>CACCCCCTTACCTGATCCCGGGTC</td><td></td><td></td><td></td><td></td><td></td><td></td><td></td><td></td><td></td><td></td></tr>
<tr><td>Oligo rev</td><td>AAACGACCCGGGATCAGGTAAGGG</td><td></td><td></td><td></td><td></td><td></td><td></td><td></td><td></td><td></td><td></td></tr>
<tr><td>Chromosome</td><td>start</td><td>end</td><td>strand</td><td>MM</td><td>target_seq</td><td>PAM</td><td>alignment</td><td>distance</td><td>position</td><td>gene name</td><td>gene id</td></tr>
<tr><td>GU305914.1</td><td>119202</td><td>119224</td><td>-</td><td>0</td><td>CCCTTACCTGATCCCGGGTC</td><td>TGG</td><td>||||||||[||||||||||||]PAM</td><td>NA</td><td>Intergenic</td><td>NA</td><td>NA</td></tr>
<tr><td></td><td></td><td></td><td></td><td></td><td></td><td></td><td></td><td></td><td></td><td></td><td></td></tr>
<tr><td>T25022</td><td>TGGACATCCACCCGAGCGTGTGG</td><td>1000</td><td></td><td></td><td></td><td></td><td></td><td></td><td></td><td></td><td></td></tr>
<tr><td>Oligo fwd</td><td>CACCTGGACATCCACCCGAGCGTG</td><td></td><td></td><td></td><td></td><td></td><td></td><td></td><td></td><td></td><td></td></tr>
<tr><td>Oligo rev</td><td>AAACCACGCTCGGGTGGATGTCCA</td><td></td><td></td><td></td><td></td><td></td><td></td><td></td><td></td><td></td><td></td></tr>
<tr><td>Chromosome</td><td>start</td><td>end</td><td>strand</td><td>MM</td><td>target_seq</td><td>PAM</td><td>alignment</td><td>distance</td><td>position</td><td>gene name</td><td>gene id</td></tr>
<tr><td>GU305914.1</td><td>119182</td><td>119204</td><td>-</td><td>0</td><td>TGGACATCCACCCGAGCGTG</td><td>TGG</td><td>||||||||[||||||||||||]PAM</td><td>NA</td><td>Intergenic</td><td>NA</td><td>NA</td></tr>
<tr><td></td><td></td><td></td><td></td><td></td><td></td><td></td><td></td><td></td><td></td><td></td><td></td></tr>
<tr><td>T25023</td><td>AACCTGTCGTTCACGATCACCGG</td><td>1000</td><td></td><td></td><td></td><td></td><td></td><td></td><td></td><td></td><td></td></tr>
<tr><td>Oligo fwd</td><td>CACCAACCTGTCGTTCACGATCAC</td><td></td><td></td><td></td><td></td><td></td><td></td><td></td><td></td><td></td><td></td></tr>
<tr><td>Oligo rev</td><td>AAACGTGATCGTGAACGACAGGTT</td><td></td><td></td><td></td><td></td><td></td><td></td><td></td><td></td><td></td><td></td></tr>
<tr><td>Chromosome</td><td>start</td><td>end</td><td>strand</td><td>MM</td><td>target_seq</td><td>PAM</td><td>alignment</td><td>distance</td><td>position</td><td>gene name</td><td>gene id</td></tr>
<tr><td>GU305914.1</td><td>119147</td><td>119169</td><td>-</td><td>0</td><td>AACCTGTCGTTCACGATCAC</td><td>CGG</td><td>||||||||[||||||||||||]PAM</td><td>NA</td><td>Intergenic</td><td>NA</td><td>NA</td></tr>
<tr><td>GU305914.1</td><td>46608</td><td>46630</td><td>-</td><td>4</td><td>AGCATGTCGTTCAGGATCTC</td><td>GGG</td><td>|-|-||||[|||||-||||-|]PAM</td><td>NA</td><td>Intergenic</td><td>NA</td><td>NA</td></tr>
<tr><td></td><td></td><td></td><td></td><td></td><td></td><td></td><td></td><td></td><td></td><td></td><td></td></tr>
<tr><td>T25024</td><td>GTTCACGATCACCGGCATGAAGG</td><td>1000</td><td></td><td></td><td></td><td></td><td></td><td></td><td></td><td></td><td></td></tr>
<tr><td>Oligo fwd</td><td>CACCGTTCACGATCACCGGCATGA</td><td></td><td></td><td></td><td></td><td></td><td></td><td></td><td></td><td></td><td></td></tr>
<tr><td>Oligo rev</td><td>AAACTCATGCCGGTGATCGTGAAC</td><td></td><td></td><td></td><td></td><td></td><td></td><td></td><td></td><td></td><td></td></tr>
<tr><td>Chromosome</td><td>start</td><td>end</td><td>strand</td><td>MM</td><td>target_seq</td><td>PAM</td><td>alignment</td><td>distance</td><td>position</td><td>gene name</td><td>gene id</td></tr>
<tr><td>GU305914.1</td><td>119139</td><td>119161</td><td>-</td><td>0</td><td>GTTCACGATCACCGGCATGA</td><td>AGG</td><td>||||||||[||||||||||||]PAM</td><td>NA</td><td>Intergenic</td><td>NA</td><td>NA</td></tr>
<tr><td></td><td></td><td></td><td></td><td></td><td></td><td></td><td></td><td></td><td></td><td></td><td></td></tr>
<tr><td>T25025</td><td>GATCACCGGCATGAAGGACATGG</td><td>1000</td><td></td><td></td><td></td><td></td><td></td><td></td><td></td><td></td><td></td></tr>
<tr><td>Oligo fwd</td><td>CACCGATCACCGGCATGAAGGACA</td><td></td><td></td><td></td><td></td><td></td><td></td><td></td><td></td><td></td><td></td></tr>
<tr><td>Oligo rev</td><td>AAACTGTCCTTCATGCCGGTGATC</td><td></td><td></td><td></td><td></td><td></td><td></td><td></td><td></td><td></td><td></td></tr>
<tr><td>Chromosome</td><td>start</td><td>end</td><td>strand</td><td>MM</td><td>target_seq</td><td>PAM</td><td>alignment</td><td>distance</td><td>position</td><td>gene name</td><td>gene id</td></tr>
<tr><td>GU305914.1</td><td>119133</td><td>119155</td><td>-</td><td>0</td><td>GATCACCGGCATGAAGGACA</td><td>TGG</td><td>||||||||[||||||||||||]PAM</td><td>NA</td><td>Intergenic</td><td>NA</td><td>NA</td></tr>
<tr><td></td><td></td><td></td><td></td><td></td><td></td><td></td><td></td><td></td><td></td><td></td><td></td></tr>
<tr><td>T25026</td><td>CACCGGCATGAAGGACATGGAGG</td><td>1000</td><td></td><td></td><td></td><td></td><td></td><td></td><td></td><td></td><td></td></tr>
<tr><td>Oligo fwd</td><td>CACCCACCGGCATGAAGGACATGG</td><td></td><td></td><td></td><td></td><td></td><td></td><td></td><td></td><td></td><td></td></tr>
<tr><td>Oligo rev</td><td>AAACCCATGTCCTTCATGCCGGTG</td><td></td><td></td><td></td><td></td><td></td><td></td><td></td><td></td><td></td><td></td></tr>
<tr><td>Chromosome</td><td>start</td><td>end</td><td>strand</td><td>MM</td><td>target_seq</td><td>PAM</td><td>alignment</td><td>distance</td><td>position</td><td>gene name</td><td>gene id</td></tr>
<tr><td>GU305914.1</td><td>119130</td><td>119152</td><td>-</td><td>0</td><td>CACCGGCATGAAGGACATGG</td><td>AGG</td><td>||||||||[||||||||||||]PAM</td><td>NA</td><td>Intergenic</td><td>NA</td><td>NA</td></tr>
<tr><td></td><td></td><td></td><td></td><td></td><td></td><td></td><td></td><td></td><td></td><td></td><td></td></tr>
<tr><td>T25027</td><td>CATGAAGGACATGGAGGTCTCGG</td><td>1000</td><td></td><td></td><td></td><td></td><td></td><td></td><td></td><td></td><td></td></tr>
<tr><td>Oligo fwd</td><td>CACCCATGAAGGACATGGAGGTCT</td><td></td><td></td><td></td><td></td><td></td><td></td><td></td><td></td><td></td><td></td></tr>
<tr><td>Oligo rev</td><td>AAACAGACCTCCATGTCCTTCATG</td><td></td><td></td><td></td><td></td><td></td><td></td><td></td><td></td><td></td><td></td></tr>
<tr><td>Chromosome</td><td>start</td><td>end</td><td>strand</td><td>MM</td><td>target_seq</td><td>PAM</td><td>alignment</td><td>distance</td><td>position</td><td>gene name</td><td>gene id</td></tr>
<tr><td>GU305914.1</td><td>119124</td><td>119146</td><td>-</td><td>0</td><td>CATGAAGGACATGGAGGTCT</td><td>CGG</td><td>||||||||[||||||||||||]PAM</td><td>NA</td><td>Intergenic</td><td>NA</td><td>NA</td></tr>
<tr><td></td><td></td><td></td><td></td><td></td><td></td><td></td><td></td><td></td><td></td><td></td><td></td></tr>
<tr><td>T25028</td><td>GAAGGACATGGAGGTCTCGGTGG</td><td>1000</td><td></td><td></td><td></td><td></td><td></td><td></td><td></td><td></td><td></td></tr>
<tr><td>Oligo fwd</td><td>CACCGAAGGACATGGAGGTCTCGG</td><td></td><td></td><td></td><td></td><td></td><td></td><td></td><td></td><td></td><td></td></tr>
<tr><td>Oligo rev</td><td>AAACCCGAGACCTCCATGTCCTTC</td><td></td><td></td><td></td><td></td><td></td><td></td><td></td><td></td><td></td><td></td></tr>
<tr><td>Chromosome</td><td>start</td><td>end</td><td>strand</td><td>MM</td><td>target_seq</td><td>PAM</td><td>alignment</td><td>distance</td><td>position</td><td>gene name</td><td>gene id</td></tr>
<tr><td>GU305914.1</td><td>119121</td><td>119143</td><td>-</td><td>0</td><td>GAAGGACATGGAGGTCTCGG</td><td>TGG</td><td>||||||||[||||||||||||]PAM</td><td>NA</td><td>Intergenic</td><td>NA</td><td>NA</td></tr>
<tr><td></td><td></td><td></td><td></td><td></td><td></td><td></td><td></td><td></td><td></td><td></td><td></td></tr>
<tr><td>T25029</td><td>GTCTCGGTGGAGACCGCCATCGG</td><td>1000</td><td></td><td></td><td></td><td></td><td></td><td></td><td></td><td></td><td></td></tr>
<tr><td>Oligo fwd</td><td>CACCGTCTCGGTGGAGACCGCCAT</td><td></td><td></td><td></td><td></td><td></td><td></td><td></td><td></td><td></td><td></td></tr>
<tr><td>Oligo rev</td><td>AAACATGGCGGTCTCCACCGAGAC</td><td></td><td></td><td></td><td></td><td></td><td></td><td></td><td></td><td></td><td></td></tr>
<tr><td>Chromosome</td><td>start</td><td>end</td><td>strand</td><td>MM</td><td>target_seq</td><td>PAM</td><td>alignment</td><td>distance</td><td>position</td><td>gene name</td><td>gene id</td></tr>
<tr><td>GU305914.1</td><td>119108</td><td>119130</td><td>-</td><td>0</td><td>GTCTCGGTGGAGACCGCCAT</td><td>CGG</td><td>||||||||[||||||||||||]PAM</td><td>NA</td><td>Intergenic</td><td>NA</td><td>NA</td></tr>
<tr><td></td><td></td><td></td><td></td><td></td><td></td><td></td><td></td><td></td><td></td><td></td><td></td></tr>
<tr><td>T25030</td><td>GAGAGCTGCGCTTCGTGTCGCGG</td><td>1000</td><td></td><td></td><td></td><td></td><td></td><td></td><td></td><td></td><td></td></tr>
<tr><td>Oligo fwd</td><td>CACCGAGAGCTGCGCTTCGTGTCG</td><td></td><td></td><td></td><td></td><td></td><td></td><td></td><td></td><td></td><td></td></tr>
<tr><td>Oligo rev</td><td>AAACCGACACGAAGCGCAGCTCTC</td><td></td><td></td><td></td><td></td><td></td><td></td><td></td><td></td><td></td><td></td></tr>
<tr><td>Chromosome</td><td>start</td><td>end</td><td>strand</td><td>MM</td><td>target_seq</td><td>PAM</td><td>alignment</td><td>distance</td><td>position</td><td>gene name</td><td>gene id</td></tr>
<tr><td>GU305914.1</td><td>119086</td><td>119108</td><td>-</td><td>0</td><td>GAGAGCTGCGCTTCGTGTCG</td><td>CGG</td><td>||||||||[||||||||||||]PAM</td><td>NA</td><td>Intergenic</td><td>NA</td><td>NA</td></tr>
<tr><td></td><td></td><td></td><td></td><td></td><td></td><td></td><td></td><td></td><td></td><td></td><td></td></tr>
<tr><td>T25031</td><td>TGCGCTTCGTGTCGCGGTCGTGG</td><td>1000</td><td></td><td></td><td></td><td></td><td></td><td></td><td></td><td></td><td></td></tr>
<tr><td>Oligo fwd</td><td>CACCTGCGCTTCGTGTCGCGGTCG</td><td></td><td></td><td></td><td></td><td></td><td></td><td></td><td></td><td></td><td></td></tr>
<tr><td>Oligo rev</td><td>AAACCGACCGCGACACGAAGCGCA</td><td></td><td></td><td></td><td></td><td></td><td></td><td></td><td></td><td></td><td></td></tr>
<tr><td>Chromosome</td><td>start</td><td>end</td><td>strand</td><td>MM</td><td>target_seq</td><td>PAM</td><td>alignment</td><td>distance</td><td>position</td><td>gene name</td><td>gene id</td></tr>
<tr><td>GU305914.1</td><td>119080</td><td>119102</td><td>-</td><td>0</td><td>TGCGCTTCGTGTCGCGGTCG</td><td>TGG</td><td>||||||||[||||||||||||]PAM</td><td>NA</td><td>Intergenic</td><td>NA</td><td>NA</td></tr>
<tr><td></td><td></td><td></td><td></td><td></td><td></td><td></td><td></td><td></td><td></td><td></td><td></td></tr>
<tr><td>T25032</td><td>CGTGTCGCGGTCGTGGATGACGG</td><td>1000</td><td></td><td></td><td></td><td></td><td></td><td></td><td></td><td></td><td></td></tr>
<tr><td>Oligo fwd</td><td>CACCCGTGTCGCGGTCGTGGATGA</td><td></td><td></td><td></td><td></td><td></td><td></td><td></td><td></td><td></td><td></td></tr>
<tr><td>Oligo rev</td><td>AAACTCATCCACGACCGCGACACG</td><td></td><td></td><td></td><td></td><td></td><td></td><td></td><td></td><td></td><td></td></tr>
<tr><td>Chromosome</td><td>start</td><td>end</td><td>strand</td><td>MM</td><td>target_seq</td><td>PAM</td><td>alignment</td><td>distance</td><td>position</td><td>gene name</td><td>gene id</td></tr>
<tr><td>GU305914.1</td><td>119073</td><td>119095</td><td>-</td><td>0</td><td>CGTGTCGCGGTCGTGGATGA</td><td>CGG</td><td>||||||||[||||||||||||]PAM</td><td>NA</td><td>Intergenic</td><td>NA</td><td>NA</td></tr>
<tr><td></td><td></td><td></td><td></td><td></td><td></td><td></td><td></td><td></td><td></td><td></td><td></td></tr>
<tr><td>T25033</td><td>GTCGCGGTCGTGGATGACGGTGG</td><td>1000</td><td></td><td></td><td></td><td></td><td></td><td></td><td></td><td></td><td></td></tr>
<tr><td>Oligo fwd</td><td>CACCGTCGCGGTCGTGGATGACGG</td><td></td><td></td><td></td><td></td><td></td><td></td><td></td><td></td><td></td><td></td></tr>
<tr><td>Oligo rev</td><td>AAACCCGTCATCCACGACCGCGAC</td><td></td><td></td><td></td><td></td><td></td><td></td><td></td><td></td><td></td><td></td></tr>
<tr><td>Chromosome</td><td>start</td><td>end</td><td>strand</td><td>MM</td><td>target_seq</td><td>PAM</td><td>alignment</td><td>distance</td><td>position</td><td>gene name</td><td>gene id</td></tr>
<tr><td>GU305914.1</td><td>119070</td><td>119092</td><td>-</td><td>0</td><td>GTCGCGGTCGTGGATGACGG</td><td>TGG</td><td>||||||||[||||||||||||]PAM</td><td>NA</td><td>Intergenic</td><td>NA</td><td>NA</td></tr>
<tr><td></td><td></td><td></td><td></td><td></td><td></td><td></td><td></td><td></td><td></td><td></td><td></td></tr>
<tr><td>T25034</td><td>CCACACGTGTTCGCAGACCGAGG</td><td>1000</td><td></td><td></td><td></td><td></td><td></td><td></td><td></td><td></td><td></td></tr>
<tr><td>Oligo fwd</td><td>CACCCCACACGTGTTCGCAGACCG</td><td></td><td></td><td></td><td></td><td></td><td></td><td></td><td></td><td></td><td></td></tr>
<tr><td>Oligo rev</td><td>AAACCGGTCTGCGAACACGTGTGG</td><td></td><td></td><td></td><td></td><td></td><td></td><td></td><td></td><td></td><td></td></tr>
<tr><td>Chromosome</td><td>start</td><td>end</td><td>strand</td><td>MM</td><td>target_seq</td><td>PAM</td><td>alignment</td><td>distance</td><td>position</td><td>gene name</td><td>gene id</td></tr>
<tr><td>GU305914.1</td><td>119025</td><td>119047</td><td>-</td><td>0</td><td>CCACACGTGTTCGCAGACCG</td><td>AGG</td><td>||||||||[||||||||||||]PAM</td><td>NA</td><td>Intergenic</td><td>NA</td><td>NA</td></tr>
<tr><td></td><td></td><td></td><td></td><td></td><td></td><td></td><td></td><td></td><td></td><td></td><td></td></tr>
<tr><td>T25035</td><td>CAGACCGAGGTGATCAACTACGG</td><td>1000</td><td></td><td></td><td></td><td></td><td></td><td></td><td></td><td></td><td></td></tr>
<tr><td>Oligo fwd</td><td>CACCCAGACCGAGGTGATCAACTA</td><td></td><td></td><td></td><td></td><td></td><td></td><td></td><td></td><td></td><td></td></tr>
<tr><td>Oligo rev</td><td>AAACTAGTTGATCACCTCGGTCTG</td><td></td><td></td><td></td><td></td><td></td><td></td><td></td><td></td><td></td><td></td></tr>
<tr><td>Chromosome</td><td>start</td><td>end</td><td>strand</td><td>MM</td><td>target_seq</td><td>PAM</td><td>alignment</td><td>distance</td><td>position</td><td>gene name</td><td>gene id</td></tr>
<tr><td>GU305914.1</td><td>119012</td><td>119034</td><td>-</td><td>0</td><td>CAGACCGAGGTGATCAACTA</td><td>CGG</td><td>||||||||[||||||||||||]PAM</td><td>NA</td><td>Intergenic</td><td>NA</td><td>NA</td></tr>
</table>

<table>
<tr><td>T25036</td><td>AGACCGAGGTGATCAACTACGGG</td><td>1000</td><td></td><td></td><td></td><td></td><td></td><td></td><td></td><td></td><td></td></tr>
<tr><td>Oligo fwd</td><td>CACCAGACCGAGGTGATCAACTAC</td><td></td><td></td><td></td><td></td><td></td><td></td><td></td><td></td><td></td><td></td></tr>
<tr><td>Oligo rev</td><td>AAACGTAGTTGATCACCTCGGTCT</td><td></td><td></td><td></td><td></td><td></td><td></td><td></td><td></td><td></td><td></td></tr>
<tr><td>Chromosome</td><td>start</td><td>end</td><td>strand</td><td>MM</td><td>target_seq</td><td>PAM</td><td>alignment</td><td>distance</td><td>position</td><td>gene name</td><td>gene id</td></tr>
<tr><td>GU305914.1</td><td>119011</td><td>119033</td><td>-</td><td>0</td><td>AGACCGAGGTGATCAACTAC</td><td>GGG</td><td>||||||||[||||||||||||]PAM</td><td>NA</td><td>Intergenic</td><td>NA</td><td>NA</td></tr>
<tr><td>T25037</td><td>CGAGGTGATCAACTACGGGCCGG</td><td>1000</td><td></td><td></td><td></td><td></td><td></td><td></td><td></td><td></td><td></td></tr>
<tr><td>Oligo fwd</td><td>CACCCGAGGTGATCAACTACGGGC</td><td></td><td></td><td></td><td></td><td></td><td></td><td></td><td></td><td></td><td></td></tr>
<tr><td>Oligo rev</td><td>AAACGCCCGTAGTTGATCACCTCG</td><td></td><td></td><td></td><td></td><td></td><td></td><td></td><td></td><td></td><td></td></tr>
<tr><td>Chromosome</td><td>start</td><td>end</td><td>strand</td><td>MM</td><td>target_seq</td><td>PAM</td><td>alignment</td><td>distance</td><td>position</td><td>gene name</td><td>gene id</td></tr>
<tr><td>GU305914.1</td><td>119007</td><td>119029</td><td>-</td><td>0</td><td>CGAGGTGATCAACTACGGGC</td><td>CGG</td><td>||||||||[||||||||||||]PAM</td><td>NA</td><td>Intergenic</td><td>NA</td><td>NA</td></tr>
<tr><td>T25038</td><td>GGTGATCAACTACGGGCCGGAGG</td><td>1000</td><td></td><td></td><td></td><td></td><td></td><td></td><td></td><td></td><td></td></tr>
<tr><td>Oligo fwd</td><td>CACCGGTGATCAACTACGGGCCGG</td><td></td><td></td><td></td><td></td><td></td><td></td><td></td><td></td><td></td><td></td></tr>
<tr><td>Oligo rev</td><td>AAACCCGGCCCGTAGTTGATCACC</td><td></td><td></td><td></td><td></td><td></td><td></td><td></td><td></td><td></td><td></td></tr>
<tr><td>Chromosome</td><td>start</td><td>end</td><td>strand</td><td>MM</td><td>target_seq</td><td>PAM</td><td>alignment</td><td>distance</td><td>position</td><td>gene name</td><td>gene id</td></tr>
<tr><td>GU305914.1</td><td>119004</td><td>119026</td><td>-</td><td>0</td><td>GGTGATCAACTACGGGCCGG</td><td>AGG</td><td>||||||||[||||||||||||]PAM</td><td>NA</td><td>Intergenic</td><td>NA</td><td>NA</td></tr>
<tr><td>T25039</td><td>GAGGTCTTCAGAAAACACAAAGG</td><td>1000</td><td></td><td></td><td></td><td></td><td></td><td></td><td></td><td></td><td></td></tr>
<tr><td>Oligo fwd</td><td>CACCGAGGTCTTCAGAAAACACAA</td><td></td><td></td><td></td><td></td><td></td><td></td><td></td><td></td><td></td><td></td></tr>
<tr><td>Oligo rev</td><td>AAACTTGTGTTTTCTGAAGACCTC</td><td></td><td></td><td></td><td></td><td></td><td></td><td></td><td></td><td></td><td></td></tr>
<tr><td>Chromosome</td><td>start</td><td>end</td><td>strand</td><td>MM</td><td>target_seq</td><td>PAM</td><td>alignment</td><td>distance</td><td>position</td><td>gene name</td><td>gene id</td></tr>
<tr><td>GU305914.1</td><td>118985</td><td>119007</td><td>-</td><td>0</td><td>GAGGTCTTCAGAAAACACAA</td><td>AGG</td><td>||||||||[||||||||||||]PAM</td><td>NA</td><td>Intergenic</td><td>NA</td><td>NA</td></tr>
<tr><td>T25040</td><td>GAGCCGACTGCGAGACCTGCCGG</td><td>1000</td><td></td><td></td><td></td><td></td><td></td><td></td><td></td><td></td><td></td></tr>
<tr><td>Oligo fwd</td><td>CACCGAGCCGACTGCGAGACCTGC</td><td></td><td></td><td></td><td></td><td></td><td></td><td></td><td></td><td></td><td></td></tr>
<tr><td>Oligo rev</td><td>AAACGCAGGTCTCGCAGTCGGCTC</td><td></td><td></td><td></td><td></td><td></td><td></td><td></td><td></td><td></td><td></td></tr>
<tr><td>Chromosome</td><td>start</td><td>end</td><td>strand</td><td>MM</td><td>target_seq</td><td>PAM</td><td>alignment</td><td>distance</td><td>position</td><td>gene name</td><td>gene id</td></tr>
<tr><td>GU305914.1</td><td>118956</td><td>118978</td><td>-</td><td>0</td><td>GAGCCGACTGCGAGACCTGC</td><td>CGG</td><td>||||||||[||||||||||||]PAM</td><td>NA</td><td>Intergenic</td><td>NA</td><td>NA</td></tr>
<tr><td>T25041</td><td>GACGCTCCCCCTCCCCATACAGG</td><td>1000</td><td></td><td></td><td></td><td></td><td></td><td></td><td></td><td></td><td></td></tr>
<tr><td>Oligo fwd</td><td>CACCGACGCTCCCCCTCCCCATAC</td><td></td><td></td><td></td><td></td><td></td><td></td><td></td><td></td><td></td><td></td></tr>
<tr><td>Oligo rev</td><td>AAACGTATGGGGAGGGGGAGCGTC</td><td></td><td></td><td></td><td></td><td></td><td></td><td></td><td></td><td></td><td></td></tr>
<tr><td>Chromosome</td><td>start</td><td>end</td><td>strand</td><td>MM</td><td>target_seq</td><td>PAM</td><td>alignment</td><td>distance</td><td>position</td><td>gene name</td><td>gene id</td></tr>
<tr><td>GU305914.1</td><td>118929</td><td>118951</td><td>-</td><td>0</td><td>GACGCTCCCCCTCCCCATAC</td><td>AGG</td><td>||||||||[||||||||||||]PAM</td><td>NA</td><td>Intergenic</td><td>NA</td><td>NA</td></tr>
<tr><td>T25042</td><td>CCCCATACAGGACCTAGAGCAGG</td><td>1000</td><td></td><td></td><td></td><td></td><td></td><td></td><td></td><td></td><td></td></tr>
<tr><td>Oligo fwd</td><td>CACCCCCCATACAGGACCTAGAGC</td><td></td><td></td><td></td><td></td><td></td><td></td><td></td><td></td><td></td><td></td></tr>
<tr><td>Oligo rev</td><td>AAACGCTCTAGGTCCTGTATGGGG</td><td></td><td></td><td></td><td></td><td></td><td></td><td></td><td></td><td></td><td></td></tr>
<tr><td>Chromosome</td><td>start</td><td>end</td><td>strand</td><td>MM</td><td>target_seq</td><td>PAM</td><td>alignment</td><td>distance</td><td>position</td><td>gene name</td><td>gene id</td></tr>
<tr><td>GU305914.1</td><td>118917</td><td>118939</td><td>-</td><td>0</td><td>CCCCATACAGGACCTAGAGC</td><td>AGG</td><td>||||||||[||||||||||||]PAM</td><td>NA</td><td>Intergenic</td><td>NA</td><td>NA</td></tr>
<tr><td>T25043</td><td>GCAGGACAACGCGTCGTCCCAGG</td><td>1000</td><td></td><td></td><td></td><td></td><td></td><td></td><td></td><td></td><td></td></tr>
<tr><td>Oligo fwd</td><td>CACCGCAGGACAACGCGTCGTCCC</td><td></td><td></td><td></td><td></td><td></td><td></td><td></td><td></td><td></td><td></td></tr>
<tr><td>Oligo rev</td><td>AAACGGGACGACGCGTTGTCCTGC</td><td></td><td></td><td></td><td></td><td></td><td></td><td></td><td></td><td></td><td></td></tr>
<tr><td>Chromosome</td><td>start</td><td>end</td><td>strand</td><td>MM</td><td>target_seq</td><td>PAM</td><td>alignment</td><td>distance</td><td>position</td><td>gene name</td><td>gene id</td></tr>
<tr><td>GU305914.1</td><td>118899</td><td>118921</td><td>-</td><td>0</td><td>GCAGGACAACGCGTCGTCCC</td><td>AGG</td><td>||||||||[||||||||||||]PAM</td><td>NA</td><td>Intergenic</td><td>NA</td><td>NA</td></tr>
<tr><td>T25044</td><td>CAACGCGTCGTCCCAGGACGAGG</td><td>1000</td><td></td><td></td><td></td><td></td><td></td><td></td><td></td><td></td><td></td></tr>
<tr><td>Oligo fwd</td><td>CACCCAACGCGTCGTCCCAGGACG</td><td></td><td></td><td></td><td></td><td></td><td></td><td></td><td></td><td></td><td></td></tr>
<tr><td>Oligo rev</td><td>AAACCGTCCTGGGACGACGCGTTG</td><td></td><td></td><td></td><td></td><td></td><td></td><td></td><td></td><td></td><td></td></tr>
<tr><td>Chromosome</td><td>start</td><td>end</td><td>strand</td><td>MM</td><td>target_seq</td><td>PAM</td><td>alignment</td><td>distance</td><td>position</td><td>gene name</td><td>gene id</td></tr>
<tr><td>GU305914.1</td><td>118893</td><td>118915</td><td>-</td><td>0</td><td>CAACGCGTCGTCCCAGGACG</td><td>AGG</td><td>||||||||[||||||||||||]PAM</td><td>NA</td><td>Intergenic</td><td>NA</td><td>NA</td></tr>
<tr><td>T25045</td><td>GGCGTCGTCGTCCGACGCCGAGG</td><td>1000</td><td></td><td></td><td></td><td></td><td></td><td></td><td></td><td></td><td></td></tr>
<tr><td>Oligo fwd</td><td>CACCGGCGTCGTCGTCCGACGCCG</td><td></td><td></td><td></td><td></td><td></td><td></td><td></td><td></td><td></td><td></td></tr>
<tr><td>Oligo rev</td><td>AAACCGGCGTCGGACGACGACGCC</td><td></td><td></td><td></td><td></td><td></td><td></td><td></td><td></td><td></td><td></td></tr>
<tr><td>Chromosome</td><td>start</td><td>end</td><td>strand</td><td>MM</td><td>target_seq</td><td>PAM</td><td>alignment</td><td>distance</td><td>position</td><td>gene name</td><td>gene id</td></tr>
<tr><td>GU305914.1</td><td>118872</td><td>118894</td><td>-</td><td>0</td><td>GGCGTCGTCGTCCGACGCCG</td><td>AGG</td><td>||||||||[||||||||||||]PAM</td><td>NA</td><td>Intergenic</td><td>NA</td><td>NA</td></tr>
<tr><td>T25046</td><td>GTCGTCGTCCGACGCCGAGGAGG</td><td>1000</td><td></td><td></td><td></td><td></td><td></td><td></td><td></td><td></td><td></td></tr>
<tr><td>Oligo fwd</td><td>CACCGTCGTCGTCCGACGCCGAGG</td><td></td><td></td><td></td><td></td><td></td><td></td><td></td><td></td><td></td><td></td></tr>
<tr><td>Oligo rev</td><td>AAACCCTCGGCGTCGGACGACGAC</td><td></td><td></td><td></td><td></td><td></td><td></td><td></td><td></td><td></td><td></td></tr>
<tr><td>Chromosome</td><td>start</td><td>end</td><td>strand</td><td>MM</td><td>target_seq</td><td>PAM</td><td>alignment</td><td>distance</td><td>position</td><td>gene name</td><td>gene id</td></tr>
<tr><td>GU305914.1</td><td>118869</td><td>118891</td><td>-</td><td>0</td><td>GTCGTCGTCCGACGCCGAGG</td><td>AGG</td><td>||||||||[||||||||||||]PAM</td><td>NA</td><td>Intergenic</td><td>NA</td><td>NA</td></tr>
<tr><td>GU305914.1</td><td>198000</td><td>198022</td><td>+</td><td>4</td><td>CTCGACGTCCGACGCCGTCG</td><td>CGG</td><td>-|||-|||[|||||||||--|]PAM</td><td>NA</td><td>Intergenic</td><td>NA</td><td>NA</td></tr>
<tr><td>T25047</td><td>GTCGTCCGACGCCGAGGAGGAGG</td><td>1000</td><td></td><td></td><td></td><td></td><td></td><td></td><td></td><td></td><td></td></tr>
<tr><td>Oligo fwd</td><td>CACCGTCGTCCGACGCCGAGGAGG</td><td></td><td></td><td></td><td></td><td></td><td></td><td></td><td></td><td></td><td></td></tr>
<tr><td>Oligo rev</td><td>AAACCCTCCTCGGCGTCGGACGAC</td><td></td><td></td><td></td><td></td><td></td><td></td><td></td><td></td><td></td><td></td></tr>
<tr><td>Chromosome</td><td>start</td><td>end</td><td>strand</td><td>MM</td><td>target_seq</td><td>PAM</td><td>alignment</td><td>distance</td><td>position</td><td>gene name</td><td>gene id</td></tr>
<tr><td>GU305914.1</td><td>118866</td><td>118888</td><td>-</td><td>0</td><td>GTCGTCCGACGCCGAGGAGG</td><td>AGG</td><td>||||||||[||||||||||||]PAM</td><td>NA</td><td>Intergenic</td><td>NA</td><td>NA</td></tr>
<tr><td>T25048</td><td>GGAGGAGGAAGAAGACGAAGAGG</td><td>1000</td><td></td><td></td><td></td><td></td><td></td><td></td><td></td><td></td><td></td></tr>
<tr><td>Oligo fwd</td><td>CACCGGAGGAGGAAGAAGACGAAG</td><td></td><td></td><td></td><td></td><td></td><td></td><td></td><td></td><td></td><td></td></tr>
<tr><td>Oligo rev</td><td>AAACCTTCGTCTTCTTCCTCCTCC</td><td></td><td></td><td></td><td></td><td></td><td></td><td></td><td></td><td></td><td></td></tr>
<tr><td>Chromosome</td><td>start</td><td>end</td><td>strand</td><td>MM</td><td>target_seq</td><td>PAM</td><td>alignment</td><td>distance</td><td>position</td><td>gene name</td><td>gene id</td></tr>
<tr><td>GU305914.1</td><td>118851</td><td>118873</td><td>-</td><td>0</td><td>GGAGGAGGAAGAAGACGAAG</td><td>AGG</td><td>||||||||[||||||||||||]PAM</td><td>NA</td><td>Intergenic</td><td>NA</td><td>NA</td></tr>
<tr><td>GU305914.1</td><td>45333</td><td>45355</td><td>-</td><td>2</td><td>GGAGGAGGAAGATGATGAAG</td><td>TGG</td><td>||||||||[||||-||-||||]PAM</td><td>NA</td><td>Intergenic</td><td>NA</td><td>NA</td></tr>
<tr><td>GU305914.1</td><td>220274</td><td>220296</td><td>-</td><td>4</td><td>AGAGGAAGATGAAGAGGAAG</td><td>AGG</td><td>-|||||-|[|-|||||-||||]PAM</td><td>NA</td><td>Intergenic</td><td>NA</td><td>NA</td></tr>
<tr><td>GU305914.1</td><td>65108</td><td>65130</td><td>+</td><td>3</td><td>GGAGGAAGAAGACGACGACG</td><td>AGG</td><td>||||||-|[||||-|||||-|]PAM</td><td>NA</td><td>Intergenic</td><td>NA</td><td>NA</td></tr>
<tr><td>T25049</td><td>AGAAGACGAAGAGGACGACGAGG</td><td>1000</td><td></td><td></td><td></td><td></td><td></td><td></td><td></td><td></td><td></td></tr>
<tr><td>Oligo fwd</td><td>CACCAGAAGACGAAGAGGACGACG</td><td></td><td></td><td></td><td></td><td></td><td></td><td></td><td></td><td></td><td></td></tr>
<tr><td>Oligo rev</td><td>AAACCGTCGTCCTCTTCGTCTTCT</td><td></td><td></td><td></td><td></td><td></td><td></td><td></td><td></td><td></td><td></td></tr>
<tr><td>Chromosome</td><td>start</td><td>end</td><td>strand</td><td>MM</td><td>target_seq</td><td>PAM</td><td>alignment</td><td>distance</td><td>position</td><td>gene name</td><td>gene id</td></tr>
<tr><td>GU305914.1</td><td>118842</td><td>118864</td><td>-</td><td>0</td><td>AGAAGACGAAGAGGACGACG</td><td>AGG</td><td>||||||||[||||||||||||]PAM</td><td>NA</td><td>Intergenic</td><td>NA</td><td>NA</td></tr>
<tr><td>GU305914.1</td><td>65108</td><td>65130</td><td>+</td><td>4</td><td>GGAGGAAGAAGACGACGACG</td><td>AGG</td><td>-||-||-|[||||-|||||||]PAM</td><td>NA</td><td>Intergenic</td><td>NA</td><td>NA</td></tr>
</table>

| | | | | | | | | | | | | |
|---|---|---|---|---|---|---|---|---|---|---|---|---|
| GU305914.1 | | 118494 | 118516 | - | 4 | GGAGGACGAGGACGACGACG | AGG | -\|\|-\|\|\|\|[\|-\|\|-\|\|\|\|\|\|\|]PAM | NA | Intergenic | NA | NA |
| GU305914.1 | | 118812 | 118834 | - | 4 | TGACGACGATGATGACGACG | AGG | -\|\|-\|\|\|\|[\|-\|\|-\|\|\|\|\|\|\|]PAM | NA | Intergenic | NA | NA |
| GU305914.1 | | 118506 | 118528 | - | 4 | CGATGATGAAGAGGAGGACG | AGG | -\|\|-\|\|-\|[\|\|\|\|\|\|\|-\|\|\|\|]PAM | NA | Intergenic | NA | NA |
| T25050 | TGACGACGATGATGACGACGAGG | | 1000 | | | | | | | | | |
| Oligo fwd | CACCTGACGACGATGATGACGACG | | | | | | | | | | | |
| Oligo rev | AAACCGTCGTCATCATCGTCGTCA | | | | | | | | | | | |
| Chromosome | start | | end | strand | MM | target_seq | PAM | alignment | distance | position | gene name | gene id |
| GU305914.1 | | 118812 | 118834 | - | 0 | TGACGACGATGATGACGACG | AGG | \|\|\|\|\|\|\|\|[\|\|\|\|\|\|\|\|\|\|\|\|]PAM | NA | Intergenic | NA | NA |
| GU305914.1 | | 114978 | 115000 | + | 3 | CGACGATGACGATGACGACG | AGG | -\|\|\|\|\|-\|[\|-\|\|\|\|\|\|\|\|\|\|]PAM | NA | Intergenic | NA | NA |
| GU305914.1 | | 118494 | 118516 | - | 4 | GGAGGACGAGGACGACGACG | AGG | -\|\|-\|\|\|\|[\|-\|\|-\|\|\|\|\|\|\|]PAM | NA | Intergenic | NA | NA |
| GU305914.1 | | 118842 | 118864 | - | 4 | AGAAGACGAAGAGGACGACG | AGG | -\|\|-\|\|\|\|[\|-\|\|-\|\|\|\|\|\|\|]PAM | NA | Intergenic | NA | NA |
| GU305914.1 | | 118809 | 118831 | - | 4 | CGACGATGATGACGACGAGG | AGG | -\|\|\|\|\|-\|[\|\|\|\|-\|\|\|\|\|-\|]PAM | NA | Intergenic | NA | NA |
| GU305914.1 | | 118512 | 118534 | - | 3 | TGAGGACGATGATGAAGAGG | AGG | \|\|\|-\|\|\|\|[\|\|\|\|\|\|\|-\|\|-\|]PAM | NA | Intergenic | NA | NA |
| T25051 | CGACGATGATGACGACGAGGAGG | | 1000 | | | | | | | | | |
| Oligo fwd | CACCCGACGATGATGACGACGAGG | | | | | | | | | | | |
| Oligo rev | AAACCCTCGTCGTCATCATCGTCG | | | | | | | | | | | |
| Chromosome | start | | end | strand | MM | target_seq | PAM | alignment | distance | position | gene name | gene id |
| GU305914.1 | | 118809 | 118831 | - | 0 | CGACGATGATGACGACGAGG | AGG | \|\|\|\|\|\|\|\|[\|\|\|\|\|\|\|\|\|\|\|\|]PAM | NA | Intergenic | NA | NA |
| GU305914.1 | | 65111 | 65133 | + | 4 | GGAAGAAGACGACGACGAGG | AGG | -\|\|-\|\|-\|[\|-\|\|\|\|\|\|\|\|\|\|]PAM | NA | Intergenic | NA | NA |
| GU305914.1 | | 140726 | 140748 | + | 4 | CGAGGAGGATGAGGATGAGG | AGG | \|\|\|-\|\|-\|[\|\|\|\|-\|\|-\|\|\|\|]PAM | NA | Intergenic | NA | NA |
| GU305914.1 | | 118812 | 118834 | - | 4 | TGACGACGATGATGACGACG | AGG | -\|\|\|\|\|-\|[\|\|\|\|-\|\|\|\|\|-\|]PAM | NA | Intergenic | NA | NA |
| T25052 | GATGATGACGACGAGGAGGACGG | | 1000 | | | | | | | | | |
| Oligo fwd | CACCGATGATGACGACGAGGAGGA | | | | | | | | | | | |
| Oligo rev | AAACTCCTCCTCGTCGTCATCATC | | | | | | | | | | | |
| Chromosome | start | | end | strand | MM | target_seq | PAM | alignment | distance | position | gene name | gene id |
| GU305914.1 | | 118805 | 118827 | - | 0 | GATGATGACGACGAGGAGGA | CGG | \|\|\|\|\|\|\|\|[\|\|\|\|\|\|\|\|\|\|\|\|]PAM | NA | Intergenic | NA | NA |
| GU305914.1 | | 65115 | 65137 | + | 2 | GAAGACGACGACGAGGAGGA | GGG | \|\|-\|\|-\|\|[\|\|\|\|\|\|\|\|\|\|\|\|]PAM | NA | Intergenic | NA | NA |
| GU305914.1 | | 77033 | 77055 | - | 4 | GACGAGGATGAGGAGGAGGA | CGG | \|\|-\|\|-\|\|[-\|\|-\|\|\|\|\|\|\|\|]PAM | NA | Intergenic | NA | NA |
| GU305914.1 | | 65118 | 65140 | + | 4 | GACGACGACGAGGAGGAGGG | CGG | \|\|-\|\|-\|\|[\|\|\|-\|\|\|\|\|\|\|-]PAM | NA | Intergenic | NA | NA |
| T25053 | ATGATGACGACGAGGAGGACGGG | | 1000 | | | | | | | | | |
| Oligo fwd | CACCATGATGACGACGAGGAGGAC | | | | | | | | | | | |
| Oligo rev | AAACGTCCTCCTCGTCGTCATCAT | | | | | | | | | | | |
| Chromosome | start | | end | strand | MM | target_seq | PAM | alignment | distance | position | gene name | gene id |
| GU305914.1 | | 118804 | 118826 | - | 0 | ATGATGACGACGAGGAGGAC | GGG | \|\|\|\|\|\|\|\|[\|\|\|\|\|\|\|\|\|\|\|\|]PAM | NA | Intergenic | NA | NA |
| T25054 | GACGGGACAGAAAACCCAAACGG | | 1000 | | | | | | | | | |
| Oligo fwd | CACCGACGGGACAGAAAACCCAAA | | | | | | | | | | | |
| Oligo rev | AAACTTTGGGTTTTCTGTCCCGTC | | | | | | | | | | | |
| Chromosome | start | | end | strand | MM | target_seq | PAM | alignment | distance | position | gene name | gene id |
| GU305914.1 | | 118787 | 118809 | - | 0 | GACGGGACAGAAAACCCAAA | CGG | \|\|\|\|\|\|\|\|[\|\|\|\|\|\|\|\|\|\|\|\|]PAM | NA | Intergenic | NA | NA |
| T25055 | ACGGGACAGAAAACCCAAACGGG | | 1000 | | | | | | | | | |
| Oligo fwd | CACCACGGGACAGAAAACCCAAAC | | | | | | | | | | | |
| Oligo rev | AAACGTTTGGGTTTTCTGTCCCGT | | | | | | | | | | | |
| Chromosome | start | | end | strand | MM | target_seq | PAM | alignment | distance | position | gene name | gene id |
| GU305914.1 | | 118786 | 118808 | - | 0 | ACGGGACAGAAAACCCAAAC | GGG | \|\|\|\|\|\|\|\|[\|\|\|\|\|\|\|\|\|\|\|\|]PAM | NA | Intergenic | NA | NA |
| T25056 | GACAGAAAACCCAAACGGGATGG | | 1000 | | | | | | | | | |
| Oligo fwd | CACCGACAGAAAACCCAAACGGGA | | | | | | | | | | | |
| Oligo rev | AAACTCCCGTTTGGGTTTTCTGTC | | | | | | | | | | | |
| Chromosome | start | | end | strand | MM | target_seq | PAM | alignment | distance | position | gene name | gene id |
| GU305914.1 | | 118782 | 118804 | - | 0 | GACAGAAAACCCAAACGGGA | TGG | \|\|\|\|\|\|\|\|[\|\|\|\|\|\|\|\|\|\|\|\|]PAM | NA | Intergenic | NA | NA |
| T25057 | AGATGCCGAACCCGAGCGACAGG | | 1000 | | | | | | | | | |
| Oligo fwd | CACCAGATGCCGAACCCGAGCGAC | | | | | | | | | | | |
| Oligo rev | AAACGTCGCTCGGGTTCGGCATCT | | | | | | | | | | | |
| Chromosome | start | | end | strand | MM | target_seq | PAM | alignment | distance | position | gene name | gene id |
| GU305914.1 | | 118734 | 118756 | - | 0 | AGATGCCGAACCCGAGCGAC | AGG | \|\|\|\|\|\|\|\|[\|\|\|\|\|\|\|\|\|\|\|\|]PAM | NA | Intergenic | NA | NA |
| T25058 | AGGCCCCACCGCAGCAACAGCGG | | 1000 | | | | | | | | | |
| Oligo fwd | CACCAGGCCCCACCGCAGCAACAG | | | | | | | | | | | |
| Oligo rev | AAACCTGTTGCTGCGGTGGGGCCT | | | | | | | | | | | |
| Chromosome | start | | end | strand | MM | target_seq | PAM | alignment | distance | position | gene name | gene id |
| GU305914.1 | | 118714 | 118736 | - | 0 | AGGCCCCACCGCAGCAACAG | CGG | \|\|\|\|\|\|\|\|[\|\|\|\|\|\|\|\|\|\|\|\|]PAM | NA | Intergenic | NA | NA |
| T25059 | ATCACCGAGAGCCAACCGCCCGG | | 1000 | | | | | | | | | |
| Oligo fwd | CACCATCACCGAGAGCCAACCGCC | | | | | | | | | | | |
| Oligo rev | AAACGGCGGTTGGCTCTCGGTGAT | | | | | | | | | | | |
| Chromosome | start | | end | strand | MM | target_seq | PAM | alignment | distance | position | gene name | gene id |
| GU305914.1 | | 118686 | 118708 | - | 0 | ATCACCGAGAGCCAACCGCC | CGG | \|\|\|\|\|\|\|\|[\|\|\|\|\|\|\|\|\|\|\|\|]PAM | NA | Intergenic | NA | NA |
| T25060 | GAGAGCCAACCGCCCGGCCGAGG | | 1000 | | | | | | | | | |
| Oligo fwd | CACCGAGAGCCAACCGCCCGGCCG | | | | | | | | | | | |
| Oligo rev | AAACCGGCCGGGCGGTTGGCTCTC | | | | | | | | | | | |
| Chromosome | start | | end | strand | MM | target_seq | PAM | alignment | distance | position | gene name | gene id |
| GU305914.1 | | 118680 | 118702 | - | 0 | GAGAGCCAACCGCCCGGCCG | AGG | \|\|\|\|\|\|\|\|[\|\|\|\|\|\|\|\|\|\|\|\|]PAM | NA | Intergenic | NA | NA |
| T25061 | CCGCCCGGCCGAGGAAGACGAGG | | 1000 | | | | | | | | | |
| Oligo fwd | CACCCCGCCCGGCCGAGGAAGACG | | | | | | | | | | | |
| Oligo rev | AAACCGTCTTCCTCGGCCGGGCGG | | | | | | | | | | | |
| Chromosome | start | | end | strand | MM | target_seq | PAM | alignment | distance | position | gene name | gene id |
| GU305914.1 | | 118671 | 118693 | - | 0 | CCGCCCGGCCGAGGAAGACG | AGG | \|\|\|\|\|\|\|\|[\|\|\|\|\|\|\|\|\|\|\|\|]PAM | NA | Intergenic | NA | NA |
| T25062 | GAGGACGCCGCTGTGTTCGCCGG | | 1000 | | | | | | | | | |
| Oligo fwd | CACCGAGGACGCCGCTGTGTTCGC | | | | | | | | | | | |

<table>
<tr><td>Oligo rev</td><td>AAACGCGAACACAGCGGCGTCCTC</td><td></td><td></td><td></td><td></td><td></td><td></td><td></td><td></td><td></td><td></td></tr>
<tr><td>Chromosome</td><td>start</td><td>end</td><td>strand</td><td>MM</td><td>target_seq</td><td>PAM</td><td>alignment</td><td>distance</td><td>position</td><td>gene name</td><td>gene id</td></tr>
<tr><td>GU305914.1</td><td>118652</td><td>118674</td><td>-</td><td>0</td><td>GAGGACGCCGCTGTGTTCGC</td><td>CGG</td><td>||||||||[||||||||||||]PAM</td><td>NA</td><td>Intergenic</td><td>NA</td><td>NA</td></tr>
<tr><td>GU305914.1</td><td>118604</td><td>118626</td><td>-</td><td>2</td><td>GATGACGCCGCTGTATTCGC</td><td>AGG</td><td>||-|||||[||||||-|||||]PAM</td><td>NA</td><td>Intergenic</td><td>NA</td><td>NA</td></tr>
<tr><td></td><td></td><td></td><td></td><td></td><td></td><td></td><td></td><td></td><td></td><td></td><td></td></tr>
<tr><td>T25063</td><td>GATGACGCCGCTGTATTCGCAGG</td><td>1000</td><td></td><td></td><td></td><td></td><td></td><td></td><td></td><td></td><td></td></tr>
<tr><td>Oligo fwd</td><td>CACCGATGACGCCGCTGTATTCGC</td><td></td><td></td><td></td><td></td><td></td><td></td><td></td><td></td><td></td><td></td></tr>
<tr><td>Oligo rev</td><td>AAACGCGAATACAGCGGCGTCATC</td><td></td><td></td><td></td><td></td><td></td><td></td><td></td><td></td><td></td><td></td></tr>
<tr><td>Chromosome</td><td>start</td><td>end</td><td>strand</td><td>MM</td><td>target_seq</td><td>PAM</td><td>alignment</td><td>distance</td><td>position</td><td>gene name</td><td>gene id</td></tr>
<tr><td>GU305914.1</td><td>118604</td><td>118626</td><td>-</td><td>0</td><td>GATGACGCCGCTGTATTCGC</td><td>AGG</td><td>||||||||[||||||||||||]PAM</td><td>NA</td><td>Intergenic</td><td>NA</td><td>NA</td></tr>
<tr><td>GU305914.1</td><td>118652</td><td>118674</td><td>-</td><td>2</td><td>GAGGACGCCGCTGTGTTCGC</td><td>CGG</td><td>||-|||||[||||||-|||||]PAM</td><td>NA</td><td>Intergenic</td><td>NA</td><td>NA</td></tr>
<tr><td></td><td></td><td></td><td></td><td></td><td></td><td></td><td></td><td></td><td></td><td></td><td></td></tr>
<tr><td>T25064</td><td>AGACGCCGACGCCGACGCCATGG</td><td>1000</td><td></td><td></td><td></td><td></td><td></td><td></td><td></td><td></td><td></td></tr>
<tr><td>Oligo fwd</td><td>CACCAGACGCCGACGCCGACGCCA</td><td></td><td></td><td></td><td></td><td></td><td></td><td></td><td></td><td></td><td></td></tr>
<tr><td>Oligo rev</td><td>AAACTGGCGTCGGCGTCGGCGTCT</td><td></td><td></td><td></td><td></td><td></td><td></td><td></td><td></td><td></td><td></td></tr>
<tr><td>Chromosome</td><td>start</td><td>end</td><td>strand</td><td>MM</td><td>target_seq</td><td>PAM</td><td>alignment</td><td>distance</td><td>position</td><td>gene name</td><td>gene id</td></tr>
<tr><td>GU305914.1</td><td>118569</td><td>118591</td><td>-</td><td>0</td><td>AGACGCCGACGCCGACGCCA</td><td>TGG</td><td>||||||||[||||||||||||]PAM</td><td>NA</td><td>Intergenic</td><td>NA</td><td>NA</td></tr>
<tr><td>GU305914.1</td><td>24130</td><td>24152</td><td>-</td><td>4</td><td>CGAGGCCGACGTCGGCGCCA</td><td>TGG</td><td>-||-||||[|||-||-|||||]PAM</td><td>NA</td><td>Intergenic</td><td>NA</td><td>NA</td></tr>
<tr><td>GU305914.1</td><td>113931</td><td>113953</td><td>+</td><td>4</td><td>CGCCGCCGACGCCGACACCT</td><td>CGG</td><td>-|-|||||[||||||||-||-]PAM</td><td>NA</td><td>Intergenic</td><td>NA</td><td>NA</td></tr>
<tr><td></td><td></td><td></td><td></td><td></td><td></td><td></td><td></td><td></td><td></td><td></td><td></td></tr>
<tr><td>T25065</td><td>GACGCCGACGCCATGGACGTCGG</td><td>1000</td><td></td><td></td><td></td><td></td><td></td><td></td><td></td><td></td><td></td></tr>
<tr><td>Oligo fwd</td><td>CACCGACGCCGACGCCATGGACGT</td><td></td><td></td><td></td><td></td><td></td><td></td><td></td><td></td><td></td><td></td></tr>
<tr><td>Oligo rev</td><td>AAACACGTCCATGGCGTCGGCGTC</td><td></td><td></td><td></td><td></td><td></td><td></td><td></td><td></td><td></td><td></td></tr>
<tr><td>Chromosome</td><td>start</td><td>end</td><td>strand</td><td>MM</td><td>target_seq</td><td>PAM</td><td>alignment</td><td>distance</td><td>position</td><td>gene name</td><td>gene id</td></tr>
<tr><td>GU305914.1</td><td>118562</td><td>118584</td><td>-</td><td>0</td><td>GACGCCGACGCCATGGACGT</td><td>CGG</td><td>||||||||[||||||||||||]PAM</td><td>NA</td><td>Intergenic</td><td>NA</td><td>NA</td></tr>
<tr><td></td><td></td><td></td><td></td><td></td><td></td><td></td><td></td><td></td><td></td><td></td><td></td></tr>
<tr><td>T25066</td><td>GCCGACGCCATGGACGTCGGCGG</td><td>1000</td><td></td><td></td><td></td><td></td><td></td><td></td><td></td><td></td><td></td></tr>
<tr><td>Oligo fwd</td><td>CACCGCCGACGCCATGGACGTCGG</td><td></td><td></td><td></td><td></td><td></td><td></td><td></td><td></td><td></td><td></td></tr>
<tr><td>Oligo rev</td><td>AAACCCGACGTCCATGGCGTCGGC</td><td></td><td></td><td></td><td></td><td></td><td></td><td></td><td></td><td></td><td></td></tr>
<tr><td>Chromosome</td><td>start</td><td>end</td><td>strand</td><td>MM</td><td>target_seq</td><td>PAM</td><td>alignment</td><td>distance</td><td>position</td><td>gene name</td><td>gene id</td></tr>
<tr><td>GU305914.1</td><td>118559</td><td>118581</td><td>-</td><td>0</td><td>GCCGACGCCATGGACGTCGG</td><td>CGG</td><td>||||||||[||||||||||||]PAM</td><td>NA</td><td>Intergenic</td><td>NA</td><td>NA</td></tr>
<tr><td>GU305914.1</td><td>24120</td><td>24142</td><td>-</td><td>4</td><td>GTCGGCGCCATGGACGTCCT</td><td>CGG</td><td>|-||-|||[||||||||||--]PAM</td><td>NA</td><td>Intergenic</td><td>NA</td><td>NA</td></tr>
<tr><td></td><td></td><td></td><td></td><td></td><td></td><td></td><td></td><td></td><td></td><td></td><td></td></tr>
<tr><td>T25067</td><td>ATGGACGTCGGCGGTCCCGCTGG</td><td>1000</td><td></td><td></td><td></td><td></td><td></td><td></td><td></td><td></td><td></td></tr>
<tr><td>Oligo fwd</td><td>CACCATGGACGTCGGCGGTCCCGC</td><td></td><td></td><td></td><td></td><td></td><td></td><td></td><td></td><td></td><td></td></tr>
<tr><td>Oligo rev</td><td>AAACGCGGGACCGCCGACGTCCAT</td><td></td><td></td><td></td><td></td><td></td><td></td><td></td><td></td><td></td><td></td></tr>
<tr><td>Chromosome</td><td>start</td><td>end</td><td>strand</td><td>MM</td><td>target_seq</td><td>PAM</td><td>alignment</td><td>distance</td><td>position</td><td>gene name</td><td>gene id</td></tr>
<tr><td>GU305914.1</td><td>118550</td><td>118572</td><td>-</td><td>0</td><td>ATGGACGTCGGCGGTCCCGC</td><td>TGG</td><td>||||||||[||||||||||||]PAM</td><td>NA</td><td>Intergenic</td><td>NA</td><td>NA</td></tr>
<tr><td></td><td></td><td></td><td></td><td></td><td></td><td></td><td></td><td></td><td></td><td></td><td></td></tr>
<tr><td>T25068</td><td>TGGCGACGAGAACGCCGATGAGG</td><td>1000</td><td></td><td></td><td></td><td></td><td></td><td></td><td></td><td></td><td></td></tr>
<tr><td>Oligo fwd</td><td>CACCTGGCGACGAGAACGCCGATG</td><td></td><td></td><td></td><td></td><td></td><td></td><td></td><td></td><td></td><td></td></tr>
<tr><td>Oligo rev</td><td>AAACCATCGGCGTTCTCGTCGCCA</td><td></td><td></td><td></td><td></td><td></td><td></td><td></td><td></td><td></td><td></td></tr>
<tr><td>Chromosome</td><td>start</td><td>end</td><td>strand</td><td>MM</td><td>target_seq</td><td>PAM</td><td>alignment</td><td>distance</td><td>position</td><td>gene name</td><td>gene id</td></tr>
<tr><td>GU305914.1</td><td>118530</td><td>118552</td><td>-</td><td>0</td><td>TGGCGACGAGAACGCCGATG</td><td>AGG</td><td>||||||||[||||||||||||]PAM</td><td>NA</td><td>Intergenic</td><td>NA</td><td>NA</td></tr>
<tr><td></td><td></td><td></td><td></td><td></td><td></td><td></td><td></td><td></td><td></td><td></td><td></td></tr>
<tr><td>T25069</td><td>CGATGAGGACGATGATGAAGAGG</td><td>1000</td><td></td><td></td><td></td><td></td><td></td><td></td><td></td><td></td><td></td></tr>
<tr><td>Oligo fwd</td><td>CACCCGATGAGGACGATGATGAAG</td><td></td><td></td><td></td><td></td><td></td><td></td><td></td><td></td><td></td><td></td></tr>
<tr><td>Oligo rev</td><td>AAACCTTCATCATCGTCCTCATCG</td><td></td><td></td><td></td><td></td><td></td><td></td><td></td><td></td><td></td><td></td></tr>
<tr><td>Chromosome</td><td>start</td><td>end</td><td>strand</td><td>MM</td><td>target_seq</td><td>PAM</td><td>alignment</td><td>distance</td><td>position</td><td>gene name</td><td>gene id</td></tr>
<tr><td>GU305914.1</td><td>118515</td><td>118537</td><td>-</td><td>0</td><td>CGATGAGGACGATGATGAAG</td><td>AGG</td><td>||||||||[||||||||||||]PAM</td><td>NA</td><td>Intergenic</td><td>NA</td><td>NA</td></tr>
<tr><td>GU305914.1</td><td>45333</td><td>45355</td><td>-</td><td>3</td><td>GGAGGAGGAAGATGATGAAG</td><td>TGG</td><td>-||-||||[|-||||||||||]PAM</td><td>NA</td><td>Intergenic</td><td>NA</td><td>NA</td></tr>
<tr><td>GU305914.1</td><td>168703</td><td>168725</td><td>+</td><td>4</td><td>CGAAGAGTCCGATGACGAAG</td><td>AGG</td><td>|||-|||-[-||||||-||||]PAM</td><td>NA</td><td>Intergenic</td><td>NA</td><td>NA</td></tr>
<tr><td>GU305914.1</td><td>27345</td><td>27367</td><td>+</td><td>4</td><td>CGAAGATGATGATGAAGAAG</td><td>AGG</td><td>|||-||-|[|-|||||-||||]PAM</td><td>NA</td><td>Intergenic</td><td>NA</td><td>NA</td></tr>
<tr><td>GU305914.1</td><td>114978</td><td>115000</td><td>+</td><td>4</td><td>CGACGATGACGATGACGACG</td><td>AGG</td><td>|||-||-|[|||||||-||-|]PAM</td><td>NA</td><td>Intergenic</td><td>NA</td><td>NA</td></tr>
<tr><td></td><td></td><td></td><td></td><td></td><td></td><td></td><td></td><td></td><td></td><td></td><td></td></tr>
<tr><td>T25070</td><td>TGAGGACGATGATGAAGAGGAGG</td><td>1000</td><td></td><td></td><td></td><td></td><td></td><td></td><td></td><td></td><td></td></tr>
<tr><td>Oligo fwd</td><td>CACCTGAGGACGATGATGAAGAGG</td><td></td><td></td><td></td><td></td><td></td><td></td><td></td><td></td><td></td><td></td></tr>
<tr><td>Oligo rev</td><td>AAACCCTCTTCATCATCGTCCTCA</td><td></td><td></td><td></td><td></td><td></td><td></td><td></td><td></td><td></td><td></td></tr>
<tr><td>Chromosome</td><td>start</td><td>end</td><td>strand</td><td>MM</td><td>target_seq</td><td>PAM</td><td>alignment</td><td>distance</td><td>position</td><td>gene name</td><td>gene id</td></tr>
<tr><td>GU305914.1</td><td>118512</td><td>118534</td><td>-</td><td>0</td><td>TGAGGACGATGATGAAGAGG</td><td>AGG</td><td>||||||||[||||||||||||]PAM</td><td>NA</td><td>Intergenic</td><td>NA</td><td>NA</td></tr>
<tr><td>GU305914.1</td><td>45330</td><td>45352</td><td>-</td><td>3</td><td>GGAGGAAGATGATGAAGTGG</td><td>AGG</td><td>-|||||-|[|||||||||-||]PAM</td><td>NA</td><td>Intergenic</td><td>NA</td><td>NA</td></tr>
<tr><td>GU305914.1</td><td>140726</td><td>140748</td><td>+</td><td>4</td><td>CGAGGAGGATGAGGATGAGG</td><td>AGG</td><td>-|||||-|[||||-||-||||]PAM</td><td>NA</td><td>Intergenic</td><td>NA</td><td>NA</td></tr>
<tr><td>GU305914.1</td><td>27345</td><td>27367</td><td>+</td><td>4</td><td>CGAAGATGATGATGAAGAAG</td><td>AGG</td><td>-||-||-|[||||||||||-|]PAM</td><td>NA</td><td>Intergenic</td><td>NA</td><td>NA</td></tr>
<tr><td>GU305914.1</td><td>118812</td><td>118834</td><td>-</td><td>3</td><td>TGACGACGATGATGACGACG</td><td>AGG</td><td>|||-||||[|||||||-||-|]PAM</td><td>NA</td><td>Intergenic</td><td>NA</td><td>NA</td></tr>
<tr><td></td><td></td><td></td><td></td><td></td><td></td><td></td><td></td><td></td><td></td><td></td><td></td></tr>
<tr><td>T25071</td><td>CGATGATGAAGAGGAGGACGAGG</td><td>1000</td><td></td><td></td><td></td><td></td><td></td><td></td><td></td><td></td><td></td></tr>
<tr><td>Oligo fwd</td><td>CACCCGATGATGAAGAGGAGGACG</td><td></td><td></td><td></td><td></td><td></td><td></td><td></td><td></td><td></td><td></td></tr>
<tr><td>Oligo rev</td><td>AAACCGTCCTCCTCTTCATCATCG</td><td></td><td></td><td></td><td></td><td></td><td></td><td></td><td></td><td></td><td></td></tr>
<tr><td>Chromosome</td><td>start</td><td>end</td><td>strand</td><td>MM</td><td>target_seq</td><td>PAM</td><td>alignment</td><td>distance</td><td>position</td><td>gene name</td><td>gene id</td></tr>
<tr><td>GU305914.1</td><td>118506</td><td>118528</td><td>-</td><td>0</td><td>CGATGATGAAGAGGAGGACG</td><td>AGG</td><td>||||||||[||||||||||||]PAM</td><td>NA</td><td>Intergenic</td><td>NA</td><td>NA</td></tr>
<tr><td>GU305914.1</td><td>118842</td><td>118864</td><td>-</td><td>4</td><td>AGAAGACGAAGAGGACGACG</td><td>AGG</td><td>-||-||-|[|||||||-||||]PAM</td><td>NA</td><td>Intergenic</td><td>NA</td><td>NA</td></tr>
<tr><td>GU305914.1</td><td>45324</td><td>45346</td><td>-</td><td>3</td><td>AGATGATGAAGTGGAGGAGG</td><td>AGG</td><td>-|||||||[|||-||||||-|]PAM</td><td>NA</td><td>Intergenic</td><td>NA</td><td>NA</td></tr>
<tr><td>GU305914.1</td><td>140717</td><td>140739</td><td>+</td><td>4</td><td>AGATGAGGACGAGGAGGATG</td><td>AGG</td><td>-|||||-|[|-||||||||-|]PAM</td><td>NA</td><td>Intergenic</td><td>NA</td><td>NA</td></tr>
<tr><td></td><td></td><td></td><td></td><td></td><td></td><td></td><td></td><td></td><td></td><td></td><td></td></tr>
<tr><td>T25072</td><td>GGAGGACGAGGACGACGACGAGG</td><td>1000</td><td></td><td></td><td></td><td></td><td></td><td></td><td></td><td></td><td></td></tr>
<tr><td>Oligo fwd</td><td>CACCGGAGGACGAGGACGACGACG</td><td></td><td></td><td></td><td></td><td></td><td></td><td></td><td></td><td></td><td></td></tr>
<tr><td>Oligo rev</td><td>AAACCGTCGTCGTCCTCGTCCTCC</td><td></td><td></td><td></td><td></td><td></td><td></td><td></td><td></td><td></td><td></td></tr>
<tr><td>Chromosome</td><td>start</td><td>end</td><td>strand</td><td>MM</td><td>target_seq</td><td>PAM</td><td>alignment</td><td>distance</td><td>position</td><td>gene name</td><td>gene id</td></tr>
<tr><td>GU305914.1</td><td>118494</td><td>118516</td><td>-</td><td>0</td><td>GGAGGACGAGGACGACGACG</td><td>AGG</td><td>||||||||[||||||||||||]PAM</td><td>NA</td><td>Intergenic</td><td>NA</td><td>NA</td></tr>
<tr><td>GU305914.1</td><td>65108</td><td>65130</td><td>+</td><td>2</td><td>GGAGGAAGAAGACGACGACG</td><td>AGG</td><td>||||||-|[|-||||||||||]PAM</td><td>NA</td><td>Intergenic</td><td>NA</td><td>NA</td></tr>
<tr><td>GU305914.1</td><td>118842</td><td>118864</td><td>-</td><td>4</td><td>AGAAGACGAAGAGGACGACG</td><td>AGG</td><td>-||-||||[|-||-|||||||]PAM</td><td>NA</td><td>Intergenic</td><td>NA</td><td>NA</td></tr>
<tr><td>GU305914.1</td><td>118812</td><td>118834</td><td>-</td><td>4</td><td>TGACGACGATGATGACGACG</td><td>AGG</td><td>-||-||||[|-||-|||||||]PAM</td><td>NA</td><td>Intergenic</td><td>NA</td><td>NA</td></tr>
<tr><td>GU305914.1</td><td>65111</td><td>65133</td><td>+</td><td>4</td><td>GGAAGAAGACGACGACGAGG</td><td>AGG</td><td>|||-||-|[|-||||||||-|]PAM</td><td>NA</td><td>Intergenic</td><td>NA</td><td>NA</td></tr>
<tr><td></td><td></td><td></td><td></td><td></td><td></td><td></td><td></td><td></td><td></td><td></td><td></td></tr>
<tr><td>T25073</td><td>GATGATGATGACGACGAAGACGG</td><td>1000</td><td></td><td></td><td></td><td></td><td></td><td></td><td></td><td></td><td></td></tr>
<tr><td>Oligo fwd</td><td>CACCGATGATGATGACGACGAAGA</td><td></td><td></td><td></td><td></td><td></td><td></td><td></td><td></td><td></td><td></td></tr>
<tr><td>Oligo rev</td><td>AAACTCTTCGTCGTCATCATCATC</td><td></td><td></td><td></td><td></td><td></td><td></td><td></td><td></td><td></td><td></td></tr>
<tr><td>Chromosome</td><td>start</td><td>end</td><td>strand</td><td>MM</td><td>target_seq</td><td>PAM</td><td>alignment</td><td>distance</td><td>position</td><td>gene name</td><td>gene id</td></tr>
<tr><td>GU305914.1</td><td>118472</td><td>118494</td><td>-</td><td>0</td><td>GATGATGATGACGACGAAGA</td><td>CGG</td><td>||||||||[||||||||||||]PAM</td><td>NA</td><td>Intergenic</td><td>NA</td><td>NA</td></tr>
</table>

<table>
<tr><td>GU305914.1</td><td>40448</td><td>40470</td><td>-</td><td>3</td><td>GATGAAGAAGACGACGATGA</td><td>CGG</td><td>|||||-||[-||||||||-||]PAM</td><td>NA</td><td>Intergenic</td><td>NA</td><td>NA</td></tr>
<tr><td>GU305914.1</td><td>52533</td><td>52555</td><td>-</td><td>3</td><td>GATGAAGATGACGACTACGA</td><td>TGG</td><td>|||||-||[|||||||-|-||]PAM</td><td>NA</td><td>Intergenic</td><td>NA</td><td>NA</td></tr>
<tr><td></td><td></td><td></td><td></td><td></td><td></td><td></td><td></td><td></td><td></td><td></td><td></td></tr>
<tr><td>T25074</td><td>AACAATGAAGAAGACGACGACGG</td><td>1000</td><td></td><td></td><td></td><td></td><td></td><td></td><td></td><td></td><td></td></tr>
<tr><td>Oligo fwd</td><td>CACCAACAATGAAGAAGACGACGA</td><td></td><td></td><td></td><td></td><td></td><td></td><td></td><td></td><td></td><td></td></tr>
<tr><td>Oligo rev</td><td>AAACTCGTCGTCTTCTTCATTGTT</td><td></td><td></td><td></td><td></td><td></td><td></td><td></td><td></td><td></td><td></td></tr>
<tr><td>Chromosome</td><td>start</td><td>end</td><td>strand</td><td>MM</td><td>target_seq</td><td>PAM</td><td>alignment</td><td>distance</td><td>position</td><td>gene name</td><td>gene id</td></tr>
<tr><td>GU305914.1</td><td>118445</td><td>118467</td><td>-</td><td>0</td><td>AACAATGAAGAAGACGACGA</td><td>CGG</td><td>||||||||[||||||||||||]PAM</td><td>NA</td><td>Intergenic</td><td>NA</td><td>NA</td></tr>
<tr><td>GU305914.1</td><td>102657</td><td>102679</td><td>-</td><td>4</td><td>GAGGAGGAAGAAGACGACGA</td><td>CGG</td><td>-|--|-||[||||||||||||]PAM</td><td>NA</td><td>Intergenic</td><td>NA</td><td>NA</td></tr>
<tr><td></td><td></td><td></td><td></td><td></td><td></td><td></td><td></td><td></td><td></td><td></td><td></td></tr>
<tr><td>T25075</td><td>CGTCAGCCGCGCCAACCGAACGG</td><td>1000</td><td></td><td></td><td></td><td></td><td></td><td></td><td></td><td></td><td></td></tr>
<tr><td>Oligo fwd</td><td>CACCCGTCAGCCGCGCCAACCGAA</td><td></td><td></td><td></td><td></td><td></td><td></td><td></td><td></td><td></td><td></td></tr>
<tr><td>Oligo rev</td><td>AAACTTCGGTTGGCGCGGCTGACG</td><td></td><td></td><td></td><td></td><td></td><td></td><td></td><td></td><td></td><td></td></tr>
<tr><td>Chromosome</td><td>start</td><td>end</td><td>strand</td><td>MM</td><td>target_seq</td><td>PAM</td><td>alignment</td><td>distance</td><td>position</td><td>gene name</td><td>gene id</td></tr>
<tr><td>GU305914.1</td><td>118418</td><td>118440</td><td>-</td><td>0</td><td>CGTCAGCCGCGCCAACCGAA</td><td>CGG</td><td>||||||||[||||||||||||]PAM</td><td>NA</td><td>Intergenic</td><td>NA</td><td>NA</td></tr>
<tr><td></td><td></td><td></td><td></td><td></td><td></td><td></td><td></td><td></td><td></td><td></td><td></td></tr>
<tr><td>T25076</td><td>CAGCCGCGCCAACCGAACGGCGG</td><td>1000</td><td></td><td></td><td></td><td></td><td></td><td></td><td></td><td></td><td></td></tr>
<tr><td>Oligo fwd</td><td>CACCCAGCCGCGCCAACCGAACGG</td><td></td><td></td><td></td><td></td><td></td><td></td><td></td><td></td><td></td><td></td></tr>
<tr><td>Oligo rev</td><td>AAACCCGTTCGGTTGGCGCGGCTG</td><td></td><td></td><td></td><td></td><td></td><td></td><td></td><td></td><td></td><td></td></tr>
<tr><td>Chromosome</td><td>start</td><td>end</td><td>strand</td><td>MM</td><td>target_seq</td><td>PAM</td><td>alignment</td><td>distance</td><td>position</td><td>gene name</td><td>gene id</td></tr>
<tr><td>GU305914.1</td><td>118415</td><td>118437</td><td>-</td><td>0</td><td>CAGCCGCGCCAACCGAACGG</td><td>CGG</td><td>||||||||[||||||||||||]PAM</td><td>NA</td><td>Intergenic</td><td>NA</td><td>NA</td></tr>
<tr><td></td><td></td><td></td><td></td><td></td><td></td><td></td><td></td><td></td><td></td><td></td><td></td></tr>
<tr><td>T25077</td><td>CCGAACGGCGGCAGCCGTGACGG</td><td>1000</td><td></td><td></td><td></td><td></td><td></td><td></td><td></td><td></td><td></td></tr>
<tr><td>Oligo fwd</td><td>CACCCCGAACGGCGGCAGCCGTGA</td><td></td><td></td><td></td><td></td><td></td><td></td><td></td><td></td><td></td><td></td></tr>
<tr><td>Oligo rev</td><td>AAACTCACGGCTGCCGCCGTTCGG</td><td></td><td></td><td></td><td></td><td></td><td></td><td></td><td></td><td></td><td></td></tr>
<tr><td>Chromosome</td><td>start</td><td>end</td><td>strand</td><td>MM</td><td>target_seq</td><td>PAM</td><td>alignment</td><td>distance</td><td>position</td><td>gene name</td><td>gene id</td></tr>
<tr><td>GU305914.1</td><td>118403</td><td>118425</td><td>-</td><td>0</td><td>CCGAACGGCGGCAGCCGTGA</td><td>CGG</td><td>||||||||[||||||||||||]PAM</td><td>NA</td><td>Intergenic</td><td>NA</td><td>NA</td></tr>
<tr><td></td><td></td><td></td><td></td><td></td><td></td><td></td><td></td><td></td><td></td><td></td><td></td></tr>
<tr><td>T25078</td><td>AGCCGTGACGGCCGCCGCCGCGG</td><td>1000</td><td></td><td></td><td></td><td></td><td></td><td></td><td></td><td></td><td></td></tr>
<tr><td>Oligo fwd</td><td>CACCAGCCGTGACGGCCGCCGCCG</td><td></td><td></td><td></td><td></td><td></td><td></td><td></td><td></td><td></td><td></td></tr>
<tr><td>Oligo rev</td><td>AAACCGGCGGCGGCCGTCACGGCT</td><td></td><td></td><td></td><td></td><td></td><td></td><td></td><td></td><td></td><td></td></tr>
<tr><td>Chromosome</td><td>start</td><td>end</td><td>strand</td><td>MM</td><td>target_seq</td><td>PAM</td><td>alignment</td><td>distance</td><td>position</td><td>gene name</td><td>gene id</td></tr>
<tr><td>GU305914.1</td><td>118391</td><td>118413</td><td>-</td><td>0</td><td>AGCCGTGACGGCCGCCGCCG</td><td>CGG</td><td>||||||||[||||||||||||]PAM</td><td>NA</td><td>Intergenic</td><td>NA</td><td>NA</td></tr>
<tr><td></td><td></td><td></td><td></td><td></td><td></td><td></td><td></td><td></td><td></td><td></td><td></td></tr>
<tr><td>T25079</td><td>GACGGCCGCCGCCGCGGTCTCGG</td><td>1000</td><td></td><td></td><td></td><td></td><td></td><td></td><td></td><td></td><td></td></tr>
<tr><td>Oligo fwd</td><td>CACCGACGGCCGCCGCCGCGGTCT</td><td></td><td></td><td></td><td></td><td></td><td></td><td></td><td></td><td></td><td></td></tr>
<tr><td>Oligo rev</td><td>AAACAGACCGCGGCGGCGGCCGTC</td><td></td><td></td><td></td><td></td><td></td><td></td><td></td><td></td><td></td><td></td></tr>
<tr><td>Chromosome</td><td>start</td><td>end</td><td>strand</td><td>MM</td><td>target_seq</td><td>PAM</td><td>alignment</td><td>distance</td><td>position</td><td>gene name</td><td>gene id</td></tr>
<tr><td>GU305914.1</td><td>118385</td><td>118407</td><td>-</td><td>0</td><td>GACGGCCGCCGCCGCGGTCT</td><td>CGG</td><td>||||||||[||||||||||||]PAM</td><td>NA</td><td>Intergenic</td><td>NA</td><td>NA</td></tr>
<tr><td></td><td></td><td></td><td></td><td></td><td></td><td></td><td></td><td></td><td></td><td></td><td></td></tr>
<tr><td>T25080</td><td>CTACCGCAACCTCGTCCGCGCGG</td><td>1000</td><td></td><td></td><td></td><td></td><td></td><td></td><td></td><td></td><td></td></tr>
<tr><td>Oligo fwd</td><td>CACCCTACCGCAACCTCGTCCGCG</td><td></td><td></td><td></td><td></td><td></td><td></td><td></td><td></td><td></td><td></td></tr>
<tr><td>Oligo rev</td><td>AAACCGCGGACGAGGTTGCGGTAG</td><td></td><td></td><td></td><td></td><td></td><td></td><td></td><td></td><td></td><td></td></tr>
<tr><td>Chromosome</td><td>start</td><td>end</td><td>strand</td><td>MM</td><td>target_seq</td><td>PAM</td><td>alignment</td><td>distance</td><td>position</td><td>gene name</td><td>gene id</td></tr>
<tr><td>GU305914.1</td><td>118350</td><td>118372</td><td>-</td><td>0</td><td>CTACCGCAACCTCGTCCGCG</td><td>CGG</td><td>||||||||[||||||||||||]PAM</td><td>NA</td><td>Intergenic</td><td>NA</td><td>NA</td></tr>
<tr><td></td><td></td><td></td><td></td><td></td><td></td><td></td><td></td><td></td><td></td><td></td><td></td></tr>
<tr><td>T25081</td><td>CGCGGCGACGAGAGACGCCATGG</td><td>1000</td><td></td><td></td><td></td><td></td><td></td><td></td><td></td><td></td><td></td></tr>
<tr><td>Oligo fwd</td><td>CACCCGCGGCGACGAGAGACGCCA</td><td></td><td></td><td></td><td></td><td></td><td></td><td></td><td></td><td></td><td></td></tr>
<tr><td>Oligo rev</td><td>AAACTGGCGTCTCTCGTCGCCGCG</td><td></td><td></td><td></td><td></td><td></td><td></td><td></td><td></td><td></td><td></td></tr>
<tr><td>Chromosome</td><td>start</td><td>end</td><td>strand</td><td>MM</td><td>target_seq</td><td>PAM</td><td>alignment</td><td>distance</td><td>position</td><td>gene name</td><td>gene id</td></tr>
<tr><td>GU305914.1</td><td>118332</td><td>118354</td><td>-</td><td>0</td><td>CGCGGCGACGAGAGACGCCA</td><td>TGG</td><td>||||||||[||||||||||||]PAM</td><td>NA</td><td>Intergenic</td><td>NA</td><td>NA</td></tr>
<tr><td></td><td></td><td></td><td></td><td></td><td></td><td></td><td></td><td></td><td></td><td></td><td></td></tr>
<tr><td>T25082</td><td>GCGGCGACGAGAGACGCCATGGG</td><td>1000</td><td></td><td></td><td></td><td></td><td></td><td></td><td></td><td></td><td></td></tr>
<tr><td>Oligo fwd</td><td>CACCGCGGCGACGAGAGACGCCAT</td><td></td><td></td><td></td><td></td><td></td><td></td><td></td><td></td><td></td><td></td></tr>
<tr><td>Oligo rev</td><td>AAACATGGCGTCTCTCGTCGCCGC</td><td></td><td></td><td></td><td></td><td></td><td></td><td></td><td></td><td></td><td></td></tr>
<tr><td>Chromosome</td><td>start</td><td>end</td><td>strand</td><td>MM</td><td>target_seq</td><td>PAM</td><td>alignment</td><td>distance</td><td>position</td><td>gene name</td><td>gene id</td></tr>
<tr><td>GU305914.1</td><td>118331</td><td>118353</td><td>-</td><td>0</td><td>GCGGCGACGAGAGACGCCAT</td><td>GGG</td><td>||||||||[||||||||||||]PAM</td><td>NA</td><td>Intergenic</td><td>NA</td><td>NA</td></tr>
<tr><td></td><td></td><td></td><td></td><td></td><td></td><td></td><td></td><td></td><td></td><td></td><td></td></tr>
<tr><td>T25083</td><td>CGCCATGGGCGCCGCCGTCGCGG</td><td>1000</td><td></td><td></td><td></td><td></td><td></td><td></td><td></td><td></td><td></td></tr>
<tr><td>Oligo fwd</td><td>CACCCGCCATGGGCGCCGCCGTCG</td><td></td><td></td><td></td><td></td><td></td><td></td><td></td><td></td><td></td><td></td></tr>
<tr><td>Oligo rev</td><td>AAACCGACGGCGGCGCCCATGGCG</td><td></td><td></td><td></td><td></td><td></td><td></td><td></td><td></td><td></td><td></td></tr>
<tr><td>Chromosome</td><td>start</td><td>end</td><td>strand</td><td>MM</td><td>target_seq</td><td>PAM</td><td>alignment</td><td>distance</td><td>position</td><td>gene name</td><td>gene id</td></tr>
<tr><td>GU305914.1</td><td>118317</td><td>118339</td><td>-</td><td>0</td><td>CGCCATGGGCGCCGCCGTCG</td><td>CGG</td><td>||||||||[||||||||||||]PAM</td><td>NA</td><td>Intergenic</td><td>NA</td><td>NA</td></tr>
<tr><td></td><td></td><td></td><td></td><td></td><td></td><td></td><td></td><td></td><td></td><td></td><td></td></tr>
<tr><td>T25084</td><td>CATGGGCGCCGCCGTCGCGGTGG</td><td>1000</td><td></td><td></td><td></td><td></td><td></td><td></td><td></td><td></td><td></td></tr>
<tr><td>Oligo fwd</td><td>CACCCATGGGCGCCGCCGTCGCGG</td><td></td><td></td><td></td><td></td><td></td><td></td><td></td><td></td><td></td><td></td></tr>
<tr><td>Oligo rev</td><td>AAACCCGCGACGGCGGCGCCCATG</td><td></td><td></td><td></td><td></td><td></td><td></td><td></td><td></td><td></td><td></td></tr>
<tr><td>Chromosome</td><td>start</td><td>end</td><td>strand</td><td>MM</td><td>target_seq</td><td>PAM</td><td>alignment</td><td>distance</td><td>position</td><td>gene name</td><td>gene id</td></tr>
<tr><td>GU305914.1</td><td>118314</td><td>118336</td><td>-</td><td>0</td><td>CATGGGCGCCGCCGTCGCGG</td><td>TGG</td><td>||||||||[||||||||||||]PAM</td><td>NA</td><td>Intergenic</td><td>NA</td><td>NA</td></tr>
<tr><td></td><td></td><td></td><td></td><td></td><td></td><td></td><td></td><td></td><td></td><td></td><td></td></tr>
<tr><td>T25085</td><td>GCCGTCGCGGTGGACGATTACGG</td><td>1000</td><td></td><td></td><td></td><td></td><td></td><td></td><td></td><td></td><td></td></tr>
<tr><td>Oligo fwd</td><td>CACCGCCGTCGCGGTGGACGATTA</td><td></td><td></td><td></td><td></td><td></td><td></td><td></td><td></td><td></td><td></td></tr>
<tr><td>Oligo rev</td><td>AAACTAATCGTCCACCGCGACGGC</td><td></td><td></td><td></td><td></td><td></td><td></td><td></td><td></td><td></td><td></td></tr>
<tr><td>Chromosome</td><td>start</td><td>end</td><td>strand</td><td>MM</td><td>target_seq</td><td>PAM</td><td>alignment</td><td>distance</td><td>position</td><td>gene name</td><td>gene id</td></tr>
<tr><td>GU305914.1</td><td>118304</td><td>118326</td><td>-</td><td>0</td><td>GCCGTCGCGGTGGACGATTA</td><td>CGG</td><td>||||||||[||||||||||||]PAM</td><td>NA</td><td>Intergenic</td><td>NA</td><td>NA</td></tr>
<tr><td>GU305914.1</td><td>142164</td><td>142186</td><td>-</td><td>4</td><td>GCCGACCCGGAGGACGATCA</td><td>GGG</td><td>||||-|-|[||-|||||||-|]PAM</td><td>NA</td><td>Intergenic</td><td>NA</td><td>NA</td></tr>
<tr><td></td><td></td><td></td><td></td><td></td><td></td><td></td><td></td><td></td><td></td><td></td><td></td></tr>
<tr><td>T25086</td><td>CCGTCGCGGTGGACGATTACGGG</td><td>1000</td><td></td><td></td><td></td><td></td><td></td><td></td><td></td><td></td><td></td></tr>
<tr><td>Oligo fwd</td><td>CACCCCGTCGCGGTGGACGATTAC</td><td></td><td></td><td></td><td></td><td></td><td></td><td></td><td></td><td></td><td></td></tr>
<tr><td>Oligo rev</td><td>AAACGTAATCGTCCACCGCGACGG</td><td></td><td></td><td></td><td></td><td></td><td></td><td></td><td></td><td></td><td></td></tr>
<tr><td>Chromosome</td><td>start</td><td>end</td><td>strand</td><td>MM</td><td>target_seq</td><td>PAM</td><td>alignment</td><td>distance</td><td>position</td><td>gene name</td><td>gene id</td></tr>
<tr><td>GU305914.1</td><td>118303</td><td>118325</td><td>-</td><td>0</td><td>CCGTCGCGGTGGACGATTAC</td><td>GGG</td><td>||||||||[||||||||||||]PAM</td><td>NA</td><td>Intergenic</td><td>NA</td><td>NA</td></tr>
<tr><td></td><td></td><td></td><td></td><td></td><td></td><td></td><td></td><td></td><td></td><td></td><td></td></tr>
<tr><td>T25087</td><td>CGTCGCGGTGGACGATTACGGGG</td><td>1000</td><td></td><td></td><td></td><td></td><td></td><td></td><td></td><td></td><td></td></tr>
<tr><td>Oligo fwd</td><td>CACCCGTCGCGGTGGACGATTACG</td><td></td><td></td><td></td><td></td><td></td><td></td><td></td><td></td><td></td><td></td></tr>
<tr><td>Oligo rev</td><td>AAACCGTAATCGTCCACCGCGACG</td><td></td><td></td><td></td><td></td><td></td><td></td><td></td><td></td><td></td><td></td></tr>
<tr><td>Chromosome</td><td>start</td><td>end</td><td>strand</td><td>MM</td><td>target_seq</td><td>PAM</td><td>alignment</td><td>distance</td><td>position</td><td>gene name</td><td>gene id</td></tr>
<tr><td>GU305914.1</td><td>118302</td><td>118324</td><td>-</td><td>0</td><td>CGTCGCGGTGGACGATTACG</td><td>GGG</td><td>||||||||[||||||||||||]PAM</td><td>NA</td><td>Intergenic</td><td>NA</td><td>NA</td></tr>
</table>

<table>
<tr><td>T25088</td><td>CGCGGTGGACGATTACGGGGTGG</td><td>1000</td><td></td><td></td><td></td><td></td><td></td><td></td><td></td><td></td><td></td></tr>
<tr><td>Oligo fwd</td><td>CACCCGCGGTGGACGATTACGGGG</td><td></td><td></td><td></td><td></td><td></td><td></td><td></td><td></td><td></td><td></td></tr>
<tr><td>Oligo rev</td><td>AAACCCCCGTAATCGTCCACCGCG</td><td></td><td></td><td></td><td></td><td></td><td></td><td></td><td></td><td></td><td></td></tr>
<tr><td>Chromosome</td><td>start</td><td>end</td><td>strand</td><td>MM</td><td>target_seq</td><td>PAM</td><td>alignment</td><td>distance</td><td>position</td><td>gene name</td><td>gene id</td></tr>
<tr><td>GU305914.1</td><td>118299</td><td>118321</td><td>-</td><td>0</td><td>CGCGGTGGACGATTACGGGG</td><td>TGG</td><td>||||||||[||||||||||||]PAM</td><td>NA</td><td>Intergenic</td><td>NA</td><td>NA</td></tr>
<tr><td>T25089</td><td>GGTGGACGATTACGGGGTGGAGG</td><td>1000</td><td></td><td></td><td></td><td></td><td></td><td></td><td></td><td></td><td></td></tr>
<tr><td>Oligo fwd</td><td>CACCGGTGGACGATTACGGGGTGG</td><td></td><td></td><td></td><td></td><td></td><td></td><td></td><td></td><td></td><td></td></tr>
<tr><td>Oligo rev</td><td>AAACCCACCCCGTAATCGTCCACC</td><td></td><td></td><td></td><td></td><td></td><td></td><td></td><td></td><td></td><td></td></tr>
<tr><td>Chromosome</td><td>start</td><td>end</td><td>strand</td><td>MM</td><td>target_seq</td><td>PAM</td><td>alignment</td><td>distance</td><td>position</td><td>gene name</td><td>gene id</td></tr>
<tr><td>GU305914.1</td><td>118296</td><td>118318</td><td>-</td><td>0</td><td>GGTGGACGATTACGGGGTGG</td><td>AGG</td><td>||||||||[||||||||||||]PAM</td><td>NA</td><td>Intergenic</td><td>NA</td><td>NA</td></tr>
<tr><td>T25090</td><td>CGAAGACGTGTACGACGAGCAGG</td><td>1000</td><td></td><td></td><td></td><td></td><td></td><td></td><td></td><td></td><td></td></tr>
<tr><td>Oligo fwd</td><td>CACCCGAAGACGTGTACGACGAGC</td><td></td><td></td><td></td><td></td><td></td><td></td><td></td><td></td><td></td><td></td></tr>
<tr><td>Oligo rev</td><td>AAACGCTCGTCGTACACGTCTTCG</td><td></td><td></td><td></td><td></td><td></td><td></td><td></td><td></td><td></td><td></td></tr>
<tr><td>Chromosome</td><td>start</td><td>end</td><td>strand</td><td>MM</td><td>target_seq</td><td>PAM</td><td>alignment</td><td>distance</td><td>position</td><td>gene name</td><td>gene id</td></tr>
<tr><td>GU305914.1</td><td>118272</td><td>118294</td><td>-</td><td>0</td><td>CGAAGACGTGTACGACGAGC</td><td>AGG</td><td>||||||||[||||||||||||]PAM</td><td>NA</td><td>Intergenic</td><td>NA</td><td>NA</td></tr>
<tr><td>T25091</td><td>GAAGACGTGTACGACGAGCAGGG</td><td>1000</td><td></td><td></td><td></td><td></td><td></td><td></td><td></td><td></td><td></td></tr>
<tr><td>Oligo fwd</td><td>CACCGAAGACGTGTACGACGAGCA</td><td></td><td></td><td></td><td></td><td></td><td></td><td></td><td></td><td></td><td></td></tr>
<tr><td>Oligo rev</td><td>AAACTGCTCGTCGTACACGTCTTC</td><td></td><td></td><td></td><td></td><td></td><td></td><td></td><td></td><td></td><td></td></tr>
<tr><td>Chromosome</td><td>start</td><td>end</td><td>strand</td><td>MM</td><td>target_seq</td><td>PAM</td><td>alignment</td><td>distance</td><td>position</td><td>gene name</td><td>gene id</td></tr>
<tr><td>GU305914.1</td><td>118271</td><td>118293</td><td>-</td><td>0</td><td>GAAGACGTGTACGACGAGCA</td><td>GGG</td><td>||||||||[||||||||||||]PAM</td><td>NA</td><td>Intergenic</td><td>NA</td><td>NA</td></tr>
<tr><td>T25092</td><td>AGGGCCGCCTGATCCTCCGTCGG</td><td>1000</td><td></td><td></td><td></td><td></td><td></td><td></td><td></td><td></td><td></td></tr>
<tr><td>Oligo fwd</td><td>CACCAGGGCCGCCTGATCCTCCGT</td><td></td><td></td><td></td><td></td><td></td><td></td><td></td><td></td><td></td><td></td></tr>
<tr><td>Oligo rev</td><td>AAACACGGAGGATCAGGCGGCCCT</td><td></td><td></td><td></td><td></td><td></td><td></td><td></td><td></td><td></td><td></td></tr>
<tr><td>Chromosome</td><td>start</td><td>end</td><td>strand</td><td>MM</td><td>target_seq</td><td>PAM</td><td>alignment</td><td>distance</td><td>position</td><td>gene name</td><td>gene id</td></tr>
<tr><td>GU305914.1</td><td>118252</td><td>118274</td><td>-</td><td>0</td><td>AGGGCCGCCTGATCCTCCGT</td><td>CGG</td><td>||||||||[||||||||||||]PAM</td><td>NA</td><td>Intergenic</td><td>NA</td><td>NA</td></tr>
<tr><td>T25093</td><td>GGGCCGCCTGATCCTCCGTCGGG</td><td>1000</td><td></td><td></td><td></td><td></td><td></td><td></td><td></td><td></td><td></td></tr>
<tr><td>Oligo fwd</td><td>CACCGGGCCGCCTGATCCTCCGTC</td><td></td><td></td><td></td><td></td><td></td><td></td><td></td><td></td><td></td><td></td></tr>
<tr><td>Oligo rev</td><td>AAACGACGGAGGATCAGGCGGCCC</td><td></td><td></td><td></td><td></td><td></td><td></td><td></td><td></td><td></td><td></td></tr>
<tr><td>Chromosome</td><td>start</td><td>end</td><td>strand</td><td>MM</td><td>target_seq</td><td>PAM</td><td>alignment</td><td>distance</td><td>position</td><td>gene name</td><td>gene id</td></tr>
<tr><td>GU305914.1</td><td>118251</td><td>118273</td><td>-</td><td>0</td><td>GGGCCGCCTGATCCTCCGTC</td><td>GGG</td><td>||||||||[||||||||||||]PAM</td><td>NA</td><td>Intergenic</td><td>NA</td><td>NA</td></tr>
<tr><td>GU305914.1</td><td>96867</td><td>96889</td><td>-</td><td>4</td><td>CGCCCGCACGATCCTCCGTC</td><td>GGG</td><td>-|-||||-[-|||||||||||]PAM</td><td>NA</td><td>Intergenic</td><td>NA</td><td>NA</td></tr>
<tr><td>T25094</td><td>CCGCCTGATCCTCCGTCGGGAGG</td><td>1000</td><td></td><td></td><td></td><td></td><td></td><td></td><td></td><td></td><td></td></tr>
<tr><td>Oligo fwd</td><td>CACCCCGCCTGATCCTCCGTCGGG</td><td></td><td></td><td></td><td></td><td></td><td></td><td></td><td></td><td></td><td></td></tr>
<tr><td>Oligo rev</td><td>AAACCCCGACGGAGGATCAGGCGG</td><td></td><td></td><td></td><td></td><td></td><td></td><td></td><td></td><td></td><td></td></tr>
<tr><td>Chromosome</td><td>start</td><td>end</td><td>strand</td><td>MM</td><td>target_seq</td><td>PAM</td><td>alignment</td><td>distance</td><td>position</td><td>gene name</td><td>gene id</td></tr>
<tr><td>GU305914.1</td><td>118248</td><td>118270</td><td>-</td><td>0</td><td>CCGCCTGATCCTCCGTCGGG</td><td>AGG</td><td>||||||||[||||||||||||]PAM</td><td>NA</td><td>Intergenic</td><td>NA</td><td>NA</td></tr>
<tr><td>T25095</td><td>CCTGATCCTCCGTCGGGAGGAGG</td><td>1000</td><td></td><td></td><td></td><td></td><td></td><td></td><td></td><td></td><td></td></tr>
<tr><td>Oligo fwd</td><td>CACCCCTGATCCTCCGTCGGGAGG</td><td></td><td></td><td></td><td></td><td></td><td></td><td></td><td></td><td></td><td></td></tr>
<tr><td>Oligo rev</td><td>AAACCCTCCCGACGGAGGATCAGG</td><td></td><td></td><td></td><td></td><td></td><td></td><td></td><td></td><td></td><td></td></tr>
<tr><td>Chromosome</td><td>start</td><td>end</td><td>strand</td><td>MM</td><td>target_seq</td><td>PAM</td><td>alignment</td><td>distance</td><td>position</td><td>gene name</td><td>gene id</td></tr>
<tr><td>GU305914.1</td><td>118245</td><td>118267</td><td>-</td><td>0</td><td>CCTGATCCTCCGTCGGGAGG</td><td>AGG</td><td>||||||||[||||||||||||]PAM</td><td>NA</td><td>Intergenic</td><td>NA</td><td>NA</td></tr>
<tr><td>T25096</td><td>CCTCCGTCGGGAGGAGGACGAGG</td><td>1000</td><td></td><td></td><td></td><td></td><td></td><td></td><td></td><td></td><td></td></tr>
<tr><td>Oligo fwd</td><td>CACCCCTCCGTCGGGAGGAGGACG</td><td></td><td></td><td></td><td></td><td></td><td></td><td></td><td></td><td></td><td></td></tr>
<tr><td>Oligo rev</td><td>AAACCGTCCTCCTCCCGACGGAGG</td><td></td><td></td><td></td><td></td><td></td><td></td><td></td><td></td><td></td><td></td></tr>
<tr><td>Chromosome</td><td>start</td><td>end</td><td>strand</td><td>MM</td><td>target_seq</td><td>PAM</td><td>alignment</td><td>distance</td><td>position</td><td>gene name</td><td>gene id</td></tr>
<tr><td>GU305914.1</td><td>118239</td><td>118261</td><td>-</td><td>0</td><td>CCTCCGTCGGGAGGAGGACG</td><td>AGG</td><td>||||||||[||||||||||||]PAM</td><td>NA</td><td>Intergenic</td><td>NA</td><td>NA</td></tr>
<tr><td>T25097</td><td>AGGACGAGGAGCCCGAACCGCGG</td><td>1000</td><td></td><td></td><td></td><td></td><td></td><td></td><td></td><td></td><td></td></tr>
<tr><td>Oligo fwd</td><td>CACCAGGACGAGGAGCCCGAACCG</td><td></td><td></td><td></td><td></td><td></td><td></td><td></td><td></td><td></td><td></td></tr>
<tr><td>Oligo rev</td><td>AAACCGGTTCGGGCTCCTCGTCCT</td><td></td><td></td><td></td><td></td><td></td><td></td><td></td><td></td><td></td><td></td></tr>
<tr><td>Chromosome</td><td>start</td><td>end</td><td>strand</td><td>MM</td><td>target_seq</td><td>PAM</td><td>alignment</td><td>distance</td><td>position</td><td>gene name</td><td>gene id</td></tr>
<tr><td>GU305914.1</td><td>118225</td><td>118247</td><td>-</td><td>0</td><td>AGGACGAGGAGCCCGAACCG</td><td>CGG</td><td>||||||||[||||||||||||]PAM</td><td>NA</td><td>Intergenic</td><td>NA</td><td>NA</td></tr>
<tr><td>T25098</td><td>ACCGCGGCCGAATCAGCCGACGG</td><td>1000</td><td></td><td></td><td></td><td></td><td></td><td></td><td></td><td></td><td></td></tr>
<tr><td>Oligo fwd</td><td>CACCACCGCGGCCGAATCAGCCGA</td><td></td><td></td><td></td><td></td><td></td><td></td><td></td><td></td><td></td><td></td></tr>
<tr><td>Oligo rev</td><td>AAACTCGGCTGATTCGGCCGCGGT</td><td></td><td></td><td></td><td></td><td></td><td></td><td></td><td></td><td></td><td></td></tr>
<tr><td>Chromosome</td><td>start</td><td>end</td><td>strand</td><td>MM</td><td>target_seq</td><td>PAM</td><td>alignment</td><td>distance</td><td>position</td><td>gene name</td><td>gene id</td></tr>
<tr><td>GU305914.1</td><td>118209</td><td>118231</td><td>-</td><td>0</td><td>ACCGCGGCCGAATCAGCCGA</td><td>CGG</td><td>||||||||[||||||||||||]PAM</td><td>NA</td><td>Intergenic</td><td>NA</td><td>NA</td></tr>
<tr><td>T25099</td><td>CCGAATCAGCCGACGGAGTTCGG</td><td>1000</td><td></td><td></td><td></td><td></td><td></td><td></td><td></td><td></td><td></td></tr>
<tr><td>Oligo fwd</td><td>CACCCCGAATCAGCCGACGGAGTT</td><td></td><td></td><td></td><td></td><td></td><td></td><td></td><td></td><td></td><td></td></tr>
<tr><td>Oligo rev</td><td>AAACAACTCCGTCGGCTGATTCGG</td><td></td><td></td><td></td><td></td><td></td><td></td><td></td><td></td><td></td><td></td></tr>
<tr><td>Chromosome</td><td>start</td><td>end</td><td>strand</td><td>MM</td><td>target_seq</td><td>PAM</td><td>alignment</td><td>distance</td><td>position</td><td>gene name</td><td>gene id</td></tr>
<tr><td>GU305914.1</td><td>118202</td><td>118224</td><td>-</td><td>0</td><td>CCGAATCAGCCGACGGAGTT</td><td>CGG</td><td>||||||||[||||||||||||]PAM</td><td>NA</td><td>Intergenic</td><td>NA</td><td>NA</td></tr>
<tr><td>T25100</td><td>TAACCGCAGAGACGAAGCCGCGG</td><td>1000</td><td></td><td></td><td></td><td></td><td></td><td></td><td></td><td></td><td></td></tr>
<tr><td>Oligo fwd</td><td>CACCTAACCGCAGAGACGAAGCCG</td><td></td><td></td><td></td><td></td><td></td><td></td><td></td><td></td><td></td><td></td></tr>
<tr><td>Oligo rev</td><td>AAACCGGCTTCGTCTCTGCGGTTA</td><td></td><td></td><td></td><td></td><td></td><td></td><td></td><td></td><td></td><td></td></tr>
<tr><td>Chromosome</td><td>start</td><td>end</td><td>strand</td><td>MM</td><td>target_seq</td><td>PAM</td><td>alignment</td><td>distance</td><td>position</td><td>gene name</td><td>gene id</td></tr>
<tr><td>GU305914.1</td><td>118179</td><td>118201</td><td>-</td><td>0</td><td>TAACCGCAGAGACGAAGCCG</td><td>CGG</td><td>||||||||[||||||||||||]PAM</td><td>NA</td><td>Intergenic</td><td>NA</td><td>NA</td></tr>
<tr><td>T25101</td><td>CGCGGAAGAAGAGCGCACACCGG</td><td>1000</td><td></td><td></td><td></td><td></td><td></td><td></td><td></td><td></td><td></td></tr>
<tr><td>Oligo fwd</td><td>CACCCGCGGAAGAAGAGCGCACAC</td><td></td><td></td><td></td><td></td><td></td><td></td><td></td><td></td><td></td><td></td></tr>
<tr><td>Oligo rev</td><td>AAACGTGTGCGCTCTTCTTCCGCG</td><td></td><td></td><td></td><td></td><td></td><td></td><td></td><td></td><td></td><td></td></tr>
<tr><td>Chromosome</td><td>start</td><td>end</td><td>strand</td><td>MM</td><td>target_seq</td><td>PAM</td><td>alignment</td><td>distance</td><td>position</td><td>gene name</td><td>gene id</td></tr>
<tr><td>GU305914.1</td><td>118161</td><td>118183</td><td>-</td><td>0</td><td>CGCGGAAGAAGAGCGCACAC</td><td>CGG</td><td>||||||||[||||||||||||]PAM</td><td>NA</td><td>Intergenic</td><td>NA</td><td>NA</td></tr>
<tr><td>T25102</td><td>AGAGCGCACACCGGATCCCGAGG</td><td>1000</td><td></td><td></td><td></td><td></td><td></td><td></td><td></td><td></td><td></td></tr>
<tr><td>Oligo fwd</td><td>CACCAGAGCGCACACCGGATCCCG</td><td></td><td></td><td></td><td></td><td></td><td></td><td></td><td></td><td></td><td></td></tr>
<tr><td>Oligo rev</td><td>AAACCGGGATCCGGTGTGCGCTCT</td><td></td><td></td><td></td><td></td><td></td><td></td><td></td><td></td><td></td><td></td></tr>
<tr><td>Chromosome</td><td>start</td><td>end</td><td>strand</td><td>MM</td><td>target_seq</td><td>PAM</td><td>alignment</td><td>distance</td><td>position</td><td>gene name</td><td>gene id</td></tr>
</table>

<table>
<tr><td>GU305914.1</td><td>118152</td><td>118174</td><td>-</td><td>0</td><td>AGAGCGCACACCGGATCCCG</td><td>AGG</td><td>||||||||[||||||||||||]PAM</td><td>NA</td><td>Intergenic</td><td>NA</td><td>NA</td></tr>
<tr><td>T25103</td><td>CGATAGCCCTCTTTCTGCAGCGG</td><td>1000</td><td></td><td></td><td></td><td></td><td></td><td></td><td></td><td></td><td></td></tr>
<tr><td>Oligo fwd</td><td>CACCCGATAGCCCTCTTTCTGCAG</td><td></td><td></td><td></td><td></td><td></td><td></td><td></td><td></td><td></td><td></td></tr>
<tr><td>Oligo rev</td><td>AAACCTGCAGAAAGAGGGCTATCG</td><td></td><td></td><td></td><td></td><td></td><td></td><td></td><td></td><td></td><td></td></tr>
<tr><td>Chromosome</td><td>start</td><td>end</td><td>strand</td><td>MM</td><td>target_seq</td><td>PAM</td><td>alignment</td><td>distance</td><td>position</td><td>gene name</td><td>gene id</td></tr>
<tr><td>GU305914.1</td><td>118129</td><td>118151</td><td>-</td><td>0</td><td>CGATAGCCCTCTTTCTGCAG</td><td>CGG</td><td>||||||||[||||||||||||]PAM</td><td>NA</td><td>Intergenic</td><td>NA</td><td>NA</td></tr>
<tr><td>T25104</td><td>GATAGCCCTCTTTCTGCAGCGGG</td><td>1000</td><td></td><td></td><td></td><td></td><td></td><td></td><td></td><td></td><td></td></tr>
<tr><td>Oligo fwd</td><td>CACCGATAGCCCTCTTTCTGCAGC</td><td></td><td></td><td></td><td></td><td></td><td></td><td></td><td></td><td></td><td></td></tr>
<tr><td>Oligo rev</td><td>AAACGCTGCAGAAAGAGGGCTATC</td><td></td><td></td><td></td><td></td><td></td><td></td><td></td><td></td><td></td><td></td></tr>
<tr><td>Chromosome</td><td>start</td><td>end</td><td>strand</td><td>MM</td><td>target_seq</td><td>PAM</td><td>alignment</td><td>distance</td><td>position</td><td>gene name</td><td>gene id</td></tr>
<tr><td>GU305914.1</td><td>118128</td><td>118150</td><td>-</td><td>0</td><td>GATAGCCCTCTTTCTGCAGC</td><td>GGG</td><td>||||||||[||||||||||||]PAM</td><td>NA</td><td>Intergenic</td><td>NA</td><td>NA</td></tr>
<tr><td>T25105</td><td>CCTCTTTCTGCAGCGGGAACCGG</td><td>1000</td><td></td><td></td><td></td><td></td><td></td><td></td><td></td><td></td><td></td></tr>
<tr><td>Oligo fwd</td><td>CACCCCTCTTTCTGCAGCGGGAAC</td><td></td><td></td><td></td><td></td><td></td><td></td><td></td><td></td><td></td><td></td></tr>
<tr><td>Oligo rev</td><td>AAACGTTCCCGCTGCAGAAAGAGG</td><td></td><td></td><td></td><td></td><td></td><td></td><td></td><td></td><td></td><td></td></tr>
<tr><td>Chromosome</td><td>start</td><td>end</td><td>strand</td><td>MM</td><td>target_seq</td><td>PAM</td><td>alignment</td><td>distance</td><td>position</td><td>gene name</td><td>gene id</td></tr>
<tr><td>GU305914.1</td><td>118122</td><td>118144</td><td>-</td><td>0</td><td>CCTCTTTCTGCAGCGGGAAC</td><td>CGG</td><td>||||||||[||||||||||||]PAM</td><td>NA</td><td>Intergenic</td><td>NA</td><td>NA</td></tr>
<tr><td>T25106</td><td>TTCTGCAGCGGGAACCGGTCAGG</td><td>1000</td><td></td><td></td><td></td><td></td><td></td><td></td><td></td><td></td><td></td></tr>
<tr><td>Oligo fwd</td><td>CACCTTCTGCAGCGGGAACCGGTC</td><td></td><td></td><td></td><td></td><td></td><td></td><td></td><td></td><td></td><td></td></tr>
<tr><td>Oligo rev</td><td>AAACGACCGGTTCCCGCTGCAGAA</td><td></td><td></td><td></td><td></td><td></td><td></td><td></td><td></td><td></td><td></td></tr>
<tr><td>Chromosome</td><td>start</td><td>end</td><td>strand</td><td>MM</td><td>target_seq</td><td>PAM</td><td>alignment</td><td>distance</td><td>position</td><td>gene name</td><td>gene id</td></tr>
<tr><td>GU305914.1</td><td>118117</td><td>118139</td><td>-</td><td>0</td><td>TTCTGCAGCGGGAACCGGTC</td><td>AGG</td><td>||||||||[||||||||||||]PAM</td><td>NA</td><td>Intergenic</td><td>NA</td><td>NA</td></tr>
<tr><td>T25107</td><td>ACCGGTCAGGATCTCTTTTCAGG</td><td>1000</td><td></td><td></td><td></td><td></td><td></td><td></td><td></td><td></td><td></td></tr>
<tr><td>Oligo fwd</td><td>CACCACCGGTCAGGATCTCTTTTC</td><td></td><td></td><td></td><td></td><td></td><td></td><td></td><td></td><td></td><td></td></tr>
<tr><td>Oligo rev</td><td>AAACGAAAAGAGATCCTGACCGGT</td><td></td><td></td><td></td><td></td><td></td><td></td><td></td><td></td><td></td><td></td></tr>
<tr><td>Chromosome</td><td>start</td><td>end</td><td>strand</td><td>MM</td><td>target_seq</td><td>PAM</td><td>alignment</td><td>distance</td><td>position</td><td>gene name</td><td>gene id</td></tr>
<tr><td>GU305914.1</td><td>118104</td><td>118126</td><td>-</td><td>0</td><td>ACCGGTCAGGATCTCTTTTC</td><td>AGG</td><td>||||||||[||||||||||||]PAM</td><td>NA</td><td>Intergenic</td><td>NA</td><td>NA</td></tr>
<tr><td>T25108</td><td>CAGGAACATTATTACAGCCACGG</td><td>1000</td><td></td><td></td><td></td><td></td><td></td><td></td><td></td><td></td><td></td></tr>
<tr><td>Oligo fwd</td><td>CACCCAGGAACATTATTACAGCCA</td><td></td><td></td><td></td><td></td><td></td><td></td><td></td><td></td><td></td><td></td></tr>
<tr><td>Oligo rev</td><td>AAACTGGCTGTAATAATGTTCCTG</td><td></td><td></td><td></td><td></td><td></td><td></td><td></td><td></td><td></td><td></td></tr>
<tr><td>Chromosome</td><td>start</td><td>end</td><td>strand</td><td>MM</td><td>target_seq</td><td>PAM</td><td>alignment</td><td>distance</td><td>position</td><td>gene name</td><td>gene id</td></tr>
<tr><td>GU305914.1</td><td>118085</td><td>118107</td><td>-</td><td>0</td><td>CAGGAACATTATTACAGCCA</td><td>CGG</td><td>||||||||[||||||||||||]PAM</td><td>NA</td><td>Intergenic</td><td>NA</td><td>NA</td></tr>
<tr><td>T25109</td><td>CGGACGACAAGTGAACCCGCTGG</td><td>1000</td><td></td><td></td><td></td><td></td><td></td><td></td><td></td><td></td><td></td></tr>
<tr><td>Oligo fwd</td><td>CACCCGGACGACAAGTGAACCCGC</td><td></td><td></td><td></td><td></td><td></td><td></td><td></td><td></td><td></td><td></td></tr>
<tr><td>Oligo rev</td><td>AAACGCGGGTTCACTTGTCGTCCG</td><td></td><td></td><td></td><td></td><td></td><td></td><td></td><td></td><td></td><td></td></tr>
<tr><td>Chromosome</td><td>start</td><td>end</td><td>strand</td><td>MM</td><td>target_seq</td><td>PAM</td><td>alignment</td><td>distance</td><td>position</td><td>gene name</td><td>gene id</td></tr>
<tr><td>GU305914.1</td><td>118065</td><td>118087</td><td>-</td><td>0</td><td>CGGACGACAAGTGAACCCGC</td><td>TGG</td><td>||||||||[||||||||||||]PAM</td><td>NA</td><td>Intergenic</td><td>NA</td><td>NA</td></tr>
<tr><td>T25110</td><td>CCCGCTGGTGTCGCCCTCGACGG</td><td>1000</td><td></td><td></td><td></td><td></td><td></td><td></td><td></td><td></td><td></td></tr>
<tr><td>Oligo fwd</td><td>CACCCCCGCTGGTGTCGCCCTCGA</td><td></td><td></td><td></td><td></td><td></td><td></td><td></td><td></td><td></td><td></td></tr>
<tr><td>Oligo rev</td><td>AAACTCGAGGGCGACACCAGCGGG</td><td></td><td></td><td></td><td></td><td></td><td></td><td></td><td></td><td></td><td></td></tr>
<tr><td>Chromosome</td><td>start</td><td>end</td><td>strand</td><td>MM</td><td>target_seq</td><td>PAM</td><td>alignment</td><td>distance</td><td>position</td><td>gene name</td><td>gene id</td></tr>
<tr><td>GU305914.1</td><td>118050</td><td>118072</td><td>-</td><td>0</td><td>CCCGCTGGTGTCGCCCTCGA</td><td>CGG</td><td>||||||||[||||||||||||]PAM</td><td>NA</td><td>Intergenic</td><td>NA</td><td>NA</td></tr>
<tr><td>T25111</td><td>AACATGAACTGCACGAAAACAGG</td><td>1000</td><td></td><td></td><td></td><td></td><td></td><td></td><td></td><td></td><td></td></tr>
<tr><td>Oligo fwd</td><td>CACCAACATGAACTGCACGAAAAC</td><td></td><td></td><td></td><td></td><td></td><td></td><td></td><td></td><td></td><td></td></tr>
<tr><td>Oligo rev</td><td>AAACGTTTTCGTGCAGTTCATGTT</td><td></td><td></td><td></td><td></td><td></td><td></td><td></td><td></td><td></td><td></td></tr>
<tr><td>Chromosome</td><td>start</td><td>end</td><td>strand</td><td>MM</td><td>target_seq</td><td>PAM</td><td>alignment</td><td>distance</td><td>position</td><td>gene name</td><td>gene id</td></tr>
<tr><td>GU305914.1</td><td>118022</td><td>118044</td><td>-</td><td>0</td><td>AACATGAACTGCACGAAAAC</td><td>AGG</td><td>||||||||[||||||||||||]PAM</td><td>NA</td><td>Intergenic</td><td>NA</td><td>NA</td></tr>
<tr><td>T25112</td><td>AGGACTGTCGAAACGAGAGATGG</td><td>1000</td><td></td><td></td><td></td><td></td><td></td><td></td><td></td><td></td><td></td></tr>
<tr><td>Oligo fwd</td><td>CACCAGGACTGTCGAAACGAGAGA</td><td></td><td></td><td></td><td></td><td></td><td></td><td></td><td></td><td></td><td></td></tr>
<tr><td>Oligo rev</td><td>AAACTCTCTCGTTTCGACAGTCCT</td><td></td><td></td><td></td><td></td><td></td><td></td><td></td><td></td><td></td><td></td></tr>
<tr><td>Chromosome</td><td>start</td><td>end</td><td>strand</td><td>MM</td><td>target_seq</td><td>PAM</td><td>alignment</td><td>distance</td><td>position</td><td>gene name</td><td>gene id</td></tr>
<tr><td>GU305914.1</td><td>118002</td><td>118024</td><td>-</td><td>0</td><td>AGGACTGTCGAAACGAGAGA</td><td>TGG</td><td>||||||||[||||||||||||]PAM</td><td>NA</td><td>Intergenic</td><td>NA</td><td>NA</td></tr>
<tr><td>T25113</td><td>AGACCTGCAGCAGTTCAATCTGG</td><td>1000</td><td></td><td></td><td></td><td></td><td></td><td></td><td></td><td></td><td></td></tr>
<tr><td>Oligo fwd</td><td>CACCAGACCTGCAGCAGTTCAATC</td><td></td><td></td><td></td><td></td><td></td><td></td><td></td><td></td><td></td><td></td></tr>
<tr><td>Oligo rev</td><td>AAACGATTGAACTGCTGCAGGTCT</td><td></td><td></td><td></td><td></td><td></td><td></td><td></td><td></td><td></td><td></td></tr>
<tr><td>Chromosome</td><td>start</td><td>end</td><td>strand</td><td>MM</td><td>target_seq</td><td>PAM</td><td>alignment</td><td>distance</td><td>position</td><td>gene name</td><td>gene id</td></tr>
<tr><td>GU305914.1</td><td>117963</td><td>117985</td><td>-</td><td>0</td><td>AGACCTGCAGCAGTTCAATC</td><td>TGG</td><td>||||||||[||||||||||||]PAM</td><td>NA</td><td>Intergenic</td><td>NA</td><td>NA</td></tr>
<tr><td>T25114</td><td>ACTGATCCTCTTTCCCCTGTCGG</td><td>1000</td><td></td><td></td><td></td><td></td><td></td><td></td><td></td><td></td><td></td></tr>
<tr><td>Oligo fwd</td><td>CACCACTGATCCTCTTTCCCCTGT</td><td></td><td></td><td></td><td></td><td></td><td></td><td></td><td></td><td></td><td></td></tr>
<tr><td>Oligo rev</td><td>AAACACAGGGGAAAGAGGATCAGT</td><td></td><td></td><td></td><td></td><td></td><td></td><td></td><td></td><td></td><td></td></tr>
<tr><td>Chromosome</td><td>start</td><td>end</td><td>strand</td><td>MM</td><td>target_seq</td><td>PAM</td><td>alignment</td><td>distance</td><td>position</td><td>gene name</td><td>gene id</td></tr>
<tr><td>GU305914.1</td><td>117927</td><td>117949</td><td>-</td><td>0</td><td>ACTGATCCTCTTTCCCCTGT</td><td>CGG</td><td>||||||||[||||||||||||]PAM</td><td>NA</td><td>Intergenic</td><td>NA</td><td>NA</td></tr>
<tr><td>T25115</td><td>CTTTCCCCTGTCGGTGTGCCTGG</td><td>1000</td><td></td><td></td><td></td><td></td><td></td><td></td><td></td><td></td><td></td></tr>
<tr><td>Oligo fwd</td><td>CACCCTTTCCCCTGTCGGTGTGCC</td><td></td><td></td><td></td><td></td><td></td><td></td><td></td><td></td><td></td><td></td></tr>
<tr><td>Oligo rev</td><td>AAACGGCACACCGACAGGGGAAAG</td><td></td><td></td><td></td><td></td><td></td><td></td><td></td><td></td><td></td><td></td></tr>
<tr><td>Chromosome</td><td>start</td><td>end</td><td>strand</td><td>MM</td><td>target_seq</td><td>PAM</td><td>alignment</td><td>distance</td><td>position</td><td>gene name</td><td>gene id</td></tr>
<tr><td>GU305914.1</td><td>117918</td><td>117940</td><td>-</td><td>0</td><td>CTTTCCCCTGTCGGTGTGCC</td><td>TGG</td><td>||||||||[||||||||||||]PAM</td><td>NA</td><td>Intergenic</td><td>NA</td><td>NA</td></tr>
<tr><td>T25116</td><td>GTCGGTGTGCCTGGACGAAACGG</td><td>1000</td><td></td><td></td><td></td><td></td><td></td><td></td><td></td><td></td><td></td></tr>
<tr><td>Oligo fwd</td><td>CACCGTCGGTGTGCCTGGACGAAA</td><td></td><td></td><td></td><td></td><td></td><td></td><td></td><td></td><td></td><td></td></tr>
<tr><td>Oligo rev</td><td>AAACTTTCGTCCAGGCACACCGAC</td><td></td><td></td><td></td><td></td><td></td><td></td><td></td><td></td><td></td><td></td></tr>
<tr><td>Chromosome</td><td>start</td><td>end</td><td>strand</td><td>MM</td><td>target_seq</td><td>PAM</td><td>alignment</td><td>distance</td><td>position</td><td>gene name</td><td>gene id</td></tr>
<tr><td>GU305914.1</td><td>117909</td><td>117931</td><td>-</td><td>0</td><td>GTCGGTGTGCCTGGACGAAA</td><td>CGG</td><td>||||||||[||||||||||||]PAM</td><td>NA</td><td>Intergenic</td><td>NA</td><td>NA</td></tr>
<tr><td>GU305914.1</td><td>75601</td><td>75623</td><td>+</td><td>4</td><td>GTTGGCGTGCCTGGTCAAAA</td><td>AGG</td><td>||-||-||[||||||-|-|||]PAM</td><td>NA</td><td>Intergenic</td><td>NA</td><td>NA</td></tr>
<tr><td>T25117</td><td>GTGCCTGGACGAAACGGTTCTGG</td><td>1000</td><td></td><td></td><td></td><td></td><td></td><td></td><td></td><td></td><td></td></tr>
<tr><td>Oligo fwd</td><td>CACCGTGCCTGGACGAAACGGTTC</td><td></td><td></td><td></td><td></td><td></td><td></td><td></td><td></td><td></td><td></td></tr>
</table>

Oligo rev AAACGAACCGTTTCGTCCAGGCAC
Chromosome start end strand MM target_seq PAM alignment distance position gene name gene id
GU305914.1 117903 117925 - 0 GTGCCTGGACGAAACGGTTC TGG ||||||||[||||||||||||]PAM NA Intergenic NA NA

T25118 GTTCCTGTTCATTATCCCCCCGG 1000
Oligo fwd CACCGTTCCTGTTCATTATCCCCC
Oligo rev AAACGGGGGATAATGAACAGGAAC
Chromosome start end strand MM target_seq PAM alignment distance position gene name gene id
GU305914.1 117876 117898 - 0 GTTCCTGTTCATTATCCCCC CGG ||||||||[||||||||||||]PAM NA Intergenic NA NA

T25119 ACCGTTCAACTTCACGTTCCTGG 1000
Oligo fwd CACCACCGTTCAACTTCACGTTCC
Oligo rev AAACGGAACGTGAAGTTGAACGGT
Chromosome start end strand MM target_seq PAM alignment distance position gene name gene id
GU305914.1 117831 117853 - 0 ACCGTTCAACTTCACGTTCC TGG ||||||||[||||||||||||]PAM NA Intergenic NA NA

T25120 TTACCGCCTGAGCCAGCTGACGG 1000
Oligo fwd CACCTTACCGCCTGAGCCAGCTGA
Oligo rev AAACTCAGCTGGCTCAGGCGGTAA
Chromosome start end strand MM target_seq PAM alignment distance position gene name gene id
GU305914.1 117792 117814 - 0 TTACCGCCTGAGCCAGCTGA CGG ||||||||[||||||||||||]PAM NA Intergenic NA NA

T25121 TACCGCCTGAGCCAGCTGACGGG 1000
Oligo fwd CACCTACCGCCTGAGCCAGCTGAC
Oligo rev AAACGTCAGCTGGCTCAGGCGGTA
Chromosome start end strand MM target_seq PAM alignment distance position gene name gene id
GU305914.1 117791 117813 - 0 TACCGCCTGAGCCAGCTGAC GGG ||||||||[||||||||||||]PAM NA Intergenic NA NA

T25122 CCTGAGCCAGCTGACGGGACAGG 1000
Oligo fwd CACCCCTGAGCCAGCTGACGGGAC
Oligo rev AAACGTCCCGTCAGCTGGCTCAGG
Chromosome start end strand MM target_seq PAM alignment distance position gene name gene id
GU305914.1 117786 117808 - 0 CCTGAGCCAGCTGACGGGAC AGG ||||||||[||||||||||||]PAM NA Intergenic NA NA

T25123 CTGACGGGACAGGAGATCCCAGG 1000
Oligo fwd CACCCTGACGGGACAGGAGATCCC
Oligo rev AAACGGGATCTCCTGTCCCGTCAG
Chromosome start end strand MM target_seq PAM alignment distance position gene name gene id
GU305914.1 117776 117798 - 0 CTGACGGGACAGGAGATCCC AGG ||||||||[||||||||||||]PAM NA Intergenic NA NA

T25124 GGGACAGGAGATCCCAGGAGCGG 1000
Oligo fwd CACCGGGACAGGAGATCCCAGGAG
Oligo rev AAACCTCCTGGGATCTCCTGTCCC
Chromosome start end strand MM target_seq PAM alignment distance position gene name gene id
GU305914.1 117771 117793 - 0 GGGACAGGAGATCCCAGGAG CGG ||||||||[||||||||||||]PAM NA Intergenic NA NA

T25125 GGACAGGAGATCCCAGGAGCGGG 1000
Oligo fwd CACCGGACAGGAGATCCCAGGAGC
Oligo rev AAACGCTCCTGGGATCTCCTGTCC
Chromosome start end strand MM target_seq PAM alignment distance position gene name gene id
GU305914.1 117770 117792 - 0 GGACAGGAGATCCCAGGAGC GGG ||||||||[||||||||||||]PAM NA Intergenic NA NA

T25126 ATTCAAACACGAAGAAAAATCGG 1000
Oligo fwd CACCATTCAAACACGAAGAAAAAT
Oligo rev AAACATTTTTCTTCGTGTTTGAAT
Chromosome start end strand MM target_seq PAM alignment distance position gene name gene id
GU305914.1 117747 117769 - 0 ATTCAAACACGAAGAAAAAT CGG ||||||||[||||||||||||]PAM NA Intergenic NA NA

T25127 AAGAAAAATCGGATCGCGTCCGG 1000
Oligo fwd CACCAAGAAAAATCGGATCGCGTC
Oligo rev AAACGACGCGATCCGATTTTTCTT
Chromosome start end strand MM target_seq PAM alignment distance position gene name gene id
GU305914.1 117736 117758 - 0 AAGAAAAATCGGATCGCGTC CGG ||||||||[||||||||||||]PAM NA Intergenic NA NA

T25128 CTGCTGCGTTCATGAAACACCGG 1000
Oligo fwd CACCCTGCTGCGTTCATGAAACAC
Oligo rev AAACGTGTTTCATGAACGCAGCAG
Chromosome start end strand MM target_seq PAM alignment distance position gene name gene id
GU305914.1 117710 117732 - 0 CTGCTGCGTTCATGAAACAC CGG ||||||||[||||||||||||]PAM NA Intergenic NA NA

T25129 CTGCGTTCATGAAACACCGGTGG 1000
Oligo fwd CACCCTGCGTTCATGAAACACCGG
Oligo rev AAACCCGGTGTTTCATGAACGCAG
Chromosome start end strand MM target_seq PAM alignment distance position gene name gene id
GU305914.1 117707 117729 - 0 CTGCGTTCATGAAACACCGG TGG ||||||||[||||||||||||]PAM NA Intergenic NA NA

T25130 CGTTCATGAAACACCGGTGGCGG 1000
Oligo fwd CACCCGTTCATGAAACACCGGTGG
Oligo rev AAACCCACCGGTGTTTCATGAACG
Chromosome start end strand MM target_seq PAM alignment distance position gene name gene id
GU305914.1 117704 117726 - 0 CGTTCATGAAACACCGGTGG CGG ||||||||[||||||||||||]PAM NA Intergenic NA NA

T25131 GAAACACCGGTGGCGGCGAGCGG 1000
Oligo fwd CACCGAAACACCGGTGGCGGCGAG
Oligo rev AAACCTCGCCGCCACCGGTGTTTC
Chromosome start end strand MM target_seq PAM alignment distance position gene name gene id
GU305914.1 117697 117719 - 0 GAAACACCGGTGGCGGCGAG CGG ||||||||[||||||||||||]PAM NA Intergenic NA NA
GU305914.1 162650 162672 - 4 GAGATGCCGGTGGCGTCGAG CGG ||-|--||[|||||||-||||]PAM NA Intergenic NA NA

T25132 CCGGTGGCGGCGAGCGGCCGCGG 1000
Oligo fwd CACCCCGGTGGCGGCGAGCGGCCG
Oligo rev AAACCGGCCGCTCGCCGCCACCGG
Chromosome start end strand MM target_seq PAM alignment distance position gene name gene id
GU305914.1 117691 117713 - 0 CCGGTGGCGGCGAGCGGCCG CGG ||||||||[||||||||||||]PAM NA Intergenic NA NA

T25133 GCGGCGAGCGGCCGCGGAGCCGG 1000
Oligo fwd CACCGCGGCGAGCGGCCGCGGAGC
Oligo rev AAACGCTCCGCGGCCGCTCGCCGC
Chromosome start end strand MM target_seq PAM alignment distance position gene name gene id
GU305914.1 117685 117707 - 0 GCGGCGAGCGGCCGCGGAGC CGG ||||||||[||||||||||||]PAM NA Intergenic NA NA

T25134 CGCAGTATTTGACTCGCGATCGG 1000
Oligo fwd CACCCGCAGTATTTGACTCGCGAT
Oligo rev AAACATCGCGAGTCAAATACTGCG
Chromosome start end strand MM target_seq PAM alignment distance position gene name gene id
GU305914.1 117658 117680 - 0 CGCAGTATTTGACTCGCGAT CGG ||||||||[||||||||||||]PAM NA Intergenic NA NA

T25135 CGCCGCGAGCCCAGCGCGCGCGG 1000
Oligo fwd CACCCGCCGCGAGCCCAGCGCGCG
Oligo rev AAACCGCGCGCTGGGCTCGCGGCG
Chromosome start end strand MM target_seq PAM alignment distance position gene name gene id
GU305914.1 117624 117646 - 0 CGCCGCGAGCCCAGCGCGCG CGG ||||||||[||||||||||||]PAM NA Intergenic NA NA

T25136 CGCGCGGCCGAGCGTCGATATGG 1000
Oligo fwd CACCCGCGCGGCCGAGCGTCGATA
Oligo rev AAACTATCGACGCTCGGCCGCGCG
Chromosome start end strand MM target_seq PAM alignment distance position gene name gene id
GU305914.1 117608 117630 - 0 CGCGCGGCCGAGCGTCGATA TGG ||||||||[||||||||||||]PAM NA Intergenic NA NA

T25137 GCCGAGCGTCGATATGGCCGAGG 1000
Oligo fwd CACCGCCGAGCGTCGATATGGCCG
Oligo rev AAACCGGCCATATCGACGCTCGGC
Chromosome start end strand MM target_seq PAM alignment distance position gene name gene id
GU305914.1 117602 117624 - 0 GCCGAGCGTCGATATGGCCG AGG ||||||||[||||||||||||]PAM NA Intergenic NA NA

T25138 CGAGGAATTTAACTTCCTGACGG 1000
Oligo fwd CACCCGAGGAATTTAACTTCCTGA
Oligo rev AAACTCAGGAAGTTAAATTCCTCG
Chromosome start end strand MM target_seq PAM alignment distance position gene name gene id
GU305914.1 117584 117606 - 0 CGAGGAATTTAACTTCCTGA CGG ||||||||[||||||||||||]PAM NA Intergenic NA NA

T25139 GGAATTTAACTTCCTGACGGCGG 1000
Oligo fwd CACCGGAATTTAACTTCCTGACGG
Oligo rev AAACCCGTCAGGAAGTTAAATTCC
Chromosome start end strand MM target_seq PAM alignment distance position gene name gene id
GU305914.1 117581 117603 - 0 GGAATTTAACTTCCTGACGG CGG ||||||||[||||||||||||]PAM NA Intergenic NA NA

T25140 GACGGCGGACCCCGACAAACCGG 1000
Oligo fwd CACCGACGGCGGACCCCGACAAAC
Oligo rev AAACGTTTGTCGGGGTCCGCCGTC
Chromosome start end strand MM target_seq PAM alignment distance position gene name gene id
GU305914.1 117566 117588 - 0 GACGGCGGACCCCGACAAAC CGG ||||||||[||||||||||||]PAM NA Intergenic NA NA

T25141 ACGGCGGACCCCGACAAACCGGG 1000
Oligo fwd CACCACGGCGGACCCCGACAAACC
Oligo rev AAACGGTTTGTCGGGGTCCGCCGT
Chromosome start end strand MM target_seq PAM alignment distance position gene name gene id
GU305914.1 117565 117587 - 0 ACGGCGGACCCCGACAAACC GGG ||||||||[||||||||||||]PAM NA Intergenic NA NA

T25142 CGGCGGACCCCGACAAACCGGGG 1000
Oligo fwd CACCCGGCGGACCCCGACAAACCG
Oligo rev AAACCGGTTTGTCGGGGTCCGCCG
Chromosome start end strand MM target_seq PAM alignment distance position gene name gene id
GU305914.1 117564 117586 - 0 CGGCGGACCCCGACAAACCG GGG ||||||||[||||||||||||]PAM NA Intergenic NA NA

T25143 CCGGGGTCCACGTTCGTTCACGG 1000
Oligo fwd CACCCCGGGGTCCACGTTCGTTCA
Oligo rev AAACTGAACGAACGTGGACCCCGG
Chromosome start end strand MM target_seq PAM alignment distance position gene name gene id
GU305914.1 117547 117569 - 0 CCGGGGTCCACGTTCGTTCA CGG ||||||||[||||||||||||]PAM NA Intergenic NA NA

T25144 GGTCCACGTTCGTTCACGGATGG 1000
Oligo fwd CACCGGTCCACGTTCGTTCACGGA
Oligo rev AAACTCCGTGAACGAACGTGGACC
Chromosome start end strand MM target_seq PAM alignment distance position gene name gene id
GU305914.1 117543 117565 - 0 GGTCCACGTTCGTTCACGGA TGG ||||||||[||||||||||||]PAM NA Intergenic NA NA

T25145 GGTGCACGAACCTGAGCCTCCGG 1000
Oligo fwd CACCGGTGCACGAACCTGAGCCTC
Oligo rev AAACGAGGCTCAGGTTCGTGCACC
Chromosome start end strand MM target_seq PAM alignment distance position gene name gene id
GU305914.1 117522 117544 - 0 GGTGCACGAACCTGAGCCTC CGG ||||||||[||||||||||||]PAM NA Intergenic NA NA

T25146 GTGCACGAACCTGAGCCTCCGGG 1000
Oligo fwd CACCGTGCACGAACCTGAGCCTCC
Oligo rev AAACGGAGGCTCAGGTTCGTGCAC
Chromosome start end strand MM target_seq PAM alignment distance position gene name gene id
GU305914.1 117521 117543 - 0 GTGCACGAACCTGAGCCTCC GGG ||||||||[||||||||||||]PAM NA Intergenic NA NA

| | | | | | | | | | | | |
|---|---|---|---|---|---|---|---|---|---|---|---|
| T25147 | CCTGAGCCTCCGGGTGACCAAGG | 1000 | | | | | | | | | |
| Oligo fwd | CACCCCTGAGCCTCCGGGTGACCA | | | | | | | | | | |
| Oligo rev | AAACTGGTCACCCGGAGGCTCAGG | | | | | | | | | | |
| Chromosome | start | end | strand | MM | target_seq | PAM | alignment | distance | position | gene name | gene id |
| GU305914.1 | 117512 | 117534 | - | 0 | CCTGAGCCTCCGGGTGACCA | AGG | \|\|\|\|\|\|\|\|[\|\|\|\|\|\|\|\|\|\|\|\|]PAM | NA | Intergenic | NA | NA |
| T25148 | CTGAGCCTCCGGGTGACCAAGGG | 1000 | | | | | | | | | |
| Oligo fwd | CACCCTGAGCCTCCGGGTGACCAA | | | | | | | | | | |
| Oligo rev | AAACTTGGTCACCCGGAGGCTCAG | | | | | | | | | | |
| Chromosome | start | end | strand | MM | target_seq | PAM | alignment | distance | position | gene name | gene id |
| GU305914.1 | 117511 | 117533 | - | 0 | CTGAGCCTCCGGGTGACCAA | GGG | \|\|\|\|\|\|\|\|[\|\|\|\|\|\|\|\|\|\|\|\|]PAM | NA | Intergenic | NA | NA |
| T25149 | ACCAAGGGCCCGCTCTTCAGAGG | 1000 | | | | | | | | | |
| Oligo fwd | CACCACCAAGGGCCCGCTCTTCAG | | | | | | | | | | |
| Oligo rev | AAACCTGAAGAGCGGGCCCTTGGT | | | | | | | | | | |
| Chromosome | start | end | strand | MM | target_seq | PAM | alignment | distance | position | gene name | gene id |
| GU305914.1 | 117496 | 117518 | - | 0 | ACCAAGGGCCCGCTCTTCAG | AGG | \|\|\|\|\|\|\|\|[\|\|\|\|\|\|\|\|\|\|\|\|]PAM | NA | Intergenic | NA | NA |
| T25150 | CCAAGGGCCCGCTCTTCAGAGGG | 1000 | | | | | | | | | |
| Oligo fwd | CACCCCAAGGGCCCGCTCTTCAGA | | | | | | | | | | |
| Oligo rev | AAACTCTGAAGAGCGGGCCCTTGG | | | | | | | | | | |
| Chromosome | start | end | strand | MM | target_seq | PAM | alignment | distance | position | gene name | gene id |
| GU305914.1 | 117495 | 117517 | - | 0 | CCAAGGGCCCGCTCTTCAGA | GGG | \|\|\|\|\|\|\|\|[\|\|\|\|\|\|\|\|\|\|\|\|]PAM | NA | Intergenic | NA | NA |
| T25151 | GGGCCCGCTCTTCAGAGGGCAGG | 1000 | | | | | | | | | |
| Oligo fwd | CACCGGGCCCGCTCTTCAGAGGGC | | | | | | | | | | |
| Oligo rev | AAACGCCCTCTGAAGAGCGGGCCC | | | | | | | | | | |
| Chromosome | start | end | strand | MM | target_seq | PAM | alignment | distance | position | gene name | gene id |
| GU305914.1 | 117491 | 117513 | - | 0 | GGGCCCGCTCTTCAGAGGGC | AGG | \|\|\|\|\|\|\|\|[\|\|\|\|\|\|\|\|\|\|\|\|]PAM | NA | Intergenic | NA | NA |
| T25152 | GAGACGAAGTACCTGAACACCGG | 1000 | | | | | | | | | |
| Oligo fwd | CACCGAGACGAAGTACCTGAACAC | | | | | | | | | | |
| Oligo rev | AAACGTGTTCAGGTACTTCGTCTC | | | | | | | | | | |
| Chromosome | start | end | strand | MM | target_seq | PAM | alignment | distance | position | gene name | gene id |
| GU305914.1 | 117469 | 117491 | - | 0 | GAGACGAAGTACCTGAACAC | CGG | \|\|\|\|\|\|\|\|[\|\|\|\|\|\|\|\|\|\|\|\|]PAM | NA | Intergenic | NA | NA |
| T25153 | CGTCCCCGCCATCACCTGCCTGG | 1000 | | | | | | | | | |
| Oligo fwd | CACCCGTCCCCGCCATCACCTGCC | | | | | | | | | | |
| Oligo rev | AAACGGCAGGTGATGGCGGGGACG | | | | | | | | | | |
| Chromosome | start | end | strand | MM | target_seq | PAM | alignment | distance | position | gene name | gene id |
| GU305914.1 | 117428 | 117450 | - | 0 | CGTCCCCGCCATCACCTGCC | TGG | \|\|\|\|\|\|\|\|[\|\|\|\|\|\|\|\|\|\|\|\|]PAM | NA | Intergenic | NA | NA |
| T25154 | CCTGGCGCAACGCCTGCCCCAGG | 1000 | | | | | | | | | |
| Oligo fwd | CACCCCTGGCGCAACGCCTGCCCC | | | | | | | | | | |
| Oligo rev | AAACGGGGCAGGCGTTGCGCCAGG | | | | | | | | | | |
| Chromosome | start | end | strand | MM | target_seq | PAM | alignment | distance | position | gene name | gene id |
| GU305914.1 | 117410 | 117432 | - | 0 | CCTGGCGCAACGCCTGCCCC | AGG | \|\|\|\|\|\|\|\|[\|\|\|\|\|\|\|\|\|\|\|\|]PAM | NA | Intergenic | NA | NA |
| T25155 | CTGGCGCAACGCCTGCCCCAGGG | 1000 | | | | | | | | | |
| Oligo fwd | CACCCTGGCGCAACGCCTGCCCCA | | | | | | | | | | |
| Oligo rev | AAACTGGGGCAGGCGTTGCGCCAG | | | | | | | | | | |
| Chromosome | start | end | strand | MM | target_seq | PAM | alignment | distance | position | gene name | gene id |
| GU305914.1 | 117409 | 117431 | - | 0 | CTGGCGCAACGCCTGCCCCA | GGG | \|\|\|\|\|\|\|\|[\|\|\|\|\|\|\|\|\|\|\|\|]PAM | NA | Intergenic | NA | NA |
| T25156 | CGCAACGCCTGCCCCAGGGCCGG | 1000 | | | | | | | | | |
| Oligo fwd | CACCCGCAACGCCTGCCCCAGGGC | | | | | | | | | | |
| Oligo rev | AAACGCCCTGGGGCAGGCGTTGCG | | | | | | | | | | |
| Chromosome | start | end | strand | MM | target_seq | PAM | alignment | distance | position | gene name | gene id |
| GU305914.1 | 117405 | 117427 | - | 0 | CGCAACGCCTGCCCCAGGGC | CGG | \|\|\|\|\|\|\|\|[\|\|\|\|\|\|\|\|\|\|\|\|]PAM | NA | Intergenic | NA | NA |
| T25157 | GCCCCAGGGCCGGACCCCGTCGG | 1000 | | | | | | | | | |
| Oligo fwd | CACCGCCCCAGGGCCGGACCCCGT | | | | | | | | | | |
| Oligo rev | AAACACGGGGTCCGGCCCTGGGGC | | | | | | | | | | |
| Chromosome | start | end | strand | MM | target_seq | PAM | alignment | distance | position | gene name | gene id |
| GU305914.1 | 117395 | 117417 | - | 0 | GCCCCAGGGCCGGACCCCGT | CGG | \|\|\|\|\|\|\|\|[\|\|\|\|\|\|\|\|\|\|\|\|]PAM | NA | Intergenic | NA | NA |
| T25158 | CCGGACCCCGTCGGCGCTGTCGG | 1000 | | | | | | | | | |
| Oligo fwd | CACCCCGGACCCCGTCGGCGCTGT | | | | | | | | | | |
| Oligo rev | AAACACAGCGCCGACGGGGTCCGG | | | | | | | | | | |
| Chromosome | start | end | strand | MM | target_seq | PAM | alignment | distance | position | gene name | gene id |
| GU305914.1 | 117386 | 117408 | - | 0 | CCGGACCCCGTCGGCGCTGT | CGG | \|\|\|\|\|\|\|\|[\|\|\|\|\|\|\|\|\|\|\|\|]PAM | NA | Intergenic | NA | NA |
| T25159 | CACCGTCCTGAGCGATCCCGAGG | 1000 | | | | | | | | | |
| Oligo fwd | CACCCACCGTCCTGAGCGATCCCG | | | | | | | | | | |
| Oligo rev | AAACCGGGATCGCTCAGGACGGTG | | | | | | | | | | |
| Chromosome | start | end | strand | MM | target_seq | PAM | alignment | distance | position | gene name | gene id |
| GU305914.1 | 117356 | 117378 | - | 0 | CACCGTCCTGAGCGATCCCG | AGG | \|\|\|\|\|\|\|\|[\|\|\|\|\|\|\|\|\|\|\|\|]PAM | NA | Intergenic | NA | NA |
| T25160 | TCCCGAGGACTTCATCAACGTGG | 1000 | | | | | | | | | |
| Oligo fwd | CACCTCCCGAGGACTTCATCAACG | | | | | | | | | | |
| Oligo rev | AAACCGTTGATGAAGTCCTCGGGA | | | | | | | | | | |
| Chromosome | start | end | strand | MM | target_seq | PAM | alignment | distance | position | gene name | gene id |
| GU305914.1 | 117341 | 117363 | - | 0 | TCCCGAGGACTTCATCAACG | TGG | \|\|\|\|\|\|\|\|[\|\|\|\|\|\|\|\|\|\|\|\|]PAM | NA | Intergenic | NA | NA |
| T25161 | GTCAACCTTCACGAGAAACTCGG | 1000 | | | | | | | | | |
| Oligo fwd | CACCGTCAACCTTCACGAGAAACT | | | | | | | | | | |
| Oligo rev | AAACAGTTTCTCGTGAAGGTTGAC | | | | | | | | | | |
| Chromosome | start | end | strand | MM | target_seq | PAM | alignment | distance | position | gene name | gene id |

GU305914.1 117299 117321 - 0 GTCAACCTTCACGAGAAACT CGG ||||||||[||||||||||||]PAM NA Intergenic NA NA

T25162 GCAAGATCAGATCTTCGTGACGG 1000
Oligo fwd CACCGCAAGATCAGATCTTCGTGA
Oligo rev AAACTCACGAAGATCTGATCTTGC
Chromosome start end strand MM target_seq PAM alignment distance position gene name gene id
GU305914.1 117269 117291 - 0 GCAAGATCAGATCTTCGTGA CGG ||||||||[||||||||||||]PAM NA Intergenic NA NA

T25163 CCAGCACCCCAACTTCAAAACGG 1000
Oligo fwd CACCCCAGCACCCCAACTTCAAAA
Oligo rev AAACTTTTGAAGTTGGGGTGCTGG
Chromosome start end strand MM target_seq PAM alignment distance position gene name gene id
GU305914.1 117194 117216 - 0 CCAGCACCCCAACTTCAAAA CGG ||||||||[||||||||||||]PAM NA Intergenic NA NA

T25164 CTTCAAAACGGATTTCGAGCAGG 1000
Oligo fwd CACCCTTCAAAACGGATTTCGAGC
Oligo rev AAACGCTCGAAATCCGTTTTGAAG
Chromosome start end strand MM target_seq PAM alignment distance position gene name gene id
GU305914.1 117182 117204 - 0 CTTCAAAACGGATTTCGAGC AGG ||||||||[||||||||||||]PAM NA Intergenic NA NA

T25165 CAAAACGGATTTCGAGCAGGAGG 1000
Oligo fwd CACCCAAAACGGATTTCGAGCAGG
Oligo rev AAACCCTGCTCGAAATCCGTTTTG
Chromosome start end strand MM target_seq PAM alignment distance position gene name gene id
GU305914.1 117179 117201 - 0 CAAAACGGATTTCGAGCAGG AGG ||||||||[||||||||||||]PAM NA Intergenic NA NA

T25166 AGATCACCCACCGCCGCATGCGG 1000
Oligo fwd CACCAGATCACCCACCGCCGCATG
Oligo rev AAACCATGCGGCGGTGGGTGATCT
Chromosome start end strand MM target_seq PAM alignment distance position gene name gene id
GU305914.1 117111 117133 - 0 AGATCACCCACCGCCGCATG CGG ||||||||[||||||||||||]PAM NA Intergenic NA NA

T25167 TCACCCACCGCCGCATGCGGTGG 1000
Oligo fwd CACCTCACCCACCGCCGCATGCGG
Oligo rev AAACCCGCATGCGGCGGTGGGTGA
Chromosome start end strand MM target_seq PAM alignment distance position gene name gene id
GU305914.1 117108 117130 - 0 TCACCCACCGCCGCATGCGG TGG ||||||||[||||||||||||]PAM NA Intergenic NA NA

T25168 CACCCACCGCCGCATGCGGTGGG 1000
Oligo fwd CACCCACCCACCGCCGCATGCGGT
Oligo rev AAACACCGCATGCGGCGGTGGGTG
Chromosome start end strand MM target_seq PAM alignment distance position gene name gene id
GU305914.1 117107 117129 - 0 CACCCACCGCCGCATGCGGT GGG ||||||||[||||||||||||]PAM NA Intergenic NA NA

T25169 TGCGGTGGGTGCATAGCAAGAGG 1000
Oligo fwd CACCTGCGGTGGGTGCATAGCAAG
Oligo rev AAACCTTGCTATGCACCCACCGCA
Chromosome start end strand MM target_seq PAM alignment distance position gene name gene id
GU305914.1 117093 117115 - 0 TGCGGTGGGTGCATAGCAAG AGG ||||||||[||||||||||||]PAM NA Intergenic NA NA

T25170 CATAGCAAGAGGATCCAGTTCGG 1000
Oligo fwd CACCCATAGCAAGAGGATCCAGTT
Oligo rev AAACAACTGGATCCTCTTGCTATG
Chromosome start end strand MM target_seq PAM alignment distance position gene name gene id
GU305914.1 117082 117104 - 0 CATAGCAAGAGGATCCAGTT CGG ||||||||[||||||||||||]PAM NA Intergenic NA NA

T25171 CACCCACTGCACCACCATCACGG 1000
Oligo fwd CACCCACCCACTGCACCACCATCA
Oligo rev AAACTGATGGTGGTGCAGTGGGTG
Chromosome start end strand MM target_seq PAM alignment distance position gene name gene id
GU305914.1 117059 117081 - 0 CACCCACTGCACCACCATCA CGG ||||||||[||||||||||||]PAM NA Intergenic NA NA

T25172 CCATCACGGCCACGCTGCCCCGG 1000
Oligo fwd CACCCCATCACGGCCACGCTGCCC
Oligo rev AAACGGGCAGCGTGGCCGTGATGG
Chromosome start end strand MM target_seq PAM alignment distance position gene name gene id
GU305914.1 117045 117067 - 0 CCATCACGGCCACGCTGCCC CGG ||||||||[||||||||||||]PAM NA Intergenic NA NA

T25173 CACGGCCACGCTGCCCCGGTCGG 1000
Oligo fwd CACCCACGGCCACGCTGCCCCGGT
Oligo rev AAACACCGGGGCAGCGTGGCCGTG
Chromosome start end strand MM target_seq PAM alignment distance position gene name gene id
GU305914.1 117041 117063 - 0 CACGGCCACGCTGCCCCGGT CGG ||||||||[||||||||||||]PAM NA Intergenic NA NA

T25174 CACGCTGCCCCGGTCGGTACCGG 1000
Oligo fwd CACCCACGCTGCCCCGGTCGGTAC
Oligo rev AAACGTACCGACCGGGGCAGCGTG
Chromosome start end strand MM target_seq PAM alignment distance position gene name gene id
GU305914.1 117035 117057 - 0 CACGCTGCCCCGGTCGGTAC CGG ||||||||[||||||||||||]PAM NA Intergenic NA NA

T25175 GGTACCGGCCAACACGTACGTGG 1000
Oligo fwd CACCGGTACCGGCCAACACGTACG
Oligo rev AAACCGTACGTGTTGGCCGGTACC
Chromosome start end strand MM target_seq PAM alignment distance position gene name gene id
GU305914.1 117020 117042 - 0 GGTACCGGCCAACACGTACG TGG ||||||||[||||||||||||]PAM NA Intergenic NA NA

T25176 GTACGTGGCCGCGCAACCGAAGG 1000
Oligo fwd CACCGTACGTGGCCGCGCAACCGA
Oligo rev AAACTCGGTTGCGCGGCCACGTAC

<table>
<tr><td>Chromosome</td><td>start</td><td>end</td><td>strand</td><td>MM</td><td>target_seq</td><td>PAM</td><td>alignment</td><td>distance</td><td>position</td><td>gene name</td><td>gene id</td></tr>
<tr><td>GU305914.1</td><td>117005</td><td>117027</td><td>-</td><td>0</td><td>GTACGTGGCCGCGCAACCGA</td><td>AGG</td><td>||||||||[||||||||||||]PAM</td><td>NA</td><td>Intergenic</td><td>NA</td><td>NA</td></tr>
<tr><td></td><td></td><td></td><td></td><td></td><td></td><td></td><td></td><td></td><td></td><td></td><td></td></tr>
<tr><td>T25177</td><td>CGCGCAACCGAAGGCGATCTCGG</td><td>1000</td><td></td><td></td><td></td><td></td><td></td><td></td><td></td><td></td><td></td></tr>
<tr><td>Oligo fwd</td><td>CACCCGCGCAACCGAAGGCGATCT</td><td></td><td></td><td></td><td></td><td></td><td></td><td></td><td></td><td></td><td></td></tr>
<tr><td>Oligo rev</td><td>AAACAGATCGCCTTCGGTTGCGCG</td><td></td><td></td><td></td><td></td><td></td><td></td><td></td><td></td><td></td><td></td></tr>
<tr><td>Chromosome</td><td>start</td><td>end</td><td>strand</td><td>MM</td><td>target_seq</td><td>PAM</td><td>alignment</td><td>distance</td><td>position</td><td>gene name</td><td>gene id</td></tr>
<tr><td>GU305914.1</td><td>116996</td><td>117018</td><td>-</td><td>0</td><td>CGCGCAACCGAAGGCGATCT</td><td>CGG</td><td>||||||||[||||||||||||]PAM</td><td>NA</td><td>Intergenic</td><td>NA</td><td>NA</td></tr>
<tr><td></td><td></td><td></td><td></td><td></td><td></td><td></td><td></td><td></td><td></td><td></td><td></td></tr>
<tr><td>T25178</td><td>GAAGGCGATCTCGGACCCGATGG</td><td>1000</td><td></td><td></td><td></td><td></td><td></td><td></td><td></td><td></td><td></td></tr>
<tr><td>Oligo fwd</td><td>CACCGAAGGCGATCTCGGACCCGA</td><td></td><td></td><td></td><td></td><td></td><td></td><td></td><td></td><td></td><td></td></tr>
<tr><td>Oligo rev</td><td>AAACTCGGGTCCGAGATCGCCTTC</td><td></td><td></td><td></td><td></td><td></td><td></td><td></td><td></td><td></td><td></td></tr>
<tr><td>Chromosome</td><td>start</td><td>end</td><td>strand</td><td>MM</td><td>target_seq</td><td>PAM</td><td>alignment</td><td>distance</td><td>position</td><td>gene name</td><td>gene id</td></tr>
<tr><td>GU305914.1</td><td>116987</td><td>117009</td><td>-</td><td>0</td><td>GAAGGCGATCTCGGACCCGA</td><td>TGG</td><td>||||||||[||||||||||||]PAM</td><td>NA</td><td>Intergenic</td><td>NA</td><td>NA</td></tr>
<tr><td></td><td></td><td></td><td></td><td></td><td></td><td></td><td></td><td></td><td></td><td></td><td></td></tr>
<tr><td>T25179</td><td>CATCCCGCGACCCAGCGAGACGG</td><td>1000</td><td></td><td></td><td></td><td></td><td></td><td></td><td></td><td></td><td></td></tr>
<tr><td>Oligo fwd</td><td>CACCCATCCCGCGACCCAGCGAGA</td><td></td><td></td><td></td><td></td><td></td><td></td><td></td><td></td><td></td><td></td></tr>
<tr><td>Oligo rev</td><td>AAACTCTCGCTGGGTCGCGGGATG</td><td></td><td></td><td></td><td></td><td></td><td></td><td></td><td></td><td></td><td></td></tr>
<tr><td>Chromosome</td><td>start</td><td>end</td><td>strand</td><td>MM</td><td>target_seq</td><td>PAM</td><td>alignment</td><td>distance</td><td>position</td><td>gene name</td><td>gene id</td></tr>
<tr><td>GU305914.1</td><td>116942</td><td>116964</td><td>-</td><td>0</td><td>CATCCCGCGACCCAGCGAGA</td><td>CGG</td><td>||||||||[||||||||||||]PAM</td><td>NA</td><td>Intergenic</td><td>NA</td><td>NA</td></tr>
<tr><td></td><td></td><td></td><td></td><td></td><td></td><td></td><td></td><td></td><td></td><td></td><td></td></tr>
<tr><td>T25180</td><td>AGACGGCGCACCACAAGCTGAGG</td><td>1000</td><td></td><td></td><td></td><td></td><td></td><td></td><td></td><td></td><td></td></tr>
<tr><td>Oligo fwd</td><td>CACCAGACGGCGCACCACAAGCTG</td><td></td><td></td><td></td><td></td><td></td><td></td><td></td><td></td><td></td><td></td></tr>
<tr><td>Oligo rev</td><td>AAACCAGCTTGTGGTGCGCCGTCT</td><td></td><td></td><td></td><td></td><td></td><td></td><td></td><td></td><td></td><td></td></tr>
<tr><td>Chromosome</td><td>start</td><td>end</td><td>strand</td><td>MM</td><td>target_seq</td><td>PAM</td><td>alignment</td><td>distance</td><td>position</td><td>gene name</td><td>gene id</td></tr>
<tr><td>GU305914.1</td><td>116925</td><td>116947</td><td>-</td><td>0</td><td>AGACGGCGCACCACAAGCTG</td><td>AGG</td><td>||||||||[||||||||||||]PAM</td><td>NA</td><td>Intergenic</td><td>NA</td><td>NA</td></tr>
<tr><td></td><td></td><td></td><td></td><td></td><td></td><td></td><td></td><td></td><td></td><td></td><td></td></tr>
<tr><td>T25181</td><td>CCACAAGCTGAGGATCCAGCTGG</td><td>1000</td><td></td><td></td><td></td><td></td><td></td><td></td><td></td><td></td><td></td></tr>
<tr><td>Oligo fwd</td><td>CACCCCACAAGCTGAGGATCCAGC</td><td></td><td></td><td></td><td></td><td></td><td></td><td></td><td></td><td></td><td></td></tr>
<tr><td>Oligo rev</td><td>AAACGCTGGATCCTCAGCTTGTGG</td><td></td><td></td><td></td><td></td><td></td><td></td><td></td><td></td><td></td><td></td></tr>
<tr><td>Chromosome</td><td>start</td><td>end</td><td>strand</td><td>MM</td><td>target_seq</td><td>PAM</td><td>alignment</td><td>distance</td><td>position</td><td>gene name</td><td>gene id</td></tr>
<tr><td>GU305914.1</td><td>116915</td><td>116937</td><td>-</td><td>0</td><td>CCACAAGCTGAGGATCCAGC</td><td>TGG</td><td>||||||||[||||||||||||]PAM</td><td>NA</td><td>Intergenic</td><td>NA</td><td>NA</td></tr>
<tr><td></td><td></td><td></td><td></td><td></td><td></td><td></td><td></td><td></td><td></td><td></td><td></td></tr>
<tr><td>T25182</td><td>AGGATCCAGCTGGTGTTCGACGG</td><td>1000</td><td></td><td></td><td></td><td></td><td></td><td></td><td></td><td></td><td></td></tr>
<tr><td>Oligo fwd</td><td>CACCAGGATCCAGCTGGTGTTCGA</td><td></td><td></td><td></td><td></td><td></td><td></td><td></td><td></td><td></td><td></td></tr>
<tr><td>Oligo rev</td><td>AAACTCGAACACCAGCTGGATCCT</td><td></td><td></td><td></td><td></td><td></td><td></td><td></td><td></td><td></td><td></td></tr>
<tr><td>Chromosome</td><td>start</td><td>end</td><td>strand</td><td>MM</td><td>target_seq</td><td>PAM</td><td>alignment</td><td>distance</td><td>position</td><td>gene name</td><td>gene id</td></tr>
<tr><td>GU305914.1</td><td>116905</td><td>116927</td><td>-</td><td>0</td><td>AGGATCCAGCTGGTGTTCGA</td><td>CGG</td><td>||||||||[||||||||||||]PAM</td><td>NA</td><td>Intergenic</td><td>NA</td><td>NA</td></tr>
<tr><td></td><td></td><td></td><td></td><td></td><td></td><td></td><td></td><td></td><td></td><td></td><td></td></tr>
<tr><td>T25183</td><td>GGATCCAGCTGGTGTTCGACGGG</td><td>1000</td><td></td><td></td><td></td><td></td><td></td><td></td><td></td><td></td><td></td></tr>
<tr><td>Oligo fwd</td><td>CACCGGATCCAGCTGGTGTTCGAC</td><td></td><td></td><td></td><td></td><td></td><td></td><td></td><td></td><td></td><td></td></tr>
<tr><td>Oligo rev</td><td>AAACGTCGAACACCAGCTGGATCC</td><td></td><td></td><td></td><td></td><td></td><td></td><td></td><td></td><td></td><td></td></tr>
<tr><td>Chromosome</td><td>start</td><td>end</td><td>strand</td><td>MM</td><td>target_seq</td><td>PAM</td><td>alignment</td><td>distance</td><td>position</td><td>gene name</td><td>gene id</td></tr>
<tr><td>GU305914.1</td><td>116904</td><td>116926</td><td>-</td><td>0</td><td>GGATCCAGCTGGTGTTCGAC</td><td>GGG</td><td>||||||||[||||||||||||]PAM</td><td>NA</td><td>Intergenic</td><td>NA</td><td>NA</td></tr>
<tr><td></td><td></td><td></td><td></td><td></td><td></td><td></td><td></td><td></td><td></td><td></td><td></td></tr>
<tr><td>T25184</td><td>GCTGGTGTTCGACGGGCACGCGG</td><td>1000</td><td></td><td></td><td></td><td></td><td></td><td></td><td></td><td></td><td></td></tr>
<tr><td>Oligo fwd</td><td>CACCGCTGGTGTTCGACGGGCACG</td><td></td><td></td><td></td><td></td><td></td><td></td><td></td><td></td><td></td><td></td></tr>
<tr><td>Oligo rev</td><td>AAACCGTGCCCGTCGAACACCAGC</td><td></td><td></td><td></td><td></td><td></td><td></td><td></td><td></td><td></td><td></td></tr>
<tr><td>Chromosome</td><td>start</td><td>end</td><td>strand</td><td>MM</td><td>target_seq</td><td>PAM</td><td>alignment</td><td>distance</td><td>position</td><td>gene name</td><td>gene id</td></tr>
<tr><td>GU305914.1</td><td>116897</td><td>116919</td><td>-</td><td>0</td><td>GCTGGTGTTCGACGGGCACG</td><td>CGG</td><td>||||||||[||||||||||||]PAM</td><td>NA</td><td>Intergenic</td><td>NA</td><td>NA</td></tr>
<tr><td></td><td></td><td></td><td></td><td></td><td></td><td></td><td></td><td></td><td></td><td></td><td></td></tr>
<tr><td>T25185</td><td>GACGGGCACGCGGAGAACAACGG</td><td>1000</td><td></td><td></td><td></td><td></td><td></td><td></td><td></td><td></td><td></td></tr>
<tr><td>Oligo fwd</td><td>CACCGACGGGCACGCGGAGAACAA</td><td></td><td></td><td></td><td></td><td></td><td></td><td></td><td></td><td></td><td></td></tr>
<tr><td>Oligo rev</td><td>AAACTTGTTCTCCGCGTGCCCGTC</td><td></td><td></td><td></td><td></td><td></td><td></td><td></td><td></td><td></td><td></td></tr>
<tr><td>Chromosome</td><td>start</td><td>end</td><td>strand</td><td>MM</td><td>target_seq</td><td>PAM</td><td>alignment</td><td>distance</td><td>position</td><td>gene name</td><td>gene id</td></tr>
<tr><td>GU305914.1</td><td>116887</td><td>116909</td><td>-</td><td>0</td><td>GACGGGCACGCGGAGAACAA</td><td>CGG</td><td>||||||||[||||||||||||]PAM</td><td>NA</td><td>Intergenic</td><td>NA</td><td>NA</td></tr>
<tr><td></td><td></td><td></td><td></td><td></td><td></td><td></td><td></td><td></td><td></td><td></td><td></td></tr>
<tr><td>T25186</td><td>CAAGCCGCCGACCTCGCTGACGG</td><td>1000</td><td></td><td></td><td></td><td></td><td></td><td></td><td></td><td></td><td></td></tr>
<tr><td>Oligo fwd</td><td>CACCCAAGCCGCCGACCTCGCTGA</td><td></td><td></td><td></td><td></td><td></td><td></td><td></td><td></td><td></td><td></td></tr>
<tr><td>Oligo rev</td><td>AAACTCAGCGAGGTCGGCGGCTTG</td><td></td><td></td><td></td><td></td><td></td><td></td><td></td><td></td><td></td><td></td></tr>
<tr><td>Chromosome</td><td>start</td><td>end</td><td>strand</td><td>MM</td><td>target_seq</td><td>PAM</td><td>alignment</td><td>distance</td><td>position</td><td>gene name</td><td>gene id</td></tr>
<tr><td>GU305914.1</td><td>116852</td><td>116874</td><td>-</td><td>0</td><td>CAAGCCGCCGACCTCGCTGA</td><td>CGG</td><td>||||||||[||||||||||||]PAM</td><td>NA</td><td>Intergenic</td><td>NA</td><td>NA</td></tr>
<tr><td></td><td></td><td></td><td></td><td></td><td></td><td></td><td></td><td></td><td></td><td></td><td></td></tr>
<tr><td>T25187</td><td>CGCTTCGACGCGTTCGACAACGG</td><td>1000</td><td></td><td></td><td></td><td></td><td></td><td></td><td></td><td></td><td></td></tr>
<tr><td>Oligo fwd</td><td>CACCCGCTTCGACGCGTTCGACAA</td><td></td><td></td><td></td><td></td><td></td><td></td><td></td><td></td><td></td><td></td></tr>
<tr><td>Oligo rev</td><td>AAACTTGTCGAACGCGTCGAAGCG</td><td></td><td></td><td></td><td></td><td></td><td></td><td></td><td></td><td></td><td></td></tr>
<tr><td>Chromosome</td><td>start</td><td>end</td><td>strand</td><td>MM</td><td>target_seq</td><td>PAM</td><td>alignment</td><td>distance</td><td>position</td><td>gene name</td><td>gene id</td></tr>
<tr><td>GU305914.1</td><td>116761</td><td>116783</td><td>-</td><td>0</td><td>CGCTTCGACGCGTTCGACAA</td><td>CGG</td><td>||||||||[||||||||||||]PAM</td><td>NA</td><td>Intergenic</td><td>NA</td><td>NA</td></tr>
<tr><td></td><td></td><td></td><td></td><td></td><td></td><td></td><td></td><td></td><td></td><td></td><td></td></tr>
<tr><td>T25188</td><td>CGCGTTCGACAACGGCTACAAGG</td><td>1000</td><td></td><td></td><td></td><td></td><td></td><td></td><td></td><td></td><td></td></tr>
<tr><td>Oligo fwd</td><td>CACCCGCGTTCGACAACGGCTACA</td><td></td><td></td><td></td><td></td><td></td><td></td><td></td><td></td><td></td><td></td></tr>
<tr><td>Oligo rev</td><td>AAACTGTAGCCGTTGTCGAACGCG</td><td></td><td></td><td></td><td></td><td></td><td></td><td></td><td></td><td></td><td></td></tr>
<tr><td>Chromosome</td><td>start</td><td>end</td><td>strand</td><td>MM</td><td>target_seq</td><td>PAM</td><td>alignment</td><td>distance</td><td>position</td><td>gene name</td><td>gene id</td></tr>
<tr><td>GU305914.1</td><td>116753</td><td>116775</td><td>-</td><td>0</td><td>CGCGTTCGACAACGGCTACA</td><td>AGG</td><td>||||||||[||||||||||||]PAM</td><td>NA</td><td>Intergenic</td><td>NA</td><td>NA</td></tr>
<tr><td></td><td></td><td></td><td></td><td></td><td></td><td></td><td></td><td></td><td></td><td></td><td></td></tr>
<tr><td>T25189</td><td>GCCCGAGAAAGATCGAGAGCAGG</td><td>1000</td><td></td><td></td><td></td><td></td><td></td><td></td><td></td><td></td><td></td></tr>
<tr><td>Oligo fwd</td><td>CACCGCCCGAGAAAGATCGAGAGC</td><td></td><td></td><td></td><td></td><td></td><td></td><td></td><td></td><td></td><td></td></tr>
<tr><td>Oligo rev</td><td>AAACGCTCTCGATCTTTCTCGGGC</td><td></td><td></td><td></td><td></td><td></td><td></td><td></td><td></td><td></td><td></td></tr>
<tr><td>Chromosome</td><td>start</td><td>end</td><td>strand</td><td>MM</td><td>target_seq</td><td>PAM</td><td>alignment</td><td>distance</td><td>position</td><td>gene name</td><td>gene id</td></tr>
<tr><td>GU305914.1</td><td>116724</td><td>116746</td><td>-</td><td>0</td><td>GCCCGAGAAAGATCGAGAGC</td><td>AGG</td><td>||||||||[||||||||||||]PAM</td><td>NA</td><td>Intergenic</td><td>NA</td><td>NA</td></tr>
<tr><td></td><td></td><td></td><td></td><td></td><td></td><td></td><td></td><td></td><td></td><td></td><td></td></tr>
<tr><td>T25190</td><td>AGAAAGATCGAGAGCAGGAAAGG</td><td>1000</td><td></td><td></td><td></td><td></td><td></td><td></td><td></td><td></td><td></td></tr>
<tr><td>Oligo fwd</td><td>CACCAGAAAGATCGAGAGCAGGAA</td><td></td><td></td><td></td><td></td><td></td><td></td><td></td><td></td><td></td><td></td></tr>
<tr><td>Oligo rev</td><td>AAACTTCCTGCTCTCGATCTTTCT</td><td></td><td></td><td></td><td></td><td></td><td></td><td></td><td></td><td></td><td></td></tr>
<tr><td>Chromosome</td><td>start</td><td>end</td><td>strand</td><td>MM</td><td>target_seq</td><td>PAM</td><td>alignment</td><td>distance</td><td>position</td><td>gene name</td><td>gene id</td></tr>
<tr><td>GU305914.1</td><td>116719</td><td>116741</td><td>-</td><td>0</td><td>AGAAAGATCGAGAGCAGGAA</td><td>AGG</td><td>||||||||[||||||||||||]PAM</td><td>NA</td><td>Intergenic</td><td>NA</td><td>NA</td></tr>
<tr><td></td><td></td><td></td><td></td><td></td><td></td><td></td><td></td><td></td><td></td><td></td><td></td></tr>
<tr><td>T25191</td><td>AGAGCAGGAAAGGCCTGACCTGG</td><td>1000</td><td></td><td></td><td></td><td></td><td></td><td></td><td></td><td></td><td></td></tr>
<tr><td>Oligo fwd</td><td>CACCAGAGCAGGAAAGGCCTGACC</td><td></td><td></td><td></td><td></td><td></td><td></td><td></td><td></td><td></td><td></td></tr>
</table>

<table>
<tr><td>Oligo rev</td><td>AAACGGTCAGGCCTTTCCTGCTCT</td><td></td><td></td><td></td><td></td><td></td><td></td><td></td><td></td><td></td><td></td></tr>
<tr><td>Chromosome</td><td>start</td><td>end</td><td>strand</td><td>MM</td><td>target_seq</td><td>PAM</td><td>alignment</td><td>distance</td><td>position</td><td>gene name</td><td>gene id</td></tr>
<tr><td>GU305914.1</td><td>116709</td><td>116731</td><td>-</td><td>0</td><td>AGAGCAGGAAAGGCCTGACC</td><td>TGG</td><td>||||||||[||||||||||||]PAM</td><td>NA</td><td>Intergenic</td><td>NA</td><td>NA</td></tr>
<tr><td>T25192</td><td>TGGACGATCACCATCGACAACGG</td><td>1000</td><td></td><td></td><td></td><td></td><td></td><td></td><td></td><td></td><td></td></tr>
<tr><td>Oligo fwd</td><td>CACCTGGACGATCACCATCGACAA</td><td></td><td></td><td></td><td></td><td></td><td></td><td></td><td></td><td></td><td></td></tr>
<tr><td>Oligo rev</td><td>AAACTTGTCGATGGTGATCGTCCA</td><td></td><td></td><td></td><td></td><td></td><td></td><td></td><td></td><td></td><td></td></tr>
<tr><td>Chromosome</td><td>start</td><td>end</td><td>strand</td><td>MM</td><td>target_seq</td><td>PAM</td><td>alignment</td><td>distance</td><td>position</td><td>gene name</td><td>gene id</td></tr>
<tr><td>GU305914.1</td><td>116689</td><td>116711</td><td>-</td><td>0</td><td>TGGACGATCACCATCGACAA</td><td>CGG</td><td>||||||||[||||||||||||]PAM</td><td>NA</td><td>Intergenic</td><td>NA</td><td>NA</td></tr>
<tr><td>T25193</td><td>GACAACGGCTATTCGTGCGACGG</td><td>1000</td><td></td><td></td><td></td><td></td><td></td><td></td><td></td><td></td><td></td></tr>
<tr><td>Oligo fwd</td><td>CACCGACAACGGCTATTCGTGCGA</td><td></td><td></td><td></td><td></td><td></td><td></td><td></td><td></td><td></td><td></td></tr>
<tr><td>Oligo rev</td><td>AAACTCGCACGAATAGCCGTTGTC</td><td></td><td></td><td></td><td></td><td></td><td></td><td></td><td></td><td></td><td></td></tr>
<tr><td>Chromosome</td><td>start</td><td>end</td><td>strand</td><td>MM</td><td>target_seq</td><td>PAM</td><td>alignment</td><td>distance</td><td>position</td><td>gene name</td><td>gene id</td></tr>
<tr><td>GU305914.1</td><td>116674</td><td>116696</td><td>-</td><td>0</td><td>GACAACGGCTATTCGTGCGA</td><td>CGG</td><td>||||||||[||||||||||||]PAM</td><td>NA</td><td>Intergenic</td><td>NA</td><td>NA</td></tr>
<tr><td>T25194</td><td>ACAACGGCTATTCGTGCGACGGG</td><td>1000</td><td></td><td></td><td></td><td></td><td></td><td></td><td></td><td></td><td></td></tr>
<tr><td>Oligo fwd</td><td>CACCACAACGGCTATTCGTGCGAC</td><td></td><td></td><td></td><td></td><td></td><td></td><td></td><td></td><td></td><td></td></tr>
<tr><td>Oligo rev</td><td>AAACGTCGCACGAATAGCCGTTGT</td><td></td><td></td><td></td><td></td><td></td><td></td><td></td><td></td><td></td><td></td></tr>
<tr><td>Chromosome</td><td>start</td><td>end</td><td>strand</td><td>MM</td><td>target_seq</td><td>PAM</td><td>alignment</td><td>distance</td><td>position</td><td>gene name</td><td>gene id</td></tr>
<tr><td>GU305914.1</td><td>116673</td><td>116695</td><td>-</td><td>0</td><td>ACAACGGCTATTCGTGCGAC</td><td>GGG</td><td>||||||||[||||||||||||]PAM</td><td>NA</td><td>Intergenic</td><td>NA</td><td>NA</td></tr>
<tr><td>T25195</td><td>TCGACATCCACGTATGCCGATGG</td><td>1000</td><td></td><td></td><td></td><td></td><td></td><td></td><td></td><td></td><td></td></tr>
<tr><td>Oligo fwd</td><td>CACCTCGACATCCACGTATGCCGA</td><td></td><td></td><td></td><td></td><td></td><td></td><td></td><td></td><td></td><td></td></tr>
<tr><td>Oligo rev</td><td>AAACTCGGCATACGTGGATGTCGA</td><td></td><td></td><td></td><td></td><td></td><td></td><td></td><td></td><td></td><td></td></tr>
<tr><td>Chromosome</td><td>start</td><td>end</td><td>strand</td><td>MM</td><td>target_seq</td><td>PAM</td><td>alignment</td><td>distance</td><td>position</td><td>gene name</td><td>gene id</td></tr>
<tr><td>GU305914.1</td><td>116610</td><td>116632</td><td>-</td><td>0</td><td>TCGACATCCACGTATGCCGA</td><td>TGG</td><td>||||||||[||||||||||||]PAM</td><td>NA</td><td>Intergenic</td><td>NA</td><td>NA</td></tr>
<tr><td>T25196</td><td>TATGCCGATGGATCGAAACCAGG</td><td>1000</td><td></td><td></td><td></td><td></td><td></td><td></td><td></td><td></td><td></td></tr>
<tr><td>Oligo fwd</td><td>CACCTATGCCGATGGATCGAAACC</td><td></td><td></td><td></td><td></td><td></td><td></td><td></td><td></td><td></td><td></td></tr>
<tr><td>Oligo rev</td><td>AAACGGTTTCGATCCATCGGCATA</td><td></td><td></td><td></td><td></td><td></td><td></td><td></td><td></td><td></td><td></td></tr>
<tr><td>Chromosome</td><td>start</td><td>end</td><td>strand</td><td>MM</td><td>target_seq</td><td>PAM</td><td>alignment</td><td>distance</td><td>position</td><td>gene name</td><td>gene id</td></tr>
<tr><td>GU305914.1</td><td>116598</td><td>116620</td><td>-</td><td>0</td><td>TATGCCGATGGATCGAAACC</td><td>AGG</td><td>||||||||[||||||||||||]PAM</td><td>NA</td><td>Intergenic</td><td>NA</td><td>NA</td></tr>
<tr><td>T25197</td><td>GACGCTCAACATCTGCATCACGG</td><td>1000</td><td></td><td></td><td></td><td></td><td></td><td></td><td></td><td></td><td></td></tr>
<tr><td>Oligo fwd</td><td>CACCGACGCTCAACATCTGCATCA</td><td></td><td></td><td></td><td></td><td></td><td></td><td></td><td></td><td></td><td></td></tr>
<tr><td>Oligo rev</td><td>AAACTGATGCAGATGTTGAGCGTC</td><td></td><td></td><td></td><td></td><td></td><td></td><td></td><td></td><td></td><td></td></tr>
<tr><td>Chromosome</td><td>start</td><td>end</td><td>strand</td><td>MM</td><td>target_seq</td><td>PAM</td><td>alignment</td><td>distance</td><td>position</td><td>gene name</td><td>gene id</td></tr>
<tr><td>GU305914.1</td><td>116576</td><td>116598</td><td>-</td><td>0</td><td>GACGCTCAACATCTGCATCA</td><td>CGG</td><td>||||||||[||||||||||||]PAM</td><td>NA</td><td>Intergenic</td><td>NA</td><td>NA</td></tr>
<tr><td>T25198</td><td>CAACATCTGCATCACGGCCATGG</td><td>1000</td><td></td><td></td><td></td><td></td><td></td><td></td><td></td><td></td><td></td></tr>
<tr><td>Oligo fwd</td><td>CACCCAACATCTGCATCACGGCCA</td><td></td><td></td><td></td><td></td><td></td><td></td><td></td><td></td><td></td><td></td></tr>
<tr><td>Oligo rev</td><td>AAACTGGCCGTGATGCAGATGTTG</td><td></td><td></td><td></td><td></td><td></td><td></td><td></td><td></td><td></td><td></td></tr>
<tr><td>Chromosome</td><td>start</td><td>end</td><td>strand</td><td>MM</td><td>target_seq</td><td>PAM</td><td>alignment</td><td>distance</td><td>position</td><td>gene name</td><td>gene id</td></tr>
<tr><td>GU305914.1</td><td>116570</td><td>116592</td><td>-</td><td>0</td><td>CAACATCTGCATCACGGCCA</td><td>TGG</td><td>||||||||[||||||||||||]PAM</td><td>NA</td><td>Intergenic</td><td>NA</td><td>NA</td></tr>
<tr><td>T25199</td><td>GCCAACGTGACCATTCCCGTAGG</td><td>1000</td><td></td><td></td><td></td><td></td><td></td><td></td><td></td><td></td><td></td></tr>
<tr><td>Oligo fwd</td><td>CACCGCCAACGTGACCATTCCCGT</td><td></td><td></td><td></td><td></td><td></td><td></td><td></td><td></td><td></td><td></td></tr>
<tr><td>Oligo rev</td><td>AAACACGGGAATGGTCACGTTGGC</td><td></td><td></td><td></td><td></td><td></td><td></td><td></td><td></td><td></td><td></td></tr>
<tr><td>Chromosome</td><td>start</td><td>end</td><td>strand</td><td>MM</td><td>target_seq</td><td>PAM</td><td>alignment</td><td>distance</td><td>position</td><td>gene name</td><td>gene id</td></tr>
<tr><td>GU305914.1</td><td>116548</td><td>116570</td><td>-</td><td>0</td><td>GCCAACGTGACCATTCCCGT</td><td>AGG</td><td>||||||||[||||||||||||]PAM</td><td>NA</td><td>Intergenic</td><td>NA</td><td>NA</td></tr>
<tr><td>T25200</td><td>CGTGACCATTCCCGTAGGAATGG</td><td>1000</td><td></td><td></td><td></td><td></td><td></td><td></td><td></td><td></td><td></td></tr>
<tr><td>Oligo fwd</td><td>CACCCGTGACCATTCCCGTAGGAA</td><td></td><td></td><td></td><td></td><td></td><td></td><td></td><td></td><td></td><td></td></tr>
<tr><td>Oligo rev</td><td>AAACTTCCTACGGGAATGGTCACG</td><td></td><td></td><td></td><td></td><td></td><td></td><td></td><td></td><td></td><td></td></tr>
<tr><td>Chromosome</td><td>start</td><td>end</td><td>strand</td><td>MM</td><td>target_seq</td><td>PAM</td><td>alignment</td><td>distance</td><td>position</td><td>gene name</td><td>gene id</td></tr>
<tr><td>GU305914.1</td><td>116543</td><td>116565</td><td>-</td><td>0</td><td>CGTGACCATTCCCGTAGGAA</td><td>TGG</td><td>||||||||[||||||||||||]PAM</td><td>NA</td><td>Intergenic</td><td>NA</td><td>NA</td></tr>
<tr><td>T25201</td><td>ATTCCCGTAGGAATGGAGCTCGG</td><td>1000</td><td></td><td></td><td></td><td></td><td></td><td></td><td></td><td></td><td></td></tr>
<tr><td>Oligo fwd</td><td>CACCATTCCCGTAGGAATGGAGCT</td><td></td><td></td><td></td><td></td><td></td><td></td><td></td><td></td><td></td><td></td></tr>
<tr><td>Oligo rev</td><td>AAACAGCTCCATTCCTACGGGAAT</td><td></td><td></td><td></td><td></td><td></td><td></td><td></td><td></td><td></td><td></td></tr>
<tr><td>Chromosome</td><td>start</td><td>end</td><td>strand</td><td>MM</td><td>target_seq</td><td>PAM</td><td>alignment</td><td>distance</td><td>position</td><td>gene name</td><td>gene id</td></tr>
<tr><td>GU305914.1</td><td>116536</td><td>116558</td><td>-</td><td>0</td><td>ATTCCCGTAGGAATGGAGCT</td><td>CGG</td><td>||||||||[||||||||||||]PAM</td><td>NA</td><td>Intergenic</td><td>NA</td><td>NA</td></tr>
<tr><td>T25202</td><td>GTCGACCTGCTGACCCCCGTCGG</td><td>1000</td><td></td><td></td><td></td><td></td><td></td><td></td><td></td><td></td><td></td></tr>
<tr><td>Oligo fwd</td><td>CACCGTCGACCTGCTGACCCCCGT</td><td></td><td></td><td></td><td></td><td></td><td></td><td></td><td></td><td></td><td></td></tr>
<tr><td>Oligo rev</td><td>AAACACGGGGGTCAGCAGGTCGAC</td><td></td><td></td><td></td><td></td><td></td><td></td><td></td><td></td><td></td><td></td></tr>
<tr><td>Chromosome</td><td>start</td><td>end</td><td>strand</td><td>MM</td><td>target_seq</td><td>PAM</td><td>alignment</td><td>distance</td><td>position</td><td>gene name</td><td>gene id</td></tr>
<tr><td>GU305914.1</td><td>116473</td><td>116495</td><td>-</td><td>0</td><td>GTCGACCTGCTGACCCCCGT</td><td>CGG</td><td>||||||||[||||||||||||]PAM</td><td>NA</td><td>Intergenic</td><td>NA</td><td>NA</td></tr>
<tr><td>T25203</td><td>CATCATCAATAAAGAGTCTATGG</td><td>1000</td><td></td><td></td><td></td><td></td><td></td><td></td><td></td><td></td><td></td></tr>
<tr><td>Oligo fwd</td><td>CACCCATCATCAATAAAGAGTCTA</td><td></td><td></td><td></td><td></td><td></td><td></td><td></td><td></td><td></td><td></td></tr>
<tr><td>Oligo rev</td><td>AAACTAGACTCTTTATTGATGATG</td><td></td><td></td><td></td><td></td><td></td><td></td><td></td><td></td><td></td><td></td></tr>
<tr><td>Chromosome</td><td>start</td><td>end</td><td>strand</td><td>MM</td><td>target_seq</td><td>PAM</td><td>alignment</td><td>distance</td><td>position</td><td>gene name</td><td>gene id</td></tr>
<tr><td>GU305914.1</td><td>116450</td><td>116472</td><td>-</td><td>0</td><td>CATCATCAATAAAGAGTCTA</td><td>TGG</td><td>||||||||[||||||||||||]PAM</td><td>NA</td><td>Intergenic</td><td>NA</td><td>NA</td></tr>
<tr><td>T25204</td><td>TGCAAACTCTTCGTCCCCGTTGG</td><td>1000</td><td></td><td></td><td></td><td></td><td></td><td></td><td></td><td></td><td></td></tr>
<tr><td>Oligo fwd</td><td>CACCTGCAAACTCTTCGTCCCCGT</td><td></td><td></td><td></td><td></td><td></td><td></td><td></td><td></td><td></td><td></td></tr>
<tr><td>Oligo rev</td><td>AAACACGGGGACGAAGAGTTTGCA</td><td></td><td></td><td></td><td></td><td></td><td></td><td></td><td></td><td></td><td></td></tr>
<tr><td>Chromosome</td><td>start</td><td>end</td><td>strand</td><td>MM</td><td>target_seq</td><td>PAM</td><td>alignment</td><td>distance</td><td>position</td><td>gene name</td><td>gene id</td></tr>
<tr><td>GU305914.1</td><td>116404</td><td>116426</td><td>-</td><td>0</td><td>TGCAAACTCTTCGTCCCCGT</td><td>TGG</td><td>||||||||[||||||||||||]PAM</td><td>NA</td><td>Intergenic</td><td>NA</td><td>NA</td></tr>
<tr><td>T25205</td><td>GGATCCGAGACCTACCGACCCGG</td><td>1000</td><td></td><td></td><td></td><td></td><td></td><td></td><td></td><td></td><td></td></tr>
<tr><td>Oligo fwd</td><td>CACCGGATCCGAGACCTACCGACC</td><td></td><td></td><td></td><td></td><td></td><td></td><td></td><td></td><td></td><td></td></tr>
<tr><td>Oligo rev</td><td>AAACGGTCGGTAGGTCTCGGATCC</td><td></td><td></td><td></td><td></td><td></td><td></td><td></td><td></td><td></td><td></td></tr>
<tr><td>Chromosome</td><td>start</td><td>end</td><td>strand</td><td>MM</td><td>target_seq</td><td>PAM</td><td>alignment</td><td>distance</td><td>position</td><td>gene name</td><td>gene id</td></tr>
<tr><td>GU305914.1</td><td>116383</td><td>116405</td><td>-</td><td>0</td><td>GGATCCGAGACCTACCGACC</td><td>CGG</td><td>||||||||[||||||||||||]PAM</td><td>NA</td><td>Intergenic</td><td>NA</td><td>NA</td></tr>
<tr><td>T25206</td><td>ACCCGGCCCCGAGCCGCCGCAGG</td><td>1000</td><td></td><td></td><td></td><td></td><td></td><td></td><td></td><td></td><td></td></tr>
</table>

Oligo fwd CACCACCCGGCCCCGAGCCGCCGC
Oligo rev AAACGCGGCGGCTCGGGGCCGGGT
Chromosome start end strand MM target_seq PAM alignment distance position gene name gene id
GU305914.1 116366 116388 - 0 ACCCGGCCCCGAGCCGCCGC AGG ||||||||[||||||||||||]PAM NA Intergenic NA NA

T25207 CTCCAGCGACCACCCCTCGCCGG 1000
Oligo fwd CACCCTCCAGCGACCACCCCTCGC
Oligo rev AAACGCGAGGGGTGGTCGCTGGAG
Chromosome start end strand MM target_seq PAM alignment distance position gene name gene id
GU305914.1 116328 116350 - 0 CTCCAGCGACCACCCCTCGC CGG ||||||||[||||||||||||]PAM NA Intergenic NA NA

T25208 CACCCCTCGCCGGATGAGCTCGG 1000
Oligo fwd CACCCACCCCTCGCCGGATGAGCT
Oligo rev AAACAGCTCATCCGGCGAGGGGTG
Chromosome start end strand MM target_seq PAM alignment distance position gene name gene id
GU305914.1 116318 116340 - 0 CACCCCTCGCCGGATGAGCT CGG ||||||||[||||||||||||]PAM NA Intergenic NA NA

T25209 CGACGAGCAGCAGCTACCCGAGG 1000
Oligo fwd CACCCGACGAGCAGCAGCTACCCG
Oligo rev AAACCGGGTAGCTGCTGCTCGTCG
Chromosome start end strand MM target_seq PAM alignment distance position gene name gene id
GU305914.1 116294 116316 - 0 CGACGAGCAGCAGCTACCCG AGG ||||||||[||||||||||||]PAM NA Intergenic NA NA

T25210 TTGAACACACCGAGACCCGTCGG 1000
Oligo fwd CACCTTGAACACACCGAGACCCGT
Oligo rev AAACACGGGTCTCGGTGTGTTCAA
Chromosome start end strand MM target_seq PAM alignment distance position gene name gene id
GU305914.1 116251 116273 - 0 TTGAACACACCGAGACCCGT CGG ||||||||[||||||||||||]PAM NA Intergenic NA NA

T25211 TCGGCAGAGAGCTGCCCCCGCGG 1000
Oligo fwd CACCTCGGCAGAGAGCTGCCCCCG
Oligo rev AAACCGGGGGCAGCTCTCTGCCGA
Chromosome start end strand MM target_seq PAM alignment distance position gene name gene id
GU305914.1 116232 116254 - 0 TCGGCAGAGAGCTGCCCCCG CGG ||||||||[||||||||||||]PAM NA Intergenic NA NA

T25212 GCTGCCCCCGCGGCGCACGTCGG 1000
Oligo fwd CACCGCTGCCCCCGCGGCGCACGT
Oligo rev AAACACGTGCGCCGCGGGGGCAGC
Chromosome start end strand MM target_seq PAM alignment distance position gene name gene id
GU305914.1 116222 116244 - 0 GCTGCCCCCGCGGCGCACGT CGG ||||||||[||||||||||||]PAM NA Intergenic NA NA

T25213 GCCCCCGCGGCGCACGTCGGCGG 1000
Oligo fwd CACCGCCCCCGCGGCGCACGTCGG
Oligo rev AAACCCGACGTGCGCCGCGGGGGC
Chromosome start end strand MM target_seq PAM alignment distance position gene name gene id
GU305914.1 116219 116241 - 0 GCCCCCGCGGCGCACGTCGG CGG ||||||||[||||||||||||]PAM NA Intergenic NA NA

T25214 CGAGCGCCTCTTCGCCCAGATGG 1000
Oligo fwd CACCCGAGCGCCTCTTCGCCCAGA
Oligo rev AAACTCTGGGCGAAGAGGCGCTCG
Chromosome start end strand MM target_seq PAM alignment distance position gene name gene id
GU305914.1 116195 116217 - 0 CGAGCGCCTCTTCGCCCAGA TGG ||||||||[||||||||||||]PAM NA Intergenic NA NA

T25215 GGACGCCGACAACGTGCCGTCGG 1000
Oligo fwd CACCGGACGCCGACAACGTGCCGT
Oligo rev AAACACGGCACGTTGTCGGCGTCC
Chromosome start end strand MM target_seq PAM alignment distance position gene name gene id
GU305914.1 116174 116196 - 0 GGACGCCGACAACGTGCCGT CGG ||||||||[||||||||||||]PAM NA Intergenic NA NA
GU305914.1 125144 125166 - 4 GGTGGCCGACGACGTGCCGC TGG ||--||||[||-||||||||-]PAM NA Intergenic NA NA

T25216 CGACAACGTGCCGTCGGACCAGG 1000
Oligo fwd CACCCGACAACGTGCCGTCGGACC
Oligo rev AAACGGTCCGACGGCACGTTGTCG
Chromosome start end strand MM target_seq PAM alignment distance position gene name gene id
GU305914.1 116168 116190 - 0 CGACAACGTGCCGTCGGACC AGG ||||||||[||||||||||||]PAM NA Intergenic NA NA

T25217 CGTGCCGTCGGACCAGGAAGAGG 1000
Oligo fwd CACCCGTGCCGTCGGACCAGGAAG
Oligo rev AAACCTTCCTGGTCCGACGGCACG
Chromosome start end strand MM target_seq PAM alignment distance position gene name gene id
GU305914.1 116162 116184 - 0 CGTGCCGTCGGACCAGGAAG AGG ||||||||[||||||||||||]PAM NA Intergenic NA NA

T25218 GAAGAGGATGATAGCGACGAAGG 1000
Oligo fwd CACCGAAGAGGATGATAGCGACGA
Oligo rev AAACTCGTCGCTATCATCCTCTTC
Chromosome start end strand MM target_seq PAM alignment distance position gene name gene id
GU305914.1 116146 116168 - 0 GAAGAGGATGATAGCGACGA AGG ||||||||[||||||||||||]PAM NA Intergenic NA NA

T25219 GGATGATAGCGACGAAGGAGAGG 1000
Oligo fwd CACCGGATGATAGCGACGAAGGAG
Oligo rev AAACCTCCTTCGTCGCTATCATCC
Chromosome start end strand MM target_seq PAM alignment distance position gene name gene id
GU305914.1 116141 116163 - 0 GGATGATAGCGACGAAGGAG AGG ||||||||[||||||||||||]PAM NA Intergenic NA NA

T25220 ATTTCGACGAAGACCGAAACTGG 1000
Oligo fwd CACCATTTCGACGAAGACCGAAAC
Oligo rev AAACGTTTCGGTCTTCGTCGAAAT
Chromosome start end strand MM target_seq PAM alignment distance position gene name gene id
GU305914.1 116100 116122 - 0 ATTTCGACGAAGACCGAAAC TGG ||||||||[||||||||||||]PAM NA Intergenic NA NA

| | | | | | | | | | | | |
|---|---|---|---|---|---|---|---|---|---|---|---|
| T25221 | TTTCGACGAAGACCGAAACTGGG | 1000 | | | | | | | | | |
| Oligo fwd | CACCTTTCGACGAAGACCGAAACT | | | | | | | | | | |
| Oligo rev | AAACAGTTTCGGTCTTCGTCGAAA | | | | | | | | | | |
| Chromosome | start | end | strand | MM | target_seq | PAM | alignment | distance | position | gene name | gene id |
| GU305914.1 | 116099 | 116121 | - | 0 | TTTCGACGAAGACCGAAACT | GGG | \|\|\|\|\|\|\|\|[\|\|\|\|\|\|\|\|\|\|\|\|]PAM | NA | Intergenic | NA | NA |
| T25222 | CGCCGTCCCCGACTCGTACGAGG | 1000 | | | | | | | | | |
| Oligo fwd | CACCCGCCGTCCCCGACTCGTACG | | | | | | | | | | |
| Oligo rev | AAACCGTACGAGTCGGGGACGGCG | | | | | | | | | | |
| Chromosome | start | end | strand | MM | target_seq | PAM | alignment | distance | position | gene name | gene id |
| GU305914.1 | 116069 | 116091 | - | 0 | CGCCGTCCCCGACTCGTACG | AGG | \|\|\|\|\|\|\|\|[\|\|\|\|\|\|\|\|\|\|\|\|]PAM | NA | Intergenic | NA | NA |
| T25223 | CGTCCCCGACTCGTACGAGGAGG | 1000 | | | | | | | | | |
| Oligo fwd | CACCCGTCCCCGACTCGTACGAGG | | | | | | | | | | |
| Oligo rev | AAACCCTCGTACGAGTCGGGGACG | | | | | | | | | | |
| Chromosome | start | end | strand | MM | target_seq | PAM | alignment | distance | position | gene name | gene id |
| GU305914.1 | 116066 | 116088 | - | 0 | CGTCCCCGACTCGTACGAGG | AGG | \|\|\|\|\|\|\|\|[\|\|\|\|\|\|\|\|\|\|\|\|]PAM | NA | Intergenic | NA | NA |
| T25224 | CCCCGACTCGTACGAGGAGGAGG | 1000 | | | | | | | | | |
| Oligo fwd | CACCCCCCGACTCGTACGAGGAGG | | | | | | | | | | |
| Oligo rev | AAACCCTCCTCGTACGAGTCGGGG | | | | | | | | | | |
| Chromosome | start | end | strand | MM | target_seq | PAM | alignment | distance | position | gene name | gene id |
| GU305914.1 | 116063 | 116085 | - | 0 | CCCCGACTCGTACGAGGAGG | AGG | \|\|\|\|\|\|\|\|[\|\|\|\|\|\|\|\|\|\|\|\|]PAM | NA | Intergenic | NA | NA |
| T25225 | CGACTCGTACGAGGAGGAGGCGG | 1000 | | | | | | | | | |
| Oligo fwd | CACCCGACTCGTACGAGGAGGAGG | | | | | | | | | | |
| Oligo rev | AAACCCTCCTCCTCGTACGAGTCG | | | | | | | | | | |
| Chromosome | start | end | strand | MM | target_seq | PAM | alignment | distance | position | gene name | gene id |
| GU305914.1 | 116060 | 116082 | - | 0 | CGACTCGTACGAGGAGGAGG | CGG | \|\|\|\|\|\|\|\|[\|\|\|\|\|\|\|\|\|\|\|\|]PAM | NA | Intergenic | NA | NA |
| GU305914.1 | 123785 | 123807 | - | 3 | CGACCCGTACGACGAGGACG | CGG | \|\|\|\|-\|\|\|[\|\|\|\|-\|\|\|\|\|-\|]PAM | NA | Intergenic | NA | NA |
| GU305914.1 | 59739 | 59761 | - | 3 | CGACTCGGACGACGAGGACG | AGG | \|\|\|\|\|\|\|-[\|\|\|\|-\|\|\|\|\|-\|]PAM | NA | Intergenic | NA | NA |
| T25226 | TGCAGCCCCACGATCTGATACGG | 1000 | | | | | | | | | |
| Oligo fwd | CACCTGCAGCCCCACGATCTGATA | | | | | | | | | | |
| Oligo rev | AAACTATCAGATCGTGGGGCTGCA | | | | | | | | | | |
| Chromosome | start | end | strand | MM | target_seq | PAM | alignment | distance | position | gene name | gene id |
| GU305914.1 | 116013 | 116035 | - | 0 | TGCAGCCCCACGATCTGATA | CGG | \|\|\|\|\|\|\|\|[\|\|\|\|\|\|\|\|\|\|\|\|]PAM | NA | Intergenic | NA | NA |
| T25227 | CCCACGATCTGATACGGATCCGG | 1000 | | | | | | | | | |
| Oligo fwd | CACCCCCACGATCTGATACGGATC | | | | | | | | | | |
| Oligo rev | AAACGATCCGTATCAGATCGTGGG | | | | | | | | | | |
| Chromosome | start | end | strand | MM | target_seq | PAM | alignment | distance | position | gene name | gene id |
| GU305914.1 | 116007 | 116029 | - | 0 | CCCACGATCTGATACGGATC | CGG | \|\|\|\|\|\|\|\|[\|\|\|\|\|\|\|\|\|\|\|\|]PAM | NA | Intergenic | NA | NA |
| T25228 | CCACGATCTGATACGGATCCGGG | 1000 | | | | | | | | | |
| Oligo fwd | CACCCCACGATCTGATACGGATCC | | | | | | | | | | |
| Oligo rev | AAACGGATCCGTATCAGATCGTGG | | | | | | | | | | |
| Chromosome | start | end | strand | MM | target_seq | PAM | alignment | distance | position | gene name | gene id |
| GU305914.1 | 116006 | 116028 | - | 0 | CCACGATCTGATACGGATCC | GGG | \|\|\|\|\|\|\|\|[\|\|\|\|\|\|\|\|\|\|\|\|]PAM | NA | Intergenic | NA | NA |
| T25229 | GATCTGATACGGATCCGGGCCGG | 1000 | | | | | | | | | |
| Oligo fwd | CACCGATCTGATACGGATCCGGGC | | | | | | | | | | |
| Oligo rev | AAACGCCCGGATCCGTATCAGATC | | | | | | | | | | |
| Chromosome | start | end | strand | MM | target_seq | PAM | alignment | distance | position | gene name | gene id |
| GU305914.1 | 116002 | 116024 | - | 0 | GATCTGATACGGATCCGGGC | CGG | \|\|\|\|\|\|\|\|[\|\|\|\|\|\|\|\|\|\|\|\|]PAM | NA | Intergenic | NA | NA |
| T25230 | ATCTGATACGGATCCGGGCCGGG | 1000 | | | | | | | | | |
| Oligo fwd | CACCATCTGATACGGATCCGGGCC | | | | | | | | | | |
| Oligo rev | AAACGGCCCGGATCCGTATCAGAT | | | | | | | | | | |
| Chromosome | start | end | strand | MM | target_seq | PAM | alignment | distance | position | gene name | gene id |
| GU305914.1 | 116001 | 116023 | - | 0 | ATCTGATACGGATCCGGGCC | GGG | \|\|\|\|\|\|\|\|[\|\|\|\|\|\|\|\|\|\|\|\|]PAM | NA | Intergenic | NA | NA |
| T25231 | ACGGATCCGGGCCGGGAGACTGG | 1000 | | | | | | | | | |
| Oligo fwd | CACCACGGATCCGGGCCGGGAGAC | | | | | | | | | | |
| Oligo rev | AAACGTCTCCCGGCCCGGATCCGT | | | | | | | | | | |
| Chromosome | start | end | strand | MM | target_seq | PAM | alignment | distance | position | gene name | gene id |
| GU305914.1 | 115994 | 116016 | - | 0 | ACGGATCCGGGCCGGGAGAC | TGG | \|\|\|\|\|\|\|\|[\|\|\|\|\|\|\|\|\|\|\|\|]PAM | NA | Intergenic | NA | NA |
| T25232 | TACACTCCGTTCATCAACGAAGG | 1000 | | | | | | | | | |
| Oligo fwd | CACCTACACTCCGTTCATCAACGA | | | | | | | | | | |
| Oligo rev | AAACTCGTTGATGAACGGAGTGTA | | | | | | | | | | |
| Chromosome | start | end | strand | MM | target_seq | PAM | alignment | distance | position | gene name | gene id |
| GU305914.1 | 115951 | 115973 | - | 0 | TACACTCCGTTCATCAACGA | AGG | \|\|\|\|\|\|\|\|[\|\|\|\|\|\|\|\|\|\|\|\|]PAM | NA | Intergenic | NA | NA |
| T25233 | CGTGTATTTCAACGAAGCCATGG | 1000 | | | | | | | | | |
| Oligo fwd | CACCCGTGTATTTCAACGAAGCCA | | | | | | | | | | |
| Oligo rev | AAACTGGCTTCGTTGAAATACACG | | | | | | | | | | |
| Chromosome | start | end | strand | MM | target_seq | PAM | alignment | distance | position | gene name | gene id |
| GU305914.1 | 115928 | 115950 | - | 0 | CGTGTATTTCAACGAAGCCA | TGG | \|\|\|\|\|\|\|\|[\|\|\|\|\|\|\|\|\|\|\|\|]PAM | NA | Intergenic | NA | NA |
| T25234 | GTGTATTTCAACGAAGCCATGGG | 1000 | | | | | | | | | |
| Oligo fwd | CACCGTGTATTTCAACGAAGCCAT | | | | | | | | | | |
| Oligo rev | AAACATGGCTTCGTTGAAATACAC | | | | | | | | | | |
| Chromosome | start | end | strand | MM | target_seq | PAM | alignment | distance | position | gene name | gene id |
| GU305914.1 | 115927 | 115949 | - | 0 | GTGTATTTCAACGAAGCCAT | GGG | \|\|\|\|\|\|\|\|[\|\|\|\|\|\|\|\|\|\|\|\|]PAM | NA | Intergenic | NA | NA |
| T25235 | CGAAGCCATGGGAGTCGACAAGG | 1000 | | | | | | | | | |
| Oligo fwd | CACCCGAAGCCATGGGAGTCGACA | | | | | | | | | | |

| | | | | | | | | | | | |
|---|---|---|---|---|---|---|---|---|---|---|---|
| Oligo rev | AAACTGTCGACTCCCATGGCTTCG | | | | | | | | | | |
| Chromosome | start | end | strand | MM | target_seq | PAM | alignment | distance | position | gene name | gene id |
| GU305914.1 | 115916 | 115938 | - | 0 | CGAAGCCATGGGAGTCGACA | AGG | \|\|\|\|\|\|\|\|[\|\|\|\|\|\|\|\|\|\|\|\|]PAM | NA | Intergenic | NA | NA |
| T25236 | GCCGTGCAGTTGACCACGAAAGG | 1000 | | | | | | | | | |
| Oligo fwd | CACCGCCGTGCAGTTGACCACGAA | | | | | | | | | | |
| Oligo rev | AAACTTCGTGGTCAACTGCACGGC | | | | | | | | | | |
| Chromosome | start | end | strand | MM | target_seq | PAM | alignment | distance | position | gene name | gene id |
| GU305914.1 | 115891 | 115913 | - | 0 | GCCGTGCAGTTGACCACGAA | AGG | \|\|\|\|\|\|\|\|[\|\|\|\|\|\|\|\|\|\|\|\|]PAM | NA | Intergenic | NA | NA |
| T25237 | TTGACCACGAAAGGTAGATACGG | 1000 | | | | | | | | | |
| Oligo fwd | CACCTTGACCACGAAAGGTAGATA | | | | | | | | | | |
| Oligo rev | AAACTATCTACCTTTCGTGGTCAA | | | | | | | | | | |
| Chromosome | start | end | strand | MM | target_seq | PAM | alignment | distance | position | gene name | gene id |
| GU305914.1 | 115882 | 115904 | - | 0 | TTGACCACGAAAGGTAGATA | CGG | \|\|\|\|\|\|\|\|[\|\|\|\|\|\|\|\|\|\|\|\|]PAM | NA | Intergenic | NA | NA |
| T25238 | ACGCTCGCCCGCCCGCATCACGG | 1000 | | | | | | | | | |
| Oligo fwd | CACCACGCTCGCCCGCCCGCATCA | | | | | | | | | | |
| Oligo rev | AAACTGATGCGGGCGGGCGAGCGT | | | | | | | | | | |
| Chromosome | start | end | strand | MM | target_seq | PAM | alignment | distance | position | gene name | gene id |
| GU305914.1 | 115753 | 115775 | - | 0 | ACGCTCGCCCGCCCGCATCA | CGG | \|\|\|\|\|\|\|\|[\|\|\|\|\|\|\|\|\|\|\|\|]PAM | NA | Intergenic | NA | NA |
| T25239 | CCCGCATCACGGTGACAGACAGG | 1000 | | | | | | | | | |
| Oligo fwd | CACCCCCGCATCACGGTGACAGAC | | | | | | | | | | |
| Oligo rev | AAACGTCTGTCACCGTGATGCGGG | | | | | | | | | | |
| Chromosome | start | end | strand | MM | target_seq | PAM | alignment | distance | position | gene name | gene id |
| GU305914.1 | 115742 | 115764 | - | 0 | CCCGCATCACGGTGACAGAC | AGG | \|\|\|\|\|\|\|\|[\|\|\|\|\|\|\|\|\|\|\|\|]PAM | NA | Intergenic | NA | NA |
| T25240 | TTTTTTTATTCTTATTTTACTGG | 1000 | | | | | | | | | |
| Oligo fwd | CACCTTTTTTTATTCTTATTTTAC | | | | | | | | | | |
| Oligo rev | AAACGTAAAATAAGAATAAAAAAA | | | | | | | | | | |
| Chromosome | start | end | strand | MM | target_seq | PAM | alignment | distance | position | gene name | gene id |
| GU305914.1 | 115719 | 115741 | - | 0 | TTTTTTTATTCTTATTTTAC | TGG | \|\|\|\|\|\|\|\|[\|\|\|\|\|\|\|\|\|\|\|\|]PAM | NA | Intergenic | NA | NA |
| T25241 | ACACGTTCACTCCGTTAGTCCGG | 1000 | | | | | | | | | |
| Oligo fwd | CACCACACGTTCACTCCGTTAGTC | | | | | | | | | | |
| Oligo rev | AAACGACTAACGGAGTGAACGTGT | | | | | | | | | | |
| Chromosome | start | end | strand | MM | target_seq | PAM | alignment | distance | position | gene name | gene id |
| GU305914.1 | 115621 | 115643 | - | 0 | ACACGTTCACTCCGTTAGTC | CGG | \|\|\|\|\|\|\|\|[\|\|\|\|\|\|\|\|\|\|\|\|]PAM | NA | Intergenic | NA | NA |
| T25242 | TCACTCATTCCATTTTATTGAGG | 1000 | | | | | | | | | |
| Oligo fwd | CACCTCACTCATTCCATTTTATTG | | | | | | | | | | |
| Oligo rev | AAACCAATAAAATGGAATGAGTGA | | | | | | | | | | |
| Chromosome | start | end | strand | MM | target_seq | PAM | alignment | distance | position | gene name | gene id |
| GU305914.1 | 115590 | 115612 | - | 0 | TCACTCATTCCATTTTATTG | AGG | \|\|\|\|\|\|\|\|[\|\|\|\|\|\|\|\|\|\|\|\|]PAM | NA | Intergenic | NA | NA |
| T25243 | CTCATTCCATTTTATTGAGGAGG | 1000 | | | | | | | | | |
| Oligo fwd | CACCCTCATTCCATTTTATTGAGG | | | | | | | | | | |
| Oligo rev | AAACCCTCAATAAAATGGAATGAG | | | | | | | | | | |
| Chromosome | start | end | strand | MM | target_seq | PAM | alignment | distance | position | gene name | gene id |
| GU305914.1 | 115587 | 115609 | - | 0 | CTCATTCCATTTTATTGAGG | AGG | \|\|\|\|\|\|\|\|[\|\|\|\|\|\|\|\|\|\|\|\|]PAM | NA | Intergenic | NA | NA |
| T25244 | TCATTCCATTTTATTGAGGAGGG | 1000 | | | | | | | | | |
| Oligo fwd | CACCTCATTCCATTTTATTGAGGA | | | | | | | | | | |
| Oligo rev | AAACTCCTCAATAAAATGGAATGA | | | | | | | | | | |
| Chromosome | start | end | strand | MM | target_seq | PAM | alignment | distance | position | gene name | gene id |
| GU305914.1 | 115586 | 115608 | - | 0 | TCATTCCATTTTATTGAGGA | GGG | \|\|\|\|\|\|\|\|[\|\|\|\|\|\|\|\|\|\|\|\|]PAM | NA | Intergenic | NA | NA |
| T25245 | TTTGTTCACCTCCAACATGCTGG | 1000 | | | | | | | | | |
| Oligo fwd | CACCTTTGTTCACCTCCAACATGC | | | | | | | | | | |
| Oligo rev | AAACGCATGTTGGAGGTGAACAAA | | | | | | | | | | |
| Chromosome | start | end | strand | MM | target_seq | PAM | alignment | distance | position | gene name | gene id |
| GU305914.1 | 115550 | 115572 | - | 0 | TTTGTTCACCTCCAACATGC | TGG | \|\|\|\|\|\|\|\|[\|\|\|\|\|\|\|\|\|\|\|\|]PAM | NA | Intergenic | NA | NA |
| T25246 | GGCCTCACCTCCCTTCCTCGAGG | 1000 | | | | | | | | | |
| Oligo fwd | CACCGGCCTCACCTCCCTTCCTCG | | | | | | | | | | |
| Oligo rev | AAACCGAGGAAGGGAGGTGAGGCC | | | | | | | | | | |
| Chromosome | start | end | strand | MM | target_seq | PAM | alignment | distance | position | gene name | gene id |
| GU305914.1 | 115529 | 115551 | - | 0 | GGCCTCACCTCCCTTCCTCG | AGG | \|\|\|\|\|\|\|\|[\|\|\|\|\|\|\|\|\|\|\|\|]PAM | NA | Intergenic | NA | NA |
| T25247 | CCTCGAGGTTCCTCCTCCCGAGG | 1000 | | | | | | | | | |
| Oligo fwd | CACCCCTCGAGGTTCCTCCTCCCG | | | | | | | | | | |
| Oligo rev | AAACCGGGAGGAGGAACCTCGAGG | | | | | | | | | | |
| Chromosome | start | end | strand | MM | target_seq | PAM | alignment | distance | position | gene name | gene id |
| GU305914.1 | 115514 | 115536 | - | 0 | CCTCGAGGTTCCTCCTCCCG | AGG | \|\|\|\|\|\|\|\|[\|\|\|\|\|\|\|\|\|\|\|\|]PAM | NA | Intergenic | NA | NA |
| T25248 | CTCGAGGTTCCTCCTCCCGAGGG | 1000 | | | | | | | | | |
| Oligo fwd | CACCCTCGAGGTTCCTCCTCCCGA | | | | | | | | | | |
| Oligo rev | AAACTCGGGAGGAGGAACCTCGAG | | | | | | | | | | |
| Chromosome | start | end | strand | MM | target_seq | PAM | alignment | distance | position | gene name | gene id |
| GU305914.1 | 115513 | 115535 | - | 0 | CTCGAGGTTCCTCCTCCCGA | GGG | \|\|\|\|\|\|\|\|[\|\|\|\|\|\|\|\|\|\|\|\|]PAM | NA | Intergenic | NA | NA |
| T25249 | CGAGGGTTCTCCCACCTCCATGG | 1000 | | | | | | | | | |
| Oligo fwd | CACCCGAGGGTTCTCCCACCTCCA | | | | | | | | | | |
| Oligo rev | AAACTGGAGGTGGGAGAACCCTCG | | | | | | | | | | |
| Chromosome | start | end | strand | MM | target_seq | PAM | alignment | distance | position | gene name | gene id |
| GU305914.1 | 115496 | 115518 | - | 0 | CGAGGGTTCTCCCACCTCCA | TGG | \|\|\|\|\|\|\|\|[\|\|\|\|\|\|\|\|\|\|\|\|]PAM | NA | Intergenic | NA | NA |
| T25250 | GAGGGTTCTCCCACCTCCATGGG | 1000 | | | | | | | | | |

| | | | | | | | | | | | |
|---|---|---|---|---|---|---|---|---|---|---|---|
| Oligo fwd | CACCGAGGGTTCTCCCACCTCCAT | | | | | | | | | | |
| Oligo rev | AAACATGGAGGTGGGAGAACCCTC | | | | | | | | | | |
| Chromosome | start | end | strand | MM | target_seq | PAM | alignment | distance | position | gene name | gene id |
| GU305914.1 | 115495 | 115517 | - | 0 | GAGGGTTCTCCCACCTCCAT | GGG | \|\|\|\|\|\|\|\|[\|\|\|\|\|\|\|\|\|\|\|\|]PAM | NA | Intergenic | NA | NA |
| T25251 | TCTCCCACCTCCATGGGAGTCGG | 1000 | | | | | | | | | |
| Oligo fwd | CACCTCTCCCACCTCCATGGGAGT | | | | | | | | | | |
| Oligo rev | AAACACTCCCATGGAGGTGGGAGA | | | | | | | | | | |
| Chromosome | start | end | strand | MM | target_seq | PAM | alignment | distance | position | gene name | gene id |
| GU305914.1 | 115489 | 115511 | - | 0 | TCTCCCACCTCCATGGGAGT | CGG | \|\|\|\|\|\|\|\|[\|\|\|\|\|\|\|\|\|\|\|\|]PAM | NA | Intergenic | NA | NA |
| T25252 | CGAAGCATTCACCAATTTCCCGG | 1000 | | | | | | | | | |
| Oligo fwd | CACCCGAAGCATTCACCAATTTCC | | | | | | | | | | |
| Oligo rev | AAACGGAAATTGGTGAATGCTTCG | | | | | | | | | | |
| Chromosome | start | end | strand | MM | target_seq | PAM | alignment | distance | position | gene name | gene id |
| GU305914.1 | 115460 | 115482 | - | 0 | CGAAGCATTCACCAATTTCC | CGG | \|\|\|\|\|\|\|\|[\|\|\|\|\|\|\|\|\|\|\|\|]PAM | NA | Intergenic | NA | NA |
| T25253 | AGCATTCACCAATTTCCCGGCGG | 1000 | | | | | | | | | |
| Oligo fwd | CACCAGCATTCACCAATTTCCCGG | | | | | | | | | | |
| Oligo rev | AAACCCGGGAAATTGGTGAATGCT | | | | | | | | | | |
| Chromosome | start | end | strand | MM | target_seq | PAM | alignment | distance | position | gene name | gene id |
| GU305914.1 | 115457 | 115479 | - | 0 | AGCATTCACCAATTTCCCGG | CGG | \|\|\|\|\|\|\|\|[\|\|\|\|\|\|\|\|\|\|\|\|]PAM | NA | Intergenic | NA | NA |
| T25254 | TTTCCCGGCGGTAAGCAGCCCGG | 1000 | | | | | | | | | |
| Oligo fwd | CACCTTTCCCGGCGGTAAGCAGCC | | | | | | | | | | |
| Oligo rev | AAACGGCTGCTTACCGCCGGGAAA | | | | | | | | | | |
| Chromosome | start | end | strand | MM | target_seq | PAM | alignment | distance | position | gene name | gene id |
| GU305914.1 | 115445 | 115467 | - | 0 | TTTCCCGGCGGTAAGCAGCC | CGG | \|\|\|\|\|\|\|\|[\|\|\|\|\|\|\|\|\|\|\|\|]PAM | NA | Intergenic | NA | NA |
| T25255 | CCCGGCGGTAAGCAGCCCGGCGG | 1000 | | | | | | | | | |
| Oligo fwd | CACCCCCGGCGGTAAGCAGCCCGG | | | | | | | | | | |
| Oligo rev | AAACCCGGGCTGCTTACCGCCGGG | | | | | | | | | | |
| Chromosome | start | end | strand | MM | target_seq | PAM | alignment | distance | position | gene name | gene id |
| GU305914.1 | 115442 | 115464 | - | 0 | CCCGGCGGTAAGCAGCCCGG | CGG | \|\|\|\|\|\|\|\|[\|\|\|\|\|\|\|\|\|\|\|\|]PAM | NA | Intergenic | NA | NA |
| T25256 | CCCGGCGGTATGTCCCGTCGCGG | 1000 | | | | | | | | | |
| Oligo fwd | CACCCCCGGCGGTATGTCCCGTCG | | | | | | | | | | |
| Oligo rev | AAACCGACGGGACATACCGCCGGG | | | | | | | | | | |
| Chromosome | start | end | strand | MM | target_seq | PAM | alignment | distance | position | gene name | gene id |
| GU305914.1 | 115427 | 115449 | - | 0 | CCCGGCGGTATGTCCCGTCG | CGG | \|\|\|\|\|\|\|\|[\|\|\|\|\|\|\|\|\|\|\|\|]PAM | NA | Intergenic | NA | NA |
| T25257 | GGCGGTATGTCCCGTCGCGGCGG | 1000 | | | | | | | | | |
| Oligo fwd | CACCGGCGGTATGTCCCGTCGCGG | | | | | | | | | | |
| Oligo rev | AAACCCGCGACGGGACATACCGCC | | | | | | | | | | |
| Chromosome | start | end | strand | MM | target_seq | PAM | alignment | distance | position | gene name | gene id |
| GU305914.1 | 115424 | 115446 | - | 0 | GGCGGTATGTCCCGTCGCGG | CGG | \|\|\|\|\|\|\|\|[\|\|\|\|\|\|\|\|\|\|\|\|]PAM | NA | Intergenic | NA | NA |
| T25258 | ATGTCCCGTCGCGGCGGCCACGG | 1000 | | | | | | | | | |
| Oligo fwd | CACCATGTCCCGTCGCGGCGGCCA | | | | | | | | | | |
| Oligo rev | AAACTGGCCGCCGCGACGGGACAT | | | | | | | | | | |
| Chromosome | start | end | strand | MM | target_seq | PAM | alignment | distance | position | gene name | gene id |
| GU305914.1 | 115418 | 115440 | - | 0 | ATGTCCCGTCGCGGCGGCCA | CGG | \|\|\|\|\|\|\|\|[\|\|\|\|\|\|\|\|\|\|\|\|]PAM | NA | Intergenic | NA | NA |
| T25259 | CCCGTCGCGGCGGCCACGGCCGG | 1000 | | | | | | | | | |
| Oligo fwd | CACCCCCGTCGCGGCGGCCACGGC | | | | | | | | | | |
| Oligo rev | AAACGCCGTGGCCGCCGCGACGGG | | | | | | | | | | |
| Chromosome | start | end | strand | MM | target_seq | PAM | alignment | distance | position | gene name | gene id |
| GU305914.1 | 115414 | 115436 | - | 0 | CCCGTCGCGGCGGCCACGGC | CGG | \|\|\|\|\|\|\|\|[\|\|\|\|\|\|\|\|\|\|\|\|]PAM | NA | Intergenic | NA | NA |
| T25260 | CGGCGGCCACGGCCGGATCCAGG | 1000 | | | | | | | | | |
| Oligo fwd | CACCCGGCGGCCACGGCCGGATCC | | | | | | | | | | |
| Oligo rev | AAACGGATCCGGCCGTGGCCGCCG | | | | | | | | | | |
| Chromosome | start | end | strand | MM | target_seq | PAM | alignment | distance | position | gene name | gene id |
| GU305914.1 | 115407 | 115429 | - | 0 | CGGCGGCCACGGCCGGATCC | AGG | \|\|\|\|\|\|\|\|[\|\|\|\|\|\|\|\|\|\|\|\|]PAM | NA | Intergenic | NA | NA |
| GU305914.1 | 121800 | 121822 | + | 4 | CTGCGGACTCAGCCGGATCC | CGG | \|-\|\|\|\|-\|[-\|-\|\|\|\|\|\|\|\|\|]PAM | NA | Intergenic | NA | NA |
| T25261 | GGCGGCCACGGCCGGATCCAGGG | 1000 | | | | | | | | | |
| Oligo fwd | CACCGGCGGCCACGGCCGGATCCA | | | | | | | | | | |
| Oligo rev | AAACTGGATCCGGCCGTGGCCGCC | | | | | | | | | | |
| Chromosome | start | end | strand | MM | target_seq | PAM | alignment | distance | position | gene name | gene id |
| GU305914.1 | 115406 | 115428 | - | 0 | GGCGGCCACGGCCGGATCCA | GGG | \|\|\|\|\|\|\|\|[\|\|\|\|\|\|\|\|\|\|\|\|]PAM | NA | Intergenic | NA | NA |
| T25262 | GGCCGGATCCAGGGCCATCCTGG | 1000 | | | | | | | | | |
| Oligo fwd | CACCGGCCGGATCCAGGGCCATCC | | | | | | | | | | |
| Oligo rev | AAACGGATGGCCCTGGATCCGGCC | | | | | | | | | | |
| Chromosome | start | end | strand | MM | target_seq | PAM | alignment | distance | position | gene name | gene id |
| GU305914.1 | 115397 | 115419 | - | 0 | GGCCGGATCCAGGGCCATCC | TGG | \|\|\|\|\|\|\|\|[\|\|\|\|\|\|\|\|\|\|\|\|]PAM | NA | Intergenic | NA | NA |
| T25263 | CAGGGCCATCCTGGCGACGACGG | 1000 | | | | | | | | | |
| Oligo fwd | CACCCAGGGCCATCCTGGCGACGA | | | | | | | | | | |
| Oligo rev | AAACTCGTCGCCAGGATGGCCCTG | | | | | | | | | | |
| Chromosome | start | end | strand | MM | target_seq | PAM | alignment | distance | position | gene name | gene id |
| GU305914.1 | 115388 | 115410 | - | 0 | CAGGGCCATCCTGGCGACGA | CGG | \|\|\|\|\|\|\|\|[\|\|\|\|\|\|\|\|\|\|\|\|]PAM | NA | Intergenic | NA | NA |
| T25264 | CCTGGCGACGACGGTGCCGCCGG | 1000 | | | | | | | | | |
| Oligo fwd | CACCCCTGGCGACGACGGTGCCGC | | | | | | | | | | |
| Oligo rev | AAACGCGGCACCGTCGTCGCCAGG | | | | | | | | | | |
| Chromosome | start | end | strand | MM | target_seq | PAM | alignment | distance | position | gene name | gene id |
| GU305914.1 | 115379 | 115401 | - | 0 | CCTGGCGACGACGGTGCCGC | CGG | \|\|\|\|\|\|\|\|[\|\|\|\|\|\|\|\|\|\|\|\|]PAM | NA | Intergenic | NA | NA |

| | | | | | | | | | | | |
|---|---|---|---|---|---|---|---|---|---|---|---|
| GU305914.1 | | 76838 | 76860 | - | 4 | CCGGACGACGACGAGGCCGC | CGG | \|\|-\|-\|\|\|[\|\|\|\|\|--\|\|\|\|\|]PAM | NA | Intergenic | NA | NA |
| T25265 | GCGACGACGGTGCCGCCGGCCGG | | 1000 | | | | | | | | | |
| Oligo fwd | CACCGCGACGACGGTGCCGCCGGC | | | | | | | | | | | |
| Oligo rev | AAACGCCGGCGGCACCGTCGTCGC | | | | | | | | | | | |
| Chromosome | start | | end | strand | MM | target_seq | PAM | alignment | distance | position | gene name | gene id |
| GU305914.1 | | 115375 | 115397 | - | 0 | GCGACGACGGTGCCGCCGGC | CGG | \|\|\|\|\|\|\|\|[\|\|\|\|\|\|\|\|\|\|\|\|]PAM | NA | Intergenic | NA | NA |
| T25266 | ACGACGGTGCCGCCGGCCGGCGG | | 1000 | | | | | | | | | |
| Oligo fwd | CACCACGACGGTGCCGCCGGCCGG | | | | | | | | | | | |
| Oligo rev | AAACCCGGCCGGCGGCACCGTCGT | | | | | | | | | | | |
| Chromosome | start | | end | strand | MM | target_seq | PAM | alignment | distance | position | gene name | gene id |
| GU305914.1 | | 115372 | 115394 | - | 0 | ACGACGGTGCCGCCGGCCGG | CGG | \|\|\|\|\|\|\|\|[\|\|\|\|\|\|\|\|\|\|\|\|]PAM | NA | Intergenic | NA | NA |
| GU305914.1 | | 90202 | 90224 | + | 4 | CCGCTGGCGCCGCCGGCCGG | AGG | -\|\|--\|\|-[\|\|\|\|\|\|\|\|\|\|\|\|]PAM | NA | Intergenic | NA | NA |
| T25267 | GCCGCCGGCCGGCGGTTCCGAGG | | 1000 | | | | | | | | | |
| Oligo fwd | CACCGCCGCCGGCCGGCGGTTCCG | | | | | | | | | | | |
| Oligo rev | AAACCGGAACCGCCGGCCGGCGGC | | | | | | | | | | | |
| Chromosome | start | | end | strand | MM | target_seq | PAM | alignment | distance | position | gene name | gene id |
| GU305914.1 | | 115364 | 115386 | - | 0 | GCCGCCGGCCGGCGGTTCCG | AGG | \|\|\|\|\|\|\|\|[\|\|\|\|\|\|\|\|\|\|\|\|]PAM | NA | Intergenic | NA | NA |
| T25268 | CGGTTCCGAGGCCATCATCTCGG | | 1000 | | | | | | | | | |
| Oligo fwd | CACCCGGTTCCGAGGCCATCATCT | | | | | | | | | | | |
| Oligo rev | AAACAGATGATGGCCTCGGAACCG | | | | | | | | | | | |
| Chromosome | start | | end | strand | MM | target_seq | PAM | alignment | distance | position | gene name | gene id |
| GU305914.1 | | 115352 | 115374 | - | 0 | CGGTTCCGAGGCCATCATCT | CGG | \|\|\|\|\|\|\|\|[\|\|\|\|\|\|\|\|\|\|\|\|]PAM | NA | Intergenic | NA | NA |
| T25269 | GGCCATCATCTCGGTGACTCCGG | | 1000 | | | | | | | | | |
| Oligo fwd | CACCGGCCATCATCTCGGTGACTC | | | | | | | | | | | |
| Oligo rev | AAACGAGTCACCGAGATGATGGCC | | | | | | | | | | | |
| Chromosome | start | | end | strand | MM | target_seq | PAM | alignment | distance | position | gene name | gene id |
| GU305914.1 | | 115343 | 115365 | - | 0 | GGCCATCATCTCGGTGACTC | CGG | \|\|\|\|\|\|\|\|[\|\|\|\|\|\|\|\|\|\|\|\|]PAM | NA | Intergenic | NA | NA |
| T25270 | GCCATCATCTCGGTGACTCCGGG | | 1000 | | | | | | | | | |
| Oligo fwd | CACCGCCATCATCTCGGTGACTCC | | | | | | | | | | | |
| Oligo rev | AAACGGAGTCACCGAGATGATGGC | | | | | | | | | | | |
| Chromosome | start | | end | strand | MM | target_seq | PAM | alignment | distance | position | gene name | gene id |
| GU305914.1 | | 115342 | 115364 | - | 0 | GCCATCATCTCGGTGACTCC | GGG | \|\|\|\|\|\|\|\|[\|\|\|\|\|\|\|\|\|\|\|\|]PAM | NA | Intergenic | NA | NA |
| T25271 | CCATCATCTCGGTGACTCCGGGG | | 1000 | | | | | | | | | |
| Oligo fwd | CACCCCATCATCTCGGTGACTCCG | | | | | | | | | | | |
| Oligo rev | AAACCGGAGTCACCGAGATGATGG | | | | | | | | | | | |
| Chromosome | start | | end | strand | MM | target_seq | PAM | alignment | distance | position | gene name | gene id |
| GU305914.1 | | 115341 | 115363 | - | 0 | CCATCATCTCGGTGACTCCG | GGG | \|\|\|\|\|\|\|\|[\|\|\|\|\|\|\|\|\|\|\|\|]PAM | NA | Intergenic | NA | NA |
| T25272 | CATCATCTCGGTGACTCCGGGGG | | 1000 | | | | | | | | | |
| Oligo fwd | CACCCATCATCTCGGTGACTCCGG | | | | | | | | | | | |
| Oligo rev | AAACCCGGAGTCACCGAGATGATG | | | | | | | | | | | |
| Chromosome | start | | end | strand | MM | target_seq | PAM | alignment | distance | position | gene name | gene id |
| GU305914.1 | | 115340 | 115362 | - | 0 | CATCATCTCGGTGACTCCGG | GGG | \|\|\|\|\|\|\|\|[\|\|\|\|\|\|\|\|\|\|\|\|]PAM | NA | Intergenic | NA | NA |
| T25273 | ATCATCTCGGTGACTCCGGGGGG | | 1000 | | | | | | | | | |
| Oligo fwd | CACCATCATCTCGGTGACTCCGGG | | | | | | | | | | | |
| Oligo rev | AAACCCCGGAGTCACCGAGATGAT | | | | | | | | | | | |
| Chromosome | start | | end | strand | MM | target_seq | PAM | alignment | distance | position | gene name | gene id |
| GU305914.1 | | 115339 | 115361 | - | 0 | ATCATCTCGGTGACTCCGGG | GGG | \|\|\|\|\|\|\|\|[\|\|\|\|\|\|\|\|\|\|\|\|]PAM | NA | Intergenic | NA | NA |
| T25274 | ATCTCGGTGACTCCGGGGGGCGG | | 1000 | | | | | | | | | |
| Oligo fwd | CACCATCTCGGTGACTCCGGGGGG | | | | | | | | | | | |
| Oligo rev | AAACCCCCCCGGAGTCACCGAGAT | | | | | | | | | | | |
| Chromosome | start | | end | strand | MM | target_seq | PAM | alignment | distance | position | gene name | gene id |
| GU305914.1 | | 115336 | 115358 | - | 0 | ATCTCGGTGACTCCGGGGGG | CGG | \|\|\|\|\|\|\|\|[\|\|\|\|\|\|\|\|\|\|\|\|]PAM | NA | Intergenic | NA | NA |
| T25275 | TCTCGGTGACTCCGGGGGGCGGG | | 1000 | | | | | | | | | |
| Oligo fwd | CACCTCTCGGTGACTCCGGGGGGC | | | | | | | | | | | |
| Oligo rev | AAACGCCCCCCGGAGTCACCGAGA | | | | | | | | | | | |
| Chromosome | start | | end | strand | MM | target_seq | PAM | alignment | distance | position | gene name | gene id |
| GU305914.1 | | 115335 | 115357 | - | 0 | TCTCGGTGACTCCGGGGGGC | GGG | \|\|\|\|\|\|\|\|[\|\|\|\|\|\|\|\|\|\|\|\|]PAM | NA | Intergenic | NA | NA |
| T25276 | CTCGGTGACTCCGGGGGGCGGGG | | 1000 | | | | | | | | | |
| Oligo fwd | CACCCTCGGTGACTCCGGGGGGCG | | | | | | | | | | | |
| Oligo rev | AAACCGCCCCCCGGAGTCACCGAG | | | | | | | | | | | |
| Chromosome | start | | end | strand | MM | target_seq | PAM | alignment | distance | position | gene name | gene id |
| GU305914.1 | | 115334 | 115356 | - | 0 | CTCGGTGACTCCGGGGGGCG | GGG | \|\|\|\|\|\|\|\|[\|\|\|\|\|\|\|\|\|\|\|\|]PAM | NA | Intergenic | NA | NA |
| T25277 | GGTGACTCCGGGGGGCGGGGCGG | | 1000 | | | | | | | | | |
| Oligo fwd | CACCGGTGACTCCGGGGGGCGGGG | | | | | | | | | | | |
| Oligo rev | AAACCCCCGCCCCCCGGAGTCACC | | | | | | | | | | | |
| Chromosome | start | | end | strand | MM | target_seq | PAM | alignment | distance | position | gene name | gene id |
| GU305914.1 | | 115331 | 115353 | - | 0 | GGTGACTCCGGGGGGCGGGG | CGG | \|\|\|\|\|\|\|\|[\|\|\|\|\|\|\|\|\|\|\|\|]PAM | NA | Intergenic | NA | NA |
| T25278 | GGGGCGGGGCGGCACATTTCTGG | | 1000 | | | | | | | | | |
| Oligo fwd | CACCGGGGCGGGGCGGCACATTTC | | | | | | | | | | | |
| Oligo rev | AAACGAAATGTGCCGCCCCGCCCC | | | | | | | | | | | |
| Chromosome | start | | end | strand | MM | target_seq | PAM | alignment | distance | position | gene name | gene id |
| GU305914.1 | | 115320 | 115342 | - | 0 | GGGGCGGGGCGGCACATTTC | TGG | \|\|\|\|\|\|\|\|[\|\|\|\|\|\|\|\|\|\|\|\|]PAM | NA | Intergenic | NA | NA |
| T25279 | CTGGTCAGAAGATCCGACCGAGG | | 1000 | | | | | | | | | |
| Oligo fwd | CACCCTGGTCAGAAGATCCGACCG | | | | | | | | | | | |

| | | | | | | | | | | | |
|---|---|---|---|---|---|---|---|---|---|---|---|
| Oligo rev | AAACCGGTCGGATCTTCTGACCAG | | | | | | | | | | |
| Chromosome | start | end | strand | MM | target_seq | PAM | alignment | distance | position | gene name | gene id |
| GU305914.1 | 115301 | 115323 | - | 0 | CTGGTCAGAAGATCCGACCG | AGG | \|\|\|\|\|\|\|\|[\|\|\|\|\|\|\|\|\|\|\|\|]PAM | NA | Intergenic | NA | NA |
| T25280 | CTCGCGCGCGCCGTTGCCCTCGG | 1000 | | | | | | | | | |
| Oligo fwd | CACCCTCGCGCGCGCCGTTGCCCT | | | | | | | | | | |
| Oligo rev | AAACAGGGCAACGGCGCGCGCGAG | | | | | | | | | | |
| Chromosome | start | end | strand | MM | target_seq | PAM | alignment | distance | position | gene name | gene id |
| GU305914.1 | 115271 | 115293 | - | 0 | CTCGCGCGCGCCGTTGCCCT | CGG | \|\|\|\|\|\|\|\|[\|\|\|\|\|\|\|\|\|\|\|\|]PAM | NA | Intergenic | NA | NA |
| T25281 | CGCGCCGTTGCCCTCGGCCCTGG | 1000 | | | | | | | | | |
| Oligo fwd | CACCCGCGCCGTTGCCCTCGGCCC | | | | | | | | | | |
| Oligo rev | AAACGGGCCGAGGGCAACGGCGCG | | | | | | | | | | |
| Chromosome | start | end | strand | MM | target_seq | PAM | alignment | distance | position | gene name | gene id |
| GU305914.1 | 115265 | 115287 | - | 0 | CGCGCCGTTGCCCTCGGCCC | TGG | \|\|\|\|\|\|\|\|[\|\|\|\|\|\|\|\|\|\|\|\|]PAM | NA | Intergenic | NA | NA |
| T25282 | GCCGTTGCCCTCGGCCCTGGCGG | 1000 | | | | | | | | | |
| Oligo fwd | CACCGCCGTTGCCCTCGGCCCTGG | | | | | | | | | | |
| Oligo rev | AAACCCAGGGCCGAGGGCAACGGC | | | | | | | | | | |
| Chromosome | start | end | strand | MM | target_seq | PAM | alignment | distance | position | gene name | gene id |
| GU305914.1 | 115262 | 115284 | - | 0 | GCCGTTGCCCTCGGCCCTGG | CGG | \|\|\|\|\|\|\|\|[\|\|\|\|\|\|\|\|\|\|\|\|]PAM | NA | Intergenic | NA | NA |
| T25283 | CCTCGGCCCTGGCGGCCGCGCGG | 1000 | | | | | | | | | |
| Oligo fwd | CACCCCTCGGCCCTGGCGGCCGCG | | | | | | | | | | |
| Oligo rev | AAACCGCGGCCGCCAGGGCCGAGG | | | | | | | | | | |
| Chromosome | start | end | strand | MM | target_seq | PAM | alignment | distance | position | gene name | gene id |
| GU305914.1 | 115254 | 115276 | - | 0 | CCTCGGCCCTGGCGGCCGCG | CGG | \|\|\|\|\|\|\|\|[\|\|\|\|\|\|\|\|\|\|\|\|]PAM | NA | Intergenic | NA | NA |
| T25284 | GGATCTGCGAGATGTCTTTGCGG | 1000 | | | | | | | | | |
| Oligo fwd | CACCGGATCTGCGAGATGTCTTTG | | | | | | | | | | |
| Oligo rev | AAACCAAAGACATCTCGCAGATCC | | | | | | | | | | |
| Chromosome | start | end | strand | MM | target_seq | PAM | alignment | distance | position | gene name | gene id |
| GU305914.1 | 115233 | 115255 | - | 0 | GGATCTGCGAGATGTCTTTG | CGG | \|\|\|\|\|\|\|\|[\|\|\|\|\|\|\|\|\|\|\|\|]PAM | NA | Intergenic | NA | NA |
| T25285 | CCTCACCTCTCCGAACTCATCGG | 1000 | | | | | | | | | |
| Oligo fwd | CACCCCTCACCTCTCCGAACTCAT | | | | | | | | | | |
| Oligo rev | AAACATGAGTTCGGAGAGGTGAGG | | | | | | | | | | |
| Chromosome | start | end | strand | MM | target_seq | PAM | alignment | distance | position | gene name | gene id |
| GU305914.1 | 115193 | 115215 | - | 0 | CCTCACCTCTCCGAACTCAT | CGG | \|\|\|\|\|\|\|\|[\|\|\|\|\|\|\|\|\|\|\|\|]PAM | NA | Intergenic | NA | NA |
| T25286 | CTCTCCGAACTCATCGGTTCTGG | 1000 | | | | | | | | | |
| Oligo fwd | CACCCTCTCCGAACTCATCGGTTC | | | | | | | | | | |
| Oligo rev | AAACGAACCGATGAGTTCGGAGAG | | | | | | | | | | |
| Chromosome | start | end | strand | MM | target_seq | PAM | alignment | distance | position | gene name | gene id |
| GU305914.1 | 115187 | 115209 | - | 0 | CTCTCCGAACTCATCGGTTC | TGG | \|\|\|\|\|\|\|\|[\|\|\|\|\|\|\|\|\|\|\|\|]PAM | NA | Intergenic | NA | NA |
| T25287 | TCCGAACTCATCGGTTCTGGAGG | 1000 | | | | | | | | | |
| Oligo fwd | CACCTCCGAACTCATCGGTTCTGG | | | | | | | | | | |
| Oligo rev | AAACCCAGAACCGATGAGTTCGGA | | | | | | | | | | |
| Chromosome | start | end | strand | MM | target_seq | PAM | alignment | distance | position | gene name | gene id |
| GU305914.1 | 115184 | 115206 | - | 0 | TCCGAACTCATCGGTTCTGG | AGG | \|\|\|\|\|\|\|\|[\|\|\|\|\|\|\|\|\|\|\|\|]PAM | NA | Intergenic | NA | NA |
| T25288 | AACTCATCGGTTCTGGAGGCCGG | 1000 | | | | | | | | | |
| Oligo fwd | CACCAACTCATCGGTTCTGGAGGC | | | | | | | | | | |
| Oligo rev | AAACGCCTCCAGAACCGATGAGTT | | | | | | | | | | |
| Chromosome | start | end | strand | MM | target_seq | PAM | alignment | distance | position | gene name | gene id |
| GU305914.1 | 115180 | 115202 | - | 0 | AACTCATCGGTTCTGGAGGC | CGG | \|\|\|\|\|\|\|\|[\|\|\|\|\|\|\|\|\|\|\|\|]PAM | NA | Intergenic | NA | NA |
| T25289 | TCATCGGTTCTGGAGGCCGGCGG | 1000 | | | | | | | | | |
| Oligo fwd | CACCTCATCGGTTCTGGAGGCCGG | | | | | | | | | | |
| Oligo rev | AAACCCGGCCTCCAGAACCGATGA | | | | | | | | | | |
| Chromosome | start | end | strand | MM | target_seq | PAM | alignment | distance | position | gene name | gene id |
| GU305914.1 | 115177 | 115199 | - | 0 | TCATCGGTTCTGGAGGCCGG | CGG | \|\|\|\|\|\|\|\|[\|\|\|\|\|\|\|\|\|\|\|\|]PAM | NA | Intergenic | NA | NA |
| T25290 | TGGAGGCCGGCGGCGCCGACTGG | 1000 | | | | | | | | | |
| Oligo fwd | CACCTGGAGGCCGGCGGCGCCGAC | | | | | | | | | | |
| Oligo rev | AAACGTCGGCGCCGCCGGCCTCCA | | | | | | | | | | |
| Chromosome | start | end | strand | MM | target_seq | PAM | alignment | distance | position | gene name | gene id |
| GU305914.1 | 115167 | 115189 | - | 0 | TGGAGGCCGGCGGCGCCGAC | TGG | \|\|\|\|\|\|\|\|[\|\|\|\|\|\|\|\|\|\|\|\|]PAM | NA | Intergenic | NA | NA |
| T25291 | CGCCGACTGGTGATGATCACCGG | 1000 | | | | | | | | | |
| Oligo fwd | CACCCGCCGACTGGTGATGATCAC | | | | | | | | | | |
| Oligo rev | AAACGTGATCATCACCAGTCGGCG | | | | | | | | | | |
| Chromosome | start | end | strand | MM | target_seq | PAM | alignment | distance | position | gene name | gene id |
| GU305914.1 | 115154 | 115176 | - | 0 | CGCCGACTGGTGATGATCAC | CGG | \|\|\|\|\|\|\|\|[\|\|\|\|\|\|\|\|\|\|\|\|]PAM | NA | Intergenic | NA | NA |
| T25292 | CGACTGGTGATGATCACCGGCGG | 1000 | | | | | | | | | |
| Oligo fwd | CACCCGACTGGTGATGATCACCGG | | | | | | | | | | |
| Oligo rev | AAACCCGGTGATCATCACCAGTCG | | | | | | | | | | |
| Chromosome | start | end | strand | MM | target_seq | PAM | alignment | distance | position | gene name | gene id |
| GU305914.1 | 115151 | 115173 | - | 0 | CGACTGGTGATGATCACCGG | CGG | \|\|\|\|\|\|\|\|[\|\|\|\|\|\|\|\|\|\|\|\|]PAM | NA | Intergenic | NA | NA |
| T25293 | ATGATCACCGGCGGTGCGCTTGG | 1000 | | | | | | | | | |
| Oligo fwd | CACCATGATCACCGGCGGTGCGCT | | | | | | | | | | |
| Oligo rev | AAACAGCGCACCGCCGGTGATCAT | | | | | | | | | | |
| Chromosome | start | end | strand | MM | target_seq | PAM | alignment | distance | position | gene name | gene id |
| GU305914.1 | 115142 | 115164 | - | 0 | ATGATCACCGGCGGTGCGCT | TGG | \|\|\|\|\|\|\|\|[\|\|\|\|\|\|\|\|\|\|\|\|]PAM | NA | Intergenic | NA | NA |
| T25294 | TGATCACCGGCGGTGCGCTTGGG | 1000 | | | | | | | | | |

<table>
<tr><td>Oligo fwd</td><td>CACCTGATCACCGGCGGTGCGCTT</td><td></td><td></td><td></td><td></td><td></td><td></td><td></td><td></td><td></td><td></td></tr>
<tr><td>Oligo rev</td><td>AAACAAGCGCACCGCCGGTGATCA</td><td></td><td></td><td></td><td></td><td></td><td></td><td></td><td></td><td></td><td></td></tr>
<tr><td>Chromosome</td><td>start</td><td>end</td><td>strand</td><td>MM</td><td>target_seq</td><td>PAM</td><td>alignment</td><td>distance</td><td>position</td><td>gene name</td><td>gene id</td></tr>
<tr><td>GU305914.1</td><td>115141</td><td>115163</td><td>-</td><td>0</td><td>TGATCACCGGCGGTGCGCTT</td><td>GGG</td><td>||||||||[||||||||||||]PAM</td><td>NA</td><td>Intergenic</td><td>NA</td><td>NA</td></tr>
<tr><td></td><td></td><td></td><td></td><td></td><td></td><td></td><td></td><td></td><td></td><td></td><td></td></tr>
<tr><td>T25295</td><td>CTTGGGATATCCCCGCTCTCCGG</td><td>1000</td><td></td><td></td><td></td><td></td><td></td><td></td><td></td><td></td><td></td></tr>
<tr><td>Oligo fwd</td><td>CACCCTTGGGATATCCCCGCTCTC</td><td></td><td></td><td></td><td></td><td></td><td></td><td></td><td></td><td></td><td></td></tr>
<tr><td>Oligo rev</td><td>AAACGAGAGCGGGGATATCCCAAG</td><td></td><td></td><td></td><td></td><td></td><td></td><td></td><td></td><td></td><td></td></tr>
<tr><td>Chromosome</td><td>start</td><td>end</td><td>strand</td><td>MM</td><td>target_seq</td><td>PAM</td><td>alignment</td><td>distance</td><td>position</td><td>gene name</td><td>gene id</td></tr>
<tr><td>GU305914.1</td><td>115124</td><td>115146</td><td>-</td><td>0</td><td>CTTGGGATATCCCCGCTCTC</td><td>CGG</td><td>||||||||[||||||||||||]PAM</td><td>NA</td><td>Intergenic</td><td>NA</td><td>NA</td></tr>
<tr><td></td><td></td><td></td><td></td><td></td><td></td><td></td><td></td><td></td><td></td><td></td><td></td></tr>
<tr><td>T25296</td><td>TTGGGATATCCCCGCTCTCCGGG</td><td>1000</td><td></td><td></td><td></td><td></td><td></td><td></td><td></td><td></td><td></td></tr>
<tr><td>Oligo fwd</td><td>CACCTTGGGATATCCCCGCTCTCC</td><td></td><td></td><td></td><td></td><td></td><td></td><td></td><td></td><td></td><td></td></tr>
<tr><td>Oligo rev</td><td>AAACGGAGAGCGGGGATATCCCAA</td><td></td><td></td><td></td><td></td><td></td><td></td><td></td><td></td><td></td><td></td></tr>
<tr><td>Chromosome</td><td>start</td><td>end</td><td>strand</td><td>MM</td><td>target_seq</td><td>PAM</td><td>alignment</td><td>distance</td><td>position</td><td>gene name</td><td>gene id</td></tr>
<tr><td>GU305914.1</td><td>115123</td><td>115145</td><td>-</td><td>0</td><td>TTGGGATATCCCCGCTCTCC</td><td>GGG</td><td>||||||||[||||||||||||]PAM</td><td>NA</td><td>Intergenic</td><td>NA</td><td>NA</td></tr>
<tr><td></td><td></td><td></td><td></td><td></td><td></td><td></td><td></td><td></td><td></td><td></td><td></td></tr>
<tr><td>T25297</td><td>GATATCCCCGCTCTCCGGGTAGG</td><td>1000</td><td></td><td></td><td></td><td></td><td></td><td></td><td></td><td></td><td></td></tr>
<tr><td>Oligo fwd</td><td>CACCGATATCCCCGCTCTCCGGGT</td><td></td><td></td><td></td><td></td><td></td><td></td><td></td><td></td><td></td><td></td></tr>
<tr><td>Oligo rev</td><td>AAACACCCGGAGAGCGGGGATATC</td><td></td><td></td><td></td><td></td><td></td><td></td><td></td><td></td><td></td><td></td></tr>
<tr><td>Chromosome</td><td>start</td><td>end</td><td>strand</td><td>MM</td><td>target_seq</td><td>PAM</td><td>alignment</td><td>distance</td><td>position</td><td>gene name</td><td>gene id</td></tr>
<tr><td>GU305914.1</td><td>115119</td><td>115141</td><td>-</td><td>0</td><td>GATATCCCCGCTCTCCGGGT</td><td>AGG</td><td>||||||||[||||||||||||]PAM</td><td>NA</td><td>Intergenic</td><td>NA</td><td>NA</td></tr>
<tr><td></td><td></td><td></td><td></td><td></td><td></td><td></td><td></td><td></td><td></td><td></td><td></td></tr>
<tr><td>T25298</td><td>ATATCCCCGCTCTCCGGGTAGGG</td><td>1000</td><td></td><td></td><td></td><td></td><td></td><td></td><td></td><td></td><td></td></tr>
<tr><td>Oligo fwd</td><td>CACCATATCCCCGCTCTCCGGGTA</td><td></td><td></td><td></td><td></td><td></td><td></td><td></td><td></td><td></td><td></td></tr>
<tr><td>Oligo rev</td><td>AAACTACCCGGAGAGCGGGGATAT</td><td></td><td></td><td></td><td></td><td></td><td></td><td></td><td></td><td></td><td></td></tr>
<tr><td>Chromosome</td><td>start</td><td>end</td><td>strand</td><td>MM</td><td>target_seq</td><td>PAM</td><td>alignment</td><td>distance</td><td>position</td><td>gene name</td><td>gene id</td></tr>
<tr><td>GU305914.1</td><td>115118</td><td>115140</td><td>-</td><td>0</td><td>ATATCCCCGCTCTCCGGGTA</td><td>GGG</td><td>||||||||[||||||||||||]PAM</td><td>NA</td><td>Intergenic</td><td>NA</td><td>NA</td></tr>
<tr><td></td><td></td><td></td><td></td><td></td><td></td><td></td><td></td><td></td><td></td><td></td><td></td></tr>
<tr><td>T25299</td><td>CCCGCTCTCCGGGTAGGGAGAGG</td><td>1000</td><td></td><td></td><td></td><td></td><td></td><td></td><td></td><td></td><td></td></tr>
<tr><td>Oligo fwd</td><td>CACCCCCGCTCTCCGGGTAGGGAG</td><td></td><td></td><td></td><td></td><td></td><td></td><td></td><td></td><td></td><td></td></tr>
<tr><td>Oligo rev</td><td>AAACCTCCCTACCCGGAGAGCGGG</td><td></td><td></td><td></td><td></td><td></td><td></td><td></td><td></td><td></td><td></td></tr>
<tr><td>Chromosome</td><td>start</td><td>end</td><td>strand</td><td>MM</td><td>target_seq</td><td>PAM</td><td>alignment</td><td>distance</td><td>position</td><td>gene name</td><td>gene id</td></tr>
<tr><td>GU305914.1</td><td>115113</td><td>115135</td><td>-</td><td>0</td><td>CCCGCTCTCCGGGTAGGGAG</td><td>AGG</td><td>||||||||[||||||||||||]PAM</td><td>NA</td><td>Intergenic</td><td>NA</td><td>NA</td></tr>
<tr><td></td><td></td><td></td><td></td><td></td><td></td><td></td><td></td><td></td><td></td><td></td><td></td></tr>
<tr><td>T25300</td><td>AGGTCACCGCCATCGTCGTCAGG</td><td>1000</td><td></td><td></td><td></td><td></td><td></td><td></td><td></td><td></td><td></td></tr>
<tr><td>Oligo fwd</td><td>CACCAGGTCACCGCCATCGTCGTC</td><td></td><td></td><td></td><td></td><td></td><td></td><td></td><td></td><td></td><td></td></tr>
<tr><td>Oligo rev</td><td>AAACGACGACGATGGCGGTGACCT</td><td></td><td></td><td></td><td></td><td></td><td></td><td></td><td></td><td></td><td></td></tr>
<tr><td>Chromosome</td><td>start</td><td>end</td><td>strand</td><td>MM</td><td>target_seq</td><td>PAM</td><td>alignment</td><td>distance</td><td>position</td><td>gene name</td><td>gene id</td></tr>
<tr><td>GU305914.1</td><td>115093</td><td>115115</td><td>-</td><td>0</td><td>AGGTCACCGCCATCGTCGTC</td><td>AGG</td><td>||||||||[||||||||||||]PAM</td><td>NA</td><td>Intergenic</td><td>NA</td><td>NA</td></tr>
<tr><td></td><td></td><td></td><td></td><td></td><td></td><td></td><td></td><td></td><td></td><td></td><td></td></tr>
<tr><td>T25301</td><td>CCGACGTCTCCTCTTCCTGCCGG</td><td>1000</td><td></td><td></td><td></td><td></td><td></td><td></td><td></td><td></td><td></td></tr>
<tr><td>Oligo fwd</td><td>CACCCCGACGTCTCCTCTTCCTGC</td><td></td><td></td><td></td><td></td><td></td><td></td><td></td><td></td><td></td><td></td></tr>
<tr><td>Oligo rev</td><td>AAACGCAGGAAGAGGAGACGTCGG</td><td></td><td></td><td></td><td></td><td></td><td></td><td></td><td></td><td></td><td></td></tr>
<tr><td>Chromosome</td><td>start</td><td>end</td><td>strand</td><td>MM</td><td>target_seq</td><td>PAM</td><td>alignment</td><td>distance</td><td>position</td><td>gene name</td><td>gene id</td></tr>
<tr><td>GU305914.1</td><td>115061</td><td>115083</td><td>-</td><td>0</td><td>CCGACGTCTCCTCTTCCTGC</td><td>CGG</td><td>||||||||[||||||||||||]PAM</td><td>NA</td><td>Intergenic</td><td>NA</td><td>NA</td></tr>
<tr><td></td><td></td><td></td><td></td><td></td><td></td><td></td><td></td><td></td><td></td><td></td><td></td></tr>
<tr><td>T25302</td><td>TCTCCTCTTCCTGCCGGAGTCGG</td><td>1000</td><td></td><td></td><td></td><td></td><td></td><td></td><td></td><td></td><td></td></tr>
<tr><td>Oligo fwd</td><td>CACCTCTCCTCTTCCTGCCGGAGT</td><td></td><td></td><td></td><td></td><td></td><td></td><td></td><td></td><td></td><td></td></tr>
<tr><td>Oligo rev</td><td>AAACACTCCGGCAGGAAGAGGAGA</td><td></td><td></td><td></td><td></td><td></td><td></td><td></td><td></td><td></td><td></td></tr>
<tr><td>Chromosome</td><td>start</td><td>end</td><td>strand</td><td>MM</td><td>target_seq</td><td>PAM</td><td>alignment</td><td>distance</td><td>position</td><td>gene name</td><td>gene id</td></tr>
<tr><td>GU305914.1</td><td>115055</td><td>115077</td><td>-</td><td>0</td><td>TCTCCTCTTCCTGCCGGAGT</td><td>CGG</td><td>||||||||[||||||||||||]PAM</td><td>NA</td><td>Intergenic</td><td>NA</td><td>NA</td></tr>
<tr><td></td><td></td><td></td><td></td><td></td><td></td><td></td><td></td><td></td><td></td><td></td><td></td></tr>
<tr><td>T25303</td><td>CTTGCCCCCCGTACGCTCGCCGG</td><td>1000</td><td></td><td></td><td></td><td></td><td></td><td></td><td></td><td></td><td></td></tr>
<tr><td>Oligo fwd</td><td>CACCCTTGCCCCCCGTACGCTCGC</td><td></td><td></td><td></td><td></td><td></td><td></td><td></td><td></td><td></td><td></td></tr>
<tr><td>Oligo rev</td><td>AAACGCGAGCGTACGGGGGGCAAG</td><td></td><td></td><td></td><td></td><td></td><td></td><td></td><td></td><td></td><td></td></tr>
<tr><td>Chromosome</td><td>start</td><td>end</td><td>strand</td><td>MM</td><td>target_seq</td><td>PAM</td><td>alignment</td><td>distance</td><td>position</td><td>gene name</td><td>gene id</td></tr>
<tr><td>GU305914.1</td><td>115010</td><td>115032</td><td>-</td><td>0</td><td>CTTGCCCCCCGTACGCTCGC</td><td>CGG</td><td>||||||||[||||||||||||]PAM</td><td>NA</td><td>Intergenic</td><td>NA</td><td>NA</td></tr>
<tr><td></td><td></td><td></td><td></td><td></td><td></td><td></td><td></td><td></td><td></td><td></td><td></td></tr>
<tr><td>T25304</td><td>GTCATCGTCGTCGTCCGAATCGG</td><td>1000</td><td></td><td></td><td></td><td></td><td></td><td></td><td></td><td></td><td></td></tr>
<tr><td>Oligo fwd</td><td>CACCGTCATCGTCGTCGTCCGAAT</td><td></td><td></td><td></td><td></td><td></td><td></td><td></td><td></td><td></td><td></td></tr>
<tr><td>Oligo rev</td><td>AAACATTCGGACGACGACGATGAC</td><td></td><td></td><td></td><td></td><td></td><td></td><td></td><td></td><td></td><td></td></tr>
<tr><td>Chromosome</td><td>start</td><td>end</td><td>strand</td><td>MM</td><td>target_seq</td><td>PAM</td><td>alignment</td><td>distance</td><td>position</td><td>gene name</td><td>gene id</td></tr>
<tr><td>GU305914.1</td><td>114965</td><td>114987</td><td>-</td><td>0</td><td>GTCATCGTCGTCGTCCGAAT</td><td>CGG</td><td>||||||||[||||||||||||]PAM</td><td>NA</td><td>Intergenic</td><td>NA</td><td>NA</td></tr>
<tr><td></td><td></td><td></td><td></td><td></td><td></td><td></td><td></td><td></td><td></td><td></td><td></td></tr>
<tr><td>T25305</td><td>TCATCGTCGTCGTCCGAATCGGG</td><td>1000</td><td></td><td></td><td></td><td></td><td></td><td></td><td></td><td></td><td></td></tr>
<tr><td>Oligo fwd</td><td>CACCTCATCGTCGTCGTCCGAATC</td><td></td><td></td><td></td><td></td><td></td><td></td><td></td><td></td><td></td><td></td></tr>
<tr><td>Oligo rev</td><td>AAACGATTCGGACGACGACGATGA</td><td></td><td></td><td></td><td></td><td></td><td></td><td></td><td></td><td></td><td></td></tr>
<tr><td>Chromosome</td><td>start</td><td>end</td><td>strand</td><td>MM</td><td>target_seq</td><td>PAM</td><td>alignment</td><td>distance</td><td>position</td><td>gene name</td><td>gene id</td></tr>
<tr><td>GU305914.1</td><td>114964</td><td>114986</td><td>-</td><td>0</td><td>TCATCGTCGTCGTCCGAATC</td><td>GGG</td><td>||||||||[||||||||||||]PAM</td><td>NA</td><td>Intergenic</td><td>NA</td><td>NA</td></tr>
<tr><td>GU305914.1</td><td>77045</td><td>77067</td><td>+</td><td>3</td><td>TCATCCTCGTCGACCGACTC</td><td>CGG</td><td>|||||-||[||||-||||-||]PAM</td><td>NA</td><td>Intergenic</td><td>NA</td><td>NA</td></tr>
<tr><td></td><td></td><td></td><td></td><td></td><td></td><td></td><td></td><td></td><td></td><td></td><td></td></tr>
<tr><td>T25306</td><td>TCGTCGTCGTCCGAATCGGGAGG</td><td>1000</td><td></td><td></td><td></td><td></td><td></td><td></td><td></td><td></td><td></td></tr>
<tr><td>Oligo fwd</td><td>CACCTCGTCGTCGTCCGAATCGGG</td><td></td><td></td><td></td><td></td><td></td><td></td><td></td><td></td><td></td><td></td></tr>
<tr><td>Oligo rev</td><td>AAACCCCGATTCGGACGACGACGA</td><td></td><td></td><td></td><td></td><td></td><td></td><td></td><td></td><td></td><td></td></tr>
<tr><td>Chromosome</td><td>start</td><td>end</td><td>strand</td><td>MM</td><td>target_seq</td><td>PAM</td><td>alignment</td><td>distance</td><td>position</td><td>gene name</td><td>gene id</td></tr>
<tr><td>GU305914.1</td><td>114961</td><td>114983</td><td>-</td><td>0</td><td>TCGTCGTCGTCCGAATCGGG</td><td>AGG</td><td>||||||||[||||||||||||]PAM</td><td>NA</td><td>Intergenic</td><td>NA</td><td>NA</td></tr>
<tr><td></td><td></td><td></td><td></td><td></td><td></td><td></td><td></td><td></td><td></td><td></td><td></td></tr>
<tr><td>T25307</td><td>CGTCGTCGTCCGAATCGGGAGGG</td><td>1000</td><td></td><td></td><td></td><td></td><td></td><td></td><td></td><td></td><td></td></tr>
<tr><td>Oligo fwd</td><td>CACCCGTCGTCGTCCGAATCGGGA</td><td></td><td></td><td></td><td></td><td></td><td></td><td></td><td></td><td></td><td></td></tr>
<tr><td>Oligo rev</td><td>AAACTCCCGATTCGGACGACGACG</td><td></td><td></td><td></td><td></td><td></td><td></td><td></td><td></td><td></td><td></td></tr>
<tr><td>Chromosome</td><td>start</td><td>end</td><td>strand</td><td>MM</td><td>target_seq</td><td>PAM</td><td>alignment</td><td>distance</td><td>position</td><td>gene name</td><td>gene id</td></tr>
<tr><td>GU305914.1</td><td>114960</td><td>114982</td><td>-</td><td>0</td><td>CGTCGTCGTCCGAATCGGGA</td><td>GGG</td><td>||||||||[||||||||||||]PAM</td><td>NA</td><td>Intergenic</td><td>NA</td><td>NA</td></tr>
<tr><td></td><td></td><td></td><td></td><td></td><td></td><td></td><td></td><td></td><td></td><td></td><td></td></tr>
<tr><td>T25308</td><td>TATTCCTCTCTCCCCTGTAGAGG</td><td>1000</td><td></td><td></td><td></td><td></td><td></td><td></td><td></td><td></td><td></td></tr>
<tr><td>Oligo fwd</td><td>CACCTATTCCTCTCTCCCCTGTAG</td><td></td><td></td><td></td><td></td><td></td><td></td><td></td><td></td><td></td><td></td></tr>
<tr><td>Oligo rev</td><td>AAACCTACAGGGGAGAGAGGAATA</td><td></td><td></td><td></td><td></td><td></td><td></td><td></td><td></td><td></td><td></td></tr>
<tr><td>Chromosome</td><td>start</td><td>end</td><td>strand</td><td>MM</td><td>target_seq</td><td>PAM</td><td>alignment</td><td>distance</td><td>position</td><td>gene name</td><td>gene id</td></tr>
<tr><td>GU305914.1</td><td>114927</td><td>114949</td><td>-</td><td>0</td><td>TATTCCTCTCTCCCCTGTAG</td><td>AGG</td><td>||||||||[||||||||||||]PAM</td><td>NA</td><td>Intergenic</td><td>NA</td><td>NA</td></tr>
</table>

T25309 ATTCCTCTCTCCCCTGTAGAGGG 1000
Oligo fwd CACCATTCCTCTCTCCCCTGTAGA
Oligo rev AAACTCTACAGGGGAGAGAGGAAT
Chromosome start end strand MM target_seq PAM alignment distance position gene name gene id
GU305914.1 114926 114948 - 0 ATTCCTCTCTCCCCTGTAGA GGG ||||||||[||||||||||||]PAM NA Intergenic NA NA

T25310 TTCCTCTCTCCCCTGTAGAGGGG 1000
Oligo fwd CACCTTCCTCTCTCCCCTGTAGAG
Oligo rev AAACCTCTACAGGGGAGAGAGGAA
Chromosome start end strand MM target_seq PAM alignment distance position gene name gene id
GU305914.1 114925 114947 - 0 TTCCTCTCTCCCCTGTAGAG GGG ||||||||[||||||||||||]PAM NA Intergenic NA NA

T25311 CTGTAGAGGGGCTGCTGCTGAGG 1000
Oligo fwd CACCCTGTAGAGGGGCTGCTGCTG
Oligo rev AAACCAGCAGCAGCCCCTCTACAG
Chromosome start end strand MM target_seq PAM alignment distance position gene name gene id
GU305914.1 114913 114935 - 0 CTGTAGAGGGGCTGCTGCTG AGG ||||||||[||||||||||||]PAM NA Intergenic NA NA

T25312 TAGAGGGGCTGCTGCTGAGGCGG 1000
Oligo fwd CACCTAGAGGGGCTGCTGCTGAGG
Oligo rev AAACCCTCAGCAGCAGCCCCTCTA
Chromosome start end strand MM target_seq PAM alignment distance position gene name gene id
GU305914.1 114910 114932 - 0 TAGAGGGGCTGCTGCTGAGG CGG ||||||||[||||||||||||]PAM NA Intergenic NA NA

T25313 AGAGGGGCTGCTGCTGAGGCGGG 1000
Oligo fwd CACCAGAGGGGCTGCTGCTGAGGC
Oligo rev AAACGCCTCAGCAGCAGCCCCTCT
Chromosome start end strand MM target_seq PAM alignment distance position gene name gene id
GU305914.1 114909 114931 - 0 AGAGGGGCTGCTGCTGAGGC GGG ||||||||[||||||||||||]PAM NA Intergenic NA NA

T25314 CTGCTGCTGAGGCGGGCGATCGG 1000
Oligo fwd CACCCTGCTGCTGAGGCGGGCGAT
Oligo rev AAACATCGCCCGCCTCAGCAGCAG
Chromosome start end strand MM target_seq PAM alignment distance position gene name gene id
GU305914.1 114902 114924 - 0 CTGCTGCTGAGGCGGGCGAT CGG ||||||||[||||||||||||]PAM NA Intergenic NA NA

T25315 GCGGGCGATCGGCCCGATAGCGG 1000
Oligo fwd CACCGCGGGCGATCGGCCCGATAG
Oligo rev AAACCTATCGGGCCGATCGCCCGC
Chromosome start end strand MM target_seq PAM alignment distance position gene name gene id
GU305914.1 114891 114913 - 0 GCGGGCGATCGGCCCGATAG CGG ||||||||[||||||||||||]PAM NA Intergenic NA NA

T25316 GTCTCCCCGCCTGTTGTTGTAGG 1000
Oligo fwd CACCGTCTCCCCGCCTGTTGTTGT
Oligo rev AAACACAACAACAGGCGGGGAGAC
Chromosome start end strand MM target_seq PAM alignment distance position gene name gene id
GU305914.1 114869 114891 - 0 GTCTCCCCGCCTGTTGTTGT AGG ||||||||[||||||||||||]PAM NA Intergenic NA NA

T25317 TCCCCGCCTGTTGTTGTAGGAGG 1000
Oligo fwd CACCTCCCCGCCTGTTGTTGTAGG
Oligo rev AAACCCTACAACAACAGGCGGGGA
Chromosome start end strand MM target_seq PAM alignment distance position gene name gene id
GU305914.1 114866 114888 - 0 TCCCCGCCTGTTGTTGTAGG AGG ||||||||[||||||||||||]PAM NA Intergenic NA NA

T25318 CCCCGCCTGTTGTTGTAGGAGGG 1000
Oligo fwd CACCCCCCGCCTGTTGTTGTAGGA
Oligo rev AAACTCCTACAACAACAGGCGGGG
Chromosome start end strand MM target_seq PAM alignment distance position gene name gene id
GU305914.1 114865 114887 - 0 CCCCGCCTGTTGTTGTAGGA GGG ||||||||[||||||||||||]PAM NA Intergenic NA NA

T25319 GTAGGAGGGATGATCGTCCCCGG 1000
Oligo fwd CACCGTAGGAGGGATGATCGTCCC
Oligo rev AAACGGGACGATCATCCCTCCTAC
Chromosome start end strand MM target_seq PAM alignment distance position gene name gene id
GU305914.1 114851 114873 - 0 GTAGGAGGGATGATCGTCCC CGG ||||||||[||||||||||||]PAM NA Intergenic NA NA

T25320 GGAGGGATGATCGTCCCCGGCGG 1000
Oligo fwd CACCGGAGGGATGATCGTCCCCGG
Oligo rev AAACCCGGGGACGATCATCCCTCC
Chromosome start end strand MM target_seq PAM alignment distance position gene name gene id
GU305914.1 114848 114870 - 0 GGAGGGATGATCGTCCCCGG CGG ||||||||[||||||||||||]PAM NA Intergenic NA NA

T25321 GAGGGATGATCGTCCCCGGCGGG 1000
Oligo fwd CACCGAGGGATGATCGTCCCCGGC
Oligo rev AAACGCCGGGGACGATCATCCCTC
Chromosome start end strand MM target_seq PAM alignment distance position gene name gene id
GU305914.1 114847 114869 - 0 GAGGGATGATCGTCCCCGGC GGG ||||||||[||||||||||||]PAM NA Intergenic NA NA

T25322 AGGGATGATCGTCCCCGGCGGGG 1000
Oligo fwd CACCAGGGATGATCGTCCCCGGCG
Oligo rev AAACCGCCGGGGACGATCATCCCT
Chromosome start end strand MM target_seq PAM alignment distance position gene name gene id
GU305914.1 114846 114868 - 0 AGGGATGATCGTCCCCGGCG GGG ||||||||[||||||||||||]PAM NA Intergenic NA NA

T25323 TCCCCGGCGGGGTCGTAGCGAGG 1000
Oligo fwd CACCTCCCCGGCGGGGTCGTAGCG
Oligo rev AAACCGCTACGACCCCGCCGGGGA
Chromosome start end strand MM target_seq PAM alignment distance position gene name gene id

| | | | | | | | | | | | |
|---|---|---|---|---|---|---|---|---|---|---|---|
| GU305914.1 | 114835 | 114857 | - | 0 | TCCCCGGCGGGGTCGTAGCG | AGG | ||||||||[||||||||||||]PAM | NA | Intergenic | NA | NA |
| GU305914.1 | 221108 | 221130 | - | 4 | TCTCCGTCGGGGTCGGAGCC | TGG | ||-|||-|[||||||||-|||-]PAM | NA | Intergenic | NA | NA |
| T25324 | CCCCGGCGGGGTCGTAGCGAGGG | 1000 | | | | | | | | | |
| Oligo fwd | CACCCCCCGGCGGGGTCGTAGCGA | | | | | | | | | | |
| Oligo rev | AAACTCGCTACGACCCCGCCGGGG | | | | | | | | | | |
| Chromosome | start | end | strand | MM | target_seq | PAM | alignment | distance | position | gene name | gene id |
| GU305914.1 | 114834 | 114856 | - | 0 | CCCCGGCGGGGTCGTAGCGA | GGG | ||||||||[||||||||||||]PAM | NA | Intergenic | NA | NA |
| T25325 | AGGGCGATACCAGCCCTCGTAGG | 1000 | | | | | | | | | |
| Oligo fwd | CACCAGGGCGATACCAGCCCTCGT | | | | | | | | | | |
| Oligo rev | AAACACGAGGGCTGGTATCGCCCT | | | | | | | | | | |
| Chromosome | start | end | strand | MM | target_seq | PAM | alignment | distance | position | gene name | gene id |
| GU305914.1 | 114815 | 114837 | - | 0 | AGGGCGATACCAGCCCTCGT | AGG | ||||||||[||||||||||||]PAM | NA | Intergenic | NA | NA |
| T25326 | GGGCGATACCAGCCCTCGTAGGG | 1000 | | | | | | | | | |
| Oligo fwd | CACCGGGCGATACCAGCCCTCGTA | | | | | | | | | | |
| Oligo rev | AAACTACGAGGGCTGGTATCGCCC | | | | | | | | | | |
| Chromosome | start | end | strand | MM | target_seq | PAM | alignment | distance | position | gene name | gene id |
| GU305914.1 | 114814 | 114836 | - | 0 | GGGCGATACCAGCCCTCGTA | GGG | ||||||||[||||||||||||]PAM | NA | Intergenic | NA | NA |
| T25327 | TCGTAGGGCACTCCGCCGCGCGG | 1000 | | | | | | | | | |
| Oligo fwd | CACCTCGTAGGGCACTCCGCCGCG | | | | | | | | | | |
| Oligo rev | AAACCGCGGCGGAGTGCCCTACGA | | | | | | | | | | |
| Chromosome | start | end | strand | MM | target_seq | PAM | alignment | distance | position | gene name | gene id |
| GU305914.1 | 114799 | 114821 | - | 0 | TCGTAGGGCACTCCGCCGCG | CGG | ||||||||[||||||||||||]PAM | NA | Intergenic | NA | NA |
| T25328 | CGTAGGGCACTCCGCCGCGCGGG | 1000 | | | | | | | | | |
| Oligo fwd | CACCCGTAGGGCACTCCGCCGCGC | | | | | | | | | | |
| Oligo rev | AAACGCGCGGCGGAGTGCCCTACG | | | | | | | | | | |
| Chromosome | start | end | strand | MM | target_seq | PAM | alignment | distance | position | gene name | gene id |
| GU305914.1 | 114798 | 114820 | - | 0 | CGTAGGGCACTCCGCCGCGC | GGG | ||||||||[||||||||||||]PAM | NA | Intergenic | NA | NA |
| T25329 | CCGCCGCGCGGGTCGCGATCAGG | 1000 | | | | | | | | | |
| Oligo fwd | CACCCCGCCGCGCGGGTCGCGATC | | | | | | | | | | |
| Oligo rev | AAACGATCGCGACCCGCGCGGCGG | | | | | | | | | | |
| Chromosome | start | end | strand | MM | target_seq | PAM | alignment | distance | position | gene name | gene id |
| GU305914.1 | 114787 | 114809 | - | 0 | CCGCCGCGCGGGTCGCGATC | AGG | ||||||||[||||||||||||]PAM | NA | Intergenic | NA | NA |
| GU305914.1 | 222022 | 222044 | - | 4 | CGGCCGAACGGATCGCGATC | CGG | |-||||--[|||-||||||||]PAM | NA | Intergenic | NA | NA |
| T25330 | GCGGGTCGCGATCAGGATGATGG | 1000 | | | | | | | | | |
| Oligo fwd | CACCGCGGGTCGCGATCAGGATGA | | | | | | | | | | |
| Oligo rev | AAACTCATCCTGATCGCGACCCGC | | | | | | | | | | |
| Chromosome | start | end | strand | MM | target_seq | PAM | alignment | distance | position | gene name | gene id |
| GU305914.1 | 114780 | 114802 | - | 0 | GCGGGTCGCGATCAGGATGA | TGG | ||||||||[||||||||||||]PAM | NA | Intergenic | NA | NA |
| T25331 | ATGATGGTATCCCCCCCGATAGG | 1000 | | | | | | | | | |
| Oligo fwd | CACCATGATGGTATCCCCCCCGAT | | | | | | | | | | |
| Oligo rev | AAACATCGGGGGGGATACCATCAT | | | | | | | | | | |
| Chromosome | start | end | strand | MM | target_seq | PAM | alignment | distance | position | gene name | gene id |
| GU305914.1 | 114764 | 114786 | - | 0 | ATGATGGTATCCCCCCCGAT | AGG | ||||||||[||||||||||||]PAM | NA | Intergenic | NA | NA |
| T25332 | TATCCCCCCCGATAGGCGTCCGG | 1000 | | | | | | | | | |
| Oligo fwd | CACCTATCCCCCCCGATAGGCGTC | | | | | | | | | | |
| Oligo rev | AAACGACGCCTATCGGGGGGGATA | | | | | | | | | | |
| Chromosome | start | end | strand | MM | target_seq | PAM | alignment | distance | position | gene name | gene id |
| GU305914.1 | 114757 | 114779 | - | 0 | TATCCCCCCCGATAGGCGTC | CGG | ||||||||[||||||||||||]PAM | NA | Intergenic | NA | NA |
| T25333 | ATCCCCCCCGATAGGCGTCCGGG | 1000 | | | | | | | | | |
| Oligo fwd | CACCATCCCCCCCGATAGGCGTCC | | | | | | | | | | |
| Oligo rev | AAACGGACGCCTATCGGGGGGGAT | | | | | | | | | | |
| Chromosome | start | end | strand | MM | target_seq | PAM | alignment | distance | position | gene name | gene id |
| GU305914.1 | 114756 | 114778 | - | 0 | ATCCCCCCCGATAGGCGTCC | GGG | ||||||||[||||||||||||]PAM | NA | Intergenic | NA | NA |
| T25334 | ATAGGCGTCCGGGTGATGATAGG | 1000 | | | | | | | | | |
| Oligo fwd | CACCATAGGCGTCCGGGTGATGAT | | | | | | | | | | |
| Oligo rev | AAACATCATCACCCGGACGCCTAT | | | | | | | | | | |
| Chromosome | start | end | strand | MM | target_seq | PAM | alignment | distance | position | gene name | gene id |
| GU305914.1 | 114746 | 114768 | - | 0 | ATAGGCGTCCGGGTGATGAT | AGG | ||||||||[||||||||||||]PAM | NA | Intergenic | NA | NA |
| T25335 | TAGGCGTCCGGGTGATGATAGGG | 1000 | | | | | | | | | |
| Oligo fwd | CACCTAGGCGTCCGGGTGATGATA | | | | | | | | | | |
| Oligo rev | AAACTATCATCACCCGGACGCCTA | | | | | | | | | | |
| Chromosome | start | end | strand | MM | target_seq | PAM | alignment | distance | position | gene name | gene id |
| GU305914.1 | 114745 | 114767 | - | 0 | TAGGCGTCCGGGTGATGATA | GGG | ||||||||[||||||||||||]PAM | NA | Intergenic | NA | NA |
| T25336 | GCGTCCGGGTGATGATAGGGAGG | 1000 | | | | | | | | | |
| Oligo fwd | CACCGCGTCCGGGTGATGATAGGG | | | | | | | | | | |
| Oligo rev | AAACCCCTATCATCACCCGGACGC | | | | | | | | | | |
| Chromosome | start | end | strand | MM | target_seq | PAM | alignment | distance | position | gene name | gene id |
| GU305914.1 | 114742 | 114764 | - | 0 | GCGTCCGGGTGATGATAGGG | AGG | ||||||||[||||||||||||]PAM | NA | Intergenic | NA | NA |
| T25337 | GGAGGACCGTATCCCCCGTGTGG | 1000 | | | | | | | | | |
| Oligo fwd | CACCGGAGGACCGTATCCCCCGTG | | | | | | | | | | |
| Oligo rev | AAACCACGGGGGATACGGTCCTCC | | | | | | | | | | |
| Chromosome | start | end | strand | MM | target_seq | PAM | alignment | distance | position | gene name | gene id |
| GU305914.1 | 114724 | 114746 | - | 0 | GGAGGACCGTATCCCCCGTG | TGG | ||||||||[||||||||||||]PAM | NA | Intergenic | NA | NA |
| T25338 | GAGGACCGTATCCCCCGTGTGGG | 1000 | | | | | | | | | |

Oligo fwd CACCGAGGACCGTATCCCCCGTGT
Oligo rev AAACACACGGGGGATACGGTCCTC
Chromosome start end strand MM target_seq PAM alignment distance position gene name gene id
GU305914.1 114723 114745 - 0 GAGGACCGTATCCCCCGTGT GGG ||||||||[||||||||||||]PAM NA Intergenic NA NA

T25339 CCGTATCCCCCGTGTGGGAGAGG 1000
Oligo fwd CACCCCGTATCCCCCGTGTGGGAG
Oligo rev AAACCTCCCACACGGGGGATACGG
Chromosome start end strand MM target_seq PAM alignment distance position gene name gene id
GU305914.1 114718 114740 - 0 CCGTATCCCCCGTGTGGGAG AGG ||||||||[||||||||||||]PAM NA Intergenic NA NA

T25340 CGTATCCCCCGTGTGGGAGAGGG 1000
Oligo fwd CACCCGTATCCCCCGTGTGGGAGA
Oligo rev AAACTCTCCCACACGGGGGATACG
Chromosome start end strand MM target_seq PAM alignment distance position gene name gene id
GU305914.1 114717 114739 - 0 CGTATCCCCCGTGTGGGAGA GGG ||||||||[||||||||||||]PAM NA Intergenic NA NA

T25341 GTATCCCCCGTGTGGGAGAGGGG 1000
Oligo fwd CACCGTATCCCCCGTGTGGGAGAG
Oligo rev AAACCTCTCCCACACGGGGGATAC
Chromosome start end strand MM target_seq PAM alignment distance position gene name gene id
GU305914.1 114716 114738 - 0 GTATCCCCCGTGTGGGAGAG GGG ||||||||[||||||||||||]PAM NA Intergenic NA NA

T25342 TATCCCCCGTGTGGGAGAGGGGG 1000
Oligo fwd CACCTATCCCCCGTGTGGGAGAGG
Oligo rev AAACCCTCTCCCACACGGGGGATA
Chromosome start end strand MM target_seq PAM alignment distance position gene name gene id
GU305914.1 114715 114737 - 0 TATCCCCCGTGTGGGAGAGG GGG ||||||||[||||||||||||]PAM NA Intergenic NA NA

T25343 AAGTACCGCCACCGCCGCGATGG 1000
Oligo fwd CACCAAGTACCGCCACCGCCGCGA
Oligo rev AAACTCGCGGCGGTGGCGGTACTT
Chromosome start end strand MM target_seq PAM alignment distance position gene name gene id
GU305914.1 114687 114709 - 0 AAGTACCGCCACCGCCGCGA TGG ||||||||[||||||||||||]PAM NA Intergenic NA NA
GU305914.1 127361 127383 - 4 CGGTAGCGCCGCCGCCGCGA TGG --|||-||[||-|||||||||]PAM NA Intergenic NA NA

T25344 CCACCGCCGCGATGGTGAGACGG 1000
Oligo fwd CACCCCACCGCCGCGATGGTGAGA
Oligo rev AAACTCTCACCATCGCGGCGGTGG
Chromosome start end strand MM target_seq PAM alignment distance position gene name gene id
GU305914.1 114679 114701 - 0 CCACCGCCGCGATGGTGAGA CGG ||||||||[||||||||||||]PAM NA Intergenic NA NA

T25345 CCGCCGCGATGGTGAGACGGCGG 1000
Oligo fwd CACCCCGCCGCGATGGTGAGACGG
Oligo rev AAACCCGTCTCACCATCGCGGCGG
Chromosome start end strand MM target_seq PAM alignment distance position gene name gene id
GU305914.1 114676 114698 - 0 CCGCCGCGATGGTGAGACGG CGG ||||||||[||||||||||||]PAM NA Intergenic NA NA

T25346 CCGCGATGGTGAGACGGCGGTGG 1000
Oligo fwd CACCCCGCGATGGTGAGACGGCGG
Oligo rev AAACCCGCCGTCTCACCATCGCGG
Chromosome start end strand MM target_seq PAM alignment distance position gene name gene id
GU305914.1 114673 114695 - 0 CCGCGATGGTGAGACGGCGG TGG ||||||||[||||||||||||]PAM NA Intergenic NA NA
GU305914.1 43701 43723 - 4 TCGCGGTGATGCGACGGCGG AGG -||||-||[-||-||||||||]PAM NA Intergenic NA NA

T25347 CGATGGTGAGACGGCGGTGGTGG 1000
Oligo fwd CACCCGATGGTGAGACGGCGGTGG
Oligo rev AAACCCACCGCCGTCTCACCATCG
Chromosome start end strand MM target_seq PAM alignment distance position gene name gene id
GU305914.1 114670 114692 - 0 CGATGGTGAGACGGCGGTGG TGG ||||||||[||||||||||||]PAM NA Intergenic NA NA
GU305914.1 201599 201621 - 4 CGAGGAGGAGACCGCGGTGG CGG |||-|--|[||||-|||||||]PAM NA Intergenic NA NA

T25348 TGGTGAGACGGCGGTGGTGGTGG 1000
Oligo fwd CACCTGGTGAGACGGCGGTGGTGG
Oligo rev AAACCCACCACCGCCGTCTCACCA
Chromosome start end strand MM target_seq PAM alignment distance position gene name gene id
GU305914.1 114667 114689 - 0 TGGTGAGACGGCGGTGGTGG TGG ||||||||[||||||||||||]PAM NA Intergenic NA NA

T25349 TGAGACGGCGGTGGTGGTGGCGG 1000
Oligo fwd CACCTGAGACGGCGGTGGTGGTGG
Oligo rev AAACCCACCACCACCGCCGTCTCA
Chromosome start end strand MM target_seq PAM alignment distance position gene name gene id
GU305914.1 114664 114686 - 0 TGAGACGGCGGTGGTGGTGG CGG ||||||||[||||||||||||]PAM NA Intergenic NA NA
GU305914.1 62440 62462 + 4 GGTGGCGGCGGGGGTGGTGG CGG -|-|-|||[|||-||||||||]PAM NA Intergenic NA NA
GU305914.1 48153 48175 + 4 TGTGGCGGCGGCGGTGGCGG CGG ||-|-|||[|||-|||||-||]PAM NA Intergenic NA NA
GU305914.1 62431 62453 + 4 TTCGACGGCGGTGGCGGCGG GGG |--|||||[||||||-||-||]PAM NA Intergenic NA NA

T25350 GACGGCGGTGGTGGTGGCGGCGG 1000
Oligo fwd CACCGACGGCGGTGGTGGTGGCGG
Oligo rev AAACCCGCCACCACCACCGCCGTC
Chromosome start end strand MM target_seq PAM alignment distance position gene name gene id
GU305914.1 114661 114683 - 0 GACGGCGGTGGTGGTGGCGG CGG ||||||||[||||||||||||]PAM NA Intergenic NA NA
GU305914.1 62443 62465 + 2 GGCGGCGGGGGTGGTGGCGG CGG |-||||||[-|||||||||||]PAM NA Intergenic NA NA
GU305914.1 19936 19958 + 3 GACGGTAGCGGTGGTGGCGG TGG |||||--|[-|||||||||||]PAM NA Intergenic NA NA
GU305914.1 74710 74732 - 4 GACCCGGGTGGCGGTGGCGG CGG |||---||[|||-||||||||]PAM NA Intergenic NA NA
GU305914.1 52716 52738 + 4 GCCGCCGCTGGCGGTGGCGG TGG |-||-||-[|||-||||||||]PAM NA Intergenic NA NA
GU305914.1 62446 62468 + 3 GGCGGGGGTGGTGGCGGCGG GGG |-|||-||[||||||-|||||]PAM NA Intergenic NA NA
GU305914.1 114658 114680 - 3 GGCGGTGGTGGTGGCGGCGG CGG |-|||-||[||||||-|||||]PAM NA Intergenic NA NA
GU305914.1 48156 48178 + 3 GGCGGCGGCGGCGGTGGCGGCGG CGG |-||||||[-|||||-|||||]PAM NA Intergenic NA NA

<table>
<tr><td>GU305914.1</td><td>48159</td><td>48181</td><td>+</td><td>3</td><td>GGCGGCGGTGGCGGCGGCGG</td><td>TGG</td><td>|-||||||[|||-||-|||||]PAM</td><td>NA</td><td>Intergenic</td><td>NA</td><td>NA</td></tr>
<tr><td>GU305914.1</td><td>58129</td><td>58151</td><td>-</td><td>4</td><td>GCGGGCGGTGGCGGCGGCGG</td><td>CGG</td><td>|--|||||[|||-||-|||||]PAM</td><td>NA</td><td>Intergenic</td><td>NA</td><td>NA</td></tr>
<tr><td>GU305914.1</td><td>168814</td><td>168836</td><td>+</td><td>4</td><td>AGCGGCGGTGGTGGCAGCGG</td><td>CGG</td><td>--||||||[||||||--||||]PAM</td><td>NA</td><td>Intergenic</td><td>NA</td><td>NA</td></tr>
<tr><td></td><td></td><td></td><td></td><td></td><td></td><td></td><td></td><td></td><td></td><td></td><td></td></tr>
<tr><td>T25351</td><td>GGCGGTGGTGGTGGCGGCGGCGG</td><td>1000</td><td></td><td></td><td></td><td></td><td></td><td></td><td></td><td></td><td></td></tr>
<tr><td>Oligo fwd</td><td>CACCGGCGGTGGTGGTGGCGGCGG</td><td></td><td></td><td></td><td></td><td></td><td></td><td></td><td></td><td></td><td></td></tr>
<tr><td>Oligo rev</td><td>AAACCCGCCGCCACCACCACCGCC</td><td></td><td></td><td></td><td></td><td></td><td></td><td></td><td></td><td></td><td></td></tr>
<tr><td>Chromosome</td><td>start</td><td>end</td><td>strand</td><td>MM</td><td>target_seq</td><td>PAM</td><td>alignment</td><td>distance</td><td>position</td><td>gene name</td><td>gene id</td></tr>
<tr><td>GU305914.1</td><td>114658</td><td>114680</td><td>-</td><td>0</td><td>GGCGGTGGTGGTGGCGGCGG</td><td>CGG</td><td>||||||||[||||||||||||]PAM</td><td>NA</td><td>Intergenic</td><td>NA</td><td>NA</td></tr>
<tr><td>GU305914.1</td><td>62446</td><td>62468</td><td>+</td><td>1</td><td>GGCGGGGGTGGTGGCGGCGG</td><td>GGG</td><td>|||||-||[||||||||||||]PAM</td><td>NA</td><td>Intergenic</td><td>NA</td><td>NA</td></tr>
<tr><td>GU305914.1</td><td>48156</td><td>48178</td><td>+</td><td>2</td><td>GGCGGCGGCGGTGGCGGCGG</td><td>CGG</td><td>|||||-||[-|||||||||||]PAM</td><td>NA</td><td>Intergenic</td><td>NA</td><td>NA</td></tr>
<tr><td>GU305914.1</td><td>48159</td><td>48181</td><td>+</td><td>2</td><td>GGCGGCGGTGGCGGCGGCGG</td><td>TGG</td><td>|||||-||[|||-||||||||]PAM</td><td>NA</td><td>Intergenic</td><td>NA</td><td>NA</td></tr>
<tr><td>GU305914.1</td><td>154404</td><td>154426</td><td>+</td><td>2</td><td>GGCGGTGGCGGCGGCGGCGG</td><td>CGG</td><td>||||||||[-||-||||||||]PAM</td><td>NA</td><td>Intergenic</td><td>NA</td><td>NA</td></tr>
<tr><td>GU305914.1</td><td>58126</td><td>58148</td><td>-</td><td>2</td><td>GGCGGTGGCGGCGGCGGCGG</td><td>CGG</td><td>||||||||[-||-||||||||]PAM</td><td>NA</td><td>Intergenic</td><td>NA</td><td>NA</td></tr>
<tr><td>GU305914.1</td><td>58129</td><td>58151</td><td>-</td><td>4</td><td>GCGGGCGGTGGCGGCGGCGG</td><td>CGG</td><td>|--||-||[|||-||||||||]PAM</td><td>NA</td><td>Intergenic</td><td>NA</td><td>NA</td></tr>
<tr><td>GU305914.1</td><td>29997</td><td>30019</td><td>+</td><td>4</td><td>ACCGGTGCTGGCGGCGGCGG</td><td>CGG</td><td>--|||||-[|||-||||||||]PAM</td><td>NA</td><td>Intergenic</td><td>NA</td><td>NA</td></tr>
<tr><td>GU305914.1</td><td>30000</td><td>30022</td><td>+</td><td>4</td><td>GGTGCTGGCGGCGGCGGCGG</td><td>CGG</td><td>||-|-|||[-||-||||||||]PAM</td><td>NA</td><td>Intergenic</td><td>NA</td><td>NA</td></tr>
<tr><td>GU305914.1</td><td>154407</td><td>154429</td><td>+</td><td>4</td><td>GGTGGCGGCGGCGGCGGCGG</td><td>CGG</td><td>||-||-||[-||-||||||||]PAM</td><td>NA</td><td>Intergenic</td><td>NA</td><td>NA</td></tr>
<tr><td>GU305914.1</td><td>58123</td><td>58145</td><td>-</td><td>4</td><td>GGTGGCGGCGGCGGCGGCGG</td><td>AGG</td><td>||-||-||[-||-||||||||]PAM</td><td>NA</td><td>Intergenic</td><td>NA</td><td>NA</td></tr>
<tr><td>GU305914.1</td><td>161754</td><td>161776</td><td>-</td><td>4</td><td>GGCCGAGGAGGCGGCGGCGG</td><td>AGG</td><td>|||-|-||[-||-||||||||]PAM</td><td>NA</td><td>Intergenic</td><td>NA</td><td>NA</td></tr>
<tr><td>GU305914.1</td><td>149482</td><td>149504</td><td>-</td><td>4</td><td>GGCGACGGCGGCGGCGGCGG</td><td>CGG</td><td>||||--||[-||-||||||||]PAM</td><td>NA</td><td>Intergenic</td><td>NA</td><td>NA</td></tr>
<tr><td>GU305914.1</td><td>168817</td><td>168839</td><td>+</td><td>2</td><td>GGCGGTGGTGGCAGCGGCGG</td><td>CGG</td><td>||||||||[|||--|||||||]PAM</td><td>NA</td><td>Intergenic</td><td>NA</td><td>NA</td></tr>
<tr><td>GU305914.1</td><td>114661</td><td>114683</td><td>-</td><td>3</td><td>GACGGCGGTGGTGGTGGCGG</td><td>CGG</td><td>|-|||-||[||||||-|||||]PAM</td><td>NA</td><td>Intergenic</td><td>NA</td><td>NA</td></tr>
<tr><td>GU305914.1</td><td>74930</td><td>74952</td><td>+</td><td>4</td><td>GGCGCAGGTTGAGGCGGCGG</td><td>TGG</td><td>||||--||[|-|-||||||||]PAM</td><td>NA</td><td>Intergenic</td><td>NA</td><td>NA</td></tr>
<tr><td>GU305914.1</td><td>168814</td><td>168836</td><td>+</td><td>3</td><td>AGCGGCGGTGGTGGCAGCGG</td><td>CGG</td><td>-||||-||[|||||||-||||]PAM</td><td>NA</td><td>Intergenic</td><td>NA</td><td>NA</td></tr>
<tr><td>GU305914.1</td><td>62443</td><td>62465</td><td>+</td><td>3</td><td>GGCGGCGGGGGTGGTGGCGG</td><td>CGG</td><td>|||||-||[-|||||-|||||]PAM</td><td>NA</td><td>Intergenic</td><td>NA</td><td>NA</td></tr>
<tr><td>GU305914.1</td><td>19936</td><td>19958</td><td>+</td><td>4</td><td>GACGGTAGCGGTGGTGGCGG</td><td>TGG</td><td>|-||||-|[-|||||-|||||]PAM</td><td>NA</td><td>Intergenic</td><td>NA</td><td>NA</td></tr>
<tr><td>GU305914.1</td><td>18653</td><td>18675</td><td>-</td><td>4</td><td>GGCAGAGGTGGCGGAGGCGG</td><td>GGG</td><td>|||-|-||[|||-||-|||||]PAM</td><td>NA</td><td>Intergenic</td><td>NA</td><td>NA</td></tr>
<tr><td>GU305914.1</td><td>45255</td><td>45277</td><td>-</td><td>4</td><td>GGTGGCGGTGGCGGCAGCGG</td><td>CGG</td><td>||-||-||[|||-|||-||||]PAM</td><td>NA</td><td>Intergenic</td><td>NA</td><td>NA</td></tr>
<tr><td>GU305914.1</td><td>52719</td><td>52741</td><td>+</td><td>4</td><td>GCCGCTGGCGGTGGCGGTGG</td><td>CGG</td><td>|-||-|||[-||||||||-||]PAM</td><td>NA</td><td>Intergenic</td><td>NA</td><td>NA</td></tr>
<tr><td>GU305914.1</td><td>62449</td><td>62471</td><td>+</td><td>3</td><td>GGGGGTGGTGGCGGCGGGGG</td><td>TGG</td><td>||-|||||[|||-|||||-||]PAM</td><td>NA</td><td>Intergenic</td><td>NA</td><td>NA</td></tr>
<tr><td>GU305914.1</td><td>45258</td><td>45280</td><td>-</td><td>4</td><td>AACGGTGGCGGTGGCGGCAG</td><td>CGG</td><td>--||||||[-|||||||||-|]PAM</td><td>NA</td><td>Intergenic</td><td>NA</td><td>NA</td></tr>
<tr><td>GU305914.1</td><td>62434</td><td>62456</td><td>+</td><td>4</td><td>GACGGCGGTGGCGGCGGGGG</td><td>TGG</td><td>|-|||-||[|||-|||||-||]PAM</td><td>NA</td><td>Intergenic</td><td>NA</td><td>NA</td></tr>
<tr><td>GU305914.1</td><td>62458</td><td>62480</td><td>+</td><td>3</td><td>GGCGGCGGGGGTGGCGGCCG</td><td>CGG</td><td>|||||-||[-|||||||||-|]PAM</td><td>NA</td><td>Intergenic</td><td>NA</td><td>NA</td></tr>
<tr><td></td><td></td><td></td><td></td><td></td><td></td><td></td><td></td><td></td><td></td><td></td><td></td></tr>
<tr><td>T25352</td><td>GCGGTGGTGGTGGCGGCGGCGGG</td><td>1000</td><td></td><td></td><td></td><td></td><td></td><td></td><td></td><td></td><td></td></tr>
<tr><td>Oligo fwd</td><td>CACCGCGGTGGTGGTGGCGGCGGC</td><td></td><td></td><td></td><td></td><td></td><td></td><td></td><td></td><td></td><td></td></tr>
<tr><td>Oligo rev</td><td>AAACGCCGCCGCCACCACCACCGC</td><td></td><td></td><td></td><td></td><td></td><td></td><td></td><td></td><td></td><td></td></tr>
<tr><td>Chromosome</td><td>start</td><td>end</td><td>strand</td><td>MM</td><td>target_seq</td><td>PAM</td><td>alignment</td><td>distance</td><td>position</td><td>gene name</td><td>gene id</td></tr>
<tr><td>GU305914.1</td><td>114657</td><td>114679</td><td>-</td><td>0</td><td>GCGGTGGTGGTGGCGGCGGC</td><td>GGG</td><td>||||||||[||||||||||||]PAM</td><td>NA</td><td>Intergenic</td><td>NA</td><td>NA</td></tr>
<tr><td>GU305914.1</td><td>221599</td><td>221621</td><td>-</td><td>3</td><td>GAGGCCGTGGTGGCGGCGGC</td><td>AGG</td><td>|-||--||[||||||||||||]PAM</td><td>NA</td><td>Intergenic</td><td>NA</td><td>NA</td></tr>
<tr><td>GU305914.1</td><td>86780</td><td>86802</td><td>-</td><td>4</td><td>GAGGGGGCGGGGGCGGCGGC</td><td>AGG</td><td>|-||-||-[||-|||||||||]PAM</td><td>NA</td><td>Intergenic</td><td>NA</td><td>NA</td></tr>
<tr><td>GU305914.1</td><td>81248</td><td>81270</td><td>-</td><td>4</td><td>GCGATCGAGGAGGCGGCGGC</td><td>GGG</td><td>|||-|-|-[||-|||||||||]PAM</td><td>NA</td><td>Intergenic</td><td>NA</td><td>NA</td></tr>
<tr><td>GU305914.1</td><td>62444</td><td>62466</td><td>+</td><td>3</td><td>GCGGCGGGGGTGGTGGCGGC</td><td>GGG</td><td>||||-||-[|||||-||||||]PAM</td><td>NA</td><td>Intergenic</td><td>NA</td><td>NA</td></tr>
<tr><td>GU305914.1</td><td>149475</td><td>149497</td><td>-</td><td>4</td><td>GCGGCGGCGGCGGCGGCAGC</td><td>GGG</td><td>||||-||-[||-||||||-||]PAM</td><td>NA</td><td>Intergenic</td><td>NA</td><td>NA</td></tr>
<tr><td>GU305914.1</td><td>48160</td><td>48182</td><td>+</td><td>3</td><td>GCGGCGGTGGCGGCGGCGGT</td><td>GGG</td><td>||||-|||[||-||||||||-]PAM</td><td>NA</td><td>Intergenic</td><td>NA</td><td>NA</td></tr>
<tr><td></td><td></td><td></td><td></td><td></td><td></td><td></td><td></td><td></td><td></td><td></td><td></td></tr>
<tr><td>T25353</td><td>AGATGATACTGAACACCGTCCGG</td><td>1000</td><td></td><td></td><td></td><td></td><td></td><td></td><td></td><td></td><td></td></tr>
<tr><td>Oligo fwd</td><td>CACCAGATGATACTGAACACCGTC</td><td></td><td></td><td></td><td></td><td></td><td></td><td></td><td></td><td></td><td></td></tr>
<tr><td>Oligo rev</td><td>AAACGACGGTGTTCAGTATCATCT</td><td></td><td></td><td></td><td></td><td></td><td></td><td></td><td></td><td></td><td></td></tr>
<tr><td>Chromosome</td><td>start</td><td>end</td><td>strand</td><td>MM</td><td>target_seq</td><td>PAM</td><td>alignment</td><td>distance</td><td>position</td><td>gene name</td><td>gene id</td></tr>
<tr><td>GU305914.1</td><td>114634</td><td>114656</td><td>-</td><td>0</td><td>AGATGATACTGAACACCGTC</td><td>CGG</td><td>||||||||[||||||||||||]PAM</td><td>NA</td><td>Intergenic</td><td>NA</td><td>NA</td></tr>
<tr><td></td><td></td><td></td><td></td><td></td><td></td><td></td><td></td><td></td><td></td><td></td><td></td></tr>
<tr><td>T25354</td><td>TGATACTGAACACCGTCCGGCGG</td><td>1000</td><td></td><td></td><td></td><td></td><td></td><td></td><td></td><td></td><td></td></tr>
<tr><td>Oligo fwd</td><td>CACCTGATACTGAACACCGTCCGG</td><td></td><td></td><td></td><td></td><td></td><td></td><td></td><td></td><td></td><td></td></tr>
<tr><td>Oligo rev</td><td>AAACCCGGACGGTGTTCAGTATCA</td><td></td><td></td><td></td><td></td><td></td><td></td><td></td><td></td><td></td><td></td></tr>
<tr><td>Chromosome</td><td>start</td><td>end</td><td>strand</td><td>MM</td><td>target_seq</td><td>PAM</td><td>alignment</td><td>distance</td><td>position</td><td>gene name</td><td>gene id</td></tr>
<tr><td>GU305914.1</td><td>114631</td><td>114653</td><td>-</td><td>0</td><td>TGATACTGAACACCGTCCGG</td><td>CGG</td><td>||||||||[||||||||||||]PAM</td><td>NA</td><td>Intergenic</td><td>NA</td><td>NA</td></tr>
<tr><td></td><td></td><td></td><td></td><td></td><td></td><td></td><td></td><td></td><td></td><td></td><td></td></tr>
<tr><td>T25355</td><td>CGGCGGCATTCCGTAATAAGAGG</td><td>1000</td><td></td><td></td><td></td><td></td><td></td><td></td><td></td><td></td><td></td></tr>
<tr><td>Oligo fwd</td><td>CACCCGGCGGCATTCCGTAATAAG</td><td></td><td></td><td></td><td></td><td></td><td></td><td></td><td></td><td></td><td></td></tr>
<tr><td>Oligo rev</td><td>AAACCTTATTACGGAATGCCGCCG</td><td></td><td></td><td></td><td></td><td></td><td></td><td></td><td></td><td></td><td></td></tr>
<tr><td>Chromosome</td><td>start</td><td>end</td><td>strand</td><td>MM</td><td>target_seq</td><td>PAM</td><td>alignment</td><td>distance</td><td>position</td><td>gene name</td><td>gene id</td></tr>
<tr><td>GU305914.1</td><td>114614</td><td>114636</td><td>-</td><td>0</td><td>CGGCGGCATTCCGTAATAAG</td><td>AGG</td><td>||||||||[||||||||||||]PAM</td><td>NA</td><td>Intergenic</td><td>NA</td><td>NA</td></tr>
<tr><td></td><td></td><td></td><td></td><td></td><td></td><td></td><td></td><td></td><td></td><td></td><td></td></tr>
<tr><td>T25356</td><td>GGCGGCATTCCGTAATAAGAGGG</td><td>1000</td><td></td><td></td><td></td><td></td><td></td><td></td><td></td><td></td><td></td></tr>
<tr><td>Oligo fwd</td><td>CACCGGCGGCATTCCGTAATAAGA</td><td></td><td></td><td></td><td></td><td></td><td></td><td></td><td></td><td></td><td></td></tr>
<tr><td>Oligo rev</td><td>AAACTCTTATTACGGAATGCCGCC</td><td></td><td></td><td></td><td></td><td></td><td></td><td></td><td></td><td></td><td></td></tr>
<tr><td>Chromosome</td><td>start</td><td>end</td><td>strand</td><td>MM</td><td>target_seq</td><td>PAM</td><td>alignment</td><td>distance</td><td>position</td><td>gene name</td><td>gene id</td></tr>
<tr><td>GU305914.1</td><td>114613</td><td>114635</td><td>-</td><td>0</td><td>GGCGGCATTCCGTAATAAGA</td><td>GGG</td><td>||||||||[||||||||||||]PAM</td><td>NA</td><td>Intergenic</td><td>NA</td><td>NA</td></tr>
<tr><td></td><td></td><td></td><td></td><td></td><td></td><td></td><td></td><td></td><td></td><td></td><td></td></tr>
<tr><td>T25357</td><td>CCGTAATAAGAGGGCACCTGAGG</td><td>1000</td><td></td><td></td><td></td><td></td><td></td><td></td><td></td><td></td><td></td></tr>
<tr><td>Oligo fwd</td><td>CACCCCGTAATAAGAGGGCACCTG</td><td></td><td></td><td></td><td></td><td></td><td></td><td></td><td></td><td></td><td></td></tr>
<tr><td>Oligo rev</td><td>AAACCAGGTGCCCTCTTATTACGG</td><td></td><td></td><td></td><td></td><td></td><td></td><td></td><td></td><td></td><td></td></tr>
<tr><td>Chromosome</td><td>start</td><td>end</td><td>strand</td><td>MM</td><td>target_seq</td><td>PAM</td><td>alignment</td><td>distance</td><td>position</td><td>gene name</td><td>gene id</td></tr>
<tr><td>GU305914.1</td><td>114604</td><td>114626</td><td>-</td><td>0</td><td>CCGTAATAAGAGGGCACCTG</td><td>AGG</td><td>||||||||[||||||||||||]PAM</td><td>NA</td><td>Intergenic</td><td>NA</td><td>NA</td></tr>
<tr><td></td><td></td><td></td><td></td><td></td><td></td><td></td><td></td><td></td><td></td><td></td><td></td></tr>
<tr><td>T25358</td><td>ATAAGAGGGCACCTGAGGCAAGG</td><td>1000</td><td></td><td></td><td></td><td></td><td></td><td></td><td></td><td></td><td></td></tr>
<tr><td>Oligo fwd</td><td>CACCATAAGAGGGCACCTGAGGCA</td><td></td><td></td><td></td><td></td><td></td><td></td><td></td><td></td><td></td><td></td></tr>
<tr><td>Oligo rev</td><td>AAACTGCCTCAGGTGCCCTCTTAT</td><td></td><td></td><td></td><td></td><td></td><td></td><td></td><td></td><td></td><td></td></tr>
<tr><td>Chromosome</td><td>start</td><td>end</td><td>strand</td><td>MM</td><td>target_seq</td><td>PAM</td><td>alignment</td><td>distance</td><td>position</td><td>gene name</td><td>gene id</td></tr>
<tr><td>GU305914.1</td><td>114599</td><td>114621</td><td>-</td><td>0</td><td>ATAAGAGGGCACCTGAGGCA</td><td>AGG</td><td>||||||||[||||||||||||]PAM</td><td>NA</td><td>Intergenic</td><td>NA</td><td>NA</td></tr>
<tr><td></td><td></td><td></td><td></td><td></td><td></td><td></td><td></td><td></td><td></td><td></td><td></td></tr>
<tr><td>T25359</td><td>ACCTGAGGCAAGGCCTGCTGCGG</td><td>1000</td><td></td><td></td><td></td><td></td><td></td><td></td><td></td><td></td><td></td></tr>
<tr><td>Oligo fwd</td><td>CACCACCTGAGGCAAGGCCTGCTG</td><td></td><td></td><td></td><td></td><td></td><td></td><td></td><td></td><td></td><td></td></tr>
<tr><td>Oligo rev</td><td>AAACCAGCAGGCCTTGCCTCAGGT</td><td></td><td></td><td></td><td></td><td></td><td></td><td></td><td></td><td></td><td></td></tr>
<tr><td>Chromosome</td><td>start</td><td>end</td><td>strand</td><td>MM</td><td>target_seq</td><td>PAM</td><td>alignment</td><td>distance</td><td>position</td><td>gene name</td><td>gene id</td></tr>
<tr><td>GU305914.1</td><td>114589</td><td>114611</td><td>-</td><td>0</td><td>ACCTGAGGCAAGGCCTGCTG</td><td>CGG</td><td>||||||||[||||||||||||]PAM</td><td>NA</td><td>Intergenic</td><td>NA</td><td>NA</td></tr>
</table>

<table>
<tr><td>T25360</td><td>CTGCGGCGACGTTAAAGCGCCGG</td><td>1000</td><td></td><td></td><td></td><td></td><td></td><td></td><td></td><td></td><td></td></tr>
<tr><td>Oligo fwd</td><td>CACCCTGCGGCGACGTTAAAGCGC</td><td></td><td></td><td></td><td></td><td></td><td></td><td></td><td></td><td></td><td></td></tr>
<tr><td>Oligo rev</td><td>AAACGCGCTTTAACGTCGCCGCAG</td><td></td><td></td><td></td><td></td><td></td><td></td><td></td><td></td><td></td><td></td></tr>
<tr><td>Chromosome</td><td>start</td><td>end</td><td>strand</td><td>MM</td><td>target_seq</td><td>PAM</td><td>alignment</td><td>distance</td><td>position</td><td>gene name</td><td>gene id</td></tr>
<tr><td>GU305914.1</td><td>114572</td><td>114594</td><td>-</td><td>0</td><td>CTGCGGCGACGTTAAAGCGC</td><td>CGG</td><td>||||||||[||||||||||||]PAM</td><td>NA</td><td>Intergenic</td><td>NA</td><td>NA</td></tr>
<tr><td>T25361</td><td>TGCGGCGACGTTAAAGCGCCGGG</td><td>1000</td><td></td><td></td><td></td><td></td><td></td><td></td><td></td><td></td><td></td></tr>
<tr><td>Oligo fwd</td><td>CACCTGCGGCGACGTTAAAGCGCC</td><td></td><td></td><td></td><td></td><td></td><td></td><td></td><td></td><td></td><td></td></tr>
<tr><td>Oligo rev</td><td>AAACGGCGCTTTAACGTCGCCGCA</td><td></td><td></td><td></td><td></td><td></td><td></td><td></td><td></td><td></td><td></td></tr>
<tr><td>Chromosome</td><td>start</td><td>end</td><td>strand</td><td>MM</td><td>target_seq</td><td>PAM</td><td>alignment</td><td>distance</td><td>position</td><td>gene name</td><td>gene id</td></tr>
<tr><td>GU305914.1</td><td>114571</td><td>114593</td><td>-</td><td>0</td><td>TGCGGCGACGTTAAAGCGCC</td><td>GGG</td><td>||||||||[||||||||||||]PAM</td><td>NA</td><td>Intergenic</td><td>NA</td><td>NA</td></tr>
<tr><td>T25362</td><td>CGACGTTAAAGCGCCGGGTACGG</td><td>1000</td><td></td><td></td><td></td><td></td><td></td><td></td><td></td><td></td><td></td></tr>
<tr><td>Oligo fwd</td><td>CACCCGACGTTAAAGCGCCGGGTA</td><td></td><td></td><td></td><td></td><td></td><td></td><td></td><td></td><td></td><td></td></tr>
<tr><td>Oligo rev</td><td>AAACTACCCGGCGCTTTAACGTCG</td><td></td><td></td><td></td><td></td><td></td><td></td><td></td><td></td><td></td><td></td></tr>
<tr><td>Chromosome</td><td>start</td><td>end</td><td>strand</td><td>MM</td><td>target_seq</td><td>PAM</td><td>alignment</td><td>distance</td><td>position</td><td>gene name</td><td>gene id</td></tr>
<tr><td>GU305914.1</td><td>114566</td><td>114588</td><td>-</td><td>0</td><td>CGACGTTAAAGCGCCGGGTA</td><td>CGG</td><td>||||||||[||||||||||||]PAM</td><td>NA</td><td>Intergenic</td><td>NA</td><td>NA</td></tr>
<tr><td>T25363</td><td>GGGTACGGCGTTCTGTTTCGCGG</td><td>1000</td><td></td><td></td><td></td><td></td><td></td><td></td><td></td><td></td><td></td></tr>
<tr><td>Oligo fwd</td><td>CACCGGGTACGGCGTTCTGTTTCG</td><td></td><td></td><td></td><td></td><td></td><td></td><td></td><td></td><td></td><td></td></tr>
<tr><td>Oligo rev</td><td>AAACCGAAACAGAACGCCGTACCC</td><td></td><td></td><td></td><td></td><td></td><td></td><td></td><td></td><td></td><td></td></tr>
<tr><td>Chromosome</td><td>start</td><td>end</td><td>strand</td><td>MM</td><td>target_seq</td><td>PAM</td><td>alignment</td><td>distance</td><td>position</td><td>gene name</td><td>gene id</td></tr>
<tr><td>GU305914.1</td><td>114551</td><td>114573</td><td>-</td><td>0</td><td>GGGTACGGCGTTCTGTTTCG</td><td>CGG</td><td>||||||||[||||||||||||]PAM</td><td>NA</td><td>Intergenic</td><td>NA</td><td>NA</td></tr>
<tr><td>T25364</td><td>TACGGCGTTCTGTTTCGCGGCGG</td><td>1000</td><td></td><td></td><td></td><td></td><td></td><td></td><td></td><td></td><td></td></tr>
<tr><td>Oligo fwd</td><td>CACCTACGGCGTTCTGTTTCGCGG</td><td></td><td></td><td></td><td></td><td></td><td></td><td></td><td></td><td></td><td></td></tr>
<tr><td>Oligo rev</td><td>AAACCCGCGAAACAGAACGCCGTA</td><td></td><td></td><td></td><td></td><td></td><td></td><td></td><td></td><td></td><td></td></tr>
<tr><td>Chromosome</td><td>start</td><td>end</td><td>strand</td><td>MM</td><td>target_seq</td><td>PAM</td><td>alignment</td><td>distance</td><td>position</td><td>gene name</td><td>gene id</td></tr>
<tr><td>GU305914.1</td><td>114548</td><td>114570</td><td>-</td><td>0</td><td>TACGGCGTTCTGTTTCGCGG</td><td>CGG</td><td>||||||||[||||||||||||]PAM</td><td>NA</td><td>Intergenic</td><td>NA</td><td>NA</td></tr>
<tr><td>T25365</td><td>TCTGTTTCGCGGCGGCCGCCAGG</td><td>1000</td><td></td><td></td><td></td><td></td><td></td><td></td><td></td><td></td><td></td></tr>
<tr><td>Oligo fwd</td><td>CACCTCTGTTTCGCGGCGGCCGCC</td><td></td><td></td><td></td><td></td><td></td><td></td><td></td><td></td><td></td><td></td></tr>
<tr><td>Oligo rev</td><td>AAACGGCGGCCGCCGCGAAACAGA</td><td></td><td></td><td></td><td></td><td></td><td></td><td></td><td></td><td></td><td></td></tr>
<tr><td>Chromosome</td><td>start</td><td>end</td><td>strand</td><td>MM</td><td>target_seq</td><td>PAM</td><td>alignment</td><td>distance</td><td>position</td><td>gene name</td><td>gene id</td></tr>
<tr><td>GU305914.1</td><td>114540</td><td>114562</td><td>-</td><td>0</td><td>TCTGTTTCGCGGCGGCCGCC</td><td>AGG</td><td>||||||||[||||||||||||]PAM</td><td>NA</td><td>Intergenic</td><td>NA</td><td>NA</td></tr>
<tr><td>T25366</td><td>TTTCGCGGCGGCCGCCAGGATGG</td><td>1000</td><td></td><td></td><td></td><td></td><td></td><td></td><td></td><td></td><td></td></tr>
<tr><td>Oligo fwd</td><td>CACCTTTCGCGGCGGCCGCCAGGA</td><td></td><td></td><td></td><td></td><td></td><td></td><td></td><td></td><td></td><td></td></tr>
<tr><td>Oligo rev</td><td>AAACTCCTGGCGGCCGCCGCGAAA</td><td></td><td></td><td></td><td></td><td></td><td></td><td></td><td></td><td></td><td></td></tr>
<tr><td>Chromosome</td><td>start</td><td>end</td><td>strand</td><td>MM</td><td>target_seq</td><td>PAM</td><td>alignment</td><td>distance</td><td>position</td><td>gene name</td><td>gene id</td></tr>
<tr><td>GU305914.1</td><td>114536</td><td>114558</td><td>-</td><td>0</td><td>TTTCGCGGCGGCCGCCAGGA</td><td>TGG</td><td>||||||||[||||||||||||]PAM</td><td>NA</td><td>Intergenic</td><td>NA</td><td>NA</td></tr>
<tr><td>T25367</td><td>ACATCAGCGCATCCCGACTGAGG</td><td>1000</td><td></td><td></td><td></td><td></td><td></td><td></td><td></td><td></td><td></td></tr>
<tr><td>Oligo fwd</td><td>CACCACATCAGCGCATCCCGACTG</td><td></td><td></td><td></td><td></td><td></td><td></td><td></td><td></td><td></td><td></td></tr>
<tr><td>Oligo rev</td><td>AAACCAGTCGGGATGCGCTGATGT</td><td></td><td></td><td></td><td></td><td></td><td></td><td></td><td></td><td></td><td></td></tr>
<tr><td>Chromosome</td><td>start</td><td>end</td><td>strand</td><td>MM</td><td>target_seq</td><td>PAM</td><td>alignment</td><td>distance</td><td>position</td><td>gene name</td><td>gene id</td></tr>
<tr><td>GU305914.1</td><td>114513</td><td>114535</td><td>-</td><td>0</td><td>ACATCAGCGCATCCCGACTG</td><td>AGG</td><td>||||||||[||||||||||||]PAM</td><td>NA</td><td>Intergenic</td><td>NA</td><td>NA</td></tr>
<tr><td>T25368</td><td>CCGACTGAGGTAGACGCAGTCGG</td><td>1000</td><td></td><td></td><td></td><td></td><td></td><td></td><td></td><td></td><td></td></tr>
<tr><td>Oligo fwd</td><td>CACCCCGACTGAGGTAGACGCAGT</td><td></td><td></td><td></td><td></td><td></td><td></td><td></td><td></td><td></td><td></td></tr>
<tr><td>Oligo rev</td><td>AAACACTGCGTCTACCTCAGTCGG</td><td></td><td></td><td></td><td></td><td></td><td></td><td></td><td></td><td></td><td></td></tr>
<tr><td>Chromosome</td><td>start</td><td>end</td><td>strand</td><td>MM</td><td>target_seq</td><td>PAM</td><td>alignment</td><td>distance</td><td>position</td><td>gene name</td><td>gene id</td></tr>
<tr><td>GU305914.1</td><td>114500</td><td>114522</td><td>-</td><td>0</td><td>CCGACTGAGGTAGACGCAGT</td><td>CGG</td><td>||||||||[||||||||||||]PAM</td><td>NA</td><td>Intergenic</td><td>NA</td><td>NA</td></tr>
<tr><td>T25369</td><td>CTGAGGTAGACGCAGTCGGCCGG</td><td>1000</td><td></td><td></td><td></td><td></td><td></td><td></td><td></td><td></td><td></td></tr>
<tr><td>Oligo fwd</td><td>CACCCTGAGGTAGACGCAGTCGGC</td><td></td><td></td><td></td><td></td><td></td><td></td><td></td><td></td><td></td><td></td></tr>
<tr><td>Oligo rev</td><td>AAACGCCGACTGCGTCTACCTCAG</td><td></td><td></td><td></td><td></td><td></td><td></td><td></td><td></td><td></td><td></td></tr>
<tr><td>Chromosome</td><td>start</td><td>end</td><td>strand</td><td>MM</td><td>target_seq</td><td>PAM</td><td>alignment</td><td>distance</td><td>position</td><td>gene name</td><td>gene id</td></tr>
<tr><td>GU305914.1</td><td>114496</td><td>114518</td><td>-</td><td>0</td><td>CTGAGGTAGACGCAGTCGGC</td><td>CGG</td><td>||||||||[||||||||||||]PAM</td><td>NA</td><td>Intergenic</td><td>NA</td><td>NA</td></tr>
<tr><td>T25370</td><td>TGAGGTAGACGCAGTCGGCCGGG</td><td>1000</td><td></td><td></td><td></td><td></td><td></td><td></td><td></td><td></td><td></td></tr>
<tr><td>Oligo fwd</td><td>CACCTGAGGTAGACGCAGTCGGCC</td><td></td><td></td><td></td><td></td><td></td><td></td><td></td><td></td><td></td><td></td></tr>
<tr><td>Oligo rev</td><td>AAACGGCCGACTGCGTCTACCTCA</td><td></td><td></td><td></td><td></td><td></td><td></td><td></td><td></td><td></td><td></td></tr>
<tr><td>Chromosome</td><td>start</td><td>end</td><td>strand</td><td>MM</td><td>target_seq</td><td>PAM</td><td>alignment</td><td>distance</td><td>position</td><td>gene name</td><td>gene id</td></tr>
<tr><td>GU305914.1</td><td>114495</td><td>114517</td><td>-</td><td>0</td><td>TGAGGTAGACGCAGTCGGCC</td><td>GGG</td><td>||||||||[||||||||||||]PAM</td><td>NA</td><td>Intergenic</td><td>NA</td><td>NA</td></tr>
<tr><td>GU305914.1</td><td>151649</td><td>151671</td><td>-</td><td>4</td><td>TGAGGTGAACGCGGTCGGCG</td><td>AGG</td><td>||||||--[||||-||||||-]PAM</td><td>NA</td><td>Intergenic</td><td>NA</td><td>NA</td></tr>
<tr><td>T25371</td><td>GACGCAGTCGGCCGGGAAAGCGG</td><td>1000</td><td></td><td></td><td></td><td></td><td></td><td></td><td></td><td></td><td></td></tr>
<tr><td>Oligo fwd</td><td>CACCGACGCAGTCGGCCGGGAAAG</td><td></td><td></td><td></td><td></td><td></td><td></td><td></td><td></td><td></td><td></td></tr>
<tr><td>Oligo rev</td><td>AAACCTTTCCCGGCCGACTGCGTC</td><td></td><td></td><td></td><td></td><td></td><td></td><td></td><td></td><td></td><td></td></tr>
<tr><td>Chromosome</td><td>start</td><td>end</td><td>strand</td><td>MM</td><td>target_seq</td><td>PAM</td><td>alignment</td><td>distance</td><td>position</td><td>gene name</td><td>gene id</td></tr>
<tr><td>GU305914.1</td><td>114488</td><td>114510</td><td>-</td><td>0</td><td>GACGCAGTCGGCCGGGAAAG</td><td>CGG</td><td>||||||||[||||||||||||]PAM</td><td>NA</td><td>Intergenic</td><td>NA</td><td>NA</td></tr>
<tr><td>T25372</td><td>GCAGTCGGCCGGGAAAGCGGAGG</td><td>1000</td><td></td><td></td><td></td><td></td><td></td><td></td><td></td><td></td><td></td></tr>
<tr><td>Oligo fwd</td><td>CACCGCAGTCGGCCGGGAAAGCGG</td><td></td><td></td><td></td><td></td><td></td><td></td><td></td><td></td><td></td><td></td></tr>
<tr><td>Oligo rev</td><td>AAACCCGCTTTCCCGGCCGACTGC</td><td></td><td></td><td></td><td></td><td></td><td></td><td></td><td></td><td></td><td></td></tr>
<tr><td>Chromosome</td><td>start</td><td>end</td><td>strand</td><td>MM</td><td>target_seq</td><td>PAM</td><td>alignment</td><td>distance</td><td>position</td><td>gene name</td><td>gene id</td></tr>
<tr><td>GU305914.1</td><td>114485</td><td>114507</td><td>-</td><td>0</td><td>GCAGTCGGCCGGGAAAGCGG</td><td>AGG</td><td>||||||||[||||||||||||]PAM</td><td>NA</td><td>Intergenic</td><td>NA</td><td>NA</td></tr>
<tr><td>T25373</td><td>GTCGGCCGGGAAAGCGGAGGCGG</td><td>1000</td><td></td><td></td><td></td><td></td><td></td><td></td><td></td><td></td><td></td></tr>
<tr><td>Oligo fwd</td><td>CACCGTCGGCCGGGAAAGCGGAGG</td><td></td><td></td><td></td><td></td><td></td><td></td><td></td><td></td><td></td><td></td></tr>
<tr><td>Oligo rev</td><td>AAACCCTCCGCTTTCCCGGCCGAC</td><td></td><td></td><td></td><td></td><td></td><td></td><td></td><td></td><td></td><td></td></tr>
<tr><td>Chromosome</td><td>start</td><td>end</td><td>strand</td><td>MM</td><td>target_seq</td><td>PAM</td><td>alignment</td><td>distance</td><td>position</td><td>gene name</td><td>gene id</td></tr>
<tr><td>GU305914.1</td><td>114482</td><td>114504</td><td>-</td><td>0</td><td>GTCGGCCGGGAAAGCGGAGG</td><td>CGG</td><td>||||||||[||||||||||||]PAM</td><td>NA</td><td>Intergenic</td><td>NA</td><td>NA</td></tr>
<tr><td>T25374</td><td>GCCGGGAAAGCGGAGGCGGCAGG</td><td>1000</td><td></td><td></td><td></td><td></td><td></td><td></td><td></td><td></td><td></td></tr>
<tr><td>Oligo fwd</td><td>CACCGCCGGGAAAGCGGAGGCGGC</td><td></td><td></td><td></td><td></td><td></td><td></td><td></td><td></td><td></td><td></td></tr>
<tr><td>Oligo rev</td><td>AAACGCCGCCTCCGCTTTCCCGGC</td><td></td><td></td><td></td><td></td><td></td><td></td><td></td><td></td><td></td><td></td></tr>
<tr><td>Chromosome</td><td>start</td><td>end</td><td>strand</td><td>MM</td><td>target_seq</td><td>PAM</td><td>alignment</td><td>distance</td><td>position</td><td>gene name</td><td>gene id</td></tr>
</table>

| | | | | | | | | | | | |
|---|---|---|---|---|---|---|---|---|---|---|---|
| GU305914.1 | 114478 | 114500 | - | 0 | GCCGGGAAAGCGGAGGCGGC | AGG | \|\|\|\|\|\|\|\|[\|\|\|\|\|\|\|\|\|\|\|\|]PAM | NA | Intergenic | NA | NA |
| T25375 | CCGGGAAAGCGGAGGCGGCAGGG | 1000 | | | | | | | | | |
| Oligo fwd | CACCCCGGGAAAGCGGAGGCGGCA | | | | | | | | | | |
| Oligo rev | AAACTGCCGCCTCCGCTTTCCCGG | | | | | | | | | | |
| Chromosome | start | end | strand | MM | target_seq | PAM | alignment | distance | position | gene name | gene id |
| GU305914.1 | 114477 | 114499 | - | 0 | CCGGGAAAGCGGAGGCGGCA | GGG | \|\|\|\|\|\|\|\|[\|\|\|\|\|\|\|\|\|\|\|\|]PAM | NA | Intergenic | NA | NA |
| T25376 | GAAAGCGGAGGCGGCAGGGTTGG | 1000 | | | | | | | | | |
| Oligo fwd | CACCGAAAGCGGAGGCGGCAGGGT | | | | | | | | | | |
| Oligo rev | AAACACCCTGCCGCCTCCGCTTTC | | | | | | | | | | |
| Chromosome | start | end | strand | MM | target_seq | PAM | alignment | distance | position | gene name | gene id |
| GU305914.1 | 114473 | 114495 | - | 0 | GAAAGCGGAGGCGGCAGGGT | TGG | \|\|\|\|\|\|\|\|[\|\|\|\|\|\|\|\|\|\|\|\|]PAM | NA | Intergenic | NA | NA |
| T25377 | AGCGGAGGCGGCAGGGTTGGCGG | 1000 | | | | | | | | | |
| Oligo fwd | CACCAGCGGAGGCGGCAGGGTTGG | | | | | | | | | | |
| Oligo rev | AAACCCAACCCTGCCGCCTCCGCT | | | | | | | | | | |
| Chromosome | start | end | strand | MM | target_seq | PAM | alignment | distance | position | gene name | gene id |
| GU305914.1 | 114470 | 114492 | - | 0 | AGCGGAGGCGGCAGGGTTGG | CGG | \|\|\|\|\|\|\|\|[\|\|\|\|\|\|\|\|\|\|\|\|]PAM | NA | Intergenic | NA | NA |
| GU305914.1 | 62437 | 62459 | + | 4 | GGCGGTGGCGGCGGGGGTGG | TGG | -\|\|\|\|-\|\|[\|\|\|\|-\|\|\|-\|\|\|]PAM | NA | Intergenic | NA | NA |
| GU305914.1 | 128993 | 129015 | + | 4 | AGCGGCAGCGGCAGCGTTGA | GGG | \|\|\|\|\|--\|[\|\|\|\|\|\|-\|\|\|\|-]PAM | NA | Intergenic | NA | NA |
| T25378 | GGCGGCAGGGTTGGCGGTCCCGG | 1000 | | | | | | | | | |
| Oligo fwd | CACCGGCGGCAGGGTTGGCGGTCC | | | | | | | | | | |
| Oligo rev | AAACGGACCGCCAACCCTGCCGCC | | | | | | | | | | |
| Chromosome | start | end | strand | MM | target_seq | PAM | alignment | distance | position | gene name | gene id |
| GU305914.1 | 114464 | 114486 | - | 0 | GGCGGCAGGGTTGGCGGTCC | CGG | \|\|\|\|\|\|\|\|[\|\|\|\|\|\|\|\|\|\|\|\|]PAM | NA | Intergenic | NA | NA |
| T25379 | GCGGCAGGGTTGGCGGTCCCGGG | 1000 | | | | | | | | | |
| Oligo fwd | CACCGCGGCAGGGTTGGCGGTCCC | | | | | | | | | | |
| Oligo rev | AAACGGGACCGCCAACCCTGCCGC | | | | | | | | | | |
| Chromosome | start | end | strand | MM | target_seq | PAM | alignment | distance | position | gene name | gene id |
| GU305914.1 | 114463 | 114485 | - | 0 | GCGGCAGGGTTGGCGGTCCC | GGG | \|\|\|\|\|\|\|\|[\|\|\|\|\|\|\|\|\|\|\|\|]PAM | NA | Intergenic | NA | NA |
| T25380 | AGGGTTGGCGGTCCCGGGCACGG | 1000 | | | | | | | | | |
| Oligo fwd | CACCAGGGTTGGCGGTCCCGGGCA | | | | | | | | | | |
| Oligo rev | AAACTGCCCGGGACCGCCAACCCT | | | | | | | | | | |
| Chromosome | start | end | strand | MM | target_seq | PAM | alignment | distance | position | gene name | gene id |
| GU305914.1 | 114458 | 114480 | - | 0 | AGGGTTGGCGGTCCCGGGCA | CGG | \|\|\|\|\|\|\|\|[\|\|\|\|\|\|\|\|\|\|\|\|]PAM | NA | Intergenic | NA | NA |
| T25381 | GGTCCCGGGCACGGAAACTGCGG | 1000 | | | | | | | | | |
| Oligo fwd | CACCGGTCCCGGGCACGGAAACTG | | | | | | | | | | |
| Oligo rev | AAACCAGTTTCCGTGCCCGGGACC | | | | | | | | | | |
| Chromosome | start | end | strand | MM | target_seq | PAM | alignment | distance | position | gene name | gene id |
| GU305914.1 | 114449 | 114471 | - | 0 | GGTCCCGGGCACGGAAACTG | CGG | \|\|\|\|\|\|\|\|[\|\|\|\|\|\|\|\|\|\|\|\|]PAM | NA | Intergenic | NA | NA |
| T25382 | CCGGGCACGGAAACTGCGGCCGG | 1000 | | | | | | | | | |
| Oligo fwd | CACCCCGGGCACGGAAACTGCGGC | | | | | | | | | | |
| Oligo rev | AAACGCCGCAGTTTCCGTGCCCGG | | | | | | | | | | |
| Chromosome | start | end | strand | MM | target_seq | PAM | alignment | distance | position | gene name | gene id |
| GU305914.1 | 114445 | 114467 | - | 0 | CCGGGCACGGAAACTGCGGC | CGG | \|\|\|\|\|\|\|\|[\|\|\|\|\|\|\|\|\|\|\|\|]PAM | NA | Intergenic | NA | NA |
| T25383 | ACGACTGAGCCATCTCCTCGCGG | 1000 | | | | | | | | | |
| Oligo fwd | CACCACGACTGAGCCATCTCCTCG | | | | | | | | | | |
| Oligo rev | AAACCGAGGAGATGGCTCAGTCGT | | | | | | | | | | |
| Chromosome | start | end | strand | MM | target_seq | PAM | alignment | distance | position | gene name | gene id |
| GU305914.1 | 114408 | 114430 | - | 0 | ACGACTGAGCCATCTCCTCG | CGG | \|\|\|\|\|\|\|\|[\|\|\|\|\|\|\|\|\|\|\|\|]PAM | NA | Intergenic | NA | NA |
| T25384 | CTGAGCCATCTCCTCGCGGCCGG | 1000 | | | | | | | | | |
| Oligo fwd | CACCCTGAGCCATCTCCTCGCGGC | | | | | | | | | | |
| Oligo rev | AAACGCCGCGAGGAGATGGCTCAG | | | | | | | | | | |
| Chromosome | start | end | strand | MM | target_seq | PAM | alignment | distance | position | gene name | gene id |
| GU305914.1 | 114404 | 114426 | - | 0 | CTGAGCCATCTCCTCGCGGC | CGG | \|\|\|\|\|\|\|\|[\|\|\|\|\|\|\|\|\|\|\|\|]PAM | NA | Intergenic | NA | NA |
| T25385 | AGCCATCTCCTCGCGGCCGGCGG | 1000 | | | | | | | | | |
| Oligo fwd | CACCAGCCATCTCCTCGCGGCCGG | | | | | | | | | | |
| Oligo rev | AAACCCGGCCGCGAGGAGATGGCT | | | | | | | | | | |
| Chromosome | start | end | strand | MM | target_seq | PAM | alignment | distance | position | gene name | gene id |
| GU305914.1 | 114401 | 114423 | - | 0 | AGCCATCTCCTCGCGGCCGG | CGG | \|\|\|\|\|\|\|\|[\|\|\|\|\|\|\|\|\|\|\|\|]PAM | NA | Intergenic | NA | NA |
| T25386 | CGGCGCGCTCCTCAGAGTCCCGG | 1000 | | | | | | | | | |
| Oligo fwd | CACCCGGCGCGCTCCTCAGAGTCC | | | | | | | | | | |
| Oligo rev | AAACGGACTCTGAGGAGCGCGCCG | | | | | | | | | | |
| Chromosome | start | end | strand | MM | target_seq | PAM | alignment | distance | position | gene name | gene id |
| GU305914.1 | 114381 | 114403 | - | 0 | CGGCGCGCTCCTCAGAGTCC | CGG | \|\|\|\|\|\|\|\|[\|\|\|\|\|\|\|\|\|\|\|\|]PAM | NA | Intergenic | NA | NA |
| T25387 | GGCGCGCTCCTCAGAGTCCCGGG | 1000 | | | | | | | | | |
| Oligo fwd | CACCGGCGCGCTCCTCAGAGTCCC | | | | | | | | | | |
| Oligo rev | AAACGGGACTCTGAGGAGCGCGCC | | | | | | | | | | |
| Chromosome | start | end | strand | MM | target_seq | PAM | alignment | distance | position | gene name | gene id |
| GU305914.1 | 114380 | 114402 | - | 0 | GGCGCGCTCCTCAGAGTCCC | GGG | \|\|\|\|\|\|\|\|[\|\|\|\|\|\|\|\|\|\|\|\|]PAM | NA | Intergenic | NA | NA |
| T25388 | GCGTCGCGTCTAATACTCTGCGG | 1000 | | | | | | | | | |
| Oligo fwd | CACCGCGTCGCGTCTAATACTCTG | | | | | | | | | | |
| Oligo rev | AAACCAGAGTATTAGACGCGACGC | | | | | | | | | | |
| Chromosome | start | end | strand | MM | target_seq | PAM | alignment | distance | position | gene name | gene id |
| GU305914.1 | 114355 | 114377 | - | 0 | GCGTCGCGTCTAATACTCTG | CGG | \|\|\|\|\|\|\|\|[\|\|\|\|\|\|\|\|\|\|\|\|]PAM | NA | Intergenic | NA | NA |
| GU305914.1 | 99092 | 99114 | + | 4 | TCGACGCGTCTAATACTGCG | AGG | -\|\|-\|\|\|\|[\|\|\|\|\|\|\|\|\|--\|]PAM | NA | Intergenic | NA | NA |

<table>
<tr><td>T25389</td><td>GCGTCTAATACTCTGCGGCTCGG</td><td>1000</td><td></td><td></td><td></td><td></td><td></td><td></td><td></td><td></td><td></td></tr>
<tr><td>Oligo fwd</td><td>CACCGCGTCTAATACTCTGCGGCT</td><td></td><td></td><td></td><td></td><td></td><td></td><td></td><td></td><td></td><td></td></tr>
<tr><td>Oligo rev</td><td>AAACAGCCGCAGAGTATTAGACGC</td><td></td><td></td><td></td><td></td><td></td><td></td><td></td><td></td><td></td><td></td></tr>
<tr><td>Chromosome</td><td>start</td><td>end</td><td>strand</td><td>MM</td><td>target_seq</td><td>PAM</td><td>alignment</td><td>distance</td><td>position</td><td>gene name</td><td>gene id</td></tr>
<tr><td>GU305914.1</td><td>114350</td><td>114372</td><td>-</td><td>0</td><td>GCGTCTAATACTCTGCGGCT</td><td>CGG</td><td>||||||||[||||||||||||]PAM</td><td>NA</td><td>Intergenic</td><td>NA</td><td>NA</td></tr>
</table>

<table>
<tr><td>T25390</td><td>CGTCTAATACTCTGCGGCTCGGG</td><td>1000</td><td></td><td></td><td></td><td></td><td></td><td></td><td></td><td></td><td></td></tr>
<tr><td>Oligo fwd</td><td>CACCCGTCTAATACTCTGCGGCTC</td><td></td><td></td><td></td><td></td><td></td><td></td><td></td><td></td><td></td><td></td></tr>
<tr><td>Oligo rev</td><td>AAACGAGCCGCAGAGTATTAGACG</td><td></td><td></td><td></td><td></td><td></td><td></td><td></td><td></td><td></td><td></td></tr>
<tr><td>Chromosome</td><td>start</td><td>end</td><td>strand</td><td>MM</td><td>target_seq</td><td>PAM</td><td>alignment</td><td>distance</td><td>position</td><td>gene name</td><td>gene id</td></tr>
<tr><td>GU305914.1</td><td>114349</td><td>114371</td><td>-</td><td>0</td><td>CGTCTAATACTCTGCGGCTC</td><td>GGG</td><td>||||||||[||||||||||||]PAM</td><td>NA</td><td>Intergenic</td><td>NA</td><td>NA</td></tr>
</table>

<table>
<tr><td>T25391</td><td>CTGCGGCTCGGGACCGTCGTCGG</td><td>1000</td><td></td><td></td><td></td><td></td><td></td><td></td><td></td><td></td><td></td></tr>
<tr><td>Oligo fwd</td><td>CACCCTGCGGCTCGGGACCGTCGT</td><td></td><td></td><td></td><td></td><td></td><td></td><td></td><td></td><td></td><td></td></tr>
<tr><td>Oligo rev</td><td>AAACACGACGGTCCCGAGCCGCAG</td><td></td><td></td><td></td><td></td><td></td><td></td><td></td><td></td><td></td><td></td></tr>
<tr><td>Chromosome</td><td>start</td><td>end</td><td>strand</td><td>MM</td><td>target_seq</td><td>PAM</td><td>alignment</td><td>distance</td><td>position</td><td>gene name</td><td>gene id</td></tr>
<tr><td>GU305914.1</td><td>114338</td><td>114360</td><td>-</td><td>0</td><td>CTGCGGCTCGGGACCGTCGT</td><td>CGG</td><td>||||||||[||||||||||||]PAM</td><td>NA</td><td>Intergenic</td><td>NA</td><td>NA</td></tr>
</table>

<table>
<tr><td>T25392</td><td>CTCGGGACCGTCGTCGGCCTCGG</td><td>1000</td><td></td><td></td><td></td><td></td><td></td><td></td><td></td><td></td><td></td></tr>
<tr><td>Oligo fwd</td><td>CACCCTCGGGACCGTCGTCGGCCT</td><td></td><td></td><td></td><td></td><td></td><td></td><td></td><td></td><td></td><td></td></tr>
<tr><td>Oligo rev</td><td>AAACAGGCCGACGACGGTCCCGAG</td><td></td><td></td><td></td><td></td><td></td><td></td><td></td><td></td><td></td><td></td></tr>
<tr><td>Chromosome</td><td>start</td><td>end</td><td>strand</td><td>MM</td><td>target_seq</td><td>PAM</td><td>alignment</td><td>distance</td><td>position</td><td>gene name</td><td>gene id</td></tr>
<tr><td>GU305914.1</td><td>114332</td><td>114354</td><td>-</td><td>0</td><td>CTCGGGACCGTCGTCGGCCT</td><td>CGG</td><td>||||||||[||||||||||||]PAM</td><td>NA</td><td>Intergenic</td><td>NA</td><td>NA</td></tr>
</table>

<table>
<tr><td>T25393</td><td>GGACCGTCGTCGGCCTCGGCCGG</td><td>1000</td><td></td><td></td><td></td><td></td><td></td><td></td><td></td><td></td><td></td></tr>
<tr><td>Oligo fwd</td><td>CACCGGACCGTCGTCGGCCTCGGC</td><td></td><td></td><td></td><td></td><td></td><td></td><td></td><td></td><td></td><td></td></tr>
<tr><td>Oligo rev</td><td>AAACGCCGAGGCCGACGACGGTCC</td><td></td><td></td><td></td><td></td><td></td><td></td><td></td><td></td><td></td><td></td></tr>
<tr><td>Chromosome</td><td>start</td><td>end</td><td>strand</td><td>MM</td><td>target_seq</td><td>PAM</td><td>alignment</td><td>distance</td><td>position</td><td>gene name</td><td>gene id</td></tr>
<tr><td>GU305914.1</td><td>114328</td><td>114350</td><td>-</td><td>0</td><td>GGACCGTCGTCGGCCTCGGC</td><td>CGG</td><td>||||||||[||||||||||||]PAM</td><td>NA</td><td>Intergenic</td><td>NA</td><td>NA</td></tr>
</table>

<table>
<tr><td>T25394</td><td>GGCCTCGGCCGGCAGCTCGCTGG</td><td>1000</td><td></td><td></td><td></td><td></td><td></td><td></td><td></td><td></td><td></td></tr>
<tr><td>Oligo fwd</td><td>CACCGGCCTCGGCCGGCAGCTCGC</td><td></td><td></td><td></td><td></td><td></td><td></td><td></td><td></td><td></td><td></td></tr>
<tr><td>Oligo rev</td><td>AAACGCGAGCTGCCGGCCGAGGCC</td><td></td><td></td><td></td><td></td><td></td><td></td><td></td><td></td><td></td><td></td></tr>
<tr><td>Chromosome</td><td>start</td><td>end</td><td>strand</td><td>MM</td><td>target_seq</td><td>PAM</td><td>alignment</td><td>distance</td><td>position</td><td>gene name</td><td>gene id</td></tr>
<tr><td>GU305914.1</td><td>114317</td><td>114339</td><td>-</td><td>0</td><td>GGCCTCGGCCGGCAGCTCGC</td><td>TGG</td><td>||||||||[||||||||||||]PAM</td><td>NA</td><td>Intergenic</td><td>NA</td><td>NA</td></tr>
</table>

<table>
<tr><td>T25395</td><td>CAGCTCGCTGGCTTTGACGTAGG</td><td>1000</td><td></td><td></td><td></td><td></td><td></td><td></td><td></td><td></td><td></td></tr>
<tr><td>Oligo fwd</td><td>CACCCAGCTCGCTGGCTTTGACGT</td><td></td><td></td><td></td><td></td><td></td><td></td><td></td><td></td><td></td><td></td></tr>
<tr><td>Oligo rev</td><td>AAACACGTCAAAGCCAGCGAGCTG</td><td></td><td></td><td></td><td></td><td></td><td></td><td></td><td></td><td></td><td></td></tr>
<tr><td>Chromosome</td><td>start</td><td>end</td><td>strand</td><td>MM</td><td>target_seq</td><td>PAM</td><td>alignment</td><td>distance</td><td>position</td><td>gene name</td><td>gene id</td></tr>
<tr><td>GU305914.1</td><td>114305</td><td>114327</td><td>-</td><td>0</td><td>CAGCTCGCTGGCTTTGACGT</td><td>AGG</td><td>||||||||[||||||||||||]PAM</td><td>NA</td><td>Intergenic</td><td>NA</td><td>NA</td></tr>
</table>

<table>
<tr><td>T25396</td><td>GGCTTTGACGTAGGTCTCCTTGG</td><td>1000</td><td></td><td></td><td></td><td></td><td></td><td></td><td></td><td></td><td></td></tr>
<tr><td>Oligo fwd</td><td>CACCGGCTTTGACGTAGGTCTCCT</td><td></td><td></td><td></td><td></td><td></td><td></td><td></td><td></td><td></td><td></td></tr>
<tr><td>Oligo rev</td><td>AAACAGGAGACCTACGTCAAAGCC</td><td></td><td></td><td></td><td></td><td></td><td></td><td></td><td></td><td></td><td></td></tr>
<tr><td>Chromosome</td><td>start</td><td>end</td><td>strand</td><td>MM</td><td>target_seq</td><td>PAM</td><td>alignment</td><td>distance</td><td>position</td><td>gene name</td><td>gene id</td></tr>
<tr><td>GU305914.1</td><td>114296</td><td>114318</td><td>-</td><td>0</td><td>GGCTTTGACGTAGGTCTCCT</td><td>TGG</td><td>||||||||[||||||||||||]PAM</td><td>NA</td><td>Intergenic</td><td>NA</td><td>NA</td></tr>
</table>

<table>
<tr><td>T25397</td><td>GCAGCCGCTTGTCGTATCTGAGG</td><td>1000</td><td></td><td></td><td></td><td></td><td></td><td></td><td></td><td></td><td></td></tr>
<tr><td>Oligo fwd</td><td>CACCGCAGCCGCTTGTCGTATCTG</td><td></td><td></td><td></td><td></td><td></td><td></td><td></td><td></td><td></td><td></td></tr>
<tr><td>Oligo rev</td><td>AAACCAGATACGACAAGCGGCTGC</td><td></td><td></td><td></td><td></td><td></td><td></td><td></td><td></td><td></td><td></td></tr>
<tr><td>Chromosome</td><td>start</td><td>end</td><td>strand</td><td>MM</td><td>target_seq</td><td>PAM</td><td>alignment</td><td>distance</td><td>position</td><td>gene name</td><td>gene id</td></tr>
<tr><td>GU305914.1</td><td>114261</td><td>114283</td><td>-</td><td>0</td><td>GCAGCCGCTTGTCGTATCTG</td><td>AGG</td><td>||||||||[||||||||||||]PAM</td><td>NA</td><td>Intergenic</td><td>NA</td><td>NA</td></tr>
</table>

<table>
<tr><td>T25398</td><td>TGTCGTATCTGAGGCGACAGAGG</td><td>1000</td><td></td><td></td><td></td><td></td><td></td><td></td><td></td><td></td><td></td></tr>
<tr><td>Oligo fwd</td><td>CACCTGTCGTATCTGAGGCGACAG</td><td></td><td></td><td></td><td></td><td></td><td></td><td></td><td></td><td></td><td></td></tr>
<tr><td>Oligo rev</td><td>AAACCTGTCGCCTCAGATACGACA</td><td></td><td></td><td></td><td></td><td></td><td></td><td></td><td></td><td></td><td></td></tr>
<tr><td>Chromosome</td><td>start</td><td>end</td><td>strand</td><td>MM</td><td>target_seq</td><td>PAM</td><td>alignment</td><td>distance</td><td>position</td><td>gene name</td><td>gene id</td></tr>
<tr><td>GU305914.1</td><td>114252</td><td>114274</td><td>-</td><td>0</td><td>TGTCGTATCTGAGGCGACAG</td><td>AGG</td><td>||||||||[||||||||||||]PAM</td><td>NA</td><td>Intergenic</td><td>NA</td><td>NA</td></tr>
</table>

<table>
<tr><td>T25399</td><td>GTAACCGTCGTCGACCGTGCTGG</td><td>1000</td><td></td><td></td><td></td><td></td><td></td><td></td><td></td><td></td><td></td></tr>
<tr><td>Oligo fwd</td><td>CACCGTAACCGTCGTCGACCGTGC</td><td></td><td></td><td></td><td></td><td></td><td></td><td></td><td></td><td></td><td></td></tr>
<tr><td>Oligo rev</td><td>AAACGCACGGTCGACGACGGTTAC</td><td></td><td></td><td></td><td></td><td></td><td></td><td></td><td></td><td></td><td></td></tr>
<tr><td>Chromosome</td><td>start</td><td>end</td><td>strand</td><td>MM</td><td>target_seq</td><td>PAM</td><td>alignment</td><td>distance</td><td>position</td><td>gene name</td><td>gene id</td></tr>
<tr><td>GU305914.1</td><td>114218</td><td>114240</td><td>-</td><td>0</td><td>GTAACCGTCGTCGACCGTGC</td><td>TGG</td><td>||||||||[||||||||||||]PAM</td><td>NA</td><td>Intergenic</td><td>NA</td><td>NA</td></tr>
</table>

<table>
<tr><td>T25400</td><td>CGTCGTCGACCGTGCTGGCGAGG</td><td>1000</td><td></td><td></td><td></td><td></td><td></td><td></td><td></td><td></td><td></td></tr>
<tr><td>Oligo fwd</td><td>CACCCGTCGTCGACCGTGCTGGCG</td><td></td><td></td><td></td><td></td><td></td><td></td><td></td><td></td><td></td><td></td></tr>
<tr><td>Oligo rev</td><td>AAACCGCCAGCACGGTCGACGACG</td><td></td><td></td><td></td><td></td><td></td><td></td><td></td><td></td><td></td><td></td></tr>
<tr><td>Chromosome</td><td>start</td><td>end</td><td>strand</td><td>MM</td><td>target_seq</td><td>PAM</td><td>alignment</td><td>distance</td><td>position</td><td>gene name</td><td>gene id</td></tr>
<tr><td>GU305914.1</td><td>114213</td><td>114235</td><td>-</td><td>0</td><td>CGTCGTCGACCGTGCTGGCG</td><td>AGG</td><td>||||||||[||||||||||||]PAM</td><td>NA</td><td>Intergenic</td><td>NA</td><td>NA</td></tr>
</table>

<table>
<tr><td>T25401</td><td>CGTCGACCGTGCTGGCGAGGAGG</td><td>1000</td><td></td><td></td><td></td><td></td><td></td><td></td><td></td><td></td><td></td></tr>
<tr><td>Oligo fwd</td><td>CACCCGTCGACCGTGCTGGCGAGG</td><td></td><td></td><td></td><td></td><td></td><td></td><td></td><td></td><td></td><td></td></tr>
<tr><td>Oligo rev</td><td>AAACCCTCGCCAGCACGGTCGACG</td><td></td><td></td><td></td><td></td><td></td><td></td><td></td><td></td><td></td><td></td></tr>
<tr><td>Chromosome</td><td>start</td><td>end</td><td>strand</td><td>MM</td><td>target_seq</td><td>PAM</td><td>alignment</td><td>distance</td><td>position</td><td>gene name</td><td>gene id</td></tr>
<tr><td>GU305914.1</td><td>114210</td><td>114232</td><td>-</td><td>0</td><td>CGTCGACCGTGCTGGCGAGG</td><td>AGG</td><td>||||||||[||||||||||||]PAM</td><td>NA</td><td>Intergenic</td><td>NA</td><td>NA</td></tr>
</table>

<table>
<tr><td>T25402</td><td>GCCGTATGAGTCCGAACCGAAGG</td><td>1000</td><td></td><td></td><td></td><td></td><td></td><td></td><td></td><td></td><td></td></tr>
<tr><td>Oligo fwd</td><td>CACCGCCGTATGAGTCCGAACCGA</td><td></td><td></td><td></td><td></td><td></td><td></td><td></td><td></td><td></td><td></td></tr>
<tr><td>Oligo rev</td><td>AAACTCGGTTCGGACTCATACGGC</td><td></td><td></td><td></td><td></td><td></td><td></td><td></td><td></td><td></td><td></td></tr>
<tr><td>Chromosome</td><td>start</td><td>end</td><td>strand</td><td>MM</td><td>target_seq</td><td>PAM</td><td>alignment</td><td>distance</td><td>position</td><td>gene name</td><td>gene id</td></tr>
<tr><td>GU305914.1</td><td>114188</td><td>114210</td><td>-</td><td>0</td><td>GCCGTATGAGTCCGAACCGA</td><td>AGG</td><td>||||||||[||||||||||||]PAM</td><td>NA</td><td>Intergenic</td><td>NA</td><td>NA</td></tr>
</table>

<table>
<tr><td>T25403</td><td>CCGTATGAGTCCGAACCGAAGGG</td><td>1000</td><td></td><td></td><td></td><td></td><td></td><td></td><td></td><td></td><td></td></tr>
<tr><td>Oligo fwd</td><td>CACCCCGTATGAGTCCGAACCGAA</td><td></td><td></td><td></td><td></td><td></td><td></td><td></td><td></td><td></td><td></td></tr>
<tr><td>Oligo rev</td><td>AAACTTCGGTTCGGACTCATACGG</td><td></td><td></td><td></td><td></td><td></td><td></td><td></td><td></td><td></td><td></td></tr>
<tr><td>Chromosome</td><td>start</td><td>end</td><td>strand</td><td>MM</td><td>target_seq</td><td>PAM</td><td>alignment</td><td>distance</td><td>position</td><td>gene name</td><td>gene id</td></tr>
<tr><td>GU305914.1</td><td>114187</td><td>114209</td><td>-</td><td>0</td><td>CCGTATGAGTCCGAACCGAA</td><td>GGG</td><td>||||||||[||||||||||||]PAM</td><td>NA</td><td>Intergenic</td><td>NA</td><td>NA</td></tr>
</table>

| | | | | | | | | | | | |
|---|---|---|---|---|---|---|---|---|---|---|---|
| T25404 | TGAGTCCGAACCGAAGGGATCGG | 1000 | | | | | | | | | |
| Oligo fwd | CACCTGAGTCCGAACCGAAGGGAT | | | | | | | | | | |
| Oligo rev | AAACATCCCTTCGGTTCGGACTCA | | | | | | | | | | |
| Chromosome | start | end | strand | MM | target_seq | PAM | alignment | distance | position | gene name | gene id |
| GU305914.1 | 114182 | 114204 | - | 0 | TGAGTCCGAACCGAAGGGAT | CGG | \|\|\|\|\|\|\|\|[\|\|\|\|\|\|\|\|\|\|\|\|]PAM | NA | Intergenic | NA | NA |
| T25405 | GCGATCTCCGCCCTCTCGTCCGG | 1000 | | | | | | | | | |
| Oligo fwd | CACCGCGATCTCCGCCCTCTCGTC | | | | | | | | | | |
| Oligo rev | AAACGACGAGAGGGCGGAGATCGC | | | | | | | | | | |
| Chromosome | start | end | strand | MM | target_seq | PAM | alignment | distance | position | gene name | gene id |
| GU305914.1 | 114118 | 114140 | - | 0 | GCGATCTCCGCCCTCTCGTC | CGG | \|\|\|\|\|\|\|\|[\|\|\|\|\|\|\|\|\|\|\|\|]PAM | NA | Intergenic | NA | NA |
| T25406 | ATCTCGTCCGTAGACCGCCAAGG | 1000 | | | | | | | | | |
| Oligo fwd | CACCATCTCGTCCGTAGACCGCCA | | | | | | | | | | |
| Oligo rev | AAACTGGCGGTCTACGGACGAGAT | | | | | | | | | | |
| Chromosome | start | end | strand | MM | target_seq | PAM | alignment | distance | position | gene name | gene id |
| GU305914.1 | 114059 | 114081 | - | 0 | ATCTCGTCCGTAGACCGCCA | AGG | \|\|\|\|\|\|\|\|[\|\|\|\|\|\|\|\|\|\|\|\|]PAM | NA | Intergenic | NA | NA |
| T25407 | TCTGAAGAAGCCGCCCGAGTCGG | 1000 | | | | | | | | | |
| Oligo fwd | CACCTCTGAAGAAGCCGCCCGAGT | | | | | | | | | | |
| Oligo rev | AAACACTCGGGCGGCTTCTTCAGA | | | | | | | | | | |
| Chromosome | start | end | strand | MM | target_seq | PAM | alignment | distance | position | gene name | gene id |
| GU305914.1 | 113999 | 114021 | - | 0 | TCTGAAGAAGCCGCCCGAGT | CGG | \|\|\|\|\|\|\|\|[\|\|\|\|\|\|\|\|\|\|\|\|]PAM | NA | Intergenic | NA | NA |
| T25408 | GAGTCGGTCGTCTGCCGCCGCGG | 1000 | | | | | | | | | |
| Oligo fwd | CACCGAGTCGGTCGTCTGCCGCCG | | | | | | | | | | |
| Oligo rev | AAACCGGCGGCAGACGACCGACTC | | | | | | | | | | |
| Chromosome | start | end | strand | MM | target_seq | PAM | alignment | distance | position | gene name | gene id |
| GU305914.1 | 113983 | 114005 | - | 0 | GAGTCGGTCGTCTGCCGCCG | CGG | \|\|\|\|\|\|\|\|[\|\|\|\|\|\|\|\|\|\|\|\|]PAM | NA | Intergenic | NA | NA |
| T25409 | CTGCCGCCGCGGCTGTCCCTCGG | 1000 | | | | | | | | | |
| Oligo fwd | CACCCTGCCGCCGCGGCTGTCCCT | | | | | | | | | | |
| Oligo rev | AAACAGGGACAGCCGCGGCGGCAG | | | | | | | | | | |
| Chromosome | start | end | strand | MM | target_seq | PAM | alignment | distance | position | gene name | gene id |
| GU305914.1 | 113972 | 113994 | - | 0 | CTGCCGCCGCGGCTGTCCCT | CGG | \|\|\|\|\|\|\|\|[\|\|\|\|\|\|\|\|\|\|\|\|]PAM | NA | Intergenic | NA | NA |
| T25410 | CTGTCCCTCGGCCTCCTCGCCGG | 1000 | | | | | | | | | |
| Oligo fwd | CACCCTGTCCCTCGGCCTCCTCGC | | | | | | | | | | |
| Oligo rev | AAACGCGAGGAGGCCGAGGGACAG | | | | | | | | | | |
| Chromosome | start | end | strand | MM | target_seq | PAM | alignment | distance | position | gene name | gene id |
| GU305914.1 | 113960 | 113982 | - | 0 | CTGTCCCTCGGCCTCCTCGC | CGG | \|\|\|\|\|\|\|\|[\|\|\|\|\|\|\|\|\|\|\|\|]PAM | NA | Intergenic | NA | NA |
| T25411 | CTCCTCGCCGGAGTTCTCCGAGG | 1000 | | | | | | | | | |
| Oligo fwd | CACCCTCCTCGCCGGAGTTCTCCG | | | | | | | | | | |
| Oligo rev | AAACCGGAGAACTCCGGCGAGGAG | | | | | | | | | | |
| Chromosome | start | end | strand | MM | target_seq | PAM | alignment | distance | position | gene name | gene id |
| GU305914.1 | 113948 | 113970 | - | 0 | CTCCTCGCCGGAGTTCTCCG | AGG | \|\|\|\|\|\|\|\|[\|\|\|\|\|\|\|\|\|\|\|\|]PAM | NA | Intergenic | NA | NA |
| T25412 | GCCGGAGTTCTCCGAGGTGTCGG | 1000 | | | | | | | | | |
| Oligo fwd | CACCGCCGGAGTTCTCCGAGGTGT | | | | | | | | | | |
| Oligo rev | AAACACACCTCGGAGAACTCCGGC | | | | | | | | | | |
| Chromosome | start | end | strand | MM | target_seq | PAM | alignment | distance | position | gene name | gene id |
| GU305914.1 | 113942 | 113964 | - | 0 | GCCGGAGTTCTCCGAGGTGT | CGG | \|\|\|\|\|\|\|\|[\|\|\|\|\|\|\|\|\|\|\|\|]PAM | NA | Intergenic | NA | NA |
| T25413 | GTTCTCCGAGGTGTCGGCGTCGG | 1000 | | | | | | | | | |
| Oligo fwd | CACCGTTCTCCGAGGTGTCGGCGT | | | | | | | | | | |
| Oligo rev | AAACACGCCGACACCTCGGAGAAC | | | | | | | | | | |
| Chromosome | start | end | strand | MM | target_seq | PAM | alignment | distance | position | gene name | gene id |
| GU305914.1 | 113936 | 113958 | - | 0 | GTTCTCCGAGGTGTCGGCGT | CGG | \|\|\|\|\|\|\|\|[\|\|\|\|\|\|\|\|\|\|\|\|]PAM | NA | Intergenic | NA | NA |
| T25414 | CTCCGAGGTGTCGGCGTCGGCGG | 1000 | | | | | | | | | |
| Oligo fwd | CACCCTCCGAGGTGTCGGCGTCGG | | | | | | | | | | |
| Oligo rev | AAACCCGACGCCGACACCTCGGAG | | | | | | | | | | |
| Chromosome | start | end | strand | MM | target_seq | PAM | alignment | distance | position | gene name | gene id |
| GU305914.1 | 113933 | 113955 | - | 0 | CTCCGAGGTGTCGGCGTCGG | CGG | \|\|\|\|\|\|\|\|[\|\|\|\|\|\|\|\|\|\|\|\|]PAM | NA | Intergenic | NA | NA |
| T25415 | CGAGGTGTCGGCGTCGGCGGCGG | 1000 | | | | | | | | | |
| Oligo fwd | CACCCGAGGTGTCGGCGTCGGCGG | | | | | | | | | | |
| Oligo rev | AAACCCGCCGACGCCGACACCTCG | | | | | | | | | | |
| Chromosome | start | end | strand | MM | target_seq | PAM | alignment | distance | position | gene name | gene id |
| GU305914.1 | 113930 | 113952 | - | 0 | CGAGGTGTCGGCGTCGGCGG | CGG | \|\|\|\|\|\|\|\|[\|\|\|\|\|\|\|\|\|\|\|\|]PAM | NA | Intergenic | NA | NA |
| GU305914.1 | 154404 | 154426 | + | 4 | GGCGGTGGCGGCGGCGGCGG | CGG | -\|-\|\|\|\|-[\|\|\|\|\|-\|\|\|\|\|\|]PAM | NA | Intergenic | NA | NA |
| GU305914.1 | 58126 | 58148 | - | 4 | GGCGGTGGCGGCGGCGGCGG | CGG | -\|-\|\|\|\|-[\|\|\|\|\|-\|\|\|\|\|\|]PAM | NA | Intergenic | NA | NA |
| GU305914.1 | 90321 | 90343 | + | 4 | CAAGAGGTCGGCGTCGGCGA | AGG | \|-\|\|--\|\|[\|\|\|\|\|\|\|\|\|\|\|-]PAM | NA | Intergenic | NA | NA |
| T25416 | GTCGGCGTCGGCGGCGGCAGCGG | 1000 | | | | | | | | | |
| Oligo fwd | CACCGTCGGCGTCGGCGGCGGCAG | | | | | | | | | | |
| Oligo rev | AAACCTGCCGCCGCCGACGCCGAC | | | | | | | | | | |
| Chromosome | start | end | strand | MM | target_seq | PAM | alignment | distance | position | gene name | gene id |
| GU305914.1 | 113924 | 113946 | - | 0 | GTCGGCGTCGGCGGCGGCAG | CGG | \|\|\|\|\|\|\|\|[\|\|\|\|\|\|\|\|\|\|\|\|]PAM | NA | Intergenic | NA | NA |
| GU305914.1 | 30006 | 30028 | + | 2 | GGCGGCGGCGGCGGCGGCAG | TGG | \|-\|\|\|\|\|-[\|\|\|\|\|\|\|\|\|\|\|\|]PAM | NA | Intergenic | NA | NA |
| GU305914.1 | 149476 | 149498 | - | 2 | GGCGGCGGCGGCGGCGGCAG | CGG | \|-\|\|\|\|\|-[\|\|\|\|\|\|\|\|\|\|\|\|]PAM | NA | Intergenic | NA | NA |
| GU305914.1 | 163902 | 163924 | + | 3 | AGCGGCGGCGGCGGCGGCAG | CGG | --\|\|\|\|\|-[\|\|\|\|\|\|\|\|\|\|\|\|]PAM | NA | Intergenic | NA | NA |
| GU305914.1 | 166410 | 166432 | + | 4 | GACGTCGGCGCCGGCGGCAG | CGG | \|-\|\|-\|\|-[\|\|-\|\|\|\|\|\|\|\|\|]PAM | NA | Intergenic | NA | NA |
| GU305914.1 | 128981 | 129003 | + | 4 | GCAGCCGTCGGCAGCGGCAG | CGG | \|--\|-\|\|\|[\|\|\|\|-\|\|\|\|\|\|\|]PAM | NA | Intergenic | NA | NA |
| GU305914.1 | 163908 | 163930 | + | 3 | GGCGGCGGCGGCAGCGGCAG | CGG | \|-\|\|\|\|\|-[\|\|\|\|-\|\|\|\|\|\|\|]PAM | NA | Intergenic | NA | NA |
| GU305914.1 | 168829 | 168851 | + | 4 | AGCGGCGGCGGCGGCGACAG | CGG | --\|\|\|\|\|-[\|\|\|\|\|\|\|\|-\|\|\|]PAM | NA | Intergenic | NA | NA |
| GU305914.1 | 154407 | 154429 | + | 4 | GGTGGCGGCGGCGGCGGCGG | CGG | \|--\|\|\|\|-[\|\|\|\|\|\|\|\|\|\|-\|]PAM | NA | Intergenic | NA | NA |

<table>
<tr><td>GU305914.1</td><td>149485</td><td>149507</td><td>-</td><td>4</td><td>GCAGGCGACGGCGGCGGCGG</td><td>CGG</td><td>|--||||-[||||||||||-|]PAM</td><td>NA</td><td>Intergenic</td><td>NA</td><td>NA</td></tr>
<tr><td>GU305914.1</td><td>58123</td><td>58145</td><td>-</td><td>4</td><td>GGTGGCGGCGGCGGCGGCGG</td><td>AGG</td><td>|--||||-[||||||||||-|]PAM</td><td>NA</td><td>Intergenic</td><td>NA</td><td>NA</td></tr>
<tr><td>GU305914.1</td><td>149482</td><td>149504</td><td>-</td><td>4</td><td>GGCGACGGCGGCGGCGGCGG</td><td>CGG</td><td>|-||-||-[||||||||||-|]PAM</td><td>NA</td><td>Intergenic</td><td>NA</td><td>NA</td></tr>
<tr><td>GU305914.1</td><td>154404</td><td>154426</td><td>+</td><td>4</td><td>GGCGGTGGCGGCGGCGGCGG</td><td>CGG</td><td>|-|||-|-[||||||||||-|]PAM</td><td>NA</td><td>Intergenic</td><td>NA</td><td>NA</td></tr>
<tr><td>GU305914.1</td><td>58126</td><td>58148</td><td>-</td><td>4</td><td>GGCGGTGGCGGCGGCGGCGG</td><td>CGG</td><td>|-|||-|-[||||||||||-|]PAM</td><td>NA</td><td>Intergenic</td><td>NA</td><td>NA</td></tr>
<tr><td>GU305914.1</td><td>48159</td><td>48181</td><td>+</td><td>4</td><td>GGCGGCGGTGGCGGCGGCGG</td><td>TGG</td><td>|-|||||-[-|||||||||-|]PAM</td><td>NA</td><td>Intergenic</td><td>NA</td><td>NA</td></tr>
<tr><td>GU305914.1</td><td>48156</td><td>48178</td><td>+</td><td>4</td><td>GGCGGCGGCGGTGGCGGCGG</td><td>CGG</td><td>|-|||||-[|||-||||||-|]PAM</td><td>NA</td><td>Intergenic</td><td>NA</td><td>NA</td></tr>
<tr><td></td><td></td><td></td><td></td><td></td><td></td><td></td><td></td><td></td><td></td><td></td><td></td></tr>
<tr><td>T25417</td><td>GTCGGCGGCGGCAGCGGCGACGG</td><td>1000</td><td></td><td></td><td></td><td></td><td></td><td></td><td></td><td></td><td></td></tr>
<tr><td>Oligo fwd</td><td>CACCGTCGGCGGCGGCAGCGGCGA</td><td></td><td></td><td></td><td></td><td></td><td></td><td></td><td></td><td></td><td></td></tr>
<tr><td>Oligo rev</td><td>AAACTCGCCGCTGCCGCCGCCGAC</td><td></td><td></td><td></td><td></td><td></td><td></td><td></td><td></td><td></td><td></td></tr>
<tr><td>Chromosome</td><td>start</td><td>end</td><td>strand</td><td>MM</td><td>target_seq</td><td>PAM</td><td>alignment</td><td>distance</td><td>position</td><td>gene name</td><td>gene id</td></tr>
<tr><td>GU305914.1</td><td>113918</td><td>113940</td><td>-</td><td>0</td><td>GTCGGCGGCGGCAGCGGCGA</td><td>CGG</td><td>||||||||[||||||||||||]PAM</td><td>NA</td><td>Intergenic</td><td>NA</td><td>NA</td></tr>
<tr><td>GU305914.1</td><td>24551</td><td>24573</td><td>+</td><td>3</td><td>GATGGCGGCGGCAGCGACGA</td><td>CGG</td><td>|--|||||[||||||||-|||]PAM</td><td>NA</td><td>Intergenic</td><td>NA</td><td>NA</td></tr>
<tr><td>GU305914.1</td><td>151362</td><td>151384</td><td>+</td><td>4</td><td>GGCGACGACGGCAGCAGCGA</td><td>AGG</td><td>|-||-||-[|||||||-||||]PAM</td><td>NA</td><td>Intergenic</td><td>NA</td><td>NA</td></tr>
<tr><td>GU305914.1</td><td>168817</td><td>168839</td><td>+</td><td>4</td><td>GGCGGTGGTGGCAGCGGCGG</td><td>CGG</td><td>|-|||-||[-||||||||||-]PAM</td><td>NA</td><td>Intergenic</td><td>NA</td><td>NA</td></tr>
<tr><td>GU305914.1</td><td>109443</td><td>109465</td><td>+</td><td>3</td><td>GACGGCGGCGGTAGCGGCGT</td><td>CGG</td><td>|-||||||[|||-|||||||-]PAM</td><td>NA</td><td>Intergenic</td><td>NA</td><td>NA</td></tr>
<tr><td>GU305914.1</td><td>128721</td><td>128743</td><td>-</td><td>4</td><td>GGCGGCAGCAGCAGCGGCGC</td><td>TGG</td><td>|-||||-|[|-|||||||||-]PAM</td><td>NA</td><td>Intergenic</td><td>NA</td><td>NA</td></tr>
<tr><td>GU305914.1</td><td>154407</td><td>154429</td><td>+</td><td>4</td><td>GGTGGCGGCGGCGGCGGCGG</td><td>CGG</td><td>|--|||||[||||-||||||-]PAM</td><td>NA</td><td>Intergenic</td><td>NA</td><td>NA</td></tr>
<tr><td>GU305914.1</td><td>58123</td><td>58145</td><td>-</td><td>4</td><td>GGTGGCGGCGGCGGCGGCGG</td><td>AGG</td><td>|--|||||[||||-||||||-]PAM</td><td>NA</td><td>Intergenic</td><td>NA</td><td>NA</td></tr>
<tr><td>GU305914.1</td><td>149482</td><td>149504</td><td>-</td><td>4</td><td>GGCGACGGCGGCGGCGGCGG</td><td>CGG</td><td>|-||-|||[||||-||||||-]PAM</td><td>NA</td><td>Intergenic</td><td>NA</td><td>NA</td></tr>
<tr><td>GU305914.1</td><td>154404</td><td>154426</td><td>+</td><td>4</td><td>GGCGGTGGCGGCGGCGGCGG</td><td>CGG</td><td>|-|||-||[||||-||||||-]PAM</td><td>NA</td><td>Intergenic</td><td>NA</td><td>NA</td></tr>
<tr><td>GU305914.1</td><td>58126</td><td>58148</td><td>-</td><td>4</td><td>GGCGGTGGCGGCGGCGGCGG</td><td>CGG</td><td>|-|||-||[||||-||||||-]PAM</td><td>NA</td><td>Intergenic</td><td>NA</td><td>NA</td></tr>
<tr><td>GU305914.1</td><td>163908</td><td>163930</td><td>+</td><td>3</td><td>GGCGGCGGCGGCAGCGGCAG</td><td>CGG</td><td>|-||||||[||||||||||--]PAM</td><td>NA</td><td>Intergenic</td><td>NA</td><td>NA</td></tr>
<tr><td></td><td></td><td></td><td></td><td></td><td></td><td></td><td></td><td></td><td></td><td></td><td></td></tr>
<tr><td>T25418</td><td>GGCGGCAGCGGCGACGGCGTCGG</td><td>1000</td><td></td><td></td><td></td><td></td><td></td><td></td><td></td><td></td><td></td></tr>
<tr><td>Oligo fwd</td><td>CACCGGCGGCAGCGGCGACGGCGT</td><td></td><td></td><td></td><td></td><td></td><td></td><td></td><td></td><td></td><td></td></tr>
<tr><td>Oligo rev</td><td>AAACACGCCGTCGCCGCTGCCGCC</td><td></td><td></td><td></td><td></td><td></td><td></td><td></td><td></td><td></td><td></td></tr>
<tr><td>Chromosome</td><td>start</td><td>end</td><td>strand</td><td>MM</td><td>target_seq</td><td>PAM</td><td>alignment</td><td>distance</td><td>position</td><td>gene name</td><td>gene id</td></tr>
<tr><td>GU305914.1</td><td>113912</td><td>113934</td><td>-</td><td>0</td><td>GGCGGCAGCGGCGACGGCGT</td><td>CGG</td><td>||||||||[||||||||||||]PAM</td><td>NA</td><td>Intergenic</td><td>NA</td><td>NA</td></tr>
<tr><td>GU305914.1</td><td>198008</td><td>198030</td><td>-</td><td>4</td><td>CACGGCGGCCGCGACGGCGT</td><td>CGG</td><td>--||||-|[|-||||||||||]PAM</td><td>NA</td><td>Intergenic</td><td>NA</td><td>NA</td></tr>
<tr><td>GU305914.1</td><td>56773</td><td>56795</td><td>+</td><td>4</td><td>CGCCGCAGCGGCGGCGGCGG</td><td>CGG</td><td>-||-||||[|||||-|||||-]PAM</td><td>NA</td><td>Intergenic</td><td>NA</td><td>NA</td></tr>
<tr><td>GU305914.1</td><td>154407</td><td>154429</td><td>+</td><td>4</td><td>GGTGGCGGCGGCGGCGGCGG</td><td>CGG</td><td>||-|||-|[|||||-|||||-]PAM</td><td>NA</td><td>Intergenic</td><td>NA</td><td>NA</td></tr>
<tr><td>GU305914.1</td><td>58123</td><td>58145</td><td>-</td><td>4</td><td>GGTGGCGGCGGCGGCGGCGG</td><td>AGG</td><td>||-|||-|[|||||-|||||-]PAM</td><td>NA</td><td>Intergenic</td><td>NA</td><td>NA</td></tr>
<tr><td>GU305914.1</td><td>149482</td><td>149504</td><td>-</td><td>4</td><td>GGCGACGGCGGCGGCGGCGG</td><td>CGG</td><td>||||-|-|[|||||-|||||-]PAM</td><td>NA</td><td>Intergenic</td><td>NA</td><td>NA</td></tr>
<tr><td>GU305914.1</td><td>154404</td><td>154426</td><td>+</td><td>4</td><td>GGCGGTGGCGGCGGCGGCGG</td><td>CGG</td><td>|||||--|[|||||-|||||-]PAM</td><td>NA</td><td>Intergenic</td><td>NA</td><td>NA</td></tr>
<tr><td>GU305914.1</td><td>58126</td><td>58148</td><td>-</td><td>4</td><td>GGCGGTGGCGGCGGCGGCGG</td><td>CGG</td><td>|||||--|[|||||-|||||-]PAM</td><td>NA</td><td>Intergenic</td><td>NA</td><td>NA</td></tr>
<tr><td>GU305914.1</td><td>110083</td><td>110105</td><td>-</td><td>4</td><td>CGCGGCCGCGGCGACGCCGG</td><td>TGG</td><td>-|||||-|[||||||||-||-]PAM</td><td>NA</td><td>Intergenic</td><td>NA</td><td>NA</td></tr>
<tr><td></td><td></td><td></td><td></td><td></td><td></td><td></td><td></td><td></td><td></td><td></td><td></td></tr>
<tr><td>T25419</td><td>GCGGCAGCGGCGACGGCGTCGGG</td><td>1000</td><td></td><td></td><td></td><td></td><td></td><td></td><td></td><td></td><td></td></tr>
<tr><td>Oligo fwd</td><td>CACCGCGGCAGCGGCGACGGCGTC</td><td></td><td></td><td></td><td></td><td></td><td></td><td></td><td></td><td></td><td></td></tr>
<tr><td>Oligo rev</td><td>AAACGACGCCGTCGCCGCTGCCGC</td><td></td><td></td><td></td><td></td><td></td><td></td><td></td><td></td><td></td><td></td></tr>
<tr><td>Chromosome</td><td>start</td><td>end</td><td>strand</td><td>MM</td><td>target_seq</td><td>PAM</td><td>alignment</td><td>distance</td><td>position</td><td>gene name</td><td>gene id</td></tr>
<tr><td>GU305914.1</td><td>113911</td><td>113933</td><td>-</td><td>0</td><td>GCGGCAGCGGCGACGGCGTC</td><td>GGG</td><td>||||||||[||||||||||||]PAM</td><td>NA</td><td>Intergenic</td><td>NA</td><td>NA</td></tr>
<tr><td>GU305914.1</td><td>117386</td><td>117408</td><td>+</td><td>4</td><td>CCGACAGCGCCGACGGGGTC</td><td>CGG</td><td>-||-||||[|-||||||-|||]PAM</td><td>NA</td><td>Intergenic</td><td>NA</td><td>NA</td></tr>
<tr><td></td><td></td><td></td><td></td><td></td><td></td><td></td><td></td><td></td><td></td><td></td><td></td></tr>
<tr><td>T25420</td><td>GACGGCGTCGGGAGAGAAGCTGG</td><td>1000</td><td></td><td></td><td></td><td></td><td></td><td></td><td></td><td></td><td></td></tr>
<tr><td>Oligo fwd</td><td>CACCGACGGCGTCGGGAGAGAAGC</td><td></td><td></td><td></td><td></td><td></td><td></td><td></td><td></td><td></td><td></td></tr>
<tr><td>Oligo rev</td><td>AAACGCTTCTCTCCCGACGCCGTC</td><td></td><td></td><td></td><td></td><td></td><td></td><td></td><td></td><td></td><td></td></tr>
<tr><td>Chromosome</td><td>start</td><td>end</td><td>strand</td><td>MM</td><td>target_seq</td><td>PAM</td><td>alignment</td><td>distance</td><td>position</td><td>gene name</td><td>gene id</td></tr>
<tr><td>GU305914.1</td><td>113900</td><td>113922</td><td>-</td><td>0</td><td>GACGGCGTCGGGAGAGAAGC</td><td>TGG</td><td>||||||||[||||||||||||]PAM</td><td>NA</td><td>Intergenic</td><td>NA</td><td>NA</td></tr>
<tr><td></td><td></td><td></td><td></td><td></td><td></td><td></td><td></td><td></td><td></td><td></td><td></td></tr>
<tr><td>T25421</td><td>GAGAGAAGCTGGAGAGCGAGAGG</td><td>1000</td><td></td><td></td><td></td><td></td><td></td><td></td><td></td><td></td><td></td></tr>
<tr><td>Oligo fwd</td><td>CACCGAGAGAAGCTGGAGAGCGAG</td><td></td><td></td><td></td><td></td><td></td><td></td><td></td><td></td><td></td><td></td></tr>
<tr><td>Oligo rev</td><td>AAACCTCGCTCTCCAGCTTCTCTC</td><td></td><td></td><td></td><td></td><td></td><td></td><td></td><td></td><td></td><td></td></tr>
<tr><td>Chromosome</td><td>start</td><td>end</td><td>strand</td><td>MM</td><td>target_seq</td><td>PAM</td><td>alignment</td><td>distance</td><td>position</td><td>gene name</td><td>gene id</td></tr>
<tr><td>GU305914.1</td><td>113889</td><td>113911</td><td>-</td><td>0</td><td>GAGAGAAGCTGGAGAGCGAG</td><td>AGG</td><td>||||||||[||||||||||||]PAM</td><td>NA</td><td>Intergenic</td><td>NA</td><td>NA</td></tr>
<tr><td></td><td></td><td></td><td></td><td></td><td></td><td></td><td></td><td></td><td></td><td></td><td></td></tr>
<tr><td>T25422</td><td>AGAGAAGCTGGAGAGCGAGAGGG</td><td>1000</td><td></td><td></td><td></td><td></td><td></td><td></td><td></td><td></td><td></td></tr>
<tr><td>Oligo fwd</td><td>CACCAGAGAAGCTGGAGAGCGAGA</td><td></td><td></td><td></td><td></td><td></td><td></td><td></td><td></td><td></td><td></td></tr>
<tr><td>Oligo rev</td><td>AAACTCTCGCTCTCCAGCTTCTCT</td><td></td><td></td><td></td><td></td><td></td><td></td><td></td><td></td><td></td><td></td></tr>
<tr><td>Chromosome</td><td>start</td><td>end</td><td>strand</td><td>MM</td><td>target_seq</td><td>PAM</td><td>alignment</td><td>distance</td><td>position</td><td>gene name</td><td>gene id</td></tr>
<tr><td>GU305914.1</td><td>113888</td><td>113910</td><td>-</td><td>0</td><td>AGAGAAGCTGGAGAGCGAGA</td><td>GGG</td><td>||||||||[||||||||||||]PAM</td><td>NA</td><td>Intergenic</td><td>NA</td><td>NA</td></tr>
<tr><td></td><td></td><td></td><td></td><td></td><td></td><td></td><td></td><td></td><td></td><td></td><td></td></tr>
<tr><td>T25423</td><td>GAAGCTGGAGAGCGAGAGGGCGG</td><td>1000</td><td></td><td></td><td></td><td></td><td></td><td></td><td></td><td></td><td></td></tr>
<tr><td>Oligo fwd</td><td>CACCGAAGCTGGAGAGCGAGAGGG</td><td></td><td></td><td></td><td></td><td></td><td></td><td></td><td></td><td></td><td></td></tr>
<tr><td>Oligo rev</td><td>AAACCCCTCTCGCTCTCCAGCTTC</td><td></td><td></td><td></td><td></td><td></td><td></td><td></td><td></td><td></td><td></td></tr>
<tr><td>Chromosome</td><td>start</td><td>end</td><td>strand</td><td>MM</td><td>target_seq</td><td>PAM</td><td>alignment</td><td>distance</td><td>position</td><td>gene name</td><td>gene id</td></tr>
<tr><td>GU305914.1</td><td>113885</td><td>113907</td><td>-</td><td>0</td><td>GAAGCTGGAGAGCGAGAGGG</td><td>CGG</td><td>||||||||[||||||||||||]PAM</td><td>NA</td><td>Intergenic</td><td>NA</td><td>NA</td></tr>
<tr><td></td><td></td><td></td><td></td><td></td><td></td><td></td><td></td><td></td><td></td><td></td><td></td></tr>
<tr><td>T25424</td><td>AAGCTGGAGAGCGAGAGGGCGGG</td><td>1000</td><td></td><td></td><td></td><td></td><td></td><td></td><td></td><td></td><td></td></tr>
<tr><td>Oligo fwd</td><td>CACCAAGCTGGAGAGCGAGAGGGC</td><td></td><td></td><td></td><td></td><td></td><td></td><td></td><td></td><td></td><td></td></tr>
<tr><td>Oligo rev</td><td>AAACGCCCTCTCGCTCTCCAGCTT</td><td></td><td></td><td></td><td></td><td></td><td></td><td></td><td></td><td></td><td></td></tr>
<tr><td>Chromosome</td><td>start</td><td>end</td><td>strand</td><td>MM</td><td>target_seq</td><td>PAM</td><td>alignment</td><td>distance</td><td>position</td><td>gene name</td><td>gene id</td></tr>
<tr><td>GU305914.1</td><td>113884</td><td>113906</td><td>-</td><td>0</td><td>AAGCTGGAGAGCGAGAGGGC</td><td>GGG</td><td>||||||||[||||||||||||]PAM</td><td>NA</td><td>Intergenic</td><td>NA</td><td>NA</td></tr>
<tr><td></td><td></td><td></td><td></td><td></td><td></td><td></td><td></td><td></td><td></td><td></td><td></td></tr>
<tr><td>T25425</td><td>AGCTGGAGAGCGAGAGGGCGGGG</td><td>1000</td><td></td><td></td><td></td><td></td><td></td><td></td><td></td><td></td><td></td></tr>
<tr><td>Oligo fwd</td><td>CACCAGCTGGAGAGCGAGAGGGCG</td><td></td><td></td><td></td><td></td><td></td><td></td><td></td><td></td><td></td><td></td></tr>
<tr><td>Oligo rev</td><td>AAACCGCCCTCTCGCTCTCCAGCT</td><td></td><td></td><td></td><td></td><td></td><td></td><td></td><td></td><td></td><td></td></tr>
<tr><td>Chromosome</td><td>start</td><td>end</td><td>strand</td><td>MM</td><td>target_seq</td><td>PAM</td><td>alignment</td><td>distance</td><td>position</td><td>gene name</td><td>gene id</td></tr>
<tr><td>GU305914.1</td><td>113883</td><td>113905</td><td>-</td><td>0</td><td>AGCTGGAGAGCGAGAGGGCG</td><td>GGG</td><td>||||||||[||||||||||||]PAM</td><td>NA</td><td>Intergenic</td><td>NA</td><td>NA</td></tr>
<tr><td></td><td></td><td></td><td></td><td></td><td></td><td></td><td></td><td></td><td></td><td></td><td></td></tr>
<tr><td>T25426</td><td>GAGCGAGAGGGCGGGGAACCCGG</td><td>1000</td><td></td><td></td><td></td><td></td><td></td><td></td><td></td><td></td><td></td></tr>
<tr><td>Oligo fwd</td><td>CACCGAGCGAGAGGGCGGGGAACC</td><td></td><td></td><td></td><td></td><td></td><td></td><td></td><td></td><td></td><td></td></tr>
<tr><td>Oligo rev</td><td>AAACGGTTCCCCGCCCTCTCGCTC</td><td></td><td></td><td></td><td></td><td></td><td></td><td></td><td></td><td></td><td></td></tr>
<tr><td>Chromosome</td><td>start</td><td>end</td><td>strand</td><td>MM</td><td>target_seq</td><td>PAM</td><td>alignment</td><td>distance</td><td>position</td><td>gene name</td><td>gene id</td></tr>
<tr><td>GU305914.1</td><td>113876</td><td>113898</td><td>-</td><td>0</td><td>GAGCGAGAGGGCGGGGAACC</td><td>CGG</td><td>||||||||[||||||||||||]PAM</td><td>NA</td><td>Intergenic</td><td>NA</td><td>NA</td></tr>
<tr><td></td><td></td><td></td><td></td><td></td><td></td><td></td><td></td><td></td><td></td><td></td><td></td></tr>
<tr><td>T25427</td><td>GGCGCTGAGATACTCGACGACGG</td><td>1000</td><td></td><td></td><td></td><td></td><td></td><td></td><td></td><td></td><td></td></tr>
</table>

<table>
<tr><td>Oligo fwd</td><td>CACCGGCGCTGAGATACTCGACGA</td><td></td><td></td><td></td><td></td><td></td><td></td><td></td><td></td><td></td><td></td></tr>
<tr><td>Oligo rev</td><td>AAACTCGTCGAGTATCTCAGCGCC</td><td></td><td></td><td></td><td></td><td></td><td></td><td></td><td></td><td></td><td></td></tr>
<tr><td>Chromosome</td><td>start</td><td>end</td><td>strand</td><td>MM</td><td>target_seq</td><td>PAM</td><td>alignment</td><td>distance</td><td>position</td><td>gene name</td><td>gene id</td></tr>
<tr><td>GU305914.1</td><td>113855</td><td>113877</td><td>-</td><td>0</td><td>GGCGCTGAGATACTCGACGA</td><td>CGG</td><td>||||||||[||||||||||||]PAM</td><td>NA</td><td>Intergenic</td><td>NA</td><td>NA</td></tr>
<tr><td></td><td></td><td></td><td></td><td></td><td></td><td></td><td></td><td></td><td></td><td></td><td></td></tr>
<tr><td>T25428</td><td>GCGCTGAGATACTCGACGACGGG</td><td>1000</td><td></td><td></td><td></td><td></td><td></td><td></td><td></td><td></td><td></td></tr>
<tr><td>Oligo fwd</td><td>CACCGCGCTGAGATACTCGACGAC</td><td></td><td></td><td></td><td></td><td></td><td></td><td></td><td></td><td></td><td></td></tr>
<tr><td>Oligo rev</td><td>AAACGTCGTCGAGTATCTCAGCGC</td><td></td><td></td><td></td><td></td><td></td><td></td><td></td><td></td><td></td><td></td></tr>
<tr><td>Chromosome</td><td>start</td><td>end</td><td>strand</td><td>MM</td><td>target_seq</td><td>PAM</td><td>alignment</td><td>distance</td><td>position</td><td>gene name</td><td>gene id</td></tr>
<tr><td>GU305914.1</td><td>113854</td><td>113876</td><td>-</td><td>0</td><td>GCGCTGAGATACTCGACGAC</td><td>GGG</td><td>||||||||[||||||||||||]PAM</td><td>NA</td><td>Intergenic</td><td>NA</td><td>NA</td></tr>
<tr><td></td><td></td><td></td><td></td><td></td><td></td><td></td><td></td><td></td><td></td><td></td><td></td></tr>
<tr><td>T25429</td><td>CGCTGAGATACTCGACGACGGGG</td><td>1000</td><td></td><td></td><td></td><td></td><td></td><td></td><td></td><td></td><td></td></tr>
<tr><td>Oligo fwd</td><td>CACCCGCTGAGATACTCGACGACG</td><td></td><td></td><td></td><td></td><td></td><td></td><td></td><td></td><td></td><td></td></tr>
<tr><td>Oligo rev</td><td>AAACCGTCGTCGAGTATCTCAGCG</td><td></td><td></td><td></td><td></td><td></td><td></td><td></td><td></td><td></td><td></td></tr>
<tr><td>Chromosome</td><td>start</td><td>end</td><td>strand</td><td>MM</td><td>target_seq</td><td>PAM</td><td>alignment</td><td>distance</td><td>position</td><td>gene name</td><td>gene id</td></tr>
<tr><td>GU305914.1</td><td>113853</td><td>113875</td><td>-</td><td>0</td><td>CGCTGAGATACTCGACGACG</td><td>GGG</td><td>||||||||[||||||||||||]PAM</td><td>NA</td><td>Intergenic</td><td>NA</td><td>NA</td></tr>
<tr><td></td><td></td><td></td><td></td><td></td><td></td><td></td><td></td><td></td><td></td><td></td><td></td></tr>
<tr><td>T25430</td><td>GGGGTCCGCTTCCAGCCCCTTGG</td><td>1000</td><td></td><td></td><td></td><td></td><td></td><td></td><td></td><td></td><td></td></tr>
<tr><td>Oligo fwd</td><td>CACCGGGGTCCGCTTCCAGCCCCT</td><td></td><td></td><td></td><td></td><td></td><td></td><td></td><td></td><td></td><td></td></tr>
<tr><td>Oligo rev</td><td>AAACAGGGGCTGGAAGCGGACCCC</td><td></td><td></td><td></td><td></td><td></td><td></td><td></td><td></td><td></td><td></td></tr>
<tr><td>Chromosome</td><td>start</td><td>end</td><td>strand</td><td>MM</td><td>target_seq</td><td>PAM</td><td>alignment</td><td>distance</td><td>position</td><td>gene name</td><td>gene id</td></tr>
<tr><td>GU305914.1</td><td>113834</td><td>113856</td><td>-</td><td>0</td><td>GGGGTCCGCTTCCAGCCCCT</td><td>TGG</td><td>||||||||[||||||||||||]PAM</td><td>NA</td><td>Intergenic</td><td>NA</td><td>NA</td></tr>
<tr><td>GU305914.1</td><td>70764</td><td>70786</td><td>-</td><td>4</td><td>GGTGTCCACGTCCAGGCCCT</td><td>CGG</td><td>||-||||-[|-|||||-||||]PAM</td><td>NA</td><td>Intergenic</td><td>NA</td><td>NA</td></tr>
<tr><td></td><td></td><td></td><td></td><td></td><td></td><td></td><td></td><td></td><td></td><td></td><td></td></tr>
<tr><td>T25431</td><td>GTCCGCTTCCAGCCCCTTGGCGG</td><td>1000</td><td></td><td></td><td></td><td></td><td></td><td></td><td></td><td></td><td></td></tr>
<tr><td>Oligo fwd</td><td>CACCGTCCGCTTCCAGCCCCTTGG</td><td></td><td></td><td></td><td></td><td></td><td></td><td></td><td></td><td></td><td></td></tr>
<tr><td>Oligo rev</td><td>AAACCCAAGGGGCTGGAAGCGGAC</td><td></td><td></td><td></td><td></td><td></td><td></td><td></td><td></td><td></td><td></td></tr>
<tr><td>Chromosome</td><td>start</td><td>end</td><td>strand</td><td>MM</td><td>target_seq</td><td>PAM</td><td>alignment</td><td>distance</td><td>position</td><td>gene name</td><td>gene id</td></tr>
<tr><td>GU305914.1</td><td>113831</td><td>113853</td><td>-</td><td>0</td><td>GTCCGCTTCCAGCCCCTTGG</td><td>CGG</td><td>||||||||[||||||||||||]PAM</td><td>NA</td><td>Intergenic</td><td>NA</td><td>NA</td></tr>
<tr><td></td><td></td><td></td><td></td><td></td><td></td><td></td><td></td><td></td><td></td><td></td><td></td></tr>
<tr><td>T25432</td><td>TCCGCTTCCAGCCCCTTGGCGGG</td><td>1000</td><td></td><td></td><td></td><td></td><td></td><td></td><td></td><td></td><td></td></tr>
<tr><td>Oligo fwd</td><td>CACCTCCGCTTCCAGCCCCTTGGC</td><td></td><td></td><td></td><td></td><td></td><td></td><td></td><td></td><td></td><td></td></tr>
<tr><td>Oligo rev</td><td>AAACGCCAAGGGGCTGGAAGCGGA</td><td></td><td></td><td></td><td></td><td></td><td></td><td></td><td></td><td></td><td></td></tr>
<tr><td>Chromosome</td><td>start</td><td>end</td><td>strand</td><td>MM</td><td>target_seq</td><td>PAM</td><td>alignment</td><td>distance</td><td>position</td><td>gene name</td><td>gene id</td></tr>
<tr><td>GU305914.1</td><td>113830</td><td>113852</td><td>-</td><td>0</td><td>TCCGCTTCCAGCCCCTTGGC</td><td>GGG</td><td>||||||||[||||||||||||]PAM</td><td>NA</td><td>Intergenic</td><td>NA</td><td>NA</td></tr>
<tr><td></td><td></td><td></td><td></td><td></td><td></td><td></td><td></td><td></td><td></td><td></td><td></td></tr>
<tr><td>T25433</td><td>CCAGCCCCTTGGCGGGACCGCGG</td><td>1000</td><td></td><td></td><td></td><td></td><td></td><td></td><td></td><td></td><td></td></tr>
<tr><td>Oligo fwd</td><td>CACCCCAGCCCCTTGGCGGGACCG</td><td></td><td></td><td></td><td></td><td></td><td></td><td></td><td></td><td></td><td></td></tr>
<tr><td>Oligo rev</td><td>AAACCGGTCCCGCCAAGGGGCTGG</td><td></td><td></td><td></td><td></td><td></td><td></td><td></td><td></td><td></td><td></td></tr>
<tr><td>Chromosome</td><td>start</td><td>end</td><td>strand</td><td>MM</td><td>target_seq</td><td>PAM</td><td>alignment</td><td>distance</td><td>position</td><td>gene name</td><td>gene id</td></tr>
<tr><td>GU305914.1</td><td>113823</td><td>113845</td><td>-</td><td>0</td><td>CCAGCCCCTTGGCGGGACCG</td><td>CGG</td><td>||||||||[||||||||||||]PAM</td><td>NA</td><td>Intergenic</td><td>NA</td><td>NA</td></tr>
<tr><td></td><td></td><td></td><td></td><td></td><td></td><td></td><td></td><td></td><td></td><td></td><td></td></tr>
<tr><td>T25434</td><td>CAGCCCCTTGGCGGGACCGCGGG</td><td>1000</td><td></td><td></td><td></td><td></td><td></td><td></td><td></td><td></td><td></td></tr>
<tr><td>Oligo fwd</td><td>CACCCAGCCCCTTGGCGGGACCGC</td><td></td><td></td><td></td><td></td><td></td><td></td><td></td><td></td><td></td><td></td></tr>
<tr><td>Oligo rev</td><td>AAACGCGGTCCCGCCAAGGGGCTG</td><td></td><td></td><td></td><td></td><td></td><td></td><td></td><td></td><td></td><td></td></tr>
<tr><td>Chromosome</td><td>start</td><td>end</td><td>strand</td><td>MM</td><td>target_seq</td><td>PAM</td><td>alignment</td><td>distance</td><td>position</td><td>gene name</td><td>gene id</td></tr>
<tr><td>GU305914.1</td><td>113822</td><td>113844</td><td>-</td><td>0</td><td>CAGCCCCTTGGCGGGACCGC</td><td>GGG</td><td>||||||||[||||||||||||]PAM</td><td>NA</td><td>Intergenic</td><td>NA</td><td>NA</td></tr>
<tr><td></td><td></td><td></td><td></td><td></td><td></td><td></td><td></td><td></td><td></td><td></td><td></td></tr>
<tr><td>T25435</td><td>GGGACCGCGGGCGACGAGTTTGG</td><td>1000</td><td></td><td></td><td></td><td></td><td></td><td></td><td></td><td></td><td></td></tr>
<tr><td>Oligo fwd</td><td>CACCGGGACCGCGGGCGACGAGTT</td><td></td><td></td><td></td><td></td><td></td><td></td><td></td><td></td><td></td><td></td></tr>
<tr><td>Oligo rev</td><td>AAACAACTCGTCGCCCGCGGTCCC</td><td></td><td></td><td></td><td></td><td></td><td></td><td></td><td></td><td></td><td></td></tr>
<tr><td>Chromosome</td><td>start</td><td>end</td><td>strand</td><td>MM</td><td>target_seq</td><td>PAM</td><td>alignment</td><td>distance</td><td>position</td><td>gene name</td><td>gene id</td></tr>
<tr><td>GU305914.1</td><td>113810</td><td>113832</td><td>-</td><td>0</td><td>GGGACCGCGGGCGACGAGTT</td><td>TGG</td><td>||||||||[||||||||||||]PAM</td><td>NA</td><td>Intergenic</td><td>NA</td><td>NA</td></tr>
<tr><td></td><td></td><td></td><td></td><td></td><td></td><td></td><td></td><td></td><td></td><td></td><td></td></tr>
<tr><td>T25436</td><td>GACGAGTTTGGATTTGCCCGCGG</td><td>1000</td><td></td><td></td><td></td><td></td><td></td><td></td><td></td><td></td><td></td></tr>
<tr><td>Oligo fwd</td><td>CACCGACGAGTTTGGATTTGCCCG</td><td></td><td></td><td></td><td></td><td></td><td></td><td></td><td></td><td></td><td></td></tr>
<tr><td>Oligo rev</td><td>AAACCGGGCAAATCCAAACTCGTC</td><td></td><td></td><td></td><td></td><td></td><td></td><td></td><td></td><td></td><td></td></tr>
<tr><td>Chromosome</td><td>start</td><td>end</td><td>strand</td><td>MM</td><td>target_seq</td><td>PAM</td><td>alignment</td><td>distance</td><td>position</td><td>gene name</td><td>gene id</td></tr>
<tr><td>GU305914.1</td><td>113798</td><td>113820</td><td>-</td><td>0</td><td>GACGAGTTTGGATTTGCCCG</td><td>CGG</td><td>||||||||[||||||||||||]PAM</td><td>NA</td><td>Intergenic</td><td>NA</td><td>NA</td></tr>
<tr><td></td><td></td><td></td><td></td><td></td><td></td><td></td><td></td><td></td><td></td><td></td><td></td></tr>
<tr><td>T25437</td><td>CTTCTCGACGATCCCCAGAAAGG</td><td>1000</td><td></td><td></td><td></td><td></td><td></td><td></td><td></td><td></td><td></td></tr>
<tr><td>Oligo fwd</td><td>CACCCTTCTCGACGATCCCCAGAA</td><td></td><td></td><td></td><td></td><td></td><td></td><td></td><td></td><td></td><td></td></tr>
<tr><td>Oligo rev</td><td>AAACTTCTGGGGATCGTCGAGAAG</td><td></td><td></td><td></td><td></td><td></td><td></td><td></td><td></td><td></td><td></td></tr>
<tr><td>Chromosome</td><td>start</td><td>end</td><td>strand</td><td>MM</td><td>target_seq</td><td>PAM</td><td>alignment</td><td>distance</td><td>position</td><td>gene name</td><td>gene id</td></tr>
<tr><td>GU305914.1</td><td>113774</td><td>113796</td><td>-</td><td>0</td><td>CTTCTCGACGATCCCCAGAA</td><td>AGG</td><td>||||||||[||||||||||||]PAM</td><td>NA</td><td>Intergenic</td><td>NA</td><td>NA</td></tr>
<tr><td></td><td></td><td></td><td></td><td></td><td></td><td></td><td></td><td></td><td></td><td></td><td></td></tr>
<tr><td>T25438</td><td>CTCGACGATCCCCAGAAAGGCGG</td><td>1000</td><td></td><td></td><td></td><td></td><td></td><td></td><td></td><td></td><td></td></tr>
<tr><td>Oligo fwd</td><td>CACCCTCGACGATCCCCAGAAAGG</td><td></td><td></td><td></td><td></td><td></td><td></td><td></td><td></td><td></td><td></td></tr>
<tr><td>Oligo rev</td><td>AAACCCTTTCTGGGGATCGTCGAG</td><td></td><td></td><td></td><td></td><td></td><td></td><td></td><td></td><td></td><td></td></tr>
<tr><td>Chromosome</td><td>start</td><td>end</td><td>strand</td><td>MM</td><td>target_seq</td><td>PAM</td><td>alignment</td><td>distance</td><td>position</td><td>gene name</td><td>gene id</td></tr>
<tr><td>GU305914.1</td><td>113771</td><td>113793</td><td>-</td><td>0</td><td>CTCGACGATCCCCAGAAAGG</td><td>CGG</td><td>||||||||[||||||||||||]PAM</td><td>NA</td><td>Intergenic</td><td>NA</td><td>NA</td></tr>
<tr><td></td><td></td><td></td><td></td><td></td><td></td><td></td><td></td><td></td><td></td><td></td><td></td></tr>
<tr><td>T25439</td><td>TCGACGATCCCCAGAAAGGCGGG</td><td>1000</td><td></td><td></td><td></td><td></td><td></td><td></td><td></td><td></td><td></td></tr>
<tr><td>Oligo fwd</td><td>CACCTCGACGATCCCCAGAAAGGC</td><td></td><td></td><td></td><td></td><td></td><td></td><td></td><td></td><td></td><td></td></tr>
<tr><td>Oligo rev</td><td>AAACGCCTTTCTGGGGATCGTCGA</td><td></td><td></td><td></td><td></td><td></td><td></td><td></td><td></td><td></td><td></td></tr>
<tr><td>Chromosome</td><td>start</td><td>end</td><td>strand</td><td>MM</td><td>target_seq</td><td>PAM</td><td>alignment</td><td>distance</td><td>position</td><td>gene name</td><td>gene id</td></tr>
<tr><td>GU305914.1</td><td>113770</td><td>113792</td><td>-</td><td>0</td><td>TCGACGATCCCCAGAAAGGC</td><td>GGG</td><td>||||||||[||||||||||||]PAM</td><td>NA</td><td>Intergenic</td><td>NA</td><td>NA</td></tr>
<tr><td></td><td></td><td></td><td></td><td></td><td></td><td></td><td></td><td></td><td></td><td></td><td></td></tr>
<tr><td>T25440</td><td>GAAAGGCGGGCGAGCTGAGCCGG</td><td>1000</td><td></td><td></td><td></td><td></td><td></td><td></td><td></td><td></td><td></td></tr>
<tr><td>Oligo fwd</td><td>CACCGAAAGGCGGGCGAGCTGAGC</td><td></td><td></td><td></td><td></td><td></td><td></td><td></td><td></td><td></td><td></td></tr>
<tr><td>Oligo rev</td><td>AAACGCTCAGCTCGCCCGCCTTTC</td><td></td><td></td><td></td><td></td><td></td><td></td><td></td><td></td><td></td><td></td></tr>
<tr><td>Chromosome</td><td>start</td><td>end</td><td>strand</td><td>MM</td><td>target_seq</td><td>PAM</td><td>alignment</td><td>distance</td><td>position</td><td>gene name</td><td>gene id</td></tr>
<tr><td>GU305914.1</td><td>113757</td><td>113779</td><td>-</td><td>0</td><td>GAAAGGCGGGCGAGCTGAGC</td><td>CGG</td><td>||||||||[||||||||||||]PAM</td><td>NA</td><td>Intergenic</td><td>NA</td><td>NA</td></tr>
<tr><td></td><td></td><td></td><td></td><td></td><td></td><td></td><td></td><td></td><td></td><td></td><td></td></tr>
<tr><td>T25441</td><td>CCGGCCCAGACAGAAGAGTCCGG</td><td>1000</td><td></td><td></td><td></td><td></td><td></td><td></td><td></td><td></td><td></td></tr>
<tr><td>Oligo fwd</td><td>CACCCCGGCCCAGACAGAAGAGTC</td><td></td><td></td><td></td><td></td><td></td><td></td><td></td><td></td><td></td><td></td></tr>
<tr><td>Oligo rev</td><td>AAACGACTCTTCTGTCTGGGCCGG</td><td></td><td></td><td></td><td></td><td></td><td></td><td></td><td></td><td></td><td></td></tr>
<tr><td>Chromosome</td><td>start</td><td>end</td><td>strand</td><td>MM</td><td>target_seq</td><td>PAM</td><td>alignment</td><td>distance</td><td>position</td><td>gene name</td><td>gene id</td></tr>
<tr><td>GU305914.1</td><td>113738</td><td>113760</td><td>-</td><td>0</td><td>CCGGCCCAGACAGAAGAGTC</td><td>CGG</td><td>||||||||[||||||||||||]PAM</td><td>NA</td><td>Intergenic</td><td>NA</td><td>NA</td></tr>
</table>

<table>
<tr><td>T25442</td><td>CAGACAGAAGAGTCCGGCCTCGG</td><td>1000</td><td></td><td></td><td></td><td></td><td></td><td></td><td></td><td></td><td></td></tr>
<tr><td>Oligo fwd</td><td>CACCCAGACAGAAGAGTCCGGCCT</td><td></td><td></td><td></td><td></td><td></td><td></td><td></td><td></td><td></td><td></td></tr>
<tr><td>Oligo rev</td><td>AAACAGGCCGGACTCTTCTGTCTG</td><td></td><td></td><td></td><td></td><td></td><td></td><td></td><td></td><td></td><td></td></tr>
<tr><td>Chromosome</td><td>start</td><td>end</td><td>strand</td><td>MM</td><td>target_seq</td><td>PAM</td><td>alignment</td><td>distance</td><td>position</td><td>gene name</td><td>gene id</td></tr>
<tr><td>GU305914.1</td><td>113732</td><td>113754</td><td>-</td><td>0</td><td>CAGACAGAAGAGTCCGGCCT</td><td>CGG</td><td>||||||||[||||||||||||]PAM</td><td>NA</td><td>Intergenic</td><td>NA</td><td>NA</td></tr>
<tr><td>T25443</td><td>CTCGGCGTCAAAGAGACGTACGG</td><td>1000</td><td></td><td></td><td></td><td></td><td></td><td></td><td></td><td></td><td></td></tr>
<tr><td>Oligo fwd</td><td>CACCCTCGGCGTCAAAGAGACGTA</td><td></td><td></td><td></td><td></td><td></td><td></td><td></td><td></td><td></td><td></td></tr>
<tr><td>Oligo rev</td><td>AAACTACGTCTCTTTGACGCCGAG</td><td></td><td></td><td></td><td></td><td></td><td></td><td></td><td></td><td></td><td></td></tr>
<tr><td>Chromosome</td><td>start</td><td>end</td><td>strand</td><td>MM</td><td>target_seq</td><td>PAM</td><td>alignment</td><td>distance</td><td>position</td><td>gene name</td><td>gene id</td></tr>
<tr><td>GU305914.1</td><td>113714</td><td>113736</td><td>-</td><td>0</td><td>CTCGGCGTCAAAGAGACGTA</td><td>CGG</td><td>||||||||[||||||||||||]PAM</td><td>NA</td><td>Intergenic</td><td>NA</td><td>NA</td></tr>
<tr><td>T25444</td><td>GAGACGTACGGTGCCGACCGTGG</td><td>1000</td><td></td><td></td><td></td><td></td><td></td><td></td><td></td><td></td><td></td></tr>
<tr><td>Oligo fwd</td><td>CACCGAGACGTACGGTGCCGACCG</td><td></td><td></td><td></td><td></td><td></td><td></td><td></td><td></td><td></td><td></td></tr>
<tr><td>Oligo rev</td><td>AAACCGGTCGGCACCGTACGTCTC</td><td></td><td></td><td></td><td></td><td></td><td></td><td></td><td></td><td></td><td></td></tr>
<tr><td>Chromosome</td><td>start</td><td>end</td><td>strand</td><td>MM</td><td>target_seq</td><td>PAM</td><td>alignment</td><td>distance</td><td>position</td><td>gene name</td><td>gene id</td></tr>
<tr><td>GU305914.1</td><td>113702</td><td>113724</td><td>-</td><td>0</td><td>GAGACGTACGGTGCCGACCG</td><td>TGG</td><td>||||||||[||||||||||||]PAM</td><td>NA</td><td>Intergenic</td><td>NA</td><td>NA</td></tr>
<tr><td>T25445</td><td>ACGTACGGTGCCGACCGTGGAGG</td><td>1000</td><td></td><td></td><td></td><td></td><td></td><td></td><td></td><td></td><td></td></tr>
<tr><td>Oligo fwd</td><td>CACCACGTACGGTGCCGACCGTGG</td><td></td><td></td><td></td><td></td><td></td><td></td><td></td><td></td><td></td><td></td></tr>
<tr><td>Oligo rev</td><td>AAACCCACGGTCGGCACCGTACGT</td><td></td><td></td><td></td><td></td><td></td><td></td><td></td><td></td><td></td><td></td></tr>
<tr><td>Chromosome</td><td>start</td><td>end</td><td>strand</td><td>MM</td><td>target_seq</td><td>PAM</td><td>alignment</td><td>distance</td><td>position</td><td>gene name</td><td>gene id</td></tr>
<tr><td>GU305914.1</td><td>113699</td><td>113721</td><td>-</td><td>0</td><td>ACGTACGGTGCCGACCGTGG</td><td>AGG</td><td>||||||||[||||||||||||]PAM</td><td>NA</td><td>Intergenic</td><td>NA</td><td>NA</td></tr>
<tr><td>T25446</td><td>ATTCGTCGTGATTGATATTGAGG</td><td>1000</td><td></td><td></td><td></td><td></td><td></td><td></td><td></td><td></td><td></td></tr>
<tr><td>Oligo fwd</td><td>CACCATTCGTCGTGATTGATATTG</td><td></td><td></td><td></td><td></td><td></td><td></td><td></td><td></td><td></td><td></td></tr>
<tr><td>Oligo rev</td><td>AAACCAATATCAATCACGACGAAT</td><td></td><td></td><td></td><td></td><td></td><td></td><td></td><td></td><td></td><td></td></tr>
<tr><td>Chromosome</td><td>start</td><td>end</td><td>strand</td><td>MM</td><td>target_seq</td><td>PAM</td><td>alignment</td><td>distance</td><td>position</td><td>gene name</td><td>gene id</td></tr>
<tr><td>GU305914.1</td><td>113676</td><td>113698</td><td>-</td><td>0</td><td>ATTCGTCGTGATTGATATTG</td><td>AGG</td><td>||||||||[||||||||||||]PAM</td><td>NA</td><td>Intergenic</td><td>NA</td><td>NA</td></tr>
<tr><td>T25447</td><td>TTCGTCGTGATTGATATTGAGGG</td><td>1000</td><td></td><td></td><td></td><td></td><td></td><td></td><td></td><td></td><td></td></tr>
<tr><td>Oligo fwd</td><td>CACCTTCGTCGTGATTGATATTGA</td><td></td><td></td><td></td><td></td><td></td><td></td><td></td><td></td><td></td><td></td></tr>
<tr><td>Oligo rev</td><td>AAACTCAATATCAATCACGACGAA</td><td></td><td></td><td></td><td></td><td></td><td></td><td></td><td></td><td></td><td></td></tr>
<tr><td>Chromosome</td><td>start</td><td>end</td><td>strand</td><td>MM</td><td>target_seq</td><td>PAM</td><td>alignment</td><td>distance</td><td>position</td><td>gene name</td><td>gene id</td></tr>
<tr><td>GU305914.1</td><td>113675</td><td>113697</td><td>-</td><td>0</td><td>TTCGTCGTGATTGATATTGA</td><td>GGG</td><td>||||||||[||||||||||||]PAM</td><td>NA</td><td>Intergenic</td><td>NA</td><td>NA</td></tr>
<tr><td>T25448</td><td>TCGTCGTGATTGATATTGAGGGG</td><td>1000</td><td></td><td></td><td></td><td></td><td></td><td></td><td></td><td></td><td></td></tr>
<tr><td>Oligo fwd</td><td>CACCTCGTCGTGATTGATATTGAG</td><td></td><td></td><td></td><td></td><td></td><td></td><td></td><td></td><td></td><td></td></tr>
<tr><td>Oligo rev</td><td>AAACCTCAATATCAATCACGACGA</td><td></td><td></td><td></td><td></td><td></td><td></td><td></td><td></td><td></td><td></td></tr>
<tr><td>Chromosome</td><td>start</td><td>end</td><td>strand</td><td>MM</td><td>target_seq</td><td>PAM</td><td>alignment</td><td>distance</td><td>position</td><td>gene name</td><td>gene id</td></tr>
<tr><td>GU305914.1</td><td>113674</td><td>113696</td><td>-</td><td>0</td><td>TCGTCGTGATTGATATTGAG</td><td>GGG</td><td>||||||||[||||||||||||]PAM</td><td>NA</td><td>Intergenic</td><td>NA</td><td>NA</td></tr>
<tr><td>T25449</td><td>GTGATTGATATTGAGGGGCACGG</td><td>1000</td><td></td><td></td><td></td><td></td><td></td><td></td><td></td><td></td><td></td></tr>
<tr><td>Oligo fwd</td><td>CACCGTGATTGATATTGAGGGGCA</td><td></td><td></td><td></td><td></td><td></td><td></td><td></td><td></td><td></td><td></td></tr>
<tr><td>Oligo rev</td><td>AAACTGCCCCTCAATATCAATCAC</td><td></td><td></td><td></td><td></td><td></td><td></td><td></td><td></td><td></td><td></td></tr>
<tr><td>Chromosome</td><td>start</td><td>end</td><td>strand</td><td>MM</td><td>target_seq</td><td>PAM</td><td>alignment</td><td>distance</td><td>position</td><td>gene name</td><td>gene id</td></tr>
<tr><td>GU305914.1</td><td>113669</td><td>113691</td><td>-</td><td>0</td><td>GTGATTGATATTGAGGGGCA</td><td>CGG</td><td>||||||||[||||||||||||]PAM</td><td>NA</td><td>Intergenic</td><td>NA</td><td>NA</td></tr>
<tr><td>T25450</td><td>ATTGATATTGAGGGGCACGGCGG</td><td>1000</td><td></td><td></td><td></td><td></td><td></td><td></td><td></td><td></td><td></td></tr>
<tr><td>Oligo fwd</td><td>CACCATTGATATTGAGGGGCACGG</td><td></td><td></td><td></td><td></td><td></td><td></td><td></td><td></td><td></td><td></td></tr>
<tr><td>Oligo rev</td><td>AAACCCGTGCCCCTCAATATCAAT</td><td></td><td></td><td></td><td></td><td></td><td></td><td></td><td></td><td></td><td></td></tr>
<tr><td>Chromosome</td><td>start</td><td>end</td><td>strand</td><td>MM</td><td>target_seq</td><td>PAM</td><td>alignment</td><td>distance</td><td>position</td><td>gene name</td><td>gene id</td></tr>
<tr><td>GU305914.1</td><td>113666</td><td>113688</td><td>-</td><td>0</td><td>ATTGATATTGAGGGGCACGG</td><td>CGG</td><td>||||||||[||||||||||||]PAM</td><td>NA</td><td>Intergenic</td><td>NA</td><td>NA</td></tr>
<tr><td>T25451</td><td>TTGATATTGAGGGGCACGGCGGG</td><td>1000</td><td></td><td></td><td></td><td></td><td></td><td></td><td></td><td></td><td></td></tr>
<tr><td>Oligo fwd</td><td>CACCTTGATATTGAGGGGCACGGC</td><td></td><td></td><td></td><td></td><td></td><td></td><td></td><td></td><td></td><td></td></tr>
<tr><td>Oligo rev</td><td>AAACGCCGTGCCCCTCAATATCAA</td><td></td><td></td><td></td><td></td><td></td><td></td><td></td><td></td><td></td><td></td></tr>
<tr><td>Chromosome</td><td>start</td><td>end</td><td>strand</td><td>MM</td><td>target_seq</td><td>PAM</td><td>alignment</td><td>distance</td><td>position</td><td>gene name</td><td>gene id</td></tr>
<tr><td>GU305914.1</td><td>113665</td><td>113687</td><td>-</td><td>0</td><td>TTGATATTGAGGGGCACGGC</td><td>GGG</td><td>||||||||[||||||||||||]PAM</td><td>NA</td><td>Intergenic</td><td>NA</td><td>NA</td></tr>
<tr><td>T25452</td><td>GAGGGGCACGGCGGGCCCCCCGG</td><td>1000</td><td></td><td></td><td></td><td></td><td></td><td></td><td></td><td></td><td></td></tr>
<tr><td>Oligo fwd</td><td>CACCGAGGGGCACGGCGGGCCCCC</td><td></td><td></td><td></td><td></td><td></td><td></td><td></td><td></td><td></td><td></td></tr>
<tr><td>Oligo rev</td><td>AAACGGGGGCCCGCCGTGCCCCTC</td><td></td><td></td><td></td><td></td><td></td><td></td><td></td><td></td><td></td><td></td></tr>
<tr><td>Chromosome</td><td>start</td><td>end</td><td>strand</td><td>MM</td><td>target_seq</td><td>PAM</td><td>alignment</td><td>distance</td><td>position</td><td>gene name</td><td>gene id</td></tr>
<tr><td>GU305914.1</td><td>113657</td><td>113679</td><td>-</td><td>0</td><td>GAGGGGCACGGCGGGCCCCC</td><td>CGG</td><td>||||||||[||||||||||||]PAM</td><td>NA</td><td>Intergenic</td><td>NA</td><td>NA</td></tr>
<tr><td>GU305914.1</td><td>52596</td><td>52618</td><td>-</td><td>4</td><td>GGGGCGCACGGCGGGGCCGC</td><td>CGG</td><td>|-||-|||[|||||||-||-|]PAM</td><td>NA</td><td>Intergenic</td><td>NA</td><td>NA</td></tr>
<tr><td>T25453</td><td>CGCCGCTCTGCGCAGCTCCCTGG</td><td>1000</td><td></td><td></td><td></td><td></td><td></td><td></td><td></td><td></td><td></td></tr>
<tr><td>Oligo fwd</td><td>CACCCGCCGCTCTGCGCAGCTCCC</td><td></td><td></td><td></td><td></td><td></td><td></td><td></td><td></td><td></td><td></td></tr>
<tr><td>Oligo rev</td><td>AAACGGGAGCTGCGCAGAGCGGCG</td><td></td><td></td><td></td><td></td><td></td><td></td><td></td><td></td><td></td><td></td></tr>
<tr><td>Chromosome</td><td>start</td><td>end</td><td>strand</td><td>MM</td><td>target_seq</td><td>PAM</td><td>alignment</td><td>distance</td><td>position</td><td>gene name</td><td>gene id</td></tr>
<tr><td>GU305914.1</td><td>113633</td><td>113655</td><td>-</td><td>0</td><td>CGCCGCTCTGCGCAGCTCCC</td><td>TGG</td><td>||||||||[||||||||||||]PAM</td><td>NA</td><td>Intergenic</td><td>NA</td><td>NA</td></tr>
<tr><td>T25454</td><td>TCCCTGGCCACGACGTCGCGAGG</td><td>1000</td><td></td><td></td><td></td><td></td><td></td><td></td><td></td><td></td><td></td></tr>
<tr><td>Oligo fwd</td><td>CACCTCCCTGGCCACGACGTCGCG</td><td></td><td></td><td></td><td></td><td></td><td></td><td></td><td></td><td></td><td></td></tr>
<tr><td>Oligo rev</td><td>AAACCGCGACGTCGTGGCCAGGGA</td><td></td><td></td><td></td><td></td><td></td><td></td><td></td><td></td><td></td><td></td></tr>
<tr><td>Chromosome</td><td>start</td><td>end</td><td>strand</td><td>MM</td><td>target_seq</td><td>PAM</td><td>alignment</td><td>distance</td><td>position</td><td>gene name</td><td>gene id</td></tr>
<tr><td>GU305914.1</td><td>113617</td><td>113639</td><td>-</td><td>0</td><td>TCCCTGGCCACGACGTCGCG</td><td>AGG</td><td>||||||||[||||||||||||]PAM</td><td>NA</td><td>Intergenic</td><td>NA</td><td>NA</td></tr>
<tr><td>T25455</td><td>CCCTGGCCACGACGTCGCGAGGG</td><td>1000</td><td></td><td></td><td></td><td></td><td></td><td></td><td></td><td></td><td></td></tr>
<tr><td>Oligo fwd</td><td>CACCCCCTGGCCACGACGTCGCGA</td><td></td><td></td><td></td><td></td><td></td><td></td><td></td><td></td><td></td><td></td></tr>
<tr><td>Oligo rev</td><td>AAACTCGCGACGTCGTGGCCAGGG</td><td></td><td></td><td></td><td></td><td></td><td></td><td></td><td></td><td></td><td></td></tr>
<tr><td>Chromosome</td><td>start</td><td>end</td><td>strand</td><td>MM</td><td>target_seq</td><td>PAM</td><td>alignment</td><td>distance</td><td>position</td><td>gene name</td><td>gene id</td></tr>
<tr><td>GU305914.1</td><td>113616</td><td>113638</td><td>-</td><td>0</td><td>CCCTGGCCACGACGTCGCGA</td><td>GGG</td><td>||||||||[||||||||||||]PAM</td><td>NA</td><td>Intergenic</td><td>NA</td><td>NA</td></tr>
<tr><td>T25456</td><td>CGACGTCGCGAGGGAGCCGCAGG</td><td>1000</td><td></td><td></td><td></td><td></td><td></td><td></td><td></td><td></td><td></td></tr>
<tr><td>Oligo fwd</td><td>CACCCGACGTCGCGAGGGAGCCGC</td><td></td><td></td><td></td><td></td><td></td><td></td><td></td><td></td><td></td><td></td></tr>
<tr><td>Oligo rev</td><td>AAACGCGGCTCCCTCGCGACGTCG</td><td></td><td></td><td></td><td></td><td></td><td></td><td></td><td></td><td></td><td></td></tr>
</table>

| | | | | | | | | | | | |
|---|---|---|---|---|---|---|---|---|---|---|---|
| Chromosome | start | end | strand | MM | target_seq | PAM | alignment | distance | position | gene name | gene id |
| GU305914.1 | 113607 | 113629 | - | 0 | CGACGTCGCGAGGGAGCCGC | AGG | \|\|\|\|\|\|\|\|[\|\|\|\|\|\|\|\|\|\|\|\|]PAM | NA | Intergenic | NA | NA |
| GU305914.1 | 75704 | 75726 | + | 4 | TGATCCCGCGAGGGAGCCGC | CGG | -\|\|---\|\|[\|\|\|\|\|\|\|\|\|\|\|\|]PAM | NA | Intergenic | NA | NA |
| GU305914.1 | 26064 | 26086 | - | 3 | CGACGCCGCGTGCGAGCCGC | TGG | \|\|\|\|\|-\|\|[\|\|-\|-\|\|\|\|\|\|\|]PAM | NA | Intergenic | NA | NA |
| T25457 | AGCCGCAGGCGTTCGTCCTGAGG | 1000 | | | | | | | | | |
| Oligo fwd | CACCAGCCGCAGGCGTTCGTCCTG | | | | | | | | | | |
| Oligo rev | AAACCAGGACGAACGCCTGCGGCT | | | | | | | | | | |
| Chromosome | start | end | strand | MM | target_seq | PAM | alignment | distance | position | gene name | gene id |
| GU305914.1 | 113593 | 113615 | - | 0 | AGCCGCAGGCGTTCGTCCTG | AGG | \|\|\|\|\|\|\|\|[\|\|\|\|\|\|\|\|\|\|\|\|]PAM | NA | Intergenic | NA | NA |
| T25458 | GCCGCAGGCGTTCGTCCTGAGGG | 1000 | | | | | | | | | |
| Oligo fwd | CACCGCCGCAGGCGTTCGTCCTGA | | | | | | | | | | |
| Oligo rev | AAACTCAGGACGAACGCCTGCGGC | | | | | | | | | | |
| Chromosome | start | end | strand | MM | target_seq | PAM | alignment | distance | position | gene name | gene id |
| GU305914.1 | 113592 | 113614 | - | 0 | GCCGCAGGCGTTCGTCCTGA | GGG | \|\|\|\|\|\|\|\|[\|\|\|\|\|\|\|\|\|\|\|\|]PAM | NA | Intergenic | NA | NA |
| T25459 | GGTCCTCGTCATACAGAGTGAGG | 1000 | | | | | | | | | |
| Oligo fwd | CACCGGTCCTCGTCATACAGAGTG | | | | | | | | | | |
| Oligo rev | AAACCACTCTGTATGACGAGGACC | | | | | | | | | | |
| Chromosome | start | end | strand | MM | target_seq | PAM | alignment | distance | position | gene name | gene id |
| GU305914.1 | 113571 | 113593 | - | 0 | GGTCCTCGTCATACAGAGTG | AGG | \|\|\|\|\|\|\|\|[\|\|\|\|\|\|\|\|\|\|\|\|]PAM | NA | Intergenic | NA | NA |
| T25460 | TAGATCATCGATCCCGCGTCCGG | 1000 | | | | | | | | | |
| Oligo fwd | CACCTAGATCATCGATCCCGCGTC | | | | | | | | | | |
| Oligo rev | AAACGACGCGGGATCGATGATCTA | | | | | | | | | | |
| Chromosome | start | end | strand | MM | target_seq | PAM | alignment | distance | position | gene name | gene id |
| GU305914.1 | 113536 | 113558 | - | 0 | TAGATCATCGATCCCGCGTC | CGG | \|\|\|\|\|\|\|\|[\|\|\|\|\|\|\|\|\|\|\|\|]PAM | NA | Intergenic | NA | NA |
| T25461 | TCCCGCGTCCGGCGCGCGAGCGG | 1000 | | | | | | | | | |
| Oligo fwd | CACCTCCCGCGTCCGGCGCGCGAG | | | | | | | | | | |
| Oligo rev | AAACCTCGCGCGCCGGACGCGGGA | | | | | | | | | | |
| Chromosome | start | end | strand | MM | target_seq | PAM | alignment | distance | position | gene name | gene id |
| GU305914.1 | 113525 | 113547 | - | 0 | TCCCGCGTCCGGCGCGCGAG | CGG | \|\|\|\|\|\|\|\|[\|\|\|\|\|\|\|\|\|\|\|\|]PAM | NA | Intergenic | NA | NA |
| GU305914.1 | 224568 | 224590 | + | 4 | TCCTGCGCCCGGCGAGCGGG | AGG | \|\|\|-\|\|\|-[\|\|\|\|\|\|-\|\|\|-\|]PAM | NA | Intergenic | NA | NA |
| T25462 | GCGAGCGGCGTCGCCCGTCATGG | 1000 | | | | | | | | | |
| Oligo fwd | CACCGCGAGCGGCGTCGCCCGTCA | | | | | | | | | | |
| Oligo rev | AAACTGACGGGCGACGCCGCTCGC | | | | | | | | | | |
| Chromosome | start | end | strand | MM | target_seq | PAM | alignment | distance | position | gene name | gene id |
| GU305914.1 | 113510 | 113532 | - | 0 | GCGAGCGGCGTCGCCCGTCA | TGG | \|\|\|\|\|\|\|\|[\|\|\|\|\|\|\|\|\|\|\|\|]PAM | NA | Intergenic | NA | NA |
| T25463 | ATGGTGAAGACGATCCGCGTCGG | 1000 | | | | | | | | | |
| Oligo fwd | CACCATGGTGAAGACGATCCGCGT | | | | | | | | | | |
| Oligo rev | AAACACGCGGATCGTCTTCACCAT | | | | | | | | | | |
| Chromosome | start | end | strand | MM | target_seq | PAM | alignment | distance | position | gene name | gene id |
| GU305914.1 | 113491 | 113513 | - | 0 | ATGGTGAAGACGATCCGCGT | CGG | \|\|\|\|\|\|\|\|[\|\|\|\|\|\|\|\|\|\|\|\|]PAM | NA | Intergenic | NA | NA |
| T25464 | TGGTGAAGACGATCCGCGTCGGG | 1000 | | | | | | | | | |
| Oligo fwd | CACCTGGTGAAGACGATCCGCGTC | | | | | | | | | | |
| Oligo rev | AAACGACGCGGATCGTCTTCACCA | | | | | | | | | | |
| Chromosome | start | end | strand | MM | target_seq | PAM | alignment | distance | position | gene name | gene id |
| GU305914.1 | 113490 | 113512 | - | 0 | TGGTGAAGACGATCCGCGTC | GGG | \|\|\|\|\|\|\|\|[\|\|\|\|\|\|\|\|\|\|\|\|]PAM | NA | Intergenic | NA | NA |
| T25465 | TGAAGACGATCCGCGTCGGGCGG | 1000 | | | | | | | | | |
| Oligo fwd | CACCTGAAGACGATCCGCGTCGGG | | | | | | | | | | |
| Oligo rev | AAACCCCGACGCGGATCGTCTTCA | | | | | | | | | | |
| Chromosome | start | end | strand | MM | target_seq | PAM | alignment | distance | position | gene name | gene id |
| GU305914.1 | 113487 | 113509 | - | 0 | TGAAGACGATCCGCGTCGGG | CGG | \|\|\|\|\|\|\|\|[\|\|\|\|\|\|\|\|\|\|\|\|]PAM | NA | Intergenic | NA | NA |
| GU305914.1 | 47368 | 47390 | + | 3 | TGAGGCCGATCCGCGGCGGG | GGG | \|\|\|-\|-\|\|[\|\|\|\|\|\|\|-\|\|\|\|]PAM | NA | Intergenic | NA | NA |
| T25466 | CGGGCGGTTTCTGCACCTGTCGG | 1000 | | | | | | | | | |
| Oligo fwd | CACCCGGGCGGTTTCTGCACCTGT | | | | | | | | | | |
| Oligo rev | AAACACAGGTGCAGAAACCGCCCG | | | | | | | | | | |
| Chromosome | start | end | strand | MM | target_seq | PAM | alignment | distance | position | gene name | gene id |
| GU305914.1 | 113471 | 113493 | - | 0 | CGGGCGGTTTCTGCACCTGT | CGG | \|\|\|\|\|\|\|\|[\|\|\|\|\|\|\|\|\|\|\|\|]PAM | NA | Intergenic | NA | NA |
| T25467 | CATCACGACGAAGCTGCTGTCGG | 1000 | | | | | | | | | |
| Oligo fwd | CACCCATCACGACGAAGCTGCTGT | | | | | | | | | | |
| Oligo rev | AAACACAGCAGCTTCGTCGTGATG | | | | | | | | | | |
| Chromosome | start | end | strand | MM | target_seq | PAM | alignment | distance | position | gene name | gene id |
| GU305914.1 | 113426 | 113448 | - | 0 | CATCACGACGAAGCTGCTGT | CGG | \|\|\|\|\|\|\|\|[\|\|\|\|\|\|\|\|\|\|\|\|]PAM | NA | Intergenic | NA | NA |
| T25468 | ATCACGACGAAGCTGCTGTCGGG | 1000 | | | | | | | | | |
| Oligo fwd | CACCATCACGACGAAGCTGCTGTC | | | | | | | | | | |
| Oligo rev | AAACGACAGCAGCTTCGTCGTGAT | | | | | | | | | | |
| Chromosome | start | end | strand | MM | target_seq | PAM | alignment | distance | position | gene name | gene id |
| GU305914.1 | 113425 | 113447 | - | 0 | ATCACGACGAAGCTGCTGTC | GGG | \|\|\|\|\|\|\|\|[\|\|\|\|\|\|\|\|\|\|\|\|]PAM | NA | Intergenic | NA | NA |
| T25469 | TCACGACGAAGCTGCTGTCGGGG | 1000 | | | | | | | | | |
| Oligo fwd | CACCTCACGACGAAGCTGCTGTCG | | | | | | | | | | |
| Oligo rev | AAACCGACAGCAGCTTCGTCGTGA | | | | | | | | | | |
| Chromosome | start | end | strand | MM | target_seq | PAM | alignment | distance | position | gene name | gene id |
| GU305914.1 | 113424 | 113446 | - | 0 | TCACGACGAAGCTGCTGTCG | GGG | \|\|\|\|\|\|\|\|[\|\|\|\|\|\|\|\|\|\|\|\|]PAM | NA | Intergenic | NA | NA |
| T25470 | GCCGCTGTCGTCCATGCGCCTGG | 1000 | | | | | | | | | |
| Oligo fwd | CACCGCCGCTGTCGTCCATGCGCC | | | | | | | | | | |
| Oligo rev | AAACGGCGCATGGACGACAGCGGC | | | | | | | | | | |
| Chromosome | start | end | strand | MM | target_seq | PAM | alignment | distance | position | gene name | gene id |

GU305914.1 113399 113421 - 0 GCCGCTGTCGTCCATGCGCC TGG ||||||||[||||||||||||]PAM NA Intergenic NA NA

T25471 GCTGTCGTCCATGCGCCTGGAGG 1000
Oligo fwd CACCGCTGTCGTCCATGCGCCTGG
Oligo rev AAACCCAGGCGCATGGACGACAGC
Chromosome start end strand MM target_seq PAM alignment distance position gene name gene id
GU305914.1 113396 113418 - 0 GCTGTCGTCCATGCGCCTGG AGG ||||||||[||||||||||||]PAM NA Intergenic NA NA

T25472 TGGAGGAGTTGAAGATCATCAGG 1000
Oligo fwd CACCTGGAGGAGTTGAAGATCATC
Oligo rev AAACGATGATCTTCAACTCCTCCA
Chromosome start end strand MM target_seq PAM alignment distance position gene name gene id
GU305914.1 113379 113401 - 0 TGGAGGAGTTGAAGATCATC AGG ||||||||[||||||||||||]PAM NA Intergenic NA NA

T25473 GGAGTTGAAGATCATCAGGCTGG 1000
Oligo fwd CACCGGAGTTGAAGATCATCAGGC
Oligo rev AAACGCCTGATGATCTTCAACTCC
Chromosome start end strand MM target_seq PAM alignment distance position gene name gene id
GU305914.1 113375 113397 - 0 GGAGTTGAAGATCATCAGGC TGG ||||||||[||||||||||||]PAM NA Intergenic NA NA

T25474 GCTGGCGTGCCTGCTGACGCTGG 1000
Oligo fwd CACCGCTGGCGTGCCTGCTGACGC
Oligo rev AAACGCGTCAGCAGGCACGCCAGC
Chromosome start end strand MM target_seq PAM alignment distance position gene name gene id
GU305914.1 113357 113379 - 0 GCTGGCGTGCCTGCTGACGC TGG ||||||||[||||||||||||]PAM NA Intergenic NA NA
GU305914.1 62684 62706 - 4 GCTGCCGCGGCTGCTGCCGC CGG ||||-||-[|-||||||-|||]PAM NA Intergenic NA NA

T25475 CTGGCGTGCCTGCTGACGCTGGG 1000
Oligo fwd CACCCTGGCGTGCCTGCTGACGCT
Oligo rev AAACAGCGTCAGCAGGCACGCCAG
Chromosome start end strand MM target_seq PAM alignment distance position gene name gene id
GU305914.1 113356 113378 - 0 CTGGCGTGCCTGCTGACGCT GGG ||||||||[||||||||||||]PAM NA Intergenic NA NA

T25476 TGGCGTGCCTGCTGACGCTGGGG 1000
Oligo fwd CACCTGGCGTGCCTGCTGACGCTG
Oligo rev AAACCAGCGTCAGCAGGCACGCCA
Chromosome start end strand MM target_seq PAM alignment distance position gene name gene id
GU305914.1 113355 113377 - 0 TGGCGTGCCTGCTGACGCTG GGG ||||||||[||||||||||||]PAM NA Intergenic NA NA

T25477 TGCCTGCTGACGCTGGGGCGCGG 1000
Oligo fwd CACCTGCCTGCTGACGCTGGGGCG
Oligo rev AAACCGCCCCAGCGTCAGCAGGCA
Chromosome start end strand MM target_seq PAM alignment distance position gene name gene id
GU305914.1 113350 113372 - 0 TGCCTGCTGACGCTGGGGCG CGG ||||||||[||||||||||||]PAM NA Intergenic NA NA

T25478 GCCTGCTGACGCTGGGGCGCGGG 1000
Oligo fwd CACCGCCTGCTGACGCTGGGGCGC
Oligo rev AAACGCGCCCCAGCGTCAGCAGGC
Chromosome start end strand MM target_seq PAM alignment distance position gene name gene id
GU305914.1 113349 113371 - 0 GCCTGCTGACGCTGGGGCGC GGG ||||||||[||||||||||||]PAM NA Intergenic NA NA

T25479 GGATCGAGCTGCTGATCCTGAGG 1000
Oligo fwd CACCGGATCGAGCTGCTGATCCTG
Oligo rev AAACCAGGATCAGCAGCTCGATCC
Chromosome start end strand MM target_seq PAM alignment distance position gene name gene id
GU305914.1 113328 113350 - 0 GGATCGAGCTGCTGATCCTG AGG ||||||||[||||||||||||]PAM NA Intergenic NA NA
GU305914.1 203215 203237 - 4 GGCTCGAACTGCAGATCGTG AGG ||-||||-[||||-||||-||]PAM NA Intergenic NA NA

T25480 GATCGAGCTGCTGATCCTGAGGG 1000
Oligo fwd CACCGATCGAGCTGCTGATCCTGA
Oligo rev AAACTCAGGATCAGCAGCTCGATC
Chromosome start end strand MM target_seq PAM alignment distance position gene name gene id
GU305914.1 113327 113349 - 0 GATCGAGCTGCTGATCCTGA GGG ||||||||[||||||||||||]PAM NA Intergenic NA NA

T25481 GCTGCTGATCCTGAGGGAGACGG 1000
Oligo fwd CACCGCTGCTGATCCTGAGGGAGA
Oligo rev AAACTCTCCCTCAGGATCAGCAGC
Chromosome start end strand MM target_seq PAM alignment distance position gene name gene id
GU305914.1 113321 113343 - 0 GCTGCTGATCCTGAGGGAGA CGG ||||||||[||||||||||||]PAM NA Intergenic NA NA
GU305914.1 113108 113130 - 4 GCACCTGATCCGGACGGAGA CGG ||--||||[|||-||-|||||]PAM NA Intergenic NA NA

T25482 GCTGATCCTGAGGGAGACGGTGG 1000
Oligo fwd CACCGCTGATCCTGAGGGAGACGG
Oligo rev AAACCCGTCTCCCTCAGGATCAGC
Chromosome start end strand MM target_seq PAM alignment distance position gene name gene id
GU305914.1 113318 113340 - 0 GCTGATCCTGAGGGAGACGG TGG ||||||||[||||||||||||]PAM NA Intergenic NA NA

T25483 AGGGAGACGGTGGCGAACAACGG 1000
Oligo fwd CACCAGGGAGACGGTGGCGAACAA
Oligo rev AAACTTGTTCGCCACCGTCTCCCT
Chromosome start end strand MM target_seq PAM alignment distance position gene name gene id
GU305914.1 113308 113330 - 0 AGGGAGACGGTGGCGAACAA CGG ||||||||[||||||||||||]PAM NA Intergenic NA NA

T25484 GGGAGACGGTGGCGAACAACGGG 1000
Oligo fwd CACCGGGAGACGGTGGCGAACAAC
Oligo rev AAACGTTGTTCGCCACCGTCTCCC
Chromosome start end strand MM target_seq PAM alignment distance position gene name gene id
GU305914.1 113307 113329 - 0 GGGAGACGGTGGCGAACAAC GGG ||||||||[||||||||||||]PAM NA Intergenic NA NA

T25485 GGAGACGGTGGCGAACAACGGGG 1000
Oligo fwd CACCGGAGACGGTGGCGAACAACG
Oligo rev AAACCGTTGTTCGCCACCGTCTCC
Chromosome start end strand MM target_seq PAM alignment distance position gene name gene id
GU305914.1 113306 113328 - 0 GGAGACGGTGGCGAACAACG GGG ||||||||[||||||||||||]PAM NA Intergenic NA NA

T25486 GGTGGCGAACAACGGGGTGTCGG 1000
Oligo fwd CACCGGTGGCGAACAACGGGGTGT
Oligo rev AAACACACCCCGTTGTTCGCCACC
Chromosome start end strand MM target_seq PAM alignment distance position gene name gene id
GU305914.1 113300 113322 - 0 GGTGGCGAACAACGGGGTGT CGG ||||||||[||||||||||||]PAM NA Intergenic NA NA

T25487 GCAAGATCTCGCCCGAGTTCTGG 1000
Oligo fwd CACCGCAAGATCTCGCCCGAGTTC
Oligo rev AAACGAACTCGGGCGAGATCTTGC
Chromosome start end strand MM target_seq PAM alignment distance position gene name gene id
GU305914.1 113259 113281 - 0 GCAAGATCTCGCCCGAGTTC TGG ||||||||[||||||||||||]PAM NA Intergenic NA NA

T25488 GTTCTGGCGCAAGATGTACGAGG 1000
Oligo fwd CACCGTTCTGGCGCAAGATGTACG
Oligo rev AAACCGTACATCTTGCGCCAGAAC
Chromosome start end strand MM target_seq PAM alignment distance position gene name gene id
GU305914.1 113243 113265 - 0 GTTCTGGCGCAAGATGTACG AGG ||||||||[||||||||||||]PAM NA Intergenic NA NA

T25489 CCGCGCCTTCAGCGAGCGTTCGG 1000
Oligo fwd CACCCCGCGCCTTCAGCGAGCGTT
Oligo rev AAACAACGCTCGCTGAAGGCGCGG
Chromosome start end strand MM target_seq PAM alignment distance position gene name gene id
GU305914.1 113186 113208 - 0 CCGCGCCTTCAGCGAGCGTT CGG ||||||||[||||||||||||]PAM NA Intergenic NA NA

T25490 CTTCAGCGAGCGTTCGGCCGCGG 1000
Oligo fwd CACCCTTCAGCGAGCGTTCGGCCG
Oligo rev AAACCGGCCGAACGCTCGCTGAAG
Chromosome start end strand MM target_seq PAM alignment distance position gene name gene id
GU305914.1 113180 113202 - 0 CTTCAGCGAGCGTTCGGCCG CGG ||||||||[||||||||||||]PAM NA Intergenic NA NA

T25491 GCGTTCGGCCGCGGCGCTGTCGG 1000
Oligo fwd CACCGCGTTCGGCCGCGGCGCTGT
Oligo rev AAACACAGCGCCGCGGCCGAACGC
Chromosome start end strand MM target_seq PAM alignment distance position gene name gene id
GU305914.1 113171 113193 - 0 GCGTTCGGCCGCGGCGCTGT CGG ||||||||[||||||||||||]PAM NA Intergenic NA NA

T25492 TTCGGCCGCGGCGCTGTCGGTGG 1000
Oligo fwd CACCTTCGGCCGCGGCGCTGTCGG
Oligo rev AAACCCGACAGCGCCGCGGCCGAA
Chromosome start end strand MM target_seq PAM alignment distance position gene name gene id
GU305914.1 113168 113190 - 0 TTCGGCCGCGGCGCTGTCGG TGG ||||||||[||||||||||||]PAM NA Intergenic NA NA
GU305914.1 152553 152575 + 4 TGCGGCGGCGGCGGTGCCGG CGG |-||||-|[||||||-||-|||]PAM NA Intergenic NA NA

T25493 GGCGCTGTCGGTGGAGATGACGG 1000
Oligo fwd CACCGGCGCTGTCGGTGGAGATGA
Oligo rev AAACTCATCTCCACCGACAGCGCC
Chromosome start end strand MM target_seq PAM alignment distance position gene name gene id
GU305914.1 113159 113181 - 0 GGCGCTGTCGGTGGAGATGA CGG ||||||||[||||||||||||]PAM NA Intergenic NA NA

T25494 GCGCTGTCGGTGGAGATGACGGG 1000
Oligo fwd CACCGCGCTGTCGGTGGAGATGAC
Oligo rev AAACGTCATCTCCACCGACAGCGC
Chromosome start end strand MM target_seq PAM alignment distance position gene name gene id
GU305914.1 113158 113180 - 0 GCGCTGTCGGTGGAGATGAC GGG ||||||||[||||||||||||]PAM NA Intergenic NA NA

T25495 TTCGCGAGCGTGCCGCGCCCTGG 1000
Oligo fwd CACCTTCGCGAGCGTGCCGCGCCC
Oligo rev AAACGGGCGCGGCACGCTCGCGAA
Chromosome start end strand MM target_seq PAM alignment distance position gene name gene id
GU305914.1 113135 113157 - 0 TTCGCGAGCGTGCCGCGCCC TGG ||||||||[||||||||||||]PAM NA Intergenic NA NA

T25496 CCCTGGTGTCGCACCTGATCCGG 1000
Oligo fwd CACCCCCTGGTGTCGCACCTGATC
Oligo rev AAACGATCAGGTGCGACACCAGGG
Chromosome start end strand MM target_seq PAM alignment distance position gene name gene id
GU305914.1 113118 113140 - 0 CCCTGGTGTCGCACCTGATC CGG ||||||||[||||||||||||]PAM NA Intergenic NA NA

T25497 GGTGTCGCACCTGATCCGGACGG 1000
Oligo fwd CACCGGTGTCGCACCTGATCCGGA
Oligo rev AAACTCCGGATCAGGTGCGACACC
Chromosome start end strand MM target_seq PAM alignment distance position gene name gene id
GU305914.1 113114 113136 - 0 GGTGTCGCACCTGATCCGGA CGG ||||||||[||||||||||||]PAM NA Intergenic NA NA

T25498 GCACCTGATCCGGACGGAGACGG 1000
Oligo fwd CACCGCACCTGATCCGGACGGAGA
Oligo rev AAACTCTCCGTCCGGATCAGGTGC
Chromosome start end strand MM target_seq PAM alignment distance position gene name gene id
GU305914.1 113108 113130 - 0 GCACCTGATCCGGACGGAGA CGG ||||||||[||||||||||||]PAM NA Intergenic NA NA
GU305914.1 113321 113343 - 4 GCTGCTGATCCTGAGGGAGA CGG ||--||||[|||-||-|||||]PAM NA Intergenic NA NA

T25499 CACCTGATCCGGACGGAGACGGG 1000
Oligo fwd CACCCACCTGATCCGGACGGAGAC
Oligo rev AAACGTCTCCGTCCGGATCAGGTG

Chromosome start end strand MM target_seq PAM alignment distance position gene name gene id
GU305914.1 113107 113129 - 0 CACCTGATCCGGACGGAGAC GGG ||||||||[||||||||||||]PAM NA Intergenic NA NA

T25500 GATCCGGACGGAGACGGGCCTGG 1000
Oligo fwd CACCGATCCGGACGGAGACGGGCC
Oligo rev AAACGGCCCGTCTCCGTCCGGATC
Chromosome start end strand MM target_seq PAM alignment distance position gene name gene id
GU305914.1 113102 113124 - 0 GATCCGGACGGAGACGGGCC TGG ||||||||[||||||||||||]PAM NA Intergenic NA NA

T25501 GGGCCTGGCCCTGAGTCTGCCGG 1000
Oligo fwd CACCGGGCCTGGCCCTGAGTCTGC
Oligo rev AAACGCAGACTCAGGGCCAGGCCC
Chromosome start end strand MM target_seq PAM alignment distance position gene name gene id
GU305914.1 113087 113109 - 0 GGGCCTGGCCCTGAGTCTGC CGG ||||||||[||||||||||||]PAM NA Intergenic NA NA

T25502 CCGGACGAGCTGCTGTCCGACGG 1000
Oligo fwd CACCCCGGACGAGCTGCTGTCCGA
Oligo rev AAACTCGGACAGCAGCTCGTCCGG
Chromosome start end strand MM target_seq PAM alignment distance position gene name gene id
GU305914.1 113068 113090 - 0 CCGGACGAGCTGCTGTCCGA CGG ||||||||[||||||||||||]PAM NA Intergenic NA NA

T25503 CGGCAACATCTTCTTCTCGCTGG 1000
Oligo fwd CACCCGGCAACATCTTCTTCTCGC
Oligo rev AAACGCGAGAAGAAGATGTTGCCG
Chromosome start end strand MM target_seq PAM alignment distance position gene name gene id
GU305914.1 113048 113070 - 0 CGGCAACATCTTCTTCTCGC TGG ||||||||[||||||||||||]PAM NA Intergenic NA NA

T25504 GGCAACATCTTCTTCTCGCTGGG 1000
Oligo fwd CACCGGCAACATCTTCTTCTCGCT
Oligo rev AAACAGCGAGAAGAAGATGTTGCC
Chromosome start end strand MM target_seq PAM alignment distance position gene name gene id
GU305914.1 113047 113069 - 0 GGCAACATCTTCTTCTCGCT GGG ||||||||[||||||||||||]PAM NA Intergenic NA NA

T25505 CATCTTCTTCTCGCTGGGAACGG 1000
Oligo fwd CACCCATCTTCTTCTCGCTGGGAA
Oligo rev AAACTTCCCAGCGAGAAGAAGATG
Chromosome start end strand MM target_seq PAM alignment distance position gene name gene id
GU305914.1 113042 113064 - 0 CATCTTCTTCTCGCTGGGAA CGG ||||||||[||||||||||||]PAM NA Intergenic NA NA

T25506 TTCTCGCTGGGAACGGTGTACGG 1000
Oligo fwd CACCTTCTCGCTGGGAACGGTGTA
Oligo rev AAACTACACCGTTCCCAGCGAGAA
Chromosome start end strand MM target_seq PAM alignment distance position gene name gene id
GU305914.1 113035 113057 - 0 TTCTCGCTGGGAACGGTGTA CGG ||||||||[||||||||||||]PAM NA Intergenic NA NA

T25507 TCTCGCTGGGAACGGTGTACGGG 1000
Oligo fwd CACCTCTCGCTGGGAACGGTGTAC
Oligo rev AAACGTACACCGTTCCCAGCGAGA
Chromosome start end strand MM target_seq PAM alignment distance position gene name gene id
GU305914.1 113034 113056 - 0 TCTCGCTGGGAACGGTGTAC GGG ||||||||[||||||||||||]PAM NA Intergenic NA NA

T25508 TGTACGGGCACCGTCTGTTCCGG 1000
Oligo fwd CACCTGTACGGGCACCGTCTGTTC
Oligo rev AAACGAACAGACGGTGCCCGTACA
Chromosome start end strand MM target_seq PAM alignment distance position gene name gene id
GU305914.1 113019 113041 - 0 TGTACGGGCACCGTCTGTTC CGG ||||||||[||||||||||||]PAM NA Intergenic NA NA

T25509 GGCTGCTGCGCTTCTTCAACCGG 1000
Oligo fwd CACCGGCTGCTGCGCTTCTTCAAC
Oligo rev AAACGTTGAAGAAGCGCAGCAGCC
Chromosome start end strand MM target_seq PAM alignment distance position gene name gene id
GU305914.1 112998 113020 - 0 GGCTGCTGCGCTTCTTCAAC CGG ||||||||[||||||||||||]PAM NA Intergenic NA NA

T25510 TGCGCTTCTTCAACCGGCACTGG 1000
Oligo fwd CACCTGCGCTTCTTCAACCGGCAC
Oligo rev AAACGTGCCGGTTGAAGAAGCGCA
Chromosome start end strand MM target_seq PAM alignment distance position gene name gene id
GU305914.1 112992 113014 - 0 TGCGCTTCTTCAACCGGCAC TGG ||||||||[||||||||||||]PAM NA Intergenic NA NA

T25511 GCGCTTCTTCAACCGGCACTGGG 1000
Oligo fwd CACCGCGCTTCTTCAACCGGCACT
Oligo rev AAACAGTGCCGGTTGAAGAAGCGC
Chromosome start end strand MM target_seq PAM alignment distance position gene name gene id
GU305914.1 112991 113013 - 0 GCGCTTCTTCAACCGGCACT GGG ||||||||[||||||||||||]PAM NA Intergenic NA NA

T25512 CGCTTCTTCAACCGGCACTGGGG 1000
Oligo fwd CACCCGCTTCTTCAACCGGCACTG
Oligo rev AAACCAGTGCCGGTTGAAGAAGCG
Chromosome start end strand MM target_seq PAM alignment distance position gene name gene id
GU305914.1 112990 113012 - 0 CGCTTCTTCAACCGGCACTG GGG ||||||||[||||||||||||]PAM NA Intergenic NA NA

T25513 CTTCAACCGGCACTGGGGCAAGG 1000
Oligo fwd CACCCTTCAACCGGCACTGGGGCA
Oligo rev AAACTGCCCCAGTGCCGGTTGAAG
Chromosome start end strand MM target_seq PAM alignment distance position gene name gene id
GU305914.1 112985 113007 - 0 CTTCAACCGGCACTGGGGCA AGG ||||||||[||||||||||||]PAM NA Intergenic NA NA

T25514 CAACCGGCACTGGGGCAAGGAGG 1000
Oligo fwd CACCCAACCGGCACTGGGGCAAGG

<table>
<tr><td>Oligo rev</td><td>AAACCCTTGCCCCAGTGCCGGTTG</td><td></td><td></td><td></td><td></td><td></td><td></td><td></td><td></td><td></td><td></td></tr>
<tr><td>Chromosome</td><td>start</td><td>end</td><td>strand</td><td>MM</td><td>target_seq</td><td>PAM</td><td>alignment</td><td>distance</td><td>position</td><td>gene name</td><td>gene id</td></tr>
<tr><td>GU305914.1</td><td>112982</td><td>113004</td><td>-</td><td>0</td><td>CAACCGGCACTGGGGCAAGG</td><td>AGG</td><td>||||||||[||||||||||||]PAM</td><td>NA</td><td>Intergenic</td><td>NA</td><td>NA</td></tr>
<tr><td>T25515</td><td>GGGCAAGGAGGCGCACGAGCCGG</td><td>1000</td><td></td><td></td><td></td><td></td><td></td><td></td><td></td><td></td><td></td></tr>
<tr><td>Oligo fwd</td><td>CACCGGGCAAGGAGGCGCACGAGC</td><td></td><td></td><td></td><td></td><td></td><td></td><td></td><td></td><td></td><td></td></tr>
<tr><td>Oligo rev</td><td>AAACGCTCGTGCGCCTCCTTGCCC</td><td></td><td></td><td></td><td></td><td></td><td></td><td></td><td></td><td></td><td></td></tr>
<tr><td>Chromosome</td><td>start</td><td>end</td><td>strand</td><td>MM</td><td>target_seq</td><td>PAM</td><td>alignment</td><td>distance</td><td>position</td><td>gene name</td><td>gene id</td></tr>
<tr><td>GU305914.1</td><td>112970</td><td>112992</td><td>-</td><td>0</td><td>GGGCAAGGAGGCGCACGAGC</td><td>CGG</td><td>||||||||[||||||||||||]PAM</td><td>NA</td><td>Intergenic</td><td>NA</td><td>NA</td></tr>
<tr><td>T25516</td><td>AGGCGCACGAGCCGGCGATCAGG</td><td>1000</td><td></td><td></td><td></td><td></td><td></td><td></td><td></td><td></td><td></td></tr>
<tr><td>Oligo fwd</td><td>CACCAGGCGCACGAGCCGGCGATC</td><td></td><td></td><td></td><td></td><td></td><td></td><td></td><td></td><td></td><td></td></tr>
<tr><td>Oligo rev</td><td>AAACGATCGCCGGCTCGTGCGCCT</td><td></td><td></td><td></td><td></td><td></td><td></td><td></td><td></td><td></td><td></td></tr>
<tr><td>Chromosome</td><td>start</td><td>end</td><td>strand</td><td>MM</td><td>target_seq</td><td>PAM</td><td>alignment</td><td>distance</td><td>position</td><td>gene name</td><td>gene id</td></tr>
<tr><td>GU305914.1</td><td>112962</td><td>112984</td><td>-</td><td>0</td><td>AGGCGCACGAGCCGGCGATC</td><td>AGG</td><td>||||||||[||||||||||||]PAM</td><td>NA</td><td>Intergenic</td><td>NA</td><td>NA</td></tr>
<tr><td>T25517</td><td>GATCAGGACGATCTGTCAGAAGG</td><td>1000</td><td></td><td></td><td></td><td></td><td></td><td></td><td></td><td></td><td></td></tr>
<tr><td>Oligo fwd</td><td>CACCGATCAGGACGATCTGTCAGA</td><td></td><td></td><td></td><td></td><td></td><td></td><td></td><td></td><td></td><td></td></tr>
<tr><td>Oligo rev</td><td>AAACTCTGACAGATCGTCCTGATC</td><td></td><td></td><td></td><td></td><td></td><td></td><td></td><td></td><td></td><td></td></tr>
<tr><td>Chromosome</td><td>start</td><td>end</td><td>strand</td><td>MM</td><td>target_seq</td><td>PAM</td><td>alignment</td><td>distance</td><td>position</td><td>gene name</td><td>gene id</td></tr>
<tr><td>GU305914.1</td><td>112946</td><td>112968</td><td>-</td><td>0</td><td>GATCAGGACGATCTGTCAGA</td><td>AGG</td><td>||||||||[||||||||||||]PAM</td><td>NA</td><td>Intergenic</td><td>NA</td><td>NA</td></tr>
<tr><td>T25518</td><td>GGACGATCTGTCAGAAGGTGTGG</td><td>1000</td><td></td><td></td><td></td><td></td><td></td><td></td><td></td><td></td><td></td></tr>
<tr><td>Oligo fwd</td><td>CACCGGACGATCTGTCAGAAGGTG</td><td></td><td></td><td></td><td></td><td></td><td></td><td></td><td></td><td></td><td></td></tr>
<tr><td>Oligo rev</td><td>AAACCACCTTCTGACAGATCGTCC</td><td></td><td></td><td></td><td></td><td></td><td></td><td></td><td></td><td></td><td></td></tr>
<tr><td>Chromosome</td><td>start</td><td>end</td><td>strand</td><td>MM</td><td>target_seq</td><td>PAM</td><td>alignment</td><td>distance</td><td>position</td><td>gene name</td><td>gene id</td></tr>
<tr><td>GU305914.1</td><td>112941</td><td>112963</td><td>-</td><td>0</td><td>GGACGATCTGTCAGAAGGTG</td><td>TGG</td><td>||||||||[||||||||||||]PAM</td><td>NA</td><td>Intergenic</td><td>NA</td><td>NA</td></tr>
<tr><td>T25519</td><td>GGTTCTTCTACCTGATCGCCTGG</td><td>1000</td><td></td><td></td><td></td><td></td><td></td><td></td><td></td><td></td><td></td></tr>
<tr><td>Oligo fwd</td><td>CACCGGTTCTTCTACCTGATCGCC</td><td></td><td></td><td></td><td></td><td></td><td></td><td></td><td></td><td></td><td></td></tr>
<tr><td>Oligo rev</td><td>AAACGGCGATCAGGTAGAAGAACC</td><td></td><td></td><td></td><td></td><td></td><td></td><td></td><td></td><td></td><td></td></tr>
<tr><td>Chromosome</td><td>start</td><td>end</td><td>strand</td><td>MM</td><td>target_seq</td><td>PAM</td><td>alignment</td><td>distance</td><td>position</td><td>gene name</td><td>gene id</td></tr>
<tr><td>GU305914.1</td><td>112920</td><td>112942</td><td>-</td><td>0</td><td>GGTTCTTCTACCTGATCGCC</td><td>TGG</td><td>||||||||[||||||||||||]PAM</td><td>NA</td><td>Intergenic</td><td>NA</td><td>NA</td></tr>
<tr><td>T25520</td><td>GATCGCCTGGAAAAAGCTGACGG</td><td>1000</td><td></td><td></td><td></td><td></td><td></td><td></td><td></td><td></td><td></td></tr>
<tr><td>Oligo fwd</td><td>CACCGATCGCCTGGAAAAAGCTGA</td><td></td><td></td><td></td><td></td><td></td><td></td><td></td><td></td><td></td><td></td></tr>
<tr><td>Oligo rev</td><td>AAACTCAGCTTTTTCCAGGCGATC</td><td></td><td></td><td></td><td></td><td></td><td></td><td></td><td></td><td></td><td></td></tr>
<tr><td>Chromosome</td><td>start</td><td>end</td><td>strand</td><td>MM</td><td>target_seq</td><td>PAM</td><td>alignment</td><td>distance</td><td>position</td><td>gene name</td><td>gene id</td></tr>
<tr><td>GU305914.1</td><td>112907</td><td>112929</td><td>-</td><td>0</td><td>GATCGCCTGGAAAAAGCTGA</td><td>CGG</td><td>||||||||[||||||||||||]PAM</td><td>NA</td><td>Intergenic</td><td>NA</td><td>NA</td></tr>
<tr><td>T25521</td><td>AAAGCTGACGGTGTCGCCCGAGG</td><td>1000</td><td></td><td></td><td></td><td></td><td></td><td></td><td></td><td></td><td></td></tr>
<tr><td>Oligo fwd</td><td>CACCAAAGCTGACGGTGTCGCCCG</td><td></td><td></td><td></td><td></td><td></td><td></td><td></td><td></td><td></td><td></td></tr>
<tr><td>Oligo rev</td><td>AAACCGGGCGACACCGTCAGCTTT</td><td></td><td></td><td></td><td></td><td></td><td></td><td></td><td></td><td></td><td></td></tr>
<tr><td>Chromosome</td><td>start</td><td>end</td><td>strand</td><td>MM</td><td>target_seq</td><td>PAM</td><td>alignment</td><td>distance</td><td>position</td><td>gene name</td><td>gene id</td></tr>
<tr><td>GU305914.1</td><td>112895</td><td>112917</td><td>-</td><td>0</td><td>AAAGCTGACGGTGTCGCCCG</td><td>AGG</td><td>||||||||[||||||||||||]PAM</td><td>NA</td><td>Intergenic</td><td>NA</td><td>NA</td></tr>
<tr><td>T25522</td><td>GCAGCGCTCTGACCACGAGCTGG</td><td>1000</td><td></td><td></td><td></td><td></td><td></td><td></td><td></td><td></td><td></td></tr>
<tr><td>Oligo fwd</td><td>CACCGCAGCGCTCTGACCACGAGC</td><td></td><td></td><td></td><td></td><td></td><td></td><td></td><td></td><td></td><td></td></tr>
<tr><td>Oligo rev</td><td>AAACGCTCGTGGTCAGAGCGCTGC</td><td></td><td></td><td></td><td></td><td></td><td></td><td></td><td></td><td></td><td></td></tr>
<tr><td>Chromosome</td><td>start</td><td>end</td><td>strand</td><td>MM</td><td>target_seq</td><td>PAM</td><td>alignment</td><td>distance</td><td>position</td><td>gene name</td><td>gene id</td></tr>
<tr><td>GU305914.1</td><td>112862</td><td>112884</td><td>-</td><td>0</td><td>GCAGCGCTCTGACCACGAGC</td><td>TGG</td><td>||||||||[||||||||||||]PAM</td><td>NA</td><td>Intergenic</td><td>NA</td><td>NA</td></tr>
<tr><td>T25523</td><td>CAGCGCTCTGACCACGAGCTGGG</td><td>1000</td><td></td><td></td><td></td><td></td><td></td><td></td><td></td><td></td><td></td></tr>
<tr><td>Oligo fwd</td><td>CACCCAGCGCTCTGACCACGAGCT</td><td></td><td></td><td></td><td></td><td></td><td></td><td></td><td></td><td></td><td></td></tr>
<tr><td>Oligo rev</td><td>AAACAGCTCGTGGTCAGAGCGCTG</td><td></td><td></td><td></td><td></td><td></td><td></td><td></td><td></td><td></td><td></td></tr>
<tr><td>Chromosome</td><td>start</td><td>end</td><td>strand</td><td>MM</td><td>target_seq</td><td>PAM</td><td>alignment</td><td>distance</td><td>position</td><td>gene name</td><td>gene id</td></tr>
<tr><td>GU305914.1</td><td>112861</td><td>112883</td><td>-</td><td>0</td><td>CAGCGCTCTGACCACGAGCT</td><td>GGG</td><td>||||||||[||||||||||||]PAM</td><td>NA</td><td>Intergenic</td><td>NA</td><td>NA</td></tr>
<tr><td>T25524</td><td>AGCGCTCTGACCACGAGCTGGGG</td><td>1000</td><td></td><td></td><td></td><td></td><td></td><td></td><td></td><td></td><td></td></tr>
<tr><td>Oligo fwd</td><td>CACCAGCGCTCTGACCACGAGCTG</td><td></td><td></td><td></td><td></td><td></td><td></td><td></td><td></td><td></td><td></td></tr>
<tr><td>Oligo rev</td><td>AAACCAGCTCGTGGTCAGAGCGCT</td><td></td><td></td><td></td><td></td><td></td><td></td><td></td><td></td><td></td><td></td></tr>
<tr><td>Chromosome</td><td>start</td><td>end</td><td>strand</td><td>MM</td><td>target_seq</td><td>PAM</td><td>alignment</td><td>distance</td><td>position</td><td>gene name</td><td>gene id</td></tr>
<tr><td>GU305914.1</td><td>112860</td><td>112882</td><td>-</td><td>0</td><td>AGCGCTCTGACCACGAGCTG</td><td>GGG</td><td>||||||||[||||||||||||]PAM</td><td>NA</td><td>Intergenic</td><td>NA</td><td>NA</td></tr>
<tr><td>T25525</td><td>GATCTTTTCCTTCCTGATCCAGG</td><td>1000</td><td></td><td></td><td></td><td></td><td></td><td></td><td></td><td></td><td></td></tr>
<tr><td>Oligo fwd</td><td>CACCGATCTTTTCCTTCCTGATCC</td><td></td><td></td><td></td><td></td><td></td><td></td><td></td><td></td><td></td><td></td></tr>
<tr><td>Oligo rev</td><td>AAACGGATCAGGAAGGAAAAGATC</td><td></td><td></td><td></td><td></td><td></td><td></td><td></td><td></td><td></td><td></td></tr>
<tr><td>Chromosome</td><td>start</td><td>end</td><td>strand</td><td>MM</td><td>target_seq</td><td>PAM</td><td>alignment</td><td>distance</td><td>position</td><td>gene name</td><td>gene id</td></tr>
<tr><td>GU305914.1</td><td>112838</td><td>112860</td><td>-</td><td>0</td><td>GATCTTTTCCTTCCTGATCC</td><td>AGG</td><td>||||||||[||||||||||||]PAM</td><td>NA</td><td>Intergenic</td><td>NA</td><td>NA</td></tr>
<tr><td>T25526</td><td>CCAGGACTACCTGACCTTCACGG</td><td>1000</td><td></td><td></td><td></td><td></td><td></td><td></td><td></td><td></td><td></td></tr>
<tr><td>Oligo fwd</td><td>CACCCCAGGACTACCTGACCTTCA</td><td></td><td></td><td></td><td></td><td></td><td></td><td></td><td></td><td></td><td></td></tr>
<tr><td>Oligo rev</td><td>AAACTGAAGGTCAGGTAGTCCTGG</td><td></td><td></td><td></td><td></td><td></td><td></td><td></td><td></td><td></td><td></td></tr>
<tr><td>Chromosome</td><td>start</td><td>end</td><td>strand</td><td>MM</td><td>target_seq</td><td>PAM</td><td>alignment</td><td>distance</td><td>position</td><td>gene name</td><td>gene id</td></tr>
<tr><td>GU305914.1</td><td>112820</td><td>112842</td><td>-</td><td>0</td><td>CCAGGACTACCTGACCTTCA</td><td>CGG</td><td>||||||||[||||||||||||]PAM</td><td>NA</td><td>Intergenic</td><td>NA</td><td>NA</td></tr>
<tr><td>T25527</td><td>CAGGACTACCTGACCTTCACGGG</td><td>1000</td><td></td><td></td><td></td><td></td><td></td><td></td><td></td><td></td><td></td></tr>
<tr><td>Oligo fwd</td><td>CACCCAGGACTACCTGACCTTCAC</td><td></td><td></td><td></td><td></td><td></td><td></td><td></td><td></td><td></td><td></td></tr>
<tr><td>Oligo rev</td><td>AAACGTGAAGGTCAGGTAGTCCTG</td><td></td><td></td><td></td><td></td><td></td><td></td><td></td><td></td><td></td><td></td></tr>
<tr><td>Chromosome</td><td>start</td><td>end</td><td>strand</td><td>MM</td><td>target_seq</td><td>PAM</td><td>alignment</td><td>distance</td><td>position</td><td>gene name</td><td>gene id</td></tr>
<tr><td>GU305914.1</td><td>112819</td><td>112841</td><td>-</td><td>0</td><td>CAGGACTACCTGACCTTCAC</td><td>GGG</td><td>||||||||[||||||||||||]PAM</td><td>NA</td><td>Intergenic</td><td>NA</td><td>NA</td></tr>
<tr><td>T25528</td><td>GCGACGCAGCACGCCGCCCATGG</td><td>1000</td><td></td><td></td><td></td><td></td><td></td><td></td><td></td><td></td><td></td></tr>
<tr><td>Oligo fwd</td><td>CACCGCGACGCAGCACGCCGCCCA</td><td></td><td></td><td></td><td></td><td></td><td></td><td></td><td></td><td></td><td></td></tr>
<tr><td>Oligo rev</td><td>AAACTGGGCGGCGTGCTGCGTCGC</td><td></td><td></td><td></td><td></td><td></td><td></td><td></td><td></td><td></td><td></td></tr>
<tr><td>Chromosome</td><td>start</td><td>end</td><td>strand</td><td>MM</td><td>target_seq</td><td>PAM</td><td>alignment</td><td>distance</td><td>position</td><td>gene name</td><td>gene id</td></tr>
<tr><td>GU305914.1</td><td>112790</td><td>112812</td><td>-</td><td>0</td><td>GCGACGCAGCACGCCGCCCA</td><td>TGG</td><td>||||||||[||||||||||||]PAM</td><td>NA</td><td>Intergenic</td><td>NA</td><td>NA</td></tr>
<tr><td>T25529</td><td>CACGCCGCCCATGGACAAGAAGG</td><td>1000</td><td></td><td></td><td></td><td></td><td></td><td></td><td></td><td></td><td></td></tr>
</table>

<table>
<tr><td>Oligo fwd</td><td>CACCCACGCCGCCCATGGACAAGA</td><td></td><td></td><td></td><td></td><td></td><td></td><td></td><td></td><td></td><td></td></tr>
<tr><td>Oligo rev</td><td>AAACTCTTGTCCATGGGCGGCGTG</td><td></td><td></td><td></td><td></td><td></td><td></td><td></td><td></td><td></td><td></td></tr>
<tr><td>Chromosome</td><td>start</td><td>end</td><td>strand</td><td>MM</td><td>target_seq</td><td>PAM</td><td>alignment</td><td>distance</td><td>position</td><td>gene name</td><td>gene id</td></tr>
<tr><td>GU305914.1</td><td>112781</td><td>112803</td><td>-</td><td>0</td><td>CACGCCGCCCATGGACAAGA</td><td>AGG</td><td>||||||||[||||||||||||]PAM</td><td>NA</td><td>Intergenic</td><td>NA</td><td>NA</td></tr>
<tr><td></td><td></td><td></td><td></td><td></td><td></td><td></td><td></td><td></td><td></td><td></td><td></td></tr>
<tr><td>T25530</td><td>GCCGCCCATGGACAAGAAGGAGG</td><td>1000</td><td></td><td></td><td></td><td></td><td></td><td></td><td></td><td></td><td></td></tr>
<tr><td>Oligo fwd</td><td>CACCGCCGCCCATGGACAAGAAGG</td><td></td><td></td><td></td><td></td><td></td><td></td><td></td><td></td><td></td><td></td></tr>
<tr><td>Oligo rev</td><td>AAACCCTTCTTGTCCATGGGCGGC</td><td></td><td></td><td></td><td></td><td></td><td></td><td></td><td></td><td></td><td></td></tr>
<tr><td>Chromosome</td><td>start</td><td>end</td><td>strand</td><td>MM</td><td>target_seq</td><td>PAM</td><td>alignment</td><td>distance</td><td>position</td><td>gene name</td><td>gene id</td></tr>
<tr><td>GU305914.1</td><td>112778</td><td>112800</td><td>-</td><td>0</td><td>GCCGCCCATGGACAAGAAGG</td><td>AGG</td><td>||||||||[||||||||||||]PAM</td><td>NA</td><td>Intergenic</td><td>NA</td><td>NA</td></tr>
<tr><td></td><td></td><td></td><td></td><td></td><td></td><td></td><td></td><td></td><td></td><td></td><td></td></tr>
<tr><td>T25531</td><td>CCCATGGACAAGAAGGAGGAAGG</td><td>1000</td><td></td><td></td><td></td><td></td><td></td><td></td><td></td><td></td><td></td></tr>
<tr><td>Oligo fwd</td><td>CACCCCCATGGACAAGAAGGAGGA</td><td></td><td></td><td></td><td></td><td></td><td></td><td></td><td></td><td></td><td></td></tr>
<tr><td>Oligo rev</td><td>AAACTCCTCCTTCTTGTCCATGGG</td><td></td><td></td><td></td><td></td><td></td><td></td><td></td><td></td><td></td><td></td></tr>
<tr><td>Chromosome</td><td>start</td><td>end</td><td>strand</td><td>MM</td><td>target_seq</td><td>PAM</td><td>alignment</td><td>distance</td><td>position</td><td>gene name</td><td>gene id</td></tr>
<tr><td>GU305914.1</td><td>112774</td><td>112796</td><td>-</td><td>0</td><td>CCCATGGACAAGAAGGAGGA</td><td>AGG</td><td>||||||||[||||||||||||]PAM</td><td>NA</td><td>Intergenic</td><td>NA</td><td>NA</td></tr>
<tr><td></td><td></td><td></td><td></td><td></td><td></td><td></td><td></td><td></td><td></td><td></td><td></td></tr>
<tr><td>T25532</td><td>GTGATCGCCGACCTGCTGAGCGG</td><td>1000</td><td></td><td></td><td></td><td></td><td></td><td></td><td></td><td></td><td></td></tr>
<tr><td>Oligo fwd</td><td>CACCGTGATCGCCGACCTGCTGAG</td><td></td><td></td><td></td><td></td><td></td><td></td><td></td><td></td><td></td><td></td></tr>
<tr><td>Oligo rev</td><td>AAACCTCAGCAGGTCGGCGATCAC</td><td></td><td></td><td></td><td></td><td></td><td></td><td></td><td></td><td></td><td></td></tr>
<tr><td>Chromosome</td><td>start</td><td>end</td><td>strand</td><td>MM</td><td>target_seq</td><td>PAM</td><td>alignment</td><td>distance</td><td>position</td><td>gene name</td><td>gene id</td></tr>
<tr><td>GU305914.1</td><td>112750</td><td>112772</td><td>-</td><td>0</td><td>GTGATCGCCGACCTGCTGAG</td><td>CGG</td><td>||||||||[||||||||||||]PAM</td><td>NA</td><td>Intergenic</td><td>NA</td><td>NA</td></tr>
<tr><td></td><td></td><td></td><td></td><td></td><td></td><td></td><td></td><td></td><td></td><td></td><td></td></tr>
<tr><td>T25533</td><td>AAAAAAAAACGACCCACACACGG</td><td>1000</td><td></td><td></td><td></td><td></td><td></td><td></td><td></td><td></td><td></td></tr>
<tr><td>Oligo fwd</td><td>CACCAAAAAAAAACGACCCACACA</td><td></td><td></td><td></td><td></td><td></td><td></td><td></td><td></td><td></td><td></td></tr>
<tr><td>Oligo rev</td><td>AAACTGTGTGGGTCGTTTTTTTTT</td><td></td><td></td><td></td><td></td><td></td><td></td><td></td><td></td><td></td><td></td></tr>
<tr><td>Chromosome</td><td>start</td><td>end</td><td>strand</td><td>MM</td><td>target_seq</td><td>PAM</td><td>alignment</td><td>distance</td><td>position</td><td>gene name</td><td>gene id</td></tr>
<tr><td>GU305914.1</td><td>112695</td><td>112717</td><td>-</td><td>0</td><td>AAAAAAAAACGACCCACACA</td><td>CGG</td><td>||||||||[||||||||||||]PAM</td><td>NA</td><td>Intergenic</td><td>NA</td><td>NA</td></tr>
<tr><td></td><td></td><td></td><td></td><td></td><td></td><td></td><td></td><td></td><td></td><td></td><td></td></tr>
<tr><td>T25534</td><td>ATCATCCACCGTCGTTACGACGG</td><td>1000</td><td></td><td></td><td></td><td></td><td></td><td></td><td></td><td></td><td></td></tr>
<tr><td>Oligo fwd</td><td>CACCATCATCCACCGTCGTTACGA</td><td></td><td></td><td></td><td></td><td></td><td></td><td></td><td></td><td></td><td></td></tr>
<tr><td>Oligo rev</td><td>AAACTCGTAACGACGGTGGATGAT</td><td></td><td></td><td></td><td></td><td></td><td></td><td></td><td></td><td></td><td></td></tr>
<tr><td>Chromosome</td><td>start</td><td>end</td><td>strand</td><td>MM</td><td>target_seq</td><td>PAM</td><td>alignment</td><td>distance</td><td>position</td><td>gene name</td><td>gene id</td></tr>
<tr><td>GU305914.1</td><td>112600</td><td>112622</td><td>-</td><td>0</td><td>ATCATCCACCGTCGTTACGA</td><td>CGG</td><td>||||||||[||||||||||||]PAM</td><td>NA</td><td>Intergenic</td><td>NA</td><td>NA</td></tr>
<tr><td></td><td></td><td></td><td></td><td></td><td></td><td></td><td></td><td></td><td></td><td></td><td></td></tr>
<tr><td>T25535</td><td>GTTTTTATTTCAGACAACAGAGG</td><td>1000</td><td></td><td></td><td></td><td></td><td></td><td></td><td></td><td></td><td></td></tr>
<tr><td>Oligo fwd</td><td>CACCGTTTTTATTTCAGACAACAG</td><td></td><td></td><td></td><td></td><td></td><td></td><td></td><td></td><td></td><td></td></tr>
<tr><td>Oligo rev</td><td>AAACCTGTTGTCTGAAATAAAAAC</td><td></td><td></td><td></td><td></td><td></td><td></td><td></td><td></td><td></td><td></td></tr>
<tr><td>Chromosome</td><td>start</td><td>end</td><td>strand</td><td>MM</td><td>target_seq</td><td>PAM</td><td>alignment</td><td>distance</td><td>position</td><td>gene name</td><td>gene id</td></tr>
<tr><td>GU305914.1</td><td>112486</td><td>112508</td><td>-</td><td>0</td><td>GTTTTTATTTCAGACAACAG</td><td>AGG</td><td>||||||||[||||||||||||]PAM</td><td>NA</td><td>Intergenic</td><td>NA</td><td>NA</td></tr>
<tr><td></td><td></td><td></td><td></td><td></td><td></td><td></td><td></td><td></td><td></td><td></td><td></td></tr>
<tr><td>T25536</td><td>TTTATTTCAGACAACAGAGGAGG</td><td>1000</td><td></td><td></td><td></td><td></td><td></td><td></td><td></td><td></td><td></td></tr>
<tr><td>Oligo fwd</td><td>CACCTTTATTTCAGACAACAGAGG</td><td></td><td></td><td></td><td></td><td></td><td></td><td></td><td></td><td></td><td></td></tr>
<tr><td>Oligo rev</td><td>AAACCCTCTGTTGTCTGAAATAAA</td><td></td><td></td><td></td><td></td><td></td><td></td><td></td><td></td><td></td><td></td></tr>
<tr><td>Chromosome</td><td>start</td><td>end</td><td>strand</td><td>MM</td><td>target_seq</td><td>PAM</td><td>alignment</td><td>distance</td><td>position</td><td>gene name</td><td>gene id</td></tr>
<tr><td>GU305914.1</td><td>112483</td><td>112505</td><td>-</td><td>0</td><td>TTTATTTCAGACAACAGAGG</td><td>AGG</td><td>||||||||[||||||||||||]PAM</td><td>NA</td><td>Intergenic</td><td>NA</td><td>NA</td></tr>
<tr><td></td><td></td><td></td><td></td><td></td><td></td><td></td><td></td><td></td><td></td><td></td><td></td></tr>
<tr><td>T25537</td><td>ATTTCAGACAACAGAGGAGGAGG</td><td>1000</td><td></td><td></td><td></td><td></td><td></td><td></td><td></td><td></td><td></td></tr>
<tr><td>Oligo fwd</td><td>CACCATTTCAGACAACAGAGGAGG</td><td></td><td></td><td></td><td></td><td></td><td></td><td></td><td></td><td></td><td></td></tr>
<tr><td>Oligo rev</td><td>AAACCCTCCTCTGTTGTCTGAAAT</td><td></td><td></td><td></td><td></td><td></td><td></td><td></td><td></td><td></td><td></td></tr>
<tr><td>Chromosome</td><td>start</td><td>end</td><td>strand</td><td>MM</td><td>target_seq</td><td>PAM</td><td>alignment</td><td>distance</td><td>position</td><td>gene name</td><td>gene id</td></tr>
<tr><td>GU305914.1</td><td>112480</td><td>112502</td><td>-</td><td>0</td><td>ATTTCAGACAACAGAGGAGG</td><td>AGG</td><td>||||||||[||||||||||||]PAM</td><td>NA</td><td>Intergenic</td><td>NA</td><td>NA</td></tr>
<tr><td></td><td></td><td></td><td></td><td></td><td></td><td></td><td></td><td></td><td></td><td></td><td></td></tr>
<tr><td>T25538</td><td>CAGACAACAGAGGAGGAGGTAGG</td><td>1000</td><td></td><td></td><td></td><td></td><td></td><td></td><td></td><td></td><td></td></tr>
<tr><td>Oligo fwd</td><td>CACCCAGACAACAGAGGAGGAGGT</td><td></td><td></td><td></td><td></td><td></td><td></td><td></td><td></td><td></td><td></td></tr>
<tr><td>Oligo rev</td><td>AAACACCTCCTCCTCTGTTGTCTG</td><td></td><td></td><td></td><td></td><td></td><td></td><td></td><td></td><td></td><td></td></tr>
<tr><td>Chromosome</td><td>start</td><td>end</td><td>strand</td><td>MM</td><td>target_seq</td><td>PAM</td><td>alignment</td><td>distance</td><td>position</td><td>gene name</td><td>gene id</td></tr>
<tr><td>GU305914.1</td><td>112476</td><td>112498</td><td>-</td><td>0</td><td>CAGACAACAGAGGAGGAGGT</td><td>AGG</td><td>||||||||[||||||||||||]PAM</td><td>NA</td><td>Intergenic</td><td>NA</td><td>NA</td></tr>
<tr><td></td><td></td><td></td><td></td><td></td><td></td><td></td><td></td><td></td><td></td><td></td><td></td></tr>
<tr><td>T25539</td><td>CAGAGGAGGAGGTAGGCAGTAGG</td><td>1000</td><td></td><td></td><td></td><td></td><td></td><td></td><td></td><td></td><td></td></tr>
<tr><td>Oligo fwd</td><td>CACCCAGAGGAGGAGGTAGGCAGT</td><td></td><td></td><td></td><td></td><td></td><td></td><td></td><td></td><td></td><td></td></tr>
<tr><td>Oligo rev</td><td>AAACACTGCCTACCTCCTCCTCTG</td><td></td><td></td><td></td><td></td><td></td><td></td><td></td><td></td><td></td><td></td></tr>
<tr><td>Chromosome</td><td>start</td><td>end</td><td>strand</td><td>MM</td><td>target_seq</td><td>PAM</td><td>alignment</td><td>distance</td><td>position</td><td>gene name</td><td>gene id</td></tr>
<tr><td>GU305914.1</td><td>112469</td><td>112491</td><td>-</td><td>0</td><td>CAGAGGAGGAGGTAGGCAGT</td><td>AGG</td><td>||||||||[||||||||||||]PAM</td><td>NA</td><td>Intergenic</td><td>NA</td><td>NA</td></tr>
<tr><td></td><td></td><td></td><td></td><td></td><td></td><td></td><td></td><td></td><td></td><td></td><td></td></tr>
<tr><td>T25540</td><td>AGAGGAGGAGGTAGGCAGTAGGG</td><td>1000</td><td></td><td></td><td></td><td></td><td></td><td></td><td></td><td></td><td></td></tr>
<tr><td>Oligo fwd</td><td>CACCAGAGGAGGAGGTAGGCAGTA</td><td></td><td></td><td></td><td></td><td></td><td></td><td></td><td></td><td></td><td></td></tr>
<tr><td>Oligo rev</td><td>AAACTACTGCCTACCTCCTCCTCT</td><td></td><td></td><td></td><td></td><td></td><td></td><td></td><td></td><td></td><td></td></tr>
<tr><td>Chromosome</td><td>start</td><td>end</td><td>strand</td><td>MM</td><td>target_seq</td><td>PAM</td><td>alignment</td><td>distance</td><td>position</td><td>gene name</td><td>gene id</td></tr>
<tr><td>GU305914.1</td><td>112468</td><td>112490</td><td>-</td><td>0</td><td>AGAGGAGGAGGTAGGCAGTA</td><td>GGG</td><td>||||||||[||||||||||||]PAM</td><td>NA</td><td>Intergenic</td><td>NA</td><td>NA</td></tr>
<tr><td></td><td></td><td></td><td></td><td></td><td></td><td></td><td></td><td></td><td></td><td></td><td></td></tr>
<tr><td>T25541</td><td>CAGTAGGGCCGAGACATCATCGG</td><td>1000</td><td></td><td></td><td></td><td></td><td></td><td></td><td></td><td></td><td></td></tr>
<tr><td>Oligo fwd</td><td>CACCCAGTAGGGCCGAGACATCAT</td><td></td><td></td><td></td><td></td><td></td><td></td><td></td><td></td><td></td><td></td></tr>
<tr><td>Oligo rev</td><td>AAACATGATGTCTCGGCCCTACTG</td><td></td><td></td><td></td><td></td><td></td><td></td><td></td><td></td><td></td><td></td></tr>
<tr><td>Chromosome</td><td>start</td><td>end</td><td>strand</td><td>MM</td><td>target_seq</td><td>PAM</td><td>alignment</td><td>distance</td><td>position</td><td>gene name</td><td>gene id</td></tr>
<tr><td>GU305914.1</td><td>112453</td><td>112475</td><td>-</td><td>0</td><td>CAGTAGGGCCGAGACATCAT</td><td>CGG</td><td>||||||||[||||||||||||]PAM</td><td>NA</td><td>Intergenic</td><td>NA</td><td>NA</td></tr>
<tr><td></td><td></td><td></td><td></td><td></td><td></td><td></td><td></td><td></td><td></td><td></td><td></td></tr>
<tr><td>T25542</td><td>GACATCATCGGTCACCCCGTCGG</td><td>1000</td><td></td><td></td><td></td><td></td><td></td><td></td><td></td><td></td><td></td></tr>
<tr><td>Oligo fwd</td><td>CACCGACATCATCGGTCACCCCGT</td><td></td><td></td><td></td><td></td><td></td><td></td><td></td><td></td><td></td><td></td></tr>
<tr><td>Oligo rev</td><td>AAACACGGGGTGACCGATGATGTC</td><td></td><td></td><td></td><td></td><td></td><td></td><td></td><td></td><td></td><td></td></tr>
<tr><td>Chromosome</td><td>start</td><td>end</td><td>strand</td><td>MM</td><td>target_seq</td><td>PAM</td><td>alignment</td><td>distance</td><td>position</td><td>gene name</td><td>gene id</td></tr>
<tr><td>GU305914.1</td><td>112441</td><td>112463</td><td>-</td><td>0</td><td>GACATCATCGGTCACCCCGT</td><td>CGG</td><td>||||||||[||||||||||||]PAM</td><td>NA</td><td>Intergenic</td><td>NA</td><td>NA</td></tr>
<tr><td></td><td></td><td></td><td></td><td></td><td></td><td></td><td></td><td></td><td></td><td></td><td></td></tr>
<tr><td>T25543</td><td>TCATCGGTCACCCCGTCGGACGG</td><td>1000</td><td></td><td></td><td></td><td></td><td></td><td></td><td></td><td></td><td></td></tr>
<tr><td>Oligo fwd</td><td>CACCTCATCGGTCACCCCGTCGGA</td><td></td><td></td><td></td><td></td><td></td><td></td><td></td><td></td><td></td><td></td></tr>
<tr><td>Oligo rev</td><td>AAACTCCGACGGGGTGACCGATGA</td><td></td><td></td><td></td><td></td><td></td><td></td><td></td><td></td><td></td><td></td></tr>
<tr><td>Chromosome</td><td>start</td><td>end</td><td>strand</td><td>MM</td><td>target_seq</td><td>PAM</td><td>alignment</td><td>distance</td><td>position</td><td>gene name</td><td>gene id</td></tr>
<tr><td>GU305914.1</td><td>112437</td><td>112459</td><td>-</td><td>0</td><td>TCATCGGTCACCCCGTCGGA</td><td>CGG</td><td>||||||||[||||||||||||]PAM</td><td>NA</td><td>Intergenic</td><td>NA</td><td>NA</td></tr>
</table>

| | | | | | | | | | | | | |
|---|---|---|---|---|---|---|---|---|---|---|---|---|
| T25544 | TCGGTCACCCCGTCGGACGGCGG | | 1000 | | | | | | | | | |
| Oligo fwd | CACCTCGGTCACCCCGTCGGACGG | | | | | | | | | | | |
| Oligo rev | AAACCCGTCCGACGGGGTGACCGA | | | | | | | | | | | |
| Chromosome | start | | end | strand | MM | target_seq | PAM | alignment | distance | position | gene name | gene id |
| GU305914.1 | | 112434 | 112456 | - | 0 | TCGGTCACCCCGTCGGACGG | CGG | \|\|\|\|\|\|\|\|[\|\|\|\|\|\|\|\|\|\|\|\|]PAM | NA | Intergenic | NA | NA |
| T25545 | ACCCCGTCGGACGGCGGTATCGG | | 1000 | | | | | | | | | |
| Oligo fwd | CACCACCCCGTCGGACGGCGGTAT | | | | | | | | | | | |
| Oligo rev | AAACATACCGCCGTCCGACGGGGT | | | | | | | | | | | |
| Chromosome | start | | end | strand | MM | target_seq | PAM | alignment | distance | position | gene name | gene id |
| GU305914.1 | | 112428 | 112450 | - | 0 | ACCCCGTCGGACGGCGGTAT | CGG | \|\|\|\|\|\|\|\|[\|\|\|\|\|\|\|\|\|\|\|\|]PAM | NA | Intergenic | NA | NA |
| T25546 | GTCCGAGTAAAAAACAGACATGG | | 1000 | | | | | | | | | |
| Oligo fwd | CACCGTCCGAGTAAAAAACAGACA | | | | | | | | | | | |
| Oligo rev | AAACTGTCTGTTTTTTACTCGGAC | | | | | | | | | | | |
| Chromosome | start | | end | strand | MM | target_seq | PAM | alignment | distance | position | gene name | gene id |
| GU305914.1 | | 112357 | 112379 | - | 0 | GTCCGAGTAAAAAACAGACA | TGG | \|\|\|\|\|\|\|\|[\|\|\|\|\|\|\|\|\|\|\|\|]PAM | NA | Intergenic | NA | NA |
| T25547 | TCCGAGTAAAAAACAGACATGGG | | 1000 | | | | | | | | | |
| Oligo fwd | CACCTCCGAGTAAAAAACAGACAT | | | | | | | | | | | |
| Oligo rev | AAACATGTCTGTTTTTTACTCGGA | | | | | | | | | | | |
| Chromosome | start | | end | strand | MM | target_seq | PAM | alignment | distance | position | gene name | gene id |
| GU305914.1 | | 112356 | 112378 | - | 0 | TCCGAGTAAAAAACAGACAT | GGG | \|\|\|\|\|\|\|\|[\|\|\|\|\|\|\|\|\|\|\|\|]PAM | NA | Intergenic | NA | NA |
| T25548 | AGTAAAAAACAGACATGGGTAGG | | 1000 | | | | | | | | | |
| Oligo fwd | CACCAGTAAAAAACAGACATGGGT | | | | | | | | | | | |
| Oligo rev | AAACACCCATGTCTGTTTTTTACT | | | | | | | | | | | |
| Chromosome | start | | end | strand | MM | target_seq | PAM | alignment | distance | position | gene name | gene id |
| GU305914.1 | | 112352 | 112374 | - | 0 | AGTAAAAAACAGACATGGGT | AGG | \|\|\|\|\|\|\|\|[\|\|\|\|\|\|\|\|\|\|\|\|]PAM | NA | Intergenic | NA | NA |
| T25549 | GTAAAAAACAGACATGGGTAGGG | | 1000 | | | | | | | | | |
| Oligo fwd | CACCGTAAAAAACAGACATGGGTA | | | | | | | | | | | |
| Oligo rev | AAACTACCCATGTCTGTTTTTTAC | | | | | | | | | | | |
| Chromosome | start | | end | strand | MM | target_seq | PAM | alignment | distance | position | gene name | gene id |
| GU305914.1 | | 112351 | 112373 | - | 0 | GTAAAAAACAGACATGGGTA | GGG | \|\|\|\|\|\|\|\|[\|\|\|\|\|\|\|\|\|\|\|\|]PAM | NA | Intergenic | NA | NA |
| T25550 | GTAGGGACGTCATCTTCTGCTGG | | 1000 | | | | | | | | | |
| Oligo fwd | CACCGTAGGGACGTCATCTTCTGC | | | | | | | | | | | |
| Oligo rev | AAACGCAGAAGATGACGTCCCTAC | | | | | | | | | | | |
| Chromosome | start | | end | strand | MM | target_seq | PAM | alignment | distance | position | gene name | gene id |
| GU305914.1 | | 112334 | 112356 | - | 0 | GTAGGGACGTCATCTTCTGC | TGG | \|\|\|\|\|\|\|\|[\|\|\|\|\|\|\|\|\|\|\|\|]PAM | NA | Intergenic | NA | NA |
| GU305914.1 | | 111835 | 111857 | + | 4 | GTCGGGTCGTCATCTCCTTC | GGG | \|\|-\|\|\|-\|[\|\|\|\|\|\|\|-\|\|-\|]PAM | NA | Intergenic | NA | NA |
| T25551 | TTCTGCTGGCTGTCCACCACAGG | | 1000 | | | | | | | | | |
| Oligo fwd | CACCTTCTGCTGGCTGTCCACCAC | | | | | | | | | | | |
| Oligo rev | AAACGTGGTGGACAGCCAGCAGAA | | | | | | | | | | | |
| Chromosome | start | | end | strand | MM | target_seq | PAM | alignment | distance | position | gene name | gene id |
| GU305914.1 | | 112320 | 112342 | - | 0 | TTCTGCTGGCTGTCCACCAC | AGG | \|\|\|\|\|\|\|\|[\|\|\|\|\|\|\|\|\|\|\|\|]PAM | NA | Intergenic | NA | NA |
| T25552 | TCTGCTGGCTGTCCACCACAGGG | | 1000 | | | | | | | | | |
| Oligo fwd | CACCTCTGCTGGCTGTCCACCACA | | | | | | | | | | | |
| Oligo rev | AAACTGTGGTGGACAGCCAGCAGA | | | | | | | | | | | |
| Chromosome | start | | end | strand | MM | target_seq | PAM | alignment | distance | position | gene name | gene id |
| GU305914.1 | | 112319 | 112341 | - | 0 | TCTGCTGGCTGTCCACCACA | GGG | \|\|\|\|\|\|\|\|[\|\|\|\|\|\|\|\|\|\|\|\|]PAM | NA | Intergenic | NA | NA |
| T25553 | CTGCTGGCTGTCCACCACAGGGG | | 1000 | | | | | | | | | |
| Oligo fwd | CACCCTGCTGGCTGTCCACCACAG | | | | | | | | | | | |
| Oligo rev | AAACCTGTGGTGGACAGCCAGCAG | | | | | | | | | | | |
| Chromosome | start | | end | strand | MM | target_seq | PAM | alignment | distance | position | gene name | gene id |
| GU305914.1 | | 112318 | 112340 | - | 0 | CTGCTGGCTGTCCACCACAG | GGG | \|\|\|\|\|\|\|\|[\|\|\|\|\|\|\|\|\|\|\|\|]PAM | NA | Intergenic | NA | NA |
| T25554 | GCTGTCCACCACAGGGGTCATGG | | 1000 | | | | | | | | | |
| Oligo fwd | CACCGCTGTCCACCACAGGGGTCA | | | | | | | | | | | |
| Oligo rev | AAACTGACCCCTGTGGTGGACAGC | | | | | | | | | | | |
| Chromosome | start | | end | strand | MM | target_seq | PAM | alignment | distance | position | gene name | gene id |
| GU305914.1 | | 112312 | 112334 | - | 0 | GCTGTCCACCACAGGGGTCA | TGG | \|\|\|\|\|\|\|\|[\|\|\|\|\|\|\|\|\|\|\|\|]PAM | NA | Intergenic | NA | NA |
| T25555 | CACAGGGGTCATGGCGATCTCGG | | 1000 | | | | | | | | | |
| Oligo fwd | CACCCACAGGGGTCATGGCGATCT | | | | | | | | | | | |
| Oligo rev | AAACAGATCGCCATGACCCCTGTG | | | | | | | | | | | |
| Chromosome | start | | end | strand | MM | target_seq | PAM | alignment | distance | position | gene name | gene id |
| GU305914.1 | | 112303 | 112325 | - | 0 | CACAGGGGTCATGGCGATCT | CGG | \|\|\|\|\|\|\|\|[\|\|\|\|\|\|\|\|\|\|\|\|]PAM | NA | Intergenic | NA | NA |
| T25556 | GGGTCATGGCGATCTCGGTGAGG | | 1000 | | | | | | | | | |
| Oligo fwd | CACCGGGTCATGGCGATCTCGGTG | | | | | | | | | | | |
| Oligo rev | AAACCACCGAGATCGCCATGACCC | | | | | | | | | | | |
| Chromosome | start | | end | strand | MM | target_seq | PAM | alignment | distance | position | gene name | gene id |
| GU305914.1 | | 112298 | 112320 | - | 0 | GGGTCATGGCGATCTCGGTG | AGG | \|\|\|\|\|\|\|\|[\|\|\|\|\|\|\|\|\|\|\|\|]PAM | NA | Intergenic | NA | NA |
| T25557 | GGTCATGGCGATCTCGGTGAGGG | | 1000 | | | | | | | | | |
| Oligo fwd | CACCGGTCATGGCGATCTCGGTGA | | | | | | | | | | | |
| Oligo rev | AAACTCACCGAGATCGCCATGACC | | | | | | | | | | | |
| Chromosome | start | | end | strand | MM | target_seq | PAM | alignment | distance | position | gene name | gene id |
| GU305914.1 | | 112297 | 112319 | - | 0 | GGTCATGGCGATCTCGGTGA | GGG | \|\|\|\|\|\|\|\|[\|\|\|\|\|\|\|\|\|\|\|\|]PAM | NA | Intergenic | NA | NA |
| T25558 | ATGGCGATCTCGGTGAGGGATGG | | 1000 | | | | | | | | | |
| Oligo fwd | CACCATGGCGATCTCGGTGAGGGA | | | | | | | | | | | |
| Oligo rev | AAACTCCCTCACCGAGATCGCCAT | | | | | | | | | | | |
| Chromosome | start | | end | strand | MM | target_seq | PAM | alignment | distance | position | gene name | gene id |

<table>
<tr><td>GU305914.1</td><td>112293</td><td>112315</td><td>-</td><td>0</td><td>ATGGCGATCTCGGTGAGGGA</td><td>TGG</td><td>||||||||[||||||||||||]PAM</td><td>NA</td><td>Intergenic</td><td>NA</td><td>NA</td></tr>
<tr><td colspan="12"></td></tr>
<tr><td>T25559</td><td>GCGATCTCGGTGAGGGATGGAGG</td><td>1000</td><td></td><td></td><td></td><td></td><td></td><td></td><td></td><td></td><td></td></tr>
<tr><td>Oligo fwd</td><td>CACCGCGATCTCGGTGAGGGATGG</td><td colspan="10"></td></tr>
<tr><td>Oligo rev</td><td>AAACCCATCCCTCACCGAGATCGC</td><td colspan="10"></td></tr>
<tr><td>Chromosome</td><td>start</td><td>end</td><td>strand</td><td>MM</td><td>target_seq</td><td>PAM</td><td>alignment</td><td>distance</td><td>position</td><td>gene name</td><td>gene id</td></tr>
<tr><td>GU305914.1</td><td>112290</td><td>112312</td><td>-</td><td>0</td><td>GCGATCTCGGTGAGGGATGG</td><td>AGG</td><td>||||||||[||||||||||||]PAM</td><td>NA</td><td>Intergenic</td><td>NA</td><td>NA</td></tr>
<tr><td colspan="12"></td></tr>
<tr><td>T25560</td><td>CGATCTCGGTGAGGGATGGAGGG</td><td>1000</td><td colspan="9"></td></tr>
<tr><td>Oligo fwd</td><td>CACCCGATCTCGGTGAGGGATGGA</td><td colspan="10"></td></tr>
<tr><td>Oligo rev</td><td>AAACTCCATCCCTCACCGAGATCG</td><td colspan="10"></td></tr>
<tr><td>Chromosome</td><td>start</td><td>end</td><td>strand</td><td>MM</td><td>target_seq</td><td>PAM</td><td>alignment</td><td>distance</td><td>position</td><td>gene name</td><td>gene id</td></tr>
<tr><td>GU305914.1</td><td>112289</td><td>112311</td><td>-</td><td>0</td><td>CGATCTCGGTGAGGGATGGA</td><td>GGG</td><td>||||||||[||||||||||||]PAM</td><td>NA</td><td>Intergenic</td><td>NA</td><td>NA</td></tr>
<tr><td colspan="12"></td></tr>
<tr><td>T25561</td><td>GATCTCGGTGAGGGATGGAGGGG</td><td>1000</td><td colspan="9"></td></tr>
<tr><td>Oligo fwd</td><td>CACCGATCTCGGTGAGGGATGGAG</td><td colspan="10"></td></tr>
<tr><td>Oligo rev</td><td>AAACCTCCATCCCTCACCGAGATC</td><td colspan="10"></td></tr>
<tr><td>Chromosome</td><td>start</td><td>end</td><td>strand</td><td>MM</td><td>target_seq</td><td>PAM</td><td>alignment</td><td>distance</td><td>position</td><td>gene name</td><td>gene id</td></tr>
<tr><td>GU305914.1</td><td>112288</td><td>112310</td><td>-</td><td>0</td><td>GATCTCGGTGAGGGATGGAG</td><td>GGG</td><td>||||||||[||||||||||||]PAM</td><td>NA</td><td>Intergenic</td><td>NA</td><td>NA</td></tr>
<tr><td colspan="12"></td></tr>
<tr><td>T25562</td><td>ATCTCGGTGAGGGATGGAGGGGG</td><td>1000</td><td colspan="9"></td></tr>
<tr><td>Oligo fwd</td><td>CACCATCTCGGTGAGGGATGGAGG</td><td colspan="10"></td></tr>
<tr><td>Oligo rev</td><td>AAACCCTCCATCCCTCACCGAGAT</td><td colspan="10"></td></tr>
<tr><td>Chromosome</td><td>start</td><td>end</td><td>strand</td><td>MM</td><td>target_seq</td><td>PAM</td><td>alignment</td><td>distance</td><td>position</td><td>gene name</td><td>gene id</td></tr>
<tr><td>GU305914.1</td><td>112287</td><td>112309</td><td>-</td><td>0</td><td>ATCTCGGTGAGGGATGGAGG</td><td>GGG</td><td>||||||||[||||||||||||]PAM</td><td>NA</td><td>Intergenic</td><td>NA</td><td>NA</td></tr>
<tr><td colspan="12"></td></tr>
<tr><td>T25563</td><td>GGTGAGGGATGGAGGGGGTCCGG</td><td>1000</td><td colspan="9"></td></tr>
<tr><td>Oligo fwd</td><td>CACCGGTGAGGGATGGAGGGGGTC</td><td colspan="10"></td></tr>
<tr><td>Oligo rev</td><td>AAACGACCCCCTCCATCCCTCACC</td><td colspan="10"></td></tr>
<tr><td>Chromosome</td><td>start</td><td>end</td><td>strand</td><td>MM</td><td>target_seq</td><td>PAM</td><td>alignment</td><td>distance</td><td>position</td><td>gene name</td><td>gene id</td></tr>
<tr><td>GU305914.1</td><td>112282</td><td>112304</td><td>-</td><td>0</td><td>GGTGAGGGATGGAGGGGGTC</td><td>CGG</td><td>||||||||[||||||||||||]PAM</td><td>NA</td><td>Intergenic</td><td>NA</td><td>NA</td></tr>
<tr><td colspan="12"></td></tr>
<tr><td>T25564</td><td>GGGTCCGGTCTTCTCACAGACGG</td><td>1000</td><td colspan="9"></td></tr>
<tr><td>Oligo fwd</td><td>CACCGGGTCCGGTCTTCTCACAGA</td><td colspan="10"></td></tr>
<tr><td>Oligo rev</td><td>AAACTCTGTGAGAAGACCGGACCC</td><td colspan="10"></td></tr>
<tr><td>Chromosome</td><td>start</td><td>end</td><td>strand</td><td>MM</td><td>target_seq</td><td>PAM</td><td>alignment</td><td>distance</td><td>position</td><td>gene name</td><td>gene id</td></tr>
<tr><td>GU305914.1</td><td>112267</td><td>112289</td><td>-</td><td>0</td><td>GGGTCCGGTCTTCTCACAGA</td><td>CGG</td><td>||||||||[||||||||||||]PAM</td><td>NA</td><td>Intergenic</td><td>NA</td><td>NA</td></tr>
<tr><td colspan="12"></td></tr>
<tr><td>T25565</td><td>TCCGGTCTTCTCACAGACGGTGG</td><td>1000</td><td colspan="9"></td></tr>
<tr><td>Oligo fwd</td><td>CACCTCCGGTCTTCTCACAGACGG</td><td colspan="10"></td></tr>
<tr><td>Oligo rev</td><td>AAACCCGTCTGTGAGAAGACCGGA</td><td colspan="10"></td></tr>
<tr><td>Chromosome</td><td>start</td><td>end</td><td>strand</td><td>MM</td><td>target_seq</td><td>PAM</td><td>alignment</td><td>distance</td><td>position</td><td>gene name</td><td>gene id</td></tr>
<tr><td>GU305914.1</td><td>112264</td><td>112286</td><td>-</td><td>0</td><td>TCCGGTCTTCTCACAGACGG</td><td>TGG</td><td>||||||||[||||||||||||]PAM</td><td>NA</td><td>Intergenic</td><td>NA</td><td>NA</td></tr>
<tr><td colspan="12"></td></tr>
<tr><td>T25566</td><td>CCGGTCTTCTCACAGACGGTGGG</td><td>1000</td><td colspan="9"></td></tr>
<tr><td>Oligo fwd</td><td>CACCCCGGTCTTCTCACAGACGGT</td><td colspan="10"></td></tr>
<tr><td>Oligo rev</td><td>AAACACCGTCTGTGAGAAGACCGG</td><td colspan="10"></td></tr>
<tr><td>Chromosome</td><td>start</td><td>end</td><td>strand</td><td>MM</td><td>target_seq</td><td>PAM</td><td>alignment</td><td>distance</td><td>position</td><td>gene name</td><td>gene id</td></tr>
<tr><td>GU305914.1</td><td>112263</td><td>112285</td><td>-</td><td>0</td><td>CCGGTCTTCTCACAGACGGT</td><td>GGG</td><td>||||||||[||||||||||||]PAM</td><td>NA</td><td>Intergenic</td><td>NA</td><td>NA</td></tr>
<tr><td colspan="12"></td></tr>
<tr><td>T25567</td><td>CGGTCTTCTCACAGACGGTGGGG</td><td>1000</td><td colspan="9"></td></tr>
<tr><td>Oligo fwd</td><td>CACCCGGTCTTCTCACAGACGGTG</td><td colspan="10"></td></tr>
<tr><td>Oligo rev</td><td>AAACCACCGTCTGTGAGAAGACCG</td><td colspan="10"></td></tr>
<tr><td>Chromosome</td><td>start</td><td>end</td><td>strand</td><td>MM</td><td>target_seq</td><td>PAM</td><td>alignment</td><td>distance</td><td>position</td><td>gene name</td><td>gene id</td></tr>
<tr><td>GU305914.1</td><td>112262</td><td>112284</td><td>-</td><td>0</td><td>CGGTCTTCTCACAGACGGTG</td><td>GGG</td><td>||||||||[||||||||||||]PAM</td><td>NA</td><td>Intergenic</td><td>NA</td><td>NA</td></tr>
<tr><td colspan="12"></td></tr>
<tr><td>T25568</td><td>CTTGTCGTCGTCGTCCTCGTCGG</td><td>1000</td><td colspan="9"></td></tr>
<tr><td>Oligo fwd</td><td>CACCCTTGTCGTCGTCGTCCTCGT</td><td colspan="10"></td></tr>
<tr><td>Oligo rev</td><td>AAACACGAGGACGACGACGACAAG</td><td colspan="10"></td></tr>
<tr><td>Chromosome</td><td>start</td><td>end</td><td>strand</td><td>MM</td><td>target_seq</td><td>PAM</td><td>alignment</td><td>distance</td><td>position</td><td>gene name</td><td>gene id</td></tr>
<tr><td>GU305914.1</td><td>112237</td><td>112259</td><td>-</td><td>0</td><td>CTTGTCGTCGTCGTCCTCGT</td><td>CGG</td><td>||||||||[||||||||||||]PAM</td><td>NA</td><td>Intergenic</td><td>NA</td><td>NA</td></tr>
<tr><td>GU305914.1</td><td>73071</td><td>73093</td><td>-</td><td>4</td><td>ATCGTCGCCCTCGTCCTCGT</td><td>GGG</td><td>-|-||||-[|-||||||||||]PAM</td><td>NA</td><td>Intergenic</td><td>NA</td><td>NA</td></tr>
<tr><td>GU305914.1</td><td>27140</td><td>27162</td><td>-</td><td>4</td><td>ATCGTCGTCATCGTCGTCGT</td><td>TGG</td><td>-|-|||||[|-|||||-||||]PAM</td><td>NA</td><td>Intergenic</td><td>NA</td><td>NA</td></tr>
<tr><td colspan="12"></td></tr>
<tr><td>T25569</td><td>TTGTCGTCGTCGTCCTCGTCGGG</td><td>1000</td><td colspan="9"></td></tr>
<tr><td>Oligo fwd</td><td>CACCTTGTCGTCGTCGTCCTCGTC</td><td colspan="10"></td></tr>
<tr><td>Oligo rev</td><td>AAACGACGAGGACGACGACGACAA</td><td colspan="10"></td></tr>
<tr><td>Chromosome</td><td>start</td><td>end</td><td>strand</td><td>MM</td><td>target_seq</td><td>PAM</td><td>alignment</td><td>distance</td><td>position</td><td>gene name</td><td>gene id</td></tr>
<tr><td>GU305914.1</td><td>112236</td><td>112258</td><td>-</td><td>0</td><td>TTGTCGTCGTCGTCCTCGTC</td><td>GGG</td><td>||||||||[||||||||||||]PAM</td><td>NA</td><td>Intergenic</td><td>NA</td><td>NA</td></tr>
<tr><td>GU305914.1</td><td>73070</td><td>73092</td><td>-</td><td>4</td><td>TCGTCGCCCTCGTCCTCGTG</td><td>GGG</td><td>|-||||-|[-||||||||||-]PAM</td><td>NA</td><td>Intergenic</td><td>NA</td><td>NA</td></tr>
<tr><td colspan="12"></td></tr>
<tr><td>T25570</td><td>CGAGAAACTCTTGTTGTCGTAGG</td><td>1000</td><td colspan="9"></td></tr>
<tr><td>Oligo fwd</td><td>CACCCGAGAAACTCTTGTTGTCGT</td><td colspan="10"></td></tr>
<tr><td>Oligo rev</td><td>AAACACGACAACAAGAGTTTCTCG</td><td colspan="10"></td></tr>
<tr><td>Chromosome</td><td>start</td><td>end</td><td>strand</td><td>MM</td><td>target_seq</td><td>PAM</td><td>alignment</td><td>distance</td><td>position</td><td>gene name</td><td>gene id</td></tr>
<tr><td>GU305914.1</td><td>112213</td><td>112235</td><td>-</td><td>0</td><td>CGAGAAACTCTTGTTGTCGT</td><td>AGG</td><td>||||||||[||||||||||||]PAM</td><td>NA</td><td>Intergenic</td><td>NA</td><td>NA</td></tr>
<tr><td colspan="12"></td></tr>
<tr><td>T25571</td><td>GAAACTCTTGTTGTCGTAGGCGG</td><td>1000</td><td colspan="9"></td></tr>
<tr><td>Oligo fwd</td><td>CACCGAAACTCTTGTTGTCGTAGG</td><td colspan="10"></td></tr>
<tr><td>Oligo rev</td><td>AAACCCTACGACAACAAGAGTTTC</td><td colspan="10"></td></tr>
<tr><td>Chromosome</td><td>start</td><td>end</td><td>strand</td><td>MM</td><td>target_seq</td><td>PAM</td><td>alignment</td><td>distance</td><td>position</td><td>gene name</td><td>gene id</td></tr>
<tr><td>GU305914.1</td><td>112210</td><td>112232</td><td>-</td><td>0</td><td>GAAACTCTTGTTGTCGTAGG</td><td>CGG</td><td>||||||||[||||||||||||]PAM</td><td>NA</td><td>Intergenic</td><td>NA</td><td>NA</td></tr>
<tr><td colspan="12"></td></tr>
<tr><td>T25572</td><td>CCCCTCGAGCGCCTTCATCGCGG</td><td>1000</td><td colspan="9"></td></tr>
<tr><td>Oligo fwd</td><td>CACCCCCCTCGAGCGCCTTCATCG</td><td colspan="10"></td></tr>
<tr><td>Oligo rev</td><td>AAACCGATGAAGGCGCTCGAGGGG</td><td colspan="10"></td></tr>
<tr><td>Chromosome</td><td>start</td><td>end</td><td>strand</td><td>MM</td><td>target_seq</td><td>PAM</td><td>alignment</td><td>distance</td><td>position</td><td>gene name</td><td>gene id</td></tr>
<tr><td>GU305914.1</td><td>112186</td><td>112208</td><td>-</td><td>0</td><td>CCCCTCGAGCGCCTTCATCG</td><td>CGG</td><td>||||||||[||||||||||||]PAM</td><td>NA</td><td>Intergenic</td><td>NA</td><td>NA</td></tr>
</table>

<table>
<tr><td>T25573</td><td>TCGAGCGCCTTCATCGCGGCCGG</td><td>1000</td><td></td><td></td><td></td><td></td><td></td><td></td><td></td><td></td><td></td></tr>
<tr><td>Oligo fwd</td><td>CACCTCGAGCGCCTTCATCGCGGC</td><td></td><td></td><td></td><td></td><td></td><td></td><td></td><td></td><td></td><td></td></tr>
<tr><td>Oligo rev</td><td>AAACGCCGCGATGAAGGCGCTCGA</td><td></td><td></td><td></td><td></td><td></td><td></td><td></td><td></td><td></td><td></td></tr>
<tr><td>Chromosome</td><td>start</td><td>end</td><td>strand</td><td>MM</td><td>target_seq</td><td>PAM</td><td>alignment</td><td>distance</td><td>position</td><td>gene name</td><td>gene id</td></tr>
<tr><td>GU305914.1</td><td>112182</td><td>112204</td><td>-</td><td>0</td><td>TCGAGCGCCTTCATCGCGGC</td><td>CGG</td><td>||||||||[||||||||||||]PAM</td><td>NA</td><td>Intergenic</td><td>NA</td><td>NA</td></tr>
<tr><td>T25574</td><td>TCATCGCGGCCGGTGACTTGAGG</td><td>1000</td><td></td><td></td><td></td><td></td><td></td><td></td><td></td><td></td><td></td></tr>
<tr><td>Oligo fwd</td><td>CACCTCATCGCGGCCGGTGACTTG</td><td></td><td></td><td></td><td></td><td></td><td></td><td></td><td></td><td></td><td></td></tr>
<tr><td>Oligo rev</td><td>AAACCAAGTCACCGGCCGCGATGA</td><td></td><td></td><td></td><td></td><td></td><td></td><td></td><td></td><td></td><td></td></tr>
<tr><td>Chromosome</td><td>start</td><td>end</td><td>strand</td><td>MM</td><td>target_seq</td><td>PAM</td><td>alignment</td><td>distance</td><td>position</td><td>gene name</td><td>gene id</td></tr>
<tr><td>GU305914.1</td><td>112172</td><td>112194</td><td>-</td><td>0</td><td>TCATCGCGGCCGGTGACTTG</td><td>AGG</td><td>||||||||[||||||||||||]PAM</td><td>NA</td><td>Intergenic</td><td>NA</td><td>NA</td></tr>
<tr><td>T25575</td><td>CATCGCGGCCGGTGACTTGAGGG</td><td>1000</td><td></td><td></td><td></td><td></td><td></td><td></td><td></td><td></td><td></td></tr>
<tr><td>Oligo fwd</td><td>CACCCATCGCGGCCGGTGACTTGA</td><td></td><td></td><td></td><td></td><td></td><td></td><td></td><td></td><td></td><td></td></tr>
<tr><td>Oligo rev</td><td>AAACTCAAGTCACCGGCCGCGATG</td><td></td><td></td><td></td><td></td><td></td><td></td><td></td><td></td><td></td><td></td></tr>
<tr><td>Chromosome</td><td>start</td><td>end</td><td>strand</td><td>MM</td><td>target_seq</td><td>PAM</td><td>alignment</td><td>distance</td><td>position</td><td>gene name</td><td>gene id</td></tr>
<tr><td>GU305914.1</td><td>112171</td><td>112193</td><td>-</td><td>0</td><td>CATCGCGGCCGGTGACTTGA</td><td>GGG</td><td>||||||||[||||||||||||]PAM</td><td>NA</td><td>Intergenic</td><td>NA</td><td>NA</td></tr>
<tr><td>T25576</td><td>GTGACTTGAGGGCGTCCTTCTGG</td><td>1000</td><td></td><td></td><td></td><td></td><td></td><td></td><td></td><td></td><td></td></tr>
<tr><td>Oligo fwd</td><td>CACCGTGACTTGAGGGCGTCCTTC</td><td></td><td></td><td></td><td></td><td></td><td></td><td></td><td></td><td></td><td></td></tr>
<tr><td>Oligo rev</td><td>AAACGAAGGACGCCCTCAAGTCAC</td><td></td><td></td><td></td><td></td><td></td><td></td><td></td><td></td><td></td><td></td></tr>
<tr><td>Chromosome</td><td>start</td><td>end</td><td>strand</td><td>MM</td><td>target_seq</td><td>PAM</td><td>alignment</td><td>distance</td><td>position</td><td>gene name</td><td>gene id</td></tr>
<tr><td>GU305914.1</td><td>112160</td><td>112182</td><td>-</td><td>0</td><td>GTGACTTGAGGGCGTCCTTC</td><td>TGG</td><td>||||||||[||||||||||||]PAM</td><td>NA</td><td>Intergenic</td><td>NA</td><td>NA</td></tr>
<tr><td>T25577</td><td>GCGTCCTTCTGGATCCCGAAAGG</td><td>1000</td><td></td><td></td><td></td><td></td><td></td><td></td><td></td><td></td><td></td></tr>
<tr><td>Oligo fwd</td><td>CACCGCGTCCTTCTGGATCCCGAA</td><td></td><td></td><td></td><td></td><td></td><td></td><td></td><td></td><td></td><td></td></tr>
<tr><td>Oligo rev</td><td>AAACTTCGGGATCCAGAAGGACGC</td><td></td><td></td><td></td><td></td><td></td><td></td><td></td><td></td><td></td><td></td></tr>
<tr><td>Chromosome</td><td>start</td><td>end</td><td>strand</td><td>MM</td><td>target_seq</td><td>PAM</td><td>alignment</td><td>distance</td><td>position</td><td>gene name</td><td>gene id</td></tr>
<tr><td>GU305914.1</td><td>112149</td><td>112171</td><td>-</td><td>0</td><td>GCGTCCTTCTGGATCCCGAA</td><td>AGG</td><td>||||||||[||||||||||||]PAM</td><td>NA</td><td>Intergenic</td><td>NA</td><td>NA</td></tr>
<tr><td>T25578</td><td>ACACAGCCTCAGCATCGTCTTGG</td><td>1000</td><td></td><td></td><td></td><td></td><td></td><td></td><td></td><td></td><td></td></tr>
<tr><td>Oligo fwd</td><td>CACCACACAGCCTCAGCATCGTCT</td><td></td><td></td><td></td><td></td><td></td><td></td><td></td><td></td><td></td><td></td></tr>
<tr><td>Oligo rev</td><td>AAACAGACGATGCTGAGGCTGTGT</td><td></td><td></td><td></td><td></td><td></td><td></td><td></td><td></td><td></td><td></td></tr>
<tr><td>Chromosome</td><td>start</td><td>end</td><td>strand</td><td>MM</td><td>target_seq</td><td>PAM</td><td>alignment</td><td>distance</td><td>position</td><td>gene name</td><td>gene id</td></tr>
<tr><td>GU305914.1</td><td>112114</td><td>112136</td><td>-</td><td>0</td><td>ACACAGCCTCAGCATCGTCT</td><td>TGG</td><td>||||||||[||||||||||||]PAM</td><td>NA</td><td>Intergenic</td><td>NA</td><td>NA</td></tr>
<tr><td>T25579</td><td>GCATCGTCTTGGCGCCCTTGAGG</td><td>1000</td><td></td><td></td><td></td><td></td><td></td><td></td><td></td><td></td><td></td></tr>
<tr><td>Oligo fwd</td><td>CACCGCATCGTCTTGGCGCCCTTG</td><td></td><td></td><td></td><td></td><td></td><td></td><td></td><td></td><td></td><td></td></tr>
<tr><td>Oligo rev</td><td>AAACCAAGGGCGCCAAGACGATGC</td><td></td><td></td><td></td><td></td><td></td><td></td><td></td><td></td><td></td><td></td></tr>
<tr><td>Chromosome</td><td>start</td><td>end</td><td>strand</td><td>MM</td><td>target_seq</td><td>PAM</td><td>alignment</td><td>distance</td><td>position</td><td>gene name</td><td>gene id</td></tr>
<tr><td>GU305914.1</td><td>112103</td><td>112125</td><td>-</td><td>0</td><td>GCATCGTCTTGGCGCCCTTG</td><td>AGG</td><td>||||||||[||||||||||||]PAM</td><td>NA</td><td>Intergenic</td><td>NA</td><td>NA</td></tr>
<tr><td>T25580</td><td>AACCGAAAGTGCTCTTGACGAGG</td><td>1000</td><td></td><td></td><td></td><td></td><td></td><td></td><td></td><td></td><td></td></tr>
<tr><td>Oligo fwd</td><td>CACCAACCGAAAGTGCTCTTGACG</td><td></td><td></td><td></td><td></td><td></td><td></td><td></td><td></td><td></td><td></td></tr>
<tr><td>Oligo rev</td><td>AAACCGTCAAGAGCACTTTCGGTT</td><td></td><td></td><td></td><td></td><td></td><td></td><td></td><td></td><td></td><td></td></tr>
<tr><td>Chromosome</td><td>start</td><td>end</td><td>strand</td><td>MM</td><td>target_seq</td><td>PAM</td><td>alignment</td><td>distance</td><td>position</td><td>gene name</td><td>gene id</td></tr>
<tr><td>GU305914.1</td><td>112076</td><td>112098</td><td>-</td><td>0</td><td>AACCGAAAGTGCTCTTGACG</td><td>AGG</td><td>||||||||[||||||||||||]PAM</td><td>NA</td><td>Intergenic</td><td>NA</td><td>NA</td></tr>
<tr><td>T25581</td><td>ACCGAAAGTGCTCTTGACGAGGG</td><td>1000</td><td></td><td></td><td></td><td></td><td></td><td></td><td></td><td></td><td></td></tr>
<tr><td>Oligo fwd</td><td>CACCACCGAAAGTGCTCTTGACGA</td><td></td><td></td><td></td><td></td><td></td><td></td><td></td><td></td><td></td><td></td></tr>
<tr><td>Oligo rev</td><td>AAACTCGTCAAGAGCACTTTCGGT</td><td></td><td></td><td></td><td></td><td></td><td></td><td></td><td></td><td></td><td></td></tr>
<tr><td>Chromosome</td><td>start</td><td>end</td><td>strand</td><td>MM</td><td>target_seq</td><td>PAM</td><td>alignment</td><td>distance</td><td>position</td><td>gene name</td><td>gene id</td></tr>
<tr><td>GU305914.1</td><td>112075</td><td>112097</td><td>-</td><td>0</td><td>ACCGAAAGTGCTCTTGACGA</td><td>GGG</td><td>||||||||[||||||||||||]PAM</td><td>NA</td><td>Intergenic</td><td>NA</td><td>NA</td></tr>
<tr><td>T25582</td><td>GAAAGTGCTCTTGACGAGGGTGG</td><td>1000</td><td></td><td></td><td></td><td></td><td></td><td></td><td></td><td></td><td></td></tr>
<tr><td>Oligo fwd</td><td>CACCGAAAGTGCTCTTGACGAGGG</td><td></td><td></td><td></td><td></td><td></td><td></td><td></td><td></td><td></td><td></td></tr>
<tr><td>Oligo rev</td><td>AAACCCCTCGTCAAGAGCACTTTC</td><td></td><td></td><td></td><td></td><td></td><td></td><td></td><td></td><td></td><td></td></tr>
<tr><td>Chromosome</td><td>start</td><td>end</td><td>strand</td><td>MM</td><td>target_seq</td><td>PAM</td><td>alignment</td><td>distance</td><td>position</td><td>gene name</td><td>gene id</td></tr>
<tr><td>GU305914.1</td><td>112072</td><td>112094</td><td>-</td><td>0</td><td>GAAAGTGCTCTTGACGAGGG</td><td>TGG</td><td>||||||||[||||||||||||]PAM</td><td>NA</td><td>Intergenic</td><td>NA</td><td>NA</td></tr>
<tr><td>T25583</td><td>TTGACGAGGGTGGTTCCGACCGG</td><td>1000</td><td></td><td></td><td></td><td></td><td></td><td></td><td></td><td></td><td></td></tr>
<tr><td>Oligo fwd</td><td>CACCTTGACGAGGGTGGTTCCGAC</td><td></td><td></td><td></td><td></td><td></td><td></td><td></td><td></td><td></td><td></td></tr>
<tr><td>Oligo rev</td><td>AAACGTCGGAACCACCCTCGTCAA</td><td></td><td></td><td></td><td></td><td></td><td></td><td></td><td></td><td></td><td></td></tr>
<tr><td>Chromosome</td><td>start</td><td>end</td><td>strand</td><td>MM</td><td>target_seq</td><td>PAM</td><td>alignment</td><td>distance</td><td>position</td><td>gene name</td><td>gene id</td></tr>
<tr><td>GU305914.1</td><td>112062</td><td>112084</td><td>-</td><td>0</td><td>TTGACGAGGGTGGTTCCGAC</td><td>CGG</td><td>||||||||[||||||||||||]PAM</td><td>NA</td><td>Intergenic</td><td>NA</td><td>NA</td></tr>
<tr><td>T25584</td><td>TGACGAGGGTGGTTCCGACCGGG</td><td>1000</td><td></td><td></td><td></td><td></td><td></td><td></td><td></td><td></td><td></td></tr>
<tr><td>Oligo fwd</td><td>CACCTGACGAGGGTGGTTCCGACC</td><td></td><td></td><td></td><td></td><td></td><td></td><td></td><td></td><td></td><td></td></tr>
<tr><td>Oligo rev</td><td>AAACGGTCGGAACCACCCTCGTCA</td><td></td><td></td><td></td><td></td><td></td><td></td><td></td><td></td><td></td><td></td></tr>
<tr><td>Chromosome</td><td>start</td><td>end</td><td>strand</td><td>MM</td><td>target_seq</td><td>PAM</td><td>alignment</td><td>distance</td><td>position</td><td>gene name</td><td>gene id</td></tr>
<tr><td>GU305914.1</td><td>112061</td><td>112083</td><td>-</td><td>0</td><td>TGACGAGGGTGGTTCCGACC</td><td>GGG</td><td>||||||||[||||||||||||]PAM</td><td>NA</td><td>Intergenic</td><td>NA</td><td>NA</td></tr>
<tr><td>T25585</td><td>GGTTCCGACCGGGTAGATGCAGG</td><td>1000</td><td></td><td></td><td></td><td></td><td></td><td></td><td></td><td></td><td></td></tr>
<tr><td>Oligo fwd</td><td>CACCGGTTCCGACCGGGTAGATGC</td><td></td><td></td><td></td><td></td><td></td><td></td><td></td><td></td><td></td><td></td></tr>
<tr><td>Oligo rev</td><td>AAACGCATCTACCCGGTCGGAACC</td><td></td><td></td><td></td><td></td><td></td><td></td><td></td><td></td><td></td><td></td></tr>
<tr><td>Chromosome</td><td>start</td><td>end</td><td>strand</td><td>MM</td><td>target_seq</td><td>PAM</td><td>alignment</td><td>distance</td><td>position</td><td>gene name</td><td>gene id</td></tr>
<tr><td>GU305914.1</td><td>112051</td><td>112073</td><td>-</td><td>0</td><td>GGTTCCGACCGGGTAGATGC</td><td>AGG</td><td>||||||||[||||||||||||]PAM</td><td>NA</td><td>Intergenic</td><td>NA</td><td>NA</td></tr>
<tr><td>T25586</td><td>TCCGACCGGGTAGATGCAGGAGG</td><td>1000</td><td></td><td></td><td></td><td></td><td></td><td></td><td></td><td></td><td></td></tr>
<tr><td>Oligo fwd</td><td>CACCTCCGACCGGGTAGATGCAGG</td><td></td><td></td><td></td><td></td><td></td><td></td><td></td><td></td><td></td><td></td></tr>
<tr><td>Oligo rev</td><td>AAACCCTGCATCTACCCGGTCGGA</td><td></td><td></td><td></td><td></td><td></td><td></td><td></td><td></td><td></td><td></td></tr>
<tr><td>Chromosome</td><td>start</td><td>end</td><td>strand</td><td>MM</td><td>target_seq</td><td>PAM</td><td>alignment</td><td>distance</td><td>position</td><td>gene name</td><td>gene id</td></tr>
<tr><td>GU305914.1</td><td>112048</td><td>112070</td><td>-</td><td>0</td><td>TCCGACCGGGTAGATGCAGG</td><td>AGG</td><td>||||||||[||||||||||||]PAM</td><td>NA</td><td>Intergenic</td><td>NA</td><td>NA</td></tr>
<tr><td>T25587</td><td>CCGACCGGGTAGATGCAGGAGGG</td><td>1000</td><td></td><td></td><td></td><td></td><td></td><td></td><td></td><td></td><td></td></tr>
<tr><td>Oligo fwd</td><td>CACCCCGACCGGGTAGATGCAGGA</td><td></td><td></td><td></td><td></td><td></td><td></td><td></td><td></td><td></td><td></td></tr>
<tr><td>Oligo rev</td><td>AAACTCCTGCATCTACCCGGTCGG</td><td></td><td></td><td></td><td></td><td></td><td></td><td></td><td></td><td></td><td></td></tr>
<tr><td>Chromosome</td><td>start</td><td>end</td><td>strand</td><td>MM</td><td>target_seq</td><td>PAM</td><td>alignment</td><td>distance</td><td>position</td><td>gene name</td><td>gene id</td></tr>
<tr><td>GU305914.1</td><td>112047</td><td>112069</td><td>-</td><td>0</td><td>CCGACCGGGTAGATGCAGGA</td><td>GGG</td><td>||||||||[||||||||||||]PAM</td><td>NA</td><td>Intergenic</td><td>NA</td><td>NA</td></tr>
</table>

<table>
<tr><td>T25588</td><td>CGACCGGGTAGATGCAGGAGGGG</td><td>1000</td><td></td><td></td><td></td><td></td><td></td><td></td><td></td><td></td><td></td></tr>
<tr><td>Oligo fwd</td><td>CACCCGACCGGGTAGATGCAGGAG</td><td></td><td></td><td></td><td></td><td></td><td></td><td></td><td></td><td></td><td></td></tr>
<tr><td>Oligo rev</td><td>AAACCTCCTGCATCTACCCGGTCG</td><td></td><td></td><td></td><td></td><td></td><td></td><td></td><td></td><td></td><td></td></tr>
<tr><td>Chromosome</td><td>start</td><td>end</td><td>strand</td><td>MM</td><td>target_seq</td><td>PAM</td><td>alignment</td><td>distance</td><td>position</td><td>gene name</td><td>gene id</td></tr>
<tr><td>GU305914.1</td><td>112046</td><td>112068</td><td>-</td><td>0</td><td>CGACCGGGTAGATGCAGGAG</td><td>GGG</td><td>||||||||[||||||||||||]PAM</td><td>NA</td><td>Intergenic</td><td>NA</td><td>NA</td></tr>
<tr><td>T25589</td><td>GGAGATCCGCGCACAAAGACCGG</td><td>1000</td><td></td><td></td><td></td><td></td><td></td><td></td><td></td><td></td><td></td></tr>
<tr><td>Oligo fwd</td><td>CACCGGAGATCCGCGCACAAAGAC</td><td></td><td></td><td></td><td></td><td></td><td></td><td></td><td></td><td></td><td></td></tr>
<tr><td>Oligo rev</td><td>AAACGTCTTTGTGCGCGGATCTCC</td><td></td><td></td><td></td><td></td><td></td><td></td><td></td><td></td><td></td><td></td></tr>
<tr><td>Chromosome</td><td>start</td><td>end</td><td>strand</td><td>MM</td><td>target_seq</td><td>PAM</td><td>alignment</td><td>distance</td><td>position</td><td>gene name</td><td>gene id</td></tr>
<tr><td>GU305914.1</td><td>112025</td><td>112047</td><td>-</td><td>0</td><td>GGAGATCCGCGCACAAAGAC</td><td>CGG</td><td>||||||||[||||||||||||]PAM</td><td>NA</td><td>Intergenic</td><td>NA</td><td>NA</td></tr>
<tr><td>T25590</td><td>GATCCGCGCACAAAGACCGGCGG</td><td>1000</td><td></td><td></td><td></td><td></td><td></td><td></td><td></td><td></td><td></td></tr>
<tr><td>Oligo fwd</td><td>CACCGATCCGCGCACAAAGACCGG</td><td></td><td></td><td></td><td></td><td></td><td></td><td></td><td></td><td></td><td></td></tr>
<tr><td>Oligo rev</td><td>AAACCCGGTCTTTGTGCGCGGATC</td><td></td><td></td><td></td><td></td><td></td><td></td><td></td><td></td><td></td><td></td></tr>
<tr><td>Chromosome</td><td>start</td><td>end</td><td>strand</td><td>MM</td><td>target_seq</td><td>PAM</td><td>alignment</td><td>distance</td><td>position</td><td>gene name</td><td>gene id</td></tr>
<tr><td>GU305914.1</td><td>112022</td><td>112044</td><td>-</td><td>0</td><td>GATCCGCGCACAAAGACCGG</td><td>CGG</td><td>||||||||[||||||||||||]PAM</td><td>NA</td><td>Intergenic</td><td>NA</td><td>NA</td></tr>
<tr><td>GU305914.1</td><td>209951</td><td>209973</td><td>-</td><td>4</td><td>GATGCGGTCACAAAGACCGT</td><td>CGG</td><td>|||-||--[|||||||||||-]PAM</td><td>NA</td><td>Intergenic</td><td>NA</td><td>NA</td></tr>
<tr><td>T25591</td><td>GCACAAAGACCGGCGGTCATCGG</td><td>1000</td><td></td><td></td><td></td><td></td><td></td><td></td><td></td><td></td><td></td></tr>
<tr><td>Oligo fwd</td><td>CACCGCACAAAGACCGGCGGTCAT</td><td></td><td></td><td></td><td></td><td></td><td></td><td></td><td></td><td></td><td></td></tr>
<tr><td>Oligo rev</td><td>AAACATGACCGCCGGTCTTTGTGC</td><td></td><td></td><td></td><td></td><td></td><td></td><td></td><td></td><td></td><td></td></tr>
<tr><td>Chromosome</td><td>start</td><td>end</td><td>strand</td><td>MM</td><td>target_seq</td><td>PAM</td><td>alignment</td><td>distance</td><td>position</td><td>gene name</td><td>gene id</td></tr>
<tr><td>GU305914.1</td><td>112015</td><td>112037</td><td>-</td><td>0</td><td>GCACAAAGACCGGCGGTCAT</td><td>CGG</td><td>||||||||[||||||||||||]PAM</td><td>NA</td><td>Intergenic</td><td>NA</td><td>NA</td></tr>
<tr><td>T25592</td><td>GCGGTCATCGGAGCCGTCGACGG</td><td>1000</td><td></td><td></td><td></td><td></td><td></td><td></td><td></td><td></td><td></td></tr>
<tr><td>Oligo fwd</td><td>CACCGCGGTCATCGGAGCCGTCGA</td><td></td><td></td><td></td><td></td><td></td><td></td><td></td><td></td><td></td><td></td></tr>
<tr><td>Oligo rev</td><td>AAACTCGACGGCTCCGATGACCGC</td><td></td><td></td><td></td><td></td><td></td><td></td><td></td><td></td><td></td><td></td></tr>
<tr><td>Chromosome</td><td>start</td><td>end</td><td>strand</td><td>MM</td><td>target_seq</td><td>PAM</td><td>alignment</td><td>distance</td><td>position</td><td>gene name</td><td>gene id</td></tr>
<tr><td>GU305914.1</td><td>112003</td><td>112025</td><td>-</td><td>0</td><td>GCGGTCATCGGAGCCGTCGA</td><td>CGG</td><td>||||||||[||||||||||||]PAM</td><td>NA</td><td>Intergenic</td><td>NA</td><td>NA</td></tr>
<tr><td>GU305914.1</td><td>83512</td><td>83534</td><td>-</td><td>4</td><td>GTGGCCATCGGGGCCGTCGG</td><td>CGG</td><td>|-||-|||[|||-|||||||-]PAM</td><td>NA</td><td>Intergenic</td><td>NA</td><td>NA</td></tr>
<tr><td>T25593</td><td>CGGAGCCGTCGACGGACGCATGG</td><td>1000</td><td></td><td></td><td></td><td></td><td></td><td></td><td></td><td></td><td></td></tr>
<tr><td>Oligo fwd</td><td>CACCCGGAGCCGTCGACGGACGCA</td><td></td><td></td><td></td><td></td><td></td><td></td><td></td><td></td><td></td><td></td></tr>
<tr><td>Oligo rev</td><td>AAACTGCGTCCGTCGACGGCTCCG</td><td></td><td></td><td></td><td></td><td></td><td></td><td></td><td></td><td></td><td></td></tr>
<tr><td>Chromosome</td><td>start</td><td>end</td><td>strand</td><td>MM</td><td>target_seq</td><td>PAM</td><td>alignment</td><td>distance</td><td>position</td><td>gene name</td><td>gene id</td></tr>
<tr><td>GU305914.1</td><td>111995</td><td>112017</td><td>-</td><td>0</td><td>CGGAGCCGTCGACGGACGCA</td><td>TGG</td><td>||||||||[||||||||||||]PAM</td><td>NA</td><td>Intergenic</td><td>NA</td><td>NA</td></tr>
<tr><td>T25594</td><td>GACGGACGCATGGAGCTCGCCGG</td><td>1000</td><td></td><td></td><td></td><td></td><td></td><td></td><td></td><td></td><td></td></tr>
<tr><td>Oligo fwd</td><td>CACCGACGGACGCATGGAGCTCGC</td><td></td><td></td><td></td><td></td><td></td><td></td><td></td><td></td><td></td><td></td></tr>
<tr><td>Oligo rev</td><td>AAACGCGAGCTCCATGCGTCCGTC</td><td></td><td></td><td></td><td></td><td></td><td></td><td></td><td></td><td></td><td></td></tr>
<tr><td>Chromosome</td><td>start</td><td>end</td><td>strand</td><td>MM</td><td>target_seq</td><td>PAM</td><td>alignment</td><td>distance</td><td>position</td><td>gene name</td><td>gene id</td></tr>
<tr><td>GU305914.1</td><td>111985</td><td>112007</td><td>-</td><td>0</td><td>GACGGACGCATGGAGCTCGC</td><td>CGG</td><td>||||||||[||||||||||||]PAM</td><td>NA</td><td>Intergenic</td><td>NA</td><td>NA</td></tr>
<tr><td>T25595</td><td>GGACGCATGGAGCTCGCCGGCGG</td><td>1000</td><td></td><td></td><td></td><td></td><td></td><td></td><td></td><td></td><td></td></tr>
<tr><td>Oligo fwd</td><td>CACCGGACGCATGGAGCTCGCCGG</td><td></td><td></td><td></td><td></td><td></td><td></td><td></td><td></td><td></td><td></td></tr>
<tr><td>Oligo rev</td><td>AAACCCGGCGAGCTCCATGCGTCC</td><td></td><td></td><td></td><td></td><td></td><td></td><td></td><td></td><td></td><td></td></tr>
<tr><td>Chromosome</td><td>start</td><td>end</td><td>strand</td><td>MM</td><td>target_seq</td><td>PAM</td><td>alignment</td><td>distance</td><td>position</td><td>gene name</td><td>gene id</td></tr>
<tr><td>GU305914.1</td><td>111982</td><td>112004</td><td>-</td><td>0</td><td>GGACGCATGGAGCTCGCCGG</td><td>CGG</td><td>||||||||[||||||||||||]PAM</td><td>NA</td><td>Intergenic</td><td>NA</td><td>NA</td></tr>
<tr><td>T25596</td><td>GTACTTGACGACCACGTCGCAGG</td><td>1000</td><td></td><td></td><td></td><td></td><td></td><td></td><td></td><td></td><td></td></tr>
<tr><td>Oligo fwd</td><td>CACCGTACTTGACGACCACGTCGC</td><td></td><td></td><td></td><td></td><td></td><td></td><td></td><td></td><td></td><td></td></tr>
<tr><td>Oligo rev</td><td>AAACGCGACGTGGTCGTCAAGTAC</td><td></td><td></td><td></td><td></td><td></td><td></td><td></td><td></td><td></td><td></td></tr>
<tr><td>Chromosome</td><td>start</td><td>end</td><td>strand</td><td>MM</td><td>target_seq</td><td>PAM</td><td>alignment</td><td>distance</td><td>position</td><td>gene name</td><td>gene id</td></tr>
<tr><td>GU305914.1</td><td>111952</td><td>111974</td><td>-</td><td>0</td><td>GTACTTGACGACCACGTCGC</td><td>AGG</td><td>||||||||[||||||||||||]PAM</td><td>NA</td><td>Intergenic</td><td>NA</td><td>NA</td></tr>
<tr><td>T25597</td><td>GACGACCACGTCGCAGGCGAAGG</td><td>1000</td><td></td><td></td><td></td><td></td><td></td><td></td><td></td><td></td><td></td></tr>
<tr><td>Oligo fwd</td><td>CACCGACGACCACGTCGCAGGCGA</td><td></td><td></td><td></td><td></td><td></td><td></td><td></td><td></td><td></td><td></td></tr>
<tr><td>Oligo rev</td><td>AAACTCGCCTGCGACGTGGTCGTC</td><td></td><td></td><td></td><td></td><td></td><td></td><td></td><td></td><td></td><td></td></tr>
<tr><td>Chromosome</td><td>start</td><td>end</td><td>strand</td><td>MM</td><td>target_seq</td><td>PAM</td><td>alignment</td><td>distance</td><td>position</td><td>gene name</td><td>gene id</td></tr>
<tr><td>GU305914.1</td><td>111946</td><td>111968</td><td>-</td><td>0</td><td>GACGACCACGTCGCAGGCGA</td><td>AGG</td><td>||||||||[||||||||||||]PAM</td><td>NA</td><td>Intergenic</td><td>NA</td><td>NA</td></tr>
<tr><td>T25598</td><td>CACGTCGCAGGCGAAGGCGACGG</td><td>1000</td><td></td><td></td><td></td><td></td><td></td><td></td><td></td><td></td><td></td></tr>
<tr><td>Oligo fwd</td><td>CACCCACGTCGCAGGCGAAGGCGA</td><td></td><td></td><td></td><td></td><td></td><td></td><td></td><td></td><td></td><td></td></tr>
<tr><td>Oligo rev</td><td>AAACTCGCCTTCGCCTGCGACGTG</td><td></td><td></td><td></td><td></td><td></td><td></td><td></td><td></td><td></td><td></td></tr>
<tr><td>Chromosome</td><td>start</td><td>end</td><td>strand</td><td>MM</td><td>target_seq</td><td>PAM</td><td>alignment</td><td>distance</td><td>position</td><td>gene name</td><td>gene id</td></tr>
<tr><td>GU305914.1</td><td>111940</td><td>111962</td><td>-</td><td>0</td><td>CACGTCGCAGGCGAAGGCGA</td><td>CGG</td><td>||||||||[||||||||||||]PAM</td><td>NA</td><td>Intergenic</td><td>NA</td><td>NA</td></tr>
<tr><td>GU305914.1</td><td>69244</td><td>69266</td><td>+</td><td>4</td><td>CGAGGCGGAGGCGAAGGCGA</td><td>CGG</td><td>|--|-||-[||||||||||||]PAM</td><td>NA</td><td>Intergenic</td><td>NA</td><td>NA</td></tr>
<tr><td>T25599</td><td>GATGCCCCTTCCCTGCCAGCCGG</td><td>1000</td><td></td><td></td><td></td><td></td><td></td><td></td><td></td><td></td><td></td></tr>
<tr><td>Oligo fwd</td><td>CACCGATGCCCCTTCCCTGCCAGC</td><td></td><td></td><td></td><td></td><td></td><td></td><td></td><td></td><td></td><td></td></tr>
<tr><td>Oligo rev</td><td>AAACGCTGGCAGGGAAGGGGCATC</td><td></td><td></td><td></td><td></td><td></td><td></td><td></td><td></td><td></td><td></td></tr>
<tr><td>Chromosome</td><td>start</td><td>end</td><td>strand</td><td>MM</td><td>target_seq</td><td>PAM</td><td>alignment</td><td>distance</td><td>position</td><td>gene name</td><td>gene id</td></tr>
<tr><td>GU305914.1</td><td>111916</td><td>111938</td><td>-</td><td>0</td><td>GATGCCCCTTCCCTGCCAGC</td><td>CGG</td><td>||||||||[||||||||||||]PAM</td><td>NA</td><td>Intergenic</td><td>NA</td><td>NA</td></tr>
<tr><td>T25600</td><td>GCCCCTTCCCTGCCAGCCGGCGG</td><td>1000</td><td></td><td></td><td></td><td></td><td></td><td></td><td></td><td></td><td></td></tr>
<tr><td>Oligo fwd</td><td>CACCGCCCCTTCCCTGCCAGCCGG</td><td></td><td></td><td></td><td></td><td></td><td></td><td></td><td></td><td></td><td></td></tr>
<tr><td>Oligo rev</td><td>AAACCCGGCTGGCAGGGAAGGGGC</td><td></td><td></td><td></td><td></td><td></td><td></td><td></td><td></td><td></td><td></td></tr>
<tr><td>Chromosome</td><td>start</td><td>end</td><td>strand</td><td>MM</td><td>target_seq</td><td>PAM</td><td>alignment</td><td>distance</td><td>position</td><td>gene name</td><td>gene id</td></tr>
<tr><td>GU305914.1</td><td>111913</td><td>111935</td><td>-</td><td>0</td><td>GCCCCTTCCCTGCCAGCCGG</td><td>CGG</td><td>||||||||[||||||||||||]PAM</td><td>NA</td><td>Intergenic</td><td>NA</td><td>NA</td></tr>
<tr><td>T25601</td><td>GAGATCGACGTACTGCAGCGTGG</td><td>1000</td><td></td><td></td><td></td><td></td><td></td><td></td><td></td><td></td><td></td></tr>
<tr><td>Oligo fwd</td><td>CACCGAGATCGACGTACTGCAGCG</td><td></td><td></td><td></td><td></td><td></td><td></td><td></td><td></td><td></td><td></td></tr>
<tr><td>Oligo rev</td><td>AAACCGCTGCAGTACGTCGATCTC</td><td></td><td></td><td></td><td></td><td></td><td></td><td></td><td></td><td></td><td></td></tr>
<tr><td>Chromosome</td><td>start</td><td>end</td><td>strand</td><td>MM</td><td>target_seq</td><td>PAM</td><td>alignment</td><td>distance</td><td>position</td><td>gene name</td><td>gene id</td></tr>
<tr><td>GU305914.1</td><td>111883</td><td>111905</td><td>-</td><td>0</td><td>GAGATCGACGTACTGCAGCG</td><td>TGG</td><td>||||||||[||||||||||||]PAM</td><td>NA</td><td>Intergenic</td><td>NA</td><td>NA</td></tr>
<tr><td>T25602</td><td>CGACGTACTGCAGCGTGGTCTGG</td><td>1000</td><td></td><td></td><td></td><td></td><td></td><td></td><td></td><td></td><td></td></tr>
</table>

| | | | | | | | | | | | |
|---|---|---|---|---|---|---|---|---|---|---|---|
| Oligo fwd | CACCCGACGTACTGCAGCGTGGTC | | | | | | | | | | |
| Oligo rev | AAACGACCACGCTGCAGTACGTCG | | | | | | | | | | |
| Chromosome | start | end | strand | MM | target_seq | PAM | alignment | distance | position | gene name | gene id |
| GU305914.1 | 111878 | 111900 | - | 0 | CGACGTACTGCAGCGTGGTC | TGG | \|\|\|\|\|\|\|\|[\|\|\|\|\|\|\|\|\|\|\|\|]PAM | NA | Intergenic | NA | NA |
| T25603 | CGAGTGTCCGATCTCCCCGAAGG | 1000 | | | | | | | | | |
| Oligo fwd | CACCCGAGTGTCCGATCTCCCCGA | | | | | | | | | | |
| Oligo rev | AAACTCGGGGAGATCGGACACTCG | | | | | | | | | | |
| Chromosome | start | end | strand | MM | target_seq | PAM | alignment | distance | position | gene name | gene id |
| GU305914.1 | 111850 | 111872 | - | 0 | CGAGTGTCCGATCTCCCCGA | AGG | \|\|\|\|\|\|\|\|[\|\|\|\|\|\|\|\|\|\|\|\|]PAM | NA | Intergenic | NA | NA |
| T25604 | GACGACCCGACAGATGATGATGG | 1000 | | | | | | | | | |
| Oligo fwd | CACCGACGACCCGACAGATGATGA | | | | | | | | | | |
| Oligo rev | AAACTCATCATCTGTCGGGTCGTC | | | | | | | | | | |
| Chromosome | start | end | strand | MM | target_seq | PAM | alignment | distance | position | gene name | gene id |
| GU305914.1 | 111823 | 111845 | - | 0 | GACGACCCGACAGATGATGA | TGG | \|\|\|\|\|\|\|\|[\|\|\|\|\|\|\|\|\|\|\|\|]PAM | NA | Intergenic | NA | NA |
| T25605 | ACGACCCGACAGATGATGATGGG | 1000 | | | | | | | | | |
| Oligo fwd | CACCACGACCCGACAGATGATGAT | | | | | | | | | | |
| Oligo rev | AAACATCATCATCTGTCGGGTCGT | | | | | | | | | | |
| Chromosome | start | end | strand | MM | target_seq | PAM | alignment | distance | position | gene name | gene id |
| GU305914.1 | 111822 | 111844 | - | 0 | ACGACCCGACAGATGATGAT | GGG | \|\|\|\|\|\|\|\|[\|\|\|\|\|\|\|\|\|\|\|\|]PAM | NA | Intergenic | NA | NA |
| T25606 | CATGTACATGACGTAGTAGATGG | 1000 | | | | | | | | | |
| Oligo fwd | CACCCATGTACATGACGTAGTAGA | | | | | | | | | | |
| Oligo rev | AAACTCTACTACGTCATGTACATG | | | | | | | | | | |
| Chromosome | start | end | strand | MM | target_seq | PAM | alignment | distance | position | gene name | gene id |
| GU305914.1 | 111787 | 111809 | - | 0 | CATGTACATGACGTAGTAGA | TGG | \|\|\|\|\|\|\|\|[\|\|\|\|\|\|\|\|\|\|\|\|]PAM | NA | Intergenic | NA | NA |
| T25607 | GTACATGACGTAGTAGATGGAGG | 1000 | | | | | | | | | |
| Oligo fwd | CACCGTACATGACGTAGTAGATGG | | | | | | | | | | |
| Oligo rev | AAACCCATCTACTACGTCATGTAC | | | | | | | | | | |
| Chromosome | start | end | strand | MM | target_seq | PAM | alignment | distance | position | gene name | gene id |
| GU305914.1 | 111784 | 111806 | - | 0 | GTACATGACGTAGTAGATGG | AGG | \|\|\|\|\|\|\|\|[\|\|\|\|\|\|\|\|\|\|\|\|]PAM | NA | Intergenic | NA | NA |
| T25608 | GGCGCCGCTCACGAAGTGCCAGG | 1000 | | | | | | | | | |
| Oligo fwd | CACCGGCGCCGCTCACGAAGTGCC | | | | | | | | | | |
| Oligo rev | AAACGGCACTTCGTGAGCGGCGCC | | | | | | | | | | |
| Chromosome | start | end | strand | MM | target_seq | PAM | alignment | distance | position | gene name | gene id |
| GU305914.1 | 111763 | 111785 | - | 0 | GGCGCCGCTCACGAAGTGCC | AGG | \|\|\|\|\|\|\|\|[\|\|\|\|\|\|\|\|\|\|\|\|]PAM | NA | Intergenic | NA | NA |
| T25609 | CGAAGTGCCAGGTGCCGTCCCGG | 1000 | | | | | | | | | |
| Oligo fwd | CACCCGAAGTGCCAGGTGCCGTCC | | | | | | | | | | |
| Oligo rev | AAACGGACGGCACCTGGCACTTCG | | | | | | | | | | |
| Chromosome | start | end | strand | MM | target_seq | PAM | alignment | distance | position | gene name | gene id |
| GU305914.1 | 111752 | 111774 | - | 0 | CGAAGTGCCAGGTGCCGTCC | CGG | \|\|\|\|\|\|\|\|[\|\|\|\|\|\|\|\|\|\|\|\|]PAM | NA | Intergenic | NA | NA |
| T25610 | GAAGTGCCAGGTGCCGTCCCGGG | 1000 | | | | | | | | | |
| Oligo fwd | CACCGAAGTGCCAGGTGCCGTCCC | | | | | | | | | | |
| Oligo rev | AAACGGGACGGCACCTGGCACTTC | | | | | | | | | | |
| Chromosome | start | end | strand | MM | target_seq | PAM | alignment | distance | position | gene name | gene id |
| GU305914.1 | 111751 | 111773 | - | 0 | GAAGTGCCAGGTGCCGTCCC | GGG | \|\|\|\|\|\|\|\|[\|\|\|\|\|\|\|\|\|\|\|\|]PAM | NA | Intergenic | NA | NA |
| T25611 | GTACCGCATCTCGAAGAGCCTGG | 1000 | | | | | | | | | |
| Oligo fwd | CACCGTACCGCATCTCGAAGAGCC | | | | | | | | | | |
| Oligo rev | AAACGGCTCTTCGAGATGCGGTAC | | | | | | | | | | |
| Chromosome | start | end | strand | MM | target_seq | PAM | alignment | distance | position | gene name | gene id |
| GU305914.1 | 111724 | 111746 | - | 0 | GTACCGCATCTCGAAGAGCC | TGG | \|\|\|\|\|\|\|\|[\|\|\|\|\|\|\|\|\|\|\|\|]PAM | NA | Intergenic | NA | NA |
| T25612 | AGAGCCTGGAGATGATGATGAGG | 1000 | | | | | | | | | |
| Oligo fwd | CACCAGAGCCTGGAGATGATGATG | | | | | | | | | | |
| Oligo rev | AAACCATCATCATCTCCAGGCTCT | | | | | | | | | | |
| Chromosome | start | end | strand | MM | target_seq | PAM | alignment | distance | position | gene name | gene id |
| GU305914.1 | 111710 | 111732 | - | 0 | AGAGCCTGGAGATGATGATG | AGG | \|\|\|\|\|\|\|\|[\|\|\|\|\|\|\|\|\|\|\|\|]PAM | NA | Intergenic | NA | NA |
| GU305914.1 | 180051 | 180073 | - | 4 | TGATTCTGAAGATGATGATG | AGG | -\|\|--\|\|\|[-\|\|\|\|\|\|\|\|\|\|\|]PAM | NA | Intergenic | NA | NA |
| T25613 | CTGGAGATGATGATGAGGAAAGG | 1000 | | | | | | | | | |
| Oligo fwd | CACCCTGGAGATGATGATGAGGAA | | | | | | | | | | |
| Oligo rev | AAACTTCCTCATCATCATCTCCAG | | | | | | | | | | |
| Chromosome | start | end | strand | MM | target_seq | PAM | alignment | distance | position | gene name | gene id |
| GU305914.1 | 111705 | 111727 | - | 0 | CTGGAGATGATGATGAGGAA | AGG | \|\|\|\|\|\|\|\|[\|\|\|\|\|\|\|\|\|\|\|\|]PAM | NA | Intergenic | NA | NA |
| T25614 | GAGGAAAGGCACGACGTAGACGG | 1000 | | | | | | | | | |
| Oligo fwd | CACCGAGGAAAGGCACGACGTAGA | | | | | | | | | | |
| Oligo rev | AAACTCTACGTCGTGCCTTTCCTC | | | | | | | | | | |
| Chromosome | start | end | strand | MM | target_seq | PAM | alignment | distance | position | gene name | gene id |
| GU305914.1 | 111691 | 111713 | - | 0 | GAGGAAAGGCACGACGTAGA | CGG | \|\|\|\|\|\|\|\|[\|\|\|\|\|\|\|\|\|\|\|\|]PAM | NA | Intergenic | NA | NA |
| T25615 | GAAAGGCACGACGTAGACGGCGG | 1000 | | | | | | | | | |
| Oligo fwd | CACCGAAAGGCACGACGTAGACGG | | | | | | | | | | |
| Oligo rev | AAACCCGTCTACGTCGTGCCTTTC | | | | | | | | | | |
| Chromosome | start | end | strand | MM | target_seq | PAM | alignment | distance | position | gene name | gene id |
| GU305914.1 | 111688 | 111710 | - | 0 | GAAAGGCACGACGTAGACGG | CGG | \|\|\|\|\|\|\|\|[\|\|\|\|\|\|\|\|\|\|\|\|]PAM | NA | Intergenic | NA | NA |
| T25616 | GGTGACATGCATGAGCACGTAGG | 1000 | | | | | | | | | |
| Oligo fwd | CACCGGTGACATGCATGAGCACGT | | | | | | | | | | |
| Oligo rev | AAACACGTGCTCATGCATGTCACC | | | | | | | | | | |
| Chromosome | start | end | strand | MM | target_seq | PAM | alignment | distance | position | gene name | gene id |
| GU305914.1 | 111667 | 111689 | - | 0 | GGTGACATGCATGAGCACGT | AGG | \|\|\|\|\|\|\|\|[\|\|\|\|\|\|\|\|\|\|\|\|]PAM | NA | Intergenic | NA | NA |

T25617 GTGACATGCATGAGCACGTAGGG 1000
Oligo fwd CACCGTGACATGCATGAGCACGTA
Oligo rev AAACTACGTGCTCATGCATGTCAC
Chromosome start end strand MM target_seq PAM alignment distance position gene name gene id
GU305914.1 111666 111688 - 0 GTGACATGCATGAGCACGTA GGG ||||||||[||||||||||||]PAM NA Intergenic NA NA

T25618 GAGCACGTAGGGCATATCGTCGG 1000
Oligo fwd CACCGAGCACGTAGGGCATATCGT
Oligo rev AAACACGATATGCCCTACGTGCTC
Chromosome start end strand MM target_seq PAM alignment distance position gene name gene id
GU305914.1 111655 111677 - 0 GAGCACGTAGGGCATATCGT CGG ||||||||[||||||||||||]PAM NA Intergenic NA NA

T25619 AGCACGTAGGGCATATCGTCGGG 1000
Oligo fwd CACCAGCACGTAGGGCATATCGTC
Oligo rev AAACGACGATATGCCCTACGTGCT
Chromosome start end strand MM target_seq PAM alignment distance position gene name gene id
GU305914.1 111654 111676 - 0 AGCACGTAGGGCATATCGTC GGG ||||||||[||||||||||||]PAM NA Intergenic NA NA

T25620 GCACGTAGGGCATATCGTCGGGG 1000
Oligo fwd CACCGCACGTAGGGCATATCGTCG
Oligo rev AAACCGACGATATGCCCTACGTGC
Chromosome start end strand MM target_seq PAM alignment distance position gene name gene id
GU305914.1 111653 111675 - 0 GCACGTAGGGCATATCGTCG GGG ||||||||[||||||||||||]PAM NA Intergenic NA NA

T25621 GGGGCCCTGCATCTCACAGACGG 1000
Oligo fwd CACCGGGGCCCTGCATCTCACAGA
Oligo rev AAACTCTGTGAGATGCAGGGCCCC
Chromosome start end strand MM target_seq PAM alignment distance position gene name gene id
GU305914.1 111634 111656 - 0 GGGGCCCTGCATCTCACAGA CGG ||||||||[||||||||||||]PAM NA Intergenic NA NA

T25622 GGGCCCTGCATCTCACAGACGGG 1000
Oligo fwd CACCGGGCCCTGCATCTCACAGAC
Oligo rev AAACGTCTGTGAGATGCAGGGCCC
Chromosome start end strand MM target_seq PAM alignment distance position gene name gene id
GU305914.1 111633 111655 - 0 GGGCCCTGCATCTCACAGAC GGG ||||||||[||||||||||||]PAM NA Intergenic NA NA

T25623 GGCCCTGCATCTCACAGACGGGG 1000
Oligo fwd CACCGGCCCTGCATCTCACAGACG
Oligo rev AAACCGTCTGTGAGATGCAGGGCC
Chromosome start end strand MM target_seq PAM alignment distance position gene name gene id
GU305914.1 111632 111654 - 0 GGCCCTGCATCTCACAGACG GGG ||||||||[||||||||||||]PAM NA Intergenic NA NA

T25624 CCTGCATCTCACAGACGGGGAGG 1000
Oligo fwd CACCCCTGCATCTCACAGACGGGG
Oligo rev AAACCCCCGTCTGTGAGATGCAGG
Chromosome start end strand MM target_seq PAM alignment distance position gene name gene id
GU305914.1 111629 111651 - 0 CCTGCATCTCACAGACGGGG AGG ||||||||[||||||||||||]PAM NA Intergenic NA NA

T25625 CTCACAGACGGGGAGGAATCCGG 1000
Oligo fwd CACCCTCACAGACGGGGAGGAATC
Oligo rev AAACGATTCCTCCCCGTCTGTGAG
Chromosome start end strand MM target_seq PAM alignment distance position gene name gene id
GU305914.1 111622 111644 - 0 CTCACAGACGGGGAGGAATC CGG ||||||||[||||||||||||]PAM NA Intergenic NA NA

T25626 AGACGGGGAGGAATCCGGCCCGG 1000
Oligo fwd CACCAGACGGGGAGGAATCCGGCC
Oligo rev AAACGGCCGGATTCCTCCCCGTCT
Chromosome start end strand MM target_seq PAM alignment distance position gene name gene id
GU305914.1 111617 111639 - 0 AGACGGGGAGGAATCCGGCC CGG ||||||||[||||||||||||]PAM NA Intergenic NA NA

T25627 CGGCCCGGTCGCGCATGACCTGG 1000
Oligo fwd CACCCGGCCCGGTCGCGCATGACC
Oligo rev AAACGGTCATGCGCGACCGGGCCG
Chromosome start end strand MM target_seq PAM alignment distance position gene name gene id
GU305914.1 111602 111624 - 0 CGGCCCGGTCGCGCATGACC TGG ||||||||[||||||||||||]PAM NA Intergenic NA NA

T25628 CCGGTCGCGCATGACCTGGCTGG 1000
Oligo fwd CACCCCGGTCGCGCATGACCTGGC
Oligo rev AAACGCCAGGTCATGCGCGACCGG
Chromosome start end strand MM target_seq PAM alignment distance position gene name gene id
GU305914.1 111598 111620 - 0 CCGGTCGCGCATGACCTGGC TGG ||||||||[||||||||||||]PAM NA Intergenic NA NA

T25629 CATGACCTGGCTGGTGAGAGCGG 1000
Oligo fwd CACCCATGACCTGGCTGGTGAGAG
Oligo rev AAACCTCTCACCAGCCAGGTCATG
Chromosome start end strand MM target_seq PAM alignment distance position gene name gene id
GU305914.1 111589 111611 - 0 CATGACCTGGCTGGTGAGAG CGG ||||||||[||||||||||||]PAM NA Intergenic NA NA

T25630 ATGACCTGGCTGGTGAGAGCGGG 1000
Oligo fwd CACCATGACCTGGCTGGTGAGAGC
Oligo rev AAACGCTCTCACCAGCCAGGTCAT
Chromosome start end strand MM target_seq PAM alignment distance position gene name gene id
GU305914.1 111588 111610 - 0 ATGACCTGGCTGGTGAGAGC GGG ||||||||[||||||||||||]PAM NA Intergenic NA NA

T25631 AGCGGGAGCGCCGACGATGAAGG 1000
Oligo fwd CACCAGCGGGAGCGCCGACGATGA
Oligo rev AAACTCATCGTCGGCGCTCCCGCT
Chromosome start end strand MM target_seq PAM alignment distance position gene name gene id

<table>
<tr><td>GU305914.1</td><td>111571</td><td>111593</td><td>-</td><td>0</td><td>AGCGGGAGCGCCGACGATGA</td><td>AGG</td><td>||||||||[||||||||||||]PAM</td><td>NA</td><td>Intergenic</td><td>NA</td><td>NA</td></tr>
<tr><td>GU305914.1</td><td>37591</td><td>37613</td><td>-</td><td>4</td><td>ACTGGGTGCCCCGACGATGA</td><td>TGG</td><td>|--|||-|[|-||||||||||]PAM</td><td>NA</td><td>Intergenic</td><td>NA</td><td>NA</td></tr>
<tr><td></td><td></td><td></td><td></td><td></td><td></td><td></td><td></td><td></td><td></td><td></td><td></td></tr>
<tr><td>T25632</td><td>GGGAGCGCCGACGATGAAGGCGG</td><td>1000</td><td></td><td></td><td></td><td></td><td></td><td></td><td></td><td></td><td></td></tr>
<tr><td>Oligo fwd</td><td>CACCGGGAGCGCCGACGATGAAGG</td><td></td><td></td><td></td><td></td><td></td><td></td><td></td><td></td><td></td><td></td></tr>
<tr><td>Oligo rev</td><td>AAACCCTTCATCGTCGGCGCTCCC</td><td></td><td></td><td></td><td></td><td></td><td></td><td></td><td></td><td></td><td></td></tr>
<tr><td>Chromosome</td><td>start</td><td>end</td><td>strand</td><td>MM</td><td>target_seq</td><td>PAM</td><td>alignment</td><td>distance</td><td>position</td><td>gene name</td><td>gene id</td></tr>
<tr><td>GU305914.1</td><td>111568</td><td>111590</td><td>-</td><td>0</td><td>GGGAGCGCCGACGATGAAGG</td><td>CGG</td><td>||||||||[||||||||||||]PAM</td><td>NA</td><td>Intergenic</td><td>NA</td><td>NA</td></tr>
<tr><td>GU305914.1</td><td>37588</td><td>37610</td><td>-</td><td>3</td><td>GGGTGCCCCGACGATGATGG</td><td>AGG</td><td>|||-||-|[|||||||||-||]PAM</td><td>NA</td><td>Intergenic</td><td>NA</td><td>NA</td></tr>
<tr><td></td><td></td><td></td><td></td><td></td><td></td><td></td><td></td><td></td><td></td><td></td><td></td></tr>
<tr><td>T25633</td><td>GACGATGAAGGCGGAGATCCAGG</td><td>1000</td><td></td><td></td><td></td><td></td><td></td><td></td><td></td><td></td><td></td></tr>
<tr><td>Oligo fwd</td><td>CACCGACGATGAAGGCGGAGATCC</td><td></td><td></td><td></td><td></td><td></td><td></td><td></td><td></td><td></td><td></td></tr>
<tr><td>Oligo rev</td><td>AAACGGATCTCCGCCTTCATCGTC</td><td></td><td></td><td></td><td></td><td></td><td></td><td></td><td></td><td></td><td></td></tr>
<tr><td>Chromosome</td><td>start</td><td>end</td><td>strand</td><td>MM</td><td>target_seq</td><td>PAM</td><td>alignment</td><td>distance</td><td>position</td><td>gene name</td><td>gene id</td></tr>
<tr><td>GU305914.1</td><td>111559</td><td>111581</td><td>-</td><td>0</td><td>GACGATGAAGGCGGAGATCC</td><td>AGG</td><td>||||||||[||||||||||||]PAM</td><td>NA</td><td>Intergenic</td><td>NA</td><td>NA</td></tr>
<tr><td></td><td></td><td></td><td></td><td></td><td></td><td></td><td></td><td></td><td></td><td></td><td></td></tr>
<tr><td>T25634</td><td>GGCGGAGATCCAGGCGACCACGG</td><td>1000</td><td></td><td></td><td></td><td></td><td></td><td></td><td></td><td></td><td></td></tr>
<tr><td>Oligo fwd</td><td>CACCGGCGGAGATCCAGGCGACCA</td><td></td><td></td><td></td><td></td><td></td><td></td><td></td><td></td><td></td><td></td></tr>
<tr><td>Oligo rev</td><td>AAACTGGTCGCCTGGATCTCCGCC</td><td></td><td></td><td></td><td></td><td></td><td></td><td></td><td></td><td></td><td></td></tr>
<tr><td>Chromosome</td><td>start</td><td>end</td><td>strand</td><td>MM</td><td>target_seq</td><td>PAM</td><td>alignment</td><td>distance</td><td>position</td><td>gene name</td><td>gene id</td></tr>
<tr><td>GU305914.1</td><td>111550</td><td>111572</td><td>-</td><td>0</td><td>GGCGGAGATCCAGGCGACCA</td><td>CGG</td><td>||||||||[||||||||||||]PAM</td><td>NA</td><td>Intergenic</td><td>NA</td><td>NA</td></tr>
<tr><td></td><td></td><td></td><td></td><td></td><td></td><td></td><td></td><td></td><td></td><td></td><td></td></tr>
<tr><td>T25635</td><td>GACGCCGACGAGATCCTTGCCGG</td><td>1000</td><td></td><td></td><td></td><td></td><td></td><td></td><td></td><td></td><td></td></tr>
<tr><td>Oligo fwd</td><td>CACCGACGCCGACGAGATCCTTGC</td><td></td><td></td><td></td><td></td><td></td><td></td><td></td><td></td><td></td><td></td></tr>
<tr><td>Oligo rev</td><td>AAACGCAAGGATCTCGTCGGCGTC</td><td></td><td></td><td></td><td></td><td></td><td></td><td></td><td></td><td></td><td></td></tr>
<tr><td>Chromosome</td><td>start</td><td>end</td><td>strand</td><td>MM</td><td>target_seq</td><td>PAM</td><td>alignment</td><td>distance</td><td>position</td><td>gene name</td><td>gene id</td></tr>
<tr><td>GU305914.1</td><td>111517</td><td>111539</td><td>-</td><td>0</td><td>GACGCCGACGAGATCCTTGC</td><td>CGG</td><td>||||||||[||||||||||||]PAM</td><td>NA</td><td>Intergenic</td><td>NA</td><td>NA</td></tr>
<tr><td></td><td></td><td></td><td></td><td></td><td></td><td></td><td></td><td></td><td></td><td></td><td></td></tr>
<tr><td>T25636</td><td>GCCGACGAGATCCTTGCCGGCGG</td><td>1000</td><td></td><td></td><td></td><td></td><td></td><td></td><td></td><td></td><td></td></tr>
<tr><td>Oligo fwd</td><td>CACCGCCGACGAGATCCTTGCCGG</td><td></td><td></td><td></td><td></td><td></td><td></td><td></td><td></td><td></td><td></td></tr>
<tr><td>Oligo rev</td><td>AAACCCGGCAAGGATCTCGTCGGC</td><td></td><td></td><td></td><td></td><td></td><td></td><td></td><td></td><td></td><td></td></tr>
<tr><td>Chromosome</td><td>start</td><td>end</td><td>strand</td><td>MM</td><td>target_seq</td><td>PAM</td><td>alignment</td><td>distance</td><td>position</td><td>gene name</td><td>gene id</td></tr>
<tr><td>GU305914.1</td><td>111514</td><td>111536</td><td>-</td><td>0</td><td>GCCGACGAGATCCTTGCCGG</td><td>CGG</td><td>||||||||[||||||||||||]PAM</td><td>NA</td><td>Intergenic</td><td>NA</td><td>NA</td></tr>
<tr><td></td><td></td><td></td><td></td><td></td><td></td><td></td><td></td><td></td><td></td><td></td><td></td></tr>
<tr><td>T25637</td><td>GACGAGATCCTTGCCGGCGGAGG</td><td>1000</td><td></td><td></td><td></td><td></td><td></td><td></td><td></td><td></td><td></td></tr>
<tr><td>Oligo fwd</td><td>CACCGACGAGATCCTTGCCGGCGG</td><td></td><td></td><td></td><td></td><td></td><td></td><td></td><td></td><td></td><td></td></tr>
<tr><td>Oligo rev</td><td>AAACCCGCCGGCAAGGATCTCGTC</td><td></td><td></td><td></td><td></td><td></td><td></td><td></td><td></td><td></td><td></td></tr>
<tr><td>Chromosome</td><td>start</td><td>end</td><td>strand</td><td>MM</td><td>target_seq</td><td>PAM</td><td>alignment</td><td>distance</td><td>position</td><td>gene name</td><td>gene id</td></tr>
<tr><td>GU305914.1</td><td>111511</td><td>111533</td><td>-</td><td>0</td><td>GACGAGATCCTTGCCGGCGG</td><td>AGG</td><td>||||||||[||||||||||||]PAM</td><td>NA</td><td>Intergenic</td><td>NA</td><td>NA</td></tr>
<tr><td></td><td></td><td></td><td></td><td></td><td></td><td></td><td></td><td></td><td></td><td></td><td></td></tr>
<tr><td>T25638</td><td>GCCGGCGGAGGTGTTGAAGATGG</td><td>1000</td><td></td><td></td><td></td><td></td><td></td><td></td><td></td><td></td><td></td></tr>
<tr><td>Oligo fwd</td><td>CACCGCCGGCGGAGGTGTTGAAGA</td><td></td><td></td><td></td><td></td><td></td><td></td><td></td><td></td><td></td><td></td></tr>
<tr><td>Oligo rev</td><td>AAACTCTTCAACACCTCCGCCGGC</td><td></td><td></td><td></td><td></td><td></td><td></td><td></td><td></td><td></td><td></td></tr>
<tr><td>Chromosome</td><td>start</td><td>end</td><td>strand</td><td>MM</td><td>target_seq</td><td>PAM</td><td>alignment</td><td>distance</td><td>position</td><td>gene name</td><td>gene id</td></tr>
<tr><td>GU305914.1</td><td>111499</td><td>111521</td><td>-</td><td>0</td><td>GCCGGCGGAGGTGTTGAAGA</td><td>TGG</td><td>||||||||[||||||||||||]PAM</td><td>NA</td><td>Intergenic</td><td>NA</td><td>NA</td></tr>
<tr><td>GU305914.1</td><td>71026</td><td>71048</td><td>+</td><td>4</td><td>GGAGGCGGAGATGTTGGAGA</td><td>CGG</td><td>|--|||||[||-|||||-|||]PAM</td><td>NA</td><td>Intergenic</td><td>NA</td><td>NA</td></tr>
<tr><td></td><td></td><td></td><td></td><td></td><td></td><td></td><td></td><td></td><td></td><td></td><td></td></tr>
<tr><td>T25639</td><td>GTTGAAGATGGTCTTCTTCTCGG</td><td>1000</td><td></td><td></td><td></td><td></td><td></td><td></td><td></td><td></td><td></td></tr>
<tr><td>Oligo fwd</td><td>CACCGTTGAAGATGGTCTTCTTCT</td><td></td><td></td><td></td><td></td><td></td><td></td><td></td><td></td><td></td><td></td></tr>
<tr><td>Oligo rev</td><td>AAACAGAAGAAGACCATCTTCAAC</td><td></td><td></td><td></td><td></td><td></td><td></td><td></td><td></td><td></td><td></td></tr>
<tr><td>Chromosome</td><td>start</td><td>end</td><td>strand</td><td>MM</td><td>target_seq</td><td>PAM</td><td>alignment</td><td>distance</td><td>position</td><td>gene name</td><td>gene id</td></tr>
<tr><td>GU305914.1</td><td>111487</td><td>111509</td><td>-</td><td>0</td><td>GTTGAAGATGGTCTTCTTCT</td><td>CGG</td><td>||||||||[||||||||||||]PAM</td><td>NA</td><td>Intergenic</td><td>NA</td><td>NA</td></tr>
<tr><td></td><td></td><td></td><td></td><td></td><td></td><td></td><td></td><td></td><td></td><td></td><td></td></tr>
<tr><td>T25640</td><td>GATGGTCTTCTTCTCGGTCACGG</td><td>1000</td><td></td><td></td><td></td><td></td><td></td><td></td><td></td><td></td><td></td></tr>
<tr><td>Oligo fwd</td><td>CACCGATGGTCTTCTTCTCGGTCA</td><td></td><td></td><td></td><td></td><td></td><td></td><td></td><td></td><td></td><td></td></tr>
<tr><td>Oligo rev</td><td>AAACTGACCGAGAAGAAGACCATC</td><td></td><td></td><td></td><td></td><td></td><td></td><td></td><td></td><td></td><td></td></tr>
<tr><td>Chromosome</td><td>start</td><td>end</td><td>strand</td><td>MM</td><td>target_seq</td><td>PAM</td><td>alignment</td><td>distance</td><td>position</td><td>gene name</td><td>gene id</td></tr>
<tr><td>GU305914.1</td><td>111481</td><td>111503</td><td>-</td><td>0</td><td>GATGGTCTTCTTCTCGGTCA</td><td>CGG</td><td>||||||||[||||||||||||]PAM</td><td>NA</td><td>Intergenic</td><td>NA</td><td>NA</td></tr>
<tr><td></td><td></td><td></td><td></td><td></td><td></td><td></td><td></td><td></td><td></td><td></td><td></td></tr>
<tr><td>T25641</td><td>TCTCGGTCACGGCGCCGACGAGG</td><td>1000</td><td></td><td></td><td></td><td></td><td></td><td></td><td></td><td></td><td></td></tr>
<tr><td>Oligo fwd</td><td>CACCTCTCGGTCACGGCGCCGACG</td><td></td><td></td><td></td><td></td><td></td><td></td><td></td><td></td><td></td><td></td></tr>
<tr><td>Oligo rev</td><td>AAACCGTCGGCGCCGTGACCGAGA</td><td></td><td></td><td></td><td></td><td></td><td></td><td></td><td></td><td></td><td></td></tr>
<tr><td>Chromosome</td><td>start</td><td>end</td><td>strand</td><td>MM</td><td>target_seq</td><td>PAM</td><td>alignment</td><td>distance</td><td>position</td><td>gene name</td><td>gene id</td></tr>
<tr><td>GU305914.1</td><td>111470</td><td>111492</td><td>-</td><td>0</td><td>TCTCGGTCACGGCGCCGACG</td><td>AGG</td><td>||||||||[||||||||||||]PAM</td><td>NA</td><td>Intergenic</td><td>NA</td><td>NA</td></tr>
<tr><td></td><td></td><td></td><td></td><td></td><td></td><td></td><td></td><td></td><td></td><td></td><td></td></tr>
<tr><td>T25642</td><td>TCACGGCGCCGACGAGGCAGAGG</td><td>1000</td><td></td><td></td><td></td><td></td><td></td><td></td><td></td><td></td><td></td></tr>
<tr><td>Oligo fwd</td><td>CACCTCACGGCGCCGACGAGGCAG</td><td></td><td></td><td></td><td></td><td></td><td></td><td></td><td></td><td></td><td></td></tr>
<tr><td>Oligo rev</td><td>AAACCTGCCTCGTCGGCGCCGTGA</td><td></td><td></td><td></td><td></td><td></td><td></td><td></td><td></td><td></td><td></td></tr>
<tr><td>Chromosome</td><td>start</td><td>end</td><td>strand</td><td>MM</td><td>target_seq</td><td>PAM</td><td>alignment</td><td>distance</td><td>position</td><td>gene name</td><td>gene id</td></tr>
<tr><td>GU305914.1</td><td>111464</td><td>111486</td><td>-</td><td>0</td><td>TCACGGCGCCGACGAGGCAG</td><td>AGG</td><td>||||||||[||||||||||||]PAM</td><td>NA</td><td>Intergenic</td><td>NA</td><td>NA</td></tr>
<tr><td></td><td></td><td></td><td></td><td></td><td></td><td></td><td></td><td></td><td></td><td></td><td></td></tr>
<tr><td>T25643</td><td>GGCAGAGGCGATCGAGCGTGAGG</td><td>1000</td><td></td><td></td><td></td><td></td><td></td><td></td><td></td><td></td><td></td></tr>
<tr><td>Oligo fwd</td><td>CACCGGCAGAGGCGATCGAGCGTG</td><td></td><td></td><td></td><td></td><td></td><td></td><td></td><td></td><td></td><td></td></tr>
<tr><td>Oligo rev</td><td>AAACCACGCTCGATCGCCTCTGCC</td><td></td><td></td><td></td><td></td><td></td><td></td><td></td><td></td><td></td><td></td></tr>
<tr><td>Chromosome</td><td>start</td><td>end</td><td>strand</td><td>MM</td><td>target_seq</td><td>PAM</td><td>alignment</td><td>distance</td><td>position</td><td>gene name</td><td>gene id</td></tr>
<tr><td>GU305914.1</td><td>111449</td><td>111471</td><td>-</td><td>0</td><td>GGCAGAGGCGATCGAGCGTG</td><td>AGG</td><td>||||||||[||||||||||||]PAM</td><td>NA</td><td>Intergenic</td><td>NA</td><td>NA</td></tr>
<tr><td></td><td></td><td></td><td></td><td></td><td></td><td></td><td></td><td></td><td></td><td></td><td></td></tr>
<tr><td>T25644</td><td>GGCGATCGAGCGTGAGGTACAGG</td><td>1000</td><td></td><td></td><td></td><td></td><td></td><td></td><td></td><td></td><td></td></tr>
<tr><td>Oligo fwd</td><td>CACCGGCGATCGAGCGTGAGGTAC</td><td></td><td></td><td></td><td></td><td></td><td></td><td></td><td></td><td></td><td></td></tr>
<tr><td>Oligo rev</td><td>AAACGTACCTCACGCTCGATCGCC</td><td></td><td></td><td></td><td></td><td></td><td></td><td></td><td></td><td></td><td></td></tr>
<tr><td>Chromosome</td><td>start</td><td>end</td><td>strand</td><td>MM</td><td>target_seq</td><td>PAM</td><td>alignment</td><td>distance</td><td>position</td><td>gene name</td><td>gene id</td></tr>
<tr><td>GU305914.1</td><td>111443</td><td>111465</td><td>-</td><td>0</td><td>GGCGATCGAGCGTGAGGTAC</td><td>AGG</td><td>||||||||[||||||||||||]PAM</td><td>NA</td><td>Intergenic</td><td>NA</td><td>NA</td></tr>
<tr><td></td><td></td><td></td><td></td><td></td><td></td><td></td><td></td><td></td><td></td><td></td><td></td></tr>
<tr><td>T25645</td><td>CGTGAGGTACAGGTAGAAGAAGG</td><td>1000</td><td></td><td></td><td></td><td></td><td></td><td></td><td></td><td></td><td></td></tr>
<tr><td>Oligo fwd</td><td>CACCCGTGAGGTACAGGTAGAAGA</td><td></td><td></td><td></td><td></td><td></td><td></td><td></td><td></td><td></td><td></td></tr>
<tr><td>Oligo rev</td><td>AAACTCTTCTACCTGTACCTCACG</td><td></td><td></td><td></td><td></td><td></td><td></td><td></td><td></td><td></td><td></td></tr>
<tr><td>Chromosome</td><td>start</td><td>end</td><td>strand</td><td>MM</td><td>target_seq</td><td>PAM</td><td>alignment</td><td>distance</td><td>position</td><td>gene name</td><td>gene id</td></tr>
<tr><td>GU305914.1</td><td>111433</td><td>111455</td><td>-</td><td>0</td><td>CGTGAGGTACAGGTAGAAGA</td><td>AGG</td><td>||||||||[||||||||||||]PAM</td><td>NA</td><td>Intergenic</td><td>NA</td><td>NA</td></tr>
</table>

<table>
<tr><td>T25646</td><td>GTACAGGTAGAAGAAGGAGACGG</td><td>1000</td><td></td><td></td><td></td><td></td><td></td><td></td><td></td><td></td><td></td></tr>
<tr><td>Oligo fwd</td><td>CACCGTACAGGTAGAAGAAGGAGA</td><td></td><td></td><td></td><td></td><td></td><td></td><td></td><td></td><td></td><td></td></tr>
<tr><td>Oligo rev</td><td>AAACTCTCCTTCTTCTACCTGTAC</td><td></td><td></td><td></td><td></td><td></td><td></td><td></td><td></td><td></td><td></td></tr>
<tr><td>Chromosome</td><td>start</td><td>end</td><td>strand</td><td>MM</td><td>target_seq</td><td>PAM</td><td>alignment</td><td>distance</td><td>position</td><td>gene name</td><td>gene id</td></tr>
<tr><td>GU305914.1</td><td>111427</td><td>111449</td><td>-</td><td>0</td><td>GTACAGGTAGAAGAAGGAGA</td><td>CGG</td><td>||||||||[||||||||||||]PAM</td><td>NA</td><td>Intergenic</td><td>NA</td><td>NA</td></tr>
<tr><td>GU305914.1</td><td>66965</td><td>66987</td><td>+</td><td>4</td><td>GTACGTGGAGACGAAGGAGA</td><td>AGG</td><td>||||--|-[|||-||||||||]PAM</td><td>NA</td><td>Intergenic</td><td>NA</td><td>NA</td></tr>
<tr><td>T25647</td><td>GAAGAAGGAGACGGCGCCTTCGG</td><td>1000</td><td></td><td></td><td></td><td></td><td></td><td></td><td></td><td></td><td></td></tr>
<tr><td>Oligo fwd</td><td>CACCGAAGAAGGAGACGGCGCCTT</td><td></td><td></td><td></td><td></td><td></td><td></td><td></td><td></td><td></td><td></td></tr>
<tr><td>Oligo rev</td><td>AAACAAGGCGCCGTCTCCTTCTTC</td><td></td><td></td><td></td><td></td><td></td><td></td><td></td><td></td><td></td><td></td></tr>
<tr><td>Chromosome</td><td>start</td><td>end</td><td>strand</td><td>MM</td><td>target_seq</td><td>PAM</td><td>alignment</td><td>distance</td><td>position</td><td>gene name</td><td>gene id</td></tr>
<tr><td>GU305914.1</td><td>111418</td><td>111440</td><td>-</td><td>0</td><td>GAAGAAGGAGACGGCGCCTT</td><td>CGG</td><td>||||||||[||||||||||||]PAM</td><td>NA</td><td>Intergenic</td><td>NA</td><td>NA</td></tr>
<tr><td>T25648</td><td>GGACATGCCGTTGACCATGATGG</td><td>1000</td><td></td><td></td><td></td><td></td><td></td><td></td><td></td><td></td><td></td></tr>
<tr><td>Oligo fwd</td><td>CACCGGACATGCCGTTGACCATGA</td><td></td><td></td><td></td><td></td><td></td><td></td><td></td><td></td><td></td><td></td></tr>
<tr><td>Oligo rev</td><td>AAACTCATGGTCAACGGCATGTCC</td><td></td><td></td><td></td><td></td><td></td><td></td><td></td><td></td><td></td><td></td></tr>
<tr><td>Chromosome</td><td>start</td><td>end</td><td>strand</td><td>MM</td><td>target_seq</td><td>PAM</td><td>alignment</td><td>distance</td><td>position</td><td>gene name</td><td>gene id</td></tr>
<tr><td>GU305914.1</td><td>111397</td><td>111419</td><td>-</td><td>0</td><td>GGACATGCCGTTGACCATGA</td><td>TGG</td><td>||||||||[||||||||||||]PAM</td><td>NA</td><td>Intergenic</td><td>NA</td><td>NA</td></tr>
<tr><td>T25649</td><td>TGCCGTTGACCATGATGGCCAGG</td><td>1000</td><td></td><td></td><td></td><td></td><td></td><td></td><td></td><td></td><td></td></tr>
<tr><td>Oligo fwd</td><td>CACCTGCCGTTGACCATGATGGCC</td><td></td><td></td><td></td><td></td><td></td><td></td><td></td><td></td><td></td><td></td></tr>
<tr><td>Oligo rev</td><td>AAACGGCCATCATGGTCAACGGCA</td><td></td><td></td><td></td><td></td><td></td><td></td><td></td><td></td><td></td><td></td></tr>
<tr><td>Chromosome</td><td>start</td><td>end</td><td>strand</td><td>MM</td><td>target_seq</td><td>PAM</td><td>alignment</td><td>distance</td><td>position</td><td>gene name</td><td>gene id</td></tr>
<tr><td>GU305914.1</td><td>111392</td><td>111414</td><td>-</td><td>0</td><td>TGCCGTTGACCATGATGGCC</td><td>AGG</td><td>||||||||[||||||||||||]PAM</td><td>NA</td><td>Intergenic</td><td>NA</td><td>NA</td></tr>
<tr><td>T25650</td><td>CGTTGACCATGATGGCCAGGCGG</td><td>1000</td><td></td><td></td><td></td><td></td><td></td><td></td><td></td><td></td><td></td></tr>
<tr><td>Oligo fwd</td><td>CACCCGTTGACCATGATGGCCAGG</td><td></td><td></td><td></td><td></td><td></td><td></td><td></td><td></td><td></td><td></td></tr>
<tr><td>Oligo rev</td><td>AAACCCTGGCCATCATGGTCAACG</td><td></td><td></td><td></td><td></td><td></td><td></td><td></td><td></td><td></td><td></td></tr>
<tr><td>Chromosome</td><td>start</td><td>end</td><td>strand</td><td>MM</td><td>target_seq</td><td>PAM</td><td>alignment</td><td>distance</td><td>position</td><td>gene name</td><td>gene id</td></tr>
<tr><td>GU305914.1</td><td>111389</td><td>111411</td><td>-</td><td>0</td><td>CGTTGACCATGATGGCCAGG</td><td>CGG</td><td>||||||||[||||||||||||]PAM</td><td>NA</td><td>Intergenic</td><td>NA</td><td>NA</td></tr>
<tr><td>T25651</td><td>CATGATGGCCAGGCGGCAGTAGG</td><td>1000</td><td></td><td></td><td></td><td></td><td></td><td></td><td></td><td></td><td></td></tr>
<tr><td>Oligo fwd</td><td>CACCCATGATGGCCAGGCGGCAGT</td><td></td><td></td><td></td><td></td><td></td><td></td><td></td><td></td><td></td><td></td></tr>
<tr><td>Oligo rev</td><td>AAACACTGCCGCCTGGCCATCATG</td><td></td><td></td><td></td><td></td><td></td><td></td><td></td><td></td><td></td><td></td></tr>
<tr><td>Chromosome</td><td>start</td><td>end</td><td>strand</td><td>MM</td><td>target_seq</td><td>PAM</td><td>alignment</td><td>distance</td><td>position</td><td>gene name</td><td>gene id</td></tr>
<tr><td>GU305914.1</td><td>111382</td><td>111404</td><td>-</td><td>0</td><td>CATGATGGCCAGGCGGCAGT</td><td>AGG</td><td>||||||||[||||||||||||]PAM</td><td>NA</td><td>Intergenic</td><td>NA</td><td>NA</td></tr>
<tr><td>T25652</td><td>ATGATGGCCAGGCGGCAGTAGGG</td><td>1000</td><td></td><td></td><td></td><td></td><td></td><td></td><td></td><td></td><td></td></tr>
<tr><td>Oligo fwd</td><td>CACCATGATGGCCAGGCGGCAGTA</td><td></td><td></td><td></td><td></td><td></td><td></td><td></td><td></td><td></td><td></td></tr>
<tr><td>Oligo rev</td><td>AAACTACTGCCGCCTGGCCATCAT</td><td></td><td></td><td></td><td></td><td></td><td></td><td></td><td></td><td></td><td></td></tr>
<tr><td>Chromosome</td><td>start</td><td>end</td><td>strand</td><td>MM</td><td>target_seq</td><td>PAM</td><td>alignment</td><td>distance</td><td>position</td><td>gene name</td><td>gene id</td></tr>
<tr><td>GU305914.1</td><td>111381</td><td>111403</td><td>-</td><td>0</td><td>ATGATGGCCAGGCGGCAGTA</td><td>GGG</td><td>||||||||[||||||||||||]PAM</td><td>NA</td><td>Intergenic</td><td>NA</td><td>NA</td></tr>
<tr><td>T25653</td><td>GGCCAGGCGGCAGTAGGGTTCGG</td><td>1000</td><td></td><td></td><td></td><td></td><td></td><td></td><td></td><td></td><td></td></tr>
<tr><td>Oligo fwd</td><td>CACCGGCCAGGCGGCAGTAGGGTT</td><td></td><td></td><td></td><td></td><td></td><td></td><td></td><td></td><td></td><td></td></tr>
<tr><td>Oligo rev</td><td>AAACAACCCTACTGCCGCCTGGCC</td><td></td><td></td><td></td><td></td><td></td><td></td><td></td><td></td><td></td><td></td></tr>
<tr><td>Chromosome</td><td>start</td><td>end</td><td>strand</td><td>MM</td><td>target_seq</td><td>PAM</td><td>alignment</td><td>distance</td><td>position</td><td>gene name</td><td>gene id</td></tr>
<tr><td>GU305914.1</td><td>111376</td><td>111398</td><td>-</td><td>0</td><td>GGCCAGGCGGCAGTAGGGTT</td><td>CGG</td><td>||||||||[||||||||||||]PAM</td><td>NA</td><td>Intergenic</td><td>NA</td><td>NA</td></tr>
<tr><td>T25654</td><td>GTAGGGTTCGGTGACCTTAGAGG</td><td>1000</td><td></td><td></td><td></td><td></td><td></td><td></td><td></td><td></td><td></td></tr>
<tr><td>Oligo fwd</td><td>CACCGTAGGGTTCGGTGACCTTAG</td><td></td><td></td><td></td><td></td><td></td><td></td><td></td><td></td><td></td><td></td></tr>
<tr><td>Oligo rev</td><td>AAACCTAAGGTCACCGAACCCTAC</td><td></td><td></td><td></td><td></td><td></td><td></td><td></td><td></td><td></td><td></td></tr>
<tr><td>Chromosome</td><td>start</td><td>end</td><td>strand</td><td>MM</td><td>target_seq</td><td>PAM</td><td>alignment</td><td>distance</td><td>position</td><td>gene name</td><td>gene id</td></tr>
<tr><td>GU305914.1</td><td>111364</td><td>111386</td><td>-</td><td>0</td><td>GTAGGGTTCGGTGACCTTAG</td><td>AGG</td><td>||||||||[||||||||||||]PAM</td><td>NA</td><td>Intergenic</td><td>NA</td><td>NA</td></tr>
<tr><td>GU305914.1</td><td>82477</td><td>82499</td><td>+</td><td>4</td><td>GCAGGGCTCGGGGACCTTAT</td><td>CGG</td><td>|-||||-|[|||-|||||||-]PAM</td><td>NA</td><td>Intergenic</td><td>NA</td><td>NA</td></tr>
<tr><td>T25655</td><td>TAGGGTTCGGTGACCTTAGAGGG</td><td>1000</td><td></td><td></td><td></td><td></td><td></td><td></td><td></td><td></td><td></td></tr>
<tr><td>Oligo fwd</td><td>CACCTAGGGTTCGGTGACCTTAGA</td><td></td><td></td><td></td><td></td><td></td><td></td><td></td><td></td><td></td><td></td></tr>
<tr><td>Oligo rev</td><td>AAACTCTAAGGTCACCGAACCCTA</td><td></td><td></td><td></td><td></td><td></td><td></td><td></td><td></td><td></td><td></td></tr>
<tr><td>Chromosome</td><td>start</td><td>end</td><td>strand</td><td>MM</td><td>target_seq</td><td>PAM</td><td>alignment</td><td>distance</td><td>position</td><td>gene name</td><td>gene id</td></tr>
<tr><td>GU305914.1</td><td>111363</td><td>111385</td><td>-</td><td>0</td><td>TAGGGTTCGGTGACCTTAGA</td><td>GGG</td><td>||||||||[||||||||||||]PAM</td><td>NA</td><td>Intergenic</td><td>NA</td><td>NA</td></tr>
<tr><td>T25656</td><td>AGGGTTCGGTGACCTTAGAGGGG</td><td>1000</td><td></td><td></td><td></td><td></td><td></td><td></td><td></td><td></td><td></td></tr>
<tr><td>Oligo fwd</td><td>CACCAGGGTTCGGTGACCTTAGAG</td><td></td><td></td><td></td><td></td><td></td><td></td><td></td><td></td><td></td><td></td></tr>
<tr><td>Oligo rev</td><td>AAACCTCTAAGGTCACCGAACCCT</td><td></td><td></td><td></td><td></td><td></td><td></td><td></td><td></td><td></td><td></td></tr>
<tr><td>Chromosome</td><td>start</td><td>end</td><td>strand</td><td>MM</td><td>target_seq</td><td>PAM</td><td>alignment</td><td>distance</td><td>position</td><td>gene name</td><td>gene id</td></tr>
<tr><td>GU305914.1</td><td>111362</td><td>111384</td><td>-</td><td>0</td><td>AGGGTTCGGTGACCTTAGAG</td><td>GGG</td><td>||||||||[||||||||||||]PAM</td><td>NA</td><td>Intergenic</td><td>NA</td><td>NA</td></tr>
<tr><td>T25657</td><td>GACGACGCGCACGAGCCAGATGG</td><td>1000</td><td></td><td></td><td></td><td></td><td></td><td></td><td></td><td></td><td></td></tr>
<tr><td>Oligo fwd</td><td>CACCGACGACGCGCACGAGCCAGA</td><td></td><td></td><td></td><td></td><td></td><td></td><td></td><td></td><td></td><td></td></tr>
<tr><td>Oligo rev</td><td>AAACTCTGGCTCGTGCGCGTCGTC</td><td></td><td></td><td></td><td></td><td></td><td></td><td></td><td></td><td></td><td></td></tr>
<tr><td>Chromosome</td><td>start</td><td>end</td><td>strand</td><td>MM</td><td>target_seq</td><td>PAM</td><td>alignment</td><td>distance</td><td>position</td><td>gene name</td><td>gene id</td></tr>
<tr><td>GU305914.1</td><td>111337</td><td>111359</td><td>-</td><td>0</td><td>GACGACGCGCACGAGCCAGA</td><td>TGG</td><td>||||||||[||||||||||||]PAM</td><td>NA</td><td>Intergenic</td><td>NA</td><td>NA</td></tr>
<tr><td>GU305914.1</td><td>187957</td><td>187979</td><td>+</td><td>4</td><td>GTCCACGGGCACGAGCCTGA</td><td>TGG</td><td>|-|-|||-[|||||||||-||]PAM</td><td>NA</td><td>Intergenic</td><td>NA</td><td>NA</td></tr>
<tr><td>T25658</td><td>GAGCCAGATGGCGACGAAGCTGG</td><td>1000</td><td></td><td></td><td></td><td></td><td></td><td></td><td></td><td></td><td></td></tr>
<tr><td>Oligo fwd</td><td>CACCGAGCCAGATGGCGACGAAGC</td><td></td><td></td><td></td><td></td><td></td><td></td><td></td><td></td><td></td><td></td></tr>
<tr><td>Oligo rev</td><td>AAACGCTTCGTCGCCATCTGGCTC</td><td></td><td></td><td></td><td></td><td></td><td></td><td></td><td></td><td></td><td></td></tr>
<tr><td>Chromosome</td><td>start</td><td>end</td><td>strand</td><td>MM</td><td>target_seq</td><td>PAM</td><td>alignment</td><td>distance</td><td>position</td><td>gene name</td><td>gene id</td></tr>
<tr><td>GU305914.1</td><td>111325</td><td>111347</td><td>-</td><td>0</td><td>GAGCCAGATGGCGACGAAGC</td><td>TGG</td><td>||||||||[||||||||||||]PAM</td><td>NA</td><td>Intergenic</td><td>NA</td><td>NA</td></tr>
<tr><td>T25659</td><td>GCTGGCGACGTACATGAACTCGG</td><td>1000</td><td></td><td></td><td></td><td></td><td></td><td></td><td></td><td></td><td></td></tr>
<tr><td>Oligo fwd</td><td>CACCGCTGGCGACGTACATGAACT</td><td></td><td></td><td></td><td></td><td></td><td></td><td></td><td></td><td></td><td></td></tr>
<tr><td>Oligo rev</td><td>AAACAGTTCATGTACGTCGCCAGC</td><td></td><td></td><td></td><td></td><td></td><td></td><td></td><td></td><td></td><td></td></tr>
<tr><td>Chromosome</td><td>start</td><td>end</td><td>strand</td><td>MM</td><td>target_seq</td><td>PAM</td><td>alignment</td><td>distance</td><td>position</td><td>gene name</td><td>gene id</td></tr>
<tr><td>GU305914.1</td><td>111307</td><td>111329</td><td>-</td><td>0</td><td>GCTGGCGACGTACATGAACT</td><td>CGG</td><td>||||||||[||||||||||||]PAM</td><td>NA</td><td>Intergenic</td><td>NA</td><td>NA</td></tr>
<tr><td>T25660</td><td>ACATGAACTCGGCCAAGAAGAGG</td><td>1000</td><td></td><td></td><td></td><td></td><td></td><td></td><td></td><td></td><td></td></tr>
<tr><td>Oligo fwd</td><td>CACCACATGAACTCGGCCAAGAAG</td><td></td><td></td><td></td><td></td><td></td><td></td><td></td><td></td><td></td><td></td></tr>
</table>

<table>
<tr><td>Oligo rev</td><td>AAACCTTCTTGGCCGAGTTCATGT</td><td></td><td></td><td></td><td></td><td></td><td></td><td></td><td></td><td></td><td></td></tr>
<tr><td>Chromosome</td><td>start</td><td>end</td><td>strand</td><td>MM</td><td>target_seq</td><td>PAM</td><td>alignment</td><td>distance</td><td>position</td><td>gene name</td><td>gene id</td></tr>
<tr><td>GU305914.1</td><td>111296</td><td>111318</td><td>-</td><td>0</td><td>ACATGAACTCGGCCAAGAAG</td><td>AGG</td><td>||||||||[||||||||||||]PAM</td><td>NA</td><td>Intergenic</td><td>NA</td><td>NA</td></tr>
<tr><td>T25661</td><td>AGATACGCGCGCCTCCGTTACGG</td><td>1000</td><td></td><td></td><td></td><td></td><td></td><td></td><td></td><td></td><td></td></tr>
<tr><td>Oligo fwd</td><td>CACCAGATACGCGCGCCTCCGTTA</td><td></td><td></td><td></td><td></td><td></td><td></td><td></td><td></td><td></td><td></td></tr>
<tr><td>Oligo rev</td><td>AAACTAACGGAGGCGCGCGTATCT</td><td></td><td></td><td></td><td></td><td></td><td></td><td></td><td></td><td></td><td></td></tr>
<tr><td>Chromosome</td><td>start</td><td>end</td><td>strand</td><td>MM</td><td>target_seq</td><td>PAM</td><td>alignment</td><td>distance</td><td>position</td><td>gene name</td><td>gene id</td></tr>
<tr><td>GU305914.1</td><td>111263</td><td>111285</td><td>-</td><td>0</td><td>AGATACGCGCGCCTCCGTTA</td><td>CGG</td><td>||||||||[||||||||||||]PAM</td><td>NA</td><td>Intergenic</td><td>NA</td><td>NA</td></tr>
<tr><td>T25662</td><td>TACGCGCGCCTCCGTTACGGCGG</td><td>1000</td><td></td><td></td><td></td><td></td><td></td><td></td><td></td><td></td><td></td></tr>
<tr><td>Oligo fwd</td><td>CACCTACGCGCGCCTCCGTTACGG</td><td></td><td></td><td></td><td></td><td></td><td></td><td></td><td></td><td></td><td></td></tr>
<tr><td>Oligo rev</td><td>AAACCCGTAACGGAGGCGCGCGTA</td><td></td><td></td><td></td><td></td><td></td><td></td><td></td><td></td><td></td><td></td></tr>
<tr><td>Chromosome</td><td>start</td><td>end</td><td>strand</td><td>MM</td><td>target_seq</td><td>PAM</td><td>alignment</td><td>distance</td><td>position</td><td>gene name</td><td>gene id</td></tr>
<tr><td>GU305914.1</td><td>111260</td><td>111282</td><td>-</td><td>0</td><td>TACGCGCGCCTCCGTTACGG</td><td>CGG</td><td>||||||||[||||||||||||]PAM</td><td>NA</td><td>Intergenic</td><td>NA</td><td>NA</td></tr>
<tr><td>T25663</td><td>AGACCTTCCTCCACATCGAGAGG</td><td>1000</td><td></td><td></td><td></td><td></td><td></td><td></td><td></td><td></td><td></td></tr>
<tr><td>Oligo fwd</td><td>CACCAGACCTTCCTCCACATCGAG</td><td></td><td></td><td></td><td></td><td></td><td></td><td></td><td></td><td></td><td></td></tr>
<tr><td>Oligo rev</td><td>AAACCTCGATGTGGAGGAAGGTCT</td><td></td><td></td><td></td><td></td><td></td><td></td><td></td><td></td><td></td><td></td></tr>
<tr><td>Chromosome</td><td>start</td><td>end</td><td>strand</td><td>MM</td><td>target_seq</td><td>PAM</td><td>alignment</td><td>distance</td><td>position</td><td>gene name</td><td>gene id</td></tr>
<tr><td>GU305914.1</td><td>111233</td><td>111255</td><td>-</td><td>0</td><td>AGACCTTCCTCCACATCGAG</td><td>AGG</td><td>||||||||[||||||||||||]PAM</td><td>NA</td><td>Intergenic</td><td>NA</td><td>NA</td></tr>
<tr><td>T25664</td><td>TCCTCCACATCGAGAGGATGAGG</td><td>1000</td><td></td><td></td><td></td><td></td><td></td><td></td><td></td><td></td><td></td></tr>
<tr><td>Oligo fwd</td><td>CACCTCCTCCACATCGAGAGGATG</td><td></td><td></td><td></td><td></td><td></td><td></td><td></td><td></td><td></td><td></td></tr>
<tr><td>Oligo rev</td><td>AAACCATCCTCTCGATGTGGAGGA</td><td></td><td></td><td></td><td></td><td></td><td></td><td></td><td></td><td></td><td></td></tr>
<tr><td>Chromosome</td><td>start</td><td>end</td><td>strand</td><td>MM</td><td>target_seq</td><td>PAM</td><td>alignment</td><td>distance</td><td>position</td><td>gene name</td><td>gene id</td></tr>
<tr><td>GU305914.1</td><td>111227</td><td>111249</td><td>-</td><td>0</td><td>TCCTCCACATCGAGAGGATG</td><td>AGG</td><td>||||||||[||||||||||||]PAM</td><td>NA</td><td>Intergenic</td><td>NA</td><td>NA</td></tr>
<tr><td>T25665</td><td>TCGAGAGGATGAGGCTCCCTAGG</td><td>1000</td><td></td><td></td><td></td><td></td><td></td><td></td><td></td><td></td><td></td></tr>
<tr><td>Oligo fwd</td><td>CACCTCGAGAGGATGAGGCTCCCT</td><td></td><td></td><td></td><td></td><td></td><td></td><td></td><td></td><td></td><td></td></tr>
<tr><td>Oligo rev</td><td>AAACAGGGAGCCTCATCCTCTCGA</td><td></td><td></td><td></td><td></td><td></td><td></td><td></td><td></td><td></td><td></td></tr>
<tr><td>Chromosome</td><td>start</td><td>end</td><td>strand</td><td>MM</td><td>target_seq</td><td>PAM</td><td>alignment</td><td>distance</td><td>position</td><td>gene name</td><td>gene id</td></tr>
<tr><td>GU305914.1</td><td>111218</td><td>111240</td><td>-</td><td>0</td><td>TCGAGAGGATGAGGCTCCCT</td><td>AGG</td><td>||||||||[||||||||||||]PAM</td><td>NA</td><td>Intergenic</td><td>NA</td><td>NA</td></tr>
<tr><td>T25666</td><td>GGATGAGAACGATCGAGATCAGG</td><td>1000</td><td></td><td></td><td></td><td></td><td></td><td></td><td></td><td></td><td></td></tr>
<tr><td>Oligo fwd</td><td>CACCGGATGAGAACGATCGAGATC</td><td></td><td></td><td></td><td></td><td></td><td></td><td></td><td></td><td></td><td></td></tr>
<tr><td>Oligo rev</td><td>AAACGATCTCGATCGTTCTCATCC</td><td></td><td></td><td></td><td></td><td></td><td></td><td></td><td></td><td></td><td></td></tr>
<tr><td>Chromosome</td><td>start</td><td>end</td><td>strand</td><td>MM</td><td>target_seq</td><td>PAM</td><td>alignment</td><td>distance</td><td>position</td><td>gene name</td><td>gene id</td></tr>
<tr><td>GU305914.1</td><td>111197</td><td>111219</td><td>-</td><td>0</td><td>GGATGAGAACGATCGAGATC</td><td>AGG</td><td>||||||||[||||||||||||]PAM</td><td>NA</td><td>Intergenic</td><td>NA</td><td>NA</td></tr>
<tr><td>T25667</td><td>GATCGAGATCAGGCCGACGACGG</td><td>1000</td><td></td><td></td><td></td><td></td><td></td><td></td><td></td><td></td><td></td></tr>
<tr><td>Oligo fwd</td><td>CACCGATCGAGATCAGGCCGACGA</td><td></td><td></td><td></td><td></td><td></td><td></td><td></td><td></td><td></td><td></td></tr>
<tr><td>Oligo rev</td><td>AAACTCGTCGGCCTGATCTCGATC</td><td></td><td></td><td></td><td></td><td></td><td></td><td></td><td></td><td></td><td></td></tr>
<tr><td>Chromosome</td><td>start</td><td>end</td><td>strand</td><td>MM</td><td>target_seq</td><td>PAM</td><td>alignment</td><td>distance</td><td>position</td><td>gene name</td><td>gene id</td></tr>
<tr><td>GU305914.1</td><td>111187</td><td>111209</td><td>-</td><td>0</td><td>GATCGAGATCAGGCCGACGA</td><td>CGG</td><td>||||||||[||||||||||||]PAM</td><td>NA</td><td>Intergenic</td><td>NA</td><td>NA</td></tr>
<tr><td>T25668</td><td>GATCAGGCCGACGACGGCGAAGG</td><td>1000</td><td></td><td></td><td></td><td></td><td></td><td></td><td></td><td></td><td></td></tr>
<tr><td>Oligo fwd</td><td>CACCGATCAGGCCGACGACGGCGA</td><td></td><td></td><td></td><td></td><td></td><td></td><td></td><td></td><td></td><td></td></tr>
<tr><td>Oligo rev</td><td>AAACTCGCCGTCGTCGGCCTGATC</td><td></td><td></td><td></td><td></td><td></td><td></td><td></td><td></td><td></td><td></td></tr>
<tr><td>Chromosome</td><td>start</td><td>end</td><td>strand</td><td>MM</td><td>target_seq</td><td>PAM</td><td>alignment</td><td>distance</td><td>position</td><td>gene name</td><td>gene id</td></tr>
<tr><td>GU305914.1</td><td>111181</td><td>111203</td><td>-</td><td>0</td><td>GATCAGGCCGACGACGGCGA</td><td>AGG</td><td>||||||||[||||||||||||]PAM</td><td>NA</td><td>Intergenic</td><td>NA</td><td>NA</td></tr>
<tr><td>GU305914.1</td><td>182975</td><td>182997</td><td>+</td><td>4</td><td>GGTCAGACCGAAGACTGCGA</td><td>CGG</td><td>|-||||-|[|||-|||-||||]PAM</td><td>NA</td><td>Intergenic</td><td>NA</td><td>NA</td></tr>
<tr><td>T25669</td><td>CAGGCCGACGACGGCGAAGGTGG</td><td>1000</td><td></td><td></td><td></td><td></td><td></td><td></td><td></td><td></td><td></td></tr>
<tr><td>Oligo fwd</td><td>CACCCAGGCCGACGACGGCGAAGG</td><td></td><td></td><td></td><td></td><td></td><td></td><td></td><td></td><td></td><td></td></tr>
<tr><td>Oligo rev</td><td>AAACCCTTCGCCGTCGTCGGCCTG</td><td></td><td></td><td></td><td></td><td></td><td></td><td></td><td></td><td></td><td></td></tr>
<tr><td>Chromosome</td><td>start</td><td>end</td><td>strand</td><td>MM</td><td>target_seq</td><td>PAM</td><td>alignment</td><td>distance</td><td>position</td><td>gene name</td><td>gene id</td></tr>
<tr><td>GU305914.1</td><td>111178</td><td>111200</td><td>-</td><td>0</td><td>CAGGCCGACGACGGCGAAGG</td><td>TGG</td><td>||||||||[||||||||||||]PAM</td><td>NA</td><td>Intergenic</td><td>NA</td><td>NA</td></tr>
<tr><td>GU305914.1</td><td>90262</td><td>90284</td><td>-</td><td>4</td><td>GAGGCGGGCGACGGCGGAGG</td><td>CGG</td><td>-||||-|-[||||||||-|||]PAM</td><td>NA</td><td>Intergenic</td><td>NA</td><td>NA</td></tr>
<tr><td>T25670</td><td>GCCGACGACGGCGAAGGTGGAGG</td><td>1000</td><td></td><td></td><td></td><td></td><td></td><td></td><td></td><td></td><td></td></tr>
<tr><td>Oligo fwd</td><td>CACCGCCGACGACGGCGAAGGTGG</td><td></td><td></td><td></td><td></td><td></td><td></td><td></td><td></td><td></td><td></td></tr>
<tr><td>Oligo rev</td><td>AAACCCACCTTCGCCGTCGTCGGC</td><td></td><td></td><td></td><td></td><td></td><td></td><td></td><td></td><td></td><td></td></tr>
<tr><td>Chromosome</td><td>start</td><td>end</td><td>strand</td><td>MM</td><td>target_seq</td><td>PAM</td><td>alignment</td><td>distance</td><td>position</td><td>gene name</td><td>gene id</td></tr>
<tr><td>GU305914.1</td><td>111175</td><td>111197</td><td>-</td><td>0</td><td>GCCGACGACGGCGAAGGTGG</td><td>AGG</td><td>||||||||[||||||||||||]PAM</td><td>NA</td><td>Intergenic</td><td>NA</td><td>NA</td></tr>
<tr><td>GU305914.1</td><td>225498</td><td>225520</td><td>+</td><td>4</td><td>GCCGAGGGCGGCGGTGGTGG</td><td>TGG</td><td>|||||-|-[|||||--|||||]PAM</td><td>NA</td><td>Intergenic</td><td>NA</td><td>NA</td></tr>
<tr><td>GU305914.1</td><td>90259</td><td>90281</td><td>-</td><td>4</td><td>GCGGGCGACGGCGGAGGCGG</td><td>CGG</td><td>||-|-|||[|||||-|||-||]PAM</td><td>NA</td><td>Intergenic</td><td>NA</td><td>NA</td></tr>
<tr><td>GU305914.1</td><td>19436</td><td>19458</td><td>+</td><td>4</td><td>GCCGGCGTCGGCGGAGGTGC</td><td>CGG</td><td>||||-||-[|||||-|||||-]PAM</td><td>NA</td><td>Intergenic</td><td>NA</td><td>NA</td></tr>
<tr><td>T25671</td><td>CGGCGAAGGTGGAGGTGACGAGG</td><td>1000</td><td></td><td></td><td></td><td></td><td></td><td></td><td></td><td></td><td></td></tr>
<tr><td>Oligo fwd</td><td>CACCCGGCGAAGGTGGAGGTGACG</td><td></td><td></td><td></td><td></td><td></td><td></td><td></td><td></td><td></td><td></td></tr>
<tr><td>Oligo rev</td><td>AAACCGTCACCTCCACCTTCGCCG</td><td></td><td></td><td></td><td></td><td></td><td></td><td></td><td></td><td></td><td></td></tr>
<tr><td>Chromosome</td><td>start</td><td>end</td><td>strand</td><td>MM</td><td>target_seq</td><td>PAM</td><td>alignment</td><td>distance</td><td>position</td><td>gene name</td><td>gene id</td></tr>
<tr><td>GU305914.1</td><td>111167</td><td>111189</td><td>-</td><td>0</td><td>CGGCGAAGGTGGAGGTGACG</td><td>AGG</td><td>||||||||[||||||||||||]PAM</td><td>NA</td><td>Intergenic</td><td>NA</td><td>NA</td></tr>
<tr><td>T25672</td><td>GGAGGTGACGAGGATGTTGTAGG</td><td>1000</td><td></td><td></td><td></td><td></td><td></td><td></td><td></td><td></td><td></td></tr>
<tr><td>Oligo fwd</td><td>CACCGGAGGTGACGAGGATGTTGT</td><td></td><td></td><td></td><td></td><td></td><td></td><td></td><td></td><td></td><td></td></tr>
<tr><td>Oligo rev</td><td>AAACACAACATCCTCGTCACCTCC</td><td></td><td></td><td></td><td></td><td></td><td></td><td></td><td></td><td></td><td></td></tr>
<tr><td>Chromosome</td><td>start</td><td>end</td><td>strand</td><td>MM</td><td>target_seq</td><td>PAM</td><td>alignment</td><td>distance</td><td>position</td><td>gene name</td><td>gene id</td></tr>
<tr><td>GU305914.1</td><td>111157</td><td>111179</td><td>-</td><td>0</td><td>GGAGGTGACGAGGATGTTGT</td><td>AGG</td><td>||||||||[||||||||||||]PAM</td><td>NA</td><td>Intergenic</td><td>NA</td><td>NA</td></tr>
<tr><td>T25673</td><td>TGACGAGGATGTTGTAGGTCTGG</td><td>1000</td><td></td><td></td><td></td><td></td><td></td><td></td><td></td><td></td><td></td></tr>
<tr><td>Oligo fwd</td><td>CACCTGACGAGGATGTTGTAGGTC</td><td></td><td></td><td></td><td></td><td></td><td></td><td></td><td></td><td></td><td></td></tr>
<tr><td>Oligo rev</td><td>AAACGACCTACAACATCCTCGTCA</td><td></td><td></td><td></td><td></td><td></td><td></td><td></td><td></td><td></td><td></td></tr>
<tr><td>Chromosome</td><td>start</td><td>end</td><td>strand</td><td>MM</td><td>target_seq</td><td>PAM</td><td>alignment</td><td>distance</td><td>position</td><td>gene name</td><td>gene id</td></tr>
<tr><td>GU305914.1</td><td>111152</td><td>111174</td><td>-</td><td>0</td><td>TGACGAGGATGTTGTAGGTC</td><td>TGG</td><td>||||||||[||||||||||||]PAM</td><td>NA</td><td>Intergenic</td><td>NA</td><td>NA</td></tr>
<tr><td>T25674</td><td>CTGGATGAAGCTGAGATGCTCGG</td><td>1000</td><td></td><td></td><td></td><td></td><td></td><td></td><td></td><td></td><td></td></tr>
<tr><td>Oligo fwd</td><td>CACCCTGGATGAAGCTGAGATGCT</td><td></td><td></td><td></td><td></td><td></td><td></td><td></td><td></td><td></td><td></td></tr>
</table>

| | | | | | | | | | | | |
|---|---|---|---|---|---|---|---|---|---|---|---|
| Oligo rev | AAACAGCATCTCAGCTTCATCCAG | | | | | | | | | | |
| Chromosome | start | end | strand | MM | target_seq | PAM | alignment | distance | position | gene name | gene id |
| GU305914.1 | 111133 | 111155 | - | 0 | CTGGATGAAGCTGAGATGCT | CGG | \|\|\|\|\|\|\|\|[\|\|\|\|\|\|\|\|\|\|\|\|]PAM | NA | Intergenic | NA | NA |
| T25675 | AGATGCTCGGCGTCGACCTCCGG | 1000 | | | | | | | | | |
| Oligo fwd | CACCAGATGCTCGGCGTCGACCTC | | | | | | | | | | |
| Oligo rev | AAACGAGGTCGACGCCGAGCATCT | | | | | | | | | | |
| Chromosome | start | end | strand | MM | target_seq | PAM | alignment | distance | position | gene name | gene id |
| GU305914.1 | 111120 | 111142 | - | 0 | AGATGCTCGGCGTCGACCTC | CGG | \|\|\|\|\|\|\|\|[\|\|\|\|\|\|\|\|\|\|\|\|]PAM | NA | Intergenic | NA | NA |
| T25676 | GATGCTCGGCGTCGACCTCCGGG | 1000 | | | | | | | | | |
| Oligo fwd | CACCGATGCTCGGCGTCGACCTCC | | | | | | | | | | |
| Oligo rev | AAACGGAGGTCGACGCCGAGCATC | | | | | | | | | | |
| Chromosome | start | end | strand | MM | target_seq | PAM | alignment | distance | position | gene name | gene id |
| GU305914.1 | 111119 | 111141 | - | 0 | GATGCTCGGCGTCGACCTCC | GGG | \|\|\|\|\|\|\|\|[\|\|\|\|\|\|\|\|\|\|\|\|]PAM | NA | Intergenic | NA | NA |
| T25677 | CTCGGCGTCGACCTCCGGGTAGG | 1000 | | | | | | | | | |
| Oligo fwd | CACCCTCGGCGTCGACCTCCGGGT | | | | | | | | | | |
| Oligo rev | AAACACCCGGAGGTCGACGCCGAG | | | | | | | | | | |
| Chromosome | start | end | strand | MM | target_seq | PAM | alignment | distance | position | gene name | gene id |
| GU305914.1 | 111115 | 111137 | - | 0 | CTCGGCGTCGACCTCCGGGT | AGG | \|\|\|\|\|\|\|\|[\|\|\|\|\|\|\|\|\|\|\|\|]PAM | NA | Intergenic | NA | NA |
| T25678 | CTCCGGGTAGGAGTAATCGACGG | 1000 | | | | | | | | | |
| Oligo fwd | CACCCTCCGGGTAGGAGTAATCGA | | | | | | | | | | |
| Oligo rev | AAACTCGATTACTCCTACCCGGAG | | | | | | | | | | |
| Chromosome | start | end | strand | MM | target_seq | PAM | alignment | distance | position | gene name | gene id |
| GU305914.1 | 111103 | 111125 | - | 0 | CTCCGGGTAGGAGTAATCGA | CGG | \|\|\|\|\|\|\|\|[\|\|\|\|\|\|\|\|\|\|\|\|]PAM | NA | Intergenic | NA | NA |
| T25679 | TCGACGGCGCACGATGAAGTCGG | 1000 | | | | | | | | | |
| Oligo fwd | CACCTCGACGGCGCACGATGAAGT | | | | | | | | | | |
| Oligo rev | AAACACTTCATCGTGCGCCGTCGA | | | | | | | | | | |
| Chromosome | start | end | strand | MM | target_seq | PAM | alignment | distance | position | gene name | gene id |
| GU305914.1 | 111087 | 111109 | - | 0 | TCGACGGCGCACGATGAAGT | CGG | \|\|\|\|\|\|\|\|[\|\|\|\|\|\|\|\|\|\|\|\|]PAM | NA | Intergenic | NA | NA |
| T25680 | GGCGCACGATGAAGTCGGCATGG | 1000 | | | | | | | | | |
| Oligo fwd | CACCGGCGCACGATGAAGTCGGCA | | | | | | | | | | |
| Oligo rev | AAACTGCCGACTTCATCGTGCGCC | | | | | | | | | | |
| Chromosome | start | end | strand | MM | target_seq | PAM | alignment | distance | position | gene name | gene id |
| GU305914.1 | 111082 | 111104 | - | 0 | GGCGCACGATGAAGTCGGCA | TGG | \|\|\|\|\|\|\|\|[\|\|\|\|\|\|\|\|\|\|\|\|]PAM | NA | Intergenic | NA | NA |
| T25681 | CGATGAAGTCGGCATGGCGACGG | 1000 | | | | | | | | | |
| Oligo fwd | CACCCGATGAAGTCGGCATGGCGA | | | | | | | | | | |
| Oligo rev | AAACTCGCCATGCCGACTTCATCG | | | | | | | | | | |
| Chromosome | start | end | strand | MM | target_seq | PAM | alignment | distance | position | gene name | gene id |
| GU305914.1 | 111076 | 111098 | - | 0 | CGATGAAGTCGGCATGGCGA | CGG | \|\|\|\|\|\|\|\|[\|\|\|\|\|\|\|\|\|\|\|\|]PAM | NA | Intergenic | NA | NA |
| T25682 | GGCATGGCGACGGTTCGACGCGG | 1000 | | | | | | | | | |
| Oligo fwd | CACCGGCATGGCGACGGTTCGACG | | | | | | | | | | |
| Oligo rev | AAACCGTCGAACCGTCGCCATGCC | | | | | | | | | | |
| Chromosome | start | end | strand | MM | target_seq | PAM | alignment | distance | position | gene name | gene id |
| GU305914.1 | 111066 | 111088 | - | 0 | GGCATGGCGACGGTTCGACG | CGG | \|\|\|\|\|\|\|\|[\|\|\|\|\|\|\|\|\|\|\|\|]PAM | NA | Intergenic | NA | NA |
| T25683 | GCATGGCGACGGTTCGACGCGGG | 1000 | | | | | | | | | |
| Oligo fwd | CACCGCATGGCGACGGTTCGACGC | | | | | | | | | | |
| Oligo rev | AAACGCGTCGAACCGTCGCCATGC | | | | | | | | | | |
| Chromosome | start | end | strand | MM | target_seq | PAM | alignment | distance | position | gene name | gene id |
| GU305914.1 | 111065 | 111087 | - | 0 | GCATGGCGACGGTTCGACGC | GGG | \|\|\|\|\|\|\|\|[\|\|\|\|\|\|\|\|\|\|\|\|]PAM | NA | Intergenic | NA | NA |
| T25684 | TGGCGACGGTTCGACGCGGGCGG | 1000 | | | | | | | | | |
| Oligo fwd | CACCTGGCGACGGTTCGACGCGGG | | | | | | | | | | |
| Oligo rev | AAACCCCGCGTCGAACCGTCGCCA | | | | | | | | | | |
| Chromosome | start | end | strand | MM | target_seq | PAM | alignment | distance | position | gene name | gene id |
| GU305914.1 | 111062 | 111084 | - | 0 | TGGCGACGGTTCGACGCGGG | CGG | \|\|\|\|\|\|\|\|[\|\|\|\|\|\|\|\|\|\|\|\|]PAM | NA | Intergenic | NA | NA |
| T25685 | GGGCGGCCCGCGCCCCGCACTGG | 1000 | | | | | | | | | |
| Oligo fwd | CACCGGGCGGCCCGCGCCCCGCAC | | | | | | | | | | |
| Oligo rev | AAACGTGCGGGGCGCGGGCCGCCC | | | | | | | | | | |
| Chromosome | start | end | strand | MM | target_seq | PAM | alignment | distance | position | gene name | gene id |
| GU305914.1 | 111045 | 111067 | - | 0 | GGGCGGCCCGCGCCCCGCAC | TGG | \|\|\|\|\|\|\|\|[\|\|\|\|\|\|\|\|\|\|\|\|]PAM | NA | Intergenic | NA | NA |
| T25686 | CCGCGCCCCGCACTGGTGCCTGG | 1000 | | | | | | | | | |
| Oligo fwd | CACCCCGCGCCCCGCACTGGTGCC | | | | | | | | | | |
| Oligo rev | AAACGGCACCAGTGCGGGGCGCGG | | | | | | | | | | |
| Chromosome | start | end | strand | MM | target_seq | PAM | alignment | distance | position | gene name | gene id |
| GU305914.1 | 111038 | 111060 | - | 0 | CCGCGCCCCGCACTGGTGCC | TGG | \|\|\|\|\|\|\|\|[\|\|\|\|\|\|\|\|\|\|\|\|]PAM | NA | Intergenic | NA | NA |
| T25687 | TGGTGCCTGGCACTGTGTGAAGG | 1000 | | | | | | | | | |
| Oligo fwd | CACCTGGTGCCTGGCACTGTGTGA | | | | | | | | | | |
| Oligo rev | AAACTCACACAGTGCCAGGCACCA | | | | | | | | | | |
| Chromosome | start | end | strand | MM | target_seq | PAM | alignment | distance | position | gene name | gene id |
| GU305914.1 | 111025 | 111047 | - | 0 | TGGTGCCTGGCACTGTGTGA | AGG | \|\|\|\|\|\|\|\|[\|\|\|\|\|\|\|\|\|\|\|\|]PAM | NA | Intergenic | NA | NA |
| T25688 | TGAGACTGTGCGCTCCGAAACGG | 1000 | | | | | | | | | |
| Oligo fwd | CACCTGAGACTGTGCGCTCCGAAA | | | | | | | | | | |
| Oligo rev | AAACTTTCGGAGCGCACAGTCTCA | | | | | | | | | | |
| Chromosome | start | end | strand | MM | target_seq | PAM | alignment | distance | position | gene name | gene id |
| GU305914.1 | 110995 | 111017 | - | 0 | TGAGACTGTGCGCTCCGAAA | CGG | \|\|\|\|\|\|\|\|[\|\|\|\|\|\|\|\|\|\|\|\|]PAM | NA | Intergenic | NA | NA |
| T25689 | GAGACTGTGCGCTCCGAAACGGG | 1000 | | | | | | | | | |

<table>
<tr><td>Oligo fwd</td><td>CACCGAGACTGTGCGCTCCGAAAC</td><td></td><td></td><td></td><td></td><td></td><td></td><td></td><td></td><td></td><td></td></tr>
<tr><td>Oligo rev</td><td>AAACGTTTCGGAGCGCACAGTCTC</td><td></td><td></td><td></td><td></td><td></td><td></td><td></td><td></td><td></td><td></td></tr>
<tr><td>Chromosome</td><td>start</td><td>end</td><td>strand</td><td>MM</td><td>target_seq</td><td>PAM</td><td>alignment</td><td>distance</td><td>position</td><td>gene name</td><td>gene id</td></tr>
<tr><td>GU305914.1</td><td>110994</td><td>111016</td><td>-</td><td>0</td><td>GAGACTGTGCGCTCCGAAAC</td><td>GGG</td><td>||||||||[||||||||||||]PAM</td><td>NA</td><td>Intergenic</td><td>NA</td><td>NA</td></tr>
<tr><td>T25690</td><td>GGCCGCGCGCCTTATATGTCAGG</td><td>1000</td><td></td><td></td><td></td><td></td><td></td><td></td><td></td><td></td><td></td></tr>
<tr><td>Oligo fwd</td><td>CACCGGCCGCGCGCCTTATATGTC</td><td></td><td></td><td></td><td></td><td></td><td></td><td></td><td></td><td></td><td></td></tr>
<tr><td>Oligo rev</td><td>AAACGACATATAAGGCGCGCGGCC</td><td></td><td></td><td></td><td></td><td></td><td></td><td></td><td></td><td></td><td></td></tr>
<tr><td>Chromosome</td><td>start</td><td>end</td><td>strand</td><td>MM</td><td>target_seq</td><td>PAM</td><td>alignment</td><td>distance</td><td>position</td><td>gene name</td><td>gene id</td></tr>
<tr><td>GU305914.1</td><td>110973</td><td>110995</td><td>-</td><td>0</td><td>GGCCGCGCGCCTTATATGTC</td><td>AGG</td><td>||||||||[||||||||||||]PAM</td><td>NA</td><td>Intergenic</td><td>NA</td><td>NA</td></tr>
<tr><td>T25691</td><td>CGCGCGCCTTATATGTCAGGCGG</td><td>1000</td><td></td><td></td><td></td><td></td><td></td><td></td><td></td><td></td><td></td></tr>
<tr><td>Oligo fwd</td><td>CACCCGCGCGCCTTATATGTCAGG</td><td></td><td></td><td></td><td></td><td></td><td></td><td></td><td></td><td></td><td></td></tr>
<tr><td>Oligo rev</td><td>AAACCCTGACATATAAGGCGCGCG</td><td></td><td></td><td></td><td></td><td></td><td></td><td></td><td></td><td></td><td></td></tr>
<tr><td>Chromosome</td><td>start</td><td>end</td><td>strand</td><td>MM</td><td>target_seq</td><td>PAM</td><td>alignment</td><td>distance</td><td>position</td><td>gene name</td><td>gene id</td></tr>
<tr><td>GU305914.1</td><td>110970</td><td>110992</td><td>-</td><td>0</td><td>CGCGCGCCTTATATGTCAGG</td><td>CGG</td><td>||||||||[||||||||||||]PAM</td><td>NA</td><td>Intergenic</td><td>NA</td><td>NA</td></tr>
<tr><td>T25692</td><td>ATATGTCAGGCGGCGCCAGAAGG</td><td>1000</td><td></td><td></td><td></td><td></td><td></td><td></td><td></td><td></td><td></td></tr>
<tr><td>Oligo fwd</td><td>CACCATATGTCAGGCGGCGCCAGA</td><td></td><td></td><td></td><td></td><td></td><td></td><td></td><td></td><td></td><td></td></tr>
<tr><td>Oligo rev</td><td>AAACTCTGGCGCCGCCTGACATAT</td><td></td><td></td><td></td><td></td><td></td><td></td><td></td><td></td><td></td><td></td></tr>
<tr><td>Chromosome</td><td>start</td><td>end</td><td>strand</td><td>MM</td><td>target_seq</td><td>PAM</td><td>alignment</td><td>distance</td><td>position</td><td>gene name</td><td>gene id</td></tr>
<tr><td>GU305914.1</td><td>110960</td><td>110982</td><td>-</td><td>0</td><td>ATATGTCAGGCGGCGCCAGA</td><td>AGG</td><td>||||||||[||||||||||||]PAM</td><td>NA</td><td>Intergenic</td><td>NA</td><td>NA</td></tr>
<tr><td>T25693</td><td>TATGTCAGGCGGCGCCAGAAGGG</td><td>1000</td><td></td><td></td><td></td><td></td><td></td><td></td><td></td><td></td><td></td></tr>
<tr><td>Oligo fwd</td><td>CACCTATGTCAGGCGGCGCCAGAA</td><td></td><td></td><td></td><td></td><td></td><td></td><td></td><td></td><td></td><td></td></tr>
<tr><td>Oligo rev</td><td>AAACTTCTGGCGCCGCCTGACATA</td><td></td><td></td><td></td><td></td><td></td><td></td><td></td><td></td><td></td><td></td></tr>
<tr><td>Chromosome</td><td>start</td><td>end</td><td>strand</td><td>MM</td><td>target_seq</td><td>PAM</td><td>alignment</td><td>distance</td><td>position</td><td>gene name</td><td>gene id</td></tr>
<tr><td>GU305914.1</td><td>110959</td><td>110981</td><td>-</td><td>0</td><td>TATGTCAGGCGGCGCCAGAA</td><td>GGG</td><td>||||||||[||||||||||||]PAM</td><td>NA</td><td>Intergenic</td><td>NA</td><td>NA</td></tr>
<tr><td>T25694</td><td>ATGTCAGGCGGCGCCAGAAGGGG</td><td>1000</td><td></td><td></td><td></td><td></td><td></td><td></td><td></td><td></td><td></td></tr>
<tr><td>Oligo fwd</td><td>CACCATGTCAGGCGGCGCCAGAAG</td><td></td><td></td><td></td><td></td><td></td><td></td><td></td><td></td><td></td><td></td></tr>
<tr><td>Oligo rev</td><td>AAACCTTCTGGCGCCGCCTGACAT</td><td></td><td></td><td></td><td></td><td></td><td></td><td></td><td></td><td></td><td></td></tr>
<tr><td>Chromosome</td><td>start</td><td>end</td><td>strand</td><td>MM</td><td>target_seq</td><td>PAM</td><td>alignment</td><td>distance</td><td>position</td><td>gene name</td><td>gene id</td></tr>
<tr><td>GU305914.1</td><td>110958</td><td>110980</td><td>-</td><td>0</td><td>ATGTCAGGCGGCGCCAGAAG</td><td>GGG</td><td>||||||||[||||||||||||]PAM</td><td>NA</td><td>Intergenic</td><td>NA</td><td>NA</td></tr>
<tr><td>T25695</td><td>CCGCCAATCCATAAAATACTCGG</td><td>1000</td><td></td><td></td><td></td><td></td><td></td><td></td><td></td><td></td><td></td></tr>
<tr><td>Oligo fwd</td><td>CACCCCGCCAATCCATAAAATACT</td><td></td><td></td><td></td><td></td><td></td><td></td><td></td><td></td><td></td><td></td></tr>
<tr><td>Oligo rev</td><td>AAACAGTATTTTATGGATTGGCGG</td><td></td><td></td><td></td><td></td><td></td><td></td><td></td><td></td><td></td><td></td></tr>
<tr><td>Chromosome</td><td>start</td><td>end</td><td>strand</td><td>MM</td><td>target_seq</td><td>PAM</td><td>alignment</td><td>distance</td><td>position</td><td>gene name</td><td>gene id</td></tr>
<tr><td>GU305914.1</td><td>110931</td><td>110953</td><td>-</td><td>0</td><td>CCGCCAATCCATAAAATACT</td><td>CGG</td><td>||||||||[||||||||||||]PAM</td><td>NA</td><td>Intergenic</td><td>NA</td><td>NA</td></tr>
<tr><td>T25696</td><td>TAAAATACTCGGCCGAGTCGCGG</td><td>1000</td><td></td><td></td><td></td><td></td><td></td><td></td><td></td><td></td><td></td></tr>
<tr><td>Oligo fwd</td><td>CACCTAAAATACTCGGCCGAGTCG</td><td></td><td></td><td></td><td></td><td></td><td></td><td></td><td></td><td></td><td></td></tr>
<tr><td>Oligo rev</td><td>AAACCGACTCGGCCGAGTATTTTA</td><td></td><td></td><td></td><td></td><td></td><td></td><td></td><td></td><td></td><td></td></tr>
<tr><td>Chromosome</td><td>start</td><td>end</td><td>strand</td><td>MM</td><td>target_seq</td><td>PAM</td><td>alignment</td><td>distance</td><td>position</td><td>gene name</td><td>gene id</td></tr>
<tr><td>GU305914.1</td><td>110920</td><td>110942</td><td>-</td><td>0</td><td>TAAAATACTCGGCCGAGTCG</td><td>CGG</td><td>||||||||[||||||||||||]PAM</td><td>NA</td><td>Intergenic</td><td>NA</td><td>NA</td></tr>
<tr><td>T25697</td><td>AATACTCGGCCGAGTCGCGGCGG</td><td>1000</td><td></td><td></td><td></td><td></td><td></td><td></td><td></td><td></td><td></td></tr>
<tr><td>Oligo fwd</td><td>CACCAATACTCGGCCGAGTCGCGG</td><td></td><td></td><td></td><td></td><td></td><td></td><td></td><td></td><td></td><td></td></tr>
<tr><td>Oligo rev</td><td>AAACCCGCGACTCGGCCGAGTATT</td><td></td><td></td><td></td><td></td><td></td><td></td><td></td><td></td><td></td><td></td></tr>
<tr><td>Chromosome</td><td>start</td><td>end</td><td>strand</td><td>MM</td><td>target_seq</td><td>PAM</td><td>alignment</td><td>distance</td><td>position</td><td>gene name</td><td>gene id</td></tr>
<tr><td>GU305914.1</td><td>110917</td><td>110939</td><td>-</td><td>0</td><td>AATACTCGGCCGAGTCGCGG</td><td>CGG</td><td>||||||||[||||||||||||]PAM</td><td>NA</td><td>Intergenic</td><td>NA</td><td>NA</td></tr>
<tr><td>T25698</td><td>ACTCGGCCGAGTCGCGGCGGCGG</td><td>1000</td><td></td><td></td><td></td><td></td><td></td><td></td><td></td><td></td><td></td></tr>
<tr><td>Oligo fwd</td><td>CACCACTCGGCCGAGTCGCGGCGG</td><td></td><td></td><td></td><td></td><td></td><td></td><td></td><td></td><td></td><td></td></tr>
<tr><td>Oligo rev</td><td>AAACCCGCCGCGACTCGGCCGAGT</td><td></td><td></td><td></td><td></td><td></td><td></td><td></td><td></td><td></td><td></td></tr>
<tr><td>Chromosome</td><td>start</td><td>end</td><td>strand</td><td>MM</td><td>target_seq</td><td>PAM</td><td>alignment</td><td>distance</td><td>position</td><td>gene name</td><td>gene id</td></tr>
<tr><td>GU305914.1</td><td>110914</td><td>110936</td><td>-</td><td>0</td><td>ACTCGGCCGAGTCGCGGCGG</td><td>CGG</td><td>||||||||[||||||||||||]PAM</td><td>NA</td><td>Intergenic</td><td>NA</td><td>NA</td></tr>
<tr><td>T25699</td><td>GCTCACGTCACACTCACCACAGG</td><td>1000</td><td></td><td></td><td></td><td></td><td></td><td></td><td></td><td></td><td></td></tr>
<tr><td>Oligo fwd</td><td>CACCGCTCACGTCACACTCACCAC</td><td></td><td></td><td></td><td></td><td></td><td></td><td></td><td></td><td></td><td></td></tr>
<tr><td>Oligo rev</td><td>AAACGTGGTGAGTGTGACGTGAGC</td><td></td><td></td><td></td><td></td><td></td><td></td><td></td><td></td><td></td><td></td></tr>
<tr><td>Chromosome</td><td>start</td><td>end</td><td>strand</td><td>MM</td><td>target_seq</td><td>PAM</td><td>alignment</td><td>distance</td><td>position</td><td>gene name</td><td>gene id</td></tr>
<tr><td>GU305914.1</td><td>110892</td><td>110914</td><td>-</td><td>0</td><td>GCTCACGTCACACTCACCAC</td><td>AGG</td><td>||||||||[||||||||||||]PAM</td><td>NA</td><td>Intergenic</td><td>NA</td><td>NA</td></tr>
<tr><td>T25700</td><td>ACCGAGCACAAAAAAATACATGG</td><td>1000</td><td></td><td></td><td></td><td></td><td></td><td></td><td></td><td></td><td></td></tr>
<tr><td>Oligo fwd</td><td>CACCACCGAGCACAAAAAAATACA</td><td></td><td></td><td></td><td></td><td></td><td></td><td></td><td></td><td></td><td></td></tr>
<tr><td>Oligo rev</td><td>AAACTGTATTTTTTTGTGCTCGGT</td><td></td><td></td><td></td><td></td><td></td><td></td><td></td><td></td><td></td><td></td></tr>
<tr><td>Chromosome</td><td>start</td><td>end</td><td>strand</td><td>MM</td><td>target_seq</td><td>PAM</td><td>alignment</td><td>distance</td><td>position</td><td>gene name</td><td>gene id</td></tr>
<tr><td>GU305914.1</td><td>110863</td><td>110885</td><td>-</td><td>0</td><td>ACCGAGCACAAAAAAATACA</td><td>TGG</td><td>||||||||[||||||||||||]PAM</td><td>NA</td><td>Intergenic</td><td>NA</td><td>NA</td></tr>
<tr><td>T25701</td><td>TGGAATCGTACACGTTCTGCAGG</td><td>1000</td><td></td><td></td><td></td><td></td><td></td><td></td><td></td><td></td><td></td></tr>
<tr><td>Oligo fwd</td><td>CACCTGGAATCGTACACGTTCTGC</td><td></td><td></td><td></td><td></td><td></td><td></td><td></td><td></td><td></td><td></td></tr>
<tr><td>Oligo rev</td><td>AAACGCAGAACGTGTACGATTCCA</td><td></td><td></td><td></td><td></td><td></td><td></td><td></td><td></td><td></td><td></td></tr>
<tr><td>Chromosome</td><td>start</td><td>end</td><td>strand</td><td>MM</td><td>target_seq</td><td>PAM</td><td>alignment</td><td>distance</td><td>position</td><td>gene name</td><td>gene id</td></tr>
<tr><td>GU305914.1</td><td>110843</td><td>110865</td><td>-</td><td>0</td><td>TGGAATCGTACACGTTCTGC</td><td>AGG</td><td>||||||||[||||||||||||]PAM</td><td>NA</td><td>Intergenic</td><td>NA</td><td>NA</td></tr>
<tr><td>T25702</td><td>GGTTGAAAAACTGCTGACCGAGG</td><td>1000</td><td></td><td></td><td></td><td></td><td></td><td></td><td></td><td></td><td></td></tr>
<tr><td>Oligo fwd</td><td>CACCGGTTGAAAAACTGCTGACCG</td><td></td><td></td><td></td><td></td><td></td><td></td><td></td><td></td><td></td><td></td></tr>
<tr><td>Oligo rev</td><td>AAACCGGTCAGCAGTTTTTCAACC</td><td></td><td></td><td></td><td></td><td></td><td></td><td></td><td></td><td></td><td></td></tr>
<tr><td>Chromosome</td><td>start</td><td>end</td><td>strand</td><td>MM</td><td>target_seq</td><td>PAM</td><td>alignment</td><td>distance</td><td>position</td><td>gene name</td><td>gene id</td></tr>
<tr><td>GU305914.1</td><td>110822</td><td>110844</td><td>-</td><td>0</td><td>GGTTGAAAAACTGCTGACCG</td><td>AGG</td><td>||||||||[||||||||||||]PAM</td><td>NA</td><td>Intergenic</td><td>NA</td><td>NA</td></tr>
<tr><td>T25703</td><td>GTTGAAAAACTGCTGACCGAGGG</td><td>1000</td><td></td><td></td><td></td><td></td><td></td><td></td><td></td><td></td><td></td></tr>
<tr><td>Oligo fwd</td><td>CACCGTTGAAAAACTGCTGACCGA</td><td></td><td></td><td></td><td></td><td></td><td></td><td></td><td></td><td></td><td></td></tr>
<tr><td>Oligo rev</td><td>AAACTCGGTCAGCAGTTTTTCAAC</td><td></td><td></td><td></td><td></td><td></td><td></td><td></td><td></td><td></td><td></td></tr>
<tr><td>Chromosome</td><td>start</td><td>end</td><td>strand</td><td>MM</td><td>target_seq</td><td>PAM</td><td>alignment</td><td>distance</td><td>position</td><td>gene name</td><td>gene id</td></tr>
<tr><td>GU305914.1</td><td>110821</td><td>110843</td><td>-</td><td>0</td><td>GTTGAAAAACTGCTGACCGA</td><td>GGG</td><td>||||||||[||||||||||||]PAM</td><td>NA</td><td>Intergenic</td><td>NA</td><td>NA</td></tr>
</table>

T25704 AAAACTGCTGACCGAGGGTCAGG 1000
Oligo fwd CACCAAAACTGCTGACCGAGGGTC
Oligo rev AAACGACCCTCGGTCAGCAGTTTT
Chromosome start end strand MM target_seq PAM alignment distance position gene name gene id
GU305914.1 110816 110838 - 0 AAAACTGCTGACCGAGGGTC AGG ||||||||[||||||||||||]PAM NA Intergenic NA NA

T25705 GAGGGTCAGGACCCTGTGCTTGG 1000
Oligo fwd CACCGAGGGTCAGGACCCTGTGCT
Oligo rev AAACAGCACAGGGTCCTGACCCTC
Chromosome start end strand MM target_seq PAM alignment distance position gene name gene id
GU305914.1 110803 110825 - 0 GAGGGTCAGGACCCTGTGCT TGG ||||||||[||||||||||||]PAM NA Intergenic NA NA

T25706 AGGGTCAGGACCCTGTGCTTGGG 1000
Oligo fwd CACCAGGGTCAGGACCCTGTGCTT
Oligo rev AAACAAGCACAGGGTCCTGACCCT
Chromosome start end strand MM target_seq PAM alignment distance position gene name gene id
GU305914.1 110802 110824 - 0 AGGGTCAGGACCCTGTGCTT GGG ||||||||[||||||||||||]PAM NA Intergenic NA NA

T25707 CCTGTGCTTGGGCAGCGTGATGG 1000
Oligo fwd CACCCCTGTGCTTGGGCAGCGTGA
Oligo rev AAACTCACGCTGCCCAAGCACAGG
Chromosome start end strand MM target_seq PAM alignment distance position gene name gene id
GU305914.1 110791 110813 - 0 CCTGTGCTTGGGCAGCGTGA TGG ||||||||[||||||||||||]PAM NA Intergenic NA NA
GU305914.1 138632 138654 - 4 CCGGTCCTTGGGCAGCGCGT TGG ||-||-||[|||||||||-|-]PAM NA Intergenic NA NA

T25708 TGCCGATCCCGTTCTCGCAGTGG 1000
Oligo fwd CACCTGCCGATCCCGTTCTCGCAG
Oligo rev AAACCTGCGAGAACGGGATCGGCA
Chromosome start end strand MM target_seq PAM alignment distance position gene name gene id
GU305914.1 110759 110781 - 0 TGCCGATCCCGTTCTCGCAG TGG ||||||||[||||||||||||]PAM NA Intergenic NA NA

T25709 GAAGAGAGCGAAGTCGTGCGCGG 1000
Oligo fwd CACCGAAGAGAGCGAAGTCGTGCG
Oligo rev AAACCGCACGACTTCGCTCTCTTC
Chromosome start end strand MM target_seq PAM alignment distance position gene name gene id
GU305914.1 110737 110759 - 0 GAAGAGAGCGAAGTCGTGCG CGG ||||||||[||||||||||||]PAM NA Intergenic NA NA

T25710 AGCCTGTCGACATCCTTGTCCGG 1000
Oligo fwd CACCAGCCTGTCGACATCCTTGTC
Oligo rev AAACGACAAGGATGTCGACAGGCT
Chromosome start end strand MM target_seq PAM alignment distance position gene name gene id
GU305914.1 110712 110734 - 0 AGCCTGTCGACATCCTTGTC CGG ||||||||[||||||||||||]PAM NA Intergenic NA NA

T25711 GCCTGTCGACATCCTTGTCCGGG 1000
Oligo fwd CACCGCCTGTCGACATCCTTGTCC
Oligo rev AAACGGACAAGGATGTCGACAGGC
Chromosome start end strand MM target_seq PAM alignment distance position gene name gene id
GU305914.1 110711 110733 - 0 GCCTGTCGACATCCTTGTCC GGG ||||||||[||||||||||||]PAM NA Intergenic NA NA

T25712 CCTGTCGACATCCTTGTCCGGGG 1000
Oligo fwd CACCCCTGTCGACATCCTTGTCCG
Oligo rev AAACCGGACAAGGATGTCGACAGG
Chromosome start end strand MM target_seq PAM alignment distance position gene name gene id
GU305914.1 110710 110732 - 0 CCTGTCGACATCCTTGTCCG GGG ||||||||[||||||||||||]PAM NA Intergenic NA NA

T25713 GTCGACATCCTTGTCCGGGGCGG 1000
Oligo fwd CACCGTCGACATCCTTGTCCGGGG
Oligo rev AAACCCCCGGACAAGGATGTCGAC
Chromosome start end strand MM target_seq PAM alignment distance position gene name gene id
GU305914.1 110707 110729 - 0 GTCGACATCCTTGTCCGGGG CGG ||||||||[||||||||||||]PAM NA Intergenic NA NA

T25714 GCAGCAGCATGTAGTCGACGAGG 1000
Oligo fwd CACCGCAGCAGCATGTAGTCGACG
Oligo rev AAACCGTCGACTACATGCTGCTGC
Chromosome start end strand MM target_seq PAM alignment distance position gene name gene id
GU305914.1 110675 110697 - 0 GCAGCAGCATGTAGTCGACG AGG ||||||||[||||||||||||]PAM NA Intergenic NA NA

T25715 CAGCAGCATGTAGTCGACGAGGG 1000
Oligo fwd CACCCAGCAGCATGTAGTCGACGA
Oligo rev AAACTCGTCGACTACATGCTGCTG
Chromosome start end strand MM target_seq PAM alignment distance position gene name gene id
GU305914.1 110674 110696 - 0 CAGCAGCATGTAGTCGACGA GGG ||||||||[||||||||||||]PAM NA Intergenic NA NA

T25716 AGCAGCATGTAGTCGACGAGGGG 1000
Oligo fwd CACCAGCAGCATGTAGTCGACGAG
Oligo rev AAACCTCGTCGACTACATGCTGCT
Chromosome start end strand MM target_seq PAM alignment distance position gene name gene id
GU305914.1 110673 110695 - 0 AGCAGCATGTAGTCGACGAG GGG ||||||||[||||||||||||]PAM NA Intergenic NA NA

T25717 GCAGCATGTAGTCGACGAGGGGG 1000
Oligo fwd CACCGCAGCATGTAGTCGACGAGG
Oligo rev AAACCCTCGTCGACTACATGCTGC
Chromosome start end strand MM target_seq PAM alignment distance position gene name gene id
GU305914.1 110672 110694 - 0 GCAGCATGTAGTCGACGAGG GGG ||||||||[||||||||||||]PAM NA Intergenic NA NA

T25718 AGTCGACGAGGGGGTTGAAACGG 1000
Oligo fwd CACCAGTCGACGAGGGGGTTGAAA
Oligo rev AAACTTTCAACCCCCTCGTCGACT
Chromosome start end strand MM target_seq PAM alignment distance position gene name gene id

<table>
<tr><td>GU305914.1</td><td>110663</td><td>110685</td><td>-</td><td>0</td><td>AGTCGACGAGGGGGTTGAAA</td><td>CGG</td><td>||||||||[||||||||||||]PAM</td><td>NA</td><td>Intergenic</td><td>NA</td><td>NA</td></tr>
<tr><td></td><td></td><td></td><td></td><td></td><td></td><td></td><td></td><td></td><td></td><td></td><td></td></tr>
<tr><td>T25719</td><td>GACGAGGGGGTTGAAACGGCTGG</td><td>1000</td><td></td><td></td><td></td><td></td><td></td><td></td><td></td><td></td><td></td></tr>
<tr><td>Oligo fwd</td><td>CACCGACGAGGGGGTTGAAACGGC</td><td></td><td></td><td></td><td></td><td></td><td></td><td></td><td></td><td></td><td></td></tr>
<tr><td>Oligo rev</td><td>AAACGCCGTTTCAACCCCCTCGTC</td><td></td><td></td><td></td><td></td><td></td><td></td><td></td><td></td><td></td><td></td></tr>
<tr><td>Chromosome</td><td>start</td><td>end</td><td>strand</td><td>MM</td><td>target_seq</td><td>PAM</td><td>alignment</td><td>distance</td><td>position</td><td>gene name</td><td>gene id</td></tr>
<tr><td>GU305914.1</td><td>110659</td><td>110681</td><td>-</td><td>0</td><td>GACGAGGGGGTTGAAACGGC</td><td>TGG</td><td>||||||||[||||||||||||]PAM</td><td>NA</td><td>Intergenic</td><td>NA</td><td>NA</td></tr>
<tr><td></td><td></td><td></td><td></td><td></td><td></td><td></td><td></td><td></td><td></td><td></td><td></td></tr>
<tr><td>T25720</td><td>GGGGGTTGAAACGGCTGGCCTGG</td><td>1000</td><td></td><td></td><td></td><td></td><td></td><td></td><td></td><td></td><td></td></tr>
<tr><td>Oligo fwd</td><td>CACCGGGGGTTGAAACGGCTGGCC</td><td></td><td></td><td></td><td></td><td></td><td></td><td></td><td></td><td></td><td></td></tr>
<tr><td>Oligo rev</td><td>AAACGGCCAGCCGTTTCAACCCCC</td><td></td><td></td><td></td><td></td><td></td><td></td><td></td><td></td><td></td><td></td></tr>
<tr><td>Chromosome</td><td>start</td><td>end</td><td>strand</td><td>MM</td><td>target_seq</td><td>PAM</td><td>alignment</td><td>distance</td><td>position</td><td>gene name</td><td>gene id</td></tr>
<tr><td>GU305914.1</td><td>110654</td><td>110676</td><td>-</td><td>0</td><td>GGGGGTTGAAACGGCTGGCC</td><td>TGG</td><td>||||||||[||||||||||||]PAM</td><td>NA</td><td>Intergenic</td><td>NA</td><td>NA</td></tr>
<tr><td></td><td></td><td></td><td></td><td></td><td></td><td></td><td></td><td></td><td></td><td></td><td></td></tr>
<tr><td>T25721</td><td>AACGGCTGGCCTGGACGTGCAGG</td><td>1000</td><td></td><td></td><td></td><td></td><td></td><td></td><td></td><td></td><td></td></tr>
<tr><td>Oligo fwd</td><td>CACCAACGGCTGGCCTGGACGTGC</td><td></td><td></td><td></td><td></td><td></td><td></td><td></td><td></td><td></td><td></td></tr>
<tr><td>Oligo rev</td><td>AAACGCACGTCCAGGCCAGCCGTT</td><td></td><td></td><td></td><td></td><td></td><td></td><td></td><td></td><td></td><td></td></tr>
<tr><td>Chromosome</td><td>start</td><td>end</td><td>strand</td><td>MM</td><td>target_seq</td><td>PAM</td><td>alignment</td><td>distance</td><td>position</td><td>gene name</td><td>gene id</td></tr>
<tr><td>GU305914.1</td><td>110645</td><td>110667</td><td>-</td><td>0</td><td>AACGGCTGGCCTGGACGTGC</td><td>AGG</td><td>||||||||[||||||||||||]PAM</td><td>NA</td><td>Intergenic</td><td>NA</td><td>NA</td></tr>
<tr><td></td><td></td><td></td><td></td><td></td><td></td><td></td><td></td><td></td><td></td><td></td><td></td></tr>
<tr><td>T25722</td><td>GGCCTGGACGTGCAGGACGCCGG</td><td>1000</td><td></td><td></td><td></td><td></td><td></td><td></td><td></td><td></td><td></td></tr>
<tr><td>Oligo fwd</td><td>CACCGGCCTGGACGTGCAGGACGC</td><td></td><td></td><td></td><td></td><td></td><td></td><td></td><td></td><td></td><td></td></tr>
<tr><td>Oligo rev</td><td>AAACGCGTCCTGCACGTCCAGGCC</td><td></td><td></td><td></td><td></td><td></td><td></td><td></td><td></td><td></td><td></td></tr>
<tr><td>Chromosome</td><td>start</td><td>end</td><td>strand</td><td>MM</td><td>target_seq</td><td>PAM</td><td>alignment</td><td>distance</td><td>position</td><td>gene name</td><td>gene id</td></tr>
<tr><td>GU305914.1</td><td>110638</td><td>110660</td><td>-</td><td>0</td><td>GGCCTGGACGTGCAGGACGC</td><td>CGG</td><td>||||||||[||||||||||||]PAM</td><td>NA</td><td>Intergenic</td><td>NA</td><td>NA</td></tr>
<tr><td></td><td></td><td></td><td></td><td></td><td></td><td></td><td></td><td></td><td></td><td></td><td></td></tr>
<tr><td>T25723</td><td>CTGGACGTGCAGGACGCCGGCGG</td><td>1000</td><td></td><td></td><td></td><td></td><td></td><td></td><td></td><td></td><td></td></tr>
<tr><td>Oligo fwd</td><td>CACCCTGGACGTGCAGGACGCCGG</td><td></td><td></td><td></td><td></td><td></td><td></td><td></td><td></td><td></td><td></td></tr>
<tr><td>Oligo rev</td><td>AAACCCGGCGTCCTGCACGTCCAG</td><td></td><td></td><td></td><td></td><td></td><td></td><td></td><td></td><td></td><td></td></tr>
<tr><td>Chromosome</td><td>start</td><td>end</td><td>strand</td><td>MM</td><td>target_seq</td><td>PAM</td><td>alignment</td><td>distance</td><td>position</td><td>gene name</td><td>gene id</td></tr>
<tr><td>GU305914.1</td><td>110635</td><td>110657</td><td>-</td><td>0</td><td>CTGGACGTGCAGGACGCCGG</td><td>CGG</td><td>||||||||[||||||||||||]PAM</td><td>NA</td><td>Intergenic</td><td>NA</td><td>NA</td></tr>
<tr><td></td><td></td><td></td><td></td><td></td><td></td><td></td><td></td><td></td><td></td><td></td><td></td></tr>
<tr><td>T25724</td><td>GACGTGCAGGACGCCGGCGGCGG</td><td>1000</td><td></td><td></td><td></td><td></td><td></td><td></td><td></td><td></td><td></td></tr>
<tr><td>Oligo fwd</td><td>CACCGACGTGCAGGACGCCGGCGG</td><td></td><td></td><td></td><td></td><td></td><td></td><td></td><td></td><td></td><td></td></tr>
<tr><td>Oligo rev</td><td>AAACCCGCCGGCGTCCTGCACGTC</td><td></td><td></td><td></td><td></td><td></td><td></td><td></td><td></td><td></td><td></td></tr>
<tr><td>Chromosome</td><td>start</td><td>end</td><td>strand</td><td>MM</td><td>target_seq</td><td>PAM</td><td>alignment</td><td>distance</td><td>position</td><td>gene name</td><td>gene id</td></tr>
<tr><td>GU305914.1</td><td>110632</td><td>110654</td><td>-</td><td>0</td><td>GACGTGCAGGACGCCGGCGG</td><td>CGG</td><td>||||||||[||||||||||||]PAM</td><td>NA</td><td>Intergenic</td><td>NA</td><td>NA</td></tr>
<tr><td>GU305914.1</td><td>75661</td><td>75683</td><td>+</td><td>4</td><td>GACGTTCGTGTCGCCGGCGG</td><td>GGG</td><td>|||||-|-[-|-|||||||||]PAM</td><td>NA</td><td>Intergenic</td><td>NA</td><td>NA</td></tr>
<tr><td>GU305914.1</td><td>96942</td><td>96964</td><td>+</td><td>4</td><td>GGCGAGCAGGGCGACGGCGG</td><td>AGG</td><td>|-||-|||[||-||-||||||]PAM</td><td>NA</td><td>Intergenic</td><td>NA</td><td>NA</td></tr>
<tr><td></td><td></td><td></td><td></td><td></td><td></td><td></td><td></td><td></td><td></td><td></td><td></td></tr>
<tr><td>T25725</td><td>GTGCAGGACGCCGGCGGCGGAGG</td><td>1000</td><td></td><td></td><td></td><td></td><td></td><td></td><td></td><td></td><td></td></tr>
<tr><td>Oligo fwd</td><td>CACCGTGCAGGACGCCGGCGGCGG</td><td></td><td></td><td></td><td></td><td></td><td></td><td></td><td></td><td></td><td></td></tr>
<tr><td>Oligo rev</td><td>AAACCCGCCGCCGGCGTCCTGCAC</td><td></td><td></td><td></td><td></td><td></td><td></td><td></td><td></td><td></td><td></td></tr>
<tr><td>Chromosome</td><td>start</td><td>end</td><td>strand</td><td>MM</td><td>target_seq</td><td>PAM</td><td>alignment</td><td>distance</td><td>position</td><td>gene name</td><td>gene id</td></tr>
<tr><td>GU305914.1</td><td>110629</td><td>110651</td><td>-</td><td>0</td><td>GTGCAGGACGCCGGCGGCGG</td><td>AGG</td><td>||||||||[||||||||||||]PAM</td><td>NA</td><td>Intergenic</td><td>NA</td><td>NA</td></tr>
<tr><td>GU305914.1</td><td>19054</td><td>19076</td><td>+</td><td>4</td><td>GGGTTGGACGACGGCGGCGG</td><td>TGG</td><td>|-|--|||[||-|||||||||]PAM</td><td>NA</td><td>Intergenic</td><td>NA</td><td>NA</td></tr>
<tr><td>GU305914.1</td><td>225984</td><td>226006</td><td>-</td><td>4</td><td>GTGGAGCGCGACGGCGGCGG</td><td>CGG</td><td>|||-||--[||-|||||||||]PAM</td><td>NA</td><td>Intergenic</td><td>NA</td><td>NA</td></tr>
<tr><td>GU305914.1</td><td>96945</td><td>96967</td><td>+</td><td>4</td><td>GAGCAGGGCGACGGCGGAGG</td><td>CGG</td><td>|-|||||-[||-||||||-||]PAM</td><td>NA</td><td>Intergenic</td><td>NA</td><td>NA</td></tr>
<tr><td></td><td></td><td></td><td></td><td></td><td></td><td></td><td></td><td></td><td></td><td></td><td></td></tr>
<tr><td>T25726</td><td>GGCGGCGGAGGAGACGCCGTCGG</td><td>1000</td><td></td><td></td><td></td><td></td><td></td><td></td><td></td><td></td><td></td></tr>
<tr><td>Oligo fwd</td><td>CACCGGCGGCGGAGGAGACGCCGT</td><td></td><td></td><td></td><td></td><td></td><td></td><td></td><td></td><td></td><td></td></tr>
<tr><td>Oligo rev</td><td>AAACACGGCGTCTCCTCCGCCGCC</td><td></td><td></td><td></td><td></td><td></td><td></td><td></td><td></td><td></td><td></td></tr>
<tr><td>Chromosome</td><td>start</td><td>end</td><td>strand</td><td>MM</td><td>target_seq</td><td>PAM</td><td>alignment</td><td>distance</td><td>position</td><td>gene name</td><td>gene id</td></tr>
<tr><td>GU305914.1</td><td>110617</td><td>110639</td><td>-</td><td>0</td><td>GGCGGCGGAGGAGACGCCGT</td><td>CGG</td><td>||||||||[||||||||||||]PAM</td><td>NA</td><td>Intergenic</td><td>NA</td><td>NA</td></tr>
<tr><td></td><td></td><td></td><td></td><td></td><td></td><td></td><td></td><td></td><td></td><td></td><td></td></tr>
<tr><td>T25727</td><td>GGAGGAGACGCCGTCGGCCGTGG</td><td>1000</td><td></td><td></td><td></td><td></td><td></td><td></td><td></td><td></td><td></td></tr>
<tr><td>Oligo fwd</td><td>CACCGGAGGAGACGCCGTCGGCCG</td><td></td><td></td><td></td><td></td><td></td><td></td><td></td><td></td><td></td><td></td></tr>
<tr><td>Oligo rev</td><td>AAACCGGCCGACGGCGTCTCCTCC</td><td></td><td></td><td></td><td></td><td></td><td></td><td></td><td></td><td></td><td></td></tr>
<tr><td>Chromosome</td><td>start</td><td>end</td><td>strand</td><td>MM</td><td>target_seq</td><td>PAM</td><td>alignment</td><td>distance</td><td>position</td><td>gene name</td><td>gene id</td></tr>
<tr><td>GU305914.1</td><td>110611</td><td>110633</td><td>-</td><td>0</td><td>GGAGGAGACGCCGTCGGCCG</td><td>TGG</td><td>||||||||[||||||||||||]PAM</td><td>NA</td><td>Intergenic</td><td>NA</td><td>NA</td></tr>
<tr><td></td><td></td><td></td><td></td><td></td><td></td><td></td><td></td><td></td><td></td><td></td><td></td></tr>
<tr><td>T25728</td><td>TCGGCCGTGGCGTGTTCGAACGG</td><td>1000</td><td></td><td></td><td></td><td></td><td></td><td></td><td></td><td></td><td></td></tr>
<tr><td>Oligo fwd</td><td>CACCTCGGCCGTGGCGTGTTCGAA</td><td></td><td></td><td></td><td></td><td></td><td></td><td></td><td></td><td></td><td></td></tr>
<tr><td>Oligo rev</td><td>AAACTTCGAACACGCCACGGCCGA</td><td></td><td></td><td></td><td></td><td></td><td></td><td></td><td></td><td></td><td></td></tr>
<tr><td>Chromosome</td><td>start</td><td>end</td><td>strand</td><td>MM</td><td>target_seq</td><td>PAM</td><td>alignment</td><td>distance</td><td>position</td><td>gene name</td><td>gene id</td></tr>
<tr><td>GU305914.1</td><td>110598</td><td>110620</td><td>-</td><td>0</td><td>TCGGCCGTGGCGTGTTCGAA</td><td>CGG</td><td>||||||||[||||||||||||]PAM</td><td>NA</td><td>Intergenic</td><td>NA</td><td>NA</td></tr>
<tr><td></td><td></td><td></td><td></td><td></td><td></td><td></td><td></td><td></td><td></td><td></td><td></td></tr>
<tr><td>T25729</td><td>GTTCGAACGGCCTGCGCGAGAGG</td><td>1000</td><td></td><td></td><td></td><td></td><td></td><td></td><td></td><td></td><td></td></tr>
<tr><td>Oligo fwd</td><td>CACCGTTCGAACGGCCTGCGCGAG</td><td></td><td></td><td></td><td></td><td></td><td></td><td></td><td></td><td></td><td></td></tr>
<tr><td>Oligo rev</td><td>AAACCTCGCGCAGGCCGTTCGAAC</td><td></td><td></td><td></td><td></td><td></td><td></td><td></td><td></td><td></td><td></td></tr>
<tr><td>Chromosome</td><td>start</td><td>end</td><td>strand</td><td>MM</td><td>target_seq</td><td>PAM</td><td>alignment</td><td>distance</td><td>position</td><td>gene name</td><td>gene id</td></tr>
<tr><td>GU305914.1</td><td>110585</td><td>110607</td><td>-</td><td>0</td><td>GTTCGAACGGCCTGCGCGAG</td><td>AGG</td><td>||||||||[||||||||||||]PAM</td><td>NA</td><td>Intergenic</td><td>NA</td><td>NA</td></tr>
<tr><td>GU305914.1</td><td>66943</td><td>66965</td><td>+</td><td>4</td><td>CGCCGAACGACCTGCGCGAG</td><td>TGG</td><td>---|||||[|-||||||||||]PAM</td><td>NA</td><td>Intergenic</td><td>NA</td><td>NA</td></tr>
<tr><td></td><td></td><td></td><td></td><td></td><td></td><td></td><td></td><td></td><td></td><td></td><td></td></tr>
<tr><td>T25730</td><td>GCGAGAGGTCCCAGCCGCTGCGG</td><td>1000</td><td></td><td></td><td></td><td></td><td></td><td></td><td></td><td></td><td></td></tr>
<tr><td>Oligo fwd</td><td>CACCGCGAGAGGTCCCAGCCGCTG</td><td></td><td></td><td></td><td></td><td></td><td></td><td></td><td></td><td></td><td></td></tr>
<tr><td>Oligo rev</td><td>AAACCAGCGGCTGGGACCTCTCGC</td><td></td><td></td><td></td><td></td><td></td><td></td><td></td><td></td><td></td><td></td></tr>
<tr><td>Chromosome</td><td>start</td><td>end</td><td>strand</td><td>MM</td><td>target_seq</td><td>PAM</td><td>alignment</td><td>distance</td><td>position</td><td>gene name</td><td>gene id</td></tr>
<tr><td>GU305914.1</td><td>110570</td><td>110592</td><td>-</td><td>0</td><td>GCGAGAGGTCCCAGCCGCTG</td><td>CGG</td><td>||||||||[||||||||||||]PAM</td><td>NA</td><td>Intergenic</td><td>NA</td><td>NA</td></tr>
<tr><td></td><td></td><td></td><td></td><td></td><td></td><td></td><td></td><td></td><td></td><td></td><td></td></tr>
<tr><td>T25731</td><td>AGAGGTCCCAGCCGCTGCGGTGG</td><td>1000</td><td></td><td></td><td></td><td></td><td></td><td></td><td></td><td></td><td></td></tr>
<tr><td>Oligo fwd</td><td>CACCAGAGGTCCCAGCCGCTGCGG</td><td></td><td></td><td></td><td></td><td></td><td></td><td></td><td></td><td></td><td></td></tr>
<tr><td>Oligo rev</td><td>AAACCCGCAGCGGCTGGGACCTCT</td><td></td><td></td><td></td><td></td><td></td><td></td><td></td><td></td><td></td><td></td></tr>
<tr><td>Chromosome</td><td>start</td><td>end</td><td>strand</td><td>MM</td><td>target_seq</td><td>PAM</td><td>alignment</td><td>distance</td><td>position</td><td>gene name</td><td>gene id</td></tr>
<tr><td>GU305914.1</td><td>110567</td><td>110589</td><td>-</td><td>0</td><td>AGAGGTCCCAGCCGCTGCGG</td><td>TGG</td><td>||||||||[||||||||||||]PAM</td><td>NA</td><td>Intergenic</td><td>NA</td><td>NA</td></tr>
<tr><td></td><td></td><td></td><td></td><td></td><td></td><td></td><td></td><td></td><td></td><td></td><td></td></tr>
<tr><td>T25732</td><td>TGCGGTGGCTCTCGCAGTAGAGG</td><td>1000</td><td></td><td></td><td></td><td></td><td></td><td></td><td></td><td></td><td></td></tr>
<tr><td>Oligo fwd</td><td>CACCTGCGGTGGCTCTCGCAGTAG</td><td></td><td></td><td></td><td></td><td></td><td></td><td></td><td></td><td></td><td></td></tr>
<tr><td>Oligo rev</td><td>AAACCTACTGCGAGAGCCACCGCA</td><td></td><td></td><td></td><td></td><td></td><td></td><td></td><td></td><td></td><td></td></tr>
</table>

| | | | | | | | | | | | |
|---|---|---|---|---|---|---|---|---|---|---|---|
| Chromosome | start | end | strand | MM | target_seq | PAM | alignment | distance | position | gene name | gene id |
| GU305914.1 | 110552 | 110574 | - | 0 | TGCGGTGGCTCTCGCAGTAG | AGG | \|\|\|\|\|\|\|\|[\|\|\|\|\|\|\|\|\|\|\|\|]PAM | NA | Intergenic | NA | NA |
| T25733 | GGTGGCTCTCGCAGTAGAGGAGG | 1000 | | | | | | | | | |
| Oligo fwd | CACCGGTGGCTCTCGCAGTAGAGG | | | | | | | | | | |
| Oligo rev | AAACCCTCTACTGCGAGAGCCACC | | | | | | | | | | |
| Chromosome | start | end | strand | MM | target_seq | PAM | alignment | distance | position | gene name | gene id |
| GU305914.1 | 110549 | 110571 | - | 0 | GGTGGCTCTCGCAGTAGAGG | AGG | \|\|\|\|\|\|\|\|[\|\|\|\|\|\|\|\|\|\|\|\|]PAM | NA | Intergenic | NA | NA |
| T25734 | TAGAGGAGGAGCACGAGCCCCGG | 1000 | | | | | | | | | |
| Oligo fwd | CACCTAGAGGAGGAGCACGAGCCC | | | | | | | | | | |
| Oligo rev | AAACGGGCTCGTGCTCCTCCTCTA | | | | | | | | | | |
| Chromosome | start | end | strand | MM | target_seq | PAM | alignment | distance | position | gene name | gene id |
| GU305914.1 | 110535 | 110557 | - | 0 | TAGAGGAGGAGCACGAGCCC | CGG | \|\|\|\|\|\|\|\|[\|\|\|\|\|\|\|\|\|\|\|\|]PAM | NA | Intergenic | NA | NA |
| T25735 | CCAGAGCTGCGAGACCGTGACGG | 1000 | | | | | | | | | |
| Oligo fwd | CACCCCAGAGCTGCGAGACCGTGA | | | | | | | | | | |
| Oligo rev | AAACTCACGGTCTCGCAGCTCTGG | | | | | | | | | | |
| Chromosome | start | end | strand | MM | target_seq | PAM | alignment | distance | position | gene name | gene id |
| GU305914.1 | 110512 | 110534 | - | 0 | CCAGAGCTGCGAGACCGTGA | CGG | \|\|\|\|\|\|\|\|[\|\|\|\|\|\|\|\|\|\|\|\|]PAM | NA | Intergenic | NA | NA |
| T25736 | AGCTGCGAGACCGTGACGGCCGG | 1000 | | | | | | | | | |
| Oligo fwd | CACCAGCTGCGAGACCGTGACGGC | | | | | | | | | | |
| Oligo rev | AAACGCCGTCACGGTCTCGCAGCT | | | | | | | | | | |
| Chromosome | start | end | strand | MM | target_seq | PAM | alignment | distance | position | gene name | gene id |
| GU305914.1 | 110508 | 110530 | - | 0 | AGCTGCGAGACCGTGACGGC | CGG | \|\|\|\|\|\|\|\|[\|\|\|\|\|\|\|\|\|\|\|\|]PAM | NA | Intergenic | NA | NA |
| T25737 | GCTGCGAGACCGTGACGGCCGGG | 1000 | | | | | | | | | |
| Oligo fwd | CACCGCTGCGAGACCGTGACGGCC | | | | | | | | | | |
| Oligo rev | AAACGGCCGTCACGGTCTCGCAGC | | | | | | | | | | |
| Chromosome | start | end | strand | MM | target_seq | PAM | alignment | distance | position | gene name | gene id |
| GU305914.1 | 110507 | 110529 | - | 0 | GCTGCGAGACCGTGACGGCC | GGG | \|\|\|\|\|\|\|\|[\|\|\|\|\|\|\|\|\|\|\|\|]PAM | NA | Intergenic | NA | NA |
| T25738 | GCCGGGTCGCGCGAGTACCACGG | 1000 | | | | | | | | | |
| Oligo fwd | CACCGCCGGGTCGCGCGAGTACCA | | | | | | | | | | |
| Oligo rev | AAACTGGTACTCGCGCGACCCGGC | | | | | | | | | | |
| Chromosome | start | end | strand | MM | target_seq | PAM | alignment | distance | position | gene name | gene id |
| GU305914.1 | 110490 | 110512 | - | 0 | GCCGGGTCGCGCGAGTACCA | CGG | \|\|\|\|\|\|\|\|[\|\|\|\|\|\|\|\|\|\|\|\|]PAM | NA | Intergenic | NA | NA |
| T25739 | CCGGGTCGCGCGAGTACCACGGG | 1000 | | | | | | | | | |
| Oligo fwd | CACCCCGGGTCGCGCGAGTACCAC | | | | | | | | | | |
| Oligo rev | AAACGTGGTACTCGCGCGACCCGG | | | | | | | | | | |
| Chromosome | start | end | strand | MM | target_seq | PAM | alignment | distance | position | gene name | gene id |
| GU305914.1 | 110489 | 110511 | - | 0 | CCGGGTCGCGCGAGTACCAC | GGG | \|\|\|\|\|\|\|\|[\|\|\|\|\|\|\|\|\|\|\|\|]PAM | NA | Intergenic | NA | NA |
| T25740 | GAAGTTCTTGACGCAGCCGCCGG | 1000 | | | | | | | | | |
| Oligo fwd | CACCGAAGTTCTTGACGCAGCCGC | | | | | | | | | | |
| Oligo rev | AAACGCGGCTGCGTCAAGAACTTC | | | | | | | | | | |
| Chromosome | start | end | strand | MM | target_seq | PAM | alignment | distance | position | gene name | gene id |
| GU305914.1 | 110440 | 110462 | - | 0 | GAAGTTCTTGACGCAGCCGC | CGG | \|\|\|\|\|\|\|\|[\|\|\|\|\|\|\|\|\|\|\|\|]PAM | NA | Intergenic | NA | NA |
| GU305914.1 | 69574 | 69596 | + | 4 | GAAGAACTCGACGCAGTCGC | TGG | \|\|\|\|--\|\|[-\|\|\|\|\|\|\|-\|\|\|]PAM | NA | Intergenic | NA | NA |
| T25741 | GAAGCCCTCTTCCCTCTCGACGG | 1000 | | | | | | | | | |
| Oligo fwd | CACCGAAGCCCTCTTCCCTCTCGA | | | | | | | | | | |
| Oligo rev | AAACTCGAGAGGGAAGAGGGCTTC | | | | | | | | | | |
| Chromosome | start | end | strand | MM | target_seq | PAM | alignment | distance | position | gene name | gene id |
| GU305914.1 | 110416 | 110438 | - | 0 | GAAGCCCTCTTCCCTCTCGA | CGG | \|\|\|\|\|\|\|\|[\|\|\|\|\|\|\|\|\|\|\|\|]PAM | NA | Intergenic | NA | NA |
| T25742 | AAGCCCTCTTCCCTCTCGACGGG | 1000 | | | | | | | | | |
| Oligo fwd | CACCAAGCCCTCTTCCCTCTCGAC | | | | | | | | | | |
| Oligo rev | AAACGTCGAGAGGGAAGAGGGCTT | | | | | | | | | | |
| Chromosome | start | end | strand | MM | target_seq | PAM | alignment | distance | position | gene name | gene id |
| GU305914.1 | 110415 | 110437 | - | 0 | AAGCCCTCTTCCCTCTCGAC | GGG | \|\|\|\|\|\|\|\|[\|\|\|\|\|\|\|\|\|\|\|\|]PAM | NA | Intergenic | NA | NA |
| T25743 | CCCTCTTCCCTCTCGACGGGCGG | 1000 | | | | | | | | | |
| Oligo fwd | CACCCCCTCTTCCCTCTCGACGGG | | | | | | | | | | |
| Oligo rev | AAACCCCGTCGAGAGGGAAGAGGG | | | | | | | | | | |
| Chromosome | start | end | strand | MM | target_seq | PAM | alignment | distance | position | gene name | gene id |
| GU305914.1 | 110412 | 110434 | - | 0 | CCCTCTTCCCTCTCGACGGG | CGG | \|\|\|\|\|\|\|\|[\|\|\|\|\|\|\|\|\|\|\|\|]PAM | NA | Intergenic | NA | NA |
| T25744 | TCTTCCCTCTCGACGGGCGGCGG | 1000 | | | | | | | | | |
| Oligo fwd | CACCTCTTCCCTCTCGACGGGCGG | | | | | | | | | | |
| Oligo rev | AAACCCGCCCGTCGAGAGGGAAGA | | | | | | | | | | |
| Chromosome | start | end | strand | MM | target_seq | PAM | alignment | distance | position | gene name | gene id |
| GU305914.1 | 110409 | 110431 | - | 0 | TCTTCCCTCTCGACGGGCGG | CGG | \|\|\|\|\|\|\|\|[\|\|\|\|\|\|\|\|\|\|\|\|]PAM | NA | Intergenic | NA | NA |
| T25745 | CGACGGGCGGCGGCGCGTCGCGG | 1000 | | | | | | | | | |
| Oligo fwd | CACCCGACGGGCGGCGGCGCGTCG | | | | | | | | | | |
| Oligo rev | AAACCGACGCGCCGCCGCCCGTCG | | | | | | | | | | |
| Chromosome | start | end | strand | MM | target_seq | PAM | alignment | distance | position | gene name | gene id |
| GU305914.1 | 110399 | 110421 | - | 0 | CGACGGGCGGCGGCGCGTCG | CGG | \|\|\|\|\|\|\|\|[\|\|\|\|\|\|\|\|\|\|\|\|]PAM | NA | Intergenic | NA | NA |
| T25746 | GCGCGTCGCGGCCGCCGCAGAGG | 1000 | | | | | | | | | |
| Oligo fwd | CACCGCGCGTCGCGGCCGCCGCAG | | | | | | | | | | |
| Oligo rev | AAACCTGCGGCGGCCGCGACGCGC | | | | | | | | | | |
| Chromosome | start | end | strand | MM | target_seq | PAM | alignment | distance | position | gene name | gene id |
| GU305914.1 | 110387 | 110409 | - | 0 | GCGCGTCGCGGCCGCCGCAG | AGG | \|\|\|\|\|\|\|\|[\|\|\|\|\|\|\|\|\|\|\|\|]PAM | NA | Intergenic | NA | NA |
| GU305914.1 | 26975 | 26997 | - | 3 | GCCCGCCGCGGCCGCCGGAG | CGG | \|\|-\|\|-\|\|[\|\|\|\|\|\|\|\|\|-\|\|]PAM | NA | Intergenic | NA | NA |
| GU305914.1 | 199810 | 199832 | - | 4 | GCACGTGGCGGCCGCCGCGT | GGG | \|\|-\|\|\|-\|[\|\|\|\|\|\|\|\|\|\|--]PAM | NA | Intergenic | NA | NA |

T25747 GTCGCGGCCGCCGCAGAGGTCGG 1000
Oligo fwd CACCGTCGCGGCCGCCGCAGAGGT
Oligo rev AAACACCTCTGCGGCGGCCGCGAC
Chromosome start end strand MM target_seq PAM alignment distance position gene name gene id
GU305914.1 110383 110405 - 0 GTCGCGGCCGCCGCAGAGGT CGG ||||||||[||||||||||||]PAM NA Intergenic NA NA

T25748 TCGCGGCCGCCGCAGAGGTCGGG 1000
Oligo fwd CACCTCGCGGCCGCCGCAGAGGTC
Oligo rev AAACGACCTCTGCGGCGGCCGCGA
Chromosome start end strand MM target_seq PAM alignment distance position gene name gene id
GU305914.1 110382 110404 - 0 TCGCGGCCGCCGCAGAGGTC GGG ||||||||[||||||||||||]PAM NA Intergenic NA NA

T25749 CGCGGCCGCCGCAGAGGTCGGGG 1000
Oligo fwd CACCCGCGGCCGCCGCAGAGGTCG
Oligo rev AAACCGACCTCTGCGGCGGCCGCG
Chromosome start end strand MM target_seq PAM alignment distance position gene name gene id
GU305914.1 110381 110403 - 0 CGCGGCCGCCGCAGAGGTCG GGG ||||||||[||||||||||||]PAM NA Intergenic NA NA

T25750 GTCCCTGAGCGAGAACTTGCCGG 1000
Oligo fwd CACCGTCCCTGAGCGAGAACTTGC
Oligo rev AAACGCAAGTTCTCGCTCAGGGAC
Chromosome start end strand MM target_seq PAM alignment distance position gene name gene id
GU305914.1 110353 110375 - 0 GTCCCTGAGCGAGAACTTGC CGG ||||||||[||||||||||||]PAM NA Intergenic NA NA

T25751 TGAGCGAGAACTTGCCGGCGCGG 1000
Oligo fwd CACCTGAGCGAGAACTTGCCGGCG
Oligo rev AAACCGCCGGCAAGTTCTCGCTCA
Chromosome start end strand MM target_seq PAM alignment distance position gene name gene id
GU305914.1 110348 110370 - 0 TGAGCGAGAACTTGCCGGCG CGG ||||||||[||||||||||||]PAM NA Intergenic NA NA

T25752 CGAGAACTTGCCGGCGCGGCCGG 1000
Oligo fwd CACCCGAGAACTTGCCGGCGCGGC
Oligo rev AAACGCCGCGCCGGCAAGTTCTCG
Chromosome start end strand MM target_seq PAM alignment distance position gene name gene id
GU305914.1 110344 110366 - 0 CGAGAACTTGCCGGCGCGGC CGG ||||||||[||||||||||||]PAM NA Intergenic NA NA

T25753 GCGGCCGGAGAAGACGCTCTCGG 1000
Oligo fwd CACCGCGGCCGGAGAAGACGCTCT
Oligo rev AAACAGAGCGTCTTCTCCGGCCGC
Chromosome start end strand MM target_seq PAM alignment distance position gene name gene id
GU305914.1 110329 110351 - 0 GCGGCCGGAGAAGACGCTCT CGG ||||||||[||||||||||||]PAM NA Intergenic NA NA

T25754 AGAAGACGCTCTCGGAGTTGAGG 1000
Oligo fwd CACCAGAAGACGCTCTCGGAGTTG
Oligo rev AAACCAACTCCGAGAGCGTCTTCT
Chromosome start end strand MM target_seq PAM alignment distance position gene name gene id
GU305914.1 110321 110343 - 0 AGAAGACGCTCTCGGAGTTG AGG ||||||||[||||||||||||]PAM NA Intergenic NA NA

T25755 CGGAGTTGAGGAGTTTCCAGAGG 1000
Oligo fwd CACCCGGAGTTGAGGAGTTTCCAG
Oligo rev AAACCTGGAAACTCCTCAACTCCG
Chromosome start end strand MM target_seq PAM alignment distance position gene name gene id
GU305914.1 110309 110331 - 0 CGGAGTTGAGGAGTTTCCAG AGG ||||||||[||||||||||||]PAM NA Intergenic NA NA

T25756 GGAGTTTCCAGAGGAACAGCAGG 1000
Oligo fwd CACCGGAGTTTCCAGAGGAACAGC
Oligo rev AAACGCTGTTCCTCTGGAAACTCC
Chromosome start end strand MM target_seq PAM alignment distance position gene name gene id
GU305914.1 110300 110322 - 0 GGAGTTTCCAGAGGAACAGC AGG ||||||||[||||||||||||]PAM NA Intergenic NA NA

T25757 TCCAGAGGAACAGCAGGCAGAGG 1000
Oligo fwd CACCTCCAGAGGAACAGCAGGCAG
Oligo rev AAACCTGCCTGCTGTTCCTCTGGA
Chromosome start end strand MM target_seq PAM alignment distance position gene name gene id
GU305914.1 110294 110316 - 0 TCCAGAGGAACAGCAGGCAG AGG ||||||||[||||||||||||]PAM NA Intergenic NA NA

T25758 GAGGAACAGCAGGCAGAGGACGG 1000
Oligo fwd CACCGAGGAACAGCAGGCAGAGGA
Oligo rev AAACTCCTCTGCCTGCTGTTCCTC
Chromosome start end strand MM target_seq PAM alignment distance position gene name gene id
GU305914.1 110290 110312 - 0 GAGGAACAGCAGGCAGAGGA CGG ||||||||[||||||||||||]PAM NA Intergenic NA NA
GU305914.1 139019 139041 - 4 GCAGAACAGCAGCCAGAGCA CGG |--|||||[||||-|||||-|]PAM NA Intergenic NA NA

T25759 CGGCCGTGAGCCCCGACCTGAGG 1000
Oligo fwd CACCCGGCCGTGAGCCCCGACCTG
Oligo rev AAACCAGGTCGGGGCTCACGGCCG
Chromosome start end strand MM target_seq PAM alignment distance position gene name gene id
GU305914.1 110270 110292 - 0 CGGCCGTGAGCCCCGACCTG AGG ||||||||[||||||||||||]PAM NA Intergenic NA NA

T25760 ACCTGAGGTAGTGCTGCTCCTGG 1000
Oligo fwd CACCACCTGAGGTAGTGCTGCTCC
Oligo rev AAACGGAGCAGCACTACCTCAGGT
Chromosome start end strand MM target_seq PAM alignment distance position gene name gene id
GU305914.1 110255 110277 - 0 ACCTGAGGTAGTGCTGCTCC TGG ||||||||[||||||||||||]PAM NA Intergenic NA NA

T25761 TGAGGTAGTGCTGCTCCTGGAGG 1000
Oligo fwd CACCTGAGGTAGTGCTGCTCCTGG
Oligo rev AAACCCAGGAGCAGCACTACCTCA

<table>
<tr><td>Chromosome</td><td>start</td><td>end</td><td>strand</td><td>MM</td><td>target_seq</td><td>PAM</td><td>alignment</td><td>distance</td><td>position</td><td>gene name</td><td>gene id</td></tr>
<tr><td>GU305914.1</td><td>110252</td><td>110274</td><td>-</td><td>0</td><td>TGAGGTAGTGCTGCTCCTGG</td><td>AGG</td><td>||||||||[||||||||||||]PAM</td><td>NA</td><td>Intergenic</td><td>NA</td><td>NA</td></tr>
<tr><td>T25762</td><td>GGTAGTGCTGCTCCTGGAGGAGG</td><td>1000</td><td></td><td></td><td></td><td></td><td></td><td></td><td></td><td></td><td></td></tr>
<tr><td>Oligo fwd</td><td>CACCGGTAGTGCTGCTCCTGGAGG</td><td></td><td></td><td></td><td></td><td></td><td></td><td></td><td></td><td></td><td></td></tr>
<tr><td>Oligo rev</td><td>AAACCCTCCAGGAGCAGCACTACC</td><td></td><td></td><td></td><td></td><td></td><td></td><td></td><td></td><td></td><td></td></tr>
<tr><td>Chromosome</td><td>start</td><td>end</td><td>strand</td><td>MM</td><td>target_seq</td><td>PAM</td><td>alignment</td><td>distance</td><td>position</td><td>gene name</td><td>gene id</td></tr>
<tr><td>GU305914.1</td><td>110249</td><td>110271</td><td>-</td><td>0</td><td>GGTAGTGCTGCTCCTGGAGG</td><td>AGG</td><td>||||||||[||||||||||||]PAM</td><td>NA</td><td>Intergenic</td><td>NA</td><td>NA</td></tr>
<tr><td>GU305914.1</td><td>168320</td><td>168342</td><td>+</td><td>4</td><td>GCTGGTGCTGCTCCTGGTTG</td><td>CGG</td><td>|-|-||||[|||||||||--|]PAM</td><td>NA</td><td>Intergenic</td><td>NA</td><td>NA</td></tr>
<tr><td>T25763</td><td>GTGCTGCTCCTGGAGGAGGTTGG</td><td>1000</td><td></td><td></td><td></td><td></td><td></td><td></td><td></td><td></td><td></td></tr>
<tr><td>Oligo fwd</td><td>CACCGTGCTGCTCCTGGAGGAGGT</td><td></td><td></td><td></td><td></td><td></td><td></td><td></td><td></td><td></td><td></td></tr>
<tr><td>Oligo rev</td><td>AAACACCTCCTCCAGGAGCAGCAC</td><td></td><td></td><td></td><td></td><td></td><td></td><td></td><td></td><td></td><td></td></tr>
<tr><td>Chromosome</td><td>start</td><td>end</td><td>strand</td><td>MM</td><td>target_seq</td><td>PAM</td><td>alignment</td><td>distance</td><td>position</td><td>gene name</td><td>gene id</td></tr>
<tr><td>GU305914.1</td><td>110245</td><td>110267</td><td>-</td><td>0</td><td>GTGCTGCTCCTGGAGGAGGT</td><td>TGG</td><td>||||||||[||||||||||||]PAM</td><td>NA</td><td>Intergenic</td><td>NA</td><td>NA</td></tr>
<tr><td>T25764</td><td>TGCTGCTCCTGGAGGAGGTTGGG</td><td>1000</td><td></td><td></td><td></td><td></td><td></td><td></td><td></td><td></td><td></td></tr>
<tr><td>Oligo fwd</td><td>CACCTGCTGCTCCTGGAGGAGGTT</td><td></td><td></td><td></td><td></td><td></td><td></td><td></td><td></td><td></td><td></td></tr>
<tr><td>Oligo rev</td><td>AAACAACCTCCTCCAGGAGCAGCA</td><td></td><td></td><td></td><td></td><td></td><td></td><td></td><td></td><td></td><td></td></tr>
<tr><td>Chromosome</td><td>start</td><td>end</td><td>strand</td><td>MM</td><td>target_seq</td><td>PAM</td><td>alignment</td><td>distance</td><td>position</td><td>gene name</td><td>gene id</td></tr>
<tr><td>GU305914.1</td><td>110244</td><td>110266</td><td>-</td><td>0</td><td>TGCTGCTCCTGGAGGAGGTT</td><td>GGG</td><td>||||||||[||||||||||||]PAM</td><td>NA</td><td>Intergenic</td><td>NA</td><td>NA</td></tr>
<tr><td>T25765</td><td>GCTGCTCCTGGAGGAGGTTGGGG</td><td>1000</td><td></td><td></td><td></td><td></td><td></td><td></td><td></td><td></td><td></td></tr>
<tr><td>Oligo fwd</td><td>CACCGCTGCTCCTGGAGGAGGTTG</td><td></td><td></td><td></td><td></td><td></td><td></td><td></td><td></td><td></td><td></td></tr>
<tr><td>Oligo rev</td><td>AAACCAACCTCCTCCAGGAGCAGC</td><td></td><td></td><td></td><td></td><td></td><td></td><td></td><td></td><td></td><td></td></tr>
<tr><td>Chromosome</td><td>start</td><td>end</td><td>strand</td><td>MM</td><td>target_seq</td><td>PAM</td><td>alignment</td><td>distance</td><td>position</td><td>gene name</td><td>gene id</td></tr>
<tr><td>GU305914.1</td><td>110243</td><td>110265</td><td>-</td><td>0</td><td>GCTGCTCCTGGAGGAGGTTG</td><td>GGG</td><td>||||||||[||||||||||||]PAM</td><td>NA</td><td>Intergenic</td><td>NA</td><td>NA</td></tr>
<tr><td>T25766</td><td>GGTTGGGGACGACGCCGCTGAGG</td><td>1000</td><td></td><td></td><td></td><td></td><td></td><td></td><td></td><td></td><td></td></tr>
<tr><td>Oligo fwd</td><td>CACCGGTTGGGGACGACGCCGCTG</td><td></td><td></td><td></td><td></td><td></td><td></td><td></td><td></td><td></td><td></td></tr>
<tr><td>Oligo rev</td><td>AAACCAGCGGCGTCGTCCCCAACC</td><td></td><td></td><td></td><td></td><td></td><td></td><td></td><td></td><td></td><td></td></tr>
<tr><td>Chromosome</td><td>start</td><td>end</td><td>strand</td><td>MM</td><td>target_seq</td><td>PAM</td><td>alignment</td><td>distance</td><td>position</td><td>gene name</td><td>gene id</td></tr>
<tr><td>GU305914.1</td><td>110228</td><td>110250</td><td>-</td><td>0</td><td>GGTTGGGGACGACGCCGCTG</td><td>AGG</td><td>||||||||[||||||||||||]PAM</td><td>NA</td><td>Intergenic</td><td>NA</td><td>NA</td></tr>
<tr><td>T25767</td><td>CGCTGAGGCGCTTGCCGAAGAGG</td><td>1000</td><td></td><td></td><td></td><td></td><td></td><td></td><td></td><td></td><td></td></tr>
<tr><td>Oligo fwd</td><td>CACCCGCTGAGGCGCTTGCCGAAG</td><td></td><td></td><td></td><td></td><td></td><td></td><td></td><td></td><td></td><td></td></tr>
<tr><td>Oligo rev</td><td>AAACCTTCGGCAAGCGCCTCAGCG</td><td></td><td></td><td></td><td></td><td></td><td></td><td></td><td></td><td></td><td></td></tr>
<tr><td>Chromosome</td><td>start</td><td>end</td><td>strand</td><td>MM</td><td>target_seq</td><td>PAM</td><td>alignment</td><td>distance</td><td>position</td><td>gene name</td><td>gene id</td></tr>
<tr><td>GU305914.1</td><td>110213</td><td>110235</td><td>-</td><td>0</td><td>CGCTGAGGCGCTTGCCGAAG</td><td>AGG</td><td>||||||||[||||||||||||]PAM</td><td>NA</td><td>Intergenic</td><td>NA</td><td>NA</td></tr>
<tr><td>T25768</td><td>GGCGCAGCGTCCAGAGCGCCAGG</td><td>1000</td><td></td><td></td><td></td><td></td><td></td><td></td><td></td><td></td><td></td></tr>
<tr><td>Oligo fwd</td><td>CACCGGCGCAGCGTCCAGAGCGCC</td><td></td><td></td><td></td><td></td><td></td><td></td><td></td><td></td><td></td><td></td></tr>
<tr><td>Oligo rev</td><td>AAACGGCGCTCTGGACGCTGCGCC</td><td></td><td></td><td></td><td></td><td></td><td></td><td></td><td></td><td></td><td></td></tr>
<tr><td>Chromosome</td><td>start</td><td>end</td><td>strand</td><td>MM</td><td>target_seq</td><td>PAM</td><td>alignment</td><td>distance</td><td>position</td><td>gene name</td><td>gene id</td></tr>
<tr><td>GU305914.1</td><td>110192</td><td>110214</td><td>-</td><td>0</td><td>GGCGCAGCGTCCAGAGCGCC</td><td>AGG</td><td>||||||||[||||||||||||]PAM</td><td>NA</td><td>Intergenic</td><td>NA</td><td>NA</td></tr>
<tr><td>T25769</td><td>CCAGAGCGCCAGGACGTCGTCGG</td><td>1000</td><td></td><td></td><td></td><td></td><td></td><td></td><td></td><td></td><td></td></tr>
<tr><td>Oligo fwd</td><td>CACCCCAGAGCGCCAGGACGTCGT</td><td></td><td></td><td></td><td></td><td></td><td></td><td></td><td></td><td></td><td></td></tr>
<tr><td>Oligo rev</td><td>AAACACGACGTCCTGGCGCTCTGG</td><td></td><td></td><td></td><td></td><td></td><td></td><td></td><td></td><td></td><td></td></tr>
<tr><td>Chromosome</td><td>start</td><td>end</td><td>strand</td><td>MM</td><td>target_seq</td><td>PAM</td><td>alignment</td><td>distance</td><td>position</td><td>gene name</td><td>gene id</td></tr>
<tr><td>GU305914.1</td><td>110182</td><td>110204</td><td>-</td><td>0</td><td>CCAGAGCGCCAGGACGTCGT</td><td>CGG</td><td>||||||||[||||||||||||]PAM</td><td>NA</td><td>Intergenic</td><td>NA</td><td>NA</td></tr>
<tr><td>T25770</td><td>GACGTCGTCGGAGACCGCGCCGG</td><td>1000</td><td></td><td></td><td></td><td></td><td></td><td></td><td></td><td></td><td></td></tr>
<tr><td>Oligo fwd</td><td>CACCGACGTCGTCGGAGACCGCGC</td><td></td><td></td><td></td><td></td><td></td><td></td><td></td><td></td><td></td><td></td></tr>
<tr><td>Oligo rev</td><td>AAACGCGCGGTCTCCGACGACGTC</td><td></td><td></td><td></td><td></td><td></td><td></td><td></td><td></td><td></td><td></td></tr>
<tr><td>Chromosome</td><td>start</td><td>end</td><td>strand</td><td>MM</td><td>target_seq</td><td>PAM</td><td>alignment</td><td>distance</td><td>position</td><td>gene name</td><td>gene id</td></tr>
<tr><td>GU305914.1</td><td>110170</td><td>110192</td><td>-</td><td>0</td><td>GACGTCGTCGGAGACCGCGC</td><td>CGG</td><td>||||||||[||||||||||||]PAM</td><td>NA</td><td>Intergenic</td><td>NA</td><td>NA</td></tr>
<tr><td>T25771</td><td>CGTCGGAGACCGCGCCGGAGAGG</td><td>1000</td><td></td><td></td><td></td><td></td><td></td><td></td><td></td><td></td><td></td></tr>
<tr><td>Oligo fwd</td><td>CACCCGTCGGAGACCGCGCCGGAG</td><td></td><td></td><td></td><td></td><td></td><td></td><td></td><td></td><td></td><td></td></tr>
<tr><td>Oligo rev</td><td>AAACCTCCGGCGCGGTCTCCGACG</td><td></td><td></td><td></td><td></td><td></td><td></td><td></td><td></td><td></td><td></td></tr>
<tr><td>Chromosome</td><td>start</td><td>end</td><td>strand</td><td>MM</td><td>target_seq</td><td>PAM</td><td>alignment</td><td>distance</td><td>position</td><td>gene name</td><td>gene id</td></tr>
<tr><td>GU305914.1</td><td>110165</td><td>110187</td><td>-</td><td>0</td><td>CGTCGGAGACCGCGCCGGAG</td><td>AGG</td><td>||||||||[||||||||||||]PAM</td><td>NA</td><td>Intergenic</td><td>NA</td><td>NA</td></tr>
<tr><td>T25772</td><td>AGAGGCGACTCATGACCTCCCGG</td><td>1000</td><td></td><td></td><td></td><td></td><td></td><td></td><td></td><td></td><td></td></tr>
<tr><td>Oligo fwd</td><td>CACCAGAGGCGACTCATGACCTCC</td><td></td><td></td><td></td><td></td><td></td><td></td><td></td><td></td><td></td><td></td></tr>
<tr><td>Oligo rev</td><td>AAACGGAGGTCATGAGTCGCCTCT</td><td></td><td></td><td></td><td></td><td></td><td></td><td></td><td></td><td></td><td></td></tr>
<tr><td>Chromosome</td><td>start</td><td>end</td><td>strand</td><td>MM</td><td>target_seq</td><td>PAM</td><td>alignment</td><td>distance</td><td>position</td><td>gene name</td><td>gene id</td></tr>
<tr><td>GU305914.1</td><td>110147</td><td>110169</td><td>-</td><td>0</td><td>AGAGGCGACTCATGACCTCC</td><td>CGG</td><td>||||||||[||||||||||||]PAM</td><td>NA</td><td>Intergenic</td><td>NA</td><td>NA</td></tr>
<tr><td>T25773</td><td>GAGGCGACTCATGACCTCCCGGG</td><td>1000</td><td></td><td></td><td></td><td></td><td></td><td></td><td></td><td></td><td></td></tr>
<tr><td>Oligo fwd</td><td>CACCGAGGCGACTCATGACCTCCC</td><td></td><td></td><td></td><td></td><td></td><td></td><td></td><td></td><td></td><td></td></tr>
<tr><td>Oligo rev</td><td>AAACGGGAGGTCATGAGTCGCCTC</td><td></td><td></td><td></td><td></td><td></td><td></td><td></td><td></td><td></td><td></td></tr>
<tr><td>Chromosome</td><td>start</td><td>end</td><td>strand</td><td>MM</td><td>target_seq</td><td>PAM</td><td>alignment</td><td>distance</td><td>position</td><td>gene name</td><td>gene id</td></tr>
<tr><td>GU305914.1</td><td>110146</td><td>110168</td><td>-</td><td>0</td><td>GAGGCGACTCATGACCTCCC</td><td>GGG</td><td>||||||||[||||||||||||]PAM</td><td>NA</td><td>Intergenic</td><td>NA</td><td>NA</td></tr>
<tr><td>T25774</td><td>CATGACCTCCCGGGACGCGACGG</td><td>1000</td><td></td><td></td><td></td><td></td><td></td><td></td><td></td><td></td><td></td></tr>
<tr><td>Oligo fwd</td><td>CACCCATGACCTCCCGGGACGCGA</td><td></td><td></td><td></td><td></td><td></td><td></td><td></td><td></td><td></td><td></td></tr>
<tr><td>Oligo rev</td><td>AAACTCGCGTCCCGGGAGGTCATG</td><td></td><td></td><td></td><td></td><td></td><td></td><td></td><td></td><td></td><td></td></tr>
<tr><td>Chromosome</td><td>start</td><td>end</td><td>strand</td><td>MM</td><td>target_seq</td><td>PAM</td><td>alignment</td><td>distance</td><td>position</td><td>gene name</td><td>gene id</td></tr>
<tr><td>GU305914.1</td><td>110137</td><td>110159</td><td>-</td><td>0</td><td>CATGACCTCCCGGGACGCGA</td><td>CGG</td><td>||||||||[||||||||||||]PAM</td><td>NA</td><td>Intergenic</td><td>NA</td><td>NA</td></tr>
<tr><td>T25775</td><td>GACCTCCCGGGACGCGACGGCGG</td><td>1000</td><td></td><td></td><td></td><td></td><td></td><td></td><td></td><td></td><td></td></tr>
<tr><td>Oligo fwd</td><td>CACCGACCTCCCGGGACGCGACGG</td><td></td><td></td><td></td><td></td><td></td><td></td><td></td><td></td><td></td><td></td></tr>
<tr><td>Oligo rev</td><td>AAACCCGTCGCGTCCCGGGAGGTC</td><td></td><td></td><td></td><td></td><td></td><td></td><td></td><td></td><td></td><td></td></tr>
<tr><td>Chromosome</td><td>start</td><td>end</td><td>strand</td><td>MM</td><td>target_seq</td><td>PAM</td><td>alignment</td><td>distance</td><td>position</td><td>gene name</td><td>gene id</td></tr>
<tr><td>GU305914.1</td><td>110134</td><td>110156</td><td>-</td><td>0</td><td>GACCTCCCGGGACGCGACGG</td><td>CGG</td><td>||||||||[||||||||||||]PAM</td><td>NA</td><td>Intergenic</td><td>NA</td><td>NA</td></tr>
<tr><td>T25776</td><td>CCGGGACGCGACGGCGGCCGCGG</td><td>1000</td><td></td><td></td><td></td><td></td><td></td><td></td><td></td><td></td><td></td></tr>
</table>

Oligo fwd CACCCCGGGACGCGACGGCGGCCG
Oligo rev AAACCGGCCGCCGTCGCGTCCCGG
Chromosome start end strand MM target_seq PAM alignment distance position gene name gene id
GU305914.1 110128 110150 - 0 CCGGGACGCGACGGCGGCCG CGG ||||||||[||||||||||||]PAM NA Intergenic NA NA

T25777 GGACGCGACGGCGGCCGCGGCGG 1000
Oligo fwd CACCGGACGCGACGGCGGCCGCGG
Oligo rev AAACCCGCGGCCGCCGTCGCGTCC
Chromosome start end strand MM target_seq PAM alignment distance position gene name gene id
GU305914.1 110125 110147 - 0 GGACGCGACGGCGGCCGCGG CGG ||||||||[||||||||||||]PAM NA Intergenic NA NA
GU305914.1 26973 26995 + 4 GACCGCTCCGGCGGCCGCGG CGG |--|||--[||||||||||||]PAM NA Intergenic NA NA
GU305914.1 149485 149507 - 3 GCAGGCGACGGCGGCGGCGG CGG |-|-||||[|||||||-||||]PAM NA Intergenic NA NA
GU305914.1 67742 67764 + 4 GGGAGCGGCGGCGGCGGCGG CGG ||--|||-[|||||||-||||]PAM NA Intergenic NA NA
GU305914.1 154407 154429 + 4 GGTGGCGGCGGCGGCGGCGG CGG ||--|||-[|||||||-||||]PAM NA Intergenic NA NA
GU305914.1 58123 58145 - 4 GGTGGCGGCGGCGGCGGCGG AGG ||--|||-[|||||||-||||]PAM NA Intergenic NA NA
GU305914.1 24884 24906 + 4 GGCGGCGACGGCGGCCTCGA CGG ||--||||[||||||||-||-]PAM NA Intergenic NA NA

T25778 GGCGGCCGCGGCGGACGAGCCGG 1000
Oligo fwd CACCGGCGGCCGCGGCGGACGAGC
Oligo rev AAACGCTCGTCCGCCGCGGCCGCC
Chromosome start end strand MM target_seq PAM alignment distance position gene name gene id
GU305914.1 110116 110138 - 0 GGCGGCCGCGGCGGACGAGC CGG ||||||||[||||||||||||]PAM NA Intergenic NA NA
GU305914.1 55854 55876 + 4 GGCGATCGCGGCTGACGACC AGG ||||--||[||||-|||||-|]PAM NA Intergenic NA NA

T25779 GCGGCCGCGGCGGACGAGCCGGG 1000
Oligo fwd CACCGCGGCCGCGGCGGACGAGCC
Oligo rev AAACGGCTCGTCCGCCGCGGCCGC
Chromosome start end strand MM target_seq PAM alignment distance position gene name gene id
GU305914.1 110115 110137 - 0 GCGGCCGCGGCGGACGAGCC GGG ||||||||[||||||||||||]PAM NA Intergenic NA NA

T25780 CGAGCCGGGACCCCCCGACGCGG 1000
Oligo fwd CACCCGAGCCGGGACCCCCCGACG
Oligo rev AAACCGTCGGGGGGTCCCGGCTCG
Chromosome start end strand MM target_seq PAM alignment distance position gene name gene id
GU305914.1 110101 110123 - 0 CGAGCCGGGACCCCCCGACG CGG ||||||||[||||||||||||]PAM NA Intergenic NA NA

T25781 GGGACCCCCCGACGCGGCCGCGG 1000
Oligo fwd CACCGGGACCCCCCGACGCGGCCG
Oligo rev AAACCGGCCGCGTCGGGGGGTCCC
Chromosome start end strand MM target_seq PAM alignment distance position gene name gene id
GU305914.1 110095 110117 - 0 GGGACCCCCCGACGCGGCCG CGG ||||||||[||||||||||||]PAM NA Intergenic NA NA

T25782 CGACGCGGCCGCGGCGACGCCGG 1000
Oligo fwd CACCCGACGCGGCCGCGGCGACGC
Oligo rev AAACGCGTCGCCGCGGCCGCGTCG
Chromosome start end strand MM target_seq PAM alignment distance position gene name gene id
GU305914.1 110086 110108 - 0 CGACGCGGCCGCGGCGACGC CGG ||||||||[||||||||||||]PAM NA Intergenic NA NA
GU305914.1 198749 198771 - 4 GAACGCGGACGCGGCGACGT CGG --||||||[-||||||||||-]PAM NA Intergenic NA NA

T25783 CGCGGCCGCGGCGACGCCGGTGG 1000
Oligo fwd CACCCGCGGCCGCGGCGACGCCGG
Oligo rev AAACCCGGCGTCGCCGCGGCCGCG
Chromosome start end strand MM target_seq PAM alignment distance position gene name gene id
GU305914.1 110083 110105 - 0 CGCGGCCGCGGCGACGCCGG TGG ||||||||[||||||||||||]PAM NA Intergenic NA NA
GU305914.1 198746 198768 - 2 CGCGGACGCGGCGACGTCGG TGG |||||-||[||||||||-|||]PAM NA Intergenic NA NA
GU305914.1 152553 152575 + 4 TGCGGCGGCGGCGGTGCCGG CGG -|||||-|[|||||--|||||]PAM NA Intergenic NA NA
GU305914.1 128803 128825 + 4 CGCAGCGGCAGCGACGTCGG CGG |||-||-|[|-||||||-|||]PAM NA Intergenic NA NA
GU305914.1 56773 56795 + 4 CGCCGCAGCGGCGGCGGCGG CGG |||-||-|[|||||-||-|||]PAM NA Intergenic NA NA
GU305914.1 113912 113934 - 4 GGCGGCAGCGGCGACGGCGT CGG -|||||-|[||||||||-||-]PAM NA Intergenic NA NA

T25784 GCGACGCCGGTGGCACCGCCCGG 1000
Oligo fwd CACCGCGACGCCGGTGGCACCGCC
Oligo rev AAACGGCGGTGCCACCGGCGTCGC
Chromosome start end strand MM target_seq PAM alignment distance position gene name gene id
GU305914.1 110073 110095 - 0 GCGACGCCGGTGGCACCGCC CGG ||||||||[||||||||||||]PAM NA Intergenic NA NA

T25785 ACCGCCCGGCACCTGCGCGATGG 1000
Oligo fwd CACCACCGCCCGGCACCTGCGCGA
Oligo rev AAACTCGCGCAGGTGCCGGGCGGT
Chromosome start end strand MM target_seq PAM alignment distance position gene name gene id
GU305914.1 110059 110081 - 0 ACCGCCCGGCACCTGCGCGA TGG ||||||||[||||||||||||]PAM NA Intergenic NA NA

T25786 CGATGGCCCCCCGCCGCAGCAGG 1000
Oligo fwd CACCCGATGGCCCCCCGCCGCAGC
Oligo rev AAACGCTGCGGCGGGGGGCCATCG
Chromosome start end strand MM target_seq PAM alignment distance position gene name gene id
GU305914.1 110042 110064 - 0 CGATGGCCCCCCGCCGCAGC AGG ||||||||[||||||||||||]PAM NA Intergenic NA NA

T25787 GGCCCCCCGCCGCAGCAGGACGG 1000
Oligo fwd CACCGGCCCCCCGCCGCAGCAGGA
Oligo rev AAACTCCTGCTGCGGCGGGGGGCC
Chromosome start end strand MM target_seq PAM alignment distance position gene name gene id
GU305914.1 110038 110060 - 0 GGCCCCCCGCCGCAGCAGGA CGG ||||||||[||||||||||||]PAM NA Intergenic NA NA

T25788 GCCGCAGCAGGACGGCGACGAGG 1000
Oligo fwd CACCGCCGCAGCAGGACGGCGACG
Oligo rev AAACCGTCGCCGTCCTGCTGCGGC
Chromosome start end strand MM target_seq PAM alignment distance position gene name gene id
GU305914.1 110030 110052 - 0 GCCGCAGCAGGACGGCGACG AGG ||||||||[||||||||||||]PAM NA Intergenic NA NA

T25789 CCGCAGCAGGACGGCGACGAGGG 1000
Oligo fwd CACCCCGCAGCAGGACGGCGACGA
Oligo rev AAACTCGTCGCCGTCCTGCTGCGG
Chromosome start end strand MM target_seq PAM alignment distance position gene name gene id
GU305914.1 110029 110051 - 0 CCGCAGCAGGACGGCGACGA GGG ||||||||[||||||||||||]PAM NA Intergenic NA NA

T25790 GTTCTCGTCGAACGTGCCCGCGG 1000
Oligo fwd CACCGTTCTCGTCGAACGTGCCCG
Oligo rev AAACCGGGCACGTTCGACGAGAAC
Chromosome start end strand MM target_seq PAM alignment distance position gene name gene id
GU305914.1 110005 110027 - 0 GTTCTCGTCGAACGTGCCCG CGG ||||||||[||||||||||||]PAM NA Intergenic NA NA

T25791 CGGCGTAGCGCCTGCCCTGCTGG 1000
Oligo fwd CACCCGGCGTAGCGCCTGCCCTGC
Oligo rev AAACGCAGGGCAGGCGCTACGCCG
Chromosome start end strand MM target_seq PAM alignment distance position gene name gene id
GU305914.1 109985 110007 - 0 CGGCGTAGCGCCTGCCCTGC TGG ||||||||[||||||||||||]PAM NA Intergenic NA NA

T25792 TAGCGCCTGCCCTGCTGGACCGG 1000
Oligo fwd CACCTAGCGCCTGCCCTGCTGGAC
Oligo rev AAACGTCCAGCAGGGCAGGCGCTA
Chromosome start end strand MM target_seq PAM alignment distance position gene name gene id
GU305914.1 109980 110002 - 0 TAGCGCCTGCCCTGCTGGAC CGG ||||||||[||||||||||||]PAM NA Intergenic NA NA

T25793 GGCGCGTAGAAACGCAAGCCCGG 1000
Oligo fwd CACCGGCGCGTAGAAACGCAAGCC
Oligo rev AAACGGCTTGCGTTTCTACGCGCC
Chromosome start end strand MM target_seq PAM alignment distance position gene name gene id
GU305914.1 109959 109981 - 0 GGCGCGTAGAAACGCAAGCC CGG ||||||||[||||||||||||]PAM NA Intergenic NA NA

T25794 GCGTAGAAACGCAAGCCCGGCGG 1000
Oligo fwd CACCGCGTAGAAACGCAAGCCCGG
Oligo rev AAACCCGGGCTTGCGTTTCTACGC
Chromosome start end strand MM target_seq PAM alignment distance position gene name gene id
GU305914.1 109956 109978 - 0 GCGTAGAAACGCAAGCCCGG CGG ||||||||[||||||||||||]PAM NA Intergenic NA NA

T25795 CGTAGAAACGCAAGCCCGGCGGG 1000
Oligo fwd CACCCGTAGAAACGCAAGCCCGGC
Oligo rev AAACGCCGGGCTTGCGTTTCTACG
Chromosome start end strand MM target_seq PAM alignment distance position gene name gene id
GU305914.1 109955 109977 - 0 CGTAGAAACGCAAGCCCGGC GGG ||||||||[||||||||||||]PAM NA Intergenic NA NA

T25796 CAAGCCCGGCGGGTCCTTGTAGG 1000
Oligo fwd CACCCAAGCCCGGCGGGTCCTTGT
Oligo rev AAACACAAGGACCCGCCGGGCTTG
Chromosome start end strand MM target_seq PAM alignment distance position gene name gene id
GU305914.1 109945 109967 - 0 CAAGCCCGGCGGGTCCTTGT AGG ||||||||[||||||||||||]PAM NA Intergenic NA NA

T25797 GGCGAAAGCGTTGCCCCCCGAGG 1000
Oligo fwd CACCGGCGAAAGCGTTGCCCCCCG
Oligo rev AAACCGGGGGGCAACGCTTTCGCC
Chromosome start end strand MM target_seq PAM alignment distance position gene name gene id
GU305914.1 109924 109946 - 0 GGCGAAAGCGTTGCCCCCCG AGG ||||||||[||||||||||||]PAM NA Intergenic NA NA

T25798 GCGAAAGCGTTGCCCCCCGAGGG 1000
Oligo fwd CACCGCGAAAGCGTTGCCCCCCGA
Oligo rev AAACTCGGGGGGCAACGCTTTCGC
Chromosome start end strand MM target_seq PAM alignment distance position gene name gene id
GU305914.1 109923 109945 - 0 GCGAAAGCGTTGCCCCCCGA GGG ||||||||[||||||||||||]PAM NA Intergenic NA NA

T25799 CGAGGGTCGCCGCGCGATCTCGG 1000
Oligo fwd CACCCGAGGGTCGCCGCGCGATCT
Oligo rev AAACAGATCGCGCGGCGACCCTCG
Chromosome start end strand MM target_seq PAM alignment distance position gene name gene id
GU305914.1 109906 109928 - 0 CGAGGGTCGCCGCGCGATCT CGG ||||||||[||||||||||||]PAM NA Intergenic NA NA

T25800 GGGTCGCCGCGCGATCTCGGCGG 1000
Oligo fwd CACCGGGTCGCCGCGCGATCTCGG
Oligo rev AAACCCGAGATCGCGCGGCGACCC
Chromosome start end strand MM target_seq PAM alignment distance position gene name gene id
GU305914.1 109903 109925 - 0 GGGTCGCCGCGCGATCTCGG CGG ||||||||[||||||||||||]PAM NA Intergenic NA NA

T25801 GATCTCGGCGGCGATGTCGTCGG 1000
Oligo fwd CACCGATCTCGGCGGCGATGTCGT
Oligo rev AAACACGACATCGCCGCCGAGATC
Chromosome start end strand MM target_seq PAM alignment distance position gene name gene id
GU305914.1 109891 109913 - 0 GATCTCGGCGGCGATGTCGT CGG ||||||||[||||||||||||]PAM NA Intergenic NA NA

T25802 CGATGTCGTCGGAGAGCGCCCGG 1000
Oligo fwd CACCCGATGTCGTCGGAGAGCGCC
Oligo rev AAACGGCGCTCTCCGACGACATCG
Chromosome start end strand MM target_seq PAM alignment distance position gene name gene id
GU305914.1 109880 109902 - 0 CGATGTCGTCGGAGAGCGCC CGG ||||||||[||||||||||||]PAM NA Intergenic NA NA

T25803 TGTCGTCGGAGAGCGCCCGGAGG 1000
Oligo fwd CACCTGTCGTCGGAGAGCGCCCGG
Oligo rev AAACCCGGGCGCTCTCCGACGACA
Chromosome start end strand MM target_seq PAM alignment distance position gene name gene id

<table>
<tr><td>GU305914.1</td><td>109877</td><td>109899</td><td>-</td><td>0</td><td>TGTCGTCGGAGAGCGCCCGG</td><td>AGG</td><td>||||||||[||||||||||||]PAM</td><td>NA</td><td>Intergenic</td><td>NA</td><td>NA</td></tr>
<tr><td></td><td></td><td></td><td></td><td></td><td></td><td></td><td></td><td></td><td></td><td></td><td></td></tr>
<tr><td>T25804</td><td>GAGCGCCCGGAGGACCCTGCCGG</td><td>1000</td><td></td><td></td><td></td><td></td><td></td><td></td><td></td><td></td><td></td></tr>
<tr><td>Oligo fwd</td><td>CACCGAGCGCCCGGAGGACCCTGC</td><td></td><td></td><td></td><td></td><td></td><td></td><td></td><td></td><td></td><td></td></tr>
<tr><td>Oligo rev</td><td>AAACGCAGGGTCCTCCGGGCGCTC</td><td></td><td></td><td></td><td></td><td></td><td></td><td></td><td></td><td></td><td></td></tr>
<tr><td>Chromosome</td><td>start</td><td>end</td><td>strand</td><td>MM</td><td>target_seq</td><td>PAM</td><td>alignment</td><td>distance</td><td>position</td><td>gene name</td><td>gene id</td></tr>
<tr><td>GU305914.1</td><td>109867</td><td>109889</td><td>-</td><td>0</td><td>GAGCGCCCGGAGGACCCTGC</td><td>CGG</td><td>||||||||[||||||||||||]PAM</td><td>NA</td><td>Intergenic</td><td>NA</td><td>NA</td></tr>
<tr><td></td><td></td><td></td><td></td><td></td><td></td><td></td><td></td><td></td><td></td><td></td><td></td></tr>
<tr><td>T25805</td><td>GGACCCTGCCGGAGTCGCGCAGG</td><td>1000</td><td></td><td></td><td></td><td></td><td></td><td></td><td></td><td></td><td></td></tr>
<tr><td>Oligo fwd</td><td>CACCGGACCCTGCCGGAGTCGCGC</td><td></td><td></td><td></td><td></td><td></td><td></td><td></td><td></td><td></td><td></td></tr>
<tr><td>Oligo rev</td><td>AAACGCGCGACTCCGGCAGGGTCC</td><td></td><td></td><td></td><td></td><td></td><td></td><td></td><td></td><td></td><td></td></tr>
<tr><td>Chromosome</td><td>start</td><td>end</td><td>strand</td><td>MM</td><td>target_seq</td><td>PAM</td><td>alignment</td><td>distance</td><td>position</td><td>gene name</td><td>gene id</td></tr>
<tr><td>GU305914.1</td><td>109856</td><td>109878</td><td>-</td><td>0</td><td>GGACCCTGCCGGAGTCGCGC</td><td>AGG</td><td>||||||||[||||||||||||]PAM</td><td>NA</td><td>Intergenic</td><td>NA</td><td>NA</td></tr>
<tr><td></td><td></td><td></td><td></td><td></td><td></td><td></td><td></td><td></td><td></td><td></td><td></td></tr>
<tr><td>T25806</td><td>GTCGCGCAGGACGCCCTCGACGG</td><td>1000</td><td></td><td></td><td></td><td></td><td></td><td></td><td></td><td></td><td></td></tr>
<tr><td>Oligo fwd</td><td>CACCGTCGCGCAGGACGCCCTCGA</td><td></td><td></td><td></td><td></td><td></td><td></td><td></td><td></td><td></td><td></td></tr>
<tr><td>Oligo rev</td><td>AAACTCGAGGGCGTCCTGCGCGAC</td><td></td><td></td><td></td><td></td><td></td><td></td><td></td><td></td><td></td><td></td></tr>
<tr><td>Chromosome</td><td>start</td><td>end</td><td>strand</td><td>MM</td><td>target_seq</td><td>PAM</td><td>alignment</td><td>distance</td><td>position</td><td>gene name</td><td>gene id</td></tr>
<tr><td>GU305914.1</td><td>109843</td><td>109865</td><td>-</td><td>0</td><td>GTCGCGCAGGACGCCCTCGA</td><td>CGG</td><td>||||||||[||||||||||||]PAM</td><td>NA</td><td>Intergenic</td><td>NA</td><td>NA</td></tr>
<tr><td></td><td></td><td></td><td></td><td></td><td></td><td></td><td></td><td></td><td></td><td></td><td></td></tr>
<tr><td>T25807</td><td>GCGCAGGACGCCCTCGACGGTGG</td><td>1000</td><td></td><td></td><td></td><td></td><td></td><td></td><td></td><td></td><td></td></tr>
<tr><td>Oligo fwd</td><td>CACCGCGCAGGACGCCCTCGACGG</td><td></td><td></td><td></td><td></td><td></td><td></td><td></td><td></td><td></td><td></td></tr>
<tr><td>Oligo rev</td><td>AAACCCGTCGAGGGCGTCCTGCGC</td><td></td><td></td><td></td><td></td><td></td><td></td><td></td><td></td><td></td><td></td></tr>
<tr><td>Chromosome</td><td>start</td><td>end</td><td>strand</td><td>MM</td><td>target_seq</td><td>PAM</td><td>alignment</td><td>distance</td><td>position</td><td>gene name</td><td>gene id</td></tr>
<tr><td>GU305914.1</td><td>109840</td><td>109862</td><td>-</td><td>0</td><td>GCGCAGGACGCCCTCGACGG</td><td>TGG</td><td>||||||||[||||||||||||]PAM</td><td>NA</td><td>Intergenic</td><td>NA</td><td>NA</td></tr>
<tr><td></td><td></td><td></td><td></td><td></td><td></td><td></td><td></td><td></td><td></td><td></td><td></td></tr>
<tr><td>T25808</td><td>CAGGACGCCCTCGACGGTGGTGG</td><td>1000</td><td></td><td></td><td></td><td></td><td></td><td></td><td></td><td></td><td></td></tr>
<tr><td>Oligo fwd</td><td>CACCCAGGACGCCCTCGACGGTGG</td><td></td><td></td><td></td><td></td><td></td><td></td><td></td><td></td><td></td><td></td></tr>
<tr><td>Oligo rev</td><td>AAACCCACCGTCGAGGGCGTCCTG</td><td></td><td></td><td></td><td></td><td></td><td></td><td></td><td></td><td></td><td></td></tr>
<tr><td>Chromosome</td><td>start</td><td>end</td><td>strand</td><td>MM</td><td>target_seq</td><td>PAM</td><td>alignment</td><td>distance</td><td>position</td><td>gene name</td><td>gene id</td></tr>
<tr><td>GU305914.1</td><td>109837</td><td>109859</td><td>-</td><td>0</td><td>CAGGACGCCCTCGACGGTGG</td><td>TGG</td><td>||||||||[||||||||||||]PAM</td><td>NA</td><td>Intergenic</td><td>NA</td><td>NA</td></tr>
<tr><td></td><td></td><td></td><td></td><td></td><td></td><td></td><td></td><td></td><td></td><td></td><td></td></tr>
<tr><td>T25809</td><td>AGGACGCCCTCGACGGTGGTGGG</td><td>1000</td><td></td><td></td><td></td><td></td><td></td><td></td><td></td><td></td><td></td></tr>
<tr><td>Oligo fwd</td><td>CACCAGGACGCCCTCGACGGTGGT</td><td></td><td></td><td></td><td></td><td></td><td></td><td></td><td></td><td></td><td></td></tr>
<tr><td>Oligo rev</td><td>AAACACCACCGTCGAGGGCGTCCT</td><td></td><td></td><td></td><td></td><td></td><td></td><td></td><td></td><td></td><td></td></tr>
<tr><td>Chromosome</td><td>start</td><td>end</td><td>strand</td><td>MM</td><td>target_seq</td><td>PAM</td><td>alignment</td><td>distance</td><td>position</td><td>gene name</td><td>gene id</td></tr>
<tr><td>GU305914.1</td><td>109836</td><td>109858</td><td>-</td><td>0</td><td>AGGACGCCCTCGACGGTGGT</td><td>GGG</td><td>||||||||[||||||||||||]PAM</td><td>NA</td><td>Intergenic</td><td>NA</td><td>NA</td></tr>
<tr><td></td><td></td><td></td><td></td><td></td><td></td><td></td><td></td><td></td><td></td><td></td><td></td></tr>
<tr><td>T25810</td><td>CTCGACGGTGGTGGGCACCGCGG</td><td>1000</td><td></td><td></td><td></td><td></td><td></td><td></td><td></td><td></td><td></td></tr>
<tr><td>Oligo fwd</td><td>CACCCTCGACGGTGGTGGGCACCG</td><td></td><td></td><td></td><td></td><td></td><td></td><td></td><td></td><td></td><td></td></tr>
<tr><td>Oligo rev</td><td>AAACCGGTGCCCACCACCGTCGAG</td><td></td><td></td><td></td><td></td><td></td><td></td><td></td><td></td><td></td><td></td></tr>
<tr><td>Chromosome</td><td>start</td><td>end</td><td>strand</td><td>MM</td><td>target_seq</td><td>PAM</td><td>alignment</td><td>distance</td><td>position</td><td>gene name</td><td>gene id</td></tr>
<tr><td>GU305914.1</td><td>109828</td><td>109850</td><td>-</td><td>0</td><td>CTCGACGGTGGTGGGCACCG</td><td>CGG</td><td>||||||||[||||||||||||]PAM</td><td>NA</td><td>Intergenic</td><td>NA</td><td>NA</td></tr>
<tr><td></td><td></td><td></td><td></td><td></td><td></td><td></td><td></td><td></td><td></td><td></td><td></td></tr>
<tr><td>T25811</td><td>CGGTGGTGGGCACCGCGGCCCGG</td><td>1000</td><td></td><td></td><td></td><td></td><td></td><td></td><td></td><td></td><td></td></tr>
<tr><td>Oligo fwd</td><td>CACCCGGTGGTGGGCACCGCGGCC</td><td></td><td></td><td></td><td></td><td></td><td></td><td></td><td></td><td></td><td></td></tr>
<tr><td>Oligo rev</td><td>AAACGGCCGCGGTGCCCACCACCG</td><td></td><td></td><td></td><td></td><td></td><td></td><td></td><td></td><td></td><td></td></tr>
<tr><td>Chromosome</td><td>start</td><td>end</td><td>strand</td><td>MM</td><td>target_seq</td><td>PAM</td><td>alignment</td><td>distance</td><td>position</td><td>gene name</td><td>gene id</td></tr>
<tr><td>GU305914.1</td><td>109823</td><td>109845</td><td>-</td><td>0</td><td>CGGTGGTGGGCACCGCGGCC</td><td>CGG</td><td>||||||||[||||||||||||]PAM</td><td>NA</td><td>Intergenic</td><td>NA</td><td>NA</td></tr>
<tr><td></td><td></td><td></td><td></td><td></td><td></td><td></td><td></td><td></td><td></td><td></td><td></td></tr>
<tr><td>T25812</td><td>GGTGGTGGGCACCGCGGCCCGGG</td><td>1000</td><td></td><td></td><td></td><td></td><td></td><td></td><td></td><td></td><td></td></tr>
<tr><td>Oligo fwd</td><td>CACCGGTGGTGGGCACCGCGGCCC</td><td></td><td></td><td></td><td></td><td></td><td></td><td></td><td></td><td></td><td></td></tr>
<tr><td>Oligo rev</td><td>AAACGGGCCGCGGTGCCCACCACC</td><td></td><td></td><td></td><td></td><td></td><td></td><td></td><td></td><td></td><td></td></tr>
<tr><td>Chromosome</td><td>start</td><td>end</td><td>strand</td><td>MM</td><td>target_seq</td><td>PAM</td><td>alignment</td><td>distance</td><td>position</td><td>gene name</td><td>gene id</td></tr>
<tr><td>GU305914.1</td><td>109822</td><td>109844</td><td>-</td><td>0</td><td>GGTGGTGGGCACCGCGGCCC</td><td>GGG</td><td>||||||||[||||||||||||]PAM</td><td>NA</td><td>Intergenic</td><td>NA</td><td>NA</td></tr>
<tr><td></td><td></td><td></td><td></td><td></td><td></td><td></td><td></td><td></td><td></td><td></td><td></td></tr>
<tr><td>T25813</td><td>GTGGTGGGCACCGCGGCCCGGGG</td><td>1000</td><td></td><td></td><td></td><td></td><td></td><td></td><td></td><td></td><td></td></tr>
<tr><td>Oligo fwd</td><td>CACCGTGGTGGGCACCGCGGCCCG</td><td></td><td></td><td></td><td></td><td></td><td></td><td></td><td></td><td></td><td></td></tr>
<tr><td>Oligo rev</td><td>AAACCGGGCCGCGGTGCCCACCAC</td><td></td><td></td><td></td><td></td><td></td><td></td><td></td><td></td><td></td><td></td></tr>
<tr><td>Chromosome</td><td>start</td><td>end</td><td>strand</td><td>MM</td><td>target_seq</td><td>PAM</td><td>alignment</td><td>distance</td><td>position</td><td>gene name</td><td>gene id</td></tr>
<tr><td>GU305914.1</td><td>109821</td><td>109843</td><td>-</td><td>0</td><td>GTGGTGGGCACCGCGGCCCG</td><td>GGG</td><td>||||||||[||||||||||||]PAM</td><td>NA</td><td>Intergenic</td><td>NA</td><td>NA</td></tr>
<tr><td></td><td></td><td></td><td></td><td></td><td></td><td></td><td></td><td></td><td></td><td></td><td></td></tr>
<tr><td>T25814</td><td>TGGTGGGCACCGCGGCCCGGGGG</td><td>1000</td><td></td><td></td><td></td><td></td><td></td><td></td><td></td><td></td><td></td></tr>
<tr><td>Oligo fwd</td><td>CACCTGGTGGGCACCGCGGCCCGG</td><td></td><td></td><td></td><td></td><td></td><td></td><td></td><td></td><td></td><td></td></tr>
<tr><td>Oligo rev</td><td>AAACCCGGGCCGCGGTGCCCACCA</td><td></td><td></td><td></td><td></td><td></td><td></td><td></td><td></td><td></td><td></td></tr>
<tr><td>Chromosome</td><td>start</td><td>end</td><td>strand</td><td>MM</td><td>target_seq</td><td>PAM</td><td>alignment</td><td>distance</td><td>position</td><td>gene name</td><td>gene id</td></tr>
<tr><td>GU305914.1</td><td>109820</td><td>109842</td><td>-</td><td>0</td><td>TGGTGGGCACCGCGGCCCGG</td><td>GGG</td><td>||||||||[||||||||||||]PAM</td><td>NA</td><td>Intergenic</td><td>NA</td><td>NA</td></tr>
<tr><td></td><td></td><td></td><td></td><td></td><td></td><td></td><td></td><td></td><td></td><td></td><td></td></tr>
<tr><td>T25815</td><td>CGCGGCCCGGGGGTCGAGCGCGG</td><td>1000</td><td></td><td></td><td></td><td></td><td></td><td></td><td></td><td></td><td></td></tr>
<tr><td>Oligo fwd</td><td>CACCCGCGGCCCGGGGGTCGAGCG</td><td></td><td></td><td></td><td></td><td></td><td></td><td></td><td></td><td></td><td></td></tr>
<tr><td>Oligo rev</td><td>AAACCGCTCGACCCCCGGGCCGCG</td><td></td><td></td><td></td><td></td><td></td><td></td><td></td><td></td><td></td><td></td></tr>
<tr><td>Chromosome</td><td>start</td><td>end</td><td>strand</td><td>MM</td><td>target_seq</td><td>PAM</td><td>alignment</td><td>distance</td><td>position</td><td>gene name</td><td>gene id</td></tr>
<tr><td>GU305914.1</td><td>109810</td><td>109832</td><td>-</td><td>0</td><td>CGCGGCCCGGGGGTCGAGCG</td><td>CGG</td><td>||||||||[||||||||||||]PAM</td><td>NA</td><td>Intergenic</td><td>NA</td><td>NA</td></tr>
<tr><td></td><td></td><td></td><td></td><td></td><td></td><td></td><td></td><td></td><td></td><td></td><td></td></tr>
<tr><td>T25816</td><td>GCGCGGCCTGATACGCCGCCAGG</td><td>1000</td><td></td><td></td><td></td><td></td><td></td><td></td><td></td><td></td><td></td></tr>
<tr><td>Oligo fwd</td><td>CACCGCGCGGCCTGATACGCCGCC</td><td></td><td></td><td></td><td></td><td></td><td></td><td></td><td></td><td></td><td></td></tr>
<tr><td>Oligo rev</td><td>AAACGGCGGCGTATCAGGCCGCGC</td><td></td><td></td><td></td><td></td><td></td><td></td><td></td><td></td><td></td><td></td></tr>
<tr><td>Chromosome</td><td>start</td><td>end</td><td>strand</td><td>MM</td><td>target_seq</td><td>PAM</td><td>alignment</td><td>distance</td><td>position</td><td>gene name</td><td>gene id</td></tr>
<tr><td>GU305914.1</td><td>109793</td><td>109815</td><td>-</td><td>0</td><td>GCGCGGCCTGATACGCCGCC</td><td>AGG</td><td>||||||||[||||||||||||]PAM</td><td>NA</td><td>Intergenic</td><td>NA</td><td>NA</td></tr>
<tr><td></td><td></td><td></td><td></td><td></td><td></td><td></td><td></td><td></td><td></td><td></td><td></td></tr>
<tr><td>T25817</td><td>CTGATACGCCGCCAGGAGACAGG</td><td>1000</td><td></td><td></td><td></td><td></td><td></td><td></td><td></td><td></td><td></td></tr>
<tr><td>Oligo fwd</td><td>CACCCTGATACGCCGCCAGGAGAC</td><td></td><td></td><td></td><td></td><td></td><td></td><td></td><td></td><td></td><td></td></tr>
<tr><td>Oligo rev</td><td>AAACGTCTCCTGGCGGCGTATCAG</td><td></td><td></td><td></td><td></td><td></td><td></td><td></td><td></td><td></td><td></td></tr>
<tr><td>Chromosome</td><td>start</td><td>end</td><td>strand</td><td>MM</td><td>target_seq</td><td>PAM</td><td>alignment</td><td>distance</td><td>position</td><td>gene name</td><td>gene id</td></tr>
<tr><td>GU305914.1</td><td>109786</td><td>109808</td><td>-</td><td>0</td><td>CTGATACGCCGCCAGGAGAC</td><td>AGG</td><td>||||||||[||||||||||||]PAM</td><td>NA</td><td>Intergenic</td><td>NA</td><td>NA</td></tr>
<tr><td></td><td></td><td></td><td></td><td></td><td></td><td></td><td></td><td></td><td></td><td></td><td></td></tr>
<tr><td>T25818</td><td>ATACGCCGCCAGGAGACAGGCGG</td><td>1000</td><td></td><td></td><td></td><td></td><td></td><td></td><td></td><td></td><td></td></tr>
<tr><td>Oligo fwd</td><td>CACCATACGCCGCCAGGAGACAGG</td><td></td><td></td><td></td><td></td><td></td><td></td><td></td><td></td><td></td><td></td></tr>
<tr><td>Oligo rev</td><td>AAACCCTGTCTCCTGGCGGCGTAT</td><td></td><td></td><td></td><td></td><td></td><td></td><td></td><td></td><td></td><td></td></tr>
</table>

<table>
<tr><td>Chromosome</td><td>start</td><td>end</td><td>strand</td><td>MM</td><td>target_seq</td><td>PAM</td><td>alignment</td><td>distance</td><td>position</td><td>gene name</td><td>gene id</td></tr>
<tr><td>GU305914.1</td><td>109783</td><td>109805</td><td>-</td><td>0</td><td>ATACGCCGCCAGGAGACAGG</td><td>CGG</td><td>||||||||[||||||||||||]PAM</td><td>NA</td><td>Intergenic</td><td>NA</td><td>NA</td></tr>
<tr><td>T25819</td><td>GGAGACAGGCGGCCACCTCGAGG</td><td>1000</td><td></td><td></td><td></td><td></td><td></td><td></td><td></td><td></td><td></td></tr>
<tr><td>Oligo fwd</td><td>CACCGGAGACAGGCGGCCACCTCG</td><td></td><td></td><td></td><td></td><td></td><td></td><td></td><td></td><td></td><td></td></tr>
<tr><td>Oligo rev</td><td>AAACCGAGGTGGCCGCCTGTCTCC</td><td></td><td></td><td></td><td></td><td></td><td></td><td></td><td></td><td></td><td></td></tr>
<tr><td>Chromosome</td><td>start</td><td>end</td><td>strand</td><td>MM</td><td>target_seq</td><td>PAM</td><td>alignment</td><td>distance</td><td>position</td><td>gene name</td><td>gene id</td></tr>
<tr><td>GU305914.1</td><td>109772</td><td>109794</td><td>-</td><td>0</td><td>GGAGACAGGCGGCCACCTCG</td><td>AGG</td><td>||||||||[||||||||||||]PAM</td><td>NA</td><td>Intergenic</td><td>NA</td><td>NA</td></tr>
<tr><td>T25820</td><td>CACCTCGAGGTCGCTGACGACGG</td><td>1000</td><td></td><td></td><td></td><td></td><td></td><td></td><td></td><td></td><td></td></tr>
<tr><td>Oligo fwd</td><td>CACCCACCTCGAGGTCGCTGACGA</td><td></td><td></td><td></td><td></td><td></td><td></td><td></td><td></td><td></td><td></td></tr>
<tr><td>Oligo rev</td><td>AAACTCGTCAGCGACCTCGAGGTG</td><td></td><td></td><td></td><td></td><td></td><td></td><td></td><td></td><td></td><td></td></tr>
<tr><td>Chromosome</td><td>start</td><td>end</td><td>strand</td><td>MM</td><td>target_seq</td><td>PAM</td><td>alignment</td><td>distance</td><td>position</td><td>gene name</td><td>gene id</td></tr>
<tr><td>GU305914.1</td><td>109759</td><td>109781</td><td>-</td><td>0</td><td>CACCTCGAGGTCGCTGACGA</td><td>CGG</td><td>||||||||[||||||||||||]PAM</td><td>NA</td><td>Intergenic</td><td>NA</td><td>NA</td></tr>
<tr><td>T25821</td><td>GCTGACGACGGTGTCGTGCCCGG</td><td>1000</td><td></td><td></td><td></td><td></td><td></td><td></td><td></td><td></td><td></td></tr>
<tr><td>Oligo fwd</td><td>CACCGCTGACGACGGTGTCGTGCC</td><td></td><td></td><td></td><td></td><td></td><td></td><td></td><td></td><td></td><td></td></tr>
<tr><td>Oligo rev</td><td>AAACGGCACGACACCGTCGTCAGC</td><td></td><td></td><td></td><td></td><td></td><td></td><td></td><td></td><td></td><td></td></tr>
<tr><td>Chromosome</td><td>start</td><td>end</td><td>strand</td><td>MM</td><td>target_seq</td><td>PAM</td><td>alignment</td><td>distance</td><td>position</td><td>gene name</td><td>gene id</td></tr>
<tr><td>GU305914.1</td><td>109747</td><td>109769</td><td>-</td><td>0</td><td>GCTGACGACGGTGTCGTGCC</td><td>CGG</td><td>||||||||[||||||||||||]PAM</td><td>NA</td><td>Intergenic</td><td>NA</td><td>NA</td></tr>
<tr><td>T25822</td><td>CTGACGACGGTGTCGTGCCCGGG</td><td>1000</td><td></td><td></td><td></td><td></td><td></td><td></td><td></td><td></td><td></td></tr>
<tr><td>Oligo fwd</td><td>CACCCTGACGACGGTGTCGTGCCC</td><td></td><td></td><td></td><td></td><td></td><td></td><td></td><td></td><td></td><td></td></tr>
<tr><td>Oligo rev</td><td>AAACGGGCACGACACCGTCGTCAG</td><td></td><td></td><td></td><td></td><td></td><td></td><td></td><td></td><td></td><td></td></tr>
<tr><td>Chromosome</td><td>start</td><td>end</td><td>strand</td><td>MM</td><td>target_seq</td><td>PAM</td><td>alignment</td><td>distance</td><td>position</td><td>gene name</td><td>gene id</td></tr>
<tr><td>GU305914.1</td><td>109746</td><td>109768</td><td>-</td><td>0</td><td>CTGACGACGGTGTCGTGCCC</td><td>GGG</td><td>||||||||[||||||||||||]PAM</td><td>NA</td><td>Intergenic</td><td>NA</td><td>NA</td></tr>
<tr><td>T25823</td><td>TGACGACGGTGTCGTGCCCGGGG</td><td>1000</td><td></td><td></td><td></td><td></td><td></td><td></td><td></td><td></td><td></td></tr>
<tr><td>Oligo fwd</td><td>CACCTGACGACGGTGTCGTGCCCG</td><td></td><td></td><td></td><td></td><td></td><td></td><td></td><td></td><td></td><td></td></tr>
<tr><td>Oligo rev</td><td>AAACCGGGCACGACACCGTCGTCA</td><td></td><td></td><td></td><td></td><td></td><td></td><td></td><td></td><td></td><td></td></tr>
<tr><td>Chromosome</td><td>start</td><td>end</td><td>strand</td><td>MM</td><td>target_seq</td><td>PAM</td><td>alignment</td><td>distance</td><td>position</td><td>gene name</td><td>gene id</td></tr>
<tr><td>GU305914.1</td><td>109745</td><td>109767</td><td>-</td><td>0</td><td>TGACGACGGTGTCGTGCCCG</td><td>GGG</td><td>||||||||[||||||||||||]PAM</td><td>NA</td><td>Intergenic</td><td>NA</td><td>NA</td></tr>
<tr><td>T25824</td><td>GGTGTCGTGCCCGGGGTAGACGG</td><td>1000</td><td></td><td></td><td></td><td></td><td></td><td></td><td></td><td></td><td></td></tr>
<tr><td>Oligo fwd</td><td>CACCGGTGTCGTGCCCGGGGTAGA</td><td></td><td></td><td></td><td></td><td></td><td></td><td></td><td></td><td></td><td></td></tr>
<tr><td>Oligo rev</td><td>AAACTCTACCCCGGGCACGACACC</td><td></td><td></td><td></td><td></td><td></td><td></td><td></td><td></td><td></td><td></td></tr>
<tr><td>Chromosome</td><td>start</td><td>end</td><td>strand</td><td>MM</td><td>target_seq</td><td>PAM</td><td>alignment</td><td>distance</td><td>position</td><td>gene name</td><td>gene id</td></tr>
<tr><td>GU305914.1</td><td>109738</td><td>109760</td><td>-</td><td>0</td><td>GGTGTCGTGCCCGGGGTAGA</td><td>CGG</td><td>||||||||[||||||||||||]PAM</td><td>NA</td><td>Intergenic</td><td>NA</td><td>NA</td></tr>
<tr><td>T25825</td><td>GTAGACGGTGACCTGCTCCACGG</td><td>1000</td><td></td><td></td><td></td><td></td><td></td><td></td><td></td><td></td><td></td></tr>
<tr><td>Oligo fwd</td><td>CACCGTAGACGGTGACCTGCTCCA</td><td></td><td></td><td></td><td></td><td></td><td></td><td></td><td></td><td></td><td></td></tr>
<tr><td>Oligo rev</td><td>AAACTGGAGCAGGTCACCGTCTAC</td><td></td><td></td><td></td><td></td><td></td><td></td><td></td><td></td><td></td><td></td></tr>
<tr><td>Chromosome</td><td>start</td><td>end</td><td>strand</td><td>MM</td><td>target_seq</td><td>PAM</td><td>alignment</td><td>distance</td><td>position</td><td>gene name</td><td>gene id</td></tr>
<tr><td>GU305914.1</td><td>109723</td><td>109745</td><td>-</td><td>0</td><td>GTAGACGGTGACCTGCTCCA</td><td>CGG</td><td>||||||||[||||||||||||]PAM</td><td>NA</td><td>Intergenic</td><td>NA</td><td>NA</td></tr>
<tr><td>T25826</td><td>TGACCTGCTCCACGGCGTTGAGG</td><td>1000</td><td></td><td></td><td></td><td></td><td></td><td></td><td></td><td></td><td></td></tr>
<tr><td>Oligo fwd</td><td>CACCTGACCTGCTCCACGGCGTTG</td><td></td><td></td><td></td><td></td><td></td><td></td><td></td><td></td><td></td><td></td></tr>
<tr><td>Oligo rev</td><td>AAACCAACGCCGTGGAGCAGGTCA</td><td></td><td></td><td></td><td></td><td></td><td></td><td></td><td></td><td></td><td></td></tr>
<tr><td>Chromosome</td><td>start</td><td>end</td><td>strand</td><td>MM</td><td>target_seq</td><td>PAM</td><td>alignment</td><td>distance</td><td>position</td><td>gene name</td><td>gene id</td></tr>
<tr><td>GU305914.1</td><td>109715</td><td>109737</td><td>-</td><td>0</td><td>TGACCTGCTCCACGGCGTTG</td><td>AGG</td><td>||||||||[||||||||||||]PAM</td><td>NA</td><td>Intergenic</td><td>NA</td><td>NA</td></tr>
<tr><td>T25827</td><td>CGGCGTTGAGGACGCCCATGAGG</td><td>1000</td><td></td><td></td><td></td><td></td><td></td><td></td><td></td><td></td><td></td></tr>
<tr><td>Oligo fwd</td><td>CACCCGGCGTTGAGGACGCCCATG</td><td></td><td></td><td></td><td></td><td></td><td></td><td></td><td></td><td></td><td></td></tr>
<tr><td>Oligo rev</td><td>AAACCATGGGCGTCCTCAACGCCG</td><td></td><td></td><td></td><td></td><td></td><td></td><td></td><td></td><td></td><td></td></tr>
<tr><td>Chromosome</td><td>start</td><td>end</td><td>strand</td><td>MM</td><td>target_seq</td><td>PAM</td><td>alignment</td><td>distance</td><td>position</td><td>gene name</td><td>gene id</td></tr>
<tr><td>GU305914.1</td><td>109703</td><td>109725</td><td>-</td><td>0</td><td>CGGCGTTGAGGACGCCCATG</td><td>AGG</td><td>||||||||[||||||||||||]PAM</td><td>NA</td><td>Intergenic</td><td>NA</td><td>NA</td></tr>
<tr><td>T25828</td><td>CATGAGGAGCTCCTGCGAGACGG</td><td>1000</td><td></td><td></td><td></td><td></td><td></td><td></td><td></td><td></td><td></td></tr>
<tr><td>Oligo fwd</td><td>CACCCATGAGGAGCTCCTGCGAGA</td><td></td><td></td><td></td><td></td><td></td><td></td><td></td><td></td><td></td><td></td></tr>
<tr><td>Oligo rev</td><td>AAACTCTCGCAGGAGCTCCTCATG</td><td></td><td></td><td></td><td></td><td></td><td></td><td></td><td></td><td></td><td></td></tr>
<tr><td>Chromosome</td><td>start</td><td>end</td><td>strand</td><td>MM</td><td>target_seq</td><td>PAM</td><td>alignment</td><td>distance</td><td>position</td><td>gene name</td><td>gene id</td></tr>
<tr><td>GU305914.1</td><td>109687</td><td>109709</td><td>-</td><td>0</td><td>CATGAGGAGCTCCTGCGAGA</td><td>CGG</td><td>||||||||[||||||||||||]PAM</td><td>NA</td><td>Intergenic</td><td>NA</td><td>NA</td></tr>
<tr><td>T25829</td><td>GGAGCTCCTGCGAGACGGACAGG</td><td>1000</td><td></td><td></td><td></td><td></td><td></td><td></td><td></td><td></td><td></td></tr>
<tr><td>Oligo fwd</td><td>CACCGGAGCTCCTGCGAGACGGAC</td><td></td><td></td><td></td><td></td><td></td><td></td><td></td><td></td><td></td><td></td></tr>
<tr><td>Oligo rev</td><td>AAACGTCCGTCTCGCAGGAGCTCC</td><td></td><td></td><td></td><td></td><td></td><td></td><td></td><td></td><td></td><td></td></tr>
<tr><td>Chromosome</td><td>start</td><td>end</td><td>strand</td><td>MM</td><td>target_seq</td><td>PAM</td><td>alignment</td><td>distance</td><td>position</td><td>gene name</td><td>gene id</td></tr>
<tr><td>GU305914.1</td><td>109682</td><td>109704</td><td>-</td><td>0</td><td>GGAGCTCCTGCGAGACGGAC</td><td>AGG</td><td>||||||||[||||||||||||]PAM</td><td>NA</td><td>Intergenic</td><td>NA</td><td>NA</td></tr>
<tr><td>T25830</td><td>ACAGGCTCTCGATCTGCGTGAGG</td><td>1000</td><td></td><td></td><td></td><td></td><td></td><td></td><td></td><td></td><td></td></tr>
<tr><td>Oligo fwd</td><td>CACCACAGGCTCTCGATCTGCGTG</td><td></td><td></td><td></td><td></td><td></td><td></td><td></td><td></td><td></td><td></td></tr>
<tr><td>Oligo rev</td><td>AAACCACGCAGATCGAGAGCCTGT</td><td></td><td></td><td></td><td></td><td></td><td></td><td></td><td></td><td></td><td></td></tr>
<tr><td>Chromosome</td><td>start</td><td>end</td><td>strand</td><td>MM</td><td>target_seq</td><td>PAM</td><td>alignment</td><td>distance</td><td>position</td><td>gene name</td><td>gene id</td></tr>
<tr><td>GU305914.1</td><td>109664</td><td>109686</td><td>-</td><td>0</td><td>ACAGGCTCTCGATCTGCGTG</td><td>AGG</td><td>||||||||[||||||||||||]PAM</td><td>NA</td><td>Intergenic</td><td>NA</td><td>NA</td></tr>
<tr><td>T25831</td><td>CGATACACGCGCCGCCAGCGCGG</td><td>1000</td><td></td><td></td><td></td><td></td><td></td><td></td><td></td><td></td><td></td></tr>
<tr><td>Oligo fwd</td><td>CACCCGATACACGCGCCGCCAGCG</td><td></td><td></td><td></td><td></td><td></td><td></td><td></td><td></td><td></td><td></td></tr>
<tr><td>Oligo rev</td><td>AAACCGCTGGCGGCGCGTGTATCG</td><td></td><td></td><td></td><td></td><td></td><td></td><td></td><td></td><td></td><td></td></tr>
<tr><td>Chromosome</td><td>start</td><td>end</td><td>strand</td><td>MM</td><td>target_seq</td><td>PAM</td><td>alignment</td><td>distance</td><td>position</td><td>gene name</td><td>gene id</td></tr>
<tr><td>GU305914.1</td><td>109641</td><td>109663</td><td>-</td><td>0</td><td>CGATACACGCGCCGCCAGCG</td><td>CGG</td><td>||||||||[||||||||||||]PAM</td><td>NA</td><td>Intergenic</td><td>NA</td><td>NA</td></tr>
<tr><td>T25832</td><td>ACACGCGCCGCCAGCGCGGCTGG</td><td>1000</td><td></td><td></td><td></td><td></td><td></td><td></td><td></td><td></td><td></td></tr>
<tr><td>Oligo fwd</td><td>CACCACACGCGCCGCCAGCGCGGC</td><td></td><td></td><td></td><td></td><td></td><td></td><td></td><td></td><td></td><td></td></tr>
<tr><td>Oligo rev</td><td>AAACGCCGCGCTGGCGGCGCGTGT</td><td></td><td></td><td></td><td></td><td></td><td></td><td></td><td></td><td></td><td></td></tr>
<tr><td>Chromosome</td><td>start</td><td>end</td><td>strand</td><td>MM</td><td>target_seq</td><td>PAM</td><td>alignment</td><td>distance</td><td>position</td><td>gene name</td><td>gene id</td></tr>
<tr><td>GU305914.1</td><td>109637</td><td>109659</td><td>-</td><td>0</td><td>ACACGCGCCGCCAGCGCGGC</td><td>TGG</td><td>||||||||[||||||||||||]PAM</td><td>NA</td><td>Intergenic</td><td>NA</td><td>NA</td></tr>
<tr><td>T25833</td><td>GGTTGTAGAGCCACCTCTTGAGG</td><td>1000</td><td></td><td></td><td></td><td></td><td></td><td></td><td></td><td></td><td></td></tr>
<tr><td>Oligo fwd</td><td>CACCGGTTGTAGAGCCACCTCTTG</td><td></td><td></td><td></td><td></td><td></td><td></td><td></td><td></td><td></td><td></td></tr>
</table>

| | | | | | | | | | | | | |
|---|---|---|---|---|---|---|---|---|---|---|---|---|
| Oligo rev | AAACCAAGAGGTGGCTCTACAACC | | | | | | | | | | | |
| Chromosome | start | | end | strand | MM | target_seq | PAM | alignment | distance | position | gene name | gene id |
| GU305914.1 | | 109616 | 109638 | - | 0 | GGTTGTAGAGCCACCTCTTG | AGG | \|\|\|\|\|\|\|\|[\|\|\|\|\|\|\|\|\|\|\|\|]PAM | NA | Intergenic | NA | NA |
| T25834 | TGTAGAGCCACCTCTTGAGGCGG | | 1000 | | | | | | | | | |
| Oligo fwd | CACCTGTAGAGCCACCTCTTGAGG | | | | | | | | | | | |
| Oligo rev | AAACCCTCAAGAGGTGGCTCTACA | | | | | | | | | | | |
| Chromosome | start | | end | strand | MM | target_seq | PAM | alignment | distance | position | gene name | gene id |
| GU305914.1 | | 109613 | 109635 | - | 0 | TGTAGAGCCACCTCTTGAGG | CGG | \|\|\|\|\|\|\|\|[\|\|\|\|\|\|\|\|\|\|\|\|]PAM | NA | Intergenic | NA | NA |
| T25835 | CCTCTTGAGGCGGTAGTACCAGG | | 1000 | | | | | | | | | |
| Oligo fwd | CACCCCTCTTGAGGCGGTAGTACC | | | | | | | | | | | |
| Oligo rev | AAACGGTACTACCGCCTCAAGAGG | | | | | | | | | | | |
| Chromosome | start | | end | strand | MM | target_seq | PAM | alignment | distance | position | gene name | gene id |
| GU305914.1 | | 109603 | 109625 | - | 0 | CCTCTTGAGGCGGTAGTACC | AGG | \|\|\|\|\|\|\|\|[\|\|\|\|\|\|\|\|\|\|\|\|]PAM | NA | Intergenic | NA | NA |
| T25836 | GCGGTAGTACCAGGAGCCGAAGG | | 1000 | | | | | | | | | |
| Oligo fwd | CACCGCGGTAGTACCAGGAGCCGA | | | | | | | | | | | |
| Oligo rev | AAACTCGGCTCCTGGTACTACCGC | | | | | | | | | | | |
| Chromosome | start | | end | strand | MM | target_seq | PAM | alignment | distance | position | gene name | gene id |
| GU305914.1 | | 109594 | 109616 | - | 0 | GCGGTAGTACCAGGAGCCGA | AGG | \|\|\|\|\|\|\|\|[\|\|\|\|\|\|\|\|\|\|\|\|]PAM | NA | Intergenic | NA | NA |
| T25837 | GGAGCCGAAGGAGAAAGTCCAGG | | 1000 | | | | | | | | | |
| Oligo fwd | CACCGGAGCCGAAGGAGAAAGTCC | | | | | | | | | | | |
| Oligo rev | AAACGGACTTTCTCCTTCGGCTCC | | | | | | | | | | | |
| Chromosome | start | | end | strand | MM | target_seq | PAM | alignment | distance | position | gene name | gene id |
| GU305914.1 | | 109582 | 109604 | - | 0 | GGAGCCGAAGGAGAAAGTCC | AGG | \|\|\|\|\|\|\|\|[\|\|\|\|\|\|\|\|\|\|\|\|]PAM | NA | Intergenic | NA | NA |
| T25838 | GGCCTGACTCACGTTGAAGATGG | | 1000 | | | | | | | | | |
| Oligo fwd | CACCGGCCTGACTCACGTTGAAGA | | | | | | | | | | | |
| Oligo rev | AAACTCTTCAACGTGAGTCAGGCC | | | | | | | | | | | |
| Chromosome | start | | end | strand | MM | target_seq | PAM | alignment | distance | position | gene name | gene id |
| GU305914.1 | | 109561 | 109583 | - | 0 | GGCCTGACTCACGTTGAAGA | TGG | \|\|\|\|\|\|\|\|[\|\|\|\|\|\|\|\|\|\|\|\|]PAM | NA | Intergenic | NA | NA |
| T25839 | TGGTCTCGACCAGCTCCCCGCGG | | 1000 | | | | | | | | | |
| Oligo fwd | CACCTGGTCTCGACCAGCTCCCCG | | | | | | | | | | | |
| Oligo rev | AAACCGGGGAGCTGGTCGAGACCA | | | | | | | | | | | |
| Chromosome | start | | end | strand | MM | target_seq | PAM | alignment | distance | position | gene name | gene id |
| GU305914.1 | | 109541 | 109563 | - | 0 | TGGTCTCGACCAGCTCCCCG | CGG | \|\|\|\|\|\|\|\|[\|\|\|\|\|\|\|\|\|\|\|\|]PAM | NA | Intergenic | NA | NA |
| T25840 | GACCAGCTCCCCGCGGAAATCGG | | 1000 | | | | | | | | | |
| Oligo fwd | CACCGACCAGCTCCCCGCGGAAAT | | | | | | | | | | | |
| Oligo rev | AAACATTTCCGCGGGGAGCTGGTC | | | | | | | | | | | |
| Chromosome | start | | end | strand | MM | target_seq | PAM | alignment | distance | position | gene name | gene id |
| GU305914.1 | | 109534 | 109556 | - | 0 | GACCAGCTCCCCGCGGAAAT | CGG | \|\|\|\|\|\|\|\|[\|\|\|\|\|\|\|\|\|\|\|\|]PAM | NA | Intergenic | NA | NA |
| T25841 | CAGCTCCCCGCGGAAATCGGTGG | | 1000 | | | | | | | | | |
| Oligo fwd | CACCCAGCTCCCCGCGGAAATCGG | | | | | | | | | | | |
| Oligo rev | AAACCCGATTTCCGCGGGGAGCTG | | | | | | | | | | | |
| Chromosome | start | | end | strand | MM | target_seq | PAM | alignment | distance | position | gene name | gene id |
| GU305914.1 | | 109531 | 109553 | - | 0 | CAGCTCCCCGCGGAAATCGG | TGG | \|\|\|\|\|\|\|\|[\|\|\|\|\|\|\|\|\|\|\|\|]PAM | NA | Intergenic | NA | NA |
| T25842 | CCCGCGGAAATCGGTGGAGTAGG | | 1000 | | | | | | | | | |
| Oligo fwd | CACCCCCGCGGAAATCGGTGGAGT | | | | | | | | | | | |
| Oligo rev | AAACACTCCACCGATTTCCGCGGG | | | | | | | | | | | |
| Chromosome | start | | end | strand | MM | target_seq | PAM | alignment | distance | position | gene name | gene id |
| GU305914.1 | | 109525 | 109547 | - | 0 | CCCGCGGAAATCGGTGGAGT | AGG | \|\|\|\|\|\|\|\|[\|\|\|\|\|\|\|\|\|\|\|\|]PAM | NA | Intergenic | NA | NA |
| T25843 | GAAATCGGTGGAGTAGGAGATGG | | 1000 | | | | | | | | | |
| Oligo fwd | CACCGAAATCGGTGGAGTAGGAGA | | | | | | | | | | | |
| Oligo rev | AAACTCTCCTACTCCACCGATTTC | | | | | | | | | | | |
| Chromosome | start | | end | strand | MM | target_seq | PAM | alignment | distance | position | gene name | gene id |
| GU305914.1 | | 109519 | 109541 | - | 0 | GAAATCGGTGGAGTAGGAGA | TGG | \|\|\|\|\|\|\|\|[\|\|\|\|\|\|\|\|\|\|\|\|]PAM | NA | Intergenic | NA | NA |
| T25844 | AAATCGGTGGAGTAGGAGATGGG | | 1000 | | | | | | | | | |
| Oligo fwd | CACCAAATCGGTGGAGTAGGAGAT | | | | | | | | | | | |
| Oligo rev | AAACATCTCCTACTCCACCGATTT | | | | | | | | | | | |
| Chromosome | start | | end | strand | MM | target_seq | PAM | alignment | distance | position | gene name | gene id |
| GU305914.1 | | 109518 | 109540 | - | 0 | AAATCGGTGGAGTAGGAGAT | GGG | \|\|\|\|\|\|\|\|[\|\|\|\|\|\|\|\|\|\|\|\|]PAM | NA | Intergenic | NA | NA |
| T25845 | AATCGGTGGAGTAGGAGATGGGG | | 1000 | | | | | | | | | |
| Oligo fwd | CACCAATCGGTGGAGTAGGAGATG | | | | | | | | | | | |
| Oligo rev | AAACCATCTCCTACTCCACCGATT | | | | | | | | | | | |
| Chromosome | start | | end | strand | MM | target_seq | PAM | alignment | distance | position | gene name | gene id |
| GU305914.1 | | 109517 | 109539 | - | 0 | AATCGGTGGAGTAGGAGATG | GGG | \|\|\|\|\|\|\|\|[\|\|\|\|\|\|\|\|\|\|\|\|]PAM | NA | Intergenic | NA | NA |
| GU305914.1 | | 87385 | 87407 | - | 4 | AGACGGTGGAGCAGGATATG | GGG | \|--\|\|\|\|\|[\|\|\|-\|\|\|\|-\|\|\|]PAM | NA | Intergenic | NA | NA |
| T25846 | ATCGGTGGAGTAGGAGATGGGGG | | 1000 | | | | | | | | | |
| Oligo fwd | CACCATCGGTGGAGTAGGAGATGG | | | | | | | | | | | |
| Oligo rev | AAACCCATCTCCTACTCCACCGAT | | | | | | | | | | | |
| Chromosome | start | | end | strand | MM | target_seq | PAM | alignment | distance | position | gene name | gene id |
| GU305914.1 | | 109516 | 109538 | - | 0 | ATCGGTGGAGTAGGAGATGG | GGG | \|\|\|\|\|\|\|\|[\|\|\|\|\|\|\|\|\|\|\|\|]PAM | NA | Intergenic | NA | NA |
| GU305914.1 | | 87384 | 87406 | - | 4 | GACGGTGGAGCAGGATATGG | GGG | --\|\|\|\|\|\|[\|\|-\|\|\|\|-\|\|\|\|]PAM | NA | Intergenic | NA | NA |
| T25847 | TCGGTGGAGTAGGAGATGGGGGG | | 1000 | | | | | | | | | |
| Oligo fwd | CACCTCGGTGGAGTAGGAGATGGG | | | | | | | | | | | |
| Oligo rev | AAACCCCATCTCCTACTCCACCGA | | | | | | | | | | | |
| Chromosome | start | | end | strand | MM | target_seq | PAM | alignment | distance | position | gene name | gene id |
| GU305914.1 | | 109515 | 109537 | - | 0 | TCGGTGGAGTAGGAGATGGG | GGG | \|\|\|\|\|\|\|\|[\|\|\|\|\|\|\|\|\|\|\|\|]PAM | NA | Intergenic | NA | NA |

<table>
<tr><td>T25848</td><td>TAGGAGATGGGGGGATCGTTCGG</td><td>1000</td><td></td><td></td><td></td><td></td><td></td><td></td><td></td><td></td><td></td></tr>
<tr><td>Oligo fwd</td><td>CACCTAGGAGATGGGGGGATCGTT</td><td></td><td></td><td></td><td></td><td></td><td></td><td></td><td></td><td></td><td></td></tr>
<tr><td>Oligo rev</td><td>AAACAACGATCCCCCCATCTCCTA</td><td></td><td></td><td></td><td></td><td></td><td></td><td></td><td></td><td></td><td></td></tr>
<tr><td>Chromosome</td><td>start</td><td>end</td><td>strand</td><td>MM</td><td>target_seq</td><td>PAM</td><td>alignment</td><td>distance</td><td>position</td><td>gene name</td><td>gene id</td></tr>
<tr><td>GU305914.1</td><td>109506</td><td>109528</td><td>-</td><td>0</td><td>TAGGAGATGGGGGGATCGTT</td><td>CGG</td><td>||||||||[||||||||||||]PAM</td><td>NA</td><td>Intergenic</td><td>NA</td><td>NA</td></tr>
<tr><td>T25849</td><td>GCCGTCTCCCGTCGCCGAGCAGG</td><td>1000</td><td></td><td></td><td></td><td></td><td></td><td></td><td></td><td></td><td></td></tr>
<tr><td>Oligo fwd</td><td>CACCGCCGTCTCCCGTCGCCGAGC</td><td></td><td></td><td></td><td></td><td></td><td></td><td></td><td></td><td></td><td></td></tr>
<tr><td>Oligo rev</td><td>AAACGCTCGGCGACGGGAGACGGC</td><td></td><td></td><td></td><td></td><td></td><td></td><td></td><td></td><td></td><td></td></tr>
<tr><td>Chromosome</td><td>start</td><td>end</td><td>strand</td><td>MM</td><td>target_seq</td><td>PAM</td><td>alignment</td><td>distance</td><td>position</td><td>gene name</td><td>gene id</td></tr>
<tr><td>GU305914.1</td><td>109426</td><td>109448</td><td>-</td><td>0</td><td>GCCGTCTCCCGTCGCCGAGC</td><td>AGG</td><td>||||||||[||||||||||||]PAM</td><td>NA</td><td>Intergenic</td><td>NA</td><td>NA</td></tr>
<tr><td>T25850</td><td>TCCCGTCGCCGAGCAGGAAGAGG</td><td>1000</td><td></td><td></td><td></td><td></td><td></td><td></td><td></td><td></td><td></td></tr>
<tr><td>Oligo fwd</td><td>CACCTCCCGTCGCCGAGCAGGAAG</td><td></td><td></td><td></td><td></td><td></td><td></td><td></td><td></td><td></td><td></td></tr>
<tr><td>Oligo rev</td><td>AAACCTTCCTGCTCGGCGACGGGA</td><td></td><td></td><td></td><td></td><td></td><td></td><td></td><td></td><td></td><td></td></tr>
<tr><td>Chromosome</td><td>start</td><td>end</td><td>strand</td><td>MM</td><td>target_seq</td><td>PAM</td><td>alignment</td><td>distance</td><td>position</td><td>gene name</td><td>gene id</td></tr>
<tr><td>GU305914.1</td><td>109420</td><td>109442</td><td>-</td><td>0</td><td>TCCCGTCGCCGAGCAGGAAG</td><td>AGG</td><td>||||||||[||||||||||||]PAM</td><td>NA</td><td>Intergenic</td><td>NA</td><td>NA</td></tr>
<tr><td>T25851</td><td>CGCCGAGCAGGAAGAGGCCGAGG</td><td>1000</td><td></td><td></td><td></td><td></td><td></td><td></td><td></td><td></td><td></td></tr>
<tr><td>Oligo fwd</td><td>CACCCGCCGAGCAGGAAGAGGCCG</td><td></td><td></td><td></td><td></td><td></td><td></td><td></td><td></td><td></td><td></td></tr>
<tr><td>Oligo rev</td><td>AAACCGGCCTCTTCCTGCTCGGCG</td><td></td><td></td><td></td><td></td><td></td><td></td><td></td><td></td><td></td><td></td></tr>
<tr><td>Chromosome</td><td>start</td><td>end</td><td>strand</td><td>MM</td><td>target_seq</td><td>PAM</td><td>alignment</td><td>distance</td><td>position</td><td>gene name</td><td>gene id</td></tr>
<tr><td>GU305914.1</td><td>109414</td><td>109436</td><td>-</td><td>0</td><td>CGCCGAGCAGGAAGAGGCCG</td><td>AGG</td><td>||||||||[||||||||||||]PAM</td><td>NA</td><td>Intergenic</td><td>NA</td><td>NA</td></tr>
<tr><td>T25852</td><td>AGAGGCCGAGGCGCCGCCGCGG</td><td>1000</td><td></td><td></td><td></td><td></td><td></td><td></td><td></td><td></td><td></td></tr>
<tr><td>Oligo fwd</td><td>CACCAGAGGCCGAGGCGCCGCCGC</td><td></td><td></td><td></td><td></td><td></td><td></td><td></td><td></td><td></td><td></td></tr>
<tr><td>Oligo rev</td><td>AAACGCGGCGGCGCCTCGGCCTCT</td><td></td><td></td><td></td><td></td><td></td><td></td><td></td><td></td><td></td><td></td></tr>
<tr><td>Chromosome</td><td>start</td><td>end</td><td>strand</td><td>MM</td><td>target_seq</td><td>PAM</td><td>alignment</td><td>distance</td><td>position</td><td>gene name</td><td>gene id</td></tr>
<tr><td>GU305914.1</td><td>109402</td><td>109424</td><td>-</td><td>0</td><td>AGAGGCCGAGGCGCCGCCGC</td><td>CGG</td><td>||||||||[||||||||||||]PAM</td><td>NA</td><td>Intergenic</td><td>NA</td><td>NA</td></tr>
<tr><td>T25853</td><td>CCTTCTTTCTCTTGTTCTTCTGG</td><td>1000</td><td></td><td></td><td></td><td></td><td></td><td></td><td></td><td></td><td></td></tr>
<tr><td>Oligo fwd</td><td>CACCCCTTCTTTCTCTTGTTCTTC</td><td></td><td></td><td></td><td></td><td></td><td></td><td></td><td></td><td></td><td></td></tr>
<tr><td>Oligo rev</td><td>AAACGAAGAACAAGAGAAAGAAGG</td><td></td><td></td><td></td><td></td><td></td><td></td><td></td><td></td><td></td><td></td></tr>
<tr><td>Chromosome</td><td>start</td><td>end</td><td>strand</td><td>MM</td><td>target_seq</td><td>PAM</td><td>alignment</td><td>distance</td><td>position</td><td>gene name</td><td>gene id</td></tr>
<tr><td>GU305914.1</td><td>109379</td><td>109401</td><td>-</td><td>0</td><td>CCTTCTTTCTCTTGTTCTTC</td><td>TGG</td><td>||||||||[||||||||||||]PAM</td><td>NA</td><td>Intergenic</td><td>NA</td><td>NA</td></tr>
<tr><td>T25854</td><td>CTTCTGGCTAGCGTTGTCGACGG</td><td>1000</td><td></td><td></td><td></td><td></td><td></td><td></td><td></td><td></td><td></td></tr>
<tr><td>Oligo fwd</td><td>CACCCTTCTGGCTAGCGTTGTCGA</td><td></td><td></td><td></td><td></td><td></td><td></td><td></td><td></td><td></td><td></td></tr>
<tr><td>Oligo rev</td><td>AAACTCGACAACGCTAGCCAGAAG</td><td></td><td></td><td></td><td></td><td></td><td></td><td></td><td></td><td></td><td></td></tr>
<tr><td>Chromosome</td><td>start</td><td>end</td><td>strand</td><td>MM</td><td>target_seq</td><td>PAM</td><td>alignment</td><td>distance</td><td>position</td><td>gene name</td><td>gene id</td></tr>
<tr><td>GU305914.1</td><td>109363</td><td>109385</td><td>-</td><td>0</td><td>CTTCTGGCTAGCGTTGTCGA</td><td>CGG</td><td>||||||||[||||||||||||]PAM</td><td>NA</td><td>Intergenic</td><td>NA</td><td>NA</td></tr>
<tr><td>T25855</td><td>TCCGCCCAGAATGTCCGACATGG</td><td>1000</td><td></td><td></td><td></td><td></td><td></td><td></td><td></td><td></td><td></td></tr>
<tr><td>Oligo fwd</td><td>CACCTCCGCCCAGAATGTCCGACA</td><td></td><td></td><td></td><td></td><td></td><td></td><td></td><td></td><td></td><td></td></tr>
<tr><td>Oligo rev</td><td>AAACTGTCGGACATTCTGGGCGGA</td><td></td><td></td><td></td><td></td><td></td><td></td><td></td><td></td><td></td><td></td></tr>
<tr><td>Chromosome</td><td>start</td><td>end</td><td>strand</td><td>MM</td><td>target_seq</td><td>PAM</td><td>alignment</td><td>distance</td><td>position</td><td>gene name</td><td>gene id</td></tr>
<tr><td>GU305914.1</td><td>109297</td><td>109319</td><td>-</td><td>0</td><td>TCCGCCCAGAATGTCCGACA</td><td>TGG</td><td>||||||||[||||||||||||]PAM</td><td>NA</td><td>Intergenic</td><td>NA</td><td>NA</td></tr>
<tr><td>T25856</td><td>CCGCCCAGAATGTCCGACATGGG</td><td>1000</td><td></td><td></td><td></td><td></td><td></td><td></td><td></td><td></td><td></td></tr>
<tr><td>Oligo fwd</td><td>CACCCCGCCCAGAATGTCCGACAT</td><td></td><td></td><td></td><td></td><td></td><td></td><td></td><td></td><td></td><td></td></tr>
<tr><td>Oligo rev</td><td>AAACATGTCGGACATTCTGGGCGG</td><td></td><td></td><td></td><td></td><td></td><td></td><td></td><td></td><td></td><td></td></tr>
<tr><td>Chromosome</td><td>start</td><td>end</td><td>strand</td><td>MM</td><td>target_seq</td><td>PAM</td><td>alignment</td><td>distance</td><td>position</td><td>gene name</td><td>gene id</td></tr>
<tr><td>GU305914.1</td><td>109296</td><td>109318</td><td>-</td><td>0</td><td>CCGCCCAGAATGTCCGACAT</td><td>GGG</td><td>||||||||[||||||||||||]PAM</td><td>NA</td><td>Intergenic</td><td>NA</td><td>NA</td></tr>
<tr><td>T25857</td><td>CAGAATGTCCGACATGGGCCCGG</td><td>1000</td><td></td><td></td><td></td><td></td><td></td><td></td><td></td><td></td><td></td></tr>
<tr><td>Oligo fwd</td><td>CACCCAGAATGTCCGACATGGGCC</td><td></td><td></td><td></td><td></td><td></td><td></td><td></td><td></td><td></td><td></td></tr>
<tr><td>Oligo rev</td><td>AAACGGCCCATGTCGGACATTCTG</td><td></td><td></td><td></td><td></td><td></td><td></td><td></td><td></td><td></td><td></td></tr>
<tr><td>Chromosome</td><td>start</td><td>end</td><td>strand</td><td>MM</td><td>target_seq</td><td>PAM</td><td>alignment</td><td>distance</td><td>position</td><td>gene name</td><td>gene id</td></tr>
<tr><td>GU305914.1</td><td>109291</td><td>109313</td><td>-</td><td>0</td><td>CAGAATGTCCGACATGGGCC</td><td>CGG</td><td>||||||||[||||||||||||]PAM</td><td>NA</td><td>Intergenic</td><td>NA</td><td>NA</td></tr>
<tr><td>T25858</td><td>CCGCCTGATCGATCCTGCCGCGG</td><td>1000</td><td></td><td></td><td></td><td></td><td></td><td></td><td></td><td></td><td></td></tr>
<tr><td>Oligo fwd</td><td>CACCCCGCCTGATCGATCCTGCCG</td><td></td><td></td><td></td><td></td><td></td><td></td><td></td><td></td><td></td><td></td></tr>
<tr><td>Oligo rev</td><td>AAACCGGCAGGATCGATCAGGCGG</td><td></td><td></td><td></td><td></td><td></td><td></td><td></td><td></td><td></td><td></td></tr>
<tr><td>Chromosome</td><td>start</td><td>end</td><td>strand</td><td>MM</td><td>target_seq</td><td>PAM</td><td>alignment</td><td>distance</td><td>position</td><td>gene name</td><td>gene id</td></tr>
<tr><td>GU305914.1</td><td>109256</td><td>109278</td><td>-</td><td>0</td><td>CCGCCTGATCGATCCTGCCG</td><td>CGG</td><td>||||||||[||||||||||||]PAM</td><td>NA</td><td>Intergenic</td><td>NA</td><td>NA</td></tr>
<tr><td>T25859</td><td>GCTCTTTTCCGAGAGACTCTAGG</td><td>1000</td><td></td><td></td><td></td><td></td><td></td><td></td><td></td><td></td><td></td></tr>
<tr><td>Oligo fwd</td><td>CACCGCTCTTTTCCGAGAGACTCT</td><td></td><td></td><td></td><td></td><td></td><td></td><td></td><td></td><td></td><td></td></tr>
<tr><td>Oligo rev</td><td>AAACAGAGTCTCTCGGAAAAGAGC</td><td></td><td></td><td></td><td></td><td></td><td></td><td></td><td></td><td></td><td></td></tr>
<tr><td>Chromosome</td><td>start</td><td>end</td><td>strand</td><td>MM</td><td>target_seq</td><td>PAM</td><td>alignment</td><td>distance</td><td>position</td><td>gene name</td><td>gene id</td></tr>
<tr><td>GU305914.1</td><td>109211</td><td>109233</td><td>-</td><td>0</td><td>GCTCTTTTCCGAGAGACTCT</td><td>AGG</td><td>||||||||[||||||||||||]PAM</td><td>NA</td><td>Intergenic</td><td>NA</td><td>NA</td></tr>
<tr><td>T25860</td><td>GGTCTTCGCCCGCCCTTCGTCGG</td><td>1000</td><td></td><td></td><td></td><td></td><td></td><td></td><td></td><td></td><td></td></tr>
<tr><td>Oligo fwd</td><td>CACCGGTCTTCGCCCGCCCTTCGT</td><td></td><td></td><td></td><td></td><td></td><td></td><td></td><td></td><td></td><td></td></tr>
<tr><td>Oligo rev</td><td>AAACACGAAGGGCGGGCGAAGACC</td><td></td><td></td><td></td><td></td><td></td><td></td><td></td><td></td><td></td><td></td></tr>
<tr><td>Chromosome</td><td>start</td><td>end</td><td>strand</td><td>MM</td><td>target_seq</td><td>PAM</td><td>alignment</td><td>distance</td><td>position</td><td>gene name</td><td>gene id</td></tr>
<tr><td>GU305914.1</td><td>109190</td><td>109212</td><td>-</td><td>0</td><td>GGTCTTCGCCCGCCCTTCGT</td><td>CGG</td><td>||||||||[||||||||||||]PAM</td><td>NA</td><td>Intergenic</td><td>NA</td><td>NA</td></tr>
<tr><td>T25861</td><td>GCCCGCCCTTCGTCGGAGCCTGG</td><td>1000</td><td></td><td></td><td></td><td></td><td></td><td></td><td></td><td></td><td></td></tr>
<tr><td>Oligo fwd</td><td>CACCGCCCGCCCTTCGTCGGAGCC</td><td></td><td></td><td></td><td></td><td></td><td></td><td></td><td></td><td></td><td></td></tr>
<tr><td>Oligo rev</td><td>AAACGGCTCCGACGAAGGGCGGGC</td><td></td><td></td><td></td><td></td><td></td><td></td><td></td><td></td><td></td><td></td></tr>
<tr><td>Chromosome</td><td>start</td><td>end</td><td>strand</td><td>MM</td><td>target_seq</td><td>PAM</td><td>alignment</td><td>distance</td><td>position</td><td>gene name</td><td>gene id</td></tr>
<tr><td>GU305914.1</td><td>109183</td><td>109205</td><td>-</td><td>0</td><td>GCCCGCCCTTCGTCGGAGCC</td><td>TGG</td><td>||||||||[||||||||||||]PAM</td><td>NA</td><td>Intergenic</td><td>NA</td><td>NA</td></tr>
<tr><td>T25862</td><td>GGAGCCTGGCGTCGTGCCCCAGG</td><td>1000</td><td></td><td></td><td></td><td></td><td></td><td></td><td></td><td></td><td></td></tr>
<tr><td>Oligo fwd</td><td>CACCGGAGCCTGGCGTCGTGCCCC</td><td></td><td></td><td></td><td></td><td></td><td></td><td></td><td></td><td></td><td></td></tr>
<tr><td>Oligo rev</td><td>AAACGGGGCACGACGCCAGGCTCC</td><td></td><td></td><td></td><td></td><td></td><td></td><td></td><td></td><td></td><td></td></tr>
<tr><td>Chromosome</td><td>start</td><td>end</td><td>strand</td><td>MM</td><td>target_seq</td><td>PAM</td><td>alignment</td><td>distance</td><td>position</td><td>gene name</td><td>gene id</td></tr>
</table>

<table>
<tr><td>GU305914.1</td><td>109169</td><td>109191</td><td>-</td><td>0</td><td>GGAGCCTGGCGTCGTGCCCC</td><td>AGG</td><td>||||||||[||||||||||||]PAM</td><td>NA</td><td>Intergenic</td><td>NA</td><td>NA</td></tr>
<tr><td>T25863</td><td>GGCGTCGTGCCCCAGGTCGTCGG</td><td>1000</td><td></td><td></td><td></td><td></td><td></td><td></td><td></td><td></td><td></td></tr>
<tr><td>Oligo fwd</td><td>CACCGGCGTCGTGCCCCAGGTCGT</td><td></td><td></td><td></td><td></td><td></td><td></td><td></td><td></td><td></td><td></td></tr>
<tr><td>Oligo rev</td><td>AAACACGACCTGGGGCACGACGCC</td><td></td><td></td><td></td><td></td><td></td><td></td><td></td><td></td><td></td><td></td></tr>
<tr><td>Chromosome</td><td>start</td><td>end</td><td>strand</td><td>MM</td><td>target_seq</td><td>PAM</td><td>alignment</td><td>distance</td><td>position</td><td>gene name</td><td>gene id</td></tr>
<tr><td>GU305914.1</td><td>109162</td><td>109184</td><td>-</td><td>0</td><td>GGCGTCGTGCCCCAGGTCGT</td><td>CGG</td><td>||||||||[||||||||||||]PAM</td><td>NA</td><td>Intergenic</td><td>NA</td><td>NA</td></tr>
<tr><td>T25864</td><td>GGTCGTCGGCGTACTGCCGCAGG</td><td>1000</td><td></td><td></td><td></td><td></td><td></td><td></td><td></td><td></td><td></td></tr>
<tr><td>Oligo fwd</td><td>CACCGGTCGTCGGCGTACTGCCGC</td><td></td><td></td><td></td><td></td><td></td><td></td><td></td><td></td><td></td><td></td></tr>
<tr><td>Oligo rev</td><td>AAACGCGGCAGTACGCCGACGACC</td><td></td><td></td><td></td><td></td><td></td><td></td><td></td><td></td><td></td><td></td></tr>
<tr><td>Chromosome</td><td>start</td><td>end</td><td>strand</td><td>MM</td><td>target_seq</td><td>PAM</td><td>alignment</td><td>distance</td><td>position</td><td>gene name</td><td>gene id</td></tr>
<tr><td>GU305914.1</td><td>109148</td><td>109170</td><td>-</td><td>0</td><td>GGTCGTCGGCGTACTGCCGC</td><td>AGG</td><td>||||||||[||||||||||||]PAM</td><td>NA</td><td>Intergenic</td><td>NA</td><td>NA</td></tr>
<tr><td>T25865</td><td>GGCGCAGCGCGCTTTCGTCGCGG</td><td>1000</td><td></td><td></td><td></td><td></td><td></td><td></td><td></td><td></td><td></td></tr>
<tr><td>Oligo fwd</td><td>CACCGGCGCAGCGCGCTTTCGTCG</td><td></td><td></td><td></td><td></td><td></td><td></td><td></td><td></td><td></td><td></td></tr>
<tr><td>Oligo rev</td><td>AAACCGACGAAAGCGCGCTGCGCC</td><td></td><td></td><td></td><td></td><td></td><td></td><td></td><td></td><td></td><td></td></tr>
<tr><td>Chromosome</td><td>start</td><td>end</td><td>strand</td><td>MM</td><td>target_seq</td><td>PAM</td><td>alignment</td><td>distance</td><td>position</td><td>gene name</td><td>gene id</td></tr>
<tr><td>GU305914.1</td><td>109127</td><td>109149</td><td>-</td><td>0</td><td>GGCGCAGCGCGCTTTCGTCG</td><td>CGG</td><td>||||||||[||||||||||||]PAM</td><td>NA</td><td>Intergenic</td><td>NA</td><td>NA</td></tr>
<tr><td>T25866</td><td>GCAGCGCGCTTTCGTCGCGGAGG</td><td>1000</td><td></td><td></td><td></td><td></td><td></td><td></td><td></td><td></td><td></td></tr>
<tr><td>Oligo fwd</td><td>CACCGCAGCGCGCTTTCGTCGCGG</td><td></td><td></td><td></td><td></td><td></td><td></td><td></td><td></td><td></td><td></td></tr>
<tr><td>Oligo rev</td><td>AAACCCGCGACGAAAGCGCGCTGC</td><td></td><td></td><td></td><td></td><td></td><td></td><td></td><td></td><td></td><td></td></tr>
<tr><td>Chromosome</td><td>start</td><td>end</td><td>strand</td><td>MM</td><td>target_seq</td><td>PAM</td><td>alignment</td><td>distance</td><td>position</td><td>gene name</td><td>gene id</td></tr>
<tr><td>GU305914.1</td><td>109124</td><td>109146</td><td>-</td><td>0</td><td>GCAGCGCGCTTTCGTCGCGG</td><td>AGG</td><td>||||||||[||||||||||||]PAM</td><td>NA</td><td>Intergenic</td><td>NA</td><td>NA</td></tr>
<tr><td>T25867</td><td>GGAGGCGCGCGAGTTTTTCTCGG</td><td>1000</td><td></td><td></td><td></td><td></td><td></td><td></td><td></td><td></td><td></td></tr>
<tr><td>Oligo fwd</td><td>CACCGGAGGCGCGCGAGTTTTTCT</td><td></td><td></td><td></td><td></td><td></td><td></td><td></td><td></td><td></td><td></td></tr>
<tr><td>Oligo rev</td><td>AAACAGAAAAACTCGCGCGCCTCC</td><td></td><td></td><td></td><td></td><td></td><td></td><td></td><td></td><td></td><td></td></tr>
<tr><td>Chromosome</td><td>start</td><td>end</td><td>strand</td><td>MM</td><td>target_seq</td><td>PAM</td><td>alignment</td><td>distance</td><td>position</td><td>gene name</td><td>gene id</td></tr>
<tr><td>GU305914.1</td><td>109106</td><td>109128</td><td>-</td><td>0</td><td>GGAGGCGCGCGAGTTTTTCT</td><td>CGG</td><td>||||||||[||||||||||||]PAM</td><td>NA</td><td>Intergenic</td><td>NA</td><td>NA</td></tr>
<tr><td>T25868</td><td>GAGGCGCGCGAGTTTTTCTCGGG</td><td>1000</td><td></td><td></td><td></td><td></td><td></td><td></td><td></td><td></td><td></td></tr>
<tr><td>Oligo fwd</td><td>CACCGAGGCGCGCGAGTTTTTCTC</td><td></td><td></td><td></td><td></td><td></td><td></td><td></td><td></td><td></td><td></td></tr>
<tr><td>Oligo rev</td><td>AAACGAGAAAAACTCGCGCGCCTC</td><td></td><td></td><td></td><td></td><td></td><td></td><td></td><td></td><td></td><td></td></tr>
<tr><td>Chromosome</td><td>start</td><td>end</td><td>strand</td><td>MM</td><td>target_seq</td><td>PAM</td><td>alignment</td><td>distance</td><td>position</td><td>gene name</td><td>gene id</td></tr>
<tr><td>GU305914.1</td><td>109105</td><td>109127</td><td>-</td><td>0</td><td>GAGGCGCGCGAGTTTTTCTC</td><td>GGG</td><td>||||||||[||||||||||||]PAM</td><td>NA</td><td>Intergenic</td><td>NA</td><td>NA</td></tr>
<tr><td>T25869</td><td>AGGCGCGCGAGTTTTTCTCGGGG</td><td>1000</td><td></td><td></td><td></td><td></td><td></td><td></td><td></td><td></td><td></td></tr>
<tr><td>Oligo fwd</td><td>CACCAGGCGCGCGAGTTTTTCTCG</td><td></td><td></td><td></td><td></td><td></td><td></td><td></td><td></td><td></td><td></td></tr>
<tr><td>Oligo rev</td><td>AAACCGAGAAAAACTCGCGCGCCT</td><td></td><td></td><td></td><td></td><td></td><td></td><td></td><td></td><td></td><td></td></tr>
<tr><td>Chromosome</td><td>start</td><td>end</td><td>strand</td><td>MM</td><td>target_seq</td><td>PAM</td><td>alignment</td><td>distance</td><td>position</td><td>gene name</td><td>gene id</td></tr>
<tr><td>GU305914.1</td><td>109104</td><td>109126</td><td>-</td><td>0</td><td>AGGCGCGCGAGTTTTTCTCG</td><td>GGG</td><td>||||||||[||||||||||||]PAM</td><td>NA</td><td>Intergenic</td><td>NA</td><td>NA</td></tr>
<tr><td>T25870</td><td>GGCGCGCGAGTTTTTCTCGGGGG</td><td>1000</td><td></td><td></td><td></td><td></td><td></td><td></td><td></td><td></td><td></td></tr>
<tr><td>Oligo fwd</td><td>CACCGGCGCGCGAGTTTTTCTCGG</td><td></td><td></td><td></td><td></td><td></td><td></td><td></td><td></td><td></td><td></td></tr>
<tr><td>Oligo rev</td><td>AAACCCGAGAAAAACTCGCGCGCC</td><td></td><td></td><td></td><td></td><td></td><td></td><td></td><td></td><td></td><td></td></tr>
<tr><td>Chromosome</td><td>start</td><td>end</td><td>strand</td><td>MM</td><td>target_seq</td><td>PAM</td><td>alignment</td><td>distance</td><td>position</td><td>gene name</td><td>gene id</td></tr>
<tr><td>GU305914.1</td><td>109103</td><td>109125</td><td>-</td><td>0</td><td>GGCGCGCGAGTTTTTCTCGG</td><td>GGG</td><td>||||||||[||||||||||||]PAM</td><td>NA</td><td>Intergenic</td><td>NA</td><td>NA</td></tr>
<tr><td>T25871</td><td>GTTTTTCTCGGGGGAAGACCAGG</td><td>1000</td><td></td><td></td><td></td><td></td><td></td><td></td><td></td><td></td><td></td></tr>
<tr><td>Oligo fwd</td><td>CACCGTTTTTCTCGGGGGAAGACC</td><td></td><td></td><td></td><td></td><td></td><td></td><td></td><td></td><td></td><td></td></tr>
<tr><td>Oligo rev</td><td>AAACGGTCTTCCCCCGAGAAAAAC</td><td></td><td></td><td></td><td></td><td></td><td></td><td></td><td></td><td></td><td></td></tr>
<tr><td>Chromosome</td><td>start</td><td>end</td><td>strand</td><td>MM</td><td>target_seq</td><td>PAM</td><td>alignment</td><td>distance</td><td>position</td><td>gene name</td><td>gene id</td></tr>
<tr><td>GU305914.1</td><td>109094</td><td>109116</td><td>-</td><td>0</td><td>GTTTTTCTCGGGGGAAGACC</td><td>AGG</td><td>||||||||[||||||||||||]PAM</td><td>NA</td><td>Intergenic</td><td>NA</td><td>NA</td></tr>
<tr><td>T25872</td><td>GTTTTCCCGATGCGCCTCGAAGG</td><td>1000</td><td></td><td></td><td></td><td></td><td></td><td></td><td></td><td></td><td></td></tr>
<tr><td>Oligo fwd</td><td>CACCGTTTTCCCGATGCGCCTCGA</td><td></td><td></td><td></td><td></td><td></td><td></td><td></td><td></td><td></td><td></td></tr>
<tr><td>Oligo rev</td><td>AAACTCGAGGCGCATCGGGAAAAC</td><td></td><td></td><td></td><td></td><td></td><td></td><td></td><td></td><td></td><td></td></tr>
<tr><td>Chromosome</td><td>start</td><td>end</td><td>strand</td><td>MM</td><td>target_seq</td><td>PAM</td><td>alignment</td><td>distance</td><td>position</td><td>gene name</td><td>gene id</td></tr>
<tr><td>GU305914.1</td><td>109069</td><td>109091</td><td>-</td><td>0</td><td>GTTTTCCCGATGCGCCTCGA</td><td>AGG</td><td>||||||||[||||||||||||]PAM</td><td>NA</td><td>Intergenic</td><td>NA</td><td>NA</td></tr>
<tr><td>T25873</td><td>GCCTCGAAGGCCACTATACACGG</td><td>1000</td><td></td><td></td><td></td><td></td><td></td><td></td><td></td><td></td><td></td></tr>
<tr><td>Oligo fwd</td><td>CACCGCCTCGAAGGCCACTATACA</td><td></td><td></td><td></td><td></td><td></td><td></td><td></td><td></td><td></td><td></td></tr>
<tr><td>Oligo rev</td><td>AAACTGTATAGTGGCCTTCGAGGC</td><td></td><td></td><td></td><td></td><td></td><td></td><td></td><td></td><td></td><td></td></tr>
<tr><td>Chromosome</td><td>start</td><td>end</td><td>strand</td><td>MM</td><td>target_seq</td><td>PAM</td><td>alignment</td><td>distance</td><td>position</td><td>gene name</td><td>gene id</td></tr>
<tr><td>GU305914.1</td><td>109056</td><td>109078</td><td>-</td><td>0</td><td>GCCTCGAAGGCCACTATACA</td><td>CGG</td><td>||||||||[||||||||||||]PAM</td><td>NA</td><td>Intergenic</td><td>NA</td><td>NA</td></tr>
<tr><td>T25874</td><td>TATACACGGCACATCGTGAAAGG</td><td>1000</td><td></td><td></td><td></td><td></td><td></td><td></td><td></td><td></td><td></td></tr>
<tr><td>Oligo fwd</td><td>CACCTATACACGGCACATCGTGAA</td><td></td><td></td><td></td><td></td><td></td><td></td><td></td><td></td><td></td><td></td></tr>
<tr><td>Oligo rev</td><td>AAACTTCACGATGTGCCGTGTATA</td><td></td><td></td><td></td><td></td><td></td><td></td><td></td><td></td><td></td><td></td></tr>
<tr><td>Chromosome</td><td>start</td><td>end</td><td>strand</td><td>MM</td><td>target_seq</td><td>PAM</td><td>alignment</td><td>distance</td><td>position</td><td>gene name</td><td>gene id</td></tr>
<tr><td>GU305914.1</td><td>109042</td><td>109064</td><td>-</td><td>0</td><td>TATACACGGCACATCGTGAA</td><td>AGG</td><td>||||||||[||||||||||||]PAM</td><td>NA</td><td>Intergenic</td><td>NA</td><td>NA</td></tr>
<tr><td>T25875</td><td>GGTTTTTAACCGACTCATCCTGG</td><td>1000</td><td></td><td></td><td></td><td></td><td></td><td></td><td></td><td></td><td></td></tr>
<tr><td>Oligo fwd</td><td>CACCGGTTTTTAACCGACTCATCC</td><td></td><td></td><td></td><td></td><td></td><td></td><td></td><td></td><td></td><td></td></tr>
<tr><td>Oligo rev</td><td>AAACGGATGAGTCGGTTAAAAACC</td><td></td><td></td><td></td><td></td><td></td><td></td><td></td><td></td><td></td><td></td></tr>
<tr><td>Chromosome</td><td>start</td><td>end</td><td>strand</td><td>MM</td><td>target_seq</td><td>PAM</td><td>alignment</td><td>distance</td><td>position</td><td>gene name</td><td>gene id</td></tr>
<tr><td>GU305914.1</td><td>109021</td><td>109043</td><td>-</td><td>0</td><td>GGTTTTTAACCGACTCATCC</td><td>TGG</td><td>||||||||[||||||||||||]PAM</td><td>NA</td><td>Intergenic</td><td>NA</td><td>NA</td></tr>
<tr><td>T25876</td><td>TAACCGACTCATCCTGGTGCAGG</td><td>1000</td><td></td><td></td><td></td><td></td><td></td><td></td><td></td><td></td><td></td></tr>
<tr><td>Oligo fwd</td><td>CACCTAACCGACTCATCCTGGTGC</td><td></td><td></td><td></td><td></td><td></td><td></td><td></td><td></td><td></td><td></td></tr>
<tr><td>Oligo rev</td><td>AAACGCACCAGGATGAGTCGGTTA</td><td></td><td></td><td></td><td></td><td></td><td></td><td></td><td></td><td></td><td></td></tr>
<tr><td>Chromosome</td><td>start</td><td>end</td><td>strand</td><td>MM</td><td>target_seq</td><td>PAM</td><td>alignment</td><td>distance</td><td>position</td><td>gene name</td><td>gene id</td></tr>
<tr><td>GU305914.1</td><td>109015</td><td>109037</td><td>-</td><td>0</td><td>TAACCGACTCATCCTGGTGC</td><td>AGG</td><td>||||||||[||||||||||||]PAM</td><td>NA</td><td>Intergenic</td><td>NA</td><td>NA</td></tr>
<tr><td>T25877</td><td>ACTCATCCTGGTGCAGGCGCAGG</td><td>1000</td><td></td><td></td><td></td><td></td><td></td><td></td><td></td><td></td><td></td></tr>
<tr><td>Oligo fwd</td><td>CACCACTCATCCTGGTGCAGGCGC</td><td></td><td></td><td></td><td></td><td></td><td></td><td></td><td></td><td></td><td></td></tr>
<tr><td>Oligo rev</td><td>AAACGCGCCTGCACCAGGATGAGT</td><td></td><td></td><td></td><td></td><td></td><td></td><td></td><td></td><td></td><td></td></tr>
</table>

<table>
<tr><td>Chromosome</td><td>start</td><td>end</td><td>strand</td><td>MM</td><td>target_seq</td><td>PAM</td><td>alignment</td><td>distance</td><td>position</td><td>gene name</td><td>gene id</td></tr>
<tr><td>GU305914.1</td><td>109009</td><td>109031</td><td>-</td><td>0</td><td>ACTCATCCTGGTGCAGGCGC</td><td>AGG</td><td>||||||||[||||||||||||]PAM</td><td>NA</td><td>Intergenic</td><td>NA</td><td>NA</td></tr>
<tr><td>T25878</td><td>GCGCAGGAAATCCAAGACGACGG</td><td>1000</td></tr>
<tr><td>Oligo fwd</td><td>CACCGCGCAGGAAATCCAAGACGA</td></tr>
<tr><td>Oligo rev</td><td>AAACTCGTCTTGGATTTCCTGCGC</td></tr>
<tr><td>Chromosome</td><td>start</td><td>end</td><td>strand</td><td>MM</td><td>target_seq</td><td>PAM</td><td>alignment</td><td>distance</td><td>position</td><td>gene name</td><td>gene id</td></tr>
<tr><td>GU305914.1</td><td>108993</td><td>109015</td><td>-</td><td>0</td><td>GCGCAGGAAATCCAAGACGA</td><td>CGG</td><td>||||||||[||||||||||||]PAM</td><td>NA</td><td>Intergenic</td><td>NA</td><td>NA</td></tr>
<tr><td>T25879</td><td>CAGGAAATCCAAGACGACGGAGG</td><td>1000</td></tr>
<tr><td>Oligo fwd</td><td>CACCCAGGAAATCCAAGACGACGG</td></tr>
<tr><td>Oligo rev</td><td>AAACCCGTCGTCTTGGATTTCCTG</td></tr>
<tr><td>Chromosome</td><td>start</td><td>end</td><td>strand</td><td>MM</td><td>target_seq</td><td>PAM</td><td>alignment</td><td>distance</td><td>position</td><td>gene name</td><td>gene id</td></tr>
<tr><td>GU305914.1</td><td>108990</td><td>109012</td><td>-</td><td>0</td><td>CAGGAAATCCAAGACGACGG</td><td>AGG</td><td>||||||||[||||||||||||]PAM</td><td>NA</td><td>Intergenic</td><td>NA</td><td>NA</td></tr>
<tr><td>T25880</td><td>AATCCAAGACGACGGAGGAGAGG</td><td>1000</td></tr>
<tr><td>Oligo fwd</td><td>CACCAATCCAAGACGACGGAGGAG</td></tr>
<tr><td>Oligo rev</td><td>AAACCTCCTCCGTCGTCTTGGATT</td></tr>
<tr><td>Chromosome</td><td>start</td><td>end</td><td>strand</td><td>MM</td><td>target_seq</td><td>PAM</td><td>alignment</td><td>distance</td><td>position</td><td>gene name</td><td>gene id</td></tr>
<tr><td>GU305914.1</td><td>108985</td><td>109007</td><td>-</td><td>0</td><td>AATCCAAGACGACGGAGGAG</td><td>AGG</td><td>||||||||[||||||||||||]PAM</td><td>NA</td><td>Intergenic</td><td>NA</td><td>NA</td></tr>
<tr><td>T25881</td><td>GGAGAGGACGCGATACTCCCTGG</td><td>1000</td></tr>
<tr><td>Oligo fwd</td><td>CACCGGAGAGGACGCGATACTCCC</td></tr>
<tr><td>Oligo rev</td><td>AAACGGGAGTATCGCGTCCTCTCC</td></tr>
<tr><td>Chromosome</td><td>start</td><td>end</td><td>strand</td><td>MM</td><td>target_seq</td><td>PAM</td><td>alignment</td><td>distance</td><td>position</td><td>gene name</td><td>gene id</td></tr>
<tr><td>GU305914.1</td><td>108969</td><td>108991</td><td>-</td><td>0</td><td>GGAGAGGACGCGATACTCCC</td><td>TGG</td><td>||||||||[||||||||||||]PAM</td><td>NA</td><td>Intergenic</td><td>NA</td><td>NA</td></tr>
<tr><td>T25882</td><td>GAGAGGACGCGATACTCCCTGGG</td><td>1000</td></tr>
<tr><td>Oligo fwd</td><td>CACCGAGAGGACGCGATACTCCCT</td></tr>
<tr><td>Oligo rev</td><td>AAACAGGGAGTATCGCGTCCTCTC</td></tr>
<tr><td>Chromosome</td><td>start</td><td>end</td><td>strand</td><td>MM</td><td>target_seq</td><td>PAM</td><td>alignment</td><td>distance</td><td>position</td><td>gene name</td><td>gene id</td></tr>
<tr><td>GU305914.1</td><td>108968</td><td>108990</td><td>-</td><td>0</td><td>GAGAGGACGCGATACTCCCT</td><td>GGG</td><td>||||||||[||||||||||||]PAM</td><td>NA</td><td>Intergenic</td><td>NA</td><td>NA</td></tr>
<tr><td>T25883</td><td>TGGGCTTGAAGACCCGCGTGAGG</td><td>1000</td></tr>
<tr><td>Oligo fwd</td><td>CACCTGGGCTTGAAGACCCGCGTG</td></tr>
<tr><td>Oligo rev</td><td>AAACCACGCGGGTCTTCAAGCCCA</td></tr>
<tr><td>Chromosome</td><td>start</td><td>end</td><td>strand</td><td>MM</td><td>target_seq</td><td>PAM</td><td>alignment</td><td>distance</td><td>position</td><td>gene name</td><td>gene id</td></tr>
<tr><td>GU305914.1</td><td>108949</td><td>108971</td><td>-</td><td>0</td><td>TGGGCTTGAAGACCCGCGTG</td><td>AGG</td><td>||||||||[||||||||||||]PAM</td><td>NA</td><td>Intergenic</td><td>NA</td><td>NA</td></tr>
<tr><td>T25884</td><td>GGGCTTGAAGACCCGCGTGAGGG</td><td>1000</td></tr>
<tr><td>Oligo fwd</td><td>CACCGGGCTTGAAGACCCGCGTGA</td></tr>
<tr><td>Oligo rev</td><td>AAACTCACGCGGGTCTTCAAGCCC</td></tr>
<tr><td>Chromosome</td><td>start</td><td>end</td><td>strand</td><td>MM</td><td>target_seq</td><td>PAM</td><td>alignment</td><td>distance</td><td>position</td><td>gene name</td><td>gene id</td></tr>
<tr><td>GU305914.1</td><td>108948</td><td>108970</td><td>-</td><td>0</td><td>GGGCTTGAAGACCCGCGTGA</td><td>GGG</td><td>||||||||[||||||||||||]PAM</td><td>NA</td><td>Intergenic</td><td>NA</td><td>NA</td></tr>
<tr><td>T25885</td><td>GGGCGACGCGCGCCCCCTTCTGG</td><td>1000</td></tr>
<tr><td>Oligo fwd</td><td>CACCGGGCGACGCGCGCCCCCTTC</td></tr>
<tr><td>Oligo rev</td><td>AAACGAAGGGGGCGCGCGTCGCCC</td></tr>
<tr><td>Chromosome</td><td>start</td><td>end</td><td>strand</td><td>MM</td><td>target_seq</td><td>PAM</td><td>alignment</td><td>distance</td><td>position</td><td>gene name</td><td>gene id</td></tr>
<tr><td>GU305914.1</td><td>108928</td><td>108950</td><td>-</td><td>0</td><td>GGGCGACGCGCGCCCCCTTC</td><td>TGG</td><td>||||||||[||||||||||||]PAM</td><td>NA</td><td>Intergenic</td><td>NA</td><td>NA</td></tr>
<tr><td>T25886</td><td>CTGGCGAAAGAAAGAGATGACGG</td><td>1000</td></tr>
<tr><td>Oligo fwd</td><td>CACCCTGGCGAAAGAAAGAGATGA</td></tr>
<tr><td>Oligo rev</td><td>AAACTCATCTCTTTCTTTCGCCAG</td></tr>
<tr><td>Chromosome</td><td>start</td><td>end</td><td>strand</td><td>MM</td><td>target_seq</td><td>PAM</td><td>alignment</td><td>distance</td><td>position</td><td>gene name</td><td>gene id</td></tr>
<tr><td>GU305914.1</td><td>108909</td><td>108931</td><td>-</td><td>0</td><td>CTGGCGAAAGAAAGAGATGA</td><td>CGG</td><td>||||||||[||||||||||||]PAM</td><td>NA</td><td>Intergenic</td><td>NA</td><td>NA</td></tr>
<tr><td>T25887</td><td>TGGCGAAAGAAAGAGATGACGGG</td><td>1000</td></tr>
<tr><td>Oligo fwd</td><td>CACCTGGCGAAAGAAAGAGATGAC</td></tr>
<tr><td>Oligo rev</td><td>AAACGTCATCTCTTTCTTTCGCCA</td></tr>
<tr><td>Chromosome</td><td>start</td><td>end</td><td>strand</td><td>MM</td><td>target_seq</td><td>PAM</td><td>alignment</td><td>distance</td><td>position</td><td>gene name</td><td>gene id</td></tr>
<tr><td>GU305914.1</td><td>108908</td><td>108930</td><td>-</td><td>0</td><td>TGGCGAAAGAAAGAGATGAC</td><td>GGG</td><td>||||||||[||||||||||||]PAM</td><td>NA</td><td>Intergenic</td><td>NA</td><td>NA</td></tr>
<tr><td>T25888</td><td>GACCTTCCAACAGACACCGTCGG</td><td>1000</td></tr>
<tr><td>Oligo fwd</td><td>CACCGACCTTCCAACAGACACCGT</td></tr>
<tr><td>Oligo rev</td><td>AAACACGGTGTCTGTTGGAAGGTC</td></tr>
<tr><td>Chromosome</td><td>start</td><td>end</td><td>strand</td><td>MM</td><td>target_seq</td><td>PAM</td><td>alignment</td><td>distance</td><td>position</td><td>gene name</td><td>gene id</td></tr>
<tr><td>GU305914.1</td><td>108882</td><td>108904</td><td>-</td><td>0</td><td>GACCTTCCAACAGACACCGT</td><td>CGG</td><td>||||||||[||||||||||||]PAM</td><td>NA</td><td>Intergenic</td><td>NA</td><td>NA</td></tr>
<tr><td>T25889</td><td>GAACTGCGAGAAGATGCGCGTGG</td><td>1000</td></tr>
<tr><td>Oligo fwd</td><td>CACCGAACTGCGAGAAGATGCGCG</td></tr>
<tr><td>Oligo rev</td><td>AAACCGCGCATCTTCTCGCAGTTC</td></tr>
<tr><td>Chromosome</td><td>start</td><td>end</td><td>strand</td><td>MM</td><td>target_seq</td><td>PAM</td><td>alignment</td><td>distance</td><td>position</td><td>gene name</td><td>gene id</td></tr>
<tr><td>GU305914.1</td><td>108852</td><td>108874</td><td>-</td><td>0</td><td>GAACTGCGAGAAGATGCGCG</td><td>TGG</td><td>||||||||[||||||||||||]PAM</td><td>NA</td><td>Intergenic</td><td>NA</td><td>NA</td></tr>
<tr><td>T25890</td><td>TGCGCGTGGAATCGCGCAGCTGG</td><td>1000</td></tr>
<tr><td>Oligo fwd</td><td>CACCTGCGCGTGGAATCGCGCAGC</td></tr>
<tr><td>Oligo rev</td><td>AAACGCTGCGCGATTCCACGCGCA</td></tr>
<tr><td>Chromosome</td><td>start</td><td>end</td><td>strand</td><td>MM</td><td>target_seq</td><td>PAM</td><td>alignment</td><td>distance</td><td>position</td><td>gene name</td><td>gene id</td></tr>
<tr><td>GU305914.1</td><td>108838</td><td>108860</td><td>-</td><td>0</td><td>TGCGCGTGGAATCGCGCAGC</td><td>TGG</td><td>||||||||[||||||||||||]PAM</td><td>NA</td><td>Intergenic</td><td>NA</td><td>NA</td></tr>
<tr><td>T25891</td><td>GCAGCTGGCGCAGACCCTGCAGG</td><td>1000</td></tr>
<tr><td>Oligo fwd</td><td>CACCGCAGCTGGCGCAGACCCTGC</td></tr>
<tr><td>Oligo rev</td><td>AAACGCAGGGTCTGCGCCAGCTGC</td></tr>
<tr><td>Chromosome</td><td>start</td><td>end</td><td>strand</td><td>MM</td><td>target_seq</td><td>PAM</td><td>alignment</td><td>distance</td><td>position</td><td>gene name</td><td>gene id</td></tr>
<tr><td>GU305914.1</td><td>108823</td><td>108845</td><td>-</td><td>0</td><td>GCAGCTGGCGCAGACCCTGC</td><td>AGG</td><td>||||||||[||||||||||||]PAM</td><td>NA</td><td>Intergenic</td><td>NA</td><td>NA</td></tr>
<tr><td>T25892</td><td>GGCGCAGACCCTGCAGGTACTGG</td><td>1000</td></tr>
<tr><td>Oligo fwd</td><td>CACCGGCGCAGACCCTGCAGGTAC</td></tr>
</table>

Oligo rev AAACGTACCTGCAGGGTCTGCGCC
Chromosome start end strand MM target_seq PAM alignment distance position gene name gene id
GU305914.1 108817 108839 - 0 GGCGCAGACCCTGCAGGTAC TGG ||||||||[||||||||||||]PAM NA Intergenic NA NA

T25893 GCAGACCCTGCAGGTACTGGAGG 1000
Oligo fwd CACCGCAGACCCTGCAGGTACTGG
Oligo rev AAACCCAGTACCTGCAGGGTCTGC
Chromosome start end strand MM target_seq PAM alignment distance position gene name gene id
GU305914.1 108814 108836 - 0 GCAGACCCTGCAGGTACTGG AGG ||||||||[||||||||||||]PAM NA Intergenic NA NA

T25894 GGTACTGGAGGCGATGCGTGCGG 1000
Oligo fwd CACCGGTACTGGAGGCGATGCGTG
Oligo rev AAACCACGCATCGCCTCCAGTACC
Chromosome start end strand MM target_seq PAM alignment distance position gene name gene id
GU305914.1 108802 108824 - 0 GGTACTGGAGGCGATGCGTG CGG ||||||||[||||||||||||]PAM NA Intergenic NA NA

T25895 CTGGAGGCGATGCGTGCGGTTGG 1000
Oligo fwd CACCCTGGAGGCGATGCGTGCGGT
Oligo rev AAACACCGCACGCATCGCCTCCAG
Chromosome start end strand MM target_seq PAM alignment distance position gene name gene id
GU305914.1 108798 108820 - 0 CTGGAGGCGATGCGTGCGGT TGG ||||||||[||||||||||||]PAM NA Intergenic NA NA

T25896 GGCGATGCGTGCGGTTGGAGCGG 1000
Oligo fwd CACCGGCGATGCGTGCGGTTGGAG
Oligo rev AAACCTCCAACCGCACGCATCGCC
Chromosome start end strand MM target_seq PAM alignment distance position gene name gene id
GU305914.1 108793 108815 - 0 GGCGATGCGTGCGGTTGGAG CGG ||||||||[||||||||||||]PAM NA Intergenic NA NA

T25897 ATGCGTGCGGTTGGAGCGGATGG 1000
Oligo fwd CACCATGCGTGCGGTTGGAGCGGA
Oligo rev AAACTCCGCTCCAACCGCACGCAT
Chromosome start end strand MM target_seq PAM alignment distance position gene name gene id
GU305914.1 108789 108811 - 0 ATGCGTGCGGTTGGAGCGGA TGG ||||||||[||||||||||||]PAM NA Intergenic NA NA

T25898 GCGGTTGGAGCGGATGGTGCCGG 1000
Oligo fwd CACCGCGGTTGGAGCGGATGGTGC
Oligo rev AAACGCACCATCCGCTCCAACCGC
Chromosome start end strand MM target_seq PAM alignment distance position gene name gene id
GU305914.1 108783 108805 - 0 GCGGTTGGAGCGGATGGTGC CGG ||||||||[||||||||||||]PAM NA Intergenic NA NA

T25899 GGAGCGGATGGTGCCGGTGTCGG 1000
Oligo fwd CACCGGAGCGGATGGTGCCGGTGT
Oligo rev AAACACACCGGCACCATCCGCTCC
Chromosome start end strand MM target_seq PAM alignment distance position gene name gene id
GU305914.1 108777 108799 - 0 GGAGCGGATGGTGCCGGTGT CGG ||||||||[||||||||||||]PAM NA Intergenic NA NA
GU305914.1 29462 29484 + 4 GGAGAAGACGGTGCCGGGGT AGG ||||--||[-||||||||-||]PAM NA Intergenic NA NA

T25900 GGTGTCGGCGCAGCGCATAGTGG 1000
Oligo fwd CACCGGTGTCGGCGCAGCGCATAG
Oligo rev AAACCTATGCGCTGCGCCGACACC
Chromosome start end strand MM target_seq PAM alignment distance position gene name gene id
GU305914.1 108762 108784 - 0 GGTGTCGGCGCAGCGCATAG TGG ||||||||[||||||||||||]PAM NA Intergenic NA NA

T25901 GAACTCGATCACATAGCACTCGG 1000
Oligo fwd CACCGAACTCGATCACATAGCACT
Oligo rev AAACAGTGCTATGTGATCGAGTTC
Chromosome start end strand MM target_seq PAM alignment distance position gene name gene id
GU305914.1 108735 108757 - 0 GAACTCGATCACATAGCACT CGG ||||||||[||||||||||||]PAM NA Intergenic NA NA

T25902 AGCACTCGGCCGAGCCGTCGCGG 1000
Oligo fwd CACCAGCACTCGGCCGAGCCGTCG
Oligo rev AAACCGACGGCTCGGCCGAGTGCT
Chromosome start end strand MM target_seq PAM alignment distance position gene name gene id
GU305914.1 108721 108743 - 0 AGCACTCGGCCGAGCCGTCG CGG ||||||||[||||||||||||]PAM NA Intergenic NA NA

T25903 AGCCGTCGCGGTCGTCGACGAGG 1000
Oligo fwd CACCAGCCGTCGCGGTCGTCGACG
Oligo rev AAACCGTCGACGACCGCGACGGCT
Chromosome start end strand MM target_seq PAM alignment distance position gene name gene id
GU305914.1 108709 108731 - 0 AGCCGTCGCGGTCGTCGACG AGG ||||||||[||||||||||||]PAM NA Intergenic NA NA

T25904 GACGAGGACCACGATGCAGTCGG 1000
Oligo fwd CACCGACGAGGACCACGATGCAGT
Oligo rev AAACACTGCATCGTGGTCCTCGTC
Chromosome start end strand MM target_seq PAM alignment distance position gene name gene id
GU305914.1 108693 108715 - 0 GACGAGGACCACGATGCAGT CGG ||||||||[||||||||||||]PAM NA Intergenic NA NA

T25905 ACGAGGACCACGATGCAGTCGGG 1000
Oligo fwd CACCACGAGGACCACGATGCAGTC
Oligo rev AAACGACTGCATCGTGGTCCTCGT
Chromosome start end strand MM target_seq PAM alignment distance position gene name gene id
GU305914.1 108692 108714 - 0 ACGAGGACCACGATGCAGTC GGG ||||||||[||||||||||||]PAM NA Intergenic NA NA

T25906 CGAGGACCACGATGCAGTCGGGG 1000
Oligo fwd CACCCGAGGACCACGATGCAGTCG
Oligo rev AAACCGACTGCATCGTGGTCCTCG
Chromosome start end strand MM target_seq PAM alignment distance position gene name gene id
GU305914.1 108691 108713 - 0 CGAGGACCACGATGCAGTCG GGG ||||||||[||||||||||||]PAM NA Intergenic NA NA

<table>
<tr><td>T25907</td><td>TCTCGTAGAAGACGTATCGACGG</td><td>1000</td><td></td><td></td><td></td><td></td><td></td><td></td><td></td><td></td><td></td></tr>
<tr><td>Oligo fwd</td><td>CACCTCTCGTAGAAGACGTATCGA</td><td></td><td></td><td></td><td></td><td></td><td></td><td></td><td></td><td></td><td></td></tr>
<tr><td>Oligo rev</td><td>AAACTCGATACGTCTTCTACGAGA</td><td></td><td></td><td></td><td></td><td></td><td></td><td></td><td></td><td></td><td></td></tr>
<tr><td>Chromosome</td><td>start</td><td>end</td><td>strand</td><td>MM</td><td>target_seq</td><td>PAM</td><td>alignment</td><td>distance</td><td>position</td><td>gene name</td><td>gene id</td></tr>
<tr><td>GU305914.1</td><td>108649</td><td>108671</td><td>-</td><td>0</td><td>TCTCGTAGAAGACGTATCGA</td><td>CGG</td><td>||||||||[||||||||||||]PAM</td><td>NA</td><td>Intergenic</td><td>NA</td><td>NA</td></tr>
<tr><td>T25908</td><td>TAGAAGACGTATCGACGGCCCGG</td><td>1000</td><td></td><td></td><td></td><td></td><td></td><td></td><td></td><td></td><td></td></tr>
<tr><td>Oligo fwd</td><td>CACCTAGAAGACGTATCGACGGCC</td><td></td><td></td><td></td><td></td><td></td><td></td><td></td><td></td><td></td><td></td></tr>
<tr><td>Oligo rev</td><td>AAACGGCCGTCGATACGTCTTCTA</td><td></td><td></td><td></td><td></td><td></td><td></td><td></td><td></td><td></td><td></td></tr>
<tr><td>Chromosome</td><td>start</td><td>end</td><td>strand</td><td>MM</td><td>target_seq</td><td>PAM</td><td>alignment</td><td>distance</td><td>position</td><td>gene name</td><td>gene id</td></tr>
<tr><td>GU305914.1</td><td>108644</td><td>108666</td><td>-</td><td>0</td><td>TAGAAGACGTATCGACGGCC</td><td>CGG</td><td>||||||||[||||||||||||]PAM</td><td>NA</td><td>Intergenic</td><td>NA</td><td>NA</td></tr>
<tr><td>T25909</td><td>AGAAGACGTATCGACGGCCCGGG</td><td>1000</td><td></td><td></td><td></td><td></td><td></td><td></td><td></td><td></td><td></td></tr>
<tr><td>Oligo fwd</td><td>CACCAGAAGACGTATCGACGGCCC</td><td></td><td></td><td></td><td></td><td></td><td></td><td></td><td></td><td></td><td></td></tr>
<tr><td>Oligo rev</td><td>AAACGGGCCGTCGATACGTCTTCT</td><td></td><td></td><td></td><td></td><td></td><td></td><td></td><td></td><td></td><td></td></tr>
<tr><td>Chromosome</td><td>start</td><td>end</td><td>strand</td><td>MM</td><td>target_seq</td><td>PAM</td><td>alignment</td><td>distance</td><td>position</td><td>gene name</td><td>gene id</td></tr>
<tr><td>GU305914.1</td><td>108643</td><td>108665</td><td>-</td><td>0</td><td>AGAAGACGTATCGACGGCCC</td><td>GGG</td><td>||||||||[||||||||||||]PAM</td><td>NA</td><td>Intergenic</td><td>NA</td><td>NA</td></tr>
<tr><td>T25910</td><td>ATCGACGGCCCGGGACCCCCCGG</td><td>1000</td><td></td><td></td><td></td><td></td><td></td><td></td><td></td><td></td><td></td></tr>
<tr><td>Oligo fwd</td><td>CACCATCGACGGCCCGGGACCCCC</td><td></td><td></td><td></td><td></td><td></td><td></td><td></td><td></td><td></td><td></td></tr>
<tr><td>Oligo rev</td><td>AAACGGGGGTCCCGGGCCGTCGAT</td><td></td><td></td><td></td><td></td><td></td><td></td><td></td><td></td><td></td><td></td></tr>
<tr><td>Chromosome</td><td>start</td><td>end</td><td>strand</td><td>MM</td><td>target_seq</td><td>PAM</td><td>alignment</td><td>distance</td><td>position</td><td>gene name</td><td>gene id</td></tr>
<tr><td>GU305914.1</td><td>108634</td><td>108656</td><td>-</td><td>0</td><td>ATCGACGGCCCGGGACCCCC</td><td>CGG</td><td>||||||||[||||||||||||]PAM</td><td>NA</td><td>Intergenic</td><td>NA</td><td>NA</td></tr>
<tr><td>T25911</td><td>ACGGCCCGGGACCCCCCGGTAGG</td><td>1000</td><td></td><td></td><td></td><td></td><td></td><td></td><td></td><td></td><td></td></tr>
<tr><td>Oligo fwd</td><td>CACCACGGCCCGGGACCCCCCGGT</td><td></td><td></td><td></td><td></td><td></td><td></td><td></td><td></td><td></td><td></td></tr>
<tr><td>Oligo rev</td><td>AAACACCGGGGGGTCCCGGGCCGT</td><td></td><td></td><td></td><td></td><td></td><td></td><td></td><td></td><td></td><td></td></tr>
<tr><td>Chromosome</td><td>start</td><td>end</td><td>strand</td><td>MM</td><td>target_seq</td><td>PAM</td><td>alignment</td><td>distance</td><td>position</td><td>gene name</td><td>gene id</td></tr>
<tr><td>GU305914.1</td><td>108630</td><td>108652</td><td>-</td><td>0</td><td>ACGGCCCGGGACCCCCCGGT</td><td>AGG</td><td>||||||||[||||||||||||]PAM</td><td>NA</td><td>Intergenic</td><td>NA</td><td>NA</td></tr>
<tr><td>T25912</td><td>CGGCCCGGGACCCCCCGGTAGGG</td><td>1000</td><td></td><td></td><td></td><td></td><td></td><td></td><td></td><td></td><td></td></tr>
<tr><td>Oligo fwd</td><td>CACCCGGCCCGGGACCCCCCGGTA</td><td></td><td></td><td></td><td></td><td></td><td></td><td></td><td></td><td></td><td></td></tr>
<tr><td>Oligo rev</td><td>AAACTACCGGGGGGTCCCGGGCCG</td><td></td><td></td><td></td><td></td><td></td><td></td><td></td><td></td><td></td><td></td></tr>
<tr><td>Chromosome</td><td>start</td><td>end</td><td>strand</td><td>MM</td><td>target_seq</td><td>PAM</td><td>alignment</td><td>distance</td><td>position</td><td>gene name</td><td>gene id</td></tr>
<tr><td>GU305914.1</td><td>108629</td><td>108651</td><td>-</td><td>0</td><td>CGGCCCGGGACCCCCCGGTA</td><td>GGG</td><td>||||||||[||||||||||||]PAM</td><td>NA</td><td>Intergenic</td><td>NA</td><td>NA</td></tr>
<tr><td>T25913</td><td>CCGGGACCCCCCGGTAGGGCAGG</td><td>1000</td><td></td><td></td><td></td><td></td><td></td><td></td><td></td><td></td><td></td></tr>
<tr><td>Oligo fwd</td><td>CACCCCGGGACCCCCCGGTAGGGC</td><td></td><td></td><td></td><td></td><td></td><td></td><td></td><td></td><td></td><td></td></tr>
<tr><td>Oligo rev</td><td>AAACGCCCTACCGGGGGGTCCCGG</td><td></td><td></td><td></td><td></td><td></td><td></td><td></td><td></td><td></td><td></td></tr>
<tr><td>Chromosome</td><td>start</td><td>end</td><td>strand</td><td>MM</td><td>target_seq</td><td>PAM</td><td>alignment</td><td>distance</td><td>position</td><td>gene name</td><td>gene id</td></tr>
<tr><td>GU305914.1</td><td>108625</td><td>108647</td><td>-</td><td>0</td><td>CCGGGACCCCCCGGTAGGGC</td><td>AGG</td><td>||||||||[||||||||||||]PAM</td><td>NA</td><td>Intergenic</td><td>NA</td><td>NA</td></tr>
<tr><td>T25914</td><td>GGTAGGGCAGGATGCCCCCGAGG</td><td>1000</td><td></td><td></td><td></td><td></td><td></td><td></td><td></td><td></td><td></td></tr>
<tr><td>Oligo fwd</td><td>CACCGGTAGGGCAGGATGCCCCCG</td><td></td><td></td><td></td><td></td><td></td><td></td><td></td><td></td><td></td><td></td></tr>
<tr><td>Oligo rev</td><td>AAACCGGGGGCATCCTGCCCTACC</td><td></td><td></td><td></td><td></td><td></td><td></td><td></td><td></td><td></td><td></td></tr>
<tr><td>Chromosome</td><td>start</td><td>end</td><td>strand</td><td>MM</td><td>target_seq</td><td>PAM</td><td>alignment</td><td>distance</td><td>position</td><td>gene name</td><td>gene id</td></tr>
<tr><td>GU305914.1</td><td>108613</td><td>108635</td><td>-</td><td>0</td><td>GGTAGGGCAGGATGCCCCCG</td><td>AGG</td><td>||||||||[||||||||||||]PAM</td><td>NA</td><td>Intergenic</td><td>NA</td><td>NA</td></tr>
<tr><td>T25915</td><td>GGGCAGGATGCCCCCGAGGATGG</td><td>1000</td><td></td><td></td><td></td><td></td><td></td><td></td><td></td><td></td><td></td></tr>
<tr><td>Oligo fwd</td><td>CACCGGGCAGGATGCCCCCGAGGA</td><td></td><td></td><td></td><td></td><td></td><td></td><td></td><td></td><td></td><td></td></tr>
<tr><td>Oligo rev</td><td>AAACTCCTCGGGGGCATCCTGCCC</td><td></td><td></td><td></td><td></td><td></td><td></td><td></td><td></td><td></td><td></td></tr>
<tr><td>Chromosome</td><td>start</td><td>end</td><td>strand</td><td>MM</td><td>target_seq</td><td>PAM</td><td>alignment</td><td>distance</td><td>position</td><td>gene name</td><td>gene id</td></tr>
<tr><td>GU305914.1</td><td>108609</td><td>108631</td><td>-</td><td>0</td><td>GGGCAGGATGCCCCCGAGGA</td><td>TGG</td><td>||||||||[||||||||||||]PAM</td><td>NA</td><td>Intergenic</td><td>NA</td><td>NA</td></tr>
<tr><td>T25916</td><td>CGCGTCGTCGTCCTCGAACACGG</td><td>1000</td><td></td><td></td><td></td><td></td><td></td><td></td><td></td><td></td><td></td></tr>
<tr><td>Oligo fwd</td><td>CACCCGCGTCGTCGTCCTCGAACA</td><td></td><td></td><td></td><td></td><td></td><td></td><td></td><td></td><td></td><td></td></tr>
<tr><td>Oligo rev</td><td>AAACTGTTCGAGGACGACGACGCG</td><td></td><td></td><td></td><td></td><td></td><td></td><td></td><td></td><td></td><td></td></tr>
<tr><td>Chromosome</td><td>start</td><td>end</td><td>strand</td><td>MM</td><td>target_seq</td><td>PAM</td><td>alignment</td><td>distance</td><td>position</td><td>gene name</td><td>gene id</td></tr>
<tr><td>GU305914.1</td><td>108579</td><td>108601</td><td>-</td><td>0</td><td>CGCGTCGTCGTCCTCGAACA</td><td>CGG</td><td>||||||||[||||||||||||]PAM</td><td>NA</td><td>Intergenic</td><td>NA</td><td>NA</td></tr>
<tr><td>GU305914.1</td><td>106460</td><td>106482</td><td>+</td><td>4</td><td>CACCTCGTCGTACTCGACCA</td><td>GGG</td><td>|-|-||||[|||-|||||-||]PAM</td><td>NA</td><td>Intergenic</td><td>NA</td><td>NA</td></tr>
<tr><td>T25917</td><td>CCCGTCGCCCCGCACGCTTGCGG</td><td>1000</td><td></td><td></td><td></td><td></td><td></td><td></td><td></td><td></td><td></td></tr>
<tr><td>Oligo fwd</td><td>CACCCCCGTCGCCCCGCACGCTTG</td><td></td><td></td><td></td><td></td><td></td><td></td><td></td><td></td><td></td><td></td></tr>
<tr><td>Oligo rev</td><td>AAACCAAGCGTGCGGGGCGACGGG</td><td></td><td></td><td></td><td></td><td></td><td></td><td></td><td></td><td></td><td></td></tr>
<tr><td>Chromosome</td><td>start</td><td>end</td><td>strand</td><td>MM</td><td>target_seq</td><td>PAM</td><td>alignment</td><td>distance</td><td>position</td><td>gene name</td><td>gene id</td></tr>
<tr><td>GU305914.1</td><td>108526</td><td>108548</td><td>-</td><td>0</td><td>CCCGTCGCCCCGCACGCTTG</td><td>CGG</td><td>||||||||[||||||||||||]PAM</td><td>NA</td><td>Intergenic</td><td>NA</td><td>NA</td></tr>
<tr><td>T25918</td><td>CCGTCGCCCCGCACGCTTGCGGG</td><td>1000</td><td></td><td></td><td></td><td></td><td></td><td></td><td></td><td></td><td></td></tr>
<tr><td>Oligo fwd</td><td>CACCCCGTCGCCCCGCACGCTTGC</td><td></td><td></td><td></td><td></td><td></td><td></td><td></td><td></td><td></td><td></td></tr>
<tr><td>Oligo rev</td><td>AAACGCAAGCGTGCGGGGCGACGG</td><td></td><td></td><td></td><td></td><td></td><td></td><td></td><td></td><td></td><td></td></tr>
<tr><td>Chromosome</td><td>start</td><td>end</td><td>strand</td><td>MM</td><td>target_seq</td><td>PAM</td><td>alignment</td><td>distance</td><td>position</td><td>gene name</td><td>gene id</td></tr>
<tr><td>GU305914.1</td><td>108525</td><td>108547</td><td>-</td><td>0</td><td>CCGTCGCCCCGCACGCTTGC</td><td>GGG</td><td>||||||||[||||||||||||]PAM</td><td>NA</td><td>Intergenic</td><td>NA</td><td>NA</td></tr>
<tr><td>T25919</td><td>CCCGCACGCTTGCGGGCGTCAGG</td><td>1000</td><td></td><td></td><td></td><td></td><td></td><td></td><td></td><td></td><td></td></tr>
<tr><td>Oligo fwd</td><td>CACCCCCGCACGCTTGCGGGCGTC</td><td></td><td></td><td></td><td></td><td></td><td></td><td></td><td></td><td></td><td></td></tr>
<tr><td>Oligo rev</td><td>AAACGACGCCCGCAAGCGTGCGGG</td><td></td><td></td><td></td><td></td><td></td><td></td><td></td><td></td><td></td><td></td></tr>
<tr><td>Chromosome</td><td>start</td><td>end</td><td>strand</td><td>MM</td><td>target_seq</td><td>PAM</td><td>alignment</td><td>distance</td><td>position</td><td>gene name</td><td>gene id</td></tr>
<tr><td>GU305914.1</td><td>108518</td><td>108540</td><td>-</td><td>0</td><td>CCCGCACGCTTGCGGGCGTC</td><td>AGG</td><td>||||||||[||||||||||||]PAM</td><td>NA</td><td>Intergenic</td><td>NA</td><td>NA</td></tr>
<tr><td>T25920</td><td>CGATAACAAATCCGTGTCCGCGG</td><td>1000</td><td></td><td></td><td></td><td></td><td></td><td></td><td></td><td></td><td></td></tr>
<tr><td>Oligo fwd</td><td>CACCCGATAACAAATCCGTGTCCG</td><td></td><td></td><td></td><td></td><td></td><td></td><td></td><td></td><td></td><td></td></tr>
<tr><td>Oligo rev</td><td>AAACCGGACACGGATTTGTTATCG</td><td></td><td></td><td></td><td></td><td></td><td></td><td></td><td></td><td></td><td></td></tr>
<tr><td>Chromosome</td><td>start</td><td>end</td><td>strand</td><td>MM</td><td>target_seq</td><td>PAM</td><td>alignment</td><td>distance</td><td>position</td><td>gene name</td><td>gene id</td></tr>
<tr><td>GU305914.1</td><td>108489</td><td>108511</td><td>-</td><td>0</td><td>CGATAACAAATCCGTGTCCG</td><td>CGG</td><td>||||||||[||||||||||||]PAM</td><td>NA</td><td>Intergenic</td><td>NA</td><td>NA</td></tr>
<tr><td>T25921</td><td>GTCCGCGGACGTCCCCATGACGG</td><td>1000</td><td></td><td></td><td></td><td></td><td></td><td></td><td></td><td></td><td></td></tr>
<tr><td>Oligo fwd</td><td>CACCGTCCGCGGACGTCCCCATGA</td><td></td><td></td><td></td><td></td><td></td><td></td><td></td><td></td><td></td><td></td></tr>
<tr><td>Oligo rev</td><td>AAACTCATGGGGACGTCCGCGGAC</td><td></td><td></td><td></td><td></td><td></td><td></td><td></td><td></td><td></td><td></td></tr>
<tr><td>Chromosome</td><td>start</td><td>end</td><td>strand</td><td>MM</td><td>target_seq</td><td>PAM</td><td>alignment</td><td>distance</td><td>position</td><td>gene name</td><td>gene id</td></tr>
</table>

<table>
<tr><td>GU305914.1</td><td>108474</td><td>108496</td><td>-</td><td>0</td><td>GTCCGCGGACGTCCCCATGA</td><td>CGG</td><td>||||||||[||||||||||||]PAM</td><td>NA</td><td>Intergenic</td><td>NA</td><td>NA</td></tr>
<tr><td>T25922</td><td>TCCGCGGACGTCCCCATGACGGG</td><td>1000</td><td></td><td></td><td></td><td></td><td></td><td></td><td></td><td></td><td></td></tr>
<tr><td>Oligo fwd</td><td>CACCTCCGCGGACGTCCCCATGAC</td><td></td><td></td><td></td><td></td><td></td><td></td><td></td><td></td><td></td><td></td></tr>
<tr><td>Oligo rev</td><td>AAACGTCATGGGGACGTCCGCGGA</td><td></td><td></td><td></td><td></td><td></td><td></td><td></td><td></td><td></td><td></td></tr>
<tr><td>Chromosome</td><td>start</td><td>end</td><td>strand</td><td>MM</td><td>target_seq</td><td>PAM</td><td>alignment</td><td>distance</td><td>position</td><td>gene name</td><td>gene id</td></tr>
<tr><td>GU305914.1</td><td>108473</td><td>108495</td><td>-</td><td>0</td><td>TCCGCGGACGTCCCCATGAC</td><td>GGG</td><td>||||||||[||||||||||||]PAM</td><td>NA</td><td>Intergenic</td><td>NA</td><td>NA</td></tr>
<tr><td>T25923</td><td>CGGACGTCCCCATGACGGGTCGG</td><td>1000</td><td></td><td></td><td></td><td></td><td></td><td></td><td></td><td></td><td></td></tr>
<tr><td>Oligo fwd</td><td>CACCCGGACGTCCCCATGACGGGT</td><td></td><td></td><td></td><td></td><td></td><td></td><td></td><td></td><td></td><td></td></tr>
<tr><td>Oligo rev</td><td>AAACACCCGTCATGGGGACGTCCG</td><td></td><td></td><td></td><td></td><td></td><td></td><td></td><td></td><td></td><td></td></tr>
<tr><td>Chromosome</td><td>start</td><td>end</td><td>strand</td><td>MM</td><td>target_seq</td><td>PAM</td><td>alignment</td><td>distance</td><td>position</td><td>gene name</td><td>gene id</td></tr>
<tr><td>GU305914.1</td><td>108469</td><td>108491</td><td>-</td><td>0</td><td>CGGACGTCCCCATGACGGGT</td><td>CGG</td><td>||||||||[||||||||||||]PAM</td><td>NA</td><td>Intergenic</td><td>NA</td><td>NA</td></tr>
<tr><td>T25924</td><td>CGTCCCCATGACGGGTCGGACGG</td><td>1000</td><td></td><td></td><td></td><td></td><td></td><td></td><td></td><td></td><td></td></tr>
<tr><td>Oligo fwd</td><td>CACCCGTCCCCATGACGGGTCGGA</td><td></td><td></td><td></td><td></td><td></td><td></td><td></td><td></td><td></td><td></td></tr>
<tr><td>Oligo rev</td><td>AAACTCCGACCCGTCATGGGGACG</td><td></td><td></td><td></td><td></td><td></td><td></td><td></td><td></td><td></td><td></td></tr>
<tr><td>Chromosome</td><td>start</td><td>end</td><td>strand</td><td>MM</td><td>target_seq</td><td>PAM</td><td>alignment</td><td>distance</td><td>position</td><td>gene name</td><td>gene id</td></tr>
<tr><td>GU305914.1</td><td>108465</td><td>108487</td><td>-</td><td>0</td><td>CGTCCCCATGACGGGTCGGA</td><td>CGG</td><td>||||||||[||||||||||||]PAM</td><td>NA</td><td>Intergenic</td><td>NA</td><td>NA</td></tr>
<tr><td>T25925</td><td>CGTAATTAAGATATTTAAACAGG</td><td>1000</td><td></td><td></td><td></td><td></td><td></td><td></td><td></td><td></td><td></td></tr>
<tr><td>Oligo fwd</td><td>CACCCGTAATTAAGATATTTAAAC</td><td></td><td></td><td></td><td></td><td></td><td></td><td></td><td></td><td></td><td></td></tr>
<tr><td>Oligo rev</td><td>AAACGTTTAAATATCTTAATTACG</td><td></td><td></td><td></td><td></td><td></td><td></td><td></td><td></td><td></td><td></td></tr>
<tr><td>Chromosome</td><td>start</td><td>end</td><td>strand</td><td>MM</td><td>target_seq</td><td>PAM</td><td>alignment</td><td>distance</td><td>position</td><td>gene name</td><td>gene id</td></tr>
<tr><td>GU305914.1</td><td>108428</td><td>108450</td><td>-</td><td>0</td><td>CGTAATTAAGATATTTAAAC</td><td>AGG</td><td>||||||||[||||||||||||]PAM</td><td>NA</td><td>Intergenic</td><td>NA</td><td>NA</td></tr>
<tr><td>T25926</td><td>TTAAACAGGATCGACGAGATAGG</td><td>1000</td><td></td><td></td><td></td><td></td><td></td><td></td><td></td><td></td><td></td></tr>
<tr><td>Oligo fwd</td><td>CACCTTAAACAGGATCGACGAGAT</td><td></td><td></td><td></td><td></td><td></td><td></td><td></td><td></td><td></td><td></td></tr>
<tr><td>Oligo rev</td><td>AAACATCTCGTCGATCCTGTTTAA</td><td></td><td></td><td></td><td></td><td></td><td></td><td></td><td></td><td></td><td></td></tr>
<tr><td>Chromosome</td><td>start</td><td>end</td><td>strand</td><td>MM</td><td>target_seq</td><td>PAM</td><td>alignment</td><td>distance</td><td>position</td><td>gene name</td><td>gene id</td></tr>
<tr><td>GU305914.1</td><td>108414</td><td>108436</td><td>-</td><td>0</td><td>TTAAACAGGATCGACGAGAT</td><td>AGG</td><td>||||||||[||||||||||||]PAM</td><td>NA</td><td>Intergenic</td><td>NA</td><td>NA</td></tr>
<tr><td>T25927</td><td>TAAACAGGATCGACGAGATAGGG</td><td>1000</td><td></td><td></td><td></td><td></td><td></td><td></td><td></td><td></td><td></td></tr>
<tr><td>Oligo fwd</td><td>CACCTAAACAGGATCGACGAGATA</td><td></td><td></td><td></td><td></td><td></td><td></td><td></td><td></td><td></td><td></td></tr>
<tr><td>Oligo rev</td><td>AAACTATCTCGTCGATCCTGTTTA</td><td></td><td></td><td></td><td></td><td></td><td></td><td></td><td></td><td></td><td></td></tr>
<tr><td>Chromosome</td><td>start</td><td>end</td><td>strand</td><td>MM</td><td>target_seq</td><td>PAM</td><td>alignment</td><td>distance</td><td>position</td><td>gene name</td><td>gene id</td></tr>
<tr><td>GU305914.1</td><td>108413</td><td>108435</td><td>-</td><td>0</td><td>TAAACAGGATCGACGAGATA</td><td>GGG</td><td>||||||||[||||||||||||]PAM</td><td>NA</td><td>Intergenic</td><td>NA</td><td>NA</td></tr>
<tr><td>T25928</td><td>TTCGACGCGCGTCGTTTACGCGG</td><td>1000</td><td></td><td></td><td></td><td></td><td></td><td></td><td></td><td></td><td></td></tr>
<tr><td>Oligo fwd</td><td>CACCTTCGACGCGCGTCGTTTACG</td><td></td><td></td><td></td><td></td><td></td><td></td><td></td><td></td><td></td><td></td></tr>
<tr><td>Oligo rev</td><td>AAACCGTAAACGACGCGCGTCGAA</td><td></td><td></td><td></td><td></td><td></td><td></td><td></td><td></td><td></td><td></td></tr>
<tr><td>Chromosome</td><td>start</td><td>end</td><td>strand</td><td>MM</td><td>target_seq</td><td>PAM</td><td>alignment</td><td>distance</td><td>position</td><td>gene name</td><td>gene id</td></tr>
<tr><td>GU305914.1</td><td>108325</td><td>108347</td><td>-</td><td>0</td><td>TTCGACGCGCGTCGTTTACG</td><td>CGG</td><td>||||||||[||||||||||||]PAM</td><td>NA</td><td>Intergenic</td><td>NA</td><td>NA</td></tr>
<tr><td>T25929</td><td>ACGCGCGTCGTTTACGCGGCTGG</td><td>1000</td><td></td><td></td><td></td><td></td><td></td><td></td><td></td><td></td><td></td></tr>
<tr><td>Oligo fwd</td><td>CACCACGCGCGTCGTTTACGCGGC</td><td></td><td></td><td></td><td></td><td></td><td></td><td></td><td></td><td></td><td></td></tr>
<tr><td>Oligo rev</td><td>AAACGCCGCGTAAACGACGCGCGT</td><td></td><td></td><td></td><td></td><td></td><td></td><td></td><td></td><td></td><td></td></tr>
<tr><td>Chromosome</td><td>start</td><td>end</td><td>strand</td><td>MM</td><td>target_seq</td><td>PAM</td><td>alignment</td><td>distance</td><td>position</td><td>gene name</td><td>gene id</td></tr>
<tr><td>GU305914.1</td><td>108321</td><td>108343</td><td>-</td><td>0</td><td>ACGCGCGTCGTTTACGCGGC</td><td>TGG</td><td>||||||||[||||||||||||]PAM</td><td>NA</td><td>Intergenic</td><td>NA</td><td>NA</td></tr>
<tr><td>T25930</td><td>CGCGCGTCGTTTACGCGGCTGGG</td><td>1000</td><td></td><td></td><td></td><td></td><td></td><td></td><td></td><td></td><td></td></tr>
<tr><td>Oligo fwd</td><td>CACCCGCGCGTCGTTTACGCGGCT</td><td></td><td></td><td></td><td></td><td></td><td></td><td></td><td></td><td></td><td></td></tr>
<tr><td>Oligo rev</td><td>AAACAGCCGCGTAAACGACGCGCG</td><td></td><td></td><td></td><td></td><td></td><td></td><td></td><td></td><td></td><td></td></tr>
<tr><td>Chromosome</td><td>start</td><td>end</td><td>strand</td><td>MM</td><td>target_seq</td><td>PAM</td><td>alignment</td><td>distance</td><td>position</td><td>gene name</td><td>gene id</td></tr>
<tr><td>GU305914.1</td><td>108320</td><td>108342</td><td>-</td><td>0</td><td>CGCGCGTCGTTTACGCGGCT</td><td>GGG</td><td>||||||||[||||||||||||]PAM</td><td>NA</td><td>Intergenic</td><td>NA</td><td>NA</td></tr>
<tr><td>T25931</td><td>GCGCGTCGTTTACGCGGCTGGGG</td><td>1000</td><td></td><td></td><td></td><td></td><td></td><td></td><td></td><td></td><td></td></tr>
<tr><td>Oligo fwd</td><td>CACCGCGCGTCGTTTACGCGGCTG</td><td></td><td></td><td></td><td></td><td></td><td></td><td></td><td></td><td></td><td></td></tr>
<tr><td>Oligo rev</td><td>AAACCAGCCGCGTAAACGACGCGC</td><td></td><td></td><td></td><td></td><td></td><td></td><td></td><td></td><td></td><td></td></tr>
<tr><td>Chromosome</td><td>start</td><td>end</td><td>strand</td><td>MM</td><td>target_seq</td><td>PAM</td><td>alignment</td><td>distance</td><td>position</td><td>gene name</td><td>gene id</td></tr>
<tr><td>GU305914.1</td><td>108319</td><td>108341</td><td>-</td><td>0</td><td>GCGCGTCGTTTACGCGGCTG</td><td>GGG</td><td>||||||||[||||||||||||]PAM</td><td>NA</td><td>Intergenic</td><td>NA</td><td>NA</td></tr>
<tr><td>T25932</td><td>CGTTTACGCGGCTGGGGCCGAGG</td><td>1000</td><td></td><td></td><td></td><td></td><td></td><td></td><td></td><td></td><td></td></tr>
<tr><td>Oligo fwd</td><td>CACCCGTTTACGCGGCTGGGGCCG</td><td></td><td></td><td></td><td></td><td></td><td></td><td></td><td></td><td></td><td></td></tr>
<tr><td>Oligo rev</td><td>AAACCGGCCCCAGCCGCGTAAACG</td><td></td><td></td><td></td><td></td><td></td><td></td><td></td><td></td><td></td><td></td></tr>
<tr><td>Chromosome</td><td>start</td><td>end</td><td>strand</td><td>MM</td><td>target_seq</td><td>PAM</td><td>alignment</td><td>distance</td><td>position</td><td>gene name</td><td>gene id</td></tr>
<tr><td>GU305914.1</td><td>108313</td><td>108335</td><td>-</td><td>0</td><td>CGTTTACGCGGCTGGGGCCG</td><td>AGG</td><td>||||||||[||||||||||||]PAM</td><td>NA</td><td>Intergenic</td><td>NA</td><td>NA</td></tr>
<tr><td>T25933</td><td>CGGCTGGGGCCGAGGCGCCGCGG</td><td>1000</td><td></td><td></td><td></td><td></td><td></td><td></td><td></td><td></td><td></td></tr>
<tr><td>Oligo fwd</td><td>CACCCGGCTGGGGCCGAGGCGCCG</td><td></td><td></td><td></td><td></td><td></td><td></td><td></td><td></td><td></td><td></td></tr>
<tr><td>Oligo rev</td><td>AAACCGGCGCCTCGGCCCCAGCCG</td><td></td><td></td><td></td><td></td><td></td><td></td><td></td><td></td><td></td><td></td></tr>
<tr><td>Chromosome</td><td>start</td><td>end</td><td>strand</td><td>MM</td><td>target_seq</td><td>PAM</td><td>alignment</td><td>distance</td><td>position</td><td>gene name</td><td>gene id</td></tr>
<tr><td>GU305914.1</td><td>108305</td><td>108327</td><td>-</td><td>0</td><td>CGGCTGGGGCCGAGGCGCCG</td><td>CGG</td><td>||||||||[||||||||||||]PAM</td><td>NA</td><td>Intergenic</td><td>NA</td><td>NA</td></tr>
<tr><td>T25934</td><td>TGCACTCGTACAACGAGAGCCGG</td><td>1000</td><td></td><td></td><td></td><td></td><td></td><td></td><td></td><td></td><td></td></tr>
<tr><td>Oligo fwd</td><td>CACCTGCACTCGTACAACGAGAGC</td><td></td><td></td><td></td><td></td><td></td><td></td><td></td><td></td><td></td><td></td></tr>
<tr><td>Oligo rev</td><td>AAACGCTCTCGTTGTACGAGTGCA</td><td></td><td></td><td></td><td></td><td></td><td></td><td></td><td></td><td></td><td></td></tr>
<tr><td>Chromosome</td><td>start</td><td>end</td><td>strand</td><td>MM</td><td>target_seq</td><td>PAM</td><td>alignment</td><td>distance</td><td>position</td><td>gene name</td><td>gene id</td></tr>
<tr><td>GU305914.1</td><td>108260</td><td>108282</td><td>-</td><td>0</td><td>TGCACTCGTACAACGAGAGC</td><td>CGG</td><td>||||||||[||||||||||||]PAM</td><td>NA</td><td>Intergenic</td><td>NA</td><td>NA</td></tr>
<tr><td>T25935</td><td>GCACTCGTACAACGAGAGCCGGG</td><td>1000</td><td></td><td></td><td></td><td></td><td></td><td></td><td></td><td></td><td></td></tr>
<tr><td>Oligo fwd</td><td>CACCGCACTCGTACAACGAGAGCC</td><td></td><td></td><td></td><td></td><td></td><td></td><td></td><td></td><td></td><td></td></tr>
<tr><td>Oligo rev</td><td>AAACGGCTCTCGTTGTACGAGTGC</td><td></td><td></td><td></td><td></td><td></td><td></td><td></td><td></td><td></td><td></td></tr>
<tr><td>Chromosome</td><td>start</td><td>end</td><td>strand</td><td>MM</td><td>target_seq</td><td>PAM</td><td>alignment</td><td>distance</td><td>position</td><td>gene name</td><td>gene id</td></tr>
<tr><td>GU305914.1</td><td>108259</td><td>108281</td><td>-</td><td>0</td><td>GCACTCGTACAACGAGAGCC</td><td>GGG</td><td>||||||||[||||||||||||]PAM</td><td>NA</td><td>Intergenic</td><td>NA</td><td>NA</td></tr>
<tr><td>T25936</td><td>ACAACGAGAGCCGGGTGTGCCGG</td><td>1000</td><td></td><td></td><td></td><td></td><td></td><td></td><td></td><td></td><td></td></tr>
<tr><td>Oligo fwd</td><td>CACCACAACGAGAGCCGGGTGTGC</td><td></td><td></td><td></td><td></td><td></td><td></td><td></td><td></td><td></td><td></td></tr>
<tr><td>Oligo rev</td><td>AAACGCACACCCGGCTCTCGTTGT</td><td></td><td></td><td></td><td></td><td></td><td></td><td></td><td></td><td></td><td></td></tr>
</table>

| Chromosome | start | end | strand | MM | target_seq | PAM | alignment | distance | position | gene name | gene id |
|---|---|---|---|---|---|---|---|---|---|---|---|
| GU305914.1 | 108251 | 108273 | - | 0 | ACAACGAGAGCCGGGTGTGC | CGG | \|\|\|\|\|\|\|\|[\|\|\|\|\|\|\|\|\|\|\|\|]PAM | NA | Intergenic | NA | NA |
| T25937 | CGACGAATCCTCTAACCAGACGG | 1000 | | | | | | | | | |
| Oligo fwd | CACCCGACGAATCCTCTAACCAGA | | | | | | | | | | |
| Oligo rev | AAACTCTGGTTAGAGGATTCGTCG | | | | | | | | | | |
| Chromosome | start | end | strand | MM | target_seq | PAM | alignment | distance | position | gene name | gene id |
| GU305914.1 | 108226 | 108248 | - | 0 | CGACGAATCCTCTAACCAGA | CGG | \|\|\|\|\|\|\|\|[\|\|\|\|\|\|\|\|\|\|\|\|]PAM | NA | Intergenic | NA | NA |
| T25938 | TAACCAGACGGTGTCTCACGCGG | 1000 | | | | | | | | | |
| Oligo fwd | CACCTAACCAGACGGTGTCTCACG | | | | | | | | | | |
| Oligo rev | AAACCGTGAGACACCGTCTGGTTA | | | | | | | | | | |
| Chromosome | start | end | strand | MM | target_seq | PAM | alignment | distance | position | gene name | gene id |
| GU305914.1 | 108214 | 108236 | - | 0 | TAACCAGACGGTGTCTCACG | CGG | \|\|\|\|\|\|\|\|[\|\|\|\|\|\|\|\|\|\|\|\|]PAM | NA | Intergenic | NA | NA |
| T25939 | CATGTTCACGTTCAACTTCCAGG | 1000 | | | | | | | | | |
| Oligo fwd | CACCCATGTTCACGTTCAACTTCC | | | | | | | | | | |
| Oligo rev | AAACGGAAGTTGAACGTGAACATG | | | | | | | | | | |
| Chromosome | start | end | strand | MM | target_seq | PAM | alignment | distance | position | gene name | gene id |
| GU305914.1 | 108190 | 108212 | - | 0 | CATGTTCACGTTCAACTTCC | AGG | \|\|\|\|\|\|\|\|[\|\|\|\|\|\|\|\|\|\|\|\|]PAM | NA | Intergenic | NA | NA |
| T25940 | TTCACGTTCAACTTCCAGGACGG | 1000 | | | | | | | | | |
| Oligo fwd | CACCTTCACGTTCAACTTCCAGGA | | | | | | | | | | |
| Oligo rev | AAACTCCTGGAAGTTGAACGTGAA | | | | | | | | | | |
| Chromosome | start | end | strand | MM | target_seq | PAM | alignment | distance | position | gene name | gene id |
| GU305914.1 | 108186 | 108208 | - | 0 | TTCACGTTCAACTTCCAGGA | CGG | \|\|\|\|\|\|\|\|[\|\|\|\|\|\|\|\|\|\|\|\|]PAM | NA | Intergenic | NA | NA |
| T25941 | TCACGTTCAACTTCCAGGACGGG | 1000 | | | | | | | | | |
| Oligo fwd | CACCTCACGTTCAACTTCCAGGAC | | | | | | | | | | |
| Oligo rev | AAACGTCCTGGAAGTTGAACGTGA | | | | | | | | | | |
| Chromosome | start | end | strand | MM | target_seq | PAM | alignment | distance | position | gene name | gene id |
| GU305914.1 | 108185 | 108207 | - | 0 | TCACGTTCAACTTCCAGGAC | GGG | \|\|\|\|\|\|\|\|[\|\|\|\|\|\|\|\|\|\|\|\|]PAM | NA | Intergenic | NA | NA |
| T25942 | CACGTTCAACTTCCAGGACGGGG | 1000 | | | | | | | | | |
| Oligo fwd | CACCCACGTTCAACTTCCAGGACG | | | | | | | | | | |
| Oligo rev | AAACCGTCCTGGAAGTTGAACGTG | | | | | | | | | | |
| Chromosome | start | end | strand | MM | target_seq | PAM | alignment | distance | position | gene name | gene id |
| GU305914.1 | 108184 | 108206 | - | 0 | CACGTTCAACTTCCAGGACG | GGG | \|\|\|\|\|\|\|\|[\|\|\|\|\|\|\|\|\|\|\|\|]PAM | NA | Intergenic | NA | NA |
| T25943 | TTCAACTTCCAGGACGGGGACGG | 1000 | | | | | | | | | |
| Oligo fwd | CACCTTCAACTTCCAGGACGGGGA | | | | | | | | | | |
| Oligo rev | AAACTCCCCGTCCTGGAAGTTGAA | | | | | | | | | | |
| Chromosome | start | end | strand | MM | target_seq | PAM | alignment | distance | position | gene name | gene id |
| GU305914.1 | 108180 | 108202 | - | 0 | TTCAACTTCCAGGACGGGGA | CGG | \|\|\|\|\|\|\|\|[\|\|\|\|\|\|\|\|\|\|\|\|]PAM | NA | Intergenic | NA | NA |
| T25944 | GGACGGCTACCGCGTCTACCAGG | 1000 | | | | | | | | | |
| Oligo fwd | CACCGGACGGCTACCGCGTCTACC | | | | | | | | | | |
| Oligo rev | AAACGGTAGACGCGGTAGCCGTCC | | | | | | | | | | |
| Chromosome | start | end | strand | MM | target_seq | PAM | alignment | distance | position | gene name | gene id |
| GU305914.1 | 108163 | 108185 | - | 0 | GGACGGCTACCGCGTCTACC | AGG | \|\|\|\|\|\|\|\|[\|\|\|\|\|\|\|\|\|\|\|\|]PAM | NA | Intergenic | NA | NA |
| T25945 | CTGTCTCTTCAACACGCACGCGG | 1000 | | | | | | | | | |
| Oligo fwd | CACCCTGTCTCTTCAACACGCACG | | | | | | | | | | |
| Oligo rev | AAACCGTGCGTGTTGAAGAGACAG | | | | | | | | | | |
| Chromosome | start | end | strand | MM | target_seq | PAM | alignment | distance | position | gene name | gene id |
| GU305914.1 | 108133 | 108155 | - | 0 | CTGTCTCTTCAACACGCACG | CGG | \|\|\|\|\|\|\|\|[\|\|\|\|\|\|\|\|\|\|\|\|]PAM | NA | Intergenic | NA | NA |
| GU305914.1 | 13219 | 13241 | - | 4 | CTTTCTCGTCGACACGTACG | CGG | \|\|-\|\|\|\|-[\|\|-\|\|\|\|\|-\|\|\|]PAM | NA | Intergenic | NA | NA |
| T25946 | CAACACGCACGCGGCGAGAGAGG | 1000 | | | | | | | | | |
| Oligo fwd | CACCCAACACGCACGCGGCGAGAG | | | | | | | | | | |
| Oligo rev | AAACCTCTCGCCGCGTGCGTGTTG | | | | | | | | | | |
| Chromosome | start | end | strand | MM | target_seq | PAM | alignment | distance | position | gene name | gene id |
| GU305914.1 | 108124 | 108146 | - | 0 | CAACACGCACGCGGCGAGAG | AGG | \|\|\|\|\|\|\|\|[\|\|\|\|\|\|\|\|\|\|\|\|]PAM | NA | Intergenic | NA | NA |
| T25947 | CGCTAGAGTCGTATCGCAAACGG | 1000 | | | | | | | | | |
| Oligo fwd | CACCCGCTAGAGTCGTATCGCAAA | | | | | | | | | | |
| Oligo rev | AAACTTTGCGATACGACTCTAGCG | | | | | | | | | | |
| Chromosome | start | end | strand | MM | target_seq | PAM | alignment | distance | position | gene name | gene id |
| GU305914.1 | 108074 | 108096 | - | 0 | CGCTAGAGTCGTATCGCAAA | CGG | \|\|\|\|\|\|\|\|[\|\|\|\|\|\|\|\|\|\|\|\|]PAM | NA | Intergenic | NA | NA |
| T25948 | AGTCGTATCGCAAACGGTTTCGG | 1000 | | | | | | | | | |
| Oligo fwd | CACCAGTCGTATCGCAAACGGTTT | | | | | | | | | | |
| Oligo rev | AAACAAACCGTTTGCGATACGACT | | | | | | | | | | |
| Chromosome | start | end | strand | MM | target_seq | PAM | alignment | distance | position | gene name | gene id |
| GU305914.1 | 108068 | 108090 | - | 0 | AGTCGTATCGCAAACGGTTT | CGG | \|\|\|\|\|\|\|\|[\|\|\|\|\|\|\|\|\|\|\|\|]PAM | NA | Intergenic | NA | NA |
| T25949 | GTCGTATCGCAAACGGTTTCGGG | 1000 | | | | | | | | | |
| Oligo fwd | CACCGTCGTATCGCAAACGGTTTC | | | | | | | | | | |
| Oligo rev | AAACGAAACCGTTTGCGATACGAC | | | | | | | | | | |
| Chromosome | start | end | strand | MM | target_seq | PAM | alignment | distance | position | gene name | gene id |
| GU305914.1 | 108067 | 108089 | - | 0 | GTCGTATCGCAAACGGTTTC | GGG | \|\|\|\|\|\|\|\|[\|\|\|\|\|\|\|\|\|\|\|\|]PAM | NA | Intergenic | NA | NA |
| T25950 | TACTTCGTCGTGCCCATCTACGG | 1000 | | | | | | | | | |
| Oligo fwd | CACCTACTTCGTCGTGCCCATCTA | | | | | | | | | | |
| Oligo rev | AAACTAGATGGGCACGACGAAGTA | | | | | | | | | | |
| Chromosome | start | end | strand | MM | target_seq | PAM | alignment | distance | position | gene name | gene id |
| GU305914.1 | 108042 | 108064 | - | 0 | TACTTCGTCGTGCCCATCTA | CGG | \|\|\|\|\|\|\|\|[\|\|\|\|\|\|\|\|\|\|\|\|]PAM | NA | Intergenic | NA | NA |
| T25951 | TGCCCATCTACGGCGCCTACAGG | 1000 | | | | | | | | | |

Oligo fwd CACCTGCCCATCTACGGCGCCTAC
Oligo rev AAACGTAGGCGCCGTAGATGGGCA
Chromosome start end strand MM target_seq PAM alignment distance position gene name gene id
GU305914.1 108032 108054 - 0 TGCCCATCTACGGCGCCTAC AGG ||||||||[||||||||||||]PAM NA Intergenic NA NA

T25952 CATCTACGGCGCCTACAGGCTGG 1000
Oligo fwd CACCCATCTACGGCGCCTACAGGC
Oligo rev AAACGCCTGTAGGCGCCGTAGATG
Chromosome start end strand MM target_seq PAM alignment distance position gene name gene id
GU305914.1 108028 108050 - 0 CATCTACGGCGCCTACAGGC TGG ||||||||[||||||||||||]PAM NA Intergenic NA NA

T25953 CTACGGCGCCTACAGGCTGGTGG 1000
Oligo fwd CACCCTACGGCGCCTACAGGCTGG
Oligo rev AAACCCAGCCTGTAGGCGCCGTAG
Chromosome start end strand MM target_seq PAM alignment distance position gene name gene id
GU305914.1 108025 108047 - 0 CTACGGCGCCTACAGGCTGG TGG ||||||||[||||||||||||]PAM NA Intergenic NA NA

T25954 GCTGGTGGCCCGCTCGCCCACGG 1000
Oligo fwd CACCGCTGGTGGCCCGCTCGCCCA
Oligo rev AAACTGGGCGAGCGGGCCACCAGC
Chromosome start end strand MM target_seq PAM alignment distance position gene name gene id
GU305914.1 108010 108032 - 0 GCTGGTGGCCCGCTCGCCCA CGG ||||||||[||||||||||||]PAM NA Intergenic NA NA

T25955 CTCGCCCACGGCCAAGTATCCGG 1000
Oligo fwd CACCCTCGCCCACGGCCAAGTATC
Oligo rev AAACGATACTTGGCCGTGGGCGAG
Chromosome start end strand MM target_seq PAM alignment distance position gene name gene id
GU305914.1 107998 108020 - 0 CTCGCCCACGGCCAAGTATC CGG ||||||||[||||||||||||]PAM NA Intergenic NA NA

T25956 TCGCCCACGGCCAAGTATCCGGG 1000
Oligo fwd CACCTCGCCCACGGCCAAGTATCC
Oligo rev AAACGGATACTTGGCCGTGGGCGA
Chromosome start end strand MM target_seq PAM alignment distance position gene name gene id
GU305914.1 107997 108019 - 0 TCGCCCACGGCCAAGTATCC GGG ||||||||[||||||||||||]PAM NA Intergenic NA NA

T25957 CCCACGGCCAAGTATCCGGGCGG 1000
Oligo fwd CACCCCCACGGCCAAGTATCCGGG
Oligo rev AAACCCCGGATACTTGGCCGTGGG
Chromosome start end strand MM target_seq PAM alignment distance position gene name gene id
GU305914.1 107994 108016 - 0 CCCACGGCCAAGTATCCGGG CGG ||||||||[||||||||||||]PAM NA Intergenic NA NA

T25958 CGGCGTCCTGAACCCCCCGCCGG 1000
Oligo fwd CACCCGGCGTCCTGAACCCCCCGC
Oligo rev AAACGCGGGGGGTTCAGGACGCCG
Chromosome start end strand MM target_seq PAM alignment distance position gene name gene id
GU305914.1 107974 107996 - 0 CGGCGTCCTGAACCCCCCGC CGG ||||||||[||||||||||||]PAM NA Intergenic NA NA

T25959 GGCCAGCTCAGTGACCATGCAGG 1000
Oligo fwd CACCGGCCAGCTCAGTGACCATGC
Oligo rev AAACGCATGGTCACTGAGCTGGCC
Chromosome start end strand MM target_seq PAM alignment distance position gene name gene id
GU305914.1 107953 107975 - 0 GGCCAGCTCAGTGACCATGC AGG ||||||||[||||||||||||]PAM NA Intergenic NA NA

T25960 CGACGCGACGAACATCCACACGG 1000
Oligo fwd CACCCGACGCGACGAACATCCACA
Oligo rev AAACTGTGGATGTTCGTCGCGTCG
Chromosome start end strand MM target_seq PAM alignment distance position gene name gene id
GU305914.1 107920 107942 - 0 CGACGCGACGAACATCCACA CGG ||||||||[||||||||||||]PAM NA Intergenic NA NA

T25961 GAACATCCACACGGTCGTTCCGG 1000
Oligo fwd CACCGAACATCCACACGGTCGTTC
Oligo rev AAACGAACGACCGTGTGGATGTTC
Chromosome start end strand MM target_seq PAM alignment distance position gene name gene id
GU305914.1 107911 107933 - 0 GAACATCCACACGGTCGTTC CGG ||||||||[||||||||||||]PAM NA Intergenic NA NA

T25962 CTGCGTCATCACCGAGCACCCGG 1000
Oligo fwd CACCCTGCGTCATCACCGAGCACC
Oligo rev AAACGGTGCTCGGTGATGACGCAG
Chromosome start end strand MM target_seq PAM alignment distance position gene name gene id
GU305914.1 107881 107903 - 0 CTGCGTCATCACCGAGCACC CGG ||||||||[||||||||||||]PAM NA Intergenic NA NA

T25963 CCCGGTGATCTTCTCCATGAAGG 1000
Oligo fwd CACCCCCGGTGATCTTCTCCATGA
Oligo rev AAACTCATGGAGAAGATCACCGGG
Chromosome start end strand MM target_seq PAM alignment distance position gene name gene id
GU305914.1 107863 107885 - 0 CCCGGTGATCTTCTCCATGA AGG ||||||||[||||||||||||]PAM NA Intergenic NA NA

T25964 GAAGGTACCGTGCAGCCATCAGG 1000
Oligo fwd CACCGAAGGTACCGTGCAGCCATC
Oligo rev AAACGATGGCTGCACGGTACCTTC
Chromosome start end strand MM target_seq PAM alignment distance position gene name gene id
GU305914.1 107845 107867 - 0 GAAGGTACCGTGCAGCCATC AGG ||||||||[||||||||||||]PAM NA Intergenic NA NA

T25965 GCAGCCATCAGGTCATCACCTGG 1000
Oligo fwd CACCGCAGCCATCAGGTCATCACC
Oligo rev AAACGGTGATGACCTGATGGCTGC
Chromosome start end strand MM target_seq PAM alignment distance position gene name gene id
GU305914.1 107834 107856 - 0 GCAGCCATCAGGTCATCACC TGG ||||||||[||||||||||||]PAM NA Intergenic NA NA

| | | | | | | | | | | | |
|---|---|---|---|---|---|---|---|---|---|---|---|
| T25966 | CCATCAGGTCATCACCTGGACGG | 1000 | | | | | | | | | |
| Oligo fwd | CACCCCATCAGGTCATCACCTGGA | | | | | | | | | | |
| Oligo rev | AAACTCCAGGTGATGACCTGATGG | | | | | | | | | | |
| Chromosome | start | end | strand | MM | target_seq | PAM | alignment | distance | position | gene name | gene id |
| GU305914.1 | 107830 | 107852 | - | 0 | CCATCAGGTCATCACCTGGA | CGG | \|\|\|\|\|\|\|\|[\|\|\|\|\|\|\|\|\|\|\|\|]PAM | NA | Intergenic | NA | NA |
| T25967 | CATCAGGTCATCACCTGGACGGG | 1000 | | | | | | | | | |
| Oligo fwd | CACCCATCAGGTCATCACCTGGAC | | | | | | | | | | |
| Oligo rev | AAACGTCCAGGTGATGACCTGATG | | | | | | | | | | |
| Chromosome | start | end | strand | MM | target_seq | PAM | alignment | distance | position | gene name | gene id |
| GU305914.1 | 107829 | 107851 | - | 0 | CATCAGGTCATCACCTGGAC | GGG | \|\|\|\|\|\|\|\|[\|\|\|\|\|\|\|\|\|\|\|\|]PAM | NA | Intergenic | NA | NA |
| T25968 | CTGGACGGGATATACCGTGACGG | 1000 | | | | | | | | | |
| Oligo fwd | CACCCTGGACGGGATATACCGTGA | | | | | | | | | | |
| Oligo rev | AAACTCACGGTATATCCCGTCCAG | | | | | | | | | | |
| Chromosome | start | end | strand | MM | target_seq | PAM | alignment | distance | position | gene name | gene id |
| GU305914.1 | 107815 | 107837 | - | 0 | CTGGACGGGATATACCGTGA | CGG | \|\|\|\|\|\|\|\|[\|\|\|\|\|\|\|\|\|\|\|\|]PAM | NA | Intergenic | NA | NA |
| T25969 | ACCTACCCGCGACCACACCTCGG | 1000 | | | | | | | | | |
| Oligo fwd | CACCACCTACCCGCGACCACACCT | | | | | | | | | | |
| Oligo rev | AAACAGGTGTGGTCGCGGGTAGGT | | | | | | | | | | |
| Chromosome | start | end | strand | MM | target_seq | PAM | alignment | distance | position | gene name | gene id |
| GU305914.1 | 107755 | 107777 | - | 0 | ACCTACCCGCGACCACACCT | CGG | \|\|\|\|\|\|\|\|[\|\|\|\|\|\|\|\|\|\|\|\|]PAM | NA | Intergenic | NA | NA |
| T25970 | CCACACCTCGGAGAACACGTTGG | 1000 | | | | | | | | | |
| Oligo fwd | CACCCCACACCTCGGAGAACACGT | | | | | | | | | | |
| Oligo rev | AAACACGTGTTCTCCGAGGTGTGG | | | | | | | | | | |
| Chromosome | start | end | strand | MM | target_seq | PAM | alignment | distance | position | gene name | gene id |
| GU305914.1 | 107743 | 107765 | - | 0 | CCACACCTCGGAGAACACGT | TGG | \|\|\|\|\|\|\|\|[\|\|\|\|\|\|\|\|\|\|\|\|]PAM | NA | Intergenic | NA | NA |
| T25971 | AACACGTTGGCCATGTTCTTCGG | 1000 | | | | | | | | | |
| Oligo fwd | CACCAACACGTTGGCCATGTTCTT | | | | | | | | | | |
| Oligo rev | AAACAAGAACATGGCCAACGTGTT | | | | | | | | | | |
| Chromosome | start | end | strand | MM | target_seq | PAM | alignment | distance | position | gene name | gene id |
| GU305914.1 | 107730 | 107752 | - | 0 | AACACGTTGGCCATGTTCTT | CGG | \|\|\|\|\|\|\|\|[\|\|\|\|\|\|\|\|\|\|\|\|]PAM | NA | Intergenic | NA | NA |
| T25972 | GTTCTTCGGAGACGTGAGAGAGG | 1000 | | | | | | | | | |
| Oligo fwd | CACCGTTCTTCGGAGACGTGAGAG | | | | | | | | | | |
| Oligo rev | AAACCTCTCACGTCTCCGAAGAAC | | | | | | | | | | |
| Chromosome | start | end | strand | MM | target_seq | PAM | alignment | distance | position | gene name | gene id |
| GU305914.1 | 107716 | 107738 | - | 0 | GTTCTTCGGAGACGTGAGAG | AGG | \|\|\|\|\|\|\|\|[\|\|\|\|\|\|\|\|\|\|\|\|]PAM | NA | Intergenic | NA | NA |
| T25973 | CTTCGGAGACGTGAGAGAGGTGG | 1000 | | | | | | | | | |
| Oligo fwd | CACCCTTCGGAGACGTGAGAGAGG | | | | | | | | | | |
| Oligo rev | AAACCCTCTCTCACGTCTCCGAAG | | | | | | | | | | |
| Chromosome | start | end | strand | MM | target_seq | PAM | alignment | distance | position | gene name | gene id |
| GU305914.1 | 107713 | 107735 | - | 0 | CTTCGGAGACGTGAGAGAGG | TGG | \|\|\|\|\|\|\|\|[\|\|\|\|\|\|\|\|\|\|\|\|]PAM | NA | Intergenic | NA | NA |
| T25974 | CGTGAGAGAGGTGGATCTGAAGG | 1000 | | | | | | | | | |
| Oligo fwd | CACCCGTGAGAGAGGTGGATCTGA | | | | | | | | | | |
| Oligo rev | AAACTCAGATCCACCTCTCTCACG | | | | | | | | | | |
| Chromosome | start | end | strand | MM | target_seq | PAM | alignment | distance | position | gene name | gene id |
| GU305914.1 | 107704 | 107726 | - | 0 | CGTGAGAGAGGTGGATCTGA | AGG | \|\|\|\|\|\|\|\|[\|\|\|\|\|\|\|\|\|\|\|\|]PAM | NA | Intergenic | NA | NA |
| T25975 | CTGAAGGCGCCGTACACCGTCGG | 1000 | | | | | | | | | |
| Oligo fwd | CACCCTGAAGGCGCCGTACACCGT | | | | | | | | | | |
| Oligo rev | AAACACGGTGTACGGCGCCTTCAG | | | | | | | | | | |
| Chromosome | start | end | strand | MM | target_seq | PAM | alignment | distance | position | gene name | gene id |
| GU305914.1 | 107688 | 107710 | - | 0 | CTGAAGGCGCCGTACACCGT | CGG | \|\|\|\|\|\|\|\|[\|\|\|\|\|\|\|\|\|\|\|\|]PAM | NA | Intergenic | NA | NA |
| T25976 | CCGTCGGAGCCTTCCTCCTGCGG | 1000 | | | | | | | | | |
| Oligo fwd | CACCCCGTCGGAGCCTTCCTCCTG | | | | | | | | | | |
| Oligo rev | AAACCAGGAGGAAGGCTCCGACGG | | | | | | | | | | |
| Chromosome | start | end | strand | MM | target_seq | PAM | alignment | distance | position | gene name | gene id |
| GU305914.1 | 107672 | 107694 | - | 0 | CCGTCGGAGCCTTCCTCCTG | CGG | \|\|\|\|\|\|\|\|[\|\|\|\|\|\|\|\|\|\|\|\|]PAM | NA | Intergenic | NA | NA |
| T25977 | GACGCCCGACCACGACCTCCTGG | 1000 | | | | | | | | | |
| Oligo fwd | CACCGACGCCCGACCACGACCTCC | | | | | | | | | | |
| Oligo rev | AAACGGAGGTCGTGGTCGGGCGTC | | | | | | | | | | |
| Chromosome | start | end | strand | MM | target_seq | PAM | alignment | distance | position | gene name | gene id |
| GU305914.1 | 107647 | 107669 | - | 0 | GACGCCCGACCACGACCTCC | TGG | \|\|\|\|\|\|\|\|[\|\|\|\|\|\|\|\|\|\|\|\|]PAM | NA | Intergenic | NA | NA |
| T25978 | GCCCGACCACGACCTCCTGGTGG | 1000 | | | | | | | | | |
| Oligo fwd | CACCGCCCGACCACGACCTCCTGG | | | | | | | | | | |
| Oligo rev | AAACCCAGGAGGTCGTGGTCGGGC | | | | | | | | | | |
| Chromosome | start | end | strand | MM | target_seq | PAM | alignment | distance | position | gene name | gene id |
| GU305914.1 | 107644 | 107666 | - | 0 | GCCCGACCACGACCTCCTGG | TGG | \|\|\|\|\|\|\|\|[\|\|\|\|\|\|\|\|\|\|\|\|]PAM | NA | Intergenic | NA | NA |
| GU305914.1 | 106378 | 106400 | - | 3 | CCCCGACAACGACATCCTGG | TGG | -\|\|\|\|\|\|-[\|\|\|\|\|-\|\|\|\|\|\|]PAM | NA | Intergenic | NA | NA |
| T25979 | CGACCACGACCTCCTGGTGGTGG | 1000 | | | | | | | | | |
| Oligo fwd | CACCCGACCACGACCTCCTGGTGG | | | | | | | | | | |
| Oligo rev | AAACCCACCAGGAGGTCGTGGTCG | | | | | | | | | | |
| Chromosome | start | end | strand | MM | target_seq | PAM | alignment | distance | position | gene name | gene id |
| GU305914.1 | 107641 | 107663 | - | 0 | CGACCACGACCTCCTGGTGG | TGG | \|\|\|\|\|\|\|\|[\|\|\|\|\|\|\|\|\|\|\|\|]PAM | NA | Intergenic | NA | NA |
| GU305914.1 | 143010 | 143032 | + | 3 | CGACGACGGCCTGCTGGTGG | TGG | \|\|\|\|-\|\|\|[-\|\|\|-\|\|\|\|\|\|\|]PAM | NA | Intergenic | NA | NA |
| T25980 | CTACATGTTCCTCACCGATGTGG | 1000 | | | | | | | | | |
| Oligo fwd | CACCCTACATGTTCCTCACCGATG | | | | | | | | | | |
| Oligo rev | AAACCATCGGTGAGGAACATGTAG | | | | | | | | | | |

Chromosome start end strand MM target_seq PAM alignment distance position gene name gene id
GU305914.1 107593 107615 - 0 CTACATGTTCCTCACCGATG TGG ||||||||[||||||||||||]PAM NA Intergenic NA NA

T25981 GTTCCTGCAACGCACGCTGTCGG 1000
Oligo fwd CACCGTTCCTGCAACGCACGCTGT
Oligo rev AAACACAGCGTGCGTTGCAGGAAC
Chromosome start end strand MM target_seq PAM alignment distance position gene name gene id
GU305914.1 107569 107591 - 0 GTTCCTGCAACGCACGCTGT CGG ||||||||[||||||||||||]PAM NA Intergenic NA NA

T25982 CACGCTGTCGGCCGACTACGCGG 1000
Oligo fwd CACCCACGCTGTCGGCCGACTACG
Oligo rev AAACCGTAGTCGGCCGACAGCGTG
Chromosome start end strand MM target_seq PAM alignment distance position gene name gene id
GU305914.1 107557 107579 - 0 CACGCTGTCGGCCGACTACG CGG ||||||||[||||||||||||]PAM NA Intergenic NA NA

T25983 CGGACACGAGCGTGTGTCTGCGG 1000
Oligo fwd CACCCGGACACGAGCGTGTGTCTG
Oligo rev AAACCAGACACACGCTCGTGTCCG
Chromosome start end strand MM target_seq PAM alignment distance position gene name gene id
GU305914.1 107537 107559 - 0 CGGACACGAGCGTGTGTCTG CGG ||||||||[||||||||||||]PAM NA Intergenic NA NA

T25984 GGACACGAGCGTGTGTCTGCGGG 1000
Oligo fwd CACCGGACACGAGCGTGTGTCTGC
Oligo rev AAACGCAGACACACGCTCGTGTCC
Chromosome start end strand MM target_seq PAM alignment distance position gene name gene id
GU305914.1 107536 107558 - 0 GGACACGAGCGTGTGTCTGC GGG ||||||||[||||||||||||]PAM NA Intergenic NA NA

T25985 TCTGCGGGTGCTGAGCGTTCTGG 1000
Oligo fwd CACCTCTGCGGGTGCTGAGCGTTC
Oligo rev AAACGAACGCTCAGCACCCGCAGA
Chromosome start end strand MM target_seq PAM alignment distance position gene name gene id
GU305914.1 107521 107543 - 0 TCTGCGGGTGCTGAGCGTTC TGG ||||||||[||||||||||||]PAM NA Intergenic NA NA

T25986 GGTGCTGAGCGTTCTGGCGTCGG 1000
Oligo fwd CACCGGTGCTGAGCGTTCTGGCGT
Oligo rev AAACACGCCAGAACGCTCAGCACC
Chromosome start end strand MM target_seq PAM alignment distance position gene name gene id
GU305914.1 107515 107537 - 0 GGTGCTGAGCGTTCTGGCGT CGG ||||||||[||||||||||||]PAM NA Intergenic NA NA

T25987 GAGCGTTCTGGCGTCGGTAGTGG 1000
Oligo fwd CACCGAGCGTTCTGGCGTCGGTAG
Oligo rev AAACCTACCGACGCCAGAACGCTC
Chromosome start end strand MM target_seq PAM alignment distance position gene name gene id
GU305914.1 107509 107531 - 0 GAGCGTTCTGGCGTCGGTAG TGG ||||||||[||||||||||||]PAM NA Intergenic NA NA
GU305914.1 203130 203152 - 4 GAGCGTCATGGCGGCGGTCG TGG ||||||--[|||||-||||-|]PAM NA Intergenic NA NA

T25988 CTGGCGTCGGTAGTGGCCAGAGG 1000
Oligo fwd CACCCTGGCGTCGGTAGTGGCCAG
Oligo rev AAACCTGGCCACTACCGACGCCAG
Chromosome start end strand MM target_seq PAM alignment distance position gene name gene id
GU305914.1 107502 107524 - 0 CTGGCGTCGGTAGTGGCCAG AGG ||||||||[||||||||||||]PAM NA Intergenic NA NA

T25989 GTGGCCAGAGGCAAGCAGTGCGG 1000
Oligo fwd CACCGTGGCCAGAGGCAAGCAGTG
Oligo rev AAACCACTGCTTGCCTCTGGCCAC
Chromosome start end strand MM target_seq PAM alignment distance position gene name gene id
GU305914.1 107490 107512 - 0 GTGGCCAGAGGCAAGCAGTG CGG ||||||||[||||||||||||]PAM NA Intergenic NA NA

T25990 CGGCCTCATCACGCGCGACACGG 1000
Oligo fwd CACCCGGCCTCATCACGCGCGACA
Oligo rev AAACTGTCGCGCGTGATGAGGCCG
Chromosome start end strand MM target_seq PAM alignment distance position gene name gene id
GU305914.1 107470 107492 - 0 CGGCCTCATCACGCGCGACA CGG ||||||||[||||||||||||]PAM NA Intergenic NA NA

T25991 CTATTCTCTGTGTCAGCTCATGG 1000
Oligo fwd CACCCTATTCTCTGTGTCAGCTCA
Oligo rev AAACTGAGCTGACACAGAGAATAG
Chromosome start end strand MM target_seq PAM alignment distance position gene name gene id
GU305914.1 107431 107453 - 0 CTATTCTCTGTGTCAGCTCA TGG ||||||||[||||||||||||]PAM NA Intergenic NA NA

T25992 CTGTGTCAGCTCATGGCGAACGG 1000
Oligo fwd CACCCTGTGTCAGCTCATGGCGAA
Oligo rev AAACTTCGCCATGAGCTGACACAG
Chromosome start end strand MM target_seq PAM alignment distance position gene name gene id
GU305914.1 107424 107446 - 0 CTGTGTCAGCTCATGGCGAA CGG ||||||||[||||||||||||]PAM NA Intergenic NA NA

T25993 TGTGTCAGCTCATGGCGAACGGG 1000
Oligo fwd CACCTGTGTCAGCTCATGGCGAAC
Oligo rev AAACGTTCGCCATGAGCTGACACA
Chromosome start end strand MM target_seq PAM alignment distance position gene name gene id
GU305914.1 107423 107445 - 0 TGTGTCAGCTCATGGCGAAC GGG ||||||||[||||||||||||]PAM NA Intergenic NA NA

T25994 AGCTCATGGCGAACGGGACGAGG 1000
Oligo fwd CACCAGCTCATGGCGAACGGGACG
Oligo rev AAACCGTCCCGTTCGCCATGAGCT
Chromosome start end strand MM target_seq PAM alignment distance position gene name gene id
GU305914.1 107417 107439 - 0 AGCTCATGGCGAACGGGACG AGG ||||||||[||||||||||||]PAM NA Intergenic NA NA

T25995 CGGGACGAGGTATCAGTCGACGG 1000

<table>
<tr><td>Oligo fwd</td><td>CACCCGGGACGAGGTATCAGTCGA</td><td></td><td></td><td></td><td></td><td></td><td></td><td></td><td></td><td></td><td></td></tr>
<tr><td>Oligo rev</td><td>AAACTCGACTGATACCTCGTCCCG</td><td></td><td></td><td></td><td></td><td></td><td></td><td></td><td></td><td></td><td></td></tr>
<tr><td>Chromosome</td><td>start</td><td>end</td><td>strand</td><td>MM</td><td>target_seq</td><td>PAM</td><td>alignment</td><td>distance</td><td>position</td><td>gene name</td><td>gene id</td></tr>
<tr><td>GU305914.1</td><td>107404</td><td>107426</td><td>-</td><td>0</td><td>CGGGACGAGGTATCAGTCGA</td><td>CGG</td><td>||||||||[||||||||||||]PAM</td><td>NA</td><td>Intergenic</td><td>NA</td><td>NA</td></tr>
<tr><td>T25996</td><td>CTCCCGTCTCGACCGCCCTCTGG</td><td>1000</td><td></td><td></td><td></td><td></td><td></td><td></td><td></td><td></td><td></td></tr>
<tr><td>Oligo fwd</td><td>CACCCTCCCGTCTCGACCGCCCTC</td><td></td><td></td><td></td><td></td><td></td><td></td><td></td><td></td><td></td><td></td></tr>
<tr><td>Oligo rev</td><td>AAACGAGGGCGGTCGAGACGGGAG</td><td></td><td></td><td></td><td></td><td></td><td></td><td></td><td></td><td></td><td></td></tr>
<tr><td>Chromosome</td><td>start</td><td>end</td><td>strand</td><td>MM</td><td>target_seq</td><td>PAM</td><td>alignment</td><td>distance</td><td>position</td><td>gene name</td><td>gene id</td></tr>
<tr><td>GU305914.1</td><td>107381</td><td>107403</td><td>-</td><td>0</td><td>CTCCCGTCTCGACCGCCCTC</td><td>TGG</td><td>||||||||[||||||||||||]PAM</td><td>NA</td><td>Intergenic</td><td>NA</td><td>NA</td></tr>
<tr><td>T25997</td><td>CCGTCTCGACCGCCCTCTGGCGG</td><td>1000</td><td></td><td></td><td></td><td></td><td></td><td></td><td></td><td></td><td></td></tr>
<tr><td>Oligo fwd</td><td>CACCCCGTCTCGACCGCCCTCTGG</td><td></td><td></td><td></td><td></td><td></td><td></td><td></td><td></td><td></td><td></td></tr>
<tr><td>Oligo rev</td><td>AAACCCAGAGGGCGGTCGAGACGG</td><td></td><td></td><td></td><td></td><td></td><td></td><td></td><td></td><td></td><td></td></tr>
<tr><td>Chromosome</td><td>start</td><td>end</td><td>strand</td><td>MM</td><td>target_seq</td><td>PAM</td><td>alignment</td><td>distance</td><td>position</td><td>gene name</td><td>gene id</td></tr>
<tr><td>GU305914.1</td><td>107378</td><td>107400</td><td>-</td><td>0</td><td>CCGTCTCGACCGCCCTCTGG</td><td>CGG</td><td>||||||||[||||||||||||]PAM</td><td>NA</td><td>Intergenic</td><td>NA</td><td>NA</td></tr>
<tr><td>T25998</td><td>CCTCTGGCGGCAGAGCGAATTGG</td><td>1000</td><td></td><td></td><td></td><td></td><td></td><td></td><td></td><td></td><td></td></tr>
<tr><td>Oligo fwd</td><td>CACCCCTCTGGCGGCAGAGCGAAT</td><td></td><td></td><td></td><td></td><td></td><td></td><td></td><td></td><td></td><td></td></tr>
<tr><td>Oligo rev</td><td>AAACATTCGCTCTGCCGCCAGAGG</td><td></td><td></td><td></td><td></td><td></td><td></td><td></td><td></td><td></td><td></td></tr>
<tr><td>Chromosome</td><td>start</td><td>end</td><td>strand</td><td>MM</td><td>target_seq</td><td>PAM</td><td>alignment</td><td>distance</td><td>position</td><td>gene name</td><td>gene id</td></tr>
<tr><td>GU305914.1</td><td>107365</td><td>107387</td><td>-</td><td>0</td><td>CCTCTGGCGGCAGAGCGAAT</td><td>TGG</td><td>||||||||[||||||||||||]PAM</td><td>NA</td><td>Intergenic</td><td>NA</td><td>NA</td></tr>
<tr><td>T25999</td><td>CAGAGCGAATTGGAGCTGTTCGG</td><td>1000</td><td></td><td></td><td></td><td></td><td></td><td></td><td></td><td></td><td></td></tr>
<tr><td>Oligo fwd</td><td>CACCCAGAGCGAATTGGAGCTGTT</td><td></td><td></td><td></td><td></td><td></td><td></td><td></td><td></td><td></td><td></td></tr>
<tr><td>Oligo rev</td><td>AAACAACAGCTCCAATTCGCTCTG</td><td></td><td></td><td></td><td></td><td></td><td></td><td></td><td></td><td></td><td></td></tr>
<tr><td>Chromosome</td><td>start</td><td>end</td><td>strand</td><td>MM</td><td>target_seq</td><td>PAM</td><td>alignment</td><td>distance</td><td>position</td><td>gene name</td><td>gene id</td></tr>
<tr><td>GU305914.1</td><td>107355</td><td>107377</td><td>-</td><td>0</td><td>CAGAGCGAATTGGAGCTGTT</td><td>CGG</td><td>||||||||[||||||||||||]PAM</td><td>NA</td><td>Intergenic</td><td>NA</td><td>NA</td></tr>
<tr><td>T26000</td><td>CACGACACCGAACCCGACGCCGG</td><td>1000</td><td></td><td></td><td></td><td></td><td></td><td></td><td></td><td></td><td></td></tr>
<tr><td>Oligo fwd</td><td>CACCCACGACACCGAACCCGACGC</td><td></td><td></td><td></td><td></td><td></td><td></td><td></td><td></td><td></td><td></td></tr>
<tr><td>Oligo rev</td><td>AAACGCGTCGGGTTCGGTGTCGTG</td><td></td><td></td><td></td><td></td><td></td><td></td><td></td><td></td><td></td><td></td></tr>
<tr><td>Chromosome</td><td>start</td><td>end</td><td>strand</td><td>MM</td><td>target_seq</td><td>PAM</td><td>alignment</td><td>distance</td><td>position</td><td>gene name</td><td>gene id</td></tr>
<tr><td>GU305914.1</td><td>107305</td><td>107327</td><td>-</td><td>0</td><td>CACGACACCGAACCCGACGC</td><td>CGG</td><td>||||||||[||||||||||||]PAM</td><td>NA</td><td>Intergenic</td><td>NA</td><td>NA</td></tr>
<tr><td>T26001</td><td>CGACGCCGGCGTTTCAGACGAGG</td><td>1000</td><td></td><td></td><td></td><td></td><td></td><td></td><td></td><td></td><td></td></tr>
<tr><td>Oligo fwd</td><td>CACCCGACGCCGGCGTTTCAGACG</td><td></td><td></td><td></td><td></td><td></td><td></td><td></td><td></td><td></td><td></td></tr>
<tr><td>Oligo rev</td><td>AAACCGTCTGAAACGCCGGCGTCG</td><td></td><td></td><td></td><td></td><td></td><td></td><td></td><td></td><td></td><td></td></tr>
<tr><td>Chromosome</td><td>start</td><td>end</td><td>strand</td><td>MM</td><td>target_seq</td><td>PAM</td><td>alignment</td><td>distance</td><td>position</td><td>gene name</td><td>gene id</td></tr>
<tr><td>GU305914.1</td><td>107291</td><td>107313</td><td>-</td><td>0</td><td>CGACGCCGGCGTTTCAGACG</td><td>AGG</td><td>||||||||[||||||||||||]PAM</td><td>NA</td><td>Intergenic</td><td>NA</td><td>NA</td></tr>
<tr><td>T26002</td><td>CAAGCAACTCGATCGACGTCCGG</td><td>1000</td><td></td><td></td><td></td><td></td><td></td><td></td><td></td><td></td><td></td></tr>
<tr><td>Oligo fwd</td><td>CACCCAAGCAACTCGATCGACGTC</td><td></td><td></td><td></td><td></td><td></td><td></td><td></td><td></td><td></td><td></td></tr>
<tr><td>Oligo rev</td><td>AAACGACGTCGATCGAGTTGCTTG</td><td></td><td></td><td></td><td></td><td></td><td></td><td></td><td></td><td></td><td></td></tr>
<tr><td>Chromosome</td><td>start</td><td>end</td><td>strand</td><td>MM</td><td>target_seq</td><td>PAM</td><td>alignment</td><td>distance</td><td>position</td><td>gene name</td><td>gene id</td></tr>
<tr><td>GU305914.1</td><td>107234</td><td>107256</td><td>-</td><td>0</td><td>CAAGCAACTCGATCGACGTC</td><td>CGG</td><td>||||||||[||||||||||||]PAM</td><td>NA</td><td>Intergenic</td><td>NA</td><td>NA</td></tr>
<tr><td>T26003</td><td>AAGCAACTCGATCGACGTCCGGG</td><td>1000</td><td></td><td></td><td></td><td></td><td></td><td></td><td></td><td></td><td></td></tr>
<tr><td>Oligo fwd</td><td>CACCAAGCAACTCGATCGACGTCC</td><td></td><td></td><td></td><td></td><td></td><td></td><td></td><td></td><td></td><td></td></tr>
<tr><td>Oligo rev</td><td>AAACGGACGTCGATCGAGTTGCTT</td><td></td><td></td><td></td><td></td><td></td><td></td><td></td><td></td><td></td><td></td></tr>
<tr><td>Chromosome</td><td>start</td><td>end</td><td>strand</td><td>MM</td><td>target_seq</td><td>PAM</td><td>alignment</td><td>distance</td><td>position</td><td>gene name</td><td>gene id</td></tr>
<tr><td>GU305914.1</td><td>107233</td><td>107255</td><td>-</td><td>0</td><td>AAGCAACTCGATCGACGTCC</td><td>GGG</td><td>||||||||[||||||||||||]PAM</td><td>NA</td><td>Intergenic</td><td>NA</td><td>NA</td></tr>
<tr><td>T26004</td><td>CTCGATCGACGTCCGGGTATTGG</td><td>1000</td><td></td><td></td><td></td><td></td><td></td><td></td><td></td><td></td><td></td></tr>
<tr><td>Oligo fwd</td><td>CACCCTCGATCGACGTCCGGGTAT</td><td></td><td></td><td></td><td></td><td></td><td></td><td></td><td></td><td></td><td></td></tr>
<tr><td>Oligo rev</td><td>AAACATACCCGGACGTCGATCGAG</td><td></td><td></td><td></td><td></td><td></td><td></td><td></td><td></td><td></td><td></td></tr>
<tr><td>Chromosome</td><td>start</td><td>end</td><td>strand</td><td>MM</td><td>target_seq</td><td>PAM</td><td>alignment</td><td>distance</td><td>position</td><td>gene name</td><td>gene id</td></tr>
<tr><td>GU305914.1</td><td>107227</td><td>107249</td><td>-</td><td>0</td><td>CTCGATCGACGTCCGGGTAT</td><td>TGG</td><td>||||||||[||||||||||||]PAM</td><td>NA</td><td>Intergenic</td><td>NA</td><td>NA</td></tr>
<tr><td>T26005</td><td>GATCGACGTCCGGGTATTGGCGG</td><td>1000</td><td></td><td></td><td></td><td></td><td></td><td></td><td></td><td></td><td></td></tr>
<tr><td>Oligo fwd</td><td>CACCGATCGACGTCCGGGTATTGG</td><td></td><td></td><td></td><td></td><td></td><td></td><td></td><td></td><td></td><td></td></tr>
<tr><td>Oligo rev</td><td>AAACCCAATACCCGGACGTCGATC</td><td></td><td></td><td></td><td></td><td></td><td></td><td></td><td></td><td></td><td></td></tr>
<tr><td>Chromosome</td><td>start</td><td>end</td><td>strand</td><td>MM</td><td>target_seq</td><td>PAM</td><td>alignment</td><td>distance</td><td>position</td><td>gene name</td><td>gene id</td></tr>
<tr><td>GU305914.1</td><td>107224</td><td>107246</td><td>-</td><td>0</td><td>GATCGACGTCCGGGTATTGG</td><td>CGG</td><td>||||||||[||||||||||||]PAM</td><td>NA</td><td>Intergenic</td><td>NA</td><td>NA</td></tr>
<tr><td>T26006</td><td>TTGGCGGCCACGTACTCGTCAGG</td><td>1000</td><td></td><td></td><td></td><td></td><td></td><td></td><td></td><td></td><td></td></tr>
<tr><td>Oligo fwd</td><td>CACCTTGGCGGCCACGTACTCGTC</td><td></td><td></td><td></td><td></td><td></td><td></td><td></td><td></td><td></td><td></td></tr>
<tr><td>Oligo rev</td><td>AAACGACGAGTACGTGGCCGCCAA</td><td></td><td></td><td></td><td></td><td></td><td></td><td></td><td></td><td></td><td></td></tr>
<tr><td>Chromosome</td><td>start</td><td>end</td><td>strand</td><td>MM</td><td>target_seq</td><td>PAM</td><td>alignment</td><td>distance</td><td>position</td><td>gene name</td><td>gene id</td></tr>
<tr><td>GU305914.1</td><td>107208</td><td>107230</td><td>-</td><td>0</td><td>TTGGCGGCCACGTACTCGTC</td><td>AGG</td><td>||||||||[||||||||||||]PAM</td><td>NA</td><td>Intergenic</td><td>NA</td><td>NA</td></tr>
<tr><td>T26007</td><td>TGGCGGCCACGTACTCGTCAGGG</td><td>1000</td><td></td><td></td><td></td><td></td><td></td><td></td><td></td><td></td><td></td></tr>
<tr><td>Oligo fwd</td><td>CACCTGGCGGCCACGTACTCGTCA</td><td></td><td></td><td></td><td></td><td></td><td></td><td></td><td></td><td></td><td></td></tr>
<tr><td>Oligo rev</td><td>AAACTGACGAGTACGTGGCCGCCA</td><td></td><td></td><td></td><td></td><td></td><td></td><td></td><td></td><td></td><td></td></tr>
<tr><td>Chromosome</td><td>start</td><td>end</td><td>strand</td><td>MM</td><td>target_seq</td><td>PAM</td><td>alignment</td><td>distance</td><td>position</td><td>gene name</td><td>gene id</td></tr>
<tr><td>GU305914.1</td><td>107207</td><td>107229</td><td>-</td><td>0</td><td>TGGCGGCCACGTACTCGTCA</td><td>GGG</td><td>||||||||[||||||||||||]PAM</td><td>NA</td><td>Intergenic</td><td>NA</td><td>NA</td></tr>
<tr><td>T26008</td><td>GTACTCGTCAGGGATGCACGCGG</td><td>1000</td><td></td><td></td><td></td><td></td><td></td><td></td><td></td><td></td><td></td></tr>
<tr><td>Oligo fwd</td><td>CACCGTACTCGTCAGGGATGCACG</td><td></td><td></td><td></td><td></td><td></td><td></td><td></td><td></td><td></td><td></td></tr>
<tr><td>Oligo rev</td><td>AAACCGTGCATCCCTGACGAGTAC</td><td></td><td></td><td></td><td></td><td></td><td></td><td></td><td></td><td></td><td></td></tr>
<tr><td>Chromosome</td><td>start</td><td>end</td><td>strand</td><td>MM</td><td>target_seq</td><td>PAM</td><td>alignment</td><td>distance</td><td>position</td><td>gene name</td><td>gene id</td></tr>
<tr><td>GU305914.1</td><td>107197</td><td>107219</td><td>-</td><td>0</td><td>GTACTCGTCAGGGATGCACG</td><td>CGG</td><td>||||||||[||||||||||||]PAM</td><td>NA</td><td>Intergenic</td><td>NA</td><td>NA</td></tr>
<tr><td>T26009</td><td>AGGGATGCACGCGGCGAGCATGG</td><td>1000</td><td></td><td></td><td></td><td></td><td></td><td></td><td></td><td></td><td></td></tr>
<tr><td>Oligo fwd</td><td>CACCAGGGATGCACGCGGCGAGCA</td><td></td><td></td><td></td><td></td><td></td><td></td><td></td><td></td><td></td><td></td></tr>
<tr><td>Oligo rev</td><td>AAACTGCTCGCCGCGTGCATCCCT</td><td></td><td></td><td></td><td></td><td></td><td></td><td></td><td></td><td></td><td></td></tr>
<tr><td>Chromosome</td><td>start</td><td>end</td><td>strand</td><td>MM</td><td>target_seq</td><td>PAM</td><td>alignment</td><td>distance</td><td>position</td><td>gene name</td><td>gene id</td></tr>
<tr><td>GU305914.1</td><td>107188</td><td>107210</td><td>-</td><td>0</td><td>AGGGATGCACGCGGCGAGCA</td><td>TGG</td><td>||||||||[||||||||||||]PAM</td><td>NA</td><td>Intergenic</td><td>NA</td><td>NA</td></tr>
</table>

<table>
<tr><td>T26010</td><td>CATCCCGCCGAATGTCAACACGG</td><td>1000</td><td></td><td></td><td></td><td></td><td></td><td></td><td></td><td></td><td></td></tr>
<tr><td>Oligo fwd</td><td>CACCCATCCCGCCGAATGTCAACA</td><td></td><td></td><td></td><td></td><td></td><td></td><td></td><td></td><td></td><td></td></tr>
<tr><td>Oligo rev</td><td>AAACTGTTGACATTCGGCGGGATG</td><td></td><td></td><td></td><td></td><td></td><td></td><td></td><td></td><td></td><td></td></tr>
<tr><td>Chromosome</td><td>start</td><td>end</td><td>strand</td><td>MM</td><td>target_seq</td><td>PAM</td><td>alignment</td><td>distance</td><td>position</td><td>gene name</td><td>gene id</td></tr>
<tr><td>GU305914.1</td><td>107134</td><td>107156</td><td>-</td><td>0</td><td>CATCCCGCCGAATGTCAACA</td><td>CGG</td><td>||||||||[||||||||||||]PAM</td><td>NA</td><td>Intergenic</td><td>NA</td><td>NA</td></tr>
<tr><td>T26011</td><td>ACAGAAGTTACTGTTCACCACGG</td><td>1000</td><td></td><td></td><td></td><td></td><td></td><td></td><td></td><td></td><td></td></tr>
<tr><td>Oligo fwd</td><td>CACCACAGAAGTTACTGTTCACCA</td><td></td><td></td><td></td><td></td><td></td><td></td><td></td><td></td><td></td><td></td></tr>
<tr><td>Oligo rev</td><td>AAACTGGTGAACAGTAACTTCTGT</td><td></td><td></td><td></td><td></td><td></td><td></td><td></td><td></td><td></td><td></td></tr>
<tr><td>Chromosome</td><td>start</td><td>end</td><td>strand</td><td>MM</td><td>target_seq</td><td>PAM</td><td>alignment</td><td>distance</td><td>position</td><td>gene name</td><td>gene id</td></tr>
<tr><td>GU305914.1</td><td>107101</td><td>107123</td><td>-</td><td>0</td><td>ACAGAAGTTACTGTTCACCA</td><td>CGG</td><td>||||||||[||||||||||||]PAM</td><td>NA</td><td>Intergenic</td><td>NA</td><td>NA</td></tr>
<tr><td>T26012</td><td>AGCTTGAAGATCCCGCTATCCGG</td><td>1000</td><td></td><td></td><td></td><td></td><td></td><td></td><td></td><td></td><td></td></tr>
<tr><td>Oligo fwd</td><td>CACCAGCTTGAAGATCCCGCTATC</td><td></td><td></td><td></td><td></td><td></td><td></td><td></td><td></td><td></td><td></td></tr>
<tr><td>Oligo rev</td><td>AAACGATAGCGGGATCTTCAAGCT</td><td></td><td></td><td></td><td></td><td></td><td></td><td></td><td></td><td></td><td></td></tr>
<tr><td>Chromosome</td><td>start</td><td>end</td><td>strand</td><td>MM</td><td>target_seq</td><td>PAM</td><td>alignment</td><td>distance</td><td>position</td><td>gene name</td><td>gene id</td></tr>
<tr><td>GU305914.1</td><td>107061</td><td>107083</td><td>-</td><td>0</td><td>AGCTTGAAGATCCCGCTATC</td><td>CGG</td><td>||||||||[||||||||||||]PAM</td><td>NA</td><td>Intergenic</td><td>NA</td><td>NA</td></tr>
<tr><td>T26013</td><td>CTTGATCCGCGTCGACCTAACGG</td><td>1000</td><td></td><td></td><td></td><td></td><td></td><td></td><td></td><td></td><td></td></tr>
<tr><td>Oligo fwd</td><td>CACCCTTGATCCGCGTCGACCTAA</td><td></td><td></td><td></td><td></td><td></td><td></td><td></td><td></td><td></td><td></td></tr>
<tr><td>Oligo rev</td><td>AAACTTAGGTCGACGCGGATCAAG</td><td></td><td></td><td></td><td></td><td></td><td></td><td></td><td></td><td></td><td></td></tr>
<tr><td>Chromosome</td><td>start</td><td>end</td><td>strand</td><td>MM</td><td>target_seq</td><td>PAM</td><td>alignment</td><td>distance</td><td>position</td><td>gene name</td><td>gene id</td></tr>
<tr><td>GU305914.1</td><td>107023</td><td>107045</td><td>-</td><td>0</td><td>CTTGATCCGCGTCGACCTAA</td><td>CGG</td><td>||||||||[||||||||||||]PAM</td><td>NA</td><td>Intergenic</td><td>NA</td><td>NA</td></tr>
<tr><td>T26014</td><td>ACAACTGAACGAGAGCAGCGTGG</td><td>1000</td><td></td><td></td><td></td><td></td><td></td><td></td><td></td><td></td><td></td></tr>
<tr><td>Oligo fwd</td><td>CACCACAACTGAACGAGAGCAGCG</td><td></td><td></td><td></td><td></td><td></td><td></td><td></td><td></td><td></td><td></td></tr>
<tr><td>Oligo rev</td><td>AAACCGCTGCTCTCGTTCAGTTGT</td><td></td><td></td><td></td><td></td><td></td><td></td><td></td><td></td><td></td><td></td></tr>
<tr><td>Chromosome</td><td>start</td><td>end</td><td>strand</td><td>MM</td><td>target_seq</td><td>PAM</td><td>alignment</td><td>distance</td><td>position</td><td>gene name</td><td>gene id</td></tr>
<tr><td>GU305914.1</td><td>106993</td><td>107015</td><td>-</td><td>0</td><td>ACAACTGAACGAGAGCAGCG</td><td>TGG</td><td>||||||||[||||||||||||]PAM</td><td>NA</td><td>Intergenic</td><td>NA</td><td>NA</td></tr>
<tr><td>T26015</td><td>ACGAGAGCAGCGTGGCGAGACGG</td><td>1000</td><td></td><td></td><td></td><td></td><td></td><td></td><td></td><td></td><td></td></tr>
<tr><td>Oligo fwd</td><td>CACCACGAGAGCAGCGTGGCGAGA</td><td></td><td></td><td></td><td></td><td></td><td></td><td></td><td></td><td></td><td></td></tr>
<tr><td>Oligo rev</td><td>AAACTCTCGCCACGCTGCTCTCGT</td><td></td><td></td><td></td><td></td><td></td><td></td><td></td><td></td><td></td><td></td></tr>
<tr><td>Chromosome</td><td>start</td><td>end</td><td>strand</td><td>MM</td><td>target_seq</td><td>PAM</td><td>alignment</td><td>distance</td><td>position</td><td>gene name</td><td>gene id</td></tr>
<tr><td>GU305914.1</td><td>106985</td><td>107007</td><td>-</td><td>0</td><td>ACGAGAGCAGCGTGGCGAGA</td><td>CGG</td><td>||||||||[||||||||||||]PAM</td><td>NA</td><td>Intergenic</td><td>NA</td><td>NA</td></tr>
<tr><td>T26016</td><td>CGCCGCGAGAACAGATCTCATGG</td><td>1000</td><td></td><td></td><td></td><td></td><td></td><td></td><td></td><td></td><td></td></tr>
<tr><td>Oligo fwd</td><td>CACCCGCCGCGAGAACAGATCTCA</td><td></td><td></td><td></td><td></td><td></td><td></td><td></td><td></td><td></td><td></td></tr>
<tr><td>Oligo rev</td><td>AAACTGAGATCTGTTCTCGCGGCG</td><td></td><td></td><td></td><td></td><td></td><td></td><td></td><td></td><td></td><td></td></tr>
<tr><td>Chromosome</td><td>start</td><td>end</td><td>strand</td><td>MM</td><td>target_seq</td><td>PAM</td><td>alignment</td><td>distance</td><td>position</td><td>gene name</td><td>gene id</td></tr>
<tr><td>GU305914.1</td><td>106934</td><td>106956</td><td>-</td><td>0</td><td>CGCCGCGAGAACAGATCTCA</td><td>TGG</td><td>||||||||[||||||||||||]PAM</td><td>NA</td><td>Intergenic</td><td>NA</td><td>NA</td></tr>
<tr><td>T26017</td><td>GCCGCGAGAACAGATCTCATGGG</td><td>1000</td><td></td><td></td><td></td><td></td><td></td><td></td><td></td><td></td><td></td></tr>
<tr><td>Oligo fwd</td><td>CACCGCCGCGAGAACAGATCTCAT</td><td></td><td></td><td></td><td></td><td></td><td></td><td></td><td></td><td></td><td></td></tr>
<tr><td>Oligo rev</td><td>AAACATGAGATCTGTTCTCGCGGC</td><td></td><td></td><td></td><td></td><td></td><td></td><td></td><td></td><td></td><td></td></tr>
<tr><td>Chromosome</td><td>start</td><td>end</td><td>strand</td><td>MM</td><td>target_seq</td><td>PAM</td><td>alignment</td><td>distance</td><td>position</td><td>gene name</td><td>gene id</td></tr>
<tr><td>GU305914.1</td><td>106933</td><td>106955</td><td>-</td><td>0</td><td>GCCGCGAGAACAGATCTCAT</td><td>GGG</td><td>||||||||[||||||||||||]PAM</td><td>NA</td><td>Intergenic</td><td>NA</td><td>NA</td></tr>
<tr><td>T26018</td><td>CCGCGAGAACAGATCTCATGGGG</td><td>1000</td><td></td><td></td><td></td><td></td><td></td><td></td><td></td><td></td><td></td></tr>
<tr><td>Oligo fwd</td><td>CACCCCGCGAGAACAGATCTCATG</td><td></td><td></td><td></td><td></td><td></td><td></td><td></td><td></td><td></td><td></td></tr>
<tr><td>Oligo rev</td><td>AAACCATGAGATCTGTTCTCGCGG</td><td></td><td></td><td></td><td></td><td></td><td></td><td></td><td></td><td></td><td></td></tr>
<tr><td>Chromosome</td><td>start</td><td>end</td><td>strand</td><td>MM</td><td>target_seq</td><td>PAM</td><td>alignment</td><td>distance</td><td>position</td><td>gene name</td><td>gene id</td></tr>
<tr><td>GU305914.1</td><td>106932</td><td>106954</td><td>-</td><td>0</td><td>CCGCGAGAACAGATCTCATG</td><td>GGG</td><td>||||||||[||||||||||||]PAM</td><td>NA</td><td>Intergenic</td><td>NA</td><td>NA</td></tr>
<tr><td>T26019</td><td>CGCGAGAACAGATCTCATGGGGG</td><td>1000</td><td></td><td></td><td></td><td></td><td></td><td></td><td></td><td></td><td></td></tr>
<tr><td>Oligo fwd</td><td>CACCCGCGAGAACAGATCTCATGG</td><td></td><td></td><td></td><td></td><td></td><td></td><td></td><td></td><td></td><td></td></tr>
<tr><td>Oligo rev</td><td>AAACCCATGAGATCTGTTCTCGCG</td><td></td><td></td><td></td><td></td><td></td><td></td><td></td><td></td><td></td><td></td></tr>
<tr><td>Chromosome</td><td>start</td><td>end</td><td>strand</td><td>MM</td><td>target_seq</td><td>PAM</td><td>alignment</td><td>distance</td><td>position</td><td>gene name</td><td>gene id</td></tr>
<tr><td>GU305914.1</td><td>106931</td><td>106953</td><td>-</td><td>0</td><td>CGCGAGAACAGATCTCATGG</td><td>GGG</td><td>||||||||[||||||||||||]PAM</td><td>NA</td><td>Intergenic</td><td>NA</td><td>NA</td></tr>
<tr><td>T26020</td><td>GCGAGAACAGATCTCATGGGGGG</td><td>1000</td><td></td><td></td><td></td><td></td><td></td><td></td><td></td><td></td><td></td></tr>
<tr><td>Oligo fwd</td><td>CACCGCGAGAACAGATCTCATGGG</td><td></td><td></td><td></td><td></td><td></td><td></td><td></td><td></td><td></td><td></td></tr>
<tr><td>Oligo rev</td><td>AAACCCCATGAGATCTGTTCTCGC</td><td></td><td></td><td></td><td></td><td></td><td></td><td></td><td></td><td></td><td></td></tr>
<tr><td>Chromosome</td><td>start</td><td>end</td><td>strand</td><td>MM</td><td>target_seq</td><td>PAM</td><td>alignment</td><td>distance</td><td>position</td><td>gene name</td><td>gene id</td></tr>
<tr><td>GU305914.1</td><td>106930</td><td>106952</td><td>-</td><td>0</td><td>GCGAGAACAGATCTCATGGG</td><td>GGG</td><td>||||||||[||||||||||||]PAM</td><td>NA</td><td>Intergenic</td><td>NA</td><td>NA</td></tr>
<tr><td>T26021</td><td>AACTGCTCATGAACCCACAACGG</td><td>1000</td><td></td><td></td><td></td><td></td><td></td><td></td><td></td><td></td><td></td></tr>
<tr><td>Oligo fwd</td><td>CACCAACTGCTCATGAACCCACAA</td><td></td><td></td><td></td><td></td><td></td><td></td><td></td><td></td><td></td><td></td></tr>
<tr><td>Oligo rev</td><td>AAACTTGTGGGTTCATGAGCAGTT</td><td></td><td></td><td></td><td></td><td></td><td></td><td></td><td></td><td></td><td></td></tr>
<tr><td>Chromosome</td><td>start</td><td>end</td><td>strand</td><td>MM</td><td>target_seq</td><td>PAM</td><td>alignment</td><td>distance</td><td>position</td><td>gene name</td><td>gene id</td></tr>
<tr><td>GU305914.1</td><td>106907</td><td>106929</td><td>-</td><td>0</td><td>AACTGCTCATGAACCCACAA</td><td>CGG</td><td>||||||||[||||||||||||]PAM</td><td>NA</td><td>Intergenic</td><td>NA</td><td>NA</td></tr>
<tr><td>T26022</td><td>ACTGCTCATGAACCCACAACGGG</td><td>1000</td><td></td><td></td><td></td><td></td><td></td><td></td><td></td><td></td><td></td></tr>
<tr><td>Oligo fwd</td><td>CACCACTGCTCATGAACCCACAAC</td><td></td><td></td><td></td><td></td><td></td><td></td><td></td><td></td><td></td><td></td></tr>
<tr><td>Oligo rev</td><td>AAACGTTGTGGGTTCATGAGCAGT</td><td></td><td></td><td></td><td></td><td></td><td></td><td></td><td></td><td></td><td></td></tr>
<tr><td>Chromosome</td><td>start</td><td>end</td><td>strand</td><td>MM</td><td>target_seq</td><td>PAM</td><td>alignment</td><td>distance</td><td>position</td><td>gene name</td><td>gene id</td></tr>
<tr><td>GU305914.1</td><td>106906</td><td>106928</td><td>-</td><td>0</td><td>ACTGCTCATGAACCCACAAC</td><td>GGG</td><td>||||||||[||||||||||||]PAM</td><td>NA</td><td>Intergenic</td><td>NA</td><td>NA</td></tr>
<tr><td>T26023</td><td>CTGCTCATGAACCCACAACGGGG</td><td>1000</td><td></td><td></td><td></td><td></td><td></td><td></td><td></td><td></td><td></td></tr>
<tr><td>Oligo fwd</td><td>CACCCTGCTCATGAACCCACAACG</td><td></td><td></td><td></td><td></td><td></td><td></td><td></td><td></td><td></td><td></td></tr>
<tr><td>Oligo rev</td><td>AAACCGTTGTGGGTTCATGAGCAG</td><td></td><td></td><td></td><td></td><td></td><td></td><td></td><td></td><td></td><td></td></tr>
<tr><td>Chromosome</td><td>start</td><td>end</td><td>strand</td><td>MM</td><td>target_seq</td><td>PAM</td><td>alignment</td><td>distance</td><td>position</td><td>gene name</td><td>gene id</td></tr>
<tr><td>GU305914.1</td><td>106905</td><td>106927</td><td>-</td><td>0</td><td>CTGCTCATGAACCCACAACG</td><td>GGG</td><td>||||||||[||||||||||||]PAM</td><td>NA</td><td>Intergenic</td><td>NA</td><td>NA</td></tr>
<tr><td>T26024</td><td>CCCACAACGGGGCGCACCGTCGG</td><td>1000</td><td></td><td></td><td></td><td></td><td></td><td></td><td></td><td></td><td></td></tr>
<tr><td>Oligo fwd</td><td>CACCCCCACAACGGGGCGCACCGT</td><td></td><td></td><td></td><td></td><td></td><td></td><td></td><td></td><td></td><td></td></tr>
<tr><td>Oligo rev</td><td>AAACACGGTGCGCCCCGTTGTGGG</td><td></td><td></td><td></td><td></td><td></td><td></td><td></td><td></td><td></td><td></td></tr>
<tr><td>Chromosome</td><td>start</td><td>end</td><td>strand</td><td>MM</td><td>target_seq</td><td>PAM</td><td>alignment</td><td>distance</td><td>position</td><td>gene name</td><td>gene id</td></tr>
<tr><td>GU305914.1</td><td>106894</td><td>106916</td><td>-</td><td>0</td><td>CCCACAACGGGGCGCACCGT</td><td>CGG</td><td>||||||||[||||||||||||]PAM</td><td>NA</td><td>Intergenic</td><td>NA</td><td>NA</td></tr>
</table>

T26025 GATCTACAGCCCGTGCGTATCGG 1000
Oligo fwd CACCGATCTACAGCCCGTGCGTAT
Oligo rev AAACATACGCACGGGCTGTAGATC
Chromosome start end strand MM target_seq PAM alignment distance position gene name gene id
GU305914.1 106870 106892 - 0 GATCTACAGCCCGTGCGTAT CGG ||||||||[||||||||||||]PAM NA Intergenic NA NA

T26026 ATCTACAGCCCGTGCGTATCGGG 1000
Oligo fwd CACCATCTACAGCCCGTGCGTATC
Oligo rev AAACGATACGCACGGGCTGTAGAT
Chromosome start end strand MM target_seq PAM alignment distance position gene name gene id
GU305914.1 106869 106891 - 0 ATCTACAGCCCGTGCGTATC GGG ||||||||[||||||||||||]PAM NA Intergenic NA NA

T26027 TACAGCCCGTGCGTATCGGGAGG 1000
Oligo fwd CACCTACAGCCCGTGCGTATCGGG
Oligo rev AAACCCCGATACGCACGGGCTGTA
Chromosome start end strand MM target_seq PAM alignment distance position gene name gene id
GU305914.1 106866 106888 - 0 TACAGCCCGTGCGTATCGGG AGG ||||||||[||||||||||||]PAM NA Intergenic NA NA

T26028 GCCCGTGCGTATCGGGAGGCAGG 1000
Oligo fwd CACCGCCCGTGCGTATCGGGAGGC
Oligo rev AAACGCCTCCCGATACGCACGGGC
Chromosome start end strand MM target_seq PAM alignment distance position gene name gene id
GU305914.1 106862 106884 - 0 GCCCGTGCGTATCGGGAGGC AGG ||||||||[||||||||||||]PAM NA Intergenic NA NA

T26029 GGGAGGCAGGCGCGACTATACGG 1000
Oligo fwd CACCGGGAGGCAGGCGCGACTATA
Oligo rev AAACTATAGTCGCGCCTGCCTCCC
Chromosome start end strand MM target_seq PAM alignment distance position gene name gene id
GU305914.1 106849 106871 - 0 GGGAGGCAGGCGCGACTATA CGG ||||||||[||||||||||||]PAM NA Intergenic NA NA

T26030 GGAGGCAGGCGCGACTATACGGG 1000
Oligo fwd CACCGGAGGCAGGCGCGACTATAC
Oligo rev AAACGTATAGTCGCGCCTGCCTCC
Chromosome start end strand MM target_seq PAM alignment distance position gene name gene id
GU305914.1 106848 106870 - 0 GGAGGCAGGCGCGACTATAC GGG ||||||||[||||||||||||]PAM NA Intergenic NA NA

T26031 GCGCGACTATACGGGTCCATCGG 1000
Oligo fwd CACCGCGCGACTATACGGGTCCAT
Oligo rev AAACATGGACCCGTATAGTCGCGC
Chromosome start end strand MM target_seq PAM alignment distance position gene name gene id
GU305914.1 106840 106862 - 0 GCGCGACTATACGGGTCCAT CGG ||||||||[||||||||||||]PAM NA Intergenic NA NA

T26032 ACTATACGGGTCCATCGGTCAGG 1000
Oligo fwd CACCACTATACGGGTCCATCGGTC
Oligo rev AAACGACCGATGGACCCGTATAGT
Chromosome start end strand MM target_seq PAM alignment distance position gene name gene id
GU305914.1 106835 106857 - 0 ACTATACGGGTCCATCGGTC AGG ||||||||[||||||||||||]PAM NA Intergenic NA NA

T26033 CTATACGGGTCCATCGGTCAGGG 1000
Oligo fwd CACCCTATACGGGTCCATCGGTCA
Oligo rev AAACTGACCGATGGACCCGTATAG
Chromosome start end strand MM target_seq PAM alignment distance position gene name gene id
GU305914.1 106834 106856 - 0 CTATACGGGTCCATCGGTCA GGG ||||||||[||||||||||||]PAM NA Intergenic NA NA

T26034 TCCATCGGTCAGGGCCCTGATGG 1000
Oligo fwd CACCTCCATCGGTCAGGGCCCTGA
Oligo rev AAACTCAGGGCCCTGACCGATGGA
Chromosome start end strand MM target_seq PAM alignment distance position gene name gene id
GU305914.1 106825 106847 - 0 TCCATCGGTCAGGGCCCTGA TGG ||||||||[||||||||||||]PAM NA Intergenic NA NA

T26035 GGTCAGGGCCCTGATGGAGTCGG 1000
Oligo fwd CACCGGTCAGGGCCCTGATGGAGT
Oligo rev AAACACTCCATCAGGGCCCTGACC
Chromosome start end strand MM target_seq PAM alignment distance position gene name gene id
GU305914.1 106819 106841 - 0 GGTCAGGGCCCTGATGGAGT CGG ||||||||[||||||||||||]PAM NA Intergenic NA NA

T26036 GATGGAGTCGGCGCACCGTCCGG 1000
Oligo fwd CACCGATGGAGTCGGCGCACCGTC
Oligo rev AAACGACGGTGCGCCGACTCCATC
Chromosome start end strand MM target_seq PAM alignment distance position gene name gene id
GU305914.1 106807 106829 - 0 GATGGAGTCGGCGCACCGTC CGG ||||||||[||||||||||||]PAM NA Intergenic NA NA

T26037 CGGCGCACCGTCCGGAGAGACGG 1000
Oligo fwd CACCCGGCGCACCGTCCGGAGAGA
Oligo rev AAACTCTCTCCGGACGGTGCGCCG
Chromosome start end strand MM target_seq PAM alignment distance position gene name gene id
GU305914.1 106799 106821 - 0 CGGCGCACCGTCCGGAGAGA CGG ||||||||[||||||||||||]PAM NA Intergenic NA NA

T26038 GGCGCACCGTCCGGAGAGACGGG 1000
Oligo fwd CACCGGCGCACCGTCCGGAGAGAC
Oligo rev AAACGTCTCTCCGGACGGTGCGCC
Chromosome start end strand MM target_seq PAM alignment distance position gene name gene id
GU305914.1 106798 106820 - 0 GGCGCACCGTCCGGAGAGAC GGG ||||||||[||||||||||||]PAM NA Intergenic NA NA

T26039 TCCGGAGAGACGGGCCGAACAGG 1000
Oligo fwd CACCTCCGGAGAGACGGGCCGAAC
Oligo rev AAACGTTCGGCCCGTCTCTCCGGA
Chromosome start end strand MM target_seq PAM alignment distance position gene name gene id

| | | | | | | | | | | | |
|---|---|---|---|---|---|---|---|---|---|---|---|
| GU305914.1 | 106789 | 106811 | - | 0 | TCCGGAGAGACGGGCCGAAC | AGG | \|\|\|\|\|\|\|\|[\|\|\|\|\|\|\|\|\|\|\|\|]PAM | NA | Intergenic | NA | NA |
| T26040 | ACGGGCCGAACAGGTGATGTCGG | 1000 | | | | | | | | | |
| Oligo fwd | CACCACGGGCCGAACAGGTGATGT | | | | | | | | | | |
| Oligo rev | AAACACATCACCTGTTCGGCCCGT | | | | | | | | | | |
| Chromosome | start | end | strand | MM | target_seq | PAM | alignment | distance | position | gene name | gene id |
| GU305914.1 | 106780 | 106802 | - | 0 | ACGGGCCGAACAGGTGATGT | CGG | \|\|\|\|\|\|\|\|[\|\|\|\|\|\|\|\|\|\|\|\|]PAM | NA | Intergenic | NA | NA |
| T26041 | CGAACAGGTGATGTCGGTGACGG | 1000 | | | | | | | | | |
| Oligo fwd | CACCCGAACAGGTGATGTCGGTGA | | | | | | | | | | |
| Oligo rev | AAACTCACCGACATCACCTGTTCG | | | | | | | | | | |
| Chromosome | start | end | strand | MM | target_seq | PAM | alignment | distance | position | gene name | gene id |
| GU305914.1 | 106774 | 106796 | - | 0 | CGAACAGGTGATGTCGGTGA | CGG | \|\|\|\|\|\|\|\|[\|\|\|\|\|\|\|\|\|\|\|\|]PAM | NA | Intergenic | NA | NA |
| T26042 | TCCGAAGCGCTCGCAGATGTCGG | 1000 | | | | | | | | | |
| Oligo fwd | CACCTCCGAAGCGCTCGCAGATGT | | | | | | | | | | |
| Oligo rev | AAACACATCTGCGAGCGCTTCGGA | | | | | | | | | | |
| Chromosome | start | end | strand | MM | target_seq | PAM | alignment | distance | position | gene name | gene id |
| GU305914.1 | 106741 | 106763 | - | 0 | TCCGAAGCGCTCGCAGATGT | CGG | \|\|\|\|\|\|\|\|[\|\|\|\|\|\|\|\|\|\|\|\|]PAM | NA | Intergenic | NA | NA |
| T26043 | GCGCTCGCAGATGTCGGACGAGG | 1000 | | | | | | | | | |
| Oligo fwd | CACCGCGCTCGCAGATGTCGGACG | | | | | | | | | | |
| Oligo rev | AAACCGTCCGACATCTGCGAGCGC | | | | | | | | | | |
| Chromosome | start | end | strand | MM | target_seq | PAM | alignment | distance | position | gene name | gene id |
| GU305914.1 | 106735 | 106757 | - | 0 | GCGCTCGCAGATGTCGGACG | AGG | \|\|\|\|\|\|\|\|[\|\|\|\|\|\|\|\|\|\|\|\|]PAM | NA | Intergenic | NA | NA |
| GU305914.1 | 69037 | 69059 | + | 3 | GCGCAAGCAGATCTCGGACG | AGG | \|\|\|\|--\|\|[\|\|\|\|-\|\|\|\|\|\|\|]PAM | NA | Intergenic | NA | NA |
| T26044 | GGACGAGGCTAACTGCGTGCCGG | 1000 | | | | | | | | | |
| Oligo fwd | CACCGGACGAGGCTAACTGCGTGC | | | | | | | | | | |
| Oligo rev | AAACGCACGCAGTTAGCCTCGTCC | | | | | | | | | | |
| Chromosome | start | end | strand | MM | target_seq | PAM | alignment | distance | position | gene name | gene id |
| GU305914.1 | 106720 | 106742 | - | 0 | GGACGAGGCTAACTGCGTGC | CGG | \|\|\|\|\|\|\|\|[\|\|\|\|\|\|\|\|\|\|\|\|]PAM | NA | Intergenic | NA | NA |
| T26045 | CTGCGTGCCGGATTCGACTCAGG | 1000 | | | | | | | | | |
| Oligo fwd | CACCCTGCGTGCCGGATTCGACTC | | | | | | | | | | |
| Oligo rev | AAACGAGTCGAATCCGGCACGCAG | | | | | | | | | | |
| Chromosome | start | end | strand | MM | target_seq | PAM | alignment | distance | position | gene name | gene id |
| GU305914.1 | 106708 | 106730 | - | 0 | CTGCGTGCCGGATTCGACTC | AGG | \|\|\|\|\|\|\|\|[\|\|\|\|\|\|\|\|\|\|\|\|]PAM | NA | Intergenic | NA | NA |
| T26046 | TGCGTGCCGGATTCGACTCAGGG | 1000 | | | | | | | | | |
| Oligo fwd | CACCTGCGTGCCGGATTCGACTCA | | | | | | | | | | |
| Oligo rev | AAACTGAGTCGAATCCGGCACGCA | | | | | | | | | | |
| Chromosome | start | end | strand | MM | target_seq | PAM | alignment | distance | position | gene name | gene id |
| GU305914.1 | 106707 | 106729 | - | 0 | TGCGTGCCGGATTCGACTCA | GGG | \|\|\|\|\|\|\|\|[\|\|\|\|\|\|\|\|\|\|\|\|]PAM | NA | Intergenic | NA | NA |
| T26047 | GCCGGATTCGACTCAGGGCGCGG | 1000 | | | | | | | | | |
| Oligo fwd | CACCGCCGGATTCGACTCAGGGCG | | | | | | | | | | |
| Oligo rev | AAACCGCCCTGAGTCGAATCCGGC | | | | | | | | | | |
| Chromosome | start | end | strand | MM | target_seq | PAM | alignment | distance | position | gene name | gene id |
| GU305914.1 | 106702 | 106724 | - | 0 | GCCGGATTCGACTCAGGGCG | CGG | \|\|\|\|\|\|\|\|[\|\|\|\|\|\|\|\|\|\|\|\|]PAM | NA | Intergenic | NA | NA |
| T26048 | GACTCAGGGCGCGGTGATAACGG | 1000 | | | | | | | | | |
| Oligo fwd | CACCGACTCAGGGCGCGGTGATAA | | | | | | | | | | |
| Oligo rev | AAACTTATCACCGCGCCCTGAGTC | | | | | | | | | | |
| Chromosome | start | end | strand | MM | target_seq | PAM | alignment | distance | position | gene name | gene id |
| GU305914.1 | 106693 | 106715 | - | 0 | GACTCAGGGCGCGGTGATAA | CGG | \|\|\|\|\|\|\|\|[\|\|\|\|\|\|\|\|\|\|\|\|]PAM | NA | Intergenic | NA | NA |
| T26049 | CTCGTCCGATTTTATCGTCAAGG | 1000 | | | | | | | | | |
| Oligo fwd | CACCCTCGTCCGATTTTATCGTCA | | | | | | | | | | |
| Oligo rev | AAACTGACGATAAAATCGGACGAG | | | | | | | | | | |
| Chromosome | start | end | strand | MM | target_seq | PAM | alignment | distance | position | gene name | gene id |
| GU305914.1 | 106648 | 106670 | - | 0 | CTCGTCCGATTTTATCGTCA | AGG | \|\|\|\|\|\|\|\|[\|\|\|\|\|\|\|\|\|\|\|\|]PAM | NA | Intergenic | NA | NA |
| T26050 | TCGTCCGATTTTATCGTCAAGGG | 1000 | | | | | | | | | |
| Oligo fwd | CACCTCGTCCGATTTTATCGTCAA | | | | | | | | | | |
| Oligo rev | AAACTTGACGATAAAATCGGACGA | | | | | | | | | | |
| Chromosome | start | end | strand | MM | target_seq | PAM | alignment | distance | position | gene name | gene id |
| GU305914.1 | 106647 | 106669 | - | 0 | TCGTCCGATTTTATCGTCAA | GGG | \|\|\|\|\|\|\|\|[\|\|\|\|\|\|\|\|\|\|\|\|]PAM | NA | Intergenic | NA | NA |
| T26051 | CGATTTTATCGTCAAGGGCCTGG | 1000 | | | | | | | | | |
| Oligo fwd | CACCCGATTTTATCGTCAAGGGCC | | | | | | | | | | |
| Oligo rev | AAACGGCCCTTGACGATAAAATCG | | | | | | | | | | |
| Chromosome | start | end | strand | MM | target_seq | PAM | alignment | distance | position | gene name | gene id |
| GU305914.1 | 106642 | 106664 | - | 0 | CGATTTTATCGTCAAGGGCC | TGG | \|\|\|\|\|\|\|\|[\|\|\|\|\|\|\|\|\|\|\|\|]PAM | NA | Intergenic | NA | NA |
| T26052 | GATCCCCGTGAGCAACACCGTGG | 1000 | | | | | | | | | |
| Oligo fwd | CACCGATCCCCGTGAGCAACACCG | | | | | | | | | | |
| Oligo rev | AAACCGGTGTTGCTCACGGGGATC | | | | | | | | | | |
| Chromosome | start | end | strand | MM | target_seq | PAM | alignment | distance | position | gene name | gene id |
| GU305914.1 | 106618 | 106640 | - | 0 | GATCCCCGTGAGCAACACCG | TGG | \|\|\|\|\|\|\|\|[\|\|\|\|\|\|\|\|\|\|\|\|]PAM | NA | Intergenic | NA | NA |
| T26053 | CCCCGTGAGCAACACCGTGGTGG | 1000 | | | | | | | | | |
| Oligo fwd | CACCCCCCGTGAGCAACACCGTGG | | | | | | | | | | |
| Oligo rev | AAACCCACGGTGTTGCTCACGGGG | | | | | | | | | | |
| Chromosome | start | end | strand | MM | target_seq | PAM | alignment | distance | position | gene name | gene id |
| GU305914.1 | 106615 | 106637 | - | 0 | CCCCGTGAGCAACACCGTGG | TGG | \|\|\|\|\|\|\|\|[\|\|\|\|\|\|\|\|\|\|\|\|]PAM | NA | Intergenic | NA | NA |
| T26054 | TGAGCAACACCGTGGTGGACAGG | 1000 | | | | | | | | | |
| Oligo fwd | CACCTGAGCAACACCGTGGTGGAC | | | | | | | | | | |

| | | | | | | | | | | | |
|---|---|---|---|---|---|---|---|---|---|---|---|
| Oligo rev | AAACGTCCACCACGGTGTTGCTCA | | | | | | | | | | |
| Chromosome | start | end | strand | MM | target_seq | PAM | alignment | distance | position | gene name | gene id |
| GU305914.1 | 106610 | 106632 | - | 0 | TGAGCAACACCGTGGTGGAC | AGG | \|\|\|\|\|\|\|\|[\|\|\|\|\|\|\|\|\|\|\|\|]PAM | NA | Intergenic | NA | NA |
| T26055 | GGACAGGAACCTGATGATCACGG | 1000 | | | | | | | | | |
| Oligo fwd | CACCGGACAGGAACCTGATGATCA | | | | | | | | | | |
| Oligo rev | AAACTGATCATCAGGTTCCTGTCC | | | | | | | | | | |
| Chromosome | start | end | strand | MM | target_seq | PAM | alignment | distance | position | gene name | gene id |
| GU305914.1 | 106594 | 106616 | - | 0 | GGACAGGAACCTGATGATCA | CGG | \|\|\|\|\|\|\|\|[\|\|\|\|\|\|\|\|\|\|\|\|]PAM | NA | Intergenic | NA | NA |
| T26056 | GAACCTGATGATCACGGTCCTGG | 1000 | | | | | | | | | |
| Oligo fwd | CACCGAACCTGATGATCACGGTCC | | | | | | | | | | |
| Oligo rev | AAACGGACCGTGATCATCAGGTTC | | | | | | | | | | |
| Chromosome | start | end | strand | MM | target_seq | PAM | alignment | distance | position | gene name | gene id |
| GU305914.1 | 106588 | 106610 | - | 0 | GAACCTGATGATCACGGTCC | TGG | \|\|\|\|\|\|\|\|[\|\|\|\|\|\|\|\|\|\|\|\|]PAM | NA | Intergenic | NA | NA |
| T26057 | TGATCACGGTCCTGGACCGCAGG | 1000 | | | | | | | | | |
| Oligo fwd | CACCTGATCACGGTCCTGGACCGC | | | | | | | | | | |
| Oligo rev | AAACGCGGTCCAGGACCGTGATCA | | | | | | | | | | |
| Chromosome | start | end | strand | MM | target_seq | PAM | alignment | distance | position | gene name | gene id |
| GU305914.1 | 106580 | 106602 | - | 0 | TGATCACGGTCCTGGACCGC | AGG | \|\|\|\|\|\|\|\|[\|\|\|\|\|\|\|\|\|\|\|\|]PAM | NA | Intergenic | NA | NA |
| T26058 | GGAGCCCCTGCGTCCTGTCGAGG | 1000 | | | | | | | | | |
| Oligo fwd | CACCGGAGCCCCTGCGTCCTGTCG | | | | | | | | | | |
| Oligo rev | AAACCGACAGGACGCAGGGGCTCC | | | | | | | | | | |
| Chromosome | start | end | strand | MM | target_seq | PAM | alignment | distance | position | gene name | gene id |
| GU305914.1 | 106559 | 106581 | - | 0 | GGAGCCCCTGCGTCCTGTCG | AGG | \|\|\|\|\|\|\|\|[\|\|\|\|\|\|\|\|\|\|\|\|]PAM | NA | Intergenic | NA | NA |
| T26059 | TCGAGGAGCTACCGCGAACGCGG | 1000 | | | | | | | | | |
| Oligo fwd | CACCTCGAGGAGCTACCGCGAACG | | | | | | | | | | |
| Oligo rev | AAACCGTTCGCGGTAGCTCCTCGA | | | | | | | | | | |
| Chromosome | start | end | strand | MM | target_seq | PAM | alignment | distance | position | gene name | gene id |
| GU305914.1 | 106542 | 106564 | - | 0 | TCGAGGAGCTACCGCGAACG | CGG | \|\|\|\|\|\|\|\|[\|\|\|\|\|\|\|\|\|\|\|\|]PAM | NA | Intergenic | NA | NA |
| T26060 | GAGCTACCGCGAACGCGGATCGG | 1000 | | | | | | | | | |
| Oligo fwd | CACCGAGCTACCGCGAACGCGGAT | | | | | | | | | | |
| Oligo rev | AAACATCCGCGTTCGCGGTAGCTC | | | | | | | | | | |
| Chromosome | start | end | strand | MM | target_seq | PAM | alignment | distance | position | gene name | gene id |
| GU305914.1 | 106537 | 106559 | - | 0 | GAGCTACCGCGAACGCGGAT | CGG | \|\|\|\|\|\|\|\|[\|\|\|\|\|\|\|\|\|\|\|\|]PAM | NA | Intergenic | NA | NA |
| T26061 | ATAACATCACGTTCACCGAACGG | 1000 | | | | | | | | | |
| Oligo fwd | CACCATAACATCACGTTCACCGAA | | | | | | | | | | |
| Oligo rev | AAACTTCGGTGAACGTGATGTTAT | | | | | | | | | | |
| Chromosome | start | end | strand | MM | target_seq | PAM | alignment | distance | position | gene name | gene id |
| GU305914.1 | 106502 | 106524 | - | 0 | ATAACATCACGTTCACCGAA | CGG | \|\|\|\|\|\|\|\|[\|\|\|\|\|\|\|\|\|\|\|\|]PAM | NA | Intergenic | NA | NA |
| T26062 | CGAGTTCTGCGCCAGCACCCTGG | 1000 | | | | | | | | | |
| Oligo fwd | CACCCGAGTTCTGCGCCAGCACCC | | | | | | | | | | |
| Oligo rev | AAACGGGTGCTGGCGCAGAACTCG | | | | | | | | | | |
| Chromosome | start | end | strand | MM | target_seq | PAM | alignment | distance | position | gene name | gene id |
| GU305914.1 | 106477 | 106499 | - | 0 | CGAGTTCTGCGCCAGCACCC | TGG | \|\|\|\|\|\|\|\|[\|\|\|\|\|\|\|\|\|\|\|\|]PAM | NA | Intergenic | NA | NA |
| T26063 | CACCCTGGTCGAGTACGACGAGG | 1000 | | | | | | | | | |
| Oligo fwd | CACCCACCCTGGTCGAGTACGACG | | | | | | | | | | |
| Oligo rev | AAACCGTCGTACTCGACCAGGGTG | | | | | | | | | | |
| Chromosome | start | end | strand | MM | target_seq | PAM | alignment | distance | position | gene name | gene id |
| GU305914.1 | 106462 | 106484 | - | 0 | CACCCTGGTCGAGTACGACG | AGG | \|\|\|\|\|\|\|\|[\|\|\|\|\|\|\|\|\|\|\|\|]PAM | NA | Intergenic | NA | NA |
| T26064 | CCTGGTCGAGTACGACGAGGTGG | 1000 | | | | | | | | | |
| Oligo fwd | CACCCCTGGTCGAGTACGACGAGG | | | | | | | | | | |
| Oligo rev | AAACCCTCGTCGTACTCGACCAGG | | | | | | | | | | |
| Chromosome | start | end | strand | MM | target_seq | PAM | alignment | distance | position | gene name | gene id |
| GU305914.1 | 106459 | 106481 | - | 0 | CCTGGTCGAGTACGACGAGG | TGG | \|\|\|\|\|\|\|\|[\|\|\|\|\|\|\|\|\|\|\|\|]PAM | NA | Intergenic | NA | NA |
| T26065 | GTCGAGTACGACGAGGTGGACGG | 1000 | | | | | | | | | |
| Oligo fwd | CACCGTCGAGTACGACGAGGTGGA | | | | | | | | | | |
| Oligo rev | AAACTCCACCTCGTCGTACTCGAC | | | | | | | | | | |
| Chromosome | start | end | strand | MM | target_seq | PAM | alignment | distance | position | gene name | gene id |
| GU305914.1 | 106455 | 106477 | - | 0 | GTCGAGTACGACGAGGTGGA | CGG | \|\|\|\|\|\|\|\|[\|\|\|\|\|\|\|\|\|\|\|\|]PAM | NA | Intergenic | NA | NA |
| T26066 | CATGCACATCCCGAGCATAGAGG | 1000 | | | | | | | | | |
| Oligo fwd | CACCCATGCACATCCCGAGCATAG | | | | | | | | | | |
| Oligo rev | AAACCTATGCTCGGGATGTGCATG | | | | | | | | | | |
| Chromosome | start | end | strand | MM | target_seq | PAM | alignment | distance | position | gene name | gene id |
| GU305914.1 | 106420 | 106442 | - | 0 | CATGCACATCCCGAGCATAG | AGG | \|\|\|\|\|\|\|\|[\|\|\|\|\|\|\|\|\|\|\|\|]PAM | NA | Intergenic | NA | NA |
| T26067 | AGAGGTCCTGAAATACCTGACGG | 1000 | | | | | | | | | |
| Oligo fwd | CACCAGAGGTCCTGAAATACCTGA | | | | | | | | | | |
| Oligo rev | AAACTCAGGTATTTCAGGACCTCT | | | | | | | | | | |
| Chromosome | start | end | strand | MM | target_seq | PAM | alignment | distance | position | gene name | gene id |
| GU305914.1 | 106402 | 106424 | - | 0 | AGAGGTCCTGAAATACCTGA | CGG | \|\|\|\|\|\|\|\|[\|\|\|\|\|\|\|\|\|\|\|\|]PAM | NA | Intergenic | NA | NA |
| T26068 | GGACCCCGACAACGACATCCTGG | 1000 | | | | | | | | | |
| Oligo fwd | CACCGGACCCCGACAACGACATCC | | | | | | | | | | |
| Oligo rev | AAACGGATGTCGTTGTCGGGGTCC | | | | | | | | | | |
| Chromosome | start | end | strand | MM | target_seq | PAM | alignment | distance | position | gene name | gene id |
| GU305914.1 | 106381 | 106403 | - | 0 | GGACCCCGACAACGACATCC | TGG | \|\|\|\|\|\|\|\|[\|\|\|\|\|\|\|\|\|\|\|\|]PAM | NA | Intergenic | NA | NA |
| T26069 | CCCCGACAACGACATCCTGGTGG | 1000 | | | | | | | | | |

| | | | | | | | | | | | | |
|---|---|---|---|---|---|---|---|---|---|---|---|---|
| Oligo fwd | CACCCCCCGACAACGACATCCTGG | | | | | | | | | | | |
| Oligo rev | AAACCCAGGATGTCGTTGTCGGGG | | | | | | | | | | | |
| Chromosome | start | | end | strand | MM | target_seq | PAM | alignment | distance | position | gene name | gene id |
| GU305914.1 | | 106378 | 106400 | - | 0 | CCCCGACAACGACATCCTGG | TGG | \|\|\|\|\|\|\|\|[\|\|\|\|\|\|\|\|\|\|\|\|]PAM | NA | Intergenic | NA | NA |
| GU305914.1 | | 107644 | 107666 | - | 3 | GCCCGACCACGACCTCCTGG | TGG | -\|\|\|\|\|\|-[\|\|\|\|\|-\|\|\|\|\|\|]PAM | NA | Intergenic | NA | NA |
| T26070 | ACATCCTGGTGGCGACGCCTAGG | | 1000 | | | | | | | | | |
| Oligo fwd | CACCACATCCTGGTGGCGACGCCT | | | | | | | | | | | |
| Oligo rev | AAACAGGCGTCGCCACCAGGATGT | | | | | | | | | | | |
| Chromosome | start | | end | strand | MM | target_seq | PAM | alignment | distance | position | gene name | gene id |
| GU305914.1 | | 106367 | 106389 | - | 0 | ACATCCTGGTGGCGACGCCT | AGG | \|\|\|\|\|\|\|\|[\|\|\|\|\|\|\|\|\|\|\|\|]PAM | NA | Intergenic | NA | NA |
| T26071 | CATCCTGGTGGCGACGCCTAGGG | | 1000 | | | | | | | | | |
| Oligo fwd | CACCCATCCTGGTGGCGACGCCTA | | | | | | | | | | | |
| Oligo rev | AAACTAGGCGTCGCCACCAGGATG | | | | | | | | | | | |
| Chromosome | start | | end | strand | MM | target_seq | PAM | alignment | distance | position | gene name | gene id |
| GU305914.1 | | 106366 | 106388 | - | 0 | CATCCTGGTGGCGACGCCTA | GGG | \|\|\|\|\|\|\|\|[\|\|\|\|\|\|\|\|\|\|\|\|]PAM | NA | Intergenic | NA | NA |
| T26072 | TACCTATTACTAACCGCGAACGG | | 1000 | | | | | | | | | |
| Oligo fwd | CACCTACCTATTACTAACCGCGAA | | | | | | | | | | | |
| Oligo rev | AAACTTCGCGGTTAGTAATAGGTA | | | | | | | | | | | |
| Chromosome | start | | end | strand | MM | target_seq | PAM | alignment | distance | position | gene name | gene id |
| GU305914.1 | | 106338 | 106360 | - | 0 | TACCTATTACTAACCGCGAA | CGG | \|\|\|\|\|\|\|\|[\|\|\|\|\|\|\|\|\|\|\|\|]PAM | NA | Intergenic | NA | NA |
| T26073 | ACCTATTACTAACCGCGAACGGG | | 1000 | | | | | | | | | |
| Oligo fwd | CACCACCTATTACTAACCGCGAAC | | | | | | | | | | | |
| Oligo rev | AAACGTTCGCGGTTAGTAATAGGT | | | | | | | | | | | |
| Chromosome | start | | end | strand | MM | target_seq | PAM | alignment | distance | position | gene name | gene id |
| GU305914.1 | | 106337 | 106359 | - | 0 | ACCTATTACTAACCGCGAAC | GGG | \|\|\|\|\|\|\|\|[\|\|\|\|\|\|\|\|\|\|\|\|]PAM | NA | Intergenic | NA | NA |
| T26074 | ATTACTAACCGCGAACGGGACGG | | 1000 | | | | | | | | | |
| Oligo fwd | CACCATTACTAACCGCGAACGGGA | | | | | | | | | | | |
| Oligo rev | AAACTCCCGTTCGCGGTTAGTAAT | | | | | | | | | | | |
| Chromosome | start | | end | strand | MM | target_seq | PAM | alignment | distance | position | gene name | gene id |
| GU305914.1 | | 106333 | 106355 | - | 0 | ATTACTAACCGCGAACGGGA | CGG | \|\|\|\|\|\|\|\|[\|\|\|\|\|\|\|\|\|\|\|\|]PAM | NA | Intergenic | NA | NA |
| T26075 | CGCGAACGGGACGGTGTTCGAGG | | 1000 | | | | | | | | | |
| Oligo fwd | CACCCGCGAACGGGACGGTGTTCG | | | | | | | | | | | |
| Oligo rev | AAACCGAACACCGTCCCGTTCGCG | | | | | | | | | | | |
| Chromosome | start | | end | strand | MM | target_seq | PAM | alignment | distance | position | gene name | gene id |
| GU305914.1 | | 106324 | 106346 | - | 0 | CGCGAACGGGACGGTGTTCG | AGG | \|\|\|\|\|\|\|\|[\|\|\|\|\|\|\|\|\|\|\|\|]PAM | NA | Intergenic | NA | NA |
| T26076 | ACCATCGATGCCCTACTCCGTGG | | 1000 | | | | | | | | | |
| Oligo fwd | CACCACCATCGATGCCCTACTCCG | | | | | | | | | | | |
| Oligo rev | AAACCGGAGTAGGGCATCGATGGT | | | | | | | | | | | |
| Chromosome | start | | end | strand | MM | target_seq | PAM | alignment | distance | position | gene name | gene id |
| GU305914.1 | | 106276 | 106298 | - | 0 | ACCATCGATGCCCTACTCCG | TGG | \|\|\|\|\|\|\|\|[\|\|\|\|\|\|\|\|\|\|\|\|]PAM | NA | Intergenic | NA | NA |
| T26077 | CTACTCCGTGGTCGTCGCCTTGG | | 1000 | | | | | | | | | |
| Oligo fwd | CACCCTACTCCGTGGTCGTCGCCT | | | | | | | | | | | |
| Oligo rev | AAACAGGCGACGACCACGGAGTAG | | | | | | | | | | | |
| Chromosome | start | | end | strand | MM | target_seq | PAM | alignment | distance | position | gene name | gene id |
| GU305914.1 | | 106264 | 106286 | - | 0 | CTACTCCGTGGTCGTCGCCT | TGG | \|\|\|\|\|\|\|\|[\|\|\|\|\|\|\|\|\|\|\|\|]PAM | NA | Intergenic | NA | NA |
| T26078 | GGTGATCATAGCGATCCTCATGG | | 1000 | | | | | | | | | |
| Oligo fwd | CACCGGTGATCATAGCGATCCTCA | | | | | | | | | | | |
| Oligo rev | AAACTGAGGATCGCTATGATCACC | | | | | | | | | | | |
| Chromosome | start | | end | strand | MM | target_seq | PAM | alignment | distance | position | gene name | gene id |
| GU305914.1 | | 106243 | 106265 | - | 0 | GGTGATCATAGCGATCCTCA | TGG | \|\|\|\|\|\|\|\|[\|\|\|\|\|\|\|\|\|\|\|\|]PAM | NA | Intergenic | NA | NA |
| T26079 | ATAGCGATCCTCATGGCGCTAGG | | 1000 | | | | | | | | | |
| Oligo fwd | CACCATAGCGATCCTCATGGCGCT | | | | | | | | | | | |
| Oligo rev | AAACAGCGCCATGAGGATCGCTAT | | | | | | | | | | | |
| Chromosome | start | | end | strand | MM | target_seq | PAM | alignment | distance | position | gene name | gene id |
| GU305914.1 | | 106236 | 106258 | - | 0 | ATAGCGATCCTCATGGCGCT | AGG | \|\|\|\|\|\|\|\|[\|\|\|\|\|\|\|\|\|\|\|\|]PAM | NA | Intergenic | NA | NA |
| T26080 | TAGCGATCCTCATGGCGCTAGGG | | 1000 | | | | | | | | | |
| Oligo fwd | CACCTAGCGATCCTCATGGCGCTA | | | | | | | | | | | |
| Oligo rev | AAACTAGCGCCATGAGGATCGCTA | | | | | | | | | | | |
| Chromosome | start | | end | strand | MM | target_seq | PAM | alignment | distance | position | gene name | gene id |
| GU305914.1 | | 106235 | 106257 | - | 0 | TAGCGATCCTCATGGCGCTA | GGG | \|\|\|\|\|\|\|\|[\|\|\|\|\|\|\|\|\|\|\|\|]PAM | NA | Intergenic | NA | NA |
| T26081 | TCATGGCGCTAGGGCTGTACCGG | | 1000 | | | | | | | | | |
| Oligo fwd | CACCTCATGGCGCTAGGGCTGTAC | | | | | | | | | | | |
| Oligo rev | AAACGTACAGCCCTAGCGCCATGA | | | | | | | | | | | |
| Chromosome | start | | end | strand | MM | target_seq | PAM | alignment | distance | position | gene name | gene id |
| GU305914.1 | | 106226 | 106248 | - | 0 | TCATGGCGCTAGGGCTGTAC | CGG | \|\|\|\|\|\|\|\|[\|\|\|\|\|\|\|\|\|\|\|\|]PAM | NA | Intergenic | NA | NA |
| T26082 | TAGGGCTGTACCGGTTGTGCCGG | | 1000 | | | | | | | | | |
| Oligo fwd | CACCTAGGGCTGTACCGGTTGTGC | | | | | | | | | | | |
| Oligo rev | AAACGCACAACCGGTACAGCCCTA | | | | | | | | | | | |
| Chromosome | start | | end | strand | MM | target_seq | PAM | alignment | distance | position | gene name | gene id |
| GU305914.1 | | 106217 | 106239 | - | 0 | TAGGGCTGTACCGGTTGTGC | CGG | \|\|\|\|\|\|\|\|[\|\|\|\|\|\|\|\|\|\|\|\|]PAM | NA | Intergenic | NA | NA |
| T26083 | TTGTGCCGGCAAAAAAGATAAGG | | 1000 | | | | | | | | | |
| Oligo fwd | CACCTTGTGCCGGCAAAAAAGATA | | | | | | | | | | | |
| Oligo rev | AAACTATCTTTTTTGCCGGCACAA | | | | | | | | | | | |
| Chromosome | start | | end | strand | MM | target_seq | PAM | alignment | distance | position | gene name | gene id |
| GU305914.1 | | 106203 | 106225 | - | 0 | TTGTGCCGGCAAAAAAGATA | AGG | \|\|\|\|\|\|\|\|[\|\|\|\|\|\|\|\|\|\|\|\|]PAM | NA | Intergenic | NA | NA |

<table>
<tr><td>T26084</td><td>GCCGGCAAAAAAGATAAGGAAGG</td><td>1000</td><td></td><td></td><td></td><td></td><td></td><td></td><td></td><td></td><td></td></tr>
<tr><td>Oligo fwd</td><td>CACCGCCGGCAAAAAAGATAAGGA</td><td></td><td></td><td></td><td></td><td></td><td></td><td></td><td></td><td></td><td></td></tr>
<tr><td>Oligo rev</td><td>AAACTCCTTATCTTTTTTGCCGGC</td><td></td><td></td><td></td><td></td><td></td><td></td><td></td><td></td><td></td><td></td></tr>
<tr><td>Chromosome</td><td>start</td><td>end</td><td>strand</td><td>MM</td><td>target_seq</td><td>PAM</td><td>alignment</td><td>distance</td><td>position</td><td>gene name</td><td>gene id</td></tr>
<tr><td>GU305914.1</td><td>106199</td><td>106221</td><td>-</td><td>0</td><td>GCCGGCAAAAAAGATAAGGA</td><td>AGG</td><td>||||||||[||||||||||||]PAM</td><td>NA</td><td>Intergenic</td><td>NA</td><td>NA</td></tr>
<tr><td>T26085</td><td>GATAAGGAAGGACACACGCGAGG</td><td>1000</td><td></td><td></td><td></td><td></td><td></td><td></td><td></td><td></td><td></td></tr>
<tr><td>Oligo fwd</td><td>CACCGATAAGGAAGGACACACGCG</td><td></td><td></td><td></td><td></td><td></td><td></td><td></td><td></td><td></td><td></td></tr>
<tr><td>Oligo rev</td><td>AAACCGCGTGTGTCCTTCCTTATC</td><td></td><td></td><td></td><td></td><td></td><td></td><td></td><td></td><td></td><td></td></tr>
<tr><td>Chromosome</td><td>start</td><td>end</td><td>strand</td><td>MM</td><td>target_seq</td><td>PAM</td><td>alignment</td><td>distance</td><td>position</td><td>gene name</td><td>gene id</td></tr>
<tr><td>GU305914.1</td><td>106187</td><td>106209</td><td>-</td><td>0</td><td>GATAAGGAAGGACACACGCG</td><td>AGG</td><td>||||||||[||||||||||||]PAM</td><td>NA</td><td>Intergenic</td><td>NA</td><td>NA</td></tr>
<tr><td>T26086</td><td>CGTCGAGCACCCCCCGCACGCGG</td><td>1000</td><td></td><td></td><td></td><td></td><td></td><td></td><td></td><td></td><td></td></tr>
<tr><td>Oligo fwd</td><td>CACCCGTCGAGCACCCCCCGCACG</td><td></td><td></td><td></td><td></td><td></td><td></td><td></td><td></td><td></td><td></td></tr>
<tr><td>Oligo rev</td><td>AAACCGTGCGGGGGGTGCTCGACG</td><td></td><td></td><td></td><td></td><td></td><td></td><td></td><td></td><td></td><td></td></tr>
<tr><td>Chromosome</td><td>start</td><td>end</td><td>strand</td><td>MM</td><td>target_seq</td><td>PAM</td><td>alignment</td><td>distance</td><td>position</td><td>gene name</td><td>gene id</td></tr>
<tr><td>GU305914.1</td><td>106135</td><td>106157</td><td>-</td><td>0</td><td>CGTCGAGCACCCCCCGCACG</td><td>CGG</td><td>||||||||[||||||||||||]PAM</td><td>NA</td><td>Intergenic</td><td>NA</td><td>NA</td></tr>
<tr><td>GU305914.1</td><td>119659</td><td>119681</td><td>-</td><td>4</td><td>CGGCGACCACCACGCGCACG</td><td>GGG</td><td>||-|||-|[|||-|-||||||]PAM</td><td>NA</td><td>Intergenic</td><td>NA</td><td>NA</td></tr>
<tr><td>T26087</td><td>CGAGCACCCCCCGCACGCGGTGG</td><td>1000</td><td></td><td></td><td></td><td></td><td></td><td></td><td></td><td></td><td></td></tr>
<tr><td>Oligo fwd</td><td>CACCCGAGCACCCCCCGCACGCGG</td><td></td><td></td><td></td><td></td><td></td><td></td><td></td><td></td><td></td><td></td></tr>
<tr><td>Oligo rev</td><td>AAACCCGCGTGCGGGGGGTGCTCG</td><td></td><td></td><td></td><td></td><td></td><td></td><td></td><td></td><td></td><td></td></tr>
<tr><td>Chromosome</td><td>start</td><td>end</td><td>strand</td><td>MM</td><td>target_seq</td><td>PAM</td><td>alignment</td><td>distance</td><td>position</td><td>gene name</td><td>gene id</td></tr>
<tr><td>GU305914.1</td><td>106132</td><td>106154</td><td>-</td><td>0</td><td>CGAGCACCCCCCGCACGCGG</td><td>TGG</td><td>||||||||[||||||||||||]PAM</td><td>NA</td><td>Intergenic</td><td>NA</td><td>NA</td></tr>
<tr><td>T26088</td><td>CGCGGTGGCGTTGCTCTTTCCGG</td><td>1000</td><td></td><td></td><td></td><td></td><td></td><td></td><td></td><td></td><td></td></tr>
<tr><td>Oligo fwd</td><td>CACCCGCGGTGGCGTTGCTCTTTC</td><td></td><td></td><td></td><td></td><td></td><td></td><td></td><td></td><td></td><td></td></tr>
<tr><td>Oligo rev</td><td>AAACGAAAGAGCAACGCCACCGCG</td><td></td><td></td><td></td><td></td><td></td><td></td><td></td><td></td><td></td><td></td></tr>
<tr><td>Chromosome</td><td>start</td><td>end</td><td>strand</td><td>MM</td><td>target_seq</td><td>PAM</td><td>alignment</td><td>distance</td><td>position</td><td>gene name</td><td>gene id</td></tr>
<tr><td>GU305914.1</td><td>106117</td><td>106139</td><td>-</td><td>0</td><td>CGCGGTGGCGTTGCTCTTTC</td><td>CGG</td><td>||||||||[||||||||||||]PAM</td><td>NA</td><td>Intergenic</td><td>NA</td><td>NA</td></tr>
<tr><td>T26089</td><td>TGAAAGCCTCCGTGCTTCTCAGG</td><td>1000</td><td></td><td></td><td></td><td></td><td></td><td></td><td></td><td></td><td></td></tr>
<tr><td>Oligo fwd</td><td>CACCTGAAAGCCTCCGTGCTTCTC</td><td></td><td></td><td></td><td></td><td></td><td></td><td></td><td></td><td></td><td></td></tr>
<tr><td>Oligo rev</td><td>AAACGAGAAGCACGGAGGCTTTCA</td><td></td><td></td><td></td><td></td><td></td><td></td><td></td><td></td><td></td><td></td></tr>
<tr><td>Chromosome</td><td>start</td><td>end</td><td>strand</td><td>MM</td><td>target_seq</td><td>PAM</td><td>alignment</td><td>distance</td><td>position</td><td>gene name</td><td>gene id</td></tr>
<tr><td>GU305914.1</td><td>106082</td><td>106104</td><td>-</td><td>0</td><td>TGAAAGCCTCCGTGCTTCTC</td><td>AGG</td><td>||||||||[||||||||||||]PAM</td><td>NA</td><td>Intergenic</td><td>NA</td><td>NA</td></tr>
<tr><td>T26090</td><td>TTCTCAGGACGATGTCATCTTGG</td><td>1000</td><td></td><td></td><td></td><td></td><td></td><td></td><td></td><td></td><td></td></tr>
<tr><td>Oligo fwd</td><td>CACCTTCTCAGGACGATGTCATCT</td><td></td><td></td><td></td><td></td><td></td><td></td><td></td><td></td><td></td><td></td></tr>
<tr><td>Oligo rev</td><td>AAACAGATGACATCGTCCTGAGAA</td><td></td><td></td><td></td><td></td><td></td><td></td><td></td><td></td><td></td><td></td></tr>
<tr><td>Chromosome</td><td>start</td><td>end</td><td>strand</td><td>MM</td><td>target_seq</td><td>PAM</td><td>alignment</td><td>distance</td><td>position</td><td>gene name</td><td>gene id</td></tr>
<tr><td>GU305914.1</td><td>106067</td><td>106089</td><td>-</td><td>0</td><td>TTCTCAGGACGATGTCATCT</td><td>TGG</td><td>||||||||[||||||||||||]PAM</td><td>NA</td><td>Intergenic</td><td>NA</td><td>NA</td></tr>
<tr><td>T26091</td><td>GTCATCTTGGAATCCGACGATGG</td><td>1000</td><td></td><td></td><td></td><td></td><td></td><td></td><td></td><td></td><td></td></tr>
<tr><td>Oligo fwd</td><td>CACCGTCATCTTGGAATCCGACGA</td><td></td><td></td><td></td><td></td><td></td><td></td><td></td><td></td><td></td><td></td></tr>
<tr><td>Oligo rev</td><td>AAACTCGTCGGATTCCAAGATGAC</td><td></td><td></td><td></td><td></td><td></td><td></td><td></td><td></td><td></td><td></td></tr>
<tr><td>Chromosome</td><td>start</td><td>end</td><td>strand</td><td>MM</td><td>target_seq</td><td>PAM</td><td>alignment</td><td>distance</td><td>position</td><td>gene name</td><td>gene id</td></tr>
<tr><td>GU305914.1</td><td>106054</td><td>106076</td><td>-</td><td>0</td><td>GTCATCTTGGAATCCGACGA</td><td>TGG</td><td>||||||||[||||||||||||]PAM</td><td>NA</td><td>Intergenic</td><td>NA</td><td>NA</td></tr>
<tr><td>T26092</td><td>CTTGGAATCCGACGATGGCTCGG</td><td>1000</td><td></td><td></td><td></td><td></td><td></td><td></td><td></td><td></td><td></td></tr>
<tr><td>Oligo fwd</td><td>CACCCTTGGAATCCGACGATGGCT</td><td></td><td></td><td></td><td></td><td></td><td></td><td></td><td></td><td></td><td></td></tr>
<tr><td>Oligo rev</td><td>AAACAGCCATCGTCGGATTCCAAG</td><td></td><td></td><td></td><td></td><td></td><td></td><td></td><td></td><td></td><td></td></tr>
<tr><td>Chromosome</td><td>start</td><td>end</td><td>strand</td><td>MM</td><td>target_seq</td><td>PAM</td><td>alignment</td><td>distance</td><td>position</td><td>gene name</td><td>gene id</td></tr>
<tr><td>GU305914.1</td><td>106049</td><td>106071</td><td>-</td><td>0</td><td>CTTGGAATCCGACGATGGCT</td><td>CGG</td><td>||||||||[||||||||||||]PAM</td><td>NA</td><td>Intergenic</td><td>NA</td><td>NA</td></tr>
<tr><td>T26093</td><td>CATCATCGCCACCATGATCACGG</td><td>1000</td><td></td><td></td><td></td><td></td><td></td><td></td><td></td><td></td><td></td></tr>
<tr><td>Oligo fwd</td><td>CACCCATCATCGCCACCATGATCA</td><td></td><td></td><td></td><td></td><td></td><td></td><td></td><td></td><td></td><td></td></tr>
<tr><td>Oligo rev</td><td>AAACTGATCATGGTGGCGATGATG</td><td></td><td></td><td></td><td></td><td></td><td></td><td></td><td></td><td></td><td></td></tr>
<tr><td>Chromosome</td><td>start</td><td>end</td><td>strand</td><td>MM</td><td>target_seq</td><td>PAM</td><td>alignment</td><td>distance</td><td>position</td><td>gene name</td><td>gene id</td></tr>
<tr><td>GU305914.1</td><td>106015</td><td>106037</td><td>-</td><td>0</td><td>CATCATCGCCACCATGATCA</td><td>CGG</td><td>||||||||[||||||||||||]PAM</td><td>NA</td><td>Intergenic</td><td>NA</td><td>NA</td></tr>
<tr><td>T26094</td><td>GATCACGGTGTTGACTATCGCGG</td><td>1000</td><td></td><td></td><td></td><td></td><td></td><td></td><td></td><td></td><td></td></tr>
<tr><td>Oligo fwd</td><td>CACCGATCACGGTGTTGACTATCG</td><td></td><td></td><td></td><td></td><td></td><td></td><td></td><td></td><td></td><td></td></tr>
<tr><td>Oligo rev</td><td>AAACCGATAGTCAACACCGTGATC</td><td></td><td></td><td></td><td></td><td></td><td></td><td></td><td></td><td></td><td></td></tr>
<tr><td>Chromosome</td><td>start</td><td>end</td><td>strand</td><td>MM</td><td>target_seq</td><td>PAM</td><td>alignment</td><td>distance</td><td>position</td><td>gene name</td><td>gene id</td></tr>
<tr><td>GU305914.1</td><td>106000</td><td>106022</td><td>-</td><td>0</td><td>GATCACGGTGTTGACTATCG</td><td>CGG</td><td>||||||||[||||||||||||]PAM</td><td>NA</td><td>Intergenic</td><td>NA</td><td>NA</td></tr>
<tr><td>T26095</td><td>AATGTCCGCTGTATTCAACGCGG</td><td>1000</td><td></td><td></td><td></td><td></td><td></td><td></td><td></td><td></td><td></td></tr>
<tr><td>Oligo fwd</td><td>CACCAATGTCCGCTGTATTCAACG</td><td></td><td></td><td></td><td></td><td></td><td></td><td></td><td></td><td></td><td></td></tr>
<tr><td>Oligo rev</td><td>AAACCGTTGAATACAGCGGACATT</td><td></td><td></td><td></td><td></td><td></td><td></td><td></td><td></td><td></td><td></td></tr>
<tr><td>Chromosome</td><td>start</td><td>end</td><td>strand</td><td>MM</td><td>target_seq</td><td>PAM</td><td>alignment</td><td>distance</td><td>position</td><td>gene name</td><td>gene id</td></tr>
<tr><td>GU305914.1</td><td>105919</td><td>105941</td><td>-</td><td>0</td><td>AATGTCCGCTGTATTCAACG</td><td>CGG</td><td>||||||||[||||||||||||]PAM</td><td>NA</td><td>Intergenic</td><td>NA</td><td>NA</td></tr>
<tr><td>T26096</td><td>ACTCATGTCGATCGTCACCGCGG</td><td>1000</td><td></td><td></td><td></td><td></td><td></td><td></td><td></td><td></td><td></td></tr>
<tr><td>Oligo fwd</td><td>CACCACTCATGTCGATCGTCACCG</td><td></td><td></td><td></td><td></td><td></td><td></td><td></td><td></td><td></td><td></td></tr>
<tr><td>Oligo rev</td><td>AAACCGGTGACGATCGACATGAGT</td><td></td><td></td><td></td><td></td><td></td><td></td><td></td><td></td><td></td><td></td></tr>
<tr><td>Chromosome</td><td>start</td><td>end</td><td>strand</td><td>MM</td><td>target_seq</td><td>PAM</td><td>alignment</td><td>distance</td><td>position</td><td>gene name</td><td>gene id</td></tr>
<tr><td>GU305914.1</td><td>105867</td><td>105889</td><td>-</td><td>0</td><td>ACTCATGTCGATCGTCACCG</td><td>CGG</td><td>||||||||[||||||||||||]PAM</td><td>NA</td><td>Intergenic</td><td>NA</td><td>NA</td></tr>
<tr><td>T26097</td><td>CGAAAGTGAACAGCTCACTACGG</td><td>1000</td><td></td><td></td><td></td><td></td><td></td><td></td><td></td><td></td><td></td></tr>
<tr><td>Oligo fwd</td><td>CACCCGAAAGTGAACAGCTCACTA</td><td></td><td></td><td></td><td></td><td></td><td></td><td></td><td></td><td></td><td></td></tr>
<tr><td>Oligo rev</td><td>AAACTAGTGAGCTGTTCACTTTCG</td><td></td><td></td><td></td><td></td><td></td><td></td><td></td><td></td><td></td><td></td></tr>
<tr><td>Chromosome</td><td>start</td><td>end</td><td>strand</td><td>MM</td><td>target_seq</td><td>PAM</td><td>alignment</td><td>distance</td><td>position</td><td>gene name</td><td>gene id</td></tr>
<tr><td>GU305914.1</td><td>105835</td><td>105857</td><td>-</td><td>0</td><td>CGAAAGTGAACAGCTCACTA</td><td>CGG</td><td>||||||||[||||||||||||]PAM</td><td>NA</td><td>Intergenic</td><td>NA</td><td>NA</td></tr>
<tr><td>T26098</td><td>GAAAAGTGTAGCATACAACCCGG</td><td>1000</td><td></td><td></td><td></td><td></td><td></td><td></td><td></td><td></td><td></td></tr>
<tr><td>Oligo fwd</td><td>CACCGAAAAGTGTAGCATACAACC</td><td></td><td></td><td></td><td></td><td></td><td></td><td></td><td></td><td></td><td></td></tr>
<tr><td>Oligo rev</td><td>AAACGGTTGTATGCTACACTTTTC</td><td></td><td></td><td></td><td></td><td></td><td></td><td></td><td></td><td></td><td></td></tr>
</table>

| | | | | | | | | | | | | |
|---|---|---|---|---|---|---|---|---|---|---|---|---|
| Chromosome | start | | end | strand | MM | target_seq | PAM | alignment | distance | position | gene name | gene id |
| GU305914.1 | | 105813 | 105835 | - | 0 | GAAAAGTGTAGCATACAACC | CGG | \|\|\|\|\|\|\|\|[\|\|\|\|\|\|\|\|\|\|\|\|]PAM | NA | Intergenic | NA | NA |
| T26099 | GCATACAACCCGGCCGTTACCGG | | 1000 | | | | | | | | | |
| Oligo fwd | CACCGCATACAACCCGGCCGTTAC | | | | | | | | | | | |
| Oligo rev | AAACGTAACGGCCGGGTTGTATGC | | | | | | | | | | | |
| Chromosome | start | | end | strand | MM | target_seq | PAM | alignment | distance | position | gene name | gene id |
| GU305914.1 | | 105803 | 105825 | - | 0 | GCATACAACCCGGCCGTTAC | CGG | \|\|\|\|\|\|\|\|[\|\|\|\|\|\|\|\|\|\|\|\|]PAM | NA | Intergenic | NA | NA |
| T26100 | AGCGAAGATGATTATGAAGACGG | | 1000 | | | | | | | | | |
| Oligo fwd | CACCAGCGAAGATGATTATGAAGA | | | | | | | | | | | |
| Oligo rev | AAACTCTTCATAATCATCTTCGCT | | | | | | | | | | | |
| Chromosome | start | | end | strand | MM | target_seq | PAM | alignment | distance | position | gene name | gene id |
| GU305914.1 | | 105767 | 105789 | - | 0 | AGCGAAGATGATTATGAAGA | CGG | \|\|\|\|\|\|\|\|[\|\|\|\|\|\|\|\|\|\|\|\|]PAM | NA | Intergenic | NA | NA |
| T26101 | GAAGATGATTATGAAGACGGAGG | | 1000 | | | | | | | | | |
| Oligo fwd | CACCGAAGATGATTATGAAGACGG | | | | | | | | | | | |
| Oligo rev | AAACCCGTCTTCATAATCATCTTC | | | | | | | | | | | |
| Chromosome | start | | end | strand | MM | target_seq | PAM | alignment | distance | position | gene name | gene id |
| GU305914.1 | | 105764 | 105786 | - | 0 | GAAGATGATTATGAAGACGG | AGG | \|\|\|\|\|\|\|\|[\|\|\|\|\|\|\|\|\|\|\|\|]PAM | NA | Intergenic | NA | NA |
| T26102 | ATCGATGCGACCGAACCTCCCGG | | 1000 | | | | | | | | | |
| Oligo fwd | CACCATCGATGCGACCGAACCTCC | | | | | | | | | | | |
| Oligo rev | AAACGGAGGTTCGGTCGCATCGAT | | | | | | | | | | | |
| Chromosome | start | | end | strand | MM | target_seq | PAM | alignment | distance | position | gene name | gene id |
| GU305914.1 | | 105731 | 105753 | - | 0 | ATCGATGCGACCGAACCTCC | CGG | \|\|\|\|\|\|\|\|[\|\|\|\|\|\|\|\|\|\|\|\|]PAM | NA | Intergenic | NA | NA |
| T26103 | CCTCCCGGCGAAATACACATAGG | | 1000 | | | | | | | | | |
| Oligo fwd | CACCCCTCCCGGCGAAATACACAT | | | | | | | | | | | |
| Oligo rev | AAACATGTGTATTTCGCCGGGAGG | | | | | | | | | | | |
| Chromosome | start | | end | strand | MM | target_seq | PAM | alignment | distance | position | gene name | gene id |
| GU305914.1 | | 105716 | 105738 | - | 0 | CCTCCCGGCGAAATACACAT | AGG | \|\|\|\|\|\|\|\|[\|\|\|\|\|\|\|\|\|\|\|\|]PAM | NA | Intergenic | NA | NA |
| T26104 | AATACACATAGGTTCGTCTCTGG | | 1000 | | | | | | | | | |
| Oligo fwd | CACCAATACACATAGGTTCGTCTC | | | | | | | | | | | |
| Oligo rev | AAACGAGACGAACCTATGTGTATT | | | | | | | | | | | |
| Chromosome | start | | end | strand | MM | target_seq | PAM | alignment | distance | position | gene name | gene id |
| GU305914.1 | | 105705 | 105727 | - | 0 | AATACACATAGGTTCGTCTC | TGG | \|\|\|\|\|\|\|\|[\|\|\|\|\|\|\|\|\|\|\|\|]PAM | NA | Intergenic | NA | NA |
| T26105 | CAAAGACTATCTCTCATTCACGG | | 1000 | | | | | | | | | |
| Oligo fwd | CACCCAAAGACTATCTCTCATTCA | | | | | | | | | | | |
| Oligo rev | AAACTGAATGAGAGATAGTCTTTG | | | | | | | | | | | |
| Chromosome | start | | end | strand | MM | target_seq | PAM | alignment | distance | position | gene name | gene id |
| GU305914.1 | | 105621 | 105643 | - | 0 | CAAAGACTATCTCTCATTCA | CGG | \|\|\|\|\|\|\|\|[\|\|\|\|\|\|\|\|\|\|\|\|]PAM | NA | Intergenic | NA | NA |
| T26106 | CAGTCAAATTTTCAAAAACATGG | | 1000 | | | | | | | | | |
| Oligo fwd | CACCCAGTCAAATTTTCAAAAACA | | | | | | | | | | | |
| Oligo rev | AAACTGTTTTTGAAAATTTGACTG | | | | | | | | | | | |
| Chromosome | start | | end | strand | MM | target_seq | PAM | alignment | distance | position | gene name | gene id |
| GU305914.1 | | 105592 | 105614 | - | 0 | CAGTCAAATTTTCAAAAACA | TGG | \|\|\|\|\|\|\|\|[\|\|\|\|\|\|\|\|\|\|\|\|]PAM | NA | Intergenic | NA | NA |
| T26107 | TTTCAAAAACATGGATCCGAAGG | | 1000 | | | | | | | | | |
| Oligo fwd | CACCTTTCAAAAACATGGATCCGA | | | | | | | | | | | |
| Oligo rev | AAACTCGGATCCATGTTTTTGAAA | | | | | | | | | | | |
| Chromosome | start | | end | strand | MM | target_seq | PAM | alignment | distance | position | gene name | gene id |
| GU305914.1 | | 105583 | 105605 | - | 0 | TTTCAAAAACATGGATCCGA | AGG | \|\|\|\|\|\|\|\|[\|\|\|\|\|\|\|\|\|\|\|\|]PAM | NA | Intergenic | NA | NA |
| T26108 | CCGAAGGAGAAATCGACGTCCGG | | 1000 | | | | | | | | | |
| Oligo fwd | CACCCCGAAGGAGAAATCGACGTC | | | | | | | | | | | |
| Oligo rev | AAACGACGTCGATTTCTCCTTCGG | | | | | | | | | | | |
| Chromosome | start | | end | strand | MM | target_seq | PAM | alignment | distance | position | gene name | gene id |
| GU305914.1 | | 105567 | 105589 | - | 0 | CCGAAGGAGAAATCGACGTC | CGG | \|\|\|\|\|\|\|\|[\|\|\|\|\|\|\|\|\|\|\|\|]PAM | NA | Intergenic | NA | NA |
| T26109 | CGGACAACGTCTTTCCCATCTGG | | 1000 | | | | | | | | | |
| Oligo fwd | CACCCGGACAACGTCTTTCCCATC | | | | | | | | | | | |
| Oligo rev | AAACGATGGGAAAGACGTTGTCCG | | | | | | | | | | | |
| Chromosome | start | | end | strand | MM | target_seq | PAM | alignment | distance | position | gene name | gene id |
| GU305914.1 | | 105547 | 105569 | - | 0 | CGGACAACGTCTTTCCCATC | TGG | \|\|\|\|\|\|\|\|[\|\|\|\|\|\|\|\|\|\|\|\|]PAM | NA | Intergenic | NA | NA |
| T26110 | TCCCATCTGGTATCTCATCGAGG | | 1000 | | | | | | | | | |
| Oligo fwd | CACCTCCCATCTGGTATCTCATCG | | | | | | | | | | | |
| Oligo rev | AAACCGATGAGATACCAGATGGGA | | | | | | | | | | | |
| Chromosome | start | | end | strand | MM | target_seq | PAM | alignment | distance | position | gene name | gene id |
| GU305914.1 | | 105534 | 105556 | - | 0 | TCCCATCTGGTATCTCATCG | AGG | \|\|\|\|\|\|\|\|[\|\|\|\|\|\|\|\|\|\|\|\|]PAM | NA | Intergenic | NA | NA |
| T26111 | CCCATCTGGTATCTCATCGAGGG | | 1000 | | | | | | | | | |
| Oligo fwd | CACCCCCATCTGGTATCTCATCGA | | | | | | | | | | | |
| Oligo rev | AAACTCGATGAGATACCAGATGGG | | | | | | | | | | | |
| Chromosome | start | | end | strand | MM | target_seq | PAM | alignment | distance | position | gene name | gene id |
| GU305914.1 | | 105533 | 105555 | - | 0 | CCCATCTGGTATCTCATCGA | GGG | \|\|\|\|\|\|\|\|[\|\|\|\|\|\|\|\|\|\|\|\|]PAM | NA | Intergenic | NA | NA |
| T26112 | GTATCTCATCGAGGGAACAGAGG | | 1000 | | | | | | | | | |
| Oligo fwd | CACCGTATCTCATCGAGGGAACAG | | | | | | | | | | | |
| Oligo rev | AAACCTGTTCCCTCGATGAGATAC | | | | | | | | | | | |
| Chromosome | start | | end | strand | MM | target_seq | PAM | alignment | distance | position | gene name | gene id |
| GU305914.1 | | 105525 | 105547 | - | 0 | GTATCTCATCGAGGGAACAG | AGG | \|\|\|\|\|\|\|\|[\|\|\|\|\|\|\|\|\|\|\|\|]PAM | NA | Intergenic | NA | NA |
| T26113 | TCTCATCGAGGGAACAGAGGTGG | | 1000 | | | | | | | | | |
| Oligo fwd | CACCTCTCATCGAGGGAACAGAGG | | | | | | | | | | | |

<table>
<tr><td>Oligo rev</td><td>AAACCCTCTGTTCCCTCGATGAGA</td><td></td><td></td><td></td><td></td><td></td><td></td><td></td><td></td><td></td><td></td></tr>
<tr><td>Chromosome</td><td>start</td><td>end</td><td>strand</td><td>MM</td><td>target_seq</td><td>PAM</td><td>alignment</td><td>distance</td><td>position</td><td>gene name</td><td>gene id</td></tr>
<tr><td>GU305914.1</td><td>105522</td><td>105544</td><td>-</td><td>0</td><td>TCTCATCGAGGGAACAGAGG</td><td>TGG</td><td>||||||||[||||||||||||]PAM</td><td>NA</td><td>Intergenic</td><td>NA</td><td>NA</td></tr>
<tr><td>T26114</td><td>GGCGAGCATCAAAGCCTACGCGG</td><td>1000</td><td></td><td></td><td></td><td></td><td></td><td></td><td></td><td></td><td></td></tr>
<tr><td>Oligo fwd</td><td>CACCGGCGAGCATCAAAGCCTACG</td><td></td><td></td><td></td><td></td><td></td><td></td><td></td><td></td><td></td><td></td></tr>
<tr><td>Oligo rev</td><td>AAACCGTAGGCTTTGATGCTCGCC</td><td></td><td></td><td></td><td></td><td></td><td></td><td></td><td></td><td></td><td></td></tr>
<tr><td>Chromosome</td><td>start</td><td>end</td><td>strand</td><td>MM</td><td>target_seq</td><td>PAM</td><td>alignment</td><td>distance</td><td>position</td><td>gene name</td><td>gene id</td></tr>
<tr><td>GU305914.1</td><td>105501</td><td>105523</td><td>-</td><td>0</td><td>GGCGAGCATCAAAGCCTACG</td><td>CGG</td><td>||||||||[||||||||||||]PAM</td><td>NA</td><td>Intergenic</td><td>NA</td><td>NA</td></tr>
<tr><td>T26115</td><td>GCGAGCATCAAAGCCTACGCGGG</td><td>1000</td><td></td><td></td><td></td><td></td><td></td><td></td><td></td><td></td><td></td></tr>
<tr><td>Oligo fwd</td><td>CACCGCGAGCATCAAAGCCTACGC</td><td></td><td></td><td></td><td></td><td></td><td></td><td></td><td></td><td></td><td></td></tr>
<tr><td>Oligo rev</td><td>AAACGCGTAGGCTTTGATGCTCGC</td><td></td><td></td><td></td><td></td><td></td><td></td><td></td><td></td><td></td><td></td></tr>
<tr><td>Chromosome</td><td>start</td><td>end</td><td>strand</td><td>MM</td><td>target_seq</td><td>PAM</td><td>alignment</td><td>distance</td><td>position</td><td>gene name</td><td>gene id</td></tr>
<tr><td>GU305914.1</td><td>105500</td><td>105522</td><td>-</td><td>0</td><td>GCGAGCATCAAAGCCTACGC</td><td>GGG</td><td>||||||||[||||||||||||]PAM</td><td>NA</td><td>Intergenic</td><td>NA</td><td>NA</td></tr>
<tr><td>T26116</td><td>AATAATAAAACTAGCACAATCGG</td><td>1000</td><td></td><td></td><td></td><td></td><td></td><td></td><td></td><td></td><td></td></tr>
<tr><td>Oligo fwd</td><td>CACCAATAATAAAACTAGCACAAT</td><td></td><td></td><td></td><td></td><td></td><td></td><td></td><td></td><td></td><td></td></tr>
<tr><td>Oligo rev</td><td>AAACATTGTGCTAGTTTTATTATT</td><td></td><td></td><td></td><td></td><td></td><td></td><td></td><td></td><td></td><td></td></tr>
<tr><td>Chromosome</td><td>start</td><td>end</td><td>strand</td><td>MM</td><td>target_seq</td><td>PAM</td><td>alignment</td><td>distance</td><td>position</td><td>gene name</td><td>gene id</td></tr>
<tr><td>GU305914.1</td><td>105437</td><td>105459</td><td>-</td><td>0</td><td>AATAATAAAACTAGCACAAT</td><td>CGG</td><td>||||||||[||||||||||||]PAM</td><td>NA</td><td>Intergenic</td><td>NA</td><td>NA</td></tr>
<tr><td>T26117</td><td>ATAATAAAACTAGCACAATCGGG</td><td>1000</td><td></td><td></td><td></td><td></td><td></td><td></td><td></td><td></td><td></td></tr>
<tr><td>Oligo fwd</td><td>CACCATAATAAAACTAGCACAATC</td><td></td><td></td><td></td><td></td><td></td><td></td><td></td><td></td><td></td><td></td></tr>
<tr><td>Oligo rev</td><td>AAACGATTGTGCTAGTTTTATTAT</td><td></td><td></td><td></td><td></td><td></td><td></td><td></td><td></td><td></td><td></td></tr>
<tr><td>Chromosome</td><td>start</td><td>end</td><td>strand</td><td>MM</td><td>target_seq</td><td>PAM</td><td>alignment</td><td>distance</td><td>position</td><td>gene name</td><td>gene id</td></tr>
<tr><td>GU305914.1</td><td>105436</td><td>105458</td><td>-</td><td>0</td><td>ATAATAAAACTAGCACAATC</td><td>GGG</td><td>||||||||[||||||||||||]PAM</td><td>NA</td><td>Intergenic</td><td>NA</td><td>NA</td></tr>
<tr><td>T26118</td><td>ATCGGGACCGTAATCGTGAGCGG</td><td>1000</td><td></td><td></td><td></td><td></td><td></td><td></td><td></td><td></td><td></td></tr>
<tr><td>Oligo fwd</td><td>CACCATCGGGACCGTAATCGTGAG</td><td></td><td></td><td></td><td></td><td></td><td></td><td></td><td></td><td></td><td></td></tr>
<tr><td>Oligo rev</td><td>AAACCTCACGATTACGGTCCCGAT</td><td></td><td></td><td></td><td></td><td></td><td></td><td></td><td></td><td></td><td></td></tr>
<tr><td>Chromosome</td><td>start</td><td>end</td><td>strand</td><td>MM</td><td>target_seq</td><td>PAM</td><td>alignment</td><td>distance</td><td>position</td><td>gene name</td><td>gene id</td></tr>
<tr><td>GU305914.1</td><td>105419</td><td>105441</td><td>-</td><td>0</td><td>ATCGGGACCGTAATCGTGAG</td><td>CGG</td><td>||||||||[||||||||||||]PAM</td><td>NA</td><td>Intergenic</td><td>NA</td><td>NA</td></tr>
<tr><td>T26119</td><td>ATCGTGAGCGGAGTACCTTGCGG</td><td>1000</td><td></td><td></td><td></td><td></td><td></td><td></td><td></td><td></td><td></td></tr>
<tr><td>Oligo fwd</td><td>CACCATCGTGAGCGGAGTACCTTG</td><td></td><td></td><td></td><td></td><td></td><td></td><td></td><td></td><td></td><td></td></tr>
<tr><td>Oligo rev</td><td>AAACCAAGGTACTCCGCTCACGAT</td><td></td><td></td><td></td><td></td><td></td><td></td><td></td><td></td><td></td><td></td></tr>
<tr><td>Chromosome</td><td>start</td><td>end</td><td>strand</td><td>MM</td><td>target_seq</td><td>PAM</td><td>alignment</td><td>distance</td><td>position</td><td>gene name</td><td>gene id</td></tr>
<tr><td>GU305914.1</td><td>105407</td><td>105429</td><td>-</td><td>0</td><td>ATCGTGAGCGGAGTACCTTG</td><td>CGG</td><td>||||||||[||||||||||||]PAM</td><td>NA</td><td>Intergenic</td><td>NA</td><td>NA</td></tr>
<tr><td>T26120</td><td>GCTTTCGTGTCACCGATCCATGG</td><td>1000</td><td></td><td></td><td></td><td></td><td></td><td></td><td></td><td></td><td></td></tr>
<tr><td>Oligo fwd</td><td>CACCGCTTTCGTGTCACCGATCCA</td><td></td><td></td><td></td><td></td><td></td><td></td><td></td><td></td><td></td><td></td></tr>
<tr><td>Oligo rev</td><td>AAACTGGATCGGTGACACGAAAGC</td><td></td><td></td><td></td><td></td><td></td><td></td><td></td><td></td><td></td><td></td></tr>
<tr><td>Chromosome</td><td>start</td><td>end</td><td>strand</td><td>MM</td><td>target_seq</td><td>PAM</td><td>alignment</td><td>distance</td><td>position</td><td>gene name</td><td>gene id</td></tr>
<tr><td>GU305914.1</td><td>105372</td><td>105394</td><td>-</td><td>0</td><td>GCTTTCGTGTCACCGATCCA</td><td>TGG</td><td>||||||||[||||||||||||]PAM</td><td>NA</td><td>Intergenic</td><td>NA</td><td>NA</td></tr>
<tr><td>T26121</td><td>CTTTCGTGTCACCGATCCATGGG</td><td>1000</td><td></td><td></td><td></td><td></td><td></td><td></td><td></td><td></td><td></td></tr>
<tr><td>Oligo fwd</td><td>CACCCTTTCGTGTCACCGATCCAT</td><td></td><td></td><td></td><td></td><td></td><td></td><td></td><td></td><td></td><td></td></tr>
<tr><td>Oligo rev</td><td>AAACATGGATCGGTGACACGAAAG</td><td></td><td></td><td></td><td></td><td></td><td></td><td></td><td></td><td></td><td></td></tr>
<tr><td>Chromosome</td><td>start</td><td>end</td><td>strand</td><td>MM</td><td>target_seq</td><td>PAM</td><td>alignment</td><td>distance</td><td>position</td><td>gene name</td><td>gene id</td></tr>
<tr><td>GU305914.1</td><td>105371</td><td>105393</td><td>-</td><td>0</td><td>CTTTCGTGTCACCGATCCAT</td><td>GGG</td><td>||||||||[||||||||||||]PAM</td><td>NA</td><td>Intergenic</td><td>NA</td><td>NA</td></tr>
<tr><td>T26122</td><td>TTTCGTGTCACCGATCCATGGGG</td><td>1000</td><td></td><td></td><td></td><td></td><td></td><td></td><td></td><td></td><td></td></tr>
<tr><td>Oligo fwd</td><td>CACCTTTCGTGTCACCGATCCATG</td><td></td><td></td><td></td><td></td><td></td><td></td><td></td><td></td><td></td><td></td></tr>
<tr><td>Oligo rev</td><td>AAACCATGGATCGGTGACACGAAA</td><td></td><td></td><td></td><td></td><td></td><td></td><td></td><td></td><td></td><td></td></tr>
<tr><td>Chromosome</td><td>start</td><td>end</td><td>strand</td><td>MM</td><td>target_seq</td><td>PAM</td><td>alignment</td><td>distance</td><td>position</td><td>gene name</td><td>gene id</td></tr>
<tr><td>GU305914.1</td><td>105370</td><td>105392</td><td>-</td><td>0</td><td>TTTCGTGTCACCGATCCATG</td><td>GGG</td><td>||||||||[||||||||||||]PAM</td><td>NA</td><td>Intergenic</td><td>NA</td><td>NA</td></tr>
<tr><td>T26123</td><td>TCCTGTACGACCTGCCCCGCTGG</td><td>1000</td><td></td><td></td><td></td><td></td><td></td><td></td><td></td><td></td><td></td></tr>
<tr><td>Oligo fwd</td><td>CACCTCCTGTACGACCTGCCCCGC</td><td></td><td></td><td></td><td></td><td></td><td></td><td></td><td></td><td></td><td></td></tr>
<tr><td>Oligo rev</td><td>AAACGCGGGGCAGGTCGTACAGGA</td><td></td><td></td><td></td><td></td><td></td><td></td><td></td><td></td><td></td><td></td></tr>
<tr><td>Chromosome</td><td>start</td><td>end</td><td>strand</td><td>MM</td><td>target_seq</td><td>PAM</td><td>alignment</td><td>distance</td><td>position</td><td>gene name</td><td>gene id</td></tr>
<tr><td>GU305914.1</td><td>105271</td><td>105293</td><td>-</td><td>0</td><td>TCCTGTACGACCTGCCCCGC</td><td>TGG</td><td>||||||||[||||||||||||]PAM</td><td>NA</td><td>Intergenic</td><td>NA</td><td>NA</td></tr>
<tr><td>T26124</td><td>CGACCTGCCCCGCTGGCAAGTGG</td><td>1000</td><td></td><td></td><td></td><td></td><td></td><td></td><td></td><td></td><td></td></tr>
<tr><td>Oligo fwd</td><td>CACCCGACCTGCCCCGCTGGCAAG</td><td></td><td></td><td></td><td></td><td></td><td></td><td></td><td></td><td></td><td></td></tr>
<tr><td>Oligo rev</td><td>AAACCTTGCCAGCGGGGCAGGTCG</td><td></td><td></td><td></td><td></td><td></td><td></td><td></td><td></td><td></td><td></td></tr>
<tr><td>Chromosome</td><td>start</td><td>end</td><td>strand</td><td>MM</td><td>target_seq</td><td>PAM</td><td>alignment</td><td>distance</td><td>position</td><td>gene name</td><td>gene id</td></tr>
<tr><td>GU305914.1</td><td>105264</td><td>105286</td><td>-</td><td>0</td><td>CGACCTGCCCCGCTGGCAAG</td><td>TGG</td><td>||||||||[||||||||||||]PAM</td><td>NA</td><td>Intergenic</td><td>NA</td><td>NA</td></tr>
<tr><td>T26125</td><td>CCCGCTGGCAAGTGGAACTGAGG</td><td>1000</td><td></td><td></td><td></td><td></td><td></td><td></td><td></td><td></td><td></td></tr>
<tr><td>Oligo fwd</td><td>CACCCCCGCTGGCAAGTGGAACTG</td><td></td><td></td><td></td><td></td><td></td><td></td><td></td><td></td><td></td><td></td></tr>
<tr><td>Oligo rev</td><td>AAACCAGTTCCACTTGCCAGCGGG</td><td></td><td></td><td></td><td></td><td></td><td></td><td></td><td></td><td></td><td></td></tr>
<tr><td>Chromosome</td><td>start</td><td>end</td><td>strand</td><td>MM</td><td>target_seq</td><td>PAM</td><td>alignment</td><td>distance</td><td>position</td><td>gene name</td><td>gene id</td></tr>
<tr><td>GU305914.1</td><td>105256</td><td>105278</td><td>-</td><td>0</td><td>CCCGCTGGCAAGTGGAACTG</td><td>AGG</td><td>||||||||[||||||||||||]PAM</td><td>NA</td><td>Intergenic</td><td>NA</td><td>NA</td></tr>
<tr><td>T26126</td><td>GCAAGTGGAACTGAGGACCCCGG</td><td>1000</td><td></td><td></td><td></td><td></td><td></td><td></td><td></td><td></td><td></td></tr>
<tr><td>Oligo fwd</td><td>CACCGCAAGTGGAACTGAGGACCC</td><td></td><td></td><td></td><td></td><td></td><td></td><td></td><td></td><td></td><td></td></tr>
<tr><td>Oligo rev</td><td>AAACGGGTCCTCAGTTCCACTTGC</td><td></td><td></td><td></td><td></td><td></td><td></td><td></td><td></td><td></td><td></td></tr>
<tr><td>Chromosome</td><td>start</td><td>end</td><td>strand</td><td>MM</td><td>target_seq</td><td>PAM</td><td>alignment</td><td>distance</td><td>position</td><td>gene name</td><td>gene id</td></tr>
<tr><td>GU305914.1</td><td>105249</td><td>105271</td><td>-</td><td>0</td><td>GCAAGTGGAACTGAGGACCC</td><td>CGG</td><td>||||||||[||||||||||||]PAM</td><td>NA</td><td>Intergenic</td><td>NA</td><td>NA</td></tr>
<tr><td>T26127</td><td>GACCCCGGACAGACGCGTGTTGG</td><td>1000</td><td></td><td></td><td></td><td></td><td></td><td></td><td></td><td></td><td></td></tr>
<tr><td>Oligo fwd</td><td>CACCGACCCCGGACAGACGCGTGT</td><td></td><td></td><td></td><td></td><td></td><td></td><td></td><td></td><td></td><td></td></tr>
<tr><td>Oligo rev</td><td>AAACACACGCGTCTGTCCGGGGTC</td><td></td><td></td><td></td><td></td><td></td><td></td><td></td><td></td><td></td><td></td></tr>
<tr><td>Chromosome</td><td>start</td><td>end</td><td>strand</td><td>MM</td><td>target_seq</td><td>PAM</td><td>alignment</td><td>distance</td><td>position</td><td>gene name</td><td>gene id</td></tr>
<tr><td>GU305914.1</td><td>105234</td><td>105256</td><td>-</td><td>0</td><td>GACCCCGGACAGACGCGTGT</td><td>TGG</td><td>||||||||[||||||||||||]PAM</td><td>NA</td><td>Intergenic</td><td>NA</td><td>NA</td></tr>
<tr><td>T26128</td><td>TTTTCAAAGTCACAGCCTCCTGG</td><td>1000</td><td></td><td></td><td></td><td></td><td></td><td></td><td></td><td></td><td></td></tr>
</table>

<table>
<tr><td>Oligo fwd</td><td>CACCTTTTCAAAGTCACAGCCTCC</td><td></td><td></td><td></td><td></td><td></td><td></td><td></td><td></td><td></td><td></td></tr>
<tr><td>Oligo rev</td><td>AAACGGAGGCTGTGACTTTGAAAA</td><td></td><td></td><td></td><td></td><td></td><td></td><td></td><td></td><td></td><td></td></tr>
<tr><td>Chromosome</td><td>start</td><td>end</td><td>strand</td><td>MM</td><td>target_seq</td><td>PAM</td><td>alignment</td><td>distance</td><td>position</td><td>gene name</td><td>gene id</td></tr>
<tr><td>GU305914.1</td><td>105091</td><td>105113</td><td>-</td><td>0</td><td>TTTTCAAAGTCACAGCCTCC</td><td>TGG</td><td>||||||||[||||||||||||]PAM</td><td>NA</td><td>Intergenic</td><td>NA</td><td>NA</td></tr>
<tr><td>T26129</td><td>CAAAGTCACAGCCTCCTGGCCGG</td><td>1000</td><td></td><td></td><td></td><td></td><td></td><td></td><td></td><td></td><td></td></tr>
<tr><td>Oligo fwd</td><td>CACCCAAAGTCACAGCCTCCTGGC</td><td></td><td></td><td></td><td></td><td></td><td></td><td></td><td></td><td></td><td></td></tr>
<tr><td>Oligo rev</td><td>AAACGCCAGGAGGCTGTGACTTTG</td><td></td><td></td><td></td><td></td><td></td><td></td><td></td><td></td><td></td><td></td></tr>
<tr><td>Chromosome</td><td>start</td><td>end</td><td>strand</td><td>MM</td><td>target_seq</td><td>PAM</td><td>alignment</td><td>distance</td><td>position</td><td>gene name</td><td>gene id</td></tr>
<tr><td>GU305914.1</td><td>105087</td><td>105109</td><td>-</td><td>0</td><td>CAAAGTCACAGCCTCCTGGC</td><td>CGG</td><td>||||||||[||||||||||||]PAM</td><td>NA</td><td>Intergenic</td><td>NA</td><td>NA</td></tr>
<tr><td>T26130</td><td>ACATCAAAACGTTTTTCGATAGG</td><td>1000</td><td></td><td></td><td></td><td></td><td></td><td></td><td></td><td></td><td></td></tr>
<tr><td>Oligo fwd</td><td>CACCACATCAAAACGTTTTTCGAT</td><td></td><td></td><td></td><td></td><td></td><td></td><td></td><td></td><td></td><td></td></tr>
<tr><td>Oligo rev</td><td>AAACATCGAAAAACGTTTTGATGT</td><td></td><td></td><td></td><td></td><td></td><td></td><td></td><td></td><td></td><td></td></tr>
<tr><td>Chromosome</td><td>start</td><td>end</td><td>strand</td><td>MM</td><td>target_seq</td><td>PAM</td><td>alignment</td><td>distance</td><td>position</td><td>gene name</td><td>gene id</td></tr>
<tr><td>GU305914.1</td><td>105064</td><td>105086</td><td>-</td><td>0</td><td>ACATCAAAACGTTTTTCGAT</td><td>AGG</td><td>||||||||[||||||||||||]PAM</td><td>NA</td><td>Intergenic</td><td>NA</td><td>NA</td></tr>
<tr><td>T26131</td><td>ATCTCAAACACTCGCACCCGCGG</td><td>1000</td><td></td><td></td><td></td><td></td><td></td><td></td><td></td><td></td><td></td></tr>
<tr><td>Oligo fwd</td><td>CACCATCTCAAACACTCGCACCCG</td><td></td><td></td><td></td><td></td><td></td><td></td><td></td><td></td><td></td><td></td></tr>
<tr><td>Oligo rev</td><td>AAACCGGGTGCGAGTGTTTGAGAT</td><td></td><td></td><td></td><td></td><td></td><td></td><td></td><td></td><td></td><td></td></tr>
<tr><td>Chromosome</td><td>start</td><td>end</td><td>strand</td><td>MM</td><td>target_seq</td><td>PAM</td><td>alignment</td><td>distance</td><td>position</td><td>gene name</td><td>gene id</td></tr>
<tr><td>GU305914.1</td><td>105027</td><td>105049</td><td>-</td><td>0</td><td>ATCTCAAACACTCGCACCCG</td><td>CGG</td><td>||||||||[||||||||||||]PAM</td><td>NA</td><td>Intergenic</td><td>NA</td><td>NA</td></tr>
<tr><td>T26132</td><td>CCGCGGTCACCTTACCCCCGAGG</td><td>1000</td><td></td><td></td><td></td><td></td><td></td><td></td><td></td><td></td><td></td></tr>
<tr><td>Oligo fwd</td><td>CACCCCGCGGTCACCTTACCCCCG</td><td></td><td></td><td></td><td></td><td></td><td></td><td></td><td></td><td></td><td></td></tr>
<tr><td>Oligo rev</td><td>AAACCGGGGGTAAGGTGACCGCGG</td><td></td><td></td><td></td><td></td><td></td><td></td><td></td><td></td><td></td><td></td></tr>
<tr><td>Chromosome</td><td>start</td><td>end</td><td>strand</td><td>MM</td><td>target_seq</td><td>PAM</td><td>alignment</td><td>distance</td><td>position</td><td>gene name</td><td>gene id</td></tr>
<tr><td>GU305914.1</td><td>105010</td><td>105032</td><td>-</td><td>0</td><td>CCGCGGTCACCTTACCCCCG</td><td>AGG</td><td>||||||||[||||||||||||]PAM</td><td>NA</td><td>Intergenic</td><td>NA</td><td>NA</td></tr>
<tr><td>T26133</td><td>CATACCGAAAAACATGCACCCGG</td><td>1000</td><td></td><td></td><td></td><td></td><td></td><td></td><td></td><td></td><td></td></tr>
<tr><td>Oligo fwd</td><td>CACCCATACCGAAAAACATGCACC</td><td></td><td></td><td></td><td></td><td></td><td></td><td></td><td></td><td></td><td></td></tr>
<tr><td>Oligo rev</td><td>AAACGGTGCATGTTTTTCGGTATG</td><td></td><td></td><td></td><td></td><td></td><td></td><td></td><td></td><td></td><td></td></tr>
<tr><td>Chromosome</td><td>start</td><td>end</td><td>strand</td><td>MM</td><td>target_seq</td><td>PAM</td><td>alignment</td><td>distance</td><td>position</td><td>gene name</td><td>gene id</td></tr>
<tr><td>GU305914.1</td><td>104949</td><td>104971</td><td>-</td><td>0</td><td>CATACCGAAAAACATGCACC</td><td>CGG</td><td>||||||||[||||||||||||]PAM</td><td>NA</td><td>Intergenic</td><td>NA</td><td>NA</td></tr>
<tr><td>T26134</td><td>ATCGCAACCGTACTAACGAACGG</td><td>1000</td><td></td><td></td><td></td><td></td><td></td><td></td><td></td><td></td><td></td></tr>
<tr><td>Oligo fwd</td><td>CACCATCGCAACCGTACTAACGAA</td><td></td><td></td><td></td><td></td><td></td><td></td><td></td><td></td><td></td><td></td></tr>
<tr><td>Oligo rev</td><td>AAACTTCGTTAGTACGGTTGCGAT</td><td></td><td></td><td></td><td></td><td></td><td></td><td></td><td></td><td></td><td></td></tr>
<tr><td>Chromosome</td><td>start</td><td>end</td><td>strand</td><td>MM</td><td>target_seq</td><td>PAM</td><td>alignment</td><td>distance</td><td>position</td><td>gene name</td><td>gene id</td></tr>
<tr><td>GU305914.1</td><td>104843</td><td>104865</td><td>-</td><td>0</td><td>ATCGCAACCGTACTAACGAA</td><td>CGG</td><td>||||||||[||||||||||||]PAM</td><td>NA</td><td>Intergenic</td><td>NA</td><td>NA</td></tr>
<tr><td>T26135</td><td>AACCGTACTAACGAACGGTACGG</td><td>1000</td><td></td><td></td><td></td><td></td><td></td><td></td><td></td><td></td><td></td></tr>
<tr><td>Oligo fwd</td><td>CACCAACCGTACTAACGAACGGTA</td><td></td><td></td><td></td><td></td><td></td><td></td><td></td><td></td><td></td><td></td></tr>
<tr><td>Oligo rev</td><td>AAACTACCGTTCGTTAGTACGGTT</td><td></td><td></td><td></td><td></td><td></td><td></td><td></td><td></td><td></td><td></td></tr>
<tr><td>Chromosome</td><td>start</td><td>end</td><td>strand</td><td>MM</td><td>target_seq</td><td>PAM</td><td>alignment</td><td>distance</td><td>position</td><td>gene name</td><td>gene id</td></tr>
<tr><td>GU305914.1</td><td>104838</td><td>104860</td><td>-</td><td>0</td><td>AACCGTACTAACGAACGGTA</td><td>CGG</td><td>||||||||[||||||||||||]PAM</td><td>NA</td><td>Intergenic</td><td>NA</td><td>NA</td></tr>
<tr><td>T26136</td><td>CGTACTAACGAACGGTACGGTGG</td><td>1000</td><td></td><td></td><td></td><td></td><td></td><td></td><td></td><td></td><td></td></tr>
<tr><td>Oligo fwd</td><td>CACCCGTACTAACGAACGGTACGG</td><td></td><td></td><td></td><td></td><td></td><td></td><td></td><td></td><td></td><td></td></tr>
<tr><td>Oligo rev</td><td>AAACCCGTACCGTTCGTTAGTACG</td><td></td><td></td><td></td><td></td><td></td><td></td><td></td><td></td><td></td><td></td></tr>
<tr><td>Chromosome</td><td>start</td><td>end</td><td>strand</td><td>MM</td><td>target_seq</td><td>PAM</td><td>alignment</td><td>distance</td><td>position</td><td>gene name</td><td>gene id</td></tr>
<tr><td>GU305914.1</td><td>104835</td><td>104857</td><td>-</td><td>0</td><td>CGTACTAACGAACGGTACGG</td><td>TGG</td><td>||||||||[||||||||||||]PAM</td><td>NA</td><td>Intergenic</td><td>NA</td><td>NA</td></tr>
<tr><td>T26137</td><td>AACGAACGGTACGGTGGAGACGG</td><td>1000</td><td></td><td></td><td></td><td></td><td></td><td></td><td></td><td></td><td></td></tr>
<tr><td>Oligo fwd</td><td>CACCAACGAACGGTACGGTGGAGA</td><td></td><td></td><td></td><td></td><td></td><td></td><td></td><td></td><td></td><td></td></tr>
<tr><td>Oligo rev</td><td>AAACTCTCCACCGTACCGTTCGTT</td><td></td><td></td><td></td><td></td><td></td><td></td><td></td><td></td><td></td><td></td></tr>
<tr><td>Chromosome</td><td>start</td><td>end</td><td>strand</td><td>MM</td><td>target_seq</td><td>PAM</td><td>alignment</td><td>distance</td><td>position</td><td>gene name</td><td>gene id</td></tr>
<tr><td>GU305914.1</td><td>104829</td><td>104851</td><td>-</td><td>0</td><td>AACGAACGGTACGGTGGAGA</td><td>CGG</td><td>||||||||[||||||||||||]PAM</td><td>NA</td><td>Intergenic</td><td>NA</td><td>NA</td></tr>
<tr><td>T26138</td><td>GGTACGGTGGAGACGGTAGTAGG</td><td>1000</td><td></td><td></td><td></td><td></td><td></td><td></td><td></td><td></td><td></td></tr>
<tr><td>Oligo fwd</td><td>CACCGGTACGGTGGAGACGGTAGT</td><td></td><td></td><td></td><td></td><td></td><td></td><td></td><td></td><td></td><td></td></tr>
<tr><td>Oligo rev</td><td>AAACACTACCGTCTCCACCGTACC</td><td></td><td></td><td></td><td></td><td></td><td></td><td></td><td></td><td></td><td></td></tr>
<tr><td>Chromosome</td><td>start</td><td>end</td><td>strand</td><td>MM</td><td>target_seq</td><td>PAM</td><td>alignment</td><td>distance</td><td>position</td><td>gene name</td><td>gene id</td></tr>
<tr><td>GU305914.1</td><td>104822</td><td>104844</td><td>-</td><td>0</td><td>GGTACGGTGGAGACGGTAGT</td><td>AGG</td><td>||||||||[||||||||||||]PAM</td><td>NA</td><td>Intergenic</td><td>NA</td><td>NA</td></tr>
<tr><td>T26139</td><td>GGTGGAGACGGTAGTAGGACCGG</td><td>1000</td><td></td><td></td><td></td><td></td><td></td><td></td><td></td><td></td><td></td></tr>
<tr><td>Oligo fwd</td><td>CACCGGTGGAGACGGTAGTAGGAC</td><td></td><td></td><td></td><td></td><td></td><td></td><td></td><td></td><td></td><td></td></tr>
<tr><td>Oligo rev</td><td>AAACGTCCTACTACCGTCTCCACC</td><td></td><td></td><td></td><td></td><td></td><td></td><td></td><td></td><td></td><td></td></tr>
<tr><td>Chromosome</td><td>start</td><td>end</td><td>strand</td><td>MM</td><td>target_seq</td><td>PAM</td><td>alignment</td><td>distance</td><td>position</td><td>gene name</td><td>gene id</td></tr>
<tr><td>GU305914.1</td><td>104817</td><td>104839</td><td>-</td><td>0</td><td>GGTGGAGACGGTAGTAGGAC</td><td>CGG</td><td>||||||||[||||||||||||]PAM</td><td>NA</td><td>Intergenic</td><td>NA</td><td>NA</td></tr>
<tr><td>T26140</td><td>CGTTACAACGTTATACAAAACGG</td><td>1000</td><td></td><td></td><td></td><td></td><td></td><td></td><td></td><td></td><td></td></tr>
<tr><td>Oligo fwd</td><td>CACCCGTTACAACGTTATACAAAA</td><td></td><td></td><td></td><td></td><td></td><td></td><td></td><td></td><td></td><td></td></tr>
<tr><td>Oligo rev</td><td>AAACTTTTGTATAACGTTGTAACG</td><td></td><td></td><td></td><td></td><td></td><td></td><td></td><td></td><td></td><td></td></tr>
<tr><td>Chromosome</td><td>start</td><td>end</td><td>strand</td><td>MM</td><td>target_seq</td><td>PAM</td><td>alignment</td><td>distance</td><td>position</td><td>gene name</td><td>gene id</td></tr>
<tr><td>GU305914.1</td><td>104790</td><td>104812</td><td>-</td><td>0</td><td>CGTTACAACGTTATACAAAA</td><td>CGG</td><td>||||||||[||||||||||||]PAM</td><td>NA</td><td>Intergenic</td><td>NA</td><td>NA</td></tr>
<tr><td>T26141</td><td>CAAAAATAATAAGATCAACCTGG</td><td>1000</td><td></td><td></td><td></td><td></td><td></td><td></td><td></td><td></td><td></td></tr>
<tr><td>Oligo fwd</td><td>CACCCAAAAATAATAAGATCAACC</td><td></td><td></td><td></td><td></td><td></td><td></td><td></td><td></td><td></td><td></td></tr>
<tr><td>Oligo rev</td><td>AAACGGTTGATCTTATTATTTTTG</td><td></td><td></td><td></td><td></td><td></td><td></td><td></td><td></td><td></td><td></td></tr>
<tr><td>Chromosome</td><td>start</td><td>end</td><td>strand</td><td>MM</td><td>target_seq</td><td>PAM</td><td>alignment</td><td>distance</td><td>position</td><td>gene name</td><td>gene id</td></tr>
<tr><td>GU305914.1</td><td>104745</td><td>104767</td><td>-</td><td>0</td><td>CAAAAATAATAAGATCAACC</td><td>TGG</td><td>||||||||[||||||||||||]PAM</td><td>NA</td><td>Intergenic</td><td>NA</td><td>NA</td></tr>
<tr><td>T26142</td><td>AAATAATAAGATCAACCTGGAGG</td><td>1000</td><td></td><td></td><td></td><td></td><td></td><td></td><td></td><td></td><td></td></tr>
<tr><td>Oligo fwd</td><td>CACCAAATAATAAGATCAACCTGG</td><td></td><td></td><td></td><td></td><td></td><td></td><td></td><td></td><td></td><td></td></tr>
<tr><td>Oligo rev</td><td>AAACCCAGGTTGATCTTATTATTT</td><td></td><td></td><td></td><td></td><td></td><td></td><td></td><td></td><td></td><td></td></tr>
<tr><td>Chromosome</td><td>start</td><td>end</td><td>strand</td><td>MM</td><td>target_seq</td><td>PAM</td><td>alignment</td><td>distance</td><td>position</td><td>gene name</td><td>gene id</td></tr>
<tr><td>GU305914.1</td><td>104742</td><td>104764</td><td>-</td><td>0</td><td>AAATAATAAGATCAACCTGG</td><td>AGG</td><td>||||||||[||||||||||||]PAM</td><td>NA</td><td>Intergenic</td><td>NA</td><td>NA</td></tr>
</table>

<table>
<tr><td>T26143</td><td>AACTCTAGACGACGTACTCATGG</td><td>1000</td><td></td><td></td><td></td><td></td><td></td><td></td><td></td><td></td><td></td></tr>
<tr><td>Oligo fwd</td><td>CACCAACTCTAGACGACGTACTCA</td><td></td><td></td><td></td><td></td><td></td><td></td><td></td><td></td><td></td><td></td></tr>
<tr><td>Oligo rev</td><td>AAACTGAGTACGTCGTCTAGAGTT</td><td></td><td></td><td></td><td></td><td></td><td></td><td></td><td></td><td></td><td></td></tr>
<tr><td>Chromosome</td><td>start</td><td>end</td><td>strand</td><td>MM</td><td>target_seq</td><td>PAM</td><td>alignment</td><td>distance</td><td>position</td><td>gene name</td><td>gene id</td></tr>
<tr><td>GU305914.1</td><td>104712</td><td>104734</td><td>-</td><td>0</td><td>AACTCTAGACGACGTACTCA</td><td>TGG</td><td>||||||||[||||||||||||]PAM</td><td>NA</td><td>Intergenic</td><td>NA</td><td>NA</td></tr>
<tr><td>T26144</td><td>CGACGTACTCATGGATTATCTGG</td><td>1000</td><td></td><td></td><td></td><td></td><td></td><td></td><td></td><td></td><td></td></tr>
<tr><td>Oligo fwd</td><td>CACCCGACGTACTCATGGATTATC</td><td></td><td></td><td></td><td></td><td></td><td></td><td></td><td></td><td></td><td></td></tr>
<tr><td>Oligo rev</td><td>AAACGATAATCCATGAGTACGTCG</td><td></td><td></td><td></td><td></td><td></td><td></td><td></td><td></td><td></td><td></td></tr>
<tr><td>Chromosome</td><td>start</td><td>end</td><td>strand</td><td>MM</td><td>target_seq</td><td>PAM</td><td>alignment</td><td>distance</td><td>position</td><td>gene name</td><td>gene id</td></tr>
<tr><td>GU305914.1</td><td>104703</td><td>104725</td><td>-</td><td>0</td><td>CGACGTACTCATGGATTATC</td><td>TGG</td><td>||||||||[||||||||||||]PAM</td><td>NA</td><td>Intergenic</td><td>NA</td><td>NA</td></tr>
<tr><td>T26145</td><td>TGACGTTGACCGCAAGCCCTTGG</td><td>1000</td><td></td><td></td><td></td><td></td><td></td><td></td><td></td><td></td><td></td></tr>
<tr><td>Oligo fwd</td><td>CACCTGACGTTGACCGCAAGCCCT</td><td></td><td></td><td></td><td></td><td></td><td></td><td></td><td></td><td></td><td></td></tr>
<tr><td>Oligo rev</td><td>AAACAGGGCTTGCGGTCAACGTCA</td><td></td><td></td><td></td><td></td><td></td><td></td><td></td><td></td><td></td><td></td></tr>
<tr><td>Chromosome</td><td>start</td><td>end</td><td>strand</td><td>MM</td><td>target_seq</td><td>PAM</td><td>alignment</td><td>distance</td><td>position</td><td>gene name</td><td>gene id</td></tr>
<tr><td>GU305914.1</td><td>104674</td><td>104696</td><td>-</td><td>0</td><td>TGACGTTGACCGCAAGCCCT</td><td>TGG</td><td>||||||||[||||||||||||]PAM</td><td>NA</td><td>Intergenic</td><td>NA</td><td>NA</td></tr>
<tr><td>T26146</td><td>CGTTGACCGCAAGCCCTTGGAGG</td><td>1000</td><td></td><td></td><td></td><td></td><td></td><td></td><td></td><td></td><td></td></tr>
<tr><td>Oligo fwd</td><td>CACCCGTTGACCGCAAGCCCTTGG</td><td></td><td></td><td></td><td></td><td></td><td></td><td></td><td></td><td></td><td></td></tr>
<tr><td>Oligo rev</td><td>AAACCCAAGGGCTTGCGGTCAACG</td><td></td><td></td><td></td><td></td><td></td><td></td><td></td><td></td><td></td><td></td></tr>
<tr><td>Chromosome</td><td>start</td><td>end</td><td>strand</td><td>MM</td><td>target_seq</td><td>PAM</td><td>alignment</td><td>distance</td><td>position</td><td>gene name</td><td>gene id</td></tr>
<tr><td>GU305914.1</td><td>104671</td><td>104693</td><td>-</td><td>0</td><td>CGTTGACCGCAAGCCCTTGG</td><td>AGG</td><td>||||||||[||||||||||||]PAM</td><td>NA</td><td>Intergenic</td><td>NA</td><td>NA</td></tr>
<tr><td>T26147</td><td>TACCGTCTCTGCCCCACAAAAGG</td><td>1000</td><td></td><td></td><td></td><td></td><td></td><td></td><td></td><td></td><td></td></tr>
<tr><td>Oligo fwd</td><td>CACCTACCGTCTCTGCCCCACAAA</td><td></td><td></td><td></td><td></td><td></td><td></td><td></td><td></td><td></td><td></td></tr>
<tr><td>Oligo rev</td><td>AAACTTTGTGGGGCAGAGACGGTA</td><td></td><td></td><td></td><td></td><td></td><td></td><td></td><td></td><td></td><td></td></tr>
<tr><td>Chromosome</td><td>start</td><td>end</td><td>strand</td><td>MM</td><td>target_seq</td><td>PAM</td><td>alignment</td><td>distance</td><td>position</td><td>gene name</td><td>gene id</td></tr>
<tr><td>GU305914.1</td><td>104620</td><td>104642</td><td>-</td><td>0</td><td>TACCGTCTCTGCCCCACAAA</td><td>AGG</td><td>||||||||[||||||||||||]PAM</td><td>NA</td><td>Intergenic</td><td>NA</td><td>NA</td></tr>
<tr><td>T26148</td><td>CTGCCCCACAAAAGGCGCACCGG</td><td>1000</td><td></td><td></td><td></td><td></td><td></td><td></td><td></td><td></td><td></td></tr>
<tr><td>Oligo fwd</td><td>CACCCTGCCCCACAAAAGGCGCAC</td><td></td><td></td><td></td><td></td><td></td><td></td><td></td><td></td><td></td><td></td></tr>
<tr><td>Oligo rev</td><td>AAACGTGCGCCTTTTGTGGGGCAG</td><td></td><td></td><td></td><td></td><td></td><td></td><td></td><td></td><td></td><td></td></tr>
<tr><td>Chromosome</td><td>start</td><td>end</td><td>strand</td><td>MM</td><td>target_seq</td><td>PAM</td><td>alignment</td><td>distance</td><td>position</td><td>gene name</td><td>gene id</td></tr>
<tr><td>GU305914.1</td><td>104612</td><td>104634</td><td>-</td><td>0</td><td>CTGCCCCACAAAAGGCGCAC</td><td>CGG</td><td>||||||||[||||||||||||]PAM</td><td>NA</td><td>Intergenic</td><td>NA</td><td>NA</td></tr>
<tr><td>T26149</td><td>ATATCCTTTAGCCGTGTCTGAGG</td><td>1000</td><td></td><td></td><td></td><td></td><td></td><td></td><td></td><td></td><td></td></tr>
<tr><td>Oligo fwd</td><td>CACCATATCCTTTAGCCGTGTCTG</td><td></td><td></td><td></td><td></td><td></td><td></td><td></td><td></td><td></td><td></td></tr>
<tr><td>Oligo rev</td><td>AAACCAGACACGGCTAAAGGATAT</td><td></td><td></td><td></td><td></td><td></td><td></td><td></td><td></td><td></td><td></td></tr>
<tr><td>Chromosome</td><td>start</td><td>end</td><td>strand</td><td>MM</td><td>target_seq</td><td>PAM</td><td>alignment</td><td>distance</td><td>position</td><td>gene name</td><td>gene id</td></tr>
<tr><td>GU305914.1</td><td>104585</td><td>104607</td><td>-</td><td>0</td><td>ATATCCTTTAGCCGTGTCTG</td><td>AGG</td><td>||||||||[||||||||||||]PAM</td><td>NA</td><td>Intergenic</td><td>NA</td><td>NA</td></tr>
<tr><td>T26150</td><td>TTTAGCCGTGTCTGAGGTGAAGG</td><td>1000</td><td></td><td></td><td></td><td></td><td></td><td></td><td></td><td></td><td></td></tr>
<tr><td>Oligo fwd</td><td>CACCTTTAGCCGTGTCTGAGGTGA</td><td></td><td></td><td></td><td></td><td></td><td></td><td></td><td></td><td></td><td></td></tr>
<tr><td>Oligo rev</td><td>AAACTCACCTCAGACACGGCTAAA</td><td></td><td></td><td></td><td></td><td></td><td></td><td></td><td></td><td></td><td></td></tr>
<tr><td>Chromosome</td><td>start</td><td>end</td><td>strand</td><td>MM</td><td>target_seq</td><td>PAM</td><td>alignment</td><td>distance</td><td>position</td><td>gene name</td><td>gene id</td></tr>
<tr><td>GU305914.1</td><td>104579</td><td>104601</td><td>-</td><td>0</td><td>TTTAGCCGTGTCTGAGGTGA</td><td>AGG</td><td>||||||||[||||||||||||]PAM</td><td>NA</td><td>Intergenic</td><td>NA</td><td>NA</td></tr>
<tr><td>T26151</td><td>GAACGCACAAATAAAGACGATGG</td><td>1000</td><td></td><td></td><td></td><td></td><td></td><td></td><td></td><td></td><td></td></tr>
<tr><td>Oligo fwd</td><td>CACCGAACGCACAAATAAAGACGA</td><td></td><td></td><td></td><td></td><td></td><td></td><td></td><td></td><td></td><td></td></tr>
<tr><td>Oligo rev</td><td>AAACTCGTCTTTATTTGTGCGTTC</td><td></td><td></td><td></td><td></td><td></td><td></td><td></td><td></td><td></td><td></td></tr>
<tr><td>Chromosome</td><td>start</td><td>end</td><td>strand</td><td>MM</td><td>target_seq</td><td>PAM</td><td>alignment</td><td>distance</td><td>position</td><td>gene name</td><td>gene id</td></tr>
<tr><td>GU305914.1</td><td>104525</td><td>104547</td><td>-</td><td>0</td><td>GAACGCACAAATAAAGACGA</td><td>TGG</td><td>||||||||[||||||||||||]PAM</td><td>NA</td><td>Intergenic</td><td>NA</td><td>NA</td></tr>
<tr><td>T26152</td><td>GGCGTTCACCATCATCCAGATGG</td><td>1000</td><td></td><td></td><td></td><td></td><td></td><td></td><td></td><td></td><td></td></tr>
<tr><td>Oligo fwd</td><td>CACCGGCGTTCACCATCATCCAGA</td><td></td><td></td><td></td><td></td><td></td><td></td><td></td><td></td><td></td><td></td></tr>
<tr><td>Oligo rev</td><td>AAACTCTGGATGATGGTGAACGCC</td><td></td><td></td><td></td><td></td><td></td><td></td><td></td><td></td><td></td><td></td></tr>
<tr><td>Chromosome</td><td>start</td><td>end</td><td>strand</td><td>MM</td><td>target_seq</td><td>PAM</td><td>alignment</td><td>distance</td><td>position</td><td>gene name</td><td>gene id</td></tr>
<tr><td>GU305914.1</td><td>104504</td><td>104526</td><td>-</td><td>0</td><td>GGCGTTCACCATCATCCAGA</td><td>TGG</td><td>||||||||[||||||||||||]PAM</td><td>NA</td><td>Intergenic</td><td>NA</td><td>NA</td></tr>
<tr><td>T26153</td><td>GTTCACCATCATCCAGATGGAGG</td><td>1000</td><td></td><td></td><td></td><td></td><td></td><td></td><td></td><td></td><td></td></tr>
<tr><td>Oligo fwd</td><td>CACCGTTCACCATCATCCAGATGG</td><td></td><td></td><td></td><td></td><td></td><td></td><td></td><td></td><td></td><td></td></tr>
<tr><td>Oligo rev</td><td>AAACCCATCTGGATGATGGTGAAC</td><td></td><td></td><td></td><td></td><td></td><td></td><td></td><td></td><td></td><td></td></tr>
<tr><td>Chromosome</td><td>start</td><td>end</td><td>strand</td><td>MM</td><td>target_seq</td><td>PAM</td><td>alignment</td><td>distance</td><td>position</td><td>gene name</td><td>gene id</td></tr>
<tr><td>GU305914.1</td><td>104501</td><td>104523</td><td>-</td><td>0</td><td>GTTCACCATCATCCAGATGG</td><td>AGG</td><td>||||||||[||||||||||||]PAM</td><td>NA</td><td>Intergenic</td><td>NA</td><td>NA</td></tr>
<tr><td>T26154</td><td>GATGGAGGCGAACGACGAGACGG</td><td>1000</td><td></td><td></td><td></td><td></td><td></td><td></td><td></td><td></td><td></td></tr>
<tr><td>Oligo fwd</td><td>CACCGATGGAGGCGAACGACGAGA</td><td></td><td></td><td></td><td></td><td></td><td></td><td></td><td></td><td></td><td></td></tr>
<tr><td>Oligo rev</td><td>AAACTCTCGTCGTTCGCCTCCATC</td><td></td><td></td><td></td><td></td><td></td><td></td><td></td><td></td><td></td><td></td></tr>
<tr><td>Chromosome</td><td>start</td><td>end</td><td>strand</td><td>MM</td><td>target_seq</td><td>PAM</td><td>alignment</td><td>distance</td><td>position</td><td>gene name</td><td>gene id</td></tr>
<tr><td>GU305914.1</td><td>104486</td><td>104508</td><td>-</td><td>0</td><td>GATGGAGGCGAACGACGAGA</td><td>CGG</td><td>||||||||[||||||||||||]PAM</td><td>NA</td><td>Intergenic</td><td>NA</td><td>NA</td></tr>
<tr><td>T26155</td><td>CTCGTTGATATTGTGCGCCTCGG</td><td>1000</td><td></td><td></td><td></td><td></td><td></td><td></td><td></td><td></td><td></td></tr>
<tr><td>Oligo fwd</td><td>CACCCTCGTTGATATTGTGCGCCT</td><td></td><td></td><td></td><td></td><td></td><td></td><td></td><td></td><td></td><td></td></tr>
<tr><td>Oligo rev</td><td>AAACAGGCGCACAATATCAACGAG</td><td></td><td></td><td></td><td></td><td></td><td></td><td></td><td></td><td></td><td></td></tr>
<tr><td>Chromosome</td><td>start</td><td>end</td><td>strand</td><td>MM</td><td>target_seq</td><td>PAM</td><td>alignment</td><td>distance</td><td>position</td><td>gene name</td><td>gene id</td></tr>
<tr><td>GU305914.1</td><td>104423</td><td>104445</td><td>-</td><td>0</td><td>CTCGTTGATATTGTGCGCCT</td><td>CGG</td><td>||||||||[||||||||||||]PAM</td><td>NA</td><td>Intergenic</td><td>NA</td><td>NA</td></tr>
<tr><td>T26156</td><td>GCTGATCACGATCTTGTTGACGG</td><td>1000</td><td></td><td></td><td></td><td></td><td></td><td></td><td></td><td></td><td></td></tr>
<tr><td>Oligo fwd</td><td>CACCGCTGATCACGATCTTGTTGA</td><td></td><td></td><td></td><td></td><td></td><td></td><td></td><td></td><td></td><td></td></tr>
<tr><td>Oligo rev</td><td>AAACTCAACAAGATCGTGATCAGC</td><td></td><td></td><td></td><td></td><td></td><td></td><td></td><td></td><td></td><td></td></tr>
<tr><td>Chromosome</td><td>start</td><td>end</td><td>strand</td><td>MM</td><td>target_seq</td><td>PAM</td><td>alignment</td><td>distance</td><td>position</td><td>gene name</td><td>gene id</td></tr>
<tr><td>GU305914.1</td><td>104393</td><td>104415</td><td>-</td><td>0</td><td>GCTGATCACGATCTTGTTGA</td><td>CGG</td><td>||||||||[||||||||||||]PAM</td><td>NA</td><td>Intergenic</td><td>NA</td><td>NA</td></tr>
<tr><td>T26157</td><td>ATCTTGTTGACGGTCGTATCCGG</td><td>1000</td><td></td><td></td><td></td><td></td><td></td><td></td><td></td><td></td><td></td></tr>
<tr><td>Oligo fwd</td><td>CACCATCTTGTTGACGGTCGTATC</td><td></td><td></td><td></td><td></td><td></td><td></td><td></td><td></td><td></td><td></td></tr>
<tr><td>Oligo rev</td><td>AAACGATACGACCGTCAACAAGAT</td><td></td><td></td><td></td><td></td><td></td><td></td><td></td><td></td><td></td><td></td></tr>
<tr><td>Chromosome</td><td>start</td><td>end</td><td>strand</td><td>MM</td><td>target_seq</td><td>PAM</td><td>alignment</td><td>distance</td><td>position</td><td>gene name</td><td>gene id</td></tr>
<tr><td>GU305914.1</td><td>104383</td><td>104405</td><td>-</td><td>0</td><td>ATCTTGTTGACGGTCGTATC</td><td>CGG</td><td>||||||||[||||||||||||]PAM</td><td>NA</td><td>Intergenic</td><td>NA</td><td>NA</td></tr>
</table>

T26158 GGTCGTATCCGGCACCGAAGCGG 1000
Oligo fwd CACCGGTCGTATCCGGCACCGAAG
Oligo rev AAACCTTCGGTGCCGGATACGACC
Chromosome start end strand MM target_seq PAM alignment distance position gene name gene id
GU305914.1 104372 104394 - 0 GGTCGTATCCGGCACCGAAG CGG ||||||||[||||||||||||]PAM NA Intergenic NA NA

T26159 ACCGAAGCGGCGACCGACACCGG 1000
Oligo fwd CACCACCGAAGCGGCGACCGACAC
Oligo rev AAACGTGTCGGTCGCCGCTTCGGT
Chromosome start end strand MM target_seq PAM alignment distance position gene name gene id
GU305914.1 104359 104381 - 0 ACCGAAGCGGCGACCGACAC CGG ||||||||[||||||||||||]PAM NA Intergenic NA NA

T26160 AGCGGCGACCGACACCGGTACGG 1000
Oligo fwd CACCAGCGGCGACCGACACCGGTA
Oligo rev AAACTACCGGTGTCGGTCGCCGCT
Chromosome start end strand MM target_seq PAM alignment distance position gene name gene id
GU305914.1 104354 104376 - 0 AGCGGCGACCGACACCGGTA CGG ||||||||[||||||||||||]PAM NA Intergenic NA NA

T26161 GGCGACCGACACCGGTACGGAGG 1000
Oligo fwd CACCGGCGACCGACACCGGTACGG
Oligo rev AAACCCGTACCGGTGTCGGTCGCC
Chromosome start end strand MM target_seq PAM alignment distance position gene name gene id
GU305914.1 104351 104373 - 0 GGCGACCGACACCGGTACGG AGG ||||||||[||||||||||||]PAM NA Intergenic NA NA

T26162 ACCGACACCGGTACGGAGGCCGG 1000
Oligo fwd CACCACCGACACCGGTACGGAGGC
Oligo rev AAACGCCTCCGTACCGGTGTCGGT
Chromosome start end strand MM target_seq PAM alignment distance position gene name gene id
GU305914.1 104347 104369 - 0 ACCGACACCGGTACGGAGGC CGG ||||||||[||||||||||||]PAM NA Intergenic NA NA

T26163 CGGTACGGAGGCCGGCGACGAGG 1000
Oligo fwd CACCCGGTACGGAGGCCGGCGACG
Oligo rev AAACCGTCGCCGGCCTCCGTACCG
Chromosome start end strand MM target_seq PAM alignment distance position gene name gene id
GU305914.1 104339 104361 - 0 CGGTACGGAGGCCGGCGACG AGG ||||||||[||||||||||||]PAM NA Intergenic NA NA

T26164 CGACGAGGATCCGCTCGCCATGG 1000
Oligo fwd CACCCGACGAGGATCCGCTCGCCA
Oligo rev AAACTGGCGAGCGGATCCTCGTCG
Chromosome start end strand MM target_seq PAM alignment distance position gene name gene id
GU305914.1 104324 104346 - 0 CGACGAGGATCCGCTCGCCA TGG ||||||||[||||||||||||]PAM NA Intergenic NA NA

T26165 CCATGGCGAAGCACAACAGAAGG 1000
Oligo fwd CACCCCATGGCGAAGCACAACAGA
Oligo rev AAACTCTGTTGTGCTTCGCCATGG
Chromosome start end strand MM target_seq PAM alignment distance position gene name gene id
GU305914.1 104307 104329 - 0 CCATGGCGAAGCACAACAGA AGG ||||||||[||||||||||||]PAM NA Intergenic NA NA

T26166 ACAGAAGGAAAAAGCTCAAACGG 1000
Oligo fwd CACCACAGAAGGAAAAAGCTCAAA
Oligo rev AAACTTTGAGCTTTTTCCTTCTGT
Chromosome start end strand MM target_seq PAM alignment distance position gene name gene id
GU305914.1 104292 104314 - 0 ACAGAAGGAAAAAGCTCAAA CGG ||||||||[||||||||||||]PAM NA Intergenic NA NA

T26167 CGGAAAATCCCTACAAAGCCTGG 1000
Oligo fwd CACCCGGAAAATCCCTACAAAGCC
Oligo rev AAACGGCTTTGTAGGGATTTTCCG
Chromosome start end strand MM target_seq PAM alignment distance position gene name gene id
GU305914.1 104272 104294 - 0 CGGAAAATCCCTACAAAGCC TGG ||||||||[||||||||||||]PAM NA Intergenic NA NA

T26168 CATGATAAAACCAAACATACCGG 1000
Oligo fwd CACCCATGATAAAACCAAACATAC
Oligo rev AAACGTATGTTTGGTTTTATCATG
Chromosome start end strand MM target_seq PAM alignment distance position gene name gene id
GU305914.1 104249 104271 - 0 CATGATAAAACCAAACATAC CGG ||||||||[||||||||||||]PAM NA Intergenic NA NA

T26169 AAAACCAAACATACCGGCTCCGG 1000
Oligo fwd CACCAAAACCAAACATACCGGCTC
Oligo rev AAACGAGCCGGTATGTTTGGTTTT
Chromosome start end strand MM target_seq PAM alignment distance position gene name gene id
GU305914.1 104243 104265 - 0 AAAACCAAACATACCGGCTC CGG ||||||||[||||||||||||]PAM NA Intergenic NA NA

T26170 GCCAGACTCAGTCTTGCCGCAGG 1000
Oligo fwd CACCGCCAGACTCAGTCTTGCCGC
Oligo rev AAACGCGGCAAGACTGAGTCTGGC
Chromosome start end strand MM target_seq PAM alignment distance position gene name gene id
GU305914.1 104195 104217 - 0 GCCAGACTCAGTCTTGCCGC AGG ||||||||[||||||||||||]PAM NA Intergenic NA NA

T26171 AGTCTTGCCGCAGGCCATGAAGG 1000
Oligo fwd CACCAGTCTTGCCGCAGGCCATGA
Oligo rev AAACTCATGGCCTGCGGCAAGACT
Chromosome start end strand MM target_seq PAM alignment distance position gene name gene id
GU305914.1 104186 104208 - 0 AGTCTTGCCGCAGGCCATGA AGG ||||||||[||||||||||||]PAM NA Intergenic NA NA

T26172 CGAACTCGTCGTTCTTAAATAGG 1000
Oligo fwd CACCCGAACTCGTCGTTCTTAAAT
Oligo rev AAACATTTAAGAACGACGAGTTCG
Chromosome start end strand MM target_seq PAM alignment distance position gene name gene id

GU305914.1 104043 104065 - 0 CGAACTCGTCGTTCTTAAAT AGG ||||||||[||||||||||||]PAM NA Intergenic NA NA

T26173 GTTCTTAAATAGGTTCTTTCCGG 1000
Oligo fwd CACCGTTCTTAAATAGGTTCTTTC
Oligo rev AAACGAAAGAACCTATTTAAGAAC
Chromosome start end strand MM target_seq PAM alignment distance position gene name gene id
GU305914.1 104033 104055 - 0 GTTCTTAAATAGGTTCTTTC CGG ||||||||[||||||||||||]PAM NA Intergenic NA NA

T26174 CTTAAATAGGTTCTTTCCGGAGG 1000
Oligo fwd CACCCTTAAATAGGTTCTTTCCGG
Oligo rev AAACCCGGAAAGAACCTATTTAAG
Chromosome start end strand MM target_seq PAM alignment distance position gene name gene id
GU305914.1 104030 104052 - 0 CTTAAATAGGTTCTTTCCGG AGG ||||||||[||||||||||||]PAM NA Intergenic NA NA

T26175 GACGACGAATCCAGCGAGAACGG 1000
Oligo fwd CACCGACGACGAATCCAGCGAGAA
Oligo rev AAACTTCTCGCTGGATTCGTCGTC
Chromosome start end strand MM target_seq PAM alignment distance position gene name gene id
GU305914.1 104008 104030 - 0 GACGACGAATCCAGCGAGAA CGG ||||||||[||||||||||||]PAM NA Intergenic NA NA

T26176 CGAATCCAGCGAGAACGGCGCGG 1000
Oligo fwd CACCCGAATCCAGCGAGAACGGCG
Oligo rev AAACCGCCGTTCTCGCTGGATTCG
Chromosome start end strand MM target_seq PAM alignment distance position gene name gene id
GU305914.1 104003 104025 - 0 CGAATCCAGCGAGAACGGCG CGG ||||||||[||||||||||||]PAM NA Intergenic NA NA

T26177 CGAGAACGGCGCGGCGCTCGCGG 1000
Oligo fwd CACCCGAGAACGGCGCGGCGCTCG
Oligo rev AAACCGAGCGCCGCGCCGTTCTCG
Chromosome start end strand MM target_seq PAM alignment distance position gene name gene id
GU305914.1 103994 104016 - 0 CGAGAACGGCGCGGCGCTCG CGG ||||||||[||||||||||||]PAM NA Intergenic NA NA

T26178 ATCAACCAGCTGCTGCGCGACGG 1000
Oligo fwd CACCATCAACCAGCTGCTGCGCGA
Oligo rev AAACTCGCGCAGCAGCTGGTTGAT
Chromosome start end strand MM target_seq PAM alignment distance position gene name gene id
GU305914.1 103969 103991 - 0 ATCAACCAGCTGCTGCGCGA CGG ||||||||[||||||||||||]PAM NA Intergenic NA NA

T26179 CCAGCTGCTGCGCGACGGTCCGG 1000
Oligo fwd CACCCCAGCTGCTGCGCGACGGTC
Oligo rev AAACGACCGTCGCGCAGCAGCTGG
Chromosome start end strand MM target_seq PAM alignment distance position gene name gene id
GU305914.1 103964 103986 - 0 CCAGCTGCTGCGCGACGGTC CGG ||||||||[||||||||||||]PAM NA Intergenic NA NA

T26180 GCTGCTGCGCGACGGTCCGGAGG 1000
Oligo fwd CACCGCTGCTGCGCGACGGTCCGG
Oligo rev AAACCCGGACCGTCGCGCAGCAGC
Chromosome start end strand MM target_seq PAM alignment distance position gene name gene id
GU305914.1 103961 103983 - 0 GCTGCTGCGCGACGGTCCGG AGG ||||||||[||||||||||||]PAM NA Intergenic NA NA
GU305914.1 125360 125382 - 4 GCGGCTGGGCGACGGCGCGG TGG ||-||||-[|||||||--|||]PAM NA Intergenic NA NA

T26181 GCCGCGCGAACCGACGCCGAAGG 1000
Oligo fwd CACCGCCGCGCGAACCGACGCCGA
Oligo rev AAACTCGGCGTCGGTTCGCGCGGC
Chromosome start end strand MM target_seq PAM alignment distance position gene name gene id
GU305914.1 103934 103956 - 0 GCCGCGCGAACCGACGCCGA AGG ||||||||[||||||||||||]PAM NA Intergenic NA NA

T26182 AAGATCTTCTTCCAGACGTACGG 1000
Oligo fwd CACCAAGATCTTCTTCCAGACGTA
Oligo rev AAACTACGTCTGGAAGAAGATCTT
Chromosome start end strand MM target_seq PAM alignment distance position gene name gene id
GU305914.1 103885 103907 - 0 AAGATCTTCTTCCAGACGTA CGG ||||||||[||||||||||||]PAM NA Intergenic NA NA

T26183 GTACGGCAACGATTTCAAAGTGG 1000
Oligo fwd CACCGTACGGCAACGATTTCAAAG
Oligo rev AAACCTTTGAAATCGTTGCCGTAC
Chromosome start end strand MM target_seq PAM alignment distance position gene name gene id
GU305914.1 103868 103890 - 0 GTACGGCAACGATTTCAAAG TGG ||||||||[||||||||||||]PAM NA Intergenic NA NA

T26184 GATTTCAAAGTGGAAGCGCACGG 1000
Oligo fwd CACCGATTTCAAAGTGGAAGCGCA
Oligo rev AAACTGCGCTTCCACTTTGAAATC
Chromosome start end strand MM target_seq PAM alignment distance position gene name gene id
GU305914.1 103858 103880 - 0 GATTTCAAAGTGGAAGCGCA CGG ||||||||[||||||||||||]PAM NA Intergenic NA NA

T26185 GCTCGCGCAGAATCAGCGCGTGG 1000
Oligo fwd CACCGCTCGCGCAGAATCAGCGCG
Oligo rev AAACCGCGCTGATTCTGCGCGAGC
Chromosome start end strand MM target_seq PAM alignment distance position gene name gene id
GU305914.1 103826 103848 - 0 GCTCGCGCAGAATCAGCGCG TGG ||||||||[||||||||||||]PAM NA Intergenic NA NA

T26186 GCGTGGTACGACTGTTCGAGCGG 1000
Oligo fwd CACCGCGTGGTACGACTGTTCGAG
Oligo rev AAACCTCGAACAGTCGTACCACGC
Chromosome start end strand MM target_seq PAM alignment distance position gene name gene id
GU305914.1 103809 103831 - 0 GCGTGGTACGACTGTTCGAG CGG ||||||||[||||||||||||]PAM NA Intergenic NA NA

T26187 CGTGGTACGACTGTTCGAGCGGG 1000
Oligo fwd CACCCGTGGTACGACTGTTCGAGC

<table>
<tr><td>Oligo rev</td><td>AAACGCTCGAACAGTCGTACCACG</td><td></td><td></td><td></td><td></td><td></td><td></td><td></td><td></td><td></td><td></td></tr>
<tr><td>Chromosome</td><td>start</td><td>end</td><td>strand</td><td>MM</td><td>target_seq</td><td>PAM</td><td>alignment</td><td>distance</td><td>position</td><td>gene name</td><td>gene id</td></tr>
<tr><td>GU305914.1</td><td>103808</td><td>103830</td><td>-</td><td>0</td><td>CGTGGTACGACTGTTCGAGC</td><td>GGG</td><td>||||||||[||||||||||||]PAM</td><td>NA</td><td>Intergenic</td><td>NA</td><td>NA</td></tr>
<tr><td>T26188</td><td>GTGGTACGACTGTTCGAGCGGGG</td><td>1000</td><td></td><td></td><td></td><td></td><td></td><td></td><td></td><td></td><td></td></tr>
<tr><td>Oligo fwd</td><td>CACCGTGGTACGACTGTTCGAGCG</td><td></td><td></td><td></td><td></td><td></td><td></td><td></td><td></td><td></td><td></td></tr>
<tr><td>Oligo rev</td><td>AAACCGCTCGAACAGTCGTACCAC</td><td></td><td></td><td></td><td></td><td></td><td></td><td></td><td></td><td></td><td></td></tr>
<tr><td>Chromosome</td><td>start</td><td>end</td><td>strand</td><td>MM</td><td>target_seq</td><td>PAM</td><td>alignment</td><td>distance</td><td>position</td><td>gene name</td><td>gene id</td></tr>
<tr><td>GU305914.1</td><td>103807</td><td>103829</td><td>-</td><td>0</td><td>GTGGTACGACTGTTCGAGCG</td><td>GGG</td><td>||||||||[||||||||||||]PAM</td><td>NA</td><td>Intergenic</td><td>NA</td><td>NA</td></tr>
<tr><td>T26189</td><td>TTCGAGCGGGGCCGCATCCTCGG</td><td>1000</td><td></td><td></td><td></td><td></td><td></td><td></td><td></td><td></td><td></td></tr>
<tr><td>Oligo fwd</td><td>CACCTTCGAGCGGGGCCGCATCCT</td><td></td><td></td><td></td><td></td><td></td><td></td><td></td><td></td><td></td><td></td></tr>
<tr><td>Oligo rev</td><td>AAACAGGATGCGGCCCCGCTCGAA</td><td></td><td></td><td></td><td></td><td></td><td></td><td></td><td></td><td></td><td></td></tr>
<tr><td>Chromosome</td><td>start</td><td>end</td><td>strand</td><td>MM</td><td>target_seq</td><td>PAM</td><td>alignment</td><td>distance</td><td>position</td><td>gene name</td><td>gene id</td></tr>
<tr><td>GU305914.1</td><td>103795</td><td>103817</td><td>-</td><td>0</td><td>TTCGAGCGGGGCCGCATCCT</td><td>CGG</td><td>||||||||[||||||||||||]PAM</td><td>NA</td><td>Intergenic</td><td>NA</td><td>NA</td></tr>
<tr><td>T26190</td><td>GCGGGGCCGCATCCTCGGTCTGG</td><td>1000</td><td></td><td></td><td></td><td></td><td></td><td></td><td></td><td></td><td></td></tr>
<tr><td>Oligo fwd</td><td>CACCGCGGGGCCGCATCCTCGGTC</td><td></td><td></td><td></td><td></td><td></td><td></td><td></td><td></td><td></td><td></td></tr>
<tr><td>Oligo rev</td><td>AAACGACCGAGGATGCGGCCCCGC</td><td></td><td></td><td></td><td></td><td></td><td></td><td></td><td></td><td></td><td></td></tr>
<tr><td>Chromosome</td><td>start</td><td>end</td><td>strand</td><td>MM</td><td>target_seq</td><td>PAM</td><td>alignment</td><td>distance</td><td>position</td><td>gene name</td><td>gene id</td></tr>
<tr><td>GU305914.1</td><td>103790</td><td>103812</td><td>-</td><td>0</td><td>GCGGGGCCGCATCCTCGGTC</td><td>TGG</td><td>||||||||[||||||||||||]PAM</td><td>NA</td><td>Intergenic</td><td>NA</td><td>NA</td></tr>
<tr><td>T26191</td><td>CGGGGCCGCATCCTCGGTCTGGG</td><td>1000</td><td></td><td></td><td></td><td></td><td></td><td></td><td></td><td></td><td></td></tr>
<tr><td>Oligo fwd</td><td>CACCCGGGGCCGCATCCTCGGTCT</td><td></td><td></td><td></td><td></td><td></td><td></td><td></td><td></td><td></td><td></td></tr>
<tr><td>Oligo rev</td><td>AAACAGACCGAGGATGCGGCCCCG</td><td></td><td></td><td></td><td></td><td></td><td></td><td></td><td></td><td></td><td></td></tr>
<tr><td>Chromosome</td><td>start</td><td>end</td><td>strand</td><td>MM</td><td>target_seq</td><td>PAM</td><td>alignment</td><td>distance</td><td>position</td><td>gene name</td><td>gene id</td></tr>
<tr><td>GU305914.1</td><td>103789</td><td>103811</td><td>-</td><td>0</td><td>CGGGGCCGCATCCTCGGTCT</td><td>GGG</td><td>||||||||[||||||||||||]PAM</td><td>NA</td><td>Intergenic</td><td>NA</td><td>NA</td></tr>
<tr><td>T26192</td><td>GGGGCCGCATCCTCGGTCTGGGG</td><td>1000</td><td></td><td></td><td></td><td></td><td></td><td></td><td></td><td></td><td></td></tr>
<tr><td>Oligo fwd</td><td>CACCGGGGCCGCATCCTCGGTCTG</td><td></td><td></td><td></td><td></td><td></td><td></td><td></td><td></td><td></td><td></td></tr>
<tr><td>Oligo rev</td><td>AAACCAGACCGAGGATGCGGCCCC</td><td></td><td></td><td></td><td></td><td></td><td></td><td></td><td></td><td></td><td></td></tr>
<tr><td>Chromosome</td><td>start</td><td>end</td><td>strand</td><td>MM</td><td>target_seq</td><td>PAM</td><td>alignment</td><td>distance</td><td>position</td><td>gene name</td><td>gene id</td></tr>
<tr><td>GU305914.1</td><td>103788</td><td>103810</td><td>-</td><td>0</td><td>GGGGCCGCATCCTCGGTCTG</td><td>GGG</td><td>||||||||[||||||||||||]PAM</td><td>NA</td><td>Intergenic</td><td>NA</td><td>NA</td></tr>
<tr><td>T26193</td><td>GGGCCGCATCCTCGGTCTGGGGG</td><td>1000</td><td></td><td></td><td></td><td></td><td></td><td></td><td></td><td></td><td></td></tr>
<tr><td>Oligo fwd</td><td>CACCGGGCCGCATCCTCGGTCTGG</td><td></td><td></td><td></td><td></td><td></td><td></td><td></td><td></td><td></td><td></td></tr>
<tr><td>Oligo rev</td><td>AAACCCAGACCGAGGATGCGGCCC</td><td></td><td></td><td></td><td></td><td></td><td></td><td></td><td></td><td></td><td></td></tr>
<tr><td>Chromosome</td><td>start</td><td>end</td><td>strand</td><td>MM</td><td>target_seq</td><td>PAM</td><td>alignment</td><td>distance</td><td>position</td><td>gene name</td><td>gene id</td></tr>
<tr><td>GU305914.1</td><td>103787</td><td>103809</td><td>-</td><td>0</td><td>GGGCCGCATCCTCGGTCTGG</td><td>GGG</td><td>||||||||[||||||||||||]PAM</td><td>NA</td><td>Intergenic</td><td>NA</td><td>NA</td></tr>
<tr><td>T26194</td><td>GGTGAAGATGAACATCCCGATGG</td><td>1000</td><td></td><td></td><td></td><td></td><td></td><td></td><td></td><td></td><td></td></tr>
<tr><td>Oligo fwd</td><td>CACCGGTGAAGATGAACATCCCGA</td><td></td><td></td><td></td><td></td><td></td><td></td><td></td><td></td><td></td><td></td></tr>
<tr><td>Oligo rev</td><td>AAACTCGGGATGTTCATCTTCACC</td><td></td><td></td><td></td><td></td><td></td><td></td><td></td><td></td><td></td><td></td></tr>
<tr><td>Chromosome</td><td>start</td><td>end</td><td>strand</td><td>MM</td><td>target_seq</td><td>PAM</td><td>alignment</td><td>distance</td><td>position</td><td>gene name</td><td>gene id</td></tr>
<tr><td>GU305914.1</td><td>103766</td><td>103788</td><td>-</td><td>0</td><td>GGTGAAGATGAACATCCCGA</td><td>TGG</td><td>||||||||[||||||||||||]PAM</td><td>NA</td><td>Intergenic</td><td>NA</td><td>NA</td></tr>
<tr><td>T26195</td><td>GTGAAGATGAACATCCCGATGGG</td><td>1000</td><td></td><td></td><td></td><td></td><td></td><td></td><td></td><td></td><td></td></tr>
<tr><td>Oligo fwd</td><td>CACCGTGAAGATGAACATCCCGAT</td><td></td><td></td><td></td><td></td><td></td><td></td><td></td><td></td><td></td><td></td></tr>
<tr><td>Oligo rev</td><td>AAACATCGGGATGTTCATCTTCAC</td><td></td><td></td><td></td><td></td><td></td><td></td><td></td><td></td><td></td><td></td></tr>
<tr><td>Chromosome</td><td>start</td><td>end</td><td>strand</td><td>MM</td><td>target_seq</td><td>PAM</td><td>alignment</td><td>distance</td><td>position</td><td>gene name</td><td>gene id</td></tr>
<tr><td>GU305914.1</td><td>103765</td><td>103787</td><td>-</td><td>0</td><td>GTGAAGATGAACATCCCGAT</td><td>GGG</td><td>||||||||[||||||||||||]PAM</td><td>NA</td><td>Intergenic</td><td>NA</td><td>NA</td></tr>
<tr><td>T26196</td><td>CACGAACTGTTTCCACCCACCGG</td><td>1000</td><td></td><td></td><td></td><td></td><td></td><td></td><td></td><td></td><td></td></tr>
<tr><td>Oligo fwd</td><td>CACCCACGAACTGTTTCCACCCAC</td><td></td><td></td><td></td><td></td><td></td><td></td><td></td><td></td><td></td><td></td></tr>
<tr><td>Oligo rev</td><td>AAACGTGGGTGGAAACAGTTCGTG</td><td></td><td></td><td></td><td></td><td></td><td></td><td></td><td></td><td></td><td></td></tr>
<tr><td>Chromosome</td><td>start</td><td>end</td><td>strand</td><td>MM</td><td>target_seq</td><td>PAM</td><td>alignment</td><td>distance</td><td>position</td><td>gene name</td><td>gene id</td></tr>
<tr><td>GU305914.1</td><td>103730</td><td>103752</td><td>-</td><td>0</td><td>CACGAACTGTTTCCACCCAC</td><td>CGG</td><td>||||||||[||||||||||||]PAM</td><td>NA</td><td>Intergenic</td><td>NA</td><td>NA</td></tr>
<tr><td>T26197</td><td>ACGAACTGTTTCCACCCACCGGG</td><td>1000</td><td></td><td></td><td></td><td></td><td></td><td></td><td></td><td></td><td></td></tr>
<tr><td>Oligo fwd</td><td>CACCACGAACTGTTTCCACCCACC</td><td></td><td></td><td></td><td></td><td></td><td></td><td></td><td></td><td></td><td></td></tr>
<tr><td>Oligo rev</td><td>AAACGGTGGGTGGAAACAGTTCGT</td><td></td><td></td><td></td><td></td><td></td><td></td><td></td><td></td><td></td><td></td></tr>
<tr><td>Chromosome</td><td>start</td><td>end</td><td>strand</td><td>MM</td><td>target_seq</td><td>PAM</td><td>alignment</td><td>distance</td><td>position</td><td>gene name</td><td>gene id</td></tr>
<tr><td>GU305914.1</td><td>103729</td><td>103751</td><td>-</td><td>0</td><td>ACGAACTGTTTCCACCCACC</td><td>GGG</td><td>||||||||[||||||||||||]PAM</td><td>NA</td><td>Intergenic</td><td>NA</td><td>NA</td></tr>
<tr><td>T26198</td><td>CGAACTGTTTCCACCCACCGGGG</td><td>1000</td><td></td><td></td><td></td><td></td><td></td><td></td><td></td><td></td><td></td></tr>
<tr><td>Oligo fwd</td><td>CACCCGAACTGTTTCCACCCACCG</td><td></td><td></td><td></td><td></td><td></td><td></td><td></td><td></td><td></td><td></td></tr>
<tr><td>Oligo rev</td><td>AAACCGGTGGGTGGAAACAGTTCG</td><td></td><td></td><td></td><td></td><td></td><td></td><td></td><td></td><td></td><td></td></tr>
<tr><td>Chromosome</td><td>start</td><td>end</td><td>strand</td><td>MM</td><td>target_seq</td><td>PAM</td><td>alignment</td><td>distance</td><td>position</td><td>gene name</td><td>gene id</td></tr>
<tr><td>GU305914.1</td><td>103728</td><td>103750</td><td>-</td><td>0</td><td>CGAACTGTTTCCACCCACCG</td><td>GGG</td><td>||||||||[||||||||||||]PAM</td><td>NA</td><td>Intergenic</td><td>NA</td><td>NA</td></tr>
<tr><td>T26199</td><td>ACCGGGGTGCGTCTGCGTGATGG</td><td>1000</td><td></td><td></td><td></td><td></td><td></td><td></td><td></td><td></td><td></td></tr>
<tr><td>Oligo fwd</td><td>CACCACCGGGGTGCGTCTGCGTGA</td><td></td><td></td><td></td><td></td><td></td><td></td><td></td><td></td><td></td><td></td></tr>
<tr><td>Oligo rev</td><td>AAACTCACGCAGACGCACCCCGGT</td><td></td><td></td><td></td><td></td><td></td><td></td><td></td><td></td><td></td><td></td></tr>
<tr><td>Chromosome</td><td>start</td><td>end</td><td>strand</td><td>MM</td><td>target_seq</td><td>PAM</td><td>alignment</td><td>distance</td><td>position</td><td>gene name</td><td>gene id</td></tr>
<tr><td>GU305914.1</td><td>103712</td><td>103734</td><td>-</td><td>0</td><td>ACCGGGGTGCGTCTGCGTGA</td><td>TGG</td><td>||||||||[||||||||||||]PAM</td><td>NA</td><td>Intergenic</td><td>NA</td><td>NA</td></tr>
<tr><td>T26200</td><td>GTGCGTCTGCGTGATGGACATGG</td><td>1000</td><td></td><td></td><td></td><td></td><td></td><td></td><td></td><td></td><td></td></tr>
<tr><td>Oligo fwd</td><td>CACCGTGCGTCTGCGTGATGGACA</td><td></td><td></td><td></td><td></td><td></td><td></td><td></td><td></td><td></td><td></td></tr>
<tr><td>Oligo rev</td><td>AAACTGTCCATCACGCAGACGCAC</td><td></td><td></td><td></td><td></td><td></td><td></td><td></td><td></td><td></td><td></td></tr>
<tr><td>Chromosome</td><td>start</td><td>end</td><td>strand</td><td>MM</td><td>target_seq</td><td>PAM</td><td>alignment</td><td>distance</td><td>position</td><td>gene name</td><td>gene id</td></tr>
<tr><td>GU305914.1</td><td>103706</td><td>103728</td><td>-</td><td>0</td><td>GTGCGTCTGCGTGATGGACA</td><td>TGG</td><td>||||||||[||||||||||||]PAM</td><td>NA</td><td>Intergenic</td><td>NA</td><td>NA</td></tr>
<tr><td>T26201</td><td>CGTCTGCGTGATGGACATGGCGG</td><td>1000</td><td></td><td></td><td></td><td></td><td></td><td></td><td></td><td></td><td></td></tr>
<tr><td>Oligo fwd</td><td>CACCCGTCTGCGTGATGGACATGG</td><td></td><td></td><td></td><td></td><td></td><td></td><td></td><td></td><td></td><td></td></tr>
<tr><td>Oligo rev</td><td>AAACCCATGTCCATCACGCAGACG</td><td></td><td></td><td></td><td></td><td></td><td></td><td></td><td></td><td></td><td></td></tr>
<tr><td>Chromosome</td><td>start</td><td>end</td><td>strand</td><td>MM</td><td>target_seq</td><td>PAM</td><td>alignment</td><td>distance</td><td>position</td><td>gene name</td><td>gene id</td></tr>
<tr><td>GU305914.1</td><td>103703</td><td>103725</td><td>-</td><td>0</td><td>CGTCTGCGTGATGGACATGG</td><td>CGG</td><td>||||||||[||||||||||||]PAM</td><td>NA</td><td>Intergenic</td><td>NA</td><td>NA</td></tr>
<tr><td>T26202</td><td>GTCTGCGTGATGGACATGGCGGG</td><td>1000</td><td></td><td></td><td></td><td></td><td></td><td></td><td></td><td></td><td></td></tr>
</table>

<table>
<tr><td>Oligo fwd</td><td>CACCGTCTGCGTGATGGACATGGC</td><td></td><td></td><td></td><td></td><td></td><td></td><td></td><td></td><td></td><td></td></tr>
<tr><td>Oligo rev</td><td>AAACGCCATGTCCATCACGCAGAC</td><td></td><td></td><td></td><td></td><td></td><td></td><td></td><td></td><td></td><td></td></tr>
<tr><td>Chromosome</td><td>start</td><td>end</td><td>strand</td><td>MM</td><td>target_seq</td><td>PAM</td><td>alignment</td><td>distance</td><td>position</td><td>gene name</td><td>gene id</td></tr>
<tr><td>GU305914.1</td><td>103702</td><td>103724</td><td>-</td><td>0</td><td>GTCTGCGTGATGGACATGGC</td><td>GGG</td><td>||||||||[||||||||||||]PAM</td><td>NA</td><td>Intergenic</td><td>NA</td><td>NA</td></tr>
<tr><td></td><td></td><td></td><td></td><td></td><td></td><td></td><td></td><td></td><td></td><td></td><td></td></tr>
<tr><td>T26203</td><td>GTGATGGACATGGCGGGTTACGG</td><td>1000</td><td></td><td></td><td></td><td></td><td></td><td></td><td></td><td></td><td></td></tr>
<tr><td>Oligo fwd</td><td>CACCGTGATGGACATGGCGGGTTA</td><td></td><td></td><td></td><td></td><td></td><td></td><td></td><td></td><td></td><td></td></tr>
<tr><td>Oligo rev</td><td>AAACTAACCCGCCATGTCCATCAC</td><td></td><td></td><td></td><td></td><td></td><td></td><td></td><td></td><td></td><td></td></tr>
<tr><td>Chromosome</td><td>start</td><td>end</td><td>strand</td><td>MM</td><td>target_seq</td><td>PAM</td><td>alignment</td><td>distance</td><td>position</td><td>gene name</td><td>gene id</td></tr>
<tr><td>GU305914.1</td><td>103696</td><td>103718</td><td>-</td><td>0</td><td>GTGATGGACATGGCGGGTTA</td><td>CGG</td><td>||||||||[||||||||||||]PAM</td><td>NA</td><td>Intergenic</td><td>NA</td><td>NA</td></tr>
<tr><td></td><td></td><td></td><td></td><td></td><td></td><td></td><td></td><td></td><td></td><td></td><td></td></tr>
<tr><td>T26204</td><td>GCAGATCGTGAACATCAGCTCGG</td><td>1000</td><td></td><td></td><td></td><td></td><td></td><td></td><td></td><td></td><td></td></tr>
<tr><td>Oligo fwd</td><td>CACCGCAGATCGTGAACATCAGCT</td><td></td><td></td><td></td><td></td><td></td><td></td><td></td><td></td><td></td><td></td></tr>
<tr><td>Oligo rev</td><td>AAACAGCTGATGTTCACGATCTGC</td><td></td><td></td><td></td><td></td><td></td><td></td><td></td><td></td><td></td><td></td></tr>
<tr><td>Chromosome</td><td>start</td><td>end</td><td>strand</td><td>MM</td><td>target_seq</td><td>PAM</td><td>alignment</td><td>distance</td><td>position</td><td>gene name</td><td>gene id</td></tr>
<tr><td>GU305914.1</td><td>103655</td><td>103677</td><td>-</td><td>0</td><td>GCAGATCGTGAACATCAGCT</td><td>CGG</td><td>||||||||[||||||||||||]PAM</td><td>NA</td><td>Intergenic</td><td>NA</td><td>NA</td></tr>
<tr><td></td><td></td><td></td><td></td><td></td><td></td><td></td><td></td><td></td><td></td><td></td><td></td></tr>
<tr><td>T26205</td><td>CAGATCGTGAACATCAGCTCGGG</td><td>1000</td><td></td><td></td><td></td><td></td><td></td><td></td><td></td><td></td><td></td></tr>
<tr><td>Oligo fwd</td><td>CACCCAGATCGTGAACATCAGCTC</td><td></td><td></td><td></td><td></td><td></td><td></td><td></td><td></td><td></td><td></td></tr>
<tr><td>Oligo rev</td><td>AAACGAGCTGATGTTCACGATCTG</td><td></td><td></td><td></td><td></td><td></td><td></td><td></td><td></td><td></td><td></td></tr>
<tr><td>Chromosome</td><td>start</td><td>end</td><td>strand</td><td>MM</td><td>target_seq</td><td>PAM</td><td>alignment</td><td>distance</td><td>position</td><td>gene name</td><td>gene id</td></tr>
<tr><td>GU305914.1</td><td>103654</td><td>103676</td><td>-</td><td>0</td><td>CAGATCGTGAACATCAGCTC</td><td>GGG</td><td>||||||||[||||||||||||]PAM</td><td>NA</td><td>Intergenic</td><td>NA</td><td>NA</td></tr>
<tr><td></td><td></td><td></td><td></td><td></td><td></td><td></td><td></td><td></td><td></td><td></td><td></td></tr>
<tr><td>T26206</td><td>GAACATCAGCTCGGGTCCGCTGG</td><td>1000</td><td></td><td></td><td></td><td></td><td></td><td></td><td></td><td></td><td></td></tr>
<tr><td>Oligo fwd</td><td>CACCGAACATCAGCTCGGGTCCGC</td><td></td><td></td><td></td><td></td><td></td><td></td><td></td><td></td><td></td><td></td></tr>
<tr><td>Oligo rev</td><td>AAACGCGGACCCGAGCTGATGTTC</td><td></td><td></td><td></td><td></td><td></td><td></td><td></td><td></td><td></td><td></td></tr>
<tr><td>Chromosome</td><td>start</td><td>end</td><td>strand</td><td>MM</td><td>target_seq</td><td>PAM</td><td>alignment</td><td>distance</td><td>position</td><td>gene name</td><td>gene id</td></tr>
<tr><td>GU305914.1</td><td>103646</td><td>103668</td><td>-</td><td>0</td><td>GAACATCAGCTCGGGTCCGC</td><td>TGG</td><td>||||||||[||||||||||||]PAM</td><td>NA</td><td>Intergenic</td><td>NA</td><td>NA</td></tr>
<tr><td></td><td></td><td></td><td></td><td></td><td></td><td></td><td></td><td></td><td></td><td></td><td></td></tr>
<tr><td>T26207</td><td>CATGTTGCCGATGCTGCTGCCGG</td><td>1000</td><td></td><td></td><td></td><td></td><td></td><td></td><td></td><td></td><td></td></tr>
<tr><td>Oligo fwd</td><td>CACCCATGTTGCCGATGCTGCTGC</td><td></td><td></td><td></td><td></td><td></td><td></td><td></td><td></td><td></td><td></td></tr>
<tr><td>Oligo rev</td><td>AAACGCAGCAGCATCGGCAACATG</td><td></td><td></td><td></td><td></td><td></td><td></td><td></td><td></td><td></td><td></td></tr>
<tr><td>Chromosome</td><td>start</td><td>end</td><td>strand</td><td>MM</td><td>target_seq</td><td>PAM</td><td>alignment</td><td>distance</td><td>position</td><td>gene name</td><td>gene id</td></tr>
<tr><td>GU305914.1</td><td>103589</td><td>103611</td><td>-</td><td>0</td><td>CATGTTGCCGATGCTGCTGC</td><td>CGG</td><td>||||||||[||||||||||||]PAM</td><td>NA</td><td>Intergenic</td><td>NA</td><td>NA</td></tr>
<tr><td></td><td></td><td></td><td></td><td></td><td></td><td></td><td></td><td></td><td></td><td></td><td></td></tr>
<tr><td>T26208</td><td>CGATGCTGCTGCCGGAGCCGTGG</td><td>1000</td><td></td><td></td><td></td><td></td><td></td><td></td><td></td><td></td><td></td></tr>
<tr><td>Oligo fwd</td><td>CACCCGATGCTGCTGCCGGAGCCG</td><td></td><td></td><td></td><td></td><td></td><td></td><td></td><td></td><td></td><td></td></tr>
<tr><td>Oligo rev</td><td>AAACCGGCTCCGGCAGCAGCATCG</td><td></td><td></td><td></td><td></td><td></td><td></td><td></td><td></td><td></td><td></td></tr>
<tr><td>Chromosome</td><td>start</td><td>end</td><td>strand</td><td>MM</td><td>target_seq</td><td>PAM</td><td>alignment</td><td>distance</td><td>position</td><td>gene name</td><td>gene id</td></tr>
<tr><td>GU305914.1</td><td>103581</td><td>103603</td><td>-</td><td>0</td><td>CGATGCTGCTGCCGGAGCCG</td><td>TGG</td><td>||||||||[||||||||||||]PAM</td><td>NA</td><td>Intergenic</td><td>NA</td><td>NA</td></tr>
<tr><td></td><td></td><td></td><td></td><td></td><td></td><td></td><td></td><td></td><td></td><td></td><td></td></tr>
<tr><td>T26209</td><td>GTGGCAGACCATCAACCTGATGG</td><td>1000</td><td></td><td></td><td></td><td></td><td></td><td></td><td></td><td></td><td></td></tr>
<tr><td>Oligo fwd</td><td>CACCGTGGCAGACCATCAACCTGA</td><td></td><td></td><td></td><td></td><td></td><td></td><td></td><td></td><td></td><td></td></tr>
<tr><td>Oligo rev</td><td>AAACTCAGGTTGATGGTCTGCCAC</td><td></td><td></td><td></td><td></td><td></td><td></td><td></td><td></td><td></td><td></td></tr>
<tr><td>Chromosome</td><td>start</td><td>end</td><td>strand</td><td>MM</td><td>target_seq</td><td>PAM</td><td>alignment</td><td>distance</td><td>position</td><td>gene name</td><td>gene id</td></tr>
<tr><td>GU305914.1</td><td>103562</td><td>103584</td><td>-</td><td>0</td><td>GTGGCAGACCATCAACCTGA</td><td>TGG</td><td>||||||||[||||||||||||]PAM</td><td>NA</td><td>Intergenic</td><td>NA</td><td>NA</td></tr>
<tr><td></td><td></td><td></td><td></td><td></td><td></td><td></td><td></td><td></td><td></td><td></td><td></td></tr>
<tr><td>T26210</td><td>CAACCTGATGGCGCCGCACCAGG</td><td>1000</td><td></td><td></td><td></td><td></td><td></td><td></td><td></td><td></td><td></td></tr>
<tr><td>Oligo fwd</td><td>CACCCAACCTGATGGCGCCGCACC</td><td></td><td></td><td></td><td></td><td></td><td></td><td></td><td></td><td></td><td></td></tr>
<tr><td>Oligo rev</td><td>AAACGGTGCGGCGCCATCAGGTTG</td><td></td><td></td><td></td><td></td><td></td><td></td><td></td><td></td><td></td><td></td></tr>
<tr><td>Chromosome</td><td>start</td><td>end</td><td>strand</td><td>MM</td><td>target_seq</td><td>PAM</td><td>alignment</td><td>distance</td><td>position</td><td>gene name</td><td>gene id</td></tr>
<tr><td>GU305914.1</td><td>103550</td><td>103572</td><td>-</td><td>0</td><td>CAACCTGATGGCGCCGCACC</td><td>AGG</td><td>||||||||[||||||||||||]PAM</td><td>NA</td><td>Intergenic</td><td>NA</td><td>NA</td></tr>
<tr><td></td><td></td><td></td><td></td><td></td><td></td><td></td><td></td><td></td><td></td><td></td><td></td></tr>
<tr><td>T26211</td><td>AACCTGATGGCGCCGCACCAGGG</td><td>1000</td><td></td><td></td><td></td><td></td><td></td><td></td><td></td><td></td><td></td></tr>
<tr><td>Oligo fwd</td><td>CACCAACCTGATGGCGCCGCACCA</td><td></td><td></td><td></td><td></td><td></td><td></td><td></td><td></td><td></td><td></td></tr>
<tr><td>Oligo rev</td><td>AAACTGGTGCGGCGCCATCAGGTT</td><td></td><td></td><td></td><td></td><td></td><td></td><td></td><td></td><td></td><td></td></tr>
<tr><td>Chromosome</td><td>start</td><td>end</td><td>strand</td><td>MM</td><td>target_seq</td><td>PAM</td><td>alignment</td><td>distance</td><td>position</td><td>gene name</td><td>gene id</td></tr>
<tr><td>GU305914.1</td><td>103549</td><td>103571</td><td>-</td><td>0</td><td>AACCTGATGGCGCCGCACCA</td><td>GGG</td><td>||||||||[||||||||||||]PAM</td><td>NA</td><td>Intergenic</td><td>NA</td><td>NA</td></tr>
<tr><td></td><td></td><td></td><td></td><td></td><td></td><td></td><td></td><td></td><td></td><td></td><td></td></tr>
<tr><td>T26212</td><td>CGTACTTCGACCTACGCCTGCGG</td><td>1000</td><td></td><td></td><td></td><td></td><td></td><td></td><td></td><td></td><td></td></tr>
<tr><td>Oligo fwd</td><td>CACCCGTACTTCGACCTACGCCTG</td><td></td><td></td><td></td><td></td><td></td><td></td><td></td><td></td><td></td><td></td></tr>
<tr><td>Oligo rev</td><td>AAACCAGGCGTAGGTCGAAGTACG</td><td></td><td></td><td></td><td></td><td></td><td></td><td></td><td></td><td></td><td></td></tr>
<tr><td>Chromosome</td><td>start</td><td>end</td><td>strand</td><td>MM</td><td>target_seq</td><td>PAM</td><td>alignment</td><td>distance</td><td>position</td><td>gene name</td><td>gene id</td></tr>
<tr><td>GU305914.1</td><td>103521</td><td>103543</td><td>-</td><td>0</td><td>CGTACTTCGACCTACGCCTG</td><td>CGG</td><td>||||||||[||||||||||||]PAM</td><td>NA</td><td>Intergenic</td><td>NA</td><td>NA</td></tr>
<tr><td></td><td></td><td></td><td></td><td></td><td></td><td></td><td></td><td></td><td></td><td></td><td></td></tr>
<tr><td>T26213</td><td>GACCGCTGAGCCTGCCGCCGCGG</td><td>1000</td><td></td><td></td><td></td><td></td><td></td><td></td><td></td><td></td><td></td></tr>
<tr><td>Oligo fwd</td><td>CACCGACCGCTGAGCCTGCCGCCG</td><td></td><td></td><td></td><td></td><td></td><td></td><td></td><td></td><td></td><td></td></tr>
<tr><td>Oligo rev</td><td>AAACCGGCGGCAGGCTCAGCGGTC</td><td></td><td></td><td></td><td></td><td></td><td></td><td></td><td></td><td></td><td></td></tr>
<tr><td>Chromosome</td><td>start</td><td>end</td><td>strand</td><td>MM</td><td>target_seq</td><td>PAM</td><td>alignment</td><td>distance</td><td>position</td><td>gene name</td><td>gene id</td></tr>
<tr><td>GU305914.1</td><td>103497</td><td>103519</td><td>-</td><td>0</td><td>GACCGCTGAGCCTGCCGCCG</td><td>CGG</td><td>||||||||[||||||||||||]PAM</td><td>NA</td><td>Intergenic</td><td>NA</td><td>NA</td></tr>
<tr><td></td><td></td><td></td><td></td><td></td><td></td><td></td><td></td><td></td><td></td><td></td><td></td></tr>
<tr><td>T26214</td><td>ACCGCTGAGCCTGCCGCCGCGGG</td><td>1000</td><td></td><td></td><td></td><td></td><td></td><td></td><td></td><td></td><td></td></tr>
<tr><td>Oligo fwd</td><td>CACCACCGCTGAGCCTGCCGCCGC</td><td></td><td></td><td></td><td></td><td></td><td></td><td></td><td></td><td></td><td></td></tr>
<tr><td>Oligo rev</td><td>AAACGCGGCGGCAGGCTCAGCGGT</td><td></td><td></td><td></td><td></td><td></td><td></td><td></td><td></td><td></td><td></td></tr>
<tr><td>Chromosome</td><td>start</td><td>end</td><td>strand</td><td>MM</td><td>target_seq</td><td>PAM</td><td>alignment</td><td>distance</td><td>position</td><td>gene name</td><td>gene id</td></tr>
<tr><td>GU305914.1</td><td>103496</td><td>103518</td><td>-</td><td>0</td><td>ACCGCTGAGCCTGCCGCCGC</td><td>GGG</td><td>||||||||[||||||||||||]PAM</td><td>NA</td><td>Intergenic</td><td>NA</td><td>NA</td></tr>
<tr><td>GU305914.1</td><td>65720</td><td>65742</td><td>+</td><td>4</td><td>CCCGCTGCCGCTGCCGCCGC</td><td>AGG</td><td>-||||||-[--||||||||||]PAM</td><td>NA</td><td>Intergenic</td><td>NA</td><td>NA</td></tr>
<tr><td></td><td></td><td></td><td></td><td></td><td></td><td></td><td></td><td></td><td></td><td></td><td></td></tr>
<tr><td>T26215</td><td>CTCGAAGAACCTGAAGTTCGAGG</td><td>1000</td><td></td><td></td><td></td><td></td><td></td><td></td><td></td><td></td><td></td></tr>
<tr><td>Oligo fwd</td><td>CACCCTCGAAGAACCTGAAGTTCG</td><td></td><td></td><td></td><td></td><td></td><td></td><td></td><td></td><td></td><td></td></tr>
<tr><td>Oligo rev</td><td>AAACCGAACTTCAGGTTCTTCGAG</td><td></td><td></td><td></td><td></td><td></td><td></td><td></td><td></td><td></td><td></td></tr>
<tr><td>Chromosome</td><td>start</td><td>end</td><td>strand</td><td>MM</td><td>target_seq</td><td>PAM</td><td>alignment</td><td>distance</td><td>position</td><td>gene name</td><td>gene id</td></tr>
<tr><td>GU305914.1</td><td>103472</td><td>103494</td><td>-</td><td>0</td><td>CTCGAAGAACCTGAAGTTCG</td><td>AGG</td><td>||||||||[||||||||||||]PAM</td><td>NA</td><td>Intergenic</td><td>NA</td><td>NA</td></tr>
<tr><td></td><td></td><td></td><td></td><td></td><td></td><td></td><td></td><td></td><td></td><td></td><td></td></tr>
<tr><td>T26216</td><td>AAGAACCTGAAGTTCGAGGCCGG</td><td>1000</td><td></td><td></td><td></td><td></td><td></td><td></td><td></td><td></td><td></td></tr>
<tr><td>Oligo fwd</td><td>CACCAAGAACCTGAAGTTCGAGGC</td><td></td><td></td><td></td><td></td><td></td><td></td><td></td><td></td><td></td><td></td></tr>
<tr><td>Oligo rev</td><td>AAACGCCTCGAACTTCAGGTTCTT</td><td></td><td></td><td></td><td></td><td></td><td></td><td></td><td></td><td></td><td></td></tr>
<tr><td>Chromosome</td><td>start</td><td>end</td><td>strand</td><td>MM</td><td>target_seq</td><td>PAM</td><td>alignment</td><td>distance</td><td>position</td><td>gene name</td><td>gene id</td></tr>
<tr><td>GU305914.1</td><td>103468</td><td>103490</td><td>-</td><td>0</td><td>AAGAACCTGAAGTTCGAGGC</td><td>CGG</td><td>||||||||[||||||||||||]PAM</td><td>NA</td><td>Intergenic</td><td>NA</td><td>NA</td></tr>
</table>

<table>
<tr><td>T26217</td><td>GGCCGGACACCTGTGTGAAGAGG</td><td>1000</td></tr>
<tr><td>Oligo fwd</td><td>CACCGGCCGGACACCTGTGTGAAG</td></tr>
<tr><td>Oligo rev</td><td>AAACCTTCACACAGGTGTCCGGCC</td></tr>
<tr><td>Chromosome</td><td>start</td><td>end</td><td>strand</td><td>MM</td><td>target_seq</td><td>PAM</td><td>alignment</td><td>distance</td><td>position</td><td>gene name</td><td>gene id</td></tr>
<tr><td>GU305914.1</td><td>103451</td><td>103473</td><td>-</td><td>0</td><td>GGCCGGACACCTGTGTGAAG</td><td>AGG</td><td>||||||||[||||||||||||]PAM</td><td>NA</td><td>Intergenic</td><td>NA</td><td>NA</td></tr>
<tr><td>T26218</td><td>AGAGGACGCGAGCAGCTGCCTGG</td><td>1000</td></tr>
<tr><td>Oligo fwd</td><td>CACCAGAGGACGCGAGCAGCTGCC</td></tr>
<tr><td>Oligo rev</td><td>AAACGGCAGCTGCTCGCGTCCTCT</td></tr>
<tr><td>Chromosome</td><td>start</td><td>end</td><td>strand</td><td>MM</td><td>target_seq</td><td>PAM</td><td>alignment</td><td>distance</td><td>position</td><td>gene name</td><td>gene id</td></tr>
<tr><td>GU305914.1</td><td>103433</td><td>103455</td><td>-</td><td>0</td><td>AGAGGACGCGAGCAGCTGCC</td><td>TGG</td><td>||||||||[||||||||||||]PAM</td><td>NA</td><td>Intergenic</td><td>NA</td><td>NA</td></tr>
<tr><td>T26219</td><td>AGCGACAAAGCGCGTGCTCCTGG</td><td>1000</td></tr>
<tr><td>Oligo fwd</td><td>CACCAGCGACAAAGCGCGTGCTCC</td></tr>
<tr><td>Oligo rev</td><td>AAACGGAGCACGCGCTTTGTCGCT</td></tr>
<tr><td>Chromosome</td><td>start</td><td>end</td><td>strand</td><td>MM</td><td>target_seq</td><td>PAM</td><td>alignment</td><td>distance</td><td>position</td><td>gene name</td><td>gene id</td></tr>
<tr><td>GU305914.1</td><td>103391</td><td>103413</td><td>-</td><td>0</td><td>AGCGACAAAGCGCGTGCTCC</td><td>TGG</td><td>||||||||[||||||||||||]PAM</td><td>NA</td><td>Intergenic</td><td>NA</td><td>NA</td></tr>
<tr><td>T26220</td><td>TGCTCCTGGACCCGACCGTGTGG</td><td>1000</td></tr>
<tr><td>Oligo fwd</td><td>CACCTGCTCCTGGACCCGACCGTG</td></tr>
<tr><td>Oligo rev</td><td>AAACCACGGTCGGGTCCAGGAGCA</td></tr>
<tr><td>Chromosome</td><td>start</td><td>end</td><td>strand</td><td>MM</td><td>target_seq</td><td>PAM</td><td>alignment</td><td>distance</td><td>position</td><td>gene name</td><td>gene id</td></tr>
<tr><td>GU305914.1</td><td>103377</td><td>103399</td><td>-</td><td>0</td><td>TGCTCCTGGACCCGACCGTG</td><td>TGG</td><td>||||||||[||||||||||||]PAM</td><td>NA</td><td>Intergenic</td><td>NA</td><td>NA</td></tr>
<tr><td>T26221</td><td>CGTGTGGCGCCCCAACTCACTGG</td><td>1000</td></tr>
<tr><td>Oligo fwd</td><td>CACCCGTGTGGCGCCCCAACTCAC</td></tr>
<tr><td>Oligo rev</td><td>AAACGTGAGTTGGGGCGCCACACG</td></tr>
<tr><td>Chromosome</td><td>start</td><td>end</td><td>strand</td><td>MM</td><td>target_seq</td><td>PAM</td><td>alignment</td><td>distance</td><td>position</td><td>gene name</td><td>gene id</td></tr>
<tr><td>GU305914.1</td><td>103361</td><td>103383</td><td>-</td><td>0</td><td>CGTGTGGCGCCCCAACTCAC</td><td>TGG</td><td>||||||||[||||||||||||]PAM</td><td>NA</td><td>Intergenic</td><td>NA</td><td>NA</td></tr>
<tr><td>T26222</td><td>GTGGCGCCCCAACTCACTGGCGG</td><td>1000</td></tr>
<tr><td>Oligo fwd</td><td>CACCGTGGCGCCCCAACTCACTGG</td></tr>
<tr><td>Oligo rev</td><td>AAACCCAGTGAGTTGGGGCGCCAC</td></tr>
<tr><td>Chromosome</td><td>start</td><td>end</td><td>strand</td><td>MM</td><td>target_seq</td><td>PAM</td><td>alignment</td><td>distance</td><td>position</td><td>gene name</td><td>gene id</td></tr>
<tr><td>GU305914.1</td><td>103358</td><td>103380</td><td>-</td><td>0</td><td>GTGGCGCCCCAACTCACTGG</td><td>CGG</td><td>||||||||[||||||||||||]PAM</td><td>NA</td><td>Intergenic</td><td>NA</td><td>NA</td></tr>
<tr><td>T26223</td><td>GCTACGCGTGCTGAACGCCTCGG</td><td>1000</td></tr>
<tr><td>Oligo fwd</td><td>CACCGCTACGCGTGCTGAACGCCT</td></tr>
<tr><td>Oligo rev</td><td>AAACAGGCGTTCAGCACGCGTAGC</td></tr>
<tr><td>Chromosome</td><td>start</td><td>end</td><td>strand</td><td>MM</td><td>target_seq</td><td>PAM</td><td>alignment</td><td>distance</td><td>position</td><td>gene name</td><td>gene id</td></tr>
<tr><td>GU305914.1</td><td>103334</td><td>103356</td><td>-</td><td>0</td><td>GCTACGCGTGCTGAACGCCT</td><td>CGG</td><td>||||||||[||||||||||||]PAM</td><td>NA</td><td>Intergenic</td><td>NA</td><td>NA</td></tr>
<tr><td>T26224</td><td>CTCGGACGAGCACGTCGATCTGG</td><td>1000</td></tr>
<tr><td>Oligo fwd</td><td>CACCCTCGGACGAGCACGTCGATC</td></tr>
<tr><td>Oligo rev</td><td>AAACGATCGACGTGCTCGTCCGAG</td></tr>
<tr><td>Chromosome</td><td>start</td><td>end</td><td>strand</td><td>MM</td><td>target_seq</td><td>PAM</td><td>alignment</td><td>distance</td><td>position</td><td>gene name</td><td>gene id</td></tr>
<tr><td>GU305914.1</td><td>103316</td><td>103338</td><td>-</td><td>0</td><td>CTCGGACGAGCACGTCGATC</td><td>TGG</td><td>||||||||[||||||||||||]PAM</td><td>NA</td><td>Intergenic</td><td>NA</td><td>NA</td></tr>
<tr><td>T26225</td><td>GGACGAGCACGTCGATCTGGAGG</td><td>1000</td></tr>
<tr><td>Oligo fwd</td><td>CACCGGACGAGCACGTCGATCTGG</td></tr>
<tr><td>Oligo rev</td><td>AAACCCAGATCGACGTGCTCGTCC</td></tr>
<tr><td>Chromosome</td><td>start</td><td>end</td><td>strand</td><td>MM</td><td>target_seq</td><td>PAM</td><td>alignment</td><td>distance</td><td>position</td><td>gene name</td><td>gene id</td></tr>
<tr><td>GU305914.1</td><td>103313</td><td>103335</td><td>-</td><td>0</td><td>GGACGAGCACGTCGATCTGG</td><td>AGG</td><td>||||||||[||||||||||||]PAM</td><td>NA</td><td>Intergenic</td><td>NA</td><td>NA</td></tr>
<tr><td>T26226</td><td>GAGCACGTCGATCTGGAGGCCGG</td><td>1000</td></tr>
<tr><td>Oligo fwd</td><td>CACCGAGCACGTCGATCTGGAGGC</td></tr>
<tr><td>Oligo rev</td><td>AAACGCCTCCAGATCGACGTGCTC</td></tr>
<tr><td>Chromosome</td><td>start</td><td>end</td><td>strand</td><td>MM</td><td>target_seq</td><td>PAM</td><td>alignment</td><td>distance</td><td>position</td><td>gene name</td><td>gene id</td></tr>
<tr><td>GU305914.1</td><td>103309</td><td>103331</td><td>-</td><td>0</td><td>GAGCACGTCGATCTGGAGGC</td><td>CGG</td><td>||||||||[||||||||||||]PAM</td><td>NA</td><td>Intergenic</td><td>NA</td><td>NA</td></tr>
<tr><td>T26227</td><td>AGCACGTCGATCTGGAGGCCGGG</td><td>1000</td></tr>
<tr><td>Oligo fwd</td><td>CACCAGCACGTCGATCTGGAGGCC</td></tr>
<tr><td>Oligo rev</td><td>AAACGGCCTCCAGATCGACGTGCT</td></tr>
<tr><td>Chromosome</td><td>start</td><td>end</td><td>strand</td><td>MM</td><td>target_seq</td><td>PAM</td><td>alignment</td><td>distance</td><td>position</td><td>gene name</td><td>gene id</td></tr>
<tr><td>GU305914.1</td><td>103308</td><td>103330</td><td>-</td><td>0</td><td>AGCACGTCGATCTGGAGGCC</td><td>GGG</td><td>||||||||[||||||||||||]PAM</td><td>NA</td><td>Intergenic</td><td>NA</td><td>NA</td></tr>
<tr><td>T26228</td><td>CGTCGATCTGGAGGCCGGGATGG</td><td>1000</td></tr>
<tr><td>Oligo fwd</td><td>CACCCGTCGATCTGGAGGCCGGGA</td></tr>
<tr><td>Oligo rev</td><td>AAACTCCCGGCCTCCAGATCGACG</td></tr>
<tr><td>Chromosome</td><td>start</td><td>end</td><td>strand</td><td>MM</td><td>target_seq</td><td>PAM</td><td>alignment</td><td>distance</td><td>position</td><td>gene name</td><td>gene id</td></tr>
<tr><td>GU305914.1</td><td>103304</td><td>103326</td><td>-</td><td>0</td><td>CGTCGATCTGGAGGCCGGGA</td><td>TGG</td><td>||||||||[||||||||||||]PAM</td><td>NA</td><td>Intergenic</td><td>NA</td><td>NA</td></tr>
<tr><td>T26229</td><td>TCTGGAGGCCGGGATGGCGATGG</td><td>1000</td></tr>
<tr><td>Oligo fwd</td><td>CACCTCTGGAGGCCGGGATGGCGA</td></tr>
<tr><td>Oligo rev</td><td>AAACTCGCCATCCCGGCCTCCAGA</td></tr>
<tr><td>Chromosome</td><td>start</td><td>end</td><td>strand</td><td>MM</td><td>target_seq</td><td>PAM</td><td>alignment</td><td>distance</td><td>position</td><td>gene name</td><td>gene id</td></tr>
<tr><td>GU305914.1</td><td>103298</td><td>103320</td><td>-</td><td>0</td><td>TCTGGAGGCCGGGATGGCGA</td><td>TGG</td><td>||||||||[||||||||||||]PAM</td><td>NA</td><td>Intergenic</td><td>NA</td><td>NA</td></tr>
<tr><td>T26230</td><td>CAAGATCATCTTCACGACGCCGG</td><td>1000</td></tr>
<tr><td>Oligo fwd</td><td>CACCCAAGATCATCTTCACGACGC</td></tr>
<tr><td>Oligo rev</td><td>AAACGCGTCGTGAAGATGATCTTG</td></tr>
<tr><td>Chromosome</td><td>start</td><td>end</td><td>strand</td><td>MM</td><td>target_seq</td><td>PAM</td><td>alignment</td><td>distance</td><td>position</td><td>gene name</td><td>gene id</td></tr>
<tr><td>GU305914.1</td><td>103274</td><td>103296</td><td>-</td><td>0</td><td>CAAGATCATCTTCACGACGC</td><td>CGG</td><td>||||||||[||||||||||||]PAM</td><td>NA</td><td>Intergenic</td><td>NA</td><td>NA</td></tr>
<tr><td>T26231</td><td>AAGATCATCTTCACGACGCCGGG</td><td>1000</td></tr>
<tr><td>Oligo fwd</td><td>CACCAAGATCATCTTCACGACGCC</td></tr>
<tr><td>Oligo rev</td><td>AAACGGCGTCGTGAAGATGATCTT</td></tr>
<tr><td>Chromosome</td><td>start</td><td>end</td><td>strand</td><td>MM</td><td>target_seq</td><td>PAM</td><td>alignment</td><td>distance</td><td>position</td><td>gene name</td><td>gene id</td></tr>
</table>

GU305914.1 103273 103295 - 0 AAGATCATCTTCACGACGCC GGG ||||||||[||||||||||||]PAM NA Intergenic NA NA

T26232 GTCTTTCGACGTGCCGCGCTCGG 1000
Oligo fwd CACCGTCTTTCGACGTGCCGCGCT
Oligo rev AAACAGCGCGGCACGTCGAAAGAC
Chromosome start end strand MM target_seq PAM alignment distance position gene name gene id
GU305914.1 103211 103233 - 0 GTCTTTCGACGTGCCGCGCT CGG ||||||||[||||||||||||]PAM NA Intergenic NA NA

T26233 CTCGGACCTAAAGCTCGTCAAGG 1000
Oligo fwd CACCCTCGGACCTAAAGCTCGTCA
Oligo rev AAACTGACGAGCTTTAGGTCCGAG
Chromosome start end strand MM target_seq PAM alignment distance position gene name gene id
GU305914.1 103193 103215 - 0 CTCGGACCTAAAGCTCGTCA AGG ||||||||[||||||||||||]PAM NA Intergenic NA NA

T26234 TCGGACCTAAAGCTCGTCAAGGG 1000
Oligo fwd CACCTCGGACCTAAAGCTCGTCAA
Oligo rev AAACTTGACGAGCTTTAGGTCCGA
Chromosome start end strand MM target_seq PAM alignment distance position gene name gene id
GU305914.1 103192 103214 - 0 TCGGACCTAAAGCTCGTCAA GGG ||||||||[||||||||||||]PAM NA Intergenic NA NA

T26235 GACCTAAAGCTCGTCAAGGGCGG 1000
Oligo fwd CACCGACCTAAAGCTCGTCAAGGG
Oligo rev AAACCCCTTGACGAGCTTTAGGTC
Chromosome start end strand MM target_seq PAM alignment distance position gene name gene id
GU305914.1 103189 103211 - 0 GACCTAAAGCTCGTCAAGGG CGG ||||||||[||||||||||||]PAM NA Intergenic NA NA

T26236 ACCTAAAGCTCGTCAAGGGCGGG 1000
Oligo fwd CACCACCTAAAGCTCGTCAAGGGC
Oligo rev AAACGCCCTTGACGAGCTTTAGGT
Chromosome start end strand MM target_seq PAM alignment distance position gene name gene id
GU305914.1 103188 103210 - 0 ACCTAAAGCTCGTCAAGGGC GGG ||||||||[||||||||||||]PAM NA Intergenic NA NA

T26237 GCGCCGCGACTACTACCGCGCGG 1000
Oligo fwd CACCGCGCCGCGACTACTACCGCG
Oligo rev AAACCGCGGTAGTAGTCGCGGCGC
Chromosome start end strand MM target_seq PAM alignment distance position gene name gene id
GU305914.1 103166 103188 - 0 GCGCCGCGACTACTACCGCG CGG ||||||||[||||||||||||]PAM NA Intergenic NA NA

T26238 GCGGAAGAGAGAATGCGAAGCGG 1000
Oligo fwd CACCGCGGAAGAGAGAATGCGAAG
Oligo rev AAACCTTCGCATTCTCTCTTCCGC
Chromosome start end strand MM target_seq PAM alignment distance position gene name gene id
GU305914.1 103147 103169 - 0 GCGGAAGAGAGAATGCGAAG CGG ||||||||[||||||||||||]PAM NA Intergenic NA NA

T26239 CGGTCGAGACTCGTCGAACGAGG 1000
Oligo fwd CACCCGGTCGAGACTCGTCGAACG
Oligo rev AAACCGTTCGACGAGTCTCGACCG
Chromosome start end strand MM target_seq PAM alignment distance position gene name gene id
GU305914.1 103127 103149 - 0 CGGTCGAGACTCGTCGAACG AGG ||||||||[||||||||||||]PAM NA Intergenic NA NA

T26240 GGTCGAGACTCGTCGAACGAGGG 1000
Oligo fwd CACCGGTCGAGACTCGTCGAACGA
Oligo rev AAACTCGTTCGACGAGTCTCGACC
Chromosome start end strand MM target_seq PAM alignment distance position gene name gene id
GU305914.1 103126 103148 - 0 GGTCGAGACTCGTCGAACGA GGG ||||||||[||||||||||||]PAM NA Intergenic NA NA

T26241 ACCTCTTGTCTAGTGTACTCAGG 1000
Oligo fwd CACCACCTCTTGTCTAGTGTACTC
Oligo rev AAACGAGTACACTAGACAAGAGGT
Chromosome start end strand MM target_seq PAM alignment distance position gene name gene id
GU305914.1 103087 103109 - 0 ACCTCTTGTCTAGTGTACTC AGG ||||||||[||||||||||||]PAM NA Intergenic NA NA

T26242 ACATTATATTTTTCTAAAAACGG 1000
Oligo fwd CACCACATTATATTTTTCTAAAAA
Oligo rev AAACTTTTTAGAAAAATATAATGT
Chromosome start end strand MM target_seq PAM alignment distance position gene name gene id
GU305914.1 102992 103014 - 0 ACATTATATTTTTCTAAAAA CGG ||||||||[||||||||||||]PAM NA Intergenic NA NA

T26243 TTTTTCTAAAAACGGTAAAGTGG 1000
Oligo fwd CACCTTTTTCTAAAAACGGTAAAG
Oligo rev AAACCTTTACCGTTTTTAGAAAAA
Chromosome start end strand MM target_seq PAM alignment distance position gene name gene id
GU305914.1 102984 103006 - 0 TTTTTCTAAAAACGGTAAAG TGG ||||||||[||||||||||||]PAM NA Intergenic NA NA

T26244 GTTAACCAATAAACACGCCCCGG 1000
Oligo fwd CACCGTTAACCAATAAACACGCCC
Oligo rev AAACGGGCGTGTTTATTGGTTAAC
Chromosome start end strand MM target_seq PAM alignment distance position gene name gene id
GU305914.1 102947 102969 - 0 GTTAACCAATAAACACGCCC CGG ||||||||[||||||||||||]PAM NA Intergenic NA NA

T26245 GAAAAGCCTCGATCACAGAGCGG 1000
Oligo fwd CACCGAAAAGCCTCGATCACAGAG
Oligo rev AAACCTCTGTGATCGAGGCTTTTC
Chromosome start end strand MM target_seq PAM alignment distance position gene name gene id
GU305914.1 102883 102905 - 0 GAAAAGCCTCGATCACAGAG CGG ||||||||[||||||||||||]PAM NA Intergenic NA NA

T26246 ATCACAGAGCGGCCATCTGCTGG 1000
Oligo fwd CACCATCACAGAGCGGCCATCTGC
Oligo rev AAACGCAGATGGCCGCTCTGTGAT

<table>
<tr><td>Chromosome</td><td>start</td><td>end</td><td>strand</td><td>MM</td><td>target_seq</td><td>PAM</td><td>alignment</td><td>distance</td><td>position</td><td>gene name</td><td>gene id</td></tr>
<tr><td>GU305914.1</td><td>102872</td><td>102894</td><td>-</td><td>0</td><td>ATCACAGAGCGGCCATCTGC</td><td>TGG</td><td>||||||||[||||||||||||]PAM</td><td>NA</td><td>Intergenic</td><td>NA</td><td>NA</td></tr>
<tr><td>T26247</td><td>GCTGCTGCTTTTGTTCCTCGAGG</td><td>1000</td><td></td><td></td><td></td><td></td><td></td><td></td><td></td><td></td><td></td></tr>
<tr><td>Oligo fwd</td><td>CACCGCTGCTGCTTTTGTTCCTCG</td><td></td><td></td><td></td><td></td><td></td><td></td><td></td><td></td><td></td><td></td></tr>
<tr><td>Oligo rev</td><td>AAACCGAGGAACAAAAGCAGCAGC</td><td></td><td></td><td></td><td></td><td></td><td></td><td></td><td></td><td></td><td></td></tr>
<tr><td>Chromosome</td><td>start</td><td>end</td><td>strand</td><td>MM</td><td>target_seq</td><td>PAM</td><td>alignment</td><td>distance</td><td>position</td><td>gene name</td><td>gene id</td></tr>
<tr><td>GU305914.1</td><td>102848</td><td>102870</td><td>-</td><td>0</td><td>GCTGCTGCTTTTGTTCCTCG</td><td>AGG</td><td>||||||||[||||||||||||]PAM</td><td>NA</td><td>Intergenic</td><td>NA</td><td>NA</td></tr>
<tr><td>T26248</td><td>GACGCCTTCCGACGTTGTGCAGG</td><td>1000</td><td></td><td></td><td></td><td></td><td></td><td></td><td></td><td></td><td></td></tr>
<tr><td>Oligo fwd</td><td>CACCGACGCCTTCCGACGTTGTGC</td><td></td><td></td><td></td><td></td><td></td><td></td><td></td><td></td><td></td><td></td></tr>
<tr><td>Oligo rev</td><td>AAACGCACAACGTCGGAAGGCGTC</td><td></td><td></td><td></td><td></td><td></td><td></td><td></td><td></td><td></td><td></td></tr>
<tr><td>Chromosome</td><td>start</td><td>end</td><td>strand</td><td>MM</td><td>target_seq</td><td>PAM</td><td>alignment</td><td>distance</td><td>position</td><td>gene name</td><td>gene id</td></tr>
<tr><td>GU305914.1</td><td>102821</td><td>102843</td><td>-</td><td>0</td><td>GACGCCTTCCGACGTTGTGC</td><td>AGG</td><td>||||||||[||||||||||||]PAM</td><td>NA</td><td>Intergenic</td><td>NA</td><td>NA</td></tr>
<tr><td>T26249</td><td>TCCCAGACGTCGCTTCGTCGCGG</td><td>1000</td><td></td><td></td><td></td><td></td><td></td><td></td><td></td><td></td><td></td></tr>
<tr><td>Oligo fwd</td><td>CACCTCCCAGACGTCGCTTCGTCG</td><td></td><td></td><td></td><td></td><td></td><td></td><td></td><td></td><td></td><td></td></tr>
<tr><td>Oligo rev</td><td>AAACCGACGAAGCGACGTCTGGGA</td><td></td><td></td><td></td><td></td><td></td><td></td><td></td><td></td><td></td><td></td></tr>
<tr><td>Chromosome</td><td>start</td><td>end</td><td>strand</td><td>MM</td><td>target_seq</td><td>PAM</td><td>alignment</td><td>distance</td><td>position</td><td>gene name</td><td>gene id</td></tr>
<tr><td>GU305914.1</td><td>102791</td><td>102813</td><td>-</td><td>0</td><td>TCCCAGACGTCGCTTCGTCG</td><td>CGG</td><td>||||||||[||||||||||||]PAM</td><td>NA</td><td>Intergenic</td><td>NA</td><td>NA</td></tr>
<tr><td>GU305914.1</td><td>209236</td><td>209258</td><td>+</td><td>4</td><td>TGCCACACGTCTCTCCGTCG</td><td>TGG</td><td>|-|||-||[|||-||-|||||]PAM</td><td>NA</td><td>Intergenic</td><td>NA</td><td>NA</td></tr>
<tr><td>T26250</td><td>CAAAATCTCGTCGATCTCCCCGG</td><td>1000</td><td></td><td></td><td></td><td></td><td></td><td></td><td></td><td></td><td></td></tr>
<tr><td>Oligo fwd</td><td>CACCCAAAATCTCGTCGATCTCCC</td><td></td><td></td><td></td><td></td><td></td><td></td><td></td><td></td><td></td><td></td></tr>
<tr><td>Oligo rev</td><td>AAACGGGAGATCGACGAGATTTTG</td><td></td><td></td><td></td><td></td><td></td><td></td><td></td><td></td><td></td><td></td></tr>
<tr><td>Chromosome</td><td>start</td><td>end</td><td>strand</td><td>MM</td><td>target_seq</td><td>PAM</td><td>alignment</td><td>distance</td><td>position</td><td>gene name</td><td>gene id</td></tr>
<tr><td>GU305914.1</td><td>102766</td><td>102788</td><td>-</td><td>0</td><td>CAAAATCTCGTCGATCTCCC</td><td>CGG</td><td>||||||||[||||||||||||]PAM</td><td>NA</td><td>Intergenic</td><td>NA</td><td>NA</td></tr>
<tr><td>T26251</td><td>AAAATCTCGTCGATCTCCCCGGG</td><td>1000</td><td></td><td></td><td></td><td></td><td></td><td></td><td></td><td></td><td></td></tr>
<tr><td>Oligo fwd</td><td>CACCAAAATCTCGTCGATCTCCCC</td><td></td><td></td><td></td><td></td><td></td><td></td><td></td><td></td><td></td><td></td></tr>
<tr><td>Oligo rev</td><td>AAACGGGGAGATCGACGAGATTTT</td><td></td><td></td><td></td><td></td><td></td><td></td><td></td><td></td><td></td><td></td></tr>
<tr><td>Chromosome</td><td>start</td><td>end</td><td>strand</td><td>MM</td><td>target_seq</td><td>PAM</td><td>alignment</td><td>distance</td><td>position</td><td>gene name</td><td>gene id</td></tr>
<tr><td>GU305914.1</td><td>102765</td><td>102787</td><td>-</td><td>0</td><td>AAAATCTCGTCGATCTCCCC</td><td>GGG</td><td>||||||||[||||||||||||]PAM</td><td>NA</td><td>Intergenic</td><td>NA</td><td>NA</td></tr>
<tr><td>T26252</td><td>AAATCTCGTCGATCTCCCCGGGG</td><td>1000</td><td></td><td></td><td></td><td></td><td></td><td></td><td></td><td></td><td></td></tr>
<tr><td>Oligo fwd</td><td>CACCAAATCTCGTCGATCTCCCCG</td><td></td><td></td><td></td><td></td><td></td><td></td><td></td><td></td><td></td><td></td></tr>
<tr><td>Oligo rev</td><td>AAACCGGGGAGATCGACGAGATTT</td><td></td><td></td><td></td><td></td><td></td><td></td><td></td><td></td><td></td><td></td></tr>
<tr><td>Chromosome</td><td>start</td><td>end</td><td>strand</td><td>MM</td><td>target_seq</td><td>PAM</td><td>alignment</td><td>distance</td><td>position</td><td>gene name</td><td>gene id</td></tr>
<tr><td>GU305914.1</td><td>102764</td><td>102786</td><td>-</td><td>0</td><td>AAATCTCGTCGATCTCCCCG</td><td>GGG</td><td>||||||||[||||||||||||]PAM</td><td>NA</td><td>Intergenic</td><td>NA</td><td>NA</td></tr>
<tr><td>T26253</td><td>TCTCGTCGATCTCCCCGGGGAGG</td><td>1000</td><td></td><td></td><td></td><td></td><td></td><td></td><td></td><td></td><td></td></tr>
<tr><td>Oligo fwd</td><td>CACCTCTCGTCGATCTCCCCGGGG</td><td></td><td></td><td></td><td></td><td></td><td></td><td></td><td></td><td></td><td></td></tr>
<tr><td>Oligo rev</td><td>AAACCCCCGGGGAGATCGACGAGA</td><td></td><td></td><td></td><td></td><td></td><td></td><td></td><td></td><td></td><td></td></tr>
<tr><td>Chromosome</td><td>start</td><td>end</td><td>strand</td><td>MM</td><td>target_seq</td><td>PAM</td><td>alignment</td><td>distance</td><td>position</td><td>gene name</td><td>gene id</td></tr>
<tr><td>GU305914.1</td><td>102761</td><td>102783</td><td>-</td><td>0</td><td>TCTCGTCGATCTCCCCGGGG</td><td>AGG</td><td>||||||||[||||||||||||]PAM</td><td>NA</td><td>Intergenic</td><td>NA</td><td>NA</td></tr>
<tr><td>T26254</td><td>CGTCGATCTCCCCGGGGAGGTGG</td><td>1000</td><td></td><td></td><td></td><td></td><td></td><td></td><td></td><td></td><td></td></tr>
<tr><td>Oligo fwd</td><td>CACCCGTCGATCTCCCCGGGGAGG</td><td></td><td></td><td></td><td></td><td></td><td></td><td></td><td></td><td></td><td></td></tr>
<tr><td>Oligo rev</td><td>AAACCCTCCCCGGGGAGATCGACG</td><td></td><td></td><td></td><td></td><td></td><td></td><td></td><td></td><td></td><td></td></tr>
<tr><td>Chromosome</td><td>start</td><td>end</td><td>strand</td><td>MM</td><td>target_seq</td><td>PAM</td><td>alignment</td><td>distance</td><td>position</td><td>gene name</td><td>gene id</td></tr>
<tr><td>GU305914.1</td><td>102758</td><td>102780</td><td>-</td><td>0</td><td>CGTCGATCTCCCCGGGGAGG</td><td>TGG</td><td>||||||||[||||||||||||]PAM</td><td>NA</td><td>Intergenic</td><td>NA</td><td>NA</td></tr>
<tr><td>T26255</td><td>GATCTCCCCGGGGAGGTGGTCGG</td><td>1000</td><td></td><td></td><td></td><td></td><td></td><td></td><td></td><td></td><td></td></tr>
<tr><td>Oligo fwd</td><td>CACCGATCTCCCCGGGGAGGTGGT</td><td></td><td></td><td></td><td></td><td></td><td></td><td></td><td></td><td></td><td></td></tr>
<tr><td>Oligo rev</td><td>AAACACCACCTCCCCGGGGAGATC</td><td></td><td></td><td></td><td></td><td></td><td></td><td></td><td></td><td></td><td></td></tr>
<tr><td>Chromosome</td><td>start</td><td>end</td><td>strand</td><td>MM</td><td>target_seq</td><td>PAM</td><td>alignment</td><td>distance</td><td>position</td><td>gene name</td><td>gene id</td></tr>
<tr><td>GU305914.1</td><td>102754</td><td>102776</td><td>-</td><td>0</td><td>GATCTCCCCGGGGAGGTGGT</td><td>CGG</td><td>||||||||[||||||||||||]PAM</td><td>NA</td><td>Intergenic</td><td>NA</td><td>NA</td></tr>
<tr><td>T26256</td><td>ATCTCCCCGGGGAGGTGGTCGGG</td><td>1000</td><td></td><td></td><td></td><td></td><td></td><td></td><td></td><td></td><td></td></tr>
<tr><td>Oligo fwd</td><td>CACCATCTCCCCGGGGAGGTGGTC</td><td></td><td></td><td></td><td></td><td></td><td></td><td></td><td></td><td></td><td></td></tr>
<tr><td>Oligo rev</td><td>AAACGACCACCTCCCCGGGGAGAT</td><td></td><td></td><td></td><td></td><td></td><td></td><td></td><td></td><td></td><td></td></tr>
<tr><td>Chromosome</td><td>start</td><td>end</td><td>strand</td><td>MM</td><td>target_seq</td><td>PAM</td><td>alignment</td><td>distance</td><td>position</td><td>gene name</td><td>gene id</td></tr>
<tr><td>GU305914.1</td><td>102753</td><td>102775</td><td>-</td><td>0</td><td>ATCTCCCCGGGGAGGTGGTC</td><td>GGG</td><td>||||||||[||||||||||||]PAM</td><td>NA</td><td>Intergenic</td><td>NA</td><td>NA</td></tr>
<tr><td>T26257</td><td>CCCGGGGAGGTGGTCGGGCGCGG</td><td>1000</td><td></td><td></td><td></td><td></td><td></td><td></td><td></td><td></td><td></td></tr>
<tr><td>Oligo fwd</td><td>CACCCCCGGGGAGGTGGTCGGGCG</td><td></td><td></td><td></td><td></td><td></td><td></td><td></td><td></td><td></td><td></td></tr>
<tr><td>Oligo rev</td><td>AAACCGCCCGACCACCTCCCCGGG</td><td></td><td></td><td></td><td></td><td></td><td></td><td></td><td></td><td></td><td></td></tr>
<tr><td>Chromosome</td><td>start</td><td>end</td><td>strand</td><td>MM</td><td>target_seq</td><td>PAM</td><td>alignment</td><td>distance</td><td>position</td><td>gene name</td><td>gene id</td></tr>
<tr><td>GU305914.1</td><td>102748</td><td>102770</td><td>-</td><td>0</td><td>CCCGGGGAGGTGGTCGGGCG</td><td>CGG</td><td>||||||||[||||||||||||]PAM</td><td>NA</td><td>Intergenic</td><td>NA</td><td>NA</td></tr>
<tr><td>T26258</td><td>GGGGAGGTGGTCGGGCGCGGCGG</td><td>1000</td><td></td><td></td><td></td><td></td><td></td><td></td><td></td><td></td><td></td></tr>
<tr><td>Oligo fwd</td><td>CACCGGGGAGGTGGTCGGGCGCGG</td><td></td><td></td><td></td><td></td><td></td><td></td><td></td><td></td><td></td><td></td></tr>
<tr><td>Oligo rev</td><td>AAACCCGCGCCCGACCACCTCCCC</td><td></td><td></td><td></td><td></td><td></td><td></td><td></td><td></td><td></td><td></td></tr>
<tr><td>Chromosome</td><td>start</td><td>end</td><td>strand</td><td>MM</td><td>target_seq</td><td>PAM</td><td>alignment</td><td>distance</td><td>position</td><td>gene name</td><td>gene id</td></tr>
<tr><td>GU305914.1</td><td>102745</td><td>102767</td><td>-</td><td>0</td><td>GGGGAGGTGGTCGGGCGCGG</td><td>CGG</td><td>||||||||[||||||||||||]PAM</td><td>NA</td><td>Intergenic</td><td>NA</td><td>NA</td></tr>
<tr><td>T26259</td><td>GGGAGGTGGTCGGGCGCGGCGGG</td><td>1000</td><td></td><td></td><td></td><td></td><td></td><td></td><td></td><td></td><td></td></tr>
<tr><td>Oligo fwd</td><td>CACCGGGAGGTGGTCGGGCGCGGC</td><td></td><td></td><td></td><td></td><td></td><td></td><td></td><td></td><td></td><td></td></tr>
<tr><td>Oligo rev</td><td>AAACGCCGCGCCCGACCACCTCCC</td><td></td><td></td><td></td><td></td><td></td><td></td><td></td><td></td><td></td><td></td></tr>
<tr><td>Chromosome</td><td>start</td><td>end</td><td>strand</td><td>MM</td><td>target_seq</td><td>PAM</td><td>alignment</td><td>distance</td><td>position</td><td>gene name</td><td>gene id</td></tr>
<tr><td>GU305914.1</td><td>102744</td><td>102766</td><td>-</td><td>0</td><td>GGGAGGTGGTCGGGCGCGGC</td><td>GGG</td><td>||||||||[||||||||||||]PAM</td><td>NA</td><td>Intergenic</td><td>NA</td><td>NA</td></tr>
<tr><td>GU305914.1</td><td>135634</td><td>135656</td><td>-</td><td>4</td><td>CTGACGGGGTCGGGCGCGGC</td><td>CGG</td><td>--||-|-|[||||||||||||]PAM</td><td>NA</td><td>Intergenic</td><td>NA</td><td>NA</td></tr>
<tr><td>T26260</td><td>TGTCCAGCTCCCTGAGTAAGAGG</td><td>1000</td><td></td><td></td><td></td><td></td><td></td><td></td><td></td><td></td><td></td></tr>
<tr><td>Oligo fwd</td><td>CACCTGTCCAGCTCCCTGAGTAAG</td><td></td><td></td><td></td><td></td><td></td><td></td><td></td><td></td><td></td><td></td></tr>
<tr><td>Oligo rev</td><td>AAACCTTACTCAGGGAGCTGGACA</td><td></td><td></td><td></td><td></td><td></td><td></td><td></td><td></td><td></td><td></td></tr>
<tr><td>Chromosome</td><td>start</td><td>end</td><td>strand</td><td>MM</td><td>target_seq</td><td>PAM</td><td>alignment</td><td>distance</td><td>position</td><td>gene name</td><td>gene id</td></tr>
<tr><td>GU305914.1</td><td>102716</td><td>102738</td><td>-</td><td>0</td><td>TGTCCAGCTCCCTGAGTAAG</td><td>AGG</td><td>||||||||[||||||||||||]PAM</td><td>NA</td><td>Intergenic</td><td>NA</td><td>NA</td></tr>
</table>

| | | | | | | | | | | | | |
|---|---|---|---|---|---|---|---|---|---|---|---|---|
| T26261 | CAGCTCCCTGAGTAAGAGGTCGG | | 1000 | | | | | | | | | |
| Oligo fwd | CACCCAGCTCCCTGAGTAAGAGGT | | | | | | | | | | | |
| Oligo rev | AAACACCTCTTACTCAGGGAGCTG | | | | | | | | | | | |
| Chromosome | start | | end | strand | MM | target_seq | PAM | alignment | distance | position | gene name | gene id |
| GU305914.1 | | 102712 | 102734 | - | 0 | CAGCTCCCTGAGTAAGAGGT | CGG | \|\|\|\|\|\|\|\|[\|\|\|\|\|\|\|\|\|\|\|\|]PAM | NA | Intergenic | NA | NA |
| T26262 | CTCCCTGAGTAAGAGGTCGGAGG | | 1000 | | | | | | | | | |
| Oligo fwd | CACCCTCCCTGAGTAAGAGGTCGG | | | | | | | | | | | |
| Oligo rev | AAACCCGACCTCTTACTCAGGGAG | | | | | | | | | | | |
| Chromosome | start | | end | strand | MM | target_seq | PAM | alignment | distance | position | gene name | gene id |
| GU305914.1 | | 102709 | 102731 | - | 0 | CTCCCTGAGTAAGAGGTCGG | AGG | \|\|\|\|\|\|\|\|[\|\|\|\|\|\|\|\|\|\|\|\|]PAM | NA | Intergenic | NA | NA |
| T26263 | GAGTAAGAGGTCGGAGGCAGAGG | | 1000 | | | | | | | | | |
| Oligo fwd | CACCGAGTAAGAGGTCGGAGGCAG | | | | | | | | | | | |
| Oligo rev | AAACCTGCCTCCGACCTCTTACTC | | | | | | | | | | | |
| Chromosome | start | | end | strand | MM | target_seq | PAM | alignment | distance | position | gene name | gene id |
| GU305914.1 | | 102703 | 102725 | - | 0 | GAGTAAGAGGTCGGAGGCAG | AGG | \|\|\|\|\|\|\|\|[\|\|\|\|\|\|\|\|\|\|\|\|]PAM | NA | Intergenic | NA | NA |
| T26264 | GAGGTCGGAGGCAGAGGAGATGG | | 1000 | | | | | | | | | |
| Oligo fwd | CACCGAGGTCGGAGGCAGAGGAGA | | | | | | | | | | | |
| Oligo rev | AAACTCTCCTCTGCCTCCGACCTC | | | | | | | | | | | |
| Chromosome | start | | end | strand | MM | target_seq | PAM | alignment | distance | position | gene name | gene id |
| GU305914.1 | | 102697 | 102719 | - | 0 | GAGGTCGGAGGCAGAGGAGA | TGG | \|\|\|\|\|\|\|\|[\|\|\|\|\|\|\|\|\|\|\|\|]PAM | NA | Intergenic | NA | NA |
| T26265 | GGCAGAGGAGATGGTAGTAGTGG | | 1000 | | | | | | | | | |
| Oligo fwd | CACCGGCAGAGGAGATGGTAGTAG | | | | | | | | | | | |
| Oligo rev | AAACCTACTACCATCTCCTCTGCC | | | | | | | | | | | |
| Chromosome | start | | end | strand | MM | target_seq | PAM | alignment | distance | position | gene name | gene id |
| GU305914.1 | | 102688 | 102710 | - | 0 | GGCAGAGGAGATGGTAGTAG | TGG | \|\|\|\|\|\|\|\|[\|\|\|\|\|\|\|\|\|\|\|\|]PAM | NA | Intergenic | NA | NA |
| GU305914.1 | | 174636 | 174658 | + | 3 | GCCGGAGGAGATGGTGGTAG | TGG | \|-\|-\|\|\|\|[\|\|\|\|\|\|\|-\|\|\|\|]PAM | NA | Intergenic | NA | NA |
| T26266 | GATGGTAGTAGTGGCCGCGTAGG | | 1000 | | | | | | | | | |
| Oligo fwd | CACCGATGGTAGTAGTGGCCGCGT | | | | | | | | | | | |
| Oligo rev | AAACACGCGGCCACTACTACCATC | | | | | | | | | | | |
| Chromosome | start | | end | strand | MM | target_seq | PAM | alignment | distance | position | gene name | gene id |
| GU305914.1 | | 102679 | 102701 | - | 0 | GATGGTAGTAGTGGCCGCGT | AGG | \|\|\|\|\|\|\|\|[\|\|\|\|\|\|\|\|\|\|\|\|]PAM | NA | Intergenic | NA | NA |
| T26267 | GGTAGTAGTGGCCGCGTAGGAGG | | 1000 | | | | | | | | | |
| Oligo fwd | CACCGGTAGTAGTGGCCGCGTAGG | | | | | | | | | | | |
| Oligo rev | AAACCCTACGCGGCCACTACTACC | | | | | | | | | | | |
| Chromosome | start | | end | strand | MM | target_seq | PAM | alignment | distance | position | gene name | gene id |
| GU305914.1 | | 102676 | 102698 | - | 0 | GGTAGTAGTGGCCGCGTAGG | AGG | \|\|\|\|\|\|\|\|[\|\|\|\|\|\|\|\|\|\|\|\|]PAM | NA | Intergenic | NA | NA |
| T26268 | AGTAGTGGCCGCGTAGGAGGAGG | | 1000 | | | | | | | | | |
| Oligo fwd | CACCAGTAGTGGCCGCGTAGGAGG | | | | | | | | | | | |
| Oligo rev | AAACCCTCCTACGCGGCCACTACT | | | | | | | | | | | |
| Chromosome | start | | end | strand | MM | target_seq | PAM | alignment | distance | position | gene name | gene id |
| GU305914.1 | | 102673 | 102695 | - | 0 | AGTAGTGGCCGCGTAGGAGG | AGG | \|\|\|\|\|\|\|\|[\|\|\|\|\|\|\|\|\|\|\|\|]PAM | NA | Intergenic | NA | NA |
| GU305914.1 | | 73786 | 73808 | - | 4 | AGCAGCGGCGGCGGAGGAGG | AGG | \|\|-\|\|-\|\|[\|-\|\|\|-\|\|\|\|\|\|]PAM | NA | Intergenic | NA | NA |
| T26269 | GAGGAGGAAGAAGACGACGACGG | | 1000 | | | | | | | | | |
| Oligo fwd | CACCGAGGAGGAAGAAGACGACGA | | | | | | | | | | | |
| Oligo rev | AAACTCGTCGTCTTCTTCCTCCTC | | | | | | | | | | | |
| Chromosome | start | | end | strand | MM | target_seq | PAM | alignment | distance | position | gene name | gene id |
| GU305914.1 | | 102657 | 102679 | - | 0 | GAGGAGGAAGAAGACGACGA | CGG | \|\|\|\|\|\|\|\|[\|\|\|\|\|\|\|\|\|\|\|\|]PAM | NA | Intergenic | NA | NA |
| GU305914.1 | | 118445 | 118467 | - | 4 | AACAATGAAGAAGACGACGA | CGG | -\|--\|-\|\|[\|\|\|\|\|\|\|\|\|\|\|\|]PAM | NA | Intergenic | NA | NA |
| GU305914.1 | | 40448 | 40470 | - | 4 | GATGAAGAAGACGACGATGA | CGG | \|\|-\|\|-\|\|[\|\|\|-\|\|\|\|\|-\|\|]PAM | NA | Intergenic | NA | NA |
| T26270 | ACGACGACGGCACGCCGTTCAGG | | 1000 | | | | | | | | | |
| Oligo fwd | CACCACGACGACGGCACGCCGTTC | | | | | | | | | | | |
| Oligo rev | AAACGAACGGCGTGCCGTCGTCGT | | | | | | | | | | | |
| Chromosome | start | | end | strand | MM | target_seq | PAM | alignment | distance | position | gene name | gene id |
| GU305914.1 | | 102644 | 102666 | - | 0 | ACGACGACGGCACGCCGTTC | AGG | \|\|\|\|\|\|\|\|[\|\|\|\|\|\|\|\|\|\|\|\|]PAM | NA | Intergenic | NA | NA |
| GU305914.1 | | 149305 | 149327 | + | 4 | ACGGCGGCGGCAGGCCGATC | GGG | \|\|\|-\|\|-\|[\|\|\|\|-\|\|\|\|-\|\|]PAM | NA | Intergenic | NA | NA |
| T26271 | GCACGCCGTTCAGGTCGCAGAGG | | 1000 | | | | | | | | | |
| Oligo fwd | CACCGCACGCCGTTCAGGTCGCAG | | | | | | | | | | | |
| Oligo rev | AAACCTGCGACCTGAACGGCGTGC | | | | | | | | | | | |
| Chromosome | start | | end | strand | MM | target_seq | PAM | alignment | distance | position | gene name | gene id |
| GU305914.1 | | 102635 | 102657 | - | 0 | GCACGCCGTTCAGGTCGCAG | AGG | \|\|\|\|\|\|\|\|[\|\|\|\|\|\|\|\|\|\|\|\|]PAM | NA | Intergenic | NA | NA |
| T26272 | CACGCCGTTCAGGTCGCAGAGGG | | 1000 | | | | | | | | | |
| Oligo fwd | CACCCACGCCGTTCAGGTCGCAGA | | | | | | | | | | | |
| Oligo rev | AAACTCTGCGACCTGAACGGCGTG | | | | | | | | | | | |
| Chromosome | start | | end | strand | MM | target_seq | PAM | alignment | distance | position | gene name | gene id |
| GU305914.1 | | 102634 | 102656 | - | 0 | CACGCCGTTCAGGTCGCAGA | GGG | \|\|\|\|\|\|\|\|[\|\|\|\|\|\|\|\|\|\|\|\|]PAM | NA | Intergenic | NA | NA |
| T26273 | ACGCCGTTCAGGTCGCAGAGGGG | | 1000 | | | | | | | | | |
| Oligo fwd | CACCACGCCGTTCAGGTCGCAGAG | | | | | | | | | | | |
| Oligo rev | AAACCTCTGCGACCTGAACGGCGT | | | | | | | | | | | |
| Chromosome | start | | end | strand | MM | target_seq | PAM | alignment | distance | position | gene name | gene id |
| GU305914.1 | | 102633 | 102655 | - | 0 | ACGCCGTTCAGGTCGCAGAG | GGG | \|\|\|\|\|\|\|\|[\|\|\|\|\|\|\|\|\|\|\|\|]PAM | NA | Intergenic | NA | NA |
| T26274 | CCGTTCAGGTCGCAGAGGGGAGG | | 1000 | | | | | | | | | |
| Oligo fwd | CACCCCGTTCAGGTCGCAGAGGGG | | | | | | | | | | | |
| Oligo rev | AAACCCCCTCTGCGACCTGAACGG | | | | | | | | | | | |
| Chromosome | start | | end | strand | MM | target_seq | PAM | alignment | distance | position | gene name | gene id |
| GU305914.1 | | 102630 | 102652 | - | 0 | CCGTTCAGGTCGCAGAGGGG | AGG | \|\|\|\|\|\|\|\|[\|\|\|\|\|\|\|\|\|\|\|\|]PAM | NA | Intergenic | NA | NA |

T26275 AGGTCGCAGAGGGGAGGCGCCGG 1000
Oligo fwd CACCAGGTCGCAGAGGGGAGGCGC
Oligo rev AAACGCGCCTCCCCTCTGCGACCT
Chromosome start end strand MM target_seq PAM alignment distance position gene name gene id
GU305914.1 102624 102646 - 0 AGGTCGCAGAGGGGAGGCGC CGG ||||||||[||||||||||||]PAM NA Intergenic NA NA

T26276 GCTCCTCCCGTCCGTCGACGAGG 1000
Oligo fwd CACCGCTCCTCCCGTCCGTCGACG
Oligo rev AAACCGTCGACGGACGGGAGGAGC
Chromosome start end strand MM target_seq PAM alignment distance position gene name gene id
GU305914.1 102602 102624 - 0 GCTCCTCCCGTCCGTCGACG AGG ||||||||[||||||||||||]PAM NA Intergenic NA NA

T26277 CGAGGCGCTGAGTGTTCTCGCGG 1000
Oligo fwd CACCCGAGGCGCTGAGTGTTCTCG
Oligo rev AAACCGAGAACACTCAGCGCCTCG
Chromosome start end strand MM target_seq PAM alignment distance position gene name gene id
GU305914.1 102584 102606 - 0 CGAGGCGCTGAGTGTTCTCG CGG ||||||||[||||||||||||]PAM NA Intergenic NA NA

T26278 TCTCGCGGTCCATGTCTATGAGG 1000
Oligo fwd CACCTCTCGCGGTCCATGTCTATG
Oligo rev AAACCATAGACATGGACCGCGAGA
Chromosome start end strand MM target_seq PAM alignment distance position gene name gene id
GU305914.1 102569 102591 - 0 TCTCGCGGTCCATGTCTATG AGG ||||||||[||||||||||||]PAM NA Intergenic NA NA

T26279 CGATCCTGCCGACCTCGCAGAGG 1000
Oligo fwd CACCCGATCCTGCCGACCTCGCAG
Oligo rev AAACCTGCGAGGTCGGCAGGATCG
Chromosome start end strand MM target_seq PAM alignment distance position gene name gene id
GU305914.1 102542 102564 - 0 CGATCCTGCCGACCTCGCAG AGG ||||||||[||||||||||||]PAM NA Intergenic NA NA

T26280 GATCCTGCCGACCTCGCAGAGGG 1000
Oligo fwd CACCGATCCTGCCGACCTCGCAGA
Oligo rev AAACTCTGCGAGGTCGGCAGGATC
Chromosome start end strand MM target_seq PAM alignment distance position gene name gene id
GU305914.1 102541 102563 - 0 GATCCTGCCGACCTCGCAGA GGG ||||||||[||||||||||||]PAM NA Intergenic NA NA
GU305914.1 126347 126369 - 4 CATCCAGCAGAGCTCGCAGA AGG -||||-||[-||-||||||||]PAM NA Intergenic NA NA

T26281 GCAGAGGGTGCTCTGATCGACGG 1000
Oligo fwd CACCGCAGAGGGTGCTCTGATCGA
Oligo rev AAACTCGATCAGAGCACCCTCTGC
Chromosome start end strand MM target_seq PAM alignment distance position gene name gene id
GU305914.1 102526 102548 - 0 GCAGAGGGTGCTCTGATCGA CGG ||||||||[||||||||||||]PAM NA Intergenic NA NA

T26282 CGACGCTGTCCTCGTCCGAGAGG 1000
Oligo fwd CACCCGACGCTGTCCTCGTCCGAG
Oligo rev AAACCTCGGACGAGGACAGCGTCG
Chromosome start end strand MM target_seq PAM alignment distance position gene name gene id
GU305914.1 102482 102504 - 0 CGACGCTGTCCTCGTCCGAG AGG ||||||||[||||||||||||]PAM NA Intergenic NA NA

T26283 GACGCTGTCCTCGTCCGAGAGGG 1000
Oligo fwd CACCGACGCTGTCCTCGTCCGAGA
Oligo rev AAACTCTCGGACGAGGACAGCGTC
Chromosome start end strand MM target_seq PAM alignment distance position gene name gene id
GU305914.1 102481 102503 - 0 GACGCTGTCCTCGTCCGAGA GGG ||||||||[||||||||||||]PAM NA Intergenic NA NA

T26284 AGAGGGTGAACTCCCCGACGAGG 1000
Oligo fwd CACCAGAGGGTGAACTCCCCGACG
Oligo rev AAACCGTCGGGGAGTTCACCCTCT
Chromosome start end strand MM target_seq PAM alignment distance position gene name gene id
GU305914.1 102464 102486 - 0 AGAGGGTGAACTCCCCGACG AGG ||||||||[||||||||||||]PAM NA Intergenic NA NA

T26285 ACTCCCCGACGAGGAGCTTGAGG 1000
Oligo fwd CACCACTCCCCGACGAGGAGCTTG
Oligo rev AAACCAAGCTCCTCGTCGGGGAGT
Chromosome start end strand MM target_seq PAM alignment distance position gene name gene id
GU305914.1 102455 102477 - 0 ACTCCCCGACGAGGAGCTTG AGG ||||||||[||||||||||||]PAM NA Intergenic NA NA

T26286 GAGCGCCTGCATGACACGTTTGG 1000
Oligo fwd CACCGAGCGCCTGCATGACACGTT
Oligo rev AAACAACGTGTCATGCAGGCGCTC
Chromosome start end strand MM target_seq PAM alignment distance position gene name gene id
GU305914.1 102421 102443 - 0 GAGCGCCTGCATGACACGTT TGG ||||||||[||||||||||||]PAM NA Intergenic NA NA

T26287 GGATTTCTGACACTCGATCTCGG 1000
Oligo fwd CACCGGATTTCTGACACTCGATCT
Oligo rev AAACAGATCGAGTGTCAGAAATCC
Chromosome start end strand MM target_seq PAM alignment distance position gene name gene id
GU305914.1 102400 102422 - 0 GGATTTCTGACACTCGATCT CGG ||||||||[||||||||||||]PAM NA Intergenic NA NA

T26288 TTTCTGACACTCGATCTCGGCGG 1000
Oligo fwd CACCTTTCTGACACTCGATCTCGG
Oligo rev AAACCCGAGATCGAGTGTCAGAAA
Chromosome start end strand MM target_seq PAM alignment distance position gene name gene id
GU305914.1 102397 102419 - 0 TTTCTGACACTCGATCTCGG CGG ||||||||[||||||||||||]PAM NA Intergenic NA NA

T26289 ACTCGATCTCGGCGGCGTTGCGG 1000
Oligo fwd CACCACTCGATCTCGGCGGCGTTG
Oligo rev AAACCAACGCCGCCGAGATCGAGT
Chromosome start end strand MM target_seq PAM alignment distance position gene name gene id

GU305914.1 102389 102411 - 0 ACTCGATCTCGGCGGCGTTG CGG ||||||||[||||||||||||]PAM NA Intergenic NA NA

T26290 CGATCTCGGCGGCGTTGCGGAGG 1000
Oligo fwd CACCCGATCTCGGCGGCGTTGCGG
Oligo rev AAACCCGCAACGCCGCCGAGATCG
Chromosome start end strand MM target_seq PAM alignment distance position gene name gene id
GU305914.1 102386 102408 - 0 CGATCTCGGCGGCGTTGCGG AGG ||||||||[||||||||||||]PAM NA Intergenic NA NA

T26291 TCGGCGGCGTTGCGGAGGTGCGG 1000
Oligo fwd CACCTCGGCGGCGTTGCGGAGGTG
Oligo rev AAACCACCTCCGCAACGCCGCCGA
Chromosome start end strand MM target_seq PAM alignment distance position gene name gene id
GU305914.1 102381 102403 - 0 TCGGCGGCGTTGCGGAGGTG CGG ||||||||[||||||||||||]PAM NA Intergenic NA NA

T26292 TGCGGAGGTGCGGTCGACATTGG 1000
Oligo fwd CACCTGCGGAGGTGCGGTCGACAT
Oligo rev AAACATGTCGACCGCACCTCCGCA
Chromosome start end strand MM target_seq PAM alignment distance position gene name gene id
GU305914.1 102371 102393 - 0 TGCGGAGGTGCGGTCGACAT TGG ||||||||[||||||||||||]PAM NA Intergenic NA NA

T26293 GACATTGGTTAGCCAGAAGACGG 1000
Oligo fwd CACCGACATTGGTTAGCCAGAAGA
Oligo rev AAACTCTTCTGGCTAACCAATGTC
Chromosome start end strand MM target_seq PAM alignment distance position gene name gene id
GU305914.1 102356 102378 - 0 GACATTGGTTAGCCAGAAGA CGG ||||||||[||||||||||||]PAM NA Intergenic NA NA

T26294 TAGCCAGAAGACGGTCGAGCCGG 1000
Oligo fwd CACCTAGCCAGAAGACGGTCGAGC
Oligo rev AAACGCTCGACCGTCTTCTGGCTA
Chromosome start end strand MM target_seq PAM alignment distance position gene name gene id
GU305914.1 102347 102369 - 0 TAGCCAGAAGACGGTCGAGC CGG ||||||||[||||||||||||]PAM NA Intergenic NA NA

T26295 AGCCAGAAGACGGTCGAGCCGGG 1000
Oligo fwd CACCAGCCAGAAGACGGTCGAGCC
Oligo rev AAACGGCTCGACCGTCTTCTGGCT
Chromosome start end strand MM target_seq PAM alignment distance position gene name gene id
GU305914.1 102346 102368 - 0 AGCCAGAAGACGGTCGAGCC GGG ||||||||[||||||||||||]PAM NA Intergenic NA NA
GU305914.1 107379 107401 + 4 CGCCAGAGGGCGGTCGAGAC GGG -||||||-[|-||||||||-|]PAM NA Intergenic NA NA

T26296 CGCACGCAGCGCTTATAGACCGG 1000
Oligo fwd CACCCGCACGCAGCGCTTATAGAC
Oligo rev AAACGTCTATAAGCGCTGCGTGCG
Chromosome start end strand MM target_seq PAM alignment distance position gene name gene id
GU305914.1 102315 102337 - 0 CGCACGCAGCGCTTATAGAC CGG ||||||||[||||||||||||]PAM NA Intergenic NA NA

T26297 CGCAGCGCTTATAGACCGGAAGG 1000
Oligo fwd CACCCGCAGCGCTTATAGACCGGA
Oligo rev AAACTCCGGTCTATAAGCGCTGCG
Chromosome start end strand MM target_seq PAM alignment distance position gene name gene id
GU305914.1 102311 102333 - 0 CGCAGCGCTTATAGACCGGA AGG ||||||||[||||||||||||]PAM NA Intergenic NA NA

T26298 AGACCGGAAGGAGCCTGAGCAGG 1000
Oligo fwd CACCAGACCGGAAGGAGCCTGAGC
Oligo rev AAACGCTCAGGCTCCTTCCGGTCT
Chromosome start end strand MM target_seq PAM alignment distance position gene name gene id
GU305914.1 102299 102321 - 0 AGACCGGAAGGAGCCTGAGC AGG ||||||||[||||||||||||]PAM NA Intergenic NA NA

T26299 AAGGAGCCTGAGCAGGTGTTTGG 1000
Oligo fwd CACCAAGGAGCCTGAGCAGGTGTT
Oligo rev AAACAACACCTGCTCAGGCTCCTT
Chromosome start end strand MM target_seq PAM alignment distance position gene name gene id
GU305914.1 102292 102314 - 0 AAGGAGCCTGAGCAGGTGTT TGG ||||||||[||||||||||||]PAM NA Intergenic NA NA

T26300 ACTCCCTATCTATTTCAGACAGG 1000
Oligo fwd CACCACTCCCTATCTATTTCAGAC
Oligo rev AAACGTCTGAAATAGATAGGGAGT
Chromosome start end strand MM target_seq PAM alignment distance position gene name gene id
GU305914.1 102260 102282 - 0 ACTCCCTATCTATTTCAGAC AGG ||||||||[||||||||||||]PAM NA Intergenic NA NA

T26301 GATTCCCAAAATTATCTTTGAGG 1000
Oligo fwd CACCGATTCCCAAAATTATCTTTG
Oligo rev AAACCAAAGATAATTTTGGGAATC
Chromosome start end strand MM target_seq PAM alignment distance position gene name gene id
GU305914.1 102218 102240 - 0 GATTCCCAAAATTATCTTTG AGG ||||||||[||||||||||||]PAM NA Intergenic NA NA

T26302 TTTGAGGAACCCTTCTAATTCGG 1000
Oligo fwd CACCTTTGAGGAACCCTTCTAATT
Oligo rev AAACAATTAGAAGGGTTCCTCAAA
Chromosome start end strand MM target_seq PAM alignment distance position gene name gene id
GU305914.1 102202 102224 - 0 TTTGAGGAACCCTTCTAATT CGG ||||||||[||||||||||||]PAM NA Intergenic NA NA

T26303 GGCCATATCGACGTCCTCGTCGG 1000
Oligo fwd CACCGGCCATATCGACGTCCTCGT
Oligo rev AAACACGAGGACGTCGATATGGCC
Chromosome start end strand MM target_seq PAM alignment distance position gene name gene id
GU305914.1 102181 102203 - 0 GGCCATATCGACGTCCTCGT CGG ||||||||[||||||||||||]PAM NA Intergenic NA NA

T26304 CGACGTCCTCGTCGGCCCTGAGG 1000
Oligo fwd CACCCGACGTCCTCGTCGGCCCTG

<table>
<tr><td>Oligo rev</td><td>AAACCAGGGCCGACGAGGACGTCG</td><td></td><td></td><td></td><td></td><td></td><td></td><td></td><td></td><td></td><td></td></tr>
<tr><td>Chromosome</td><td>start</td><td>end</td><td>strand</td><td>MM</td><td>target_seq</td><td>PAM</td><td>alignment</td><td>distance</td><td>position</td><td>gene name</td><td>gene id</td></tr>
<tr><td>GU305914.1</td><td>102173</td><td>102195</td><td>-</td><td>0</td><td>CGACGTCCTCGTCGGCCCTG</td><td>AGG</td><td>||||||||[||||||||||||]PAM</td><td>NA</td><td>Intergenic</td><td>NA</td><td>NA</td></tr>
<tr><td></td><td></td><td></td><td></td><td></td><td></td><td></td><td></td><td></td><td></td><td></td><td></td></tr>
<tr><td>T26305</td><td>GGCCCTGAGGACGATATATTCGG</td><td>1000</td><td></td><td></td><td></td><td></td><td></td><td></td><td></td><td></td><td></td></tr>
<tr><td>Oligo fwd</td><td>CACCGGCCCTGAGGACGATATATT</td><td></td><td></td><td></td><td></td><td></td><td></td><td></td><td></td><td></td><td></td></tr>
<tr><td>Oligo rev</td><td>AAACAATATATCGTCCTCAGGGCC</td><td></td><td></td><td></td><td></td><td></td><td></td><td></td><td></td><td></td><td></td></tr>
<tr><td>Chromosome</td><td>start</td><td>end</td><td>strand</td><td>MM</td><td>target_seq</td><td>PAM</td><td>alignment</td><td>distance</td><td>position</td><td>gene name</td><td>gene id</td></tr>
<tr><td>GU305914.1</td><td>102160</td><td>102182</td><td>-</td><td>0</td><td>GGCCCTGAGGACGATATATT</td><td>CGG</td><td>||||||||[||||||||||||]PAM</td><td>NA</td><td>Intergenic</td><td>NA</td><td>NA</td></tr>
<tr><td></td><td></td><td></td><td></td><td></td><td></td><td></td><td></td><td></td><td></td><td></td><td></td></tr>
<tr><td>T26306</td><td>CCCGAATCGTGCGTCGACACAGG</td><td>1000</td><td></td><td></td><td></td><td></td><td></td><td></td><td></td><td></td><td></td></tr>
<tr><td>Oligo fwd</td><td>CACCCCCGAATCGTGCGTCGACAC</td><td></td><td></td><td></td><td></td><td></td><td></td><td></td><td></td><td></td><td></td></tr>
<tr><td>Oligo rev</td><td>AAACGTGTCGACGCACGATTCGGG</td><td></td><td></td><td></td><td></td><td></td><td></td><td></td><td></td><td></td><td></td></tr>
<tr><td>Chromosome</td><td>start</td><td>end</td><td>strand</td><td>MM</td><td>target_seq</td><td>PAM</td><td>alignment</td><td>distance</td><td>position</td><td>gene name</td><td>gene id</td></tr>
<tr><td>GU305914.1</td><td>102131</td><td>102153</td><td>-</td><td>0</td><td>CCCGAATCGTGCGTCGACAC</td><td>AGG</td><td>||||||||[||||||||||||]PAM</td><td>NA</td><td>Intergenic</td><td>NA</td><td>NA</td></tr>
<tr><td></td><td></td><td></td><td></td><td></td><td></td><td></td><td></td><td></td><td></td><td></td><td></td></tr>
<tr><td>T26307</td><td>CCGAATCGTGCGTCGACACAGGG</td><td>1000</td><td></td><td></td><td></td><td></td><td></td><td></td><td></td><td></td><td></td></tr>
<tr><td>Oligo fwd</td><td>CACCCCGAATCGTGCGTCGACACA</td><td></td><td></td><td></td><td></td><td></td><td></td><td></td><td></td><td></td><td></td></tr>
<tr><td>Oligo rev</td><td>AAACTGTGTCGACGCACGATTCGG</td><td></td><td></td><td></td><td></td><td></td><td></td><td></td><td></td><td></td><td></td></tr>
<tr><td>Chromosome</td><td>start</td><td>end</td><td>strand</td><td>MM</td><td>target_seq</td><td>PAM</td><td>alignment</td><td>distance</td><td>position</td><td>gene name</td><td>gene id</td></tr>
<tr><td>GU305914.1</td><td>102130</td><td>102152</td><td>-</td><td>0</td><td>CCGAATCGTGCGTCGACACA</td><td>GGG</td><td>||||||||[||||||||||||]PAM</td><td>NA</td><td>Intergenic</td><td>NA</td><td>NA</td></tr>
<tr><td></td><td></td><td></td><td></td><td></td><td></td><td></td><td></td><td></td><td></td><td></td><td></td></tr>
<tr><td>T26308</td><td>AGCATCCTCCTCCAGCAGCTCGG</td><td>1000</td><td></td><td></td><td></td><td></td><td></td><td></td><td></td><td></td><td></td></tr>
<tr><td>Oligo fwd</td><td>CACCAGCATCCTCCTCCAGCAGCT</td><td></td><td></td><td></td><td></td><td></td><td></td><td></td><td></td><td></td><td></td></tr>
<tr><td>Oligo rev</td><td>AAACAGCTGCTGGAGGAGGATGCT</td><td></td><td></td><td></td><td></td><td></td><td></td><td></td><td></td><td></td><td></td></tr>
<tr><td>Chromosome</td><td>start</td><td>end</td><td>strand</td><td>MM</td><td>target_seq</td><td>PAM</td><td>alignment</td><td>distance</td><td>position</td><td>gene name</td><td>gene id</td></tr>
<tr><td>GU305914.1</td><td>102099</td><td>102121</td><td>-</td><td>0</td><td>AGCATCCTCCTCCAGCAGCT</td><td>CGG</td><td>||||||||[||||||||||||]PAM</td><td>NA</td><td>Intergenic</td><td>NA</td><td>NA</td></tr>
<tr><td></td><td></td><td></td><td></td><td></td><td></td><td></td><td></td><td></td><td></td><td></td><td></td></tr>
<tr><td>T26309</td><td>ATCCTCCTCCAGCAGCTCGGAGG</td><td>1000</td><td></td><td></td><td></td><td></td><td></td><td></td><td></td><td></td><td></td></tr>
<tr><td>Oligo fwd</td><td>CACCATCCTCCTCCAGCAGCTCGG</td><td></td><td></td><td></td><td></td><td></td><td></td><td></td><td></td><td></td><td></td></tr>
<tr><td>Oligo rev</td><td>AAACCCGAGCTGCTGGAGGAGGAT</td><td></td><td></td><td></td><td></td><td></td><td></td><td></td><td></td><td></td><td></td></tr>
<tr><td>Chromosome</td><td>start</td><td>end</td><td>strand</td><td>MM</td><td>target_seq</td><td>PAM</td><td>alignment</td><td>distance</td><td>position</td><td>gene name</td><td>gene id</td></tr>
<tr><td>GU305914.1</td><td>102096</td><td>102118</td><td>-</td><td>0</td><td>ATCCTCCTCCAGCAGCTCGG</td><td>AGG</td><td>||||||||[||||||||||||]PAM</td><td>NA</td><td>Intergenic</td><td>NA</td><td>NA</td></tr>
<tr><td></td><td></td><td></td><td></td><td></td><td></td><td></td><td></td><td></td><td></td><td></td><td></td></tr>
<tr><td>T26310</td><td>GCAGCTCGGAGGAGCCGCCCCGG</td><td>1000</td><td></td><td></td><td></td><td></td><td></td><td></td><td></td><td></td><td></td></tr>
<tr><td>Oligo fwd</td><td>CACCGCAGCTCGGAGGAGCCGCCC</td><td></td><td></td><td></td><td></td><td></td><td></td><td></td><td></td><td></td><td></td></tr>
<tr><td>Oligo rev</td><td>AAACGGGCGGCTCCTCCGAGCTGC</td><td></td><td></td><td></td><td></td><td></td><td></td><td></td><td></td><td></td><td></td></tr>
<tr><td>Chromosome</td><td>start</td><td>end</td><td>strand</td><td>MM</td><td>target_seq</td><td>PAM</td><td>alignment</td><td>distance</td><td>position</td><td>gene name</td><td>gene id</td></tr>
<tr><td>GU305914.1</td><td>102085</td><td>102107</td><td>-</td><td>0</td><td>GCAGCTCGGAGGAGCCGCCC</td><td>CGG</td><td>||||||||[||||||||||||]PAM</td><td>NA</td><td>Intergenic</td><td>NA</td><td>NA</td></tr>
<tr><td></td><td></td><td></td><td></td><td></td><td></td><td></td><td></td><td></td><td></td><td></td><td></td></tr>
<tr><td>T26311</td><td>CGCCCCGGCGTCGTCGTCTGCGG</td><td>1000</td><td></td><td></td><td></td><td></td><td></td><td></td><td></td><td></td><td></td></tr>
<tr><td>Oligo fwd</td><td>CACCCGCCCCGGCGTCGTCGTCTG</td><td></td><td></td><td></td><td></td><td></td><td></td><td></td><td></td><td></td><td></td></tr>
<tr><td>Oligo rev</td><td>AAACCAGACGACGACGCCGGGGCG</td><td></td><td></td><td></td><td></td><td></td><td></td><td></td><td></td><td></td><td></td></tr>
<tr><td>Chromosome</td><td>start</td><td>end</td><td>strand</td><td>MM</td><td>target_seq</td><td>PAM</td><td>alignment</td><td>distance</td><td>position</td><td>gene name</td><td>gene id</td></tr>
<tr><td>GU305914.1</td><td>102070</td><td>102092</td><td>-</td><td>0</td><td>CGCCCCGGCGTCGTCGTCTG</td><td>CGG</td><td>||||||||[||||||||||||]PAM</td><td>NA</td><td>Intergenic</td><td>NA</td><td>NA</td></tr>
<tr><td></td><td></td><td></td><td></td><td></td><td></td><td></td><td></td><td></td><td></td><td></td><td></td></tr>
<tr><td>T26312</td><td>CCCGGCGTCGTCGTCTGCGGCGG</td><td>1000</td><td></td><td></td><td></td><td></td><td></td><td></td><td></td><td></td><td></td></tr>
<tr><td>Oligo fwd</td><td>CACCCCCGGCGTCGTCGTCTGCGG</td><td></td><td></td><td></td><td></td><td></td><td></td><td></td><td></td><td></td><td></td></tr>
<tr><td>Oligo rev</td><td>AAACCCGCAGACGACGACGCCGGG</td><td></td><td></td><td></td><td></td><td></td><td></td><td></td><td></td><td></td><td></td></tr>
<tr><td>Chromosome</td><td>start</td><td>end</td><td>strand</td><td>MM</td><td>target_seq</td><td>PAM</td><td>alignment</td><td>distance</td><td>position</td><td>gene name</td><td>gene id</td></tr>
<tr><td>GU305914.1</td><td>102067</td><td>102089</td><td>-</td><td>0</td><td>CCCGGCGTCGTCGTCTGCGG</td><td>CGG</td><td>||||||||[||||||||||||]PAM</td><td>NA</td><td>Intergenic</td><td>NA</td><td>NA</td></tr>
<tr><td></td><td></td><td></td><td></td><td></td><td></td><td></td><td></td><td></td><td></td><td></td><td></td></tr>
<tr><td>T26313</td><td>AGTCCCGCCACCGCCGAAGTCGG</td><td>1000</td><td></td><td></td><td></td><td></td><td></td><td></td><td></td><td></td><td></td></tr>
<tr><td>Oligo fwd</td><td>CACCAGTCCCGCCACCGCCGAAGT</td><td></td><td></td><td></td><td></td><td></td><td></td><td></td><td></td><td></td><td></td></tr>
<tr><td>Oligo rev</td><td>AAACACTTCGGCGGTGGCGGGACT</td><td></td><td></td><td></td><td></td><td></td><td></td><td></td><td></td><td></td><td></td></tr>
<tr><td>Chromosome</td><td>start</td><td>end</td><td>strand</td><td>MM</td><td>target_seq</td><td>PAM</td><td>alignment</td><td>distance</td><td>position</td><td>gene name</td><td>gene id</td></tr>
<tr><td>GU305914.1</td><td>102004</td><td>102026</td><td>-</td><td>0</td><td>AGTCCCGCCACCGCCGAAGT</td><td>CGG</td><td>||||||||[||||||||||||]PAM</td><td>NA</td><td>Intergenic</td><td>NA</td><td>NA</td></tr>
<tr><td></td><td></td><td></td><td></td><td></td><td></td><td></td><td></td><td></td><td></td><td></td><td></td></tr>
<tr><td>T26314</td><td>TATCGTAGTCAATATGCTGTCGG</td><td>1000</td><td></td><td></td><td></td><td></td><td></td><td></td><td></td><td></td><td></td></tr>
<tr><td>Oligo fwd</td><td>CACCTATCGTAGTCAATATGCTGT</td><td></td><td></td><td></td><td></td><td></td><td></td><td></td><td></td><td></td><td></td></tr>
<tr><td>Oligo rev</td><td>AAACACAGCATATTGACTACGATA</td><td></td><td></td><td></td><td></td><td></td><td></td><td></td><td></td><td></td><td></td></tr>
<tr><td>Chromosome</td><td>start</td><td>end</td><td>strand</td><td>MM</td><td>target_seq</td><td>PAM</td><td>alignment</td><td>distance</td><td>position</td><td>gene name</td><td>gene id</td></tr>
<tr><td>GU305914.1</td><td>101932</td><td>101954</td><td>-</td><td>0</td><td>TATCGTAGTCAATATGCTGT</td><td>CGG</td><td>||||||||[||||||||||||]PAM</td><td>NA</td><td>Intergenic</td><td>NA</td><td>NA</td></tr>
<tr><td></td><td></td><td></td><td></td><td></td><td></td><td></td><td></td><td></td><td></td><td></td><td></td></tr>
<tr><td>T26315</td><td>TATGCTGTCGGAGACACCGACGG</td><td>1000</td><td></td><td></td><td></td><td></td><td></td><td></td><td></td><td></td><td></td></tr>
<tr><td>Oligo fwd</td><td>CACCTATGCTGTCGGAGACACCGA</td><td></td><td></td><td></td><td></td><td></td><td></td><td></td><td></td><td></td><td></td></tr>
<tr><td>Oligo rev</td><td>AAACTCGGTGTCTCCGACAGCATA</td><td></td><td></td><td></td><td></td><td></td><td></td><td></td><td></td><td></td><td></td></tr>
<tr><td>Chromosome</td><td>start</td><td>end</td><td>strand</td><td>MM</td><td>target_seq</td><td>PAM</td><td>alignment</td><td>distance</td><td>position</td><td>gene name</td><td>gene id</td></tr>
<tr><td>GU305914.1</td><td>101920</td><td>101942</td><td>-</td><td>0</td><td>TATGCTGTCGGAGACACCGA</td><td>CGG</td><td>||||||||[||||||||||||]PAM</td><td>NA</td><td>Intergenic</td><td>NA</td><td>NA</td></tr>
<tr><td></td><td></td><td></td><td></td><td></td><td></td><td></td><td></td><td></td><td></td><td></td><td></td></tr>
<tr><td>T26316</td><td>AATTCGTACTTCAGACCCAGAGG</td><td>1000</td><td></td><td></td><td></td><td></td><td></td><td></td><td></td><td></td><td></td></tr>
<tr><td>Oligo fwd</td><td>CACCAATTCGTACTTCAGACCCAG</td><td></td><td></td><td></td><td></td><td></td><td></td><td></td><td></td><td></td><td></td></tr>
<tr><td>Oligo rev</td><td>AAACCTGGGTCTGAAGTACGAATT</td><td></td><td></td><td></td><td></td><td></td><td></td><td></td><td></td><td></td><td></td></tr>
<tr><td>Chromosome</td><td>start</td><td>end</td><td>strand</td><td>MM</td><td>target_seq</td><td>PAM</td><td>alignment</td><td>distance</td><td>position</td><td>gene name</td><td>gene id</td></tr>
<tr><td>GU305914.1</td><td>101846</td><td>101868</td><td>-</td><td>0</td><td>AATTCGTACTTCAGACCCAG</td><td>AGG</td><td>||||||||[||||||||||||]PAM</td><td>NA</td><td>Intergenic</td><td>NA</td><td>NA</td></tr>
<tr><td></td><td></td><td></td><td></td><td></td><td></td><td></td><td></td><td></td><td></td><td></td><td></td></tr>
<tr><td>T26317</td><td>ACTTCAGACCCAGAGGTGCCCGG</td><td>1000</td><td></td><td></td><td></td><td></td><td></td><td></td><td></td><td></td><td></td></tr>
<tr><td>Oligo fwd</td><td>CACCACTTCAGACCCAGAGGTGCC</td><td></td><td></td><td></td><td></td><td></td><td></td><td></td><td></td><td></td><td></td></tr>
<tr><td>Oligo rev</td><td>AAACGGCACCTCTGGGTCTGAAGT</td><td></td><td></td><td></td><td></td><td></td><td></td><td></td><td></td><td></td><td></td></tr>
<tr><td>Chromosome</td><td>start</td><td>end</td><td>strand</td><td>MM</td><td>target_seq</td><td>PAM</td><td>alignment</td><td>distance</td><td>position</td><td>gene name</td><td>gene id</td></tr>
<tr><td>GU305914.1</td><td>101839</td><td>101861</td><td>-</td><td>0</td><td>ACTTCAGACCCAGAGGTGCC</td><td>CGG</td><td>||||||||[||||||||||||]PAM</td><td>NA</td><td>Intergenic</td><td>NA</td><td>NA</td></tr>
<tr><td></td><td></td><td></td><td></td><td></td><td></td><td></td><td></td><td></td><td></td><td></td><td></td></tr>
<tr><td>T26318</td><td>AGAGGTGCCCGGATTCGACCCGG</td><td>1000</td><td></td><td></td><td></td><td></td><td></td><td></td><td></td><td></td><td></td></tr>
<tr><td>Oligo fwd</td><td>CACCAGAGGTGCCCGGATTCGACC</td><td></td><td></td><td></td><td></td><td></td><td></td><td></td><td></td><td></td><td></td></tr>
<tr><td>Oligo rev</td><td>AAACGGTCGAATCCGGGCACCTCT</td><td></td><td></td><td></td><td></td><td></td><td></td><td></td><td></td><td></td><td></td></tr>
<tr><td>Chromosome</td><td>start</td><td>end</td><td>strand</td><td>MM</td><td>target_seq</td><td>PAM</td><td>alignment</td><td>distance</td><td>position</td><td>gene name</td><td>gene id</td></tr>
<tr><td>GU305914.1</td><td>101828</td><td>101850</td><td>-</td><td>0</td><td>AGAGGTGCCCGGATTCGACC</td><td>CGG</td><td>||||||||[||||||||||||]PAM</td><td>NA</td><td>Intergenic</td><td>NA</td><td>NA</td></tr>
<tr><td></td><td></td><td></td><td></td><td></td><td></td><td></td><td></td><td></td><td></td><td></td><td></td></tr>
<tr><td>T26319</td><td>GAGGTGCCCGGATTCGACCCGGG</td><td>1000</td><td></td><td></td><td></td><td></td><td></td><td></td><td></td><td></td><td></td></tr>
</table>

<table>
<tr><td>Oligo fwd</td><td>CACCGAGGTGCCCGGATTCGACCC</td></tr>
<tr><td>Oligo rev</td><td>AAACGGGTCGAATCCGGGCACCTC</td></tr>
<tr><td>Chromosome</td><td>start</td><td>end</td><td>strand</td><td>MM</td><td>target_seq</td><td>PAM</td><td>alignment</td><td>distance</td><td>position</td><td>gene name</td><td>gene id</td></tr>
<tr><td>GU305914.1</td><td>101827</td><td>101849</td><td>-</td><td>0</td><td>GAGGTGCCCGGATTCGACCC</td><td>GGG</td><td>||||||||[||||||||||||]PAM</td><td>NA</td><td>Intergenic</td><td>NA</td><td>NA</td></tr>
<tr><td></td></tr>
<tr><td>T26320</td><td>CCCGCGCGCGTTATCTGCGCCGG</td><td>1000</td></tr>
<tr><td>Oligo fwd</td><td>CACCCCCGCGCGCGTTATCTGCGC</td></tr>
<tr><td>Oligo rev</td><td>AAACGCGCAGATAACGCGCGCGGG</td></tr>
<tr><td>Chromosome</td><td>start</td><td>end</td><td>strand</td><td>MM</td><td>target_seq</td><td>PAM</td><td>alignment</td><td>distance</td><td>position</td><td>gene name</td><td>gene id</td></tr>
<tr><td>GU305914.1</td><td>101733</td><td>101755</td><td>-</td><td>0</td><td>CCCGCGCGCGTTATCTGCGC</td><td>CGG</td><td>||||||||[||||||||||||]PAM</td><td>NA</td><td>Intergenic</td><td>NA</td><td>NA</td></tr>
<tr><td></td></tr>
<tr><td>T26321</td><td>CGATCGTAAAAAAGACGAGACGG</td><td>1000</td></tr>
<tr><td>Oligo fwd</td><td>CACCCGATCGTAAAAAAGACGAGA</td></tr>
<tr><td>Oligo rev</td><td>AAACTCTCGTCTTTTTTACGATCG</td></tr>
<tr><td>Chromosome</td><td>start</td><td>end</td><td>strand</td><td>MM</td><td>target_seq</td><td>PAM</td><td>alignment</td><td>distance</td><td>position</td><td>gene name</td><td>gene id</td></tr>
<tr><td>GU305914.1</td><td>101668</td><td>101690</td><td>-</td><td>0</td><td>CGATCGTAAAAAAGACGAGA</td><td>CGG</td><td>||||||||[||||||||||||]PAM</td><td>NA</td><td>Intergenic</td><td>NA</td><td>NA</td></tr>
<tr><td></td></tr>
<tr><td>T26322</td><td>GATCGTAAAAAAGACGAGACGGG</td><td>1000</td></tr>
<tr><td>Oligo fwd</td><td>CACCGATCGTAAAAAAGACGAGAC</td></tr>
<tr><td>Oligo rev</td><td>AAACGTCTCGTCTTTTTTACGATC</td></tr>
<tr><td>Chromosome</td><td>start</td><td>end</td><td>strand</td><td>MM</td><td>target_seq</td><td>PAM</td><td>alignment</td><td>distance</td><td>position</td><td>gene name</td><td>gene id</td></tr>
<tr><td>GU305914.1</td><td>101667</td><td>101689</td><td>-</td><td>0</td><td>GATCGTAAAAAAGACGAGAC</td><td>GGG</td><td>||||||||[||||||||||||]PAM</td><td>NA</td><td>Intergenic</td><td>NA</td><td>NA</td></tr>
<tr><td></td></tr>
<tr><td>T26323</td><td>AGACGAGACGGGTCATGACGAGG</td><td>1000</td></tr>
<tr><td>Oligo fwd</td><td>CACCAGACGAGACGGGTCATGACG</td></tr>
<tr><td>Oligo rev</td><td>AAACCGTCATGACCCGTCTCGTCT</td></tr>
<tr><td>Chromosome</td><td>start</td><td>end</td><td>strand</td><td>MM</td><td>target_seq</td><td>PAM</td><td>alignment</td><td>distance</td><td>position</td><td>gene name</td><td>gene id</td></tr>
<tr><td>GU305914.1</td><td>101656</td><td>101678</td><td>-</td><td>0</td><td>AGACGAGACGGGTCATGACG</td><td>AGG</td><td>||||||||[||||||||||||]PAM</td><td>NA</td><td>Intergenic</td><td>NA</td><td>NA</td></tr>
<tr><td></td></tr>
<tr><td>T26324</td><td>ACGGGTCATGACGAGGAGAACGG</td><td>1000</td></tr>
<tr><td>Oligo fwd</td><td>CACCACGGGTCATGACGAGGAGAA</td></tr>
<tr><td>Oligo rev</td><td>AAACTTCTCCTCGTCATGACCCGT</td></tr>
<tr><td>Chromosome</td><td>start</td><td>end</td><td>strand</td><td>MM</td><td>target_seq</td><td>PAM</td><td>alignment</td><td>distance</td><td>position</td><td>gene name</td><td>gene id</td></tr>
<tr><td>GU305914.1</td><td>101649</td><td>101671</td><td>-</td><td>0</td><td>ACGGGTCATGACGAGGAGAA</td><td>CGG</td><td>||||||||[||||||||||||]PAM</td><td>NA</td><td>Intergenic</td><td>NA</td><td>NA</td></tr>
<tr><td></td></tr>
<tr><td>T26325</td><td>CGGGTCATGACGAGGAGAACGGG</td><td>1000</td></tr>
<tr><td>Oligo fwd</td><td>CACCCGGGTCATGACGAGGAGAAC</td></tr>
<tr><td>Oligo rev</td><td>AAACGTTCTCCTCGTCATGACCCG</td></tr>
<tr><td>Chromosome</td><td>start</td><td>end</td><td>strand</td><td>MM</td><td>target_seq</td><td>PAM</td><td>alignment</td><td>distance</td><td>position</td><td>gene name</td><td>gene id</td></tr>
<tr><td>GU305914.1</td><td>101648</td><td>101670</td><td>-</td><td>0</td><td>CGGGTCATGACGAGGAGAAC</td><td>GGG</td><td>||||||||[||||||||||||]PAM</td><td>NA</td><td>Intergenic</td><td>NA</td><td>NA</td></tr>
<tr><td></td></tr>
<tr><td>T26326</td><td>GACGAGGAGAACGGGTCCGCCGG</td><td>1000</td></tr>
<tr><td>Oligo fwd</td><td>CACCGACGAGGAGAACGGGTCCGC</td></tr>
<tr><td>Oligo rev</td><td>AAACGCGGACCCGTTCTCCTCGTC</td></tr>
<tr><td>Chromosome</td><td>start</td><td>end</td><td>strand</td><td>MM</td><td>target_seq</td><td>PAM</td><td>alignment</td><td>distance</td><td>position</td><td>gene name</td><td>gene id</td></tr>
<tr><td>GU305914.1</td><td>101640</td><td>101662</td><td>-</td><td>0</td><td>GACGAGGAGAACGGGTCCGC</td><td>CGG</td><td>||||||||[||||||||||||]PAM</td><td>NA</td><td>Intergenic</td><td>NA</td><td>NA</td></tr>
<tr><td></td></tr>
<tr><td>T26327</td><td>GAGGAGAACGGGTCCGCCGGCGG</td><td>1000</td></tr>
<tr><td>Oligo fwd</td><td>CACCGAGGAGAACGGGTCCGCCGG</td></tr>
<tr><td>Oligo rev</td><td>AAACCCGGCGGACCCGTTCTCCTC</td></tr>
<tr><td>Chromosome</td><td>start</td><td>end</td><td>strand</td><td>MM</td><td>target_seq</td><td>PAM</td><td>alignment</td><td>distance</td><td>position</td><td>gene name</td><td>gene id</td></tr>
<tr><td>GU305914.1</td><td>101637</td><td>101659</td><td>-</td><td>0</td><td>GAGGAGAACGGGTCCGCCGG</td><td>CGG</td><td>||||||||[||||||||||||]PAM</td><td>NA</td><td>Intergenic</td><td>NA</td><td>NA</td></tr>
<tr><td></td></tr>
<tr><td>T26328</td><td>CACCAGCACCCTCAAATGCCTGG</td><td>1000</td></tr>
<tr><td>Oligo fwd</td><td>CACCCACCAGCACCCTCAAATGCC</td></tr>
<tr><td>Oligo rev</td><td>AAACGGCATTTGAGGGTGCTGGTG</td></tr>
<tr><td>Chromosome</td><td>start</td><td>end</td><td>strand</td><td>MM</td><td>target_seq</td><td>PAM</td><td>alignment</td><td>distance</td><td>position</td><td>gene name</td><td>gene id</td></tr>
<tr><td>GU305914.1</td><td>101596</td><td>101618</td><td>-</td><td>0</td><td>CACCAGCACCCTCAAATGCC</td><td>TGG</td><td>||||||||[||||||||||||]PAM</td><td>NA</td><td>Intergenic</td><td>NA</td><td>NA</td></tr>
<tr><td></td></tr>
<tr><td>T26329</td><td>TGTCCCCCACGTCCCCCTCCTGG</td><td>1000</td></tr>
<tr><td>Oligo fwd</td><td>CACCTGTCCCCCACGTCCCCCTCC</td></tr>
<tr><td>Oligo rev</td><td>AAACGGAGGGGGACGTGGGGGACA</td></tr>
<tr><td>Chromosome</td><td>start</td><td>end</td><td>strand</td><td>MM</td><td>target_seq</td><td>PAM</td><td>alignment</td><td>distance</td><td>position</td><td>gene name</td><td>gene id</td></tr>
<tr><td>GU305914.1</td><td>101525</td><td>101547</td><td>-</td><td>0</td><td>TGTCCCCCACGTCCCCCTCC</td><td>TGG</td><td>||||||||[||||||||||||]PAM</td><td>NA</td><td>Intergenic</td><td>NA</td><td>NA</td></tr>
<tr><td></td></tr>
<tr><td>T26330</td><td>TCCTGGTTCCTGAGCACGTACGG</td><td>1000</td></tr>
<tr><td>Oligo fwd</td><td>CACCTCCTGGTTCCTGAGCACGTA</td></tr>
<tr><td>Oligo rev</td><td>AAACTACGTGCTCAGGAACCAGGA</td></tr>
<tr><td>Chromosome</td><td>start</td><td>end</td><td>strand</td><td>MM</td><td>target_seq</td><td>PAM</td><td>alignment</td><td>distance</td><td>position</td><td>gene name</td><td>gene id</td></tr>
<tr><td>GU305914.1</td><td>101508</td><td>101530</td><td>-</td><td>0</td><td>TCCTGGTTCCTGAGCACGTA</td><td>CGG</td><td>||||||||[||||||||||||]PAM</td><td>NA</td><td>Intergenic</td><td>NA</td><td>NA</td></tr>
<tr><td></td></tr>
<tr><td>T26331</td><td>CCTGGTTCCTGAGCACGTACGGG</td><td>1000</td></tr>
<tr><td>Oligo fwd</td><td>CACCCCTGGTTCCTGAGCACGTAC</td></tr>
<tr><td>Oligo rev</td><td>AAACGTACGTGCTCAGGAACCAGG</td></tr>
<tr><td>Chromosome</td><td>start</td><td>end</td><td>strand</td><td>MM</td><td>target_seq</td><td>PAM</td><td>alignment</td><td>distance</td><td>position</td><td>gene name</td><td>gene id</td></tr>
<tr><td>GU305914.1</td><td>101507</td><td>101529</td><td>-</td><td>0</td><td>CCTGGTTCCTGAGCACGTAC</td><td>GGG</td><td>||||||||[||||||||||||]PAM</td><td>NA</td><td>Intergenic</td><td>NA</td><td>NA</td></tr>
<tr><td></td></tr>
<tr><td>T26332</td><td>GAGCACGTACGGGACCCACGAGG</td><td>1000</td></tr>
<tr><td>Oligo fwd</td><td>CACCGAGCACGTACGGGACCCACG</td></tr>
<tr><td>Oligo rev</td><td>AAACCGTGGGTCCCGTACGTGCTC</td></tr>
<tr><td>Chromosome</td><td>start</td><td>end</td><td>strand</td><td>MM</td><td>target_seq</td><td>PAM</td><td>alignment</td><td>distance</td><td>position</td><td>gene name</td><td>gene id</td></tr>
<tr><td>GU305914.1</td><td>101497</td><td>101519</td><td>-</td><td>0</td><td>GAGCACGTACGGGACCCACG</td><td>AGG</td><td>||||||||[||||||||||||]PAM</td><td>NA</td><td>Intergenic</td><td>NA</td><td>NA</td></tr>
<tr><td></td></tr>
<tr><td>T26333</td><td>AGCACGTACGGGACCCACGAGGG</td><td>1000</td></tr>
<tr><td>Oligo fwd</td><td>CACCAGCACGTACGGGACCCACGA</td></tr>
<tr><td>Oligo rev</td><td>AAACTCGTGGGTCCCGTACGTGCT</td></tr>
<tr><td>Chromosome</td><td>start</td><td>end</td><td>strand</td><td>MM</td><td>target_seq</td><td>PAM</td><td>alignment</td><td>distance</td><td>position</td><td>gene name</td><td>gene id</td></tr>
<tr><td>GU305914.1</td><td>101496</td><td>101518</td><td>-</td><td>0</td><td>AGCACGTACGGGACCCACGA</td><td>GGG</td><td>||||||||[||||||||||||]PAM</td><td>NA</td><td>Intergenic</td><td>NA</td><td>NA</td></tr>
</table>

<table>
<tr><td>T26334</td><td>TGTCCTACTATCTGTTCGAGAGG</td><td>1000</td><td></td><td></td><td></td><td></td><td></td><td></td><td></td><td></td><td></td></tr>
<tr><td>Oligo fwd</td><td>CACCTGTCCTACTATCTGTTCGAG</td><td></td><td></td><td></td><td></td><td></td><td></td><td></td><td></td><td></td><td></td></tr>
<tr><td>Oligo rev</td><td>AAACCTCGAACAGATAGTAGGACA</td><td></td><td></td><td></td><td></td><td></td><td></td><td></td><td></td><td></td><td></td></tr>
<tr><td>Chromosome</td><td>start</td><td>end</td><td>strand</td><td>MM</td><td>target_seq</td><td>PAM</td><td>alignment</td><td>distance</td><td>position</td><td>gene name</td><td>gene id</td></tr>
<tr><td>GU305914.1</td><td>101456</td><td>101478</td><td>-</td><td>0</td><td>TGTCCTACTATCTGTTCGAG</td><td>AGG</td><td>||||||||[||||||||||||]PAM</td><td>NA</td><td>Intergenic</td><td>NA</td><td>NA</td></tr>
<tr><td>T26335</td><td>AAATGTTTCACCGCCGACCCCGG</td><td>1000</td><td></td><td></td><td></td><td></td><td></td><td></td><td></td><td></td><td></td></tr>
<tr><td>Oligo fwd</td><td>CACCAAATGTTTCACCGCCGACCC</td><td></td><td></td><td></td><td></td><td></td><td></td><td></td><td></td><td></td><td></td></tr>
<tr><td>Oligo rev</td><td>AAACGGGTCGGCGGTGAAACATTT</td><td></td><td></td><td></td><td></td><td></td><td></td><td></td><td></td><td></td><td></td></tr>
<tr><td>Chromosome</td><td>start</td><td>end</td><td>strand</td><td>MM</td><td>target_seq</td><td>PAM</td><td>alignment</td><td>distance</td><td>position</td><td>gene name</td><td>gene id</td></tr>
<tr><td>GU305914.1</td><td>101397</td><td>101419</td><td>-</td><td>0</td><td>AAATGTTTCACCGCCGACCC</td><td>CGG</td><td>||||||||[||||||||||||]PAM</td><td>NA</td><td>Intergenic</td><td>NA</td><td>NA</td></tr>
<tr><td>T26336</td><td>CATCAACATGCGCGATTTCATGG</td><td>1000</td><td></td><td></td><td></td><td></td><td></td><td></td><td></td><td></td><td></td></tr>
<tr><td>Oligo fwd</td><td>CACCCATCAACATGCGCGATTTCA</td><td></td><td></td><td></td><td></td><td></td><td></td><td></td><td></td><td></td><td></td></tr>
<tr><td>Oligo rev</td><td>AAACTGAAATCGCGCATGTTGATG</td><td></td><td></td><td></td><td></td><td></td><td></td><td></td><td></td><td></td><td></td></tr>
<tr><td>Chromosome</td><td>start</td><td>end</td><td>strand</td><td>MM</td><td>target_seq</td><td>PAM</td><td>alignment</td><td>distance</td><td>position</td><td>gene name</td><td>gene id</td></tr>
<tr><td>GU305914.1</td><td>101356</td><td>101378</td><td>-</td><td>0</td><td>CATCAACATGCGCGATTTCA</td><td>TGG</td><td>||||||||[||||||||||||]PAM</td><td>NA</td><td>Intergenic</td><td>NA</td><td>NA</td></tr>
<tr><td>T26337</td><td>GATTTCATGGCGACCATGAACGG</td><td>1000</td><td></td><td></td><td></td><td></td><td></td><td></td><td></td><td></td><td></td></tr>
<tr><td>Oligo fwd</td><td>CACCGATTTCATGGCGACCATGAA</td><td></td><td></td><td></td><td></td><td></td><td></td><td></td><td></td><td></td><td></td></tr>
<tr><td>Oligo rev</td><td>AAACTTCATGGTCGCCATGAAATC</td><td></td><td></td><td></td><td></td><td></td><td></td><td></td><td></td><td></td><td></td></tr>
<tr><td>Chromosome</td><td>start</td><td>end</td><td>strand</td><td>MM</td><td>target_seq</td><td>PAM</td><td>alignment</td><td>distance</td><td>position</td><td>gene name</td><td>gene id</td></tr>
<tr><td>GU305914.1</td><td>101343</td><td>101365</td><td>-</td><td>0</td><td>GATTTCATGGCGACCATGAA</td><td>CGG</td><td>||||||||[||||||||||||]PAM</td><td>NA</td><td>Intergenic</td><td>NA</td><td>NA</td></tr>
<tr><td>T26338</td><td>TGGCGACCATGAACGGCTCGCGG</td><td>1000</td><td></td><td></td><td></td><td></td><td></td><td></td><td></td><td></td><td></td></tr>
<tr><td>Oligo fwd</td><td>CACCTGGCGACCATGAACGGCTCG</td><td></td><td></td><td></td><td></td><td></td><td></td><td></td><td></td><td></td><td></td></tr>
<tr><td>Oligo rev</td><td>AAACCGAGCCGTTCATGGTCGCCA</td><td></td><td></td><td></td><td></td><td></td><td></td><td></td><td></td><td></td><td></td></tr>
<tr><td>Chromosome</td><td>start</td><td>end</td><td>strand</td><td>MM</td><td>target_seq</td><td>PAM</td><td>alignment</td><td>distance</td><td>position</td><td>gene name</td><td>gene id</td></tr>
<tr><td>GU305914.1</td><td>101336</td><td>101358</td><td>-</td><td>0</td><td>TGGCGACCATGAACGGCTCG</td><td>CGG</td><td>||||||||[||||||||||||]PAM</td><td>NA</td><td>Intergenic</td><td>NA</td><td>NA</td></tr>
<tr><td>T26339</td><td>GGTTCCGCAAGCAGACCGCGCGG</td><td>1000</td><td></td><td></td><td></td><td></td><td></td><td></td><td></td><td></td><td></td></tr>
<tr><td>Oligo fwd</td><td>CACCGGTTCCGCAAGCAGACCGCG</td><td></td><td></td><td></td><td></td><td></td><td></td><td></td><td></td><td></td><td></td></tr>
<tr><td>Oligo rev</td><td>AAACCGCGGTCTGCTTGCGGAACC</td><td></td><td></td><td></td><td></td><td></td><td></td><td></td><td></td><td></td><td></td></tr>
<tr><td>Chromosome</td><td>start</td><td>end</td><td>strand</td><td>MM</td><td>target_seq</td><td>PAM</td><td>alignment</td><td>distance</td><td>position</td><td>gene name</td><td>gene id</td></tr>
<tr><td>GU305914.1</td><td>101315</td><td>101337</td><td>-</td><td>0</td><td>GGTTCCGCAAGCAGACCGCG</td><td>CGG</td><td>||||||||[||||||||||||]PAM</td><td>NA</td><td>Intergenic</td><td>NA</td><td>NA</td></tr>
<tr><td>T26340</td><td>CAAGCAGACCGCGCGGTTCGCGG</td><td>1000</td><td></td><td></td><td></td><td></td><td></td><td></td><td></td><td></td><td></td></tr>
<tr><td>Oligo fwd</td><td>CACCCAAGCAGACCGCGCGGTTCG</td><td></td><td></td><td></td><td></td><td></td><td></td><td></td><td></td><td></td><td></td></tr>
<tr><td>Oligo rev</td><td>AAACCGAACCGCGCGGTCTGCTTG</td><td></td><td></td><td></td><td></td><td></td><td></td><td></td><td></td><td></td><td></td></tr>
<tr><td>Chromosome</td><td>start</td><td>end</td><td>strand</td><td>MM</td><td>target_seq</td><td>PAM</td><td>alignment</td><td>distance</td><td>position</td><td>gene name</td><td>gene id</td></tr>
<tr><td>GU305914.1</td><td>101308</td><td>101330</td><td>-</td><td>0</td><td>CAAGCAGACCGCGCGGTTCG</td><td>CGG</td><td>||||||||[||||||||||||]PAM</td><td>NA</td><td>Intergenic</td><td>NA</td><td>NA</td></tr>
<tr><td>GU305914.1</td><td>66230</td><td>66252</td><td>+</td><td>4</td><td>CCAGCTGGCCGAGCGGTTCG</td><td>CGG</td><td>|-|||-|-[|||-||||||||]PAM</td><td>NA</td><td>Intergenic</td><td>NA</td><td>NA</td></tr>
<tr><td>T26341</td><td>GCGGTTCGCGGCGTTCGCCAAGG</td><td>1000</td><td></td><td></td><td></td><td></td><td></td><td></td><td></td><td></td><td></td></tr>
<tr><td>Oligo fwd</td><td>CACCGCGGTTCGCGGCGTTCGCCA</td><td></td><td></td><td></td><td></td><td></td><td></td><td></td><td></td><td></td><td></td></tr>
<tr><td>Oligo rev</td><td>AAACTGGCGAACGCCGCGAACCGC</td><td></td><td></td><td></td><td></td><td></td><td></td><td></td><td></td><td></td><td></td></tr>
<tr><td>Chromosome</td><td>start</td><td>end</td><td>strand</td><td>MM</td><td>target_seq</td><td>PAM</td><td>alignment</td><td>distance</td><td>position</td><td>gene name</td><td>gene id</td></tr>
<tr><td>GU305914.1</td><td>101296</td><td>101318</td><td>-</td><td>0</td><td>GCGGTTCGCGGCGTTCGCCA</td><td>AGG</td><td>||||||||[||||||||||||]PAM</td><td>NA</td><td>Intergenic</td><td>NA</td><td>NA</td></tr>
<tr><td>T26342</td><td>AGGCGAGAAACGCCCGAGACCGG</td><td>1000</td><td></td><td></td><td></td><td></td><td></td><td></td><td></td><td></td><td></td></tr>
<tr><td>Oligo fwd</td><td>CACCAGGCGAGAAACGCCCGAGAC</td><td></td><td></td><td></td><td></td><td></td><td></td><td></td><td></td><td></td><td></td></tr>
<tr><td>Oligo rev</td><td>AAACGTCTCGGGCGTTTCTCGCCT</td><td></td><td></td><td></td><td></td><td></td><td></td><td></td><td></td><td></td><td></td></tr>
<tr><td>Chromosome</td><td>start</td><td>end</td><td>strand</td><td>MM</td><td>target_seq</td><td>PAM</td><td>alignment</td><td>distance</td><td>position</td><td>gene name</td><td>gene id</td></tr>
<tr><td>GU305914.1</td><td>101276</td><td>101298</td><td>-</td><td>0</td><td>AGGCGAGAAACGCCCGAGAC</td><td>CGG</td><td>||||||||[||||||||||||]PAM</td><td>NA</td><td>Intergenic</td><td>NA</td><td>NA</td></tr>
<tr><td>T26343</td><td>CGCCCGAGACCGGCGCGAGCTGG</td><td>1000</td><td></td><td></td><td></td><td></td><td></td><td></td><td></td><td></td><td></td></tr>
<tr><td>Oligo fwd</td><td>CACCCGCCCGAGACCGGCGCGAGC</td><td></td><td></td><td></td><td></td><td></td><td></td><td></td><td></td><td></td><td></td></tr>
<tr><td>Oligo rev</td><td>AAACGCTCGCGCCGGTCTCGGGCG</td><td></td><td></td><td></td><td></td><td></td><td></td><td></td><td></td><td></td><td></td></tr>
<tr><td>Chromosome</td><td>start</td><td>end</td><td>strand</td><td>MM</td><td>target_seq</td><td>PAM</td><td>alignment</td><td>distance</td><td>position</td><td>gene name</td><td>gene id</td></tr>
<tr><td>GU305914.1</td><td>101266</td><td>101288</td><td>-</td><td>0</td><td>CGCCCGAGACCGGCGCGAGC</td><td>TGG</td><td>||||||||[||||||||||||]PAM</td><td>NA</td><td>Intergenic</td><td>NA</td><td>NA</td></tr>
<tr><td>T26344</td><td>CCGGCGCGAGCTGGAATACGTGG</td><td>1000</td><td></td><td></td><td></td><td></td><td></td><td></td><td></td><td></td><td></td></tr>
<tr><td>Oligo fwd</td><td>CACCCCGGCGCGAGCTGGAATACG</td><td></td><td></td><td></td><td></td><td></td><td></td><td></td><td></td><td></td><td></td></tr>
<tr><td>Oligo rev</td><td>AAACCGTATTCCAGCTCGCGCCGG</td><td></td><td></td><td></td><td></td><td></td><td></td><td></td><td></td><td></td><td></td></tr>
<tr><td>Chromosome</td><td>start</td><td>end</td><td>strand</td><td>MM</td><td>target_seq</td><td>PAM</td><td>alignment</td><td>distance</td><td>position</td><td>gene name</td><td>gene id</td></tr>
<tr><td>GU305914.1</td><td>101257</td><td>101279</td><td>-</td><td>0</td><td>CCGGCGCGAGCTGGAATACG</td><td>TGG</td><td>||||||||[||||||||||||]PAM</td><td>NA</td><td>Intergenic</td><td>NA</td><td>NA</td></tr>
<tr><td>T26345</td><td>CAAGATCAACGCGTTCCGCGAGG</td><td>1000</td><td></td><td></td><td></td><td></td><td></td><td></td><td></td><td></td><td></td></tr>
<tr><td>Oligo fwd</td><td>CACCCAAGATCAACGCGTTCCGCG</td><td></td><td></td><td></td><td></td><td></td><td></td><td></td><td></td><td></td><td></td></tr>
<tr><td>Oligo rev</td><td>AAACCGCGGAACGCGTTGATCTTG</td><td></td><td></td><td></td><td></td><td></td><td></td><td></td><td></td><td></td><td></td></tr>
<tr><td>Chromosome</td><td>start</td><td>end</td><td>strand</td><td>MM</td><td>target_seq</td><td>PAM</td><td>alignment</td><td>distance</td><td>position</td><td>gene name</td><td>gene id</td></tr>
<tr><td>GU305914.1</td><td>101230</td><td>101252</td><td>-</td><td>0</td><td>CAAGATCAACGCGTTCCGCG</td><td>AGG</td><td>||||||||[||||||||||||]PAM</td><td>NA</td><td>Intergenic</td><td>NA</td><td>NA</td></tr>
<tr><td>T26346</td><td>GTTCCGCGAGGAATCGCGCCTGG</td><td>1000</td><td></td><td></td><td></td><td></td><td></td><td></td><td></td><td></td><td></td></tr>
<tr><td>Oligo fwd</td><td>CACCGTTCCGCGAGGAATCGCGCC</td><td></td><td></td><td></td><td></td><td></td><td></td><td></td><td></td><td></td><td></td></tr>
<tr><td>Oligo rev</td><td>AAACGGCGCGATTCCTCGCGGAAC</td><td></td><td></td><td></td><td></td><td></td><td></td><td></td><td></td><td></td><td></td></tr>
<tr><td>Chromosome</td><td>start</td><td>end</td><td>strand</td><td>MM</td><td>target_seq</td><td>PAM</td><td>alignment</td><td>distance</td><td>position</td><td>gene name</td><td>gene id</td></tr>
<tr><td>GU305914.1</td><td>101218</td><td>101240</td><td>-</td><td>0</td><td>GTTCCGCGAGGAATCGCGCC</td><td>TGG</td><td>||||||||[||||||||||||]PAM</td><td>NA</td><td>Intergenic</td><td>NA</td><td>NA</td></tr>
<tr><td>T26347</td><td>CCGCGAGGAATCGCGCCTGGCGG</td><td>1000</td><td></td><td></td><td></td><td></td><td></td><td></td><td></td><td></td><td></td></tr>
<tr><td>Oligo fwd</td><td>CACCCCGCGAGGAATCGCGCCTGG</td><td></td><td></td><td></td><td></td><td></td><td></td><td></td><td></td><td></td><td></td></tr>
<tr><td>Oligo rev</td><td>AAACCCAGGCGCGATTCCTCGCGG</td><td></td><td></td><td></td><td></td><td></td><td></td><td></td><td></td><td></td><td></td></tr>
<tr><td>Chromosome</td><td>start</td><td>end</td><td>strand</td><td>MM</td><td>target_seq</td><td>PAM</td><td>alignment</td><td>distance</td><td>position</td><td>gene name</td><td>gene id</td></tr>
<tr><td>GU305914.1</td><td>101215</td><td>101237</td><td>-</td><td>0</td><td>CCGCGAGGAATCGCGCCTGG</td><td>CGG</td><td>||||||||[||||||||||||]PAM</td><td>NA</td><td>Intergenic</td><td>NA</td><td>NA</td></tr>
<tr><td>T26348</td><td>GTGCGTGTACTACGTGTACCTGG</td><td>1000</td><td></td><td></td><td></td><td></td><td></td><td></td><td></td><td></td><td></td></tr>
<tr><td>Oligo fwd</td><td>CACCGTGCGTGTACTACGTGTACC</td><td></td><td></td><td></td><td></td><td></td><td></td><td></td><td></td><td></td><td></td></tr>
<tr><td>Oligo rev</td><td>AAACGGTACACGTAGTACACGCAC</td><td></td><td></td><td></td><td></td><td></td><td></td><td></td><td></td><td></td><td></td></tr>
<tr><td>Chromosome</td><td>start</td><td>end</td><td>strand</td><td>MM</td><td>target_seq</td><td>PAM</td><td>alignment</td><td>distance</td><td>position</td><td>gene name</td><td>gene id</td></tr>
</table>

| | | | | | | | | | | | |
|---|---|---|---|---|---|---|---|---|---|---|---|
| GU305914.1 | 101185 | 101207 | - | 0 | GTGCGTGTACTACGTGTACC | TGG | \|\|\|\|\|\|\|\|[\|\|\|\|\|\|\|\|\|\|\|\|]PAM | NA | Intergenic | NA | NA |
| T26349 | ACTACGTGTACCTGGCCTACCGG | 1000 | | | | | | | | | |
| Oligo fwd | CACCACTACGTGTACCTGGCCTAC | | | | | | | | | | |
| Oligo rev | AAACGTAGGCCAGGTACACGTAGT | | | | | | | | | | |
| Chromosome | start | end | strand | MM | target_seq | PAM | alignment | distance | position | gene name | gene id |
| GU305914.1 | 101177 | 101199 | - | 0 | ACTACGTGTACCTGGCCTAC | CGG | \|\|\|\|\|\|\|\|[\|\|\|\|\|\|\|\|\|\|\|\|]PAM | NA | Intergenic | NA | NA |
| T26350 | CGTGTACCTGGCCTACCGGACGG | 1000 | | | | | | | | | |
| Oligo fwd | CACCCGTGTACCTGGCCTACCGGA | | | | | | | | | | |
| Oligo rev | AAACTCCGGTAGGCCAGGTACACG | | | | | | | | | | |
| Chromosome | start | end | strand | MM | target_seq | PAM | alignment | distance | position | gene name | gene id |
| GU305914.1 | 101173 | 101195 | - | 0 | CGTGTACCTGGCCTACCGGA | CGG | \|\|\|\|\|\|\|\|[\|\|\|\|\|\|\|\|\|\|\|\|]PAM | NA | Intergenic | NA | NA |
| T26351 | CCTACCGGACGGCGCTGTGCCGG | 1000 | | | | | | | | | |
| Oligo fwd | CACCCCTACCGGACGGCGCTGTGC | | | | | | | | | | |
| Oligo rev | AAACGCACAGCGCCGTCCGGTAGG | | | | | | | | | | |
| Chromosome | start | end | strand | MM | target_seq | PAM | alignment | distance | position | gene name | gene id |
| GU305914.1 | 101162 | 101184 | - | 0 | CCTACCGGACGGCGCTGTGC | CGG | \|\|\|\|\|\|\|\|[\|\|\|\|\|\|\|\|\|\|\|\|]PAM | NA | Intergenic | NA | NA |
| T26352 | CTACCGGACGGCGCTGTGCCGGG | 1000 | | | | | | | | | |
| Oligo fwd | CACCCTACCGGACGGCGCTGTGCC | | | | | | | | | | |
| Oligo rev | AAACGGCACAGCGCCGTCCGGTAG | | | | | | | | | | |
| Chromosome | start | end | strand | MM | target_seq | PAM | alignment | distance | position | gene name | gene id |
| GU305914.1 | 101161 | 101183 | - | 0 | CTACCGGACGGCGCTGTGCC | GGG | \|\|\|\|\|\|\|\|[\|\|\|\|\|\|\|\|\|\|\|\|]PAM | NA | Intergenic | NA | NA |
| T26353 | CCTGCAGTACTGCGAGCACACGG | 1000 | | | | | | | | | |
| Oligo fwd | CACCCCTGCAGTACTGCGAGCACA | | | | | | | | | | |
| Oligo rev | AAACTGTGCTCGCAGTACTGCAGG | | | | | | | | | | |
| Chromosome | start | end | strand | MM | target_seq | PAM | alignment | distance | position | gene name | gene id |
| GU305914.1 | 101131 | 101153 | - | 0 | CCTGCAGTACTGCGAGCACA | CGG | \|\|\|\|\|\|\|\|[\|\|\|\|\|\|\|\|\|\|\|\|]PAM | NA | Intergenic | NA | NA |
| T26354 | GGCGTACGACAAGAACCTGCCGG | 1000 | | | | | | | | | |
| Oligo fwd | CACCGGCGTACGACAAGAACCTGC | | | | | | | | | | |
| Oligo rev | AAACGCAGGTTCTTGTCGTACGCC | | | | | | | | | | |
| Chromosome | start | end | strand | MM | target_seq | PAM | alignment | distance | position | gene name | gene id |
| GU305914.1 | 101110 | 101132 | - | 0 | GGCGTACGACAAGAACCTGC | CGG | \|\|\|\|\|\|\|\|[\|\|\|\|\|\|\|\|\|\|\|\|]PAM | NA | Intergenic | NA | NA |
| T26355 | GCCGGACGATCAGCAGTGCGCGG | 1000 | | | | | | | | | |
| Oligo fwd | CACCGCCGGACGATCAGCAGTGCG | | | | | | | | | | |
| Oligo rev | AAACCGCACTGCTGATCGTCCGGC | | | | | | | | | | |
| Chromosome | start | end | strand | MM | target_seq | PAM | alignment | distance | position | gene name | gene id |
| GU305914.1 | 101092 | 101114 | - | 0 | GCCGGACGATCAGCAGTGCG | CGG | \|\|\|\|\|\|\|\|[\|\|\|\|\|\|\|\|\|\|\|\|]PAM | NA | Intergenic | NA | NA |
| T26356 | GGACGATCAGCAGTGCGCGGCGG | 1000 | | | | | | | | | |
| Oligo fwd | CACCGGACGATCAGCAGTGCGCGG | | | | | | | | | | |
| Oligo rev | AAACCCGCGCACTGCTGATCGTCC | | | | | | | | | | |
| Chromosome | start | end | strand | MM | target_seq | PAM | alignment | distance | position | gene name | gene id |
| GU305914.1 | 101089 | 101111 | - | 0 | GGACGATCAGCAGTGCGCGG | CGG | \|\|\|\|\|\|\|\|[\|\|\|\|\|\|\|\|\|\|\|\|]PAM | NA | Intergenic | NA | NA |
| T26357 | CGATCAGCAGTGCGCGGCGGAGG | 1000 | | | | | | | | | |
| Oligo fwd | CACCCGATCAGCAGTGCGCGGCGG | | | | | | | | | | |
| Oligo rev | AAACCCGCCGCGCACTGCTGATCG | | | | | | | | | | |
| Chromosome | start | end | strand | MM | target_seq | PAM | alignment | distance | position | gene name | gene id |
| GU305914.1 | 101086 | 101108 | - | 0 | CGATCAGCAGTGCGCGGCGG | AGG | \|\|\|\|\|\|\|\|[\|\|\|\|\|\|\|\|\|\|\|\|]PAM | NA | Intergenic | NA | NA |
| T26358 | CGCGGCGGAGGAGAACTACCTGG | 1000 | | | | | | | | | |
| Oligo fwd | CACCCGCGGCGGAGGAGAACTACC | | | | | | | | | | |
| Oligo rev | AAACGGTAGTTCTCCTCCGCCGCG | | | | | | | | | | |
| Chromosome | start | end | strand | MM | target_seq | PAM | alignment | distance | position | gene name | gene id |
| GU305914.1 | 101074 | 101096 | - | 0 | CGCGGCGGAGGAGAACTACC | TGG | \|\|\|\|\|\|\|\|[\|\|\|\|\|\|\|\|\|\|\|\|]PAM | NA | Intergenic | NA | NA |
| T26359 | GCGGCGGAGGAGAACTACCTGGG | 1000 | | | | | | | | | |
| Oligo fwd | CACCGCGGCGGAGGAGAACTACCT | | | | | | | | | | |
| Oligo rev | AAACAGGTAGTTCTCCTCCGCCGC | | | | | | | | | | |
| Chromosome | start | end | strand | MM | target_seq | PAM | alignment | distance | position | gene name | gene id |
| GU305914.1 | 101073 | 101095 | - | 0 | GCGGCGGAGGAGAACTACCT | GGG | \|\|\|\|\|\|\|\|[\|\|\|\|\|\|\|\|\|\|\|\|]PAM | NA | Intergenic | NA | NA |
| GU305914.1 | 195536 | 195558 | - | 4 | GCCGCGGTGGTGCACTACCT | CGG | \|\|-\|\|\|\|-[\|\|-\|-\|\|\|\|\|\|\|]PAM | NA | Intergenic | NA | NA |
| T26360 | CGGCGGAGGAGAACTACCTGGGG | 1000 | | | | | | | | | |
| Oligo fwd | CACCCGGCGGAGGAGAACTACCTG | | | | | | | | | | |
| Oligo rev | AAACCAGGTAGTTCTCCTCCGCCG | | | | | | | | | | |
| Chromosome | start | end | strand | MM | target_seq | PAM | alignment | distance | position | gene name | gene id |
| GU305914.1 | 101072 | 101094 | - | 0 | CGGCGGAGGAGAACTACCTG | GGG | \|\|\|\|\|\|\|\|[\|\|\|\|\|\|\|\|\|\|\|\|]PAM | NA | Intergenic | NA | NA |
| T26361 | GAACTACCTGGGGCGCAGCCTGG | 1000 | | | | | | | | | |
| Oligo fwd | CACCGAACTACCTGGGGCGCAGCC | | | | | | | | | | |
| Oligo rev | AAACGGCTGCGCCCCAGGTAGTTC | | | | | | | | | | |
| Chromosome | start | end | strand | MM | target_seq | PAM | alignment | distance | position | gene name | gene id |
| GU305914.1 | 101062 | 101084 | - | 0 | GAACTACCTGGGGCGCAGCC | TGG | \|\|\|\|\|\|\|\|[\|\|\|\|\|\|\|\|\|\|\|\|]PAM | NA | Intergenic | NA | NA |
| T26362 | CATCATGAATACCTACTTTTCGG | 1000 | | | | | | | | | |
| Oligo fwd | CACCCATCATGAATACCTACTTTT | | | | | | | | | | |
| Oligo rev | AAACAAAAGTAGGTATTCATGATG | | | | | | | | | | |
| Chromosome | start | end | strand | MM | target_seq | PAM | alignment | distance | position | gene name | gene id |
| GU305914.1 | 101023 | 101045 | - | 0 | CATCATGAATACCTACTTTT | CGG | \|\|\|\|\|\|\|\|[\|\|\|\|\|\|\|\|\|\|\|\|]PAM | NA | Intergenic | NA | NA |
| T26363 | CATGAATACCTACTTTTCGGTGG | 1000 | | | | | | | | | |
| Oligo fwd | CACCCATGAATACCTACTTTTCGG | | | | | | | | | | |

<table>
<tr><td>Oligo rev</td><td>AAACCCGAAAAGTAGGTATTCATG</td><td></td><td></td><td></td><td></td><td></td><td></td><td></td><td></td><td></td><td></td></tr>
<tr><td>Chromosome</td><td>start</td><td>end</td><td>strand</td><td>MM</td><td>target_seq</td><td>PAM</td><td>alignment</td><td>distance</td><td>position</td><td>gene name</td><td>gene id</td></tr>
<tr><td>GU305914.1</td><td>101020</td><td>101042</td><td>-</td><td>0</td><td>CATGAATACCTACTTTTCGG</td><td>TGG</td><td>||||||||[||||||||||||]PAM</td><td>NA</td><td>Intergenic</td><td>NA</td><td>NA</td></tr>
<tr><td></td><td></td><td></td><td></td><td></td><td></td><td></td><td></td><td></td><td></td><td></td><td></td></tr>
<tr><td>T26364</td><td>GAATACCTACTTTTCGGTGGAGG</td><td>1000</td><td></td><td></td><td></td><td></td><td></td><td></td><td></td><td></td><td></td></tr>
<tr><td>Oligo fwd</td><td>CACCGAATACCTACTTTTCGGTGG</td><td></td><td></td><td></td><td></td><td></td><td></td><td></td><td></td><td></td><td></td></tr>
<tr><td>Oligo rev</td><td>AAACCCACCGAAAAGTAGGTATTC</td><td></td><td></td><td></td><td></td><td></td><td></td><td></td><td></td><td></td><td></td></tr>
<tr><td>Chromosome</td><td>start</td><td>end</td><td>strand</td><td>MM</td><td>target_seq</td><td>PAM</td><td>alignment</td><td>distance</td><td>position</td><td>gene name</td><td>gene id</td></tr>
<tr><td>GU305914.1</td><td>101017</td><td>101039</td><td>-</td><td>0</td><td>GAATACCTACTTTTCGGTGG</td><td>AGG</td><td>||||||||[||||||||||||]PAM</td><td>NA</td><td>Intergenic</td><td>NA</td><td>NA</td></tr>
<tr><td></td><td></td><td></td><td></td><td></td><td></td><td></td><td></td><td></td><td></td><td></td><td></td></tr>
<tr><td>T26365</td><td>AATACCTACTTTTCGGTGGAGGG</td><td>1000</td><td></td><td></td><td></td><td></td><td></td><td></td><td></td><td></td><td></td></tr>
<tr><td>Oligo fwd</td><td>CACCAATACCTACTTTTCGGTGGA</td><td></td><td></td><td></td><td></td><td></td><td></td><td></td><td></td><td></td><td></td></tr>
<tr><td>Oligo rev</td><td>AAACTCCACCGAAAAGTAGGTATT</td><td></td><td></td><td></td><td></td><td></td><td></td><td></td><td></td><td></td><td></td></tr>
<tr><td>Chromosome</td><td>start</td><td>end</td><td>strand</td><td>MM</td><td>target_seq</td><td>PAM</td><td>alignment</td><td>distance</td><td>position</td><td>gene name</td><td>gene id</td></tr>
<tr><td>GU305914.1</td><td>101016</td><td>101038</td><td>-</td><td>0</td><td>AATACCTACTTTTCGGTGGA</td><td>GGG</td><td>||||||||[||||||||||||]PAM</td><td>NA</td><td>Intergenic</td><td>NA</td><td>NA</td></tr>
<tr><td></td><td></td><td></td><td></td><td></td><td></td><td></td><td></td><td></td><td></td><td></td><td></td></tr>
<tr><td>T26366</td><td>TTTTCGGTGGAGGGCTATTTTGG</td><td>1000</td><td></td><td></td><td></td><td></td><td></td><td></td><td></td><td></td><td></td></tr>
<tr><td>Oligo fwd</td><td>CACCTTTTCGGTGGAGGGCTATTT</td><td></td><td></td><td></td><td></td><td></td><td></td><td></td><td></td><td></td><td></td></tr>
<tr><td>Oligo rev</td><td>AAACAAATAGCCCTCCACCGAAAA</td><td></td><td></td><td></td><td></td><td></td><td></td><td></td><td></td><td></td><td></td></tr>
<tr><td>Chromosome</td><td>start</td><td>end</td><td>strand</td><td>MM</td><td>target_seq</td><td>PAM</td><td>alignment</td><td>distance</td><td>position</td><td>gene name</td><td>gene id</td></tr>
<tr><td>GU305914.1</td><td>101007</td><td>101029</td><td>-</td><td>0</td><td>TTTTCGGTGGAGGGCTATTT</td><td>TGG</td><td>||||||||[||||||||||||]PAM</td><td>NA</td><td>Intergenic</td><td>NA</td><td>NA</td></tr>
<tr><td></td><td></td><td></td><td></td><td></td><td></td><td></td><td></td><td></td><td></td><td></td><td></td></tr>
<tr><td>T26367</td><td>AGCCCGCCCCACAGCTACCGCGG</td><td>1000</td><td></td><td></td><td></td><td></td><td></td><td></td><td></td><td></td><td></td></tr>
<tr><td>Oligo fwd</td><td>CACCAGCCCGCCCCACAGCTACCG</td><td></td><td></td><td></td><td></td><td></td><td></td><td></td><td></td><td></td><td></td></tr>
<tr><td>Oligo rev</td><td>AAACCGGTAGCTGTGGGGCGGGCT</td><td></td><td></td><td></td><td></td><td></td><td></td><td></td><td></td><td></td><td></td></tr>
<tr><td>Chromosome</td><td>start</td><td>end</td><td>strand</td><td>MM</td><td>target_seq</td><td>PAM</td><td>alignment</td><td>distance</td><td>position</td><td>gene name</td><td>gene id</td></tr>
<tr><td>GU305914.1</td><td>100953</td><td>100975</td><td>-</td><td>0</td><td>AGCCCGCCCCACAGCTACCG</td><td>CGG</td><td>||||||||[||||||||||||]PAM</td><td>NA</td><td>Intergenic</td><td>NA</td><td>NA</td></tr>
<tr><td></td><td></td><td></td><td></td><td></td><td></td><td></td><td></td><td></td><td></td><td></td><td></td></tr>
<tr><td>T26368</td><td>ACAGCTACCGCGGCTACGACTGG</td><td>1000</td><td></td><td></td><td></td><td></td><td></td><td></td><td></td><td></td><td></td></tr>
<tr><td>Oligo fwd</td><td>CACCACAGCTACCGCGGCTACGAC</td><td></td><td></td><td></td><td></td><td></td><td></td><td></td><td></td><td></td><td></td></tr>
<tr><td>Oligo rev</td><td>AAACGTCGTAGCCGCGGTAGCTGT</td><td></td><td></td><td></td><td></td><td></td><td></td><td></td><td></td><td></td><td></td></tr>
<tr><td>Chromosome</td><td>start</td><td>end</td><td>strand</td><td>MM</td><td>target_seq</td><td>PAM</td><td>alignment</td><td>distance</td><td>position</td><td>gene name</td><td>gene id</td></tr>
<tr><td>GU305914.1</td><td>100943</td><td>100965</td><td>-</td><td>0</td><td>ACAGCTACCGCGGCTACGAC</td><td>TGG</td><td>||||||||[||||||||||||]PAM</td><td>NA</td><td>Intergenic</td><td>NA</td><td>NA</td></tr>
<tr><td></td><td></td><td></td><td></td><td></td><td></td><td></td><td></td><td></td><td></td><td></td><td></td></tr>
<tr><td>T26369</td><td>CCGCGGCTACGACTGGAACACGG</td><td>1000</td><td></td><td></td><td></td><td></td><td></td><td></td><td></td><td></td><td></td></tr>
<tr><td>Oligo fwd</td><td>CACCCCGCGGCTACGACTGGAACA</td><td></td><td></td><td></td><td></td><td></td><td></td><td></td><td></td><td></td><td></td></tr>
<tr><td>Oligo rev</td><td>AAACTGTTCCAGTCGTAGCCGCGG</td><td></td><td></td><td></td><td></td><td></td><td></td><td></td><td></td><td></td><td></td></tr>
<tr><td>Chromosome</td><td>start</td><td>end</td><td>strand</td><td>MM</td><td>target_seq</td><td>PAM</td><td>alignment</td><td>distance</td><td>position</td><td>gene name</td><td>gene id</td></tr>
<tr><td>GU305914.1</td><td>100936</td><td>100958</td><td>-</td><td>0</td><td>CCGCGGCTACGACTGGAACA</td><td>CGG</td><td>||||||||[||||||||||||]PAM</td><td>NA</td><td>Intergenic</td><td>NA</td><td>NA</td></tr>
<tr><td></td><td></td><td></td><td></td><td></td><td></td><td></td><td></td><td></td><td></td><td></td><td></td></tr>
<tr><td>T26370</td><td>GAACACGGAAGCCGACACGATGG</td><td>1000</td><td></td><td></td><td></td><td></td><td></td><td></td><td></td><td></td><td></td></tr>
<tr><td>Oligo fwd</td><td>CACCGAACACGGAAGCCGACACGA</td><td></td><td></td><td></td><td></td><td></td><td></td><td></td><td></td><td></td><td></td></tr>
<tr><td>Oligo rev</td><td>AAACTCGTGTCGGCTTCCGTGTTC</td><td></td><td></td><td></td><td></td><td></td><td></td><td></td><td></td><td></td><td></td></tr>
<tr><td>Chromosome</td><td>start</td><td>end</td><td>strand</td><td>MM</td><td>target_seq</td><td>PAM</td><td>alignment</td><td>distance</td><td>position</td><td>gene name</td><td>gene id</td></tr>
<tr><td>GU305914.1</td><td>100921</td><td>100943</td><td>-</td><td>0</td><td>GAACACGGAAGCCGACACGA</td><td>TGG</td><td>||||||||[||||||||||||]PAM</td><td>NA</td><td>Intergenic</td><td>NA</td><td>NA</td></tr>
<tr><td></td><td></td><td></td><td></td><td></td><td></td><td></td><td></td><td></td><td></td><td></td><td></td></tr>
<tr><td>T26371</td><td>CACGGAAGCCGACACGATGGTGG</td><td>1000</td><td></td><td></td><td></td><td></td><td></td><td></td><td></td><td></td><td></td></tr>
<tr><td>Oligo fwd</td><td>CACCCACGGAAGCCGACACGATGG</td><td></td><td></td><td></td><td></td><td></td><td></td><td></td><td></td><td></td><td></td></tr>
<tr><td>Oligo rev</td><td>AAACCCATCGTGTCGGCTTCCGTG</td><td></td><td></td><td></td><td></td><td></td><td></td><td></td><td></td><td></td><td></td></tr>
<tr><td>Chromosome</td><td>start</td><td>end</td><td>strand</td><td>MM</td><td>target_seq</td><td>PAM</td><td>alignment</td><td>distance</td><td>position</td><td>gene name</td><td>gene id</td></tr>
<tr><td>GU305914.1</td><td>100918</td><td>100940</td><td>-</td><td>0</td><td>CACGGAAGCCGACACGATGG</td><td>TGG</td><td>||||||||[||||||||||||]PAM</td><td>NA</td><td>Intergenic</td><td>NA</td><td>NA</td></tr>
<tr><td></td><td></td><td></td><td></td><td></td><td></td><td></td><td></td><td></td><td></td><td></td><td></td></tr>
<tr><td>T26372</td><td>ACGGAAGCCGACACGATGGTGGG</td><td>1000</td><td></td><td></td><td></td><td></td><td></td><td></td><td></td><td></td><td></td></tr>
<tr><td>Oligo fwd</td><td>CACCACGGAAGCCGACACGATGGT</td><td></td><td></td><td></td><td></td><td></td><td></td><td></td><td></td><td></td><td></td></tr>
<tr><td>Oligo rev</td><td>AAACACCATCGTGTCGGCTTCCGT</td><td></td><td></td><td></td><td></td><td></td><td></td><td></td><td></td><td></td><td></td></tr>
<tr><td>Chromosome</td><td>start</td><td>end</td><td>strand</td><td>MM</td><td>target_seq</td><td>PAM</td><td>alignment</td><td>distance</td><td>position</td><td>gene name</td><td>gene id</td></tr>
<tr><td>GU305914.1</td><td>100917</td><td>100939</td><td>-</td><td>0</td><td>ACGGAAGCCGACACGATGGT</td><td>GGG</td><td>||||||||[||||||||||||]PAM</td><td>NA</td><td>Intergenic</td><td>NA</td><td>NA</td></tr>
<tr><td></td><td></td><td></td><td></td><td></td><td></td><td></td><td></td><td></td><td></td><td></td><td></td></tr>
<tr><td>T26373</td><td>CTCTTCCACCGCGACGAACCTGG</td><td>1000</td><td></td><td></td><td></td><td></td><td></td><td></td><td></td><td></td><td></td></tr>
<tr><td>Oligo fwd</td><td>CACCCTCTTCCACCGCGACGAACC</td><td></td><td></td><td></td><td></td><td></td><td></td><td></td><td></td><td></td><td></td></tr>
<tr><td>Oligo rev</td><td>AAACGGTTCGTCGCGGTGGAAGAG</td><td></td><td></td><td></td><td></td><td></td><td></td><td></td><td></td><td></td><td></td></tr>
<tr><td>Chromosome</td><td>start</td><td>end</td><td>strand</td><td>MM</td><td>target_seq</td><td>PAM</td><td>alignment</td><td>distance</td><td>position</td><td>gene name</td><td>gene id</td></tr>
<tr><td>GU305914.1</td><td>100891</td><td>100913</td><td>-</td><td>0</td><td>CTCTTCCACCGCGACGAACC</td><td>TGG</td><td>||||||||[||||||||||||]PAM</td><td>NA</td><td>Intergenic</td><td>NA</td><td>NA</td></tr>
<tr><td></td><td></td><td></td><td></td><td></td><td></td><td></td><td></td><td></td><td></td><td></td><td></td></tr>
<tr><td>T26374</td><td>GCCGCTGCCGCCGACGCTGATGG</td><td>1000</td><td></td><td></td><td></td><td></td><td></td><td></td><td></td><td></td><td></td></tr>
<tr><td>Oligo fwd</td><td>CACCGCCGCTGCCGCCGACGCTGA</td><td></td><td></td><td></td><td></td><td></td><td></td><td></td><td></td><td></td><td></td></tr>
<tr><td>Oligo rev</td><td>AAACTCAGCGTCGGCGGCAGCGGC</td><td></td><td></td><td></td><td></td><td></td><td></td><td></td><td></td><td></td><td></td></tr>
<tr><td>Chromosome</td><td>start</td><td>end</td><td>strand</td><td>MM</td><td>target_seq</td><td>PAM</td><td>alignment</td><td>distance</td><td>position</td><td>gene name</td><td>gene id</td></tr>
<tr><td>GU305914.1</td><td>100822</td><td>100844</td><td>-</td><td>0</td><td>GCCGCTGCCGCCGACGCTGA</td><td>TGG</td><td>||||||||[||||||||||||]PAM</td><td>NA</td><td>Intergenic</td><td>NA</td><td>NA</td></tr>
<tr><td>GU305914.1</td><td>70188</td><td>70210</td><td>-</td><td>4</td><td>GGCGCCGCCGCCGGCGCTGC</td><td>CGG</td><td>|-|||-||[|||||-|||||-]PAM</td><td>NA</td><td>Intergenic</td><td>NA</td><td>NA</td></tr>
<tr><td></td><td></td><td></td><td></td><td></td><td></td><td></td><td></td><td></td><td></td><td></td><td></td></tr>
<tr><td>T26375</td><td>CCGCTGCCGCCGACGCTGATGGG</td><td>1000</td><td></td><td></td><td></td><td></td><td></td><td></td><td></td><td></td><td></td></tr>
<tr><td>Oligo fwd</td><td>CACCCCGCTGCCGCCGACGCTGAT</td><td></td><td></td><td></td><td></td><td></td><td></td><td></td><td></td><td></td><td></td></tr>
<tr><td>Oligo rev</td><td>AAACATCAGCGTCGGCGGCAGCGG</td><td></td><td></td><td></td><td></td><td></td><td></td><td></td><td></td><td></td><td></td></tr>
<tr><td>Chromosome</td><td>start</td><td>end</td><td>strand</td><td>MM</td><td>target_seq</td><td>PAM</td><td>alignment</td><td>distance</td><td>position</td><td>gene name</td><td>gene id</td></tr>
<tr><td>GU305914.1</td><td>100821</td><td>100843</td><td>-</td><td>0</td><td>CCGCTGCCGCCGACGCTGAT</td><td>GGG</td><td>||||||||[||||||||||||]PAM</td><td>NA</td><td>Intergenic</td><td>NA</td><td>NA</td></tr>
<tr><td>GU305914.1</td><td>100326</td><td>100348</td><td>-</td><td>4</td><td>GCGCTGCGGCGGACCCTGAT</td><td>CGG</td><td>-||||||-[||-|||-|||||]PAM</td><td>NA</td><td>Intergenic</td><td>NA</td><td>NA</td></tr>
<tr><td></td><td></td><td></td><td></td><td></td><td></td><td></td><td></td><td></td><td></td><td></td><td></td></tr>
<tr><td>T26376</td><td>CGCTGCCGCCGACGCTGATGGGG</td><td>1000</td><td></td><td></td><td></td><td></td><td></td><td></td><td></td><td></td><td></td></tr>
<tr><td>Oligo fwd</td><td>CACCCGCTGCCGCCGACGCTGATG</td><td></td><td></td><td></td><td></td><td></td><td></td><td></td><td></td><td></td><td></td></tr>
<tr><td>Oligo rev</td><td>AAACCATCAGCGTCGGCGGCAGCG</td><td></td><td></td><td></td><td></td><td></td><td></td><td></td><td></td><td></td><td></td></tr>
<tr><td>Chromosome</td><td>start</td><td>end</td><td>strand</td><td>MM</td><td>target_seq</td><td>PAM</td><td>alignment</td><td>distance</td><td>position</td><td>gene name</td><td>gene id</td></tr>
<tr><td>GU305914.1</td><td>100820</td><td>100842</td><td>-</td><td>0</td><td>CGCTGCCGCCGACGCTGATG</td><td>GGG</td><td>||||||||[||||||||||||]PAM</td><td>NA</td><td>Intergenic</td><td>NA</td><td>NA</td></tr>
<tr><td></td><td></td><td></td><td></td><td></td><td></td><td></td><td></td><td></td><td></td><td></td><td></td></tr>
<tr><td>T26377</td><td>GCTGCTGAAGCTGTGCGCCTCGG</td><td>1000</td><td></td><td></td><td></td><td></td><td></td><td></td><td></td><td></td><td></td></tr>
<tr><td>Oligo fwd</td><td>CACCGCTGCTGAAGCTGTGCGCCT</td><td></td><td></td><td></td><td></td><td></td><td></td><td></td><td></td><td></td><td></td></tr>
<tr><td>Oligo rev</td><td>AAACAGGCGCACAGCTTCAGCAGC</td><td></td><td></td><td></td><td></td><td></td><td></td><td></td><td></td><td></td><td></td></tr>
<tr><td>Chromosome</td><td>start</td><td>end</td><td>strand</td><td>MM</td><td>target_seq</td><td>PAM</td><td>alignment</td><td>distance</td><td>position</td><td>gene name</td><td>gene id</td></tr>
<tr><td>GU305914.1</td><td>100798</td><td>100820</td><td>-</td><td>0</td><td>GCTGCTGAAGCTGTGCGCCT</td><td>CGG</td><td>||||||||[||||||||||||]PAM</td><td>NA</td><td>Intergenic</td><td>NA</td><td>NA</td></tr>
</table>

| | | | | | | | | | | | |
|---|---|---|---|---|---|---|---|---|---|---|---|
| T26378 | TGAAGCTGTGCGCCTCGGACCGG | 1000 | | | | | | | | | |
| Oligo fwd | CACCTGAAGCTGTGCGCCTCGGAC | | | | | | | | | | |
| Oligo rev | AAACGTCCGAGGCGCACAGCTTCA | | | | | | | | | | |
| Chromosome | start | end | strand | MM | target_seq | PAM | alignment | distance | position | gene name | gene id |
| GU305914.1 | 100793 | 100815 | - | 0 | TGAAGCTGTGCGCCTCGGAC | CGG | \|\|\|\|\|\|\|\|[\|\|\|\|\|\|\|\|\|\|\|\|]PAM | NA | Intergenic | NA | NA |

| | | | | | | | | | | | |
|---|---|---|---|---|---|---|---|---|---|---|---|
| T26379 | CCTCGGACCGGTACGTGCCGCGG | 1000 | | | | | | | | | |
| Oligo fwd | CACCCCTCGGACCGGTACGTGCCG | | | | | | | | | | |
| Oligo rev | AAACCGGCACGTACCGGTCCGAGG | | | | | | | | | | |
| Chromosome | start | end | strand | MM | target_seq | PAM | alignment | distance | position | gene name | gene id |
| GU305914.1 | 100781 | 100803 | - | 0 | CCTCGGACCGGTACGTGCCG | CGG | \|\|\|\|\|\|\|\|[\|\|\|\|\|\|\|\|\|\|\|\|]PAM | NA | Intergenic | NA | NA |

| | | | | | | | | | | | |
|---|---|---|---|---|---|---|---|---|---|---|---|
| T26380 | CTCGGACCGGTACGTGCCGCGGG | 1000 | | | | | | | | | |
| Oligo fwd | CACCCTCGGACCGGTACGTGCCGC | | | | | | | | | | |
| Oligo rev | AAACGCGGCACGTACCGGTCCGAG | | | | | | | | | | |
| Chromosome | start | end | strand | MM | target_seq | PAM | alignment | distance | position | gene name | gene id |
| GU305914.1 | 100780 | 100802 | - | 0 | CTCGGACCGGTACGTGCCGC | GGG | \|\|\|\|\|\|\|\|[\|\|\|\|\|\|\|\|\|\|\|\|]PAM | NA | Intergenic | NA | NA |

| | | | | | | | | | | | |
|---|---|---|---|---|---|---|---|---|---|---|---|
| T26381 | GGACCGGTACGTGCCGCGGGCGG | 1000 | | | | | | | | | |
| Oligo fwd | CACCGGACCGGTACGTGCCGCGGG | | | | | | | | | | |
| Oligo rev | AAACCCCGCGGCACGTACCGGTCC | | | | | | | | | | |
| Chromosome | start | end | strand | MM | target_seq | PAM | alignment | distance | position | gene name | gene id |
| GU305914.1 | 100777 | 100799 | - | 0 | GGACCGGTACGTGCCGCGGG | CGG | \|\|\|\|\|\|\|\|[\|\|\|\|\|\|\|\|\|\|\|\|]PAM | NA | Intergenic | NA | NA |
| GU305914.1 | 198857 | 198879 | - | 4 | GCAGCGGTCCGTGCCGCGCG | CGG | \|-\|-\|\|\|\|[-\|\|\|\|\|\|\|\|\|-\|]PAM | NA | Intergenic | NA | NA |

| | | | | | | | | | | | |
|---|---|---|---|---|---|---|---|---|---|---|---|
| T26382 | AAAAGCCGCAAGCGCACCAGCGG | 1000 | | | | | | | | | |
| Oligo fwd | CACCAAAAGCCGCAAGCGCACCAG | | | | | | | | | | |
| Oligo rev | AAACCTGGTGCGCTTGCGGCTTTT | | | | | | | | | | |
| Chromosome | start | end | strand | MM | target_seq | PAM | alignment | distance | position | gene name | gene id |
| GU305914.1 | 100752 | 100774 | - | 0 | AAAAGCCGCAAGCGCACCAG | CGG | \|\|\|\|\|\|\|\|[\|\|\|\|\|\|\|\|\|\|\|\|]PAM | NA | Intergenic | NA | NA |

| | | | | | | | | | | | |
|---|---|---|---|---|---|---|---|---|---|---|---|
| T26383 | AGCCGCAAGCGCACCAGCGGCGG | 1000 | | | | | | | | | |
| Oligo fwd | CACCAGCCGCAAGCGCACCAGCGG | | | | | | | | | | |
| Oligo rev | AAACCCGCTGGTGCGCTTGCGGCT | | | | | | | | | | |
| Chromosome | start | end | strand | MM | target_seq | PAM | alignment | distance | position | gene name | gene id |
| GU305914.1 | 100749 | 100771 | - | 0 | AGCCGCAAGCGCACCAGCGG | CGG | \|\|\|\|\|\|\|\|[\|\|\|\|\|\|\|\|\|\|\|\|]PAM | NA | Intergenic | NA | NA |

| | | | | | | | | | | | |
|---|---|---|---|---|---|---|---|---|---|---|---|
| T26384 | AAGACGAGACGCGCGTGTGCCGG | 1000 | | | | | | | | | |
| Oligo fwd | CACCAAGACGAGACGCGCGTGTGC | | | | | | | | | | |
| Oligo rev | AAACGCACACGCGCGTCTCGTCTT | | | | | | | | | | |
| Chromosome | start | end | strand | MM | target_seq | PAM | alignment | distance | position | gene name | gene id |
| GU305914.1 | 100715 | 100737 | - | 0 | AAGACGAGACGCGCGTGTGC | CGG | \|\|\|\|\|\|\|\|[\|\|\|\|\|\|\|\|\|\|\|\|]PAM | NA | Intergenic | NA | NA |

| | | | | | | | | | | | |
|---|---|---|---|---|---|---|---|---|---|---|---|
| T26385 | AACGACACGAGCCGTCCGATCGG | 1000 | | | | | | | | | |
| Oligo fwd | CACCAACGACACGAGCCGTCCGAT | | | | | | | | | | |
| Oligo rev | AAACATCGGACGGCTCGTGTCGTT | | | | | | | | | | |
| Chromosome | start | end | strand | MM | target_seq | PAM | alignment | distance | position | gene name | gene id |
| GU305914.1 | 100677 | 100699 | - | 0 | AACGACACGAGCCGTCCGAT | CGG | \|\|\|\|\|\|\|\|[\|\|\|\|\|\|\|\|\|\|\|\|]PAM | NA | Intergenic | NA | NA |

| | | | | | | | | | | | |
|---|---|---|---|---|---|---|---|---|---|---|---|
| T26386 | ACGACACGAGCCGTCCGATCGGG | 1000 | | | | | | | | | |
| Oligo fwd | CACCACGACACGAGCCGTCCGATC | | | | | | | | | | |
| Oligo rev | AAACGATCGGACGGCTCGTGTCGT | | | | | | | | | | |
| Chromosome | start | end | strand | MM | target_seq | PAM | alignment | distance | position | gene name | gene id |
| GU305914.1 | 100676 | 100698 | - | 0 | ACGACACGAGCCGTCCGATC | GGG | \|\|\|\|\|\|\|\|[\|\|\|\|\|\|\|\|\|\|\|\|]PAM | NA | Intergenic | NA | NA |

| | | | | | | | | | | | |
|---|---|---|---|---|---|---|---|---|---|---|---|
| T26387 | CCGTCCGATCGGGCCGATGCCGG | 1000 | | | | | | | | | |
| Oligo fwd | CACCCCGTCCGATCGGGCCGATGC | | | | | | | | | | |
| Oligo rev | AAACGCATCGGCCCGATCGGACGG | | | | | | | | | | |
| Chromosome | start | end | strand | MM | target_seq | PAM | alignment | distance | position | gene name | gene id |
| GU305914.1 | 100666 | 100688 | - | 0 | CCGTCCGATCGGGCCGATGC | CGG | \|\|\|\|\|\|\|\|[\|\|\|\|\|\|\|\|\|\|\|\|]PAM | NA | Intergenic | NA | NA |

| | | | | | | | | | | | |
|---|---|---|---|---|---|---|---|---|---|---|---|
| T26388 | TCGGGCCGATGCCGGTGTTCCGG | 1000 | | | | | | | | | |
| Oligo fwd | CACCTCGGGCCGATGCCGGTGTTC | | | | | | | | | | |
| Oligo rev | AAACGAACACCGGCATCGGCCCGA | | | | | | | | | | |
| Chromosome | start | end | strand | MM | target_seq | PAM | alignment | distance | position | gene name | gene id |
| GU305914.1 | 100658 | 100680 | - | 0 | TCGGGCCGATGCCGGTGTTC | CGG | \|\|\|\|\|\|\|\|[\|\|\|\|\|\|\|\|\|\|\|\|]PAM | NA | Intergenic | NA | NA |

| | | | | | | | | | | | |
|---|---|---|---|---|---|---|---|---|---|---|---|
| T26389 | CGGGCCGATGCCGGTGTTCCGGG | 1000 | | | | | | | | | |
| Oligo fwd | CACCCGGGCCGATGCCGGTGTTCC | | | | | | | | | | |
| Oligo rev | AAACGGAACACCGGCATCGGCCCG | | | | | | | | | | |
| Chromosome | start | end | strand | MM | target_seq | PAM | alignment | distance | position | gene name | gene id |
| GU305914.1 | 100657 | 100679 | - | 0 | CGGGCCGATGCCGGTGTTCC | GGG | \|\|\|\|\|\|\|\|[\|\|\|\|\|\|\|\|\|\|\|\|]PAM | NA | Intergenic | NA | NA |

| | | | | | | | | | | | |
|---|---|---|---|---|---|---|---|---|---|---|---|
| T26390 | GCCGATGCCGGTGTTCCGGGTGG | 1000 | | | | | | | | | |
| Oligo fwd | CACCGCCGATGCCGGTGTTCCGGG | | | | | | | | | | |
| Oligo rev | AAACCCCGGAACACCGGCATCGGC | | | | | | | | | | |
| Chromosome | start | end | strand | MM | target_seq | PAM | alignment | distance | position | gene name | gene id |
| GU305914.1 | 100654 | 100676 | - | 0 | GCCGATGCCGGTGTTCCGGG | TGG | \|\|\|\|\|\|\|\|[\|\|\|\|\|\|\|\|\|\|\|\|]PAM | NA | Intergenic | NA | NA |

| | | | | | | | | | | | |
|---|---|---|---|---|---|---|---|---|---|---|---|
| T26391 | GGTGTTCCGGGTGGAGATGCCGG | 1000 | | | | | | | | | |
| Oligo fwd | CACCGGTGTTCCGGGTGGAGATGC | | | | | | | | | | |
| Oligo rev | AAACGCATCTCCACCCGGAACACC | | | | | | | | | | |
| Chromosome | start | end | strand | MM | target_seq | PAM | alignment | distance | position | gene name | gene id |
| GU305914.1 | 100645 | 100667 | - | 0 | GGTGTTCCGGGTGGAGATGC | CGG | \|\|\|\|\|\|\|\|[\|\|\|\|\|\|\|\|\|\|\|\|]PAM | NA | Intergenic | NA | NA |

| | | |
|---|---|---|
| T26392 | GGGTGGAGATGCCGGAGAAGCGG | 1000 |
| Oligo fwd | CACCGGGTGGAGATGCCGGAGAAG | |
| Oligo rev | AAACCTTCTCCGGCATCTCCACCC | |

Chromosome start end strand MM target_seq PAM alignment distance position gene name gene id
GU305914.1 100637 100659 - 0 GGGTGGAGATGCCGGAGAAG CGG ||||||||[||||||||||||]PAM NA Intergenic NA NA

T26393 GAAGCGGCACGTGTTCTGCGCGG 1000
Oligo fwd CACCGAAGCGGCACGTGTTCTGCG
Oligo rev AAACCGCAGAACACGTGCCGCTTC
Chromosome start end strand MM target_seq PAM alignment distance position gene name gene id
GU305914.1 100621 100643 - 0 GAAGCGGCACGTGTTCTGCG CGG ||||||||[||||||||||||]PAM NA Intergenic NA NA

T26394 CGTGTTCTGCGCGGTCAGCGCGG 1000
Oligo fwd CACCCGTGTTCTGCGCGGTCAGCG
Oligo rev AAACCGCTGACCGCGCAGAACACG
Chromosome start end strand MM target_seq PAM alignment distance position gene name gene id
GU305914.1 100612 100634 - 0 CGTGTTCTGCGCGGTCAGCG CGG ||||||||[||||||||||||]PAM NA Intergenic NA NA

T26395 GCGCGGTCAGCGCGGAGAACTGG 1000
Oligo fwd CACCGCGCGGTCAGCGCGGAGAAC
Oligo rev AAACGTTCTCCGCGCTGACCGCGC
Chromosome start end strand MM target_seq PAM alignment distance position gene name gene id
GU305914.1 100604 100626 - 0 GCGCGGTCAGCGCGGAGAAC TGG ||||||||[||||||||||||]PAM NA Intergenic NA NA

T26396 GCGCGGAGAACTGGACCCGCCGG 1000
Oligo fwd CACCGCGCGGAGAACTGGACCCGC
Oligo rev AAACGCGGGTCCAGTTCTCCGCGC
Chromosome start end strand MM target_seq PAM alignment distance position gene name gene id
GU305914.1 100595 100617 - 0 GCGCGGAGAACTGGACCCGC CGG ||||||||[||||||||||||]PAM NA Intergenic NA NA

T26397 GACCCGCCGGCTGCTGCCCAAGG 1000
Oligo fwd CACCGACCCGCCGGCTGCTGCCCA
Oligo rev AAACTGGGCAGCAGCCGGCGGGTC
Chromosome start end strand MM target_seq PAM alignment distance position gene name gene id
GU305914.1 100582 100604 - 0 GACCCGCCGGCTGCTGCCCA AGG ||||||||[||||||||||||]PAM NA Intergenic NA NA

T26398 CCTGCCGTCCGAGTACGTCTCGG 1000
Oligo fwd CACCCCTGCCGTCCGAGTACGTCT
Oligo rev AAACAGACGTACTCGGACGGCAGG
Chromosome start end strand MM target_seq PAM alignment distance position gene name gene id
GU305914.1 100546 100568 - 0 CCTGCCGTCCGAGTACGTCT CGG ||||||||[||||||||||||]PAM NA Intergenic NA NA

T26399 CGTCTCGGACGAGTGTCTGACGG 1000
Oligo fwd CACCCGTCTCGGACGAGTGTCTGA
Oligo rev AAACTCAGACACTCGTCCGAGACG
Chromosome start end strand MM target_seq PAM alignment distance position gene name gene id
GU305914.1 100531 100553 - 0 CGTCTCGGACGAGTGTCTGA CGG ||||||||[||||||||||||]PAM NA Intergenic NA NA

T26400 GGACGAGTGTCTGACGGACGCGG 1000
Oligo fwd CACCGGACGAGTGTCTGACGGACG
Oligo rev AAACCGTCCGTCAGACACTCGTCC
Chromosome start end strand MM target_seq PAM alignment distance position gene name gene id
GU305914.1 100525 100547 - 0 GGACGAGTGTCTGACGGACG CGG ||||||||[||||||||||||]PAM NA Intergenic NA NA

T26401 AGTGTCTGACGGACGCGGTGTGG 1000
Oligo fwd CACCAGTGTCTGACGGACGCGGTG
Oligo rev AAACCACCGCGTCCGTCAGACACT
Chromosome start end strand MM target_seq PAM alignment distance position gene name gene id
GU305914.1 100520 100542 - 0 AGTGTCTGACGGACGCGGTG TGG ||||||||[||||||||||||]PAM NA Intergenic NA NA

T26402 GTGGCTGCGCGAAGACATCGCGG 1000
Oligo fwd CACCGTGGCTGCGCGAAGACATCG
Oligo rev AAACCGATGTCTTCGCGCAGCCAC
Chromosome start end strand MM target_seq PAM alignment distance position gene name gene id
GU305914.1 100501 100523 - 0 GTGGCTGCGCGAAGACATCG CGG ||||||||[||||||||||||]PAM NA Intergenic NA NA

T26403 AGACATCGCGGCCTCGTGCGAGG 1000
Oligo fwd CACCAGACATCGCGGCCTCGTGCG
Oligo rev AAACCGCACGAGGCCGCGATGTCT
Chromosome start end strand MM target_seq PAM alignment distance position gene name gene id
GU305914.1 100489 100511 - 0 AGACATCGCGGCCTCGTGCG AGG ||||||||[||||||||||||]PAM NA Intergenic NA NA

T26404 CATCGCGGCCTCGTGCGAGGTGG 1000
Oligo fwd CACCCATCGCGGCCTCGTGCGAGG
Oligo rev AAACCCTCGCACGAGGCCGCGATG
Chromosome start end strand MM target_seq PAM alignment distance position gene name gene id
GU305914.1 100486 100508 - 0 CATCGCGGCCTCGTGCGAGG TGG ||||||||[||||||||||||]PAM NA Intergenic NA NA

T26405 ATCGCGGCCTCGTGCGAGGTGGG 1000
Oligo fwd CACCATCGCGGCCTCGTGCGAGGT
Oligo rev AAACACCTCGCACGAGGCCGCGAT
Chromosome start end strand MM target_seq PAM alignment distance position gene name gene id
GU305914.1 100485 100507 - 0 ATCGCGGCCTCGTGCGAGGT GGG ||||||||[||||||||||||]PAM NA Intergenic NA NA

T26406 CCTGCCCGTGTTCAACTTCGTGG 1000
Oligo fwd CACCCCTGCCCGTGTTCAACTTCG
Oligo rev AAACCGAAGTTGAACACGGGCAGG
Chromosome start end strand MM target_seq PAM alignment distance position gene name gene id
GU305914.1 100420 100442 - 0 CCTGCCCGTGTTCAACTTCG TGG ||||||||[||||||||||||]PAM NA Intergenic NA NA

T26407 CTGCCCGTGTTCAACTTCGTGGG 1000
Oligo fwd CACCCTGCCCGTGTTCAACTTCGT

<table>
<tr><td>Oligo rev</td><td>AAACACGAAGTTGAACACGGGCAG</td><td></td><td></td><td></td><td></td><td></td><td></td><td></td><td></td><td></td><td></td></tr>
<tr><td>Chromosome</td><td>start</td><td>end</td><td>strand</td><td>MM</td><td>target_seq</td><td>PAM</td><td>alignment</td><td>distance</td><td>position</td><td>gene name</td><td>gene id</td></tr>
<tr><td>GU305914.1</td><td>100419</td><td>100441</td><td>-</td><td>0</td><td>CTGCCCGTGTTCAACTTCGT</td><td>GGG</td><td>||||||||[||||||||||||]PAM</td><td>NA</td><td>Intergenic</td><td>NA</td><td>NA</td></tr>
<tr><td></td><td></td><td></td><td></td><td></td><td></td><td></td><td></td><td></td><td></td><td></td><td></td></tr>
<tr><td>T26408</td><td>GTTCAACTTCGTGGGCGACGTGG</td><td>1000</td><td></td><td></td><td></td><td></td><td></td><td></td><td></td><td></td><td></td></tr>
<tr><td>Oligo fwd</td><td>CACCGTTCAACTTCGTGGGCGACG</td><td></td><td></td><td></td><td></td><td></td><td></td><td></td><td></td><td></td><td></td></tr>
<tr><td>Oligo rev</td><td>AAACCGTCGCCCACGAAGTTGAAC</td><td></td><td></td><td></td><td></td><td></td><td></td><td></td><td></td><td></td><td></td></tr>
<tr><td>Chromosome</td><td>start</td><td>end</td><td>strand</td><td>MM</td><td>target_seq</td><td>PAM</td><td>alignment</td><td>distance</td><td>position</td><td>gene name</td><td>gene id</td></tr>
<tr><td>GU305914.1</td><td>100411</td><td>100433</td><td>-</td><td>0</td><td>GTTCAACTTCGTGGGCGACG</td><td>TGG</td><td>||||||||[||||||||||||]PAM</td><td>NA</td><td>Intergenic</td><td>NA</td><td>NA</td></tr>
<tr><td>GU305914.1</td><td>145061</td><td>145083</td><td>-</td><td>4</td><td>CATCAACTTCATGGGCGACT</td><td>CGG</td><td>--||||||[||-||||||||-]PAM</td><td>NA</td><td>Intergenic</td><td>NA</td><td>NA</td></tr>
<tr><td></td><td></td><td></td><td></td><td></td><td></td><td></td><td></td><td></td><td></td><td></td><td></td></tr>
<tr><td>T26409</td><td>CGTGGACCTCAAGCTGCGCGAGG</td><td>1000</td><td></td><td></td><td></td><td></td><td></td><td></td><td></td><td></td><td></td></tr>
<tr><td>Oligo fwd</td><td>CACCCGTGGACCTCAAGCTGCGCG</td><td></td><td></td><td></td><td></td><td></td><td></td><td></td><td></td><td></td><td></td></tr>
<tr><td>Oligo rev</td><td>AAACCGCGCAGCTTGAGGTCCACG</td><td></td><td></td><td></td><td></td><td></td><td></td><td></td><td></td><td></td><td></td></tr>
<tr><td>Chromosome</td><td>start</td><td>end</td><td>strand</td><td>MM</td><td>target_seq</td><td>PAM</td><td>alignment</td><td>distance</td><td>position</td><td>gene name</td><td>gene id</td></tr>
<tr><td>GU305914.1</td><td>100393</td><td>100415</td><td>-</td><td>0</td><td>CGTGGACCTCAAGCTGCGCG</td><td>AGG</td><td>||||||||[||||||||||||]PAM</td><td>NA</td><td>Intergenic</td><td>NA</td><td>NA</td></tr>
<tr><td></td><td></td><td></td><td></td><td></td><td></td><td></td><td></td><td></td><td></td><td></td><td></td></tr>
<tr><td>T26410</td><td>CAAGCTGCGCGAGGACCTGCAGG</td><td>1000</td><td></td><td></td><td></td><td></td><td></td><td></td><td></td><td></td><td></td></tr>
<tr><td>Oligo fwd</td><td>CACCCAAGCTGCGCGAGGACCTGC</td><td></td><td></td><td></td><td></td><td></td><td></td><td></td><td></td><td></td><td></td></tr>
<tr><td>Oligo rev</td><td>AAACGCAGGTCCTCGCGCAGCTTG</td><td></td><td></td><td></td><td></td><td></td><td></td><td></td><td></td><td></td><td></td></tr>
<tr><td>Chromosome</td><td>start</td><td>end</td><td>strand</td><td>MM</td><td>target_seq</td><td>PAM</td><td>alignment</td><td>distance</td><td>position</td><td>gene name</td><td>gene id</td></tr>
<tr><td>GU305914.1</td><td>100384</td><td>100406</td><td>-</td><td>0</td><td>CAAGCTGCGCGAGGACCTGC</td><td>AGG</td><td>||||||||[||||||||||||]PAM</td><td>NA</td><td>Intergenic</td><td>NA</td><td>NA</td></tr>
<tr><td>GU305914.1</td><td>69496</td><td>69518</td><td>+</td><td>4</td><td>CCAGGTGCGCGCCGACCTGC</td><td>TGG</td><td>|-||-|||[|||--|||||||]PAM</td><td>NA</td><td>Intergenic</td><td>NA</td><td>NA</td></tr>
<tr><td></td><td></td><td></td><td></td><td></td><td></td><td></td><td></td><td></td><td></td><td></td><td></td></tr>
<tr><td>T26411</td><td>AAGCTGCGCGAGGACCTGCAGGG</td><td>1000</td><td></td><td></td><td></td><td></td><td></td><td></td><td></td><td></td><td></td></tr>
<tr><td>Oligo fwd</td><td>CACCAAGCTGCGCGAGGACCTGCA</td><td></td><td></td><td></td><td></td><td></td><td></td><td></td><td></td><td></td><td></td></tr>
<tr><td>Oligo rev</td><td>AAACTGCAGGTCCTCGCGCAGCTT</td><td></td><td></td><td></td><td></td><td></td><td></td><td></td><td></td><td></td><td></td></tr>
<tr><td>Chromosome</td><td>start</td><td>end</td><td>strand</td><td>MM</td><td>target_seq</td><td>PAM</td><td>alignment</td><td>distance</td><td>position</td><td>gene name</td><td>gene id</td></tr>
<tr><td>GU305914.1</td><td>100383</td><td>100405</td><td>-</td><td>0</td><td>AAGCTGCGCGAGGACCTGCA</td><td>GGG</td><td>||||||||[||||||||||||]PAM</td><td>NA</td><td>Intergenic</td><td>NA</td><td>NA</td></tr>
<tr><td></td><td></td><td></td><td></td><td></td><td></td><td></td><td></td><td></td><td></td><td></td><td></td></tr>
<tr><td>T26412</td><td>CCTGCAGGGTCTGAGCCGTCAGG</td><td>1000</td><td></td><td></td><td></td><td></td><td></td><td></td><td></td><td></td><td></td></tr>
<tr><td>Oligo fwd</td><td>CACCCCTGCAGGGTCTGAGCCGTC</td><td></td><td></td><td></td><td></td><td></td><td></td><td></td><td></td><td></td><td></td></tr>
<tr><td>Oligo rev</td><td>AAACGACGGCTCAGACCCTGCAGG</td><td></td><td></td><td></td><td></td><td></td><td></td><td></td><td></td><td></td><td></td></tr>
<tr><td>Chromosome</td><td>start</td><td>end</td><td>strand</td><td>MM</td><td>target_seq</td><td>PAM</td><td>alignment</td><td>distance</td><td>position</td><td>gene name</td><td>gene id</td></tr>
<tr><td>GU305914.1</td><td>100369</td><td>100391</td><td>-</td><td>0</td><td>CCTGCAGGGTCTGAGCCGTC</td><td>AGG</td><td>||||||||[||||||||||||]PAM</td><td>NA</td><td>Intergenic</td><td>NA</td><td>NA</td></tr>
<tr><td></td><td></td><td></td><td></td><td></td><td></td><td></td><td></td><td></td><td></td><td></td><td></td></tr>
<tr><td>T26413</td><td>GCAGGGTCTGAGCCGTCAGGAGG</td><td>1000</td><td></td><td></td><td></td><td></td><td></td><td></td><td></td><td></td><td></td></tr>
<tr><td>Oligo fwd</td><td>CACCGCAGGGTCTGAGCCGTCAGG</td><td></td><td></td><td></td><td></td><td></td><td></td><td></td><td></td><td></td><td></td></tr>
<tr><td>Oligo rev</td><td>AAACCCTGACGGCTCAGACCCTGC</td><td></td><td></td><td></td><td></td><td></td><td></td><td></td><td></td><td></td><td></td></tr>
<tr><td>Chromosome</td><td>start</td><td>end</td><td>strand</td><td>MM</td><td>target_seq</td><td>PAM</td><td>alignment</td><td>distance</td><td>position</td><td>gene name</td><td>gene id</td></tr>
<tr><td>GU305914.1</td><td>100366</td><td>100388</td><td>-</td><td>0</td><td>GCAGGGTCTGAGCCGTCAGG</td><td>AGG</td><td>||||||||[||||||||||||]PAM</td><td>NA</td><td>Intergenic</td><td>NA</td><td>NA</td></tr>
<tr><td></td><td></td><td></td><td></td><td></td><td></td><td></td><td></td><td></td><td></td><td></td><td></td></tr>
<tr><td>T26414</td><td>TCGACCTGTGCCGCGCGCTGCGG</td><td>1000</td><td></td><td></td><td></td><td></td><td></td><td></td><td></td><td></td><td></td></tr>
<tr><td>Oligo fwd</td><td>CACCTCGACCTGTGCCGCGCGCTG</td><td></td><td></td><td></td><td></td><td></td><td></td><td></td><td></td><td></td><td></td></tr>
<tr><td>Oligo rev</td><td>AAACCAGCGCGCGGCACAGGTCGA</td><td></td><td></td><td></td><td></td><td></td><td></td><td></td><td></td><td></td><td></td></tr>
<tr><td>Chromosome</td><td>start</td><td>end</td><td>strand</td><td>MM</td><td>target_seq</td><td>PAM</td><td>alignment</td><td>distance</td><td>position</td><td>gene name</td><td>gene id</td></tr>
<tr><td>GU305914.1</td><td>100340</td><td>100362</td><td>-</td><td>0</td><td>TCGACCTGTGCCGCGCGCTG</td><td>CGG</td><td>||||||||[||||||||||||]PAM</td><td>NA</td><td>Intergenic</td><td>NA</td><td>NA</td></tr>
<tr><td></td><td></td><td></td><td></td><td></td><td></td><td></td><td></td><td></td><td></td><td></td><td></td></tr>
<tr><td>T26415</td><td>ACCTGTGCCGCGCGCTGCGGCGG</td><td>1000</td><td></td><td></td><td></td><td></td><td></td><td></td><td></td><td></td><td></td></tr>
<tr><td>Oligo fwd</td><td>CACCACCTGTGCCGCGCGCTGCGG</td><td></td><td></td><td></td><td></td><td></td><td></td><td></td><td></td><td></td><td></td></tr>
<tr><td>Oligo rev</td><td>AAACCCGCAGCGCGCGGCACAGGT</td><td></td><td></td><td></td><td></td><td></td><td></td><td></td><td></td><td></td><td></td></tr>
<tr><td>Chromosome</td><td>start</td><td>end</td><td>strand</td><td>MM</td><td>target_seq</td><td>PAM</td><td>alignment</td><td>distance</td><td>position</td><td>gene name</td><td>gene id</td></tr>
<tr><td>GU305914.1</td><td>100337</td><td>100359</td><td>-</td><td>0</td><td>ACCTGTGCCGCGCGCTGCGG</td><td>CGG</td><td>||||||||[||||||||||||]PAM</td><td>NA</td><td>Intergenic</td><td>NA</td><td>NA</td></tr>
<tr><td></td><td></td><td></td><td></td><td></td><td></td><td></td><td></td><td></td><td></td><td></td><td></td></tr>
<tr><td>T26416</td><td>GCGCTGCGGCGGACCCTGATCGG</td><td>1000</td><td></td><td></td><td></td><td></td><td></td><td></td><td></td><td></td><td></td></tr>
<tr><td>Oligo fwd</td><td>CACCGCGCTGCGGCGGACCCTGAT</td><td></td><td></td><td></td><td></td><td></td><td></td><td></td><td></td><td></td><td></td></tr>
<tr><td>Oligo rev</td><td>AAACATCAGGGTCCGCCGCAGCGC</td><td></td><td></td><td></td><td></td><td></td><td></td><td></td><td></td><td></td><td></td></tr>
<tr><td>Chromosome</td><td>start</td><td>end</td><td>strand</td><td>MM</td><td>target_seq</td><td>PAM</td><td>alignment</td><td>distance</td><td>position</td><td>gene name</td><td>gene id</td></tr>
<tr><td>GU305914.1</td><td>100326</td><td>100348</td><td>-</td><td>0</td><td>GCGCTGCGGCGGACCCTGAT</td><td>CGG</td><td>||||||||[||||||||||||]PAM</td><td>NA</td><td>Intergenic</td><td>NA</td><td>NA</td></tr>
<tr><td>GU305914.1</td><td>100821</td><td>100843</td><td>-</td><td>4</td><td>CCGCTGCCGCCGACGCTGAT</td><td>GGG</td><td>-||||||-[||-|||-|||||]PAM</td><td>NA</td><td>Intergenic</td><td>NA</td><td>NA</td></tr>
<tr><td></td><td></td><td></td><td></td><td></td><td></td><td></td><td></td><td></td><td></td><td></td><td></td></tr>
<tr><td>T26417</td><td>CGCTGCGGCGGACCCTGATCGGG</td><td>1000</td><td></td><td></td><td></td><td></td><td></td><td></td><td></td><td></td><td></td></tr>
<tr><td>Oligo fwd</td><td>CACCCGCTGCGGCGGACCCTGATC</td><td></td><td></td><td></td><td></td><td></td><td></td><td></td><td></td><td></td><td></td></tr>
<tr><td>Oligo rev</td><td>AAACGATCAGGGTCCGCCGCAGCG</td><td></td><td></td><td></td><td></td><td></td><td></td><td></td><td></td><td></td><td></td></tr>
<tr><td>Chromosome</td><td>start</td><td>end</td><td>strand</td><td>MM</td><td>target_seq</td><td>PAM</td><td>alignment</td><td>distance</td><td>position</td><td>gene name</td><td>gene id</td></tr>
<tr><td>GU305914.1</td><td>100325</td><td>100347</td><td>-</td><td>0</td><td>CGCTGCGGCGGACCCTGATC</td><td>GGG</td><td>||||||||[||||||||||||]PAM</td><td>NA</td><td>Intergenic</td><td>NA</td><td>NA</td></tr>
<tr><td></td><td></td><td></td><td></td><td></td><td></td><td></td><td></td><td></td><td></td><td></td><td></td></tr>
<tr><td>T26418</td><td>GCTGCGGCGGACCCTGATCGGGG</td><td>1000</td><td></td><td></td><td></td><td></td><td></td><td></td><td></td><td></td><td></td></tr>
<tr><td>Oligo fwd</td><td>CACCGCTGCGGCGGACCCTGATCG</td><td></td><td></td><td></td><td></td><td></td><td></td><td></td><td></td><td></td><td></td></tr>
<tr><td>Oligo rev</td><td>AAACCGATCAGGGTCCGCCGCAGC</td><td></td><td></td><td></td><td></td><td></td><td></td><td></td><td></td><td></td><td></td></tr>
<tr><td>Chromosome</td><td>start</td><td>end</td><td>strand</td><td>MM</td><td>target_seq</td><td>PAM</td><td>alignment</td><td>distance</td><td>position</td><td>gene name</td><td>gene id</td></tr>
<tr><td>GU305914.1</td><td>100324</td><td>100346</td><td>-</td><td>0</td><td>GCTGCGGCGGACCCTGATCG</td><td>GGG</td><td>||||||||[||||||||||||]PAM</td><td>NA</td><td>Intergenic</td><td>NA</td><td>NA</td></tr>
<tr><td></td><td></td><td></td><td></td><td></td><td></td><td></td><td></td><td></td><td></td><td></td><td></td></tr>
<tr><td>T26419</td><td>GGCGGACCCTGATCGGGGCCTGG</td><td>1000</td><td></td><td></td><td></td><td></td><td></td><td></td><td></td><td></td><td></td></tr>
<tr><td>Oligo fwd</td><td>CACCGGCGGACCCTGATCGGGGCC</td><td></td><td></td><td></td><td></td><td></td><td></td><td></td><td></td><td></td><td></td></tr>
<tr><td>Oligo rev</td><td>AAACGGCCCCGATCAGGGTCCGCC</td><td></td><td></td><td></td><td></td><td></td><td></td><td></td><td></td><td></td><td></td></tr>
<tr><td>Chromosome</td><td>start</td><td>end</td><td>strand</td><td>MM</td><td>target_seq</td><td>PAM</td><td>alignment</td><td>distance</td><td>position</td><td>gene name</td><td>gene id</td></tr>
<tr><td>GU305914.1</td><td>100319</td><td>100341</td><td>-</td><td>0</td><td>GGCGGACCCTGATCGGGGCC</td><td>TGG</td><td>||||||||[||||||||||||]PAM</td><td>NA</td><td>Intergenic</td><td>NA</td><td>NA</td></tr>
<tr><td></td><td></td><td></td><td></td><td></td><td></td><td></td><td></td><td></td><td></td><td></td><td></td></tr>
<tr><td>T26420</td><td>GGACCCTGATCGGGGCCTGGCGG</td><td>1000</td><td></td><td></td><td></td><td></td><td></td><td></td><td></td><td></td><td></td></tr>
<tr><td>Oligo fwd</td><td>CACCGGACCCTGATCGGGGCCTGG</td><td></td><td></td><td></td><td></td><td></td><td></td><td></td><td></td><td></td><td></td></tr>
<tr><td>Oligo rev</td><td>AAACCCAGGCCCCGATCAGGGTCC</td><td></td><td></td><td></td><td></td><td></td><td></td><td></td><td></td><td></td><td></td></tr>
<tr><td>Chromosome</td><td>start</td><td>end</td><td>strand</td><td>MM</td><td>target_seq</td><td>PAM</td><td>alignment</td><td>distance</td><td>position</td><td>gene name</td><td>gene id</td></tr>
<tr><td>GU305914.1</td><td>100316</td><td>100338</td><td>-</td><td>0</td><td>GGACCCTGATCGGGGCCTGG</td><td>CGG</td><td>||||||||[||||||||||||]PAM</td><td>NA</td><td>Intergenic</td><td>NA</td><td>NA</td></tr>
<tr><td></td><td></td><td></td><td></td><td></td><td></td><td></td><td></td><td></td><td></td><td></td><td></td></tr>
<tr><td>T26421</td><td>CTGGCGGCACCTGTTTCCCGAGG</td><td>1000</td><td></td><td></td><td></td><td></td><td></td><td></td><td></td><td></td><td></td></tr>
<tr><td>Oligo fwd</td><td>CACCCTGGCGGCACCTGTTTCCCG</td><td></td><td></td><td></td><td></td><td></td><td></td><td></td><td></td><td></td><td></td></tr>
<tr><td>Oligo rev</td><td>AAACCGGGAAACAGGTGCCGCCAG</td><td></td><td></td><td></td><td></td><td></td><td></td><td></td><td></td><td></td><td></td></tr>
<tr><td>Chromosome</td><td>start</td><td>end</td><td>strand</td><td>MM</td><td>target_seq</td><td>PAM</td><td>alignment</td><td>distance</td><td>position</td><td>gene name</td><td>gene id</td></tr>
</table>

<table>
<tr><td>GU305914.1</td><td>100300</td><td>100322</td><td>-</td><td>0</td><td>CTGGCGGCACCTGTTTCCCG</td><td>AGG</td><td>||||||||[||||||||||||]PAM</td><td>NA</td><td>Intergenic</td><td>NA</td><td>NA</td></tr>
<tr><td>T26422</td><td>CCTGTTTCCCGAGGTAGACCCGG</td><td>1000</td><td></td><td></td><td></td><td></td><td></td><td></td><td></td><td></td><td></td></tr>
<tr><td>Oligo fwd</td><td>CACCCCTGTTTCCCGAGGTAGACC</td><td></td><td></td><td></td><td></td><td></td><td></td><td></td><td></td><td></td><td></td></tr>
<tr><td>Oligo rev</td><td>AAACGGTCTACCTCGGGAAACAGG</td><td></td><td></td><td></td><td></td><td></td><td></td><td></td><td></td><td></td><td></td></tr>
<tr><td>Chromosome</td><td>start</td><td>end</td><td>strand</td><td>MM</td><td>target_seq</td><td>PAM</td><td>alignment</td><td>distance</td><td>position</td><td>gene name</td><td>gene id</td></tr>
<tr><td>GU305914.1</td><td>100291</td><td>100313</td><td>-</td><td>0</td><td>CCTGTTTCCCGAGGTAGACC</td><td>CGG</td><td>||||||||[||||||||||||]PAM</td><td>NA</td><td>Intergenic</td><td>NA</td><td>NA</td></tr>
<tr><td>T26423</td><td>CGCGTGTCCGCAGAACGCCGCGG</td><td>1000</td><td></td><td></td><td></td><td></td><td></td><td></td><td></td><td></td><td></td></tr>
<tr><td>Oligo fwd</td><td>CACCCGCGTGTCCGCAGAACGCCG</td><td></td><td></td><td></td><td></td><td></td><td></td><td></td><td></td><td></td><td></td></tr>
<tr><td>Oligo rev</td><td>AAACCGGCGTTCTGCGGACACGCG</td><td></td><td></td><td></td><td></td><td></td><td></td><td></td><td></td><td></td><td></td></tr>
<tr><td>Chromosome</td><td>start</td><td>end</td><td>strand</td><td>MM</td><td>target_seq</td><td>PAM</td><td>alignment</td><td>distance</td><td>position</td><td>gene name</td><td>gene id</td></tr>
<tr><td>GU305914.1</td><td>100240</td><td>100262</td><td>-</td><td>0</td><td>CGCGTGTCCGCAGAACGCCG</td><td>CGG</td><td>||||||||[||||||||||||]PAM</td><td>NA</td><td>Intergenic</td><td>NA</td><td>NA</td></tr>
<tr><td>T26424</td><td>GCGTGTCCGCAGAACGCCGCGGG</td><td>1000</td><td></td><td></td><td></td><td></td><td></td><td></td><td></td><td></td><td></td></tr>
<tr><td>Oligo fwd</td><td>CACCGCGTGTCCGCAGAACGCCGC</td><td></td><td></td><td></td><td></td><td></td><td></td><td></td><td></td><td></td><td></td></tr>
<tr><td>Oligo rev</td><td>AAACGCGGCGTTCTGCGGACACGC</td><td></td><td></td><td></td><td></td><td></td><td></td><td></td><td></td><td></td><td></td></tr>
<tr><td>Chromosome</td><td>start</td><td>end</td><td>strand</td><td>MM</td><td>target_seq</td><td>PAM</td><td>alignment</td><td>distance</td><td>position</td><td>gene name</td><td>gene id</td></tr>
<tr><td>GU305914.1</td><td>100239</td><td>100261</td><td>-</td><td>0</td><td>GCGTGTCCGCAGAACGCCGC</td><td>GGG</td><td>||||||||[||||||||||||]PAM</td><td>NA</td><td>Intergenic</td><td>NA</td><td>NA</td></tr>
<tr><td>T26425</td><td>CGTGTCCGCAGAACGCCGCGGGG</td><td>1000</td><td></td><td></td><td></td><td></td><td></td><td></td><td></td><td></td><td></td></tr>
<tr><td>Oligo fwd</td><td>CACCCGTGTCCGCAGAACGCCGCG</td><td></td><td></td><td></td><td></td><td></td><td></td><td></td><td></td><td></td><td></td></tr>
<tr><td>Oligo rev</td><td>AAACCGCGGCGTTCTGCGGACACG</td><td></td><td></td><td></td><td></td><td></td><td></td><td></td><td></td><td></td><td></td></tr>
<tr><td>Chromosome</td><td>start</td><td>end</td><td>strand</td><td>MM</td><td>target_seq</td><td>PAM</td><td>alignment</td><td>distance</td><td>position</td><td>gene name</td><td>gene id</td></tr>
<tr><td>GU305914.1</td><td>100238</td><td>100260</td><td>-</td><td>0</td><td>CGTGTCCGCAGAACGCCGCG</td><td>GGG</td><td>||||||||[||||||||||||]PAM</td><td>NA</td><td>Intergenic</td><td>NA</td><td>NA</td></tr>
<tr><td>T26426</td><td>GTGTCCGCAGAACGCCGCGGGGG</td><td>1000</td><td></td><td></td><td></td><td></td><td></td><td></td><td></td><td></td><td></td></tr>
<tr><td>Oligo fwd</td><td>CACCGTGTCCGCAGAACGCCGCGG</td><td></td><td></td><td></td><td></td><td></td><td></td><td></td><td></td><td></td><td></td></tr>
<tr><td>Oligo rev</td><td>AAACCCGCGGCGTTCTGCGGACAC</td><td></td><td></td><td></td><td></td><td></td><td></td><td></td><td></td><td></td><td></td></tr>
<tr><td>Chromosome</td><td>start</td><td>end</td><td>strand</td><td>MM</td><td>target_seq</td><td>PAM</td><td>alignment</td><td>distance</td><td>position</td><td>gene name</td><td>gene id</td></tr>
<tr><td>GU305914.1</td><td>100237</td><td>100259</td><td>-</td><td>0</td><td>GTGTCCGCAGAACGCCGCGG</td><td>GGG</td><td>||||||||[||||||||||||]PAM</td><td>NA</td><td>Intergenic</td><td>NA</td><td>NA</td></tr>
<tr><td>T26427</td><td>TCCGCAGAACGCCGCGGGGGCGG</td><td>1000</td><td></td><td></td><td></td><td></td><td></td><td></td><td></td><td></td><td></td></tr>
<tr><td>Oligo fwd</td><td>CACCTCCGCAGAACGCCGCGGGGG</td><td></td><td></td><td></td><td></td><td></td><td></td><td></td><td></td><td></td><td></td></tr>
<tr><td>Oligo rev</td><td>AAACCCCCCGCGGCGTTCTGCGGA</td><td></td><td></td><td></td><td></td><td></td><td></td><td></td><td></td><td></td><td></td></tr>
<tr><td>Chromosome</td><td>start</td><td>end</td><td>strand</td><td>MM</td><td>target_seq</td><td>PAM</td><td>alignment</td><td>distance</td><td>position</td><td>gene name</td><td>gene id</td></tr>
<tr><td>GU305914.1</td><td>100234</td><td>100256</td><td>-</td><td>0</td><td>TCCGCAGAACGCCGCGGGGG</td><td>CGG</td><td>||||||||[||||||||||||]PAM</td><td>NA</td><td>Intergenic</td><td>NA</td><td>NA</td></tr>
<tr><td>T26428</td><td>GCAGAACGCCGCGGGGGCGGCGG</td><td>1000</td><td></td><td></td><td></td><td></td><td></td><td></td><td></td><td></td><td></td></tr>
<tr><td>Oligo fwd</td><td>CACCGCAGAACGCCGCGGGGGCGG</td><td></td><td></td><td></td><td></td><td></td><td></td><td></td><td></td><td></td><td></td></tr>
<tr><td>Oligo rev</td><td>AAACCCGCCCCCGCGGCGTTCTGC</td><td></td><td></td><td></td><td></td><td></td><td></td><td></td><td></td><td></td><td></td></tr>
<tr><td>Chromosome</td><td>start</td><td>end</td><td>strand</td><td>MM</td><td>target_seq</td><td>PAM</td><td>alignment</td><td>distance</td><td>position</td><td>gene name</td><td>gene id</td></tr>
<tr><td>GU305914.1</td><td>100231</td><td>100253</td><td>-</td><td>0</td><td>GCAGAACGCCGCGGGGGCGG</td><td>CGG</td><td>||||||||[||||||||||||]PAM</td><td>NA</td><td>Intergenic</td><td>NA</td><td>NA</td></tr>
<tr><td>T26429</td><td>CGCCGCGGGGGCGGCGGACGAGG</td><td>1000</td><td></td><td></td><td></td><td></td><td></td><td></td><td></td><td></td><td></td></tr>
<tr><td>Oligo fwd</td><td>CACCCGCCGCGGGGGCGGCGGACG</td><td></td><td></td><td></td><td></td><td></td><td></td><td></td><td></td><td></td><td></td></tr>
<tr><td>Oligo rev</td><td>AAACCGTCCGCCGCCCCCGCGGCG</td><td></td><td></td><td></td><td></td><td></td><td></td><td></td><td></td><td></td><td></td></tr>
<tr><td>Chromosome</td><td>start</td><td>end</td><td>strand</td><td>MM</td><td>target_seq</td><td>PAM</td><td>alignment</td><td>distance</td><td>position</td><td>gene name</td><td>gene id</td></tr>
<tr><td>GU305914.1</td><td>100225</td><td>100247</td><td>-</td><td>0</td><td>CGCCGCGGGGGCGGCGGACG</td><td>AGG</td><td>||||||||[||||||||||||]PAM</td><td>NA</td><td>Intergenic</td><td>NA</td><td>NA</td></tr>
<tr><td>GU305914.1</td><td>113523</td><td>113545</td><td>+</td><td>4</td><td>CGCCGCTCGCGCGCCGGACG</td><td>CGG</td><td>||||||--[|-|||-||||||]PAM</td><td>NA</td><td>Intergenic</td><td>NA</td><td>NA</td></tr>
<tr><td>GU305914.1</td><td>62458</td><td>62480</td><td>+</td><td>4</td><td>GGCGGCGGGGGTGGCGGCCG</td><td>CGG</td><td>-||-||||[|||-|||||-||]PAM</td><td>NA</td><td>Intergenic</td><td>NA</td><td>NA</td></tr>
<tr><td>T26430</td><td>GCGGACGAGGCGATGCTGTACGG</td><td>1000</td><td></td><td></td><td></td><td></td><td></td><td></td><td></td><td></td><td></td></tr>
<tr><td>Oligo fwd</td><td>CACCGCGGACGAGGCGATGCTGTA</td><td></td><td></td><td></td><td></td><td></td><td></td><td></td><td></td><td></td><td></td></tr>
<tr><td>Oligo rev</td><td>AAACTACAGCATCGCCTCGTCCGC</td><td></td><td></td><td></td><td></td><td></td><td></td><td></td><td></td><td></td><td></td></tr>
<tr><td>Chromosome</td><td>start</td><td>end</td><td>strand</td><td>MM</td><td>target_seq</td><td>PAM</td><td>alignment</td><td>distance</td><td>position</td><td>gene name</td><td>gene id</td></tr>
<tr><td>GU305914.1</td><td>100212</td><td>100234</td><td>-</td><td>0</td><td>GCGGACGAGGCGATGCTGTA</td><td>CGG</td><td>||||||||[||||||||||||]PAM</td><td>NA</td><td>Intergenic</td><td>NA</td><td>NA</td></tr>
<tr><td>T26431</td><td>GACGAGGCGATGCTGTACGGCGG</td><td>1000</td><td></td><td></td><td></td><td></td><td></td><td></td><td></td><td></td><td></td></tr>
<tr><td>Oligo fwd</td><td>CACCGACGAGGCGATGCTGTACGG</td><td></td><td></td><td></td><td></td><td></td><td></td><td></td><td></td><td></td><td></td></tr>
<tr><td>Oligo rev</td><td>AAACCCGTACAGCATCGCCTCGTC</td><td></td><td></td><td></td><td></td><td></td><td></td><td></td><td></td><td></td><td></td></tr>
<tr><td>Chromosome</td><td>start</td><td>end</td><td>strand</td><td>MM</td><td>target_seq</td><td>PAM</td><td>alignment</td><td>distance</td><td>position</td><td>gene name</td><td>gene id</td></tr>
<tr><td>GU305914.1</td><td>100209</td><td>100231</td><td>-</td><td>0</td><td>GACGAGGCGATGCTGTACGG</td><td>CGG</td><td>||||||||[||||||||||||]PAM</td><td>NA</td><td>Intergenic</td><td>NA</td><td>NA</td></tr>
<tr><td>T26432</td><td>GAGGCGATGCTGTACGGCGGCGG</td><td>1000</td><td></td><td></td><td></td><td></td><td></td><td></td><td></td><td></td><td></td></tr>
<tr><td>Oligo fwd</td><td>CACCGAGGCGATGCTGTACGGCGG</td><td></td><td></td><td></td><td></td><td></td><td></td><td></td><td></td><td></td><td></td></tr>
<tr><td>Oligo rev</td><td>AAACCCGCCGTACAGCATCGCCTC</td><td></td><td></td><td></td><td></td><td></td><td></td><td></td><td></td><td></td><td></td></tr>
<tr><td>Chromosome</td><td>start</td><td>end</td><td>strand</td><td>MM</td><td>target_seq</td><td>PAM</td><td>alignment</td><td>distance</td><td>position</td><td>gene name</td><td>gene id</td></tr>
<tr><td>GU305914.1</td><td>100206</td><td>100228</td><td>-</td><td>0</td><td>GAGGCGATGCTGTACGGCGG</td><td>CGG</td><td>||||||||[||||||||||||]PAM</td><td>NA</td><td>Intergenic</td><td>NA</td><td>NA</td></tr>
<tr><td>T26433</td><td>GCGATGCTGTACGGCGGCGGCGG</td><td>1000</td><td></td><td></td><td></td><td></td><td></td><td></td><td></td><td></td><td></td></tr>
<tr><td>Oligo fwd</td><td>CACCGCGATGCTGTACGGCGGCGG</td><td></td><td></td><td></td><td></td><td></td><td></td><td></td><td></td><td></td><td></td></tr>
<tr><td>Oligo rev</td><td>AAACCCGCCGCCGTACAGCATCGC</td><td></td><td></td><td></td><td></td><td></td><td></td><td></td><td></td><td></td><td></td></tr>
<tr><td>Chromosome</td><td>start</td><td>end</td><td>strand</td><td>MM</td><td>target_seq</td><td>PAM</td><td>alignment</td><td>distance</td><td>position</td><td>gene name</td><td>gene id</td></tr>
<tr><td>GU305914.1</td><td>100203</td><td>100225</td><td>-</td><td>0</td><td>GCGATGCTGTACGGCGGCGG</td><td>CGG</td><td>||||||||[||||||||||||]PAM</td><td>NA</td><td>Intergenic</td><td>NA</td><td>NA</td></tr>
<tr><td>T26434</td><td>ACGACGAAGACGACGACCCGCGG</td><td>1000</td><td></td><td></td><td></td><td></td><td></td><td></td><td></td><td></td><td></td></tr>
<tr><td>Oligo fwd</td><td>CACCACGACGAAGACGACGACCCG</td><td></td><td></td><td></td><td></td><td></td><td></td><td></td><td></td><td></td><td></td></tr>
<tr><td>Oligo rev</td><td>AAACCGGGTCGTCGTCTTCGTCGT</td><td></td><td></td><td></td><td></td><td></td><td></td><td></td><td></td><td></td><td></td></tr>
<tr><td>Chromosome</td><td>start</td><td>end</td><td>strand</td><td>MM</td><td>target_seq</td><td>PAM</td><td>alignment</td><td>distance</td><td>position</td><td>gene name</td><td>gene id</td></tr>
<tr><td>GU305914.1</td><td>100178</td><td>100200</td><td>-</td><td>0</td><td>ACGACGAAGACGACGACCCG</td><td>CGG</td><td>||||||||[||||||||||||]PAM</td><td>NA</td><td>Intergenic</td><td>NA</td><td>NA</td></tr>
<tr><td>T26435</td><td>CCCGCGGCCCGAGCACGCCGCGG</td><td>1000</td><td></td><td></td><td></td><td></td><td></td><td></td><td></td><td></td><td></td></tr>
<tr><td>Oligo fwd</td><td>CACCCCCGCGGCCCGAGCACGCCG</td><td></td><td></td><td></td><td></td><td></td><td></td><td></td><td></td><td></td><td></td></tr>
<tr><td>Oligo rev</td><td>AAACCGGCGTGCTCGGGCCGCGGG</td><td></td><td></td><td></td><td></td><td></td><td></td><td></td><td></td><td></td><td></td></tr>
<tr><td>Chromosome</td><td>start</td><td>end</td><td>strand</td><td>MM</td><td>target_seq</td><td>PAM</td><td>alignment</td><td>distance</td><td>position</td><td>gene name</td><td>gene id</td></tr>
<tr><td>GU305914.1</td><td>100162</td><td>100184</td><td>-</td><td>0</td><td>CCCGCGGCCCGAGCACGCCG</td><td>CGG</td><td>||||||||[||||||||||||]PAM</td><td>NA</td><td>Intergenic</td><td>NA</td><td>NA</td></tr>
<tr><td>T26436</td><td>GCCCGAGCACGCCGCGGCGATGG</td><td>1000</td><td></td><td></td><td></td><td></td><td></td><td></td><td></td><td></td><td></td></tr>
</table>

<table>
<tr><td>Oligo fwd</td><td>CACCGCCCGAGCACGCCGCGGCGA</td><td></td><td></td><td></td><td></td><td></td><td></td><td></td><td></td><td></td><td></td></tr>
<tr><td>Oligo rev</td><td>AAACTCGCCGCGGCGTGCTCGGGC</td><td></td><td></td><td></td><td></td><td></td><td></td><td></td><td></td><td></td><td></td></tr>
<tr><td>Chromosome</td><td>start</td><td>end</td><td>strand</td><td>MM</td><td>target_seq</td><td>PAM</td><td>alignment</td><td>distance</td><td>position</td><td>gene name</td><td>gene id</td></tr>
<tr><td>GU305914.1</td><td>100156</td><td>100178</td><td>-</td><td>0</td><td>GCCCGAGCACGCCGCGGCGA</td><td>TGG</td><td>||||||||[||||||||||||]PAM</td><td>NA</td><td>Intergenic</td><td>NA</td><td>NA</td></tr>
<tr><td>T26437</td><td>GCCGCGGCGATGGTCGACTACGG</td><td>1000</td><td></td><td></td><td></td><td></td><td></td><td></td><td></td><td></td><td></td></tr>
<tr><td>Oligo fwd</td><td>CACCGCCGCGGCGATGGTCGACTA</td><td></td><td></td><td></td><td></td><td></td><td></td><td></td><td></td><td></td><td></td></tr>
<tr><td>Oligo rev</td><td>AAACTAGTCGACCATCGCCGCGGC</td><td></td><td></td><td></td><td></td><td></td><td></td><td></td><td></td><td></td><td></td></tr>
<tr><td>Chromosome</td><td>start</td><td>end</td><td>strand</td><td>MM</td><td>target_seq</td><td>PAM</td><td>alignment</td><td>distance</td><td>position</td><td>gene name</td><td>gene id</td></tr>
<tr><td>GU305914.1</td><td>100146</td><td>100168</td><td>-</td><td>0</td><td>GCCGCGGCGATGGTCGACTA</td><td>CGG</td><td>||||||||[||||||||||||]PAM</td><td>NA</td><td>Intergenic</td><td>NA</td><td>NA</td></tr>
<tr><td>T26438</td><td>TCGACTACGGCGATGCCGTTCGG</td><td>1000</td><td></td><td></td><td></td><td></td><td></td><td></td><td></td><td></td><td></td></tr>
<tr><td>Oligo fwd</td><td>CACCTCGACTACGGCGATGCCGTT</td><td></td><td></td><td></td><td></td><td></td><td></td><td></td><td></td><td></td><td></td></tr>
<tr><td>Oligo rev</td><td>AAACAACGGCATCGCCGTAGTCGA</td><td></td><td></td><td></td><td></td><td></td><td></td><td></td><td></td><td></td><td></td></tr>
<tr><td>Chromosome</td><td>start</td><td>end</td><td>strand</td><td>MM</td><td>target_seq</td><td>PAM</td><td>alignment</td><td>distance</td><td>position</td><td>gene name</td><td>gene id</td></tr>
<tr><td>GU305914.1</td><td>100133</td><td>100155</td><td>-</td><td>0</td><td>TCGACTACGGCGATGCCGTT</td><td>CGG</td><td>||||||||[||||||||||||]PAM</td><td>NA</td><td>Intergenic</td><td>NA</td><td>NA</td></tr>
<tr><td>T26439</td><td>GCCCGCCCTTCTGTGTCTGTCGG</td><td>1000</td><td></td><td></td><td></td><td></td><td></td><td></td><td></td><td></td><td></td></tr>
<tr><td>Oligo fwd</td><td>CACCGCCCGCCCTTCTGTGTCTGT</td><td></td><td></td><td></td><td></td><td></td><td></td><td></td><td></td><td></td><td></td></tr>
<tr><td>Oligo rev</td><td>AAACACAGACACAGAAGGGCGGGC</td><td></td><td></td><td></td><td></td><td></td><td></td><td></td><td></td><td></td><td></td></tr>
<tr><td>Chromosome</td><td>start</td><td>end</td><td>strand</td><td>MM</td><td>target_seq</td><td>PAM</td><td>alignment</td><td>distance</td><td>position</td><td>gene name</td><td>gene id</td></tr>
<tr><td>GU305914.1</td><td>100109</td><td>100131</td><td>-</td><td>0</td><td>GCCCGCCCTTCTGTGTCTGT</td><td>CGG</td><td>||||||||[||||||||||||]PAM</td><td>NA</td><td>Intergenic</td><td>NA</td><td>NA</td></tr>
<tr><td>T26440</td><td>CTGTGTCTGTCGGCGCAAGCTGG</td><td>1000</td><td></td><td></td><td></td><td></td><td></td><td></td><td></td><td></td><td></td></tr>
<tr><td>Oligo fwd</td><td>CACCCTGTGTCTGTCGGCGCAAGC</td><td></td><td></td><td></td><td></td><td></td><td></td><td></td><td></td><td></td><td></td></tr>
<tr><td>Oligo rev</td><td>AAACGCTTGCGCCGACAGACACAG</td><td></td><td></td><td></td><td></td><td></td><td></td><td></td><td></td><td></td><td></td></tr>
<tr><td>Chromosome</td><td>start</td><td>end</td><td>strand</td><td>MM</td><td>target_seq</td><td>PAM</td><td>alignment</td><td>distance</td><td>position</td><td>gene name</td><td>gene id</td></tr>
<tr><td>GU305914.1</td><td>100099</td><td>100121</td><td>-</td><td>0</td><td>CTGTGTCTGTCGGCGCAAGC</td><td>TGG</td><td>||||||||[||||||||||||]PAM</td><td>NA</td><td>Intergenic</td><td>NA</td><td>NA</td></tr>
<tr><td>T26441</td><td>TGTGTCTGTCGGCGCAAGCTGGG</td><td>1000</td><td></td><td></td><td></td><td></td><td></td><td></td><td></td><td></td><td></td></tr>
<tr><td>Oligo fwd</td><td>CACCTGTGTCTGTCGGCGCAAGCT</td><td></td><td></td><td></td><td></td><td></td><td></td><td></td><td></td><td></td><td></td></tr>
<tr><td>Oligo rev</td><td>AAACAGCTTGCGCCGACAGACACA</td><td></td><td></td><td></td><td></td><td></td><td></td><td></td><td></td><td></td><td></td></tr>
<tr><td>Chromosome</td><td>start</td><td>end</td><td>strand</td><td>MM</td><td>target_seq</td><td>PAM</td><td>alignment</td><td>distance</td><td>position</td><td>gene name</td><td>gene id</td></tr>
<tr><td>GU305914.1</td><td>100098</td><td>100120</td><td>-</td><td>0</td><td>TGTGTCTGTCGGCGCAAGCT</td><td>GGG</td><td>||||||||[||||||||||||]PAM</td><td>NA</td><td>Intergenic</td><td>NA</td><td>NA</td></tr>
<tr><td>T26442</td><td>GTCGGCGCAAGCTGGGCCTGAGG</td><td>1000</td><td></td><td></td><td></td><td></td><td></td><td></td><td></td><td></td><td></td></tr>
<tr><td>Oligo fwd</td><td>CACCGTCGGCGCAAGCTGGGCCTG</td><td></td><td></td><td></td><td></td><td></td><td></td><td></td><td></td><td></td><td></td></tr>
<tr><td>Oligo rev</td><td>AAACCAGGCCCAGCTTGCGCCGAC</td><td></td><td></td><td></td><td></td><td></td><td></td><td></td><td></td><td></td><td></td></tr>
<tr><td>Chromosome</td><td>start</td><td>end</td><td>strand</td><td>MM</td><td>target_seq</td><td>PAM</td><td>alignment</td><td>distance</td><td>position</td><td>gene name</td><td>gene id</td></tr>
<tr><td>GU305914.1</td><td>100091</td><td>100113</td><td>-</td><td>0</td><td>GTCGGCGCAAGCTGGGCCTG</td><td>AGG</td><td>||||||||[||||||||||||]PAM</td><td>NA</td><td>Intergenic</td><td>NA</td><td>NA</td></tr>
<tr><td>T26443</td><td>TCGGCGCAAGCTGGGCCTGAGGG</td><td>1000</td><td></td><td></td><td></td><td></td><td></td><td></td><td></td><td></td><td></td></tr>
<tr><td>Oligo fwd</td><td>CACCTCGGCGCAAGCTGGGCCTGA</td><td></td><td></td><td></td><td></td><td></td><td></td><td></td><td></td><td></td><td></td></tr>
<tr><td>Oligo rev</td><td>AAACTCAGGCCCAGCTTGCGCCGA</td><td></td><td></td><td></td><td></td><td></td><td></td><td></td><td></td><td></td><td></td></tr>
<tr><td>Chromosome</td><td>start</td><td>end</td><td>strand</td><td>MM</td><td>target_seq</td><td>PAM</td><td>alignment</td><td>distance</td><td>position</td><td>gene name</td><td>gene id</td></tr>
<tr><td>GU305914.1</td><td>100090</td><td>100112</td><td>-</td><td>0</td><td>TCGGCGCAAGCTGGGCCTGA</td><td>GGG</td><td>||||||||[||||||||||||]PAM</td><td>NA</td><td>Intergenic</td><td>NA</td><td>NA</td></tr>
<tr><td>T26444</td><td>CATCCCTTTCCCCCCTCGCACGG</td><td>1000</td><td></td><td></td><td></td><td></td><td></td><td></td><td></td><td></td><td></td></tr>
<tr><td>Oligo fwd</td><td>CACCCATCCCTTTCCCCCCTCGCA</td><td></td><td></td><td></td><td></td><td></td><td></td><td></td><td></td><td></td><td></td></tr>
<tr><td>Oligo rev</td><td>AAACTGCGAGGGGGGAAAGGGATG</td><td></td><td></td><td></td><td></td><td></td><td></td><td></td><td></td><td></td><td></td></tr>
<tr><td>Chromosome</td><td>start</td><td>end</td><td>strand</td><td>MM</td><td>target_seq</td><td>PAM</td><td>alignment</td><td>distance</td><td>position</td><td>gene name</td><td>gene id</td></tr>
<tr><td>GU305914.1</td><td>100063</td><td>100085</td><td>-</td><td>0</td><td>CATCCCTTTCCCCCCTCGCA</td><td>CGG</td><td>||||||||[||||||||||||]PAM</td><td>NA</td><td>Intergenic</td><td>NA</td><td>NA</td></tr>
<tr><td>T26445</td><td>CCCTTTCCCCCCTCGCACGGCGG</td><td>1000</td><td></td><td></td><td></td><td></td><td></td><td></td><td></td><td></td><td></td></tr>
<tr><td>Oligo fwd</td><td>CACCCCCTTTCCCCCCTCGCACGG</td><td></td><td></td><td></td><td></td><td></td><td></td><td></td><td></td><td></td><td></td></tr>
<tr><td>Oligo rev</td><td>AAACCCGTGCGAGGGGGGAAAGGG</td><td></td><td></td><td></td><td></td><td></td><td></td><td></td><td></td><td></td><td></td></tr>
<tr><td>Chromosome</td><td>start</td><td>end</td><td>strand</td><td>MM</td><td>target_seq</td><td>PAM</td><td>alignment</td><td>distance</td><td>position</td><td>gene name</td><td>gene id</td></tr>
<tr><td>GU305914.1</td><td>100060</td><td>100082</td><td>-</td><td>0</td><td>CCCTTTCCCCCCTCGCACGG</td><td>CGG</td><td>||||||||[||||||||||||]PAM</td><td>NA</td><td>Intergenic</td><td>NA</td><td>NA</td></tr>
<tr><td>T26446</td><td>CCCCCTCGCACGGCGGCGATCGG</td><td>1000</td><td></td><td></td><td></td><td></td><td></td><td></td><td></td><td></td><td></td></tr>
<tr><td>Oligo fwd</td><td>CACCCCCCCTCGCACGGCGGCGAT</td><td></td><td></td><td></td><td></td><td></td><td></td><td></td><td></td><td></td><td></td></tr>
<tr><td>Oligo rev</td><td>AAACATCGCCGCCGTGCGAGGGGG</td><td></td><td></td><td></td><td></td><td></td><td></td><td></td><td></td><td></td><td></td></tr>
<tr><td>Chromosome</td><td>start</td><td>end</td><td>strand</td><td>MM</td><td>target_seq</td><td>PAM</td><td>alignment</td><td>distance</td><td>position</td><td>gene name</td><td>gene id</td></tr>
<tr><td>GU305914.1</td><td>100053</td><td>100075</td><td>-</td><td>0</td><td>CCCCCTCGCACGGCGGCGAT</td><td>CGG</td><td>||||||||[||||||||||||]PAM</td><td>NA</td><td>Intergenic</td><td>NA</td><td>NA</td></tr>
<tr><td>T26447</td><td>CCCCTCGCACGGCGGCGATCGGG</td><td>1000</td><td></td><td></td><td></td><td></td><td></td><td></td><td></td><td></td><td></td></tr>
<tr><td>Oligo fwd</td><td>CACCCCCCTCGCACGGCGGCGATC</td><td></td><td></td><td></td><td></td><td></td><td></td><td></td><td></td><td></td><td></td></tr>
<tr><td>Oligo rev</td><td>AAACGATCGCCGCCGTGCGAGGGG</td><td></td><td></td><td></td><td></td><td></td><td></td><td></td><td></td><td></td><td></td></tr>
<tr><td>Chromosome</td><td>start</td><td>end</td><td>strand</td><td>MM</td><td>target_seq</td><td>PAM</td><td>alignment</td><td>distance</td><td>position</td><td>gene name</td><td>gene id</td></tr>
<tr><td>GU305914.1</td><td>100052</td><td>100074</td><td>-</td><td>0</td><td>CCCCTCGCACGGCGGCGATC</td><td>GGG</td><td>||||||||[||||||||||||]PAM</td><td>NA</td><td>Intergenic</td><td>NA</td><td>NA</td></tr>
<tr><td>T26448</td><td>CCCTCGCACGGCGGCGATCGGGG</td><td>1000</td><td></td><td></td><td></td><td></td><td></td><td></td><td></td><td></td><td></td></tr>
<tr><td>Oligo fwd</td><td>CACCCCCTCGCACGGCGGCGATCG</td><td></td><td></td><td></td><td></td><td></td><td></td><td></td><td></td><td></td><td></td></tr>
<tr><td>Oligo rev</td><td>AAACCGATCGCCGCCGTGCGAGGG</td><td></td><td></td><td></td><td></td><td></td><td></td><td></td><td></td><td></td><td></td></tr>
<tr><td>Chromosome</td><td>start</td><td>end</td><td>strand</td><td>MM</td><td>target_seq</td><td>PAM</td><td>alignment</td><td>distance</td><td>position</td><td>gene name</td><td>gene id</td></tr>
<tr><td>GU305914.1</td><td>100051</td><td>100073</td><td>-</td><td>0</td><td>CCCTCGCACGGCGGCGATCG</td><td>GGG</td><td>||||||||[||||||||||||]PAM</td><td>NA</td><td>Intergenic</td><td>NA</td><td>NA</td></tr>
<tr><td>T26449</td><td>GGCGCAGACGCTGAAACGCCTGG</td><td>1000</td><td></td><td></td><td></td><td></td><td></td><td></td><td></td><td></td><td></td></tr>
<tr><td>Oligo fwd</td><td>CACCGGCGCAGACGCTGAAACGCC</td><td></td><td></td><td></td><td></td><td></td><td></td><td></td><td></td><td></td><td></td></tr>
<tr><td>Oligo rev</td><td>AAACGGCGTTTCAGCGTCTGCGCC</td><td></td><td></td><td></td><td></td><td></td><td></td><td></td><td></td><td></td><td></td></tr>
<tr><td>Chromosome</td><td>start</td><td>end</td><td>strand</td><td>MM</td><td>target_seq</td><td>PAM</td><td>alignment</td><td>distance</td><td>position</td><td>gene name</td><td>gene id</td></tr>
<tr><td>GU305914.1</td><td>100030</td><td>100052</td><td>-</td><td>0</td><td>GGCGCAGACGCTGAAACGCC</td><td>TGG</td><td>||||||||[||||||||||||]PAM</td><td>NA</td><td>Intergenic</td><td>NA</td><td>NA</td></tr>
<tr><td>T26450</td><td>CAGACGCTGAAACGCCTGGCCGG</td><td>1000</td><td></td><td></td><td></td><td></td><td></td><td></td><td></td><td></td><td></td></tr>
<tr><td>Oligo fwd</td><td>CACCCAGACGCTGAAACGCCTGGC</td><td></td><td></td><td></td><td></td><td></td><td></td><td></td><td></td><td></td><td></td></tr>
<tr><td>Oligo rev</td><td>AAACGCCAGGCGTTTCAGCGTCTG</td><td></td><td></td><td></td><td></td><td></td><td></td><td></td><td></td><td></td><td></td></tr>
<tr><td>Chromosome</td><td>start</td><td>end</td><td>strand</td><td>MM</td><td>target_seq</td><td>PAM</td><td>alignment</td><td>distance</td><td>position</td><td>gene name</td><td>gene id</td></tr>
<tr><td>GU305914.1</td><td>100026</td><td>100048</td><td>-</td><td>0</td><td>CAGACGCTGAAACGCCTGGC</td><td>CGG</td><td>||||||||[||||||||||||]PAM</td><td>NA</td><td>Intergenic</td><td>NA</td><td>NA</td></tr>
</table>

<table>
<tr><td>T26451</td><td>TCTGTGCCTCGATCGAGACCTGG</td><td>1000</td><td></td><td></td><td></td><td></td><td></td><td></td><td></td><td></td><td></td></tr>
<tr><td>Oligo fwd</td><td>CACCTCTGTGCCTCGATCGAGACC</td><td></td><td></td><td></td><td></td><td></td><td></td><td></td><td></td><td></td><td></td></tr>
<tr><td>Oligo rev</td><td>AAACGGTCTCGATCGAGGCACAGA</td><td></td><td></td><td></td><td></td><td></td><td></td><td></td><td></td><td></td><td></td></tr>
<tr><td>Chromosome</td><td>start</td><td>end</td><td>strand</td><td>MM</td><td>target_seq</td><td>PAM</td><td>alignment</td><td>distance</td><td>position</td><td>gene name</td><td>gene id</td></tr>
<tr><td>GU305914.1</td><td>99988</td><td>100010</td><td>-</td><td>0</td><td>TCTGTGCCTCGATCGAGACC</td><td>TGG</td><td>||||||||[||||||||||||]PAM</td><td>NA</td><td>Intergenic</td><td>NA</td><td>NA</td></tr>
<tr><td>GU305914.1</td><td>124343</td><td>124365</td><td>+</td><td>4</td><td>TCAGCGCCGCGATCGACACC</td><td>GGG</td><td>||-|-|||[-|||||||-|||]PAM</td><td>NA</td><td>Intergenic</td><td>NA</td><td>NA</td></tr>
<tr><td>T26452</td><td>CTCAACGCCATCTCGCACCCCGG</td><td>1000</td><td></td><td></td><td></td><td></td><td></td><td></td><td></td><td></td><td></td></tr>
<tr><td>Oligo fwd</td><td>CACCCTCAACGCCATCTCGCACCC</td><td></td><td></td><td></td><td></td><td></td><td></td><td></td><td></td><td></td><td></td></tr>
<tr><td>Oligo rev</td><td>AAACGGGTGCGAGATGGCGTTGAG</td><td></td><td></td><td></td><td></td><td></td><td></td><td></td><td></td><td></td><td></td></tr>
<tr><td>Chromosome</td><td>start</td><td>end</td><td>strand</td><td>MM</td><td>target_seq</td><td>PAM</td><td>alignment</td><td>distance</td><td>position</td><td>gene name</td><td>gene id</td></tr>
<tr><td>GU305914.1</td><td>99957</td><td>99979</td><td>-</td><td>0</td><td>CTCAACGCCATCTCGCACCC</td><td>CGG</td><td>||||||||[||||||||||||]PAM</td><td>NA</td><td>Intergenic</td><td>NA</td><td>NA</td></tr>
<tr><td>T26453</td><td>CCCCGGCGAGTGTTTCGACACGG</td><td>1000</td><td></td><td></td><td></td><td></td><td></td><td></td><td></td><td></td><td></td></tr>
<tr><td>Oligo fwd</td><td>CACCCCCCGGCGAGTGTTTCGACA</td><td></td><td></td><td></td><td></td><td></td><td></td><td></td><td></td><td></td><td></td></tr>
<tr><td>Oligo rev</td><td>AAACTGTCGAAACACTCGCCGGGG</td><td></td><td></td><td></td><td></td><td></td><td></td><td></td><td></td><td></td><td></td></tr>
<tr><td>Chromosome</td><td>start</td><td>end</td><td>strand</td><td>MM</td><td>target_seq</td><td>PAM</td><td>alignment</td><td>distance</td><td>position</td><td>gene name</td><td>gene id</td></tr>
<tr><td>GU305914.1</td><td>99940</td><td>99962</td><td>-</td><td>0</td><td>CCCCGGCGAGTGTTTCGACA</td><td>CGG</td><td>||||||||[||||||||||||]PAM</td><td>NA</td><td>Intergenic</td><td>NA</td><td>NA</td></tr>
<tr><td>T26454</td><td>CCCGGCGAGTGTTTCGACACGGG</td><td>1000</td><td></td><td></td><td></td><td></td><td></td><td></td><td></td><td></td><td></td></tr>
<tr><td>Oligo fwd</td><td>CACCCCCGGCGAGTGTTTCGACAC</td><td></td><td></td><td></td><td></td><td></td><td></td><td></td><td></td><td></td><td></td></tr>
<tr><td>Oligo rev</td><td>AAACGTGTCGAAACACTCGCCGGG</td><td></td><td></td><td></td><td></td><td></td><td></td><td></td><td></td><td></td><td></td></tr>
<tr><td>Chromosome</td><td>start</td><td>end</td><td>strand</td><td>MM</td><td>target_seq</td><td>PAM</td><td>alignment</td><td>distance</td><td>position</td><td>gene name</td><td>gene id</td></tr>
<tr><td>GU305914.1</td><td>99939</td><td>99961</td><td>-</td><td>0</td><td>CCCGGCGAGTGTTTCGACAC</td><td>GGG</td><td>||||||||[||||||||||||]PAM</td><td>NA</td><td>Intergenic</td><td>NA</td><td>NA</td></tr>
<tr><td>T26455</td><td>CCGGCGAGTGTTTCGACACGGGG</td><td>1000</td><td></td><td></td><td></td><td></td><td></td><td></td><td></td><td></td><td></td></tr>
<tr><td>Oligo fwd</td><td>CACCCCGGCGAGTGTTTCGACACG</td><td></td><td></td><td></td><td></td><td></td><td></td><td></td><td></td><td></td><td></td></tr>
<tr><td>Oligo rev</td><td>AAACCGTGTCGAAACACTCGCCGG</td><td></td><td></td><td></td><td></td><td></td><td></td><td></td><td></td><td></td><td></td></tr>
<tr><td>Chromosome</td><td>start</td><td>end</td><td>strand</td><td>MM</td><td>target_seq</td><td>PAM</td><td>alignment</td><td>distance</td><td>position</td><td>gene name</td><td>gene id</td></tr>
<tr><td>GU305914.1</td><td>99938</td><td>99960</td><td>-</td><td>0</td><td>CCGGCGAGTGTTTCGACACG</td><td>GGG</td><td>||||||||[||||||||||||]PAM</td><td>NA</td><td>Intergenic</td><td>NA</td><td>NA</td></tr>
<tr><td>T26456</td><td>GACACGGGGATCTACAGCCACGG</td><td>1000</td><td></td><td></td><td></td><td></td><td></td><td></td><td></td><td></td><td></td></tr>
<tr><td>Oligo fwd</td><td>CACCGACACGGGGATCTACAGCCA</td><td></td><td></td><td></td><td></td><td></td><td></td><td></td><td></td><td></td><td></td></tr>
<tr><td>Oligo rev</td><td>AAACTGGCTGTAGATCCCCGTGTC</td><td></td><td></td><td></td><td></td><td></td><td></td><td></td><td></td><td></td><td></td></tr>
<tr><td>Chromosome</td><td>start</td><td>end</td><td>strand</td><td>MM</td><td>target_seq</td><td>PAM</td><td>alignment</td><td>distance</td><td>position</td><td>gene name</td><td>gene id</td></tr>
<tr><td>GU305914.1</td><td>99924</td><td>99946</td><td>-</td><td>0</td><td>GACACGGGGATCTACAGCCA</td><td>CGG</td><td>||||||||[||||||||||||]PAM</td><td>NA</td><td>Intergenic</td><td>NA</td><td>NA</td></tr>
<tr><td>T26457</td><td>ACACGGGGATCTACAGCCACGGG</td><td>1000</td><td></td><td></td><td></td><td></td><td></td><td></td><td></td><td></td><td></td></tr>
<tr><td>Oligo fwd</td><td>CACCACACGGGGATCTACAGCCAC</td><td></td><td></td><td></td><td></td><td></td><td></td><td></td><td></td><td></td><td></td></tr>
<tr><td>Oligo rev</td><td>AAACGTGGCTGTAGATCCCCGTGT</td><td></td><td></td><td></td><td></td><td></td><td></td><td></td><td></td><td></td><td></td></tr>
<tr><td>Chromosome</td><td>start</td><td>end</td><td>strand</td><td>MM</td><td>target_seq</td><td>PAM</td><td>alignment</td><td>distance</td><td>position</td><td>gene name</td><td>gene id</td></tr>
<tr><td>GU305914.1</td><td>99923</td><td>99945</td><td>-</td><td>0</td><td>ACACGGGGATCTACAGCCAC</td><td>GGG</td><td>||||||||[||||||||||||]PAM</td><td>NA</td><td>Intergenic</td><td>NA</td><td>NA</td></tr>
<tr><td>T26458</td><td>CATGCCGCTGATGTACAAGCTGG</td><td>1000</td><td></td><td></td><td></td><td></td><td></td><td></td><td></td><td></td><td></td></tr>
<tr><td>Oligo fwd</td><td>CACCCATGCCGCTGATGTACAAGC</td><td></td><td></td><td></td><td></td><td></td><td></td><td></td><td></td><td></td><td></td></tr>
<tr><td>Oligo rev</td><td>AAACGCTTGTACATCAGCGGCATG</td><td></td><td></td><td></td><td></td><td></td><td></td><td></td><td></td><td></td><td></td></tr>
<tr><td>Chromosome</td><td>start</td><td>end</td><td>strand</td><td>MM</td><td>target_seq</td><td>PAM</td><td>alignment</td><td>distance</td><td>position</td><td>gene name</td><td>gene id</td></tr>
<tr><td>GU305914.1</td><td>99889</td><td>99911</td><td>-</td><td>0</td><td>CATGCCGCTGATGTACAAGC</td><td>TGG</td><td>||||||||[||||||||||||]PAM</td><td>NA</td><td>Intergenic</td><td>NA</td><td>NA</td></tr>
<tr><td>T26459</td><td>GCTGATGTACAAGCTGGACGAGG</td><td>1000</td><td></td><td></td><td></td><td></td><td></td><td></td><td></td><td></td><td></td></tr>
<tr><td>Oligo fwd</td><td>CACCGCTGATGTACAAGCTGGACG</td><td></td><td></td><td></td><td></td><td></td><td></td><td></td><td></td><td></td><td></td></tr>
<tr><td>Oligo rev</td><td>AAACCGTCCAGCTTGTACATCAGC</td><td></td><td></td><td></td><td></td><td></td><td></td><td></td><td></td><td></td><td></td></tr>
<tr><td>Chromosome</td><td>start</td><td>end</td><td>strand</td><td>MM</td><td>target_seq</td><td>PAM</td><td>alignment</td><td>distance</td><td>position</td><td>gene name</td><td>gene id</td></tr>
<tr><td>GU305914.1</td><td>99883</td><td>99905</td><td>-</td><td>0</td><td>GCTGATGTACAAGCTGGACG</td><td>AGG</td><td>||||||||[||||||||||||]PAM</td><td>NA</td><td>Intergenic</td><td>NA</td><td>NA</td></tr>
<tr><td>T26460</td><td>TACAAGCTGGACGAGGCCAGCGG</td><td>1000</td><td></td><td></td><td></td><td></td><td></td><td></td><td></td><td></td><td></td></tr>
<tr><td>Oligo fwd</td><td>CACCTACAAGCTGGACGAGGCCAG</td><td></td><td></td><td></td><td></td><td></td><td></td><td></td><td></td><td></td><td></td></tr>
<tr><td>Oligo rev</td><td>AAACCTGGCCTCGTCCAGCTTGTA</td><td></td><td></td><td></td><td></td><td></td><td></td><td></td><td></td><td></td><td></td></tr>
<tr><td>Chromosome</td><td>start</td><td>end</td><td>strand</td><td>MM</td><td>target_seq</td><td>PAM</td><td>alignment</td><td>distance</td><td>position</td><td>gene name</td><td>gene id</td></tr>
<tr><td>GU305914.1</td><td>99876</td><td>99898</td><td>-</td><td>0</td><td>TACAAGCTGGACGAGGCCAG</td><td>CGG</td><td>||||||||[||||||||||||]PAM</td><td>NA</td><td>Intergenic</td><td>NA</td><td>NA</td></tr>
<tr><td>T26461</td><td>ACAAGCTGGACGAGGCCAGCGGG</td><td>1000</td><td></td><td></td><td></td><td></td><td></td><td></td><td></td><td></td><td></td></tr>
<tr><td>Oligo fwd</td><td>CACCACAAGCTGGACGAGGCCAGC</td><td></td><td></td><td></td><td></td><td></td><td></td><td></td><td></td><td></td><td></td></tr>
<tr><td>Oligo rev</td><td>AAACGCTGGCCTCGTCCAGCTTGT</td><td></td><td></td><td></td><td></td><td></td><td></td><td></td><td></td><td></td><td></td></tr>
<tr><td>Chromosome</td><td>start</td><td>end</td><td>strand</td><td>MM</td><td>target_seq</td><td>PAM</td><td>alignment</td><td>distance</td><td>position</td><td>gene name</td><td>gene id</td></tr>
<tr><td>GU305914.1</td><td>99875</td><td>99897</td><td>-</td><td>0</td><td>ACAAGCTGGACGAGGCCAGC</td><td>GGG</td><td>||||||||[||||||||||||]PAM</td><td>NA</td><td>Intergenic</td><td>NA</td><td>NA</td></tr>
<tr><td>T26462</td><td>CCCATCTTCATCGTCCCCGCCGG</td><td>1000</td><td></td><td></td><td></td><td></td><td></td><td></td><td></td><td></td><td></td></tr>
<tr><td>Oligo fwd</td><td>CACCCCCATCTTCATCGTCCCCGC</td><td></td><td></td><td></td><td></td><td></td><td></td><td></td><td></td><td></td><td></td></tr>
<tr><td>Oligo rev</td><td>AAACGCGGGGACGATGAAGATGGG</td><td></td><td></td><td></td><td></td><td></td><td></td><td></td><td></td><td></td><td></td></tr>
<tr><td>Chromosome</td><td>start</td><td>end</td><td>strand</td><td>MM</td><td>target_seq</td><td>PAM</td><td>alignment</td><td>distance</td><td>position</td><td>gene name</td><td>gene id</td></tr>
<tr><td>GU305914.1</td><td>99828</td><td>99850</td><td>-</td><td>0</td><td>CCCATCTTCATCGTCCCCGC</td><td>CGG</td><td>||||||||[||||||||||||]PAM</td><td>NA</td><td>Intergenic</td><td>NA</td><td>NA</td></tr>
<tr><td>T26463</td><td>CCATCTTCATCGTCCCCGCCGGG</td><td>1000</td><td></td><td></td><td></td><td></td><td></td><td></td><td></td><td></td><td></td></tr>
<tr><td>Oligo fwd</td><td>CACCCCATCTTCATCGTCCCCGCC</td><td></td><td></td><td></td><td></td><td></td><td></td><td></td><td></td><td></td><td></td></tr>
<tr><td>Oligo rev</td><td>AAACGGCGGGGACGATGAAGATGG</td><td></td><td></td><td></td><td></td><td></td><td></td><td></td><td></td><td></td><td></td></tr>
<tr><td>Chromosome</td><td>start</td><td>end</td><td>strand</td><td>MM</td><td>target_seq</td><td>PAM</td><td>alignment</td><td>distance</td><td>position</td><td>gene name</td><td>gene id</td></tr>
<tr><td>GU305914.1</td><td>99827</td><td>99849</td><td>-</td><td>0</td><td>CCATCTTCATCGTCCCCGCC</td><td>GGG</td><td>||||||||[||||||||||||]PAM</td><td>NA</td><td>Intergenic</td><td>NA</td><td>NA</td></tr>
<tr><td>GU305914.1</td><td>39446</td><td>39468</td><td>+</td><td>4</td><td>CCACCTCCATCCTCCCCGAC</td><td>GGG</td><td>|||-||-|[|||-||||||-|]PAM</td><td>NA</td><td>Intergenic</td><td>NA</td><td>NA</td></tr>
<tr><td>T26464</td><td>CGCCGGGTATCGCGACCGTCCGG</td><td>1000</td><td></td><td></td><td></td><td></td><td></td><td></td><td></td><td></td><td></td></tr>
<tr><td>Oligo fwd</td><td>CACCCGCCGGGTATCGCGACCGTC</td><td></td><td></td><td></td><td></td><td></td><td></td><td></td><td></td><td></td><td></td></tr>
<tr><td>Oligo rev</td><td>AAACGACGGTCGCGATACCCGGCG</td><td></td><td></td><td></td><td></td><td></td><td></td><td></td><td></td><td></td><td></td></tr>
<tr><td>Chromosome</td><td>start</td><td>end</td><td>strand</td><td>MM</td><td>target_seq</td><td>PAM</td><td>alignment</td><td>distance</td><td>position</td><td>gene name</td><td>gene id</td></tr>
<tr><td>GU305914.1</td><td>99811</td><td>99833</td><td>-</td><td>0</td><td>CGCCGGGTATCGCGACCGTC</td><td>CGG</td><td>||||||||[||||||||||||]PAM</td><td>NA</td><td>Intergenic</td><td>NA</td><td>NA</td></tr>
<tr><td>T26465</td><td>CGGGTATCGCGACCGTCCGGCGG</td><td>1000</td><td></td><td></td><td></td><td></td><td></td><td></td><td></td><td></td><td></td></tr>
<tr><td>Oligo fwd</td><td>CACCCGGGTATCGCGACCGTCCGG</td><td></td><td></td><td></td><td></td><td></td><td></td><td></td><td></td><td></td><td></td></tr>
<tr><td>Oligo rev</td><td>AAACCCGGACGGTCGCGATACCCG</td><td></td><td></td><td></td><td></td><td></td><td></td><td></td><td></td><td></td><td></td></tr>
</table>

<table>
<tr><td>Chromosome</td><td>start</td><td>end</td><td>strand</td><td>MM</td><td>target_seq</td><td>PAM</td><td>alignment</td><td>distance</td><td>position</td><td>gene name</td><td>gene id</td></tr>
<tr><td>GU305914.1</td><td>99808</td><td>99830</td><td>-</td><td>0</td><td>CGGGTATCGCGACCGTCCGG</td><td>CGG</td><td>||||||||[||||||||||||]PAM</td><td>NA</td><td>Intergenic</td><td>NA</td><td>NA</td></tr>
<tr><td></td><td></td><td></td><td></td><td></td><td></td><td></td><td></td><td></td><td></td><td></td><td></td></tr>
<tr><td>T26466</td><td>CCCCAGAACCTGACGCACCACGG</td><td>1000</td><td></td><td></td><td></td><td></td><td></td><td></td><td></td><td></td><td></td></tr>
<tr><td>Oligo fwd</td><td>CACCCCCCAGAACCTGACGCACCA</td><td></td><td></td><td></td><td></td><td></td><td></td><td></td><td></td><td></td><td></td></tr>
<tr><td>Oligo rev</td><td>AAACTGGTGCGTCAGGTTCTGGGG</td><td></td><td></td><td></td><td></td><td></td><td></td><td></td><td></td><td></td><td></td></tr>
<tr><td>Chromosome</td><td>start</td><td>end</td><td>strand</td><td>MM</td><td>target_seq</td><td>PAM</td><td>alignment</td><td>distance</td><td>position</td><td>gene name</td><td>gene id</td></tr>
<tr><td>GU305914.1</td><td>99762</td><td>99784</td><td>-</td><td>0</td><td>CCCCAGAACCTGACGCACCA</td><td>CGG</td><td>||||||||[||||||||||||]PAM</td><td>NA</td><td>Intergenic</td><td>NA</td><td>NA</td></tr>
<tr><td></td><td></td><td></td><td></td><td></td><td></td><td></td><td></td><td></td><td></td><td></td><td></td></tr>
<tr><td>T26467</td><td>CGCACCACGGCCGTCCGCCGCGG</td><td>1000</td><td></td><td></td><td></td><td></td><td></td><td></td><td></td><td></td><td></td></tr>
<tr><td>Oligo fwd</td><td>CACCCGCACCACGGCCGTCCGCCG</td><td></td><td></td><td></td><td></td><td></td><td></td><td></td><td></td><td></td><td></td></tr>
<tr><td>Oligo rev</td><td>AAACCGGCGGACGGCCGTGGTGCG</td><td></td><td></td><td></td><td></td><td></td><td></td><td></td><td></td><td></td><td></td></tr>
<tr><td>Chromosome</td><td>start</td><td>end</td><td>strand</td><td>MM</td><td>target_seq</td><td>PAM</td><td>alignment</td><td>distance</td><td>position</td><td>gene name</td><td>gene id</td></tr>
<tr><td>GU305914.1</td><td>99749</td><td>99771</td><td>-</td><td>0</td><td>CGCACCACGGCCGTCCGCCG</td><td>CGG</td><td>||||||||[||||||||||||]PAM</td><td>NA</td><td>Intergenic</td><td>NA</td><td>NA</td></tr>
<tr><td></td><td></td><td></td><td></td><td></td><td></td><td></td><td></td><td></td><td></td><td></td><td></td></tr>
<tr><td>T26468</td><td>GGCCGTCCGCCGCGGCGAGACGG</td><td>1000</td><td></td><td></td><td></td><td></td><td></td><td></td><td></td><td></td><td></td></tr>
<tr><td>Oligo fwd</td><td>CACCGGCCGTCCGCCGCGGCGAGA</td><td></td><td></td><td></td><td></td><td></td><td></td><td></td><td></td><td></td><td></td></tr>
<tr><td>Oligo rev</td><td>AAACTCTCGCCGCGGCGGACGGCC</td><td></td><td></td><td></td><td></td><td></td><td></td><td></td><td></td><td></td><td></td></tr>
<tr><td>Chromosome</td><td>start</td><td>end</td><td>strand</td><td>MM</td><td>target_seq</td><td>PAM</td><td>alignment</td><td>distance</td><td>position</td><td>gene name</td><td>gene id</td></tr>
<tr><td>GU305914.1</td><td>99741</td><td>99763</td><td>-</td><td>0</td><td>GGCCGTCCGCCGCGGCGAGA</td><td>CGG</td><td>||||||||[||||||||||||]PAM</td><td>NA</td><td>Intergenic</td><td>NA</td><td>NA</td></tr>
<tr><td></td><td></td><td></td><td></td><td></td><td></td><td></td><td></td><td></td><td></td><td></td><td></td></tr>
<tr><td>T26469</td><td>GCCGTCCGCCGCGGCGAGACGGG</td><td>1000</td><td></td><td></td><td></td><td></td><td></td><td></td><td></td><td></td><td></td></tr>
<tr><td>Oligo fwd</td><td>CACCGCCGTCCGCCGCGGCGAGAC</td><td></td><td></td><td></td><td></td><td></td><td></td><td></td><td></td><td></td><td></td></tr>
<tr><td>Oligo rev</td><td>AAACGTCTCGCCGCGGCGGACGGC</td><td></td><td></td><td></td><td></td><td></td><td></td><td></td><td></td><td></td><td></td></tr>
<tr><td>Chromosome</td><td>start</td><td>end</td><td>strand</td><td>MM</td><td>target_seq</td><td>PAM</td><td>alignment</td><td>distance</td><td>position</td><td>gene name</td><td>gene id</td></tr>
<tr><td>GU305914.1</td><td>99740</td><td>99762</td><td>-</td><td>0</td><td>GCCGTCCGCCGCGGCGAGAC</td><td>GGG</td><td>||||||||[||||||||||||]PAM</td><td>NA</td><td>Intergenic</td><td>NA</td><td>NA</td></tr>
<tr><td>GU305914.1</td><td>83369</td><td>83391</td><td>-</td><td>4</td><td>GCGGTCGGCCGCCGCGAGGC</td><td>CGG</td><td>||-|||-|[||||-|||||-|]PAM</td><td>NA</td><td>Intergenic</td><td>NA</td><td>NA</td></tr>
<tr><td></td><td></td><td></td><td></td><td></td><td></td><td></td><td></td><td></td><td></td><td></td><td></td></tr>
<tr><td>T26470</td><td>CGGGAGCGCCGACCAGCTGACGG</td><td>1000</td><td></td><td></td><td></td><td></td><td></td><td></td><td></td><td></td><td></td></tr>
<tr><td>Oligo fwd</td><td>CACCCGGGAGCGCCGACCAGCTGA</td><td></td><td></td><td></td><td></td><td></td><td></td><td></td><td></td><td></td><td></td></tr>
<tr><td>Oligo rev</td><td>AAACTCAGCTGGTCGGCGCTCCCG</td><td></td><td></td><td></td><td></td><td></td><td></td><td></td><td></td><td></td><td></td></tr>
<tr><td>Chromosome</td><td>start</td><td>end</td><td>strand</td><td>MM</td><td>target_seq</td><td>PAM</td><td>alignment</td><td>distance</td><td>position</td><td>gene name</td><td>gene id</td></tr>
<tr><td>GU305914.1</td><td>99721</td><td>99743</td><td>-</td><td>0</td><td>CGGGAGCGCCGACCAGCTGA</td><td>CGG</td><td>||||||||[||||||||||||]PAM</td><td>NA</td><td>Intergenic</td><td>NA</td><td>NA</td></tr>
<tr><td></td><td></td><td></td><td></td><td></td><td></td><td></td><td></td><td></td><td></td><td></td><td></td></tr>
<tr><td>T26471</td><td>GAGCGCCGACCAGCTGACGGAGG</td><td>1000</td><td></td><td></td><td></td><td></td><td></td><td></td><td></td><td></td><td></td></tr>
<tr><td>Oligo fwd</td><td>CACCGAGCGCCGACCAGCTGACGG</td><td></td><td></td><td></td><td></td><td></td><td></td><td></td><td></td><td></td><td></td></tr>
<tr><td>Oligo rev</td><td>AAACCCGTCAGCTGGTCGGCGCTC</td><td></td><td></td><td></td><td></td><td></td><td></td><td></td><td></td><td></td><td></td></tr>
<tr><td>Chromosome</td><td>start</td><td>end</td><td>strand</td><td>MM</td><td>target_seq</td><td>PAM</td><td>alignment</td><td>distance</td><td>position</td><td>gene name</td><td>gene id</td></tr>
<tr><td>GU305914.1</td><td>99718</td><td>99740</td><td>-</td><td>0</td><td>GAGCGCCGACCAGCTGACGG</td><td>AGG</td><td>||||||||[||||||||||||]PAM</td><td>NA</td><td>Intergenic</td><td>NA</td><td>NA</td></tr>
<tr><td></td><td></td><td></td><td></td><td></td><td></td><td></td><td></td><td></td><td></td><td></td><td></td></tr>
<tr><td>T26472</td><td>CGCCGACCAGCTGACGGAGGTGG</td><td>1000</td><td></td><td></td><td></td><td></td><td></td><td></td><td></td><td></td><td></td></tr>
<tr><td>Oligo fwd</td><td>CACCCGCCGACCAGCTGACGGAGG</td><td></td><td></td><td></td><td></td><td></td><td></td><td></td><td></td><td></td><td></td></tr>
<tr><td>Oligo rev</td><td>AAACCCTCCGTCAGCTGGTCGGCG</td><td></td><td></td><td></td><td></td><td></td><td></td><td></td><td></td><td></td><td></td></tr>
<tr><td>Chromosome</td><td>start</td><td>end</td><td>strand</td><td>MM</td><td>target_seq</td><td>PAM</td><td>alignment</td><td>distance</td><td>position</td><td>gene name</td><td>gene id</td></tr>
<tr><td>GU305914.1</td><td>99715</td><td>99737</td><td>-</td><td>0</td><td>CGCCGACCAGCTGACGGAGG</td><td>TGG</td><td>||||||||[||||||||||||]PAM</td><td>NA</td><td>Intergenic</td><td>NA</td><td>NA</td></tr>
<tr><td></td><td></td><td></td><td></td><td></td><td></td><td></td><td></td><td></td><td></td><td></td><td></td></tr>
<tr><td>T26473</td><td>CGCGCATGCGCCGACAGCGACGG</td><td>1000</td><td></td><td></td><td></td><td></td><td></td><td></td><td></td><td></td><td></td></tr>
<tr><td>Oligo fwd</td><td>CACCCGCGCATGCGCCGACAGCGA</td><td></td><td></td><td></td><td></td><td></td><td></td><td></td><td></td><td></td><td></td></tr>
<tr><td>Oligo rev</td><td>AAACTCGCTGTCGGCGCATGCGCG</td><td></td><td></td><td></td><td></td><td></td><td></td><td></td><td></td><td></td><td></td></tr>
<tr><td>Chromosome</td><td>start</td><td>end</td><td>strand</td><td>MM</td><td>target_seq</td><td>PAM</td><td>alignment</td><td>distance</td><td>position</td><td>gene name</td><td>gene id</td></tr>
<tr><td>GU305914.1</td><td>99675</td><td>99697</td><td>-</td><td>0</td><td>CGCGCATGCGCCGACAGCGA</td><td>CGG</td><td>||||||||[||||||||||||]PAM</td><td>NA</td><td>Intergenic</td><td>NA</td><td>NA</td></tr>
<tr><td></td><td></td><td></td><td></td><td></td><td></td><td></td><td></td><td></td><td></td><td></td><td></td></tr>
<tr><td>T26474</td><td>ACGGCAACTTTCTGCAATCGCGG</td><td>1000</td><td></td><td></td><td></td><td></td><td></td><td></td><td></td><td></td><td></td></tr>
<tr><td>Oligo fwd</td><td>CACCACGGCAACTTTCTGCAATCG</td><td></td><td></td><td></td><td></td><td></td><td></td><td></td><td></td><td></td><td></td></tr>
<tr><td>Oligo rev</td><td>AAACCGATTGCAGAAAGTTGCCGT</td><td></td><td></td><td></td><td></td><td></td><td></td><td></td><td></td><td></td><td></td></tr>
<tr><td>Chromosome</td><td>start</td><td>end</td><td>strand</td><td>MM</td><td>target_seq</td><td>PAM</td><td>alignment</td><td>distance</td><td>position</td><td>gene name</td><td>gene id</td></tr>
<tr><td>GU305914.1</td><td>99656</td><td>99678</td><td>-</td><td>0</td><td>ACGGCAACTTTCTGCAATCG</td><td>CGG</td><td>||||||||[||||||||||||]PAM</td><td>NA</td><td>Intergenic</td><td>NA</td><td>NA</td></tr>
<tr><td></td><td></td><td></td><td></td><td></td><td></td><td></td><td></td><td></td><td></td><td></td><td></td></tr>
<tr><td>T26475</td><td>CGGCAACTTTCTGCAATCGCGGG</td><td>1000</td><td></td><td></td><td></td><td></td><td></td><td></td><td></td><td></td><td></td></tr>
<tr><td>Oligo fwd</td><td>CACCCGGCAACTTTCTGCAATCGC</td><td></td><td></td><td></td><td></td><td></td><td></td><td></td><td></td><td></td><td></td></tr>
<tr><td>Oligo rev</td><td>AAACGCGATTGCAGAAAGTTGCCG</td><td></td><td></td><td></td><td></td><td></td><td></td><td></td><td></td><td></td><td></td></tr>
<tr><td>Chromosome</td><td>start</td><td>end</td><td>strand</td><td>MM</td><td>target_seq</td><td>PAM</td><td>alignment</td><td>distance</td><td>position</td><td>gene name</td><td>gene id</td></tr>
<tr><td>GU305914.1</td><td>99655</td><td>99677</td><td>-</td><td>0</td><td>CGGCAACTTTCTGCAATCGC</td><td>GGG</td><td>||||||||[||||||||||||]PAM</td><td>NA</td><td>Intergenic</td><td>NA</td><td>NA</td></tr>
<tr><td></td><td></td><td></td><td></td><td></td><td></td><td></td><td></td><td></td><td></td><td></td><td></td></tr>
<tr><td>T26476</td><td>ACTTTCTGCAATCGCGGGCGAGG</td><td>1000</td><td></td><td></td><td></td><td></td><td></td><td></td><td></td><td></td><td></td></tr>
<tr><td>Oligo fwd</td><td>CACCACTTTCTGCAATCGCGGGCG</td><td></td><td></td><td></td><td></td><td></td><td></td><td></td><td></td><td></td><td></td></tr>
<tr><td>Oligo rev</td><td>AAACCGCCCGCGATTGCAGAAAGT</td><td></td><td></td><td></td><td></td><td></td><td></td><td></td><td></td><td></td><td></td></tr>
<tr><td>Chromosome</td><td>start</td><td>end</td><td>strand</td><td>MM</td><td>target_seq</td><td>PAM</td><td>alignment</td><td>distance</td><td>position</td><td>gene name</td><td>gene id</td></tr>
<tr><td>GU305914.1</td><td>99650</td><td>99672</td><td>-</td><td>0</td><td>ACTTTCTGCAATCGCGGGCG</td><td>AGG</td><td>||||||||[||||||||||||]PAM</td><td>NA</td><td>Intergenic</td><td>NA</td><td>NA</td></tr>
<tr><td></td><td></td><td></td><td></td><td></td><td></td><td></td><td></td><td></td><td></td><td></td><td></td></tr>
<tr><td>T26477</td><td>GGGCGAGGCGCGCGATGTCGAGG</td><td>1000</td><td></td><td></td><td></td><td></td><td></td><td></td><td></td><td></td><td></td></tr>
<tr><td>Oligo fwd</td><td>CACCGGGCGAGGCGCGCGATGTCG</td><td></td><td></td><td></td><td></td><td></td><td></td><td></td><td></td><td></td><td></td></tr>
<tr><td>Oligo rev</td><td>AAACCGACATCGCGCGCCTCGCCC</td><td></td><td></td><td></td><td></td><td></td><td></td><td></td><td></td><td></td><td></td></tr>
<tr><td>Chromosome</td><td>start</td><td>end</td><td>strand</td><td>MM</td><td>target_seq</td><td>PAM</td><td>alignment</td><td>distance</td><td>position</td><td>gene name</td><td>gene id</td></tr>
<tr><td>GU305914.1</td><td>99635</td><td>99657</td><td>-</td><td>0</td><td>GGGCGAGGCGCGCGATGTCG</td><td>AGG</td><td>||||||||[||||||||||||]PAM</td><td>NA</td><td>Intergenic</td><td>NA</td><td>NA</td></tr>
<tr><td></td><td></td><td></td><td></td><td></td><td></td><td></td><td></td><td></td><td></td><td></td><td></td></tr>
<tr><td>T26478</td><td>CGAGGCGCGCGATGTCGAGGCGG</td><td>1000</td><td></td><td></td><td></td><td></td><td></td><td></td><td></td><td></td><td></td></tr>
<tr><td>Oligo fwd</td><td>CACCCGAGGCGCGCGATGTCGAGG</td><td></td><td></td><td></td><td></td><td></td><td></td><td></td><td></td><td></td><td></td></tr>
<tr><td>Oligo rev</td><td>AAACCCTCGACATCGCGCGCCTCG</td><td></td><td></td><td></td><td></td><td></td><td></td><td></td><td></td><td></td><td></td></tr>
<tr><td>Chromosome</td><td>start</td><td>end</td><td>strand</td><td>MM</td><td>target_seq</td><td>PAM</td><td>alignment</td><td>distance</td><td>position</td><td>gene name</td><td>gene id</td></tr>
<tr><td>GU305914.1</td><td>99632</td><td>99654</td><td>-</td><td>0</td><td>CGAGGCGCGCGATGTCGAGG</td><td>CGG</td><td>||||||||[||||||||||||]PAM</td><td>NA</td><td>Intergenic</td><td>NA</td><td>NA</td></tr>
<tr><td></td><td></td><td></td><td></td><td></td><td></td><td></td><td></td><td></td><td></td><td></td><td></td></tr>
<tr><td>T26479</td><td>TCGAGGCGGCGTCTGCCGCTCGG</td><td>1000</td><td></td><td></td><td></td><td></td><td></td><td></td><td></td><td></td><td></td></tr>
<tr><td>Oligo fwd</td><td>CACCTCGAGGCGGCGTCTGCCGCT</td><td></td><td></td><td></td><td></td><td></td><td></td><td></td><td></td><td></td><td></td></tr>
<tr><td>Oligo rev</td><td>AAACAGCGGCAGACGCCGCCTCGA</td><td></td><td></td><td></td><td></td><td></td><td></td><td></td><td></td><td></td><td></td></tr>
<tr><td>Chromosome</td><td>start</td><td>end</td><td>strand</td><td>MM</td><td>target_seq</td><td>PAM</td><td>alignment</td><td>distance</td><td>position</td><td>gene name</td><td>gene id</td></tr>
<tr><td>GU305914.1</td><td>99618</td><td>99640</td><td>-</td><td>0</td><td>TCGAGGCGGCGTCTGCCGCT</td><td>CGG</td><td>||||||||[||||||||||||]PAM</td><td>NA</td><td>Intergenic</td><td>NA</td><td>NA</td></tr>
<tr><td></td><td></td><td></td><td></td><td></td><td></td><td></td><td></td><td></td><td></td><td></td><td></td></tr>
<tr><td>T26480</td><td>CGAGGCGGCGTCTGCCGCTCGGG</td><td>1000</td><td></td><td></td><td></td><td></td><td></td><td></td><td></td><td></td><td></td></tr>
</table>

Oligo fwd CACCCGAGGCGGCGTCTGCCGCTC
Oligo rev AAACGAGCGGCAGACGCCGCCTCG
Chromosome start end strand MM target_seq PAM alignment distance position gene name gene id
GU305914.1 99617 99639 - 0 CGAGGCGGCGTCTGCCGCTC GGG ||||||||[||||||||||||]PAM NA Intergenic NA NA

T26481 AGCGCACCTGTCGCTCGAGTCGG 1000
Oligo fwd CACCAGCGCACCTGTCGCTCGAGT
Oligo rev AAACACTCGAGCGACAGGTGCGCT
Chromosome start end strand MM target_seq PAM alignment distance position gene name gene id
GU305914.1 99583 99605 - 0 AGCGCACCTGTCGCTCGAGT CGG ||||||||[||||||||||||]PAM NA Intergenic NA NA

T26482 GCGCACCTGTCGCTCGAGTCGGG 1000
Oligo fwd CACCGCGCACCTGTCGCTCGAGTC
Oligo rev AAACGACTCGAGCGACAGGTGCGC
Chromosome start end strand MM target_seq PAM alignment distance position gene name gene id
GU305914.1 99582 99604 - 0 GCGCACCTGTCGCTCGAGTC GGG ||||||||[||||||||||||]PAM NA Intergenic NA NA

T26483 GTCGCTCGAGTCGGGACAGTCGG 1000
Oligo fwd CACCGTCGCTCGAGTCGGGACAGT
Oligo rev AAACACTGTCCCGACTCGAGCGAC
Chromosome start end strand MM target_seq PAM alignment distance position gene name gene id
GU305914.1 99574 99596 - 0 GTCGCTCGAGTCGGGACAGT CGG ||||||||[||||||||||||]PAM NA Intergenic NA NA
GU305914.1 77746 77768 + 4 GGCGCTCAAGACGCGACAGT CGG |-|||||-[||-||-||||||]PAM NA Intergenic NA NA
GU305914.1 83930 83952 - 4 GAAGCTCGAGTCGGGCCAGC TGG |--|||||[|||||||-|||-]PAM NA Intergenic NA NA

T26484 CCGTCGCTGCCAACGCTCGTCGG 1000
Oligo fwd CACCCCGTCGCTGCCAACGCTCGT
Oligo rev AAACACGAGCGTTGGCAGCGACGG
Chromosome start end strand MM target_seq PAM alignment distance position gene name gene id
GU305914.1 99549 99571 - 0 CCGTCGCTGCCAACGCTCGT CGG ||||||||[||||||||||||]PAM NA Intergenic NA NA

T26485 CGTCGCTGCCAACGCTCGTCGGG 1000
Oligo fwd CACCCGTCGCTGCCAACGCTCGTC
Oligo rev AAACGACGAGCGTTGGCAGCGACG
Chromosome start end strand MM target_seq PAM alignment distance position gene name gene id
GU305914.1 99548 99570 - 0 CGTCGCTGCCAACGCTCGTC GGG ||||||||[||||||||||||]PAM NA Intergenic NA NA

T26486 CTGCCAACGCTCGTCGGGCGAGG 1000
Oligo fwd CACCCTGCCAACGCTCGTCGGGCG
Oligo rev AAACCGCCCGACGAGCGTTGGCAG
Chromosome start end strand MM target_seq PAM alignment distance position gene name gene id
GU305914.1 99543 99565 - 0 CTGCCAACGCTCGTCGGGCG AGG ||||||||[||||||||||||]PAM NA Intergenic NA NA

T26487 CCAACGCTCGTCGGGCGAGGAGG 1000
Oligo fwd CACCCCAACGCTCGTCGGGCGAGG
Oligo rev AAACCCTCGCCCGACGAGCGTTGG
Chromosome start end strand MM target_seq PAM alignment distance position gene name gene id
GU305914.1 99540 99562 - 0 CCAACGCTCGTCGGGCGAGG AGG ||||||||[||||||||||||]PAM NA Intergenic NA NA
GU305914.1 88246 88268 - 4 CCGCCGCTCGTCCGGCGAGC GGG ||--||||[||||-||||||-]PAM NA Intergenic NA NA

T26488 ACGCTCGTCGGGCGAGGAGGCGG 1000
Oligo fwd CACCACGCTCGTCGGGCGAGGAGG
Oligo rev AAACCCTCCTCGCCCGACGAGCGT
Chromosome start end strand MM target_seq PAM alignment distance position gene name gene id
GU305914.1 99537 99559 - 0 ACGCTCGTCGGGCGAGGAGG CGG ||||||||[||||||||||||]PAM NA Intergenic NA NA
GU305914.1 71332 71354 + 4 CCCCTCGACGGGGGAGGAGG TGG -|-||||-[||||-|||||||]PAM NA Intergenic NA NA

T26489 CTCGTCGGGCGAGGAGGCGGAGG 1000
Oligo fwd CACCCTCGTCGGGCGAGGAGGCGG
Oligo rev AAACCCGCCTCCTCGCCCGACGAG
Chromosome start end strand MM target_seq PAM alignment distance position gene name gene id
GU305914.1 99534 99556 - 0 CTCGTCGGGCGAGGAGGCGG AGG ||||||||[||||||||||||]PAM NA Intergenic NA NA
GU305914.1 161760 161782 - 4 CCCGGAGGCCGAGGAGGCGG CGG |-||--||[-|||||||||||]PAM NA Intergenic NA NA

T26490 GGGCGAGGAGGCGGAGGAGAAGG 1000
Oligo fwd CACCGGGCGAGGAGGCGGAGGAGA
Oligo rev AAACTCTCCTCCGCCTCCTCGCCC
Chromosome start end strand MM target_seq PAM alignment distance position gene name gene id
GU305914.1 99528 99550 - 0 GGGCGAGGAGGCGGAGGAGA AGG ||||||||[||||||||||||]PAM NA Intergenic NA NA
GU305914.1 45315 45337 - 4 AGTGGAGGAGGAGGAGGAGA AGG -|--||||[|||-||||||||]PAM NA Intergenic NA NA

T26491 CGAGGAGGCGGAGGAGAAGGAGG 1000
Oligo fwd CACCCGAGGAGGCGGAGGAGAAGG
Oligo rev AAACCCTTCTCCTCCGCCTCCTCG
Chromosome start end strand MM target_seq PAM alignment distance position gene name gene id
GU305914.1 99525 99547 - 0 CGAGGAGGCGGAGGAGAAGG AGG ||||||||[||||||||||||]PAM NA Intergenic NA NA

T26492 AGGCGCGTCGAGCGATTACGAGG 1000
Oligo fwd CACCAGGCGCGTCGAGCGATTACG
Oligo rev AAACCGTAATCGCTCGACGCGCCT
Chromosome start end strand MM target_seq PAM alignment distance position gene name gene id
GU305914.1 99505 99527 - 0 AGGCGCGTCGAGCGATTACG AGG ||||||||[||||||||||||]PAM NA Intergenic NA NA

T26493 CGCGTCGAGCGATTACGAGGAGG 1000
Oligo fwd CACCCGCGTCGAGCGATTACGAGG
Oligo rev AAACCCTCGTAATCGCTCGACGCG
Chromosome start end strand MM target_seq PAM alignment distance position gene name gene id
GU305914.1 99502 99524 - 0 CGCGTCGAGCGATTACGAGG AGG ||||||||[||||||||||||]PAM NA Intergenic NA NA

| | | | | | | | | | | | |
|---|---|---|---|---|---|---|---|---|---|---|---|
| T26494 | CGAGCGATTACGAGGAGGAGAGG | 1000 | | | | | | | | | |
| Oligo fwd | CACCCGAGCGATTACGAGGAGGAG | | | | | | | | | | |
| Oligo rev | AAACCTCCTCCTCGTAATCGCTCG | | | | | | | | | | |
| Chromosome | start | end | strand | MM | target_seq | PAM | alignment | distance | position | gene name | gene id |
| GU305914.1 | 99497 | 99519 | - | 0 | CGAGCGATTACGAGGAGGAG | AGG | \|\|\|\|\|\|\|\|[\|\|\|\|\|\|\|\|\|\|\|\|]PAM | NA | Intergenic | NA | NA |
| T26495 | GAGCGATTACGAGGAGGAGAGGG | 1000 | | | | | | | | | |
| Oligo fwd | CACCGAGCGATTACGAGGAGGAGA | | | | | | | | | | |
| Oligo rev | AAACTCTCCTCCTCGTAATCGCTC | | | | | | | | | | |
| Chromosome | start | end | strand | MM | target_seq | PAM | alignment | distance | position | gene name | gene id |
| GU305914.1 | 99496 | 99518 | - | 0 | GAGCGATTACGAGGAGGAGA | GGG | \|\|\|\|\|\|\|\|[\|\|\|\|\|\|\|\|\|\|\|\|]PAM | NA | Intergenic | NA | NA |
| T26496 | CGATTACGAGGAGGAGAGGGCGG | 1000 | | | | | | | | | |
| Oligo fwd | CACCCGATTACGAGGAGGAGAGGG | | | | | | | | | | |
| Oligo rev | AAACCCCTCTCCTCCTCGTAATCG | | | | | | | | | | |
| Chromosome | start | end | strand | MM | target_seq | PAM | alignment | distance | position | gene name | gene id |
| GU305914.1 | 99493 | 99515 | - | 0 | CGATTACGAGGAGGAGAGGG | CGG | \|\|\|\|\|\|\|\|[\|\|\|\|\|\|\|\|\|\|\|\|]PAM | NA | Intergenic | NA | NA |
| T26497 | TTACGAGGAGGAGAGGGCGGTGG | 1000 | | | | | | | | | |
| Oligo fwd | CACCTTACGAGGAGGAGAGGGCGG | | | | | | | | | | |
| Oligo rev | AAACCCGCCCTCTCCTCCTCGTAA | | | | | | | | | | |
| Chromosome | start | end | strand | MM | target_seq | PAM | alignment | distance | position | gene name | gene id |
| GU305914.1 | 99490 | 99512 | - | 0 | TTACGAGGAGGAGAGGGCGG | TGG | \|\|\|\|\|\|\|\|[\|\|\|\|\|\|\|\|\|\|\|\|]PAM | NA | Intergenic | NA | NA |
| GU305914.1 | 6142 | 6164 | - | 4 | TGACGAGTAGGAGCTGGCGG | CGG | \|-\|\|\|\|\|-[\|\|\|\|\|--\|\|\|\|\|]PAM | NA | Intergenic | NA | NA |
| GU305914.1 | 201602 | 201624 | - | 4 | GTCCGAGGAGGAGACCGCGG | TGG | -\|-\|\|\|\|\|[\|\|\|\|\|\|--\|\|\|\|]PAM | NA | Intergenic | NA | NA |
| T26498 | TACGAGGAGGAGAGGGCGGTGGG | 1000 | | | | | | | | | |
| Oligo fwd | CACCTACGAGGAGGAGAGGGCGGT | | | | | | | | | | |
| Oligo rev | AAACACCGCCCTCTCCTCCTCGTA | | | | | | | | | | |
| Chromosome | start | end | strand | MM | target_seq | PAM | alignment | distance | position | gene name | gene id |
| GU305914.1 | 99489 | 99511 | - | 0 | TACGAGGAGGAGAGGGCGGT | GGG | \|\|\|\|\|\|\|\|[\|\|\|\|\|\|\|\|\|\|\|\|]PAM | NA | Intergenic | NA | NA |
| T26499 | GGAGGAGAGGGCGGTGGGATCGG | 1000 | | | | | | | | | |
| Oligo fwd | CACCGGAGGAGAGGGCGGTGGGAT | | | | | | | | | | |
| Oligo rev | AAACATCCCACCGCCCTCTCCTCC | | | | | | | | | | |
| Chromosome | start | end | strand | MM | target_seq | PAM | alignment | distance | position | gene name | gene id |
| GU305914.1 | 99484 | 99506 | - | 0 | GGAGGAGAGGGCGGTGGGAT | CGG | \|\|\|\|\|\|\|\|[\|\|\|\|\|\|\|\|\|\|\|\|]PAM | NA | Intergenic | NA | NA |
| T26500 | GAGGGCGGTGGGATCGGACGAGG | 1000 | | | | | | | | | |
| Oligo fwd | CACCGAGGGCGGTGGGATCGGACG | | | | | | | | | | |
| Oligo rev | AAACCGTCCGATCCCACCGCCCTC | | | | | | | | | | |
| Chromosome | start | end | strand | MM | target_seq | PAM | alignment | distance | position | gene name | gene id |
| GU305914.1 | 99478 | 99500 | - | 0 | GAGGGCGGTGGGATCGGACG | AGG | \|\|\|\|\|\|\|\|[\|\|\|\|\|\|\|\|\|\|\|\|]PAM | NA | Intergenic | NA | NA |
| T26501 | GGCGGTGGGATCGGACGAGGAGG | 1000 | | | | | | | | | |
| Oligo fwd | CACCGGCGGTGGGATCGGACGAGG | | | | | | | | | | |
| Oligo rev | AAACCCTCGTCCGATCCCACCGCC | | | | | | | | | | |
| Chromosome | start | end | strand | MM | target_seq | PAM | alignment | distance | position | gene name | gene id |
| GU305914.1 | 99475 | 99497 | - | 0 | GGCGGTGGGATCGGACGAGG | AGG | \|\|\|\|\|\|\|\|[\|\|\|\|\|\|\|\|\|\|\|\|]PAM | NA | Intergenic | NA | NA |
| T26502 | GGTGGGATCGGACGAGGAGGAGG | 1000 | | | | | | | | | |
| Oligo fwd | CACCGGTGGGATCGGACGAGGAGG | | | | | | | | | | |
| Oligo rev | AAACCCTCCTCGTCCGATCCCACC | | | | | | | | | | |
| Chromosome | start | end | strand | MM | target_seq | PAM | alignment | distance | position | gene name | gene id |
| GU305914.1 | 99472 | 99494 | - | 0 | GGTGGGATCGGACGAGGAGG | AGG | \|\|\|\|\|\|\|\|[\|\|\|\|\|\|\|\|\|\|\|\|]PAM | NA | Intergenic | NA | NA |
| T26503 | CGACGACGTCGAGAACCTGCAGG | 1000 | | | | | | | | | |
| Oligo fwd | CACCCGACGACGTCGAGAACCTGC | | | | | | | | | | |
| Oligo rev | AAACGCAGGTTCTCGACGTCGTCG | | | | | | | | | | |
| Chromosome | start | end | strand | MM | target_seq | PAM | alignment | distance | position | gene name | gene id |
| GU305914.1 | 99445 | 99467 | - | 0 | CGACGACGTCGAGAACCTGC | AGG | \|\|\|\|\|\|\|\|[\|\|\|\|\|\|\|\|\|\|\|\|]PAM | NA | Intergenic | NA | NA |
| T26504 | AGAACCTGCAGGCGTTCGCGAGG | 1000 | | | | | | | | | |
| Oligo fwd | CACCAGAACCTGCAGGCGTTCGCG | | | | | | | | | | |
| Oligo rev | AAACCGCGAACGCCTGCAGGTTCT | | | | | | | | | | |
| Chromosome | start | end | strand | MM | target_seq | PAM | alignment | distance | position | gene name | gene id |
| GU305914.1 | 99434 | 99456 | - | 0 | AGAACCTGCAGGCGTTCGCG | AGG | \|\|\|\|\|\|\|\|[\|\|\|\|\|\|\|\|\|\|\|\|]PAM | NA | Intergenic | NA | NA |
| T26505 | ACCTGCAGGCGTTCGCGAGGCGG | 1000 | | | | | | | | | |
| Oligo fwd | CACCACCTGCAGGCGTTCGCGAGG | | | | | | | | | | |
| Oligo rev | AAACCCTCGCGAACGCCTGCAGGT | | | | | | | | | | |
| Chromosome | start | end | strand | MM | target_seq | PAM | alignment | distance | position | gene name | gene id |
| GU305914.1 | 99431 | 99453 | - | 0 | ACCTGCAGGCGTTCGCGAGG | CGG | \|\|\|\|\|\|\|\|[\|\|\|\|\|\|\|\|\|\|\|\|]PAM | NA | Intergenic | NA | NA |
| T26506 | CGTTCGCGAGGCGGATCGCCTGG | 1000 | | | | | | | | | |
| Oligo fwd | CACCCGTTCGCGAGGCGGATCGCC | | | | | | | | | | |
| Oligo rev | AAACGGCGATCCGCCTCGCGAACG | | | | | | | | | | |
| Chromosome | start | end | strand | MM | target_seq | PAM | alignment | distance | position | gene name | gene id |
| GU305914.1 | 99422 | 99444 | - | 0 | CGTTCGCGAGGCGGATCGCC | TGG | \|\|\|\|\|\|\|\|[\|\|\|\|\|\|\|\|\|\|\|\|]PAM | NA | Intergenic | NA | NA |
| T26507 | CGCGAGGCGGATCGCCTGGCCGG | 1000 | | | | | | | | | |
| Oligo fwd | CACCCGCGAGGCGGATCGCCTGGC | | | | | | | | | | |
| Oligo rev | AAACGCCAGGCGATCCGCCTCGCG | | | | | | | | | | |
| Chromosome | start | end | strand | MM | target_seq | PAM | alignment | distance | position | gene name | gene id |
| GU305914.1 | 99418 | 99440 | - | 0 | CGCGAGGCGGATCGCCTGGC | CGG | \|\|\|\|\|\|\|\|[\|\|\|\|\|\|\|\|\|\|\|\|]PAM | NA | Intergenic | NA | NA |
| T26508 | CACGCGTAACCACTACCGCGAGG | 1000 | | | | | | | | | |
| Oligo fwd | CACCCACGCGTAACCACTACCGCG | | | | | | | | | | |
| Oligo rev | AAACCGCGGTAGTGGTTACGCGTG | | | | | | | | | | |

Chromosome start end strand MM target_seq PAM alignment distance position gene name gene id
GU305914.1 99382 99404 - 0 CACGCGTAACCACTACCGCG AGG ||||||||[||||||||||||]PAM NA Intergenic NA NA

T26509 GCGTAACCACTACCGCGAGGAGG 1000
Oligo fwd CACCGCGTAACCACTACCGCGAGG
Oligo rev AAACCCTCGCGGTAGTGGTTACGC
Chromosome start end strand MM target_seq PAM alignment distance position gene name gene id
GU305914.1 99379 99401 - 0 GCGTAACCACTACCGCGAGG AGG ||||||||[||||||||||||]PAM NA Intergenic NA NA

T26510 CGAGGAGGTGCAGCAGCAGCTGG 1000
Oligo fwd CACCCGAGGAGGTGCAGCAGCAGC
Oligo rev AAACGCTGCTGCTGCACCTCCTCG
Chromosome start end strand MM target_seq PAM alignment distance position gene name gene id
GU305914.1 99364 99386 - 0 CGAGGAGGTGCAGCAGCAGC TGG ||||||||[||||||||||||]PAM NA Intergenic NA NA

T26511 GGAGGTGCAGCAGCAGCTGGAGG 1000
Oligo fwd CACCGGAGGTGCAGCAGCAGCTGG
Oligo rev AAACCCAGCTGCTGCTGCACCTCC
Chromosome start end strand MM target_seq PAM alignment distance position gene name gene id
GU305914.1 99361 99383 - 0 GGAGGTGCAGCAGCAGCTGG AGG ||||||||[||||||||||||]PAM NA Intergenic NA NA

T26512 GGTGCAGCAGCAGCTGGAGGCGG 1000
Oligo fwd CACCGGTGCAGCAGCAGCTGGAGG
Oligo rev AAACCCTCCAGCTGCTGCTGCACC
Chromosome start end strand MM target_seq PAM alignment distance position gene name gene id
GU305914.1 99358 99380 - 0 GGTGCAGCAGCAGCTGGAGG CGG ||||||||[||||||||||||]PAM NA Intergenic NA NA

T26513 GCAGCAGCTGGAGGCGGCCACGG 1000
Oligo fwd CACCGCAGCAGCTGGAGGCGGCCA
Oligo rev AAACTGGCCGCCTCCAGCTGCTGC
Chromosome start end strand MM target_seq PAM alignment distance position gene name gene id
GU305914.1 99352 99374 - 0 GCAGCAGCTGGAGGCGGCCA CGG ||||||||[||||||||||||]PAM NA Intergenic NA NA

T26514 GCCACGGTGTTTACCGCCGTCGG 1000
Oligo fwd CACCGCCACGGTGTTTACCGCCGT
Oligo rev AAACACGGCGGTAAACACCGTGGC
Chromosome start end strand MM target_seq PAM alignment distance position gene name gene id
GU305914.1 99336 99358 - 0 GCCACGGTGTTTACCGCCGT CGG ||||||||[||||||||||||]PAM NA Intergenic NA NA
GU305914.1 138274 138296 - 4 ACCACGGAGTTTCCCGACGT GGG -||||||-[||||-|||-|||]PAM NA Intergenic NA NA

T26515 GCACCTGCGTTGCCGTCAAACGG 1000
Oligo fwd CACCGCACCTGCGTTGCCGTCAAA
Oligo rev AAACTTTGACGGCAACGCAGGTGC
Chromosome start end strand MM target_seq PAM alignment distance position gene name gene id
GU305914.1 99311 99333 - 0 GCACCTGCGTTGCCGTCAAA CGG ||||||||[||||||||||||]PAM NA Intergenic NA NA

T26516 CACCTGCGTTGCCGTCAAACGGG 1000
Oligo fwd CACCCACCTGCGTTGCCGTCAAAC
Oligo rev AAACGTTTGACGGCAACGCAGGTG
Chromosome start end strand MM target_seq PAM alignment distance position gene name gene id
GU305914.1 99310 99332 - 0 CACCTGCGTTGCCGTCAAAC GGG ||||||||[||||||||||||]PAM NA Intergenic NA NA

T26517 ACCTGCGTTGCCGTCAAACGGGG 1000
Oligo fwd CACCACCTGCGTTGCCGTCAAACG
Oligo rev AAACCGTTTGACGGCAACGCAGGT
Chromosome start end strand MM target_seq PAM alignment distance position gene name gene id
GU305914.1 99309 99331 - 0 ACCTGCGTTGCCGTCAAACG GGG ||||||||[||||||||||||]PAM NA Intergenic NA NA

T26518 GCCGTCAAACGGGGTTTATACGG 1000
Oligo fwd CACCGCCGTCAAACGGGGTTTATA
Oligo rev AAACTATAAACCCCGTTTGACGGC
Chromosome start end strand MM target_seq PAM alignment distance position gene name gene id
GU305914.1 99300 99322 - 0 GCCGTCAAACGGGGTTTATA CGG ||||||||[||||||||||||]PAM NA Intergenic NA NA

T26519 GCGCGAGCACTACACTCGCCAGG 1000
Oligo fwd CACCGCGCGAGCACTACACTCGCC
Oligo rev AAACGGCGAGTGTAGTGCTCGCGC
Chromosome start end strand MM target_seq PAM alignment distance position gene name gene id
GU305914.1 99250 99272 - 0 GCGCGAGCACTACACTCGCC AGG ||||||||[||||||||||||]PAM NA Intergenic NA NA

T26520 GCACTACACTCGCCAGGAGACGG 1000
Oligo fwd CACCGCACTACACTCGCCAGGAGA
Oligo rev AAACTCTCCTGGCGAGTGTAGTGC
Chromosome start end strand MM target_seq PAM alignment distance position gene name gene id
GU305914.1 99244 99266 - 0 GCACTACACTCGCCAGGAGA CGG ||||||||[||||||||||||]PAM NA Intergenic NA NA

T26521 CACTCGCCAGGAGACGGTGCAGG 1000
Oligo fwd CACCCACTCGCCAGGAGACGGTGC
Oligo rev AAACGCACCGTCTCCTGGCGAGTG
Chromosome start end strand MM target_seq PAM alignment distance position gene name gene id
GU305914.1 99238 99260 - 0 CACTCGCCAGGAGACGGTGC AGG ||||||||[||||||||||||]PAM NA Intergenic NA NA

T26522 GGAGACGGTGCAGGTATTCCTGG 1000
Oligo fwd CACCGGAGACGGTGCAGGTATTCC
Oligo rev AAACGGAATACCTGCACCGTCTCC
Chromosome start end strand MM target_seq PAM alignment distance position gene name gene id
GU305914.1 99229 99251 - 0 GGAGACGGTGCAGGTATTCC TGG ||||||||[||||||||||||]PAM NA Intergenic NA NA

T26523 CAGGTATTCCTGGACATCCGCGG 1000

<table>
<tr><td>Oligo fwd</td><td>CACCCAGGTATTCCTGGACATCCG</td><td></td><td></td><td></td><td></td><td></td><td></td><td></td><td></td><td></td><td></td></tr>
<tr><td>Oligo rev</td><td>AAACCGGATGTCCAGGAATACCTG</td><td></td><td></td><td></td><td></td><td></td><td></td><td></td><td></td><td></td><td></td></tr>
<tr><td>Chromosome</td><td>start</td><td>end</td><td>strand</td><td>MM</td><td>target_seq</td><td>PAM</td><td>alignment</td><td>distance</td><td>position</td><td>gene name</td><td>gene id</td></tr>
<tr><td>GU305914.1</td><td>99219</td><td>99241</td><td>-</td><td>0</td><td>CAGGTATTCCTGGACATCCG</td><td>CGG</td><td>||||||||[||||||||||||]PAM</td><td>NA</td><td>Intergenic</td><td>NA</td><td>NA</td></tr>
<tr><td></td><td></td><td></td><td></td><td></td><td></td><td></td><td></td><td></td><td></td><td></td><td></td></tr>
<tr><td>T26524</td><td>TGGACATCCGCGGAGACCAGCGG</td><td>1000</td><td></td><td></td><td></td><td></td><td></td><td></td><td></td><td></td><td></td></tr>
<tr><td>Oligo fwd</td><td>CACCTGGACATCCGCGGAGACCAG</td><td></td><td></td><td></td><td></td><td></td><td></td><td></td><td></td><td></td><td></td></tr>
<tr><td>Oligo rev</td><td>AAACCTGGTCTCCGCGGATGTCCA</td><td></td><td></td><td></td><td></td><td></td><td></td><td></td><td></td><td></td><td></td></tr>
<tr><td>Chromosome</td><td>start</td><td>end</td><td>strand</td><td>MM</td><td>target_seq</td><td>PAM</td><td>alignment</td><td>distance</td><td>position</td><td>gene name</td><td>gene id</td></tr>
<tr><td>GU305914.1</td><td>99209</td><td>99231</td><td>-</td><td>0</td><td>TGGACATCCGCGGAGACCAG</td><td>CGG</td><td>||||||||[||||||||||||]PAM</td><td>NA</td><td>Intergenic</td><td>NA</td><td>NA</td></tr>
<tr><td></td><td></td><td></td><td></td><td></td><td></td><td></td><td></td><td></td><td></td><td></td><td></td></tr>
<tr><td>T26525</td><td>ACATCCGCGGAGACCAGCGGCGG</td><td>1000</td><td></td><td></td><td></td><td></td><td></td><td></td><td></td><td></td><td></td></tr>
<tr><td>Oligo fwd</td><td>CACCACATCCGCGGAGACCAGCGG</td><td></td><td></td><td></td><td></td><td></td><td></td><td></td><td></td><td></td><td></td></tr>
<tr><td>Oligo rev</td><td>AAACCCGCTGGTCTCCGCGGATGT</td><td></td><td></td><td></td><td></td><td></td><td></td><td></td><td></td><td></td><td></td></tr>
<tr><td>Chromosome</td><td>start</td><td>end</td><td>strand</td><td>MM</td><td>target_seq</td><td>PAM</td><td>alignment</td><td>distance</td><td>position</td><td>gene name</td><td>gene id</td></tr>
<tr><td>GU305914.1</td><td>99206</td><td>99228</td><td>-</td><td>0</td><td>ACATCCGCGGAGACCAGCGG</td><td>CGG</td><td>||||||||[||||||||||||]PAM</td><td>NA</td><td>Intergenic</td><td>NA</td><td>NA</td></tr>
<tr><td></td><td></td><td></td><td></td><td></td><td></td><td></td><td></td><td></td><td></td><td></td><td></td></tr>
<tr><td>T26526</td><td>GAGACCAGCGGCGGAACGTGTGG</td><td>1000</td><td></td><td></td><td></td><td></td><td></td><td></td><td></td><td></td><td></td></tr>
<tr><td>Oligo fwd</td><td>CACCGAGACCAGCGGCGGAACGTG</td><td></td><td></td><td></td><td></td><td></td><td></td><td></td><td></td><td></td><td></td></tr>
<tr><td>Oligo rev</td><td>AAACCACGTTCCGCCGCTGGTCTC</td><td></td><td></td><td></td><td></td><td></td><td></td><td></td><td></td><td></td><td></td></tr>
<tr><td>Chromosome</td><td>start</td><td>end</td><td>strand</td><td>MM</td><td>target_seq</td><td>PAM</td><td>alignment</td><td>distance</td><td>position</td><td>gene name</td><td>gene id</td></tr>
<tr><td>GU305914.1</td><td>99197</td><td>99219</td><td>-</td><td>0</td><td>GAGACCAGCGGCGGAACGTG</td><td>TGG</td><td>||||||||[||||||||||||]PAM</td><td>NA</td><td>Intergenic</td><td>NA</td><td>NA</td></tr>
<tr><td></td><td></td><td></td><td></td><td></td><td></td><td></td><td></td><td></td><td></td><td></td><td></td></tr>
<tr><td>T26527</td><td>AGACCAGCGGCGGAACGTGTGGG</td><td>1000</td><td></td><td></td><td></td><td></td><td></td><td></td><td></td><td></td><td></td></tr>
<tr><td>Oligo fwd</td><td>CACCAGACCAGCGGCGGAACGTGT</td><td></td><td></td><td></td><td></td><td></td><td></td><td></td><td></td><td></td><td></td></tr>
<tr><td>Oligo rev</td><td>AAACACACGTTCCGCCGCTGGTCT</td><td></td><td></td><td></td><td></td><td></td><td></td><td></td><td></td><td></td><td></td></tr>
<tr><td>Chromosome</td><td>start</td><td>end</td><td>strand</td><td>MM</td><td>target_seq</td><td>PAM</td><td>alignment</td><td>distance</td><td>position</td><td>gene name</td><td>gene id</td></tr>
<tr><td>GU305914.1</td><td>99196</td><td>99218</td><td>-</td><td>0</td><td>AGACCAGCGGCGGAACGTGT</td><td>GGG</td><td>||||||||[||||||||||||]PAM</td><td>NA</td><td>Intergenic</td><td>NA</td><td>NA</td></tr>
<tr><td></td><td></td><td></td><td></td><td></td><td></td><td></td><td></td><td></td><td></td><td></td><td></td></tr>
<tr><td>T26528</td><td>GGAACGTGTGGGCGACCCTGTGG</td><td>1000</td><td></td><td></td><td></td><td></td><td></td><td></td><td></td><td></td><td></td></tr>
<tr><td>Oligo fwd</td><td>CACCGGAACGTGTGGGCGACCCTG</td><td></td><td></td><td></td><td></td><td></td><td></td><td></td><td></td><td></td><td></td></tr>
<tr><td>Oligo rev</td><td>AAACCAGGGTCGCCCACACGTTCC</td><td></td><td></td><td></td><td></td><td></td><td></td><td></td><td></td><td></td><td></td></tr>
<tr><td>Chromosome</td><td>start</td><td>end</td><td>strand</td><td>MM</td><td>target_seq</td><td>PAM</td><td>alignment</td><td>distance</td><td>position</td><td>gene name</td><td>gene id</td></tr>
<tr><td>GU305914.1</td><td>99185</td><td>99207</td><td>-</td><td>0</td><td>GGAACGTGTGGGCGACCCTG</td><td>TGG</td><td>||||||||[||||||||||||]PAM</td><td>NA</td><td>Intergenic</td><td>NA</td><td>NA</td></tr>
<tr><td></td><td></td><td></td><td></td><td></td><td></td><td></td><td></td><td></td><td></td><td></td><td></td></tr>
<tr><td>T26529</td><td>TGTGGGCGACCCTGTGGAGCAGG</td><td>1000</td><td></td><td></td><td></td><td></td><td></td><td></td><td></td><td></td><td></td></tr>
<tr><td>Oligo fwd</td><td>CACCTGTGGGCGACCCTGTGGAGC</td><td></td><td></td><td></td><td></td><td></td><td></td><td></td><td></td><td></td><td></td></tr>
<tr><td>Oligo rev</td><td>AAACGCTCCACAGGGTCGCCCACA</td><td></td><td></td><td></td><td></td><td></td><td></td><td></td><td></td><td></td><td></td></tr>
<tr><td>Chromosome</td><td>start</td><td>end</td><td>strand</td><td>MM</td><td>target_seq</td><td>PAM</td><td>alignment</td><td>distance</td><td>position</td><td>gene name</td><td>gene id</td></tr>
<tr><td>GU305914.1</td><td>99179</td><td>99201</td><td>-</td><td>0</td><td>TGTGGGCGACCCTGTGGAGC</td><td>AGG</td><td>||||||||[||||||||||||]PAM</td><td>NA</td><td>Intergenic</td><td>NA</td><td>NA</td></tr>
<tr><td></td><td></td><td></td><td></td><td></td><td></td><td></td><td></td><td></td><td></td><td></td><td></td></tr>
<tr><td>T26530</td><td>TGTGGAGCAGGTGTTTCACCCGG</td><td>1000</td><td></td><td></td><td></td><td></td><td></td><td></td><td></td><td></td><td></td></tr>
<tr><td>Oligo fwd</td><td>CACCTGTGGAGCAGGTGTTTCACC</td><td></td><td></td><td></td><td></td><td></td><td></td><td></td><td></td><td></td><td></td></tr>
<tr><td>Oligo rev</td><td>AAACGGTGAAACACCTGCTCCACA</td><td></td><td></td><td></td><td></td><td></td><td></td><td></td><td></td><td></td><td></td></tr>
<tr><td>Chromosome</td><td>start</td><td>end</td><td>strand</td><td>MM</td><td>target_seq</td><td>PAM</td><td>alignment</td><td>distance</td><td>position</td><td>gene name</td><td>gene id</td></tr>
<tr><td>GU305914.1</td><td>99167</td><td>99189</td><td>-</td><td>0</td><td>TGTGGAGCAGGTGTTTCACC</td><td>CGG</td><td>||||||||[||||||||||||]PAM</td><td>NA</td><td>Intergenic</td><td>NA</td><td>NA</td></tr>
<tr><td></td><td></td><td></td><td></td><td></td><td></td><td></td><td></td><td></td><td></td><td></td><td></td></tr>
<tr><td>T26531</td><td>TAGACGCGTCGACGAGCGCGTGG</td><td>1000</td><td></td><td></td><td></td><td></td><td></td><td></td><td></td><td></td><td></td></tr>
<tr><td>Oligo fwd</td><td>CACCTAGACGCGTCGACGAGCGCG</td><td></td><td></td><td></td><td></td><td></td><td></td><td></td><td></td><td></td><td></td></tr>
<tr><td>Oligo rev</td><td>AAACCGCGCTCGTCGACGCGTCTA</td><td></td><td></td><td></td><td></td><td></td><td></td><td></td><td></td><td></td><td></td></tr>
<tr><td>Chromosome</td><td>start</td><td>end</td><td>strand</td><td>MM</td><td>target_seq</td><td>PAM</td><td>alignment</td><td>distance</td><td>position</td><td>gene name</td><td>gene id</td></tr>
<tr><td>GU305914.1</td><td>99081</td><td>99103</td><td>-</td><td>0</td><td>TAGACGCGTCGACGAGCGCG</td><td>TGG</td><td>||||||||[||||||||||||]PAM</td><td>NA</td><td>Intergenic</td><td>NA</td><td>NA</td></tr>
<tr><td></td><td></td><td></td><td></td><td></td><td></td><td></td><td></td><td></td><td></td><td></td><td></td></tr>
<tr><td>T26532</td><td>AGACGCGTCGACGAGCGCGTGGG</td><td>1000</td><td></td><td></td><td></td><td></td><td></td><td></td><td></td><td></td><td></td></tr>
<tr><td>Oligo fwd</td><td>CACCAGACGCGTCGACGAGCGCGT</td><td></td><td></td><td></td><td></td><td></td><td></td><td></td><td></td><td></td><td></td></tr>
<tr><td>Oligo rev</td><td>AAACACGCGCTCGTCGACGCGTCT</td><td></td><td></td><td></td><td></td><td></td><td></td><td></td><td></td><td></td><td></td></tr>
<tr><td>Chromosome</td><td>start</td><td>end</td><td>strand</td><td>MM</td><td>target_seq</td><td>PAM</td><td>alignment</td><td>distance</td><td>position</td><td>gene name</td><td>gene id</td></tr>
<tr><td>GU305914.1</td><td>99080</td><td>99102</td><td>-</td><td>0</td><td>AGACGCGTCGACGAGCGCGT</td><td>GGG</td><td>||||||||[||||||||||||]PAM</td><td>NA</td><td>Intergenic</td><td>NA</td><td>NA</td></tr>
<tr><td></td><td></td><td></td><td></td><td></td><td></td><td></td><td></td><td></td><td></td><td></td><td></td></tr>
<tr><td>T26533</td><td>GGCGACGACGACCCCCGTTCCGG</td><td>1000</td><td></td><td></td><td></td><td></td><td></td><td></td><td></td><td></td><td></td></tr>
<tr><td>Oligo fwd</td><td>CACCGGCGACGACGACCCCCGTTC</td><td></td><td></td><td></td><td></td><td></td><td></td><td></td><td></td><td></td><td></td></tr>
<tr><td>Oligo rev</td><td>AAACGAACGGGGGTCGTCGTCGCC</td><td></td><td></td><td></td><td></td><td></td><td></td><td></td><td></td><td></td><td></td></tr>
<tr><td>Chromosome</td><td>start</td><td>end</td><td>strand</td><td>MM</td><td>target_seq</td><td>PAM</td><td>alignment</td><td>distance</td><td>position</td><td>gene name</td><td>gene id</td></tr>
<tr><td>GU305914.1</td><td>99059</td><td>99081</td><td>-</td><td>0</td><td>GGCGACGACGACCCCCGTTC</td><td>CGG</td><td>||||||||[||||||||||||]PAM</td><td>NA</td><td>Intergenic</td><td>NA</td><td>NA</td></tr>
<tr><td></td><td></td><td></td><td></td><td></td><td></td><td></td><td></td><td></td><td></td><td></td><td></td></tr>
<tr><td>T26534</td><td>CACGCATCCTGCTTGCGTCAAGG</td><td>1000</td><td></td><td></td><td></td><td></td><td></td><td></td><td></td><td></td><td></td></tr>
<tr><td>Oligo fwd</td><td>CACCCACGCATCCTGCTTGCGTCA</td><td></td><td></td><td></td><td></td><td></td><td></td><td></td><td></td><td></td><td></td></tr>
<tr><td>Oligo rev</td><td>AAACTGACGCAAGCAGGATGCGTG</td><td></td><td></td><td></td><td></td><td></td><td></td><td></td><td></td><td></td><td></td></tr>
<tr><td>Chromosome</td><td>start</td><td>end</td><td>strand</td><td>MM</td><td>target_seq</td><td>PAM</td><td>alignment</td><td>distance</td><td>position</td><td>gene name</td><td>gene id</td></tr>
<tr><td>GU305914.1</td><td>99027</td><td>99049</td><td>-</td><td>0</td><td>CACGCATCCTGCTTGCGTCA</td><td>AGG</td><td>||||||||[||||||||||||]PAM</td><td>NA</td><td>Intergenic</td><td>NA</td><td>NA</td></tr>
<tr><td></td><td></td><td></td><td></td><td></td><td></td><td></td><td></td><td></td><td></td><td></td><td></td></tr>
<tr><td>T26535</td><td>CGTCAAGGAATAAATCCGTCCGG</td><td>1000</td><td></td><td></td><td></td><td></td><td></td><td></td><td></td><td></td><td></td></tr>
<tr><td>Oligo fwd</td><td>CACCCGTCAAGGAATAAATCCGTC</td><td></td><td></td><td></td><td></td><td></td><td></td><td></td><td></td><td></td><td></td></tr>
<tr><td>Oligo rev</td><td>AAACGACGGATTTATTCCTTGACG</td><td></td><td></td><td></td><td></td><td></td><td></td><td></td><td></td><td></td><td></td></tr>
<tr><td>Chromosome</td><td>start</td><td>end</td><td>strand</td><td>MM</td><td>target_seq</td><td>PAM</td><td>alignment</td><td>distance</td><td>position</td><td>gene name</td><td>gene id</td></tr>
<tr><td>GU305914.1</td><td>99012</td><td>99034</td><td>-</td><td>0</td><td>CGTCAAGGAATAAATCCGTC</td><td>CGG</td><td>||||||||[||||||||||||]PAM</td><td>NA</td><td>Intergenic</td><td>NA</td><td>NA</td></tr>
<tr><td></td><td></td><td></td><td></td><td></td><td></td><td></td><td></td><td></td><td></td><td></td><td></td></tr>
<tr><td>T26536</td><td>TAAATCCGTCCGGCCCACGCTGG</td><td>1000</td><td></td><td></td><td></td><td></td><td></td><td></td><td></td><td></td><td></td></tr>
<tr><td>Oligo fwd</td><td>CACCTAAATCCGTCCGGCCCACGC</td><td></td><td></td><td></td><td></td><td></td><td></td><td></td><td></td><td></td><td></td></tr>
<tr><td>Oligo rev</td><td>AAACGCGTGGGCCGGACGGATTTA</td><td></td><td></td><td></td><td></td><td></td><td></td><td></td><td></td><td></td><td></td></tr>
<tr><td>Chromosome</td><td>start</td><td>end</td><td>strand</td><td>MM</td><td>target_seq</td><td>PAM</td><td>alignment</td><td>distance</td><td>position</td><td>gene name</td><td>gene id</td></tr>
<tr><td>GU305914.1</td><td>99002</td><td>99024</td><td>-</td><td>0</td><td>TAAATCCGTCCGGCCCACGC</td><td>TGG</td><td>||||||||[||||||||||||]PAM</td><td>NA</td><td>Intergenic</td><td>NA</td><td>NA</td></tr>
<tr><td></td><td></td><td></td><td></td><td></td><td></td><td></td><td></td><td></td><td></td><td></td><td></td></tr>
<tr><td>T26537</td><td>GTCCGGCCCACGCTGGCACCCGG</td><td>1000</td><td></td><td></td><td></td><td></td><td></td><td></td><td></td><td></td><td></td></tr>
<tr><td>Oligo fwd</td><td>CACCGTCCGGCCCACGCTGGCACC</td><td></td><td></td><td></td><td></td><td></td><td></td><td></td><td></td><td></td><td></td></tr>
<tr><td>Oligo rev</td><td>AAACGGTGCCAGCGTGGGCCGGAC</td><td></td><td></td><td></td><td></td><td></td><td></td><td></td><td></td><td></td><td></td></tr>
<tr><td>Chromosome</td><td>start</td><td>end</td><td>strand</td><td>MM</td><td>target_seq</td><td>PAM</td><td>alignment</td><td>distance</td><td>position</td><td>gene name</td><td>gene id</td></tr>
<tr><td>GU305914.1</td><td>98995</td><td>99017</td><td>-</td><td>0</td><td>GTCCGGCCCACGCTGGCACC</td><td>CGG</td><td>||||||||[||||||||||||]PAM</td><td>NA</td><td>Intergenic</td><td>NA</td><td>NA</td></tr>
</table>

<table>
<tr><td>T26538</td><td>CCTCCCGCGATATAAAAAAGCGG</td><td></td><td>1000</td><td></td><td></td><td></td><td></td><td></td><td></td><td></td><td></td><td></td></tr>
<tr><td>Oligo fwd</td><td>CACCCCTCCCGCGATATAAAAAAG</td><td></td><td></td><td></td><td></td><td></td><td></td><td></td><td></td><td></td><td></td><td></td></tr>
<tr><td>Oligo rev</td><td>AAACCTTTTTTATATCGCGGGAGG</td><td></td><td></td><td></td><td></td><td></td><td></td><td></td><td></td><td></td><td></td><td></td></tr>
<tr><td>Chromosome</td><td>start</td><td></td><td>end</td><td>strand</td><td>MM</td><td>target_seq</td><td>PAM</td><td>alignment</td><td>distance</td><td>position</td><td>gene name</td><td>gene id</td></tr>
<tr><td>GU305914.1</td><td></td><td>98944</td><td>98966</td><td>-</td><td>0</td><td>CCTCCCGCGATATAAAAAAG</td><td>CGG</td><td>||||||||[||||||||||||]PAM</td><td>NA</td><td>Intergenic</td><td>NA</td><td>NA</td></tr>
<tr><td>T26539</td><td>CTCCCGCGATATAAAAAAGCGGG</td><td></td><td>1000</td><td></td><td></td><td></td><td></td><td></td><td></td><td></td><td></td><td></td></tr>
<tr><td>Oligo fwd</td><td>CACCCTCCCGCGATATAAAAAAGC</td><td></td><td></td><td></td><td></td><td></td><td></td><td></td><td></td><td></td><td></td><td></td></tr>
<tr><td>Oligo rev</td><td>AAACGCTTTTTTATATCGCGGGAG</td><td></td><td></td><td></td><td></td><td></td><td></td><td></td><td></td><td></td><td></td><td></td></tr>
<tr><td>Chromosome</td><td>start</td><td></td><td>end</td><td>strand</td><td>MM</td><td>target_seq</td><td>PAM</td><td>alignment</td><td>distance</td><td>position</td><td>gene name</td><td>gene id</td></tr>
<tr><td>GU305914.1</td><td></td><td>98943</td><td>98965</td><td>-</td><td>0</td><td>CTCCCGCGATATAAAAAAGC</td><td>GGG</td><td>||||||||[||||||||||||]PAM</td><td>NA</td><td>Intergenic</td><td>NA</td><td>NA</td></tr>
<tr><td>T26540</td><td>TCCCGCGATATAAAAAAGCGGGG</td><td></td><td>1000</td><td></td><td></td><td></td><td></td><td></td><td></td><td></td><td></td><td></td></tr>
<tr><td>Oligo fwd</td><td>CACCTCCCGCGATATAAAAAAGCG</td><td></td><td></td><td></td><td></td><td></td><td></td><td></td><td></td><td></td><td></td><td></td></tr>
<tr><td>Oligo rev</td><td>AAACCGCTTTTTTATATCGCGGGA</td><td></td><td></td><td></td><td></td><td></td><td></td><td></td><td></td><td></td><td></td><td></td></tr>
<tr><td>Chromosome</td><td>start</td><td></td><td>end</td><td>strand</td><td>MM</td><td>target_seq</td><td>PAM</td><td>alignment</td><td>distance</td><td>position</td><td>gene name</td><td>gene id</td></tr>
<tr><td>GU305914.1</td><td></td><td>98942</td><td>98964</td><td>-</td><td>0</td><td>TCCCGCGATATAAAAAAGCG</td><td>GGG</td><td>||||||||[||||||||||||]PAM</td><td>NA</td><td>Intergenic</td><td>NA</td><td>NA</td></tr>
<tr><td>T26541</td><td>CCCGCGATATAAAAAAGCGGGGG</td><td></td><td>1000</td><td></td><td></td><td></td><td></td><td></td><td></td><td></td><td></td><td></td></tr>
<tr><td>Oligo fwd</td><td>CACCCCCGCGATATAAAAAAGCGG</td><td></td><td></td><td></td><td></td><td></td><td></td><td></td><td></td><td></td><td></td><td></td></tr>
<tr><td>Oligo rev</td><td>AAACCCGCTTTTTTATATCGCGGG</td><td></td><td></td><td></td><td></td><td></td><td></td><td></td><td></td><td></td><td></td><td></td></tr>
<tr><td>Chromosome</td><td>start</td><td></td><td>end</td><td>strand</td><td>MM</td><td>target_seq</td><td>PAM</td><td>alignment</td><td>distance</td><td>position</td><td>gene name</td><td>gene id</td></tr>
<tr><td>GU305914.1</td><td></td><td>98941</td><td>98963</td><td>-</td><td>0</td><td>CCCGCGATATAAAAAAGCGG</td><td>GGG</td><td>||||||||[||||||||||||]PAM</td><td>NA</td><td>Intergenic</td><td>NA</td><td>NA</td></tr>
<tr><td>T26542</td><td>CCGCGATATAAAAAAGCGGGGGG</td><td></td><td>1000</td><td></td><td></td><td></td><td></td><td></td><td></td><td></td><td></td><td></td></tr>
<tr><td>Oligo fwd</td><td>CACCCCGCGATATAAAAAAGCGGG</td><td></td><td></td><td></td><td></td><td></td><td></td><td></td><td></td><td></td><td></td><td></td></tr>
<tr><td>Oligo rev</td><td>AAACCCCGCTTTTTTATATCGCGG</td><td></td><td></td><td></td><td></td><td></td><td></td><td></td><td></td><td></td><td></td><td></td></tr>
<tr><td>Chromosome</td><td>start</td><td></td><td>end</td><td>strand</td><td>MM</td><td>target_seq</td><td>PAM</td><td>alignment</td><td>distance</td><td>position</td><td>gene name</td><td>gene id</td></tr>
<tr><td>GU305914.1</td><td></td><td>98940</td><td>98962</td><td>-</td><td>0</td><td>CCGCGATATAAAAAAGCGGG</td><td>GGG</td><td>||||||||[||||||||||||]PAM</td><td>NA</td><td>Intergenic</td><td>NA</td><td>NA</td></tr>
<tr><td>T26543</td><td>AAAGCGGGGGGTTTCCTGCGCGG</td><td></td><td>1000</td><td></td><td></td><td></td><td></td><td></td><td></td><td></td><td></td><td></td></tr>
<tr><td>Oligo fwd</td><td>CACCAAAGCGGGGGGTTTCCTGCG</td><td></td><td></td><td></td><td></td><td></td><td></td><td></td><td></td><td></td><td></td><td></td></tr>
<tr><td>Oligo rev</td><td>AAACCGCAGGAAACCCCCCGCTTT</td><td></td><td></td><td></td><td></td><td></td><td></td><td></td><td></td><td></td><td></td><td></td></tr>
<tr><td>Chromosome</td><td>start</td><td></td><td>end</td><td>strand</td><td>MM</td><td>target_seq</td><td>PAM</td><td>alignment</td><td>distance</td><td>position</td><td>gene name</td><td>gene id</td></tr>
<tr><td>GU305914.1</td><td></td><td>98928</td><td>98950</td><td>-</td><td>0</td><td>AAAGCGGGGGGTTTCCTGCG</td><td>CGG</td><td>||||||||[||||||||||||]PAM</td><td>NA</td><td>Intergenic</td><td>NA</td><td>NA</td></tr>
<tr><td>T26544</td><td>CGGGGGGTTTCCTGCGCGGAAGG</td><td></td><td>1000</td><td></td><td></td><td></td><td></td><td></td><td></td><td></td><td></td><td></td></tr>
<tr><td>Oligo fwd</td><td>CACCCGGGGGGTTTCCTGCGCGGA</td><td></td><td></td><td></td><td></td><td></td><td></td><td></td><td></td><td></td><td></td><td></td></tr>
<tr><td>Oligo rev</td><td>AAACTCCGCGCAGGAAACCCCCCG</td><td></td><td></td><td></td><td></td><td></td><td></td><td></td><td></td><td></td><td></td><td></td></tr>
<tr><td>Chromosome</td><td>start</td><td></td><td>end</td><td>strand</td><td>MM</td><td>target_seq</td><td>PAM</td><td>alignment</td><td>distance</td><td>position</td><td>gene name</td><td>gene id</td></tr>
<tr><td>GU305914.1</td><td></td><td>98924</td><td>98946</td><td>-</td><td>0</td><td>CGGGGGGTTTCCTGCGCGGA</td><td>AGG</td><td>||||||||[||||||||||||]PAM</td><td>NA</td><td>Intergenic</td><td>NA</td><td>NA</td></tr>
<tr><td>T26545</td><td>CACCTAGCCACATTTCCGCGCGG</td><td></td><td>1000</td><td></td><td></td><td></td><td></td><td></td><td></td><td></td><td></td><td></td></tr>
<tr><td>Oligo fwd</td><td>CACCCACCTAGCCACATTTCCGCG</td><td></td><td></td><td></td><td></td><td></td><td></td><td></td><td></td><td></td><td></td><td></td></tr>
<tr><td>Oligo rev</td><td>AAACCGCGGAAATGTGGCTAGGTG</td><td></td><td></td><td></td><td></td><td></td><td></td><td></td><td></td><td></td><td></td><td></td></tr>
<tr><td>Chromosome</td><td>start</td><td></td><td>end</td><td>strand</td><td>MM</td><td>target_seq</td><td>PAM</td><td>alignment</td><td>distance</td><td>position</td><td>gene name</td><td>gene id</td></tr>
<tr><td>GU305914.1</td><td></td><td>98878</td><td>98900</td><td>-</td><td>0</td><td>CACCTAGCCACATTTCCGCG</td><td>CGG</td><td>||||||||[||||||||||||]PAM</td><td>NA</td><td>Intergenic</td><td>NA</td><td>NA</td></tr>
<tr><td>T26546</td><td>ACCTAGCCACATTTCCGCGCGGG</td><td></td><td>1000</td><td></td><td></td><td></td><td></td><td></td><td></td><td></td><td></td><td></td></tr>
<tr><td>Oligo fwd</td><td>CACCACCTAGCCACATTTCCGCGC</td><td></td><td></td><td></td><td></td><td></td><td></td><td></td><td></td><td></td><td></td><td></td></tr>
<tr><td>Oligo rev</td><td>AAACGCGCGGAAATGTGGCTAGGT</td><td></td><td></td><td></td><td></td><td></td><td></td><td></td><td></td><td></td><td></td><td></td></tr>
<tr><td>Chromosome</td><td>start</td><td></td><td>end</td><td>strand</td><td>MM</td><td>target_seq</td><td>PAM</td><td>alignment</td><td>distance</td><td>position</td><td>gene name</td><td>gene id</td></tr>
<tr><td>GU305914.1</td><td></td><td>98877</td><td>98899</td><td>-</td><td>0</td><td>ACCTAGCCACATTTCCGCGC</td><td>GGG</td><td>||||||||[||||||||||||]PAM</td><td>NA</td><td>Intergenic</td><td>NA</td><td>NA</td></tr>
<tr><td>T26547</td><td>CGAGCGACCGAGCGCACCGAAGG</td><td></td><td>1000</td><td></td><td></td><td></td><td></td><td></td><td></td><td></td><td></td><td></td></tr>
<tr><td>Oligo fwd</td><td>CACCCGAGCGACCGAGCGCACCGA</td><td></td><td></td><td></td><td></td><td></td><td></td><td></td><td></td><td></td><td></td><td></td></tr>
<tr><td>Oligo rev</td><td>AAACTCGGTGCGCTCGGTCGCTCG</td><td></td><td></td><td></td><td></td><td></td><td></td><td></td><td></td><td></td><td></td><td></td></tr>
<tr><td>Chromosome</td><td>start</td><td></td><td>end</td><td>strand</td><td>MM</td><td>target_seq</td><td>PAM</td><td>alignment</td><td>distance</td><td>position</td><td>gene name</td><td>gene id</td></tr>
<tr><td>GU305914.1</td><td></td><td>98823</td><td>98845</td><td>-</td><td>0</td><td>CGAGCGACCGAGCGCACCGA</td><td>AGG</td><td>||||||||[||||||||||||]PAM</td><td>NA</td><td>Intergenic</td><td>NA</td><td>NA</td></tr>
<tr><td>T26548</td><td>GCGACCGAGCGCACCGAAGGCGG</td><td></td><td>1000</td><td></td><td></td><td></td><td></td><td></td><td></td><td></td><td></td><td></td></tr>
<tr><td>Oligo fwd</td><td>CACCGCGACCGAGCGCACCGAAGG</td><td></td><td></td><td></td><td></td><td></td><td></td><td></td><td></td><td></td><td></td><td></td></tr>
<tr><td>Oligo rev</td><td>AAACCCTTCGGTGCGCTCGGTCGC</td><td></td><td></td><td></td><td></td><td></td><td></td><td></td><td></td><td></td><td></td><td></td></tr>
<tr><td>Chromosome</td><td>start</td><td></td><td>end</td><td>strand</td><td>MM</td><td>target_seq</td><td>PAM</td><td>alignment</td><td>distance</td><td>position</td><td>gene name</td><td>gene id</td></tr>
<tr><td>GU305914.1</td><td></td><td>98820</td><td>98842</td><td>-</td><td>0</td><td>GCGACCGAGCGCACCGAAGG</td><td>CGG</td><td>||||||||[||||||||||||]PAM</td><td>NA</td><td>Intergenic</td><td>NA</td><td>NA</td></tr>
<tr><td>T26549</td><td>AAGGCGGCGAAACCATGCTGCGG</td><td></td><td>1000</td><td></td><td></td><td></td><td></td><td></td><td></td><td></td><td></td><td></td></tr>
<tr><td>Oligo fwd</td><td>CACCAAGGCGGCGAAACCATGCTG</td><td></td><td></td><td></td><td></td><td></td><td></td><td></td><td></td><td></td><td></td><td></td></tr>
<tr><td>Oligo rev</td><td>AAACCAGCATGGTTTCGCCGCCTT</td><td></td><td></td><td></td><td></td><td></td><td></td><td></td><td></td><td></td><td></td><td></td></tr>
<tr><td>Chromosome</td><td>start</td><td></td><td>end</td><td>strand</td><td>MM</td><td>target_seq</td><td>PAM</td><td>alignment</td><td>distance</td><td>position</td><td>gene name</td><td>gene id</td></tr>
<tr><td>GU305914.1</td><td></td><td>98804</td><td>98826</td><td>-</td><td>0</td><td>AAGGCGGCGAAACCATGCTG</td><td>CGG</td><td>||||||||[||||||||||||]PAM</td><td>NA</td><td>Intergenic</td><td>NA</td><td>NA</td></tr>
<tr><td>T26550</td><td>GGCGAAACCATGCTGCGGACCGG</td><td></td><td>1000</td><td></td><td></td><td></td><td></td><td></td><td></td><td></td><td></td><td></td></tr>
<tr><td>Oligo fwd</td><td>CACCGGCGAAACCATGCTGCGGAC</td><td></td><td></td><td></td><td></td><td></td><td></td><td></td><td></td><td></td><td></td><td></td></tr>
<tr><td>Oligo rev</td><td>AAACGTCCGCAGCATGGTTTCGCC</td><td></td><td></td><td></td><td></td><td></td><td></td><td></td><td></td><td></td><td></td><td></td></tr>
<tr><td>Chromosome</td><td>start</td><td></td><td>end</td><td>strand</td><td>MM</td><td>target_seq</td><td>PAM</td><td>alignment</td><td>distance</td><td>position</td><td>gene name</td><td>gene id</td></tr>
<tr><td>GU305914.1</td><td></td><td>98799</td><td>98821</td><td>-</td><td>0</td><td>GGCGAAACCATGCTGCGGAC</td><td>CGG</td><td>||||||||[||||||||||||]PAM</td><td>NA</td><td>Intergenic</td><td>NA</td><td>NA</td></tr>
<tr><td>T26551</td><td>TGCGGACCGGTGTCAAGAGACGG</td><td></td><td>1000</td><td></td><td></td><td></td><td></td><td></td><td></td><td></td><td></td><td></td></tr>
<tr><td>Oligo fwd</td><td>CACCTGCGGACCGGTGTCAAGAGA</td><td></td><td></td><td></td><td></td><td></td><td></td><td></td><td></td><td></td><td></td><td></td></tr>
<tr><td>Oligo rev</td><td>AAACTCTCTTGACACCGGTCCGCA</td><td></td><td></td><td></td><td></td><td></td><td></td><td></td><td></td><td></td><td></td><td></td></tr>
<tr><td>Chromosome</td><td>start</td><td></td><td>end</td><td>strand</td><td>MM</td><td>target_seq</td><td>PAM</td><td>alignment</td><td>distance</td><td>position</td><td>gene name</td><td>gene id</td></tr>
<tr><td>GU305914.1</td><td></td><td>98786</td><td>98808</td><td>-</td><td>0</td><td>TGCGGACCGGTGTCAAGAGA</td><td>CGG</td><td>||||||||[||||||||||||]PAM</td><td>NA</td><td>Intergenic</td><td>NA</td><td>NA</td></tr>
<tr><td>T26552</td><td>GACCGGTGTCAAGAGACGGCTGG</td><td></td><td>1000</td><td></td><td></td><td></td><td></td><td></td><td></td><td></td><td></td><td></td></tr>
<tr><td>Oligo fwd</td><td>CACCGACCGGTGTCAAGAGACGGC</td><td></td><td></td><td></td><td></td><td></td><td></td><td></td><td></td><td></td><td></td><td></td></tr>
<tr><td>Oligo rev</td><td>AAACGCCGTCTCTTGACACCGGTC</td><td></td><td></td><td></td><td></td><td></td><td></td><td></td><td></td><td></td><td></td><td></td></tr>
<tr><td>Chromosome</td><td>start</td><td></td><td>end</td><td>strand</td><td>MM</td><td>target_seq</td><td>PAM</td><td>alignment</td><td>distance</td><td>position</td><td>gene name</td><td>gene id</td></tr>
<tr><td>GU305914.1</td><td></td><td>98782</td><td>98804</td><td>-</td><td>0</td><td>GACCGGTGTCAAGAGACGGC</td><td>TGG</td><td>||||||||[||||||||||||]PAM</td><td>NA</td><td>Intergenic</td><td>NA</td><td>NA</td></tr>
</table>

<table>
<tr><td>T26553</td><td>ACCGGTGTCAAGAGACGGCTGGG</td><td>1000</td><td></td><td></td><td></td><td></td><td></td><td></td><td></td><td></td><td></td></tr>
<tr><td>Oligo fwd</td><td>CACCACCGGTGTCAAGAGACGGCT</td><td></td><td></td><td></td><td></td><td></td><td></td><td></td><td></td><td></td><td></td></tr>
<tr><td>Oligo rev</td><td>AAACAGCCGTCTCTTGACACCGGT</td><td></td><td></td><td></td><td></td><td></td><td></td><td></td><td></td><td></td><td></td></tr>
<tr><td>Chromosome</td><td>start</td><td>end</td><td>strand</td><td>MM</td><td>target_seq</td><td>PAM</td><td>alignment</td><td>distance</td><td>position</td><td>gene name</td><td>gene id</td></tr>
<tr><td>GU305914.1</td><td>98781</td><td>98803</td><td>-</td><td>0</td><td>ACCGGTGTCAAGAGACGGCT</td><td>GGG</td><td>||||||||[||||||||||||]PAM</td><td>NA</td><td>Intergenic</td><td>NA</td><td>NA</td></tr>
<tr><td>T26554</td><td>AGACGGCTGGGCCCGTTCGCCGG</td><td>1000</td><td></td><td></td><td></td><td></td><td></td><td></td><td></td><td></td><td></td></tr>
<tr><td>Oligo fwd</td><td>CACCAGACGGCTGGGCCCGTTCGC</td><td></td><td></td><td></td><td></td><td></td><td></td><td></td><td></td><td></td><td></td></tr>
<tr><td>Oligo rev</td><td>AAACGCGAACGGGCCCAGCCGTCT</td><td></td><td></td><td></td><td></td><td></td><td></td><td></td><td></td><td></td><td></td></tr>
<tr><td>Chromosome</td><td>start</td><td>end</td><td>strand</td><td>MM</td><td>target_seq</td><td>PAM</td><td>alignment</td><td>distance</td><td>position</td><td>gene name</td><td>gene id</td></tr>
<tr><td>GU305914.1</td><td>98769</td><td>98791</td><td>-</td><td>0</td><td>AGACGGCTGGGCCCGTTCGC</td><td>CGG</td><td>||||||||[||||||||||||]PAM</td><td>NA</td><td>Intergenic</td><td>NA</td><td>NA</td></tr>
<tr><td>T26555</td><td>CCCGTTCGCCGGATACGACGAGG</td><td>1000</td><td></td><td></td><td></td><td></td><td></td><td></td><td></td><td></td><td></td></tr>
<tr><td>Oligo fwd</td><td>CACCCCCGTTCGCCGGATACGACG</td><td></td><td></td><td></td><td></td><td></td><td></td><td></td><td></td><td></td><td></td></tr>
<tr><td>Oligo rev</td><td>AAACCGTCGTATCCGGCGAACGGG</td><td></td><td></td><td></td><td></td><td></td><td></td><td></td><td></td><td></td><td></td></tr>
<tr><td>Chromosome</td><td>start</td><td>end</td><td>strand</td><td>MM</td><td>target_seq</td><td>PAM</td><td>alignment</td><td>distance</td><td>position</td><td>gene name</td><td>gene id</td></tr>
<tr><td>GU305914.1</td><td>98758</td><td>98780</td><td>-</td><td>0</td><td>CCCGTTCGCCGGATACGACG</td><td>AGG</td><td>||||||||[||||||||||||]PAM</td><td>NA</td><td>Intergenic</td><td>NA</td><td>NA</td></tr>
<tr><td>T26556</td><td>CGACGAGGACGATGCCGCGACGG</td><td>1000</td><td></td><td></td><td></td><td></td><td></td><td></td><td></td><td></td><td></td></tr>
<tr><td>Oligo fwd</td><td>CACCCGACGAGGACGATGCCGCGA</td><td></td><td></td><td></td><td></td><td></td><td></td><td></td><td></td><td></td><td></td></tr>
<tr><td>Oligo rev</td><td>AAACTCGCGGCATCGTCCTCGTCG</td><td></td><td></td><td></td><td></td><td></td><td></td><td></td><td></td><td></td><td></td></tr>
<tr><td>Chromosome</td><td>start</td><td>end</td><td>strand</td><td>MM</td><td>target_seq</td><td>PAM</td><td>alignment</td><td>distance</td><td>position</td><td>gene name</td><td>gene id</td></tr>
<tr><td>GU305914.1</td><td>98743</td><td>98765</td><td>-</td><td>0</td><td>CGACGAGGACGATGCCGCGA</td><td>CGG</td><td>||||||||[||||||||||||]PAM</td><td>NA</td><td>Intergenic</td><td>NA</td><td>NA</td></tr>
<tr><td>GU305914.1</td><td>142232</td><td>142254</td><td>+</td><td>4</td><td>CGATGACGACGATACCGCGG</td><td>CGG</td><td>|||-||-|[|||||-|||||-]PAM</td><td>NA</td><td>Intergenic</td><td>NA</td><td>NA</td></tr>
<tr><td>T26557</td><td>GACGAGGACGATGCCGCGACGGG</td><td>1000</td><td></td><td></td><td></td><td></td><td></td><td></td><td></td><td></td><td></td></tr>
<tr><td>Oligo fwd</td><td>CACCGACGAGGACGATGCCGCGAC</td><td></td><td></td><td></td><td></td><td></td><td></td><td></td><td></td><td></td><td></td></tr>
<tr><td>Oligo rev</td><td>AAACGTCGCGGCATCGTCCTCGTC</td><td></td><td></td><td></td><td></td><td></td><td></td><td></td><td></td><td></td><td></td></tr>
<tr><td>Chromosome</td><td>start</td><td>end</td><td>strand</td><td>MM</td><td>target_seq</td><td>PAM</td><td>alignment</td><td>distance</td><td>position</td><td>gene name</td><td>gene id</td></tr>
<tr><td>GU305914.1</td><td>98742</td><td>98764</td><td>-</td><td>0</td><td>GACGAGGACGATGCCGCGAC</td><td>GGG</td><td>||||||||[||||||||||||]PAM</td><td>NA</td><td>Intergenic</td><td>NA</td><td>NA</td></tr>
<tr><td>T26558</td><td>GAGGACGATGCCGCGACGGGCGG</td><td>1000</td><td></td><td></td><td></td><td></td><td></td><td></td><td></td><td></td><td></td></tr>
<tr><td>Oligo fwd</td><td>CACCGAGGACGATGCCGCGACGGG</td><td></td><td></td><td></td><td></td><td></td><td></td><td></td><td></td><td></td><td></td></tr>
<tr><td>Oligo rev</td><td>AAACCCCGTCGCGGCATCGTCCTC</td><td></td><td></td><td></td><td></td><td></td><td></td><td></td><td></td><td></td><td></td></tr>
<tr><td>Chromosome</td><td>start</td><td>end</td><td>strand</td><td>MM</td><td>target_seq</td><td>PAM</td><td>alignment</td><td>distance</td><td>position</td><td>gene name</td><td>gene id</td></tr>
<tr><td>GU305914.1</td><td>98739</td><td>98761</td><td>-</td><td>0</td><td>GAGGACGATGCCGCGACGGG</td><td>CGG</td><td>||||||||[||||||||||||]PAM</td><td>NA</td><td>Intergenic</td><td>NA</td><td>NA</td></tr>
<tr><td>T26559</td><td>CAGTCTCAGCACTACTACTACGG</td><td>1000</td><td></td><td></td><td></td><td></td><td></td><td></td><td></td><td></td><td></td></tr>
<tr><td>Oligo fwd</td><td>CACCCAGTCTCAGCACTACTACTA</td><td></td><td></td><td></td><td></td><td></td><td></td><td></td><td></td><td></td><td></td></tr>
<tr><td>Oligo rev</td><td>AAACTAGTAGTAGTGCTGAGACTG</td><td></td><td></td><td></td><td></td><td></td><td></td><td></td><td></td><td></td><td></td></tr>
<tr><td>Chromosome</td><td>start</td><td>end</td><td>strand</td><td>MM</td><td>target_seq</td><td>PAM</td><td>alignment</td><td>distance</td><td>position</td><td>gene name</td><td>gene id</td></tr>
<tr><td>GU305914.1</td><td>98685</td><td>98707</td><td>-</td><td>0</td><td>CAGTCTCAGCACTACTACTA</td><td>CGG</td><td>||||||||[||||||||||||]PAM</td><td>NA</td><td>Intergenic</td><td>NA</td><td>NA</td></tr>
<tr><td>T26560</td><td>CCAGTCGTCTTACCGCGATTCGG</td><td>1000</td><td></td><td></td><td></td><td></td><td></td><td></td><td></td><td></td><td></td></tr>
<tr><td>Oligo fwd</td><td>CACCCCAGTCGTCTTACCGCGATT</td><td></td><td></td><td></td><td></td><td></td><td></td><td></td><td></td><td></td><td></td></tr>
<tr><td>Oligo rev</td><td>AAACAATCGCGGTAAGACGACTGG</td><td></td><td></td><td></td><td></td><td></td><td></td><td></td><td></td><td></td><td></td></tr>
<tr><td>Chromosome</td><td>start</td><td>end</td><td>strand</td><td>MM</td><td>target_seq</td><td>PAM</td><td>alignment</td><td>distance</td><td>position</td><td>gene name</td><td>gene id</td></tr>
<tr><td>GU305914.1</td><td>98656</td><td>98678</td><td>-</td><td>0</td><td>CCAGTCGTCTTACCGCGATT</td><td>CGG</td><td>||||||||[||||||||||||]PAM</td><td>NA</td><td>Intergenic</td><td>NA</td><td>NA</td></tr>
<tr><td>T26561</td><td>CAGTCGTCTTACCGCGATTCGGG</td><td>1000</td><td></td><td></td><td></td><td></td><td></td><td></td><td></td><td></td><td></td></tr>
<tr><td>Oligo fwd</td><td>CACCCAGTCGTCTTACCGCGATTC</td><td></td><td></td><td></td><td></td><td></td><td></td><td></td><td></td><td></td><td></td></tr>
<tr><td>Oligo rev</td><td>AAACGAATCGCGGTAAGACGACTG</td><td></td><td></td><td></td><td></td><td></td><td></td><td></td><td></td><td></td><td></td></tr>
<tr><td>Chromosome</td><td>start</td><td>end</td><td>strand</td><td>MM</td><td>target_seq</td><td>PAM</td><td>alignment</td><td>distance</td><td>position</td><td>gene name</td><td>gene id</td></tr>
<tr><td>GU305914.1</td><td>98655</td><td>98677</td><td>-</td><td>0</td><td>CAGTCGTCTTACCGCGATTC</td><td>GGG</td><td>||||||||[||||||||||||]PAM</td><td>NA</td><td>Intergenic</td><td>NA</td><td>NA</td></tr>
<tr><td>T26562</td><td>CGGGCGCCTCTCATCCCAACTGG</td><td>1000</td><td></td><td></td><td></td><td></td><td></td><td></td><td></td><td></td><td></td></tr>
<tr><td>Oligo fwd</td><td>CACCCGGGCGCCTCTCATCCCAAC</td><td></td><td></td><td></td><td></td><td></td><td></td><td></td><td></td><td></td><td></td></tr>
<tr><td>Oligo rev</td><td>AAACGTTGGGATGAGAGGCGCCCG</td><td></td><td></td><td></td><td></td><td></td><td></td><td></td><td></td><td></td><td></td></tr>
<tr><td>Chromosome</td><td>start</td><td>end</td><td>strand</td><td>MM</td><td>target_seq</td><td>PAM</td><td>alignment</td><td>distance</td><td>position</td><td>gene name</td><td>gene id</td></tr>
<tr><td>GU305914.1</td><td>98636</td><td>98658</td><td>-</td><td>0</td><td>CGGGCGCCTCTCATCCCAAC</td><td>TGG</td><td>||||||||[||||||||||||]PAM</td><td>NA</td><td>Intergenic</td><td>NA</td><td>NA</td></tr>
<tr><td>T26563</td><td>CCAGTATGACAAAAACATCGCGG</td><td>1000</td><td></td><td></td><td></td><td></td><td></td><td></td><td></td><td></td><td></td></tr>
<tr><td>Oligo fwd</td><td>CACCCCAGTATGACAAAAACATCG</td><td></td><td></td><td></td><td></td><td></td><td></td><td></td><td></td><td></td><td></td></tr>
<tr><td>Oligo rev</td><td>AAACCGATGTTTTTGTCATACTGG</td><td></td><td></td><td></td><td></td><td></td><td></td><td></td><td></td><td></td><td></td></tr>
<tr><td>Chromosome</td><td>start</td><td>end</td><td>strand</td><td>MM</td><td>target_seq</td><td>PAM</td><td>alignment</td><td>distance</td><td>position</td><td>gene name</td><td>gene id</td></tr>
<tr><td>GU305914.1</td><td>98557</td><td>98579</td><td>-</td><td>0</td><td>CCAGTATGACAAAAACATCG</td><td>CGG</td><td>||||||||[||||||||||||]PAM</td><td>NA</td><td>Intergenic</td><td>NA</td><td>NA</td></tr>
<tr><td>T26564</td><td>GAATAAAAAACTAGACTGTCTGG</td><td>1000</td><td></td><td></td><td></td><td></td><td></td><td></td><td></td><td></td><td></td></tr>
<tr><td>Oligo fwd</td><td>CACCGAATAAAAAACTAGACTGTC</td><td></td><td></td><td></td><td></td><td></td><td></td><td></td><td></td><td></td><td></td></tr>
<tr><td>Oligo rev</td><td>AAACGACAGTCTAGTTTTTTATTC</td><td></td><td></td><td></td><td></td><td></td><td></td><td></td><td></td><td></td><td></td></tr>
<tr><td>Chromosome</td><td>start</td><td>end</td><td>strand</td><td>MM</td><td>target_seq</td><td>PAM</td><td>alignment</td><td>distance</td><td>position</td><td>gene name</td><td>gene id</td></tr>
<tr><td>GU305914.1</td><td>98521</td><td>98543</td><td>-</td><td>0</td><td>GAATAAAAAACTAGACTGTC</td><td>TGG</td><td>||||||||[||||||||||||]PAM</td><td>NA</td><td>Intergenic</td><td>NA</td><td>NA</td></tr>
<tr><td>T26565</td><td>AATAAAAAACTAGACTGTCTGGG</td><td>1000</td><td></td><td></td><td></td><td></td><td></td><td></td><td></td><td></td><td></td></tr>
<tr><td>Oligo fwd</td><td>CACCAATAAAAAACTAGACTGTCT</td><td></td><td></td><td></td><td></td><td></td><td></td><td></td><td></td><td></td><td></td></tr>
<tr><td>Oligo rev</td><td>AAACAGACAGTCTAGTTTTTTATT</td><td></td><td></td><td></td><td></td><td></td><td></td><td></td><td></td><td></td><td></td></tr>
<tr><td>Chromosome</td><td>start</td><td>end</td><td>strand</td><td>MM</td><td>target_seq</td><td>PAM</td><td>alignment</td><td>distance</td><td>position</td><td>gene name</td><td>gene id</td></tr>
<tr><td>GU305914.1</td><td>98520</td><td>98542</td><td>-</td><td>0</td><td>AATAAAAAACTAGACTGTCT</td><td>GGG</td><td>||||||||[||||||||||||]PAM</td><td>NA</td><td>Intergenic</td><td>NA</td><td>NA</td></tr>
<tr><td>T26566</td><td>CTGTCTGGGACCCGACGACCTGG</td><td>1000</td><td></td><td></td><td></td><td></td><td></td><td></td><td></td><td></td><td></td></tr>
<tr><td>Oligo fwd</td><td>CACCCTGTCTGGGACCCGACGACC</td><td></td><td></td><td></td><td></td><td></td><td></td><td></td><td></td><td></td><td></td></tr>
<tr><td>Oligo rev</td><td>AAACGGTCGTCGGGTCCCAGACAG</td><td></td><td></td><td></td><td></td><td></td><td></td><td></td><td></td><td></td><td></td></tr>
<tr><td>Chromosome</td><td>start</td><td>end</td><td>strand</td><td>MM</td><td>target_seq</td><td>PAM</td><td>alignment</td><td>distance</td><td>position</td><td>gene name</td><td>gene id</td></tr>
<tr><td>GU305914.1</td><td>98506</td><td>98528</td><td>-</td><td>0</td><td>CTGTCTGGGACCCGACGACC</td><td>TGG</td><td>||||||||[||||||||||||]PAM</td><td>NA</td><td>Intergenic</td><td>NA</td><td>NA</td></tr>
<tr><td>T26567</td><td>AATGCTTGAAAGCCATGATCCGG</td><td>1000</td><td></td><td></td><td></td><td></td><td></td><td></td><td></td><td></td><td></td></tr>
<tr><td>Oligo fwd</td><td>CACCAATGCTTGAAAGCCATGATC</td><td></td><td></td><td></td><td></td><td></td><td></td><td></td><td></td><td></td><td></td></tr>
<tr><td>Oligo rev</td><td>AAACGATCATGGCTTTCAAGCATT</td><td></td><td></td><td></td><td></td><td></td><td></td><td></td><td></td><td></td><td></td></tr>
</table>

<table>
<tr><td>Chromosome</td><td>start</td><td>end</td><td>strand</td><td>MM</td><td>target_seq</td><td>PAM</td><td>alignment</td><td>distance</td><td>position</td><td>gene name</td><td>gene id</td></tr>
<tr><td>GU305914.1</td><td>98483</td><td>98505</td><td>-</td><td>0</td><td>AATGCTTGAAAGCCATGATC</td><td>CGG</td><td>||||||||[||||||||||||]PAM</td><td>NA</td><td>Intergenic</td><td>NA</td><td>NA</td></tr>
<tr><td></td><td></td><td></td><td></td><td></td><td></td><td></td><td></td><td></td><td></td><td></td><td></td></tr>
<tr><td>T26568</td><td>AGCCATGATCCGGATCCGCGAGG</td><td>1000</td><td></td><td></td><td></td><td></td><td></td><td></td><td></td><td></td><td></td></tr>
<tr><td>Oligo fwd</td><td>CACCAGCCATGATCCGGATCCGCG</td><td></td><td></td><td></td><td></td><td></td><td></td><td></td><td></td><td></td><td></td></tr>
<tr><td>Oligo rev</td><td>AAACCGCGGATCCGGATCATGGCT</td><td></td><td></td><td></td><td></td><td></td><td></td><td></td><td></td><td></td><td></td></tr>
<tr><td>Chromosome</td><td>start</td><td>end</td><td>strand</td><td>MM</td><td>target_seq</td><td>PAM</td><td>alignment</td><td>distance</td><td>position</td><td>gene name</td><td>gene id</td></tr>
<tr><td>GU305914.1</td><td>98473</td><td>98495</td><td>-</td><td>0</td><td>AGCCATGATCCGGATCCGCG</td><td>AGG</td><td>||||||||[||||||||||||]PAM</td><td>NA</td><td>Intergenic</td><td>NA</td><td>NA</td></tr>
<tr><td></td><td></td><td></td><td></td><td></td><td></td><td></td><td></td><td></td><td></td><td></td><td></td></tr>
<tr><td>T26569</td><td>GATCCGCGAGGCGCGCGCCCAGG</td><td>1000</td><td></td><td></td><td></td><td></td><td></td><td></td><td></td><td></td><td></td></tr>
<tr><td>Oligo fwd</td><td>CACCGATCCGCGAGGCGCGCGCCC</td><td></td><td></td><td></td><td></td><td></td><td></td><td></td><td></td><td></td><td></td></tr>
<tr><td>Oligo rev</td><td>AAACGGGCGCGCGCCTCGCGGATC</td><td></td><td></td><td></td><td></td><td></td><td></td><td></td><td></td><td></td><td></td></tr>
<tr><td>Chromosome</td><td>start</td><td>end</td><td>strand</td><td>MM</td><td>target_seq</td><td>PAM</td><td>alignment</td><td>distance</td><td>position</td><td>gene name</td><td>gene id</td></tr>
<tr><td>GU305914.1</td><td>98461</td><td>98483</td><td>-</td><td>0</td><td>GATCCGCGAGGCGCGCGCCC</td><td>AGG</td><td>||||||||[||||||||||||]PAM</td><td>NA</td><td>Intergenic</td><td>NA</td><td>NA</td></tr>
<tr><td></td><td></td><td></td><td></td><td></td><td></td><td></td><td></td><td></td><td></td><td></td><td></td></tr>
<tr><td>T26570</td><td>ATCCGCGAGGCGCGCGCCCAGGG</td><td>1000</td><td></td><td></td><td></td><td></td><td></td><td></td><td></td><td></td><td></td></tr>
<tr><td>Oligo fwd</td><td>CACCATCCGCGAGGCGCGCGCCCA</td><td></td><td></td><td></td><td></td><td></td><td></td><td></td><td></td><td></td><td></td></tr>
<tr><td>Oligo rev</td><td>AAACTGGGCGCGCGCCTCGCGGAT</td><td></td><td></td><td></td><td></td><td></td><td></td><td></td><td></td><td></td><td></td></tr>
<tr><td>Chromosome</td><td>start</td><td>end</td><td>strand</td><td>MM</td><td>target_seq</td><td>PAM</td><td>alignment</td><td>distance</td><td>position</td><td>gene name</td><td>gene id</td></tr>
<tr><td>GU305914.1</td><td>98460</td><td>98482</td><td>-</td><td>0</td><td>ATCCGCGAGGCGCGCGCCCA</td><td>GGG</td><td>||||||||[||||||||||||]PAM</td><td>NA</td><td>Intergenic</td><td>NA</td><td>NA</td></tr>
<tr><td></td><td></td><td></td><td></td><td></td><td></td><td></td><td></td><td></td><td></td><td></td><td></td></tr>
<tr><td>T26571</td><td>CCGCCGTCCCGAGCCGTCGTCGG</td><td>1000</td><td></td><td></td><td></td><td></td><td></td><td></td><td></td><td></td><td></td></tr>
<tr><td>Oligo fwd</td><td>CACCCCGCCGTCCCGAGCCGTCGT</td><td></td><td></td><td></td><td></td><td></td><td></td><td></td><td></td><td></td><td></td></tr>
<tr><td>Oligo rev</td><td>AAACACGACGGCTCGGGACGGCGG</td><td></td><td></td><td></td><td></td><td></td><td></td><td></td><td></td><td></td><td></td></tr>
<tr><td>Chromosome</td><td>start</td><td>end</td><td>strand</td><td>MM</td><td>target_seq</td><td>PAM</td><td>alignment</td><td>distance</td><td>position</td><td>gene name</td><td>gene id</td></tr>
<tr><td>GU305914.1</td><td>98437</td><td>98459</td><td>-</td><td>0</td><td>CCGCCGTCCCGAGCCGTCGT</td><td>CGG</td><td>||||||||[||||||||||||]PAM</td><td>NA</td><td>Intergenic</td><td>NA</td><td>NA</td></tr>
<tr><td></td><td></td><td></td><td></td><td></td><td></td><td></td><td></td><td></td><td></td><td></td><td></td></tr>
<tr><td>T26572</td><td>AACAATACCACGCTGAGCCTAGG</td><td>1000</td><td></td><td></td><td></td><td></td><td></td><td></td><td></td><td></td><td></td></tr>
<tr><td>Oligo fwd</td><td>CACCAACAATACCACGCTGAGCCT</td><td></td><td></td><td></td><td></td><td></td><td></td><td></td><td></td><td></td><td></td></tr>
<tr><td>Oligo rev</td><td>AAACAGGCTCAGCGTGGTATTGTT</td><td></td><td></td><td></td><td></td><td></td><td></td><td></td><td></td><td></td><td></td></tr>
<tr><td>Chromosome</td><td>start</td><td>end</td><td>strand</td><td>MM</td><td>target_seq</td><td>PAM</td><td>alignment</td><td>distance</td><td>position</td><td>gene name</td><td>gene id</td></tr>
<tr><td>GU305914.1</td><td>98367</td><td>98389</td><td>-</td><td>0</td><td>AACAATACCACGCTGAGCCT</td><td>AGG</td><td>||||||||[||||||||||||]PAM</td><td>NA</td><td>Intergenic</td><td>NA</td><td>NA</td></tr>
<tr><td></td><td></td><td></td><td></td><td></td><td></td><td></td><td></td><td></td><td></td><td></td><td></td></tr>
<tr><td>T26573</td><td>AATACCACGCTGAGCCTAGGCGG</td><td>1000</td><td></td><td></td><td></td><td></td><td></td><td></td><td></td><td></td><td></td></tr>
<tr><td>Oligo fwd</td><td>CACCAATACCACGCTGAGCCTAGG</td><td></td><td></td><td></td><td></td><td></td><td></td><td></td><td></td><td></td><td></td></tr>
<tr><td>Oligo rev</td><td>AAACCCTAGGCTCAGCGTGGTATT</td><td></td><td></td><td></td><td></td><td></td><td></td><td></td><td></td><td></td><td></td></tr>
<tr><td>Chromosome</td><td>start</td><td>end</td><td>strand</td><td>MM</td><td>target_seq</td><td>PAM</td><td>alignment</td><td>distance</td><td>position</td><td>gene name</td><td>gene id</td></tr>
<tr><td>GU305914.1</td><td>98364</td><td>98386</td><td>-</td><td>0</td><td>AATACCACGCTGAGCCTAGG</td><td>CGG</td><td>||||||||[||||||||||||]PAM</td><td>NA</td><td>Intergenic</td><td>NA</td><td>NA</td></tr>
<tr><td></td><td></td><td></td><td></td><td></td><td></td><td></td><td></td><td></td><td></td><td></td><td></td></tr>
<tr><td>T26574</td><td>ACCACGCTGAGCCTAGGCGGCGG</td><td>1000</td><td></td><td></td><td></td><td></td><td></td><td></td><td></td><td></td><td></td></tr>
<tr><td>Oligo fwd</td><td>CACCACCACGCTGAGCCTAGGCGG</td><td></td><td></td><td></td><td></td><td></td><td></td><td></td><td></td><td></td><td></td></tr>
<tr><td>Oligo rev</td><td>AAACCCGCCTAGGCTCAGCGTGGT</td><td></td><td></td><td></td><td></td><td></td><td></td><td></td><td></td><td></td><td></td></tr>
<tr><td>Chromosome</td><td>start</td><td>end</td><td>strand</td><td>MM</td><td>target_seq</td><td>PAM</td><td>alignment</td><td>distance</td><td>position</td><td>gene name</td><td>gene id</td></tr>
<tr><td>GU305914.1</td><td>98361</td><td>98383</td><td>-</td><td>0</td><td>ACCACGCTGAGCCTAGGCGG</td><td>CGG</td><td>||||||||[||||||||||||]PAM</td><td>NA</td><td>Intergenic</td><td>NA</td><td>NA</td></tr>
<tr><td></td><td></td><td></td><td></td><td></td><td></td><td></td><td></td><td></td><td></td><td></td><td></td></tr>
<tr><td>T26575</td><td>ACGCTGAGCCTAGGCGGCGGTGG</td><td>1000</td><td></td><td></td><td></td><td></td><td></td><td></td><td></td><td></td><td></td></tr>
<tr><td>Oligo fwd</td><td>CACCACGCTGAGCCTAGGCGGCGG</td><td></td><td></td><td></td><td></td><td></td><td></td><td></td><td></td><td></td><td></td></tr>
<tr><td>Oligo rev</td><td>AAACCCGCCGCCTAGGCTCAGCGT</td><td></td><td></td><td></td><td></td><td></td><td></td><td></td><td></td><td></td><td></td></tr>
<tr><td>Chromosome</td><td>start</td><td>end</td><td>strand</td><td>MM</td><td>target_seq</td><td>PAM</td><td>alignment</td><td>distance</td><td>position</td><td>gene name</td><td>gene id</td></tr>
<tr><td>GU305914.1</td><td>98358</td><td>98380</td><td>-</td><td>0</td><td>ACGCTGAGCCTAGGCGGCGG</td><td>TGG</td><td>||||||||[||||||||||||]PAM</td><td>NA</td><td>Intergenic</td><td>NA</td><td>NA</td></tr>
<tr><td></td><td></td><td></td><td></td><td></td><td></td><td></td><td></td><td></td><td></td><td></td><td></td></tr>
<tr><td>T26576</td><td>CTGAGCCTAGGCGGCGGTGGCGG</td><td>1000</td><td></td><td></td><td></td><td></td><td></td><td></td><td></td><td></td><td></td></tr>
<tr><td>Oligo fwd</td><td>CACCCTGAGCCTAGGCGGCGGTGG</td><td></td><td></td><td></td><td></td><td></td><td></td><td></td><td></td><td></td><td></td></tr>
<tr><td>Oligo rev</td><td>AAACCCACCGCCGCCTAGGCTCAG</td><td></td><td></td><td></td><td></td><td></td><td></td><td></td><td></td><td></td><td></td></tr>
<tr><td>Chromosome</td><td>start</td><td>end</td><td>strand</td><td>MM</td><td>target_seq</td><td>PAM</td><td>alignment</td><td>distance</td><td>position</td><td>gene name</td><td>gene id</td></tr>
<tr><td>GU305914.1</td><td>98355</td><td>98377</td><td>-</td><td>0</td><td>CTGAGCCTAGGCGGCGGTGG</td><td>CGG</td><td>||||||||[||||||||||||]PAM</td><td>NA</td><td>Intergenic</td><td>NA</td><td>NA</td></tr>
<tr><td></td><td></td><td></td><td></td><td></td><td></td><td></td><td></td><td></td><td></td><td></td><td></td></tr>
<tr><td>T26577</td><td>AGCCTAGGCGGCGGTGGCGGCGG</td><td>1000</td><td></td><td></td><td></td><td></td><td></td><td></td><td></td><td></td><td></td></tr>
<tr><td>Oligo fwd</td><td>CACCAGCCTAGGCGGCGGTGGCGG</td><td></td><td></td><td></td><td></td><td></td><td></td><td></td><td></td><td></td><td></td></tr>
<tr><td>Oligo rev</td><td>AAACCCGCCACCGCCGCCTAGGCT</td><td></td><td></td><td></td><td></td><td></td><td></td><td></td><td></td><td></td><td></td></tr>
<tr><td>Chromosome</td><td>start</td><td>end</td><td>strand</td><td>MM</td><td>target_seq</td><td>PAM</td><td>alignment</td><td>distance</td><td>position</td><td>gene name</td><td>gene id</td></tr>
<tr><td>GU305914.1</td><td>98352</td><td>98374</td><td>-</td><td>0</td><td>AGCCTAGGCGGCGGTGGCGG</td><td>CGG</td><td>||||||||[||||||||||||]PAM</td><td>NA</td><td>Intergenic</td><td>NA</td><td>NA</td></tr>
<tr><td>GU305914.1</td><td>35653</td><td>35675</td><td>+</td><td>4</td><td>ATCCACGGCGGCGGCGGCGG</td><td>CGG</td><td>|-||--||[||||||-|||||]PAM</td><td>NA</td><td>Intergenic</td><td>NA</td><td>NA</td></tr>
<tr><td>GU305914.1</td><td>161754</td><td>161776</td><td>-</td><td>4</td><td>GGCCGAGGAGGCGGCGGCGG</td><td>AGG</td><td>-|||-|||[-|||||-|||||]PAM</td><td>NA</td><td>Intergenic</td><td>NA</td><td>NA</td></tr>
<tr><td></td><td></td><td></td><td></td><td></td><td></td><td></td><td></td><td></td><td></td><td></td><td></td></tr>
<tr><td>T26578</td><td>CTATCACCGCCAGACCTCTCCGG</td><td>1000</td><td></td><td></td><td></td><td></td><td></td><td></td><td></td><td></td><td></td></tr>
<tr><td>Oligo fwd</td><td>CACCCTATCACCGCCAGACCTCTC</td><td></td><td></td><td></td><td></td><td></td><td></td><td></td><td></td><td></td><td></td></tr>
<tr><td>Oligo rev</td><td>AAACGAGAGGTCTGGCGGTGATAG</td><td></td><td></td><td></td><td></td><td></td><td></td><td></td><td></td><td></td><td></td></tr>
<tr><td>Chromosome</td><td>start</td><td>end</td><td>strand</td><td>MM</td><td>target_seq</td><td>PAM</td><td>alignment</td><td>distance</td><td>position</td><td>gene name</td><td>gene id</td></tr>
<tr><td>GU305914.1</td><td>98326</td><td>98348</td><td>-</td><td>0</td><td>CTATCACCGCCAGACCTCTC</td><td>CGG</td><td>||||||||[||||||||||||]PAM</td><td>NA</td><td>Intergenic</td><td>NA</td><td>NA</td></tr>
<tr><td></td><td></td><td></td><td></td><td></td><td></td><td></td><td></td><td></td><td></td><td></td><td></td></tr>
<tr><td>T26579</td><td>GGACATCCGCGACTACACGACGG</td><td>1000</td><td></td><td></td><td></td><td></td><td></td><td></td><td></td><td></td><td></td></tr>
<tr><td>Oligo fwd</td><td>CACCGGACATCCGCGACTACACGA</td><td></td><td></td><td></td><td></td><td></td><td></td><td></td><td></td><td></td><td></td></tr>
<tr><td>Oligo rev</td><td>AAACTCGTGTAGTCGCGGATGTCC</td><td></td><td></td><td></td><td></td><td></td><td></td><td></td><td></td><td></td><td></td></tr>
<tr><td>Chromosome</td><td>start</td><td>end</td><td>strand</td><td>MM</td><td>target_seq</td><td>PAM</td><td>alignment</td><td>distance</td><td>position</td><td>gene name</td><td>gene id</td></tr>
<tr><td>GU305914.1</td><td>98305</td><td>98327</td><td>-</td><td>0</td><td>GGACATCCGCGACTACACGA</td><td>CGG</td><td>||||||||[||||||||||||]PAM</td><td>NA</td><td>Intergenic</td><td>NA</td><td>NA</td></tr>
<tr><td></td><td></td><td></td><td></td><td></td><td></td><td></td><td></td><td></td><td></td><td></td><td></td></tr>
<tr><td>T26580</td><td>GACATCCGCGACTACACGACGGG</td><td>1000</td><td></td><td></td><td></td><td></td><td></td><td></td><td></td><td></td><td></td></tr>
<tr><td>Oligo fwd</td><td>CACCGACATCCGCGACTACACGAC</td><td></td><td></td><td></td><td></td><td></td><td></td><td></td><td></td><td></td><td></td></tr>
<tr><td>Oligo rev</td><td>AAACGTCGTGTAGTCGCGGATGTC</td><td></td><td></td><td></td><td></td><td></td><td></td><td></td><td></td><td></td><td></td></tr>
<tr><td>Chromosome</td><td>start</td><td>end</td><td>strand</td><td>MM</td><td>target_seq</td><td>PAM</td><td>alignment</td><td>distance</td><td>position</td><td>gene name</td><td>gene id</td></tr>
<tr><td>GU305914.1</td><td>98304</td><td>98326</td><td>-</td><td>0</td><td>GACATCCGCGACTACACGAC</td><td>GGG</td><td>||||||||[||||||||||||]PAM</td><td>NA</td><td>Intergenic</td><td>NA</td><td>NA</td></tr>
<tr><td></td><td></td><td></td><td></td><td></td><td></td><td></td><td></td><td></td><td></td><td></td><td></td></tr>
<tr><td>T26581</td><td>CGACTACACGACGGGTTCCTTGG</td><td>1000</td><td></td><td></td><td></td><td></td><td></td><td></td><td></td><td></td><td></td></tr>
<tr><td>Oligo fwd</td><td>CACCCGACTACACGACGGGTTCCT</td><td></td><td></td><td></td><td></td><td></td><td></td><td></td><td></td><td></td><td></td></tr>
<tr><td>Oligo rev</td><td>AAACAGGAACCCGTCGTGTAGTCG</td><td></td><td></td><td></td><td></td><td></td><td></td><td></td><td></td><td></td><td></td></tr>
<tr><td>Chromosome</td><td>start</td><td>end</td><td>strand</td><td>MM</td><td>target_seq</td><td>PAM</td><td>alignment</td><td>distance</td><td>position</td><td>gene name</td><td>gene id</td></tr>
<tr><td>GU305914.1</td><td>98296</td><td>98318</td><td>-</td><td>0</td><td>CGACTACACGACGGGTTCCT</td><td>TGG</td><td>||||||||[||||||||||||]PAM</td><td>NA</td><td>Intergenic</td><td>NA</td><td>NA</td></tr>
</table>

<table>
<tr><td>T26582</td><td>GACTACACGACGGGTTCCTTGGG</td><td>1000</td><td></td><td></td><td></td><td></td><td></td><td></td><td></td><td></td><td></td></tr>
<tr><td>Oligo fwd</td><td>CACCGACTACACGACGGGTTCCTT</td><td></td><td></td><td></td><td></td><td></td><td></td><td></td><td></td><td></td><td></td></tr>
<tr><td>Oligo rev</td><td>AAACAAGGAACCCGTCGTGTAGTC</td><td></td><td></td><td></td><td></td><td></td><td></td><td></td><td></td><td></td><td></td></tr>
<tr><td>Chromosome</td><td>start</td><td>end</td><td>strand</td><td>MM</td><td>target_seq</td><td>PAM</td><td>alignment</td><td>distance</td><td>position</td><td>gene name</td><td>gene id</td></tr>
<tr><td>GU305914.1</td><td>98295</td><td>98317</td><td>-</td><td>0</td><td>GACTACACGACGGGTTCCTT</td><td>GGG</td><td>||||||||[||||||||||||]PAM</td><td>NA</td><td>Intergenic</td><td>NA</td><td>NA</td></tr>
<tr><td>T26583</td><td>GGGCCTGTGCATGTTCCCCATGG</td><td>1000</td><td></td><td></td><td></td><td></td><td></td><td></td><td></td><td></td><td></td></tr>
<tr><td>Oligo fwd</td><td>CACCGGGCCTGTGCATGTTCCCCA</td><td></td><td></td><td></td><td></td><td></td><td></td><td></td><td></td><td></td><td></td></tr>
<tr><td>Oligo rev</td><td>AAACTGGGGAACATGCACAGGCCC</td><td></td><td></td><td></td><td></td><td></td><td></td><td></td><td></td><td></td><td></td></tr>
<tr><td>Chromosome</td><td>start</td><td>end</td><td>strand</td><td>MM</td><td>target_seq</td><td>PAM</td><td>alignment</td><td>distance</td><td>position</td><td>gene name</td><td>gene id</td></tr>
<tr><td>GU305914.1</td><td>98275</td><td>98297</td><td>-</td><td>0</td><td>GGGCCTGTGCATGTTCCCCA</td><td>TGG</td><td>||||||||[||||||||||||]PAM</td><td>NA</td><td>Intergenic</td><td>NA</td><td>NA</td></tr>
<tr><td>T26584</td><td>CCATCAAGCTGCTAGAGAACAGG</td><td>1000</td><td></td><td></td><td></td><td></td><td></td><td></td><td></td><td></td><td></td></tr>
<tr><td>Oligo fwd</td><td>CACCCCATCAAGCTGCTAGAGAAC</td><td></td><td></td><td></td><td></td><td></td><td></td><td></td><td></td><td></td><td></td></tr>
<tr><td>Oligo rev</td><td>AAACGTTCTCTAGCAGCTTGATGG</td><td></td><td></td><td></td><td></td><td></td><td></td><td></td><td></td><td></td><td></td></tr>
<tr><td>Chromosome</td><td>start</td><td>end</td><td>strand</td><td>MM</td><td>target_seq</td><td>PAM</td><td>alignment</td><td>distance</td><td>position</td><td>gene name</td><td>gene id</td></tr>
<tr><td>GU305914.1</td><td>98240</td><td>98262</td><td>-</td><td>0</td><td>CCATCAAGCTGCTAGAGAAC</td><td>AGG</td><td>||||||||[||||||||||||]PAM</td><td>NA</td><td>Intergenic</td><td>NA</td><td>NA</td></tr>
<tr><td>T26585</td><td>CGACAACGACCGCCACGCGCCGG</td><td>1000</td><td></td><td></td><td></td><td></td><td></td><td></td><td></td><td></td><td></td></tr>
<tr><td>Oligo fwd</td><td>CACCCGACAACGACCGCCACGCGC</td><td></td><td></td><td></td><td></td><td></td><td></td><td></td><td></td><td></td><td></td></tr>
<tr><td>Oligo rev</td><td>AAACGCGCGTGGCGGTCGTTGTCG</td><td></td><td></td><td></td><td></td><td></td><td></td><td></td><td></td><td></td><td></td></tr>
<tr><td>Chromosome</td><td>start</td><td>end</td><td>strand</td><td>MM</td><td>target_seq</td><td>PAM</td><td>alignment</td><td>distance</td><td>position</td><td>gene name</td><td>gene id</td></tr>
<tr><td>GU305914.1</td><td>98212</td><td>98234</td><td>-</td><td>0</td><td>CGACAACGACCGCCACGCGC</td><td>CGG</td><td>||||||||[||||||||||||]PAM</td><td>NA</td><td>Intergenic</td><td>NA</td><td>NA</td></tr>
<tr><td>T26586</td><td>CAACGACCGCCACGCGCCGGCGG</td><td>1000</td><td></td><td></td><td></td><td></td><td></td><td></td><td></td><td></td><td></td></tr>
<tr><td>Oligo fwd</td><td>CACCCAACGACCGCCACGCGCCGG</td><td></td><td></td><td></td><td></td><td></td><td></td><td></td><td></td><td></td><td></td></tr>
<tr><td>Oligo rev</td><td>AAACCCGGCGCGTGGCGGTCGTTG</td><td></td><td></td><td></td><td></td><td></td><td></td><td></td><td></td><td></td><td></td></tr>
<tr><td>Chromosome</td><td>start</td><td>end</td><td>strand</td><td>MM</td><td>target_seq</td><td>PAM</td><td>alignment</td><td>distance</td><td>position</td><td>gene name</td><td>gene id</td></tr>
<tr><td>GU305914.1</td><td>98209</td><td>98231</td><td>-</td><td>0</td><td>CAACGACCGCCACGCGCCGG</td><td>CGG</td><td>||||||||[||||||||||||]PAM</td><td>NA</td><td>Intergenic</td><td>NA</td><td>NA</td></tr>
<tr><td>T26587</td><td>CGACCGCCACGCGCCGGCGGTGG</td><td>1000</td><td></td><td></td><td></td><td></td><td></td><td></td><td></td><td></td><td></td></tr>
<tr><td>Oligo fwd</td><td>CACCCGACCGCCACGCGCCGGCGG</td><td></td><td></td><td></td><td></td><td></td><td></td><td></td><td></td><td></td><td></td></tr>
<tr><td>Oligo rev</td><td>AAACCCGCCGGCGCGTGGCGGTCG</td><td></td><td></td><td></td><td></td><td></td><td></td><td></td><td></td><td></td><td></td></tr>
<tr><td>Chromosome</td><td>start</td><td>end</td><td>strand</td><td>MM</td><td>target_seq</td><td>PAM</td><td>alignment</td><td>distance</td><td>position</td><td>gene name</td><td>gene id</td></tr>
<tr><td>GU305914.1</td><td>98206</td><td>98228</td><td>-</td><td>0</td><td>CGACCGCCACGCGCCGGCGG</td><td>TGG</td><td>||||||||[||||||||||||]PAM</td><td>NA</td><td>Intergenic</td><td>NA</td><td>NA</td></tr>
<tr><td>T26588</td><td>GCACTTCGACGCGCTGCCGCCGG</td><td>1000</td><td></td><td></td><td></td><td></td><td></td><td></td><td></td><td></td><td></td></tr>
<tr><td>Oligo fwd</td><td>CACCGCACTTCGACGCGCTGCCGC</td><td></td><td></td><td></td><td></td><td></td><td></td><td></td><td></td><td></td><td></td></tr>
<tr><td>Oligo rev</td><td>AAACGCGGCAGCGCGTCGAAGTGC</td><td></td><td></td><td></td><td></td><td></td><td></td><td></td><td></td><td></td><td></td></tr>
<tr><td>Chromosome</td><td>start</td><td>end</td><td>strand</td><td>MM</td><td>target_seq</td><td>PAM</td><td>alignment</td><td>distance</td><td>position</td><td>gene name</td><td>gene id</td></tr>
<tr><td>GU305914.1</td><td>98134</td><td>98156</td><td>-</td><td>0</td><td>GCACTTCGACGCGCTGCCGC</td><td>CGG</td><td>||||||||[||||||||||||]PAM</td><td>NA</td><td>Intergenic</td><td>NA</td><td>NA</td></tr>
<tr><td>T26589</td><td>CTTCGACGCGCTGCCGCCGGAGG</td><td>1000</td><td></td><td></td><td></td><td></td><td></td><td></td><td></td><td></td><td></td></tr>
<tr><td>Oligo fwd</td><td>CACCCTTCGACGCGCTGCCGCCGG</td><td></td><td></td><td></td><td></td><td></td><td></td><td></td><td></td><td></td><td></td></tr>
<tr><td>Oligo rev</td><td>AAACCCGGCGGCAGCGCGTCGAAG</td><td></td><td></td><td></td><td></td><td></td><td></td><td></td><td></td><td></td><td></td></tr>
<tr><td>Chromosome</td><td>start</td><td>end</td><td>strand</td><td>MM</td><td>target_seq</td><td>PAM</td><td>alignment</td><td>distance</td><td>position</td><td>gene name</td><td>gene id</td></tr>
<tr><td>GU305914.1</td><td>98131</td><td>98153</td><td>-</td><td>0</td><td>CTTCGACGCGCTGCCGCCGG</td><td>AGG</td><td>||||||||[||||||||||||]PAM</td><td>NA</td><td>Intergenic</td><td>NA</td><td>NA</td></tr>
<tr><td>T26590</td><td>CGCTGCCGCCGGAGGAACTGCGG</td><td>1000</td><td></td><td></td><td></td><td></td><td></td><td></td><td></td><td></td><td></td></tr>
<tr><td>Oligo fwd</td><td>CACCCGCTGCCGCCGGAGGAACTG</td><td></td><td></td><td></td><td></td><td></td><td></td><td></td><td></td><td></td><td></td></tr>
<tr><td>Oligo rev</td><td>AAACCAGTTCCTCCGGCGGCAGCG</td><td></td><td></td><td></td><td></td><td></td><td></td><td></td><td></td><td></td><td></td></tr>
<tr><td>Chromosome</td><td>start</td><td>end</td><td>strand</td><td>MM</td><td>target_seq</td><td>PAM</td><td>alignment</td><td>distance</td><td>position</td><td>gene name</td><td>gene id</td></tr>
<tr><td>GU305914.1</td><td>98123</td><td>98145</td><td>-</td><td>0</td><td>CGCTGCCGCCGGAGGAACTG</td><td>CGG</td><td>||||||||[||||||||||||]PAM</td><td>NA</td><td>Intergenic</td><td>NA</td><td>NA</td></tr>
<tr><td>T26591</td><td>GCTGCCGCCGGAGGAACTGCGGG</td><td>1000</td><td></td><td></td><td></td><td></td><td></td><td></td><td></td><td></td><td></td></tr>
<tr><td>Oligo fwd</td><td>CACCGCTGCCGCCGGAGGAACTGC</td><td></td><td></td><td></td><td></td><td></td><td></td><td></td><td></td><td></td><td></td></tr>
<tr><td>Oligo rev</td><td>AAACGCAGTTCCTCCGGCGGCAGC</td><td></td><td></td><td></td><td></td><td></td><td></td><td></td><td></td><td></td><td></td></tr>
<tr><td>Chromosome</td><td>start</td><td>end</td><td>strand</td><td>MM</td><td>target_seq</td><td>PAM</td><td>alignment</td><td>distance</td><td>position</td><td>gene name</td><td>gene id</td></tr>
<tr><td>GU305914.1</td><td>98122</td><td>98144</td><td>-</td><td>0</td><td>GCTGCCGCCGGAGGAACTGC</td><td>GGG</td><td>||||||||[||||||||||||]PAM</td><td>NA</td><td>Intergenic</td><td>NA</td><td>NA</td></tr>
<tr><td>T26592</td><td>GCCGGAGGAACTGCGGGTCCTGG</td><td>1000</td><td></td><td></td><td></td><td></td><td></td><td></td><td></td><td></td><td></td></tr>
<tr><td>Oligo fwd</td><td>CACCGCCGGAGGAACTGCGGGTCC</td><td></td><td></td><td></td><td></td><td></td><td></td><td></td><td></td><td></td><td></td></tr>
<tr><td>Oligo rev</td><td>AAACGGACCCGCAGTTCCTCCGGC</td><td></td><td></td><td></td><td></td><td></td><td></td><td></td><td></td><td></td><td></td></tr>
<tr><td>Chromosome</td><td>start</td><td>end</td><td>strand</td><td>MM</td><td>target_seq</td><td>PAM</td><td>alignment</td><td>distance</td><td>position</td><td>gene name</td><td>gene id</td></tr>
<tr><td>GU305914.1</td><td>98116</td><td>98138</td><td>-</td><td>0</td><td>GCCGGAGGAACTGCGGGTCC</td><td>TGG</td><td>||||||||[||||||||||||]PAM</td><td>NA</td><td>Intergenic</td><td>NA</td><td>NA</td></tr>
<tr><td>T26593</td><td>CATCAACAACGCGCCGAGCCTGG</td><td>1000</td><td></td><td></td><td></td><td></td><td></td><td></td><td></td><td></td><td></td></tr>
<tr><td>Oligo fwd</td><td>CACCCATCAACAACGCGCCGAGCC</td><td></td><td></td><td></td><td></td><td></td><td></td><td></td><td></td><td></td><td></td></tr>
<tr><td>Oligo rev</td><td>AAACGGCTCGGCGCGTTGTTGATG</td><td></td><td></td><td></td><td></td><td></td><td></td><td></td><td></td><td></td><td></td></tr>
<tr><td>Chromosome</td><td>start</td><td>end</td><td>strand</td><td>MM</td><td>target_seq</td><td>PAM</td><td>alignment</td><td>distance</td><td>position</td><td>gene name</td><td>gene id</td></tr>
<tr><td>GU305914.1</td><td>98074</td><td>98096</td><td>-</td><td>0</td><td>CATCAACAACGCGCCGAGCC</td><td>TGG</td><td>||||||||[||||||||||||]PAM</td><td>NA</td><td>Intergenic</td><td>NA</td><td>NA</td></tr>
<tr><td>T26594</td><td>CAACGCGCCGAGCCTGGACGTGG</td><td>1000</td><td></td><td></td><td></td><td></td><td></td><td></td><td></td><td></td><td></td></tr>
<tr><td>Oligo fwd</td><td>CACCCAACGCGCCGAGCCTGGACG</td><td></td><td></td><td></td><td></td><td></td><td></td><td></td><td></td><td></td><td></td></tr>
<tr><td>Oligo rev</td><td>AAACCGTCCAGGCTCGGCGCGTTG</td><td></td><td></td><td></td><td></td><td></td><td></td><td></td><td></td><td></td><td></td></tr>
<tr><td>Chromosome</td><td>start</td><td>end</td><td>strand</td><td>MM</td><td>target_seq</td><td>PAM</td><td>alignment</td><td>distance</td><td>position</td><td>gene name</td><td>gene id</td></tr>
<tr><td>GU305914.1</td><td>98068</td><td>98090</td><td>-</td><td>0</td><td>CAACGCGCCGAGCCTGGACG</td><td>TGG</td><td>||||||||[||||||||||||]PAM</td><td>NA</td><td>Intergenic</td><td>NA</td><td>NA</td></tr>
<tr><td>T26595</td><td>CGCGCCGAGCCTGGACGTGGTGG</td><td>1000</td><td></td><td></td><td></td><td></td><td></td><td></td><td></td><td></td><td></td></tr>
<tr><td>Oligo fwd</td><td>CACCCGCGCCGAGCCTGGACGTGG</td><td></td><td></td><td></td><td></td><td></td><td></td><td></td><td></td><td></td><td></td></tr>
<tr><td>Oligo rev</td><td>AAACCCACGTCCAGGCTCGGCGCG</td><td></td><td></td><td></td><td></td><td></td><td></td><td></td><td></td><td></td><td></td></tr>
<tr><td>Chromosome</td><td>start</td><td>end</td><td>strand</td><td>MM</td><td>target_seq</td><td>PAM</td><td>alignment</td><td>distance</td><td>position</td><td>gene name</td><td>gene id</td></tr>
<tr><td>GU305914.1</td><td>98065</td><td>98087</td><td>-</td><td>0</td><td>CGCGCCGAGCCTGGACGTGG</td><td>TGG</td><td>||||||||[||||||||||||]PAM</td><td>NA</td><td>Intergenic</td><td>NA</td><td>NA</td></tr>
<tr><td>T26596</td><td>GCCGAGCCTGGACGTGGTGGCGG</td><td>1000</td><td></td><td></td><td></td><td></td><td></td><td></td><td></td><td></td><td></td></tr>
<tr><td>Oligo fwd</td><td>CACCGCCGAGCCTGGACGTGGTGG</td><td></td><td></td><td></td><td></td><td></td><td></td><td></td><td></td><td></td><td></td></tr>
<tr><td>Oligo rev</td><td>AAACCCACCACGTCCAGGCTCGGC</td><td></td><td></td><td></td><td></td><td></td><td></td><td></td><td></td><td></td><td></td></tr>
<tr><td>Chromosome</td><td>start</td><td>end</td><td>strand</td><td>MM</td><td>target_seq</td><td>PAM</td><td>alignment</td><td>distance</td><td>position</td><td>gene name</td><td>gene id</td></tr>
<tr><td>GU305914.1</td><td>98062</td><td>98084</td><td>-</td><td>0</td><td>GCCGAGCCTGGACGTGGTGG</td><td>CGG</td><td>||||||||[||||||||||||]PAM</td><td>NA</td><td>Intergenic</td><td>NA</td><td>NA</td></tr>
</table>

T26597 CCTGGACGTGGTGGCGGCGATGG 1000
Oligo fwd CACCCCTGGACGTGGTGGCGGCGA
Oligo rev AAACTCGCCGCCACCACGTCCAGG
Chromosome start end strand MM target_seq PAM alignment distance position gene name gene id
GU305914.1 98056 98078 - 0 CCTGGACGTGGTGGCGGCGA TGG ||||||||[||||||||||||]PAM NA Intergenic NA NA

T26598 GGACGTGGTGGCGGCGATGGCGG 1000
Oligo fwd CACCGGACGTGGTGGCGGCGATGG
Oligo rev AAACCCATCGCCGCCACCACGTCC
Chromosome start end strand MM target_seq PAM alignment distance position gene name gene id
GU305914.1 98053 98075 - 0 GGACGTGGTGGCGGCGATGG CGG ||||||||[||||||||||||]PAM NA Intergenic NA NA
GU305914.1 62449 62471 + 4 GGGGGTGGTGGCGGCGGGGG TGG ||--||||[||||||||--||]PAM NA Intergenic NA NA

T26599 CCACCGCGTACACAACCTGCCGG 1000
Oligo fwd CACCCCACCGCGTACACAACCTGC
Oligo rev AAACGCAGGTTGTGTACGCGGTGG
Chromosome start end strand MM target_seq PAM alignment distance position gene name gene id
GU305914.1 98005 98027 - 0 CCACCGCGTACACAACCTGC CGG ||||||||[||||||||||||]PAM NA Intergenic NA NA

T26600 AGACCTGTACATGAGCACCCTGG 1000
Oligo fwd CACCAGACCTGTACATGAGCACCC
Oligo rev AAACGGGTGCTCATGTACAGGTCT
Chromosome start end strand MM target_seq PAM alignment distance position gene name gene id
GU305914.1 97972 97994 - 0 AGACCTGTACATGAGCACCC TGG ||||||||[||||||||||||]PAM NA Intergenic NA NA

T26601 GACCTGTACATGAGCACCCTGGG 1000
Oligo fwd CACCGACCTGTACATGAGCACCCT
Oligo rev AAACAGGGTGCTCATGTACAGGTC
Chromosome start end strand MM target_seq PAM alignment distance position gene name gene id
GU305914.1 97971 97993 - 0 GACCTGTACATGAGCACCCT GGG ||||||||[||||||||||||]PAM NA Intergenic NA NA

T26602 ACGTTCAACAAACTGAACCTCGG 1000
Oligo fwd CACCACGTTCAACAAACTGAACCT
Oligo rev AAACAGGTTCAGTTTGTTGAACGT
Chromosome start end strand MM target_seq PAM alignment distance position gene name gene id
GU305914.1 97932 97954 - 0 ACGTTCAACAAACTGAACCT CGG ||||||||[||||||||||||]PAM NA Intergenic NA NA

T26603 CGGCGAGCTGTCGTGCCTCCTGG 1000
Oligo fwd CACCCGGCGAGCTGTCGTGCCTCC
Oligo rev AAACGGAGGCACGACAGCTCGCCG
Chromosome start end strand MM target_seq PAM alignment distance position gene name gene id
GU305914.1 97912 97934 - 0 CGGCGAGCTGTCGTGCCTCC TGG ||||||||[||||||||||||]PAM NA Intergenic NA NA

T26604 GTCGTGCCTCCTGGACAGCCCGG 1000
Oligo fwd CACCGTCGTGCCTCCTGGACAGCC
Oligo rev AAACGGCTGTCCAGGAGGCACGAC
Chromosome start end strand MM target_seq PAM alignment distance position gene name gene id
GU305914.1 97903 97925 - 0 GTCGTGCCTCCTGGACAGCC CGG ||||||||[||||||||||||]PAM NA Intergenic NA NA

T26605 TCGTGCCTCCTGGACAGCCCGGG 1000
Oligo fwd CACCTCGTGCCTCCTGGACAGCCC
Oligo rev AAACGGGCTGTCCAGGAGGCACGA
Chromosome start end strand MM target_seq PAM alignment distance position gene name gene id
GU305914.1 97902 97924 - 0 TCGTGCCTCCTGGACAGCCC GGG ||||||||[||||||||||||]PAM NA Intergenic NA NA

T26606 TGCCTCCTGGACAGCCCGGGCGG 1000
Oligo fwd CACCTGCCTCCTGGACAGCCCGGG
Oligo rev AAACCCCGGGCTGTCCAGGAGGCA
Chromosome start end strand MM target_seq PAM alignment distance position gene name gene id
GU305914.1 97899 97921 - 0 TGCCTCCTGGACAGCCCGGG CGG ||||||||[||||||||||||]PAM NA Intergenic NA NA

T26607 CTCCTGGACAGCCCGGGCGGCGG 1000
Oligo fwd CACCCTCCTGGACAGCCCGGGCGG
Oligo rev AAACCCGCCCGGGCTGTCCAGGAG
Chromosome start end strand MM target_seq PAM alignment distance position gene name gene id
GU305914.1 97896 97918 - 0 CTCCTGGACAGCCCGGGCGG CGG ||||||||[||||||||||||]PAM NA Intergenic NA NA

T26608 CTGGACAGCCCGGGCGGCGGCGG 1000
Oligo fwd CACCCTGGACAGCCCGGGCGGCGG
Oligo rev AAACCCGCCGCCCGGGCTGTCCAG
Chromosome start end strand MM target_seq PAM alignment distance position gene name gene id
GU305914.1 97893 97915 - 0 CTGGACAGCCCGGGCGGCGG CGG ||||||||[||||||||||||]PAM NA Intergenic NA NA

T26609 GCGCCAGATCGCGAACAAACCGG 1000
Oligo fwd CACCGCGCCAGATCGCGAACAAAC
Oligo rev AAACGTTTGTTCGCGATCTGGCGC
Chromosome start end strand MM target_seq PAM alignment distance position gene name gene id
GU305914.1 97846 97868 - 0 GCGCCAGATCGCGAACAAAC CGG ||||||||[||||||||||||]PAM NA Intergenic NA NA

T26610 ACCGGCGTCTCCGTGCCGCAAGG 1000
Oligo fwd CACCACCGGCGTCTCCGTGCCGCA
Oligo rev AAACTGCGGCACGGAGACGCCGGT
Chromosome start end strand MM target_seq PAM alignment distance position gene name gene id
GU305914.1 97828 97850 - 0 ACCGGCGTCTCCGTGCCGCA AGG ||||||||[||||||||||||]PAM NA Intergenic NA NA

T26611 CCGGCGTCTCCGTGCCGCAAGGG 1000
Oligo fwd CACCCCGGCGTCTCCGTGCCGCAA
Oligo rev AAACTTGCGGCACGGAGACGCCGG

Chromosome start end strand MM target_seq PAM alignment distance position gene name gene id
GU305914.1 97827 97849 - 0 CCGGCGTCTCCGTGCCGCAA GGG ||||||||[||||||||||||]PAM NA Intergenic NA NA

T26612 GCGTCTCCGTGCCGCAAGGGCGG 1000
Oligo fwd CACCGCGTCTCCGTGCCGCAAGGG
Oligo rev AAACCCCTTGCGGCACGGAGACGC
Chromosome start end strand MM target_seq PAM alignment distance position gene name gene id
GU305914.1 97824 97846 - 0 GCGTCTCCGTGCCGCAAGGG CGG ||||||||[||||||||||||]PAM NA Intergenic NA NA

T26613 GTGCCGCAAGGGCGGCTCTTCGG 1000
Oligo fwd CACCGTGCCGCAAGGGCGGCTCTT
Oligo rev AAACAAGAGCCGCCCTTGCGGCAC
Chromosome start end strand MM target_seq PAM alignment distance position gene name gene id
GU305914.1 97816 97838 - 0 GTGCCGCAAGGGCGGCTCTT CGG ||||||||[||||||||||||]PAM NA Intergenic NA NA

T26614 CAAGGGCGGCTCTTCGGCAGCGG 1000
Oligo fwd CACCCAAGGGCGGCTCTTCGGCAG
Oligo rev AAACCTGCCGAAGAGCCGCCCTTG
Chromosome start end strand MM target_seq PAM alignment distance position gene name gene id
GU305914.1 97810 97832 - 0 CAAGGGCGGCTCTTCGGCAG CGG ||||||||[||||||||||||]PAM NA Intergenic NA NA

T26615 GGGCGGCTCTTCGGCAGCGGCGG 1000
Oligo fwd CACCGGGCGGCTCTTCGGCAGCGG
Oligo rev AAACCCGCTGCCGAAGAGCCGCCC
Chromosome start end strand MM target_seq PAM alignment distance position gene name gene id
GU305914.1 97807 97829 - 0 GGGCGGCTCTTCGGCAGCGG CGG ||||||||[||||||||||||]PAM NA Intergenic NA NA

T26616 CGGCTCTTCGGCAGCGGCGGCGG 1000
Oligo fwd CACCCGGCTCTTCGGCAGCGGCGG
Oligo rev AAACCCGCCGCTGCCGAAGAGCCG
Chromosome start end strand MM target_seq PAM alignment distance position gene name gene id
GU305914.1 97804 97826 - 0 CGGCTCTTCGGCAGCGGCGG CGG ||||||||[||||||||||||]PAM NA Intergenic NA NA

T26617 TTTCAAGCACCCGCTGCAGCAGG 1000
Oligo fwd CACCTTTCAAGCACCCGCTGCAGC
Oligo rev AAACGCTGCAGCGGGTGCTTGAAA
Chromosome start end strand MM target_seq PAM alignment distance position gene name gene id
GU305914.1 97747 97769 - 0 TTTCAAGCACCCGCTGCAGC AGG ||||||||[||||||||||||]PAM NA Intergenic NA NA

T26618 GCACCCGCTGCAGCAGGCGCTGG 1000
Oligo fwd CACCGCACCCGCTGCAGCAGGCGC
Oligo rev AAACGCGCCTGCTGCAGCGGGTGC
Chromosome start end strand MM target_seq PAM alignment distance position gene name gene id
GU305914.1 97741 97763 - 0 GCACCCGCTGCAGCAGGCGC TGG ||||||||[||||||||||||]PAM NA Intergenic NA NA

T26619 GCGTCGTTCGCGCGCATCGTCGG 1000
Oligo fwd CACCGCGTCGTTCGCGCGCATCGT
Oligo rev AAACACGATGCGCGCGAACGACGC
Chromosome start end strand MM target_seq PAM alignment distance position gene name gene id
GU305914.1 97710 97732 - 0 GCGTCGTTCGCGCGCATCGT CGG ||||||||[||||||||||||]PAM NA Intergenic NA NA

T26620 CGTCGTTCGCGCGCATCGTCGGG 1000
Oligo fwd CACCCGTCGTTCGCGCGCATCGTC
Oligo rev AAACGACGATGCGCGCGAACGACG
Chromosome start end strand MM target_seq PAM alignment distance position gene name gene id
GU305914.1 97709 97731 - 0 CGTCGTTCGCGCGCATCGTC GGG ||||||||[||||||||||||]PAM NA Intergenic NA NA

T26621 GTCGTTCGCGCGCATCGTCGGGG 1000
Oligo fwd CACCGTCGTTCGCGCGCATCGTCG
Oligo rev AAACCGACGATGCGCGCGAACGAC
Chromosome start end strand MM target_seq PAM alignment distance position gene name gene id
GU305914.1 97708 97730 - 0 GTCGTTCGCGCGCATCGTCG GGG ||||||||[||||||||||||]PAM NA Intergenic NA NA

T26622 TGATCCGCCGCAGATCGCTGCGG 1000
Oligo fwd CACCTGATCCGCCGCAGATCGCTG
Oligo rev AAACCAGCGATCTGCGGCGGATCA
Chromosome start end strand MM target_seq PAM alignment distance position gene name gene id
GU305914.1 97685 97707 - 0 TGATCCGCCGCAGATCGCTG CGG ||||||||[||||||||||||]PAM NA Intergenic NA NA

T26623 CGCAGATCGCTGCGGCACTCCGG 1000
Oligo fwd CACCCGCAGATCGCTGCGGCACTC
Oligo rev AAACGAGTGCCGCAGCGATCTGCG
Chromosome start end strand MM target_seq PAM alignment distance position gene name gene id
GU305914.1 97677 97699 - 0 CGCAGATCGCTGCGGCACTC CGG ||||||||[||||||||||||]PAM NA Intergenic NA NA

T26624 GCAGATCGCTGCGGCACTCCGGG 1000
Oligo fwd CACCGCAGATCGCTGCGGCACTCC
Oligo rev AAACGGAGTGCCGCAGCGATCTGC
Chromosome start end strand MM target_seq PAM alignment distance position gene name gene id
GU305914.1 97676 97698 - 0 GCAGATCGCTGCGGCACTCC GGG ||||||||[||||||||||||]PAM NA Intergenic NA NA

T26625 CATCCGCGACTTCGACGACACGG 1000
Oligo fwd CACCCATCCGCGACTTCGACGACA
Oligo rev AAACTGTCGTCGAAGTCGCGGATG
Chromosome start end strand MM target_seq PAM alignment distance position gene name gene id
GU305914.1 97645 97667 - 0 CATCCGCGACTTCGACGACA CGG ||||||||[||||||||||||]PAM NA Intergenic NA NA

T26626 ATCCGCGACTTCGACGACACGGG 1000
Oligo fwd CACCATCCGCGACTTCGACGACAC

Oligo rev AAACGTGTCGTCGAAGTCGCGGAT
Chromosome start end strand MM target_seq PAM alignment distance position gene name gene id
GU305914.1 97644 97666 - 0 ATCCGCGACTTCGACGACAC GGG ||||||||[||||||||||||]PAM NA Intergenic NA NA

T26627 CTTCGACGACACGGGCGCCACGG 1000
Oligo fwd CACCCTTCGACGACACGGGCGCCA
Oligo rev AAACTGGCGCCCGTGTCGTCGAAG
Chromosome start end strand MM target_seq PAM alignment distance position gene name gene id
GU305914.1 97636 97658 - 0 CTTCGACGACACGGGCGCCA CGG ||||||||[||||||||||||]PAM NA Intergenic NA NA

T26628 GCCACGGATTCCTACCGCTGCGG 1000
Oligo fwd CACCGCCACGGATTCCTACCGCTG
Oligo rev AAACCAGCGGTAGGAATCCGTGGC
Chromosome start end strand MM target_seq PAM alignment distance position gene name gene id
GU305914.1 97620 97642 - 0 GCCACGGATTCCTACCGCTG CGG ||||||||[||||||||||||]PAM NA Intergenic NA NA

T26629 CTGATCTTCGACTACCTGCGCGG 1000
Oligo fwd CACCCTGATCTTCGACTACCTGCG
Oligo rev AAACCGCAGGTAGTCGAAGATCAG
Chromosome start end strand MM target_seq PAM alignment distance position gene name gene id
GU305914.1 97584 97606 - 0 CTGATCTTCGACTACCTGCG CGG ||||||||[||||||||||||]PAM NA Intergenic NA NA

T26630 TCTGCGCGTACAACAACACGAGG 1000
Oligo fwd CACCTCTGCGCGTACAACAACACG
Oligo rev AAACCGTGTTGTTGTACGCGCAGA
Chromosome start end strand MM target_seq PAM alignment distance position gene name gene id
GU305914.1 97487 97509 - 0 TCTGCGCGTACAACAACACG AGG ||||||||[||||||||||||]PAM NA Intergenic NA NA

T26631 AACACGAGGAAGCACCCCAACGG 1000
Oligo fwd CACCAACACGAGGAAGCACCCCAA
Oligo rev AAACTTGGGGTGCTTCCTCGTGTT
Chromosome start end strand MM target_seq PAM alignment distance position gene name gene id
GU305914.1 97473 97495 - 0 AACACGAGGAAGCACCCCAA CGG ||||||||[||||||||||||]PAM NA Intergenic NA NA

T26632 AGCACCCCAACGGCACCTACCGG 1000
Oligo fwd CACCAGCACCCCAACGGCACCTAC
Oligo rev AAACGTAGGTGCCGTTGGGGTGCT
Chromosome start end strand MM target_seq PAM alignment distance position gene name gene id
GU305914.1 97463 97485 - 0 AGCACCCCAACGGCACCTAC CGG ||||||||[||||||||||||]PAM NA Intergenic NA NA

T26633 AACGGCACCTACCGGCGCGAAGG 1000
Oligo fwd CACCAACGGCACCTACCGGCGCGA
Oligo rev AAACTCGCGCCGGTAGGTGCCGTT
Chromosome start end strand MM target_seq PAM alignment distance position gene name gene id
GU305914.1 97455 97477 - 0 AACGGCACCTACCGGCGCGA AGG ||||||||[||||||||||||]PAM NA Intergenic NA NA

T26634 TCGCCAGAAGCGCCGTGCGCCGG 1000
Oligo fwd CACCTCGCCAGAAGCGCCGTGCGC
Oligo rev AAACGCGCACGGCGCTTCTGGCGA
Chromosome start end strand MM target_seq PAM alignment distance position gene name gene id
GU305914.1 97432 97454 - 0 TCGCCAGAAGCGCCGTGCGC CGG ||||||||[||||||||||||]PAM NA Intergenic NA NA

T26635 ACGCGACCCCGAACATCCCCAGG 1000
Oligo fwd CACCACGCGACCCCGAACATCCCC
Oligo rev AAACGGGGATGTTCGGGGTCGCGT
Chromosome start end strand MM target_seq PAM alignment distance position gene name gene id
GU305914.1 97409 97431 - 0 ACGCGACCCCGAACATCCCC AGG ||||||||[||||||||||||]PAM NA Intergenic NA NA

T26636 GACCCCGAACATCCCCAGGCTGG 1000
Oligo fwd CACCGACCCCGAACATCCCCAGGC
Oligo rev AAACGCCTGGGGATGTTCGGGGTC
Chromosome start end strand MM target_seq PAM alignment distance position gene name gene id
GU305914.1 97405 97427 - 0 GACCCCGAACATCCCCAGGC TGG ||||||||[||||||||||||]PAM NA Intergenic NA NA

T26637 ACATCCCCAGGCTGGCCTACAGG 1000
Oligo fwd CACCACATCCCCAGGCTGGCCTAC
Oligo rev AAACGTAGGCCAGCCTGGGGATGT
Chromosome start end strand MM target_seq PAM alignment distance position gene name gene id
GU305914.1 97397 97419 - 0 ACATCCCCAGGCTGGCCTAC AGG ||||||||[||||||||||||]PAM NA Intergenic NA NA

T26638 CGCGACGACGAGCCCCGAAGTGG 1000
Oligo fwd CACCCGCGACGACGAGCCCCGAAG
Oligo rev AAACCTTCGGGGCTCGTCGTCGCG
Chromosome start end strand MM target_seq PAM alignment distance position gene name gene id
GU305914.1 97369 97391 - 0 CGCGACGACGAGCCCCGAAG TGG ||||||||[||||||||||||]PAM NA Intergenic NA NA

T26639 TGGAACCCGCCCCTCCGAGTCGG 1000
Oligo fwd CACCTGGAACCCGCCCCTCCGAGT
Oligo rev AAACACTCGGAGGGGCGGGTTCCA
Chromosome start end strand MM target_seq PAM alignment distance position gene name gene id
GU305914.1 97349 97371 - 0 TGGAACCCGCCCCTCCGAGT CGG ||||||||[||||||||||||]PAM NA Intergenic NA NA

T26640 GACCAGCAGCAGTCCTCGTGTGG 1000
Oligo fwd CACCGACCAGCAGCAGTCCTCGTG
Oligo rev AAACCACGAGGACTGCTGCTGGTC
Chromosome start end strand MM target_seq PAM alignment distance position gene name gene id
GU305914.1 97324 97346 - 0 GACCAGCAGCAGTCCTCGTG TGG ||||||||[||||||||||||]PAM NA Intergenic NA NA

T26641 AGTCCTCGTGTGGACAGCAGAGG 1000

| | | | | | | | | | | | | |
|---|---|---|---|---|---|---|---|---|---|---|---|---|
| Oligo fwd | CACCAGTCCTCGTGTGGACAGCAG | | | | | | | | | | | |
| Oligo rev | AAACCTGCTGTCCACACGAGGACT | | | | | | | | | | | |
| Chromosome | start | | end | strand | MM | target_seq | PAM | alignment | distance | position | gene name | gene id |
| GU305914.1 | | 97314 | 97336 | - | 0 | AGTCCTCGTGTGGACAGCAG | AGG | \|\|\|\|\|\|\|\|[\|\|\|\|\|\|\|\|\|\|\|\|]PAM | NA | Intergenic | NA | NA |
| T26642 | CCTCGTGTGGACAGCAGAGGAGG | 1000 | | | | | | | | | | |
| Oligo fwd | CACCCCTCGTGTGGACAGCAGAGG | | | | | | | | | | | |
| Oligo rev | AAACCCTCTGCTGTCCACACGAGG | | | | | | | | | | | |
| Chromosome | start | | end | strand | MM | target_seq | PAM | alignment | distance | position | gene name | gene id |
| GU305914.1 | | 97311 | 97333 | - | 0 | CCTCGTGTGGACAGCAGAGG | AGG | \|\|\|\|\|\|\|\|[\|\|\|\|\|\|\|\|\|\|\|\|]PAM | NA | Intergenic | NA | NA |
| T26643 | CGTGTGGACAGCAGAGGAGGAGG | 1000 | | | | | | | | | | |
| Oligo fwd | CACCCGTGTGGACAGCAGAGGAGG | | | | | | | | | | | |
| Oligo rev | AAACCCTCCTCTGCTGTCCACACG | | | | | | | | | | | |
| Chromosome | start | | end | strand | MM | target_seq | PAM | alignment | distance | position | gene name | gene id |
| GU305914.1 | | 97308 | 97330 | - | 0 | CGTGTGGACAGCAGAGGAGG | AGG | \|\|\|\|\|\|\|\|[\|\|\|\|\|\|\|\|\|\|\|\|]PAM | NA | Intergenic | NA | NA |
| T26644 | GTGGACAGCAGAGGAGGAGGAGG | 1000 | | | | | | | | | | |
| Oligo fwd | CACCGTGGACAGCAGAGGAGGAGG | | | | | | | | | | | |
| Oligo rev | AAACCCTCCTCCTCTGCTGTCCAC | | | | | | | | | | | |
| Chromosome | start | | end | strand | MM | target_seq | PAM | alignment | distance | position | gene name | gene id |
| GU305914.1 | | 97305 | 97327 | - | 0 | GTGGACAGCAGAGGAGGAGG | AGG | \|\|\|\|\|\|\|\|[\|\|\|\|\|\|\|\|\|\|\|\|]PAM | NA | Intergenic | NA | NA |
| T26645 | GGAGGAGGAGGAGACCGTCGCGG | 1000 | | | | | | | | | | |
| Oligo fwd | CACCGGAGGAGGAGGAGACCGTCG | | | | | | | | | | | |
| Oligo rev | AAACCGACGGTCTCCTCCTCCTCC | | | | | | | | | | | |
| Chromosome | start | | end | strand | MM | target_seq | PAM | alignment | distance | position | gene name | gene id |
| GU305914.1 | | 97293 | 97315 | - | 0 | GGAGGAGGAGGAGACCGTCG | CGG | \|\|\|\|\|\|\|\|[\|\|\|\|\|\|\|\|\|\|\|\|]PAM | NA | Intergenic | NA | NA |
| T26646 | ATCACCATCGCCACCACACCAGG | 1000 | | | | | | | | | | |
| Oligo fwd | CACCATCACCATCGCCACCACACC | | | | | | | | | | | |
| Oligo rev | AAACGGTGTGGTGGCGATGGTGAT | | | | | | | | | | | |
| Chromosome | start | | end | strand | MM | target_seq | PAM | alignment | distance | position | gene name | gene id |
| GU305914.1 | | 97247 | 97269 | - | 0 | ATCACCATCGCCACCACACC | AGG | \|\|\|\|\|\|\|\|[\|\|\|\|\|\|\|\|\|\|\|\|]PAM | NA | Intergenic | NA | NA |
| T26647 | ACCCACGACAGTAGCAGCAGCGG | 1000 | | | | | | | | | | |
| Oligo fwd | CACCACCCACGACAGTAGCAGCAG | | | | | | | | | | | |
| Oligo rev | AAACCTGCTGCTACTGTCGTGGGT | | | | | | | | | | | |
| Chromosome | start | | end | strand | MM | target_seq | PAM | alignment | distance | position | gene name | gene id |
| GU305914.1 | | 97206 | 97228 | - | 0 | ACCCACGACAGTAGCAGCAG | CGG | \|\|\|\|\|\|\|\|[\|\|\|\|\|\|\|\|\|\|\|\|]PAM | NA | Intergenic | NA | NA |
| T26648 | CAGACGTCGCAGCAGCGCGACGG | 1000 | | | | | | | | | | |
| Oligo fwd | CACCCAGACGTCGCAGCAGCGCGA | | | | | | | | | | | |
| Oligo rev | AAACTCGCGCTGCTGCGACGTCTG | | | | | | | | | | | |
| Chromosome | start | | end | strand | MM | target_seq | PAM | alignment | distance | position | gene name | gene id |
| GU305914.1 | | 97180 | 97202 | - | 0 | CAGACGTCGCAGCAGCGCGA | CGG | \|\|\|\|\|\|\|\|[\|\|\|\|\|\|\|\|\|\|\|\|]PAM | NA | Intergenic | NA | NA |
| T26649 | CGTCGCAGCAGCGCGACGGACGG | 1000 | | | | | | | | | | |
| Oligo fwd | CACCCGTCGCAGCAGCGCGACGGA | | | | | | | | | | | |
| Oligo rev | AAACTCCGTCGCGCTGCTGCGACG | | | | | | | | | | | |
| Chromosome | start | | end | strand | MM | target_seq | PAM | alignment | distance | position | gene name | gene id |
| GU305914.1 | | 97176 | 97198 | - | 0 | CGTCGCAGCAGCGCGACGGA | CGG | \|\|\|\|\|\|\|\|[\|\|\|\|\|\|\|\|\|\|\|\|]PAM | NA | Intergenic | NA | NA |
| T26650 | GCAGCAGCGCGACGGACGGCCGG | 1000 | | | | | | | | | | |
| Oligo fwd | CACCGCAGCAGCGCGACGGACGGC | | | | | | | | | | | |
| Oligo rev | AAACGCCGTCCGTCGCGCTGCTGC | | | | | | | | | | | |
| Chromosome | start | | end | strand | MM | target_seq | PAM | alignment | distance | position | gene name | gene id |
| GU305914.1 | | 97172 | 97194 | - | 0 | GCAGCAGCGCGACGGACGGC | CGG | \|\|\|\|\|\|\|\|[\|\|\|\|\|\|\|\|\|\|\|\|]PAM | NA | Intergenic | NA | NA |
| T26651 | CGACGGACGGCCGGCGCAGCAGG | 1000 | | | | | | | | | | |
| Oligo fwd | CACCCGACGGACGGCCGGCGCAGC | | | | | | | | | | | |
| Oligo rev | AAACGCTGCGCCGGCCGTCCGTCG | | | | | | | | | | | |
| Chromosome | start | | end | strand | MM | target_seq | PAM | alignment | distance | position | gene name | gene id |
| GU305914.1 | | 97163 | 97185 | - | 0 | CGACGGACGGCCGGCGCAGC | AGG | \|\|\|\|\|\|\|\|[\|\|\|\|\|\|\|\|\|\|\|\|]PAM | NA | Intergenic | NA | NA |
| T26652 | GACGGCCGGCGCAGCAGGCGCGG | 1000 | | | | | | | | | | |
| Oligo fwd | CACCGACGGCCGGCGCAGCAGGCG | | | | | | | | | | | |
| Oligo rev | AAACCGCCTGCTGCGCCGGCCGTC | | | | | | | | | | | |
| Chromosome | start | | end | strand | MM | target_seq | PAM | alignment | distance | position | gene name | gene id |
| GU305914.1 | | 97158 | 97180 | - | 0 | GACGGCCGGCGCAGCAGGCG | CGG | \|\|\|\|\|\|\|\|[\|\|\|\|\|\|\|\|\|\|\|\|]PAM | NA | Intergenic | NA | NA |
| T26653 | GCAGCAGGCGCGGCAGCAGACGG | 1000 | | | | | | | | | | |
| Oligo fwd | CACCGCAGCAGGCGCGGCAGCAGA | | | | | | | | | | | |
| Oligo rev | AAACTCTGCTGCCGCGCCTGCTGC | | | | | | | | | | | |
| Chromosome | start | | end | strand | MM | target_seq | PAM | alignment | distance | position | gene name | gene id |
| GU305914.1 | | 97148 | 97170 | - | 0 | GCAGCAGGCGCGGCAGCAGA | CGG | \|\|\|\|\|\|\|\|[\|\|\|\|\|\|\|\|\|\|\|\|]PAM | NA | Intergenic | NA | NA |
| T26654 | CAGCAGGCGCGGCAGCAGACGGG | 1000 | | | | | | | | | | |
| Oligo fwd | CACCCAGCAGGCGCGGCAGCAGAC | | | | | | | | | | | |
| Oligo rev | AAACGTCTGCTGCCGCGCCTGCTG | | | | | | | | | | | |
| Chromosome | start | | end | strand | MM | target_seq | PAM | alignment | distance | position | gene name | gene id |
| GU305914.1 | | 97147 | 97169 | - | 0 | CAGCAGGCGCGGCAGCAGAC | GGG | \|\|\|\|\|\|\|\|[\|\|\|\|\|\|\|\|\|\|\|\|]PAM | NA | Intergenic | NA | NA |
| T26655 | AGCAGGCGCGGCAGCAGACGGGG | 1000 | | | | | | | | | | |
| Oligo fwd | CACCAGCAGGCGCGGCAGCAGACG | | | | | | | | | | | |
| Oligo rev | AAACCGTCTGCTGCCGCGCCTGCT | | | | | | | | | | | |
| Chromosome | start | | end | strand | MM | target_seq | PAM | alignment | distance | position | gene name | gene id |
| GU305914.1 | | 97146 | 97168 | - | 0 | AGCAGGCGCGGCAGCAGACG | GGG | \|\|\|\|\|\|\|\|[\|\|\|\|\|\|\|\|\|\|\|\|]PAM | NA | Intergenic | NA | NA |
| GU305914.1 | | 188338 | 188360 | + | 4 | AACAGACGAGGCAGCACACG | AGG | \|-\|\|\|-\|\|[-\|\|\|\|\|\|\|-\|\|\|]PAM | NA | Intergenic | NA | NA |

| | | | | | | | | | | | |
|---|---|---|---|---|---|---|---|---|---|---|---|
| T26656 | GCGCGGCAGCAGACGGGGAGAGG | 1000 | | | | | | | | | |
| Oligo fwd | CACCGCGCGGCAGCAGACGGGGAG | | | | | | | | | | |
| Oligo rev | AAACCTCCCCGTCTGCTGCCGCGC | | | | | | | | | | |
| Chromosome | start | end | strand | MM | target_seq | PAM | alignment | distance | position | gene name | gene id |
| GU305914.1 | 97141 | 97163 | - | 0 | GCGCGGCAGCAGACGGGGAG | AGG | \|\|\|\|\|\|\|\|[\|\|\|\|\|\|\|\|\|\|\|\|]PAM | NA | Intergenic | NA | NA |

| | | | | | | | | | | | |
|---|---|---|---|---|---|---|---|---|---|---|---|
| T26657 | GAGGCGCAGCGCGAGTCTAACGG | 1000 | | | | | | | | | |
| Oligo fwd | CACCGAGGCGCAGCGCGAGTCTAA | | | | | | | | | | |
| Oligo rev | AAACTTAGACTCGCGCTGCGCCTC | | | | | | | | | | |
| Chromosome | start | end | strand | MM | target_seq | PAM | alignment | distance | position | gene name | gene id |
| GU305914.1 | 97122 | 97144 | - | 0 | GAGGCGCAGCGCGAGTCTAA | CGG | \|\|\|\|\|\|\|\|[\|\|\|\|\|\|\|\|\|\|\|\|]PAM | NA | Intergenic | NA | NA |

| | | | | | | | | | | | |
|---|---|---|---|---|---|---|---|---|---|---|---|
| T26658 | TTCGACCGTGAGCTCCACCACGG | 1000 | | | | | | | | | |
| Oligo fwd | CACCTTCGACCGTGAGCTCCACCA | | | | | | | | | | |
| Oligo rev | AAACTGGTGGAGCTCACGGTCGAA | | | | | | | | | | |
| Chromosome | start | end | strand | MM | target_seq | PAM | alignment | distance | position | gene name | gene id |
| GU305914.1 | 97078 | 97100 | - | 0 | TTCGACCGTGAGCTCCACCA | CGG | \|\|\|\|\|\|\|\|[\|\|\|\|\|\|\|\|\|\|\|\|]PAM | NA | Intergenic | NA | NA |

| | | | | | | | | | | | |
|---|---|---|---|---|---|---|---|---|---|---|---|
| T26659 | GTGAGCTCCACCACGGTCCACGG | 1000 | | | | | | | | | |
| Oligo fwd | CACCGTGAGCTCCACCACGGTCCA | | | | | | | | | | |
| Oligo rev | AAACTGGACCGTGGTGGAGCTCAC | | | | | | | | | | |
| Chromosome | start | end | strand | MM | target_seq | PAM | alignment | distance | position | gene name | gene id |
| GU305914.1 | 97071 | 97093 | - | 0 | GTGAGCTCCACCACGGTCCA | CGG | \|\|\|\|\|\|\|\|[\|\|\|\|\|\|\|\|\|\|\|\|]PAM | NA | Intergenic | NA | NA |

| | | | | | | | | | | | |
|---|---|---|---|---|---|---|---|---|---|---|---|
| T26660 | ACCACGGTCCACGGCCAGAACGG | 1000 | | | | | | | | | |
| Oligo fwd | CACCACCACGGTCCACGGCCAGAA | | | | | | | | | | |
| Oligo rev | AAACTTCTGGCCGTGGACCGTGGT | | | | | | | | | | |
| Chromosome | start | end | strand | MM | target_seq | PAM | alignment | distance | position | gene name | gene id |
| GU305914.1 | 97062 | 97084 | - | 0 | ACCACGGTCCACGGCCAGAA | CGG | \|\|\|\|\|\|\|\|[\|\|\|\|\|\|\|\|\|\|\|\|]PAM | NA | Intergenic | NA | NA |

| | | | | | | | | | | | |
|---|---|---|---|---|---|---|---|---|---|---|---|
| T26661 | CACGGCCAGAACGGCGCCCGCGG | 1000 | | | | | | | | | |
| Oligo fwd | CACCCACGGCCAGAACGGCGCCCG | | | | | | | | | | |
| Oligo rev | AAACCGGGCGCCGTTCTGGCCGTG | | | | | | | | | | |
| Chromosome | start | end | strand | MM | target_seq | PAM | alignment | distance | position | gene name | gene id |
| GU305914.1 | 97053 | 97075 | - | 0 | CACGGCCAGAACGGCGCCCG | CGG | \|\|\|\|\|\|\|\|[\|\|\|\|\|\|\|\|\|\|\|\|]PAM | NA | Intergenic | NA | NA |

| | | | | | | | | | | | |
|---|---|---|---|---|---|---|---|---|---|---|---|
| T26662 | GTCCAAAGAGTCGTCGAAGACGG | 1000 | | | | | | | | | |
| Oligo fwd | CACCGTCCAAAGAGTCGTCGAAGA | | | | | | | | | | |
| Oligo rev | AAACTCTTCGACGACTCTTTGGAC | | | | | | | | | | |
| Chromosome | start | end | strand | MM | target_seq | PAM | alignment | distance | position | gene name | gene id |
| GU305914.1 | 96976 | 96998 | - | 0 | GTCCAAAGAGTCGTCGAAGA | CGG | \|\|\|\|\|\|\|\|[\|\|\|\|\|\|\|\|\|\|\|\|]PAM | NA | Intergenic | NA | NA |

| | | | | | | | | | | | |
|---|---|---|---|---|---|---|---|---|---|---|---|
| T26663 | CCTCGCCCGCGTCCAGAGAGAGG | 1000 | | | | | | | | | |
| Oligo fwd | CACCCCTCGCCCGCGTCCAGAGAG | | | | | | | | | | |
| Oligo rev | AAACCTCTCTGGACGCGGGCGAGG | | | | | | | | | | |
| Chromosome | start | end | strand | MM | target_seq | PAM | alignment | distance | position | gene name | gene id |
| GU305914.1 | 96920 | 96942 | - | 0 | CCTCGCCCGCGTCCAGAGAG | AGG | \|\|\|\|\|\|\|\|[\|\|\|\|\|\|\|\|\|\|\|\|]PAM | NA | Intergenic | NA | NA |

| | | | | | | | | | | | |
|---|---|---|---|---|---|---|---|---|---|---|---|
| T26664 | TCGCCCGCACGATCCTCCGTCGG | 1000 | | | | | | | | | |
| Oligo fwd | CACCTCGCCCGCACGATCCTCCGT | | | | | | | | | | |
| Oligo rev | AAACACGGAGGATCGTGCGGGCGA | | | | | | | | | | |
| Chromosome | start | end | strand | MM | target_seq | PAM | alignment | distance | position | gene name | gene id |
| GU305914.1 | 96868 | 96890 | - | 0 | TCGCCCGCACGATCCTCCGT | CGG | \|\|\|\|\|\|\|\|[\|\|\|\|\|\|\|\|\|\|\|\|]PAM | NA | Intergenic | NA | NA |

| | | | | | | | | | | | |
|---|---|---|---|---|---|---|---|---|---|---|---|
| T26665 | CGCCCGCACGATCCTCCGTCGGG | 1000 | | | | | | | | | |
| Oligo fwd | CACCCGCCCGCACGATCCTCCGTC | | | | | | | | | | |
| Oligo rev | AAACGACGGAGGATCGTGCGGGCG | | | | | | | | | | |
| Chromosome | start | end | strand | MM | target_seq | PAM | alignment | distance | position | gene name | gene id |
| GU305914.1 | 96867 | 96889 | - | 0 | CGCCCGCACGATCCTCCGTC | GGG | \|\|\|\|\|\|\|\|[\|\|\|\|\|\|\|\|\|\|\|\|]PAM | NA | Intergenic | NA | NA |
| GU305914.1 | 118251 | 118273 | - | 4 | GGGCCGCCTGATCCTCCGTC | GGG | -\|-\|\|\|\|-[-\|\|\|\|\|\|\|\|\|\|\|]PAM | NA | Intergenic | NA | NA |

| | | | | | | | | | | | |
|---|---|---|---|---|---|---|---|---|---|---|---|
| T26666 | GAGAAAGAGCTCGCGACCGCCGG | 1000 | | | | | | | | | |
| Oligo fwd | CACCGAGAAAGAGCTCGCGACCGC | | | | | | | | | | |
| Oligo rev | AAACGCGGTCGCGAGCTCTTTCTC | | | | | | | | | | |
| Chromosome | start | end | strand | MM | target_seq | PAM | alignment | distance | position | gene name | gene id |
| GU305914.1 | 96828 | 96850 | - | 0 | GAGAAAGAGCTCGCGACCGC | CGG | \|\|\|\|\|\|\|\|[\|\|\|\|\|\|\|\|\|\|\|\|]PAM | NA | Intergenic | NA | NA |

| | | | | | | | | | | | |
|---|---|---|---|---|---|---|---|---|---|---|---|
| T26667 | GCTCGCGACCGCCGGCGACGAGG | 1000 | | | | | | | | | |
| Oligo fwd | CACCGCTCGCGACCGCCGGCGACG | | | | | | | | | | |
| Oligo rev | AAACCGTCGCCGGCGGTCGCGAGC | | | | | | | | | | |
| Chromosome | start | end | strand | MM | target_seq | PAM | alignment | distance | position | gene name | gene id |
| GU305914.1 | 96820 | 96842 | - | 0 | GCTCGCGACCGCCGGCGACG | AGG | \|\|\|\|\|\|\|\|[\|\|\|\|\|\|\|\|\|\|\|\|]PAM | NA | Intergenic | NA | NA |
| GU305914.1 | 139605 | 139627 | + | 4 | GCTCTCCACCACCGACGACG | CGG | \|\|\|\|-\|-\|[\|\|-\|\|\|-\|\|\|\|\|]PAM | NA | Intergenic | NA | NA |

| | | | | | | | | | | | |
|---|---|---|---|---|---|---|---|---|---|---|---|
| T26668 | ACCGCCGGCGACGAGGACGAAGG | 1000 | | | | | | | | | |
| Oligo fwd | CACCACCGCCGGCGACGAGGACGA | | | | | | | | | | |
| Oligo rev | AAACTCGTCCTCGTCGCCGGCGGT | | | | | | | | | | |
| Chromosome | start | end | strand | MM | target_seq | PAM | alignment | distance | position | gene name | gene id |
| GU305914.1 | 96813 | 96835 | - | 0 | ACCGCCGGCGACGAGGACGA | AGG | \|\|\|\|\|\|\|\|[\|\|\|\|\|\|\|\|\|\|\|\|]PAM | NA | Intergenic | NA | NA |
| GU305914.1 | 113956 | 113978 | + | 4 | AACTCCGGCGAGGAGGCCGA | GGG | \|-\|-\|\|\|\|[\|\|\|-\|\|\|\|-\|\|\|]PAM | NA | Intergenic | NA | NA |

| | | | | | | | | | | | |
|---|---|---|---|---|---|---|---|---|---|---|---|
| T26669 | CCGCCGGCGACGAGGACGAAGGG | 1000 | | | | | | | | | |
| Oligo fwd | CACCCCGCCGGCGACGAGGACGAA | | | | | | | | | | |
| Oligo rev | AAACTTCGTCCTCGTCGCCGGCGG | | | | | | | | | | |
| Chromosome | start | end | strand | MM | target_seq | PAM | alignment | distance | position | gene name | gene id |
| GU305914.1 | 96812 | 96834 | - | 0 | CCGCCGGCGACGAGGACGAA | GGG | \|\|\|\|\|\|\|\|[\|\|\|\|\|\|\|\|\|\|\|\|]PAM | NA | Intergenic | NA | NA |

| | | |
|---|---|---|
| T26670 | CGCCGGCGACGAGGACGAAGGGG | 1000 |

| | | | | | | | | | | | |
|---|---|---|---|---|---|---|---|---|---|---|---|
| Oligo fwd | CACCCGCCGGCGACGAGGACGAAG | | | | | | | | | | |
| Oligo rev | AAACCTTCGTCCTCGTCGCCGGCG | | | | | | | | | | |
| Chromosome | start | end | strand | MM | target_seq | PAM | alignment | distance | position | gene name | gene id |
| GU305914.1 | 96811 | 96833 | - | 0 | CGCCGGCGACGAGGACGAAG | GGG | \|\|\|\|\|\|\|\|[\|\|\|\|\|\|\|\|\|\|\|\|]PAM | NA | Intergenic | NA | NA |
| GU305914.1 | 59736 | 59758 | - | 4 | CTCGGACGACGAGGACGAGG | TGG | \|-\|-\|-\|\|[\|\|\|\|\|\|\|\|\|\|-\|]PAM | NA | Intergenic | NA | NA |
| GU305914.1 | 174261 | 174283 | + | 3 | CGACGGCGACGATGACGAAC | AGG | \|\|-\|\|\|\|\|[\|\|\|\|-\|\|\|\|\|\|-]PAM | NA | Intergenic | NA | NA |
| GU305914.1 | 68313 | 68335 | + | 4 | CGGCGGTGACGAGGACGGAC | CGG | \|\|-\|\|\|-\|[\|\|\|\|\|\|\|\|\|-\|-]PAM | NA | Intergenic | NA | NA |
| T26671 | GACGAAGGGGTCAGATCTCCCGG | 1000 | | | | | | | | | |
| Oligo fwd | CACCGACGAAGGGGTCAGATCTCC | | | | | | | | | | |
| Oligo rev | AAACGGAGATCTGACCCCTTCGTC | | | | | | | | | | |
| Chromosome | start | end | strand | MM | target_seq | PAM | alignment | distance | position | gene name | gene id |
| GU305914.1 | 96798 | 96820 | - | 0 | GACGAAGGGGTCAGATCTCC | CGG | \|\|\|\|\|\|\|\|[\|\|\|\|\|\|\|\|\|\|\|\|]PAM | NA | Intergenic | NA | NA |
| T26672 | ACGAAGGGGTCAGATCTCCCGGG | 1000 | | | | | | | | | |
| Oligo fwd | CACCACGAAGGGGTCAGATCTCCC | | | | | | | | | | |
| Oligo rev | AAACGGGAGATCTGACCCCTTCGT | | | | | | | | | | |
| Chromosome | start | end | strand | MM | target_seq | PAM | alignment | distance | position | gene name | gene id |
| GU305914.1 | 96797 | 96819 | - | 0 | ACGAAGGGGTCAGATCTCCC | GGG | \|\|\|\|\|\|\|\|[\|\|\|\|\|\|\|\|\|\|\|\|]PAM | NA | Intergenic | NA | NA |
| T26673 | CGAAGGGGTCAGATCTCCCGGGG | 1000 | | | | | | | | | |
| Oligo fwd | CACCCGAAGGGGTCAGATCTCCCG | | | | | | | | | | |
| Oligo rev | AAACCGGGAGATCTGACCCCTTCG | | | | | | | | | | |
| Chromosome | start | end | strand | MM | target_seq | PAM | alignment | distance | position | gene name | gene id |
| GU305914.1 | 96796 | 96818 | - | 0 | CGAAGGGGTCAGATCTCCCG | GGG | \|\|\|\|\|\|\|\|[\|\|\|\|\|\|\|\|\|\|\|\|]PAM | NA | Intergenic | NA | NA |
| T26674 | TGCAGCGTCGCCACGCGTCGCGG | 1000 | | | | | | | | | |
| Oligo fwd | CACCTGCAGCGTCGCCACGCGTCG | | | | | | | | | | |
| Oligo rev | AAACCGACGCGTGGCGACGCTGCA | | | | | | | | | | |
| Chromosome | start | end | strand | MM | target_seq | PAM | alignment | distance | position | gene name | gene id |
| GU305914.1 | 96771 | 96793 | - | 0 | TGCAGCGTCGCCACGCGTCG | CGG | \|\|\|\|\|\|\|\|[\|\|\|\|\|\|\|\|\|\|\|\|]PAM | NA | Intergenic | NA | NA |
| T26675 | CGCCACGCGTCGCGGCTCGTCGG | 1000 | | | | | | | | | |
| Oligo fwd | CACCCGCCACGCGTCGCGGCTCGT | | | | | | | | | | |
| Oligo rev | AAACACGAGCCGCGACGCGTGGCG | | | | | | | | | | |
| Chromosome | start | end | strand | MM | target_seq | PAM | alignment | distance | position | gene name | gene id |
| GU305914.1 | 96763 | 96785 | - | 0 | CGCCACGCGTCGCGGCTCGT | CGG | \|\|\|\|\|\|\|\|[\|\|\|\|\|\|\|\|\|\|\|\|]PAM | NA | Intergenic | NA | NA |
| T26676 | CACGCGTCGCGGCTCGTCGGCGG | 1000 | | | | | | | | | |
| Oligo fwd | CACCCACGCGTCGCGGCTCGTCGG | | | | | | | | | | |
| Oligo rev | AAACCCGACGAGCCGCGACGCGTG | | | | | | | | | | |
| Chromosome | start | end | strand | MM | target_seq | PAM | alignment | distance | position | gene name | gene id |
| GU305914.1 | 96760 | 96782 | - | 0 | CACGCGTCGCGGCTCGTCGG | CGG | \|\|\|\|\|\|\|\|[\|\|\|\|\|\|\|\|\|\|\|\|]PAM | NA | Intergenic | NA | NA |
| T26677 | CGACTCCTCGTCCAGCTCCTCGG | 1000 | | | | | | | | | |
| Oligo fwd | CACCCGACTCCTCGTCCAGCTCCT | | | | | | | | | | |
| Oligo rev | AAACAGGAGCTGGACGAGGAGTCG | | | | | | | | | | |
| Chromosome | start | end | strand | MM | target_seq | PAM | alignment | distance | position | gene name | gene id |
| GU305914.1 | 96727 | 96749 | - | 0 | CGACTCCTCGTCCAGCTCCT | CGG | \|\|\|\|\|\|\|\|[\|\|\|\|\|\|\|\|\|\|\|\|]PAM | NA | Intergenic | NA | NA |
| T26678 | CTCGTCGTCGTCCTCCGACGAGG | 1000 | | | | | | | | | |
| Oligo fwd | CACCCTCGTCGTCGTCCTCCGACG | | | | | | | | | | |
| Oligo rev | AAACCGTCGGAGGACGACGACGAG | | | | | | | | | | |
| Chromosome | start | end | strand | MM | target_seq | PAM | alignment | distance | position | gene name | gene id |
| GU305914.1 | 96703 | 96725 | - | 0 | CTCGTCGTCGTCCTCCGACG | AGG | \|\|\|\|\|\|\|\|[\|\|\|\|\|\|\|\|\|\|\|\|]PAM | NA | Intergenic | NA | NA |
| T26679 | GTCGTCGTCCTCCGACGAGGAGG | 1000 | | | | | | | | | |
| Oligo fwd | CACCGTCGTCGTCCTCCGACGAGG | | | | | | | | | | |
| Oligo rev | AAACCCTCGTCGGAGGACGACGAC | | | | | | | | | | |
| Chromosome | start | end | strand | MM | target_seq | PAM | alignment | distance | position | gene name | gene id |
| GU305914.1 | 96700 | 96722 | - | 0 | GTCGTCGTCCTCCGACGAGG | AGG | \|\|\|\|\|\|\|\|[\|\|\|\|\|\|\|\|\|\|\|\|]PAM | NA | Intergenic | NA | NA |
| T26680 | CGAGGAGGAATCTGACGTCGAGG | 1000 | | | | | | | | | |
| Oligo fwd | CACCCGAGGAGGAATCTGACGTCG | | | | | | | | | | |
| Oligo rev | AAACCGACGTCAGATTCCTCCTCG | | | | | | | | | | |
| Chromosome | start | end | strand | MM | target_seq | PAM | alignment | distance | position | gene name | gene id |
| GU305914.1 | 96685 | 96707 | - | 0 | CGAGGAGGAATCTGACGTCG | AGG | \|\|\|\|\|\|\|\|[\|\|\|\|\|\|\|\|\|\|\|\|]PAM | NA | Intergenic | NA | NA |
| T26681 | GCAGTCGAAACGACTCGAAGAGG | 1000 | | | | | | | | | |
| Oligo fwd | CACCGCAGTCGAAACGACTCGAAG | | | | | | | | | | |
| Oligo rev | AAACCTTCGAGTCGTTTCGACTGC | | | | | | | | | | |
| Chromosome | start | end | strand | MM | target_seq | PAM | alignment | distance | position | gene name | gene id |
| GU305914.1 | 96643 | 96665 | - | 0 | GCAGTCGAAACGACTCGAAG | AGG | \|\|\|\|\|\|\|\|[\|\|\|\|\|\|\|\|\|\|\|\|]PAM | NA | Intergenic | NA | NA |
| T26682 | ACGACTCGAAGAGGCGCTCGAGG | 1000 | | | | | | | | | |
| Oligo fwd | CACCACGACTCGAAGAGGCGCTCG | | | | | | | | | | |
| Oligo rev | AAACCGAGCGCCTCTTCGAGTCGT | | | | | | | | | | |
| Chromosome | start | end | strand | MM | target_seq | PAM | alignment | distance | position | gene name | gene id |
| GU305914.1 | 96634 | 96656 | - | 0 | ACGACTCGAAGAGGCGCTCG | AGG | \|\|\|\|\|\|\|\|[\|\|\|\|\|\|\|\|\|\|\|\|]PAM | NA | Intergenic | NA | NA |
| T26683 | GCGTTGCGAGCGCGATTTCGAGG | 1000 | | | | | | | | | |
| Oligo fwd | CACCGCGTTGCGAGCGCGATTTCG | | | | | | | | | | |
| Oligo rev | AAACCGAAATCGCGCTCGCAACGC | | | | | | | | | | |
| Chromosome | start | end | strand | MM | target_seq | PAM | alignment | distance | position | gene name | gene id |
| GU305914.1 | 96610 | 96632 | - | 0 | GCGTTGCGAGCGCGATTTCG | AGG | \|\|\|\|\|\|\|\|[\|\|\|\|\|\|\|\|\|\|\|\|]PAM | NA | Intergenic | NA | NA |
| T26684 | TTGCGAGCGCGATTTCGAGGCGG | 1000 | | | | | | | | | |
| Oligo fwd | CACCTTGCGAGCGCGATTTCGAGG | | | | | | | | | | |
| Oligo rev | AAACCCTCGAAATCGCGCTCGCAA | | | | | | | | | | |

| | | | | | | | | | | | |
|---|---|---|---|---|---|---|---|---|---|---|---|
| Chromosome | start | end | strand | MM | target_seq | PAM | alignment | distance | position | gene name | gene id |
| GU305914.1 | 96607 | 96629 | - | 0 | TTGCGAGCGCGATTTCGAGG | CGG | \|\|\|\|\|\|\|\|[\|\|\|\|\|\|\|\|\|\|\|\|]PAM | NA | Intergenic | NA | NA |
| T26685 | CGATTTCGAGGCGGACGACGAGG | 1000 | | | | | | | | | |
| Oligo fwd | CACCCGATTTCGAGGCGGACGACG | | | | | | | | | | |
| Oligo rev | AAACCGTCGTCCGCCTCGAAATCG | | | | | | | | | | |
| Chromosome | start | end | strand | MM | target_seq | PAM | alignment | distance | position | gene name | gene id |
| GU305914.1 | 96598 | 96620 | - | 0 | CGATTTCGAGGCGGACGACG | AGG | \|\|\|\|\|\|\|\|[\|\|\|\|\|\|\|\|\|\|\|\|]PAM | NA | Intergenic | NA | NA |
| T26686 | GGAGTTCGCCGAGCCGATCGAGG | 1000 | | | | | | | | | |
| Oligo fwd | CACCGGAGTTCGCCGAGCCGATCG | | | | | | | | | | |
| Oligo rev | AAACCGATCGGCTCGGCGAACTCC | | | | | | | | | | |
| Chromosome | start | end | strand | MM | target_seq | PAM | alignment | distance | position | gene name | gene id |
| GU305914.1 | 96577 | 96599 | - | 0 | GGAGTTCGCCGAGCCGATCG | AGG | \|\|\|\|\|\|\|\|[\|\|\|\|\|\|\|\|\|\|\|\|]PAM | NA | Intergenic | NA | NA |
| GU305914.1 | 198764 | 198786 | - | 4 | GGTGTTCTCCGACCCGAACG | CGG | \|\|-\|\|\|\|-[\|\|\|\|-\|\|\|\|-\|\|]PAM | NA | Intergenic | NA | NA |
| T26687 | CCTACACTGCTCGCTAGACATGG | 1000 | | | | | | | | | |
| Oligo fwd | CACCCCTACACTGCTCGCTAGACA | | | | | | | | | | |
| Oligo rev | AAACTGTCTAGCGAGCAGTGTAGG | | | | | | | | | | |
| Chromosome | start | end | strand | MM | target_seq | PAM | alignment | distance | position | gene name | gene id |
| GU305914.1 | 96547 | 96569 | - | 0 | CCTACACTGCTCGCTAGACA | TGG | \|\|\|\|\|\|\|\|[\|\|\|\|\|\|\|\|\|\|\|\|]PAM | NA | Intergenic | NA | NA |
| T26688 | AGACATGGAAGAAGACATCGAGG | 1000 | | | | | | | | | |
| Oligo fwd | CACCAGACATGGAAGAAGACATCG | | | | | | | | | | |
| Oligo rev | AAACCGATGTCTTCTTCCATGTCT | | | | | | | | | | |
| Chromosome | start | end | strand | MM | target_seq | PAM | alignment | distance | position | gene name | gene id |
| GU305914.1 | 96532 | 96554 | - | 0 | AGACATGGAAGAAGACATCG | AGG | \|\|\|\|\|\|\|\|[\|\|\|\|\|\|\|\|\|\|\|\|]PAM | NA | Intergenic | NA | NA |
| T26689 | AGACATCGAGGACGAGCCCTTGG | 1000 | | | | | | | | | |
| Oligo fwd | CACCAGACATCGAGGACGAGCCCT | | | | | | | | | | |
| Oligo rev | AAACAGGGCTCGTCCTCGATGTCT | | | | | | | | | | |
| Chromosome | start | end | strand | MM | target_seq | PAM | alignment | distance | position | gene name | gene id |
| GU305914.1 | 96520 | 96542 | - | 0 | AGACATCGAGGACGAGCCCT | TGG | \|\|\|\|\|\|\|\|[\|\|\|\|\|\|\|\|\|\|\|\|]PAM | NA | Intergenic | NA | NA |
| T26690 | CGAGCCCTTGGACCCCGAGACGG | 1000 | | | | | | | | | |
| Oligo fwd | CACCCGAGCCCTTGGACCCCGAGA | | | | | | | | | | |
| Oligo rev | AAACTCTCGGGGTCCAAGGGCTCG | | | | | | | | | | |
| Chromosome | start | end | strand | MM | target_seq | PAM | alignment | distance | position | gene name | gene id |
| GU305914.1 | 96508 | 96530 | - | 0 | CGAGCCCTTGGACCCCGAGA | CGG | \|\|\|\|\|\|\|\|[\|\|\|\|\|\|\|\|\|\|\|\|]PAM | NA | Intergenic | NA | NA |
| T26691 | ACCCCGAGACGGAGTCCGTGTGG | 1000 | | | | | | | | | |
| Oligo fwd | CACCACCCCGAGACGGAGTCCGTG | | | | | | | | | | |
| Oligo rev | AAACCACGGACTCCGTCTCGGGGT | | | | | | | | | | |
| Chromosome | start | end | strand | MM | target_seq | PAM | alignment | distance | position | gene name | gene id |
| GU305914.1 | 96497 | 96519 | - | 0 | ACCCCGAGACGGAGTCCGTG | TGG | \|\|\|\|\|\|\|\|[\|\|\|\|\|\|\|\|\|\|\|\|]PAM | NA | Intergenic | NA | NA |
| T26692 | CGAGACGGAGTCCGTGTGGACGG | 1000 | | | | | | | | | |
| Oligo fwd | CACCCGAGACGGAGTCCGTGTGGA | | | | | | | | | | |
| Oligo rev | AAACTCCACACGGACTCCGTCTCG | | | | | | | | | | |
| Chromosome | start | end | strand | MM | target_seq | PAM | alignment | distance | position | gene name | gene id |
| GU305914.1 | 96493 | 96515 | - | 0 | CGAGACGGAGTCCGTGTGGA | CGG | \|\|\|\|\|\|\|\|[\|\|\|\|\|\|\|\|\|\|\|\|]PAM | NA | Intergenic | NA | NA |
| T26693 | GACGGCGTCCGTGACGCCCCTGG | 1000 | | | | | | | | | |
| Oligo fwd | CACCGACGGCGTCCGTGACGCCCC | | | | | | | | | | |
| Oligo rev | AAACGGGGCGTCACGGACGCCGTC | | | | | | | | | | |
| Chromosome | start | end | strand | MM | target_seq | PAM | alignment | distance | position | gene name | gene id |
| GU305914.1 | 96475 | 96497 | - | 0 | GACGGCGTCCGTGACGCCCC | TGG | \|\|\|\|\|\|\|\|[\|\|\|\|\|\|\|\|\|\|\|\|]PAM | NA | Intergenic | NA | NA |
| T26694 | CCGCATCCTCGACCACGAGCCGG | 1000 | | | | | | | | | |
| Oligo fwd | CACCCCGCATCCTCGACCACGAGC | | | | | | | | | | |
| Oligo rev | AAACGCTCGTGGTCGAGGATGCGG | | | | | | | | | | |
| Chromosome | start | end | strand | MM | target_seq | PAM | alignment | distance | position | gene name | gene id |
| GU305914.1 | 96436 | 96458 | - | 0 | CCGCATCCTCGACCACGAGC | CGG | \|\|\|\|\|\|\|\|[\|\|\|\|\|\|\|\|\|\|\|\|]PAM | NA | Intergenic | NA | NA |
| T26695 | CGCATCCTCGACCACGAGCCGGG | 1000 | | | | | | | | | |
| Oligo fwd | CACCCGCATCCTCGACCACGAGCC | | | | | | | | | | |
| Oligo rev | AAACGGCTCGTGGTCGAGGATGCG | | | | | | | | | | |
| Chromosome | start | end | strand | MM | target_seq | PAM | alignment | distance | position | gene name | gene id |
| GU305914.1 | 96435 | 96457 | - | 0 | CGCATCCTCGACCACGAGCC | GGG | \|\|\|\|\|\|\|\|[\|\|\|\|\|\|\|\|\|\|\|\|]PAM | NA | Intergenic | NA | NA |
| T26696 | CGACCACGAGCCGGGAGACGCGG | 1000 | | | | | | | | | |
| Oligo fwd | CACCCGACCACGAGCCGGGAGACG | | | | | | | | | | |
| Oligo rev | AAACCGTCTCCCGGCTCGTGGTCG | | | | | | | | | | |
| Chromosome | start | end | strand | MM | target_seq | PAM | alignment | distance | position | gene name | gene id |
| GU305914.1 | 96427 | 96449 | - | 0 | CGACCACGAGCCGGGAGACG | CGG | \|\|\|\|\|\|\|\|[\|\|\|\|\|\|\|\|\|\|\|\|]PAM | NA | Intergenic | NA | NA |
| T26697 | CCACGAGCCGGGAGACGCGGAGG | 1000 | | | | | | | | | |
| Oligo fwd | CACCCCACGAGCCGGGAGACGCGG | | | | | | | | | | |
| Oligo rev | AAACCCGCGTCTCCCGGCTCGTGG | | | | | | | | | | |
| Chromosome | start | end | strand | MM | target_seq | PAM | alignment | distance | position | gene name | gene id |
| GU305914.1 | 96424 | 96446 | - | 0 | CCACGAGCCGGGAGACGCGG | AGG | \|\|\|\|\|\|\|\|[\|\|\|\|\|\|\|\|\|\|\|\|]PAM | NA | Intergenic | NA | NA |
| T26698 | GGGAGACGCGGAGGAAGAAGAGG | 1000 | | | | | | | | | |
| Oligo fwd | CACCGGGAGACGCGGAGGAAGAAG | | | | | | | | | | |
| Oligo rev | AAACCTTCTTCCTCCGCGTCTCCC | | | | | | | | | | |
| Chromosome | start | end | strand | MM | target_seq | PAM | alignment | distance | position | gene name | gene id |
| GU305914.1 | 96415 | 96437 | - | 0 | GGGAGACGCGGAGGAAGAAG | AGG | \|\|\|\|\|\|\|\|[\|\|\|\|\|\|\|\|\|\|\|\|]PAM | NA | Intergenic | NA | NA |
| GU305914.1 | 45065 | 45087 | + | 4 | AGGAGAAGAGGAGGAGGAAG | AGG | -\|\|\|\|\|-\|[-\|\|\|\|\|\|-\|\|\|\|]PAM | NA | Intergenic | NA | NA |

<table>
<tr><td>T26699</td><td>GGAAGAAGAGGAGTCCGACACGG</td><td>1000</td><td></td><td></td><td></td><td></td><td></td><td></td><td></td><td></td><td></td></tr>
<tr><td>Oligo fwd</td><td>CACCGGAAGAAGAGGAGTCCGACA</td><td></td><td></td><td></td><td></td><td></td><td></td><td></td><td></td><td></td><td></td></tr>
<tr><td>Oligo rev</td><td>AAACTGTCGGACTCCTCTTCTTCC</td><td></td><td></td><td></td><td></td><td></td><td></td><td></td><td></td><td></td><td></td></tr>
<tr><td>Chromosome</td><td>start</td><td>end</td><td>strand</td><td>MM</td><td>target_seq</td><td>PAM</td><td>alignment</td><td>distance</td><td>position</td><td>gene name</td><td>gene id</td></tr>
<tr><td>GU305914.1</td><td>96403</td><td>96425</td><td>-</td><td>0</td><td>GGAAGAAGAGGAGTCCGACA</td><td>CGG</td><td>||||||||[||||||||||||]PAM</td><td>NA</td><td>Intergenic</td><td>NA</td><td>NA</td></tr>
<tr><td>T26700</td><td>CACGGACTTCTACGACGAGACGG</td><td>1000</td><td></td><td></td><td></td><td></td><td></td><td></td><td></td><td></td><td></td></tr>
<tr><td>Oligo fwd</td><td>CACCCACGGACTTCTACGACGAGA</td><td></td><td></td><td></td><td></td><td></td><td></td><td></td><td></td><td></td><td></td></tr>
<tr><td>Oligo rev</td><td>AAACTCTCGTCGTAGAAGTCCGTG</td><td></td><td></td><td></td><td></td><td></td><td></td><td></td><td></td><td></td><td></td></tr>
<tr><td>Chromosome</td><td>start</td><td>end</td><td>strand</td><td>MM</td><td>target_seq</td><td>PAM</td><td>alignment</td><td>distance</td><td>position</td><td>gene name</td><td>gene id</td></tr>
<tr><td>GU305914.1</td><td>96385</td><td>96407</td><td>-</td><td>0</td><td>CACGGACTTCTACGACGAGA</td><td>CGG</td><td>||||||||[||||||||||||]PAM</td><td>NA</td><td>Intergenic</td><td>NA</td><td>NA</td></tr>
<tr><td>T26701</td><td>TCAACAAACGCATCCACTTGAGG</td><td>1000</td><td></td><td></td><td></td><td></td><td></td><td></td><td></td><td></td><td></td></tr>
<tr><td>Oligo fwd</td><td>CACCTCAACAAACGCATCCACTTG</td><td></td><td></td><td></td><td></td><td></td><td></td><td></td><td></td><td></td><td></td></tr>
<tr><td>Oligo rev</td><td>AAACCAAGTGGATGCGTTTGTTGA</td><td></td><td></td><td></td><td></td><td></td><td></td><td></td><td></td><td></td><td></td></tr>
<tr><td>Chromosome</td><td>start</td><td>end</td><td>strand</td><td>MM</td><td>target_seq</td><td>PAM</td><td>alignment</td><td>distance</td><td>position</td><td>gene name</td><td>gene id</td></tr>
<tr><td>GU305914.1</td><td>96353</td><td>96375</td><td>-</td><td>0</td><td>TCAACAAACGCATCCACTTG</td><td>AGG</td><td>||||||||[||||||||||||]PAM</td><td>NA</td><td>Intergenic</td><td>NA</td><td>NA</td></tr>
<tr><td>T26702</td><td>CTTGAGGAGCGCGACCCCCACGG</td><td>1000</td><td></td><td></td><td></td><td></td><td></td><td></td><td></td><td></td><td></td></tr>
<tr><td>Oligo fwd</td><td>CACCCTTGAGGAGCGCGACCCCCA</td><td></td><td></td><td></td><td></td><td></td><td></td><td></td><td></td><td></td><td></td></tr>
<tr><td>Oligo rev</td><td>AAACTGGGGGTCGCGCTCCTCAAG</td><td></td><td></td><td></td><td></td><td></td><td></td><td></td><td></td><td></td><td></td></tr>
<tr><td>Chromosome</td><td>start</td><td>end</td><td>strand</td><td>MM</td><td>target_seq</td><td>PAM</td><td>alignment</td><td>distance</td><td>position</td><td>gene name</td><td>gene id</td></tr>
<tr><td>GU305914.1</td><td>96337</td><td>96359</td><td>-</td><td>0</td><td>CTTGAGGAGCGCGACCCCCA</td><td>CGG</td><td>||||||||[||||||||||||]PAM</td><td>NA</td><td>Intergenic</td><td>NA</td><td>NA</td></tr>
<tr><td>T26703</td><td>CCCCACGGACGACGTGATCATGG</td><td>1000</td><td></td><td></td><td></td><td></td><td></td><td></td><td></td><td></td><td></td></tr>
<tr><td>Oligo fwd</td><td>CACCCCCCACGGACGACGTGATCA</td><td></td><td></td><td></td><td></td><td></td><td></td><td></td><td></td><td></td><td></td></tr>
<tr><td>Oligo rev</td><td>AAACTGATCACGTCGTCCGTGGGG</td><td></td><td></td><td></td><td></td><td></td><td></td><td></td><td></td><td></td><td></td></tr>
<tr><td>Chromosome</td><td>start</td><td>end</td><td>strand</td><td>MM</td><td>target_seq</td><td>PAM</td><td>alignment</td><td>distance</td><td>position</td><td>gene name</td><td>gene id</td></tr>
<tr><td>GU305914.1</td><td>96322</td><td>96344</td><td>-</td><td>0</td><td>CCCCACGGACGACGTGATCA</td><td>TGG</td><td>||||||||[||||||||||||]PAM</td><td>NA</td><td>Intergenic</td><td>NA</td><td>NA</td></tr>
<tr><td>T26704</td><td>CGACCTCTCCTACAGCGAGATGG</td><td>1000</td><td></td><td></td><td></td><td></td><td></td><td></td><td></td><td></td><td></td></tr>
<tr><td>Oligo fwd</td><td>CACCCGACCTCTCCTACAGCGAGA</td><td></td><td></td><td></td><td></td><td></td><td></td><td></td><td></td><td></td><td></td></tr>
<tr><td>Oligo rev</td><td>AAACTCTCGCTGTAGGAGAGGTCG</td><td></td><td></td><td></td><td></td><td></td><td></td><td></td><td></td><td></td><td></td></tr>
<tr><td>Chromosome</td><td>start</td><td>end</td><td>strand</td><td>MM</td><td>target_seq</td><td>PAM</td><td>alignment</td><td>distance</td><td>position</td><td>gene name</td><td>gene id</td></tr>
<tr><td>GU305914.1</td><td>96295</td><td>96317</td><td>-</td><td>0</td><td>CGACCTCTCCTACAGCGAGA</td><td>TGG</td><td>||||||||[||||||||||||]PAM</td><td>NA</td><td>Intergenic</td><td>NA</td><td>NA</td></tr>
<tr><td>T26705</td><td>TGGACTCTGACTGATGAAATCGG</td><td>1000</td><td></td><td></td><td></td><td></td><td></td><td></td><td></td><td></td><td></td></tr>
<tr><td>Oligo fwd</td><td>CACCTGGACTCTGACTGATGAAAT</td><td></td><td></td><td></td><td></td><td></td><td></td><td></td><td></td><td></td><td></td></tr>
<tr><td>Oligo rev</td><td>AAACATTTCATCAGTCAGAGTCCA</td><td></td><td></td><td></td><td></td><td></td><td></td><td></td><td></td><td></td><td></td></tr>
<tr><td>Chromosome</td><td>start</td><td>end</td><td>strand</td><td>MM</td><td>target_seq</td><td>PAM</td><td>alignment</td><td>distance</td><td>position</td><td>gene name</td><td>gene id</td></tr>
<tr><td>GU305914.1</td><td>96275</td><td>96297</td><td>-</td><td>0</td><td>TGGACTCTGACTGATGAAAT</td><td>CGG</td><td>||||||||[||||||||||||]PAM</td><td>NA</td><td>Intergenic</td><td>NA</td><td>NA</td></tr>
<tr><td>T26706</td><td>TCCCCTATCCTCTCAGACGTTGG</td><td>1000</td><td></td><td></td><td></td><td></td><td></td><td></td><td></td><td></td><td></td></tr>
<tr><td>Oligo fwd</td><td>CACCTCCCCTATCCTCTCAGACGT</td><td></td><td></td><td></td><td></td><td></td><td></td><td></td><td></td><td></td><td></td></tr>
<tr><td>Oligo rev</td><td>AAACACGTCTGAGAGGATAGGGGA</td><td></td><td></td><td></td><td></td><td></td><td></td><td></td><td></td><td></td><td></td></tr>
<tr><td>Chromosome</td><td>start</td><td>end</td><td>strand</td><td>MM</td><td>target_seq</td><td>PAM</td><td>alignment</td><td>distance</td><td>position</td><td>gene name</td><td>gene id</td></tr>
<tr><td>GU305914.1</td><td>96218</td><td>96240</td><td>-</td><td>0</td><td>TCCCCTATCCTCTCAGACGT</td><td>TGG</td><td>||||||||[||||||||||||]PAM</td><td>NA</td><td>Intergenic</td><td>NA</td><td>NA</td></tr>
<tr><td>T26707</td><td>GTCAAACGAGCGTCATTCTGTGG</td><td>1000</td><td></td><td></td><td></td><td></td><td></td><td></td><td></td><td></td><td></td></tr>
<tr><td>Oligo fwd</td><td>CACCGTCAAACGAGCGTCATTCTG</td><td></td><td></td><td></td><td></td><td></td><td></td><td></td><td></td><td></td><td></td></tr>
<tr><td>Oligo rev</td><td>AAACCAGAATGACGCTCGTTTGAC</td><td></td><td></td><td></td><td></td><td></td><td></td><td></td><td></td><td></td><td></td></tr>
<tr><td>Chromosome</td><td>start</td><td>end</td><td>strand</td><td>MM</td><td>target_seq</td><td>PAM</td><td>alignment</td><td>distance</td><td>position</td><td>gene name</td><td>gene id</td></tr>
<tr><td>GU305914.1</td><td>96087</td><td>96109</td><td>-</td><td>0</td><td>GTCAAACGAGCGTCATTCTG</td><td>TGG</td><td>||||||||[||||||||||||]PAM</td><td>NA</td><td>Intergenic</td><td>NA</td><td>NA</td></tr>
<tr><td>T26708</td><td>TCAAACGAGCGTCATTCTGTGGG</td><td>1000</td><td></td><td></td><td></td><td></td><td></td><td></td><td></td><td></td><td></td></tr>
<tr><td>Oligo fwd</td><td>CACCTCAAACGAGCGTCATTCTGT</td><td></td><td></td><td></td><td></td><td></td><td></td><td></td><td></td><td></td><td></td></tr>
<tr><td>Oligo rev</td><td>AAACACAGAATGACGCTCGTTTGA</td><td></td><td></td><td></td><td></td><td></td><td></td><td></td><td></td><td></td><td></td></tr>
<tr><td>Chromosome</td><td>start</td><td>end</td><td>strand</td><td>MM</td><td>target_seq</td><td>PAM</td><td>alignment</td><td>distance</td><td>position</td><td>gene name</td><td>gene id</td></tr>
<tr><td>GU305914.1</td><td>96086</td><td>96108</td><td>-</td><td>0</td><td>TCAAACGAGCGTCATTCTGT</td><td>GGG</td><td>||||||||[||||||||||||]PAM</td><td>NA</td><td>Intergenic</td><td>NA</td><td>NA</td></tr>
<tr><td>T26709</td><td>GTGGGTCTGCGACTCAGAGACGG</td><td>1000</td><td></td><td></td><td></td><td></td><td></td><td></td><td></td><td></td><td></td></tr>
<tr><td>Oligo fwd</td><td>CACCGTGGGTCTGCGACTCAGAGA</td><td></td><td></td><td></td><td></td><td></td><td></td><td></td><td></td><td></td><td></td></tr>
<tr><td>Oligo rev</td><td>AAACTCTCTGAGTCGCAGACCCAC</td><td></td><td></td><td></td><td></td><td></td><td></td><td></td><td></td><td></td><td></td></tr>
<tr><td>Chromosome</td><td>start</td><td>end</td><td>strand</td><td>MM</td><td>target_seq</td><td>PAM</td><td>alignment</td><td>distance</td><td>position</td><td>gene name</td><td>gene id</td></tr>
<tr><td>GU305914.1</td><td>96068</td><td>96090</td><td>-</td><td>0</td><td>GTGGGTCTGCGACTCAGAGA</td><td>CGG</td><td>||||||||[||||||||||||]PAM</td><td>NA</td><td>Intergenic</td><td>NA</td><td>NA</td></tr>
<tr><td>T26710</td><td>TGGGTCTGCGACTCAGAGACGGG</td><td>1000</td><td></td><td></td><td></td><td></td><td></td><td></td><td></td><td></td><td></td></tr>
<tr><td>Oligo fwd</td><td>CACCTGGGTCTGCGACTCAGAGAC</td><td></td><td></td><td></td><td></td><td></td><td></td><td></td><td></td><td></td><td></td></tr>
<tr><td>Oligo rev</td><td>AAACGTCTCTGAGTCGCAGACCCA</td><td></td><td></td><td></td><td></td><td></td><td></td><td></td><td></td><td></td><td></td></tr>
<tr><td>Chromosome</td><td>start</td><td>end</td><td>strand</td><td>MM</td><td>target_seq</td><td>PAM</td><td>alignment</td><td>distance</td><td>position</td><td>gene name</td><td>gene id</td></tr>
<tr><td>GU305914.1</td><td>96067</td><td>96089</td><td>-</td><td>0</td><td>TGGGTCTGCGACTCAGAGAC</td><td>GGG</td><td>||||||||[||||||||||||]PAM</td><td>NA</td><td>Intergenic</td><td>NA</td><td>NA</td></tr>
<tr><td>T26711</td><td>GTGCACCGTCGATGACTAGATGG</td><td>1000</td><td></td><td></td><td></td><td></td><td></td><td></td><td></td><td></td><td></td></tr>
<tr><td>Oligo fwd</td><td>CACCGTGCACCGTCGATGACTAGA</td><td></td><td></td><td></td><td></td><td></td><td></td><td></td><td></td><td></td><td></td></tr>
<tr><td>Oligo rev</td><td>AAACTCTAGTCATCGACGGTGCAC</td><td></td><td></td><td></td><td></td><td></td><td></td><td></td><td></td><td></td><td></td></tr>
<tr><td>Chromosome</td><td>start</td><td>end</td><td>strand</td><td>MM</td><td>target_seq</td><td>PAM</td><td>alignment</td><td>distance</td><td>position</td><td>gene name</td><td>gene id</td></tr>
<tr><td>GU305914.1</td><td>96039</td><td>96061</td><td>-</td><td>0</td><td>GTGCACCGTCGATGACTAGA</td><td>TGG</td><td>||||||||[||||||||||||]PAM</td><td>NA</td><td>Intergenic</td><td>NA</td><td>NA</td></tr>
<tr><td>T26712</td><td>TGAACCGTTAGAAATCTTGCCGG</td><td>1000</td><td></td><td></td><td></td><td></td><td></td><td></td><td></td><td></td><td></td></tr>
<tr><td>Oligo fwd</td><td>CACCTGAACCGTTAGAAATCTTGC</td><td></td><td></td><td></td><td></td><td></td><td></td><td></td><td></td><td></td><td></td></tr>
<tr><td>Oligo rev</td><td>AAACGCAAGATTTCTAACGGTTCA</td><td></td><td></td><td></td><td></td><td></td><td></td><td></td><td></td><td></td><td></td></tr>
<tr><td>Chromosome</td><td>start</td><td>end</td><td>strand</td><td>MM</td><td>target_seq</td><td>PAM</td><td>alignment</td><td>distance</td><td>position</td><td>gene name</td><td>gene id</td></tr>
<tr><td>GU305914.1</td><td>96010</td><td>96032</td><td>-</td><td>0</td><td>TGAACCGTTAGAAATCTTGC</td><td>CGG</td><td>||||||||[||||||||||||]PAM</td><td>NA</td><td>Intergenic</td><td>NA</td><td>NA</td></tr>
<tr><td>T26713</td><td>GAACCGTTAGAAATCTTGCCGGG</td><td>1000</td><td></td><td></td><td></td><td></td><td></td><td></td><td></td><td></td><td></td></tr>
<tr><td>Oligo fwd</td><td>CACCGAACCGTTAGAAATCTTGCC</td><td></td><td></td><td></td><td></td><td></td><td></td><td></td><td></td><td></td><td></td></tr>
<tr><td>Oligo rev</td><td>AAACGGCAAGATTTCTAACGGTTC</td><td></td><td></td><td></td><td></td><td></td><td></td><td></td><td></td><td></td><td></td></tr>
<tr><td>Chromosome</td><td>start</td><td>end</td><td>strand</td><td>MM</td><td>target_seq</td><td>PAM</td><td>alignment</td><td>distance</td><td>position</td><td>gene name</td><td>gene id</td></tr>
<tr><td>GU305914.1</td><td>96009</td><td>96031</td><td>-</td><td>0</td><td>GAACCGTTAGAAATCTTGCC</td><td>GGG</td><td>||||||||[||||||||||||]PAM</td><td>NA</td><td>Intergenic</td><td>NA</td><td>NA</td></tr>
</table>

<table>
<tr><td>T26714</td><td>GACCTGTCACTCAACGTGCACGG</td><td>1000</td><td></td><td></td><td></td><td></td><td></td><td></td><td></td><td></td><td></td></tr>
<tr><td>Oligo fwd</td><td>CACCGACCTGTCACTCAACGTGCA</td><td></td><td></td><td></td><td></td><td></td><td></td><td></td><td></td><td></td><td></td></tr>
<tr><td>Oligo rev</td><td>AAACTGCACGTTGAGTGACAGGTC</td><td></td><td></td><td></td><td></td><td></td><td></td><td></td><td></td><td></td><td></td></tr>
<tr><td>Chromosome</td><td>start</td><td>end</td><td>strand</td><td>MM</td><td>target_seq</td><td>PAM</td><td>alignment</td><td>distance</td><td>position</td><td>gene name</td><td>gene id</td></tr>
<tr><td>GU305914.1</td><td>95983</td><td>96005</td><td>-</td><td>0</td><td>GACCTGTCACTCAACGTGCA</td><td>CGG</td><td>||||||||[||||||||||||]PAM</td><td>NA</td><td>Intergenic</td><td>NA</td><td>NA</td></tr>
<tr><td>T26715</td><td>ACCTGTCACTCAACGTGCACGGG</td><td>1000</td><td></td><td></td><td></td><td></td><td></td><td></td><td></td><td></td><td></td></tr>
<tr><td>Oligo fwd</td><td>CACCACCTGTCACTCAACGTGCAC</td><td></td><td></td><td></td><td></td><td></td><td></td><td></td><td></td><td></td><td></td></tr>
<tr><td>Oligo rev</td><td>AAACGTGCACGTTGAGTGACAGGT</td><td></td><td></td><td></td><td></td><td></td><td></td><td></td><td></td><td></td><td></td></tr>
<tr><td>Chromosome</td><td>start</td><td>end</td><td>strand</td><td>MM</td><td>target_seq</td><td>PAM</td><td>alignment</td><td>distance</td><td>position</td><td>gene name</td><td>gene id</td></tr>
<tr><td>GU305914.1</td><td>95982</td><td>96004</td><td>-</td><td>0</td><td>ACCTGTCACTCAACGTGCAC</td><td>GGG</td><td>||||||||[||||||||||||]PAM</td><td>NA</td><td>Intergenic</td><td>NA</td><td>NA</td></tr>
<tr><td>T26716</td><td>CCTGTCACTCAACGTGCACGGGG</td><td>1000</td><td></td><td></td><td></td><td></td><td></td><td></td><td></td><td></td><td></td></tr>
<tr><td>Oligo fwd</td><td>CACCCCTGTCACTCAACGTGCACG</td><td></td><td></td><td></td><td></td><td></td><td></td><td></td><td></td><td></td><td></td></tr>
<tr><td>Oligo rev</td><td>AAACCGTGCACGTTGAGTGACAGG</td><td></td><td></td><td></td><td></td><td></td><td></td><td></td><td></td><td></td><td></td></tr>
<tr><td>Chromosome</td><td>start</td><td>end</td><td>strand</td><td>MM</td><td>target_seq</td><td>PAM</td><td>alignment</td><td>distance</td><td>position</td><td>gene name</td><td>gene id</td></tr>
<tr><td>GU305914.1</td><td>95981</td><td>96003</td><td>-</td><td>0</td><td>CCTGTCACTCAACGTGCACG</td><td>GGG</td><td>||||||||[||||||||||||]PAM</td><td>NA</td><td>Intergenic</td><td>NA</td><td>NA</td></tr>
<tr><td>T26717</td><td>GGACCGTACGAGCTTTCGATTGG</td><td>1000</td><td></td><td></td><td></td><td></td><td></td><td></td><td></td><td></td><td></td></tr>
<tr><td>Oligo fwd</td><td>CACCGGACCGTACGAGCTTTCGAT</td><td></td><td></td><td></td><td></td><td></td><td></td><td></td><td></td><td></td><td></td></tr>
<tr><td>Oligo rev</td><td>AAACATCGAAAGCTCGTACGGTCC</td><td></td><td></td><td></td><td></td><td></td><td></td><td></td><td></td><td></td><td></td></tr>
<tr><td>Chromosome</td><td>start</td><td>end</td><td>strand</td><td>MM</td><td>target_seq</td><td>PAM</td><td>alignment</td><td>distance</td><td>position</td><td>gene name</td><td>gene id</td></tr>
<tr><td>GU305914.1</td><td>95960</td><td>95982</td><td>-</td><td>0</td><td>GGACCGTACGAGCTTTCGAT</td><td>TGG</td><td>||||||||[||||||||||||]PAM</td><td>NA</td><td>Intergenic</td><td>NA</td><td>NA</td></tr>
<tr><td>T26718</td><td>CCGTACGAGCTTTCGATTGGTGG</td><td>1000</td><td></td><td></td><td></td><td></td><td></td><td></td><td></td><td></td><td></td></tr>
<tr><td>Oligo fwd</td><td>CACCCCGTACGAGCTTTCGATTGG</td><td></td><td></td><td></td><td></td><td></td><td></td><td></td><td></td><td></td><td></td></tr>
<tr><td>Oligo rev</td><td>AAACCCAATCGAAAGCTCGTACGG</td><td></td><td></td><td></td><td></td><td></td><td></td><td></td><td></td><td></td><td></td></tr>
<tr><td>Chromosome</td><td>start</td><td>end</td><td>strand</td><td>MM</td><td>target_seq</td><td>PAM</td><td>alignment</td><td>distance</td><td>position</td><td>gene name</td><td>gene id</td></tr>
<tr><td>GU305914.1</td><td>95957</td><td>95979</td><td>-</td><td>0</td><td>CCGTACGAGCTTTCGATTGG</td><td>TGG</td><td>||||||||[||||||||||||]PAM</td><td>NA</td><td>Intergenic</td><td>NA</td><td>NA</td></tr>
<tr><td>T26719</td><td>ACTGTCCATCAACGATAAGCTGG</td><td>1000</td><td></td><td></td><td></td><td></td><td></td><td></td><td></td><td></td><td></td></tr>
<tr><td>Oligo fwd</td><td>CACCACTGTCCATCAACGATAAGC</td><td></td><td></td><td></td><td></td><td></td><td></td><td></td><td></td><td></td><td></td></tr>
<tr><td>Oligo rev</td><td>AAACGCTTATCGTTGATGGACAGT</td><td></td><td></td><td></td><td></td><td></td><td></td><td></td><td></td><td></td><td></td></tr>
<tr><td>Chromosome</td><td>start</td><td>end</td><td>strand</td><td>MM</td><td>target_seq</td><td>PAM</td><td>alignment</td><td>distance</td><td>position</td><td>gene name</td><td>gene id</td></tr>
<tr><td>GU305914.1</td><td>95929</td><td>95951</td><td>-</td><td>0</td><td>ACTGTCCATCAACGATAAGC</td><td>TGG</td><td>||||||||[||||||||||||]PAM</td><td>NA</td><td>Intergenic</td><td>NA</td><td>NA</td></tr>
<tr><td>T26720</td><td>GCTGGCTACCATCCGAATATCGG</td><td>1000</td><td></td><td></td><td></td><td></td><td></td><td></td><td></td><td></td><td></td></tr>
<tr><td>Oligo fwd</td><td>CACCGCTGGCTACCATCCGAATAT</td><td></td><td></td><td></td><td></td><td></td><td></td><td></td><td></td><td></td><td></td></tr>
<tr><td>Oligo rev</td><td>AAACATATTCGGATGGTAGCCAGC</td><td></td><td></td><td></td><td></td><td></td><td></td><td></td><td></td><td></td><td></td></tr>
<tr><td>Chromosome</td><td>start</td><td>end</td><td>strand</td><td>MM</td><td>target_seq</td><td>PAM</td><td>alignment</td><td>distance</td><td>position</td><td>gene name</td><td>gene id</td></tr>
<tr><td>GU305914.1</td><td>95911</td><td>95933</td><td>-</td><td>0</td><td>GCTGGCTACCATCCGAATAT</td><td>CGG</td><td>||||||||[||||||||||||]PAM</td><td>NA</td><td>Intergenic</td><td>NA</td><td>NA</td></tr>
<tr><td>T26721</td><td>CCATCCGAATATCGGAGTTCTGG</td><td>1000</td><td></td><td></td><td></td><td></td><td></td><td></td><td></td><td></td><td></td></tr>
<tr><td>Oligo fwd</td><td>CACCCCATCCGAATATCGGAGTTC</td><td></td><td></td><td></td><td></td><td></td><td></td><td></td><td></td><td></td><td></td></tr>
<tr><td>Oligo rev</td><td>AAACGAACTCCGATATTCGGATGG</td><td></td><td></td><td></td><td></td><td></td><td></td><td></td><td></td><td></td><td></td></tr>
<tr><td>Chromosome</td><td>start</td><td>end</td><td>strand</td><td>MM</td><td>target_seq</td><td>PAM</td><td>alignment</td><td>distance</td><td>position</td><td>gene name</td><td>gene id</td></tr>
<tr><td>GU305914.1</td><td>95903</td><td>95925</td><td>-</td><td>0</td><td>CCATCCGAATATCGGAGTTC</td><td>TGG</td><td>||||||||[||||||||||||]PAM</td><td>NA</td><td>Intergenic</td><td>NA</td><td>NA</td></tr>
<tr><td>T26722</td><td>CATCCGAATATCGGAGTTCTGGG</td><td>1000</td><td></td><td></td><td></td><td></td><td></td><td></td><td></td><td></td><td></td></tr>
<tr><td>Oligo fwd</td><td>CACCCATCCGAATATCGGAGTTCT</td><td></td><td></td><td></td><td></td><td></td><td></td><td></td><td></td><td></td><td></td></tr>
<tr><td>Oligo rev</td><td>AAACAGAACTCCGATATTCGGATG</td><td></td><td></td><td></td><td></td><td></td><td></td><td></td><td></td><td></td><td></td></tr>
<tr><td>Chromosome</td><td>start</td><td>end</td><td>strand</td><td>MM</td><td>target_seq</td><td>PAM</td><td>alignment</td><td>distance</td><td>position</td><td>gene name</td><td>gene id</td></tr>
<tr><td>GU305914.1</td><td>95902</td><td>95924</td><td>-</td><td>0</td><td>CATCCGAATATCGGAGTTCT</td><td>GGG</td><td>||||||||[||||||||||||]PAM</td><td>NA</td><td>Intergenic</td><td>NA</td><td>NA</td></tr>
<tr><td>T26723</td><td>GCGCATGCGCATACACATGATGG</td><td>1000</td><td></td><td></td><td></td><td></td><td></td><td></td><td></td><td></td><td></td></tr>
<tr><td>Oligo fwd</td><td>CACCGCGCATGCGCATACACATGA</td><td></td><td></td><td></td><td></td><td></td><td></td><td></td><td></td><td></td><td></td></tr>
<tr><td>Oligo rev</td><td>AAACTCATGTGTATGCGCATGCGC</td><td></td><td></td><td></td><td></td><td></td><td></td><td></td><td></td><td></td><td></td></tr>
<tr><td>Chromosome</td><td>start</td><td>end</td><td>strand</td><td>MM</td><td>target_seq</td><td>PAM</td><td>alignment</td><td>distance</td><td>position</td><td>gene name</td><td>gene id</td></tr>
<tr><td>GU305914.1</td><td>95869</td><td>95891</td><td>-</td><td>0</td><td>GCGCATGCGCATACACATGA</td><td>TGG</td><td>||||||||[||||||||||||]PAM</td><td>NA</td><td>Intergenic</td><td>NA</td><td>NA</td></tr>
<tr><td>T26724</td><td>TCCTAGCGACCGATACACATAGG</td><td>1000</td><td></td><td></td><td></td><td></td><td></td><td></td><td></td><td></td><td></td></tr>
<tr><td>Oligo fwd</td><td>CACCTCCTAGCGACCGATACACAT</td><td></td><td></td><td></td><td></td><td></td><td></td><td></td><td></td><td></td><td></td></tr>
<tr><td>Oligo rev</td><td>AAACATGTGTATCGGTCGCTAGGA</td><td></td><td></td><td></td><td></td><td></td><td></td><td></td><td></td><td></td><td></td></tr>
<tr><td>Chromosome</td><td>start</td><td>end</td><td>strand</td><td>MM</td><td>target_seq</td><td>PAM</td><td>alignment</td><td>distance</td><td>position</td><td>gene name</td><td>gene id</td></tr>
<tr><td>GU305914.1</td><td>95825</td><td>95847</td><td>-</td><td>0</td><td>TCCTAGCGACCGATACACAT</td><td>AGG</td><td>||||||||[||||||||||||]PAM</td><td>NA</td><td>Intergenic</td><td>NA</td><td>NA</td></tr>
<tr><td>T26725</td><td>ACCGATACACATAGGCCCACTGG</td><td>1000</td><td></td><td></td><td></td><td></td><td></td><td></td><td></td><td></td><td></td></tr>
<tr><td>Oligo fwd</td><td>CACCACCGATACACATAGGCCCAC</td><td></td><td></td><td></td><td></td><td></td><td></td><td></td><td></td><td></td><td></td></tr>
<tr><td>Oligo rev</td><td>AAACGTGGGCCTATGTGTATCGGT</td><td></td><td></td><td></td><td></td><td></td><td></td><td></td><td></td><td></td><td></td></tr>
<tr><td>Chromosome</td><td>start</td><td>end</td><td>strand</td><td>MM</td><td>target_seq</td><td>PAM</td><td>alignment</td><td>distance</td><td>position</td><td>gene name</td><td>gene id</td></tr>
<tr><td>GU305914.1</td><td>95817</td><td>95839</td><td>-</td><td>0</td><td>ACCGATACACATAGGCCCAC</td><td>TGG</td><td>||||||||[||||||||||||]PAM</td><td>NA</td><td>Intergenic</td><td>NA</td><td>NA</td></tr>
<tr><td>T26726</td><td>AGGCCCACTGGCACACTTCCCGG</td><td>1000</td><td></td><td></td><td></td><td></td><td></td><td></td><td></td><td></td><td></td></tr>
<tr><td>Oligo fwd</td><td>CACCAGGCCCACTGGCACACTTCC</td><td></td><td></td><td></td><td></td><td></td><td></td><td></td><td></td><td></td><td></td></tr>
<tr><td>Oligo rev</td><td>AAACGGAAGTGTGCCAGTGGGCCT</td><td></td><td></td><td></td><td></td><td></td><td></td><td></td><td></td><td></td><td></td></tr>
<tr><td>Chromosome</td><td>start</td><td>end</td><td>strand</td><td>MM</td><td>target_seq</td><td>PAM</td><td>alignment</td><td>distance</td><td>position</td><td>gene name</td><td>gene id</td></tr>
<tr><td>GU305914.1</td><td>95805</td><td>95827</td><td>-</td><td>0</td><td>AGGCCCACTGGCACACTTCC</td><td>CGG</td><td>||||||||[||||||||||||]PAM</td><td>NA</td><td>Intergenic</td><td>NA</td><td>NA</td></tr>
<tr><td>T26727</td><td>GGCCCACTGGCACACTTCCCGGG</td><td>1000</td><td></td><td></td><td></td><td></td><td></td><td></td><td></td><td></td><td></td></tr>
<tr><td>Oligo fwd</td><td>CACCGGCCCACTGGCACACTTCCC</td><td></td><td></td><td></td><td></td><td></td><td></td><td></td><td></td><td></td><td></td></tr>
<tr><td>Oligo rev</td><td>AAACGGGAAGTGTGCCAGTGGGCC</td><td></td><td></td><td></td><td></td><td></td><td></td><td></td><td></td><td></td><td></td></tr>
<tr><td>Chromosome</td><td>start</td><td>end</td><td>strand</td><td>MM</td><td>target_seq</td><td>PAM</td><td>alignment</td><td>distance</td><td>position</td><td>gene name</td><td>gene id</td></tr>
<tr><td>GU305914.1</td><td>95804</td><td>95826</td><td>-</td><td>0</td><td>GGCCCACTGGCACACTTCCC</td><td>GGG</td><td>||||||||[||||||||||||]PAM</td><td>NA</td><td>Intergenic</td><td>NA</td><td>NA</td></tr>
<tr><td>T26728</td><td>GCATACACGCATATTAATTATGG</td><td>1000</td><td></td><td></td><td></td><td></td><td></td><td></td><td></td><td></td><td></td></tr>
<tr><td>Oligo fwd</td><td>CACCGCATACACGCATATTAATTA</td><td></td><td></td><td></td><td></td><td></td><td></td><td></td><td></td><td></td><td></td></tr>
<tr><td>Oligo rev</td><td>AAACTAATTAATATGCGTGTATGC</td><td></td><td></td><td></td><td></td><td></td><td></td><td></td><td></td><td></td><td></td></tr>
<tr><td>Chromosome</td><td>start</td><td>end</td><td>strand</td><td>MM</td><td>target_seq</td><td>PAM</td><td>alignment</td><td>distance</td><td>position</td><td>gene name</td><td>gene id</td></tr>
</table>

GU305914.1 95756 95778 - 0 GCATACACGCATATTAATTA TGG ||||||||[||||||||||||]PAM NA Intergenic NA NA

T26729 TACACGCATATTAATTATGGAGG 1000
Oligo fwd CACCTACACGCATATTAATTATGG
Oligo rev AAACCCATAATTAATATGCGTGTA
Chromosome start end strand MM target_seq PAM alignment distance position gene name gene id
GU305914.1 95753 95775 - 0 TACACGCATATTAATTATGG AGG ||||||||[||||||||||||]PAM NA Intergenic NA NA

T26730 CACATGCATCTGATCGCTATCGG 1000
Oligo fwd CACCCACATGCATCTGATCGCTAT
Oligo rev AAACATAGCGATCAGATGCATGTG
Chromosome start end strand MM target_seq PAM alignment distance position gene name gene id
GU305914.1 95722 95744 - 0 CACATGCATCTGATCGCTAT CGG ||||||||[||||||||||||]PAM NA Intergenic NA NA

T26731 TCGACAGTGACTCAGCAACTAGG 1000
Oligo fwd CACCTCGACAGTGACTCAGCAACT
Oligo rev AAACAGTTGCTGAGTCACTGTCGA
Chromosome start end strand MM target_seq PAM alignment distance position gene name gene id
GU305914.1 95689 95711 - 0 TCGACAGTGACTCAGCAACT AGG ||||||||[||||||||||||]PAM NA Intergenic NA NA

T26732 CGACAGTGACTCAGCAACTAGGG 1000
Oligo fwd CACCCGACAGTGACTCAGCAACTA
Oligo rev AAACTAGTTGCTGAGTCACTGTCG
Chromosome start end strand MM target_seq PAM alignment distance position gene name gene id
GU305914.1 95688 95710 - 0 CGACAGTGACTCAGCAACTA GGG ||||||||[||||||||||||]PAM NA Intergenic NA NA

T26733 TCATGCTATAGATGAACGATCGG 1000
Oligo fwd CACCTCATGCTATAGATGAACGAT
Oligo rev AAACATCGTTCATCTATAGCATGA
Chromosome start end strand MM target_seq PAM alignment distance position gene name gene id
GU305914.1 95633 95655 - 0 TCATGCTATAGATGAACGAT CGG ||||||||[||||||||||||]PAM NA Intergenic NA NA

T26734 ATGAACGATCGGTTCTATATTGG 1000
Oligo fwd CACCATGAACGATCGGTTCTATAT
Oligo rev AAACATATAGAACCGATCGTTCAT
Chromosome start end strand MM target_seq PAM alignment distance position gene name gene id
GU305914.1 95622 95644 - 0 ATGAACGATCGGTTCTATAT TGG ||||||||[||||||||||||]PAM NA Intergenic NA NA

T26735 TGAACGATCGGTTCTATATTGGG 1000
Oligo fwd CACCTGAACGATCGGTTCTATATT
Oligo rev AAACAATATAGAACCGATCGTTCA
Chromosome start end strand MM target_seq PAM alignment distance position gene name gene id
GU305914.1 95621 95643 - 0 TGAACGATCGGTTCTATATT GGG ||||||||[||||||||||||]PAM NA Intergenic NA NA

T26736 CTCCTCACACGCATGCGCAGTGG 1000
Oligo fwd CACCCTCCTCACACGCATGCGCAG
Oligo rev AAACCTGCGCATGCGTGTGAGGAG
Chromosome start end strand MM target_seq PAM alignment distance position gene name gene id
GU305914.1 95592 95614 - 0 CTCCTCACACGCATGCGCAG TGG ||||||||[||||||||||||]PAM NA Intergenic NA NA

T26737 GCGCAGTGGACTTTTGTGTTCGG 1000
Oligo fwd CACCGCGCAGTGGACTTTTGTGTT
Oligo rev AAACAACACAAAAGTCCACTGCGC
Chromosome start end strand MM target_seq PAM alignment distance position gene name gene id
GU305914.1 95578 95600 - 0 GCGCAGTGGACTTTTGTGTT CGG ||||||||[||||||||||||]PAM NA Intergenic NA NA

T26738 CCTACACCGACACACCCCCTCGG 1000
Oligo fwd CACCCCTACACCGACACACCCCCT
Oligo rev AAACAGGGGGTGTGTCGGTGTAGG
Chromosome start end strand MM target_seq PAM alignment distance position gene name gene id
GU305914.1 95541 95563 - 0 CCTACACCGACACACCCCCT CGG ||||||||[||||||||||||]PAM NA Intergenic NA NA

T26739 CTACACCGACACACCCCCTCGGG 1000
Oligo fwd CACCCTACACCGACACACCCCCTC
Oligo rev AAACGAGGGGGTGTGTCGGTGTAG
Chromosome start end strand MM target_seq PAM alignment distance position gene name gene id
GU305914.1 95540 95562 - 0 CTACACCGACACACCCCCTC GGG ||||||||[||||||||||||]PAM NA Intergenic NA NA

T26740 TAAGTCTGAATCACGTGATCAGG 1000
Oligo fwd CACCTAAGTCTGAATCACGTGATC
Oligo rev AAACGATCACGTGATTCAGACTTA
Chromosome start end strand MM target_seq PAM alignment distance position gene name gene id
GU305914.1 95409 95431 - 0 TAAGTCTGAATCACGTGATC AGG ||||||||[||||||||||||]PAM NA Intergenic NA NA

T26741 AAGTCTGAATCACGTGATCAGGG 1000
Oligo fwd CACCAAGTCTGAATCACGTGATCA
Oligo rev AAACTGATCACGTGATTCAGACTT
Chromosome start end strand MM target_seq PAM alignment distance position gene name gene id
GU305914.1 95408 95430 - 0 AAGTCTGAATCACGTGATCA GGG ||||||||[||||||||||||]PAM NA Intergenic NA NA

T26742 TTATTACCCCCCACCCAACGAGG 1000
Oligo fwd CACCTTATTACCCCCCACCCAACG
Oligo rev AAACCGTTGGGTGGGGGGTAATAA
Chromosome start end strand MM target_seq PAM alignment distance position gene name gene id
GU305914.1 95350 95372 - 0 TTATTACCCCCCACCCAACG AGG ||||||||[||||||||||||]PAM NA Intergenic NA NA

T26743 ACCCAACGAGGTCGTCCGCTAGG 1000
Oligo fwd CACCACCCAACGAGGTCGTCCGCT
Oligo rev AAACAGCGGACGACCTCGTTGGGT

<table>
<tr><td>Chromosome</td><td>start</td><td>end</td><td>strand</td><td>MM</td><td>target_seq</td><td>PAM</td><td>alignment</td><td>distance</td><td>position</td><td>gene name</td><td>gene id</td></tr>
<tr><td>GU305914.1</td><td>95338</td><td>95360</td><td>-</td><td>0</td><td>ACCCAACGAGGTCGTCCGCT</td><td>AGG</td><td>||||||||[||||||||||||]PAM</td><td>NA</td><td>Intergenic</td><td>NA</td><td>NA</td></tr>
<tr><td>T26744</td><td>CAACGAGGTCGTCCGCTAGGTGG</td><td>1000</td><td></td><td></td><td></td><td></td><td></td><td></td><td></td><td></td><td></td></tr>
<tr><td>Oligo fwd</td><td>CACCCAACGAGGTCGTCCGCTAGG</td><td></td><td></td><td></td><td></td><td></td><td></td><td></td><td></td><td></td><td></td></tr>
<tr><td>Oligo rev</td><td>AAACCCTAGCGGACGACCTCGTTG</td><td></td><td></td><td></td><td></td><td></td><td></td><td></td><td></td><td></td><td></td></tr>
<tr><td>Chromosome</td><td>start</td><td>end</td><td>strand</td><td>MM</td><td>target_seq</td><td>PAM</td><td>alignment</td><td>distance</td><td>position</td><td>gene name</td><td>gene id</td></tr>
<tr><td>GU305914.1</td><td>95335</td><td>95357</td><td>-</td><td>0</td><td>CAACGAGGTCGTCCGCTAGG</td><td>TGG</td><td>||||||||[||||||||||||]PAM</td><td>NA</td><td>Intergenic</td><td>NA</td><td>NA</td></tr>
<tr><td>T26745</td><td>CGAGGTCGTCCGCTAGGTGGTGG</td><td>1000</td><td></td><td></td><td></td><td></td><td></td><td></td><td></td><td></td><td></td></tr>
<tr><td>Oligo fwd</td><td>CACCCGAGGTCGTCCGCTAGGTGG</td><td></td><td></td><td></td><td></td><td></td><td></td><td></td><td></td><td></td><td></td></tr>
<tr><td>Oligo rev</td><td>AAACCCACCTAGCGGACGACCTCG</td><td></td><td></td><td></td><td></td><td></td><td></td><td></td><td></td><td></td><td></td></tr>
<tr><td>Chromosome</td><td>start</td><td>end</td><td>strand</td><td>MM</td><td>target_seq</td><td>PAM</td><td>alignment</td><td>distance</td><td>position</td><td>gene name</td><td>gene id</td></tr>
<tr><td>GU305914.1</td><td>95332</td><td>95354</td><td>-</td><td>0</td><td>CGAGGTCGTCCGCTAGGTGG</td><td>TGG</td><td>||||||||[||||||||||||]PAM</td><td>NA</td><td>Intergenic</td><td>NA</td><td>NA</td></tr>
<tr><td>T26746</td><td>TCCGCTAGGTGGTGGTCGAGAGG</td><td>1000</td><td></td><td></td><td></td><td></td><td></td><td></td><td></td><td></td><td></td></tr>
<tr><td>Oligo fwd</td><td>CACCTCCGCTAGGTGGTGGTCGAG</td><td></td><td></td><td></td><td></td><td></td><td></td><td></td><td></td><td></td><td></td></tr>
<tr><td>Oligo rev</td><td>AAACCTCGACCACCACCTAGCGGA</td><td></td><td></td><td></td><td></td><td></td><td></td><td></td><td></td><td></td><td></td></tr>
<tr><td>Chromosome</td><td>start</td><td>end</td><td>strand</td><td>MM</td><td>target_seq</td><td>PAM</td><td>alignment</td><td>distance</td><td>position</td><td>gene name</td><td>gene id</td></tr>
<tr><td>GU305914.1</td><td>95324</td><td>95346</td><td>-</td><td>0</td><td>TCCGCTAGGTGGTGGTCGAG</td><td>AGG</td><td>||||||||[||||||||||||]PAM</td><td>NA</td><td>Intergenic</td><td>NA</td><td>NA</td></tr>
<tr><td>T26747</td><td>CCGCTAGGTGGTGGTCGAGAGGG</td><td>1000</td><td></td><td></td><td></td><td></td><td></td><td></td><td></td><td></td><td></td></tr>
<tr><td>Oligo fwd</td><td>CACCCCGCTAGGTGGTGGTCGAGA</td><td></td><td></td><td></td><td></td><td></td><td></td><td></td><td></td><td></td><td></td></tr>
<tr><td>Oligo rev</td><td>AAACTCTCGACCACCACCTAGCGG</td><td></td><td></td><td></td><td></td><td></td><td></td><td></td><td></td><td></td><td></td></tr>
<tr><td>Chromosome</td><td>start</td><td>end</td><td>strand</td><td>MM</td><td>target_seq</td><td>PAM</td><td>alignment</td><td>distance</td><td>position</td><td>gene name</td><td>gene id</td></tr>
<tr><td>GU305914.1</td><td>95323</td><td>95345</td><td>-</td><td>0</td><td>CCGCTAGGTGGTGGTCGAGA</td><td>GGG</td><td>||||||||[||||||||||||]PAM</td><td>NA</td><td>Intergenic</td><td>NA</td><td>NA</td></tr>
<tr><td>GU305914.1</td><td>107378</td><td>107400</td><td>+</td><td>4</td><td>CCGCCAGAGGGCGGTCGAGA</td><td>CGG</td><td>||||-||-[-||-||||||||]PAM</td><td>NA</td><td>Intergenic</td><td>NA</td><td>NA</td></tr>
<tr><td>T26748</td><td>GGTGGTCGAGAGGGAGATCTAGG</td><td>1000</td><td></td><td></td><td></td><td></td><td></td><td></td><td></td><td></td><td></td></tr>
<tr><td>Oligo fwd</td><td>CACCGGTGGTCGAGAGGGAGATCT</td><td></td><td></td><td></td><td></td><td></td><td></td><td></td><td></td><td></td><td></td></tr>
<tr><td>Oligo rev</td><td>AAACAGATCTCCCTCTCGACCACC</td><td></td><td></td><td></td><td></td><td></td><td></td><td></td><td></td><td></td><td></td></tr>
<tr><td>Chromosome</td><td>start</td><td>end</td><td>strand</td><td>MM</td><td>target_seq</td><td>PAM</td><td>alignment</td><td>distance</td><td>position</td><td>gene name</td><td>gene id</td></tr>
<tr><td>GU305914.1</td><td>95314</td><td>95336</td><td>-</td><td>0</td><td>GGTGGTCGAGAGGGAGATCT</td><td>AGG</td><td>||||||||[||||||||||||]PAM</td><td>NA</td><td>Intergenic</td><td>NA</td><td>NA</td></tr>
<tr><td>T26749</td><td>GACCACCTTGTGTCCTTATAAGG</td><td>1000</td><td></td><td></td><td></td><td></td><td></td><td></td><td></td><td></td><td></td></tr>
<tr><td>Oligo fwd</td><td>CACCGACCACCTTGTGTCCTTATA</td><td></td><td></td><td></td><td></td><td></td><td></td><td></td><td></td><td></td><td></td></tr>
<tr><td>Oligo rev</td><td>AAACTATAAGGACACAAGGTGGTC</td><td></td><td></td><td></td><td></td><td></td><td></td><td></td><td></td><td></td><td></td></tr>
<tr><td>Chromosome</td><td>start</td><td>end</td><td>strand</td><td>MM</td><td>target_seq</td><td>PAM</td><td>alignment</td><td>distance</td><td>position</td><td>gene name</td><td>gene id</td></tr>
<tr><td>GU305914.1</td><td>95292</td><td>95314</td><td>-</td><td>0</td><td>GACCACCTTGTGTCCTTATA</td><td>AGG</td><td>||||||||[||||||||||||]PAM</td><td>NA</td><td>Intergenic</td><td>NA</td><td>NA</td></tr>
<tr><td>T26750</td><td>AGATTATTCAAGCGATTTCTTGG</td><td>1000</td><td></td><td></td><td></td><td></td><td></td><td></td><td></td><td></td><td></td></tr>
<tr><td>Oligo fwd</td><td>CACCAGATTATTCAAGCGATTTCT</td><td></td><td></td><td></td><td></td><td></td><td></td><td></td><td></td><td></td><td></td></tr>
<tr><td>Oligo rev</td><td>AAACAGAAATCGCTTGAATAATCT</td><td></td><td></td><td></td><td></td><td></td><td></td><td></td><td></td><td></td><td></td></tr>
<tr><td>Chromosome</td><td>start</td><td>end</td><td>strand</td><td>MM</td><td>target_seq</td><td>PAM</td><td>alignment</td><td>distance</td><td>position</td><td>gene name</td><td>gene id</td></tr>
<tr><td>GU305914.1</td><td>95269</td><td>95291</td><td>-</td><td>0</td><td>AGATTATTCAAGCGATTTCT</td><td>TGG</td><td>||||||||[||||||||||||]PAM</td><td>NA</td><td>Intergenic</td><td>NA</td><td>NA</td></tr>
<tr><td>T26751</td><td>ACCATGTGACGCCCCATATACGG</td><td>1000</td><td></td><td></td><td></td><td></td><td></td><td></td><td></td><td></td><td></td></tr>
<tr><td>Oligo fwd</td><td>CACCACCATGTGACGCCCCATATA</td><td></td><td></td><td></td><td></td><td></td><td></td><td></td><td></td><td></td><td></td></tr>
<tr><td>Oligo rev</td><td>AAACTATATGGGGCGTCACATGGT</td><td></td><td></td><td></td><td></td><td></td><td></td><td></td><td></td><td></td><td></td></tr>
<tr><td>Chromosome</td><td>start</td><td>end</td><td>strand</td><td>MM</td><td>target_seq</td><td>PAM</td><td>alignment</td><td>distance</td><td>position</td><td>gene name</td><td>gene id</td></tr>
<tr><td>GU305914.1</td><td>95134</td><td>95156</td><td>-</td><td>0</td><td>ACCATGTGACGCCCCATATA</td><td>CGG</td><td>||||||||[||||||||||||]PAM</td><td>NA</td><td>Intergenic</td><td>NA</td><td>NA</td></tr>
<tr><td>T26752</td><td>GACGCCCCATATACGGTCATAGG</td><td>1000</td><td></td><td></td><td></td><td></td><td></td><td></td><td></td><td></td><td></td></tr>
<tr><td>Oligo fwd</td><td>CACCGACGCCCCATATACGGTCAT</td><td></td><td></td><td></td><td></td><td></td><td></td><td></td><td></td><td></td><td></td></tr>
<tr><td>Oligo rev</td><td>AAACATGACCGTATATGGGGCGTC</td><td></td><td></td><td></td><td></td><td></td><td></td><td></td><td></td><td></td><td></td></tr>
<tr><td>Chromosome</td><td>start</td><td>end</td><td>strand</td><td>MM</td><td>target_seq</td><td>PAM</td><td>alignment</td><td>distance</td><td>position</td><td>gene name</td><td>gene id</td></tr>
<tr><td>GU305914.1</td><td>95127</td><td>95149</td><td>-</td><td>0</td><td>GACGCCCCATATACGGTCAT</td><td>AGG</td><td>||||||||[||||||||||||]PAM</td><td>NA</td><td>Intergenic</td><td>NA</td><td>NA</td></tr>
<tr><td>T26753</td><td>GTCATAGGAACGCCTTTCCGAGG</td><td>1000</td><td></td><td></td><td></td><td></td><td></td><td></td><td></td><td></td><td></td></tr>
<tr><td>Oligo fwd</td><td>CACCGTCATAGGAACGCCTTTCCG</td><td></td><td></td><td></td><td></td><td></td><td></td><td></td><td></td><td></td><td></td></tr>
<tr><td>Oligo rev</td><td>AAACCGGAAAGGCGTTCCTATGAC</td><td></td><td></td><td></td><td></td><td></td><td></td><td></td><td></td><td></td><td></td></tr>
<tr><td>Chromosome</td><td>start</td><td>end</td><td>strand</td><td>MM</td><td>target_seq</td><td>PAM</td><td>alignment</td><td>distance</td><td>position</td><td>gene name</td><td>gene id</td></tr>
<tr><td>GU305914.1</td><td>95112</td><td>95134</td><td>-</td><td>0</td><td>GTCATAGGAACGCCTTTCCG</td><td>AGG</td><td>||||||||[||||||||||||]PAM</td><td>NA</td><td>Intergenic</td><td>NA</td><td>NA</td></tr>
<tr><td>T26754</td><td>CGCCTTTCCGAGGCCGACAGCGG</td><td>1000</td><td></td><td></td><td></td><td></td><td></td><td></td><td></td><td></td><td></td></tr>
<tr><td>Oligo fwd</td><td>CACCCGCCTTTCCGAGGCCGACAG</td><td></td><td></td><td></td><td></td><td></td><td></td><td></td><td></td><td></td><td></td></tr>
<tr><td>Oligo rev</td><td>AAACCTGTCGGCCTCGGAAAGGCG</td><td></td><td></td><td></td><td></td><td></td><td></td><td></td><td></td><td></td><td></td></tr>
<tr><td>Chromosome</td><td>start</td><td>end</td><td>strand</td><td>MM</td><td>target_seq</td><td>PAM</td><td>alignment</td><td>distance</td><td>position</td><td>gene name</td><td>gene id</td></tr>
<tr><td>GU305914.1</td><td>95102</td><td>95124</td><td>-</td><td>0</td><td>CGCCTTTCCGAGGCCGACAG</td><td>CGG</td><td>||||||||[||||||||||||]PAM</td><td>NA</td><td>Intergenic</td><td>NA</td><td>NA</td></tr>
<tr><td>T26755</td><td>CCGACAGCGGCTAAGTCCGATGG</td><td>1000</td><td></td><td></td><td></td><td></td><td></td><td></td><td></td><td></td><td></td></tr>
<tr><td>Oligo fwd</td><td>CACCCCGACAGCGGCTAAGTCCGA</td><td></td><td></td><td></td><td></td><td></td><td></td><td></td><td></td><td></td><td></td></tr>
<tr><td>Oligo rev</td><td>AAACTCGGACTTAGCCGCTGTCGG</td><td></td><td></td><td></td><td></td><td></td><td></td><td></td><td></td><td></td><td></td></tr>
<tr><td>Chromosome</td><td>start</td><td>end</td><td>strand</td><td>MM</td><td>target_seq</td><td>PAM</td><td>alignment</td><td>distance</td><td>position</td><td>gene name</td><td>gene id</td></tr>
<tr><td>GU305914.1</td><td>95089</td><td>95111</td><td>-</td><td>0</td><td>CCGACAGCGGCTAAGTCCGA</td><td>TGG</td><td>||||||||[||||||||||||]PAM</td><td>NA</td><td>Intergenic</td><td>NA</td><td>NA</td></tr>
<tr><td>T26756</td><td>GCTAAGTCCGATGGACACTTAGG</td><td>1000</td><td></td><td></td><td></td><td></td><td></td><td></td><td></td><td></td><td></td></tr>
<tr><td>Oligo fwd</td><td>CACCGCTAAGTCCGATGGACACTT</td><td></td><td></td><td></td><td></td><td></td><td></td><td></td><td></td><td></td><td></td></tr>
<tr><td>Oligo rev</td><td>AAACAAGTGTCCATCGGACTTAGC</td><td></td><td></td><td></td><td></td><td></td><td></td><td></td><td></td><td></td><td></td></tr>
<tr><td>Chromosome</td><td>start</td><td>end</td><td>strand</td><td>MM</td><td>target_seq</td><td>PAM</td><td>alignment</td><td>distance</td><td>position</td><td>gene name</td><td>gene id</td></tr>
<tr><td>GU305914.1</td><td>95080</td><td>95102</td><td>-</td><td>0</td><td>GCTAAGTCCGATGGACACTT</td><td>AGG</td><td>||||||||[||||||||||||]PAM</td><td>NA</td><td>Intergenic</td><td>NA</td><td>NA</td></tr>
<tr><td>T26757</td><td>ATGGACACTTAGGCTCTGTGAGG</td><td>1000</td><td></td><td></td><td></td><td></td><td></td><td></td><td></td><td></td><td></td></tr>
<tr><td>Oligo fwd</td><td>CACCATGGACACTTAGGCTCTGTG</td><td></td><td></td><td></td><td></td><td></td><td></td><td></td><td></td><td></td><td></td></tr>
<tr><td>Oligo rev</td><td>AAACCACAGAGCCTAAGTGTCCAT</td><td></td><td></td><td></td><td></td><td></td><td></td><td></td><td></td><td></td><td></td></tr>
<tr><td>Chromosome</td><td>start</td><td>end</td><td>strand</td><td>MM</td><td>target_seq</td><td>PAM</td><td>alignment</td><td>distance</td><td>position</td><td>gene name</td><td>gene id</td></tr>
<tr><td>GU305914.1</td><td>95070</td><td>95092</td><td>-</td><td>0</td><td>ATGGACACTTAGGCTCTGTG</td><td>AGG</td><td>||||||||[||||||||||||]PAM</td><td>NA</td><td>Intergenic</td><td>NA</td><td>NA</td></tr>
<tr><td>T26758</td><td>TGGACACTTAGGCTCTGTGAGGG</td><td>1000</td><td></td><td></td><td></td><td></td><td></td><td></td><td></td><td></td><td></td></tr>
</table>

<table>
<tr><td>Oligo fwd</td><td>CACCTGGACACTTAGGCTCTGTGA</td><td></td><td></td><td></td><td></td><td></td><td></td><td></td><td></td><td></td><td></td></tr>
<tr><td>Oligo rev</td><td>AAACTCACAGAGCCTAAGTGTCCA</td><td></td><td></td><td></td><td></td><td></td><td></td><td></td><td></td><td></td><td></td></tr>
<tr><td>Chromosome</td><td>start</td><td>end</td><td>strand</td><td>MM</td><td>target_seq</td><td>PAM</td><td>alignment</td><td>distance</td><td>position</td><td>gene name</td><td>gene id</td></tr>
<tr><td>GU305914.1</td><td>95069</td><td>95091</td><td>-</td><td>0</td><td>TGGACACTTAGGCTCTGTGA</td><td>GGG</td><td>||||||||[||||||||||||]PAM</td><td>NA</td><td>Intergenic</td><td>NA</td><td>NA</td></tr>
<tr><td>T26759</td><td>TAGGCTCTGTGAGGGCTCTTCGG</td><td>1000</td><td></td><td></td><td></td><td></td><td></td><td></td><td></td><td></td><td></td></tr>
<tr><td>Oligo fwd</td><td>CACCTAGGCTCTGTGAGGGCTCTT</td><td></td><td></td><td></td><td></td><td></td><td></td><td></td><td></td><td></td><td></td></tr>
<tr><td>Oligo rev</td><td>AAACAAGAGCCCTCACAGAGCCTA</td><td></td><td></td><td></td><td></td><td></td><td></td><td></td><td></td><td></td><td></td></tr>
<tr><td>Chromosome</td><td>start</td><td>end</td><td>strand</td><td>MM</td><td>target_seq</td><td>PAM</td><td>alignment</td><td>distance</td><td>position</td><td>gene name</td><td>gene id</td></tr>
<tr><td>GU305914.1</td><td>95061</td><td>95083</td><td>-</td><td>0</td><td>TAGGCTCTGTGAGGGCTCTT</td><td>CGG</td><td>||||||||[||||||||||||]PAM</td><td>NA</td><td>Intergenic</td><td>NA</td><td>NA</td></tr>
<tr><td>T26760</td><td>AGGCTCTGTGAGGGCTCTTCGGG</td><td>1000</td><td></td><td></td><td></td><td></td><td></td><td></td><td></td><td></td><td></td></tr>
<tr><td>Oligo fwd</td><td>CACCAGGCTCTGTGAGGGCTCTTC</td><td></td><td></td><td></td><td></td><td></td><td></td><td></td><td></td><td></td><td></td></tr>
<tr><td>Oligo rev</td><td>AAACGAAGAGCCCTCACAGAGCCT</td><td></td><td></td><td></td><td></td><td></td><td></td><td></td><td></td><td></td><td></td></tr>
<tr><td>Chromosome</td><td>start</td><td>end</td><td>strand</td><td>MM</td><td>target_seq</td><td>PAM</td><td>alignment</td><td>distance</td><td>position</td><td>gene name</td><td>gene id</td></tr>
<tr><td>GU305914.1</td><td>95060</td><td>95082</td><td>-</td><td>0</td><td>AGGCTCTGTGAGGGCTCTTC</td><td>GGG</td><td>||||||||[||||||||||||]PAM</td><td>NA</td><td>Intergenic</td><td>NA</td><td>NA</td></tr>
<tr><td>T26761</td><td>GATGATTGACAGCCCCTAAGCGG</td><td>1000</td><td></td><td></td><td></td><td></td><td></td><td></td><td></td><td></td><td></td></tr>
<tr><td>Oligo fwd</td><td>CACCGATGATTGACAGCCCCTAAG</td><td></td><td></td><td></td><td></td><td></td><td></td><td></td><td></td><td></td><td></td></tr>
<tr><td>Oligo rev</td><td>AAACCTTAGGGGCTGTCAATCATC</td><td></td><td></td><td></td><td></td><td></td><td></td><td></td><td></td><td></td><td></td></tr>
<tr><td>Chromosome</td><td>start</td><td>end</td><td>strand</td><td>MM</td><td>target_seq</td><td>PAM</td><td>alignment</td><td>distance</td><td>position</td><td>gene name</td><td>gene id</td></tr>
<tr><td>GU305914.1</td><td>95031</td><td>95053</td><td>-</td><td>0</td><td>GATGATTGACAGCCCCTAAG</td><td>CGG</td><td>||||||||[||||||||||||]PAM</td><td>NA</td><td>Intergenic</td><td>NA</td><td>NA</td></tr>
<tr><td>T26762</td><td>ATTGACAGCCCCTAAGCGGAAGG</td><td>1000</td><td></td><td></td><td></td><td></td><td></td><td></td><td></td><td></td><td></td></tr>
<tr><td>Oligo fwd</td><td>CACCATTGACAGCCCCTAAGCGGA</td><td></td><td></td><td></td><td></td><td></td><td></td><td></td><td></td><td></td><td></td></tr>
<tr><td>Oligo rev</td><td>AAACTCCGCTTAGGGGCTGTCAAT</td><td></td><td></td><td></td><td></td><td></td><td></td><td></td><td></td><td></td><td></td></tr>
<tr><td>Chromosome</td><td>start</td><td>end</td><td>strand</td><td>MM</td><td>target_seq</td><td>PAM</td><td>alignment</td><td>distance</td><td>position</td><td>gene name</td><td>gene id</td></tr>
<tr><td>GU305914.1</td><td>95027</td><td>95049</td><td>-</td><td>0</td><td>ATTGACAGCCCCTAAGCGGA</td><td>AGG</td><td>||||||||[||||||||||||]PAM</td><td>NA</td><td>Intergenic</td><td>NA</td><td>NA</td></tr>
<tr><td>T26763</td><td>TTGACAGCCCCTAAGCGGAAGGG</td><td>1000</td><td></td><td></td><td></td><td></td><td></td><td></td><td></td><td></td><td></td></tr>
<tr><td>Oligo fwd</td><td>CACCTTGACAGCCCCTAAGCGGAA</td><td></td><td></td><td></td><td></td><td></td><td></td><td></td><td></td><td></td><td></td></tr>
<tr><td>Oligo rev</td><td>AAACTTCCGCTTAGGGGCTGTCAA</td><td></td><td></td><td></td><td></td><td></td><td></td><td></td><td></td><td></td><td></td></tr>
<tr><td>Chromosome</td><td>start</td><td>end</td><td>strand</td><td>MM</td><td>target_seq</td><td>PAM</td><td>alignment</td><td>distance</td><td>position</td><td>gene name</td><td>gene id</td></tr>
<tr><td>GU305914.1</td><td>95026</td><td>95048</td><td>-</td><td>0</td><td>TTGACAGCCCCTAAGCGGAA</td><td>GGG</td><td>||||||||[||||||||||||]PAM</td><td>NA</td><td>Intergenic</td><td>NA</td><td>NA</td></tr>
<tr><td>T26764</td><td>TGACAGCCCCTAAGCGGAAGGGG</td><td>1000</td><td></td><td></td><td></td><td></td><td></td><td></td><td></td><td></td><td></td></tr>
<tr><td>Oligo fwd</td><td>CACCTGACAGCCCCTAAGCGGAAG</td><td></td><td></td><td></td><td></td><td></td><td></td><td></td><td></td><td></td><td></td></tr>
<tr><td>Oligo rev</td><td>AAACCTTCCGCTTAGGGGCTGTCA</td><td></td><td></td><td></td><td></td><td></td><td></td><td></td><td></td><td></td><td></td></tr>
<tr><td>Chromosome</td><td>start</td><td>end</td><td>strand</td><td>MM</td><td>target_seq</td><td>PAM</td><td>alignment</td><td>distance</td><td>position</td><td>gene name</td><td>gene id</td></tr>
<tr><td>GU305914.1</td><td>95025</td><td>95047</td><td>-</td><td>0</td><td>TGACAGCCCCTAAGCGGAAG</td><td>GGG</td><td>||||||||[||||||||||||]PAM</td><td>NA</td><td>Intergenic</td><td>NA</td><td>NA</td></tr>
<tr><td>T26765</td><td>CAGCCCCTAAGCGGAAGGGGCGG</td><td>1000</td><td></td><td></td><td></td><td></td><td></td><td></td><td></td><td></td><td></td></tr>
<tr><td>Oligo fwd</td><td>CACCCAGCCCCTAAGCGGAAGGGG</td><td></td><td></td><td></td><td></td><td></td><td></td><td></td><td></td><td></td><td></td></tr>
<tr><td>Oligo rev</td><td>AAACCCCCTTCCGCTTAGGGGCTG</td><td></td><td></td><td></td><td></td><td></td><td></td><td></td><td></td><td></td><td></td></tr>
<tr><td>Chromosome</td><td>start</td><td>end</td><td>strand</td><td>MM</td><td>target_seq</td><td>PAM</td><td>alignment</td><td>distance</td><td>position</td><td>gene name</td><td>gene id</td></tr>
<tr><td>GU305914.1</td><td>95022</td><td>95044</td><td>-</td><td>0</td><td>CAGCCCCTAAGCGGAAGGGG</td><td>CGG</td><td>||||||||[||||||||||||]PAM</td><td>NA</td><td>Intergenic</td><td>NA</td><td>NA</td></tr>
<tr><td>T26766</td><td>AGCCCCTAAGCGGAAGGGGCGGG</td><td>1000</td><td></td><td></td><td></td><td></td><td></td><td></td><td></td><td></td><td></td></tr>
<tr><td>Oligo fwd</td><td>CACCAGCCCCTAAGCGGAAGGGGC</td><td></td><td></td><td></td><td></td><td></td><td></td><td></td><td></td><td></td><td></td></tr>
<tr><td>Oligo rev</td><td>AAACGCCCCTTCCGCTTAGGGGCT</td><td></td><td></td><td></td><td></td><td></td><td></td><td></td><td></td><td></td><td></td></tr>
<tr><td>Chromosome</td><td>start</td><td>end</td><td>strand</td><td>MM</td><td>target_seq</td><td>PAM</td><td>alignment</td><td>distance</td><td>position</td><td>gene name</td><td>gene id</td></tr>
<tr><td>GU305914.1</td><td>95021</td><td>95043</td><td>-</td><td>0</td><td>AGCCCCTAAGCGGAAGGGGC</td><td>GGG</td><td>||||||||[||||||||||||]PAM</td><td>NA</td><td>Intergenic</td><td>NA</td><td>NA</td></tr>
<tr><td>T26767</td><td>GGAAGGGGCGGGCATATGACCGG</td><td>1000</td><td></td><td></td><td></td><td></td><td></td><td></td><td></td><td></td><td></td></tr>
<tr><td>Oligo fwd</td><td>CACCGGAAGGGGCGGGCATATGAC</td><td></td><td></td><td></td><td></td><td></td><td></td><td></td><td></td><td></td><td></td></tr>
<tr><td>Oligo rev</td><td>AAACGTCATATGCCCGCCCCTTCC</td><td></td><td></td><td></td><td></td><td></td><td></td><td></td><td></td><td></td><td></td></tr>
<tr><td>Chromosome</td><td>start</td><td>end</td><td>strand</td><td>MM</td><td>target_seq</td><td>PAM</td><td>alignment</td><td>distance</td><td>position</td><td>gene name</td><td>gene id</td></tr>
<tr><td>GU305914.1</td><td>95010</td><td>95032</td><td>-</td><td>0</td><td>GGAAGGGGCGGGCATATGAC</td><td>CGG</td><td>||||||||[||||||||||||]PAM</td><td>NA</td><td>Intergenic</td><td>NA</td><td>NA</td></tr>
<tr><td>T26768</td><td>AACCAAGAAGAAACCGCAAACGG</td><td>1000</td><td></td><td></td><td></td><td></td><td></td><td></td><td></td><td></td><td></td></tr>
<tr><td>Oligo fwd</td><td>CACCAACCAAGAAGAAACCGCAAA</td><td></td><td></td><td></td><td></td><td></td><td></td><td></td><td></td><td></td><td></td></tr>
<tr><td>Oligo rev</td><td>AAACTTTGCGGTTTCTTCTTGGTT</td><td></td><td></td><td></td><td></td><td></td><td></td><td></td><td></td><td></td><td></td></tr>
<tr><td>Chromosome</td><td>start</td><td>end</td><td>strand</td><td>MM</td><td>target_seq</td><td>PAM</td><td>alignment</td><td>distance</td><td>position</td><td>gene name</td><td>gene id</td></tr>
<tr><td>GU305914.1</td><td>94924</td><td>94946</td><td>-</td><td>0</td><td>AACCAAGAAGAAACCGCAAA</td><td>CGG</td><td>||||||||[||||||||||||]PAM</td><td>NA</td><td>Intergenic</td><td>NA</td><td>NA</td></tr>
<tr><td>T26769</td><td>GCCATCTTAGATAAAGCACATGG</td><td>1000</td><td></td><td></td><td></td><td></td><td></td><td></td><td></td><td></td><td></td></tr>
<tr><td>Oligo fwd</td><td>CACCGCCATCTTAGATAAAGCACA</td><td></td><td></td><td></td><td></td><td></td><td></td><td></td><td></td><td></td><td></td></tr>
<tr><td>Oligo rev</td><td>AAACTGTGCTTTATCTAAGATGGC</td><td></td><td></td><td></td><td></td><td></td><td></td><td></td><td></td><td></td><td></td></tr>
<tr><td>Chromosome</td><td>start</td><td>end</td><td>strand</td><td>MM</td><td>target_seq</td><td>PAM</td><td>alignment</td><td>distance</td><td>position</td><td>gene name</td><td>gene id</td></tr>
<tr><td>GU305914.1</td><td>94886</td><td>94908</td><td>-</td><td>0</td><td>GCCATCTTAGATAAAGCACA</td><td>TGG</td><td>||||||||[||||||||||||]PAM</td><td>NA</td><td>Intergenic</td><td>NA</td><td>NA</td></tr>
<tr><td>T26770</td><td>TCTTAGATAAAGCACATGGCAGG</td><td>1000</td><td></td><td></td><td></td><td></td><td></td><td></td><td></td><td></td><td></td></tr>
<tr><td>Oligo fwd</td><td>CACCTCTTAGATAAAGCACATGGC</td><td></td><td></td><td></td><td></td><td></td><td></td><td></td><td></td><td></td><td></td></tr>
<tr><td>Oligo rev</td><td>AAACGCCATGTGCTTTATCTAAGA</td><td></td><td></td><td></td><td></td><td></td><td></td><td></td><td></td><td></td><td></td></tr>
<tr><td>Chromosome</td><td>start</td><td>end</td><td>strand</td><td>MM</td><td>target_seq</td><td>PAM</td><td>alignment</td><td>distance</td><td>position</td><td>gene name</td><td>gene id</td></tr>
<tr><td>GU305914.1</td><td>94882</td><td>94904</td><td>-</td><td>0</td><td>TCTTAGATAAAGCACATGGC</td><td>AGG</td><td>||||||||[||||||||||||]PAM</td><td>NA</td><td>Intergenic</td><td>NA</td><td>NA</td></tr>
<tr><td>T26771</td><td>ATAATTCCAATGTTTGCTTTAGG</td><td>1000</td><td></td><td></td><td></td><td></td><td></td><td></td><td></td><td></td><td></td></tr>
<tr><td>Oligo fwd</td><td>CACCATAATTCCAATGTTTGCTTT</td><td></td><td></td><td></td><td></td><td></td><td></td><td></td><td></td><td></td><td></td></tr>
<tr><td>Oligo rev</td><td>AAACAAAGCAAACATTGGAATTAT</td><td></td><td></td><td></td><td></td><td></td><td></td><td></td><td></td><td></td><td></td></tr>
<tr><td>Chromosome</td><td>start</td><td>end</td><td>strand</td><td>MM</td><td>target_seq</td><td>PAM</td><td>alignment</td><td>distance</td><td>position</td><td>gene name</td><td>gene id</td></tr>
<tr><td>GU305914.1</td><td>94838</td><td>94860</td><td>-</td><td>0</td><td>ATAATTCCAATGTTTGCTTT</td><td>AGG</td><td>||||||||[||||||||||||]PAM</td><td>NA</td><td>Intergenic</td><td>NA</td><td>NA</td></tr>
<tr><td>T26772</td><td>GTTTCAGAAATTCCTATCAACGG</td><td>1000</td><td></td><td></td><td></td><td></td><td></td><td></td><td></td><td></td><td></td></tr>
<tr><td>Oligo fwd</td><td>CACCGTTTCAGAAATTCCTATCAA</td><td></td><td></td><td></td><td></td><td></td><td></td><td></td><td></td><td></td><td></td></tr>
<tr><td>Oligo rev</td><td>AAACTTGATAGGAATTTCTGAAAC</td><td></td><td></td><td></td><td></td><td></td><td></td><td></td><td></td><td></td><td></td></tr>
<tr><td>Chromosome</td><td>start</td><td>end</td><td>strand</td><td>MM</td><td>target_seq</td><td>PAM</td><td>alignment</td><td>distance</td><td>position</td><td>gene name</td><td>gene id</td></tr>
<tr><td>GU305914.1</td><td>94797</td><td>94819</td><td>-</td><td>0</td><td>GTTTCAGAAATTCCTATCAA</td><td>CGG</td><td>||||||||[||||||||||||]PAM</td><td>NA</td><td>Intergenic</td><td>NA</td><td>NA</td></tr>
</table>

<table>
<tr><td>T26773</td><td>TTTCAGAAATTCCTATCAACGGG</td><td>1000</td><td></td><td></td><td></td><td></td><td></td><td></td><td></td><td></td><td></td></tr>
<tr><td>Oligo fwd</td><td>CACCTTTCAGAAATTCCTATCAAC</td><td></td><td></td><td></td><td></td><td></td><td></td><td></td><td></td><td></td><td></td></tr>
<tr><td>Oligo rev</td><td>AAACGTTGATAGGAATTTCTGAAA</td><td></td><td></td><td></td><td></td><td></td><td></td><td></td><td></td><td></td><td></td></tr>
<tr><td>Chromosome</td><td>start</td><td>end</td><td>strand</td><td>MM</td><td>target_seq</td><td>PAM</td><td>alignment</td><td>distance</td><td>position</td><td>gene name</td><td>gene id</td></tr>
<tr><td>GU305914.1</td><td>94796</td><td>94818</td><td>-</td><td>0</td><td>TTTCAGAAATTCCTATCAAC</td><td>GGG</td><td>||||||||[||||||||||||]PAM</td><td>NA</td><td>Intergenic</td><td>NA</td><td>NA</td></tr>
<tr><td>T26774</td><td>TCATTTTTAAGCCTTTTTCGAGG</td><td>1000</td><td></td><td></td><td></td><td></td><td></td><td></td><td></td><td></td><td></td></tr>
<tr><td>Oligo fwd</td><td>CACCTCATTTTTAAGCCTTTTTCG</td><td></td><td></td><td></td><td></td><td></td><td></td><td></td><td></td><td></td><td></td></tr>
<tr><td>Oligo rev</td><td>AAACCGAAAAAGGCTTAAAAATGA</td><td></td><td></td><td></td><td></td><td></td><td></td><td></td><td></td><td></td><td></td></tr>
<tr><td>Chromosome</td><td>start</td><td>end</td><td>strand</td><td>MM</td><td>target_seq</td><td>PAM</td><td>alignment</td><td>distance</td><td>position</td><td>gene name</td><td>gene id</td></tr>
<tr><td>GU305914.1</td><td>94744</td><td>94766</td><td>-</td><td>0</td><td>TCATTTTTAAGCCTTTTTCG</td><td>AGG</td><td>||||||||[||||||||||||]PAM</td><td>NA</td><td>Intergenic</td><td>NA</td><td>NA</td></tr>
<tr><td>T26775</td><td>TTTAAGCCTTTTTCGAGGTCTGG</td><td>1000</td><td></td><td></td><td></td><td></td><td></td><td></td><td></td><td></td><td></td></tr>
<tr><td>Oligo fwd</td><td>CACCTTTAAGCCTTTTTCGAGGTC</td><td></td><td></td><td></td><td></td><td></td><td></td><td></td><td></td><td></td><td></td></tr>
<tr><td>Oligo rev</td><td>AAACGACCTCGAAAAAGGCTTAAA</td><td></td><td></td><td></td><td></td><td></td><td></td><td></td><td></td><td></td><td></td></tr>
<tr><td>Chromosome</td><td>start</td><td>end</td><td>strand</td><td>MM</td><td>target_seq</td><td>PAM</td><td>alignment</td><td>distance</td><td>position</td><td>gene name</td><td>gene id</td></tr>
<tr><td>GU305914.1</td><td>94739</td><td>94761</td><td>-</td><td>0</td><td>TTTAAGCCTTTTTCGAGGTC</td><td>TGG</td><td>||||||||[||||||||||||]PAM</td><td>NA</td><td>Intergenic</td><td>NA</td><td>NA</td></tr>
<tr><td>GU305914.1</td><td>26949</td><td>26971</td><td>-</td><td>4</td><td>CTTGAGCCTTCTTCGGGGTC</td><td>CGG</td><td>-||-||||[||-||||-||||]PAM</td><td>NA</td><td>Intergenic</td><td>NA</td><td>NA</td></tr>
<tr><td>T26776</td><td>TAAAAAAAGTTGTTTGTTCGAGG</td><td>1000</td><td></td><td></td><td></td><td></td><td></td><td></td><td></td><td></td><td></td></tr>
<tr><td>Oligo fwd</td><td>CACCTAAAAAAAGTTGTTTGTTCG</td><td></td><td></td><td></td><td></td><td></td><td></td><td></td><td></td><td></td><td></td></tr>
<tr><td>Oligo rev</td><td>AAACCGAACAAACAACTTTTTTTA</td><td></td><td></td><td></td><td></td><td></td><td></td><td></td><td></td><td></td><td></td></tr>
<tr><td>Chromosome</td><td>start</td><td>end</td><td>strand</td><td>MM</td><td>target_seq</td><td>PAM</td><td>alignment</td><td>distance</td><td>position</td><td>gene name</td><td>gene id</td></tr>
<tr><td>GU305914.1</td><td>94711</td><td>94733</td><td>-</td><td>0</td><td>TAAAAAAAGTTGTTTGTTCG</td><td>AGG</td><td>||||||||[||||||||||||]PAM</td><td>NA</td><td>Intergenic</td><td>NA</td><td>NA</td></tr>
<tr><td>T26777</td><td>AAAAAAAGTTGTTTGTTCGAGGG</td><td>1000</td><td></td><td></td><td></td><td></td><td></td><td></td><td></td><td></td><td></td></tr>
<tr><td>Oligo fwd</td><td>CACCAAAAAAAGTTGTTTGTTCGA</td><td></td><td></td><td></td><td></td><td></td><td></td><td></td><td></td><td></td><td></td></tr>
<tr><td>Oligo rev</td><td>AAACTCGAACAAACAACTTTTTTT</td><td></td><td></td><td></td><td></td><td></td><td></td><td></td><td></td><td></td><td></td></tr>
<tr><td>Chromosome</td><td>start</td><td>end</td><td>strand</td><td>MM</td><td>target_seq</td><td>PAM</td><td>alignment</td><td>distance</td><td>position</td><td>gene name</td><td>gene id</td></tr>
<tr><td>GU305914.1</td><td>94710</td><td>94732</td><td>-</td><td>0</td><td>AAAAAAAGTTGTTTGTTCGA</td><td>GGG</td><td>||||||||[||||||||||||]PAM</td><td>NA</td><td>Intergenic</td><td>NA</td><td>NA</td></tr>
<tr><td>T26778</td><td>TTTGTTCGAGGGAGACCGAACGG</td><td>1000</td><td></td><td></td><td></td><td></td><td></td><td></td><td></td><td></td><td></td></tr>
<tr><td>Oligo fwd</td><td>CACCTTTGTTCGAGGGAGACCGAA</td><td></td><td></td><td></td><td></td><td></td><td></td><td></td><td></td><td></td><td></td></tr>
<tr><td>Oligo rev</td><td>AAACTTCGGTCTCCCTCGAACAAA</td><td></td><td></td><td></td><td></td><td></td><td></td><td></td><td></td><td></td><td></td></tr>
<tr><td>Chromosome</td><td>start</td><td>end</td><td>strand</td><td>MM</td><td>target_seq</td><td>PAM</td><td>alignment</td><td>distance</td><td>position</td><td>gene name</td><td>gene id</td></tr>
<tr><td>GU305914.1</td><td>94699</td><td>94721</td><td>-</td><td>0</td><td>TTTGTTCGAGGGAGACCGAA</td><td>CGG</td><td>||||||||[||||||||||||]PAM</td><td>NA</td><td>Intergenic</td><td>NA</td><td>NA</td></tr>
<tr><td>T26779</td><td>CCTGATCTTTTCAAAGCCACAGG</td><td>1000</td><td></td><td></td><td></td><td></td><td></td><td></td><td></td><td></td><td></td></tr>
<tr><td>Oligo fwd</td><td>CACCCCTGATCTTTTCAAAGCCAC</td><td></td><td></td><td></td><td></td><td></td><td></td><td></td><td></td><td></td><td></td></tr>
<tr><td>Oligo rev</td><td>AAACGTGGCTTTGAAAAGATCAGG</td><td></td><td></td><td></td><td></td><td></td><td></td><td></td><td></td><td></td><td></td></tr>
<tr><td>Chromosome</td><td>start</td><td>end</td><td>strand</td><td>MM</td><td>target_seq</td><td>PAM</td><td>alignment</td><td>distance</td><td>position</td><td>gene name</td><td>gene id</td></tr>
<tr><td>GU305914.1</td><td>94667</td><td>94689</td><td>-</td><td>0</td><td>CCTGATCTTTTCAAAGCCAC</td><td>AGG</td><td>||||||||[||||||||||||]PAM</td><td>NA</td><td>Intergenic</td><td>NA</td><td>NA</td></tr>
<tr><td>T26780</td><td>CTGATCTTTTCAAAGCCACAGGG</td><td>1000</td><td></td><td></td><td></td><td></td><td></td><td></td><td></td><td></td><td></td></tr>
<tr><td>Oligo fwd</td><td>CACCCTGATCTTTTCAAAGCCACA</td><td></td><td></td><td></td><td></td><td></td><td></td><td></td><td></td><td></td><td></td></tr>
<tr><td>Oligo rev</td><td>AAACTGTGGCTTTGAAAAGATCAG</td><td></td><td></td><td></td><td></td><td></td><td></td><td></td><td></td><td></td><td></td></tr>
<tr><td>Chromosome</td><td>start</td><td>end</td><td>strand</td><td>MM</td><td>target_seq</td><td>PAM</td><td>alignment</td><td>distance</td><td>position</td><td>gene name</td><td>gene id</td></tr>
<tr><td>GU305914.1</td><td>94666</td><td>94688</td><td>-</td><td>0</td><td>CTGATCTTTTCAAAGCCACA</td><td>GGG</td><td>||||||||[||||||||||||]PAM</td><td>NA</td><td>Intergenic</td><td>NA</td><td>NA</td></tr>
<tr><td>T26781</td><td>ATGTCATCTTTTTCAACTGACGG</td><td>1000</td><td></td><td></td><td></td><td></td><td></td><td></td><td></td><td></td><td></td></tr>
<tr><td>Oligo fwd</td><td>CACCATGTCATCTTTTTCAACTGA</td><td></td><td></td><td></td><td></td><td></td><td></td><td></td><td></td><td></td><td></td></tr>
<tr><td>Oligo rev</td><td>AAACTCAGTTGAAAAAGATGACAT</td><td></td><td></td><td></td><td></td><td></td><td></td><td></td><td></td><td></td><td></td></tr>
<tr><td>Chromosome</td><td>start</td><td>end</td><td>strand</td><td>MM</td><td>target_seq</td><td>PAM</td><td>alignment</td><td>distance</td><td>position</td><td>gene name</td><td>gene id</td></tr>
<tr><td>GU305914.1</td><td>94638</td><td>94660</td><td>-</td><td>0</td><td>ATGTCATCTTTTTCAACTGA</td><td>CGG</td><td>||||||||[||||||||||||]PAM</td><td>NA</td><td>Intergenic</td><td>NA</td><td>NA</td></tr>
<tr><td>T26782</td><td>TCTTTTTCAACTGACGGTCGAGG</td><td>1000</td><td></td><td></td><td></td><td></td><td></td><td></td><td></td><td></td><td></td></tr>
<tr><td>Oligo fwd</td><td>CACCTCTTTTTCAACTGACGGTCG</td><td></td><td></td><td></td><td></td><td></td><td></td><td></td><td></td><td></td><td></td></tr>
<tr><td>Oligo rev</td><td>AAACCGACCGTCAGTTGAAAAAGA</td><td></td><td></td><td></td><td></td><td></td><td></td><td></td><td></td><td></td><td></td></tr>
<tr><td>Chromosome</td><td>start</td><td>end</td><td>strand</td><td>MM</td><td>target_seq</td><td>PAM</td><td>alignment</td><td>distance</td><td>position</td><td>gene name</td><td>gene id</td></tr>
<tr><td>GU305914.1</td><td>94632</td><td>94654</td><td>-</td><td>0</td><td>TCTTTTTCAACTGACGGTCG</td><td>AGG</td><td>||||||||[||||||||||||]PAM</td><td>NA</td><td>Intergenic</td><td>NA</td><td>NA</td></tr>
<tr><td>T26783</td><td>ATTTTGCACTACCGCTCAAAAGG</td><td>1000</td><td></td><td></td><td></td><td></td><td></td><td></td><td></td><td></td><td></td></tr>
<tr><td>Oligo fwd</td><td>CACCATTTTGCACTACCGCTCAAA</td><td></td><td></td><td></td><td></td><td></td><td></td><td></td><td></td><td></td><td></td></tr>
<tr><td>Oligo rev</td><td>AAACTTTGAGCGGTAGTGCAAAAT</td><td></td><td></td><td></td><td></td><td></td><td></td><td></td><td></td><td></td><td></td></tr>
<tr><td>Chromosome</td><td>start</td><td>end</td><td>strand</td><td>MM</td><td>target_seq</td><td>PAM</td><td>alignment</td><td>distance</td><td>position</td><td>gene name</td><td>gene id</td></tr>
<tr><td>GU305914.1</td><td>94600</td><td>94622</td><td>-</td><td>0</td><td>ATTTTGCACTACCGCTCAAA</td><td>AGG</td><td>||||||||[||||||||||||]PAM</td><td>NA</td><td>Intergenic</td><td>NA</td><td>NA</td></tr>
<tr><td>T26784</td><td>GCTCAAAAGGAGCAGTGCCTCGG</td><td>1000</td><td></td><td></td><td></td><td></td><td></td><td></td><td></td><td></td><td></td></tr>
<tr><td>Oligo fwd</td><td>CACCGCTCAAAAGGAGCAGTGCCT</td><td></td><td></td><td></td><td></td><td></td><td></td><td></td><td></td><td></td><td></td></tr>
<tr><td>Oligo rev</td><td>AAACAGGCACTGCTCCTTTTGAGC</td><td></td><td></td><td></td><td></td><td></td><td></td><td></td><td></td><td></td><td></td></tr>
<tr><td>Chromosome</td><td>start</td><td>end</td><td>strand</td><td>MM</td><td>target_seq</td><td>PAM</td><td>alignment</td><td>distance</td><td>position</td><td>gene name</td><td>gene id</td></tr>
<tr><td>GU305914.1</td><td>94587</td><td>94609</td><td>-</td><td>0</td><td>GCTCAAAAGGAGCAGTGCCT</td><td>CGG</td><td>||||||||[||||||||||||]PAM</td><td>NA</td><td>Intergenic</td><td>NA</td><td>NA</td></tr>
<tr><td>T26785</td><td>ACAGCGACCTAGACCCTCTGCGG</td><td>1000</td><td></td><td></td><td></td><td></td><td></td><td></td><td></td><td></td><td></td></tr>
<tr><td>Oligo fwd</td><td>CACCACAGCGACCTAGACCCTCTG</td><td></td><td></td><td></td><td></td><td></td><td></td><td></td><td></td><td></td><td></td></tr>
<tr><td>Oligo rev</td><td>AAACCAGAGGGTCTAGGTCGCTGT</td><td></td><td></td><td></td><td></td><td></td><td></td><td></td><td></td><td></td><td></td></tr>
<tr><td>Chromosome</td><td>start</td><td>end</td><td>strand</td><td>MM</td><td>target_seq</td><td>PAM</td><td>alignment</td><td>distance</td><td>position</td><td>gene name</td><td>gene id</td></tr>
<tr><td>GU305914.1</td><td>94561</td><td>94583</td><td>-</td><td>0</td><td>ACAGCGACCTAGACCCTCTG</td><td>CGG</td><td>||||||||[||||||||||||]PAM</td><td>NA</td><td>Intergenic</td><td>NA</td><td>NA</td></tr>
<tr><td>T26786</td><td>TGCGGACCCCCCTTTGAAAGAGG</td><td>1000</td><td></td><td></td><td></td><td></td><td></td><td></td><td></td><td></td><td></td></tr>
<tr><td>Oligo fwd</td><td>CACCTGCGGACCCCCCTTTGAAAG</td><td></td><td></td><td></td><td></td><td></td><td></td><td></td><td></td><td></td><td></td></tr>
<tr><td>Oligo rev</td><td>AAACCTTTCAAAGGGGGGTCCGCA</td><td></td><td></td><td></td><td></td><td></td><td></td><td></td><td></td><td></td><td></td></tr>
<tr><td>Chromosome</td><td>start</td><td>end</td><td>strand</td><td>MM</td><td>target_seq</td><td>PAM</td><td>alignment</td><td>distance</td><td>position</td><td>gene name</td><td>gene id</td></tr>
<tr><td>GU305914.1</td><td>94543</td><td>94565</td><td>-</td><td>0</td><td>TGCGGACCCCCCTTTGAAAG</td><td>AGG</td><td>||||||||[||||||||||||]PAM</td><td>NA</td><td>Intergenic</td><td>NA</td><td>NA</td></tr>
<tr><td>T26787</td><td>AAAGAGGAAAGTGAGATTTATGG</td><td>1000</td><td></td><td></td><td></td><td></td><td></td><td></td><td></td><td></td><td></td></tr>
<tr><td>Oligo fwd</td><td>CACCAAAGAGGAAAGTGAGATTTA</td><td></td><td></td><td></td><td></td><td></td><td></td><td></td><td></td><td></td><td></td></tr>
<tr><td>Oligo rev</td><td>AAACTAAATCTCACTTTCCTCTTT</td><td></td><td></td><td></td><td></td><td></td><td></td><td></td><td></td><td></td><td></td></tr>
<tr><td>Chromosome</td><td>start</td><td>end</td><td>strand</td><td>MM</td><td>target_seq</td><td>PAM</td><td>alignment</td><td>distance</td><td>position</td><td>gene name</td><td>gene id</td></tr>
</table>

GU305914.1 94527 94549 - 0 AAAGAGGAAAGTGAGATTTA TGG ||||||||[||||||||||||]PAM NA Intergenic NA NA

T26788 AGTGAGATTTATGGAGATCTCGG 1000
Oligo fwd CACCAGTGAGATTTATGGAGATCT
Oligo rev AAACAGATCTCCATAAATCTCACT
Chromosome start end strand MM target_seq PAM alignment distance position gene name gene id
GU305914.1 94518 94540 - 0 AGTGAGATTTATGGAGATCT CGG ||||||||[||||||||||||]PAM NA Intergenic NA NA

T26789 CGTGCTTAGTGAACGTGACGAGG 1000
Oligo fwd CACCCGTGCTTAGTGAACGTGACG
Oligo rev AAACCGTCACGTTCACTAAGCACG
Chromosome start end strand MM target_seq PAM alignment distance position gene name gene id
GU305914.1 94485 94507 - 0 CGTGCTTAGTGAACGTGACG AGG ||||||||[||||||||||||]PAM NA Intergenic NA NA

T26790 GTGCTTAGTGAACGTGACGAGGG 1000
Oligo fwd CACCGTGCTTAGTGAACGTGACGA
Oligo rev AAACTCGTCACGTTCACTAAGCAC
Chromosome start end strand MM target_seq PAM alignment distance position gene name gene id
GU305914.1 94484 94506 - 0 GTGCTTAGTGAACGTGACGA GGG ||||||||[||||||||||||]PAM NA Intergenic NA NA

T26791 GTGACGAGGGTTTGTGTGTCAGG 1000
Oligo fwd CACCGTGACGAGGGTTTGTGTGTC
Oligo rev AAACGACACACAAACCCTCGTCAC
Chromosome start end strand MM target_seq PAM alignment distance position gene name gene id
GU305914.1 94471 94493 - 0 GTGACGAGGGTTTGTGTGTC AGG ||||||||[||||||||||||]PAM NA Intergenic NA NA

T26792 TGACGAGGGTTTGTGTGTCAGGG 1000
Oligo fwd CACCTGACGAGGGTTTGTGTGTCA
Oligo rev AAACTGACACACAAACCCTCGTCA
Chromosome start end strand MM target_seq PAM alignment distance position gene name gene id
GU305914.1 94470 94492 - 0 TGACGAGGGTTTGTGTGTCA GGG ||||||||[||||||||||||]PAM NA Intergenic NA NA

T26793 GCGCTTGTGCGAACGTTAAATGG 1000
Oligo fwd CACCGCGCTTGTGCGAACGTTAAA
Oligo rev AAACTTTAACGTTCGCACAAGCGC
Chromosome start end strand MM target_seq PAM alignment distance position gene name gene id
GU305914.1 94443 94465 - 0 GCGCTTGTGCGAACGTTAAA TGG ||||||||[||||||||||||]PAM NA Intergenic NA NA

T26794 CTTGTGCGAACGTTAAATGGCGG 1000
Oligo fwd CACCCTTGTGCGAACGTTAAATGG
Oligo rev AAACCCATTTAACGTTCGCACAAG
Chromosome start end strand MM target_seq PAM alignment distance position gene name gene id
GU305914.1 94440 94462 - 0 CTTGTGCGAACGTTAAATGG CGG ||||||||[||||||||||||]PAM NA Intergenic NA NA

T26795 GTGCGAACGTTAAATGGCGGAGG 1000
Oligo fwd CACCGTGCGAACGTTAAATGGCGG
Oligo rev AAACCCGCCATTTAACGTTCGCAC
Chromosome start end strand MM target_seq PAM alignment distance position gene name gene id
GU305914.1 94437 94459 - 0 GTGCGAACGTTAAATGGCGG AGG ||||||||[||||||||||||]PAM NA Intergenic NA NA

T26796 AATGGCGGAGGCCACCAAAATGG 1000
Oligo fwd CACCAATGGCGGAGGCCACCAAAA
Oligo rev AAACTTTTGGTGGCCTCCGCCATT
Chromosome start end strand MM target_seq PAM alignment distance position gene name gene id
GU305914.1 94425 94447 - 0 AATGGCGGAGGCCACCAAAA TGG ||||||||[||||||||||||]PAM NA Intergenic NA NA

T26797 GGAGGCCACCAAAATGGCGTCGG 1000
Oligo fwd CACCGGAGGCCACCAAAATGGCGT
Oligo rev AAACACGCCATTTTGGTGGCCTCC
Chromosome start end strand MM target_seq PAM alignment distance position gene name gene id
GU305914.1 94419 94441 - 0 GGAGGCCACCAAAATGGCGT CGG ||||||||[||||||||||||]PAM NA Intergenic NA NA

T26798 GAGGCCACCAAAATGGCGTCGGG 1000
Oligo fwd CACCGAGGCCACCAAAATGGCGTC
Oligo rev AAACGACGCCATTTTGGTGGCCTC
Chromosome start end strand MM target_seq PAM alignment distance position gene name gene id
GU305914.1 94418 94440 - 0 GAGGCCACCAAAATGGCGTC GGG ||||||||[||||||||||||]PAM NA Intergenic NA NA

T26799 AAAATGGCGTCGGGACCAAATGG 1000
Oligo fwd CACCAAAATGGCGTCGGGACCAAA
Oligo rev AAACTTTGGTCCCGACGCCATTTT
Chromosome start end strand MM target_seq PAM alignment distance position gene name gene id
GU305914.1 94409 94431 - 0 AAAATGGCGTCGGGACCAAA TGG ||||||||[||||||||||||]PAM NA Intergenic NA NA

T26800 TAAACGAAAACTCGACAAAATGG 1000
Oligo fwd CACCTAAACGAAAACTCGACAAAA
Oligo rev AAACTTTTGTCGAGTTTTCGTTTA
Chromosome start end strand MM target_seq PAM alignment distance position gene name gene id
GU305914.1 94370 94392 - 0 TAAACGAAAACTCGACAAAA TGG ||||||||[||||||||||||]PAM NA Intergenic NA NA

T26801 CTCGACAAAATGGCCGCCAACGG 1000
Oligo fwd CACCCTCGACAAAATGGCCGCCAA
Oligo rev AAACTTGGCGGCCATTTTGTCGAG
Chromosome start end strand MM target_seq PAM alignment distance position gene name gene id
GU305914.1 94360 94382 - 0 CTCGACAAAATGGCCGCCAA CGG ||||||||[||||||||||||]PAM NA Intergenic NA NA

T26802 GGCAAGCTAAAACACAAAAAAGG 1000
Oligo fwd CACCGGCAAGCTAAAACACAAAAA
Oligo rev AAACTTTTTGTGTTTTAGCTTGCC

<table>
<tr><td>Chromosome</td><td>start</td><td></td><td>end</td><td>strand</td><td>MM</td><td>target_seq</td><td>PAM</td><td>alignment</td><td>distance</td><td>position</td><td>gene name</td><td>gene id</td></tr>
<tr><td>GU305914.1</td><td></td><td>94339</td><td>94361</td><td>-</td><td>0</td><td>GGCAAGCTAAAACACAAAAA</td><td>AGG</td><td>||||||||[||||||||||||]PAM</td><td>NA</td><td>Intergenic</td><td>NA</td><td>NA</td></tr>
<tr><td>T26803</td><td>GCAAGCTAAAACACAAAAAAGGG</td><td></td><td>1000</td><td></td><td></td><td></td><td></td><td></td><td></td><td></td><td></td><td></td></tr>
<tr><td>Oligo fwd</td><td>CACCGCAAGCTAAAACACAAAAAA</td><td></td><td></td><td></td><td></td><td></td><td></td><td></td><td></td><td></td><td></td><td></td></tr>
<tr><td>Oligo rev</td><td>AAACTTTTTTGTGTTTTAGCTTGC</td><td></td><td></td><td></td><td></td><td></td><td></td><td></td><td></td><td></td><td></td><td></td></tr>
<tr><td>Chromosome</td><td>start</td><td></td><td>end</td><td>strand</td><td>MM</td><td>target_seq</td><td>PAM</td><td>alignment</td><td>distance</td><td>position</td><td>gene name</td><td>gene id</td></tr>
<tr><td>GU305914.1</td><td></td><td>94338</td><td>94360</td><td>-</td><td>0</td><td>GCAAGCTAAAACACAAAAAA</td><td>GGG</td><td>||||||||[||||||||||||]PAM</td><td>NA</td><td>Intergenic</td><td>NA</td><td>NA</td></tr>
<tr><td>T26804</td><td>CAAGCTAAAACACAAAAAAGGGG</td><td></td><td>1000</td><td></td><td></td><td></td><td></td><td></td><td></td><td></td><td></td><td></td></tr>
<tr><td>Oligo fwd</td><td>CACCCAAGCTAAAACACAAAAAAG</td><td></td><td></td><td></td><td></td><td></td><td></td><td></td><td></td><td></td><td></td><td></td></tr>
<tr><td>Oligo rev</td><td>AAACCTTTTTTGTGTTTTAGCTTG</td><td></td><td></td><td></td><td></td><td></td><td></td><td></td><td></td><td></td><td></td><td></td></tr>
<tr><td>Chromosome</td><td>start</td><td></td><td>end</td><td>strand</td><td>MM</td><td>target_seq</td><td>PAM</td><td>alignment</td><td>distance</td><td>position</td><td>gene name</td><td>gene id</td></tr>
<tr><td>GU305914.1</td><td></td><td>94337</td><td>94359</td><td>-</td><td>0</td><td>CAAGCTAAAACACAAAAAAG</td><td>GGG</td><td>||||||||[||||||||||||]PAM</td><td>NA</td><td>Intergenic</td><td>NA</td><td>NA</td></tr>
<tr><td>T26805</td><td>AAAAAAGGGGCATTTTGCAGCGG</td><td></td><td>1000</td><td></td><td></td><td></td><td></td><td></td><td></td><td></td><td></td><td></td></tr>
<tr><td>Oligo fwd</td><td>CACCAAAAAAGGGGCATTTTGCAG</td><td></td><td></td><td></td><td></td><td></td><td></td><td></td><td></td><td></td><td></td><td></td></tr>
<tr><td>Oligo rev</td><td>AAACCTGCAAAATGCCCCTTTTTT</td><td></td><td></td><td></td><td></td><td></td><td></td><td></td><td></td><td></td><td></td><td></td></tr>
<tr><td>Chromosome</td><td>start</td><td></td><td>end</td><td>strand</td><td>MM</td><td>target_seq</td><td>PAM</td><td>alignment</td><td>distance</td><td>position</td><td>gene name</td><td>gene id</td></tr>
<tr><td>GU305914.1</td><td></td><td>94324</td><td>94346</td><td>-</td><td>0</td><td>AAAAAAGGGGCATTTTGCAG</td><td>CGG</td><td>||||||||[||||||||||||]PAM</td><td>NA</td><td>Intergenic</td><td>NA</td><td>NA</td></tr>
<tr><td>T26806</td><td>AAGGGGCATTTTGCAGCGGCTGG</td><td></td><td>1000</td><td></td><td></td><td></td><td></td><td></td><td></td><td></td><td></td><td></td></tr>
<tr><td>Oligo fwd</td><td>CACCAAGGGGCATTTTGCAGCGGC</td><td></td><td></td><td></td><td></td><td></td><td></td><td></td><td></td><td></td><td></td><td></td></tr>
<tr><td>Oligo rev</td><td>AAACGCCGCTGCAAAATGCCCCTT</td><td></td><td></td><td></td><td></td><td></td><td></td><td></td><td></td><td></td><td></td><td></td></tr>
<tr><td>Chromosome</td><td>start</td><td></td><td>end</td><td>strand</td><td>MM</td><td>target_seq</td><td>PAM</td><td>alignment</td><td>distance</td><td>position</td><td>gene name</td><td>gene id</td></tr>
<tr><td>GU305914.1</td><td></td><td>94320</td><td>94342</td><td>-</td><td>0</td><td>AAGGGGCATTTTGCAGCGGC</td><td>TGG</td><td>||||||||[||||||||||||]PAM</td><td>NA</td><td>Intergenic</td><td>NA</td><td>NA</td></tr>
<tr><td>T26807</td><td>AGGGGCATTTTGCAGCGGCTGGG</td><td></td><td>1000</td><td></td><td></td><td></td><td></td><td></td><td></td><td></td><td></td><td></td></tr>
<tr><td>Oligo fwd</td><td>CACCAGGGGCATTTTGCAGCGGCT</td><td></td><td></td><td></td><td></td><td></td><td></td><td></td><td></td><td></td><td></td><td></td></tr>
<tr><td>Oligo rev</td><td>AAACAGCCGCTGCAAAATGCCCCT</td><td></td><td></td><td></td><td></td><td></td><td></td><td></td><td></td><td></td><td></td><td></td></tr>
<tr><td>Chromosome</td><td>start</td><td></td><td>end</td><td>strand</td><td>MM</td><td>target_seq</td><td>PAM</td><td>alignment</td><td>distance</td><td>position</td><td>gene name</td><td>gene id</td></tr>
<tr><td>GU305914.1</td><td></td><td>94319</td><td>94341</td><td>-</td><td>0</td><td>AGGGGCATTTTGCAGCGGCT</td><td>GGG</td><td>||||||||[||||||||||||]PAM</td><td>NA</td><td>Intergenic</td><td>NA</td><td>NA</td></tr>
<tr><td>T26808</td><td>ATTTTGCAGCGGCTGGGACAAGG</td><td></td><td>1000</td><td></td><td></td><td></td><td></td><td></td><td></td><td></td><td></td><td></td></tr>
<tr><td>Oligo fwd</td><td>CACCATTTTGCAGCGGCTGGGACA</td><td></td><td></td><td></td><td></td><td></td><td></td><td></td><td></td><td></td><td></td><td></td></tr>
<tr><td>Oligo rev</td><td>AAACTGTCCCAGCCGCTGCAAAAT</td><td></td><td></td><td></td><td></td><td></td><td></td><td></td><td></td><td></td><td></td><td></td></tr>
<tr><td>Chromosome</td><td>start</td><td></td><td>end</td><td>strand</td><td>MM</td><td>target_seq</td><td>PAM</td><td>alignment</td><td>distance</td><td>position</td><td>gene name</td><td>gene id</td></tr>
<tr><td>GU305914.1</td><td></td><td>94313</td><td>94335</td><td>-</td><td>0</td><td>ATTTTGCAGCGGCTGGGACA</td><td>AGG</td><td>||||||||[||||||||||||]PAM</td><td>NA</td><td>Intergenic</td><td>NA</td><td>NA</td></tr>
<tr><td>T26809</td><td>AAAACGCAGCGCCGAGCGAAAGG</td><td></td><td>1000</td><td></td><td></td><td></td><td></td><td></td><td></td><td></td><td></td><td></td></tr>
<tr><td>Oligo fwd</td><td>CACCAAAACGCAGCGCCGAGCGAA</td><td></td><td></td><td></td><td></td><td></td><td></td><td></td><td></td><td></td><td></td><td></td></tr>
<tr><td>Oligo rev</td><td>AAACTTCGCTCGGCGCTGCGTTTT</td><td></td><td></td><td></td><td></td><td></td><td></td><td></td><td></td><td></td><td></td><td></td></tr>
<tr><td>Chromosome</td><td>start</td><td></td><td>end</td><td>strand</td><td>MM</td><td>target_seq</td><td>PAM</td><td>alignment</td><td>distance</td><td>position</td><td>gene name</td><td>gene id</td></tr>
<tr><td>GU305914.1</td><td></td><td>94281</td><td>94303</td><td>-</td><td>0</td><td>AAAACGCAGCGCCGAGCGAA</td><td>AGG</td><td>||||||||[||||||||||||]PAM</td><td>NA</td><td>Intergenic</td><td>NA</td><td>NA</td></tr>
<tr><td>GU305914.1</td><td></td><td>36286</td><td>36308</td><td>-</td><td>4</td><td>AAAGCGCCGCTCCGAGCGGA</td><td>AGG</td><td>|||-|||-[||-|||||||-|]PAM</td><td>NA</td><td>Intergenic</td><td>NA</td><td>NA</td></tr>
<tr><td>T26810</td><td>AGCGCCGAGCGAAAGGCGAGAGG</td><td></td><td>1000</td><td></td><td></td><td></td><td></td><td></td><td></td><td></td><td></td><td></td></tr>
<tr><td>Oligo fwd</td><td>CACCAGCGCCGAGCGAAAGGCGAG</td><td></td><td></td><td></td><td></td><td></td><td></td><td></td><td></td><td></td><td></td><td></td></tr>
<tr><td>Oligo rev</td><td>AAACCTCGCCTTTCGCTCGGCGCT</td><td></td><td></td><td></td><td></td><td></td><td></td><td></td><td></td><td></td><td></td><td></td></tr>
<tr><td>Chromosome</td><td>start</td><td></td><td>end</td><td>strand</td><td>MM</td><td>target_seq</td><td>PAM</td><td>alignment</td><td>distance</td><td>position</td><td>gene name</td><td>gene id</td></tr>
<tr><td>GU305914.1</td><td></td><td>94274</td><td>94296</td><td>-</td><td>0</td><td>AGCGCCGAGCGAAAGGCGAG</td><td>AGG</td><td>||||||||[||||||||||||]PAM</td><td>NA</td><td>Intergenic</td><td>NA</td><td>NA</td></tr>
<tr><td>T26811</td><td>GGCGAGAGGCAGAGCAAAAGCGG</td><td></td><td>1000</td><td></td><td></td><td></td><td></td><td></td><td></td><td></td><td></td><td></td></tr>
<tr><td>Oligo fwd</td><td>CACCGGCGAGAGGCAGAGCAAAAG</td><td></td><td></td><td></td><td></td><td></td><td></td><td></td><td></td><td></td><td></td><td></td></tr>
<tr><td>Oligo rev</td><td>AAACCTTTTGCTCTGCCTCTCGCC</td><td></td><td></td><td></td><td></td><td></td><td></td><td></td><td></td><td></td><td></td><td></td></tr>
<tr><td>Chromosome</td><td>start</td><td></td><td>end</td><td>strand</td><td>MM</td><td>target_seq</td><td>PAM</td><td>alignment</td><td>distance</td><td>position</td><td>gene name</td><td>gene id</td></tr>
<tr><td>GU305914.1</td><td></td><td>94260</td><td>94282</td><td>-</td><td>0</td><td>GGCGAGAGGCAGAGCAAAAG</td><td>CGG</td><td>||||||||[||||||||||||]PAM</td><td>NA</td><td>Intergenic</td><td>NA</td><td>NA</td></tr>
<tr><td>T26812</td><td>AGCAAAAGCGGAGCGTCGAGCGG</td><td></td><td>1000</td><td></td><td></td><td></td><td></td><td></td><td></td><td></td><td></td><td></td></tr>
<tr><td>Oligo fwd</td><td>CACCAGCAAAAGCGGAGCGTCGAG</td><td></td><td></td><td></td><td></td><td></td><td></td><td></td><td></td><td></td><td></td><td></td></tr>
<tr><td>Oligo rev</td><td>AAACCTCGACGCTCCGCTTTTGCT</td><td></td><td></td><td></td><td></td><td></td><td></td><td></td><td></td><td></td><td></td><td></td></tr>
<tr><td>Chromosome</td><td>start</td><td></td><td>end</td><td>strand</td><td>MM</td><td>target_seq</td><td>PAM</td><td>alignment</td><td>distance</td><td>position</td><td>gene name</td><td>gene id</td></tr>
<tr><td>GU305914.1</td><td></td><td>94248</td><td>94270</td><td>-</td><td>0</td><td>AGCAAAAGCGGAGCGTCGAG</td><td>CGG</td><td>||||||||[||||||||||||]PAM</td><td>NA</td><td>Intergenic</td><td>NA</td><td>NA</td></tr>
<tr><td>T26813</td><td>GCAAAAGCGGAGCGTCGAGCGGG</td><td></td><td>1000</td><td></td><td></td><td></td><td></td><td></td><td></td><td></td><td></td><td></td></tr>
<tr><td>Oligo fwd</td><td>CACCGCAAAAGCGGAGCGTCGAGC</td><td></td><td></td><td></td><td></td><td></td><td></td><td></td><td></td><td></td><td></td><td></td></tr>
<tr><td>Oligo rev</td><td>AAACGCTCGACGCTCCGCTTTTGC</td><td></td><td></td><td></td><td></td><td></td><td></td><td></td><td></td><td></td><td></td><td></td></tr>
<tr><td>Chromosome</td><td>start</td><td></td><td>end</td><td>strand</td><td>MM</td><td>target_seq</td><td>PAM</td><td>alignment</td><td>distance</td><td>position</td><td>gene name</td><td>gene id</td></tr>
<tr><td>GU305914.1</td><td></td><td>94247</td><td>94269</td><td>-</td><td>0</td><td>GCAAAAGCGGAGCGTCGAGC</td><td>GGG</td><td>||||||||[||||||||||||]PAM</td><td>NA</td><td>Intergenic</td><td>NA</td><td>NA</td></tr>
<tr><td>T26814</td><td>GCGGAGCGTCGAGCGGGATTCGG</td><td></td><td>1000</td><td></td><td></td><td></td><td></td><td></td><td></td><td></td><td></td><td></td></tr>
<tr><td>Oligo fwd</td><td>CACCGCGGAGCGTCGAGCGGGATT</td><td></td><td></td><td></td><td></td><td></td><td></td><td></td><td></td><td></td><td></td><td></td></tr>
<tr><td>Oligo rev</td><td>AAACAATCCCGCTCGACGCTCCGC</td><td></td><td></td><td></td><td></td><td></td><td></td><td></td><td></td><td></td><td></td><td></td></tr>
<tr><td>Chromosome</td><td>start</td><td></td><td>end</td><td>strand</td><td>MM</td><td>target_seq</td><td>PAM</td><td>alignment</td><td>distance</td><td>position</td><td>gene name</td><td>gene id</td></tr>
<tr><td>GU305914.1</td><td></td><td>94241</td><td>94263</td><td>-</td><td>0</td><td>GCGGAGCGTCGAGCGGGATT</td><td>CGG</td><td>||||||||[||||||||||||]PAM</td><td>NA</td><td>Intergenic</td><td>NA</td><td>NA</td></tr>
<tr><td>T26815</td><td>CGGAGCGTCGAGCGGGATTCGGG</td><td></td><td>1000</td><td></td><td></td><td></td><td></td><td></td><td></td><td></td><td></td><td></td></tr>
<tr><td>Oligo fwd</td><td>CACCCGGAGCGTCGAGCGGGATTC</td><td></td><td></td><td></td><td></td><td></td><td></td><td></td><td></td><td></td><td></td><td></td></tr>
<tr><td>Oligo rev</td><td>AAACGAATCCCGCTCGACGCTCCG</td><td></td><td></td><td></td><td></td><td></td><td></td><td></td><td></td><td></td><td></td><td></td></tr>
<tr><td>Chromosome</td><td>start</td><td></td><td>end</td><td>strand</td><td>MM</td><td>target_seq</td><td>PAM</td><td>alignment</td><td>distance</td><td>position</td><td>gene name</td><td>gene id</td></tr>
<tr><td>GU305914.1</td><td></td><td>94240</td><td>94262</td><td>-</td><td>0</td><td>CGGAGCGTCGAGCGGGATTC</td><td>GGG</td><td>||||||||[||||||||||||]PAM</td><td>NA</td><td>Intergenic</td><td>NA</td><td>NA</td></tr>
<tr><td>T26816</td><td>GTCGAGCGGGATTCGGGAACCGG</td><td></td><td>1000</td><td></td><td></td><td></td><td></td><td></td><td></td><td></td><td></td><td></td></tr>
<tr><td>Oligo fwd</td><td>CACCGTCGAGCGGGATTCGGGAAC</td><td></td><td></td><td></td><td></td><td></td><td></td><td></td><td></td><td></td><td></td><td></td></tr>
<tr><td>Oligo rev</td><td>AAACGTTCCCGAATCCCGCTCGAC</td><td></td><td></td><td></td><td></td><td></td><td></td><td></td><td></td><td></td><td></td><td></td></tr>
<tr><td>Chromosome</td><td>start</td><td></td><td>end</td><td>strand</td><td>MM</td><td>target_seq</td><td>PAM</td><td>alignment</td><td>distance</td><td>position</td><td>gene name</td><td>gene id</td></tr>
<tr><td>GU305914.1</td><td></td><td>94234</td><td>94256</td><td>-</td><td>0</td><td>GTCGAGCGGGATTCGGGAAC</td><td>CGG</td><td>||||||||[||||||||||||]PAM</td><td>NA</td><td>Intergenic</td><td>NA</td><td>NA</td></tr>
<tr><td>T26817</td><td>TCGAGCGGGATTCGGGAACCGGG</td><td></td><td>1000</td><td></td><td></td><td></td><td></td><td></td><td></td><td></td><td></td><td></td></tr>
</table>

<table>
<tr><td>Oligo fwd</td><td>CACCTCGAGCGGGATTCGGGAACC</td></tr>
<tr><td>Oligo rev</td><td>AAACGGTTCCCGAATCCCGCTCGA</td></tr>
<tr><td>Chromosome</td><td>start</td><td>end</td><td>strand</td><td>MM</td><td>target_seq</td><td>PAM</td><td>alignment</td><td>distance</td><td>position</td><td>gene name</td><td>gene id</td></tr>
<tr><td>GU305914.1</td><td>94233</td><td>94255</td><td>-</td><td>0</td><td>TCGAGCGGGATTCGGGAACC</td><td>GGG</td><td>||||||||[||||||||||||]PAM</td><td>NA</td><td>Intergenic</td><td>NA</td><td>NA</td></tr>
<tr><td>T26818</td><td>GGGATTCGGGAACCGGGACCTGG</td><td>1000</td></tr>
<tr><td>Oligo fwd</td><td>CACCGGGATTCGGGAACCGGGACC</td></tr>
<tr><td>Oligo rev</td><td>AAACGGTCCCGGTTCCCGAATCCC</td></tr>
<tr><td>Chromosome</td><td>start</td><td>end</td><td>strand</td><td>MM</td><td>target_seq</td><td>PAM</td><td>alignment</td><td>distance</td><td>position</td><td>gene name</td><td>gene id</td></tr>
<tr><td>GU305914.1</td><td>94227</td><td>94249</td><td>-</td><td>0</td><td>GGGATTCGGGAACCGGGACC</td><td>TGG</td><td>||||||||[||||||||||||]PAM</td><td>NA</td><td>Intergenic</td><td>NA</td><td>NA</td></tr>
<tr><td>T26819</td><td>GGATTCGGGAACCGGGACCTGGG</td><td>1000</td></tr>
<tr><td>Oligo fwd</td><td>CACCGGATTCGGGAACCGGGACCT</td></tr>
<tr><td>Oligo rev</td><td>AAACAGGTCCCGGTTCCCGAATCC</td></tr>
<tr><td>Chromosome</td><td>start</td><td>end</td><td>strand</td><td>MM</td><td>target_seq</td><td>PAM</td><td>alignment</td><td>distance</td><td>position</td><td>gene name</td><td>gene id</td></tr>
<tr><td>GU305914.1</td><td>94226</td><td>94248</td><td>-</td><td>0</td><td>GGATTCGGGAACCGGGACCT</td><td>GGG</td><td>||||||||[||||||||||||]PAM</td><td>NA</td><td>Intergenic</td><td>NA</td><td>NA</td></tr>
<tr><td>T26820</td><td>TCGGGAACCGGGACCTGGGCAGG</td><td>1000</td></tr>
<tr><td>Oligo fwd</td><td>CACCTCGGGAACCGGGACCTGGGC</td></tr>
<tr><td>Oligo rev</td><td>AAACGCCCAGGTCCCGGTTCCCGA</td></tr>
<tr><td>Chromosome</td><td>start</td><td>end</td><td>strand</td><td>MM</td><td>target_seq</td><td>PAM</td><td>alignment</td><td>distance</td><td>position</td><td>gene name</td><td>gene id</td></tr>
<tr><td>GU305914.1</td><td>94222</td><td>94244</td><td>-</td><td>0</td><td>TCGGGAACCGGGACCTGGGC</td><td>AGG</td><td>||||||||[||||||||||||]PAM</td><td>NA</td><td>Intergenic</td><td>NA</td><td>NA</td></tr>
<tr><td>GU305914.1</td><td>94126</td><td>94148</td><td>-</td><td>3</td><td>ACGGCAATCGGGACCTGGGC</td><td>AGG</td><td>-|||-||-[||||||||||||]PAM</td><td>NA</td><td>Intergenic</td><td>NA</td><td>NA</td></tr>
<tr><td>T26821</td><td>CGGGAACCGGGACCTGGGCAGGG</td><td>1000</td></tr>
<tr><td>Oligo fwd</td><td>CACCCGGGAACCGGGACCTGGGCA</td></tr>
<tr><td>Oligo rev</td><td>AAACTGCCCAGGTCCCGGTTCCCG</td></tr>
<tr><td>Chromosome</td><td>start</td><td>end</td><td>strand</td><td>MM</td><td>target_seq</td><td>PAM</td><td>alignment</td><td>distance</td><td>position</td><td>gene name</td><td>gene id</td></tr>
<tr><td>GU305914.1</td><td>94221</td><td>94243</td><td>-</td><td>0</td><td>CGGGAACCGGGACCTGGGCA</td><td>GGG</td><td>||||||||[||||||||||||]PAM</td><td>NA</td><td>Intergenic</td><td>NA</td><td>NA</td></tr>
<tr><td>GU305914.1</td><td>94125</td><td>94147</td><td>-</td><td>2</td><td>CGGCAATCGGGACCTGGGCA</td><td>GGG</td><td>|||-||-|[||||||||||||]PAM</td><td>NA</td><td>Intergenic</td><td>NA</td><td>NA</td></tr>
<tr><td>T26822</td><td>TGGGCAGGGAGCAACGAAGCCGG</td><td>1000</td></tr>
<tr><td>Oligo fwd</td><td>CACCTGGGCAGGGAGCAACGAAGC</td></tr>
<tr><td>Oligo rev</td><td>AAACGCTTCGTTGCTCCCTGCCCA</td></tr>
<tr><td>Chromosome</td><td>start</td><td>end</td><td>strand</td><td>MM</td><td>target_seq</td><td>PAM</td><td>alignment</td><td>distance</td><td>position</td><td>gene name</td><td>gene id</td></tr>
<tr><td>GU305914.1</td><td>94207</td><td>94229</td><td>-</td><td>0</td><td>TGGGCAGGGAGCAACGAAGC</td><td>CGG</td><td>||||||||[||||||||||||]PAM</td><td>NA</td><td>Intergenic</td><td>NA</td><td>NA</td></tr>
<tr><td>GU305914.1</td><td>94111</td><td>94133</td><td>-</td><td>2</td><td>TGGGCAGGGTGCAATGAAGC</td><td>GGG</td><td>||||||||[|-||||-|||||]PAM</td><td>NA</td><td>Intergenic</td><td>NA</td><td>NA</td></tr>
<tr><td>T26823</td><td>GGGAGCAACGAAGCCGGAACCGG</td><td>1000</td></tr>
<tr><td>Oligo fwd</td><td>CACCGGGAGCAACGAAGCCGGAAC</td></tr>
<tr><td>Oligo rev</td><td>AAACGTTCCGGCTTCGTTGCTCCC</td></tr>
<tr><td>Chromosome</td><td>start</td><td>end</td><td>strand</td><td>MM</td><td>target_seq</td><td>PAM</td><td>alignment</td><td>distance</td><td>position</td><td>gene name</td><td>gene id</td></tr>
<tr><td>GU305914.1</td><td>94201</td><td>94223</td><td>-</td><td>0</td><td>GGGAGCAACGAAGCCGGAAC</td><td>CGG</td><td>||||||||[||||||||||||]PAM</td><td>NA</td><td>Intergenic</td><td>NA</td><td>NA</td></tr>
<tr><td>T26824</td><td>GGAGCAACGAAGCCGGAACCGGG</td><td>1000</td></tr>
<tr><td>Oligo fwd</td><td>CACCGGAGCAACGAAGCCGGAACC</td></tr>
<tr><td>Oligo rev</td><td>AAACGGTTCCGGCTTCGTTGCTCC</td></tr>
<tr><td>Chromosome</td><td>start</td><td>end</td><td>strand</td><td>MM</td><td>target_seq</td><td>PAM</td><td>alignment</td><td>distance</td><td>position</td><td>gene name</td><td>gene id</td></tr>
<tr><td>GU305914.1</td><td>94200</td><td>94222</td><td>-</td><td>0</td><td>GGAGCAACGAAGCCGGAACC</td><td>GGG</td><td>||||||||[||||||||||||]PAM</td><td>NA</td><td>Intergenic</td><td>NA</td><td>NA</td></tr>
<tr><td>GU305914.1</td><td>94104</td><td>94126</td><td>-</td><td>4</td><td>GGTGCAATGAAGCGGGGACC</td><td>GGG</td><td>||-||||-[|||||-||-|||]PAM</td><td>NA</td><td>Intergenic</td><td>NA</td><td>NA</td></tr>
<tr><td>T26825</td><td>GAACCGGGACCCCGAAAAGCCGG</td><td>1000</td></tr>
<tr><td>Oligo fwd</td><td>CACCGAACCGGGACCCCGAAAAGC</td></tr>
<tr><td>Oligo rev</td><td>AAACGCTTTTCGGGGTCCCGGTTC</td></tr>
<tr><td>Chromosome</td><td>start</td><td>end</td><td>strand</td><td>MM</td><td>target_seq</td><td>PAM</td><td>alignment</td><td>distance</td><td>position</td><td>gene name</td><td>gene id</td></tr>
<tr><td>GU305914.1</td><td>94185</td><td>94207</td><td>-</td><td>0</td><td>GAACCGGGACCCCGAAAAGC</td><td>CGG</td><td>||||||||[||||||||||||]PAM</td><td>NA</td><td>Intergenic</td><td>NA</td><td>NA</td></tr>
<tr><td>T26826</td><td>GGGACCCCGAAAAGCCGGCCAGG</td><td>1000</td></tr>
<tr><td>Oligo fwd</td><td>CACCGGGACCCCGAAAAGCCGGCC</td></tr>
<tr><td>Oligo rev</td><td>AAACGGCCGGCTTTTCGGGGTCCC</td></tr>
<tr><td>Chromosome</td><td>start</td><td>end</td><td>strand</td><td>MM</td><td>target_seq</td><td>PAM</td><td>alignment</td><td>distance</td><td>position</td><td>gene name</td><td>gene id</td></tr>
<tr><td>GU305914.1</td><td>94180</td><td>94202</td><td>-</td><td>0</td><td>GGGACCCCGAAAAGCCGGCC</td><td>AGG</td><td>||||||||[||||||||||||]PAM</td><td>NA</td><td>Intergenic</td><td>NA</td><td>NA</td></tr>
<tr><td>T26827</td><td>GGACCCCGAAAAGCCGGCCAGGG</td><td>1000</td></tr>
<tr><td>Oligo fwd</td><td>CACCGGACCCCGAAAAGCCGGCCA</td></tr>
<tr><td>Oligo rev</td><td>AAACTGGCCGGCTTTTCGGGGTCC</td></tr>
<tr><td>Chromosome</td><td>start</td><td>end</td><td>strand</td><td>MM</td><td>target_seq</td><td>PAM</td><td>alignment</td><td>distance</td><td>position</td><td>gene name</td><td>gene id</td></tr>
<tr><td>GU305914.1</td><td>94179</td><td>94201</td><td>-</td><td>0</td><td>GGACCCCGAAAAGCCGGCCA</td><td>GGG</td><td>||||||||[||||||||||||]PAM</td><td>NA</td><td>Intergenic</td><td>NA</td><td>NA</td></tr>
<tr><td>T26828</td><td>AAGCCGGCCAGGGTGCCGAACGG</td><td>1000</td></tr>
<tr><td>Oligo fwd</td><td>CACCAAGCCGGCCAGGGTGCCGAA</td></tr>
<tr><td>Oligo rev</td><td>AAACTTCGGCACCCTGGCCGGCTT</td></tr>
<tr><td>Chromosome</td><td>start</td><td>end</td><td>strand</td><td>MM</td><td>target_seq</td><td>PAM</td><td>alignment</td><td>distance</td><td>position</td><td>gene name</td><td>gene id</td></tr>
<tr><td>GU305914.1</td><td>94169</td><td>94191</td><td>-</td><td>0</td><td>AAGCCGGCCAGGGTGCCGAA</td><td>CGG</td><td>||||||||[||||||||||||]PAM</td><td>NA</td><td>Intergenic</td><td>NA</td><td>NA</td></tr>
<tr><td>T26829</td><td>GAACGGAGCATCGAGCCGCTCGG</td><td>1000</td></tr>
<tr><td>Oligo fwd</td><td>CACCGAACGGAGCATCGAGCCGCT</td></tr>
<tr><td>Oligo rev</td><td>AAACAGCGGCTCGATGCTCCGTTC</td></tr>
<tr><td>Chromosome</td><td>start</td><td>end</td><td>strand</td><td>MM</td><td>target_seq</td><td>PAM</td><td>alignment</td><td>distance</td><td>position</td><td>gene name</td><td>gene id</td></tr>
<tr><td>GU305914.1</td><td>94152</td><td>94174</td><td>-</td><td>0</td><td>GAACGGAGCATCGAGCCGCT</td><td>CGG</td><td>||||||||[||||||||||||]PAM</td><td>NA</td><td>Intergenic</td><td>NA</td><td>NA</td></tr>
<tr><td>T26830</td><td>GCATCGAGCCGCTCGGAGAACGG</td><td>1000</td></tr>
<tr><td>Oligo fwd</td><td>CACCGCATCGAGCCGCTCGGAGAA</td></tr>
<tr><td>Oligo rev</td><td>AAACTTCTCCGAGCGGCTCGATGC</td></tr>
<tr><td>Chromosome</td><td>start</td><td>end</td><td>strand</td><td>MM</td><td>target_seq</td><td>PAM</td><td>alignment</td><td>distance</td><td>position</td><td>gene name</td><td>gene id</td></tr>
<tr><td>GU305914.1</td><td>94145</td><td>94167</td><td>-</td><td>0</td><td>GCATCGAGCCGCTCGGAGAA</td><td>CGG</td><td>||||||||[||||||||||||]PAM</td><td>NA</td><td>Intergenic</td><td>NA</td><td>NA</td></tr>
<tr><td>T26831</td><td>GCCGCTCGGAGAACGGCAATCGG</td><td>1000</td></tr>
<tr><td>Oligo fwd</td><td>CACCGCCGCTCGGAGAACGGCAAT</td></tr>
</table>

<table>
<tr><td>Oligo rev</td><td>AAACATTGCCGTTCTCCGAGCGGC</td><td></td><td></td><td></td><td></td><td></td><td></td><td></td><td></td><td></td><td></td></tr>
<tr><td>Chromosome</td><td>start</td><td>end</td><td>strand</td><td>MM</td><td>target_seq</td><td>PAM</td><td>alignment</td><td>distance</td><td>position</td><td>gene name</td><td>gene id</td></tr>
<tr><td>GU305914.1</td><td>94138</td><td>94160</td><td>-</td><td>0</td><td>GCCGCTCGGAGAACGGCAAT</td><td>CGG</td><td>||||||||[||||||||||||]PAM</td><td>NA</td><td>Intergenic</td><td>NA</td><td>NA</td></tr>
<tr><td>T26832</td><td>CCGCTCGGAGAACGGCAATCGGG</td><td>1000</td><td></td><td></td><td></td><td></td><td></td><td></td><td></td><td></td><td></td></tr>
<tr><td>Oligo fwd</td><td>CACCCCGCTCGGAGAACGGCAATC</td><td></td><td></td><td></td><td></td><td></td><td></td><td></td><td></td><td></td><td></td></tr>
<tr><td>Oligo rev</td><td>AAACGATTGCCGTTCTCCGAGCGG</td><td></td><td></td><td></td><td></td><td></td><td></td><td></td><td></td><td></td><td></td></tr>
<tr><td>Chromosome</td><td>start</td><td>end</td><td>strand</td><td>MM</td><td>target_seq</td><td>PAM</td><td>alignment</td><td>distance</td><td>position</td><td>gene name</td><td>gene id</td></tr>
<tr><td>GU305914.1</td><td>94137</td><td>94159</td><td>-</td><td>0</td><td>CCGCTCGGAGAACGGCAATC</td><td>GGG</td><td>||||||||[||||||||||||]PAM</td><td>NA</td><td>Intergenic</td><td>NA</td><td>NA</td></tr>
<tr><td>T26833</td><td>GGAGAACGGCAATCGGGACCTGG</td><td>1000</td><td></td><td></td><td></td><td></td><td></td><td></td><td></td><td></td><td></td></tr>
<tr><td>Oligo fwd</td><td>CACCGGAGAACGGCAATCGGGACC</td><td></td><td></td><td></td><td></td><td></td><td></td><td></td><td></td><td></td><td></td></tr>
<tr><td>Oligo rev</td><td>AAACGGTCCCGATTGCCGTTCTCC</td><td></td><td></td><td></td><td></td><td></td><td></td><td></td><td></td><td></td><td></td></tr>
<tr><td>Chromosome</td><td>start</td><td>end</td><td>strand</td><td>MM</td><td>target_seq</td><td>PAM</td><td>alignment</td><td>distance</td><td>position</td><td>gene name</td><td>gene id</td></tr>
<tr><td>GU305914.1</td><td>94131</td><td>94153</td><td>-</td><td>0</td><td>GGAGAACGGCAATCGGGACC</td><td>TGG</td><td>||||||||[||||||||||||]PAM</td><td>NA</td><td>Intergenic</td><td>NA</td><td>NA</td></tr>
<tr><td>T26834</td><td>GAGAACGGCAATCGGGACCTGGG</td><td>1000</td><td></td><td></td><td></td><td></td><td></td><td></td><td></td><td></td><td></td></tr>
<tr><td>Oligo fwd</td><td>CACCGAGAACGGCAATCGGGACCT</td><td></td><td></td><td></td><td></td><td></td><td></td><td></td><td></td><td></td><td></td></tr>
<tr><td>Oligo rev</td><td>AAACAGGTCCCGATTGCCGTTCTC</td><td></td><td></td><td></td><td></td><td></td><td></td><td></td><td></td><td></td><td></td></tr>
<tr><td>Chromosome</td><td>start</td><td>end</td><td>strand</td><td>MM</td><td>target_seq</td><td>PAM</td><td>alignment</td><td>distance</td><td>position</td><td>gene name</td><td>gene id</td></tr>
<tr><td>GU305914.1</td><td>94130</td><td>94152</td><td>-</td><td>0</td><td>GAGAACGGCAATCGGGACCT</td><td>GGG</td><td>||||||||[||||||||||||]PAM</td><td>NA</td><td>Intergenic</td><td>NA</td><td>NA</td></tr>
<tr><td>T26835</td><td>ACGGCAATCGGGACCTGGGCAGG</td><td>1000</td><td></td><td></td><td></td><td></td><td></td><td></td><td></td><td></td><td></td></tr>
<tr><td>Oligo fwd</td><td>CACCACGGCAATCGGGACCTGGGC</td><td></td><td></td><td></td><td></td><td></td><td></td><td></td><td></td><td></td><td></td></tr>
<tr><td>Oligo rev</td><td>AAACGCCCAGGTCCCGATTGCCGT</td><td></td><td></td><td></td><td></td><td></td><td></td><td></td><td></td><td></td><td></td></tr>
<tr><td>Chromosome</td><td>start</td><td>end</td><td>strand</td><td>MM</td><td>target_seq</td><td>PAM</td><td>alignment</td><td>distance</td><td>position</td><td>gene name</td><td>gene id</td></tr>
<tr><td>GU305914.1</td><td>94126</td><td>94148</td><td>-</td><td>0</td><td>ACGGCAATCGGGACCTGGGC</td><td>AGG</td><td>||||||||[||||||||||||]PAM</td><td>NA</td><td>Intergenic</td><td>NA</td><td>NA</td></tr>
<tr><td>GU305914.1</td><td>94222</td><td>94244</td><td>-</td><td>3</td><td>TCGGGAACCGGGACCTGGGC</td><td>AGG</td><td>-|||-||-[||||||||||||]PAM</td><td>NA</td><td>Intergenic</td><td>NA</td><td>NA</td></tr>
<tr><td>T26836</td><td>CGGCAATCGGGACCTGGGCAGGG</td><td>1000</td><td></td><td></td><td></td><td></td><td></td><td></td><td></td><td></td><td></td></tr>
<tr><td>Oligo fwd</td><td>CACCCGGCAATCGGGACCTGGGCA</td><td></td><td></td><td></td><td></td><td></td><td></td><td></td><td></td><td></td><td></td></tr>
<tr><td>Oligo rev</td><td>AAACTGCCCAGGTCCCGATTGCCG</td><td></td><td></td><td></td><td></td><td></td><td></td><td></td><td></td><td></td><td></td></tr>
<tr><td>Chromosome</td><td>start</td><td>end</td><td>strand</td><td>MM</td><td>target_seq</td><td>PAM</td><td>alignment</td><td>distance</td><td>position</td><td>gene name</td><td>gene id</td></tr>
<tr><td>GU305914.1</td><td>94125</td><td>94147</td><td>-</td><td>0</td><td>CGGCAATCGGGACCTGGGCA</td><td>GGG</td><td>||||||||[||||||||||||]PAM</td><td>NA</td><td>Intergenic</td><td>NA</td><td>NA</td></tr>
<tr><td>GU305914.1</td><td>94221</td><td>94243</td><td>-</td><td>2</td><td>CGGGAACCGGGACCTGGGCA</td><td>GGG</td><td>|||-||-|[||||||||||||]PAM</td><td>NA</td><td>Intergenic</td><td>NA</td><td>NA</td></tr>
<tr><td>T26837</td><td>CTGGGCAGGGTGCAATGAAGCGG</td><td>1000</td><td></td><td></td><td></td><td></td><td></td><td></td><td></td><td></td><td></td></tr>
<tr><td>Oligo fwd</td><td>CACCCTGGGCAGGGTGCAATGAAG</td><td></td><td></td><td></td><td></td><td></td><td></td><td></td><td></td><td></td><td></td></tr>
<tr><td>Oligo rev</td><td>AAACCTTCATTGCACCCTGCCCAG</td><td></td><td></td><td></td><td></td><td></td><td></td><td></td><td></td><td></td><td></td></tr>
<tr><td>Chromosome</td><td>start</td><td>end</td><td>strand</td><td>MM</td><td>target_seq</td><td>PAM</td><td>alignment</td><td>distance</td><td>position</td><td>gene name</td><td>gene id</td></tr>
<tr><td>GU305914.1</td><td>94112</td><td>94134</td><td>-</td><td>0</td><td>CTGGGCAGGGTGCAATGAAG</td><td>CGG</td><td>||||||||[||||||||||||]PAM</td><td>NA</td><td>Intergenic</td><td>NA</td><td>NA</td></tr>
<tr><td>T26838</td><td>TGGGCAGGGTGCAATGAAGCGGG</td><td>1000</td><td></td><td></td><td></td><td></td><td></td><td></td><td></td><td></td><td></td></tr>
<tr><td>Oligo fwd</td><td>CACCTGGGCAGGGTGCAATGAAGC</td><td></td><td></td><td></td><td></td><td></td><td></td><td></td><td></td><td></td><td></td></tr>
<tr><td>Oligo rev</td><td>AAACGCTTCATTGCACCCTGCCCA</td><td></td><td></td><td></td><td></td><td></td><td></td><td></td><td></td><td></td><td></td></tr>
<tr><td>Chromosome</td><td>start</td><td>end</td><td>strand</td><td>MM</td><td>target_seq</td><td>PAM</td><td>alignment</td><td>distance</td><td>position</td><td>gene name</td><td>gene id</td></tr>
<tr><td>GU305914.1</td><td>94111</td><td>94133</td><td>-</td><td>0</td><td>TGGGCAGGGTGCAATGAAGC</td><td>GGG</td><td>||||||||[||||||||||||]PAM</td><td>NA</td><td>Intergenic</td><td>NA</td><td>NA</td></tr>
<tr><td>GU305914.1</td><td>94207</td><td>94229</td><td>-</td><td>2</td><td>TGGGCAGGGAGCAACGAAGC</td><td>CGG</td><td>||||||||[|-||||-|||||]PAM</td><td>NA</td><td>Intergenic</td><td>NA</td><td>NA</td></tr>
<tr><td>T26839</td><td>GGGCAGGGTGCAATGAAGCGGGG</td><td>1000</td><td></td><td></td><td></td><td></td><td></td><td></td><td></td><td></td><td></td></tr>
<tr><td>Oligo fwd</td><td>CACCGGGCAGGGTGCAATGAAGCG</td><td></td><td></td><td></td><td></td><td></td><td></td><td></td><td></td><td></td><td></td></tr>
<tr><td>Oligo rev</td><td>AAACCGCTTCATTGCACCCTGCCC</td><td></td><td></td><td></td><td></td><td></td><td></td><td></td><td></td><td></td><td></td></tr>
<tr><td>Chromosome</td><td>start</td><td>end</td><td>strand</td><td>MM</td><td>target_seq</td><td>PAM</td><td>alignment</td><td>distance</td><td>position</td><td>gene name</td><td>gene id</td></tr>
<tr><td>GU305914.1</td><td>94110</td><td>94132</td><td>-</td><td>0</td><td>GGGCAGGGTGCAATGAAGCG</td><td>GGG</td><td>||||||||[||||||||||||]PAM</td><td>NA</td><td>Intergenic</td><td>NA</td><td>NA</td></tr>
<tr><td>T26840</td><td>GGGTGCAATGAAGCGGGGACCGG</td><td>1000</td><td></td><td></td><td></td><td></td><td></td><td></td><td></td><td></td><td></td></tr>
<tr><td>Oligo fwd</td><td>CACCGGGTGCAATGAAGCGGGGAC</td><td></td><td></td><td></td><td></td><td></td><td></td><td></td><td></td><td></td><td></td></tr>
<tr><td>Oligo rev</td><td>AAACGTCCCCGCTTCATTGCACCC</td><td></td><td></td><td></td><td></td><td></td><td></td><td></td><td></td><td></td><td></td></tr>
<tr><td>Chromosome</td><td>start</td><td>end</td><td>strand</td><td>MM</td><td>target_seq</td><td>PAM</td><td>alignment</td><td>distance</td><td>position</td><td>gene name</td><td>gene id</td></tr>
<tr><td>GU305914.1</td><td>94105</td><td>94127</td><td>-</td><td>0</td><td>GGGTGCAATGAAGCGGGGAC</td><td>CGG</td><td>||||||||[||||||||||||]PAM</td><td>NA</td><td>Intergenic</td><td>NA</td><td>NA</td></tr>
<tr><td>T26841</td><td>GGTGCAATGAAGCGGGGACCGGG</td><td>1000</td><td></td><td></td><td></td><td></td><td></td><td></td><td></td><td></td><td></td></tr>
<tr><td>Oligo fwd</td><td>CACCGGTGCAATGAAGCGGGGACC</td><td></td><td></td><td></td><td></td><td></td><td></td><td></td><td></td><td></td><td></td></tr>
<tr><td>Oligo rev</td><td>AAACGGTCCCCGCTTCATTGCACC</td><td></td><td></td><td></td><td></td><td></td><td></td><td></td><td></td><td></td><td></td></tr>
<tr><td>Chromosome</td><td>start</td><td>end</td><td>strand</td><td>MM</td><td>target_seq</td><td>PAM</td><td>alignment</td><td>distance</td><td>position</td><td>gene name</td><td>gene id</td></tr>
<tr><td>GU305914.1</td><td>94104</td><td>94126</td><td>-</td><td>0</td><td>GGTGCAATGAAGCGGGGACC</td><td>GGG</td><td>||||||||[||||||||||||]PAM</td><td>NA</td><td>Intergenic</td><td>NA</td><td>NA</td></tr>
<tr><td>GU305914.1</td><td>94200</td><td>94222</td><td>-</td><td>4</td><td>GGAGCAACGAAGCCGGAACC</td><td>GGG</td><td>||-||||-[|||||-||-|||]PAM</td><td>NA</td><td>Intergenic</td><td>NA</td><td>NA</td></tr>
<tr><td>T26842</td><td>TGAAGCGGGGACCGGGAGCTCGG</td><td>1000</td><td></td><td></td><td></td><td></td><td></td><td></td><td></td><td></td><td></td></tr>
<tr><td>Oligo fwd</td><td>CACCTGAAGCGGGGACCGGGAGCT</td><td></td><td></td><td></td><td></td><td></td><td></td><td></td><td></td><td></td><td></td></tr>
<tr><td>Oligo rev</td><td>AAACAGCTCCCGGTCCCCGCTTCA</td><td></td><td></td><td></td><td></td><td></td><td></td><td></td><td></td><td></td><td></td></tr>
<tr><td>Chromosome</td><td>start</td><td>end</td><td>strand</td><td>MM</td><td>target_seq</td><td>PAM</td><td>alignment</td><td>distance</td><td>position</td><td>gene name</td><td>gene id</td></tr>
<tr><td>GU305914.1</td><td>94097</td><td>94119</td><td>-</td><td>0</td><td>TGAAGCGGGGACCGGGAGCT</td><td>CGG</td><td>||||||||[||||||||||||]PAM</td><td>NA</td><td>Intergenic</td><td>NA</td><td>NA</td></tr>
<tr><td>T26843</td><td>GAAGCGGGGACCGGGAGCTCGGG</td><td>1000</td><td></td><td></td><td></td><td></td><td></td><td></td><td></td><td></td><td></td></tr>
<tr><td>Oligo fwd</td><td>CACCGAAGCGGGGACCGGGAGCTC</td><td></td><td></td><td></td><td></td><td></td><td></td><td></td><td></td><td></td><td></td></tr>
<tr><td>Oligo rev</td><td>AAACGAGCTCCCGGTCCCCGCTTC</td><td></td><td></td><td></td><td></td><td></td><td></td><td></td><td></td><td></td><td></td></tr>
<tr><td>Chromosome</td><td>start</td><td>end</td><td>strand</td><td>MM</td><td>target_seq</td><td>PAM</td><td>alignment</td><td>distance</td><td>position</td><td>gene name</td><td>gene id</td></tr>
<tr><td>GU305914.1</td><td>94096</td><td>94118</td><td>-</td><td>0</td><td>GAAGCGGGGACCGGGAGCTC</td><td>GGG</td><td>||||||||[||||||||||||]PAM</td><td>NA</td><td>Intergenic</td><td>NA</td><td>NA</td></tr>
<tr><td>T26844</td><td>AAGCGGGGACCGGGAGCTCGGGG</td><td>1000</td><td></td><td></td><td></td><td></td><td></td><td></td><td></td><td></td><td></td></tr>
<tr><td>Oligo fwd</td><td>CACCAAGCGGGGACCGGGAGCTCG</td><td></td><td></td><td></td><td></td><td></td><td></td><td></td><td></td><td></td><td></td></tr>
<tr><td>Oligo rev</td><td>AAACCGAGCTCCCGGTCCCCGCTT</td><td></td><td></td><td></td><td></td><td></td><td></td><td></td><td></td><td></td><td></td></tr>
<tr><td>Chromosome</td><td>start</td><td>end</td><td>strand</td><td>MM</td><td>target_seq</td><td>PAM</td><td>alignment</td><td>distance</td><td>position</td><td>gene name</td><td>gene id</td></tr>
<tr><td>GU305914.1</td><td>94095</td><td>94117</td><td>-</td><td>0</td><td>AAGCGGGGACCGGGAGCTCG</td><td>GGG</td><td>||||||||[||||||||||||]PAM</td><td>NA</td><td>Intergenic</td><td>NA</td><td>NA</td></tr>
<tr><td>GU305914.1</td><td>94031</td><td>94053</td><td>-</td><td>4</td><td>GGACAGGGACCGGGAGCTCG</td><td>GGG</td><td>---|-|||[||||||||||||]PAM</td><td>NA</td><td>Intergenic</td><td>NA</td><td>NA</td></tr>
<tr><td>GU305914.1</td><td>93999</td><td>94021</td><td>-</td><td>4</td><td>GGACAGGGACCGGGAGCTCG</td><td>GGG</td><td>---|-|||[||||||||||||]PAM</td><td>NA</td><td>Intergenic</td><td>NA</td><td>NA</td></tr>
<tr><td>GU305914.1</td><td>93967</td><td>93989</td><td>-</td><td>4</td><td>GGACAGGGACCGGGAGCTCG</td><td>GGG</td><td>---|-|||[||||||||||||]PAM</td><td>NA</td><td>Intergenic</td><td>NA</td><td>NA</td></tr>
<tr><td>GU305914.1</td><td>93935</td><td>93957</td><td>-</td><td>4</td><td>GGACAGGGACCGGGAGCTCG</td><td>GGG</td><td>---|-|||[||||||||||||]PAM</td><td>NA</td><td>Intergenic</td><td>NA</td><td>NA</td></tr>
</table>

<table>
<tr><td>GU305914.1</td><td>93903</td><td>93925</td><td>-</td><td>4</td><td>GGACAGGGACCGGGAGCTCG</td><td>GGG</td><td>---|-|||[||||||||||||]PAM</td><td>NA</td><td>Intergenic</td><td>NA</td><td>NA</td></tr>
<tr><td>GU305914.1</td><td>94063</td><td>94085</td><td>-</td><td>4</td><td>GGACAGGGACCGGGAGCTCG</td><td>GGG</td><td>---|-|||[||||||||||||]PAM</td><td>NA</td><td>Intergenic</td><td>NA</td><td>NA</td></tr>
<tr><td></td><td></td><td></td><td></td><td></td><td></td><td></td><td></td><td></td><td></td><td></td><td></td></tr>
<tr><td>T26845</td><td>CGGGGACCGGGAGCTCGGGGAGG</td><td>1000</td><td></td><td></td><td></td><td></td><td></td><td></td><td></td><td></td><td></td></tr>
<tr><td>Oligo fwd</td><td>CACCCGGGGACCGGGAGCTCGGGG</td><td></td><td></td><td></td><td></td><td></td><td></td><td></td><td></td><td></td><td></td></tr>
<tr><td>Oligo rev</td><td>AAACCCCCGAGCTCCCGGTCCCCG</td><td></td><td></td><td></td><td></td><td></td><td></td><td></td><td></td><td></td><td></td></tr>
<tr><td>Chromosome</td><td>start</td><td>end</td><td>strand</td><td>MM</td><td>target_seq</td><td>PAM</td><td>alignment</td><td>distance</td><td>position</td><td>gene name</td><td>gene id</td></tr>
<tr><td>GU305914.1</td><td>94092</td><td>94114</td><td>-</td><td>0</td><td>CGGGGACCGGGAGCTCGGGG</td><td>AGG</td><td>||||||||[||||||||||||]PAM</td><td>NA</td><td>Intergenic</td><td>NA</td><td>NA</td></tr>
<tr><td>GU305914.1</td><td>93932</td><td>93954</td><td>-</td><td>1</td><td>CAGGGACCGGGAGCTCGGGG</td><td>AGG</td><td>|-||||||[||||||||||||]PAM</td><td>NA</td><td>Intergenic</td><td>NA</td><td>NA</td></tr>
<tr><td>GU305914.1</td><td>94060</td><td>94082</td><td>-</td><td>1</td><td>CAGGGACCGGGAGCTCGGGG</td><td>AGG</td><td>|-||||||[||||||||||||]PAM</td><td>NA</td><td>Intergenic</td><td>NA</td><td>NA</td></tr>
<tr><td>GU305914.1</td><td>94028</td><td>94050</td><td>-</td><td>1</td><td>CAGGGACCGGGAGCTCGGGG</td><td>AGG</td><td>|-||||||[||||||||||||]PAM</td><td>NA</td><td>Intergenic</td><td>NA</td><td>NA</td></tr>
<tr><td>GU305914.1</td><td>93996</td><td>94018</td><td>-</td><td>1</td><td>CAGGGACCGGGAGCTCGGGG</td><td>AGG</td><td>|-||||||[||||||||||||]PAM</td><td>NA</td><td>Intergenic</td><td>NA</td><td>NA</td></tr>
<tr><td>GU305914.1</td><td>93964</td><td>93986</td><td>-</td><td>1</td><td>CAGGGACCGGGAGCTCGGGG</td><td>AGG</td><td>|-||||||[||||||||||||]PAM</td><td>NA</td><td>Intergenic</td><td>NA</td><td>NA</td></tr>
<tr><td></td><td></td><td></td><td></td><td></td><td></td><td></td><td></td><td></td><td></td><td></td><td></td></tr>
<tr><td>T26846</td><td>GGGGACCGGGAGCTCGGGGAGGG</td><td>1000</td><td></td><td></td><td></td><td></td><td></td><td></td><td></td><td></td><td></td></tr>
<tr><td>Oligo fwd</td><td>CACCGGGGACCGGGAGCTCGGGGA</td><td></td><td></td><td></td><td></td><td></td><td></td><td></td><td></td><td></td><td></td></tr>
<tr><td>Oligo rev</td><td>AAACTCCCCGAGCTCCCGGTCCCC</td><td></td><td></td><td></td><td></td><td></td><td></td><td></td><td></td><td></td><td></td></tr>
<tr><td>Chromosome</td><td>start</td><td>end</td><td>strand</td><td>MM</td><td>target_seq</td><td>PAM</td><td>alignment</td><td>distance</td><td>position</td><td>gene name</td><td>gene id</td></tr>
<tr><td>GU305914.1</td><td>94091</td><td>94113</td><td>-</td><td>0</td><td>GGGGACCGGGAGCTCGGGGA</td><td>GGG</td><td>||||||||[||||||||||||]PAM</td><td>NA</td><td>Intergenic</td><td>NA</td><td>NA</td></tr>
<tr><td>GU305914.1</td><td>94027</td><td>94049</td><td>-</td><td>1</td><td>AGGGACCGGGAGCTCGGGGA</td><td>GGG</td><td>-|||||||[||||||||||||]PAM</td><td>NA</td><td>Intergenic</td><td>NA</td><td>NA</td></tr>
<tr><td>GU305914.1</td><td>93995</td><td>94017</td><td>-</td><td>1</td><td>AGGGACCGGGAGCTCGGGGA</td><td>GGG</td><td>-|||||||[||||||||||||]PAM</td><td>NA</td><td>Intergenic</td><td>NA</td><td>NA</td></tr>
<tr><td>GU305914.1</td><td>93963</td><td>93985</td><td>-</td><td>1</td><td>AGGGACCGGGAGCTCGGGGA</td><td>GGG</td><td>-|||||||[||||||||||||]PAM</td><td>NA</td><td>Intergenic</td><td>NA</td><td>NA</td></tr>
<tr><td>GU305914.1</td><td>93931</td><td>93953</td><td>-</td><td>1</td><td>AGGGACCGGGAGCTCGGGGA</td><td>GGG</td><td>-|||||||[||||||||||||]PAM</td><td>NA</td><td>Intergenic</td><td>NA</td><td>NA</td></tr>
<tr><td>GU305914.1</td><td>94059</td><td>94081</td><td>-</td><td>1</td><td>AGGGACCGGGAGCTCGGGGA</td><td>GGG</td><td>-|||||||[||||||||||||]PAM</td><td>NA</td><td>Intergenic</td><td>NA</td><td>NA</td></tr>
<tr><td></td><td></td><td></td><td></td><td></td><td></td><td></td><td></td><td></td><td></td><td></td><td></td></tr>
<tr><td>T26847</td><td>GGGAGCTCGGGGAGGGAAGCCGG</td><td>1000</td><td></td><td></td><td></td><td></td><td></td><td></td><td></td><td></td><td></td></tr>
<tr><td>Oligo fwd</td><td>CACCGGGAGCTCGGGGAGGGAAGC</td><td></td><td></td><td></td><td></td><td></td><td></td><td></td><td></td><td></td><td></td></tr>
<tr><td>Oligo rev</td><td>AAACGCTTCCCTCCCCGAGCTCCC</td><td></td><td></td><td></td><td></td><td></td><td></td><td></td><td></td><td></td><td></td></tr>
<tr><td>Chromosome</td><td>start</td><td>end</td><td>strand</td><td>MM</td><td>target_seq</td><td>PAM</td><td>alignment</td><td>distance</td><td>position</td><td>gene name</td><td>gene id</td></tr>
<tr><td>GU305914.1</td><td>94052</td><td>94074</td><td>-</td><td>0</td><td>GGGAGCTCGGGGAGGGAAGC</td><td>CGG</td><td>||||||||[||||||||||||]PAM</td><td>NA</td><td>Intergenic</td><td>NA</td><td>NA</td></tr>
<tr><td>GU305914.1</td><td>94020</td><td>94042</td><td>-</td><td>0</td><td>GGGAGCTCGGGGAGGGAAGC</td><td>CGG</td><td>||||||||[||||||||||||]PAM</td><td>NA</td><td>Intergenic</td><td>NA</td><td>NA</td></tr>
<tr><td>GU305914.1</td><td>93988</td><td>94010</td><td>-</td><td>0</td><td>GGGAGCTCGGGGAGGGAAGC</td><td>CGG</td><td>||||||||[||||||||||||]PAM</td><td>NA</td><td>Intergenic</td><td>NA</td><td>NA</td></tr>
<tr><td>GU305914.1</td><td>93956</td><td>93978</td><td>-</td><td>0</td><td>GGGAGCTCGGGGAGGGAAGC</td><td>CGG</td><td>||||||||[||||||||||||]PAM</td><td>NA</td><td>Intergenic</td><td>NA</td><td>NA</td></tr>
<tr><td>GU305914.1</td><td>93924</td><td>93946</td><td>-</td><td>0</td><td>GGGAGCTCGGGGAGGGAAGC</td><td>CGG</td><td>||||||||[||||||||||||]PAM</td><td>NA</td><td>Intergenic</td><td>NA</td><td>NA</td></tr>
<tr><td>GU305914.1</td><td>94084</td><td>94106</td><td>-</td><td>0</td><td>GGGAGCTCGGGGAGGGAAGC</td><td>CGG</td><td>||||||||[||||||||||||]PAM</td><td>NA</td><td>Intergenic</td><td>NA</td><td>NA</td></tr>
<tr><td>GU305914.1</td><td>93892</td><td>93914</td><td>-</td><td>2</td><td>GGGAGCTCGGGGAGAGGAGC</td><td>CGG</td><td>||||||||[||||||-|-|||]PAM</td><td>NA</td><td>Intergenic</td><td>NA</td><td>NA</td></tr>
<tr><td></td><td></td><td></td><td></td><td></td><td></td><td></td><td></td><td></td><td></td><td></td><td></td></tr>
<tr><td>T26848</td><td>CTCGGGGAGGGAAGCCGGACAGG</td><td>1000</td><td></td><td></td><td></td><td></td><td></td><td></td><td></td><td></td><td></td></tr>
<tr><td>Oligo fwd</td><td>CACCCTCGGGGAGGGAAGCCGGAC</td><td></td><td></td><td></td><td></td><td></td><td></td><td></td><td></td><td></td><td></td></tr>
<tr><td>Oligo rev</td><td>AAACGTCCGGCTTCCCTCCCCGAG</td><td></td><td></td><td></td><td></td><td></td><td></td><td></td><td></td><td></td><td></td></tr>
<tr><td>Chromosome</td><td>start</td><td>end</td><td>strand</td><td>MM</td><td>target_seq</td><td>PAM</td><td>alignment</td><td>distance</td><td>position</td><td>gene name</td><td>gene id</td></tr>
<tr><td>GU305914.1</td><td>93951</td><td>93973</td><td>-</td><td>0</td><td>CTCGGGGAGGGAAGCCGGAC</td><td>AGG</td><td>||||||||[||||||||||||]PAM</td><td>NA</td><td>Intergenic</td><td>NA</td><td>NA</td></tr>
<tr><td>GU305914.1</td><td>93919</td><td>93941</td><td>-</td><td>0</td><td>CTCGGGGAGGGAAGCCGGAC</td><td>AGG</td><td>||||||||[||||||||||||]PAM</td><td>NA</td><td>Intergenic</td><td>NA</td><td>NA</td></tr>
<tr><td>GU305914.1</td><td>94079</td><td>94101</td><td>-</td><td>0</td><td>CTCGGGGAGGGAAGCCGGAC</td><td>AGG</td><td>||||||||[||||||||||||]PAM</td><td>NA</td><td>Intergenic</td><td>NA</td><td>NA</td></tr>
<tr><td>GU305914.1</td><td>94047</td><td>94069</td><td>-</td><td>0</td><td>CTCGGGGAGGGAAGCCGGAC</td><td>AGG</td><td>||||||||[||||||||||||]PAM</td><td>NA</td><td>Intergenic</td><td>NA</td><td>NA</td></tr>
<tr><td>GU305914.1</td><td>94015</td><td>94037</td><td>-</td><td>0</td><td>CTCGGGGAGGGAAGCCGGAC</td><td>AGG</td><td>||||||||[||||||||||||]PAM</td><td>NA</td><td>Intergenic</td><td>NA</td><td>NA</td></tr>
<tr><td>GU305914.1</td><td>93983</td><td>94005</td><td>-</td><td>0</td><td>CTCGGGGAGGGAAGCCGGAC</td><td>AGG</td><td>||||||||[||||||||||||]PAM</td><td>NA</td><td>Intergenic</td><td>NA</td><td>NA</td></tr>
<tr><td>GU305914.1</td><td>93887</td><td>93909</td><td>-</td><td>2</td><td>CTCGGGGAGAGGAGCCGGAC</td><td>AGG</td><td>||||||||[|-|-||||||||]PAM</td><td>NA</td><td>Intergenic</td><td>NA</td><td>NA</td></tr>
<tr><td></td><td></td><td></td><td></td><td></td><td></td><td></td><td></td><td></td><td></td><td></td><td></td></tr>
<tr><td>T26849</td><td>TCGGGGAGGGAAGCCGGACAGGG</td><td>1000</td><td></td><td></td><td></td><td></td><td></td><td></td><td></td><td></td><td></td></tr>
<tr><td>Oligo fwd</td><td>CACCTCGGGGAGGGAAGCCGGACA</td><td></td><td></td><td></td><td></td><td></td><td></td><td></td><td></td><td></td><td></td></tr>
<tr><td>Oligo rev</td><td>AAACTGTCCGGCTTCCCTCCCCGA</td><td></td><td></td><td></td><td></td><td></td><td></td><td></td><td></td><td></td><td></td></tr>
<tr><td>Chromosome</td><td>start</td><td>end</td><td>strand</td><td>MM</td><td>target_seq</td><td>PAM</td><td>alignment</td><td>distance</td><td>position</td><td>gene name</td><td>gene id</td></tr>
<tr><td>GU305914.1</td><td>94078</td><td>94100</td><td>-</td><td>0</td><td>TCGGGGAGGGAAGCCGGACA</td><td>GGG</td><td>||||||||[||||||||||||]PAM</td><td>NA</td><td>Intergenic</td><td>NA</td><td>NA</td></tr>
<tr><td>GU305914.1</td><td>94046</td><td>94068</td><td>-</td><td>0</td><td>TCGGGGAGGGAAGCCGGACA</td><td>GGG</td><td>||||||||[||||||||||||]PAM</td><td>NA</td><td>Intergenic</td><td>NA</td><td>NA</td></tr>
<tr><td>GU305914.1</td><td>94014</td><td>94036</td><td>-</td><td>0</td><td>TCGGGGAGGGAAGCCGGACA</td><td>GGG</td><td>||||||||[||||||||||||]PAM</td><td>NA</td><td>Intergenic</td><td>NA</td><td>NA</td></tr>
<tr><td>GU305914.1</td><td>93982</td><td>94004</td><td>-</td><td>0</td><td>TCGGGGAGGGAAGCCGGACA</td><td>GGG</td><td>||||||||[||||||||||||]PAM</td><td>NA</td><td>Intergenic</td><td>NA</td><td>NA</td></tr>
<tr><td>GU305914.1</td><td>93950</td><td>93972</td><td>-</td><td>0</td><td>TCGGGGAGGGAAGCCGGACA</td><td>GGG</td><td>||||||||[||||||||||||]PAM</td><td>NA</td><td>Intergenic</td><td>NA</td><td>NA</td></tr>
<tr><td>GU305914.1</td><td>93918</td><td>93940</td><td>-</td><td>0</td><td>TCGGGGAGGGAAGCCGGACA</td><td>GGG</td><td>||||||||[||||||||||||]PAM</td><td>NA</td><td>Intergenic</td><td>NA</td><td>NA</td></tr>
<tr><td></td><td></td><td></td><td></td><td></td><td></td><td></td><td></td><td></td><td></td><td></td><td></td></tr>
<tr><td>T26850</td><td>GAGGGAAGCCGGACAGGGACCGG</td><td>1000</td><td></td><td></td><td></td><td></td><td></td><td></td><td></td><td></td><td></td></tr>
<tr><td>Oligo fwd</td><td>CACCGAGGGAAGCCGGACAGGGAC</td><td></td><td></td><td></td><td></td><td></td><td></td><td></td><td></td><td></td><td></td></tr>
<tr><td>Oligo rev</td><td>AAACGTCCCTGTCCGGCTTCCCTC</td><td></td><td></td><td></td><td></td><td></td><td></td><td></td><td></td><td></td><td></td></tr>
<tr><td>Chromosome</td><td>start</td><td>end</td><td>strand</td><td>MM</td><td>target_seq</td><td>PAM</td><td>alignment</td><td>distance</td><td>position</td><td>gene name</td><td>gene id</td></tr>
<tr><td>GU305914.1</td><td>93913</td><td>93935</td><td>-</td><td>0</td><td>GAGGGAAGCCGGACAGGGAC</td><td>CGG</td><td>||||||||[||||||||||||]PAM</td><td>NA</td><td>Intergenic</td><td>NA</td><td>NA</td></tr>
<tr><td>GU305914.1</td><td>94073</td><td>94095</td><td>-</td><td>0</td><td>GAGGGAAGCCGGACAGGGAC</td><td>CGG</td><td>||||||||[||||||||||||]PAM</td><td>NA</td><td>Intergenic</td><td>NA</td><td>NA</td></tr>
<tr><td>GU305914.1</td><td>94041</td><td>94063</td><td>-</td><td>0</td><td>GAGGGAAGCCGGACAGGGAC</td><td>CGG</td><td>||||||||[||||||||||||]PAM</td><td>NA</td><td>Intergenic</td><td>NA</td><td>NA</td></tr>
<tr><td>GU305914.1</td><td>94009</td><td>94031</td><td>-</td><td>0</td><td>GAGGGAAGCCGGACAGGGAC</td><td>CGG</td><td>||||||||[||||||||||||]PAM</td><td>NA</td><td>Intergenic</td><td>NA</td><td>NA</td></tr>
<tr><td>GU305914.1</td><td>93977</td><td>93999</td><td>-</td><td>0</td><td>GAGGGAAGCCGGACAGGGAC</td><td>CGG</td><td>||||||||[||||||||||||]PAM</td><td>NA</td><td>Intergenic</td><td>NA</td><td>NA</td></tr>
<tr><td>GU305914.1</td><td>93945</td><td>93967</td><td>-</td><td>0</td><td>GAGGGAAGCCGGACAGGGAC</td><td>CGG</td><td>||||||||[||||||||||||]PAM</td><td>NA</td><td>Intergenic</td><td>NA</td><td>NA</td></tr>
<tr><td>GU305914.1</td><td>93881</td><td>93903</td><td>-</td><td>3</td><td>GAGAGGAGCCGGACAGGAAC</td><td>CGG</td><td>|||-|-||[|||||||||-||]PAM</td><td>NA</td><td>Intergenic</td><td>NA</td><td>NA</td></tr>
<tr><td></td><td></td><td></td><td></td><td></td><td></td><td></td><td></td><td></td><td></td><td></td><td></td></tr>
<tr><td>T26851</td><td>AGGGAAGCCGGACAGGGACCGGG</td><td>1000</td><td></td><td></td><td></td><td></td><td></td><td></td><td></td><td></td><td></td></tr>
<tr><td>Oligo fwd</td><td>CACCAGGGAAGCCGGACAGGGACC</td><td></td><td></td><td></td><td></td><td></td><td></td><td></td><td></td><td></td><td></td></tr>
<tr><td>Oligo rev</td><td>AAACGGTCCCTGTCCGGCTTCCCT</td><td></td><td></td><td></td><td></td><td></td><td></td><td></td><td></td><td></td><td></td></tr>
<tr><td>Chromosome</td><td>start</td><td>end</td><td>strand</td><td>MM</td><td>target_seq</td><td>PAM</td><td>alignment</td><td>distance</td><td>position</td><td>gene name</td><td>gene id</td></tr>
<tr><td>GU305914.1</td><td>93976</td><td>93998</td><td>-</td><td>0</td><td>AGGGAAGCCGGACAGGGACC</td><td>GGG</td><td>||||||||[||||||||||||]PAM</td><td>NA</td><td>Intergenic</td><td>NA</td><td>NA</td></tr>
<tr><td>GU305914.1</td><td>93944</td><td>93966</td><td>-</td><td>0</td><td>AGGGAAGCCGGACAGGGACC</td><td>GGG</td><td>||||||||[||||||||||||]PAM</td><td>NA</td><td>Intergenic</td><td>NA</td><td>NA</td></tr>
<tr><td>GU305914.1</td><td>93912</td><td>93934</td><td>-</td><td>0</td><td>AGGGAAGCCGGACAGGGACC</td><td>GGG</td><td>||||||||[||||||||||||]PAM</td><td>NA</td><td>Intergenic</td><td>NA</td><td>NA</td></tr>
<tr><td>GU305914.1</td><td>94072</td><td>94094</td><td>-</td><td>0</td><td>AGGGAAGCCGGACAGGGACC</td><td>GGG</td><td>||||||||[||||||||||||]PAM</td><td>NA</td><td>Intergenic</td><td>NA</td><td>NA</td></tr>
<tr><td>GU305914.1</td><td>94040</td><td>94062</td><td>-</td><td>0</td><td>AGGGAAGCCGGACAGGGACC</td><td>GGG</td><td>||||||||[||||||||||||]PAM</td><td>NA</td><td>Intergenic</td><td>NA</td><td>NA</td></tr>
<tr><td>GU305914.1</td><td>94008</td><td>94030</td><td>-</td><td>0</td><td>AGGGAAGCCGGACAGGGACC</td><td>GGG</td><td>||||||||[||||||||||||]PAM</td><td>NA</td><td>Intergenic</td><td>NA</td><td>NA</td></tr>
<tr><td>GU305914.1</td><td>93880</td><td>93902</td><td>-</td><td>3</td><td>AGAGGAGCCGGACAGGAACC</td><td>GGG</td><td>||-|-|||[||||||||-|||]PAM</td><td>NA</td><td>Intergenic</td><td>NA</td><td>NA</td></tr>
<tr><td></td><td></td><td></td><td></td><td></td><td></td><td></td><td></td><td></td><td></td><td></td><td></td></tr>
<tr><td>T26852</td><td>CCGGACAGGGACCGGGAGCTCGG</td><td>1000</td><td></td><td></td><td></td><td></td><td></td><td></td><td></td><td></td><td></td></tr>
<tr><td>Oligo fwd</td><td>CACCCCGGACAGGGACCGGGAGCT</td><td></td><td></td><td></td><td></td><td></td><td></td><td></td><td></td><td></td><td></td></tr>
<tr><td>Oligo rev</td><td>AAACAGCTCCCGGTCCCTGTCCGG</td><td></td><td></td><td></td><td></td><td></td><td></td><td></td><td></td><td></td><td></td></tr>
<tr><td>Chromosome</td><td>start</td><td>end</td><td>strand</td><td>MM</td><td>target_seq</td><td>PAM</td><td>alignment</td><td>distance</td><td>position</td><td>gene name</td><td>gene id</td></tr>
<tr><td>GU305914.1</td><td>94001</td><td>94023</td><td>-</td><td>0</td><td>CCGGACAGGGACCGGGAGCT</td><td>CGG</td><td>||||||||[||||||||||||]PAM</td><td>NA</td><td>Intergenic</td><td>NA</td><td>NA</td></tr>
</table>

<table>
<tr><td>GU305914.1</td><td>93969</td><td>93991</td><td>-</td><td>0</td><td>CCGGACAGGGACCGGGAGCT</td><td>CGG</td><td>||||||||[||||||||||||]PAM</td><td>NA</td><td>Intergenic</td><td>NA</td><td>NA</td></tr>
<tr><td>GU305914.1</td><td>93937</td><td>93959</td><td>-</td><td>0</td><td>CCGGACAGGGACCGGGAGCT</td><td>CGG</td><td>||||||||[||||||||||||]PAM</td><td>NA</td><td>Intergenic</td><td>NA</td><td>NA</td></tr>
<tr><td>GU305914.1</td><td>93905</td><td>93927</td><td>-</td><td>0</td><td>CCGGACAGGGACCGGGAGCT</td><td>CGG</td><td>||||||||[||||||||||||]PAM</td><td>NA</td><td>Intergenic</td><td>NA</td><td>NA</td></tr>
<tr><td>GU305914.1</td><td>94065</td><td>94087</td><td>-</td><td>0</td><td>CCGGACAGGGACCGGGAGCT</td><td>CGG</td><td>||||||||[||||||||||||]PAM</td><td>NA</td><td>Intergenic</td><td>NA</td><td>NA</td></tr>
<tr><td>GU305914.1</td><td>94033</td><td>94055</td><td>-</td><td>0</td><td>CCGGACAGGGACCGGGAGCT</td><td>CGG</td><td>||||||||[||||||||||||]PAM</td><td>NA</td><td>Intergenic</td><td>NA</td><td>NA</td></tr>
<tr><td>GU305914.1</td><td>93873</td><td>93895</td><td>-</td><td>2</td><td>CCGGACAGGAACCGGGAGCG</td><td>GGG</td><td>||||||||[|-|||||||||-]PAM</td><td>NA</td><td>Intergenic</td><td>NA</td><td>NA</td></tr>
<tr><td></td><td></td><td></td><td></td><td></td><td></td><td></td><td></td><td></td><td></td><td></td><td></td></tr>
<tr><td>T26853</td><td>CGGACAGGGACCGGGAGCTCGGG</td><td>1000</td><td></td><td></td><td></td><td></td><td></td><td></td><td></td><td></td><td></td></tr>
<tr><td>Oligo fwd</td><td>CACCCGGACAGGGACCGGGAGCTC</td><td></td><td></td><td></td><td></td><td></td><td></td><td></td><td></td><td></td><td></td></tr>
<tr><td>Oligo rev</td><td>AAACGAGCTCCCGGTCCCTGTCCG</td><td></td><td></td><td></td><td></td><td></td><td></td><td></td><td></td><td></td><td></td></tr>
<tr><td>Chromosome</td><td>start</td><td>end</td><td>strand</td><td>MM</td><td>target_seq</td><td>PAM</td><td>alignment</td><td>distance</td><td>position</td><td>gene name</td><td>gene id</td></tr>
<tr><td>GU305914.1</td><td>93968</td><td>93990</td><td>-</td><td>0</td><td>CGGACAGGGACCGGGAGCTC</td><td>GGG</td><td>||||||||[||||||||||||]PAM</td><td>NA</td><td>Intergenic</td><td>NA</td><td>NA</td></tr>
<tr><td>GU305914.1</td><td>93936</td><td>93958</td><td>-</td><td>0</td><td>CGGACAGGGACCGGGAGCTC</td><td>GGG</td><td>||||||||[||||||||||||]PAM</td><td>NA</td><td>Intergenic</td><td>NA</td><td>NA</td></tr>
<tr><td>GU305914.1</td><td>93904</td><td>93926</td><td>-</td><td>0</td><td>CGGACAGGGACCGGGAGCTC</td><td>GGG</td><td>||||||||[||||||||||||]PAM</td><td>NA</td><td>Intergenic</td><td>NA</td><td>NA</td></tr>
<tr><td>GU305914.1</td><td>94064</td><td>94086</td><td>-</td><td>0</td><td>CGGACAGGGACCGGGAGCTC</td><td>GGG</td><td>||||||||[||||||||||||]PAM</td><td>NA</td><td>Intergenic</td><td>NA</td><td>NA</td></tr>
<tr><td>GU305914.1</td><td>94032</td><td>94054</td><td>-</td><td>0</td><td>CGGACAGGGACCGGGAGCTC</td><td>GGG</td><td>||||||||[||||||||||||]PAM</td><td>NA</td><td>Intergenic</td><td>NA</td><td>NA</td></tr>
<tr><td>GU305914.1</td><td>94000</td><td>94022</td><td>-</td><td>0</td><td>CGGACAGGGACCGGGAGCTC</td><td>GGG</td><td>||||||||[||||||||||||]PAM</td><td>NA</td><td>Intergenic</td><td>NA</td><td>NA</td></tr>
<tr><td></td><td></td><td></td><td></td><td></td><td></td><td></td><td></td><td></td><td></td><td></td><td></td></tr>
<tr><td>T26854</td><td>GGACAGGGACCGGGAGCTCGGGG</td><td>1000</td><td></td><td></td><td></td><td></td><td></td><td></td><td></td><td></td><td></td></tr>
<tr><td>Oligo fwd</td><td>CACCGGACAGGGACCGGGAGCTCG</td><td></td><td></td><td></td><td></td><td></td><td></td><td></td><td></td><td></td><td></td></tr>
<tr><td>Oligo rev</td><td>AAACCGAGCTCCCGGTCCCTGTCC</td><td></td><td></td><td></td><td></td><td></td><td></td><td></td><td></td><td></td><td></td></tr>
<tr><td>Chromosome</td><td>start</td><td>end</td><td>strand</td><td>MM</td><td>target_seq</td><td>PAM</td><td>alignment</td><td>distance</td><td>position</td><td>gene name</td><td>gene id</td></tr>
<tr><td>GU305914.1</td><td>93903</td><td>93925</td><td>-</td><td>0</td><td>GGACAGGGACCGGGAGCTCG</td><td>GGG</td><td>||||||||[||||||||||||]PAM</td><td>NA</td><td>Intergenic</td><td>NA</td><td>NA</td></tr>
<tr><td>GU305914.1</td><td>94063</td><td>94085</td><td>-</td><td>0</td><td>GGACAGGGACCGGGAGCTCG</td><td>GGG</td><td>||||||||[||||||||||||]PAM</td><td>NA</td><td>Intergenic</td><td>NA</td><td>NA</td></tr>
<tr><td>GU305914.1</td><td>94031</td><td>94053</td><td>-</td><td>0</td><td>GGACAGGGACCGGGAGCTCG</td><td>GGG</td><td>||||||||[||||||||||||]PAM</td><td>NA</td><td>Intergenic</td><td>NA</td><td>NA</td></tr>
<tr><td>GU305914.1</td><td>93999</td><td>94021</td><td>-</td><td>0</td><td>GGACAGGGACCGGGAGCTCG</td><td>GGG</td><td>||||||||[||||||||||||]PAM</td><td>NA</td><td>Intergenic</td><td>NA</td><td>NA</td></tr>
<tr><td>GU305914.1</td><td>93967</td><td>93989</td><td>-</td><td>0</td><td>GGACAGGGACCGGGAGCTCG</td><td>GGG</td><td>||||||||[||||||||||||]PAM</td><td>NA</td><td>Intergenic</td><td>NA</td><td>NA</td></tr>
<tr><td>GU305914.1</td><td>93935</td><td>93957</td><td>-</td><td>0</td><td>GGACAGGGACCGGGAGCTCG</td><td>GGG</td><td>||||||||[||||||||||||]PAM</td><td>NA</td><td>Intergenic</td><td>NA</td><td>NA</td></tr>
<tr><td>GU305914.1</td><td>94095</td><td>94117</td><td>-</td><td>4</td><td>AAGCGGGGACCGGGAGCTCG</td><td>GGG</td><td>---|-|||[||||||||||||]PAM</td><td>NA</td><td>Intergenic</td><td>NA</td><td>NA</td></tr>
<tr><td></td><td></td><td></td><td></td><td></td><td></td><td></td><td></td><td></td><td></td><td></td><td></td></tr>
<tr><td>T26855</td><td>CAGGGACCGGGAGCTCGGGGAGG</td><td>1000</td><td></td><td></td><td></td><td></td><td></td><td></td><td></td><td></td><td></td></tr>
<tr><td>Oligo fwd</td><td>CACCCAGGGACCGGGAGCTCGGGG</td><td></td><td></td><td></td><td></td><td></td><td></td><td></td><td></td><td></td><td></td></tr>
<tr><td>Oligo rev</td><td>AAACCCCCGAGCTCCCGGTCCCTG</td><td></td><td></td><td></td><td></td><td></td><td></td><td></td><td></td><td></td><td></td></tr>
<tr><td>Chromosome</td><td>start</td><td>end</td><td>strand</td><td>MM</td><td>target_seq</td><td>PAM</td><td>alignment</td><td>distance</td><td>position</td><td>gene name</td><td>gene id</td></tr>
<tr><td>GU305914.1</td><td>93996</td><td>94018</td><td>-</td><td>0</td><td>CAGGGACCGGGAGCTCGGGG</td><td>AGG</td><td>||||||||[||||||||||||]PAM</td><td>NA</td><td>Intergenic</td><td>NA</td><td>NA</td></tr>
<tr><td>GU305914.1</td><td>93964</td><td>93986</td><td>-</td><td>0</td><td>CAGGGACCGGGAGCTCGGGG</td><td>AGG</td><td>||||||||[||||||||||||]PAM</td><td>NA</td><td>Intergenic</td><td>NA</td><td>NA</td></tr>
<tr><td>GU305914.1</td><td>93932</td><td>93954</td><td>-</td><td>0</td><td>CAGGGACCGGGAGCTCGGGG</td><td>AGG</td><td>||||||||[||||||||||||]PAM</td><td>NA</td><td>Intergenic</td><td>NA</td><td>NA</td></tr>
<tr><td>GU305914.1</td><td>94060</td><td>94082</td><td>-</td><td>0</td><td>CAGGGACCGGGAGCTCGGGG</td><td>AGG</td><td>||||||||[||||||||||||]PAM</td><td>NA</td><td>Intergenic</td><td>NA</td><td>NA</td></tr>
<tr><td>GU305914.1</td><td>94028</td><td>94050</td><td>-</td><td>0</td><td>CAGGGACCGGGAGCTCGGGG</td><td>AGG</td><td>||||||||[||||||||||||]PAM</td><td>NA</td><td>Intergenic</td><td>NA</td><td>NA</td></tr>
<tr><td>GU305914.1</td><td>94092</td><td>94114</td><td>-</td><td>1</td><td>CGGGGACCGGGAGCTCGGGG</td><td>AGG</td><td>|-||||||[||||||||||||]PAM</td><td>NA</td><td>Intergenic</td><td>NA</td><td>NA</td></tr>
<tr><td></td><td></td><td></td><td></td><td></td><td></td><td></td><td></td><td></td><td></td><td></td><td></td></tr>
<tr><td>T26856</td><td>AGGGACCGGGAGCTCGGGGAGGG</td><td>1000</td><td></td><td></td><td></td><td></td><td></td><td></td><td></td><td></td><td></td></tr>
<tr><td>Oligo fwd</td><td>CACCAGGGACCGGGAGCTCGGGGA</td><td></td><td></td><td></td><td></td><td></td><td></td><td></td><td></td><td></td><td></td></tr>
<tr><td>Oligo rev</td><td>AAACTCCCCGAGCTCCCGGTCCCT</td><td></td><td></td><td></td><td></td><td></td><td></td><td></td><td></td><td></td><td></td></tr>
<tr><td>Chromosome</td><td>start</td><td>end</td><td>strand</td><td>MM</td><td>target_seq</td><td>PAM</td><td>alignment</td><td>distance</td><td>position</td><td>gene name</td><td>gene id</td></tr>
<tr><td>GU305914.1</td><td>93995</td><td>94017</td><td>-</td><td>0</td><td>AGGGACCGGGAGCTCGGGGA</td><td>GGG</td><td>||||||||[||||||||||||]PAM</td><td>NA</td><td>Intergenic</td><td>NA</td><td>NA</td></tr>
<tr><td>GU305914.1</td><td>93963</td><td>93985</td><td>-</td><td>0</td><td>AGGGACCGGGAGCTCGGGGA</td><td>GGG</td><td>||||||||[||||||||||||]PAM</td><td>NA</td><td>Intergenic</td><td>NA</td><td>NA</td></tr>
<tr><td>GU305914.1</td><td>93931</td><td>93953</td><td>-</td><td>0</td><td>AGGGACCGGGAGCTCGGGGA</td><td>GGG</td><td>||||||||[||||||||||||]PAM</td><td>NA</td><td>Intergenic</td><td>NA</td><td>NA</td></tr>
<tr><td>GU305914.1</td><td>94059</td><td>94081</td><td>-</td><td>0</td><td>AGGGACCGGGAGCTCGGGGA</td><td>GGG</td><td>||||||||[||||||||||||]PAM</td><td>NA</td><td>Intergenic</td><td>NA</td><td>NA</td></tr>
<tr><td>GU305914.1</td><td>94027</td><td>94049</td><td>-</td><td>0</td><td>AGGGACCGGGAGCTCGGGGA</td><td>GGG</td><td>||||||||[||||||||||||]PAM</td><td>NA</td><td>Intergenic</td><td>NA</td><td>NA</td></tr>
<tr><td>GU305914.1</td><td>94091</td><td>94113</td><td>-</td><td>1</td><td>GGGGACCGGGAGCTCGGGGA</td><td>GGG</td><td>-|||||||[||||||||||||]PAM</td><td>NA</td><td>Intergenic</td><td>NA</td><td>NA</td></tr>
<tr><td></td><td></td><td></td><td></td><td></td><td></td><td></td><td></td><td></td><td></td><td></td><td></td></tr>
<tr><td>T26857</td><td>GGGAGCTCGGGGAGGGAAGCCGG</td><td>1000</td><td></td><td></td><td></td><td></td><td></td><td></td><td></td><td></td><td></td></tr>
<tr><td>Oligo fwd</td><td>CACCGGGAGCTCGGGGAGGGAAGC</td><td></td><td></td><td></td><td></td><td></td><td></td><td></td><td></td><td></td><td></td></tr>
<tr><td>Oligo rev</td><td>AAACGCTTCCCTCCCCGAGCTCCC</td><td></td><td></td><td></td><td></td><td></td><td></td><td></td><td></td><td></td><td></td></tr>
<tr><td>Chromosome</td><td>start</td><td>end</td><td>strand</td><td>MM</td><td>target_seq</td><td>PAM</td><td>alignment</td><td>distance</td><td>position</td><td>gene name</td><td>gene id</td></tr>
<tr><td>GU305914.1</td><td>94052</td><td>94074</td><td>-</td><td>0</td><td>GGGAGCTCGGGGAGGGAAGC</td><td>CGG</td><td>||||||||[||||||||||||]PAM</td><td>NA</td><td>Intergenic</td><td>NA</td><td>NA</td></tr>
<tr><td>GU305914.1</td><td>94020</td><td>94042</td><td>-</td><td>0</td><td>GGGAGCTCGGGGAGGGAAGC</td><td>CGG</td><td>||||||||[||||||||||||]PAM</td><td>NA</td><td>Intergenic</td><td>NA</td><td>NA</td></tr>
<tr><td>GU305914.1</td><td>93988</td><td>94010</td><td>-</td><td>0</td><td>GGGAGCTCGGGGAGGGAAGC</td><td>CGG</td><td>||||||||[||||||||||||]PAM</td><td>NA</td><td>Intergenic</td><td>NA</td><td>NA</td></tr>
<tr><td>GU305914.1</td><td>93956</td><td>93978</td><td>-</td><td>0</td><td>GGGAGCTCGGGGAGGGAAGC</td><td>CGG</td><td>||||||||[||||||||||||]PAM</td><td>NA</td><td>Intergenic</td><td>NA</td><td>NA</td></tr>
<tr><td>GU305914.1</td><td>93924</td><td>93946</td><td>-</td><td>0</td><td>GGGAGCTCGGGGAGGGAAGC</td><td>CGG</td><td>||||||||[||||||||||||]PAM</td><td>NA</td><td>Intergenic</td><td>NA</td><td>NA</td></tr>
<tr><td>GU305914.1</td><td>94084</td><td>94106</td><td>-</td><td>0</td><td>GGGAGCTCGGGGAGGGAAGC</td><td>CGG</td><td>||||||||[||||||||||||]PAM</td><td>NA</td><td>Intergenic</td><td>NA</td><td>NA</td></tr>
<tr><td>GU305914.1</td><td>93892</td><td>93914</td><td>-</td><td>2</td><td>GGGAGCTCGGGGAGAGGAGC</td><td>CGG</td><td>||||||||[||||||-|-|||]PAM</td><td>NA</td><td>Intergenic</td><td>NA</td><td>NA</td></tr>
<tr><td></td><td></td><td></td><td></td><td></td><td></td><td></td><td></td><td></td><td></td><td></td><td></td></tr>
<tr><td>T26858</td><td>CTCGGGGAGGGAAGCCGGACAGG</td><td>1000</td><td></td><td></td><td></td><td></td><td></td><td></td><td></td><td></td><td></td></tr>
<tr><td>Oligo fwd</td><td>CACCCTCGGGGAGGGAAGCCGGAC</td><td></td><td></td><td></td><td></td><td></td><td></td><td></td><td></td><td></td><td></td></tr>
<tr><td>Oligo rev</td><td>AAACGTCCGGCTTCCCTCCCCGAG</td><td></td><td></td><td></td><td></td><td></td><td></td><td></td><td></td><td></td><td></td></tr>
<tr><td>Chromosome</td><td>start</td><td>end</td><td>strand</td><td>MM</td><td>target_seq</td><td>PAM</td><td>alignment</td><td>distance</td><td>position</td><td>gene name</td><td>gene id</td></tr>
<tr><td>GU305914.1</td><td>93951</td><td>93973</td><td>-</td><td>0</td><td>CTCGGGGAGGGAAGCCGGAC</td><td>AGG</td><td>||||||||[||||||||||||]PAM</td><td>NA</td><td>Intergenic</td><td>NA</td><td>NA</td></tr>
<tr><td>GU305914.1</td><td>93919</td><td>93941</td><td>-</td><td>0</td><td>CTCGGGGAGGGAAGCCGGAC</td><td>AGG</td><td>||||||||[||||||||||||]PAM</td><td>NA</td><td>Intergenic</td><td>NA</td><td>NA</td></tr>
<tr><td>GU305914.1</td><td>94079</td><td>94101</td><td>-</td><td>0</td><td>CTCGGGGAGGGAAGCCGGAC</td><td>AGG</td><td>||||||||[||||||||||||]PAM</td><td>NA</td><td>Intergenic</td><td>NA</td><td>NA</td></tr>
<tr><td>GU305914.1</td><td>94047</td><td>94069</td><td>-</td><td>0</td><td>CTCGGGGAGGGAAGCCGGAC</td><td>AGG</td><td>||||||||[||||||||||||]PAM</td><td>NA</td><td>Intergenic</td><td>NA</td><td>NA</td></tr>
<tr><td>GU305914.1</td><td>94015</td><td>94037</td><td>-</td><td>0</td><td>CTCGGGGAGGGAAGCCGGAC</td><td>AGG</td><td>||||||||[||||||||||||]PAM</td><td>NA</td><td>Intergenic</td><td>NA</td><td>NA</td></tr>
<tr><td>GU305914.1</td><td>93983</td><td>94005</td><td>-</td><td>0</td><td>CTCGGGGAGGGAAGCCGGAC</td><td>AGG</td><td>||||||||[||||||||||||]PAM</td><td>NA</td><td>Intergenic</td><td>NA</td><td>NA</td></tr>
<tr><td>GU305914.1</td><td>93887</td><td>93909</td><td>-</td><td>2</td><td>CTCGGGGAGAGGAGCCGGAC</td><td>AGG</td><td>||||||||[|-|-||||||||]PAM</td><td>NA</td><td>Intergenic</td><td>NA</td><td>NA</td></tr>
<tr><td></td><td></td><td></td><td></td><td></td><td></td><td></td><td></td><td></td><td></td><td></td><td></td></tr>
<tr><td>T26859</td><td>TCGGGGAGGGAAGCCGGACAGGG</td><td>1000</td><td></td><td></td><td></td><td></td><td></td><td></td><td></td><td></td><td></td></tr>
<tr><td>Oligo fwd</td><td>CACCTCGGGGAGGGAAGCCGGACA</td><td></td><td></td><td></td><td></td><td></td><td></td><td></td><td></td><td></td><td></td></tr>
<tr><td>Oligo rev</td><td>AAACTGTCCGGCTTCCCTCCCCGA</td><td></td><td></td><td></td><td></td><td></td><td></td><td></td><td></td><td></td><td></td></tr>
<tr><td>Chromosome</td><td>start</td><td>end</td><td>strand</td><td>MM</td><td>target_seq</td><td>PAM</td><td>alignment</td><td>distance</td><td>position</td><td>gene name</td><td>gene id</td></tr>
<tr><td>GU305914.1</td><td>94078</td><td>94100</td><td>-</td><td>0</td><td>TCGGGGAGGGAAGCCGGACA</td><td>GGG</td><td>||||||||[||||||||||||]PAM</td><td>NA</td><td>Intergenic</td><td>NA</td><td>NA</td></tr>
<tr><td>GU305914.1</td><td>94046</td><td>94068</td><td>-</td><td>0</td><td>TCGGGGAGGGAAGCCGGACA</td><td>GGG</td><td>||||||||[||||||||||||]PAM</td><td>NA</td><td>Intergenic</td><td>NA</td><td>NA</td></tr>
<tr><td>GU305914.1</td><td>94014</td><td>94036</td><td>-</td><td>0</td><td>TCGGGGAGGGAAGCCGGACA</td><td>GGG</td><td>||||||||[||||||||||||]PAM</td><td>NA</td><td>Intergenic</td><td>NA</td><td>NA</td></tr>
<tr><td>GU305914.1</td><td>93982</td><td>94004</td><td>-</td><td>0</td><td>TCGGGGAGGGAAGCCGGACA</td><td>GGG</td><td>||||||||[||||||||||||]PAM</td><td>NA</td><td>Intergenic</td><td>NA</td><td>NA</td></tr>
<tr><td>GU305914.1</td><td>93950</td><td>93972</td><td>-</td><td>0</td><td>TCGGGGAGGGAAGCCGGACA</td><td>GGG</td><td>||||||||[||||||||||||]PAM</td><td>NA</td><td>Intergenic</td><td>NA</td><td>NA</td></tr>
<tr><td>GU305914.1</td><td>93918</td><td>93940</td><td>-</td><td>0</td><td>TCGGGGAGGGAAGCCGGACA</td><td>GGG</td><td>||||||||[||||||||||||]PAM</td><td>NA</td><td>Intergenic</td><td>NA</td><td>NA</td></tr>
<tr><td></td><td></td><td></td><td></td><td></td><td></td><td></td><td></td><td></td><td></td><td></td><td></td></tr>
<tr><td>T26860</td><td>GAGGGAAGCCGGACAGGGACCGG</td><td>1000</td><td></td><td></td><td></td><td></td><td></td><td></td><td></td><td></td><td></td></tr>
<tr><td>Oligo fwd</td><td>CACCGAGGGAAGCCGGACAGGGAC</td><td></td><td></td><td></td><td></td><td></td><td></td><td></td><td></td><td></td><td></td></tr>
</table>

<table>
<tr><td>Oligo rev</td><td>AAACGTCCCTGTCCGGCTTCCCTC</td><td></td><td></td><td></td><td></td><td></td><td></td><td></td><td></td><td></td><td></td></tr>
<tr><td>Chromosome</td><td>start</td><td>end</td><td>strand</td><td>MM</td><td>target_seq</td><td>PAM</td><td>alignment</td><td>distance</td><td>position</td><td>gene name</td><td>gene id</td></tr>
<tr><td>GU305914.1</td><td>93913</td><td>93935</td><td>-</td><td>0</td><td>GAGGGAAGCCGGACAGGGAC</td><td>CGG</td><td>||||||||[||||||||||||]PAM</td><td>NA</td><td>Intergenic</td><td>NA</td><td>NA</td></tr>
<tr><td>GU305914.1</td><td>94073</td><td>94095</td><td>-</td><td>0</td><td>GAGGGAAGCCGGACAGGGAC</td><td>CGG</td><td>||||||||[||||||||||||]PAM</td><td>NA</td><td>Intergenic</td><td>NA</td><td>NA</td></tr>
<tr><td>GU305914.1</td><td>94041</td><td>94063</td><td>-</td><td>0</td><td>GAGGGAAGCCGGACAGGGAC</td><td>CGG</td><td>||||||||[||||||||||||]PAM</td><td>NA</td><td>Intergenic</td><td>NA</td><td>NA</td></tr>
<tr><td>GU305914.1</td><td>94009</td><td>94031</td><td>-</td><td>0</td><td>GAGGGAAGCCGGACAGGGAC</td><td>CGG</td><td>||||||||[||||||||||||]PAM</td><td>NA</td><td>Intergenic</td><td>NA</td><td>NA</td></tr>
<tr><td>GU305914.1</td><td>93977</td><td>93999</td><td>-</td><td>0</td><td>GAGGGAAGCCGGACAGGGAC</td><td>CGG</td><td>||||||||[||||||||||||]PAM</td><td>NA</td><td>Intergenic</td><td>NA</td><td>NA</td></tr>
<tr><td>GU305914.1</td><td>93945</td><td>93967</td><td>-</td><td>0</td><td>GAGGGAAGCCGGACAGGGAC</td><td>CGG</td><td>||||||||[||||||||||||]PAM</td><td>NA</td><td>Intergenic</td><td>NA</td><td>NA</td></tr>
<tr><td>GU305914.1</td><td>93881</td><td>93903</td><td>-</td><td>3</td><td>GAGAGGAGCCGGACAGGAAC</td><td>CGG</td><td>|||-|-||[|||||||||-||]PAM</td><td>NA</td><td>Intergenic</td><td>NA</td><td>NA</td></tr>
<tr><td></td><td></td><td></td><td></td><td></td><td></td><td></td><td></td><td></td><td></td><td></td><td></td></tr>
<tr><td>T26861</td><td>AGGGAAGCCGGACAGGGACCGGG</td><td>1000</td><td></td><td></td><td></td><td></td><td></td><td></td><td></td><td></td><td></td></tr>
<tr><td>Oligo fwd</td><td>CACCAGGGAAGCCGGACAGGGACC</td><td></td><td></td><td></td><td></td><td></td><td></td><td></td><td></td><td></td><td></td></tr>
<tr><td>Oligo rev</td><td>AAACGGTCCCTGTCCGGCTTCCCT</td><td></td><td></td><td></td><td></td><td></td><td></td><td></td><td></td><td></td><td></td></tr>
<tr><td>Chromosome</td><td>start</td><td>end</td><td>strand</td><td>MM</td><td>target_seq</td><td>PAM</td><td>alignment</td><td>distance</td><td>position</td><td>gene name</td><td>gene id</td></tr>
<tr><td>GU305914.1</td><td>93976</td><td>93998</td><td>-</td><td>0</td><td>AGGGAAGCCGGACAGGGACC</td><td>GGG</td><td>||||||||[||||||||||||]PAM</td><td>NA</td><td>Intergenic</td><td>NA</td><td>NA</td></tr>
<tr><td>GU305914.1</td><td>93944</td><td>93966</td><td>-</td><td>0</td><td>AGGGAAGCCGGACAGGGACC</td><td>GGG</td><td>||||||||[||||||||||||]PAM</td><td>NA</td><td>Intergenic</td><td>NA</td><td>NA</td></tr>
<tr><td>GU305914.1</td><td>93912</td><td>93934</td><td>-</td><td>0</td><td>AGGGAAGCCGGACAGGGACC</td><td>GGG</td><td>||||||||[||||||||||||]PAM</td><td>NA</td><td>Intergenic</td><td>NA</td><td>NA</td></tr>
<tr><td>GU305914.1</td><td>94072</td><td>94094</td><td>-</td><td>0</td><td>AGGGAAGCCGGACAGGGACC</td><td>GGG</td><td>||||||||[||||||||||||]PAM</td><td>NA</td><td>Intergenic</td><td>NA</td><td>NA</td></tr>
<tr><td>GU305914.1</td><td>94040</td><td>94062</td><td>-</td><td>0</td><td>AGGGAAGCCGGACAGGGACC</td><td>GGG</td><td>||||||||[||||||||||||]PAM</td><td>NA</td><td>Intergenic</td><td>NA</td><td>NA</td></tr>
<tr><td>GU305914.1</td><td>94008</td><td>94030</td><td>-</td><td>0</td><td>AGGGAAGCCGGACAGGGACC</td><td>GGG</td><td>||||||||[||||||||||||]PAM</td><td>NA</td><td>Intergenic</td><td>NA</td><td>NA</td></tr>
<tr><td>GU305914.1</td><td>93880</td><td>93902</td><td>-</td><td>3</td><td>AGAGGAGCCGGACAGGAACC</td><td>GGG</td><td>||-|-|||[||||||||-|||]PAM</td><td>NA</td><td>Intergenic</td><td>NA</td><td>NA</td></tr>
<tr><td></td><td></td><td></td><td></td><td></td><td></td><td></td><td></td><td></td><td></td><td></td><td></td></tr>
<tr><td>T26862</td><td>CCGGACAGGGACCGGGAGCTCGG</td><td>1000</td><td></td><td></td><td></td><td></td><td></td><td></td><td></td><td></td><td></td></tr>
<tr><td>Oligo fwd</td><td>CACCCCGGACAGGGACCGGGAGCT</td><td></td><td></td><td></td><td></td><td></td><td></td><td></td><td></td><td></td><td></td></tr>
<tr><td>Oligo rev</td><td>AAACAGCTCCCGGTCCCTGTCCGG</td><td></td><td></td><td></td><td></td><td></td><td></td><td></td><td></td><td></td><td></td></tr>
<tr><td>Chromosome</td><td>start</td><td>end</td><td>strand</td><td>MM</td><td>target_seq</td><td>PAM</td><td>alignment</td><td>distance</td><td>position</td><td>gene name</td><td>gene id</td></tr>
<tr><td>GU305914.1</td><td>94001</td><td>94023</td><td>-</td><td>0</td><td>CCGGACAGGGACCGGGAGCT</td><td>CGG</td><td>||||||||[||||||||||||]PAM</td><td>NA</td><td>Intergenic</td><td>NA</td><td>NA</td></tr>
<tr><td>GU305914.1</td><td>93969</td><td>93991</td><td>-</td><td>0</td><td>CCGGACAGGGACCGGGAGCT</td><td>CGG</td><td>||||||||[||||||||||||]PAM</td><td>NA</td><td>Intergenic</td><td>NA</td><td>NA</td></tr>
<tr><td>GU305914.1</td><td>93937</td><td>93959</td><td>-</td><td>0</td><td>CCGGACAGGGACCGGGAGCT</td><td>CGG</td><td>||||||||[||||||||||||]PAM</td><td>NA</td><td>Intergenic</td><td>NA</td><td>NA</td></tr>
<tr><td>GU305914.1</td><td>93905</td><td>93927</td><td>-</td><td>0</td><td>CCGGACAGGGACCGGGAGCT</td><td>CGG</td><td>||||||||[||||||||||||]PAM</td><td>NA</td><td>Intergenic</td><td>NA</td><td>NA</td></tr>
<tr><td>GU305914.1</td><td>94065</td><td>94087</td><td>-</td><td>0</td><td>CCGGACAGGGACCGGGAGCT</td><td>CGG</td><td>||||||||[||||||||||||]PAM</td><td>NA</td><td>Intergenic</td><td>NA</td><td>NA</td></tr>
<tr><td>GU305914.1</td><td>94033</td><td>94055</td><td>-</td><td>0</td><td>CCGGACAGGGACCGGGAGCT</td><td>CGG</td><td>||||||||[||||||||||||]PAM</td><td>NA</td><td>Intergenic</td><td>NA</td><td>NA</td></tr>
<tr><td>GU305914.1</td><td>93873</td><td>93895</td><td>-</td><td>2</td><td>CCGGACAGGAACCGGGAGCG</td><td>GGG</td><td>||||||||[|-|||||||||-]PAM</td><td>NA</td><td>Intergenic</td><td>NA</td><td>NA</td></tr>
<tr><td></td><td></td><td></td><td></td><td></td><td></td><td></td><td></td><td></td><td></td><td></td><td></td></tr>
<tr><td>T26863</td><td>CGGACAGGGACCGGGAGCTCGGG</td><td>1000</td><td></td><td></td><td></td><td></td><td></td><td></td><td></td><td></td><td></td></tr>
<tr><td>Oligo fwd</td><td>CACCCGGACAGGGACCGGGAGCTC</td><td></td><td></td><td></td><td></td><td></td><td></td><td></td><td></td><td></td><td></td></tr>
<tr><td>Oligo rev</td><td>AAACGAGCTCCCGGTCCCTGTCCG</td><td></td><td></td><td></td><td></td><td></td><td></td><td></td><td></td><td></td><td></td></tr>
<tr><td>Chromosome</td><td>start</td><td>end</td><td>strand</td><td>MM</td><td>target_seq</td><td>PAM</td><td>alignment</td><td>distance</td><td>position</td><td>gene name</td><td>gene id</td></tr>
<tr><td>GU305914.1</td><td>93968</td><td>93990</td><td>-</td><td>0</td><td>CGGACAGGGACCGGGAGCTC</td><td>GGG</td><td>||||||||[||||||||||||]PAM</td><td>NA</td><td>Intergenic</td><td>NA</td><td>NA</td></tr>
<tr><td>GU305914.1</td><td>93936</td><td>93958</td><td>-</td><td>0</td><td>CGGACAGGGACCGGGAGCTC</td><td>GGG</td><td>||||||||[||||||||||||]PAM</td><td>NA</td><td>Intergenic</td><td>NA</td><td>NA</td></tr>
<tr><td>GU305914.1</td><td>93904</td><td>93926</td><td>-</td><td>0</td><td>CGGACAGGGACCGGGAGCTC</td><td>GGG</td><td>||||||||[||||||||||||]PAM</td><td>NA</td><td>Intergenic</td><td>NA</td><td>NA</td></tr>
<tr><td>GU305914.1</td><td>94064</td><td>94086</td><td>-</td><td>0</td><td>CGGACAGGGACCGGGAGCTC</td><td>GGG</td><td>||||||||[||||||||||||]PAM</td><td>NA</td><td>Intergenic</td><td>NA</td><td>NA</td></tr>
<tr><td>GU305914.1</td><td>94032</td><td>94054</td><td>-</td><td>0</td><td>CGGACAGGGACCGGGAGCTC</td><td>GGG</td><td>||||||||[||||||||||||]PAM</td><td>NA</td><td>Intergenic</td><td>NA</td><td>NA</td></tr>
<tr><td>GU305914.1</td><td>94000</td><td>94022</td><td>-</td><td>0</td><td>CGGACAGGGACCGGGAGCTC</td><td>GGG</td><td>||||||||[||||||||||||]PAM</td><td>NA</td><td>Intergenic</td><td>NA</td><td>NA</td></tr>
<tr><td></td><td></td><td></td><td></td><td></td><td></td><td></td><td></td><td></td><td></td><td></td><td></td></tr>
<tr><td>T26864</td><td>GGACAGGGACCGGGAGCTCGGGG</td><td>1000</td><td></td><td></td><td></td><td></td><td></td><td></td><td></td><td></td><td></td></tr>
<tr><td>Oligo fwd</td><td>CACCGGACAGGGACCGGGAGCTCG</td><td></td><td></td><td></td><td></td><td></td><td></td><td></td><td></td><td></td><td></td></tr>
<tr><td>Oligo rev</td><td>AAACCGAGCTCCCGGTCCCTGTCC</td><td></td><td></td><td></td><td></td><td></td><td></td><td></td><td></td><td></td><td></td></tr>
<tr><td>Chromosome</td><td>start</td><td>end</td><td>strand</td><td>MM</td><td>target_seq</td><td>PAM</td><td>alignment</td><td>distance</td><td>position</td><td>gene name</td><td>gene id</td></tr>
<tr><td>GU305914.1</td><td>93903</td><td>93925</td><td>-</td><td>0</td><td>GGACAGGGACCGGGAGCTCG</td><td>GGG</td><td>||||||||[||||||||||||]PAM</td><td>NA</td><td>Intergenic</td><td>NA</td><td>NA</td></tr>
<tr><td>GU305914.1</td><td>94063</td><td>94085</td><td>-</td><td>0</td><td>GGACAGGGACCGGGAGCTCG</td><td>GGG</td><td>||||||||[||||||||||||]PAM</td><td>NA</td><td>Intergenic</td><td>NA</td><td>NA</td></tr>
<tr><td>GU305914.1</td><td>94031</td><td>94053</td><td>-</td><td>0</td><td>GGACAGGGACCGGGAGCTCG</td><td>GGG</td><td>||||||||[||||||||||||]PAM</td><td>NA</td><td>Intergenic</td><td>NA</td><td>NA</td></tr>
<tr><td>GU305914.1</td><td>93999</td><td>94021</td><td>-</td><td>0</td><td>GGACAGGGACCGGGAGCTCG</td><td>GGG</td><td>||||||||[||||||||||||]PAM</td><td>NA</td><td>Intergenic</td><td>NA</td><td>NA</td></tr>
<tr><td>GU305914.1</td><td>93967</td><td>93989</td><td>-</td><td>0</td><td>GGACAGGGACCGGGAGCTCG</td><td>GGG</td><td>||||||||[||||||||||||]PAM</td><td>NA</td><td>Intergenic</td><td>NA</td><td>NA</td></tr>
<tr><td>GU305914.1</td><td>93935</td><td>93957</td><td>-</td><td>0</td><td>GGACAGGGACCGGGAGCTCG</td><td>GGG</td><td>||||||||[||||||||||||]PAM</td><td>NA</td><td>Intergenic</td><td>NA</td><td>NA</td></tr>
<tr><td>GU305914.1</td><td>94095</td><td>94117</td><td>-</td><td>4</td><td>AAGCGGGGACCGGGAGCTCG</td><td>GGG</td><td>---|-|||[||||||||||||]PAM</td><td>NA</td><td>Intergenic</td><td>NA</td><td>NA</td></tr>
<tr><td></td><td></td><td></td><td></td><td></td><td></td><td></td><td></td><td></td><td></td><td></td><td></td></tr>
<tr><td>T26865</td><td>CAGGGACCGGGAGCTCGGGGAGG</td><td>1000</td><td></td><td></td><td></td><td></td><td></td><td></td><td></td><td></td><td></td></tr>
<tr><td>Oligo fwd</td><td>CACCCAGGGACCGGGAGCTCGGGG</td><td></td><td></td><td></td><td></td><td></td><td></td><td></td><td></td><td></td><td></td></tr>
<tr><td>Oligo rev</td><td>AAACCCCCGAGCTCCCGGTCCCTG</td><td></td><td></td><td></td><td></td><td></td><td></td><td></td><td></td><td></td><td></td></tr>
<tr><td>Chromosome</td><td>start</td><td>end</td><td>strand</td><td>MM</td><td>target_seq</td><td>PAM</td><td>alignment</td><td>distance</td><td>position</td><td>gene name</td><td>gene id</td></tr>
<tr><td>GU305914.1</td><td>93996</td><td>94018</td><td>-</td><td>0</td><td>CAGGGACCGGGAGCTCGGGG</td><td>AGG</td><td>||||||||[||||||||||||]PAM</td><td>NA</td><td>Intergenic</td><td>NA</td><td>NA</td></tr>
<tr><td>GU305914.1</td><td>93964</td><td>93986</td><td>-</td><td>0</td><td>CAGGGACCGGGAGCTCGGGG</td><td>AGG</td><td>||||||||[||||||||||||]PAM</td><td>NA</td><td>Intergenic</td><td>NA</td><td>NA</td></tr>
<tr><td>GU305914.1</td><td>93932</td><td>93954</td><td>-</td><td>0</td><td>CAGGGACCGGGAGCTCGGGG</td><td>AGG</td><td>||||||||[||||||||||||]PAM</td><td>NA</td><td>Intergenic</td><td>NA</td><td>NA</td></tr>
<tr><td>GU305914.1</td><td>94060</td><td>94082</td><td>-</td><td>0</td><td>CAGGGACCGGGAGCTCGGGG</td><td>AGG</td><td>||||||||[||||||||||||]PAM</td><td>NA</td><td>Intergenic</td><td>NA</td><td>NA</td></tr>
<tr><td>GU305914.1</td><td>94028</td><td>94050</td><td>-</td><td>0</td><td>CAGGGACCGGGAGCTCGGGG</td><td>AGG</td><td>||||||||[||||||||||||]PAM</td><td>NA</td><td>Intergenic</td><td>NA</td><td>NA</td></tr>
<tr><td>GU305914.1</td><td>94092</td><td>94114</td><td>-</td><td>1</td><td>CGGGGACCGGGAGCTCGGGG</td><td>AGG</td><td>|-||||||[||||||||||||]PAM</td><td>NA</td><td>Intergenic</td><td>NA</td><td>NA</td></tr>
<tr><td></td><td></td><td></td><td></td><td></td><td></td><td></td><td></td><td></td><td></td><td></td><td></td></tr>
<tr><td>T26866</td><td>AGGGACCGGGAGCTCGGGGAGGG</td><td>1000</td><td></td><td></td><td></td><td></td><td></td><td></td><td></td><td></td><td></td></tr>
<tr><td>Oligo fwd</td><td>CACCAGGGACCGGGAGCTCGGGGA</td><td></td><td></td><td></td><td></td><td></td><td></td><td></td><td></td><td></td><td></td></tr>
<tr><td>Oligo rev</td><td>AAACTCCCCGAGCTCCCGGTCCCT</td><td></td><td></td><td></td><td></td><td></td><td></td><td></td><td></td><td></td><td></td></tr>
<tr><td>Chromosome</td><td>start</td><td>end</td><td>strand</td><td>MM</td><td>target_seq</td><td>PAM</td><td>alignment</td><td>distance</td><td>position</td><td>gene name</td><td>gene id</td></tr>
<tr><td>GU305914.1</td><td>93995</td><td>94017</td><td>-</td><td>0</td><td>AGGGACCGGGAGCTCGGGGA</td><td>GGG</td><td>||||||||[||||||||||||]PAM</td><td>NA</td><td>Intergenic</td><td>NA</td><td>NA</td></tr>
<tr><td>GU305914.1</td><td>93963</td><td>93985</td><td>-</td><td>0</td><td>AGGGACCGGGAGCTCGGGGA</td><td>GGG</td><td>||||||||[||||||||||||]PAM</td><td>NA</td><td>Intergenic</td><td>NA</td><td>NA</td></tr>
<tr><td>GU305914.1</td><td>93931</td><td>93953</td><td>-</td><td>0</td><td>AGGGACCGGGAGCTCGGGGA</td><td>GGG</td><td>||||||||[||||||||||||]PAM</td><td>NA</td><td>Intergenic</td><td>NA</td><td>NA</td></tr>
<tr><td>GU305914.1</td><td>94059</td><td>94081</td><td>-</td><td>0</td><td>AGGGACCGGGAGCTCGGGGA</td><td>GGG</td><td>||||||||[||||||||||||]PAM</td><td>NA</td><td>Intergenic</td><td>NA</td><td>NA</td></tr>
<tr><td>GU305914.1</td><td>94027</td><td>94049</td><td>-</td><td>0</td><td>AGGGACCGGGAGCTCGGGGA</td><td>GGG</td><td>||||||||[||||||||||||]PAM</td><td>NA</td><td>Intergenic</td><td>NA</td><td>NA</td></tr>
<tr><td>GU305914.1</td><td>94091</td><td>94113</td><td>-</td><td>1</td><td>GGGGACCGGGAGCTCGGGGA</td><td>GGG</td><td>-|||||||[||||||||||||]PAM</td><td>NA</td><td>Intergenic</td><td>NA</td><td>NA</td></tr>
<tr><td></td><td></td><td></td><td></td><td></td><td></td><td></td><td></td><td></td><td></td><td></td><td></td></tr>
<tr><td>T26867</td><td>GGGAGCTCGGGGAGGGAAGCCGG</td><td>1000</td><td></td><td></td><td></td><td></td><td></td><td></td><td></td><td></td><td></td></tr>
<tr><td>Oligo fwd</td><td>CACCGGGAGCTCGGGGAGGGAAGC</td><td></td><td></td><td></td><td></td><td></td><td></td><td></td><td></td><td></td><td></td></tr>
<tr><td>Oligo rev</td><td>AAACGCTTCCCTCCCCGAGCTCCC</td><td></td><td></td><td></td><td></td><td></td><td></td><td></td><td></td><td></td><td></td></tr>
<tr><td>Chromosome</td><td>start</td><td>end</td><td>strand</td><td>MM</td><td>target_seq</td><td>PAM</td><td>alignment</td><td>distance</td><td>position</td><td>gene name</td><td>gene id</td></tr>
<tr><td>GU305914.1</td><td>94052</td><td>94074</td><td>-</td><td>0</td><td>GGGAGCTCGGGGAGGGAAGC</td><td>CGG</td><td>||||||||[||||||||||||]PAM</td><td>NA</td><td>Intergenic</td><td>NA</td><td>NA</td></tr>
<tr><td>GU305914.1</td><td>94020</td><td>94042</td><td>-</td><td>0</td><td>GGGAGCTCGGGGAGGGAAGC</td><td>CGG</td><td>||||||||[||||||||||||]PAM</td><td>NA</td><td>Intergenic</td><td>NA</td><td>NA</td></tr>
<tr><td>GU305914.1</td><td>93988</td><td>94010</td><td>-</td><td>0</td><td>GGGAGCTCGGGGAGGGAAGC</td><td>CGG</td><td>||||||||[||||||||||||]PAM</td><td>NA</td><td>Intergenic</td><td>NA</td><td>NA</td></tr>
<tr><td>GU305914.1</td><td>93956</td><td>93978</td><td>-</td><td>0</td><td>GGGAGCTCGGGGAGGGAAGC</td><td>CGG</td><td>||||||||[||||||||||||]PAM</td><td>NA</td><td>Intergenic</td><td>NA</td><td>NA</td></tr>
<tr><td>GU305914.1</td><td>93924</td><td>93946</td><td>-</td><td>0</td><td>GGGAGCTCGGGGAGGGAAGC</td><td>CGG</td><td>||||||||[||||||||||||]PAM</td><td>NA</td><td>Intergenic</td><td>NA</td><td>NA</td></tr>
<tr><td>GU305914.1</td><td>94084</td><td>94106</td><td>-</td><td>0</td><td>GGGAGCTCGGGGAGGGAAGC</td><td>CGG</td><td>||||||||[||||||||||||]PAM</td><td>NA</td><td>Intergenic</td><td>NA</td><td>NA</td></tr>
</table>

<table>
<tr><td>GU305914.1</td><td></td><td>93892</td><td>93914</td><td>-</td><td>2</td><td>GGGAGCTCGGGGAGAGGAGC</td><td>CGG</td><td>||||||||[||||||-|-|||]PAM</td><td>NA</td><td>Intergenic</td><td>NA</td><td>NA</td></tr>
<tr><td></td><td></td><td></td><td></td><td></td><td></td><td></td><td></td><td></td><td></td><td></td><td></td><td></td></tr>
<tr><td>T26868</td><td>CTCGGGGAGGGAAGCCGGACAGG</td><td></td><td>1000</td><td></td><td></td><td></td><td></td><td></td><td></td><td></td><td></td><td></td></tr>
<tr><td>Oligo fwd</td><td>CACCCTCGGGGAGGGAAGCCGGAC</td><td></td><td></td><td></td><td></td><td></td><td></td><td></td><td></td><td></td><td></td><td></td></tr>
<tr><td>Oligo rev</td><td>AAACGTCCGGCTTCCCTCCCCGAG</td><td></td><td></td><td></td><td></td><td></td><td></td><td></td><td></td><td></td><td></td><td></td></tr>
<tr><td>Chromosome</td><td>start</td><td></td><td>end</td><td>strand</td><td>MM</td><td>target_seq</td><td>PAM</td><td>alignment</td><td>distance</td><td>position</td><td>gene name</td><td>gene id</td></tr>
<tr><td>GU305914.1</td><td></td><td>93951</td><td>93973</td><td>-</td><td>0</td><td>CTCGGGGAGGGAAGCCGGAC</td><td>AGG</td><td>||||||||[||||||||||||]PAM</td><td>NA</td><td>Intergenic</td><td>NA</td><td>NA</td></tr>
<tr><td>GU305914.1</td><td></td><td>93919</td><td>93941</td><td>-</td><td>0</td><td>CTCGGGGAGGGAAGCCGGAC</td><td>AGG</td><td>||||||||[||||||||||||]PAM</td><td>NA</td><td>Intergenic</td><td>NA</td><td>NA</td></tr>
<tr><td>GU305914.1</td><td></td><td>94079</td><td>94101</td><td>-</td><td>0</td><td>CTCGGGGAGGGAAGCCGGAC</td><td>AGG</td><td>||||||||[||||||||||||]PAM</td><td>NA</td><td>Intergenic</td><td>NA</td><td>NA</td></tr>
<tr><td>GU305914.1</td><td></td><td>94047</td><td>94069</td><td>-</td><td>0</td><td>CTCGGGGAGGGAAGCCGGAC</td><td>AGG</td><td>||||||||[||||||||||||]PAM</td><td>NA</td><td>Intergenic</td><td>NA</td><td>NA</td></tr>
<tr><td>GU305914.1</td><td></td><td>94015</td><td>94037</td><td>-</td><td>0</td><td>CTCGGGGAGGGAAGCCGGAC</td><td>AGG</td><td>||||||||[||||||||||||]PAM</td><td>NA</td><td>Intergenic</td><td>NA</td><td>NA</td></tr>
<tr><td>GU305914.1</td><td></td><td>93983</td><td>94005</td><td>-</td><td>0</td><td>CTCGGGGAGGGAAGCCGGAC</td><td>AGG</td><td>||||||||[||||||||||||]PAM</td><td>NA</td><td>Intergenic</td><td>NA</td><td>NA</td></tr>
<tr><td>GU305914.1</td><td></td><td>93887</td><td>93909</td><td>-</td><td>2</td><td>CTCGGGGAGAGGAGCCGGAC</td><td>AGG</td><td>||||||||[|-|-||||||||]PAM</td><td>NA</td><td>Intergenic</td><td>NA</td><td>NA</td></tr>
<tr><td></td><td></td><td></td><td></td><td></td><td></td><td></td><td></td><td></td><td></td><td></td><td></td><td></td></tr>
<tr><td>T26869</td><td>TCGGGGAGGGAAGCCGGACAGGG</td><td></td><td>1000</td><td></td><td></td><td></td><td></td><td></td><td></td><td></td><td></td><td></td></tr>
<tr><td>Oligo fwd</td><td>CACCTCGGGGAGGGAAGCCGGACA</td><td></td><td></td><td></td><td></td><td></td><td></td><td></td><td></td><td></td><td></td><td></td></tr>
<tr><td>Oligo rev</td><td>AAACTGTCCGGCTTCCCTCCCCGA</td><td></td><td></td><td></td><td></td><td></td><td></td><td></td><td></td><td></td><td></td><td></td></tr>
<tr><td>Chromosome</td><td>start</td><td></td><td>end</td><td>strand</td><td>MM</td><td>target_seq</td><td>PAM</td><td>alignment</td><td>distance</td><td>position</td><td>gene name</td><td>gene id</td></tr>
<tr><td>GU305914.1</td><td></td><td>94078</td><td>94100</td><td>-</td><td>0</td><td>TCGGGGAGGGAAGCCGGACA</td><td>GGG</td><td>||||||||[||||||||||||]PAM</td><td>NA</td><td>Intergenic</td><td>NA</td><td>NA</td></tr>
<tr><td>GU305914.1</td><td></td><td>94046</td><td>94068</td><td>-</td><td>0</td><td>TCGGGGAGGGAAGCCGGACA</td><td>GGG</td><td>||||||||[||||||||||||]PAM</td><td>NA</td><td>Intergenic</td><td>NA</td><td>NA</td></tr>
<tr><td>GU305914.1</td><td></td><td>94014</td><td>94036</td><td>-</td><td>0</td><td>TCGGGGAGGGAAGCCGGACA</td><td>GGG</td><td>||||||||[||||||||||||]PAM</td><td>NA</td><td>Intergenic</td><td>NA</td><td>NA</td></tr>
<tr><td>GU305914.1</td><td></td><td>93982</td><td>94004</td><td>-</td><td>0</td><td>TCGGGGAGGGAAGCCGGACA</td><td>GGG</td><td>||||||||[||||||||||||]PAM</td><td>NA</td><td>Intergenic</td><td>NA</td><td>NA</td></tr>
<tr><td>GU305914.1</td><td></td><td>93950</td><td>93972</td><td>-</td><td>0</td><td>TCGGGGAGGGAAGCCGGACA</td><td>GGG</td><td>||||||||[||||||||||||]PAM</td><td>NA</td><td>Intergenic</td><td>NA</td><td>NA</td></tr>
<tr><td>GU305914.1</td><td></td><td>93918</td><td>93940</td><td>-</td><td>0</td><td>TCGGGGAGGGAAGCCGGACA</td><td>GGG</td><td>||||||||[||||||||||||]PAM</td><td>NA</td><td>Intergenic</td><td>NA</td><td>NA</td></tr>
<tr><td></td><td></td><td></td><td></td><td></td><td></td><td></td><td></td><td></td><td></td><td></td><td></td><td></td></tr>
<tr><td>T26870</td><td>GAGGGAAGCCGGACAGGGACCGG</td><td></td><td>1000</td><td></td><td></td><td></td><td></td><td></td><td></td><td></td><td></td><td></td></tr>
<tr><td>Oligo fwd</td><td>CACCGAGGGAAGCCGGACAGGGAC</td><td></td><td></td><td></td><td></td><td></td><td></td><td></td><td></td><td></td><td></td><td></td></tr>
<tr><td>Oligo rev</td><td>AAACGTCCCTGTCCGGCTTCCCTC</td><td></td><td></td><td></td><td></td><td></td><td></td><td></td><td></td><td></td><td></td><td></td></tr>
<tr><td>Chromosome</td><td>start</td><td></td><td>end</td><td>strand</td><td>MM</td><td>target_seq</td><td>PAM</td><td>alignment</td><td>distance</td><td>position</td><td>gene name</td><td>gene id</td></tr>
<tr><td>GU305914.1</td><td></td><td>93913</td><td>93935</td><td>-</td><td>0</td><td>GAGGGAAGCCGGACAGGGAC</td><td>CGG</td><td>||||||||[||||||||||||]PAM</td><td>NA</td><td>Intergenic</td><td>NA</td><td>NA</td></tr>
<tr><td>GU305914.1</td><td></td><td>94073</td><td>94095</td><td>-</td><td>0</td><td>GAGGGAAGCCGGACAGGGAC</td><td>CGG</td><td>||||||||[||||||||||||]PAM</td><td>NA</td><td>Intergenic</td><td>NA</td><td>NA</td></tr>
<tr><td>GU305914.1</td><td></td><td>94041</td><td>94063</td><td>-</td><td>0</td><td>GAGGGAAGCCGGACAGGGAC</td><td>CGG</td><td>||||||||[||||||||||||]PAM</td><td>NA</td><td>Intergenic</td><td>NA</td><td>NA</td></tr>
<tr><td>GU305914.1</td><td></td><td>94009</td><td>94031</td><td>-</td><td>0</td><td>GAGGGAAGCCGGACAGGGAC</td><td>CGG</td><td>||||||||[||||||||||||]PAM</td><td>NA</td><td>Intergenic</td><td>NA</td><td>NA</td></tr>
<tr><td>GU305914.1</td><td></td><td>93977</td><td>93999</td><td>-</td><td>0</td><td>GAGGGAAGCCGGACAGGGAC</td><td>CGG</td><td>||||||||[||||||||||||]PAM</td><td>NA</td><td>Intergenic</td><td>NA</td><td>NA</td></tr>
<tr><td>GU305914.1</td><td></td><td>93945</td><td>93967</td><td>-</td><td>0</td><td>GAGGGAAGCCGGACAGGGAC</td><td>CGG</td><td>||||||||[||||||||||||]PAM</td><td>NA</td><td>Intergenic</td><td>NA</td><td>NA</td></tr>
<tr><td>GU305914.1</td><td></td><td>93881</td><td>93903</td><td>-</td><td>3</td><td>GAGAGGAGCCGGACAGGAAC</td><td>CGG</td><td>|||-|-||[|||||||||-||]PAM</td><td>NA</td><td>Intergenic</td><td>NA</td><td>NA</td></tr>
<tr><td></td><td></td><td></td><td></td><td></td><td></td><td></td><td></td><td></td><td></td><td></td><td></td><td></td></tr>
<tr><td>T26871</td><td>AGGGAAGCCGGACAGGGACCGGG</td><td></td><td>1000</td><td></td><td></td><td></td><td></td><td></td><td></td><td></td><td></td><td></td></tr>
<tr><td>Oligo fwd</td><td>CACCAGGGAAGCCGGACAGGGACC</td><td></td><td></td><td></td><td></td><td></td><td></td><td></td><td></td><td></td><td></td><td></td></tr>
<tr><td>Oligo rev</td><td>AAACGGTCCCTGTCCGGCTTCCCT</td><td></td><td></td><td></td><td></td><td></td><td></td><td></td><td></td><td></td><td></td><td></td></tr>
<tr><td>Chromosome</td><td>start</td><td></td><td>end</td><td>strand</td><td>MM</td><td>target_seq</td><td>PAM</td><td>alignment</td><td>distance</td><td>position</td><td>gene name</td><td>gene id</td></tr>
<tr><td>GU305914.1</td><td></td><td>93976</td><td>93998</td><td>-</td><td>0</td><td>AGGGAAGCCGGACAGGGACC</td><td>GGG</td><td>||||||||[||||||||||||]PAM</td><td>NA</td><td>Intergenic</td><td>NA</td><td>NA</td></tr>
<tr><td>GU305914.1</td><td></td><td>93944</td><td>93966</td><td>-</td><td>0</td><td>AGGGAAGCCGGACAGGGACC</td><td>GGG</td><td>||||||||[||||||||||||]PAM</td><td>NA</td><td>Intergenic</td><td>NA</td><td>NA</td></tr>
<tr><td>GU305914.1</td><td></td><td>93912</td><td>93934</td><td>-</td><td>0</td><td>AGGGAAGCCGGACAGGGACC</td><td>GGG</td><td>||||||||[||||||||||||]PAM</td><td>NA</td><td>Intergenic</td><td>NA</td><td>NA</td></tr>
<tr><td>GU305914.1</td><td></td><td>94072</td><td>94094</td><td>-</td><td>0</td><td>AGGGAAGCCGGACAGGGACC</td><td>GGG</td><td>||||||||[||||||||||||]PAM</td><td>NA</td><td>Intergenic</td><td>NA</td><td>NA</td></tr>
<tr><td>GU305914.1</td><td></td><td>94040</td><td>94062</td><td>-</td><td>0</td><td>AGGGAAGCCGGACAGGGACC</td><td>GGG</td><td>||||||||[||||||||||||]PAM</td><td>NA</td><td>Intergenic</td><td>NA</td><td>NA</td></tr>
<tr><td>GU305914.1</td><td></td><td>94008</td><td>94030</td><td>-</td><td>0</td><td>AGGGAAGCCGGACAGGGACC</td><td>GGG</td><td>||||||||[||||||||||||]PAM</td><td>NA</td><td>Intergenic</td><td>NA</td><td>NA</td></tr>
<tr><td>GU305914.1</td><td></td><td>93880</td><td>93902</td><td>-</td><td>3</td><td>AGAGGAGCCGGACAGGAACC</td><td>GGG</td><td>||-|-|||[||||||||-|||]PAM</td><td>NA</td><td>Intergenic</td><td>NA</td><td>NA</td></tr>
<tr><td></td><td></td><td></td><td></td><td></td><td></td><td></td><td></td><td></td><td></td><td></td><td></td><td></td></tr>
<tr><td>T26872</td><td>CCGGACAGGGACCGGGAGCTCGG</td><td></td><td>1000</td><td></td><td></td><td></td><td></td><td></td><td></td><td></td><td></td><td></td></tr>
<tr><td>Oligo fwd</td><td>CACCCCGGACAGGGACCGGGAGCT</td><td></td><td></td><td></td><td></td><td></td><td></td><td></td><td></td><td></td><td></td><td></td></tr>
<tr><td>Oligo rev</td><td>AAACAGCTCCCGGTCCCTGTCCGG</td><td></td><td></td><td></td><td></td><td></td><td></td><td></td><td></td><td></td><td></td><td></td></tr>
<tr><td>Chromosome</td><td>start</td><td></td><td>end</td><td>strand</td><td>MM</td><td>target_seq</td><td>PAM</td><td>alignment</td><td>distance</td><td>position</td><td>gene name</td><td>gene id</td></tr>
<tr><td>GU305914.1</td><td></td><td>94001</td><td>94023</td><td>-</td><td>0</td><td>CCGGACAGGGACCGGGAGCT</td><td>CGG</td><td>||||||||[||||||||||||]PAM</td><td>NA</td><td>Intergenic</td><td>NA</td><td>NA</td></tr>
<tr><td>GU305914.1</td><td></td><td>93969</td><td>93991</td><td>-</td><td>0</td><td>CCGGACAGGGACCGGGAGCT</td><td>CGG</td><td>||||||||[||||||||||||]PAM</td><td>NA</td><td>Intergenic</td><td>NA</td><td>NA</td></tr>
<tr><td>GU305914.1</td><td></td><td>93937</td><td>93959</td><td>-</td><td>0</td><td>CCGGACAGGGACCGGGAGCT</td><td>CGG</td><td>||||||||[||||||||||||]PAM</td><td>NA</td><td>Intergenic</td><td>NA</td><td>NA</td></tr>
<tr><td>GU305914.1</td><td></td><td>93905</td><td>93927</td><td>-</td><td>0</td><td>CCGGACAGGGACCGGGAGCT</td><td>CGG</td><td>||||||||[||||||||||||]PAM</td><td>NA</td><td>Intergenic</td><td>NA</td><td>NA</td></tr>
<tr><td>GU305914.1</td><td></td><td>94065</td><td>94087</td><td>-</td><td>0</td><td>CCGGACAGGGACCGGGAGCT</td><td>CGG</td><td>||||||||[||||||||||||]PAM</td><td>NA</td><td>Intergenic</td><td>NA</td><td>NA</td></tr>
<tr><td>GU305914.1</td><td></td><td>94033</td><td>94055</td><td>-</td><td>0</td><td>CCGGACAGGGACCGGGAGCT</td><td>CGG</td><td>||||||||[||||||||||||]PAM</td><td>NA</td><td>Intergenic</td><td>NA</td><td>NA</td></tr>
<tr><td>GU305914.1</td><td></td><td>93873</td><td>93895</td><td>-</td><td>2</td><td>CCGGACAGGAACCGGGAGCG</td><td>GGG</td><td>||||||||[|-|||||||||-]PAM</td><td>NA</td><td>Intergenic</td><td>NA</td><td>NA</td></tr>
<tr><td></td><td></td><td></td><td></td><td></td><td></td><td></td><td></td><td></td><td></td><td></td><td></td><td></td></tr>
<tr><td>T26873</td><td>CGGACAGGGACCGGGAGCTCGGG</td><td></td><td>1000</td><td></td><td></td><td></td><td></td><td></td><td></td><td></td><td></td><td></td></tr>
<tr><td>Oligo fwd</td><td>CACCCGGACAGGGACCGGGAGCTC</td><td></td><td></td><td></td><td></td><td></td><td></td><td></td><td></td><td></td><td></td><td></td></tr>
<tr><td>Oligo rev</td><td>AAACGAGCTCCCGGTCCCTGTCCG</td><td></td><td></td><td></td><td></td><td></td><td></td><td></td><td></td><td></td><td></td><td></td></tr>
<tr><td>Chromosome</td><td>start</td><td></td><td>end</td><td>strand</td><td>MM</td><td>target_seq</td><td>PAM</td><td>alignment</td><td>distance</td><td>position</td><td>gene name</td><td>gene id</td></tr>
<tr><td>GU305914.1</td><td></td><td>93968</td><td>93990</td><td>-</td><td>0</td><td>CGGACAGGGACCGGGAGCTC</td><td>GGG</td><td>||||||||[||||||||||||]PAM</td><td>NA</td><td>Intergenic</td><td>NA</td><td>NA</td></tr>
<tr><td>GU305914.1</td><td></td><td>93936</td><td>93958</td><td>-</td><td>0</td><td>CGGACAGGGACCGGGAGCTC</td><td>GGG</td><td>||||||||[||||||||||||]PAM</td><td>NA</td><td>Intergenic</td><td>NA</td><td>NA</td></tr>
<tr><td>GU305914.1</td><td></td><td>93904</td><td>93926</td><td>-</td><td>0</td><td>CGGACAGGGACCGGGAGCTC</td><td>GGG</td><td>||||||||[||||||||||||]PAM</td><td>NA</td><td>Intergenic</td><td>NA</td><td>NA</td></tr>
<tr><td>GU305914.1</td><td></td><td>94064</td><td>94086</td><td>-</td><td>0</td><td>CGGACAGGGACCGGGAGCTC</td><td>GGG</td><td>||||||||[||||||||||||]PAM</td><td>NA</td><td>Intergenic</td><td>NA</td><td>NA</td></tr>
<tr><td>GU305914.1</td><td></td><td>94032</td><td>94054</td><td>-</td><td>0</td><td>CGGACAGGGACCGGGAGCTC</td><td>GGG</td><td>||||||||[||||||||||||]PAM</td><td>NA</td><td>Intergenic</td><td>NA</td><td>NA</td></tr>
<tr><td>GU305914.1</td><td></td><td>94000</td><td>94022</td><td>-</td><td>0</td><td>CGGACAGGGACCGGGAGCTC</td><td>GGG</td><td>||||||||[||||||||||||]PAM</td><td>NA</td><td>Intergenic</td><td>NA</td><td>NA</td></tr>
<tr><td></td><td></td><td></td><td></td><td></td><td></td><td></td><td></td><td></td><td></td><td></td><td></td><td></td></tr>
<tr><td>T26874</td><td>GGACAGGGACCGGGAGCTCGGGG</td><td></td><td>1000</td><td></td><td></td><td></td><td></td><td></td><td></td><td></td><td></td><td></td></tr>
<tr><td>Oligo fwd</td><td>CACCGGACAGGGACCGGGAGCTCG</td><td></td><td></td><td></td><td></td><td></td><td></td><td></td><td></td><td></td><td></td><td></td></tr>
<tr><td>Oligo rev</td><td>AAACCGAGCTCCCGGTCCCTGTCC</td><td></td><td></td><td></td><td></td><td></td><td></td><td></td><td></td><td></td><td></td><td></td></tr>
<tr><td>Chromosome</td><td>start</td><td></td><td>end</td><td>strand</td><td>MM</td><td>target_seq</td><td>PAM</td><td>alignment</td><td>distance</td><td>position</td><td>gene name</td><td>gene id</td></tr>
<tr><td>GU305914.1</td><td></td><td>93903</td><td>93925</td><td>-</td><td>0</td><td>GGACAGGGACCGGGAGCTCG</td><td>GGG</td><td>||||||||[||||||||||||]PAM</td><td>NA</td><td>Intergenic</td><td>NA</td><td>NA</td></tr>
<tr><td>GU305914.1</td><td></td><td>94063</td><td>94085</td><td>-</td><td>0</td><td>GGACAGGGACCGGGAGCTCG</td><td>GGG</td><td>||||||||[||||||||||||]PAM</td><td>NA</td><td>Intergenic</td><td>NA</td><td>NA</td></tr>
<tr><td>GU305914.1</td><td></td><td>94031</td><td>94053</td><td>-</td><td>0</td><td>GGACAGGGACCGGGAGCTCG</td><td>GGG</td><td>||||||||[||||||||||||]PAM</td><td>NA</td><td>Intergenic</td><td>NA</td><td>NA</td></tr>
<tr><td>GU305914.1</td><td></td><td>93999</td><td>94021</td><td>-</td><td>0</td><td>GGACAGGGACCGGGAGCTCG</td><td>GGG</td><td>||||||||[||||||||||||]PAM</td><td>NA</td><td>Intergenic</td><td>NA</td><td>NA</td></tr>
<tr><td>GU305914.1</td><td></td><td>93967</td><td>93989</td><td>-</td><td>0</td><td>GGACAGGGACCGGGAGCTCG</td><td>GGG</td><td>||||||||[||||||||||||]PAM</td><td>NA</td><td>Intergenic</td><td>NA</td><td>NA</td></tr>
<tr><td>GU305914.1</td><td></td><td>93935</td><td>93957</td><td>-</td><td>0</td><td>GGACAGGGACCGGGAGCTCG</td><td>GGG</td><td>||||||||[||||||||||||]PAM</td><td>NA</td><td>Intergenic</td><td>NA</td><td>NA</td></tr>
<tr><td>GU305914.1</td><td></td><td>94095</td><td>94117</td><td>-</td><td>4</td><td>AAGCGGGGACCGGGAGCTCG</td><td>GGG</td><td>---|-|||[||||||||||||]PAM</td><td>NA</td><td>Intergenic</td><td>NA</td><td>NA</td></tr>
<tr><td></td><td></td><td></td><td></td><td></td><td></td><td></td><td></td><td></td><td></td><td></td><td></td><td></td></tr>
<tr><td>T26875</td><td>CAGGGACCGGGAGCTCGGGGAGG</td><td></td><td>1000</td><td></td><td></td><td></td><td></td><td></td><td></td><td></td><td></td><td></td></tr>
<tr><td>Oligo fwd</td><td>CACCCAGGGACCGGGAGCTCGGGG</td><td></td><td></td><td></td><td></td><td></td><td></td><td></td><td></td><td></td><td></td><td></td></tr>
<tr><td>Oligo rev</td><td>AAACCCCCGAGCTCCCGGTCCCTG</td><td></td><td></td><td></td><td></td><td></td><td></td><td></td><td></td><td></td><td></td><td></td></tr>
<tr><td>Chromosome</td><td>start</td><td></td><td>end</td><td>strand</td><td>MM</td><td>target_seq</td><td>PAM</td><td>alignment</td><td>distance</td><td>position</td><td>gene name</td><td>gene id</td></tr>
<tr><td>GU305914.1</td><td></td><td>93996</td><td>94018</td><td>-</td><td>0</td><td>CAGGGACCGGGAGCTCGGGG</td><td>AGG</td><td>||||||||[||||||||||||]PAM</td><td>NA</td><td>Intergenic</td><td>NA</td><td>NA</td></tr>
</table>

<table>
<tr><td>GU305914.1</td><td>93964</td><td>93986</td><td>-</td><td>0</td><td>CAGGGACCGGGAGCTCGGGG</td><td>AGG</td><td>||||||||[||||||||||||]PAM</td><td>NA</td><td>Intergenic</td><td>NA</td><td>NA</td></tr>
<tr><td>GU305914.1</td><td>93932</td><td>93954</td><td>-</td><td>0</td><td>CAGGGACCGGGAGCTCGGGG</td><td>AGG</td><td>||||||||[||||||||||||]PAM</td><td>NA</td><td>Intergenic</td><td>NA</td><td>NA</td></tr>
<tr><td>GU305914.1</td><td>94060</td><td>94082</td><td>-</td><td>0</td><td>CAGGGACCGGGAGCTCGGGG</td><td>AGG</td><td>||||||||[||||||||||||]PAM</td><td>NA</td><td>Intergenic</td><td>NA</td><td>NA</td></tr>
<tr><td>GU305914.1</td><td>94028</td><td>94050</td><td>-</td><td>0</td><td>CAGGGACCGGGAGCTCGGGG</td><td>AGG</td><td>||||||||[||||||||||||]PAM</td><td>NA</td><td>Intergenic</td><td>NA</td><td>NA</td></tr>
<tr><td>GU305914.1</td><td>94092</td><td>94114</td><td>-</td><td>1</td><td>CGGGGACCGGGAGCTCGGGG</td><td>AGG</td><td>|-||||||[||||||||||||]PAM</td><td>NA</td><td>Intergenic</td><td>NA</td><td>NA</td></tr>
<tr><td></td></tr>
<tr><td>T26876</td><td>AGGGACCGGGAGCTCGGGGAGGG</td><td>1000</td></tr>
<tr><td>Oligo fwd</td><td>CACCAGGGACCGGGAGCTCGGGGA</td></tr>
<tr><td>Oligo rev</td><td>AAACTCCCCGAGCTCCCGGTCCCT</td></tr>
<tr><td>Chromosome</td><td>start</td><td>end</td><td>strand</td><td>MM</td><td>target_seq</td><td>PAM</td><td>alignment</td><td>distance</td><td>position</td><td>gene name</td><td>gene id</td></tr>
<tr><td>GU305914.1</td><td>93995</td><td>94017</td><td>-</td><td>0</td><td>AGGGACCGGGAGCTCGGGGA</td><td>GGG</td><td>||||||||[||||||||||||]PAM</td><td>NA</td><td>Intergenic</td><td>NA</td><td>NA</td></tr>
<tr><td>GU305914.1</td><td>93963</td><td>93985</td><td>-</td><td>0</td><td>AGGGACCGGGAGCTCGGGGA</td><td>GGG</td><td>||||||||[||||||||||||]PAM</td><td>NA</td><td>Intergenic</td><td>NA</td><td>NA</td></tr>
<tr><td>GU305914.1</td><td>93931</td><td>93953</td><td>-</td><td>0</td><td>AGGGACCGGGAGCTCGGGGA</td><td>GGG</td><td>||||||||[||||||||||||]PAM</td><td>NA</td><td>Intergenic</td><td>NA</td><td>NA</td></tr>
<tr><td>GU305914.1</td><td>94059</td><td>94081</td><td>-</td><td>0</td><td>AGGGACCGGGAGCTCGGGGA</td><td>GGG</td><td>||||||||[||||||||||||]PAM</td><td>NA</td><td>Intergenic</td><td>NA</td><td>NA</td></tr>
<tr><td>GU305914.1</td><td>94027</td><td>94049</td><td>-</td><td>0</td><td>AGGGACCGGGAGCTCGGGGA</td><td>GGG</td><td>||||||||[||||||||||||]PAM</td><td>NA</td><td>Intergenic</td><td>NA</td><td>NA</td></tr>
<tr><td>GU305914.1</td><td>94091</td><td>94113</td><td>-</td><td>1</td><td>GGGGACCGGGAGCTCGGGGA</td><td>GGG</td><td>-|||||||[||||||||||||]PAM</td><td>NA</td><td>Intergenic</td><td>NA</td><td>NA</td></tr>
<tr><td></td></tr>
<tr><td>T26877</td><td>GGGAGCTCGGGGAGGGAAGCCGG</td><td>1000</td></tr>
<tr><td>Oligo fwd</td><td>CACCGGGAGCTCGGGGAGGGAAGC</td></tr>
<tr><td>Oligo rev</td><td>AAACGCTTCCCTCCCCGAGCTCCC</td></tr>
<tr><td>Chromosome</td><td>start</td><td>end</td><td>strand</td><td>MM</td><td>target_seq</td><td>PAM</td><td>alignment</td><td>distance</td><td>position</td><td>gene name</td><td>gene id</td></tr>
<tr><td>GU305914.1</td><td>94052</td><td>94074</td><td>-</td><td>0</td><td>GGGAGCTCGGGGAGGGAAGC</td><td>CGG</td><td>||||||||[||||||||||||]PAM</td><td>NA</td><td>Intergenic</td><td>NA</td><td>NA</td></tr>
<tr><td>GU305914.1</td><td>94020</td><td>94042</td><td>-</td><td>0</td><td>GGGAGCTCGGGGAGGGAAGC</td><td>CGG</td><td>||||||||[||||||||||||]PAM</td><td>NA</td><td>Intergenic</td><td>NA</td><td>NA</td></tr>
<tr><td>GU305914.1</td><td>93988</td><td>94010</td><td>-</td><td>0</td><td>GGGAGCTCGGGGAGGGAAGC</td><td>CGG</td><td>||||||||[||||||||||||]PAM</td><td>NA</td><td>Intergenic</td><td>NA</td><td>NA</td></tr>
<tr><td>GU305914.1</td><td>93956</td><td>93978</td><td>-</td><td>0</td><td>GGGAGCTCGGGGAGGGAAGC</td><td>CGG</td><td>||||||||[||||||||||||]PAM</td><td>NA</td><td>Intergenic</td><td>NA</td><td>NA</td></tr>
<tr><td>GU305914.1</td><td>93924</td><td>93946</td><td>-</td><td>0</td><td>GGGAGCTCGGGGAGGGAAGC</td><td>CGG</td><td>||||||||[||||||||||||]PAM</td><td>NA</td><td>Intergenic</td><td>NA</td><td>NA</td></tr>
<tr><td>GU305914.1</td><td>94084</td><td>94106</td><td>-</td><td>0</td><td>GGGAGCTCGGGGAGGGAAGC</td><td>CGG</td><td>||||||||[||||||||||||]PAM</td><td>NA</td><td>Intergenic</td><td>NA</td><td>NA</td></tr>
<tr><td>GU305914.1</td><td>93892</td><td>93914</td><td>-</td><td>2</td><td>GGGAGCTCGGGGAGAGGAGC</td><td>CGG</td><td>||||||||[||||||-|-|||]PAM</td><td>NA</td><td>Intergenic</td><td>NA</td><td>NA</td></tr>
<tr><td></td></tr>
<tr><td>T26878</td><td>CTCGGGGAGGGAAGCCGGACAGG</td><td>1000</td></tr>
<tr><td>Oligo fwd</td><td>CACCCTCGGGGAGGGAAGCCGGAC</td></tr>
<tr><td>Oligo rev</td><td>AAACGTCCGGCTTCCCTCCCCGAG</td></tr>
<tr><td>Chromosome</td><td>start</td><td>end</td><td>strand</td><td>MM</td><td>target_seq</td><td>PAM</td><td>alignment</td><td>distance</td><td>position</td><td>gene name</td><td>gene id</td></tr>
<tr><td>GU305914.1</td><td>93951</td><td>93973</td><td>-</td><td>0</td><td>CTCGGGGAGGGAAGCCGGAC</td><td>AGG</td><td>||||||||[||||||||||||]PAM</td><td>NA</td><td>Intergenic</td><td>NA</td><td>NA</td></tr>
<tr><td>GU305914.1</td><td>93919</td><td>93941</td><td>-</td><td>0</td><td>CTCGGGGAGGGAAGCCGGAC</td><td>AGG</td><td>||||||||[||||||||||||]PAM</td><td>NA</td><td>Intergenic</td><td>NA</td><td>NA</td></tr>
<tr><td>GU305914.1</td><td>94079</td><td>94101</td><td>-</td><td>0</td><td>CTCGGGGAGGGAAGCCGGAC</td><td>AGG</td><td>||||||||[||||||||||||]PAM</td><td>NA</td><td>Intergenic</td><td>NA</td><td>NA</td></tr>
<tr><td>GU305914.1</td><td>94047</td><td>94069</td><td>-</td><td>0</td><td>CTCGGGGAGGGAAGCCGGAC</td><td>AGG</td><td>||||||||[||||||||||||]PAM</td><td>NA</td><td>Intergenic</td><td>NA</td><td>NA</td></tr>
<tr><td>GU305914.1</td><td>94015</td><td>94037</td><td>-</td><td>0</td><td>CTCGGGGAGGGAAGCCGGAC</td><td>AGG</td><td>||||||||[||||||||||||]PAM</td><td>NA</td><td>Intergenic</td><td>NA</td><td>NA</td></tr>
<tr><td>GU305914.1</td><td>93983</td><td>94005</td><td>-</td><td>0</td><td>CTCGGGGAGGGAAGCCGGAC</td><td>AGG</td><td>||||||||[||||||||||||]PAM</td><td>NA</td><td>Intergenic</td><td>NA</td><td>NA</td></tr>
<tr><td>GU305914.1</td><td>93887</td><td>93909</td><td>-</td><td>2</td><td>CTCGGGGAGAGGAGCCGGAC</td><td>AGG</td><td>||||||||[|-|-||||||||]PAM</td><td>NA</td><td>Intergenic</td><td>NA</td><td>NA</td></tr>
<tr><td></td></tr>
<tr><td>T26879</td><td>TCGGGGAGGGAAGCCGGACAGGG</td><td>1000</td></tr>
<tr><td>Oligo fwd</td><td>CACCTCGGGGAGGGAAGCCGGACA</td></tr>
<tr><td>Oligo rev</td><td>AAACTGTCCGGCTTCCCTCCCCGA</td></tr>
<tr><td>Chromosome</td><td>start</td><td>end</td><td>strand</td><td>MM</td><td>target_seq</td><td>PAM</td><td>alignment</td><td>distance</td><td>position</td><td>gene name</td><td>gene id</td></tr>
<tr><td>GU305914.1</td><td>94078</td><td>94100</td><td>-</td><td>0</td><td>TCGGGGAGGGAAGCCGGACA</td><td>GGG</td><td>||||||||[||||||||||||]PAM</td><td>NA</td><td>Intergenic</td><td>NA</td><td>NA</td></tr>
<tr><td>GU305914.1</td><td>94046</td><td>94068</td><td>-</td><td>0</td><td>TCGGGGAGGGAAGCCGGACA</td><td>GGG</td><td>||||||||[||||||||||||]PAM</td><td>NA</td><td>Intergenic</td><td>NA</td><td>NA</td></tr>
<tr><td>GU305914.1</td><td>94014</td><td>94036</td><td>-</td><td>0</td><td>TCGGGGAGGGAAGCCGGACA</td><td>GGG</td><td>||||||||[||||||||||||]PAM</td><td>NA</td><td>Intergenic</td><td>NA</td><td>NA</td></tr>
<tr><td>GU305914.1</td><td>93982</td><td>94004</td><td>-</td><td>0</td><td>TCGGGGAGGGAAGCCGGACA</td><td>GGG</td><td>||||||||[||||||||||||]PAM</td><td>NA</td><td>Intergenic</td><td>NA</td><td>NA</td></tr>
<tr><td>GU305914.1</td><td>93950</td><td>93972</td><td>-</td><td>0</td><td>TCGGGGAGGGAAGCCGGACA</td><td>GGG</td><td>||||||||[||||||||||||]PAM</td><td>NA</td><td>Intergenic</td><td>NA</td><td>NA</td></tr>
<tr><td>GU305914.1</td><td>93918</td><td>93940</td><td>-</td><td>0</td><td>TCGGGGAGGGAAGCCGGACA</td><td>GGG</td><td>||||||||[||||||||||||]PAM</td><td>NA</td><td>Intergenic</td><td>NA</td><td>NA</td></tr>
<tr><td></td></tr>
<tr><td>T26880</td><td>GAGGGAAGCCGGACAGGGACCGG</td><td>1000</td></tr>
<tr><td>Oligo fwd</td><td>CACCGAGGGAAGCCGGACAGGGAC</td></tr>
<tr><td>Oligo rev</td><td>AAACGTCCCTGTCCGGCTTCCCTC</td></tr>
<tr><td>Chromosome</td><td>start</td><td>end</td><td>strand</td><td>MM</td><td>target_seq</td><td>PAM</td><td>alignment</td><td>distance</td><td>position</td><td>gene name</td><td>gene id</td></tr>
<tr><td>GU305914.1</td><td>93913</td><td>93935</td><td>-</td><td>0</td><td>GAGGGAAGCCGGACAGGGAC</td><td>CGG</td><td>||||||||[||||||||||||]PAM</td><td>NA</td><td>Intergenic</td><td>NA</td><td>NA</td></tr>
<tr><td>GU305914.1</td><td>94073</td><td>94095</td><td>-</td><td>0</td><td>GAGGGAAGCCGGACAGGGAC</td><td>CGG</td><td>||||||||[||||||||||||]PAM</td><td>NA</td><td>Intergenic</td><td>NA</td><td>NA</td></tr>
<tr><td>GU305914.1</td><td>94041</td><td>94063</td><td>-</td><td>0</td><td>GAGGGAAGCCGGACAGGGAC</td><td>CGG</td><td>||||||||[||||||||||||]PAM</td><td>NA</td><td>Intergenic</td><td>NA</td><td>NA</td></tr>
<tr><td>GU305914.1</td><td>94009</td><td>94031</td><td>-</td><td>0</td><td>GAGGGAAGCCGGACAGGGAC</td><td>CGG</td><td>||||||||[||||||||||||]PAM</td><td>NA</td><td>Intergenic</td><td>NA</td><td>NA</td></tr>
<tr><td>GU305914.1</td><td>93977</td><td>93999</td><td>-</td><td>0</td><td>GAGGGAAGCCGGACAGGGAC</td><td>CGG</td><td>||||||||[||||||||||||]PAM</td><td>NA</td><td>Intergenic</td><td>NA</td><td>NA</td></tr>
<tr><td>GU305914.1</td><td>93945</td><td>93967</td><td>-</td><td>0</td><td>GAGGGAAGCCGGACAGGGAC</td><td>CGG</td><td>||||||||[||||||||||||]PAM</td><td>NA</td><td>Intergenic</td><td>NA</td><td>NA</td></tr>
<tr><td>GU305914.1</td><td>93881</td><td>93903</td><td>-</td><td>3</td><td>GAGAGGAGCCGGACAGGAAC</td><td>CGG</td><td>|||-|-||[|||||||||-||]PAM</td><td>NA</td><td>Intergenic</td><td>NA</td><td>NA</td></tr>
<tr><td></td></tr>
<tr><td>T26881</td><td>AGGGAAGCCGGACAGGGACCGGG</td><td>1000</td></tr>
<tr><td>Oligo fwd</td><td>CACCAGGGAAGCCGGACAGGGACC</td></tr>
<tr><td>Oligo rev</td><td>AAACGGTCCCTGTCCGGCTTCCCT</td></tr>
<tr><td>Chromosome</td><td>start</td><td>end</td><td>strand</td><td>MM</td><td>target_seq</td><td>PAM</td><td>alignment</td><td>distance</td><td>position</td><td>gene name</td><td>gene id</td></tr>
<tr><td>GU305914.1</td><td>93976</td><td>93998</td><td>-</td><td>0</td><td>AGGGAAGCCGGACAGGGACC</td><td>GGG</td><td>||||||||[||||||||||||]PAM</td><td>NA</td><td>Intergenic</td><td>NA</td><td>NA</td></tr>
<tr><td>GU305914.1</td><td>93944</td><td>93966</td><td>-</td><td>0</td><td>AGGGAAGCCGGACAGGGACC</td><td>GGG</td><td>||||||||[||||||||||||]PAM</td><td>NA</td><td>Intergenic</td><td>NA</td><td>NA</td></tr>
<tr><td>GU305914.1</td><td>93912</td><td>93934</td><td>-</td><td>0</td><td>AGGGAAGCCGGACAGGGACC</td><td>GGG</td><td>||||||||[||||||||||||]PAM</td><td>NA</td><td>Intergenic</td><td>NA</td><td>NA</td></tr>
<tr><td>GU305914.1</td><td>94072</td><td>94094</td><td>-</td><td>0</td><td>AGGGAAGCCGGACAGGGACC</td><td>GGG</td><td>||||||||[||||||||||||]PAM</td><td>NA</td><td>Intergenic</td><td>NA</td><td>NA</td></tr>
<tr><td>GU305914.1</td><td>94040</td><td>94062</td><td>-</td><td>0</td><td>AGGGAAGCCGGACAGGGACC</td><td>GGG</td><td>||||||||[||||||||||||]PAM</td><td>NA</td><td>Intergenic</td><td>NA</td><td>NA</td></tr>
<tr><td>GU305914.1</td><td>94008</td><td>94030</td><td>-</td><td>0</td><td>AGGGAAGCCGGACAGGGACC</td><td>GGG</td><td>||||||||[||||||||||||]PAM</td><td>NA</td><td>Intergenic</td><td>NA</td><td>NA</td></tr>
<tr><td>GU305914.1</td><td>93880</td><td>93902</td><td>-</td><td>3</td><td>AGAGGAGCCGGACAGGAACC</td><td>GGG</td><td>||-|-|||[||||||||-|||]PAM</td><td>NA</td><td>Intergenic</td><td>NA</td><td>NA</td></tr>
<tr><td></td></tr>
<tr><td>T26882</td><td>CCGGACAGGGACCGGGAGCTCGG</td><td>1000</td></tr>
<tr><td>Oligo fwd</td><td>CACCCCGGACAGGGACCGGGAGCT</td></tr>
<tr><td>Oligo rev</td><td>AAACAGCTCCCGGTCCCTGTCCGG</td></tr>
<tr><td>Chromosome</td><td>start</td><td>end</td><td>strand</td><td>MM</td><td>target_seq</td><td>PAM</td><td>alignment</td><td>distance</td><td>position</td><td>gene name</td><td>gene id</td></tr>
<tr><td>GU305914.1</td><td>94001</td><td>94023</td><td>-</td><td>0</td><td>CCGGACAGGGACCGGGAGCT</td><td>CGG</td><td>||||||||[||||||||||||]PAM</td><td>NA</td><td>Intergenic</td><td>NA</td><td>NA</td></tr>
<tr><td>GU305914.1</td><td>93969</td><td>93991</td><td>-</td><td>0</td><td>CCGGACAGGGACCGGGAGCT</td><td>CGG</td><td>||||||||[||||||||||||]PAM</td><td>NA</td><td>Intergenic</td><td>NA</td><td>NA</td></tr>
<tr><td>GU305914.1</td><td>93937</td><td>93959</td><td>-</td><td>0</td><td>CCGGACAGGGACCGGGAGCT</td><td>CGG</td><td>||||||||[||||||||||||]PAM</td><td>NA</td><td>Intergenic</td><td>NA</td><td>NA</td></tr>
<tr><td>GU305914.1</td><td>93905</td><td>93927</td><td>-</td><td>0</td><td>CCGGACAGGGACCGGGAGCT</td><td>CGG</td><td>||||||||[||||||||||||]PAM</td><td>NA</td><td>Intergenic</td><td>NA</td><td>NA</td></tr>
<tr><td>GU305914.1</td><td>94065</td><td>94087</td><td>-</td><td>0</td><td>CCGGACAGGGACCGGGAGCT</td><td>CGG</td><td>||||||||[||||||||||||]PAM</td><td>NA</td><td>Intergenic</td><td>NA</td><td>NA</td></tr>
<tr><td>GU305914.1</td><td>94033</td><td>94055</td><td>-</td><td>0</td><td>CCGGACAGGGACCGGGAGCT</td><td>CGG</td><td>||||||||[||||||||||||]PAM</td><td>NA</td><td>Intergenic</td><td>NA</td><td>NA</td></tr>
<tr><td>GU305914.1</td><td>93873</td><td>93895</td><td>-</td><td>2</td><td>CCGGACAGGAACCGGGAGCG</td><td>GGG</td><td>||||||||[|-|||||||||-]PAM</td><td>NA</td><td>Intergenic</td><td>NA</td><td>NA</td></tr>
<tr><td></td></tr>
<tr><td>T26883</td><td>CGGACAGGGACCGGGAGCTCGGG</td><td>1000</td></tr>
</table>

| | | | | | | | | | | | |
|---|---|---|---|---|---|---|---|---|---|---|---|
| Oligo fwd | CACCCGGACAGGGACCGGGAGCTC | | | | | | | | | | |
| Oligo rev | AAACGAGCTCCCGGTCCCTGTCCG | | | | | | | | | | |
| Chromosome | start | end | strand | MM | target_seq | PAM | alignment | distance | position | gene name | gene id |
| GU305914.1 | 93968 | 93990 | - | 0 | CGGACAGGGACCGGGAGCTC | GGG | \|\|\|\|\|\|\|\|[\|\|\|\|\|\|\|\|\|\|\|\|]PAM | NA | Intergenic | NA | NA |
| GU305914.1 | 93936 | 93958 | - | 0 | CGGACAGGGACCGGGAGCTC | GGG | \|\|\|\|\|\|\|\|[\|\|\|\|\|\|\|\|\|\|\|\|]PAM | NA | Intergenic | NA | NA |
| GU305914.1 | 93904 | 93926 | - | 0 | CGGACAGGGACCGGGAGCTC | GGG | \|\|\|\|\|\|\|\|[\|\|\|\|\|\|\|\|\|\|\|\|]PAM | NA | Intergenic | NA | NA |
| GU305914.1 | 94064 | 94086 | - | 0 | CGGACAGGGACCGGGAGCTC | GGG | \|\|\|\|\|\|\|\|[\|\|\|\|\|\|\|\|\|\|\|\|]PAM | NA | Intergenic | NA | NA |
| GU305914.1 | 94032 | 94054 | - | 0 | CGGACAGGGACCGGGAGCTC | GGG | \|\|\|\|\|\|\|\|[\|\|\|\|\|\|\|\|\|\|\|\|]PAM | NA | Intergenic | NA | NA |
| GU305914.1 | 94000 | 94022 | - | 0 | CGGACAGGGACCGGGAGCTC | GGG | \|\|\|\|\|\|\|\|[\|\|\|\|\|\|\|\|\|\|\|\|]PAM | NA | Intergenic | NA | NA |
| T26884 | GGACAGGGACCGGGAGCTCGGGG | 1000 | | | | | | | | | |
| Oligo fwd | CACCGGACAGGGACCGGGAGCTCG | | | | | | | | | | |
| Oligo rev | AAACCGAGCTCCCGGTCCCTGTCC | | | | | | | | | | |
| Chromosome | start | end | strand | MM | target_seq | PAM | alignment | distance | position | gene name | gene id |
| GU305914.1 | 93903 | 93925 | - | 0 | GGACAGGGACCGGGAGCTCG | GGG | \|\|\|\|\|\|\|\|[\|\|\|\|\|\|\|\|\|\|\|\|]PAM | NA | Intergenic | NA | NA |
| GU305914.1 | 94063 | 94085 | - | 0 | GGACAGGGACCGGGAGCTCG | GGG | \|\|\|\|\|\|\|\|[\|\|\|\|\|\|\|\|\|\|\|\|]PAM | NA | Intergenic | NA | NA |
| GU305914.1 | 94031 | 94053 | - | 0 | GGACAGGGACCGGGAGCTCG | GGG | \|\|\|\|\|\|\|\|[\|\|\|\|\|\|\|\|\|\|\|\|]PAM | NA | Intergenic | NA | NA |
| GU305914.1 | 93999 | 94021 | - | 0 | GGACAGGGACCGGGAGCTCG | GGG | \|\|\|\|\|\|\|\|[\|\|\|\|\|\|\|\|\|\|\|\|]PAM | NA | Intergenic | NA | NA |
| GU305914.1 | 93967 | 93989 | - | 0 | GGACAGGGACCGGGAGCTCG | GGG | \|\|\|\|\|\|\|\|[\|\|\|\|\|\|\|\|\|\|\|\|]PAM | NA | Intergenic | NA | NA |
| GU305914.1 | 93935 | 93957 | - | 0 | GGACAGGGACCGGGAGCTCG | GGG | \|\|\|\|\|\|\|\|[\|\|\|\|\|\|\|\|\|\|\|\|]PAM | NA | Intergenic | NA | NA |
| GU305914.1 | 94095 | 94117 | - | 4 | AAGCGGGGACCGGGAGCTCG | GGG | ---\|-\|\|\|[\|\|\|\|\|\|\|\|\|\|\|\|]PAM | NA | Intergenic | NA | NA |
| T26885 | CAGGGACCGGGAGCTCGGGGAGG | 1000 | | | | | | | | | |
| Oligo fwd | CACCCAGGGACCGGGAGCTCGGGG | | | | | | | | | | |
| Oligo rev | AAACCCCCGAGCTCCCGGTCCCTG | | | | | | | | | | |
| Chromosome | start | end | strand | MM | target_seq | PAM | alignment | distance | position | gene name | gene id |
| GU305914.1 | 93996 | 94018 | - | 0 | CAGGGACCGGGAGCTCGGGG | AGG | \|\|\|\|\|\|\|\|[\|\|\|\|\|\|\|\|\|\|\|\|]PAM | NA | Intergenic | NA | NA |
| GU305914.1 | 93964 | 93986 | - | 0 | CAGGGACCGGGAGCTCGGGG | AGG | \|\|\|\|\|\|\|\|[\|\|\|\|\|\|\|\|\|\|\|\|]PAM | NA | Intergenic | NA | NA |
| GU305914.1 | 93932 | 93954 | - | 0 | CAGGGACCGGGAGCTCGGGG | AGG | \|\|\|\|\|\|\|\|[\|\|\|\|\|\|\|\|\|\|\|\|]PAM | NA | Intergenic | NA | NA |
| GU305914.1 | 94060 | 94082 | - | 0 | CAGGGACCGGGAGCTCGGGG | AGG | \|\|\|\|\|\|\|\|[\|\|\|\|\|\|\|\|\|\|\|\|]PAM | NA | Intergenic | NA | NA |
| GU305914.1 | 94028 | 94050 | - | 0 | CAGGGACCGGGAGCTCGGGG | AGG | \|\|\|\|\|\|\|\|[\|\|\|\|\|\|\|\|\|\|\|\|]PAM | NA | Intergenic | NA | NA |
| GU305914.1 | 94092 | 94114 | - | 1 | CGGGGACCGGGAGCTCGGGG | AGG | \|-\|\|\|\|\|\|[\|\|\|\|\|\|\|\|\|\|\|\|]PAM | NA | Intergenic | NA | NA |
| T26886 | AGGGACCGGGAGCTCGGGGAGGG | 1000 | | | | | | | | | |
| Oligo fwd | CACCAGGGACCGGGAGCTCGGGGA | | | | | | | | | | |
| Oligo rev | AAACTCCCCGAGCTCCCGGTCCCT | | | | | | | | | | |
| Chromosome | start | end | strand | MM | target_seq | PAM | alignment | distance | position | gene name | gene id |
| GU305914.1 | 93995 | 94017 | - | 0 | AGGGACCGGGAGCTCGGGGA | GGG | \|\|\|\|\|\|\|\|[\|\|\|\|\|\|\|\|\|\|\|\|]PAM | NA | Intergenic | NA | NA |
| GU305914.1 | 93963 | 93985 | - | 0 | AGGGACCGGGAGCTCGGGGA | GGG | \|\|\|\|\|\|\|\|[\|\|\|\|\|\|\|\|\|\|\|\|]PAM | NA | Intergenic | NA | NA |
| GU305914.1 | 93931 | 93953 | - | 0 | AGGGACCGGGAGCTCGGGGA | GGG | \|\|\|\|\|\|\|\|[\|\|\|\|\|\|\|\|\|\|\|\|]PAM | NA | Intergenic | NA | NA |
| GU305914.1 | 94059 | 94081 | - | 0 | AGGGACCGGGAGCTCGGGGA | GGG | \|\|\|\|\|\|\|\|[\|\|\|\|\|\|\|\|\|\|\|\|]PAM | NA | Intergenic | NA | NA |
| GU305914.1 | 94027 | 94049 | - | 0 | AGGGACCGGGAGCTCGGGGA | GGG | \|\|\|\|\|\|\|\|[\|\|\|\|\|\|\|\|\|\|\|\|]PAM | NA | Intergenic | NA | NA |
| GU305914.1 | 94091 | 94113 | - | 1 | GGGGACCGGGAGCTCGGGGA | GGG | -\|\|\|\|\|\|\|[\|\|\|\|\|\|\|\|\|\|\|\|]PAM | NA | Intergenic | NA | NA |
| T26887 | GGGAGCTCGGGGAGGGAAGCCGG | 1000 | | | | | | | | | |
| Oligo fwd | CACCGGGAGCTCGGGGAGGGAAGC | | | | | | | | | | |
| Oligo rev | AAACGCTTCCCTCCCCGAGCTCCC | | | | | | | | | | |
| Chromosome | start | end | strand | MM | target_seq | PAM | alignment | distance | position | gene name | gene id |
| GU305914.1 | 94052 | 94074 | - | 0 | GGGAGCTCGGGGAGGGAAGC | CGG | \|\|\|\|\|\|\|\|[\|\|\|\|\|\|\|\|\|\|\|\|]PAM | NA | Intergenic | NA | NA |
| GU305914.1 | 94020 | 94042 | - | 0 | GGGAGCTCGGGGAGGGAAGC | CGG | \|\|\|\|\|\|\|\|[\|\|\|\|\|\|\|\|\|\|\|\|]PAM | NA | Intergenic | NA | NA |
| GU305914.1 | 93988 | 94010 | - | 0 | GGGAGCTCGGGGAGGGAAGC | CGG | \|\|\|\|\|\|\|\|[\|\|\|\|\|\|\|\|\|\|\|\|]PAM | NA | Intergenic | NA | NA |
| GU305914.1 | 93956 | 93978 | - | 0 | GGGAGCTCGGGGAGGGAAGC | CGG | \|\|\|\|\|\|\|\|[\|\|\|\|\|\|\|\|\|\|\|\|]PAM | NA | Intergenic | NA | NA |
| GU305914.1 | 93924 | 93946 | - | 0 | GGGAGCTCGGGGAGGGAAGC | CGG | \|\|\|\|\|\|\|\|[\|\|\|\|\|\|\|\|\|\|\|\|]PAM | NA | Intergenic | NA | NA |
| GU305914.1 | 94084 | 94106 | - | 0 | GGGAGCTCGGGGAGGGAAGC | CGG | \|\|\|\|\|\|\|\|[\|\|\|\|\|\|\|\|\|\|\|\|]PAM | NA | Intergenic | NA | NA |
| GU305914.1 | 93892 | 93914 | - | 2 | GGGAGCTCGGGGAGAGGAGC | CGG | \|\|\|\|\|\|\|\|[\|\|\|\|\|\|-\|-\|\|\|]PAM | NA | Intergenic | NA | NA |
| T26888 | CTCGGGGAGGGAAGCCGGACAGG | 1000 | | | | | | | | | |
| Oligo fwd | CACCCTCGGGGAGGGAAGCCGGAC | | | | | | | | | | |
| Oligo rev | AAACGTCCGGCTTCCCTCCCCGAG | | | | | | | | | | |
| Chromosome | start | end | strand | MM | target_seq | PAM | alignment | distance | position | gene name | gene id |
| GU305914.1 | 93951 | 93973 | - | 0 | CTCGGGGAGGGAAGCCGGAC | AGG | \|\|\|\|\|\|\|\|[\|\|\|\|\|\|\|\|\|\|\|\|]PAM | NA | Intergenic | NA | NA |
| GU305914.1 | 93919 | 93941 | - | 0 | CTCGGGGAGGGAAGCCGGAC | AGG | \|\|\|\|\|\|\|\|[\|\|\|\|\|\|\|\|\|\|\|\|]PAM | NA | Intergenic | NA | NA |
| GU305914.1 | 94079 | 94101 | - | 0 | CTCGGGGAGGGAAGCCGGAC | AGG | \|\|\|\|\|\|\|\|[\|\|\|\|\|\|\|\|\|\|\|\|]PAM | NA | Intergenic | NA | NA |
| GU305914.1 | 94047 | 94069 | - | 0 | CTCGGGGAGGGAAGCCGGAC | AGG | \|\|\|\|\|\|\|\|[\|\|\|\|\|\|\|\|\|\|\|\|]PAM | NA | Intergenic | NA | NA |
| GU305914.1 | 94015 | 94037 | - | 0 | CTCGGGGAGGGAAGCCGGAC | AGG | \|\|\|\|\|\|\|\|[\|\|\|\|\|\|\|\|\|\|\|\|]PAM | NA | Intergenic | NA | NA |
| GU305914.1 | 93983 | 94005 | - | 0 | CTCGGGGAGGGAAGCCGGAC | AGG | \|\|\|\|\|\|\|\|[\|\|\|\|\|\|\|\|\|\|\|\|]PAM | NA | Intergenic | NA | NA |
| GU305914.1 | 93887 | 93909 | - | 2 | CTCGGGGAGAGGAGCCGGAC | AGG | \|\|\|\|\|\|\|\|[\|-\|-\|\|\|\|\|\|\|\|]PAM | NA | Intergenic | NA | NA |
| T26889 | TCGGGGAGGGAAGCCGGACAGGG | 1000 | | | | | | | | | |
| Oligo fwd | CACCTCGGGGAGGGAAGCCGGACA | | | | | | | | | | |
| Oligo rev | AAACTGTCCGGCTTCCCTCCCCGA | | | | | | | | | | |
| Chromosome | start | end | strand | MM | target_seq | PAM | alignment | distance | position | gene name | gene id |
| GU305914.1 | 94078 | 94100 | - | 0 | TCGGGGAGGGAAGCCGGACA | GGG | \|\|\|\|\|\|\|\|[\|\|\|\|\|\|\|\|\|\|\|\|]PAM | NA | Intergenic | NA | NA |
| GU305914.1 | 94046 | 94068 | - | 0 | TCGGGGAGGGAAGCCGGACA | GGG | \|\|\|\|\|\|\|\|[\|\|\|\|\|\|\|\|\|\|\|\|]PAM | NA | Intergenic | NA | NA |
| GU305914.1 | 94014 | 94036 | - | 0 | TCGGGGAGGGAAGCCGGACA | GGG | \|\|\|\|\|\|\|\|[\|\|\|\|\|\|\|\|\|\|\|\|]PAM | NA | Intergenic | NA | NA |
| GU305914.1 | 93982 | 94004 | - | 0 | TCGGGGAGGGAAGCCGGACA | GGG | \|\|\|\|\|\|\|\|[\|\|\|\|\|\|\|\|\|\|\|\|]PAM | NA | Intergenic | NA | NA |
| GU305914.1 | 93950 | 93972 | - | 0 | TCGGGGAGGGAAGCCGGACA | GGG | \|\|\|\|\|\|\|\|[\|\|\|\|\|\|\|\|\|\|\|\|]PAM | NA | Intergenic | NA | NA |
| GU305914.1 | 93918 | 93940 | - | 0 | TCGGGGAGGGAAGCCGGACA | GGG | \|\|\|\|\|\|\|\|[\|\|\|\|\|\|\|\|\|\|\|\|]PAM | NA | Intergenic | NA | NA |
| T26890 | GAGGGAAGCCGGACAGGGACCGG | 1000 | | | | | | | | | |
| Oligo fwd | CACCGAGGGAAGCCGGACAGGGAC | | | | | | | | | | |
| Oligo rev | AAACGTCCCTGTCCGGCTTCCCTC | | | | | | | | | | |
| Chromosome | start | end | strand | MM | target_seq | PAM | alignment | distance | position | gene name | gene id |
| GU305914.1 | 93913 | 93935 | - | 0 | GAGGGAAGCCGGACAGGGAC | CGG | \|\|\|\|\|\|\|\|[\|\|\|\|\|\|\|\|\|\|\|\|]PAM | NA | Intergenic | NA | NA |
| GU305914.1 | 94073 | 94095 | - | 0 | GAGGGAAGCCGGACAGGGAC | CGG | \|\|\|\|\|\|\|\|[\|\|\|\|\|\|\|\|\|\|\|\|]PAM | NA | Intergenic | NA | NA |
| GU305914.1 | 94041 | 94063 | - | 0 | GAGGGAAGCCGGACAGGGAC | CGG | \|\|\|\|\|\|\|\|[\|\|\|\|\|\|\|\|\|\|\|\|]PAM | NA | Intergenic | NA | NA |
| GU305914.1 | 94009 | 94031 | - | 0 | GAGGGAAGCCGGACAGGGAC | CGG | \|\|\|\|\|\|\|\|[\|\|\|\|\|\|\|\|\|\|\|\|]PAM | NA | Intergenic | NA | NA |
| GU305914.1 | 93977 | 93999 | - | 0 | GAGGGAAGCCGGACAGGGAC | CGG | \|\|\|\|\|\|\|\|[\|\|\|\|\|\|\|\|\|\|\|\|]PAM | NA | Intergenic | NA | NA |
| GU305914.1 | 93945 | 93967 | - | 0 | GAGGGAAGCCGGACAGGGAC | CGG | \|\|\|\|\|\|\|\|[\|\|\|\|\|\|\|\|\|\|\|\|]PAM | NA | Intergenic | NA | NA |

| | | | | | | | | | | | |
|---|---|---|---|---|---|---|---|---|---|---|---|
| GU305914.1 | 93881 | 93903 | - | 3 | GAGAGGAGCCGGACAGGAAC | CGG | |||-|-||[|||||||||-||]PAM | NA | Intergenic | NA | NA |
| T26891 | AGGGAAGCCGGACAGGGACCGGG | 1000 | | | | | | | | | |
| Oligo fwd | CACCAGGGAAGCCGGACAGGGACC | | | | | | | | | | |
| Oligo rev | AAACGGTCCCTGTCCGGCTTCCCT | | | | | | | | | | |
| Chromosome | start | end | strand | MM | target_seq | PAM | alignment | distance | position | gene name | gene id |
| GU305914.1 | 93976 | 93998 | - | 0 | AGGGAAGCCGGACAGGGACC | GGG | ||||||||[||||||||||||]PAM | NA | Intergenic | NA | NA |
| GU305914.1 | 93944 | 93966 | - | 0 | AGGGAAGCCGGACAGGGACC | GGG | ||||||||[||||||||||||]PAM | NA | Intergenic | NA | NA |
| GU305914.1 | 93912 | 93934 | - | 0 | AGGGAAGCCGGACAGGGACC | GGG | ||||||||[||||||||||||]PAM | NA | Intergenic | NA | NA |
| GU305914.1 | 94072 | 94094 | - | 0 | AGGGAAGCCGGACAGGGACC | GGG | ||||||||[||||||||||||]PAM | NA | Intergenic | NA | NA |
| GU305914.1 | 94040 | 94062 | - | 0 | AGGGAAGCCGGACAGGGACC | GGG | ||||||||[||||||||||||]PAM | NA | Intergenic | NA | NA |
| GU305914.1 | 94008 | 94030 | - | 0 | AGGGAAGCCGGACAGGGACC | GGG | ||||||||[||||||||||||]PAM | NA | Intergenic | NA | NA |
| GU305914.1 | 93880 | 93902 | - | 3 | AGAGGAGCCGGACAGGAACC | GGG | ||-|-|||[||||||||-|||]PAM | NA | Intergenic | NA | NA |
| T26892 | CCGGACAGGGACCGGGAGCTCGG | 1000 | | | | | | | | | |
| Oligo fwd | CACCCCGGACAGGGACCGGGAGCT | | | | | | | | | | |
| Oligo rev | AAACAGCTCCCGGTCCCTGTCCGG | | | | | | | | | | |
| Chromosome | start | end | strand | MM | target_seq | PAM | alignment | distance | position | gene name | gene id |
| GU305914.1 | 94001 | 94023 | - | 0 | CCGGACAGGGACCGGGAGCT | CGG | ||||||||[||||||||||||]PAM | NA | Intergenic | NA | NA |
| GU305914.1 | 93969 | 93991 | - | 0 | CCGGACAGGGACCGGGAGCT | CGG | ||||||||[||||||||||||]PAM | NA | Intergenic | NA | NA |
| GU305914.1 | 93937 | 93959 | - | 0 | CCGGACAGGGACCGGGAGCT | CGG | ||||||||[||||||||||||]PAM | NA | Intergenic | NA | NA |
| GU305914.1 | 93905 | 93927 | - | 0 | CCGGACAGGGACCGGGAGCT | CGG | ||||||||[||||||||||||]PAM | NA | Intergenic | NA | NA |
| GU305914.1 | 94065 | 94087 | - | 0 | CCGGACAGGGACCGGGAGCT | CGG | ||||||||[||||||||||||]PAM | NA | Intergenic | NA | NA |
| GU305914.1 | 94033 | 94055 | - | 0 | CCGGACAGGGACCGGGAGCT | CGG | ||||||||[||||||||||||]PAM | NA | Intergenic | NA | NA |
| GU305914.1 | 93873 | 93895 | - | 2 | CCGGACAGGAACCGGGAGCG | GGG | ||||||||[|-|||||||||-]PAM | NA | Intergenic | NA | NA |
| T26893 | CGGACAGGGACCGGGAGCTCGGG | 1000 | | | | | | | | | |
| Oligo fwd | CACCCGGACAGGGACCGGGAGCTC | | | | | | | | | | |
| Oligo rev | AAACGAGCTCCCGGTCCCTGTCCG | | | | | | | | | | |
| Chromosome | start | end | strand | MM | target_seq | PAM | alignment | distance | position | gene name | gene id |
| GU305914.1 | 93968 | 93990 | - | 0 | CGGACAGGGACCGGGAGCTC | GGG | ||||||||[||||||||||||]PAM | NA | Intergenic | NA | NA |
| GU305914.1 | 93936 | 93958 | - | 0 | CGGACAGGGACCGGGAGCTC | GGG | ||||||||[||||||||||||]PAM | NA | Intergenic | NA | NA |
| GU305914.1 | 93904 | 93926 | - | 0 | CGGACAGGGACCGGGAGCTC | GGG | ||||||||[||||||||||||]PAM | NA | Intergenic | NA | NA |
| GU305914.1 | 94064 | 94086 | - | 0 | CGGACAGGGACCGGGAGCTC | GGG | ||||||||[||||||||||||]PAM | NA | Intergenic | NA | NA |
| GU305914.1 | 94032 | 94054 | - | 0 | CGGACAGGGACCGGGAGCTC | GGG | ||||||||[||||||||||||]PAM | NA | Intergenic | NA | NA |
| GU305914.1 | 94000 | 94022 | - | 0 | CGGACAGGGACCGGGAGCTC | GGG | ||||||||[||||||||||||]PAM | NA | Intergenic | NA | NA |
| T26894 | GGACAGGGACCGGGAGCTCGGGG | 1000 | | | | | | | | | |
| Oligo fwd | CACCGGACAGGGACCGGGAGCTCG | | | | | | | | | | |
| Oligo rev | AAACCGAGCTCCCGGTCCCTGTCC | | | | | | | | | | |
| Chromosome | start | end | strand | MM | target_seq | PAM | alignment | distance | position | gene name | gene id |
| GU305914.1 | 93903 | 93925 | - | 0 | GGACAGGGACCGGGAGCTCG | GGG | ||||||||[||||||||||||]PAM | NA | Intergenic | NA | NA |
| GU305914.1 | 94063 | 94085 | - | 0 | GGACAGGGACCGGGAGCTCG | GGG | ||||||||[||||||||||||]PAM | NA | Intergenic | NA | NA |
| GU305914.1 | 94031 | 94053 | - | 0 | GGACAGGGACCGGGAGCTCG | GGG | ||||||||[||||||||||||]PAM | NA | Intergenic | NA | NA |
| GU305914.1 | 93999 | 94021 | - | 0 | GGACAGGGACCGGGAGCTCG | GGG | ||||||||[||||||||||||]PAM | NA | Intergenic | NA | NA |
| GU305914.1 | 93967 | 93989 | - | 0 | GGACAGGGACCGGGAGCTCG | GGG | ||||||||[||||||||||||]PAM | NA | Intergenic | NA | NA |
| GU305914.1 | 93935 | 93957 | - | 0 | GGACAGGGACCGGGAGCTCG | GGG | ||||||||[||||||||||||]PAM | NA | Intergenic | NA | NA |
| GU305914.1 | 94095 | 94117 | - | 4 | AAGCGGGGACCGGGAGCTCG | GGG | ---|-|||[||||||||||||]PAM | NA | Intergenic | NA | NA |
| T26895 | CAGGGACCGGGAGCTCGGGGAGG | 1000 | | | | | | | | | |
| Oligo fwd | CACCCAGGGACCGGGAGCTCGGGG | | | | | | | | | | |
| Oligo rev | AAACCCCCGAGCTCCCGGTCCCTG | | | | | | | | | | |
| Chromosome | start | end | strand | MM | target_seq | PAM | alignment | distance | position | gene name | gene id |
| GU305914.1 | 93996 | 94018 | - | 0 | CAGGGACCGGGAGCTCGGGG | AGG | ||||||||[||||||||||||]PAM | NA | Intergenic | NA | NA |
| GU305914.1 | 93964 | 93986 | - | 0 | CAGGGACCGGGAGCTCGGGG | AGG | ||||||||[||||||||||||]PAM | NA | Intergenic | NA | NA |
| GU305914.1 | 93932 | 93954 | - | 0 | CAGGGACCGGGAGCTCGGGG | AGG | ||||||||[||||||||||||]PAM | NA | Intergenic | NA | NA |
| GU305914.1 | 94060 | 94082 | - | 0 | CAGGGACCGGGAGCTCGGGG | AGG | ||||||||[||||||||||||]PAM | NA | Intergenic | NA | NA |
| GU305914.1 | 94028 | 94050 | - | 0 | CAGGGACCGGGAGCTCGGGG | AGG | ||||||||[||||||||||||]PAM | NA | Intergenic | NA | NA |
| GU305914.1 | 94092 | 94114 | - | 1 | CGGGGACCGGGAGCTCGGGG | AGG | |-||||||[||||||||||||]PAM | NA | Intergenic | NA | NA |
| T26896 | AGGGACCGGGAGCTCGGGGAGGG | 1000 | | | | | | | | | |
| Oligo fwd | CACCAGGGACCGGGAGCTCGGGGA | | | | | | | | | | |
| Oligo rev | AAACTCCCCGAGCTCCCGGTCCCT | | | | | | | | | | |
| Chromosome | start | end | strand | MM | target_seq | PAM | alignment | distance | position | gene name | gene id |
| GU305914.1 | 93995 | 94017 | - | 0 | AGGGACCGGGAGCTCGGGGA | GGG | ||||||||[||||||||||||]PAM | NA | Intergenic | NA | NA |
| GU305914.1 | 93963 | 93985 | - | 0 | AGGGACCGGGAGCTCGGGGA | GGG | ||||||||[||||||||||||]PAM | NA | Intergenic | NA | NA |
| GU305914.1 | 93931 | 93953 | - | 0 | AGGGACCGGGAGCTCGGGGA | GGG | ||||||||[||||||||||||]PAM | NA | Intergenic | NA | NA |
| GU305914.1 | 94059 | 94081 | - | 0 | AGGGACCGGGAGCTCGGGGA | GGG | ||||||||[||||||||||||]PAM | NA | Intergenic | NA | NA |
| GU305914.1 | 94027 | 94049 | - | 0 | AGGGACCGGGAGCTCGGGGA | GGG | ||||||||[||||||||||||]PAM | NA | Intergenic | NA | NA |
| GU305914.1 | 94091 | 94113 | - | 1 | GGGGACCGGGAGCTCGGGGA | GGG | -|||||||[||||||||||||]PAM | NA | Intergenic | NA | NA |
| T26897 | GGGAGCTCGGGGAGGGAAGCCGG | 1000 | | | | | | | | | |
| Oligo fwd | CACCGGGAGCTCGGGGAGGGAAGC | | | | | | | | | | |
| Oligo rev | AAACGCTTCCCTCCCCGAGCTCCC | | | | | | | | | | |
| Chromosome | start | end | strand | MM | target_seq | PAM | alignment | distance | position | gene name | gene id |
| GU305914.1 | 94052 | 94074 | - | 0 | GGGAGCTCGGGGAGGGAAGC | CGG | ||||||||[||||||||||||]PAM | NA | Intergenic | NA | NA |
| GU305914.1 | 94020 | 94042 | - | 0 | GGGAGCTCGGGGAGGGAAGC | CGG | ||||||||[||||||||||||]PAM | NA | Intergenic | NA | NA |
| GU305914.1 | 93988 | 94010 | - | 0 | GGGAGCTCGGGGAGGGAAGC | CGG | ||||||||[||||||||||||]PAM | NA | Intergenic | NA | NA |
| GU305914.1 | 93956 | 93978 | - | 0 | GGGAGCTCGGGGAGGGAAGC | CGG | ||||||||[||||||||||||]PAM | NA | Intergenic | NA | NA |
| GU305914.1 | 93924 | 93946 | - | 0 | GGGAGCTCGGGGAGGGAAGC | CGG | ||||||||[||||||||||||]PAM | NA | Intergenic | NA | NA |
| GU305914.1 | 94084 | 94106 | - | 0 | GGGAGCTCGGGGAGGGAAGC | CGG | ||||||||[||||||||||||]PAM | NA | Intergenic | NA | NA |
| GU305914.1 | 93892 | 93914 | - | 2 | GGGAGCTCGGGGAGAGGAGC | CGG | ||||||||[||||||-|-|||]PAM | NA | Intergenic | NA | NA |
| T26898 | CTCGGGGAGGGAAGCCGGACAGG | 1000 | | | | | | | | | |
| Oligo fwd | CACCCTCGGGGAGGGAAGCCGGAC | | | | | | | | | | |
| Oligo rev | AAACGTCCGGCTTCCCTCCCCGAG | | | | | | | | | | |
| Chromosome | start | end | strand | MM | target_seq | PAM | alignment | distance | position | gene name | gene id |
| GU305914.1 | 93951 | 93973 | - | 0 | CTCGGGGAGGGAAGCCGGAC | AGG | ||||||||[||||||||||||]PAM | NA | Intergenic | NA | NA |
| GU305914.1 | 93919 | 93941 | - | 0 | CTCGGGGAGGGAAGCCGGAC | AGG | ||||||||[||||||||||||]PAM | NA | Intergenic | NA | NA |

| | | | | | | | | | | | |
|---|---|---|---|---|---|---|---|---|---|---|---|
| GU305914.1 | 94079 | 94101 | - | 0 | CTCGGGGAGGGAAGCCGGAC | AGG | \|\|\|\|\|\|\|\|[\|\|\|\|\|\|\|\|\|\|\|\|]PAM | NA | Intergenic | NA | NA |
| GU305914.1 | 94047 | 94069 | - | 0 | CTCGGGGAGGGAAGCCGGAC | AGG | \|\|\|\|\|\|\|\|[\|\|\|\|\|\|\|\|\|\|\|\|]PAM | NA | Intergenic | NA | NA |
| GU305914.1 | 94015 | 94037 | - | 0 | CTCGGGGAGGGAAGCCGGAC | AGG | \|\|\|\|\|\|\|\|[\|\|\|\|\|\|\|\|\|\|\|\|]PAM | NA | Intergenic | NA | NA |
| GU305914.1 | 93983 | 94005 | - | 0 | CTCGGGGAGGGAAGCCGGAC | AGG | \|\|\|\|\|\|\|\|[\|\|\|\|\|\|\|\|\|\|\|\|]PAM | NA | Intergenic | NA | NA |
| GU305914.1 | 93887 | 93909 | - | 2 | CTCGGGGAGAGGAGCCGGAC | AGG | \|\|\|\|\|\|\|\|[\|-\|-\|\|\|\|\|\|\|\|]PAM | NA | Intergenic | NA | NA |
| T26899 | TCGGGGAGGGAAGCCGGACAGGG | 1000 | | | | | | | | | |
| Oligo fwd | CACCTCGGGGAGGGAAGCCGGACA | | | | | | | | | | |
| Oligo rev | AAACTGTCCGGCTTCCCTCCCCGA | | | | | | | | | | |
| Chromosome | start | end | strand | MM | target_seq | PAM | alignment | distance | position | gene name | gene id |
| GU305914.1 | 94078 | 94100 | - | 0 | TCGGGGAGGGAAGCCGGACA | GGG | \|\|\|\|\|\|\|\|[\|\|\|\|\|\|\|\|\|\|\|\|]PAM | NA | Intergenic | NA | NA |
| GU305914.1 | 94046 | 94068 | - | 0 | TCGGGGAGGGAAGCCGGACA | GGG | \|\|\|\|\|\|\|\|[\|\|\|\|\|\|\|\|\|\|\|\|]PAM | NA | Intergenic | NA | NA |
| GU305914.1 | 94014 | 94036 | - | 0 | TCGGGGAGGGAAGCCGGACA | GGG | \|\|\|\|\|\|\|\|[\|\|\|\|\|\|\|\|\|\|\|\|]PAM | NA | Intergenic | NA | NA |
| GU305914.1 | 93982 | 94004 | - | 0 | TCGGGGAGGGAAGCCGGACA | GGG | \|\|\|\|\|\|\|\|[\|\|\|\|\|\|\|\|\|\|\|\|]PAM | NA | Intergenic | NA | NA |
| GU305914.1 | 93950 | 93972 | - | 0 | TCGGGGAGGGAAGCCGGACA | GGG | \|\|\|\|\|\|\|\|[\|\|\|\|\|\|\|\|\|\|\|\|]PAM | NA | Intergenic | NA | NA |
| GU305914.1 | 93918 | 93940 | - | 0 | TCGGGGAGGGAAGCCGGACA | GGG | \|\|\|\|\|\|\|\|[\|\|\|\|\|\|\|\|\|\|\|\|]PAM | NA | Intergenic | NA | NA |
| T26900 | GAGGGAAGCCGGACAGGGACCGG | 1000 | | | | | | | | | |
| Oligo fwd | CACCGAGGGAAGCCGGACAGGGAC | | | | | | | | | | |
| Oligo rev | AAACGTCCCTGTCCGGCTTCCCTC | | | | | | | | | | |
| Chromosome | start | end | strand | MM | target_seq | PAM | alignment | distance | position | gene name | gene id |
| GU305914.1 | 93913 | 93935 | - | 0 | GAGGGAAGCCGGACAGGGAC | CGG | \|\|\|\|\|\|\|\|[\|\|\|\|\|\|\|\|\|\|\|\|]PAM | NA | Intergenic | NA | NA |
| GU305914.1 | 94073 | 94095 | - | 0 | GAGGGAAGCCGGACAGGGAC | CGG | \|\|\|\|\|\|\|\|[\|\|\|\|\|\|\|\|\|\|\|\|]PAM | NA | Intergenic | NA | NA |
| GU305914.1 | 94041 | 94063 | - | 0 | GAGGGAAGCCGGACAGGGAC | CGG | \|\|\|\|\|\|\|\|[\|\|\|\|\|\|\|\|\|\|\|\|]PAM | NA | Intergenic | NA | NA |
| GU305914.1 | 94009 | 94031 | - | 0 | GAGGGAAGCCGGACAGGGAC | CGG | \|\|\|\|\|\|\|\|[\|\|\|\|\|\|\|\|\|\|\|\|]PAM | NA | Intergenic | NA | NA |
| GU305914.1 | 93977 | 93999 | - | 0 | GAGGGAAGCCGGACAGGGAC | CGG | \|\|\|\|\|\|\|\|[\|\|\|\|\|\|\|\|\|\|\|\|]PAM | NA | Intergenic | NA | NA |
| GU305914.1 | 93945 | 93967 | - | 0 | GAGGGAAGCCGGACAGGGAC | CGG | \|\|\|\|\|\|\|\|[\|\|\|\|\|\|\|\|\|\|\|\|]PAM | NA | Intergenic | NA | NA |
| GU305914.1 | 93881 | 93903 | - | 3 | GAGAGGAGCCGGACAGGAAC | CGG | \|\|\|-\|-\|\|[\|\|\|\|\|\|\|\|\|-\|\|]PAM | NA | Intergenic | NA | NA |
| T26901 | AGGGAAGCCGGACAGGGACCGGG | 1000 | | | | | | | | | |
| Oligo fwd | CACCAGGGAAGCCGGACAGGGACC | | | | | | | | | | |
| Oligo rev | AAACGGTCCCTGTCCGGCTTCCCT | | | | | | | | | | |
| Chromosome | start | end | strand | MM | target_seq | PAM | alignment | distance | position | gene name | gene id |
| GU305914.1 | 93976 | 93998 | - | 0 | AGGGAAGCCGGACAGGGACC | GGG | \|\|\|\|\|\|\|\|[\|\|\|\|\|\|\|\|\|\|\|\|]PAM | NA | Intergenic | NA | NA |
| GU305914.1 | 93944 | 93966 | - | 0 | AGGGAAGCCGGACAGGGACC | GGG | \|\|\|\|\|\|\|\|[\|\|\|\|\|\|\|\|\|\|\|\|]PAM | NA | Intergenic | NA | NA |
| GU305914.1 | 93912 | 93934 | - | 0 | AGGGAAGCCGGACAGGGACC | GGG | \|\|\|\|\|\|\|\|[\|\|\|\|\|\|\|\|\|\|\|\|]PAM | NA | Intergenic | NA | NA |
| GU305914.1 | 94072 | 94094 | - | 0 | AGGGAAGCCGGACAGGGACC | GGG | \|\|\|\|\|\|\|\|[\|\|\|\|\|\|\|\|\|\|\|\|]PAM | NA | Intergenic | NA | NA |
| GU305914.1 | 94040 | 94062 | - | 0 | AGGGAAGCCGGACAGGGACC | GGG | \|\|\|\|\|\|\|\|[\|\|\|\|\|\|\|\|\|\|\|\|]PAM | NA | Intergenic | NA | NA |
| GU305914.1 | 94008 | 94030 | - | 0 | AGGGAAGCCGGACAGGGACC | GGG | \|\|\|\|\|\|\|\|[\|\|\|\|\|\|\|\|\|\|\|\|]PAM | NA | Intergenic | NA | NA |
| GU305914.1 | 93880 | 93902 | - | 3 | AGAGGAGCCGGACAGGAACC | GGG | \|\|-\|-\|\|\|[\|\|\|\|\|\|\|\|-\|\|\|]PAM | NA | Intergenic | NA | NA |
| T26902 | CCGGACAGGGACCGGGAGCTCGG | 1000 | | | | | | | | | |
| Oligo fwd | CACCCCGGACAGGGACCGGGAGCT | | | | | | | | | | |
| Oligo rev | AAACAGCTCCCGGTCCCTGTCCGG | | | | | | | | | | |
| Chromosome | start | end | strand | MM | target_seq | PAM | alignment | distance | position | gene name | gene id |
| GU305914.1 | 94001 | 94023 | - | 0 | CCGGACAGGGACCGGGAGCT | CGG | \|\|\|\|\|\|\|\|[\|\|\|\|\|\|\|\|\|\|\|\|]PAM | NA | Intergenic | NA | NA |
| GU305914.1 | 93969 | 93991 | - | 0 | CCGGACAGGGACCGGGAGCT | CGG | \|\|\|\|\|\|\|\|[\|\|\|\|\|\|\|\|\|\|\|\|]PAM | NA | Intergenic | NA | NA |
| GU305914.1 | 93937 | 93959 | - | 0 | CCGGACAGGGACCGGGAGCT | CGG | \|\|\|\|\|\|\|\|[\|\|\|\|\|\|\|\|\|\|\|\|]PAM | NA | Intergenic | NA | NA |
| GU305914.1 | 93905 | 93927 | - | 0 | CCGGACAGGGACCGGGAGCT | CGG | \|\|\|\|\|\|\|\|[\|\|\|\|\|\|\|\|\|\|\|\|]PAM | NA | Intergenic | NA | NA |
| GU305914.1 | 94065 | 94087 | - | 0 | CCGGACAGGGACCGGGAGCT | CGG | \|\|\|\|\|\|\|\|[\|\|\|\|\|\|\|\|\|\|\|\|]PAM | NA | Intergenic | NA | NA |
| GU305914.1 | 94033 | 94055 | - | 0 | CCGGACAGGGACCGGGAGCT | CGG | \|\|\|\|\|\|\|\|[\|\|\|\|\|\|\|\|\|\|\|\|]PAM | NA | Intergenic | NA | NA |
| GU305914.1 | 93873 | 93895 | - | 2 | CCGGACAGGAACCGGGAGCG | GGG | \|\|\|\|\|\|\|\|[\|-\|\|\|\|\|\|\|\|\|-]PAM | NA | Intergenic | NA | NA |
| T26903 | CGGACAGGGACCGGGAGCTCGGG | 1000 | | | | | | | | | |
| Oligo fwd | CACCCGGACAGGGACCGGGAGCTC | | | | | | | | | | |
| Oligo rev | AAACGAGCTCCCGGTCCCTGTCCG | | | | | | | | | | |
| Chromosome | start | end | strand | MM | target_seq | PAM | alignment | distance | position | gene name | gene id |
| GU305914.1 | 93968 | 93990 | - | 0 | CGGACAGGGACCGGGAGCTC | GGG | \|\|\|\|\|\|\|\|[\|\|\|\|\|\|\|\|\|\|\|\|]PAM | NA | Intergenic | NA | NA |
| GU305914.1 | 93936 | 93958 | - | 0 | CGGACAGGGACCGGGAGCTC | GGG | \|\|\|\|\|\|\|\|[\|\|\|\|\|\|\|\|\|\|\|\|]PAM | NA | Intergenic | NA | NA |
| GU305914.1 | 93904 | 93926 | - | 0 | CGGACAGGGACCGGGAGCTC | GGG | \|\|\|\|\|\|\|\|[\|\|\|\|\|\|\|\|\|\|\|\|]PAM | NA | Intergenic | NA | NA |
| GU305914.1 | 94064 | 94086 | - | 0 | CGGACAGGGACCGGGAGCTC | GGG | \|\|\|\|\|\|\|\|[\|\|\|\|\|\|\|\|\|\|\|\|]PAM | NA | Intergenic | NA | NA |
| GU305914.1 | 94032 | 94054 | - | 0 | CGGACAGGGACCGGGAGCTC | GGG | \|\|\|\|\|\|\|\|[\|\|\|\|\|\|\|\|\|\|\|\|]PAM | NA | Intergenic | NA | NA |
| GU305914.1 | 94000 | 94022 | - | 0 | CGGACAGGGACCGGGAGCTC | GGG | \|\|\|\|\|\|\|\|[\|\|\|\|\|\|\|\|\|\|\|\|]PAM | NA | Intergenic | NA | NA |
| T26904 | GGACAGGGACCGGGAGCTCGGGG | 1000 | | | | | | | | | |
| Oligo fwd | CACCGGACAGGGACCGGGAGCTCG | | | | | | | | | | |
| Oligo rev | AAACCGAGCTCCCGGTCCCTGTCC | | | | | | | | | | |
| Chromosome | start | end | strand | MM | target_seq | PAM | alignment | distance | position | gene name | gene id |
| GU305914.1 | 93903 | 93925 | - | 0 | GGACAGGGACCGGGAGCTCG | GGG | \|\|\|\|\|\|\|\|[\|\|\|\|\|\|\|\|\|\|\|\|]PAM | NA | Intergenic | NA | NA |
| GU305914.1 | 94063 | 94085 | - | 0 | GGACAGGGACCGGGAGCTCG | GGG | \|\|\|\|\|\|\|\|[\|\|\|\|\|\|\|\|\|\|\|\|]PAM | NA | Intergenic | NA | NA |
| GU305914.1 | 94031 | 94053 | - | 0 | GGACAGGGACCGGGAGCTCG | GGG | \|\|\|\|\|\|\|\|[\|\|\|\|\|\|\|\|\|\|\|\|]PAM | NA | Intergenic | NA | NA |
| GU305914.1 | 93999 | 94021 | - | 0 | GGACAGGGACCGGGAGCTCG | GGG | \|\|\|\|\|\|\|\|[\|\|\|\|\|\|\|\|\|\|\|\|]PAM | NA | Intergenic | NA | NA |
| GU305914.1 | 93967 | 93989 | - | 0 | GGACAGGGACCGGGAGCTCG | GGG | \|\|\|\|\|\|\|\|[\|\|\|\|\|\|\|\|\|\|\|\|]PAM | NA | Intergenic | NA | NA |
| GU305914.1 | 93935 | 93957 | - | 0 | GGACAGGGACCGGGAGCTCG | GGG | \|\|\|\|\|\|\|\|[\|\|\|\|\|\|\|\|\|\|\|\|]PAM | NA | Intergenic | NA | NA |
| GU305914.1 | 94095 | 94117 | - | 4 | AAGCGGGGACCGGGAGCTCG | GGG | ---\|-\|\|\|[\|\|\|\|\|\|\|\|\|\|\|\|]PAM | NA | Intergenic | NA | NA |
| T26905 | GGGACCGGGAGCTCGGGGAGAGG | 1000 | | | | | | | | | |
| Oligo fwd | CACCGGGACCGGGAGCTCGGGGAG | | | | | | | | | | |
| Oligo rev | AAACCTCCCCGAGCTCCCGGTCCC | | | | | | | | | | |
| Chromosome | start | end | strand | MM | target_seq | PAM | alignment | distance | position | gene name | gene id |
| GU305914.1 | 93898 | 93920 | - | 0 | GGGACCGGGAGCTCGGGGAG | AGG | \|\|\|\|\|\|\|\|[\|\|\|\|\|\|\|\|\|\|\|\|]PAM | NA | Intergenic | NA | NA |
| T26906 | GGGAGCTCGGGGAGAGGAGCCGG | 1000 | | | | | | | | | |
| Oligo fwd | CACCGGGAGCTCGGGGAGAGGAGC | | | | | | | | | | |
| Oligo rev | AAACGCTCCTCTCCCCGAGCTCCC | | | | | | | | | | |
| Chromosome | start | end | strand | MM | target_seq | PAM | alignment | distance | position | gene name | gene id |
| GU305914.1 | 93892 | 93914 | - | 0 | GGGAGCTCGGGGAGAGGAGC | CGG | \|\|\|\|\|\|\|\|[\|\|\|\|\|\|\|\|\|\|\|\|]PAM | NA | Intergenic | NA | NA |
| GU305914.1 | 93956 | 93978 | - | 2 | GGGAGCTCGGGGAGGGAAGC | CGG | \|\|\|\|\|\|\|\|[\|\|\|\|\|\|-\|-\|\|\|]PAM | NA | Intergenic | NA | NA |
| GU305914.1 | 93924 | 93946 | - | 2 | GGGAGCTCGGGGAGGGAAGC | CGG | \|\|\|\|\|\|\|\|[\|\|\|\|\|\|-\|-\|\|\|]PAM | NA | Intergenic | NA | NA |

<table>
<tr><td>GU305914.1</td><td>94084</td><td>94106</td><td>-</td><td>2</td><td>GGGAGCTCGGGGAGGGAAGC</td><td>CGG</td><td>||||||||[||||||-|-|||]PAM</td><td>NA</td><td>Intergenic</td><td>NA</td><td>NA</td></tr>
<tr><td>GU305914.1</td><td>94052</td><td>94074</td><td>-</td><td>2</td><td>GGGAGCTCGGGGAGGGAAGC</td><td>CGG</td><td>||||||||[||||||-|-|||]PAM</td><td>NA</td><td>Intergenic</td><td>NA</td><td>NA</td></tr>
<tr><td>GU305914.1</td><td>94020</td><td>94042</td><td>-</td><td>2</td><td>GGGAGCTCGGGGAGGGAAGC</td><td>CGG</td><td>||||||||[||||||-|-|||]PAM</td><td>NA</td><td>Intergenic</td><td>NA</td><td>NA</td></tr>
<tr><td>GU305914.1</td><td>93988</td><td>94010</td><td>-</td><td>2</td><td>GGGAGCTCGGGGAGGGAAGC</td><td>CGG</td><td>||||||||[||||||-|-|||]PAM</td><td>NA</td><td>Intergenic</td><td>NA</td><td>NA</td></tr>
<tr><td></td><td></td><td></td><td></td><td></td><td></td><td></td><td></td><td></td><td></td><td></td><td></td></tr>
<tr><td>T26907</td><td>CTCGGGGAGAGGAGCCGGACAGG</td><td>1000</td><td></td><td></td><td></td><td></td><td></td><td></td><td></td><td></td><td></td></tr>
<tr><td>Oligo fwd</td><td>CACCCTCGGGGAGAGGAGCCGGAC</td><td></td><td></td><td></td><td></td><td></td><td></td><td></td><td></td><td></td><td></td></tr>
<tr><td>Oligo rev</td><td>AAACGTCCGGCTCCTCTCCCCGAG</td><td></td><td></td><td></td><td></td><td></td><td></td><td></td><td></td><td></td><td></td></tr>
<tr><td>Chromosome</td><td>start</td><td>end</td><td>strand</td><td>MM</td><td>target_seq</td><td>PAM</td><td>alignment</td><td>distance</td><td>position</td><td>gene name</td><td>gene id</td></tr>
<tr><td>GU305914.1</td><td>93887</td><td>93909</td><td>-</td><td>0</td><td>CTCGGGGAGAGGAGCCGGAC</td><td>AGG</td><td>||||||||[||||||||||||]PAM</td><td>NA</td><td>Intergenic</td><td>NA</td><td>NA</td></tr>
<tr><td>GU305914.1</td><td>94015</td><td>94037</td><td>-</td><td>2</td><td>CTCGGGGAGGGAAGCCGGAC</td><td>AGG</td><td>||||||||[|-|-||||||||]PAM</td><td>NA</td><td>Intergenic</td><td>NA</td><td>NA</td></tr>
<tr><td>GU305914.1</td><td>93983</td><td>94005</td><td>-</td><td>2</td><td>CTCGGGGAGGGAAGCCGGAC</td><td>AGG</td><td>||||||||[|-|-||||||||]PAM</td><td>NA</td><td>Intergenic</td><td>NA</td><td>NA</td></tr>
<tr><td>GU305914.1</td><td>93951</td><td>93973</td><td>-</td><td>2</td><td>CTCGGGGAGGGAAGCCGGAC</td><td>AGG</td><td>||||||||[|-|-||||||||]PAM</td><td>NA</td><td>Intergenic</td><td>NA</td><td>NA</td></tr>
<tr><td>GU305914.1</td><td>93919</td><td>93941</td><td>-</td><td>2</td><td>CTCGGGGAGGGAAGCCGGAC</td><td>AGG</td><td>||||||||[|-|-||||||||]PAM</td><td>NA</td><td>Intergenic</td><td>NA</td><td>NA</td></tr>
<tr><td>GU305914.1</td><td>94079</td><td>94101</td><td>-</td><td>2</td><td>CTCGGGGAGGGAAGCCGGAC</td><td>AGG</td><td>||||||||[|-|-||||||||]PAM</td><td>NA</td><td>Intergenic</td><td>NA</td><td>NA</td></tr>
<tr><td>GU305914.1</td><td>94047</td><td>94069</td><td>-</td><td>2</td><td>CTCGGGGAGGGAAGCCGGAC</td><td>AGG</td><td>||||||||[|-|-||||||||]PAM</td><td>NA</td><td>Intergenic</td><td>NA</td><td>NA</td></tr>
<tr><td></td><td></td><td></td><td></td><td></td><td></td><td></td><td></td><td></td><td></td><td></td><td></td></tr>
<tr><td>T26908</td><td>GAGAGGAGCCGGACAGGAACCGG</td><td>1000</td><td></td><td></td><td></td><td></td><td></td><td></td><td></td><td></td><td></td></tr>
<tr><td>Oligo fwd</td><td>CACCGAGAGGAGCCGGACAGGAAC</td><td></td><td></td><td></td><td></td><td></td><td></td><td></td><td></td><td></td><td></td></tr>
<tr><td>Oligo rev</td><td>AAACGTTCCTGTCCGGCTCCTCTC</td><td></td><td></td><td></td><td></td><td></td><td></td><td></td><td></td><td></td><td></td></tr>
<tr><td>Chromosome</td><td>start</td><td>end</td><td>strand</td><td>MM</td><td>target_seq</td><td>PAM</td><td>alignment</td><td>distance</td><td>position</td><td>gene name</td><td>gene id</td></tr>
<tr><td>GU305914.1</td><td>93881</td><td>93903</td><td>-</td><td>0</td><td>GAGAGGAGCCGGACAGGAAC</td><td>CGG</td><td>||||||||[||||||||||||]PAM</td><td>NA</td><td>Intergenic</td><td>NA</td><td>NA</td></tr>
<tr><td>GU305914.1</td><td>93913</td><td>93935</td><td>-</td><td>3</td><td>GAGGGAAGCCGGACAGGGAC</td><td>CGG</td><td>|||-|-||[|||||||||-||]PAM</td><td>NA</td><td>Intergenic</td><td>NA</td><td>NA</td></tr>
<tr><td>GU305914.1</td><td>94073</td><td>94095</td><td>-</td><td>3</td><td>GAGGGAAGCCGGACAGGGAC</td><td>CGG</td><td>|||-|-||[|||||||||-||]PAM</td><td>NA</td><td>Intergenic</td><td>NA</td><td>NA</td></tr>
<tr><td>GU305914.1</td><td>94041</td><td>94063</td><td>-</td><td>3</td><td>GAGGGAAGCCGGACAGGGAC</td><td>CGG</td><td>|||-|-||[|||||||||-||]PAM</td><td>NA</td><td>Intergenic</td><td>NA</td><td>NA</td></tr>
<tr><td>GU305914.1</td><td>94009</td><td>94031</td><td>-</td><td>3</td><td>GAGGGAAGCCGGACAGGGAC</td><td>CGG</td><td>|||-|-||[|||||||||-||]PAM</td><td>NA</td><td>Intergenic</td><td>NA</td><td>NA</td></tr>
<tr><td>GU305914.1</td><td>93977</td><td>93999</td><td>-</td><td>3</td><td>GAGGGAAGCCGGACAGGGAC</td><td>CGG</td><td>|||-|-||[|||||||||-||]PAM</td><td>NA</td><td>Intergenic</td><td>NA</td><td>NA</td></tr>
<tr><td>GU305914.1</td><td>93945</td><td>93967</td><td>-</td><td>3</td><td>GAGGGAAGCCGGACAGGGAC</td><td>CGG</td><td>|||-|-||[|||||||||-||]PAM</td><td>NA</td><td>Intergenic</td><td>NA</td><td>NA</td></tr>
<tr><td></td><td></td><td></td><td></td><td></td><td></td><td></td><td></td><td></td><td></td><td></td><td></td></tr>
<tr><td>T26909</td><td>AGAGGAGCCGGACAGGAACCGGG</td><td>1000</td><td></td><td></td><td></td><td></td><td></td><td></td><td></td><td></td><td></td></tr>
<tr><td>Oligo fwd</td><td>CACCAGAGGAGCCGGACAGGAACC</td><td></td><td></td><td></td><td></td><td></td><td></td><td></td><td></td><td></td><td></td></tr>
<tr><td>Oligo rev</td><td>AAACGGTTCCTGTCCGGCTCCTCT</td><td></td><td></td><td></td><td></td><td></td><td></td><td></td><td></td><td></td><td></td></tr>
<tr><td>Chromosome</td><td>start</td><td>end</td><td>strand</td><td>MM</td><td>target_seq</td><td>PAM</td><td>alignment</td><td>distance</td><td>position</td><td>gene name</td><td>gene id</td></tr>
<tr><td>GU305914.1</td><td>93880</td><td>93902</td><td>-</td><td>0</td><td>AGAGGAGCCGGACAGGAACC</td><td>GGG</td><td>||||||||[||||||||||||]PAM</td><td>NA</td><td>Intergenic</td><td>NA</td><td>NA</td></tr>
<tr><td>GU305914.1</td><td>94072</td><td>94094</td><td>-</td><td>3</td><td>AGGGAAGCCGGACAGGGACC</td><td>GGG</td><td>||-|-|||[||||||||-|||]PAM</td><td>NA</td><td>Intergenic</td><td>NA</td><td>NA</td></tr>
<tr><td>GU305914.1</td><td>94040</td><td>94062</td><td>-</td><td>3</td><td>AGGGAAGCCGGACAGGGACC</td><td>GGG</td><td>||-|-|||[||||||||-|||]PAM</td><td>NA</td><td>Intergenic</td><td>NA</td><td>NA</td></tr>
<tr><td>GU305914.1</td><td>94008</td><td>94030</td><td>-</td><td>3</td><td>AGGGAAGCCGGACAGGGACC</td><td>GGG</td><td>||-|-|||[||||||||-|||]PAM</td><td>NA</td><td>Intergenic</td><td>NA</td><td>NA</td></tr>
<tr><td>GU305914.1</td><td>93976</td><td>93998</td><td>-</td><td>3</td><td>AGGGAAGCCGGACAGGGACC</td><td>GGG</td><td>||-|-|||[||||||||-|||]PAM</td><td>NA</td><td>Intergenic</td><td>NA</td><td>NA</td></tr>
<tr><td>GU305914.1</td><td>93944</td><td>93966</td><td>-</td><td>3</td><td>AGGGAAGCCGGACAGGGACC</td><td>GGG</td><td>||-|-|||[||||||||-|||]PAM</td><td>NA</td><td>Intergenic</td><td>NA</td><td>NA</td></tr>
<tr><td>GU305914.1</td><td>93912</td><td>93934</td><td>-</td><td>3</td><td>AGGGAAGCCGGACAGGGACC</td><td>GGG</td><td>||-|-|||[||||||||-|||]PAM</td><td>NA</td><td>Intergenic</td><td>NA</td><td>NA</td></tr>
<tr><td></td><td></td><td></td><td></td><td></td><td></td><td></td><td></td><td></td><td></td><td></td><td></td></tr>
<tr><td>T26910</td><td>AGCCGGACAGGAACCGGGAGCGG</td><td>1000</td><td></td><td></td><td></td><td></td><td></td><td></td><td></td><td></td><td></td></tr>
<tr><td>Oligo fwd</td><td>CACCAGCCGGACAGGAACCGGGAG</td><td></td><td></td><td></td><td></td><td></td><td></td><td></td><td></td><td></td><td></td></tr>
<tr><td>Oligo rev</td><td>AAACCTCCCGGTTCCTGTCCGGCT</td><td></td><td></td><td></td><td></td><td></td><td></td><td></td><td></td><td></td><td></td></tr>
<tr><td>Chromosome</td><td>start</td><td>end</td><td>strand</td><td>MM</td><td>target_seq</td><td>PAM</td><td>alignment</td><td>distance</td><td>position</td><td>gene name</td><td>gene id</td></tr>
<tr><td>GU305914.1</td><td>93875</td><td>93897</td><td>-</td><td>0</td><td>AGCCGGACAGGAACCGGGAG</td><td>CGG</td><td>||||||||[||||||||||||]PAM</td><td>NA</td><td>Intergenic</td><td>NA</td><td>NA</td></tr>
<tr><td></td><td></td><td></td><td></td><td></td><td></td><td></td><td></td><td></td><td></td><td></td><td></td></tr>
<tr><td>T26911</td><td>GCCGGACAGGAACCGGGAGCGGG</td><td>1000</td><td></td><td></td><td></td><td></td><td></td><td></td><td></td><td></td><td></td></tr>
<tr><td>Oligo fwd</td><td>CACCGCCGGACAGGAACCGGGAGC</td><td></td><td></td><td></td><td></td><td></td><td></td><td></td><td></td><td></td><td></td></tr>
<tr><td>Oligo rev</td><td>AAACGCTCCCGGTTCCTGTCCGGC</td><td></td><td></td><td></td><td></td><td></td><td></td><td></td><td></td><td></td><td></td></tr>
<tr><td>Chromosome</td><td>start</td><td>end</td><td>strand</td><td>MM</td><td>target_seq</td><td>PAM</td><td>alignment</td><td>distance</td><td>position</td><td>gene name</td><td>gene id</td></tr>
<tr><td>GU305914.1</td><td>93874</td><td>93896</td><td>-</td><td>0</td><td>GCCGGACAGGAACCGGGAGC</td><td>GGG</td><td>||||||||[||||||||||||]PAM</td><td>NA</td><td>Intergenic</td><td>NA</td><td>NA</td></tr>
<tr><td></td><td></td><td></td><td></td><td></td><td></td><td></td><td></td><td></td><td></td><td></td><td></td></tr>
<tr><td>T26912</td><td>CCGGACAGGAACCGGGAGCGGGG</td><td>1000</td><td></td><td></td><td></td><td></td><td></td><td></td><td></td><td></td><td></td></tr>
<tr><td>Oligo fwd</td><td>CACCCCGGACAGGAACCGGGAGCG</td><td></td><td></td><td></td><td></td><td></td><td></td><td></td><td></td><td></td><td></td></tr>
<tr><td>Oligo rev</td><td>AAACCGCTCCCGGTTCCTGTCCGG</td><td></td><td></td><td></td><td></td><td></td><td></td><td></td><td></td><td></td><td></td></tr>
<tr><td>Chromosome</td><td>start</td><td>end</td><td>strand</td><td>MM</td><td>target_seq</td><td>PAM</td><td>alignment</td><td>distance</td><td>position</td><td>gene name</td><td>gene id</td></tr>
<tr><td>GU305914.1</td><td>93873</td><td>93895</td><td>-</td><td>0</td><td>CCGGACAGGAACCGGGAGCG</td><td>GGG</td><td>||||||||[||||||||||||]PAM</td><td>NA</td><td>Intergenic</td><td>NA</td><td>NA</td></tr>
<tr><td>GU305914.1</td><td>93905</td><td>93927</td><td>-</td><td>2</td><td>CCGGACAGGGACCGGGAGCT</td><td>CGG</td><td>||||||||[|-|||||||||-]PAM</td><td>NA</td><td>Intergenic</td><td>NA</td><td>NA</td></tr>
<tr><td>GU305914.1</td><td>94065</td><td>94087</td><td>-</td><td>2</td><td>CCGGACAGGGACCGGGAGCT</td><td>CGG</td><td>||||||||[|-|||||||||-]PAM</td><td>NA</td><td>Intergenic</td><td>NA</td><td>NA</td></tr>
<tr><td>GU305914.1</td><td>94033</td><td>94055</td><td>-</td><td>2</td><td>CCGGACAGGGACCGGGAGCT</td><td>CGG</td><td>||||||||[|-|||||||||-]PAM</td><td>NA</td><td>Intergenic</td><td>NA</td><td>NA</td></tr>
<tr><td>GU305914.1</td><td>94001</td><td>94023</td><td>-</td><td>2</td><td>CCGGACAGGGACCGGGAGCT</td><td>CGG</td><td>||||||||[|-|||||||||-]PAM</td><td>NA</td><td>Intergenic</td><td>NA</td><td>NA</td></tr>
<tr><td>GU305914.1</td><td>93969</td><td>93991</td><td>-</td><td>2</td><td>CCGGACAGGGACCGGGAGCT</td><td>CGG</td><td>||||||||[|-|||||||||-]PAM</td><td>NA</td><td>Intergenic</td><td>NA</td><td>NA</td></tr>
<tr><td>GU305914.1</td><td>93937</td><td>93959</td><td>-</td><td>2</td><td>CCGGACAGGGACCGGGAGCT</td><td>CGG</td><td>||||||||[|-|||||||||-]PAM</td><td>NA</td><td>Intergenic</td><td>NA</td><td>NA</td></tr>
<tr><td></td><td></td><td></td><td></td><td></td><td></td><td></td><td></td><td></td><td></td><td></td><td></td></tr>
<tr><td>T26913</td><td>AGGAACCGGGAGCGGGGCGCTGG</td><td>1000</td><td></td><td></td><td></td><td></td><td></td><td></td><td></td><td></td><td></td></tr>
<tr><td>Oligo fwd</td><td>CACCAGGAACCGGGAGCGGGGCGC</td><td></td><td></td><td></td><td></td><td></td><td></td><td></td><td></td><td></td><td></td></tr>
<tr><td>Oligo rev</td><td>AAACGCGCCCCGCTCCCGGTTCCT</td><td></td><td></td><td></td><td></td><td></td><td></td><td></td><td></td><td></td><td></td></tr>
<tr><td>Chromosome</td><td>start</td><td>end</td><td>strand</td><td>MM</td><td>target_seq</td><td>PAM</td><td>alignment</td><td>distance</td><td>position</td><td>gene name</td><td>gene id</td></tr>
<tr><td>GU305914.1</td><td>93867</td><td>93889</td><td>-</td><td>0</td><td>AGGAACCGGGAGCGGGGCGC</td><td>TGG</td><td>||||||||[||||||||||||]PAM</td><td>NA</td><td>Intergenic</td><td>NA</td><td>NA</td></tr>
<tr><td></td><td></td><td></td><td></td><td></td><td></td><td></td><td></td><td></td><td></td><td></td><td></td></tr>
<tr><td>T26914</td><td>CCGGGAGCGGGGCGCTGGTCTGG</td><td>1000</td><td></td><td></td><td></td><td></td><td></td><td></td><td></td><td></td><td></td></tr>
<tr><td>Oligo fwd</td><td>CACCCCGGGAGCGGGGCGCTGGTC</td><td></td><td></td><td></td><td></td><td></td><td></td><td></td><td></td><td></td><td></td></tr>
<tr><td>Oligo rev</td><td>AAACGACCAGCGCCCCGCTCCCGG</td><td></td><td></td><td></td><td></td><td></td><td></td><td></td><td></td><td></td><td></td></tr>
<tr><td>Chromosome</td><td>start</td><td>end</td><td>strand</td><td>MM</td><td>target_seq</td><td>PAM</td><td>alignment</td><td>distance</td><td>position</td><td>gene name</td><td>gene id</td></tr>
<tr><td>GU305914.1</td><td>93862</td><td>93884</td><td>-</td><td>0</td><td>CCGGGAGCGGGGCGCTGGTC</td><td>TGG</td><td>||||||||[||||||||||||]PAM</td><td>NA</td><td>Intergenic</td><td>NA</td><td>NA</td></tr>
<tr><td></td><td></td><td></td><td></td><td></td><td></td><td></td><td></td><td></td><td></td><td></td><td></td></tr>
<tr><td>T26915</td><td>CGGGAGCGGGGCGCTGGTCTGGG</td><td>1000</td><td></td><td></td><td></td><td></td><td></td><td></td><td></td><td></td><td></td></tr>
<tr><td>Oligo fwd</td><td>CACCCGGGAGCGGGGCGCTGGTCT</td><td></td><td></td><td></td><td></td><td></td><td></td><td></td><td></td><td></td><td></td></tr>
<tr><td>Oligo rev</td><td>AAACAGACCAGCGCCCCGCTCCCG</td><td></td><td></td><td></td><td></td><td></td><td></td><td></td><td></td><td></td><td></td></tr>
<tr><td>Chromosome</td><td>start</td><td>end</td><td>strand</td><td>MM</td><td>target_seq</td><td>PAM</td><td>alignment</td><td>distance</td><td>position</td><td>gene name</td><td>gene id</td></tr>
<tr><td>GU305914.1</td><td>93861</td><td>93883</td><td>-</td><td>0</td><td>CGGGAGCGGGGCGCTGGTCT</td><td>GGG</td><td>||||||||[||||||||||||]PAM</td><td>NA</td><td>Intergenic</td><td>NA</td><td>NA</td></tr>
<tr><td>GU305914.1</td><td>93588</td><td>93610</td><td>-</td><td>4</td><td>CGGGTCCGGGGTGCTGGTCC</td><td>TGG</td><td>||||--||[|||-|||||||-]PAM</td><td>NA</td><td>Intergenic</td><td>NA</td><td>NA</td></tr>
<tr><td></td><td></td><td></td><td></td><td></td><td></td><td></td><td></td><td></td><td></td><td></td><td></td></tr>
<tr><td>T26916</td><td>GGGAGCGGGGCGCTGGTCTGGGG</td><td>1000</td><td></td><td></td><td></td><td></td><td></td><td></td><td></td><td></td><td></td></tr>
<tr><td>Oligo fwd</td><td>CACCGGGAGCGGGGCGCTGGTCTG</td><td></td><td></td><td></td><td></td><td></td><td></td><td></td><td></td><td></td><td></td></tr>
<tr><td>Oligo rev</td><td>AAACCAGACCAGCGCCCCGCTCCC</td><td></td><td></td><td></td><td></td><td></td><td></td><td></td><td></td><td></td><td></td></tr>
<tr><td>Chromosome</td><td>start</td><td>end</td><td>strand</td><td>MM</td><td>target_seq</td><td>PAM</td><td>alignment</td><td>distance</td><td>position</td><td>gene name</td><td>gene id</td></tr>
<tr><td>GU305914.1</td><td>93860</td><td>93882</td><td>-</td><td>0</td><td>GGGAGCGGGGCGCTGGTCTG</td><td>GGG</td><td>||||||||[||||||||||||]PAM</td><td>NA</td><td>Intergenic</td><td>NA</td><td>NA</td></tr>
</table>

GU305914.1 93505 93527 - 2 GGGTGCGGGGCGCTGGTCCG AGG |||-||||[||||||||||-|]PAM NA Intergenic NA NA

T26917 CGCTGGTCTGGGGAGCCAAGCGG 1000
Oligo fwd CACCCGCTGGTCTGGGGAGCCAAG
Oligo rev AAACCTTGGCTCCCCAGACCAGCG
Chromosome start end strand MM target_seq PAM alignment distance position gene name gene id
GU305914.1 93850 93872 - 0 CGCTGGTCTGGGGAGCCAAG CGG ||||||||[||||||||||||]PAM NA Intergenic NA NA

T26918 GCTGGTCTGGGGAGCCAAGCGGG 1000
Oligo fwd CACCGCTGGTCTGGGGAGCCAAGC
Oligo rev AAACGCTTGGCTCCCCAGACCAGC
Chromosome start end strand MM target_seq PAM alignment distance position gene name gene id
GU305914.1 93849 93871 - 0 GCTGGTCTGGGGAGCCAAGC GGG ||||||||[||||||||||||]PAM NA Intergenic NA NA
GU305914.1 93607 93629 - 3 GCTGGTCCGGGAAGCCAGGC GGG |||||||-[|||-|||||-||]PAM NA Intergenic NA NA

T26919 CTGGTCTGGGGAGCCAAGCGGGG 1000
Oligo fwd CACCCTGGTCTGGGGAGCCAAGCG
Oligo rev AAACCGCTTGGCTCCCCAGACCAG
Chromosome start end strand MM target_seq PAM alignment distance position gene name gene id
GU305914.1 93848 93870 - 0 CTGGTCTGGGGAGCCAAGCG GGG ||||||||[||||||||||||]PAM NA Intergenic NA NA

T26920 CTGGGGAGCCAAGCGGGGTGCGG 1000
Oligo fwd CACCCTGGGGAGCCAAGCGGGGTG
Oligo rev AAACCACCCCGCTTGGCTCCCCAG
Chromosome start end strand MM target_seq PAM alignment distance position gene name gene id
GU305914.1 93843 93865 - 0 CTGGGGAGCCAAGCGGGGTG CGG ||||||||[||||||||||||]PAM NA Intergenic NA NA

T26921 TGGGGAGCCAAGCGGGGTGCGGG 1000
Oligo fwd CACCTGGGGAGCCAAGCGGGGTGC
Oligo rev AAACGCACCCCGCTTGGCTCCCCA
Chromosome start end strand MM target_seq PAM alignment distance position gene name gene id
GU305914.1 93842 93864 - 0 TGGGGAGCCAAGCGGGGTGC GGG ||||||||[||||||||||||]PAM NA Intergenic NA NA

T26922 GGGGAGCCAAGCGGGGTGCGGGG 1000
Oligo fwd CACCGGGGAGCCAAGCGGGGTGCG
Oligo rev AAACCGCACCCCGCTTGGCTCCCC
Chromosome start end strand MM target_seq PAM alignment distance position gene name gene id
GU305914.1 93841 93863 - 0 GGGGAGCCAAGCGGGGTGCG GGG ||||||||[||||||||||||]PAM NA Intergenic NA NA

T26923 CGGGGTGCGGGGAATCGAGCCGG 1000
Oligo fwd CACCCGGGGTGCGGGGAATCGAGC
Oligo rev AAACGCTCGATTCCCCGCACCCCG
Chromosome start end strand MM target_seq PAM alignment distance position gene name gene id
GU305914.1 93830 93852 - 0 CGGGGTGCGGGGAATCGAGC CGG ||||||||[||||||||||||]PAM NA Intergenic NA NA

T26924 GGGGTGCGGGGAATCGAGCCGGG 1000
Oligo fwd CACCGGGGTGCGGGGAATCGAGCC
Oligo rev AAACGGCTCGATTCCCCGCACCCC
Chromosome start end strand MM target_seq PAM alignment distance position gene name gene id
GU305914.1 93829 93851 - 0 GGGGTGCGGGGAATCGAGCC GGG ||||||||[||||||||||||]PAM NA Intergenic NA NA

T26925 GGGTGCGGGGAATCGAGCCGGGG 1000
Oligo fwd CACCGGGTGCGGGGAATCGAGCCG
Oligo rev AAACCGGCTCGATTCCCCGCACCC
Chromosome start end strand MM target_seq PAM alignment distance position gene name gene id
GU305914.1 93828 93850 - 0 GGGTGCGGGGAATCGAGCCG GGG ||||||||[||||||||||||]PAM NA Intergenic NA NA

T26926 GGGGAATCGAGCCGGGGAGCTGG 1000
Oligo fwd CACCGGGGAATCGAGCCGGGGAGC
Oligo rev AAACGCTCCCCGGCTCGATTCCCC
Chromosome start end strand MM target_seq PAM alignment distance position gene name gene id
GU305914.1 93822 93844 - 0 GGGGAATCGAGCCGGGGAGC TGG ||||||||[||||||||||||]PAM NA Intergenic NA NA
GU305914.1 93758 93780 - 4 GGGGAGCCGAGCCGAGAAGC CGG |||||--|[||||||-|-|||]PAM NA Intergenic NA NA
GU305914.1 93654 93676 - 4 GGGGAGCCGAGCCGGAAAGC CGG |||||--|[|||||||--|||]PAM NA Intergenic NA NA

T26927 TCGAGCCGGGGAGCTGGTCCTGG 1000
Oligo fwd CACCTCGAGCCGGGGAGCTGGTCC
Oligo rev AAACGGACCAGCTCCCCGGCTCGA
Chromosome start end strand MM target_seq PAM alignment distance position gene name gene id
GU305914.1 93816 93838 - 0 TCGAGCCGGGGAGCTGGTCC TGG ||||||||[||||||||||||]PAM NA Intergenic NA NA
GU305914.1 93712 93734 - 3 GGTAGCCGGGGAGCTGGTCC TGG ---|||||[||||||||||||]PAM NA Intergenic NA NA
GU305914.1 93619 93641 - 3 GGTAGCCGGGGAGCTGGTCC GGG ---|||||[||||||||||||]PAM NA Intergenic NA NA

T26928 AGCCGGGGAGCTGGTCCTGGAGG 1000
Oligo fwd CACCAGCCGGGGAGCTGGTCCTGG
Oligo rev AAACCCAGGACCAGCTCCCCGGCT
Chromosome start end strand MM target_seq PAM alignment distance position gene name gene id
GU305914.1 93709 93731 - 0 AGCCGGGGAGCTGGTCCTGG GGG ||||||||[||||||||||||]PAM NA Intergenic NA NA
GU305914.1 93813 93835 - 0 AGCCGGGGAGCTGGTCCTGG AGG ||||||||[||||||||||||]PAM NA Intergenic NA NA

T26929 GGGAGCTGGTCCTGGAGGTCCGG 1000
Oligo fwd CACCGGGAGCTGGTCCTGGAGGTC
Oligo rev AAACGACCTCCAGGACCAGCTCCC
Chromosome start end strand MM target_seq PAM alignment distance position gene name gene id
GU305914.1 93808 93830 - 0 GGGAGCTGGTCCTGGAGGTC CGG ||||||||[||||||||||||]PAM NA Intergenic NA NA
GU305914.1 93704 93726 - 2 GGGAGCTGGTCCTGGGGGCC CGG ||||||||[|||||||-||-|]PAM NA Intergenic NA NA

T26930 CCTGGAGGTCCGGTGAGCCGAGG 1000
Oligo fwd CACCCCTGGAGGTCCGGTGAGCCG

Oligo rev AAACCGGCTCACCGGACCTCCAGG
Chromosome start end strand MM target_seq PAM alignment distance position gene name gene id
GU305914.1 93798 93820 - 0 CCTGGAGGTCCGGTGAGCCG AGG ||||||||[||||||||||||]PAM NA Intergenic NA NA
GU305914.1 93694 93716 - 2 CCTGGGGGCCCGGTGAGCCG AGG |||||-||[-|||||||||||]PAM NA Intergenic NA NA

T26931 GGAGGTCCGGTGAGCCGAGGAGG 1000
Oligo fwd CACCGGAGGTCCGGTGAGCCGAGG
Oligo rev AAACCCTCGGCTCACCGGACCTCC
Chromosome start end strand MM target_seq PAM alignment distance position gene name gene id
GU305914.1 93795 93817 - 0 GGAGGTCCGGTGAGCCGAGG AGG ||||||||[||||||||||||]PAM NA Intergenic NA NA

T26932 GAGGTCCGGTGAGCCGAGGAGGG 1000
Oligo fwd CACCGAGGTCCGGTGAGCCGAGGA
Oligo rev AAACTCCTCGGCTCACCGGACCTC
Chromosome start end strand MM target_seq PAM alignment distance position gene name gene id
GU305914.1 93794 93816 - 0 GAGGTCCGGTGAGCCGAGGA GGG ||||||||[||||||||||||]PAM NA Intergenic NA NA
GU305914.1 93690 93712 - 3 GGGGCCCGGTGAGCCGAGGT CGG |-||-|||[|||||||||||-]PAM NA Intergenic NA NA
GU305914.1 93567 93589 - 3 GGGGTCCGGGGAGCCGAGGC CGG |-||||||[|-|||||||||-]PAM NA Intergenic NA NA

T26933 GGTGAGCCGAGGAGGGTAGCCGG 1000
Oligo fwd CACCGGTGAGCCGAGGAGGGTAGC
Oligo rev AAACGCTACCCTCCTCGGCTCACC
Chromosome start end strand MM target_seq PAM alignment distance position gene name gene id
GU305914.1 93787 93809 - 0 GGTGAGCCGAGGAGGGTAGC CGG ||||||||[||||||||||||]PAM NA Intergenic NA NA
GU305914.1 93633 93655 - 3 GGGGAGCCGAGGCCGGTAGC CGG ||-|||||[||||--||||||]PAM NA Intergenic NA NA
GU305914.1 93560 93582 - 3 GGGGAGCCGAGGCCGGTAGC CGG ||-|||||[||||--||||||]PAM NA Intergenic NA NA
GU305914.1 93726 93748 - 4 GGGGATCCGAGGTCGGTAGC CGG ||-||-||[||||--||||||]PAM NA Intergenic NA NA

T26934 GTGAGCCGAGGAGGGTAGCCGGG 1000
Oligo fwd CACCGTGAGCCGAGGAGGGTAGCC
Oligo rev AAACGGCTACCCTCCTCGGCTCAC
Chromosome start end strand MM target_seq PAM alignment distance position gene name gene id
GU305914.1 93786 93808 - 0 GTGAGCCGAGGAGGGTAGCC GGG ||||||||[||||||||||||]PAM NA Intergenic NA NA
GU305914.1 93632 93654 - 3 GGGAGCCGAGGCCGGTAGCC GGG |-||||||[|||--|||||||]PAM NA Intergenic NA NA
GU305914.1 93559 93581 - 3 GGGAGCCGAGGCCGGTAGCC GGG |-||||||[|||--|||||||]PAM NA Intergenic NA NA
GU305914.1 93725 93747 - 4 GGGATCCGAGGTCGGTAGCC GGG |-||-|||[|||--|||||||]PAM NA Intergenic NA NA

T26935 GAGGAGGGTAGCCGGGAAGCTGG 1000
Oligo fwd CACCGAGGAGGGTAGCCGGGAAGC
Oligo rev AAACGCTTCCCGGCTACCCTCCTC
Chromosome start end strand MM target_seq PAM alignment distance position gene name gene id
GU305914.1 93779 93801 - 0 GAGGAGGGTAGCCGGGAAGC TGG ||||||||[||||||||||||]PAM NA Intergenic NA NA
GU305914.1 93675 93697 - 3 GAGGTCGGTAGGCGGGAAGC TGG ||||--||[|||-||||||||]PAM NA Intergenic NA NA
GU305914.1 93718 93740 - 3 GAGGTCGGTAGCCGGGGAGC TGG ||||--||[||||||||-|||]PAM NA Intergenic NA NA
GU305914.1 93625 93647 - 3 GAGGCCGGTAGCCGGGGAGC TGG ||||--||[||||||||-|||]PAM NA Intergenic NA NA
GU305914.1 93552 93574 - 3 GAGGCCGGTAGCCGGGGAGC TGG ||||--||[||||||||-|||]PAM NA Intergenic NA NA

T26936 AGGAGGGTAGCCGGGAAGCTGGG 1000
Oligo fwd CACCAGGAGGGTAGCCGGGAAGCT
Oligo rev AAACAGCTTCCCGGCTACCCTCCT
Chromosome start end strand MM target_seq PAM alignment distance position gene name gene id
GU305914.1 93778 93800 - 0 AGGAGGGTAGCCGGGAAGCT GGG ||||||||[||||||||||||]PAM NA Intergenic NA NA
GU305914.1 93674 93696 - 3 AGGTCGGTAGGCGGGAAGCT GGG |||--|||[||-|||||||||]PAM NA Intergenic NA NA

T26937 GGAGGGTAGCCGGGAAGCTGGGG 1000
Oligo fwd CACCGGAGGGTAGCCGGGAAGCTG
Oligo rev AAACCAGCTTCCCGGCTACCCTCC
Chromosome start end strand MM target_seq PAM alignment distance position gene name gene id
GU305914.1 93777 93799 - 0 GGAGGGTAGCCGGGAAGCTG GGG ||||||||[||||||||||||]PAM NA Intergenic NA NA
GU305914.1 93673 93695 - 3 GGTCGGTAGGCGGGAAGCTG GGG ||--||||[|-||||||||||]PAM NA Intergenic NA NA

T26938 GGGGAGCCGAGCCGAGAAGCCGG 1000
Oligo fwd CACCGGGGAGCCGAGCCGAGAAGC
Oligo rev AAACGCTTCTCGGCTCGGCTCCCC
Chromosome start end strand MM target_seq PAM alignment distance position gene name gene id
GU305914.1 93758 93780 - 0 GGGGAGCCGAGCCGAGAAGC CGG ||||||||[||||||||||||]PAM NA Intergenic NA NA
GU305914.1 93654 93676 - 2 GGGGAGCCGAGCCGGAAAGC CGG ||||||||[||||||--||||]PAM NA Intergenic NA NA
GU305914.1 93822 93844 - 4 GGGGAATCGAGCCGGGGAGC TGG |||||--|[||||||-|-|||]PAM NA Intergenic NA NA

T26939 GGGAGCCGAGCCGAGAAGCCGGG 1000
Oligo fwd CACCGGGAGCCGAGCCGAGAAGCC
Oligo rev AAACGGCTTCTCGGCTCGGCTCCC
Chromosome start end strand MM target_seq PAM alignment distance position gene name gene id
GU305914.1 93757 93779 - 0 GGGAGCCGAGCCGAGAAGCC GGG ||||||||[||||||||||||]PAM NA Intergenic NA NA
GU305914.1 93653 93675 - 2 GGGAGCCGAGCCGGAAAGCC GGG ||||||||[|||||--|||||]PAM NA Intergenic NA NA

T26940 GCCGAGCCGAGAAGCCGGGAAGG 1000
Oligo fwd CACCGCCGAGCCGAGAAGCCGGGA
Oligo rev AAACTCCCGGCTTCTCGGCTCGGC
Chromosome start end strand MM target_seq PAM alignment distance position gene name gene id
GU305914.1 93753 93775 - 0 GCCGAGCCGAGAAGCCGGGA AGG ||||||||[||||||||||||]PAM NA Intergenic NA NA

T26941 CCGAGCCGAGAAGCCGGGAAGGG 1000
Oligo fwd CACCCCGAGCCGAGAAGCCGGGAA
Oligo rev AAACTTCCCGGCTTCTCGGCTCGG
Chromosome start end strand MM target_seq PAM alignment distance position gene name gene id
GU305914.1 93752 93774 - 0 CCGAGCCGAGAAGCCGGGAA GGG ||||||||[||||||||||||]PAM NA Intergenic NA NA

T26942 CCGAGAAGCCGGGAAGGGTGCGG 1000

| | | | | | | | | | | | |
|---|---|---|---|---|---|---|---|---|---|---|---|
| Oligo fwd | CACCCCGAGAAGCCGGGAAGGGTG | | | | | | | | | | |
| Oligo rev | AAACCACCCTTCCCGGCTTCTCGG | | | | | | | | | | |
| Chromosome | start | end | strand | MM | target_seq | PAM | alignment | distance | position | gene name | gene id |
| GU305914.1 | 93747 | 93769 | - | 0 | CCGAGAAGCCGGGAAGGGTG | CGG | \|\|\|\|\|\|\|\|[\|\|\|\|\|\|\|\|\|\|\|\|]PAM | NA | Intergenic | NA | NA |

| | | | | | | | | | | | |
|---|---|---|---|---|---|---|---|---|---|---|---|
| T26943 | CGAGAAGCCGGGAAGGGTGCGGG | 1000 | | | | | | | | | |
| Oligo fwd | CACCCGAGAAGCCGGGAAGGGTGC | | | | | | | | | | |
| Oligo rev | AAACGCACCCTTCCCGGCTTCTCG | | | | | | | | | | |
| Chromosome | start | end | strand | MM | target_seq | PAM | alignment | distance | position | gene name | gene id |
| GU305914.1 | 93746 | 93768 | - | 0 | CGAGAAGCCGGGAAGGGTGC | GGG | \|\|\|\|\|\|\|\|[\|\|\|\|\|\|\|\|\|\|\|\|]PAM | NA | Intergenic | NA | NA |

| | | | | | | | | | | | |
|---|---|---|---|---|---|---|---|---|---|---|---|
| T26944 | GAGAAGCCGGGAAGGGTGCGGGG | 1000 | | | | | | | | | |
| Oligo fwd | CACCGAGAAGCCGGGAAGGGTGCG | | | | | | | | | | |
| Oligo rev | AAACCGCACCCTTCCCGGCTTCTC | | | | | | | | | | |
| Chromosome | start | end | strand | MM | target_seq | PAM | alignment | distance | position | gene name | gene id |
| GU305914.1 | 93745 | 93767 | - | 0 | GAGAAGCCGGGAAGGGTGCG | GGG | \|\|\|\|\|\|\|\|[\|\|\|\|\|\|\|\|\|\|\|\|]PAM | NA | Intergenic | NA | NA |

| | | | | | | | | | | | |
|---|---|---|---|---|---|---|---|---|---|---|---|
| T26945 | GGGAAGGGTGCGGGGATCCGAGG | 1000 | | | | | | | | | |
| Oligo fwd | CACCGGGAAGGGTGCGGGGATCCG | | | | | | | | | | |
| Oligo rev | AAACCGGATCCCCGCACCCTTCCC | | | | | | | | | | |
| Chromosome | start | end | strand | MM | target_seq | PAM | alignment | distance | position | gene name | gene id |
| GU305914.1 | 93737 | 93759 | - | 0 | GGGAAGGGTGCGGGGATCCG | AGG | \|\|\|\|\|\|\|\|[\|\|\|\|\|\|\|\|\|\|\|\|]PAM | NA | Intergenic | NA | NA |
| GU305914.1 | 170441 | 170463 | + | 4 | GGGAAATGTGAGGGGATCGG | AGG | \|\|\|\|\|--\|[\|\|-\|\|\|\|\|\|\|-\|]PAM | NA | Intergenic | NA | NA |

| | | | | | | | | | | | |
|---|---|---|---|---|---|---|---|---|---|---|---|
| T26946 | AGGGTGCGGGGATCCGAGGTCGG | 1000 | | | | | | | | | |
| Oligo fwd | CACCAGGGTGCGGGGATCCGAGGT | | | | | | | | | | |
| Oligo rev | AAACACCTCGGATCCCCGCACCCT | | | | | | | | | | |
| Chromosome | start | end | strand | MM | target_seq | PAM | alignment | distance | position | gene name | gene id |
| GU305914.1 | 93733 | 93755 | - | 0 | AGGGTGCGGGGATCCGAGGT | CGG | \|\|\|\|\|\|\|\|[\|\|\|\|\|\|\|\|\|\|\|\|]PAM | NA | Intergenic | NA | NA |
| GU305914.1 | 93567 | 93589 | - | 4 | GGGGTCCGGGGAGCCGAGGC | CGG | -\|\|\|\|-\|\|[\|\|\|\|-\|\|\|\|\|\|-]PAM | NA | Intergenic | NA | NA |

| | | | | | | | | | | | |
|---|---|---|---|---|---|---|---|---|---|---|---|
| T26947 | GGGGATCCGAGGTCGGTAGCCGG | 1000 | | | | | | | | | |
| Oligo fwd | CACCGGGGATCCGAGGTCGGTAGC | | | | | | | | | | |
| Oligo rev | AAACGCTACCGACCTCGGATCCCC | | | | | | | | | | |
| Chromosome | start | end | strand | MM | target_seq | PAM | alignment | distance | position | gene name | gene id |
| GU305914.1 | 93726 | 93748 | - | 0 | GGGGATCCGAGGTCGGTAGC | CGG | \|\|\|\|\|\|\|\|[\|\|\|\|\|\|\|\|\|\|\|\|]PAM | NA | Intergenic | NA | NA |
| GU305914.1 | 93560 | 93582 | - | 2 | GGGGAGCCGAGGCCGGTAGC | CGG | \|\|\|\|\|-\|\|[\|\|\|\|-\|\|\|\|\|\|\|]PAM | NA | Intergenic | NA | NA |
| GU305914.1 | 93633 | 93655 | - | 2 | GGGGAGCCGAGGCCGGTAGC | CGG | \|\|\|\|\|-\|\|[\|\|\|\|-\|\|\|\|\|\|\|]PAM | NA | Intergenic | NA | NA |
| GU305914.1 | 93787 | 93809 | - | 4 | GGTGAGCCGAGGAGGGTAGC | CGG | \|\|-\|\|-\|\|[\|\|\|\|--\|\|\|\|\|\|]PAM | NA | Intergenic | NA | NA |
| GU305914.1 | 93683 | 93705 | - | 3 | GGTGAGCCGAGGTCGGTAGG | CGG | \|\|-\|\|-\|\|[\|\|\|\|\|\|\|\|\|\|\|-]PAM | NA | Intergenic | NA | NA |

| | | | | | | | | | | | |
|---|---|---|---|---|---|---|---|---|---|---|---|
| T26948 | GGGATCCGAGGTCGGTAGCCGGG | 1000 | | | | | | | | | |
| Oligo fwd | CACCGGGATCCGAGGTCGGTAGCC | | | | | | | | | | |
| Oligo rev | AAACGGCTACCGACCTCGGATCCC | | | | | | | | | | |
| Chromosome | start | end | strand | MM | target_seq | PAM | alignment | distance | position | gene name | gene id |
| GU305914.1 | 93725 | 93747 | - | 0 | GGGATCCGAGGTCGGTAGCC | GGG | \|\|\|\|\|\|\|\|[\|\|\|\|\|\|\|\|\|\|\|\|]PAM | NA | Intergenic | NA | NA |
| GU305914.1 | 93559 | 93581 | - | 2 | GGGAGCCGAGGCCGGTAGCC | GGG | \|\|\|\|-\|\|\|[\|\|\|-\|\|\|\|\|\|\|\|]PAM | NA | Intergenic | NA | NA |
| GU305914.1 | 93632 | 93654 | - | 2 | GGGAGCCGAGGCCGGTAGCC | GGG | \|\|\|\|-\|\|\|[\|\|\|-\|\|\|\|\|\|\|\|]PAM | NA | Intergenic | NA | NA |
| GU305914.1 | 93786 | 93808 | - | 4 | GTGAGCCGAGGAGGGTAGCC | GGG | \|-\|\|-\|\|\|[\|\|\|--\|\|\|\|\|\|\|]PAM | NA | Intergenic | NA | NA |
| GU305914.1 | 93682 | 93704 | - | 3 | GTGAGCCGAGGTCGGTAGGC | GGG | \|-\|\|-\|\|\|[\|\|\|\|\|\|\|\|\|\|-\|]PAM | NA | Intergenic | NA | NA |

| | | | | | | | | | | | |
|---|---|---|---|---|---|---|---|---|---|---|---|
| T26949 | GGATCCGAGGTCGGTAGCCGGGG | 1000 | | | | | | | | | |
| Oligo fwd | CACCGGATCCGAGGTCGGTAGCCG | | | | | | | | | | |
| Oligo rev | AAACCGGCTACCGACCTCGGATCC | | | | | | | | | | |
| Chromosome | start | end | strand | MM | target_seq | PAM | alignment | distance | position | gene name | gene id |
| GU305914.1 | 93724 | 93746 | - | 0 | GGATCCGAGGTCGGTAGCCG | GGG | \|\|\|\|\|\|\|\|[\|\|\|\|\|\|\|\|\|\|\|\|]PAM | NA | Intergenic | NA | NA |
| GU305914.1 | 93631 | 93653 | - | 2 | GGAGCCGAGGCCGGTAGCCG | GGG | \|\|\|-\|\|\|\|[\|\|-\|\|\|\|\|\|\|\|\|]PAM | NA | Intergenic | NA | NA |
| GU305914.1 | 93558 | 93580 | - | 2 | GGAGCCGAGGCCGGTAGCCG | GGG | \|\|\|-\|\|\|\|[\|\|-\|\|\|\|\|\|\|\|\|]PAM | NA | Intergenic | NA | NA |

| | | | | | | | | | | | |
|---|---|---|---|---|---|---|---|---|---|---|---|
| T26950 | GAGGTCGGTAGCCGGGGAGCTGG | 1000 | | | | | | | | | |
| Oligo fwd | CACCGAGGTCGGTAGCCGGGGAGC | | | | | | | | | | |
| Oligo rev | AAACGCTCCCCGGCTACCGACCTC | | | | | | | | | | |
| Chromosome | start | end | strand | MM | target_seq | PAM | alignment | distance | position | gene name | gene id |
| GU305914.1 | 93718 | 93740 | - | 0 | GAGGTCGGTAGCCGGGGAGC | TGG | \|\|\|\|\|\|\|\|[\|\|\|\|\|\|\|\|\|\|\|\|]PAM | NA | Intergenic | NA | NA |
| GU305914.1 | 93552 | 93574 | - | 1 | GAGGCCGGTAGCCGGGGAGC | TGG | \|\|\|\|-\|\|\|[\|\|\|\|\|\|\|\|\|\|\|\|]PAM | NA | Intergenic | NA | NA |
| GU305914.1 | 93625 | 93647 | - | 1 | GAGGCCGGTAGCCGGGGAGC | TGG | \|\|\|\|-\|\|\|[\|\|\|\|\|\|\|\|\|\|\|\|]PAM | NA | Intergenic | NA | NA |
| GU305914.1 | 93779 | 93801 | - | 3 | GAGGAGGGTAGCCGGGAAGC | TGG | \|\|\|\|--\|\|[\|\|\|\|\|\|\|\|-\|\|\|]PAM | NA | Intergenic | NA | NA |
| GU305914.1 | 93675 | 93697 | - | 2 | GAGGTCGGTAGGCGGGAAGC | TGG | \|\|\|\|\|\|\|\|[\|\|\|-\|\|\|\|-\|\|\|]PAM | NA | Intergenic | NA | NA |

| | | | | | | | | | | | |
|---|---|---|---|---|---|---|---|---|---|---|---|
| T26951 | GGTAGCCGGGGAGCTGGTCCTGG | 1000 | | | | | | | | | |
| Oligo fwd | CACCGGTAGCCGGGGAGCTGGTCC | | | | | | | | | | |
| Oligo rev | AAACGGACCAGCTCCCCGGCTACC | | | | | | | | | | |
| Chromosome | start | end | strand | MM | target_seq | PAM | alignment | distance | position | gene name | gene id |
| GU305914.1 | 93619 | 93641 | - | 0 | GGTAGCCGGGGAGCTGGTCC | GGG | \|\|\|\|\|\|\|\|[\|\|\|\|\|\|\|\|\|\|\|\|]PAM | NA | Intergenic | NA | NA |
| GU305914.1 | 93712 | 93734 | - | 0 | GGTAGCCGGGGAGCTGGTCC | TGG | \|\|\|\|\|\|\|\|[\|\|\|\|\|\|\|\|\|\|\|\|]PAM | NA | Intergenic | NA | NA |
| GU305914.1 | 93816 | 93838 | - | 3 | TCGAGCCGGGGAGCTGGTCC | TGG | ---\|\|\|\|\|[\|\|\|\|\|\|\|\|\|\|\|\|]PAM | NA | Intergenic | NA | NA |

| | | | | | | | | | | | |
|---|---|---|---|---|---|---|---|---|---|---|---|
| T26952 | GTAGCCGGGGAGCTGGTCCTGGG | 1000 | | | | | | | | | |
| Oligo fwd | CACCGTAGCCGGGGAGCTGGTCCT | | | | | | | | | | |
| Oligo rev | AAACAGGACCAGCTCCCCGGCTAC | | | | | | | | | | |
| Chromosome | start | end | strand | MM | target_seq | PAM | alignment | distance | position | gene name | gene id |
| GU305914.1 | 93711 | 93733 | - | 0 | GTAGCCGGGGAGCTGGTCCT | GGG | \|\|\|\|\|\|\|\|[\|\|\|\|\|\|\|\|\|\|\|\|]PAM | NA | Intergenic | NA | NA |
| GU305914.1 | 93587 | 93609 | - | 4 | GGGTCCGGGGTGCTGGTCCT | GGG | \|---\|\|\|\|[\|\|-\|\|\|\|\|\|\|\|\|]PAM | NA | Intergenic | NA | NA |

| | | | | | | | | | | | |
|---|---|---|---|---|---|---|---|---|---|---|---|
| T26953 | TAGCCGGGGAGCTGGTCCTGGGG | 1000 | | | | | | | | | |
| Oligo fwd | CACCTAGCCGGGGAGCTGGTCCTG | | | | | | | | | | |
| Oligo rev | AAACCAGGACCAGCTCCCCGGCTA | | | | | | | | | | |
| Chromosome | start | end | strand | MM | target_seq | PAM | alignment | distance | position | gene name | gene id |
| GU305914.1 | 93544 | 93566 | - | 0 | TAGCCGGGGAGCTGGTCCTG | CGG | \|\|\|\|\|\|\|\|[\|\|\|\|\|\|\|\|\|\|\|\|]PAM | NA | Intergenic | NA | NA |

| | | | | | | | | | | | |
|---|---|---|---|---|---|---|---|---|---|---|---|
| GU305914.1 | 93710 | 93732 | - | 0 | TAGCCGGGGAGCTGGTCCTG | GGG | \|\|\|\|\|\|\|\|[\|\|\|\|\|\|\|\|\|\|\|\|]PAM | NA | Intergenic | NA | NA |
| GU305914.1 | 93586 | 93608 | - | 4 | GGTCCGGGGTGCTGGTCCTG | GGG | ---\|\|\|\|\|[\|-\|\|\|\|\|\|\|\|\|\|]PAM | NA | Intergenic | NA | NA |
| T26954 | AGCCGGGGAGCTGGTCCTGGGGG | 1000 | | | | | | | | | |
| Oligo fwd | CACCAGCCGGGGAGCTGGTCCTGG | | | | | | | | | | |
| Oligo rev | AAACCCAGGACCAGCTCCCCGGCT | | | | | | | | | | |
| Chromosome | start | end | strand | MM | target_seq | PAM | alignment | distance | position | gene name | gene id |
| GU305914.1 | 93709 | 93731 | - | 0 | AGCCGGGGAGCTGGTCCTGG | GGG | \|\|\|\|\|\|\|\|[\|\|\|\|\|\|\|\|\|\|\|\|]PAM | NA | Intergenic | NA | NA |
| GU305914.1 | 93813 | 93835 | - | 0 | AGCCGGGGAGCTGGTCCTGG | AGG | \|\|\|\|\|\|\|\|[\|\|\|\|\|\|\|\|\|\|\|\|]PAM | NA | Intergenic | NA | NA |
| T26955 | GGGAGCTGGTCCTGGGGGCCCGG | 1000 | | | | | | | | | |
| Oligo fwd | CACCGGGAGCTGGTCCTGGGGGCC | | | | | | | | | | |
| Oligo rev | AAACGGCCCCCAGGACCAGCTCCC | | | | | | | | | | |
| Chromosome | start | end | strand | MM | target_seq | PAM | alignment | distance | position | gene name | gene id |
| GU305914.1 | 93704 | 93726 | - | 0 | GGGAGCTGGTCCTGGGGGCC | CGG | \|\|\|\|\|\|\|\|[\|\|\|\|\|\|\|\|\|\|\|\|]PAM | NA | Intergenic | NA | NA |
| GU305914.1 | 93580 | 93602 | - | 2 | GGGTGCTGGTCCTGGGGTCC | GGG | \|\|\|-\|\|\|\|[\|\|\|\|\|\|\|\|\|-\|\|]PAM | NA | Intergenic | NA | NA |
| GU305914.1 | 93538 | 93560 | - | 2 | GGGAGCTGGTCCTGCGGTCC | GGG | \|\|\|\|\|\|\|\|[\|\|\|\|\|\|-\|\|-\|\|]PAM | NA | Intergenic | NA | NA |
| GU305914.1 | 93808 | 93830 | - | 2 | GGGAGCTGGTCCTGGAGGTC | CGG | \|\|\|\|\|\|\|\|[\|\|\|\|\|\|\|-\|\|-\|]PAM | NA | Intergenic | NA | NA |
| T26956 | CCTGGGGGCCCGGTGAGCCGAGG | 1000 | | | | | | | | | |
| Oligo fwd | CACCCCTGGGGGCCCGGTGAGCCG | | | | | | | | | | |
| Oligo rev | AAACCGGCTCACCGGGCCCCCAGG | | | | | | | | | | |
| Chromosome | start | end | strand | MM | target_seq | PAM | alignment | distance | position | gene name | gene id |
| GU305914.1 | 93694 | 93716 | - | 0 | CCTGGGGGCCCGGTGAGCCG | AGG | \|\|\|\|\|\|\|\|[\|\|\|\|\|\|\|\|\|\|\|\|]PAM | NA | Intergenic | NA | NA |
| GU305914.1 | 93798 | 93820 | - | 2 | CCTGGAGGTCCGGTGAGCCG | AGG | \|\|\|\|\|-\|\|[-\|\|\|\|\|\|\|\|\|\|\|]PAM | NA | Intergenic | NA | NA |
| T26957 | GGGGCCCGGTGAGCCGAGGTCGG | 1000 | | | | | | | | | |
| Oligo fwd | CACCGGGGCCCGGTGAGCCGAGGT | | | | | | | | | | |
| Oligo rev | AAACACCTCGGCTCACCGGGCCCC | | | | | | | | | | |
| Chromosome | start | end | strand | MM | target_seq | PAM | alignment | distance | position | gene name | gene id |
| GU305914.1 | 93690 | 93712 | - | 0 | GGGGCCCGGTGAGCCGAGGT | CGG | \|\|\|\|\|\|\|\|[\|\|\|\|\|\|\|\|\|\|\|\|]PAM | NA | Intergenic | NA | NA |
| GU305914.1 | 93794 | 93816 | - | 3 | GAGGTCCGGTGAGCCGAGGA | GGG | \|-\|\|-\|\|\|[\|\|\|\|\|\|\|\|\|\|\|-]PAM | NA | Intergenic | NA | NA |
| GU305914.1 | 93567 | 93589 | - | 3 | GGGGTCCGGGGAGCCGAGGC | CGG | \|\|\|\|-\|\|\|[\|-\|\|\|\|\|\|\|\|\|-]PAM | NA | Intergenic | NA | NA |
| T26958 | CCCGGTGAGCCGAGGTCGGTAGG | 1000 | | | | | | | | | |
| Oligo fwd | CACCCCCGGTGAGCCGAGGTCGGT | | | | | | | | | | |
| Oligo rev | AAACACCGACCTCGGCTCACCGGG | | | | | | | | | | |
| Chromosome | start | end | strand | MM | target_seq | PAM | alignment | distance | position | gene name | gene id |
| GU305914.1 | 93686 | 93708 | - | 0 | CCCGGTGAGCCGAGGTCGGT | AGG | \|\|\|\|\|\|\|\|[\|\|\|\|\|\|\|\|\|\|\|\|]PAM | NA | Intergenic | NA | NA |
| T26959 | GGTGAGCCGAGGTCGGTAGGCGG | 1000 | | | | | | | | | |
| Oligo fwd | CACCGGTGAGCCGAGGTCGGTAGG | | | | | | | | | | |
| Oligo rev | AAACCCTACCGACCTCGGCTCACC | | | | | | | | | | |
| Chromosome | start | end | strand | MM | target_seq | PAM | alignment | distance | position | gene name | gene id |
| GU305914.1 | 93683 | 93705 | - | 0 | GGTGAGCCGAGGTCGGTAGG | CGG | \|\|\|\|\|\|\|\|[\|\|\|\|\|\|\|\|\|\|\|\|]PAM | NA | Intergenic | NA | NA |
| GU305914.1 | 93726 | 93748 | - | 3 | GGGGATCCGAGGTCGGTAGC | CGG | \|\|-\|\|-\|\|[\|\|\|\|\|\|\|\|\|\|\|-]PAM | NA | Intergenic | NA | NA |
| GU305914.1 | 93560 | 93582 | - | 3 | GGGGAGCCGAGGCCGGTAGC | CGG | \|\|-\|\|\|\|\|[\|\|\|\|-\|\|\|\|\|\|-]PAM | NA | Intergenic | NA | NA |
| GU305914.1 | 93633 | 93655 | - | 3 | GGGGAGCCGAGGCCGGTAGC | CGG | \|\|-\|\|\|\|\|[\|\|\|\|-\|\|\|\|\|\|-]PAM | NA | Intergenic | NA | NA |
| T26960 | GTGAGCCGAGGTCGGTAGGCGGG | 1000 | | | | | | | | | |
| Oligo fwd | CACCGTGAGCCGAGGTCGGTAGGC | | | | | | | | | | |
| Oligo rev | AAACGCCTACCGACCTCGGCTCAC | | | | | | | | | | |
| Chromosome | start | end | strand | MM | target_seq | PAM | alignment | distance | position | gene name | gene id |
| GU305914.1 | 93682 | 93704 | - | 0 | GTGAGCCGAGGTCGGTAGGC | GGG | \|\|\|\|\|\|\|\|[\|\|\|\|\|\|\|\|\|\|\|\|]PAM | NA | Intergenic | NA | NA |
| GU305914.1 | 93725 | 93747 | - | 3 | GGGATCCGAGGTCGGTAGCC | GGG | \|-\|\|-\|\|\|[\|\|\|\|\|\|\|\|\|\|-\|]PAM | NA | Intergenic | NA | NA |
| GU305914.1 | 93632 | 93654 | - | 3 | GGGAGCCGAGGCCGGTAGCC | GGG | \|-\|\|\|\|\|\|[\|\|\|-\|\|\|\|\|\|-\|]PAM | NA | Intergenic | NA | NA |
| GU305914.1 | 93559 | 93581 | - | 3 | GGGAGCCGAGGCCGGTAGCC | GGG | \|-\|\|\|\|\|\|[\|\|\|-\|\|\|\|\|\|-\|]PAM | NA | Intergenic | NA | NA |
| T26961 | GAGGTCGGTAGGCGGGAAGCTGG | 1000 | | | | | | | | | |
| Oligo fwd | CACCGAGGTCGGTAGGCGGGAAGC | | | | | | | | | | |
| Oligo rev | AAACGCTTCCCGCCTACCGACCTC | | | | | | | | | | |
| Chromosome | start | end | strand | MM | target_seq | PAM | alignment | distance | position | gene name | gene id |
| GU305914.1 | 93675 | 93697 | - | 0 | GAGGTCGGTAGGCGGGAAGC | TGG | \|\|\|\|\|\|\|\|[\|\|\|\|\|\|\|\|\|\|\|\|]PAM | NA | Intergenic | NA | NA |
| GU305914.1 | 93779 | 93801 | - | 3 | GAGGAGGGTAGCCGGGAAGC | TGG | \|\|\|\|--\|\|[\|\|\|-\|\|\|\|\|\|\|\|]PAM | NA | Intergenic | NA | NA |
| GU305914.1 | 93718 | 93740 | - | 2 | GAGGTCGGTAGCCGGGGAGC | TGG | \|\|\|\|\|\|\|\|[\|\|\|-\|\|\|\|-\|\|\|]PAM | NA | Intergenic | NA | NA |
| GU305914.1 | 93625 | 93647 | - | 3 | GAGGCCGGTAGCCGGGGAGC | TGG | \|\|\|\|-\|\|\|[\|\|\|-\|\|\|\|-\|\|\|]PAM | NA | Intergenic | NA | NA |
| GU305914.1 | 93552 | 93574 | - | 3 | GAGGCCGGTAGCCGGGGAGC | TGG | \|\|\|\|-\|\|\|[\|\|\|-\|\|\|\|-\|\|\|]PAM | NA | Intergenic | NA | NA |
| T26962 | AGGTCGGTAGGCGGGAAGCTGGG | 1000 | | | | | | | | | |
| Oligo fwd | CACCAGGTCGGTAGGCGGGAAGCT | | | | | | | | | | |
| Oligo rev | AAACAGCTTCCCGCCTACCGACCT | | | | | | | | | | |
| Chromosome | start | end | strand | MM | target_seq | PAM | alignment | distance | position | gene name | gene id |
| GU305914.1 | 93674 | 93696 | - | 0 | AGGTCGGTAGGCGGGAAGCT | GGG | \|\|\|\|\|\|\|\|[\|\|\|\|\|\|\|\|\|\|\|\|]PAM | NA | Intergenic | NA | NA |
| GU305914.1 | 93778 | 93800 | - | 3 | AGGAGGGTAGCCGGGAAGCT | GGG | \|\|\|--\|\|\|[\|\|-\|\|\|\|\|\|\|\|\|]PAM | NA | Intergenic | NA | NA |
| T26963 | GGTCGGTAGGCGGGAAGCTGGGG | 1000 | | | | | | | | | |
| Oligo fwd | CACCGGTCGGTAGGCGGGAAGCTG | | | | | | | | | | |
| Oligo rev | AAACCAGCTTCCCGCCTACCGACC | | | | | | | | | | |
| Chromosome | start | end | strand | MM | target_seq | PAM | alignment | distance | position | gene name | gene id |
| GU305914.1 | 93673 | 93695 | - | 0 | GGTCGGTAGGCGGGAAGCTG | GGG | \|\|\|\|\|\|\|\|[\|\|\|\|\|\|\|\|\|\|\|\|]PAM | NA | Intergenic | NA | NA |
| GU305914.1 | 93777 | 93799 | - | 3 | GGAGGGTAGCCGGGAAGCTG | GGG | \|\|--\|\|\|\|[\|-\|\|\|\|\|\|\|\|\|\|]PAM | NA | Intergenic | NA | NA |
| T26964 | GGGAAGCTGGGGAGCCGAGCCGG | 1000 | | | | | | | | | |
| Oligo fwd | CACCGGGAAGCTGGGGAGCCGAGC | | | | | | | | | | |
| Oligo rev | AAACGCTCGGCTCCCCAGCTTCCC | | | | | | | | | | |
| Chromosome | start | end | strand | MM | target_seq | PAM | alignment | distance | position | gene name | gene id |
| GU305914.1 | 93662 | 93684 | - | 0 | GGGAAGCTGGGGAGCCGAGC | CGG | \|\|\|\|\|\|\|\|[\|\|\|\|\|\|\|\|\|\|\|\|]PAM | NA | Intergenic | NA | NA |
| T26965 | GGGGAGCCGAGCCGGAAAGCCGG | 1000 | | | | | | | | | |

| | | | | | | | | | | | |
|---|---|---|---|---|---|---|---|---|---|---|---|
| Oligo fwd | CACCGGGGAGCCGAGCCGGAAAGC | | | | | | | | | | |
| Oligo rev | AAACGCTTTCCGGCTCGGCTCCCC | | | | | | | | | | |
| Chromosome | start | end | strand | MM | target_seq | PAM | alignment | distance | position | gene name | gene id |
| GU305914.1 | 93654 | 93676 | - | 0 | GGGGAGCCGAGCCGGAAAGC | CGG | \|\|\|\|\|\|\|\|[\|\|\|\|\|\|\|\|\|\|\|\|]PAM | NA | Intergenic | NA | NA |
| GU305914.1 | 93758 | 93780 | - | 2 | GGGGAGCCGAGCCGAGAAGC | CGG | \|\|\|\|\|\|\|\|[\|\|\|\|\|\|--\|\|\|\|]PAM | NA | Intergenic | NA | NA |
| GU305914.1 | 93822 | 93844 | - | 4 | GGGGAATCGAGCCGGGGAGC | TGG | \|\|\|\|\|--\|[\|\|\|\|\|\|\|--\|\|\|]PAM | NA | Intergenic | NA | NA |
| T26966 | GGGAGCCGAGCCGGAAAGCCGGG | 1000 | | | | | | | | | |
| Oligo fwd | CACCGGGAGCCGAGCCGGAAAGCC | | | | | | | | | | |
| Oligo rev | AAACGGCTTTCCGGCTCGGCTCCC | | | | | | | | | | |
| Chromosome | start | end | strand | MM | target_seq | PAM | alignment | distance | position | gene name | gene id |
| GU305914.1 | 93653 | 93675 | - | 0 | GGGAGCCGAGCCGGAAAGCC | GGG | \|\|\|\|\|\|\|\|[\|\|\|\|\|\|\|\|\|\|\|\|]PAM | NA | Intergenic | NA | NA |
| GU305914.1 | 93757 | 93779 | - | 2 | GGGAGCCGAGCCGAGAAGCC | GGG | \|\|\|\|\|\|\|\|[\|\|\|\|\|--\|\|\|\|\|]PAM | NA | Intergenic | NA | NA |
| T26967 | GGAGCCGAGCCGGAAAGCCGGGG | 1000 | | | | | | | | | |
| Oligo fwd | CACCGGAGCCGAGCCGGAAAGCCG | | | | | | | | | | |
| Oligo rev | AAACCGGCTTTCCGGCTCGGCTCC | | | | | | | | | | |
| Chromosome | start | end | strand | MM | target_seq | PAM | alignment | distance | position | gene name | gene id |
| GU305914.1 | 93652 | 93674 | - | 0 | GGAGCCGAGCCGGAAAGCCG | GGG | \|\|\|\|\|\|\|\|[\|\|\|\|\|\|\|\|\|\|\|\|]PAM | NA | Intergenic | NA | NA |
| T26968 | GCCGGAAAGCCGGGGAGCCGAGG | 1000 | | | | | | | | | |
| Oligo fwd | CACCGCCGGAAAGCCGGGGAGCCG | | | | | | | | | | |
| Oligo rev | AAACCGGCTCCCCGGCTTTCCGGC | | | | | | | | | | |
| Chromosome | start | end | strand | MM | target_seq | PAM | alignment | distance | position | gene name | gene id |
| GU305914.1 | 93644 | 93666 | - | 0 | GCCGGAAAGCCGGGGAGCCG | AGG | \|\|\|\|\|\|\|\|[\|\|\|\|\|\|\|\|\|\|\|\|]PAM | NA | Intergenic | NA | NA |
| T26969 | GAAAGCCGGGGAGCCGAGGCCGG | 1000 | | | | | | | | | |
| Oligo fwd | CACCGAAAGCCGGGGAGCCGAGGC | | | | | | | | | | |
| Oligo rev | AAACGCCTCGGCTCCCCGGCTTTC | | | | | | | | | | |
| Chromosome | start | end | strand | MM | target_seq | PAM | alignment | distance | position | gene name | gene id |
| GU305914.1 | 93640 | 93662 | - | 0 | GAAAGCCGGGGAGCCGAGGC | CGG | \|\|\|\|\|\|\|\|[\|\|\|\|\|\|\|\|\|\|\|\|]PAM | NA | Intergenic | NA | NA |
| GU305914.1 | 93567 | 93589 | - | 4 | GGGGTCCGGGGAGCCGAGGC | CGG | \|----\|\|\|[\|\|\|\|\|\|\|\|\|\|\|\|]PAM | NA | Intergenic | NA | NA |
| T26970 | GGGGAGCCGAGGCCGGTAGCCGG | 1000 | | | | | | | | | |
| Oligo fwd | CACCGGGGAGCCGAGGCCGGTAGC | | | | | | | | | | |
| Oligo rev | AAACGCTACCGGCCTCGGCTCCCC | | | | | | | | | | |
| Chromosome | start | end | strand | MM | target_seq | PAM | alignment | distance | position | gene name | gene id |
| GU305914.1 | 93560 | 93582 | - | 0 | GGGGAGCCGAGGCCGGTAGC | CGG | \|\|\|\|\|\|\|\|[\|\|\|\|\|\|\|\|\|\|\|\|]PAM | NA | Intergenic | NA | NA |
| GU305914.1 | 93633 | 93655 | - | 0 | GGGGAGCCGAGGCCGGTAGC | CGG | \|\|\|\|\|\|\|\|[\|\|\|\|\|\|\|\|\|\|\|\|]PAM | NA | Intergenic | NA | NA |
| GU305914.1 | 93726 | 93748 | - | 2 | GGGGATCCGAGGTCGGTAGC | CGG | \|\|\|\|\|-\|\|[\|\|\|\|-\|\|\|\|\|\|\|]PAM | NA | Intergenic | NA | NA |
| GU305914.1 | 93787 | 93809 | - | 3 | GGTGAGCCGAGGAGGGTAGC | CGG | \|\|-\|\|\|\|\|[\|\|\|\|--\|\|\|\|\|\|]PAM | NA | Intergenic | NA | NA |
| GU305914.1 | 93683 | 93705 | - | 3 | GGTGAGCCGAGGTCGGTAGG | CGG | \|\|-\|\|\|\|\|[\|\|\|\|-\|\|\|\|\|\|-]PAM | NA | Intergenic | NA | NA |
| T26971 | GGGAGCCGAGGCCGGTAGCCGGG | 1000 | | | | | | | | | |
| Oligo fwd | CACCGGGAGCCGAGGCCGGTAGCC | | | | | | | | | | |
| Oligo rev | AAACGGCTACCGGCCTCGGCTCCC | | | | | | | | | | |
| Chromosome | start | end | strand | MM | target_seq | PAM | alignment | distance | position | gene name | gene id |
| GU305914.1 | 93559 | 93581 | - | 0 | GGGAGCCGAGGCCGGTAGCC | GGG | \|\|\|\|\|\|\|\|[\|\|\|\|\|\|\|\|\|\|\|\|]PAM | NA | Intergenic | NA | NA |
| GU305914.1 | 93632 | 93654 | - | 0 | GGGAGCCGAGGCCGGTAGCC | GGG | \|\|\|\|\|\|\|\|[\|\|\|\|\|\|\|\|\|\|\|\|]PAM | NA | Intergenic | NA | NA |
| GU305914.1 | 93725 | 93747 | - | 2 | GGGATCCGAGGTCGGTAGCC | GGG | \|\|\|\|-\|\|\|[\|\|\|-\|\|\|\|\|\|\|\|]PAM | NA | Intergenic | NA | NA |
| GU305914.1 | 93786 | 93808 | - | 3 | GTGAGCCGAGGAGGGTAGCC | GGG | \|-\|\|\|\|\|\|[\|\|\|--\|\|\|\|\|\|\|]PAM | NA | Intergenic | NA | NA |
| GU305914.1 | 93682 | 93704 | - | 3 | GTGAGCCGAGGTCGGTAGGC | GGG | \|-\|\|\|\|\|\|[\|\|\|-\|\|\|\|\|\|-\|]PAM | NA | Intergenic | NA | NA |
| T26972 | GGAGCCGAGGCCGGTAGCCGGGG | 1000 | | | | | | | | | |
| Oligo fwd | CACCGGAGCCGAGGCCGGTAGCCG | | | | | | | | | | |
| Oligo rev | AAACCGGCTACCGGCCTCGGCTCC | | | | | | | | | | |
| Chromosome | start | end | strand | MM | target_seq | PAM | alignment | distance | position | gene name | gene id |
| GU305914.1 | 93558 | 93580 | - | 0 | GGAGCCGAGGCCGGTAGCCG | GGG | \|\|\|\|\|\|\|\|[\|\|\|\|\|\|\|\|\|\|\|\|]PAM | NA | Intergenic | NA | NA |
| GU305914.1 | 93631 | 93653 | - | 0 | GGAGCCGAGGCCGGTAGCCG | GGG | \|\|\|\|\|\|\|\|[\|\|\|\|\|\|\|\|\|\|\|\|]PAM | NA | Intergenic | NA | NA |
| GU305914.1 | 93724 | 93746 | - | 2 | GGATCCGAGGTCGGTAGCCG | GGG | \|\|\|-\|\|\|\|[\|\|-\|\|\|\|\|\|\|\|\|]PAM | NA | Intergenic | NA | NA |
| T26973 | GAGGCCGGTAGCCGGGGAGCTGG | 1000 | | | | | | | | | |
| Oligo fwd | CACCGAGGCCGGTAGCCGGGGAGC | | | | | | | | | | |
| Oligo rev | AAACGCTCCCCGGCTACCGGCCTC | | | | | | | | | | |
| Chromosome | start | end | strand | MM | target_seq | PAM | alignment | distance | position | gene name | gene id |
| GU305914.1 | 93625 | 93647 | - | 0 | GAGGCCGGTAGCCGGGGAGC | TGG | \|\|\|\|\|\|\|\|[\|\|\|\|\|\|\|\|\|\|\|\|]PAM | NA | Intergenic | NA | NA |
| GU305914.1 | 93552 | 93574 | - | 0 | GAGGCCGGTAGCCGGGGAGC | TGG | \|\|\|\|\|\|\|\|[\|\|\|\|\|\|\|\|\|\|\|\|]PAM | NA | Intergenic | NA | NA |
| GU305914.1 | 93718 | 93740 | - | 1 | GAGGTCGGTAGCCGGGGAGC | TGG | \|\|\|\|-\|\|\|[\|\|\|\|\|\|\|\|\|\|\|\|]PAM | NA | Intergenic | NA | NA |
| GU305914.1 | 93779 | 93801 | - | 3 | GAGGAGGGTAGCCGGGAAGC | TGG | \|\|\|\|--\|\|[\|\|\|\|\|\|\|\|-\|\|\|]PAM | NA | Intergenic | NA | NA |
| GU305914.1 | 93675 | 93697 | - | 3 | GAGGTCGGTAGGCGGGAAGC | TGG | \|\|\|\|-\|\|\|[\|\|\|-\|\|\|\|-\|\|\|]PAM | NA | Intergenic | NA | NA |
| T26974 | CGGTAGCCGGGGAGCTGGTCCGG | 1000 | | | | | | | | | |
| Oligo fwd | CACCCGGTAGCCGGGGAGCTGGTC | | | | | | | | | | |
| Oligo rev | AAACGACCAGCTCCCCGGCTACCG | | | | | | | | | | |
| Chromosome | start | end | strand | MM | target_seq | PAM | alignment | distance | position | gene name | gene id |
| GU305914.1 | 93620 | 93642 | - | 0 | CGGTAGCCGGGGAGCTGGTC | CGG | \|\|\|\|\|\|\|\|[\|\|\|\|\|\|\|\|\|\|\|\|]PAM | NA | Intergenic | NA | NA |
| T26975 | GGTAGCCGGGGAGCTGGTCCGGG | 1000 | | | | | | | | | |
| Oligo fwd | CACCGGTAGCCGGGGAGCTGGTCC | | | | | | | | | | |
| Oligo rev | AAACGGACCAGCTCCCCGGCTACC | | | | | | | | | | |
| Chromosome | start | end | strand | MM | target_seq | PAM | alignment | distance | position | gene name | gene id |
| GU305914.1 | 93619 | 93641 | - | 0 | GGTAGCCGGGGAGCTGGTCC | GGG | \|\|\|\|\|\|\|\|[\|\|\|\|\|\|\|\|\|\|\|\|]PAM | NA | Intergenic | NA | NA |
| GU305914.1 | 93712 | 93734 | - | 0 | GGTAGCCGGGGAGCTGGTCC | TGG | \|\|\|\|\|\|\|\|[\|\|\|\|\|\|\|\|\|\|\|\|]PAM | NA | Intergenic | NA | NA |
| GU305914.1 | 93816 | 93838 | - | 3 | TCGAGCCGGGGAGCTGGTCC | TGG | ---\|\|\|\|\|[\|\|\|\|\|\|\|\|\|\|\|\|]PAM | NA | Intergenic | NA | NA |
| T26976 | GGGAGCTGGTCCGGGAAGCCAGG | 1000 | | | | | | | | | |
| Oligo fwd | CACCGGGAGCTGGTCCGGGAAGCC | | | | | | | | | | |
| Oligo rev | AAACGGCTTCCCGGACCAGCTCCC | | | | | | | | | | |
| Chromosome | start | end | strand | MM | target_seq | PAM | alignment | distance | position | gene name | gene id |

<table>
<tr><td>GU305914.1</td><td>93611</td><td>93633</td><td>-</td><td>0</td><td>GGGAGCTGGTCCGGGAAGCC</td><td>AGG</td><td>||||||||[||||||||||||]PAM</td><td>NA</td><td>Intergenic</td><td>NA</td><td>NA</td></tr>
<tr><td></td><td></td><td></td><td></td><td></td><td></td><td></td><td></td><td></td><td></td><td></td><td></td></tr>
<tr><td>T26977</td><td>AGCTGGTCCGGGAAGCCAGGCGG</td><td>1000</td><td></td><td></td><td></td><td></td><td></td><td></td><td></td><td></td><td></td></tr>
<tr><td>Oligo fwd</td><td>CACCAGCTGGTCCGGGAAGCCAGG</td><td></td><td></td><td></td><td></td><td></td><td></td><td></td><td></td><td></td><td></td></tr>
<tr><td>Oligo rev</td><td>AAACCCTGGCTTCCCGGACCAGCT</td><td></td><td></td><td></td><td></td><td></td><td></td><td></td><td></td><td></td><td></td></tr>
<tr><td>Chromosome</td><td>start</td><td>end</td><td>strand</td><td>MM</td><td>target_seq</td><td>PAM</td><td>alignment</td><td>distance</td><td>position</td><td>gene name</td><td>gene id</td></tr>
<tr><td>GU305914.1</td><td>93608</td><td>93630</td><td>-</td><td>0</td><td>AGCTGGTCCGGGAAGCCAGG</td><td>CGG</td><td>||||||||[||||||||||||]PAM</td><td>NA</td><td>Intergenic</td><td>NA</td><td>NA</td></tr>
<tr><td></td><td></td><td></td><td></td><td></td><td></td><td></td><td></td><td></td><td></td><td></td><td></td></tr>
<tr><td>T26978</td><td>GCTGGTCCGGGAAGCCAGGCGGG</td><td>1000</td><td></td><td></td><td></td><td></td><td></td><td></td><td></td><td></td><td></td></tr>
<tr><td>Oligo fwd</td><td>CACCGCTGGTCCGGGAAGCCAGGC</td><td></td><td></td><td></td><td></td><td></td><td></td><td></td><td></td><td></td><td></td></tr>
<tr><td>Oligo rev</td><td>AAACGCCTGGCTTCCCGGACCAGC</td><td></td><td></td><td></td><td></td><td></td><td></td><td></td><td></td><td></td><td></td></tr>
<tr><td>Chromosome</td><td>start</td><td>end</td><td>strand</td><td>MM</td><td>target_seq</td><td>PAM</td><td>alignment</td><td>distance</td><td>position</td><td>gene name</td><td>gene id</td></tr>
<tr><td>GU305914.1</td><td>93607</td><td>93629</td><td>-</td><td>0</td><td>GCTGGTCCGGGAAGCCAGGC</td><td>GGG</td><td>||||||||[||||||||||||]PAM</td><td>NA</td><td>Intergenic</td><td>NA</td><td>NA</td></tr>
<tr><td>GU305914.1</td><td>93849</td><td>93871</td><td>-</td><td>3</td><td>GCTGGTCTGGGGAGCCAAGC</td><td>GGG</td><td>|||||||-[|||-|||||-||]PAM</td><td>NA</td><td>Intergenic</td><td>NA</td><td>NA</td></tr>
<tr><td></td><td></td><td></td><td></td><td></td><td></td><td></td><td></td><td></td><td></td><td></td><td></td></tr>
<tr><td>T26979</td><td>TCCGGGAAGCCAGGCGGGTCCGG</td><td>1000</td><td></td><td></td><td></td><td></td><td></td><td></td><td></td><td></td><td></td></tr>
<tr><td>Oligo fwd</td><td>CACCTCCGGGAAGCCAGGCGGGTC</td><td></td><td></td><td></td><td></td><td></td><td></td><td></td><td></td><td></td><td></td></tr>
<tr><td>Oligo rev</td><td>AAACGACCCGCCTGGCTTCCCGGA</td><td></td><td></td><td></td><td></td><td></td><td></td><td></td><td></td><td></td><td></td></tr>
<tr><td>Chromosome</td><td>start</td><td>end</td><td>strand</td><td>MM</td><td>target_seq</td><td>PAM</td><td>alignment</td><td>distance</td><td>position</td><td>gene name</td><td>gene id</td></tr>
<tr><td>GU305914.1</td><td>93602</td><td>93624</td><td>-</td><td>0</td><td>TCCGGGAAGCCAGGCGGGTC</td><td>CGG</td><td>||||||||[||||||||||||]PAM</td><td>NA</td><td>Intergenic</td><td>NA</td><td>NA</td></tr>
<tr><td></td><td></td><td></td><td></td><td></td><td></td><td></td><td></td><td></td><td></td><td></td><td></td></tr>
<tr><td>T26980</td><td>CCGGGAAGCCAGGCGGGTCCGGG</td><td>1000</td><td></td><td></td><td></td><td></td><td></td><td></td><td></td><td></td><td></td></tr>
<tr><td>Oligo fwd</td><td>CACCCCGGGAAGCCAGGCGGGTCC</td><td></td><td></td><td></td><td></td><td></td><td></td><td></td><td></td><td></td><td></td></tr>
<tr><td>Oligo rev</td><td>AAACGGACCCGCCTGGCTTCCCGG</td><td></td><td></td><td></td><td></td><td></td><td></td><td></td><td></td><td></td><td></td></tr>
<tr><td>Chromosome</td><td>start</td><td>end</td><td>strand</td><td>MM</td><td>target_seq</td><td>PAM</td><td>alignment</td><td>distance</td><td>position</td><td>gene name</td><td>gene id</td></tr>
<tr><td>GU305914.1</td><td>93601</td><td>93623</td><td>-</td><td>0</td><td>CCGGGAAGCCAGGCGGGTCC</td><td>GGG</td><td>||||||||[||||||||||||]PAM</td><td>NA</td><td>Intergenic</td><td>NA</td><td>NA</td></tr>
<tr><td></td><td></td><td></td><td></td><td></td><td></td><td></td><td></td><td></td><td></td><td></td><td></td></tr>
<tr><td>T26981</td><td>CGGGAAGCCAGGCGGGTCCGGGG</td><td>1000</td><td></td><td></td><td></td><td></td><td></td><td></td><td></td><td></td><td></td></tr>
<tr><td>Oligo fwd</td><td>CACCCGGGAAGCCAGGCGGGTCCG</td><td></td><td></td><td></td><td></td><td></td><td></td><td></td><td></td><td></td><td></td></tr>
<tr><td>Oligo rev</td><td>AAACCGGACCCGCCTGGCTTCCCG</td><td></td><td></td><td></td><td></td><td></td><td></td><td></td><td></td><td></td><td></td></tr>
<tr><td>Chromosome</td><td>start</td><td>end</td><td>strand</td><td>MM</td><td>target_seq</td><td>PAM</td><td>alignment</td><td>distance</td><td>position</td><td>gene name</td><td>gene id</td></tr>
<tr><td>GU305914.1</td><td>93600</td><td>93622</td><td>-</td><td>0</td><td>CGGGAAGCCAGGCGGGTCCG</td><td>GGG</td><td>||||||||[||||||||||||]PAM</td><td>NA</td><td>Intergenic</td><td>NA</td><td>NA</td></tr>
<tr><td></td><td></td><td></td><td></td><td></td><td></td><td></td><td></td><td></td><td></td><td></td><td></td></tr>
<tr><td>T26982</td><td>GCCAGGCGGGTCCGGGGTGCTGG</td><td>1000</td><td></td><td></td><td></td><td></td><td></td><td></td><td></td><td></td><td></td></tr>
<tr><td>Oligo fwd</td><td>CACCGCCAGGCGGGTCCGGGGTGC</td><td></td><td></td><td></td><td></td><td></td><td></td><td></td><td></td><td></td><td></td></tr>
<tr><td>Oligo rev</td><td>AAACGCACCCCGGACCCGCCTGGC</td><td></td><td></td><td></td><td></td><td></td><td></td><td></td><td></td><td></td><td></td></tr>
<tr><td>Chromosome</td><td>start</td><td>end</td><td>strand</td><td>MM</td><td>target_seq</td><td>PAM</td><td>alignment</td><td>distance</td><td>position</td><td>gene name</td><td>gene id</td></tr>
<tr><td>GU305914.1</td><td>93594</td><td>93616</td><td>-</td><td>0</td><td>GCCAGGCGGGTCCGGGGTGC</td><td>TGG</td><td>||||||||[||||||||||||]PAM</td><td>NA</td><td>Intergenic</td><td>NA</td><td>NA</td></tr>
<tr><td></td><td></td><td></td><td></td><td></td><td></td><td></td><td></td><td></td><td></td><td></td><td></td></tr>
<tr><td>T26983</td><td>CGGGTCCGGGGTGCTGGTCCTGG</td><td>1000</td><td></td><td></td><td></td><td></td><td></td><td></td><td></td><td></td><td></td></tr>
<tr><td>Oligo fwd</td><td>CACCCGGGTCCGGGGTGCTGGTCC</td><td></td><td></td><td></td><td></td><td></td><td></td><td></td><td></td><td></td><td></td></tr>
<tr><td>Oligo rev</td><td>AAACGGACCAGCACCCCGGACCCG</td><td></td><td></td><td></td><td></td><td></td><td></td><td></td><td></td><td></td><td></td></tr>
<tr><td>Chromosome</td><td>start</td><td>end</td><td>strand</td><td>MM</td><td>target_seq</td><td>PAM</td><td>alignment</td><td>distance</td><td>position</td><td>gene name</td><td>gene id</td></tr>
<tr><td>GU305914.1</td><td>93588</td><td>93610</td><td>-</td><td>0</td><td>CGGGTCCGGGGTGCTGGTCC</td><td>TGG</td><td>||||||||[||||||||||||]PAM</td><td>NA</td><td>Intergenic</td><td>NA</td><td>NA</td></tr>
<tr><td>GU305914.1</td><td>93861</td><td>93883</td><td>-</td><td>4</td><td>CGGGAGCGGGGCGCTGGTCT</td><td>GGG</td><td>||||--||[|||-|||||||-]PAM</td><td>NA</td><td>Intergenic</td><td>NA</td><td>NA</td></tr>
<tr><td></td><td></td><td></td><td></td><td></td><td></td><td></td><td></td><td></td><td></td><td></td><td></td></tr>
<tr><td>T26984</td><td>GGGTCCGGGGTGCTGGTCCTGGG</td><td>1000</td><td></td><td></td><td></td><td></td><td></td><td></td><td></td><td></td><td></td></tr>
<tr><td>Oligo fwd</td><td>CACCGGGTCCGGGGTGCTGGTCCT</td><td></td><td></td><td></td><td></td><td></td><td></td><td></td><td></td><td></td><td></td></tr>
<tr><td>Oligo rev</td><td>AAACAGGACCAGCACCCCGGACCC</td><td></td><td></td><td></td><td></td><td></td><td></td><td></td><td></td><td></td><td></td></tr>
<tr><td>Chromosome</td><td>start</td><td>end</td><td>strand</td><td>MM</td><td>target_seq</td><td>PAM</td><td>alignment</td><td>distance</td><td>position</td><td>gene name</td><td>gene id</td></tr>
<tr><td>GU305914.1</td><td>93587</td><td>93609</td><td>-</td><td>0</td><td>GGGTCCGGGGTGCTGGTCCT</td><td>GGG</td><td>||||||||[||||||||||||]PAM</td><td>NA</td><td>Intergenic</td><td>NA</td><td>NA</td></tr>
<tr><td>GU305914.1</td><td>93711</td><td>93733</td><td>-</td><td>4</td><td>GTAGCCGGGGAGCTGGTCCT</td><td>GGG</td><td>|---||||[||-|||||||||]PAM</td><td>NA</td><td>Intergenic</td><td>NA</td><td>NA</td></tr>
<tr><td>GU305914.1</td><td>93505</td><td>93527</td><td>-</td><td>3</td><td>GGGTGCGGGGCGCTGGTCCG</td><td>AGG</td><td>||||-|||[||-||||||||-]PAM</td><td>NA</td><td>Intergenic</td><td>NA</td><td>NA</td></tr>
<tr><td></td><td></td><td></td><td></td><td></td><td></td><td></td><td></td><td></td><td></td><td></td><td></td></tr>
<tr><td>T26985</td><td>GGTCCGGGGTGCTGGTCCTGGGG</td><td>1000</td><td></td><td></td><td></td><td></td><td></td><td></td><td></td><td></td><td></td></tr>
<tr><td>Oligo fwd</td><td>CACCGGTCCGGGGTGCTGGTCCTG</td><td></td><td></td><td></td><td></td><td></td><td></td><td></td><td></td><td></td><td></td></tr>
<tr><td>Oligo rev</td><td>AAACCAGGACCAGCACCCCGGACC</td><td></td><td></td><td></td><td></td><td></td><td></td><td></td><td></td><td></td><td></td></tr>
<tr><td>Chromosome</td><td>start</td><td>end</td><td>strand</td><td>MM</td><td>target_seq</td><td>PAM</td><td>alignment</td><td>distance</td><td>position</td><td>gene name</td><td>gene id</td></tr>
<tr><td>GU305914.1</td><td>93586</td><td>93608</td><td>-</td><td>0</td><td>GGTCCGGGGTGCTGGTCCTG</td><td>GGG</td><td>||||||||[||||||||||||]PAM</td><td>NA</td><td>Intergenic</td><td>NA</td><td>NA</td></tr>
<tr><td>GU305914.1</td><td>93710</td><td>93732</td><td>-</td><td>4</td><td>TAGCCGGGGAGCTGGTCCTG</td><td>GGG</td><td>---|||||[|-||||||||||]PAM</td><td>NA</td><td>Intergenic</td><td>NA</td><td>NA</td></tr>
<tr><td>GU305914.1</td><td>93544</td><td>93566</td><td>-</td><td>4</td><td>TAGCCGGGGAGCTGGTCCTG</td><td>CGG</td><td>---|||||[|-||||||||||]PAM</td><td>NA</td><td>Intergenic</td><td>NA</td><td>NA</td></tr>
<tr><td></td><td></td><td></td><td></td><td></td><td></td><td></td><td></td><td></td><td></td><td></td><td></td></tr>
<tr><td>T26986</td><td>GGGGTGCTGGTCCTGGGGTCCGG</td><td>1000</td><td></td><td></td><td></td><td></td><td></td><td></td><td></td><td></td><td></td></tr>
<tr><td>Oligo fwd</td><td>CACCGGGGTGCTGGTCCTGGGGTC</td><td></td><td></td><td></td><td></td><td></td><td></td><td></td><td></td><td></td><td></td></tr>
<tr><td>Oligo rev</td><td>AAACGACCCCAGGACCAGCACCCC</td><td></td><td></td><td></td><td></td><td></td><td></td><td></td><td></td><td></td><td></td></tr>
<tr><td>Chromosome</td><td>start</td><td>end</td><td>strand</td><td>MM</td><td>target_seq</td><td>PAM</td><td>alignment</td><td>distance</td><td>position</td><td>gene name</td><td>gene id</td></tr>
<tr><td>GU305914.1</td><td>93581</td><td>93603</td><td>-</td><td>0</td><td>GGGGTGCTGGTCCTGGGGTC</td><td>CGG</td><td>||||||||[||||||||||||]PAM</td><td>NA</td><td>Intergenic</td><td>NA</td><td>NA</td></tr>
<tr><td>GU305914.1</td><td>93539</td><td>93561</td><td>-</td><td>2</td><td>GGGGAGCTGGTCCTGCGGTC</td><td>CGG</td><td>||||-|||[|||||||-||||]PAM</td><td>NA</td><td>Intergenic</td><td>NA</td><td>NA</td></tr>
<tr><td></td><td></td><td></td><td></td><td></td><td></td><td></td><td></td><td></td><td></td><td></td><td></td></tr>
<tr><td>T26987</td><td>GGGTGCTGGTCCTGGGGTCCGGG</td><td>1000</td><td></td><td></td><td></td><td></td><td></td><td></td><td></td><td></td><td></td></tr>
<tr><td>Oligo fwd</td><td>CACCGGGTGCTGGTCCTGGGGTCC</td><td></td><td></td><td></td><td></td><td></td><td></td><td></td><td></td><td></td><td></td></tr>
<tr><td>Oligo rev</td><td>AAACGGACCCCAGGACCAGCACCC</td><td></td><td></td><td></td><td></td><td></td><td></td><td></td><td></td><td></td><td></td></tr>
<tr><td>Chromosome</td><td>start</td><td>end</td><td>strand</td><td>MM</td><td>target_seq</td><td>PAM</td><td>alignment</td><td>distance</td><td>position</td><td>gene name</td><td>gene id</td></tr>
<tr><td>GU305914.1</td><td>93580</td><td>93602</td><td>-</td><td>0</td><td>GGGTGCTGGTCCTGGGGTCC</td><td>GGG</td><td>||||||||[||||||||||||]PAM</td><td>NA</td><td>Intergenic</td><td>NA</td><td>NA</td></tr>
<tr><td>GU305914.1</td><td>93538</td><td>93560</td><td>-</td><td>2</td><td>GGGAGCTGGTCCTGCGGTCC</td><td>GGG</td><td>|||-||||[||||||-|||||]PAM</td><td>NA</td><td>Intergenic</td><td>NA</td><td>NA</td></tr>
<tr><td>GU305914.1</td><td>93704</td><td>93726</td><td>-</td><td>2</td><td>GGGAGCTGGTCCTGGGGGCC</td><td>CGG</td><td>|||-||||[|||||||||-||]PAM</td><td>NA</td><td>Intergenic</td><td>NA</td><td>NA</td></tr>
<tr><td></td><td></td><td></td><td></td><td></td><td></td><td></td><td></td><td></td><td></td><td></td><td></td></tr>
<tr><td>T26988</td><td>GGTGCTGGTCCTGGGGTCCGGGG</td><td>1000</td><td></td><td></td><td></td><td></td><td></td><td></td><td></td><td></td><td></td></tr>
<tr><td>Oligo fwd</td><td>CACCGGTGCTGGTCCTGGGGTCCG</td><td></td><td></td><td></td><td></td><td></td><td></td><td></td><td></td><td></td><td></td></tr>
<tr><td>Oligo rev</td><td>AAACCGGACCCCAGGACCAGCACC</td><td></td><td></td><td></td><td></td><td></td><td></td><td></td><td></td><td></td><td></td></tr>
<tr><td>Chromosome</td><td>start</td><td>end</td><td>strand</td><td>MM</td><td>target_seq</td><td>PAM</td><td>alignment</td><td>distance</td><td>position</td><td>gene name</td><td>gene id</td></tr>
<tr><td>GU305914.1</td><td>93579</td><td>93601</td><td>-</td><td>0</td><td>GGTGCTGGTCCTGGGGTCCG</td><td>GGG</td><td>||||||||[||||||||||||]PAM</td><td>NA</td><td>Intergenic</td><td>NA</td><td>NA</td></tr>
<tr><td>GU305914.1</td><td>93537</td><td>93559</td><td>-</td><td>2</td><td>GGAGCTGGTCCTGCGGTCCG</td><td>GGG</td><td>||-|||||[|||||-||||||]PAM</td><td>NA</td><td>Intergenic</td><td>NA</td><td>NA</td></tr>
<tr><td>GU305914.1</td><td>124139</td><td>124161</td><td>-</td><td>4</td><td>GGAGATGGGCCTGGGGTACG</td><td>GGG</td><td>||-|-|||[-||||||||-||]PAM</td><td>NA</td><td>Intergenic</td><td>NA</td><td>NA</td></tr>
<tr><td></td><td></td><td></td><td></td><td></td><td></td><td></td><td></td><td></td><td></td><td></td><td></td></tr>
<tr><td>T26989</td><td>TCCTGGGGTCCGGGGAGCCGAGG</td><td>1000</td><td></td><td></td><td></td><td></td><td></td><td></td><td></td><td></td><td></td></tr>
<tr><td>Oligo fwd</td><td>CACCTCCTGGGGTCCGGGGAGCCG</td><td></td><td></td><td></td><td></td><td></td><td></td><td></td><td></td><td></td><td></td></tr>
<tr><td>Oligo rev</td><td>AAACCGGCTCCCCGGACCCCAGGA</td><td></td><td></td><td></td><td></td><td></td><td></td><td></td><td></td><td></td><td></td></tr>
<tr><td>Chromosome</td><td>start</td><td>end</td><td>strand</td><td>MM</td><td>target_seq</td><td>PAM</td><td>alignment</td><td>distance</td><td>position</td><td>gene name</td><td>gene id</td></tr>
</table>

<table>
<tr><td>GU305914.1</td><td></td><td>93571</td><td>93593</td><td>-</td><td>0</td><td>TCCTGGGGTCCGGGGAGCCG</td><td>AGG</td><td>||||||||[||||||||||||]PAM</td><td>NA</td><td>Intergenic</td><td>NA</td><td>NA</td></tr>
<tr><td>T26990</td><td>GGGGTCCGGGGAGCCGAGGCCGG</td><td></td><td>1000</td><td></td><td></td><td></td><td></td><td></td><td></td><td></td><td></td><td></td></tr>
<tr><td>Oligo fwd</td><td>CACCGGGGTCCGGGGAGCCGAGGC</td><td></td><td></td><td></td><td></td><td></td><td></td><td></td><td></td><td></td><td></td><td></td></tr>
<tr><td>Oligo rev</td><td>AAACGCCTCGGCTCCCCGGACCCC</td><td></td><td></td><td></td><td></td><td></td><td></td><td></td><td></td><td></td><td></td><td></td></tr>
<tr><td>Chromosome</td><td>start</td><td></td><td>end</td><td>strand</td><td>MM</td><td>target_seq</td><td>PAM</td><td>alignment</td><td>distance</td><td>position</td><td>gene name</td><td>gene id</td></tr>
<tr><td>GU305914.1</td><td></td><td>93567</td><td>93589</td><td>-</td><td>0</td><td>GGGGTCCGGGGAGCCGAGGC</td><td>CGG</td><td>||||||||[||||||||||||]PAM</td><td>NA</td><td>Intergenic</td><td>NA</td><td>NA</td></tr>
<tr><td>GU305914.1</td><td></td><td>93640</td><td>93662</td><td>-</td><td>4</td><td>GAAAGCCGGGGAGCCGAGGC</td><td>CGG</td><td>|----|||[||||||||||||]PAM</td><td>NA</td><td>Intergenic</td><td>NA</td><td>NA</td></tr>
<tr><td>GU305914.1</td><td></td><td>93794</td><td>93816</td><td>-</td><td>3</td><td>GAGGTCCGGTGAGCCGAGGA</td><td>GGG</td><td>|-||||||[|-|||||||||-]PAM</td><td>NA</td><td>Intergenic</td><td>NA</td><td>NA</td></tr>
<tr><td>GU305914.1</td><td></td><td>93690</td><td>93712</td><td>-</td><td>3</td><td>GGGGCCCGGTGAGCCGAGGT</td><td>CGG</td><td>||||-|||[|-|||||||||-]PAM</td><td>NA</td><td>Intergenic</td><td>NA</td><td>NA</td></tr>
<tr><td>GU305914.1</td><td></td><td>93733</td><td>93755</td><td>-</td><td>4</td><td>AGGGTGCGGGGATCCGAGGT</td><td>CGG</td><td>-||||-||[||||-||||||-]PAM</td><td>NA</td><td>Intergenic</td><td>NA</td><td>NA</td></tr>
<tr><td>T26991</td><td>GGGGAGCCGAGGCCGGTAGCCGG</td><td></td><td>1000</td><td></td><td></td><td></td><td></td><td></td><td></td><td></td><td></td><td></td></tr>
<tr><td>Oligo fwd</td><td>CACCGGGGAGCCGAGGCCGGTAGC</td><td></td><td></td><td></td><td></td><td></td><td></td><td></td><td></td><td></td><td></td><td></td></tr>
<tr><td>Oligo rev</td><td>AAACGCTACCGGCCTCGGCTCCCC</td><td></td><td></td><td></td><td></td><td></td><td></td><td></td><td></td><td></td><td></td><td></td></tr>
<tr><td>Chromosome</td><td>start</td><td></td><td>end</td><td>strand</td><td>MM</td><td>target_seq</td><td>PAM</td><td>alignment</td><td>distance</td><td>position</td><td>gene name</td><td>gene id</td></tr>
<tr><td>GU305914.1</td><td></td><td>93560</td><td>93582</td><td>-</td><td>0</td><td>GGGGAGCCGAGGCCGGTAGC</td><td>CGG</td><td>||||||||[||||||||||||]PAM</td><td>NA</td><td>Intergenic</td><td>NA</td><td>NA</td></tr>
<tr><td>GU305914.1</td><td></td><td>93633</td><td>93655</td><td>-</td><td>0</td><td>GGGGAGCCGAGGCCGGTAGC</td><td>CGG</td><td>||||||||[||||||||||||]PAM</td><td>NA</td><td>Intergenic</td><td>NA</td><td>NA</td></tr>
<tr><td>GU305914.1</td><td></td><td>93726</td><td>93748</td><td>-</td><td>2</td><td>GGGGATCCGAGGTCGGTAGC</td><td>CGG</td><td>|||||-||[||||-|||||||]PAM</td><td>NA</td><td>Intergenic</td><td>NA</td><td>NA</td></tr>
<tr><td>GU305914.1</td><td></td><td>93787</td><td>93809</td><td>-</td><td>3</td><td>GGTGAGCCGAGGAGGGTAGC</td><td>CGG</td><td>||-|||||[||||--||||||]PAM</td><td>NA</td><td>Intergenic</td><td>NA</td><td>NA</td></tr>
<tr><td>GU305914.1</td><td></td><td>93683</td><td>93705</td><td>-</td><td>3</td><td>GGTGAGCCGAGGTCGGTAGG</td><td>CGG</td><td>||-|||||[||||-||||||-]PAM</td><td>NA</td><td>Intergenic</td><td>NA</td><td>NA</td></tr>
<tr><td>T26992</td><td>GGGAGCCGAGGCCGGTAGCCGGG</td><td></td><td>1000</td><td></td><td></td><td></td><td></td><td></td><td></td><td></td><td></td><td></td></tr>
<tr><td>Oligo fwd</td><td>CACCGGGAGCCGAGGCCGGTAGCC</td><td></td><td></td><td></td><td></td><td></td><td></td><td></td><td></td><td></td><td></td><td></td></tr>
<tr><td>Oligo rev</td><td>AAACGGCTACCGGCCTCGGCTCCC</td><td></td><td></td><td></td><td></td><td></td><td></td><td></td><td></td><td></td><td></td><td></td></tr>
<tr><td>Chromosome</td><td>start</td><td></td><td>end</td><td>strand</td><td>MM</td><td>target_seq</td><td>PAM</td><td>alignment</td><td>distance</td><td>position</td><td>gene name</td><td>gene id</td></tr>
<tr><td>GU305914.1</td><td></td><td>93559</td><td>93581</td><td>-</td><td>0</td><td>GGGAGCCGAGGCCGGTAGCC</td><td>GGG</td><td>||||||||[||||||||||||]PAM</td><td>NA</td><td>Intergenic</td><td>NA</td><td>NA</td></tr>
<tr><td>GU305914.1</td><td></td><td>93632</td><td>93654</td><td>-</td><td>0</td><td>GGGAGCCGAGGCCGGTAGCC</td><td>GGG</td><td>||||||||[||||||||||||]PAM</td><td>NA</td><td>Intergenic</td><td>NA</td><td>NA</td></tr>
<tr><td>GU305914.1</td><td></td><td>93725</td><td>93747</td><td>-</td><td>2</td><td>GGGATCCGAGGTCGGTAGCC</td><td>GGG</td><td>||||-|||[|||-||||||||]PAM</td><td>NA</td><td>Intergenic</td><td>NA</td><td>NA</td></tr>
<tr><td>GU305914.1</td><td></td><td>93786</td><td>93808</td><td>-</td><td>3</td><td>GTGAGCCGAGGAGGGTAGCC</td><td>GGG</td><td>|-||||||[|||--|||||||]PAM</td><td>NA</td><td>Intergenic</td><td>NA</td><td>NA</td></tr>
<tr><td>GU305914.1</td><td></td><td>93682</td><td>93704</td><td>-</td><td>3</td><td>GTGAGCCGAGGTCGGTAGGC</td><td>GGG</td><td>|-||||||[|||-||||||-|]PAM</td><td>NA</td><td>Intergenic</td><td>NA</td><td>NA</td></tr>
<tr><td>T26993</td><td>GGAGCCGAGGCCGGTAGCCGGGG</td><td></td><td>1000</td><td></td><td></td><td></td><td></td><td></td><td></td><td></td><td></td><td></td></tr>
<tr><td>Oligo fwd</td><td>CACCGGAGCCGAGGCCGGTAGCCG</td><td></td><td></td><td></td><td></td><td></td><td></td><td></td><td></td><td></td><td></td><td></td></tr>
<tr><td>Oligo rev</td><td>AAACCGGCTACCGGCCTCGGCTCC</td><td></td><td></td><td></td><td></td><td></td><td></td><td></td><td></td><td></td><td></td><td></td></tr>
<tr><td>Chromosome</td><td>start</td><td></td><td>end</td><td>strand</td><td>MM</td><td>target_seq</td><td>PAM</td><td>alignment</td><td>distance</td><td>position</td><td>gene name</td><td>gene id</td></tr>
<tr><td>GU305914.1</td><td></td><td>93558</td><td>93580</td><td>-</td><td>0</td><td>GGAGCCGAGGCCGGTAGCCG</td><td>GGG</td><td>||||||||[||||||||||||]PAM</td><td>NA</td><td>Intergenic</td><td>NA</td><td>NA</td></tr>
<tr><td>GU305914.1</td><td></td><td>93631</td><td>93653</td><td>-</td><td>0</td><td>GGAGCCGAGGCCGGTAGCCG</td><td>GGG</td><td>||||||||[||||||||||||]PAM</td><td>NA</td><td>Intergenic</td><td>NA</td><td>NA</td></tr>
<tr><td>GU305914.1</td><td></td><td>93724</td><td>93746</td><td>-</td><td>2</td><td>GGATCCGAGGTCGGTAGCCG</td><td>GGG</td><td>|||-||||[||-|||||||||]PAM</td><td>NA</td><td>Intergenic</td><td>NA</td><td>NA</td></tr>
<tr><td>T26994</td><td>GAGGCCGGTAGCCGGGGAGCTGG</td><td></td><td>1000</td><td></td><td></td><td></td><td></td><td></td><td></td><td></td><td></td><td></td></tr>
<tr><td>Oligo fwd</td><td>CACCGAGGCCGGTAGCCGGGGAGC</td><td></td><td></td><td></td><td></td><td></td><td></td><td></td><td></td><td></td><td></td><td></td></tr>
<tr><td>Oligo rev</td><td>AAACGCTCCCCGGCTACCGGCCTC</td><td></td><td></td><td></td><td></td><td></td><td></td><td></td><td></td><td></td><td></td><td></td></tr>
<tr><td>Chromosome</td><td>start</td><td></td><td>end</td><td>strand</td><td>MM</td><td>target_seq</td><td>PAM</td><td>alignment</td><td>distance</td><td>position</td><td>gene name</td><td>gene id</td></tr>
<tr><td>GU305914.1</td><td></td><td>93625</td><td>93647</td><td>-</td><td>0</td><td>GAGGCCGGTAGCCGGGGAGC</td><td>TGG</td><td>||||||||[||||||||||||]PAM</td><td>NA</td><td>Intergenic</td><td>NA</td><td>NA</td></tr>
<tr><td>GU305914.1</td><td></td><td>93552</td><td>93574</td><td>-</td><td>0</td><td>GAGGCCGGTAGCCGGGGAGC</td><td>TGG</td><td>||||||||[||||||||||||]PAM</td><td>NA</td><td>Intergenic</td><td>NA</td><td>NA</td></tr>
<tr><td>GU305914.1</td><td></td><td>93718</td><td>93740</td><td>-</td><td>1</td><td>GAGGTCGGTAGCCGGGGAGC</td><td>TGG</td><td>||||-|||[||||||||||||]PAM</td><td>NA</td><td>Intergenic</td><td>NA</td><td>NA</td></tr>
<tr><td>GU305914.1</td><td></td><td>93779</td><td>93801</td><td>-</td><td>3</td><td>GAGGAGGGTAGCCGGGAAGC</td><td>TGG</td><td>||||--||[||||||||-|||]PAM</td><td>NA</td><td>Intergenic</td><td>NA</td><td>NA</td></tr>
<tr><td>GU305914.1</td><td></td><td>93675</td><td>93697</td><td>-</td><td>3</td><td>GAGGTCGGTAGGCGGGAAGC</td><td>TGG</td><td>||||-|||[|||-||||-|||]PAM</td><td>NA</td><td>Intergenic</td><td>NA</td><td>NA</td></tr>
<tr><td>T26995</td><td>TAGCCGGGGAGCTGGTCCTGCGG</td><td></td><td>1000</td><td></td><td></td><td></td><td></td><td></td><td></td><td></td><td></td><td></td></tr>
<tr><td>Oligo fwd</td><td>CACCTAGCCGGGGAGCTGGTCCTG</td><td></td><td></td><td></td><td></td><td></td><td></td><td></td><td></td><td></td><td></td><td></td></tr>
<tr><td>Oligo rev</td><td>AAACCAGGACCAGCTCCCCGGCTA</td><td></td><td></td><td></td><td></td><td></td><td></td><td></td><td></td><td></td><td></td><td></td></tr>
<tr><td>Chromosome</td><td>start</td><td></td><td>end</td><td>strand</td><td>MM</td><td>target_seq</td><td>PAM</td><td>alignment</td><td>distance</td><td>position</td><td>gene name</td><td>gene id</td></tr>
<tr><td>GU305914.1</td><td></td><td>93544</td><td>93566</td><td>-</td><td>0</td><td>TAGCCGGGGAGCTGGTCCTG</td><td>CGG</td><td>||||||||[||||||||||||]PAM</td><td>NA</td><td>Intergenic</td><td>NA</td><td>NA</td></tr>
<tr><td>GU305914.1</td><td></td><td>93710</td><td>93732</td><td>-</td><td>0</td><td>TAGCCGGGGAGCTGGTCCTG</td><td>GGG</td><td>||||||||[||||||||||||]PAM</td><td>NA</td><td>Intergenic</td><td>NA</td><td>NA</td></tr>
<tr><td>GU305914.1</td><td></td><td>93586</td><td>93608</td><td>-</td><td>4</td><td>GGTCCGGGGTGCTGGTCCTG</td><td>GGG</td><td>---|||||[|-||||||||||]PAM</td><td>NA</td><td>Intergenic</td><td>NA</td><td>NA</td></tr>
<tr><td>T26996</td><td>GGGGAGCTGGTCCTGCGGTCCGG</td><td></td><td>1000</td><td></td><td></td><td></td><td></td><td></td><td></td><td></td><td></td><td></td></tr>
<tr><td>Oligo fwd</td><td>CACCGGGGAGCTGGTCCTGCGGTC</td><td></td><td></td><td></td><td></td><td></td><td></td><td></td><td></td><td></td><td></td><td></td></tr>
<tr><td>Oligo rev</td><td>AAACGACCGCAGGACCAGCTCCCC</td><td></td><td></td><td></td><td></td><td></td><td></td><td></td><td></td><td></td><td></td><td></td></tr>
<tr><td>Chromosome</td><td>start</td><td></td><td>end</td><td>strand</td><td>MM</td><td>target_seq</td><td>PAM</td><td>alignment</td><td>distance</td><td>position</td><td>gene name</td><td>gene id</td></tr>
<tr><td>GU305914.1</td><td></td><td>93539</td><td>93561</td><td>-</td><td>0</td><td>GGGGAGCTGGTCCTGCGGTC</td><td>CGG</td><td>||||||||[||||||||||||]PAM</td><td>NA</td><td>Intergenic</td><td>NA</td><td>NA</td></tr>
<tr><td>GU305914.1</td><td></td><td>93581</td><td>93603</td><td>-</td><td>2</td><td>GGGGTGCTGGTCCTGGGGTC</td><td>CGG</td><td>||||-|||[|||||||-||||]PAM</td><td>NA</td><td>Intergenic</td><td>NA</td><td>NA</td></tr>
<tr><td>T26997</td><td>GGGAGCTGGTCCTGCGGTCCGGG</td><td></td><td>1000</td><td></td><td></td><td></td><td></td><td></td><td></td><td></td><td></td><td></td></tr>
<tr><td>Oligo fwd</td><td>CACCGGGAGCTGGTCCTGCGGTCC</td><td></td><td></td><td></td><td></td><td></td><td></td><td></td><td></td><td></td><td></td><td></td></tr>
<tr><td>Oligo rev</td><td>AAACGGACCGCAGGACCAGCTCCC</td><td></td><td></td><td></td><td></td><td></td><td></td><td></td><td></td><td></td><td></td><td></td></tr>
<tr><td>Chromosome</td><td>start</td><td></td><td>end</td><td>strand</td><td>MM</td><td>target_seq</td><td>PAM</td><td>alignment</td><td>distance</td><td>position</td><td>gene name</td><td>gene id</td></tr>
<tr><td>GU305914.1</td><td></td><td>93538</td><td>93560</td><td>-</td><td>0</td><td>GGGAGCTGGTCCTGCGGTCC</td><td>GGG</td><td>||||||||[||||||||||||]PAM</td><td>NA</td><td>Intergenic</td><td>NA</td><td>NA</td></tr>
<tr><td>GU305914.1</td><td></td><td>93580</td><td>93602</td><td>-</td><td>2</td><td>GGGTGCTGGTCCTGGGGTCC</td><td>GGG</td><td>|||-||||[||||||-|||||]PAM</td><td>NA</td><td>Intergenic</td><td>NA</td><td>NA</td></tr>
<tr><td>GU305914.1</td><td></td><td>93704</td><td>93726</td><td>-</td><td>2</td><td>GGGAGCTGGTCCTGGGGGCC</td><td>CGG</td><td>||||||||[||||||-||-||]PAM</td><td>NA</td><td>Intergenic</td><td>NA</td><td>NA</td></tr>
<tr><td>T26998</td><td>GGAGCTGGTCCTGCGGTCCGGGG</td><td></td><td>1000</td><td></td><td></td><td></td><td></td><td></td><td></td><td></td><td></td><td></td></tr>
<tr><td>Oligo fwd</td><td>CACCGGAGCTGGTCCTGCGGTCCG</td><td></td><td></td><td></td><td></td><td></td><td></td><td></td><td></td><td></td><td></td><td></td></tr>
<tr><td>Oligo rev</td><td>AAACCGGACCGCAGGACCAGCTCC</td><td></td><td></td><td></td><td></td><td></td><td></td><td></td><td></td><td></td><td></td><td></td></tr>
<tr><td>Chromosome</td><td>start</td><td></td><td>end</td><td>strand</td><td>MM</td><td>target_seq</td><td>PAM</td><td>alignment</td><td>distance</td><td>position</td><td>gene name</td><td>gene id</td></tr>
<tr><td>GU305914.1</td><td></td><td>93537</td><td>93559</td><td>-</td><td>0</td><td>GGAGCTGGTCCTGCGGTCCG</td><td>GGG</td><td>||||||||[||||||||||||]PAM</td><td>NA</td><td>Intergenic</td><td>NA</td><td>NA</td></tr>
<tr><td>GU305914.1</td><td></td><td>93579</td><td>93601</td><td>-</td><td>2</td><td>GGTGCTGGTCCTGGGGTCCG</td><td>GGG</td><td>||-|||||[|||||-||||||]PAM</td><td>NA</td><td>Intergenic</td><td>NA</td><td>NA</td></tr>
<tr><td>T26999</td><td>GGTCCTGCGGTCCGGGGATCCGG</td><td></td><td>1000</td><td></td><td></td><td></td><td></td><td></td><td></td><td></td><td></td><td></td></tr>
<tr><td>Oligo fwd</td><td>CACCGGTCCTGCGGTCCGGGGATC</td><td></td><td></td><td></td><td></td><td></td><td></td><td></td><td></td><td></td><td></td><td></td></tr>
<tr><td>Oligo rev</td><td>AAACGATCCCCGGACCGCAGGACC</td><td></td><td></td><td></td><td></td><td></td><td></td><td></td><td></td><td></td><td></td><td></td></tr>
<tr><td>Chromosome</td><td>start</td><td></td><td>end</td><td>strand</td><td>MM</td><td>target_seq</td><td>PAM</td><td>alignment</td><td>distance</td><td>position</td><td>gene name</td><td>gene id</td></tr>
<tr><td>GU305914.1</td><td></td><td>93531</td><td>93553</td><td>-</td><td>0</td><td>GGTCCTGCGGTCCGGGGATC</td><td>CGG</td><td>||||||||[||||||||||||]PAM</td><td>NA</td><td>Intergenic</td><td>NA</td><td>NA</td></tr>
<tr><td>T27000</td><td>GTCCTGCGGTCCGGGGATCCGGG</td><td></td><td>1000</td><td></td><td></td><td></td><td></td><td></td><td></td><td></td><td></td><td></td></tr>
<tr><td>Oligo fwd</td><td>CACCGTCCTGCGGTCCGGGGATCC</td><td></td><td></td><td></td><td></td><td></td><td></td><td></td><td></td><td></td><td></td><td></td></tr>
<tr><td>Oligo rev</td><td>AAACGGATCCCCGGACCGCAGGAC</td><td></td><td></td><td></td><td></td><td></td><td></td><td></td><td></td><td></td><td></td><td></td></tr>
</table>

<table>
<tr><td>Chromosome</td><td>start</td><td>end</td><td>strand</td><td>MM</td><td>target_seq</td><td>PAM</td><td>alignment</td><td>distance</td><td>position</td><td>gene name</td><td>gene id</td></tr>
<tr><td>GU305914.1</td><td>93530</td><td>93552</td><td>-</td><td>0</td><td>GTCCTGCGGTCCGGGGATCC</td><td>GGG</td><td>||||||||[||||||||||||]PAM</td><td>NA</td><td>Intergenic</td><td>NA</td><td>NA</td></tr>
<tr><td></td><td></td><td></td><td></td><td></td><td></td><td></td><td></td><td></td><td></td><td></td><td></td></tr>
<tr><td>T27001</td><td>CTGCGGTCCGGGGATCCGGGTGG</td><td>1000</td><td></td><td></td><td></td><td></td><td></td><td></td><td></td><td></td><td></td></tr>
<tr><td>Oligo fwd</td><td>CACCCTGCGGTCCGGGGATCCGGG</td><td></td><td></td><td></td><td></td><td></td><td></td><td></td><td></td><td></td><td></td></tr>
<tr><td>Oligo rev</td><td>AAACCCCGGATCCCCGGACCGCAG</td><td></td><td></td><td></td><td></td><td></td><td></td><td></td><td></td><td></td><td></td></tr>
<tr><td>Chromosome</td><td>start</td><td>end</td><td>strand</td><td>MM</td><td>target_seq</td><td>PAM</td><td>alignment</td><td>distance</td><td>position</td><td>gene name</td><td>gene id</td></tr>
<tr><td>GU305914.1</td><td>93527</td><td>93549</td><td>-</td><td>0</td><td>CTGCGGTCCGGGGATCCGGG</td><td>TGG</td><td>||||||||[||||||||||||]PAM</td><td>NA</td><td>Intergenic</td><td>NA</td><td>NA</td></tr>
<tr><td></td><td></td><td></td><td></td><td></td><td></td><td></td><td></td><td></td><td></td><td></td><td></td></tr>
<tr><td>T27002</td><td>TGCGGTCCGGGGATCCGGGTGGG</td><td>1000</td><td></td><td></td><td></td><td></td><td></td><td></td><td></td><td></td><td></td></tr>
<tr><td>Oligo fwd</td><td>CACCTGCGGTCCGGGGATCCGGGT</td><td></td><td></td><td></td><td></td><td></td><td></td><td></td><td></td><td></td><td></td></tr>
<tr><td>Oligo rev</td><td>AAACACCCGGATCCCCGGACCGCA</td><td></td><td></td><td></td><td></td><td></td><td></td><td></td><td></td><td></td><td></td></tr>
<tr><td>Chromosome</td><td>start</td><td>end</td><td>strand</td><td>MM</td><td>target_seq</td><td>PAM</td><td>alignment</td><td>distance</td><td>position</td><td>gene name</td><td>gene id</td></tr>
<tr><td>GU305914.1</td><td>93526</td><td>93548</td><td>-</td><td>0</td><td>TGCGGTCCGGGGATCCGGGT</td><td>GGG</td><td>||||||||[||||||||||||]PAM</td><td>NA</td><td>Intergenic</td><td>NA</td><td>NA</td></tr>
<tr><td></td><td></td><td></td><td></td><td></td><td></td><td></td><td></td><td></td><td></td><td></td><td></td></tr>
<tr><td>T27003</td><td>GCGGTCCGGGGATCCGGGTGGGG</td><td>1000</td><td></td><td></td><td></td><td></td><td></td><td></td><td></td><td></td><td></td></tr>
<tr><td>Oligo fwd</td><td>CACCGCGGTCCGGGGATCCGGGTG</td><td></td><td></td><td></td><td></td><td></td><td></td><td></td><td></td><td></td><td></td></tr>
<tr><td>Oligo rev</td><td>AAACCACCCGGATCCCCGGACCGC</td><td></td><td></td><td></td><td></td><td></td><td></td><td></td><td></td><td></td><td></td></tr>
<tr><td>Chromosome</td><td>start</td><td>end</td><td>strand</td><td>MM</td><td>target_seq</td><td>PAM</td><td>alignment</td><td>distance</td><td>position</td><td>gene name</td><td>gene id</td></tr>
<tr><td>GU305914.1</td><td>93525</td><td>93547</td><td>-</td><td>0</td><td>GCGGTCCGGGGATCCGGGTG</td><td>GGG</td><td>||||||||[||||||||||||]PAM</td><td>NA</td><td>Intergenic</td><td>NA</td><td>NA</td></tr>
<tr><td></td><td></td><td></td><td></td><td></td><td></td><td></td><td></td><td></td><td></td><td></td><td></td></tr>
<tr><td>T27004</td><td>CCGGGGATCCGGGTGGGGTGCGG</td><td>1000</td><td></td><td></td><td></td><td></td><td></td><td></td><td></td><td></td><td></td></tr>
<tr><td>Oligo fwd</td><td>CACCCCGGGGATCCGGGTGGGGTG</td><td></td><td></td><td></td><td></td><td></td><td></td><td></td><td></td><td></td><td></td></tr>
<tr><td>Oligo rev</td><td>AAACCACCCCACCCGGATCCCCGG</td><td></td><td></td><td></td><td></td><td></td><td></td><td></td><td></td><td></td><td></td></tr>
<tr><td>Chromosome</td><td>start</td><td>end</td><td>strand</td><td>MM</td><td>target_seq</td><td>PAM</td><td>alignment</td><td>distance</td><td>position</td><td>gene name</td><td>gene id</td></tr>
<tr><td>GU305914.1</td><td>93520</td><td>93542</td><td>-</td><td>0</td><td>CCGGGGATCCGGGTGGGGTG</td><td>CGG</td><td>||||||||[||||||||||||]PAM</td><td>NA</td><td>Intergenic</td><td>NA</td><td>NA</td></tr>
<tr><td></td><td></td><td></td><td></td><td></td><td></td><td></td><td></td><td></td><td></td><td></td><td></td></tr>
<tr><td>T27005</td><td>CGGGGATCCGGGTGGGGTGCGGG</td><td>1000</td><td></td><td></td><td></td><td></td><td></td><td></td><td></td><td></td><td></td></tr>
<tr><td>Oligo fwd</td><td>CACCCGGGGATCCGGGTGGGGTGC</td><td></td><td></td><td></td><td></td><td></td><td></td><td></td><td></td><td></td><td></td></tr>
<tr><td>Oligo rev</td><td>AAACGCACCCCACCCGGATCCCCG</td><td></td><td></td><td></td><td></td><td></td><td></td><td></td><td></td><td></td><td></td></tr>
<tr><td>Chromosome</td><td>start</td><td>end</td><td>strand</td><td>MM</td><td>target_seq</td><td>PAM</td><td>alignment</td><td>distance</td><td>position</td><td>gene name</td><td>gene id</td></tr>
<tr><td>GU305914.1</td><td>93519</td><td>93541</td><td>-</td><td>0</td><td>CGGGGATCCGGGTGGGGTGC</td><td>GGG</td><td>||||||||[||||||||||||]PAM</td><td>NA</td><td>Intergenic</td><td>NA</td><td>NA</td></tr>
<tr><td></td><td></td><td></td><td></td><td></td><td></td><td></td><td></td><td></td><td></td><td></td><td></td></tr>
<tr><td>T27006</td><td>GGGGATCCGGGTGGGGTGCGGGG</td><td>1000</td><td></td><td></td><td></td><td></td><td></td><td></td><td></td><td></td><td></td></tr>
<tr><td>Oligo fwd</td><td>CACCGGGGATCCGGGTGGGGTGCG</td><td></td><td></td><td></td><td></td><td></td><td></td><td></td><td></td><td></td><td></td></tr>
<tr><td>Oligo rev</td><td>AAACCGCACCCCACCCGGATCCCC</td><td></td><td></td><td></td><td></td><td></td><td></td><td></td><td></td><td></td><td></td></tr>
<tr><td>Chromosome</td><td>start</td><td>end</td><td>strand</td><td>MM</td><td>target_seq</td><td>PAM</td><td>alignment</td><td>distance</td><td>position</td><td>gene name</td><td>gene id</td></tr>
<tr><td>GU305914.1</td><td>93518</td><td>93540</td><td>-</td><td>0</td><td>GGGGATCCGGGTGGGGTGCG</td><td>GGG</td><td>||||||||[||||||||||||]PAM</td><td>NA</td><td>Intergenic</td><td>NA</td><td>NA</td></tr>
<tr><td></td><td></td><td></td><td></td><td></td><td></td><td></td><td></td><td></td><td></td><td></td><td></td></tr>
<tr><td>T27007</td><td>CCGGGTGGGGTGCGGGGCGCTGG</td><td>1000</td><td></td><td></td><td></td><td></td><td></td><td></td><td></td><td></td><td></td></tr>
<tr><td>Oligo fwd</td><td>CACCCCGGGTGGGGTGCGGGGCGC</td><td></td><td></td><td></td><td></td><td></td><td></td><td></td><td></td><td></td><td></td></tr>
<tr><td>Oligo rev</td><td>AAACGCGCCCCGCACCCCACCCGG</td><td></td><td></td><td></td><td></td><td></td><td></td><td></td><td></td><td></td><td></td></tr>
<tr><td>Chromosome</td><td>start</td><td>end</td><td>strand</td><td>MM</td><td>target_seq</td><td>PAM</td><td>alignment</td><td>distance</td><td>position</td><td>gene name</td><td>gene id</td></tr>
<tr><td>GU305914.1</td><td>93512</td><td>93534</td><td>-</td><td>0</td><td>CCGGGTGGGGTGCGGGGCGC</td><td>TGG</td><td>||||||||[||||||||||||]PAM</td><td>NA</td><td>Intergenic</td><td>NA</td><td>NA</td></tr>
<tr><td></td><td></td><td></td><td></td><td></td><td></td><td></td><td></td><td></td><td></td><td></td><td></td></tr>
<tr><td>T27008</td><td>GGGTGCGGGGCGCTGGTCCGAGG</td><td>1000</td><td></td><td></td><td></td><td></td><td></td><td></td><td></td><td></td><td></td></tr>
<tr><td>Oligo fwd</td><td>CACCGGGTGCGGGGCGCTGGTCCG</td><td></td><td></td><td></td><td></td><td></td><td></td><td></td><td></td><td></td><td></td></tr>
<tr><td>Oligo rev</td><td>AAACCGGACCAGCGCCCCGCACCC</td><td></td><td></td><td></td><td></td><td></td><td></td><td></td><td></td><td></td><td></td></tr>
<tr><td>Chromosome</td><td>start</td><td>end</td><td>strand</td><td>MM</td><td>target_seq</td><td>PAM</td><td>alignment</td><td>distance</td><td>position</td><td>gene name</td><td>gene id</td></tr>
<tr><td>GU305914.1</td><td>93505</td><td>93527</td><td>-</td><td>0</td><td>GGGTGCGGGGCGCTGGTCCG</td><td>AGG</td><td>||||||||[||||||||||||]PAM</td><td>NA</td><td>Intergenic</td><td>NA</td><td>NA</td></tr>
<tr><td>GU305914.1</td><td>93860</td><td>93882</td><td>-</td><td>2</td><td>GGGAGCGGGGCGCTGGTCTG</td><td>GGG</td><td>|||-||||[||||||||||-|]PAM</td><td>NA</td><td>Intergenic</td><td>NA</td><td>NA</td></tr>
<tr><td>GU305914.1</td><td>93587</td><td>93609</td><td>-</td><td>3</td><td>GGGTCCGGGGTGCTGGTCCT</td><td>GGG</td><td>||||-|||[||-||||||||-]PAM</td><td>NA</td><td>Intergenic</td><td>NA</td><td>NA</td></tr>
<tr><td></td><td></td><td></td><td></td><td></td><td></td><td></td><td></td><td></td><td></td><td></td><td></td></tr>
<tr><td>T27009</td><td>TGCGGGGCGCTGGTCCGAGGCGG</td><td>1000</td><td></td><td></td><td></td><td></td><td></td><td></td><td></td><td></td><td></td></tr>
<tr><td>Oligo fwd</td><td>CACCTGCGGGGCGCTGGTCCGAGG</td><td></td><td></td><td></td><td></td><td></td><td></td><td></td><td></td><td></td><td></td></tr>
<tr><td>Oligo rev</td><td>AAACCCTCGGACCAGCGCCCCGCA</td><td></td><td></td><td></td><td></td><td></td><td></td><td></td><td></td><td></td><td></td></tr>
<tr><td>Chromosome</td><td>start</td><td>end</td><td>strand</td><td>MM</td><td>target_seq</td><td>PAM</td><td>alignment</td><td>distance</td><td>position</td><td>gene name</td><td>gene id</td></tr>
<tr><td>GU305914.1</td><td>93502</td><td>93524</td><td>-</td><td>0</td><td>TGCGGGGCGCTGGTCCGAGG</td><td>CGG</td><td>||||||||[||||||||||||]PAM</td><td>NA</td><td>Intergenic</td><td>NA</td><td>NA</td></tr>
<tr><td></td><td></td><td></td><td></td><td></td><td></td><td></td><td></td><td></td><td></td><td></td><td></td></tr>
<tr><td>T27010</td><td>GGGGCGCTGGTCCGAGGCGGCGG</td><td>1000</td><td></td><td></td><td></td><td></td><td></td><td></td><td></td><td></td><td></td></tr>
<tr><td>Oligo fwd</td><td>CACCGGGGCGCTGGTCCGAGGCGG</td><td></td><td></td><td></td><td></td><td></td><td></td><td></td><td></td><td></td><td></td></tr>
<tr><td>Oligo rev</td><td>AAACCCGCCTCGGACCAGCGCCCC</td><td></td><td></td><td></td><td></td><td></td><td></td><td></td><td></td><td></td><td></td></tr>
<tr><td>Chromosome</td><td>start</td><td>end</td><td>strand</td><td>MM</td><td>target_seq</td><td>PAM</td><td>alignment</td><td>distance</td><td>position</td><td>gene name</td><td>gene id</td></tr>
<tr><td>GU305914.1</td><td>93499</td><td>93521</td><td>-</td><td>0</td><td>GGGGCGCTGGTCCGAGGCGG</td><td>CGG</td><td>||||||||[||||||||||||]PAM</td><td>NA</td><td>Intergenic</td><td>NA</td><td>NA</td></tr>
<tr><td></td><td></td><td></td><td></td><td></td><td></td><td></td><td></td><td></td><td></td><td></td><td></td></tr>
<tr><td>T27011</td><td>GCTGGTCCGAGGCGGCGGTCCGG</td><td>1000</td><td></td><td></td><td></td><td></td><td></td><td></td><td></td><td></td><td></td></tr>
<tr><td>Oligo fwd</td><td>CACCGCTGGTCCGAGGCGGCGGTC</td><td></td><td></td><td></td><td></td><td></td><td></td><td></td><td></td><td></td><td></td></tr>
<tr><td>Oligo rev</td><td>AAACGACCGCCGCCTCGGACCAGC</td><td></td><td></td><td></td><td></td><td></td><td></td><td></td><td></td><td></td><td></td></tr>
<tr><td>Chromosome</td><td>start</td><td>end</td><td>strand</td><td>MM</td><td>target_seq</td><td>PAM</td><td>alignment</td><td>distance</td><td>position</td><td>gene name</td><td>gene id</td></tr>
<tr><td>GU305914.1</td><td>93494</td><td>93516</td><td>-</td><td>0</td><td>GCTGGTCCGAGGCGGCGGTC</td><td>CGG</td><td>||||||||[||||||||||||]PAM</td><td>NA</td><td>Intergenic</td><td>NA</td><td>NA</td></tr>
<tr><td></td><td></td><td></td><td></td><td></td><td></td><td></td><td></td><td></td><td></td><td></td><td></td></tr>
<tr><td>T27012</td><td>TCCGAGGCGGCGGTCCGGAGCGG</td><td>1000</td><td></td><td></td><td></td><td></td><td></td><td></td><td></td><td></td><td></td></tr>
<tr><td>Oligo fwd</td><td>CACCTCCGAGGCGGCGGTCCGGAG</td><td></td><td></td><td></td><td></td><td></td><td></td><td></td><td></td><td></td><td></td></tr>
<tr><td>Oligo rev</td><td>AAACCTCCGGACCGCCGCCTCGGA</td><td></td><td></td><td></td><td></td><td></td><td></td><td></td><td></td><td></td><td></td></tr>
<tr><td>Chromosome</td><td>start</td><td>end</td><td>strand</td><td>MM</td><td>target_seq</td><td>PAM</td><td>alignment</td><td>distance</td><td>position</td><td>gene name</td><td>gene id</td></tr>
<tr><td>GU305914.1</td><td>93489</td><td>93511</td><td>-</td><td>0</td><td>TCCGAGGCGGCGGTCCGGAG</td><td>CGG</td><td>||||||||[||||||||||||]PAM</td><td>NA</td><td>Intergenic</td><td>NA</td><td>NA</td></tr>
<tr><td></td><td></td><td></td><td></td><td></td><td></td><td></td><td></td><td></td><td></td><td></td><td></td></tr>
<tr><td>T27013</td><td>GGCGGCGGTCCGGAGCGGCGTGG</td><td>1000</td><td></td><td></td><td></td><td></td><td></td><td></td><td></td><td></td><td></td></tr>
<tr><td>Oligo fwd</td><td>CACCGGCGGCGGTCCGGAGCGGCG</td><td></td><td></td><td></td><td></td><td></td><td></td><td></td><td></td><td></td><td></td></tr>
<tr><td>Oligo rev</td><td>AAACCGCCGCTCCGGACCGCCGCC</td><td></td><td></td><td></td><td></td><td></td><td></td><td></td><td></td><td></td><td></td></tr>
<tr><td>Chromosome</td><td>start</td><td>end</td><td>strand</td><td>MM</td><td>target_seq</td><td>PAM</td><td>alignment</td><td>distance</td><td>position</td><td>gene name</td><td>gene id</td></tr>
<tr><td>GU305914.1</td><td>93484</td><td>93506</td><td>-</td><td>0</td><td>GGCGGCGGTCCGGAGCGGCG</td><td>TGG</td><td>||||||||[||||||||||||]PAM</td><td>NA</td><td>Intergenic</td><td>NA</td><td>NA</td></tr>
<tr><td></td><td></td><td></td><td></td><td></td><td></td><td></td><td></td><td></td><td></td><td></td><td></td></tr>
<tr><td>T27014</td><td>GTCCGGAGCGGCGTGGATCCTGG</td><td>1000</td><td></td><td></td><td></td><td></td><td></td><td></td><td></td><td></td><td></td></tr>
<tr><td>Oligo fwd</td><td>CACCGTCCGGAGCGGCGTGGATCC</td><td></td><td></td><td></td><td></td><td></td><td></td><td></td><td></td><td></td><td></td></tr>
<tr><td>Oligo rev</td><td>AAACGGATCCACGCCGCTCCGGAC</td><td></td><td></td><td></td><td></td><td></td><td></td><td></td><td></td><td></td><td></td></tr>
<tr><td>Chromosome</td><td>start</td><td>end</td><td>strand</td><td>MM</td><td>target_seq</td><td>PAM</td><td>alignment</td><td>distance</td><td>position</td><td>gene name</td><td>gene id</td></tr>
<tr><td>GU305914.1</td><td>93477</td><td>93499</td><td>-</td><td>0</td><td>GTCCGGAGCGGCGTGGATCC</td><td>TGG</td><td>||||||||[||||||||||||]PAM</td><td>NA</td><td>Intergenic</td><td>NA</td><td>NA</td></tr>
</table>

| | | | | | | | | | | | | |
|---|---|---|---|---|---|---|---|---|---|---|---|---|
| T27015 | CGGAGCGGCGTGGATCCTGGTGG | | 1000 | | | | | | | | | |
| Oligo fwd | CACCCGGAGCGGCGTGGATCCTGG | | | | | | | | | | | |
| Oligo rev | AAACCCAGGATCCACGCCGCTCCG | | | | | | | | | | | |
| Chromosome | start | | end | strand | MM | target_seq | PAM | alignment | distance | position | gene name | gene id |
| GU305914.1 | | 93474 | 93496 | - | 0 | CGGAGCGGCGTGGATCCTGG | TGG | \|\|\|\|\|\|\|\|[\|\|\|\|\|\|\|\|\|\|\|\|]PAM | NA | Intergenic | NA | NA |
| T27016 | TGGATCCTGGTGGCGAGCGACGG | | 1000 | | | | | | | | | |
| Oligo fwd | CACCTGGATCCTGGTGGCGAGCGA | | | | | | | | | | | |
| Oligo rev | AAACTCGCTCGCCACCAGGATCCA | | | | | | | | | | | |
| Chromosome | start | | end | strand | MM | target_seq | PAM | alignment | distance | position | gene name | gene id |
| GU305914.1 | | 93464 | 93486 | - | 0 | TGGATCCTGGTGGCGAGCGA | CGG | \|\|\|\|\|\|\|\|[\|\|\|\|\|\|\|\|\|\|\|\|]PAM | NA | Intergenic | NA | NA |
| T27017 | GGATCCTGGTGGCGAGCGACGGG | | 1000 | | | | | | | | | |
| Oligo fwd | CACCGGATCCTGGTGGCGAGCGAC | | | | | | | | | | | |
| Oligo rev | AAACGTCGCTCGCCACCAGGATCC | | | | | | | | | | | |
| Chromosome | start | | end | strand | MM | target_seq | PAM | alignment | distance | position | gene name | gene id |
| GU305914.1 | | 93463 | 93485 | - | 0 | GGATCCTGGTGGCGAGCGAC | GGG | \|\|\|\|\|\|\|\|[\|\|\|\|\|\|\|\|\|\|\|\|]PAM | NA | Intergenic | NA | NA |
| T27018 | TCCTGGTGGCGAGCGACGGGCGG | | 1000 | | | | | | | | | |
| Oligo fwd | CACCTCCTGGTGGCGAGCGACGGG | | | | | | | | | | | |
| Oligo rev | AAACCCCGTCGCTCGCCACCAGGA | | | | | | | | | | | |
| Chromosome | start | | end | strand | MM | target_seq | PAM | alignment | distance | position | gene name | gene id |
| GU305914.1 | | 93460 | 93482 | - | 0 | TCCTGGTGGCGAGCGACGGG | CGG | \|\|\|\|\|\|\|\|[\|\|\|\|\|\|\|\|\|\|\|\|]PAM | NA | Intergenic | NA | NA |
| T27019 | CCTGGTGGCGAGCGACGGGCGGG | | 1000 | | | | | | | | | |
| Oligo fwd | CACCCCTGGTGGCGAGCGACGGGC | | | | | | | | | | | |
| Oligo rev | AAACGCCCGTCGCTCGCCACCAGG | | | | | | | | | | | |
| Chromosome | start | | end | strand | MM | target_seq | PAM | alignment | distance | position | gene name | gene id |
| GU305914.1 | | 93459 | 93481 | - | 0 | CCTGGTGGCGAGCGACGGGC | GGG | \|\|\|\|\|\|\|\|[\|\|\|\|\|\|\|\|\|\|\|\|]PAM | NA | Intergenic | NA | NA |
| T27020 | TGGCGAGCGACGGGCGGGTCTGG | | 1000 | | | | | | | | | |
| Oligo fwd | CACCTGGCGAGCGACGGGCGGGTC | | | | | | | | | | | |
| Oligo rev | AAACGACCCGCCCGTCGCTCGCCA | | | | | | | | | | | |
| Chromosome | start | | end | strand | MM | target_seq | PAM | alignment | distance | position | gene name | gene id |
| GU305914.1 | | 93454 | 93476 | - | 0 | TGGCGAGCGACGGGCGGGTC | TGG | \|\|\|\|\|\|\|\|[\|\|\|\|\|\|\|\|\|\|\|\|]PAM | NA | Intergenic | NA | NA |
| T27021 | GAGCGACGGGCGGGTCTGGCCGG | | 1000 | | | | | | | | | |
| Oligo fwd | CACCGAGCGACGGGCGGGTCTGGC | | | | | | | | | | | |
| Oligo rev | AAACGCCAGACCCGCCCGTCGCTC | | | | | | | | | | | |
| Chromosome | start | | end | strand | MM | target_seq | PAM | alignment | distance | position | gene name | gene id |
| GU305914.1 | | 93450 | 93472 | - | 0 | GAGCGACGGGCGGGTCTGGC | CGG | \|\|\|\|\|\|\|\|[\|\|\|\|\|\|\|\|\|\|\|\|]PAM | NA | Intergenic | NA | NA |
| T27022 | CGACGGGCGGGTCTGGCCGGCGG | | 1000 | | | | | | | | | |
| Oligo fwd | CACCCGACGGGCGGGTCTGGCCGG | | | | | | | | | | | |
| Oligo rev | AAACCCGGCCAGACCCGCCCGTCG | | | | | | | | | | | |
| Chromosome | start | | end | strand | MM | target_seq | PAM | alignment | distance | position | gene name | gene id |
| GU305914.1 | | 93447 | 93469 | - | 0 | CGACGGGCGGGTCTGGCCGG | CGG | \|\|\|\|\|\|\|\|[\|\|\|\|\|\|\|\|\|\|\|\|]PAM | NA | Intergenic | NA | NA |
| T27023 | CGGGCGGGTCTGGCCGGCGGCGG | | 1000 | | | | | | | | | |
| Oligo fwd | CACCCGGGCGGGTCTGGCCGGCGG | | | | | | | | | | | |
| Oligo rev | AAACCCGCCGGCCAGACCCGCCCG | | | | | | | | | | | |
| Chromosome | start | | end | strand | MM | target_seq | PAM | alignment | distance | position | gene name | gene id |
| GU305914.1 | | 93444 | 93466 | - | 0 | CGGGCGGGTCTGGCCGGCGG | CGG | \|\|\|\|\|\|\|\|[\|\|\|\|\|\|\|\|\|\|\|\|]PAM | NA | Intergenic | NA | NA |
| T27024 | GGGCGGGTCTGGCCGGCGGCGGG | | 1000 | | | | | | | | | |
| Oligo fwd | CACCGGGCGGGTCTGGCCGGCGGC | | | | | | | | | | | |
| Oligo rev | AAACGCCGCCGGCCAGACCCGCCC | | | | | | | | | | | |
| Chromosome | start | | end | strand | MM | target_seq | PAM | alignment | distance | position | gene name | gene id |
| GU305914.1 | | 93443 | 93465 | - | 0 | GGGCGGGTCTGGCCGGCGGC | GGG | \|\|\|\|\|\|\|\|[\|\|\|\|\|\|\|\|\|\|\|\|]PAM | NA | Intergenic | NA | NA |
| T27025 | GGGTCTGGCCGGCGGCGGGCCGG | | 1000 | | | | | | | | | |
| Oligo fwd | CACCGGGTCTGGCCGGCGGCGGGC | | | | | | | | | | | |
| Oligo rev | AAACGCCCGCCGCCGGCCAGACCC | | | | | | | | | | | |
| Chromosome | start | | end | strand | MM | target_seq | PAM | alignment | distance | position | gene name | gene id |
| GU305914.1 | | 93439 | 93461 | - | 0 | GGGTCTGGCCGGCGGCGGGC | CGG | \|\|\|\|\|\|\|\|[\|\|\|\|\|\|\|\|\|\|\|\|]PAM | NA | Intergenic | NA | NA |
| T27026 | TATCCGCCACGATGACGCATAGG | | 1000 | | | | | | | | | |
| Oligo fwd | CACCTATCCGCCACGATGACGCAT | | | | | | | | | | | |
| Oligo rev | AAACATGCGTCATCGTGGCGGATA | | | | | | | | | | | |
| Chromosome | start | | end | strand | MM | target_seq | PAM | alignment | distance | position | gene name | gene id |
| GU305914.1 | | 93406 | 93428 | - | 0 | TATCCGCCACGATGACGCAT | AGG | \|\|\|\|\|\|\|\|[\|\|\|\|\|\|\|\|\|\|\|\|]PAM | NA | Intergenic | NA | NA |
| T27027 | CATAGGTCAGCTGACCAAAATGG | | 1000 | | | | | | | | | |
| Oligo fwd | CACCCATAGGTCAGCTGACCAAAA | | | | | | | | | | | |
| Oligo rev | AAACTTTTGGTCAGCTGACCTATG | | | | | | | | | | | |
| Chromosome | start | | end | strand | MM | target_seq | PAM | alignment | distance | position | gene name | gene id |
| GU305914.1 | | 93389 | 93411 | - | 0 | CATAGGTCAGCTGACCAAAA | TGG | \|\|\|\|\|\|\|\|[\|\|\|\|\|\|\|\|\|\|\|\|]PAM | NA | Intergenic | NA | NA |
| T27028 | ATAGGTCAGCTGACCAAAATGGG | | 1000 | | | | | | | | | |
| Oligo fwd | CACCATAGGTCAGCTGACCAAAAT | | | | | | | | | | | |
| Oligo rev | AAACATTTTGGTCAGCTGACCTAT | | | | | | | | | | | |
| Chromosome | start | | end | strand | MM | target_seq | PAM | alignment | distance | position | gene name | gene id |
| GU305914.1 | | 93388 | 93410 | - | 0 | ATAGGTCAGCTGACCAAAAT | GGG | \|\|\|\|\|\|\|\|[\|\|\|\|\|\|\|\|\|\|\|\|]PAM | NA | Intergenic | NA | NA |
| T27029 | TCAGCTGACCAAAATGGGCGCGG | | 1000 | | | | | | | | | |
| Oligo fwd | CACCTCAGCTGACCAAAATGGGCG | | | | | | | | | | | |
| Oligo rev | AAACCGCCCATTTTGGTCAGCTGA | | | | | | | | | | | |
| Chromosome | start | | end | strand | MM | target_seq | PAM | alignment | distance | position | gene name | gene id |
| GU305914.1 | | 93383 | 93405 | - | 0 | TCAGCTGACCAAAATGGGCG | CGG | \|\|\|\|\|\|\|\|[\|\|\|\|\|\|\|\|\|\|\|\|]PAM | NA | Intergenic | NA | NA |

<table>
<tr><td>T27030</td><td>CCAAAATGGGCGCGGTTTCGCGG</td><td>1000</td></tr>
<tr><td>Oligo fwd</td><td>CACCCCAAAATGGGCGCGGTTTCG</td><td></td></tr>
<tr><td>Oligo rev</td><td>AAACCGAAACCGCGCCCATTTTGG</td><td></td></tr>
</table>
<table>
<tr><td>Chromosome</td><td>start</td><td>end</td><td>strand</td><td>MM</td><td>target_seq</td><td>PAM</td><td>alignment</td><td>distance</td><td>position</td><td>gene name</td><td>gene id</td></tr>
<tr><td>GU305914.1</td><td>93375</td><td>93397</td><td>-</td><td>0</td><td>CCAAAATGGGCGCGGTTTCG</td><td>CGG</td><td>||||||||[||||||||||||]PAM</td><td>NA</td><td>Intergenic</td><td>NA</td><td>NA</td></tr>
</table>

<table>
<tr><td>T27031</td><td>CAAAATGGGCGCGGTTTCGCGGG</td><td>1000</td></tr>
<tr><td>Oligo fwd</td><td>CACCCAAAATGGGCGCGGTTTCGC</td><td></td></tr>
<tr><td>Oligo rev</td><td>AAACGCGAAACCGCGCCCATTTTG</td><td></td></tr>
</table>
<table>
<tr><td>Chromosome</td><td>start</td><td>end</td><td>strand</td><td>MM</td><td>target_seq</td><td>PAM</td><td>alignment</td><td>distance</td><td>position</td><td>gene name</td><td>gene id</td></tr>
<tr><td>GU305914.1</td><td>93374</td><td>93396</td><td>-</td><td>0</td><td>CAAAATGGGCGCGGTTTCGC</td><td>GGG</td><td>||||||||[||||||||||||]PAM</td><td>NA</td><td>Intergenic</td><td>NA</td><td>NA</td></tr>
</table>

<table>
<tr><td>T27032</td><td>AAAATGGGCGCGGTTTCGCGGGG</td><td>1000</td></tr>
<tr><td>Oligo fwd</td><td>CACCAAAATGGGCGCGGTTTCGCG</td><td></td></tr>
<tr><td>Oligo rev</td><td>AAACCGCGAAACCGCGCCCATTTT</td><td></td></tr>
</table>
<table>
<tr><td>Chromosome</td><td>start</td><td>end</td><td>strand</td><td>MM</td><td>target_seq</td><td>PAM</td><td>alignment</td><td>distance</td><td>position</td><td>gene name</td><td>gene id</td></tr>
<tr><td>GU305914.1</td><td>93373</td><td>93395</td><td>-</td><td>0</td><td>AAAATGGGCGCGGTTTCGCG</td><td>GGG</td><td>||||||||[||||||||||||]PAM</td><td>NA</td><td>Intergenic</td><td>NA</td><td>NA</td></tr>
</table>

<table>
<tr><td>T27033</td><td>AAATGGGCGCGGTTTCGCGGGGG</td><td>1000</td></tr>
<tr><td>Oligo fwd</td><td>CACCAAATGGGCGCGGTTTCGCGG</td><td></td></tr>
<tr><td>Oligo rev</td><td>AAACCCGCGAAACCGCGCCCATTT</td><td></td></tr>
</table>
<table>
<tr><td>Chromosome</td><td>start</td><td>end</td><td>strand</td><td>MM</td><td>target_seq</td><td>PAM</td><td>alignment</td><td>distance</td><td>position</td><td>gene name</td><td>gene id</td></tr>
<tr><td>GU305914.1</td><td>93372</td><td>93394</td><td>-</td><td>0</td><td>AAATGGGCGCGGTTTCGCGG</td><td>GGG</td><td>||||||||[||||||||||||]PAM</td><td>NA</td><td>Intergenic</td><td>NA</td><td>NA</td></tr>
</table>

<table>
<tr><td>T27034</td><td>GCGCGGTTTCGCGGGGGATGCGG</td><td>1000</td></tr>
<tr><td>Oligo fwd</td><td>CACCGCGCGGTTTCGCGGGGGATG</td><td></td></tr>
<tr><td>Oligo rev</td><td>AAACCATCCCCCGCGAAACCGCGC</td><td></td></tr>
</table>
<table>
<tr><td>Chromosome</td><td>start</td><td>end</td><td>strand</td><td>MM</td><td>target_seq</td><td>PAM</td><td>alignment</td><td>distance</td><td>position</td><td>gene name</td><td>gene id</td></tr>
<tr><td>GU305914.1</td><td>93366</td><td>93388</td><td>-</td><td>0</td><td>GCGCGGTTTCGCGGGGGATG</td><td>CGG</td><td>||||||||[||||||||||||]PAM</td><td>NA</td><td>Intergenic</td><td>NA</td><td>NA</td></tr>
</table>

<table>
<tr><td>T27035</td><td>CGCGGTTTCGCGGGGGATGCGGG</td><td>1000</td></tr>
<tr><td>Oligo fwd</td><td>CACCCGCGGTTTCGCGGGGGATGC</td><td></td></tr>
<tr><td>Oligo rev</td><td>AAACGCATCCCCCGCGAAACCGCG</td><td></td></tr>
</table>
<table>
<tr><td>Chromosome</td><td>start</td><td>end</td><td>strand</td><td>MM</td><td>target_seq</td><td>PAM</td><td>alignment</td><td>distance</td><td>position</td><td>gene name</td><td>gene id</td></tr>
<tr><td>GU305914.1</td><td>93365</td><td>93387</td><td>-</td><td>0</td><td>CGCGGTTTCGCGGGGGATGC</td><td>GGG</td><td>||||||||[||||||||||||]PAM</td><td>NA</td><td>Intergenic</td><td>NA</td><td>NA</td></tr>
</table>

<table>
<tr><td>T27036</td><td>GCGGTTTCGCGGGGGATGCGGGG</td><td>1000</td></tr>
<tr><td>Oligo fwd</td><td>CACCGCGGTTTCGCGGGGGATGCG</td><td></td></tr>
<tr><td>Oligo rev</td><td>AAACCGCATCCCCCGCGAAACCGC</td><td></td></tr>
</table>
<table>
<tr><td>Chromosome</td><td>start</td><td>end</td><td>strand</td><td>MM</td><td>target_seq</td><td>PAM</td><td>alignment</td><td>distance</td><td>position</td><td>gene name</td><td>gene id</td></tr>
<tr><td>GU305914.1</td><td>93364</td><td>93386</td><td>-</td><td>0</td><td>GCGGTTTCGCGGGGGATGCG</td><td>GGG</td><td>||||||||[||||||||||||]PAM</td><td>NA</td><td>Intergenic</td><td>NA</td><td>NA</td></tr>
</table>

<table>
<tr><td>T27037</td><td>CGGTTTCGCGGGGGATGCGGGGG</td><td>1000</td></tr>
<tr><td>Oligo fwd</td><td>CACCCGGTTTCGCGGGGGATGCGG</td><td></td></tr>
<tr><td>Oligo rev</td><td>AAACCCGCATCCCCCGCGAAACCG</td><td></td></tr>
</table>
<table>
<tr><td>Chromosome</td><td>start</td><td>end</td><td>strand</td><td>MM</td><td>target_seq</td><td>PAM</td><td>alignment</td><td>distance</td><td>position</td><td>gene name</td><td>gene id</td></tr>
<tr><td>GU305914.1</td><td>93363</td><td>93385</td><td>-</td><td>0</td><td>CGGTTTCGCGGGGGATGCGG</td><td>GGG</td><td>||||||||[||||||||||||]PAM</td><td>NA</td><td>Intergenic</td><td>NA</td><td>NA</td></tr>
<tr><td>GU305914.1</td><td>105850</td><td>105872</td><td>+</td><td>4</td><td>CACTTTCGCGGCGGATGCCG</td><td>CGG</td><td>|--|||||[|||-||||||-|]PAM</td><td>NA</td><td>Intergenic</td><td>NA</td><td>NA</td></tr>
</table>

<table>
<tr><td>T27038</td><td>GGTTTCGCGGGGGATGCGGGGGG</td><td>1000</td></tr>
<tr><td>Oligo fwd</td><td>CACCGGTTTCGCGGGGGATGCGGG</td><td></td></tr>
<tr><td>Oligo rev</td><td>AAACCCCGCATCCCCCGCGAAACC</td><td></td></tr>
</table>
<table>
<tr><td>Chromosome</td><td>start</td><td>end</td><td>strand</td><td>MM</td><td>target_seq</td><td>PAM</td><td>alignment</td><td>distance</td><td>position</td><td>gene name</td><td>gene id</td></tr>
<tr><td>GU305914.1</td><td>93362</td><td>93384</td><td>-</td><td>0</td><td>GGTTTCGCGGGGGATGCGGG</td><td>GGG</td><td>||||||||[||||||||||||]PAM</td><td>NA</td><td>Intergenic</td><td>NA</td><td>NA</td></tr>
</table>

<table>
<tr><td>T27039</td><td>GGGGATGCGGGGGGAGAGCGTGG</td><td>1000</td></tr>
<tr><td>Oligo fwd</td><td>CACCGGGGATGCGGGGGGAGAGCG</td><td></td></tr>
<tr><td>Oligo rev</td><td>AAACCGCTCTCCCCCCGCATCCCC</td><td></td></tr>
</table>
<table>
<tr><td>Chromosome</td><td>start</td><td>end</td><td>strand</td><td>MM</td><td>target_seq</td><td>PAM</td><td>alignment</td><td>distance</td><td>position</td><td>gene name</td><td>gene id</td></tr>
<tr><td>GU305914.1</td><td>93353</td><td>93375</td><td>-</td><td>0</td><td>GGGGATGCGGGGGGAGAGCG</td><td>TGG</td><td>||||||||[||||||||||||]PAM</td><td>NA</td><td>Intergenic</td><td>NA</td><td>NA</td></tr>
</table>

<table>
<tr><td>T27040</td><td>GGGATGCGGGGGGAGAGCGTGGG</td><td>1000</td></tr>
<tr><td>Oligo fwd</td><td>CACCGGGATGCGGGGGGAGAGCGT</td><td></td></tr>
<tr><td>Oligo rev</td><td>AAACACGCTCTCCCCCCGCATCCC</td><td></td></tr>
</table>
<table>
<tr><td>Chromosome</td><td>start</td><td>end</td><td>strand</td><td>MM</td><td>target_seq</td><td>PAM</td><td>alignment</td><td>distance</td><td>position</td><td>gene name</td><td>gene id</td></tr>
<tr><td>GU305914.1</td><td>93352</td><td>93374</td><td>-</td><td>0</td><td>GGGATGCGGGGGGAGAGCGT</td><td>GGG</td><td>||||||||[||||||||||||]PAM</td><td>NA</td><td>Intergenic</td><td>NA</td><td>NA</td></tr>
</table>

<table>
<tr><td>T27041</td><td>GGATGCGGGGGGAGAGCGTGGGG</td><td>1000</td></tr>
<tr><td>Oligo fwd</td><td>CACCGGATGCGGGGGGAGAGCGTG</td><td></td></tr>
<tr><td>Oligo rev</td><td>AAACCACGCTCTCCCCCCGCATCC</td><td></td></tr>
</table>
<table>
<tr><td>Chromosome</td><td>start</td><td>end</td><td>strand</td><td>MM</td><td>target_seq</td><td>PAM</td><td>alignment</td><td>distance</td><td>position</td><td>gene name</td><td>gene id</td></tr>
<tr><td>GU305914.1</td><td>93351</td><td>93373</td><td>-</td><td>0</td><td>GGATGCGGGGGGAGAGCGTG</td><td>GGG</td><td>||||||||[||||||||||||]PAM</td><td>NA</td><td>Intergenic</td><td>NA</td><td>NA</td></tr>
</table>

<table>
<tr><td>T27042</td><td>GATGCGGGGGGAGAGCGTGGGGG</td><td>1000</td></tr>
<tr><td>Oligo fwd</td><td>CACCGATGCGGGGGGAGAGCGTGG</td><td></td></tr>
<tr><td>Oligo rev</td><td>AAACCCACGCTCTCCCCCCGCATC</td><td></td></tr>
</table>
<table>
<tr><td>Chromosome</td><td>start</td><td>end</td><td>strand</td><td>MM</td><td>target_seq</td><td>PAM</td><td>alignment</td><td>distance</td><td>position</td><td>gene name</td><td>gene id</td></tr>
<tr><td>GU305914.1</td><td>93350</td><td>93372</td><td>-</td><td>0</td><td>GATGCGGGGGGAGAGCGTGG</td><td>GGG</td><td>||||||||[||||||||||||]PAM</td><td>NA</td><td>Intergenic</td><td>NA</td><td>NA</td></tr>
</table>

<table>
<tr><td>T27043</td><td>GGGGGAGAGCGTGGGGGATGCGG</td><td>1000</td></tr>
<tr><td>Oligo fwd</td><td>CACCGGGGGAGAGCGTGGGGGATG</td><td></td></tr>
<tr><td>Oligo rev</td><td>AAACCATCCCCCACGCTCTCCCCC</td><td></td></tr>
</table>
<table>
<tr><td>Chromosome</td><td>start</td><td>end</td><td>strand</td><td>MM</td><td>target_seq</td><td>PAM</td><td>alignment</td><td>distance</td><td>position</td><td>gene name</td><td>gene id</td></tr>
<tr><td>GU305914.1</td><td>93344</td><td>93366</td><td>-</td><td>0</td><td>GGGGGAGAGCGTGGGGGATG</td><td>CGG</td><td>||||||||[||||||||||||]PAM</td><td>NA</td><td>Intergenic</td><td>NA</td><td>NA</td></tr>
</table>

<table>
<tr><td>T27044</td><td>GGGGAGAGCGTGGGGGATGCGGG</td><td>1000</td></tr>
<tr><td>Oligo fwd</td><td>CACCGGGGAGAGCGTGGGGGATGC</td><td></td></tr>
<tr><td>Oligo rev</td><td>AAACGCATCCCCCACGCTCTCCCC</td><td></td></tr>
</table>

| | | | | | | | | | | | |
|---|---|---|---|---|---|---|---|---|---|---|---|
| Chromosome | start | end | strand | MM | target_seq | PAM | alignment | distance | position | gene name | gene id |
| GU305914.1 | 93343 | 93365 | - | 0 | GGGGAGAGCGTGGGGGATGC | GGG | \|\|\|\|\|\|\|\|[\|\|\|\|\|\|\|\|\|\|\|\|]PAM | NA | Intergenic | NA | NA |
| T27045 | GGGAGAGCGTGGGGGATGCGGGG | 1000 | | | | | | | | | |
| Oligo fwd | CACCGGGAGAGCGTGGGGGATGCG | | | | | | | | | | |
| Oligo rev | AAACCGCATCCCCCACGCTCTCCC | | | | | | | | | | |
| Chromosome | start | end | strand | MM | target_seq | PAM | alignment | distance | position | gene name | gene id |
| GU305914.1 | 93342 | 93364 | - | 0 | GGGAGAGCGTGGGGGATGCG | GGG | \|\|\|\|\|\|\|\|[\|\|\|\|\|\|\|\|\|\|\|\|]PAM | NA | Intergenic | NA | NA |
| T27046 | GGAGAGCGTGGGGGATGCGGGGG | 1000 | | | | | | | | | |
| Oligo fwd | CACCGGAGAGCGTGGGGGATGCGG | | | | | | | | | | |
| Oligo rev | AAACCCGCATCCCCCACGCTCTCC | | | | | | | | | | |
| Chromosome | start | end | strand | MM | target_seq | PAM | alignment | distance | position | gene name | gene id |
| GU305914.1 | 93341 | 93363 | - | 0 | GGAGAGCGTGGGGGATGCGG | GGG | \|\|\|\|\|\|\|\|[\|\|\|\|\|\|\|\|\|\|\|\|]PAM | NA | Intergenic | NA | NA |
| T27047 | GAGAGCGTGGGGGATGCGGGGGG | 1000 | | | | | | | | | |
| Oligo fwd | CACCGAGAGCGTGGGGGATGCGGG | | | | | | | | | | |
| Oligo rev | AAACCCCGCATCCCCCACGCTCTC | | | | | | | | | | |
| Chromosome | start | end | strand | MM | target_seq | PAM | alignment | distance | position | gene name | gene id |
| GU305914.1 | 93340 | 93362 | - | 0 | GAGAGCGTGGGGGATGCGGG | GGG | \|\|\|\|\|\|\|\|[\|\|\|\|\|\|\|\|\|\|\|\|]PAM | NA | Intergenic | NA | NA |
| T27048 | AGAGCGTGGGGGATGCGGGGGGG | 1000 | | | | | | | | | |
| Oligo fwd | CACCAGAGCGTGGGGGATGCGGGG | | | | | | | | | | |
| Oligo rev | AAACCCCCGCATCCCCCACGCTCT | | | | | | | | | | |
| Chromosome | start | end | strand | MM | target_seq | PAM | alignment | distance | position | gene name | gene id |
| GU305914.1 | 93339 | 93361 | - | 0 | AGAGCGTGGGGGATGCGGGG | GGG | \|\|\|\|\|\|\|\|[\|\|\|\|\|\|\|\|\|\|\|\|]PAM | NA | Intergenic | NA | NA |
| T27049 | GCGGGGGGGCCAGCTCGAGCTGG | 1000 | | | | | | | | | |
| Oligo fwd | CACCGCGGGGGGGCCAGCTCGAGC | | | | | | | | | | |
| Oligo rev | AAACGCTCGAGCTGGCCCCCCCGC | | | | | | | | | | |
| Chromosome | start | end | strand | MM | target_seq | PAM | alignment | distance | position | gene name | gene id |
| GU305914.1 | 93325 | 93347 | - | 0 | GCGGGGGGGCCAGCTCGAGC | TGG | \|\|\|\|\|\|\|\|[\|\|\|\|\|\|\|\|\|\|\|\|]PAM | NA | Intergenic | NA | NA |
| T27050 | CGGGGGGGCCAGCTCGAGCTGGG | 1000 | | | | | | | | | |
| Oligo fwd | CACCCGGGGGGGCCAGCTCGAGCT | | | | | | | | | | |
| Oligo rev | AAACAGCTCGAGCTGGCCCCCCCG | | | | | | | | | | |
| Chromosome | start | end | strand | MM | target_seq | PAM | alignment | distance | position | gene name | gene id |
| GU305914.1 | 93324 | 93346 | - | 0 | CGGGGGGGCCAGCTCGAGCT | GGG | \|\|\|\|\|\|\|\|[\|\|\|\|\|\|\|\|\|\|\|\|]PAM | NA | Intergenic | NA | NA |
| T27051 | GGGGGCCAGCTCGAGCTGGGAGG | 1000 | | | | | | | | | |
| Oligo fwd | CACCGGGGGCCAGCTCGAGCTGGG | | | | | | | | | | |
| Oligo rev | AAACCCCAGCTCGAGCTGGCCCCC | | | | | | | | | | |
| Chromosome | start | end | strand | MM | target_seq | PAM | alignment | distance | position | gene name | gene id |
| GU305914.1 | 93321 | 93343 | - | 0 | GGGGGCCAGCTCGAGCTGGG | AGG | \|\|\|\|\|\|\|\|[\|\|\|\|\|\|\|\|\|\|\|\|]PAM | NA | Intergenic | NA | NA |
| T27052 | GGGGCCAGCTCGAGCTGGGAGGG | 1000 | | | | | | | | | |
| Oligo fwd | CACCGGGGCCAGCTCGAGCTGGGA | | | | | | | | | | |
| Oligo rev | AAACTCCCAGCTCGAGCTGGCCCC | | | | | | | | | | |
| Chromosome | start | end | strand | MM | target_seq | PAM | alignment | distance | position | gene name | gene id |
| GU305914.1 | 93320 | 93342 | - | 0 | GGGGCCAGCTCGAGCTGGGA | GGG | \|\|\|\|\|\|\|\|[\|\|\|\|\|\|\|\|\|\|\|\|]PAM | NA | Intergenic | NA | NA |
| T27053 | GCCAGCTCGAGCTGGGAGGGCGG | 1000 | | | | | | | | | |
| Oligo fwd | CACCGCCAGCTCGAGCTGGGAGGG | | | | | | | | | | |
| Oligo rev | AAACCCCTCCCAGCTCGAGCTGGC | | | | | | | | | | |
| Chromosome | start | end | strand | MM | target_seq | PAM | alignment | distance | position | gene name | gene id |
| GU305914.1 | 93317 | 93339 | - | 0 | GCCAGCTCGAGCTGGGAGGG | CGG | \|\|\|\|\|\|\|\|[\|\|\|\|\|\|\|\|\|\|\|\|]PAM | NA | Intergenic | NA | NA |
| T27054 | CCAGCTCGAGCTGGGAGGGCGGG | 1000 | | | | | | | | | |
| Oligo fwd | CACCCCAGCTCGAGCTGGGAGGGC | | | | | | | | | | |
| Oligo rev | AAACGCCCTCCCAGCTCGAGCTGG | | | | | | | | | | |
| Chromosome | start | end | strand | MM | target_seq | PAM | alignment | distance | position | gene name | gene id |
| GU305914.1 | 93316 | 93338 | - | 0 | CCAGCTCGAGCTGGGAGGGC | GGG | \|\|\|\|\|\|\|\|[\|\|\|\|\|\|\|\|\|\|\|\|]PAM | NA | Intergenic | NA | NA |
| T27055 | CAGCTCGAGCTGGGAGGGCGGGG | 1000 | | | | | | | | | |
| Oligo fwd | CACCCAGCTCGAGCTGGGAGGGCG | | | | | | | | | | |
| Oligo rev | AAACCGCCCTCCCAGCTCGAGCTG | | | | | | | | | | |
| Chromosome | start | end | strand | MM | target_seq | PAM | alignment | distance | position | gene name | gene id |
| GU305914.1 | 93315 | 93337 | - | 0 | CAGCTCGAGCTGGGAGGGCG | GGG | \|\|\|\|\|\|\|\|[\|\|\|\|\|\|\|\|\|\|\|\|]PAM | NA | Intergenic | NA | NA |
| T27056 | TCGAGCTGGGAGGGCGGGGAAGG | 1000 | | | | | | | | | |
| Oligo fwd | CACCTCGAGCTGGGAGGGCGGGGA | | | | | | | | | | |
| Oligo rev | AAACTCCCCGCCCTCCCAGCTCGA | | | | | | | | | | |
| Chromosome | start | end | strand | MM | target_seq | PAM | alignment | distance | position | gene name | gene id |
| GU305914.1 | 93311 | 93333 | - | 0 | TCGAGCTGGGAGGGCGGGGA | AGG | \|\|\|\|\|\|\|\|[\|\|\|\|\|\|\|\|\|\|\|\|]PAM | NA | Intergenic | NA | NA |
| T27057 | CGAGCTGGGAGGGCGGGGAAGGG | 1000 | | | | | | | | | |
| Oligo fwd | CACCCGAGCTGGGAGGGCGGGGAA | | | | | | | | | | |
| Oligo rev | AAACTTCCCCGCCCTCCCAGCTCG | | | | | | | | | | |
| Chromosome | start | end | strand | MM | target_seq | PAM | alignment | distance | position | gene name | gene id |
| GU305914.1 | 93310 | 93332 | - | 0 | CGAGCTGGGAGGGCGGGGAA | GGG | \|\|\|\|\|\|\|\|[\|\|\|\|\|\|\|\|\|\|\|\|]PAM | NA | Intergenic | NA | NA |
| T27058 | GCTGGGAGGGCGGGGAAGGGAGG | 1000 | | | | | | | | | |
| Oligo fwd | CACCGCTGGGAGGGCGGGGAAGGG | | | | | | | | | | |
| Oligo rev | AAACCCCTTCCCCGCCCTCCCAGC | | | | | | | | | | |
| Chromosome | start | end | strand | MM | target_seq | PAM | alignment | distance | position | gene name | gene id |
| GU305914.1 | 93307 | 93329 | - | 0 | GCTGGGAGGGCGGGGAAGGG | AGG | \|\|\|\|\|\|\|\|[\|\|\|\|\|\|\|\|\|\|\|\|]PAM | NA | Intergenic | NA | NA |
| T27059 | CTGGGAGGGCGGGGAAGGGAGGG | 1000 | | | | | | | | | |
| Oligo fwd | CACCCTGGGAGGGCGGGGAAGGGA | | | | | | | | | | |

| | | | | | | | | | | | |
|---|---|---|---|---|---|---|---|---|---|---|---|
| Oligo rev | AAACTCCCTTCCCCGCCCTCCCAG | | | | | | | | | | |
| Chromosome | start | end | strand | MM | target_seq | PAM | alignment | distance | position | gene name | gene id |
| GU305914.1 | 93306 | 93328 | - | 0 | CTGGGAGGGCGGGGAAGGGA | GGG | \|\|\|\|\|\|\|\|[\|\|\|\|\|\|\|\|\|\|\|\|]PAM | NA | Intergenic | NA | NA |
| T27060 | TGGGAGGGCGGGGAAGGGAGGGG | 1000 | | | | | | | | | |
| Oligo fwd | CACCTGGGAGGGCGGGGAAGGGAG | | | | | | | | | | |
| Oligo rev | AAACCTCCCTTCCCCGCCCTCCCA | | | | | | | | | | |
| Chromosome | start | end | strand | MM | target_seq | PAM | alignment | distance | position | gene name | gene id |
| GU305914.1 | 93305 | 93327 | - | 0 | TGGGAGGGCGGGGAAGGGAG | GGG | \|\|\|\|\|\|\|\|[\|\|\|\|\|\|\|\|\|\|\|\|]PAM | NA | Intergenic | NA | NA |
| T27061 | GGGAGGGCGGGGAAGGGAGGGGG | 1000 | | | | | | | | | |
| Oligo fwd | CACCGGGAGGGCGGGGAAGGGAGG | | | | | | | | | | |
| Oligo rev | AAACCCTCCCTTCCCCGCCCTCCC | | | | | | | | | | |
| Chromosome | start | end | strand | MM | target_seq | PAM | alignment | distance | position | gene name | gene id |
| GU305914.1 | 93304 | 93326 | - | 0 | GGGAGGGCGGGGAAGGGAGG | GGG | \|\|\|\|\|\|\|\|[\|\|\|\|\|\|\|\|\|\|\|\|]PAM | NA | Intergenic | NA | NA |
| T27062 | GGAGGGCGGGGAAGGGAGGGGGG | 1000 | | | | | | | | | |
| Oligo fwd | CACCGGAGGGCGGGGAAGGGAGGG | | | | | | | | | | |
| Oligo rev | AAACCCCTCCCTTCCCCGCCCTCC | | | | | | | | | | |
| Chromosome | start | end | strand | MM | target_seq | PAM | alignment | distance | position | gene name | gene id |
| GU305914.1 | 93303 | 93325 | - | 0 | GGAGGGCGGGGAAGGGAGGG | GGG | \|\|\|\|\|\|\|\|[\|\|\|\|\|\|\|\|\|\|\|\|]PAM | NA | Intergenic | NA | NA |
| T27063 | GAGGGCGGGGAAGGGAGGGGGGG | 1000 | | | | | | | | | |
| Oligo fwd | CACCGAGGGCGGGGAAGGGAGGGG | | | | | | | | | | |
| Oligo rev | AAACCCCCTCCCTTCCCCGCCCTC | | | | | | | | | | |
| Chromosome | start | end | strand | MM | target_seq | PAM | alignment | distance | position | gene name | gene id |
| GU305914.1 | 93302 | 93324 | - | 0 | GAGGGCGGGGAAGGGAGGGG | GGG | \|\|\|\|\|\|\|\|[\|\|\|\|\|\|\|\|\|\|\|\|]PAM | NA | Intergenic | NA | NA |
| T27064 | AGGGCGGGGAAGGGAGGGGGGGG | 1000 | | | | | | | | | |
| Oligo fwd | CACCAGGGCGGGGAAGGGAGGGGG | | | | | | | | | | |
| Oligo rev | AAACCCCCCTCCCTTCCCCGCCCT | | | | | | | | | | |
| Chromosome | start | end | strand | MM | target_seq | PAM | alignment | distance | position | gene name | gene id |
| GU305914.1 | 93301 | 93323 | - | 0 | AGGGCGGGGAAGGGAGGGGG | GGG | \|\|\|\|\|\|\|\|[\|\|\|\|\|\|\|\|\|\|\|\|]PAM | NA | Intergenic | NA | NA |
| T27065 | GGGCGGGGAAGGGAGGGGGGGGG | 1000 | | | | | | | | | |
| Oligo fwd | CACCGGGCGGGGAAGGGAGGGGGG | | | | | | | | | | |
| Oligo rev | AAACCCCCCCTCCCTTCCCCGCCC | | | | | | | | | | |
| Chromosome | start | end | strand | MM | target_seq | PAM | alignment | distance | position | gene name | gene id |
| GU305914.1 | 93300 | 93322 | - | 0 | GGGCGGGGAAGGGAGGGGGG | GGG | \|\|\|\|\|\|\|\|[\|\|\|\|\|\|\|\|\|\|\|\|]PAM | NA | Intergenic | NA | NA |
| T27066 | GGGAAGGGAGGGGGGGGGTTTGG | 1000 | | | | | | | | | |
| Oligo fwd | CACCGGGAAGGGAGGGGGGGGGTT | | | | | | | | | | |
| Oligo rev | AAACAACCCCCCCCCTCCCTTCCC | | | | | | | | | | |
| Chromosome | start | end | strand | MM | target_seq | PAM | alignment | distance | position | gene name | gene id |
| GU305914.1 | 93295 | 93317 | - | 0 | GGGAAGGGAGGGGGGGGGTT | TGG | \|\|\|\|\|\|\|\|[\|\|\|\|\|\|\|\|\|\|\|\|]PAM | NA | Intergenic | NA | NA |
| T27067 | GGAAGGGAGGGGGGGGGTTTGGG | 1000 | | | | | | | | | |
| Oligo fwd | CACCGGAAGGGAGGGGGGGGGTTT | | | | | | | | | | |
| Oligo rev | AAACAAACCCCCCCCCTCCCTTCC | | | | | | | | | | |
| Chromosome | start | end | strand | MM | target_seq | PAM | alignment | distance | position | gene name | gene id |
| GU305914.1 | 93294 | 93316 | - | 0 | GGAAGGGAGGGGGGGGGTTT | GGG | \|\|\|\|\|\|\|\|[\|\|\|\|\|\|\|\|\|\|\|\|]PAM | NA | Intergenic | NA | NA |
| T27068 | GAAGGGAGGGGGGGGGTTTGGGG | 1000 | | | | | | | | | |
| Oligo fwd | CACCGAAGGGAGGGGGGGGGTTTG | | | | | | | | | | |
| Oligo rev | AAACCAAACCCCCCCCCTCCCTTC | | | | | | | | | | |
| Chromosome | start | end | strand | MM | target_seq | PAM | alignment | distance | position | gene name | gene id |
| GU305914.1 | 93293 | 93315 | - | 0 | GAAGGGAGGGGGGGGGTTTG | GGG | \|\|\|\|\|\|\|\|[\|\|\|\|\|\|\|\|\|\|\|\|]PAM | NA | Intergenic | NA | NA |
| T27069 | AAGGGAGGGGGGGGGTTTGGGGG | 1000 | | | | | | | | | |
| Oligo fwd | CACCAAGGGAGGGGGGGGGTTTGG | | | | | | | | | | |
| Oligo rev | AAACCCAAACCCCCCCCCTCCCTT | | | | | | | | | | |
| Chromosome | start | end | strand | MM | target_seq | PAM | alignment | distance | position | gene name | gene id |
| GU305914.1 | 93292 | 93314 | - | 0 | AAGGGAGGGGGGGGGTTTGG | GGG | \|\|\|\|\|\|\|\|[\|\|\|\|\|\|\|\|\|\|\|\|]PAM | NA | Intergenic | NA | NA |
| T27070 | AGGGAGGGGGGGGGTTTGGGGGG | 1000 | | | | | | | | | |
| Oligo fwd | CACCAGGGAGGGGGGGGGTTTGGG | | | | | | | | | | |
| Oligo rev | AAACCCCAAACCCCCCCCCTCCCT | | | | | | | | | | |
| Chromosome | start | end | strand | MM | target_seq | PAM | alignment | distance | position | gene name | gene id |
| GU305914.1 | 93291 | 93313 | - | 0 | AGGGAGGGGGGGGGTTTGGG | GGG | \|\|\|\|\|\|\|\|[\|\|\|\|\|\|\|\|\|\|\|\|]PAM | NA | Intergenic | NA | NA |
| T27071 | GGGAGGGGGGGGGTTTGGGGGGG | 1000 | | | | | | | | | |
| Oligo fwd | CACCGGGAGGGGGGGGGTTTGGGG | | | | | | | | | | |
| Oligo rev | AAACCCCCAAACCCCCCCCCTCCC | | | | | | | | | | |
| Chromosome | start | end | strand | MM | target_seq | PAM | alignment | distance | position | gene name | gene id |
| GU305914.1 | 93290 | 93312 | - | 0 | GGGAGGGGGGGGGTTTGGGG | GGG | \|\|\|\|\|\|\|\|[\|\|\|\|\|\|\|\|\|\|\|\|]PAM | NA | Intergenic | NA | NA |
| GU305914.1 | 93289 | 93311 | - | 4 | GGAGGGGGGGGGTTTGGGGG | GGG | \|\|--\|\|\|\|[\|\|\|\|-\|\|-\|\|\|\|]PAM | NA | Intergenic | NA | NA |
| T27072 | GGAGGGGGGGGGTTTGGGGGGGG | 1000 | | | | | | | | | |
| Oligo fwd | CACCGGAGGGGGGGGGTTTGGGGG | | | | | | | | | | |
| Oligo rev | AAACCCCCCAAACCCCCCCCCTCC | | | | | | | | | | |
| Chromosome | start | end | strand | MM | target_seq | PAM | alignment | distance | position | gene name | gene id |
| GU305914.1 | 93289 | 93311 | - | 0 | GGAGGGGGGGGGTTTGGGGG | GGG | \|\|\|\|\|\|\|\|[\|\|\|\|\|\|\|\|\|\|\|\|]PAM | NA | Intergenic | NA | NA |
| GU305914.1 | 93288 | 93310 | - | 4 | GAGGGGGGGGGTTTGGGGGG | GGG | \|--\|\|\|\|\|[\|\|\|-\|\|-\|\|\|\|\|]PAM | NA | Intergenic | NA | NA |
| GU305914.1 | 93290 | 93312 | - | 4 | GGGAGGGGGGGGGTTTGGGG | GGG | \|\|--\|\|\|\|[\|\|\|\|-\|\|-\|\|\|\|]PAM | NA | Intergenic | NA | NA |
| T27073 | GAGGGGGGGGGTTTGGGGGGGGG | 1000 | | | | | | | | | |
| Oligo fwd | CACCGAGGGGGGGGGTTTGGGGGG | | | | | | | | | | |
| Oligo rev | AAACCCCCCCAAACCCCCCCCCTC | | | | | | | | | | |
| Chromosome | start | end | strand | MM | target_seq | PAM | alignment | distance | position | gene name | gene id |

| | | | | | | | | | | | |
|---|---|---|---|---|---|---|---|---|---|---|---|
| GU305914.1 | 93288 | 93310 | - | 0 | GAGGGGGGGGGTTTGGGGGG | GGG | ||||||||[||||||||||||]PAM | NA | Intergenic | NA | NA |
| GU305914.1 | 93289 | 93311 | - | 4 | GGAGGGGGGGGGTTTGGGGG | GGG | |--|||||[|||-||-|||||]PAM | NA | Intergenic | NA | NA |
| T27074 | GGGTTTGGGGGGGGGTGAGTTGG | 1000 | | | | | | | | | |
| Oligo fwd | CACCGGGTTTGGGGGGGGGTGAGT | | | | | | | | | | |
| Oligo rev | AAACACTCACCCCCCCCCAAACCC | | | | | | | | | | |
| Chromosome | start | end | strand | MM | target_seq | PAM | alignment | distance | position | gene name | gene id |
| GU305914.1 | 93280 | 93302 | - | 0 | GGGTTTGGGGGGGGGTGAGT | TGG | ||||||||[||||||||||||]PAM | NA | Intergenic | NA | NA |
| T27075 | GGTTTGGGGGGGGGTGAGTTGGG | 1000 | | | | | | | | | |
| Oligo fwd | CACCGGTTTGGGGGGGGGTGAGTT | | | | | | | | | | |
| Oligo rev | AAACAACTCACCCCCCCCCAAACC | | | | | | | | | | |
| Chromosome | start | end | strand | MM | target_seq | PAM | alignment | distance | position | gene name | gene id |
| GU305914.1 | 93279 | 93301 | - | 0 | GGTTTGGGGGGGGGTGAGTT | GGG | ||||||||[||||||||||||]PAM | NA | Intergenic | NA | NA |
| T27076 | GTTTGGGGGGGGGTGAGTTGGGG | 1000 | | | | | | | | | |
| Oligo fwd | CACCGTTTGGGGGGGGGTGAGTTG | | | | | | | | | | |
| Oligo rev | AAACCAACTCACCCCCCCCCAAAC | | | | | | | | | | |
| Chromosome | start | end | strand | MM | target_seq | PAM | alignment | distance | position | gene name | gene id |
| GU305914.1 | 93278 | 93300 | - | 0 | GTTTGGGGGGGGGTGAGTTG | GGG | ||||||||[||||||||||||]PAM | NA | Intergenic | NA | NA |
| T27077 | TGGGGGGGGGTGAGTTGGGGTGG | 1000 | | | | | | | | | |
| Oligo fwd | CACCTGGGGGGGGGTGAGTTGGGG | | | | | | | | | | |
| Oligo rev | AAACCCCCAACTCACCCCCCCCCA | | | | | | | | | | |
| Chromosome | start | end | strand | MM | target_seq | PAM | alignment | distance | position | gene name | gene id |
| GU305914.1 | 93275 | 93297 | - | 0 | TGGGGGGGGGTGAGTTGGGG | TGG | ||||||||[||||||||||||]PAM | NA | Intergenic | NA | NA |
| T27078 | GGGGGGTGAGTTGGGGTGGAAGG | 1000 | | | | | | | | | |
| Oligo fwd | CACCGGGGGGTGAGTTGGGGTGGA | | | | | | | | | | |
| Oligo rev | AAACTCCACCCCAACTCACCCCCC | | | | | | | | | | |
| Chromosome | start | end | strand | MM | target_seq | PAM | alignment | distance | position | gene name | gene id |
| GU305914.1 | 93271 | 93293 | - | 0 | GGGGGGTGAGTTGGGGTGGA | AGG | ||||||||[||||||||||||]PAM | NA | Intergenic | NA | NA |
| T27079 | GGGGGTGAGTTGGGGTGGAAGGG | 1000 | | | | | | | | | |
| Oligo fwd | CACCGGGGGTGAGTTGGGGTGGAA | | | | | | | | | | |
| Oligo rev | AAACTTCCACCCCAACTCACCCCC | | | | | | | | | | |
| Chromosome | start | end | strand | MM | target_seq | PAM | alignment | distance | position | gene name | gene id |
| GU305914.1 | 93270 | 93292 | - | 0 | GGGGGTGAGTTGGGGTGGAA | GGG | ||||||||[||||||||||||]PAM | NA | Intergenic | NA | NA |
| T27080 | TATTATTTTATCCCTTAAGTAGG | 1000 | | | | | | | | | |
| Oligo fwd | CACCTATTATTTTATCCCTTAAGT | | | | | | | | | | |
| Oligo rev | AAACACTTAAGGGATAAAATAATA | | | | | | | | | | |
| Chromosome | start | end | strand | MM | target_seq | PAM | alignment | distance | position | gene name | gene id |
| GU305914.1 | 93212 | 93234 | - | 0 | TATTATTTTATCCCTTAAGT | AGG | ||||||||[||||||||||||]PAM | NA | Intergenic | NA | NA |
| T27081 | ATCCCTTAAGTAGGATCCATAGG | 1000 | | | | | | | | | |
| Oligo fwd | CACCATCCCTTAAGTAGGATCCAT | | | | | | | | | | |
| Oligo rev | AAACATGGATCCTACTTAAGGGAT | | | | | | | | | | |
| Chromosome | start | end | strand | MM | target_seq | PAM | alignment | distance | position | gene name | gene id |
| GU305914.1 | 93203 | 93225 | - | 0 | ATCCCTTAAGTAGGATCCAT | AGG | ||||||||[||||||||||||]PAM | NA | Intergenic | NA | NA |
| T27082 | TTAAGTAGGATCCATAGGATAGG | 1000 | | | | | | | | | |
| Oligo fwd | CACCTTAAGTAGGATCCATAGGAT | | | | | | | | | | |
| Oligo rev | AAACATCCTATGGATCCTACTTAA | | | | | | | | | | |
| Chromosome | start | end | strand | MM | target_seq | PAM | alignment | distance | position | gene name | gene id |
| GU305914.1 | 93198 | 93220 | - | 0 | TTAAGTAGGATCCATAGGAT | AGG | ||||||||[||||||||||||]PAM | NA | Intergenic | NA | NA |
| T27083 | TAAGTAGGATCCATAGGATAGGG | 1000 | | | | | | | | | |
| Oligo fwd | CACCTAAGTAGGATCCATAGGATA | | | | | | | | | | |
| Oligo rev | AAACTATCCTATGGATCCTACTTA | | | | | | | | | | |
| Chromosome | start | end | strand | MM | target_seq | PAM | alignment | distance | position | gene name | gene id |
| GU305914.1 | 93197 | 93219 | - | 0 | TAAGTAGGATCCATAGGATA | GGG | ||||||||[||||||||||||]PAM | NA | Intergenic | NA | NA |
| T27084 | ATAGTAGACCCTATGTATATAGG | 1000 | | | | | | | | | |
| Oligo fwd | CACCATAGTAGACCCTATGTATAT | | | | | | | | | | |
| Oligo rev | AAACATATACATAGGGTCTACTAT | | | | | | | | | | |
| Chromosome | start | end | strand | MM | target_seq | PAM | alignment | distance | position | gene name | gene id |
| GU305914.1 | 93168 | 93190 | - | 0 | ATAGTAGACCCTATGTATAT | AGG | ||||||||[||||||||||||]PAM | NA | Intergenic | NA | NA |
| T27085 | CCAGACGTATACTATTCCACAGG | 1000 | | | | | | | | | |
| Oligo fwd | CACCCCAGACGTATACTATTCCAC | | | | | | | | | | |
| Oligo rev | AAACGTGGAATAGTATACGTCTGG | | | | | | | | | | |
| Chromosome | start | end | strand | MM | target_seq | PAM | alignment | distance | position | gene name | gene id |
| GU305914.1 | 93128 | 93150 | - | 0 | CCAGACGTATACTATTCCAC | AGG | ||||||||[||||||||||||]PAM | NA | Intergenic | NA | NA |
| T27086 | CAGACGTATACTATTCCACAGGG | 1000 | | | | | | | | | |
| Oligo fwd | CACCCAGACGTATACTATTCCACA | | | | | | | | | | |
| Oligo rev | AAACTGTGGAATAGTATACGTCTG | | | | | | | | | | |
| Chromosome | start | end | strand | MM | target_seq | PAM | alignment | distance | position | gene name | gene id |
| GU305914.1 | 93127 | 93149 | - | 0 | CAGACGTATACTATTCCACA | GGG | ||||||||[||||||||||||]PAM | NA | Intergenic | NA | NA |
| T27087 | TTCGTTATCCAACGTATACATGG | 1000 | | | | | | | | | |
| Oligo fwd | CACCTTCGTTATCCAACGTATACA | | | | | | | | | | |
| Oligo rev | AAACTGTATACGTTGGATAACGAA | | | | | | | | | | |
| Chromosome | start | end | strand | MM | target_seq | PAM | alignment | distance | position | gene name | gene id |
| GU305914.1 | 93094 | 93116 | - | 0 | TTCGTTATCCAACGTATACA | TGG | ||||||||[||||||||||||]PAM | NA | Intergenic | NA | NA |
| T27088 | GTATACATGGCTTAATGTATAGG | 1000 | | | | | | | | | |
| Oligo fwd | CACCGTATACATGGCTTAATGTAT | | | | | | | | | | |

<table>
<tr><td>Oligo rev</td><td>AAACATACATTAAGCCATGTATAC</td><td></td><td></td><td></td><td></td><td></td><td></td><td></td><td></td><td></td><td></td><td></td></tr>
<tr><td>Chromosome</td><td>start</td><td></td><td>end</td><td>strand</td><td>MM</td><td>target_seq</td><td>PAM</td><td>alignment</td><td>distance</td><td>position</td><td>gene name</td><td>gene id</td></tr>
<tr><td>GU305914.1</td><td></td><td>93081</td><td>93103</td><td>-</td><td>0</td><td>GTATACATGGCTTAATGTAT</td><td>AGG</td><td>||||||||[||||||||||||]PAM</td><td>NA</td><td>Intergenic</td><td>NA</td><td>NA</td></tr>
<tr><td>T27089</td><td>TATACATGGCTTAATGTATAGGG</td><td></td><td>1000</td><td></td><td></td><td></td><td></td><td></td><td></td><td></td><td></td><td></td></tr>
<tr><td>Oligo fwd</td><td>CACCTATACATGGCTTAATGTATA</td><td></td><td></td><td></td><td></td><td></td><td></td><td></td><td></td><td></td><td></td><td></td></tr>
<tr><td>Oligo rev</td><td>AAACTATACATTAAGCCATGTATA</td><td></td><td></td><td></td><td></td><td></td><td></td><td></td><td></td><td></td><td></td><td></td></tr>
<tr><td>Chromosome</td><td>start</td><td></td><td>end</td><td>strand</td><td>MM</td><td>target_seq</td><td>PAM</td><td>alignment</td><td>distance</td><td>position</td><td>gene name</td><td>gene id</td></tr>
<tr><td>GU305914.1</td><td></td><td>93080</td><td>93102</td><td>-</td><td>0</td><td>TATACATGGCTTAATGTATA</td><td>GGG</td><td>||||||||[||||||||||||]PAM</td><td>NA</td><td>Intergenic</td><td>NA</td><td>NA</td></tr>
<tr><td>T27090</td><td>ATCGATAACCTTTCAACATCCGG</td><td></td><td>1000</td><td></td><td></td><td></td><td></td><td></td><td></td><td></td><td></td><td></td></tr>
<tr><td>Oligo fwd</td><td>CACCATCGATAACCTTTCAACATC</td><td></td><td></td><td></td><td></td><td></td><td></td><td></td><td></td><td></td><td></td><td></td></tr>
<tr><td>Oligo rev</td><td>AAACGATGTTGAAAGGTTATCGAT</td><td></td><td></td><td></td><td></td><td></td><td></td><td></td><td></td><td></td><td></td><td></td></tr>
<tr><td>Chromosome</td><td>start</td><td></td><td>end</td><td>strand</td><td>MM</td><td>target_seq</td><td>PAM</td><td>alignment</td><td>distance</td><td>position</td><td>gene name</td><td>gene id</td></tr>
<tr><td>GU305914.1</td><td></td><td>93057</td><td>93079</td><td>-</td><td>0</td><td>ATCGATAACCTTTCAACATC</td><td>CGG</td><td>||||||||[||||||||||||]PAM</td><td>NA</td><td>Intergenic</td><td>NA</td><td>NA</td></tr>
<tr><td>T27091</td><td>TTCAACATCCGGTTGACCTTCGG</td><td></td><td>1000</td><td></td><td></td><td></td><td></td><td></td><td></td><td></td><td></td><td></td></tr>
<tr><td>Oligo fwd</td><td>CACCTTCAACATCCGGTTGACCTT</td><td></td><td></td><td></td><td></td><td></td><td></td><td></td><td></td><td></td><td></td><td></td></tr>
<tr><td>Oligo rev</td><td>AAACAAGGTCAACCGGATGTTGAA</td><td></td><td></td><td></td><td></td><td></td><td></td><td></td><td></td><td></td><td></td><td></td></tr>
<tr><td>Chromosome</td><td>start</td><td></td><td>end</td><td>strand</td><td>MM</td><td>target_seq</td><td>PAM</td><td>alignment</td><td>distance</td><td>position</td><td>gene name</td><td>gene id</td></tr>
<tr><td>GU305914.1</td><td></td><td>93046</td><td>93068</td><td>-</td><td>0</td><td>TTCAACATCCGGTTGACCTT</td><td>CGG</td><td>||||||||[||||||||||||]PAM</td><td>NA</td><td>Intergenic</td><td>NA</td><td>NA</td></tr>
<tr><td>GU305914.1</td><td></td><td>93030</td><td>93052</td><td>+</td><td>3</td><td>TTCGACATCCGGTTGACCGA</td><td>AGG</td><td>|||-||||[||||||||||--]PAM</td><td>NA</td><td>Intergenic</td><td>NA</td><td>NA</td></tr>
<tr><td>T27092</td><td>CCGGTTGACCTTCGGTCAACCGG</td><td></td><td>1000</td><td></td><td></td><td></td><td></td><td></td><td></td><td></td><td></td><td></td></tr>
<tr><td>Oligo fwd</td><td>CACCCCGGTTGACCTTCGGTCAAC</td><td></td><td></td><td></td><td></td><td></td><td></td><td></td><td></td><td></td><td></td><td></td></tr>
<tr><td>Oligo rev</td><td>AAACGTTGACCGAAGGTCAACCGG</td><td></td><td></td><td></td><td></td><td></td><td></td><td></td><td></td><td></td><td></td><td></td></tr>
<tr><td>Chromosome</td><td>start</td><td></td><td>end</td><td>strand</td><td>MM</td><td>target_seq</td><td>PAM</td><td>alignment</td><td>distance</td><td>position</td><td>gene name</td><td>gene id</td></tr>
<tr><td>GU305914.1</td><td></td><td>93038</td><td>93060</td><td>-</td><td>0</td><td>CCGGTTGACCTTCGGTCAAC</td><td>CGG</td><td>||||||||[||||||||||||]PAM</td><td>NA</td><td>Intergenic</td><td>NA</td><td>NA</td></tr>
<tr><td>T27093</td><td>AGTAATCAATAATATACGTTAGG</td><td></td><td>1000</td><td></td><td></td><td></td><td></td><td></td><td></td><td></td><td></td><td></td></tr>
<tr><td>Oligo fwd</td><td>CACCAGTAATCAATAATATACGTT</td><td></td><td></td><td></td><td></td><td></td><td></td><td></td><td></td><td></td><td></td><td></td></tr>
<tr><td>Oligo rev</td><td>AAACAACGTATATTATTGATTACT</td><td></td><td></td><td></td><td></td><td></td><td></td><td></td><td></td><td></td><td></td><td></td></tr>
<tr><td>Chromosome</td><td>start</td><td></td><td>end</td><td>strand</td><td>MM</td><td>target_seq</td><td>PAM</td><td>alignment</td><td>distance</td><td>position</td><td>gene name</td><td>gene id</td></tr>
<tr><td>GU305914.1</td><td></td><td>92986</td><td>93008</td><td>-</td><td>0</td><td>AGTAATCAATAATATACGTT</td><td>AGG</td><td>||||||||[||||||||||||]PAM</td><td>NA</td><td>Intergenic</td><td>NA</td><td>NA</td></tr>
<tr><td>T27094</td><td>GTAATCAATAATATACGTTAGGG</td><td></td><td>1000</td><td></td><td></td><td></td><td></td><td></td><td></td><td></td><td></td><td></td></tr>
<tr><td>Oligo fwd</td><td>CACCGTAATCAATAATATACGTTA</td><td></td><td></td><td></td><td></td><td></td><td></td><td></td><td></td><td></td><td></td><td></td></tr>
<tr><td>Oligo rev</td><td>AAACTAACGTATATTATTGATTAC</td><td></td><td></td><td></td><td></td><td></td><td></td><td></td><td></td><td></td><td></td><td></td></tr>
<tr><td>Chromosome</td><td>start</td><td></td><td>end</td><td>strand</td><td>MM</td><td>target_seq</td><td>PAM</td><td>alignment</td><td>distance</td><td>position</td><td>gene name</td><td>gene id</td></tr>
<tr><td>GU305914.1</td><td></td><td>92985</td><td>93007</td><td>-</td><td>0</td><td>GTAATCAATAATATACGTTA</td><td>GGG</td><td>||||||||[||||||||||||]PAM</td><td>NA</td><td>Intergenic</td><td>NA</td><td>NA</td></tr>
<tr><td>T27095</td><td>TAATCAATAATATACGTTAGGGG</td><td></td><td>1000</td><td></td><td></td><td></td><td></td><td></td><td></td><td></td><td></td><td></td></tr>
<tr><td>Oligo fwd</td><td>CACCTAATCAATAATATACGTTAG</td><td></td><td></td><td></td><td></td><td></td><td></td><td></td><td></td><td></td><td></td><td></td></tr>
<tr><td>Oligo rev</td><td>AAACCTAACGTATATTATTGATTA</td><td></td><td></td><td></td><td></td><td></td><td></td><td></td><td></td><td></td><td></td><td></td></tr>
<tr><td>Chromosome</td><td>start</td><td></td><td>end</td><td>strand</td><td>MM</td><td>target_seq</td><td>PAM</td><td>alignment</td><td>distance</td><td>position</td><td>gene name</td><td>gene id</td></tr>
<tr><td>GU305914.1</td><td></td><td>92984</td><td>93006</td><td>-</td><td>0</td><td>TAATCAATAATATACGTTAG</td><td>GGG</td><td>||||||||[||||||||||||]PAM</td><td>NA</td><td>Intergenic</td><td>NA</td><td>NA</td></tr>
<tr><td>T27096</td><td>AATCAATAATATACGTTAGGGGG</td><td></td><td>1000</td><td></td><td></td><td></td><td></td><td></td><td></td><td></td><td></td><td></td></tr>
<tr><td>Oligo fwd</td><td>CACCAATCAATAATATACGTTAGG</td><td></td><td></td><td></td><td></td><td></td><td></td><td></td><td></td><td></td><td></td><td></td></tr>
<tr><td>Oligo rev</td><td>AAACCCTAACGTATATTATTGATT</td><td></td><td></td><td></td><td></td><td></td><td></td><td></td><td></td><td></td><td></td><td></td></tr>
<tr><td>Chromosome</td><td>start</td><td></td><td>end</td><td>strand</td><td>MM</td><td>target_seq</td><td>PAM</td><td>alignment</td><td>distance</td><td>position</td><td>gene name</td><td>gene id</td></tr>
<tr><td>GU305914.1</td><td></td><td>92983</td><td>93005</td><td>-</td><td>0</td><td>AATCAATAATATACGTTAGG</td><td>GGG</td><td>||||||||[||||||||||||]PAM</td><td>NA</td><td>Intergenic</td><td>NA</td><td>NA</td></tr>
<tr><td>T27097</td><td>TAATGAGCATTACGTCACCTAGG</td><td></td><td>1000</td><td></td><td></td><td></td><td></td><td></td><td></td><td></td><td></td><td></td></tr>
<tr><td>Oligo fwd</td><td>CACCTAATGAGCATTACGTCACCT</td><td></td><td></td><td></td><td></td><td></td><td></td><td></td><td></td><td></td><td></td><td></td></tr>
<tr><td>Oligo rev</td><td>AAACAGGTGACGTAATGCTCATTA</td><td></td><td></td><td></td><td></td><td></td><td></td><td></td><td></td><td></td><td></td><td></td></tr>
<tr><td>Chromosome</td><td>start</td><td></td><td>end</td><td>strand</td><td>MM</td><td>target_seq</td><td>PAM</td><td>alignment</td><td>distance</td><td>position</td><td>gene name</td><td>gene id</td></tr>
<tr><td>GU305914.1</td><td></td><td>92948</td><td>92970</td><td>-</td><td>0</td><td>TAATGAGCATTACGTCACCT</td><td>AGG</td><td>||||||||[||||||||||||]PAM</td><td>NA</td><td>Intergenic</td><td>NA</td><td>NA</td></tr>
<tr><td>T27098</td><td>GCATTACGTCACCTAGGTGCTGG</td><td></td><td>1000</td><td></td><td></td><td></td><td></td><td></td><td></td><td></td><td></td><td></td></tr>
<tr><td>Oligo fwd</td><td>CACCGCATTACGTCACCTAGGTGC</td><td></td><td></td><td></td><td></td><td></td><td></td><td></td><td></td><td></td><td></td><td></td></tr>
<tr><td>Oligo rev</td><td>AAACGCACCTAGGTGACGTAATGC</td><td></td><td></td><td></td><td></td><td></td><td></td><td></td><td></td><td></td><td></td><td></td></tr>
<tr><td>Chromosome</td><td>start</td><td></td><td>end</td><td>strand</td><td>MM</td><td>target_seq</td><td>PAM</td><td>alignment</td><td>distance</td><td>position</td><td>gene name</td><td>gene id</td></tr>
<tr><td>GU305914.1</td><td></td><td>92942</td><td>92964</td><td>-</td><td>0</td><td>GCATTACGTCACCTAGGTGC</td><td>TGG</td><td>||||||||[||||||||||||]PAM</td><td>NA</td><td>Intergenic</td><td>NA</td><td>NA</td></tr>
<tr><td>T27099</td><td>CGTCACCTAGGTGCTGGTTACGG</td><td></td><td>1000</td><td></td><td></td><td></td><td></td><td></td><td></td><td></td><td></td><td></td></tr>
<tr><td>Oligo fwd</td><td>CACCCGTCACCTAGGTGCTGGTTA</td><td></td><td></td><td></td><td></td><td></td><td></td><td></td><td></td><td></td><td></td><td></td></tr>
<tr><td>Oligo rev</td><td>AAACTAACCAGCACCTAGGTGACG</td><td></td><td></td><td></td><td></td><td></td><td></td><td></td><td></td><td></td><td></td><td></td></tr>
<tr><td>Chromosome</td><td>start</td><td></td><td>end</td><td>strand</td><td>MM</td><td>target_seq</td><td>PAM</td><td>alignment</td><td>distance</td><td>position</td><td>gene name</td><td>gene id</td></tr>
<tr><td>GU305914.1</td><td></td><td>92936</td><td>92958</td><td>-</td><td>0</td><td>CGTCACCTAGGTGCTGGTTA</td><td>CGG</td><td>||||||||[||||||||||||]PAM</td><td>NA</td><td>Intergenic</td><td>NA</td><td>NA</td></tr>
<tr><td>T27100</td><td>GTCACCTAGGTGCTGGTTACGGG</td><td></td><td>1000</td><td></td><td></td><td></td><td></td><td></td><td></td><td></td><td></td><td></td></tr>
<tr><td>Oligo fwd</td><td>CACCGTCACCTAGGTGCTGGTTAC</td><td></td><td></td><td></td><td></td><td></td><td></td><td></td><td></td><td></td><td></td><td></td></tr>
<tr><td>Oligo rev</td><td>AAACGTAACCAGCACCTAGGTGAC</td><td></td><td></td><td></td><td></td><td></td><td></td><td></td><td></td><td></td><td></td><td></td></tr>
<tr><td>Chromosome</td><td>start</td><td></td><td>end</td><td>strand</td><td>MM</td><td>target_seq</td><td>PAM</td><td>alignment</td><td>distance</td><td>position</td><td>gene name</td><td>gene id</td></tr>
<tr><td>GU305914.1</td><td></td><td>92935</td><td>92957</td><td>-</td><td>0</td><td>GTCACCTAGGTGCTGGTTAC</td><td>GGG</td><td>||||||||[||||||||||||]PAM</td><td>NA</td><td>Intergenic</td><td>NA</td><td>NA</td></tr>
<tr><td>T27101</td><td>TCACCTAGGTGCTGGTTACGGGG</td><td></td><td>1000</td><td></td><td></td><td></td><td></td><td></td><td></td><td></td><td></td><td></td></tr>
<tr><td>Oligo fwd</td><td>CACCTCACCTAGGTGCTGGTTACG</td><td></td><td></td><td></td><td></td><td></td><td></td><td></td><td></td><td></td><td></td><td></td></tr>
<tr><td>Oligo rev</td><td>AAACCGTAACCAGCACCTAGGTGA</td><td></td><td></td><td></td><td></td><td></td><td></td><td></td><td></td><td></td><td></td><td></td></tr>
<tr><td>Chromosome</td><td>start</td><td></td><td>end</td><td>strand</td><td>MM</td><td>target_seq</td><td>PAM</td><td>alignment</td><td>distance</td><td>position</td><td>gene name</td><td>gene id</td></tr>
<tr><td>GU305914.1</td><td></td><td>92934</td><td>92956</td><td>-</td><td>0</td><td>TCACCTAGGTGCTGGTTACG</td><td>GGG</td><td>||||||||[||||||||||||]PAM</td><td>NA</td><td>Intergenic</td><td>NA</td><td>NA</td></tr>
<tr><td>T27102</td><td>CACCTAGGTGCTGGTTACGGGGG</td><td></td><td>1000</td><td></td><td></td><td></td><td></td><td></td><td></td><td></td><td></td><td></td></tr>
<tr><td>Oligo fwd</td><td>CACCCACCTAGGTGCTGGTTACGG</td><td></td><td></td><td></td><td></td><td></td><td></td><td></td><td></td><td></td><td></td><td></td></tr>
<tr><td>Oligo rev</td><td>AAACCCGTAACCAGCACCTAGGTG</td><td></td><td></td><td></td><td></td><td></td><td></td><td></td><td></td><td></td><td></td><td></td></tr>
<tr><td>Chromosome</td><td>start</td><td></td><td>end</td><td>strand</td><td>MM</td><td>target_seq</td><td>PAM</td><td>alignment</td><td>distance</td><td>position</td><td>gene name</td><td>gene id</td></tr>
<tr><td>GU305914.1</td><td></td><td>92933</td><td>92955</td><td>-</td><td>0</td><td>CACCTAGGTGCTGGTTACGG</td><td>GGG</td><td>||||||||[||||||||||||]PAM</td><td>NA</td><td>Intergenic</td><td>NA</td><td>NA</td></tr>
</table>

T27103 CTAGGTGCTGGTTACGGGGGAGG 1000
Oligo fwd CACCCTAGGTGCTGGTTACGGGGG
Oligo rev AAACCCCCCGTAACCAGCACCTAG
Chromosome start end strand MM target_seq PAM alignment distance position gene name gene id
GU305914.1 92930 92952 - 0 CTAGGTGCTGGTTACGGGGG AGG ||||||||[||||||||||||]PAM NA Intergenic NA NA

T27104 TAGGTGCTGGTTACGGGGGAGGG 1000
Oligo fwd CACCTAGGTGCTGGTTACGGGGGA
Oligo rev AAACTCCCCCGTAACCAGCACCTA
Chromosome start end strand MM target_seq PAM alignment distance position gene name gene id
GU305914.1 92929 92951 - 0 TAGGTGCTGGTTACGGGGGA GGG ||||||||[||||||||||||]PAM NA Intergenic NA NA

T27105 AGGTGCTGGTTACGGGGGAGGGG 1000
Oligo fwd CACCAGGTGCTGGTTACGGGGGAG
Oligo rev AAACCTCCCCCGTAACCAGCACCT
Chromosome start end strand MM target_seq PAM alignment distance position gene name gene id
GU305914.1 92928 92950 - 0 AGGTGCTGGTTACGGGGGAG GGG ||||||||[||||||||||||]PAM NA Intergenic NA NA

T27106 TACGGGGGAGGGGCTATGAATGG 1000
Oligo fwd CACCTACGGGGGAGGGGCTATGAA
Oligo rev AAACTTCATAGCCCCTCCCCCGTA
Chromosome start end strand MM target_seq PAM alignment distance position gene name gene id
GU305914.1 92918 92940 - 0 TACGGGGGAGGGGCTATGAA TGG ||||||||[||||||||||||]PAM NA Intergenic NA NA

T27107 ACGGGGGAGGGGCTATGAATGGG 1000
Oligo fwd CACCACGGGGGAGGGGCTATGAAT
Oligo rev AAACATTCATAGCCCCTCCCCCGT
Chromosome start end strand MM target_seq PAM alignment distance position gene name gene id
GU305914.1 92917 92939 - 0 ACGGGGGAGGGGCTATGAAT GGG ||||||||[||||||||||||]PAM NA Intergenic NA NA

T27108 GGAGGGGCTATGAATGGGCGTGG 1000
Oligo fwd CACCGGAGGGGCTATGAATGGGCG
Oligo rev AAACCGCCCATTCATAGCCCCTCC
Chromosome start end strand MM target_seq PAM alignment distance position gene name gene id
GU305914.1 92912 92934 - 0 GGAGGGGCTATGAATGGGCG TGG ||||||||[||||||||||||]PAM NA Intergenic NA NA

T27109 ATGAATGGGCGTGGTTACCTAGG 1000
Oligo fwd CACCATGAATGGGCGTGGTTACCT
Oligo rev AAACAGGTAACCACGCCCATTCAT
Chromosome start end strand MM target_seq PAM alignment distance position gene name gene id
GU305914.1 92903 92925 - 0 ATGAATGGGCGTGGTTACCT AGG ||||||||[||||||||||||]PAM NA Intergenic NA NA

T27110 CGTGGTTACCTAGGTTTCGATGG 1000
Oligo fwd CACCCGTGGTTACCTAGGTTTCGA
Oligo rev AAACTCGAAACCTAGGTAACCACG
Chromosome start end strand MM target_seq PAM alignment distance position gene name gene id
GU305914.1 92894 92916 - 0 CGTGGTTACCTAGGTTTCGA TGG ||||||||[||||||||||||]PAM NA Intergenic NA NA

T27111 CCTAGGTTTCGATGGTTCACCGG 1000
Oligo fwd CACCCCTAGGTTTCGATGGTTCAC
Oligo rev AAACGTGAACCATCGAAACCTAGG
Chromosome start end strand MM target_seq PAM alignment distance position gene name gene id
GU305914.1 92886 92908 - 0 CCTAGGTTTCGATGGTTCAC CGG ||||||||[||||||||||||]PAM NA Intergenic NA NA

T27112 TTCGATGGTTCACCGGCCGTAGG 1000
Oligo fwd CACCTTCGATGGTTCACCGGCCGT
Oligo rev AAACACGGCCGGTGAACCATCGAA
Chromosome start end strand MM target_seq PAM alignment distance position gene name gene id
GU305914.1 92879 92901 - 0 TTCGATGGTTCACCGGCCGT AGG ||||||||[||||||||||||]PAM NA Intergenic NA NA

T27113 ATGGTTCACCGGCCGTAGGCCGG 1000
Oligo fwd CACCATGGTTCACCGGCCGTAGGC
Oligo rev AAACGCCTACGGCCGGTGAACCAT
Chromosome start end strand MM target_seq PAM alignment distance position gene name gene id
GU305914.1 92875 92897 - 0 ATGGTTCACCGGCCGTAGGC CGG ||||||||[||||||||||||]PAM NA Intergenic NA NA

T27114 AGGCCGGTGAACCAGTCGACTGG 1000
Oligo fwd CACCAGGCCGGTGAACCAGTCGAC
Oligo rev AAACGTCGACTGGTTCACCGGCCT
Chromosome start end strand MM target_seq PAM alignment distance position gene name gene id
GU305914.1 92859 92881 - 0 AGGCCGGTGAACCAGTCGAC TGG ||||||||[||||||||||||]PAM NA Intergenic NA NA

T27115 TGGTCCATAATTAGTTGTACAGG 1000
Oligo fwd CACCTGGTCCATAATTAGTTGTAC
Oligo rev AAACGTACAACTAATTATGGACCA
Chromosome start end strand MM target_seq PAM alignment distance position gene name gene id
GU305914.1 92839 92861 - 0 TGGTCCATAATTAGTTGTAC AGG ||||||||[||||||||||||]PAM NA Intergenic NA NA

T27116 GGTCCATAATTAGTTGTACAGGG 1000
Oligo fwd CACCGGTCCATAATTAGTTGTACA
Oligo rev AAACTGTACAACTAATTATGGACC
Chromosome start end strand MM target_seq PAM alignment distance position gene name gene id
GU305914.1 92838 92860 - 0 GGTCCATAATTAGTTGTACA GGG ||||||||[||||||||||||]PAM NA Intergenic NA NA

T27117 GTCCATAATTAGTTGTACAGGGG 1000
Oligo fwd CACCGTCCATAATTAGTTGTACAG
Oligo rev AAACCTGTACAACTAATTATGGAC
Chromosome start end strand MM target_seq PAM alignment distance position gene name gene id
GU305914.1 92837 92859 - 0 GTCCATAATTAGTTGTACAG GGG ||||||||[||||||||||||]PAM NA Intergenic NA NA

<table>
<tr><td>T27118</td><td>AATTAGCATAAATTAATTAACGG</td><td>1000</td><td></td><td></td><td></td><td></td><td></td><td></td><td></td><td></td><td></td></tr>
<tr><td>Oligo fwd</td><td>CACCAATTAGCATAAATTAATTAA</td><td></td><td></td><td></td><td></td><td></td><td></td><td></td><td></td><td></td><td></td></tr>
<tr><td>Oligo rev</td><td>AAACTTAATTAATTTATGCTAATT</td><td></td><td></td><td></td><td></td><td></td><td></td><td></td><td></td><td></td><td></td></tr>
<tr><td>Chromosome</td><td>start</td><td>end</td><td>strand</td><td>MM</td><td>target_seq</td><td>PAM</td><td>alignment</td><td>distance</td><td>position</td><td>gene name</td><td>gene id</td></tr>
<tr><td>GU305914.1</td><td>92812</td><td>92834</td><td>-</td><td>0</td><td>AATTAGCATAAATTAATTAA</td><td>CGG</td><td>||||||||[||||||||||||]PAM</td><td>NA</td><td>Intergenic</td><td>NA</td><td>NA</td></tr>
<tr><td>T27119</td><td>TCTGCTAATTTGCATAATTCAGG</td><td>1000</td><td></td><td></td><td></td><td></td><td></td><td></td><td></td><td></td><td></td></tr>
<tr><td>Oligo fwd</td><td>CACCTCTGCTAATTTGCATAATTC</td><td></td><td></td><td></td><td></td><td></td><td></td><td></td><td></td><td></td><td></td></tr>
<tr><td>Oligo rev</td><td>AAACGAATTATGCAAATTAGCAGA</td><td></td><td></td><td></td><td></td><td></td><td></td><td></td><td></td><td></td><td></td></tr>
<tr><td>Chromosome</td><td>start</td><td>end</td><td>strand</td><td>MM</td><td>target_seq</td><td>PAM</td><td>alignment</td><td>distance</td><td>position</td><td>gene name</td><td>gene id</td></tr>
<tr><td>GU305914.1</td><td>92748</td><td>92770</td><td>-</td><td>0</td><td>TCTGCTAATTTGCATAATTC</td><td>AGG</td><td>||||||||[||||||||||||]PAM</td><td>NA</td><td>Intergenic</td><td>NA</td><td>NA</td></tr>
<tr><td>T27120</td><td>GCTAATTTGCATAATTCAGGAGG</td><td>1000</td><td></td><td></td><td></td><td></td><td></td><td></td><td></td><td></td><td></td></tr>
<tr><td>Oligo fwd</td><td>CACCGCTAATTTGCATAATTCAGG</td><td></td><td></td><td></td><td></td><td></td><td></td><td></td><td></td><td></td><td></td></tr>
<tr><td>Oligo rev</td><td>AAACCCTGAATTATGCAAATTAGC</td><td></td><td></td><td></td><td></td><td></td><td></td><td></td><td></td><td></td><td></td></tr>
<tr><td>Chromosome</td><td>start</td><td>end</td><td>strand</td><td>MM</td><td>target_seq</td><td>PAM</td><td>alignment</td><td>distance</td><td>position</td><td>gene name</td><td>gene id</td></tr>
<tr><td>GU305914.1</td><td>92745</td><td>92767</td><td>-</td><td>0</td><td>GCTAATTTGCATAATTCAGG</td><td>AGG</td><td>||||||||[||||||||||||]PAM</td><td>NA</td><td>Intergenic</td><td>NA</td><td>NA</td></tr>
<tr><td>T27121</td><td>GGAGGTTACGCCCATTGTTTAGG</td><td>1000</td><td></td><td></td><td></td><td></td><td></td><td></td><td></td><td></td><td></td></tr>
<tr><td>Oligo fwd</td><td>CACCGGAGGTTACGCCCATTGTTT</td><td></td><td></td><td></td><td></td><td></td><td></td><td></td><td></td><td></td><td></td></tr>
<tr><td>Oligo rev</td><td>AAACAAACAATGGGCGTAACCTCC</td><td></td><td></td><td></td><td></td><td></td><td></td><td></td><td></td><td></td><td></td></tr>
<tr><td>Chromosome</td><td>start</td><td>end</td><td>strand</td><td>MM</td><td>target_seq</td><td>PAM</td><td>alignment</td><td>distance</td><td>position</td><td>gene name</td><td>gene id</td></tr>
<tr><td>GU305914.1</td><td>92727</td><td>92749</td><td>-</td><td>0</td><td>GGAGGTTACGCCCATTGTTT</td><td>AGG</td><td>||||||||[||||||||||||]PAM</td><td>NA</td><td>Intergenic</td><td>NA</td><td>NA</td></tr>
<tr><td>T27122</td><td>GGTTACGCCCATTGTTTAGGTGG</td><td>1000</td><td></td><td></td><td></td><td></td><td></td><td></td><td></td><td></td><td></td></tr>
<tr><td>Oligo fwd</td><td>CACCGGTTACGCCCATTGTTTAGG</td><td></td><td></td><td></td><td></td><td></td><td></td><td></td><td></td><td></td><td></td></tr>
<tr><td>Oligo rev</td><td>AAACCCTAAACAATGGGCGTAACC</td><td></td><td></td><td></td><td></td><td></td><td></td><td></td><td></td><td></td><td></td></tr>
<tr><td>Chromosome</td><td>start</td><td>end</td><td>strand</td><td>MM</td><td>target_seq</td><td>PAM</td><td>alignment</td><td>distance</td><td>position</td><td>gene name</td><td>gene id</td></tr>
<tr><td>GU305914.1</td><td>92724</td><td>92746</td><td>-</td><td>0</td><td>GGTTACGCCCATTGTTTAGG</td><td>TGG</td><td>||||||||[||||||||||||]PAM</td><td>NA</td><td>Intergenic</td><td>NA</td><td>NA</td></tr>
<tr><td>T27123</td><td>GTTACGCCCATTGTTTAGGTGGG</td><td>1000</td><td></td><td></td><td></td><td></td><td></td><td></td><td></td><td></td><td></td></tr>
<tr><td>Oligo fwd</td><td>CACCGTTACGCCCATTGTTTAGGT</td><td></td><td></td><td></td><td></td><td></td><td></td><td></td><td></td><td></td><td></td></tr>
<tr><td>Oligo rev</td><td>AAACACCTAAACAATGGGCGTAAC</td><td></td><td></td><td></td><td></td><td></td><td></td><td></td><td></td><td></td><td></td></tr>
<tr><td>Chromosome</td><td>start</td><td>end</td><td>strand</td><td>MM</td><td>target_seq</td><td>PAM</td><td>alignment</td><td>distance</td><td>position</td><td>gene name</td><td>gene id</td></tr>
<tr><td>GU305914.1</td><td>92723</td><td>92745</td><td>-</td><td>0</td><td>GTTACGCCCATTGTTTAGGT</td><td>GGG</td><td>||||||||[||||||||||||]PAM</td><td>NA</td><td>Intergenic</td><td>NA</td><td>NA</td></tr>
<tr><td>T27124</td><td>TAGGTGGGCGTACCTAGTTTAGG</td><td>1000</td><td></td><td></td><td></td><td></td><td></td><td></td><td></td><td></td><td></td></tr>
<tr><td>Oligo fwd</td><td>CACCTAGGTGGGCGTACCTAGTTT</td><td></td><td></td><td></td><td></td><td></td><td></td><td></td><td></td><td></td><td></td></tr>
<tr><td>Oligo rev</td><td>AAACAAACTAGGTACGCCCACCTA</td><td></td><td></td><td></td><td></td><td></td><td></td><td></td><td></td><td></td><td></td></tr>
<tr><td>Chromosome</td><td>start</td><td>end</td><td>strand</td><td>MM</td><td>target_seq</td><td>PAM</td><td>alignment</td><td>distance</td><td>position</td><td>gene name</td><td>gene id</td></tr>
<tr><td>GU305914.1</td><td>92708</td><td>92730</td><td>-</td><td>0</td><td>TAGGTGGGCGTACCTAGTTT</td><td>AGG</td><td>||||||||[||||||||||||]PAM</td><td>NA</td><td>Intergenic</td><td>NA</td><td>NA</td></tr>
<tr><td>T27125</td><td>AGGTGGGCGTACCTAGTTTAGGG</td><td>1000</td><td></td><td></td><td></td><td></td><td></td><td></td><td></td><td></td><td></td></tr>
<tr><td>Oligo fwd</td><td>CACCAGGTGGGCGTACCTAGTTTA</td><td></td><td></td><td></td><td></td><td></td><td></td><td></td><td></td><td></td><td></td></tr>
<tr><td>Oligo rev</td><td>AAACTAAACTAGGTACGCCCACCT</td><td></td><td></td><td></td><td></td><td></td><td></td><td></td><td></td><td></td><td></td></tr>
<tr><td>Chromosome</td><td>start</td><td>end</td><td>strand</td><td>MM</td><td>target_seq</td><td>PAM</td><td>alignment</td><td>distance</td><td>position</td><td>gene name</td><td>gene id</td></tr>
<tr><td>GU305914.1</td><td>92707</td><td>92729</td><td>-</td><td>0</td><td>AGGTGGGCGTACCTAGTTTA</td><td>GGG</td><td>||||||||[||||||||||||]PAM</td><td>NA</td><td>Intergenic</td><td>NA</td><td>NA</td></tr>
<tr><td>T27126</td><td>GGTGGGCGTACCTAGTTTAGGGG</td><td>1000</td><td></td><td></td><td></td><td></td><td></td><td></td><td></td><td></td><td></td></tr>
<tr><td>Oligo fwd</td><td>CACCGGTGGGCGTACCTAGTTTAG</td><td></td><td></td><td></td><td></td><td></td><td></td><td></td><td></td><td></td><td></td></tr>
<tr><td>Oligo rev</td><td>AAACCTAAACTAGGTACGCCCACC</td><td></td><td></td><td></td><td></td><td></td><td></td><td></td><td></td><td></td><td></td></tr>
<tr><td>Chromosome</td><td>start</td><td>end</td><td>strand</td><td>MM</td><td>target_seq</td><td>PAM</td><td>alignment</td><td>distance</td><td>position</td><td>gene name</td><td>gene id</td></tr>
<tr><td>GU305914.1</td><td>92706</td><td>92728</td><td>-</td><td>0</td><td>GGTGGGCGTACCTAGTTTAG</td><td>GGG</td><td>||||||||[||||||||||||]PAM</td><td>NA</td><td>Intergenic</td><td>NA</td><td>NA</td></tr>
<tr><td>T27127</td><td>GTGGGCGTACCTAGTTTAGGGGG</td><td>1000</td><td></td><td></td><td></td><td></td><td></td><td></td><td></td><td></td><td></td></tr>
<tr><td>Oligo fwd</td><td>CACCGTGGGCGTACCTAGTTTAGG</td><td></td><td></td><td></td><td></td><td></td><td></td><td></td><td></td><td></td><td></td></tr>
<tr><td>Oligo rev</td><td>AAACCCTAAACTAGGTACGCCCAC</td><td></td><td></td><td></td><td></td><td></td><td></td><td></td><td></td><td></td><td></td></tr>
<tr><td>Chromosome</td><td>start</td><td>end</td><td>strand</td><td>MM</td><td>target_seq</td><td>PAM</td><td>alignment</td><td>distance</td><td>position</td><td>gene name</td><td>gene id</td></tr>
<tr><td>GU305914.1</td><td>92705</td><td>92727</td><td>-</td><td>0</td><td>GTGGGCGTACCTAGTTTAGG</td><td>GGG</td><td>||||||||[||||||||||||]PAM</td><td>NA</td><td>Intergenic</td><td>NA</td><td>NA</td></tr>
<tr><td>T27128</td><td>CGTACCTAGTTTAGGGGGCGTGG</td><td>1000</td><td></td><td></td><td></td><td></td><td></td><td></td><td></td><td></td><td></td></tr>
<tr><td>Oligo fwd</td><td>CACCCGTACCTAGTTTAGGGGGCG</td><td></td><td></td><td></td><td></td><td></td><td></td><td></td><td></td><td></td><td></td></tr>
<tr><td>Oligo rev</td><td>AAACCGCCCCCTAAACTAGGTACG</td><td></td><td></td><td></td><td></td><td></td><td></td><td></td><td></td><td></td><td></td></tr>
<tr><td>Chromosome</td><td>start</td><td>end</td><td>strand</td><td>MM</td><td>target_seq</td><td>PAM</td><td>alignment</td><td>distance</td><td>position</td><td>gene name</td><td>gene id</td></tr>
<tr><td>GU305914.1</td><td>92700</td><td>92722</td><td>-</td><td>0</td><td>CGTACCTAGTTTAGGGGGCG</td><td>TGG</td><td>||||||||[||||||||||||]PAM</td><td>NA</td><td>Intergenic</td><td>NA</td><td>NA</td></tr>
<tr><td>T27129</td><td>TAGTTTAGGGGGCGTGGTTCCGG</td><td>1000</td><td></td><td></td><td></td><td></td><td></td><td></td><td></td><td></td><td></td></tr>
<tr><td>Oligo fwd</td><td>CACCTAGTTTAGGGGGCGTGGTTC</td><td></td><td></td><td></td><td></td><td></td><td></td><td></td><td></td><td></td><td></td></tr>
<tr><td>Oligo rev</td><td>AAACGAACCACGCCCCCTAAACTA</td><td></td><td></td><td></td><td></td><td></td><td></td><td></td><td></td><td></td><td></td></tr>
<tr><td>Chromosome</td><td>start</td><td>end</td><td>strand</td><td>MM</td><td>target_seq</td><td>PAM</td><td>alignment</td><td>distance</td><td>position</td><td>gene name</td><td>gene id</td></tr>
<tr><td>GU305914.1</td><td>92694</td><td>92716</td><td>-</td><td>0</td><td>TAGTTTAGGGGGCGTGGTTC</td><td>CGG</td><td>||||||||[||||||||||||]PAM</td><td>NA</td><td>Intergenic</td><td>NA</td><td>NA</td></tr>
<tr><td>T27130</td><td>AGTTTAGGGGGCGTGGTTCCGGG</td><td>1000</td><td></td><td></td><td></td><td></td><td></td><td></td><td></td><td></td><td></td></tr>
<tr><td>Oligo fwd</td><td>CACCAGTTTAGGGGGCGTGGTTCC</td><td></td><td></td><td></td><td></td><td></td><td></td><td></td><td></td><td></td><td></td></tr>
<tr><td>Oligo rev</td><td>AAACGGAACCACGCCCCCTAAACT</td><td></td><td></td><td></td><td></td><td></td><td></td><td></td><td></td><td></td><td></td></tr>
<tr><td>Chromosome</td><td>start</td><td>end</td><td>strand</td><td>MM</td><td>target_seq</td><td>PAM</td><td>alignment</td><td>distance</td><td>position</td><td>gene name</td><td>gene id</td></tr>
<tr><td>GU305914.1</td><td>92693</td><td>92715</td><td>-</td><td>0</td><td>AGTTTAGGGGGCGTGGTTCC</td><td>GGG</td><td>||||||||[||||||||||||]PAM</td><td>NA</td><td>Intergenic</td><td>NA</td><td>NA</td></tr>
<tr><td>T27131</td><td>GTTTAGGGGGCGTGGTTCCGGGG</td><td>1000</td><td></td><td></td><td></td><td></td><td></td><td></td><td></td><td></td><td></td></tr>
<tr><td>Oligo fwd</td><td>CACCGTTTAGGGGGCGTGGTTCCG</td><td></td><td></td><td></td><td></td><td></td><td></td><td></td><td></td><td></td><td></td></tr>
<tr><td>Oligo rev</td><td>AAACCGGAACCACGCCCCCTAAAC</td><td></td><td></td><td></td><td></td><td></td><td></td><td></td><td></td><td></td><td></td></tr>
<tr><td>Chromosome</td><td>start</td><td>end</td><td>strand</td><td>MM</td><td>target_seq</td><td>PAM</td><td>alignment</td><td>distance</td><td>position</td><td>gene name</td><td>gene id</td></tr>
<tr><td>GU305914.1</td><td>92692</td><td>92714</td><td>-</td><td>0</td><td>GTTTAGGGGGCGTGGTTCCG</td><td>GGG</td><td>||||||||[||||||||||||]PAM</td><td>NA</td><td>Intergenic</td><td>NA</td><td>NA</td></tr>
<tr><td>T27132</td><td>TTTAGGGGGCGTGGTTCCGGGGG</td><td>1000</td><td></td><td></td><td></td><td></td><td></td><td></td><td></td><td></td><td></td></tr>
<tr><td>Oligo fwd</td><td>CACCTTTAGGGGGCGTGGTTCCGG</td><td></td><td></td><td></td><td></td><td></td><td></td><td></td><td></td><td></td><td></td></tr>
<tr><td>Oligo rev</td><td>AAACCCGGAACCACGCCCCCTAAA</td><td></td><td></td><td></td><td></td><td></td><td></td><td></td><td></td><td></td><td></td></tr>
<tr><td>Chromosome</td><td>start</td><td>end</td><td>strand</td><td>MM</td><td>target_seq</td><td>PAM</td><td>alignment</td><td>distance</td><td>position</td><td>gene name</td><td>gene id</td></tr>
</table>

GU305914.1 92691 92713 - 0 TTTAGGGGGCGTGGTTCCGG GGG ||||||||[||||||||||||]PAM NA Intergenic NA NA

T27133 TGGTTCCGGGGGTTGCGCTTCGG 1000
Oligo fwd CACCTGGTTCCGGGGGTTGCGCTT
Oligo rev AAACAAGCGCAACCCCCGGAACCA
Chromosome start end strand MM target_seq PAM alignment distance position gene name gene id
GU305914.1 92680 92702 - 0 TGGTTCCGGGGGTTGCGCTT CGG ||||||||[||||||||||||]PAM NA Intergenic NA NA

T27134 GGTTCCGGGGGTTGCGCTTCGGG 1000
Oligo fwd CACCGGTTCCGGGGGTTGCGCTTC
Oligo rev AAACGAAGCGCAACCCCCGGAACC
Chromosome start end strand MM target_seq PAM alignment distance position gene name gene id
GU305914.1 92679 92701 - 0 GGTTCCGGGGGTTGCGCTTC GGG ||||||||[||||||||||||]PAM NA Intergenic NA NA

T27135 CCCGCCCGTACGCGCGCGCGAGG 1000
Oligo fwd CACCCCCGCCCGTACGCGCGCGCG
Oligo rev AAACCGCGCGCGCGTACGGGCGGG
Chromosome start end strand MM target_seq PAM alignment distance position gene name gene id
GU305914.1 92614 92636 - 0 CCCGCCCGTACGCGCGCGCG AGG ||||||||[||||||||||||]PAM NA Intergenic NA NA

T27136 CGTACGCGCGCGCGAGGCACCGG 1000
Oligo fwd CACCCGTACGCGCGCGCGAGGCAC
Oligo rev AAACGTGCCTCGCGCGCGCGTACG
Chromosome start end strand MM target_seq PAM alignment distance position gene name gene id
GU305914.1 92608 92630 - 0 CGTACGCGCGCGCGAGGCAC CGG ||||||||[||||||||||||]PAM NA Intergenic NA NA

T27137 GTACGCGCGCGCGAGGCACCGGG 1000
Oligo fwd CACCGTACGCGCGCGCGAGGCACC
Oligo rev AAACGGTGCCTCGCGCGCGCGTAC
Chromosome start end strand MM target_seq PAM alignment distance position gene name gene id
GU305914.1 92607 92629 - 0 GTACGCGCGCGCGAGGCACC GGG ||||||||[||||||||||||]PAM NA Intergenic NA NA

T27138 GCGCGAGGCACCGGGCCAAAAGG 1000
Oligo fwd CACCGCGCGAGGCACCGGGCCAAA
Oligo rev AAACTTTGGCCCGGTGCCTCGCGC
Chromosome start end strand MM target_seq PAM alignment distance position gene name gene id
GU305914.1 92599 92621 - 0 GCGCGAGGCACCGGGCCAAA AGG ||||||||[||||||||||||]PAM NA Intergenic NA NA

T27139 CGAGGCACCGGGCCAAAAGGCGG 1000
Oligo fwd CACCCGAGGCACCGGGCCAAAAGG
Oligo rev AAACCCTTTTGGCCCGGTGCCTCG
Chromosome start end strand MM target_seq PAM alignment distance position gene name gene id
GU305914.1 92596 92618 - 0 CGAGGCACCGGGCCAAAAGG CGG ||||||||[||||||||||||]PAM NA Intergenic NA NA

T27140 GAGGCACCGGGCCAAAAGGCGGG 1000
Oligo fwd CACCGAGGCACCGGGCCAAAAGGC
Oligo rev AAACGCCTTTTGGCCCGGTGCCTC
Chromosome start end strand MM target_seq PAM alignment distance position gene name gene id
GU305914.1 92595 92617 - 0 GAGGCACCGGGCCAAAAGGC GGG ||||||||[||||||||||||]PAM NA Intergenic NA NA

T27141 GGGCCAAAAGGCGGGACTTCCGG 1000
Oligo fwd CACCGGGCCAAAAGGCGGGACTTC
Oligo rev AAACGAAGTCCCGCCTTTTGGCCC
Chromosome start end strand MM target_seq PAM alignment distance position gene name gene id
GU305914.1 92587 92609 - 0 GGGCCAAAAGGCGGGACTTC CGG ||||||||[||||||||||||]PAM NA Intergenic NA NA

T27142 CGGGACTTCCGGTCAGCCTCTGG 1000
Oligo fwd CACCCGGGACTTCCGGTCAGCCTC
Oligo rev AAACGAGGCTGACCGGAAGTCCCG
Chromosome start end strand MM target_seq PAM alignment distance position gene name gene id
GU305914.1 92576 92598 - 0 CGGGACTTCCGGTCAGCCTC TGG ||||||||[||||||||||||]PAM NA Intergenic NA NA

T27143 CTTCCGGTCAGCCTCTGGCCCGG 1000
Oligo fwd CACCCTTCCGGTCAGCCTCTGGCC
Oligo rev AAACGGCCAGAGGCTGACCGGAAG
Chromosome start end strand MM target_seq PAM alignment distance position gene name gene id
GU305914.1 92571 92593 - 0 CTTCCGGTCAGCCTCTGGCC CGG ||||||||[||||||||||||]PAM NA Intergenic NA NA

T27144 TCTGGCCCGGCACATCGCCGCGG 1000
Oligo fwd CACCTCTGGCCCGGCACATCGCCG
Oligo rev AAACCGGCGATGTGCCGGGCCAGA
Chromosome start end strand MM target_seq PAM alignment distance position gene name gene id
GU305914.1 92558 92580 - 0 TCTGGCCCGGCACATCGCCG CGG ||||||||[||||||||||||]PAM NA Intergenic NA NA

T27145 GCCCGGCACATCGCCGCGGAAGG 1000
Oligo fwd CACCGCCCGGCACATCGCCGCGGA
Oligo rev AAACTCCGCGGCGATGTGCCGGGC
Chromosome start end strand MM target_seq PAM alignment distance position gene name gene id
GU305914.1 92554 92576 - 0 GCCCGGCACATCGCCGCGGA AGG ||||||||[||||||||||||]PAM NA Intergenic NA NA

T27146 CGGCACATCGCCGCGGAAGGCGG 1000
Oligo fwd CACCCGGCACATCGCCGCGGAAGG
Oligo rev AAACCCTTCCGCGGCGATGTGCCG
Chromosome start end strand MM target_seq PAM alignment distance position gene name gene id
GU305914.1 92551 92573 - 0 CGGCACATCGCCGCGGAAGG CGG ||||||||[||||||||||||]PAM NA Intergenic NA NA

T27147 GGCACATCGCCGCGGAAGGCGGG 1000
Oligo fwd CACCGGCACATCGCCGCGGAAGGC
Oligo rev AAACGCCTTCCGCGGCGATGTGCC

| | | | | | | | | | | | | |
|---|---|---|---|---|---|---|---|---|---|---|---|---|
| Chromosome | start | | end | strand | MM | target_seq | PAM | alignment | distance | position | gene name | gene id |
| GU305914.1 | | 92550 | 92572 | - | 0 | GGCACATCGCCGCGGAAGGC | GGG | ||||||||[||||||||||||]PAM | NA | Intergenic | NA | NA |
| T27148 | GAAGGCGGGACATGTAACCATGG | | 1000 | | | | | | | | | |
| Oligo fwd | CACCGAAGGCGGGACATGTAACCA | | | | | | | | | | | |
| Oligo rev | AAACTGGTTACATGTCCCGCCTTC | | | | | | | | | | | |
| Chromosome | start | | end | strand | MM | target_seq | PAM | alignment | distance | position | gene name | gene id |
| GU305914.1 | | 92536 | 92558 | - | 0 | GAAGGCGGGACATGTAACCA | TGG | ||||||||[||||||||||||]PAM | NA | Intergenic | NA | NA |
| T27149 | CCTGCCCGCCCGATAAGCGTTGG | | 1000 | | | | | | | | | |
| Oligo fwd | CACCCCTGCCCGCCCGATAAGCGT | | | | | | | | | | | |
| Oligo rev | AAACACGCTTATCGGGCGGGCAGG | | | | | | | | | | | |
| Chromosome | start | | end | strand | MM | target_seq | PAM | alignment | distance | position | gene name | gene id |
| GU305914.1 | | 92507 | 92529 | - | 0 | CCTGCCCGCCCGATAAGCGT | TGG | ||||||||[||||||||||||]PAM | NA | Intergenic | NA | NA |
| GU305914.1 | | 99540 | 99562 | + | 4 | CCTCCTCGCCCGACGAGCGT | TGG | |||-|-||[|||||--|||||]PAM | NA | Intergenic | NA | NA |
| T27150 | CATGTGCTGCGTCTTCCGCTAGG | | 1000 | | | | | | | | | |
| Oligo fwd | CACCCATGTGCTGCGTCTTCCGCT | | | | | | | | | | | |
| Oligo rev | AAACAGCGGAAGACGCAGCACATG | | | | | | | | | | | |
| Chromosome | start | | end | strand | MM | target_seq | PAM | alignment | distance | position | gene name | gene id |
| GU305914.1 | | 92466 | 92488 | - | 0 | CATGTGCTGCGTCTTCCGCT | AGG | ||||||||[||||||||||||]PAM | NA | Intergenic | NA | NA |
| T27151 | ATGTGCTGCGTCTTCCGCTAGGG | | 1000 | | | | | | | | | |
| Oligo fwd | CACCATGTGCTGCGTCTTCCGCTA | | | | | | | | | | | |
| Oligo rev | AAACTAGCGGAAGACGCAGCACAT | | | | | | | | | | | |
| Chromosome | start | | end | strand | MM | target_seq | PAM | alignment | distance | position | gene name | gene id |
| GU305914.1 | | 92465 | 92487 | - | 0 | ATGTGCTGCGTCTTCCGCTA | GGG | ||||||||[||||||||||||]PAM | NA | Intergenic | NA | NA |
| T27152 | TGTGCTGCGTCTTCCGCTAGGGG | | 1000 | | | | | | | | | |
| Oligo fwd | CACCTGTGCTGCGTCTTCCGCTAG | | | | | | | | | | | |
| Oligo rev | AAACCTAGCGGAAGACGCAGCACA | | | | | | | | | | | |
| Chromosome | start | | end | strand | MM | target_seq | PAM | alignment | distance | position | gene name | gene id |
| GU305914.1 | | 92464 | 92486 | - | 0 | TGTGCTGCGTCTTCCGCTAG | GGG | ||||||||[||||||||||||]PAM | NA | Intergenic | NA | NA |
| T27153 | GTGCTGCGTCTTCCGCTAGGGGG | | 1000 | | | | | | | | | |
| Oligo fwd | CACCGTGCTGCGTCTTCCGCTAGG | | | | | | | | | | | |
| Oligo rev | AAACCCTAGCGGAAGACGCAGCAC | | | | | | | | | | | |
| Chromosome | start | | end | strand | MM | target_seq | PAM | alignment | distance | position | gene name | gene id |
| GU305914.1 | | 92463 | 92485 | - | 0 | GTGCTGCGTCTTCCGCTAGG | GGG | ||||||||[||||||||||||]PAM | NA | Intergenic | NA | NA |
| T27154 | GGTGCTACGAGACCACAAATCGG | | 1000 | | | | | | | | | |
| Oligo fwd | CACCGGTGCTACGAGACCACAAAT | | | | | | | | | | | |
| Oligo rev | AAACATTTGTGGTCTCGTAGCACC | | | | | | | | | | | |
| Chromosome | start | | end | strand | MM | target_seq | PAM | alignment | distance | position | gene name | gene id |
| GU305914.1 | | 92442 | 92464 | - | 0 | GGTGCTACGAGACCACAAAT | CGG | ||||||||[||||||||||||]PAM | NA | Intergenic | NA | NA |
| T27155 | ACGAGACCACAAATCGGTTGCGG | | 1000 | | | | | | | | | |
| Oligo fwd | CACCACGAGACCACAAATCGGTTG | | | | | | | | | | | |
| Oligo rev | AAACCAACCGATTTGTGGTCTCGT | | | | | | | | | | | |
| Chromosome | start | | end | strand | MM | target_seq | PAM | alignment | distance | position | gene name | gene id |
| GU305914.1 | | 92436 | 92458 | - | 0 | ACGAGACCACAAATCGGTTG | CGG | ||||||||[||||||||||||]PAM | NA | Intergenic | NA | NA |
| T27156 | CGAGACCACAAATCGGTTGCGGG | | 1000 | | | | | | | | | |
| Oligo fwd | CACCCGAGACCACAAATCGGTTGC | | | | | | | | | | | |
| Oligo rev | AAACGCAACCGATTTGTGGTCTCG | | | | | | | | | | | |
| Chromosome | start | | end | strand | MM | target_seq | PAM | alignment | distance | position | gene name | gene id |
| GU305914.1 | | 92435 | 92457 | - | 0 | CGAGACCACAAATCGGTTGC | GGG | ||||||||[||||||||||||]PAM | NA | Intergenic | NA | NA |
| T27157 | GAGACCACAAATCGGTTGCGGGG | | 1000 | | | | | | | | | |
| Oligo fwd | CACCGAGACCACAAATCGGTTGCG | | | | | | | | | | | |
| Oligo rev | AAACCGCAACCGATTTGTGGTCTC | | | | | | | | | | | |
| Chromosome | start | | end | strand | MM | target_seq | PAM | alignment | distance | position | gene name | gene id |
| GU305914.1 | | 92434 | 92456 | - | 0 | GAGACCACAAATCGGTTGCG | GGG | ||||||||[||||||||||||]PAM | NA | Intergenic | NA | NA |
| T27158 | ACCACAAATCGGTTGCGGGGCGG | | 1000 | | | | | | | | | |
| Oligo fwd | CACCACCACAAATCGGTTGCGGGG | | | | | | | | | | | |
| Oligo rev | AAACCCCCGCAACCGATTTGTGGT | | | | | | | | | | | |
| Chromosome | start | | end | strand | MM | target_seq | PAM | alignment | distance | position | gene name | gene id |
| GU305914.1 | | 92431 | 92453 | - | 0 | ACCACAAATCGGTTGCGGGG | CGG | ||||||||[||||||||||||]PAM | NA | Intergenic | NA | NA |
| T27159 | CCACAAATCGGTTGCGGGGCGGG | | 1000 | | | | | | | | | |
| Oligo fwd | CACCCCACAAATCGGTTGCGGGGC | | | | | | | | | | | |
| Oligo rev | AAACGCCCCGCAACCGATTTGTGG | | | | | | | | | | | |
| Chromosome | start | | end | strand | MM | target_seq | PAM | alignment | distance | position | gene name | gene id |
| GU305914.1 | | 92430 | 92452 | - | 0 | CCACAAATCGGTTGCGGGGC | GGG | ||||||||[||||||||||||]PAM | NA | Intergenic | NA | NA |
| T27160 | TTTGAGTCAGTACCGTCTAACGG | | 1000 | | | | | | | | | |
| Oligo fwd | CACCTTTGAGTCAGTACCGTCTAA | | | | | | | | | | | |
| Oligo rev | AAACTTAGACGGTACTGACTCAAA | | | | | | | | | | | |
| Chromosome | start | | end | strand | MM | target_seq | PAM | alignment | distance | position | gene name | gene id |
| GU305914.1 | | 92404 | 92426 | - | 0 | TTTGAGTCAGTACCGTCTAA | CGG | ||||||||[||||||||||||]PAM | NA | Intergenic | NA | NA |
| T27161 | TCTAACGGTGTGTCAAACACCGG | | 1000 | | | | | | | | | |
| Oligo fwd | CACCTCTAACGGTGTGTCAAACAC | | | | | | | | | | | |
| Oligo rev | AAACGTGTTTGACACACCGTTAGA | | | | | | | | | | | |
| Chromosome | start | | end | strand | MM | target_seq | PAM | alignment | distance | position | gene name | gene id |
| GU305914.1 | | 92389 | 92411 | - | 0 | TCTAACGGTGTGTCAAACAC | CGG | ||||||||[||||||||||||]PAM | NA | Intergenic | NA | NA |
| T27162 | CTCGTCTGCGTAGTGTGTGTCGG | | 1000 | | | | | | | | | |

<table>
<tr><td>Oligo fwd</td><td>CACCCTCGTCTGCGTAGTGTGTGT</td><td></td><td></td><td></td><td></td><td></td><td></td><td></td><td></td><td></td><td></td></tr>
<tr><td>Oligo rev</td><td>AAACACACACACTACGCAGACGAG</td><td></td><td></td><td></td><td></td><td></td><td></td><td></td><td></td><td></td><td></td></tr>
<tr><td>Chromosome</td><td>start</td><td>end</td><td>strand</td><td>MM</td><td>target_seq</td><td>PAM</td><td>alignment</td><td>distance</td><td>position</td><td>gene name</td><td>gene id</td></tr>
<tr><td>GU305914.1</td><td>92323</td><td>92345</td><td>-</td><td>0</td><td>CTCGTCTGCGTAGTGTGTGT</td><td>CGG</td><td>||||||||[||||||||||||]PAM</td><td>NA</td><td>Intergenic</td><td>NA</td><td>NA</td></tr>
<tr><td>GU305914.1</td><td>1148</td><td>1170</td><td>+</td><td>4</td><td>CTCGTCCCCGTGGTGTCTGT</td><td>CGG</td><td>||||||--[|||-||||-|||]PAM</td><td>NA</td><td>Intergenic</td><td>NA</td><td>NA</td></tr>
<tr><td></td><td></td><td></td><td></td><td></td><td></td><td></td><td></td><td></td><td></td><td></td><td></td></tr>
<tr><td>T27163</td><td>TCGTCTGCGTAGTGTGTGTCGGG</td><td>1000</td><td></td><td></td><td></td><td></td><td></td><td></td><td></td><td></td><td></td></tr>
<tr><td>Oligo fwd</td><td>CACCTCGTCTGCGTAGTGTGTGTC</td><td></td><td></td><td></td><td></td><td></td><td></td><td></td><td></td><td></td><td></td></tr>
<tr><td>Oligo rev</td><td>AAACGACACACACTACGCAGACGA</td><td></td><td></td><td></td><td></td><td></td><td></td><td></td><td></td><td></td><td></td></tr>
<tr><td>Chromosome</td><td>start</td><td>end</td><td>strand</td><td>MM</td><td>target_seq</td><td>PAM</td><td>alignment</td><td>distance</td><td>position</td><td>gene name</td><td>gene id</td></tr>
<tr><td>GU305914.1</td><td>92322</td><td>92344</td><td>-</td><td>0</td><td>TCGTCTGCGTAGTGTGTGTC</td><td>GGG</td><td>||||||||[||||||||||||]PAM</td><td>NA</td><td>Intergenic</td><td>NA</td><td>NA</td></tr>
<tr><td></td><td></td><td></td><td></td><td></td><td></td><td></td><td></td><td></td><td></td><td></td><td></td></tr>
<tr><td>T27164</td><td>CGTCTGCGTAGTGTGTGTCGGGG</td><td>1000</td><td></td><td></td><td></td><td></td><td></td><td></td><td></td><td></td><td></td></tr>
<tr><td>Oligo fwd</td><td>CACCCGTCTGCGTAGTGTGTGTCG</td><td></td><td></td><td></td><td></td><td></td><td></td><td></td><td></td><td></td><td></td></tr>
<tr><td>Oligo rev</td><td>AAACCGACACACACTACGCAGACG</td><td></td><td></td><td></td><td></td><td></td><td></td><td></td><td></td><td></td><td></td></tr>
<tr><td>Chromosome</td><td>start</td><td>end</td><td>strand</td><td>MM</td><td>target_seq</td><td>PAM</td><td>alignment</td><td>distance</td><td>position</td><td>gene name</td><td>gene id</td></tr>
<tr><td>GU305914.1</td><td>92321</td><td>92343</td><td>-</td><td>0</td><td>CGTCTGCGTAGTGTGTGTCG</td><td>GGG</td><td>||||||||[||||||||||||]PAM</td><td>NA</td><td>Intergenic</td><td>NA</td><td>NA</td></tr>
<tr><td></td><td></td><td></td><td></td><td></td><td></td><td></td><td></td><td></td><td></td><td></td><td></td></tr>
<tr><td>T27165</td><td>CGTAGTGTGTGTCGGGGATGTGG</td><td>1000</td><td></td><td></td><td></td><td></td><td></td><td></td><td></td><td></td><td></td></tr>
<tr><td>Oligo fwd</td><td>CACCCGTAGTGTGTGTCGGGGATG</td><td></td><td></td><td></td><td></td><td></td><td></td><td></td><td></td><td></td><td></td></tr>
<tr><td>Oligo rev</td><td>AAACCATCCCCGACACACACTACG</td><td></td><td></td><td></td><td></td><td></td><td></td><td></td><td></td><td></td><td></td></tr>
<tr><td>Chromosome</td><td>start</td><td>end</td><td>strand</td><td>MM</td><td>target_seq</td><td>PAM</td><td>alignment</td><td>distance</td><td>position</td><td>gene name</td><td>gene id</td></tr>
<tr><td>GU305914.1</td><td>92315</td><td>92337</td><td>-</td><td>0</td><td>CGTAGTGTGTGTCGGGGATG</td><td>TGG</td><td>||||||||[||||||||||||]PAM</td><td>NA</td><td>Intergenic</td><td>NA</td><td>NA</td></tr>
<tr><td></td><td></td><td></td><td></td><td></td><td></td><td></td><td></td><td></td><td></td><td></td><td></td></tr>
<tr><td>T27166</td><td>GTAGTGTGTGTCGGGGATGTGGG</td><td>1000</td><td></td><td></td><td></td><td></td><td></td><td></td><td></td><td></td><td></td></tr>
<tr><td>Oligo fwd</td><td>CACCGTAGTGTGTGTCGGGGATGT</td><td></td><td></td><td></td><td></td><td></td><td></td><td></td><td></td><td></td><td></td></tr>
<tr><td>Oligo rev</td><td>AAACACATCCCCGACACACACTAC</td><td></td><td></td><td></td><td></td><td></td><td></td><td></td><td></td><td></td><td></td></tr>
<tr><td>Chromosome</td><td>start</td><td>end</td><td>strand</td><td>MM</td><td>target_seq</td><td>PAM</td><td>alignment</td><td>distance</td><td>position</td><td>gene name</td><td>gene id</td></tr>
<tr><td>GU305914.1</td><td>92314</td><td>92336</td><td>-</td><td>0</td><td>GTAGTGTGTGTCGGGGATGT</td><td>GGG</td><td>||||||||[||||||||||||]PAM</td><td>NA</td><td>Intergenic</td><td>NA</td><td>NA</td></tr>
<tr><td></td><td></td><td></td><td></td><td></td><td></td><td></td><td></td><td></td><td></td><td></td><td></td></tr>
<tr><td>T27167</td><td>CGGGGATGTGGGCCTGTTCGCGG</td><td>1000</td><td></td><td></td><td></td><td></td><td></td><td></td><td></td><td></td><td></td></tr>
<tr><td>Oligo fwd</td><td>CACCCGGGGATGTGGGCCTGTTCG</td><td></td><td></td><td></td><td></td><td></td><td></td><td></td><td></td><td></td><td></td></tr>
<tr><td>Oligo rev</td><td>AAACCGAACAGGCCCACATCCCCG</td><td></td><td></td><td></td><td></td><td></td><td></td><td></td><td></td><td></td><td></td></tr>
<tr><td>Chromosome</td><td>start</td><td>end</td><td>strand</td><td>MM</td><td>target_seq</td><td>PAM</td><td>alignment</td><td>distance</td><td>position</td><td>gene name</td><td>gene id</td></tr>
<tr><td>GU305914.1</td><td>92303</td><td>92325</td><td>-</td><td>0</td><td>CGGGGATGTGGGCCTGTTCG</td><td>CGG</td><td>||||||||[||||||||||||]PAM</td><td>NA</td><td>Intergenic</td><td>NA</td><td>NA</td></tr>
<tr><td></td><td></td><td></td><td></td><td></td><td></td><td></td><td></td><td></td><td></td><td></td><td></td></tr>
<tr><td>T27168</td><td>CGTTTCGCGTGTGTGTTTTGTGG</td><td>1000</td><td></td><td></td><td></td><td></td><td></td><td></td><td></td><td></td><td></td></tr>
<tr><td>Oligo fwd</td><td>CACCCGTTTCGCGTGTGTGTTTTG</td><td></td><td></td><td></td><td></td><td></td><td></td><td></td><td></td><td></td><td></td></tr>
<tr><td>Oligo rev</td><td>AAACCAAAACACACACGCGAAACG</td><td></td><td></td><td></td><td></td><td></td><td></td><td></td><td></td><td></td><td></td></tr>
<tr><td>Chromosome</td><td>start</td><td>end</td><td>strand</td><td>MM</td><td>target_seq</td><td>PAM</td><td>alignment</td><td>distance</td><td>position</td><td>gene name</td><td>gene id</td></tr>
<tr><td>GU305914.1</td><td>92271</td><td>92293</td><td>-</td><td>0</td><td>CGTTTCGCGTGTGTGTTTTG</td><td>TGG</td><td>||||||||[||||||||||||]PAM</td><td>NA</td><td>Intergenic</td><td>NA</td><td>NA</td></tr>
<tr><td></td><td></td><td></td><td></td><td></td><td></td><td></td><td></td><td></td><td></td><td></td><td></td></tr>
<tr><td>T27169</td><td>GTTTCGCGTGTGTGTTTTGTGGG</td><td>1000</td><td></td><td></td><td></td><td></td><td></td><td></td><td></td><td></td><td></td></tr>
<tr><td>Oligo fwd</td><td>CACCGTTTCGCGTGTGTGTTTTGT</td><td></td><td></td><td></td><td></td><td></td><td></td><td></td><td></td><td></td><td></td></tr>
<tr><td>Oligo rev</td><td>AAACACAAAACACACACGCGAAAC</td><td></td><td></td><td></td><td></td><td></td><td></td><td></td><td></td><td></td><td></td></tr>
<tr><td>Chromosome</td><td>start</td><td>end</td><td>strand</td><td>MM</td><td>target_seq</td><td>PAM</td><td>alignment</td><td>distance</td><td>position</td><td>gene name</td><td>gene id</td></tr>
<tr><td>GU305914.1</td><td>92270</td><td>92292</td><td>-</td><td>0</td><td>GTTTCGCGTGTGTGTTTTGT</td><td>GGG</td><td>||||||||[||||||||||||]PAM</td><td>NA</td><td>Intergenic</td><td>NA</td><td>NA</td></tr>
<tr><td></td><td></td><td></td><td></td><td></td><td></td><td></td><td></td><td></td><td></td><td></td><td></td></tr>
<tr><td>T27170</td><td>CGTGTGTGTTTTGTGGGACACGG</td><td>1000</td><td></td><td></td><td></td><td></td><td></td><td></td><td></td><td></td><td></td></tr>
<tr><td>Oligo fwd</td><td>CACCCGTGTGTGTTTTGTGGGACA</td><td></td><td></td><td></td><td></td><td></td><td></td><td></td><td></td><td></td><td></td></tr>
<tr><td>Oligo rev</td><td>AAACTGTCCCACAAAACACACACG</td><td></td><td></td><td></td><td></td><td></td><td></td><td></td><td></td><td></td><td></td></tr>
<tr><td>Chromosome</td><td>start</td><td>end</td><td>strand</td><td>MM</td><td>target_seq</td><td>PAM</td><td>alignment</td><td>distance</td><td>position</td><td>gene name</td><td>gene id</td></tr>
<tr><td>GU305914.1</td><td>92264</td><td>92286</td><td>-</td><td>0</td><td>CGTGTGTGTTTTGTGGGACA</td><td>CGG</td><td>||||||||[||||||||||||]PAM</td><td>NA</td><td>Intergenic</td><td>NA</td><td>NA</td></tr>
<tr><td></td><td></td><td></td><td></td><td></td><td></td><td></td><td></td><td></td><td></td><td></td><td></td></tr>
<tr><td>T27171</td><td>GTGTGTTTTGTGGGACACGGTGG</td><td>1000</td><td></td><td></td><td></td><td></td><td></td><td></td><td></td><td></td><td></td></tr>
<tr><td>Oligo fwd</td><td>CACCGTGTGTTTTGTGGGACACGG</td><td></td><td></td><td></td><td></td><td></td><td></td><td></td><td></td><td></td><td></td></tr>
<tr><td>Oligo rev</td><td>AAACCCGTGTCCCACAAAACACAC</td><td></td><td></td><td></td><td></td><td></td><td></td><td></td><td></td><td></td><td></td></tr>
<tr><td>Chromosome</td><td>start</td><td>end</td><td>strand</td><td>MM</td><td>target_seq</td><td>PAM</td><td>alignment</td><td>distance</td><td>position</td><td>gene name</td><td>gene id</td></tr>
<tr><td>GU305914.1</td><td>92261</td><td>92283</td><td>-</td><td>0</td><td>GTGTGTTTTGTGGGACACGG</td><td>TGG</td><td>||||||||[||||||||||||]PAM</td><td>NA</td><td>Intergenic</td><td>NA</td><td>NA</td></tr>
<tr><td></td><td></td><td></td><td></td><td></td><td></td><td></td><td></td><td></td><td></td><td></td><td></td></tr>
<tr><td>T27172</td><td>TGTGTTTTGTGGGACACGGTGGG</td><td>1000</td><td></td><td></td><td></td><td></td><td></td><td></td><td></td><td></td><td></td></tr>
<tr><td>Oligo fwd</td><td>CACCTGTGTTTTGTGGGACACGGT</td><td></td><td></td><td></td><td></td><td></td><td></td><td></td><td></td><td></td><td></td></tr>
<tr><td>Oligo rev</td><td>AAACACCGTGTCCCACAAAACACA</td><td></td><td></td><td></td><td></td><td></td><td></td><td></td><td></td><td></td><td></td></tr>
<tr><td>Chromosome</td><td>start</td><td>end</td><td>strand</td><td>MM</td><td>target_seq</td><td>PAM</td><td>alignment</td><td>distance</td><td>position</td><td>gene name</td><td>gene id</td></tr>
<tr><td>GU305914.1</td><td>92260</td><td>92282</td><td>-</td><td>0</td><td>TGTGTTTTGTGGGACACGGT</td><td>GGG</td><td>||||||||[||||||||||||]PAM</td><td>NA</td><td>Intergenic</td><td>NA</td><td>NA</td></tr>
<tr><td></td><td></td><td></td><td></td><td></td><td></td><td></td><td></td><td></td><td></td><td></td><td></td></tr>
<tr><td>T27173</td><td>GTGTTTTGTGGGACACGGTGGGG</td><td>1000</td><td></td><td></td><td></td><td></td><td></td><td></td><td></td><td></td><td></td></tr>
<tr><td>Oligo fwd</td><td>CACCGTGTTTTGTGGGACACGGTG</td><td></td><td></td><td></td><td></td><td></td><td></td><td></td><td></td><td></td><td></td></tr>
<tr><td>Oligo rev</td><td>AAACCACCGTGTCCCACAAAACAC</td><td></td><td></td><td></td><td></td><td></td><td></td><td></td><td></td><td></td><td></td></tr>
<tr><td>Chromosome</td><td>start</td><td>end</td><td>strand</td><td>MM</td><td>target_seq</td><td>PAM</td><td>alignment</td><td>distance</td><td>position</td><td>gene name</td><td>gene id</td></tr>
<tr><td>GU305914.1</td><td>92259</td><td>92281</td><td>-</td><td>0</td><td>GTGTTTTGTGGGACACGGTG</td><td>GGG</td><td>||||||||[||||||||||||]PAM</td><td>NA</td><td>Intergenic</td><td>NA</td><td>NA</td></tr>
<tr><td></td><td></td><td></td><td></td><td></td><td></td><td></td><td></td><td></td><td></td><td></td><td></td></tr>
<tr><td>T27174</td><td>GTGGGACACGGTGGGGATTTCGG</td><td>1000</td><td></td><td></td><td></td><td></td><td></td><td></td><td></td><td></td><td></td></tr>
<tr><td>Oligo fwd</td><td>CACCGTGGGACACGGTGGGGATTT</td><td></td><td></td><td></td><td></td><td></td><td></td><td></td><td></td><td></td><td></td></tr>
<tr><td>Oligo rev</td><td>AAACAAATCCCCACCGTGTCCCAC</td><td></td><td></td><td></td><td></td><td></td><td></td><td></td><td></td><td></td><td></td></tr>
<tr><td>Chromosome</td><td>start</td><td>end</td><td>strand</td><td>MM</td><td>target_seq</td><td>PAM</td><td>alignment</td><td>distance</td><td>position</td><td>gene name</td><td>gene id</td></tr>
<tr><td>GU305914.1</td><td>92252</td><td>92274</td><td>-</td><td>0</td><td>GTGGGACACGGTGGGGATTT</td><td>CGG</td><td>||||||||[||||||||||||]PAM</td><td>NA</td><td>Intergenic</td><td>NA</td><td>NA</td></tr>
<tr><td></td><td></td><td></td><td></td><td></td><td></td><td></td><td></td><td></td><td></td><td></td><td></td></tr>
<tr><td>T27175</td><td>ACGGTGGGGATTTCGGTGTTTGG</td><td>1000</td><td></td><td></td><td></td><td></td><td></td><td></td><td></td><td></td><td></td></tr>
<tr><td>Oligo fwd</td><td>CACCACGGTGGGGATTTCGGTGTT</td><td></td><td></td><td></td><td></td><td></td><td></td><td></td><td></td><td></td><td></td></tr>
<tr><td>Oligo rev</td><td>AAACAACACCGAAATCCCCACCGT</td><td></td><td></td><td></td><td></td><td></td><td></td><td></td><td></td><td></td><td></td></tr>
<tr><td>Chromosome</td><td>start</td><td>end</td><td>strand</td><td>MM</td><td>target_seq</td><td>PAM</td><td>alignment</td><td>distance</td><td>position</td><td>gene name</td><td>gene id</td></tr>
<tr><td>GU305914.1</td><td>92245</td><td>92267</td><td>-</td><td>0</td><td>ACGGTGGGGATTTCGGTGTT</td><td>TGG</td><td>||||||||[||||||||||||]PAM</td><td>NA</td><td>Intergenic</td><td>NA</td><td>NA</td></tr>
<tr><td></td><td></td><td></td><td></td><td></td><td></td><td></td><td></td><td></td><td></td><td></td><td></td></tr>
<tr><td>T27176</td><td>TTTTCGTCAATGCCGTCTGACGG</td><td>1000</td><td></td><td></td><td></td><td></td><td></td><td></td><td></td><td></td><td></td></tr>
<tr><td>Oligo fwd</td><td>CACCTTTTCGTCAATGCCGTCTGA</td><td></td><td></td><td></td><td></td><td></td><td></td><td></td><td></td><td></td><td></td></tr>
<tr><td>Oligo rev</td><td>AAACTCAGACGGCATTGACGAAAA</td><td></td><td></td><td></td><td></td><td></td><td></td><td></td><td></td><td></td><td></td></tr>
<tr><td>Chromosome</td><td>start</td><td>end</td><td>strand</td><td>MM</td><td>target_seq</td><td>PAM</td><td>alignment</td><td>distance</td><td>position</td><td>gene name</td><td>gene id</td></tr>
<tr><td>GU305914.1</td><td>92203</td><td>92225</td><td>-</td><td>0</td><td>TTTTCGTCAATGCCGTCTGA</td><td>CGG</td><td>||||||||[||||||||||||]PAM</td><td>NA</td><td>Intergenic</td><td>NA</td><td>NA</td></tr>
</table>

<table>
<tr><td>T27177</td><td>GTCAATGCCGTCTGACGGCCCGG</td><td>1000</td></tr>
<tr><td>Oligo fwd</td><td>CACCGTCAATGCCGTCTGACGGCC</td></tr>
<tr><td>Oligo rev</td><td>AAACGGCCGTCAGACGGCATTGAC</td></tr>
<tr><td>Chromosome</td><td>start</td><td>end</td><td>strand</td><td>MM</td><td>target_seq</td><td>PAM</td><td>alignment</td><td>distance</td><td>position</td><td>gene name</td><td>gene id</td></tr>
<tr><td>GU305914.1</td><td>92198</td><td>92220</td><td>-</td><td>0</td><td>GTCAATGCCGTCTGACGGCC</td><td>CGG</td><td>||||||||[||||||||||||]PAM</td><td>NA</td><td>Intergenic</td><td>NA</td><td>NA</td></tr>
<tr><td>T27178</td><td>TCAATGCCGTCTGACGGCCCGGG</td><td>1000</td></tr>
<tr><td>Oligo fwd</td><td>CACCTCAATGCCGTCTGACGGCCC</td></tr>
<tr><td>Oligo rev</td><td>AAACGGGCCGTCAGACGGCATTGA</td></tr>
<tr><td>Chromosome</td><td>start</td><td>end</td><td>strand</td><td>MM</td><td>target_seq</td><td>PAM</td><td>alignment</td><td>distance</td><td>position</td><td>gene name</td><td>gene id</td></tr>
<tr><td>GU305914.1</td><td>92197</td><td>92219</td><td>-</td><td>0</td><td>TCAATGCCGTCTGACGGCCC</td><td>GGG</td><td>||||||||[||||||||||||]PAM</td><td>NA</td><td>Intergenic</td><td>NA</td><td>NA</td></tr>
<tr><td>T27179</td><td>GCCGTCTGACGGCCCGGGCTCGG</td><td>1000</td></tr>
<tr><td>Oligo fwd</td><td>CACCGCCGTCTGACGGCCCGGGCT</td></tr>
<tr><td>Oligo rev</td><td>AAACAGCCCGGGCCGTCAGACGGC</td></tr>
<tr><td>Chromosome</td><td>start</td><td>end</td><td>strand</td><td>MM</td><td>target_seq</td><td>PAM</td><td>alignment</td><td>distance</td><td>position</td><td>gene name</td><td>gene id</td></tr>
<tr><td>GU305914.1</td><td>92192</td><td>92214</td><td>-</td><td>0</td><td>GCCGTCTGACGGCCCGGGCT</td><td>CGG</td><td>||||||||[||||||||||||]PAM</td><td>NA</td><td>Intergenic</td><td>NA</td><td>NA</td></tr>
<tr><td>T27180</td><td>CGGCCCGGGCTCGGAGCTGCTGG</td><td>1000</td></tr>
<tr><td>Oligo fwd</td><td>CACCCGGCCCGGGCTCGGAGCTGC</td></tr>
<tr><td>Oligo rev</td><td>AAACGCAGCTCCGAGCCCGGGCCG</td></tr>
<tr><td>Chromosome</td><td>start</td><td>end</td><td>strand</td><td>MM</td><td>target_seq</td><td>PAM</td><td>alignment</td><td>distance</td><td>position</td><td>gene name</td><td>gene id</td></tr>
<tr><td>GU305914.1</td><td>92183</td><td>92205</td><td>-</td><td>0</td><td>CGGCCCGGGCTCGGAGCTGC</td><td>TGG</td><td>||||||||[||||||||||||]PAM</td><td>NA</td><td>Intergenic</td><td>NA</td><td>NA</td></tr>
<tr><td>T27181</td><td>AGCTGCTGGCGCAACGCCGCCGG</td><td>1000</td></tr>
<tr><td>Oligo fwd</td><td>CACCAGCTGCTGGCGCAACGCCGC</td></tr>
<tr><td>Oligo rev</td><td>AAACGCGGCGTTGCGCCAGCAGCT</td></tr>
<tr><td>Chromosome</td><td>start</td><td>end</td><td>strand</td><td>MM</td><td>target_seq</td><td>PAM</td><td>alignment</td><td>distance</td><td>position</td><td>gene name</td><td>gene id</td></tr>
<tr><td>GU305914.1</td><td>92169</td><td>92191</td><td>-</td><td>0</td><td>AGCTGCTGGCGCAACGCCGC</td><td>CGG</td><td>||||||||[||||||||||||]PAM</td><td>NA</td><td>Intergenic</td><td>NA</td><td>NA</td></tr>
<tr><td>T27182</td><td>TCGAGCGCGCCAAGAGCCCGCGG</td><td>1000</td></tr>
<tr><td>Oligo fwd</td><td>CACCTCGAGCGCGCCAAGAGCCCG</td></tr>
<tr><td>Oligo rev</td><td>AAACCGGGCTCTTGGCGCGCTCGA</td></tr>
<tr><td>Chromosome</td><td>start</td><td>end</td><td>strand</td><td>MM</td><td>target_seq</td><td>PAM</td><td>alignment</td><td>distance</td><td>position</td><td>gene name</td><td>gene id</td></tr>
<tr><td>GU305914.1</td><td>92122</td><td>92144</td><td>-</td><td>0</td><td>TCGAGCGCGCCAAGAGCCCG</td><td>CGG</td><td>||||||||[||||||||||||]PAM</td><td>NA</td><td>Intergenic</td><td>NA</td><td>NA</td></tr>
<tr><td>T27183</td><td>CCCGCGGTCCTCTGCGATTGCGG</td><td>1000</td></tr>
<tr><td>Oligo fwd</td><td>CACCCCCGCGGTCCTCTGCGATTG</td></tr>
<tr><td>Oligo rev</td><td>AAACCAATCGCAGAGGACCGCGGG</td></tr>
<tr><td>Chromosome</td><td>start</td><td>end</td><td>strand</td><td>MM</td><td>target_seq</td><td>PAM</td><td>alignment</td><td>distance</td><td>position</td><td>gene name</td><td>gene id</td></tr>
<tr><td>GU305914.1</td><td>92106</td><td>92128</td><td>-</td><td>0</td><td>CCCGCGGTCCTCTGCGATTG</td><td>CGG</td><td>||||||||[||||||||||||]PAM</td><td>NA</td><td>Intergenic</td><td>NA</td><td>NA</td></tr>
<tr><td>T27184</td><td>TCCTCTGCGATTGCGGCGCACGG</td><td>1000</td></tr>
<tr><td>Oligo fwd</td><td>CACCTCCTCTGCGATTGCGGCGCA</td></tr>
<tr><td>Oligo rev</td><td>AAACTGCGCCGCAATCGCAGAGGA</td></tr>
<tr><td>Chromosome</td><td>start</td><td>end</td><td>strand</td><td>MM</td><td>target_seq</td><td>PAM</td><td>alignment</td><td>distance</td><td>position</td><td>gene name</td><td>gene id</td></tr>
<tr><td>GU305914.1</td><td>92099</td><td>92121</td><td>-</td><td>0</td><td>TCCTCTGCGATTGCGGCGCA</td><td>CGG</td><td>||||||||[||||||||||||]PAM</td><td>NA</td><td>Intergenic</td><td>NA</td><td>NA</td></tr>
<tr><td>T27185</td><td>TTGCGGCGCACGGAGCCCTTCGG</td><td>1000</td></tr>
<tr><td>Oligo fwd</td><td>CACCTTGCGGCGCACGGAGCCCTT</td></tr>
<tr><td>Oligo rev</td><td>AAACAAGGGCTCCGTGCGCCGCAA</td></tr>
<tr><td>Chromosome</td><td>start</td><td>end</td><td>strand</td><td>MM</td><td>target_seq</td><td>PAM</td><td>alignment</td><td>distance</td><td>position</td><td>gene name</td><td>gene id</td></tr>
<tr><td>GU305914.1</td><td>92089</td><td>92111</td><td>-</td><td>0</td><td>TTGCGGCGCACGGAGCCCTT</td><td>CGG</td><td>||||||||[||||||||||||]PAM</td><td>NA</td><td>Intergenic</td><td>NA</td><td>NA</td></tr>
<tr><td>T27186</td><td>TGCGGCGCACGGAGCCCTTCGGG</td><td>1000</td></tr>
<tr><td>Oligo fwd</td><td>CACCTGCGGCGCACGGAGCCCTTC</td></tr>
<tr><td>Oligo rev</td><td>AAACGAAGGGCTCCGTGCGCCGCA</td></tr>
<tr><td>Chromosome</td><td>start</td><td>end</td><td>strand</td><td>MM</td><td>target_seq</td><td>PAM</td><td>alignment</td><td>distance</td><td>position</td><td>gene name</td><td>gene id</td></tr>
<tr><td>GU305914.1</td><td>92088</td><td>92110</td><td>-</td><td>0</td><td>TGCGGCGCACGGAGCCCTTC</td><td>GGG</td><td>||||||||[||||||||||||]PAM</td><td>NA</td><td>Intergenic</td><td>NA</td><td>NA</td></tr>
<tr><td>T27187</td><td>GCGGCGCACGGAGCCCTTCGGGG</td><td>1000</td></tr>
<tr><td>Oligo fwd</td><td>CACCGCGGCGCACGGAGCCCTTCG</td></tr>
<tr><td>Oligo rev</td><td>AAACCGAAGGGCTCCGTGCGCCGC</td></tr>
<tr><td>Chromosome</td><td>start</td><td>end</td><td>strand</td><td>MM</td><td>target_seq</td><td>PAM</td><td>alignment</td><td>distance</td><td>position</td><td>gene name</td><td>gene id</td></tr>
<tr><td>GU305914.1</td><td>92087</td><td>92109</td><td>-</td><td>0</td><td>GCGGCGCACGGAGCCCTTCG</td><td>GGG</td><td>||||||||[||||||||||||]PAM</td><td>NA</td><td>Intergenic</td><td>NA</td><td>NA</td></tr>
<tr><td>T27188</td><td>GCGCACGGAGCCCTTCGGGGCGG</td><td>1000</td></tr>
<tr><td>Oligo fwd</td><td>CACCGCGCACGGAGCCCTTCGGGG</td></tr>
<tr><td>Oligo rev</td><td>AAACCCCCGAAGGGCTCCGTGCGC</td></tr>
<tr><td>Chromosome</td><td>start</td><td>end</td><td>strand</td><td>MM</td><td>target_seq</td><td>PAM</td><td>alignment</td><td>distance</td><td>position</td><td>gene name</td><td>gene id</td></tr>
<tr><td>GU305914.1</td><td>92084</td><td>92106</td><td>-</td><td>0</td><td>GCGCACGGAGCCCTTCGGGG</td><td>CGG</td><td>||||||||[||||||||||||]PAM</td><td>NA</td><td>Intergenic</td><td>NA</td><td>NA</td></tr>
<tr><td>T27189</td><td>TGTCCAGTACGTTGTGACAAAGG</td><td>1000</td></tr>
<tr><td>Oligo fwd</td><td>CACCTGTCCAGTACGTTGTGACAA</td></tr>
<tr><td>Oligo rev</td><td>AAACTTGTCACAACGTACTGGACA</td></tr>
<tr><td>Chromosome</td><td>start</td><td>end</td><td>strand</td><td>MM</td><td>target_seq</td><td>PAM</td><td>alignment</td><td>distance</td><td>position</td><td>gene name</td><td>gene id</td></tr>
<tr><td>GU305914.1</td><td>92031</td><td>92053</td><td>-</td><td>0</td><td>TGTCCAGTACGTTGTGACAA</td><td>AGG</td><td>||||||||[||||||||||||]PAM</td><td>NA</td><td>Intergenic</td><td>NA</td><td>NA</td></tr>
<tr><td>T27190</td><td>ACGTTGTGACAAAGGCTTCGAGG</td><td>1000</td></tr>
<tr><td>Oligo fwd</td><td>CACCACGTTGTGACAAAGGCTTCG</td></tr>
<tr><td>Oligo rev</td><td>AAACCGAAGCCTTTGTCACAACGT</td></tr>
<tr><td>Chromosome</td><td>start</td><td>end</td><td>strand</td><td>MM</td><td>target_seq</td><td>PAM</td><td>alignment</td><td>distance</td><td>position</td><td>gene name</td><td>gene id</td></tr>
<tr><td>GU305914.1</td><td>92023</td><td>92045</td><td>-</td><td>0</td><td>ACGTTGTGACAAAGGCTTCG</td><td>AGG</td><td>||||||||[||||||||||||]PAM</td><td>NA</td><td>Intergenic</td><td>NA</td><td>NA</td></tr>
<tr><td>T27191</td><td>CGTTGTGACAAAGGCTTCGAGGG</td><td>1000</td></tr>
<tr><td>Oligo fwd</td><td>CACCCGTTGTGACAAAGGCTTCGA</td></tr>
<tr><td>Oligo rev</td><td>AAACTCGAAGCCTTTGTCACAACG</td></tr>
<tr><td>Chromosome</td><td>start</td><td>end</td><td>strand</td><td>MM</td><td>target_seq</td><td>PAM</td><td>alignment</td><td>distance</td><td>position</td><td>gene name</td><td>gene id</td></tr>
</table>

<table>
<tr><td>GU305914.1</td><td>92022</td><td>92044</td><td>-</td><td>0</td><td>CGTTGTGACAAAGGCTTCGA</td><td>GGG</td><td>||||||||[||||||||||||]PAM</td><td>NA</td><td>Intergenic</td><td>NA</td><td>NA</td></tr>
<tr><td></td><td></td><td></td><td></td><td></td><td></td><td></td><td></td><td></td><td></td><td></td><td></td></tr>
<tr><td>T27192</td><td>GCTTCGAGGGCGCTTGCGTGAGG</td><td>1000</td><td></td><td></td><td></td><td></td><td></td><td></td><td></td><td></td><td></td></tr>
<tr><td>Oligo fwd</td><td>CACCGCTTCGAGGGCGCTTGCGTG</td><td></td><td></td><td></td><td></td><td></td><td></td><td></td><td></td><td></td><td></td></tr>
<tr><td>Oligo rev</td><td>AAACCACGCAAGCGCCCTCGAAGC</td><td></td><td></td><td></td><td></td><td></td><td></td><td></td><td></td><td></td><td></td></tr>
<tr><td>Chromosome</td><td>start</td><td>end</td><td>strand</td><td>MM</td><td>target_seq</td><td>PAM</td><td>alignment</td><td>distance</td><td>position</td><td>gene name</td><td>gene id</td></tr>
<tr><td>GU305914.1</td><td>92009</td><td>92031</td><td>-</td><td>0</td><td>GCTTCGAGGGCGCTTGCGTG</td><td>AGG</td><td>||||||||[||||||||||||]PAM</td><td>NA</td><td>Intergenic</td><td>NA</td><td>NA</td></tr>
<tr><td></td><td></td><td></td><td></td><td></td><td></td><td></td><td></td><td></td><td></td><td></td><td></td></tr>
<tr><td>T27193</td><td>GGATTGTGACGTTTTCGTCGAGG</td><td>1000</td><td></td><td></td><td></td><td></td><td></td><td></td><td></td><td></td><td></td></tr>
<tr><td>Oligo fwd</td><td>CACCGGATTGTGACGTTTTCGTCG</td><td></td><td></td><td></td><td></td><td></td><td></td><td></td><td></td><td></td><td></td></tr>
<tr><td>Oligo rev</td><td>AAACCGACGAAAACGTCACAATCC</td><td></td><td></td><td></td><td></td><td></td><td></td><td></td><td></td><td></td><td></td></tr>
<tr><td>Chromosome</td><td>start</td><td>end</td><td>strand</td><td>MM</td><td>target_seq</td><td>PAM</td><td>alignment</td><td>distance</td><td>position</td><td>gene name</td><td>gene id</td></tr>
<tr><td>GU305914.1</td><td>91988</td><td>92010</td><td>-</td><td>0</td><td>GGATTGTGACGTTTTCGTCG</td><td>AGG</td><td>||||||||[||||||||||||]PAM</td><td>NA</td><td>Intergenic</td><td>NA</td><td>NA</td></tr>
<tr><td></td><td></td><td></td><td></td><td></td><td></td><td></td><td></td><td></td><td></td><td></td><td></td></tr>
<tr><td>T27194</td><td>TTTCGTCGAGGAGAACCCGCCGG</td><td>1000</td><td></td><td></td><td></td><td></td><td></td><td></td><td></td><td></td><td></td></tr>
<tr><td>Oligo fwd</td><td>CACCTTTCGTCGAGGAGAACCCGC</td><td></td><td></td><td></td><td></td><td></td><td></td><td></td><td></td><td></td><td></td></tr>
<tr><td>Oligo rev</td><td>AAACGCGGGTTCTCCTCGACGAAA</td><td></td><td></td><td></td><td></td><td></td><td></td><td></td><td></td><td></td><td></td></tr>
<tr><td>Chromosome</td><td>start</td><td>end</td><td>strand</td><td>MM</td><td>target_seq</td><td>PAM</td><td>alignment</td><td>distance</td><td>position</td><td>gene name</td><td>gene id</td></tr>
<tr><td>GU305914.1</td><td>91976</td><td>91998</td><td>-</td><td>0</td><td>TTTCGTCGAGGAGAACCCGC</td><td>CGG</td><td>||||||||[||||||||||||]PAM</td><td>NA</td><td>Intergenic</td><td>NA</td><td>NA</td></tr>
<tr><td></td><td></td><td></td><td></td><td></td><td></td><td></td><td></td><td></td><td></td><td></td><td></td></tr>
<tr><td>T27195</td><td>TTCGTCGAGGAGAACCCGCCGGG</td><td>1000</td><td></td><td></td><td></td><td></td><td></td><td></td><td></td><td></td><td></td></tr>
<tr><td>Oligo fwd</td><td>CACCTTCGTCGAGGAGAACCCGCC</td><td></td><td></td><td></td><td></td><td></td><td></td><td></td><td></td><td></td><td></td></tr>
<tr><td>Oligo rev</td><td>AAACGGCGGGTTCTCCTCGACGAA</td><td></td><td></td><td></td><td></td><td></td><td></td><td></td><td></td><td></td><td></td></tr>
<tr><td>Chromosome</td><td>start</td><td>end</td><td>strand</td><td>MM</td><td>target_seq</td><td>PAM</td><td>alignment</td><td>distance</td><td>position</td><td>gene name</td><td>gene id</td></tr>
<tr><td>GU305914.1</td><td>91975</td><td>91997</td><td>-</td><td>0</td><td>TTCGTCGAGGAGAACCCGCC</td><td>GGG</td><td>||||||||[||||||||||||]PAM</td><td>NA</td><td>Intergenic</td><td>NA</td><td>NA</td></tr>
<tr><td></td><td></td><td></td><td></td><td></td><td></td><td></td><td></td><td></td><td></td><td></td><td></td></tr>
<tr><td>T27196</td><td>TCGTCGAGGAGAACCCGCCGGGG</td><td>1000</td><td></td><td></td><td></td><td></td><td></td><td></td><td></td><td></td><td></td></tr>
<tr><td>Oligo fwd</td><td>CACCTCGTCGAGGAGAACCCGCCG</td><td></td><td></td><td></td><td></td><td></td><td></td><td></td><td></td><td></td><td></td></tr>
<tr><td>Oligo rev</td><td>AAACCGGCGGGTTCTCCTCGACGA</td><td></td><td></td><td></td><td></td><td></td><td></td><td></td><td></td><td></td><td></td></tr>
<tr><td>Chromosome</td><td>start</td><td>end</td><td>strand</td><td>MM</td><td>target_seq</td><td>PAM</td><td>alignment</td><td>distance</td><td>position</td><td>gene name</td><td>gene id</td></tr>
<tr><td>GU305914.1</td><td>91974</td><td>91996</td><td>-</td><td>0</td><td>TCGTCGAGGAGAACCCGCCG</td><td>GGG</td><td>||||||||[||||||||||||]PAM</td><td>NA</td><td>Intergenic</td><td>NA</td><td>NA</td></tr>
<tr><td></td><td></td><td></td><td></td><td></td><td></td><td></td><td></td><td></td><td></td><td></td><td></td></tr>
<tr><td>T27197</td><td>CGTCGAGGAGAACCCGCCGGGGG</td><td>1000</td><td></td><td></td><td></td><td></td><td></td><td></td><td></td><td></td><td></td></tr>
<tr><td>Oligo fwd</td><td>CACCCGTCGAGGAGAACCCGCCGG</td><td></td><td></td><td></td><td></td><td></td><td></td><td></td><td></td><td></td><td></td></tr>
<tr><td>Oligo rev</td><td>AAACCCGGCGGGTTCTCCTCGACG</td><td></td><td></td><td></td><td></td><td></td><td></td><td></td><td></td><td></td><td></td></tr>
<tr><td>Chromosome</td><td>start</td><td>end</td><td>strand</td><td>MM</td><td>target_seq</td><td>PAM</td><td>alignment</td><td>distance</td><td>position</td><td>gene name</td><td>gene id</td></tr>
<tr><td>GU305914.1</td><td>91973</td><td>91995</td><td>-</td><td>0</td><td>CGTCGAGGAGAACCCGCCGG</td><td>GGG</td><td>||||||||[||||||||||||]PAM</td><td>NA</td><td>Intergenic</td><td>NA</td><td>NA</td></tr>
<tr><td></td><td></td><td></td><td></td><td></td><td></td><td></td><td></td><td></td><td></td><td></td><td></td></tr>
<tr><td>T27198</td><td>CGGGGGCTTCACGTATATAAAGG</td><td>1000</td><td></td><td></td><td></td><td></td><td></td><td></td><td></td><td></td><td></td></tr>
<tr><td>Oligo fwd</td><td>CACCCGGGGGCTTCACGTATATAA</td><td></td><td></td><td></td><td></td><td></td><td></td><td></td><td></td><td></td><td></td></tr>
<tr><td>Oligo rev</td><td>AAACTTATATACGTGAAGCCCCCG</td><td></td><td></td><td></td><td></td><td></td><td></td><td></td><td></td><td></td><td></td></tr>
<tr><td>Chromosome</td><td>start</td><td>end</td><td>strand</td><td>MM</td><td>target_seq</td><td>PAM</td><td>alignment</td><td>distance</td><td>position</td><td>gene name</td><td>gene id</td></tr>
<tr><td>GU305914.1</td><td>91956</td><td>91978</td><td>-</td><td>0</td><td>CGGGGGCTTCACGTATATAA</td><td>AGG</td><td>||||||||[||||||||||||]PAM</td><td>NA</td><td>Intergenic</td><td>NA</td><td>NA</td></tr>
<tr><td></td><td></td><td></td><td></td><td></td><td></td><td></td><td></td><td></td><td></td><td></td><td></td></tr>
<tr><td>T27199</td><td>CTTCACGTATATAAAGGTTCTGG</td><td>1000</td><td></td><td></td><td></td><td></td><td></td><td></td><td></td><td></td><td></td></tr>
<tr><td>Oligo fwd</td><td>CACCCTTCACGTATATAAAGGTTC</td><td></td><td></td><td></td><td></td><td></td><td></td><td></td><td></td><td></td><td></td></tr>
<tr><td>Oligo rev</td><td>AAACGAACCTTTATATACGTGAAG</td><td></td><td></td><td></td><td></td><td></td><td></td><td></td><td></td><td></td><td></td></tr>
<tr><td>Chromosome</td><td>start</td><td>end</td><td>strand</td><td>MM</td><td>target_seq</td><td>PAM</td><td>alignment</td><td>distance</td><td>position</td><td>gene name</td><td>gene id</td></tr>
<tr><td>GU305914.1</td><td>91950</td><td>91972</td><td>-</td><td>0</td><td>CTTCACGTATATAAAGGTTC</td><td>TGG</td><td>||||||||[||||||||||||]PAM</td><td>NA</td><td>Intergenic</td><td>NA</td><td>NA</td></tr>
<tr><td></td><td></td><td></td><td></td><td></td><td></td><td></td><td></td><td></td><td></td><td></td><td></td></tr>
<tr><td>T27200</td><td>ATATAAAGGTTCTGGTGCCGCGG</td><td>1000</td><td></td><td></td><td></td><td></td><td></td><td></td><td></td><td></td><td></td></tr>
<tr><td>Oligo fwd</td><td>CACCATATAAAGGTTCTGGTGCCG</td><td></td><td></td><td></td><td></td><td></td><td></td><td></td><td></td><td></td><td></td></tr>
<tr><td>Oligo rev</td><td>AAACCGGCACCAGAACCTTTATAT</td><td></td><td></td><td></td><td></td><td></td><td></td><td></td><td></td><td></td><td></td></tr>
<tr><td>Chromosome</td><td>start</td><td>end</td><td>strand</td><td>MM</td><td>target_seq</td><td>PAM</td><td>alignment</td><td>distance</td><td>position</td><td>gene name</td><td>gene id</td></tr>
<tr><td>GU305914.1</td><td>91942</td><td>91964</td><td>-</td><td>0</td><td>ATATAAAGGTTCTGGTGCCG</td><td>CGG</td><td>||||||||[||||||||||||]PAM</td><td>NA</td><td>Intergenic</td><td>NA</td><td>NA</td></tr>
<tr><td></td><td></td><td></td><td></td><td></td><td></td><td></td><td></td><td></td><td></td><td></td><td></td></tr>
<tr><td>T27201</td><td>TATAAAGGTTCTGGTGCCGCGGG</td><td>1000</td><td></td><td></td><td></td><td></td><td></td><td></td><td></td><td></td><td></td></tr>
<tr><td>Oligo fwd</td><td>CACCTATAAAGGTTCTGGTGCCGC</td><td></td><td></td><td></td><td></td><td></td><td></td><td></td><td></td><td></td><td></td></tr>
<tr><td>Oligo rev</td><td>AAACGCGGCACCAGAACCTTTATA</td><td></td><td></td><td></td><td></td><td></td><td></td><td></td><td></td><td></td><td></td></tr>
<tr><td>Chromosome</td><td>start</td><td>end</td><td>strand</td><td>MM</td><td>target_seq</td><td>PAM</td><td>alignment</td><td>distance</td><td>position</td><td>gene name</td><td>gene id</td></tr>
<tr><td>GU305914.1</td><td>91941</td><td>91963</td><td>-</td><td>0</td><td>TATAAAGGTTCTGGTGCCGC</td><td>GGG</td><td>||||||||[||||||||||||]PAM</td><td>NA</td><td>Intergenic</td><td>NA</td><td>NA</td></tr>
<tr><td></td><td></td><td></td><td></td><td></td><td></td><td></td><td></td><td></td><td></td><td></td><td></td></tr>
<tr><td>T27202</td><td>CTCCTGTATAATTTCGATGCCGG</td><td>1000</td><td></td><td></td><td></td><td></td><td></td><td></td><td></td><td></td><td></td></tr>
<tr><td>Oligo fwd</td><td>CACCCTCCTGTATAATTTCGATGC</td><td></td><td></td><td></td><td></td><td></td><td></td><td></td><td></td><td></td><td></td></tr>
<tr><td>Oligo rev</td><td>AAACGCATCGAAATTATACAGGAG</td><td></td><td></td><td></td><td></td><td></td><td></td><td></td><td></td><td></td><td></td></tr>
<tr><td>Chromosome</td><td>start</td><td>end</td><td>strand</td><td>MM</td><td>target_seq</td><td>PAM</td><td>alignment</td><td>distance</td><td>position</td><td>gene name</td><td>gene id</td></tr>
<tr><td>GU305914.1</td><td>91900</td><td>91922</td><td>-</td><td>0</td><td>CTCCTGTATAATTTCGATGC</td><td>CGG</td><td>||||||||[||||||||||||]PAM</td><td>NA</td><td>Intergenic</td><td>NA</td><td>NA</td></tr>
<tr><td></td><td></td><td></td><td></td><td></td><td></td><td></td><td></td><td></td><td></td><td></td><td></td></tr>
<tr><td>T27203</td><td>TCCTGTATAATTTCGATGCCGGG</td><td>1000</td><td></td><td></td><td></td><td></td><td></td><td></td><td></td><td></td><td></td></tr>
<tr><td>Oligo fwd</td><td>CACCTCCTGTATAATTTCGATGCC</td><td></td><td></td><td></td><td></td><td></td><td></td><td></td><td></td><td></td><td></td></tr>
<tr><td>Oligo rev</td><td>AAACGGCATCGAAATTATACAGGA</td><td></td><td></td><td></td><td></td><td></td><td></td><td></td><td></td><td></td><td></td></tr>
<tr><td>Chromosome</td><td>start</td><td>end</td><td>strand</td><td>MM</td><td>target_seq</td><td>PAM</td><td>alignment</td><td>distance</td><td>position</td><td>gene name</td><td>gene id</td></tr>
<tr><td>GU305914.1</td><td>91899</td><td>91921</td><td>-</td><td>0</td><td>TCCTGTATAATTTCGATGCC</td><td>GGG</td><td>||||||||[||||||||||||]PAM</td><td>NA</td><td>Intergenic</td><td>NA</td><td>NA</td></tr>
<tr><td></td><td></td><td></td><td></td><td></td><td></td><td></td><td></td><td></td><td></td><td></td><td></td></tr>
<tr><td>T27204</td><td>CCTGTATAATTTCGATGCCGGGG</td><td>1000</td><td></td><td></td><td></td><td></td><td></td><td></td><td></td><td></td><td></td></tr>
<tr><td>Oligo fwd</td><td>CACCCCTGTATAATTTCGATGCCG</td><td></td><td></td><td></td><td></td><td></td><td></td><td></td><td></td><td></td><td></td></tr>
<tr><td>Oligo rev</td><td>AAACCGGCATCGAAATTATACAGG</td><td></td><td></td><td></td><td></td><td></td><td></td><td></td><td></td><td></td><td></td></tr>
<tr><td>Chromosome</td><td>start</td><td>end</td><td>strand</td><td>MM</td><td>target_seq</td><td>PAM</td><td>alignment</td><td>distance</td><td>position</td><td>gene name</td><td>gene id</td></tr>
<tr><td>GU305914.1</td><td>91898</td><td>91920</td><td>-</td><td>0</td><td>CCTGTATAATTTCGATGCCG</td><td>GGG</td><td>||||||||[||||||||||||]PAM</td><td>NA</td><td>Intergenic</td><td>NA</td><td>NA</td></tr>
<tr><td></td><td></td><td></td><td></td><td></td><td></td><td></td><td></td><td></td><td></td><td></td><td></td></tr>
<tr><td>T27205</td><td>GTATAATTTCGATGCCGGGGAGG</td><td>1000</td><td></td><td></td><td></td><td></td><td></td><td></td><td></td><td></td><td></td></tr>
<tr><td>Oligo fwd</td><td>CACCGTATAATTTCGATGCCGGGG</td><td></td><td></td><td></td><td></td><td></td><td></td><td></td><td></td><td></td><td></td></tr>
<tr><td>Oligo rev</td><td>AAACCCCCGGCATCGAAATTATAC</td><td></td><td></td><td></td><td></td><td></td><td></td><td></td><td></td><td></td><td></td></tr>
<tr><td>Chromosome</td><td>start</td><td>end</td><td>strand</td><td>MM</td><td>target_seq</td><td>PAM</td><td>alignment</td><td>distance</td><td>position</td><td>gene name</td><td>gene id</td></tr>
<tr><td>GU305914.1</td><td>91895</td><td>91917</td><td>-</td><td>0</td><td>GTATAATTTCGATGCCGGGG</td><td>AGG</td><td>||||||||[||||||||||||]PAM</td><td>NA</td><td>Intergenic</td><td>NA</td><td>NA</td></tr>
<tr><td></td><td></td><td></td><td></td><td></td><td></td><td></td><td></td><td></td><td></td><td></td><td></td></tr>
<tr><td>T27206</td><td>CGATGCCGGGGAGGTCGCCATGG</td><td>1000</td><td></td><td></td><td></td><td></td><td></td><td></td><td></td><td></td><td></td></tr>
<tr><td>Oligo fwd</td><td>CACCCGATGCCGGGGAGGTCGCCA</td><td></td><td></td><td></td><td></td><td></td><td></td><td></td><td></td><td></td><td></td></tr>
<tr><td>Oligo rev</td><td>AAACTGGCGACCTCCCCGGCATCG</td><td></td><td></td><td></td><td></td><td></td><td></td><td></td><td></td><td></td><td></td></tr>
</table>

Chromosome start end strand MM target_seq PAM alignment distance position gene name gene id
GU305914.1 91886 91908 - 0 CGATGCCGGGGAGGTCGCCA TGG ||||||||[||||||||||||]PAM NA Intergenic NA NA

T27207 TGCCGGGGAGGTCGCCATGGCGG 1000
Oligo fwd CACCTGCCGGGGAGGTCGCCATGG
Oligo rev AAACCCATGGCGACCTCCCCGGCA
Chromosome start end strand MM target_seq PAM alignment distance position gene name gene id
GU305914.1 91883 91905 - 0 TGCCGGGGAGGTCGCCATGG CGG ||||||||[||||||||||||]PAM NA Intergenic NA NA

T27208 CCCCCGTCGCTCCCGCCGTCTGG 1000
Oligo fwd CACCCCCCCGTCGCTCCCGCCGTC
Oligo rev AAACGACGGCGGGAGCGACGGGGG
Chromosome start end strand MM target_seq PAM alignment distance position gene name gene id
GU305914.1 91839 91861 - 0 CCCCCGTCGCTCCCGCCGTC TGG ||||||||[||||||||||||]PAM NA Intergenic NA NA

T27209 TGTTCTTCCTCAAAAAGACGCGG 1000
Oligo fwd CACCTGTTCTTCCTCAAAAAGACG
Oligo rev AAACCGTCTTTTTGAGGAAGAACA
Chromosome start end strand MM target_seq PAM alignment distance position gene name gene id
GU305914.1 91815 91837 - 0 TGTTCTTCCTCAAAAAGACG CGG ||||||||[||||||||||||]PAM NA Intergenic NA NA

T27210 GTTCTTCCTCAAAAAGACGCGGG 1000
Oligo fwd CACCGTTCTTCCTCAAAAAGACGC
Oligo rev AAACGCGTCTTTTTGAGGAAGAAC
Chromosome start end strand MM target_seq PAM alignment distance position gene name gene id
GU305914.1 91814 91836 - 0 GTTCTTCCTCAAAAAGACGC GGG ||||||||[||||||||||||]PAM NA Intergenic NA NA

T27211 CAAAAAGACGCGGGAGCTCGCGG 1000
Oligo fwd CACCCAAAAAGACGCGGGAGCTCG
Oligo rev AAACCGAGCTCCCGCGTCTTTTTG
Chromosome start end strand MM target_seq PAM alignment distance position gene name gene id
GU305914.1 91805 91827 - 0 CAAAAAGACGCGGGAGCTCG CGG ||||||||[||||||||||||]PAM NA Intergenic NA NA

T27212 GCGGGAGCTCGCGGACATCGTGG 1000
Oligo fwd CACCGCGGGAGCTCGCGGACATCG
Oligo rev AAACCGATGTCCGCGAGCTCCCGC
Chromosome start end strand MM target_seq PAM alignment distance position gene name gene id
GU305914.1 91796 91818 - 0 GCGGGAGCTCGCGGACATCG TGG ||||||||[||||||||||||]PAM NA Intergenic NA NA

T27213 CGCGATGTCGCTCTGCGACAAGG 1000
Oligo fwd CACCCGCGATGTCGCTCTGCGACA
Oligo rev AAACTGTCGCAGAGCGACATCGCG
Chromosome start end strand MM target_seq PAM alignment distance position gene name gene id
GU305914.1 91772 91794 - 0 CGCGATGTCGCTCTGCGACA AGG ||||||||[||||||||||||]PAM NA Intergenic NA NA

T27214 ACCGTCGACCGCGACTTCTGCGG 1000
Oligo fwd CACCACCGTCGACCGCGACTTCTG
Oligo rev AAACCAGAAGTCGCGGTCGACGGT
Chromosome start end strand MM target_seq PAM alignment distance position gene name gene id
GU305914.1 91711 91733 - 0 ACCGTCGACCGCGACTTCTG CGG ||||||||[||||||||||||]PAM NA Intergenic NA NA

T27215 CGACCGCGACTTCTGCGGCGCGG 1000
Oligo fwd CACCCGACCGCGACTTCTGCGGCG
Oligo rev AAACCGCCGCAGAAGTCGCGGTCG
Chromosome start end strand MM target_seq PAM alignment distance position gene name gene id
GU305914.1 91706 91728 - 0 CGACCGCGACTTCTGCGGCG CGG ||||||||[||||||||||||]PAM NA Intergenic NA NA

T27216 GCGACTTCTGCGGCGCGGTCAGG 1000
Oligo fwd CACCGCGACTTCTGCGGCGCGGTC
Oligo rev AAACGACCGCGCCGCAGAAGTCGC
Chromosome start end strand MM target_seq PAM alignment distance position gene name gene id
GU305914.1 91701 91723 - 0 GCGACTTCTGCGGCGCGGTC AGG ||||||||[||||||||||||]PAM NA Intergenic NA NA

T27217 GACGCCGATGAGCACATACGAGG 1000
Oligo fwd CACCGACGCCGATGAGCACATACG
Oligo rev AAACCGTATGTGCTCATCGGCGTC
Chromosome start end strand MM target_seq PAM alignment distance position gene name gene id
GU305914.1 91679 91701 - 0 GACGCCGATGAGCACATACG AGG ||||||||[||||||||||||]PAM NA Intergenic NA NA

T27218 ACGCCGATGAGCACATACGAGGG 1000
Oligo fwd CACCACGCCGATGAGCACATACGA
Oligo rev AAACTCGTATGTGCTCATCGGCGT
Chromosome start end strand MM target_seq PAM alignment distance position gene name gene id
GU305914.1 91678 91700 - 0 ACGCCGATGAGCACATACGA GGG ||||||||[||||||||||||]PAM NA Intergenic NA NA

T27219 CGCCGATGAGCACATACGAGGGG 1000
Oligo fwd CACCCGCCGATGAGCACATACGAG
Oligo rev AAACCTCGTATGTGCTCATCGGCG
Chromosome start end strand MM target_seq PAM alignment distance position gene name gene id
GU305914.1 91677 91699 - 0 CGCCGATGAGCACATACGAG GGG ||||||||[||||||||||||]PAM NA Intergenic NA NA

T27220 GCCGATGAGCACATACGAGGGGG 1000
Oligo fwd CACCGCCGATGAGCACATACGAGG
Oligo rev AAACCCTCGTATGTGCTCATCGGC
Chromosome start end strand MM target_seq PAM alignment distance position gene name gene id
GU305914.1 91676 91698 - 0 GCCGATGAGCACATACGAGG GGG ||||||||[||||||||||||]PAM NA Intergenic NA NA

T27221 CCGATGAGCACATACGAGGGGGG 1000
Oligo fwd CACCCCGATGAGCACATACGAGGG

<table>
<tr><td>Oligo rev</td><td>AAACCCCTCGTATGTGCTCATCGG</td><td></td><td></td><td></td><td></td><td></td><td></td><td></td><td></td><td></td><td></td><td></td></tr>
<tr><td>Chromosome</td><td>start</td><td></td><td>end</td><td>strand</td><td>MM</td><td>target_seq</td><td>PAM</td><td>alignment</td><td>distance</td><td>position</td><td>gene name</td><td>gene id</td></tr>
<tr><td>GU305914.1</td><td></td><td>91675</td><td>91697</td><td>-</td><td>0</td><td>CCGATGAGCACATACGAGGG</td><td>GGG</td><td>||||||||[||||||||||||]PAM</td><td>NA</td><td>Intergenic</td><td>NA</td><td>NA</td></tr>
<tr><td>T27222</td><td>CGAGATCCTCGACGTCGTCGAGG</td><td></td><td>1000</td><td></td><td></td><td></td><td></td><td></td><td></td><td></td><td></td><td></td></tr>
<tr><td>Oligo fwd</td><td>CACCCGAGATCCTCGACGTCGTCG</td><td></td><td></td><td></td><td></td><td></td><td></td><td></td><td></td><td></td><td></td><td></td></tr>
<tr><td>Oligo rev</td><td>AAACCGACGACGTCGAGGATCTCG</td><td></td><td></td><td></td><td></td><td></td><td></td><td></td><td></td><td></td><td></td><td></td></tr>
<tr><td>Chromosome</td><td>start</td><td></td><td>end</td><td>strand</td><td>MM</td><td>target_seq</td><td>PAM</td><td>alignment</td><td>distance</td><td>position</td><td>gene name</td><td>gene id</td></tr>
<tr><td>GU305914.1</td><td></td><td>91595</td><td>91617</td><td>-</td><td>0</td><td>CGAGATCCTCGACGTCGTCG</td><td>AGG</td><td>||||||||[||||||||||||]PAM</td><td>NA</td><td>Intergenic</td><td>NA</td><td>NA</td></tr>
<tr><td>GU305914.1</td><td></td><td>91589</td><td>91611</td><td>+</td><td>2</td><td>CGTGATCCTCGACGACGTCG</td><td>AGG</td><td>||-|||||[||||||-|||||]PAM</td><td>NA</td><td>Intergenic</td><td>NA</td><td>NA</td></tr>
<tr><td>T27223</td><td>CTCGACGTCGTCGAGGATCACGG</td><td></td><td>1000</td><td></td><td></td><td></td><td></td><td></td><td></td><td></td><td></td><td></td></tr>
<tr><td>Oligo fwd</td><td>CACCCTCGACGTCGTCGAGGATCA</td><td></td><td></td><td></td><td></td><td></td><td></td><td></td><td></td><td></td><td></td><td></td></tr>
<tr><td>Oligo rev</td><td>AAACTGATCCTCGACGACGTCGAG</td><td></td><td></td><td></td><td></td><td></td><td></td><td></td><td></td><td></td><td></td><td></td></tr>
<tr><td>Chromosome</td><td>start</td><td></td><td>end</td><td>strand</td><td>MM</td><td>target_seq</td><td>PAM</td><td>alignment</td><td>distance</td><td>position</td><td>gene name</td><td>gene id</td></tr>
<tr><td>GU305914.1</td><td></td><td>91588</td><td>91610</td><td>-</td><td>0</td><td>CTCGACGTCGTCGAGGATCA</td><td>CGG</td><td>||||||||[||||||||||||]PAM</td><td>NA</td><td>Intergenic</td><td>NA</td><td>NA</td></tr>
<tr><td>T27224</td><td>TGCACCTCTGTGACGATGCCAGG</td><td></td><td>1000</td><td></td><td></td><td></td><td></td><td></td><td></td><td></td><td></td><td></td></tr>
<tr><td>Oligo fwd</td><td>CACCTGCACCTCTGTGACGATGCC</td><td></td><td></td><td></td><td></td><td></td><td></td><td></td><td></td><td></td><td></td><td></td></tr>
<tr><td>Oligo rev</td><td>AAACGGCATCGTCACAGAGGTGCA</td><td></td><td></td><td></td><td></td><td></td><td></td><td></td><td></td><td></td><td></td><td></td></tr>
<tr><td>Chromosome</td><td>start</td><td></td><td>end</td><td>strand</td><td>MM</td><td>target_seq</td><td>PAM</td><td>alignment</td><td>distance</td><td>position</td><td>gene name</td><td>gene id</td></tr>
<tr><td>GU305914.1</td><td></td><td>91557</td><td>91579</td><td>-</td><td>0</td><td>TGCACCTCTGTGACGATGCC</td><td>AGG</td><td>||||||||[||||||||||||]PAM</td><td>NA</td><td>Intergenic</td><td>NA</td><td>NA</td></tr>
<tr><td>T27225</td><td>GACGATGCCAGGCGCCGCTTCGG</td><td></td><td>1000</td><td></td><td></td><td></td><td></td><td></td><td></td><td></td><td></td><td></td></tr>
<tr><td>Oligo fwd</td><td>CACCGACGATGCCAGGCGCCGCTT</td><td></td><td></td><td></td><td></td><td></td><td></td><td></td><td></td><td></td><td></td><td></td></tr>
<tr><td>Oligo rev</td><td>AAACAAGCGGCGCCTGGCATCGTC</td><td></td><td></td><td></td><td></td><td></td><td></td><td></td><td></td><td></td><td></td><td></td></tr>
<tr><td>Chromosome</td><td>start</td><td></td><td>end</td><td>strand</td><td>MM</td><td>target_seq</td><td>PAM</td><td>alignment</td><td>distance</td><td>position</td><td>gene name</td><td>gene id</td></tr>
<tr><td>GU305914.1</td><td></td><td>91546</td><td>91568</td><td>-</td><td>0</td><td>GACGATGCCAGGCGCCGCTT</td><td>CGG</td><td>||||||||[||||||||||||]PAM</td><td>NA</td><td>Intergenic</td><td>NA</td><td>NA</td></tr>
<tr><td>T27226</td><td>CAGGCGCCGCTTCGGCGTCCAGG</td><td></td><td>1000</td><td></td><td></td><td></td><td></td><td></td><td></td><td></td><td></td><td></td></tr>
<tr><td>Oligo fwd</td><td>CACCCAGGCGCCGCTTCGGCGTCC</td><td></td><td></td><td></td><td></td><td></td><td></td><td></td><td></td><td></td><td></td><td></td></tr>
<tr><td>Oligo rev</td><td>AAACGGACGCCGAAGCGGCGCCTG</td><td></td><td></td><td></td><td></td><td></td><td></td><td></td><td></td><td></td><td></td><td></td></tr>
<tr><td>Chromosome</td><td>start</td><td></td><td>end</td><td>strand</td><td>MM</td><td>target_seq</td><td>PAM</td><td>alignment</td><td>distance</td><td>position</td><td>gene name</td><td>gene id</td></tr>
<tr><td>GU305914.1</td><td></td><td>91538</td><td>91560</td><td>-</td><td>0</td><td>CAGGCGCCGCTTCGGCGTCC</td><td>AGG</td><td>||||||||[||||||||||||]PAM</td><td>NA</td><td>Intergenic</td><td>NA</td><td>NA</td></tr>
<tr><td>T27227</td><td>GCTCGCTAATCGAGATCGCACGG</td><td></td><td>1000</td><td></td><td></td><td></td><td></td><td></td><td></td><td></td><td></td><td></td></tr>
<tr><td>Oligo fwd</td><td>CACCGCTCGCTAATCGAGATCGCA</td><td></td><td></td><td></td><td></td><td></td><td></td><td></td><td></td><td></td><td></td><td></td></tr>
<tr><td>Oligo rev</td><td>AAACTGCGATCTCGATTAGCGAGC</td><td></td><td></td><td></td><td></td><td></td><td></td><td></td><td></td><td></td><td></td><td></td></tr>
<tr><td>Chromosome</td><td>start</td><td></td><td>end</td><td>strand</td><td>MM</td><td>target_seq</td><td>PAM</td><td>alignment</td><td>distance</td><td>position</td><td>gene name</td><td>gene id</td></tr>
<tr><td>GU305914.1</td><td></td><td>91505</td><td>91527</td><td>-</td><td>0</td><td>GCTCGCTAATCGAGATCGCA</td><td>CGG</td><td>||||||||[||||||||||||]PAM</td><td>NA</td><td>Intergenic</td><td>NA</td><td>NA</td></tr>
<tr><td>T27228</td><td>GACGTCCTCTGCGACGAGCTAGG</td><td></td><td>1000</td><td></td><td></td><td></td><td></td><td></td><td></td><td></td><td></td><td></td></tr>
<tr><td>Oligo fwd</td><td>CACCGACGTCCTCTGCGACGAGCT</td><td></td><td></td><td></td><td></td><td></td><td></td><td></td><td></td><td></td><td></td><td></td></tr>
<tr><td>Oligo rev</td><td>AAACAGCTCGTCGCAGAGGACGTC</td><td></td><td></td><td></td><td></td><td></td><td></td><td></td><td></td><td></td><td></td><td></td></tr>
<tr><td>Chromosome</td><td>start</td><td></td><td>end</td><td>strand</td><td>MM</td><td>target_seq</td><td>PAM</td><td>alignment</td><td>distance</td><td>position</td><td>gene name</td><td>gene id</td></tr>
<tr><td>GU305914.1</td><td></td><td>91477</td><td>91499</td><td>-</td><td>0</td><td>GACGTCCTCTGCGACGAGCT</td><td>AGG</td><td>||||||||[||||||||||||]PAM</td><td>NA</td><td>Intergenic</td><td>NA</td><td>NA</td></tr>
<tr><td>T27229</td><td>TATCGCCCCCGCCGAGTACACGG</td><td></td><td>1000</td><td></td><td></td><td></td><td></td><td></td><td></td><td></td><td></td><td></td></tr>
<tr><td>Oligo fwd</td><td>CACCTATCGCCCCCGCCGAGTACA</td><td></td><td></td><td></td><td></td><td></td><td></td><td></td><td></td><td></td><td></td><td></td></tr>
<tr><td>Oligo rev</td><td>AAACTGTACTCGGCGGGGGCGATA</td><td></td><td></td><td></td><td></td><td></td><td></td><td></td><td></td><td></td><td></td><td></td></tr>
<tr><td>Chromosome</td><td>start</td><td></td><td>end</td><td>strand</td><td>MM</td><td>target_seq</td><td>PAM</td><td>alignment</td><td>distance</td><td>position</td><td>gene name</td><td>gene id</td></tr>
<tr><td>GU305914.1</td><td></td><td>91454</td><td>91476</td><td>-</td><td>0</td><td>TATCGCCCCCGCCGAGTACA</td><td>CGG</td><td>||||||||[||||||||||||]PAM</td><td>NA</td><td>Intergenic</td><td>NA</td><td>NA</td></tr>
<tr><td>T27230</td><td>ATCGCCCCCGCCGAGTACACGGG</td><td></td><td>1000</td><td></td><td></td><td></td><td></td><td></td><td></td><td></td><td></td><td></td></tr>
<tr><td>Oligo fwd</td><td>CACCATCGCCCCCGCCGAGTACAC</td><td></td><td></td><td></td><td></td><td></td><td></td><td></td><td></td><td></td><td></td><td></td></tr>
<tr><td>Oligo rev</td><td>AAACGTGTACTCGGCGGGGGCGAT</td><td></td><td></td><td></td><td></td><td></td><td></td><td></td><td></td><td></td><td></td><td></td></tr>
<tr><td>Chromosome</td><td>start</td><td></td><td>end</td><td>strand</td><td>MM</td><td>target_seq</td><td>PAM</td><td>alignment</td><td>distance</td><td>position</td><td>gene name</td><td>gene id</td></tr>
<tr><td>GU305914.1</td><td></td><td>91453</td><td>91475</td><td>-</td><td>0</td><td>ATCGCCCCCGCCGAGTACAC</td><td>GGG</td><td>||||||||[||||||||||||]PAM</td><td>NA</td><td>Intergenic</td><td>NA</td><td>NA</td></tr>
<tr><td>T27231</td><td>TCGCCCCCGCCGAGTACACGGGG</td><td></td><td>1000</td><td></td><td></td><td></td><td></td><td></td><td></td><td></td><td></td><td></td></tr>
<tr><td>Oligo fwd</td><td>CACCTCGCCCCCGCCGAGTACACG</td><td></td><td></td><td></td><td></td><td></td><td></td><td></td><td></td><td></td><td></td><td></td></tr>
<tr><td>Oligo rev</td><td>AAACCGTGTACTCGGCGGGGGCGA</td><td></td><td></td><td></td><td></td><td></td><td></td><td></td><td></td><td></td><td></td><td></td></tr>
<tr><td>Chromosome</td><td>start</td><td></td><td>end</td><td>strand</td><td>MM</td><td>target_seq</td><td>PAM</td><td>alignment</td><td>distance</td><td>position</td><td>gene name</td><td>gene id</td></tr>
<tr><td>GU305914.1</td><td></td><td>91452</td><td>91474</td><td>-</td><td>0</td><td>TCGCCCCCGCCGAGTACACG</td><td>GGG</td><td>||||||||[||||||||||||]PAM</td><td>NA</td><td>Intergenic</td><td>NA</td><td>NA</td></tr>
<tr><td>T27232</td><td>TACACGGGGCACGTCGTGTGCGG</td><td></td><td>1000</td><td></td><td></td><td></td><td></td><td></td><td></td><td></td><td></td><td></td></tr>
<tr><td>Oligo fwd</td><td>CACCTACACGGGGCACGTCGTGTG</td><td></td><td></td><td></td><td></td><td></td><td></td><td></td><td></td><td></td><td></td><td></td></tr>
<tr><td>Oligo rev</td><td>AAACCACACGACGTGCCCCGTGTA</td><td></td><td></td><td></td><td></td><td></td><td></td><td></td><td></td><td></td><td></td><td></td></tr>
<tr><td>Chromosome</td><td>start</td><td></td><td>end</td><td>strand</td><td>MM</td><td>target_seq</td><td>PAM</td><td>alignment</td><td>distance</td><td>position</td><td>gene name</td><td>gene id</td></tr>
<tr><td>GU305914.1</td><td></td><td>91438</td><td>91460</td><td>-</td><td>0</td><td>TACACGGGGCACGTCGTGTG</td><td>CGG</td><td>||||||||[||||||||||||]PAM</td><td>NA</td><td>Intergenic</td><td>NA</td><td>NA</td></tr>
<tr><td>GU305914.1</td><td></td><td>144721</td><td>144743</td><td>-</td><td>4</td><td>AAGACGGCGCACGTCGTGTT</td><td>CGG</td><td>-|-||||-[|||||||||||-]PAM</td><td>NA</td><td>Intergenic</td><td>NA</td><td>NA</td></tr>
<tr><td>T27233</td><td>GGGCACGTCGTGTGCGGAAACGG</td><td></td><td>1000</td><td></td><td></td><td></td><td></td><td></td><td></td><td></td><td></td><td></td></tr>
<tr><td>Oligo fwd</td><td>CACCGGGCACGTCGTGTGCGGAAA</td><td></td><td></td><td></td><td></td><td></td><td></td><td></td><td></td><td></td><td></td><td></td></tr>
<tr><td>Oligo rev</td><td>AAACTTTCCGCACACGACGTGCCC</td><td></td><td></td><td></td><td></td><td></td><td></td><td></td><td></td><td></td><td></td><td></td></tr>
<tr><td>Chromosome</td><td>start</td><td></td><td>end</td><td>strand</td><td>MM</td><td>target_seq</td><td>PAM</td><td>alignment</td><td>distance</td><td>position</td><td>gene name</td><td>gene id</td></tr>
<tr><td>GU305914.1</td><td></td><td>91432</td><td>91454</td><td>-</td><td>0</td><td>GGGCACGTCGTGTGCGGAAA</td><td>CGG</td><td>||||||||[||||||||||||]PAM</td><td>NA</td><td>Intergenic</td><td>NA</td><td>NA</td></tr>
<tr><td>T27234</td><td>CGTGTGCGGAAACGGCCTCAAGG</td><td></td><td>1000</td><td></td><td></td><td></td><td></td><td></td><td></td><td></td><td></td><td></td></tr>
<tr><td>Oligo fwd</td><td>CACCCGTGTGCGGAAACGGCCTCA</td><td></td><td></td><td></td><td></td><td></td><td></td><td></td><td></td><td></td><td></td><td></td></tr>
<tr><td>Oligo rev</td><td>AAACTGAGGCCGTTTCCGCACACG</td><td></td><td></td><td></td><td></td><td></td><td></td><td></td><td></td><td></td><td></td><td></td></tr>
<tr><td>Chromosome</td><td>start</td><td></td><td>end</td><td>strand</td><td>MM</td><td>target_seq</td><td>PAM</td><td>alignment</td><td>distance</td><td>position</td><td>gene name</td><td>gene id</td></tr>
<tr><td>GU305914.1</td><td></td><td>91424</td><td>91446</td><td>-</td><td>0</td><td>CGTGTGCGGAAACGGCCTCA</td><td>AGG</td><td>||||||||[||||||||||||]PAM</td><td>NA</td><td>Intergenic</td><td>NA</td><td>NA</td></tr>
<tr><td>T27235</td><td>CCTCAAGGAGCTCCTCTACGCGG</td><td></td><td>1000</td><td></td><td></td><td></td><td></td><td></td><td></td><td></td><td></td><td></td></tr>
<tr><td>Oligo fwd</td><td>CACCCCTCAAGGAGCTCCTCTACG</td><td></td><td></td><td></td><td></td><td></td><td></td><td></td><td></td><td></td><td></td><td></td></tr>
<tr><td>Oligo rev</td><td>AAACCGTAGAGGAGCTCCTTGAGG</td><td></td><td></td><td></td><td></td><td></td><td></td><td></td><td></td><td></td><td></td><td></td></tr>
<tr><td>Chromosome</td><td>start</td><td></td><td>end</td><td>strand</td><td>MM</td><td>target_seq</td><td>PAM</td><td>alignment</td><td>distance</td><td>position</td><td>gene name</td><td>gene id</td></tr>
<tr><td>GU305914.1</td><td></td><td>91409</td><td>91431</td><td>-</td><td>0</td><td>CCTCAAGGAGCTCCTCTACG</td><td>CGG</td><td>||||||||[||||||||||||]PAM</td><td>NA</td><td>Intergenic</td><td>NA</td><td>NA</td></tr>
</table>

T27236 CTCAAGGAGCTCCTCTACGCGGG 1000
Oligo fwd CACCCTCAAGGAGCTCCTCTACGC
Oligo rev AAACGCGTAGAGGAGCTCCTTGAG
Chromosome start end strand MM target_seq PAM alignment distance position gene name gene id
GU305914.1 91408 91430 - 0 CTCAAGGAGCTCCTCTACGC GGG ||||||||[||||||||||||]PAM NA Intergenic NA NA

T27237 TCAAGGAGCTCCTCTACGCGGGG 1000
Oligo fwd CACCTCAAGGAGCTCCTCTACGCG
Oligo rev AAACCGCGTAGAGGAGCTCCTTGA
Chromosome start end strand MM target_seq PAM alignment distance position gene name gene id
GU305914.1 91407 91429 - 0 TCAAGGAGCTCCTCTACGCG GGG ||||||||[||||||||||||]PAM NA Intergenic NA NA

T27238 GCTCATCCCCTGTCCCGAAGAGG 1000
Oligo fwd CACCGCTCATCCCCTGTCCCGAAG
Oligo rev AAACCTTCGGGACAGGGGATGAGC
Chromosome start end strand MM target_seq PAM alignment distance position gene name gene id
GU305914.1 91382 91404 - 0 GCTCATCCCCTGTCCCGAAG AGG ||||||||[||||||||||||]PAM NA Intergenic NA NA

T27239 CTGTCCCGAAGAGGCCGTGAAGG 1000
Oligo fwd CACCCTGTCCCGAAGAGGCCGTGA
Oligo rev AAACTCACGGCCTCTTCGGGACAG
Chromosome start end strand MM target_seq PAM alignment distance position gene name gene id
GU305914.1 91373 91395 - 0 CTGTCCCGAAGAGGCCGTGA AGG ||||||||[||||||||||||]PAM NA Intergenic NA NA

T27240 CGAAGAGGCCGTGAAGGTGCAGG 1000
Oligo fwd CACCCGAAGAGGCCGTGAAGGTGC
Oligo rev AAACGCACCTTCACGGCCTCTTCG
Chromosome start end strand MM target_seq PAM alignment distance position gene name gene id
GU305914.1 91367 91389 - 0 CGAAGAGGCCGTGAAGGTGC AGG ||||||||[||||||||||||]PAM NA Intergenic NA NA

T27241 GAGGCCGTGAAGGTGCAGGTCGG 1000
Oligo fwd CACCGAGGCCGTGAAGGTGCAGGT
Oligo rev AAACACCTGCACCTTCACGGCCTC
Chromosome start end strand MM target_seq PAM alignment distance position gene name gene id
GU305914.1 91363 91385 - 0 GAGGCCGTGAAGGTGCAGGT CGG ||||||||[||||||||||||]PAM NA Intergenic NA NA

T27242 CGTGAAGGTGCAGGTCGGCGCGG 1000
Oligo fwd CACCCGTGAAGGTGCAGGTCGGCG
Oligo rev AAACCGCCGACCTGCACCTTCACG
Chromosome start end strand MM target_seq PAM alignment distance position gene name gene id
GU305914.1 91358 91380 - 0 CGTGAAGGTGCAGGTCGGCG CGG ||||||||[||||||||||||]PAM NA Intergenic NA NA
GU305914.1 75733 75755 + 4 GGTGGAAGTGGAGGTCGGCG AGG -|||-|-|[||-|||||||||]PAM NA Intergenic NA NA

T27243 GTGCAGGTCGGCGCGGTCGATGG 1000
Oligo fwd CACCGTGCAGGTCGGCGCGGTCGA
Oligo rev AAACTCGACCGCGCCGACCTGCAC
Chromosome start end strand MM target_seq PAM alignment distance position gene name gene id
GU305914.1 91351 91373 - 0 GTGCAGGTCGGCGCGGTCGA TGG ||||||||[||||||||||||]PAM NA Intergenic NA NA

T27244 CGGCGCGGTCGATGGCGTCAAGG 1000
Oligo fwd CACCCGGCGCGGTCGATGGCGTCA
Oligo rev AAACTGACGCCATCGACCGCGCCG
Chromosome start end strand MM target_seq PAM alignment distance position gene name gene id
GU305914.1 91343 91365 - 0 CGGCGCGGTCGATGGCGTCA AGG ||||||||[||||||||||||]PAM NA Intergenic NA NA

T27245 TATCCCTACACTCTCTTCTCCGG 1000
Oligo fwd CACCTATCCCTACACTCTCTTCTC
Oligo rev AAACGAGAAGAGAGTGTAGGGATA
Chromosome start end strand MM target_seq PAM alignment distance position gene name gene id
GU305914.1 91312 91334 - 0 TATCCCTACACTCTCTTCTC CGG ||||||||[||||||||||||]PAM NA Intergenic NA NA

T27246 CCCTACACTCTCTTCTCCGGCGG 1000
Oligo fwd CACCCCCTACACTCTCTTCTCCGG
Oligo rev AAACCCGGAGAAGAGAGTGTAGGG
Chromosome start end strand MM target_seq PAM alignment distance position gene name gene id
GU305914.1 91309 91331 - 0 CCCTACACTCTCTTCTCCGG CGG ||||||||[||||||||||||]PAM NA Intergenic NA NA

T27247 GCCGATGCCGCGCACGCCGACGG 1000
Oligo fwd CACCGCCGATGCCGCGCACGCCGA
Oligo rev AAACTCGGCGTGCGCGGCATCGGC
Chromosome start end strand MM target_seq PAM alignment distance position gene name gene id
GU305914.1 91285 91307 - 0 GCCGATGCCGCGCACGCCGA CGG ||||||||[||||||||||||]PAM NA Intergenic NA NA

T27248 GCACGCCGACGGCGCTTCCGCGG 1000
Oligo fwd CACCGCACGCCGACGGCGCTTCCG
Oligo rev AAACCGGAAGCGCCGTCGGCGTGC
Chromosome start end strand MM target_seq PAM alignment distance position gene name gene id
GU305914.1 91274 91296 - 0 GCACGCCGACGGCGCTTCCG CGG ||||||||[||||||||||||]PAM NA Intergenic NA NA

T27249 CCGTCGCCTGCGACGATCCGTGG 1000
Oligo fwd CACCCCGTCGCCTGCGACGATCCG
Oligo rev AAACCGGATCGTCGCAGGCGACGG
Chromosome start end strand MM target_seq PAM alignment distance position gene name gene id
GU305914.1 91251 91273 - 0 CCGTCGCCTGCGACGATCCG TGG ||||||||[||||||||||||]PAM NA Intergenic NA NA

T27250 CGTCGCCTGCGACGATCCGTGGG 1000
Oligo fwd CACCCGTCGCCTGCGACGATCCGT
Oligo rev AAACACGGATCGTCGCAGGCGACG

Chromosome start end strand MM target_seq PAM alignment distance position gene name gene id
GU305914.1 91250 91272 - 0 CGTCGCCTGCGACGATCCGT GGG ||||||||[||||||||||||]PAM NA Intergenic NA NA

T27251 GATCCGTGGGTCCTCGAGCACGG 1000
Oligo fwd CACCGATCCGTGGGTCCTCGAGCA
Oligo rev AAACTGCTCGAGGACCCACGGATC
Chromosome start end strand MM target_seq PAM alignment distance position gene name gene id
GU305914.1 91237 91259 - 0 GATCCGTGGGTCCTCGAGCA CGG ||||||||[||||||||||||]PAM NA Intergenic NA NA

T27252 ATCCGTGGGTCCTCGAGCACGGG 1000
Oligo fwd CACCATCCGTGGGTCCTCGAGCAC
Oligo rev AAACGTGCTCGAGGACCCACGGAT
Chromosome start end strand MM target_seq PAM alignment distance position gene name gene id
GU305914.1 91236 91258 - 0 ATCCGTGGGTCCTCGAGCAC GGG ||||||||[||||||||||||]PAM NA Intergenic NA NA

T27253 CTACGACCCCGCGCTCAGCGAGG 1000
Oligo fwd CACCCTACGACCCCGCGCTCAGCG
Oligo rev AAACCGCTGAGCGCGGGGTCGTAG
Chromosome start end strand MM target_seq PAM alignment distance position gene name gene id
GU305914.1 91211 91233 - 0 CTACGACCCCGCGCTCAGCG AGG ||||||||[||||||||||||]PAM NA Intergenic NA NA

T27254 TCTACTTCATGTTCACCTCCTGG 1000
Oligo fwd CACCTCTACTTCATGTTCACCTCC
Oligo rev AAACGGAGGTGAACATGAAGTAGA
Chromosome start end strand MM target_seq PAM alignment distance position gene name gene id
GU305914.1 91182 91204 - 0 TCTACTTCATGTTCACCTCC TGG ||||||||[||||||||||||]PAM NA Intergenic NA NA

T27255 CTACTTCATGTTCACCTCCTGGG 1000
Oligo fwd CACCCTACTTCATGTTCACCTCCT
Oligo rev AAACAGGAGGTGAACATGAAGTAG
Chromosome start end strand MM target_seq PAM alignment distance position gene name gene id
GU305914.1 91181 91203 - 0 CTACTTCATGTTCACCTCCT GGG ||||||||[||||||||||||]PAM NA Intergenic NA NA

T27256 TACTTCATGTTCACCTCCTGGGG 1000
Oligo fwd CACCTACTTCATGTTCACCTCCTG
Oligo rev AAACCAGGAGGTGAACATGAAGTA
Chromosome start end strand MM target_seq PAM alignment distance position gene name gene id
GU305914.1 91180 91202 - 0 TACTTCATGTTCACCTCCTG GGG ||||||||[||||||||||||]PAM NA Intergenic NA NA

T27257 CGAGACCAGTCGCCTCATCGAGG 1000
Oligo fwd CACCCGAGACCAGTCGCCTCATCG
Oligo rev AAACCGATGAGGCGACTGGTCTCG
Chromosome start end strand MM target_seq PAM alignment distance position gene name gene id
GU305914.1 91139 91161 - 0 CGAGACCAGTCGCCTCATCG AGG ||||||||[||||||||||||]PAM NA Intergenic NA NA

T27258 ACCAGTCGCCTCATCGAGGCCGG 1000
Oligo fwd CACCACCAGTCGCCTCATCGAGGC
Oligo rev AAACGCCTCGATGAGGCGACTGGT
Chromosome start end strand MM target_seq PAM alignment distance position gene name gene id
GU305914.1 91135 91157 - 0 ACCAGTCGCCTCATCGAGGC CGG ||||||||[||||||||||||]PAM NA Intergenic NA NA

T27259 CGGTCTGCAGCAGTTCGTCGAGG 1000
Oligo fwd CACCCGGTCTGCAGCAGTTCGTCG
Oligo rev AAACCGACGAACTGCTGCAGACCG
Chromosome start end strand MM target_seq PAM alignment distance position gene name gene id
GU305914.1 91115 91137 - 0 CGGTCTGCAGCAGTTCGTCG AGG ||||||||[||||||||||||]PAM NA Intergenic NA NA

T27260 CGTCGAGGACACGCAGCAGACGG 1000
Oligo fwd CACCCGTCGAGGACACGCAGCAGA
Oligo rev AAACTCTGCTGCGTGTCCTCGACG
Chromosome start end strand MM target_seq PAM alignment distance position gene name gene id
GU305914.1 91100 91122 - 0 CGTCGAGGACACGCAGCAGA CGG ||||||||[||||||||||||]PAM NA Intergenic NA NA

T27261 ACGCCGTTCAAGAAGTACCACGG 1000
Oligo fwd CACCACGCCGTTCAAGAAGTACCA
Oligo rev AAACTGGTACTTCTTGAACGGCGT
Chromosome start end strand MM target_seq PAM alignment distance position gene name gene id
GU305914.1 91069 91091 - 0 ACGCCGTTCAAGAAGTACCA CGG ||||||||[||||||||||||]PAM NA Intergenic NA NA

T27262 AGAAGCTCACCGCCGTCGAGAGG 1000
Oligo fwd CACCAGAAGCTCACCGCCGTCGAG
Oligo rev AAACCTCGACGGCGGTGAGCTTCT
Chromosome start end strand MM target_seq PAM alignment distance position gene name gene id
GU305914.1 91035 91057 - 0 AGAAGCTCACCGCCGTCGAG AGG ||||||||[||||||||||||]PAM NA Intergenic NA NA

T27263 GAAGCTCACCGCCGTCGAGAGGG 1000
Oligo fwd CACCGAAGCTCACCGCCGTCGAGA
Oligo rev AAACTCTCGACGGCGGTGAGCTTC
Chromosome start end strand MM target_seq PAM alignment distance position gene name gene id
GU305914.1 91034 91056 - 0 GAAGCTCACCGCCGTCGAGA GGG ||||||||[||||||||||||]PAM NA Intergenic NA NA
GU305914.1 138477 138499 + 4 GGCGCTCGCCGCCGTCGAGG AGG |--||||-[|||||||||||-]PAM NA Intergenic NA NA

T27264 CGAGAGGGACCAGCTCATGACGG 1000
Oligo fwd CACCCGAGAGGGACCAGCTCATGA
Oligo rev AAACTCATGAGCTGGTCCCTCTCG
Chromosome start end strand MM target_seq PAM alignment distance position gene name gene id
GU305914.1 91019 91041 - 0 CGAGAGGGACCAGCTCATGA CGG ||||||||[||||||||||||]PAM NA Intergenic NA NA

T27265 CCAGCTCATGACGGTCGACGCGG 1000

<table>
<tr><td>Oligo fwd</td><td>CACCCCAGCTCATGACGGTCGACG</td><td></td><td></td><td></td><td></td><td></td><td></td><td></td><td></td><td></td><td></td><td></td></tr>
<tr><td>Oligo rev</td><td>AAACCGTCGACCGTCATGAGCTGG</td><td></td><td></td><td></td><td></td><td></td><td></td><td></td><td></td><td></td><td></td><td></td></tr>
<tr><td>Chromosome</td><td>start</td><td></td><td>end</td><td>strand</td><td>MM</td><td>target_seq</td><td>PAM</td><td>alignment</td><td>distance</td><td>position</td><td>gene name</td><td>gene id</td></tr>
<tr><td>GU305914.1</td><td></td><td>91010</td><td>91032</td><td>-</td><td>0</td><td>CCAGCTCATGACGGTCGACG</td><td>CGG</td><td>||||||||[||||||||||||]PAM</td><td>NA</td><td>Intergenic</td><td>NA</td><td>NA</td></tr>
<tr><td></td><td></td><td></td><td></td><td></td><td></td><td></td><td></td><td></td><td></td><td></td><td></td><td></td></tr>
<tr><td>T27266</td><td>TTCCGTCTACGAATTCAGCACGG</td><td></td><td>1000</td><td></td><td></td><td></td><td></td><td></td><td></td><td></td><td></td><td></td></tr>
<tr><td>Oligo fwd</td><td>CACCTTCCGTCTACGAATTCAGCA</td><td></td><td></td><td></td><td></td><td></td><td></td><td></td><td></td><td></td><td></td><td></td></tr>
<tr><td>Oligo rev</td><td>AAACTGCTGAATTCGTAGACGGAA</td><td></td><td></td><td></td><td></td><td></td><td></td><td></td><td></td><td></td><td></td><td></td></tr>
<tr><td>Chromosome</td><td>start</td><td></td><td>end</td><td>strand</td><td>MM</td><td>target_seq</td><td>PAM</td><td>alignment</td><td>distance</td><td>position</td><td>gene name</td><td>gene id</td></tr>
<tr><td>GU305914.1</td><td></td><td>90944</td><td>90966</td><td>-</td><td>0</td><td>TTCCGTCTACGAATTCAGCA</td><td>CGG</td><td>||||||||[||||||||||||]PAM</td><td>NA</td><td>Intergenic</td><td>NA</td><td>NA</td></tr>
<tr><td></td><td></td><td></td><td></td><td></td><td></td><td></td><td></td><td></td><td></td><td></td><td></td><td></td></tr>
<tr><td>T27267</td><td>CGGCCTCCAACTTTCTCGAATGG</td><td></td><td>1000</td><td></td><td></td><td></td><td></td><td></td><td></td><td></td><td></td><td></td></tr>
<tr><td>Oligo fwd</td><td>CACCCGGCCTCCAACTTTCTCGAA</td><td></td><td></td><td></td><td></td><td></td><td></td><td></td><td></td><td></td><td></td><td></td></tr>
<tr><td>Oligo rev</td><td>AAACTTCGAGAAAGTTGGAGGCCG</td><td></td><td></td><td></td><td></td><td></td><td></td><td></td><td></td><td></td><td></td><td></td></tr>
<tr><td>Chromosome</td><td>start</td><td></td><td>end</td><td>strand</td><td>MM</td><td>target_seq</td><td>PAM</td><td>alignment</td><td>distance</td><td>position</td><td>gene name</td><td>gene id</td></tr>
<tr><td>GU305914.1</td><td></td><td>90924</td><td>90946</td><td>-</td><td>0</td><td>CGGCCTCCAACTTTCTCGAA</td><td>TGG</td><td>||||||||[||||||||||||]PAM</td><td>NA</td><td>Intergenic</td><td>NA</td><td>NA</td></tr>
<tr><td></td><td></td><td></td><td></td><td></td><td></td><td></td><td></td><td></td><td></td><td></td><td></td><td></td></tr>
<tr><td>T27268</td><td>CAAGACCCACGCCGATCTGCTGG</td><td></td><td>1000</td><td></td><td></td><td></td><td></td><td></td><td></td><td></td><td></td><td></td></tr>
<tr><td>Oligo fwd</td><td>CACCCAAGACCCACGCCGATCTGC</td><td></td><td></td><td></td><td></td><td></td><td></td><td></td><td></td><td></td><td></td><td></td></tr>
<tr><td>Oligo rev</td><td>AAACGCAGATCGGCGTGGGTCTTG</td><td></td><td></td><td></td><td></td><td></td><td></td><td></td><td></td><td></td><td></td><td></td></tr>
<tr><td>Chromosome</td><td>start</td><td></td><td>end</td><td>strand</td><td>MM</td><td>target_seq</td><td>PAM</td><td>alignment</td><td>distance</td><td>position</td><td>gene name</td><td>gene id</td></tr>
<tr><td>GU305914.1</td><td></td><td>90884</td><td>90906</td><td>-</td><td>0</td><td>CAAGACCCACGCCGATCTGC</td><td>TGG</td><td>||||||||[||||||||||||]PAM</td><td>NA</td><td>Intergenic</td><td>NA</td><td>NA</td></tr>
<tr><td></td><td></td><td></td><td></td><td></td><td></td><td></td><td></td><td></td><td></td><td></td><td></td><td></td></tr>
<tr><td>T27269</td><td>CCGATCTGCTGGACAACCTGCGG</td><td></td><td>1000</td><td></td><td></td><td></td><td></td><td></td><td></td><td></td><td></td><td></td></tr>
<tr><td>Oligo fwd</td><td>CACCCCGATCTGCTGGACAACCTG</td><td></td><td></td><td></td><td></td><td></td><td></td><td></td><td></td><td></td><td></td><td></td></tr>
<tr><td>Oligo rev</td><td>AAACCAGGTTGTCCAGCAGATCGG</td><td></td><td></td><td></td><td></td><td></td><td></td><td></td><td></td><td></td><td></td><td></td></tr>
<tr><td>Chromosome</td><td>start</td><td></td><td>end</td><td>strand</td><td>MM</td><td>target_seq</td><td>PAM</td><td>alignment</td><td>distance</td><td>position</td><td>gene name</td><td>gene id</td></tr>
<tr><td>GU305914.1</td><td></td><td>90873</td><td>90895</td><td>-</td><td>0</td><td>CCGATCTGCTGGACAACCTG</td><td>CGG</td><td>||||||||[||||||||||||]PAM</td><td>NA</td><td>Intergenic</td><td>NA</td><td>NA</td></tr>
<tr><td></td><td></td><td></td><td></td><td></td><td></td><td></td><td></td><td></td><td></td><td></td><td></td><td></td></tr>
<tr><td>T27270</td><td>CGATCTGCTGGACAACCTGCGGG</td><td></td><td>1000</td><td></td><td></td><td></td><td></td><td></td><td></td><td></td><td></td><td></td></tr>
<tr><td>Oligo fwd</td><td>CACCCGATCTGCTGGACAACCTGC</td><td></td><td></td><td></td><td></td><td></td><td></td><td></td><td></td><td></td><td></td><td></td></tr>
<tr><td>Oligo rev</td><td>AAACGCAGGTTGTCCAGCAGATCG</td><td></td><td></td><td></td><td></td><td></td><td></td><td></td><td></td><td></td><td></td><td></td></tr>
<tr><td>Chromosome</td><td>start</td><td></td><td>end</td><td>strand</td><td>MM</td><td>target_seq</td><td>PAM</td><td>alignment</td><td>distance</td><td>position</td><td>gene name</td><td>gene id</td></tr>
<tr><td>GU305914.1</td><td></td><td>90872</td><td>90894</td><td>-</td><td>0</td><td>CGATCTGCTGGACAACCTGC</td><td>GGG</td><td>||||||||[||||||||||||]PAM</td><td>NA</td><td>Intergenic</td><td>NA</td><td>NA</td></tr>
<tr><td>GU305914.1</td><td></td><td>22673</td><td>22695</td><td>+</td><td>4</td><td>CATTCTGCTGTACGACCTGC</td><td>CGG</td><td>|--|||||[||-||-||||||]PAM</td><td>NA</td><td>Intergenic</td><td>NA</td><td>NA</td></tr>
<tr><td></td><td></td><td></td><td></td><td></td><td></td><td></td><td></td><td></td><td></td><td></td><td></td><td></td></tr>
<tr><td>T27271</td><td>ACTCGATCCTCTACCAGACGCGG</td><td></td><td>1000</td><td></td><td></td><td></td><td></td><td></td><td></td><td></td><td></td><td></td></tr>
<tr><td>Oligo fwd</td><td>CACCACTCGATCCTCTACCAGACG</td><td></td><td></td><td></td><td></td><td></td><td></td><td></td><td></td><td></td><td></td><td></td></tr>
<tr><td>Oligo rev</td><td>AAACCGTCTGGTAGAGGATCGAGT</td><td></td><td></td><td></td><td></td><td></td><td></td><td></td><td></td><td></td><td></td><td></td></tr>
<tr><td>Chromosome</td><td>start</td><td></td><td>end</td><td>strand</td><td>MM</td><td>target_seq</td><td>PAM</td><td>alignment</td><td>distance</td><td>position</td><td>gene name</td><td>gene id</td></tr>
<tr><td>GU305914.1</td><td></td><td>90798</td><td>90820</td><td>-</td><td>0</td><td>ACTCGATCCTCTACCAGACG</td><td>CGG</td><td>||||||||[||||||||||||]PAM</td><td>NA</td><td>Intergenic</td><td>NA</td><td>NA</td></tr>
<tr><td></td><td></td><td></td><td></td><td></td><td></td><td></td><td></td><td></td><td></td><td></td><td></td><td></td></tr>
<tr><td>T27272</td><td>ATCGTGTTCGTGCCCAGCGCCGG</td><td></td><td>1000</td><td></td><td></td><td></td><td></td><td></td><td></td><td></td><td></td><td></td></tr>
<tr><td>Oligo fwd</td><td>CACCATCGTGTTCGTGCCCAGCGC</td><td></td><td></td><td></td><td></td><td></td><td></td><td></td><td></td><td></td><td></td><td></td></tr>
<tr><td>Oligo rev</td><td>AAACGCGCTGGGCACGAACACGAT</td><td></td><td></td><td></td><td></td><td></td><td></td><td></td><td></td><td></td><td></td><td></td></tr>
<tr><td>Chromosome</td><td>start</td><td></td><td>end</td><td>strand</td><td>MM</td><td>target_seq</td><td>PAM</td><td>alignment</td><td>distance</td><td>position</td><td>gene name</td><td>gene id</td></tr>
<tr><td>GU305914.1</td><td></td><td>90775</td><td>90797</td><td>-</td><td>0</td><td>ATCGTGTTCGTGCCCAGCGC</td><td>CGG</td><td>||||||||[||||||||||||]PAM</td><td>NA</td><td>Intergenic</td><td>NA</td><td>NA</td></tr>
<tr><td></td><td></td><td></td><td></td><td></td><td></td><td></td><td></td><td></td><td></td><td></td><td></td><td></td></tr>
<tr><td>T27273</td><td>GTTCGTGCCCAGCGCCGGCAAGG</td><td></td><td>1000</td><td></td><td></td><td></td><td></td><td></td><td></td><td></td><td></td><td></td></tr>
<tr><td>Oligo fwd</td><td>CACCGTTCGTGCCCAGCGCCGGCA</td><td></td><td></td><td></td><td></td><td></td><td></td><td></td><td></td><td></td><td></td><td></td></tr>
<tr><td>Oligo rev</td><td>AAACTGCCGGCGCTGGGCACGAAC</td><td></td><td></td><td></td><td></td><td></td><td></td><td></td><td></td><td></td><td></td><td></td></tr>
<tr><td>Chromosome</td><td>start</td><td></td><td>end</td><td>strand</td><td>MM</td><td>target_seq</td><td>PAM</td><td>alignment</td><td>distance</td><td>position</td><td>gene name</td><td>gene id</td></tr>
<tr><td>GU305914.1</td><td></td><td>90770</td><td>90792</td><td>-</td><td>0</td><td>GTTCGTGCCCAGCGCCGGCA</td><td>AGG</td><td>||||||||[||||||||||||]PAM</td><td>NA</td><td>Intergenic</td><td>NA</td><td>NA</td></tr>
<tr><td></td><td></td><td></td><td></td><td></td><td></td><td></td><td></td><td></td><td></td><td></td><td></td><td></td></tr>
<tr><td>T27274</td><td>TTCGTGCCCAGCGCCGGCAAGGG</td><td></td><td>1000</td><td></td><td></td><td></td><td></td><td></td><td></td><td></td><td></td><td></td></tr>
<tr><td>Oligo fwd</td><td>CACCTTCGTGCCCAGCGCCGGCAA</td><td></td><td></td><td></td><td></td><td></td><td></td><td></td><td></td><td></td><td></td><td></td></tr>
<tr><td>Oligo rev</td><td>AAACTTGCCGGCGCTGGGCACGAA</td><td></td><td></td><td></td><td></td><td></td><td></td><td></td><td></td><td></td><td></td><td></td></tr>
<tr><td>Chromosome</td><td>start</td><td></td><td>end</td><td>strand</td><td>MM</td><td>target_seq</td><td>PAM</td><td>alignment</td><td>distance</td><td>position</td><td>gene name</td><td>gene id</td></tr>
<tr><td>GU305914.1</td><td></td><td>90769</td><td>90791</td><td>-</td><td>0</td><td>TTCGTGCCCAGCGCCGGCAA</td><td>GGG</td><td>||||||||[||||||||||||]PAM</td><td>NA</td><td>Intergenic</td><td>NA</td><td>NA</td></tr>
<tr><td></td><td></td><td></td><td></td><td></td><td></td><td></td><td></td><td></td><td></td><td></td><td></td><td></td></tr>
<tr><td>T27275</td><td>TCGTGCCCAGCGCCGGCAAGGGG</td><td></td><td>1000</td><td></td><td></td><td></td><td></td><td></td><td></td><td></td><td></td><td></td></tr>
<tr><td>Oligo fwd</td><td>CACCTCGTGCCCAGCGCCGGCAAG</td><td></td><td></td><td></td><td></td><td></td><td></td><td></td><td></td><td></td><td></td><td></td></tr>
<tr><td>Oligo rev</td><td>AAACCTTGCCGGCGCTGGGCACGA</td><td></td><td></td><td></td><td></td><td></td><td></td><td></td><td></td><td></td><td></td><td></td></tr>
<tr><td>Chromosome</td><td>start</td><td></td><td>end</td><td>strand</td><td>MM</td><td>target_seq</td><td>PAM</td><td>alignment</td><td>distance</td><td>position</td><td>gene name</td><td>gene id</td></tr>
<tr><td>GU305914.1</td><td></td><td>90768</td><td>90790</td><td>-</td><td>0</td><td>TCGTGCCCAGCGCCGGCAAG</td><td>GGG</td><td>||||||||[||||||||||||]PAM</td><td>NA</td><td>Intergenic</td><td>NA</td><td>NA</td></tr>
<tr><td></td><td></td><td></td><td></td><td></td><td></td><td></td><td></td><td></td><td></td><td></td><td></td><td></td></tr>
<tr><td>T27276</td><td>CGTGCCCAGCGCCGGCAAGGGGG</td><td></td><td>1000</td><td></td><td></td><td></td><td></td><td></td><td></td><td></td><td></td><td></td></tr>
<tr><td>Oligo fwd</td><td>CACCCGTGCCCAGCGCCGGCAAGG</td><td></td><td></td><td></td><td></td><td></td><td></td><td></td><td></td><td></td><td></td><td></td></tr>
<tr><td>Oligo rev</td><td>AAACCCTTGCCGGCGCTGGGCACG</td><td></td><td></td><td></td><td></td><td></td><td></td><td></td><td></td><td></td><td></td><td></td></tr>
<tr><td>Chromosome</td><td>start</td><td></td><td>end</td><td>strand</td><td>MM</td><td>target_seq</td><td>PAM</td><td>alignment</td><td>distance</td><td>position</td><td>gene name</td><td>gene id</td></tr>
<tr><td>GU305914.1</td><td></td><td>90767</td><td>90789</td><td>-</td><td>0</td><td>CGTGCCCAGCGCCGGCAAGG</td><td>GGG</td><td>||||||||[||||||||||||]PAM</td><td>NA</td><td>Intergenic</td><td>NA</td><td>NA</td></tr>
<tr><td></td><td></td><td></td><td></td><td></td><td></td><td></td><td></td><td></td><td></td><td></td><td></td><td></td></tr>
<tr><td>T27277</td><td>GGGGGCCAACAGCAACCCGTCGG</td><td></td><td>1000</td><td></td><td></td><td></td><td></td><td></td><td></td><td></td><td></td><td></td></tr>
<tr><td>Oligo fwd</td><td>CACCGGGGGCCAACAGCAACCCGT</td><td></td><td></td><td></td><td></td><td></td><td></td><td></td><td></td><td></td><td></td><td></td></tr>
<tr><td>Oligo rev</td><td>AAACACGGGTTGCTGTTGGCCCCC</td><td></td><td></td><td></td><td></td><td></td><td></td><td></td><td></td><td></td><td></td><td></td></tr>
<tr><td>Chromosome</td><td>start</td><td></td><td>end</td><td>strand</td><td>MM</td><td>target_seq</td><td>PAM</td><td>alignment</td><td>distance</td><td>position</td><td>gene name</td><td>gene id</td></tr>
<tr><td>GU305914.1</td><td></td><td>90749</td><td>90771</td><td>-</td><td>0</td><td>GGGGGCCAACAGCAACCCGT</td><td>CGG</td><td>||||||||[||||||||||||]PAM</td><td>NA</td><td>Intergenic</td><td>NA</td><td>NA</td></tr>
<tr><td></td><td></td><td></td><td></td><td></td><td></td><td></td><td></td><td></td><td></td><td></td><td></td><td></td></tr>
<tr><td>T27278</td><td>CAACAGCAACCCGTCGGCGCAGG</td><td></td><td>1000</td><td></td><td></td><td></td><td></td><td></td><td></td><td></td><td></td><td></td></tr>
<tr><td>Oligo fwd</td><td>CACCCAACAGCAACCCGTCGGCGC</td><td></td><td></td><td></td><td></td><td></td><td></td><td></td><td></td><td></td><td></td><td></td></tr>
<tr><td>Oligo rev</td><td>AAACGCGCCGACGGGTTGCTGTTG</td><td></td><td></td><td></td><td></td><td></td><td></td><td></td><td></td><td></td><td></td><td></td></tr>
<tr><td>Chromosome</td><td>start</td><td></td><td>end</td><td>strand</td><td>MM</td><td>target_seq</td><td>PAM</td><td>alignment</td><td>distance</td><td>position</td><td>gene name</td><td>gene id</td></tr>
<tr><td>GU305914.1</td><td></td><td>90743</td><td>90765</td><td>-</td><td>0</td><td>CAACAGCAACCCGTCGGCGC</td><td>AGG</td><td>||||||||[||||||||||||]PAM</td><td>NA</td><td>Intergenic</td><td>NA</td><td>NA</td></tr>
<tr><td></td><td></td><td></td><td></td><td></td><td></td><td></td><td></td><td></td><td></td><td></td><td></td><td></td></tr>
<tr><td>T27279</td><td>AAGTCCATCCGCTTCTTCAACGG</td><td></td><td>1000</td><td></td><td></td><td></td><td></td><td></td><td></td><td></td><td></td><td></td></tr>
<tr><td>Oligo fwd</td><td>CACCAAGTCCATCCGCTTCTTCAA</td><td></td><td></td><td></td><td></td><td></td><td></td><td></td><td></td><td></td><td></td><td></td></tr>
<tr><td>Oligo rev</td><td>AAACTTGAAGAAGCGGATGGACTT</td><td></td><td></td><td></td><td></td><td></td><td></td><td></td><td></td><td></td><td></td><td></td></tr>
<tr><td>Chromosome</td><td>start</td><td></td><td>end</td><td>strand</td><td>MM</td><td>target_seq</td><td>PAM</td><td>alignment</td><td>distance</td><td>position</td><td>gene name</td><td>gene id</td></tr>
<tr><td>GU305914.1</td><td></td><td>90709</td><td>90731</td><td>-</td><td>0</td><td>AAGTCCATCCGCTTCTTCAA</td><td>CGG</td><td>||||||||[||||||||||||]PAM</td><td>NA</td><td>Intergenic</td><td>NA</td><td>NA</td></tr>
</table>

| | | | | | | | | | | | |
|---|---|---|---|---|---|---|---|---|---|---|---|
| T27280 | CCGCTTCTTCAACGGCCTCACGG | 1000 | | | | | | | | | |
| Oligo fwd | CACCCCGCTTCTTCAACGGCCTCA | | | | | | | | | | |
| Oligo rev | AAACTGAGGCCGTTGAAGAAGCGG | | | | | | | | | | |
| Chromosome | start | end | strand | MM | target_seq | PAM | alignment | distance | position | gene name | gene id |
| GU305914.1 | 90701 | 90723 | - | 0 | CCGCTTCTTCAACGGCCTCA | CGG | \|\|\|\|\|\|\|\|[\|\|\|\|\|\|\|\|\|\|\|\|]PAM | NA | Intergenic | NA | NA |

| | | | | | | | | | | | |
|---|---|---|---|---|---|---|---|---|---|---|---|
| T27281 | CGCTTCTTCAACGGCCTCACGGG | 1000 | | | | | | | | | |
| Oligo fwd | CACCCGCTTCTTCAACGGCCTCAC | | | | | | | | | | |
| Oligo rev | AAACGTGAGGCCGTTGAAGAAGCG | | | | | | | | | | |
| Chromosome | start | end | strand | MM | target_seq | PAM | alignment | distance | position | gene name | gene id |
| GU305914.1 | 90700 | 90722 | - | 0 | CGCTTCTTCAACGGCCTCAC | GGG | \|\|\|\|\|\|\|\|[\|\|\|\|\|\|\|\|\|\|\|\|]PAM | NA | Intergenic | NA | NA |

| | | | | | | | | | | | |
|---|---|---|---|---|---|---|---|---|---|---|---|
| T27282 | ACGACGCCAAGAAGACCATCAGG | 1000 | | | | | | | | | |
| Oligo fwd | CACCACGACGCCAAGAAGACCATC | | | | | | | | | | |
| Oligo rev | AAACGATGGTCTTCTTGGCGTCGT | | | | | | | | | | |
| Chromosome | start | end | strand | MM | target_seq | PAM | alignment | distance | position | gene name | gene id |
| GU305914.1 | 90657 | 90679 | - | 0 | ACGACGCCAAGAAGACCATC | AGG | \|\|\|\|\|\|\|\|[\|\|\|\|\|\|\|\|\|\|\|\|]PAM | NA | Intergenic | NA | NA |

| | | | | | | | | | | | |
|---|---|---|---|---|---|---|---|---|---|---|---|
| T27283 | CAAGAAGACCATCAGGTTCGAGG | 1000 | | | | | | | | | |
| Oligo fwd | CACCCAAGAAGACCATCAGGTTCG | | | | | | | | | | |
| Oligo rev | AAACCGAACCTGATGGTCTTCTTG | | | | | | | | | | |
| Chromosome | start | end | strand | MM | target_seq | PAM | alignment | distance | position | gene name | gene id |
| GU305914.1 | 90650 | 90672 | - | 0 | CAAGAAGACCATCAGGTTCG | AGG | \|\|\|\|\|\|\|\|[\|\|\|\|\|\|\|\|\|\|\|\|]PAM | NA | Intergenic | NA | NA |

| | | | | | | | | | | | |
|---|---|---|---|---|---|---|---|---|---|---|---|
| T27284 | AAGAAGACCATCAGGTTCGAGGG | 1000 | | | | | | | | | |
| Oligo fwd | CACCAAGAAGACCATCAGGTTCGA | | | | | | | | | | |
| Oligo rev | AAACTCGAACCTGATGGTCTTCTT | | | | | | | | | | |
| Chromosome | start | end | strand | MM | target_seq | PAM | alignment | distance | position | gene name | gene id |
| GU305914.1 | 90649 | 90671 | - | 0 | AAGAAGACCATCAGGTTCGA | GGG | \|\|\|\|\|\|\|\|[\|\|\|\|\|\|\|\|\|\|\|\|]PAM | NA | Intergenic | NA | NA |
| GU305914.1 | 196913 | 196935 | + | 4 | AGGCAGGCCTTCAGGTTCGA | CGG | \|-\|-\|\|-\|[\|-\|\|\|\|\|\|\|\|\|\|]PAM | NA | Intergenic | NA | NA |

| | | | | | | | | | | | |
|---|---|---|---|---|---|---|---|---|---|---|---|
| T27285 | AGAAGACCATCAGGTTCGAGGGG | 1000 | | | | | | | | | |
| Oligo fwd | CACCAGAAGACCATCAGGTTCGAG | | | | | | | | | | |
| Oligo rev | AAACCTCGAACCTGATGGTCTTCT | | | | | | | | | | |
| Chromosome | start | end | strand | MM | target_seq | PAM | alignment | distance | position | gene name | gene id |
| GU305914.1 | 90648 | 90670 | - | 0 | AGAAGACCATCAGGTTCGAG | GGG | \|\|\|\|\|\|\|\|[\|\|\|\|\|\|\|\|\|\|\|\|]PAM | NA | Intergenic | NA | NA |

| | | | | | | | | | | | |
|---|---|---|---|---|---|---|---|---|---|---|---|
| T27286 | GAAGACCATCAGGTTCGAGGGGG | 1000 | | | | | | | | | |
| Oligo fwd | CACCGAAGACCATCAGGTTCGAGG | | | | | | | | | | |
| Oligo rev | AAACCCTCGAACCTGATGGTCTTC | | | | | | | | | | |
| Chromosome | start | end | strand | MM | target_seq | PAM | alignment | distance | position | gene name | gene id |
| GU305914.1 | 90647 | 90669 | - | 0 | GAAGACCATCAGGTTCGAGG | GGG | \|\|\|\|\|\|\|\|[\|\|\|\|\|\|\|\|\|\|\|\|]PAM | NA | Intergenic | NA | NA |

| | | | | | | | | | | | |
|---|---|---|---|---|---|---|---|---|---|---|---|
| T27287 | ATCAGGTTCGAGGGGGCCGTCGG | 1000 | | | | | | | | | |
| Oligo fwd | CACCATCAGGTTCGAGGGGGCCGT | | | | | | | | | | |
| Oligo rev | AAACACGGCCCCCTCGAACCTGAT | | | | | | | | | | |
| Chromosome | start | end | strand | MM | target_seq | PAM | alignment | distance | position | gene name | gene id |
| GU305914.1 | 90640 | 90662 | - | 0 | ATCAGGTTCGAGGGGGCCGT | CGG | \|\|\|\|\|\|\|\|[\|\|\|\|\|\|\|\|\|\|\|\|]PAM | NA | Intergenic | NA | NA |

| | | | | | | | | | | | |
|---|---|---|---|---|---|---|---|---|---|---|---|
| T27288 | TCAGGTTCGAGGGGGCCGTCGGG | 1000 | | | | | | | | | |
| Oligo fwd | CACCTCAGGTTCGAGGGGGCCGTC | | | | | | | | | | |
| Oligo rev | AAACGACGGCCCCCTCGAACCTGA | | | | | | | | | | |
| Chromosome | start | end | strand | MM | target_seq | PAM | alignment | distance | position | gene name | gene id |
| GU305914.1 | 90639 | 90661 | - | 0 | TCAGGTTCGAGGGGGCCGTC | GGG | \|\|\|\|\|\|\|\|[\|\|\|\|\|\|\|\|\|\|\|\|]PAM | NA | Intergenic | NA | NA |

| | | | | | | | | | | | |
|---|---|---|---|---|---|---|---|---|---|---|---|
| T27289 | GGTTCGAGGGGGCCGTCGGGCGG | 1000 | | | | | | | | | |
| Oligo fwd | CACCGGTTCGAGGGGGCCGTCGGG | | | | | | | | | | |
| Oligo rev | AAACCCCGACGGCCCCCTCGAACC | | | | | | | | | | |
| Chromosome | start | end | strand | MM | target_seq | PAM | alignment | distance | position | gene name | gene id |
| GU305914.1 | 90636 | 90658 | - | 0 | GGTTCGAGGGGGCCGTCGGG | CGG | \|\|\|\|\|\|\|\|[\|\|\|\|\|\|\|\|\|\|\|\|]PAM | NA | Intergenic | NA | NA |

| | | | | | | | | | | | |
|---|---|---|---|---|---|---|---|---|---|---|---|
| T27290 | GTTCGAGGGGGCCGTCGGGCGGG | 1000 | | | | | | | | | |
| Oligo fwd | CACCGTTCGAGGGGGCCGTCGGGC | | | | | | | | | | |
| Oligo rev | AAACGCCCGACGGCCCCCTCGAAC | | | | | | | | | | |
| Chromosome | start | end | strand | MM | target_seq | PAM | alignment | distance | position | gene name | gene id |
| GU305914.1 | 90635 | 90657 | - | 0 | GTTCGAGGGGGCCGTCGGGC | GGG | \|\|\|\|\|\|\|\|[\|\|\|\|\|\|\|\|\|\|\|\|]PAM | NA | Intergenic | NA | NA |

| | | | | | | | | | | | |
|---|---|---|---|---|---|---|---|---|---|---|---|
| T27291 | GAAGTACTCCCCGCACCACCTGG | 1000 | | | | | | | | | |
| Oligo fwd | CACCGAAGTACTCCCCGCACCACC | | | | | | | | | | |
| Oligo rev | AAACGGTGGTGCGGGGAGTACTTC | | | | | | | | | | |
| Chromosome | start | end | strand | MM | target_seq | PAM | alignment | distance | position | gene name | gene id |
| GU305914.1 | 90608 | 90630 | - | 0 | GAAGTACTCCCCGCACCACC | TGG | \|\|\|\|\|\|\|\|[\|\|\|\|\|\|\|\|\|\|\|\|]PAM | NA | Intergenic | NA | NA |

| | | | | | | | | | | | |
|---|---|---|---|---|---|---|---|---|---|---|---|
| T27292 | CACCACCTGGCCTACTTCTGCGG | 1000 | | | | | | | | | |
| Oligo fwd | CACCCACCACCTGGCCTACTTCTG | | | | | | | | | | |
| Oligo rev | AAACCAGAAGTAGGCCAGGTGGTG | | | | | | | | | | |
| Chromosome | start | end | strand | MM | target_seq | PAM | alignment | distance | position | gene name | gene id |
| GU305914.1 | 90595 | 90617 | - | 0 | CACCACCTGGCCTACTTCTG | CGG | \|\|\|\|\|\|\|\|[\|\|\|\|\|\|\|\|\|\|\|\|]PAM | NA | Intergenic | NA | NA |

| | | | | | | | | | | | |
|---|---|---|---|---|---|---|---|---|---|---|---|
| T27293 | AGCTCTTCTCCACCCTCATGTGG | 1000 | | | | | | | | | |
| Oligo fwd | CACCAGCTCTTCTCCACCCTCATG | | | | | | | | | | |
| Oligo rev | AAACCATGAGGGTGGAGAAGAGCT | | | | | | | | | | |
| Chromosome | start | end | strand | MM | target_seq | PAM | alignment | distance | position | gene name | gene id |
| GU305914.1 | 90561 | 90583 | - | 0 | AGCTCTTCTCCACCCTCATG | TGG | \|\|\|\|\|\|\|\|[\|\|\|\|\|\|\|\|\|\|\|\|]PAM | NA | Intergenic | NA | NA |

| | | |
|---|---|---|
| T27294 | CGCATGTCCATCTACTCGACCGG | 1000 |
| Oligo fwd | CACCCGCATGTCCATCTACTCGAC | |
| Oligo rev | AAACGTCGAGTAGATGGACATGCG | |

| | | | | | | | | | | | | |
|---|---|---|---|---|---|---|---|---|---|---|---|---|
| Chromosome | start | | end | strand | MM | target_seq | PAM | alignment | distance | position | gene name | gene id |
| GU305914.1 | | 90529 | 90551 | - | 0 | CGCATGTCCATCTACTCGAC | CGG | \|\|\|\|\|\|\|\|[\|\|\|\|\|\|\|\|\|\|\|\|]PAM | NA | Intergenic | NA | NA |
| T27295 | GCATGTCCATCTACTCGACCGGG | 1000 | | | | | | | | | | |
| Oligo fwd | CACCGCATGTCCATCTACTCGACC | | | | | | | | | | | |
| Oligo rev | AAACGGTCGAGTAGATGGACATGC | | | | | | | | | | | |
| Chromosome | start | | end | strand | MM | target_seq | PAM | alignment | distance | position | gene name | gene id |
| GU305914.1 | | 90528 | 90550 | - | 0 | GCATGTCCATCTACTCGACC | GGG | \|\|\|\|\|\|\|\|[\|\|\|\|\|\|\|\|\|\|\|\|]PAM | NA | Intergenic | NA | NA |
| T27296 | CATGTCCATCTACTCGACCGGGG | 1000 | | | | | | | | | | |
| Oligo fwd | CACCCATGTCCATCTACTCGACCG | | | | | | | | | | | |
| Oligo rev | AAACCGGTCGAGTAGATGGACATG | | | | | | | | | | | |
| Chromosome | start | | end | strand | MM | target_seq | PAM | alignment | distance | position | gene name | gene id |
| GU305914.1 | | 90527 | 90549 | - | 0 | CATGTCCATCTACTCGACCG | GGG | \|\|\|\|\|\|\|\|[\|\|\|\|\|\|\|\|\|\|\|\|]PAM | NA | Intergenic | NA | NA |
| T27297 | TACTCGACCGGGGTCACGAGCGG | 1000 | | | | | | | | | | |
| Oligo fwd | CACCTACTCGACCGGGGTCACGAG | | | | | | | | | | | |
| Oligo rev | AAACCTCGTGACCCCGGTCGAGTA | | | | | | | | | | | |
| Chromosome | start | | end | strand | MM | target_seq | PAM | alignment | distance | position | gene name | gene id |
| GU305914.1 | | 90517 | 90539 | - | 0 | TACTCGACCGGGGTCACGAG | CGG | \|\|\|\|\|\|\|\|[\|\|\|\|\|\|\|\|\|\|\|\|]PAM | NA | Intergenic | NA | NA |
| T27298 | CGGGGTCACGAGCGGCGACACGG | 1000 | | | | | | | | | | |
| Oligo fwd | CACCCGGGGTCACGAGCGGCGACA | | | | | | | | | | | |
| Oligo rev | AAACTGTCGCCGCTCGTGACCCCG | | | | | | | | | | | |
| Chromosome | start | | end | strand | MM | target_seq | PAM | alignment | distance | position | gene name | gene id |
| GU305914.1 | | 90509 | 90531 | - | 0 | CGGGGTCACGAGCGGCGACA | CGG | \|\|\|\|\|\|\|\|[\|\|\|\|\|\|\|\|\|\|\|\|]PAM | NA | Intergenic | NA | NA |
| GU305914.1 | | 122664 | 122686 | - | 4 | CGAGATCACGATCGACGACA | CGG | \|\|-\|-\|\|\|[\|\|\|-\|\|-\|\|\|\|\|]PAM | NA | Intergenic | NA | NA |
| T27299 | TTCAGTCACATCGTGAACGCCGG | 1000 | | | | | | | | | | |
| Oligo fwd | CACCTTCAGTCACATCGTGAACGC | | | | | | | | | | | |
| Oligo rev | AAACGCGTTCACGATGTGACTGAA | | | | | | | | | | | |
| Chromosome | start | | end | strand | MM | target_seq | PAM | alignment | distance | position | gene name | gene id |
| GU305914.1 | | 90484 | 90506 | - | 0 | TTCAGTCACATCGTGAACGC | CGG | \|\|\|\|\|\|\|\|[\|\|\|\|\|\|\|\|\|\|\|\|]PAM | NA | Intergenic | NA | NA |
| T27300 | AACGCCGGCTCCAAGCTGTGCGG | 1000 | | | | | | | | | | |
| Oligo fwd | CACCAACGCCGGCTCCAAGCTGTG | | | | | | | | | | | |
| Oligo rev | AAACCACAGCTTGGAGCCGGCGTT | | | | | | | | | | | |
| Chromosome | start | | end | strand | MM | target_seq | PAM | alignment | distance | position | gene name | gene id |
| GU305914.1 | | 90469 | 90491 | - | 0 | AACGCCGGCTCCAAGCTGTG | CGG | \|\|\|\|\|\|\|\|[\|\|\|\|\|\|\|\|\|\|\|\|]PAM | NA | Intergenic | NA | NA |
| T27301 | TCCAAGCTGTGCGGCGCGTGCGG | 1000 | | | | | | | | | | |
| Oligo fwd | CACCTCCAAGCTGTGCGGCGCGTG | | | | | | | | | | | |
| Oligo rev | AAACCACGCGCCGCACAGCTTGGA | | | | | | | | | | | |
| Chromosome | start | | end | strand | MM | target_seq | PAM | alignment | distance | position | gene name | gene id |
| GU305914.1 | | 90460 | 90482 | - | 0 | TCCAAGCTGTGCGGCGCGTG | CGG | \|\|\|\|\|\|\|\|[\|\|\|\|\|\|\|\|\|\|\|\|]PAM | NA | Intergenic | NA | NA |
| T27302 | AAGCTGTGCGGCGCGTGCGGCGG | 1000 | | | | | | | | | | |
| Oligo fwd | CACCAAGCTGTGCGGCGCGTGCGG | | | | | | | | | | | |
| Oligo rev | AAACCCGCACGCGCCGCACAGCTT | | | | | | | | | | | |
| Chromosome | start | | end | strand | MM | target_seq | PAM | alignment | distance | position | gene name | gene id |
| GU305914.1 | | 90457 | 90479 | - | 0 | AAGCTGTGCGGCGCGTGCGG | CGG | \|\|\|\|\|\|\|\|[\|\|\|\|\|\|\|\|\|\|\|\|]PAM | NA | Intergenic | NA | NA |
| T27303 | GCTACGCCACCTCCTTCATCAGG | 1000 | | | | | | | | | | |
| Oligo fwd | CACCGCTACGCCACCTCCTTCATC | | | | | | | | | | | |
| Oligo rev | AAACGATGAAGGAGGTGGCGTAGC | | | | | | | | | | | |
| Chromosome | start | | end | strand | MM | target_seq | PAM | alignment | distance | position | gene name | gene id |
| GU305914.1 | | 90417 | 90439 | - | 0 | GCTACGCCACCTCCTTCATC | AGG | \|\|\|\|\|\|\|\|[\|\|\|\|\|\|\|\|\|\|\|\|]PAM | NA | Intergenic | NA | NA |
| GU305914.1 | | 99709 | 99731 | + | 4 | GCAACACCACCTCCGTCAGC | TGG | \|\|-\|\|-\|\|[\|\|\|\|\|\|-\|\|\|-\|]PAM | NA | Intergenic | NA | NA |
| T27304 | CTACGCCACCTCCTTCATCAGGG | 1000 | | | | | | | | | | |
| Oligo fwd | CACCCTACGCCACCTCCTTCATCA | | | | | | | | | | | |
| Oligo rev | AAACTGATGAAGGAGGTGGCGTAG | | | | | | | | | | | |
| Chromosome | start | | end | strand | MM | target_seq | PAM | alignment | distance | position | gene name | gene id |
| GU305914.1 | | 90416 | 90438 | - | 0 | CTACGCCACCTCCTTCATCA | GGG | \|\|\|\|\|\|\|\|[\|\|\|\|\|\|\|\|\|\|\|\|]PAM | NA | Intergenic | NA | NA |
| T27305 | CCTTCATCAGGGTGAACACGCGG | 1000 | | | | | | | | | | |
| Oligo fwd | CACCCCTTCATCAGGGTGAACACG | | | | | | | | | | | |
| Oligo rev | AAACCGTGTTCACCCTGATGAAGG | | | | | | | | | | | |
| Chromosome | start | | end | strand | MM | target_seq | PAM | alignment | distance | position | gene name | gene id |
| GU305914.1 | | 90405 | 90427 | - | 0 | CCTTCATCAGGGTGAACACG | CGG | \|\|\|\|\|\|\|\|[\|\|\|\|\|\|\|\|\|\|\|\|]PAM | NA | Intergenic | NA | NA |
| T27306 | CAGGGTGAACACGCGGCTGCCGG | 1000 | | | | | | | | | | |
| Oligo fwd | CACCCAGGGTGAACACGCGGCTGC | | | | | | | | | | | |
| Oligo rev | AAACGCAGCCGCGTGTTCACCCTG | | | | | | | | | | | |
| Chromosome | start | | end | strand | MM | target_seq | PAM | alignment | distance | position | gene name | gene id |
| GU305914.1 | | 90398 | 90420 | - | 0 | CAGGGTGAACACGCGGCTGC | CGG | \|\|\|\|\|\|\|\|[\|\|\|\|\|\|\|\|\|\|\|\|]PAM | NA | Intergenic | NA | NA |
| T27307 | AGGGTGAACACGCGGCTGCCGGG | 1000 | | | | | | | | | | |
| Oligo fwd | CACCAGGGTGAACACGCGGCTGCC | | | | | | | | | | | |
| Oligo rev | AAACGGCAGCCGCGTGTTCACCCT | | | | | | | | | | | |
| Chromosome | start | | end | strand | MM | target_seq | PAM | alignment | distance | position | gene name | gene id |
| GU305914.1 | | 90397 | 90419 | - | 0 | AGGGTGAACACGCGGCTGCC | GGG | \|\|\|\|\|\|\|\|[\|\|\|\|\|\|\|\|\|\|\|\|]PAM | NA | Intergenic | NA | NA |
| T27308 | CATCCCCAAGCAGATCAAGAAGG | 1000 | | | | | | | | | | |
| Oligo fwd | CACCCATCCCCAAGCAGATCAAGA | | | | | | | | | | | |
| Oligo rev | AAACTCTTGATCTGCTTGGGGATG | | | | | | | | | | | |
| Chromosome | start | | end | strand | MM | target_seq | PAM | alignment | distance | position | gene name | gene id |
| GU305914.1 | | 90374 | 90396 | - | 0 | CATCCCCAAGCAGATCAAGA | AGG | \|\|\|\|\|\|\|\|[\|\|\|\|\|\|\|\|\|\|\|\|]PAM | NA | Intergenic | NA | NA |

<table>
<tr><td>T27309</td><td>GATCAAGAAGGAGCCCGTCGTGG</td><td>1000</td><td></td><td></td><td></td><td></td><td></td><td></td><td></td><td></td><td></td></tr>
<tr><td>Oligo fwd</td><td>CACCGATCAAGAAGGAGCCCGTCG</td><td></td><td></td><td></td><td></td><td></td><td></td><td></td><td></td><td></td><td></td></tr>
<tr><td>Oligo rev</td><td>AAACCGACGGGCTCCTTCTTGATC</td><td></td><td></td><td></td><td></td><td></td><td></td><td></td><td></td><td></td><td></td></tr>
<tr><td>Chromosome</td><td>start</td><td>end</td><td>strand</td><td>MM</td><td>target_seq</td><td>PAM</td><td>alignment</td><td>distance</td><td>position</td><td>gene name</td><td>gene id</td></tr>
<tr><td>GU305914.1</td><td>90362</td><td>90384</td><td>-</td><td>0</td><td>GATCAAGAAGGAGCCCGTCG</td><td>TGG</td><td>||||||||[||||||||||||]PAM</td><td>NA</td><td>Intergenic</td><td>NA</td><td>NA</td></tr>
<tr><td>T27310</td><td>TCGTGGTCACTCTGCTCTCGCGG</td><td>1000</td><td></td><td></td><td></td><td></td><td></td><td></td><td></td><td></td><td></td></tr>
<tr><td>Oligo fwd</td><td>CACCTCGTGGTCACTCTGCTCTCG</td><td></td><td></td><td></td><td></td><td></td><td></td><td></td><td></td><td></td><td></td></tr>
<tr><td>Oligo rev</td><td>AAACCGAGAGCAGAGTGACCACGA</td><td></td><td></td><td></td><td></td><td></td><td></td><td></td><td></td><td></td><td></td></tr>
<tr><td>Chromosome</td><td>start</td><td>end</td><td>strand</td><td>MM</td><td>target_seq</td><td>PAM</td><td>alignment</td><td>distance</td><td>position</td><td>gene name</td><td>gene id</td></tr>
<tr><td>GU305914.1</td><td>90345</td><td>90367</td><td>-</td><td>0</td><td>TCGTGGTCACTCTGCTCTCG</td><td>CGG</td><td>||||||||[||||||||||||]PAM</td><td>NA</td><td>Intergenic</td><td>NA</td><td>NA</td></tr>
<tr><td>T27311</td><td>CGTGGTCACTCTGCTCTCGCGGG</td><td>1000</td><td></td><td></td><td></td><td></td><td></td><td></td><td></td><td></td><td></td></tr>
<tr><td>Oligo fwd</td><td>CACCCGTGGTCACTCTGCTCTCGC</td><td></td><td></td><td></td><td></td><td></td><td></td><td></td><td></td><td></td><td></td></tr>
<tr><td>Oligo rev</td><td>AAACGCGAGAGCAGAGTGACCACG</td><td></td><td></td><td></td><td></td><td></td><td></td><td></td><td></td><td></td><td></td></tr>
<tr><td>Chromosome</td><td>start</td><td>end</td><td>strand</td><td>MM</td><td>target_seq</td><td>PAM</td><td>alignment</td><td>distance</td><td>position</td><td>gene name</td><td>gene id</td></tr>
<tr><td>GU305914.1</td><td>90344</td><td>90366</td><td>-</td><td>0</td><td>CGTGGTCACTCTGCTCTCGC</td><td>GGG</td><td>||||||||[||||||||||||]PAM</td><td>NA</td><td>Intergenic</td><td>NA</td><td>NA</td></tr>
<tr><td>T27312</td><td>CTTCGCCGACGCCGACCTCTTGG</td><td>1000</td><td></td><td></td><td></td><td></td><td></td><td></td><td></td><td></td><td></td></tr>
<tr><td>Oligo fwd</td><td>CACCCTTCGCCGACGCCGACCTCT</td><td></td><td></td><td></td><td></td><td></td><td></td><td></td><td></td><td></td><td></td></tr>
<tr><td>Oligo rev</td><td>AAACAGAGGTCGGCGTCGGCGAAG</td><td></td><td></td><td></td><td></td><td></td><td></td><td></td><td></td><td></td><td></td></tr>
<tr><td>Chromosome</td><td>start</td><td>end</td><td>strand</td><td>MM</td><td>target_seq</td><td>PAM</td><td>alignment</td><td>distance</td><td>position</td><td>gene name</td><td>gene id</td></tr>
<tr><td>GU305914.1</td><td>90320</td><td>90342</td><td>-</td><td>0</td><td>CTTCGCCGACGCCGACCTCT</td><td>TGG</td><td>||||||||[||||||||||||]PAM</td><td>NA</td><td>Intergenic</td><td>NA</td><td>NA</td></tr>
<tr><td>GU305914.1</td><td>113931</td><td>113953</td><td>+</td><td>4</td><td>CGCCGCCGACGCCGACACCT</td><td>CGG</td><td>|--|||||[||||||||--||]PAM</td><td>NA</td><td>Intergenic</td><td>NA</td><td>NA</td></tr>
<tr><td>T27313</td><td>TTCGCCGACGCCGACCTCTTGGG</td><td>1000</td><td></td><td></td><td></td><td></td><td></td><td></td><td></td><td></td><td></td></tr>
<tr><td>Oligo fwd</td><td>CACCTTCGCCGACGCCGACCTCTT</td><td></td><td></td><td></td><td></td><td></td><td></td><td></td><td></td><td></td><td></td></tr>
<tr><td>Oligo rev</td><td>AAACAAGAGGTCGGCGTCGGCGAA</td><td></td><td></td><td></td><td></td><td></td><td></td><td></td><td></td><td></td><td></td></tr>
<tr><td>Chromosome</td><td>start</td><td>end</td><td>strand</td><td>MM</td><td>target_seq</td><td>PAM</td><td>alignment</td><td>distance</td><td>position</td><td>gene name</td><td>gene id</td></tr>
<tr><td>GU305914.1</td><td>90319</td><td>90341</td><td>-</td><td>0</td><td>TTCGCCGACGCCGACCTCTT</td><td>GGG</td><td>||||||||[||||||||||||]PAM</td><td>NA</td><td>Intergenic</td><td>NA</td><td>NA</td></tr>
<tr><td>GU305914.1</td><td>77576</td><td>77598</td><td>-</td><td>4</td><td>TCCGCCGGCGCCGGCTTCTT</td><td>CGG</td><td>|-|||||-[|||||-|-||||]PAM</td><td>NA</td><td>Intergenic</td><td>NA</td><td>NA</td></tr>
<tr><td>T27314</td><td>GCCGACCTCTTGGGAAACTATGG</td><td>1000</td><td></td><td></td><td></td><td></td><td></td><td></td><td></td><td></td><td></td></tr>
<tr><td>Oligo fwd</td><td>CACCGCCGACCTCTTGGGAAACTA</td><td></td><td></td><td></td><td></td><td></td><td></td><td></td><td></td><td></td><td></td></tr>
<tr><td>Oligo rev</td><td>AAACTAGTTTCCCAAGAGGTCGGC</td><td></td><td></td><td></td><td></td><td></td><td></td><td></td><td></td><td></td><td></td></tr>
<tr><td>Chromosome</td><td>start</td><td>end</td><td>strand</td><td>MM</td><td>target_seq</td><td>PAM</td><td>alignment</td><td>distance</td><td>position</td><td>gene name</td><td>gene id</td></tr>
<tr><td>GU305914.1</td><td>90310</td><td>90332</td><td>-</td><td>0</td><td>GCCGACCTCTTGGGAAACTA</td><td>TGG</td><td>||||||||[||||||||||||]PAM</td><td>NA</td><td>Intergenic</td><td>NA</td><td>NA</td></tr>
<tr><td>T27315</td><td>GGAAACTATGGTAAGAGATACGG</td><td>1000</td><td></td><td></td><td></td><td></td><td></td><td></td><td></td><td></td><td></td></tr>
<tr><td>Oligo fwd</td><td>CACCGGAAACTATGGTAAGAGATA</td><td></td><td></td><td></td><td></td><td></td><td></td><td></td><td></td><td></td><td></td></tr>
<tr><td>Oligo rev</td><td>AAACTATCTCTTACCATAGTTTCC</td><td></td><td></td><td></td><td></td><td></td><td></td><td></td><td></td><td></td><td></td></tr>
<tr><td>Chromosome</td><td>start</td><td>end</td><td>strand</td><td>MM</td><td>target_seq</td><td>PAM</td><td>alignment</td><td>distance</td><td>position</td><td>gene name</td><td>gene id</td></tr>
<tr><td>GU305914.1</td><td>90298</td><td>90320</td><td>-</td><td>0</td><td>GGAAACTATGGTAAGAGATA</td><td>CGG</td><td>||||||||[||||||||||||]PAM</td><td>NA</td><td>Intergenic</td><td>NA</td><td>NA</td></tr>
<tr><td>T27316</td><td>CTATGGTAAGAGATACGGCCTGG</td><td>1000</td><td></td><td></td><td></td><td></td><td></td><td></td><td></td><td></td><td></td></tr>
<tr><td>Oligo fwd</td><td>CACCCTATGGTAAGAGATACGGCC</td><td></td><td></td><td></td><td></td><td></td><td></td><td></td><td></td><td></td><td></td></tr>
<tr><td>Oligo rev</td><td>AAACGGCCGTATCTCTTACCATAG</td><td></td><td></td><td></td><td></td><td></td><td></td><td></td><td></td><td></td><td></td></tr>
<tr><td>Chromosome</td><td>start</td><td>end</td><td>strand</td><td>MM</td><td>target_seq</td><td>PAM</td><td>alignment</td><td>distance</td><td>position</td><td>gene name</td><td>gene id</td></tr>
<tr><td>GU305914.1</td><td>90293</td><td>90315</td><td>-</td><td>0</td><td>CTATGGTAAGAGATACGGCC</td><td>TGG</td><td>||||||||[||||||||||||]PAM</td><td>NA</td><td>Intergenic</td><td>NA</td><td>NA</td></tr>
<tr><td>T27317</td><td>AGAGATACGGCCTGGAGTCCCGG</td><td>1000</td><td></td><td></td><td></td><td></td><td></td><td></td><td></td><td></td><td></td></tr>
<tr><td>Oligo fwd</td><td>CACCAGAGATACGGCCTGGAGTCC</td><td></td><td></td><td></td><td></td><td></td><td></td><td></td><td></td><td></td><td></td></tr>
<tr><td>Oligo rev</td><td>AAACGGACTCCAGGCCGTATCTCT</td><td></td><td></td><td></td><td></td><td></td><td></td><td></td><td></td><td></td><td></td></tr>
<tr><td>Chromosome</td><td>start</td><td>end</td><td>strand</td><td>MM</td><td>target_seq</td><td>PAM</td><td>alignment</td><td>distance</td><td>position</td><td>gene name</td><td>gene id</td></tr>
<tr><td>GU305914.1</td><td>90285</td><td>90307</td><td>-</td><td>0</td><td>AGAGATACGGCCTGGAGTCC</td><td>CGG</td><td>||||||||[||||||||||||]PAM</td><td>NA</td><td>Intergenic</td><td>NA</td><td>NA</td></tr>
<tr><td>T27318</td><td>GAGATACGGCCTGGAGTCCCGGG</td><td>1000</td><td></td><td></td><td></td><td></td><td></td><td></td><td></td><td></td><td></td></tr>
<tr><td>Oligo fwd</td><td>CACCGAGATACGGCCTGGAGTCCC</td><td></td><td></td><td></td><td></td><td></td><td></td><td></td><td></td><td></td><td></td></tr>
<tr><td>Oligo rev</td><td>AAACGGGACTCCAGGCCGTATCTC</td><td></td><td></td><td></td><td></td><td></td><td></td><td></td><td></td><td></td><td></td></tr>
<tr><td>Chromosome</td><td>start</td><td>end</td><td>strand</td><td>MM</td><td>target_seq</td><td>PAM</td><td>alignment</td><td>distance</td><td>position</td><td>gene name</td><td>gene id</td></tr>
<tr><td>GU305914.1</td><td>90284</td><td>90306</td><td>-</td><td>0</td><td>GAGATACGGCCTGGAGTCCC</td><td>GGG</td><td>||||||||[||||||||||||]PAM</td><td>NA</td><td>Intergenic</td><td>NA</td><td>NA</td></tr>
<tr><td>T27319</td><td>ATACGGCCTGGAGTCCCGGGAGG</td><td>1000</td><td></td><td></td><td></td><td></td><td></td><td></td><td></td><td></td><td></td></tr>
<tr><td>Oligo fwd</td><td>CACCATACGGCCTGGAGTCCCGGG</td><td></td><td></td><td></td><td></td><td></td><td></td><td></td><td></td><td></td><td></td></tr>
<tr><td>Oligo rev</td><td>AAACCCCGGGACTCCAGGCCGTAT</td><td></td><td></td><td></td><td></td><td></td><td></td><td></td><td></td><td></td><td></td></tr>
<tr><td>Chromosome</td><td>start</td><td>end</td><td>strand</td><td>MM</td><td>target_seq</td><td>PAM</td><td>alignment</td><td>distance</td><td>position</td><td>gene name</td><td>gene id</td></tr>
<tr><td>GU305914.1</td><td>90281</td><td>90303</td><td>-</td><td>0</td><td>ATACGGCCTGGAGTCCCGGG</td><td>AGG</td><td>||||||||[||||||||||||]PAM</td><td>NA</td><td>Intergenic</td><td>NA</td><td>NA</td></tr>
<tr><td>T27320</td><td>CGGCCTGGAGTCCCGGGAGGCGG</td><td>1000</td><td></td><td></td><td></td><td></td><td></td><td></td><td></td><td></td><td></td></tr>
<tr><td>Oligo fwd</td><td>CACCCGGCCTGGAGTCCCGGGAGG</td><td></td><td></td><td></td><td></td><td></td><td></td><td></td><td></td><td></td><td></td></tr>
<tr><td>Oligo rev</td><td>AAACCCTCCCGGGACTCCAGGCCG</td><td></td><td></td><td></td><td></td><td></td><td></td><td></td><td></td><td></td><td></td></tr>
<tr><td>Chromosome</td><td>start</td><td>end</td><td>strand</td><td>MM</td><td>target_seq</td><td>PAM</td><td>alignment</td><td>distance</td><td>position</td><td>gene name</td><td>gene id</td></tr>
<tr><td>GU305914.1</td><td>90278</td><td>90300</td><td>-</td><td>0</td><td>CGGCCTGGAGTCCCGGGAGG</td><td>CGG</td><td>||||||||[||||||||||||]PAM</td><td>NA</td><td>Intergenic</td><td>NA</td><td>NA</td></tr>
<tr><td>T27321</td><td>GGCCTGGAGTCCCGGGAGGCGGG</td><td>1000</td><td></td><td></td><td></td><td></td><td></td><td></td><td></td><td></td><td></td></tr>
<tr><td>Oligo fwd</td><td>CACCGGCCTGGAGTCCCGGGAGGC</td><td></td><td></td><td></td><td></td><td></td><td></td><td></td><td></td><td></td><td></td></tr>
<tr><td>Oligo rev</td><td>AAACGCCTCCCGGGACTCCAGGCC</td><td></td><td></td><td></td><td></td><td></td><td></td><td></td><td></td><td></td><td></td></tr>
<tr><td>Chromosome</td><td>start</td><td>end</td><td>strand</td><td>MM</td><td>target_seq</td><td>PAM</td><td>alignment</td><td>distance</td><td>position</td><td>gene name</td><td>gene id</td></tr>
<tr><td>GU305914.1</td><td>90277</td><td>90299</td><td>-</td><td>0</td><td>GGCCTGGAGTCCCGGGAGGC</td><td>GGG</td><td>||||||||[||||||||||||]PAM</td><td>NA</td><td>Intergenic</td><td>NA</td><td>NA</td></tr>
<tr><td>T27322</td><td>GAGTCCCGGGAGGCGGGCGACGG</td><td>1000</td><td></td><td></td><td></td><td></td><td></td><td></td><td></td><td></td><td></td></tr>
<tr><td>Oligo fwd</td><td>CACCGAGTCCCGGGAGGCGGGCGA</td><td></td><td></td><td></td><td></td><td></td><td></td><td></td><td></td><td></td><td></td></tr>
<tr><td>Oligo rev</td><td>AAACTCGCCCGCCTCCCGGGACTC</td><td></td><td></td><td></td><td></td><td></td><td></td><td></td><td></td><td></td><td></td></tr>
<tr><td>Chromosome</td><td>start</td><td>end</td><td>strand</td><td>MM</td><td>target_seq</td><td>PAM</td><td>alignment</td><td>distance</td><td>position</td><td>gene name</td><td>gene id</td></tr>
<tr><td>GU305914.1</td><td>90271</td><td>90293</td><td>-</td><td>0</td><td>GAGTCCCGGGAGGCGGGCGA</td><td>CGG</td><td>||||||||[||||||||||||]PAM</td><td>NA</td><td>Intergenic</td><td>NA</td><td>NA</td></tr>
<tr><td>T27323</td><td>TCCCGGGAGGCGGGCGACGGCGG</td><td>1000</td><td></td><td></td><td></td><td></td><td></td><td></td><td></td><td></td><td></td></tr>
<tr><td>Oligo fwd</td><td>CACCTCCCGGGAGGCGGGCGACGG</td><td></td><td></td><td></td><td></td><td></td><td></td><td></td><td></td><td></td><td></td></tr>
<tr><td>Oligo rev</td><td>AAACCCGTCGCCCGCCTCCCGGGA</td><td></td><td></td><td></td><td></td><td></td><td></td><td></td><td></td><td></td><td></td></tr>
</table>

<table>
<tr><td>Chromosome</td><td>start</td><td>end</td><td>strand</td><td>MM</td><td>target_seq</td><td>PAM</td><td>alignment</td><td>distance</td><td>position</td><td>gene name</td><td>gene id</td></tr>
<tr><td>GU305914.1</td><td>90268</td><td>90290</td><td>-</td><td>0</td><td>TCCCGGGAGGCGGGCGACGG</td><td>CGG</td><td>||||||||[||||||||||||]PAM</td><td>NA</td><td>Intergenic</td><td>NA</td><td>NA</td></tr>
<tr><td></td><td></td><td></td><td></td><td></td><td></td><td></td><td></td><td></td><td></td><td></td><td></td></tr>
<tr><td>T27324</td><td>CGGGAGGCGGGCGACGGCGGAGG</td><td>1000</td><td></td><td></td><td></td><td></td><td></td><td></td><td></td><td></td><td></td></tr>
<tr><td>Oligo fwd</td><td>CACCCGGGAGGCGGGCGACGGCGG</td><td></td><td></td><td></td><td></td><td></td><td></td><td></td><td></td><td></td><td></td></tr>
<tr><td>Oligo rev</td><td>AAACCCGCCGTCGCCCGCCTCCCG</td><td></td><td></td><td></td><td></td><td></td><td></td><td></td><td></td><td></td><td></td></tr>
<tr><td>Chromosome</td><td>start</td><td>end</td><td>strand</td><td>MM</td><td>target_seq</td><td>PAM</td><td>alignment</td><td>distance</td><td>position</td><td>gene name</td><td>gene id</td></tr>
<tr><td>GU305914.1</td><td>90265</td><td>90287</td><td>-</td><td>0</td><td>CGGGAGGCGGGCGACGGCGG</td><td>AGG</td><td>||||||||[||||||||||||]PAM</td><td>NA</td><td>Intergenic</td><td>NA</td><td>NA</td></tr>
<tr><td>GU305914.1</td><td>96942</td><td>96964</td><td>+</td><td>4</td><td>GGCGAGCAGGGCGACGGCGG</td><td>AGG</td><td>-|-|||--[||||||||||||]PAM</td><td>NA</td><td>Intergenic</td><td>NA</td><td>NA</td></tr>
<tr><td>GU305914.1</td><td>225987</td><td>226009</td><td>-</td><td>4</td><td>CGCGTGGAGCGCGACGGCGG</td><td>CGG</td><td>||-|-||-[|-||||||||||]PAM</td><td>NA</td><td>Intergenic</td><td>NA</td><td>NA</td></tr>
<tr><td>GU305914.1</td><td>75553</td><td>75575</td><td>+</td><td>4</td><td>CAGGACGAGGGCGATGGCGG</td><td>CGG</td><td>|-|||-|-[||||||-|||||]PAM</td><td>NA</td><td>Intergenic</td><td>NA</td><td>NA</td></tr>
<tr><td>GU305914.1</td><td>58135</td><td>58157</td><td>-</td><td>4</td><td>CGCAAGGCGGGCGGTGGCGG</td><td>CGG</td><td>||--||||[|||||--|||||]PAM</td><td>NA</td><td>Intergenic</td><td>NA</td><td>NA</td></tr>
<tr><td></td><td></td><td></td><td></td><td></td><td></td><td></td><td></td><td></td><td></td><td></td><td></td></tr>
<tr><td>T27325</td><td>GAGGCGGGCGACGGCGGAGGCGG</td><td>1000</td><td></td><td></td><td></td><td></td><td></td><td></td><td></td><td></td><td></td></tr>
<tr><td>Oligo fwd</td><td>CACCGAGGCGGGCGACGGCGGAGG</td><td></td><td></td><td></td><td></td><td></td><td></td><td></td><td></td><td></td><td></td></tr>
<tr><td>Oligo rev</td><td>AAACCCTCCGCCGTCGCCCGCCTC</td><td></td><td></td><td></td><td></td><td></td><td></td><td></td><td></td><td></td><td></td></tr>
<tr><td>Chromosome</td><td>start</td><td>end</td><td>strand</td><td>MM</td><td>target_seq</td><td>PAM</td><td>alignment</td><td>distance</td><td>position</td><td>gene name</td><td>gene id</td></tr>
<tr><td>GU305914.1</td><td>90262</td><td>90284</td><td>-</td><td>0</td><td>GAGGCGGGCGACGGCGGAGG</td><td>CGG</td><td>||||||||[||||||||||||]PAM</td><td>NA</td><td>Intergenic</td><td>NA</td><td>NA</td></tr>
<tr><td>GU305914.1</td><td>96945</td><td>96967</td><td>+</td><td>2</td><td>GAGCAGGGCGACGGCGGAGG</td><td>CGG</td><td>|||--|||[||||||||||||]PAM</td><td>NA</td><td>Intergenic</td><td>NA</td><td>NA</td></tr>
<tr><td>GU305914.1</td><td>111178</td><td>111200</td><td>-</td><td>4</td><td>CAGGCCGACGACGGCGAAGG</td><td>TGG</td><td>-||||-|-[||||||||-|||]PAM</td><td>NA</td><td>Intergenic</td><td>NA</td><td>NA</td></tr>
<tr><td>GU305914.1</td><td>149488</td><td>149510</td><td>-</td><td>4</td><td>GGAGCAGGCGACGGCGGCGG</td><td>CGG</td><td>|--||-||[|||||||||-||]PAM</td><td>NA</td><td>Intergenic</td><td>NA</td><td>NA</td></tr>
<tr><td>GU305914.1</td><td>176454</td><td>176476</td><td>+</td><td>4</td><td>GACGATGGCGACGGCGGCGG</td><td>AGG</td><td>||-|--||[|||||||||-||]PAM</td><td>NA</td><td>Intergenic</td><td>NA</td><td>NA</td></tr>
<tr><td>GU305914.1</td><td>225984</td><td>226006</td><td>-</td><td>4</td><td>GTGGAGCGCGACGGCGGCGG</td><td>CGG</td><td>|-||-|-|[|||||||||-||]PAM</td><td>NA</td><td>Intergenic</td><td>NA</td><td>NA</td></tr>
<tr><td>GU305914.1</td><td>149292</td><td>149314</td><td>+</td><td>4</td><td>GGAGCGGGCGACGACGGCGG</td><td>CGG</td><td>|--|||||[|||||-|||-||]PAM</td><td>NA</td><td>Intergenic</td><td>NA</td><td>NA</td></tr>
<tr><td></td><td></td><td></td><td></td><td></td><td></td><td></td><td></td><td></td><td></td><td></td><td></td></tr>
<tr><td>T27326</td><td>GCGGGCGACGGCGGAGGCGGCGG</td><td>1000</td><td></td><td></td><td></td><td></td><td></td><td></td><td></td><td></td><td></td></tr>
<tr><td>Oligo fwd</td><td>CACCGCGGGCGACGGCGGAGGCGG</td><td></td><td></td><td></td><td></td><td></td><td></td><td></td><td></td><td></td><td></td></tr>
<tr><td>Oligo rev</td><td>AAACCCGCCTCCGCCGTCGCCCGC</td><td></td><td></td><td></td><td></td><td></td><td></td><td></td><td></td><td></td><td></td></tr>
<tr><td>Chromosome</td><td>start</td><td>end</td><td>strand</td><td>MM</td><td>target_seq</td><td>PAM</td><td>alignment</td><td>distance</td><td>position</td><td>gene name</td><td>gene id</td></tr>
<tr><td>GU305914.1</td><td>90259</td><td>90281</td><td>-</td><td>0</td><td>GCGGGCGACGGCGGAGGCGG</td><td>CGG</td><td>||||||||[||||||||||||]PAM</td><td>NA</td><td>Intergenic</td><td>NA</td><td>NA</td></tr>
<tr><td>GU305914.1</td><td>149485</td><td>149507</td><td>-</td><td>2</td><td>GCAGGCGACGGCGGCGGCGG</td><td>CGG</td><td>||-|||||[||||||-|||||]PAM</td><td>NA</td><td>Intergenic</td><td>NA</td><td>NA</td></tr>
<tr><td>GU305914.1</td><td>154407</td><td>154429</td><td>+</td><td>4</td><td>GGTGGCGGCGGCGGCGGCGG</td><td>CGG</td><td>|--||||-[||||||-|||||]PAM</td><td>NA</td><td>Intergenic</td><td>NA</td><td>NA</td></tr>
<tr><td>GU305914.1</td><td>58123</td><td>58145</td><td>-</td><td>4</td><td>GGTGGCGGCGGCGGCGGCGG</td><td>AGG</td><td>|--||||-[||||||-|||||]PAM</td><td>NA</td><td>Intergenic</td><td>NA</td><td>NA</td></tr>
<tr><td>GU305914.1</td><td>67742</td><td>67764</td><td>+</td><td>4</td><td>GGGAGCGGCGGCGGCGGCGG</td><td>CGG</td><td>|-|-|||-[||||||-|||||]PAM</td><td>NA</td><td>Intergenic</td><td>NA</td><td>NA</td></tr>
<tr><td>GU305914.1</td><td>62672</td><td>62694</td><td>-</td><td>4</td><td>GCTGCCGCCGGCGGGGGCGG</td><td>CGG</td><td>||-|-||-[||||||-|||||]PAM</td><td>NA</td><td>Intergenic</td><td>NA</td><td>NA</td></tr>
<tr><td>GU305914.1</td><td>58129</td><td>58151</td><td>-</td><td>3</td><td>GCGGGCGGTGGCGGCGGCGG</td><td>CGG</td><td>|||||||-[-|||||-|||||]PAM</td><td>NA</td><td>Intergenic</td><td>NA</td><td>NA</td></tr>
<tr><td>GU305914.1</td><td>154401</td><td>154423</td><td>+</td><td>4</td><td>CCGGGCGGTGGCGGCGGCGG</td><td>CGG</td><td>-||||||-[-|||||-|||||]PAM</td><td>NA</td><td>Intergenic</td><td>NA</td><td>NA</td></tr>
<tr><td>GU305914.1</td><td>111175</td><td>111197</td><td>-</td><td>4</td><td>GCCGACGACGGCGAAGGTGG</td><td>AGG</td><td>||-|-|||[|||||-|||-||]PAM</td><td>NA</td><td>Intergenic</td><td>NA</td><td>NA</td></tr>
<tr><td>GU305914.1</td><td>77584</td><td>77606</td><td>+</td><td>4</td><td>GCCGGCGCCGGCGGAGCCCG</td><td>AGG</td><td>||-||||-[||||||||-|-|]PAM</td><td>NA</td><td>Intergenic</td><td>NA</td><td>NA</td></tr>
<tr><td>GU305914.1</td><td>19436</td><td>19458</td><td>+</td><td>4</td><td>GCCGGCGTCGGCGGAGGTGC</td><td>CGG</td><td>||-||||-[|||||||||-|-]PAM</td><td>NA</td><td>Intergenic</td><td>NA</td><td>NA</td></tr>
<tr><td></td><td></td><td></td><td></td><td></td><td></td><td></td><td></td><td></td><td></td><td></td><td></td></tr>
<tr><td>T27327</td><td>GACGGCGGAGGCGGCGGAGCCGG</td><td>1000</td><td></td><td></td><td></td><td></td><td></td><td></td><td></td><td></td><td></td></tr>
<tr><td>Oligo fwd</td><td>CACCGACGGCGGAGGCGGCGGAGC</td><td></td><td></td><td></td><td></td><td></td><td></td><td></td><td></td><td></td><td></td></tr>
<tr><td>Oligo rev</td><td>AAACGCTCCGCCGCCTCCGCCGTC</td><td></td><td></td><td></td><td></td><td></td><td></td><td></td><td></td><td></td><td></td></tr>
<tr><td>Chromosome</td><td>start</td><td>end</td><td>strand</td><td>MM</td><td>target_seq</td><td>PAM</td><td>alignment</td><td>distance</td><td>position</td><td>gene name</td><td>gene id</td></tr>
<tr><td>GU305914.1</td><td>90253</td><td>90275</td><td>-</td><td>0</td><td>GACGGCGGAGGCGGCGGAGC</td><td>CGG</td><td>||||||||[||||||||||||]PAM</td><td>NA</td><td>Intergenic</td><td>NA</td><td>NA</td></tr>
<tr><td>GU305914.1</td><td>18656</td><td>18678</td><td>-</td><td>3</td><td>GACGGCAGAGGTGGCGGAGG</td><td>CGG</td><td>||||||-|[|||-|||||||-]PAM</td><td>NA</td><td>Intergenic</td><td>NA</td><td>NA</td></tr>
<tr><td></td><td></td><td></td><td></td><td></td><td></td><td></td><td></td><td></td><td></td><td></td><td></td></tr>
<tr><td>T27328</td><td>GGCGGAGGCGGCGGAGCCGGCGG</td><td>1000</td><td></td><td></td><td></td><td></td><td></td><td></td><td></td><td></td><td></td></tr>
<tr><td>Oligo fwd</td><td>CACCGGCGGAGGCGGCGGAGCCGG</td><td></td><td></td><td></td><td></td><td></td><td></td><td></td><td></td><td></td><td></td></tr>
<tr><td>Oligo rev</td><td>AAACCCGGCTCCGCCGCCTCCGCC</td><td></td><td></td><td></td><td></td><td></td><td></td><td></td><td></td><td></td><td></td></tr>
<tr><td>Chromosome</td><td>start</td><td>end</td><td>strand</td><td>MM</td><td>target_seq</td><td>PAM</td><td>alignment</td><td>distance</td><td>position</td><td>gene name</td><td>gene id</td></tr>
<tr><td>GU305914.1</td><td>90250</td><td>90272</td><td>-</td><td>0</td><td>GGCGGAGGCGGCGGAGCCGG</td><td>CGG</td><td>||||||||[||||||||||||]PAM</td><td>NA</td><td>Intergenic</td><td>NA</td><td>NA</td></tr>
<tr><td>GU305914.1</td><td>152553</td><td>152575</td><td>+</td><td>3</td><td>TGCGGCGGCGGCGGTGCCGG</td><td>CGG</td><td>-||||-||[||||||-|||||]PAM</td><td>NA</td><td>Intergenic</td><td>NA</td><td>NA</td></tr>
<tr><td>GU305914.1</td><td>166404</td><td>166426</td><td>+</td><td>4</td><td>GGCAGAGACGTCGGCGCCGG</td><td>CGG</td><td>|||-|||-[||-|||-|||||]PAM</td><td>NA</td><td>Intergenic</td><td>NA</td><td>NA</td></tr>
<tr><td>GU305914.1</td><td>18653</td><td>18675</td><td>-</td><td>3</td><td>GGCAGAGGTGGCGGAGGCGG</td><td>GGG</td><td>|||-||||[-|||||||-|||]PAM</td><td>NA</td><td>Intergenic</td><td>NA</td><td>NA</td></tr>
<tr><td>GU305914.1</td><td>19439</td><td>19461</td><td>+</td><td>4</td><td>GGCGTCGGCGGAGGTGCCGG</td><td>AGG</td><td>||||--||[|||-||-|||||]PAM</td><td>NA</td><td>Intergenic</td><td>NA</td><td>NA</td></tr>
<tr><td>GU305914.1</td><td>154404</td><td>154426</td><td>+</td><td>3</td><td>GGCGGTGGCGGCGGCGGCGG</td><td>CGG</td><td>|||||-||[||||||-|-|||]PAM</td><td>NA</td><td>Intergenic</td><td>NA</td><td>NA</td></tr>
<tr><td>GU305914.1</td><td>58126</td><td>58148</td><td>-</td><td>3</td><td>GGCGGTGGCGGCGGCGGCGG</td><td>CGG</td><td>|||||-||[||||||-|-|||]PAM</td><td>NA</td><td>Intergenic</td><td>NA</td><td>NA</td></tr>
<tr><td>GU305914.1</td><td>77584</td><td>77606</td><td>+</td><td>4</td><td>GCCGGCGCCGGCGGAGCCCG</td><td>AGG</td><td>|-|||-|-[||||||||||-|]PAM</td><td>NA</td><td>Intergenic</td><td>NA</td><td>NA</td></tr>
<tr><td>GU305914.1</td><td>154407</td><td>154429</td><td>+</td><td>4</td><td>GGTGGCGGCGGCGGCGGCGG</td><td>CGG</td><td>||-||-||[||||||-|-|||]PAM</td><td>NA</td><td>Intergenic</td><td>NA</td><td>NA</td></tr>
<tr><td>GU305914.1</td><td>58123</td><td>58145</td><td>-</td><td>4</td><td>GGTGGCGGCGGCGGCGGCGG</td><td>AGG</td><td>||-||-||[||||||-|-|||]PAM</td><td>NA</td><td>Intergenic</td><td>NA</td><td>NA</td></tr>
<tr><td>GU305914.1</td><td>149482</td><td>149504</td><td>-</td><td>4</td><td>GGCGACGGCGGCGGCGGCGG</td><td>CGG</td><td>||||--||[||||||-|-|||]PAM</td><td>NA</td><td>Intergenic</td><td>NA</td><td>NA</td></tr>
<tr><td></td><td></td><td></td><td></td><td></td><td></td><td></td><td></td><td></td><td></td><td></td><td></td></tr>
<tr><td>T27329</td><td>GAGGCGGCGGAGCCGGCGGCAGG</td><td>1000</td><td></td><td></td><td></td><td></td><td></td><td></td><td></td><td></td><td></td></tr>
<tr><td>Oligo fwd</td><td>CACCGAGGCGGCGGAGCCGGCGGC</td><td></td><td></td><td></td><td></td><td></td><td></td><td></td><td></td><td></td><td></td></tr>
<tr><td>Oligo rev</td><td>AAACGCCGCCGGCTCCGCCGCCTC</td><td></td><td></td><td></td><td></td><td></td><td></td><td></td><td></td><td></td><td></td></tr>
<tr><td>Chromosome</td><td>start</td><td>end</td><td>strand</td><td>MM</td><td>target_seq</td><td>PAM</td><td>alignment</td><td>distance</td><td>position</td><td>gene name</td><td>gene id</td></tr>
<tr><td>GU305914.1</td><td>90246</td><td>90268</td><td>-</td><td>0</td><td>GAGGCGGCGGAGCCGGCGGC</td><td>AGG</td><td>||||||||[||||||||||||]PAM</td><td>NA</td><td>Intergenic</td><td>NA</td><td>NA</td></tr>
<tr><td>GU305914.1</td><td>86780</td><td>86802</td><td>-</td><td>3</td><td>GAGGGGGCGGGGGCGGCGGC</td><td>AGG</td><td>||||-|||[||-|-|||||||]PAM</td><td>NA</td><td>Intergenic</td><td>NA</td><td>NA</td></tr>
<tr><td>GU305914.1</td><td>67743</td><td>67765</td><td>+</td><td>4</td><td>GGAGCGGCGGCGGCGGCGGC</td><td>GGG</td><td>|--|||||[||-|-|||||||]PAM</td><td>NA</td><td>Intergenic</td><td>NA</td><td>NA</td></tr>
<tr><td>GU305914.1</td><td>221599</td><td>221621</td><td>-</td><td>4</td><td>GAGGCCGTGGTGGCGGCGGC</td><td>AGG</td><td>|||||-|-[||-|-|||||||]PAM</td><td>NA</td><td>Intergenic</td><td>NA</td><td>NA</td></tr>
<tr><td>GU305914.1</td><td>75852</td><td>75874</td><td>+</td><td>4</td><td>GGGGAGGCGGAGGCGGCGGT</td><td>CGG</td><td>|-||-|||[||||-||||||-]PAM</td><td>NA</td><td>Intergenic</td><td>NA</td><td>NA</td></tr>
<tr><td></td><td></td><td></td><td></td><td></td><td></td><td></td><td></td><td></td><td></td><td></td><td></td></tr>
<tr><td>T27330</td><td>CGGCGGAGCCGGCGGCAGGACGG</td><td>1000</td><td></td><td></td><td></td><td></td><td></td><td></td><td></td><td></td><td></td></tr>
<tr><td>Oligo fwd</td><td>CACCCGGCGGAGCCGGCGGCAGGA</td><td></td><td></td><td></td><td></td><td></td><td></td><td></td><td></td><td></td><td></td></tr>
<tr><td>Oligo rev</td><td>AAACTCCTGCCGCCGGCTCCGCCG</td><td></td><td></td><td></td><td></td><td></td><td></td><td></td><td></td><td></td><td></td></tr>
<tr><td>Chromosome</td><td>start</td><td>end</td><td>strand</td><td>MM</td><td>target_seq</td><td>PAM</td><td>alignment</td><td>distance</td><td>position</td><td>gene name</td><td>gene id</td></tr>
<tr><td>GU305914.1</td><td>90242</td><td>90264</td><td>-</td><td>0</td><td>CGGCGGAGCCGGCGGCAGGA</td><td>CGG</td><td>||||||||[||||||||||||]PAM</td><td>NA</td><td>Intergenic</td><td>NA</td><td>NA</td></tr>
<tr><td></td><td></td><td></td><td></td><td></td><td></td><td></td><td></td><td></td><td></td><td></td><td></td></tr>
<tr><td>T27331</td><td>AGCCGGCGGCAGGACGGACGAGG</td><td>1000</td><td></td><td></td><td></td><td></td><td></td><td></td><td></td><td></td><td></td></tr>
<tr><td>Oligo fwd</td><td>CACCAGCCGGCGGCAGGACGGACG</td><td></td><td></td><td></td><td></td><td></td><td></td><td></td><td></td><td></td><td></td></tr>
<tr><td>Oligo rev</td><td>AAACCGTCCGTCCTGCCGCCGGCT</td><td></td><td></td><td></td><td></td><td></td><td></td><td></td><td></td><td></td><td></td></tr>
<tr><td>Chromosome</td><td>start</td><td>end</td><td>strand</td><td>MM</td><td>target_seq</td><td>PAM</td><td>alignment</td><td>distance</td><td>position</td><td>gene name</td><td>gene id</td></tr>
<tr><td>GU305914.1</td><td>90236</td><td>90258</td><td>-</td><td>0</td><td>AGCCGGCGGCAGGACGGACG</td><td>AGG</td><td>||||||||[||||||||||||]PAM</td><td>NA</td><td>Intergenic</td><td>NA</td><td>NA</td></tr>
<tr><td></td><td></td><td></td><td></td><td></td><td></td><td></td><td></td><td></td><td></td><td></td><td></td></tr>
<tr><td>T27332</td><td>CGGCGGCAGGACGGACGAGGTGG</td><td>1000</td><td></td><td></td><td></td><td></td><td></td><td></td><td></td><td></td><td></td></tr>
<tr><td>Oligo fwd</td><td>CACCCGGCGGCAGGACGGACGAGG</td><td></td><td></td><td></td><td></td><td></td><td></td><td></td><td></td><td></td><td></td></tr>
<tr><td>Oligo rev</td><td>AAACCCTCGTCCGTCCTGCCGCCG</td><td></td><td></td><td></td><td></td><td></td><td></td><td></td><td></td><td></td><td></td></tr>
</table>

| | | | | | | | | | | | |
|---|---|---|---|---|---|---|---|---|---|---|---|
| Chromosome | start | end | strand | MM | target_seq | PAM | alignment | distance | position | gene name | gene id |
| GU305914.1 | 90233 | 90255 | - | 0 | CGGCGGCAGGACGGACGAGG | TGG | \|\|\|\|\|\|\|\|[\|\|\|\|\|\|\|\|\|\|\|\|]PAM | NA | Intergenic | NA | NA |
| T27333 | CAGGACGGACGAGGTGGCCGCGG | 1000 | | | | | | | | | |
| Oligo fwd | CACCCAGGACGGACGAGGTGGCCG | | | | | | | | | | |
| Oligo rev | AAACCGGCCACCTCGTCCGTCCTG | | | | | | | | | | |
| Chromosome | start | end | strand | MM | target_seq | PAM | alignment | distance | position | gene name | gene id |
| GU305914.1 | 90227 | 90249 | - | 0 | CAGGACGGACGAGGTGGCCG | CGG | \|\|\|\|\|\|\|\|[\|\|\|\|\|\|\|\|\|\|\|\|]PAM | NA | Intergenic | NA | NA |
| T27334 | AGGACGGACGAGGTGGCCGCGGG | 1000 | | | | | | | | | |
| Oligo fwd | CACCAGGACGGACGAGGTGGCCGC | | | | | | | | | | |
| Oligo rev | AAACGCGGCCACCTCGTCCGTCCT | | | | | | | | | | |
| Chromosome | start | end | strand | MM | target_seq | PAM | alignment | distance | position | gene name | gene id |
| GU305914.1 | 90226 | 90248 | - | 0 | AGGACGGACGAGGTGGCCGC | GGG | \|\|\|\|\|\|\|\|[\|\|\|\|\|\|\|\|\|\|\|\|]PAM | NA | Intergenic | NA | NA |
| T27335 | GGACGGACGAGGTGGCCGCGGGG | 1000 | | | | | | | | | |
| Oligo fwd | CACCGGACGGACGAGGTGGCCGCG | | | | | | | | | | |
| Oligo rev | AAACCGCGGCCACCTCGTCCGTCC | | | | | | | | | | |
| Chromosome | start | end | strand | MM | target_seq | PAM | alignment | distance | position | gene name | gene id |
| GU305914.1 | 90225 | 90247 | - | 0 | GGACGGACGAGGTGGCCGCG | GGG | \|\|\|\|\|\|\|\|[\|\|\|\|\|\|\|\|\|\|\|\|]PAM | NA | Intergenic | NA | NA |
| T27336 | CGAGGTGGCCGCGGGGCCTCCGG | 1000 | | | | | | | | | |
| Oligo fwd | CACCCGAGGTGGCCGCGGGGCCTC | | | | | | | | | | |
| Oligo rev | AAACGAGGCCCCGCGGCCACCTCG | | | | | | | | | | |
| Chromosome | start | end | strand | MM | target_seq | PAM | alignment | distance | position | gene name | gene id |
| GU305914.1 | 90218 | 90240 | - | 0 | CGAGGTGGCCGCGGGGCCTC | CGG | \|\|\|\|\|\|\|\|[\|\|\|\|\|\|\|\|\|\|\|\|]PAM | NA | Intergenic | NA | NA |
| T27337 | GTGGCCGCGGGGCCTCCGGCCGG | 1000 | | | | | | | | | |
| Oligo fwd | CACCGTGGCCGCGGGGCCTCCGGC | | | | | | | | | | |
| Oligo rev | AAACGCCGGAGGCCCCGCGGCCAC | | | | | | | | | | |
| Chromosome | start | end | strand | MM | target_seq | PAM | alignment | distance | position | gene name | gene id |
| GU305914.1 | 90214 | 90236 | - | 0 | GTGGCCGCGGGGCCTCCGGC | CGG | \|\|\|\|\|\|\|\|[\|\|\|\|\|\|\|\|\|\|\|\|]PAM | NA | Intergenic | NA | NA |
| T27338 | GCCGCGGGGCCTCCGGCCGGCGG | 1000 | | | | | | | | | |
| Oligo fwd | CACCGCCGCGGGGCCTCCGGCCGG | | | | | | | | | | |
| Oligo rev | AAACCCGGCCGGAGGCCCCGCGGC | | | | | | | | | | |
| Chromosome | start | end | strand | MM | target_seq | PAM | alignment | distance | position | gene name | gene id |
| GU305914.1 | 90211 | 90233 | - | 0 | GCCGCGGGGCCTCCGGCCGG | CGG | \|\|\|\|\|\|\|\|[\|\|\|\|\|\|\|\|\|\|\|\|]PAM | NA | Intergenic | NA | NA |
| GU305914.1 | 115363 | 115385 | + | 4 | GCCTCGGAACCGCCGGCCGG | CGG | \|\|\|-\|\|\|-[-\|\|-\|\|\|\|\|\|\|\|]PAM | NA | Intergenic | NA | NA |
| T27339 | CCTCCGGCCGGCGGCGCCAGCGG | 1000 | | | | | | | | | |
| Oligo fwd | CACCCCTCCGGCCGGCGGCGCCAG | | | | | | | | | | |
| Oligo rev | AAACCTGGCGCCGCCGGCCGGAGG | | | | | | | | | | |
| Chromosome | start | end | strand | MM | target_seq | PAM | alignment | distance | position | gene name | gene id |
| GU305914.1 | 90202 | 90224 | - | 0 | CCTCCGGCCGGCGGCGCCAG | CGG | \|\|\|\|\|\|\|\|[\|\|\|\|\|\|\|\|\|\|\|\|]PAM | NA | Intergenic | NA | NA |
| GU305914.1 | 121642 | 121664 | + | 4 | CCGCCGGACGGCGGCTTCAG | CGG | \|\|-\|\|\|\|-[\|\|\|\|\|\|\|--\|\|\|]PAM | NA | Intergenic | NA | NA |
| T27340 | CTCCGGCCGGCGGCGCCAGCGGG | 1000 | | | | | | | | | |
| Oligo fwd | CACCCTCCGGCCGGCGGCGCCAGC | | | | | | | | | | |
| Oligo rev | AAACGCTGGCGCCGCCGGCCGGAG | | | | | | | | | | |
| Chromosome | start | end | strand | MM | target_seq | PAM | alignment | distance | position | gene name | gene id |
| GU305914.1 | 90201 | 90223 | - | 0 | CTCCGGCCGGCGGCGCCAGC | GGG | \|\|\|\|\|\|\|\|[\|\|\|\|\|\|\|\|\|\|\|\|]PAM | NA | Intergenic | NA | NA |
| T27341 | TGAACTTCGTCTCCGTCGACAGG | 1000 | | | | | | | | | |
| Oligo fwd | CACCTGAACTTCGTCTCCGTCGAC | | | | | | | | | | |
| Oligo rev | AAACGTCGACGGAGACGAAGTTCA | | | | | | | | | | |
| Chromosome | start | end | strand | MM | target_seq | PAM | alignment | distance | position | gene name | gene id |
| GU305914.1 | 90177 | 90199 | - | 0 | TGAACTTCGTCTCCGTCGAC | AGG | \|\|\|\|\|\|\|\|[\|\|\|\|\|\|\|\|\|\|\|\|]PAM | NA | Intergenic | NA | NA |
| T27342 | GTCGACAGGATGAAGTATCTCGG | 1000 | | | | | | | | | |
| Oligo fwd | CACCGTCGACAGGATGAAGTATCT | | | | | | | | | | |
| Oligo rev | AAACAGATACTTCATCCTGTCGAC | | | | | | | | | | |
| Chromosome | start | end | strand | MM | target_seq | PAM | alignment | distance | position | gene name | gene id |
| GU305914.1 | 90163 | 90185 | - | 0 | GTCGACAGGATGAAGTATCT | CGG | \|\|\|\|\|\|\|\|[\|\|\|\|\|\|\|\|\|\|\|\|]PAM | NA | Intergenic | NA | NA |
| T27343 | TCGACAGGATGAAGTATCTCGGG | 1000 | | | | | | | | | |
| Oligo fwd | CACCTCGACAGGATGAAGTATCTC | | | | | | | | | | |
| Oligo rev | AAACGAGATACTTCATCCTGTCGA | | | | | | | | | | |
| Chromosome | start | end | strand | MM | target_seq | PAM | alignment | distance | position | gene name | gene id |
| GU305914.1 | 90162 | 90184 | - | 0 | TCGACAGGATGAAGTATCTC | GGG | \|\|\|\|\|\|\|\|[\|\|\|\|\|\|\|\|\|\|\|\|]PAM | NA | Intergenic | NA | NA |
| T27344 | CAGGATGAAGTATCTCGGGCAGG | 1000 | | | | | | | | | |
| Oligo fwd | CACCCAGGATGAAGTATCTCGGGC | | | | | | | | | | |
| Oligo rev | AAACGCCCGAGATACTTCATCCTG | | | | | | | | | | |
| Chromosome | start | end | strand | MM | target_seq | PAM | alignment | distance | position | gene name | gene id |
| GU305914.1 | 90158 | 90180 | - | 0 | CAGGATGAAGTATCTCGGGC | AGG | \|\|\|\|\|\|\|\|[\|\|\|\|\|\|\|\|\|\|\|\|]PAM | NA | Intergenic | NA | NA |
| T27345 | GAAGTATCTCGGGCAGGTCCTGG | 1000 | | | | | | | | | |
| Oligo fwd | CACCGAAGTATCTCGGGCAGGTCC | | | | | | | | | | |
| Oligo rev | AAACGGACCTGCCCGAGATACTTC | | | | | | | | | | |
| Chromosome | start | end | strand | MM | target_seq | PAM | alignment | distance | position | gene name | gene id |
| GU305914.1 | 90152 | 90174 | - | 0 | GAAGTATCTCGGGCAGGTCC | TGG | \|\|\|\|\|\|\|\|[\|\|\|\|\|\|\|\|\|\|\|\|]PAM | NA | Intergenic | NA | NA |
| T27346 | TCGCTGATCGACGCGATCACCGG | 1000 | | | | | | | | | |
| Oligo fwd | CACCTCGCTGATCGACGCGATCAC | | | | | | | | | | |
| Oligo rev | AAACGTGATCGCGTCGATCAGCGA | | | | | | | | | | |
| Chromosome | start | end | strand | MM | target_seq | PAM | alignment | distance | position | gene name | gene id |
| GU305914.1 | 90112 | 90134 | - | 0 | TCGCTGATCGACGCGATCAC | CGG | \|\|\|\|\|\|\|\|[\|\|\|\|\|\|\|\|\|\|\|\|]PAM | NA | Intergenic | NA | NA |

| | | | | | | | | | | | |
|---|---|---|---|---|---|---|---|---|---|---|---|
| T27347 | GATCGACGCGATCACCGGCGAGG | 1000 | | | | | | | | | |
| Oligo fwd | CACCGATCGACGCGATCACCGGCG | | | | | | | | | | |
| Oligo rev | AAACCGCCGGTGATCGCGTCGATC | | | | | | | | | | |
| Chromosome | start | end | strand | MM | target_seq | PAM | alignment | distance | position | gene name | gene id |
| GU305914.1 | 90107 | 90129 | - | 0 | GATCGACGCGATCACCGGCG | AGG | \|\|\|\|\|\|\|\|[\|\|\|\|\|\|\|\|\|\|\|\|]PAM | NA | Intergenic | NA | NA |
| GU305914.1 | 138353 | 138375 | - | 4 | CACCGACGCGATCGCCGTCG | CGG | -\|-\|\|\|\|\|[\|\|\|\|\|-\|\|\|-\|\|]PAM | NA | Intergenic | NA | NA |
| T27348 | CAAGAGAGACTTCGTCGCGACGG | 1000 | | | | | | | | | |
| Oligo fwd | CACCCAAGAGAGACTTCGTCGCGA | | | | | | | | | | |
| Oligo rev | AAACTCGCGACGAAGTCTCTCTTG | | | | | | | | | | |
| Chromosome | start | end | strand | MM | target_seq | PAM | alignment | distance | position | gene name | gene id |
| GU305914.1 | 90065 | 90087 | - | 0 | CAAGAGAGACTTCGTCGCGA | CGG | \|\|\|\|\|\|\|\|[\|\|\|\|\|\|\|\|\|\|\|\|]PAM | NA | Intergenic | NA | NA |
| T27349 | AGACTTCGTCGCGACGGTCACGG | 1000 | | | | | | | | | |
| Oligo fwd | CACCAGACTTCGTCGCGACGGTCA | | | | | | | | | | |
| Oligo rev | AAACTGACCGTCGCGACGAAGTCT | | | | | | | | | | |
| Chromosome | start | end | strand | MM | target_seq | PAM | alignment | distance | position | gene name | gene id |
| GU305914.1 | 90059 | 90081 | - | 0 | AGACTTCGTCGCGACGGTCA | CGG | \|\|\|\|\|\|\|\|[\|\|\|\|\|\|\|\|\|\|\|\|]PAM | NA | Intergenic | NA | NA |
| T27350 | CGATCGACGATGCCGTGTGCAGG | 1000 | | | | | | | | | |
| Oligo fwd | CACCCGATCGACGATGCCGTGTGC | | | | | | | | | | |
| Oligo rev | AAACGCACACGGCATCGTCGATCG | | | | | | | | | | |
| Chromosome | start | end | strand | MM | target_seq | PAM | alignment | distance | position | gene name | gene id |
| GU305914.1 | 90024 | 90046 | - | 0 | CGATCGACGATGCCGTGTGC | AGG | \|\|\|\|\|\|\|\|[\|\|\|\|\|\|\|\|\|\|\|\|]PAM | NA | Intergenic | NA | NA |
| T27351 | TGCCGTGTGCAGGTTCGCCATGG | 1000 | | | | | | | | | |
| Oligo fwd | CACCTGCCGTGTGCAGGTTCGCCA | | | | | | | | | | |
| Oligo rev | AAACTGGCGAACCTGCACACGGCA | | | | | | | | | | |
| Chromosome | start | end | strand | MM | target_seq | PAM | alignment | distance | position | gene name | gene id |
| GU305914.1 | 90014 | 90036 | - | 0 | TGCCGTGTGCAGGTTCGCCA | TGG | \|\|\|\|\|\|\|\|[\|\|\|\|\|\|\|\|\|\|\|\|]PAM | NA | Intergenic | NA | NA |
| T27352 | GCAGGTTCGCCATGGACGTGCGG | 1000 | | | | | | | | | |
| Oligo fwd | CACCGCAGGTTCGCCATGGACGTG | | | | | | | | | | |
| Oligo rev | AAACCACGTCCATGGCGAACCTGC | | | | | | | | | | |
| Chromosome | start | end | strand | MM | target_seq | PAM | alignment | distance | position | gene name | gene id |
| GU305914.1 | 90006 | 90028 | - | 0 | GCAGGTTCGCCATGGACGTG | CGG | \|\|\|\|\|\|\|\|[\|\|\|\|\|\|\|\|\|\|\|\|]PAM | NA | Intergenic | NA | NA |
| T27353 | GGTTCGCCATGGACGTGCGGAGG | 1000 | | | | | | | | | |
| Oligo fwd | CACCGGTTCGCCATGGACGTGCGG | | | | | | | | | | |
| Oligo rev | AAACCCGCACGTCCATGGCGAACC | | | | | | | | | | |
| Chromosome | start | end | strand | MM | target_seq | PAM | alignment | distance | position | gene name | gene id |
| GU305914.1 | 90003 | 90025 | - | 0 | GGTTCGCCATGGACGTGCGG | AGG | \|\|\|\|\|\|\|\|[\|\|\|\|\|\|\|\|\|\|\|\|]PAM | NA | Intergenic | NA | NA |
| T27354 | GCCATGGACGTGCGGAGGTCCGG | 1000 | | | | | | | | | |
| Oligo fwd | CACCGCCATGGACGTGCGGAGGTC | | | | | | | | | | |
| Oligo rev | AAACGACCTCCGCACGTCCATGGC | | | | | | | | | | |
| Chromosome | start | end | strand | MM | target_seq | PAM | alignment | distance | position | gene name | gene id |
| GU305914.1 | 89998 | 90020 | - | 0 | GCCATGGACGTGCGGAGGTC | CGG | \|\|\|\|\|\|\|\|[\|\|\|\|\|\|\|\|\|\|\|\|]PAM | NA | Intergenic | NA | NA |
| T27355 | CCATGGACGTGCGGAGGTCCGGG | 1000 | | | | | | | | | |
| Oligo fwd | CACCCCATGGACGTGCGGAGGTCC | | | | | | | | | | |
| Oligo rev | AAACGGACCTCCGCACGTCCATGG | | | | | | | | | | |
| Chromosome | start | end | strand | MM | target_seq | PAM | alignment | distance | position | gene name | gene id |
| GU305914.1 | 89997 | 90019 | - | 0 | CCATGGACGTGCGGAGGTCC | GGG | \|\|\|\|\|\|\|\|[\|\|\|\|\|\|\|\|\|\|\|\|]PAM | NA | Intergenic | NA | NA |
| T27356 | GACGTGCGGAGGTCCGGGCACGG | 1000 | | | | | | | | | |
| Oligo fwd | CACCGACGTGCGGAGGTCCGGGCA | | | | | | | | | | |
| Oligo rev | AAACTGCCCGGACCTCCGCACGTC | | | | | | | | | | |
| Chromosome | start | end | strand | MM | target_seq | PAM | alignment | distance | position | gene name | gene id |
| GU305914.1 | 89992 | 90014 | - | 0 | GACGTGCGGAGGTCCGGGCA | CGG | \|\|\|\|\|\|\|\|[\|\|\|\|\|\|\|\|\|\|\|\|]PAM | NA | Intergenic | NA | NA |
| GU305914.1 | 226196 | 226218 | + | 4 | CACGGGCGGAGGTCCGGACT | TGG | -\|\|\|-\|\|\|[\|\|\|\|\|\|\|\|\|-\|-]PAM | NA | Intergenic | NA | NA |
| T27357 | ACGTGCGGAGGTCCGGGCACGGG | 1000 | | | | | | | | | |
| Oligo fwd | CACCACGTGCGGAGGTCCGGGCAC | | | | | | | | | | |
| Oligo rev | AAACGTGCCCGGACCTCCGCACGT | | | | | | | | | | |
| Chromosome | start | end | strand | MM | target_seq | PAM | alignment | distance | position | gene name | gene id |
| GU305914.1 | 89991 | 90013 | - | 0 | ACGTGCGGAGGTCCGGGCAC | GGG | \|\|\|\|\|\|\|\|[\|\|\|\|\|\|\|\|\|\|\|\|]PAM | NA | Intergenic | NA | NA |
| T27358 | TGCGGAGGTCCGGGCACGGGAGG | 1000 | | | | | | | | | |
| Oligo fwd | CACCTGCGGAGGTCCGGGCACGGG | | | | | | | | | | |
| Oligo rev | AAACCCCGTGCCCGGACCTCCGCA | | | | | | | | | | |
| Chromosome | start | end | strand | MM | target_seq | PAM | alignment | distance | position | gene name | gene id |
| GU305914.1 | 89988 | 90010 | - | 0 | TGCGGAGGTCCGGGCACGGG | AGG | \|\|\|\|\|\|\|\|[\|\|\|\|\|\|\|\|\|\|\|\|]PAM | NA | Intergenic | NA | NA |
| T27359 | GCGGAGGTCCGGGCACGGGAGGG | 1000 | | | | | | | | | |
| Oligo fwd | CACCGCGGAGGTCCGGGCACGGGA | | | | | | | | | | |
| Oligo rev | AAACTCCCGTGCCCGGACCTCCGC | | | | | | | | | | |
| Chromosome | start | end | strand | MM | target_seq | PAM | alignment | distance | position | gene name | gene id |
| GU305914.1 | 89987 | 90009 | - | 0 | GCGGAGGTCCGGGCACGGGA | GGG | \|\|\|\|\|\|\|\|[\|\|\|\|\|\|\|\|\|\|\|\|]PAM | NA | Intergenic | NA | NA |
| T27360 | CACGGGAGGGACGAGATCAGCGG | 1000 | | | | | | | | | |
| Oligo fwd | CACCCACGGGAGGGACGAGATCAG | | | | | | | | | | |
| Oligo rev | AAACCTGATCTCGTCCCTCCCGTG | | | | | | | | | | |
| Chromosome | start | end | strand | MM | target_seq | PAM | alignment | distance | position | gene name | gene id |
| GU305914.1 | 89974 | 89996 | - | 0 | CACGGGAGGGACGAGATCAG | CGG | \|\|\|\|\|\|\|\|[\|\|\|\|\|\|\|\|\|\|\|\|]PAM | NA | Intergenic | NA | NA |
| T27361 | ACGGGAGGGACGAGATCAGCGGG | 1000 | | | | | | | | | |
| Oligo fwd | CACCACGGGAGGGACGAGATCAGC | | | | | | | | | | |
| Oligo rev | AAACGCTGATCTCGTCCCTCCCGT | | | | | | | | | | |

<table>
<tr><td>Chromosome</td><td>start</td><td>end</td><td>strand</td><td>MM</td><td>target_seq</td><td>PAM</td><td>alignment</td><td>distance</td><td>position</td><td>gene name</td><td>gene id</td></tr>
<tr><td>GU305914.1</td><td>89973</td><td>89995</td><td>-</td><td>0</td><td>ACGGGAGGGACGAGATCAGC</td><td>GGG</td><td>||||||||[||||||||||||]PAM</td><td>NA</td><td>Intergenic</td><td>NA</td><td>NA</td></tr>
<tr><td>T27362</td><td>GAGCACGCAGTCCTTCAACCTGG</td><td>1000</td><td></td><td></td><td></td><td></td><td></td><td></td><td></td><td></td><td></td></tr>
<tr><td>Oligo fwd</td><td>CACCGAGCACGCAGTCCTTCAACC</td><td></td><td></td><td></td><td></td><td></td><td></td><td></td><td></td><td></td><td></td></tr>
<tr><td>Oligo rev</td><td>AAACGGTTGAAGGACTGCGTGCTC</td><td></td><td></td><td></td><td></td><td></td><td></td><td></td><td></td><td></td><td></td></tr>
<tr><td>Chromosome</td><td>start</td><td>end</td><td>strand</td><td>MM</td><td>target_seq</td><td>PAM</td><td>alignment</td><td>distance</td><td>position</td><td>gene name</td><td>gene id</td></tr>
<tr><td>GU305914.1</td><td>89951</td><td>89973</td><td>-</td><td>0</td><td>GAGCACGCAGTCCTTCAACC</td><td>TGG</td><td>||||||||[||||||||||||]PAM</td><td>NA</td><td>Intergenic</td><td>NA</td><td>NA</td></tr>
<tr><td>T27363</td><td>GGACCTCAGCCCCTACGCCACGG</td><td>1000</td><td></td><td></td><td></td><td></td><td></td><td></td><td></td><td></td><td></td></tr>
<tr><td>Oligo fwd</td><td>CACCGGACCTCAGCCCCTACGCCA</td><td></td><td></td><td></td><td></td><td></td><td></td><td></td><td></td><td></td><td></td></tr>
<tr><td>Oligo rev</td><td>AAACTGGCGTAGGGGCTGAGGTCC</td><td></td><td></td><td></td><td></td><td></td><td></td><td></td><td></td><td></td><td></td></tr>
<tr><td>Chromosome</td><td>start</td><td>end</td><td>strand</td><td>MM</td><td>target_seq</td><td>PAM</td><td>alignment</td><td>distance</td><td>position</td><td>gene name</td><td>gene id</td></tr>
<tr><td>GU305914.1</td><td>89930</td><td>89952</td><td>-</td><td>0</td><td>GGACCTCAGCCCCTACGCCA</td><td>CGG</td><td>||||||||[||||||||||||]PAM</td><td>NA</td><td>Intergenic</td><td>NA</td><td>NA</td></tr>
<tr><td>T27364</td><td>TCCTGTCGTTCCAGTATTACCGG</td><td>1000</td><td></td><td></td><td></td><td></td><td></td><td></td><td></td><td></td><td></td></tr>
<tr><td>Oligo fwd</td><td>CACCTCCTGTCGTTCCAGTATTAC</td><td></td><td></td><td></td><td></td><td></td><td></td><td></td><td></td><td></td><td></td></tr>
<tr><td>Oligo rev</td><td>AAACGTAATACTGGAACGACAGGA</td><td></td><td></td><td></td><td></td><td></td><td></td><td></td><td></td><td></td><td></td></tr>
<tr><td>Chromosome</td><td>start</td><td>end</td><td>strand</td><td>MM</td><td>target_seq</td><td>PAM</td><td>alignment</td><td>distance</td><td>position</td><td>gene name</td><td>gene id</td></tr>
<tr><td>GU305914.1</td><td>89895</td><td>89917</td><td>-</td><td>0</td><td>TCCTGTCGTTCCAGTATTAC</td><td>CGG</td><td>||||||||[||||||||||||]PAM</td><td>NA</td><td>Intergenic</td><td>NA</td><td>NA</td></tr>
<tr><td>T27365</td><td>GTTCTCCATCATCCAGAACCTGG</td><td>1000</td><td></td><td></td><td></td><td></td><td></td><td></td><td></td><td></td><td></td></tr>
<tr><td>Oligo fwd</td><td>CACCGTTCTCCATCATCCAGAACC</td><td></td><td></td><td></td><td></td><td></td><td></td><td></td><td></td><td></td><td></td></tr>
<tr><td>Oligo rev</td><td>AAACGGTTCTGGATGATGGAGAAC</td><td></td><td></td><td></td><td></td><td></td><td></td><td></td><td></td><td></td><td></td></tr>
<tr><td>Chromosome</td><td>start</td><td>end</td><td>strand</td><td>MM</td><td>target_seq</td><td>PAM</td><td>alignment</td><td>distance</td><td>position</td><td>gene name</td><td>gene id</td></tr>
<tr><td>GU305914.1</td><td>89867</td><td>89889</td><td>-</td><td>0</td><td>GTTCTCCATCATCCAGAACC</td><td>TGG</td><td>||||||||[||||||||||||]PAM</td><td>NA</td><td>Intergenic</td><td>NA</td><td>NA</td></tr>
<tr><td>T27366</td><td>CATCAACGCCGCGTCGTACGTGG</td><td>1000</td><td></td><td></td><td></td><td></td><td></td><td></td><td></td><td></td><td></td></tr>
<tr><td>Oligo fwd</td><td>CACCCATCAACGCCGCGTCGTACG</td><td></td><td></td><td></td><td></td><td></td><td></td><td></td><td></td><td></td><td></td></tr>
<tr><td>Oligo rev</td><td>AAACCGTACGACGCGGCGTTGATG</td><td></td><td></td><td></td><td></td><td></td><td></td><td></td><td></td><td></td><td></td></tr>
<tr><td>Chromosome</td><td>start</td><td>end</td><td>strand</td><td>MM</td><td>target_seq</td><td>PAM</td><td>alignment</td><td>distance</td><td>position</td><td>gene name</td><td>gene id</td></tr>
<tr><td>GU305914.1</td><td>89840</td><td>89862</td><td>-</td><td>0</td><td>CATCAACGCCGCGTCGTACG</td><td>TGG</td><td>||||||||[||||||||||||]PAM</td><td>NA</td><td>Intergenic</td><td>NA</td><td>NA</td></tr>
<tr><td>T27367</td><td>GGTCGACAACCCCCTGACGACGG</td><td>1000</td><td></td><td></td><td></td><td></td><td></td><td></td><td></td><td></td><td></td></tr>
<tr><td>Oligo fwd</td><td>CACCGGTCGACAACCCCCTGACGA</td><td></td><td></td><td></td><td></td><td></td><td></td><td></td><td></td><td></td><td></td></tr>
<tr><td>Oligo rev</td><td>AAACTCGTCAGGGGGTTGTCGACC</td><td></td><td></td><td></td><td></td><td></td><td></td><td></td><td></td><td></td><td></td></tr>
<tr><td>Chromosome</td><td>start</td><td>end</td><td>strand</td><td>MM</td><td>target_seq</td><td>PAM</td><td>alignment</td><td>distance</td><td>position</td><td>gene name</td><td>gene id</td></tr>
<tr><td>GU305914.1</td><td>89819</td><td>89841</td><td>-</td><td>0</td><td>GGTCGACAACCCCCTGACGA</td><td>CGG</td><td>||||||||[||||||||||||]PAM</td><td>NA</td><td>Intergenic</td><td>NA</td><td>NA</td></tr>
<tr><td>T27368</td><td>CGACGGCGCAGATCTCCAAGTGG</td><td>1000</td><td></td><td></td><td></td><td></td><td></td><td></td><td></td><td></td><td></td></tr>
<tr><td>Oligo fwd</td><td>CACCCGACGGCGCAGATCTCCAAG</td><td></td><td></td><td></td><td></td><td></td><td></td><td></td><td></td><td></td><td></td></tr>
<tr><td>Oligo rev</td><td>AAACCTTGGAGATCTGCGCCGTCG</td><td></td><td></td><td></td><td></td><td></td><td></td><td></td><td></td><td></td><td></td></tr>
<tr><td>Chromosome</td><td>start</td><td>end</td><td>strand</td><td>MM</td><td>target_seq</td><td>PAM</td><td>alignment</td><td>distance</td><td>position</td><td>gene name</td><td>gene id</td></tr>
<tr><td>GU305914.1</td><td>89802</td><td>89824</td><td>-</td><td>0</td><td>CGACGGCGCAGATCTCCAAG</td><td>TGG</td><td>||||||||[||||||||||||]PAM</td><td>NA</td><td>Intergenic</td><td>NA</td><td>NA</td></tr>
<tr><td>T27369</td><td>GACGGCGCAGATCTCCAAGTGGG</td><td>1000</td><td></td><td></td><td></td><td></td><td></td><td></td><td></td><td></td><td></td></tr>
<tr><td>Oligo fwd</td><td>CACCGACGGCGCAGATCTCCAAGT</td><td></td><td></td><td></td><td></td><td></td><td></td><td></td><td></td><td></td><td></td></tr>
<tr><td>Oligo rev</td><td>AAACACTTGGAGATCTGCGCCGTC</td><td></td><td></td><td></td><td></td><td></td><td></td><td></td><td></td><td></td><td></td></tr>
<tr><td>Chromosome</td><td>start</td><td>end</td><td>strand</td><td>MM</td><td>target_seq</td><td>PAM</td><td>alignment</td><td>distance</td><td>position</td><td>gene name</td><td>gene id</td></tr>
<tr><td>GU305914.1</td><td>89801</td><td>89823</td><td>-</td><td>0</td><td>GACGGCGCAGATCTCCAAGT</td><td>GGG</td><td>||||||||[||||||||||||]PAM</td><td>NA</td><td>Intergenic</td><td>NA</td><td>NA</td></tr>
<tr><td>T27370</td><td>CTGCACTTCCAGTCCATCTGCGG</td><td>1000</td><td></td><td></td><td></td><td></td><td></td><td></td><td></td><td></td><td></td></tr>
<tr><td>Oligo fwd</td><td>CACCCTGCACTTCCAGTCCATCTG</td><td></td><td></td><td></td><td></td><td></td><td></td><td></td><td></td><td></td><td></td></tr>
<tr><td>Oligo rev</td><td>AAACCAGATGGACTGGAAGTGCAG</td><td></td><td></td><td></td><td></td><td></td><td></td><td></td><td></td><td></td><td></td></tr>
<tr><td>Chromosome</td><td>start</td><td>end</td><td>strand</td><td>MM</td><td>target_seq</td><td>PAM</td><td>alignment</td><td>distance</td><td>position</td><td>gene name</td><td>gene id</td></tr>
<tr><td>GU305914.1</td><td>89773</td><td>89795</td><td>-</td><td>0</td><td>CTGCACTTCCAGTCCATCTG</td><td>CGG</td><td>||||||||[||||||||||||]PAM</td><td>NA</td><td>Intergenic</td><td>NA</td><td>NA</td></tr>
<tr><td>T27371</td><td>CAGTCCATCTGCGGCGCGTTCGG</td><td>1000</td><td></td><td></td><td></td><td></td><td></td><td></td><td></td><td></td><td></td></tr>
<tr><td>Oligo fwd</td><td>CACCCAGTCCATCTGCGGCGCGTT</td><td></td><td></td><td></td><td></td><td></td><td></td><td></td><td></td><td></td><td></td></tr>
<tr><td>Oligo rev</td><td>AAACAACGCGCCGCAGATGGACTG</td><td></td><td></td><td></td><td></td><td></td><td></td><td></td><td></td><td></td><td></td></tr>
<tr><td>Chromosome</td><td>start</td><td>end</td><td>strand</td><td>MM</td><td>target_seq</td><td>PAM</td><td>alignment</td><td>distance</td><td>position</td><td>gene name</td><td>gene id</td></tr>
<tr><td>GU305914.1</td><td>89764</td><td>89786</td><td>-</td><td>0</td><td>CAGTCCATCTGCGGCGCGTT</td><td>CGG</td><td>||||||||[||||||||||||]PAM</td><td>NA</td><td>Intergenic</td><td>NA</td><td>NA</td></tr>
<tr><td>T27372</td><td>AGTCCATCTGCGGCGCGTTCGGG</td><td>1000</td><td></td><td></td><td></td><td></td><td></td><td></td><td></td><td></td><td></td></tr>
<tr><td>Oligo fwd</td><td>CACCAGTCCATCTGCGGCGCGTTC</td><td></td><td></td><td></td><td></td><td></td><td></td><td></td><td></td><td></td><td></td></tr>
<tr><td>Oligo rev</td><td>AAACGAACGCGCCGCAGATGGACT</td><td></td><td></td><td></td><td></td><td></td><td></td><td></td><td></td><td></td><td></td></tr>
<tr><td>Chromosome</td><td>start</td><td>end</td><td>strand</td><td>MM</td><td>target_seq</td><td>PAM</td><td>alignment</td><td>distance</td><td>position</td><td>gene name</td><td>gene id</td></tr>
<tr><td>GU305914.1</td><td>89763</td><td>89785</td><td>-</td><td>0</td><td>AGTCCATCTGCGGCGCGTTC</td><td>GGG</td><td>||||||||[||||||||||||]PAM</td><td>NA</td><td>Intergenic</td><td>NA</td><td>NA</td></tr>
<tr><td>T27373</td><td>CGGGACGACGCCCCTCAAGAAGG</td><td>1000</td><td></td><td></td><td></td><td></td><td></td><td></td><td></td><td></td><td></td></tr>
<tr><td>Oligo fwd</td><td>CACCCGGGACGACGCCCCTCAAGA</td><td></td><td></td><td></td><td></td><td></td><td></td><td></td><td></td><td></td><td></td></tr>
<tr><td>Oligo rev</td><td>AAACTCTTGAGGGGCGTCGTCCCG</td><td></td><td></td><td></td><td></td><td></td><td></td><td></td><td></td><td></td><td></td></tr>
<tr><td>Chromosome</td><td>start</td><td>end</td><td>strand</td><td>MM</td><td>target_seq</td><td>PAM</td><td>alignment</td><td>distance</td><td>position</td><td>gene name</td><td>gene id</td></tr>
<tr><td>GU305914.1</td><td>89744</td><td>89766</td><td>-</td><td>0</td><td>CGGGACGACGCCCCTCAAGA</td><td>AGG</td><td>||||||||[||||||||||||]PAM</td><td>NA</td><td>Intergenic</td><td>NA</td><td>NA</td></tr>
<tr><td>T27374</td><td>GGGACGACGCCCCTCAAGAAGGG</td><td>1000</td><td></td><td></td><td></td><td></td><td></td><td></td><td></td><td></td><td></td></tr>
<tr><td>Oligo fwd</td><td>CACCGGGACGACGCCCCTCAAGAA</td><td></td><td></td><td></td><td></td><td></td><td></td><td></td><td></td><td></td><td></td></tr>
<tr><td>Oligo rev</td><td>AAACTTCTTGAGGGGCGTCGTCCC</td><td></td><td></td><td></td><td></td><td></td><td></td><td></td><td></td><td></td><td></td></tr>
<tr><td>Chromosome</td><td>start</td><td>end</td><td>strand</td><td>MM</td><td>target_seq</td><td>PAM</td><td>alignment</td><td>distance</td><td>position</td><td>gene name</td><td>gene id</td></tr>
<tr><td>GU305914.1</td><td>89743</td><td>89765</td><td>-</td><td>0</td><td>GGGACGACGCCCCTCAAGAA</td><td>GGG</td><td>||||||||[||||||||||||]PAM</td><td>NA</td><td>Intergenic</td><td>NA</td><td>NA</td></tr>
<tr><td>T27375</td><td>GGACGACGCCCCTCAAGAAGGGG</td><td>1000</td><td></td><td></td><td></td><td></td><td></td><td></td><td></td><td></td><td></td></tr>
<tr><td>Oligo fwd</td><td>CACCGGACGACGCCCCTCAAGAAG</td><td></td><td></td><td></td><td></td><td></td><td></td><td></td><td></td><td></td><td></td></tr>
<tr><td>Oligo rev</td><td>AAACCTTCTTGAGGGGCGTCGTCC</td><td></td><td></td><td></td><td></td><td></td><td></td><td></td><td></td><td></td><td></td></tr>
<tr><td>Chromosome</td><td>start</td><td>end</td><td>strand</td><td>MM</td><td>target_seq</td><td>PAM</td><td>alignment</td><td>distance</td><td>position</td><td>gene name</td><td>gene id</td></tr>
<tr><td>GU305914.1</td><td>89742</td><td>89764</td><td>-</td><td>0</td><td>GGACGACGCCCCTCAAGAAG</td><td>GGG</td><td>||||||||[||||||||||||]PAM</td><td>NA</td><td>Intergenic</td><td>NA</td><td>NA</td></tr>
<tr><td>T27376</td><td>GAAGGGGTTCCTGAACGTCAAGG</td><td>1000</td><td></td><td></td><td></td><td></td><td></td><td></td><td></td><td></td><td></td></tr>
<tr><td>Oligo fwd</td><td>CACCGAAGGGGTTCCTGAACGTCA</td><td></td><td></td><td></td><td></td><td></td><td></td><td></td><td></td><td></td><td></td></tr>
</table>

Oligo rev AAACTGACGTTCAGGAACCCCTTC
Chromosome start end strand MM target_seq PAM alignment distance position gene name gene id
GU305914.1 89726 89748 - 0 GAAGGGGTTCCTGAACGTCA AGG ||||||||[||||||||||||]PAM NA Intergenic NA NA

T27377 CGTCGAGTTCGAGCGCATCATGG 1000
Oligo fwd CACCCGTCGAGTTCGAGCGCATCA
Oligo rev AAACTGATGCGCTCGAACTCGACG
Chromosome start end strand MM target_seq PAM alignment distance position gene name gene id
GU305914.1 89684 89706 - 0 CGTCGAGTTCGAGCGCATCA TGG ||||||||[||||||||||||]PAM NA Intergenic NA NA

T27378 TCGAGCGCATCATGGACTTCAGG 1000
Oligo fwd CACCTCGAGCGCATCATGGACTTC
Oligo rev AAACGAAGTCCATGATGCGCTCGA
Chromosome start end strand MM target_seq PAM alignment distance position gene name gene id
GU305914.1 89676 89698 - 0 TCGAGCGCATCATGGACTTC AGG ||||||||[||||||||||||]PAM NA Intergenic NA NA

T27379 CATGGACTTCAGGTCCTTCCAGG 1000
Oligo fwd CACCCATGGACTTCAGGTCCTTCC
Oligo rev AAACGGAAGGACCTGAAGTCCATG
Chromosome start end strand MM target_seq PAM alignment distance position gene name gene id
GU305914.1 89666 89688 - 0 CATGGACTTCAGGTCCTTCC AGG ||||||||[||||||||||||]PAM NA Intergenic NA NA

T27380 CTTCAGGTCCTTCCAGGAGACGG 1000
Oligo fwd CACCCTTCAGGTCCTTCCAGGAGA
Oligo rev AAACTCTCCTGGAAGGACCTGAAG
Chromosome start end strand MM target_seq PAM alignment distance position gene name gene id
GU305914.1 89660 89682 - 0 CTTCAGGTCCTTCCAGGAGA CGG ||||||||[||||||||||||]PAM NA Intergenic NA NA

T27381 TTCAGGTCCTTCCAGGAGACGGG 1000
Oligo fwd CACCTTCAGGTCCTTCCAGGAGAC
Oligo rev AAACGTCTCCTGGAAGGACCTGAA
Chromosome start end strand MM target_seq PAM alignment distance position gene name gene id
GU305914.1 89659 89681 - 0 TTCAGGTCCTTCCAGGAGAC GGG ||||||||[||||||||||||]PAM NA Intergenic NA NA

T27382 GATCAAGTCGTGTAAGATGTCGG 1000
Oligo fwd CACCGATCAAGTCGTGTAAGATGT
Oligo rev AAACACATCTTACACGACTTGATC
Chromosome start end strand MM target_seq PAM alignment distance position gene name gene id
GU305914.1 89612 89634 - 0 GATCAAGTCGTGTAAGATGT CGG ||||||||[||||||||||||]PAM NA Intergenic NA NA

T27383 AGAGCTGCCGCATCAAGAATCGG 1000
Oligo fwd CACCAGAGCTGCCGCATCAAGAAT
Oligo rev AAACATTCTTGATGCGGCAGCTCT
Chromosome start end strand MM target_seq PAM alignment distance position gene name gene id
GU305914.1 89577 89599 - 0 AGAGCTGCCGCATCAAGAAT CGG ||||||||[||||||||||||]PAM NA Intergenic NA NA

T27384 GAGCAGCGTCTTCTTCAAGAAGG 1000
Oligo fwd CACCGAGCAGCGTCTTCTTCAAGA
Oligo rev AAACTCTTGAAGAAGACGCTGCTC
Chromosome start end strand MM target_seq PAM alignment distance position gene name gene id
GU305914.1 89534 89556 - 0 GAGCAGCGTCTTCTTCAAGA AGG ||||||||[||||||||||||]PAM NA Intergenic NA NA

T27385 AGCAGCGTCTTCTTCAAGAAGGG 1000
Oligo fwd CACCAGCAGCGTCTTCTTCAAGAA
Oligo rev AAACTTCTTGAAGAAGACGCTGCT
Chromosome start end strand MM target_seq PAM alignment distance position gene name gene id
GU305914.1 89533 89555 - 0 AGCAGCGTCTTCTTCAAGAA GGG ||||||||[||||||||||||]PAM NA Intergenic NA NA

T27386 TCAAGAAGGGCGCCCTGCAGAGG 1000
Oligo fwd CACCTCAAGAAGGGCGCCCTGCAG
Oligo rev AAACCTGCAGGGCGCCCTTCTTGA
Chromosome start end strand MM target_seq PAM alignment distance position gene name gene id
GU305914.1 89520 89542 - 0 TCAAGAAGGGCGCCCTGCAG AGG ||||||||[||||||||||||]PAM NA Intergenic NA NA

T27387 GCAGAGGAAGAATCCCATCAAGG 1000
Oligo fwd CACCGCAGAGGAAGAATCCCATCA
Oligo rev AAACTGATGGGATTCTTCCTCTGC
Chromosome start end strand MM target_seq PAM alignment distance position gene name gene id
GU305914.1 89504 89526 - 0 GCAGAGGAAGAATCCCATCA AGG ||||||||[||||||||||||]PAM NA Intergenic NA NA

T27388 CAGAGGAAGAATCCCATCAAGGG 1000
Oligo fwd CACCCAGAGGAAGAATCCCATCAA
Oligo rev AAACTTGATGGGATTCTTCCTCTG
Chromosome start end strand MM target_seq PAM alignment distance position gene name gene id
GU305914.1 89503 89525 - 0 CAGAGGAAGAATCCCATCAA GGG ||||||||[||||||||||||]PAM NA Intergenic NA NA

T27389 CTGCCACGAGAAGCTCTTCCCGG 1000
Oligo fwd CACCCTGCCACGAGAAGCTCTTCC
Oligo rev AAACGGAAGAGCTTCTCGTGGCAG
Chromosome start end strand MM target_seq PAM alignment distance position gene name gene id
GU305914.1 89456 89478 - 0 CTGCCACGAGAAGCTCTTCC CGG ||||||||[||||||||||||]PAM NA Intergenic NA NA

T27390 GAGAAGCTCTTCCCGGCCTGCGG 1000
Oligo fwd CACCGAGAAGCTCTTCCCGGCCTG
Oligo rev AAACCAGGCCGGGAAGAGCTTCTC
Chromosome start end strand MM target_seq PAM alignment distance position gene name gene id
GU305914.1 89449 89471 - 0 GAGAAGCTCTTCCCGGCCTG CGG ||||||||[||||||||||||]PAM NA Intergenic NA NA

T27391 GCCTGTCGTGCCTCGAGTTCTGG 1000

| | | | | | | | | | | | |
|---|---|---|---|---|---|---|---|---|---|---|---|
| Oligo fwd | CACCGCCTGTCGTGCCTCGAGTTC | | | | | | | | | | |
| Oligo rev | AAACGAACTCGAGGCACGACAGGC | | | | | | | | | | |
| Chromosome | start | end | strand | MM | target_seq | PAM | alignment | distance | position | gene name | gene id |
| GU305914.1 | 89427 | 89449 | - | 0 | GCCTGTCGTGCCTCGAGTTC | TGG | \|\|\|\|\|\|\|\|[\|\|\|\|\|\|\|\|\|\|\|\|]PAM | NA | Intergenic | NA | NA |
| T27392 | GCAGAACTCGCTGCCGCGCTCGG | 1000 | | | | | | | | | |
| Oligo fwd | CACCGCAGAACTCGCTGCCGCGCT | | | | | | | | | | |
| Oligo rev | AAACAGCGCGGCAGCGAGTTCTGC | | | | | | | | | | |
| Chromosome | start | end | strand | MM | target_seq | PAM | alignment | distance | position | gene name | gene id |
| GU305914.1 | 89393 | 89415 | - | 0 | GCAGAACTCGCTGCCGCGCT | CGG | \|\|\|\|\|\|\|\|[\|\|\|\|\|\|\|\|\|\|\|\|]PAM | NA | Intergenic | NA | NA |
| T27393 | GCTGCCGCGCTCGGTCAACGTGG | 1000 | | | | | | | | | |
| Oligo fwd | CACCGCTGCCGCGCTCGGTCAACG | | | | | | | | | | |
| Oligo rev | AAACCGTTGACCGAGCGCGGCAGC | | | | | | | | | | |
| Chromosome | start | end | strand | MM | target_seq | PAM | alignment | distance | position | gene name | gene id |
| GU305914.1 | 89384 | 89406 | - | 0 | GCTGCCGCGCTCGGTCAACG | TGG | \|\|\|\|\|\|\|\|[\|\|\|\|\|\|\|\|\|\|\|\|]PAM | NA | Intergenic | NA | NA |
| GU305914.1 | 125825 | 125847 | - | 4 | CCCGCCGCGCACGGCCAACG | AGG | -\|-\|\|\|\|\|[\|\|-\|\|\|-\|\|\|\|\|]PAM | NA | Intergenic | NA | NA |
| T27394 | CTGCCGCGCTCGGTCAACGTGGG | 1000 | | | | | | | | | |
| Oligo fwd | CACCCTGCCGCGCTCGGTCAACGT | | | | | | | | | | |
| Oligo rev | AAACACGTTGACCGAGCGCGGCAG | | | | | | | | | | |
| Chromosome | start | end | strand | MM | target_seq | PAM | alignment | distance | position | gene name | gene id |
| GU305914.1 | 89383 | 89405 | - | 0 | CTGCCGCGCTCGGTCAACGT | GGG | \|\|\|\|\|\|\|\|[\|\|\|\|\|\|\|\|\|\|\|\|]PAM | NA | Intergenic | NA | NA |
| GU305914.1 | 153088 | 153110 | - | 4 | CTGCCGACCTCGATCGACGT | AGG | \|\|\|\|\|\|--[\|\|\|\|-\|\|-\|\|\|\|]PAM | NA | Intergenic | NA | NA |
| T27395 | GCGCTCGGTCAACGTGGGCAAGG | 1000 | | | | | | | | | |
| Oligo fwd | CACCGCGCTCGGTCAACGTGGGCA | | | | | | | | | | |
| Oligo rev | AAACTGCCCACGTTGACCGAGCGC | | | | | | | | | | |
| Chromosome | start | end | strand | MM | target_seq | PAM | alignment | distance | position | gene name | gene id |
| GU305914.1 | 89378 | 89400 | - | 0 | GCGCTCGGTCAACGTGGGCA | AGG | \|\|\|\|\|\|\|\|[\|\|\|\|\|\|\|\|\|\|\|\|]PAM | NA | Intergenic | NA | NA |
| T27396 | CTCGGTCAACGTGGGCAAGGTGG | 1000 | | | | | | | | | |
| Oligo fwd | CACCCTCGGTCAACGTGGGCAAGG | | | | | | | | | | |
| Oligo rev | AAACCCTTGCCCACGTTGACCGAG | | | | | | | | | | |
| Chromosome | start | end | strand | MM | target_seq | PAM | alignment | distance | position | gene name | gene id |
| GU305914.1 | 89375 | 89397 | - | 0 | CTCGGTCAACGTGGGCAAGG | TGG | \|\|\|\|\|\|\|\|[\|\|\|\|\|\|\|\|\|\|\|\|]PAM | NA | Intergenic | NA | NA |
| T27397 | GGTCAACGTGGGCAAGGTGGAGG | 1000 | | | | | | | | | |
| Oligo fwd | CACCGGTCAACGTGGGCAAGGTGG | | | | | | | | | | |
| Oligo rev | AAACCCACCTTGCCCACGTTGACC | | | | | | | | | | |
| Chromosome | start | end | strand | MM | target_seq | PAM | alignment | distance | position | gene name | gene id |
| GU305914.1 | 89372 | 89394 | - | 0 | GGTCAACGTGGGCAAGGTGG | AGG | \|\|\|\|\|\|\|\|[\|\|\|\|\|\|\|\|\|\|\|\|]PAM | NA | Intergenic | NA | NA |
| T27398 | GGTGGAGGACTTCGACAACCTGG | 1000 | | | | | | | | | |
| Oligo fwd | CACCGGTGGAGGACTTCGACAACC | | | | | | | | | | |
| Oligo rev | AAACGGTTGTCGAAGTCCTCCACC | | | | | | | | | | |
| Chromosome | start | end | strand | MM | target_seq | PAM | alignment | distance | position | gene name | gene id |
| GU305914.1 | 89357 | 89379 | - | 0 | GGTGGAGGACTTCGACAACC | TGG | \|\|\|\|\|\|\|\|[\|\|\|\|\|\|\|\|\|\|\|\|]PAM | NA | Intergenic | NA | NA |
| T27399 | CGACTACGACGAGAGCGACGTGG | 1000 | | | | | | | | | |
| Oligo fwd | CACCCGACTACGACGAGAGCGACG | | | | | | | | | | |
| Oligo rev | AAACCGTCGCTCTCGTCGTAGTCG | | | | | | | | | | |
| Chromosome | start | end | strand | MM | target_seq | PAM | alignment | distance | position | gene name | gene id |
| GU305914.1 | 89309 | 89331 | - | 0 | CGACTACGACGAGAGCGACG | TGG | \|\|\|\|\|\|\|\|[\|\|\|\|\|\|\|\|\|\|\|\|]PAM | NA | Intergenic | NA | NA |
| T27400 | CTACGACGAGAGCGACGTGGTGG | 1000 | | | | | | | | | |
| Oligo fwd | CACCCTACGACGAGAGCGACGTGG | | | | | | | | | | |
| Oligo rev | AAACCCACGTCGCTCTCGTCGTAG | | | | | | | | | | |
| Chromosome | start | end | strand | MM | target_seq | PAM | alignment | distance | position | gene name | gene id |
| GU305914.1 | 89306 | 89328 | - | 0 | CTACGACGAGAGCGACGTGG | TGG | \|\|\|\|\|\|\|\|[\|\|\|\|\|\|\|\|\|\|\|\|]PAM | NA | Intergenic | NA | NA |
| T27401 | CGACGTGGTGGACATCCAGCCGG | 1000 | | | | | | | | | |
| Oligo fwd | CACCCGACGTGGTGGACATCCAGC | | | | | | | | | | |
| Oligo rev | AAACGCTGGATGTCCACCACGTCG | | | | | | | | | | |
| Chromosome | start | end | strand | MM | target_seq | PAM | alignment | distance | position | gene name | gene id |
| GU305914.1 | 89294 | 89316 | - | 0 | CGACGTGGTGGACATCCAGC | CGG | \|\|\|\|\|\|\|\|[\|\|\|\|\|\|\|\|\|\|\|\|]PAM | NA | Intergenic | NA | NA |
| T27402 | GGACTGCCTGCTCAGCTACGTGG | 1000 | | | | | | | | | |
| Oligo fwd | CACCGGACTGCCTGCTCAGCTACG | | | | | | | | | | |
| Oligo rev | AAACCGTAGCTGAGCAGGCAGTCC | | | | | | | | | | |
| Chromosome | start | end | strand | MM | target_seq | PAM | alignment | distance | position | gene name | gene id |
| GU305914.1 | 89273 | 89295 | - | 0 | GGACTGCCTGCTCAGCTACG | TGG | \|\|\|\|\|\|\|\|[\|\|\|\|\|\|\|\|\|\|\|\|]PAM | NA | Intergenic | NA | NA |
| T27403 | AACAAGTTCCTCTACATGTTCGG | 1000 | | | | | | | | | |
| Oligo fwd | CACCAACAAGTTCCTCTACATGTT | | | | | | | | | | |
| Oligo rev | AAACAACATGTAGAGGAACTTGTT | | | | | | | | | | |
| Chromosome | start | end | strand | MM | target_seq | PAM | alignment | distance | position | gene name | gene id |
| GU305914.1 | 89236 | 89258 | - | 0 | AACAAGTTCCTCTACATGTT | CGG | \|\|\|\|\|\|\|\|[\|\|\|\|\|\|\|\|\|\|\|\|]PAM | NA | Intergenic | NA | NA |
| T27404 | TCCTCTACATGTTCGGCTTCAGG | 1000 | | | | | | | | | |
| Oligo fwd | CACCTCCTCTACATGTTCGGCTTC | | | | | | | | | | |
| Oligo rev | AAACGAAGCCGAACATGTAGAGGA | | | | | | | | | | |
| Chromosome | start | end | strand | MM | target_seq | PAM | alignment | distance | position | gene name | gene id |
| GU305914.1 | 89229 | 89251 | - | 0 | TCCTCTACATGTTCGGCTTC | AGG | \|\|\|\|\|\|\|\|[\|\|\|\|\|\|\|\|\|\|\|\|]PAM | NA | Intergenic | NA | NA |
| T27405 | CCTCTACATGTTCGGCTTCAGGG | 1000 | | | | | | | | | |
| Oligo fwd | CACCCCTCTACATGTTCGGCTTCA | | | | | | | | | | |
| Oligo rev | AAACTGAAGCCGAACATGTAGAGG | | | | | | | | | | |
| Chromosome | start | end | strand | MM | target_seq | PAM | alignment | distance | position | gene name | gene id |

GU305914.1 89228 89250 - 0 CCTCTACATGTTCGGCTTCA GGG ||||||||[||||||||||||]PAM NA Intergenic NA NA

T27406 GGACTACATGAGCACCATCCAGG 1000
Oligo fwd CACCGGACTACATGAGCACCATCC
Oligo rev AAACGGATGGTGCTCATGTAGTCC
Chromosome start end strand MM target_seq PAM alignment distance position gene name gene id
GU305914.1 89207 89229 - 0 GGACTACATGAGCACCATCC AGG ||||||||[||||||||||||]PAM NA Intergenic NA NA

T27407 GACTACATGAGCACCATCCAGGG 1000
Oligo fwd CACCGACTACATGAGCACCATCCA
Oligo rev AAACTGGATGGTGCTCATGTAGTC
Chromosome start end strand MM target_seq PAM alignment distance position gene name gene id
GU305914.1 89206 89228 - 0 GACTACATGAGCACCATCCA GGG ||||||||[||||||||||||]PAM NA Intergenic NA NA

T27408 GCAGTTCCCCTGTCTCCTCAAGG 1000
Oligo fwd CACCGCAGTTCCCCTGTCTCCTCA
Oligo rev AAACTGAGGAGACAGGGGAACTGC
Chromosome start end strand MM target_seq PAM alignment distance position gene name gene id
GU305914.1 89150 89172 - 0 GCAGTTCCCCTGTCTCCTCA AGG ||||||||[||||||||||||]PAM NA Intergenic NA NA

T27409 GCACTTCAAGAAGATGAAGCTGG 1000
Oligo fwd CACCGCACTTCAAGAAGATGAAGC
Oligo rev AAACGCTTCATCTTCTTGAAGTGC
Chromosome start end strand MM target_seq PAM alignment distance position gene name gene id
GU305914.1 89090 89112 - 0 GCACTTCAAGAAGATGAAGC TGG ||||||||[||||||||||||]PAM NA Intergenic NA NA
GU305914.1 42212 42234 - 4 GTAGTTGAAGAAGATGATGC AGG |-|-||-|[|||||||||-||]PAM NA Intergenic NA NA

T27410 TTCAAGAAGATGAAGCTGGACGG 1000
Oligo fwd CACCTTCAAGAAGATGAAGCTGGA
Oligo rev AAACTCCAGCTTCATCTTCTTGAA
Chromosome start end strand MM target_seq PAM alignment distance position gene name gene id
GU305914.1 89086 89108 - 0 TTCAAGAAGATGAAGCTGGA CGG ||||||||[||||||||||||]PAM NA Intergenic NA NA

T27411 GATGAAGCTGGACGGCGTCAAGG 1000
Oligo fwd CACCGATGAAGCTGGACGGCGTCA
Oligo rev AAACTGACGCCGTCCAGCTTCATC
Chromosome start end strand MM target_seq PAM alignment distance position gene name gene id
GU305914.1 89078 89100 - 0 GATGAAGCTGGACGGCGTCA AGG ||||||||[||||||||||||]PAM NA Intergenic NA NA

T27412 GGACGGCGTCAAGGCGCCCCAGG 1000
Oligo fwd CACCGGACGGCGTCAAGGCGCCCC
Oligo rev AAACGGGGCGCCTTGACGCCGTCC
Chromosome start end strand MM target_seq PAM alignment distance position gene name gene id
GU305914.1 89069 89091 - 0 GGACGGCGTCAAGGCGCCCC AGG ||||||||[||||||||||||]PAM NA Intergenic NA NA

T27413 CGGCGTCAAGGCGCCCCAGGTGG 1000
Oligo fwd CACCCGGCGTCAAGGCGCCCCAGG
Oligo rev AAACCCTGGGGCGCCTTGACGCCG
Chromosome start end strand MM target_seq PAM alignment distance position gene name gene id
GU305914.1 89066 89088 - 0 CGGCGTCAAGGCGCCCCAGG TGG ||||||||[||||||||||||]PAM NA Intergenic NA NA

T27414 GTGCTCAAGAACGTCTTCGACGG 1000
Oligo fwd CACCGTGCTCAAGAACGTCTTCGA
Oligo rev AAACTCGAAGACGTTCTTGAGCAC
Chromosome start end strand MM target_seq PAM alignment distance position gene name gene id
GU305914.1 89023 89045 - 0 GTGCTCAAGAACGTCTTCGA CGG ||||||||[||||||||||||]PAM NA Intergenic NA NA

T27415 CGAGAAGTACTCCAGCTCCATGG 1000
Oligo fwd CACCCGAGAAGTACTCCAGCTCCA
Oligo rev AAACTGGAGCTGGAGTACTTCTCG
Chromosome start end strand MM target_seq PAM alignment distance position gene name gene id
GU305914.1 88970 88992 - 0 CGAGAAGTACTCCAGCTCCA TGG ||||||||[||||||||||||]PAM NA Intergenic NA NA

T27416 GAGAAGTACTCCAGCTCCATGGG 1000
Oligo fwd CACCGAGAAGTACTCCAGCTCCAT
Oligo rev AAACATGGAGCTGGAGTACTTCTC
Chromosome start end strand MM target_seq PAM alignment distance position gene name gene id
GU305914.1 88969 88991 - 0 GAGAAGTACTCCAGCTCCAT GGG ||||||||[||||||||||||]PAM NA Intergenic NA NA

T27417 ACTCCAGCTCCATGGGCACCCGG 1000
Oligo fwd CACCACTCCAGCTCCATGGGCACC
Oligo rev AAACGGTGCCCATGGAGCTGGAGT
Chromosome start end strand MM target_seq PAM alignment distance position gene name gene id
GU305914.1 88962 88984 - 0 ACTCCAGCTCCATGGGCACC CGG ||||||||[||||||||||||]PAM NA Intergenic NA NA

T27418 CTCCAGCTCCATGGGCACCCGGG 1000
Oligo fwd CACCCTCCAGCTCCATGGGCACCC
Oligo rev AAACGGGTGCCCATGGAGCTGGAG
Chromosome start end strand MM target_seq PAM alignment distance position gene name gene id
GU305914.1 88961 88983 - 0 CTCCAGCTCCATGGGCACCC GGG ||||||||[||||||||||||]PAM NA Intergenic NA NA
GU305914.1 88877 88899 - 4 CACGGGCTCCATGGGCACGC AGG |-|--|||[||||||||||-|]PAM NA Intergenic NA NA

T27419 ACCCGGGACGTGTTTCAGTTCGG 1000
Oligo fwd CACCACCCGGGACGTGTTTCAGTT
Oligo rev AAACAACTGAAACACGTCCCGGGT
Chromosome start end strand MM target_seq PAM alignment distance position gene name gene id
GU305914.1 88945 88967 - 0 ACCCGGGACGTGTTTCAGTT CGG ||||||||[||||||||||||]PAM NA Intergenic NA NA

T27420 GTGTTTCAGTTCGGACAGATCGG 1000

<table>
<tr><td>Oligo fwd</td><td>CACCGTGTTTCAGTTCGGACAGAT</td><td></td><td></td><td></td><td></td><td></td><td></td><td></td><td></td><td></td><td></td></tr>
<tr><td>Oligo rev</td><td>AAACATCTGTCCGAACTGAAACAC</td><td></td><td></td><td></td><td></td><td></td><td></td><td></td><td></td><td></td><td></td></tr>
<tr><td>Chromosome</td><td>start</td><td>end</td><td>strand</td><td>MM</td><td>target_seq</td><td>PAM</td><td>alignment</td><td>distance</td><td>position</td><td>gene name</td><td>gene id</td></tr>
<tr><td>GU305914.1</td><td>88936</td><td>88958</td><td>-</td><td>0</td><td>GTGTTTCAGTTCGGACAGAT</td><td>CGG</td><td>||||||||[||||||||||||]PAM</td><td>NA</td><td>Intergenic</td><td>NA</td><td>NA</td></tr>
<tr><td>T27421</td><td>TGTTTCAGTTCGGACAGATCGGG</td><td>1000</td><td></td><td></td><td></td><td></td><td></td><td></td><td></td><td></td><td></td></tr>
<tr><td>Oligo fwd</td><td>CACCTGTTTCAGTTCGGACAGATC</td><td></td><td></td><td></td><td></td><td></td><td></td><td></td><td></td><td></td><td></td></tr>
<tr><td>Oligo rev</td><td>AAACGATCTGTCCGAACTGAAACA</td><td></td><td></td><td></td><td></td><td></td><td></td><td></td><td></td><td></td><td></td></tr>
<tr><td>Chromosome</td><td>start</td><td>end</td><td>strand</td><td>MM</td><td>target_seq</td><td>PAM</td><td>alignment</td><td>distance</td><td>position</td><td>gene name</td><td>gene id</td></tr>
<tr><td>GU305914.1</td><td>88935</td><td>88957</td><td>-</td><td>0</td><td>TGTTTCAGTTCGGACAGATC</td><td>GGG</td><td>||||||||[||||||||||||]PAM</td><td>NA</td><td>Intergenic</td><td>NA</td><td>NA</td></tr>
<tr><td>T27422</td><td>CGGACAGATCGGGTACTACGTGG</td><td>1000</td><td></td><td></td><td></td><td></td><td></td><td></td><td></td><td></td><td></td></tr>
<tr><td>Oligo fwd</td><td>CACCCGGACAGATCGGGTACTACG</td><td></td><td></td><td></td><td></td><td></td><td></td><td></td><td></td><td></td><td></td></tr>
<tr><td>Oligo rev</td><td>AAACCGTAGTACCCGATCTGTCCG</td><td></td><td></td><td></td><td></td><td></td><td></td><td></td><td></td><td></td><td></td></tr>
<tr><td>Chromosome</td><td>start</td><td>end</td><td>strand</td><td>MM</td><td>target_seq</td><td>PAM</td><td>alignment</td><td>distance</td><td>position</td><td>gene name</td><td>gene id</td></tr>
<tr><td>GU305914.1</td><td>88925</td><td>88947</td><td>-</td><td>0</td><td>CGGACAGATCGGGTACTACG</td><td>TGG</td><td>||||||||[||||||||||||]PAM</td><td>NA</td><td>Intergenic</td><td>NA</td><td>NA</td></tr>
<tr><td>T27423</td><td>GGACAGATCGGGTACTACGTGGG</td><td>1000</td><td></td><td></td><td></td><td></td><td></td><td></td><td></td><td></td><td></td></tr>
<tr><td>Oligo fwd</td><td>CACCGGACAGATCGGGTACTACGT</td><td></td><td></td><td></td><td></td><td></td><td></td><td></td><td></td><td></td><td></td></tr>
<tr><td>Oligo rev</td><td>AAACACGTAGTACCCGATCTGTCC</td><td></td><td></td><td></td><td></td><td></td><td></td><td></td><td></td><td></td><td></td></tr>
<tr><td>Chromosome</td><td>start</td><td>end</td><td>strand</td><td>MM</td><td>target_seq</td><td>PAM</td><td>alignment</td><td>distance</td><td>position</td><td>gene name</td><td>gene id</td></tr>
<tr><td>GU305914.1</td><td>88924</td><td>88946</td><td>-</td><td>0</td><td>GGACAGATCGGGTACTACGT</td><td>GGG</td><td>||||||||[||||||||||||]PAM</td><td>NA</td><td>Intergenic</td><td>NA</td><td>NA</td></tr>
<tr><td>T27424</td><td>GACAGATCGGGTACTACGTGGGG</td><td>1000</td><td></td><td></td><td></td><td></td><td></td><td></td><td></td><td></td><td></td></tr>
<tr><td>Oligo fwd</td><td>CACCGACAGATCGGGTACTACGTG</td><td></td><td></td><td></td><td></td><td></td><td></td><td></td><td></td><td></td><td></td></tr>
<tr><td>Oligo rev</td><td>AAACCACGTAGTACCCGATCTGTC</td><td></td><td></td><td></td><td></td><td></td><td></td><td></td><td></td><td></td><td></td></tr>
<tr><td>Chromosome</td><td>start</td><td>end</td><td>strand</td><td>MM</td><td>target_seq</td><td>PAM</td><td>alignment</td><td>distance</td><td>position</td><td>gene name</td><td>gene id</td></tr>
<tr><td>GU305914.1</td><td>88923</td><td>88945</td><td>-</td><td>0</td><td>GACAGATCGGGTACTACGTG</td><td>GGG</td><td>||||||||[||||||||||||]PAM</td><td>NA</td><td>Intergenic</td><td>NA</td><td>NA</td></tr>
<tr><td>T27425</td><td>GATCGGGTACTACGTGGGGTCGG</td><td>1000</td><td></td><td></td><td></td><td></td><td></td><td></td><td></td><td></td><td></td></tr>
<tr><td>Oligo fwd</td><td>CACCGATCGGGTACTACGTGGGGT</td><td></td><td></td><td></td><td></td><td></td><td></td><td></td><td></td><td></td><td></td></tr>
<tr><td>Oligo rev</td><td>AAACACCCCACGTAGTACCCGATC</td><td></td><td></td><td></td><td></td><td></td><td></td><td></td><td></td><td></td><td></td></tr>
<tr><td>Chromosome</td><td>start</td><td>end</td><td>strand</td><td>MM</td><td>target_seq</td><td>PAM</td><td>alignment</td><td>distance</td><td>position</td><td>gene name</td><td>gene id</td></tr>
<tr><td>GU305914.1</td><td>88919</td><td>88941</td><td>-</td><td>0</td><td>GATCGGGTACTACGTGGGGT</td><td>CGG</td><td>||||||||[||||||||||||]PAM</td><td>NA</td><td>Intergenic</td><td>NA</td><td>NA</td></tr>
<tr><td>T27426</td><td>ATCGGGTACTACGTGGGGTCGGG</td><td>1000</td><td></td><td></td><td></td><td></td><td></td><td></td><td></td><td></td><td></td></tr>
<tr><td>Oligo fwd</td><td>CACCATCGGGTACTACGTGGGGTC</td><td></td><td></td><td></td><td></td><td></td><td></td><td></td><td></td><td></td><td></td></tr>
<tr><td>Oligo rev</td><td>AAACGACCCCACGTAGTACCCGAT</td><td></td><td></td><td></td><td></td><td></td><td></td><td></td><td></td><td></td><td></td></tr>
<tr><td>Chromosome</td><td>start</td><td>end</td><td>strand</td><td>MM</td><td>target_seq</td><td>PAM</td><td>alignment</td><td>distance</td><td>position</td><td>gene name</td><td>gene id</td></tr>
<tr><td>GU305914.1</td><td>88918</td><td>88940</td><td>-</td><td>0</td><td>ATCGGGTACTACGTGGGGTC</td><td>GGG</td><td>||||||||[||||||||||||]PAM</td><td>NA</td><td>Intergenic</td><td>NA</td><td>NA</td></tr>
<tr><td>T27427</td><td>TCGGGTACTACGTGGGGTCGGGG</td><td>1000</td><td></td><td></td><td></td><td></td><td></td><td></td><td></td><td></td><td></td></tr>
<tr><td>Oligo fwd</td><td>CACCTCGGGTACTACGTGGGGTCG</td><td></td><td></td><td></td><td></td><td></td><td></td><td></td><td></td><td></td><td></td></tr>
<tr><td>Oligo rev</td><td>AAACCGACCCCACGTAGTACCCGA</td><td></td><td></td><td></td><td></td><td></td><td></td><td></td><td></td><td></td><td></td></tr>
<tr><td>Chromosome</td><td>start</td><td>end</td><td>strand</td><td>MM</td><td>target_seq</td><td>PAM</td><td>alignment</td><td>distance</td><td>position</td><td>gene name</td><td>gene id</td></tr>
<tr><td>GU305914.1</td><td>88917</td><td>88939</td><td>-</td><td>0</td><td>TCGGGTACTACGTGGGGTCG</td><td>GGG</td><td>||||||||[||||||||||||]PAM</td><td>NA</td><td>Intergenic</td><td>NA</td><td>NA</td></tr>
<tr><td>T27428</td><td>CGGGTACTACGTGGGGTCGGGGG</td><td>1000</td><td></td><td></td><td></td><td></td><td></td><td></td><td></td><td></td><td></td></tr>
<tr><td>Oligo fwd</td><td>CACCCGGGTACTACGTGGGGTCGG</td><td></td><td></td><td></td><td></td><td></td><td></td><td></td><td></td><td></td><td></td></tr>
<tr><td>Oligo rev</td><td>AAACCCGACCCCACGTAGTACCCG</td><td></td><td></td><td></td><td></td><td></td><td></td><td></td><td></td><td></td><td></td></tr>
<tr><td>Chromosome</td><td>start</td><td>end</td><td>strand</td><td>MM</td><td>target_seq</td><td>PAM</td><td>alignment</td><td>distance</td><td>position</td><td>gene name</td><td>gene id</td></tr>
<tr><td>GU305914.1</td><td>88916</td><td>88938</td><td>-</td><td>0</td><td>CGGGTACTACGTGGGGTCGG</td><td>GGG</td><td>||||||||[||||||||||||]PAM</td><td>NA</td><td>Intergenic</td><td>NA</td><td>NA</td></tr>
<tr><td>T27429</td><td>GTACTACGTGGGGTCGGGGGTGG</td><td>1000</td><td></td><td></td><td></td><td></td><td></td><td></td><td></td><td></td><td></td></tr>
<tr><td>Oligo fwd</td><td>CACCGTACTACGTGGGGTCGGGGG</td><td></td><td></td><td></td><td></td><td></td><td></td><td></td><td></td><td></td><td></td></tr>
<tr><td>Oligo rev</td><td>AAACCCCCCGACCCCACGTAGTAC</td><td></td><td></td><td></td><td></td><td></td><td></td><td></td><td></td><td></td><td></td></tr>
<tr><td>Chromosome</td><td>start</td><td>end</td><td>strand</td><td>MM</td><td>target_seq</td><td>PAM</td><td>alignment</td><td>distance</td><td>position</td><td>gene name</td><td>gene id</td></tr>
<tr><td>GU305914.1</td><td>88913</td><td>88935</td><td>-</td><td>0</td><td>GTACTACGTGGGGTCGGGGG</td><td>TGG</td><td>||||||||[||||||||||||]PAM</td><td>NA</td><td>Intergenic</td><td>NA</td><td>NA</td></tr>
<tr><td>T27430</td><td>GGTGGACCGCAGCCTCAACACGG</td><td>1000</td><td></td><td></td><td></td><td></td><td></td><td></td><td></td><td></td><td></td></tr>
<tr><td>Oligo fwd</td><td>CACCGGTGGACCGCAGCCTCAACA</td><td></td><td></td><td></td><td></td><td></td><td></td><td></td><td></td><td></td><td></td></tr>
<tr><td>Oligo rev</td><td>AAACTGTTGAGGCTGCGGTCCACC</td><td></td><td></td><td></td><td></td><td></td><td></td><td></td><td></td><td></td><td></td></tr>
<tr><td>Chromosome</td><td>start</td><td>end</td><td>strand</td><td>MM</td><td>target_seq</td><td>PAM</td><td>alignment</td><td>distance</td><td>position</td><td>gene name</td><td>gene id</td></tr>
<tr><td>GU305914.1</td><td>88895</td><td>88917</td><td>-</td><td>0</td><td>GGTGGACCGCAGCCTCAACA</td><td>CGG</td><td>||||||||[||||||||||||]PAM</td><td>NA</td><td>Intergenic</td><td>NA</td><td>NA</td></tr>
<tr><td>T27431</td><td>GTGGACCGCAGCCTCAACACGGG</td><td>1000</td><td></td><td></td><td></td><td></td><td></td><td></td><td></td><td></td><td></td></tr>
<tr><td>Oligo fwd</td><td>CACCGTGGACCGCAGCCTCAACAC</td><td></td><td></td><td></td><td></td><td></td><td></td><td></td><td></td><td></td><td></td></tr>
<tr><td>Oligo rev</td><td>AAACGTGTTGAGGCTGCGGTCCAC</td><td></td><td></td><td></td><td></td><td></td><td></td><td></td><td></td><td></td><td></td></tr>
<tr><td>Chromosome</td><td>start</td><td>end</td><td>strand</td><td>MM</td><td>target_seq</td><td>PAM</td><td>alignment</td><td>distance</td><td>position</td><td>gene name</td><td>gene id</td></tr>
<tr><td>GU305914.1</td><td>88894</td><td>88916</td><td>-</td><td>0</td><td>GTGGACCGCAGCCTCAACAC</td><td>GGG</td><td>||||||||[||||||||||||]PAM</td><td>NA</td><td>Intergenic</td><td>NA</td><td>NA</td></tr>
<tr><td>T27432</td><td>CAGCCTCAACACGGGCTCCATGG</td><td>1000</td><td></td><td></td><td></td><td></td><td></td><td></td><td></td><td></td><td></td></tr>
<tr><td>Oligo fwd</td><td>CACCCAGCCTCAACACGGGCTCCA</td><td></td><td></td><td></td><td></td><td></td><td></td><td></td><td></td><td></td><td></td></tr>
<tr><td>Oligo rev</td><td>AAACTGGAGCCCGTGTTGAGGCTG</td><td></td><td></td><td></td><td></td><td></td><td></td><td></td><td></td><td></td><td></td></tr>
<tr><td>Chromosome</td><td>start</td><td>end</td><td>strand</td><td>MM</td><td>target_seq</td><td>PAM</td><td>alignment</td><td>distance</td><td>position</td><td>gene name</td><td>gene id</td></tr>
<tr><td>GU305914.1</td><td>88886</td><td>88908</td><td>-</td><td>0</td><td>CAGCCTCAACACGGGCTCCA</td><td>TGG</td><td>||||||||[||||||||||||]PAM</td><td>NA</td><td>Intergenic</td><td>NA</td><td>NA</td></tr>
<tr><td>GU305914.1</td><td>124389</td><td>124411</td><td>+</td><td>3</td><td>CAGCGTCAACACCGACTCCA</td><td>GGG</td><td>||||-|||[||||-|-|||||]PAM</td><td>NA</td><td>Intergenic</td><td>NA</td><td>NA</td></tr>
<tr><td>T27433</td><td>AGCCTCAACACGGGCTCCATGGG</td><td>1000</td><td></td><td></td><td></td><td></td><td></td><td></td><td></td><td></td><td></td></tr>
<tr><td>Oligo fwd</td><td>CACCAGCCTCAACACGGGCTCCAT</td><td></td><td></td><td></td><td></td><td></td><td></td><td></td><td></td><td></td><td></td></tr>
<tr><td>Oligo rev</td><td>AAACATGGAGCCCGTGTTGAGGCT</td><td></td><td></td><td></td><td></td><td></td><td></td><td></td><td></td><td></td><td></td></tr>
<tr><td>Chromosome</td><td>start</td><td>end</td><td>strand</td><td>MM</td><td>target_seq</td><td>PAM</td><td>alignment</td><td>distance</td><td>position</td><td>gene name</td><td>gene id</td></tr>
<tr><td>GU305914.1</td><td>88885</td><td>88907</td><td>-</td><td>0</td><td>AGCCTCAACACGGGCTCCAT</td><td>GGG</td><td>||||||||[||||||||||||]PAM</td><td>NA</td><td>Intergenic</td><td>NA</td><td>NA</td></tr>
<tr><td>T27434</td><td>CACGGGCTCCATGGGCACGCAGG</td><td>1000</td><td></td><td></td><td></td><td></td><td></td><td></td><td></td><td></td><td></td></tr>
<tr><td>Oligo fwd</td><td>CACCCACGGGCTCCATGGGCACGC</td><td></td><td></td><td></td><td></td><td></td><td></td><td></td><td></td><td></td><td></td></tr>
<tr><td>Oligo rev</td><td>AAACGCGTGCCCATGGAGCCCGTG</td><td></td><td></td><td></td><td></td><td></td><td></td><td></td><td></td><td></td><td></td></tr>
<tr><td>Chromosome</td><td>start</td><td>end</td><td>strand</td><td>MM</td><td>target_seq</td><td>PAM</td><td>alignment</td><td>distance</td><td>position</td><td>gene name</td><td>gene id</td></tr>
<tr><td>GU305914.1</td><td>88877</td><td>88899</td><td>-</td><td>0</td><td>CACGGGCTCCATGGGCACGC</td><td>AGG</td><td>||||||||[||||||||||||]PAM</td><td>NA</td><td>Intergenic</td><td>NA</td><td>NA</td></tr>
</table>

| | | | | | | | | | | | |
|---|---|---|---|---|---|---|---|---|---|---|---|
| GU305914.1 | 88961 | 88983 | - | 4 | CTCCAGCTCCATGGGCACCC | GGG | \|-\|--\|\|\|[\|\|\|\|\|\|\|\|\|\|-\|]PAM | NA | Intergenic | NA | NA |
| T27435 | ACTACCGCTTCATGCGCTACAGG | 1000 | | | | | | | | | |
| Oligo fwd | CACCACTACCGCTTCATGCGCTAC | | | | | | | | | | |
| Oligo rev | AAACGTAGCGCATGAAGCGGTAGT | | | | | | | | | | |
| Chromosome | start | end | strand | MM | target_seq | PAM | alignment | distance | position | gene name | gene id |
| GU305914.1 | 88854 | 88876 | - | 0 | ACTACCGCTTCATGCGCTAC | AGG | \|\|\|\|\|\|\|\|[\|\|\|\|\|\|\|\|\|\|\|\|]PAM | NA | Intergenic | NA | NA |
| T27436 | GTACATCATCGCCACCAAGCTGG | 1000 | | | | | | | | | |
| Oligo fwd | CACCGTACATCATCGCCACCAAGC | | | | | | | | | | |
| Oligo rev | AAACGCTTGGTGGCGATGATGTAC | | | | | | | | | | |
| Chromosome | start | end | strand | MM | target_seq | PAM | alignment | distance | position | gene name | gene id |
| GU305914.1 | 88832 | 88854 | - | 0 | GTACATCATCGCCACCAAGC | TGG | \|\|\|\|\|\|\|\|[\|\|\|\|\|\|\|\|\|\|\|\|]PAM | NA | Intergenic | NA | NA |
| T27437 | CATCATCGCCACCAAGCTGGTGG | 1000 | | | | | | | | | |
| Oligo fwd | CACCCATCATCGCCACCAAGCTGG | | | | | | | | | | |
| Oligo rev | AAACCCAGCTTGGTGGCGATGATG | | | | | | | | | | |
| Chromosome | start | end | strand | MM | target_seq | PAM | alignment | distance | position | gene name | gene id |
| GU305914.1 | 88829 | 88851 | - | 0 | CATCATCGCCACCAAGCTGG | TGG | \|\|\|\|\|\|\|\|[\|\|\|\|\|\|\|\|\|\|\|\|]PAM | NA | Intergenic | NA | NA |
| T27438 | CGAGAACGTCATGTACGACGCGG | 1000 | | | | | | | | | |
| Oligo fwd | CACCCGAGAACGTCATGTACGACG | | | | | | | | | | |
| Oligo rev | AAACCGTCGTACATGACGTTCTCG | | | | | | | | | | |
| Chromosome | start | end | strand | MM | target_seq | PAM | alignment | distance | position | gene name | gene id |
| GU305914.1 | 88781 | 88803 | - | 0 | CGAGAACGTCATGTACGACG | CGG | \|\|\|\|\|\|\|\|[\|\|\|\|\|\|\|\|\|\|\|\|]PAM | NA | Intergenic | NA | NA |
| T27439 | ACGCGGACGTCGTCCGCTCGCGG | 1000 | | | | | | | | | |
| Oligo fwd | CACCACGCGGACGTCGTCCGCTCG | | | | | | | | | | |
| Oligo rev | AAACCGAGCGGACGACGTCCGCGT | | | | | | | | | | |
| Chromosome | start | end | strand | MM | target_seq | PAM | alignment | distance | position | gene name | gene id |
| GU305914.1 | 88764 | 88786 | - | 0 | ACGCGGACGTCGTCCGCTCG | CGG | \|\|\|\|\|\|\|\|[\|\|\|\|\|\|\|\|\|\|\|\|]PAM | NA | Intergenic | NA | NA |
| T27440 | CGCGGACGTCGTCCGCTCGCGGG | 1000 | | | | | | | | | |
| Oligo fwd | CACCCGCGGACGTCGTCCGCTCGC | | | | | | | | | | |
| Oligo rev | AAACGCGAGCGGACGACGTCCGCG | | | | | | | | | | |
| Chromosome | start | end | strand | MM | target_seq | PAM | alignment | distance | position | gene name | gene id |
| GU305914.1 | 88763 | 88785 | - | 0 | CGCGGACGTCGTCCGCTCGC | GGG | \|\|\|\|\|\|\|\|[\|\|\|\|\|\|\|\|\|\|\|\|]PAM | NA | Intergenic | NA | NA |
| T27441 | CGTCGTCCGCTCGCGGGTGCTGG | 1000 | | | | | | | | | |
| Oligo fwd | CACCCGTCGTCCGCTCGCGGGTGC | | | | | | | | | | |
| Oligo rev | AAACGCACCCGCGAGCGGACGACG | | | | | | | | | | |
| Chromosome | start | end | strand | MM | target_seq | PAM | alignment | distance | position | gene name | gene id |
| GU305914.1 | 88757 | 88779 | - | 0 | CGTCGTCCGCTCGCGGGTGC | TGG | \|\|\|\|\|\|\|\|[\|\|\|\|\|\|\|\|\|\|\|\|]PAM | NA | Intergenic | NA | NA |
| T27442 | CGTCCGCTCGCGGGTGCTGGCGG | 1000 | | | | | | | | | |
| Oligo fwd | CACCCGTCCGCTCGCGGGTGCTGG | | | | | | | | | | |
| Oligo rev | AAACCCAGCACCCGCGAGCGGACG | | | | | | | | | | |
| Chromosome | start | end | strand | MM | target_seq | PAM | alignment | distance | position | gene name | gene id |
| GU305914.1 | 88754 | 88776 | - | 0 | CGTCCGCTCGCGGGTGCTGG | CGG | \|\|\|\|\|\|\|\|[\|\|\|\|\|\|\|\|\|\|\|\|]PAM | NA | Intergenic | NA | NA |
| T27443 | GCTGGCGGCGCTCGACTCGACGG | 1000 | | | | | | | | | |
| Oligo fwd | CACCGCTGGCGGCGCTCGACTCGA | | | | | | | | | | |
| Oligo rev | AAACTCGAGTCGAGCGCCGCCAGC | | | | | | | | | | |
| Chromosome | start | end | strand | MM | target_seq | PAM | alignment | distance | position | gene name | gene id |
| GU305914.1 | 88739 | 88761 | - | 0 | GCTGGCGGCGCTCGACTCGA | CGG | \|\|\|\|\|\|\|\|[\|\|\|\|\|\|\|\|\|\|\|\|]PAM | NA | Intergenic | NA | NA |
| T27444 | CTGGCGGCGCTCGACTCGACGGG | 1000 | | | | | | | | | |
| Oligo fwd | CACCCTGGCGGCGCTCGACTCGAC | | | | | | | | | | |
| Oligo rev | AAACGTCGAGTCGAGCGCCGCCAG | | | | | | | | | | |
| Chromosome | start | end | strand | MM | target_seq | PAM | alignment | distance | position | gene name | gene id |
| GU305914.1 | 88738 | 88760 | - | 0 | CTGGCGGCGCTCGACTCGAC | GGG | \|\|\|\|\|\|\|\|[\|\|\|\|\|\|\|\|\|\|\|\|]PAM | NA | Intergenic | NA | NA |
| T27445 | TGGCGGCGCTCGACTCGACGGGG | 1000 | | | | | | | | | |
| Oligo fwd | CACCTGGCGGCGCTCGACTCGACG | | | | | | | | | | |
| Oligo rev | AAACCGTCGAGTCGAGCGCCGCCA | | | | | | | | | | |
| Chromosome | start | end | strand | MM | target_seq | PAM | alignment | distance | position | gene name | gene id |
| GU305914.1 | 88737 | 88759 | - | 0 | TGGCGGCGCTCGACTCGACG | GGG | \|\|\|\|\|\|\|\|[\|\|\|\|\|\|\|\|\|\|\|\|]PAM | NA | Intergenic | NA | NA |
| T27446 | CGACTCGACGGGGCTCGACGTGG | 1000 | | | | | | | | | |
| Oligo fwd | CACCCGACTCGACGGGGCTCGACG | | | | | | | | | | |
| Oligo rev | AAACCGTCGAGCCCCGTCGAGTCG | | | | | | | | | | |
| Chromosome | start | end | strand | MM | target_seq | PAM | alignment | distance | position | gene name | gene id |
| GU305914.1 | 88727 | 88749 | - | 0 | CGACTCGACGGGGCTCGACG | TGG | \|\|\|\|\|\|\|\|[\|\|\|\|\|\|\|\|\|\|\|\|]PAM | NA | Intergenic | NA | NA |
| T27447 | GCTCGACGTGGATCCCGAGCTGG | 1000 | | | | | | | | | |
| Oligo fwd | CACCGCTCGACGTGGATCCCGAGC | | | | | | | | | | |
| Oligo rev | AAACGCTCGGGATCCACGTCGAGC | | | | | | | | | | |
| Chromosome | start | end | strand | MM | target_seq | PAM | alignment | distance | position | gene name | gene id |
| GU305914.1 | 88715 | 88737 | - | 0 | GCTCGACGTGGATCCCGAGC | TGG | \|\|\|\|\|\|\|\|[\|\|\|\|\|\|\|\|\|\|\|\|]PAM | NA | Intergenic | NA | NA |
| T27448 | CGACGTGGATCCCGAGCTGGCGG | 1000 | | | | | | | | | |
| Oligo fwd | CACCCGACGTGGATCCCGAGCTGG | | | | | | | | | | |
| Oligo rev | AAACCCAGCTCGGGATCCACGTCG | | | | | | | | | | |
| Chromosome | start | end | strand | MM | target_seq | PAM | alignment | distance | position | gene name | gene id |
| GU305914.1 | 88712 | 88734 | - | 0 | CGACGTGGATCCCGAGCTGG | CGG | \|\|\|\|\|\|\|\|[\|\|\|\|\|\|\|\|\|\|\|\|]PAM | NA | Intergenic | NA | NA |
| T27449 | GGCGGCCATCGCCGAGCTCATGG | 1000 | | | | | | | | | |
| Oligo fwd | CACCGGCGGCCATCGCCGAGCTCA | | | | | | | | | | |
| Oligo rev | AAACTGAGCTCGGCGATGGCCGCC | | | | | | | | | | |

Chromosome start end strand MM target_seq PAM alignment distance position gene name gene id
GU305914.1 88694 88716 - 0 GGCGGCCATCGCCGAGCTCA TGG ||||||||[||||||||||||]PAM NA Intergenic NA NA

T27450 GGCCATCGCCGAGCTCATGGAGG 1000
Oligo fwd CACCGGCCATCGCCGAGCTCATGG
Oligo rev AAACCCATGAGCTCGGCGATGGCC
Chromosome start end strand MM target_seq PAM alignment distance position gene name gene id
GU305914.1 88691 88713 - 0 GGCCATCGCCGAGCTCATGG AGG ||||||||[||||||||||||]PAM NA Intergenic NA NA

T27451 GCCATCGCCGAGCTCATGGAGGG 1000
Oligo fwd CACCGCCATCGCCGAGCTCATGGA
Oligo rev AAACTCCATGAGCTCGGCGATGGC
Chromosome start end strand MM target_seq PAM alignment distance position gene name gene id
GU305914.1 88690 88712 - 0 GCCATCGCCGAGCTCATGGA GGG ||||||||[||||||||||||]PAM NA Intergenic NA NA

T27452 GCTCATGGAGGGCCGCGACGAGG 1000
Oligo fwd CACCGCTCATGGAGGGCCGCGACG
Oligo rev AAACCGTCGCGGCCCTCCATGAGC
Chromosome start end strand MM target_seq PAM alignment distance position gene name gene id
GU305914.1 88679 88701 - 0 GCTCATGGAGGGCCGCGACG AGG ||||||||[||||||||||||]PAM NA Intergenic NA NA
GU305914.1 172332 172354 - 4 GGCCACGGAGGACCGCGACG AGG |--||-||[|||-||||||||]PAM NA Intergenic NA NA

T27453 CTCATGGAGGGCCGCGACGAGGG 1000
Oligo fwd CACCCTCATGGAGGGCCGCGACGA
Oligo rev AAACTCGTCGCGGCCCTCCATGAG
Chromosome start end strand MM target_seq PAM alignment distance position gene name gene id
GU305914.1 88678 88700 - 0 CTCATGGAGGGCCGCGACGA GGG ||||||||[||||||||||||]PAM NA Intergenic NA NA

T27454 CGACGACATCCTCTTCTACGTGG 1000
Oligo fwd CACCCGACGACATCCTCTTCTACG
Oligo rev AAACCGTAGAAGAGGATGTCGTCG
Chromosome start end strand MM target_seq PAM alignment distance position gene name gene id
GU305914.1 88640 88662 - 0 CGACGACATCCTCTTCTACG TGG ||||||||[||||||||||||]PAM NA Intergenic NA NA

T27455 CCTCTTCTACGTGGACCAGCAGG 1000
Oligo fwd CACCCCTCTTCTACGTGGACCAGC
Oligo rev AAACGCTGGTCCACGTAGAAGAGG
Chromosome start end strand MM target_seq PAM alignment distance position gene name gene id
GU305914.1 88631 88653 - 0 CCTCTTCTACGTGGACCAGC AGG ||||||||[||||||||||||]PAM NA Intergenic NA NA

T27456 ACCAGCAGGAGTACATCGCCAGG 1000
Oligo fwd CACCACCAGCAGGAGTACATCGCC
Oligo rev AAACGGCGATGTACTCCTGCTGGT
Chromosome start end strand MM target_seq PAM alignment distance position gene name gene id
GU305914.1 88617 88639 - 0 ACCAGCAGGAGTACATCGCC AGG ||||||||[||||||||||||]PAM NA Intergenic NA NA

T27457 GTATCGCAAGATGCGCAGTCTGG 1000
Oligo fwd CACCGTATCGCAAGATGCGCAGTC
Oligo rev AAACGACTGCGCATCTTGCGATAC
Chromosome start end strand MM target_seq PAM alignment distance position gene name gene id
GU305914.1 88589 88611 - 0 GTATCGCAAGATGCGCAGTC TGG ||||||||[||||||||||||]PAM NA Intergenic NA NA

T27458 AGATGCGCAGTCTGGCCGAGAGG 1000
Oligo fwd CACCAGATGCGCAGTCTGGCCGAG
Oligo rev AAACCTCGGCCAGACTGCGCATCT
Chromosome start end strand MM target_seq PAM alignment distance position gene name gene id
GU305914.1 88581 88603 - 0 AGATGCGCAGTCTGGCCGAG AGG ||||||||[||||||||||||]PAM NA Intergenic NA NA

T27459 GATGCGCAGTCTGGCCGAGAGGG 1000
Oligo fwd CACCGATGCGCAGTCTGGCCGAGA
Oligo rev AAACTCTCGGCCAGACTGCGCATC
Chromosome start end strand MM target_seq PAM alignment distance position gene name gene id
GU305914.1 88580 88602 - 0 GATGCGCAGTCTGGCCGAGA GGG ||||||||[||||||||||||]PAM NA Intergenic NA NA

T27460 ATGCGCAGTCTGGCCGAGAGGGG 1000
Oligo fwd CACCATGCGCAGTCTGGCCGAGAG
Oligo rev AAACCTCTCGGCCAGACTGCGCAT
Chromosome start end strand MM target_seq PAM alignment distance position gene name gene id
GU305914.1 88579 88601 - 0 ATGCGCAGTCTGGCCGAGAG GGG ||||||||[||||||||||||]PAM NA Intergenic NA NA

T27461 TGCGCAGTCTGGCCGAGAGGGGG 1000
Oligo fwd CACCTGCGCAGTCTGGCCGAGAGG
Oligo rev AAACCCTCTCGGCCAGACTGCGCA
Chromosome start end strand MM target_seq PAM alignment distance position gene name gene id
GU305914.1 88578 88600 - 0 TGCGCAGTCTGGCCGAGAGG GGG ||||||||[||||||||||||]PAM NA Intergenic NA NA

T27462 GCGCAGTCTGGCCGAGAGGGGGG 1000
Oligo fwd CACCGCGCAGTCTGGCCGAGAGGG
Oligo rev AAACCCCTCTCGGCCAGACTGCGC
Chromosome start end strand MM target_seq PAM alignment distance position gene name gene id
GU305914.1 88577 88599 - 0 GCGCAGTCTGGCCGAGAGGG GGG ||||||||[||||||||||||]PAM NA Intergenic NA NA

T27463 TCTGGCCGAGAGGGGGGTGACGG 1000
Oligo fwd CACCTCTGGCCGAGAGGGGGGTGA
Oligo rev AAACTCACCCCCCTCTCGGCCAGA
Chromosome start end strand MM target_seq PAM alignment distance position gene name gene id
GU305914.1 88571 88593 - 0 TCTGGCCGAGAGGGGGGTGA CGG ||||||||[||||||||||||]PAM NA Intergenic NA NA

T27464 GGGGGTGACGGACTTTTCGCTGG 1000

<table>
<tr><td>Oligo fwd</td><td>CACCGGGGGTGACGGACTTTTCGC</td><td></td><td></td><td></td><td></td><td></td><td></td><td></td><td></td><td></td><td></td></tr>
<tr><td>Oligo rev</td><td>AAACGCGAAAAGTCCGTCACCCCC</td><td></td><td></td><td></td><td></td><td></td><td></td><td></td><td></td><td></td><td></td></tr>
<tr><td>Chromosome</td><td>start</td><td>end</td><td>strand</td><td>MM</td><td>target_seq</td><td>PAM</td><td>alignment</td><td>distance</td><td>position</td><td>gene name</td><td>gene id</td></tr>
<tr><td>GU305914.1</td><td>88559</td><td>88581</td><td>-</td><td>0</td><td>GGGGGTGACGGACTTTTCGC</td><td>TGG</td><td>||||||||[||||||||||||]PAM</td><td>NA</td><td>Intergenic</td><td>NA</td><td>NA</td></tr>
<tr><td></td><td></td><td></td><td></td><td></td><td></td><td></td><td></td><td></td><td></td><td></td><td></td></tr>
<tr><td>T27465</td><td>TTCGCTGGCCTCTCTGCGCGAGG</td><td>1000</td><td></td><td></td><td></td><td></td><td></td><td></td><td></td><td></td><td></td></tr>
<tr><td>Oligo fwd</td><td>CACCTTCGCTGGCCTCTCTGCGCG</td><td></td><td></td><td></td><td></td><td></td><td></td><td></td><td></td><td></td><td></td></tr>
<tr><td>Oligo rev</td><td>AAACCGCGCAGAGAGGCCAGCGAA</td><td></td><td></td><td></td><td></td><td></td><td></td><td></td><td></td><td></td><td></td></tr>
<tr><td>Chromosome</td><td>start</td><td>end</td><td>strand</td><td>MM</td><td>target_seq</td><td>PAM</td><td>alignment</td><td>distance</td><td>position</td><td>gene name</td><td>gene id</td></tr>
<tr><td>GU305914.1</td><td>88544</td><td>88566</td><td>-</td><td>0</td><td>TTCGCTGGCCTCTCTGCGCG</td><td>AGG</td><td>||||||||[||||||||||||]PAM</td><td>NA</td><td>Intergenic</td><td>NA</td><td>NA</td></tr>
<tr><td></td><td></td><td></td><td></td><td></td><td></td><td></td><td></td><td></td><td></td><td></td><td></td></tr>
<tr><td>T27466</td><td>GGCGACCGCCACCAACGCGACGG</td><td>1000</td><td></td><td></td><td></td><td></td><td></td><td></td><td></td><td></td><td></td></tr>
<tr><td>Oligo fwd</td><td>CACCGGCGACCGCCACCAACGCGA</td><td></td><td></td><td></td><td></td><td></td><td></td><td></td><td></td><td></td><td></td></tr>
<tr><td>Oligo rev</td><td>AAACTCGCGTTGGTGGCGGTCGCC</td><td></td><td></td><td></td><td></td><td></td><td></td><td></td><td></td><td></td><td></td></tr>
<tr><td>Chromosome</td><td>start</td><td>end</td><td>strand</td><td>MM</td><td>target_seq</td><td>PAM</td><td>alignment</td><td>distance</td><td>position</td><td>gene name</td><td>gene id</td></tr>
<tr><td>GU305914.1</td><td>88523</td><td>88545</td><td>-</td><td>0</td><td>GGCGACCGCCACCAACGCGA</td><td>CGG</td><td>||||||||[||||||||||||]PAM</td><td>NA</td><td>Intergenic</td><td>NA</td><td>NA</td></tr>
<tr><td></td><td></td><td></td><td></td><td></td><td></td><td></td><td></td><td></td><td></td><td></td><td></td></tr>
<tr><td>T27467</td><td>GACCGCCACCAACGCGACGGCGG</td><td>1000</td><td></td><td></td><td></td><td></td><td></td><td></td><td></td><td></td><td></td></tr>
<tr><td>Oligo fwd</td><td>CACCGACCGCCACCAACGCGACGG</td><td></td><td></td><td></td><td></td><td></td><td></td><td></td><td></td><td></td><td></td></tr>
<tr><td>Oligo rev</td><td>AAACCCGTCGCGTTGGTGGCGGTC</td><td></td><td></td><td></td><td></td><td></td><td></td><td></td><td></td><td></td><td></td></tr>
<tr><td>Chromosome</td><td>start</td><td>end</td><td>strand</td><td>MM</td><td>target_seq</td><td>PAM</td><td>alignment</td><td>distance</td><td>position</td><td>gene name</td><td>gene id</td></tr>
<tr><td>GU305914.1</td><td>88520</td><td>88542</td><td>-</td><td>0</td><td>GACCGCCACCAACGCGACGG</td><td>CGG</td><td>||||||||[||||||||||||]PAM</td><td>NA</td><td>Intergenic</td><td>NA</td><td>NA</td></tr>
<tr><td></td><td></td><td></td><td></td><td></td><td></td><td></td><td></td><td></td><td></td><td></td><td></td></tr>
<tr><td>T27468</td><td>CGCCACCAACGCGACGGCGGCGG</td><td>1000</td><td></td><td></td><td></td><td></td><td></td><td></td><td></td><td></td><td></td></tr>
<tr><td>Oligo fwd</td><td>CACCCGCCACCAACGCGACGGCGG</td><td></td><td></td><td></td><td></td><td></td><td></td><td></td><td></td><td></td><td></td></tr>
<tr><td>Oligo rev</td><td>AAACCCGCCGTCGCGTTGGTGGCG</td><td></td><td></td><td></td><td></td><td></td><td></td><td></td><td></td><td></td><td></td></tr>
<tr><td>Chromosome</td><td>start</td><td>end</td><td>strand</td><td>MM</td><td>target_seq</td><td>PAM</td><td>alignment</td><td>distance</td><td>position</td><td>gene name</td><td>gene id</td></tr>
<tr><td>GU305914.1</td><td>88517</td><td>88539</td><td>-</td><td>0</td><td>CGCCACCAACGCGACGGCGG</td><td>CGG</td><td>||||||||[||||||||||||]PAM</td><td>NA</td><td>Intergenic</td><td>NA</td><td>NA</td></tr>
<tr><td></td><td></td><td></td><td></td><td></td><td></td><td></td><td></td><td></td><td></td><td></td><td></td></tr>
<tr><td>T27469</td><td>GCCACCAACGCGACGGCGGCGGG</td><td>1000</td><td></td><td></td><td></td><td></td><td></td><td></td><td></td><td></td><td></td></tr>
<tr><td>Oligo fwd</td><td>CACCGCCACCAACGCGACGGCGGC</td><td></td><td></td><td></td><td></td><td></td><td></td><td></td><td></td><td></td><td></td></tr>
<tr><td>Oligo rev</td><td>AAACGCCGCCGTCGCGTTGGTGGC</td><td></td><td></td><td></td><td></td><td></td><td></td><td></td><td></td><td></td><td></td></tr>
<tr><td>Chromosome</td><td>start</td><td>end</td><td>strand</td><td>MM</td><td>target_seq</td><td>PAM</td><td>alignment</td><td>distance</td><td>position</td><td>gene name</td><td>gene id</td></tr>
<tr><td>GU305914.1</td><td>88516</td><td>88538</td><td>-</td><td>0</td><td>GCCACCAACGCGACGGCGGC</td><td>GGG</td><td>||||||||[||||||||||||]PAM</td><td>NA</td><td>Intergenic</td><td>NA</td><td>NA</td></tr>
<tr><td></td><td></td><td></td><td></td><td></td><td></td><td></td><td></td><td></td><td></td><td></td><td></td></tr>
<tr><td>T27470</td><td>ACGGCGGCGGGCTCCGCTGCTGG</td><td>1000</td><td></td><td></td><td></td><td></td><td></td><td></td><td></td><td></td><td></td></tr>
<tr><td>Oligo fwd</td><td>CACCACGGCGGCGGGCTCCGCTGC</td><td></td><td></td><td></td><td></td><td></td><td></td><td></td><td></td><td></td><td></td></tr>
<tr><td>Oligo rev</td><td>AAACGCAGCGGAGCCCGCCGCCGT</td><td></td><td></td><td></td><td></td><td></td><td></td><td></td><td></td><td></td><td></td></tr>
<tr><td>Chromosome</td><td>start</td><td>end</td><td>strand</td><td>MM</td><td>target_seq</td><td>PAM</td><td>alignment</td><td>distance</td><td>position</td><td>gene name</td><td>gene id</td></tr>
<tr><td>GU305914.1</td><td>88504</td><td>88526</td><td>-</td><td>0</td><td>ACGGCGGCGGGCTCCGCTGC</td><td>TGG</td><td>||||||||[||||||||||||]PAM</td><td>NA</td><td>Intergenic</td><td>NA</td><td>NA</td></tr>
<tr><td></td><td></td><td></td><td></td><td></td><td></td><td></td><td></td><td></td><td></td><td></td><td></td></tr>
<tr><td>T27471</td><td>GCGGCGGGCTCCGCTGCTGGCGG</td><td>1000</td><td></td><td></td><td></td><td></td><td></td><td></td><td></td><td></td><td></td></tr>
<tr><td>Oligo fwd</td><td>CACCGCGGCGGGCTCCGCTGCTGG</td><td></td><td></td><td></td><td></td><td></td><td></td><td></td><td></td><td></td><td></td></tr>
<tr><td>Oligo rev</td><td>AAACCCAGCAGCGGAGCCCGCCGC</td><td></td><td></td><td></td><td></td><td></td><td></td><td></td><td></td><td></td><td></td></tr>
<tr><td>Chromosome</td><td>start</td><td>end</td><td>strand</td><td>MM</td><td>target_seq</td><td>PAM</td><td>alignment</td><td>distance</td><td>position</td><td>gene name</td><td>gene id</td></tr>
<tr><td>GU305914.1</td><td>88501</td><td>88523</td><td>-</td><td>0</td><td>GCGGCGGGCTCCGCTGCTGG</td><td>CGG</td><td>||||||||[||||||||||||]PAM</td><td>NA</td><td>Intergenic</td><td>NA</td><td>NA</td></tr>
<tr><td></td><td></td><td></td><td></td><td></td><td></td><td></td><td></td><td></td><td></td><td></td><td></td></tr>
<tr><td>T27472</td><td>GCGGGCTCCGCTGCTGGCGGTGG</td><td>1000</td><td></td><td></td><td></td><td></td><td></td><td></td><td></td><td></td><td></td></tr>
<tr><td>Oligo fwd</td><td>CACCGCGGGCTCCGCTGCTGGCGG</td><td></td><td></td><td></td><td></td><td></td><td></td><td></td><td></td><td></td><td></td></tr>
<tr><td>Oligo rev</td><td>AAACCCGCCAGCAGCGGAGCCCGC</td><td></td><td></td><td></td><td></td><td></td><td></td><td></td><td></td><td></td><td></td></tr>
<tr><td>Chromosome</td><td>start</td><td>end</td><td>strand</td><td>MM</td><td>target_seq</td><td>PAM</td><td>alignment</td><td>distance</td><td>position</td><td>gene name</td><td>gene id</td></tr>
<tr><td>GU305914.1</td><td>88498</td><td>88520</td><td>-</td><td>0</td><td>GCGGGCTCCGCTGCTGGCGG</td><td>TGG</td><td>||||||||[||||||||||||]PAM</td><td>NA</td><td>Intergenic</td><td>NA</td><td>NA</td></tr>
<tr><td>GU305914.1</td><td>29991</td><td>30013</td><td>+</td><td>4</td><td>TCGGGGACCGGTGCTGGCGG</td><td>CGG</td><td>-||||--|[||-|||||||||]PAM</td><td>NA</td><td>Intergenic</td><td>NA</td><td>NA</td></tr>
<tr><td></td><td></td><td></td><td></td><td></td><td></td><td></td><td></td><td></td><td></td><td></td><td></td></tr>
<tr><td>T27473</td><td>GGCTCCGCTGCTGGCGGTGGCGG</td><td>1000</td><td></td><td></td><td></td><td></td><td></td><td></td><td></td><td></td><td></td></tr>
<tr><td>Oligo fwd</td><td>CACCGGCTCCGCTGCTGGCGGTGG</td><td></td><td></td><td></td><td></td><td></td><td></td><td></td><td></td><td></td><td></td></tr>
<tr><td>Oligo rev</td><td>AAACCCACCGCCAGCAGCGGAGCC</td><td></td><td></td><td></td><td></td><td></td><td></td><td></td><td></td><td></td><td></td></tr>
<tr><td>Chromosome</td><td>start</td><td>end</td><td>strand</td><td>MM</td><td>target_seq</td><td>PAM</td><td>alignment</td><td>distance</td><td>position</td><td>gene name</td><td>gene id</td></tr>
<tr><td>GU305914.1</td><td>88495</td><td>88517</td><td>-</td><td>0</td><td>GGCTCCGCTGCTGGCGGTGG</td><td>CGG</td><td>||||||||[||||||||||||]PAM</td><td>NA</td><td>Intergenic</td><td>NA</td><td>NA</td></tr>
<tr><td>GU305914.1</td><td>29994</td><td>30016</td><td>+</td><td>4</td><td>GGGACCGGTGCTGGCGGCGG</td><td>CGG</td><td>||--|||-[|||||||||-||]PAM</td><td>NA</td><td>Intergenic</td><td>NA</td><td>NA</td></tr>
<tr><td></td><td></td><td></td><td></td><td></td><td></td><td></td><td></td><td></td><td></td><td></td><td></td></tr>
<tr><td>T27474</td><td>CGCTGCTGGCGGTGGCGGCTCGG</td><td>1000</td><td></td><td></td><td></td><td></td><td></td><td></td><td></td><td></td><td></td></tr>
<tr><td>Oligo fwd</td><td>CACCCGCTGCTGGCGGTGGCGGCT</td><td></td><td></td><td></td><td></td><td></td><td></td><td></td><td></td><td></td><td></td></tr>
<tr><td>Oligo rev</td><td>AAACAGCCGCCACCGCCAGCAGCG</td><td></td><td></td><td></td><td></td><td></td><td></td><td></td><td></td><td></td><td></td></tr>
<tr><td>Chromosome</td><td>start</td><td>end</td><td>strand</td><td>MM</td><td>target_seq</td><td>PAM</td><td>alignment</td><td>distance</td><td>position</td><td>gene name</td><td>gene id</td></tr>
<tr><td>GU305914.1</td><td>88490</td><td>88512</td><td>-</td><td>0</td><td>CGCTGCTGGCGGTGGCGGCT</td><td>CGG</td><td>||||||||[||||||||||||]PAM</td><td>NA</td><td>Intergenic</td><td>NA</td><td>NA</td></tr>
<tr><td></td><td></td><td></td><td></td><td></td><td></td><td></td><td></td><td></td><td></td><td></td><td></td></tr>
<tr><td>T27475</td><td>TGGCGGTGGCGGCTCGGCGACGG</td><td>1000</td><td></td><td></td><td></td><td></td><td></td><td></td><td></td><td></td><td></td></tr>
<tr><td>Oligo fwd</td><td>CACCTGGCGGTGGCGGCTCGGCGA</td><td></td><td></td><td></td><td></td><td></td><td></td><td></td><td></td><td></td><td></td></tr>
<tr><td>Oligo rev</td><td>AAACTCGCCGAGCCGCCACCGCCA</td><td></td><td></td><td></td><td></td><td></td><td></td><td></td><td></td><td></td><td></td></tr>
<tr><td>Chromosome</td><td>start</td><td>end</td><td>strand</td><td>MM</td><td>target_seq</td><td>PAM</td><td>alignment</td><td>distance</td><td>position</td><td>gene name</td><td>gene id</td></tr>
<tr><td>GU305914.1</td><td>88484</td><td>88506</td><td>-</td><td>0</td><td>TGGCGGTGGCGGCTCGGCGA</td><td>CGG</td><td>||||||||[||||||||||||]PAM</td><td>NA</td><td>Intergenic</td><td>NA</td><td>NA</td></tr>
<tr><td></td><td></td><td></td><td></td><td></td><td></td><td></td><td></td><td></td><td></td><td></td><td></td></tr>
<tr><td>T27476</td><td>CGGTGGCGGCTCGGCGACGGAGG</td><td>1000</td><td></td><td></td><td></td><td></td><td></td><td></td><td></td><td></td><td></td></tr>
<tr><td>Oligo fwd</td><td>CACCCGGTGGCGGCTCGGCGACGG</td><td></td><td></td><td></td><td></td><td></td><td></td><td></td><td></td><td></td><td></td></tr>
<tr><td>Oligo rev</td><td>AAACCCGTCGCCGAGCCGCCACCG</td><td></td><td></td><td></td><td></td><td></td><td></td><td></td><td></td><td></td><td></td></tr>
<tr><td>Chromosome</td><td>start</td><td>end</td><td>strand</td><td>MM</td><td>target_seq</td><td>PAM</td><td>alignment</td><td>distance</td><td>position</td><td>gene name</td><td>gene id</td></tr>
<tr><td>GU305914.1</td><td>88481</td><td>88503</td><td>-</td><td>0</td><td>CGGTGGCGGCTCGGCGACGG</td><td>AGG</td><td>||||||||[||||||||||||]PAM</td><td>NA</td><td>Intergenic</td><td>NA</td><td>NA</td></tr>
<tr><td></td><td></td><td></td><td></td><td></td><td></td><td></td><td></td><td></td><td></td><td></td><td></td></tr>
<tr><td>T27477</td><td>GGTGGCGGCTCGGCGACGGAGGG</td><td>1000</td><td></td><td></td><td></td><td></td><td></td><td></td><td></td><td></td><td></td></tr>
<tr><td>Oligo fwd</td><td>CACCGGTGGCGGCTCGGCGACGGA</td><td></td><td></td><td></td><td></td><td></td><td></td><td></td><td></td><td></td><td></td></tr>
<tr><td>Oligo rev</td><td>AAACTCCGTCGCCGAGCCGCCACC</td><td></td><td></td><td></td><td></td><td></td><td></td><td></td><td></td><td></td><td></td></tr>
<tr><td>Chromosome</td><td>start</td><td>end</td><td>strand</td><td>MM</td><td>target_seq</td><td>PAM</td><td>alignment</td><td>distance</td><td>position</td><td>gene name</td><td>gene id</td></tr>
<tr><td>GU305914.1</td><td>88480</td><td>88502</td><td>-</td><td>0</td><td>GGTGGCGGCTCGGCGACGGA</td><td>GGG</td><td>||||||||[||||||||||||]PAM</td><td>NA</td><td>Intergenic</td><td>NA</td><td>NA</td></tr>
<tr><td></td><td></td><td></td><td></td><td></td><td></td><td></td><td></td><td></td><td></td><td></td><td></td></tr>
<tr><td>T27478</td><td>GGCGGCTCGGCGACGGAGGGCGG</td><td>1000</td><td></td><td></td><td></td><td></td><td></td><td></td><td></td><td></td><td></td></tr>
<tr><td>Oligo fwd</td><td>CACCGGCGGCTCGGCGACGGAGGG</td><td></td><td></td><td></td><td></td><td></td><td></td><td></td><td></td><td></td><td></td></tr>
<tr><td>Oligo rev</td><td>AAACCCCTCCGTCGCCGAGCCGCC</td><td></td><td></td><td></td><td></td><td></td><td></td><td></td><td></td><td></td><td></td></tr>
<tr><td>Chromosome</td><td>start</td><td>end</td><td>strand</td><td>MM</td><td>target_seq</td><td>PAM</td><td>alignment</td><td>distance</td><td>position</td><td>gene name</td><td>gene id</td></tr>
</table>

<table>
<tr><td>GU305914.1</td><td>88477</td><td>88499</td><td>-</td><td>0</td><td>GGCGGCTCGGCGACGGAGGG</td><td>CGG</td><td>||||||||[||||||||||||]PAM</td><td>NA</td><td>Intergenic</td><td>NA</td><td>NA</td></tr>
<tr><td>T27479</td><td>GGCTCGGCGACGGAGGGCGGTGG</td><td>1000</td><td></td><td></td><td></td><td></td><td></td><td></td><td></td><td></td><td></td></tr>
<tr><td>Oligo fwd</td><td>CACCGGCTCGGCGACGGAGGGCGG</td><td></td><td></td><td></td><td></td><td></td><td></td><td></td><td></td><td></td><td></td></tr>
<tr><td>Oligo rev</td><td>AAACCCGCCCTCCGTCGCCGAGCC</td><td></td><td></td><td></td><td></td><td></td><td></td><td></td><td></td><td></td><td></td></tr>
<tr><td>Chromosome</td><td>start</td><td>end</td><td>strand</td><td>MM</td><td>target_seq</td><td>PAM</td><td>alignment</td><td>distance</td><td>position</td><td>gene name</td><td>gene id</td></tr>
<tr><td>GU305914.1</td><td>88474</td><td>88496</td><td>-</td><td>0</td><td>GGCTCGGCGACGGAGGGCGG</td><td>TGG</td><td>||||||||[||||||||||||]PAM</td><td>NA</td><td>Intergenic</td><td>NA</td><td>NA</td></tr>
<tr><td>T27480</td><td>TCGGCGACGGAGGGCGGTGGCGG</td><td>1000</td><td></td><td></td><td></td><td></td><td></td><td></td><td></td><td></td><td></td></tr>
<tr><td>Oligo fwd</td><td>CACCTCGGCGACGGAGGGCGGTGG</td><td></td><td></td><td></td><td></td><td></td><td></td><td></td><td></td><td></td><td></td></tr>
<tr><td>Oligo rev</td><td>AAACCCACCGCCCTCCGTCGCCGA</td><td></td><td></td><td></td><td></td><td></td><td></td><td></td><td></td><td></td><td></td></tr>
<tr><td>Chromosome</td><td>start</td><td>end</td><td>strand</td><td>MM</td><td>target_seq</td><td>PAM</td><td>alignment</td><td>distance</td><td>position</td><td>gene name</td><td>gene id</td></tr>
<tr><td>GU305914.1</td><td>88471</td><td>88493</td><td>-</td><td>0</td><td>TCGGCGACGGAGGGCGGTGG</td><td>CGG</td><td>||||||||[||||||||||||]PAM</td><td>NA</td><td>Intergenic</td><td>NA</td><td>NA</td></tr>
<tr><td>T27481</td><td>GCGACGGAGGGCGGTGGCGGTGG</td><td>1000</td><td></td><td></td><td></td><td></td><td></td><td></td><td></td><td></td><td></td></tr>
<tr><td>Oligo fwd</td><td>CACCGCGACGGAGGGCGGTGGCGG</td><td></td><td></td><td></td><td></td><td></td><td></td><td></td><td></td><td></td><td></td></tr>
<tr><td>Oligo rev</td><td>AAACCCGCCACCGCCCTCCGTCGC</td><td></td><td></td><td></td><td></td><td></td><td></td><td></td><td></td><td></td><td></td></tr>
<tr><td>Chromosome</td><td>start</td><td>end</td><td>strand</td><td>MM</td><td>target_seq</td><td>PAM</td><td>alignment</td><td>distance</td><td>position</td><td>gene name</td><td>gene id</td></tr>
<tr><td>GU305914.1</td><td>88468</td><td>88490</td><td>-</td><td>0</td><td>GCGACGGAGGGCGGTGGCGG</td><td>TGG</td><td>||||||||[||||||||||||]PAM</td><td>NA</td><td>Intergenic</td><td>NA</td><td>NA</td></tr>
<tr><td>T27482</td><td>CGACGGAGGGCGGTGGCGGTGGG</td><td>1000</td><td></td><td></td><td></td><td></td><td></td><td></td><td></td><td></td><td></td></tr>
<tr><td>Oligo fwd</td><td>CACCCGACGGAGGGCGGTGGCGGT</td><td></td><td></td><td></td><td></td><td></td><td></td><td></td><td></td><td></td><td></td></tr>
<tr><td>Oligo rev</td><td>AAACACCGCCACCGCCCTCCGTCG</td><td></td><td></td><td></td><td></td><td></td><td></td><td></td><td></td><td></td><td></td></tr>
<tr><td>Chromosome</td><td>start</td><td>end</td><td>strand</td><td>MM</td><td>target_seq</td><td>PAM</td><td>alignment</td><td>distance</td><td>position</td><td>gene name</td><td>gene id</td></tr>
<tr><td>GU305914.1</td><td>88467</td><td>88489</td><td>-</td><td>0</td><td>CGACGGAGGGCGGTGGCGGT</td><td>GGG</td><td>||||||||[||||||||||||]PAM</td><td>NA</td><td>Intergenic</td><td>NA</td><td>NA</td></tr>
<tr><td>T27483</td><td>GACGGAGGGCGGTGGCGGTGGGG</td><td>1000</td><td></td><td></td><td></td><td></td><td></td><td></td><td></td><td></td><td></td></tr>
<tr><td>Oligo fwd</td><td>CACCGACGGAGGGCGGTGGCGGTG</td><td></td><td></td><td></td><td></td><td></td><td></td><td></td><td></td><td></td><td></td></tr>
<tr><td>Oligo rev</td><td>AAACCACCGCCACCGCCCTCCGTC</td><td></td><td></td><td></td><td></td><td></td><td></td><td></td><td></td><td></td><td></td></tr>
<tr><td>Chromosome</td><td>start</td><td>end</td><td>strand</td><td>MM</td><td>target_seq</td><td>PAM</td><td>alignment</td><td>distance</td><td>position</td><td>gene name</td><td>gene id</td></tr>
<tr><td>GU305914.1</td><td>88466</td><td>88488</td><td>-</td><td>0</td><td>GACGGAGGGCGGTGGCGGTG</td><td>GGG</td><td>||||||||[||||||||||||]PAM</td><td>NA</td><td>Intergenic</td><td>NA</td><td>NA</td></tr>
<tr><td>T27484</td><td>GGAGGGCGGTGGCGGTGGGGCGG</td><td>1000</td><td></td><td></td><td></td><td></td><td></td><td></td><td></td><td></td><td></td></tr>
<tr><td>Oligo fwd</td><td>CACCGGAGGGCGGTGGCGGTGGGG</td><td></td><td></td><td></td><td></td><td></td><td></td><td></td><td></td><td></td><td></td></tr>
<tr><td>Oligo rev</td><td>AAACCCCCACCGCCACCGCCCTCC</td><td></td><td></td><td></td><td></td><td></td><td></td><td></td><td></td><td></td><td></td></tr>
<tr><td>Chromosome</td><td>start</td><td>end</td><td>strand</td><td>MM</td><td>target_seq</td><td>PAM</td><td>alignment</td><td>distance</td><td>position</td><td>gene name</td><td>gene id</td></tr>
<tr><td>GU305914.1</td><td>88463</td><td>88485</td><td>-</td><td>0</td><td>GGAGGGCGGTGGCGGTGGGG</td><td>CGG</td><td>||||||||[||||||||||||]PAM</td><td>NA</td><td>Intergenic</td><td>NA</td><td>NA</td></tr>
<tr><td>T27485</td><td>CGGTGGCGGTGGGGCGGCCGCGG</td><td>1000</td><td></td><td></td><td></td><td></td><td></td><td></td><td></td><td></td><td></td></tr>
<tr><td>Oligo fwd</td><td>CACCCGGTGGCGGTGGGGCGGCCG</td><td></td><td></td><td></td><td></td><td></td><td></td><td></td><td></td><td></td><td></td></tr>
<tr><td>Oligo rev</td><td>AAACCGGCCGCCCCACCGCCACCG</td><td></td><td></td><td></td><td></td><td></td><td></td><td></td><td></td><td></td><td></td></tr>
<tr><td>Chromosome</td><td>start</td><td>end</td><td>strand</td><td>MM</td><td>target_seq</td><td>PAM</td><td>alignment</td><td>distance</td><td>position</td><td>gene name</td><td>gene id</td></tr>
<tr><td>GU305914.1</td><td>88457</td><td>88479</td><td>-</td><td>0</td><td>CGGTGGCGGTGGGGCGGCCG</td><td>CGG</td><td>||||||||[||||||||||||]PAM</td><td>NA</td><td>Intergenic</td><td>NA</td><td>NA</td></tr>
<tr><td>T27486</td><td>GGGGCGGCCGCGGACGAGAGCGG</td><td>1000</td><td></td><td></td><td></td><td></td><td></td><td></td><td></td><td></td><td></td></tr>
<tr><td>Oligo fwd</td><td>CACCGGGGCGGCCGCGGACGAGAG</td><td></td><td></td><td></td><td></td><td></td><td></td><td></td><td></td><td></td><td></td></tr>
<tr><td>Oligo rev</td><td>AAACCTCTCGTCCGCGGCCGCCCC</td><td></td><td></td><td></td><td></td><td></td><td></td><td></td><td></td><td></td><td></td></tr>
<tr><td>Chromosome</td><td>start</td><td>end</td><td>strand</td><td>MM</td><td>target_seq</td><td>PAM</td><td>alignment</td><td>distance</td><td>position</td><td>gene name</td><td>gene id</td></tr>
<tr><td>GU305914.1</td><td>88447</td><td>88469</td><td>-</td><td>0</td><td>GGGGCGGCCGCGGACGAGAG</td><td>CGG</td><td>||||||||[||||||||||||]PAM</td><td>NA</td><td>Intergenic</td><td>NA</td><td>NA</td></tr>
<tr><td>T27487</td><td>ATTTCTCCGCTCTGTTTTCGAGG</td><td>1000</td><td></td><td></td><td></td><td></td><td></td><td></td><td></td><td></td><td></td></tr>
<tr><td>Oligo fwd</td><td>CACCATTTCTCCGCTCTGTTTTCG</td><td></td><td></td><td></td><td></td><td></td><td></td><td></td><td></td><td></td><td></td></tr>
<tr><td>Oligo rev</td><td>AAACCGAAAACAGAGCGGAGAAAT</td><td></td><td></td><td></td><td></td><td></td><td></td><td></td><td></td><td></td><td></td></tr>
<tr><td>Chromosome</td><td>start</td><td>end</td><td>strand</td><td>MM</td><td>target_seq</td><td>PAM</td><td>alignment</td><td>distance</td><td>position</td><td>gene name</td><td>gene id</td></tr>
<tr><td>GU305914.1</td><td>88413</td><td>88435</td><td>-</td><td>0</td><td>ATTTCTCCGCTCTGTTTTCG</td><td>AGG</td><td>||||||||[||||||||||||]PAM</td><td>NA</td><td>Intergenic</td><td>NA</td><td>NA</td></tr>
<tr><td>T27488</td><td>TCTGTTTTCGAGGCGCGACGAGG</td><td>1000</td><td></td><td></td><td></td><td></td><td></td><td></td><td></td><td></td><td></td></tr>
<tr><td>Oligo fwd</td><td>CACCTCTGTTTTCGAGGCGCGACG</td><td></td><td></td><td></td><td></td><td></td><td></td><td></td><td></td><td></td><td></td></tr>
<tr><td>Oligo rev</td><td>AAACCGTCGCGCCTCGAAAACAGA</td><td></td><td></td><td></td><td></td><td></td><td></td><td></td><td></td><td></td><td></td></tr>
<tr><td>Chromosome</td><td>start</td><td>end</td><td>strand</td><td>MM</td><td>target_seq</td><td>PAM</td><td>alignment</td><td>distance</td><td>position</td><td>gene name</td><td>gene id</td></tr>
<tr><td>GU305914.1</td><td>88403</td><td>88425</td><td>-</td><td>0</td><td>TCTGTTTTCGAGGCGCGACG</td><td>AGG</td><td>||||||||[||||||||||||]PAM</td><td>NA</td><td>Intergenic</td><td>NA</td><td>NA</td></tr>
<tr><td>T27489</td><td>TTCGAGGCGCGACGAGGCCGAGG</td><td>1000</td><td></td><td></td><td></td><td></td><td></td><td></td><td></td><td></td><td></td></tr>
<tr><td>Oligo fwd</td><td>CACCTTCGAGGCGCGACGAGGCCG</td><td></td><td></td><td></td><td></td><td></td><td></td><td></td><td></td><td></td><td></td></tr>
<tr><td>Oligo rev</td><td>AAACCGGCCTCGTCGCGCCTCGAA</td><td></td><td></td><td></td><td></td><td></td><td></td><td></td><td></td><td></td><td></td></tr>
<tr><td>Chromosome</td><td>start</td><td>end</td><td>strand</td><td>MM</td><td>target_seq</td><td>PAM</td><td>alignment</td><td>distance</td><td>position</td><td>gene name</td><td>gene id</td></tr>
<tr><td>GU305914.1</td><td>88397</td><td>88419</td><td>-</td><td>0</td><td>TTCGAGGCGCGACGAGGCCG</td><td>AGG</td><td>||||||||[||||||||||||]PAM</td><td>NA</td><td>Intergenic</td><td>NA</td><td>NA</td></tr>
<tr><td>T27490</td><td>GAGGCCGAGGACGTGAACGCCGG</td><td>1000</td><td></td><td></td><td></td><td></td><td></td><td></td><td></td><td></td><td></td></tr>
<tr><td>Oligo fwd</td><td>CACCGAGGCCGAGGACGTGAACGC</td><td></td><td></td><td></td><td></td><td></td><td></td><td></td><td></td><td></td><td></td></tr>
<tr><td>Oligo rev</td><td>AAACGCGTTCACGTCCTCGGCCTC</td><td></td><td></td><td></td><td></td><td></td><td></td><td></td><td></td><td></td><td></td></tr>
<tr><td>Chromosome</td><td>start</td><td>end</td><td>strand</td><td>MM</td><td>target_seq</td><td>PAM</td><td>alignment</td><td>distance</td><td>position</td><td>gene name</td><td>gene id</td></tr>
<tr><td>GU305914.1</td><td>88384</td><td>88406</td><td>-</td><td>0</td><td>GAGGCCGAGGACGTGAACGC</td><td>CGG</td><td>||||||||[||||||||||||]PAM</td><td>NA</td><td>Intergenic</td><td>NA</td><td>NA</td></tr>
<tr><td>T27491</td><td>GTGAACGCCGGCCTCATCAACGG</td><td>1000</td><td></td><td></td><td></td><td></td><td></td><td></td><td></td><td></td><td></td></tr>
<tr><td>Oligo fwd</td><td>CACCGTGAACGCCGGCCTCATCAA</td><td></td><td></td><td></td><td></td><td></td><td></td><td></td><td></td><td></td><td></td></tr>
<tr><td>Oligo rev</td><td>AAACTTGATGAGGCCGGCGTTCAC</td><td></td><td></td><td></td><td></td><td></td><td></td><td></td><td></td><td></td><td></td></tr>
<tr><td>Chromosome</td><td>start</td><td>end</td><td>strand</td><td>MM</td><td>target_seq</td><td>PAM</td><td>alignment</td><td>distance</td><td>position</td><td>gene name</td><td>gene id</td></tr>
<tr><td>GU305914.1</td><td>88372</td><td>88394</td><td>-</td><td>0</td><td>GTGAACGCCGGCCTCATCAA</td><td>CGG</td><td>||||||||[||||||||||||]PAM</td><td>NA</td><td>Intergenic</td><td>NA</td><td>NA</td></tr>
<tr><td>T27492</td><td>TCATCAACGGCGACGACGTGCGG</td><td>1000</td><td></td><td></td><td></td><td></td><td></td><td></td><td></td><td></td><td></td></tr>
<tr><td>Oligo fwd</td><td>CACCTCATCAACGGCGACGACGTG</td><td></td><td></td><td></td><td></td><td></td><td></td><td></td><td></td><td></td><td></td></tr>
<tr><td>Oligo rev</td><td>AAACCACGTCGTCGCCGTTGATGA</td><td></td><td></td><td></td><td></td><td></td><td></td><td></td><td></td><td></td><td></td></tr>
<tr><td>Chromosome</td><td>start</td><td>end</td><td>strand</td><td>MM</td><td>target_seq</td><td>PAM</td><td>alignment</td><td>distance</td><td>position</td><td>gene name</td><td>gene id</td></tr>
<tr><td>GU305914.1</td><td>88359</td><td>88381</td><td>-</td><td>0</td><td>TCATCAACGGCGACGACGTG</td><td>CGG</td><td>||||||||[||||||||||||]PAM</td><td>NA</td><td>Intergenic</td><td>NA</td><td>NA</td></tr>
<tr><td>T27493</td><td>CATCAACGGCGACGACGTGCGGG</td><td>1000</td><td></td><td></td><td></td><td></td><td></td><td></td><td></td><td></td><td></td></tr>
<tr><td>Oligo fwd</td><td>CACCCATCAACGGCGACGACGTGC</td><td></td><td></td><td></td><td></td><td></td><td></td><td></td><td></td><td></td><td></td></tr>
<tr><td>Oligo rev</td><td>AAACGCACGTCGTCGCCGTTGATG</td><td></td><td></td><td></td><td></td><td></td><td></td><td></td><td></td><td></td><td></td></tr>
</table>

| | | | | | | | | | | | |
|---|---|---|---|---|---|---|---|---|---|---|---|
| Chromosome | start | end | strand | MM | target_seq | PAM | alignment | distance | position | gene name | gene id |
| GU305914.1 | 88358 | 88380 | - | 0 | CATCAACGGCGACGACGTGC | GGG | \|\|\|\|\|\|\|\|[\|\|\|\|\|\|\|\|\|\|\|\|]PAM | NA | Intergenic | NA | NA |
| T27494 | ATCAACGGCGACGACGTGCGGGG | 1000 | | | | | | | | | |
| Oligo fwd | CACCATCAACGGCGACGACGTGCG | | | | | | | | | | |
| Oligo rev | AAACCGCACGTCGTCGCCGTTGAT | | | | | | | | | | |
| Chromosome | start | end | strand | MM | target_seq | PAM | alignment | distance | position | gene name | gene id |
| GU305914.1 | 88357 | 88379 | - | 0 | ATCAACGGCGACGACGTGCG | GGG | \|\|\|\|\|\|\|\|[\|\|\|\|\|\|\|\|\|\|\|\|]PAM | NA | Intergenic | NA | NA |
| T27495 | TCAACGGCGACGACGTGCGGGGG | 1000 | | | | | | | | | |
| Oligo fwd | CACCTCAACGGCGACGACGTGCGG | | | | | | | | | | |
| Oligo rev | AAACCCGCACGTCGTCGCCGTTGA | | | | | | | | | | |
| Chromosome | start | end | strand | MM | target_seq | PAM | alignment | distance | position | gene name | gene id |
| GU305914.1 | 88356 | 88378 | - | 0 | TCAACGGCGACGACGTGCGG | GGG | \|\|\|\|\|\|\|\|[\|\|\|\|\|\|\|\|\|\|\|\|]PAM | NA | Intergenic | NA | NA |
| T27496 | CAACGGCGACGACGTGCGGGGGG | 1000 | | | | | | | | | |
| Oligo fwd | CACCCAACGGCGACGACGTGCGGG | | | | | | | | | | |
| Oligo rev | AAACCCCGCACGTCGTCGCCGTTG | | | | | | | | | | |
| Chromosome | start | end | strand | MM | target_seq | PAM | alignment | distance | position | gene name | gene id |
| GU305914.1 | 88355 | 88377 | - | 0 | CAACGGCGACGACGTGCGGG | GGG | \|\|\|\|\|\|\|\|[\|\|\|\|\|\|\|\|\|\|\|\|]PAM | NA | Intergenic | NA | NA |
| T27497 | AGTTCGAACTTCCCAGCAAACGG | 1000 | | | | | | | | | |
| Oligo fwd | CACCAGTTCGAACTTCCCAGCAAA | | | | | | | | | | |
| Oligo rev | AAACTTTGCTGGGAAGTTCGAACT | | | | | | | | | | |
| Chromosome | start | end | strand | MM | target_seq | PAM | alignment | distance | position | gene name | gene id |
| GU305914.1 | 88326 | 88348 | - | 0 | AGTTCGAACTTCCCAGCAAA | CGG | \|\|\|\|\|\|\|\|[\|\|\|\|\|\|\|\|\|\|\|\|]PAM | NA | Intergenic | NA | NA |
| T27498 | AACTTCCCAGCAAACGGTCTCGG | 1000 | | | | | | | | | |
| Oligo fwd | CACCAACTTCCCAGCAAACGGTCT | | | | | | | | | | |
| Oligo rev | AAACAGACCGTTTGCTGGGAAGTT | | | | | | | | | | |
| Chromosome | start | end | strand | MM | target_seq | PAM | alignment | distance | position | gene name | gene id |
| GU305914.1 | 88320 | 88342 | - | 0 | AACTTCCCAGCAAACGGTCT | CGG | \|\|\|\|\|\|\|\|[\|\|\|\|\|\|\|\|\|\|\|\|]PAM | NA | Intergenic | NA | NA |
| T27499 | CGGTCTCGGTTGTGAGTATTCGG | 1000 | | | | | | | | | |
| Oligo fwd | CACCCGGTCTCGGTTGTGAGTATT | | | | | | | | | | |
| Oligo rev | AAACAATACTCACAACCGAGACCG | | | | | | | | | | |
| Chromosome | start | end | strand | MM | target_seq | PAM | alignment | distance | position | gene name | gene id |
| GU305914.1 | 88306 | 88328 | - | 0 | CGGTCTCGGTTGTGAGTATT | CGG | \|\|\|\|\|\|\|\|[\|\|\|\|\|\|\|\|\|\|\|\|]PAM | NA | Intergenic | NA | NA |
| T27500 | GGTCTCGGTTGTGAGTATTCGGG | 1000 | | | | | | | | | |
| Oligo fwd | CACCGGTCTCGGTTGTGAGTATTC | | | | | | | | | | |
| Oligo rev | AAACGAATACTCACAACCGAGACC | | | | | | | | | | |
| Chromosome | start | end | strand | MM | target_seq | PAM | alignment | distance | position | gene name | gene id |
| GU305914.1 | 88305 | 88327 | - | 0 | GGTCTCGGTTGTGAGTATTC | GGG | \|\|\|\|\|\|\|\|[\|\|\|\|\|\|\|\|\|\|\|\|]PAM | NA | Intergenic | NA | NA |
| T27501 | TTTCCCGTCCGCCGCTCGTCCGG | 1000 | | | | | | | | | |
| Oligo fwd | CACCTTTCCCGTCCGCCGCTCGTC | | | | | | | | | | |
| Oligo rev | AAACGACGAGCGGCGGACGGGAAA | | | | | | | | | | |
| Chromosome | start | end | strand | MM | target_seq | PAM | alignment | distance | position | gene name | gene id |
| GU305914.1 | 88254 | 88276 | - | 0 | TTTCCCGTCCGCCGCTCGTC | CGG | \|\|\|\|\|\|\|\|[\|\|\|\|\|\|\|\|\|\|\|\|]PAM | NA | Intergenic | NA | NA |
| T27502 | TCCGCCGCTCGTCCGGCGAGCGG | 1000 | | | | | | | | | |
| Oligo fwd | CACCTCCGCCGCTCGTCCGGCGAG | | | | | | | | | | |
| Oligo rev | AAACCTCGCCGGACGAGCGGCGGA | | | | | | | | | | |
| Chromosome | start | end | strand | MM | target_seq | PAM | alignment | distance | position | gene name | gene id |
| GU305914.1 | 88247 | 88269 | - | 0 | TCCGCCGCTCGTCCGGCGAG | CGG | \|\|\|\|\|\|\|\|[\|\|\|\|\|\|\|\|\|\|\|\|]PAM | NA | Intergenic | NA | NA |
| T27503 | CCGCCGCTCGTCCGGCGAGCGGG | 1000 | | | | | | | | | |
| Oligo fwd | CACCCCGCCGCTCGTCCGGCGAGC | | | | | | | | | | |
| Oligo rev | AAACGCTCGCCGGACGAGCGGCGG | | | | | | | | | | |
| Chromosome | start | end | strand | MM | target_seq | PAM | alignment | distance | position | gene name | gene id |
| GU305914.1 | 88246 | 88268 | - | 0 | CCGCCGCTCGTCCGGCGAGC | GGG | \|\|\|\|\|\|\|\|[\|\|\|\|\|\|\|\|\|\|\|\|]PAM | NA | Intergenic | NA | NA |
| GU305914.1 | 229680 | 229702 | + | 4 | CGGTCGCTCGACCAGCGAGC | GGG | \|-\|-\|\|\|\|[\|\|-\|\|-\|\|\|\|\|\|]PAM | NA | Intergenic | NA | NA |
| GU305914.1 | 99540 | 99562 | - | 4 | CCAACGCTCGTCGGGCGAGG | AGG | \|\|--\|\|\|\|[\|\|\|\|-\|\|\|\|\|\|-]PAM | NA | Intergenic | NA | NA |
| T27504 | ATTAAAGCTCGTTGAAATCACGG | 1000 | | | | | | | | | |
| Oligo fwd | CACCATTAAAGCTCGTTGAAATCA | | | | | | | | | | |
| Oligo rev | AAACTGATTTCAACGAGCTTTAAT | | | | | | | | | | |
| Chromosome | start | end | strand | MM | target_seq | PAM | alignment | distance | position | gene name | gene id |
| GU305914.1 | 88212 | 88234 | - | 0 | ATTAAAGCTCGTTGAAATCA | CGG | \|\|\|\|\|\|\|\|[\|\|\|\|\|\|\|\|\|\|\|\|]PAM | NA | Intergenic | NA | NA |
| T27505 | GAAATCACGGTGTAACGTAGCGG | 1000 | | | | | | | | | |
| Oligo fwd | CACCGAAATCACGGTGTAACGTAG | | | | | | | | | | |
| Oligo rev | AAACCTACGTTACACCGTGATTTC | | | | | | | | | | |
| Chromosome | start | end | strand | MM | target_seq | PAM | alignment | distance | position | gene name | gene id |
| GU305914.1 | 88199 | 88221 | - | 0 | GAAATCACGGTGTAACGTAG | CGG | \|\|\|\|\|\|\|\|[\|\|\|\|\|\|\|\|\|\|\|\|]PAM | NA | Intergenic | NA | NA |
| T27506 | GCGGTGTCTGTATCGCCGTTCGG | 1000 | | | | | | | | | |
| Oligo fwd | CACCGCGGTGTCTGTATCGCCGTT | | | | | | | | | | |
| Oligo rev | AAACAACGGCGATACAGACACCGC | | | | | | | | | | |
| Chromosome | start | end | strand | MM | target_seq | PAM | alignment | distance | position | gene name | gene id |
| GU305914.1 | 88180 | 88202 | - | 0 | GCGGTGTCTGTATCGCCGTT | CGG | \|\|\|\|\|\|\|\|[\|\|\|\|\|\|\|\|\|\|\|\|]PAM | NA | Intergenic | NA | NA |
| T27507 | CGGTGTCTGTATCGCCGTTCGGG | 1000 | | | | | | | | | |
| Oligo fwd | CACCCGGTGTCTGTATCGCCGTTC | | | | | | | | | | |
| Oligo rev | AAACGAACGGCGATACAGACACCG | | | | | | | | | | |
| Chromosome | start | end | strand | MM | target_seq | PAM | alignment | distance | position | gene name | gene id |
| GU305914.1 | 88179 | 88201 | - | 0 | CGGTGTCTGTATCGCCGTTC | GGG | \|\|\|\|\|\|\|\|[\|\|\|\|\|\|\|\|\|\|\|\|]PAM | NA | Intergenic | NA | NA |

<table>
<tr><td>T27508</td><td>GGTGTCTGTATCGCCGTTCGGGG</td><td></td><td>1000</td><td></td><td></td><td></td><td></td><td></td><td></td><td></td><td></td><td></td></tr>
<tr><td>Oligo fwd</td><td>CACCGGTGTCTGTATCGCCGTTCG</td><td></td><td></td><td></td><td></td><td></td><td></td><td></td><td></td><td></td><td></td><td></td></tr>
<tr><td>Oligo rev</td><td>AAACCGAACGGCGATACAGACACC</td><td></td><td></td><td></td><td></td><td></td><td></td><td></td><td></td><td></td><td></td><td></td></tr>
<tr><td>Chromosome</td><td>start</td><td></td><td>end</td><td>strand</td><td>MM</td><td>target_seq</td><td>PAM</td><td>alignment</td><td>distance</td><td>position</td><td>gene name</td><td>gene id</td></tr>
<tr><td>GU305914.1</td><td></td><td>88178</td><td>88200</td><td>-</td><td>0</td><td>GGTGTCTGTATCGCCGTTCG</td><td>GGG</td><td>||||||||[||||||||||||]PAM</td><td>NA</td><td>Intergenic</td><td>NA</td><td>NA</td></tr>
<tr><td>T27509</td><td>CGTTCGGGGTTGTCTGTGCGCGG</td><td></td><td>1000</td><td></td><td></td><td></td><td></td><td></td><td></td><td></td><td></td><td></td></tr>
<tr><td>Oligo fwd</td><td>CACCGTTCGGGGTTGTCTGTGCG</td><td></td><td></td><td></td><td></td><td></td><td></td><td></td><td></td><td></td><td></td><td></td></tr>
<tr><td>Oligo rev</td><td>AAACCGCACAGACAACCCCGAACG</td><td></td><td></td><td></td><td></td><td></td><td></td><td></td><td></td><td></td><td></td><td></td></tr>
<tr><td>Chromosome</td><td>start</td><td></td><td>end</td><td>strand</td><td>MM</td><td>target_seq</td><td>PAM</td><td>alignment</td><td>distance</td><td>position</td><td>gene name</td><td>gene id</td></tr>
<tr><td>GU305914.1</td><td></td><td>88164</td><td>88186</td><td>-</td><td>0</td><td>CGTTCGGGGTTGTCTGTGCG</td><td>CGG</td><td>||||||||[||||||||||||]PAM</td><td>NA</td><td>Intergenic</td><td>NA</td><td>NA</td></tr>
<tr><td>T27510</td><td>CTGTGCGCGGTATGTATAAGCGG</td><td></td><td>1000</td><td></td><td></td><td></td><td></td><td></td><td></td><td></td><td></td><td></td></tr>
<tr><td>Oligo fwd</td><td>CACCCTGTGCGCGGTATGTATAAG</td><td></td><td></td><td></td><td></td><td></td><td></td><td></td><td></td><td></td><td></td><td></td></tr>
<tr><td>Oligo rev</td><td>AAACCTTATACATACCGCGCACAG</td><td></td><td></td><td></td><td></td><td></td><td></td><td></td><td></td><td></td><td></td><td></td></tr>
<tr><td>Chromosome</td><td>start</td><td></td><td>end</td><td>strand</td><td>MM</td><td>target_seq</td><td>PAM</td><td>alignment</td><td>distance</td><td>position</td><td>gene name</td><td>gene id</td></tr>
<tr><td>GU305914.1</td><td></td><td>88151</td><td>88173</td><td>-</td><td>0</td><td>CTGTGCGCGGTATGTATAAG</td><td>CGG</td><td>||||||||[||||||||||||]PAM</td><td>NA</td><td>Intergenic</td><td>NA</td><td>NA</td></tr>
<tr><td>T27511</td><td>TGCGCGGTATGTATAAGCGGAGG</td><td></td><td>1000</td><td></td><td></td><td></td><td></td><td></td><td></td><td></td><td></td><td></td></tr>
<tr><td>Oligo fwd</td><td>CACCTGCGCGGTATGTATAAGCGG</td><td></td><td></td><td></td><td></td><td></td><td></td><td></td><td></td><td></td><td></td><td></td></tr>
<tr><td>Oligo rev</td><td>AAACCCGCTTATACATACCGCGCA</td><td></td><td></td><td></td><td></td><td></td><td></td><td></td><td></td><td></td><td></td><td></td></tr>
<tr><td>Chromosome</td><td>start</td><td></td><td>end</td><td>strand</td><td>MM</td><td>target_seq</td><td>PAM</td><td>alignment</td><td>distance</td><td>position</td><td>gene name</td><td>gene id</td></tr>
<tr><td>GU305914.1</td><td></td><td>88148</td><td>88170</td><td>-</td><td>0</td><td>TGCGCGGTATGTATAAGCGG</td><td>AGG</td><td>||||||||[||||||||||||]PAM</td><td>NA</td><td>Intergenic</td><td>NA</td><td>NA</td></tr>
<tr><td>T27512</td><td>GCGCGGTATGTATAAGCGGAGGG</td><td></td><td>1000</td><td></td><td></td><td></td><td></td><td></td><td></td><td></td><td></td><td></td></tr>
<tr><td>Oligo fwd</td><td>CACCGCGCGGTATGTATAAGCGGA</td><td></td><td></td><td></td><td></td><td></td><td></td><td></td><td></td><td></td><td></td><td></td></tr>
<tr><td>Oligo rev</td><td>AAACTCCGCTTATACATACCGCGC</td><td></td><td></td><td></td><td></td><td></td><td></td><td></td><td></td><td></td><td></td><td></td></tr>
<tr><td>Chromosome</td><td>start</td><td></td><td>end</td><td>strand</td><td>MM</td><td>target_seq</td><td>PAM</td><td>alignment</td><td>distance</td><td>position</td><td>gene name</td><td>gene id</td></tr>
<tr><td>GU305914.1</td><td></td><td>88147</td><td>88169</td><td>-</td><td>0</td><td>GCGCGGTATGTATAAGCGGA</td><td>GGG</td><td>||||||||[||||||||||||]PAM</td><td>NA</td><td>Intergenic</td><td>NA</td><td>NA</td></tr>
<tr><td>T27513</td><td>CGCGGTATGTATAAGCGGAGGGG</td><td></td><td>1000</td><td></td><td></td><td></td><td></td><td></td><td></td><td></td><td></td><td></td></tr>
<tr><td>Oligo fwd</td><td>CACCCGCGGTATGTATAAGCGGAG</td><td></td><td></td><td></td><td></td><td></td><td></td><td></td><td></td><td></td><td></td><td></td></tr>
<tr><td>Oligo rev</td><td>AAACCTCCGCTTATACATACCGCG</td><td></td><td></td><td></td><td></td><td></td><td></td><td></td><td></td><td></td><td></td><td></td></tr>
<tr><td>Chromosome</td><td>start</td><td></td><td>end</td><td>strand</td><td>MM</td><td>target_seq</td><td>PAM</td><td>alignment</td><td>distance</td><td>position</td><td>gene name</td><td>gene id</td></tr>
<tr><td>GU305914.1</td><td></td><td>88146</td><td>88168</td><td>-</td><td>0</td><td>CGCGGTATGTATAAGCGGAG</td><td>GGG</td><td>||||||||[||||||||||||]PAM</td><td>NA</td><td>Intergenic</td><td>NA</td><td>NA</td></tr>
<tr><td>T27514</td><td>GTATGTATAAGCGGAGGGGTAGG</td><td></td><td>1000</td><td></td><td></td><td></td><td></td><td></td><td></td><td></td><td></td><td></td></tr>
<tr><td>Oligo fwd</td><td>CACCGTATGTATAAGCGGAGGGGT</td><td></td><td></td><td></td><td></td><td></td><td></td><td></td><td></td><td></td><td></td><td></td></tr>
<tr><td>Oligo rev</td><td>AAACACCCCTCCGCTTATACATAC</td><td></td><td></td><td></td><td></td><td></td><td></td><td></td><td></td><td></td><td></td><td></td></tr>
<tr><td>Chromosome</td><td>start</td><td></td><td>end</td><td>strand</td><td>MM</td><td>target_seq</td><td>PAM</td><td>alignment</td><td>distance</td><td>position</td><td>gene name</td><td>gene id</td></tr>
<tr><td>GU305914.1</td><td></td><td>88142</td><td>88164</td><td>-</td><td>0</td><td>GTATGTATAAGCGGAGGGGT</td><td>AGG</td><td>||||||||[||||||||||||]PAM</td><td>NA</td><td>Intergenic</td><td>NA</td><td>NA</td></tr>
<tr><td>T27515</td><td>TATGTATAAGCGGAGGGGTAGGG</td><td></td><td>1000</td><td></td><td></td><td></td><td></td><td></td><td></td><td></td><td></td><td></td></tr>
<tr><td>Oligo fwd</td><td>CACCTATGTATAAGCGGAGGGGTA</td><td></td><td></td><td></td><td></td><td></td><td></td><td></td><td></td><td></td><td></td><td></td></tr>
<tr><td>Oligo rev</td><td>AAACTACCCCTCCGCTTATACATA</td><td></td><td></td><td></td><td></td><td></td><td></td><td></td><td></td><td></td><td></td><td></td></tr>
<tr><td>Chromosome</td><td>start</td><td></td><td>end</td><td>strand</td><td>MM</td><td>target_seq</td><td>PAM</td><td>alignment</td><td>distance</td><td>position</td><td>gene name</td><td>gene id</td></tr>
<tr><td>GU305914.1</td><td></td><td>88141</td><td>88163</td><td>-</td><td>0</td><td>TATGTATAAGCGGAGGGGTA</td><td>GGG</td><td>||||||||[||||||||||||]PAM</td><td>NA</td><td>Intergenic</td><td>NA</td><td>NA</td></tr>
<tr><td>T27516</td><td>ATGTATAAGCGGAGGGGTAGGGG</td><td></td><td>1000</td><td></td><td></td><td></td><td></td><td></td><td></td><td></td><td></td><td></td></tr>
<tr><td>Oligo fwd</td><td>CACCATGTATAAGCGGAGGGGTAG</td><td></td><td></td><td></td><td></td><td></td><td></td><td></td><td></td><td></td><td></td><td></td></tr>
<tr><td>Oligo rev</td><td>AAACCTACCCCTCCGCTTATACAT</td><td></td><td></td><td></td><td></td><td></td><td></td><td></td><td></td><td></td><td></td><td></td></tr>
<tr><td>Chromosome</td><td>start</td><td></td><td>end</td><td>strand</td><td>MM</td><td>target_seq</td><td>PAM</td><td>alignment</td><td>distance</td><td>position</td><td>gene name</td><td>gene id</td></tr>
<tr><td>GU305914.1</td><td></td><td>88140</td><td>88162</td><td>-</td><td>0</td><td>ATGTATAAGCGGAGGGGTAG</td><td>GGG</td><td>||||||||[||||||||||||]PAM</td><td>NA</td><td>Intergenic</td><td>NA</td><td>NA</td></tr>
<tr><td>T27517</td><td>GACCTGCTCGCGCGCAACGTCGG</td><td></td><td>1000</td><td></td><td></td><td></td><td></td><td></td><td></td><td></td><td></td><td></td></tr>
<tr><td>Oligo fwd</td><td>CACCGACCTGCTCGCGCGCAACGT</td><td></td><td></td><td></td><td></td><td></td><td></td><td></td><td></td><td></td><td></td><td></td></tr>
<tr><td>Oligo rev</td><td>AAACACGTTGCGCGCGAGCAGGTC</td><td></td><td></td><td></td><td></td><td></td><td></td><td></td><td></td><td></td><td></td><td></td></tr>
<tr><td>Chromosome</td><td>start</td><td></td><td>end</td><td>strand</td><td>MM</td><td>target_seq</td><td>PAM</td><td>alignment</td><td>distance</td><td>position</td><td>gene name</td><td>gene id</td></tr>
<tr><td>GU305914.1</td><td></td><td>88113</td><td>88135</td><td>-</td><td>0</td><td>GACCTGCTCGCGCGCAACGT</td><td>CGG</td><td>||||||||[||||||||||||]PAM</td><td>NA</td><td>Intergenic</td><td>NA</td><td>NA</td></tr>
<tr><td>T27518</td><td>GCTCGCGCGCAACGTCGGCAAGG</td><td></td><td>1000</td><td></td><td></td><td></td><td></td><td></td><td></td><td></td><td></td><td></td></tr>
<tr><td>Oligo fwd</td><td>CACCGCTCGCGCGCAACGTCGGCA</td><td></td><td></td><td></td><td></td><td></td><td></td><td></td><td></td><td></td><td></td><td></td></tr>
<tr><td>Oligo rev</td><td>AAACTGCCGACGTTGCGCGCGAGC</td><td></td><td></td><td></td><td></td><td></td><td></td><td></td><td></td><td></td><td></td><td></td></tr>
<tr><td>Chromosome</td><td>start</td><td></td><td>end</td><td>strand</td><td>MM</td><td>target_seq</td><td>PAM</td><td>alignment</td><td>distance</td><td>position</td><td>gene name</td><td>gene id</td></tr>
<tr><td>GU305914.1</td><td></td><td>88108</td><td>88130</td><td>-</td><td>0</td><td>GCTCGCGCGCAACGTCGGCA</td><td>AGG</td><td>||||||||[||||||||||||]PAM</td><td>NA</td><td>Intergenic</td><td>NA</td><td>NA</td></tr>
<tr><td>T27519</td><td>GCGCGCAACGTCGGCAAGGATGG</td><td></td><td>1000</td><td></td><td></td><td></td><td></td><td></td><td></td><td></td><td></td><td></td></tr>
<tr><td>Oligo fwd</td><td>CACCGCGCGCAACGTCGGCAAGGA</td><td></td><td></td><td></td><td></td><td></td><td></td><td></td><td></td><td></td><td></td><td></td></tr>
<tr><td>Oligo rev</td><td>AAACTCCTTGCCGACGTTGCGCGC</td><td></td><td></td><td></td><td></td><td></td><td></td><td></td><td></td><td></td><td></td><td></td></tr>
<tr><td>Chromosome</td><td>start</td><td></td><td>end</td><td>strand</td><td>MM</td><td>target_seq</td><td>PAM</td><td>alignment</td><td>distance</td><td>position</td><td>gene name</td><td>gene id</td></tr>
<tr><td>GU305914.1</td><td></td><td>88104</td><td>88126</td><td>-</td><td>0</td><td>GCGCGCAACGTCGGCAAGGA</td><td>TGG</td><td>||||||||[||||||||||||]PAM</td><td>NA</td><td>Intergenic</td><td>NA</td><td>NA</td></tr>
<tr><td>T27520</td><td>CAAATGCAACGAGTGCGCGATGG</td><td></td><td>1000</td><td></td><td></td><td></td><td></td><td></td><td></td><td></td><td></td><td></td></tr>
<tr><td>Oligo fwd</td><td>CACCCAAATGCAACGAGTGCGCGA</td><td></td><td></td><td></td><td></td><td></td><td></td><td></td><td></td><td></td><td></td><td></td></tr>
<tr><td>Oligo rev</td><td>AAACTCGCGCACTCGTTGCATTTG</td><td></td><td></td><td></td><td></td><td></td><td></td><td></td><td></td><td></td><td></td><td></td></tr>
<tr><td>Chromosome</td><td>start</td><td></td><td>end</td><td>strand</td><td>MM</td><td>target_seq</td><td>PAM</td><td>alignment</td><td>distance</td><td>position</td><td>gene name</td><td>gene id</td></tr>
<tr><td>GU305914.1</td><td></td><td>88044</td><td>88066</td><td>-</td><td>0</td><td>CAAATGCAACGAGTGCGCGA</td><td>TGG</td><td>||||||||[||||||||||||]PAM</td><td>NA</td><td>Intergenic</td><td>NA</td><td>NA</td></tr>
<tr><td>T27521</td><td>CAACGAGTGCGCGATGGATGTGG</td><td></td><td>1000</td><td></td><td></td><td></td><td></td><td></td><td></td><td></td><td></td><td></td></tr>
<tr><td>Oligo fwd</td><td>CACCCAACGAGTGCGCGATGGATG</td><td></td><td></td><td></td><td></td><td></td><td></td><td></td><td></td><td></td><td></td><td></td></tr>
<tr><td>Oligo rev</td><td>AAACCATCCATCGCGCACTCGTTG</td><td></td><td></td><td></td><td></td><td></td><td></td><td></td><td></td><td></td><td></td><td></td></tr>
<tr><td>Chromosome</td><td>start</td><td></td><td>end</td><td>strand</td><td>MM</td><td>target_seq</td><td>PAM</td><td>alignment</td><td>distance</td><td>position</td><td>gene name</td><td>gene id</td></tr>
<tr><td>GU305914.1</td><td></td><td>88038</td><td>88060</td><td>-</td><td>0</td><td>CAACGAGTGCGCGATGGATG</td><td>TGG</td><td>||||||||[||||||||||||]PAM</td><td>NA</td><td>Intergenic</td><td>NA</td><td>NA</td></tr>
<tr><td>T27522</td><td>CGATCCGAATATTGTGTCGATGG</td><td></td><td>1000</td><td></td><td></td><td></td><td></td><td></td><td></td><td></td><td></td><td></td></tr>
<tr><td>Oligo fwd</td><td>CACCCGATCCGAATATTGTGTCGA</td><td></td><td></td><td></td><td></td><td></td><td></td><td></td><td></td><td></td><td></td><td></td></tr>
<tr><td>Oligo rev</td><td>AAACTCGACACAATATTCGGATCG</td><td></td><td></td><td></td><td></td><td></td><td></td><td></td><td></td><td></td><td></td><td></td></tr>
<tr><td>Chromosome</td><td>start</td><td></td><td>end</td><td>strand</td><td>MM</td><td>target_seq</td><td>PAM</td><td>alignment</td><td>distance</td><td>position</td><td>gene name</td><td>gene id</td></tr>
<tr><td>GU305914.1</td><td></td><td>87999</td><td>88021</td><td>-</td><td>0</td><td>CGATCCGAATATTGTGTCGA</td><td>TGG</td><td>||||||||[||||||||||||]PAM</td><td>NA</td><td>Intergenic</td><td>NA</td><td>NA</td></tr>
</table>

<table>
<tr><td>T27523</td><td>TATTGTGTCGATGGATTCGACGG</td><td>1000</td><td></td><td></td><td></td><td></td><td></td><td></td><td></td><td></td><td></td></tr>
<tr><td>Oligo fwd</td><td>CACCTATTGTGTCGATGGATTCGA</td><td></td><td></td><td></td><td></td><td></td><td></td><td></td><td></td><td></td><td></td></tr>
<tr><td>Oligo rev</td><td>AAACTCGAATCCATCGACACAATA</td><td></td><td></td><td></td><td></td><td></td><td></td><td></td><td></td><td></td><td></td></tr>
<tr><td>Chromosome</td><td>start</td><td>end</td><td>strand</td><td>MM</td><td>target_seq</td><td>PAM</td><td>alignment</td><td>distance</td><td>position</td><td>gene name</td><td>gene id</td></tr>
<tr><td>GU305914.1</td><td>87990</td><td>88012</td><td>-</td><td>0</td><td>TATTGTGTCGATGGATTCGA</td><td>CGG</td><td>||||||||[||||||||||||]PAM</td><td>NA</td><td>Intergenic</td><td>NA</td><td>NA</td></tr>
<tr><td>T27524</td><td>CGATGGATTCGACGGCCTTCAGG</td><td>1000</td><td></td><td></td><td></td><td></td><td></td><td></td><td></td><td></td><td></td></tr>
<tr><td>Oligo fwd</td><td>CACCCGATGGATTCGACGGCCTTC</td><td></td><td></td><td></td><td></td><td></td><td></td><td></td><td></td><td></td><td></td></tr>
<tr><td>Oligo rev</td><td>AAACGAAGGCCGTCGAATCCATCG</td><td></td><td></td><td></td><td></td><td></td><td></td><td></td><td></td><td></td><td></td></tr>
<tr><td>Chromosome</td><td>start</td><td>end</td><td>strand</td><td>MM</td><td>target_seq</td><td>PAM</td><td>alignment</td><td>distance</td><td>position</td><td>gene name</td><td>gene id</td></tr>
<tr><td>GU305914.1</td><td>87982</td><td>88004</td><td>-</td><td>0</td><td>CGATGGATTCGACGGCCTTC</td><td>AGG</td><td>||||||||[||||||||||||]PAM</td><td>NA</td><td>Intergenic</td><td>NA</td><td>NA</td></tr>
<tr><td>T27525</td><td>TGGATTCGACGGCCTTCAGGAGG</td><td>1000</td><td></td><td></td><td></td><td></td><td></td><td></td><td></td><td></td><td></td></tr>
<tr><td>Oligo fwd</td><td>CACCTGGATTCGACGGCCTTCAGG</td><td></td><td></td><td></td><td></td><td></td><td></td><td></td><td></td><td></td><td></td></tr>
<tr><td>Oligo rev</td><td>AAACCCTGAAGGCCGTCGAATCCA</td><td></td><td></td><td></td><td></td><td></td><td></td><td></td><td></td><td></td><td></td></tr>
<tr><td>Chromosome</td><td>start</td><td>end</td><td>strand</td><td>MM</td><td>target_seq</td><td>PAM</td><td>alignment</td><td>distance</td><td>position</td><td>gene name</td><td>gene id</td></tr>
<tr><td>GU305914.1</td><td>87979</td><td>88001</td><td>-</td><td>0</td><td>TGGATTCGACGGCCTTCAGG</td><td>AGG</td><td>||||||||[||||||||||||]PAM</td><td>NA</td><td>Intergenic</td><td>NA</td><td>NA</td></tr>
<tr><td>T27526</td><td>TCGACGGCCTTCAGGAGGAACGG</td><td>1000</td><td></td><td></td><td></td><td></td><td></td><td></td><td></td><td></td><td></td></tr>
<tr><td>Oligo fwd</td><td>CACCTCGACGGCCTTCAGGAGGAA</td><td></td><td></td><td></td><td></td><td></td><td></td><td></td><td></td><td></td><td></td></tr>
<tr><td>Oligo rev</td><td>AAACTTCCTCCTGAAGGCCGTCGA</td><td></td><td></td><td></td><td></td><td></td><td></td><td></td><td></td><td></td><td></td></tr>
<tr><td>Chromosome</td><td>start</td><td>end</td><td>strand</td><td>MM</td><td>target_seq</td><td>PAM</td><td>alignment</td><td>distance</td><td>position</td><td>gene name</td><td>gene id</td></tr>
<tr><td>GU305914.1</td><td>87974</td><td>87996</td><td>-</td><td>0</td><td>TCGACGGCCTTCAGGAGGAA</td><td>CGG</td><td>||||||||[||||||||||||]PAM</td><td>NA</td><td>Intergenic</td><td>NA</td><td>NA</td></tr>
<tr><td>T27527</td><td>CGACGGCCTTCAGGAGGAACGGG</td><td>1000</td><td></td><td></td><td></td><td></td><td></td><td></td><td></td><td></td><td></td></tr>
<tr><td>Oligo fwd</td><td>CACCCGACGGCCTTCAGGAGGAAC</td><td></td><td></td><td></td><td></td><td></td><td></td><td></td><td></td><td></td><td></td></tr>
<tr><td>Oligo rev</td><td>AAACGTTCCTCCTGAAGGCCGTCG</td><td></td><td></td><td></td><td></td><td></td><td></td><td></td><td></td><td></td><td></td></tr>
<tr><td>Chromosome</td><td>start</td><td>end</td><td>strand</td><td>MM</td><td>target_seq</td><td>PAM</td><td>alignment</td><td>distance</td><td>position</td><td>gene name</td><td>gene id</td></tr>
<tr><td>GU305914.1</td><td>87973</td><td>87995</td><td>-</td><td>0</td><td>CGACGGCCTTCAGGAGGAAC</td><td>GGG</td><td>||||||||[||||||||||||]PAM</td><td>NA</td><td>Intergenic</td><td>NA</td><td>NA</td></tr>
<tr><td>T27528</td><td>GACGGCCTTCAGGAGGAACGGGG</td><td>1000</td><td></td><td></td><td></td><td></td><td></td><td></td><td></td><td></td><td></td></tr>
<tr><td>Oligo fwd</td><td>CACCGACGGCCTTCAGGAGGAACG</td><td></td><td></td><td></td><td></td><td></td><td></td><td></td><td></td><td></td><td></td></tr>
<tr><td>Oligo rev</td><td>AAACCGTTCCTCCTGAAGGCCGTC</td><td></td><td></td><td></td><td></td><td></td><td></td><td></td><td></td><td></td><td></td></tr>
<tr><td>Chromosome</td><td>start</td><td>end</td><td>strand</td><td>MM</td><td>target_seq</td><td>PAM</td><td>alignment</td><td>distance</td><td>position</td><td>gene name</td><td>gene id</td></tr>
<tr><td>GU305914.1</td><td>87972</td><td>87994</td><td>-</td><td>0</td><td>GACGGCCTTCAGGAGGAACG</td><td>GGG</td><td>||||||||[||||||||||||]PAM</td><td>NA</td><td>Intergenic</td><td>NA</td><td>NA</td></tr>
<tr><td>T27529</td><td>CTTCAGGAGGAACGGGGTCATGG</td><td>1000</td><td></td><td></td><td></td><td></td><td></td><td></td><td></td><td></td><td></td></tr>
<tr><td>Oligo fwd</td><td>CACCCTTCAGGAGGAACGGGGTCA</td><td></td><td></td><td></td><td></td><td></td><td></td><td></td><td></td><td></td><td></td></tr>
<tr><td>Oligo rev</td><td>AAACTGACCCCGTTCCTCCTGAAG</td><td></td><td></td><td></td><td></td><td></td><td></td><td></td><td></td><td></td><td></td></tr>
<tr><td>Chromosome</td><td>start</td><td>end</td><td>strand</td><td>MM</td><td>target_seq</td><td>PAM</td><td>alignment</td><td>distance</td><td>position</td><td>gene name</td><td>gene id</td></tr>
<tr><td>GU305914.1</td><td>87966</td><td>87988</td><td>-</td><td>0</td><td>CTTCAGGAGGAACGGGGTCA</td><td>TGG</td><td>||||||||[||||||||||||]PAM</td><td>NA</td><td>Intergenic</td><td>NA</td><td>NA</td></tr>
<tr><td>T27530</td><td>TCATGGTGATACATTTGTATCGG</td><td>1000</td><td></td><td></td><td></td><td></td><td></td><td></td><td></td><td></td><td></td></tr>
<tr><td>Oligo fwd</td><td>CACCTCATGGTGATACATTTGTAT</td><td></td><td></td><td></td><td></td><td></td><td></td><td></td><td></td><td></td><td></td></tr>
<tr><td>Oligo rev</td><td>AAACATACAAATGTATCACCATGA</td><td></td><td></td><td></td><td></td><td></td><td></td><td></td><td></td><td></td><td></td></tr>
<tr><td>Chromosome</td><td>start</td><td>end</td><td>strand</td><td>MM</td><td>target_seq</td><td>PAM</td><td>alignment</td><td>distance</td><td>position</td><td>gene name</td><td>gene id</td></tr>
<tr><td>GU305914.1</td><td>87949</td><td>87971</td><td>-</td><td>0</td><td>TCATGGTGATACATTTGTAT</td><td>CGG</td><td>||||||||[||||||||||||]PAM</td><td>NA</td><td>Intergenic</td><td>NA</td><td>NA</td></tr>
<tr><td>T27531</td><td>TTTGTATCGGACCCTCTATCCGG</td><td>1000</td><td></td><td></td><td></td><td></td><td></td><td></td><td></td><td></td><td></td></tr>
<tr><td>Oligo fwd</td><td>CACCTTTGTATCGGACCCTCTATC</td><td></td><td></td><td></td><td></td><td></td><td></td><td></td><td></td><td></td><td></td></tr>
<tr><td>Oligo rev</td><td>AAACGATAGAGGGTCCGATACAAA</td><td></td><td></td><td></td><td></td><td></td><td></td><td></td><td></td><td></td><td></td></tr>
<tr><td>Chromosome</td><td>start</td><td>end</td><td>strand</td><td>MM</td><td>target_seq</td><td>PAM</td><td>alignment</td><td>distance</td><td>position</td><td>gene name</td><td>gene id</td></tr>
<tr><td>GU305914.1</td><td>87936</td><td>87958</td><td>-</td><td>0</td><td>TTTGTATCGGACCCTCTATC</td><td>CGG</td><td>||||||||[||||||||||||]PAM</td><td>NA</td><td>Intergenic</td><td>NA</td><td>NA</td></tr>
<tr><td>T27532</td><td>TCGGACCCTCTATCCGGCGCTGG</td><td>1000</td><td></td><td></td><td></td><td></td><td></td><td></td><td></td><td></td><td></td></tr>
<tr><td>Oligo fwd</td><td>CACCTCGGACCCTCTATCCGGCGC</td><td></td><td></td><td></td><td></td><td></td><td></td><td></td><td></td><td></td><td></td></tr>
<tr><td>Oligo rev</td><td>AAACGCGCCGGATAGAGGGTCCGA</td><td></td><td></td><td></td><td></td><td></td><td></td><td></td><td></td><td></td><td></td></tr>
<tr><td>Chromosome</td><td>start</td><td>end</td><td>strand</td><td>MM</td><td>target_seq</td><td>PAM</td><td>alignment</td><td>distance</td><td>position</td><td>gene name</td><td>gene id</td></tr>
<tr><td>GU305914.1</td><td>87930</td><td>87952</td><td>-</td><td>0</td><td>TCGGACCCTCTATCCGGCGC</td><td>TGG</td><td>||||||||[||||||||||||]PAM</td><td>NA</td><td>Intergenic</td><td>NA</td><td>NA</td></tr>
<tr><td>T27533</td><td>GGCGCTGGTCTCGCAGAACGCGG</td><td>1000</td><td></td><td></td><td></td><td></td><td></td><td></td><td></td><td></td><td></td></tr>
<tr><td>Oligo fwd</td><td>CACCGGCGCTGGTCTCGCAGAACG</td><td></td><td></td><td></td><td></td><td></td><td></td><td></td><td></td><td></td><td></td></tr>
<tr><td>Oligo rev</td><td>AAACCGTTCTGCGAGACCAGCGCC</td><td></td><td></td><td></td><td></td><td></td><td></td><td></td><td></td><td></td><td></td></tr>
<tr><td>Chromosome</td><td>start</td><td>end</td><td>strand</td><td>MM</td><td>target_seq</td><td>PAM</td><td>alignment</td><td>distance</td><td>position</td><td>gene name</td><td>gene id</td></tr>
<tr><td>GU305914.1</td><td>87915</td><td>87937</td><td>-</td><td>0</td><td>GGCGCTGGTCTCGCAGAACG</td><td>CGG</td><td>||||||||[||||||||||||]PAM</td><td>NA</td><td>Intergenic</td><td>NA</td><td>NA</td></tr>
<tr><td>T27534</td><td>GCAGAACGCGGTGCAGACCTCGG</td><td>1000</td><td></td><td></td><td></td><td></td><td></td><td></td><td></td><td></td><td></td></tr>
<tr><td>Oligo fwd</td><td>CACCGCAGAACGCGGTGCAGACCT</td><td></td><td></td><td></td><td></td><td></td><td></td><td></td><td></td><td></td><td></td></tr>
<tr><td>Oligo rev</td><td>AAACAGGTCTGCACCGCGTTCTGC</td><td></td><td></td><td></td><td></td><td></td><td></td><td></td><td></td><td></td><td></td></tr>
<tr><td>Chromosome</td><td>start</td><td>end</td><td>strand</td><td>MM</td><td>target_seq</td><td>PAM</td><td>alignment</td><td>distance</td><td>position</td><td>gene name</td><td>gene id</td></tr>
<tr><td>GU305914.1</td><td>87903</td><td>87925</td><td>-</td><td>0</td><td>GCAGAACGCGGTGCAGACCT</td><td>CGG</td><td>||||||||[||||||||||||]PAM</td><td>NA</td><td>Intergenic</td><td>NA</td><td>NA</td></tr>
<tr><td>T27535</td><td>CTCGGTGCTGACCCTCTACATGG</td><td>1000</td><td></td><td></td><td></td><td></td><td></td><td></td><td></td><td></td><td></td></tr>
<tr><td>Oligo fwd</td><td>CACCCTCGGTGCTGACCCTCTACA</td><td></td><td></td><td></td><td></td><td></td><td></td><td></td><td></td><td></td><td></td></tr>
<tr><td>Oligo rev</td><td>AAACTGTAGAGGGTCAGCACCGAG</td><td></td><td></td><td></td><td></td><td></td><td></td><td></td><td></td><td></td><td></td></tr>
<tr><td>Chromosome</td><td>start</td><td>end</td><td>strand</td><td>MM</td><td>target_seq</td><td>PAM</td><td>alignment</td><td>distance</td><td>position</td><td>gene name</td><td>gene id</td></tr>
<tr><td>GU305914.1</td><td>87885</td><td>87907</td><td>-</td><td>0</td><td>CTCGGTGCTGACCCTCTACA</td><td>TGG</td><td>||||||||[||||||||||||]PAM</td><td>NA</td><td>Intergenic</td><td>NA</td><td>NA</td></tr>
<tr><td>T27536</td><td>CTACATGGAGATGCTGCTGCAGG</td><td>1000</td><td></td><td></td><td></td><td></td><td></td><td></td><td></td><td></td><td></td></tr>
<tr><td>Oligo fwd</td><td>CACCCTACATGGAGATGCTGCTGC</td><td></td><td></td><td></td><td></td><td></td><td></td><td></td><td></td><td></td><td></td></tr>
<tr><td>Oligo rev</td><td>AAACGCAGCAGCATCTCCATGTAG</td><td></td><td></td><td></td><td></td><td></td><td></td><td></td><td></td><td></td><td></td></tr>
<tr><td>Chromosome</td><td>start</td><td>end</td><td>strand</td><td>MM</td><td>target_seq</td><td>PAM</td><td>alignment</td><td>distance</td><td>position</td><td>gene name</td><td>gene id</td></tr>
<tr><td>GU305914.1</td><td>87870</td><td>87892</td><td>-</td><td>0</td><td>CTACATGGAGATGCTGCTGC</td><td>AGG</td><td>||||||||[||||||||||||]PAM</td><td>NA</td><td>Intergenic</td><td>NA</td><td>NA</td></tr>
<tr><td>T27537</td><td>TACATGGAGATGCTGCTGCAGGG</td><td>1000</td><td></td><td></td><td></td><td></td><td></td><td></td><td></td><td></td><td></td></tr>
<tr><td>Oligo fwd</td><td>CACCTACATGGAGATGCTGCTGCA</td><td></td><td></td><td></td><td></td><td></td><td></td><td></td><td></td><td></td><td></td></tr>
<tr><td>Oligo rev</td><td>AAACTGCAGCAGCATCTCCATGTA</td><td></td><td></td><td></td><td></td><td></td><td></td><td></td><td></td><td></td><td></td></tr>
<tr><td>Chromosome</td><td>start</td><td>end</td><td>strand</td><td>MM</td><td>target_seq</td><td>PAM</td><td>alignment</td><td>distance</td><td>position</td><td>gene name</td><td>gene id</td></tr>
</table>

GU305914.1 87869 87891 - 0 TACATGGAGATGCTGCTGCA GGG ||||||||[||||||||||||]PAM NA Intergenic NA NA

T27538 AGGGCTTGTACGATACGATGAGG 1000
Oligo fwd CACCAGGGCTTGTACGATACGATG
Oligo rev AAACCATCGTATCGTACAAGCCCT
Chromosome start end strand MM target_seq PAM alignment distance position gene name gene id
GU305914.1 87850 87872 - 0 AGGGCTTGTACGATACGATG AGG ||||||||[||||||||||||]PAM NA Intergenic NA NA

T27539 GGGCTTGTACGATACGATGAGGG 1000
Oligo fwd CACCGGGCTTGTACGATACGATGA
Oligo rev AAACTCATCGTATCGTACAAGCCC
Chromosome start end strand MM target_seq PAM alignment distance position gene name gene id
GU305914.1 87849 87871 - 0 GGGCTTGTACGATACGATGA GGG ||||||||[||||||||||||]PAM NA Intergenic NA NA

T27540 TACGATGAGGGAGATCGACATGG 1000
Oligo fwd CACCTACGATGAGGGAGATCGACA
Oligo rev AAACTGTCGATCTCCCTCATCGTA
Chromosome start end strand MM target_seq PAM alignment distance position gene name gene id
GU305914.1 87837 87859 - 0 TACGATGAGGGAGATCGACA TGG ||||||||[||||||||||||]PAM NA Intergenic NA NA

T27541 GGAGATCGACATGGCTTTGACGG 1000
Oligo fwd CACCGGAGATCGACATGGCTTTGA
Oligo rev AAACTCAAAGCCATGTCGATCTCC
Chromosome start end strand MM target_seq PAM alignment distance position gene name gene id
GU305914.1 87828 87850 - 0 GGAGATCGACATGGCTTTGA CGG ||||||||[||||||||||||]PAM NA Intergenic NA NA

T27542 GACATGGCTTTGACGGACTTTGG 1000
Oligo fwd CACCGACATGGCTTTGACGGACTT
Oligo rev AAACAAGTCCGTCAAAGCCATGTC
Chromosome start end strand MM target_seq PAM alignment distance position gene name gene id
GU305914.1 87821 87843 - 0 GACATGGCTTTGACGGACTT TGG ||||||||[||||||||||||]PAM NA Intergenic NA NA

T27543 ACATGGCTTTGACGGACTTTGGG 1000
Oligo fwd CACCACATGGCTTTGACGGACTTT
Oligo rev AAACAAAGTCCGTCAAAGCCATGT
Chromosome start end strand MM target_seq PAM alignment distance position gene name gene id
GU305914.1 87820 87842 - 0 ACATGGCTTTGACGGACTTT GGG ||||||||[||||||||||||]PAM NA Intergenic NA NA

T27544 TGACGGACTTTGGGACGCATCGG 1000
Oligo fwd CACCTGACGGACTTTGGGACGCAT
Oligo rev AAACATGCGTCCCAAAGTCCGTCA
Chromosome start end strand MM target_seq PAM alignment distance position gene name gene id
GU305914.1 87811 87833 - 0 TGACGGACTTTGGGACGCAT CGG ||||||||[||||||||||||]PAM NA Intergenic NA NA

T27545 GACGGACTTTGGGACGCATCGGG 1000
Oligo fwd CACCGACGGACTTTGGGACGCATC
Oligo rev AAACGATGCGTCCCAAAGTCCGTC
Chromosome start end strand MM target_seq PAM alignment distance position gene name gene id
GU305914.1 87810 87832 - 0 GACGGACTTTGGGACGCATC GGG ||||||||[||||||||||||]PAM NA Intergenic NA NA

T27546 ACTTTGGGACGCATCGGGACCGG 1000
Oligo fwd CACCACTTTGGGACGCATCGGGAC
Oligo rev AAACGTCCCGATGCGTCCCAAAGT
Chromosome start end strand MM target_seq PAM alignment distance position gene name gene id
GU305914.1 87805 87827 - 0 ACTTTGGGACGCATCGGGAC CGG ||||||||[||||||||||||]PAM NA Intergenic NA NA

T27547 GGGACCGGCAGCGATACTACCGG 1000
Oligo fwd CACCGGGACCGGCAGCGATACTAC
Oligo rev AAACGTAGTATCGCTGCCGGTCCC
Chromosome start end strand MM target_seq PAM alignment distance position gene name gene id
GU305914.1 87790 87812 - 0 GGGACCGGCAGCGATACTAC CGG ||||||||[||||||||||||]PAM NA Intergenic NA NA

T27548 ACCGGCAGCGATACTACCGGCGG 1000
Oligo fwd CACCACCGGCAGCGATACTACCGG
Oligo rev AAACCCGGTAGTATCGCTGCCGGT
Chromosome start end strand MM target_seq PAM alignment distance position gene name gene id
GU305914.1 87787 87809 - 0 ACCGGCAGCGATACTACCGG CGG ||||||||[||||||||||||]PAM NA Intergenic NA NA

T27549 CCGGCAGCGATACTACCGGCGGG 1000
Oligo fwd CACCCCGGCAGCGATACTACCGGC
Oligo rev AAACGCCGGTAGTATCGCTGCCGG
Chromosome start end strand MM target_seq PAM alignment distance position gene name gene id
GU305914.1 87786 87808 - 0 CCGGCAGCGATACTACCGGC GGG ||||||||[||||||||||||]PAM NA Intergenic NA NA

T27550 GATAACTTTTGCGCCCGAGCTGG 1000
Oligo fwd CACCGATAACTTTTGCGCCCGAGC
Oligo rev AAACGCTCGGGCGCAAAAGTTATC
Chromosome start end strand MM target_seq PAM alignment distance position gene name gene id
GU305914.1 87723 87745 - 0 GATAACTTTTGCGCCCGAGC TGG ||||||||[||||||||||||]PAM NA Intergenic NA NA

T27551 GCTGGCGCTGACGATCGACCTGG 1000
Oligo fwd CACCGCTGGCGCTGACGATCGACC
Oligo rev AAACGGTCGATCGTCAGCGCCAGC
Chromosome start end strand MM target_seq PAM alignment distance position gene name gene id
GU305914.1 87705 87727 - 0 GCTGGCGCTGACGATCGACC TGG ||||||||[||||||||||||]PAM NA Intergenic NA NA

T27552 CCTGGCGACGCTGAACGACGTGG 1000
Oligo fwd CACCCCTGGCGACGCTGAACGACG
Oligo rev AAACCGTCGTTCAGCGTCGCCAGG

<table>
<tr><td>Chromosome</td><td>start</td><td>end</td><td>strand</td><td>MM</td><td>target_seq</td><td>PAM</td><td>alignment</td><td>distance</td><td>position</td><td>gene name</td><td>gene id</td></tr>
<tr><td>GU305914.1</td><td>87687</td><td>87709</td><td>-</td><td>0</td><td>CCTGGCGACGCTGAACGACG</td><td>TGG</td><td>||||||||[||||||||||||]PAM</td><td>NA</td><td>Intergenic</td><td>NA</td><td>NA</td></tr>
<tr><td>GU305914.1</td><td>69229</td><td>69251</td><td>+</td><td>4</td><td>GCTGTCGACGCTGGACGAGG</td><td>CGG</td><td>-|||-|||[|||||-||||-|]PAM</td><td>NA</td><td>Intergenic</td><td>NA</td><td>NA</td></tr>
<tr><td>GU305914.1</td><td>213222</td><td>213244</td><td>-</td><td>4</td><td>CCTCACGACGCTGATCGACC</td><td>GGG</td><td>|||--|||[||||||-||||-]PAM</td><td>NA</td><td>Intergenic</td><td>NA</td><td>NA</td></tr>
<tr><td>T27553</td><td>TGCAAAATCAACTGTGTGTATGG</td><td>1000</td><td></td><td></td><td></td><td></td><td></td><td></td><td></td><td></td><td></td></tr>
<tr><td>Oligo fwd</td><td>CACCTGCAAAATCAACTGTGTGTA</td><td></td><td></td><td></td><td></td><td></td><td></td><td></td><td></td><td></td><td></td></tr>
<tr><td>Oligo rev</td><td>AAACTACACACAGTTGATTTTGCA</td><td></td><td></td><td></td><td></td><td></td><td></td><td></td><td></td><td></td><td></td></tr>
<tr><td>Chromosome</td><td>start</td><td>end</td><td>strand</td><td>MM</td><td>target_seq</td><td>PAM</td><td>alignment</td><td>distance</td><td>position</td><td>gene name</td><td>gene id</td></tr>
<tr><td>GU305914.1</td><td>87653</td><td>87675</td><td>-</td><td>0</td><td>TGCAAAATCAACTGTGTGTA</td><td>TGG</td><td>||||||||[||||||||||||]PAM</td><td>NA</td><td>Intergenic</td><td>NA</td><td>NA</td></tr>
<tr><td>T27554</td><td>GCAAAATCAACTGTGTGTATGGG</td><td>1000</td><td></td><td></td><td></td><td></td><td></td><td></td><td></td><td></td><td></td></tr>
<tr><td>Oligo fwd</td><td>CACCGCAAAATCAACTGTGTGTAT</td><td></td><td></td><td></td><td></td><td></td><td></td><td></td><td></td><td></td><td></td></tr>
<tr><td>Oligo rev</td><td>AAACATACACACAGTTGATTTTGC</td><td></td><td></td><td></td><td></td><td></td><td></td><td></td><td></td><td></td><td></td></tr>
<tr><td>Chromosome</td><td>start</td><td>end</td><td>strand</td><td>MM</td><td>target_seq</td><td>PAM</td><td>alignment</td><td>distance</td><td>position</td><td>gene name</td><td>gene id</td></tr>
<tr><td>GU305914.1</td><td>87652</td><td>87674</td><td>-</td><td>0</td><td>GCAAAATCAACTGTGTGTAT</td><td>GGG</td><td>||||||||[||||||||||||]PAM</td><td>NA</td><td>Intergenic</td><td>NA</td><td>NA</td></tr>
<tr><td>T27555</td><td>CAAAATCAACTGTGTGTATGGGG</td><td>1000</td><td></td><td></td><td></td><td></td><td></td><td></td><td></td><td></td><td></td></tr>
<tr><td>Oligo fwd</td><td>CACCCAAAATCAACTGTGTGTATG</td><td></td><td></td><td></td><td></td><td></td><td></td><td></td><td></td><td></td><td></td></tr>
<tr><td>Oligo rev</td><td>AAACCATACACACAGTTGATTTTG</td><td></td><td></td><td></td><td></td><td></td><td></td><td></td><td></td><td></td><td></td></tr>
<tr><td>Chromosome</td><td>start</td><td>end</td><td>strand</td><td>MM</td><td>target_seq</td><td>PAM</td><td>alignment</td><td>distance</td><td>position</td><td>gene name</td><td>gene id</td></tr>
<tr><td>GU305914.1</td><td>87651</td><td>87673</td><td>-</td><td>0</td><td>CAAAATCAACTGTGTGTATG</td><td>GGG</td><td>||||||||[||||||||||||]PAM</td><td>NA</td><td>Intergenic</td><td>NA</td><td>NA</td></tr>
<tr><td>T27556</td><td>AATCAACTGTGTGTATGGGGCGG</td><td>1000</td><td></td><td></td><td></td><td></td><td></td><td></td><td></td><td></td><td></td></tr>
<tr><td>Oligo fwd</td><td>CACCAATCAACTGTGTGTATGGGG</td><td></td><td></td><td></td><td></td><td></td><td></td><td></td><td></td><td></td><td></td></tr>
<tr><td>Oligo rev</td><td>AAACCCCCATACACACAGTTGATT</td><td></td><td></td><td></td><td></td><td></td><td></td><td></td><td></td><td></td><td></td></tr>
<tr><td>Chromosome</td><td>start</td><td>end</td><td>strand</td><td>MM</td><td>target_seq</td><td>PAM</td><td>alignment</td><td>distance</td><td>position</td><td>gene name</td><td>gene id</td></tr>
<tr><td>GU305914.1</td><td>87648</td><td>87670</td><td>-</td><td>0</td><td>AATCAACTGTGTGTATGGGG</td><td>CGG</td><td>||||||||[||||||||||||]PAM</td><td>NA</td><td>Intergenic</td><td>NA</td><td>NA</td></tr>
<tr><td>T27557</td><td>CAACTGTGTGTATGGGGCGGTGG</td><td>1000</td><td></td><td></td><td></td><td></td><td></td><td></td><td></td><td></td><td></td></tr>
<tr><td>Oligo fwd</td><td>CACCCAACTGTGTGTATGGGGCGG</td><td></td><td></td><td></td><td></td><td></td><td></td><td></td><td></td><td></td><td></td></tr>
<tr><td>Oligo rev</td><td>AAACCCGCCCCATACACACAGTTG</td><td></td><td></td><td></td><td></td><td></td><td></td><td></td><td></td><td></td><td></td></tr>
<tr><td>Chromosome</td><td>start</td><td>end</td><td>strand</td><td>MM</td><td>target_seq</td><td>PAM</td><td>alignment</td><td>distance</td><td>position</td><td>gene name</td><td>gene id</td></tr>
<tr><td>GU305914.1</td><td>87645</td><td>87667</td><td>-</td><td>0</td><td>CAACTGTGTGTATGGGGCGG</td><td>TGG</td><td>||||||||[||||||||||||]PAM</td><td>NA</td><td>Intergenic</td><td>NA</td><td>NA</td></tr>
<tr><td>T27558</td><td>TGGGGCGGTGGACGCGAGCCAGG</td><td>1000</td><td></td><td></td><td></td><td></td><td></td><td></td><td></td><td></td><td></td></tr>
<tr><td>Oligo fwd</td><td>CACCTGGGGCGGTGGACGCGAGCC</td><td></td><td></td><td></td><td></td><td></td><td></td><td></td><td></td><td></td><td></td></tr>
<tr><td>Oligo rev</td><td>AAACGGCTCGCGTCCACCGCCCCA</td><td></td><td></td><td></td><td></td><td></td><td></td><td></td><td></td><td></td><td></td></tr>
<tr><td>Chromosome</td><td>start</td><td>end</td><td>strand</td><td>MM</td><td>target_seq</td><td>PAM</td><td>alignment</td><td>distance</td><td>position</td><td>gene name</td><td>gene id</td></tr>
<tr><td>GU305914.1</td><td>87633</td><td>87655</td><td>-</td><td>0</td><td>TGGGGCGGTGGACGCGAGCC</td><td>AGG</td><td>||||||||[||||||||||||]PAM</td><td>NA</td><td>Intergenic</td><td>NA</td><td>NA</td></tr>
<tr><td>T27559</td><td>GGGGCGGTGGACGCGAGCCAGGG</td><td>1000</td><td></td><td></td><td></td><td></td><td></td><td></td><td></td><td></td><td></td></tr>
<tr><td>Oligo fwd</td><td>CACCGGGGCGGTGGACGCGAGCCA</td><td></td><td></td><td></td><td></td><td></td><td></td><td></td><td></td><td></td><td></td></tr>
<tr><td>Oligo rev</td><td>AAACTGGCTCGCGTCCACCGCCCC</td><td></td><td></td><td></td><td></td><td></td><td></td><td></td><td></td><td></td><td></td></tr>
<tr><td>Chromosome</td><td>start</td><td>end</td><td>strand</td><td>MM</td><td>target_seq</td><td>PAM</td><td>alignment</td><td>distance</td><td>position</td><td>gene name</td><td>gene id</td></tr>
<tr><td>GU305914.1</td><td>87632</td><td>87654</td><td>-</td><td>0</td><td>GGGGCGGTGGACGCGAGCCA</td><td>GGG</td><td>||||||||[||||||||||||]PAM</td><td>NA</td><td>Intergenic</td><td>NA</td><td>NA</td></tr>
<tr><td>T27560</td><td>GGGCGGTGGACGCGAGCCAGGGG</td><td>1000</td><td></td><td></td><td></td><td></td><td></td><td></td><td></td><td></td><td></td></tr>
<tr><td>Oligo fwd</td><td>CACCGGGCGGTGGACGCGAGCCAG</td><td></td><td></td><td></td><td></td><td></td><td></td><td></td><td></td><td></td><td></td></tr>
<tr><td>Oligo rev</td><td>AAACCTGGCTCGCGTCCACCGCCC</td><td></td><td></td><td></td><td></td><td></td><td></td><td></td><td></td><td></td><td></td></tr>
<tr><td>Chromosome</td><td>start</td><td>end</td><td>strand</td><td>MM</td><td>target_seq</td><td>PAM</td><td>alignment</td><td>distance</td><td>position</td><td>gene name</td><td>gene id</td></tr>
<tr><td>GU305914.1</td><td>87631</td><td>87653</td><td>-</td><td>0</td><td>GGGCGGTGGACGCGAGCCAG</td><td>GGG</td><td>||||||||[||||||||||||]PAM</td><td>NA</td><td>Intergenic</td><td>NA</td><td>NA</td></tr>
<tr><td>T27561</td><td>GGCGGTGGACGCGAGCCAGGGGG</td><td>1000</td><td></td><td></td><td></td><td></td><td></td><td></td><td></td><td></td><td></td></tr>
<tr><td>Oligo fwd</td><td>CACCGGCGGTGGACGCGAGCCAGG</td><td></td><td></td><td></td><td></td><td></td><td></td><td></td><td></td><td></td><td></td></tr>
<tr><td>Oligo rev</td><td>AAACCCTGGCTCGCGTCCACCGCC</td><td></td><td></td><td></td><td></td><td></td><td></td><td></td><td></td><td></td><td></td></tr>
<tr><td>Chromosome</td><td>start</td><td>end</td><td>strand</td><td>MM</td><td>target_seq</td><td>PAM</td><td>alignment</td><td>distance</td><td>position</td><td>gene name</td><td>gene id</td></tr>
<tr><td>GU305914.1</td><td>87630</td><td>87652</td><td>-</td><td>0</td><td>GGCGGTGGACGCGAGCCAGG</td><td>GGG</td><td>||||||||[||||||||||||]PAM</td><td>NA</td><td>Intergenic</td><td>NA</td><td>NA</td></tr>
<tr><td>T27562</td><td>GGTGGACGCGAGCCAGGGGGTGG</td><td>1000</td><td></td><td></td><td></td><td></td><td></td><td></td><td></td><td></td><td></td></tr>
<tr><td>Oligo fwd</td><td>CACCGGTGGACGCGAGCCAGGGGG</td><td></td><td></td><td></td><td></td><td></td><td></td><td></td><td></td><td></td><td></td></tr>
<tr><td>Oligo rev</td><td>AAACCCCCCTGGCTCGCGTCCACC</td><td></td><td></td><td></td><td></td><td></td><td></td><td></td><td></td><td></td><td></td></tr>
<tr><td>Chromosome</td><td>start</td><td>end</td><td>strand</td><td>MM</td><td>target_seq</td><td>PAM</td><td>alignment</td><td>distance</td><td>position</td><td>gene name</td><td>gene id</td></tr>
<tr><td>GU305914.1</td><td>87627</td><td>87649</td><td>-</td><td>0</td><td>GGTGGACGCGAGCCAGGGGG</td><td>TGG</td><td>||||||||[||||||||||||]PAM</td><td>NA</td><td>Intergenic</td><td>NA</td><td>NA</td></tr>
<tr><td>T27563</td><td>GGACGCGAGCCAGGGGGTGGCGG</td><td>1000</td><td></td><td></td><td></td><td></td><td></td><td></td><td></td><td></td><td></td></tr>
<tr><td>Oligo fwd</td><td>CACCGGACGCGAGCCAGGGGGTGG</td><td></td><td></td><td></td><td></td><td></td><td></td><td></td><td></td><td></td><td></td></tr>
<tr><td>Oligo rev</td><td>AAACCCACCCCCTGGCTCGCGTCC</td><td></td><td></td><td></td><td></td><td></td><td></td><td></td><td></td><td></td><td></td></tr>
<tr><td>Chromosome</td><td>start</td><td>end</td><td>strand</td><td>MM</td><td>target_seq</td><td>PAM</td><td>alignment</td><td>distance</td><td>position</td><td>gene name</td><td>gene id</td></tr>
<tr><td>GU305914.1</td><td>87624</td><td>87646</td><td>-</td><td>0</td><td>GGACGCGAGCCAGGGGGTGG</td><td>CGG</td><td>||||||||[||||||||||||]PAM</td><td>NA</td><td>Intergenic</td><td>NA</td><td>NA</td></tr>
<tr><td>T27564</td><td>AGGGGGTGGCGGTCTGTCGACGG</td><td>1000</td><td></td><td></td><td></td><td></td><td></td><td></td><td></td><td></td><td></td></tr>
<tr><td>Oligo fwd</td><td>CACCAGGGGGTGGCGGTCTGTCGA</td><td></td><td></td><td></td><td></td><td></td><td></td><td></td><td></td><td></td><td></td></tr>
<tr><td>Oligo rev</td><td>AAACTCGACAGACCGCCACCCCCT</td><td></td><td></td><td></td><td></td><td></td><td></td><td></td><td></td><td></td><td></td></tr>
<tr><td>Chromosome</td><td>start</td><td>end</td><td>strand</td><td>MM</td><td>target_seq</td><td>PAM</td><td>alignment</td><td>distance</td><td>position</td><td>gene name</td><td>gene id</td></tr>
<tr><td>GU305914.1</td><td>87613</td><td>87635</td><td>-</td><td>0</td><td>AGGGGGTGGCGGTCTGTCGA</td><td>CGG</td><td>||||||||[||||||||||||]PAM</td><td>NA</td><td>Intergenic</td><td>NA</td><td>NA</td></tr>
<tr><td>T27565</td><td>TCGACGGCTCTTGTCCCTGCTGG</td><td>1000</td><td></td><td></td><td></td><td></td><td></td><td></td><td></td><td></td><td></td></tr>
<tr><td>Oligo fwd</td><td>CACCTCGACGGCTCTTGTCCCTGC</td><td></td><td></td><td></td><td></td><td></td><td></td><td></td><td></td><td></td><td></td></tr>
<tr><td>Oligo rev</td><td>AAACGCAGGGACAAGAGCCGTCGA</td><td></td><td></td><td></td><td></td><td></td><td></td><td></td><td></td><td></td><td></td></tr>
<tr><td>Chromosome</td><td>start</td><td>end</td><td>strand</td><td>MM</td><td>target_seq</td><td>PAM</td><td>alignment</td><td>distance</td><td>position</td><td>gene name</td><td>gene id</td></tr>
<tr><td>GU305914.1</td><td>87597</td><td>87619</td><td>-</td><td>0</td><td>TCGACGGCTCTTGTCCCTGC</td><td>TGG</td><td>||||||||[||||||||||||]PAM</td><td>NA</td><td>Intergenic</td><td>NA</td><td>NA</td></tr>
<tr><td>T27566</td><td>GGCTCTTGTCCCTGCTGGCGAGG</td><td>1000</td><td></td><td></td><td></td><td></td><td></td><td></td><td></td><td></td><td></td></tr>
<tr><td>Oligo fwd</td><td>CACCGGCTCTTGTCCCTGCTGGCG</td><td></td><td></td><td></td><td></td><td></td><td></td><td></td><td></td><td></td><td></td></tr>
<tr><td>Oligo rev</td><td>AAACCGCCAGCAGGGACAAGAGCC</td><td></td><td></td><td></td><td></td><td></td><td></td><td></td><td></td><td></td><td></td></tr>
<tr><td>Chromosome</td><td>start</td><td>end</td><td>strand</td><td>MM</td><td>target_seq</td><td>PAM</td><td>alignment</td><td>distance</td><td>position</td><td>gene name</td><td>gene id</td></tr>
<tr><td>GU305914.1</td><td>87592</td><td>87614</td><td>-</td><td>0</td><td>GGCTCTTGTCCCTGCTGGCG</td><td>AGG</td><td>||||||||[||||||||||||]PAM</td><td>NA</td><td>Intergenic</td><td>NA</td><td>NA</td></tr>
</table>

<table>
<tr><td>T27567</td><td>GAGGTTGTGCGATATCTGCCCGG</td><td>1000</td><td></td><td></td><td></td><td></td><td></td><td></td><td></td><td></td><td></td></tr>
<tr><td>Oligo fwd</td><td>CACCGAGGTTGTGCGATATCTGCC</td><td></td><td></td><td></td><td></td><td></td><td></td><td></td><td></td><td></td><td></td></tr>
<tr><td>Oligo rev</td><td>AAACGGCAGATATCGCACAACCTC</td><td></td><td></td><td></td><td></td><td></td><td></td><td></td><td></td><td></td><td></td></tr>
<tr><td>Chromosome</td><td>start</td><td>end</td><td>strand</td><td>MM</td><td>target_seq</td><td>PAM</td><td>alignment</td><td>distance</td><td>position</td><td>gene name</td><td>gene id</td></tr>
<tr><td>GU305914.1</td><td>87573</td><td>87595</td><td>-</td><td>0</td><td>GAGGTTGTGCGATATCTGCC</td><td>CGG</td><td>||||||||[||||||||||||]PAM</td><td>NA</td><td>Intergenic</td><td>NA</td><td>NA</td></tr>
<tr><td>T27568</td><td>GTTGTGCGATATCTGCCCGGTGG</td><td>1000</td><td></td><td></td><td></td><td></td><td></td><td></td><td></td><td></td><td></td></tr>
<tr><td>Oligo fwd</td><td>CACCGTTGTGCGATATCTGCCCGG</td><td></td><td></td><td></td><td></td><td></td><td></td><td></td><td></td><td></td><td></td></tr>
<tr><td>Oligo rev</td><td>AAACCCGGGCAGATATCGCACAAC</td><td></td><td></td><td></td><td></td><td></td><td></td><td></td><td></td><td></td><td></td></tr>
<tr><td>Chromosome</td><td>start</td><td>end</td><td>strand</td><td>MM</td><td>target_seq</td><td>PAM</td><td>alignment</td><td>distance</td><td>position</td><td>gene name</td><td>gene id</td></tr>
<tr><td>GU305914.1</td><td>87570</td><td>87592</td><td>-</td><td>0</td><td>GTTGTGCGATATCTGCCCGG</td><td>TGG</td><td>||||||||[||||||||||||]PAM</td><td>NA</td><td>Intergenic</td><td>NA</td><td>NA</td></tr>
<tr><td>T27569</td><td>GTGCGATATCTGCCCGGTGGCGG</td><td>1000</td><td></td><td></td><td></td><td></td><td></td><td></td><td></td><td></td><td></td></tr>
<tr><td>Oligo fwd</td><td>CACCGTGCGATATCTGCCCGGTGG</td><td></td><td></td><td></td><td></td><td></td><td></td><td></td><td></td><td></td><td></td></tr>
<tr><td>Oligo rev</td><td>AAACCCACCGGGCAGATATCGCAC</td><td></td><td></td><td></td><td></td><td></td><td></td><td></td><td></td><td></td><td></td></tr>
<tr><td>Chromosome</td><td>start</td><td>end</td><td>strand</td><td>MM</td><td>target_seq</td><td>PAM</td><td>alignment</td><td>distance</td><td>position</td><td>gene name</td><td>gene id</td></tr>
<tr><td>GU305914.1</td><td>87567</td><td>87589</td><td>-</td><td>0</td><td>GTGCGATATCTGCCCGGTGG</td><td>CGG</td><td>||||||||[||||||||||||]PAM</td><td>NA</td><td>Intergenic</td><td>NA</td><td>NA</td></tr>
<tr><td>T27570</td><td>TGCGATATCTGCCCGGTGGCGGG</td><td>1000</td><td></td><td></td><td></td><td></td><td></td><td></td><td></td><td></td><td></td></tr>
<tr><td>Oligo fwd</td><td>CACCTGCGATATCTGCCCGGTGGC</td><td></td><td></td><td></td><td></td><td></td><td></td><td></td><td></td><td></td><td></td></tr>
<tr><td>Oligo rev</td><td>AAACGCCACCGGGCAGATATCGCA</td><td></td><td></td><td></td><td></td><td></td><td></td><td></td><td></td><td></td><td></td></tr>
<tr><td>Chromosome</td><td>start</td><td>end</td><td>strand</td><td>MM</td><td>target_seq</td><td>PAM</td><td>alignment</td><td>distance</td><td>position</td><td>gene name</td><td>gene id</td></tr>
<tr><td>GU305914.1</td><td>87566</td><td>87588</td><td>-</td><td>0</td><td>TGCGATATCTGCCCGGTGGC</td><td>GGG</td><td>||||||||[||||||||||||]PAM</td><td>NA</td><td>Intergenic</td><td>NA</td><td>NA</td></tr>
<tr><td>T27571</td><td>TATCTGCCCGGTGGCGGGTCCGG</td><td>1000</td><td></td><td></td><td></td><td></td><td></td><td></td><td></td><td></td><td></td></tr>
<tr><td>Oligo fwd</td><td>CACCTATCTGCCCGGTGGCGGGTC</td><td></td><td></td><td></td><td></td><td></td><td></td><td></td><td></td><td></td><td></td></tr>
<tr><td>Oligo rev</td><td>AAACGACCCGCCACCGGGCAGATA</td><td></td><td></td><td></td><td></td><td></td><td></td><td></td><td></td><td></td><td></td></tr>
<tr><td>Chromosome</td><td>start</td><td>end</td><td>strand</td><td>MM</td><td>target_seq</td><td>PAM</td><td>alignment</td><td>distance</td><td>position</td><td>gene name</td><td>gene id</td></tr>
<tr><td>GU305914.1</td><td>87561</td><td>87583</td><td>-</td><td>0</td><td>TATCTGCCCGGTGGCGGGTC</td><td>CGG</td><td>||||||||[||||||||||||]PAM</td><td>NA</td><td>Intergenic</td><td>NA</td><td>NA</td></tr>
<tr><td>T27572</td><td>TGGCGGGTCCGGAGATCTATCGG</td><td>1000</td><td></td><td></td><td></td><td></td><td></td><td></td><td></td><td></td><td></td></tr>
<tr><td>Oligo fwd</td><td>CACCTGGCGGGTCCGGAGATCTAT</td><td></td><td></td><td></td><td></td><td></td><td></td><td></td><td></td><td></td><td></td></tr>
<tr><td>Oligo rev</td><td>AAACATAGATCTCCGGACCCGCCA</td><td></td><td></td><td></td><td></td><td></td><td></td><td></td><td></td><td></td><td></td></tr>
<tr><td>Chromosome</td><td>start</td><td>end</td><td>strand</td><td>MM</td><td>target_seq</td><td>PAM</td><td>alignment</td><td>distance</td><td>position</td><td>gene name</td><td>gene id</td></tr>
<tr><td>GU305914.1</td><td>87550</td><td>87572</td><td>-</td><td>0</td><td>TGGCGGGTCCGGAGATCTAT</td><td>CGG</td><td>||||||||[||||||||||||]PAM</td><td>NA</td><td>Intergenic</td><td>NA</td><td>NA</td></tr>
<tr><td>T27573</td><td>GGCGGGTCCGGAGATCTATCGGG</td><td>1000</td><td></td><td></td><td></td><td></td><td></td><td></td><td></td><td></td><td></td></tr>
<tr><td>Oligo fwd</td><td>CACCGGCGGGTCCGGAGATCTATC</td><td></td><td></td><td></td><td></td><td></td><td></td><td></td><td></td><td></td><td></td></tr>
<tr><td>Oligo rev</td><td>AAACGATAGATCTCCGGACCCGCC</td><td></td><td></td><td></td><td></td><td></td><td></td><td></td><td></td><td></td><td></td></tr>
<tr><td>Chromosome</td><td>start</td><td>end</td><td>strand</td><td>MM</td><td>target_seq</td><td>PAM</td><td>alignment</td><td>distance</td><td>position</td><td>gene name</td><td>gene id</td></tr>
<tr><td>GU305914.1</td><td>87549</td><td>87571</td><td>-</td><td>0</td><td>GGCGGGTCCGGAGATCTATC</td><td>GGG</td><td>||||||||[||||||||||||]PAM</td><td>NA</td><td>Intergenic</td><td>NA</td><td>NA</td></tr>
<tr><td>T27574</td><td>TCCGGAGATCTATCGGGAGACGG</td><td>1000</td><td></td><td></td><td></td><td></td><td></td><td></td><td></td><td></td><td></td></tr>
<tr><td>Oligo fwd</td><td>CACCTCCGGAGATCTATCGGGAGA</td><td></td><td></td><td></td><td></td><td></td><td></td><td></td><td></td><td></td><td></td></tr>
<tr><td>Oligo rev</td><td>AAACTCTCCCGATAGATCTCCGGA</td><td></td><td></td><td></td><td></td><td></td><td></td><td></td><td></td><td></td><td></td></tr>
<tr><td>Chromosome</td><td>start</td><td>end</td><td>strand</td><td>MM</td><td>target_seq</td><td>PAM</td><td>alignment</td><td>distance</td><td>position</td><td>gene name</td><td>gene id</td></tr>
<tr><td>GU305914.1</td><td>87543</td><td>87565</td><td>-</td><td>0</td><td>TCCGGAGATCTATCGGGAGA</td><td>CGG</td><td>||||||||[||||||||||||]PAM</td><td>NA</td><td>Intergenic</td><td>NA</td><td>NA</td></tr>
<tr><td>T27575</td><td>CACCTGTTTTCAGTGCTACGAGG</td><td>1000</td><td></td><td></td><td></td><td></td><td></td><td></td><td></td><td></td><td></td></tr>
<tr><td>Oligo fwd</td><td>CACCCACCTGTTTTCAGTGCTACG</td><td></td><td></td><td></td><td></td><td></td><td></td><td></td><td></td><td></td><td></td></tr>
<tr><td>Oligo rev</td><td>AAACCGTAGCACTGAAAACAGGTG</td><td></td><td></td><td></td><td></td><td></td><td></td><td></td><td></td><td></td><td></td></tr>
<tr><td>Chromosome</td><td>start</td><td>end</td><td>strand</td><td>MM</td><td>target_seq</td><td>PAM</td><td>alignment</td><td>distance</td><td>position</td><td>gene name</td><td>gene id</td></tr>
<tr><td>GU305914.1</td><td>87519</td><td>87541</td><td>-</td><td>0</td><td>CACCTGTTTTCAGTGCTACG</td><td>AGG</td><td>||||||||[||||||||||||]PAM</td><td>NA</td><td>Intergenic</td><td>NA</td><td>NA</td></tr>
<tr><td>T27576</td><td>TCAGTGCTACGAGGAGCTGATGG</td><td>1000</td><td></td><td></td><td></td><td></td><td></td><td></td><td></td><td></td><td></td></tr>
<tr><td>Oligo fwd</td><td>CACCTCAGTGCTACGAGGAGCTGA</td><td></td><td></td><td></td><td></td><td></td><td></td><td></td><td></td><td></td><td></td></tr>
<tr><td>Oligo rev</td><td>AAACTCAGCTCCTCGTAGCACTGA</td><td></td><td></td><td></td><td></td><td></td><td></td><td></td><td></td><td></td><td></td></tr>
<tr><td>Chromosome</td><td>start</td><td>end</td><td>strand</td><td>MM</td><td>target_seq</td><td>PAM</td><td>alignment</td><td>distance</td><td>position</td><td>gene name</td><td>gene id</td></tr>
<tr><td>GU305914.1</td><td>87510</td><td>87532</td><td>-</td><td>0</td><td>TCAGTGCTACGAGGAGCTGA</td><td>TGG</td><td>||||||||[||||||||||||]PAM</td><td>NA</td><td>Intergenic</td><td>NA</td><td>NA</td></tr>
<tr><td>T27577</td><td>GTGCTACGAGGAGCTGATGGCGG</td><td>1000</td><td></td><td></td><td></td><td></td><td></td><td></td><td></td><td></td><td></td></tr>
<tr><td>Oligo fwd</td><td>CACCGTGCTACGAGGAGCTGATGG</td><td></td><td></td><td></td><td></td><td></td><td></td><td></td><td></td><td></td><td></td></tr>
<tr><td>Oligo rev</td><td>AAACCCATCAGCTCCTCGTAGCAC</td><td></td><td></td><td></td><td></td><td></td><td></td><td></td><td></td><td></td><td></td></tr>
<tr><td>Chromosome</td><td>start</td><td>end</td><td>strand</td><td>MM</td><td>target_seq</td><td>PAM</td><td>alignment</td><td>distance</td><td>position</td><td>gene name</td><td>gene id</td></tr>
<tr><td>GU305914.1</td><td>87507</td><td>87529</td><td>-</td><td>0</td><td>GTGCTACGAGGAGCTGATGG</td><td>CGG</td><td>||||||||[||||||||||||]PAM</td><td>NA</td><td>Intergenic</td><td>NA</td><td>NA</td></tr>
<tr><td>T27578</td><td>CTGATGGCGGTGCCGAACCAAGG</td><td>1000</td><td></td><td></td><td></td><td></td><td></td><td></td><td></td><td></td><td></td></tr>
<tr><td>Oligo fwd</td><td>CACCCTGATGGCGGTGCCGAACCA</td><td></td><td></td><td></td><td></td><td></td><td></td><td></td><td></td><td></td><td></td></tr>
<tr><td>Oligo rev</td><td>AAACTGGTTCGGCACCGCCATCAG</td><td></td><td></td><td></td><td></td><td></td><td></td><td></td><td></td><td></td><td></td></tr>
<tr><td>Chromosome</td><td>start</td><td>end</td><td>strand</td><td>MM</td><td>target_seq</td><td>PAM</td><td>alignment</td><td>distance</td><td>position</td><td>gene name</td><td>gene id</td></tr>
<tr><td>GU305914.1</td><td>87494</td><td>87516</td><td>-</td><td>0</td><td>CTGATGGCGGTGCCGAACCA</td><td>AGG</td><td>||||||||[||||||||||||]PAM</td><td>NA</td><td>Intergenic</td><td>NA</td><td>NA</td></tr>
<tr><td>T27579</td><td>ACCAAGGCCGCTCGATCAACCGG</td><td>1000</td><td></td><td></td><td></td><td></td><td></td><td></td><td></td><td></td><td></td></tr>
<tr><td>Oligo fwd</td><td>CACCACCAAGGCCGCTCGATCAAC</td><td></td><td></td><td></td><td></td><td></td><td></td><td></td><td></td><td></td><td></td></tr>
<tr><td>Oligo rev</td><td>AAACGTTGATCGAGCGGCCTTGGT</td><td></td><td></td><td></td><td></td><td></td><td></td><td></td><td></td><td></td><td></td></tr>
<tr><td>Chromosome</td><td>start</td><td>end</td><td>strand</td><td>MM</td><td>target_seq</td><td>PAM</td><td>alignment</td><td>distance</td><td>position</td><td>gene name</td><td>gene id</td></tr>
<tr><td>GU305914.1</td><td>87478</td><td>87500</td><td>-</td><td>0</td><td>ACCAAGGCCGCTCGATCAAC</td><td>CGG</td><td>||||||||[||||||||||||]PAM</td><td>NA</td><td>Intergenic</td><td>NA</td><td>NA</td></tr>
<tr><td>T27580</td><td>AAGGCCGCTCGATCAACCGGCGG</td><td>1000</td><td></td><td></td><td></td><td></td><td></td><td></td><td></td><td></td><td></td></tr>
<tr><td>Oligo fwd</td><td>CACCAAGGCCGCTCGATCAACCGG</td><td></td><td></td><td></td><td></td><td></td><td></td><td></td><td></td><td></td><td></td></tr>
<tr><td>Oligo rev</td><td>AAACCCGGTTGATCGAGCGGCCTT</td><td></td><td></td><td></td><td></td><td></td><td></td><td></td><td></td><td></td><td></td></tr>
<tr><td>Chromosome</td><td>start</td><td>end</td><td>strand</td><td>MM</td><td>target_seq</td><td>PAM</td><td>alignment</td><td>distance</td><td>position</td><td>gene name</td><td>gene id</td></tr>
<tr><td>GU305914.1</td><td>87475</td><td>87497</td><td>-</td><td>0</td><td>AAGGCCGCTCGATCAACCGG</td><td>CGG</td><td>||||||||[||||||||||||]PAM</td><td>NA</td><td>Intergenic</td><td>NA</td><td>NA</td></tr>
<tr><td>T27581</td><td>CTCGATCAACCGGCGGATGCAGG</td><td>1000</td><td></td><td></td><td></td><td></td><td></td><td></td><td></td><td></td><td></td></tr>
<tr><td>Oligo fwd</td><td>CACCCTCGATCAACCGGCGGATGC</td><td></td><td></td><td></td><td></td><td></td><td></td><td></td><td></td><td></td><td></td></tr>
<tr><td>Oligo rev</td><td>AAACGCATCCGCCGGTTGATCGAG</td><td></td><td></td><td></td><td></td><td></td><td></td><td></td><td></td><td></td><td></td></tr>
<tr><td>Chromosome</td><td>start</td><td>end</td><td>strand</td><td>MM</td><td>target_seq</td><td>PAM</td><td>alignment</td><td>distance</td><td>position</td><td>gene name</td><td>gene id</td></tr>
<tr><td>GU305914.1</td><td>87468</td><td>87490</td><td>-</td><td>0</td><td>CTCGATCAACCGGCGGATGC</td><td>AGG</td><td>||||||||[||||||||||||]PAM</td><td>NA</td><td>Intergenic</td><td>NA</td><td>NA</td></tr>
</table>

<table>
<tr><td>T27582</td><td>TCGATCAACCGGCGGATGCAGGG</td><td></td><td>1000</td><td></td><td></td><td></td><td></td><td></td><td></td><td></td><td></td><td></td></tr>
<tr><td>Oligo fwd</td><td>CACCTCGATCAACCGGCGGATGCA</td><td></td><td></td><td></td><td></td><td></td><td></td><td></td><td></td><td></td><td></td><td></td></tr>
<tr><td>Oligo rev</td><td>AAACTGCATCCGCCGGTTGATCGA</td><td></td><td></td><td></td><td></td><td></td><td></td><td></td><td></td><td></td><td></td><td></td></tr>
<tr><td>Chromosome</td><td>start</td><td></td><td>end</td><td>strand</td><td>MM</td><td>target_seq</td><td>PAM</td><td>alignment</td><td>distance</td><td>position</td><td>gene name</td><td>gene id</td></tr>
<tr><td>GU305914.1</td><td></td><td>87467</td><td>87489</td><td>-</td><td>0</td><td>TCGATCAACCGGCGGATGCA</td><td>GGG</td><td>||||||||[||||||||||||]PAM</td><td>NA</td><td>Intergenic</td><td>NA</td><td>NA</td></tr>
<tr><td>T27583</td><td>CGACCATATCACTATCAAAAAGG</td><td></td><td>1000</td><td></td><td></td><td></td><td></td><td></td><td></td><td></td><td></td><td></td></tr>
<tr><td>Oligo fwd</td><td>CACCCGACCATATCACTATCAAAA</td><td></td><td></td><td></td><td></td><td></td><td></td><td></td><td></td><td></td><td></td><td></td></tr>
<tr><td>Oligo rev</td><td>AAACTTTTGATAGTGATATGGTCG</td><td></td><td></td><td></td><td></td><td></td><td></td><td></td><td></td><td></td><td></td><td></td></tr>
<tr><td>Chromosome</td><td>start</td><td></td><td>end</td><td>strand</td><td>MM</td><td>target_seq</td><td>PAM</td><td>alignment</td><td>distance</td><td>position</td><td>gene name</td><td>gene id</td></tr>
<tr><td>GU305914.1</td><td></td><td>87435</td><td>87457</td><td>-</td><td>0</td><td>CGACCATATCACTATCAAAA</td><td>AGG</td><td>||||||||[||||||||||||]PAM</td><td>NA</td><td>Intergenic</td><td>NA</td><td>NA</td></tr>
<tr><td>T27584</td><td>TATCACTATCAAAAAGGTCTTGG</td><td></td><td>1000</td><td></td><td></td><td></td><td></td><td></td><td></td><td></td><td></td><td></td></tr>
<tr><td>Oligo fwd</td><td>CACCTATCACTATCAAAAAGGTCT</td><td></td><td></td><td></td><td></td><td></td><td></td><td></td><td></td><td></td><td></td><td></td></tr>
<tr><td>Oligo rev</td><td>AAACAGACCTTTTTGATAGTGATA</td><td></td><td></td><td></td><td></td><td></td><td></td><td></td><td></td><td></td><td></td><td></td></tr>
<tr><td>Chromosome</td><td>start</td><td></td><td>end</td><td>strand</td><td>MM</td><td>target_seq</td><td>PAM</td><td>alignment</td><td>distance</td><td>position</td><td>gene name</td><td>gene id</td></tr>
<tr><td>GU305914.1</td><td></td><td>87429</td><td>87451</td><td>-</td><td>0</td><td>TATCACTATCAAAAAGGTCT</td><td>TGG</td><td>||||||||[||||||||||||]PAM</td><td>NA</td><td>Intergenic</td><td>NA</td><td>NA</td></tr>
<tr><td>T27585</td><td>CAAAAAGGTCTTGGTGCAGCTGG</td><td></td><td>1000</td><td></td><td></td><td></td><td></td><td></td><td></td><td></td><td></td><td></td></tr>
<tr><td>Oligo fwd</td><td>CACCCAAAAAGGTCTTGGTGCAGC</td><td></td><td></td><td></td><td></td><td></td><td></td><td></td><td></td><td></td><td></td><td></td></tr>
<tr><td>Oligo rev</td><td>AAACGCTGCACCAAGACCTTTTTG</td><td></td><td></td><td></td><td></td><td></td><td></td><td></td><td></td><td></td><td></td><td></td></tr>
<tr><td>Chromosome</td><td>start</td><td></td><td>end</td><td>strand</td><td>MM</td><td>target_seq</td><td>PAM</td><td>alignment</td><td>distance</td><td>position</td><td>gene name</td><td>gene id</td></tr>
<tr><td>GU305914.1</td><td></td><td>87420</td><td>87442</td><td>-</td><td>0</td><td>CAAAAAGGTCTTGGTGCAGC</td><td>TGG</td><td>||||||||[||||||||||||]PAM</td><td>NA</td><td>Intergenic</td><td>NA</td><td>NA</td></tr>
<tr><td>T27586</td><td>GGTCTTGGTGCAGCTGGATATGG</td><td></td><td>1000</td><td></td><td></td><td></td><td></td><td></td><td></td><td></td><td></td><td></td></tr>
<tr><td>Oligo fwd</td><td>CACCGGTCTTGGTGCAGCTGGATA</td><td></td><td></td><td></td><td></td><td></td><td></td><td></td><td></td><td></td><td></td><td></td></tr>
<tr><td>Oligo rev</td><td>AAACTATCCAGCTGCACCAAGACC</td><td></td><td></td><td></td><td></td><td></td><td></td><td></td><td></td><td></td><td></td><td></td></tr>
<tr><td>Chromosome</td><td>start</td><td></td><td>end</td><td>strand</td><td>MM</td><td>target_seq</td><td>PAM</td><td>alignment</td><td>distance</td><td>position</td><td>gene name</td><td>gene id</td></tr>
<tr><td>GU305914.1</td><td></td><td>87414</td><td>87436</td><td>-</td><td>0</td><td>GGTCTTGGTGCAGCTGGATA</td><td>TGG</td><td>||||||||[||||||||||||]PAM</td><td>NA</td><td>Intergenic</td><td>NA</td><td>NA</td></tr>
<tr><td>T27587</td><td>GCTGGATATGGATGCCCAGACGG</td><td></td><td>1000</td><td></td><td></td><td></td><td></td><td></td><td></td><td></td><td></td><td></td></tr>
<tr><td>Oligo fwd</td><td>CACCGCTGGATATGGATGCCCAGA</td><td></td><td></td><td></td><td></td><td></td><td></td><td></td><td></td><td></td><td></td><td></td></tr>
<tr><td>Oligo rev</td><td>AAACTCTGGGCATCCATATCCAGC</td><td></td><td></td><td></td><td></td><td></td><td></td><td></td><td></td><td></td><td></td><td></td></tr>
<tr><td>Chromosome</td><td>start</td><td></td><td>end</td><td>strand</td><td>MM</td><td>target_seq</td><td>PAM</td><td>alignment</td><td>distance</td><td>position</td><td>gene name</td><td>gene id</td></tr>
<tr><td>GU305914.1</td><td></td><td>87402</td><td>87424</td><td>-</td><td>0</td><td>GCTGGATATGGATGCCCAGA</td><td>CGG</td><td>||||||||[||||||||||||]PAM</td><td>NA</td><td>Intergenic</td><td>NA</td><td>NA</td></tr>
<tr><td>T27588</td><td>GGATATGGATGCCCAGACGGTGG</td><td></td><td>1000</td><td></td><td></td><td></td><td></td><td></td><td></td><td></td><td></td><td></td></tr>
<tr><td>Oligo fwd</td><td>CACCGGATATGGATGCCCAGACGG</td><td></td><td></td><td></td><td></td><td></td><td></td><td></td><td></td><td></td><td></td><td></td></tr>
<tr><td>Oligo rev</td><td>AAACCCGTCTGGGCATCCATATCC</td><td></td><td></td><td></td><td></td><td></td><td></td><td></td><td></td><td></td><td></td><td></td></tr>
<tr><td>Chromosome</td><td>start</td><td></td><td>end</td><td>strand</td><td>MM</td><td>target_seq</td><td>PAM</td><td>alignment</td><td>distance</td><td>position</td><td>gene name</td><td>gene id</td></tr>
<tr><td>GU305914.1</td><td></td><td>87399</td><td>87421</td><td>-</td><td>0</td><td>GGATATGGATGCCCAGACGG</td><td>TGG</td><td>||||||||[||||||||||||]PAM</td><td>NA</td><td>Intergenic</td><td>NA</td><td>NA</td></tr>
<tr><td>T27589</td><td>GGATGCCCAGACGGTGGAGCAGG</td><td></td><td>1000</td><td></td><td></td><td></td><td></td><td></td><td></td><td></td><td></td><td></td></tr>
<tr><td>Oligo fwd</td><td>CACCGGATGCCCAGACGGTGGAGC</td><td></td><td></td><td></td><td></td><td></td><td></td><td></td><td></td><td></td><td></td><td></td></tr>
<tr><td>Oligo rev</td><td>AAACGCTCCACCGTCTGGGCATCC</td><td></td><td></td><td></td><td></td><td></td><td></td><td></td><td></td><td></td><td></td><td></td></tr>
<tr><td>Chromosome</td><td>start</td><td></td><td>end</td><td>strand</td><td>MM</td><td>target_seq</td><td>PAM</td><td>alignment</td><td>distance</td><td>position</td><td>gene name</td><td>gene id</td></tr>
<tr><td>GU305914.1</td><td></td><td>87393</td><td>87415</td><td>-</td><td>0</td><td>GGATGCCCAGACGGTGGAGC</td><td>AGG</td><td>||||||||[||||||||||||]PAM</td><td>NA</td><td>Intergenic</td><td>NA</td><td>NA</td></tr>
<tr><td>T27590</td><td>CCAGACGGTGGAGCAGGATATGG</td><td></td><td>1000</td><td></td><td></td><td></td><td></td><td></td><td></td><td></td><td></td><td></td></tr>
<tr><td>Oligo fwd</td><td>CACCCCAGACGGTGGAGCAGGATA</td><td></td><td></td><td></td><td></td><td></td><td></td><td></td><td></td><td></td><td></td><td></td></tr>
<tr><td>Oligo rev</td><td>AAACTATCCTGCTCCACCGTCTGG</td><td></td><td></td><td></td><td></td><td></td><td></td><td></td><td></td><td></td><td></td><td></td></tr>
<tr><td>Chromosome</td><td>start</td><td></td><td>end</td><td>strand</td><td>MM</td><td>target_seq</td><td>PAM</td><td>alignment</td><td>distance</td><td>position</td><td>gene name</td><td>gene id</td></tr>
<tr><td>GU305914.1</td><td></td><td>87387</td><td>87409</td><td>-</td><td>0</td><td>CCAGACGGTGGAGCAGGATA</td><td>TGG</td><td>||||||||[||||||||||||]PAM</td><td>NA</td><td>Intergenic</td><td>NA</td><td>NA</td></tr>
<tr><td>T27591</td><td>CAGACGGTGGAGCAGGATATGGG</td><td></td><td>1000</td><td></td><td></td><td></td><td></td><td></td><td></td><td></td><td></td><td></td></tr>
<tr><td>Oligo fwd</td><td>CACCCAGACGGTGGAGCAGGATAT</td><td></td><td></td><td></td><td></td><td></td><td></td><td></td><td></td><td></td><td></td><td></td></tr>
<tr><td>Oligo rev</td><td>AAACATATCCTGCTCCACCGTCTG</td><td></td><td></td><td></td><td></td><td></td><td></td><td></td><td></td><td></td><td></td><td></td></tr>
<tr><td>Chromosome</td><td>start</td><td></td><td>end</td><td>strand</td><td>MM</td><td>target_seq</td><td>PAM</td><td>alignment</td><td>distance</td><td>position</td><td>gene name</td><td>gene id</td></tr>
<tr><td>GU305914.1</td><td></td><td>87386</td><td>87408</td><td>-</td><td>0</td><td>CAGACGGTGGAGCAGGATAT</td><td>GGG</td><td>||||||||[||||||||||||]PAM</td><td>NA</td><td>Intergenic</td><td>NA</td><td>NA</td></tr>
<tr><td>T27592</td><td>AGACGGTGGAGCAGGATATGGGG</td><td></td><td>1000</td><td></td><td></td><td></td><td></td><td></td><td></td><td></td><td></td><td></td></tr>
<tr><td>Oligo fwd</td><td>CACCAGACGGTGGAGCAGGATATG</td><td></td><td></td><td></td><td></td><td></td><td></td><td></td><td></td><td></td><td></td><td></td></tr>
<tr><td>Oligo rev</td><td>AAACCATATCCTGCTCCACCGTCT</td><td></td><td></td><td></td><td></td><td></td><td></td><td></td><td></td><td></td><td></td><td></td></tr>
<tr><td>Chromosome</td><td>start</td><td></td><td>end</td><td>strand</td><td>MM</td><td>target_seq</td><td>PAM</td><td>alignment</td><td>distance</td><td>position</td><td>gene name</td><td>gene id</td></tr>
<tr><td>GU305914.1</td><td></td><td>87385</td><td>87407</td><td>-</td><td>0</td><td>AGACGGTGGAGCAGGATATG</td><td>GGG</td><td>||||||||[||||||||||||]PAM</td><td>NA</td><td>Intergenic</td><td>NA</td><td>NA</td></tr>
<tr><td>GU305914.1</td><td></td><td>109517</td><td>109539</td><td>-</td><td>4</td><td>AATCGGTGGAGTAGGAGATG</td><td>GGG</td><td>|--|||||[|||-||||-|||]PAM</td><td>NA</td><td>Intergenic</td><td>NA</td><td>NA</td></tr>
<tr><td>T27593</td><td>GACGGTGGAGCAGGATATGGGGG</td><td></td><td>1000</td><td></td><td></td><td></td><td></td><td></td><td></td><td></td><td></td><td></td></tr>
<tr><td>Oligo fwd</td><td>CACCGACGGTGGAGCAGGATATGG</td><td></td><td></td><td></td><td></td><td></td><td></td><td></td><td></td><td></td><td></td><td></td></tr>
<tr><td>Oligo rev</td><td>AAACCCATATCCTGCTCCACCGTC</td><td></td><td></td><td></td><td></td><td></td><td></td><td></td><td></td><td></td><td></td><td></td></tr>
<tr><td>Chromosome</td><td>start</td><td></td><td>end</td><td>strand</td><td>MM</td><td>target_seq</td><td>PAM</td><td>alignment</td><td>distance</td><td>position</td><td>gene name</td><td>gene id</td></tr>
<tr><td>GU305914.1</td><td></td><td>87384</td><td>87406</td><td>-</td><td>0</td><td>GACGGTGGAGCAGGATATGG</td><td>GGG</td><td>||||||||[||||||||||||]PAM</td><td>NA</td><td>Intergenic</td><td>NA</td><td>NA</td></tr>
<tr><td>GU305914.1</td><td></td><td>109516</td><td>109538</td><td>-</td><td>4</td><td>ATCGGTGGAGTAGGAGATGG</td><td>GGG</td><td>--||||||[||-||||-||||]PAM</td><td>NA</td><td>Intergenic</td><td>NA</td><td>NA</td></tr>
<tr><td>T27594</td><td>ATATGGGGGATATTGCGATCAGG</td><td></td><td>1000</td><td></td><td></td><td></td><td></td><td></td><td></td><td></td><td></td><td></td></tr>
<tr><td>Oligo fwd</td><td>CACCATATGGGGGATATTGCGATC</td><td></td><td></td><td></td><td></td><td></td><td></td><td></td><td></td><td></td><td></td><td></td></tr>
<tr><td>Oligo rev</td><td>AAACGATCGCAATATCCCCCATAT</td><td></td><td></td><td></td><td></td><td></td><td></td><td></td><td></td><td></td><td></td><td></td></tr>
<tr><td>Chromosome</td><td>start</td><td></td><td>end</td><td>strand</td><td>MM</td><td>target_seq</td><td>PAM</td><td>alignment</td><td>distance</td><td>position</td><td>gene name</td><td>gene id</td></tr>
<tr><td>GU305914.1</td><td></td><td>87370</td><td>87392</td><td>-</td><td>0</td><td>ATATGGGGGATATTGCGATC</td><td>AGG</td><td>||||||||[||||||||||||]PAM</td><td>NA</td><td>Intergenic</td><td>NA</td><td>NA</td></tr>
<tr><td>T27595</td><td>TATGGGGGATATTGCGATCAGGG</td><td></td><td>1000</td><td></td><td></td><td></td><td></td><td></td><td></td><td></td><td></td><td></td></tr>
<tr><td>Oligo fwd</td><td>CACCTATGGGGGATATTGCGATCA</td><td></td><td></td><td></td><td></td><td></td><td></td><td></td><td></td><td></td><td></td><td></td></tr>
<tr><td>Oligo rev</td><td>AAACTGATCGCAATATCCCCCATA</td><td></td><td></td><td></td><td></td><td></td><td></td><td></td><td></td><td></td><td></td><td></td></tr>
<tr><td>Chromosome</td><td>start</td><td></td><td>end</td><td>strand</td><td>MM</td><td>target_seq</td><td>PAM</td><td>alignment</td><td>distance</td><td>position</td><td>gene name</td><td>gene id</td></tr>
<tr><td>GU305914.1</td><td></td><td>87369</td><td>87391</td><td>-</td><td>0</td><td>TATGGGGGATATTGCGATCA</td><td>GGG</td><td>||||||||[||||||||||||]PAM</td><td>NA</td><td>Intergenic</td><td>NA</td><td>NA</td></tr>
<tr><td>T27596</td><td>ATCAGGGCGCCCTCTGTGAAAGG</td><td></td><td>1000</td><td></td><td></td><td></td><td></td><td></td><td></td><td></td><td></td><td></td></tr>
<tr><td>Oligo fwd</td><td>CACCATCAGGGCGCCCTCTGTGAA</td><td></td><td></td><td></td><td></td><td></td><td></td><td></td><td></td><td></td><td></td><td></td></tr>
</table>

<table>
<tr><td>Oligo rev</td><td>AAACTTCACAGAGGGCGCCCTGAT</td><td></td><td></td><td></td><td></td><td></td><td></td><td></td><td></td><td></td><td></td><td></td></tr>
<tr><td>Chromosome</td><td>start</td><td></td><td>end</td><td>strand</td><td>MM</td><td>target_seq</td><td>PAM</td><td>alignment</td><td>distance</td><td>position</td><td>gene name</td><td>gene id</td></tr>
<tr><td>GU305914.1</td><td></td><td>87353</td><td>87375</td><td>-</td><td>0</td><td>ATCAGGGCGCCCTCTGTGAA</td><td>AGG</td><td>||||||||[||||||||||||]PAM</td><td>NA</td><td>Intergenic</td><td>NA</td><td>NA</td></tr>
<tr><td>T27597</td><td>CCTCTGTGAAAGGTATCATCCGG</td><td></td><td>1000</td><td></td><td></td><td></td><td></td><td></td><td></td><td></td><td></td><td></td></tr>
<tr><td>Oligo fwd</td><td>CACCCCTCTGTGAAAGGTATCATC</td><td></td><td></td><td></td><td></td><td></td><td></td><td></td><td></td><td></td><td></td><td></td></tr>
<tr><td>Oligo rev</td><td>AAACGATGATACCTTTCACAGAGG</td><td></td><td></td><td></td><td></td><td></td><td></td><td></td><td></td><td></td><td></td><td></td></tr>
<tr><td>Chromosome</td><td>start</td><td></td><td>end</td><td>strand</td><td>MM</td><td>target_seq</td><td>PAM</td><td>alignment</td><td>distance</td><td>position</td><td>gene name</td><td>gene id</td></tr>
<tr><td>GU305914.1</td><td></td><td>87343</td><td>87365</td><td>-</td><td>0</td><td>CCTCTGTGAAAGGTATCATC</td><td>CGG</td><td>||||||||[||||||||||||]PAM</td><td>NA</td><td>Intergenic</td><td>NA</td><td>NA</td></tr>
<tr><td>T27598</td><td>CTCTGTGAAAGGTATCATCCGGG</td><td></td><td>1000</td><td></td><td></td><td></td><td></td><td></td><td></td><td></td><td></td><td></td></tr>
<tr><td>Oligo fwd</td><td>CACCCTCTGTGAAAGGTATCATCC</td><td></td><td></td><td></td><td></td><td></td><td></td><td></td><td></td><td></td><td></td><td></td></tr>
<tr><td>Oligo rev</td><td>AAACGGATGATACCTTTCACAGAG</td><td></td><td></td><td></td><td></td><td></td><td></td><td></td><td></td><td></td><td></td><td></td></tr>
<tr><td>Chromosome</td><td>start</td><td></td><td>end</td><td>strand</td><td>MM</td><td>target_seq</td><td>PAM</td><td>alignment</td><td>distance</td><td>position</td><td>gene name</td><td>gene id</td></tr>
<tr><td>GU305914.1</td><td></td><td>87342</td><td>87364</td><td>-</td><td>0</td><td>CTCTGTGAAAGGTATCATCC</td><td>GGG</td><td>||||||||[||||||||||||]PAM</td><td>NA</td><td>Intergenic</td><td>NA</td><td>NA</td></tr>
<tr><td>T27599</td><td>GAGTCTCGCGTCGTTCTCACCGG</td><td></td><td>1000</td><td></td><td></td><td></td><td></td><td></td><td></td><td></td><td></td><td></td></tr>
<tr><td>Oligo fwd</td><td>CACCGAGTCTCGCGTCGTTCTCAC</td><td></td><td></td><td></td><td></td><td></td><td></td><td></td><td></td><td></td><td></td><td></td></tr>
<tr><td>Oligo rev</td><td>AAACGTGAGAACGACGCGAGACTC</td><td></td><td></td><td></td><td></td><td></td><td></td><td></td><td></td><td></td><td></td><td></td></tr>
<tr><td>Chromosome</td><td>start</td><td></td><td>end</td><td>strand</td><td>MM</td><td>target_seq</td><td>PAM</td><td>alignment</td><td>distance</td><td>position</td><td>gene name</td><td>gene id</td></tr>
<tr><td>GU305914.1</td><td></td><td>87312</td><td>87334</td><td>-</td><td>0</td><td>GAGTCTCGCGTCGTTCTCAC</td><td>CGG</td><td>||||||||[||||||||||||]PAM</td><td>NA</td><td>Intergenic</td><td>NA</td><td>NA</td></tr>
<tr><td>T27600</td><td>GTATGCGTACATCAACGACGCGG</td><td></td><td>1000</td><td></td><td></td><td></td><td></td><td></td><td></td><td></td><td></td><td></td></tr>
<tr><td>Oligo fwd</td><td>CACCGTATGCGTACATCAACGACG</td><td></td><td></td><td></td><td></td><td></td><td></td><td></td><td></td><td></td><td></td><td></td></tr>
<tr><td>Oligo rev</td><td>AAACCGTCGTTGATGTACGCATAC</td><td></td><td></td><td></td><td></td><td></td><td></td><td></td><td></td><td></td><td></td><td></td></tr>
<tr><td>Chromosome</td><td>start</td><td></td><td>end</td><td>strand</td><td>MM</td><td>target_seq</td><td>PAM</td><td>alignment</td><td>distance</td><td>position</td><td>gene name</td><td>gene id</td></tr>
<tr><td>GU305914.1</td><td></td><td>87285</td><td>87307</td><td>-</td><td>0</td><td>GTATGCGTACATCAACGACG</td><td>CGG</td><td>||||||||[||||||||||||]PAM</td><td>NA</td><td>Intergenic</td><td>NA</td><td>NA</td></tr>
<tr><td>T27601</td><td>TGCGTACATCAACGACGCGGAGG</td><td></td><td>1000</td><td></td><td></td><td></td><td></td><td></td><td></td><td></td><td></td><td></td></tr>
<tr><td>Oligo fwd</td><td>CACCTGCGTACATCAACGACGCGG</td><td></td><td></td><td></td><td></td><td></td><td></td><td></td><td></td><td></td><td></td><td></td></tr>
<tr><td>Oligo rev</td><td>AAACCCGCGTCGTTGATGTACGCA</td><td></td><td></td><td></td><td></td><td></td><td></td><td></td><td></td><td></td><td></td><td></td></tr>
<tr><td>Chromosome</td><td>start</td><td></td><td>end</td><td>strand</td><td>MM</td><td>target_seq</td><td>PAM</td><td>alignment</td><td>distance</td><td>position</td><td>gene name</td><td>gene id</td></tr>
<tr><td>GU305914.1</td><td></td><td>87282</td><td>87304</td><td>-</td><td>0</td><td>TGCGTACATCAACGACGCGG</td><td>AGG</td><td>||||||||[||||||||||||]PAM</td><td>NA</td><td>Intergenic</td><td>NA</td><td>NA</td></tr>
<tr><td>T27602</td><td>GTACATCAACGACGCGGAGGAGG</td><td></td><td>1000</td><td></td><td></td><td></td><td></td><td></td><td></td><td></td><td></td><td></td></tr>
<tr><td>Oligo fwd</td><td>CACCGTACATCAACGACGCGGAGG</td><td></td><td></td><td></td><td></td><td></td><td></td><td></td><td></td><td></td><td></td><td></td></tr>
<tr><td>Oligo rev</td><td>AAACCCTCCGCGTCGTTGATGTAC</td><td></td><td></td><td></td><td></td><td></td><td></td><td></td><td></td><td></td><td></td><td></td></tr>
<tr><td>Chromosome</td><td>start</td><td></td><td>end</td><td>strand</td><td>MM</td><td>target_seq</td><td>PAM</td><td>alignment</td><td>distance</td><td>position</td><td>gene name</td><td>gene id</td></tr>
<tr><td>GU305914.1</td><td></td><td>87279</td><td>87301</td><td>-</td><td>0</td><td>GTACATCAACGACGCGGAGG</td><td>AGG</td><td>||||||||[||||||||||||]PAM</td><td>NA</td><td>Intergenic</td><td>NA</td><td>NA</td></tr>
<tr><td>T27603</td><td>ACGACGCGGAGGAGGCGTTGAGG</td><td></td><td>1000</td><td></td><td></td><td></td><td></td><td></td><td></td><td></td><td></td><td></td></tr>
<tr><td>Oligo fwd</td><td>CACCACGACGCGGAGGAGGCGTTG</td><td></td><td></td><td></td><td></td><td></td><td></td><td></td><td></td><td></td><td></td><td></td></tr>
<tr><td>Oligo rev</td><td>AAACCAACGCCTCCTCCGCGTCGT</td><td></td><td></td><td></td><td></td><td></td><td></td><td></td><td></td><td></td><td></td><td></td></tr>
<tr><td>Chromosome</td><td>start</td><td></td><td>end</td><td>strand</td><td>MM</td><td>target_seq</td><td>PAM</td><td>alignment</td><td>distance</td><td>position</td><td>gene name</td><td>gene id</td></tr>
<tr><td>GU305914.1</td><td></td><td>87271</td><td>87293</td><td>-</td><td>0</td><td>ACGACGCGGAGGAGGCGTTG</td><td>AGG</td><td>||||||||[||||||||||||]PAM</td><td>NA</td><td>Intergenic</td><td>NA</td><td>NA</td></tr>
<tr><td>T27604</td><td>CGACGCGGAGGAGGCGTTGAGGG</td><td></td><td>1000</td><td></td><td></td><td></td><td></td><td></td><td></td><td></td><td></td><td></td></tr>
<tr><td>Oligo fwd</td><td>CACCCGACGCGGAGGAGGCGTTGA</td><td></td><td></td><td></td><td></td><td></td><td></td><td></td><td></td><td></td><td></td><td></td></tr>
<tr><td>Oligo rev</td><td>AAACTCAACGCCTCCTCCGCGTCG</td><td></td><td></td><td></td><td></td><td></td><td></td><td></td><td></td><td></td><td></td><td></td></tr>
<tr><td>Chromosome</td><td>start</td><td></td><td>end</td><td>strand</td><td>MM</td><td>target_seq</td><td>PAM</td><td>alignment</td><td>distance</td><td>position</td><td>gene name</td><td>gene id</td></tr>
<tr><td>GU305914.1</td><td></td><td>87270</td><td>87292</td><td>-</td><td>0</td><td>CGACGCGGAGGAGGCGTTGA</td><td>GGG</td><td>||||||||[||||||||||||]PAM</td><td>NA</td><td>Intergenic</td><td>NA</td><td>NA</td></tr>
<tr><td>T27605</td><td>GACGCGGAGGAGGCGTTGAGGGG</td><td></td><td>1000</td><td></td><td></td><td></td><td></td><td></td><td></td><td></td><td></td><td></td></tr>
<tr><td>Oligo fwd</td><td>CACCGACGCGGAGGAGGCGTTGAG</td><td></td><td></td><td></td><td></td><td></td><td></td><td></td><td></td><td></td><td></td><td></td></tr>
<tr><td>Oligo rev</td><td>AAACCTCAACGCCTCCTCCGCGTC</td><td></td><td></td><td></td><td></td><td></td><td></td><td></td><td></td><td></td><td></td><td></td></tr>
<tr><td>Chromosome</td><td>start</td><td></td><td>end</td><td>strand</td><td>MM</td><td>target_seq</td><td>PAM</td><td>alignment</td><td>distance</td><td>position</td><td>gene name</td><td>gene id</td></tr>
<tr><td>GU305914.1</td><td></td><td>87269</td><td>87291</td><td>-</td><td>0</td><td>GACGCGGAGGAGGCGTTGAG</td><td>GGG</td><td>||||||||[||||||||||||]PAM</td><td>NA</td><td>Intergenic</td><td>NA</td><td>NA</td></tr>
<tr><td>T27606</td><td>ACGCGGAGGAGGCGTTGAGGGGG</td><td></td><td>1000</td><td></td><td></td><td></td><td></td><td></td><td></td><td></td><td></td><td></td></tr>
<tr><td>Oligo fwd</td><td>CACCACGCGGAGGAGGCGTTGAGG</td><td></td><td></td><td></td><td></td><td></td><td></td><td></td><td></td><td></td><td></td><td></td></tr>
<tr><td>Oligo rev</td><td>AAACCCTCAACGCCTCCTCCGCGT</td><td></td><td></td><td></td><td></td><td></td><td></td><td></td><td></td><td></td><td></td><td></td></tr>
<tr><td>Chromosome</td><td>start</td><td></td><td>end</td><td>strand</td><td>MM</td><td>target_seq</td><td>PAM</td><td>alignment</td><td>distance</td><td>position</td><td>gene name</td><td>gene id</td></tr>
<tr><td>GU305914.1</td><td></td><td>87268</td><td>87290</td><td>-</td><td>0</td><td>ACGCGGAGGAGGCGTTGAGG</td><td>GGG</td><td>||||||||[||||||||||||]PAM</td><td>NA</td><td>Intergenic</td><td>NA</td><td>NA</td></tr>
<tr><td>T27607</td><td>CAACTTGTTCAGCGAGATCCCGG</td><td></td><td>1000</td><td></td><td></td><td></td><td></td><td></td><td></td><td></td><td></td><td></td></tr>
<tr><td>Oligo fwd</td><td>CACCCAACTTGTTCAGCGAGATCC</td><td></td><td></td><td></td><td></td><td></td><td></td><td></td><td></td><td></td><td></td><td></td></tr>
<tr><td>Oligo rev</td><td>AAACGGATCTCGCTGAACAAGTTG</td><td></td><td></td><td></td><td></td><td></td><td></td><td></td><td></td><td></td><td></td><td></td></tr>
<tr><td>Chromosome</td><td>start</td><td></td><td>end</td><td>strand</td><td>MM</td><td>target_seq</td><td>PAM</td><td>alignment</td><td>distance</td><td>position</td><td>gene name</td><td>gene id</td></tr>
<tr><td>GU305914.1</td><td></td><td>87243</td><td>87265</td><td>-</td><td>0</td><td>CAACTTGTTCAGCGAGATCC</td><td>CGG</td><td>||||||||[||||||||||||]PAM</td><td>NA</td><td>Intergenic</td><td>NA</td><td>NA</td></tr>
<tr><td>T27608</td><td>TGTTCAGCGAGATCCCGGATCGG</td><td></td><td>1000</td><td></td><td></td><td></td><td></td><td></td><td></td><td></td><td></td><td></td></tr>
<tr><td>Oligo fwd</td><td>CACCTGTTCAGCGAGATCCCGGAT</td><td></td><td></td><td></td><td></td><td></td><td></td><td></td><td></td><td></td><td></td><td></td></tr>
<tr><td>Oligo rev</td><td>AAACATCCGGGATCTCGCTGAACA</td><td></td><td></td><td></td><td></td><td></td><td></td><td></td><td></td><td></td><td></td><td></td></tr>
<tr><td>Chromosome</td><td>start</td><td></td><td>end</td><td>strand</td><td>MM</td><td>target_seq</td><td>PAM</td><td>alignment</td><td>distance</td><td>position</td><td>gene name</td><td>gene id</td></tr>
<tr><td>GU305914.1</td><td></td><td>87238</td><td>87260</td><td>-</td><td>0</td><td>TGTTCAGCGAGATCCCGGAT</td><td>CGG</td><td>||||||||[||||||||||||]PAM</td><td>NA</td><td>Intergenic</td><td>NA</td><td>NA</td></tr>
<tr><td>T27609</td><td>GGATCGGATCTACTCACTGTCGG</td><td></td><td>1000</td><td></td><td></td><td></td><td></td><td></td><td></td><td></td><td></td><td></td></tr>
<tr><td>Oligo fwd</td><td>CACCGGATCGGATCTACTCACTGT</td><td></td><td></td><td></td><td></td><td></td><td></td><td></td><td></td><td></td><td></td><td></td></tr>
<tr><td>Oligo rev</td><td>AAACACAGTGAGTAGATCCGATCC</td><td></td><td></td><td></td><td></td><td></td><td></td><td></td><td></td><td></td><td></td><td></td></tr>
<tr><td>Chromosome</td><td>start</td><td></td><td>end</td><td>strand</td><td>MM</td><td>target_seq</td><td>PAM</td><td>alignment</td><td>distance</td><td>position</td><td>gene name</td><td>gene id</td></tr>
<tr><td>GU305914.1</td><td></td><td>87222</td><td>87244</td><td>-</td><td>0</td><td>GGATCGGATCTACTCACTGT</td><td>CGG</td><td>||||||||[||||||||||||]PAM</td><td>NA</td><td>Intergenic</td><td>NA</td><td>NA</td></tr>
<tr><td>T27610</td><td>CACTGTCGGACTACACATACTGG</td><td></td><td>1000</td><td></td><td></td><td></td><td></td><td></td><td></td><td></td><td></td><td></td></tr>
<tr><td>Oligo fwd</td><td>CACCCACTGTCGGACTACACATAC</td><td></td><td></td><td></td><td></td><td></td><td></td><td></td><td></td><td></td><td></td><td></td></tr>
<tr><td>Oligo rev</td><td>AAACGTATGTGTAGTCCGACAGTG</td><td></td><td></td><td></td><td></td><td></td><td></td><td></td><td></td><td></td><td></td><td></td></tr>
<tr><td>Chromosome</td><td>start</td><td></td><td>end</td><td>strand</td><td>MM</td><td>target_seq</td><td>PAM</td><td>alignment</td><td>distance</td><td>position</td><td>gene name</td><td>gene id</td></tr>
<tr><td>GU305914.1</td><td></td><td>87208</td><td>87230</td><td>-</td><td>0</td><td>CACTGTCGGACTACACATAC</td><td>TGG</td><td>||||||||[||||||||||||]PAM</td><td>NA</td><td>Intergenic</td><td>NA</td><td>NA</td></tr>
<tr><td>T27611</td><td>CACATACTGGTCGAAAACGTCGG</td><td></td><td>1000</td><td></td><td></td><td></td><td></td><td></td><td></td><td></td><td></td><td></td></tr>
</table>

<table>
<tr><td>Oligo fwd</td><td>CACCCACATACTGGTCGAAAACGT</td><td></td><td></td><td></td><td></td><td></td><td></td><td></td><td></td><td></td><td></td></tr>
<tr><td>Oligo rev</td><td>AAACACGTTTTCGACCAGTATGTG</td><td></td><td></td><td></td><td></td><td></td><td></td><td></td><td></td><td></td><td></td></tr>
<tr><td>Chromosome</td><td>start</td><td>end</td><td>strand</td><td>MM</td><td>target_seq</td><td>PAM</td><td>alignment</td><td>distance</td><td>position</td><td>gene name</td><td>gene id</td></tr>
<tr><td>GU305914.1</td><td>87195</td><td>87217</td><td>-</td><td>0</td><td>CACATACTGGTCGAAAACGT</td><td>CGG</td><td>||||||||[||||||||||||]PAM</td><td>NA</td><td>Intergenic</td><td>NA</td><td>NA</td></tr>
<tr><td></td><td></td><td></td><td></td><td></td><td></td><td></td><td></td><td></td><td></td><td></td><td></td></tr>
<tr><td>T27612</td><td>ATACTGGTCGAAAACGTCGGAGG</td><td>1000</td><td></td><td></td><td></td><td></td><td></td><td></td><td></td><td></td><td></td></tr>
<tr><td>Oligo fwd</td><td>CACCATACTGGTCGAAAACGTCGG</td><td></td><td></td><td></td><td></td><td></td><td></td><td></td><td></td><td></td><td></td></tr>
<tr><td>Oligo rev</td><td>AAACCCGACGTTTTCGACCAGTAT</td><td></td><td></td><td></td><td></td><td></td><td></td><td></td><td></td><td></td><td></td></tr>
<tr><td>Chromosome</td><td>start</td><td>end</td><td>strand</td><td>MM</td><td>target_seq</td><td>PAM</td><td>alignment</td><td>distance</td><td>position</td><td>gene name</td><td>gene id</td></tr>
<tr><td>GU305914.1</td><td>87192</td><td>87214</td><td>-</td><td>0</td><td>ATACTGGTCGAAAACGTCGG</td><td>AGG</td><td>||||||||[||||||||||||]PAM</td><td>NA</td><td>Intergenic</td><td>NA</td><td>NA</td></tr>
<tr><td></td><td></td><td></td><td></td><td></td><td></td><td></td><td></td><td></td><td></td><td></td><td></td></tr>
<tr><td>T27613</td><td>GGAGGCGATCGTCCGCCACGTGG</td><td>1000</td><td></td><td></td><td></td><td></td><td></td><td></td><td></td><td></td><td></td></tr>
<tr><td>Oligo fwd</td><td>CACCGGAGGCGATCGTCCGCCACG</td><td></td><td></td><td></td><td></td><td></td><td></td><td></td><td></td><td></td><td></td></tr>
<tr><td>Oligo rev</td><td>AAACCGTGGCGGACGATCGCCTCC</td><td></td><td></td><td></td><td></td><td></td><td></td><td></td><td></td><td></td><td></td></tr>
<tr><td>Chromosome</td><td>start</td><td>end</td><td>strand</td><td>MM</td><td>target_seq</td><td>PAM</td><td>alignment</td><td>distance</td><td>position</td><td>gene name</td><td>gene id</td></tr>
<tr><td>GU305914.1</td><td>87174</td><td>87196</td><td>-</td><td>0</td><td>GGAGGCGATCGTCCGCCACG</td><td>TGG</td><td>||||||||[||||||||||||]PAM</td><td>NA</td><td>Intergenic</td><td>NA</td><td>NA</td></tr>
<tr><td></td><td></td><td></td><td></td><td></td><td></td><td></td><td></td><td></td><td></td><td></td><td></td></tr>
<tr><td>T27614</td><td>GAGGCGATCGTCCGCCACGTGGG</td><td>1000</td><td></td><td></td><td></td><td></td><td></td><td></td><td></td><td></td><td></td></tr>
<tr><td>Oligo fwd</td><td>CACCGAGGCGATCGTCCGCCACGT</td><td></td><td></td><td></td><td></td><td></td><td></td><td></td><td></td><td></td><td></td></tr>
<tr><td>Oligo rev</td><td>AAACACGTGGCGGACGATCGCCTC</td><td></td><td></td><td></td><td></td><td></td><td></td><td></td><td></td><td></td><td></td></tr>
<tr><td>Chromosome</td><td>start</td><td>end</td><td>strand</td><td>MM</td><td>target_seq</td><td>PAM</td><td>alignment</td><td>distance</td><td>position</td><td>gene name</td><td>gene id</td></tr>
<tr><td>GU305914.1</td><td>87173</td><td>87195</td><td>-</td><td>0</td><td>GAGGCGATCGTCCGCCACGT</td><td>GGG</td><td>||||||||[||||||||||||]PAM</td><td>NA</td><td>Intergenic</td><td>NA</td><td>NA</td></tr>
<tr><td></td><td></td><td></td><td></td><td></td><td></td><td></td><td></td><td></td><td></td><td></td><td></td></tr>
<tr><td>T27615</td><td>AGGCGATCGTCCGCCACGTGGGG</td><td>1000</td><td></td><td></td><td></td><td></td><td></td><td></td><td></td><td></td><td></td></tr>
<tr><td>Oligo fwd</td><td>CACCAGGCGATCGTCCGCCACGTG</td><td></td><td></td><td></td><td></td><td></td><td></td><td></td><td></td><td></td><td></td></tr>
<tr><td>Oligo rev</td><td>AAACCACGTGGCGGACGATCGCCT</td><td></td><td></td><td></td><td></td><td></td><td></td><td></td><td></td><td></td><td></td></tr>
<tr><td>Chromosome</td><td>start</td><td>end</td><td>strand</td><td>MM</td><td>target_seq</td><td>PAM</td><td>alignment</td><td>distance</td><td>position</td><td>gene name</td><td>gene id</td></tr>
<tr><td>GU305914.1</td><td>87172</td><td>87194</td><td>-</td><td>0</td><td>AGGCGATCGTCCGCCACGTG</td><td>GGG</td><td>||||||||[||||||||||||]PAM</td><td>NA</td><td>Intergenic</td><td>NA</td><td>NA</td></tr>
<tr><td></td><td></td><td></td><td></td><td></td><td></td><td></td><td></td><td></td><td></td><td></td><td></td></tr>
<tr><td>T27616</td><td>GCCACGTGGGGATTACGATGAGG</td><td>1000</td><td></td><td></td><td></td><td></td><td></td><td></td><td></td><td></td><td></td></tr>
<tr><td>Oligo fwd</td><td>CACCGCCACGTGGGGATTACGATG</td><td></td><td></td><td></td><td></td><td></td><td></td><td></td><td></td><td></td><td></td></tr>
<tr><td>Oligo rev</td><td>AAACCATCGTAATCCCCACGTGGC</td><td></td><td></td><td></td><td></td><td></td><td></td><td></td><td></td><td></td><td></td></tr>
<tr><td>Chromosome</td><td>start</td><td>end</td><td>strand</td><td>MM</td><td>target_seq</td><td>PAM</td><td>alignment</td><td>distance</td><td>position</td><td>gene name</td><td>gene id</td></tr>
<tr><td>GU305914.1</td><td>87160</td><td>87182</td><td>-</td><td>0</td><td>GCCACGTGGGGATTACGATG</td><td>AGG</td><td>||||||||[||||||||||||]PAM</td><td>NA</td><td>Intergenic</td><td>NA</td><td>NA</td></tr>
<tr><td></td><td></td><td></td><td></td><td></td><td></td><td></td><td></td><td></td><td></td><td></td><td></td></tr>
<tr><td>T27617</td><td>TCAACGTCTCCCACAGCCTCTGG</td><td>1000</td><td></td><td></td><td></td><td></td><td></td><td></td><td></td><td></td><td></td></tr>
<tr><td>Oligo fwd</td><td>CACCTCAACGTCTCCCACAGCCTC</td><td></td><td></td><td></td><td></td><td></td><td></td><td></td><td></td><td></td><td></td></tr>
<tr><td>Oligo rev</td><td>AAACGAGGCTGTGGGAGACGTTGA</td><td></td><td></td><td></td><td></td><td></td><td></td><td></td><td></td><td></td><td></td></tr>
<tr><td>Chromosome</td><td>start</td><td>end</td><td>strand</td><td>MM</td><td>target_seq</td><td>PAM</td><td>alignment</td><td>distance</td><td>position</td><td>gene name</td><td>gene id</td></tr>
<tr><td>GU305914.1</td><td>87133</td><td>87155</td><td>-</td><td>0</td><td>TCAACGTCTCCCACAGCCTC</td><td>TGG</td><td>||||||||[||||||||||||]PAM</td><td>NA</td><td>Intergenic</td><td>NA</td><td>NA</td></tr>
<tr><td></td><td></td><td></td><td></td><td></td><td></td><td></td><td></td><td></td><td></td><td></td><td></td></tr>
<tr><td>T27618</td><td>ACAGCCTCTGGAAAACCTTGCGG</td><td>1000</td><td></td><td></td><td></td><td></td><td></td><td></td><td></td><td></td><td></td></tr>
<tr><td>Oligo fwd</td><td>CACCACAGCCTCTGGAAAACCTTG</td><td></td><td></td><td></td><td></td><td></td><td></td><td></td><td></td><td></td><td></td></tr>
<tr><td>Oligo rev</td><td>AAACCAAGGTTTTCCAGAGGCTGT</td><td></td><td></td><td></td><td></td><td></td><td></td><td></td><td></td><td></td><td></td></tr>
<tr><td>Chromosome</td><td>start</td><td>end</td><td>strand</td><td>MM</td><td>target_seq</td><td>PAM</td><td>alignment</td><td>distance</td><td>position</td><td>gene name</td><td>gene id</td></tr>
<tr><td>GU305914.1</td><td>87121</td><td>87143</td><td>-</td><td>0</td><td>ACAGCCTCTGGAAAACCTTG</td><td>CGG</td><td>||||||||[||||||||||||]PAM</td><td>NA</td><td>Intergenic</td><td>NA</td><td>NA</td></tr>
<tr><td></td><td></td><td></td><td></td><td></td><td></td><td></td><td></td><td></td><td></td><td></td><td></td></tr>
<tr><td>T27619</td><td>CCTCTGGAAAACCTTGCGGACGG</td><td>1000</td><td></td><td></td><td></td><td></td><td></td><td></td><td></td><td></td><td></td></tr>
<tr><td>Oligo fwd</td><td>CACCCCTCTGGAAAACCTTGCGGA</td><td></td><td></td><td></td><td></td><td></td><td></td><td></td><td></td><td></td><td></td></tr>
<tr><td>Oligo rev</td><td>AAACTCCGCAAGGTTTTCCAGAGG</td><td></td><td></td><td></td><td></td><td></td><td></td><td></td><td></td><td></td><td></td></tr>
<tr><td>Chromosome</td><td>start</td><td>end</td><td>strand</td><td>MM</td><td>target_seq</td><td>PAM</td><td>alignment</td><td>distance</td><td>position</td><td>gene name</td><td>gene id</td></tr>
<tr><td>GU305914.1</td><td>87117</td><td>87139</td><td>-</td><td>0</td><td>CCTCTGGAAAACCTTGCGGA</td><td>CGG</td><td>||||||||[||||||||||||]PAM</td><td>NA</td><td>Intergenic</td><td>NA</td><td>NA</td></tr>
<tr><td></td><td></td><td></td><td></td><td></td><td></td><td></td><td></td><td></td><td></td><td></td><td></td></tr>
<tr><td>T27620</td><td>GAATTGAGTCGCTATCACTACGG</td><td>1000</td><td></td><td></td><td></td><td></td><td></td><td></td><td></td><td></td><td></td></tr>
<tr><td>Oligo fwd</td><td>CACCGAATTGAGTCGCTATCACTA</td><td></td><td></td><td></td><td></td><td></td><td></td><td></td><td></td><td></td><td></td></tr>
<tr><td>Oligo rev</td><td>AAACTAGTGATAGCGACTCAATTC</td><td></td><td></td><td></td><td></td><td></td><td></td><td></td><td></td><td></td><td></td></tr>
<tr><td>Chromosome</td><td>start</td><td>end</td><td>strand</td><td>MM</td><td>target_seq</td><td>PAM</td><td>alignment</td><td>distance</td><td>position</td><td>gene name</td><td>gene id</td></tr>
<tr><td>GU305914.1</td><td>87095</td><td>87117</td><td>-</td><td>0</td><td>GAATTGAGTCGCTATCACTA</td><td>CGG</td><td>||||||||[||||||||||||]PAM</td><td>NA</td><td>Intergenic</td><td>NA</td><td>NA</td></tr>
<tr><td></td><td></td><td></td><td></td><td></td><td></td><td></td><td></td><td></td><td></td><td></td><td></td></tr>
<tr><td>T27621</td><td>AATTGAGTCGCTATCACTACGGG</td><td>1000</td><td></td><td></td><td></td><td></td><td></td><td></td><td></td><td></td><td></td></tr>
<tr><td>Oligo fwd</td><td>CACCAATTGAGTCGCTATCACTAC</td><td></td><td></td><td></td><td></td><td></td><td></td><td></td><td></td><td></td><td></td></tr>
<tr><td>Oligo rev</td><td>AAACGTAGTGATAGCGACTCAATT</td><td></td><td></td><td></td><td></td><td></td><td></td><td></td><td></td><td></td><td></td></tr>
<tr><td>Chromosome</td><td>start</td><td>end</td><td>strand</td><td>MM</td><td>target_seq</td><td>PAM</td><td>alignment</td><td>distance</td><td>position</td><td>gene name</td><td>gene id</td></tr>
<tr><td>GU305914.1</td><td>87094</td><td>87116</td><td>-</td><td>0</td><td>AATTGAGTCGCTATCACTAC</td><td>GGG</td><td>||||||||[||||||||||||]PAM</td><td>NA</td><td>Intergenic</td><td>NA</td><td>NA</td></tr>
<tr><td></td><td></td><td></td><td></td><td></td><td></td><td></td><td></td><td></td><td></td><td></td><td></td></tr>
<tr><td>T27622</td><td>ATTGAGTCGCTATCACTACGGGG</td><td>1000</td><td></td><td></td><td></td><td></td><td></td><td></td><td></td><td></td><td></td></tr>
<tr><td>Oligo fwd</td><td>CACCATTGAGTCGCTATCACTACG</td><td></td><td></td><td></td><td></td><td></td><td></td><td></td><td></td><td></td><td></td></tr>
<tr><td>Oligo rev</td><td>AAACCGTAGTGATAGCGACTCAAT</td><td></td><td></td><td></td><td></td><td></td><td></td><td></td><td></td><td></td><td></td></tr>
<tr><td>Chromosome</td><td>start</td><td>end</td><td>strand</td><td>MM</td><td>target_seq</td><td>PAM</td><td>alignment</td><td>distance</td><td>position</td><td>gene name</td><td>gene id</td></tr>
<tr><td>GU305914.1</td><td>87093</td><td>87115</td><td>-</td><td>0</td><td>ATTGAGTCGCTATCACTACG</td><td>GGG</td><td>||||||||[||||||||||||]PAM</td><td>NA</td><td>Intergenic</td><td>NA</td><td>NA</td></tr>
<tr><td></td><td></td><td></td><td></td><td></td><td></td><td></td><td></td><td></td><td></td><td></td><td></td></tr>
<tr><td>T27623</td><td>TCACTACGGGGAAGACCTCGAGG</td><td>1000</td><td></td><td></td><td></td><td></td><td></td><td></td><td></td><td></td><td></td></tr>
<tr><td>Oligo fwd</td><td>CACCTCACTACGGGGAAGACCTCG</td><td></td><td></td><td></td><td></td><td></td><td></td><td></td><td></td><td></td><td></td></tr>
<tr><td>Oligo rev</td><td>AAACCGAGGTCTTCCCCGTAGTGA</td><td></td><td></td><td></td><td></td><td></td><td></td><td></td><td></td><td></td><td></td></tr>
<tr><td>Chromosome</td><td>start</td><td>end</td><td>strand</td><td>MM</td><td>target_seq</td><td>PAM</td><td>alignment</td><td>distance</td><td>position</td><td>gene name</td><td>gene id</td></tr>
<tr><td>GU305914.1</td><td>87081</td><td>87103</td><td>-</td><td>0</td><td>TCACTACGGGGAAGACCTCG</td><td>AGG</td><td>||||||||[||||||||||||]PAM</td><td>NA</td><td>Intergenic</td><td>NA</td><td>NA</td></tr>
<tr><td></td><td></td><td></td><td></td><td></td><td></td><td></td><td></td><td></td><td></td><td></td><td></td></tr>
<tr><td>T27624</td><td>CCTCGAGGACGTTTTCACTCTGG</td><td>1000</td><td></td><td></td><td></td><td></td><td></td><td></td><td></td><td></td><td></td></tr>
<tr><td>Oligo fwd</td><td>CACCCCTCGAGGACGTTTTCACTC</td><td></td><td></td><td></td><td></td><td></td><td></td><td></td><td></td><td></td><td></td></tr>
<tr><td>Oligo rev</td><td>AAACGAGTGAAAACGTCCTCGAGG</td><td></td><td></td><td></td><td></td><td></td><td></td><td></td><td></td><td></td><td></td></tr>
<tr><td>Chromosome</td><td>start</td><td>end</td><td>strand</td><td>MM</td><td>target_seq</td><td>PAM</td><td>alignment</td><td>distance</td><td>position</td><td>gene name</td><td>gene id</td></tr>
<tr><td>GU305914.1</td><td>87066</td><td>87088</td><td>-</td><td>0</td><td>CCTCGAGGACGTTTTCACTC</td><td>TGG</td><td>||||||||[||||||||||||]PAM</td><td>NA</td><td>Intergenic</td><td>NA</td><td>NA</td></tr>
<tr><td></td><td></td><td></td><td></td><td></td><td></td><td></td><td></td><td></td><td></td><td></td><td></td></tr>
<tr><td>T27625</td><td>CTCGAGGACGTTTTCACTCTGGG</td><td>1000</td><td></td><td></td><td></td><td></td><td></td><td></td><td></td><td></td><td></td></tr>
<tr><td>Oligo fwd</td><td>CACCCTCGAGGACGTTTTCACTCT</td><td></td><td></td><td></td><td></td><td></td><td></td><td></td><td></td><td></td><td></td></tr>
<tr><td>Oligo rev</td><td>AAACAGAGTGAAAACGTCCTCGAG</td><td></td><td></td><td></td><td></td><td></td><td></td><td></td><td></td><td></td><td></td></tr>
<tr><td>Chromosome</td><td>start</td><td>end</td><td>strand</td><td>MM</td><td>target_seq</td><td>PAM</td><td>alignment</td><td>distance</td><td>position</td><td>gene name</td><td>gene id</td></tr>
<tr><td>GU305914.1</td><td>87065</td><td>87087</td><td>-</td><td>0</td><td>CTCGAGGACGTTTTCACTCT</td><td>GGG</td><td>||||||||[||||||||||||]PAM</td><td>NA</td><td>Intergenic</td><td>NA</td><td>NA</td></tr>
</table>

T27626 GGACGTTTTCACTCTGGGCGAGG 1000
Oligo fwd CACCGGACGTTTTCACTCTGGGCG
Oligo rev AAACCGCCCAGAGTGAAAACGTCC
Chromosome start end strand MM target_seq PAM alignment distance position gene name gene id
GU305914.1 87060 87082 - 0 GGACGTTTTCACTCTGGGCG AGG ||||||||[||||||||||||]PAM NA Intergenic NA NA

T27627 GACGTTTTCACTCTGGGCGAGGG 1000
Oligo fwd CACCGACGTTTTCACTCTGGGCGA
Oligo rev AAACTCGCCCAGAGTGAAAACGTC
Chromosome start end strand MM target_seq PAM alignment distance position gene name gene id
GU305914.1 87059 87081 - 0 GACGTTTTCACTCTGGGCGA GGG ||||||||[||||||||||||]PAM NA Intergenic NA NA

T27628 ACGTTTTCACTCTGGGCGAGGGG 1000
Oligo fwd CACCACGTTTTCACTCTGGGCGAG
Oligo rev AAACCTCGCCCAGAGTGAAAACGT
Chromosome start end strand MM target_seq PAM alignment distance position gene name gene id
GU305914.1 87058 87080 - 0 ACGTTTTCACTCTGGGCGAG GGG ||||||||[||||||||||||]PAM NA Intergenic NA NA

T27629 ACTCTGGGCGAGGGGCGCTTCGG 1000
Oligo fwd CACCACTCTGGGCGAGGGGCGCTT
Oligo rev AAACAAGCGCCCCTCGCCCAGAGT
Chromosome start end strand MM target_seq PAM alignment distance position gene name gene id
GU305914.1 87050 87072 - 0 ACTCTGGGCGAGGGGCGCTT CGG ||||||||[||||||||||||]PAM NA Intergenic NA NA

T27630 CTCTGGGCGAGGGGCGCTTCGGG 1000
Oligo fwd CACCCTCTGGGCGAGGGGCGCTTC
Oligo rev AAACGAAGCGCCCCTCGCCCAGAG
Chromosome start end strand MM target_seq PAM alignment distance position gene name gene id
GU305914.1 87049 87071 - 0 CTCTGGGCGAGGGGCGCTTC GGG ||||||||[||||||||||||]PAM NA Intergenic NA NA

T27631 TCTGGGCGAGGGGCGCTTCGGGG 1000
Oligo fwd CACCTCTGGGCGAGGGGCGCTTCG
Oligo rev AAACCGAAGCGCCCCTCGCCCAGA
Chromosome start end strand MM target_seq PAM alignment distance position gene name gene id
GU305914.1 87048 87070 - 0 TCTGGGCGAGGGGCGCTTCG GGG ||||||||[||||||||||||]PAM NA Intergenic NA NA

T27632 CTGGGCGAGGGGCGCTTCGGGGG 1000
Oligo fwd CACCCTGGGCGAGGGGCGCTTCGG
Oligo rev AAACCCGAAGCGCCCCTCGCCCAG
Chromosome start end strand MM target_seq PAM alignment distance position gene name gene id
GU305914.1 87047 87069 - 0 CTGGGCGAGGGGCGCTTCGG GGG ||||||||[||||||||||||]PAM NA Intergenic NA NA

T27633 GGGCGACGAGCGAATCTACGTGG 1000
Oligo fwd CACCGGGCGACGAGCGAATCTACG
Oligo rev AAACCGTAGATTCGCTCGTCGCCC
Chromosome start end strand MM target_seq PAM alignment distance position gene name gene id
GU305914.1 87027 87049 - 0 GGGCGACGAGCGAATCTACG TGG ||||||||[||||||||||||]PAM NA Intergenic NA NA

T27634 GGCGACGAGCGAATCTACGTGGG 1000
Oligo fwd CACCGGCGACGAGCGAATCTACGT
Oligo rev AAACACGTAGATTCGCTCGTCGCC
Chromosome start end strand MM target_seq PAM alignment distance position gene name gene id
GU305914.1 87026 87048 - 0 GGCGACGAGCGAATCTACGT GGG ||||||||[||||||||||||]PAM NA Intergenic NA NA

T27635 GCGACGAGCGAATCTACGTGGGG 1000
Oligo fwd CACCGCGACGAGCGAATCTACGTG
Oligo rev AAACCACGTAGATTCGCTCGTCGC
Chromosome start end strand MM target_seq PAM alignment distance position gene name gene id
GU305914.1 87025 87047 - 0 GCGACGAGCGAATCTACGTG GGG ||||||||[||||||||||||]PAM NA Intergenic NA NA

T27636 CTACGTGGGGTCCATCTTCGCGG 1000
Oligo fwd CACCCTACGTGGGGTCCATCTTCG
Oligo rev AAACCGAAGATGGACCCCACGTAG
Chromosome start end strand MM target_seq PAM alignment distance position gene name gene id
GU305914.1 87012 87034 - 0 CTACGTGGGGTCCATCTTCG CGG ||||||||[||||||||||||]PAM NA Intergenic NA NA

T27637 GGGGTCCATCTTCGCGGCGCCGG 1000
Oligo fwd CACCGGGGTCCATCTTCGCGGCGC
Oligo rev AAACGCGCCGCGAAGATGGACCCC
Chromosome start end strand MM target_seq PAM alignment distance position gene name gene id
GU305914.1 87006 87028 - 0 GGGGTCCATCTTCGCGGCGC CGG ||||||||[||||||||||||]PAM NA Intergenic NA NA

T27638 GGGTCCATCTTCGCGGCGCCGGG 1000
Oligo fwd CACCGGGTCCATCTTCGCGGCGCC
Oligo rev AAACGGCGCCGCGAAGATGGACCC
Chromosome start end strand MM target_seq PAM alignment distance position gene name gene id
GU305914.1 87005 87027 - 0 GGGTCCATCTTCGCGGCGCC GGG ||||||||[||||||||||||]PAM NA Intergenic NA NA

T27639 CATCTTCGCGGCGCCGGGAAAGG 1000
Oligo fwd CACCCATCTTCGCGGCGCCGGGAA
Oligo rev AAACTTCCCGGCGCCGCGAAGATG
Chromosome start end strand MM target_seq PAM alignment distance position gene name gene id
GU305914.1 87000 87022 - 0 CATCTTCGCGGCGCCGGGAA AGG ||||||||[||||||||||||]PAM NA Intergenic NA NA

T27640 CAAGATCAAGTCGCTGATCGAGG 1000
Oligo fwd CACCCAAGATCAAGTCGCTGATCG
Oligo rev AAACCGATCAGCGACTTGATCTTG
Chromosome start end strand MM target_seq PAM alignment distance position gene name gene id
GU305914.1 86889 86911 - 0 CAAGATCAAGTCGCTGATCG AGG ||||||||[||||||||||||]PAM NA Intergenic NA NA

| | | | | | | | | | | | |
|---|---|---|---|---|---|---|---|---|---|---|---|
| T27641 | AGTCGCTGATCGAGGAAATCCGG | 1000 | | | | | | | | | |
| Oligo fwd | CACCAGTCGCTGATCGAGGAAATC | | | | | | | | | | |
| Oligo rev | AAACGATTTCCTCGATCAGCGACT | | | | | | | | | | |
| Chromosome | start | end | strand | MM | target_seq | PAM | alignment | distance | position | gene name | gene id |
| GU305914.1 | 86881 | 86903 | - | 0 | AGTCGCTGATCGAGGAAATC | CGG | \|\|\|\|\|\|\|\|[\|\|\|\|\|\|\|\|\|\|\|\|]PAM | NA | Intergenic | NA | NA |
| T27642 | GTCGCTGATCGAGGAAATCCGGG | 1000 | | | | | | | | | |
| Oligo fwd | CACCGTCGCTGATCGAGGAAATCC | | | | | | | | | | |
| Oligo rev | AAACGGATTTCCTCGATCAGCGAC | | | | | | | | | | |
| Chromosome | start | end | strand | MM | target_seq | PAM | alignment | distance | position | gene name | gene id |
| GU305914.1 | 86880 | 86902 | - | 0 | GTCGCTGATCGAGGAAATCC | GGG | \|\|\|\|\|\|\|\|[\|\|\|\|\|\|\|\|\|\|\|\|]PAM | NA | Intergenic | NA | NA |
| T27643 | TCGCTGATCGAGGAAATCCGGGG | 1000 | | | | | | | | | |
| Oligo fwd | CACCTCGCTGATCGAGGAAATCCG | | | | | | | | | | |
| Oligo rev | AAACCGGATTTCCTCGATCAGCGA | | | | | | | | | | |
| Chromosome | start | end | strand | MM | target_seq | PAM | alignment | distance | position | gene name | gene id |
| GU305914.1 | 86879 | 86901 | - | 0 | TCGCTGATCGAGGAAATCCG | GGG | \|\|\|\|\|\|\|\|[\|\|\|\|\|\|\|\|\|\|\|\|]PAM | NA | Intergenic | NA | NA |
| T27644 | CGCTGATCGAGGAAATCCGGGGG | 1000 | | | | | | | | | |
| Oligo fwd | CACCCGCTGATCGAGGAAATCCGG | | | | | | | | | | |
| Oligo rev | AAACCCGGATTTCCTCGATCAGCG | | | | | | | | | | |
| Chromosome | start | end | strand | MM | target_seq | PAM | alignment | distance | position | gene name | gene id |
| GU305914.1 | 86878 | 86900 | - | 0 | CGCTGATCGAGGAAATCCGG | GGG | \|\|\|\|\|\|\|\|[\|\|\|\|\|\|\|\|\|\|\|\|]PAM | NA | Intergenic | NA | NA |
| T27645 | GCTGATCGAGGAAATCCGGGGGG | 1000 | | | | | | | | | |
| Oligo fwd | CACCGCTGATCGAGGAAATCCGGG | | | | | | | | | | |
| Oligo rev | AAACCCCGGATTTCCTCGATCAGC | | | | | | | | | | |
| Chromosome | start | end | strand | MM | target_seq | PAM | alignment | distance | position | gene name | gene id |
| GU305914.1 | 86877 | 86899 | - | 0 | GCTGATCGAGGAAATCCGGG | GGG | \|\|\|\|\|\|\|\|[\|\|\|\|\|\|\|\|\|\|\|\|]PAM | NA | Intergenic | NA | NA |
| T27646 | GATCGAGGAAATCCGGGGGGTGG | 1000 | | | | | | | | | |
| Oligo fwd | CACCGATCGAGGAAATCCGGGGGG | | | | | | | | | | |
| Oligo rev | AAACCCCCCCGGATTTCCTCGATC | | | | | | | | | | |
| Chromosome | start | end | strand | MM | target_seq | PAM | alignment | distance | position | gene name | gene id |
| GU305914.1 | 86874 | 86896 | - | 0 | GATCGAGGAAATCCGGGGGG | TGG | \|\|\|\|\|\|\|\|[\|\|\|\|\|\|\|\|\|\|\|\|]PAM | NA | Intergenic | NA | NA |
| T27647 | ATCGAGGAAATCCGGGGGGTGGG | 1000 | | | | | | | | | |
| Oligo fwd | CACCATCGAGGAAATCCGGGGGGT | | | | | | | | | | |
| Oligo rev | AAACACCCCCCGGATTTCCTCGAT | | | | | | | | | | |
| Chromosome | start | end | strand | MM | target_seq | PAM | alignment | distance | position | gene name | gene id |
| GU305914.1 | 86873 | 86895 | - | 0 | ATCGAGGAAATCCGGGGGGT | GGG | \|\|\|\|\|\|\|\|[\|\|\|\|\|\|\|\|\|\|\|\|]PAM | NA | Intergenic | NA | NA |
| T27648 | GAAATCCGGGGGGTGGGAGATGG | 1000 | | | | | | | | | |
| Oligo fwd | CACCGAAATCCGGGGGGTGGGAGA | | | | | | | | | | |
| Oligo rev | AAACTCTCCCACCCCCCGGATTTC | | | | | | | | | | |
| Chromosome | start | end | strand | MM | target_seq | PAM | alignment | distance | position | gene name | gene id |
| GU305914.1 | 86867 | 86889 | - | 0 | GAAATCCGGGGGGTGGGAGA | TGG | \|\|\|\|\|\|\|\|[\|\|\|\|\|\|\|\|\|\|\|\|]PAM | NA | Intergenic | NA | NA |
| T27649 | GGGGGTGGGAGATGGACCCGCGG | 1000 | | | | | | | | | |
| Oligo fwd | CACCGGGGGTGGGAGATGGACCCG | | | | | | | | | | |
| Oligo rev | AAACCGGGTCCATCTCCCACCCCC | | | | | | | | | | |
| Chromosome | start | end | strand | MM | target_seq | PAM | alignment | distance | position | gene name | gene id |
| GU305914.1 | 86859 | 86881 | - | 0 | GGGGGTGGGAGATGGACCCG | CGG | \|\|\|\|\|\|\|\|[\|\|\|\|\|\|\|\|\|\|\|\|]PAM | NA | Intergenic | NA | NA |
| T27650 | GGTGGGAGATGGACCCGCGGCGG | 1000 | | | | | | | | | |
| Oligo fwd | CACCGGTGGGAGATGGACCCGCGG | | | | | | | | | | |
| Oligo rev | AAACCCGCGGGTCCATCTCCCACC | | | | | | | | | | |
| Chromosome | start | end | strand | MM | target_seq | PAM | alignment | distance | position | gene name | gene id |
| GU305914.1 | 86856 | 86878 | - | 0 | GGTGGGAGATGGACCCGCGG | CGG | \|\|\|\|\|\|\|\|[\|\|\|\|\|\|\|\|\|\|\|\|]PAM | NA | Intergenic | NA | NA |
| T27651 | GTGGGAGATGGACCCGCGGCGGG | 1000 | | | | | | | | | |
| Oligo fwd | CACCGTGGGAGATGGACCCGCGGC | | | | | | | | | | |
| Oligo rev | AAACGCCGCGGGTCCATCTCCCAC | | | | | | | | | | |
| Chromosome | start | end | strand | MM | target_seq | PAM | alignment | distance | position | gene name | gene id |
| GU305914.1 | 86855 | 86877 | - | 0 | GTGGGAGATGGACCCGCGGC | GGG | \|\|\|\|\|\|\|\|[\|\|\|\|\|\|\|\|\|\|\|\|]PAM | NA | Intergenic | NA | NA |
| T27652 | TGGGAGATGGACCCGCGGCGGGG | 1000 | | | | | | | | | |
| Oligo fwd | CACCTGGGAGATGGACCCGCGGCG | | | | | | | | | | |
| Oligo rev | AAACCGCCGCGGGTCCATCTCCCA | | | | | | | | | | |
| Chromosome | start | end | strand | MM | target_seq | PAM | alignment | distance | position | gene name | gene id |
| GU305914.1 | 86854 | 86876 | - | 0 | TGGGAGATGGACCCGCGGCG | GGG | \|\|\|\|\|\|\|\|[\|\|\|\|\|\|\|\|\|\|\|\|]PAM | NA | Intergenic | NA | NA |
| T27653 | GGGAGATGGACCCGCGGCGGGGG | 1000 | | | | | | | | | |
| Oligo fwd | CACCGGGAGATGGACCCGCGGCGG | | | | | | | | | | |
| Oligo rev | AAACCCGCCGCGGGTCCATCTCCC | | | | | | | | | | |
| Chromosome | start | end | strand | MM | target_seq | PAM | alignment | distance | position | gene name | gene id |
| GU305914.1 | 86853 | 86875 | - | 0 | GGGAGATGGACCCGCGGCGG | GGG | \|\|\|\|\|\|\|\|[\|\|\|\|\|\|\|\|\|\|\|\|]PAM | NA | Intergenic | NA | NA |
| T27654 | AGATGGACCCGCGGCGGGGGCGG | 1000 | | | | | | | | | |
| Oligo fwd | CACCAGATGGACCCGCGGCGGGGG | | | | | | | | | | |
| Oligo rev | AAACCCCCCGCCGCGGGTCCATCT | | | | | | | | | | |
| Chromosome | start | end | strand | MM | target_seq | PAM | alignment | distance | position | gene name | gene id |
| GU305914.1 | 86850 | 86872 | - | 0 | AGATGGACCCGCGGCGGGGG | CGG | \|\|\|\|\|\|\|\|[\|\|\|\|\|\|\|\|\|\|\|\|]PAM | NA | Intergenic | NA | NA |
| T27655 | CGGCGGGGGCGGCAGCTTCAAGG | 1000 | | | | | | | | | |
| Oligo fwd | CACCCGGCGGGGGCGGCAGCTTCA | | | | | | | | | | |
| Oligo rev | AAACTGAAGCTGCCGCCCCCGCCG | | | | | | | | | | |
| Chromosome | start | end | strand | MM | target_seq | PAM | alignment | distance | position | gene name | gene id |

<table>
<tr><td>GU305914.1</td><td>86839</td><td>86861</td><td>-</td><td>0</td><td>CGGCGGGGGCGGCAGCTTCA</td><td>AGG</td><td>||||||||[||||||||||||]PAM</td><td>NA</td><td>Intergenic</td><td>NA</td><td>NA</td></tr>
<tr><td>T27656</td><td>GGCGGGGGCGGCAGCTTCAAGGG</td><td>1000</td><td></td><td></td><td></td><td></td><td></td><td></td><td></td><td></td><td></td></tr>
<tr><td>Oligo fwd</td><td>CACCGGCGGGGGCGGCAGCTTCAA</td><td></td><td></td><td></td><td></td><td></td><td></td><td></td><td></td><td></td><td></td></tr>
<tr><td>Oligo rev</td><td>AAACTTGAAGCTGCCGCCCCCGCC</td><td></td><td></td><td></td><td></td><td></td><td></td><td></td><td></td><td></td><td></td></tr>
<tr><td>Chromosome</td><td>start</td><td>end</td><td>strand</td><td>MM</td><td>target_seq</td><td>PAM</td><td>alignment</td><td>distance</td><td>position</td><td>gene name</td><td>gene id</td></tr>
<tr><td>GU305914.1</td><td>86838</td><td>86860</td><td>-</td><td>0</td><td>GGCGGGGGCGGCAGCTTCAA</td><td>GGG</td><td>||||||||[||||||||||||]PAM</td><td>NA</td><td>Intergenic</td><td>NA</td><td>NA</td></tr>
<tr><td>T27657</td><td>GGCGGCAGCTTCAAGGGCCGAGG</td><td>1000</td><td></td><td></td><td></td><td></td><td></td><td></td><td></td><td></td><td></td></tr>
<tr><td>Oligo fwd</td><td>CACCGGCGGCAGCTTCAAGGGCCG</td><td></td><td></td><td></td><td></td><td></td><td></td><td></td><td></td><td></td><td></td></tr>
<tr><td>Oligo rev</td><td>AAACCGGCCCTTGAAGCTGCCGCC</td><td></td><td></td><td></td><td></td><td></td><td></td><td></td><td></td><td></td><td></td></tr>
<tr><td>Chromosome</td><td>start</td><td>end</td><td>strand</td><td>MM</td><td>target_seq</td><td>PAM</td><td>alignment</td><td>distance</td><td>position</td><td>gene name</td><td>gene id</td></tr>
<tr><td>GU305914.1</td><td>86832</td><td>86854</td><td>-</td><td>0</td><td>GGCGGCAGCTTCAAGGGCCG</td><td>AGG</td><td>||||||||[||||||||||||]PAM</td><td>NA</td><td>Intergenic</td><td>NA</td><td>NA</td></tr>
<tr><td>T27658</td><td>GGCAGCTTCAAGGGCCGAGGCGG</td><td>1000</td><td></td><td></td><td></td><td></td><td></td><td></td><td></td><td></td><td></td></tr>
<tr><td>Oligo fwd</td><td>CACCGGCAGCTTCAAGGGCCGAGG</td><td></td><td></td><td></td><td></td><td></td><td></td><td></td><td></td><td></td><td></td></tr>
<tr><td>Oligo rev</td><td>AAACCCTCGGCCCTTGAAGCTGCC</td><td></td><td></td><td></td><td></td><td></td><td></td><td></td><td></td><td></td><td></td></tr>
<tr><td>Chromosome</td><td>start</td><td>end</td><td>strand</td><td>MM</td><td>target_seq</td><td>PAM</td><td>alignment</td><td>distance</td><td>position</td><td>gene name</td><td>gene id</td></tr>
<tr><td>GU305914.1</td><td>86829</td><td>86851</td><td>-</td><td>0</td><td>GGCAGCTTCAAGGGCCGAGG</td><td>CGG</td><td>||||||||[||||||||||||]PAM</td><td>NA</td><td>Intergenic</td><td>NA</td><td>NA</td></tr>
<tr><td>T27659</td><td>TTCAAGGGCCGAGGCGGCGTCGG</td><td>1000</td><td></td><td></td><td></td><td></td><td></td><td></td><td></td><td></td><td></td></tr>
<tr><td>Oligo fwd</td><td>CACCTTCAAGGGCCGAGGCGGCGT</td><td></td><td></td><td></td><td></td><td></td><td></td><td></td><td></td><td></td><td></td></tr>
<tr><td>Oligo rev</td><td>AAACACGCCGCCTCGGCCCTTGAA</td><td></td><td></td><td></td><td></td><td></td><td></td><td></td><td></td><td></td><td></td></tr>
<tr><td>Chromosome</td><td>start</td><td>end</td><td>strand</td><td>MM</td><td>target_seq</td><td>PAM</td><td>alignment</td><td>distance</td><td>position</td><td>gene name</td><td>gene id</td></tr>
<tr><td>GU305914.1</td><td>86823</td><td>86845</td><td>-</td><td>0</td><td>TTCAAGGGCCGAGGCGGCGT</td><td>CGG</td><td>||||||||[||||||||||||]PAM</td><td>NA</td><td>Intergenic</td><td>NA</td><td>NA</td></tr>
<tr><td>T27660</td><td>TCAAGGGCCGAGGCGGCGTCGGG</td><td>1000</td><td></td><td></td><td></td><td></td><td></td><td></td><td></td><td></td><td></td></tr>
<tr><td>Oligo fwd</td><td>CACCTCAAGGGCCGAGGCGGCGTC</td><td></td><td></td><td></td><td></td><td></td><td></td><td></td><td></td><td></td><td></td></tr>
<tr><td>Oligo rev</td><td>AAACGACGCCGCCTCGGCCCTTGA</td><td></td><td></td><td></td><td></td><td></td><td></td><td></td><td></td><td></td><td></td></tr>
<tr><td>Chromosome</td><td>start</td><td>end</td><td>strand</td><td>MM</td><td>target_seq</td><td>PAM</td><td>alignment</td><td>distance</td><td>position</td><td>gene name</td><td>gene id</td></tr>
<tr><td>GU305914.1</td><td>86822</td><td>86844</td><td>-</td><td>0</td><td>TCAAGGGCCGAGGCGGCGTC</td><td>GGG</td><td>||||||||[||||||||||||]PAM</td><td>NA</td><td>Intergenic</td><td>NA</td><td>NA</td></tr>
<tr><td>T27661</td><td>CAAGGGCCGAGGCGGCGTCGGGG</td><td>1000</td><td></td><td></td><td></td><td></td><td></td><td></td><td></td><td></td><td></td></tr>
<tr><td>Oligo fwd</td><td>CACCCAAGGGCCGAGGCGGCGTCG</td><td></td><td></td><td></td><td></td><td></td><td></td><td></td><td></td><td></td><td></td></tr>
<tr><td>Oligo rev</td><td>AAACCGACGCCGCCTCGGCCCTTG</td><td></td><td></td><td></td><td></td><td></td><td></td><td></td><td></td><td></td><td></td></tr>
<tr><td>Chromosome</td><td>start</td><td>end</td><td>strand</td><td>MM</td><td>target_seq</td><td>PAM</td><td>alignment</td><td>distance</td><td>position</td><td>gene name</td><td>gene id</td></tr>
<tr><td>GU305914.1</td><td>86821</td><td>86843</td><td>-</td><td>0</td><td>CAAGGGCCGAGGCGGCGTCG</td><td>GGG</td><td>||||||||[||||||||||||]PAM</td><td>NA</td><td>Intergenic</td><td>NA</td><td>NA</td></tr>
<tr><td>GU305914.1</td><td>96953</td><td>96975</td><td>+</td><td>4</td><td>CGACGGCGGAGGCGGCATCG</td><td>CGG</td><td>|-|-|||-[||||||||-|||]PAM</td><td>NA</td><td>Intergenic</td><td>NA</td><td>NA</td></tr>
<tr><td>T27662</td><td>AAGGGCCGAGGCGGCGTCGGGGG</td><td>1000</td><td></td><td></td><td></td><td></td><td></td><td></td><td></td><td></td><td></td></tr>
<tr><td>Oligo fwd</td><td>CACCAAGGGCCGAGGCGGCGTCGG</td><td></td><td></td><td></td><td></td><td></td><td></td><td></td><td></td><td></td><td></td></tr>
<tr><td>Oligo rev</td><td>AAACCCGACGCCGCCTCGGCCCTT</td><td></td><td></td><td></td><td></td><td></td><td></td><td></td><td></td><td></td><td></td></tr>
<tr><td>Chromosome</td><td>start</td><td>end</td><td>strand</td><td>MM</td><td>target_seq</td><td>PAM</td><td>alignment</td><td>distance</td><td>position</td><td>gene name</td><td>gene id</td></tr>
<tr><td>GU305914.1</td><td>86820</td><td>86842</td><td>-</td><td>0</td><td>AAGGGCCGAGGCGGCGTCGG</td><td>GGG</td><td>||||||||[||||||||||||]PAM</td><td>NA</td><td>Intergenic</td><td>NA</td><td>NA</td></tr>
<tr><td>T27663</td><td>GCCGAGGCGGCGTCGGGGGCCGG</td><td>1000</td><td></td><td></td><td></td><td></td><td></td><td></td><td></td><td></td><td></td></tr>
<tr><td>Oligo fwd</td><td>CACCGCCGAGGCGGCGTCGGGGGC</td><td></td><td></td><td></td><td></td><td></td><td></td><td></td><td></td><td></td><td></td></tr>
<tr><td>Oligo rev</td><td>AAACGCCCCCGACGCCGCCTCGGC</td><td></td><td></td><td></td><td></td><td></td><td></td><td></td><td></td><td></td><td></td></tr>
<tr><td>Chromosome</td><td>start</td><td>end</td><td>strand</td><td>MM</td><td>target_seq</td><td>PAM</td><td>alignment</td><td>distance</td><td>position</td><td>gene name</td><td>gene id</td></tr>
<tr><td>GU305914.1</td><td>86816</td><td>86838</td><td>-</td><td>0</td><td>GCCGAGGCGGCGTCGGGGGC</td><td>CGG</td><td>||||||||[||||||||||||]PAM</td><td>NA</td><td>Intergenic</td><td>NA</td><td>NA</td></tr>
<tr><td>T27664</td><td>CCGAGGCGGCGTCGGGGGCCGGG</td><td>1000</td><td></td><td></td><td></td><td></td><td></td><td></td><td></td><td></td><td></td></tr>
<tr><td>Oligo fwd</td><td>CACCCCGAGGCGGCGTCGGGGGCC</td><td></td><td></td><td></td><td></td><td></td><td></td><td></td><td></td><td></td><td></td></tr>
<tr><td>Oligo rev</td><td>AAACGGCCCCCGACGCCGCCTCGG</td><td></td><td></td><td></td><td></td><td></td><td></td><td></td><td></td><td></td><td></td></tr>
<tr><td>Chromosome</td><td>start</td><td>end</td><td>strand</td><td>MM</td><td>target_seq</td><td>PAM</td><td>alignment</td><td>distance</td><td>position</td><td>gene name</td><td>gene id</td></tr>
<tr><td>GU305914.1</td><td>86815</td><td>86837</td><td>-</td><td>0</td><td>CCGAGGCGGCGTCGGGGGCC</td><td>GGG</td><td>||||||||[||||||||||||]PAM</td><td>NA</td><td>Intergenic</td><td>NA</td><td>NA</td></tr>
<tr><td>T27665</td><td>CGAGGCGGCGTCGGGGGCCGGGG</td><td>1000</td><td></td><td></td><td></td><td></td><td></td><td></td><td></td><td></td><td></td></tr>
<tr><td>Oligo fwd</td><td>CACCCGAGGCGGCGTCGGGGGCCG</td><td></td><td></td><td></td><td></td><td></td><td></td><td></td><td></td><td></td><td></td></tr>
<tr><td>Oligo rev</td><td>AAACCGGCCCCCGACGCCGCCTCG</td><td></td><td></td><td></td><td></td><td></td><td></td><td></td><td></td><td></td><td></td></tr>
<tr><td>Chromosome</td><td>start</td><td>end</td><td>strand</td><td>MM</td><td>target_seq</td><td>PAM</td><td>alignment</td><td>distance</td><td>position</td><td>gene name</td><td>gene id</td></tr>
<tr><td>GU305914.1</td><td>86814</td><td>86836</td><td>-</td><td>0</td><td>CGAGGCGGCGTCGGGGGCCG</td><td>GGG</td><td>||||||||[||||||||||||]PAM</td><td>NA</td><td>Intergenic</td><td>NA</td><td>NA</td></tr>
<tr><td>T27666</td><td>GCGGCGTCGGGGGCCGGGGCAGG</td><td>1000</td><td></td><td></td><td></td><td></td><td></td><td></td><td></td><td></td><td></td></tr>
<tr><td>Oligo fwd</td><td>CACCGCGGCGTCGGGGGCCGGGGC</td><td></td><td></td><td></td><td></td><td></td><td></td><td></td><td></td><td></td><td></td></tr>
<tr><td>Oligo rev</td><td>AAACGCCCCGGCCCCCGACGCCGC</td><td></td><td></td><td></td><td></td><td></td><td></td><td></td><td></td><td></td><td></td></tr>
<tr><td>Chromosome</td><td>start</td><td>end</td><td>strand</td><td>MM</td><td>target_seq</td><td>PAM</td><td>alignment</td><td>distance</td><td>position</td><td>gene name</td><td>gene id</td></tr>
<tr><td>GU305914.1</td><td>86810</td><td>86832</td><td>-</td><td>0</td><td>GCGGCGTCGGGGGCCGGGGC</td><td>AGG</td><td>||||||||[||||||||||||]PAM</td><td>NA</td><td>Intergenic</td><td>NA</td><td>NA</td></tr>
<tr><td>T27667</td><td>GCGTCGGGGGCCGGGGCAGGAGG</td><td>1000</td><td></td><td></td><td></td><td></td><td></td><td></td><td></td><td></td><td></td></tr>
<tr><td>Oligo fwd</td><td>CACCGCGTCGGGGGCCGGGGCAGG</td><td></td><td></td><td></td><td></td><td></td><td></td><td></td><td></td><td></td><td></td></tr>
<tr><td>Oligo rev</td><td>AAACCCTGCCCCGGCCCCCGACGC</td><td></td><td></td><td></td><td></td><td></td><td></td><td></td><td></td><td></td><td></td></tr>
<tr><td>Chromosome</td><td>start</td><td>end</td><td>strand</td><td>MM</td><td>target_seq</td><td>PAM</td><td>alignment</td><td>distance</td><td>position</td><td>gene name</td><td>gene id</td></tr>
<tr><td>GU305914.1</td><td>86807</td><td>86829</td><td>-</td><td>0</td><td>GCGTCGGGGGCCGGGGCAGG</td><td>AGG</td><td>||||||||[||||||||||||]PAM</td><td>NA</td><td>Intergenic</td><td>NA</td><td>NA</td></tr>
<tr><td>T27668</td><td>CGTCGGGGGCCGGGGCAGGAGGG</td><td>1000</td><td></td><td></td><td></td><td></td><td></td><td></td><td></td><td></td><td></td></tr>
<tr><td>Oligo fwd</td><td>CACCCGTCGGGGGCCGGGGCAGGA</td><td></td><td></td><td></td><td></td><td></td><td></td><td></td><td></td><td></td><td></td></tr>
<tr><td>Oligo rev</td><td>AAACTCCTGCCCCGGCCCCCGACG</td><td></td><td></td><td></td><td></td><td></td><td></td><td></td><td></td><td></td><td></td></tr>
<tr><td>Chromosome</td><td>start</td><td>end</td><td>strand</td><td>MM</td><td>target_seq</td><td>PAM</td><td>alignment</td><td>distance</td><td>position</td><td>gene name</td><td>gene id</td></tr>
<tr><td>GU305914.1</td><td>86806</td><td>86828</td><td>-</td><td>0</td><td>CGTCGGGGGCCGGGGCAGGA</td><td>GGG</td><td>||||||||[||||||||||||]PAM</td><td>NA</td><td>Intergenic</td><td>NA</td><td>NA</td></tr>
<tr><td>T27669</td><td>GTCGGGGGCCGGGGCAGGAGGGG</td><td>1000</td><td></td><td></td><td></td><td></td><td></td><td></td><td></td><td></td><td></td></tr>
<tr><td>Oligo fwd</td><td>CACCGTCGGGGGCCGGGGCAGGAG</td><td></td><td></td><td></td><td></td><td></td><td></td><td></td><td></td><td></td><td></td></tr>
<tr><td>Oligo rev</td><td>AAACCTCCTGCCCCGGCCCCCGAC</td><td></td><td></td><td></td><td></td><td></td><td></td><td></td><td></td><td></td><td></td></tr>
<tr><td>Chromosome</td><td>start</td><td>end</td><td>strand</td><td>MM</td><td>target_seq</td><td>PAM</td><td>alignment</td><td>distance</td><td>position</td><td>gene name</td><td>gene id</td></tr>
<tr><td>GU305914.1</td><td>86805</td><td>86827</td><td>-</td><td>0</td><td>GTCGGGGGCCGGGGCAGGAG</td><td>GGG</td><td>||||||||[||||||||||||]PAM</td><td>NA</td><td>Intergenic</td><td>NA</td><td>NA</td></tr>
<tr><td>T27670</td><td>GGCCGGGGCAGGAGGGGAAGAGG</td><td>1000</td><td></td><td></td><td></td><td></td><td></td><td></td><td></td><td></td><td></td></tr>
<tr><td>Oligo fwd</td><td>CACCGGCCGGGGCAGGAGGGGAAG</td><td></td><td></td><td></td><td></td><td></td><td></td><td></td><td></td><td></td><td></td></tr>
</table>

Oligo rev AAACCTTCCCCTCCTGCCCCGGCC
Chromosome start end strand MM target_seq PAM alignment distance position gene name gene id
GU305914.1 86799 86821 - 0 GGCCGGGGCAGGAGGGGAAG AGG ||||||||[||||||||||||]PAM NA Intergenic NA NA

T27671 GCCGGGGCAGGAGGGGAAGAGGG 1000
Oligo fwd CACCGCCGGGGCAGGAGGGGAAGA
Oligo rev AAACTCTTCCCCTCCTGCCCCGGC
Chromosome start end strand MM target_seq PAM alignment distance position gene name gene id
GU305914.1 86798 86820 - 0 GCCGGGGCAGGAGGGGAAGA GGG ||||||||[||||||||||||]PAM NA Intergenic NA NA
GU305914.1 18851 18873 - 3 GGTGGGGGAGGAGGGGAAGA CGG |--||||-[||||||||||||]PAM NA Intergenic NA NA

T27672 CCGGGGCAGGAGGGGAAGAGGGG 1000
Oligo fwd CACCCCGGGGCAGGAGGGGAAGAG
Oligo rev AAACCTCTTCCCCTCCTGCCCCGG
Chromosome start end strand MM target_seq PAM alignment distance position gene name gene id
GU305914.1 86797 86819 - 0 CCGGGGCAGGAGGGGAAGAG GGG ||||||||[||||||||||||]PAM NA Intergenic NA NA

T27673 CGGGGCAGGAGGGGAAGAGGGGG 1000
Oligo fwd CACCCGGGGCAGGAGGGGAAGAGG
Oligo rev AAACCCTCTTCCCCTCCTGCCCCG
Chromosome start end strand MM target_seq PAM alignment distance position gene name gene id
GU305914.1 86796 86818 - 0 CGGGGCAGGAGGGGAAGAGG GGG ||||||||[||||||||||||]PAM NA Intergenic NA NA
GU305914.1 165009 165031 + 4 AGGAGTAGGAGGGGAAGACG AGG -||-|-||[||||||||||-|]PAM NA Intergenic NA NA

T27674 GGCAGGAGGGGAAGAGGGGGCGG 1000
Oligo fwd CACCGGCAGGAGGGGAAGAGGGGG
Oligo rev AAACCCCCCTCTTCCCCTCCTGCC
Chromosome start end strand MM target_seq PAM alignment distance position gene name gene id
GU305914.1 86793 86815 - 0 GGCAGGAGGGGAAGAGGGGG CGG ||||||||[||||||||||||]PAM NA Intergenic NA NA
GU305914.1 45059 45081 + 4 AGTAGGAGGAGAAGAGGAGG AGG -|-|||||[|-|||||||-||]PAM NA Intergenic NA NA

T27675 GCAGGAGGGGAAGAGGGGGCGGG 1000
Oligo fwd CACCGCAGGAGGGGAAGAGGGGGC
Oligo rev AAACGCCCCCTCTTCCCCTCCTGC
Chromosome start end strand MM target_seq PAM alignment distance position gene name gene id
GU305914.1 86792 86814 - 0 GCAGGAGGGGAAGAGGGGGC GGG ||||||||[||||||||||||]PAM NA Intergenic NA NA

T27676 CAGGAGGGGAAGAGGGGGCGGGG 1000
Oligo fwd CACCCAGGAGGGGAAGAGGGGGCG
Oligo rev AAACCGCCCCCTCTTCCCCTCCTG
Chromosome start end strand MM target_seq PAM alignment distance position gene name gene id
GU305914.1 86791 86813 - 0 CAGGAGGGGAAGAGGGGGCG GGG ||||||||[||||||||||||]PAM NA Intergenic NA NA

T27677 AGGAGGGGAAGAGGGGGCGGGGG 1000
Oligo fwd CACCAGGAGGGGAAGAGGGGGCGG
Oligo rev AAACCCGCCCCCTCTTCCCCTCCT
Chromosome start end strand MM target_seq PAM alignment distance position gene name gene id
GU305914.1 86790 86812 - 0 AGGAGGGGAAGAGGGGGCGG GGG ||||||||[||||||||||||]PAM NA Intergenic NA NA

T27678 AGGGGAAGAGGGGGCGGGGGCGG 1000
Oligo fwd CACCAGGGGAAGAGGGGGCGGGGG
Oligo rev AAACCCCCCGCCCCCTCTTCCCCT
Chromosome start end strand MM target_seq PAM alignment distance position gene name gene id
GU305914.1 86787 86809 - 0 AGGGGAAGAGGGGGCGGGGG CGG ||||||||[||||||||||||]PAM NA Intergenic NA NA

T27679 GGAAGAGGGGGCGGGGGCGGCGG 1000
Oligo fwd CACCGGAAGAGGGGGCGGGGGCGG
Oligo rev AAACCCGCCCCCGCCCCCTCTTCC
Chromosome start end strand MM target_seq PAM alignment distance position gene name gene id
GU305914.1 86784 86806 - 0 GGAAGAGGGGGCGGGGGCGG CGG ||||||||[||||||||||||]PAM NA Intergenic NA NA
GU305914.1 18653 18675 - 3 GGCAGAGGTGGCGGAGGCGG GGG ||-|||||[-|||||-|||||]PAM NA Intergenic NA NA
GU305914.1 161754 161776 - 4 GGCCGAGGAGGCGGCGGCGG AGG ||--||||[-|||||-|||||]PAM NA Intergenic NA NA
GU305914.1 67742 67764 + 4 GGGAGCGGCGGCGGCGGCGG CGG ||-||-||[-|||||-|||||]PAM NA Intergenic NA NA
GU305914.1 57997 58019 - 4 GGTGGAGGGGCCGGGGGAGG CGG ||--||||[||-||||||-||]PAM NA Intergenic NA NA

T27680 GAGGGGGCGGGGGCGGCGGCAGG 1000
Oligo fwd CACCGAGGGGGCGGGGGCGGCGGC
Oligo rev AAACGCCGCCGCCCCCGCCCCCTC
Chromosome start end strand MM target_seq PAM alignment distance position gene name gene id
GU305914.1 86780 86802 - 0 GAGGGGGCGGGGGCGGCGGC AGG ||||||||[||||||||||||]PAM NA Intergenic NA NA
GU305914.1 67743 67765 + 4 GGAGCGGCGGCGGCGGCGGC GGG |--|-|||[||-|||||||||]PAM NA Intergenic NA NA
GU305914.1 114657 114679 - 4 GCGGTGGTGGTGGCGGCGGC GGG |-||-||-[||-|||||||||]PAM NA Intergenic NA NA
GU305914.1 221599 221621 - 4 GAGGCCGTGGTGGCGGCGGC AGG ||||--|-[||-|||||||||]PAM NA Intergenic NA NA
GU305914.1 90246 90268 - 3 GAGGCGGCGGAGCCGGCGGC AGG ||||-|||[||-|-|||||||]PAM NA Intergenic NA NA
GU305914.1 149475 149497 - 4 GCGGCGGCGGCGGCGGCAGC GGG |-||-|||[||-||||||-||]PAM NA Intergenic NA NA
GU305914.1 75852 75874 + 4 GGGGAGGCGGAGGCGGCGGT CGG |-||-|||[||-||||||||-]PAM NA Intergenic NA NA
GU305914.1 58122 58144 - 4 GTGGCGGCGGCGGCGGCGGA GGG |-||-|||[||-||||||||-]PAM NA Intergenic NA NA

T27681 AGGGGGCGGGGGCGGCGGCAGGG 1000
Oligo fwd CACCAGGGGGCGGGGGCGGCGGCA
Oligo rev AAACTGCCGCCGCCCCCGCCCCCT
Chromosome start end strand MM target_seq PAM alignment distance position gene name gene id
GU305914.1 86779 86801 - 0 AGGGGGCGGGGGCGGCGGCA GGG ||||||||[||||||||||||]PAM NA Intergenic NA NA
GU305914.1 221598 221620 - 4 AGGCCGTGGTGGCGGCGGCA GGG |||--|-|[|-||||||||||]PAM NA Intergenic NA NA

T27682 GCGGGGGCGGCGGCAGGGCGAGG 1000
Oligo fwd CACCGCGGGGGCGGCGGCAGGGCG
Oligo rev AAACCGCCCTGCCGCCGCCCCCGC
Chromosome start end strand MM target_seq PAM alignment distance position gene name gene id

GU305914.1 86774 86796 - 0 GCGGGGGCGGCGGCAGGGCG AGG ||||||||[||||||||||||]PAM NA Intergenic NA NA

T27683 CGGGGGCGGCGGCAGGGCGAGGG 1000
Oligo fwd CACCCGGGGGCGGCGGCAGGGCGA
Oligo rev AAACTCGCCCTGCCGCCGCCCCCG
Chromosome start end strand MM target_seq PAM alignment distance position gene name gene id
GU305914.1 86773 86795 - 0 CGGGGGCGGCGGCAGGGCGA GGG ||||||||[||||||||||||]PAM NA Intergenic NA NA

T27684 GCGGCAGGGCGAGGGAACACTGG 1000
Oligo fwd CACCGCGGCAGGGCGAGGGAACAC
Oligo rev AAACGTGTTCCCTCGCCCTGCCGC
Chromosome start end strand MM target_seq PAM alignment distance position gene name gene id
GU305914.1 86765 86787 - 0 GCGGCAGGGCGAGGGAACAC TGG ||||||||[||||||||||||]PAM NA Intergenic NA NA

T27685 CGGCAGGGCGAGGGAACACTGGG 1000
Oligo fwd CACCCGGCAGGGCGAGGGAACACT
Oligo rev AAACAGTGTTCCCTCGCCCTGCCG
Chromosome start end strand MM target_seq PAM alignment distance position gene name gene id
GU305914.1 86764 86786 - 0 CGGCAGGGCGAGGGAACACT GGG ||||||||[||||||||||||]PAM NA Intergenic NA NA

T27686 GGCAGGGCGAGGGAACACTGGGG 1000
Oligo fwd CACCGGCAGGGCGAGGGAACACTG
Oligo rev AAACCAGTGTTCCCTCGCCCTGCC
Chromosome start end strand MM target_seq PAM alignment distance position gene name gene id
GU305914.1 86763 86785 - 0 GGCAGGGCGAGGGAACACTG GGG ||||||||[||||||||||||]PAM NA Intergenic NA NA

T27687 GCAGGGCGAGGGAACACTGGGGG 1000
Oligo fwd CACCGCAGGGCGAGGGAACACTGG
Oligo rev AAACCCAGTGTTCCCTCGCCCTGC
Chromosome start end strand MM target_seq PAM alignment distance position gene name gene id
GU305914.1 86762 86784 - 0 GCAGGGCGAGGGAACACTGG GGG ||||||||[||||||||||||]PAM NA Intergenic NA NA

T27688 CAGGGCGAGGGAACACTGGGGGG 1000
Oligo fwd CACCCAGGGCGAGGGAACACTGGG
Oligo rev AAACCCCAGTGTTCCCTCGCCCTG
Chromosome start end strand MM target_seq PAM alignment distance position gene name gene id
GU305914.1 86761 86783 - 0 CAGGGCGAGGGAACACTGGG GGG ||||||||[||||||||||||]PAM NA Intergenic NA NA

T27689 AGGGCGAGGGAACACTGGGGGGG 1000
Oligo fwd CACCAGGGCGAGGGAACACTGGGG
Oligo rev AAACCCCCAGTGTTCCCTCGCCCT
Chromosome start end strand MM target_seq PAM alignment distance position gene name gene id
GU305914.1 86760 86782 - 0 AGGGCGAGGGAACACTGGGG GGG ||||||||[||||||||||||]PAM NA Intergenic NA NA

T27690 AGGGAACACTGGGGGGGACGAGG 1000
Oligo fwd CACCAGGGAACACTGGGGGGGACG
Oligo rev AAACCGTCCCCCCCAGTGTTCCCT
Chromosome start end strand MM target_seq PAM alignment distance position gene name gene id
GU305914.1 86754 86776 - 0 AGGGAACACTGGGGGGGACG AGG ||||||||[||||||||||||]PAM NA Intergenic NA NA

T27691 GGGAACACTGGGGGGGACGAGGG 1000
Oligo fwd CACCGGGAACACTGGGGGGGACGA
Oligo rev AAACTCGTCCCCCCCAGTGTTCCC
Chromosome start end strand MM target_seq PAM alignment distance position gene name gene id
GU305914.1 86753 86775 - 0 GGGAACACTGGGGGGGACGA GGG ||||||||[||||||||||||]PAM NA Intergenic NA NA

T27692 CACTGGGGGGGACGAGGGCGCGG 1000
Oligo fwd CACCCACTGGGGGGGACGAGGGCG
Oligo rev AAACCGCCCTCGTCCCCCCCAGTG
Chromosome start end strand MM target_seq PAM alignment distance position gene name gene id
GU305914.1 86748 86770 - 0 CACTGGGGGGGACGAGGGCG CGG ||||||||[||||||||||||]PAM NA Intergenic NA NA

T27693 ACTGGGGGGGACGAGGGCGCGGG 1000
Oligo fwd CACCACTGGGGGGGACGAGGGCGC
Oligo rev AAACGCGCCCTCGTCCCCCCCAGT
Chromosome start end strand MM target_seq PAM alignment distance position gene name gene id
GU305914.1 86747 86769 - 0 ACTGGGGGGGACGAGGGCGC GGG ||||||||[||||||||||||]PAM NA Intergenic NA NA

T27694 CGAGGGCGCGGGAACAACAACGG 1000
Oligo fwd CACCCGAGGGCGCGGGAACAACAA
Oligo rev AAACTTGTTGTTCCCGCGCCCTCG
Chromosome start end strand MM target_seq PAM alignment distance position gene name gene id
GU305914.1 86736 86758 - 0 CGAGGGCGCGGGAACAACAA CGG ||||||||[||||||||||||]PAM NA Intergenic NA NA

T27695 AACGGCAATGAGTTCCGCGTTGG 1000
Oligo fwd CACCAACGGCAATGAGTTCCGCGT
Oligo rev AAACACGCGGAACTCATTGCCGTT
Chromosome start end strand MM target_seq PAM alignment distance position gene name gene id
GU305914.1 86718 86740 - 0 AACGGCAATGAGTTCCGCGT TGG ||||||||[||||||||||||]PAM NA Intergenic NA NA

T27696 ATGAGTTCCGCGTTGGAGTGTGG 1000
Oligo fwd CACCATGAGTTCCGCGTTGGAGTG
Oligo rev AAACCACTCCAACGCGGAACTCAT
Chromosome start end strand MM target_seq PAM alignment distance position gene name gene id
GU305914.1 86711 86733 - 0 ATGAGTTCCGCGTTGGAGTG TGG ||||||||[||||||||||||]PAM NA Intergenic NA NA

T27697 AGTGTGGAGATCCGCTGCTGAGG 1000
Oligo fwd CACCAGTGTGGAGATCCGCTGCTG
Oligo rev AAACCAGCAGCGGATCTCCACACT

Chromosome start end strand MM target_seq PAM alignment distance position gene name gene id
GU305914.1 86695 86717 - 0 AGTGTGGAGATCCGCTGCTG AGG ||||||||[||||||||||||]PAM NA Intergenic NA NA

T27698 GTGTGGAGATCCGCTGCTGAGGG 1000
Oligo fwd CACCGTGTGGAGATCCGCTGCTGA
Oligo rev AAACTCAGCAGCGGATCTCCACAC
Chromosome start end strand MM target_seq PAM alignment distance position gene name gene id
GU305914.1 86694 86716 - 0 GTGTGGAGATCCGCTGCTGA GGG ||||||||[||||||||||||]PAM NA Intergenic NA NA

T27699 GAGGGTGCACGATGTGAACAAGG 1000
Oligo fwd CACCGAGGGTGCACGATGTGAACA
Oligo rev AAACTGTTCACATCGTGCACCCTC
Chromosome start end strand MM target_seq PAM alignment distance position gene name gene id
GU305914.1 86676 86698 - 0 GAGGGTGCACGATGTGAACA AGG ||||||||[||||||||||||]PAM NA Intergenic NA NA

T27700 GGTGCACGATGTGAACAAGGAGG 1000
Oligo fwd CACCGGTGCACGATGTGAACAAGG
Oligo rev AAACCCTTGTTCACATCGTGCACC
Chromosome start end strand MM target_seq PAM alignment distance position gene name gene id
GU305914.1 86673 86695 - 0 GGTGCACGATGTGAACAAGG AGG ||||||||[||||||||||||]PAM NA Intergenic NA NA

T27701 AGGAGGTGAATGTCCGAAAAAGG 1000
Oligo fwd CACCAGGAGGTGAATGTCCGAAAA
Oligo rev AAACTTTTCGGACATTCACCTCCT
Chromosome start end strand MM target_seq PAM alignment distance position gene name gene id
GU305914.1 86656 86678 - 0 AGGAGGTGAATGTCCGAAAA AGG ||||||||[||||||||||||]PAM NA Intergenic NA NA

T27702 GGAGGTGAATGTCCGAAAAAGGG 1000
Oligo fwd CACCGGAGGTGAATGTCCGAAAAA
Oligo rev AAACTTTTTCGGACATTCACCTCC
Chromosome start end strand MM target_seq PAM alignment distance position gene name gene id
GU305914.1 86655 86677 - 0 GGAGGTGAATGTCCGAAAAA GGG ||||||||[||||||||||||]PAM NA Intergenic NA NA

T27703 AAAAAGGGCATACTTAAAAAAGG 1000
Oligo fwd CACCAAAAAGGGCATACTTAAAAA
Oligo rev AAACTTTTTAAGTATGCCCTTTTT
Chromosome start end strand MM target_seq PAM alignment distance position gene name gene id
GU305914.1 86640 86662 - 0 AAAAAGGGCATACTTAAAAA AGG ||||||||[||||||||||||]PAM NA Intergenic NA NA

T27704 CTTAAAAAAGGTTTCCGAAATGG 1000
Oligo fwd CACCCTTAAAAAAGGTTTCCGAAA
Oligo rev AAACTTTCGGAAACCTTTTTTAAG
Chromosome start end strand MM target_seq PAM alignment distance position gene name gene id
GU305914.1 86628 86650 - 0 CTTAAAAAAGGTTTCCGAAA TGG ||||||||[||||||||||||]PAM NA Intergenic NA NA

T27705 TTAAAAAAGGTTTCCGAAATGGG 1000
Oligo fwd CACCTTAAAAAAGGTTTCCGAAAT
Oligo rev AAACATTTCGGAAACCTTTTTTAA
Chromosome start end strand MM target_seq PAM alignment distance position gene name gene id
GU305914.1 86627 86649 - 0 TTAAAAAAGGTTTCCGAAAT GGG ||||||||[||||||||||||]PAM NA Intergenic NA NA

T27706 TTCCGAAATGGGCTACAACAAGG 1000
Oligo fwd CACCTTCCGAAATGGGCTACAACA
Oligo rev AAACTGTTGTAGCCCATTTCGGAA
Chromosome start end strand MM target_seq PAM alignment distance position gene name gene id
GU305914.1 86616 86638 - 0 TTCCGAAATGGGCTACAACA AGG ||||||||[||||||||||||]PAM NA Intergenic NA NA

T27707 AATGGGCTACAACAAGGTTATGG 1000
Oligo fwd CACCAATGGGCTACAACAAGGTTA
Oligo rev AAACTAACCTTGTTGTAGCCCATT
Chromosome start end strand MM target_seq PAM alignment distance position gene name gene id
GU305914.1 86610 86632 - 0 AATGGGCTACAACAAGGTTA TGG ||||||||[||||||||||||]PAM NA Intergenic NA NA

T27708 TATGGCCTGTATTCGAAACCAGG 1000
Oligo fwd CACCTATGGCCTGTATTCGAAACC
Oligo rev AAACGGTTTCGAATACAGGCCATA
Chromosome start end strand MM target_seq PAM alignment distance position gene name gene id
GU305914.1 86592 86614 - 0 TATGGCCTGTATTCGAAACC AGG ||||||||[||||||||||||]PAM NA Intergenic NA NA

T27709 TATTCGAAACCAGGAGCATCTGG 1000
Oligo fwd CACCTATTCGAAACCAGGAGCATC
Oligo rev AAACGATGCTCCTGGTTTCGAATA
Chromosome start end strand MM target_seq PAM alignment distance position gene name gene id
GU305914.1 86583 86605 - 0 TATTCGAAACCAGGAGCATC TGG ||||||||[||||||||||||]PAM NA Intergenic NA NA

T27710 GGAGCATCTGGTTACGAAACTGG 1000
Oligo fwd CACCGGAGCATCTGGTTACGAAAC
Oligo rev AAACGTTTCGTAACCAGATGCTCC
Chromosome start end strand MM target_seq PAM alignment distance position gene name gene id
GU305914.1 86571 86593 - 0 GGAGCATCTGGTTACGAAAC TGG ||||||||[||||||||||||]PAM NA Intergenic NA NA

T27711 GAAACTGGTGAATGTCAACCTGG 1000
Oligo fwd CACCGAAACTGGTGAATGTCAACC
Oligo rev AAACGGTTGACATTCACCAGTTTC
Chromosome start end strand MM target_seq PAM alignment distance position gene name gene id
GU305914.1 86556 86578 - 0 GAAACTGGTGAATGTCAACC TGG ||||||||[||||||||||||]PAM NA Intergenic NA NA

T27712 ACTGGTGAATGTCAACCTGGTGG 1000
Oligo fwd CACCACTGGTGAATGTCAACCTGG

| | | | | | | | | | | | | |
|---|---|---|---|---|---|---|---|---|---|---|---|---|
| Oligo rev | AAACCCAGGTTGACATTCACCAGT | | | | | | | | | | | |
| Chromosome | start | | end | strand | MM | target_seq | PAM | alignment | distance | position | gene name | gene id |
| GU305914.1 | | 86553 | 86575 | - | 0 | ACTGGTGAATGTCAACCTGG | TGG | \|\|\|\|\|\|\|\|[\|\|\|\|\|\|\|\|\|\|\|\|]PAM | NA | Intergenic | NA | NA |
| T27713 | CTGGTGAATGTCAACCTGGTGGG | | 1000 | | | | | | | | | |
| Oligo fwd | CACCCTGGTGAATGTCAACCTGGT | | | | | | | | | | | |
| Oligo rev | AAACACCAGGTTGACATTCACCAG | | | | | | | | | | | |
| Chromosome | start | | end | strand | MM | target_seq | PAM | alignment | distance | position | gene name | gene id |
| GU305914.1 | | 86552 | 86574 | - | 0 | CTGGTGAATGTCAACCTGGT | GGG | \|\|\|\|\|\|\|\|[\|\|\|\|\|\|\|\|\|\|\|\|]PAM | NA | Intergenic | NA | NA |
| T27714 | GAATGTCAACCTGGTGGGAACGG | | 1000 | | | | | | | | | |
| Oligo fwd | CACCGAATGTCAACCTGGTGGGAA | | | | | | | | | | | |
| Oligo rev | AAACTTCCCACCAGGTTGACATTC | | | | | | | | | | | |
| Chromosome | start | | end | strand | MM | target_seq | PAM | alignment | distance | position | gene name | gene id |
| GU305914.1 | | 86547 | 86569 | - | 0 | GAATGTCAACCTGGTGGGAA | CGG | \|\|\|\|\|\|\|\|[\|\|\|\|\|\|\|\|\|\|\|\|]PAM | NA | Intergenic | NA | NA |
| T27715 | CCTGGTGGGAACGGTGTGTTTGG | | 1000 | | | | | | | | | |
| Oligo fwd | CACCCCTGGTGGGAACGGTGTGTT | | | | | | | | | | | |
| Oligo rev | AAACAACACACCGTTCCCACCAGG | | | | | | | | | | | |
| Chromosome | start | | end | strand | MM | target_seq | PAM | alignment | distance | position | gene name | gene id |
| GU305914.1 | | 86538 | 86560 | - | 0 | CCTGGTGGGAACGGTGTGTT | TGG | \|\|\|\|\|\|\|\|[\|\|\|\|\|\|\|\|\|\|\|\|]PAM | NA | Intergenic | NA | NA |
| T27716 | GGTGGGAACGGTGTGTTTGGAGG | | 1000 | | | | | | | | | |
| Oligo fwd | CACCGGTGGGAACGGTGTGTTTGG | | | | | | | | | | | |
| Oligo rev | AAACCCAAACACACCGTTCCCACC | | | | | | | | | | | |
| Chromosome | start | | end | strand | MM | target_seq | PAM | alignment | distance | position | gene name | gene id |
| GU305914.1 | | 86535 | 86557 | - | 0 | GGTGGGAACGGTGTGTTTGG | AGG | \|\|\|\|\|\|\|\|[\|\|\|\|\|\|\|\|\|\|\|\|]PAM | NA | Intergenic | NA | NA |
| T27717 | GGGAACGGTGTGTTTGGAGGCGG | | 1000 | | | | | | | | | |
| Oligo fwd | CACCGGGAACGGTGTGTTTGGAGG | | | | | | | | | | | |
| Oligo rev | AAACCCTCCAAACACACCGTTCCC | | | | | | | | | | | |
| Chromosome | start | | end | strand | MM | target_seq | PAM | alignment | distance | position | gene name | gene id |
| GU305914.1 | | 86532 | 86554 | - | 0 | GGGAACGGTGTGTTTGGAGG | CGG | \|\|\|\|\|\|\|\|[\|\|\|\|\|\|\|\|\|\|\|\|]PAM | NA | Intergenic | NA | NA |
| T27718 | GCGGTTTCCAAAATTATGAACGG | | 1000 | | | | | | | | | |
| Oligo fwd | CACCGCGGTTTCCAAAATTATGAA | | | | | | | | | | | |
| Oligo rev | AAACTTCATAATTTTGGAAACCGC | | | | | | | | | | | |
| Chromosome | start | | end | strand | MM | target_seq | PAM | alignment | distance | position | gene name | gene id |
| GU305914.1 | | 86513 | 86535 | - | 0 | GCGGTTTCCAAAATTATGAA | CGG | \|\|\|\|\|\|\|\|[\|\|\|\|\|\|\|\|\|\|\|\|]PAM | NA | Intergenic | NA | NA |
| T27719 | ACGGTTTTCTCTCCCGACAGCGG | | 1000 | | | | | | | | | |
| Oligo fwd | CACCACGGTTTTCTCTCCCGACAG | | | | | | | | | | | |
| Oligo rev | AAACCTGTCGGGAGAGAAAACCGT | | | | | | | | | | | |
| Chromosome | start | | end | strand | MM | target_seq | PAM | alignment | distance | position | gene name | gene id |
| GU305914.1 | | 86494 | 86516 | - | 0 | ACGGTTTTCTCTCCCGACAG | CGG | \|\|\|\|\|\|\|\|[\|\|\|\|\|\|\|\|\|\|\|\|]PAM | NA | Intergenic | NA | NA |
| T27720 | CCGACAGCGGTCTATAACTGAGG | | 1000 | | | | | | | | | |
| Oligo fwd | CACCCCGACAGCGGTCTATAACTG | | | | | | | | | | | |
| Oligo rev | AAACCAGTTATAGACCGCTGTCGG | | | | | | | | | | | |
| Chromosome | start | | end | strand | MM | target_seq | PAM | alignment | distance | position | gene name | gene id |
| GU305914.1 | | 86481 | 86503 | - | 0 | CCGACAGCGGTCTATAACTG | AGG | \|\|\|\|\|\|\|\|[\|\|\|\|\|\|\|\|\|\|\|\|]PAM | NA | Intergenic | NA | NA |
| T27721 | GGCCGAGACCTACCCCGACGTGG | | 1000 | | | | | | | | | |
| Oligo fwd | CACCGGCCGAGACCTACCCCGACG | | | | | | | | | | | |
| Oligo rev | AAACCGTCGGGGTAGGTCTCGGCC | | | | | | | | | | | |
| Chromosome | start | | end | strand | MM | target_seq | PAM | alignment | distance | position | gene name | gene id |
| GU305914.1 | | 86460 | 86482 | - | 0 | GGCCGAGACCTACCCCGACG | TGG | \|\|\|\|\|\|\|\|[\|\|\|\|\|\|\|\|\|\|\|\|]PAM | NA | Intergenic | NA | NA |
| T27722 | CCCGACGTGGCCGAGAGCCTCGG | | 1000 | | | | | | | | | |
| Oligo fwd | CACCCCCGACGTGGCCGAGAGCCT | | | | | | | | | | | |
| Oligo rev | AAACAGGCTCTCGGCCACGTCGGG | | | | | | | | | | | |
| Chromosome | start | | end | strand | MM | target_seq | PAM | alignment | distance | position | gene name | gene id |
| GU305914.1 | | 86447 | 86469 | - | 0 | CCCGACGTGGCCGAGAGCCT | CGG | \|\|\|\|\|\|\|\|[\|\|\|\|\|\|\|\|\|\|\|\|]PAM | NA | Intergenic | NA | NA |
| T27723 | TCAATAATCTCGTGCACAAGAGG | | 1000 | | | | | | | | | |
| Oligo fwd | CACCTCAATAATCTCGTGCACAAG | | | | | | | | | | | |
| Oligo rev | AAACCTTGTGCACGAGATTATTGA | | | | | | | | | | | |
| Chromosome | start | | end | strand | MM | target_seq | PAM | alignment | distance | position | gene name | gene id |
| GU305914.1 | | 86401 | 86423 | - | 0 | TCAATAATCTCGTGCACAAG | AGG | \|\|\|\|\|\|\|\|[\|\|\|\|\|\|\|\|\|\|\|\|]PAM | NA | Intergenic | NA | NA |
| T27724 | TCCGAGCTGCTGCCCCAGCTCGG | | 1000 | | | | | | | | | |
| Oligo fwd | CACCTCCGAGCTGCTGCCCCAGCT | | | | | | | | | | | |
| Oligo rev | AAACAGCTGGGGCAGCAGCTCGGA | | | | | | | | | | | |
| Chromosome | start | | end | strand | MM | target_seq | PAM | alignment | distance | position | gene name | gene id |
| GU305914.1 | | 86372 | 86394 | - | 0 | TCCGAGCTGCTGCCCCAGCT | CGG | \|\|\|\|\|\|\|\|[\|\|\|\|\|\|\|\|\|\|\|\|]PAM | NA | Intergenic | NA | NA |
| T27725 | CCGAGCTGCTGCCCCAGCTCGGG | | 1000 | | | | | | | | | |
| Oligo fwd | CACCCCGAGCTGCTGCCCCAGCTC | | | | | | | | | | | |
| Oligo rev | AAACGAGCTGGGGCAGCAGCTCGG | | | | | | | | | | | |
| Chromosome | start | | end | strand | MM | target_seq | PAM | alignment | distance | position | gene name | gene id |
| GU305914.1 | | 86371 | 86393 | - | 0 | CCGAGCTGCTGCCCCAGCTC | GGG | \|\|\|\|\|\|\|\|[\|\|\|\|\|\|\|\|\|\|\|\|]PAM | NA | Intergenic | NA | NA |
| T27726 | CAGATCTACCGTTTCATCAACGG | | 1000 | | | | | | | | | |
| Oligo fwd | CACCCAGATCTACCGTTTCATCAA | | | | | | | | | | | |
| Oligo rev | AAACTTGATGAAACGGTAGATCTG | | | | | | | | | | | |
| Chromosome | start | | end | strand | MM | target_seq | PAM | alignment | distance | position | gene name | gene id |
| GU305914.1 | | 86345 | 86367 | - | 0 | CAGATCTACCGTTTCATCAA | CGG | \|\|\|\|\|\|\|\|[\|\|\|\|\|\|\|\|\|\|\|\|]PAM | NA | Intergenic | NA | NA |
| T27727 | CCTGCCCTATAACGTTAACATGG | | 1000 | | | | | | | | | |

<table>
<tr><td>Oligo fwd</td><td>CACCCCTGCCCTATAACGTTAACA</td><td></td><td></td><td></td><td></td><td></td><td></td><td></td><td></td><td></td><td></td></tr>
<tr><td>Oligo rev</td><td>AAACTGTTAACGTTATAGGGCAGG</td><td></td><td></td><td></td><td></td><td></td><td></td><td></td><td></td><td></td><td></td></tr>
<tr><td>Chromosome</td><td>start</td><td>end</td><td>strand</td><td>MM</td><td>target_seq</td><td>PAM</td><td>alignment</td><td>distance</td><td>position</td><td>gene name</td><td>gene id</td></tr>
<tr><td>GU305914.1</td><td>86289</td><td>86311</td><td>-</td><td>0</td><td>CCTGCCCTATAACGTTAACA</td><td>TGG</td><td>||||||||[||||||||||||]PAM</td><td>NA</td><td>Intergenic</td><td>NA</td><td>NA</td></tr>
<tr><td>T27728</td><td>GCCTACGCCTGCGATAACGCCGG</td><td>1000</td><td></td><td></td><td></td><td></td><td></td><td></td><td></td><td></td><td></td></tr>
<tr><td>Oligo fwd</td><td>CACCGCCTACGCCTGCGATAACGC</td><td></td><td></td><td></td><td></td><td></td><td></td><td></td><td></td><td></td><td></td></tr>
<tr><td>Oligo rev</td><td>AAACGCGTTATCGCAGGCGTAGGC</td><td></td><td></td><td></td><td></td><td></td><td></td><td></td><td></td><td></td><td></td></tr>
<tr><td>Chromosome</td><td>start</td><td>end</td><td>strand</td><td>MM</td><td>target_seq</td><td>PAM</td><td>alignment</td><td>distance</td><td>position</td><td>gene name</td><td>gene id</td></tr>
<tr><td>GU305914.1</td><td>86267</td><td>86289</td><td>-</td><td>0</td><td>GCCTACGCCTGCGATAACGC</td><td>CGG</td><td>||||||||[||||||||||||]PAM</td><td>NA</td><td>Intergenic</td><td>NA</td><td>NA</td></tr>
<tr><td>T27729</td><td>CATCCTGCCGCACGTCAAAGAGG</td><td>1000</td><td></td><td></td><td></td><td></td><td></td><td></td><td></td><td></td><td></td></tr>
<tr><td>Oligo fwd</td><td>CACCCATCCTGCCGCACGTCAAAG</td><td></td><td></td><td></td><td></td><td></td><td></td><td></td><td></td><td></td><td></td></tr>
<tr><td>Oligo rev</td><td>AAACCTTTGACGTGCGGCAGGATG</td><td></td><td></td><td></td><td></td><td></td><td></td><td></td><td></td><td></td><td></td></tr>
<tr><td>Chromosome</td><td>start</td><td>end</td><td>strand</td><td>MM</td><td>target_seq</td><td>PAM</td><td>alignment</td><td>distance</td><td>position</td><td>gene name</td><td>gene id</td></tr>
<tr><td>GU305914.1</td><td>86244</td><td>86266</td><td>-</td><td>0</td><td>CATCCTGCCGCACGTCAAAG</td><td>AGG</td><td>||||||||[||||||||||||]PAM</td><td>NA</td><td>Intergenic</td><td>NA</td><td>NA</td></tr>
<tr><td>T27730</td><td>ACGTCAAAGAGGATCTCGTCAGG</td><td>1000</td><td></td><td></td><td></td><td></td><td></td><td></td><td></td><td></td><td></td></tr>
<tr><td>Oligo fwd</td><td>CACCACGTCAAAGAGGATCTCGTC</td><td></td><td></td><td></td><td></td><td></td><td></td><td></td><td></td><td></td><td></td></tr>
<tr><td>Oligo rev</td><td>AAACGACGAGATCCTCTTTGACGT</td><td></td><td></td><td></td><td></td><td></td><td></td><td></td><td></td><td></td><td></td></tr>
<tr><td>Chromosome</td><td>start</td><td>end</td><td>strand</td><td>MM</td><td>target_seq</td><td>PAM</td><td>alignment</td><td>distance</td><td>position</td><td>gene name</td><td>gene id</td></tr>
<tr><td>GU305914.1</td><td>86233</td><td>86255</td><td>-</td><td>0</td><td>ACGTCAAAGAGGATCTCGTC</td><td>AGG</td><td>||||||||[||||||||||||]PAM</td><td>NA</td><td>Intergenic</td><td>NA</td><td>NA</td></tr>
<tr><td>T27731</td><td>AGAGGATCTCGTCAGGTGCGCGG</td><td>1000</td><td></td><td></td><td></td><td></td><td></td><td></td><td></td><td></td><td></td></tr>
<tr><td>Oligo fwd</td><td>CACCAGAGGATCTCGTCAGGTGCG</td><td></td><td></td><td></td><td></td><td></td><td></td><td></td><td></td><td></td><td></td></tr>
<tr><td>Oligo rev</td><td>AAACCGCACCTGACGAGATCCTCT</td><td></td><td></td><td></td><td></td><td></td><td></td><td></td><td></td><td></td><td></td></tr>
<tr><td>Chromosome</td><td>start</td><td>end</td><td>strand</td><td>MM</td><td>target_seq</td><td>PAM</td><td>alignment</td><td>distance</td><td>position</td><td>gene name</td><td>gene id</td></tr>
<tr><td>GU305914.1</td><td>86226</td><td>86248</td><td>-</td><td>0</td><td>AGAGGATCTCGTCAGGTGCG</td><td>CGG</td><td>||||||||[||||||||||||]PAM</td><td>NA</td><td>Intergenic</td><td>NA</td><td>NA</td></tr>
<tr><td>T27732</td><td>GATCTCGTCAGGTGCGCGGACGG</td><td>1000</td><td></td><td></td><td></td><td></td><td></td><td></td><td></td><td></td><td></td></tr>
<tr><td>Oligo fwd</td><td>CACCGATCTCGTCAGGTGCGCGGA</td><td></td><td></td><td></td><td></td><td></td><td></td><td></td><td></td><td></td><td></td></tr>
<tr><td>Oligo rev</td><td>AAACTCCGCGCACCTGACGAGATC</td><td></td><td></td><td></td><td></td><td></td><td></td><td></td><td></td><td></td><td></td></tr>
<tr><td>Chromosome</td><td>start</td><td>end</td><td>strand</td><td>MM</td><td>target_seq</td><td>PAM</td><td>alignment</td><td>distance</td><td>position</td><td>gene name</td><td>gene id</td></tr>
<tr><td>GU305914.1</td><td>86222</td><td>86244</td><td>-</td><td>0</td><td>GATCTCGTCAGGTGCGCGGA</td><td>CGG</td><td>||||||||[||||||||||||]PAM</td><td>NA</td><td>Intergenic</td><td>NA</td><td>NA</td></tr>
<tr><td>T27733</td><td>ATCTCGTCAGGTGCGCGGACGGG</td><td>1000</td><td></td><td></td><td></td><td></td><td></td><td></td><td></td><td></td><td></td></tr>
<tr><td>Oligo fwd</td><td>CACCATCTCGTCAGGTGCGCGGAC</td><td></td><td></td><td></td><td></td><td></td><td></td><td></td><td></td><td></td><td></td></tr>
<tr><td>Oligo rev</td><td>AAACGTCCGCGCACCTGACGAGAT</td><td></td><td></td><td></td><td></td><td></td><td></td><td></td><td></td><td></td><td></td></tr>
<tr><td>Chromosome</td><td>start</td><td>end</td><td>strand</td><td>MM</td><td>target_seq</td><td>PAM</td><td>alignment</td><td>distance</td><td>position</td><td>gene name</td><td>gene id</td></tr>
<tr><td>GU305914.1</td><td>86221</td><td>86243</td><td>-</td><td>0</td><td>ATCTCGTCAGGTGCGCGGAC</td><td>GGG</td><td>||||||||[||||||||||||]PAM</td><td>NA</td><td>Intergenic</td><td>NA</td><td>NA</td></tr>
<tr><td>T27734</td><td>CGTCAGGTGCGCGGACGGGACGG</td><td>1000</td><td></td><td></td><td></td><td></td><td></td><td></td><td></td><td></td><td></td></tr>
<tr><td>Oligo fwd</td><td>CACCCGTCAGGTGCGCGGACGGGA</td><td></td><td></td><td></td><td></td><td></td><td></td><td></td><td></td><td></td><td></td></tr>
<tr><td>Oligo rev</td><td>AAACTCCCGTCCGCGCACCTGACG</td><td></td><td></td><td></td><td></td><td></td><td></td><td></td><td></td><td></td><td></td></tr>
<tr><td>Chromosome</td><td>start</td><td>end</td><td>strand</td><td>MM</td><td>target_seq</td><td>PAM</td><td>alignment</td><td>distance</td><td>position</td><td>gene name</td><td>gene id</td></tr>
<tr><td>GU305914.1</td><td>86217</td><td>86239</td><td>-</td><td>0</td><td>CGTCAGGTGCGCGGACGGGA</td><td>CGG</td><td>||||||||[||||||||||||]PAM</td><td>NA</td><td>Intergenic</td><td>NA</td><td>NA</td></tr>
<tr><td>T27735</td><td>CAGGTGCGCGGACGGGACGGTGG</td><td>1000</td><td></td><td></td><td></td><td></td><td></td><td></td><td></td><td></td><td></td></tr>
<tr><td>Oligo fwd</td><td>CACCCAGGTGCGCGGACGGGACGG</td><td></td><td></td><td></td><td></td><td></td><td></td><td></td><td></td><td></td><td></td></tr>
<tr><td>Oligo rev</td><td>AAACCCGTCCCGTCCGCGCACCTG</td><td></td><td></td><td></td><td></td><td></td><td></td><td></td><td></td><td></td><td></td></tr>
<tr><td>Chromosome</td><td>start</td><td>end</td><td>strand</td><td>MM</td><td>target_seq</td><td>PAM</td><td>alignment</td><td>distance</td><td>position</td><td>gene name</td><td>gene id</td></tr>
<tr><td>GU305914.1</td><td>86214</td><td>86236</td><td>-</td><td>0</td><td>CAGGTGCGCGGACGGGACGG</td><td>TGG</td><td>||||||||[||||||||||||]PAM</td><td>NA</td><td>Intergenic</td><td>NA</td><td>NA</td></tr>
<tr><td>T27736</td><td>GGACGGTGGTGCCCAGCGACTGG</td><td>1000</td><td></td><td></td><td></td><td></td><td></td><td></td><td></td><td></td><td></td></tr>
<tr><td>Oligo fwd</td><td>CACCGGACGGTGGTGCCCAGCGAC</td><td></td><td></td><td></td><td></td><td></td><td></td><td></td><td></td><td></td><td></td></tr>
<tr><td>Oligo rev</td><td>AAACGTCGCTGGGCACCACCGTCC</td><td></td><td></td><td></td><td></td><td></td><td></td><td></td><td></td><td></td><td></td></tr>
<tr><td>Chromosome</td><td>start</td><td>end</td><td>strand</td><td>MM</td><td>target_seq</td><td>PAM</td><td>alignment</td><td>distance</td><td>position</td><td>gene name</td><td>gene id</td></tr>
<tr><td>GU305914.1</td><td>86200</td><td>86222</td><td>-</td><td>0</td><td>GGACGGTGGTGCCCAGCGAC</td><td>TGG</td><td>||||||||[||||||||||||]PAM</td><td>NA</td><td>Intergenic</td><td>NA</td><td>NA</td></tr>
<tr><td>T27737</td><td>CCCAGCGACTGGATGACAGTCGG</td><td>1000</td><td></td><td></td><td></td><td></td><td></td><td></td><td></td><td></td><td></td></tr>
<tr><td>Oligo fwd</td><td>CACCCCCAGCGACTGGATGACAGT</td><td></td><td></td><td></td><td></td><td></td><td></td><td></td><td></td><td></td><td></td></tr>
<tr><td>Oligo rev</td><td>AAACACTGTCATCCAGTCGCTGGG</td><td></td><td></td><td></td><td></td><td></td><td></td><td></td><td></td><td></td><td></td></tr>
<tr><td>Chromosome</td><td>start</td><td>end</td><td>strand</td><td>MM</td><td>target_seq</td><td>PAM</td><td>alignment</td><td>distance</td><td>position</td><td>gene name</td><td>gene id</td></tr>
<tr><td>GU305914.1</td><td>86189</td><td>86211</td><td>-</td><td>0</td><td>CCCAGCGACTGGATGACAGT</td><td>CGG</td><td>||||||||[||||||||||||]PAM</td><td>NA</td><td>Intergenic</td><td>NA</td><td>NA</td></tr>
<tr><td>T27738</td><td>CTGGATGACAGTCGGATACATGG</td><td>1000</td><td></td><td></td><td></td><td></td><td></td><td></td><td></td><td></td><td></td></tr>
<tr><td>Oligo fwd</td><td>CACCCTGGATGACAGTCGGATACA</td><td></td><td></td><td></td><td></td><td></td><td></td><td></td><td></td><td></td><td></td></tr>
<tr><td>Oligo rev</td><td>AAACTGTATCCGACTGTCATCCAG</td><td></td><td></td><td></td><td></td><td></td><td></td><td></td><td></td><td></td><td></td></tr>
<tr><td>Chromosome</td><td>start</td><td>end</td><td>strand</td><td>MM</td><td>target_seq</td><td>PAM</td><td>alignment</td><td>distance</td><td>position</td><td>gene name</td><td>gene id</td></tr>
<tr><td>GU305914.1</td><td>86181</td><td>86203</td><td>-</td><td>0</td><td>CTGGATGACAGTCGGATACA</td><td>TGG</td><td>||||||||[||||||||||||]PAM</td><td>NA</td><td>Intergenic</td><td>NA</td><td>NA</td></tr>
<tr><td>T27739</td><td>TGGATGACAGTCGGATACATGGG</td><td>1000</td><td></td><td></td><td></td><td></td><td></td><td></td><td></td><td></td><td></td></tr>
<tr><td>Oligo fwd</td><td>CACCTGGATGACAGTCGGATACAT</td><td></td><td></td><td></td><td></td><td></td><td></td><td></td><td></td><td></td><td></td></tr>
<tr><td>Oligo rev</td><td>AAACATGTATCCGACTGTCATCCA</td><td></td><td></td><td></td><td></td><td></td><td></td><td></td><td></td><td></td><td></td></tr>
<tr><td>Chromosome</td><td>start</td><td>end</td><td>strand</td><td>MM</td><td>target_seq</td><td>PAM</td><td>alignment</td><td>distance</td><td>position</td><td>gene name</td><td>gene id</td></tr>
<tr><td>GU305914.1</td><td>86180</td><td>86202</td><td>-</td><td>0</td><td>TGGATGACAGTCGGATACAT</td><td>GGG</td><td>||||||||[||||||||||||]PAM</td><td>NA</td><td>Intergenic</td><td>NA</td><td>NA</td></tr>
<tr><td>T27740</td><td>TCGGATACATGGGCTTCTTCAGG</td><td>1000</td><td></td><td></td><td></td><td></td><td></td><td></td><td></td><td></td><td></td></tr>
<tr><td>Oligo fwd</td><td>CACCTCGGATACATGGGCTTCTTC</td><td></td><td></td><td></td><td></td><td></td><td></td><td></td><td></td><td></td><td></td></tr>
<tr><td>Oligo rev</td><td>AAACGAAGAAGCCCATGTATCCGA</td><td></td><td></td><td></td><td></td><td></td><td></td><td></td><td></td><td></td><td></td></tr>
<tr><td>Chromosome</td><td>start</td><td>end</td><td>strand</td><td>MM</td><td>target_seq</td><td>PAM</td><td>alignment</td><td>distance</td><td>position</td><td>gene name</td><td>gene id</td></tr>
<tr><td>GU305914.1</td><td>86170</td><td>86192</td><td>-</td><td>0</td><td>TCGGATACATGGGCTTCTTC</td><td>AGG</td><td>||||||||[||||||||||||]PAM</td><td>NA</td><td>Intergenic</td><td>NA</td><td>NA</td></tr>
<tr><td>T27741</td><td>GCTCAACGACCTGCAGAAGATGG</td><td>1000</td><td></td><td></td><td></td><td></td><td></td><td></td><td></td><td></td><td></td></tr>
<tr><td>Oligo fwd</td><td>CACCGCTCAACGACCTGCAGAAGA</td><td></td><td></td><td></td><td></td><td></td><td></td><td></td><td></td><td></td><td></td></tr>
<tr><td>Oligo rev</td><td>AAACTCTTCTGCAGGTCGTTGAGC</td><td></td><td></td><td></td><td></td><td></td><td></td><td></td><td></td><td></td><td></td></tr>
<tr><td>Chromosome</td><td>start</td><td>end</td><td>strand</td><td>MM</td><td>target_seq</td><td>PAM</td><td>alignment</td><td>distance</td><td>position</td><td>gene name</td><td>gene id</td></tr>
<tr><td>GU305914.1</td><td>86130</td><td>86152</td><td>-</td><td>0</td><td>GCTCAACGACCTGCAGAAGA</td><td>TGG</td><td>||||||||[||||||||||||]PAM</td><td>NA</td><td>Intergenic</td><td>NA</td><td>NA</td></tr>
</table>

<table>
<tr><td>T27742</td><td>ACGACCTGCAGAAGATGGTCTGG</td><td>1000</td><td></td><td></td><td></td><td></td><td></td><td></td><td></td><td></td><td></td></tr>
<tr><td>Oligo fwd</td><td>CACCACGACCTGCAGAAGATGGTC</td><td></td><td></td><td></td><td></td><td></td><td></td><td></td><td></td><td></td><td></td></tr>
<tr><td>Oligo rev</td><td>AAACGACCATCTTCTGCAGGTCGT</td><td></td><td></td><td></td><td></td><td></td><td></td><td></td><td></td><td></td><td></td></tr>
<tr><td>Chromosome</td><td>start</td><td>end</td><td>strand</td><td>MM</td><td>target_seq</td><td>PAM</td><td>alignment</td><td>distance</td><td>position</td><td>gene name</td><td>gene id</td></tr>
<tr><td>GU305914.1</td><td>86125</td><td>86147</td><td>-</td><td>0</td><td>ACGACCTGCAGAAGATGGTC</td><td>TGG</td><td>||||||||[||||||||||||]PAM</td><td>NA</td><td>Intergenic</td><td>NA</td><td>NA</td></tr>
<tr><td>T27743</td><td>CGACCTGCAGAAGATGGTCTGGG</td><td>1000</td><td></td><td></td><td></td><td></td><td></td><td></td><td></td><td></td><td></td></tr>
<tr><td>Oligo fwd</td><td>CACCCGACCTGCAGAAGATGGTCT</td><td></td><td></td><td></td><td></td><td></td><td></td><td></td><td></td><td></td><td></td></tr>
<tr><td>Oligo rev</td><td>AAACAGACCATCTTCTGCAGGTCG</td><td></td><td></td><td></td><td></td><td></td><td></td><td></td><td></td><td></td><td></td></tr>
<tr><td>Chromosome</td><td>start</td><td>end</td><td>strand</td><td>MM</td><td>target_seq</td><td>PAM</td><td>alignment</td><td>distance</td><td>position</td><td>gene name</td><td>gene id</td></tr>
<tr><td>GU305914.1</td><td>86124</td><td>86146</td><td>-</td><td>0</td><td>CGACCTGCAGAAGATGGTCT</td><td>GGG</td><td>||||||||[||||||||||||]PAM</td><td>NA</td><td>Intergenic</td><td>NA</td><td>NA</td></tr>
<tr><td>T27744</td><td>GCACTATATAACGAGACGTTCGG</td><td>1000</td><td></td><td></td><td></td><td></td><td></td><td></td><td></td><td></td><td></td></tr>
<tr><td>Oligo fwd</td><td>CACCGCACTATATAACGAGACGTT</td><td></td><td></td><td></td><td></td><td></td><td></td><td></td><td></td><td></td><td></td></tr>
<tr><td>Oligo rev</td><td>AAACAACGTCTCGTTATATAGTGC</td><td></td><td></td><td></td><td></td><td></td><td></td><td></td><td></td><td></td><td></td></tr>
<tr><td>Chromosome</td><td>start</td><td>end</td><td>strand</td><td>MM</td><td>target_seq</td><td>PAM</td><td>alignment</td><td>distance</td><td>position</td><td>gene name</td><td>gene id</td></tr>
<tr><td>GU305914.1</td><td>86072</td><td>86094</td><td>-</td><td>0</td><td>GCACTATATAACGAGACGTT</td><td>CGG</td><td>||||||||[||||||||||||]PAM</td><td>NA</td><td>Intergenic</td><td>NA</td><td>NA</td></tr>
<tr><td>T27745</td><td>CGAGACGTTCGGCAAGCAGTTGG</td><td>1000</td><td></td><td></td><td></td><td></td><td></td><td></td><td></td><td></td><td></td></tr>
<tr><td>Oligo fwd</td><td>CACCCGAGACGTTCGGCAAGCAGT</td><td></td><td></td><td></td><td></td><td></td><td></td><td></td><td></td><td></td><td></td></tr>
<tr><td>Oligo rev</td><td>AAACACTGCTTGCCGAACGTCTCG</td><td></td><td></td><td></td><td></td><td></td><td></td><td></td><td></td><td></td><td></td></tr>
<tr><td>Chromosome</td><td>start</td><td>end</td><td>strand</td><td>MM</td><td>target_seq</td><td>PAM</td><td>alignment</td><td>distance</td><td>position</td><td>gene name</td><td>gene id</td></tr>
<tr><td>GU305914.1</td><td>86061</td><td>86083</td><td>-</td><td>0</td><td>CGAGACGTTCGGCAAGCAGT</td><td>TGG</td><td>||||||||[||||||||||||]PAM</td><td>NA</td><td>Intergenic</td><td>NA</td><td>NA</td></tr>
<tr><td>T27746</td><td>TCGGCAAGCAGTTGGCCTTGTGG</td><td>1000</td><td></td><td></td><td></td><td></td><td></td><td></td><td></td><td></td><td></td></tr>
<tr><td>Oligo fwd</td><td>CACCTCGGCAAGCAGTTGGCCTTG</td><td></td><td></td><td></td><td></td><td></td><td></td><td></td><td></td><td></td><td></td></tr>
<tr><td>Oligo rev</td><td>AAACCAAGGCCAACTGCTTGCCGA</td><td></td><td></td><td></td><td></td><td></td><td></td><td></td><td></td><td></td><td></td></tr>
<tr><td>Chromosome</td><td>start</td><td>end</td><td>strand</td><td>MM</td><td>target_seq</td><td>PAM</td><td>alignment</td><td>distance</td><td>position</td><td>gene name</td><td>gene id</td></tr>
<tr><td>GU305914.1</td><td>86053</td><td>86075</td><td>-</td><td>0</td><td>TCGGCAAGCAGTTGGCCTTG</td><td>TGG</td><td>||||||||[||||||||||||]PAM</td><td>NA</td><td>Intergenic</td><td>NA</td><td>NA</td></tr>
<tr><td>T27747</td><td>GTTGGCCTTGTGGCGCGTCGAGG</td><td>1000</td><td></td><td></td><td></td><td></td><td></td><td></td><td></td><td></td><td></td></tr>
<tr><td>Oligo fwd</td><td>CACCGTTGGCCTTGTGGCGCGTCG</td><td></td><td></td><td></td><td></td><td></td><td></td><td></td><td></td><td></td><td></td></tr>
<tr><td>Oligo rev</td><td>AAACCGACGCGCCACAAGGCCAAC</td><td></td><td></td><td></td><td></td><td></td><td></td><td></td><td></td><td></td><td></td></tr>
<tr><td>Chromosome</td><td>start</td><td>end</td><td>strand</td><td>MM</td><td>target_seq</td><td>PAM</td><td>alignment</td><td>distance</td><td>position</td><td>gene name</td><td>gene id</td></tr>
<tr><td>GU305914.1</td><td>86043</td><td>86065</td><td>-</td><td>0</td><td>GTTGGCCTTGTGGCGCGTCG</td><td>AGG</td><td>||||||||[||||||||||||]PAM</td><td>NA</td><td>Intergenic</td><td>NA</td><td>NA</td></tr>
<tr><td>T27748</td><td>GCCTTGTGGCGCGTCGAGGACGG</td><td>1000</td><td></td><td></td><td></td><td></td><td></td><td></td><td></td><td></td><td></td></tr>
<tr><td>Oligo fwd</td><td>CACCGCCTTGTGGCGCGTCGAGGA</td><td></td><td></td><td></td><td></td><td></td><td></td><td></td><td></td><td></td><td></td></tr>
<tr><td>Oligo rev</td><td>AAACTCCTCGACGCGCCACAAGGC</td><td></td><td></td><td></td><td></td><td></td><td></td><td></td><td></td><td></td><td></td></tr>
<tr><td>Chromosome</td><td>start</td><td>end</td><td>strand</td><td>MM</td><td>target_seq</td><td>PAM</td><td>alignment</td><td>distance</td><td>position</td><td>gene name</td><td>gene id</td></tr>
<tr><td>GU305914.1</td><td>86039</td><td>86061</td><td>-</td><td>0</td><td>GCCTTGTGGCGCGTCGAGGA</td><td>CGG</td><td>||||||||[||||||||||||]PAM</td><td>NA</td><td>Intergenic</td><td>NA</td><td>NA</td></tr>
<tr><td>T27749</td><td>GTCGAGGACGGCGATGAGATCGG</td><td>1000</td><td></td><td></td><td></td><td></td><td></td><td></td><td></td><td></td><td></td></tr>
<tr><td>Oligo fwd</td><td>CACCGTCGAGGACGGCGATGAGAT</td><td></td><td></td><td></td><td></td><td></td><td></td><td></td><td></td><td></td><td></td></tr>
<tr><td>Oligo rev</td><td>AAACATCTCATCGCCGTCCTCGAC</td><td></td><td></td><td></td><td></td><td></td><td></td><td></td><td></td><td></td><td></td></tr>
<tr><td>Chromosome</td><td>start</td><td>end</td><td>strand</td><td>MM</td><td>target_seq</td><td>PAM</td><td>alignment</td><td>distance</td><td>position</td><td>gene name</td><td>gene id</td></tr>
<tr><td>GU305914.1</td><td>86027</td><td>86049</td><td>-</td><td>0</td><td>GTCGAGGACGGCGATGAGAT</td><td>CGG</td><td>||||||||[||||||||||||]PAM</td><td>NA</td><td>Intergenic</td><td>NA</td><td>NA</td></tr>
<tr><td>GU305914.1</td><td>98586</td><td>98608</td><td>+</td><td>3</td><td>GGCGAGGAGGGCGATGAGAG</td><td>TGG</td><td>|-||||||[-||||||||||-]PAM</td><td>NA</td><td>Intergenic</td><td>NA</td><td>NA</td></tr>
<tr><td>T27750</td><td>GACGGCGATGAGATCGGCGACGG</td><td>1000</td><td></td><td></td><td></td><td></td><td></td><td></td><td></td><td></td><td></td></tr>
<tr><td>Oligo fwd</td><td>CACCGACGGCGATGAGATCGGCGA</td><td></td><td></td><td></td><td></td><td></td><td></td><td></td><td></td><td></td><td></td></tr>
<tr><td>Oligo rev</td><td>AAACTCGCCGATCTCATCGCCGTC</td><td></td><td></td><td></td><td></td><td></td><td></td><td></td><td></td><td></td><td></td></tr>
<tr><td>Chromosome</td><td>start</td><td>end</td><td>strand</td><td>MM</td><td>target_seq</td><td>PAM</td><td>alignment</td><td>distance</td><td>position</td><td>gene name</td><td>gene id</td></tr>
<tr><td>GU305914.1</td><td>86021</td><td>86043</td><td>-</td><td>0</td><td>GACGGCGATGAGATCGGCGA</td><td>CGG</td><td>||||||||[||||||||||||]PAM</td><td>NA</td><td>Intergenic</td><td>NA</td><td>NA</td></tr>
<tr><td>GU305914.1</td><td>27373</td><td>27395</td><td>+</td><td>4</td><td>GATTGCGAGGAGATCTGCGA</td><td>CGG</td><td>||--||||[-||||||-||||]PAM</td><td>NA</td><td>Intergenic</td><td>NA</td><td>NA</td></tr>
<tr><td>T27751</td><td>ACGGCGATGAGATCGGCGACGGG</td><td>1000</td><td></td><td></td><td></td><td></td><td></td><td></td><td></td><td></td><td></td></tr>
<tr><td>Oligo fwd</td><td>CACCACGGCGATGAGATCGGCGAC</td><td></td><td></td><td></td><td></td><td></td><td></td><td></td><td></td><td></td><td></td></tr>
<tr><td>Oligo rev</td><td>AAACGTCGCCGATCTCATCGCCGT</td><td></td><td></td><td></td><td></td><td></td><td></td><td></td><td></td><td></td><td></td></tr>
<tr><td>Chromosome</td><td>start</td><td>end</td><td>strand</td><td>MM</td><td>target_seq</td><td>PAM</td><td>alignment</td><td>distance</td><td>position</td><td>gene name</td><td>gene id</td></tr>
<tr><td>GU305914.1</td><td>86020</td><td>86042</td><td>-</td><td>0</td><td>ACGGCGATGAGATCGGCGAC</td><td>GGG</td><td>||||||||[||||||||||||]PAM</td><td>NA</td><td>Intergenic</td><td>NA</td><td>NA</td></tr>
<tr><td>GU305914.1</td><td>35672</td><td>35694</td><td>+</td><td>4</td><td>GCGGGGATGTCATCGGCGAC</td><td>AGG</td><td>-|||-|||[|--|||||||||]PAM</td><td>NA</td><td>Intergenic</td><td>NA</td><td>NA</td></tr>
<tr><td>GU305914.1</td><td>27374</td><td>27396</td><td>+</td><td>4</td><td>ATTGCGAGGAGATCTGCGAC</td><td>GGG</td><td>|--||||-[||||||-|||||]PAM</td><td>NA</td><td>Intergenic</td><td>NA</td><td>NA</td></tr>
<tr><td>T27752</td><td>TCTCCGTTGATCCTGCGTCGCGG</td><td>1000</td><td></td><td></td><td></td><td></td><td></td><td></td><td></td><td></td><td></td></tr>
<tr><td>Oligo fwd</td><td>CACCTCTCCGTTGATCCTGCGTCG</td><td></td><td></td><td></td><td></td><td></td><td></td><td></td><td></td><td></td><td></td></tr>
<tr><td>Oligo rev</td><td>AAACCGACGCAGGATCAACGGAGA</td><td></td><td></td><td></td><td></td><td></td><td></td><td></td><td></td><td></td><td></td></tr>
<tr><td>Chromosome</td><td>start</td><td>end</td><td>strand</td><td>MM</td><td>target_seq</td><td>PAM</td><td>alignment</td><td>distance</td><td>position</td><td>gene name</td><td>gene id</td></tr>
<tr><td>GU305914.1</td><td>85973</td><td>85995</td><td>-</td><td>0</td><td>TCTCCGTTGATCCTGCGTCG</td><td>CGG</td><td>||||||||[||||||||||||]PAM</td><td>NA</td><td>Intergenic</td><td>NA</td><td>NA</td></tr>
<tr><td>T27753</td><td>GAGATCGAAGTTACAGATCCAGG</td><td>1000</td><td></td><td></td><td></td><td></td><td></td><td></td><td></td><td></td><td></td></tr>
<tr><td>Oligo fwd</td><td>CACCGAGATCGAAGTTACAGATCC</td><td></td><td></td><td></td><td></td><td></td><td></td><td></td><td></td><td></td><td></td></tr>
<tr><td>Oligo rev</td><td>AAACGGATCTGTAACTTCGATCTC</td><td></td><td></td><td></td><td></td><td></td><td></td><td></td><td></td><td></td><td></td></tr>
<tr><td>Chromosome</td><td>start</td><td>end</td><td>strand</td><td>MM</td><td>target_seq</td><td>PAM</td><td>alignment</td><td>distance</td><td>position</td><td>gene name</td><td>gene id</td></tr>
<tr><td>GU305914.1</td><td>85951</td><td>85973</td><td>-</td><td>0</td><td>GAGATCGAAGTTACAGATCC</td><td>AGG</td><td>||||||||[||||||||||||]PAM</td><td>NA</td><td>Intergenic</td><td>NA</td><td>NA</td></tr>
<tr><td>T27754</td><td>AGATCGAAGTTACAGATCCAGGG</td><td>1000</td><td></td><td></td><td></td><td></td><td></td><td></td><td></td><td></td><td></td></tr>
<tr><td>Oligo fwd</td><td>CACCAGATCGAAGTTACAGATCCA</td><td></td><td></td><td></td><td></td><td></td><td></td><td></td><td></td><td></td><td></td></tr>
<tr><td>Oligo rev</td><td>AAACTGGATCTGTAACTTCGATCT</td><td></td><td></td><td></td><td></td><td></td><td></td><td></td><td></td><td></td><td></td></tr>
<tr><td>Chromosome</td><td>start</td><td>end</td><td>strand</td><td>MM</td><td>target_seq</td><td>PAM</td><td>alignment</td><td>distance</td><td>position</td><td>gene name</td><td>gene id</td></tr>
<tr><td>GU305914.1</td><td>85950</td><td>85972</td><td>-</td><td>0</td><td>AGATCGAAGTTACAGATCCA</td><td>GGG</td><td>||||||||[||||||||||||]PAM</td><td>NA</td><td>Intergenic</td><td>NA</td><td>NA</td></tr>
<tr><td>T27755</td><td>GCTGCTCTACAAGCATCTCTCGG</td><td>1000</td><td></td><td></td><td></td><td></td><td></td><td></td><td></td><td></td><td></td></tr>
<tr><td>Oligo fwd</td><td>CACCGCTGCTCTACAAGCATCTCT</td><td></td><td></td><td></td><td></td><td></td><td></td><td></td><td></td><td></td><td></td></tr>
<tr><td>Oligo rev</td><td>AAACAGAGATGCTTGTAGAGCAGC</td><td></td><td></td><td></td><td></td><td></td><td></td><td></td><td></td><td></td><td></td></tr>
<tr><td>Chromosome</td><td>start</td><td>end</td><td>strand</td><td>MM</td><td>target_seq</td><td>PAM</td><td>alignment</td><td>distance</td><td>position</td><td>gene name</td><td>gene id</td></tr>
<tr><td>GU305914.1</td><td>85917</td><td>85939</td><td>-</td><td>0</td><td>GCTGCTCTACAAGCATCTCT</td><td>CGG</td><td>||||||||[||||||||||||]PAM</td><td>NA</td><td>Intergenic</td><td>NA</td><td>NA</td></tr>
<tr><td>T27756</td><td>TCCGAGACTCTCGCCGACGCCGG</td><td>1000</td><td></td><td></td><td></td><td></td><td></td><td></td><td></td><td></td><td></td></tr>
</table>

<table>
<tr><td>Oligo fwd</td><td>CACCTCCGAGACTCTCGCCGACGC</td><td></td><td></td><td></td><td></td><td></td><td></td><td></td><td></td><td></td><td></td><td></td></tr>
<tr><td>Oligo rev</td><td>AAACGCGTCGGCGAGAGTCTCGGA</td><td></td><td></td><td></td><td></td><td></td><td></td><td></td><td></td><td></td><td></td><td></td></tr>
<tr><td>Chromosome</td><td>start</td><td></td><td>end</td><td>strand</td><td>MM</td><td>target_seq</td><td>PAM</td><td>alignment</td><td>distance</td><td>position</td><td>gene name</td><td>gene id</td></tr>
<tr><td>GU305914.1</td><td></td><td>85889</td><td>85911</td><td>-</td><td>0</td><td>TCCGAGACTCTCGCCGACGC</td><td>CGG</td><td>||||||||[||||||||||||]PAM</td><td>NA</td><td>Intergenic</td><td>NA</td><td>NA</td></tr>
<tr><td>T27757</td><td>CGACGCCGGTTCTCGCGCCTCGG</td><td></td><td>1000</td><td></td><td></td><td></td><td></td><td></td><td></td><td></td><td></td><td></td></tr>
<tr><td>Oligo fwd</td><td>CACCCGACGCCGGTTCTCGCGCCT</td><td></td><td></td><td></td><td></td><td></td><td></td><td></td><td></td><td></td><td></td><td></td></tr>
<tr><td>Oligo rev</td><td>AAACAGGCGCGAGAACCGGCGTCG</td><td></td><td></td><td></td><td></td><td></td><td></td><td></td><td></td><td></td><td></td><td></td></tr>
<tr><td>Chromosome</td><td>start</td><td></td><td>end</td><td>strand</td><td>MM</td><td>target_seq</td><td>PAM</td><td>alignment</td><td>distance</td><td>position</td><td>gene name</td><td>gene id</td></tr>
<tr><td>GU305914.1</td><td></td><td>85875</td><td>85897</td><td>-</td><td>0</td><td>CGACGCCGGTTCTCGCGCCT</td><td>CGG</td><td>||||||||[||||||||||||]PAM</td><td>NA</td><td>Intergenic</td><td>NA</td><td>NA</td></tr>
<tr><td>T27758</td><td>CTCGGTCGCCGACTTATGTCAGG</td><td></td><td>1000</td><td></td><td></td><td></td><td></td><td></td><td></td><td></td><td></td><td></td></tr>
<tr><td>Oligo fwd</td><td>CACCCTCGGTCGCCGACTTATGTC</td><td></td><td></td><td></td><td></td><td></td><td></td><td></td><td></td><td></td><td></td><td></td></tr>
<tr><td>Oligo rev</td><td>AAACGACATAAGTCGGCGACCGAG</td><td></td><td></td><td></td><td></td><td></td><td></td><td></td><td></td><td></td><td></td><td></td></tr>
<tr><td>Chromosome</td><td>start</td><td></td><td>end</td><td>strand</td><td>MM</td><td>target_seq</td><td>PAM</td><td>alignment</td><td>distance</td><td>position</td><td>gene name</td><td>gene id</td></tr>
<tr><td>GU305914.1</td><td></td><td>85857</td><td>85879</td><td>-</td><td>0</td><td>CTCGGTCGCCGACTTATGTC</td><td>AGG</td><td>||||||||[||||||||||||]PAM</td><td>NA</td><td>Intergenic</td><td>NA</td><td>NA</td></tr>
<tr><td>T27759</td><td>TTATGTCAGGTCGAGAGACCCGG</td><td></td><td>1000</td><td></td><td></td><td></td><td></td><td></td><td></td><td></td><td></td><td></td></tr>
<tr><td>Oligo fwd</td><td>CACCTTATGTCAGGTCGAGAGACC</td><td></td><td></td><td></td><td></td><td></td><td></td><td></td><td></td><td></td><td></td><td></td></tr>
<tr><td>Oligo rev</td><td>AAACGGTCTCTCGACCTGACATAA</td><td></td><td></td><td></td><td></td><td></td><td></td><td></td><td></td><td></td><td></td><td></td></tr>
<tr><td>Chromosome</td><td>start</td><td></td><td>end</td><td>strand</td><td>MM</td><td>target_seq</td><td>PAM</td><td>alignment</td><td>distance</td><td>position</td><td>gene name</td><td>gene id</td></tr>
<tr><td>GU305914.1</td><td></td><td>85844</td><td>85866</td><td>-</td><td>0</td><td>TTATGTCAGGTCGAGAGACC</td><td>CGG</td><td>||||||||[||||||||||||]PAM</td><td>NA</td><td>Intergenic</td><td>NA</td><td>NA</td></tr>
<tr><td>T27760</td><td>TATGTCAGGTCGAGAGACCCGGG</td><td></td><td>1000</td><td></td><td></td><td></td><td></td><td></td><td></td><td></td><td></td><td></td></tr>
<tr><td>Oligo fwd</td><td>CACCTATGTCAGGTCGAGAGACCC</td><td></td><td></td><td></td><td></td><td></td><td></td><td></td><td></td><td></td><td></td><td></td></tr>
<tr><td>Oligo rev</td><td>AAACGGGTCTCTCGACCTGACATA</td><td></td><td></td><td></td><td></td><td></td><td></td><td></td><td></td><td></td><td></td><td></td></tr>
<tr><td>Chromosome</td><td>start</td><td></td><td>end</td><td>strand</td><td>MM</td><td>target_seq</td><td>PAM</td><td>alignment</td><td>distance</td><td>position</td><td>gene name</td><td>gene id</td></tr>
<tr><td>GU305914.1</td><td></td><td>85843</td><td>85865</td><td>-</td><td>0</td><td>TATGTCAGGTCGAGAGACCC</td><td>GGG</td><td>||||||||[||||||||||||]PAM</td><td>NA</td><td>Intergenic</td><td>NA</td><td>NA</td></tr>
<tr><td>T27761</td><td>ATGTCAAGAAGAAACGAAAGAGG</td><td></td><td>1000</td><td></td><td></td><td></td><td></td><td></td><td></td><td></td><td></td><td></td></tr>
<tr><td>Oligo fwd</td><td>CACCATGTCAAGAAGAAACGAAAG</td><td></td><td></td><td></td><td></td><td></td><td></td><td></td><td></td><td></td><td></td><td></td></tr>
<tr><td>Oligo rev</td><td>AAACCTTTCGTTTCTTCTTGACAT</td><td></td><td></td><td></td><td></td><td></td><td></td><td></td><td></td><td></td><td></td><td></td></tr>
<tr><td>Chromosome</td><td>start</td><td></td><td>end</td><td>strand</td><td>MM</td><td>target_seq</td><td>PAM</td><td>alignment</td><td>distance</td><td>position</td><td>gene name</td><td>gene id</td></tr>
<tr><td>GU305914.1</td><td></td><td>85789</td><td>85811</td><td>-</td><td>0</td><td>ATGTCAAGAAGAAACGAAAG</td><td>AGG</td><td>||||||||[||||||||||||]PAM</td><td>NA</td><td>Intergenic</td><td>NA</td><td>NA</td></tr>
<tr><td>T27762</td><td>GAGGATGTCGCTCCTCGAGCTGG</td><td></td><td>1000</td><td></td><td></td><td></td><td></td><td></td><td></td><td></td><td></td><td></td></tr>
<tr><td>Oligo fwd</td><td>CACCGAGGATGTCGCTCCTCGAGC</td><td></td><td></td><td></td><td></td><td></td><td></td><td></td><td></td><td></td><td></td><td></td></tr>
<tr><td>Oligo rev</td><td>AAACGCTCGAGGAGCGACATCCTC</td><td></td><td></td><td></td><td></td><td></td><td></td><td></td><td></td><td></td><td></td><td></td></tr>
<tr><td>Chromosome</td><td>start</td><td></td><td>end</td><td>strand</td><td>MM</td><td>target_seq</td><td>PAM</td><td>alignment</td><td>distance</td><td>position</td><td>gene name</td><td>gene id</td></tr>
<tr><td>GU305914.1</td><td></td><td>85770</td><td>85792</td><td>-</td><td>0</td><td>GAGGATGTCGCTCCTCGAGC</td><td>TGG</td><td>||||||||[||||||||||||]PAM</td><td>NA</td><td>Intergenic</td><td>NA</td><td>NA</td></tr>
<tr><td>T27763</td><td>GAGCTGGTACGCGATGTCGACGG</td><td></td><td>1000</td><td></td><td></td><td></td><td></td><td></td><td></td><td></td><td></td><td></td></tr>
<tr><td>Oligo fwd</td><td>CACCGAGCTGGTACGCGATGTCGA</td><td></td><td></td><td></td><td></td><td></td><td></td><td></td><td></td><td></td><td></td><td></td></tr>
<tr><td>Oligo rev</td><td>AAACTCGACATCGCGTACCAGCTC</td><td></td><td></td><td></td><td></td><td></td><td></td><td></td><td></td><td></td><td></td><td></td></tr>
<tr><td>Chromosome</td><td>start</td><td></td><td>end</td><td>strand</td><td>MM</td><td>target_seq</td><td>PAM</td><td>alignment</td><td>distance</td><td>position</td><td>gene name</td><td>gene id</td></tr>
<tr><td>GU305914.1</td><td></td><td>85754</td><td>85776</td><td>-</td><td>0</td><td>GAGCTGGTACGCGATGTCGA</td><td>CGG</td><td>||||||||[||||||||||||]PAM</td><td>NA</td><td>Intergenic</td><td>NA</td><td>NA</td></tr>
<tr><td>T27764</td><td>GTACGCGATGTCGACGGCGCTGG</td><td></td><td>1000</td><td></td><td></td><td></td><td></td><td></td><td></td><td></td><td></td><td></td></tr>
<tr><td>Oligo fwd</td><td>CACCGTACGCGATGTCGACGGCGC</td><td></td><td></td><td></td><td></td><td></td><td></td><td></td><td></td><td></td><td></td><td></td></tr>
<tr><td>Oligo rev</td><td>AAACGCGCCGTCGACATCGCGTAC</td><td></td><td></td><td></td><td></td><td></td><td></td><td></td><td></td><td></td><td></td><td></td></tr>
<tr><td>Chromosome</td><td>start</td><td></td><td>end</td><td>strand</td><td>MM</td><td>target_seq</td><td>PAM</td><td>alignment</td><td>distance</td><td>position</td><td>gene name</td><td>gene id</td></tr>
<tr><td>GU305914.1</td><td></td><td>85748</td><td>85770</td><td>-</td><td>0</td><td>GTACGCGATGTCGACGGCGC</td><td>TGG</td><td>||||||||[||||||||||||]PAM</td><td>NA</td><td>Intergenic</td><td>NA</td><td>NA</td></tr>
<tr><td>T27765</td><td>CGCGATGTCGACGGCGCTGGCGG</td><td></td><td>1000</td><td></td><td></td><td></td><td></td><td></td><td></td><td></td><td></td><td></td></tr>
<tr><td>Oligo fwd</td><td>CACCCGCGATGTCGACGGCGCTGG</td><td></td><td></td><td></td><td></td><td></td><td></td><td></td><td></td><td></td><td></td><td></td></tr>
<tr><td>Oligo rev</td><td>AAACCCAGCGCCGTCGACATCGCG</td><td></td><td></td><td></td><td></td><td></td><td></td><td></td><td></td><td></td><td></td><td></td></tr>
<tr><td>Chromosome</td><td>start</td><td></td><td>end</td><td>strand</td><td>MM</td><td>target_seq</td><td>PAM</td><td>alignment</td><td>distance</td><td>position</td><td>gene name</td><td>gene id</td></tr>
<tr><td>GU305914.1</td><td></td><td>85745</td><td>85767</td><td>-</td><td>0</td><td>CGCGATGTCGACGGCGCTGG</td><td>CGG</td><td>||||||||[||||||||||||]PAM</td><td>NA</td><td>Intergenic</td><td>NA</td><td>NA</td></tr>
<tr><td>T27766</td><td>CGGCGCTGGCGGTGACGATCTGG</td><td></td><td>1000</td><td></td><td></td><td></td><td></td><td></td><td></td><td></td><td></td><td></td></tr>
<tr><td>Oligo fwd</td><td>CACCCGGCGCTGGCGGTGACGATC</td><td></td><td></td><td></td><td></td><td></td><td></td><td></td><td></td><td></td><td></td><td></td></tr>
<tr><td>Oligo rev</td><td>AAACGATCGTCACCGCCAGCGCCG</td><td></td><td></td><td></td><td></td><td></td><td></td><td></td><td></td><td></td><td></td><td></td></tr>
<tr><td>Chromosome</td><td>start</td><td></td><td>end</td><td>strand</td><td>MM</td><td>target_seq</td><td>PAM</td><td>alignment</td><td>distance</td><td>position</td><td>gene name</td><td>gene id</td></tr>
<tr><td>GU305914.1</td><td></td><td>85734</td><td>85756</td><td>-</td><td>0</td><td>CGGCGCTGGCGGTGACGATC</td><td>TGG</td><td>||||||||[||||||||||||]PAM</td><td>NA</td><td>Intergenic</td><td>NA</td><td>NA</td></tr>
<tr><td>T27767</td><td>CCTCCTTGCCTGTACAAGTGAGG</td><td></td><td>1000</td><td></td><td></td><td></td><td></td><td></td><td></td><td></td><td></td><td></td></tr>
<tr><td>Oligo fwd</td><td>CACCCCTCCTTGCCTGTACAAGTG</td><td></td><td></td><td></td><td></td><td></td><td></td><td></td><td></td><td></td><td></td><td></td></tr>
<tr><td>Oligo rev</td><td>AAACCACTTGTACAGGCAAGGAGG</td><td></td><td></td><td></td><td></td><td></td><td></td><td></td><td></td><td></td><td></td><td></td></tr>
<tr><td>Chromosome</td><td>start</td><td></td><td>end</td><td>strand</td><td>MM</td><td>target_seq</td><td>PAM</td><td>alignment</td><td>distance</td><td>position</td><td>gene name</td><td>gene id</td></tr>
<tr><td>GU305914.1</td><td></td><td>85709</td><td>85731</td><td>-</td><td>0</td><td>CCTCCTTGCCTGTACAAGTG</td><td>AGG</td><td>||||||||[||||||||||||]PAM</td><td>NA</td><td>Intergenic</td><td>NA</td><td>NA</td></tr>
<tr><td>T27768</td><td>CTGTACAAGTGAGGTAATCGCGG</td><td></td><td>1000</td><td></td><td></td><td></td><td></td><td></td><td></td><td></td><td></td><td></td></tr>
<tr><td>Oligo fwd</td><td>CACCCTGTACAAGTGAGGTAATCG</td><td></td><td></td><td></td><td></td><td></td><td></td><td></td><td></td><td></td><td></td><td></td></tr>
<tr><td>Oligo rev</td><td>AAACCGATTACCTCACTTGTACAG</td><td></td><td></td><td></td><td></td><td></td><td></td><td></td><td></td><td></td><td></td><td></td></tr>
<tr><td>Chromosome</td><td>start</td><td></td><td>end</td><td>strand</td><td>MM</td><td>target_seq</td><td>PAM</td><td>alignment</td><td>distance</td><td>position</td><td>gene name</td><td>gene id</td></tr>
<tr><td>GU305914.1</td><td></td><td>85700</td><td>85722</td><td>-</td><td>0</td><td>CTGTACAAGTGAGGTAATCG</td><td>CGG</td><td>||||||||[||||||||||||]PAM</td><td>NA</td><td>Intergenic</td><td>NA</td><td>NA</td></tr>
<tr><td>T27769</td><td>TACAAGTGAGGTAATCGCGGCGG</td><td></td><td>1000</td><td></td><td></td><td></td><td></td><td></td><td></td><td></td><td></td><td></td></tr>
<tr><td>Oligo fwd</td><td>CACCTACAAGTGAGGTAATCGCGG</td><td></td><td></td><td></td><td></td><td></td><td></td><td></td><td></td><td></td><td></td><td></td></tr>
<tr><td>Oligo rev</td><td>AAACCCGCGATTACCTCACTTGTA</td><td></td><td></td><td></td><td></td><td></td><td></td><td></td><td></td><td></td><td></td><td></td></tr>
<tr><td>Chromosome</td><td>start</td><td></td><td>end</td><td>strand</td><td>MM</td><td>target_seq</td><td>PAM</td><td>alignment</td><td>distance</td><td>position</td><td>gene name</td><td>gene id</td></tr>
<tr><td>GU305914.1</td><td></td><td>85697</td><td>85719</td><td>-</td><td>0</td><td>TACAAGTGAGGTAATCGCGG</td><td>CGG</td><td>||||||||[||||||||||||]PAM</td><td>NA</td><td>Intergenic</td><td>NA</td><td>NA</td></tr>
<tr><td>T27770</td><td>ATCGCGGCGGCGAGTACACCAGG</td><td></td><td>1000</td><td></td><td></td><td></td><td></td><td></td><td></td><td></td><td></td><td></td></tr>
<tr><td>Oligo fwd</td><td>CACCATCGCGGCGGCGAGTACACC</td><td></td><td></td><td></td><td></td><td></td><td></td><td></td><td></td><td></td><td></td><td></td></tr>
<tr><td>Oligo rev</td><td>AAACGGTGTACTCGCCGCCGCGAT</td><td></td><td></td><td></td><td></td><td></td><td></td><td></td><td></td><td></td><td></td><td></td></tr>
<tr><td>Chromosome</td><td>start</td><td></td><td>end</td><td>strand</td><td>MM</td><td>target_seq</td><td>PAM</td><td>alignment</td><td>distance</td><td>position</td><td>gene name</td><td>gene id</td></tr>
<tr><td>GU305914.1</td><td></td><td>85684</td><td>85706</td><td>-</td><td>0</td><td>ATCGCGGCGGCGAGTACACC</td><td>AGG</td><td>||||||||[||||||||||||]PAM</td><td>NA</td><td>Intergenic</td><td>NA</td><td>NA</td></tr>
</table>

<table>
<tr><td>T27771</td><td>CGCGTCCGCAGAAACAGAGACGG</td><td>1000</td><td></td><td></td><td></td><td></td><td></td><td></td><td></td><td></td><td></td></tr>
<tr><td>Oligo fwd</td><td>CACCCGCGTCCGCAGAAACAGAGA</td><td></td><td></td><td></td><td></td><td></td><td></td><td></td><td></td><td></td><td></td></tr>
<tr><td>Oligo rev</td><td>AAACTCTCTGTTTCTGCGGACGCG</td><td></td><td></td><td></td><td></td><td></td><td></td><td></td><td></td><td></td><td></td></tr>
<tr><td>Chromosome</td><td>start</td><td>end</td><td>strand</td><td>MM</td><td>target_seq</td><td>PAM</td><td>alignment</td><td>distance</td><td>position</td><td>gene name</td><td>gene id</td></tr>
<tr><td>GU305914.1</td><td>85622</td><td>85644</td><td>-</td><td>0</td><td>CGCGTCCGCAGAAACAGAGA</td><td>CGG</td><td>||||||||[||||||||||||]PAM</td><td>NA</td><td>Intergenic</td><td>NA</td><td>NA</td></tr>
<tr><td>GU305914.1</td><td>122038</td><td>122060</td><td>-</td><td>4</td><td>TCCGTCCGCAGAAGGAGAGA</td><td>GGG</td><td>--||||||[|||||--|||||]PAM</td><td>NA</td><td>Intergenic</td><td>NA</td><td>NA</td></tr>
<tr><td></td><td></td><td></td><td></td><td></td><td></td><td></td><td></td><td></td><td></td><td></td><td></td></tr>
<tr><td>T27772</td><td>AGAAACAGAGACGGCGACTTCGG</td><td>1000</td><td></td><td></td><td></td><td></td><td></td><td></td><td></td><td></td><td></td></tr>
<tr><td>Oligo fwd</td><td>CACCAGAAACAGAGACGGCGACTT</td><td></td><td></td><td></td><td></td><td></td><td></td><td></td><td></td><td></td><td></td></tr>
<tr><td>Oligo rev</td><td>AAACAAGTCGCCGTCTCTGTTTCT</td><td></td><td></td><td></td><td></td><td></td><td></td><td></td><td></td><td></td><td></td></tr>
<tr><td>Chromosome</td><td>start</td><td>end</td><td>strand</td><td>MM</td><td>target_seq</td><td>PAM</td><td>alignment</td><td>distance</td><td>position</td><td>gene name</td><td>gene id</td></tr>
<tr><td>GU305914.1</td><td>85613</td><td>85635</td><td>-</td><td>0</td><td>AGAAACAGAGACGGCGACTT</td><td>CGG</td><td>||||||||[||||||||||||]PAM</td><td>NA</td><td>Intergenic</td><td>NA</td><td>NA</td></tr>
<tr><td></td><td></td><td></td><td></td><td></td><td></td><td></td><td></td><td></td><td></td><td></td><td></td></tr>
<tr><td>T27773</td><td>GAAACAGAGACGGCGACTTCGGG</td><td>1000</td><td></td><td></td><td></td><td></td><td></td><td></td><td></td><td></td><td></td></tr>
<tr><td>Oligo fwd</td><td>CACCGAAACAGAGACGGCGACTTC</td><td></td><td></td><td></td><td></td><td></td><td></td><td></td><td></td><td></td><td></td></tr>
<tr><td>Oligo rev</td><td>AAACGAAGTCGCCGTCTCTGTTTC</td><td></td><td></td><td></td><td></td><td></td><td></td><td></td><td></td><td></td><td></td></tr>
<tr><td>Chromosome</td><td>start</td><td>end</td><td>strand</td><td>MM</td><td>target_seq</td><td>PAM</td><td>alignment</td><td>distance</td><td>position</td><td>gene name</td><td>gene id</td></tr>
<tr><td>GU305914.1</td><td>85612</td><td>85634</td><td>-</td><td>0</td><td>GAAACAGAGACGGCGACTTC</td><td>GGG</td><td>||||||||[||||||||||||]PAM</td><td>NA</td><td>Intergenic</td><td>NA</td><td>NA</td></tr>
<tr><td></td><td></td><td></td><td></td><td></td><td></td><td></td><td></td><td></td><td></td><td></td><td></td></tr>
<tr><td>T27774</td><td>AGAGACGGCGACTTCGGGCGCGG</td><td>1000</td><td></td><td></td><td></td><td></td><td></td><td></td><td></td><td></td><td></td></tr>
<tr><td>Oligo fwd</td><td>CACCAGAGACGGCGACTTCGGGCG</td><td></td><td></td><td></td><td></td><td></td><td></td><td></td><td></td><td></td><td></td></tr>
<tr><td>Oligo rev</td><td>AAACCGCCCGAAGTCGCCGTCTCT</td><td></td><td></td><td></td><td></td><td></td><td></td><td></td><td></td><td></td><td></td></tr>
<tr><td>Chromosome</td><td>start</td><td>end</td><td>strand</td><td>MM</td><td>target_seq</td><td>PAM</td><td>alignment</td><td>distance</td><td>position</td><td>gene name</td><td>gene id</td></tr>
<tr><td>GU305914.1</td><td>85607</td><td>85629</td><td>-</td><td>0</td><td>AGAGACGGCGACTTCGGGCG</td><td>CGG</td><td>||||||||[||||||||||||]PAM</td><td>NA</td><td>Intergenic</td><td>NA</td><td>NA</td></tr>
<tr><td></td><td></td><td></td><td></td><td></td><td></td><td></td><td></td><td></td><td></td><td></td><td></td></tr>
<tr><td>T27775</td><td>GCGACTTCGGGCGCGGCGACCGG</td><td>1000</td><td></td><td></td><td></td><td></td><td></td><td></td><td></td><td></td><td></td></tr>
<tr><td>Oligo fwd</td><td>CACCGCGACTTCGGGCGCGGCGAC</td><td></td><td></td><td></td><td></td><td></td><td></td><td></td><td></td><td></td><td></td></tr>
<tr><td>Oligo rev</td><td>AAACGTCGCCGCGCCCGAAGTCGC</td><td></td><td></td><td></td><td></td><td></td><td></td><td></td><td></td><td></td><td></td></tr>
<tr><td>Chromosome</td><td>start</td><td>end</td><td>strand</td><td>MM</td><td>target_seq</td><td>PAM</td><td>alignment</td><td>distance</td><td>position</td><td>gene name</td><td>gene id</td></tr>
<tr><td>GU305914.1</td><td>85600</td><td>85622</td><td>-</td><td>0</td><td>GCGACTTCGGGCGCGGCGAC</td><td>CGG</td><td>||||||||[||||||||||||]PAM</td><td>NA</td><td>Intergenic</td><td>NA</td><td>NA</td></tr>
<tr><td></td><td></td><td></td><td></td><td></td><td></td><td></td><td></td><td></td><td></td><td></td><td></td></tr>
<tr><td>T27776</td><td>GGGCGCGGCGACCGGAAAGAAGG</td><td>1000</td><td></td><td></td><td></td><td></td><td></td><td></td><td></td><td></td><td></td></tr>
<tr><td>Oligo fwd</td><td>CACCGGGCGCGGCGACCGGAAAGA</td><td></td><td></td><td></td><td></td><td></td><td></td><td></td><td></td><td></td><td></td></tr>
<tr><td>Oligo rev</td><td>AAACTCTTTCCGGTCGCCGCGCCC</td><td></td><td></td><td></td><td></td><td></td><td></td><td></td><td></td><td></td><td></td></tr>
<tr><td>Chromosome</td><td>start</td><td>end</td><td>strand</td><td>MM</td><td>target_seq</td><td>PAM</td><td>alignment</td><td>distance</td><td>position</td><td>gene name</td><td>gene id</td></tr>
<tr><td>GU305914.1</td><td>85592</td><td>85614</td><td>-</td><td>0</td><td>GGGCGCGGCGACCGGAAAGA</td><td>AGG</td><td>||||||||[||||||||||||]PAM</td><td>NA</td><td>Intergenic</td><td>NA</td><td>NA</td></tr>
<tr><td></td><td></td><td></td><td></td><td></td><td></td><td></td><td></td><td></td><td></td><td></td><td></td></tr>
<tr><td>T27777</td><td>GAAGGAAGCCACGCCGACTCAGG</td><td>1000</td><td></td><td></td><td></td><td></td><td></td><td></td><td></td><td></td><td></td></tr>
<tr><td>Oligo fwd</td><td>CACCGAAGGAAGCCACGCCGACTC</td><td></td><td></td><td></td><td></td><td></td><td></td><td></td><td></td><td></td><td></td></tr>
<tr><td>Oligo rev</td><td>AAACGAGTCGGCGTGGCTTCCTTC</td><td></td><td></td><td></td><td></td><td></td><td></td><td></td><td></td><td></td><td></td></tr>
<tr><td>Chromosome</td><td>start</td><td>end</td><td>strand</td><td>MM</td><td>target_seq</td><td>PAM</td><td>alignment</td><td>distance</td><td>position</td><td>gene name</td><td>gene id</td></tr>
<tr><td>GU305914.1</td><td>85574</td><td>85596</td><td>-</td><td>0</td><td>GAAGGAAGCCACGCCGACTC</td><td>AGG</td><td>||||||||[||||||||||||]PAM</td><td>NA</td><td>Intergenic</td><td>NA</td><td>NA</td></tr>
<tr><td></td><td></td><td></td><td></td><td></td><td></td><td></td><td></td><td></td><td></td><td></td><td></td></tr>
<tr><td>T27778</td><td>GACTCAGGCCTCGAAGATAACGG</td><td>1000</td><td></td><td></td><td></td><td></td><td></td><td></td><td></td><td></td><td></td></tr>
<tr><td>Oligo fwd</td><td>CACCGACTCAGGCCTCGAAGATAA</td><td></td><td></td><td></td><td></td><td></td><td></td><td></td><td></td><td></td><td></td></tr>
<tr><td>Oligo rev</td><td>AAACTTATCTTCGAGGCCTGAGTC</td><td></td><td></td><td></td><td></td><td></td><td></td><td></td><td></td><td></td><td></td></tr>
<tr><td>Chromosome</td><td>start</td><td>end</td><td>strand</td><td>MM</td><td>target_seq</td><td>PAM</td><td>alignment</td><td>distance</td><td>position</td><td>gene name</td><td>gene id</td></tr>
<tr><td>GU305914.1</td><td>85559</td><td>85581</td><td>-</td><td>0</td><td>GACTCAGGCCTCGAAGATAA</td><td>CGG</td><td>||||||||[||||||||||||]PAM</td><td>NA</td><td>Intergenic</td><td>NA</td><td>NA</td></tr>
<tr><td></td><td></td><td></td><td></td><td></td><td></td><td></td><td></td><td></td><td></td><td></td><td></td></tr>
<tr><td>T27779</td><td>ACTCAGGCCTCGAAGATAACGGG</td><td>1000</td><td></td><td></td><td></td><td></td><td></td><td></td><td></td><td></td><td></td></tr>
<tr><td>Oligo fwd</td><td>CACCACTCAGGCCTCGAAGATAAC</td><td></td><td></td><td></td><td></td><td></td><td></td><td></td><td></td><td></td><td></td></tr>
<tr><td>Oligo rev</td><td>AAACGTTATCTTCGAGGCCTGAGT</td><td></td><td></td><td></td><td></td><td></td><td></td><td></td><td></td><td></td><td></td></tr>
<tr><td>Chromosome</td><td>start</td><td>end</td><td>strand</td><td>MM</td><td>target_seq</td><td>PAM</td><td>alignment</td><td>distance</td><td>position</td><td>gene name</td><td>gene id</td></tr>
<tr><td>GU305914.1</td><td>85558</td><td>85580</td><td>-</td><td>0</td><td>ACTCAGGCCTCGAAGATAAC</td><td>GGG</td><td>||||||||[||||||||||||]PAM</td><td>NA</td><td>Intergenic</td><td>NA</td><td>NA</td></tr>
<tr><td></td><td></td><td></td><td></td><td></td><td></td><td></td><td></td><td></td><td></td><td></td><td></td></tr>
<tr><td>T27780</td><td>CTCAGGCCTCGAAGATAACGGGG</td><td>1000</td><td></td><td></td><td></td><td></td><td></td><td></td><td></td><td></td><td></td></tr>
<tr><td>Oligo fwd</td><td>CACCCTCAGGCCTCGAAGATAACG</td><td></td><td></td><td></td><td></td><td></td><td></td><td></td><td></td><td></td><td></td></tr>
<tr><td>Oligo rev</td><td>AAACCGTTATCTTCGAGGCCTGAG</td><td></td><td></td><td></td><td></td><td></td><td></td><td></td><td></td><td></td><td></td></tr>
<tr><td>Chromosome</td><td>start</td><td>end</td><td>strand</td><td>MM</td><td>target_seq</td><td>PAM</td><td>alignment</td><td>distance</td><td>position</td><td>gene name</td><td>gene id</td></tr>
<tr><td>GU305914.1</td><td>85557</td><td>85579</td><td>-</td><td>0</td><td>CTCAGGCCTCGAAGATAACG</td><td>GGG</td><td>||||||||[||||||||||||]PAM</td><td>NA</td><td>Intergenic</td><td>NA</td><td>NA</td></tr>
<tr><td></td><td></td><td></td><td></td><td></td><td></td><td></td><td></td><td></td><td></td><td></td><td></td></tr>
<tr><td>T27781</td><td>CGTTCCCTTCGTAAACGAGACGG</td><td>1000</td><td></td><td></td><td></td><td></td><td></td><td></td><td></td><td></td><td></td></tr>
<tr><td>Oligo fwd</td><td>CACCCGTTCCCTTCGTAAACGAGA</td><td></td><td></td><td></td><td></td><td></td><td></td><td></td><td></td><td></td><td></td></tr>
<tr><td>Oligo rev</td><td>AAACTCTCGTTTACGAAGGGAACG</td><td></td><td></td><td></td><td></td><td></td><td></td><td></td><td></td><td></td><td></td></tr>
<tr><td>Chromosome</td><td>start</td><td>end</td><td>strand</td><td>MM</td><td>target_seq</td><td>PAM</td><td>alignment</td><td>distance</td><td>position</td><td>gene name</td><td>gene id</td></tr>
<tr><td>GU305914.1</td><td>85526</td><td>85548</td><td>-</td><td>0</td><td>CGTTCCCTTCGTAAACGAGA</td><td>CGG</td><td>||||||||[||||||||||||]PAM</td><td>NA</td><td>Intergenic</td><td>NA</td><td>NA</td></tr>
<tr><td></td><td></td><td></td><td></td><td></td><td></td><td></td><td></td><td></td><td></td><td></td><td></td></tr>
<tr><td>T27782</td><td>TCCCTTCGTAAACGAGACGGAGG</td><td>1000</td><td></td><td></td><td></td><td></td><td></td><td></td><td></td><td></td><td></td></tr>
<tr><td>Oligo fwd</td><td>CACCTCCCTTCGTAAACGAGACGG</td><td></td><td></td><td></td><td></td><td></td><td></td><td></td><td></td><td></td><td></td></tr>
<tr><td>Oligo rev</td><td>AAACCCGTCTCGTTTACGAAGGGA</td><td></td><td></td><td></td><td></td><td></td><td></td><td></td><td></td><td></td><td></td></tr>
<tr><td>Chromosome</td><td>start</td><td>end</td><td>strand</td><td>MM</td><td>target_seq</td><td>PAM</td><td>alignment</td><td>distance</td><td>position</td><td>gene name</td><td>gene id</td></tr>
<tr><td>GU305914.1</td><td>85523</td><td>85545</td><td>-</td><td>0</td><td>TCCCTTCGTAAACGAGACGG</td><td>AGG</td><td>||||||||[||||||||||||]PAM</td><td>NA</td><td>Intergenic</td><td>NA</td><td>NA</td></tr>
<tr><td></td><td></td><td></td><td></td><td></td><td></td><td></td><td></td><td></td><td></td><td></td><td></td></tr>
<tr><td>T27783</td><td>CGTAAACGAGACGGAGGACATGG</td><td>1000</td><td></td><td></td><td></td><td></td><td></td><td></td><td></td><td></td><td></td></tr>
<tr><td>Oligo fwd</td><td>CACCCGTAAACGAGACGGAGGACA</td><td></td><td></td><td></td><td></td><td></td><td></td><td></td><td></td><td></td><td></td></tr>
<tr><td>Oligo rev</td><td>AAACTGTCCTCCGTCTCGTTTACG</td><td></td><td></td><td></td><td></td><td></td><td></td><td></td><td></td><td></td><td></td></tr>
<tr><td>Chromosome</td><td>start</td><td>end</td><td>strand</td><td>MM</td><td>target_seq</td><td>PAM</td><td>alignment</td><td>distance</td><td>position</td><td>gene name</td><td>gene id</td></tr>
<tr><td>GU305914.1</td><td>85517</td><td>85539</td><td>-</td><td>0</td><td>CGTAAACGAGACGGAGGACA</td><td>TGG</td><td>||||||||[||||||||||||]PAM</td><td>NA</td><td>Intergenic</td><td>NA</td><td>NA</td></tr>
<tr><td></td><td></td><td></td><td></td><td></td><td></td><td></td><td></td><td></td><td></td><td></td><td></td></tr>
<tr><td>T27784</td><td>ATATCCTTACAGAGTGTGCATGG</td><td>1000</td><td></td><td></td><td></td><td></td><td></td><td></td><td></td><td></td><td></td></tr>
<tr><td>Oligo fwd</td><td>CACCATATCCTTACAGAGTGTGCA</td><td></td><td></td><td></td><td></td><td></td><td></td><td></td><td></td><td></td><td></td></tr>
<tr><td>Oligo rev</td><td>AAACTGCACACTCTGTAAGGATAT</td><td></td><td></td><td></td><td></td><td></td><td></td><td></td><td></td><td></td><td></td></tr>
<tr><td>Chromosome</td><td>start</td><td>end</td><td>strand</td><td>MM</td><td>target_seq</td><td>PAM</td><td>alignment</td><td>distance</td><td>position</td><td>gene name</td><td>gene id</td></tr>
<tr><td>GU305914.1</td><td>85475</td><td>85497</td><td>-</td><td>0</td><td>ATATCCTTACAGAGTGTGCA</td><td>TGG</td><td>||||||||[||||||||||||]PAM</td><td>NA</td><td>Intergenic</td><td>NA</td><td>NA</td></tr>
<tr><td></td><td></td><td></td><td></td><td></td><td></td><td></td><td></td><td></td><td></td><td></td><td></td></tr>
<tr><td>T27785</td><td>CTGTGTCTACTGATCTCGTTCGG</td><td>1000</td><td></td><td></td><td></td><td></td><td></td><td></td><td></td><td></td><td></td></tr>
<tr><td>Oligo fwd</td><td>CACCCTGTGTCTACTGATCTCGTT</td><td></td><td></td><td></td><td></td><td></td><td></td><td></td><td></td><td></td><td></td></tr>
<tr><td>Oligo rev</td><td>AAACAACGAGATCAGTAGACACAG</td><td></td><td></td><td></td><td></td><td></td><td></td><td></td><td></td><td></td><td></td></tr>
<tr><td>Chromosome</td><td>start</td><td>end</td><td>strand</td><td>MM</td><td>target_seq</td><td>PAM</td><td>alignment</td><td>distance</td><td>position</td><td>gene name</td><td>gene id</td></tr>
</table>

<table>
<tr><td>GU305914.1</td><td></td><td>85452</td><td>85474</td><td>-</td><td>0</td><td>CTGTGTCTACTGATCTCGTT</td><td>CGG</td><td>||||||||[||||||||||||]PAM</td><td>NA</td><td>Intergenic</td><td>NA</td><td>NA</td></tr>
<tr><td>T27786</td><td>TCTACTGATCTCGTTCGGTTCGG</td><td></td><td>1000</td><td></td><td></td><td></td><td></td><td></td><td></td><td></td><td></td><td></td></tr>
<tr><td>Oligo fwd</td><td>CACCTCTACTGATCTCGTTCGGTT</td><td></td><td></td><td></td><td></td><td></td><td></td><td></td><td></td><td></td><td></td><td></td></tr>
<tr><td>Oligo rev</td><td>AAACAACCGAACGAGATCAGTAGA</td><td></td><td></td><td></td><td></td><td></td><td></td><td></td><td></td><td></td><td></td><td></td></tr>
<tr><td>Chromosome</td><td>start</td><td></td><td>end</td><td>strand</td><td>MM</td><td>target_seq</td><td>PAM</td><td>alignment</td><td>distance</td><td>position</td><td>gene name</td><td>gene id</td></tr>
<tr><td>GU305914.1</td><td></td><td>85447</td><td>85469</td><td>-</td><td>0</td><td>TCTACTGATCTCGTTCGGTT</td><td>CGG</td><td>||||||||[||||||||||||]PAM</td><td>NA</td><td>Intergenic</td><td>NA</td><td>NA</td></tr>
<tr><td>T27787</td><td>CTACTGATCTCGTTCGGTTCGGG</td><td></td><td>1000</td><td></td><td></td><td></td><td></td><td></td><td></td><td></td><td></td><td></td></tr>
<tr><td>Oligo fwd</td><td>CACCCTACTGATCTCGTTCGGTTC</td><td></td><td></td><td></td><td></td><td></td><td></td><td></td><td></td><td></td><td></td><td></td></tr>
<tr><td>Oligo rev</td><td>AAACGAACCGAACGAGATCAGTAG</td><td></td><td></td><td></td><td></td><td></td><td></td><td></td><td></td><td></td><td></td><td></td></tr>
<tr><td>Chromosome</td><td>start</td><td></td><td>end</td><td>strand</td><td>MM</td><td>target_seq</td><td>PAM</td><td>alignment</td><td>distance</td><td>position</td><td>gene name</td><td>gene id</td></tr>
<tr><td>GU305914.1</td><td></td><td>85446</td><td>85468</td><td>-</td><td>0</td><td>CTACTGATCTCGTTCGGTTC</td><td>GGG</td><td>||||||||[||||||||||||]PAM</td><td>NA</td><td>Intergenic</td><td>NA</td><td>NA</td></tr>
<tr><td>T27788</td><td>CCCGAAGACACCCGTACAAGAGG</td><td></td><td>1000</td><td></td><td></td><td></td><td></td><td></td><td></td><td></td><td></td><td></td></tr>
<tr><td>Oligo fwd</td><td>CACCCCCGAAGACACCCGTACAAG</td><td></td><td></td><td></td><td></td><td></td><td></td><td></td><td></td><td></td><td></td><td></td></tr>
<tr><td>Oligo rev</td><td>AAACCTTGTACGGGTGTCTTCGGG</td><td></td><td></td><td></td><td></td><td></td><td></td><td></td><td></td><td></td><td></td><td></td></tr>
<tr><td>Chromosome</td><td>start</td><td></td><td>end</td><td>strand</td><td>MM</td><td>target_seq</td><td>PAM</td><td>alignment</td><td>distance</td><td>position</td><td>gene name</td><td>gene id</td></tr>
<tr><td>GU305914.1</td><td></td><td>85397</td><td>85419</td><td>-</td><td>0</td><td>CCCGAAGACACCCGTACAAG</td><td>AGG</td><td>||||||||[||||||||||||]PAM</td><td>NA</td><td>Intergenic</td><td>NA</td><td>NA</td></tr>
<tr><td>T27789</td><td>CCGAAGACACCCGTACAAGAGGG</td><td></td><td>1000</td><td></td><td></td><td></td><td></td><td></td><td></td><td></td><td></td><td></td></tr>
<tr><td>Oligo fwd</td><td>CACCCCGAAGACACCCGTACAAGA</td><td></td><td></td><td></td><td></td><td></td><td></td><td></td><td></td><td></td><td></td><td></td></tr>
<tr><td>Oligo rev</td><td>AAACTCTTGTACGGGTGTCTTCGG</td><td></td><td></td><td></td><td></td><td></td><td></td><td></td><td></td><td></td><td></td><td></td></tr>
<tr><td>Chromosome</td><td>start</td><td></td><td>end</td><td>strand</td><td>MM</td><td>target_seq</td><td>PAM</td><td>alignment</td><td>distance</td><td>position</td><td>gene name</td><td>gene id</td></tr>
<tr><td>GU305914.1</td><td></td><td>85396</td><td>85418</td><td>-</td><td>0</td><td>CCGAAGACACCCGTACAAGA</td><td>GGG</td><td>||||||||[||||||||||||]PAM</td><td>NA</td><td>Intergenic</td><td>NA</td><td>NA</td></tr>
<tr><td>T27790</td><td>CGAAGACACCCGTACAAGAGGGG</td><td></td><td>1000</td><td></td><td></td><td></td><td></td><td></td><td></td><td></td><td></td><td></td></tr>
<tr><td>Oligo fwd</td><td>CACCCGAAGACACCCGTACAAGAG</td><td></td><td></td><td></td><td></td><td></td><td></td><td></td><td></td><td></td><td></td><td></td></tr>
<tr><td>Oligo rev</td><td>AAACCTCTTGTACGGGTGTCTTCG</td><td></td><td></td><td></td><td></td><td></td><td></td><td></td><td></td><td></td><td></td><td></td></tr>
<tr><td>Chromosome</td><td>start</td><td></td><td>end</td><td>strand</td><td>MM</td><td>target_seq</td><td>PAM</td><td>alignment</td><td>distance</td><td>position</td><td>gene name</td><td>gene id</td></tr>
<tr><td>GU305914.1</td><td></td><td>85395</td><td>85417</td><td>-</td><td>0</td><td>CGAAGACACCCGTACAAGAG</td><td>GGG</td><td>||||||||[||||||||||||]PAM</td><td>NA</td><td>Intergenic</td><td>NA</td><td>NA</td></tr>
<tr><td>T27791</td><td>ACCCGTACAAGAGGGGATCATGG</td><td></td><td>1000</td><td></td><td></td><td></td><td></td><td></td><td></td><td></td><td></td><td></td></tr>
<tr><td>Oligo fwd</td><td>CACCACCCGTACAAGAGGGGATCA</td><td></td><td></td><td></td><td></td><td></td><td></td><td></td><td></td><td></td><td></td><td></td></tr>
<tr><td>Oligo rev</td><td>AAACTGATCCCCTCTTGTACGGGT</td><td></td><td></td><td></td><td></td><td></td><td></td><td></td><td></td><td></td><td></td><td></td></tr>
<tr><td>Chromosome</td><td>start</td><td></td><td>end</td><td>strand</td><td>MM</td><td>target_seq</td><td>PAM</td><td>alignment</td><td>distance</td><td>position</td><td>gene name</td><td>gene id</td></tr>
<tr><td>GU305914.1</td><td></td><td>85388</td><td>85410</td><td>-</td><td>0</td><td>ACCCGTACAAGAGGGGATCA</td><td>TGG</td><td>||||||||[||||||||||||]PAM</td><td>NA</td><td>Intergenic</td><td>NA</td><td>NA</td></tr>
<tr><td>T27792</td><td>AGTTTACAAAGAAAATATCGTGG</td><td></td><td>1000</td><td></td><td></td><td></td><td></td><td></td><td></td><td></td><td></td><td></td></tr>
<tr><td>Oligo fwd</td><td>CACCAGTTTACAAAGAAAATATCG</td><td></td><td></td><td></td><td></td><td></td><td></td><td></td><td></td><td></td><td></td><td></td></tr>
<tr><td>Oligo rev</td><td>AAACCGATATTTTCTTTGTAAACT</td><td></td><td></td><td></td><td></td><td></td><td></td><td></td><td></td><td></td><td></td><td></td></tr>
<tr><td>Chromosome</td><td>start</td><td></td><td>end</td><td>strand</td><td>MM</td><td>target_seq</td><td>PAM</td><td>alignment</td><td>distance</td><td>position</td><td>gene name</td><td>gene id</td></tr>
<tr><td>GU305914.1</td><td></td><td>85364</td><td>85386</td><td>-</td><td>0</td><td>AGTTTACAAAGAAAATATCG</td><td>TGG</td><td>||||||||[||||||||||||]PAM</td><td>NA</td><td>Intergenic</td><td>NA</td><td>NA</td></tr>
<tr><td>T27793</td><td>TATCGTGGCTTACACGTTTGAGG</td><td></td><td>1000</td><td></td><td></td><td></td><td></td><td></td><td></td><td></td><td></td><td></td></tr>
<tr><td>Oligo fwd</td><td>CACCTATCGTGGCTTACACGTTTG</td><td></td><td></td><td></td><td></td><td></td><td></td><td></td><td></td><td></td><td></td><td></td></tr>
<tr><td>Oligo rev</td><td>AAACCAAACGTGTAAGCCACGATA</td><td></td><td></td><td></td><td></td><td></td><td></td><td></td><td></td><td></td><td></td><td></td></tr>
<tr><td>Chromosome</td><td>start</td><td></td><td>end</td><td>strand</td><td>MM</td><td>target_seq</td><td>PAM</td><td>alignment</td><td>distance</td><td>position</td><td>gene name</td><td>gene id</td></tr>
<tr><td>GU305914.1</td><td></td><td>85349</td><td>85371</td><td>-</td><td>0</td><td>TATCGTGGCTTACACGTTTG</td><td>AGG</td><td>||||||||[||||||||||||]PAM</td><td>NA</td><td>Intergenic</td><td>NA</td><td>NA</td></tr>
<tr><td>T27794</td><td>CACGACTACTAATTATTTCTTGG</td><td></td><td>1000</td><td></td><td></td><td></td><td></td><td></td><td></td><td></td><td></td><td></td></tr>
<tr><td>Oligo fwd</td><td>CACCCACGACTACTAATTATTTCT</td><td></td><td></td><td></td><td></td><td></td><td></td><td></td><td></td><td></td><td></td><td></td></tr>
<tr><td>Oligo rev</td><td>AAACAGAAATAATTAGTAGTCGTG</td><td></td><td></td><td></td><td></td><td></td><td></td><td></td><td></td><td></td><td></td><td></td></tr>
<tr><td>Chromosome</td><td>start</td><td></td><td>end</td><td>strand</td><td>MM</td><td>target_seq</td><td>PAM</td><td>alignment</td><td>distance</td><td>position</td><td>gene name</td><td>gene id</td></tr>
<tr><td>GU305914.1</td><td></td><td>85280</td><td>85302</td><td>-</td><td>0</td><td>CACGACTACTAATTATTTCT</td><td>TGG</td><td>||||||||[||||||||||||]PAM</td><td>NA</td><td>Intergenic</td><td>NA</td><td>NA</td></tr>
<tr><td>T27795</td><td>ACGACTACTAATTATTTCTTGGG</td><td></td><td>1000</td><td></td><td></td><td></td><td></td><td></td><td></td><td></td><td></td><td></td></tr>
<tr><td>Oligo fwd</td><td>CACCACGACTACTAATTATTTCTT</td><td></td><td></td><td></td><td></td><td></td><td></td><td></td><td></td><td></td><td></td><td></td></tr>
<tr><td>Oligo rev</td><td>AAACAAGAAATAATTAGTAGTCGT</td><td></td><td></td><td></td><td></td><td></td><td></td><td></td><td></td><td></td><td></td><td></td></tr>
<tr><td>Chromosome</td><td>start</td><td></td><td>end</td><td>strand</td><td>MM</td><td>target_seq</td><td>PAM</td><td>alignment</td><td>distance</td><td>position</td><td>gene name</td><td>gene id</td></tr>
<tr><td>GU305914.1</td><td></td><td>85279</td><td>85301</td><td>-</td><td>0</td><td>ACGACTACTAATTATTTCTT</td><td>GGG</td><td>||||||||[||||||||||||]PAM</td><td>NA</td><td>Intergenic</td><td>NA</td><td>NA</td></tr>
<tr><td>T27796</td><td>CGACTACTAATTATTTCTTGGGG</td><td></td><td>1000</td><td></td><td></td><td></td><td></td><td></td><td></td><td></td><td></td><td></td></tr>
<tr><td>Oligo fwd</td><td>CACCCGACTACTAATTATTTCTTG</td><td></td><td></td><td></td><td></td><td></td><td></td><td></td><td></td><td></td><td></td><td></td></tr>
<tr><td>Oligo rev</td><td>AAACCAAGAAATAATTAGTAGTCG</td><td></td><td></td><td></td><td></td><td></td><td></td><td></td><td></td><td></td><td></td><td></td></tr>
<tr><td>Chromosome</td><td>start</td><td></td><td>end</td><td>strand</td><td>MM</td><td>target_seq</td><td>PAM</td><td>alignment</td><td>distance</td><td>position</td><td>gene name</td><td>gene id</td></tr>
<tr><td>GU305914.1</td><td></td><td>85278</td><td>85300</td><td>-</td><td>0</td><td>CGACTACTAATTATTTCTTG</td><td>GGG</td><td>||||||||[||||||||||||]PAM</td><td>NA</td><td>Intergenic</td><td>NA</td><td>NA</td></tr>
<tr><td>T27797</td><td>GGGGACTAGCGTGACTAAGATGG</td><td></td><td>1000</td><td></td><td></td><td></td><td></td><td></td><td></td><td></td><td></td><td></td></tr>
<tr><td>Oligo fwd</td><td>CACCGGGGACTAGCGTGACTAAGA</td><td></td><td></td><td></td><td></td><td></td><td></td><td></td><td></td><td></td><td></td><td></td></tr>
<tr><td>Oligo rev</td><td>AAACTCTTAGTCACGCTAGTCCCC</td><td></td><td></td><td></td><td></td><td></td><td></td><td></td><td></td><td></td><td></td><td></td></tr>
<tr><td>Chromosome</td><td>start</td><td></td><td>end</td><td>strand</td><td>MM</td><td>target_seq</td><td>PAM</td><td>alignment</td><td>distance</td><td>position</td><td>gene name</td><td>gene id</td></tr>
<tr><td>GU305914.1</td><td></td><td>85259</td><td>85281</td><td>-</td><td>0</td><td>GGGGACTAGCGTGACTAAGA</td><td>TGG</td><td>||||||||[||||||||||||]PAM</td><td>NA</td><td>Intergenic</td><td>NA</td><td>NA</td></tr>
<tr><td>T27798</td><td>CTAAGATGGCCTTCCCCATCTGG</td><td></td><td>1000</td><td></td><td></td><td></td><td></td><td></td><td></td><td></td><td></td><td></td></tr>
<tr><td>Oligo fwd</td><td>CACCCTAAGATGGCCTTCCCCATC</td><td></td><td></td><td></td><td></td><td></td><td></td><td></td><td></td><td></td><td></td><td></td></tr>
<tr><td>Oligo rev</td><td>AAACGATGGGGAAGGCCATCTTAG</td><td></td><td></td><td></td><td></td><td></td><td></td><td></td><td></td><td></td><td></td><td></td></tr>
<tr><td>Chromosome</td><td>start</td><td></td><td>end</td><td>strand</td><td>MM</td><td>target_seq</td><td>PAM</td><td>alignment</td><td>distance</td><td>position</td><td>gene name</td><td>gene id</td></tr>
<tr><td>GU305914.1</td><td></td><td>85245</td><td>85267</td><td>-</td><td>0</td><td>CTAAGATGGCCTTCCCCATC</td><td>TGG</td><td>||||||||[||||||||||||]PAM</td><td>NA</td><td>Intergenic</td><td>NA</td><td>NA</td></tr>
<tr><td>T27799</td><td>TAAGATGGCCTTCCCCATCTGGG</td><td></td><td>1000</td><td></td><td></td><td></td><td></td><td></td><td></td><td></td><td></td><td></td></tr>
<tr><td>Oligo fwd</td><td>CACCTAAGATGGCCTTCCCCATCT</td><td></td><td></td><td></td><td></td><td></td><td></td><td></td><td></td><td></td><td></td><td></td></tr>
<tr><td>Oligo rev</td><td>AAACAGATGGGGAAGGCCATCTTA</td><td></td><td></td><td></td><td></td><td></td><td></td><td></td><td></td><td></td><td></td><td></td></tr>
<tr><td>Chromosome</td><td>start</td><td></td><td>end</td><td>strand</td><td>MM</td><td>target_seq</td><td>PAM</td><td>alignment</td><td>distance</td><td>position</td><td>gene name</td><td>gene id</td></tr>
<tr><td>GU305914.1</td><td></td><td>85244</td><td>85266</td><td>-</td><td>0</td><td>TAAGATGGCCTTCCCCATCT</td><td>GGG</td><td>||||||||[||||||||||||]PAM</td><td>NA</td><td>Intergenic</td><td>NA</td><td>NA</td></tr>
<tr><td>T27800</td><td>GGCCTTCCCCATCTGGGAGCTGG</td><td></td><td>1000</td><td></td><td></td><td></td><td></td><td></td><td></td><td></td><td></td><td></td></tr>
<tr><td>Oligo fwd</td><td>CACCGGCCTTCCCCATCTGGGAGC</td><td></td><td></td><td></td><td></td><td></td><td></td><td></td><td></td><td></td><td></td><td></td></tr>
<tr><td>Oligo rev</td><td>AAACGCTCCCAGATGGGGAAGGCC</td><td></td><td></td><td></td><td></td><td></td><td></td><td></td><td></td><td></td><td></td><td></td></tr>
</table>

Chromosome start end strand MM target_seq PAM alignment distance position gene name gene id
GU305914.1 85238 85260 - 0 GGCCTTCCCCATCTGGGAGC TGG ||||||||[||||||||||||]PAM NA Intergenic NA NA

T27801 CCCCATCTGGGAGCTGGATGAGG 1000
Oligo fwd CACCCCCCATCTGGGAGCTGGATG
Oligo rev AAACCATCCAGCTCCCAGATGGGG
Chromosome start end strand MM target_seq PAM alignment distance position gene name gene id
GU305914.1 85232 85254 - 0 CCCCATCTGGGAGCTGGATG AGG ||||||||[||||||||||||]PAM NA Intergenic NA NA

T27802 ATGAGGTAAACAGAAACAACAGG 1000
Oligo fwd CACCATGAGGTAAACAGAAACAAC
Oligo rev AAACGTTGTTTCTGTTTACCTCAT
Chromosome start end strand MM target_seq PAM alignment distance position gene name gene id
GU305914.1 85215 85237 - 0 ATGAGGTAAACAGAAACAAC AGG ||||||||[||||||||||||]PAM NA Intergenic NA NA

T27803 GGTGTTACTCTGCCGCGAGTAGG 1000
Oligo fwd CACCGGTGTTACTCTGCCGCGAGT
Oligo rev AAACACTCGCGGCAGAGTAACACC
Chromosome start end strand MM target_seq PAM alignment distance position gene name gene id
GU305914.1 85194 85216 - 0 GGTGTTACTCTGCCGCGAGT AGG ||||||||[||||||||||||]PAM NA Intergenic NA NA

T27804 GCCGCGAGTAGGATACTCAATGG 1000
Oligo fwd CACCGCCGCGAGTAGGATACTCAA
Oligo rev AAACTTGAGTATCCTACTCGCGGC
Chromosome start end strand MM target_seq PAM alignment distance position gene name gene id
GU305914.1 85183 85205 - 0 GCCGCGAGTAGGATACTCAA TGG ||||||||[||||||||||||]PAM NA Intergenic NA NA

T27805 CCGCGAGTAGGATACTCAATGGG 1000
Oligo fwd CACCCCGCGAGTAGGATACTCAAT
Oligo rev AAACATTGAGTATCCTACTCGCGG
Chromosome start end strand MM target_seq PAM alignment distance position gene name gene id
GU305914.1 85182 85204 - 0 CCGCGAGTAGGATACTCAAT GGG ||||||||[||||||||||||]PAM NA Intergenic NA NA

T27806 CGCGAGTAGGATACTCAATGGGG 1000
Oligo fwd CACCCGCGAGTAGGATACTCAATG
Oligo rev AAACCATTGAGTATCCTACTCGCG
Chromosome start end strand MM target_seq PAM alignment distance position gene name gene id
GU305914.1 85181 85203 - 0 CGCGAGTAGGATACTCAATG GGG ||||||||[||||||||||||]PAM NA Intergenic NA NA

T27807 CTTACCACGAAGATAGCTATAGG 1000
Oligo fwd CACCCTTACCACGAAGATAGCTAT
Oligo rev AAACATAGCTATCTTCGTGGTAAG
Chromosome start end strand MM target_seq PAM alignment distance position gene name gene id
GU305914.1 85146 85168 - 0 CTTACCACGAAGATAGCTAT AGG ||||||||[||||||||||||]PAM NA Intergenic NA NA

T27808 TAGCTATAGGAACTATACTATGG 1000
Oligo fwd CACCTAGCTATAGGAACTATACTA
Oligo rev AAACTAGTATAGTTCCTATAGCTA
Chromosome start end strand MM target_seq PAM alignment distance position gene name gene id
GU305914.1 85133 85155 - 0 TAGCTATAGGAACTATACTA TGG ||||||||[||||||||||||]PAM NA Intergenic NA NA

T27809 GAACTATACTATGGTGCTTGTGG 1000
Oligo fwd CACCGAACTATACTATGGTGCTTG
Oligo rev AAACCAAGCACCATAGTATAGTTC
Chromosome start end strand MM target_seq PAM alignment distance position gene name gene id
GU305914.1 85124 85146 - 0 GAACTATACTATGGTGCTTG TGG ||||||||[||||||||||||]PAM NA Intergenic NA NA

T27810 CTATACTATGGTGCTTGTGGAGG 1000
Oligo fwd CACCCTATACTATGGTGCTTGTGG
Oligo rev AAACCCACAAGCACCATAGTATAG
Chromosome start end strand MM target_seq PAM alignment distance position gene name gene id
GU305914.1 85121 85143 - 0 CTATACTATGGTGCTTGTGG AGG ||||||||[||||||||||||]PAM NA Intergenic NA NA

T27811 TGCTTGTGGAGGATGATTACAGG 1000
Oligo fwd CACCTGCTTGTGGAGGATGATTAC
Oligo rev AAACGTAATCATCCTCCACAAGCA
Chromosome start end strand MM target_seq PAM alignment distance position gene name gene id
GU305914.1 85110 85132 - 0 TGCTTGTGGAGGATGATTAC AGG ||||||||[||||||||||||]PAM NA Intergenic NA NA

T27812 GATACGTGACCACCAAGAGTAGG 1000
Oligo fwd CACCGATACGTGACCACCAAGAGT
Oligo rev AAACACTCTTGGTGGTCACGTATC
Chromosome start end strand MM target_seq PAM alignment distance position gene name gene id
GU305914.1 85071 85093 - 0 GATACGTGACCACCAAGAGT AGG ||||||||[||||||||||||]PAM NA Intergenic NA NA

T27813 CACCAAGAGTAGGTATCATAAGG 1000
Oligo fwd CACCCACCAAGAGTAGGTATCATA
Oligo rev AAACTATGATACCTACTCTTGGTG
Chromosome start end strand MM target_seq PAM alignment distance position gene name gene id
GU305914.1 85061 85083 - 0 CACCAAGAGTAGGTATCATA AGG ||||||||[||||||||||||]PAM NA Intergenic NA NA

T27814 ACCAAGAGTAGGTATCATAAGGG 1000
Oligo fwd CACCACCAAGAGTAGGTATCATAA
Oligo rev AAACTTATGATACCTACTCTTGGT
Chromosome start end strand MM target_seq PAM alignment distance position gene name gene id
GU305914.1 85060 85082 - 0 ACCAAGAGTAGGTATCATAA GGG ||||||||[||||||||||||]PAM NA Intergenic NA NA

T27815 GTAGGTATCATAAGGGCGCTTGG 1000
Oligo fwd CACCGTAGGTATCATAAGGGCGCT

<table>
<tr><td>Oligo rev</td><td>AAACAGCGCCCTTATGATACCTAC</td><td></td><td></td><td></td><td></td><td></td><td></td><td></td><td></td><td></td><td></td><td></td></tr>
<tr><td>Chromosome</td><td>start</td><td></td><td>end</td><td>strand</td><td>MM</td><td>target_seq</td><td>PAM</td><td>alignment</td><td>distance</td><td>position</td><td>gene name</td><td>gene id</td></tr>
<tr><td>GU305914.1</td><td></td><td>85053</td><td>85075</td><td>-</td><td>0</td><td>GTAGGTATCATAAGGGCGCT</td><td>TGG</td><td>||||||||[||||||||||||]PAM</td><td>NA</td><td>Intergenic</td><td>NA</td><td>NA</td></tr>
<tr><td></td><td></td><td></td><td></td><td></td><td></td><td></td><td></td><td></td><td></td><td></td><td></td><td></td></tr>
<tr><td>T27816</td><td>ATCATAAGGGCGCTTGGACGTGG</td><td></td><td>1000</td><td></td><td></td><td></td><td></td><td></td><td></td><td></td><td></td><td></td></tr>
<tr><td>Oligo fwd</td><td>CACCATCATAAGGGCGCTTGGACG</td><td></td><td></td><td></td><td></td><td></td><td></td><td></td><td></td><td></td><td></td><td></td></tr>
<tr><td>Oligo rev</td><td>AAACCGTCCAAGCGCCCTTATGAT</td><td></td><td></td><td></td><td></td><td></td><td></td><td></td><td></td><td></td><td></td><td></td></tr>
<tr><td>Chromosome</td><td>start</td><td></td><td>end</td><td>strand</td><td>MM</td><td>target_seq</td><td>PAM</td><td>alignment</td><td>distance</td><td>position</td><td>gene name</td><td>gene id</td></tr>
<tr><td>GU305914.1</td><td></td><td>85047</td><td>85069</td><td>-</td><td>0</td><td>ATCATAAGGGCGCTTGGACG</td><td>TGG</td><td>||||||||[||||||||||||]PAM</td><td>NA</td><td>Intergenic</td><td>NA</td><td>NA</td></tr>
<tr><td></td><td></td><td></td><td></td><td></td><td></td><td></td><td></td><td></td><td></td><td></td><td></td><td></td></tr>
<tr><td>T27817</td><td>TCGTCGTCGTCACGAAAGCTAGG</td><td></td><td>1000</td><td></td><td></td><td></td><td></td><td></td><td></td><td></td><td></td><td></td></tr>
<tr><td>Oligo fwd</td><td>CACCTCGTCGTCGTCACGAAAGCT</td><td></td><td></td><td></td><td></td><td></td><td></td><td></td><td></td><td></td><td></td><td></td></tr>
<tr><td>Oligo rev</td><td>AAACAGCTTTCGTGACGACGACGA</td><td></td><td></td><td></td><td></td><td></td><td></td><td></td><td></td><td></td><td></td><td></td></tr>
<tr><td>Chromosome</td><td>start</td><td></td><td>end</td><td>strand</td><td>MM</td><td>target_seq</td><td>PAM</td><td>alignment</td><td>distance</td><td>position</td><td>gene name</td><td>gene id</td></tr>
<tr><td>GU305914.1</td><td></td><td>84993</td><td>85015</td><td>-</td><td>0</td><td>TCGTCGTCGTCACGAAAGCT</td><td>AGG</td><td>||||||||[||||||||||||]PAM</td><td>NA</td><td>Intergenic</td><td>NA</td><td>NA</td></tr>
<tr><td></td><td></td><td></td><td></td><td></td><td></td><td></td><td></td><td></td><td></td><td></td><td></td><td></td></tr>
<tr><td>T27818</td><td>GAATTCTTCGTGCTCAGCTCCGG</td><td></td><td>1000</td><td></td><td></td><td></td><td></td><td></td><td></td><td></td><td></td><td></td></tr>
<tr><td>Oligo fwd</td><td>CACCGAATTCTTCGTGCTCAGCTC</td><td></td><td></td><td></td><td></td><td></td><td></td><td></td><td></td><td></td><td></td><td></td></tr>
<tr><td>Oligo rev</td><td>AAACGAGCTGAGCACGAAGAATTC</td><td></td><td></td><td></td><td></td><td></td><td></td><td></td><td></td><td></td><td></td><td></td></tr>
<tr><td>Chromosome</td><td>start</td><td></td><td>end</td><td>strand</td><td>MM</td><td>target_seq</td><td>PAM</td><td>alignment</td><td>distance</td><td>position</td><td>gene name</td><td>gene id</td></tr>
<tr><td>GU305914.1</td><td></td><td>84955</td><td>84977</td><td>-</td><td>0</td><td>GAATTCTTCGTGCTCAGCTC</td><td>CGG</td><td>||||||||[||||||||||||]PAM</td><td>NA</td><td>Intergenic</td><td>NA</td><td>NA</td></tr>
<tr><td></td><td></td><td></td><td></td><td></td><td></td><td></td><td></td><td></td><td></td><td></td><td></td><td></td></tr>
<tr><td>T27819</td><td>AATTCTTCGTGCTCAGCTCCGGG</td><td></td><td>1000</td><td></td><td></td><td></td><td></td><td></td><td></td><td></td><td></td><td></td></tr>
<tr><td>Oligo fwd</td><td>CACCAATTCTTCGTGCTCAGCTCC</td><td></td><td></td><td></td><td></td><td></td><td></td><td></td><td></td><td></td><td></td><td></td></tr>
<tr><td>Oligo rev</td><td>AAACGGAGCTGAGCACGAAGAATT</td><td></td><td></td><td></td><td></td><td></td><td></td><td></td><td></td><td></td><td></td><td></td></tr>
<tr><td>Chromosome</td><td>start</td><td></td><td>end</td><td>strand</td><td>MM</td><td>target_seq</td><td>PAM</td><td>alignment</td><td>distance</td><td>position</td><td>gene name</td><td>gene id</td></tr>
<tr><td>GU305914.1</td><td></td><td>84954</td><td>84976</td><td>-</td><td>0</td><td>AATTCTTCGTGCTCAGCTCC</td><td>GGG</td><td>||||||||[||||||||||||]PAM</td><td>NA</td><td>Intergenic</td><td>NA</td><td>NA</td></tr>
<tr><td></td><td></td><td></td><td></td><td></td><td></td><td></td><td></td><td></td><td></td><td></td><td></td><td></td></tr>
<tr><td>T27820</td><td>ATTCTTCGTGCTCAGCTCCGGGG</td><td></td><td>1000</td><td></td><td></td><td></td><td></td><td></td><td></td><td></td><td></td><td></td></tr>
<tr><td>Oligo fwd</td><td>CACCATTCTTCGTGCTCAGCTCCG</td><td></td><td></td><td></td><td></td><td></td><td></td><td></td><td></td><td></td><td></td><td></td></tr>
<tr><td>Oligo rev</td><td>AAACCGGAGCTGAGCACGAAGAAT</td><td></td><td></td><td></td><td></td><td></td><td></td><td></td><td></td><td></td><td></td><td></td></tr>
<tr><td>Chromosome</td><td>start</td><td></td><td>end</td><td>strand</td><td>MM</td><td>target_seq</td><td>PAM</td><td>alignment</td><td>distance</td><td>position</td><td>gene name</td><td>gene id</td></tr>
<tr><td>GU305914.1</td><td></td><td>84953</td><td>84975</td><td>-</td><td>0</td><td>ATTCTTCGTGCTCAGCTCCG</td><td>GGG</td><td>||||||||[||||||||||||]PAM</td><td>NA</td><td>Intergenic</td><td>NA</td><td>NA</td></tr>
<tr><td></td><td></td><td></td><td></td><td></td><td></td><td></td><td></td><td></td><td></td><td></td><td></td><td></td></tr>
<tr><td>T27821</td><td>GAGATATCACCGTTTTACAACGG</td><td></td><td>1000</td><td></td><td></td><td></td><td></td><td></td><td></td><td></td><td></td><td></td></tr>
<tr><td>Oligo fwd</td><td>CACCGAGATATCACCGTTTTACAA</td><td></td><td></td><td></td><td></td><td></td><td></td><td></td><td></td><td></td><td></td><td></td></tr>
<tr><td>Oligo rev</td><td>AAACTTGTAAAACGGTGATATCTC</td><td></td><td></td><td></td><td></td><td></td><td></td><td></td><td></td><td></td><td></td><td></td></tr>
<tr><td>Chromosome</td><td>start</td><td></td><td>end</td><td>strand</td><td>MM</td><td>target_seq</td><td>PAM</td><td>alignment</td><td>distance</td><td>position</td><td>gene name</td><td>gene id</td></tr>
<tr><td>GU305914.1</td><td></td><td>84922</td><td>84944</td><td>-</td><td>0</td><td>GAGATATCACCGTTTTACAA</td><td>CGG</td><td>||||||||[||||||||||||]PAM</td><td>NA</td><td>Intergenic</td><td>NA</td><td>NA</td></tr>
<tr><td>GU305914.1</td><td></td><td>84913</td><td>84935</td><td>+</td><td>4</td><td>GAGTTTTCACCGTTGTAAAA</td><td>CGG</td><td>|||-|-||[||||||-||-||]PAM</td><td>NA</td><td>Intergenic</td><td>NA</td><td>NA</td></tr>
<tr><td></td><td></td><td></td><td></td><td></td><td></td><td></td><td></td><td></td><td></td><td></td><td></td><td></td></tr>
<tr><td>T27822</td><td>AAACTCCGAGCCCTTCGAAGAGG</td><td></td><td>1000</td><td></td><td></td><td></td><td></td><td></td><td></td><td></td><td></td><td></td></tr>
<tr><td>Oligo fwd</td><td>CACCAAACTCCGAGCCCTTCGAAG</td><td></td><td></td><td></td><td></td><td></td><td></td><td></td><td></td><td></td><td></td><td></td></tr>
<tr><td>Oligo rev</td><td>AAACCTTCGAAGGGCTCGGAGTTT</td><td></td><td></td><td></td><td></td><td></td><td></td><td></td><td></td><td></td><td></td><td></td></tr>
<tr><td>Chromosome</td><td>start</td><td></td><td>end</td><td>strand</td><td>MM</td><td>target_seq</td><td>PAM</td><td>alignment</td><td>distance</td><td>position</td><td>gene name</td><td>gene id</td></tr>
<tr><td>GU305914.1</td><td></td><td>84896</td><td>84918</td><td>-</td><td>0</td><td>AAACTCCGAGCCCTTCGAAG</td><td>AGG</td><td>||||||||[||||||||||||]PAM</td><td>NA</td><td>Intergenic</td><td>NA</td><td>NA</td></tr>
<tr><td></td><td></td><td></td><td></td><td></td><td></td><td></td><td></td><td></td><td></td><td></td><td></td><td></td></tr>
<tr><td>T27823</td><td>AAGAGGACACGAGAAATTTCTGG</td><td></td><td>1000</td><td></td><td></td><td></td><td></td><td></td><td></td><td></td><td></td><td></td></tr>
<tr><td>Oligo fwd</td><td>CACCAAGAGGACACGAGAAATTTC</td><td></td><td></td><td></td><td></td><td></td><td></td><td></td><td></td><td></td><td></td><td></td></tr>
<tr><td>Oligo rev</td><td>AAACGAAATTTCTCGTGTCCTCTT</td><td></td><td></td><td></td><td></td><td></td><td></td><td></td><td></td><td></td><td></td><td></td></tr>
<tr><td>Chromosome</td><td>start</td><td></td><td>end</td><td>strand</td><td>MM</td><td>target_seq</td><td>PAM</td><td>alignment</td><td>distance</td><td>position</td><td>gene name</td><td>gene id</td></tr>
<tr><td>GU305914.1</td><td></td><td>84879</td><td>84901</td><td>-</td><td>0</td><td>AAGAGGACACGAGAAATTTC</td><td>TGG</td><td>||||||||[||||||||||||]PAM</td><td>NA</td><td>Intergenic</td><td>NA</td><td>NA</td></tr>
<tr><td></td><td></td><td></td><td></td><td></td><td></td><td></td><td></td><td></td><td></td><td></td><td></td><td></td></tr>
<tr><td>T27824</td><td>TACACCATGAAGACGTACTTCGG</td><td></td><td>1000</td><td></td><td></td><td></td><td></td><td></td><td></td><td></td><td></td><td></td></tr>
<tr><td>Oligo fwd</td><td>CACCTACACCATGAAGACGTACTT</td><td></td><td></td><td></td><td></td><td></td><td></td><td></td><td></td><td></td><td></td><td></td></tr>
<tr><td>Oligo rev</td><td>AAACAAGTACGTCTTCATGGTGTA</td><td></td><td></td><td></td><td></td><td></td><td></td><td></td><td></td><td></td><td></td><td></td></tr>
<tr><td>Chromosome</td><td>start</td><td></td><td>end</td><td>strand</td><td>MM</td><td>target_seq</td><td>PAM</td><td>alignment</td><td>distance</td><td>position</td><td>gene name</td><td>gene id</td></tr>
<tr><td>GU305914.1</td><td></td><td>84844</td><td>84866</td><td>-</td><td>0</td><td>TACACCATGAAGACGTACTT</td><td>CGG</td><td>||||||||[||||||||||||]PAM</td><td>NA</td><td>Intergenic</td><td>NA</td><td>NA</td></tr>
<tr><td></td><td></td><td></td><td></td><td></td><td></td><td></td><td></td><td></td><td></td><td></td><td></td><td></td></tr>
<tr><td>T27825</td><td>GACGTACTTCGGTGAACTCGCGG</td><td></td><td>1000</td><td></td><td></td><td></td><td></td><td></td><td></td><td></td><td></td><td></td></tr>
<tr><td>Oligo fwd</td><td>CACCGACGTACTTCGGTGAACTCG</td><td></td><td></td><td></td><td></td><td></td><td></td><td></td><td></td><td></td><td></td><td></td></tr>
<tr><td>Oligo rev</td><td>AAACCGAGTTCACCGAAGTACGTC</td><td></td><td></td><td></td><td></td><td></td><td></td><td></td><td></td><td></td><td></td><td></td></tr>
<tr><td>Chromosome</td><td>start</td><td></td><td>end</td><td>strand</td><td>MM</td><td>target_seq</td><td>PAM</td><td>alignment</td><td>distance</td><td>position</td><td>gene name</td><td>gene id</td></tr>
<tr><td>GU305914.1</td><td></td><td>84833</td><td>84855</td><td>-</td><td>0</td><td>GACGTACTTCGGTGAACTCG</td><td>CGG</td><td>||||||||[||||||||||||]PAM</td><td>NA</td><td>Intergenic</td><td>NA</td><td>NA</td></tr>
<tr><td></td><td></td><td></td><td></td><td></td><td></td><td></td><td></td><td></td><td></td><td></td><td></td><td></td></tr>
<tr><td>T27826</td><td>CAAAAAAGTAGTACCTCTGATGG</td><td></td><td>1000</td><td></td><td></td><td></td><td></td><td></td><td></td><td></td><td></td><td></td></tr>
<tr><td>Oligo fwd</td><td>CACCCAAAAAAGTAGTACCTCTGA</td><td></td><td></td><td></td><td></td><td></td><td></td><td></td><td></td><td></td><td></td><td></td></tr>
<tr><td>Oligo rev</td><td>AAACTCAGAGGTACTACTTTTTTG</td><td></td><td></td><td></td><td></td><td></td><td></td><td></td><td></td><td></td><td></td><td></td></tr>
<tr><td>Chromosome</td><td>start</td><td></td><td>end</td><td>strand</td><td>MM</td><td>target_seq</td><td>PAM</td><td>alignment</td><td>distance</td><td>position</td><td>gene name</td><td>gene id</td></tr>
<tr><td>GU305914.1</td><td></td><td>84806</td><td>84828</td><td>-</td><td>0</td><td>CAAAAAAGTAGTACCTCTGA</td><td>TGG</td><td>||||||||[||||||||||||]PAM</td><td>NA</td><td>Intergenic</td><td>NA</td><td>NA</td></tr>
<tr><td></td><td></td><td></td><td></td><td></td><td></td><td></td><td></td><td></td><td></td><td></td><td></td><td></td></tr>
<tr><td>T27827</td><td>CTCTGATGGCTTTTCTAGAGAGG</td><td></td><td>1000</td><td></td><td></td><td></td><td></td><td></td><td></td><td></td><td></td><td></td></tr>
<tr><td>Oligo fwd</td><td>CACCCTCTGATGGCTTTTCTAGAG</td><td></td><td></td><td></td><td></td><td></td><td></td><td></td><td></td><td></td><td></td><td></td></tr>
<tr><td>Oligo rev</td><td>AAACCTCTAGAAAAGCCATCAGAG</td><td></td><td></td><td></td><td></td><td></td><td></td><td></td><td></td><td></td><td></td><td></td></tr>
<tr><td>Chromosome</td><td>start</td><td></td><td>end</td><td>strand</td><td>MM</td><td>target_seq</td><td>PAM</td><td>alignment</td><td>distance</td><td>position</td><td>gene name</td><td>gene id</td></tr>
<tr><td>GU305914.1</td><td></td><td>84792</td><td>84814</td><td>-</td><td>0</td><td>CTCTGATGGCTTTTCTAGAG</td><td>AGG</td><td>||||||||[||||||||||||]PAM</td><td>NA</td><td>Intergenic</td><td>NA</td><td>NA</td></tr>
<tr><td></td><td></td><td></td><td></td><td></td><td></td><td></td><td></td><td></td><td></td><td></td><td></td><td></td></tr>
<tr><td>T27828</td><td>TCTGATGGCTTTTCTAGAGAGGG</td><td></td><td>1000</td><td></td><td></td><td></td><td></td><td></td><td></td><td></td><td></td><td></td></tr>
<tr><td>Oligo fwd</td><td>CACCTCTGATGGCTTTTCTAGAGA</td><td></td><td></td><td></td><td></td><td></td><td></td><td></td><td></td><td></td><td></td><td></td></tr>
<tr><td>Oligo rev</td><td>AAACTCTCTAGAAAAGCCATCAGA</td><td></td><td></td><td></td><td></td><td></td><td></td><td></td><td></td><td></td><td></td><td></td></tr>
<tr><td>Chromosome</td><td>start</td><td></td><td>end</td><td>strand</td><td>MM</td><td>target_seq</td><td>PAM</td><td>alignment</td><td>distance</td><td>position</td><td>gene name</td><td>gene id</td></tr>
<tr><td>GU305914.1</td><td></td><td>84791</td><td>84813</td><td>-</td><td>0</td><td>TCTGATGGCTTTTCTAGAGA</td><td>GGG</td><td>||||||||[||||||||||||]PAM</td><td>NA</td><td>Intergenic</td><td>NA</td><td>NA</td></tr>
<tr><td></td><td></td><td></td><td></td><td></td><td></td><td></td><td></td><td></td><td></td><td></td><td></td><td></td></tr>
<tr><td>T27829</td><td>GATGGCTTTTCTAGAGAGGGAGG</td><td></td><td>1000</td><td></td><td></td><td></td><td></td><td></td><td></td><td></td><td></td><td></td></tr>
<tr><td>Oligo fwd</td><td>CACCGATGGCTTTTCTAGAGAGGG</td><td></td><td></td><td></td><td></td><td></td><td></td><td></td><td></td><td></td><td></td><td></td></tr>
<tr><td>Oligo rev</td><td>AAACCCCTCTCTAGAAAAGCCATC</td><td></td><td></td><td></td><td></td><td></td><td></td><td></td><td></td><td></td><td></td><td></td></tr>
<tr><td>Chromosome</td><td>start</td><td></td><td>end</td><td>strand</td><td>MM</td><td>target_seq</td><td>PAM</td><td>alignment</td><td>distance</td><td>position</td><td>gene name</td><td>gene id</td></tr>
<tr><td>GU305914.1</td><td></td><td>84788</td><td>84810</td><td>-</td><td>0</td><td>GATGGCTTTTCTAGAGAGGG</td><td>AGG</td><td>||||||||[||||||||||||]PAM</td><td>NA</td><td>Intergenic</td><td>NA</td><td>NA</td></tr>
</table>

<table>
<tr><td>T27830</td><td>GAGAGGGAGGACATGACCATCGG</td><td>1000</td><td></td><td></td><td></td><td></td><td></td><td></td><td></td><td></td><td></td></tr>
<tr><td>Oligo fwd</td><td>CACCGAGAGGGAGGACATGACCAT</td><td></td><td></td><td></td><td></td><td></td><td></td><td></td><td></td><td></td><td></td></tr>
<tr><td>Oligo rev</td><td>AAACATGGTCATGTCCTCCCTCTC</td><td></td><td></td><td></td><td></td><td></td><td></td><td></td><td></td><td></td><td></td></tr>
<tr><td>Chromosome</td><td>start</td><td>end</td><td>strand</td><td>MM</td><td>target_seq</td><td>PAM</td><td>alignment</td><td>distance</td><td>position</td><td>gene name</td><td>gene id</td></tr>
<tr><td>GU305914.1</td><td>84775</td><td>84797</td><td>-</td><td>0</td><td>GAGAGGGAGGACATGACCAT</td><td>CGG</td><td>||||||||[||||||||||||]PAM</td><td>NA</td><td>Intergenic</td><td>NA</td><td>NA</td></tr>
<tr><td>T27831</td><td>AGAGGGAGGACATGACCATCGGG</td><td>1000</td><td></td><td></td><td></td><td></td><td></td><td></td><td></td><td></td><td></td></tr>
<tr><td>Oligo fwd</td><td>CACCAGAGGGAGGACATGACCATC</td><td></td><td></td><td></td><td></td><td></td><td></td><td></td><td></td><td></td><td></td></tr>
<tr><td>Oligo rev</td><td>AAACGATGGTCATGTCCTCCCTCT</td><td></td><td></td><td></td><td></td><td></td><td></td><td></td><td></td><td></td><td></td></tr>
<tr><td>Chromosome</td><td>start</td><td>end</td><td>strand</td><td>MM</td><td>target_seq</td><td>PAM</td><td>alignment</td><td>distance</td><td>position</td><td>gene name</td><td>gene id</td></tr>
<tr><td>GU305914.1</td><td>84774</td><td>84796</td><td>-</td><td>0</td><td>AGAGGGAGGACATGACCATC</td><td>GGG</td><td>||||||||[||||||||||||]PAM</td><td>NA</td><td>Intergenic</td><td>NA</td><td>NA</td></tr>
<tr><td>T27832</td><td>GGGAGGACATGACCATCGGGTGG</td><td>1000</td><td></td><td></td><td></td><td></td><td></td><td></td><td></td><td></td><td></td></tr>
<tr><td>Oligo fwd</td><td>CACCGGGAGGACATGACCATCGGG</td><td></td><td></td><td></td><td></td><td></td><td></td><td></td><td></td><td></td><td></td></tr>
<tr><td>Oligo rev</td><td>AAACCCCGATGGTCATGTCCTCCC</td><td></td><td></td><td></td><td></td><td></td><td></td><td></td><td></td><td></td><td></td></tr>
<tr><td>Chromosome</td><td>start</td><td>end</td><td>strand</td><td>MM</td><td>target_seq</td><td>PAM</td><td>alignment</td><td>distance</td><td>position</td><td>gene name</td><td>gene id</td></tr>
<tr><td>GU305914.1</td><td>84771</td><td>84793</td><td>-</td><td>0</td><td>GGGAGGACATGACCATCGGG</td><td>TGG</td><td>||||||||[||||||||||||]PAM</td><td>NA</td><td>Intergenic</td><td>NA</td><td>NA</td></tr>
<tr><td>T27833</td><td>GGAGGACATGACCATCGGGTGGG</td><td>1000</td><td></td><td></td><td></td><td></td><td></td><td></td><td></td><td></td><td></td></tr>
<tr><td>Oligo fwd</td><td>CACCGGAGGACATGACCATCGGGT</td><td></td><td></td><td></td><td></td><td></td><td></td><td></td><td></td><td></td><td></td></tr>
<tr><td>Oligo rev</td><td>AAACACCCGATGGTCATGTCCTCC</td><td></td><td></td><td></td><td></td><td></td><td></td><td></td><td></td><td></td><td></td></tr>
<tr><td>Chromosome</td><td>start</td><td>end</td><td>strand</td><td>MM</td><td>target_seq</td><td>PAM</td><td>alignment</td><td>distance</td><td>position</td><td>gene name</td><td>gene id</td></tr>
<tr><td>GU305914.1</td><td>84770</td><td>84792</td><td>-</td><td>0</td><td>GGAGGACATGACCATCGGGT</td><td>GGG</td><td>||||||||[||||||||||||]PAM</td><td>NA</td><td>Intergenic</td><td>NA</td><td>NA</td></tr>
<tr><td>T27834</td><td>AACAGAATGTCACGTGCGACTGG</td><td>1000</td><td></td><td></td><td></td><td></td><td></td><td></td><td></td><td></td><td></td></tr>
<tr><td>Oligo fwd</td><td>CACCAACAGAATGTCACGTGCGAC</td><td></td><td></td><td></td><td></td><td></td><td></td><td></td><td></td><td></td><td></td></tr>
<tr><td>Oligo rev</td><td>AAACGTCGCACGTGACATTCTGTT</td><td></td><td></td><td></td><td></td><td></td><td></td><td></td><td></td><td></td><td></td></tr>
<tr><td>Chromosome</td><td>start</td><td>end</td><td>strand</td><td>MM</td><td>target_seq</td><td>PAM</td><td>alignment</td><td>distance</td><td>position</td><td>gene name</td><td>gene id</td></tr>
<tr><td>GU305914.1</td><td>84735</td><td>84757</td><td>-</td><td>0</td><td>AACAGAATGTCACGTGCGAC</td><td>TGG</td><td>||||||||[||||||||||||]PAM</td><td>NA</td><td>Intergenic</td><td>NA</td><td>NA</td></tr>
<tr><td>T27835</td><td>TCACGTGCGACTGGAAGAAATGG</td><td>1000</td><td></td><td></td><td></td><td></td><td></td><td></td><td></td><td></td><td></td></tr>
<tr><td>Oligo fwd</td><td>CACCTCACGTGCGACTGGAAGAAA</td><td></td><td></td><td></td><td></td><td></td><td></td><td></td><td></td><td></td><td></td></tr>
<tr><td>Oligo rev</td><td>AAACTTTCTTCCAGTCGCACGTGA</td><td></td><td></td><td></td><td></td><td></td><td></td><td></td><td></td><td></td><td></td></tr>
<tr><td>Chromosome</td><td>start</td><td>end</td><td>strand</td><td>MM</td><td>target_seq</td><td>PAM</td><td>alignment</td><td>distance</td><td>position</td><td>gene name</td><td>gene id</td></tr>
<tr><td>GU305914.1</td><td>84726</td><td>84748</td><td>-</td><td>0</td><td>TCACGTGCGACTGGAAGAAA</td><td>TGG</td><td>||||||||[||||||||||||]PAM</td><td>NA</td><td>Intergenic</td><td>NA</td><td>NA</td></tr>
<tr><td>T27836</td><td>CGACTGGAAGAAATGGCAGACGG</td><td>1000</td><td></td><td></td><td></td><td></td><td></td><td></td><td></td><td></td><td></td></tr>
<tr><td>Oligo fwd</td><td>CACCCGACTGGAAGAAATGGCAGA</td><td></td><td></td><td></td><td></td><td></td><td></td><td></td><td></td><td></td><td></td></tr>
<tr><td>Oligo rev</td><td>AAACTCTGCCATTTCTTCCAGTCG</td><td></td><td></td><td></td><td></td><td></td><td></td><td></td><td></td><td></td><td></td></tr>
<tr><td>Chromosome</td><td>start</td><td>end</td><td>strand</td><td>MM</td><td>target_seq</td><td>PAM</td><td>alignment</td><td>distance</td><td>position</td><td>gene name</td><td>gene id</td></tr>
<tr><td>GU305914.1</td><td>84719</td><td>84741</td><td>-</td><td>0</td><td>CGACTGGAAGAAATGGCAGA</td><td>CGG</td><td>||||||||[||||||||||||]PAM</td><td>NA</td><td>Intergenic</td><td>NA</td><td>NA</td></tr>
<tr><td>T27837</td><td>AGAAATGGCAGACGGTTTCGAGG</td><td>1000</td><td></td><td></td><td></td><td></td><td></td><td></td><td></td><td></td><td></td></tr>
<tr><td>Oligo fwd</td><td>CACCAGAAATGGCAGACGGTTTCG</td><td></td><td></td><td></td><td></td><td></td><td></td><td></td><td></td><td></td><td></td></tr>
<tr><td>Oligo rev</td><td>AAACCGAAACCGTCTGCCATTTCT</td><td></td><td></td><td></td><td></td><td></td><td></td><td></td><td></td><td></td><td></td></tr>
<tr><td>Chromosome</td><td>start</td><td>end</td><td>strand</td><td>MM</td><td>target_seq</td><td>PAM</td><td>alignment</td><td>distance</td><td>position</td><td>gene name</td><td>gene id</td></tr>
<tr><td>GU305914.1</td><td>84711</td><td>84733</td><td>-</td><td>0</td><td>AGAAATGGCAGACGGTTTCG</td><td>AGG</td><td>||||||||[||||||||||||]PAM</td><td>NA</td><td>Intergenic</td><td>NA</td><td>NA</td></tr>
<tr><td>T27838</td><td>GAAATGGCAGACGGTTTCGAGGG</td><td>1000</td><td></td><td></td><td></td><td></td><td></td><td></td><td></td><td></td><td></td></tr>
<tr><td>Oligo fwd</td><td>CACCGAAATGGCAGACGGTTTCGA</td><td></td><td></td><td></td><td></td><td></td><td></td><td></td><td></td><td></td><td></td></tr>
<tr><td>Oligo rev</td><td>AAACTCGAAACCGTCTGCCATTTC</td><td></td><td></td><td></td><td></td><td></td><td></td><td></td><td></td><td></td><td></td></tr>
<tr><td>Chromosome</td><td>start</td><td>end</td><td>strand</td><td>MM</td><td>target_seq</td><td>PAM</td><td>alignment</td><td>distance</td><td>position</td><td>gene name</td><td>gene id</td></tr>
<tr><td>GU305914.1</td><td>84710</td><td>84732</td><td>-</td><td>0</td><td>GAAATGGCAGACGGTTTCGA</td><td>GGG</td><td>||||||||[||||||||||||]PAM</td><td>NA</td><td>Intergenic</td><td>NA</td><td>NA</td></tr>
<tr><td>T27839</td><td>GAGTTTAACTGCTACTTTTGTGG</td><td>1000</td><td></td><td></td><td></td><td></td><td></td><td></td><td></td><td></td><td></td></tr>
<tr><td>Oligo fwd</td><td>CACCGAGTTTAACTGCTACTTTTG</td><td></td><td></td><td></td><td></td><td></td><td></td><td></td><td></td><td></td><td></td></tr>
<tr><td>Oligo rev</td><td>AAACCAAAAGTAGCAGTTAAACTC</td><td></td><td></td><td></td><td></td><td></td><td></td><td></td><td></td><td></td><td></td></tr>
<tr><td>Chromosome</td><td>start</td><td>end</td><td>strand</td><td>MM</td><td>target_seq</td><td>PAM</td><td>alignment</td><td>distance</td><td>position</td><td>gene name</td><td>gene id</td></tr>
<tr><td>GU305914.1</td><td>84644</td><td>84666</td><td>-</td><td>0</td><td>GAGTTTAACTGCTACTTTTG</td><td>TGG</td><td>||||||||[||||||||||||]PAM</td><td>NA</td><td>Intergenic</td><td>NA</td><td>NA</td></tr>
<tr><td>T27840</td><td>GATTATAATACTACTACCGAAGG</td><td>1000</td><td></td><td></td><td></td><td></td><td></td><td></td><td></td><td></td><td></td></tr>
<tr><td>Oligo fwd</td><td>CACCGATTATAATACTACTACCGA</td><td></td><td></td><td></td><td></td><td></td><td></td><td></td><td></td><td></td><td></td></tr>
<tr><td>Oligo rev</td><td>AAACTCGGTAGTAGTATTATAATC</td><td></td><td></td><td></td><td></td><td></td><td></td><td></td><td></td><td></td><td></td></tr>
<tr><td>Chromosome</td><td>start</td><td>end</td><td>strand</td><td>MM</td><td>target_seq</td><td>PAM</td><td>alignment</td><td>distance</td><td>position</td><td>gene name</td><td>gene id</td></tr>
<tr><td>GU305914.1</td><td>84604</td><td>84626</td><td>-</td><td>0</td><td>GATTATAATACTACTACCGA</td><td>AGG</td><td>||||||||[||||||||||||]PAM</td><td>NA</td><td>Intergenic</td><td>NA</td><td>NA</td></tr>
<tr><td>T27841</td><td>AATGAGACGCATGTGAAAGACGG</td><td>1000</td><td></td><td></td><td></td><td></td><td></td><td></td><td></td><td></td><td></td></tr>
<tr><td>Oligo fwd</td><td>CACCAATGAGACGCATGTGAAAGA</td><td></td><td></td><td></td><td></td><td></td><td></td><td></td><td></td><td></td><td></td></tr>
<tr><td>Oligo rev</td><td>AAACTCTTTCACATGCGTCTCATT</td><td></td><td></td><td></td><td></td><td></td><td></td><td></td><td></td><td></td><td></td></tr>
<tr><td>Chromosome</td><td>start</td><td>end</td><td>strand</td><td>MM</td><td>target_seq</td><td>PAM</td><td>alignment</td><td>distance</td><td>position</td><td>gene name</td><td>gene id</td></tr>
<tr><td>GU305914.1</td><td>84505</td><td>84527</td><td>-</td><td>0</td><td>AATGAGACGCATGTGAAAGA</td><td>CGG</td><td>||||||||[||||||||||||]PAM</td><td>NA</td><td>Intergenic</td><td>NA</td><td>NA</td></tr>
<tr><td>T27842</td><td>ATGAGACGCATGTGAAAGACGGG</td><td>1000</td><td></td><td></td><td></td><td></td><td></td><td></td><td></td><td></td><td></td></tr>
<tr><td>Oligo fwd</td><td>CACCATGAGACGCATGTGAAAGAC</td><td></td><td></td><td></td><td></td><td></td><td></td><td></td><td></td><td></td><td></td></tr>
<tr><td>Oligo rev</td><td>AAACGTCTTTCACATGCGTCTCAT</td><td></td><td></td><td></td><td></td><td></td><td></td><td></td><td></td><td></td><td></td></tr>
<tr><td>Chromosome</td><td>start</td><td>end</td><td>strand</td><td>MM</td><td>target_seq</td><td>PAM</td><td>alignment</td><td>distance</td><td>position</td><td>gene name</td><td>gene id</td></tr>
<tr><td>GU305914.1</td><td>84504</td><td>84526</td><td>-</td><td>0</td><td>ATGAGACGCATGTGAAAGAC</td><td>GGG</td><td>||||||||[||||||||||||]PAM</td><td>NA</td><td>Intergenic</td><td>NA</td><td>NA</td></tr>
<tr><td>T27843</td><td>TGAGACGCATGTGAAAGACGGGG</td><td>1000</td><td></td><td></td><td></td><td></td><td></td><td></td><td></td><td></td><td></td></tr>
<tr><td>Oligo fwd</td><td>CACCTGAGACGCATGTGAAAGACG</td><td></td><td></td><td></td><td></td><td></td><td></td><td></td><td></td><td></td><td></td></tr>
<tr><td>Oligo rev</td><td>AAACCGTCTTTCACATGCGTCTCA</td><td></td><td></td><td></td><td></td><td></td><td></td><td></td><td></td><td></td><td></td></tr>
<tr><td>Chromosome</td><td>start</td><td>end</td><td>strand</td><td>MM</td><td>target_seq</td><td>PAM</td><td>alignment</td><td>distance</td><td>position</td><td>gene name</td><td>gene id</td></tr>
<tr><td>GU305914.1</td><td>84503</td><td>84525</td><td>-</td><td>0</td><td>TGAGACGCATGTGAAAGACG</td><td>GGG</td><td>||||||||[||||||||||||]PAM</td><td>NA</td><td>Intergenic</td><td>NA</td><td>NA</td></tr>
<tr><td>T27844</td><td>GCTTGAAATGTATAGAACAACGG</td><td>1000</td><td></td><td></td><td></td><td></td><td></td><td></td><td></td><td></td><td></td></tr>
<tr><td>Oligo fwd</td><td>CACCGCTTGAAATGTATAGAACAA</td><td></td><td></td><td></td><td></td><td></td><td></td><td></td><td></td><td></td><td></td></tr>
<tr><td>Oligo rev</td><td>AAACTTGTTCTATACATTTCAAGC</td><td></td><td></td><td></td><td></td><td></td><td></td><td></td><td></td><td></td><td></td></tr>
<tr><td>Chromosome</td><td>start</td><td>end</td><td>strand</td><td>MM</td><td>target_seq</td><td>PAM</td><td>alignment</td><td>distance</td><td>position</td><td>gene name</td><td>gene id</td></tr>
<tr><td>GU305914.1</td><td>84479</td><td>84501</td><td>-</td><td>0</td><td>GCTTGAAATGTATAGAACAA</td><td>CGG</td><td>||||||||[||||||||||||]PAM</td><td>NA</td><td>Intergenic</td><td>NA</td><td>NA</td></tr>
</table>

T27845 CTTGAAATGTATAGAACAACGGG 1000
Oligo fwd CACCCTTGAAATGTATAGAACAAC
Oligo rev AAACGTTGTTCTATACATTTCAAG
Chromosome start end strand MM target_seq PAM alignment distance position gene name gene id
GU305914.1 84478 84500 - 0 CTTGAAATGTATAGAACAAC GGG ||||||||[||||||||||||]PAM NA Intergenic NA NA

T27846 GAAATGTATAGAACAACGGGAGG 1000
Oligo fwd CACCGAAATGTATAGAACAACGGG
Oligo rev AAACCCCGTTGTTCTATACATTTC
Chromosome start end strand MM target_seq PAM alignment distance position gene name gene id
GU305914.1 84475 84497 - 0 GAAATGTATAGAACAACGGG AGG ||||||||[||||||||||||]PAM NA Intergenic NA NA

T27847 CGGGAGGTCTTATTGTATTGTGG 1000
Oligo fwd CACCCGGGAGGTCTTATTGTATTG
Oligo rev AAACCAATACAATAAGACCTCCCG
Chromosome start end strand MM target_seq PAM alignment distance position gene name gene id
GU305914.1 84459 84481 - 0 CGGGAGGTCTTATTGTATTG TGG ||||||||[||||||||||||]PAM NA Intergenic NA NA

T27848 AGGTCTTATTGTATTGTGGCAGG 1000
Oligo fwd CACCAGGTCTTATTGTATTGTGGC
Oligo rev AAACGCCACAATACAATAAGACCT
Chromosome start end strand MM target_seq PAM alignment distance position gene name gene id
GU305914.1 84455 84477 - 0 AGGTCTTATTGTATTGTGGC AGG ||||||||[||||||||||||]PAM NA Intergenic NA NA

T27849 GGTCTTATTGTATTGTGGCAGGG 1000
Oligo fwd CACCGGTCTTATTGTATTGTGGCA
Oligo rev AAACTGCCACAATACAATAAGACC
Chromosome start end strand MM target_seq PAM alignment distance position gene name gene id
GU305914.1 84454 84476 - 0 GGTCTTATTGTATTGTGGCA GGG ||||||||[||||||||||||]PAM NA Intergenic NA NA

T27850 AGCGAAATCACTGCATAATCTGG 1000
Oligo fwd CACCAGCGAAATCACTGCATAATC
Oligo rev AAACGATTATGCAGTGATTTCGCT
Chromosome start end strand MM target_seq PAM alignment distance position gene name gene id
GU305914.1 84425 84447 - 0 AGCGAAATCACTGCATAATC TGG ||||||||[||||||||||||]PAM NA Intergenic NA NA

T27851 CGCGTTGAATAACGTTTCAGTGG 1000
Oligo fwd CACCCGCGTTGAATAACGTTTCAG
Oligo rev AAACCTGAAACGTTATTCAACGCG
Chromosome start end strand MM target_seq PAM alignment distance position gene name gene id
GU305914.1 84392 84414 - 0 CGCGTTGAATAACGTTTCAG TGG ||||||||[||||||||||||]PAM NA Intergenic NA NA

T27852 ACCACTGCCGCGACTGAGAATGG 1000
Oligo fwd CACCACCACTGCCGCGACTGAGAA
Oligo rev AAACTTCTCAGTCGCGGCAGTGGT
Chromosome start end strand MM target_seq PAM alignment distance position gene name gene id
GU305914.1 84349 84371 - 0 ACCACTGCCGCGACTGAGAA TGG ||||||||[||||||||||||]PAM NA Intergenic NA NA

T27853 CTTTGACAACCTAGATGATGTGG 1000
Oligo fwd CACCCTTTGACAACCTAGATGATG
Oligo rev AAACCATCATCTAGGTTGTCAAAG
Chromosome start end strand MM target_seq PAM alignment distance position gene name gene id
GU305914.1 84296 84318 - 0 CTTTGACAACCTAGATGATG TGG ||||||||[||||||||||||]PAM NA Intergenic NA NA

T27854 CAACCTAGATGATGTGGTGACGG 1000
Oligo fwd CACCCAACCTAGATGATGTGGTGA
Oligo rev AAACTCACCACATCATCTAGGTTG
Chromosome start end strand MM target_seq PAM alignment distance position gene name gene id
GU305914.1 84290 84312 - 0 CAACCTAGATGATGTGGTGA CGG ||||||||[||||||||||||]PAM NA Intergenic NA NA

T27855 ATTTACCTACGACGTGCTCAAGG 1000
Oligo fwd CACCATTTACCTACGACGTGCTCA
Oligo rev AAACTGAGCACGTCGTAGGTAAAT
Chromosome start end strand MM target_seq PAM alignment distance position gene name gene id
GU305914.1 84245 84267 - 0 ATTTACCTACGACGTGCTCA AGG ||||||||[||||||||||||]PAM NA Intergenic NA NA

T27856 CGATGCGCTCCGAAACATCATGG 1000
Oligo fwd CACCCGATGCGCTCCGAAACATCA
Oligo rev AAACTGATGTTTCGGAGCGCATCG
Chromosome start end strand MM target_seq PAM alignment distance position gene name gene id
GU305914.1 84212 84234 - 0 CGATGCGCTCCGAAACATCA TGG ||||||||[||||||||||||]PAM NA Intergenic NA NA

T27857 TCCGAAACATCATGGACGCTTGG 1000
Oligo fwd CACCTCCGAAACATCATGGACGCT
Oligo rev AAACAGCGTCCATGATGTTTCGGA
Chromosome start end strand MM target_seq PAM alignment distance position gene name gene id
GU305914.1 84204 84226 - 0 TCCGAAACATCATGGACGCT TGG ||||||||[||||||||||||]PAM NA Intergenic NA NA

T27858 CTTGGTGTCGCGACCAGAAACGG 1000
Oligo fwd CACCCTTGGTGTCGCGACCAGAAA
Oligo rev AAACTTTCTGGTCGCGACACCAAG
Chromosome start end strand MM target_seq PAM alignment distance position gene name gene id
GU305914.1 84186 84208 - 0 CTTGGTGTCGCGACCAGAAA CGG ||||||||[||||||||||||]PAM NA Intergenic NA NA

T27859 GTGTCGCGACCAGAAACGGACGG 1000
Oligo fwd CACCGTGTCGCGACCAGAAACGGA
Oligo rev AAACTCCGTTTCTGGTCGCGACAC
Chromosome start end strand MM target_seq PAM alignment distance position gene name gene id

| | | | | | | | | | | | |
|---|---|---|---|---|---|---|---|---|---|---|---|
| GU305914.1 | 84182 | 84204 | - | 0 | GTGTCGCGACCAGAAACGGA | CGG | \|\|\|\|\|\|\|\|[\|\|\|\|\|\|\|\|\|\|\|\|]PAM | NA | Intergenic | NA | NA |
| T27860 | TCGCGACCAGAAACGGACGGCGG | 1000 | | | | | | | | | |
| Oligo fwd | CACCTCGCGACCAGAAACGGACGG | | | | | | | | | | |
| Oligo rev | AAACCCGTCCGTTTCTGGTCGCGA | | | | | | | | | | |
| Chromosome | start | end | strand | MM | target_seq | PAM | alignment | distance | position | gene name | gene id |
| GU305914.1 | 84179 | 84201 | - | 0 | TCGCGACCAGAAACGGACGG | CGG | \|\|\|\|\|\|\|\|[\|\|\|\|\|\|\|\|\|\|\|\|]PAM | NA | Intergenic | NA | NA |
| T27861 | ACGGACGGCGGAGATGCTCAAGG | 1000 | | | | | | | | | |
| Oligo fwd | CACCACGGACGGCGGAGATGCTCA | | | | | | | | | | |
| Oligo rev | AAACTGAGCATCTCCGCCGTCCGT | | | | | | | | | | |
| Chromosome | start | end | strand | MM | target_seq | PAM | alignment | distance | position | gene name | gene id |
| GU305914.1 | 84167 | 84189 | - | 0 | ACGGACGGCGGAGATGCTCA | AGG | \|\|\|\|\|\|\|\|[\|\|\|\|\|\|\|\|\|\|\|\|]PAM | NA | Intergenic | NA | NA |
| T27862 | CAATCCGTCCAACATCTTGTCGG | 1000 | | | | | | | | | |
| Oligo fwd | CACCCAATCCGTCCAACATCTTGT | | | | | | | | | | |
| Oligo rev | AAACACAAGATGTTGGACGGATTG | | | | | | | | | | |
| Chromosome | start | end | strand | MM | target_seq | PAM | alignment | distance | position | gene name | gene id |
| GU305914.1 | 84131 | 84153 | - | 0 | CAATCCGTCCAACATCTTGT | CGG | \|\|\|\|\|\|\|\|[\|\|\|\|\|\|\|\|\|\|\|\|]PAM | NA | Intergenic | NA | NA |
| T27863 | GTCGGCCATCTACGAGAGACCGG | 1000 | | | | | | | | | |
| Oligo fwd | CACCGTCGGCCATCTACGAGAGAC | | | | | | | | | | |
| Oligo rev | AAACGTCTCTCGTAGATGGCCGAC | | | | | | | | | | |
| Chromosome | start | end | strand | MM | target_seq | PAM | alignment | distance | position | gene name | gene id |
| GU305914.1 | 84113 | 84135 | - | 0 | GTCGGCCATCTACGAGAGAC | CGG | \|\|\|\|\|\|\|\|[\|\|\|\|\|\|\|\|\|\|\|\|]PAM | NA | Intergenic | NA | NA |
| T27864 | GAGACCGGTCACCGCCAAGCTGG | 1000 | | | | | | | | | |
| Oligo fwd | CACCGAGACCGGTCACCGCCAAGC | | | | | | | | | | |
| Oligo rev | AAACGCTTGGCGGTGACCGGTCTC | | | | | | | | | | |
| Chromosome | start | end | strand | MM | target_seq | PAM | alignment | distance | position | gene name | gene id |
| GU305914.1 | 84098 | 84120 | - | 0 | GAGACCGGTCACCGCCAAGC | TGG | \|\|\|\|\|\|\|\|[\|\|\|\|\|\|\|\|\|\|\|\|]PAM | NA | Intergenic | NA | NA |
| T27865 | ACCGGTCACCGCCAAGCTGGCGG | 1000 | | | | | | | | | |
| Oligo fwd | CACCACCGGTCACCGCCAAGCTGG | | | | | | | | | | |
| Oligo rev | AAACCCAGCTTGGCGGTGACCGGT | | | | | | | | | | |
| Chromosome | start | end | strand | MM | target_seq | PAM | alignment | distance | position | gene name | gene id |
| GU305914.1 | 84095 | 84117 | - | 0 | ACCGGTCACCGCCAAGCTGG | CGG | \|\|\|\|\|\|\|\|[\|\|\|\|\|\|\|\|\|\|\|\|]PAM | NA | Intergenic | NA | NA |
| T27866 | CCGGTCACCGCCAAGCTGGCGGG | 1000 | | | | | | | | | |
| Oligo fwd | CACCCCGGTCACCGCCAAGCTGGC | | | | | | | | | | |
| Oligo rev | AAACGCCAGCTTGGCGGTGACCGG | | | | | | | | | | |
| Chromosome | start | end | strand | MM | target_seq | PAM | alignment | distance | position | gene name | gene id |
| GU305914.1 | 84094 | 84116 | - | 0 | CCGGTCACCGCCAAGCTGGC | GGG | \|\|\|\|\|\|\|\|[\|\|\|\|\|\|\|\|\|\|\|\|]PAM | NA | Intergenic | NA | NA |
| T27867 | CGCGATGTCCGAGTGTGTCAAGG | 1000 | | | | | | | | | |
| Oligo fwd | CACCCGCGATGTCCGAGTGTGTCA | | | | | | | | | | |
| Oligo rev | AAACTGACACACTCGGACATCGCG | | | | | | | | | | |
| Chromosome | start | end | strand | MM | target_seq | PAM | alignment | distance | position | gene name | gene id |
| GU305914.1 | 84062 | 84084 | - | 0 | CGCGATGTCCGAGTGTGTCA | AGG | \|\|\|\|\|\|\|\|[\|\|\|\|\|\|\|\|\|\|\|\|]PAM | NA | Intergenic | NA | NA |
| T27868 | GGTCGATCAGAGTAGTGTGAAGG | 1000 | | | | | | | | | |
| Oligo fwd | CACCGGTCGATCAGAGTAGTGTGA | | | | | | | | | | |
| Oligo rev | AAACTCACACTACTCTGATCGACC | | | | | | | | | | |
| Chromosome | start | end | strand | MM | target_seq | PAM | alignment | distance | position | gene name | gene id |
| GU305914.1 | 84041 | 84063 | - | 0 | GGTCGATCAGAGTAGTGTGA | AGG | \|\|\|\|\|\|\|\|[\|\|\|\|\|\|\|\|\|\|\|\|]PAM | NA | Intergenic | NA | NA |
| T27869 | GAGTAGTGTGAAGGTCCTGAAGG | 1000 | | | | | | | | | |
| Oligo fwd | CACCGAGTAGTGTGAAGGTCCTGA | | | | | | | | | | |
| Oligo rev | AAACTCAGGACCTTCACACTACTC | | | | | | | | | | |
| Chromosome | start | end | strand | MM | target_seq | PAM | alignment | distance | position | gene name | gene id |
| GU305914.1 | 84032 | 84054 | - | 0 | GAGTAGTGTGAAGGTCCTGA | AGG | \|\|\|\|\|\|\|\|[\|\|\|\|\|\|\|\|\|\|\|\|]PAM | NA | Intergenic | NA | NA |
| T27870 | TGAAGGTCCTGAAGGACATGCGG | 1000 | | | | | | | | | |
| Oligo fwd | CACCTGAAGGTCCTGAAGGACATG | | | | | | | | | | |
| Oligo rev | AAACCATGTCCTTCAGGACCTTCA | | | | | | | | | | |
| Chromosome | start | end | strand | MM | target_seq | PAM | alignment | distance | position | gene name | gene id |
| GU305914.1 | 84024 | 84046 | - | 0 | TGAAGGTCCTGAAGGACATG | CGG | \|\|\|\|\|\|\|\|[\|\|\|\|\|\|\|\|\|\|\|\|]PAM | NA | Intergenic | NA | NA |
| T27871 | GAAGGACATGCGGATCTTCCAGG | 1000 | | | | | | | | | |
| Oligo fwd | CACCGAAGGACATGCGGATCTTCC | | | | | | | | | | |
| Oligo rev | AAACGGAAGATCCGCATGTCCTTC | | | | | | | | | | |
| Chromosome | start | end | strand | MM | target_seq | PAM | alignment | distance | position | gene name | gene id |
| GU305914.1 | 84014 | 84036 | - | 0 | GAAGGACATGCGGATCTTCC | AGG | \|\|\|\|\|\|\|\|[\|\|\|\|\|\|\|\|\|\|\|\|]PAM | NA | Intergenic | NA | NA |
| T27872 | GACATGCGGATCTTCCAGGACGG | 1000 | | | | | | | | | |
| Oligo fwd | CACCGACATGCGGATCTTCCAGGA | | | | | | | | | | |
| Oligo rev | AAACTCCTGGAAGATCCGCATGTC | | | | | | | | | | |
| Chromosome | start | end | strand | MM | target_seq | PAM | alignment | distance | position | gene name | gene id |
| GU305914.1 | 84010 | 84032 | - | 0 | GACATGCGGATCTTCCAGGA | CGG | \|\|\|\|\|\|\|\|[\|\|\|\|\|\|\|\|\|\|\|\|]PAM | NA | Intergenic | NA | NA |
| T27873 | ACATGCGGATCTTCCAGGACGGG | 1000 | | | | | | | | | |
| Oligo fwd | CACCACATGCGGATCTTCCAGGAC | | | | | | | | | | |
| Oligo rev | AAACGTCCTGGAAGATCCGCATGT | | | | | | | | | | |
| Chromosome | start | end | strand | MM | target_seq | PAM | alignment | distance | position | gene name | gene id |
| GU305914.1 | 84009 | 84031 | - | 0 | ACATGCGGATCTTCCAGGAC | GGG | \|\|\|\|\|\|\|\|[\|\|\|\|\|\|\|\|\|\|\|\|]PAM | NA | Intergenic | NA | NA |
| T27874 | GCGGATCTTCCAGGACGGGAAGG | 1000 | | | | | | | | | |
| Oligo fwd | CACCGCGGATCTTCCAGGACGGGA | | | | | | | | | | |
| Oligo rev | AAACTCCCGTCCTGGAAGATCCGC | | | | | | | | | | |

<table>
<tr><td>Chromosome</td><td>start</td><td>end</td><td>strand</td><td>MM</td><td>target_seq</td><td>PAM</td><td>alignment</td><td>distance</td><td>position</td><td>gene name</td><td>gene id</td></tr>
<tr><td>GU305914.1</td><td>84005</td><td>84027</td><td>-</td><td>0</td><td>GCGGATCTTCCAGGACGGGA</td><td>AGG</td><td>||||||||[||||||||||||]PAM</td><td>NA</td><td>Intergenic</td><td>NA</td><td>NA</td></tr>
<tr><td>T27875</td><td>CAACTGTTATTCGAGACCGCTGG</td><td>1000</td><td></td><td></td><td></td><td></td><td></td><td></td><td></td><td></td><td></td></tr>
<tr><td>Oligo fwd</td><td>CACCCAACTGTTATTCGAGACCGC</td><td></td><td></td><td></td><td></td><td></td><td></td><td></td><td></td><td></td><td></td></tr>
<tr><td>Oligo rev</td><td>AAACGCGGTCTCGAATAACAGTTG</td><td></td><td></td><td></td><td></td><td></td><td></td><td></td><td></td><td></td><td></td></tr>
<tr><td>Chromosome</td><td>start</td><td>end</td><td>strand</td><td>MM</td><td>target_seq</td><td>PAM</td><td>alignment</td><td>distance</td><td>position</td><td>gene name</td><td>gene id</td></tr>
<tr><td>GU305914.1</td><td>83978</td><td>84000</td><td>-</td><td>0</td><td>CAACTGTTATTCGAGACCGC</td><td>TGG</td><td>||||||||[||||||||||||]PAM</td><td>NA</td><td>Intergenic</td><td>NA</td><td>NA</td></tr>
<tr><td>T27876</td><td>CAACTCGACGAAGCTCGAGTCGG</td><td>1000</td><td></td><td></td><td></td><td></td><td></td><td></td><td></td><td></td><td></td></tr>
<tr><td>Oligo fwd</td><td>CACCCAACTCGACGAAGCTCGAGT</td><td></td><td></td><td></td><td></td><td></td><td></td><td></td><td></td><td></td><td></td></tr>
<tr><td>Oligo rev</td><td>AAACACTCGAGCTTCGTCGAGTTG</td><td></td><td></td><td></td><td></td><td></td><td></td><td></td><td></td><td></td><td></td></tr>
<tr><td>Chromosome</td><td>start</td><td>end</td><td>strand</td><td>MM</td><td>target_seq</td><td>PAM</td><td>alignment</td><td>distance</td><td>position</td><td>gene name</td><td>gene id</td></tr>
<tr><td>GU305914.1</td><td>83939</td><td>83961</td><td>-</td><td>0</td><td>CAACTCGACGAAGCTCGAGT</td><td>CGG</td><td>||||||||[||||||||||||]PAM</td><td>NA</td><td>Intergenic</td><td>NA</td><td>NA</td></tr>
<tr><td>T27877</td><td>AACTCGACGAAGCTCGAGTCGGG</td><td>1000</td><td></td><td></td><td></td><td></td><td></td><td></td><td></td><td></td><td></td></tr>
<tr><td>Oligo fwd</td><td>CACCAACTCGACGAAGCTCGAGTC</td><td></td><td></td><td></td><td></td><td></td><td></td><td></td><td></td><td></td><td></td></tr>
<tr><td>Oligo rev</td><td>AAACGACTCGAGCTTCGTCGAGTT</td><td></td><td></td><td></td><td></td><td></td><td></td><td></td><td></td><td></td><td></td></tr>
<tr><td>Chromosome</td><td>start</td><td>end</td><td>strand</td><td>MM</td><td>target_seq</td><td>PAM</td><td>alignment</td><td>distance</td><td>position</td><td>gene name</td><td>gene id</td></tr>
<tr><td>GU305914.1</td><td>83938</td><td>83960</td><td>-</td><td>0</td><td>AACTCGACGAAGCTCGAGTC</td><td>GGG</td><td>||||||||[||||||||||||]PAM</td><td>NA</td><td>Intergenic</td><td>NA</td><td>NA</td></tr>
<tr><td>T27878</td><td>GAAGCTCGAGTCGGGCCAGCTGG</td><td>1000</td><td></td><td></td><td></td><td></td><td></td><td></td><td></td><td></td><td></td></tr>
<tr><td>Oligo fwd</td><td>CACCGAAGCTCGAGTCGGGCCAGC</td><td></td><td></td><td></td><td></td><td></td><td></td><td></td><td></td><td></td><td></td></tr>
<tr><td>Oligo rev</td><td>AAACGCTGGCCCGACTCGAGCTTC</td><td></td><td></td><td></td><td></td><td></td><td></td><td></td><td></td><td></td><td></td></tr>
<tr><td>Chromosome</td><td>start</td><td>end</td><td>strand</td><td>MM</td><td>target_seq</td><td>PAM</td><td>alignment</td><td>distance</td><td>position</td><td>gene name</td><td>gene id</td></tr>
<tr><td>GU305914.1</td><td>83930</td><td>83952</td><td>-</td><td>0</td><td>GAAGCTCGAGTCGGGCCAGC</td><td>TGG</td><td>||||||||[||||||||||||]PAM</td><td>NA</td><td>Intergenic</td><td>NA</td><td>NA</td></tr>
<tr><td>GU305914.1</td><td>99574</td><td>99596</td><td>-</td><td>4</td><td>GTCGCTCGAGTCGGGACAGT</td><td>CGG</td><td>|--|||||[|||||||-|||-]PAM</td><td>NA</td><td>Intergenic</td><td>NA</td><td>NA</td></tr>
<tr><td>T27879</td><td>AAGCTCGAGTCGGGCCAGCTGGG</td><td>1000</td><td></td><td></td><td></td><td></td><td></td><td></td><td></td><td></td><td></td></tr>
<tr><td>Oligo fwd</td><td>CACCAAGCTCGAGTCGGGCCAGCT</td><td></td><td></td><td></td><td></td><td></td><td></td><td></td><td></td><td></td><td></td></tr>
<tr><td>Oligo rev</td><td>AAACAGCTGGCCCGACTCGAGCTT</td><td></td><td></td><td></td><td></td><td></td><td></td><td></td><td></td><td></td><td></td></tr>
<tr><td>Chromosome</td><td>start</td><td>end</td><td>strand</td><td>MM</td><td>target_seq</td><td>PAM</td><td>alignment</td><td>distance</td><td>position</td><td>gene name</td><td>gene id</td></tr>
<tr><td>GU305914.1</td><td>83929</td><td>83951</td><td>-</td><td>0</td><td>AAGCTCGAGTCGGGCCAGCT</td><td>GGG</td><td>||||||||[||||||||||||]PAM</td><td>NA</td><td>Intergenic</td><td>NA</td><td>NA</td></tr>
<tr><td>T27880</td><td>CGAGAACAACGAGATCATGCTGG</td><td>1000</td><td></td><td></td><td></td><td></td><td></td><td></td><td></td><td></td><td></td></tr>
<tr><td>Oligo fwd</td><td>CACCCGAGAACAACGAGATCATGC</td><td></td><td></td><td></td><td></td><td></td><td></td><td></td><td></td><td></td><td></td></tr>
<tr><td>Oligo rev</td><td>AAACGCATGATCTCGTTGTTCTCG</td><td></td><td></td><td></td><td></td><td></td><td></td><td></td><td></td><td></td><td></td></tr>
<tr><td>Chromosome</td><td>start</td><td>end</td><td>strand</td><td>MM</td><td>target_seq</td><td>PAM</td><td>alignment</td><td>distance</td><td>position</td><td>gene name</td><td>gene id</td></tr>
<tr><td>GU305914.1</td><td>83906</td><td>83928</td><td>-</td><td>0</td><td>CGAGAACAACGAGATCATGC</td><td>TGG</td><td>||||||||[||||||||||||]PAM</td><td>NA</td><td>Intergenic</td><td>NA</td><td>NA</td></tr>
<tr><td>T27881</td><td>GAGAACAACGAGATCATGCTGGG</td><td>1000</td><td></td><td></td><td></td><td></td><td></td><td></td><td></td><td></td><td></td></tr>
<tr><td>Oligo fwd</td><td>CACCGAGAACAACGAGATCATGCT</td><td></td><td></td><td></td><td></td><td></td><td></td><td></td><td></td><td></td><td></td></tr>
<tr><td>Oligo rev</td><td>AAACAGCATGATCTCGTTGTTCTC</td><td></td><td></td><td></td><td></td><td></td><td></td><td></td><td></td><td></td><td></td></tr>
<tr><td>Chromosome</td><td>start</td><td>end</td><td>strand</td><td>MM</td><td>target_seq</td><td>PAM</td><td>alignment</td><td>distance</td><td>position</td><td>gene name</td><td>gene id</td></tr>
<tr><td>GU305914.1</td><td>83905</td><td>83927</td><td>-</td><td>0</td><td>GAGAACAACGAGATCATGCT</td><td>GGG</td><td>||||||||[||||||||||||]PAM</td><td>NA</td><td>Intergenic</td><td>NA</td><td>NA</td></tr>
<tr><td>T27882</td><td>AGATCATGCTGGGCACCTTTCGG</td><td>1000</td><td></td><td></td><td></td><td></td><td></td><td></td><td></td><td></td><td></td></tr>
<tr><td>Oligo fwd</td><td>CACCAGATCATGCTGGGCACCTTT</td><td></td><td></td><td></td><td></td><td></td><td></td><td></td><td></td><td></td><td></td></tr>
<tr><td>Oligo rev</td><td>AAACAAAGGTGCCCAGCATGATCT</td><td></td><td></td><td></td><td></td><td></td><td></td><td></td><td></td><td></td><td></td></tr>
<tr><td>Chromosome</td><td>start</td><td>end</td><td>strand</td><td>MM</td><td>target_seq</td><td>PAM</td><td>alignment</td><td>distance</td><td>position</td><td>gene name</td><td>gene id</td></tr>
<tr><td>GU305914.1</td><td>83895</td><td>83917</td><td>-</td><td>0</td><td>AGATCATGCTGGGCACCTTT</td><td>CGG</td><td>||||||||[||||||||||||]PAM</td><td>NA</td><td>Intergenic</td><td>NA</td><td>NA</td></tr>
<tr><td>T27883</td><td>CATGCTGGGCACCTTTCGGACGG</td><td>1000</td><td></td><td></td><td></td><td></td><td></td><td></td><td></td><td></td><td></td></tr>
<tr><td>Oligo fwd</td><td>CACCCATGCTGGGCACCTTTCGGA</td><td></td><td></td><td></td><td></td><td></td><td></td><td></td><td></td><td></td><td></td></tr>
<tr><td>Oligo rev</td><td>AAACTCCGAAAGGTGCCCAGCATG</td><td></td><td></td><td></td><td></td><td></td><td></td><td></td><td></td><td></td><td></td></tr>
<tr><td>Chromosome</td><td>start</td><td>end</td><td>strand</td><td>MM</td><td>target_seq</td><td>PAM</td><td>alignment</td><td>distance</td><td>position</td><td>gene name</td><td>gene id</td></tr>
<tr><td>GU305914.1</td><td>83891</td><td>83913</td><td>-</td><td>0</td><td>CATGCTGGGCACCTTTCGGA</td><td>CGG</td><td>||||||||[||||||||||||]PAM</td><td>NA</td><td>Intergenic</td><td>NA</td><td>NA</td></tr>
<tr><td>T27884</td><td>AGAACTGCGACACGAACAGCAGG</td><td>1000</td><td></td><td></td><td></td><td></td><td></td><td></td><td></td><td></td><td></td></tr>
<tr><td>Oligo fwd</td><td>CACCAGAACTGCGACACGAACAGC</td><td></td><td></td><td></td><td></td><td></td><td></td><td></td><td></td><td></td><td></td></tr>
<tr><td>Oligo rev</td><td>AAACGCTGTTCGTGTCGCAGTTCT</td><td></td><td></td><td></td><td></td><td></td><td></td><td></td><td></td><td></td><td></td></tr>
<tr><td>Chromosome</td><td>start</td><td>end</td><td>strand</td><td>MM</td><td>target_seq</td><td>PAM</td><td>alignment</td><td>distance</td><td>position</td><td>gene name</td><td>gene id</td></tr>
<tr><td>GU305914.1</td><td>83868</td><td>83890</td><td>-</td><td>0</td><td>AGAACTGCGACACGAACAGC</td><td>AGG</td><td>||||||||[||||||||||||]PAM</td><td>NA</td><td>Intergenic</td><td>NA</td><td>NA</td></tr>
<tr><td>T27885</td><td>CAGCAGGAAGATCTTCGTCGTGG</td><td>1000</td><td></td><td></td><td></td><td></td><td></td><td></td><td></td><td></td><td></td></tr>
<tr><td>Oligo fwd</td><td>CACCCAGCAGGAAGATCTTCGTCG</td><td></td><td></td><td></td><td></td><td></td><td></td><td></td><td></td><td></td><td></td></tr>
<tr><td>Oligo rev</td><td>AAACCGACGAAGATCTTCCTGCTG</td><td></td><td></td><td></td><td></td><td></td><td></td><td></td><td></td><td></td><td></td></tr>
<tr><td>Chromosome</td><td>start</td><td>end</td><td>strand</td><td>MM</td><td>target_seq</td><td>PAM</td><td>alignment</td><td>distance</td><td>position</td><td>gene name</td><td>gene id</td></tr>
<tr><td>GU305914.1</td><td>83852</td><td>83874</td><td>-</td><td>0</td><td>CAGCAGGAAGATCTTCGTCG</td><td>TGG</td><td>||||||||[||||||||||||]PAM</td><td>NA</td><td>Intergenic</td><td>NA</td><td>NA</td></tr>
<tr><td>T27886</td><td>AGCAGGAAGATCTTCGTCGTGGG</td><td>1000</td><td></td><td></td><td></td><td></td><td></td><td></td><td></td><td></td><td></td></tr>
<tr><td>Oligo fwd</td><td>CACCAGCAGGAAGATCTTCGTCGT</td><td></td><td></td><td></td><td></td><td></td><td></td><td></td><td></td><td></td><td></td></tr>
<tr><td>Oligo rev</td><td>AAACACGACGAAGATCTTCCTGCT</td><td></td><td></td><td></td><td></td><td></td><td></td><td></td><td></td><td></td><td></td></tr>
<tr><td>Chromosome</td><td>start</td><td>end</td><td>strand</td><td>MM</td><td>target_seq</td><td>PAM</td><td>alignment</td><td>distance</td><td>position</td><td>gene name</td><td>gene id</td></tr>
<tr><td>GU305914.1</td><td>83851</td><td>83873</td><td>-</td><td>0</td><td>AGCAGGAAGATCTTCGTCGT</td><td>GGG</td><td>||||||||[||||||||||||]PAM</td><td>NA</td><td>Intergenic</td><td>NA</td><td>NA</td></tr>
<tr><td>GU305914.1</td><td>136877</td><td>136899</td><td>+</td><td>4</td><td>AACATGAAGCTCTTCGTCTT</td><td>CGG</td><td>|-||-|||[|-||||||||-|]PAM</td><td>NA</td><td>Intergenic</td><td>NA</td><td>NA</td></tr>
<tr><td>T27887</td><td>GCAGGAAGATCTTCGTCGTGGGG</td><td>1000</td><td></td><td></td><td></td><td></td><td></td><td></td><td></td><td></td><td></td></tr>
<tr><td>Oligo fwd</td><td>CACCGCAGGAAGATCTTCGTCGTG</td><td></td><td></td><td></td><td></td><td></td><td></td><td></td><td></td><td></td><td></td></tr>
<tr><td>Oligo rev</td><td>AAACCACGACGAAGATCTTCCTGC</td><td></td><td></td><td></td><td></td><td></td><td></td><td></td><td></td><td></td><td></td></tr>
<tr><td>Chromosome</td><td>start</td><td>end</td><td>strand</td><td>MM</td><td>target_seq</td><td>PAM</td><td>alignment</td><td>distance</td><td>position</td><td>gene name</td><td>gene id</td></tr>
<tr><td>GU305914.1</td><td>83850</td><td>83872</td><td>-</td><td>0</td><td>GCAGGAAGATCTTCGTCGTG</td><td>GGG</td><td>||||||||[||||||||||||]PAM</td><td>NA</td><td>Intergenic</td><td>NA</td><td>NA</td></tr>
<tr><td>T27888</td><td>GATCTTCGTCGTGGGGACTGTGG</td><td>1000</td><td></td><td></td><td></td><td></td><td></td><td></td><td></td><td></td><td></td></tr>
<tr><td>Oligo fwd</td><td>CACCGATCTTCGTCGTGGGGACTG</td><td></td><td></td><td></td><td></td><td></td><td></td><td></td><td></td><td></td><td></td></tr>
<tr><td>Oligo rev</td><td>AAACCAGTCCCCACGACGAAGATC</td><td></td><td></td><td></td><td></td><td></td><td></td><td></td><td></td><td></td><td></td></tr>
<tr><td>Chromosome</td><td>start</td><td>end</td><td>strand</td><td>MM</td><td>target_seq</td><td>PAM</td><td>alignment</td><td>distance</td><td>position</td><td>gene name</td><td>gene id</td></tr>
<tr><td>GU305914.1</td><td>83843</td><td>83865</td><td>-</td><td>0</td><td>GATCTTCGTCGTGGGGACTG</td><td>TGG</td><td>||||||||[||||||||||||]PAM</td><td>NA</td><td>Intergenic</td><td>NA</td><td>NA</td></tr>
</table>

<table>
<tr><td>T27889</td><td>ATCTTCGTCGTGGGGACTGTGGG</td><td>1000</td><td></td><td></td><td></td><td></td><td></td><td></td><td></td><td></td><td></td></tr>
<tr><td>Oligo fwd</td><td>CACCATCTTCGTCGTGGGGACTGT</td><td></td><td></td><td></td><td></td><td></td><td></td><td></td><td></td><td></td><td></td></tr>
<tr><td>Oligo rev</td><td>AAACACAGTCCCCACGACGAAGAT</td><td></td><td></td><td></td><td></td><td></td><td></td><td></td><td></td><td></td><td></td></tr>
<tr><td>Chromosome</td><td>start</td><td>end</td><td>strand</td><td>MM</td><td>target_seq</td><td>PAM</td><td>alignment</td><td>distance</td><td>position</td><td>gene name</td><td>gene id</td></tr>
<tr><td>GU305914.1</td><td>83842</td><td>83864</td><td>-</td><td>0</td><td>ATCTTCGTCGTGGGGACTGT</td><td>GGG</td><td>||||||||[||||||||||||]PAM</td><td>NA</td><td>Intergenic</td><td>NA</td><td>NA</td></tr>
<tr><td>T27890</td><td>GGACTGTGGGCTACGAGTATCGG</td><td>1000</td><td></td><td></td><td></td><td></td><td></td><td></td><td></td><td></td><td></td></tr>
<tr><td>Oligo fwd</td><td>CACCGGACTGTGGGCTACGAGTAT</td><td></td><td></td><td></td><td></td><td></td><td></td><td></td><td></td><td></td><td></td></tr>
<tr><td>Oligo rev</td><td>AAACATACTCGTAGCCCACAGTCC</td><td></td><td></td><td></td><td></td><td></td><td></td><td></td><td></td><td></td><td></td></tr>
<tr><td>Chromosome</td><td>start</td><td>end</td><td>strand</td><td>MM</td><td>target_seq</td><td>PAM</td><td>alignment</td><td>distance</td><td>position</td><td>gene name</td><td>gene id</td></tr>
<tr><td>GU305914.1</td><td>83829</td><td>83851</td><td>-</td><td>0</td><td>GGACTGTGGGCTACGAGTAT</td><td>CGG</td><td>||||||||[||||||||||||]PAM</td><td>NA</td><td>Intergenic</td><td>NA</td><td>NA</td></tr>
<tr><td>T27891</td><td>GACTGTGGGCTACGAGTATCGGG</td><td>1000</td><td></td><td></td><td></td><td></td><td></td><td></td><td></td><td></td><td></td></tr>
<tr><td>Oligo fwd</td><td>CACCGACTGTGGGCTACGAGTATC</td><td></td><td></td><td></td><td></td><td></td><td></td><td></td><td></td><td></td><td></td></tr>
<tr><td>Oligo rev</td><td>AAACGATACTCGTAGCCCACAGTC</td><td></td><td></td><td></td><td></td><td></td><td></td><td></td><td></td><td></td><td></td></tr>
<tr><td>Chromosome</td><td>start</td><td>end</td><td>strand</td><td>MM</td><td>target_seq</td><td>PAM</td><td>alignment</td><td>distance</td><td>position</td><td>gene name</td><td>gene id</td></tr>
<tr><td>GU305914.1</td><td>83828</td><td>83850</td><td>-</td><td>0</td><td>GACTGTGGGCTACGAGTATC</td><td>GGG</td><td>||||||||[||||||||||||]PAM</td><td>NA</td><td>Intergenic</td><td>NA</td><td>NA</td></tr>
<tr><td>T27892</td><td>AGTATCGGGACTACCGTTTCAGG</td><td>1000</td><td></td><td></td><td></td><td></td><td></td><td></td><td></td><td></td><td></td></tr>
<tr><td>Oligo fwd</td><td>CACCAGTATCGGGACTACCGTTTC</td><td></td><td></td><td></td><td></td><td></td><td></td><td></td><td></td><td></td><td></td></tr>
<tr><td>Oligo rev</td><td>AAACGAAACGGTAGTCCCGATACT</td><td></td><td></td><td></td><td></td><td></td><td></td><td></td><td></td><td></td><td></td></tr>
<tr><td>Chromosome</td><td>start</td><td>end</td><td>strand</td><td>MM</td><td>target_seq</td><td>PAM</td><td>alignment</td><td>distance</td><td>position</td><td>gene name</td><td>gene id</td></tr>
<tr><td>GU305914.1</td><td>83814</td><td>83836</td><td>-</td><td>0</td><td>AGTATCGGGACTACCGTTTC</td><td>AGG</td><td>||||||||[||||||||||||]PAM</td><td>NA</td><td>Intergenic</td><td>NA</td><td>NA</td></tr>
<tr><td>T27893</td><td>TTTCAGGAACGTGACCAGTCTGG</td><td>1000</td><td></td><td></td><td></td><td></td><td></td><td></td><td></td><td></td><td></td></tr>
<tr><td>Oligo fwd</td><td>CACCTTTCAGGAACGTGACCAGTC</td><td></td><td></td><td></td><td></td><td></td><td></td><td></td><td></td><td></td><td></td></tr>
<tr><td>Oligo rev</td><td>AAACGACTGGTCACGTTCCTGAAA</td><td></td><td></td><td></td><td></td><td></td><td></td><td></td><td></td><td></td><td></td></tr>
<tr><td>Chromosome</td><td>start</td><td>end</td><td>strand</td><td>MM</td><td>target_seq</td><td>PAM</td><td>alignment</td><td>distance</td><td>position</td><td>gene name</td><td>gene id</td></tr>
<tr><td>GU305914.1</td><td>83798</td><td>83820</td><td>-</td><td>0</td><td>TTTCAGGAACGTGACCAGTC</td><td>TGG</td><td>||||||||[||||||||||||]PAM</td><td>NA</td><td>Intergenic</td><td>NA</td><td>NA</td></tr>
<tr><td>T27894</td><td>GACCTCGTCGACACCCTGATCGG</td><td>1000</td><td></td><td></td><td></td><td></td><td></td><td></td><td></td><td></td><td></td></tr>
<tr><td>Oligo fwd</td><td>CACCGACCTCGTCGACACCCTGAT</td><td></td><td></td><td></td><td></td><td></td><td></td><td></td><td></td><td></td><td></td></tr>
<tr><td>Oligo rev</td><td>AAACATCAGGGTGTCGACGAGGTC</td><td></td><td></td><td></td><td></td><td></td><td></td><td></td><td></td><td></td><td></td></tr>
<tr><td>Chromosome</td><td>start</td><td>end</td><td>strand</td><td>MM</td><td>target_seq</td><td>PAM</td><td>alignment</td><td>distance</td><td>position</td><td>gene name</td><td>gene id</td></tr>
<tr><td>GU305914.1</td><td>83767</td><td>83789</td><td>-</td><td>0</td><td>GACCTCGTCGACACCCTGAT</td><td>CGG</td><td>||||||||[||||||||||||]PAM</td><td>NA</td><td>Intergenic</td><td>NA</td><td>NA</td></tr>
<tr><td>T27895</td><td>ACCTCGTCGACACCCTGATCGGG</td><td>1000</td><td></td><td></td><td></td><td></td><td></td><td></td><td></td><td></td><td></td></tr>
<tr><td>Oligo fwd</td><td>CACCACCTCGTCGACACCCTGATC</td><td></td><td></td><td></td><td></td><td></td><td></td><td></td><td></td><td></td><td></td></tr>
<tr><td>Oligo rev</td><td>AAACGATCAGGGTGTCGACGAGGT</td><td></td><td></td><td></td><td></td><td></td><td></td><td></td><td></td><td></td><td></td></tr>
<tr><td>Chromosome</td><td>start</td><td>end</td><td>strand</td><td>MM</td><td>target_seq</td><td>PAM</td><td>alignment</td><td>distance</td><td>position</td><td>gene name</td><td>gene id</td></tr>
<tr><td>GU305914.1</td><td>83766</td><td>83788</td><td>-</td><td>0</td><td>ACCTCGTCGACACCCTGATC</td><td>GGG</td><td>||||||||[||||||||||||]PAM</td><td>NA</td><td>Intergenic</td><td>NA</td><td>NA</td></tr>
<tr><td>T27896</td><td>CGTCGACACCCTGATCGGGCTGG</td><td>1000</td><td></td><td></td><td></td><td></td><td></td><td></td><td></td><td></td><td></td></tr>
<tr><td>Oligo fwd</td><td>CACCCGTCGACACCCTGATCGGGC</td><td></td><td></td><td></td><td></td><td></td><td></td><td></td><td></td><td></td><td></td></tr>
<tr><td>Oligo rev</td><td>AAACGCCCGATCAGGGTGTCGACG</td><td></td><td></td><td></td><td></td><td></td><td></td><td></td><td></td><td></td><td></td></tr>
<tr><td>Chromosome</td><td>start</td><td>end</td><td>strand</td><td>MM</td><td>target_seq</td><td>PAM</td><td>alignment</td><td>distance</td><td>position</td><td>gene name</td><td>gene id</td></tr>
<tr><td>GU305914.1</td><td>83762</td><td>83784</td><td>-</td><td>0</td><td>CGTCGACACCCTGATCGGGC</td><td>TGG</td><td>||||||||[||||||||||||]PAM</td><td>NA</td><td>Intergenic</td><td>NA</td><td>NA</td></tr>
<tr><td>T27897</td><td>CGGGCTGGACATCGAGCCCCTGG</td><td>1000</td><td></td><td></td><td></td><td></td><td></td><td></td><td></td><td></td><td></td></tr>
<tr><td>Oligo fwd</td><td>CACCCGGGCTGGACATCGAGCCCC</td><td></td><td></td><td></td><td></td><td></td><td></td><td></td><td></td><td></td><td></td></tr>
<tr><td>Oligo rev</td><td>AAACGGGGCTCGATGTCCAGCCCG</td><td></td><td></td><td></td><td></td><td></td><td></td><td></td><td></td><td></td><td></td></tr>
<tr><td>Chromosome</td><td>start</td><td>end</td><td>strand</td><td>MM</td><td>target_seq</td><td>PAM</td><td>alignment</td><td>distance</td><td>position</td><td>gene name</td><td>gene id</td></tr>
<tr><td>GU305914.1</td><td>83747</td><td>83769</td><td>-</td><td>0</td><td>CGGGCTGGACATCGAGCCCC</td><td>TGG</td><td>||||||||[||||||||||||]PAM</td><td>NA</td><td>Intergenic</td><td>NA</td><td>NA</td></tr>
<tr><td>T27898</td><td>CCTGGAGAACACCGACTTCAAGG</td><td>1000</td><td></td><td></td><td></td><td></td><td></td><td></td><td></td><td></td><td></td></tr>
<tr><td>Oligo fwd</td><td>CACCCCTGGAGAACACCGACTTCA</td><td></td><td></td><td></td><td></td><td></td><td></td><td></td><td></td><td></td><td></td></tr>
<tr><td>Oligo rev</td><td>AAACTGAAGTCGGTGTTCTCCAGG</td><td></td><td></td><td></td><td></td><td></td><td></td><td></td><td></td><td></td><td></td></tr>
<tr><td>Chromosome</td><td>start</td><td>end</td><td>strand</td><td>MM</td><td>target_seq</td><td>PAM</td><td>alignment</td><td>distance</td><td>position</td><td>gene name</td><td>gene id</td></tr>
<tr><td>GU305914.1</td><td>83729</td><td>83751</td><td>-</td><td>0</td><td>CCTGGAGAACACCGACTTCA</td><td>AGG</td><td>||||||||[||||||||||||]PAM</td><td>NA</td><td>Intergenic</td><td>NA</td><td>NA</td></tr>
<tr><td>T27899</td><td>GGTGCTCGAGCTGTACTCGAAGG</td><td>1000</td><td></td><td></td><td></td><td></td><td></td><td></td><td></td><td></td><td></td></tr>
<tr><td>Oligo fwd</td><td>CACCGGTGCTCGAGCTGTACTCGA</td><td></td><td></td><td></td><td></td><td></td><td></td><td></td><td></td><td></td><td></td></tr>
<tr><td>Oligo rev</td><td>AAACTCGAGTACAGCTCGAGCACC</td><td></td><td></td><td></td><td></td><td></td><td></td><td></td><td></td><td></td><td></td></tr>
<tr><td>Chromosome</td><td>start</td><td>end</td><td>strand</td><td>MM</td><td>target_seq</td><td>PAM</td><td>alignment</td><td>distance</td><td>position</td><td>gene name</td><td>gene id</td></tr>
<tr><td>GU305914.1</td><td>83708</td><td>83730</td><td>-</td><td>0</td><td>GGTGCTCGAGCTGTACTCGA</td><td>AGG</td><td>||||||||[||||||||||||]PAM</td><td>NA</td><td>Intergenic</td><td>NA</td><td>NA</td></tr>
<tr><td>T27900</td><td>GTGCTCGAGCTGTACTCGAAGGG</td><td>1000</td><td></td><td></td><td></td><td></td><td></td><td></td><td></td><td></td><td></td></tr>
<tr><td>Oligo fwd</td><td>CACCGTGCTCGAGCTGTACTCGAA</td><td></td><td></td><td></td><td></td><td></td><td></td><td></td><td></td><td></td><td></td></tr>
<tr><td>Oligo rev</td><td>AAACTTCGAGTACAGCTCGAGCAC</td><td></td><td></td><td></td><td></td><td></td><td></td><td></td><td></td><td></td><td></td></tr>
<tr><td>Chromosome</td><td>start</td><td>end</td><td>strand</td><td>MM</td><td>target_seq</td><td>PAM</td><td>alignment</td><td>distance</td><td>position</td><td>gene name</td><td>gene id</td></tr>
<tr><td>GU305914.1</td><td>83707</td><td>83729</td><td>-</td><td>0</td><td>GTGCTCGAGCTGTACTCGAA</td><td>GGG</td><td>||||||||[||||||||||||]PAM</td><td>NA</td><td>Intergenic</td><td>NA</td><td>NA</td></tr>
<tr><td>T27901</td><td>CGCCTCGAACGTGTTCAGCCTGG</td><td>1000</td><td></td><td></td><td></td><td></td><td></td><td></td><td></td><td></td><td></td></tr>
<tr><td>Oligo fwd</td><td>CACCCGCCTCGAACGTGTTCAGCC</td><td></td><td></td><td></td><td></td><td></td><td></td><td></td><td></td><td></td><td></td></tr>
<tr><td>Oligo rev</td><td>AAACGGCTGAACACGTTCGAGGCG</td><td></td><td></td><td></td><td></td><td></td><td></td><td></td><td></td><td></td><td></td></tr>
<tr><td>Chromosome</td><td>start</td><td>end</td><td>strand</td><td>MM</td><td>target_seq</td><td>PAM</td><td>alignment</td><td>distance</td><td>position</td><td>gene name</td><td>gene id</td></tr>
<tr><td>GU305914.1</td><td>83675</td><td>83697</td><td>-</td><td>0</td><td>CGCCTCGAACGTGTTCAGCC</td><td>TGG</td><td>||||||||[||||||||||||]PAM</td><td>NA</td><td>Intergenic</td><td>NA</td><td>NA</td></tr>
<tr><td>T27902</td><td>TCAGCCTGGACGAGATCATGCGG</td><td>1000</td><td></td><td></td><td></td><td></td><td></td><td></td><td></td><td></td><td></td></tr>
<tr><td>Oligo fwd</td><td>CACCTCAGCCTGGACGAGATCATG</td><td></td><td></td><td></td><td></td><td></td><td></td><td></td><td></td><td></td><td></td></tr>
<tr><td>Oligo rev</td><td>AAACCATGATCTCGTCCAGGCTGA</td><td></td><td></td><td></td><td></td><td></td><td></td><td></td><td></td><td></td><td></td></tr>
<tr><td>Chromosome</td><td>start</td><td>end</td><td>strand</td><td>MM</td><td>target_seq</td><td>PAM</td><td>alignment</td><td>distance</td><td>position</td><td>gene name</td><td>gene id</td></tr>
<tr><td>GU305914.1</td><td>83661</td><td>83683</td><td>-</td><td>0</td><td>TCAGCCTGGACGAGATCATG</td><td>CGG</td><td>||||||||[||||||||||||]PAM</td><td>NA</td><td>Intergenic</td><td>NA</td><td>NA</td></tr>
<tr><td>GU305914.1</td><td>196897</td><td>196919</td><td>-</td><td>3</td><td>CCTGCCTGGACGAGATCACG</td><td>GGG</td><td>-|-|||||[||||||||||-|]PAM</td><td>NA</td><td>Intergenic</td><td>NA</td><td>NA</td></tr>
<tr><td>T27903</td><td>CAGCCTGGACGAGATCATGCGGG</td><td>1000</td><td></td><td></td><td></td><td></td><td></td><td></td><td></td><td></td><td></td></tr>
<tr><td>Oligo fwd</td><td>CACCCAGCCTGGACGAGATCATGC</td><td></td><td></td><td></td><td></td><td></td><td></td><td></td><td></td><td></td><td></td></tr>
<tr><td>Oligo rev</td><td>AAACGCATGATCTCGTCCAGGCTG</td><td></td><td></td><td></td><td></td><td></td><td></td><td></td><td></td><td></td><td></td></tr>
<tr><td>Chromosome</td><td>start</td><td>end</td><td>strand</td><td>MM</td><td>target_seq</td><td>PAM</td><td>alignment</td><td>distance</td><td>position</td><td>gene name</td><td>gene id</td></tr>
</table>

| | | | | | | | | | | | |
|---|---|---|---|---|---|---|---|---|---|---|---|
| GU305914.1 | 83660 | 83682 | - | 0 | CAGCCTGGACGAGATCATGC | GGG | \|\|\|\|\|\|\|\|[\|\|\|\|\|\|\|\|\|\|\|\|]PAM | NA | Intergenic | NA | NA |
| T27904 | CAACACGCCCTCTTATCTCTTGG | 1000 | | | | | | | | | |
| Oligo fwd | CACCCAACACGCCCTCTTATCTCT | | | | | | | | | | |
| Oligo rev | AAACAGAGATAAGAGGGCGTGTTG | | | | | | | | | | |
| Chromosome | start | end | strand | MM | target_seq | PAM | alignment | distance | position | gene name | gene id |
| GU305914.1 | 83585 | 83607 | - | 0 | CAACACGCCCTCTTATCTCT | TGG | \|\|\|\|\|\|\|\|[\|\|\|\|\|\|\|\|\|\|\|\|]PAM | NA | Intergenic | NA | NA |
| T27905 | AACACGCCCTCTTATCTCTTGGG | 1000 | | | | | | | | | |
| Oligo fwd | CACCAACACGCCCTCTTATCTCTT | | | | | | | | | | |
| Oligo rev | AAACAAGAGATAAGAGGGCGTGTT | | | | | | | | | | |
| Chromosome | start | end | strand | MM | target_seq | PAM | alignment | distance | position | gene name | gene id |
| GU305914.1 | 83584 | 83606 | - | 0 | AACACGCCCTCTTATCTCTT | GGG | \|\|\|\|\|\|\|\|[\|\|\|\|\|\|\|\|\|\|\|\|]PAM | NA | Intergenic | NA | NA |
| T27906 | ACACGCCCTCTTATCTCTTGGGG | 1000 | | | | | | | | | |
| Oligo fwd | CACCACACGCCCTCTTATCTCTTG | | | | | | | | | | |
| Oligo rev | AAACCAAGAGATAAGAGGGCGTGT | | | | | | | | | | |
| Chromosome | start | end | strand | MM | target_seq | PAM | alignment | distance | position | gene name | gene id |
| GU305914.1 | 83583 | 83605 | - | 0 | ACACGCCCTCTTATCTCTTG | GGG | \|\|\|\|\|\|\|\|[\|\|\|\|\|\|\|\|\|\|\|\|]PAM | NA | Intergenic | NA | NA |
| T27907 | GCCCTCTTATCTCTTGGGGTTGG | 1000 | | | | | | | | | |
| Oligo fwd | CACCGCCCTCTTATCTCTTGGGGT | | | | | | | | | | |
| Oligo rev | AAACACCCCAAGAGATAAGAGGGC | | | | | | | | | | |
| Chromosome | start | end | strand | MM | target_seq | PAM | alignment | distance | position | gene name | gene id |
| GU305914.1 | 83579 | 83601 | - | 0 | GCCCTCTTATCTCTTGGGGT | TGG | \|\|\|\|\|\|\|\|[\|\|\|\|\|\|\|\|\|\|\|\|]PAM | NA | Intergenic | NA | NA |
| T27908 | GGGGTTGGATACTTTCATGCAGG | 1000 | | | | | | | | | |
| Oligo fwd | CACCGGGGTTGGATACTTTCATGC | | | | | | | | | | |
| Oligo rev | AAACGCATGAAAGTATCCAACCCC | | | | | | | | | | |
| Chromosome | start | end | strand | MM | target_seq | PAM | alignment | distance | position | gene name | gene id |
| GU305914.1 | 83564 | 83586 | - | 0 | GGGGTTGGATACTTTCATGC | AGG | \|\|\|\|\|\|\|\|[\|\|\|\|\|\|\|\|\|\|\|\|]PAM | NA | Intergenic | NA | NA |
| T27909 | GGGTTGGATACTTTCATGCAGGG | 1000 | | | | | | | | | |
| Oligo fwd | CACCGGGTTGGATACTTTCATGCA | | | | | | | | | | |
| Oligo rev | AAACTGCATGAAAGTATCCAACCC | | | | | | | | | | |
| Chromosome | start | end | strand | MM | target_seq | PAM | alignment | distance | position | gene name | gene id |
| GU305914.1 | 83563 | 83585 | - | 0 | GGGTTGGATACTTTCATGCA | GGG | \|\|\|\|\|\|\|\|[\|\|\|\|\|\|\|\|\|\|\|\|]PAM | NA | Intergenic | NA | NA |
| T27910 | GGTTGGATACTTTCATGCAGGGG | 1000 | | | | | | | | | |
| Oligo fwd | CACCGGTTGGATACTTTCATGCAG | | | | | | | | | | |
| Oligo rev | AAACCTGCATGAAAGTATCCAACC | | | | | | | | | | |
| Chromosome | start | end | strand | MM | target_seq | PAM | alignment | distance | position | gene name | gene id |
| GU305914.1 | 83562 | 83584 | - | 0 | GGTTGGATACTTTCATGCAG | GGG | \|\|\|\|\|\|\|\|[\|\|\|\|\|\|\|\|\|\|\|\|]PAM | NA | Intergenic | NA | NA |
| T27911 | GATACTTTCATGCAGGGGCTAGG | 1000 | | | | | | | | | |
| Oligo fwd | CACCGATACTTTCATGCAGGGGCT | | | | | | | | | | |
| Oligo rev | AAACAGCCCCTGCATGAAAGTATC | | | | | | | | | | |
| Chromosome | start | end | strand | MM | target_seq | PAM | alignment | distance | position | gene name | gene id |
| GU305914.1 | 83557 | 83579 | - | 0 | GATACTTTCATGCAGGGGCT | AGG | \|\|\|\|\|\|\|\|[\|\|\|\|\|\|\|\|\|\|\|\|]PAM | NA | Intergenic | NA | NA |
| T27912 | ATACTTTCATGCAGGGGCTAGGG | 1000 | | | | | | | | | |
| Oligo fwd | CACCATACTTTCATGCAGGGGCTA | | | | | | | | | | |
| Oligo rev | AAACTAGCCCCTGCATGAAAGTAT | | | | | | | | | | |
| Chromosome | start | end | strand | MM | target_seq | PAM | alignment | distance | position | gene name | gene id |
| GU305914.1 | 83556 | 83578 | - | 0 | ATACTTTCATGCAGGGGCTA | GGG | \|\|\|\|\|\|\|\|[\|\|\|\|\|\|\|\|\|\|\|\|]PAM | NA | Intergenic | NA | NA |
| T27913 | TACTTTCATGCAGGGGCTAGGGG | 1000 | | | | | | | | | |
| Oligo fwd | CACCTACTTTCATGCAGGGGCTAG | | | | | | | | | | |
| Oligo rev | AAACCTAGCCCCTGCATGAAAGTA | | | | | | | | | | |
| Chromosome | start | end | strand | MM | target_seq | PAM | alignment | distance | position | gene name | gene id |
| GU305914.1 | 83555 | 83577 | - | 0 | TACTTTCATGCAGGGGCTAG | GGG | \|\|\|\|\|\|\|\|[\|\|\|\|\|\|\|\|\|\|\|\|]PAM | NA | Intergenic | NA | NA |
| T27914 | CATGCAGGGGCTAGGGGTCGCGG | 1000 | | | | | | | | | |
| Oligo fwd | CACCCATGCAGGGGCTAGGGGTCG | | | | | | | | | | |
| Oligo rev | AAACCGACCCCTAGCCCCTGCATG | | | | | | | | | | |
| Chromosome | start | end | strand | MM | target_seq | PAM | alignment | distance | position | gene name | gene id |
| GU305914.1 | 83549 | 83571 | - | 0 | CATGCAGGGGCTAGGGGTCG | CGG | \|\|\|\|\|\|\|\|[\|\|\|\|\|\|\|\|\|\|\|\|]PAM | NA | Intergenic | NA | NA |
| T27915 | ATGCAGGGGCTAGGGGTCGCGGG | 1000 | | | | | | | | | |
| Oligo fwd | CACCATGCAGGGGCTAGGGGTCGC | | | | | | | | | | |
| Oligo rev | AAACGCGACCCCTAGCCCCTGCAT | | | | | | | | | | |
| Chromosome | start | end | strand | MM | target_seq | PAM | alignment | distance | position | gene name | gene id |
| GU305914.1 | 83548 | 83570 | - | 0 | ATGCAGGGGCTAGGGGTCGC | GGG | \|\|\|\|\|\|\|\|[\|\|\|\|\|\|\|\|\|\|\|\|]PAM | NA | Intergenic | NA | NA |
| T27916 | GGGGCTAGGGGTCGCGGGTAAGG | 1000 | | | | | | | | | |
| Oligo fwd | CACCGGGGCTAGGGGTCGCGGGTA | | | | | | | | | | |
| Oligo rev | AAACTACCCGCGACCCCTAGCCCC | | | | | | | | | | |
| Chromosome | start | end | strand | MM | target_seq | PAM | alignment | distance | position | gene name | gene id |
| GU305914.1 | 83543 | 83565 | - | 0 | GGGGCTAGGGGTCGCGGGTA | AGG | \|\|\|\|\|\|\|\|[\|\|\|\|\|\|\|\|\|\|\|\|]PAM | NA | Intergenic | NA | NA |
| T27917 | GGGCTAGGGGTCGCGGGTAAGGG | 1000 | | | | | | | | | |
| Oligo fwd | CACCGGGCTAGGGGTCGCGGGTAA | | | | | | | | | | |
| Oligo rev | AAACTTACCCGCGACCCCTAGCCC | | | | | | | | | | |
| Chromosome | start | end | strand | MM | target_seq | PAM | alignment | distance | position | gene name | gene id |
| GU305914.1 | 83542 | 83564 | - | 0 | GGGCTAGGGGTCGCGGGTAA | GGG | \|\|\|\|\|\|\|\|[\|\|\|\|\|\|\|\|\|\|\|\|]PAM | NA | Intergenic | NA | NA |
| T27918 | GGCTAGGGGTCGCGGGTAAGGGG | 1000 | | | | | | | | | |
| Oligo fwd | CACCGGCTAGGGGTCGCGGGTAAG | | | | | | | | | | |
| Oligo rev | AAACCTTACCCGCGACCCCTAGCC | | | | | | | | | | |

<table>
<tr><td>Chromosome</td><td>start</td><td>end</td><td>strand</td><td>MM</td><td>target_seq</td><td>PAM</td><td>alignment</td><td>distance</td><td>position</td><td>gene name</td><td>gene id</td></tr>
<tr><td>GU305914.1</td><td>83541</td><td>83563</td><td>-</td><td>0</td><td>GGCTAGGGGTCGCGGGTAAG</td><td>GGG</td><td>||||||||[||||||||||||]PAM</td><td>NA</td><td>Intergenic</td><td>NA</td><td>NA</td></tr>
<tr><td></td><td></td><td></td><td></td><td></td><td></td><td></td><td></td><td></td><td></td><td></td><td></td></tr>
<tr><td>T27919</td><td>GGGGTCGCGGGTAAGGGGATCGG</td><td>1000</td><td></td><td></td><td></td><td></td><td></td><td></td><td></td><td></td><td></td></tr>
<tr><td>Oligo fwd</td><td>CACCGGGGTCGCGGGTAAGGGGAT</td><td></td><td></td><td></td><td></td><td></td><td></td><td></td><td></td><td></td><td></td></tr>
<tr><td>Oligo rev</td><td>AAACATCCCCTTACCCGCGACCCC</td><td></td><td></td><td></td><td></td><td></td><td></td><td></td><td></td><td></td><td></td></tr>
<tr><td>Chromosome</td><td>start</td><td>end</td><td>strand</td><td>MM</td><td>target_seq</td><td>PAM</td><td>alignment</td><td>distance</td><td>position</td><td>gene name</td><td>gene id</td></tr>
<tr><td>GU305914.1</td><td>83536</td><td>83558</td><td>-</td><td>0</td><td>GGGGTCGCGGGTAAGGGGAT</td><td>CGG</td><td>||||||||[||||||||||||]PAM</td><td>NA</td><td>Intergenic</td><td>NA</td><td>NA</td></tr>
<tr><td></td><td></td><td></td><td></td><td></td><td></td><td></td><td></td><td></td><td></td><td></td><td></td></tr>
<tr><td>T27920</td><td>GGGTCGCGGGTAAGGGGATCGGG</td><td>1000</td><td></td><td></td><td></td><td></td><td></td><td></td><td></td><td></td><td></td></tr>
<tr><td>Oligo fwd</td><td>CACCGGGTCGCGGGTAAGGGGATC</td><td></td><td></td><td></td><td></td><td></td><td></td><td></td><td></td><td></td><td></td></tr>
<tr><td>Oligo rev</td><td>AAACGATCCCCTTACCCGCGACCC</td><td></td><td></td><td></td><td></td><td></td><td></td><td></td><td></td><td></td><td></td></tr>
<tr><td>Chromosome</td><td>start</td><td>end</td><td>strand</td><td>MM</td><td>target_seq</td><td>PAM</td><td>alignment</td><td>distance</td><td>position</td><td>gene name</td><td>gene id</td></tr>
<tr><td>GU305914.1</td><td>83535</td><td>83557</td><td>-</td><td>0</td><td>GGGTCGCGGGTAAGGGGATC</td><td>GGG</td><td>||||||||[||||||||||||]PAM</td><td>NA</td><td>Intergenic</td><td>NA</td><td>NA</td></tr>
<tr><td></td><td></td><td></td><td></td><td></td><td></td><td></td><td></td><td></td><td></td><td></td><td></td></tr>
<tr><td>T27921</td><td>GGTCGCGGGTAAGGGGATCGGGG</td><td>1000</td><td></td><td></td><td></td><td></td><td></td><td></td><td></td><td></td><td></td></tr>
<tr><td>Oligo fwd</td><td>CACCGGTCGCGGGTAAGGGGATCG</td><td></td><td></td><td></td><td></td><td></td><td></td><td></td><td></td><td></td><td></td></tr>
<tr><td>Oligo rev</td><td>AAACCGATCCCCTTACCCGCGACC</td><td></td><td></td><td></td><td></td><td></td><td></td><td></td><td></td><td></td><td></td></tr>
<tr><td>Chromosome</td><td>start</td><td>end</td><td>strand</td><td>MM</td><td>target_seq</td><td>PAM</td><td>alignment</td><td>distance</td><td>position</td><td>gene name</td><td>gene id</td></tr>
<tr><td>GU305914.1</td><td>83534</td><td>83556</td><td>-</td><td>0</td><td>GGTCGCGGGTAAGGGGATCG</td><td>GGG</td><td>||||||||[||||||||||||]PAM</td><td>NA</td><td>Intergenic</td><td>NA</td><td>NA</td></tr>
<tr><td></td><td></td><td></td><td></td><td></td><td></td><td></td><td></td><td></td><td></td><td></td><td></td></tr>
<tr><td>T27922</td><td>CGCGGGTAAGGGGATCGGGGTGG</td><td>1000</td><td></td><td></td><td></td><td></td><td></td><td></td><td></td><td></td><td></td></tr>
<tr><td>Oligo fwd</td><td>CACCCGCGGGTAAGGGGATCGGGG</td><td></td><td></td><td></td><td></td><td></td><td></td><td></td><td></td><td></td><td></td></tr>
<tr><td>Oligo rev</td><td>AAACCCCCGATCCCCTTACCCGCG</td><td></td><td></td><td></td><td></td><td></td><td></td><td></td><td></td><td></td><td></td></tr>
<tr><td>Chromosome</td><td>start</td><td>end</td><td>strand</td><td>MM</td><td>target_seq</td><td>PAM</td><td>alignment</td><td>distance</td><td>position</td><td>gene name</td><td>gene id</td></tr>
<tr><td>GU305914.1</td><td>83531</td><td>83553</td><td>-</td><td>0</td><td>CGCGGGTAAGGGGATCGGGG</td><td>TGG</td><td>||||||||[||||||||||||]PAM</td><td>NA</td><td>Intergenic</td><td>NA</td><td>NA</td></tr>
<tr><td></td><td></td><td></td><td></td><td></td><td></td><td></td><td></td><td></td><td></td><td></td><td></td></tr>
<tr><td>T27923</td><td>AAGGGGATCGGGGTGGCCATCGG</td><td>1000</td><td></td><td></td><td></td><td></td><td></td><td></td><td></td><td></td><td></td></tr>
<tr><td>Oligo fwd</td><td>CACCAAGGGGATCGGGGTGGCCAT</td><td></td><td></td><td></td><td></td><td></td><td></td><td></td><td></td><td></td><td></td></tr>
<tr><td>Oligo rev</td><td>AAACATGGCCACCCCGATCCCCTT</td><td></td><td></td><td></td><td></td><td></td><td></td><td></td><td></td><td></td><td></td></tr>
<tr><td>Chromosome</td><td>start</td><td>end</td><td>strand</td><td>MM</td><td>target_seq</td><td>PAM</td><td>alignment</td><td>distance</td><td>position</td><td>gene name</td><td>gene id</td></tr>
<tr><td>GU305914.1</td><td>83524</td><td>83546</td><td>-</td><td>0</td><td>AAGGGGATCGGGGTGGCCAT</td><td>CGG</td><td>||||||||[||||||||||||]PAM</td><td>NA</td><td>Intergenic</td><td>NA</td><td>NA</td></tr>
<tr><td></td><td></td><td></td><td></td><td></td><td></td><td></td><td></td><td></td><td></td><td></td><td></td></tr>
<tr><td>T27924</td><td>AGGGGATCGGGGTGGCCATCGGG</td><td>1000</td><td></td><td></td><td></td><td></td><td></td><td></td><td></td><td></td><td></td></tr>
<tr><td>Oligo fwd</td><td>CACCAGGGGATCGGGGTGGCCATC</td><td></td><td></td><td></td><td></td><td></td><td></td><td></td><td></td><td></td><td></td></tr>
<tr><td>Oligo rev</td><td>AAACGATGGCCACCCCGATCCCCT</td><td></td><td></td><td></td><td></td><td></td><td></td><td></td><td></td><td></td><td></td></tr>
<tr><td>Chromosome</td><td>start</td><td>end</td><td>strand</td><td>MM</td><td>target_seq</td><td>PAM</td><td>alignment</td><td>distance</td><td>position</td><td>gene name</td><td>gene id</td></tr>
<tr><td>GU305914.1</td><td>83523</td><td>83545</td><td>-</td><td>0</td><td>AGGGGATCGGGGTGGCCATC</td><td>GGG</td><td>||||||||[||||||||||||]PAM</td><td>NA</td><td>Intergenic</td><td>NA</td><td>NA</td></tr>
<tr><td></td><td></td><td></td><td></td><td></td><td></td><td></td><td></td><td></td><td></td><td></td><td></td></tr>
<tr><td>T27925</td><td>GGGGATCGGGGTGGCCATCGGGG</td><td>1000</td><td></td><td></td><td></td><td></td><td></td><td></td><td></td><td></td><td></td></tr>
<tr><td>Oligo fwd</td><td>CACCGGGGATCGGGGTGGCCATCG</td><td></td><td></td><td></td><td></td><td></td><td></td><td></td><td></td><td></td><td></td></tr>
<tr><td>Oligo rev</td><td>AAACCGATGGCCACCCCGATCCCC</td><td></td><td></td><td></td><td></td><td></td><td></td><td></td><td></td><td></td><td></td></tr>
<tr><td>Chromosome</td><td>start</td><td>end</td><td>strand</td><td>MM</td><td>target_seq</td><td>PAM</td><td>alignment</td><td>distance</td><td>position</td><td>gene name</td><td>gene id</td></tr>
<tr><td>GU305914.1</td><td>83522</td><td>83544</td><td>-</td><td>0</td><td>GGGGATCGGGGTGGCCATCG</td><td>GGG</td><td>||||||||[||||||||||||]PAM</td><td>NA</td><td>Intergenic</td><td>NA</td><td>NA</td></tr>
<tr><td></td><td></td><td></td><td></td><td></td><td></td><td></td><td></td><td></td><td></td><td></td><td></td></tr>
<tr><td>T27926</td><td>GGGGTGGCCATCGGGGCCGTCGG</td><td>1000</td><td></td><td></td><td></td><td></td><td></td><td></td><td></td><td></td><td></td></tr>
<tr><td>Oligo fwd</td><td>CACCGGGGTGGCCATCGGGGCCGT</td><td></td><td></td><td></td><td></td><td></td><td></td><td></td><td></td><td></td><td></td></tr>
<tr><td>Oligo rev</td><td>AAACACGGCCCCGATGGCCACCCC</td><td></td><td></td><td></td><td></td><td></td><td></td><td></td><td></td><td></td><td></td></tr>
<tr><td>Chromosome</td><td>start</td><td>end</td><td>strand</td><td>MM</td><td>target_seq</td><td>PAM</td><td>alignment</td><td>distance</td><td>position</td><td>gene name</td><td>gene id</td></tr>
<tr><td>GU305914.1</td><td>83515</td><td>83537</td><td>-</td><td>0</td><td>GGGGTGGCCATCGGGGCCGT</td><td>CGG</td><td>||||||||[||||||||||||]PAM</td><td>NA</td><td>Intergenic</td><td>NA</td><td>NA</td></tr>
<tr><td></td><td></td><td></td><td></td><td></td><td></td><td></td><td></td><td></td><td></td><td></td><td></td></tr>
<tr><td>T27927</td><td>GTGGCCATCGGGGCCGTCGGCGG</td><td>1000</td><td></td><td></td><td></td><td></td><td></td><td></td><td></td><td></td><td></td></tr>
<tr><td>Oligo fwd</td><td>CACCGTGGCCATCGGGGCCGTCGG</td><td></td><td></td><td></td><td></td><td></td><td></td><td></td><td></td><td></td><td></td></tr>
<tr><td>Oligo rev</td><td>AAACCCGACGGCCCCGATGGCCAC</td><td></td><td></td><td></td><td></td><td></td><td></td><td></td><td></td><td></td><td></td></tr>
<tr><td>Chromosome</td><td>start</td><td>end</td><td>strand</td><td>MM</td><td>target_seq</td><td>PAM</td><td>alignment</td><td>distance</td><td>position</td><td>gene name</td><td>gene id</td></tr>
<tr><td>GU305914.1</td><td>83512</td><td>83534</td><td>-</td><td>0</td><td>GTGGCCATCGGGGCCGTCGG</td><td>CGG</td><td>||||||||[||||||||||||]PAM</td><td>NA</td><td>Intergenic</td><td>NA</td><td>NA</td></tr>
<tr><td>GU305914.1</td><td>112003</td><td>112025</td><td>-</td><td>4</td><td>GCGGTCATCGGAGCCGTCGA</td><td>CGG</td><td>|-||-|||[|||-|||||||-]PAM</td><td>NA</td><td>Intergenic</td><td>NA</td><td>NA</td></tr>
<tr><td></td><td></td><td></td><td></td><td></td><td></td><td></td><td></td><td></td><td></td><td></td><td></td></tr>
<tr><td>T27928</td><td>TGGCCATCGGGGCCGTCGGCGGG</td><td>1000</td><td></td><td></td><td></td><td></td><td></td><td></td><td></td><td></td><td></td></tr>
<tr><td>Oligo fwd</td><td>CACCTGGCCATCGGGGCCGTCGGC</td><td></td><td></td><td></td><td></td><td></td><td></td><td></td><td></td><td></td><td></td></tr>
<tr><td>Oligo rev</td><td>AAACGCCGACGGCCCCGATGGCCA</td><td></td><td></td><td></td><td></td><td></td><td></td><td></td><td></td><td></td><td></td></tr>
<tr><td>Chromosome</td><td>start</td><td>end</td><td>strand</td><td>MM</td><td>target_seq</td><td>PAM</td><td>alignment</td><td>distance</td><td>position</td><td>gene name</td><td>gene id</td></tr>
<tr><td>GU305914.1</td><td>83511</td><td>83533</td><td>-</td><td>0</td><td>TGGCCATCGGGGCCGTCGGC</td><td>GGG</td><td>||||||||[||||||||||||]PAM</td><td>NA</td><td>Intergenic</td><td>NA</td><td>NA</td></tr>
<tr><td></td><td></td><td></td><td></td><td></td><td></td><td></td><td></td><td></td><td></td><td></td><td></td></tr>
<tr><td>T27929</td><td>GGCCATCGGGGCCGTCGGCGGGG</td><td>1000</td><td></td><td></td><td></td><td></td><td></td><td></td><td></td><td></td><td></td></tr>
<tr><td>Oligo fwd</td><td>CACCGGCCATCGGGGCCGTCGGCG</td><td></td><td></td><td></td><td></td><td></td><td></td><td></td><td></td><td></td><td></td></tr>
<tr><td>Oligo rev</td><td>AAACCGCCGACGGCCCCGATGGCC</td><td></td><td></td><td></td><td></td><td></td><td></td><td></td><td></td><td></td><td></td></tr>
<tr><td>Chromosome</td><td>start</td><td>end</td><td>strand</td><td>MM</td><td>target_seq</td><td>PAM</td><td>alignment</td><td>distance</td><td>position</td><td>gene name</td><td>gene id</td></tr>
<tr><td>GU305914.1</td><td>83510</td><td>83532</td><td>-</td><td>0</td><td>GGCCATCGGGGCCGTCGGCG</td><td>GGG</td><td>||||||||[||||||||||||]PAM</td><td>NA</td><td>Intergenic</td><td>NA</td><td>NA</td></tr>
<tr><td></td><td></td><td></td><td></td><td></td><td></td><td></td><td></td><td></td><td></td><td></td><td></td></tr>
<tr><td>T27930</td><td>CGTCGGCGGGGCCGTGTCTTCGG</td><td>1000</td><td></td><td></td><td></td><td></td><td></td><td></td><td></td><td></td><td></td></tr>
<tr><td>Oligo fwd</td><td>CACCCGTCGGCGGGGCCGTGTCTT</td><td></td><td></td><td></td><td></td><td></td><td></td><td></td><td></td><td></td><td></td></tr>
<tr><td>Oligo rev</td><td>AAACAAGACACGGCCCCGCCGACG</td><td></td><td></td><td></td><td></td><td></td><td></td><td></td><td></td><td></td><td></td></tr>
<tr><td>Chromosome</td><td>start</td><td>end</td><td>strand</td><td>MM</td><td>target_seq</td><td>PAM</td><td>alignment</td><td>distance</td><td>position</td><td>gene name</td><td>gene id</td></tr>
<tr><td>GU305914.1</td><td>83498</td><td>83520</td><td>-</td><td>0</td><td>CGTCGGCGGGGCCGTGTCTT</td><td>CGG</td><td>||||||||[||||||||||||]PAM</td><td>NA</td><td>Intergenic</td><td>NA</td><td>NA</td></tr>
<tr><td></td><td></td><td></td><td></td><td></td><td></td><td></td><td></td><td></td><td></td><td></td><td></td></tr>
<tr><td>T27931</td><td>CGGCGGGGCCGTGTCTTCGGTGG</td><td>1000</td><td></td><td></td><td></td><td></td><td></td><td></td><td></td><td></td><td></td></tr>
<tr><td>Oligo fwd</td><td>CACCCGGCGGGGCCGTGTCTTCGG</td><td></td><td></td><td></td><td></td><td></td><td></td><td></td><td></td><td></td><td></td></tr>
<tr><td>Oligo rev</td><td>AAACCCGAAGACACGGCCCCGCCG</td><td></td><td></td><td></td><td></td><td></td><td></td><td></td><td></td><td></td><td></td></tr>
<tr><td>Chromosome</td><td>start</td><td>end</td><td>strand</td><td>MM</td><td>target_seq</td><td>PAM</td><td>alignment</td><td>distance</td><td>position</td><td>gene name</td><td>gene id</td></tr>
<tr><td>GU305914.1</td><td>83495</td><td>83517</td><td>-</td><td>0</td><td>CGGCGGGGCCGTGTCTTCGG</td><td>TGG</td><td>||||||||[||||||||||||]PAM</td><td>NA</td><td>Intergenic</td><td>NA</td><td>NA</td></tr>
<tr><td></td><td></td><td></td><td></td><td></td><td></td><td></td><td></td><td></td><td></td><td></td><td></td></tr>
<tr><td>T27932</td><td>TTCGGTGGTGAACGCCGTCACGG</td><td>1000</td><td></td><td></td><td></td><td></td><td></td><td></td><td></td><td></td><td></td></tr>
<tr><td>Oligo fwd</td><td>CACCTTCGGTGGTGAACGCCGTCA</td><td></td><td></td><td></td><td></td><td></td><td></td><td></td><td></td><td></td><td></td></tr>
<tr><td>Oligo rev</td><td>AAACTGACGGCGTTCACCACCGAA</td><td></td><td></td><td></td><td></td><td></td><td></td><td></td><td></td><td></td><td></td></tr>
<tr><td>Chromosome</td><td>start</td><td>end</td><td>strand</td><td>MM</td><td>target_seq</td><td>PAM</td><td>alignment</td><td>distance</td><td>position</td><td>gene name</td><td>gene id</td></tr>
<tr><td>GU305914.1</td><td>83480</td><td>83502</td><td>-</td><td>0</td><td>TTCGGTGGTGAACGCCGTCA</td><td>CGG</td><td>||||||||[||||||||||||]PAM</td><td>NA</td><td>Intergenic</td><td>NA</td><td>NA</td></tr>
<tr><td></td><td></td><td></td><td></td><td></td><td></td><td></td><td></td><td></td><td></td><td></td><td></td></tr>
<tr><td>T27933</td><td>TCGGTGGTGAACGCCGTCACGGG</td><td>1000</td><td></td><td></td><td></td><td></td><td></td><td></td><td></td><td></td><td></td></tr>
</table>

<table>
<tr><td>Oligo fwd</td><td>CACCTCGGTGGTGAACGCCGTCAC</td><td></td><td></td><td></td><td></td><td></td><td></td><td></td><td></td><td></td><td></td></tr>
<tr><td>Oligo rev</td><td>AAACGTGACGGCGTTCACCACCGA</td><td></td><td></td><td></td><td></td><td></td><td></td><td></td><td></td><td></td><td></td></tr>
<tr><td>Chromosome</td><td>start</td><td>end</td><td>strand</td><td>MM</td><td>target_seq</td><td>PAM</td><td>alignment</td><td>distance</td><td>position</td><td>gene name</td><td>gene id</td></tr>
<tr><td>GU305914.1</td><td>83479</td><td>83501</td><td>-</td><td>0</td><td>TCGGTGGTGAACGCCGTCAC</td><td>GGG</td><td>||||||||[||||||||||||]PAM</td><td>NA</td><td>Intergenic</td><td>NA</td><td>NA</td></tr>
<tr><td></td><td></td><td></td><td></td><td></td><td></td><td></td><td></td><td></td><td></td><td></td><td></td></tr>
<tr><td>T27934</td><td>GGCTTCCTGACGAACCCGTTTGG</td><td>1000</td><td></td><td></td><td></td><td></td><td></td><td></td><td></td><td></td><td></td></tr>
<tr><td>Oligo fwd</td><td>CACCGGCTTCCTGACGAACCCGTT</td><td></td><td></td><td></td><td></td><td></td><td></td><td></td><td></td><td></td><td></td></tr>
<tr><td>Oligo rev</td><td>AAACAACGGGTTCGTCAGGAAGCC</td><td></td><td></td><td></td><td></td><td></td><td></td><td></td><td></td><td></td><td></td></tr>
<tr><td>Chromosome</td><td>start</td><td>end</td><td>strand</td><td>MM</td><td>target_seq</td><td>PAM</td><td>alignment</td><td>distance</td><td>position</td><td>gene name</td><td>gene id</td></tr>
<tr><td>GU305914.1</td><td>83458</td><td>83480</td><td>-</td><td>0</td><td>GGCTTCCTGACGAACCCGTT</td><td>TGG</td><td>||||||||[||||||||||||]PAM</td><td>NA</td><td>Intergenic</td><td>NA</td><td>NA</td></tr>
<tr><td></td><td></td><td></td><td></td><td></td><td></td><td></td><td></td><td></td><td></td><td></td><td></td></tr>
<tr><td>T27935</td><td>TTCCTGACGAACCCGTTTGGAGG</td><td>1000</td><td></td><td></td><td></td><td></td><td></td><td></td><td></td><td></td><td></td></tr>
<tr><td>Oligo fwd</td><td>CACCTTCCTGACGAACCCGTTTGG</td><td></td><td></td><td></td><td></td><td></td><td></td><td></td><td></td><td></td><td></td></tr>
<tr><td>Oligo rev</td><td>AAACCCAAACGGGTTCGTCAGGAA</td><td></td><td></td><td></td><td></td><td></td><td></td><td></td><td></td><td></td><td></td></tr>
<tr><td>Chromosome</td><td>start</td><td>end</td><td>strand</td><td>MM</td><td>target_seq</td><td>PAM</td><td>alignment</td><td>distance</td><td>position</td><td>gene name</td><td>gene id</td></tr>
<tr><td>GU305914.1</td><td>83455</td><td>83477</td><td>-</td><td>0</td><td>TTCCTGACGAACCCGTTTGG</td><td>AGG</td><td>||||||||[||||||||||||]PAM</td><td>NA</td><td>Intergenic</td><td>NA</td><td>NA</td></tr>
<tr><td></td><td></td><td></td><td></td><td></td><td></td><td></td><td></td><td></td><td></td><td></td><td></td></tr>
<tr><td>T27936</td><td>AGGTTTCACCACCATCCTCCTGG</td><td>1000</td><td></td><td></td><td></td><td></td><td></td><td></td><td></td><td></td><td></td></tr>
<tr><td>Oligo fwd</td><td>CACCAGGTTTCACCACCATCCTCC</td><td></td><td></td><td></td><td></td><td></td><td></td><td></td><td></td><td></td><td></td></tr>
<tr><td>Oligo rev</td><td>AAACGGAGGATGGTGGTGAAACCT</td><td></td><td></td><td></td><td></td><td></td><td></td><td></td><td></td><td></td><td></td></tr>
<tr><td>Chromosome</td><td>start</td><td>end</td><td>strand</td><td>MM</td><td>target_seq</td><td>PAM</td><td>alignment</td><td>distance</td><td>position</td><td>gene name</td><td>gene id</td></tr>
<tr><td>GU305914.1</td><td>83435</td><td>83457</td><td>-</td><td>0</td><td>AGGTTTCACCACCATCCTCC</td><td>TGG</td><td>||||||||[||||||||||||]PAM</td><td>NA</td><td>Intergenic</td><td>NA</td><td>NA</td></tr>
<tr><td></td><td></td><td></td><td></td><td></td><td></td><td></td><td></td><td></td><td></td><td></td><td></td></tr>
<tr><td>T27937</td><td>ACCACCATCCTCCTGGTGATCGG</td><td>1000</td><td></td><td></td><td></td><td></td><td></td><td></td><td></td><td></td><td></td></tr>
<tr><td>Oligo fwd</td><td>CACCACCACCATCCTCCTGGTGAT</td><td></td><td></td><td></td><td></td><td></td><td></td><td></td><td></td><td></td><td></td></tr>
<tr><td>Oligo rev</td><td>AAACATCACCAGGAGGATGGTGGT</td><td></td><td></td><td></td><td></td><td></td><td></td><td></td><td></td><td></td><td></td></tr>
<tr><td>Chromosome</td><td>start</td><td>end</td><td>strand</td><td>MM</td><td>target_seq</td><td>PAM</td><td>alignment</td><td>distance</td><td>position</td><td>gene name</td><td>gene id</td></tr>
<tr><td>GU305914.1</td><td>83428</td><td>83450</td><td>-</td><td>0</td><td>ACCACCATCCTCCTGGTGAT</td><td>CGG</td><td>||||||||[||||||||||||]PAM</td><td>NA</td><td>Intergenic</td><td>NA</td><td>NA</td></tr>
<tr><td></td><td></td><td></td><td></td><td></td><td></td><td></td><td></td><td></td><td></td><td></td><td></td></tr>
<tr><td>T27938</td><td>TCGTGTATCTCATCTTCACGAGG</td><td>1000</td><td></td><td></td><td></td><td></td><td></td><td></td><td></td><td></td><td></td></tr>
<tr><td>Oligo fwd</td><td>CACCTCGTGTATCTCATCTTCACG</td><td></td><td></td><td></td><td></td><td></td><td></td><td></td><td></td><td></td><td></td></tr>
<tr><td>Oligo rev</td><td>AAACCGTGAAGATGAGATACACGA</td><td></td><td></td><td></td><td></td><td></td><td></td><td></td><td></td><td></td><td></td></tr>
<tr><td>Chromosome</td><td>start</td><td>end</td><td>strand</td><td>MM</td><td>target_seq</td><td>PAM</td><td>alignment</td><td>distance</td><td>position</td><td>gene name</td><td>gene id</td></tr>
<tr><td>GU305914.1</td><td>83394</td><td>83416</td><td>-</td><td>0</td><td>TCGTGTATCTCATCTTCACG</td><td>AGG</td><td>||||||||[||||||||||||]PAM</td><td>NA</td><td>Intergenic</td><td>NA</td><td>NA</td></tr>
<tr><td></td><td></td><td></td><td></td><td></td><td></td><td></td><td></td><td></td><td></td><td></td><td></td></tr>
<tr><td>T27939</td><td>ATCTCATCTTCACGAGGCAGCGG</td><td>1000</td><td></td><td></td><td></td><td></td><td></td><td></td><td></td><td></td><td></td></tr>
<tr><td>Oligo fwd</td><td>CACCATCTCATCTTCACGAGGCAG</td><td></td><td></td><td></td><td></td><td></td><td></td><td></td><td></td><td></td><td></td></tr>
<tr><td>Oligo rev</td><td>AAACCTGCCTCGTGAAGATGAGAT</td><td></td><td></td><td></td><td></td><td></td><td></td><td></td><td></td><td></td><td></td></tr>
<tr><td>Chromosome</td><td>start</td><td>end</td><td>strand</td><td>MM</td><td>target_seq</td><td>PAM</td><td>alignment</td><td>distance</td><td>position</td><td>gene name</td><td>gene id</td></tr>
<tr><td>GU305914.1</td><td>83388</td><td>83410</td><td>-</td><td>0</td><td>ATCTCATCTTCACGAGGCAG</td><td>CGG</td><td>||||||||[||||||||||||]PAM</td><td>NA</td><td>Intergenic</td><td>NA</td><td>NA</td></tr>
<tr><td></td><td></td><td></td><td></td><td></td><td></td><td></td><td></td><td></td><td></td><td></td><td></td></tr>
<tr><td>T27940</td><td>CATCTTCACGAGGCAGCGGTCGG</td><td>1000</td><td></td><td></td><td></td><td></td><td></td><td></td><td></td><td></td><td></td></tr>
<tr><td>Oligo fwd</td><td>CACCCATCTTCACGAGGCAGCGGT</td><td></td><td></td><td></td><td></td><td></td><td></td><td></td><td></td><td></td><td></td></tr>
<tr><td>Oligo rev</td><td>AAACACCGCTGCCTCGTGAAGATG</td><td></td><td></td><td></td><td></td><td></td><td></td><td></td><td></td><td></td><td></td></tr>
<tr><td>Chromosome</td><td>start</td><td>end</td><td>strand</td><td>MM</td><td>target_seq</td><td>PAM</td><td>alignment</td><td>distance</td><td>position</td><td>gene name</td><td>gene id</td></tr>
<tr><td>GU305914.1</td><td>83384</td><td>83406</td><td>-</td><td>0</td><td>CATCTTCACGAGGCAGCGGT</td><td>CGG</td><td>||||||||[||||||||||||]PAM</td><td>NA</td><td>Intergenic</td><td>NA</td><td>NA</td></tr>
<tr><td></td><td></td><td></td><td></td><td></td><td></td><td></td><td></td><td></td><td></td><td></td><td></td></tr>
<tr><td>T27941</td><td>GGCAGCGGTCGGCCGCCGCGAGG</td><td>1000</td><td></td><td></td><td></td><td></td><td></td><td></td><td></td><td></td><td></td></tr>
<tr><td>Oligo fwd</td><td>CACCGGCAGCGGTCGGCCGCCGCG</td><td></td><td></td><td></td><td></td><td></td><td></td><td></td><td></td><td></td><td></td></tr>
<tr><td>Oligo rev</td><td>AAACCGCGGCGGCCGACCGCTGCC</td><td></td><td></td><td></td><td></td><td></td><td></td><td></td><td></td><td></td><td></td></tr>
<tr><td>Chromosome</td><td>start</td><td>end</td><td>strand</td><td>MM</td><td>target_seq</td><td>PAM</td><td>alignment</td><td>distance</td><td>position</td><td>gene name</td><td>gene id</td></tr>
<tr><td>GU305914.1</td><td>83373</td><td>83395</td><td>-</td><td>0</td><td>GGCAGCGGTCGGCCGCCGCG</td><td>AGG</td><td>||||||||[||||||||||||]PAM</td><td>NA</td><td>Intergenic</td><td>NA</td><td>NA</td></tr>
<tr><td></td><td></td><td></td><td></td><td></td><td></td><td></td><td></td><td></td><td></td><td></td><td></td></tr>
<tr><td>T27942</td><td>GCGGTCGGCCGCCGCGAGGCCGG</td><td>1000</td><td></td><td></td><td></td><td></td><td></td><td></td><td></td><td></td><td></td></tr>
<tr><td>Oligo fwd</td><td>CACCGCGGTCGGCCGCCGCGAGGC</td><td></td><td></td><td></td><td></td><td></td><td></td><td></td><td></td><td></td><td></td></tr>
<tr><td>Oligo rev</td><td>AAACGCCTCGCGGCGGCCGACCGC</td><td></td><td></td><td></td><td></td><td></td><td></td><td></td><td></td><td></td><td></td></tr>
<tr><td>Chromosome</td><td>start</td><td>end</td><td>strand</td><td>MM</td><td>target_seq</td><td>PAM</td><td>alignment</td><td>distance</td><td>position</td><td>gene name</td><td>gene id</td></tr>
<tr><td>GU305914.1</td><td>83369</td><td>83391</td><td>-</td><td>0</td><td>GCGGTCGGCCGCCGCGAGGC</td><td>CGG</td><td>||||||||[||||||||||||]PAM</td><td>NA</td><td>Intergenic</td><td>NA</td><td>NA</td></tr>
<tr><td>GU305914.1</td><td>99740</td><td>99762</td><td>-</td><td>4</td><td>GCCGTCCGCCGCGGCGAGAC</td><td>GGG</td><td>||-|||-|[||||-|||||-|]PAM</td><td>NA</td><td>Intergenic</td><td>NA</td><td>NA</td></tr>
<tr><td></td><td></td><td></td><td></td><td></td><td></td><td></td><td></td><td></td><td></td><td></td><td></td></tr>
<tr><td>T27943</td><td>GCCGTGCAGTATGCTCCTCCCGG</td><td>1000</td><td></td><td></td><td></td><td></td><td></td><td></td><td></td><td></td><td></td></tr>
<tr><td>Oligo fwd</td><td>CACCGCCGTGCAGTATGCTCCTCC</td><td></td><td></td><td></td><td></td><td></td><td></td><td></td><td></td><td></td><td></td></tr>
<tr><td>Oligo rev</td><td>AAACGGAGGAGCATACTGCACGGC</td><td></td><td></td><td></td><td></td><td></td><td></td><td></td><td></td><td></td><td></td></tr>
<tr><td>Chromosome</td><td>start</td><td>end</td><td>strand</td><td>MM</td><td>target_seq</td><td>PAM</td><td>alignment</td><td>distance</td><td>position</td><td>gene name</td><td>gene id</td></tr>
<tr><td>GU305914.1</td><td>83314</td><td>83336</td><td>-</td><td>0</td><td>GCCGTGCAGTATGCTCCTCC</td><td>CGG</td><td>||||||||[||||||||||||]PAM</td><td>NA</td><td>Intergenic</td><td>NA</td><td>NA</td></tr>
<tr><td></td><td></td><td></td><td></td><td></td><td></td><td></td><td></td><td></td><td></td><td></td><td></td></tr>
<tr><td>T27944</td><td>GTGCAGTATGCTCCTCCCGGCGG</td><td>1000</td><td></td><td></td><td></td><td></td><td></td><td></td><td></td><td></td><td></td></tr>
<tr><td>Oligo fwd</td><td>CACCGTGCAGTATGCTCCTCCCGG</td><td></td><td></td><td></td><td></td><td></td><td></td><td></td><td></td><td></td><td></td></tr>
<tr><td>Oligo rev</td><td>AAACCCGGGAGGAGCATACTGCAC</td><td></td><td></td><td></td><td></td><td></td><td></td><td></td><td></td><td></td><td></td></tr>
<tr><td>Chromosome</td><td>start</td><td>end</td><td>strand</td><td>MM</td><td>target_seq</td><td>PAM</td><td>alignment</td><td>distance</td><td>position</td><td>gene name</td><td>gene id</td></tr>
<tr><td>GU305914.1</td><td>83311</td><td>83333</td><td>-</td><td>0</td><td>GTGCAGTATGCTCCTCCCGG</td><td>CGG</td><td>||||||||[||||||||||||]PAM</td><td>NA</td><td>Intergenic</td><td>NA</td><td>NA</td></tr>
<tr><td></td><td></td><td></td><td></td><td></td><td></td><td></td><td></td><td></td><td></td><td></td><td></td></tr>
<tr><td>T27945</td><td>GCTCCTCCCGGCGGCGCGCACGG</td><td>1000</td><td></td><td></td><td></td><td></td><td></td><td></td><td></td><td></td><td></td></tr>
<tr><td>Oligo fwd</td><td>CACCGCTCCTCCCGGCGGCGCGCA</td><td></td><td></td><td></td><td></td><td></td><td></td><td></td><td></td><td></td><td></td></tr>
<tr><td>Oligo rev</td><td>AAACTGCGCGCCGCCGGGAGGAGC</td><td></td><td></td><td></td><td></td><td></td><td></td><td></td><td></td><td></td><td></td></tr>
<tr><td>Chromosome</td><td>start</td><td>end</td><td>strand</td><td>MM</td><td>target_seq</td><td>PAM</td><td>alignment</td><td>distance</td><td>position</td><td>gene name</td><td>gene id</td></tr>
<tr><td>GU305914.1</td><td>83302</td><td>83324</td><td>-</td><td>0</td><td>GCTCCTCCCGGCGGCGCGCA</td><td>CGG</td><td>||||||||[||||||||||||]PAM</td><td>NA</td><td>Intergenic</td><td>NA</td><td>NA</td></tr>
<tr><td></td><td></td><td></td><td></td><td></td><td></td><td></td><td></td><td></td><td></td><td></td><td></td></tr>
<tr><td>T27946</td><td>CTCCTCCCGGCGGCGCGCACGGG</td><td>1000</td><td></td><td></td><td></td><td></td><td></td><td></td><td></td><td></td><td></td></tr>
<tr><td>Oligo fwd</td><td>CACCCTCCTCCCGGCGGCGCGCAC</td><td></td><td></td><td></td><td></td><td></td><td></td><td></td><td></td><td></td><td></td></tr>
<tr><td>Oligo rev</td><td>AAACGTGCGCGCCGCCGGGAGGAG</td><td></td><td></td><td></td><td></td><td></td><td></td><td></td><td></td><td></td><td></td></tr>
<tr><td>Chromosome</td><td>start</td><td>end</td><td>strand</td><td>MM</td><td>target_seq</td><td>PAM</td><td>alignment</td><td>distance</td><td>position</td><td>gene name</td><td>gene id</td></tr>
<tr><td>GU305914.1</td><td>83301</td><td>83323</td><td>-</td><td>0</td><td>CTCCTCCCGGCGGCGCGCAC</td><td>GGG</td><td>||||||||[||||||||||||]PAM</td><td>NA</td><td>Intergenic</td><td>NA</td><td>NA</td></tr>
<tr><td></td><td></td><td></td><td></td><td></td><td></td><td></td><td></td><td></td><td></td><td></td><td></td></tr>
<tr><td>T27947</td><td>TCCTCCCGGCGGCGCGCACGGGG</td><td>1000</td><td></td><td></td><td></td><td></td><td></td><td></td><td></td><td></td><td></td></tr>
<tr><td>Oligo fwd</td><td>CACCTCCTCCCGGCGGCGCGCACG</td><td></td><td></td><td></td><td></td><td></td><td></td><td></td><td></td><td></td><td></td></tr>
<tr><td>Oligo rev</td><td>AAACCGTGCGCGCCGCCGGGAGGA</td><td></td><td></td><td></td><td></td><td></td><td></td><td></td><td></td><td></td><td></td></tr>
<tr><td>Chromosome</td><td>start</td><td>end</td><td>strand</td><td>MM</td><td>target_seq</td><td>PAM</td><td>alignment</td><td>distance</td><td>position</td><td>gene name</td><td>gene id</td></tr>
<tr><td>GU305914.1</td><td>83300</td><td>83322</td><td>-</td><td>0</td><td>TCCTCCCGGCGGCGCGCACG</td><td>GGG</td><td>||||||||[||||||||||||]PAM</td><td>NA</td><td>Intergenic</td><td>NA</td><td>NA</td></tr>
</table>

GU305914.1 199825 199847 - 4 GGCGCCCGGCTGCGCGCACG TGG --|-||||[||-|||||||||]PAM NA Intergenic NA NA

T27948 CCTCCCGGCGGCGCGCACGGGGG 1000
Oligo fwd CACCCCTCCCGGCGGCGCGCACGG
Oligo rev AAACCCGTGCGCGCCGCCGGGAGG
Chromosome start end strand MM target_seq PAM alignment distance position gene name gene id
GU305914.1 83299 83321 - 0 CCTCCCGGCGGCGCGCACGG GGG ||||||||[||||||||||||]PAM NA Intergenic NA NA

T27949 GCGCACGGGGGACTCGAGAGCGG 1000
Oligo fwd CACCGCGCACGGGGGACTCGAGAG
Oligo rev AAACCTCTCGAGTCCCCCGTGCGC
Chromosome start end strand MM target_seq PAM alignment distance position gene name gene id
GU305914.1 83287 83309 - 0 GCGCACGGGGGACTCGAGAG CGG ||||||||[||||||||||||]PAM NA Intergenic NA NA

T27950 CGCACGGGGGACTCGAGAGCGGG 1000
Oligo fwd CACCCGCACGGGGGACTCGAGAGC
Oligo rev AAACGCTCTCGAGTCCCCCGTGCG
Chromosome start end strand MM target_seq PAM alignment distance position gene name gene id
GU305914.1 83286 83308 - 0 CGCACGGGGGACTCGAGAGC GGG ||||||||[||||||||||||]PAM NA Intergenic NA NA

T27951 GGGACTCGAGAGCGGGCCGCCGG 1000
Oligo fwd CACCGGGACTCGAGAGCGGGCCGC
Oligo rev AAACGCGGCCCGCTCTCGAGTCCC
Chromosome start end strand MM target_seq PAM alignment distance position gene name gene id
GU305914.1 83279 83301 - 0 GGGACTCGAGAGCGGGCCGC CGG ||||||||[||||||||||||]PAM NA Intergenic NA NA

T27952 GGACTCGAGAGCGGGCCGCCGGG 1000
Oligo fwd CACCGGACTCGAGAGCGGGCCGCC
Oligo rev AAACGGCGGCCCGCTCTCGAGTCC
Chromosome start end strand MM target_seq PAM alignment distance position gene name gene id
GU305914.1 83278 83300 - 0 GGACTCGAGAGCGGGCCGCC GGG ||||||||[||||||||||||]PAM NA Intergenic NA NA

T27953 AGCGGGCCGCCGGGAGCGCCTGG 1000
Oligo fwd CACCAGCGGGCCGCCGGGAGCGCC
Oligo rev AAACGGCGCTCCCGGCGGCCCGCT
Chromosome start end strand MM target_seq PAM alignment distance position gene name gene id
GU305914.1 83269 83291 - 0 AGCGGGCCGCCGGGAGCGCC TGG ||||||||[||||||||||||]PAM NA Intergenic NA NA
GU305914.1 126554 126576 - 4 GGTGGGCGGCGGGGAGCGCC TGG -|-||||-[||-|||||||||]PAM NA Intergenic NA NA

T27954 CGGGAGCGCCTGGTCTGCATCGG 1000
Oligo fwd CACCCGGGAGCGCCTGGTCTGCAT
Oligo rev AAACATGCAGACCAGGCGCTCCCG
Chromosome start end strand MM target_seq PAM alignment distance position gene name gene id
GU305914.1 83259 83281 - 0 CGGGAGCGCCTGGTCTGCAT CGG ||||||||[||||||||||||]PAM NA Intergenic NA NA

T27955 TCTGCATCGGCGCGTGAACGCGG 1000
Oligo fwd CACCTCTGCATCGGCGCGTGAACG
Oligo rev AAACCGTTCACGCGCCGATGCAGA
Chromosome start end strand MM target_seq PAM alignment distance position gene name gene id
GU305914.1 83246 83268 - 0 TCTGCATCGGCGCGTGAACG CGG ||||||||[||||||||||||]PAM NA Intergenic NA NA

T27956 CTGCATCGGCGCGTGAACGCGGG 1000
Oligo fwd CACCCTGCATCGGCGCGTGAACGC
Oligo rev AAACGCGTTCACGCGCCGATGCAG
Chromosome start end strand MM target_seq PAM alignment distance position gene name gene id
GU305914.1 83245 83267 - 0 CTGCATCGGCGCGTGAACGC GGG ||||||||[||||||||||||]PAM NA Intergenic NA NA

T27957 CATCGGCGCGTGAACGCGGGCGG 1000
Oligo fwd CACCCATCGGCGCGTGAACGCGGG
Oligo rev AAACCCCGCGTTCACGCGCCGATG
Chromosome start end strand MM target_seq PAM alignment distance position gene name gene id
GU305914.1 83242 83264 - 0 CATCGGCGCGTGAACGCGGG CGG ||||||||[||||||||||||]PAM NA Intergenic NA NA

T27958 CGCGGGCGGCAGTGACGATTCGG 1000
Oligo fwd CACCCGCGGGCGGCAGTGACGATT
Oligo rev AAACAATCGTCACTGCCGCCCGCG
Chromosome start end strand MM target_seq PAM alignment distance position gene name gene id
GU305914.1 83228 83250 - 0 CGCGGGCGGCAGTGACGATT CGG ||||||||[||||||||||||]PAM NA Intergenic NA NA

T27959 GCGGGCGGCAGTGACGATTCGGG 1000
Oligo fwd CACCGCGGGCGGCAGTGACGATTC
Oligo rev AAACGAATCGTCACTGCCGCCCGC
Chromosome start end strand MM target_seq PAM alignment distance position gene name gene id
GU305914.1 83227 83249 - 0 GCGGGCGGCAGTGACGATTC GGG ||||||||[||||||||||||]PAM NA Intergenic NA NA

T27960 CGGCAGTGACGATTCGGGTAAGG 1000
Oligo fwd CACCCGGCAGTGACGATTCGGGTA
Oligo rev AAACTACCCGAATCGTCACTGCCG
Chromosome start end strand MM target_seq PAM alignment distance position gene name gene id
GU305914.1 83222 83244 - 0 CGGCAGTGACGATTCGGGTA AGG ||||||||[||||||||||||]PAM NA Intergenic NA NA

T27961 GTGACGATTCGGGTAAGGCGTGG 1000
Oligo fwd CACCGTGACGATTCGGGTAAGGCG
Oligo rev AAACCGCCTTACCCGAATCGTCAC
Chromosome start end strand MM target_seq PAM alignment distance position gene name gene id
GU305914.1 83217 83239 - 0 GTGACGATTCGGGTAAGGCG TGG ||||||||[||||||||||||]PAM NA Intergenic NA NA

T27962 GGCGTGGACTAGCGATAAAAAGG 1000
Oligo fwd CACCGGCGTGGACTAGCGATAAAA

<table>
<tr><td>Oligo rev</td><td>AAACTTTTATCGCTAGTCCACGCC</td><td></td><td></td><td></td><td></td><td></td><td></td><td></td><td></td><td></td><td></td><td></td></tr>
<tr><td>Chromosome</td><td>start</td><td></td><td>end</td><td>strand</td><td>MM</td><td>target_seq</td><td>PAM</td><td>alignment</td><td>distance</td><td>position</td><td>gene name</td><td>gene id</td></tr>
<tr><td>GU305914.1</td><td></td><td>83201</td><td>83223</td><td>-</td><td>0</td><td>GGCGTGGACTAGCGATAAAA</td><td>AGG</td><td>||||||||[||||||||||||]PAM</td><td>NA</td><td>Intergenic</td><td>NA</td><td>NA</td></tr>
<tr><td>T27963</td><td>GCGTGGACTAGCGATAAAAAGGG</td><td></td><td>1000</td><td></td><td></td><td></td><td></td><td></td><td></td><td></td><td></td><td></td></tr>
<tr><td>Oligo fwd</td><td>CACCGCGTGGACTAGCGATAAAAA</td><td></td><td></td><td></td><td></td><td></td><td></td><td></td><td></td><td></td><td></td><td></td></tr>
<tr><td>Oligo rev</td><td>AAACTTTTTATCGCTAGTCCACGC</td><td></td><td></td><td></td><td></td><td></td><td></td><td></td><td></td><td></td><td></td><td></td></tr>
<tr><td>Chromosome</td><td>start</td><td></td><td>end</td><td>strand</td><td>MM</td><td>target_seq</td><td>PAM</td><td>alignment</td><td>distance</td><td>position</td><td>gene name</td><td>gene id</td></tr>
<tr><td>GU305914.1</td><td></td><td>83200</td><td>83222</td><td>-</td><td>0</td><td>GCGTGGACTAGCGATAAAAA</td><td>GGG</td><td>||||||||[||||||||||||]PAM</td><td>NA</td><td>Intergenic</td><td>NA</td><td>NA</td></tr>
<tr><td>T27964</td><td>GACTAGCGATAAAAAGGGCTTGG</td><td></td><td>1000</td><td></td><td></td><td></td><td></td><td></td><td></td><td></td><td></td><td></td></tr>
<tr><td>Oligo fwd</td><td>CACCGACTAGCGATAAAAAGGGCT</td><td></td><td></td><td></td><td></td><td></td><td></td><td></td><td></td><td></td><td></td><td></td></tr>
<tr><td>Oligo rev</td><td>AAACAGCCCTTTTTATCGCTAGTC</td><td></td><td></td><td></td><td></td><td></td><td></td><td></td><td></td><td></td><td></td><td></td></tr>
<tr><td>Chromosome</td><td>start</td><td></td><td>end</td><td>strand</td><td>MM</td><td>target_seq</td><td>PAM</td><td>alignment</td><td>distance</td><td>position</td><td>gene name</td><td>gene id</td></tr>
<tr><td>GU305914.1</td><td></td><td>83195</td><td>83217</td><td>-</td><td>0</td><td>GACTAGCGATAAAAAGGGCT</td><td>TGG</td><td>||||||||[||||||||||||]PAM</td><td>NA</td><td>Intergenic</td><td>NA</td><td>NA</td></tr>
<tr><td>T27965</td><td>GCGATAAAAAGGGCTTGGAGAGG</td><td></td><td>1000</td><td></td><td></td><td></td><td></td><td></td><td></td><td></td><td></td><td></td></tr>
<tr><td>Oligo fwd</td><td>CACCGCGATAAAAAGGGCTTGGAG</td><td></td><td></td><td></td><td></td><td></td><td></td><td></td><td></td><td></td><td></td><td></td></tr>
<tr><td>Oligo rev</td><td>AAACCTCCAAGCCCTTTTTATCGC</td><td></td><td></td><td></td><td></td><td></td><td></td><td></td><td></td><td></td><td></td><td></td></tr>
<tr><td>Chromosome</td><td>start</td><td></td><td>end</td><td>strand</td><td>MM</td><td>target_seq</td><td>PAM</td><td>alignment</td><td>distance</td><td>position</td><td>gene name</td><td>gene id</td></tr>
<tr><td>GU305914.1</td><td></td><td>83190</td><td>83212</td><td>-</td><td>0</td><td>GCGATAAAAAGGGCTTGGAG</td><td>AGG</td><td>||||||||[||||||||||||]PAM</td><td>NA</td><td>Intergenic</td><td>NA</td><td>NA</td></tr>
<tr><td>T27966</td><td>GAAAGCGACTAGACTGCCGACGG</td><td></td><td>1000</td><td></td><td></td><td></td><td></td><td></td><td></td><td></td><td></td><td></td></tr>
<tr><td>Oligo fwd</td><td>CACCGAAAGCGACTAGACTGCCGA</td><td></td><td></td><td></td><td></td><td></td><td></td><td></td><td></td><td></td><td></td><td></td></tr>
<tr><td>Oligo rev</td><td>AAACTCGGCAGTCTAGTCGCTTTC</td><td></td><td></td><td></td><td></td><td></td><td></td><td></td><td></td><td></td><td></td><td></td></tr>
<tr><td>Chromosome</td><td>start</td><td></td><td>end</td><td>strand</td><td>MM</td><td>target_seq</td><td>PAM</td><td>alignment</td><td>distance</td><td>position</td><td>gene name</td><td>gene id</td></tr>
<tr><td>GU305914.1</td><td></td><td>83084</td><td>83106</td><td>-</td><td>0</td><td>GAAAGCGACTAGACTGCCGA</td><td>CGG</td><td>||||||||[||||||||||||]PAM</td><td>NA</td><td>Intergenic</td><td>NA</td><td>NA</td></tr>
<tr><td>GU305914.1</td><td></td><td>78992</td><td>79014</td><td>+</td><td>4</td><td>GGAGACGACGAGACTGCCGA</td><td>CGG</td><td>|-|--|||[|-||||||||||]PAM</td><td>NA</td><td>Intergenic</td><td>NA</td><td>NA</td></tr>
<tr><td>T27967</td><td>AAAGCGACTAGACTGCCGACGGG</td><td></td><td>1000</td><td></td><td></td><td></td><td></td><td></td><td></td><td></td><td></td><td></td></tr>
<tr><td>Oligo fwd</td><td>CACCAAAGCGACTAGACTGCCGAC</td><td></td><td></td><td></td><td></td><td></td><td></td><td></td><td></td><td></td><td></td><td></td></tr>
<tr><td>Oligo rev</td><td>AAACGTCGGCAGTCTAGTCGCTTT</td><td></td><td></td><td></td><td></td><td></td><td></td><td></td><td></td><td></td><td></td><td></td></tr>
<tr><td>Chromosome</td><td>start</td><td></td><td>end</td><td>strand</td><td>MM</td><td>target_seq</td><td>PAM</td><td>alignment</td><td>distance</td><td>position</td><td>gene name</td><td>gene id</td></tr>
<tr><td>GU305914.1</td><td></td><td>83083</td><td>83105</td><td>-</td><td>0</td><td>AAAGCGACTAGACTGCCGAC</td><td>GGG</td><td>||||||||[||||||||||||]PAM</td><td>NA</td><td>Intergenic</td><td>NA</td><td>NA</td></tr>
<tr><td>T27968</td><td>TAGACTGCCGACGGGCATCTTGG</td><td></td><td>1000</td><td></td><td></td><td></td><td></td><td></td><td></td><td></td><td></td><td></td></tr>
<tr><td>Oligo fwd</td><td>CACCTAGACTGCCGACGGGCATCT</td><td></td><td></td><td></td><td></td><td></td><td></td><td></td><td></td><td></td><td></td><td></td></tr>
<tr><td>Oligo rev</td><td>AAACAGATGCCCGTCGGCAGTCTA</td><td></td><td></td><td></td><td></td><td></td><td></td><td></td><td></td><td></td><td></td><td></td></tr>
<tr><td>Chromosome</td><td>start</td><td></td><td>end</td><td>strand</td><td>MM</td><td>target_seq</td><td>PAM</td><td>alignment</td><td>distance</td><td>position</td><td>gene name</td><td>gene id</td></tr>
<tr><td>GU305914.1</td><td></td><td>83075</td><td>83097</td><td>-</td><td>0</td><td>TAGACTGCCGACGGGCATCT</td><td>TGG</td><td>||||||||[||||||||||||]PAM</td><td>NA</td><td>Intergenic</td><td>NA</td><td>NA</td></tr>
<tr><td>T27969</td><td>GGCATCTTGGACCGTCTCAAAGG</td><td></td><td>1000</td><td></td><td></td><td></td><td></td><td></td><td></td><td></td><td></td><td></td></tr>
<tr><td>Oligo fwd</td><td>CACCGGCATCTTGGACCGTCTCAA</td><td></td><td></td><td></td><td></td><td></td><td></td><td></td><td></td><td></td><td></td><td></td></tr>
<tr><td>Oligo rev</td><td>AAACTTGAGACGGTCCAAGATGCC</td><td></td><td></td><td></td><td></td><td></td><td></td><td></td><td></td><td></td><td></td><td></td></tr>
<tr><td>Chromosome</td><td>start</td><td></td><td>end</td><td>strand</td><td>MM</td><td>target_seq</td><td>PAM</td><td>alignment</td><td>distance</td><td>position</td><td>gene name</td><td>gene id</td></tr>
<tr><td>GU305914.1</td><td></td><td>83062</td><td>83084</td><td>-</td><td>0</td><td>GGCATCTTGGACCGTCTCAA</td><td>AGG</td><td>||||||||[||||||||||||]PAM</td><td>NA</td><td>Intergenic</td><td>NA</td><td>NA</td></tr>
<tr><td>T27970</td><td>CTCAAAGGTAACGACACCAGTGG</td><td></td><td>1000</td><td></td><td></td><td></td><td></td><td></td><td></td><td></td><td></td><td></td></tr>
<tr><td>Oligo fwd</td><td>CACCCTCAAAGGTAACGACACCAG</td><td></td><td></td><td></td><td></td><td></td><td></td><td></td><td></td><td></td><td></td><td></td></tr>
<tr><td>Oligo rev</td><td>AAACCTGGTGTCGTTACCTTTGAG</td><td></td><td></td><td></td><td></td><td></td><td></td><td></td><td></td><td></td><td></td><td></td></tr>
<tr><td>Chromosome</td><td>start</td><td></td><td>end</td><td>strand</td><td>MM</td><td>target_seq</td><td>PAM</td><td>alignment</td><td>distance</td><td>position</td><td>gene name</td><td>gene id</td></tr>
<tr><td>GU305914.1</td><td></td><td>83047</td><td>83069</td><td>-</td><td>0</td><td>CTCAAAGGTAACGACACCAG</td><td>TGG</td><td>||||||||[||||||||||||]PAM</td><td>NA</td><td>Intergenic</td><td>NA</td><td>NA</td></tr>
<tr><td>T27971</td><td>CAGTGGCTACCAGCGATTACCGG</td><td></td><td>1000</td><td></td><td></td><td></td><td></td><td></td><td></td><td></td><td></td><td></td></tr>
<tr><td>Oligo fwd</td><td>CACCCAGTGGCTACCAGCGATTAC</td><td></td><td></td><td></td><td></td><td></td><td></td><td></td><td></td><td></td><td></td><td></td></tr>
<tr><td>Oligo rev</td><td>AAACGTAATCGCTGGTAGCCACTG</td><td></td><td></td><td></td><td></td><td></td><td></td><td></td><td></td><td></td><td></td><td></td></tr>
<tr><td>Chromosome</td><td>start</td><td></td><td>end</td><td>strand</td><td>MM</td><td>target_seq</td><td>PAM</td><td>alignment</td><td>distance</td><td>position</td><td>gene name</td><td>gene id</td></tr>
<tr><td>GU305914.1</td><td></td><td>83030</td><td>83052</td><td>-</td><td>0</td><td>CAGTGGCTACCAGCGATTAC</td><td>CGG</td><td>||||||||[||||||||||||]PAM</td><td>NA</td><td>Intergenic</td><td>NA</td><td>NA</td></tr>
<tr><td>T27972</td><td>TGGCTACCAGCGATTACCGGCGG</td><td></td><td>1000</td><td></td><td></td><td></td><td></td><td></td><td></td><td></td><td></td><td></td></tr>
<tr><td>Oligo fwd</td><td>CACCTGGCTACCAGCGATTACCGG</td><td></td><td></td><td></td><td></td><td></td><td></td><td></td><td></td><td></td><td></td><td></td></tr>
<tr><td>Oligo rev</td><td>AAACCCGGTAATCGCTGGTAGCCA</td><td></td><td></td><td></td><td></td><td></td><td></td><td></td><td></td><td></td><td></td><td></td></tr>
<tr><td>Chromosome</td><td>start</td><td></td><td>end</td><td>strand</td><td>MM</td><td>target_seq</td><td>PAM</td><td>alignment</td><td>distance</td><td>position</td><td>gene name</td><td>gene id</td></tr>
<tr><td>GU305914.1</td><td></td><td>83027</td><td>83049</td><td>-</td><td>0</td><td>TGGCTACCAGCGATTACCGG</td><td>CGG</td><td>||||||||[||||||||||||]PAM</td><td>NA</td><td>Intergenic</td><td>NA</td><td>NA</td></tr>
<tr><td>T27973</td><td>CTACCAGCGATTACCGGCGGAGG</td><td></td><td>1000</td><td></td><td></td><td></td><td></td><td></td><td></td><td></td><td></td><td></td></tr>
<tr><td>Oligo fwd</td><td>CACCCTACCAGCGATTACCGGCGG</td><td></td><td></td><td></td><td></td><td></td><td></td><td></td><td></td><td></td><td></td><td></td></tr>
<tr><td>Oligo rev</td><td>AAACCCGCCGGTAATCGCTGGTAG</td><td></td><td></td><td></td><td></td><td></td><td></td><td></td><td></td><td></td><td></td><td></td></tr>
<tr><td>Chromosome</td><td>start</td><td></td><td>end</td><td>strand</td><td>MM</td><td>target_seq</td><td>PAM</td><td>alignment</td><td>distance</td><td>position</td><td>gene name</td><td>gene id</td></tr>
<tr><td>GU305914.1</td><td></td><td>83024</td><td>83046</td><td>-</td><td>0</td><td>CTACCAGCGATTACCGGCGG</td><td>AGG</td><td>||||||||[||||||||||||]PAM</td><td>NA</td><td>Intergenic</td><td>NA</td><td>NA</td></tr>
<tr><td>T27974</td><td>ACTCCGATTTCGAGTACTGACGG</td><td></td><td>1000</td><td></td><td></td><td></td><td></td><td></td><td></td><td></td><td></td><td></td></tr>
<tr><td>Oligo fwd</td><td>CACCACTCCGATTTCGAGTACTGA</td><td></td><td></td><td></td><td></td><td></td><td></td><td></td><td></td><td></td><td></td><td></td></tr>
<tr><td>Oligo rev</td><td>AAACTCAGTACTCGAAATCGGAGT</td><td></td><td></td><td></td><td></td><td></td><td></td><td></td><td></td><td></td><td></td><td></td></tr>
<tr><td>Chromosome</td><td>start</td><td></td><td>end</td><td>strand</td><td>MM</td><td>target_seq</td><td>PAM</td><td>alignment</td><td>distance</td><td>position</td><td>gene name</td><td>gene id</td></tr>
<tr><td>GU305914.1</td><td></td><td>83001</td><td>83023</td><td>-</td><td>0</td><td>ACTCCGATTTCGAGTACTGA</td><td>CGG</td><td>||||||||[||||||||||||]PAM</td><td>NA</td><td>Intergenic</td><td>NA</td><td>NA</td></tr>
<tr><td>T27975</td><td>TCGAGTACTGACGGCCGACATGG</td><td></td><td>1000</td><td></td><td></td><td></td><td></td><td></td><td></td><td></td><td></td><td></td></tr>
<tr><td>Oligo fwd</td><td>CACCTCGAGTACTGACGGCCGACA</td><td></td><td></td><td></td><td></td><td></td><td></td><td></td><td></td><td></td><td></td><td></td></tr>
<tr><td>Oligo rev</td><td>AAACTGTCGGCCGTCAGTACTCGA</td><td></td><td></td><td></td><td></td><td></td><td></td><td></td><td></td><td></td><td></td><td></td></tr>
<tr><td>Chromosome</td><td>start</td><td></td><td>end</td><td>strand</td><td>MM</td><td>target_seq</td><td>PAM</td><td>alignment</td><td>distance</td><td>position</td><td>gene name</td><td>gene id</td></tr>
<tr><td>GU305914.1</td><td></td><td>82992</td><td>83014</td><td>-</td><td>0</td><td>TCGAGTACTGACGGCCGACA</td><td>TGG</td><td>||||||||[||||||||||||]PAM</td><td>NA</td><td>Intergenic</td><td>NA</td><td>NA</td></tr>
<tr><td>T27976</td><td>GGCCGACATGGACACTTGCGTGG</td><td></td><td>1000</td><td></td><td></td><td></td><td></td><td></td><td></td><td></td><td></td><td></td></tr>
<tr><td>Oligo fwd</td><td>CACCGGCCGACATGGACACTTGCG</td><td></td><td></td><td></td><td></td><td></td><td></td><td></td><td></td><td></td><td></td><td></td></tr>
<tr><td>Oligo rev</td><td>AAACCGCAAGTGTCCATGTCGGCC</td><td></td><td></td><td></td><td></td><td></td><td></td><td></td><td></td><td></td><td></td><td></td></tr>
<tr><td>Chromosome</td><td>start</td><td></td><td>end</td><td>strand</td><td>MM</td><td>target_seq</td><td>PAM</td><td>alignment</td><td>distance</td><td>position</td><td>gene name</td><td>gene id</td></tr>
<tr><td>GU305914.1</td><td></td><td>82980</td><td>83002</td><td>-</td><td>0</td><td>GGCCGACATGGACACTTGCG</td><td>TGG</td><td>||||||||[||||||||||||]PAM</td><td>NA</td><td>Intergenic</td><td>NA</td><td>NA</td></tr>
</table>

<table>
<tr><td>T27977</td><td>TCTTCAACCCCTACCTGCGACGG</td><td></td><td>1000</td><td colspan="9"></td></tr>
<tr><td>Oligo fwd</td><td>CACCTCTTCAACCCCTACCTGCGA</td><td colspan="11"></td></tr>
<tr><td>Oligo rev</td><td>AAACTCGCAGGTAGGGGTTGAAGA</td><td colspan="11"></td></tr>
<tr><td>Chromosome</td><td>start</td><td></td><td>end</td><td>strand</td><td>MM</td><td>target_seq</td><td>PAM</td><td>alignment</td><td>distance</td><td>position</td><td>gene name</td><td>gene id</td></tr>
<tr><td>GU305914.1</td><td></td><td>82951</td><td>82973</td><td>-</td><td>0</td><td>TCTTCAACCCCTACCTGCGA</td><td>CGG</td><td>||||||||[||||||||||||]PAM</td><td>NA</td><td>Intergenic</td><td>NA</td><td>NA</td></tr>
<tr><td>T27978</td><td>GACGGAAACCCCGTCGCGATTGG</td><td></td><td>1000</td><td colspan="9"></td></tr>
<tr><td>Oligo fwd</td><td>CACCGACGGAAACCCCGTCGCGAT</td><td colspan="11"></td></tr>
<tr><td>Oligo rev</td><td>AAACATCGCGACGGGGTTTCCGTC</td><td colspan="11"></td></tr>
<tr><td>Chromosome</td><td>start</td><td></td><td>end</td><td>strand</td><td>MM</td><td>target_seq</td><td>PAM</td><td>alignment</td><td>distance</td><td>position</td><td>gene name</td><td>gene id</td></tr>
<tr><td>GU305914.1</td><td></td><td>82933</td><td>82955</td><td>-</td><td>0</td><td>GACGGAAACCCCGTCGCGAT</td><td>TGG</td><td>||||||||[||||||||||||]PAM</td><td>NA</td><td>Intergenic</td><td>NA</td><td>NA</td></tr>
<tr><td>T27979</td><td>TCGCGATTGGAGACGCTGCGAGG</td><td></td><td>1000</td><td colspan="9"></td></tr>
<tr><td>Oligo fwd</td><td>CACCTCGCGATTGGAGACGCTGCG</td><td colspan="11"></td></tr>
<tr><td>Oligo rev</td><td>AAACCGCAGCGTCTCCAATCGCGA</td><td colspan="11"></td></tr>
<tr><td>Chromosome</td><td>start</td><td></td><td>end</td><td>strand</td><td>MM</td><td>target_seq</td><td>PAM</td><td>alignment</td><td>distance</td><td>position</td><td>gene name</td><td>gene id</td></tr>
<tr><td>GU305914.1</td><td></td><td>82920</td><td>82942</td><td>-</td><td>0</td><td>TCGCGATTGGAGACGCTGCG</td><td>AGG</td><td>||||||||[||||||||||||]PAM</td><td>NA</td><td>Intergenic</td><td>NA</td><td>NA</td></tr>
<tr><td>T27980</td><td>TTTTTACAAGTCGTTCCGCGTGG</td><td></td><td>1000</td><td colspan="9"></td></tr>
<tr><td>Oligo fwd</td><td>CACCTTTTTACAAGTCGTTCCGCG</td><td colspan="11"></td></tr>
<tr><td>Oligo rev</td><td>AAACCGCGGAACGACTTGTAAAAA</td><td colspan="11"></td></tr>
<tr><td>Chromosome</td><td>start</td><td></td><td>end</td><td>strand</td><td>MM</td><td>target_seq</td><td>PAM</td><td>alignment</td><td>distance</td><td>position</td><td>gene name</td><td>gene id</td></tr>
<tr><td>GU305914.1</td><td></td><td>82883</td><td>82905</td><td>-</td><td>0</td><td>TTTTTACAAGTCGTTCCGCG</td><td>TGG</td><td>||||||||[||||||||||||]PAM</td><td>NA</td><td>Intergenic</td><td>NA</td><td>NA</td></tr>
<tr><td>T27981</td><td>CCGCGTGGCGTGTTGTATGATGG</td><td></td><td>1000</td><td colspan="9"></td></tr>
<tr><td>Oligo fwd</td><td>CACCCCGCGTGGCGTGTTGTATGA</td><td colspan="11"></td></tr>
<tr><td>Oligo rev</td><td>AAACTCATACAACACGCCACGCGG</td><td colspan="11"></td></tr>
<tr><td>Chromosome</td><td>start</td><td></td><td>end</td><td>strand</td><td>MM</td><td>target_seq</td><td>PAM</td><td>alignment</td><td>distance</td><td>position</td><td>gene name</td><td>gene id</td></tr>
<tr><td>GU305914.1</td><td></td><td>82868</td><td>82890</td><td>-</td><td>0</td><td>CCGCGTGGCGTGTTGTATGA</td><td>TGG</td><td>||||||||[||||||||||||]PAM</td><td>NA</td><td>Intergenic</td><td>NA</td><td>NA</td></tr>
<tr><td>T27982</td><td>CGTGTTGTATGATGGTGCGACGG</td><td></td><td>1000</td><td colspan="9"></td></tr>
<tr><td>Oligo fwd</td><td>CACCCGTGTTGTATGATGGTGCGA</td><td colspan="11"></td></tr>
<tr><td>Oligo rev</td><td>AAACTCGCACCATCATACAACACG</td><td colspan="11"></td></tr>
<tr><td>Chromosome</td><td>start</td><td></td><td>end</td><td>strand</td><td>MM</td><td>target_seq</td><td>PAM</td><td>alignment</td><td>distance</td><td>position</td><td>gene name</td><td>gene id</td></tr>
<tr><td>GU305914.1</td><td></td><td>82860</td><td>82882</td><td>-</td><td>0</td><td>CGTGTTGTATGATGGTGCGA</td><td>CGG</td><td>||||||||[||||||||||||]PAM</td><td>NA</td><td>Intergenic</td><td>NA</td><td>NA</td></tr>
<tr><td>T27983</td><td>GTGTTGTATGATGGTGCGACGGG</td><td></td><td>1000</td><td colspan="9"></td></tr>
<tr><td>Oligo fwd</td><td>CACCGTGTTGTATGATGGTGCGAC</td><td colspan="11"></td></tr>
<tr><td>Oligo rev</td><td>AAACGTCGCACCATCATACAACAC</td><td colspan="11"></td></tr>
<tr><td>Chromosome</td><td>start</td><td></td><td>end</td><td>strand</td><td>MM</td><td>target_seq</td><td>PAM</td><td>alignment</td><td>distance</td><td>position</td><td>gene name</td><td>gene id</td></tr>
<tr><td>GU305914.1</td><td></td><td>82859</td><td>82881</td><td>-</td><td>0</td><td>GTGTTGTATGATGGTGCGAC</td><td>GGG</td><td>||||||||[||||||||||||]PAM</td><td>NA</td><td>Intergenic</td><td>NA</td><td>NA</td></tr>
<tr><td>T27984</td><td>TGTTGTATGATGGTGCGACGGGG</td><td></td><td>1000</td><td colspan="9"></td></tr>
<tr><td>Oligo fwd</td><td>CACCTGTTGTATGATGGTGCGACG</td><td colspan="11"></td></tr>
<tr><td>Oligo rev</td><td>AAACCGTCGCACCATCATACAACA</td><td colspan="11"></td></tr>
<tr><td>Chromosome</td><td>start</td><td></td><td>end</td><td>strand</td><td>MM</td><td>target_seq</td><td>PAM</td><td>alignment</td><td>distance</td><td>position</td><td>gene name</td><td>gene id</td></tr>
<tr><td>GU305914.1</td><td></td><td>82858</td><td>82880</td><td>-</td><td>0</td><td>TGTTGTATGATGGTGCGACG</td><td>GGG</td><td>||||||||[||||||||||||]PAM</td><td>NA</td><td>Intergenic</td><td>NA</td><td>NA</td></tr>
<tr><td>T27985</td><td>TGGTGCGACGGGGTTAATAAAGG</td><td></td><td>1000</td><td colspan="9"></td></tr>
<tr><td>Oligo fwd</td><td>CACCTGGTGCGACGGGGTTAATAA</td><td colspan="11"></td></tr>
<tr><td>Oligo rev</td><td>AAACTTATTAACCCCGTCGCACCA</td><td colspan="11"></td></tr>
<tr><td>Chromosome</td><td>start</td><td></td><td>end</td><td>strand</td><td>MM</td><td>target_seq</td><td>PAM</td><td>alignment</td><td>distance</td><td>position</td><td>gene name</td><td>gene id</td></tr>
<tr><td>GU305914.1</td><td></td><td>82848</td><td>82870</td><td>-</td><td>0</td><td>TGGTGCGACGGGGTTAATAA</td><td>AGG</td><td>||||||||[||||||||||||]PAM</td><td>NA</td><td>Intergenic</td><td>NA</td><td>NA</td></tr>
<tr><td>T27986</td><td>GGGTTAATAAAGGTACAGTCCGG</td><td></td><td>1000</td><td colspan="9"></td></tr>
<tr><td>Oligo fwd</td><td>CACCGGGTTAATAAAGGTACAGTC</td><td colspan="11"></td></tr>
<tr><td>Oligo rev</td><td>AAACGACTGTACCTTTATTAACCC</td><td colspan="11"></td></tr>
<tr><td>Chromosome</td><td>start</td><td></td><td>end</td><td>strand</td><td>MM</td><td>target_seq</td><td>PAM</td><td>alignment</td><td>distance</td><td>position</td><td>gene name</td><td>gene id</td></tr>
<tr><td>GU305914.1</td><td></td><td>82838</td><td>82860</td><td>-</td><td>0</td><td>GGGTTAATAAAGGTACAGTC</td><td>CGG</td><td>||||||||[||||||||||||]PAM</td><td>NA</td><td>Intergenic</td><td>NA</td><td>NA</td></tr>
<tr><td>T27987</td><td>AATAAAGGTACAGTCCGGCATGG</td><td></td><td>1000</td><td colspan="9"></td></tr>
<tr><td>Oligo fwd</td><td>CACCAATAAAGGTACAGTCCGGCA</td><td colspan="11"></td></tr>
<tr><td>Oligo rev</td><td>AAACTGCCGGACTGTACCTTTATT</td><td colspan="11"></td></tr>
<tr><td>Chromosome</td><td>start</td><td></td><td>end</td><td>strand</td><td>MM</td><td>target_seq</td><td>PAM</td><td>alignment</td><td>distance</td><td>position</td><td>gene name</td><td>gene id</td></tr>
<tr><td>GU305914.1</td><td></td><td>82833</td><td>82855</td><td>-</td><td>0</td><td>AATAAAGGTACAGTCCGGCA</td><td>TGG</td><td>||||||||[||||||||||||]PAM</td><td>NA</td><td>Intergenic</td><td>NA</td><td>NA</td></tr>
<tr><td>T27988</td><td>TACAGTCCGGCATGGAACCCAGG</td><td></td><td>1000</td><td colspan="9"></td></tr>
<tr><td>Oligo fwd</td><td>CACCTACAGTCCGGCATGGAACCC</td><td colspan="11"></td></tr>
<tr><td>Oligo rev</td><td>AAACGGGTTCCATGCCGGACTGTA</td><td colspan="11"></td></tr>
<tr><td>Chromosome</td><td>start</td><td></td><td>end</td><td>strand</td><td>MM</td><td>target_seq</td><td>PAM</td><td>alignment</td><td>distance</td><td>position</td><td>gene name</td><td>gene id</td></tr>
<tr><td>GU305914.1</td><td></td><td>82825</td><td>82847</td><td>-</td><td>0</td><td>TACAGTCCGGCATGGAACCC</td><td>AGG</td><td>||||||||[||||||||||||]PAM</td><td>NA</td><td>Intergenic</td><td>NA</td><td>NA</td></tr>
<tr><td>T27989</td><td>TGCTCAACCCCGACAAGCCGTGG</td><td></td><td>1000</td><td colspan="9"></td></tr>
<tr><td>Oligo fwd</td><td>CACCTGCTCAACCCCGACAAGCCG</td><td colspan="11"></td></tr>
<tr><td>Oligo rev</td><td>AAACCGGCTTGTCGGGGTTGAGCA</td><td colspan="11"></td></tr>
<tr><td>Chromosome</td><td>start</td><td></td><td>end</td><td>strand</td><td>MM</td><td>target_seq</td><td>PAM</td><td>alignment</td><td>distance</td><td>position</td><td>gene name</td><td>gene id</td></tr>
<tr><td>GU305914.1</td><td></td><td>82777</td><td>82799</td><td>-</td><td>0</td><td>TGCTCAACCCCGACAAGCCG</td><td>TGG</td><td>||||||||[||||||||||||]PAM</td><td>NA</td><td>Intergenic</td><td>NA</td><td>NA</td></tr>
<tr><td>T27990</td><td>TGGCCGACCCTGCGCACCAGAGG</td><td></td><td>1000</td><td colspan="9"></td></tr>
<tr><td>Oligo fwd</td><td>CACCTGGCCGACCCTGCGCACCAG</td><td colspan="11"></td></tr>
<tr><td>Oligo rev</td><td>AAACCTGGTGCGCAGGGTCGGCCA</td><td colspan="11"></td></tr>
<tr><td>Chromosome</td><td>start</td><td></td><td>end</td><td>strand</td><td>MM</td><td>target_seq</td><td>PAM</td><td>alignment</td><td>distance</td><td>position</td><td>gene name</td><td>gene id</td></tr>
<tr><td>GU305914.1</td><td></td><td>82757</td><td>82779</td><td>-</td><td>0</td><td>TGGCCGACCCTGCGCACCAG</td><td>AGG</td><td>||||||||[||||||||||||]PAM</td><td>NA</td><td>Intergenic</td><td>NA</td><td>NA</td></tr>
<tr><td>T27991</td><td>CGACCCTGCGCACCAGAGGTTGG</td><td></td><td>1000</td><td colspan="9"></td></tr>
<tr><td>Oligo fwd</td><td>CACCCGACCCTGCGCACCAGAGGT</td><td colspan="11"></td></tr>
<tr><td>Oligo rev</td><td>AAACACCTCTGGTGCGCAGGGTCG</td><td colspan="11"></td></tr>
<tr><td>Chromosome</td><td>start</td><td></td><td>end</td><td>strand</td><td>MM</td><td>target_seq</td><td>PAM</td><td>alignment</td><td>distance</td><td>position</td><td>gene name</td><td>gene id</td></tr>
<tr><td>GU305914.1</td><td></td><td>82753</td><td>82775</td><td>-</td><td>0</td><td>CGACCCTGCGCACCAGAGGT</td><td>TGG</td><td>||||||||[||||||||||||]PAM</td><td>NA</td><td>Intergenic</td><td>NA</td><td>NA</td></tr>
</table>

| | | | | | | | | | | | |
|---|---|---|---|---|---|---|---|---|---|---|---|
| T27992 | TGCGCACCAGAGGTTGGTGCCGG | 1000 | | | | | | | | | |
| Oligo fwd | CACCTGCGCACCAGAGGTTGGTGC | | | | | | | | | | |
| Oligo rev | AAACGCACCAACCTCTGGTGCGCA | | | | | | | | | | |
| Chromosome | start | end | strand | MM | target_seq | PAM | alignment | distance | position | gene name | gene id |
| GU305914.1 | 82747 | 82769 | - | 0 | TGCGCACCAGAGGTTGGTGC | CGG | \|\|\|\|\|\|\|\|[\|\|\|\|\|\|\|\|\|\|\|\|]PAM | NA | Intergenic | NA | NA |
| T27993 | GCGCACCAGAGGTTGGTGCCGGG | 1000 | | | | | | | | | |
| Oligo fwd | CACCGCGCACCAGAGGTTGGTGCC | | | | | | | | | | |
| Oligo rev | AAACGGCACCAACCTCTGGTGCGC | | | | | | | | | | |
| Chromosome | start | end | strand | MM | target_seq | PAM | alignment | distance | position | gene name | gene id |
| GU305914.1 | 82746 | 82768 | - | 0 | GCGCACCAGAGGTTGGTGCC | GGG | \|\|\|\|\|\|\|\|[\|\|\|\|\|\|\|\|\|\|\|\|]PAM | NA | Intergenic | NA | NA |
| T27994 | CGCACCAGAGGTTGGTGCCGGGG | 1000 | | | | | | | | | |
| Oligo fwd | CACCCGCACCAGAGGTTGGTGCCG | | | | | | | | | | |
| Oligo rev | AAACCGGCACCAACCTCTGGTGCG | | | | | | | | | | |
| Chromosome | start | end | strand | MM | target_seq | PAM | alignment | distance | position | gene name | gene id |
| GU305914.1 | 82745 | 82767 | - | 0 | CGCACCAGAGGTTGGTGCCG | GGG | \|\|\|\|\|\|\|\|[\|\|\|\|\|\|\|\|\|\|\|\|]PAM | NA | Intergenic | NA | NA |
| T27995 | GCGCTTCCACACCTACGACCAGG | 1000 | | | | | | | | | |
| Oligo fwd | CACCGCGCTTCCACACCTACGACC | | | | | | | | | | |
| Oligo rev | AAACGGTCGTAGGTGTGGAAGCGC | | | | | | | | | | |
| Chromosome | start | end | strand | MM | target_seq | PAM | alignment | distance | position | gene name | gene id |
| GU305914.1 | 82704 | 82726 | - | 0 | GCGCTTCCACACCTACGACC | AGG | \|\|\|\|\|\|\|\|[\|\|\|\|\|\|\|\|\|\|\|\|]PAM | NA | Intergenic | NA | NA |
| T27996 | CTACGACCAGGTCGTCAACCTGG | 1000 | | | | | | | | | |
| Oligo fwd | CACCCTACGACCAGGTCGTCAACC | | | | | | | | | | |
| Oligo rev | AAACGGTTGACGACCTGGTCGTAG | | | | | | | | | | |
| Chromosome | start | end | strand | MM | target_seq | PAM | alignment | distance | position | gene name | gene id |
| GU305914.1 | 82692 | 82714 | - | 0 | CTACGACCAGGTCGTCAACC | TGG | \|\|\|\|\|\|\|\|[\|\|\|\|\|\|\|\|\|\|\|\|]PAM | NA | Intergenic | NA | NA |
| T27997 | ACGAGCAGATCTCGCCCCGCTGG | 1000 | | | | | | | | | |
| Oligo fwd | CACCACGAGCAGATCTCGCCCCGC | | | | | | | | | | |
| Oligo rev | AAACGCGGGGCGAGATCTGCTCGT | | | | | | | | | | |
| Chromosome | start | end | strand | MM | target_seq | PAM | alignment | distance | position | gene name | gene id |
| GU305914.1 | 82654 | 82676 | - | 0 | ACGAGCAGATCTCGCCCCGC | TGG | \|\|\|\|\|\|\|\|[\|\|\|\|\|\|\|\|\|\|\|\|]PAM | NA | Intergenic | NA | NA |
| T27998 | GCCCCGCTGGAAGCACCATGTGG | 1000 | | | | | | | | | |
| Oligo fwd | CACCGCCCCGCTGGAAGCACCATG | | | | | | | | | | |
| Oligo rev | AAACCATGGTGCTTCCAGCGGGGC | | | | | | | | | | |
| Chromosome | start | end | strand | MM | target_seq | PAM | alignment | distance | position | gene name | gene id |
| GU305914.1 | 82641 | 82663 | - | 0 | GCCCCGCTGGAAGCACCATG | TGG | \|\|\|\|\|\|\|\|[\|\|\|\|\|\|\|\|\|\|\|\|]PAM | NA | Intergenic | NA | NA |
| T27999 | GAAGCACCATGTGGTGCCGTCGG | 1000 | | | | | | | | | |
| Oligo fwd | CACCGAAGCACCATGTGGTGCCGT | | | | | | | | | | |
| Oligo rev | AAACACGGCACCACATGGTGCTTC | | | | | | | | | | |
| Chromosome | start | end | strand | MM | target_seq | PAM | alignment | distance | position | gene name | gene id |
| GU305914.1 | 82632 | 82654 | - | 0 | GAAGCACCATGTGGTGCCGT | CGG | \|\|\|\|\|\|\|\|[\|\|\|\|\|\|\|\|\|\|\|\|]PAM | NA | Intergenic | NA | NA |
| T28000 | AAGCACCATGTGGTGCCGTCGGG | 1000 | | | | | | | | | |
| Oligo fwd | CACCAAGCACCATGTGGTGCCGTC | | | | | | | | | | |
| Oligo rev | AAACGACGGCACCACATGGTGCTT | | | | | | | | | | |
| Chromosome | start | end | strand | MM | target_seq | PAM | alignment | distance | position | gene name | gene id |
| GU305914.1 | 82631 | 82653 | - | 0 | AAGCACCATGTGGTGCCGTC | GGG | \|\|\|\|\|\|\|\|[\|\|\|\|\|\|\|\|\|\|\|\|]PAM | NA | Intergenic | NA | NA |
| T28001 | AGCACCATGTGGTGCCGTCGGGG | 1000 | | | | | | | | | |
| Oligo fwd | CACCAGCACCATGTGGTGCCGTCG | | | | | | | | | | |
| Oligo rev | AAACCGACGGCACCACATGGTGCT | | | | | | | | | | |
| Chromosome | start | end | strand | MM | target_seq | PAM | alignment | distance | position | gene name | gene id |
| GU305914.1 | 82630 | 82652 | - | 0 | AGCACCATGTGGTGCCGTCG | GGG | \|\|\|\|\|\|\|\|[\|\|\|\|\|\|\|\|\|\|\|\|]PAM | NA | Intergenic | NA | NA |
| T28002 | GGGAACGTCATCCGCATGTTCGG | 1000 | | | | | | | | | |
| Oligo fwd | CACCGGGAACGTCATCCGCATGTT | | | | | | | | | | |
| Oligo rev | AAACAACATGCGGATGACGTTCCC | | | | | | | | | | |
| Chromosome | start | end | strand | MM | target_seq | PAM | alignment | distance | position | gene name | gene id |
| GU305914.1 | 82610 | 82632 | - | 0 | GGGAACGTCATCCGCATGTT | CGG | \|\|\|\|\|\|\|\|[\|\|\|\|\|\|\|\|\|\|\|\|]PAM | NA | Intergenic | NA | NA |
| T28003 | GGAACGTCATCCGCATGTTCGGG | 1000 | | | | | | | | | |
| Oligo fwd | CACCGGAACGTCATCCGCATGTTC | | | | | | | | | | |
| Oligo rev | AAACGAACATGCGGATGACGTTCC | | | | | | | | | | |
| Chromosome | start | end | strand | MM | target_seq | PAM | alignment | distance | position | gene name | gene id |
| GU305914.1 | 82609 | 82631 | - | 0 | GGAACGTCATCCGCATGTTC | GGG | \|\|\|\|\|\|\|\|[\|\|\|\|\|\|\|\|\|\|\|\|]PAM | NA | Intergenic | NA | NA |
| T28004 | GAACGTCATCCGCATGTTCGGGG | 1000 | | | | | | | | | |
| Oligo fwd | CACCGAACGTCATCCGCATGTTCG | | | | | | | | | | |
| Oligo rev | AAACCGAACATGCGGATGACGTTC | | | | | | | | | | |
| Chromosome | start | end | strand | MM | target_seq | PAM | alignment | distance | position | gene name | gene id |
| GU305914.1 | 82608 | 82630 | - | 0 | GAACGTCATCCGCATGTTCG | GGG | \|\|\|\|\|\|\|\|[\|\|\|\|\|\|\|\|\|\|\|\|]PAM | NA | Intergenic | NA | NA |
| T28005 | CATGTTCGGGGCCACCGATGAGG | 1000 | | | | | | | | | |
| Oligo fwd | CACCCATGTTCGGGGCCACCGATG | | | | | | | | | | |
| Oligo rev | AAACCATCGGTGGCCCCGAACATG | | | | | | | | | | |
| Chromosome | start | end | strand | MM | target_seq | PAM | alignment | distance | position | gene name | gene id |
| GU305914.1 | 82596 | 82618 | - | 0 | CATGTTCGGGGCCACCGATG | AGG | \|\|\|\|\|\|\|\|[\|\|\|\|\|\|\|\|\|\|\|\|]PAM | NA | Intergenic | NA | NA |
| T28006 | ATGTTCGGGGCCACCGATGAGGG | 1000 | | | | | | | | | |
| Oligo fwd | CACCATGTTCGGGGCCACCGATGA | | | | | | | | | | |
| Oligo rev | AAACTCATCGGTGGCCCCGAACAT | | | | | | | | | | |
| Chromosome | start | end | strand | MM | target_seq | PAM | alignment | distance | position | gene name | gene id |

<table>
<tr><td>GU305914.1</td><td>82595</td><td>82617</td><td>-</td><td>0</td><td>ATGTTCGGGGCCACCGATGA</td><td>GGG</td><td>||||||||[||||||||||||]PAM</td><td>NA</td><td>Intergenic</td><td>NA</td><td>NA</td></tr>
<tr><td>T28007</td><td>TGTTCGGGGCCACCGATGAGGGG</td><td>1000</td><td></td><td></td><td></td><td></td><td></td><td></td><td></td><td></td><td></td></tr>
<tr><td>Oligo fwd</td><td>CACCTGTTCGGGGCCACCGATGAG</td><td></td><td></td><td></td><td></td><td></td><td></td><td></td><td></td><td></td><td></td></tr>
<tr><td>Oligo rev</td><td>AAACCTCATCGGTGGCCCCGAACA</td><td></td><td></td><td></td><td></td><td></td><td></td><td></td><td></td><td></td><td></td></tr>
<tr><td>Chromosome</td><td>start</td><td>end</td><td>strand</td><td>MM</td><td>target_seq</td><td>PAM</td><td>alignment</td><td>distance</td><td>position</td><td>gene name</td><td>gene id</td></tr>
<tr><td>GU305914.1</td><td>82594</td><td>82616</td><td>-</td><td>0</td><td>TGTTCGGGGCCACCGATGAG</td><td>GGG</td><td>||||||||[||||||||||||]PAM</td><td>NA</td><td>Intergenic</td><td>NA</td><td>NA</td></tr>
<tr><td>T28008</td><td>GTTCGGGGCCACCGATGAGGGGG</td><td>1000</td><td></td><td></td><td></td><td></td><td></td><td></td><td></td><td></td><td></td></tr>
<tr><td>Oligo fwd</td><td>CACCGTTCGGGGCCACCGATGAGG</td><td></td><td></td><td></td><td></td><td></td><td></td><td></td><td></td><td></td><td></td></tr>
<tr><td>Oligo rev</td><td>AAACCCTCATCGGTGGCCCCGAAC</td><td></td><td></td><td></td><td></td><td></td><td></td><td></td><td></td><td></td><td></td></tr>
<tr><td>Chromosome</td><td>start</td><td>end</td><td>strand</td><td>MM</td><td>target_seq</td><td>PAM</td><td>alignment</td><td>distance</td><td>position</td><td>gene name</td><td>gene id</td></tr>
<tr><td>GU305914.1</td><td>82593</td><td>82615</td><td>-</td><td>0</td><td>GTTCGGGGCCACCGATGAGG</td><td>GGG</td><td>||||||||[||||||||||||]PAM</td><td>NA</td><td>Intergenic</td><td>NA</td><td>NA</td></tr>
<tr><td>T28009</td><td>GGCCACCGATGAGGGGGTCTCGG</td><td>1000</td><td></td><td></td><td></td><td></td><td></td><td></td><td></td><td></td><td></td></tr>
<tr><td>Oligo fwd</td><td>CACCGGCCACCGATGAGGGGGTCT</td><td></td><td></td><td></td><td></td><td></td><td></td><td></td><td></td><td></td><td></td></tr>
<tr><td>Oligo rev</td><td>AAACAGACCCCCTCATCGGTGGCC</td><td></td><td></td><td></td><td></td><td></td><td></td><td></td><td></td><td></td><td></td></tr>
<tr><td>Chromosome</td><td>start</td><td>end</td><td>strand</td><td>MM</td><td>target_seq</td><td>PAM</td><td>alignment</td><td>distance</td><td>position</td><td>gene name</td><td>gene id</td></tr>
<tr><td>GU305914.1</td><td>82587</td><td>82609</td><td>-</td><td>0</td><td>GGCCACCGATGAGGGGGTCT</td><td>CGG</td><td>||||||||[||||||||||||]PAM</td><td>NA</td><td>Intergenic</td><td>NA</td><td>NA</td></tr>
<tr><td>T28010</td><td>TCGGTCTGCGTCAACGTGTTCGG</td><td>1000</td><td></td><td></td><td></td><td></td><td></td><td></td><td></td><td></td><td></td></tr>
<tr><td>Oligo fwd</td><td>CACCTCGGTCTGCGTCAACGTGTT</td><td></td><td></td><td></td><td></td><td></td><td></td><td></td><td></td><td></td><td></td></tr>
<tr><td>Oligo rev</td><td>AAACAACACGTTGACGCAGACCGA</td><td></td><td></td><td></td><td></td><td></td><td></td><td></td><td></td><td></td><td></td></tr>
<tr><td>Chromosome</td><td>start</td><td>end</td><td>strand</td><td>MM</td><td>target_seq</td><td>PAM</td><td>alignment</td><td>distance</td><td>position</td><td>gene name</td><td>gene id</td></tr>
<tr><td>GU305914.1</td><td>82568</td><td>82590</td><td>-</td><td>0</td><td>TCGGTCTGCGTCAACGTGTT</td><td>CGG</td><td>||||||||[||||||||||||]PAM</td><td>NA</td><td>Intergenic</td><td>NA</td><td>NA</td></tr>
<tr><td>T28011</td><td>CGTCAACGTGTTCGGCCAGAAGG</td><td>1000</td><td></td><td></td><td></td><td></td><td></td><td></td><td></td><td></td><td></td></tr>
<tr><td>Oligo fwd</td><td>CACCCGTCAACGTGTTCGGCCAGA</td><td></td><td></td><td></td><td></td><td></td><td></td><td></td><td></td><td></td><td></td></tr>
<tr><td>Oligo rev</td><td>AAACTCTGGCCGAACACGTTGACG</td><td></td><td></td><td></td><td></td><td></td><td></td><td></td><td></td><td></td><td></td></tr>
<tr><td>Chromosome</td><td>start</td><td>end</td><td>strand</td><td>MM</td><td>target_seq</td><td>PAM</td><td>alignment</td><td>distance</td><td>position</td><td>gene name</td><td>gene id</td></tr>
<tr><td>GU305914.1</td><td>82560</td><td>82582</td><td>-</td><td>0</td><td>CGTCAACGTGTTCGGCCAGA</td><td>AGG</td><td>||||||||[||||||||||||]PAM</td><td>NA</td><td>Intergenic</td><td>NA</td><td>NA</td></tr>
<tr><td>T28012</td><td>AGGCCTACTTCTACTGCGAGAGG</td><td>1000</td><td></td><td></td><td></td><td></td><td></td><td></td><td></td><td></td><td></td></tr>
<tr><td>Oligo fwd</td><td>CACCAGGCCTACTTCTACTGCGAG</td><td></td><td></td><td></td><td></td><td></td><td></td><td></td><td></td><td></td><td></td></tr>
<tr><td>Oligo rev</td><td>AAACCTCGCAGTAGAAGTAGGCCT</td><td></td><td></td><td></td><td></td><td></td><td></td><td></td><td></td><td></td><td></td></tr>
<tr><td>Chromosome</td><td>start</td><td>end</td><td>strand</td><td>MM</td><td>target_seq</td><td>PAM</td><td>alignment</td><td>distance</td><td>position</td><td>gene name</td><td>gene id</td></tr>
<tr><td>GU305914.1</td><td>82540</td><td>82562</td><td>-</td><td>0</td><td>AGGCCTACTTCTACTGCGAG</td><td>AGG</td><td>||||||||[||||||||||||]PAM</td><td>NA</td><td>Intergenic</td><td>NA</td><td>NA</td></tr>
<tr><td>T28013</td><td>CTGCGAGAGGATGCAGTCCGAGG</td><td>1000</td><td></td><td></td><td></td><td></td><td></td><td></td><td></td><td></td><td></td></tr>
<tr><td>Oligo fwd</td><td>CACCCTGCGAGAGGATGCAGTCCG</td><td></td><td></td><td></td><td></td><td></td><td></td><td></td><td></td><td></td><td></td></tr>
<tr><td>Oligo rev</td><td>AAACCGGACTGCATCCTCTCGCAG</td><td></td><td></td><td></td><td></td><td></td><td></td><td></td><td></td><td></td><td></td></tr>
<tr><td>Chromosome</td><td>start</td><td>end</td><td>strand</td><td>MM</td><td>target_seq</td><td>PAM</td><td>alignment</td><td>distance</td><td>position</td><td>gene name</td><td>gene id</td></tr>
<tr><td>GU305914.1</td><td>82527</td><td>82549</td><td>-</td><td>0</td><td>CTGCGAGAGGATGCAGTCCG</td><td>AGG</td><td>||||||||[||||||||||||]PAM</td><td>NA</td><td>Intergenic</td><td>NA</td><td>NA</td></tr>
<tr><td>T28014</td><td>CGTCTACGATATCGCCGATAAGG</td><td>1000</td><td></td><td></td><td></td><td></td><td></td><td></td><td></td><td></td><td></td></tr>
<tr><td>Oligo fwd</td><td>CACCCGTCTACGATATCGCCGATA</td><td></td><td></td><td></td><td></td><td></td><td></td><td></td><td></td><td></td><td></td></tr>
<tr><td>Oligo rev</td><td>AAACTATCGGCGATATCGTAGACG</td><td></td><td></td><td></td><td></td><td></td><td></td><td></td><td></td><td></td><td></td></tr>
<tr><td>Chromosome</td><td>start</td><td>end</td><td>strand</td><td>MM</td><td>target_seq</td><td>PAM</td><td>alignment</td><td>distance</td><td>position</td><td>gene name</td><td>gene id</td></tr>
<tr><td>GU305914.1</td><td>82491</td><td>82513</td><td>-</td><td>0</td><td>CGTCTACGATATCGCCGATA</td><td>AGG</td><td>||||||||[||||||||||||]PAM</td><td>NA</td><td>Intergenic</td><td>NA</td><td>NA</td></tr>
<tr><td>T28015</td><td>GTCACCAAATCCTCCTTCTACGG</td><td>1000</td><td></td><td></td><td></td><td></td><td></td><td></td><td></td><td></td><td></td></tr>
<tr><td>Oligo fwd</td><td>CACCGTCACCAAATCCTCCTTCTA</td><td></td><td></td><td></td><td></td><td></td><td></td><td></td><td></td><td></td><td></td></tr>
<tr><td>Oligo rev</td><td>AAACTAGAAGGAGGATTTGGTGAC</td><td></td><td></td><td></td><td></td><td></td><td></td><td></td><td></td><td></td><td></td></tr>
<tr><td>Chromosome</td><td>start</td><td>end</td><td>strand</td><td>MM</td><td>target_seq</td><td>PAM</td><td>alignment</td><td>distance</td><td>position</td><td>gene name</td><td>gene id</td></tr>
<tr><td>GU305914.1</td><td>82427</td><td>82449</td><td>-</td><td>0</td><td>GTCACCAAATCCTCCTTCTA</td><td>CGG</td><td>||||||||[||||||||||||]PAM</td><td>NA</td><td>Intergenic</td><td>NA</td><td>NA</td></tr>
<tr><td>T28016</td><td>AAATCCTCCTTCTACGGCTACGG</td><td>1000</td><td></td><td></td><td></td><td></td><td></td><td></td><td></td><td></td><td></td></tr>
<tr><td>Oligo fwd</td><td>CACCAAATCCTCCTTCTACGGCTA</td><td></td><td></td><td></td><td></td><td></td><td></td><td></td><td></td><td></td><td></td></tr>
<tr><td>Oligo rev</td><td>AAACTAGCCGTAGAAGGAGGATTT</td><td></td><td></td><td></td><td></td><td></td><td></td><td></td><td></td><td></td><td></td></tr>
<tr><td>Chromosome</td><td>start</td><td>end</td><td>strand</td><td>MM</td><td>target_seq</td><td>PAM</td><td>alignment</td><td>distance</td><td>position</td><td>gene name</td><td>gene id</td></tr>
<tr><td>GU305914.1</td><td>82421</td><td>82443</td><td>-</td><td>0</td><td>AAATCCTCCTTCTACGGCTA</td><td>CGG</td><td>||||||||[||||||||||||]PAM</td><td>NA</td><td>Intergenic</td><td>NA</td><td>NA</td></tr>
<tr><td>T28017</td><td>TCCTTCTACGGCTACGGCCTCGG</td><td>1000</td><td></td><td></td><td></td><td></td><td></td><td></td><td></td><td></td><td></td></tr>
<tr><td>Oligo fwd</td><td>CACCTCCTTCTACGGCTACGGCCT</td><td></td><td></td><td></td><td></td><td></td><td></td><td></td><td></td><td></td><td></td></tr>
<tr><td>Oligo rev</td><td>AAACAGGCCGTAGCCGTAGAAGGA</td><td></td><td></td><td></td><td></td><td></td><td></td><td></td><td></td><td></td><td></td></tr>
<tr><td>Chromosome</td><td>start</td><td>end</td><td>strand</td><td>MM</td><td>target_seq</td><td>PAM</td><td>alignment</td><td>distance</td><td>position</td><td>gene name</td><td>gene id</td></tr>
<tr><td>GU305914.1</td><td>82415</td><td>82437</td><td>-</td><td>0</td><td>TCCTTCTACGGCTACGGCCT</td><td>CGG</td><td>||||||||[||||||||||||]PAM</td><td>NA</td><td>Intergenic</td><td>NA</td><td>NA</td></tr>
<tr><td>T28018</td><td>CCTTCTACGGCTACGGCCTCGGG</td><td>1000</td><td></td><td></td><td></td><td></td><td></td><td></td><td></td><td></td><td></td></tr>
<tr><td>Oligo fwd</td><td>CACCCCTTCTACGGCTACGGCCTC</td><td></td><td></td><td></td><td></td><td></td><td></td><td></td><td></td><td></td><td></td></tr>
<tr><td>Oligo rev</td><td>AAACGAGGCCGTAGCCGTAGAAGG</td><td></td><td></td><td></td><td></td><td></td><td></td><td></td><td></td><td></td><td></td></tr>
<tr><td>Chromosome</td><td>start</td><td>end</td><td>strand</td><td>MM</td><td>target_seq</td><td>PAM</td><td>alignment</td><td>distance</td><td>position</td><td>gene name</td><td>gene id</td></tr>
<tr><td>GU305914.1</td><td>82414</td><td>82436</td><td>-</td><td>0</td><td>CCTTCTACGGCTACGGCCTC</td><td>GGG</td><td>||||||||[||||||||||||]PAM</td><td>NA</td><td>Intergenic</td><td>NA</td><td>NA</td></tr>
<tr><td>T28019</td><td>GGCACATCCCCAACCTGTACCGG</td><td>1000</td><td></td><td></td><td></td><td></td><td></td><td></td><td></td><td></td><td></td></tr>
<tr><td>Oligo fwd</td><td>CACCGGCACATCCCCAACCTGTAC</td><td></td><td></td><td></td><td></td><td></td><td></td><td></td><td></td><td></td><td></td></tr>
<tr><td>Oligo rev</td><td>AAACGTACAGGTTGGGGATGTGCC</td><td></td><td></td><td></td><td></td><td></td><td></td><td></td><td></td><td></td><td></td></tr>
<tr><td>Chromosome</td><td>start</td><td>end</td><td>strand</td><td>MM</td><td>target_seq</td><td>PAM</td><td>alignment</td><td>distance</td><td>position</td><td>gene name</td><td>gene id</td></tr>
<tr><td>GU305914.1</td><td>82393</td><td>82415</td><td>-</td><td>0</td><td>GGCACATCCCCAACCTGTAC</td><td>CGG</td><td>||||||||[||||||||||||]PAM</td><td>NA</td><td>Intergenic</td><td>NA</td><td>NA</td></tr>
<tr><td>T28020</td><td>ACCGGCTCTCCTTCAACAACTGG</td><td>1000</td><td></td><td></td><td></td><td></td><td></td><td></td><td></td><td></td><td></td></tr>
<tr><td>Oligo fwd</td><td>CACCACCGGCTCTCCTTCAACAAC</td><td></td><td></td><td></td><td></td><td></td><td></td><td></td><td></td><td></td><td></td></tr>
<tr><td>Oligo rev</td><td>AAACGTTGTTGAAGGAGAGCCGGT</td><td></td><td></td><td></td><td></td><td></td><td></td><td></td><td></td><td></td><td></td></tr>
<tr><td>Chromosome</td><td>start</td><td>end</td><td>strand</td><td>MM</td><td>target_seq</td><td>PAM</td><td>alignment</td><td>distance</td><td>position</td><td>gene name</td><td>gene id</td></tr>
<tr><td>GU305914.1</td><td>82375</td><td>82397</td><td>-</td><td>0</td><td>ACCGGCTCTCCTTCAACAAC</td><td>TGG</td><td>||||||||[||||||||||||]PAM</td><td>NA</td><td>Intergenic</td><td>NA</td><td>NA</td></tr>
<tr><td>T28021</td><td>TGGAACATGTGCCGCAAGATCGG</td><td>1000</td><td></td><td></td><td></td><td></td><td></td><td></td><td></td><td></td><td></td></tr>
<tr><td>Oligo fwd</td><td>CACCTGGAACATGTGCCGCAAGAT</td><td></td><td></td><td></td><td></td><td></td><td></td><td></td><td></td><td></td><td></td></tr>
<tr><td>Oligo rev</td><td>AAACATCTTGCGGCACATGTTCCA</td><td></td><td></td><td></td><td></td><td></td><td></td><td></td><td></td><td></td><td></td></tr>
</table>

Chromosome start end strand MM target_seq PAM alignment distance position gene name gene id
GU305914.1 82355 82377 - 0 TGGAACATGTGCCGCAAGAT CGG ||||||||[||||||||||||]PAM NA Intergenic NA NA

T28022 TGTGCCGCAAGATCGGCAAGCGG 1000
Oligo fwd CACCTGTGCCGCAAGATCGGCAAG
Oligo rev AAACCTTGCCGATCTTGCGGCACA
Chromosome start end strand MM target_seq PAM alignment distance position gene name gene id
GU305914.1 82348 82370 - 0 TGTGCCGCAAGATCGGCAAG CGG ||||||||[||||||||||||]PAM NA Intergenic NA NA

T28023 CAAGATCGGCAAGCGGATGCTGG 1000
Oligo fwd CACCCAAGATCGGCAAGCGGATGC
Oligo rev AAACGCATCCGCTTGCCGATCTTG
Chromosome start end strand MM target_seq PAM alignment distance position gene name gene id
GU305914.1 82341 82363 - 0 CAAGATCGGCAAGCGGATGC TGG ||||||||[||||||||||||]PAM NA Intergenic NA NA

T28024 GATCGGCAAGCGGATGCTGGAGG 1000
Oligo fwd CACCGATCGGCAAGCGGATGCTGG
Oligo rev AAACCCAGCATCCGCTTGCCGATC
Chromosome start end strand MM target_seq PAM alignment distance position gene name gene id
GU305914.1 82338 82360 - 0 GATCGGCAAGCGGATGCTGG AGG ||||||||[||||||||||||]PAM NA Intergenic NA NA

T28025 CGGCAAGCGGATGCTGGAGGAGG 1000
Oligo fwd CACCCGGCAAGCGGATGCTGGAGG
Oligo rev AAACCCTCCAGCATCCGCTTGCCG
Chromosome start end strand MM target_seq PAM alignment distance position gene name gene id
GU305914.1 82335 82357 - 0 CGGCAAGCGGATGCTGGAGG AGG ||||||||[||||||||||||]PAM NA Intergenic NA NA

T28026 GGCAAGCGGATGCTGGAGGAGGG 1000
Oligo fwd CACCGGCAAGCGGATGCTGGAGGA
Oligo rev AAACTCCTCCAGCATCCGCTTGCC
Chromosome start end strand MM target_seq PAM alignment distance position gene name gene id
GU305914.1 82334 82356 - 0 GGCAAGCGGATGCTGGAGGA GGG ||||||||[||||||||||||]PAM NA Intergenic NA NA

T28027 GCAAGCGGATGCTGGAGGAGGGG 1000
Oligo fwd CACCGCAAGCGGATGCTGGAGGAG
Oligo rev AAACCTCCTCCAGCATCCGCTTGC
Chromosome start end strand MM target_seq PAM alignment distance position gene name gene id
GU305914.1 82333 82355 - 0 GCAAGCGGATGCTGGAGGAG GGG ||||||||[||||||||||||]PAM NA Intergenic NA NA

T28028 AGCGGATGCTGGAGGAGGGGAGG 1000
Oligo fwd CACCAGCGGATGCTGGAGGAGGGG
Oligo rev AAACCCCCTCCTCCAGCATCCGCT
Chromosome start end strand MM target_seq PAM alignment distance position gene name gene id
GU305914.1 82330 82352 - 0 AGCGGATGCTGGAGGAGGGG AGG ||||||||[||||||||||||]PAM NA Intergenic NA NA

T28029 GATGCTGGAGGAGGGGAGGAAGG 1000
Oligo fwd CACCGATGCTGGAGGAGGGGAGGA
Oligo rev AAACTCCTCCCCTCCTCCAGCATC
Chromosome start end strand MM target_seq PAM alignment distance position gene name gene id
GU305914.1 82326 82348 - 0 GATGCTGGAGGAGGGGAGGA AGG ||||||||[||||||||||||]PAM NA Intergenic NA NA
GU305914.1 151438 151460 - 4 GAGGCAGGAGAAGGGTAGGA AGG ||-||-||[||-||||-||||]PAM NA Intergenic NA NA
GU305914.1 18851 18873 - 4 GGTGGGGGAGGAGGGGAAGA CGG |-||--||[|||||||||-||]PAM NA Intergenic NA NA

T28030 GGGGAGGAAGGTGTACGAGTTGG 1000
Oligo fwd CACCGGGGAGGAAGGTGTACGAGT
Oligo rev AAACACTCGTACACCTTCCTCCCC
Chromosome start end strand MM target_seq PAM alignment distance position gene name gene id
GU305914.1 82314 82336 - 0 GGGGAGGAAGGTGTACGAGT TGG ||||||||[||||||||||||]PAM NA Intergenic NA NA

T28031 GGGAGGAAGGTGTACGAGTTGGG 1000
Oligo fwd CACCGGGAGGAAGGTGTACGAGTT
Oligo rev AAACAACTCGTACACCTTCCTCCC
Chromosome start end strand MM target_seq PAM alignment distance position gene name gene id
GU305914.1 82313 82335 - 0 GGGAGGAAGGTGTACGAGTT GGG ||||||||[||||||||||||]PAM NA Intergenic NA NA

T28032 GAAGGTGTACGAGTTGGGAGTGG 1000
Oligo fwd CACCGAAGGTGTACGAGTTGGGAG
Oligo rev AAACCTCCCAACTCGTACACCTTC
Chromosome start end strand MM target_seq PAM alignment distance position gene name gene id
GU305914.1 82308 82330 - 0 GAAGGTGTACGAGTTGGGAG TGG ||||||||[||||||||||||]PAM NA Intergenic NA NA

T28033 CGAGTTGGGAGTGGACCCCTTGG 1000
Oligo fwd CACCCGAGTTGGGAGTGGACCCCT
Oligo rev AAACAGGGGTCCACTCCCAACTCG
Chromosome start end strand MM target_seq PAM alignment distance position gene name gene id
GU305914.1 82299 82321 - 0 CGAGTTGGGAGTGGACCCCT TGG ||||||||[||||||||||||]PAM NA Intergenic NA NA

T28034 TGGCGCGCTTCCTGATCGACAGG 1000
Oligo fwd CACCTGGCGCGCTTCCTGATCGAC
Oligo rev AAACGTCGATCAGGAAGCGCGCCA
Chromosome start end strand MM target_seq PAM alignment distance position gene name gene id
GU305914.1 82279 82301 - 0 TGGCGCGCTTCCTGATCGAC AGG ||||||||[||||||||||||]PAM NA Intergenic NA NA

T28035 GACAGGAAGATCCCCAGCTTCGG 1000
Oligo fwd CACCGACAGGAAGATCCCCAGCTT
Oligo rev AAACAAGCTGGGGATCTTCCTGTC
Chromosome start end strand MM target_seq PAM alignment distance position gene name gene id
GU305914.1 82262 82284 - 0 GACAGGAAGATCCCCAGCTT CGG ||||||||[||||||||||||]PAM NA Intergenic NA NA

| | | | | | | | | | | | |
|---|---|---|---|---|---|---|---|---|---|---|---|
| T28036 | GGAAGATCCCCAGCTTCGGCTGG | 1000 | | | | | | | | | |
| Oligo fwd | CACCGGAAGATCCCCAGCTTCGGC | | | | | | | | | | |
| Oligo rev | AAACGCCGAAGCTGGGGATCTTCC | | | | | | | | | | |
| Chromosome | start | end | strand | MM | target_seq | PAM | alignment | distance | position | gene name | gene id |
| GU305914.1 | 82258 | 82280 | - | 0 | GGAAGATCCCCAGCTTCGGC | TGG | \|\|\|\|\|\|\|\|[\|\|\|\|\|\|\|\|\|\|\|\|]PAM | NA | Intergenic | NA | NA |
| T28037 | GCTTCGGCTGGTGCCTCGCCAGG | 1000 | | | | | | | | | |
| Oligo fwd | CACCGCTTCGGCTGGTGCCTCGCC | | | | | | | | | | |
| Oligo rev | AAACGGCGAGGCACCAGCCGAAGC | | | | | | | | | | |
| Chromosome | start | end | strand | MM | target_seq | PAM | alignment | distance | position | gene name | gene id |
| GU305914.1 | 82246 | 82268 | - | 0 | GCTTCGGCTGGTGCCTCGCC | AGG | \|\|\|\|\|\|\|\|[\|\|\|\|\|\|\|\|\|\|\|\|]PAM | NA | Intergenic | NA | NA |
| T28038 | TCGGCTGGTGCCTCGCCAGGAGG | 1000 | | | | | | | | | |
| Oligo fwd | CACCTCGGCTGGTGCCTCGCCAGG | | | | | | | | | | |
| Oligo rev | AAACCCTGGCGAGGCACCAGCCGA | | | | | | | | | | |
| Chromosome | start | end | strand | MM | target_seq | PAM | alignment | distance | position | gene name | gene id |
| GU305914.1 | 82243 | 82265 | - | 0 | TCGGCTGGTGCCTCGCCAGG | AGG | \|\|\|\|\|\|\|\|[\|\|\|\|\|\|\|\|\|\|\|\|]PAM | NA | Intergenic | NA | NA |
| T28039 | AGGTACAGCGTCCGCGCCGCCGG | 1000 | | | | | | | | | |
| Oligo fwd | CACCAGGTACAGCGTCCGCGCCGC | | | | | | | | | | |
| Oligo rev | AAACGCGGCGCGGACGCTGTACCT | | | | | | | | | | |
| Chromosome | start | end | strand | MM | target_seq | PAM | alignment | distance | position | gene name | gene id |
| GU305914.1 | 82223 | 82245 | - | 0 | AGGTACAGCGTCCGCGCCGC | CGG | \|\|\|\|\|\|\|\|[\|\|\|\|\|\|\|\|\|\|\|\|]PAM | NA | Intergenic | NA | NA |
| T28040 | GCGCCGCCGGATACGTCAGCCGG | 1000 | | | | | | | | | |
| Oligo fwd | CACCGCGCCGCCGGATACGTCAGC | | | | | | | | | | |
| Oligo rev | AAACGCTGACGTATCCGGCGGCGC | | | | | | | | | | |
| Chromosome | start | end | strand | MM | target_seq | PAM | alignment | distance | position | gene name | gene id |
| GU305914.1 | 82210 | 82232 | - | 0 | GCGCCGCCGGATACGTCAGC | CGG | \|\|\|\|\|\|\|\|[\|\|\|\|\|\|\|\|\|\|\|\|]PAM | NA | Intergenic | NA | NA |
| T28041 | CGCCGCCGGATACGTCAGCCGGG | 1000 | | | | | | | | | |
| Oligo fwd | CACCCGCCGCCGGATACGTCAGCC | | | | | | | | | | |
| Oligo rev | AAACGGCTGACGTATCCGGCGGCG | | | | | | | | | | |
| Chromosome | start | end | strand | MM | target_seq | PAM | alignment | distance | position | gene name | gene id |
| GU305914.1 | 82209 | 82231 | - | 0 | CGCCGCCGGATACGTCAGCC | GGG | \|\|\|\|\|\|\|\|[\|\|\|\|\|\|\|\|\|\|\|\|]PAM | NA | Intergenic | NA | NA |
| T28042 | ATACGTCAGCCGGGCGCAGCTGG | 1000 | | | | | | | | | |
| Oligo fwd | CACCATACGTCAGCCGGGCGCAGC | | | | | | | | | | |
| Oligo rev | AAACGCTGCGCCCGGCTGACGTAT | | | | | | | | | | |
| Chromosome | start | end | strand | MM | target_seq | PAM | alignment | distance | position | gene name | gene id |
| GU305914.1 | 82200 | 82222 | - | 0 | ATACGTCAGCCGGGCGCAGC | TGG | \|\|\|\|\|\|\|\|[\|\|\|\|\|\|\|\|\|\|\|\|]PAM | NA | Intergenic | NA | NA |
| T28043 | CGATCGAAGAGCAGAGCAACTGG | 1000 | | | | | | | | | |
| Oligo fwd | CACCCGATCGAAGAGCAGAGCAAC | | | | | | | | | | |
| Oligo rev | AAACGTTGCTCTGCTCTTCGATCG | | | | | | | | | | |
| Chromosome | start | end | strand | MM | target_seq | PAM | alignment | distance | position | gene name | gene id |
| GU305914.1 | 82147 | 82169 | - | 0 | CGATCGAAGAGCAGAGCAAC | TGG | \|\|\|\|\|\|\|\|[\|\|\|\|\|\|\|\|\|\|\|\|]PAM | NA | Intergenic | NA | NA |
| T28044 | TTCGATATCGAGTGTATGAGCGG | 1000 | | | | | | | | | |
| Oligo fwd | CACCTTCGATATCGAGTGTATGAG | | | | | | | | | | |
| Oligo rev | AAACCTCATACACTCGATATCGAA | | | | | | | | | | |
| Chromosome | start | end | strand | MM | target_seq | PAM | alignment | distance | position | gene name | gene id |
| GU305914.1 | 82103 | 82125 | - | 0 | TTCGATATCGAGTGTATGAG | CGG | \|\|\|\|\|\|\|\|[\|\|\|\|\|\|\|\|\|\|\|\|]PAM | NA | Intergenic | NA | NA |
| T28045 | TATCGAGTGTATGAGCGGCACGG | 1000 | | | | | | | | | |
| Oligo fwd | CACCTATCGAGTGTATGAGCGGCA | | | | | | | | | | |
| Oligo rev | AAACTGCCGCTCATACACTCGATA | | | | | | | | | | |
| Chromosome | start | end | strand | MM | target_seq | PAM | alignment | distance | position | gene name | gene id |
| GU305914.1 | 82098 | 82120 | - | 0 | TATCGAGTGTATGAGCGGCA | CGG | \|\|\|\|\|\|\|\|[\|\|\|\|\|\|\|\|\|\|\|\|]PAM | NA | Intergenic | NA | NA |
| T28046 | ATCGAGTGTATGAGCGGCACGGG | 1000 | | | | | | | | | |
| Oligo fwd | CACCATCGAGTGTATGAGCGGCAC | | | | | | | | | | |
| Oligo rev | AAACGTGCCGCTCATACACTCGAT | | | | | | | | | | |
| Chromosome | start | end | strand | MM | target_seq | PAM | alignment | distance | position | gene name | gene id |
| GU305914.1 | 82097 | 82119 | - | 0 | ATCGAGTGTATGAGCGGCAC | GGG | \|\|\|\|\|\|\|\|[\|\|\|\|\|\|\|\|\|\|\|\|]PAM | NA | Intergenic | NA | NA |
| GU305914.1 | 175005 | 175027 | - | 4 | AGCGAGTTTATGAGAGCCAC | GGG | \|-\|\|\|\|\|-[\|\|\|\|\|\|-\|-\|\|\|]PAM | NA | Intergenic | NA | NA |
| T28047 | CAGATCTCCTGCGTCTGTTTCGG | 1000 | | | | | | | | | |
| Oligo fwd | CACCCAGATCTCCTGCGTCTGTTT | | | | | | | | | | |
| Oligo rev | AAACAAACAGACGCAGGAGATCTG | | | | | | | | | | |
| Chromosome | start | end | strand | MM | target_seq | PAM | alignment | distance | position | gene name | gene id |
| GU305914.1 | 82034 | 82056 | - | 0 | CAGATCTCCTGCGTCTGTTT | CGG | \|\|\|\|\|\|\|\|[\|\|\|\|\|\|\|\|\|\|\|\|]PAM | NA | Intergenic | NA | NA |
| GU305914.1 | 16851 | 16873 | - | 4 | CAGAACTGCGGCGTCAGTTT | CGG | \|\|\|\|-\|\|-[\|-\|\|\|\|\|-\|\|\|\|]PAM | NA | Intergenic | NA | NA |
| T28048 | TCCTGCGTCTGTTTCGGCGTCGG | 1000 | | | | | | | | | |
| Oligo fwd | CACCTCCTGCGTCTGTTTCGGCGT | | | | | | | | | | |
| Oligo rev | AAACACGCCGAAACAGACGCAGGA | | | | | | | | | | |
| Chromosome | start | end | strand | MM | target_seq | PAM | alignment | distance | position | gene name | gene id |
| GU305914.1 | 82028 | 82050 | - | 0 | TCCTGCGTCTGTTTCGGCGT | CGG | \|\|\|\|\|\|\|\|[\|\|\|\|\|\|\|\|\|\|\|\|]PAM | NA | Intergenic | NA | NA |
| T28049 | CTGTTTCGGCGTCGGAGAGATGG | 1000 | | | | | | | | | |
| Oligo fwd | CACCCTGTTTCGGCGTCGGAGAGA | | | | | | | | | | |
| Oligo rev | AAACTCTCTCCGACGCCGAAACAG | | | | | | | | | | |
| Chromosome | start | end | strand | MM | target_seq | PAM | alignment | distance | position | gene name | gene id |
| GU305914.1 | 82020 | 82042 | - | 0 | CTGTTTCGGCGTCGGAGAGA | TGG | \|\|\|\|\|\|\|\|[\|\|\|\|\|\|\|\|\|\|\|\|]PAM | NA | Intergenic | NA | NA |
| T28050 | GAGAACCACCTCTTCACCATCGG | 1000 | | | | | | | | | |
| Oligo fwd | CACCGAGAACCACCTCTTCACCAT | | | | | | | | | | |
| Oligo rev | AAACATGGTGAAGAGGTGGTTCTC | | | | | | | | | | |

<table>
<tr><td>Chromosome</td><td>start</td><td>end</td><td>strand</td><td>MM</td><td>target_seq</td><td>PAM</td><td>alignment</td><td>distance</td><td>position</td><td>gene name</td><td>gene id</td></tr>
<tr><td>GU305914.1</td><td>81947</td><td>81969</td><td>-</td><td>0</td><td>GAGAACCACCTCTTCACCAT</td><td>CGG</td><td>||||||||[||||||||||||]PAM</td><td>NA</td><td>Intergenic</td><td>NA</td><td>NA</td></tr>
<tr><td></td><td></td><td></td><td></td><td></td><td></td><td></td><td></td><td></td><td></td><td></td><td></td></tr>
<tr><td>T28051</td><td>AGAACCACCTCTTCACCATCGGG</td><td>1000</td><td></td><td></td><td></td><td></td><td></td><td></td><td></td><td></td><td></td></tr>
<tr><td>Oligo fwd</td><td>CACCAGAACCACCTCTTCACCATC</td><td></td><td></td><td></td><td></td><td></td><td></td><td></td><td></td><td></td><td></td></tr>
<tr><td>Oligo rev</td><td>AAACGATGGTGAAGAGGTGGTTCT</td><td></td><td></td><td></td><td></td><td></td><td></td><td></td><td></td><td></td><td></td></tr>
<tr><td>Chromosome</td><td>start</td><td>end</td><td>strand</td><td>MM</td><td>target_seq</td><td>PAM</td><td>alignment</td><td>distance</td><td>position</td><td>gene name</td><td>gene id</td></tr>
<tr><td>GU305914.1</td><td>81946</td><td>81968</td><td>-</td><td>0</td><td>AGAACCACCTCTTCACCATC</td><td>GGG</td><td>||||||||[||||||||||||]PAM</td><td>NA</td><td>Intergenic</td><td>NA</td><td>NA</td></tr>
<tr><td></td><td></td><td></td><td></td><td></td><td></td><td></td><td></td><td></td><td></td><td></td><td></td></tr>
<tr><td>T28052</td><td>CCTCCGAGTACGAGATGTTGCGG</td><td>1000</td><td></td><td></td><td></td><td></td><td></td><td></td><td></td><td></td><td></td></tr>
<tr><td>Oligo fwd</td><td>CACCCCTCCGAGTACGAGATGTTG</td><td></td><td></td><td></td><td></td><td></td><td></td><td></td><td></td><td></td><td></td></tr>
<tr><td>Oligo rev</td><td>AAACCAACATCTCGTACTCGGAGG</td><td></td><td></td><td></td><td></td><td></td><td></td><td></td><td></td><td></td><td></td></tr>
<tr><td>Chromosome</td><td>start</td><td>end</td><td>strand</td><td>MM</td><td>target_seq</td><td>PAM</td><td>alignment</td><td>distance</td><td>position</td><td>gene name</td><td>gene id</td></tr>
<tr><td>GU305914.1</td><td>81883</td><td>81905</td><td>-</td><td>0</td><td>CCTCCGAGTACGAGATGTTG</td><td>CGG</td><td>||||||||[||||||||||||]PAM</td><td>NA</td><td>Intergenic</td><td>NA</td><td>NA</td></tr>
<tr><td></td><td></td><td></td><td></td><td></td><td></td><td></td><td></td><td></td><td></td><td></td><td></td></tr>
<tr><td>T28053</td><td>CTCCGAGTACGAGATGTTGCGGG</td><td>1000</td><td></td><td></td><td></td><td></td><td></td><td></td><td></td><td></td><td></td></tr>
<tr><td>Oligo fwd</td><td>CACCCTCCGAGTACGAGATGTTGC</td><td></td><td></td><td></td><td></td><td></td><td></td><td></td><td></td><td></td><td></td></tr>
<tr><td>Oligo rev</td><td>AAACGCAACATCTCGTACTCGGAG</td><td></td><td></td><td></td><td></td><td></td><td></td><td></td><td></td><td></td><td></td></tr>
<tr><td>Chromosome</td><td>start</td><td>end</td><td>strand</td><td>MM</td><td>target_seq</td><td>PAM</td><td>alignment</td><td>distance</td><td>position</td><td>gene name</td><td>gene id</td></tr>
<tr><td>GU305914.1</td><td>81882</td><td>81904</td><td>-</td><td>0</td><td>CTCCGAGTACGAGATGTTGC</td><td>GGG</td><td>||||||||[||||||||||||]PAM</td><td>NA</td><td>Intergenic</td><td>NA</td><td>NA</td></tr>
<tr><td></td><td></td><td></td><td></td><td></td><td></td><td></td><td></td><td></td><td></td><td></td><td></td></tr>
<tr><td>T28054</td><td>TCCGAGTACGAGATGTTGCGGGG</td><td>1000</td><td></td><td></td><td></td><td></td><td></td><td></td><td></td><td></td><td></td></tr>
<tr><td>Oligo fwd</td><td>CACCTCCGAGTACGAGATGTTGCG</td><td></td><td></td><td></td><td></td><td></td><td></td><td></td><td></td><td></td><td></td></tr>
<tr><td>Oligo rev</td><td>AAACCGCAACATCTCGTACTCGGA</td><td></td><td></td><td></td><td></td><td></td><td></td><td></td><td></td><td></td><td></td></tr>
<tr><td>Chromosome</td><td>start</td><td>end</td><td>strand</td><td>MM</td><td>target_seq</td><td>PAM</td><td>alignment</td><td>distance</td><td>position</td><td>gene name</td><td>gene id</td></tr>
<tr><td>GU305914.1</td><td>81881</td><td>81903</td><td>-</td><td>0</td><td>TCCGAGTACGAGATGTTGCG</td><td>GGG</td><td>||||||||[||||||||||||]PAM</td><td>NA</td><td>Intergenic</td><td>NA</td><td>NA</td></tr>
<tr><td></td><td></td><td></td><td></td><td></td><td></td><td></td><td></td><td></td><td></td><td></td><td></td></tr>
<tr><td>T28055</td><td>GTTTCTTCATCTTCCTGAGCTGG</td><td>1000</td><td></td><td></td><td></td><td></td><td></td><td></td><td></td><td></td><td></td></tr>
<tr><td>Oligo fwd</td><td>CACCGTTTCTTCATCTTCCTGAGC</td><td></td><td></td><td></td><td></td><td></td><td></td><td></td><td></td><td></td><td></td></tr>
<tr><td>Oligo rev</td><td>AAACGCTCAGGAAGATGAAGAAAC</td><td></td><td></td><td></td><td></td><td></td><td></td><td></td><td></td><td></td><td></td></tr>
<tr><td>Chromosome</td><td>start</td><td>end</td><td>strand</td><td>MM</td><td>target_seq</td><td>PAM</td><td>alignment</td><td>distance</td><td>position</td><td>gene name</td><td>gene id</td></tr>
<tr><td>GU305914.1</td><td>81859</td><td>81881</td><td>-</td><td>0</td><td>GTTTCTTCATCTTCCTGAGC</td><td>TGG</td><td>||||||||[||||||||||||]PAM</td><td>NA</td><td>Intergenic</td><td>NA</td><td>NA</td></tr>
<tr><td></td><td></td><td></td><td></td><td></td><td></td><td></td><td></td><td></td><td></td><td></td><td></td></tr>
<tr><td>T28056</td><td>TACTCGCCCGAGTTCATCACCGG</td><td>1000</td><td></td><td></td><td></td><td></td><td></td><td></td><td></td><td></td><td></td></tr>
<tr><td>Oligo fwd</td><td>CACCTACTCGCCCGAGTTCATCAC</td><td></td><td></td><td></td><td></td><td></td><td></td><td></td><td></td><td></td><td></td></tr>
<tr><td>Oligo rev</td><td>AAACGTGATGAACTCGGGCGAGTA</td><td></td><td></td><td></td><td></td><td></td><td></td><td></td><td></td><td></td><td></td></tr>
<tr><td>Chromosome</td><td>start</td><td>end</td><td>strand</td><td>MM</td><td>target_seq</td><td>PAM</td><td>alignment</td><td>distance</td><td>position</td><td>gene name</td><td>gene id</td></tr>
<tr><td>GU305914.1</td><td>81836</td><td>81858</td><td>-</td><td>0</td><td>TACTCGCCCGAGTTCATCAC</td><td>CGG</td><td>||||||||[||||||||||||]PAM</td><td>NA</td><td>Intergenic</td><td>NA</td><td>NA</td></tr>
<tr><td></td><td></td><td></td><td></td><td></td><td></td><td></td><td></td><td></td><td></td><td></td><td></td></tr>
<tr><td>T28057</td><td>ATCACCGGCTACAACATCAACGG</td><td>1000</td><td></td><td></td><td></td><td></td><td></td><td></td><td></td><td></td><td></td></tr>
<tr><td>Oligo fwd</td><td>CACCATCACCGGCTACAACATCAA</td><td></td><td></td><td></td><td></td><td></td><td></td><td></td><td></td><td></td><td></td></tr>
<tr><td>Oligo rev</td><td>AAACTTGATGTTGTAGCCGGTGAT</td><td></td><td></td><td></td><td></td><td></td><td></td><td></td><td></td><td></td><td></td></tr>
<tr><td>Chromosome</td><td>start</td><td>end</td><td>strand</td><td>MM</td><td>target_seq</td><td>PAM</td><td>alignment</td><td>distance</td><td>position</td><td>gene name</td><td>gene id</td></tr>
<tr><td>GU305914.1</td><td>81821</td><td>81843</td><td>-</td><td>0</td><td>ATCACCGGCTACAACATCAA</td><td>CGG</td><td>||||||||[||||||||||||]PAM</td><td>NA</td><td>Intergenic</td><td>NA</td><td>NA</td></tr>
<tr><td></td><td></td><td></td><td></td><td></td><td></td><td></td><td></td><td></td><td></td><td></td><td></td></tr>
<tr><td>T28058</td><td>CGCCGAGAAACTCTACAAGATGG</td><td>1000</td><td></td><td></td><td></td><td></td><td></td><td></td><td></td><td></td><td></td></tr>
<tr><td>Oligo fwd</td><td>CACCCGCCGAGAAACTCTACAAGA</td><td></td><td></td><td></td><td></td><td></td><td></td><td></td><td></td><td></td><td></td></tr>
<tr><td>Oligo rev</td><td>AAACTCTTGTAGAGTTTCTCGGCG</td><td></td><td></td><td></td><td></td><td></td><td></td><td></td><td></td><td></td><td></td></tr>
<tr><td>Chromosome</td><td>start</td><td>end</td><td>strand</td><td>MM</td><td>target_seq</td><td>PAM</td><td>alignment</td><td>distance</td><td>position</td><td>gene name</td><td>gene id</td></tr>
<tr><td>GU305914.1</td><td>81771</td><td>81793</td><td>-</td><td>0</td><td>CGCCGAGAAACTCTACAAGA</td><td>TGG</td><td>||||||||[||||||||||||]PAM</td><td>NA</td><td>Intergenic</td><td>NA</td><td>NA</td></tr>
<tr><td></td><td></td><td></td><td></td><td></td><td></td><td></td><td></td><td></td><td></td><td></td><td></td></tr>
<tr><td>T28059</td><td>GAAACTCTACAAGATGGATGTGG</td><td>1000</td><td></td><td></td><td></td><td></td><td></td><td></td><td></td><td></td><td></td></tr>
<tr><td>Oligo fwd</td><td>CACCGAAACTCTACAAGATGGATG</td><td></td><td></td><td></td><td></td><td></td><td></td><td></td><td></td><td></td><td></td></tr>
<tr><td>Oligo rev</td><td>AAACCATCCATCTTGTAGAGTTTC</td><td></td><td></td><td></td><td></td><td></td><td></td><td></td><td></td><td></td><td></td></tr>
<tr><td>Chromosome</td><td>start</td><td>end</td><td>strand</td><td>MM</td><td>target_seq</td><td>PAM</td><td>alignment</td><td>distance</td><td>position</td><td>gene name</td><td>gene id</td></tr>
<tr><td>GU305914.1</td><td>81765</td><td>81787</td><td>-</td><td>0</td><td>GAAACTCTACAAGATGGATG</td><td>TGG</td><td>||||||||[||||||||||||]PAM</td><td>NA</td><td>Intergenic</td><td>NA</td><td>NA</td></tr>
<tr><td></td><td></td><td></td><td></td><td></td><td></td><td></td><td></td><td></td><td></td><td></td><td></td></tr>
<tr><td>T28060</td><td>AAACTCTACAAGATGGATGTGGG</td><td>1000</td><td></td><td></td><td></td><td></td><td></td><td></td><td></td><td></td><td></td></tr>
<tr><td>Oligo fwd</td><td>CACCAAACTCTACAAGATGGATGT</td><td></td><td></td><td></td><td></td><td></td><td></td><td></td><td></td><td></td><td></td></tr>
<tr><td>Oligo rev</td><td>AAACACATCCATCTTGTAGAGTTT</td><td></td><td></td><td></td><td></td><td></td><td></td><td></td><td></td><td></td><td></td></tr>
<tr><td>Chromosome</td><td>start</td><td>end</td><td>strand</td><td>MM</td><td>target_seq</td><td>PAM</td><td>alignment</td><td>distance</td><td>position</td><td>gene name</td><td>gene id</td></tr>
<tr><td>GU305914.1</td><td>81764</td><td>81786</td><td>-</td><td>0</td><td>AAACTCTACAAGATGGATGT</td><td>GGG</td><td>||||||||[||||||||||||]PAM</td><td>NA</td><td>Intergenic</td><td>NA</td><td>NA</td></tr>
<tr><td></td><td></td><td></td><td></td><td></td><td></td><td></td><td></td><td></td><td></td><td></td><td></td></tr>
<tr><td>T28061</td><td>TGGGCCAATTCACCAAGCTGCGG</td><td>1000</td><td></td><td></td><td></td><td></td><td></td><td></td><td></td><td></td><td></td></tr>
<tr><td>Oligo fwd</td><td>CACCTGGGCCAATTCACCAAGCTG</td><td></td><td></td><td></td><td></td><td></td><td></td><td></td><td></td><td></td><td></td></tr>
<tr><td>Oligo rev</td><td>AAACCAGCTTGGTGAATTGGCCCA</td><td></td><td></td><td></td><td></td><td></td><td></td><td></td><td></td><td></td><td></td></tr>
<tr><td>Chromosome</td><td>start</td><td>end</td><td>strand</td><td>MM</td><td>target_seq</td><td>PAM</td><td>alignment</td><td>distance</td><td>position</td><td>gene name</td><td>gene id</td></tr>
<tr><td>GU305914.1</td><td>81745</td><td>81767</td><td>-</td><td>0</td><td>TGGGCCAATTCACCAAGCTG</td><td>CGG</td><td>||||||||[||||||||||||]PAM</td><td>NA</td><td>Intergenic</td><td>NA</td><td>NA</td></tr>
<tr><td></td><td></td><td></td><td></td><td></td><td></td><td></td><td></td><td></td><td></td><td></td><td></td></tr>
<tr><td>T28062</td><td>GCCAATTCACCAAGCTGCGGAGG</td><td>1000</td><td></td><td></td><td></td><td></td><td></td><td></td><td></td><td></td><td></td></tr>
<tr><td>Oligo fwd</td><td>CACCGCCAATTCACCAAGCTGCGG</td><td></td><td></td><td></td><td></td><td></td><td></td><td></td><td></td><td></td><td></td></tr>
<tr><td>Oligo rev</td><td>AAACCCGCAGCTTGGTGAATTGGC</td><td></td><td></td><td></td><td></td><td></td><td></td><td></td><td></td><td></td><td></td></tr>
<tr><td>Chromosome</td><td>start</td><td>end</td><td>strand</td><td>MM</td><td>target_seq</td><td>PAM</td><td>alignment</td><td>distance</td><td>position</td><td>gene name</td><td>gene id</td></tr>
<tr><td>GU305914.1</td><td>81742</td><td>81764</td><td>-</td><td>0</td><td>GCCAATTCACCAAGCTGCGG</td><td>AGG</td><td>||||||||[||||||||||||]PAM</td><td>NA</td><td>Intergenic</td><td>NA</td><td>NA</td></tr>
<tr><td></td><td></td><td></td><td></td><td></td><td></td><td></td><td></td><td></td><td></td><td></td><td></td></tr>
<tr><td>T28063</td><td>CCAATTCACCAAGCTGCGGAGGG</td><td>1000</td><td></td><td></td><td></td><td></td><td></td><td></td><td></td><td></td><td></td></tr>
<tr><td>Oligo fwd</td><td>CACCCCAATTCACCAAGCTGCGGA</td><td></td><td></td><td></td><td></td><td></td><td></td><td></td><td></td><td></td><td></td></tr>
<tr><td>Oligo rev</td><td>AAACTCCGCAGCTTGGTGAATTGG</td><td></td><td></td><td></td><td></td><td></td><td></td><td></td><td></td><td></td><td></td></tr>
<tr><td>Chromosome</td><td>start</td><td>end</td><td>strand</td><td>MM</td><td>target_seq</td><td>PAM</td><td>alignment</td><td>distance</td><td>position</td><td>gene name</td><td>gene id</td></tr>
<tr><td>GU305914.1</td><td>81741</td><td>81763</td><td>-</td><td>0</td><td>CCAATTCACCAAGCTGCGGA</td><td>GGG</td><td>||||||||[||||||||||||]PAM</td><td>NA</td><td>Intergenic</td><td>NA</td><td>NA</td></tr>
<tr><td></td><td></td><td></td><td></td><td></td><td></td><td></td><td></td><td></td><td></td><td></td><td></td></tr>
<tr><td>T28064</td><td>CAATTCACCAAGCTGCGGAGGGG</td><td>1000</td><td></td><td></td><td></td><td></td><td></td><td></td><td></td><td></td><td></td></tr>
<tr><td>Oligo fwd</td><td>CACCCAATTCACCAAGCTGCGGAG</td><td></td><td></td><td></td><td></td><td></td><td></td><td></td><td></td><td></td><td></td></tr>
<tr><td>Oligo rev</td><td>AAACCTCCGCAGCTTGGTGAATTG</td><td></td><td></td><td></td><td></td><td></td><td></td><td></td><td></td><td></td><td></td></tr>
<tr><td>Chromosome</td><td>start</td><td>end</td><td>strand</td><td>MM</td><td>target_seq</td><td>PAM</td><td>alignment</td><td>distance</td><td>position</td><td>gene name</td><td>gene id</td></tr>
<tr><td>GU305914.1</td><td>81740</td><td>81762</td><td>-</td><td>0</td><td>CAATTCACCAAGCTGCGGAG</td><td>GGG</td><td>||||||||[||||||||||||]PAM</td><td>NA</td><td>Intergenic</td><td>NA</td><td>NA</td></tr>
<tr><td></td><td></td><td></td><td></td><td></td><td></td><td></td><td></td><td></td><td></td><td></td><td></td></tr>
<tr><td>T28065</td><td>AATTCACCAAGCTGCGGAGGGGG</td><td>1000</td><td></td><td></td><td></td><td></td><td></td><td></td><td></td><td></td><td></td></tr>
<tr><td>Oligo fwd</td><td>CACCAATTCACCAAGCTGCGGAGG</td><td></td><td></td><td></td><td></td><td></td><td></td><td></td><td></td><td></td><td></td></tr>
</table>

Oligo rev AAACCCTCCGCAGCTTGGTGAATT
Chromosome start end strand MM target_seq PAM alignment distance position gene name gene id
GU305914.1 81739 81761 - 0 AATTCACCAAGCTGCGGAGG GGG ||||||||[||||||||||||]PAM NA Intergenic NA NA

T28066 ATTCACCAAGCTGCGGAGGGGGG 1000
Oligo fwd CACCATTCACCAAGCTGCGGAGGG
Oligo rev AAACCCCTCCGCAGCTTGGTGAAT
Chromosome start end strand MM target_seq PAM alignment distance position gene name gene id
GU305914.1 81738 81760 - 0 ATTCACCAAGCTGCGGAGGG GGG ||||||||[||||||||||||]PAM NA Intergenic NA NA

T28067 TTCACCAAGCTGCGGAGGGGGGG 1000
Oligo fwd CACCTTCACCAAGCTGCGGAGGGG
Oligo rev AAACCCCCTCCGCAGCTTGGTGAA
Chromosome start end strand MM target_seq PAM alignment distance position gene name gene id
GU305914.1 81737 81759 - 0 TTCACCAAGCTGCGGAGGGG GGG ||||||||[||||||||||||]PAM NA Intergenic NA NA

T28068 GTTCGTCTTCAGCCCCGAGAAGG 1000
Oligo fwd CACCGTTCGTCTTCAGCCCCGAGA
Oligo rev AAACTCTCGGGGCTGAAGACGAAC
Chromosome start end strand MM target_seq PAM alignment distance position gene name gene id
GU305914.1 81708 81730 - 0 GTTCGTCTTCAGCCCCGAGA AGG ||||||||[||||||||||||]PAM NA Intergenic NA NA

T28069 TTCGTCTTCAGCCCCGAGAAGGG 1000
Oligo fwd CACCTTCGTCTTCAGCCCCGAGAA
Oligo rev AAACTTCTCGGGGCTGAAGACGAA
Chromosome start end strand MM target_seq PAM alignment distance position gene name gene id
GU305914.1 81707 81729 - 0 TTCGTCTTCAGCCCCGAGAA GGG ||||||||[||||||||||||]PAM NA Intergenic NA NA

T28070 CTTCAGCCCCGAGAAGGGCAAGG 1000
Oligo fwd CACCCTTCAGCCCCGAGAAGGGCA
Oligo rev AAACTGCCCTTCTCGGGGCTGAAG
Chromosome start end strand MM target_seq PAM alignment distance position gene name gene id
GU305914.1 81702 81724 - 0 CTTCAGCCCCGAGAAGGGCA AGG ||||||||[||||||||||||]PAM NA Intergenic NA NA

T28071 AGCCCCGAGAAGGGCAAGGCCGG 1000
Oligo fwd CACCAGCCCCGAGAAGGGCAAGGC
Oligo rev AAACGCCTTGCCCTTCTCGGGGCT
Chromosome start end strand MM target_seq PAM alignment distance position gene name gene id
GU305914.1 81698 81720 - 0 AGCCCCGAGAAGGGCAAGGC CGG ||||||||[||||||||||||]PAM NA Intergenic NA NA

T28072 GAGAAGGGCAAGGCCGGCTTCGG 1000
Oligo fwd CACCGAGAAGGGCAAGGCCGGCTT
Oligo rev AAACAAGCCGGCCTTGCCCTTCTC
Chromosome start end strand MM target_seq PAM alignment distance position gene name gene id
GU305914.1 81692 81714 - 0 GAGAAGGGCAAGGCCGGCTT CGG ||||||||[||||||||||||]PAM NA Intergenic NA NA

T28073 CGGCTTCGGCACCTCCAACACGG 1000
Oligo fwd CACCCGGCTTCGGCACCTCCAACA
Oligo rev AAACTGTTGGAGGTGCCGAAGCCG
Chromosome start end strand MM target_seq PAM alignment distance position gene name gene id
GU305914.1 81678 81700 - 0 CGGCTTCGGCACCTCCAACA CGG ||||||||[||||||||||||]PAM NA Intergenic NA NA

T28074 CGGCACCTCCAACACGGTCAAGG 1000
Oligo fwd CACCCGGCACCTCCAACACGGTCA
Oligo rev AAACTGACCGTGTTGGAGGTGCCG
Chromosome start end strand MM target_seq PAM alignment distance position gene name gene id
GU305914.1 81672 81694 - 0 CGGCACCTCCAACACGGTCA AGG ||||||||[||||||||||||]PAM NA Intergenic NA NA

T28075 CCAACACGGTCAAGGTCTTCTGG 1000
Oligo fwd CACCCCAACACGGTCAAGGTCTTC
Oligo rev AAACGAAGACCTTGACCGTGTTGG
Chromosome start end strand MM target_seq PAM alignment distance position gene name gene id
GU305914.1 81664 81686 - 0 CCAACACGGTCAAGGTCTTC TGG ||||||||[||||||||||||]PAM NA Intergenic NA NA

T28076 ACGGTCAAGGTCTTCTGGAGCGG 1000
Oligo fwd CACCACGGTCAAGGTCTTCTGGAG
Oligo rev AAACCTCCAGAAGACCTTGACCGT
Chromosome start end strand MM target_seq PAM alignment distance position gene name gene id
GU305914.1 81659 81681 - 0 ACGGTCAAGGTCTTCTGGAG CGG ||||||||[||||||||||||]PAM NA Intergenic NA NA

T28077 GTACCCCGTCTGTACCGCCAAGG 1000
Oligo fwd CACCGTACCCCGTCTGTACCGCCA
Oligo rev AAACTGGCGGTACAGACGGGGTAC
Chromosome start end strand MM target_seq PAM alignment distance position gene name gene id
GU305914.1 81618 81640 - 0 GTACCCCGTCTGTACCGCCA AGG ||||||||[||||||||||||]PAM NA Intergenic NA NA

T28078 CAACTACAAGCTCGACACCATGG 1000
Oligo fwd CACCCAACTACAAGCTCGACACCA
Oligo rev AAACTGGTGTCGAGCTTGTAGTTG
Chromosome start end strand MM target_seq PAM alignment distance position gene name gene id
GU305914.1 81585 81607 - 0 CAACTACAAGCTCGACACCA TGG ||||||||[||||||||||||]PAM NA Intergenic NA NA

T28079 ATCCCCGTCCAGTTCTCCGCCGG 1000
Oligo fwd CACCATCCCCGTCCAGTTCTCCGC
Oligo rev AAACGCGGAGAACTGGACGGGGAT
Chromosome start end strand MM target_seq PAM alignment distance position gene name gene id
GU305914.1 81515 81537 - 0 ATCCCCGTCCAGTTCTCCGC CGG ||||||||[||||||||||||]PAM NA Intergenic NA NA

T28080 TCCCCGTCCAGTTCTCCGCCGGG 1000

| | | | | | | | | | | | | |
|---|---|---|---|---|---|---|---|---|---|---|---|---|
| Oligo fwd | CACCTCCCCGTCCAGTTCTCCGCC | | | | | | | | | | | |
| Oligo rev | AAACGGCGGAGAACTGGACGGGGA | | | | | | | | | | | |
| Chromosome | start | | end | strand | MM | target_seq | PAM | alignment | distance | position | gene name | gene id |
| GU305914.1 | | 81514 | 81536 | - | 0 | TCCCCGTCCAGTTCTCCGCC | GGG | \|\|\|\|\|\|\|\|[\|\|\|\|\|\|\|\|\|\|\|\|]PAM | NA | Intergenic | NA | NA |
| T28081 | CCCCGTCCAGTTCTCCGCCGGGG | 1000 | | | | | | | | | | |
| Oligo fwd | CACCCCCCGTCCAGTTCTCCGCCG | | | | | | | | | | | |
| Oligo rev | AAACCGGCGGAGAACTGGACGGGG | | | | | | | | | | | |
| Chromosome | start | | end | strand | MM | target_seq | PAM | alignment | distance | position | gene name | gene id |
| GU305914.1 | | 81513 | 81535 | - | 0 | CCCCGTCCAGTTCTCCGCCG | GGG | \|\|\|\|\|\|\|\|[\|\|\|\|\|\|\|\|\|\|\|\|]PAM | NA | Intergenic | NA | NA |
| T28082 | CCAGTTCTCCGCCGGGGACGAGG | 1000 | | | | | | | | | | |
| Oligo fwd | CACCCCAGTTCTCCGCCGGGGACG | | | | | | | | | | | |
| Oligo rev | AAACCGTCCCCGGCGGAGAACTGG | | | | | | | | | | | |
| Chromosome | start | | end | strand | MM | target_seq | PAM | alignment | distance | position | gene name | gene id |
| GU305914.1 | | 81507 | 81529 | - | 0 | CCAGTTCTCCGCCGGGGACG | AGG | \|\|\|\|\|\|\|\|[\|\|\|\|\|\|\|\|\|\|\|\|]PAM | NA | Intergenic | NA | NA |
| T28083 | CAGTTCTCCGCCGGGGACGAGGG | 1000 | | | | | | | | | | |
| Oligo fwd | CACCCAGTTCTCCGCCGGGGACGA | | | | | | | | | | | |
| Oligo rev | AAACTCGTCCCCGGCGGAGAACTG | | | | | | | | | | | |
| Chromosome | start | | end | strand | MM | target_seq | PAM | alignment | distance | position | gene name | gene id |
| GU305914.1 | | 81506 | 81528 | - | 0 | CAGTTCTCCGCCGGGGACGA | GGG | \|\|\|\|\|\|\|\|[\|\|\|\|\|\|\|\|\|\|\|\|]PAM | NA | Intergenic | NA | NA |
| T28084 | TCTCCGCCGGGGACGAGGGACGG | 1000 | | | | | | | | | | |
| Oligo fwd | CACCTCTCCGCCGGGGACGAGGGA | | | | | | | | | | | |
| Oligo rev | AAACTCCCTCGTCCCCGGCGGAGA | | | | | | | | | | | |
| Chromosome | start | | end | strand | MM | target_seq | PAM | alignment | distance | position | gene name | gene id |
| GU305914.1 | | 81502 | 81524 | - | 0 | TCTCCGCCGGGGACGAGGGA | CGG | \|\|\|\|\|\|\|\|[\|\|\|\|\|\|\|\|\|\|\|\|]PAM | NA | Intergenic | NA | NA |
| T28085 | CTCCGCCGGGGACGAGGGACGGG | 1000 | | | | | | | | | | |
| Oligo fwd | CACCCTCCGCCGGGGACGAGGGAC | | | | | | | | | | | |
| Oligo rev | AAACGTCCCTCGTCCCCGGCGGAG | | | | | | | | | | | |
| Chromosome | start | | end | strand | MM | target_seq | PAM | alignment | distance | position | gene name | gene id |
| GU305914.1 | | 81501 | 81523 | - | 0 | CTCCGCCGGGGACGAGGGAC | GGG | \|\|\|\|\|\|\|\|[\|\|\|\|\|\|\|\|\|\|\|\|]PAM | NA | Intergenic | NA | NA |
| T28086 | CCGGGGACGAGGGACGGGCCCGG | 1000 | | | | | | | | | | |
| Oligo fwd | CACCCCGGGGACGAGGGACGGGCC | | | | | | | | | | | |
| Oligo rev | AAACGGCCCGTCCCTCGTCCCCGG | | | | | | | | | | | |
| Chromosome | start | | end | strand | MM | target_seq | PAM | alignment | distance | position | gene name | gene id |
| GU305914.1 | | 81496 | 81518 | - | 0 | CCGGGGACGAGGGACGGGCC | CGG | \|\|\|\|\|\|\|\|[\|\|\|\|\|\|\|\|\|\|\|\|]PAM | NA | Intergenic | NA | NA |
| T28087 | CGGGGACGAGGGACGGGCCCGGG | 1000 | | | | | | | | | | |
| Oligo fwd | CACCCGGGGACGAGGGACGGGCCC | | | | | | | | | | | |
| Oligo rev | AAACGGGCCCGTCCCTCGTCCCCG | | | | | | | | | | | |
| Chromosome | start | | end | strand | MM | target_seq | PAM | alignment | distance | position | gene name | gene id |
| GU305914.1 | | 81495 | 81517 | - | 0 | CGGGGACGAGGGACGGGCCC | GGG | \|\|\|\|\|\|\|\|[\|\|\|\|\|\|\|\|\|\|\|\|]PAM | NA | Intergenic | NA | NA |
| T28088 | GGACGAGGGACGGGCCCGGGTGG | 1000 | | | | | | | | | | |
| Oligo fwd | CACCGGACGAGGGACGGGCCCGGG | | | | | | | | | | | |
| Oligo rev | AAACCCCGGGCCCGTCCCTCGTCC | | | | | | | | | | | |
| Chromosome | start | | end | strand | MM | target_seq | PAM | alignment | distance | position | gene name | gene id |
| GU305914.1 | | 81492 | 81514 | - | 0 | GGACGAGGGACGGGCCCGGG | TGG | \|\|\|\|\|\|\|\|[\|\|\|\|\|\|\|\|\|\|\|\|]PAM | NA | Intergenic | NA | NA |
| T28089 | GACGAGGGACGGGCCCGGGTGGG | 1000 | | | | | | | | | | |
| Oligo fwd | CACCGACGAGGGACGGGCCCGGGT | | | | | | | | | | | |
| Oligo rev | AAACACCCGGGCCCGTCCCTCGTC | | | | | | | | | | | |
| Chromosome | start | | end | strand | MM | target_seq | PAM | alignment | distance | position | gene name | gene id |
| GU305914.1 | | 81491 | 81513 | - | 0 | GACGAGGGACGGGCCCGGGT | GGG | \|\|\|\|\|\|\|\|[\|\|\|\|\|\|\|\|\|\|\|\|]PAM | NA | Intergenic | NA | NA |
| T28090 | GGTGGGCAAGTACTGCCTGCAGG | 1000 | | | | | | | | | | |
| Oligo fwd | CACCGGTGGGCAAGTACTGCCTGC | | | | | | | | | | | |
| Oligo rev | AAACGCAGGCAGTACTTGCCCACC | | | | | | | | | | | |
| Chromosome | start | | end | strand | MM | target_seq | PAM | alignment | distance | position | gene name | gene id |
| GU305914.1 | | 81474 | 81496 | - | 0 | GGTGGGCAAGTACTGCCTGC | AGG | \|\|\|\|\|\|\|\|[\|\|\|\|\|\|\|\|\|\|\|\|]PAM | NA | Intergenic | NA | NA |
| T28091 | GATGCTCGCTTTTCACTTCGAGG | 1000 | | | | | | | | | | |
| Oligo fwd | CACCGATGCTCGCTTTTCACTTCG | | | | | | | | | | | |
| Oligo rev | AAACCGAAGTGAAAAGCGAGCATC | | | | | | | | | | | |
| Chromosome | start | | end | strand | MM | target_seq | PAM | alignment | distance | position | gene name | gene id |
| GU305914.1 | | 81423 | 81445 | - | 0 | GATGCTCGCTTTTCACTTCG | AGG | \|\|\|\|\|\|\|\|[\|\|\|\|\|\|\|\|\|\|\|\|]PAM | NA | Intergenic | NA | NA |
| T28092 | CGCTTTTCACTTCGAGGCCGCGG | 1000 | | | | | | | | | | |
| Oligo fwd | CACCCGCTTTTCACTTCGAGGCCG | | | | | | | | | | | |
| Oligo rev | AAACCGGCCTCGAAGTGAAAAGCG | | | | | | | | | | | |
| Chromosome | start | | end | strand | MM | target_seq | PAM | alignment | distance | position | gene name | gene id |
| GU305914.1 | | 81417 | 81439 | - | 0 | CGCTTTTCACTTCGAGGCCG | CGG | \|\|\|\|\|\|\|\|[\|\|\|\|\|\|\|\|\|\|\|\|]PAM | NA | Intergenic | NA | NA |
| T28093 | TCGAGGCCGCGGCCATCGCCAGG | 1000 | | | | | | | | | | |
| Oligo fwd | CACCTCGAGGCCGCGGCCATCGCC | | | | | | | | | | | |
| Oligo rev | AAACGGCGATGGCCGCGGCCTCGA | | | | | | | | | | | |
| Chromosome | start | | end | strand | MM | target_seq | PAM | alignment | distance | position | gene name | gene id |
| GU305914.1 | | 81406 | 81428 | - | 0 | TCGAGGCCGCGGCCATCGCC | AGG | \|\|\|\|\|\|\|\|[\|\|\|\|\|\|\|\|\|\|\|\|]PAM | NA | Intergenic | NA | NA |
| T28094 | GGCCGCGGCCATCGCCAGGTTGG | 1000 | | | | | | | | | | |
| Oligo fwd | CACCGGCCGCGGCCATCGCCAGGT | | | | | | | | | | | |
| Oligo rev | AAACACCTGGCGATGGCCGCGGCC | | | | | | | | | | | |
| Chromosome | start | | end | strand | MM | target_seq | PAM | alignment | distance | position | gene name | gene id |
| GU305914.1 | | 81402 | 81424 | - | 0 | GGCCGCGGCCATCGCCAGGT | TGG | \|\|\|\|\|\|\|\|[\|\|\|\|\|\|\|\|\|\|\|\|]PAM | NA | Intergenic | NA | NA |

<table>
<tr><td>T28095</td><td>CGGCCATCGCCAGGTTGGCGAGG</td><td>1000</td></tr>
<tr><td>Oligo fwd</td><td>CACCCGGCCATCGCCAGGTTGGCG</td></tr>
<tr><td>Oligo rev</td><td>AAACCGCCAACCTGGCGATGGCCG</td></tr>
<tr><td>Chromosome</td><td>start</td><td>end</td><td>strand</td><td>MM</td><td>target_seq</td><td>PAM</td><td>alignment</td><td>distance</td><td>position</td><td>gene name</td><td>gene id</td></tr>
<tr><td>GU305914.1</td><td>81397</td><td>81419</td><td>-</td><td>0</td><td>CGGCCATCGCCAGGTTGGCG</td><td>AGG</td><td>||||||||[||||||||||||]PAM</td><td>NA</td><td>Intergenic</td><td>NA</td><td>NA</td></tr>
<tr><td>T28096</td><td>GGTTGGCGAGGATCCCCCTCAGG</td><td>1000</td></tr>
<tr><td>Oligo fwd</td><td>CACCGGTTGGCGAGGATCCCCCTC</td></tr>
<tr><td>Oligo rev</td><td>AAACGAGGGGGATCCTCGCCAACC</td></tr>
<tr><td>Chromosome</td><td>start</td><td>end</td><td>strand</td><td>MM</td><td>target_seq</td><td>PAM</td><td>alignment</td><td>distance</td><td>position</td><td>gene name</td><td>gene id</td></tr>
<tr><td>GU305914.1</td><td>81385</td><td>81407</td><td>-</td><td>0</td><td>GGTTGGCGAGGATCCCCCTC</td><td>AGG</td><td>||||||||[||||||||||||]PAM</td><td>NA</td><td>Intergenic</td><td>NA</td><td>NA</td></tr>
<tr><td>T28097</td><td>GGCGAGGATCCCCCTCAGGAAGG</td><td>1000</td></tr>
<tr><td>Oligo fwd</td><td>CACCGGCGAGGATCCCCCTCAGGA</td></tr>
<tr><td>Oligo rev</td><td>AAACTCCTGAGGGGGATCCTCGCC</td></tr>
<tr><td>Chromosome</td><td>start</td><td>end</td><td>strand</td><td>MM</td><td>target_seq</td><td>PAM</td><td>alignment</td><td>distance</td><td>position</td><td>gene name</td><td>gene id</td></tr>
<tr><td>GU305914.1</td><td>81381</td><td>81403</td><td>-</td><td>0</td><td>GGCGAGGATCCCCCTCAGGA</td><td>AGG</td><td>||||||||[||||||||||||]PAM</td><td>NA</td><td>Intergenic</td><td>NA</td><td>NA</td></tr>
<tr><td>T28098</td><td>CTCAGGAAGGTCATCTTCGACGG</td><td>1000</td></tr>
<tr><td>Oligo fwd</td><td>CACCCTCAGGAAGGTCATCTTCGA</td></tr>
<tr><td>Oligo rev</td><td>AAACTCGAAGATGACCTTCCTGAG</td></tr>
<tr><td>Chromosome</td><td>start</td><td>end</td><td>strand</td><td>MM</td><td>target_seq</td><td>PAM</td><td>alignment</td><td>distance</td><td>position</td><td>gene name</td><td>gene id</td></tr>
<tr><td>GU305914.1</td><td>81368</td><td>81390</td><td>-</td><td>0</td><td>CTCAGGAAGGTCATCTTCGA</td><td>CGG</td><td>||||||||[||||||||||||]PAM</td><td>NA</td><td>Intergenic</td><td>NA</td><td>NA</td></tr>
<tr><td>T28099</td><td>TCTTCGACGGACAGCAGATCCGG</td><td>1000</td></tr>
<tr><td>Oligo fwd</td><td>CACCTCTTCGACGGACAGCAGATC</td></tr>
<tr><td>Oligo rev</td><td>AAACGATCTGCTGTCCGTCGAAGA</td></tr>
<tr><td>Chromosome</td><td>start</td><td>end</td><td>strand</td><td>MM</td><td>target_seq</td><td>PAM</td><td>alignment</td><td>distance</td><td>position</td><td>gene name</td><td>gene id</td></tr>
<tr><td>GU305914.1</td><td>81355</td><td>81377</td><td>-</td><td>0</td><td>TCTTCGACGGACAGCAGATC</td><td>CGG</td><td>||||||||[||||||||||||]PAM</td><td>NA</td><td>Intergenic</td><td>NA</td><td>NA</td></tr>
<tr><td>T28100</td><td>GATCTACACCTGTCTGCTCGAGG</td><td>1000</td></tr>
<tr><td>Oligo fwd</td><td>CACCGATCTACACCTGTCTGCTCG</td></tr>
<tr><td>Oligo rev</td><td>AAACCGAGCAGACAGGTGTAGATC</td></tr>
<tr><td>Chromosome</td><td>start</td><td>end</td><td>strand</td><td>MM</td><td>target_seq</td><td>PAM</td><td>alignment</td><td>distance</td><td>position</td><td>gene name</td><td>gene id</td></tr>
<tr><td>GU305914.1</td><td>81333</td><td>81355</td><td>-</td><td>0</td><td>GATCTACACCTGTCTGCTCG</td><td>AGG</td><td>||||||||[||||||||||||]PAM</td><td>NA</td><td>Intergenic</td><td>NA</td><td>NA</td></tr>
<tr><td>T28101</td><td>CTGTCTGCTCGAGGAGTGTTCGG</td><td>1000</td></tr>
<tr><td>Oligo fwd</td><td>CACCCTGTCTGCTCGAGGAGTGTT</td></tr>
<tr><td>Oligo rev</td><td>AAACAACACTCCTCGAGCAGACAG</td></tr>
<tr><td>Chromosome</td><td>start</td><td>end</td><td>strand</td><td>MM</td><td>target_seq</td><td>PAM</td><td>alignment</td><td>distance</td><td>position</td><td>gene name</td><td>gene id</td></tr>
<tr><td>GU305914.1</td><td>81324</td><td>81346</td><td>-</td><td>0</td><td>CTGTCTGCTCGAGGAGTGTT</td><td>CGG</td><td>||||||||[||||||||||||]PAM</td><td>NA</td><td>Intergenic</td><td>NA</td><td>NA</td></tr>
<tr><td>T28102</td><td>TGTCTGCTCGAGGAGTGTTCGGG</td><td>1000</td></tr>
<tr><td>Oligo fwd</td><td>CACCTGTCTGCTCGAGGAGTGTTC</td></tr>
<tr><td>Oligo rev</td><td>AAACGAACACTCCTCGAGCAGACA</td></tr>
<tr><td>Chromosome</td><td>start</td><td>end</td><td>strand</td><td>MM</td><td>target_seq</td><td>PAM</td><td>alignment</td><td>distance</td><td>position</td><td>gene name</td><td>gene id</td></tr>
<tr><td>GU305914.1</td><td>81323</td><td>81345</td><td>-</td><td>0</td><td>TGTCTGCTCGAGGAGTGTTC</td><td>GGG</td><td>||||||||[||||||||||||]PAM</td><td>NA</td><td>Intergenic</td><td>NA</td><td>NA</td></tr>
<tr><td>T28103</td><td>TGCTCGAGGAGTGTTCGGGTCGG</td><td>1000</td></tr>
<tr><td>Oligo fwd</td><td>CACCTGCTCGAGGAGTGTTCGGGT</td></tr>
<tr><td>Oligo rev</td><td>AAACACCCGAACACTCCTCGAGCA</td></tr>
<tr><td>Chromosome</td><td>start</td><td>end</td><td>strand</td><td>MM</td><td>target_seq</td><td>PAM</td><td>alignment</td><td>distance</td><td>position</td><td>gene name</td><td>gene id</td></tr>
<tr><td>GU305914.1</td><td>81319</td><td>81341</td><td>-</td><td>0</td><td>TGCTCGAGGAGTGTTCGGGT</td><td>CGG</td><td>||||||||[||||||||||||]PAM</td><td>NA</td><td>Intergenic</td><td>NA</td><td>NA</td></tr>
<tr><td>T28104</td><td>GCTCGAGGAGTGTTCGGGTCGGG</td><td>1000</td></tr>
<tr><td>Oligo fwd</td><td>CACCGCTCGAGGAGTGTTCGGGTC</td></tr>
<tr><td>Oligo rev</td><td>AAACGACCCGAACACTCCTCGAGC</td></tr>
<tr><td>Chromosome</td><td>start</td><td>end</td><td>strand</td><td>MM</td><td>target_seq</td><td>PAM</td><td>alignment</td><td>distance</td><td>position</td><td>gene name</td><td>gene id</td></tr>
<tr><td>GU305914.1</td><td>81318</td><td>81340</td><td>-</td><td>0</td><td>GCTCGAGGAGTGTTCGGGTC</td><td>GGG</td><td>||||||||[||||||||||||]PAM</td><td>NA</td><td>Intergenic</td><td>NA</td><td>NA</td></tr>
<tr><td>T28105</td><td>CTGCCCAACATGCCCTCGCTCGG</td><td>1000</td></tr>
<tr><td>Oligo fwd</td><td>CACCCTGCCCAACATGCCCTCGCT</td></tr>
<tr><td>Oligo rev</td><td>AAACAGCGAGGGCATGTTGGGCAG</td></tr>
<tr><td>Chromosome</td><td>start</td><td>end</td><td>strand</td><td>MM</td><td>target_seq</td><td>PAM</td><td>alignment</td><td>distance</td><td>position</td><td>gene name</td><td>gene id</td></tr>
<tr><td>GU305914.1</td><td>81287</td><td>81309</td><td>-</td><td>0</td><td>CTGCCCAACATGCCCTCGCT</td><td>CGG</td><td>||||||||[||||||||||||]PAM</td><td>NA</td><td>Intergenic</td><td>NA</td><td>NA</td></tr>
<tr><td>T28106</td><td>CATGCCCTCGCTCGGCCACGAGG</td><td>1000</td></tr>
<tr><td>Oligo fwd</td><td>CACCCATGCCCTCGCTCGGCCACG</td></tr>
<tr><td>Oligo rev</td><td>AAACCGTGGCCGAGCGAGGGCATG</td></tr>
<tr><td>Chromosome</td><td>start</td><td>end</td><td>strand</td><td>MM</td><td>target_seq</td><td>PAM</td><td>alignment</td><td>distance</td><td>position</td><td>gene name</td><td>gene id</td></tr>
<tr><td>GU305914.1</td><td>81279</td><td>81301</td><td>-</td><td>0</td><td>CATGCCCTCGCTCGGCCACG</td><td>AGG</td><td>||||||||[||||||||||||]PAM</td><td>NA</td><td>Intergenic</td><td>NA</td><td>NA</td></tr>
<tr><td>T28107</td><td>GCCCTCGCTCGGCCACGAGGCGG</td><td>1000</td></tr>
<tr><td>Oligo fwd</td><td>CACCGCCCTCGCTCGGCCACGAGG</td></tr>
<tr><td>Oligo rev</td><td>AAACCCTCGTGGCCGAGCGAGGGC</td></tr>
<tr><td>Chromosome</td><td>start</td><td>end</td><td>strand</td><td>MM</td><td>target_seq</td><td>PAM</td><td>alignment</td><td>distance</td><td>position</td><td>gene name</td><td>gene id</td></tr>
<tr><td>GU305914.1</td><td>81276</td><td>81298</td><td>-</td><td>0</td><td>GCCCTCGCTCGGCCACGAGG</td><td>CGG</td><td>||||||||[||||||||||||]PAM</td><td>NA</td><td>Intergenic</td><td>NA</td><td>NA</td></tr>
<tr><td>T28108</td><td>CGAGGCGGCCGCCGCGATCGAGG</td><td>1000</td></tr>
<tr><td>Oligo fwd</td><td>CACCCGAGGCGGCCGCCGCGATCG</td></tr>
<tr><td>Oligo rev</td><td>AAACCGATCGCGGCGGCCGCCTCG</td></tr>
<tr><td>Chromosome</td><td>start</td><td>end</td><td>strand</td><td>MM</td><td>target_seq</td><td>PAM</td><td>alignment</td><td>distance</td><td>position</td><td>gene name</td><td>gene id</td></tr>
<tr><td>GU305914.1</td><td>81261</td><td>81283</td><td>-</td><td>0</td><td>CGAGGCGGCCGCCGCGATCG</td><td>AGG</td><td>||||||||[||||||||||||]PAM</td><td>NA</td><td>Intergenic</td><td>NA</td><td>NA</td></tr>
<tr><td>GU305914.1</td><td>170603</td><td>170625</td><td>+</td><td>4</td><td>CGATGCCGCTGCCGCGAACG</td><td>TGG</td><td>|||-||-|[|-|||||||-||]PAM</td><td>NA</td><td>Intergenic</td><td>NA</td><td>NA</td></tr>
<tr><td>T28109</td><td>GGCGGCCGCCGCGATCGAGGAGG</td><td>1000</td></tr>
<tr><td>Oligo fwd</td><td>CACCGGCGGCCGCCGCGATCGAGG</td></tr>
<tr><td>Oligo rev</td><td>AAACCCTCGATCGCGGCGGCCGCC</td></tr>
<tr><td>Chromosome</td><td>start</td><td>end</td><td>strand</td><td>MM</td><td>target_seq</td><td>PAM</td><td>alignment</td><td>distance</td><td>position</td><td>gene name</td><td>gene id</td></tr>
</table>

| | | | | | | | | | | | |
|---|---|---|---|---|---|---|---|---|---|---|---|
| GU305914.1 | 81258 | 81280 | - | 0 | GGCGGCCGCCGCGATCGAGG | AGG | \|\|\|\|\|\|\|\|[\|\|\|\|\|\|\|\|\|\|\|\|]PAM | NA | Intergenic | NA | NA |
| GU305914.1 | 138477 | 138499 | + | 4 | GGCGCTCGCCGCCGTCGAGG | AGG | \|\|\|\|--\|\|[\|\|\|\|--\|\|\|\|\|\|]PAM | NA | Intergenic | NA | NA |
| T28110 | GGCCGCCGCGATCGAGGAGGCGG | 1000 | | | | | | | | | |
| Oligo fwd | CACCGGCCGCCGCGATCGAGGAGG | | | | | | | | | | |
| Oligo rev | AAACCCTCCTCGATCGCGGCGGCC | | | | | | | | | | |
| Chromosome | start | end | strand | MM | target_seq | PAM | alignment | distance | position | gene name | gene id |
| GU305914.1 | 81255 | 81277 | - | 0 | GGCCGCCGCGATCGAGGAGG | CGG | \|\|\|\|\|\|\|\|[\|\|\|\|\|\|\|\|\|\|\|\|]PAM | NA | Intergenic | NA | NA |
| T28111 | CGCCGCGATCGAGGAGGCGGCGG | 1000 | | | | | | | | | |
| Oligo fwd | CACCCGCCGCGATCGAGGAGGCGG | | | | | | | | | | |
| Oligo rev | AAACCCGCCTCCTCGATCGCGGCG | | | | | | | | | | |
| Chromosome | start | end | strand | MM | target_seq | PAM | alignment | distance | position | gene name | gene id |
| GU305914.1 | 81252 | 81274 | - | 0 | CGCCGCGATCGAGGAGGCGG | CGG | \|\|\|\|\|\|\|\|[\|\|\|\|\|\|\|\|\|\|\|\|]PAM | NA | Intergenic | NA | NA |
| GU305914.1 | 72468 | 72490 | - | 4 | GGACGCTATCGAGGAGGCGT | AGG | -\|-\|\|\|-\|[\|\|\|\|\|\|\|\|\|\|\|-]PAM | NA | Intergenic | NA | NA |
| T28112 | CGCGATCGAGGAGGCGGCGGCGG | 1000 | | | | | | | | | |
| Oligo fwd | CACCCGCGATCGAGGAGGCGGCGG | | | | | | | | | | |
| Oligo rev | AAACCCGCCGCCTCCTCGATCGCG | | | | | | | | | | |
| Chromosome | start | end | strand | MM | target_seq | PAM | alignment | distance | position | gene name | gene id |
| GU305914.1 | 81249 | 81271 | - | 0 | CGCGATCGAGGAGGCGGCGG | CGG | \|\|\|\|\|\|\|\|[\|\|\|\|\|\|\|\|\|\|\|\|]PAM | NA | Intergenic | NA | NA |
| GU305914.1 | 161757 | 161779 | - | 4 | GGAGGCCGAGGAGGCGGCGG | CGG | -\|-\|--\|\|[\|\|\|\|\|\|\|\|\|\|\|\|]PAM | NA | Intergenic | NA | NA |
| T28113 | GCGATCGAGGAGGCGGCGGCGGG | 1000 | | | | | | | | | |
| Oligo fwd | CACCGCGATCGAGGAGGCGGCGGC | | | | | | | | | | |
| Oligo rev | AAACGCCGCCGCCTCCTCGATCGC | | | | | | | | | | |
| Chromosome | start | end | strand | MM | target_seq | PAM | alignment | distance | position | gene name | gene id |
| GU305914.1 | 81248 | 81270 | - | 0 | GCGATCGAGGAGGCGGCGGC | GGG | \|\|\|\|\|\|\|\|[\|\|\|\|\|\|\|\|\|\|\|\|]PAM | NA | Intergenic | NA | NA |
| GU305914.1 | 114657 | 114679 | - | 4 | GCGGTGGTGGTGGCGGCGGC | GGG | \|\|\|-\|-\|-[\|\|-\|\|\|\|\|\|\|\|\|]PAM | NA | Intergenic | NA | NA |
| GU305914.1 | 72464 | 72486 | - | 3 | GCTATCGAGGAGGCGTAGGC | CGG | \|\|-\|\|\|\|\|[\|\|\|\|\|\|\|--\|\|\|]PAM | NA | Intergenic | NA | NA |
| T28114 | CGATCGAGGAGGCGGCGGCGGGG | 1000 | | | | | | | | | |
| Oligo fwd | CACCCGATCGAGGAGGCGGCGGCG | | | | | | | | | | |
| Oligo rev | AAACCGCCGCCGCCTCCTCGATCG | | | | | | | | | | |
| Chromosome | start | end | strand | MM | target_seq | PAM | alignment | distance | position | gene name | gene id |
| GU305914.1 | 81247 | 81269 | - | 0 | CGATCGAGGAGGCGGCGGCG | GGG | \|\|\|\|\|\|\|\|[\|\|\|\|\|\|\|\|\|\|\|\|]PAM | NA | Intergenic | NA | NA |
| GU305914.1 | 130607 | 130629 | - | 4 | AGATGGAGAAGCCGGCGGCG | CGG | -\|\|\|-\|\|\|[-\|\|-\|\|\|\|\|\|\|\|]PAM | NA | Intergenic | NA | NA |
| T28115 | GATCGAGGAGGCGGCGGCGGGGG | 1000 | | | | | | | | | |
| Oligo fwd | CACCGATCGAGGAGGCGGCGGCGG | | | | | | | | | | |
| Oligo rev | AAACCCGCCGCCGCCTCCTCGATC | | | | | | | | | | |
| Chromosome | start | end | strand | MM | target_seq | PAM | alignment | distance | position | gene name | gene id |
| GU305914.1 | 81246 | 81268 | - | 0 | GATCGAGGAGGCGGCGGCGG | GGG | \|\|\|\|\|\|\|\|[\|\|\|\|\|\|\|\|\|\|\|\|]PAM | NA | Intergenic | NA | NA |
| GU305914.1 | 161754 | 161776 | - | 2 | GGCCGAGGAGGCGGCGGCGG | AGG | \|--\|\|\|\|\|[\|\|\|\|\|\|\|\|\|\|\|\|]PAM | NA | Intergenic | NA | NA |
| GU305914.1 | 154407 | 154429 | + | 4 | GGTGGCGGCGGCGGCGGCGG | CGG | \|-\|-\|-\|\|[-\|\|\|\|\|\|\|\|\|\|\|]PAM | NA | Intergenic | NA | NA |
| GU305914.1 | 58123 | 58145 | - | 4 | GGTGGCGGCGGCGGCGGCGG | AGG | \|-\|-\|-\|\|[-\|\|\|\|\|\|\|\|\|\|\|]PAM | NA | Intergenic | NA | NA |
| GU305914.1 | 149491 | 149513 | - | 4 | GAGGGAGCAGGCGACGGCGG | CGG | \|\|--\|\|\|-[\|\|\|\|\|-\|\|\|\|\|\|]PAM | NA | Intergenic | NA | NA |
| T28116 | ATCGAGGAGGCGGCGGCGGGGGG | 1000 | | | | | | | | | |
| Oligo fwd | CACCATCGAGGAGGCGGCGGCGGG | | | | | | | | | | |
| Oligo rev | AAACCCCGCCGCCGCCTCCTCGAT | | | | | | | | | | |
| Chromosome | start | end | strand | MM | target_seq | PAM | alignment | distance | position | gene name | gene id |
| GU305914.1 | 81245 | 81267 | - | 0 | ATCGAGGAGGCGGCGGCGGG | GGG | \|\|\|\|\|\|\|\|[\|\|\|\|\|\|\|\|\|\|\|\|]PAM | NA | Intergenic | NA | NA |
| T28117 | GGAGGCGGCGGCGGGGGGTGAGG | 1000 | | | | | | | | | |
| Oligo fwd | CACCGGAGGCGGCGGCGGGGGGTG | | | | | | | | | | |
| Oligo rev | AAACCACCCCCCGCCGCCGCCTCC | | | | | | | | | | |
| Chromosome | start | end | strand | MM | target_seq | PAM | alignment | distance | position | gene name | gene id |
| GU305914.1 | 81240 | 81262 | - | 0 | GGAGGCGGCGGCGGGGGGTG | AGG | \|\|\|\|\|\|\|\|[\|\|\|\|\|\|\|\|\|\|\|\|]PAM | NA | Intergenic | NA | NA |
| GU305914.1 | 62452 | 62474 | + | 4 | GGTGGTGGCGGCGGGGGTGG | CGG | \|\|-\|\|-\|\|[\|\|\|\|\|\|\|\|\|--\|]PAM | NA | Intergenic | NA | NA |
| GU305914.1 | 62437 | 62459 | + | 4 | GGCGGTGGCGGCGGGGGTGG | TGG | \|\|-\|\|-\|\|[\|\|\|\|\|\|\|\|\|--\|]PAM | NA | Intergenic | NA | NA |
| T28118 | GAGGCGGCGGCGGGGGGTGAGGG | 1000 | | | | | | | | | |
| Oligo fwd | CACCGAGGCGGCGGCGGGGGGTGA | | | | | | | | | | |
| Oligo rev | AAACTCACCCCCCGCCGCCGCCTC | | | | | | | | | | |
| Chromosome | start | end | strand | MM | target_seq | PAM | alignment | distance | position | gene name | gene id |
| GU305914.1 | 81239 | 81261 | - | 0 | GAGGCGGCGGCGGGGGGTGA | GGG | \|\|\|\|\|\|\|\|[\|\|\|\|\|\|\|\|\|\|\|\|]PAM | NA | Intergenic | NA | NA |
| GU305914.1 | 109335 | 109357 | + | 4 | GTGGCGGGGACGGCGGGTGA | CGG | \|-\|\|\|\|\|-[\|-\|\|\|-\|\|\|\|\|\|]PAM | NA | Intergenic | NA | NA |
| T28119 | AGGCGGCGGCGGGGGGTGAGGGG | 1000 | | | | | | | | | |
| Oligo fwd | CACCAGGCGGCGGCGGGGGGTGAG | | | | | | | | | | |
| Oligo rev | AAACCTCACCCCCCGCCGCCGCCT | | | | | | | | | | |
| Chromosome | start | end | strand | MM | target_seq | PAM | alignment | distance | position | gene name | gene id |
| GU305914.1 | 81238 | 81260 | - | 0 | AGGCGGCGGCGGGGGGTGAG | GGG | \|\|\|\|\|\|\|\|[\|\|\|\|\|\|\|\|\|\|\|\|]PAM | NA | Intergenic | NA | NA |
| T28120 | GGCGGCGGCGGGGGGTGAGGGGG | 1000 | | | | | | | | | |
| Oligo fwd | CACCGGCGGCGGCGGGGGGTGAGG | | | | | | | | | | |
| Oligo rev | AAACCCTCACCCCCCGCCGCCGCC | | | | | | | | | | |
| Chromosome | start | end | strand | MM | target_seq | PAM | alignment | distance | position | gene name | gene id |
| GU305914.1 | 81237 | 81259 | - | 0 | GGCGGCGGCGGGGGGTGAGG | GGG | \|\|\|\|\|\|\|\|[\|\|\|\|\|\|\|\|\|\|\|\|]PAM | NA | Intergenic | NA | NA |
| T28121 | GGGTGAGGGGGACGAAACCTCGG | 1000 | | | | | | | | | |
| Oligo fwd | CACCGGGTGAGGGGGACGAAACCT | | | | | | | | | | |
| Oligo rev | AAACAGGTTTCGTCCCCCTCACCC | | | | | | | | | | |
| Chromosome | start | end | strand | MM | target_seq | PAM | alignment | distance | position | gene name | gene id |
| GU305914.1 | 81225 | 81247 | - | 0 | GGGTGAGGGGGACGAAACCT | CGG | \|\|\|\|\|\|\|\|[\|\|\|\|\|\|\|\|\|\|\|\|]PAM | NA | Intergenic | NA | NA |
| T28122 | TGAGGGGGACGAAACCTCGGAGG | 1000 | | | | | | | | | |
| Oligo fwd | CACCTGAGGGGGACGAAACCTCGG | | | | | | | | | | |

<table>
<tr><td>Oligo rev</td><td>AAACCCGAGGTTTCGTCCCCCTCA</td><td></td><td></td><td></td><td></td><td></td><td></td><td></td><td></td><td></td><td></td><td></td></tr>
<tr><td>Chromosome</td><td>start</td><td></td><td>end</td><td>strand</td><td>MM</td><td>target_seq</td><td>PAM</td><td>alignment</td><td>distance</td><td>position</td><td>gene name</td><td>gene id</td></tr>
<tr><td>GU305914.1</td><td></td><td>81222</td><td>81244</td><td>-</td><td>0</td><td>TGAGGGGGACGAAACCTCGG</td><td>AGG</td><td>||||||||[||||||||||||]PAM</td><td>NA</td><td>Intergenic</td><td>NA</td><td>NA</td></tr>
<tr><td>GU305914.1</td><td></td><td>12477</td><td>12499</td><td>-</td><td>4</td><td>TGAAGCGGTCGAAACCTCTG</td><td>CGG</td><td>|||-|-||[-|||||||||-|]PAM</td><td>NA</td><td>Intergenic</td><td>NA</td><td>NA</td></tr>
<tr><td></td><td></td><td></td><td></td><td></td><td></td><td></td><td></td><td></td><td></td><td></td><td></td><td></td></tr>
<tr><td>T28123</td><td>GAGGGGGACGAAACCTCGGAGGG</td><td></td><td>1000</td><td></td><td></td><td></td><td></td><td></td><td></td><td></td><td></td><td></td></tr>
<tr><td>Oligo fwd</td><td>CACCGAGGGGGACGAAACCTCGGA</td><td></td><td></td><td></td><td></td><td></td><td></td><td></td><td></td><td></td><td></td><td></td></tr>
<tr><td>Oligo rev</td><td>AAACTCCGAGGTTTCGTCCCCCTC</td><td></td><td></td><td></td><td></td><td></td><td></td><td></td><td></td><td></td><td></td><td></td></tr>
<tr><td>Chromosome</td><td>start</td><td></td><td>end</td><td>strand</td><td>MM</td><td>target_seq</td><td>PAM</td><td>alignment</td><td>distance</td><td>position</td><td>gene name</td><td>gene id</td></tr>
<tr><td>GU305914.1</td><td></td><td>81221</td><td>81243</td><td>-</td><td>0</td><td>GAGGGGGACGAAACCTCGGA</td><td>GGG</td><td>||||||||[||||||||||||]PAM</td><td>NA</td><td>Intergenic</td><td>NA</td><td>NA</td></tr>
<tr><td></td><td></td><td></td><td></td><td></td><td></td><td></td><td></td><td></td><td></td><td></td><td></td><td></td></tr>
<tr><td>T28124</td><td>GCGAGAACTCGAACAACTCGCGG</td><td></td><td>1000</td><td></td><td></td><td></td><td></td><td></td><td></td><td></td><td></td><td></td></tr>
<tr><td>Oligo fwd</td><td>CACCGCGAGAACTCGAACAACTCG</td><td></td><td></td><td></td><td></td><td></td><td></td><td></td><td></td><td></td><td></td><td></td></tr>
<tr><td>Oligo rev</td><td>AAACCGAGTTGTTCGAGTTCTCGC</td><td></td><td></td><td></td><td></td><td></td><td></td><td></td><td></td><td></td><td></td><td></td></tr>
<tr><td>Chromosome</td><td>start</td><td></td><td>end</td><td>strand</td><td>MM</td><td>target_seq</td><td>PAM</td><td>alignment</td><td>distance</td><td>position</td><td>gene name</td><td>gene id</td></tr>
<tr><td>GU305914.1</td><td></td><td>81199</td><td>81221</td><td>-</td><td>0</td><td>GCGAGAACTCGAACAACTCG</td><td>CGG</td><td>||||||||[||||||||||||]PAM</td><td>NA</td><td>Intergenic</td><td>NA</td><td>NA</td></tr>
<tr><td></td><td></td><td></td><td></td><td></td><td></td><td></td><td></td><td></td><td></td><td></td><td></td><td></td></tr>
<tr><td>T28125</td><td>GAACTCGAACAACTCGCGGACGG</td><td></td><td>1000</td><td></td><td></td><td></td><td></td><td></td><td></td><td></td><td></td><td></td></tr>
<tr><td>Oligo fwd</td><td>CACCGAACTCGAACAACTCGCGGA</td><td></td><td></td><td></td><td></td><td></td><td></td><td></td><td></td><td></td><td></td><td></td></tr>
<tr><td>Oligo rev</td><td>AAACTCCGCGAGTTGTTCGAGTTC</td><td></td><td></td><td></td><td></td><td></td><td></td><td></td><td></td><td></td><td></td><td></td></tr>
<tr><td>Chromosome</td><td>start</td><td></td><td>end</td><td>strand</td><td>MM</td><td>target_seq</td><td>PAM</td><td>alignment</td><td>distance</td><td>position</td><td>gene name</td><td>gene id</td></tr>
<tr><td>GU305914.1</td><td></td><td>81195</td><td>81217</td><td>-</td><td>0</td><td>GAACTCGAACAACTCGCGGA</td><td>CGG</td><td>||||||||[||||||||||||]PAM</td><td>NA</td><td>Intergenic</td><td>NA</td><td>NA</td></tr>
<tr><td></td><td></td><td></td><td></td><td></td><td></td><td></td><td></td><td></td><td></td><td></td><td></td><td></td></tr>
<tr><td>T28126</td><td>CTCGAACAACTCGCGGACGGTGG</td><td></td><td>1000</td><td></td><td></td><td></td><td></td><td></td><td></td><td></td><td></td><td></td></tr>
<tr><td>Oligo fwd</td><td>CACCCTCGAACAACTCGCGGACGG</td><td></td><td></td><td></td><td></td><td></td><td></td><td></td><td></td><td></td><td></td><td></td></tr>
<tr><td>Oligo rev</td><td>AAACCCGTCCGCGAGTTGTTCGAG</td><td></td><td></td><td></td><td></td><td></td><td></td><td></td><td></td><td></td><td></td><td></td></tr>
<tr><td>Chromosome</td><td>start</td><td></td><td>end</td><td>strand</td><td>MM</td><td>target_seq</td><td>PAM</td><td>alignment</td><td>distance</td><td>position</td><td>gene name</td><td>gene id</td></tr>
<tr><td>GU305914.1</td><td></td><td>81192</td><td>81214</td><td>-</td><td>0</td><td>CTCGAACAACTCGCGGACGG</td><td>TGG</td><td>||||||||[||||||||||||]PAM</td><td>NA</td><td>Intergenic</td><td>NA</td><td>NA</td></tr>
<tr><td></td><td></td><td></td><td></td><td></td><td></td><td></td><td></td><td></td><td></td><td></td><td></td><td></td></tr>
<tr><td>T28127</td><td>TCGAACAACTCGCGGACGGTGGG</td><td></td><td>1000</td><td></td><td></td><td></td><td></td><td></td><td></td><td></td><td></td><td></td></tr>
<tr><td>Oligo fwd</td><td>CACCTCGAACAACTCGCGGACGGT</td><td></td><td></td><td></td><td></td><td></td><td></td><td></td><td></td><td></td><td></td><td></td></tr>
<tr><td>Oligo rev</td><td>AAACACCGTCCGCGAGTTGTTCGA</td><td></td><td></td><td></td><td></td><td></td><td></td><td></td><td></td><td></td><td></td><td></td></tr>
<tr><td>Chromosome</td><td>start</td><td></td><td>end</td><td>strand</td><td>MM</td><td>target_seq</td><td>PAM</td><td>alignment</td><td>distance</td><td>position</td><td>gene name</td><td>gene id</td></tr>
<tr><td>GU305914.1</td><td></td><td>81191</td><td>81213</td><td>-</td><td>0</td><td>TCGAACAACTCGCGGACGGT</td><td>GGG</td><td>||||||||[||||||||||||]PAM</td><td>NA</td><td>Intergenic</td><td>NA</td><td>NA</td></tr>
<tr><td></td><td></td><td></td><td></td><td></td><td></td><td></td><td></td><td></td><td></td><td></td><td></td><td></td></tr>
<tr><td>T28128</td><td>CGAACAACTCGCGGACGGTGGGG</td><td></td><td>1000</td><td></td><td></td><td></td><td></td><td></td><td></td><td></td><td></td><td></td></tr>
<tr><td>Oligo fwd</td><td>CACCCGAACAACTCGCGGACGGTG</td><td></td><td></td><td></td><td></td><td></td><td></td><td></td><td></td><td></td><td></td><td></td></tr>
<tr><td>Oligo rev</td><td>AAACCACCGTCCGCGAGTTGTTCG</td><td></td><td></td><td></td><td></td><td></td><td></td><td></td><td></td><td></td><td></td><td></td></tr>
<tr><td>Chromosome</td><td>start</td><td></td><td>end</td><td>strand</td><td>MM</td><td>target_seq</td><td>PAM</td><td>alignment</td><td>distance</td><td>position</td><td>gene name</td><td>gene id</td></tr>
<tr><td>GU305914.1</td><td></td><td>81190</td><td>81212</td><td>-</td><td>0</td><td>CGAACAACTCGCGGACGGTG</td><td>GGG</td><td>||||||||[||||||||||||]PAM</td><td>NA</td><td>Intergenic</td><td>NA</td><td>NA</td></tr>
<tr><td></td><td></td><td></td><td></td><td></td><td></td><td></td><td></td><td></td><td></td><td></td><td></td><td></td></tr>
<tr><td>T28129</td><td>CTCGCGGACGGTGGGGTACCAGG</td><td></td><td>1000</td><td></td><td></td><td></td><td></td><td></td><td></td><td></td><td></td><td></td></tr>
<tr><td>Oligo fwd</td><td>CACCCTCGCGGACGGTGGGGTACC</td><td></td><td></td><td></td><td></td><td></td><td></td><td></td><td></td><td></td><td></td><td></td></tr>
<tr><td>Oligo rev</td><td>AAACGGTACCCCACCGTCCGCGAG</td><td></td><td></td><td></td><td></td><td></td><td></td><td></td><td></td><td></td><td></td><td></td></tr>
<tr><td>Chromosome</td><td>start</td><td></td><td>end</td><td>strand</td><td>MM</td><td>target_seq</td><td>PAM</td><td>alignment</td><td>distance</td><td>position</td><td>gene name</td><td>gene id</td></tr>
<tr><td>GU305914.1</td><td></td><td>81183</td><td>81205</td><td>-</td><td>0</td><td>CTCGCGGACGGTGGGGTACC</td><td>AGG</td><td>||||||||[||||||||||||]PAM</td><td>NA</td><td>Intergenic</td><td>NA</td><td>NA</td></tr>
<tr><td>GU305914.1</td><td></td><td>128103</td><td>128125</td><td>-</td><td>3</td><td>CTCGGGCACGGTGAGGTACC</td><td>TGG</td><td>||||-|-|[|||||-||||||]PAM</td><td>NA</td><td>Intergenic</td><td>NA</td><td>NA</td></tr>
<tr><td></td><td></td><td></td><td></td><td></td><td></td><td></td><td></td><td></td><td></td><td></td><td></td><td></td></tr>
<tr><td>T28130</td><td>TCGCGGACGGTGGGGTACCAGGG</td><td></td><td>1000</td><td></td><td></td><td></td><td></td><td></td><td></td><td></td><td></td><td></td></tr>
<tr><td>Oligo fwd</td><td>CACCTCGCGGACGGTGGGGTACCA</td><td></td><td></td><td></td><td></td><td></td><td></td><td></td><td></td><td></td><td></td><td></td></tr>
<tr><td>Oligo rev</td><td>AAACTGGTACCCCACCGTCCGCGA</td><td></td><td></td><td></td><td></td><td></td><td></td><td></td><td></td><td></td><td></td><td></td></tr>
<tr><td>Chromosome</td><td>start</td><td></td><td>end</td><td>strand</td><td>MM</td><td>target_seq</td><td>PAM</td><td>alignment</td><td>distance</td><td>position</td><td>gene name</td><td>gene id</td></tr>
<tr><td>GU305914.1</td><td></td><td>81182</td><td>81204</td><td>-</td><td>0</td><td>TCGCGGACGGTGGGGTACCA</td><td>GGG</td><td>||||||||[||||||||||||]PAM</td><td>NA</td><td>Intergenic</td><td>NA</td><td>NA</td></tr>
<tr><td></td><td></td><td></td><td></td><td></td><td></td><td></td><td></td><td></td><td></td><td></td><td></td><td></td></tr>
<tr><td>T28131</td><td>GGTGGGGTACCAGGGAGCTACGG</td><td></td><td>1000</td><td></td><td></td><td></td><td></td><td></td><td></td><td></td><td></td><td></td></tr>
<tr><td>Oligo fwd</td><td>CACCGGTGGGGTACCAGGGAGCTA</td><td></td><td></td><td></td><td></td><td></td><td></td><td></td><td></td><td></td><td></td><td></td></tr>
<tr><td>Oligo rev</td><td>AAACTAGCTCCCTGGTACCCCACC</td><td></td><td></td><td></td><td></td><td></td><td></td><td></td><td></td><td></td><td></td><td></td></tr>
<tr><td>Chromosome</td><td>start</td><td></td><td>end</td><td>strand</td><td>MM</td><td>target_seq</td><td>PAM</td><td>alignment</td><td>distance</td><td>position</td><td>gene name</td><td>gene id</td></tr>
<tr><td>GU305914.1</td><td></td><td>81174</td><td>81196</td><td>-</td><td>0</td><td>GGTGGGGTACCAGGGAGCTA</td><td>CGG</td><td>||||||||[||||||||||||]PAM</td><td>NA</td><td>Intergenic</td><td>NA</td><td>NA</td></tr>
<tr><td></td><td></td><td></td><td></td><td></td><td></td><td></td><td></td><td></td><td></td><td></td><td></td><td></td></tr>
<tr><td>T28132</td><td>GGGAGCTACGGTGCTCGAACCGG</td><td></td><td>1000</td><td></td><td></td><td></td><td></td><td></td><td></td><td></td><td></td><td></td></tr>
<tr><td>Oligo fwd</td><td>CACCGGGAGCTACGGTGCTCGAAC</td><td></td><td></td><td></td><td></td><td></td><td></td><td></td><td></td><td></td><td></td><td></td></tr>
<tr><td>Oligo rev</td><td>AAACGTTCGAGCACCGTAGCTCCC</td><td></td><td></td><td></td><td></td><td></td><td></td><td></td><td></td><td></td><td></td><td></td></tr>
<tr><td>Chromosome</td><td>start</td><td></td><td>end</td><td>strand</td><td>MM</td><td>target_seq</td><td>PAM</td><td>alignment</td><td>distance</td><td>position</td><td>gene name</td><td>gene id</td></tr>
<tr><td>GU305914.1</td><td></td><td>81162</td><td>81184</td><td>-</td><td>0</td><td>GGGAGCTACGGTGCTCGAAC</td><td>CGG</td><td>||||||||[||||||||||||]PAM</td><td>NA</td><td>Intergenic</td><td>NA</td><td>NA</td></tr>
<tr><td></td><td></td><td></td><td></td><td></td><td></td><td></td><td></td><td></td><td></td><td></td><td></td><td></td></tr>
<tr><td>T28133</td><td>ACGGTGCTCGAACCGGAGTGCGG</td><td></td><td>1000</td><td></td><td></td><td></td><td></td><td></td><td></td><td></td><td></td><td></td></tr>
<tr><td>Oligo fwd</td><td>CACCACGGTGCTCGAACCGGAGTG</td><td></td><td></td><td></td><td></td><td></td><td></td><td></td><td></td><td></td><td></td><td></td></tr>
<tr><td>Oligo rev</td><td>AAACCACTCCGGTTCGAGCACCGT</td><td></td><td></td><td></td><td></td><td></td><td></td><td></td><td></td><td></td><td></td><td></td></tr>
<tr><td>Chromosome</td><td>start</td><td></td><td>end</td><td>strand</td><td>MM</td><td>target_seq</td><td>PAM</td><td>alignment</td><td>distance</td><td>position</td><td>gene name</td><td>gene id</td></tr>
<tr><td>GU305914.1</td><td></td><td>81155</td><td>81177</td><td>-</td><td>0</td><td>ACGGTGCTCGAACCGGAGTG</td><td>CGG</td><td>||||||||[||||||||||||]PAM</td><td>NA</td><td>Intergenic</td><td>NA</td><td>NA</td></tr>
<tr><td></td><td></td><td></td><td></td><td></td><td></td><td></td><td></td><td></td><td></td><td></td><td></td><td></td></tr>
<tr><td>T28134</td><td>CGGTGCTCGAACCGGAGTGCGGG</td><td></td><td>1000</td><td></td><td></td><td></td><td></td><td></td><td></td><td></td><td></td><td></td></tr>
<tr><td>Oligo fwd</td><td>CACCCGGTGCTCGAACCGGAGTGC</td><td></td><td></td><td></td><td></td><td></td><td></td><td></td><td></td><td></td><td></td><td></td></tr>
<tr><td>Oligo rev</td><td>AAACGCACTCCGGTTCGAGCACCG</td><td></td><td></td><td></td><td></td><td></td><td></td><td></td><td></td><td></td><td></td><td></td></tr>
<tr><td>Chromosome</td><td>start</td><td></td><td>end</td><td>strand</td><td>MM</td><td>target_seq</td><td>PAM</td><td>alignment</td><td>distance</td><td>position</td><td>gene name</td><td>gene id</td></tr>
<tr><td>GU305914.1</td><td></td><td>81154</td><td>81176</td><td>-</td><td>0</td><td>CGGTGCTCGAACCGGAGTGC</td><td>GGG</td><td>||||||||[||||||||||||]PAM</td><td>NA</td><td>Intergenic</td><td>NA</td><td>NA</td></tr>
<tr><td></td><td></td><td></td><td></td><td></td><td></td><td></td><td></td><td></td><td></td><td></td><td></td><td></td></tr>
<tr><td>T28135</td><td>TCTGTGCTACTCCACCCTCTTGG</td><td></td><td>1000</td><td></td><td></td><td></td><td></td><td></td><td></td><td></td><td></td><td></td></tr>
<tr><td>Oligo fwd</td><td>CACCTCTGTGCTACTCCACCCTCT</td><td></td><td></td><td></td><td></td><td></td><td></td><td></td><td></td><td></td><td></td><td></td></tr>
<tr><td>Oligo rev</td><td>AAACAGAGGGTGGAGTAGCACAGA</td><td></td><td></td><td></td><td></td><td></td><td></td><td></td><td></td><td></td><td></td><td></td></tr>
<tr><td>Chromosome</td><td>start</td><td></td><td>end</td><td>strand</td><td>MM</td><td>target_seq</td><td>PAM</td><td>alignment</td><td>distance</td><td>position</td><td>gene name</td><td>gene id</td></tr>
<tr><td>GU305914.1</td><td></td><td>81063</td><td>81085</td><td>-</td><td>0</td><td>TCTGTGCTACTCCACCCTCT</td><td>TGG</td><td>||||||||[||||||||||||]PAM</td><td>NA</td><td>Intergenic</td><td>NA</td><td>NA</td></tr>
<tr><td></td><td></td><td></td><td></td><td></td><td></td><td></td><td></td><td></td><td></td><td></td><td></td><td></td></tr>
<tr><td>T28136</td><td>GTGCTACTCCACCCTCTTGGTGG</td><td></td><td>1000</td><td></td><td></td><td></td><td></td><td></td><td></td><td></td><td></td><td></td></tr>
<tr><td>Oligo fwd</td><td>CACCGTGCTACTCCACCCTCTTGG</td><td></td><td></td><td></td><td></td><td></td><td></td><td></td><td></td><td></td><td></td><td></td></tr>
<tr><td>Oligo rev</td><td>AAACCCAAGAGGGTGGAGTAGCAC</td><td></td><td></td><td></td><td></td><td></td><td></td><td></td><td></td><td></td><td></td><td></td></tr>
<tr><td>Chromosome</td><td>start</td><td></td><td>end</td><td>strand</td><td>MM</td><td>target_seq</td><td>PAM</td><td>alignment</td><td>distance</td><td>position</td><td>gene name</td><td>gene id</td></tr>
<tr><td>GU305914.1</td><td></td><td>81060</td><td>81082</td><td>-</td><td>0</td><td>GTGCTACTCCACCCTCTTGG</td><td>TGG</td><td>||||||||[||||||||||||]PAM</td><td>NA</td><td>Intergenic</td><td>NA</td><td>NA</td></tr>
</table>

<table>
<tr><td>T28137</td><td>CTACTCCACCCTCTTGGTGGAGG</td><td>1000</td><td></td><td></td><td></td><td></td><td></td><td></td><td></td><td></td><td></td></tr>
<tr><td>Oligo fwd</td><td>CACCCTACTCCACCCTCTTGGTGG</td><td></td><td></td><td></td><td></td><td></td><td></td><td></td><td></td><td></td><td></td></tr>
<tr><td>Oligo rev</td><td>AAACCCACCAAGAGGGTGGAGTAG</td><td></td><td></td><td></td><td></td><td></td><td></td><td></td><td></td><td></td><td></td></tr>
<tr><td>Chromosome</td><td>start</td><td>end</td><td>strand</td><td>MM</td><td>target_seq</td><td>PAM</td><td>alignment</td><td>distance</td><td>position</td><td>gene name</td><td>gene id</td></tr>
<tr><td>GU305914.1</td><td>81057</td><td>81079</td><td>-</td><td>0</td><td>CTACTCCACCCTCTTGGTGG</td><td>AGG</td><td>||||||||[||||||||||||]PAM</td><td>NA</td><td>Intergenic</td><td>NA</td><td>NA</td></tr>
<tr><td>T28138</td><td>TACTCCACCCTCTTGGTGGAGGG</td><td>1000</td><td></td><td></td><td></td><td></td><td></td><td></td><td></td><td></td><td></td></tr>
<tr><td>Oligo fwd</td><td>CACCTACTCCACCCTCTTGGTGGA</td><td></td><td></td><td></td><td></td><td></td><td></td><td></td><td></td><td></td><td></td></tr>
<tr><td>Oligo rev</td><td>AAACTCCACCAAGAGGGTGGAGTA</td><td></td><td></td><td></td><td></td><td></td><td></td><td></td><td></td><td></td><td></td></tr>
<tr><td>Chromosome</td><td>start</td><td>end</td><td>strand</td><td>MM</td><td>target_seq</td><td>PAM</td><td>alignment</td><td>distance</td><td>position</td><td>gene name</td><td>gene id</td></tr>
<tr><td>GU305914.1</td><td>81056</td><td>81078</td><td>-</td><td>0</td><td>TACTCCACCCTCTTGGTGGA</td><td>GGG</td><td>||||||||[||||||||||||]PAM</td><td>NA</td><td>Intergenic</td><td>NA</td><td>NA</td></tr>
<tr><td>T28139</td><td>CTTGGTGGAGGGCTCCCCCGAGG</td><td>1000</td><td></td><td></td><td></td><td></td><td></td><td></td><td></td><td></td><td></td></tr>
<tr><td>Oligo fwd</td><td>CACCCTTGGTGGAGGGCTCCCCCG</td><td></td><td></td><td></td><td></td><td></td><td></td><td></td><td></td><td></td><td></td></tr>
<tr><td>Oligo rev</td><td>AAACCGGGGGAGCCCTCCACCAAG</td><td></td><td></td><td></td><td></td><td></td><td></td><td></td><td></td><td></td><td></td></tr>
<tr><td>Chromosome</td><td>start</td><td>end</td><td>strand</td><td>MM</td><td>target_seq</td><td>PAM</td><td>alignment</td><td>distance</td><td>position</td><td>gene name</td><td>gene id</td></tr>
<tr><td>GU305914.1</td><td>81045</td><td>81067</td><td>-</td><td>0</td><td>CTTGGTGGAGGGCTCCCCCG</td><td>AGG</td><td>||||||||[||||||||||||]PAM</td><td>NA</td><td>Intergenic</td><td>NA</td><td>NA</td></tr>
<tr><td>T28140</td><td>CTCCCCCGAGGTGCCTGAAAAGG</td><td>1000</td><td></td><td></td><td></td><td></td><td></td><td></td><td></td><td></td><td></td></tr>
<tr><td>Oligo fwd</td><td>CACCCTCCCCCGAGGTGCCTGAAA</td><td></td><td></td><td></td><td></td><td></td><td></td><td></td><td></td><td></td><td></td></tr>
<tr><td>Oligo rev</td><td>AAACTTTCAGGCACCTCGGGGGAG</td><td></td><td></td><td></td><td></td><td></td><td></td><td></td><td></td><td></td><td></td></tr>
<tr><td>Chromosome</td><td>start</td><td>end</td><td>strand</td><td>MM</td><td>target_seq</td><td>PAM</td><td>alignment</td><td>distance</td><td>position</td><td>gene name</td><td>gene id</td></tr>
<tr><td>GU305914.1</td><td>81033</td><td>81055</td><td>-</td><td>0</td><td>CTCCCCCGAGGTGCCTGAAA</td><td>AGG</td><td>||||||||[||||||||||||]PAM</td><td>NA</td><td>Intergenic</td><td>NA</td><td>NA</td></tr>
<tr><td>T28141</td><td>TGCCTGAAAAGGACGTCCTGAGG</td><td>1000</td><td></td><td></td><td></td><td></td><td></td><td></td><td></td><td></td><td></td></tr>
<tr><td>Oligo fwd</td><td>CACCTGCCTGAAAAGGACGTCCTG</td><td></td><td></td><td></td><td></td><td></td><td></td><td></td><td></td><td></td><td></td></tr>
<tr><td>Oligo rev</td><td>AAACCAGGACGTCCTTTTCAGGCA</td><td></td><td></td><td></td><td></td><td></td><td></td><td></td><td></td><td></td><td></td></tr>
<tr><td>Chromosome</td><td>start</td><td>end</td><td>strand</td><td>MM</td><td>target_seq</td><td>PAM</td><td>alignment</td><td>distance</td><td>position</td><td>gene name</td><td>gene id</td></tr>
<tr><td>GU305914.1</td><td>81022</td><td>81044</td><td>-</td><td>0</td><td>TGCCTGAAAAGGACGTCCTG</td><td>AGG</td><td>||||||||[||||||||||||]PAM</td><td>NA</td><td>Intergenic</td><td>NA</td><td>NA</td></tr>
<tr><td>T28142</td><td>GCCTGAAAAGGACGTCCTGAGGG</td><td>1000</td><td></td><td></td><td></td><td></td><td></td><td></td><td></td><td></td><td></td></tr>
<tr><td>Oligo fwd</td><td>CACCGCCTGAAAAGGACGTCCTGA</td><td></td><td></td><td></td><td></td><td></td><td></td><td></td><td></td><td></td><td></td></tr>
<tr><td>Oligo rev</td><td>AAACTCAGGACGTCCTTTTCAGGC</td><td></td><td></td><td></td><td></td><td></td><td></td><td></td><td></td><td></td><td></td></tr>
<tr><td>Chromosome</td><td>start</td><td>end</td><td>strand</td><td>MM</td><td>target_seq</td><td>PAM</td><td>alignment</td><td>distance</td><td>position</td><td>gene name</td><td>gene id</td></tr>
<tr><td>GU305914.1</td><td>81021</td><td>81043</td><td>-</td><td>0</td><td>GCCTGAAAAGGACGTCCTGA</td><td>GGG</td><td>||||||||[||||||||||||]PAM</td><td>NA</td><td>Intergenic</td><td>NA</td><td>NA</td></tr>
<tr><td>T28143</td><td>GACGTCCTGAGGGTAGAAATCGG</td><td>1000</td><td></td><td></td><td></td><td></td><td></td><td></td><td></td><td></td><td></td></tr>
<tr><td>Oligo fwd</td><td>CACCGACGTCCTGAGGGTAGAAAT</td><td></td><td></td><td></td><td></td><td></td><td></td><td></td><td></td><td></td><td></td></tr>
<tr><td>Oligo rev</td><td>AAACATTTCTACCCTCAGGACGTC</td><td></td><td></td><td></td><td></td><td></td><td></td><td></td><td></td><td></td><td></td></tr>
<tr><td>Chromosome</td><td>start</td><td>end</td><td>strand</td><td>MM</td><td>target_seq</td><td>PAM</td><td>alignment</td><td>distance</td><td>position</td><td>gene name</td><td>gene id</td></tr>
<tr><td>GU305914.1</td><td>81011</td><td>81033</td><td>-</td><td>0</td><td>GACGTCCTGAGGGTAGAAAT</td><td>CGG</td><td>||||||||[||||||||||||]PAM</td><td>NA</td><td>Intergenic</td><td>NA</td><td>NA</td></tr>
<tr><td>T28144</td><td>AAATCGGTGATCAGTGTCACCGG</td><td>1000</td><td></td><td></td><td></td><td></td><td></td><td></td><td></td><td></td><td></td></tr>
<tr><td>Oligo fwd</td><td>CACCAAATCGGTGATCAGTGTCAC</td><td></td><td></td><td></td><td></td><td></td><td></td><td></td><td></td><td></td><td></td></tr>
<tr><td>Oligo rev</td><td>AAACGTGACACTGATCACCGATTT</td><td></td><td></td><td></td><td></td><td></td><td></td><td></td><td></td><td></td><td></td></tr>
<tr><td>Chromosome</td><td>start</td><td>end</td><td>strand</td><td>MM</td><td>target_seq</td><td>PAM</td><td>alignment</td><td>distance</td><td>position</td><td>gene name</td><td>gene id</td></tr>
<tr><td>GU305914.1</td><td>80995</td><td>81017</td><td>-</td><td>0</td><td>AAATCGGTGATCAGTGTCAC</td><td>CGG</td><td>||||||||[||||||||||||]PAM</td><td>NA</td><td>Intergenic</td><td>NA</td><td>NA</td></tr>
<tr><td>T28145</td><td>ATCAGTGTCACCGGTTTGTGCGG</td><td>1000</td><td></td><td></td><td></td><td></td><td></td><td></td><td></td><td></td><td></td></tr>
<tr><td>Oligo fwd</td><td>CACCATCAGTGTCACCGGTTTGTG</td><td></td><td></td><td></td><td></td><td></td><td></td><td></td><td></td><td></td><td></td></tr>
<tr><td>Oligo rev</td><td>AAACCACAAACCGGTGACACTGAT</td><td></td><td></td><td></td><td></td><td></td><td></td><td></td><td></td><td></td><td></td></tr>
<tr><td>Chromosome</td><td>start</td><td>end</td><td>strand</td><td>MM</td><td>target_seq</td><td>PAM</td><td>alignment</td><td>distance</td><td>position</td><td>gene name</td><td>gene id</td></tr>
<tr><td>GU305914.1</td><td>80986</td><td>81008</td><td>-</td><td>0</td><td>ATCAGTGTCACCGGTTTGTG</td><td>CGG</td><td>||||||||[||||||||||||]PAM</td><td>NA</td><td>Intergenic</td><td>NA</td><td>NA</td></tr>
<tr><td>T28146</td><td>TCAGTGTCACCGGTTTGTGCGGG</td><td>1000</td><td></td><td></td><td></td><td></td><td></td><td></td><td></td><td></td><td></td></tr>
<tr><td>Oligo fwd</td><td>CACCTCAGTGTCACCGGTTTGTGC</td><td></td><td></td><td></td><td></td><td></td><td></td><td></td><td></td><td></td><td></td></tr>
<tr><td>Oligo rev</td><td>AAACGCACAAACCGGTGACACTGA</td><td></td><td></td><td></td><td></td><td></td><td></td><td></td><td></td><td></td><td></td></tr>
<tr><td>Chromosome</td><td>start</td><td>end</td><td>strand</td><td>MM</td><td>target_seq</td><td>PAM</td><td>alignment</td><td>distance</td><td>position</td><td>gene name</td><td>gene id</td></tr>
<tr><td>GU305914.1</td><td>80985</td><td>81007</td><td>-</td><td>0</td><td>TCAGTGTCACCGGTTTGTGC</td><td>GGG</td><td>||||||||[||||||||||||]PAM</td><td>NA</td><td>Intergenic</td><td>NA</td><td>NA</td></tr>
<tr><td>T28147</td><td>GAACGTACATCGCTCTCTGCTGG</td><td>1000</td><td></td><td></td><td></td><td></td><td></td><td></td><td></td><td></td><td></td></tr>
<tr><td>Oligo fwd</td><td>CACCGAACGTACATCGCTCTCTGC</td><td></td><td></td><td></td><td></td><td></td><td></td><td></td><td></td><td></td><td></td></tr>
<tr><td>Oligo rev</td><td>AAACGCAGAGAGCGATGTACGTTC</td><td></td><td></td><td></td><td></td><td></td><td></td><td></td><td></td><td></td><td></td></tr>
<tr><td>Chromosome</td><td>start</td><td>end</td><td>strand</td><td>MM</td><td>target_seq</td><td>PAM</td><td>alignment</td><td>distance</td><td>position</td><td>gene name</td><td>gene id</td></tr>
<tr><td>GU305914.1</td><td>80961</td><td>80983</td><td>-</td><td>0</td><td>GAACGTACATCGCTCTCTGC</td><td>TGG</td><td>||||||||[||||||||||||]PAM</td><td>NA</td><td>Intergenic</td><td>NA</td><td>NA</td></tr>
<tr><td>T28148</td><td>CTCTCTGCTGGCCGAGCTGCTGG</td><td>1000</td><td></td><td></td><td></td><td></td><td></td><td></td><td></td><td></td><td></td></tr>
<tr><td>Oligo fwd</td><td>CACCCTCTCTGCTGGCCGAGCTGC</td><td></td><td></td><td></td><td></td><td></td><td></td><td></td><td></td><td></td><td></td></tr>
<tr><td>Oligo rev</td><td>AAACGCAGCTCGGCCAGCAGAGAG</td><td></td><td></td><td></td><td></td><td></td><td></td><td></td><td></td><td></td><td></td></tr>
<tr><td>Chromosome</td><td>start</td><td>end</td><td>strand</td><td>MM</td><td>target_seq</td><td>PAM</td><td>alignment</td><td>distance</td><td>position</td><td>gene name</td><td>gene id</td></tr>
<tr><td>GU305914.1</td><td>80949</td><td>80971</td><td>-</td><td>0</td><td>CTCTCTGCTGGCCGAGCTGC</td><td>TGG</td><td>||||||||[||||||||||||]PAM</td><td>NA</td><td>Intergenic</td><td>NA</td><td>NA</td></tr>
<tr><td>T28149</td><td>TGGCCGAGCTGCTGGTACGATGG</td><td>1000</td><td></td><td></td><td></td><td></td><td></td><td></td><td></td><td></td><td></td></tr>
<tr><td>Oligo fwd</td><td>CACCTGGCCGAGCTGCTGGTACGA</td><td></td><td></td><td></td><td></td><td></td><td></td><td></td><td></td><td></td><td></td></tr>
<tr><td>Oligo rev</td><td>AAACTCGTACCAGCAGCTCGGCCA</td><td></td><td></td><td></td><td></td><td></td><td></td><td></td><td></td><td></td><td></td></tr>
<tr><td>Chromosome</td><td>start</td><td>end</td><td>strand</td><td>MM</td><td>target_seq</td><td>PAM</td><td>alignment</td><td>distance</td><td>position</td><td>gene name</td><td>gene id</td></tr>
<tr><td>GU305914.1</td><td>80941</td><td>80963</td><td>-</td><td>0</td><td>TGGCCGAGCTGCTGGTACGA</td><td>TGG</td><td>||||||||[||||||||||||]PAM</td><td>NA</td><td>Intergenic</td><td>NA</td><td>NA</td></tr>
<tr><td>T28150</td><td>ATGGCTAACCCAACGCAAGCTGG</td><td>1000</td><td></td><td></td><td></td><td></td><td></td><td></td><td></td><td></td><td></td></tr>
<tr><td>Oligo fwd</td><td>CACCATGGCTAACCCAACGCAAGC</td><td></td><td></td><td></td><td></td><td></td><td></td><td></td><td></td><td></td><td></td></tr>
<tr><td>Oligo rev</td><td>AAACGCTTGCGTTGGGTTAGCCAT</td><td></td><td></td><td></td><td></td><td></td><td></td><td></td><td></td><td></td><td></td></tr>
<tr><td>Chromosome</td><td>start</td><td>end</td><td>strand</td><td>MM</td><td>target_seq</td><td>PAM</td><td>alignment</td><td>distance</td><td>position</td><td>gene name</td><td>gene id</td></tr>
<tr><td>GU305914.1</td><td>80922</td><td>80944</td><td>-</td><td>0</td><td>ATGGCTAACCCAACGCAAGC</td><td>TGG</td><td>||||||||[||||||||||||]PAM</td><td>NA</td><td>Intergenic</td><td>NA</td><td>NA</td></tr>
<tr><td>T28151</td><td>TAACCCAACGCAAGCTGGTCCGG</td><td>1000</td><td></td><td></td><td></td><td></td><td></td><td></td><td></td><td></td><td></td></tr>
<tr><td>Oligo fwd</td><td>CACCTAACCCAACGCAAGCTGGTC</td><td></td><td></td><td></td><td></td><td></td><td></td><td></td><td></td><td></td><td></td></tr>
<tr><td>Oligo rev</td><td>AAACGACCAGCTTGCGTTGGGTTA</td><td></td><td></td><td></td><td></td><td></td><td></td><td></td><td></td><td></td><td></td></tr>
<tr><td>Chromosome</td><td>start</td><td>end</td><td>strand</td><td>MM</td><td>target_seq</td><td>PAM</td><td>alignment</td><td>distance</td><td>position</td><td>gene name</td><td>gene id</td></tr>
</table>

| | | | | | | | | | | | |
|---|---|---|---|---|---|---|---|---|---|---|---|
| GU305914.1 | 80917 | 80939 | - | 0 | TAACCCAACGCAAGCTGGTC | CGG | \|\|\|\|\|\|\|\|[\|\|\|\|\|\|\|\|\|\|\|\|]PAM | NA | Intergenic | NA | NA |
| T28152 | AACCCAACGCAAGCTGGTCCGGG | 1000 | | | | | | | | | |
| Oligo fwd | CACCAACCCAACGCAAGCTGGTCC | | | | | | | | | | |
| Oligo rev | AAACGGACCAGCTTGCGTTGGGTT | | | | | | | | | | |
| Chromosome | start | end | strand | MM | target_seq | PAM | alignment | distance | position | gene name | gene id |
| GU305914.1 | 80916 | 80938 | - | 0 | AACCCAACGCAAGCTGGTCC | GGG | \|\|\|\|\|\|\|\|[\|\|\|\|\|\|\|\|\|\|\|\|]PAM | NA | Intergenic | NA | NA |
| T28153 | CCAACGCAAGCTGGTCCGGGAGG | 1000 | | | | | | | | | |
| Oligo fwd | CACCCCAACGCAAGCTGGTCCGGG | | | | | | | | | | |
| Oligo rev | AAACCCCGGACCAGCTTGCGTTGG | | | | | | | | | | |
| Chromosome | start | end | strand | MM | target_seq | PAM | alignment | distance | position | gene name | gene id |
| GU305914.1 | 80913 | 80935 | - | 0 | CCAACGCAAGCTGGTCCGGG | AGG | \|\|\|\|\|\|\|\|[\|\|\|\|\|\|\|\|\|\|\|\|]PAM | NA | Intergenic | NA | NA |
| T28154 | AGTGCACCAACGAGATGCAACGG | 1000 | | | | | | | | | |
| Oligo fwd | CACCAGTGCACCAACGAGATGCAA | | | | | | | | | | |
| Oligo rev | AAACTTGCATCTCGTTGGTGCACT | | | | | | | | | | |
| Chromosome | start | end | strand | MM | target_seq | PAM | alignment | distance | position | gene name | gene id |
| GU305914.1 | 80881 | 80903 | - | 0 | AGTGCACCAACGAGATGCAA | CGG | \|\|\|\|\|\|\|\|[\|\|\|\|\|\|\|\|\|\|\|\|]PAM | NA | Intergenic | NA | NA |
| T28155 | CGAGATGCAACGGATGATCATGG | 1000 | | | | | | | | | |
| Oligo fwd | CACCCGAGATGCAACGGATGATCA | | | | | | | | | | |
| Oligo rev | AAACTGATCATCCGTTGCATCTCG | | | | | | | | | | |
| Chromosome | start | end | strand | MM | target_seq | PAM | alignment | distance | position | gene name | gene id |
| GU305914.1 | 80871 | 80893 | - | 0 | CGAGATGCAACGGATGATCA | TGG | \|\|\|\|\|\|\|\|[\|\|\|\|\|\|\|\|\|\|\|\|]PAM | NA | Intergenic | NA | NA |
| T28156 | GATCATGGACAAGCAGCAGCTGG | 1000 | | | | | | | | | |
| Oligo fwd | CACCGATCATGGACAAGCAGCAGC | | | | | | | | | | |
| Oligo rev | AAACGCTGCTGCTTGTCCATGATC | | | | | | | | | | |
| Chromosome | start | end | strand | MM | target_seq | PAM | alignment | distance | position | gene name | gene id |
| GU305914.1 | 80856 | 80878 | - | 0 | GATCATGGACAAGCAGCAGC | TGG | \|\|\|\|\|\|\|\|[\|\|\|\|\|\|\|\|\|\|\|\|]PAM | NA | Intergenic | NA | NA |
| T28157 | GTAACGTGCAACGCTTTCTACGG | 1000 | | | | | | | | | |
| Oligo fwd | CACCGTAACGTGCAACGCTTTCTA | | | | | | | | | | |
| Oligo rev | AAACTAGAAAGCGTTGCACGTTAC | | | | | | | | | | |
| Chromosome | start | end | strand | MM | target_seq | PAM | alignment | distance | position | gene name | gene id |
| GU305914.1 | 80825 | 80847 | - | 0 | GTAACGTGCAACGCTTTCTA | CGG | \|\|\|\|\|\|\|\|[\|\|\|\|\|\|\|\|\|\|\|\|]PAM | NA | Intergenic | NA | NA |
| T28158 | CAACGCTTTCTACGGTTTCACGG | 1000 | | | | | | | | | |
| Oligo fwd | CACCCAACGCTTTCTACGGTTTCA | | | | | | | | | | |
| Oligo rev | AAACTGAAACCGTAGAAAGCGTTG | | | | | | | | | | |
| Chromosome | start | end | strand | MM | target_seq | PAM | alignment | distance | position | gene name | gene id |
| GU305914.1 | 80817 | 80839 | - | 0 | CAACGCTTTCTACGGTTTCA | CGG | \|\|\|\|\|\|\|\|[\|\|\|\|\|\|\|\|\|\|\|\|]PAM | NA | Intergenic | NA | NA |
| T28159 | AACGCTTTCTACGGTTTCACGGG | 1000 | | | | | | | | | |
| Oligo fwd | CACCAACGCTTTCTACGGTTTCAC | | | | | | | | | | |
| Oligo rev | AAACGTGAAACCGTAGAAAGCGTT | | | | | | | | | | |
| Chromosome | start | end | strand | MM | target_seq | PAM | alignment | distance | position | gene name | gene id |
| GU305914.1 | 80816 | 80838 | - | 0 | AACGCTTTCTACGGTTTCAC | GGG | \|\|\|\|\|\|\|\|[\|\|\|\|\|\|\|\|\|\|\|\|]PAM | NA | Intergenic | NA | NA |
| T28160 | ACGCTTTCTACGGTTTCACGGGG | 1000 | | | | | | | | | |
| Oligo fwd | CACCACGCTTTCTACGGTTTCACG | | | | | | | | | | |
| Oligo rev | AAACCGTGAAACCGTAGAAAGCGT | | | | | | | | | | |
| Chromosome | start | end | strand | MM | target_seq | PAM | alignment | distance | position | gene name | gene id |
| GU305914.1 | 80815 | 80837 | - | 0 | ACGCTTTCTACGGTTTCACG | GGG | \|\|\|\|\|\|\|\|[\|\|\|\|\|\|\|\|\|\|\|\|]PAM | NA | Intergenic | NA | NA |
| T28161 | CGCTTTCTACGGTTTCACGGGGG | 1000 | | | | | | | | | |
| Oligo fwd | CACCCGCTTTCTACGGTTTCACGG | | | | | | | | | | |
| Oligo rev | AAACCCGTGAAACCGTAGAAAGCG | | | | | | | | | | |
| Chromosome | start | end | strand | MM | target_seq | PAM | alignment | distance | position | gene name | gene id |
| GU305914.1 | 80814 | 80836 | - | 0 | CGCTTTCTACGGTTTCACGG | GGG | \|\|\|\|\|\|\|\|[\|\|\|\|\|\|\|\|\|\|\|\|]PAM | NA | Intergenic | NA | NA |
| T28162 | CTACGGTTTCACGGGGGTAGCGG | 1000 | | | | | | | | | |
| Oligo fwd | CACCCTACGGTTTCACGGGGGTAG | | | | | | | | | | |
| Oligo rev | AAACCTACCCCCGTGAAACCGTAG | | | | | | | | | | |
| Chromosome | start | end | strand | MM | target_seq | PAM | alignment | distance | position | gene name | gene id |
| GU305914.1 | 80808 | 80830 | - | 0 | CTACGGTTTCACGGGGGTAG | CGG | \|\|\|\|\|\|\|\|[\|\|\|\|\|\|\|\|\|\|\|\|]PAM | NA | Intergenic | NA | NA |
| GU305914.1 | 223195 | 223217 | - | 4 | CGCCGGTTTCACGGGGATGG | CGG | \|--\|\|\|\|\|[\|\|\|\|\|\|\|\|-\|-\|]PAM | NA | Intergenic | NA | NA |
| T28163 | GGTTTCACGGGGGTAGCGGCCGG | 1000 | | | | | | | | | |
| Oligo fwd | CACCGGTTTCACGGGGGTAGCGGC | | | | | | | | | | |
| Oligo rev | AAACGCCGCTACCCCCGTGAAACC | | | | | | | | | | |
| Chromosome | start | end | strand | MM | target_seq | PAM | alignment | distance | position | gene name | gene id |
| GU305914.1 | 80804 | 80826 | - | 0 | GGTTTCACGGGGGTAGCGGC | CGG | \|\|\|\|\|\|\|\|[\|\|\|\|\|\|\|\|\|\|\|\|]PAM | NA | Intergenic | NA | NA |
| T28164 | GTTTCACGGGGGTAGCGGCCGGG | 1000 | | | | | | | | | |
| Oligo fwd | CACCGTTTCACGGGGGTAGCGGCC | | | | | | | | | | |
| Oligo rev | AAACGGCCGCTACCCCCGTGAAAC | | | | | | | | | | |
| Chromosome | start | end | strand | MM | target_seq | PAM | alignment | distance | position | gene name | gene id |
| GU305914.1 | 80803 | 80825 | - | 0 | GTTTCACGGGGGTAGCGGCC | GGG | \|\|\|\|\|\|\|\|[\|\|\|\|\|\|\|\|\|\|\|\|]PAM | NA | Intergenic | NA | NA |
| T28165 | GCCGCTTCTATCACCAAGATCGG | 1000 | | | | | | | | | |
| Oligo fwd | CACCGCCGCTTCTATCACCAAGAT | | | | | | | | | | |
| Oligo rev | AAACATCTTGGTGATAGAAGCGGC | | | | | | | | | | |
| Chromosome | start | end | strand | MM | target_seq | PAM | alignment | distance | position | gene name | gene id |
| GU305914.1 | 80759 | 80781 | - | 0 | GCCGCTTCTATCACCAAGAT | CGG | \|\|\|\|\|\|\|\|[\|\|\|\|\|\|\|\|\|\|\|\|]PAM | NA | Intergenic | NA | NA |
| T28166 | CTTCTATCACCAAGATCGGCAGG | 1000 | | | | | | | | | |
| Oligo fwd | CACCCTTCTATCACCAAGATCGGC | | | | | | | | | | |

Oligo rev AAACGCCGATCTTGGTGATAGAAG
Chromosome start end strand MM target_seq PAM alignment distance position gene name gene id
GU305914.1 80755 80777 - 0 CTTCTATCACCAAGATCGGC AGG ||||||||[||||||||||||]PAM NA Intergenic NA NA

T28167 TTCTATCACCAAGATCGGCAGGG 1000
Oligo fwd CACCTTCTATCACCAAGATCGGCA
Oligo rev AAACTGCCGATCTTGGTGATAGAA
Chromosome start end strand MM target_seq PAM alignment distance position gene name gene id
GU305914.1 80754 80776 - 0 TTCTATCACCAAGATCGGCA GGG ||||||||[||||||||||||]PAM NA Intergenic NA NA

T28168 GATATGTTGCTCGCGACCGCCGG 1000
Oligo fwd CACCGATATGTTGCTCGCGACCGC
Oligo rev AAACGCGGTCGCGAGCAACATATC
Chromosome start end strand MM target_seq PAM alignment distance position gene name gene id
GU305914.1 80732 80754 - 0 GATATGTTGCTCGCGACCGC CGG ||||||||[||||||||||||]PAM NA Intergenic NA NA

T28169 CGCGACCGCCGGCCATATCGAGG 1000
Oligo fwd CACCCGCGACCGCCGGCCATATCG
Oligo rev AAACCGATATGGCCGGCGGTCGCG
Chromosome start end strand MM target_seq PAM alignment distance position gene name gene id
GU305914.1 80721 80743 - 0 CGCGACCGCCGGCCATATCG AGG ||||||||[||||||||||||]PAM NA Intergenic NA NA

T28170 CCGCCGGCCATATCGAGGACCGG 1000
Oligo fwd CACCCCGCCGGCCATATCGAGGAC
Oligo rev AAACGTCCTCGATATGGCCGGCGG
Chromosome start end strand MM target_seq PAM alignment distance position gene name gene id
GU305914.1 80716 80738 - 0 CCGCCGGCCATATCGAGGAC CGG ||||||||[||||||||||||]PAM NA Intergenic NA NA

T28171 GATTTTCTCCGTACCGTCCTCGG 1000
Oligo fwd CACCGATTTTCTCCGTACCGTCCT
Oligo rev AAACAGGACGGTACGGAGAAAATC
Chromosome start end strand MM target_seq PAM alignment distance position gene name gene id
GU305914.1 80681 80703 - 0 GATTTTCTCCGTACCGTCCT CGG ||||||||[||||||||||||]PAM NA Intergenic NA NA

T28172 CGTCCTCGGACTCCCCCCCGAGG 1000
Oligo fwd CACCCGTCCTCGGACTCCCCCCCG
Oligo rev AAACCGGGGGGGAGTCCGAGGACG
Chromosome start end strand MM target_seq PAM alignment distance position gene name gene id
GU305914.1 80667 80689 - 0 CGTCCTCGGACTCCCCCCCG AGG ||||||||[||||||||||||]PAM NA Intergenic NA NA

T28173 CTATCGACCCCGAAGCCCTCCGG 1000
Oligo fwd CACCCTATCGACCCCGAAGCCCTC
Oligo rev AAACGAGGGCTTCGGGGTCGATAG
Chromosome start end strand MM target_seq PAM alignment distance position gene name gene id
GU305914.1 80644 80666 - 0 CTATCGACCCCGAAGCCCTC CGG ||||||||[||||||||||||]PAM NA Intergenic NA NA

T28174 TATCGACCCCGAAGCCCTCCGGG 1000
Oligo fwd CACCTATCGACCCCGAAGCCCTCC
Oligo rev AAACGGAGGGCTTCGGGGTCGATA
Chromosome start end strand MM target_seq PAM alignment distance position gene name gene id
GU305914.1 80643 80665 - 0 TATCGACCCCGAAGCCCTCC GGG ||||||||[||||||||||||]PAM NA Intergenic NA NA

T28175 CTCCGGGTCAAGATCATCTACGG 1000
Oligo fwd CACCCTCCGGGTCAAGATCATCTA
Oligo rev AAACTAGATGATCTTGACCCGGAG
Chromosome start end strand MM target_seq PAM alignment distance position gene name gene id
GU305914.1 80627 80649 - 0 CTCCGGGTCAAGATCATCTA CGG ||||||||[||||||||||||]PAM NA Intergenic NA NA

T28176 CGACACCGACAGTGTGTTTGCGG 1000
Oligo fwd CACCCGACACCGACAGTGTGTTTG
Oligo rev AAACCAAACACACTGTCGGTGTCG
Chromosome start end strand MM target_seq PAM alignment distance position gene name gene id
GU305914.1 80604 80626 - 0 CGACACCGACAGTGTGTTTG CGG ||||||||[||||||||||||]PAM NA Intergenic NA NA

T28177 AGTGTGTTTGCGGCTTTCTACGG 1000
Oligo fwd CACCAGTGTGTTTGCGGCTTTCTA
Oligo rev AAACTAGAAAGCCGCAAACACACT
Chromosome start end strand MM target_seq PAM alignment distance position gene name gene id
GU305914.1 80594 80616 - 0 AGTGTGTTTGCGGCTTTCTA CGG ||||||||[||||||||||||]PAM NA Intergenic NA NA

T28178 GGCTTTCTACGGCATCGACAAGG 1000
Oligo fwd CACCGGCTTTCTACGGCATCGACA
Oligo rev AAACTGTCGATGCCGTAGAAAGCC
Chromosome start end strand MM target_seq PAM alignment distance position gene name gene id
GU305914.1 80583 80605 - 0 GGCTTTCTACGGCATCGACA AGG ||||||||[||||||||||||]PAM NA Intergenic NA NA

T28179 CGACAAGGAAGCCCTTTTGAAGG 1000
Oligo fwd CACCCGACAAGGAAGCCCTTTTGA
Oligo rev AAACTCAAAAGGGCTTCCTTGTCG
Chromosome start end strand MM target_seq PAM alignment distance position gene name gene id
GU305914.1 80568 80590 - 0 CGACAAGGAAGCCCTTTTGA AGG ||||||||[||||||||||||]PAM NA Intergenic NA NA

T28180 GAAGCCCTTTTGAAGGCCGTTGG 1000
Oligo fwd CACCGAAGCCCTTTTGAAGGCCGT
Oligo rev AAACACGGCCTTCAAAAGGGCTTC
Chromosome start end strand MM target_seq PAM alignment distance position gene name gene id
GU305914.1 80561 80583 - 0 GAAGCCCTTTTGAAGGCCGT TGG ||||||||[||||||||||||]PAM NA Intergenic NA NA

T28181 TTATGATGATATGCAAGAAGAGG 1000

<table>
<tr><td>Oligo fwd</td><td>CACCTTATGATGATATGCAAGAAG</td><td></td><td></td><td></td><td></td><td></td><td></td><td></td><td></td><td></td><td></td></tr>
<tr><td>Oligo rev</td><td>AAACCTTCTTGCATATCATCATAA</td><td></td><td></td><td></td><td></td><td></td><td></td><td></td><td></td><td></td><td></td></tr>
<tr><td>Chromosome</td><td>start</td><td>end</td><td>strand</td><td>MM</td><td>target_seq</td><td>PAM</td><td>alignment</td><td>distance</td><td>position</td><td>gene name</td><td>gene id</td></tr>
<tr><td>GU305914.1</td><td>80461</td><td>80483</td><td>-</td><td>0</td><td>TTATGATGATATGCAAGAAG</td><td>AGG</td><td>||||||||[||||||||||||]PAM</td><td>NA</td><td>Intergenic</td><td>NA</td><td>NA</td></tr>
<tr><td>T28182</td><td>ATATGCAAGAAGAGGTATATCGG</td><td>1000</td><td></td><td></td><td></td><td></td><td></td><td></td><td></td><td></td><td></td></tr>
<tr><td>Oligo fwd</td><td>CACCATATGCAAGAAGAGGTATAT</td><td></td><td></td><td></td><td></td><td></td><td></td><td></td><td></td><td></td><td></td></tr>
<tr><td>Oligo rev</td><td>AAACATATACCTCTTCTTGCATAT</td><td></td><td></td><td></td><td></td><td></td><td></td><td></td><td></td><td></td><td></td></tr>
<tr><td>Chromosome</td><td>start</td><td>end</td><td>strand</td><td>MM</td><td>target_seq</td><td>PAM</td><td>alignment</td><td>distance</td><td>position</td><td>gene name</td><td>gene id</td></tr>
<tr><td>GU305914.1</td><td>80453</td><td>80475</td><td>-</td><td>0</td><td>ATATGCAAGAAGAGGTATAT</td><td>CGG</td><td>||||||||[||||||||||||]PAM</td><td>NA</td><td>Intergenic</td><td>NA</td><td>NA</td></tr>
<tr><td>T28183</td><td>CAAGAAGAGGTATATCGGCAAGG</td><td>1000</td><td></td><td></td><td></td><td></td><td></td><td></td><td></td><td></td><td></td></tr>
<tr><td>Oligo fwd</td><td>CACCCAAGAAGAGGTATATCGGCA</td><td></td><td></td><td></td><td></td><td></td><td></td><td></td><td></td><td></td><td></td></tr>
<tr><td>Oligo rev</td><td>AAACTGCCGATATACCTCTTCTTG</td><td></td><td></td><td></td><td></td><td></td><td></td><td></td><td></td><td></td><td></td></tr>
<tr><td>Chromosome</td><td>start</td><td>end</td><td>strand</td><td>MM</td><td>target_seq</td><td>PAM</td><td>alignment</td><td>distance</td><td>position</td><td>gene name</td><td>gene id</td></tr>
<tr><td>GU305914.1</td><td>80448</td><td>80470</td><td>-</td><td>0</td><td>CAAGAAGAGGTATATCGGCA</td><td>AGG</td><td>||||||||[||||||||||||]PAM</td><td>NA</td><td>Intergenic</td><td>NA</td><td>NA</td></tr>
<tr><td>T28184</td><td>AGGTATATCGGCAAGGTCCACGG</td><td>1000</td><td></td><td></td><td></td><td></td><td></td><td></td><td></td><td></td><td></td></tr>
<tr><td>Oligo fwd</td><td>CACCAGGTATATCGGCAAGGTCCA</td><td></td><td></td><td></td><td></td><td></td><td></td><td></td><td></td><td></td><td></td></tr>
<tr><td>Oligo rev</td><td>AAACTGGACCTTGCCGATATACCT</td><td></td><td></td><td></td><td></td><td></td><td></td><td></td><td></td><td></td><td></td></tr>
<tr><td>Chromosome</td><td>start</td><td>end</td><td>strand</td><td>MM</td><td>target_seq</td><td>PAM</td><td>alignment</td><td>distance</td><td>position</td><td>gene name</td><td>gene id</td></tr>
<tr><td>GU305914.1</td><td>80441</td><td>80463</td><td>-</td><td>0</td><td>AGGTATATCGGCAAGGTCCA</td><td>CGG</td><td>||||||||[||||||||||||]PAM</td><td>NA</td><td>Intergenic</td><td>NA</td><td>NA</td></tr>
<tr><td>T28185</td><td>TCCCAGAACCTCAGTATGAAAGG</td><td>1000</td><td></td><td></td><td></td><td></td><td></td><td></td><td></td><td></td><td></td></tr>
<tr><td>Oligo fwd</td><td>CACCTCCCAGAACCTCAGTATGAA</td><td></td><td></td><td></td><td></td><td></td><td></td><td></td><td></td><td></td><td></td></tr>
<tr><td>Oligo rev</td><td>AAACTTCATACTGAGGTTCTGGGA</td><td></td><td></td><td></td><td></td><td></td><td></td><td></td><td></td><td></td><td></td></tr>
<tr><td>Chromosome</td><td>start</td><td>end</td><td>strand</td><td>MM</td><td>target_seq</td><td>PAM</td><td>alignment</td><td>distance</td><td>position</td><td>gene name</td><td>gene id</td></tr>
<tr><td>GU305914.1</td><td>80417</td><td>80439</td><td>-</td><td>0</td><td>TCCCAGAACCTCAGTATGAA</td><td>AGG</td><td>||||||||[||||||||||||]PAM</td><td>NA</td><td>Intergenic</td><td>NA</td><td>NA</td></tr>
<tr><td>T28186</td><td>CTCGTTCGCCGCACCGCATGCGG</td><td>1000</td><td></td><td></td><td></td><td></td><td></td><td></td><td></td><td></td><td></td></tr>
<tr><td>Oligo fwd</td><td>CACCCTCGTTCGCCGCACCGCATG</td><td></td><td></td><td></td><td></td><td></td><td></td><td></td><td></td><td></td><td></td></tr>
<tr><td>Oligo rev</td><td>AAACCATGCGGTGCGGCGAACGAG</td><td></td><td></td><td></td><td></td><td></td><td></td><td></td><td></td><td></td><td></td></tr>
<tr><td>Chromosome</td><td>start</td><td>end</td><td>strand</td><td>MM</td><td>target_seq</td><td>PAM</td><td>alignment</td><td>distance</td><td>position</td><td>gene name</td><td>gene id</td></tr>
<tr><td>GU305914.1</td><td>80387</td><td>80409</td><td>-</td><td>0</td><td>CTCGTTCGCCGCACCGCATG</td><td>CGG</td><td>||||||||[||||||||||||]PAM</td><td>NA</td><td>Intergenic</td><td>NA</td><td>NA</td></tr>
<tr><td>T28187</td><td>CACCGCATGCGGTTTCGTCAAGG</td><td>1000</td><td></td><td></td><td></td><td></td><td></td><td></td><td></td><td></td><td></td></tr>
<tr><td>Oligo fwd</td><td>CACCCACCGCATGCGGTTTCGTCA</td><td></td><td></td><td></td><td></td><td></td><td></td><td></td><td></td><td></td><td></td></tr>
<tr><td>Oligo rev</td><td>AAACTGACGAAACCGCATGCGGTG</td><td></td><td></td><td></td><td></td><td></td><td></td><td></td><td></td><td></td><td></td></tr>
<tr><td>Chromosome</td><td>start</td><td>end</td><td>strand</td><td>MM</td><td>target_seq</td><td>PAM</td><td>alignment</td><td>distance</td><td>position</td><td>gene name</td><td>gene id</td></tr>
<tr><td>GU305914.1</td><td>80376</td><td>80398</td><td>-</td><td>0</td><td>CACCGCATGCGGTTTCGTCA</td><td>AGG</td><td>||||||||[||||||||||||]PAM</td><td>NA</td><td>Intergenic</td><td>NA</td><td>NA</td></tr>
<tr><td>T28188</td><td>CGCATGCGGTTTCGTCAAGGCGG</td><td>1000</td><td></td><td></td><td></td><td></td><td></td><td></td><td></td><td></td><td></td></tr>
<tr><td>Oligo fwd</td><td>CACCCGCATGCGGTTTCGTCAAGG</td><td></td><td></td><td></td><td></td><td></td><td></td><td></td><td></td><td></td><td></td></tr>
<tr><td>Oligo rev</td><td>AAACCCTTGACGAAACCGCATGCG</td><td></td><td></td><td></td><td></td><td></td><td></td><td></td><td></td><td></td><td></td></tr>
<tr><td>Chromosome</td><td>start</td><td>end</td><td>strand</td><td>MM</td><td>target_seq</td><td>PAM</td><td>alignment</td><td>distance</td><td>position</td><td>gene name</td><td>gene id</td></tr>
<tr><td>GU305914.1</td><td>80373</td><td>80395</td><td>-</td><td>0</td><td>CGCATGCGGTTTCGTCAAGG</td><td>CGG</td><td>||||||||[||||||||||||]PAM</td><td>NA</td><td>Intergenic</td><td>NA</td><td>NA</td></tr>
<tr><td>T28189</td><td>ATGCGGTTTCGTCAAGGCGGTGG</td><td>1000</td><td></td><td></td><td></td><td></td><td></td><td></td><td></td><td></td><td></td></tr>
<tr><td>Oligo fwd</td><td>CACCATGCGGTTTCGTCAAGGCGG</td><td></td><td></td><td></td><td></td><td></td><td></td><td></td><td></td><td></td><td></td></tr>
<tr><td>Oligo rev</td><td>AAACCCGCCTTGACGAAACCGCAT</td><td></td><td></td><td></td><td></td><td></td><td></td><td></td><td></td><td></td><td></td></tr>
<tr><td>Chromosome</td><td>start</td><td>end</td><td>strand</td><td>MM</td><td>target_seq</td><td>PAM</td><td>alignment</td><td>distance</td><td>position</td><td>gene name</td><td>gene id</td></tr>
<tr><td>GU305914.1</td><td>80370</td><td>80392</td><td>-</td><td>0</td><td>ATGCGGTTTCGTCAAGGCGG</td><td>TGG</td><td>||||||||[||||||||||||]PAM</td><td>NA</td><td>Intergenic</td><td>NA</td><td>NA</td></tr>
<tr><td>T28190</td><td>GGTGAGCGACGTCCTCCATATGG</td><td>1000</td><td></td><td></td><td></td><td></td><td></td><td></td><td></td><td></td><td></td></tr>
<tr><td>Oligo fwd</td><td>CACCGGTGAGCGACGTCCTCCATA</td><td></td><td></td><td></td><td></td><td></td><td></td><td></td><td></td><td></td><td></td></tr>
<tr><td>Oligo rev</td><td>AAACTATGGAGGACGTCGCTCACC</td><td></td><td></td><td></td><td></td><td></td><td></td><td></td><td></td><td></td><td></td></tr>
<tr><td>Chromosome</td><td>start</td><td>end</td><td>strand</td><td>MM</td><td>target_seq</td><td>PAM</td><td>alignment</td><td>distance</td><td>position</td><td>gene name</td><td>gene id</td></tr>
<tr><td>GU305914.1</td><td>80349</td><td>80371</td><td>-</td><td>0</td><td>GGTGAGCGACGTCCTCCATA</td><td>TGG</td><td>||||||||[||||||||||||]PAM</td><td>NA</td><td>Intergenic</td><td>NA</td><td>NA</td></tr>
<tr><td>T28191</td><td>TTTCAACGACGAGACCGTTTCGG</td><td>1000</td><td></td><td></td><td></td><td></td><td></td><td></td><td></td><td></td><td></td></tr>
<tr><td>Oligo fwd</td><td>CACCTTTCAACGACGAGACCGTTT</td><td></td><td></td><td></td><td></td><td></td><td></td><td></td><td></td><td></td><td></td></tr>
<tr><td>Oligo rev</td><td>AAACAAACGGTCTCGTCGTTGAAA</td><td></td><td></td><td></td><td></td><td></td><td></td><td></td><td></td><td></td><td></td></tr>
<tr><td>Chromosome</td><td>start</td><td>end</td><td>strand</td><td>MM</td><td>target_seq</td><td>PAM</td><td>alignment</td><td>distance</td><td>position</td><td>gene name</td><td>gene id</td></tr>
<tr><td>GU305914.1</td><td>80325</td><td>80347</td><td>-</td><td>0</td><td>TTTCAACGACGAGACCGTTT</td><td>CGG</td><td>||||||||[||||||||||||]PAM</td><td>NA</td><td>Intergenic</td><td>NA</td><td>NA</td></tr>
<tr><td>T28192</td><td>CAACGACGAGACCGTTTCGGAGG</td><td>1000</td><td></td><td></td><td></td><td></td><td></td><td></td><td></td><td></td><td></td></tr>
<tr><td>Oligo fwd</td><td>CACCCAACGACGAGACCGTTTCGG</td><td></td><td></td><td></td><td></td><td></td><td></td><td></td><td></td><td></td><td></td></tr>
<tr><td>Oligo rev</td><td>AAACCCGAAACGGTCTCGTCGTTG</td><td></td><td></td><td></td><td></td><td></td><td></td><td></td><td></td><td></td><td></td></tr>
<tr><td>Chromosome</td><td>start</td><td>end</td><td>strand</td><td>MM</td><td>target_seq</td><td>PAM</td><td>alignment</td><td>distance</td><td>position</td><td>gene name</td><td>gene id</td></tr>
<tr><td>GU305914.1</td><td>80322</td><td>80344</td><td>-</td><td>0</td><td>CAACGACGAGACCGTTTCGG</td><td>AGG</td><td>||||||||[||||||||||||]PAM</td><td>NA</td><td>Intergenic</td><td>NA</td><td>NA</td></tr>
<tr><td>T28193</td><td>AACGACGAGACCGTTTCGGAGGG</td><td>1000</td><td></td><td></td><td></td><td></td><td></td><td></td><td></td><td></td><td></td></tr>
<tr><td>Oligo fwd</td><td>CACCAACGACGAGACCGTTTCGGA</td><td></td><td></td><td></td><td></td><td></td><td></td><td></td><td></td><td></td><td></td></tr>
<tr><td>Oligo rev</td><td>AAACTCCGAAACGGTCTCGTCGTT</td><td></td><td></td><td></td><td></td><td></td><td></td><td></td><td></td><td></td><td></td></tr>
<tr><td>Chromosome</td><td>start</td><td>end</td><td>strand</td><td>MM</td><td>target_seq</td><td>PAM</td><td>alignment</td><td>distance</td><td>position</td><td>gene name</td><td>gene id</td></tr>
<tr><td>GU305914.1</td><td>80321</td><td>80343</td><td>-</td><td>0</td><td>AACGACGAGACCGTTTCGGA</td><td>GGG</td><td>||||||||[||||||||||||]PAM</td><td>NA</td><td>Intergenic</td><td>NA</td><td>NA</td></tr>
<tr><td>T28194</td><td>ACGACGAGACCGTTTCGGAGGGG</td><td>1000</td><td></td><td></td><td></td><td></td><td></td><td></td><td></td><td></td><td></td></tr>
<tr><td>Oligo fwd</td><td>CACCACGACGAGACCGTTTCGGAG</td><td></td><td></td><td></td><td></td><td></td><td></td><td></td><td></td><td></td><td></td></tr>
<tr><td>Oligo rev</td><td>AAACCTCCGAAACGGTCTCGTCGT</td><td></td><td></td><td></td><td></td><td></td><td></td><td></td><td></td><td></td><td></td></tr>
<tr><td>Chromosome</td><td>start</td><td>end</td><td>strand</td><td>MM</td><td>target_seq</td><td>PAM</td><td>alignment</td><td>distance</td><td>position</td><td>gene name</td><td>gene id</td></tr>
<tr><td>GU305914.1</td><td>80320</td><td>80342</td><td>-</td><td>0</td><td>ACGACGAGACCGTTTCGGAG</td><td>GGG</td><td>||||||||[||||||||||||]PAM</td><td>NA</td><td>Intergenic</td><td>NA</td><td>NA</td></tr>
<tr><td>T28195</td><td>AGGGGACTATGAAGCTTTCCCGG</td><td>1000</td><td></td><td></td><td></td><td></td><td></td><td></td><td></td><td></td><td></td></tr>
<tr><td>Oligo fwd</td><td>CACCAGGGGACTATGAAGCTTTCC</td><td></td><td></td><td></td><td></td><td></td><td></td><td></td><td></td><td></td><td></td></tr>
<tr><td>Oligo rev</td><td>AAACGGAAAGCTTCATAGTCCCCT</td><td></td><td></td><td></td><td></td><td></td><td></td><td></td><td></td><td></td><td></td></tr>
<tr><td>Chromosome</td><td>start</td><td>end</td><td>strand</td><td>MM</td><td>target_seq</td><td>PAM</td><td>alignment</td><td>distance</td><td>position</td><td>gene name</td><td>gene id</td></tr>
<tr><td>GU305914.1</td><td>80302</td><td>80324</td><td>-</td><td>0</td><td>AGGGGACTATGAAGCTTTCC</td><td>CGG</td><td>||||||||[||||||||||||]PAM</td><td>NA</td><td>Intergenic</td><td>NA</td><td>NA</td></tr>
</table>

T28196 TTCGACGACCTCAAGAAGAACGG 1000
Oligo fwd CACCTTCGACGACCTCAAGAAGAA
Oligo rev AAACTTCTTCTTGAGGTCGTCGAA
Chromosome start end strand MM target_seq PAM alignment distance position gene name gene id
GU305914.1 80273 80295 - 0 TTCGACGACCTCAAGAAGAA CGG ||||||||[||||||||||||]PAM NA Intergenic NA NA

T28197 AACGGTATCCCATGTGAGTTCGG 1000
Oligo fwd CACCAACGGTATCCCATGTGAGTT
Oligo rev AAACAACTCACATGGGATACCGTT
Chromosome start end strand MM target_seq PAM alignment distance position gene name gene id
GU305914.1 80255 80277 - 0 AACGGTATCCCATGTGAGTT CGG ||||||||[||||||||||||]PAM NA Intergenic NA NA

T28198 ACGGTATCCCATGTGAGTTCGGG 1000
Oligo fwd CACCACGGTATCCCATGTGAGTTC
Oligo rev AAACGAACTCACATGGGATACCGT
Chromosome start end strand MM target_seq PAM alignment distance position gene name gene id
GU305914.1 80254 80276 - 0 ACGGTATCCCATGTGAGTTC GGG ||||||||[||||||||||||]PAM NA Intergenic NA NA

T28199 TCAGCCGCCTCTGCCGCGCCAGG 1000
Oligo fwd CACCTCAGCCGCCTCTGCCGCGCC
Oligo rev AAACGGCGCGGCAGAGGCGGCTGA
Chromosome start end strand MM target_seq PAM alignment distance position gene name gene id
GU305914.1 80224 80246 - 0 TCAGCCGCCTCTGCCGCGCC AGG ||||||||[||||||||||||]PAM NA Intergenic NA NA

T28200 CAGCCGCCTCTGCCGCGCCAGGG 1000
Oligo fwd CACCCAGCCGCCTCTGCCGCGCCA
Oligo rev AAACTGGCGCGGCAGAGGCGGCTG
Chromosome start end strand MM target_seq PAM alignment distance position gene name gene id
GU305914.1 80223 80245 - 0 CAGCCGCCTCTGCCGCGCCA GGG ||||||||[||||||||||||]PAM NA Intergenic NA NA

T28201 GGACGACCTGCACCTGAAGAAGG 1000
Oligo fwd CACCGGACGACCTGCACCTGAAGA
Oligo rev AAACTCTTCAGGTGCAGGTCGTCC
Chromosome start end strand MM target_seq PAM alignment distance position gene name gene id
GU305914.1 80202 80224 - 0 GGACGACCTGCACCTGAAGA AGG ||||||||[||||||||||||]PAM NA Intergenic NA NA

T28202 CCTTGCCGTCATCCGACGTCTGG 1000
Oligo fwd CACCCCTTGCCGTCATCCGACGTC
Oligo rev AAACGACGTCGGATGACGGCAAGG
Chromosome start end strand MM target_seq PAM alignment distance position gene name gene id
GU305914.1 80103 80125 - 0 CCTTGCCGTCATCCGACGTC TGG ||||||||[||||||||||||]PAM NA Intergenic NA NA

T28203 TGCCGTCATCCGACGTCTGGCGG 1000
Oligo fwd CACCTGCCGTCATCCGACGTCTGG
Oligo rev AAACCCAGACGTCGGATGACGGCA
Chromosome start end strand MM target_seq PAM alignment distance position gene name gene id
GU305914.1 80100 80122 - 0 TGCCGTCATCCGACGTCTGG CGG ||||||||[||||||||||||]PAM NA Intergenic NA NA

T28204 TCATCCGACGTCTGGCGGCTCGG 1000
Oligo fwd CACCTCATCCGACGTCTGGCGGCT
Oligo rev AAACAGCCGCCAGACGTCGGATGA
Chromosome start end strand MM target_seq PAM alignment distance position gene name gene id
GU305914.1 80095 80117 - 0 TCATCCGACGTCTGGCGGCT CGG ||||||||[||||||||||||]PAM NA Intergenic NA NA

T28205 CCGACGTCTGGCGGCTCGGAAGG 1000
Oligo fwd CACCCCGACGTCTGGCGGCTCGGA
Oligo rev AAACTCCGAGCCGCCAGACGTCGG
Chromosome start end strand MM target_seq PAM alignment distance position gene name gene id
GU305914.1 80091 80113 - 0 CCGACGTCTGGCGGCTCGGA AGG ||||||||[||||||||||||]PAM NA Intergenic NA NA

T28206 ACGTCTGGCGGCTCGGAAGGAGG 1000
Oligo fwd CACCACGTCTGGCGGCTCGGAAGG
Oligo rev AAACCCTTCCGAGCCGCCAGACGT
Chromosome start end strand MM target_seq PAM alignment distance position gene name gene id
GU305914.1 80088 80110 - 0 ACGTCTGGCGGCTCGGAAGG AGG ||||||||[||||||||||||]PAM NA Intergenic NA NA

T28207 TCGGAAGGAGGAACTGCCCGCGG 1000
Oligo fwd CACCTCGGAAGGAGGAACTGCCCG
Oligo rev AAACCGGGCAGTTCCTCCTTCCGA
Chromosome start end strand MM target_seq PAM alignment distance position gene name gene id
GU305914.1 80076 80098 - 0 TCGGAAGGAGGAACTGCCCG CGG ||||||||[||||||||||||]PAM NA Intergenic NA NA

T28208 AAGGAGGAACTGCCCGCGGTCGG 1000
Oligo fwd CACCAAGGAGGAACTGCCCGCGGT
Oligo rev AAACACCGCGGGCAGTTCCTCCTT
Chromosome start end strand MM target_seq PAM alignment distance position gene name gene id
GU305914.1 80072 80094 - 0 AAGGAGGAACTGCCCGCGGT CGG ||||||||[||||||||||||]PAM NA Intergenic NA NA

T28209 TACGTCCTCACCCTGCCCGACGG 1000
Oligo fwd CACCTACGTCCTCACCCTGCCCGA
Oligo rev AAACTCGGGCAGGGTGAGGACGTA
Chromosome start end strand MM target_seq PAM alignment distance position gene name gene id
GU305914.1 80036 80058 - 0 TACGTCCTCACCCTGCCCGA CGG ||||||||[||||||||||||]PAM NA Intergenic NA NA

T28210 TCCCAACTACGAGATCGCCGAGG 1000
Oligo fwd CACCTCCCAACTACGAGATCGCCG
Oligo rev AAACCGGCGATCTCGTAGTTGGGA
Chromosome start end strand MM target_seq PAM alignment distance position gene name gene id
GU305914.1 79998 80020 - 0 TCCCAACTACGAGATCGCCG AGG ||||||||[||||||||||||]PAM NA Intergenic NA NA

| | | | | | | | | | | | |
|---|---|---|---|---|---|---|---|---|---|---|---|
| T28211 | GGATCCGCGACACGTCGTCGAGG | 1000 | | | | | | | | | |
| Oligo fwd | CACCGGATCCGCGACACGTCGTCG | | | | | | | | | | |
| Oligo rev | AAACCGACGACGTGTCGCGGATCC | | | | | | | | | | |
| Chromosome | start | end | strand | MM | target_seq | PAM | alignment | distance | position | gene name | gene id |
| GU305914.1 | 79977 | 79999 | - | 0 | GGATCCGCGACACGTCGTCG | AGG | \|\|\|\|\|\|\|\|[\|\|\|\|\|\|\|\|\|\|\|\|]PAM | NA | Intergenic | NA | NA |
| GU305914.1 | 26766 | 26788 | + | 2 | CGATTCGCGACACGTCGTCG | TGG | -\|\|\|-\|\|\|[\|\|\|\|\|\|\|\|\|\|\|\|]PAM | NA | Intergenic | NA | NA |

| | | | | | | | | | | | |
|---|---|---|---|---|---|---|---|---|---|---|---|
| T28212 | CGCCGAAAAGTACTACGAGCAGG | 1000 | | | | | | | | | |
| Oligo fwd | CACCCGCCGAAAAGTACTACGAGC | | | | | | | | | | |
| Oligo rev | AAACGCTCGTAGTACTTTTCGGCG | | | | | | | | | | |
| Chromosome | start | end | strand | MM | target_seq | PAM | alignment | distance | position | gene name | gene id |
| GU305914.1 | 79938 | 79960 | - | 0 | CGCCGAAAAGTACTACGAGC | AGG | \|\|\|\|\|\|\|\|[\|\|\|\|\|\|\|\|\|\|\|\|]PAM | NA | Intergenic | NA | NA |

| | | | | | | | | | | | |
|---|---|---|---|---|---|---|---|---|---|---|---|
| T28213 | GTACTACGAGCAGGTCGTCAAGG | 1000 | | | | | | | | | |
| Oligo fwd | CACCGTACTACGAGCAGGTCGTCA | | | | | | | | | | |
| Oligo rev | AAACTGACGACCTGCTCGTAGTAC | | | | | | | | | | |
| Chromosome | start | end | strand | MM | target_seq | PAM | alignment | distance | position | gene name | gene id |
| GU305914.1 | 79929 | 79951 | - | 0 | GTACTACGAGCAGGTCGTCA | AGG | \|\|\|\|\|\|\|\|[\|\|\|\|\|\|\|\|\|\|\|\|]PAM | NA | Intergenic | NA | NA |

| | | | | | | | | | | | |
|---|---|---|---|---|---|---|---|---|---|---|---|
| T28214 | CGCTCATGCCCGTCTTCCCCCGG | 1000 | | | | | | | | | |
| Oligo fwd | CACCCGCTCATGCCCGTCTTCCCC | | | | | | | | | | |
| Oligo rev | AAACGGGGAAGACGGGCATGAGCG | | | | | | | | | | |
| Chromosome | start | end | strand | MM | target_seq | PAM | alignment | distance | position | gene name | gene id |
| GU305914.1 | 79894 | 79916 | - | 0 | CGCTCATGCCCGTCTTCCCC | CGG | \|\|\|\|\|\|\|\|[\|\|\|\|\|\|\|\|\|\|\|\|]PAM | NA | Intergenic | NA | NA |

| | | | | | | | | | | | |
|---|---|---|---|---|---|---|---|---|---|---|---|
| T28215 | GCTCATGCCCGTCTTCCCCCGGG | 1000 | | | | | | | | | |
| Oligo fwd | CACCGCTCATGCCCGTCTTCCCCC | | | | | | | | | | |
| Oligo rev | AAACGGGGGAAGACGGGCATGAGC | | | | | | | | | | |
| Chromosome | start | end | strand | MM | target_seq | PAM | alignment | distance | position | gene name | gene id |
| GU305914.1 | 79893 | 79915 | - | 0 | GCTCATGCCCGTCTTCCCCC | GGG | \|\|\|\|\|\|\|\|[\|\|\|\|\|\|\|\|\|\|\|\|]PAM | NA | Intergenic | NA | NA |

| | | | | | | | | | | | |
|---|---|---|---|---|---|---|---|---|---|---|---|
| T28216 | TCCCCCGGGACATGCCCAAGAGG | 1000 | | | | | | | | | |
| Oligo fwd | CACCTCCCCCGGGACATGCCCAAG | | | | | | | | | | |
| Oligo rev | AAACCTTGGGCATGTCCCGGGGGA | | | | | | | | | | |
| Chromosome | start | end | strand | MM | target_seq | PAM | alignment | distance | position | gene name | gene id |
| GU305914.1 | 79879 | 79901 | - | 0 | TCCCCCGGGACATGCCCAAG | AGG | \|\|\|\|\|\|\|\|[\|\|\|\|\|\|\|\|\|\|\|\|]PAM | NA | Intergenic | NA | NA |

| | | | | | | | | | | | |
|---|---|---|---|---|---|---|---|---|---|---|---|
| T28217 | CCCCCGGGACATGCCCAAGAGGG | 1000 | | | | | | | | | |
| Oligo fwd | CACCCCCCCGGGACATGCCCAAGA | | | | | | | | | | |
| Oligo rev | AAACTCTTGGGCATGTCCCGGGGG | | | | | | | | | | |
| Chromosome | start | end | strand | MM | target_seq | PAM | alignment | distance | position | gene name | gene id |
| GU305914.1 | 79878 | 79900 | - | 0 | CCCCCGGGACATGCCCAAGA | GGG | \|\|\|\|\|\|\|\|[\|\|\|\|\|\|\|\|\|\|\|\|]PAM | NA | Intergenic | NA | NA |

| | | | | | | | | | | | |
|---|---|---|---|---|---|---|---|---|---|---|---|
| T28218 | GGAGAAGTTCTTCTCCCTCGTGG | 1000 | | | | | | | | | |
| Oligo fwd | CACCGGAGAAGTTCTTCTCCCTCG | | | | | | | | | | |
| Oligo rev | AAACCGAGGGAGAAGAACTTCTCC | | | | | | | | | | |
| Chromosome | start | end | strand | MM | target_seq | PAM | alignment | distance | position | gene name | gene id |
| GU305914.1 | 79857 | 79879 | - | 0 | GGAGAAGTTCTTCTCCCTCG | TGG | \|\|\|\|\|\|\|\|[\|\|\|\|\|\|\|\|\|\|\|\|]PAM | NA | Intergenic | NA | NA |
| GU305914.1 | 79864 | 79886 | + | 3 | GGAGAAGAACTTCTCCCTCT | TGG | \|\|\|\|\|\|\|-[-\|\|\|\|\|\|\|\|\|\|-]PAM | NA | Intergenic | NA | NA |

| | | | | | | | | | | | |
|---|---|---|---|---|---|---|---|---|---|---|---|
| T28219 | TCTCCCTCGTGGTGCCCCAGAGG | 1000 | | | | | | | | | |
| Oligo fwd | CACCTCTCCCTCGTGGTGCCCCAG | | | | | | | | | | |
| Oligo rev | AAACCTGGGGCACCACGAGGGAGA | | | | | | | | | | |
| Chromosome | start | end | strand | MM | target_seq | PAM | alignment | distance | position | gene name | gene id |
| GU305914.1 | 79846 | 79868 | - | 0 | TCTCCCTCGTGGTGCCCCAG | AGG | \|\|\|\|\|\|\|\|[\|\|\|\|\|\|\|\|\|\|\|\|]PAM | NA | Intergenic | NA | NA |

| | | | | | | | | | | | |
|---|---|---|---|---|---|---|---|---|---|---|---|
| T28220 | GACCAGTTCCTACACCTCTGTGG | 1000 | | | | | | | | | |
| Oligo fwd | CACCGACCAGTTCCTACACCTCTG | | | | | | | | | | |
| Oligo rev | AAACCAGAGGTGTAGGAACTGGTC | | | | | | | | | | |
| Chromosome | start | end | strand | MM | target_seq | PAM | alignment | distance | position | gene name | gene id |
| GU305914.1 | 79811 | 79833 | - | 0 | GACCAGTTCCTACACCTCTG | TGG | \|\|\|\|\|\|\|\|[\|\|\|\|\|\|\|\|\|\|\|\|]PAM | NA | Intergenic | NA | NA |

| | | | | | | | | | | | |
|---|---|---|---|---|---|---|---|---|---|---|---|
| T28221 | GAAACGTGAACGAACTCGCGAGG | 1000 | | | | | | | | | |
| Oligo fwd | CACCGAAACGTGAACGAACTCGCG | | | | | | | | | | |
| Oligo rev | AAACCGCGAGTTCGTTCACGTTTC | | | | | | | | | | |
| Chromosome | start | end | strand | MM | target_seq | PAM | alignment | distance | position | gene name | gene id |
| GU305914.1 | 79789 | 79811 | - | 0 | GAAACGTGAACGAACTCGCG | AGG | \|\|\|\|\|\|\|\|[\|\|\|\|\|\|\|\|\|\|\|\|]PAM | NA | Intergenic | NA | NA |

| | | | | | | | | | | | |
|---|---|---|---|---|---|---|---|---|---|---|---|
| T28222 | AAACGTGAACGAACTCGCGAGGG | 1000 | | | | | | | | | |
| Oligo fwd | CACCAAACGTGAACGAACTCGCGA | | | | | | | | | | |
| Oligo rev | AAACTCGCGAGTTCGTTCACGTTT | | | | | | | | | | |
| Chromosome | start | end | strand | MM | target_seq | PAM | alignment | distance | position | gene name | gene id |
| GU305914.1 | 79788 | 79810 | - | 0 | AAACGTGAACGAACTCGCGA | GGG | \|\|\|\|\|\|\|\|[\|\|\|\|\|\|\|\|\|\|\|\|]PAM | NA | Intergenic | NA | NA |

| | | | | | | | | | | | |
|---|---|---|---|---|---|---|---|---|---|---|---|
| T28223 | AACGTGAACGAACTCGCGAGGGG | 1000 | | | | | | | | | |
| Oligo fwd | CACCAACGTGAACGAACTCGCGAG | | | | | | | | | | |
| Oligo rev | AAACCTCGCGAGTTCGTTCACGTT | | | | | | | | | | |
| Chromosome | start | end | strand | MM | target_seq | PAM | alignment | distance | position | gene name | gene id |
| GU305914.1 | 79787 | 79809 | - | 0 | AACGTGAACGAACTCGCGAG | GGG | \|\|\|\|\|\|\|\|[\|\|\|\|\|\|\|\|\|\|\|\|]PAM | NA | Intergenic | NA | NA |

| | | | | | | | | | | | |
|---|---|---|---|---|---|---|---|---|---|---|---|
| T28224 | GTGAACGAACTCGCGAGGGGAGG | 1000 | | | | | | | | | |
| Oligo fwd | CACCGTGAACGAACTCGCGAGGGG | | | | | | | | | | |
| Oligo rev | AAACCCCCTCGCGAGTTCGTTCAC | | | | | | | | | | |
| Chromosome | start | end | strand | MM | target_seq | PAM | alignment | distance | position | gene name | gene id |
| GU305914.1 | 79784 | 79806 | - | 0 | GTGAACGAACTCGCGAGGGG | AGG | \|\|\|\|\|\|\|\|[\|\|\|\|\|\|\|\|\|\|\|\|]PAM | NA | Intergenic | NA | NA |

| | | |
|---|---|---|
| T28225 | TGAACGAACTCGCGAGGGGAGGG | 1000 |
| Oligo fwd | CACCTGAACGAACTCGCGAGGGGA | |

| | | | | | | | | | | | |
|---|---|---|---|---|---|---|---|---|---|---|---|
| Oligo rev | AAACTCCCCTCGCGAGTTCGTTCA | | | | | | | | | | |
| Chromosome | start | end | strand | MM | target_seq | PAM | alignment | distance | position | gene name | gene id |
| GU305914.1 | 79783 | 79805 | - | 0 | TGAACGAACTCGCGAGGGGA | GGG | \|\|\|\|\|\|\|\|[\|\|\|\|\|\|\|\|\|\|\|\|]PAM | NA | Intergenic | NA | NA |
| T28226 | GAACGAACTCGCGAGGGGAGGGG | 1000 | | | | | | | | | |
| Oligo fwd | CACCGAACGAACTCGCGAGGGGAG | | | | | | | | | | |
| Oligo rev | AAACCTCCCCTCGCGAGTTCGTTC | | | | | | | | | | |
| Chromosome | start | end | strand | MM | target_seq | PAM | alignment | distance | position | gene name | gene id |
| GU305914.1 | 79782 | 79804 | - | 0 | GAACGAACTCGCGAGGGGAG | GGG | \|\|\|\|\|\|\|\|[\|\|\|\|\|\|\|\|\|\|\|\|]PAM | NA | Intergenic | NA | NA |
| T28227 | AGGGGAGGGGACGACAGCGACGG | 1000 | | | | | | | | | |
| Oligo fwd | CACCAGGGGAGGGGACGACAGCGA | | | | | | | | | | |
| Oligo rev | AAACTCGCTGTCGTCCCCTCCCCT | | | | | | | | | | |
| Chromosome | start | end | strand | MM | target_seq | PAM | alignment | distance | position | gene name | gene id |
| GU305914.1 | 79769 | 79791 | - | 0 | AGGGGAGGGGACGACAGCGA | CGG | \|\|\|\|\|\|\|\|[\|\|\|\|\|\|\|\|\|\|\|\|]PAM | NA | Intergenic | NA | NA |
| T28228 | GGGGAGGGGACGACAGCGACGGG | 1000 | | | | | | | | | |
| Oligo fwd | CACCGGGGAGGGGACGACAGCGAC | | | | | | | | | | |
| Oligo rev | AAACGTCGCTGTCGTCCCCTCCCC | | | | | | | | | | |
| Chromosome | start | end | strand | MM | target_seq | PAM | alignment | distance | position | gene name | gene id |
| GU305914.1 | 79768 | 79790 | - | 0 | GGGGAGGGGACGACAGCGAC | GGG | \|\|\|\|\|\|\|\|[\|\|\|\|\|\|\|\|\|\|\|\|]PAM | NA | Intergenic | NA | NA |
| T28229 | GGGAGGGGACGACAGCGACGGGG | 1000 | | | | | | | | | |
| Oligo fwd | CACCGGGAGGGGACGACAGCGACG | | | | | | | | | | |
| Oligo rev | AAACCGTCGCTGTCGTCCCCTCCC | | | | | | | | | | |
| Chromosome | start | end | strand | MM | target_seq | PAM | alignment | distance | position | gene name | gene id |
| GU305914.1 | 79767 | 79789 | - | 0 | GGGAGGGGACGACAGCGACG | GGG | \|\|\|\|\|\|\|\|[\|\|\|\|\|\|\|\|\|\|\|\|]PAM | NA | Intergenic | NA | NA |
| T28230 | GGAGGGGACGACAGCGACGGGGG | 1000 | | | | | | | | | |
| Oligo fwd | CACCGGAGGGGACGACAGCGACGG | | | | | | | | | | |
| Oligo rev | AAACCCGTCGCTGTCGTCCCCTCC | | | | | | | | | | |
| Chromosome | start | end | strand | MM | target_seq | PAM | alignment | distance | position | gene name | gene id |
| GU305914.1 | 79766 | 79788 | - | 0 | GGAGGGGACGACAGCGACGG | GGG | \|\|\|\|\|\|\|\|[\|\|\|\|\|\|\|\|\|\|\|\|]PAM | NA | Intergenic | NA | NA |
| T28231 | GAGGGGACGACAGCGACGGGGGG | 1000 | | | | | | | | | |
| Oligo fwd | CACCGAGGGGACGACAGCGACGGG | | | | | | | | | | |
| Oligo rev | AAACCCCGTCGCTGTCGTCCCCTC | | | | | | | | | | |
| Chromosome | start | end | strand | MM | target_seq | PAM | alignment | distance | position | gene name | gene id |
| GU305914.1 | 79765 | 79787 | - | 0 | GAGGGGACGACAGCGACGGG | GGG | \|\|\|\|\|\|\|\|[\|\|\|\|\|\|\|\|\|\|\|\|]PAM | NA | Intergenic | NA | NA |
| T28232 | AGGGGACGACAGCGACGGGGGGG | 1000 | | | | | | | | | |
| Oligo fwd | CACCAGGGGACGACAGCGACGGGG | | | | | | | | | | |
| Oligo rev | AAACCCCCGTCGCTGTCGTCCCCT | | | | | | | | | | |
| Chromosome | start | end | strand | MM | target_seq | PAM | alignment | distance | position | gene name | gene id |
| GU305914.1 | 79764 | 79786 | - | 0 | AGGGGACGACAGCGACGGGG | GGG | \|\|\|\|\|\|\|\|[\|\|\|\|\|\|\|\|\|\|\|\|]PAM | NA | Intergenic | NA | NA |
| T28233 | GGATAGTGAGAAAGAAAACATGG | 1000 | | | | | | | | | |
| Oligo fwd | CACCGGATAGTGAGAAAGAAAACA | | | | | | | | | | |
| Oligo rev | AAACTGTTTTCTTTCTCACTATCC | | | | | | | | | | |
| Chromosome | start | end | strand | MM | target_seq | PAM | alignment | distance | position | gene name | gene id |
| GU305914.1 | 79743 | 79765 | - | 0 | GGATAGTGAGAAAGAAAACA | TGG | \|\|\|\|\|\|\|\|[\|\|\|\|\|\|\|\|\|\|\|\|]PAM | NA | Intergenic | NA | NA |
| T28234 | TGAGAAAGAAAACATGGACACGG | 1000 | | | | | | | | | |
| Oligo fwd | CACCTGAGAAAGAAAACATGGACA | | | | | | | | | | |
| Oligo rev | AAACTGTCCATGTTTTCTTTCTCA | | | | | | | | | | |
| Chromosome | start | end | strand | MM | target_seq | PAM | alignment | distance | position | gene name | gene id |
| GU305914.1 | 79737 | 79759 | - | 0 | TGAGAAAGAAAACATGGACA | CGG | \|\|\|\|\|\|\|\|[\|\|\|\|\|\|\|\|\|\|\|\|]PAM | NA | Intergenic | NA | NA |
| T28235 | AAGAAAACATGGACACGGAGCGG | 1000 | | | | | | | | | |
| Oligo fwd | CACCAAGAAAACATGGACACGGAG | | | | | | | | | | |
| Oligo rev | AAACCTCCGTGTCCATGTTTTCTT | | | | | | | | | | |
| Chromosome | start | end | strand | MM | target_seq | PAM | alignment | distance | position | gene name | gene id |
| GU305914.1 | 79732 | 79754 | - | 0 | AAGAAAACATGGACACGGAG | CGG | \|\|\|\|\|\|\|\|[\|\|\|\|\|\|\|\|\|\|\|\|]PAM | NA | Intergenic | NA | NA |
| T28236 | GGAGCGGTCGTCCTCGCACGAGG | 1000 | | | | | | | | | |
| Oligo fwd | CACCGGAGCGGTCGTCCTCGCACG | | | | | | | | | | |
| Oligo rev | AAACCGTGCGAGGACGACCGCTCC | | | | | | | | | | |
| Chromosome | start | end | strand | MM | target_seq | PAM | alignment | distance | position | gene name | gene id |
| GU305914.1 | 79716 | 79738 | - | 0 | GGAGCGGTCGTCCTCGCACG | AGG | \|\|\|\|\|\|\|\|[\|\|\|\|\|\|\|\|\|\|\|\|]PAM | NA | Intergenic | NA | NA |
| GU305914.1 | 134861 | 134883 | - | 4 | GTCGCGGTCGTCCTCGCTAG | CGG | \|--\|\|\|\|\|[\|\|\|\|\|\|\|\|\|--\|]PAM | NA | Intergenic | NA | NA |
| T28237 | GTCGTCCTCGCACGAGGCGATGG | 1000 | | | | | | | | | |
| Oligo fwd | CACCGTCGTCCTCGCACGAGGCGA | | | | | | | | | | |
| Oligo rev | AAACTCGCCTCGTGCGAGGACGAC | | | | | | | | | | |
| Chromosome | start | end | strand | MM | target_seq | PAM | alignment | distance | position | gene name | gene id |
| GU305914.1 | 79710 | 79732 | - | 0 | GTCGTCCTCGCACGAGGCGA | TGG | \|\|\|\|\|\|\|\|[\|\|\|\|\|\|\|\|\|\|\|\|]PAM | NA | Intergenic | NA | NA |
| T28238 | AGGCGATGGAGACGTGATCCCGG | 1000 | | | | | | | | | |
| Oligo fwd | CACCAGGCGATGGAGACGTGATCC | | | | | | | | | | |
| Oligo rev | AAACGGATCACGTCTCCATCGCCT | | | | | | | | | | |
| Chromosome | start | end | strand | MM | target_seq | PAM | alignment | distance | position | gene name | gene id |
| GU305914.1 | 79696 | 79718 | - | 0 | AGGCGATGGAGACGTGATCC | CGG | \|\|\|\|\|\|\|\|[\|\|\|\|\|\|\|\|\|\|\|\|]PAM | NA | Intergenic | NA | NA |
| T28239 | GGCGATGGAGACGTGATCCCGGG | 1000 | | | | | | | | | |
| Oligo fwd | CACCGGCGATGGAGACGTGATCCC | | | | | | | | | | |
| Oligo rev | AAACGGGATCACGTCTCCATCGCC | | | | | | | | | | |
| Chromosome | start | end | strand | MM | target_seq | PAM | alignment | distance | position | gene name | gene id |
| GU305914.1 | 79695 | 79717 | - | 0 | GGCGATGGAGACGTGATCCC | GGG | \|\|\|\|\|\|\|\|[\|\|\|\|\|\|\|\|\|\|\|\|]PAM | NA | Intergenic | NA | NA |

<table>
<tr><td>T28240</td><td>GCGATGGAGACGTGATCCCGGGG</td><td>1000</td><td colspan="9"></td></tr>
<tr><td>Oligo fwd</td><td>CACCGCGATGGAGACGTGATCCCG</td><td colspan="10"></td></tr>
<tr><td>Oligo rev</td><td>AAACCGGGATCACGTCTCCATCGC</td><td colspan="10"></td></tr>
<tr><td>Chromosome</td><td>start</td><td>end</td><td>strand</td><td>MM</td><td>target_seq</td><td>PAM</td><td>alignment</td><td>distance</td><td>position</td><td>gene name</td><td>gene id</td></tr>
<tr><td>GU305914.1</td><td>79694</td><td>79716</td><td>-</td><td>0</td><td>GCGATGGAGACGTGATCCCG</td><td>GGG</td><td>||||||||[||||||||||||]PAM</td><td>NA</td><td>Intergenic</td><td>NA</td><td>NA</td></tr>
<tr><td>T28241</td><td>CGATGGAGACGTGATCCCGGGGG</td><td>1000</td><td colspan="9"></td></tr>
<tr><td>Oligo fwd</td><td>CACCCGATGGAGACGTGATCCCGG</td><td colspan="10"></td></tr>
<tr><td>Oligo rev</td><td>AAACCCGGGATCACGTCTCCATCG</td><td colspan="10"></td></tr>
<tr><td>Chromosome</td><td>start</td><td>end</td><td>strand</td><td>MM</td><td>target_seq</td><td>PAM</td><td>alignment</td><td>distance</td><td>position</td><td>gene name</td><td>gene id</td></tr>
<tr><td>GU305914.1</td><td>79693</td><td>79715</td><td>-</td><td>0</td><td>CGATGGAGACGTGATCCCGG</td><td>GGG</td><td>||||||||[||||||||||||]PAM</td><td>NA</td><td>Intergenic</td><td>NA</td><td>NA</td></tr>
<tr><td>T28242</td><td>TGTGACCGACAACAATGTCGCGG</td><td>1000</td><td colspan="9"></td></tr>
<tr><td>Oligo fwd</td><td>CACCTGTGACCGACAACAATGTCG</td><td colspan="10"></td></tr>
<tr><td>Oligo rev</td><td>AAACCGACATTGTTGTCGGTCACA</td><td colspan="10"></td></tr>
<tr><td>Chromosome</td><td>start</td><td>end</td><td>strand</td><td>MM</td><td>target_seq</td><td>PAM</td><td>alignment</td><td>distance</td><td>position</td><td>gene name</td><td>gene id</td></tr>
<tr><td>GU305914.1</td><td>79575</td><td>79597</td><td>-</td><td>0</td><td>TGTGACCGACAACAATGTCG</td><td>CGG</td><td>||||||||[||||||||||||]PAM</td><td>NA</td><td>Intergenic</td><td>NA</td><td>NA</td></tr>
<tr><td>T28243</td><td>GAACTTCTCGTTCGTCTCGTTGG</td><td>1000</td><td colspan="9"></td></tr>
<tr><td>Oligo fwd</td><td>CACCGAACTTCTCGTTCGTCTCGT</td><td colspan="10"></td></tr>
<tr><td>Oligo rev</td><td>AAACACGAGACGAACGAGAAGTTC</td><td colspan="10"></td></tr>
<tr><td>Chromosome</td><td>start</td><td>end</td><td>strand</td><td>MM</td><td>target_seq</td><td>PAM</td><td>alignment</td><td>distance</td><td>position</td><td>gene name</td><td>gene id</td></tr>
<tr><td>GU305914.1</td><td>79423</td><td>79445</td><td>-</td><td>0</td><td>GAACTTCTCGTTCGTCTCGT</td><td>TGG</td><td>||||||||[||||||||||||]PAM</td><td>NA</td><td>Intergenic</td><td>NA</td><td>NA</td></tr>
<tr><td>T28244</td><td>AACTTCTCGTTCGTCTCGTTGGG</td><td>1000</td><td colspan="9"></td></tr>
<tr><td>Oligo fwd</td><td>CACCAACTTCTCGTTCGTCTCGTT</td><td colspan="10"></td></tr>
<tr><td>Oligo rev</td><td>AAACAACGAGACGAACGAGAAGTT</td><td colspan="10"></td></tr>
<tr><td>Chromosome</td><td>start</td><td>end</td><td>strand</td><td>MM</td><td>target_seq</td><td>PAM</td><td>alignment</td><td>distance</td><td>position</td><td>gene name</td><td>gene id</td></tr>
<tr><td>GU305914.1</td><td>79422</td><td>79444</td><td>-</td><td>0</td><td>AACTTCTCGTTCGTCTCGTT</td><td>GGG</td><td>||||||||[||||||||||||]PAM</td><td>NA</td><td>Intergenic</td><td>NA</td><td>NA</td></tr>
<tr><td>T28245</td><td>ACTTCTCGTTCGTCTCGTTGGGG</td><td>1000</td><td colspan="9"></td></tr>
<tr><td>Oligo fwd</td><td>CACCACTTCTCGTTCGTCTCGTTG</td><td colspan="10"></td></tr>
<tr><td>Oligo rev</td><td>AAACCAACGAGACGAACGAGAAGT</td><td colspan="10"></td></tr>
<tr><td>Chromosome</td><td>start</td><td>end</td><td>strand</td><td>MM</td><td>target_seq</td><td>PAM</td><td>alignment</td><td>distance</td><td>position</td><td>gene name</td><td>gene id</td></tr>
<tr><td>GU305914.1</td><td>79421</td><td>79443</td><td>-</td><td>0</td><td>ACTTCTCGTTCGTCTCGTTG</td><td>GGG</td><td>||||||||[||||||||||||]PAM</td><td>NA</td><td>Intergenic</td><td>NA</td><td>NA</td></tr>
<tr><td>T28246</td><td>GCATCAGCACGTCGATGCGTAGG</td><td>1000</td><td colspan="9"></td></tr>
<tr><td>Oligo fwd</td><td>CACCGCATCAGCACGTCGATGCGT</td><td colspan="10"></td></tr>
<tr><td>Oligo rev</td><td>AAACACGCATCGACGTGCTGATGC</td><td colspan="10"></td></tr>
<tr><td>Chromosome</td><td>start</td><td>end</td><td>strand</td><td>MM</td><td>target_seq</td><td>PAM</td><td>alignment</td><td>distance</td><td>position</td><td>gene name</td><td>gene id</td></tr>
<tr><td>GU305914.1</td><td>79370</td><td>79392</td><td>-</td><td>0</td><td>GCATCAGCACGTCGATGCGT</td><td>AGG</td><td>||||||||[||||||||||||]PAM</td><td>NA</td><td>Intergenic</td><td>NA</td><td>NA</td></tr>
<tr><td>T28247</td><td>CATCAGCACGTCGATGCGTAGGG</td><td>1000</td><td colspan="9"></td></tr>
<tr><td>Oligo fwd</td><td>CACCCATCAGCACGTCGATGCGTA</td><td colspan="10"></td></tr>
<tr><td>Oligo rev</td><td>AAACTACGCATCGACGTGCTGATG</td><td colspan="10"></td></tr>
<tr><td>Chromosome</td><td>start</td><td>end</td><td>strand</td><td>MM</td><td>target_seq</td><td>PAM</td><td>alignment</td><td>distance</td><td>position</td><td>gene name</td><td>gene id</td></tr>
<tr><td>GU305914.1</td><td>79369</td><td>79391</td><td>-</td><td>0</td><td>CATCAGCACGTCGATGCGTA</td><td>GGG</td><td>||||||||[||||||||||||]PAM</td><td>NA</td><td>Intergenic</td><td>NA</td><td>NA</td></tr>
<tr><td>T28248</td><td>GCGCCGTCAGCGTGATCCGCTGG</td><td>1000</td><td colspan="9"></td></tr>
<tr><td>Oligo fwd</td><td>CACCGCGCCGTCAGCGTGATCCGC</td><td colspan="10"></td></tr>
<tr><td>Oligo rev</td><td>AAACGCGGATCACGCTGACGGCGC</td><td colspan="10"></td></tr>
<tr><td>Chromosome</td><td>start</td><td>end</td><td>strand</td><td>MM</td><td>target_seq</td><td>PAM</td><td>alignment</td><td>distance</td><td>position</td><td>gene name</td><td>gene id</td></tr>
<tr><td>GU305914.1</td><td>79322</td><td>79344</td><td>-</td><td>0</td><td>GCGCCGTCAGCGTGATCCGC</td><td>TGG</td><td>||||||||[||||||||||||]PAM</td><td>NA</td><td>Intergenic</td><td>NA</td><td>NA</td></tr>
<tr><td>T28249</td><td>TCCGCTGGTGCGCGATGTCCAGG</td><td>1000</td><td colspan="9"></td></tr>
<tr><td>Oligo fwd</td><td>CACCTCCGCTGGTGCGCGATGTCC</td><td colspan="10"></td></tr>
<tr><td>Oligo rev</td><td>AAACGGACATCGCGCACCAGCGGA</td><td colspan="10"></td></tr>
<tr><td>Chromosome</td><td>start</td><td>end</td><td>strand</td><td>MM</td><td>target_seq</td><td>PAM</td><td>alignment</td><td>distance</td><td>position</td><td>gene name</td><td>gene id</td></tr>
<tr><td>GU305914.1</td><td>79307</td><td>79329</td><td>-</td><td>0</td><td>TCCGCTGGTGCGCGATGTCC</td><td>AGG</td><td>||||||||[||||||||||||]PAM</td><td>NA</td><td>Intergenic</td><td>NA</td><td>NA</td></tr>
<tr><td>T28250</td><td>ACGCGTCCTTCGCCACCGTTAGG</td><td>1000</td><td colspan="9"></td></tr>
<tr><td>Oligo fwd</td><td>CACCACGCGTCCTTCGCCACCGTT</td><td colspan="10"></td></tr>
<tr><td>Oligo rev</td><td>AAACAACGGTGGCGAAGGACGCGT</td><td colspan="10"></td></tr>
<tr><td>Chromosome</td><td>start</td><td>end</td><td>strand</td><td>MM</td><td>target_seq</td><td>PAM</td><td>alignment</td><td>distance</td><td>position</td><td>gene name</td><td>gene id</td></tr>
<tr><td>GU305914.1</td><td>79274</td><td>79296</td><td>-</td><td>0</td><td>ACGCGTCCTTCGCCACCGTT</td><td>AGG</td><td>||||||||[||||||||||||]PAM</td><td>NA</td><td>Intergenic</td><td>NA</td><td>NA</td></tr>
<tr><td>T28251</td><td>TTAGGATCTGCTCGATGCACTGG</td><td>1000</td><td colspan="9"></td></tr>
<tr><td>Oligo fwd</td><td>CACCTTAGGATCTGCTCGATGCAC</td><td colspan="10"></td></tr>
<tr><td>Oligo rev</td><td>AAACGTGCATCGAGCAGATCCTAA</td><td colspan="10"></td></tr>
<tr><td>Chromosome</td><td>start</td><td>end</td><td>strand</td><td>MM</td><td>target_seq</td><td>PAM</td><td>alignment</td><td>distance</td><td>position</td><td>gene name</td><td>gene id</td></tr>
<tr><td>GU305914.1</td><td>79256</td><td>79278</td><td>-</td><td>0</td><td>TTAGGATCTGCTCGATGCAC</td><td>TGG</td><td>||||||||[||||||||||||]PAM</td><td>NA</td><td>Intergenic</td><td>NA</td><td>NA</td></tr>
<tr><td>T28252</td><td>TAGGATCTGCTCGATGCACTGGG</td><td>1000</td><td colspan="9"></td></tr>
<tr><td>Oligo fwd</td><td>CACCTAGGATCTGCTCGATGCACT</td><td colspan="10"></td></tr>
<tr><td>Oligo rev</td><td>AAACAGTGCATCGAGCAGATCCTA</td><td colspan="10"></td></tr>
<tr><td>Chromosome</td><td>start</td><td>end</td><td>strand</td><td>MM</td><td>target_seq</td><td>PAM</td><td>alignment</td><td>distance</td><td>position</td><td>gene name</td><td>gene id</td></tr>
<tr><td>GU305914.1</td><td>79255</td><td>79277</td><td>-</td><td>0</td><td>TAGGATCTGCTCGATGCACT</td><td>GGG</td><td>||||||||[||||||||||||]PAM</td><td>NA</td><td>Intergenic</td><td>NA</td><td>NA</td></tr>
<tr><td>T28253</td><td>ATCTGCTCGATGCACTGGGACGG</td><td>1000</td><td colspan="9"></td></tr>
<tr><td>Oligo fwd</td><td>CACCATCTGCTCGATGCACTGGGA</td><td colspan="10"></td></tr>
<tr><td>Oligo rev</td><td>AAACTCCCAGTGCATCGAGCAGAT</td><td colspan="10"></td></tr>
<tr><td>Chromosome</td><td>start</td><td>end</td><td>strand</td><td>MM</td><td>target_seq</td><td>PAM</td><td>alignment</td><td>distance</td><td>position</td><td>gene name</td><td>gene id</td></tr>
<tr><td>GU305914.1</td><td>79251</td><td>79273</td><td>-</td><td>0</td><td>ATCTGCTCGATGCACTGGGA</td><td>CGG</td><td>||||||||[||||||||||||]PAM</td><td>NA</td><td>Intergenic</td><td>NA</td><td>NA</td></tr>
<tr><td>T28254</td><td>TCTGCTCGATGCACTGGGACGGG</td><td>1000</td><td colspan="9"></td></tr>
<tr><td>Oligo fwd</td><td>CACCTCTGCTCGATGCACTGGGAC</td><td colspan="10"></td></tr>
<tr><td>Oligo rev</td><td>AAACGTCCCAGTGCATCGAGCAGA</td><td colspan="10"></td></tr>
<tr><td>Chromosome</td><td>start</td><td>end</td><td>strand</td><td>MM</td><td>target_seq</td><td>PAM</td><td>alignment</td><td>distance</td><td>position</td><td>gene name</td><td>gene id</td></tr>
<tr><td>GU305914.1</td><td>79250</td><td>79272</td><td>-</td><td>0</td><td>TCTGCTCGATGCACTGGGAC</td><td>GGG</td><td>||||||||[||||||||||||]PAM</td><td>NA</td><td>Intergenic</td><td>NA</td><td>NA</td></tr>
</table>

<table>
<tr><td>T28255</td><td>GTTGCATGTGCGCCATCTCTCGG</td><td>1000</td><td></td><td></td><td></td><td></td><td></td><td></td><td></td><td></td><td></td></tr>
<tr><td>Oligo fwd</td><td>CACCGTTGCATGTGCGCCATCTCT</td><td></td><td></td><td></td><td></td><td></td><td></td><td></td><td></td><td></td><td></td></tr>
<tr><td>Oligo rev</td><td>AAACAGAGATGGCGCACATGCAAC</td><td></td><td></td><td></td><td></td><td></td><td></td><td></td><td></td><td></td><td></td></tr>
<tr><td>Chromosome</td><td>start</td><td>end</td><td>strand</td><td>MM</td><td>target_seq</td><td>PAM</td><td>alignment</td><td>distance</td><td>position</td><td>gene name</td><td>gene id</td></tr>
<tr><td>GU305914.1</td><td>79199</td><td>79221</td><td>-</td><td>0</td><td>GTTGCATGTGCGCCATCTCT</td><td>CGG</td><td>||||||||[||||||||||||]PAM</td><td>NA</td><td>Intergenic</td><td>NA</td><td>NA</td></tr>
<tr><td>T28256</td><td>TCTCGGTCGTAGTAGATCAGCGG</td><td>1000</td><td></td><td></td><td></td><td></td><td></td><td></td><td></td><td></td><td></td></tr>
<tr><td>Oligo fwd</td><td>CACCTCTCGGTCGTAGTAGATCAG</td><td></td><td></td><td></td><td></td><td></td><td></td><td></td><td></td><td></td><td></td></tr>
<tr><td>Oligo rev</td><td>AAACCTGATCTACTACGACCGAGA</td><td></td><td></td><td></td><td></td><td></td><td></td><td></td><td></td><td></td><td></td></tr>
<tr><td>Chromosome</td><td>start</td><td>end</td><td>strand</td><td>MM</td><td>target_seq</td><td>PAM</td><td>alignment</td><td>distance</td><td>position</td><td>gene name</td><td>gene id</td></tr>
<tr><td>GU305914.1</td><td>79182</td><td>79204</td><td>-</td><td>0</td><td>TCTCGGTCGTAGTAGATCAG</td><td>CGG</td><td>||||||||[||||||||||||]PAM</td><td>NA</td><td>Intergenic</td><td>NA</td><td>NA</td></tr>
<tr><td>T28257</td><td>ACTGCACGATGTAGTAGATCTGG</td><td>1000</td><td></td><td></td><td></td><td></td><td></td><td></td><td></td><td></td><td></td></tr>
<tr><td>Oligo fwd</td><td>CACCACTGCACGATGTAGTAGATC</td><td></td><td></td><td></td><td></td><td></td><td></td><td></td><td></td><td></td><td></td></tr>
<tr><td>Oligo rev</td><td>AAACGATCTACTACATCGTGCAGT</td><td></td><td></td><td></td><td></td><td></td><td></td><td></td><td></td><td></td><td></td></tr>
<tr><td>Chromosome</td><td>start</td><td>end</td><td>strand</td><td>MM</td><td>target_seq</td><td>PAM</td><td>alignment</td><td>distance</td><td>position</td><td>gene name</td><td>gene id</td></tr>
<tr><td>GU305914.1</td><td>79139</td><td>79161</td><td>-</td><td>0</td><td>ACTGCACGATGTAGTAGATC</td><td>TGG</td><td>||||||||[||||||||||||]PAM</td><td>NA</td><td>Intergenic</td><td>NA</td><td>NA</td></tr>
<tr><td>T28258</td><td>TAGTAGATCTGGAACAGCTGCGG</td><td>1000</td><td></td><td></td><td></td><td></td><td></td><td></td><td></td><td></td><td></td></tr>
<tr><td>Oligo fwd</td><td>CACCTAGTAGATCTGGAACAGCTG</td><td></td><td></td><td></td><td></td><td></td><td></td><td></td><td></td><td></td><td></td></tr>
<tr><td>Oligo rev</td><td>AAACCAGCTGTTCCAGATCTACTA</td><td></td><td></td><td></td><td></td><td></td><td></td><td></td><td></td><td></td><td></td></tr>
<tr><td>Chromosome</td><td>start</td><td>end</td><td>strand</td><td>MM</td><td>target_seq</td><td>PAM</td><td>alignment</td><td>distance</td><td>position</td><td>gene name</td><td>gene id</td></tr>
<tr><td>GU305914.1</td><td>79128</td><td>79150</td><td>-</td><td>0</td><td>TAGTAGATCTGGAACAGCTG</td><td>CGG</td><td>||||||||[||||||||||||]PAM</td><td>NA</td><td>Intergenic</td><td>NA</td><td>NA</td></tr>
<tr><td>T28259</td><td>AGATCTGGAACAGCTGCGGCTGG</td><td>1000</td><td></td><td></td><td></td><td></td><td></td><td></td><td></td><td></td><td></td></tr>
<tr><td>Oligo fwd</td><td>CACCAGATCTGGAACAGCTGCGGC</td><td></td><td></td><td></td><td></td><td></td><td></td><td></td><td></td><td></td><td></td></tr>
<tr><td>Oligo rev</td><td>AAACGCCGCAGCTGTTCCAGATCT</td><td></td><td></td><td></td><td></td><td></td><td></td><td></td><td></td><td></td><td></td></tr>
<tr><td>Chromosome</td><td>start</td><td>end</td><td>strand</td><td>MM</td><td>target_seq</td><td>PAM</td><td>alignment</td><td>distance</td><td>position</td><td>gene name</td><td>gene id</td></tr>
<tr><td>GU305914.1</td><td>79124</td><td>79146</td><td>-</td><td>0</td><td>AGATCTGGAACAGCTGCGGC</td><td>TGG</td><td>||||||||[||||||||||||]PAM</td><td>NA</td><td>Intergenic</td><td>NA</td><td>NA</td></tr>
<tr><td>T28260</td><td>GAACAGCTGCGGCTGGTTAGCGG</td><td>1000</td><td></td><td></td><td></td><td></td><td></td><td></td><td></td><td></td><td></td></tr>
<tr><td>Oligo fwd</td><td>CACCGAACAGCTGCGGCTGGTTAG</td><td></td><td></td><td></td><td></td><td></td><td></td><td></td><td></td><td></td><td></td></tr>
<tr><td>Oligo rev</td><td>AAACCTAACCAGCCGCAGCTGTTC</td><td></td><td></td><td></td><td></td><td></td><td></td><td></td><td></td><td></td><td></td></tr>
<tr><td>Chromosome</td><td>start</td><td>end</td><td>strand</td><td>MM</td><td>target_seq</td><td>PAM</td><td>alignment</td><td>distance</td><td>position</td><td>gene name</td><td>gene id</td></tr>
<tr><td>GU305914.1</td><td>79117</td><td>79139</td><td>-</td><td>0</td><td>GAACAGCTGCGGCTGGTTAG</td><td>CGG</td><td>||||||||[||||||||||||]PAM</td><td>NA</td><td>Intergenic</td><td>NA</td><td>NA</td></tr>
<tr><td>T28261</td><td>TCAGCAGTTCCATGTTCTTACGG</td><td>1000</td><td></td><td></td><td></td><td></td><td></td><td></td><td></td><td></td><td></td></tr>
<tr><td>Oligo fwd</td><td>CACCTCAGCAGTTCCATGTTCTTA</td><td></td><td></td><td></td><td></td><td></td><td></td><td></td><td></td><td></td><td></td></tr>
<tr><td>Oligo rev</td><td>AAACTAAGAACATGGAACTGCTGA</td><td></td><td></td><td></td><td></td><td></td><td></td><td></td><td></td><td></td><td></td></tr>
<tr><td>Chromosome</td><td>start</td><td>end</td><td>strand</td><td>MM</td><td>target_seq</td><td>PAM</td><td>alignment</td><td>distance</td><td>position</td><td>gene name</td><td>gene id</td></tr>
<tr><td>GU305914.1</td><td>79088</td><td>79110</td><td>-</td><td>0</td><td>TCAGCAGTTCCATGTTCTTA</td><td>CGG</td><td>||||||||[||||||||||||]PAM</td><td>NA</td><td>Intergenic</td><td>NA</td><td>NA</td></tr>
<tr><td>T28262</td><td>GGAACCCGAAGTAGAACTTGCGG</td><td>1000</td><td></td><td></td><td></td><td></td><td></td><td></td><td></td><td></td><td></td></tr>
<tr><td>Oligo fwd</td><td>CACCGGAACCCGAAGTAGAACTTG</td><td></td><td></td><td></td><td></td><td></td><td></td><td></td><td></td><td></td><td></td></tr>
<tr><td>Oligo rev</td><td>AAACCAAGTTCTACTTCGGGTTCC</td><td></td><td></td><td></td><td></td><td></td><td></td><td></td><td></td><td></td><td></td></tr>
<tr><td>Chromosome</td><td>start</td><td>end</td><td>strand</td><td>MM</td><td>target_seq</td><td>PAM</td><td>alignment</td><td>distance</td><td>position</td><td>gene name</td><td>gene id</td></tr>
<tr><td>GU305914.1</td><td>79067</td><td>79089</td><td>-</td><td>0</td><td>GGAACCCGAAGTAGAACTTG</td><td>CGG</td><td>||||||||[||||||||||||]PAM</td><td>NA</td><td>Intergenic</td><td>NA</td><td>NA</td></tr>
<tr><td>T28263</td><td>GCTTCATCACGTTAGACGTCTGG</td><td>1000</td><td></td><td></td><td></td><td></td><td></td><td></td><td></td><td></td><td></td></tr>
<tr><td>Oligo fwd</td><td>CACCGCTTCATCACGTTAGACGTC</td><td></td><td></td><td></td><td></td><td></td><td></td><td></td><td></td><td></td><td></td></tr>
<tr><td>Oligo rev</td><td>AAACGACGTCTAACGTGATGAAGC</td><td></td><td></td><td></td><td></td><td></td><td></td><td></td><td></td><td></td><td></td></tr>
<tr><td>Chromosome</td><td>start</td><td>end</td><td>strand</td><td>MM</td><td>target_seq</td><td>PAM</td><td>alignment</td><td>distance</td><td>position</td><td>gene name</td><td>gene id</td></tr>
<tr><td>GU305914.1</td><td>79043</td><td>79065</td><td>-</td><td>0</td><td>GCTTCATCACGTTAGACGTC</td><td>TGG</td><td>||||||||[||||||||||||]PAM</td><td>NA</td><td>Intergenic</td><td>NA</td><td>NA</td></tr>
<tr><td>T28264</td><td>ACCATGCTGTCCGACGCCGTCGG</td><td>1000</td><td></td><td></td><td></td><td></td><td></td><td></td><td></td><td></td><td></td></tr>
<tr><td>Oligo fwd</td><td>CACCACCATGCTGTCCGACGCCGT</td><td></td><td></td><td></td><td></td><td></td><td></td><td></td><td></td><td></td><td></td></tr>
<tr><td>Oligo rev</td><td>AAACACGGCGTCGGACAGCATGGT</td><td></td><td></td><td></td><td></td><td></td><td></td><td></td><td></td><td></td><td></td></tr>
<tr><td>Chromosome</td><td>start</td><td>end</td><td>strand</td><td>MM</td><td>target_seq</td><td>PAM</td><td>alignment</td><td>distance</td><td>position</td><td>gene name</td><td>gene id</td></tr>
<tr><td>GU305914.1</td><td>79008</td><td>79030</td><td>-</td><td>0</td><td>ACCATGCTGTCCGACGCCGT</td><td>CGG</td><td>||||||||[||||||||||||]PAM</td><td>NA</td><td>Intergenic</td><td>NA</td><td>NA</td></tr>
<tr><td>T28265</td><td>CAGTCTCGTCGTCTCCTTGCAGG</td><td>1000</td><td></td><td></td><td></td><td></td><td></td><td></td><td></td><td></td><td></td></tr>
<tr><td>Oligo fwd</td><td>CACCCAGTCTCGTCGTCTCCTTGC</td><td></td><td></td><td></td><td></td><td></td><td></td><td></td><td></td><td></td><td></td></tr>
<tr><td>Oligo rev</td><td>AAACGCAAGGAGACGACGAGACTG</td><td></td><td></td><td></td><td></td><td></td><td></td><td></td><td></td><td></td><td></td></tr>
<tr><td>Chromosome</td><td>start</td><td>end</td><td>strand</td><td>MM</td><td>target_seq</td><td>PAM</td><td>alignment</td><td>distance</td><td>position</td><td>gene name</td><td>gene id</td></tr>
<tr><td>GU305914.1</td><td>78985</td><td>79007</td><td>-</td><td>0</td><td>CAGTCTCGTCGTCTCCTTGC</td><td>AGG</td><td>||||||||[||||||||||||]PAM</td><td>NA</td><td>Intergenic</td><td>NA</td><td>NA</td></tr>
<tr><td>T28266</td><td>TCTCGTCGTCTCCTTGCAGGTGG</td><td>1000</td><td></td><td></td><td></td><td></td><td></td><td></td><td></td><td></td><td></td></tr>
<tr><td>Oligo fwd</td><td>CACCTCTCGTCGTCTCCTTGCAGG</td><td></td><td></td><td></td><td></td><td></td><td></td><td></td><td></td><td></td><td></td></tr>
<tr><td>Oligo rev</td><td>AAACCCTGCAAGGAGACGACGAGA</td><td></td><td></td><td></td><td></td><td></td><td></td><td></td><td></td><td></td><td></td></tr>
<tr><td>Chromosome</td><td>start</td><td>end</td><td>strand</td><td>MM</td><td>target_seq</td><td>PAM</td><td>alignment</td><td>distance</td><td>position</td><td>gene name</td><td>gene id</td></tr>
<tr><td>GU305914.1</td><td>78982</td><td>79004</td><td>-</td><td>0</td><td>TCTCGTCGTCTCCTTGCAGG</td><td>TGG</td><td>||||||||[||||||||||||]PAM</td><td>NA</td><td>Intergenic</td><td>NA</td><td>NA</td></tr>
<tr><td>T28267</td><td>TGCACGCGCTCCGCGACACCTGG</td><td>1000</td><td></td><td></td><td></td><td></td><td></td><td></td><td></td><td></td><td></td></tr>
<tr><td>Oligo fwd</td><td>CACCTGCACGCGCTCCGCGACACC</td><td></td><td></td><td></td><td></td><td></td><td></td><td></td><td></td><td></td><td></td></tr>
<tr><td>Oligo rev</td><td>AAACGGTGTCGCGGAGCGCGTGCA</td><td></td><td></td><td></td><td></td><td></td><td></td><td></td><td></td><td></td><td></td></tr>
<tr><td>Chromosome</td><td>start</td><td>end</td><td>strand</td><td>MM</td><td>target_seq</td><td>PAM</td><td>alignment</td><td>distance</td><td>position</td><td>gene name</td><td>gene id</td></tr>
<tr><td>GU305914.1</td><td>78959</td><td>78981</td><td>-</td><td>0</td><td>TGCACGCGCTCCGCGACACC</td><td>TGG</td><td>||||||||[||||||||||||]PAM</td><td>NA</td><td>Intergenic</td><td>NA</td><td>NA</td></tr>
<tr><td>T28268</td><td>CGACACCTGGTCGTTCCCGTAGG</td><td>1000</td><td></td><td></td><td></td><td></td><td></td><td></td><td></td><td></td><td></td></tr>
<tr><td>Oligo fwd</td><td>CACCCGACACCTGGTCGTTCCCGT</td><td></td><td></td><td></td><td></td><td></td><td></td><td></td><td></td><td></td><td></td></tr>
<tr><td>Oligo rev</td><td>AAACACGGGAACGACCAGGTGTCG</td><td></td><td></td><td></td><td></td><td></td><td></td><td></td><td></td><td></td><td></td></tr>
<tr><td>Chromosome</td><td>start</td><td>end</td><td>strand</td><td>MM</td><td>target_seq</td><td>PAM</td><td>alignment</td><td>distance</td><td>position</td><td>gene name</td><td>gene id</td></tr>
<tr><td>GU305914.1</td><td>78946</td><td>78968</td><td>-</td><td>0</td><td>CGACACCTGGTCGTTCCCGT</td><td>AGG</td><td>||||||||[||||||||||||]PAM</td><td>NA</td><td>Intergenic</td><td>NA</td><td>NA</td></tr>
<tr><td>T28269</td><td>GACACCTGGTCGTTCCCGTAGGG</td><td>1000</td><td></td><td></td><td></td><td></td><td></td><td></td><td></td><td></td><td></td></tr>
<tr><td>Oligo fwd</td><td>CACCGACACCTGGTCGTTCCCGTA</td><td></td><td></td><td></td><td></td><td></td><td></td><td></td><td></td><td></td><td></td></tr>
<tr><td>Oligo rev</td><td>AAACTACGGGAACGACCAGGTGTC</td><td></td><td></td><td></td><td></td><td></td><td></td><td></td><td></td><td></td><td></td></tr>
<tr><td>Chromosome</td><td>start</td><td>end</td><td>strand</td><td>MM</td><td>target_seq</td><td>PAM</td><td>alignment</td><td>distance</td><td>position</td><td>gene name</td><td>gene id</td></tr>
</table>

GU305914.1 78945 78967 - 0 GACACCTGGTCGTTCCCGTA GGG ||||||||[||||||||||||]PAM NA Intergenic NA NA

T28270 ACACCTGGTCGTTCCCGTAGGGG 1000
Oligo fwd CACCACACCTGGTCGTTCCCGTAG
Oligo rev AAACCTACGGGAACGACCAGGTGT
Chromosome start end strand MM target_seq PAM alignment distance position gene name gene id
GU305914.1 78944 78966 - 0 ACACCTGGTCGTTCCCGTAG GGG ||||||||[||||||||||||]PAM NA Intergenic NA NA

T28271 CACCTGGTCGTTCCCGTAGGGGG 1000
Oligo fwd CACCCACCTGGTCGTTCCCGTAGG
Oligo rev AAACCCTACGGGAACGACCAGGTG
Chromosome start end strand MM target_seq PAM alignment distance position gene name gene id
GU305914.1 78943 78965 - 0 CACCTGGTCGTTCCCGTAGG GGG ||||||||[||||||||||||]PAM NA Intergenic NA NA

T28272 GGTCGTTCCCGTAGGGGGAGAGG 1000
Oligo fwd CACCGGTCGTTCCCGTAGGGGGAG
Oligo rev AAACCTCCCCCTACGGGAACGACC
Chromosome start end strand MM target_seq PAM alignment distance position gene name gene id
GU305914.1 78938 78960 - 0 GGTCGTTCCCGTAGGGGGAG AGG ||||||||[||||||||||||]PAM NA Intergenic NA NA

T28273 GGTCCAGACACGTGTGCTGCCGG 1000
Oligo fwd CACCGGTCCAGACACGTGTGCTGC
Oligo rev AAACGCAGCACACGTGTCTGGACC
Chromosome start end strand MM target_seq PAM alignment distance position gene name gene id
GU305914.1 78917 78939 - 0 GGTCCAGACACGTGTGCTGC CGG ||||||||[||||||||||||]PAM NA Intergenic NA NA

T28274 CGGTGCGACTTGAAGTCGAACGG 1000
Oligo fwd CACCCGGTGCGACTTGAAGTCGAA
Oligo rev AAACTTCGACTTCAAGTCGCACCG
Chromosome start end strand MM target_seq PAM alignment distance position gene name gene id
GU305914.1 78897 78919 - 0 CGGTGCGACTTGAAGTCGAA CGG ||||||||[||||||||||||]PAM NA Intergenic NA NA

T28275 GGTGCGACTTGAAGTCGAACGGG 1000
Oligo fwd CACCGGTGCGACTTGAAGTCGAAC
Oligo rev AAACGTTCGACTTCAAGTCGCACC
Chromosome start end strand MM target_seq PAM alignment distance position gene name gene id
GU305914.1 78896 78918 - 0 GGTGCGACTTGAAGTCGAAC GGG ||||||||[||||||||||||]PAM NA Intergenic NA NA
GU305914.1 78898 78920 + 4 CGTTCGACTTCAAGTCGCAC CGG -||-||||[||-||||||-||]PAM NA Intergenic NA NA

T28276 ACTTGAAGTCGAACGGGAGTCGG 1000
Oligo fwd CACCACTTGAAGTCGAACGGGAGT
Oligo rev AAACACTCCCGTTCGACTTCAAGT
Chromosome start end strand MM target_seq PAM alignment distance position gene name gene id
GU305914.1 78890 78912 - 0 ACTTGAAGTCGAACGGGAGT CGG ||||||||[||||||||||||]PAM NA Intergenic NA NA

T28277 ATCGACTCCTTGCCCGTGATCGG 1000
Oligo fwd CACCATCGACTCCTTGCCCGTGAT
Oligo rev AAACATCACGGGCAAGGAGTCGAT
Chromosome start end strand MM target_seq PAM alignment distance position gene name gene id
GU305914.1 78867 78889 - 0 ATCGACTCCTTGCCCGTGAT CGG ||||||||[||||||||||||]PAM NA Intergenic NA NA

T28278 TCGGTAGCTTCATGATCTTGAGG 1000
Oligo fwd CACCTCGGTAGCTTCATGATCTTG
Oligo rev AAACCAAGATCATGAAGCTACCGA
Chromosome start end strand MM target_seq PAM alignment distance position gene name gene id
GU305914.1 78848 78870 - 0 TCGGTAGCTTCATGATCTTG AGG ||||||||[||||||||||||]PAM NA Intergenic NA NA

T28279 TCTTGAGGTACTTCTGTTCGAGG 1000
Oligo fwd CACCTCTTGAGGTACTTCTGTTCG
Oligo rev AAACCGAACAGAAGTACCTCAAGA
Chromosome start end strand MM target_seq PAM alignment distance position gene name gene id
GU305914.1 78833 78855 - 0 TCTTGAGGTACTTCTGTTCG AGG ||||||||[||||||||||||]PAM NA Intergenic NA NA

T28280 GAGGTACTTCTGTTCGAGGTCGG 1000
Oligo fwd CACCGAGGTACTTCTGTTCGAGGT
Oligo rev AAACACCTCGAACAGAAGTACCTC
Chromosome start end strand MM target_seq PAM alignment distance position gene name gene id
GU305914.1 78829 78851 - 0 GAGGTACTTCTGTTCGAGGT CGG ||||||||[||||||||||||]PAM NA Intergenic NA NA

T28281 AGGTACTTCTGTTCGAGGTCGGG 1000
Oligo fwd CACCAGGTACTTCTGTTCGAGGTC
Oligo rev AAACGACCTCGAACAGAAGTACCT
Chromosome start end strand MM target_seq PAM alignment distance position gene name gene id
GU305914.1 78828 78850 - 0 AGGTACTTCTGTTCGAGGTC GGG ||||||||[||||||||||||]PAM NA Intergenic NA NA

T28282 CACGTTCAGATACTCGCTCTCGG 1000
Oligo fwd CACCCACGTTCAGATACTCGCTCT
Oligo rev AAACAGAGCGAGTATCTGAACGTG
Chromosome start end strand MM target_seq PAM alignment distance position gene name gene id
GU305914.1 78769 78791 - 0 CACGTTCAGATACTCGCTCT CGG ||||||||[||||||||||||]PAM NA Intergenic NA NA

T28283 ACGTTCAGATACTCGCTCTCGGG 1000
Oligo fwd CACCACGTTCAGATACTCGCTCTC
Oligo rev AAACGAGAGCGAGTATCTGAACGT
Chromosome start end strand MM target_seq PAM alignment distance position gene name gene id
GU305914.1 78768 78790 - 0 ACGTTCAGATACTCGCTCTC GGG ||||||||[||||||||||||]PAM NA Intergenic NA NA

T28284 CTCGCTCTCGGGCGCCGCGCCGG 1000
Oligo fwd CACCCTCGCTCTCGGGCGCCGCGC

<table>
<tr><td>Oligo rev</td><td>AAACGCGCGGCGCCCGAGAGCGAG</td><td></td><td></td><td></td><td></td><td></td><td></td><td></td><td></td><td></td><td></td></tr>
<tr><td>Chromosome</td><td>start</td><td>end</td><td>strand</td><td>MM</td><td>target_seq</td><td>PAM</td><td>alignment</td><td>distance</td><td>position</td><td>gene name</td><td>gene id</td></tr>
<tr><td>GU305914.1</td><td>78757</td><td>78779</td><td>-</td><td>0</td><td>CTCGCTCTCGGGCGCCGCGC</td><td>CGG</td><td>||||||||[||||||||||||]PAM</td><td>NA</td><td>Intergenic</td><td>NA</td><td>NA</td></tr>
<tr><td></td><td></td><td></td><td></td><td></td><td></td><td></td><td></td><td></td><td></td><td></td><td></td></tr>
<tr><td>T28285</td><td>CGCCGCGCCGGCCGCTGCGCTGG</td><td>1000</td><td></td><td></td><td></td><td></td><td></td><td></td><td></td><td></td><td></td></tr>
<tr><td>Oligo fwd</td><td>CACCCGCCGCGCCGGCCGCTGCGC</td><td></td><td></td><td></td><td></td><td></td><td></td><td></td><td></td><td></td><td></td></tr>
<tr><td>Oligo rev</td><td>AAACGCGCAGCGGCCGGCGCGGCG</td><td></td><td></td><td></td><td></td><td></td><td></td><td></td><td></td><td></td><td></td></tr>
<tr><td>Chromosome</td><td>start</td><td>end</td><td>strand</td><td>MM</td><td>target_seq</td><td>PAM</td><td>alignment</td><td>distance</td><td>position</td><td>gene name</td><td>gene id</td></tr>
<tr><td>GU305914.1</td><td>78745</td><td>78767</td><td>-</td><td>0</td><td>CGCCGCGCCGGCCGCTGCGC</td><td>TGG</td><td>||||||||[||||||||||||]PAM</td><td>NA</td><td>Intergenic</td><td>NA</td><td>NA</td></tr>
<tr><td>GU305914.1</td><td>169857</td><td>169879</td><td>+</td><td>4</td><td>CGCGGCTCCGGCCGGTTCGC</td><td>GGG</td><td>|||-||-|[||||||-|-|||]PAM</td><td>NA</td><td>Intergenic</td><td>NA</td><td>NA</td></tr>
<tr><td>GU305914.1</td><td>56772</td><td>56794</td><td>-</td><td>4</td><td>CGCCGCCGCCGCCGCTGCGG</td><td>CGG</td><td>||||||--[|-|||||||||-]PAM</td><td>NA</td><td>Intergenic</td><td>NA</td><td>NA</td></tr>
<tr><td></td><td></td><td></td><td></td><td></td><td></td><td></td><td></td><td></td><td></td><td></td><td></td></tr>
<tr><td>T28286</td><td>CGCGCCGGCCGCTGCGCTGGCGG</td><td>1000</td><td></td><td></td><td></td><td></td><td></td><td></td><td></td><td></td><td></td></tr>
<tr><td>Oligo fwd</td><td>CACCCGCGCCGGCCGCTGCGCTGG</td><td></td><td></td><td></td><td></td><td></td><td></td><td></td><td></td><td></td><td></td></tr>
<tr><td>Oligo rev</td><td>AAACCCAGCGCAGCGGCCGGCGCG</td><td></td><td></td><td></td><td></td><td></td><td></td><td></td><td></td><td></td><td></td></tr>
<tr><td>Chromosome</td><td>start</td><td>end</td><td>strand</td><td>MM</td><td>target_seq</td><td>PAM</td><td>alignment</td><td>distance</td><td>position</td><td>gene name</td><td>gene id</td></tr>
<tr><td>GU305914.1</td><td>78742</td><td>78764</td><td>-</td><td>0</td><td>CGCGCCGGCCGCTGCGCTGG</td><td>CGG</td><td>||||||||[||||||||||||]PAM</td><td>NA</td><td>Intergenic</td><td>NA</td><td>NA</td></tr>
<tr><td></td><td></td><td></td><td></td><td></td><td></td><td></td><td></td><td></td><td></td><td></td><td></td></tr>
<tr><td>T28287</td><td>TGCGCTGGCGGCCGATCTCTTGG</td><td>1000</td><td></td><td></td><td></td><td></td><td></td><td></td><td></td><td></td><td></td></tr>
<tr><td>Oligo fwd</td><td>CACCTGCGCTGGCGGCCGATCTCT</td><td></td><td></td><td></td><td></td><td></td><td></td><td></td><td></td><td></td><td></td></tr>
<tr><td>Oligo rev</td><td>AAACAGAGATCGGCCGCCAGCGCA</td><td></td><td></td><td></td><td></td><td></td><td></td><td></td><td></td><td></td><td></td></tr>
<tr><td>Chromosome</td><td>start</td><td>end</td><td>strand</td><td>MM</td><td>target_seq</td><td>PAM</td><td>alignment</td><td>distance</td><td>position</td><td>gene name</td><td>gene id</td></tr>
<tr><td>GU305914.1</td><td>78730</td><td>78752</td><td>-</td><td>0</td><td>TGCGCTGGCGGCCGATCTCT</td><td>TGG</td><td>||||||||[||||||||||||]PAM</td><td>NA</td><td>Intergenic</td><td>NA</td><td>NA</td></tr>
<tr><td></td><td></td><td></td><td></td><td></td><td></td><td></td><td></td><td></td><td></td><td></td><td></td></tr>
<tr><td>T28288</td><td>GCGCTGGCGGCCGATCTCTTGGG</td><td>1000</td><td></td><td></td><td></td><td></td><td></td><td></td><td></td><td></td><td></td></tr>
<tr><td>Oligo fwd</td><td>CACCGCGCTGGCGGCCGATCTCTT</td><td></td><td></td><td></td><td></td><td></td><td></td><td></td><td></td><td></td><td></td></tr>
<tr><td>Oligo rev</td><td>AAACAAGAGATCGGCCGCCAGCGC</td><td></td><td></td><td></td><td></td><td></td><td></td><td></td><td></td><td></td><td></td></tr>
<tr><td>Chromosome</td><td>start</td><td>end</td><td>strand</td><td>MM</td><td>target_seq</td><td>PAM</td><td>alignment</td><td>distance</td><td>position</td><td>gene name</td><td>gene id</td></tr>
<tr><td>GU305914.1</td><td>78729</td><td>78751</td><td>-</td><td>0</td><td>GCGCTGGCGGCCGATCTCTT</td><td>GGG</td><td>||||||||[||||||||||||]PAM</td><td>NA</td><td>Intergenic</td><td>NA</td><td>NA</td></tr>
<tr><td></td><td></td><td></td><td></td><td></td><td></td><td></td><td></td><td></td><td></td><td></td><td></td></tr>
<tr><td>T28289</td><td>CTGGCGGCCGATCTCTTGGGTGG</td><td>1000</td><td></td><td></td><td></td><td></td><td></td><td></td><td></td><td></td><td></td></tr>
<tr><td>Oligo fwd</td><td>CACCCTGGCGGCCGATCTCTTGGG</td><td></td><td></td><td></td><td></td><td></td><td></td><td></td><td></td><td></td><td></td></tr>
<tr><td>Oligo rev</td><td>AAACCCCAAGAGATCGGCCGCCAG</td><td></td><td></td><td></td><td></td><td></td><td></td><td></td><td></td><td></td><td></td></tr>
<tr><td>Chromosome</td><td>start</td><td>end</td><td>strand</td><td>MM</td><td>target_seq</td><td>PAM</td><td>alignment</td><td>distance</td><td>position</td><td>gene name</td><td>gene id</td></tr>
<tr><td>GU305914.1</td><td>78726</td><td>78748</td><td>-</td><td>0</td><td>CTGGCGGCCGATCTCTTGGG</td><td>TGG</td><td>||||||||[||||||||||||]PAM</td><td>NA</td><td>Intergenic</td><td>NA</td><td>NA</td></tr>
<tr><td></td><td></td><td></td><td></td><td></td><td></td><td></td><td></td><td></td><td></td><td></td><td></td></tr>
<tr><td>T28290</td><td>GCGGCCGATCTCTTGGGTGGCGG</td><td>1000</td><td></td><td></td><td></td><td></td><td></td><td></td><td></td><td></td><td></td></tr>
<tr><td>Oligo fwd</td><td>CACCGCGGCCGATCTCTTGGGTGG</td><td></td><td></td><td></td><td></td><td></td><td></td><td></td><td></td><td></td><td></td></tr>
<tr><td>Oligo rev</td><td>AAACCCACCCAAGAGATCGGCCGC</td><td></td><td></td><td></td><td></td><td></td><td></td><td></td><td></td><td></td><td></td></tr>
<tr><td>Chromosome</td><td>start</td><td>end</td><td>strand</td><td>MM</td><td>target_seq</td><td>PAM</td><td>alignment</td><td>distance</td><td>position</td><td>gene name</td><td>gene id</td></tr>
<tr><td>GU305914.1</td><td>78723</td><td>78745</td><td>-</td><td>0</td><td>GCGGCCGATCTCTTGGGTGG</td><td>CGG</td><td>||||||||[||||||||||||]PAM</td><td>NA</td><td>Intergenic</td><td>NA</td><td>NA</td></tr>
<tr><td></td><td></td><td></td><td></td><td></td><td></td><td></td><td></td><td></td><td></td><td></td><td></td></tr>
<tr><td>T28291</td><td>GCCGATCTCTTGGGTGGCGGAGG</td><td>1000</td><td></td><td></td><td></td><td></td><td></td><td></td><td></td><td></td><td></td></tr>
<tr><td>Oligo fwd</td><td>CACCGCCGATCTCTTGGGTGGCGG</td><td></td><td></td><td></td><td></td><td></td><td></td><td></td><td></td><td></td><td></td></tr>
<tr><td>Oligo rev</td><td>AAACCCGCCACCCAAGAGATCGGC</td><td></td><td></td><td></td><td></td><td></td><td></td><td></td><td></td><td></td><td></td></tr>
<tr><td>Chromosome</td><td>start</td><td>end</td><td>strand</td><td>MM</td><td>target_seq</td><td>PAM</td><td>alignment</td><td>distance</td><td>position</td><td>gene name</td><td>gene id</td></tr>
<tr><td>GU305914.1</td><td>78720</td><td>78742</td><td>-</td><td>0</td><td>GCCGATCTCTTGGGTGGCGG</td><td>AGG</td><td>||||||||[||||||||||||]PAM</td><td>NA</td><td>Intergenic</td><td>NA</td><td>NA</td></tr>
<tr><td></td><td></td><td></td><td></td><td></td><td></td><td></td><td></td><td></td><td></td><td></td><td></td></tr>
<tr><td>T28292</td><td>GATCTCTTGGGTGGCGGAGGCGG</td><td>1000</td><td></td><td></td><td></td><td></td><td></td><td></td><td></td><td></td><td></td></tr>
<tr><td>Oligo fwd</td><td>CACCGATCTCTTGGGTGGCGGAGG</td><td></td><td></td><td></td><td></td><td></td><td></td><td></td><td></td><td></td><td></td></tr>
<tr><td>Oligo rev</td><td>AAACCCTCCGCCACCCAAGAGATC</td><td></td><td></td><td></td><td></td><td></td><td></td><td></td><td></td><td></td><td></td></tr>
<tr><td>Chromosome</td><td>start</td><td>end</td><td>strand</td><td>MM</td><td>target_seq</td><td>PAM</td><td>alignment</td><td>distance</td><td>position</td><td>gene name</td><td>gene id</td></tr>
<tr><td>GU305914.1</td><td>78717</td><td>78739</td><td>-</td><td>0</td><td>GATCTCTTGGGTGGCGGAGG</td><td>CGG</td><td>||||||||[||||||||||||]PAM</td><td>NA</td><td>Intergenic</td><td>NA</td><td>NA</td></tr>
<tr><td></td><td></td><td></td><td></td><td></td><td></td><td></td><td></td><td></td><td></td><td></td><td></td></tr>
<tr><td>T28293</td><td>ATCTCTTGGGTGGCGGAGGCGGG</td><td>1000</td><td></td><td></td><td></td><td></td><td></td><td></td><td></td><td></td><td></td></tr>
<tr><td>Oligo fwd</td><td>CACCATCTCTTGGGTGGCGGAGGC</td><td></td><td></td><td></td><td></td><td></td><td></td><td></td><td></td><td></td><td></td></tr>
<tr><td>Oligo rev</td><td>AAACGCCTCCGCCACCCAAGAGAT</td><td></td><td></td><td></td><td></td><td></td><td></td><td></td><td></td><td></td><td></td></tr>
<tr><td>Chromosome</td><td>start</td><td>end</td><td>strand</td><td>MM</td><td>target_seq</td><td>PAM</td><td>alignment</td><td>distance</td><td>position</td><td>gene name</td><td>gene id</td></tr>
<tr><td>GU305914.1</td><td>78716</td><td>78738</td><td>-</td><td>0</td><td>ATCTCTTGGGTGGCGGAGGC</td><td>GGG</td><td>||||||||[||||||||||||]PAM</td><td>NA</td><td>Intergenic</td><td>NA</td><td>NA</td></tr>
<tr><td></td><td></td><td></td><td></td><td></td><td></td><td></td><td></td><td></td><td></td><td></td><td></td></tr>
<tr><td>T28294</td><td>CTTGGGTGGCGGAGGCGGGTCGG</td><td>1000</td><td></td><td></td><td></td><td></td><td></td><td></td><td></td><td></td><td></td></tr>
<tr><td>Oligo fwd</td><td>CACCCTTGGGTGGCGGAGGCGGGT</td><td></td><td></td><td></td><td></td><td></td><td></td><td></td><td></td><td></td><td></td></tr>
<tr><td>Oligo rev</td><td>AAACACCCGCCTCCGCCACCCAAG</td><td></td><td></td><td></td><td></td><td></td><td></td><td></td><td></td><td></td><td></td></tr>
<tr><td>Chromosome</td><td>start</td><td>end</td><td>strand</td><td>MM</td><td>target_seq</td><td>PAM</td><td>alignment</td><td>distance</td><td>position</td><td>gene name</td><td>gene id</td></tr>
<tr><td>GU305914.1</td><td>78712</td><td>78734</td><td>-</td><td>0</td><td>CTTGGGTGGCGGAGGCGGGT</td><td>CGG</td><td>||||||||[||||||||||||]PAM</td><td>NA</td><td>Intergenic</td><td>NA</td><td>NA</td></tr>
<tr><td></td><td></td><td></td><td></td><td></td><td></td><td></td><td></td><td></td><td></td><td></td><td></td></tr>
<tr><td>T28295</td><td>GGTGGCGGAGGCGGGTCGGTCGG</td><td>1000</td><td></td><td></td><td></td><td></td><td></td><td></td><td></td><td></td><td></td></tr>
<tr><td>Oligo fwd</td><td>CACCGGTGGCGGAGGCGGGTCGGT</td><td></td><td></td><td></td><td></td><td></td><td></td><td></td><td></td><td></td><td></td></tr>
<tr><td>Oligo rev</td><td>AAACACCGACCCGCCTCCGCCACC</td><td></td><td></td><td></td><td></td><td></td><td></td><td></td><td></td><td></td><td></td></tr>
<tr><td>Chromosome</td><td>start</td><td>end</td><td>strand</td><td>MM</td><td>target_seq</td><td>PAM</td><td>alignment</td><td>distance</td><td>position</td><td>gene name</td><td>gene id</td></tr>
<tr><td>GU305914.1</td><td>78708</td><td>78730</td><td>-</td><td>0</td><td>GGTGGCGGAGGCGGGTCGGT</td><td>CGG</td><td>||||||||[||||||||||||]PAM</td><td>NA</td><td>Intergenic</td><td>NA</td><td>NA</td></tr>
<tr><td></td><td></td><td></td><td></td><td></td><td></td><td></td><td></td><td></td><td></td><td></td><td></td></tr>
<tr><td>T28296</td><td>GTGGCGGAGGCGGGTCGGTCGGG</td><td>1000</td><td></td><td></td><td></td><td></td><td></td><td></td><td></td><td></td><td></td></tr>
<tr><td>Oligo fwd</td><td>CACCGTGGCGGAGGCGGGTCGGTC</td><td></td><td></td><td></td><td></td><td></td><td></td><td></td><td></td><td></td><td></td></tr>
<tr><td>Oligo rev</td><td>AAACGACCGACCCGCCTCCGCCAC</td><td></td><td></td><td></td><td></td><td></td><td></td><td></td><td></td><td></td><td></td></tr>
<tr><td>Chromosome</td><td>start</td><td>end</td><td>strand</td><td>MM</td><td>target_seq</td><td>PAM</td><td>alignment</td><td>distance</td><td>position</td><td>gene name</td><td>gene id</td></tr>
<tr><td>GU305914.1</td><td>78707</td><td>78729</td><td>-</td><td>0</td><td>GTGGCGGAGGCGGGTCGGTC</td><td>GGG</td><td>||||||||[||||||||||||]PAM</td><td>NA</td><td>Intergenic</td><td>NA</td><td>NA</td></tr>
<tr><td></td><td></td><td></td><td></td><td></td><td></td><td></td><td></td><td></td><td></td><td></td><td></td></tr>
<tr><td>T28297</td><td>TGGCGGAGGCGGGTCGGTCGGGG</td><td>1000</td><td></td><td></td><td></td><td></td><td></td><td></td><td></td><td></td><td></td></tr>
<tr><td>Oligo fwd</td><td>CACCTGGCGGAGGCGGGTCGGTCG</td><td></td><td></td><td></td><td></td><td></td><td></td><td></td><td></td><td></td><td></td></tr>
<tr><td>Oligo rev</td><td>AAACCGACCGACCCGCCTCCGCCA</td><td></td><td></td><td></td><td></td><td></td><td></td><td></td><td></td><td></td><td></td></tr>
<tr><td>Chromosome</td><td>start</td><td>end</td><td>strand</td><td>MM</td><td>target_seq</td><td>PAM</td><td>alignment</td><td>distance</td><td>position</td><td>gene name</td><td>gene id</td></tr>
<tr><td>GU305914.1</td><td>78706</td><td>78728</td><td>-</td><td>0</td><td>TGGCGGAGGCGGGTCGGTCG</td><td>GGG</td><td>||||||||[||||||||||||]PAM</td><td>NA</td><td>Intergenic</td><td>NA</td><td>NA</td></tr>
<tr><td></td><td></td><td></td><td></td><td></td><td></td><td></td><td></td><td></td><td></td><td></td><td></td></tr>
<tr><td>T28298</td><td>CGGAGGCGGGTCGGTCGGGGTGG</td><td>1000</td><td></td><td></td><td></td><td></td><td></td><td></td><td></td><td></td><td></td></tr>
<tr><td>Oligo fwd</td><td>CACCCGGAGGCGGGTCGGTCGGGG</td><td></td><td></td><td></td><td></td><td></td><td></td><td></td><td></td><td></td><td></td></tr>
<tr><td>Oligo rev</td><td>AAACCCCCGACCGACCCGCCTCCG</td><td></td><td></td><td></td><td></td><td></td><td></td><td></td><td></td><td></td><td></td></tr>
<tr><td>Chromosome</td><td>start</td><td>end</td><td>strand</td><td>MM</td><td>target_seq</td><td>PAM</td><td>alignment</td><td>distance</td><td>position</td><td>gene name</td><td>gene id</td></tr>
<tr><td>GU305914.1</td><td>78703</td><td>78725</td><td>-</td><td>0</td><td>CGGAGGCGGGTCGGTCGGGG</td><td>TGG</td><td>||||||||[||||||||||||]PAM</td><td>NA</td><td>Intergenic</td><td>NA</td><td>NA</td></tr>
</table>

T28299 CGCTGTTGCCACTGTCGCTCCGG 1000
Oligo fwd CACCCGCTGTTGCCACTGTCGCTC
Oligo rev AAACGAGCGACAGTGGCAACAGCG
Chromosome start end strand MM target_seq PAM alignment distance position gene name gene id
GU305914.1 78657 78679 - 0 CGCTGTTGCCACTGTCGCTC CGG ||||||||[||||||||||||]PAM NA Intergenic NA NA

T28300 TTGCCACTGTCGCTCCGGAGAGG 1000
Oligo fwd CACCTTGCCACTGTCGCTCCGGAG
Oligo rev AAACCTCCGGAGCGACAGTGGCAA
Chromosome start end strand MM target_seq PAM alignment distance position gene name gene id
GU305914.1 78652 78674 - 0 TTGCCACTGTCGCTCCGGAG AGG ||||||||[||||||||||||]PAM NA Intergenic NA NA

T28301 TGCCACTGTCGCTCCGGAGAGGG 1000
Oligo fwd CACCTGCCACTGTCGCTCCGGAGA
Oligo rev AAACTCTCCGGAGCGACAGTGGCA
Chromosome start end strand MM target_seq PAM alignment distance position gene name gene id
GU305914.1 78651 78673 - 0 TGCCACTGTCGCTCCGGAGA GGG ||||||||[||||||||||||]PAM NA Intergenic NA NA

T28302 CTGTCGCTCCGGAGAGGGAGAGG 1000
Oligo fwd CACCCTGTCGCTCCGGAGAGGGAG
Oligo rev AAACCTCCCTCTCCGGAGCGACAG
Chromosome start end strand MM target_seq PAM alignment distance position gene name gene id
GU305914.1 78646 78668 - 0 CTGTCGCTCCGGAGAGGGAG AGG ||||||||[||||||||||||]PAM NA Intergenic NA NA

T28303 CCGGAGAGGGAGAGGAGCAGCGG 1000
Oligo fwd CACCCCGGAGAGGGAGAGGAGCAG
Oligo rev AAACCTGCTCCTCTCCCTCTCCGG
Chromosome start end strand MM target_seq PAM alignment distance position gene name gene id
GU305914.1 78638 78660 - 0 CCGGAGAGGGAGAGGAGCAG CGG ||||||||[||||||||||||]PAM NA Intergenic NA NA

T28304 GGGAGAGGAGCAGCGGCACGCGG 1000
Oligo fwd CACCGGGAGAGGAGCAGCGGCACG
Oligo rev AAACCGTGCCGCTGCTCCTCTCCC
Chromosome start end strand MM target_seq PAM alignment distance position gene name gene id
GU305914.1 78631 78653 - 0 GGGAGAGGAGCAGCGGCACG CGG ||||||||[||||||||||||]PAM NA Intergenic NA NA

T28305 GAGGAGCAGCGGCACGCGGACGG 1000
Oligo fwd CACCGAGGAGCAGCGGCACGCGGA
Oligo rev AAACTCCGCGTGCCGCTGCTCCTC
Chromosome start end strand MM target_seq PAM alignment distance position gene name gene id
GU305914.1 78627 78649 - 0 GAGGAGCAGCGGCACGCGGA CGG ||||||||[||||||||||||]PAM NA Intergenic NA NA

T28306 AGGAGCAGCGGCACGCGGACGGG 1000
Oligo fwd CACCAGGAGCAGCGGCACGCGGAC
Oligo rev AAACGTCCGCGTGCCGCTGCTCCT
Chromosome start end strand MM target_seq PAM alignment distance position gene name gene id
GU305914.1 78626 78648 - 0 AGGAGCAGCGGCACGCGGAC GGG ||||||||[||||||||||||]PAM NA Intergenic NA NA

T28307 CACGCGGACGGGCTCCTCAGCGG 1000
Oligo fwd CACCCACGCGGACGGGCTCCTCAG
Oligo rev AAACCTGAGGAGCCCGTCCGCGTG
Chromosome start end strand MM target_seq PAM alignment distance position gene name gene id
GU305914.1 78615 78637 - 0 CACGCGGACGGGCTCCTCAG CGG ||||||||[||||||||||||]PAM NA Intergenic NA NA

T28308 GGCTCCTCAGCGGTGTTCTGAGG 1000
Oligo fwd CACCGGCTCCTCAGCGGTGTTCTG
Oligo rev AAACCAGAACACCGCTGAGGAGCC
Chromosome start end strand MM target_seq PAM alignment distance position gene name gene id
GU305914.1 78605 78627 - 0 GGCTCCTCAGCGGTGTTCTG AGG ||||||||[||||||||||||]PAM NA Intergenic NA NA

T28309 CGGTGTTCTGAGGCCGCTGCCGG 1000
Oligo fwd CACCCGGTGTTCTGAGGCCGCTGC
Oligo rev AAACGCAGCGGCCTCAGAACACCG
Chromosome start end strand MM target_seq PAM alignment distance position gene name gene id
GU305914.1 78595 78617 - 0 CGGTGTTCTGAGGCCGCTGC CGG ||||||||[||||||||||||]PAM NA Intergenic NA NA

T28310 GTTCTGAGGCCGCTGCCGGCCGG 1000
Oligo fwd CACCGTTCTGAGGCCGCTGCCGGC
Oligo rev AAACGCCGGCAGCGGCCTCAGAAC
Chromosome start end strand MM target_seq PAM alignment distance position gene name gene id
GU305914.1 78591 78613 - 0 GTTCTGAGGCCGCTGCCGGC CGG ||||||||[||||||||||||]PAM NA Intergenic NA NA

T28311 TTCTGAGGCCGCTGCCGGCCGGG 1000
Oligo fwd CACCTTCTGAGGCCGCTGCCGGCC
Oligo rev AAACGGCCGGCAGCGGCCTCAGAA
Chromosome start end strand MM target_seq PAM alignment distance position gene name gene id
GU305914.1 78590 78612 - 0 TTCTGAGGCCGCTGCCGGCC GGG ||||||||[||||||||||||]PAM NA Intergenic NA NA

T28312 TGAGGCCGCTGCCGGCCGGGCGG 1000
Oligo fwd CACCTGAGGCCGCTGCCGGCCGGG
Oligo rev AAACCCCGGCCGGCAGCGGCCTCA
Chromosome start end strand MM target_seq PAM alignment distance position gene name gene id
GU305914.1 78587 78609 - 0 TGAGGCCGCTGCCGGCCGGG CGG ||||||||[||||||||||||]PAM NA Intergenic NA NA

T28313 CGCTGCCGGCCGGGCGGTGTCGG 1000
Oligo fwd CACCCGCTGCCGGCCGGGCGGTGT
Oligo rev AAACACACCGCCCGGCCGGCAGCG
Chromosome start end strand MM target_seq PAM alignment distance position gene name gene id

| | | | | | | | | | | | |
|---|---|---|---|---|---|---|---|---|---|---|---|
| GU305914.1 | 78581 | 78603 | - | 0 | CGCTGCCGGCCGGGCGGTGT | CGG | ||||||||[||||||||||||]PAM | NA | Intergenic | NA | NA |

T28314 GCTGCCGGCCGGGCGGTGTCGGG 1000
Oligo fwd CACCGCTGCCGGCCGGGCGGTGTC
Oligo rev AAACGACACCGCCCGGCCGGCAGC

| Chromosome | start | end | strand | MM | target_seq | PAM | alignment | distance | position | gene name | gene id |
|---|---|---|---|---|---|---|---|---|---|---|---|
| GU305914.1 | 78580 | 78602 | - | 0 | GCTGCCGGCCGGGCGGTGTC | GGG | ||||||||[||||||||||||]PAM | NA | Intergenic | NA | NA |

T28315 GGTGTCGGGCGCGCTTGACGCGG 1000
Oligo fwd CACCGGTGTCGGGCGCGCTTGACG
Oligo rev AAACCGTCAAGCGCGCCCGACACC

| Chromosome | start | end | strand | MM | target_seq | PAM | alignment | distance | position | gene name | gene id |
|---|---|---|---|---|---|---|---|---|---|---|---|
| GU305914.1 | 78566 | 78588 | - | 0 | GGTGTCGGGCGCGCTTGACG | CGG | ||||||||[||||||||||||]PAM | NA | Intergenic | NA | NA |

T28316 CGGGCGCGCTTGACGCGGCGCGG 1000
Oligo fwd CACCCGGGCGCGCTTGACGCGGCG
Oligo rev AAACCGCCGCGTCAAGCGCGCCCG

| Chromosome | start | end | strand | MM | target_seq | PAM | alignment | distance | position | gene name | gene id |
|---|---|---|---|---|---|---|---|---|---|---|---|
| GU305914.1 | 78561 | 78583 | - | 0 | CGGGCGCGCTTGACGCGGCG | CGG | ||||||||[||||||||||||]PAM | NA | Intergenic | NA | NA |

T28317 CTCCTCCTTCCTCCTCTTCTCGG 1000
Oligo fwd CACCCTCCTCCTTCCTCCTCTTCT
Oligo rev AAACAGAAGAGGAGGAAGGAGGAG

| Chromosome | start | end | strand | MM | target_seq | PAM | alignment | distance | position | gene name | gene id |
|---|---|---|---|---|---|---|---|---|---|---|---|
| GU305914.1 | 78506 | 78528 | - | 0 | CTCCTCCTTCCTCCTCTTCT | CGG | ||||||||[||||||||||||]PAM | NA | Intergenic | NA | NA |
| GU305914.1 | 39677 | 39699 | - | 4 | GTGCTCCTTCCTCCTCCCCT | CGG | -|-|||||[||||||||--||]PAM | NA | Intergenic | NA | NA |

T28318 TCCTTCCTCCTCTTCTCGGTCGG 1000
Oligo fwd CACCTCCTTCCTCCTCTTCTCGGT
Oligo rev AAACACCGAGAAGAGGAGGAAGGA

| Chromosome | start | end | strand | MM | target_seq | PAM | alignment | distance | position | gene name | gene id |
|---|---|---|---|---|---|---|---|---|---|---|---|
| GU305914.1 | 78502 | 78524 | - | 0 | TCCTTCCTCCTCTTCTCGGT | CGG | ||||||||[||||||||||||]PAM | NA | Intergenic | NA | NA |

T28319 TTCCTCCTCTTCTCGGTCGGCGG 1000
Oligo fwd CACCTTCCTCCTCTTCTCGGTCGG
Oligo rev AAACCCGACCGAGAAGAGGAGGAA

| Chromosome | start | end | strand | MM | target_seq | PAM | alignment | distance | position | gene name | gene id |
|---|---|---|---|---|---|---|---|---|---|---|---|
| GU305914.1 | 78499 | 78521 | - | 0 | TTCCTCCTCTTCTCGGTCGG | CGG | ||||||||[||||||||||||]PAM | NA | Intergenic | NA | NA |

T28320 GCGTCCCGTTCCTCTCCCTCCGG 1000
Oligo fwd CACCGCGTCCCGTTCCTCTCCCTC
Oligo rev AAACGAGGGAGAGGAACGGGACGC

| Chromosome | start | end | strand | MM | target_seq | PAM | alignment | distance | position | gene name | gene id |
|---|---|---|---|---|---|---|---|---|---|---|---|
| GU305914.1 | 78477 | 78499 | - | 0 | GCGTCCCGTTCCTCTCCCTC | CGG | ||||||||[||||||||||||]PAM | NA | Intergenic | NA | NA |

T28321 CGTCCCGTTCCTCTCCCTCCGGG 1000
Oligo fwd CACCCGTCCCGTTCCTCTCCCTCC
Oligo rev AAACGGAGGGAGAGGAACGGGACG

| Chromosome | start | end | strand | MM | target_seq | PAM | alignment | distance | position | gene name | gene id |
|---|---|---|---|---|---|---|---|---|---|---|---|
| GU305914.1 | 78476 | 78498 | - | 0 | CGTCCCGTTCCTCTCCCTCC | GGG | ||||||||[||||||||||||]PAM | NA | Intergenic | NA | NA |

T28322 CTCCTAAACATACTTCTCGACGG 1000
Oligo fwd CACCCTCCTAAACATACTTCTCGA
Oligo rev AAACTCGAGAAGTATGTTTAGGAG

| Chromosome | start | end | strand | MM | target_seq | PAM | alignment | distance | position | gene name | gene id |
|---|---|---|---|---|---|---|---|---|---|---|---|
| GU305914.1 | 78453 | 78475 | - | 0 | CTCCTAAACATACTTCTCGA | CGG | ||||||||[||||||||||||]PAM | NA | Intergenic | NA | NA |

T28323 ATACTTCTCGACGGTGAAGCTGG 1000
Oligo fwd CACCATACTTCTCGACGGTGAAGC
Oligo rev AAACGCTTCACCGTCGAGAAGTAT

| Chromosome | start | end | strand | MM | target_seq | PAM | alignment | distance | position | gene name | gene id |
|---|---|---|---|---|---|---|---|---|---|---|---|
| GU305914.1 | 78444 | 78466 | - | 0 | ATACTTCTCGACGGTGAAGC | TGG | ||||||||[||||||||||||]PAM | NA | Intergenic | NA | NA |

T28324 TACTTCTCGACGGTGAAGCTGGG 1000
Oligo fwd CACCTACTTCTCGACGGTGAAGCT
Oligo rev AAACAGCTTCACCGTCGAGAAGTA

| Chromosome | start | end | strand | MM | target_seq | PAM | alignment | distance | position | gene name | gene id |
|---|---|---|---|---|---|---|---|---|---|---|---|
| GU305914.1 | 78443 | 78465 | - | 0 | TACTTCTCGACGGTGAAGCT | GGG | ||||||||[||||||||||||]PAM | NA | Intergenic | NA | NA |

T28325 ACTTCTCGACGGTGAAGCTGGGG 1000
Oligo fwd CACCACTTCTCGACGGTGAAGCTG
Oligo rev AAACCAGCTTCACCGTCGAGAAGT

| Chromosome | start | end | strand | MM | target_seq | PAM | alignment | distance | position | gene name | gene id |
|---|---|---|---|---|---|---|---|---|---|---|---|
| GU305914.1 | 78442 | 78464 | - | 0 | ACTTCTCGACGGTGAAGCTG | GGG | ||||||||[||||||||||||]PAM | NA | Intergenic | NA | NA |

T28326 CGGTGAAGCTGGGGTCCACGAGG 1000
Oligo fwd CACCCGGTGAAGCTGGGGTCCACG
Oligo rev AAACCGTGGACCCCAGCTTCACCG

| Chromosome | start | end | strand | MM | target_seq | PAM | alignment | distance | position | gene name | gene id |
|---|---|---|---|---|---|---|---|---|---|---|---|
| GU305914.1 | 78433 | 78455 | - | 0 | CGGTGAAGCTGGGGTCCACG | AGG | ||||||||[||||||||||||]PAM | NA | Intergenic | NA | NA |

T28327 AAATCGCGCTCGACGCGCGACGG 1000
Oligo fwd CACCAAATCGCGCTCGACGCGCGA
Oligo rev AAACTCGCGCGTCGAGCGCGATTT

| Chromosome | start | end | strand | MM | target_seq | PAM | alignment | distance | position | gene name | gene id |
|---|---|---|---|---|---|---|---|---|---|---|---|
| GU305914.1 | 78290 | 78312 | - | 0 | AAATCGCGCTCGACGCGCGA | CGG | ||||||||[||||||||||||]PAM | NA | Intergenic | NA | NA |

T28328 AATCGCGCTCGACGCGCGACGGG 1000
Oligo fwd CACCAATCGCGCTCGACGCGCGAC

<table>
<tr><td>Oligo rev</td><td>AAACGTCGCGCGTCGAGCGCGATT</td><td></td><td></td><td></td><td></td><td></td><td></td><td></td><td></td><td></td><td></td><td></td></tr>
<tr><td>Chromosome</td><td>start</td><td></td><td>end</td><td>strand</td><td>MM</td><td>target_seq</td><td>PAM</td><td>alignment</td><td>distance</td><td>position</td><td>gene name</td><td>gene id</td></tr>
<tr><td>GU305914.1</td><td></td><td>78289</td><td>78311</td><td>-</td><td>0</td><td>AATCGCGCTCGACGCGCGAC</td><td>GGG</td><td>||||||||[||||||||||||]PAM</td><td>NA</td><td>Intergenic</td><td>NA</td><td>NA</td></tr>
<tr><td></td><td></td><td></td><td></td><td></td><td></td><td></td><td></td><td></td><td></td><td></td><td></td><td></td></tr>
<tr><td>T28329</td><td>GTACGCGTTCGTCGAGCAGATGG</td><td></td><td>1000</td><td></td><td></td><td></td><td></td><td></td><td></td><td></td><td></td><td></td></tr>
<tr><td>Oligo fwd</td><td>CACCGTACGCGTTCGTCGAGCAGA</td><td></td><td></td><td></td><td></td><td></td><td></td><td></td><td></td><td></td><td></td><td></td></tr>
<tr><td>Oligo rev</td><td>AAACTCTGCTCGACGAACGCGTAC</td><td></td><td></td><td></td><td></td><td></td><td></td><td></td><td></td><td></td><td></td><td></td></tr>
<tr><td>Chromosome</td><td>start</td><td></td><td>end</td><td>strand</td><td>MM</td><td>target_seq</td><td>PAM</td><td>alignment</td><td>distance</td><td>position</td><td>gene name</td><td>gene id</td></tr>
<tr><td>GU305914.1</td><td></td><td>78267</td><td>78289</td><td>-</td><td>0</td><td>GTACGCGTTCGTCGAGCAGA</td><td>TGG</td><td>||||||||[||||||||||||]PAM</td><td>NA</td><td>Intergenic</td><td>NA</td><td>NA</td></tr>
<tr><td></td><td></td><td></td><td></td><td></td><td></td><td></td><td></td><td></td><td></td><td></td><td></td><td></td></tr>
<tr><td>T28330</td><td>CGCTTCGTTTAAATACTCGTTGG</td><td></td><td>1000</td><td></td><td></td><td></td><td></td><td></td><td></td><td></td><td></td><td></td></tr>
<tr><td>Oligo fwd</td><td>CACCCGCTTCGTTTAAATACTCGT</td><td></td><td></td><td></td><td></td><td></td><td></td><td></td><td></td><td></td><td></td><td></td></tr>
<tr><td>Oligo rev</td><td>AAACACGAGTATTTAAACGAAGCG</td><td></td><td></td><td></td><td></td><td></td><td></td><td></td><td></td><td></td><td></td><td></td></tr>
<tr><td>Chromosome</td><td>start</td><td></td><td>end</td><td>strand</td><td>MM</td><td>target_seq</td><td>PAM</td><td>alignment</td><td>distance</td><td>position</td><td>gene name</td><td>gene id</td></tr>
<tr><td>GU305914.1</td><td></td><td>78237</td><td>78259</td><td>-</td><td>0</td><td>CGCTTCGTTTAAATACTCGT</td><td>TGG</td><td>||||||||[||||||||||||]PAM</td><td>NA</td><td>Intergenic</td><td>NA</td><td>NA</td></tr>
<tr><td></td><td></td><td></td><td></td><td></td><td></td><td></td><td></td><td></td><td></td><td></td><td></td><td></td></tr>
<tr><td>T28331</td><td>GCTTCGTTTAAATACTCGTTGGG</td><td></td><td>1000</td><td></td><td></td><td></td><td></td><td></td><td></td><td></td><td></td><td></td></tr>
<tr><td>Oligo fwd</td><td>CACCGCTTCGTTTAAATACTCGTT</td><td></td><td></td><td></td><td></td><td></td><td></td><td></td><td></td><td></td><td></td><td></td></tr>
<tr><td>Oligo rev</td><td>AAACAACGAGTATTTAAACGAAGC</td><td></td><td></td><td></td><td></td><td></td><td></td><td></td><td></td><td></td><td></td><td></td></tr>
<tr><td>Chromosome</td><td>start</td><td></td><td>end</td><td>strand</td><td>MM</td><td>target_seq</td><td>PAM</td><td>alignment</td><td>distance</td><td>position</td><td>gene name</td><td>gene id</td></tr>
<tr><td>GU305914.1</td><td></td><td>78236</td><td>78258</td><td>-</td><td>0</td><td>GCTTCGTTTAAATACTCGTT</td><td>GGG</td><td>||||||||[||||||||||||]PAM</td><td>NA</td><td>Intergenic</td><td>NA</td><td>NA</td></tr>
<tr><td></td><td></td><td></td><td></td><td></td><td></td><td></td><td></td><td></td><td></td><td></td><td></td><td></td></tr>
<tr><td>T28332</td><td>CGTTGGGCACGTCGAAAAAGAGG</td><td></td><td>1000</td><td></td><td></td><td></td><td></td><td></td><td></td><td></td><td></td><td></td></tr>
<tr><td>Oligo fwd</td><td>CACCCGTTGGGCACGTCGAAAAAG</td><td></td><td></td><td></td><td></td><td></td><td></td><td></td><td></td><td></td><td></td><td></td></tr>
<tr><td>Oligo rev</td><td>AAACCTTTTTCGACGTGCCCAACG</td><td></td><td></td><td></td><td></td><td></td><td></td><td></td><td></td><td></td><td></td><td></td></tr>
<tr><td>Chromosome</td><td>start</td><td></td><td>end</td><td>strand</td><td>MM</td><td>target_seq</td><td>PAM</td><td>alignment</td><td>distance</td><td>position</td><td>gene name</td><td>gene id</td></tr>
<tr><td>GU305914.1</td><td></td><td>78220</td><td>78242</td><td>-</td><td>0</td><td>CGTTGGGCACGTCGAAAAAG</td><td>AGG</td><td>||||||||[||||||||||||]PAM</td><td>NA</td><td>Intergenic</td><td>NA</td><td>NA</td></tr>
<tr><td></td><td></td><td></td><td></td><td></td><td></td><td></td><td></td><td></td><td></td><td></td><td></td><td></td></tr>
<tr><td>T28333</td><td>ACGTCGAAAAAGAGGATGTTCGG</td><td></td><td>1000</td><td></td><td></td><td></td><td></td><td></td><td></td><td></td><td></td><td></td></tr>
<tr><td>Oligo fwd</td><td>CACCACGTCGAAAAAGAGGATGTT</td><td></td><td></td><td></td><td></td><td></td><td></td><td></td><td></td><td></td><td></td><td></td></tr>
<tr><td>Oligo rev</td><td>AAACAACATCCTCTTTTTCGACGT</td><td></td><td></td><td></td><td></td><td></td><td></td><td></td><td></td><td></td><td></td><td></td></tr>
<tr><td>Chromosome</td><td>start</td><td></td><td>end</td><td>strand</td><td>MM</td><td>target_seq</td><td>PAM</td><td>alignment</td><td>distance</td><td>position</td><td>gene name</td><td>gene id</td></tr>
<tr><td>GU305914.1</td><td></td><td>78212</td><td>78234</td><td>-</td><td>0</td><td>ACGTCGAAAAAGAGGATGTT</td><td>CGG</td><td>||||||||[||||||||||||]PAM</td><td>NA</td><td>Intergenic</td><td>NA</td><td>NA</td></tr>
<tr><td></td><td></td><td></td><td></td><td></td><td></td><td></td><td></td><td></td><td></td><td></td><td></td><td></td></tr>
<tr><td>T28334</td><td>CGCATGTTAGCCGCGCACAGCGG</td><td></td><td>1000</td><td></td><td></td><td></td><td></td><td></td><td></td><td></td><td></td><td></td></tr>
<tr><td>Oligo fwd</td><td>CACCCGCATGTTAGCCGCGCACAG</td><td></td><td></td><td></td><td></td><td></td><td></td><td></td><td></td><td></td><td></td><td></td></tr>
<tr><td>Oligo rev</td><td>AAACCTGTGCGCGGCTAACATGCG</td><td></td><td></td><td></td><td></td><td></td><td></td><td></td><td></td><td></td><td></td><td></td></tr>
<tr><td>Chromosome</td><td>start</td><td></td><td>end</td><td>strand</td><td>MM</td><td>target_seq</td><td>PAM</td><td>alignment</td><td>distance</td><td>position</td><td>gene name</td><td>gene id</td></tr>
<tr><td>GU305914.1</td><td></td><td>78179</td><td>78201</td><td>-</td><td>0</td><td>CGCATGTTAGCCGCGCACAG</td><td>CGG</td><td>||||||||[||||||||||||]PAM</td><td>NA</td><td>Intergenic</td><td>NA</td><td>NA</td></tr>
<tr><td></td><td></td><td></td><td></td><td></td><td></td><td></td><td></td><td></td><td></td><td></td><td></td><td></td></tr>
<tr><td>T28335</td><td>GCATGTTAGCCGCGCACAGCGGG</td><td></td><td>1000</td><td></td><td></td><td></td><td></td><td></td><td></td><td></td><td></td><td></td></tr>
<tr><td>Oligo fwd</td><td>CACCGCATGTTAGCCGCGCACAGC</td><td></td><td></td><td></td><td></td><td></td><td></td><td></td><td></td><td></td><td></td><td></td></tr>
<tr><td>Oligo rev</td><td>AAACGCTGTGCGCGGCTAACATGC</td><td></td><td></td><td></td><td></td><td></td><td></td><td></td><td></td><td></td><td></td><td></td></tr>
<tr><td>Chromosome</td><td>start</td><td></td><td>end</td><td>strand</td><td>MM</td><td>target_seq</td><td>PAM</td><td>alignment</td><td>distance</td><td>position</td><td>gene name</td><td>gene id</td></tr>
<tr><td>GU305914.1</td><td></td><td>78178</td><td>78200</td><td>-</td><td>0</td><td>GCATGTTAGCCGCGCACAGC</td><td>GGG</td><td>||||||||[||||||||||||]PAM</td><td>NA</td><td>Intergenic</td><td>NA</td><td>NA</td></tr>
<tr><td></td><td></td><td></td><td></td><td></td><td></td><td></td><td></td><td></td><td></td><td></td><td></td><td></td></tr>
<tr><td>T28336</td><td>CGCACAGCGGGTCCGCGAACAGG</td><td></td><td>1000</td><td></td><td></td><td></td><td></td><td></td><td></td><td></td><td></td><td></td></tr>
<tr><td>Oligo fwd</td><td>CACCCGCACAGCGGGTCCGCGAAC</td><td></td><td></td><td></td><td></td><td></td><td></td><td></td><td></td><td></td><td></td><td></td></tr>
<tr><td>Oligo rev</td><td>AAACGTTCGCGGACCCGCTGTGCG</td><td></td><td></td><td></td><td></td><td></td><td></td><td></td><td></td><td></td><td></td><td></td></tr>
<tr><td>Chromosome</td><td>start</td><td></td><td>end</td><td>strand</td><td>MM</td><td>target_seq</td><td>PAM</td><td>alignment</td><td>distance</td><td>position</td><td>gene name</td><td>gene id</td></tr>
<tr><td>GU305914.1</td><td></td><td>78166</td><td>78188</td><td>-</td><td>0</td><td>CGCACAGCGGGTCCGCGAAC</td><td>AGG</td><td>||||||||[||||||||||||]PAM</td><td>NA</td><td>Intergenic</td><td>NA</td><td>NA</td></tr>
<tr><td></td><td></td><td></td><td></td><td></td><td></td><td></td><td></td><td></td><td></td><td></td><td></td><td></td></tr>
<tr><td>T28337</td><td>CGATCACGCCGACCTGACGCAGG</td><td></td><td>1000</td><td></td><td></td><td></td><td></td><td></td><td></td><td></td><td></td><td></td></tr>
<tr><td>Oligo fwd</td><td>CACCCGATCACGCCGACCTGACGC</td><td></td><td></td><td></td><td></td><td></td><td></td><td></td><td></td><td></td><td></td><td></td></tr>
<tr><td>Oligo rev</td><td>AAACGCGTCAGGTCGGCGTGATCG</td><td></td><td></td><td></td><td></td><td></td><td></td><td></td><td></td><td></td><td></td><td></td></tr>
<tr><td>Chromosome</td><td>start</td><td></td><td>end</td><td>strand</td><td>MM</td><td>target_seq</td><td>PAM</td><td>alignment</td><td>distance</td><td>position</td><td>gene name</td><td>gene id</td></tr>
<tr><td>GU305914.1</td><td></td><td>78130</td><td>78152</td><td>-</td><td>0</td><td>CGATCACGCCGACCTGACGC</td><td>AGG</td><td>||||||||[||||||||||||]PAM</td><td>NA</td><td>Intergenic</td><td>NA</td><td>NA</td></tr>
<tr><td></td><td></td><td></td><td></td><td></td><td></td><td></td><td></td><td></td><td></td><td></td><td></td><td></td></tr>
<tr><td>T28338</td><td>CGCCGACCTGACGCAGGACCAGG</td><td></td><td>1000</td><td></td><td></td><td></td><td></td><td></td><td></td><td></td><td></td><td></td></tr>
<tr><td>Oligo fwd</td><td>CACCCGCCGACCTGACGCAGGACC</td><td></td><td></td><td></td><td></td><td></td><td></td><td></td><td></td><td></td><td></td><td></td></tr>
<tr><td>Oligo rev</td><td>AAACGGTCCTGCGTCAGGTCGGCG</td><td></td><td></td><td></td><td></td><td></td><td></td><td></td><td></td><td></td><td></td><td></td></tr>
<tr><td>Chromosome</td><td>start</td><td></td><td>end</td><td>strand</td><td>MM</td><td>target_seq</td><td>PAM</td><td>alignment</td><td>distance</td><td>position</td><td>gene name</td><td>gene id</td></tr>
<tr><td>GU305914.1</td><td></td><td>78124</td><td>78146</td><td>-</td><td>0</td><td>CGCCGACCTGACGCAGGACC</td><td>AGG</td><td>||||||||[||||||||||||]PAM</td><td>NA</td><td>Intergenic</td><td>NA</td><td>NA</td></tr>
<tr><td></td><td></td><td></td><td></td><td></td><td></td><td></td><td></td><td></td><td></td><td></td><td></td><td></td></tr>
<tr><td>T28339</td><td>GACCTGACGCAGGACCAGGTAGG</td><td></td><td>1000</td><td></td><td></td><td></td><td></td><td></td><td></td><td></td><td></td><td></td></tr>
<tr><td>Oligo fwd</td><td>CACCGACCTGACGCAGGACCAGGT</td><td></td><td></td><td></td><td></td><td></td><td></td><td></td><td></td><td></td><td></td><td></td></tr>
<tr><td>Oligo rev</td><td>AAACACCTGGTCCTGCGTCAGGTC</td><td></td><td></td><td></td><td></td><td></td><td></td><td></td><td></td><td></td><td></td><td></td></tr>
<tr><td>Chromosome</td><td>start</td><td></td><td>end</td><td>strand</td><td>MM</td><td>target_seq</td><td>PAM</td><td>alignment</td><td>distance</td><td>position</td><td>gene name</td><td>gene id</td></tr>
<tr><td>GU305914.1</td><td></td><td>78120</td><td>78142</td><td>-</td><td>0</td><td>GACCTGACGCAGGACCAGGT</td><td>AGG</td><td>||||||||[||||||||||||]PAM</td><td>NA</td><td>Intergenic</td><td>NA</td><td>NA</td></tr>
<tr><td></td><td></td><td></td><td></td><td></td><td></td><td></td><td></td><td></td><td></td><td></td><td></td><td></td></tr>
<tr><td>T28340</td><td>GTCAGCGCGCTCAGCTCGAACGG</td><td></td><td>1000</td><td></td><td></td><td></td><td></td><td></td><td></td><td></td><td></td><td></td></tr>
<tr><td>Oligo fwd</td><td>CACCGTCAGCGCGCTCAGCTCGAA</td><td></td><td></td><td></td><td></td><td></td><td></td><td></td><td></td><td></td><td></td><td></td></tr>
<tr><td>Oligo rev</td><td>AAACTTCGAGCTGAGCGCGCTGAC</td><td></td><td></td><td></td><td></td><td></td><td></td><td></td><td></td><td></td><td></td><td></td></tr>
<tr><td>Chromosome</td><td>start</td><td></td><td>end</td><td>strand</td><td>MM</td><td>target_seq</td><td>PAM</td><td>alignment</td><td>distance</td><td>position</td><td>gene name</td><td>gene id</td></tr>
<tr><td>GU305914.1</td><td></td><td>78098</td><td>78120</td><td>-</td><td>0</td><td>GTCAGCGCGCTCAGCTCGAA</td><td>CGG</td><td>||||||||[||||||||||||]PAM</td><td>NA</td><td>Intergenic</td><td>NA</td><td>NA</td></tr>
<tr><td></td><td></td><td></td><td></td><td></td><td></td><td></td><td></td><td></td><td></td><td></td><td></td><td></td></tr>
<tr><td>T28341</td><td>CTCGAACGGCTCCTCCGTCCCGG</td><td></td><td>1000</td><td></td><td></td><td></td><td></td><td></td><td></td><td></td><td></td><td></td></tr>
<tr><td>Oligo fwd</td><td>CACCCTCGAACGGCTCCTCCGTCC</td><td></td><td></td><td></td><td></td><td></td><td></td><td></td><td></td><td></td><td></td><td></td></tr>
<tr><td>Oligo rev</td><td>AAACGGACGGAGGAGCCGTTCGAG</td><td></td><td></td><td></td><td></td><td></td><td></td><td></td><td></td><td></td><td></td><td></td></tr>
<tr><td>Chromosome</td><td>start</td><td></td><td>end</td><td>strand</td><td>MM</td><td>target_seq</td><td>PAM</td><td>alignment</td><td>distance</td><td>position</td><td>gene name</td><td>gene id</td></tr>
<tr><td>GU305914.1</td><td></td><td>78084</td><td>78106</td><td>-</td><td>0</td><td>CTCGAACGGCTCCTCCGTCC</td><td>CGG</td><td>||||||||[||||||||||||]PAM</td><td>NA</td><td>Intergenic</td><td>NA</td><td>NA</td></tr>
<tr><td></td><td></td><td></td><td></td><td></td><td></td><td></td><td></td><td></td><td></td><td></td><td></td><td></td></tr>
<tr><td>T28342</td><td>TCGAACGGCTCCTCCGTCCCGGG</td><td></td><td>1000</td><td></td><td></td><td></td><td></td><td></td><td></td><td></td><td></td><td></td></tr>
<tr><td>Oligo fwd</td><td>CACCTCGAACGGCTCCTCCGTCCC</td><td></td><td></td><td></td><td></td><td></td><td></td><td></td><td></td><td></td><td></td><td></td></tr>
<tr><td>Oligo rev</td><td>AAACGGGACGGAGGAGCCGTTCGA</td><td></td><td></td><td></td><td></td><td></td><td></td><td></td><td></td><td></td><td></td><td></td></tr>
<tr><td>Chromosome</td><td>start</td><td></td><td>end</td><td>strand</td><td>MM</td><td>target_seq</td><td>PAM</td><td>alignment</td><td>distance</td><td>position</td><td>gene name</td><td>gene id</td></tr>
<tr><td>GU305914.1</td><td></td><td>78083</td><td>78105</td><td>-</td><td>0</td><td>TCGAACGGCTCCTCCGTCCC</td><td>GGG</td><td>||||||||[||||||||||||]PAM</td><td>NA</td><td>Intergenic</td><td>NA</td><td>NA</td></tr>
<tr><td></td><td></td><td></td><td></td><td></td><td></td><td></td><td></td><td></td><td></td><td></td><td></td><td></td></tr>
<tr><td>T28343</td><td>CGAACGGCTCCTCCGTCCCGGGG</td><td></td><td>1000</td><td></td><td></td><td></td><td></td><td></td><td></td><td></td><td></td><td></td></tr>
</table>

<table>
<tr><td>Oligo fwd</td><td>CACCCGAACGGCTCCTCCGTCCCG</td><td></td><td></td><td></td><td></td><td></td><td></td><td></td><td></td><td></td><td></td></tr>
<tr><td>Oligo rev</td><td>AAACCGGGACGGAGGAGCCGTTCG</td><td></td><td></td><td></td><td></td><td></td><td></td><td></td><td></td><td></td><td></td></tr>
<tr><td>Chromosome</td><td>start</td><td>end</td><td>strand</td><td>MM</td><td>target_seq</td><td>PAM</td><td>alignment</td><td>distance</td><td>position</td><td>gene name</td><td>gene id</td></tr>
<tr><td>GU305914.1</td><td>78082</td><td>78104</td><td>-</td><td>0</td><td>CGAACGGCTCCTCCGTCCCG</td><td>GGG</td><td>||||||||[||||||||||||]PAM</td><td>NA</td><td>Intergenic</td><td>NA</td><td>NA</td></tr>
<tr><td>GU305914.1</td><td>203913</td><td>203935</td><td>+</td><td>4</td><td>CGGACGAGTCCTCTGTCCCG</td><td>TGG</td><td>||-|||--[|||||-||||||]PAM</td><td>NA</td><td>Intergenic</td><td>NA</td><td>NA</td></tr>
<tr><td>T28344</td><td>ACGGCTCCTCCGTCCCGGGGCGG</td><td>1000</td><td></td><td></td><td></td><td></td><td></td><td></td><td></td><td></td><td></td></tr>
<tr><td>Oligo fwd</td><td>CACCACGGCTCCTCCGTCCCGGGG</td><td></td><td></td><td></td><td></td><td></td><td></td><td></td><td></td><td></td><td></td></tr>
<tr><td>Oligo rev</td><td>AAACCCCCGGGACGGAGGAGCCGT</td><td></td><td></td><td></td><td></td><td></td><td></td><td></td><td></td><td></td><td></td></tr>
<tr><td>Chromosome</td><td>start</td><td>end</td><td>strand</td><td>MM</td><td>target_seq</td><td>PAM</td><td>alignment</td><td>distance</td><td>position</td><td>gene name</td><td>gene id</td></tr>
<tr><td>GU305914.1</td><td>78079</td><td>78101</td><td>-</td><td>0</td><td>ACGGCTCCTCCGTCCCGGGG</td><td>CGG</td><td>||||||||[||||||||||||]PAM</td><td>NA</td><td>Intergenic</td><td>NA</td><td>NA</td></tr>
<tr><td>T28345</td><td>CGTCCCGGGGCGGCTCGCCGCGG</td><td>1000</td><td></td><td></td><td></td><td></td><td></td><td></td><td></td><td></td><td></td></tr>
<tr><td>Oligo fwd</td><td>CACCCGTCCCGGGGCGGCTCGCCG</td><td></td><td></td><td></td><td></td><td></td><td></td><td></td><td></td><td></td><td></td></tr>
<tr><td>Oligo rev</td><td>AAACCGGCGAGCCGCCCCGGGACG</td><td></td><td></td><td></td><td></td><td></td><td></td><td></td><td></td><td></td><td></td></tr>
<tr><td>Chromosome</td><td>start</td><td>end</td><td>strand</td><td>MM</td><td>target_seq</td><td>PAM</td><td>alignment</td><td>distance</td><td>position</td><td>gene name</td><td>gene id</td></tr>
<tr><td>GU305914.1</td><td>78069</td><td>78091</td><td>-</td><td>0</td><td>CGTCCCGGGGCGGCTCGCCG</td><td>CGG</td><td>||||||||[||||||||||||]PAM</td><td>NA</td><td>Intergenic</td><td>NA</td><td>NA</td></tr>
<tr><td>T28346</td><td>GCTCGCCGCGGCCAGCGTCTCGG</td><td>1000</td><td></td><td></td><td></td><td></td><td></td><td></td><td></td><td></td><td></td></tr>
<tr><td>Oligo fwd</td><td>CACCGCTCGCCGCGGCCAGCGTCT</td><td></td><td></td><td></td><td></td><td></td><td></td><td></td><td></td><td></td><td></td></tr>
<tr><td>Oligo rev</td><td>AAACAGACGCTGGCCGCGGCGAGC</td><td></td><td></td><td></td><td></td><td></td><td></td><td></td><td></td><td></td><td></td></tr>
<tr><td>Chromosome</td><td>start</td><td>end</td><td>strand</td><td>MM</td><td>target_seq</td><td>PAM</td><td>alignment</td><td>distance</td><td>position</td><td>gene name</td><td>gene id</td></tr>
<tr><td>GU305914.1</td><td>78057</td><td>78079</td><td>-</td><td>0</td><td>GCTCGCCGCGGCCAGCGTCT</td><td>CGG</td><td>||||||||[||||||||||||]PAM</td><td>NA</td><td>Intergenic</td><td>NA</td><td>NA</td></tr>
<tr><td>T28347</td><td>CTCGCCGCGGCCAGCGTCTCGGG</td><td>1000</td><td></td><td></td><td></td><td></td><td></td><td></td><td></td><td></td><td></td></tr>
<tr><td>Oligo fwd</td><td>CACCCTCGCCGCGGCCAGCGTCTC</td><td></td><td></td><td></td><td></td><td></td><td></td><td></td><td></td><td></td><td></td></tr>
<tr><td>Oligo rev</td><td>AAACGAGACGCTGGCCGCGGCGAG</td><td></td><td></td><td></td><td></td><td></td><td></td><td></td><td></td><td></td><td></td></tr>
<tr><td>Chromosome</td><td>start</td><td>end</td><td>strand</td><td>MM</td><td>target_seq</td><td>PAM</td><td>alignment</td><td>distance</td><td>position</td><td>gene name</td><td>gene id</td></tr>
<tr><td>GU305914.1</td><td>78056</td><td>78078</td><td>-</td><td>0</td><td>CTCGCCGCGGCCAGCGTCTC</td><td>GGG</td><td>||||||||[||||||||||||]PAM</td><td>NA</td><td>Intergenic</td><td>NA</td><td>NA</td></tr>
<tr><td>T28348</td><td>TCGCCGCGGCCAGCGTCTCGGGG</td><td>1000</td><td></td><td></td><td></td><td></td><td></td><td></td><td></td><td></td><td></td></tr>
<tr><td>Oligo fwd</td><td>CACCTCGCCGCGGCCAGCGTCTCG</td><td></td><td></td><td></td><td></td><td></td><td></td><td></td><td></td><td></td><td></td></tr>
<tr><td>Oligo rev</td><td>AAACCGAGACGCTGGCCGCGGCGA</td><td></td><td></td><td></td><td></td><td></td><td></td><td></td><td></td><td></td><td></td></tr>
<tr><td>Chromosome</td><td>start</td><td>end</td><td>strand</td><td>MM</td><td>target_seq</td><td>PAM</td><td>alignment</td><td>distance</td><td>position</td><td>gene name</td><td>gene id</td></tr>
<tr><td>GU305914.1</td><td>78055</td><td>78077</td><td>-</td><td>0</td><td>TCGCCGCGGCCAGCGTCTCG</td><td>GGG</td><td>||||||||[||||||||||||]PAM</td><td>NA</td><td>Intergenic</td><td>NA</td><td>NA</td></tr>
<tr><td>T28349</td><td>TCCTTGTTCTTGAGCTGCTGCGG</td><td>1000</td><td></td><td></td><td></td><td></td><td></td><td></td><td></td><td></td><td></td></tr>
<tr><td>Oligo fwd</td><td>CACCTCCTTGTTCTTGAGCTGCTG</td><td></td><td></td><td></td><td></td><td></td><td></td><td></td><td></td><td></td><td></td></tr>
<tr><td>Oligo rev</td><td>AAACCAGCAGCTCAAGAACAAGGA</td><td></td><td></td><td></td><td></td><td></td><td></td><td></td><td></td><td></td><td></td></tr>
<tr><td>Chromosome</td><td>start</td><td>end</td><td>strand</td><td>MM</td><td>target_seq</td><td>PAM</td><td>alignment</td><td>distance</td><td>position</td><td>gene name</td><td>gene id</td></tr>
<tr><td>GU305914.1</td><td>78008</td><td>78030</td><td>-</td><td>0</td><td>TCCTTGTTCTTGAGCTGCTG</td><td>CGG</td><td>||||||||[||||||||||||]PAM</td><td>NA</td><td>Intergenic</td><td>NA</td><td>NA</td></tr>
<tr><td>T28350</td><td>CCTTGTTCTTGAGCTGCTGCGGG</td><td>1000</td><td></td><td></td><td></td><td></td><td></td><td></td><td></td><td></td><td></td></tr>
<tr><td>Oligo fwd</td><td>CACCCCTTGTTCTTGAGCTGCTGC</td><td></td><td></td><td></td><td></td><td></td><td></td><td></td><td></td><td></td><td></td></tr>
<tr><td>Oligo rev</td><td>AAACGCAGCAGCTCAAGAACAAGG</td><td></td><td></td><td></td><td></td><td></td><td></td><td></td><td></td><td></td><td></td></tr>
<tr><td>Chromosome</td><td>start</td><td>end</td><td>strand</td><td>MM</td><td>target_seq</td><td>PAM</td><td>alignment</td><td>distance</td><td>position</td><td>gene name</td><td>gene id</td></tr>
<tr><td>GU305914.1</td><td>78007</td><td>78029</td><td>-</td><td>0</td><td>CCTTGTTCTTGAGCTGCTGC</td><td>GGG</td><td>||||||||[||||||||||||]PAM</td><td>NA</td><td>Intergenic</td><td>NA</td><td>NA</td></tr>
<tr><td>T28351</td><td>CCGAGTTGCGTCCGTGCGACAGG</td><td>1000</td><td></td><td></td><td></td><td></td><td></td><td></td><td></td><td></td><td></td></tr>
<tr><td>Oligo fwd</td><td>CACCCCGAGTTGCGTCCGTGCGAC</td><td></td><td></td><td></td><td></td><td></td><td></td><td></td><td></td><td></td><td></td></tr>
<tr><td>Oligo rev</td><td>AAACGTCGCACGGACGCAACTCGG</td><td></td><td></td><td></td><td></td><td></td><td></td><td></td><td></td><td></td><td></td></tr>
<tr><td>Chromosome</td><td>start</td><td>end</td><td>strand</td><td>MM</td><td>target_seq</td><td>PAM</td><td>alignment</td><td>distance</td><td>position</td><td>gene name</td><td>gene id</td></tr>
<tr><td>GU305914.1</td><td>77980</td><td>78002</td><td>-</td><td>0</td><td>CCGAGTTGCGTCCGTGCGAC</td><td>AGG</td><td>||||||||[||||||||||||]PAM</td><td>NA</td><td>Intergenic</td><td>NA</td><td>NA</td></tr>
<tr><td>T28352</td><td>GTGCGACAGGATCTCCGCCATGG</td><td>1000</td><td></td><td></td><td></td><td></td><td></td><td></td><td></td><td></td><td></td></tr>
<tr><td>Oligo fwd</td><td>CACCGTGCGACAGGATCTCCGCCA</td><td></td><td></td><td></td><td></td><td></td><td></td><td></td><td></td><td></td><td></td></tr>
<tr><td>Oligo rev</td><td>AAACTGGCGGAGATCCTGTCGCAC</td><td></td><td></td><td></td><td></td><td></td><td></td><td></td><td></td><td></td><td></td></tr>
<tr><td>Chromosome</td><td>start</td><td>end</td><td>strand</td><td>MM</td><td>target_seq</td><td>PAM</td><td>alignment</td><td>distance</td><td>position</td><td>gene name</td><td>gene id</td></tr>
<tr><td>GU305914.1</td><td>77967</td><td>77989</td><td>-</td><td>0</td><td>GTGCGACAGGATCTCCGCCA</td><td>TGG</td><td>||||||||[||||||||||||]PAM</td><td>NA</td><td>Intergenic</td><td>NA</td><td>NA</td></tr>
<tr><td>T28353</td><td>CCGCCATGGTCAGCTGCGTCCGG</td><td>1000</td><td></td><td></td><td></td><td></td><td></td><td></td><td></td><td></td><td></td></tr>
<tr><td>Oligo fwd</td><td>CACCCCGCCATGGTCAGCTGCGTC</td><td></td><td></td><td></td><td></td><td></td><td></td><td></td><td></td><td></td><td></td></tr>
<tr><td>Oligo rev</td><td>AAACGACGCAGCTGACCATGGCGG</td><td></td><td></td><td></td><td></td><td></td><td></td><td></td><td></td><td></td><td></td></tr>
<tr><td>Chromosome</td><td>start</td><td>end</td><td>strand</td><td>MM</td><td>target_seq</td><td>PAM</td><td>alignment</td><td>distance</td><td>position</td><td>gene name</td><td>gene id</td></tr>
<tr><td>GU305914.1</td><td>77953</td><td>77975</td><td>-</td><td>0</td><td>CCGCCATGGTCAGCTGCGTC</td><td>CGG</td><td>||||||||[||||||||||||]PAM</td><td>NA</td><td>Intergenic</td><td>NA</td><td>NA</td></tr>
<tr><td>GU305914.1</td><td>132693</td><td>132715</td><td>+</td><td>4</td><td>GCGCCGAGGTCAGCTTCGTC</td><td>AGG</td><td>-||||--|[|||||||-||||]PAM</td><td>NA</td><td>Intergenic</td><td>NA</td><td>NA</td></tr>
<tr><td>T28354</td><td>GCGTCCGGTCGATGATCTTCAGG</td><td>1000</td><td></td><td></td><td></td><td></td><td></td><td></td><td></td><td></td><td></td></tr>
<tr><td>Oligo fwd</td><td>CACCGCGTCCGGTCGATGATCTTC</td><td></td><td></td><td></td><td></td><td></td><td></td><td></td><td></td><td></td><td></td></tr>
<tr><td>Oligo rev</td><td>AAACGAAGATCATCGACCGGACGC</td><td></td><td></td><td></td><td></td><td></td><td></td><td></td><td></td><td></td><td></td></tr>
<tr><td>Chromosome</td><td>start</td><td>end</td><td>strand</td><td>MM</td><td>target_seq</td><td>PAM</td><td>alignment</td><td>distance</td><td>position</td><td>gene name</td><td>gene id</td></tr>
<tr><td>GU305914.1</td><td>77938</td><td>77960</td><td>-</td><td>0</td><td>GCGTCCGGTCGATGATCTTC</td><td>AGG</td><td>||||||||[||||||||||||]PAM</td><td>NA</td><td>Intergenic</td><td>NA</td><td>NA</td></tr>
<tr><td>T28355</td><td>CAGGTTGTTGCGCGAGTAGTTGG</td><td>1000</td><td></td><td></td><td></td><td></td><td></td><td></td><td></td><td></td><td></td></tr>
<tr><td>Oligo fwd</td><td>CACCCAGGTTGTTGCGCGAGTAGT</td><td></td><td></td><td></td><td></td><td></td><td></td><td></td><td></td><td></td><td></td></tr>
<tr><td>Oligo rev</td><td>AAACACTACTCGCGCAACAACCTG</td><td></td><td></td><td></td><td></td><td></td><td></td><td></td><td></td><td></td><td></td></tr>
<tr><td>Chromosome</td><td>start</td><td>end</td><td>strand</td><td>MM</td><td>target_seq</td><td>PAM</td><td>alignment</td><td>distance</td><td>position</td><td>gene name</td><td>gene id</td></tr>
<tr><td>GU305914.1</td><td>77919</td><td>77941</td><td>-</td><td>0</td><td>CAGGTTGTTGCGCGAGTAGT</td><td>TGG</td><td>||||||||[||||||||||||]PAM</td><td>NA</td><td>Intergenic</td><td>NA</td><td>NA</td></tr>
<tr><td>T28356</td><td>TGCGCTCCCGCAGCTGCTGCAGG</td><td>1000</td><td></td><td></td><td></td><td></td><td></td><td></td><td></td><td></td><td></td></tr>
<tr><td>Oligo fwd</td><td>CACCTGCGCTCCCGCAGCTGCTGC</td><td></td><td></td><td></td><td></td><td></td><td></td><td></td><td></td><td></td><td></td></tr>
<tr><td>Oligo rev</td><td>AAACGCAGCAGCTGCGGGAGCGCA</td><td></td><td></td><td></td><td></td><td></td><td></td><td></td><td></td><td></td><td></td></tr>
<tr><td>Chromosome</td><td>start</td><td>end</td><td>strand</td><td>MM</td><td>target_seq</td><td>PAM</td><td>alignment</td><td>distance</td><td>position</td><td>gene name</td><td>gene id</td></tr>
<tr><td>GU305914.1</td><td>77893</td><td>77915</td><td>-</td><td>0</td><td>TGCGCTCCCGCAGCTGCTGC</td><td>AGG</td><td>||||||||[||||||||||||]PAM</td><td>NA</td><td>Intergenic</td><td>NA</td><td>NA</td></tr>
<tr><td>T28357</td><td>GCTGCAGGAACACGTTGTCGCGG</td><td>1000</td><td></td><td></td><td></td><td></td><td></td><td></td><td></td><td></td><td></td></tr>
<tr><td>Oligo fwd</td><td>CACCGCTGCAGGAACACGTTGTCG</td><td></td><td></td><td></td><td></td><td></td><td></td><td></td><td></td><td></td><td></td></tr>
<tr><td>Oligo rev</td><td>AAACCGACAACGTGTTCCTGCAGC</td><td></td><td></td><td></td><td></td><td></td><td></td><td></td><td></td><td></td><td></td></tr>
<tr><td>Chromosome</td><td>start</td><td>end</td><td>strand</td><td>MM</td><td>target_seq</td><td>PAM</td><td>alignment</td><td>distance</td><td>position</td><td>gene name</td><td>gene id</td></tr>
</table>

| | | | | | | | | | | | |
|---|---|---|---|---|---|---|---|---|---|---|---|
| GU305914.1 | 77878 | 77900 | - | 0 | GCTGCAGGAACACGTTGTCG | CGG | \|\|\|\|\|\|\|\|[\|\|\|\|\|\|\|\|\|\|\|\|]PAM | NA | Intergenic | NA | NA |
| GU305914.1 | 59979 | 60001 | + | 4 | GCTCGAGAAAGACGTTGTCG | CGG | \|\|\|--\|\|-[\|\|-\|\|\|\|\|\|\|\|\|]PAM | NA | Intergenic | NA | NA |
| T28358 | ACGCCATCAGCTCGCACAGCAGG | 1000 | | | | | | | | | |
| Oligo fwd | CACCACGCCATCAGCTCGCACAGC | | | | | | | | | | |
| Oligo rev | AAACGCTGTGCGAGCTGATGGCGT | | | | | | | | | | |
| Chromosome | start | end | strand | MM | target_seq | PAM | alignment | distance | position | gene name | gene id |
| GU305914.1 | 77848 | 77870 | - | 0 | ACGCCATCAGCTCGCACAGC | AGG | \|\|\|\|\|\|\|\|[\|\|\|\|\|\|\|\|\|\|\|\|]PAM | NA | Intergenic | NA | NA |
| T28359 | CTCGCACAGCAGGCACACCGTGG | 1000 | | | | | | | | | |
| Oligo fwd | CACCCTCGCACAGCAGGCACACCG | | | | | | | | | | |
| Oligo rev | AAACCGGTGTGCCTGCTGTGCGAG | | | | | | | | | | |
| Chromosome | start | end | strand | MM | target_seq | PAM | alignment | distance | position | gene name | gene id |
| GU305914.1 | 77838 | 77860 | - | 0 | CTCGCACAGCAGGCACACCG | TGG | \|\|\|\|\|\|\|\|[\|\|\|\|\|\|\|\|\|\|\|\|]PAM | NA | Intergenic | NA | NA |
| T28360 | CAGCAGGCACACCGTGGTCGTGG | 1000 | | | | | | | | | |
| Oligo fwd | CACCCAGCAGGCACACCGTGGTCG | | | | | | | | | | |
| Oligo rev | AAACCGACCACGGTGTGCCTGCTG | | | | | | | | | | |
| Chromosome | start | end | strand | MM | target_seq | PAM | alignment | distance | position | gene name | gene id |
| GU305914.1 | 77832 | 77854 | - | 0 | CAGCAGGCACACCGTGGTCG | TGG | \|\|\|\|\|\|\|\|[\|\|\|\|\|\|\|\|\|\|\|\|]PAM | NA | Intergenic | NA | NA |
| GU305914.1 | 126518 | 126540 | - | 4 | GAGCACGCACACCGTGGTGC | AGG | -\|\|\|\|-\|\|[\|\|\|\|\|\|\|\|\|\|--]PAM | NA | Intergenic | NA | NA |
| T28361 | CCGTGGTCGTGGCGTTGCGCAGG | 1000 | | | | | | | | | |
| Oligo fwd | CACCCCGTGGTCGTGGCGTTGCGC | | | | | | | | | | |
| Oligo rev | AAACGCGCAACGCCACGACCACGG | | | | | | | | | | |
| Chromosome | start | end | strand | MM | target_seq | PAM | alignment | distance | position | gene name | gene id |
| GU305914.1 | 77821 | 77843 | - | 0 | CCGTGGTCGTGGCGTTGCGC | AGG | \|\|\|\|\|\|\|\|[\|\|\|\|\|\|\|\|\|\|\|\|]PAM | NA | Intergenic | NA | NA |
| T28362 | TGGTCGTGGCGTTGCGCAGGCGG | 1000 | | | | | | | | | |
| Oligo fwd | CACCTGGTCGTGGCGTTGCGCAGG | | | | | | | | | | |
| Oligo rev | AAACCCTGCGCAACGCCACGACCA | | | | | | | | | | |
| Chromosome | start | end | strand | MM | target_seq | PAM | alignment | distance | position | gene name | gene id |
| GU305914.1 | 77818 | 77840 | - | 0 | TGGTCGTGGCGTTGCGCAGG | CGG | \|\|\|\|\|\|\|\|[\|\|\|\|\|\|\|\|\|\|\|\|]PAM | NA | Intergenic | NA | NA |
| T28363 | GCGTTGCGCAGGCGGAACGTCGG | 1000 | | | | | | | | | |
| Oligo fwd | CACCGCGTTGCGCAGGCGGAACGT | | | | | | | | | | |
| Oligo rev | AAACACGTTCCGCCTGCGCAACGC | | | | | | | | | | |
| Chromosome | start | end | strand | MM | target_seq | PAM | alignment | distance | position | gene name | gene id |
| GU305914.1 | 77810 | 77832 | - | 0 | GCGTTGCGCAGGCGGAACGT | CGG | \|\|\|\|\|\|\|\|[\|\|\|\|\|\|\|\|\|\|\|\|]PAM | NA | Intergenic | NA | NA |
| T28364 | CGTTGCGCAGGCGGAACGTCGGG | 1000 | | | | | | | | | |
| Oligo fwd | CACCCGTTGCGCAGGCGGAACGTC | | | | | | | | | | |
| Oligo rev | AAACGACGTTCCGCCTGCGCAACG | | | | | | | | | | |
| Chromosome | start | end | strand | MM | target_seq | PAM | alignment | distance | position | gene name | gene id |
| GU305914.1 | 77809 | 77831 | - | 0 | CGTTGCGCAGGCGGAACGTC | GGG | \|\|\|\|\|\|\|\|[\|\|\|\|\|\|\|\|\|\|\|\|]PAM | NA | Intergenic | NA | NA |
| T28365 | CGCAGGCGGAACGTCGGGATCGG | 1000 | | | | | | | | | |
| Oligo fwd | CACCCGCAGGCGGAACGTCGGGAT | | | | | | | | | | |
| Oligo rev | AAACATCCCGACGTTCCGCCTGCG | | | | | | | | | | |
| Chromosome | start | end | strand | MM | target_seq | PAM | alignment | distance | position | gene name | gene id |
| GU305914.1 | 77804 | 77826 | - | 0 | CGCAGGCGGAACGTCGGGAT | CGG | \|\|\|\|\|\|\|\|[\|\|\|\|\|\|\|\|\|\|\|\|]PAM | NA | Intergenic | NA | NA |
| T28366 | GGAACGTCGGGATCGGCGATAGG | 1000 | | | | | | | | | |
| Oligo fwd | CACCGGAACGTCGGGATCGGCGAT | | | | | | | | | | |
| Oligo rev | AAACATCGCCGATCCCGACGTTCC | | | | | | | | | | |
| Chromosome | start | end | strand | MM | target_seq | PAM | alignment | distance | position | gene name | gene id |
| GU305914.1 | 77797 | 77819 | - | 0 | GGAACGTCGGGATCGGCGAT | AGG | \|\|\|\|\|\|\|\|[\|\|\|\|\|\|\|\|\|\|\|\|]PAM | NA | Intergenic | NA | NA |
| T28367 | CGGGATCGGCGATAGGAACACGG | 1000 | | | | | | | | | |
| Oligo fwd | CACCCGGGATCGGCGATAGGAACA | | | | | | | | | | |
| Oligo rev | AAACTGTTCCTATCGCCGATCCCG | | | | | | | | | | |
| Chromosome | start | end | strand | MM | target_seq | PAM | alignment | distance | position | gene name | gene id |
| GU305914.1 | 77790 | 77812 | - | 0 | CGGGATCGGCGATAGGAACA | CGG | \|\|\|\|\|\|\|\|[\|\|\|\|\|\|\|\|\|\|\|\|]PAM | NA | Intergenic | NA | NA |
| T28368 | GGGATCGGCGATAGGAACACGGG | 1000 | | | | | | | | | |
| Oligo fwd | CACCGGGATCGGCGATAGGAACAC | | | | | | | | | | |
| Oligo rev | AAACGTGTTCCTATCGCCGATCCC | | | | | | | | | | |
| Chromosome | start | end | strand | MM | target_seq | PAM | alignment | distance | position | gene name | gene id |
| GU305914.1 | 77789 | 77811 | - | 0 | GGGATCGGCGATAGGAACAC | GGG | \|\|\|\|\|\|\|\|[\|\|\|\|\|\|\|\|\|\|\|\|]PAM | NA | Intergenic | NA | NA |
| T28369 | GGATCGGCGATAGGAACACGGGG | 1000 | | | | | | | | | |
| Oligo fwd | CACCGGATCGGCGATAGGAACACG | | | | | | | | | | |
| Oligo rev | AAACCGTGTTCCTATCGCCGATCC | | | | | | | | | | |
| Chromosome | start | end | strand | MM | target_seq | PAM | alignment | distance | position | gene name | gene id |
| GU305914.1 | 77788 | 77810 | - | 0 | GGATCGGCGATAGGAACACG | GGG | \|\|\|\|\|\|\|\|[\|\|\|\|\|\|\|\|\|\|\|\|]PAM | NA | Intergenic | NA | NA |
| T28370 | AGGAACACGGGGCCCACGCTCGG | 1000 | | | | | | | | | |
| Oligo fwd | CACCAGGAACACGGGGCCCACGCT | | | | | | | | | | |
| Oligo rev | AAACAGCGTGGGCCCCGTGTTCCT | | | | | | | | | | |
| Chromosome | start | end | strand | MM | target_seq | PAM | alignment | distance | position | gene name | gene id |
| GU305914.1 | 77777 | 77799 | - | 0 | AGGAACACGGGGCCCACGCT | CGG | \|\|\|\|\|\|\|\|[\|\|\|\|\|\|\|\|\|\|\|\|]PAM | NA | Intergenic | NA | NA |
| T28371 | CCTTGCTCTTCTTGATCGCCAGG | 1000 | | | | | | | | | |
| Oligo fwd | CACCCCTTGCTCTTCTTGATCGCC | | | | | | | | | | |
| Oligo rev | AAACGGCGATCAAGAAGAGCAAGG | | | | | | | | | | |
| Chromosome | start | end | strand | MM | target_seq | PAM | alignment | distance | position | gene name | gene id |
| GU305914.1 | 77716 | 77738 | - | 0 | CCTTGCTCTTCTTGATCGCC | AGG | \|\|\|\|\|\|\|\|[\|\|\|\|\|\|\|\|\|\|\|\|]PAM | NA | Intergenic | NA | NA |
| T28372 | CACCCGCGCGAACGAGCCCTCGG | 1000 | | | | | | | | | |

| | | | | | | | | | | | | |
|---|---|---|---|---|---|---|---|---|---|---|---|---|
| Oligo fwd | CACCCACCCGCGCGAACGAGCCCT | | | | | | | | | | | |
| Oligo rev | AAACAGGGCTCGTTCGCGCGGGTG | | | | | | | | | | | |
| Chromosome | start | | end | strand | MM | target_seq | PAM | alignment | distance | position | gene name | gene id |
| GU305914.1 | | 77640 | 77662 | - | 0 | CACCCGCGCGAACGAGCCCT | CGG | \|\|\|\|\|\|\|\|[\|\|\|\|\|\|\|\|\|\|\|\|]PAM | NA | Intergenic | NA | NA |
| T28373 | ACGAGCCCTCGGTCGCGCGCCGG | | 1000 | | | | | | | | | |
| Oligo fwd | CACCACGAGCCCTCGGTCGCGCGC | | | | | | | | | | | |
| Oligo rev | AAACGCGCGCGACCGAGGGCTCGT | | | | | | | | | | | |
| Chromosome | start | | end | strand | MM | target_seq | PAM | alignment | distance | position | gene name | gene id |
| GU305914.1 | | 77629 | 77651 | - | 0 | ACGAGCCCTCGGTCGCGCGC | CGG | \|\|\|\|\|\|\|\|[\|\|\|\|\|\|\|\|\|\|\|\|]PAM | NA | Intergenic | NA | NA |
| T28374 | GTCCAGCAGCCGCAGCTCCTCGG | | 1000 | | | | | | | | | |
| Oligo fwd | CACCGTCCAGCAGCCGCAGCTCCT | | | | | | | | | | | |
| Oligo rev | AAACAGGAGCTGCGGCTGCTGGAC | | | | | | | | | | | |
| Chromosome | start | | end | strand | MM | target_seq | PAM | alignment | distance | position | gene name | gene id |
| GU305914.1 | | 77601 | 77623 | - | 0 | GTCCAGCAGCCGCAGCTCCT | CGG | \|\|\|\|\|\|\|\|[\|\|\|\|\|\|\|\|\|\|\|\|]PAM | NA | Intergenic | NA | NA |
| T28375 | TCCAGCAGCCGCAGCTCCTCGGG | | 1000 | | | | | | | | | |
| Oligo fwd | CACCTCCAGCAGCCGCAGCTCCTC | | | | | | | | | | | |
| Oligo rev | AAACGAGGAGCTGCGGCTGCTGGA | | | | | | | | | | | |
| Chromosome | start | | end | strand | MM | target_seq | PAM | alignment | distance | position | gene name | gene id |
| GU305914.1 | | 77600 | 77622 | - | 0 | TCCAGCAGCCGCAGCTCCTC | GGG | \|\|\|\|\|\|\|\|[\|\|\|\|\|\|\|\|\|\|\|\|]PAM | NA | Intergenic | NA | NA |
| GU305914.1 | | 98115 | 98137 | + | 4 | ACCAGGACCCGCAGTTCCTC | CGG | -\|\|\|\|-\|-[\|\|\|\|\|\|-\|\|\|\|\|]PAM | NA | Intergenic | NA | NA |
| T28376 | CGCAGCTCCTCGGGCTCCGCCGG | | 1000 | | | | | | | | | |
| Oligo fwd | CACCCGCAGCTCCTCGGGCTCCGC | | | | | | | | | | | |
| Oligo rev | AAACGCGGAGCCCGAGGAGCTGCG | | | | | | | | | | | |
| Chromosome | start | | end | strand | MM | target_seq | PAM | alignment | distance | position | gene name | gene id |
| GU305914.1 | | 77591 | 77613 | - | 0 | CGCAGCTCCTCGGGCTCCGC | CGG | \|\|\|\|\|\|\|\|[\|\|\|\|\|\|\|\|\|\|\|\|]PAM | NA | Intergenic | NA | NA |
| T28377 | TCCTCGGGCTCCGCCGGCGCCGG | | 1000 | | | | | | | | | |
| Oligo fwd | CACCTCCTCGGGCTCCGCCGGCGC | | | | | | | | | | | |
| Oligo rev | AAACGCGCCGGCGGAGCCCGAGGA | | | | | | | | | | | |
| Chromosome | start | | end | strand | MM | target_seq | PAM | alignment | distance | position | gene name | gene id |
| GU305914.1 | | 77585 | 77607 | - | 0 | TCCTCGGGCTCCGCCGGCGC | CGG | \|\|\|\|\|\|\|\|[\|\|\|\|\|\|\|\|\|\|\|\|]PAM | NA | Intergenic | NA | NA |
| T28378 | TCCGCCGGCGCCGGCTTCTTCGG | | 1000 | | | | | | | | | |
| Oligo fwd | CACCTCCGCCGGCGCCGGCTTCTT | | | | | | | | | | | |
| Oligo rev | AAACAAGAAGCCGGCGCCGGCGGA | | | | | | | | | | | |
| Chromosome | start | | end | strand | MM | target_seq | PAM | alignment | distance | position | gene name | gene id |
| GU305914.1 | | 77576 | 77598 | - | 0 | TCCGCCGGCGCCGGCTTCTT | CGG | \|\|\|\|\|\|\|\|[\|\|\|\|\|\|\|\|\|\|\|\|]PAM | NA | Intergenic | NA | NA |
| GU305914.1 | | 90319 | 90341 | - | 4 | TTCGCCGACGCCGACCTCTT | GGG | \|-\|\|\|\|\|-[\|\|\|\|\|-\|-\|\|\|\|]PAM | NA | Intergenic | NA | NA |
| T28379 | GCTTCTTCGGCAGCTTGTACTGG | | 1000 | | | | | | | | | |
| Oligo fwd | CACCGCTTCTTCGGCAGCTTGTAC | | | | | | | | | | | |
| Oligo rev | AAACGTACAAGCTGCCGAAGAAGC | | | | | | | | | | | |
| Chromosome | start | | end | strand | MM | target_seq | PAM | alignment | distance | position | gene name | gene id |
| GU305914.1 | | 77563 | 77585 | - | 0 | GCTTCTTCGGCAGCTTGTAC | TGG | \|\|\|\|\|\|\|\|[\|\|\|\|\|\|\|\|\|\|\|\|]PAM | NA | Intergenic | NA | NA |
| T28380 | GTACTGGATCTTGCGCCGCTTGG | | 1000 | | | | | | | | | |
| Oligo fwd | CACCGTACTGGATCTTGCGCCGCT | | | | | | | | | | | |
| Oligo rev | AAACAGCGGCGCAAGATCCAGTAC | | | | | | | | | | | |
| Chromosome | start | | end | strand | MM | target_seq | PAM | alignment | distance | position | gene name | gene id |
| GU305914.1 | | 77547 | 77569 | - | 0 | GTACTGGATCTTGCGCCGCT | TGG | \|\|\|\|\|\|\|\|[\|\|\|\|\|\|\|\|\|\|\|\|]PAM | NA | Intergenic | NA | NA |
| T28381 | ATCTTGCGCCGCTTGGTGTACGG | | 1000 | | | | | | | | | |
| Oligo fwd | CACCATCTTGCGCCGCTTGGTGTA | | | | | | | | | | | |
| Oligo rev | AAACTACACCAAGCGGCGCAAGAT | | | | | | | | | | | |
| Chromosome | start | | end | strand | MM | target_seq | PAM | alignment | distance | position | gene name | gene id |
| GU305914.1 | | 77540 | 77562 | - | 0 | ATCTTGCGCCGCTTGGTGTA | CGG | \|\|\|\|\|\|\|\|[\|\|\|\|\|\|\|\|\|\|\|\|]PAM | NA | Intergenic | NA | NA |
| T28382 | CGTCGTCCGTCAGCAGCGCCCGG | | 1000 | | | | | | | | | |
| Oligo fwd | CACCCGTCGTCCGTCAGCAGCGCC | | | | | | | | | | | |
| Oligo rev | AAACGGCGCTGCTGACGGACGACG | | | | | | | | | | | |
| Chromosome | start | | end | strand | MM | target_seq | PAM | alignment | distance | position | gene name | gene id |
| GU305914.1 | | 77509 | 77531 | - | 0 | CGTCGTCCGTCAGCAGCGCC | CGG | \|\|\|\|\|\|\|\|[\|\|\|\|\|\|\|\|\|\|\|\|]PAM | NA | Intergenic | NA | NA |
| T28383 | GTCGTCCGTCAGCAGCGCCCGGG | | 1000 | | | | | | | | | |
| Oligo fwd | CACCGTCGTCCGTCAGCAGCGCCC | | | | | | | | | | | |
| Oligo rev | AAACGGGCGCTGCTGACGGACGAC | | | | | | | | | | | |
| Chromosome | start | | end | strand | MM | target_seq | PAM | alignment | distance | position | gene name | gene id |
| GU305914.1 | | 77508 | 77530 | - | 0 | GTCGTCCGTCAGCAGCGCCC | GGG | \|\|\|\|\|\|\|\|[\|\|\|\|\|\|\|\|\|\|\|\|]PAM | NA | Intergenic | NA | NA |
| T28384 | GTGATGTTCTCCACCATCACCGG | | 1000 | | | | | | | | | |
| Oligo fwd | CACCGTGATGTTCTCCACCATCAC | | | | | | | | | | | |
| Oligo rev | AAACGTGATGGTGGAGAACATCAC | | | | | | | | | | | |
| Chromosome | start | | end | strand | MM | target_seq | PAM | alignment | distance | position | gene name | gene id |
| GU305914.1 | | 77465 | 77487 | - | 0 | GTGATGTTCTCCACCATCAC | CGG | \|\|\|\|\|\|\|\|[\|\|\|\|\|\|\|\|\|\|\|\|]PAM | NA | Intergenic | NA | NA |
| T28385 | TGATGTTCTCCACCATCACCGGG | | 1000 | | | | | | | | | |
| Oligo fwd | CACCTGATGTTCTCCACCATCACC | | | | | | | | | | | |
| Oligo rev | AAACGGTGATGGTGGAGAACATCA | | | | | | | | | | | |
| Chromosome | start | | end | strand | MM | target_seq | PAM | alignment | distance | position | gene name | gene id |
| GU305914.1 | | 77464 | 77486 | - | 0 | TGATGTTCTCCACCATCACC | GGG | \|\|\|\|\|\|\|\|[\|\|\|\|\|\|\|\|\|\|\|\|]PAM | NA | Intergenic | NA | NA |
| T28386 | CGTCGCGCTGCGTGTAGCACCGG | | 1000 | | | | | | | | | |
| Oligo fwd | CACCCGTCGCGCTGCGTGTAGCAC | | | | | | | | | | | |
| Oligo rev | AAACGTGCTACACGCAGCGCGACG | | | | | | | | | | | |
| Chromosome | start | | end | strand | MM | target_seq | PAM | alignment | distance | position | gene name | gene id |

<table>
<tr><td>GU305914.1</td><td>77437</td><td>77459</td><td>-</td><td>0</td><td>CGTCGCGCTGCGTGTAGCAC</td><td>CGG</td><td>||||||||[||||||||||||]PAM</td><td>NA</td><td>Intergenic</td><td>NA</td><td>NA</td></tr>
<tr><td></td><td></td><td></td><td></td><td></td><td></td><td></td><td></td><td></td><td></td><td></td><td></td></tr>
<tr><td>T28387</td><td>GGTTGATGAAGAAGTGCGCGTGG</td><td>1000</td><td></td><td></td><td></td><td></td><td></td><td></td><td></td><td></td><td></td></tr>
<tr><td>Oligo fwd</td><td>CACCGGTTGATGAAGAAGTGCGCG</td><td></td><td></td><td></td><td></td><td></td><td></td><td></td><td></td><td></td><td></td></tr>
<tr><td>Oligo rev</td><td>AAACCGCGCACTTCTTCATCAACC</td><td></td><td></td><td></td><td></td><td></td><td></td><td></td><td></td><td></td><td></td></tr>
<tr><td>Chromosome</td><td>start</td><td>end</td><td>strand</td><td>MM</td><td>target_seq</td><td>PAM</td><td>alignment</td><td>distance</td><td>position</td><td>gene name</td><td>gene id</td></tr>
<tr><td>GU305914.1</td><td>77416</td><td>77438</td><td>-</td><td>0</td><td>GGTTGATGAAGAAGTGCGCG</td><td>TGG</td><td>||||||||[||||||||||||]PAM</td><td>NA</td><td>Intergenic</td><td>NA</td><td>NA</td></tr>
<tr><td>GU305914.1</td><td>116513</td><td>116535</td><td>+</td><td>4</td><td>GGTTGGGGAAGAAGTGCACT</td><td>CGG</td><td>|||||--|[|||||||||-|-]PAM</td><td>NA</td><td>Intergenic</td><td>NA</td><td>NA</td></tr>
<tr><td></td><td></td><td></td><td></td><td></td><td></td><td></td><td></td><td></td><td></td><td></td><td></td></tr>
<tr><td>T28388</td><td>GTGCGCGTGGAAGTCGAAGATGG</td><td>1000</td><td></td><td></td><td></td><td></td><td></td><td></td><td></td><td></td><td></td></tr>
<tr><td>Oligo fwd</td><td>CACCGTGCGCGTGGAAGTCGAAGA</td><td></td><td></td><td></td><td></td><td></td><td></td><td></td><td></td><td></td><td></td></tr>
<tr><td>Oligo rev</td><td>AAACTCTTCGACTTCCACGCGCAC</td><td></td><td></td><td></td><td></td><td></td><td></td><td></td><td></td><td></td><td></td></tr>
<tr><td>Chromosome</td><td>start</td><td>end</td><td>strand</td><td>MM</td><td>target_seq</td><td>PAM</td><td>alignment</td><td>distance</td><td>position</td><td>gene name</td><td>gene id</td></tr>
<tr><td>GU305914.1</td><td>77403</td><td>77425</td><td>-</td><td>0</td><td>GTGCGCGTGGAAGTCGAAGA</td><td>TGG</td><td>||||||||[||||||||||||]PAM</td><td>NA</td><td>Intergenic</td><td>NA</td><td>NA</td></tr>
<tr><td>GU305914.1</td><td>146630</td><td>146652</td><td>-</td><td>4</td><td>GGGCCCGAAGAAGTCGAAGA</td><td>CGG</td><td>|-||-||-[-|||||||||||]PAM</td><td>NA</td><td>Intergenic</td><td>NA</td><td>NA</td></tr>
<tr><td></td><td></td><td></td><td></td><td></td><td></td><td></td><td></td><td></td><td></td><td></td><td></td></tr>
<tr><td>T28389</td><td>CTCCTTGAAGATGTTGTCGAAGG</td><td>1000</td><td></td><td></td><td></td><td></td><td></td><td></td><td></td><td></td><td></td></tr>
<tr><td>Oligo fwd</td><td>CACCCTCCTTGAAGATGTTGTCGA</td><td></td><td></td><td></td><td></td><td></td><td></td><td></td><td></td><td></td><td></td></tr>
<tr><td>Oligo rev</td><td>AAACTCGACAACATCTTCAAGGAG</td><td></td><td></td><td></td><td></td><td></td><td></td><td></td><td></td><td></td><td></td></tr>
<tr><td>Chromosome</td><td>start</td><td>end</td><td>strand</td><td>MM</td><td>target_seq</td><td>PAM</td><td>alignment</td><td>distance</td><td>position</td><td>gene name</td><td>gene id</td></tr>
<tr><td>GU305914.1</td><td>77370</td><td>77392</td><td>-</td><td>0</td><td>CTCCTTGAAGATGTTGTCGA</td><td>AGG</td><td>||||||||[||||||||||||]PAM</td><td>NA</td><td>Intergenic</td><td>NA</td><td>NA</td></tr>
<tr><td></td><td></td><td></td><td></td><td></td><td></td><td></td><td></td><td></td><td></td><td></td><td></td></tr>
<tr><td>T28390</td><td>CGCGAAAGTGCAGCTTCATCAGG</td><td>1000</td><td></td><td></td><td></td><td></td><td></td><td></td><td></td><td></td><td></td></tr>
<tr><td>Oligo fwd</td><td>CACCCGCGAAAGTGCAGCTTCATC</td><td></td><td></td><td></td><td></td><td></td><td></td><td></td><td></td><td></td><td></td></tr>
<tr><td>Oligo rev</td><td>AAACGATGAAGCTGCACTTTCGCG</td><td></td><td></td><td></td><td></td><td></td><td></td><td></td><td></td><td></td><td></td></tr>
<tr><td>Chromosome</td><td>start</td><td>end</td><td>strand</td><td>MM</td><td>target_seq</td><td>PAM</td><td>alignment</td><td>distance</td><td>position</td><td>gene name</td><td>gene id</td></tr>
<tr><td>GU305914.1</td><td>77341</td><td>77363</td><td>-</td><td>0</td><td>CGCGAAAGTGCAGCTTCATC</td><td>AGG</td><td>||||||||[||||||||||||]PAM</td><td>NA</td><td>Intergenic</td><td>NA</td><td>NA</td></tr>
<tr><td></td><td></td><td></td><td></td><td></td><td></td><td></td><td></td><td></td><td></td><td></td><td></td></tr>
<tr><td>T28391</td><td>AGTGCAGCTTCATCAGGTGCAGG</td><td>1000</td><td></td><td></td><td></td><td></td><td></td><td></td><td></td><td></td><td></td></tr>
<tr><td>Oligo fwd</td><td>CACCAGTGCAGCTTCATCAGGTGC</td><td></td><td></td><td></td><td></td><td></td><td></td><td></td><td></td><td></td><td></td></tr>
<tr><td>Oligo rev</td><td>AAACGCACCTGATGAAGCTGCACT</td><td></td><td></td><td></td><td></td><td></td><td></td><td></td><td></td><td></td><td></td></tr>
<tr><td>Chromosome</td><td>start</td><td>end</td><td>strand</td><td>MM</td><td>target_seq</td><td>PAM</td><td>alignment</td><td>distance</td><td>position</td><td>gene name</td><td>gene id</td></tr>
<tr><td>GU305914.1</td><td>77335</td><td>77357</td><td>-</td><td>0</td><td>AGTGCAGCTTCATCAGGTGC</td><td>AGG</td><td>||||||||[||||||||||||]PAM</td><td>NA</td><td>Intergenic</td><td>NA</td><td>NA</td></tr>
<tr><td>GU305914.1</td><td>80196</td><td>80218</td><td>+</td><td>4</td><td>CGGGCACCTTCTTCAGGTGC</td><td>AGG</td><td>-|-|||-|[|||-||||||||]PAM</td><td>NA</td><td>Intergenic</td><td>NA</td><td>NA</td></tr>
<tr><td></td><td></td><td></td><td></td><td></td><td></td><td></td><td></td><td></td><td></td><td></td><td></td></tr>
<tr><td>T28392</td><td>CTTCATCAGGTGCAGGAACTCGG</td><td>1000</td><td></td><td></td><td></td><td></td><td></td><td></td><td></td><td></td><td></td></tr>
<tr><td>Oligo fwd</td><td>CACCCTTCATCAGGTGCAGGAACT</td><td></td><td></td><td></td><td></td><td></td><td></td><td></td><td></td><td></td><td></td></tr>
<tr><td>Oligo rev</td><td>AAACAGTTCCTGCACCTGATGAAG</td><td></td><td></td><td></td><td></td><td></td><td></td><td></td><td></td><td></td><td></td></tr>
<tr><td>Chromosome</td><td>start</td><td>end</td><td>strand</td><td>MM</td><td>target_seq</td><td>PAM</td><td>alignment</td><td>distance</td><td>position</td><td>gene name</td><td>gene id</td></tr>
<tr><td>GU305914.1</td><td>77328</td><td>77350</td><td>-</td><td>0</td><td>CTTCATCAGGTGCAGGAACT</td><td>CGG</td><td>||||||||[||||||||||||]PAM</td><td>NA</td><td>Intergenic</td><td>NA</td><td>NA</td></tr>
<tr><td></td><td></td><td></td><td></td><td></td><td></td><td></td><td></td><td></td><td></td><td></td><td></td></tr>
<tr><td>T28393</td><td>CATCAGGTGCAGGAACTCGGTGG</td><td>1000</td><td></td><td></td><td></td><td></td><td></td><td></td><td></td><td></td><td></td></tr>
<tr><td>Oligo fwd</td><td>CACCCATCAGGTGCAGGAACTCGG</td><td></td><td></td><td></td><td></td><td></td><td></td><td></td><td></td><td></td><td></td></tr>
<tr><td>Oligo rev</td><td>AAACCCGAGTTCCTGCACCTGATG</td><td></td><td></td><td></td><td></td><td></td><td></td><td></td><td></td><td></td><td></td></tr>
<tr><td>Chromosome</td><td>start</td><td>end</td><td>strand</td><td>MM</td><td>target_seq</td><td>PAM</td><td>alignment</td><td>distance</td><td>position</td><td>gene name</td><td>gene id</td></tr>
<tr><td>GU305914.1</td><td>77325</td><td>77347</td><td>-</td><td>0</td><td>CATCAGGTGCAGGAACTCGG</td><td>TGG</td><td>||||||||[||||||||||||]PAM</td><td>NA</td><td>Intergenic</td><td>NA</td><td>NA</td></tr>
<tr><td></td><td></td><td></td><td></td><td></td><td></td><td></td><td></td><td></td><td></td><td></td><td></td></tr>
<tr><td>T28394</td><td>CTCGGTGGCCACGATCAGCGTGG</td><td>1000</td><td></td><td></td><td></td><td></td><td></td><td></td><td></td><td></td><td></td></tr>
<tr><td>Oligo fwd</td><td>CACCCTCGGTGGCCACGATCAGCG</td><td></td><td></td><td></td><td></td><td></td><td></td><td></td><td></td><td></td><td></td></tr>
<tr><td>Oligo rev</td><td>AAACCGCTGATCGTGGCCACCGAG</td><td></td><td></td><td></td><td></td><td></td><td></td><td></td><td></td><td></td><td></td></tr>
<tr><td>Chromosome</td><td>start</td><td>end</td><td>strand</td><td>MM</td><td>target_seq</td><td>PAM</td><td>alignment</td><td>distance</td><td>position</td><td>gene name</td><td>gene id</td></tr>
<tr><td>GU305914.1</td><td>77310</td><td>77332</td><td>-</td><td>0</td><td>CTCGGTGGCCACGATCAGCG</td><td>TGG</td><td>||||||||[||||||||||||]PAM</td><td>NA</td><td>Intergenic</td><td>NA</td><td>NA</td></tr>
<tr><td></td><td></td><td></td><td></td><td></td><td></td><td></td><td></td><td></td><td></td><td></td><td></td></tr>
<tr><td>T28395</td><td>TCAGCGTGGCCATCGCCGTGCGG</td><td>1000</td><td></td><td></td><td></td><td></td><td></td><td></td><td></td><td></td><td></td></tr>
<tr><td>Oligo fwd</td><td>CACCTCAGCGTGGCCATCGCCGTG</td><td></td><td></td><td></td><td></td><td></td><td></td><td></td><td></td><td></td><td></td></tr>
<tr><td>Oligo rev</td><td>AAACCACGGCGATGGCCACGCTGA</td><td></td><td></td><td></td><td></td><td></td><td></td><td></td><td></td><td></td><td></td></tr>
<tr><td>Chromosome</td><td>start</td><td>end</td><td>strand</td><td>MM</td><td>target_seq</td><td>PAM</td><td>alignment</td><td>distance</td><td>position</td><td>gene name</td><td>gene id</td></tr>
<tr><td>GU305914.1</td><td>77296</td><td>77318</td><td>-</td><td>0</td><td>TCAGCGTGGCCATCGCCGTG</td><td>CGG</td><td>||||||||[||||||||||||]PAM</td><td>NA</td><td>Intergenic</td><td>NA</td><td>NA</td></tr>
<tr><td></td><td></td><td></td><td></td><td></td><td></td><td></td><td></td><td></td><td></td><td></td><td></td></tr>
<tr><td>T28396</td><td>CAGCGTGGCCATCGCCGTGCGGG</td><td>1000</td><td></td><td></td><td></td><td></td><td></td><td></td><td></td><td></td><td></td></tr>
<tr><td>Oligo fwd</td><td>CACCCAGCGTGGCCATCGCCGTGC</td><td></td><td></td><td></td><td></td><td></td><td></td><td></td><td></td><td></td><td></td></tr>
<tr><td>Oligo rev</td><td>AAACGCACGGCGATGGCCACGCTG</td><td></td><td></td><td></td><td></td><td></td><td></td><td></td><td></td><td></td><td></td></tr>
<tr><td>Chromosome</td><td>start</td><td>end</td><td>strand</td><td>MM</td><td>target_seq</td><td>PAM</td><td>alignment</td><td>distance</td><td>position</td><td>gene name</td><td>gene id</td></tr>
<tr><td>GU305914.1</td><td>77295</td><td>77317</td><td>-</td><td>0</td><td>CAGCGTGGCCATCGCCGTGC</td><td>GGG</td><td>||||||||[||||||||||||]PAM</td><td>NA</td><td>Intergenic</td><td>NA</td><td>NA</td></tr>
<tr><td></td><td></td><td></td><td></td><td></td><td></td><td></td><td></td><td></td><td></td><td></td><td></td></tr>
<tr><td>T28397</td><td>AGCGTGGCCATCGCCGTGCGGGG</td><td>1000</td><td></td><td></td><td></td><td></td><td></td><td></td><td></td><td></td><td></td></tr>
<tr><td>Oligo fwd</td><td>CACCAGCGTGGCCATCGCCGTGCG</td><td></td><td></td><td></td><td></td><td></td><td></td><td></td><td></td><td></td><td></td></tr>
<tr><td>Oligo rev</td><td>AAACCGCACGGCGATGGCCACGCT</td><td></td><td></td><td></td><td></td><td></td><td></td><td></td><td></td><td></td><td></td></tr>
<tr><td>Chromosome</td><td>start</td><td>end</td><td>strand</td><td>MM</td><td>target_seq</td><td>PAM</td><td>alignment</td><td>distance</td><td>position</td><td>gene name</td><td>gene id</td></tr>
<tr><td>GU305914.1</td><td>77294</td><td>77316</td><td>-</td><td>0</td><td>AGCGTGGCCATCGCCGTGCG</td><td>GGG</td><td>||||||||[||||||||||||]PAM</td><td>NA</td><td>Intergenic</td><td>NA</td><td>NA</td></tr>
<tr><td></td><td></td><td></td><td></td><td></td><td></td><td></td><td></td><td></td><td></td><td></td><td></td></tr>
<tr><td>T28398</td><td>GGGGCGCCGCCGCGCACTTGTGG</td><td>1000</td><td></td><td></td><td></td><td></td><td></td><td></td><td></td><td></td><td></td></tr>
<tr><td>Oligo fwd</td><td>CACCGGGGCGCCGCCGCGCACTTG</td><td></td><td></td><td></td><td></td><td></td><td></td><td></td><td></td><td></td><td></td></tr>
<tr><td>Oligo rev</td><td>AAACCAAGTGCGCGGCGGCGCCCC</td><td></td><td></td><td></td><td></td><td></td><td></td><td></td><td></td><td></td><td></td></tr>
<tr><td>Chromosome</td><td>start</td><td>end</td><td>strand</td><td>MM</td><td>target_seq</td><td>PAM</td><td>alignment</td><td>distance</td><td>position</td><td>gene name</td><td>gene id</td></tr>
<tr><td>GU305914.1</td><td>77275</td><td>77297</td><td>-</td><td>0</td><td>GGGGCGCCGCCGCGCACTTG</td><td>TGG</td><td>||||||||[||||||||||||]PAM</td><td>NA</td><td>Intergenic</td><td>NA</td><td>NA</td></tr>
<tr><td></td><td></td><td></td><td></td><td></td><td></td><td></td><td></td><td></td><td></td><td></td><td></td></tr>
<tr><td>T28399</td><td>CCGCCGCGCACTTGTGGCACAGG</td><td>1000</td><td></td><td></td><td></td><td></td><td></td><td></td><td></td><td></td><td></td></tr>
<tr><td>Oligo fwd</td><td>CACCCCGCCGCGCACTTGTGGCAC</td><td></td><td></td><td></td><td></td><td></td><td></td><td></td><td></td><td></td><td></td></tr>
<tr><td>Oligo rev</td><td>AAACGTGCCACAAGTGCGCGGCGG</td><td></td><td></td><td></td><td></td><td></td><td></td><td></td><td></td><td></td><td></td></tr>
<tr><td>Chromosome</td><td>start</td><td>end</td><td>strand</td><td>MM</td><td>target_seq</td><td>PAM</td><td>alignment</td><td>distance</td><td>position</td><td>gene name</td><td>gene id</td></tr>
<tr><td>GU305914.1</td><td>77269</td><td>77291</td><td>-</td><td>0</td><td>CCGCCGCGCACTTGTGGCAC</td><td>AGG</td><td>||||||||[||||||||||||]PAM</td><td>NA</td><td>Intergenic</td><td>NA</td><td>NA</td></tr>
<tr><td></td><td></td><td></td><td></td><td></td><td></td><td></td><td></td><td></td><td></td><td></td><td></td></tr>
<tr><td>T28400</td><td>TCCAGCAGCCATTTCTGCTGCGG</td><td>1000</td><td></td><td></td><td></td><td></td><td></td><td></td><td></td><td></td><td></td></tr>
<tr><td>Oligo fwd</td><td>CACCTCCAGCAGCCATTTCTGCTG</td><td></td><td></td><td></td><td></td><td></td><td></td><td></td><td></td><td></td><td></td></tr>
<tr><td>Oligo rev</td><td>AAACCAGCAGAAATGGCTGCTGGA</td><td></td><td></td><td></td><td></td><td></td><td></td><td></td><td></td><td></td><td></td></tr>
<tr><td>Chromosome</td><td>start</td><td>end</td><td>strand</td><td>MM</td><td>target_seq</td><td>PAM</td><td>alignment</td><td>distance</td><td>position</td><td>gene name</td><td>gene id</td></tr>
<tr><td>GU305914.1</td><td>77237</td><td>77259</td><td>-</td><td>0</td><td>TCCAGCAGCCATTTCTGCTG</td><td>CGG</td><td>||||||||[||||||||||||]PAM</td><td>NA</td><td>Intergenic</td><td>NA</td><td>NA</td></tr>
</table>

| | | | | | | | | | | | |
|---|---|---|---|---|---|---|---|---|---|---|---|
| T28401 | CCAGCAGCCATTTCTGCTGCGGG | 1000 | | | | | | | | | |
| Oligo fwd | CACCCCAGCAGCCATTTCTGCTGC | | | | | | | | | | |
| Oligo rev | AAACGCAGCAGAAATGGCTGCTGG | | | | | | | | | | |
| Chromosome | start | end | strand | MM | target_seq | PAM | alignment | distance | position | gene name | gene id |
| GU305914.1 | 77236 | 77258 | - | 0 | CCAGCAGCCATTTCTGCTGC | GGG | \|\|\|\|\|\|\|\|[\|\|\|\|\|\|\|\|\|\|\|\|]PAM | NA | Intergenic | NA | NA |
| T28402 | TCTGCTGCGGGTCGCGCCGCAGG | 1000 | | | | | | | | | |
| Oligo fwd | CACCTCTGCTGCGGGTCGCGCCGC | | | | | | | | | | |
| Oligo rev | AAACGCGGCGCGACCCGCAGCAGA | | | | | | | | | | |
| Chromosome | start | end | strand | MM | target_seq | PAM | alignment | distance | position | gene name | gene id |
| GU305914.1 | 77224 | 77246 | - | 0 | TCTGCTGCGGGTCGCGCCGC | AGG | \|\|\|\|\|\|\|\|[\|\|\|\|\|\|\|\|\|\|\|\|]PAM | NA | Intergenic | NA | NA |
| T28403 | AGATGCTGACGATCGCGCACAGG | 1000 | | | | | | | | | |
| Oligo fwd | CACCAGATGCTGACGATCGCGCAC | | | | | | | | | | |
| Oligo rev | AAACGTGCGCGATCGTCAGCATCT | | | | | | | | | | |
| Chromosome | start | end | strand | MM | target_seq | PAM | alignment | distance | position | gene name | gene id |
| GU305914.1 | 77200 | 77222 | - | 0 | AGATGCTGACGATCGCGCAC | AGG | \|\|\|\|\|\|\|\|[\|\|\|\|\|\|\|\|\|\|\|\|]PAM | NA | Intergenic | NA | NA |
| GU305914.1 | 132378 | 132400 | + | 4 | AGACGCTCAGGATCGCGCCC | AGG | \|\|\|-\|\|\|-[\|-\|\|\|\|\|\|\|\|-\|]PAM | NA | Intergenic | NA | NA |
| T28404 | CAGCGCGAGTCGATCTCCAGCGG | 1000 | | | | | | | | | |
| Oligo fwd | CACCCAGCGCGAGTCGATCTCCAG | | | | | | | | | | |
| Oligo rev | AAACCTGGAGATCGACTCGCGCTG | | | | | | | | | | |
| Chromosome | start | end | strand | MM | target_seq | PAM | alignment | distance | position | gene name | gene id |
| GU305914.1 | 77174 | 77196 | - | 0 | CAGCGCGAGTCGATCTCCAG | CGG | \|\|\|\|\|\|\|\|[\|\|\|\|\|\|\|\|\|\|\|\|]PAM | NA | Intergenic | NA | NA |
| T28405 | GCGAGTCGATCTCCAGCGGCAGG | 1000 | | | | | | | | | |
| Oligo fwd | CACCGCGAGTCGATCTCCAGCGGC | | | | | | | | | | |
| Oligo rev | AAACGCCGCTGGAGATCGACTCGC | | | | | | | | | | |
| Chromosome | start | end | strand | MM | target_seq | PAM | alignment | distance | position | gene name | gene id |
| GU305914.1 | 77170 | 77192 | - | 0 | GCGAGTCGATCTCCAGCGGC | AGG | \|\|\|\|\|\|\|\|[\|\|\|\|\|\|\|\|\|\|\|\|]PAM | NA | Intergenic | NA | NA |
| T28406 | CCAGCGGCAGGCACAGCTCGCGG | 1000 | | | | | | | | | |
| Oligo fwd | CACCCCAGCGGCAGGCACAGCTCG | | | | | | | | | | |
| Oligo rev | AAACCGAGCTGTGCCTGCCGCTGG | | | | | | | | | | |
| Chromosome | start | end | strand | MM | target_seq | PAM | alignment | distance | position | gene name | gene id |
| GU305914.1 | 77158 | 77180 | - | 0 | CCAGCGGCAGGCACAGCTCG | CGG | \|\|\|\|\|\|\|\|[\|\|\|\|\|\|\|\|\|\|\|\|]PAM | NA | Intergenic | NA | NA |
| T28407 | CAGGCACAGCTCGCGGATCTCGG | 1000 | | | | | | | | | |
| Oligo fwd | CACCCAGGCACAGCTCGCGGATCT | | | | | | | | | | |
| Oligo rev | AAACAGATCCGCGAGCTGTGCCTG | | | | | | | | | | |
| Chromosome | start | end | strand | MM | target_seq | PAM | alignment | distance | position | gene name | gene id |
| GU305914.1 | 77151 | 77173 | - | 0 | CAGGCACAGCTCGCGGATCT | CGG | \|\|\|\|\|\|\|\|[\|\|\|\|\|\|\|\|\|\|\|\|]PAM | NA | Intergenic | NA | NA |
| T28408 | TCCGCGCCAGCTCCGAGTCGAGG | 1000 | | | | | | | | | |
| Oligo fwd | CACCTCCGCGCCAGCTCCGAGTCG | | | | | | | | | | |
| Oligo rev | AAACCGACTCGGAGCTGGCGCGGA | | | | | | | | | | |
| Chromosome | start | end | strand | MM | target_seq | PAM | alignment | distance | position | gene name | gene id |
| GU305914.1 | 77110 | 77132 | - | 0 | TCCGCGCCAGCTCCGAGTCG | AGG | \|\|\|\|\|\|\|\|[\|\|\|\|\|\|\|\|\|\|\|\|]PAM | NA | Intergenic | NA | NA |
| T28409 | CTCCAGCATCGTCTCCCGCCCGG | 1000 | | | | | | | | | |
| Oligo fwd | CACCCTCCAGCATCGTCTCCCGCC | | | | | | | | | | |
| Oligo rev | AAACGGCGGGAGACGATGCTGGAG | | | | | | | | | | |
| Chromosome | start | end | strand | MM | target_seq | PAM | alignment | distance | position | gene name | gene id |
| GU305914.1 | 77064 | 77086 | - | 0 | CTCCAGCATCGTCTCCCGCC | CGG | \|\|\|\|\|\|\|\|[\|\|\|\|\|\|\|\|\|\|\|\|]PAM | NA | Intergenic | NA | NA |
| GU305914.1 | 18262 | 18284 | + | 4 | CTCCGGAGTCGTCTCCCGAC | TGG | \|\|\|\|-\|--[\|\|\|\|\|\|\|\|\|\|-\|]PAM | NA | Intergenic | NA | NA |
| T28410 | CATCGTCTCCCGCCCGGAGTCGG | 1000 | | | | | | | | | |
| Oligo fwd | CACCCATCGTCTCCCGCCCGGAGT | | | | | | | | | | |
| Oligo rev | AAACACTCCGGGCGGGAGACGATG | | | | | | | | | | |
| Chromosome | start | end | strand | MM | target_seq | PAM | alignment | distance | position | gene name | gene id |
| GU305914.1 | 77058 | 77080 | - | 0 | CATCGTCTCCCGCCCGGAGT | CGG | \|\|\|\|\|\|\|\|[\|\|\|\|\|\|\|\|\|\|\|\|]PAM | NA | Intergenic | NA | NA |
| T28411 | CCGCCCGGAGTCGGTCGACGAGG | 1000 | | | | | | | | | |
| Oligo fwd | CACCCCGCCCGGAGTCGGTCGACG | | | | | | | | | | |
| Oligo rev | AAACCGTCGACCGACTCCGGGCGG | | | | | | | | | | |
| Chromosome | start | end | strand | MM | target_seq | PAM | alignment | distance | position | gene name | gene id |
| GU305914.1 | 77049 | 77071 | - | 0 | CCGCCCGGAGTCGGTCGACG | AGG | \|\|\|\|\|\|\|\|[\|\|\|\|\|\|\|\|\|\|\|\|]PAM | NA | Intergenic | NA | NA |
| T28412 | GGAGTCGGTCGACGAGGATGAGG | 1000 | | | | | | | | | |
| Oligo fwd | CACCGGAGTCGGTCGACGAGGATG | | | | | | | | | | |
| Oligo rev | AAACCATCCTCGTCGACCGACTCC | | | | | | | | | | |
| Chromosome | start | end | strand | MM | target_seq | PAM | alignment | distance | position | gene name | gene id |
| GU305914.1 | 77043 | 77065 | - | 0 | GGAGTCGGTCGACGAGGATG | AGG | \|\|\|\|\|\|\|\|[\|\|\|\|\|\|\|\|\|\|\|\|]PAM | NA | Intergenic | NA | NA |
| GU305914.1 | 59739 | 59761 | - | 4 | CGACTCGGACGACGAGGACG | AGG | -\|\|-\|\|\|\|[-\|\|\|\|\|\|\|\|\|-\|]PAM | NA | Intergenic | NA | NA |
| GU305914.1 | 54574 | 54596 | + | 3 | GGAGGCGGTTGACGAGGATA | CGG | \|\|\|\|-\|\|\|[\|-\|\|\|\|\|\|\|\|\|-]PAM | NA | Intergenic | NA | NA |
| T28413 | GTCGGTCGACGAGGATGAGGAGG | 1000 | | | | | | | | | |
| Oligo fwd | CACCGTCGGTCGACGAGGATGAGG | | | | | | | | | | |
| Oligo rev | AAACCCTCATCCTCGTCGACCGAC | | | | | | | | | | |
| Chromosome | start | end | strand | MM | target_seq | PAM | alignment | distance | position | gene name | gene id |
| GU305914.1 | 77040 | 77062 | - | 0 | GTCGGTCGACGAGGATGAGG | AGG | \|\|\|\|\|\|\|\|[\|\|\|\|\|\|\|\|\|\|\|\|]PAM | NA | Intergenic | NA | NA |
| GU305914.1 | 59736 | 59758 | - | 3 | CTCGGACGACGAGGACGAGG | TGG | -\|\|\|\|-\|\|[\|\|\|\|\|\|\|-\|\|\|\|]PAM | NA | Intergenic | NA | NA |
| T28414 | GGTCGACGAGGATGAGGAGGAGG | 1000 | | | | | | | | | |
| Oligo fwd | CACCGGTCGACGAGGATGAGGAGG | | | | | | | | | | |
| Oligo rev | AAACCCTCCTCATCCTCGTCGACC | | | | | | | | | | |
| Chromosome | start | end | strand | MM | target_seq | PAM | alignment | distance | position | gene name | gene id |
| GU305914.1 | 77037 | 77059 | - | 0 | GGTCGACGAGGATGAGGAGG | AGG | \|\|\|\|\|\|\|\|[\|\|\|\|\|\|\|\|\|\|\|\|]PAM | NA | Intergenic | NA | NA |
| GU305914.1 | 140723 | 140745 | + | 3 | GGACGAGGAGGATGAGGATG | AGG | \|\|-\|\|\|-\|[\|\|\|\|\|\|\|\|\|\|-\|]PAM | NA | Intergenic | NA | NA |

<table>
<tr><td>T28415</td><td>GACGAGGATGAGGAGGAGGACGG</td><td>1000</td><td colspan="9"></td></tr>
<tr><td>Oligo fwd</td><td>CACCGACGAGGATGAGGAGGAGGA</td><td colspan="10"></td></tr>
<tr><td>Oligo rev</td><td>AAACTCCTCCTCCTCATCCTCGTC</td><td colspan="10"></td></tr>
<tr><td>Chromosome</td><td>start</td><td>end</td><td>strand</td><td>MM</td><td>target_seq</td><td>PAM</td><td>alignment</td><td>distance</td><td>position</td><td>gene name</td><td>gene id</td></tr>
<tr><td>GU305914.1</td><td>77033</td><td>77055</td><td>-</td><td>0</td><td>GACGAGGATGAGGAGGAGGA</td><td>CGG</td><td>||||||||[||||||||||||]PAM</td><td>NA</td><td>Intergenic</td><td>NA</td><td>NA</td></tr>
<tr><td>GU305914.1</td><td>65115</td><td>65137</td><td>+</td><td>4</td><td>GAAGACGACGACGAGGAGGA</td><td>GGG</td><td>||-||-||[-||-||||||||]PAM</td><td>NA</td><td>Intergenic</td><td>NA</td><td>NA</td></tr>
<tr><td>GU305914.1</td><td>118805</td><td>118827</td><td>-</td><td>4</td><td>GATGATGACGACGAGGAGGA</td><td>CGG</td><td>||-||-||[-||-||||||||]PAM</td><td>NA</td><td>Intergenic</td><td>NA</td><td>NA</td></tr>
<tr><td>GU305914.1</td><td>65118</td><td>65140</td><td>+</td><td>3</td><td>GACGACGACGAGGAGGAGGG</td><td>CGG</td><td>|||||-||[-||||||||||-]PAM</td><td>NA</td><td>Intergenic</td><td>NA</td><td>NA</td></tr>
<tr><td>T28416</td><td>ATGAGGAGGAGGACGGCGTCTGG</td><td>1000</td><td colspan="9"></td></tr>
<tr><td>Oligo fwd</td><td>CACCATGAGGAGGAGGACGGCGTC</td><td colspan="10"></td></tr>
<tr><td>Oligo rev</td><td>AAACGACGCCGTCCTCCTCCTCAT</td><td colspan="10"></td></tr>
<tr><td>Chromosome</td><td>start</td><td>end</td><td>strand</td><td>MM</td><td>target_seq</td><td>PAM</td><td>alignment</td><td>distance</td><td>position</td><td>gene name</td><td>gene id</td></tr>
<tr><td>GU305914.1</td><td>77026</td><td>77048</td><td>-</td><td>0</td><td>ATGAGGAGGAGGACGGCGTC</td><td>TGG</td><td>||||||||[||||||||||||]PAM</td><td>NA</td><td>Intergenic</td><td>NA</td><td>NA</td></tr>
<tr><td>T28417</td><td>GTCTGGCCCTCCGTCACTTCTGG</td><td>1000</td><td colspan="9"></td></tr>
<tr><td>Oligo fwd</td><td>CACCGTCTGGCCCTCCGTCACTTC</td><td colspan="10"></td></tr>
<tr><td>Oligo rev</td><td>AAACGAAGTGACGGAGGGCCAGAC</td><td colspan="10"></td></tr>
<tr><td>Chromosome</td><td>start</td><td>end</td><td>strand</td><td>MM</td><td>target_seq</td><td>PAM</td><td>alignment</td><td>distance</td><td>position</td><td>gene name</td><td>gene id</td></tr>
<tr><td>GU305914.1</td><td>77009</td><td>77031</td><td>-</td><td>0</td><td>GTCTGGCCCTCCGTCACTTC</td><td>TGG</td><td>||||||||[||||||||||||]PAM</td><td>NA</td><td>Intergenic</td><td>NA</td><td>NA</td></tr>
<tr><td>T28418</td><td>TGGCCCTCCGTCACTTCTGGAGG</td><td>1000</td><td colspan="9"></td></tr>
<tr><td>Oligo fwd</td><td>CACCTGGCCCTCCGTCACTTCTGG</td><td colspan="10"></td></tr>
<tr><td>Oligo rev</td><td>AAACCCAGAAGTGACGGAGGGCCA</td><td colspan="10"></td></tr>
<tr><td>Chromosome</td><td>start</td><td>end</td><td>strand</td><td>MM</td><td>target_seq</td><td>PAM</td><td>alignment</td><td>distance</td><td>position</td><td>gene name</td><td>gene id</td></tr>
<tr><td>GU305914.1</td><td>77006</td><td>77028</td><td>-</td><td>0</td><td>TGGCCCTCCGTCACTTCTGG</td><td>AGG</td><td>||||||||[||||||||||||]PAM</td><td>NA</td><td>Intergenic</td><td>NA</td><td>NA</td></tr>
<tr><td>T28419</td><td>GGCCCTCCGTCACTTCTGGAGGG</td><td>1000</td><td colspan="9"></td></tr>
<tr><td>Oligo fwd</td><td>CACCGGCCCTCCGTCACTTCTGGA</td><td colspan="10"></td></tr>
<tr><td>Oligo rev</td><td>AAACTCCAGAAGTGACGGAGGGCC</td><td colspan="10"></td></tr>
<tr><td>Chromosome</td><td>start</td><td>end</td><td>strand</td><td>MM</td><td>target_seq</td><td>PAM</td><td>alignment</td><td>distance</td><td>position</td><td>gene name</td><td>gene id</td></tr>
<tr><td>GU305914.1</td><td>77005</td><td>77027</td><td>-</td><td>0</td><td>GGCCCTCCGTCACTTCTGGA</td><td>GGG</td><td>||||||||[||||||||||||]PAM</td><td>NA</td><td>Intergenic</td><td>NA</td><td>NA</td></tr>
<tr><td>T28420</td><td>GCCCTCCGTCACTTCTGGAGGGG</td><td>1000</td><td colspan="9"></td></tr>
<tr><td>Oligo fwd</td><td>CACCGCCCTCCGTCACTTCTGGAG</td><td colspan="10"></td></tr>
<tr><td>Oligo rev</td><td>AAACCTCCAGAAGTGACGGAGGGC</td><td colspan="10"></td></tr>
<tr><td>Chromosome</td><td>start</td><td>end</td><td>strand</td><td>MM</td><td>target_seq</td><td>PAM</td><td>alignment</td><td>distance</td><td>position</td><td>gene name</td><td>gene id</td></tr>
<tr><td>GU305914.1</td><td>77004</td><td>77026</td><td>-</td><td>0</td><td>GCCCTCCGTCACTTCTGGAG</td><td>GGG</td><td>||||||||[||||||||||||]PAM</td><td>NA</td><td>Intergenic</td><td>NA</td><td>NA</td></tr>
<tr><td>T28421</td><td>CGTCACTTCTGGAGGGGTTTTGG</td><td>1000</td><td colspan="9"></td></tr>
<tr><td>Oligo fwd</td><td>CACCCGTCACTTCTGGAGGGGTTT</td><td colspan="10"></td></tr>
<tr><td>Oligo rev</td><td>AAACAAACCCCTCCAGAAGTGACG</td><td colspan="10"></td></tr>
<tr><td>Chromosome</td><td>start</td><td>end</td><td>strand</td><td>MM</td><td>target_seq</td><td>PAM</td><td>alignment</td><td>distance</td><td>position</td><td>gene name</td><td>gene id</td></tr>
<tr><td>GU305914.1</td><td>76998</td><td>77020</td><td>-</td><td>0</td><td>CGTCACTTCTGGAGGGGTTT</td><td>TGG</td><td>||||||||[||||||||||||]PAM</td><td>NA</td><td>Intergenic</td><td>NA</td><td>NA</td></tr>
<tr><td>T28422</td><td>ACTTCTGGAGGGGTTTTGGTCGG</td><td>1000</td><td colspan="9"></td></tr>
<tr><td>Oligo fwd</td><td>CACCACTTCTGGAGGGGTTTTGGT</td><td colspan="10"></td></tr>
<tr><td>Oligo rev</td><td>AAACACCAAAACCCCTCCAGAAGT</td><td colspan="10"></td></tr>
<tr><td>Chromosome</td><td>start</td><td>end</td><td>strand</td><td>MM</td><td>target_seq</td><td>PAM</td><td>alignment</td><td>distance</td><td>position</td><td>gene name</td><td>gene id</td></tr>
<tr><td>GU305914.1</td><td>76994</td><td>77016</td><td>-</td><td>0</td><td>ACTTCTGGAGGGGTTTTGGT</td><td>CGG</td><td>||||||||[||||||||||||]PAM</td><td>NA</td><td>Intergenic</td><td>NA</td><td>NA</td></tr>
<tr><td>T28423</td><td>CTTCTGGAGGGGTTTTGGTCGGG</td><td>1000</td><td colspan="9"></td></tr>
<tr><td>Oligo fwd</td><td>CACCCTTCTGGAGGGGTTTTGGTC</td><td colspan="10"></td></tr>
<tr><td>Oligo rev</td><td>AAACGACCAAAACCCCTCCAGAAG</td><td colspan="10"></td></tr>
<tr><td>Chromosome</td><td>start</td><td>end</td><td>strand</td><td>MM</td><td>target_seq</td><td>PAM</td><td>alignment</td><td>distance</td><td>position</td><td>gene name</td><td>gene id</td></tr>
<tr><td>GU305914.1</td><td>76993</td><td>77015</td><td>-</td><td>0</td><td>CTTCTGGAGGGGTTTTGGTC</td><td>GGG</td><td>||||||||[||||||||||||]PAM</td><td>NA</td><td>Intergenic</td><td>NA</td><td>NA</td></tr>
<tr><td>T28424</td><td>GCAGCAACAGCTCGTTGACGAGG</td><td>1000</td><td colspan="9"></td></tr>
<tr><td>Oligo fwd</td><td>CACCGCAGCAACAGCTCGTTGACG</td><td colspan="10"></td></tr>
<tr><td>Oligo rev</td><td>AAACCGTCAACGAGCTGTTGCTGC</td><td colspan="10"></td></tr>
<tr><td>Chromosome</td><td>start</td><td>end</td><td>strand</td><td>MM</td><td>target_seq</td><td>PAM</td><td>alignment</td><td>distance</td><td>position</td><td>gene name</td><td>gene id</td></tr>
<tr><td>GU305914.1</td><td>76963</td><td>76985</td><td>-</td><td>0</td><td>GCAGCAACAGCTCGTTGACG</td><td>AGG</td><td>||||||||[||||||||||||]PAM</td><td>NA</td><td>Intergenic</td><td>NA</td><td>NA</td></tr>
<tr><td>T28425</td><td>CGAGTCGATCGTCGTCAGCGAGG</td><td>1000</td><td colspan="9"></td></tr>
<tr><td>Oligo fwd</td><td>CACCCGAGTCGATCGTCGTCAGCG</td><td colspan="10"></td></tr>
<tr><td>Oligo rev</td><td>AAACCGCTGACGACGATCGACTCG</td><td colspan="10"></td></tr>
<tr><td>Chromosome</td><td>start</td><td>end</td><td>strand</td><td>MM</td><td>target_seq</td><td>PAM</td><td>alignment</td><td>distance</td><td>position</td><td>gene name</td><td>gene id</td></tr>
<tr><td>GU305914.1</td><td>76938</td><td>76960</td><td>-</td><td>0</td><td>CGAGTCGATCGTCGTCAGCG</td><td>AGG</td><td>||||||||[||||||||||||]PAM</td><td>NA</td><td>Intergenic</td><td>NA</td><td>NA</td></tr>
<tr><td>GU305914.1</td><td>142554</td><td>142576</td><td>+</td><td>4</td><td>CTTGTCGATCGTCGACCGCG</td><td>AGG</td><td>|--|||||[||||||-|-|||]PAM</td><td>NA</td><td>Intergenic</td><td>NA</td><td>NA</td></tr>
<tr><td>T28426</td><td>GAGTCGATCGTCGTCAGCGAGGG</td><td>1000</td><td colspan="9"></td></tr>
<tr><td>Oligo fwd</td><td>CACCGAGTCGATCGTCGTCAGCGA</td><td colspan="10"></td></tr>
<tr><td>Oligo rev</td><td>AAACTCGCTGACGACGATCGACTC</td><td colspan="10"></td></tr>
<tr><td>Chromosome</td><td>start</td><td>end</td><td>strand</td><td>MM</td><td>target_seq</td><td>PAM</td><td>alignment</td><td>distance</td><td>position</td><td>gene name</td><td>gene id</td></tr>
<tr><td>GU305914.1</td><td>76937</td><td>76959</td><td>-</td><td>0</td><td>GAGTCGATCGTCGTCAGCGA</td><td>GGG</td><td>||||||||[||||||||||||]PAM</td><td>NA</td><td>Intergenic</td><td>NA</td><td>NA</td></tr>
<tr><td>T28427</td><td>CGTCGTCAGCGAGGGATCCCAGG</td><td>1000</td><td colspan="9"></td></tr>
<tr><td>Oligo fwd</td><td>CACCCGTCGTCAGCGAGGGATCCC</td><td colspan="10"></td></tr>
<tr><td>Oligo rev</td><td>AAACGGGATCCCTCGCTGACGACG</td><td colspan="10"></td></tr>
<tr><td>Chromosome</td><td>start</td><td>end</td><td>strand</td><td>MM</td><td>target_seq</td><td>PAM</td><td>alignment</td><td>distance</td><td>position</td><td>gene name</td><td>gene id</td></tr>
<tr><td>GU305914.1</td><td>76929</td><td>76951</td><td>-</td><td>0</td><td>CGTCGTCAGCGAGGGATCCC</td><td>AGG</td><td>||||||||[||||||||||||]PAM</td><td>NA</td><td>Intergenic</td><td>NA</td><td>NA</td></tr>
<tr><td>GU305914.1</td><td>104406</td><td>104428</td><td>+</td><td>4</td><td>CGTGATCAGCGACGGATCCG</td><td>AGG</td><td>|||--|||[||||-||||||-]PAM</td><td>NA</td><td>Intergenic</td><td>NA</td><td>NA</td></tr>
<tr><td>T28428</td><td>GTACATCGCGAGCGCCGCGACGG</td><td>1000</td><td colspan="9"></td></tr>
<tr><td>Oligo fwd</td><td>CACCGTACATCGCGAGCGCCGCGA</td><td colspan="10"></td></tr>
<tr><td>Oligo rev</td><td>AAACTCGCGGCGCTCGCGATGTAC</td><td colspan="10"></td></tr>
<tr><td>Chromosome</td><td>start</td><td>end</td><td>strand</td><td>MM</td><td>target_seq</td><td>PAM</td><td>alignment</td><td>distance</td><td>position</td><td>gene name</td><td>gene id</td></tr>
<tr><td>GU305914.1</td><td>76902</td><td>76924</td><td>-</td><td>0</td><td>GTACATCGCGAGCGCCGCGA</td><td>CGG</td><td>||||||||[||||||||||||]PAM</td><td>NA</td><td>Intergenic</td><td>NA</td><td>NA</td></tr>
</table>

T28429 AGCGCCGCGACGGCGATGAACGG 1000
Oligo fwd CACCAGCGCCGCGACGGCGATGAA
Oligo rev AAACTTCATCGCCGTCGCGGCGCT
Chromosome start end strand MM target_seq PAM alignment distance position gene name gene id
GU305914.1 76892 76914 - 0 AGCGCCGCGACGGCGATGAA CGG ||||||||[||||||||||||]PAM NA Intergenic NA NA

T28430 GGTGATAGCGTTCGCCGTCGCGG 1000
Oligo fwd CACCGGTGATAGCGTTCGCCGTCG
Oligo rev AAACCGACGGCGAACGCTATCACC
Chromosome start end strand MM target_seq PAM alignment distance position gene name gene id
GU305914.1 76871 76893 - 0 GGTGATAGCGTTCGCCGTCG CGG ||||||||[||||||||||||]PAM NA Intergenic NA NA

T28431 GATAGCGTTCGCCGTCGCGGCGG 1000
Oligo fwd CACCGATAGCGTTCGCCGTCGCGG
Oligo rev AAACCCGCGACGGCGAACGCTATC
Chromosome start end strand MM target_seq PAM alignment distance position gene name gene id
GU305914.1 76868 76890 - 0 GATAGCGTTCGCCGTCGCGG CGG ||||||||[||||||||||||]PAM NA Intergenic NA NA

T28432 CCGTCGCGGCGGCGATGCGCCGG 1000
Oligo fwd CACCCCGTCGCGGCGGCGATGCGC
Oligo rev AAACGCGCATCGCCGCCGCGACGG
Chromosome start end strand MM target_seq PAM alignment distance position gene name gene id
GU305914.1 76857 76879 - 0 CCGTCGCGGCGGCGATGCGC CGG ||||||||[||||||||||||]PAM NA Intergenic NA NA

T28433 CGATGCGCCGGACGACGACGAGG 1000
Oligo fwd CACCCGATGCGCCGGACGACGACG
Oligo rev AAACCGTCGTCGTCCGGCGCATCG
Chromosome start end strand MM target_seq PAM alignment distance position gene name gene id
GU305914.1 76845 76867 - 0 CGATGCGCCGGACGACGACG AGG ||||||||[||||||||||||]PAM NA Intergenic NA NA

T28434 CCGGACGACGACGAGGCCGCCGG 1000
Oligo fwd CACCCCGGACGACGACGAGGCCGC
Oligo rev AAACGCGGCCTCGTCGTCGTCCGG
Chromosome start end strand MM target_seq PAM alignment distance position gene name gene id
GU305914.1 76838 76860 - 0 CCGGACGACGACGAGGCCGC CGG ||||||||[||||||||||||]PAM NA Intergenic NA NA
GU305914.1 115379 115401 - 4 CCTGGCGACGACGGTGCCGC CGG ||-|-|||[|||||--|||||]PAM NA Intergenic NA NA

T28435 CGCCGCGCCCGTCCCCGCCGAGG 1000
Oligo fwd CACCCGCCGCGCCCGTCCCCGCCG
Oligo rev AAACCGGCGGGGACGGGCGCGGCG
Chromosome start end strand MM target_seq PAM alignment distance position gene name gene id
GU305914.1 76815 76837 - 0 CGCCGCGCCCGTCCCCGCCG AGG ||||||||[||||||||||||]PAM NA Intergenic NA NA

T28436 GCGCTCTTCGAACGCTTCTTCGG 1000
Oligo fwd CACCGCGCTCTTCGAACGCTTCTT
Oligo rev AAACAAGAAGCGTTCGAAGAGCGC
Chromosome start end strand MM target_seq PAM alignment distance position gene name gene id
GU305914.1 76778 76800 - 0 GCGCTCTTCGAACGCTTCTT CGG ||||||||[||||||||||||]PAM NA Intergenic NA NA

T28437 CGCTCTTCGAACGCTTCTTCGGG 1000
Oligo fwd CACCCGCTCTTCGAACGCTTCTTC
Oligo rev AAACGAAGAAGCGTTCGAAGAGCG
Chromosome start end strand MM target_seq PAM alignment distance position gene name gene id
GU305914.1 76777 76799 - 0 CGCTCTTCGAACGCTTCTTC GGG ||||||||[||||||||||||]PAM NA Intergenic NA NA

T28438 GCTCTTCGAACGCTTCTTCGGGG 1000
Oligo fwd CACCGCTCTTCGAACGCTTCTTCG
Oligo rev AAACCGAAGAAGCGTTCGAAGAGC
Chromosome start end strand MM target_seq PAM alignment distance position gene name gene id
GU305914.1 76776 76798 - 0 GCTCTTCGAACGCTTCTTCG GGG ||||||||[||||||||||||]PAM NA Intergenic NA NA

T28439 TTCGAACGCTTCTTCGGGGACGG 1000
Oligo fwd CACCTTCGAACGCTTCTTCGGGGA
Oligo rev AAACTCCCCGAAGAAGCGTTCGAA
Chromosome start end strand MM target_seq PAM alignment distance position gene name gene id
GU305914.1 76772 76794 - 0 TTCGAACGCTTCTTCGGGGA CGG ||||||||[||||||||||||]PAM NA Intergenic NA NA

T28440 GAACGCTTCTTCGGGGACGGCGG 1000
Oligo fwd CACCGAACGCTTCTTCGGGGACGG
Oligo rev AAACCCGTCCCCGAAGAAGCGTTC
Chromosome start end strand MM target_seq PAM alignment distance position gene name gene id
GU305914.1 76769 76791 - 0 GAACGCTTCTTCGGGGACGG CGG ||||||||[||||||||||||]PAM NA Intergenic NA NA

T28441 GTTTCGAGCCGATGCTGCCTAGG 1000
Oligo fwd CACCGTTTCGAGCCGATGCTGCCT
Oligo rev AAACAGGCAGCATCGGCTCGAAAC
Chromosome start end strand MM target_seq PAM alignment distance position gene name gene id
GU305914.1 76732 76754 - 0 GTTTCGAGCCGATGCTGCCT AGG ||||||||[||||||||||||]PAM NA Intergenic NA NA

T28442 TTTCGAGCCGATGCTGCCTAGGG 1000
Oligo fwd CACCTTTCGAGCCGATGCTGCCTA
Oligo rev AAACTAGGCAGCATCGGCTCGAAA
Chromosome start end strand MM target_seq PAM alignment distance position gene name gene id
GU305914.1 76731 76753 - 0 TTTCGAGCCGATGCTGCCTA GGG ||||||||[||||||||||||]PAM NA Intergenic NA NA

T28443 CGTTGCCGTCTATCGACTCGAGG 1000
Oligo fwd CACCCGTTGCCGTCTATCGACTCG
Oligo rev AAACCGAGTCGATAGACGGCAACG

Chromosome start end strand MM target_seq PAM alignment distance position gene name gene id
GU305914.1 76696 76718 - 0 CGTTGCCGTCTATCGACTCG AGG ||||||||[||||||||||||]PAM NA Intergenic NA NA

T28444 AGGTTAAATTTTATCAACGTCGG 1000
Oligo fwd CACCAGGTTAAATTTTATCAACGT
Oligo rev AAACACGTTGATAAAATTTAACCT
Chromosome start end strand MM target_seq PAM alignment distance position gene name gene id
GU305914.1 76676 76698 - 0 AGGTTAAATTTTATCAACGT CGG ||||||||[||||||||||||]PAM NA Intergenic NA NA

T28445 GACGTCACGCCGCTTTTCTGCGG 1000
Oligo fwd CACCGACGTCACGCCGCTTTTCTG
Oligo rev AAACCAGAAAAGCGGCGTGACGTC
Chromosome start end strand MM target_seq PAM alignment distance position gene name gene id
GU305914.1 76651 76673 - 0 GACGTCACGCCGCTTTTCTG CGG ||||||||[||||||||||||]PAM NA Intergenic NA NA

T28446 GCTTTTCTGCGGCATGTGTACGG 1000
Oligo fwd CACCGCTTTTCTGCGGCATGTGTA
Oligo rev AAACTACACATGCCGCAGAAAAGC
Chromosome start end strand MM target_seq PAM alignment distance position gene name gene id
GU305914.1 76640 76662 - 0 GCTTTTCTGCGGCATGTGTA CGG ||||||||[||||||||||||]PAM NA Intergenic NA NA

T28447 CTTTTCTGCGGCATGTGTACGGG 1000
Oligo fwd CACCCTTTTCTGCGGCATGTGTAC
Oligo rev AAACGTACACATGCCGCAGAAAAG
Chromosome start end strand MM target_seq PAM alignment distance position gene name gene id
GU305914.1 76639 76661 - 0 CTTTTCTGCGGCATGTGTAC GGG ||||||||[||||||||||||]PAM NA Intergenic NA NA

T28448 TTTTCTGCGGCATGTGTACGGGG 1000
Oligo fwd CACCTTTTCTGCGGCATGTGTACG
Oligo rev AAACCGTACACATGCCGCAGAAAA
Chromosome start end strand MM target_seq PAM alignment distance position gene name gene id
GU305914.1 76638 76660 - 0 TTTTCTGCGGCATGTGTACG GGG ||||||||[||||||||||||]PAM NA Intergenic NA NA

T28449 TTTCTGCGGCATGTGTACGGGGG 1000
Oligo fwd CACCTTTCTGCGGCATGTGTACGG
Oligo rev AAACCCGTACACATGCCGCAGAAA
Chromosome start end strand MM target_seq PAM alignment distance position gene name gene id
GU305914.1 76637 76659 - 0 TTTCTGCGGCATGTGTACGG GGG ||||||||[||||||||||||]PAM NA Intergenic NA NA

T28450 TTCTGCGGCATGTGTACGGGGGG 1000
Oligo fwd CACCTTCTGCGGCATGTGTACGGG
Oligo rev AAACCCCGTACACATGCCGCAGAA
Chromosome start end strand MM target_seq PAM alignment distance position gene name gene id
GU305914.1 76636 76658 - 0 TTCTGCGGCATGTGTACGGG GGG ||||||||[||||||||||||]PAM NA Intergenic NA NA

T28451 ATGTGTACGGGGGGTGCGACAGG 1000
Oligo fwd CACCATGTGTACGGGGGGTGCGAC
Oligo rev AAACGTCGCACCCCCCGTACACAT
Chromosome start end strand MM target_seq PAM alignment distance position gene name gene id
GU305914.1 76627 76649 - 0 ATGTGTACGGGGGGTGCGAC AGG ||||||||[||||||||||||]PAM NA Intergenic NA NA

T28452 CGAGAAGATGAAGCTCTTTACGG 1000
Oligo fwd CACCCGAGAAGATGAAGCTCTTTA
Oligo rev AAACTAAAGAGCTTCATCTTCTCG
Chromosome start end strand MM target_seq PAM alignment distance position gene name gene id
GU305914.1 76581 76603 - 0 CGAGAAGATGAAGCTCTTTA CGG ||||||||[||||||||||||]PAM NA Intergenic NA NA

T28453 ACTAAATTATTAGACGTGAATGG 1000
Oligo fwd CACCACTAAATTATTAGACGTGAA
Oligo rev AAACTTCACGTCTAATAATTTAGT
Chromosome start end strand MM target_seq PAM alignment distance position gene name gene id
GU305914.1 76550 76572 - 0 ACTAAATTATTAGACGTGAA TGG ||||||||[||||||||||||]PAM NA Intergenic NA NA

T28454 TGAATGGAATATTAGAACGCAGG 1000
Oligo fwd CACCTGAATGGAATATTAGAACGC
Oligo rev AAACGCGTTCTAATATTCCATTCA
Chromosome start end strand MM target_seq PAM alignment distance position gene name gene id
GU305914.1 76534 76556 - 0 TGAATGGAATATTAGAACGC AGG ||||||||[||||||||||||]PAM NA Intergenic NA NA

T28455 GAATGGAATATTAGAACGCAGGG 1000
Oligo fwd CACCGAATGGAATATTAGAACGCA
Oligo rev AAACTGCGTTCTAATATTCCATTC
Chromosome start end strand MM target_seq PAM alignment distance position gene name gene id
GU305914.1 76533 76555 - 0 GAATGGAATATTAGAACGCA GGG ||||||||[||||||||||||]PAM NA Intergenic NA NA

T28456 CAGGGAAACGACCGACTGATCGG 1000
Oligo fwd CACCCAGGGAAACGACCGACTGAT
Oligo rev AAACATCAGTCGGTCGTTTCCCTG
Chromosome start end strand MM target_seq PAM alignment distance position gene name gene id
GU305914.1 76515 76537 - 0 CAGGGAAACGACCGACTGAT CGG ||||||||[||||||||||||]PAM NA Intergenic NA NA

T28457 AGGGAAACGACCGACTGATCGGG 1000
Oligo fwd CACCAGGGAAACGACCGACTGATC
Oligo rev AAACGATCAGTCGGTCGTTTCCCT
Chromosome start end strand MM target_seq PAM alignment distance position gene name gene id
GU305914.1 76514 76536 - 0 AGGGAAACGACCGACTGATC GGG ||||||||[||||||||||||]PAM NA Intergenic NA NA

T28458 AAACGACCGACTGATCGGGTCGG 1000
Oligo fwd CACCAAACGACCGACTGATCGGGT

| | | | | | | | | | | | |
|---|---|---|---|---|---|---|---|---|---|---|---|
| Oligo rev | AAACACCCGATCAGTCGGTCGTTT | | | | | | | | | | |
| Chromosome | start | end | strand | MM | target_seq | PAM | alignment | distance | position | gene name | gene id |
| GU305914.1 | 76510 | 76532 | - | 0 | AAACGACCGACTGATCGGGT | CGG | \|\|\|\|\|\|\|\|[\|\|\|\|\|\|\|\|\|\|\|\|]PAM | NA | Intergenic | NA | NA |
| T28459 | ACCGACTGATCGGGTCGGCTCGG | 1000 | | | | | | | | | |
| Oligo fwd | CACCACCGACTGATCGGGTCGGCT | | | | | | | | | | |
| Oligo rev | AAACAGCCGACCCGATCAGTCGGT | | | | | | | | | | |
| Chromosome | start | end | strand | MM | target_seq | PAM | alignment | distance | position | gene name | gene id |
| GU305914.1 | 76505 | 76527 | - | 0 | ACCGACTGATCGGGTCGGCT | CGG | \|\|\|\|\|\|\|\|[\|\|\|\|\|\|\|\|\|\|\|\|]PAM | NA | Intergenic | NA | NA |
| T28460 | CCGACTGATCGGGTCGGCTCGGG | 1000 | | | | | | | | | |
| Oligo fwd | CACCCCGACTGATCGGGTCGGCTC | | | | | | | | | | |
| Oligo rev | AAACGAGCCGACCCGATCAGTCGG | | | | | | | | | | |
| Chromosome | start | end | strand | MM | target_seq | PAM | alignment | distance | position | gene name | gene id |
| GU305914.1 | 76504 | 76526 | - | 0 | CCGACTGATCGGGTCGGCTC | GGG | \|\|\|\|\|\|\|\|[\|\|\|\|\|\|\|\|\|\|\|\|]PAM | NA | Intergenic | NA | NA |
| T28461 | ACTGATCGGGTCGGCTCGGGCGG | 1000 | | | | | | | | | |
| Oligo fwd | CACCACTGATCGGGTCGGCTCGGG | | | | | | | | | | |
| Oligo rev | AAACCCCGAGCCGACCCGATCAGT | | | | | | | | | | |
| Chromosome | start | end | strand | MM | target_seq | PAM | alignment | distance | position | gene name | gene id |
| GU305914.1 | 76501 | 76523 | - | 0 | ACTGATCGGGTCGGCTCGGG | CGG | \|\|\|\|\|\|\|\|[\|\|\|\|\|\|\|\|\|\|\|\|]PAM | NA | Intergenic | NA | NA |
| T28462 | TCGGCTCGGGCGGCGCCACTCGG | 1000 | | | | | | | | | |
| Oligo fwd | CACCTCGGCTCGGGCGGCGCCACT | | | | | | | | | | |
| Oligo rev | AAACAGTGGCGCCGCCCGAGCCGA | | | | | | | | | | |
| Chromosome | start | end | strand | MM | target_seq | PAM | alignment | distance | position | gene name | gene id |
| GU305914.1 | 76491 | 76513 | - | 0 | TCGGCTCGGGCGGCGCCACT | CGG | \|\|\|\|\|\|\|\|[\|\|\|\|\|\|\|\|\|\|\|\|]PAM | NA | Intergenic | NA | NA |
| T28463 | CTCGGGCGGCGCCACTCGGACGG | 1000 | | | | | | | | | |
| Oligo fwd | CACCCTCGGGCGGCGCCACTCGGA | | | | | | | | | | |
| Oligo rev | AAACTCCGAGTGGCGCCGCCCGAG | | | | | | | | | | |
| Chromosome | start | end | strand | MM | target_seq | PAM | alignment | distance | position | gene name | gene id |
| GU305914.1 | 76487 | 76509 | - | 0 | CTCGGGCGGCGCCACTCGGA | CGG | \|\|\|\|\|\|\|\|[\|\|\|\|\|\|\|\|\|\|\|\|]PAM | NA | Intergenic | NA | NA |
| T28464 | GGGCGGCGCCACTCGGACGGCGG | 1000 | | | | | | | | | |
| Oligo fwd | CACCGGGCGGCGCCACTCGGACGG | | | | | | | | | | |
| Oligo rev | AAACCCGTCCGAGTGGCGCCGCCC | | | | | | | | | | |
| Chromosome | start | end | strand | MM | target_seq | PAM | alignment | distance | position | gene name | gene id |
| GU305914.1 | 76484 | 76506 | - | 0 | GGGCGGCGCCACTCGGACGG | CGG | \|\|\|\|\|\|\|\|[\|\|\|\|\|\|\|\|\|\|\|\|]PAM | NA | Intergenic | NA | NA |
| T28465 | GACGGCGGCGAGCTCATCCGCGG | 1000 | | | | | | | | | |
| Oligo fwd | CACCGACGGCGGCGAGCTCATCCG | | | | | | | | | | |
| Oligo rev | AAACCGGATGAGCTCGCCGCCGTC | | | | | | | | | | |
| Chromosome | start | end | strand | MM | target_seq | PAM | alignment | distance | position | gene name | gene id |
| GU305914.1 | 76469 | 76491 | - | 0 | GACGGCGGCGAGCTCATCCG | CGG | \|\|\|\|\|\|\|\|[\|\|\|\|\|\|\|\|\|\|\|\|]PAM | NA | Intergenic | NA | NA |
| T28466 | GGCGGCGAGCTCATCCGCGGCGG | 1000 | | | | | | | | | |
| Oligo fwd | CACCGGCGGCGAGCTCATCCGCGG | | | | | | | | | | |
| Oligo rev | AAACCCGCGGATGAGCTCGCCGCC | | | | | | | | | | |
| Chromosome | start | end | strand | MM | target_seq | PAM | alignment | distance | position | gene name | gene id |
| GU305914.1 | 76466 | 76488 | - | 0 | GGCGGCGAGCTCATCCGCGG | CGG | \|\|\|\|\|\|\|\|[\|\|\|\|\|\|\|\|\|\|\|\|]PAM | NA | Intergenic | NA | NA |
| GU305914.1 | 147953 | 147975 | - | 4 | GGCGTCCAGCTCATCCGAGC | CGG | \|\|\|\|-\|-\|[\|\|\|\|\|\|\|\|\|-\|-]PAM | NA | Intergenic | NA | NA |
| T28467 | GAGCTCATCCGCGGCGGCGCCGG | 1000 | | | | | | | | | |
| Oligo fwd | CACCGAGCTCATCCGCGGCGGCGC | | | | | | | | | | |
| Oligo rev | AAACGCGCCGCCGCGGATGAGCTC | | | | | | | | | | |
| Chromosome | start | end | strand | MM | target_seq | PAM | alignment | distance | position | gene name | gene id |
| GU305914.1 | 76460 | 76482 | - | 0 | GAGCTCATCCGCGGCGGCGC | CGG | \|\|\|\|\|\|\|\|[\|\|\|\|\|\|\|\|\|\|\|\|]PAM | NA | Intergenic | NA | NA |
| T28468 | CCGCGGCGGCGCCGGCGCGATGG | 1000 | | | | | | | | | |
| Oligo fwd | CACCCCGCGGCGGCGCCGGCGCGA | | | | | | | | | | |
| Oligo rev | AAACTCGCGCCGGCGCCGCCGCGG | | | | | | | | | | |
| Chromosome | start | end | strand | MM | target_seq | PAM | alignment | distance | position | gene name | gene id |
| GU305914.1 | 76452 | 76474 | - | 0 | CCGCGGCGGCGCCGGCGCGA | TGG | \|\|\|\|\|\|\|\|[\|\|\|\|\|\|\|\|\|\|\|\|]PAM | NA | Intergenic | NA | NA |
| GU305914.1 | 152918 | 152940 | + | 3 | CAACGGCGGCGACGGCGCGA | GGG | \|--\|\|\|\|\|[\|\|\|-\|\|\|\|\|\|\|\|]PAM | NA | Intergenic | NA | NA |
| GU305914.1 | 125187 | 125209 | + | 4 | GCGCGGCAGCGCCTGCACGA | AGG | -\|\|\|\|\|\|-[\|\|\|\|\|-\|\|-\|\|\|]PAM | NA | Intergenic | NA | NA |
| T28469 | GATGGAGATCGACAAGAATGTGG | 1000 | | | | | | | | | |
| Oligo fwd | CACCGATGGAGATCGACAAGAATG | | | | | | | | | | |
| Oligo rev | AAACCATTCTTGTCGATCTCCATC | | | | | | | | | | |
| Chromosome | start | end | strand | MM | target_seq | PAM | alignment | distance | position | gene name | gene id |
| GU305914.1 | 76434 | 76456 | - | 0 | GATGGAGATCGACAAGAATG | TGG | \|\|\|\|\|\|\|\|[\|\|\|\|\|\|\|\|\|\|\|\|]PAM | NA | Intergenic | NA | NA |
| T28470 | ATGGAGATCGACAAGAATGTGGG | 1000 | | | | | | | | | |
| Oligo fwd | CACCATGGAGATCGACAAGAATGT | | | | | | | | | | |
| Oligo rev | AAACACATTCTTGTCGATCTCCAT | | | | | | | | | | |
| Chromosome | start | end | strand | MM | target_seq | PAM | alignment | distance | position | gene name | gene id |
| GU305914.1 | 76433 | 76455 | - | 0 | ATGGAGATCGACAAGAATGT | GGG | \|\|\|\|\|\|\|\|[\|\|\|\|\|\|\|\|\|\|\|\|]PAM | NA | Intergenic | NA | NA |
| T28471 | CCGACCTGATTTCCAACACGCGG | 1000 | | | | | | | | | |
| Oligo fwd | CACCCCGACCTGATTTCCAACACG | | | | | | | | | | |
| Oligo rev | AAACCGTGTTGGAAATCAGGTCGG | | | | | | | | | | |
| Chromosome | start | end | strand | MM | target_seq | PAM | alignment | distance | position | gene name | gene id |
| GU305914.1 | 76408 | 76430 | - | 0 | CCGACCTGATTTCCAACACG | CGG | \|\|\|\|\|\|\|\|[\|\|\|\|\|\|\|\|\|\|\|\|]PAM | NA | Intergenic | NA | NA |
| T28472 | ACCTGATTTCCAACACGCGGCGG | 1000 | | | | | | | | | |
| Oligo fwd | CACCACCTGATTTCCAACACGCGG | | | | | | | | | | |
| Oligo rev | AAACCCGCGTGTTGGAAATCAGGT | | | | | | | | | | |
| Chromosome | start | end | strand | MM | target_seq | PAM | alignment | distance | position | gene name | gene id |

GU305914.1 76405 76427 - 0 ACCTGATTTCCAACACGCGG CGG ||||||||[||||||||||||]PAM NA Intergenic NA NA

T28473 CCAACACGCGGCGGATCCTCCGG 1000
Oligo fwd CACCCCAACACGCGGCGGATCCTC
Oligo rev AAACGAGGATCCGCCGCGTGTTGG
Chromosome start end strand MM target_seq PAM alignment distance position gene name gene id
GU305914.1 76396 76418 - 0 CCAACACGCGGCGGATCCTC CGG ||||||||[||||||||||||]PAM NA Intergenic NA NA

T28474 GGCTCGACGAGAACGAGCTCAGG 1000
Oligo fwd CACCGGCTCGACGAGAACGAGCTC
Oligo rev AAACGAGCTCGTTCTCGTCGAGCC
Chromosome start end strand MM target_seq PAM alignment distance position gene name gene id
GU305914.1 76375 76397 - 0 GGCTCGACGAGAACGAGCTC AGG ||||||||[||||||||||||]PAM NA Intergenic NA NA

T28475 GACCACCGATATCGTCTATCCGG 1000
Oligo fwd CACCGACCACCGATATCGTCTATC
Oligo rev AAACGATAGACGATATCGGTGGTC
Chromosome start end strand MM target_seq PAM alignment distance position gene name gene id
GU305914.1 76293 76315 - 0 GACCACCGATATCGTCTATC CGG ||||||||[||||||||||||]PAM NA Intergenic NA NA

T28476 TCGAGTATCTCCTCAGCTATTGG 1000
Oligo fwd CACCTCGAGTATCTCCTCAGCTAT
Oligo rev AAACATAGCTGAGGAGATACTCGA
Chromosome start end strand MM target_seq PAM alignment distance position gene name gene id
GU305914.1 76270 76292 - 0 TCGAGTATCTCCTCAGCTAT TGG ||||||||[||||||||||||]PAM NA Intergenic NA NA

T28477 CGAGTATCTCCTCAGCTATTGGG 1000
Oligo fwd CACCCGAGTATCTCCTCAGCTATT
Oligo rev AAACAATAGCTGAGGAGATACTCG
Chromosome start end strand MM target_seq PAM alignment distance position gene name gene id
GU305914.1 76269 76291 - 0 CGAGTATCTCCTCAGCTATT GGG ||||||||[||||||||||||]PAM NA Intergenic NA NA

T28478 TCCTCAGCTATTGGGAGTGTAGG 1000
Oligo fwd CACCTCCTCAGCTATTGGGAGTGT
Oligo rev AAACACACTCCCAATAGCTGAGGA
Chromosome start end strand MM target_seq PAM alignment distance position gene name gene id
GU305914.1 76261 76283 - 0 TCCTCAGCTATTGGGAGTGT AGG ||||||||[||||||||||||]PAM NA Intergenic NA NA

T28479 AGCTATTGGGAGTGTAGGAGCGG 1000
Oligo fwd CACCAGCTATTGGGAGTGTAGGAG
Oligo rev AAACCTCCTACACTCCCAATAGCT
Chromosome start end strand MM target_seq PAM alignment distance position gene name gene id
GU305914.1 76256 76278 - 0 AGCTATTGGGAGTGTAGGAG CGG ||||||||[||||||||||||]PAM NA Intergenic NA NA

T28480 ATTGGGAGTGTAGGAGCGGCAGG 1000
Oligo fwd CACCATTGGGAGTGTAGGAGCGGC
Oligo rev AAACGCCGCTCCTACACTCCCAAT
Chromosome start end strand MM target_seq PAM alignment distance position gene name gene id
GU305914.1 76252 76274 - 0 ATTGGGAGTGTAGGAGCGGC AGG ||||||||[||||||||||||]PAM NA Intergenic NA NA

T28481 TGCTTCGTGTTCAAAAACACCGG 1000
Oligo fwd CACCTGCTTCGTGTTCAAAAACAC
Oligo rev AAACGTGTTTTTGAACACGAAGCA
Chromosome start end strand MM target_seq PAM alignment distance position gene name gene id
GU305914.1 76223 76245 - 0 TGCTTCGTGTTCAAAAACAC CGG ||||||||[||||||||||||]PAM NA Intergenic NA NA

T28482 TGTTCAAAAACACCGGCTGTCGG 1000
Oligo fwd CACCTGTTCAAAAACACCGGCTGT
Oligo rev AAACACAGCCGGTGTTTTTGAACA
Chromosome start end strand MM target_seq PAM alignment distance position gene name gene id
GU305914.1 76216 76238 - 0 TGTTCAAAAACACCGGCTGT CGG ||||||||[||||||||||||]PAM NA Intergenic NA NA

T28483 GTTCAAAAACACCGGCTGTCGGG 1000
Oligo fwd CACCGTTCAAAAACACCGGCTGTC
Oligo rev AAACGACAGCCGGTGTTTTTGAAC
Chromosome start end strand MM target_seq PAM alignment distance position gene name gene id
GU305914.1 76215 76237 - 0 GTTCAAAAACACCGGCTGTC GGG ||||||||[||||||||||||]PAM NA Intergenic NA NA

T28484 GTCTCCCTCTCCTGCTATATAGG 1000
Oligo fwd CACCGTCTCCCTCTCCTGCTATAT
Oligo rev AAACATATAGCAGGAGAGGGAGAC
Chromosome start end strand MM target_seq PAM alignment distance position gene name gene id
GU305914.1 76193 76215 - 0 GTCTCCCTCTCCTGCTATAT AGG ||||||||[||||||||||||]PAM NA Intergenic NA NA

T28485 AGGCTTTCCCGAGCGCCTCAAGG 1000
Oligo fwd CACCAGGCTTTCCCGAGCGCCTCA
Oligo rev AAACTGAGGCGCTCGGGAAAGCCT
Chromosome start end strand MM target_seq PAM alignment distance position gene name gene id
GU305914.1 76173 76195 - 0 AGGCTTTCCCGAGCGCCTCA AGG ||||||||[||||||||||||]PAM NA Intergenic NA NA

T28486 AGCGCCTCAAGGACCTCAAGAGG 1000
Oligo fwd CACCAGCGCCTCAAGGACCTCAAG
Oligo rev AAACCTTGAGGTCCTTGAGGCGCT
Chromosome start end strand MM target_seq PAM alignment distance position gene name gene id
GU305914.1 76162 76184 - 0 AGCGCCTCAAGGACCTCAAG AGG ||||||||[||||||||||||]PAM NA Intergenic NA NA

T28487 GCGCCTCAAGGACCTCAAGAGGG 1000
Oligo fwd CACCGCGCCTCAAGGACCTCAAGA
Oligo rev AAACTCTTGAGGTCCTTGAGGCGC

Chromosome start end strand MM target_seq PAM alignment distance position gene name gene id
GU305914.1 76161 76183 - 0 GCGCCTCAAGGACCTCAAGA GGG ||||||||[||||||||||||]PAM NA Intergenic NA NA

T28488 GAGCGTCAACGAAGCCCTCGTGG 1000
Oligo fwd CACCGAGCGTCAACGAAGCCCTCG
Oligo rev AAACCGAGGGCTTCGTTGACGCTC
Chromosome start end strand MM target_seq PAM alignment distance position gene name gene id
GU305914.1 76119 76141 - 0 GAGCGTCAACGAAGCCCTCG TGG ||||||||[||||||||||||]PAM NA Intergenic NA NA

T28489 CGAAGCCCTCGTGGTCACGCTGG 1000
Oligo fwd CACCCGAAGCCCTCGTGGTCACGC
Oligo rev AAACGCGTGACCACGAGGGCTTCG
Chromosome start end strand MM target_seq PAM alignment distance position gene name gene id
GU305914.1 76110 76132 - 0 CGAAGCCCTCGTGGTCACGC TGG ||||||||[||||||||||||]PAM NA Intergenic NA NA

T28490 TCACGCTGGCCGACATCGAGAGG 1000
Oligo fwd CACCTCACGCTGGCCGACATCGAG
Oligo rev AAACCTCGATGTCGGCCAGCGTGA
Chromosome start end strand MM target_seq PAM alignment distance position gene name gene id
GU305914.1 76096 76118 - 0 TCACGCTGGCCGACATCGAG AGG ||||||||[||||||||||||]PAM NA Intergenic NA NA

T28491 GAGGATCAAGCCCTGCGACAAGG 1000
Oligo fwd CACCGAGGATCAAGCCCTGCGACA
Oligo rev AAACTGTCGCAGGGCTTGATCCTC
Chromosome start end strand MM target_seq PAM alignment distance position gene name gene id
GU305914.1 76077 76099 - 0 GAGGATCAAGCCCTGCGACA AGG ||||||||[||||||||||||]PAM NA Intergenic NA NA

T28492 AGGATCAAGCCCTGCGACAAGGG 1000
Oligo fwd CACCAGGATCAAGCCCTGCGACAA
Oligo rev AAACTTGTCGCAGGGCTTGATCCT
Chromosome start end strand MM target_seq PAM alignment distance position gene name gene id
GU305914.1 76076 76098 - 0 AGGATCAAGCCCTGCGACAA GGG ||||||||[||||||||||||]PAM NA Intergenic NA NA

T28493 GGATCAAGCCCTGCGACAAGGGG 1000
Oligo fwd CACCGGATCAAGCCCTGCGACAAG
Oligo rev AAACCTTGTCGCAGGGCTTGATCC
Chromosome start end strand MM target_seq PAM alignment distance position gene name gene id
GU305914.1 76075 76097 - 0 GGATCAAGCCCTGCGACAAG GGG ||||||||[||||||||||||]PAM NA Intergenic NA NA

T28494 GATCAAGCCCTGCGACAAGGGGG 1000
Oligo fwd CACCGATCAAGCCCTGCGACAAGG
Oligo rev AAACCCTTGTCGCAGGGCTTGATC
Chromosome start end strand MM target_seq PAM alignment distance position gene name gene id
GU305914.1 76074 76096 - 0 GATCAAGCCCTGCGACAAGG GGG ||||||||[||||||||||||]PAM NA Intergenic NA NA

T28495 TGCTCACCAACTGCGTCGTCAGG 1000
Oligo fwd CACCTGCTCACCAACTGCGTCGTC
Oligo rev AAACGACGACGCAGTTGGTGAGCA
Chromosome start end strand MM target_seq PAM alignment distance position gene name gene id
GU305914.1 76051 76073 - 0 TGCTCACCAACTGCGTCGTC AGG ||||||||[||||||||||||]PAM NA Intergenic NA NA

T28496 GTCGTCAGGAAGTCCAACAGCGG 1000
Oligo fwd CACCGTCGTCAGGAAGTCCAACAG
Oligo rev AAACCTGTTGGACTTCCTGACGAC
Chromosome start end strand MM target_seq PAM alignment distance position gene name gene id
GU305914.1 76037 76059 - 0 GTCGTCAGGAAGTCCAACAG CGG ||||||||[||||||||||||]PAM NA Intergenic NA NA

T28497 CGGCATGTCCTACAACATCGAGG 1000
Oligo fwd CACCCGGCATGTCCTACAACATCG
Oligo rev AAACCGATGTTGTAGGACATGCCG
Chromosome start end strand MM target_seq PAM alignment distance position gene name gene id
GU305914.1 76017 76039 - 0 CGGCATGTCCTACAACATCG AGG ||||||||[||||||||||||]PAM NA Intergenic NA NA

T28498 AACATCGAGGTCGTCGCCTTCGG 1000
Oligo fwd CACCAACATCGAGGTCGTCGCCTT
Oligo rev AAACAAGGCGACGACCTCGATGTT
Chromosome start end strand MM target_seq PAM alignment distance position gene name gene id
GU305914.1 76004 76026 - 0 AACATCGAGGTCGTCGCCTT CGG ||||||||[||||||||||||]PAM NA Intergenic NA NA

T28499 CGCCTTCGGACCCGACAACGAGG 1000
Oligo fwd CACCCGCCTTCGGACCCGACAACG
Oligo rev AAACCGTTGTCGGGTCCGAAGGCG
Chromosome start end strand MM target_seq PAM alignment distance position gene name gene id
GU305914.1 75990 76012 - 0 CGCCTTCGGACCCGACAACG AGG ||||||||[||||||||||||]PAM NA Intergenic NA NA

T28500 CGACAACGAGGCCGAGTACCAGG 1000
Oligo fwd CACCCGACAACGAGGCCGAGTACC
Oligo rev AAACGGTACTCGGCCTCGTTGTCG
Chromosome start end strand MM target_seq PAM alignment distance position gene name gene id
GU305914.1 75978 76000 - 0 CGACAACGAGGCCGAGTACC AGG ||||||||[||||||||||||]PAM NA Intergenic NA NA

T28501 TCCTGCGCGACATCTACGCGCGG 1000
Oligo fwd CACCTCCTGCGCGACATCTACGCG
Oligo rev AAACCGCGTAGATGTCGCGCAGGA
Chromosome start end strand MM target_seq PAM alignment distance position gene name gene id
GU305914.1 75952 75974 - 0 TCCTGCGCGACATCTACGCG CGG ||||||||[||||||||||||]PAM NA Intergenic NA NA

T28502 TGCGCGACATCTACGCGCGGAGG 1000
Oligo fwd CACCTGCGCGACATCTACGCGCGG

Oligo rev AAACCCGCGCGTAGATGTCGCGCA
Chromosome start end strand MM target_seq PAM alignment distance position gene name gene id
GU305914.1 75949 75971 - 0 TGCGCGACATCTACGCGCGG AGG ||||||||[||||||||||||]PAM NA Intergenic NA NA

T28503 ACATCCGTTCCCACCGACTGCGG 1000
Oligo fwd CACCACATCCGTTCCCACCGACTG
Oligo rev AAACCAGTCGGTGGGAACGGATGT
Chromosome start end strand MM target_seq PAM alignment distance position gene name gene id
GU305914.1 75923 75945 - 0 ACATCCGTTCCCACCGACTG CGG ||||||||[||||||||||||]PAM NA Intergenic NA NA

T28504 CATCCGTTCCCACCGACTGCGGG 1000
Oligo fwd CACCCATCCGTTCCCACCGACTGC
Oligo rev AAACGCAGTCGGTGGGAACGGATG
Chromosome start end strand MM target_seq PAM alignment distance position gene name gene id
GU305914.1 75922 75944 - 0 CATCCGTTCCCACCGACTGC GGG ||||||||[||||||||||||]PAM NA Intergenic NA NA

T28505 CCCTCATCTGTCGCCGTGCCAGG 1000
Oligo fwd CACCCCCTCATCTGTCGCCGTGCC
Oligo rev AAACGGCACGGCGACAGATGAGGG
Chromosome start end strand MM target_seq PAM alignment distance position gene name gene id
GU305914.1 75898 75920 - 0 CCCTCATCTGTCGCCGTGCC AGG ||||||||[||||||||||||]PAM NA Intergenic NA NA

T28506 ACCGCCGCCTCCGCCTCCCCCGG 1000
Oligo fwd CACCACCGCCGCCTCCGCCTCCCC
Oligo rev AAACGGGGAGGCGGAGGCGGCGGT
Chromosome start end strand MM target_seq PAM alignment distance position gene name gene id
GU305914.1 75849 75871 - 0 ACCGCCGCCTCCGCCTCCCC CGG ||||||||[||||||||||||]PAM NA Intergenic NA NA

T28507 CCGCCGCCTCCGCCTCCCCCGGG 1000
Oligo fwd CACCCCGCCGCCTCCGCCTCCCCC
Oligo rev AAACGGGGGAGGCGGAGGCGGCGG
Chromosome start end strand MM target_seq PAM alignment distance position gene name gene id
GU305914.1 75848 75870 - 0 CCGCCGCCTCCGCCTCCCCC GGG ||||||||[||||||||||||]PAM NA Intergenic NA NA

T28508 CGCCTCCCCCGGGTCAGCGCTGG 1000
Oligo fwd CACCCGCCTCCCCCGGGTCAGCGC
Oligo rev AAACGCGCTGACCCGGGGGAGGCG
Chromosome start end strand MM target_seq PAM alignment distance position gene name gene id
GU305914.1 75838 75860 - 0 CGCCTCCCCCGGGTCAGCGC TGG ||||||||[||||||||||||]PAM NA Intergenic NA NA

T28509 GCCTCCCCCGGGTCAGCGCTGGG 1000
Oligo fwd CACCGCCTCCCCCGGGTCAGCGCT
Oligo rev AAACAGCGCTGACCCGGGGGAGGC
Chromosome start end strand MM target_seq PAM alignment distance position gene name gene id
GU305914.1 75837 75859 - 0 GCCTCCCCCGGGTCAGCGCT GGG ||||||||[||||||||||||]PAM NA Intergenic NA NA

T28510 CCTCCCCCGGGTCAGCGCTGGGG 1000
Oligo fwd CACCCCTCCCCCGGGTCAGCGCTG
Oligo rev AAACCAGCGCTGACCCGGGGGAGG
Chromosome start end strand MM target_seq PAM alignment distance position gene name gene id
GU305914.1 75836 75858 - 0 CCTCCCCCGGGTCAGCGCTG GGG ||||||||[||||||||||||]PAM NA Intergenic NA NA

T28511 TGGGGCAGCCTGCGCAAACACGG 1000
Oligo fwd CACCTGGGGCAGCCTGCGCAAACA
Oligo rev AAACTGTTTGCGCAGGCTGCCCCA
Chromosome start end strand MM target_seq PAM alignment distance position gene name gene id
GU305914.1 75818 75840 - 0 TGGGGCAGCCTGCGCAAACA CGG ||||||||[||||||||||||]PAM NA Intergenic NA NA

T28512 CAGCCTGCGCAAACACGGCCCGG 1000
Oligo fwd CACCCAGCCTGCGCAAACACGGCC
Oligo rev AAACGGCCGTGTTTGCGCAGGCTG
Chromosome start end strand MM target_seq PAM alignment distance position gene name gene id
GU305914.1 75813 75835 - 0 CAGCCTGCGCAAACACGGCC CGG ||||||||[||||||||||||]PAM NA Intergenic NA NA

T28513 GTTCTCACGCGAAGATACGCAGG 1000
Oligo fwd CACCGTTCTCACGCGAAGATACGC
Oligo rev AAACGCGTATCTTCGCGTGAGAAC
Chromosome start end strand MM target_seq PAM alignment distance position gene name gene id
GU305914.1 75791 75813 - 0 GTTCTCACGCGAAGATACGC AGG ||||||||[||||||||||||]PAM NA Intergenic NA NA

T28514 CTCACGCGAAGATACGCAGGAGG 1000
Oligo fwd CACCCTCACGCGAAGATACGCAGG
Oligo rev AAACCCTGCGTATCTTCGCGTGAG
Chromosome start end strand MM target_seq PAM alignment distance position gene name gene id
GU305914.1 75788 75810 - 0 CTCACGCGAAGATACGCAGG AGG ||||||||[||||||||||||]PAM NA Intergenic NA NA

T28515 ACGCGAAGATACGCAGGAGGTGG 1000
Oligo fwd CACCACGCGAAGATACGCAGGAGG
Oligo rev AAACCCTCCTGCGTATCTTCGCGT
Chromosome start end strand MM target_seq PAM alignment distance position gene name gene id
GU305914.1 75785 75807 - 0 ACGCGAAGATACGCAGGAGG TGG ||||||||[||||||||||||]PAM NA Intergenic NA NA

T28516 CGCGAAGATACGCAGGAGGTGGG 1000
Oligo fwd CACCCGCGAAGATACGCAGGAGGT
Oligo rev AAACACCTCCTGCGTATCTTCGCG
Chromosome start end strand MM target_seq PAM alignment distance position gene name gene id
GU305914.1 75784 75806 - 0 CGCGAAGATACGCAGGAGGT GGG ||||||||[||||||||||||]PAM NA Intergenic NA NA

T28517 GCGAAGATACGCAGGAGGTGGGG 1000

<table>
<tr><td>Oligo fwd</td><td>CACCGCGAAGATACGCAGGAGGTG</td><td></td><td></td><td></td><td></td><td></td><td></td><td></td><td></td><td></td><td></td></tr>
<tr><td>Oligo rev</td><td>AAACCACCTCCTGCGTATCTTCGC</td><td></td><td></td><td></td><td></td><td></td><td></td><td></td><td></td><td></td><td></td></tr>
<tr><td>Chromosome</td><td>start</td><td>end</td><td>strand</td><td>MM</td><td>target_seq</td><td>PAM</td><td>alignment</td><td>distance</td><td>position</td><td>gene name</td><td>gene id</td></tr>
<tr><td>GU305914.1</td><td>75783</td><td>75805</td><td>-</td><td>0</td><td>GCGAAGATACGCAGGAGGTG</td><td>GGG</td><td>||||||||[||||||||||||]PAM</td><td>NA</td><td>Intergenic</td><td>NA</td><td>NA</td></tr>
<tr><td></td><td></td><td></td><td></td><td></td><td></td><td></td><td></td><td></td><td></td><td></td><td></td></tr>
<tr><td>T28518</td><td>CGAAGATACGCAGGAGGTGGGGG</td><td>1000</td><td></td><td></td><td></td><td></td><td></td><td></td><td></td><td></td><td></td></tr>
<tr><td>Oligo fwd</td><td>CACCCGAAGATACGCAGGAGGTGG</td><td></td><td></td><td></td><td></td><td></td><td></td><td></td><td></td><td></td><td></td></tr>
<tr><td>Oligo rev</td><td>AAACCCACCTCCTGCGTATCTTCG</td><td></td><td></td><td></td><td></td><td></td><td></td><td></td><td></td><td></td><td></td></tr>
<tr><td>Chromosome</td><td>start</td><td>end</td><td>strand</td><td>MM</td><td>target_seq</td><td>PAM</td><td>alignment</td><td>distance</td><td>position</td><td>gene name</td><td>gene id</td></tr>
<tr><td>GU305914.1</td><td>75782</td><td>75804</td><td>-</td><td>0</td><td>CGAAGATACGCAGGAGGTGG</td><td>GGG</td><td>||||||||[||||||||||||]PAM</td><td>NA</td><td>Intergenic</td><td>NA</td><td>NA</td></tr>
<tr><td></td><td></td><td></td><td></td><td></td><td></td><td></td><td></td><td></td><td></td><td></td><td></td></tr>
<tr><td>T28519</td><td>ATACGCAGGAGGTGGGGGCGCGG</td><td>1000</td><td></td><td></td><td></td><td></td><td></td><td></td><td></td><td></td><td></td></tr>
<tr><td>Oligo fwd</td><td>CACCATACGCAGGAGGTGGGGGCG</td><td></td><td></td><td></td><td></td><td></td><td></td><td></td><td></td><td></td><td></td></tr>
<tr><td>Oligo rev</td><td>AAACCGCCCCCACCTCCTGCGTAT</td><td></td><td></td><td></td><td></td><td></td><td></td><td></td><td></td><td></td><td></td></tr>
<tr><td>Chromosome</td><td>start</td><td>end</td><td>strand</td><td>MM</td><td>target_seq</td><td>PAM</td><td>alignment</td><td>distance</td><td>position</td><td>gene name</td><td>gene id</td></tr>
<tr><td>GU305914.1</td><td>75777</td><td>75799</td><td>-</td><td>0</td><td>ATACGCAGGAGGTGGGGGCG</td><td>CGG</td><td>||||||||[||||||||||||]PAM</td><td>NA</td><td>Intergenic</td><td>NA</td><td>NA</td></tr>
<tr><td></td><td></td><td></td><td></td><td></td><td></td><td></td><td></td><td></td><td></td><td></td><td></td></tr>
<tr><td>T28520</td><td>CGCGGCCAAGAATCAGCCCGCGG</td><td>1000</td><td></td><td></td><td></td><td></td><td></td><td></td><td></td><td></td><td></td></tr>
<tr><td>Oligo fwd</td><td>CACCCGCGGCCAAGAATCAGCCCG</td><td></td><td></td><td></td><td></td><td></td><td></td><td></td><td></td><td></td><td></td></tr>
<tr><td>Oligo rev</td><td>AAACCGGGCTGATTCTTGGCCGCG</td><td></td><td></td><td></td><td></td><td></td><td></td><td></td><td></td><td></td><td></td></tr>
<tr><td>Chromosome</td><td>start</td><td>end</td><td>strand</td><td>MM</td><td>target_seq</td><td>PAM</td><td>alignment</td><td>distance</td><td>position</td><td>gene name</td><td>gene id</td></tr>
<tr><td>GU305914.1</td><td>75759</td><td>75781</td><td>-</td><td>0</td><td>CGCGGCCAAGAATCAGCCCG</td><td>CGG</td><td>||||||||[||||||||||||]PAM</td><td>NA</td><td>Intergenic</td><td>NA</td><td>NA</td></tr>
<tr><td></td><td></td><td></td><td></td><td></td><td></td><td></td><td></td><td></td><td></td><td></td><td></td></tr>
<tr><td>T28521</td><td>CTCCACTTCCACCTCCTCCCCGG</td><td>1000</td><td></td><td></td><td></td><td></td><td></td><td></td><td></td><td></td><td></td></tr>
<tr><td>Oligo fwd</td><td>CACCCTCCACTTCCACCTCCTCCC</td><td></td><td></td><td></td><td></td><td></td><td></td><td></td><td></td><td></td><td></td></tr>
<tr><td>Oligo rev</td><td>AAACGGGAGGAGGTGGAAGTGGAG</td><td></td><td></td><td></td><td></td><td></td><td></td><td></td><td></td><td></td><td></td></tr>
<tr><td>Chromosome</td><td>start</td><td>end</td><td>strand</td><td>MM</td><td>target_seq</td><td>PAM</td><td>alignment</td><td>distance</td><td>position</td><td>gene name</td><td>gene id</td></tr>
<tr><td>GU305914.1</td><td>75723</td><td>75745</td><td>-</td><td>0</td><td>CTCCACTTCCACCTCCTCCC</td><td>CGG</td><td>||||||||[||||||||||||]PAM</td><td>NA</td><td>Intergenic</td><td>NA</td><td>NA</td></tr>
<tr><td>GU305914.1</td><td>72576</td><td>72598</td><td>-</td><td>4</td><td>CTGCCGTTCCACCTTCTCCC</td><td>CGG</td><td>||-|--||[||||||-|||||]PAM</td><td>NA</td><td>Intergenic</td><td>NA</td><td>NA</td></tr>
<tr><td></td><td></td><td></td><td></td><td></td><td></td><td></td><td></td><td></td><td></td><td></td><td></td></tr>
<tr><td>T28522</td><td>CACTTCCACCTCCTCCCCGGCGG</td><td>1000</td><td></td><td></td><td></td><td></td><td></td><td></td><td></td><td></td><td></td></tr>
<tr><td>Oligo fwd</td><td>CACCCACTTCCACCTCCTCCCCGG</td><td></td><td></td><td></td><td></td><td></td><td></td><td></td><td></td><td></td><td></td></tr>
<tr><td>Oligo rev</td><td>AAACCCGGGGAGGAGGTGGAAGTG</td><td></td><td></td><td></td><td></td><td></td><td></td><td></td><td></td><td></td><td></td></tr>
<tr><td>Chromosome</td><td>start</td><td>end</td><td>strand</td><td>MM</td><td>target_seq</td><td>PAM</td><td>alignment</td><td>distance</td><td>position</td><td>gene name</td><td>gene id</td></tr>
<tr><td>GU305914.1</td><td>75720</td><td>75742</td><td>-</td><td>0</td><td>CACTTCCACCTCCTCCCCGG</td><td>CGG</td><td>||||||||[||||||||||||]PAM</td><td>NA</td><td>Intergenic</td><td>NA</td><td>NA</td></tr>
<tr><td></td><td></td><td></td><td></td><td></td><td></td><td></td><td></td><td></td><td></td><td></td><td></td></tr>
<tr><td>T28523</td><td>CCTCCCCGGCGGCTCCCTCGCGG</td><td>1000</td><td></td><td></td><td></td><td></td><td></td><td></td><td></td><td></td><td></td></tr>
<tr><td>Oligo fwd</td><td>CACCCCTCCCCGGCGGCTCCCTCG</td><td></td><td></td><td></td><td></td><td></td><td></td><td></td><td></td><td></td><td></td></tr>
<tr><td>Oligo rev</td><td>AAACCGAGGGAGCCGCCGGGGAGG</td><td></td><td></td><td></td><td></td><td></td><td></td><td></td><td></td><td></td><td></td></tr>
<tr><td>Chromosome</td><td>start</td><td>end</td><td>strand</td><td>MM</td><td>target_seq</td><td>PAM</td><td>alignment</td><td>distance</td><td>position</td><td>gene name</td><td>gene id</td></tr>
<tr><td>GU305914.1</td><td>75709</td><td>75731</td><td>-</td><td>0</td><td>CCTCCCCGGCGGCTCCCTCG</td><td>CGG</td><td>||||||||[||||||||||||]PAM</td><td>NA</td><td>Intergenic</td><td>NA</td><td>NA</td></tr>
<tr><td></td><td></td><td></td><td></td><td></td><td></td><td></td><td></td><td></td><td></td><td></td><td></td></tr>
<tr><td>T28524</td><td>CTCCCCGGCGGCTCCCTCGCGGG</td><td>1000</td><td></td><td></td><td></td><td></td><td></td><td></td><td></td><td></td><td></td></tr>
<tr><td>Oligo fwd</td><td>CACCCTCCCCGGCGGCTCCCTCGC</td><td></td><td></td><td></td><td></td><td></td><td></td><td></td><td></td><td></td><td></td></tr>
<tr><td>Oligo rev</td><td>AAACGCGAGGGAGCCGCCGGGGAG</td><td></td><td></td><td></td><td></td><td></td><td></td><td></td><td></td><td></td><td></td></tr>
<tr><td>Chromosome</td><td>start</td><td>end</td><td>strand</td><td>MM</td><td>target_seq</td><td>PAM</td><td>alignment</td><td>distance</td><td>position</td><td>gene name</td><td>gene id</td></tr>
<tr><td>GU305914.1</td><td>75708</td><td>75730</td><td>-</td><td>0</td><td>CTCCCCGGCGGCTCCCTCGC</td><td>GGG</td><td>||||||||[||||||||||||]PAM</td><td>NA</td><td>Intergenic</td><td>NA</td><td>NA</td></tr>
<tr><td></td><td></td><td></td><td></td><td></td><td></td><td></td><td></td><td></td><td></td><td></td><td></td></tr>
<tr><td>T28525</td><td>GGCGGCTCCCTCGCGGGATCAGG</td><td>1000</td><td></td><td></td><td></td><td></td><td></td><td></td><td></td><td></td><td></td></tr>
<tr><td>Oligo fwd</td><td>CACCGGCGGCTCCCTCGCGGGATC</td><td></td><td></td><td></td><td></td><td></td><td></td><td></td><td></td><td></td><td></td></tr>
<tr><td>Oligo rev</td><td>AAACGATCCCGCGAGGGAGCCGCC</td><td></td><td></td><td></td><td></td><td></td><td></td><td></td><td></td><td></td><td></td></tr>
<tr><td>Chromosome</td><td>start</td><td>end</td><td>strand</td><td>MM</td><td>target_seq</td><td>PAM</td><td>alignment</td><td>distance</td><td>position</td><td>gene name</td><td>gene id</td></tr>
<tr><td>GU305914.1</td><td>75702</td><td>75724</td><td>-</td><td>0</td><td>GGCGGCTCCCTCGCGGGATC</td><td>AGG</td><td>||||||||[||||||||||||]PAM</td><td>NA</td><td>Intergenic</td><td>NA</td><td>NA</td></tr>
<tr><td></td><td></td><td></td><td></td><td></td><td></td><td></td><td></td><td></td><td></td><td></td><td></td></tr>
<tr><td>T28526</td><td>GGGATCAGGACCAGACCCAGCGG</td><td>1000</td><td></td><td></td><td></td><td></td><td></td><td></td><td></td><td></td><td></td></tr>
<tr><td>Oligo fwd</td><td>CACCGGGATCAGGACCAGACCCAG</td><td></td><td></td><td></td><td></td><td></td><td></td><td></td><td></td><td></td><td></td></tr>
<tr><td>Oligo rev</td><td>AAACCTGGGTCTGGTCCTGATCCC</td><td></td><td></td><td></td><td></td><td></td><td></td><td></td><td></td><td></td><td></td></tr>
<tr><td>Chromosome</td><td>start</td><td>end</td><td>strand</td><td>MM</td><td>target_seq</td><td>PAM</td><td>alignment</td><td>distance</td><td>position</td><td>gene name</td><td>gene id</td></tr>
<tr><td>GU305914.1</td><td>75688</td><td>75710</td><td>-</td><td>0</td><td>GGGATCAGGACCAGACCCAG</td><td>CGG</td><td>||||||||[||||||||||||]PAM</td><td>NA</td><td>Intergenic</td><td>NA</td><td>NA</td></tr>
<tr><td></td><td></td><td></td><td></td><td></td><td></td><td></td><td></td><td></td><td></td><td></td><td></td></tr>
<tr><td>T28527</td><td>ACCCAGCGGCCTCCCCCGCCGG</td><td>1000</td><td></td><td></td><td></td><td></td><td></td><td></td><td></td><td></td><td></td></tr>
<tr><td>Oligo fwd</td><td>CACCACCCAGCGGCCTCCCCCCGC</td><td></td><td></td><td></td><td></td><td></td><td></td><td></td><td></td><td></td><td></td></tr>
<tr><td>Oligo rev</td><td>AAACGCGGGGGGAGGCCGCTGGGT</td><td></td><td></td><td></td><td></td><td></td><td></td><td></td><td></td><td></td><td></td></tr>
<tr><td>Chromosome</td><td>start</td><td>end</td><td>strand</td><td>MM</td><td>target_seq</td><td>PAM</td><td>alignment</td><td>distance</td><td>position</td><td>gene name</td><td>gene id</td></tr>
<tr><td>GU305914.1</td><td>75674</td><td>75696</td><td>-</td><td>0</td><td>ACCCAGCGGCCTCCCCCCGC</td><td>CGG</td><td>||||||||[||||||||||||]PAM</td><td>NA</td><td>Intergenic</td><td>NA</td><td>NA</td></tr>
<tr><td></td><td></td><td></td><td></td><td></td><td></td><td></td><td></td><td></td><td></td><td></td><td></td></tr>
<tr><td>T28528</td><td>CGCCGGCGACACGAACGTCACGG</td><td>1000</td><td></td><td></td><td></td><td></td><td></td><td></td><td></td><td></td><td></td></tr>
<tr><td>Oligo fwd</td><td>CACCCGCCGGCGACACGAACGTCA</td><td></td><td></td><td></td><td></td><td></td><td></td><td></td><td></td><td></td><td></td></tr>
<tr><td>Oligo rev</td><td>AAACTGACGTTCGTGTCGCCGGCG</td><td></td><td></td><td></td><td></td><td></td><td></td><td></td><td></td><td></td><td></td></tr>
<tr><td>Chromosome</td><td>start</td><td>end</td><td>strand</td><td>MM</td><td>target_seq</td><td>PAM</td><td>alignment</td><td>distance</td><td>position</td><td>gene name</td><td>gene id</td></tr>
<tr><td>GU305914.1</td><td>75657</td><td>75679</td><td>-</td><td>0</td><td>CGCCGGCGACACGAACGTCA</td><td>CGG</td><td>||||||||[||||||||||||]PAM</td><td>NA</td><td>Intergenic</td><td>NA</td><td>NA</td></tr>
<tr><td></td><td></td><td></td><td></td><td></td><td></td><td></td><td></td><td></td><td></td><td></td><td></td></tr>
<tr><td>T28529</td><td>GAACTATTTCCTTTTTGACCAGG</td><td>1000</td><td></td><td></td><td></td><td></td><td></td><td></td><td></td><td></td><td></td></tr>
<tr><td>Oligo fwd</td><td>CACCGAACTATTTCCTTTTTGACC</td><td></td><td></td><td></td><td></td><td></td><td></td><td></td><td></td><td></td><td></td></tr>
<tr><td>Oligo rev</td><td>AAACGGTCAAAAAGGAAATAGTTC</td><td></td><td></td><td></td><td></td><td></td><td></td><td></td><td></td><td></td><td></td></tr>
<tr><td>Chromosome</td><td>start</td><td>end</td><td>strand</td><td>MM</td><td>target_seq</td><td>PAM</td><td>alignment</td><td>distance</td><td>position</td><td>gene name</td><td>gene id</td></tr>
<tr><td>GU305914.1</td><td>75610</td><td>75632</td><td>-</td><td>0</td><td>GAACTATTTCCTTTTTGACC</td><td>AGG</td><td>||||||||[||||||||||||]PAM</td><td>NA</td><td>Intergenic</td><td>NA</td><td>NA</td></tr>
<tr><td></td><td></td><td></td><td></td><td></td><td></td><td></td><td></td><td></td><td></td><td></td><td></td></tr>
<tr><td>T28530</td><td>CCATCGCCCTCGTCCTGCTCTGG</td><td>1000</td><td></td><td></td><td></td><td></td><td></td><td></td><td></td><td></td><td></td></tr>
<tr><td>Oligo fwd</td><td>CACCCCATCGCCCTCGTCCTGCTC</td><td></td><td></td><td></td><td></td><td></td><td></td><td></td><td></td><td></td><td></td></tr>
<tr><td>Oligo rev</td><td>AAACGAGCAGGACGAGGGCGATGG</td><td></td><td></td><td></td><td></td><td></td><td></td><td></td><td></td><td></td><td></td></tr>
<tr><td>Chromosome</td><td>start</td><td>end</td><td>strand</td><td>MM</td><td>target_seq</td><td>PAM</td><td>alignment</td><td>distance</td><td>position</td><td>gene name</td><td>gene id</td></tr>
<tr><td>GU305914.1</td><td>75547</td><td>75569</td><td>-</td><td>0</td><td>CCATCGCCCTCGTCCTGCTC</td><td>TGG</td><td>||||||||[||||||||||||]PAM</td><td>NA</td><td>Intergenic</td><td>NA</td><td>NA</td></tr>
<tr><td></td><td></td><td></td><td></td><td></td><td></td><td></td><td></td><td></td><td></td><td></td><td></td></tr>
<tr><td>T28531</td><td>TCCTGCTCTGGCTCCTCTACTGG</td><td>1000</td><td></td><td></td><td></td><td></td><td></td><td></td><td></td><td></td><td></td></tr>
<tr><td>Oligo fwd</td><td>CACCTCCTGCTCTGGCTCCTCTAC</td><td></td><td></td><td></td><td></td><td></td><td></td><td></td><td></td><td></td><td></td></tr>
<tr><td>Oligo rev</td><td>AAACGTAGAGGAGCCAGAGCAGGA</td><td></td><td></td><td></td><td></td><td></td><td></td><td></td><td></td><td></td><td></td></tr>
<tr><td>Chromosome</td><td>start</td><td>end</td><td>strand</td><td>MM</td><td>target_seq</td><td>PAM</td><td>alignment</td><td>distance</td><td>position</td><td>gene name</td><td>gene id</td></tr>
<tr><td>GU305914.1</td><td>75535</td><td>75557</td><td>-</td><td>0</td><td>TCCTGCTCTGGCTCCTCTAC</td><td>TGG</td><td>||||||||[||||||||||||]PAM</td><td>NA</td><td>Intergenic</td><td>NA</td><td>NA</td></tr>
</table>

T28532 CTGGCTCCTCTACTGGCATGCGG 1000
Oligo fwd CACCCTGGCTCCTCTACTGGCATG
Oligo rev AAACCATGCCAGTAGAGGAGCCAG
Chromosome start end strand MM target_seq PAM alignment distance position gene name gene id
GU305914.1 75528 75550 - 0 CTGGCTCCTCTACTGGCATG CGG ||||||||[||||||||||||]PAM NA Intergenic NA NA

T28533 CTACTGGCATGCGGCCCGTTCGG 1000
Oligo fwd CACCCTACTGGCATGCGGCCCGTT
Oligo rev AAACAACGGGCCGCATGCCAGTAG
Chromosome start end strand MM target_seq PAM alignment distance position gene name gene id
GU305914.1 75519 75541 - 0 CTACTGGCATGCGGCCCGTT CGG ||||||||[||||||||||||]PAM NA Intergenic NA NA

T28534 CTGGCATGCGGCCCGTTCGGCGG 1000
Oligo fwd CACCCTGGCATGCGGCCCGTTCGG
Oligo rev AAACCCGAACGGGCCGCATGCCAG
Chromosome start end strand MM target_seq PAM alignment distance position gene name gene id
GU305914.1 75516 75538 - 0 CTGGCATGCGGCCCGTTCGG CGG ||||||||[||||||||||||]PAM NA Intergenic NA NA

T28535 TGGCATGCGGCCCGTTCGGCGGG 1000
Oligo fwd CACCTGGCATGCGGCCCGTTCGGC
Oligo rev AAACGCCGAACGGGCCGCATGCCA
Chromosome start end strand MM target_seq PAM alignment distance position gene name gene id
GU305914.1 75515 75537 - 0 TGGCATGCGGCCCGTTCGGC GGG ||||||||[||||||||||||]PAM NA Intergenic NA NA

T28536 CGGGTCATCCGTGAGCTCTTCGG 1000
Oligo fwd CACCCGGGTCATCCGTGAGCTCTT
Oligo rev AAACAAGAGCTCACGGATGACCCG
Chromosome start end strand MM target_seq PAM alignment distance position gene name gene id
GU305914.1 75496 75518 - 0 CGGGTCATCCGTGAGCTCTT CGG ||||||||[||||||||||||]PAM NA Intergenic NA NA

T28537 GGGTCATCCGTGAGCTCTTCGGG 1000
Oligo fwd CACCGGGTCATCCGTGAGCTCTTC
Oligo rev AAACGAAGAGCTCACGGATGACCC
Chromosome start end strand MM target_seq PAM alignment distance position gene name gene id
GU305914.1 75495 75517 - 0 GGGTCATCCGTGAGCTCTTC GGG ||||||||[||||||||||||]PAM NA Intergenic NA NA
GU305914.1 150113 150135 - 4 AGGACATCCGAGAGCTCTTT AGG -||-||||[||-|||||||||-]PAM NA Intergenic NA NA

T28538 GGTCATCCGTGAGCTCTTCGGGG 1000
Oligo fwd CACCGGTCATCCGTGAGCTCTTCG
Oligo rev AAACCGAAGAGCTCACGGATGACC
Chromosome start end strand MM target_seq PAM alignment distance position gene name gene id
GU305914.1 75494 75516 - 0 GGTCATCCGTGAGCTCTTCG GGG ||||||||[||||||||||||]PAM NA Intergenic NA NA

T28539 GTCATCCGTGAGCTCTTCGGGGG 1000
Oligo fwd CACCGTCATCCGTGAGCTCTTCGG
Oligo rev AAACCCGAAGAGCTCACGGATGAC
Chromosome start end strand MM target_seq PAM alignment distance position gene name gene id
GU305914.1 75493 75515 - 0 GTCATCCGTGAGCTCTTCGG GGG ||||||||[||||||||||||]PAM NA Intergenic NA NA

T28540 TCACCTCCAGCTGCTCCTCTCGG 1000
Oligo fwd CACCTCACCTCCAGCTGCTCCTCT
Oligo rev AAACAGAGGAGCAGCTGGAGGTGA
Chromosome start end strand MM target_seq PAM alignment distance position gene name gene id
GU305914.1 75446 75468 - 0 TCACCTCCAGCTGCTCCTCT CGG ||||||||[||||||||||||]PAM NA Intergenic NA NA

T28541 CACCTCCAGCTGCTCCTCTCGGG 1000
Oligo fwd CACCCACCTCCAGCTGCTCCTCTC
Oligo rev AAACGAGAGGAGCAGCTGGAGGTG
Chromosome start end strand MM target_seq PAM alignment distance position gene name gene id
GU305914.1 75445 75467 - 0 CACCTCCAGCTGCTCCTCTC GGG ||||||||[||||||||||||]PAM NA Intergenic NA NA

T28542 GCTCCTCTCGGGCACCGTCGAGG 1000
Oligo fwd CACCGCTCCTCTCGGGCACCGTCG
Oligo rev AAACCGACGGTGCCCGAGAGGAGC
Chromosome start end strand MM target_seq PAM alignment distance position gene name gene id
GU305914.1 75434 75456 - 0 GCTCCTCTCGGGCACCGTCG AGG ||||||||[||||||||||||]PAM NA Intergenic NA NA

T28543 ACCGTCGAGGACGAACGCGACGG 1000
Oligo fwd CACCACCGTCGAGGACGAACGCGA
Oligo rev AAACTCGCGTTCGTCCTCGACGGT
Chromosome start end strand MM target_seq PAM alignment distance position gene name gene id
GU305914.1 75421 75443 - 0 ACCGTCGAGGACGAACGCGA CGG ||||||||[||||||||||||]PAM NA Intergenic NA NA

T28544 GTCCTCAACATGTTCATGAACGG 1000
Oligo fwd CACCGTCCTCAACATGTTCATGAA
Oligo rev AAACTTCATGAACATGTTGAGGAC
Chromosome start end strand MM target_seq PAM alignment distance position gene name gene id
GU305914.1 75382 75404 - 0 GTCCTCAACATGTTCATGAA CGG ||||||||[||||||||||||]PAM NA Intergenic NA NA

T28545 CGGCAAGCGCTTTCTCTGCAAGG 1000
Oligo fwd CACCCGGCAAGCGCTTTCTCTGCA
Oligo rev AAACTGCAGAGAAAGCGCTTGCCG
Chromosome start end strand MM target_seq PAM alignment distance position gene name gene id
GU305914.1 75362 75384 - 0 CGGCAAGCGCTTTCTCTGCA AGG ||||||||[||||||||||||]PAM NA Intergenic NA NA

T28546 ACTGTCGCTTCCTCGAACGCTGG 1000
Oligo fwd CACCACTGTCGCTTCCTCGAACGC
Oligo rev AAACGCGTTCGAGGAAGCGACAGT

Chromosome start end strand MM target_seq PAM alignment distance position gene name gene id
GU305914.1 75321 75343 - 0 ACTGTCGCTTCCTCGAACGC TGG ||||||||[||||||||||||]PAM NA Intergenic NA NA

T28547 CGAACGCTGGTCCTCCTGCTCGG 1000
Oligo fwd CACCCGAACGCTGGTCCTCCTGCT
Oligo rev AAACAGCAGGAGGACCAGCGTTCG
Chromosome start end strand MM target_seq PAM alignment distance position gene name gene id
GU305914.1 75308 75330 - 0 CGAACGCTGGTCCTCCTGCT CGG ||||||||[||||||||||||]PAM NA Intergenic NA NA

T28548 CCGCGAGATCTCGCAGAAAGAGG 1000
Oligo fwd CACCCCGCGAGATCTCGCAGAAAG
Oligo rev AAACCTTTCTGCGAGATCTCGCGG
Chromosome start end strand MM target_seq PAM alignment distance position gene name gene id
GU305914.1 75266 75288 - 0 CCGCGAGATCTCGCAGAAAG AGG ||||||||[||||||||||||]PAM NA Intergenic NA NA

T28549 CTTTTACGCCGTGCGCTGCATGG 1000
Oligo fwd CACCCTTTTACGCCGTGCGCTGCA
Oligo rev AAACTGCAGCGCACGGCGTAAAAG
Chromosome start end strand MM target_seq PAM alignment distance position gene name gene id
GU305914.1 75209 75231 - 0 CTTTTACGCCGTGCGCTGCA TGG ||||||||[||||||||||||]PAM NA Intergenic NA NA

T28550 ACGTCTCCTGCCTCTACCTCAGG 1000
Oligo fwd CACCACGTCTCCTGCCTCTACCTC
Oligo rev AAACGAGGTAGAGGCAGGAGACGT
Chromosome start end strand MM target_seq PAM alignment distance position gene name gene id
GU305914.1 75171 75193 - 0 ACGTCTCCTGCCTCTACCTC AGG ||||||||[||||||||||||]PAM NA Intergenic NA NA

T28551 TCCTGCCTCTACCTCAGGAGCGG 1000
Oligo fwd CACCTCCTGCCTCTACCTCAGGAG
Oligo rev AAACCTCCTGAGGTAGAGGCAGGA
Chromosome start end strand MM target_seq PAM alignment distance position gene name gene id
GU305914.1 75166 75188 - 0 TCCTGCCTCTACCTCAGGAG CGG ||||||||[||||||||||||]PAM NA Intergenic NA NA

T28552 GCCTCGAGCACGTCTTCAAGAGG 1000
Oligo fwd CACCGCCTCGAGCACGTCTTCAAG
Oligo rev AAACCTTGAAGACGTGCTCGAGGC
Chromosome start end strand MM target_seq PAM alignment distance position gene name gene id
GU305914.1 75132 75154 - 0 GCCTCGAGCACGTCTTCAAG AGG ||||||||[||||||||||||]PAM NA Intergenic NA NA

T28553 CAAGAGGTACAGCCGCTCCAAGG 1000
Oligo fwd CACCCAAGAGGTACAGCCGCTCCA
Oligo rev AAACTGGAGCGGCTGTACCTCTTG
Chromosome start end strand MM target_seq PAM alignment distance position gene name gene id
GU305914.1 75116 75138 - 0 CAAGAGGTACAGCCGCTCCA AGG ||||||||[||||||||||||]PAM NA Intergenic NA NA

T28554 CCGCTCCAAGGTCGACGCCCTGG 1000
Oligo fwd CACCCCGCTCCAAGGTCGACGCCC
Oligo rev AAACGGGCGTCGACCTTGGAGCGG
Chromosome start end strand MM target_seq PAM alignment distance position gene name gene id
GU305914.1 75104 75126 - 0 CCGCTCCAAGGTCGACGCCC TGG ||||||||[||||||||||||]PAM NA Intergenic NA NA

T28555 GGTCGACAACGCCAGCTACCAGG 1000
Oligo fwd CACCGGTCGACAACGCCAGCTACC
Oligo rev AAACGGTAGCTGGCGTTGTCGACC
Chromosome start end strand MM target_seq PAM alignment distance position gene name gene id
GU305914.1 75083 75105 - 0 GGTCGACAACGCCAGCTACC AGG ||||||||[||||||||||||]PAM NA Intergenic NA NA
GU305914.1 29641 29663 - 4 CGTCGACGACGCCATCTATC TGG -||||||-[||||||-|||-|]PAM NA Intergenic NA NA

T28556 GTCGACAACGCCAGCTACCAGGG 1000
Oligo fwd CACCGTCGACAACGCCAGCTACCA
Oligo rev AAACTGGTAGCTGGCGTTGTCGAC
Chromosome start end strand MM target_seq PAM alignment distance position gene name gene id
GU305914.1 75082 75104 - 0 GTCGACAACGCCAGCTACCA GGG ||||||||[||||||||||||]PAM NA Intergenic NA NA

T28557 TTCGTCTTCCACTTCCCCTTCGG 1000
Oligo fwd CACCTTCGTCTTCCACTTCCCCTT
Oligo rev AAACAAGGGGAAGTGGAAGACGAA
Chromosome start end strand MM target_seq PAM alignment distance position gene name gene id
GU305914.1 75040 75062 - 0 TTCGTCTTCCACTTCCCCTT CGG ||||||||[||||||||||||]PAM NA Intergenic NA NA

T28558 CGGCATCCCCATTCCCAACCAGG 1000
Oligo fwd CACCCGGCATCCCCATTCCCAACC
Oligo rev AAACGGTTGGGAATGGGGATGCCG
Chromosome start end strand MM target_seq PAM alignment distance position gene name gene id
GU305914.1 75020 75042 - 0 CGGCATCCCCATTCCCAACC AGG ||||||||[||||||||||||]PAM NA Intergenic NA NA

T28559 CCTCCGCGCCCGCGAGTACGAGG 1000
Oligo fwd CACCCCTCCGCGCCCGCGAGTACG
Oligo rev AAACCGTACTCGCGGGCGCGGAGG
Chromosome start end strand MM target_seq PAM alignment distance position gene name gene id
GU305914.1 74975 74997 - 0 CCTCCGCGCCCGCGAGTACG AGG ||||||||[||||||||||||]PAM NA Intergenic NA NA

T28560 CGCGCCCGCGAGTACGAGGTCGG 1000
Oligo fwd CACCCGCGCCCGCGAGTACGAGGT
Oligo rev AAACACCTCGTACTCGCGGGCGCG
Chromosome start end strand MM target_seq PAM alignment distance position gene name gene id
GU305914.1 74971 74993 - 0 CGCGCCCGCGAGTACGAGGT CGG ||||||||[||||||||||||]PAM NA Intergenic NA NA

T28561 CGCCTCAACCTGCGCCGCCTCGG 1000

Oligo fwd CACCCGCCTCAACCTGCGCCGCCT
Oligo rev AAACAGGCGGCGCAGGTTGAGGCG
Chromosome start end strand MM target_seq PAM alignment distance position gene name gene id
GU305914.1 74923 74945 - 0 CGCCTCAACCTGCGCCGCCT CGG ||||||||[||||||||||||]PAM NA Intergenic NA NA

T28562 GCGCCGCCTCGGTCCCGACCCGG 1000
Oligo fwd CACCGCGCCGCCTCGGTCCCGACC
Oligo rev AAACGGTCGGGACCGAGGCGGCGC
Chromosome start end strand MM target_seq PAM alignment distance position gene name gene id
GU305914.1 74912 74934 - 0 GCGCCGCCTCGGTCCCGACC CGG ||||||||[||||||||||||]PAM NA Intergenic NA NA

T28563 CGCCGCCTCGGTCCCGACCCGGG 1000
Oligo fwd CACCCGCCGCCTCGGTCCCGACCC
Oligo rev AAACGGGTCGGGACCGAGGCGGCG
Chromosome start end strand MM target_seq PAM alignment distance position gene name gene id
GU305914.1 74911 74933 - 0 CGCCGCCTCGGTCCCGACCC GGG ||||||||[||||||||||||]PAM NA Intergenic NA NA

T28564 CGCTGCCGCGAGACTCCCTGCGG 1000
Oligo fwd CACCCGCTGCCGCGAGACTCCCTG
Oligo rev AAACCAGGGAGTCTCGCGGCAGCG
Chromosome start end strand MM target_seq PAM alignment distance position gene name gene id
GU305914.1 74857 74879 - 0 CGCTGCCGCGAGACTCCCTG CGG ||||||||[||||||||||||]PAM NA Intergenic NA NA

T28565 CGGAAACCCCTTCTACGTCATGG 1000
Oligo fwd CACCCGGAAACCCCTTCTACGTCA
Oligo rev AAACTGACGTAGAAGGGGTTTCCG
Chromosome start end strand MM target_seq PAM alignment distance position gene name gene id
GU305914.1 74837 74859 - 0 CGGAAACCCCTTCTACGTCA TGG ||||||||[||||||||||||]PAM NA Intergenic NA NA

T28566 TCGTCGAACGCTACTGTCGCAGG 1000
Oligo fwd CACCTCGTCGAACGCTACTGTCGC
Oligo rev AAACGCGACAGTAGCGTTCGACGA
Chromosome start end strand MM target_seq PAM alignment distance position gene name gene id
GU305914.1 74805 74827 - 0 TCGTCGAACGCTACTGTCGC AGG ||||||||[||||||||||||]PAM NA Intergenic NA NA

T28567 CCGCGCTTCCTCATCCCCATCGG 1000
Oligo fwd CACCCCGCGCTTCCTCATCCCCAT
Oligo rev AAACATGGGGATGAGGAAGCGCGG
Chromosome start end strand MM target_seq PAM alignment distance position gene name gene id
GU305914.1 74779 74801 - 0 CCGCGCTTCCTCATCCCCAT CGG ||||||||[||||||||||||]PAM NA Intergenic NA NA

T28568 TCCTCATCCCCATCGGCAATCGG 1000
Oligo fwd CACCTCCTCATCCCCATCGGCAAT
Oligo rev AAACATTGCCGATGGGGATGAGGA
Chromosome start end strand MM target_seq PAM alignment distance position gene name gene id
GU305914.1 74772 74794 - 0 TCCTCATCCCCATCGGCAAT CGG ||||||||[||||||||||||]PAM NA Intergenic NA NA

T28569 GGCAATCGGACCCTGCGCTACGG 1000
Oligo fwd CACCGGCAATCGGACCCTGCGCTA
Oligo rev AAACTAGCGCAGGGTCCGATTGCC
Chromosome start end strand MM target_seq PAM alignment distance position gene name gene id
GU305914.1 74758 74780 - 0 GGCAATCGGACCCTGCGCTA CGG ||||||||[||||||||||||]PAM NA Intergenic NA NA

T28570 AATCGGACCCTGCGCTACGGCGG 1000
Oligo fwd CACCAATCGGACCCTGCGCTACGG
Oligo rev AAACCCGTAGCGCAGGGTCCGATT
Chromosome start end strand MM target_seq PAM alignment distance position gene name gene id
GU305914.1 74755 74777 - 0 AATCGGACCCTGCGCTACGG CGG ||||||||[||||||||||||]PAM NA Intergenic NA NA

T28571 GCGCTACGGCGGACGCGCCGCGG 1000
Oligo fwd CACCGCGCTACGGCGGACGCGCCG
Oligo rev AAACCGGCGCGTCCGCCGTAGCGC
Chromosome start end strand MM target_seq PAM alignment distance position gene name gene id
GU305914.1 74744 74766 - 0 GCGCTACGGCGGACGCGCCG CGG ||||||||[||||||||||||]PAM NA Intergenic NA NA

T28572 ACGGCGGACGCGCCGCGGTCCGG 1000
Oligo fwd CACCACGGCGGACGCGCCGCGGTC
Oligo rev AAACGACCGCGGCGCGTCCGCCGT
Chromosome start end strand MM target_seq PAM alignment distance position gene name gene id
GU305914.1 74739 74761 - 0 ACGGCGGACGCGCCGCGGTC CGG ||||||||[||||||||||||]PAM NA Intergenic NA NA

T28573 CGCGGTCCGGACCTCAGACCCGG 1000
Oligo fwd CACCCGCGGTCCGGACCTCAGACC
Oligo rev AAACGGTCTGAGGTCCGGACCGCG
Chromosome start end strand MM target_seq PAM alignment distance position gene name gene id
GU305914.1 74726 74748 - 0 CGCGGTCCGGACCTCAGACC CGG ||||||||[||||||||||||]PAM NA Intergenic NA NA

T28574 GCGGTCCGGACCTCAGACCCGGG 1000
Oligo fwd CACCGCGGTCCGGACCTCAGACCC
Oligo rev AAACGGGTCTGAGGTCCGGACCGC
Chromosome start end strand MM target_seq PAM alignment distance position gene name gene id
GU305914.1 74725 74747 - 0 GCGGTCCGGACCTCAGACCC GGG ||||||||[||||||||||||]PAM NA Intergenic NA NA

T28575 GTCCGGACCTCAGACCCGGGTGG 1000
Oligo fwd CACCGTCCGGACCTCAGACCCGGG
Oligo rev AAACCCCGGGTCTGAGGTCCGGAC
Chromosome start end strand MM target_seq PAM alignment distance position gene name gene id
GU305914.1 74722 74744 - 0 GTCCGGACCTCAGACCCGGG TGG ||||||||[||||||||||||]PAM NA Intergenic NA NA

<table>
<tr><td>T28576</td><td>CGGACCTCAGACCCGGGTGGCGG</td><td>1000</td><td></td><td></td><td></td><td></td><td></td><td></td><td></td><td></td><td></td></tr>
<tr><td>Oligo fwd</td><td>CACCCGGACCTCAGACCCGGGTGG</td><td></td><td></td><td></td><td></td><td></td><td></td><td></td><td></td><td></td><td></td></tr>
<tr><td>Oligo rev</td><td>AAACCCACCCGGGTCTGAGGTCCG</td><td></td><td></td><td></td><td></td><td></td><td></td><td></td><td></td><td></td><td></td></tr>
<tr><td>Chromosome</td><td>start</td><td>end</td><td>strand</td><td>MM</td><td>target_seq</td><td>PAM</td><td>alignment</td><td>distance</td><td>position</td><td>gene name</td><td>gene id</td></tr>
<tr><td>GU305914.1</td><td>74719</td><td>74741</td><td>-</td><td>0</td><td>CGGACCTCAGACCCGGGTGG</td><td>CGG</td><td>||||||||[||||||||||||]PAM</td><td>NA</td><td>Intergenic</td><td>NA</td><td>NA</td></tr>
<tr><td>T28577</td><td>ACCTCAGACCCGGGTGGCGGTGG</td><td>1000</td><td></td><td></td><td></td><td></td><td></td><td></td><td></td><td></td><td></td></tr>
<tr><td>Oligo fwd</td><td>CACCACCTCAGACCCGGGTGGCGG</td><td></td><td></td><td></td><td></td><td></td><td></td><td></td><td></td><td></td><td></td></tr>
<tr><td>Oligo rev</td><td>AAACCCGCCACCCGGGTCTGAGGT</td><td></td><td></td><td></td><td></td><td></td><td></td><td></td><td></td><td></td><td></td></tr>
<tr><td>Chromosome</td><td>start</td><td>end</td><td>strand</td><td>MM</td><td>target_seq</td><td>PAM</td><td>alignment</td><td>distance</td><td>position</td><td>gene name</td><td>gene id</td></tr>
<tr><td>GU305914.1</td><td>74716</td><td>74738</td><td>-</td><td>0</td><td>ACCTCAGACCCGGGTGGCGG</td><td>TGG</td><td>||||||||[||||||||||||]PAM</td><td>NA</td><td>Intergenic</td><td>NA</td><td>NA</td></tr>
<tr><td>T28578</td><td>TCAGACCCGGGTGGCGGTGGCGG</td><td>1000</td><td></td><td></td><td></td><td></td><td></td><td></td><td></td><td></td><td></td></tr>
<tr><td>Oligo fwd</td><td>CACCTCAGACCCGGGTGGCGGTGG</td><td></td><td></td><td></td><td></td><td></td><td></td><td></td><td></td><td></td><td></td></tr>
<tr><td>Oligo rev</td><td>AAACCCACCGCCACCCGGGTCTGA</td><td></td><td></td><td></td><td></td><td></td><td></td><td></td><td></td><td></td><td></td></tr>
<tr><td>Chromosome</td><td>start</td><td>end</td><td>strand</td><td>MM</td><td>target_seq</td><td>PAM</td><td>alignment</td><td>distance</td><td>position</td><td>gene name</td><td>gene id</td></tr>
<tr><td>GU305914.1</td><td>74713</td><td>74735</td><td>-</td><td>0</td><td>TCAGACCCGGGTGGCGGTGG</td><td>CGG</td><td>||||||||[||||||||||||]PAM</td><td>NA</td><td>Intergenic</td><td>NA</td><td>NA</td></tr>
<tr><td>T28579</td><td>GACCCGGGTGGCGGTGGCGGCGG</td><td>1000</td><td></td><td></td><td></td><td></td><td></td><td></td><td></td><td></td><td></td></tr>
<tr><td>Oligo fwd</td><td>CACCGACCCGGGTGGCGGTGGCGG</td><td></td><td></td><td></td><td></td><td></td><td></td><td></td><td></td><td></td><td></td></tr>
<tr><td>Oligo rev</td><td>AAACCCGCCACCGCCACCCGGGTC</td><td></td><td></td><td></td><td></td><td></td><td></td><td></td><td></td><td></td><td></td></tr>
<tr><td>Chromosome</td><td>start</td><td>end</td><td>strand</td><td>MM</td><td>target_seq</td><td>PAM</td><td>alignment</td><td>distance</td><td>position</td><td>gene name</td><td>gene id</td></tr>
<tr><td>GU305914.1</td><td>74710</td><td>74732</td><td>-</td><td>0</td><td>GACCCGGGTGGCGGTGGCGG</td><td>CGG</td><td>||||||||[||||||||||||]PAM</td><td>NA</td><td>Intergenic</td><td>NA</td><td>NA</td></tr>
<tr><td>GU305914.1</td><td>52716</td><td>52738</td><td>+</td><td>4</td><td>GCCGCCGCTGGCGGTGGCGG</td><td>TGG</td><td>|-|-|-|-[||||||||||||]PAM</td><td>NA</td><td>Intergenic</td><td>NA</td><td>NA</td></tr>
<tr><td>GU305914.1</td><td>114661</td><td>114683</td><td>-</td><td>4</td><td>GACGGCGGTGGTGGTGGCGG</td><td>CGG</td><td>|||---||[|||-||||||||]PAM</td><td>NA</td><td>Intergenic</td><td>NA</td><td>NA</td></tr>
<tr><td>T28580</td><td>GGTGGCGGTGGCGGCGGTTCCGG</td><td>1000</td><td></td><td></td><td></td><td></td><td></td><td></td><td></td><td></td><td></td></tr>
<tr><td>Oligo fwd</td><td>CACCGGTGGCGGTGGCGGCGGTTC</td><td></td><td></td><td></td><td></td><td></td><td></td><td></td><td></td><td></td><td></td></tr>
<tr><td>Oligo rev</td><td>AAACGAACCGCCGCCACCGCCACC</td><td></td><td></td><td></td><td></td><td></td><td></td><td></td><td></td><td></td><td></td></tr>
<tr><td>Chromosome</td><td>start</td><td>end</td><td>strand</td><td>MM</td><td>target_seq</td><td>PAM</td><td>alignment</td><td>distance</td><td>position</td><td>gene name</td><td>gene id</td></tr>
<tr><td>GU305914.1</td><td>74704</td><td>74726</td><td>-</td><td>0</td><td>GGTGGCGGTGGCGGCGGTTC</td><td>CGG</td><td>||||||||[||||||||||||]PAM</td><td>NA</td><td>Intergenic</td><td>NA</td><td>NA</td></tr>
<tr><td>T28581</td><td>GTGGCGGTGGCGGCGGTTCCGGG</td><td>1000</td><td></td><td></td><td></td><td></td><td></td><td></td><td></td><td></td><td></td></tr>
<tr><td>Oligo fwd</td><td>CACCGTGGCGGTGGCGGCGGTTCC</td><td></td><td></td><td></td><td></td><td></td><td></td><td></td><td></td><td></td><td></td></tr>
<tr><td>Oligo rev</td><td>AAACGGAACCGCCGCCACCGCCAC</td><td></td><td></td><td></td><td></td><td></td><td></td><td></td><td></td><td></td><td></td></tr>
<tr><td>Chromosome</td><td>start</td><td>end</td><td>strand</td><td>MM</td><td>target_seq</td><td>PAM</td><td>alignment</td><td>distance</td><td>position</td><td>gene name</td><td>gene id</td></tr>
<tr><td>GU305914.1</td><td>74703</td><td>74725</td><td>-</td><td>0</td><td>GTGGCGGTGGCGGCGGTTCC</td><td>GGG</td><td>||||||||[||||||||||||]PAM</td><td>NA</td><td>Intergenic</td><td>NA</td><td>NA</td></tr>
<tr><td>GU305914.1</td><td>154417</td><td>154439</td><td>+</td><td>4</td><td>GCGGCGGCGGCGGCTGTTGC</td><td>TGG</td><td>|-|||||-[||||||-|||-|]PAM</td><td>NA</td><td>Intergenic</td><td>NA</td><td>NA</td></tr>
<tr><td>T28582</td><td>TGGCGGCGGTTCCGGGAACTCGG</td><td>1000</td><td></td><td></td><td></td><td></td><td></td><td></td><td></td><td></td><td></td></tr>
<tr><td>Oligo fwd</td><td>CACCTGGCGGCGGTTCCGGGAACT</td><td></td><td></td><td></td><td></td><td></td><td></td><td></td><td></td><td></td><td></td></tr>
<tr><td>Oligo rev</td><td>AAACAGTTCCCGGAACCGCCGCCA</td><td></td><td></td><td></td><td></td><td></td><td></td><td></td><td></td><td></td><td></td></tr>
<tr><td>Chromosome</td><td>start</td><td>end</td><td>strand</td><td>MM</td><td>target_seq</td><td>PAM</td><td>alignment</td><td>distance</td><td>position</td><td>gene name</td><td>gene id</td></tr>
<tr><td>GU305914.1</td><td>74696</td><td>74718</td><td>-</td><td>0</td><td>TGGCGGCGGTTCCGGGAACT</td><td>CGG</td><td>||||||||[||||||||||||]PAM</td><td>NA</td><td>Intergenic</td><td>NA</td><td>NA</td></tr>
<tr><td>T28583</td><td>TCGGCCGCCACCGCCACCGACGG</td><td>1000</td><td></td><td></td><td></td><td></td><td></td><td></td><td></td><td></td><td></td></tr>
<tr><td>Oligo fwd</td><td>CACCTCGGCCGCCACCGCCACCGA</td><td></td><td></td><td></td><td></td><td></td><td></td><td></td><td></td><td></td><td></td></tr>
<tr><td>Oligo rev</td><td>AAACTCGGTGGCGGTGGCGGCCGA</td><td></td><td></td><td></td><td></td><td></td><td></td><td></td><td></td><td></td><td></td></tr>
<tr><td>Chromosome</td><td>start</td><td>end</td><td>strand</td><td>MM</td><td>target_seq</td><td>PAM</td><td>alignment</td><td>distance</td><td>position</td><td>gene name</td><td>gene id</td></tr>
<tr><td>GU305914.1</td><td>74677</td><td>74699</td><td>-</td><td>0</td><td>TCGGCCGCCACCGCCACCGA</td><td>CGG</td><td>||||||||[||||||||||||]PAM</td><td>NA</td><td>Intergenic</td><td>NA</td><td>NA</td></tr>
<tr><td>T28584</td><td>CCGACGGCACCAACTGTCCCTGG</td><td>1000</td><td></td><td></td><td></td><td></td><td></td><td></td><td></td><td></td><td></td></tr>
<tr><td>Oligo fwd</td><td>CACCCCGACGGCACCAACTGTCCC</td><td></td><td></td><td></td><td></td><td></td><td></td><td></td><td></td><td></td><td></td></tr>
<tr><td>Oligo rev</td><td>AAACGGGACAGTTGGTGCCGTCGG</td><td></td><td></td><td></td><td></td><td></td><td></td><td></td><td></td><td></td><td></td></tr>
<tr><td>Chromosome</td><td>start</td><td>end</td><td>strand</td><td>MM</td><td>target_seq</td><td>PAM</td><td>alignment</td><td>distance</td><td>position</td><td>gene name</td><td>gene id</td></tr>
<tr><td>GU305914.1</td><td>74661</td><td>74683</td><td>-</td><td>0</td><td>CCGACGGCACCAACTGTCCC</td><td>TGG</td><td>||||||||[||||||||||||]PAM</td><td>NA</td><td>Intergenic</td><td>NA</td><td>NA</td></tr>
<tr><td>T28585</td><td>GCTCTCCACCTTCGCCACCTCGG</td><td>1000</td><td></td><td></td><td></td><td></td><td></td><td></td><td></td><td></td><td></td></tr>
<tr><td>Oligo fwd</td><td>CACCGCTCTCCACCTTCGCCACCT</td><td></td><td></td><td></td><td></td><td></td><td></td><td></td><td></td><td></td><td></td></tr>
<tr><td>Oligo rev</td><td>AAACAGGTGGCGAAGGTGGAGAGC</td><td></td><td></td><td></td><td></td><td></td><td></td><td></td><td></td><td></td><td></td></tr>
<tr><td>Chromosome</td><td>start</td><td>end</td><td>strand</td><td>MM</td><td>target_seq</td><td>PAM</td><td>alignment</td><td>distance</td><td>position</td><td>gene name</td><td>gene id</td></tr>
<tr><td>GU305914.1</td><td>74627</td><td>74649</td><td>-</td><td>0</td><td>GCTCTCCACCTTCGCCACCT</td><td>CGG</td><td>||||||||[||||||||||||]PAM</td><td>NA</td><td>Intergenic</td><td>NA</td><td>NA</td></tr>
<tr><td>T28586</td><td>ACCTCGGTCGTGCTCAGAAACGG</td><td>1000</td><td></td><td></td><td></td><td></td><td></td><td></td><td></td><td></td><td></td></tr>
<tr><td>Oligo fwd</td><td>CACCACCTCGGTCGTGCTCAGAAA</td><td></td><td></td><td></td><td></td><td></td><td></td><td></td><td></td><td></td><td></td></tr>
<tr><td>Oligo rev</td><td>AAACTTTCTGAGCACGACCGAGGT</td><td></td><td></td><td></td><td></td><td></td><td></td><td></td><td></td><td></td><td></td></tr>
<tr><td>Chromosome</td><td>start</td><td>end</td><td>strand</td><td>MM</td><td>target_seq</td><td>PAM</td><td>alignment</td><td>distance</td><td>position</td><td>gene name</td><td>gene id</td></tr>
<tr><td>GU305914.1</td><td>74611</td><td>74633</td><td>-</td><td>0</td><td>ACCTCGGTCGTGCTCAGAAA</td><td>CGG</td><td>||||||||[||||||||||||]PAM</td><td>NA</td><td>Intergenic</td><td>NA</td><td>NA</td></tr>
<tr><td>T28587</td><td>GCCTCATCGACCTCCCCGTGTGG</td><td>1000</td><td></td><td></td><td></td><td></td><td></td><td></td><td></td><td></td><td></td></tr>
<tr><td>Oligo fwd</td><td>CACCGCCTCATCGACCTCCCCGTG</td><td></td><td></td><td></td><td></td><td></td><td></td><td></td><td></td><td></td><td></td></tr>
<tr><td>Oligo rev</td><td>AAACCACGGGGAGGTCGATGAGGC</td><td></td><td></td><td></td><td></td><td></td><td></td><td></td><td></td><td></td><td></td></tr>
<tr><td>Chromosome</td><td>start</td><td>end</td><td>strand</td><td>MM</td><td>target_seq</td><td>PAM</td><td>alignment</td><td>distance</td><td>position</td><td>gene name</td><td>gene id</td></tr>
<tr><td>GU305914.1</td><td>74577</td><td>74599</td><td>-</td><td>0</td><td>GCCTCATCGACCTCCCCGTG</td><td>TGG</td><td>||||||||[||||||||||||]PAM</td><td>NA</td><td>Intergenic</td><td>NA</td><td>NA</td></tr>
<tr><td>T28588</td><td>CAAGTGTCAGCGCTACGCCGAGG</td><td>1000</td><td></td><td></td><td></td><td></td><td></td><td></td><td></td><td></td><td></td></tr>
<tr><td>Oligo fwd</td><td>CACCCAAGTGTCAGCGCTACGCCG</td><td></td><td></td><td></td><td></td><td></td><td></td><td></td><td></td><td></td><td></td></tr>
<tr><td>Oligo rev</td><td>AAACCGGCGTAGCGCTGACACTTG</td><td></td><td></td><td></td><td></td><td></td><td></td><td></td><td></td><td></td><td></td></tr>
<tr><td>Chromosome</td><td>start</td><td>end</td><td>strand</td><td>MM</td><td>target_seq</td><td>PAM</td><td>alignment</td><td>distance</td><td>position</td><td>gene name</td><td>gene id</td></tr>
<tr><td>GU305914.1</td><td>74540</td><td>74562</td><td>-</td><td>0</td><td>CAAGTGTCAGCGCTACGCCG</td><td>AGG</td><td>||||||||[||||||||||||]PAM</td><td>NA</td><td>Intergenic</td><td>NA</td><td>NA</td></tr>
<tr><td>T28589</td><td>TGTCAGCGCTACGCCGAGGACGG</td><td>1000</td><td></td><td></td><td></td><td></td><td></td><td></td><td></td><td></td><td></td></tr>
<tr><td>Oligo fwd</td><td>CACCTGTCAGCGCTACGCCGAGGA</td><td></td><td></td><td></td><td></td><td></td><td></td><td></td><td></td><td></td><td></td></tr>
<tr><td>Oligo rev</td><td>AAACTCCTCGGCGTAGCGCTGACA</td><td></td><td></td><td></td><td></td><td></td><td></td><td></td><td></td><td></td><td></td></tr>
<tr><td>Chromosome</td><td>start</td><td>end</td><td>strand</td><td>MM</td><td>target_seq</td><td>PAM</td><td>alignment</td><td>distance</td><td>position</td><td>gene name</td><td>gene id</td></tr>
<tr><td>GU305914.1</td><td>74536</td><td>74558</td><td>-</td><td>0</td><td>TGTCAGCGCTACGCCGAGGA</td><td>CGG</td><td>||||||||[||||||||||||]PAM</td><td>NA</td><td>Intergenic</td><td>NA</td><td>NA</td></tr>
<tr><td>T28590</td><td>CGCCGAGGACGGCGTGCTCGAGG</td><td>1000</td><td></td><td></td><td></td><td></td><td></td><td></td><td></td><td></td><td></td></tr>
<tr><td>Oligo fwd</td><td>CACCCGCCGAGGACGGCGTGCTCG</td><td></td><td></td><td></td><td></td><td></td><td></td><td></td><td></td><td></td><td></td></tr>
</table>

<table>
<tr><td>Oligo rev</td><td>AAACCGAGCACGCCGTCCTCGGCG</td><td></td><td></td><td></td><td></td><td></td><td></td><td></td><td></td><td></td><td></td></tr>
<tr><td>Chromosome</td><td>start</td><td>end</td><td>strand</td><td>MM</td><td>target_seq</td><td>PAM</td><td>alignment</td><td>distance</td><td>position</td><td>gene name</td><td>gene id</td></tr>
<tr><td>GU305914.1</td><td>74525</td><td>74547</td><td>-</td><td>0</td><td>CGCCGAGGACGGCGTGCTCG</td><td>AGG</td><td>||||||||[||||||||||||]PAM</td><td>NA</td><td>Intergenic</td><td>NA</td><td>NA</td></tr>
<tr><td>GU305914.1</td><td>143007</td><td>143029</td><td>+</td><td>4</td><td>CGACGACGACGGCCTGCTGG</td><td>TGG</td><td>||-|||-|[|||||-||||-|]PAM</td><td>NA</td><td>Intergenic</td><td>NA</td><td>NA</td></tr>
<tr><td>T28591</td><td>GCCATCCTCTGTGACAACTGCGG</td><td>1000</td><td></td><td></td><td></td><td></td><td></td><td></td><td></td><td></td><td></td></tr>
<tr><td>Oligo fwd</td><td>CACCGCCATCCTCTGTGACAACTG</td><td></td><td></td><td></td><td></td><td></td><td></td><td></td><td></td><td></td><td></td></tr>
<tr><td>Oligo rev</td><td>AAACCAGTTGTCACAGAGGATGGC</td><td></td><td></td><td></td><td></td><td></td><td></td><td></td><td></td><td></td><td></td></tr>
<tr><td>Chromosome</td><td>start</td><td>end</td><td>strand</td><td>MM</td><td>target_seq</td><td>PAM</td><td>alignment</td><td>distance</td><td>position</td><td>gene name</td><td>gene id</td></tr>
<tr><td>GU305914.1</td><td>74503</td><td>74525</td><td>-</td><td>0</td><td>GCCATCCTCTGTGACAACTG</td><td>CGG</td><td>||||||||[||||||||||||]PAM</td><td>NA</td><td>Intergenic</td><td>NA</td><td>NA</td></tr>
<tr><td>T28592</td><td>CTGCGGACACTGTCTCAACATGG</td><td>1000</td><td></td><td></td><td></td><td></td><td></td><td></td><td></td><td></td><td></td></tr>
<tr><td>Oligo fwd</td><td>CACCCTGCGGACACTGTCTCAACA</td><td></td><td></td><td></td><td></td><td></td><td></td><td></td><td></td><td></td><td></td></tr>
<tr><td>Oligo rev</td><td>AAACTGTTGAGACAGTGTCCGCAG</td><td></td><td></td><td></td><td></td><td></td><td></td><td></td><td></td><td></td><td></td></tr>
<tr><td>Chromosome</td><td>start</td><td>end</td><td>strand</td><td>MM</td><td>target_seq</td><td>PAM</td><td>alignment</td><td>distance</td><td>position</td><td>gene name</td><td>gene id</td></tr>
<tr><td>GU305914.1</td><td>74486</td><td>74508</td><td>-</td><td>0</td><td>CTGCGGACACTGTCTCAACA</td><td>TGG</td><td>||||||||[||||||||||||]PAM</td><td>NA</td><td>Intergenic</td><td>NA</td><td>NA</td></tr>
<tr><td>T28593</td><td>TGCGGACACTGTCTCAACATGGG</td><td>1000</td><td></td><td></td><td></td><td></td><td></td><td></td><td></td><td></td><td></td></tr>
<tr><td>Oligo fwd</td><td>CACCTGCGGACACTGTCTCAACAT</td><td></td><td></td><td></td><td></td><td></td><td></td><td></td><td></td><td></td><td></td></tr>
<tr><td>Oligo rev</td><td>AAACATGTTGAGACAGTGTCCGCA</td><td></td><td></td><td></td><td></td><td></td><td></td><td></td><td></td><td></td><td></td></tr>
<tr><td>Chromosome</td><td>start</td><td>end</td><td>strand</td><td>MM</td><td>target_seq</td><td>PAM</td><td>alignment</td><td>distance</td><td>position</td><td>gene name</td><td>gene id</td></tr>
<tr><td>GU305914.1</td><td>74485</td><td>74507</td><td>-</td><td>0</td><td>TGCGGACACTGTCTCAACAT</td><td>GGG</td><td>||||||||[||||||||||||]PAM</td><td>NA</td><td>Intergenic</td><td>NA</td><td>NA</td></tr>
<tr><td>T28594</td><td>ACACTGTCTCAACATGGGCAAGG</td><td>1000</td><td></td><td></td><td></td><td></td><td></td><td></td><td></td><td></td><td></td></tr>
<tr><td>Oligo fwd</td><td>CACCACACTGTCTCAACATGGGCA</td><td></td><td></td><td></td><td></td><td></td><td></td><td></td><td></td><td></td><td></td></tr>
<tr><td>Oligo rev</td><td>AAACTGCCCATGTTGAGACAGTGT</td><td></td><td></td><td></td><td></td><td></td><td></td><td></td><td></td><td></td><td></td></tr>
<tr><td>Chromosome</td><td>start</td><td>end</td><td>strand</td><td>MM</td><td>target_seq</td><td>PAM</td><td>alignment</td><td>distance</td><td>position</td><td>gene name</td><td>gene id</td></tr>
<tr><td>GU305914.1</td><td>74480</td><td>74502</td><td>-</td><td>0</td><td>ACACTGTCTCAACATGGGCA</td><td>AGG</td><td>||||||||[||||||||||||]PAM</td><td>NA</td><td>Intergenic</td><td>NA</td><td>NA</td></tr>
<tr><td>T28595</td><td>CAACATGGGCAAGGACAAGTTGG</td><td>1000</td><td></td><td></td><td></td><td></td><td></td><td></td><td></td><td></td><td></td></tr>
<tr><td>Oligo fwd</td><td>CACCCAACATGGGCAAGGACAAGT</td><td></td><td></td><td></td><td></td><td></td><td></td><td></td><td></td><td></td><td></td></tr>
<tr><td>Oligo rev</td><td>AAACACTTGTCCTTGCCCATGTTG</td><td></td><td></td><td></td><td></td><td></td><td></td><td></td><td></td><td></td><td></td></tr>
<tr><td>Chromosome</td><td>start</td><td>end</td><td>strand</td><td>MM</td><td>target_seq</td><td>PAM</td><td>alignment</td><td>distance</td><td>position</td><td>gene name</td><td>gene id</td></tr>
<tr><td>GU305914.1</td><td>74471</td><td>74493</td><td>-</td><td>0</td><td>CAACATGGGCAAGGACAAGT</td><td>TGG</td><td>||||||||[||||||||||||]PAM</td><td>NA</td><td>Intergenic</td><td>NA</td><td>NA</td></tr>
<tr><td>T28596</td><td>CATGGGCAAGGACAAGTTGGAGG</td><td>1000</td><td></td><td></td><td></td><td></td><td></td><td></td><td></td><td></td><td></td></tr>
<tr><td>Oligo fwd</td><td>CACCCATGGGCAAGGACAAGTTGG</td><td></td><td></td><td></td><td></td><td></td><td></td><td></td><td></td><td></td><td></td></tr>
<tr><td>Oligo rev</td><td>AAACCCAACTTGTCCTTGCCCATG</td><td></td><td></td><td></td><td></td><td></td><td></td><td></td><td></td><td></td><td></td></tr>
<tr><td>Chromosome</td><td>start</td><td>end</td><td>strand</td><td>MM</td><td>target_seq</td><td>PAM</td><td>alignment</td><td>distance</td><td>position</td><td>gene name</td><td>gene id</td></tr>
<tr><td>GU305914.1</td><td>74468</td><td>74490</td><td>-</td><td>0</td><td>CATGGGCAAGGACAAGTTGG</td><td>AGG</td><td>||||||||[||||||||||||]PAM</td><td>NA</td><td>Intergenic</td><td>NA</td><td>NA</td></tr>
<tr><td>T28597</td><td>ATGGGCAAGGACAAGTTGGAGGG</td><td>1000</td><td></td><td></td><td></td><td></td><td></td><td></td><td></td><td></td><td></td></tr>
<tr><td>Oligo fwd</td><td>CACCATGGGCAAGGACAAGTTGGA</td><td></td><td></td><td></td><td></td><td></td><td></td><td></td><td></td><td></td><td></td></tr>
<tr><td>Oligo rev</td><td>AAACTCCAACTTGTCCTTGCCCAT</td><td></td><td></td><td></td><td></td><td></td><td></td><td></td><td></td><td></td><td></td></tr>
<tr><td>Chromosome</td><td>start</td><td>end</td><td>strand</td><td>MM</td><td>target_seq</td><td>PAM</td><td>alignment</td><td>distance</td><td>position</td><td>gene name</td><td>gene id</td></tr>
<tr><td>GU305914.1</td><td>74467</td><td>74489</td><td>-</td><td>0</td><td>ATGGGCAAGGACAAGTTGGA</td><td>GGG</td><td>||||||||[||||||||||||]PAM</td><td>NA</td><td>Intergenic</td><td>NA</td><td>NA</td></tr>
<tr><td>T28598</td><td>TCAACTGTATCTTTTATTACAGG</td><td>1000</td><td></td><td></td><td></td><td></td><td></td><td></td><td></td><td></td><td></td></tr>
<tr><td>Oligo fwd</td><td>CACCTCAACTGTATCTTTTATTAC</td><td></td><td></td><td></td><td></td><td></td><td></td><td></td><td></td><td></td><td></td></tr>
<tr><td>Oligo rev</td><td>AAACGTAATAAAAGATACAGTTGA</td><td></td><td></td><td></td><td></td><td></td><td></td><td></td><td></td><td></td><td></td></tr>
<tr><td>Chromosome</td><td>start</td><td>end</td><td>strand</td><td>MM</td><td>target_seq</td><td>PAM</td><td>alignment</td><td>distance</td><td>position</td><td>gene name</td><td>gene id</td></tr>
<tr><td>GU305914.1</td><td>74427</td><td>74449</td><td>-</td><td>0</td><td>TCAACTGTATCTTTTATTAC</td><td>AGG</td><td>||||||||[||||||||||||]PAM</td><td>NA</td><td>Intergenic</td><td>NA</td><td>NA</td></tr>
<tr><td>T28599</td><td>CAACTGTATCTTTTATTACAGGG</td><td>1000</td><td></td><td></td><td></td><td></td><td></td><td></td><td></td><td></td><td></td></tr>
<tr><td>Oligo fwd</td><td>CACCCAACTGTATCTTTTATTACA</td><td></td><td></td><td></td><td></td><td></td><td></td><td></td><td></td><td></td><td></td></tr>
<tr><td>Oligo rev</td><td>AAACTGTAATAAAAGATACAGTTG</td><td></td><td></td><td></td><td></td><td></td><td></td><td></td><td></td><td></td><td></td></tr>
<tr><td>Chromosome</td><td>start</td><td>end</td><td>strand</td><td>MM</td><td>target_seq</td><td>PAM</td><td>alignment</td><td>distance</td><td>position</td><td>gene name</td><td>gene id</td></tr>
<tr><td>GU305914.1</td><td>74426</td><td>74448</td><td>-</td><td>0</td><td>CAACTGTATCTTTTATTACA</td><td>GGG</td><td>||||||||[||||||||||||]PAM</td><td>NA</td><td>Intergenic</td><td>NA</td><td>NA</td></tr>
<tr><td>T28600</td><td>CTTTTATTACAGGGACCGCCAGG</td><td>1000</td><td></td><td></td><td></td><td></td><td></td><td></td><td></td><td></td><td></td></tr>
<tr><td>Oligo fwd</td><td>CACCCTTTTATTACAGGGACCGCC</td><td></td><td></td><td></td><td></td><td></td><td></td><td></td><td></td><td></td><td></td></tr>
<tr><td>Oligo rev</td><td>AAACGGCGGTCCCTGTAATAAAAG</td><td></td><td></td><td></td><td></td><td></td><td></td><td></td><td></td><td></td><td></td></tr>
<tr><td>Chromosome</td><td>start</td><td>end</td><td>strand</td><td>MM</td><td>target_seq</td><td>PAM</td><td>alignment</td><td>distance</td><td>position</td><td>gene name</td><td>gene id</td></tr>
<tr><td>GU305914.1</td><td>74417</td><td>74439</td><td>-</td><td>0</td><td>CTTTTATTACAGGGACCGCC</td><td>AGG</td><td>||||||||[||||||||||||]PAM</td><td>NA</td><td>Intergenic</td><td>NA</td><td>NA</td></tr>
<tr><td>T28601</td><td>ACCGCTCACTGTTCCCTGTGTGG</td><td>1000</td><td></td><td></td><td></td><td></td><td></td><td></td><td></td><td></td><td></td></tr>
<tr><td>Oligo fwd</td><td>CACCACCGCTCACTGTTCCCTGTG</td><td></td><td></td><td></td><td></td><td></td><td></td><td></td><td></td><td></td><td></td></tr>
<tr><td>Oligo rev</td><td>AAACCACAGGGAACAGTGAGCGGT</td><td></td><td></td><td></td><td></td><td></td><td></td><td></td><td></td><td></td><td></td></tr>
<tr><td>Chromosome</td><td>start</td><td>end</td><td>strand</td><td>MM</td><td>target_seq</td><td>PAM</td><td>alignment</td><td>distance</td><td>position</td><td>gene name</td><td>gene id</td></tr>
<tr><td>GU305914.1</td><td>74362</td><td>74384</td><td>-</td><td>0</td><td>ACCGCTCACTGTTCCCTGTG</td><td>TGG</td><td>||||||||[||||||||||||]PAM</td><td>NA</td><td>Intergenic</td><td>NA</td><td>NA</td></tr>
<tr><td>T28602</td><td>CCGCTCACTGTTCCCTGTGTGGG</td><td>1000</td><td></td><td></td><td></td><td></td><td></td><td></td><td></td><td></td><td></td></tr>
<tr><td>Oligo fwd</td><td>CACCCCGCTCACTGTTCCCTGTGT</td><td></td><td></td><td></td><td></td><td></td><td></td><td></td><td></td><td></td><td></td></tr>
<tr><td>Oligo rev</td><td>AAACACACAGGGAACAGTGAGCGG</td><td></td><td></td><td></td><td></td><td></td><td></td><td></td><td></td><td></td><td></td></tr>
<tr><td>Chromosome</td><td>start</td><td>end</td><td>strand</td><td>MM</td><td>target_seq</td><td>PAM</td><td>alignment</td><td>distance</td><td>position</td><td>gene name</td><td>gene id</td></tr>
<tr><td>GU305914.1</td><td>74361</td><td>74383</td><td>-</td><td>0</td><td>CCGCTCACTGTTCCCTGTGT</td><td>GGG</td><td>||||||||[||||||||||||]PAM</td><td>NA</td><td>Intergenic</td><td>NA</td><td>NA</td></tr>
<tr><td>T28603</td><td>ACCAGTATTTAAGCCGAGAGCGG</td><td>1000</td><td></td><td></td><td></td><td></td><td></td><td></td><td></td><td></td><td></td></tr>
<tr><td>Oligo fwd</td><td>CACCACCAGTATTTAAGCCGAGAG</td><td></td><td></td><td></td><td></td><td></td><td></td><td></td><td></td><td></td><td></td></tr>
<tr><td>Oligo rev</td><td>AAACCTCTCGGCTTAAATACTGGT</td><td></td><td></td><td></td><td></td><td></td><td></td><td></td><td></td><td></td><td></td></tr>
<tr><td>Chromosome</td><td>start</td><td>end</td><td>strand</td><td>MM</td><td>target_seq</td><td>PAM</td><td>alignment</td><td>distance</td><td>position</td><td>gene name</td><td>gene id</td></tr>
<tr><td>GU305914.1</td><td>74337</td><td>74359</td><td>-</td><td>0</td><td>ACCAGTATTTAAGCCGAGAG</td><td>CGG</td><td>||||||||[||||||||||||]PAM</td><td>NA</td><td>Intergenic</td><td>NA</td><td>NA</td></tr>
<tr><td>T28604</td><td>AAGCCGAGAGCGGATCTACGAGG</td><td>1000</td><td></td><td></td><td></td><td></td><td></td><td></td><td></td><td></td><td></td></tr>
<tr><td>Oligo fwd</td><td>CACCAAGCCGAGAGCGGATCTACG</td><td></td><td></td><td></td><td></td><td></td><td></td><td></td><td></td><td></td><td></td></tr>
<tr><td>Oligo rev</td><td>AAACCGTAGATCCGCTCTCGGCTT</td><td></td><td></td><td></td><td></td><td></td><td></td><td></td><td></td><td></td><td></td></tr>
<tr><td>Chromosome</td><td>start</td><td>end</td><td>strand</td><td>MM</td><td>target_seq</td><td>PAM</td><td>alignment</td><td>distance</td><td>position</td><td>gene name</td><td>gene id</td></tr>
<tr><td>GU305914.1</td><td>74327</td><td>74349</td><td>-</td><td>0</td><td>AAGCCGAGAGCGGATCTACG</td><td>AGG</td><td>||||||||[||||||||||||]PAM</td><td>NA</td><td>Intergenic</td><td>NA</td><td>NA</td></tr>
</table>

| | | | | | | | | | | | | |
|---|---|---|---|---|---|---|---|---|---|---|---|---|
| T28605 | ACTCGCGCCGATTACCATCACGG | | 1000 | | | | | | | | | |
| Oligo fwd | CACCACTCGCGCCGATTACCATCA | | | | | | | | | | | |
| Oligo rev | AAACTGATGGTAATCGGCGCGAGT | | | | | | | | | | | |
| Chromosome | start | | end | strand | MM | target_seq | PAM | alignment | distance | position | gene name | gene id |
| GU305914.1 | | 74302 | 74324 | - | 0 | ACTCGCGCCGATTACCATCA | CGG | \|\|\|\|\|\|\|\|[\|\|\|\|\|\|\|\|\|\|\|\|]PAM | NA | Intergenic | NA | NA |
| T28606 | CTCGCGCCGATTACCATCACGGG | | 1000 | | | | | | | | | |
| Oligo fwd | CACCCTCGCGCCGATTACCATCAC | | | | | | | | | | | |
| Oligo rev | AAACGTGATGGTAATCGGCGCGAG | | | | | | | | | | | |
| Chromosome | start | | end | strand | MM | target_seq | PAM | alignment | distance | position | gene name | gene id |
| GU305914.1 | | 74301 | 74323 | - | 0 | CTCGCGCCGATTACCATCAC | GGG | \|\|\|\|\|\|\|\|[\|\|\|\|\|\|\|\|\|\|\|\|]PAM | NA | Intergenic | NA | NA |
| T28607 | CCGATTACCATCACGGGCTCAGG | | 1000 | | | | | | | | | |
| Oligo fwd | CACCCCGATTACCATCACGGGCTC | | | | | | | | | | | |
| Oligo rev | AAACGAGCCCGTGATGGTAATCGG | | | | | | | | | | | |
| Chromosome | start | | end | strand | MM | target_seq | PAM | alignment | distance | position | gene name | gene id |
| GU305914.1 | | 74295 | 74317 | - | 0 | CCGATTACCATCACGGGCTC | AGG | \|\|\|\|\|\|\|\|[\|\|\|\|\|\|\|\|\|\|\|\|]PAM | NA | Intergenic | NA | NA |
| T28608 | TCACGGGCTCAGGTTCGCGACGG | | 1000 | | | | | | | | | |
| Oligo fwd | CACCTCACGGGCTCAGGTTCGCGA | | | | | | | | | | | |
| Oligo rev | AAACTCGCGAACCTGAGCCCGTGA | | | | | | | | | | | |
| Chromosome | start | | end | strand | MM | target_seq | PAM | alignment | distance | position | gene name | gene id |
| GU305914.1 | | 74285 | 74307 | - | 0 | TCACGGGCTCAGGTTCGCGA | CGG | \|\|\|\|\|\|\|\|[\|\|\|\|\|\|\|\|\|\|\|\|]PAM | NA | Intergenic | NA | NA |
| T28609 | TCAGGTTCGCGACGGTGCGCTGG | | 1000 | | | | | | | | | |
| Oligo fwd | CACCTCAGGTTCGCGACGGTGCGC | | | | | | | | | | | |
| Oligo rev | AAACGCGCACCGTCGCGAACCTGA | | | | | | | | | | | |
| Chromosome | start | | end | strand | MM | target_seq | PAM | alignment | distance | position | gene name | gene id |
| GU305914.1 | | 74277 | 74299 | - | 0 | TCAGGTTCGCGACGGTGCGC | TGG | \|\|\|\|\|\|\|\|[\|\|\|\|\|\|\|\|\|\|\|\|]PAM | NA | Intergenic | NA | NA |
| T28610 | GGTTCGCGACGGTGCGCTGGAGG | | 1000 | | | | | | | | | |
| Oligo fwd | CACCGGTTCGCGACGGTGCGCTGG | | | | | | | | | | | |
| Oligo rev | AAACCCAGCGCACCGTCGCGAACC | | | | | | | | | | | |
| Chromosome | start | | end | strand | MM | target_seq | PAM | alignment | distance | position | gene name | gene id |
| GU305914.1 | | 74274 | 74296 | - | 0 | GGTTCGCGACGGTGCGCTGG | AGG | \|\|\|\|\|\|\|\|[\|\|\|\|\|\|\|\|\|\|\|\|]PAM | NA | Intergenic | NA | NA |
| T28611 | GTTCGCGACGGTGCGCTGGAGGG | | 1000 | | | | | | | | | |
| Oligo fwd | CACCGTTCGCGACGGTGCGCTGGA | | | | | | | | | | | |
| Oligo rev | AAACTCCAGCGCACCGTCGCGAAC | | | | | | | | | | | |
| Chromosome | start | | end | strand | MM | target_seq | PAM | alignment | distance | position | gene name | gene id |
| GU305914.1 | | 74273 | 74295 | - | 0 | GTTCGCGACGGTGCGCTGGA | GGG | \|\|\|\|\|\|\|\|[\|\|\|\|\|\|\|\|\|\|\|\|]PAM | NA | Intergenic | NA | NA |
| T28612 | CGCGACGGTGCGCTGGAGGGCGG | | 1000 | | | | | | | | | |
| Oligo fwd | CACCCGCGACGGTGCGCTGGAGGG | | | | | | | | | | | |
| Oligo rev | AAACCCCTCCAGCGCACCGTCGCG | | | | | | | | | | | |
| Chromosome | start | | end | strand | MM | target_seq | PAM | alignment | distance | position | gene name | gene id |
| GU305914.1 | | 74270 | 74292 | - | 0 | CGCGACGGTGCGCTGGAGGG | CGG | \|\|\|\|\|\|\|\|[\|\|\|\|\|\|\|\|\|\|\|\|]PAM | NA | Intergenic | NA | NA |
| T28613 | GTGCGCTGGAGGGCGGTCATCGG | | 1000 | | | | | | | | | |
| Oligo fwd | CACCGTGCGCTGGAGGGCGGTCAT | | | | | | | | | | | |
| Oligo rev | AAACATGACCGCCCTCCAGCGCAC | | | | | | | | | | | |
| Chromosome | start | | end | strand | MM | target_seq | PAM | alignment | distance | position | gene name | gene id |
| GU305914.1 | | 74263 | 74285 | - | 0 | GTGCGCTGGAGGGCGGTCAT | CGG | \|\|\|\|\|\|\|\|[\|\|\|\|\|\|\|\|\|\|\|\|]PAM | NA | Intergenic | NA | NA |
| T28614 | ATCGGCTCGAACGCCGCCTGCGG | | 1000 | | | | | | | | | |
| Oligo fwd | CACCATCGGCTCGAACGCCGCCTG | | | | | | | | | | | |
| Oligo rev | AAACCAGGCGGCGTTCGAGCCGAT | | | | | | | | | | | |
| Chromosome | start | | end | strand | MM | target_seq | PAM | alignment | distance | position | gene name | gene id |
| GU305914.1 | | 74245 | 74267 | - | 0 | ATCGGCTCGAACGCCGCCTG | CGG | \|\|\|\|\|\|\|\|[\|\|\|\|\|\|\|\|\|\|\|\|]PAM | NA | Intergenic | NA | NA |
| T28615 | TCGGCTCGAACGCCGCCTGCGGG | | 1000 | | | | | | | | | |
| Oligo fwd | CACCTCGGCTCGAACGCCGCCTGC | | | | | | | | | | | |
| Oligo rev | AAACGCAGGCGGCGTTCGAGCCGA | | | | | | | | | | | |
| Chromosome | start | | end | strand | MM | target_seq | PAM | alignment | distance | position | gene name | gene id |
| GU305914.1 | | 74244 | 74266 | - | 0 | TCGGCTCGAACGCCGCCTGC | GGG | \|\|\|\|\|\|\|\|[\|\|\|\|\|\|\|\|\|\|\|\|]PAM | NA | Intergenic | NA | NA |
| T28616 | CGGCTCGAACGCCGCCTGCGGGG | | 1000 | | | | | | | | | |
| Oligo fwd | CACCCGGCTCGAACGCCGCCTGCG | | | | | | | | | | | |
| Oligo rev | AAACCGCAGGCGGCGTTCGAGCCG | | | | | | | | | | | |
| Chromosome | start | | end | strand | MM | target_seq | PAM | alignment | distance | position | gene name | gene id |
| GU305914.1 | | 74243 | 74265 | - | 0 | CGGCTCGAACGCCGCCTGCG | GGG | \|\|\|\|\|\|\|\|[\|\|\|\|\|\|\|\|\|\|\|\|]PAM | NA | Intergenic | NA | NA |
| T28617 | AACGCCGCCTGCGGGGTTCTCGG | | 1000 | | | | | | | | | |
| Oligo fwd | CACCAACGCCGCCTGCGGGGTTCT | | | | | | | | | | | |
| Oligo rev | AAACAGAACCCCGCAGGCGGCGTT | | | | | | | | | | | |
| Chromosome | start | | end | strand | MM | target_seq | PAM | alignment | distance | position | gene name | gene id |
| GU305914.1 | | 74236 | 74258 | - | 0 | AACGCCGCCTGCGGGGTTCT | CGG | \|\|\|\|\|\|\|\|[\|\|\|\|\|\|\|\|\|\|\|\|]PAM | NA | Intergenic | NA | NA |
| T28618 | CGCCTGCGGGGTTCTCGGCCCGG | | 1000 | | | | | | | | | |
| Oligo fwd | CACCCGCCTGCGGGGTTCTCGGCC | | | | | | | | | | | |
| Oligo rev | AAACGGCCGAGAACCCCGCAGGCG | | | | | | | | | | | |
| Chromosome | start | | end | strand | MM | target_seq | PAM | alignment | distance | position | gene name | gene id |
| GU305914.1 | | 74231 | 74253 | - | 0 | CGCCTGCGGGGTTCTCGGCC | CGG | \|\|\|\|\|\|\|\|[\|\|\|\|\|\|\|\|\|\|\|\|]PAM | NA | Intergenic | NA | NA |
| T28619 | GCCTGCGGGGTTCTCGGCCCGGG | | 1000 | | | | | | | | | |
| Oligo fwd | CACCGCCTGCGGGGTTCTCGGCCC | | | | | | | | | | | |
| Oligo rev | AAACGGGCCGAGAACCCCGCAGGC | | | | | | | | | | | |
| Chromosome | start | | end | strand | MM | target_seq | PAM | alignment | distance | position | gene name | gene id |
| GU305914.1 | | 74230 | 74252 | - | 0 | GCCTGCGGGGTTCTCGGCCC | GGG | \|\|\|\|\|\|\|\|[\|\|\|\|\|\|\|\|\|\|\|\|]PAM | NA | Intergenic | NA | NA |

T28620 TCCTCGTGCCCTGCTCCTCTAGG 1000
Oligo fwd CACCTCCTCGTGCCCTGCTCCTCT
Oligo rev AAACAGAGGAGCAGGGCACGAGGA

| Chromosome | start | end | strand | MM | target_seq | PAM | alignment | distance | position | gene name | gene id |
|---|---|---|---|---|---|---|---|---|---|---|---|
| GU305914.1 | 74193 | 74215 | - | 0 | TCCTCGTGCCCTGCTCCTCT | AGG | ||||||||[||||||||||||]PAM | NA | Intergenic | NA | NA |

T28621 TGCTCCTCTAGGACCTGTTTCGG 1000
Oligo fwd CACCTGCTCCTCTAGGACCTGTTT
Oligo rev AAACAAACAGGTCCTAGAGGAGCA

| Chromosome | start | end | strand | MM | target_seq | PAM | alignment | distance | position | gene name | gene id |
|---|---|---|---|---|---|---|---|---|---|---|---|
| GU305914.1 | 74182 | 74204 | - | 0 | TGCTCCTCTAGGACCTGTTT | CGG | ||||||||[||||||||||||]PAM | NA | Intergenic | NA | NA |
| GU305914.1 | 118916 | 118938 | + | 4 | TCCTGCTCTAGGTCCTGTAT | GGG | |-||-|||[||||-|||||-|]PAM | NA | Intergenic | NA | NA |

T28622 CTCCGCCTCGTCTCGCACTCCGG 1000
Oligo fwd CACCCTCCGCCTCGTCTCGCACTC
Oligo rev AAACGAGTGCGAGACGAGGCGGAG

| Chromosome | start | end | strand | MM | target_seq | PAM | alignment | distance | position | gene name | gene id |
|---|---|---|---|---|---|---|---|---|---|---|---|
| GU305914.1 | 74122 | 74144 | - | 0 | CTCCGCCTCGTCTCGCACTC | CGG | ||||||||[||||||||||||]PAM | NA | Intergenic | NA | NA |

T28623 AGTCCTGTCAGAACGTCTACAGG 1000
Oligo fwd CACCAGTCCTGTCAGAACGTCTAC
Oligo rev AAACGTAGACGTTCTGACAGGACT

| Chromosome | start | end | strand | MM | target_seq | PAM | alignment | distance | position | gene name | gene id |
|---|---|---|---|---|---|---|---|---|---|---|---|
| GU305914.1 | 74085 | 74107 | - | 0 | AGTCCTGTCAGAACGTCTAC | AGG | ||||||||[||||||||||||]PAM | NA | Intergenic | NA | NA |

T28624 GTCCTGTCAGAACGTCTACAGGG 1000
Oligo fwd CACCGTCCTGTCAGAACGTCTACA
Oligo rev AAACTGTAGACGTTCTGACAGGAC

| Chromosome | start | end | strand | MM | target_seq | PAM | alignment | distance | position | gene name | gene id |
|---|---|---|---|---|---|---|---|---|---|---|---|
| GU305914.1 | 74084 | 74106 | - | 0 | GTCCTGTCAGAACGTCTACA | GGG | ||||||||[||||||||||||]PAM | NA | Intergenic | NA | NA |

T28625 CGTCTACAGGGAGACCTGCCTGG 1000
Oligo fwd CACCCGTCTACAGGGAGACCTGCC
Oligo rev AAACGGCAGGTCTCCCTGTAGACG

| Chromosome | start | end | strand | MM | target_seq | PAM | alignment | distance | position | gene name | gene id |
|---|---|---|---|---|---|---|---|---|---|---|---|
| GU305914.1 | 74072 | 74094 | - | 0 | CGTCTACAGGGAGACCTGCC | TGG | ||||||||[||||||||||||]PAM | NA | Intergenic | NA | NA |

T28626 GACCTCGCCGACGCCTGTCTCGG 1000
Oligo fwd CACCGACCTCGCCGACGCCTGTCT
Oligo rev AAACAGACAGGCGTCGGCGAGGTC

| Chromosome | start | end | strand | MM | target_seq | PAM | alignment | distance | position | gene name | gene id |
|---|---|---|---|---|---|---|---|---|---|---|---|
| GU305914.1 | 74035 | 74057 | - | 0 | GACCTCGCCGACGCCTGTCT | CGG | ||||||||[||||||||||||]PAM | NA | Intergenic | NA | NA |

T28627 CCACTCTCGCTGTTCCTGTCTGG 1000
Oligo fwd CACCCCACTCTCGCTGTTCCTGTC
Oligo rev AAACGACAGGAACAGCGAGAGTGG

| Chromosome | start | end | strand | MM | target_seq | PAM | alignment | distance | position | gene name | gene id |
|---|---|---|---|---|---|---|---|---|---|---|---|
| GU305914.1 | 74003 | 74025 | - | 0 | CCACTCTCGCTGTTCCTGTC | TGG | ||||||||[||||||||||||]PAM | NA | Intergenic | NA | NA |
| GU305914.1 | 93873 | 93895 | + | 4 | CCCCGCTCCCGGTTCCTGTC | CGG | ||-|-|||[-|-|||||||||]PAM | NA | Intergenic | NA | NA |

T28628 GCTGTTCCTGTCTGGAGATCAGG 1000
Oligo fwd CACCGCTGTTCCTGTCTGGAGATC
Oligo rev AAACGATCTCCAGACAGGAACAGC

| Chromosome | start | end | strand | MM | target_seq | PAM | alignment | distance | position | gene name | gene id |
|---|---|---|---|---|---|---|---|---|---|---|---|
| GU305914.1 | 73995 | 74017 | - | 0 | GCTGTTCCTGTCTGGAGATC | AGG | ||||||||[||||||||||||]PAM | NA | Intergenic | NA | NA |

T28629 CCTGTCTGGAGATCAGGCGTAGG 1000
Oligo fwd CACCCCTGTCTGGAGATCAGGCGT
Oligo rev AAACACGCCTGATCTCCAGACAGG

| Chromosome | start | end | strand | MM | target_seq | PAM | alignment | distance | position | gene name | gene id |
|---|---|---|---|---|---|---|---|---|---|---|---|
| GU305914.1 | 73989 | 74011 | - | 0 | CCTGTCTGGAGATCAGGCGT | AGG | ||||||||[||||||||||||]PAM | NA | Intergenic | NA | NA |

T28630 AACTATAACAACAACAACACCGG 1000
Oligo fwd CACCAACTATAACAACAACAACAC
Oligo rev AAACGTGTTGTTGTTGTTATAGTT

| Chromosome | start | end | strand | MM | target_seq | PAM | alignment | distance | position | gene name | gene id |
|---|---|---|---|---|---|---|---|---|---|---|---|
| GU305914.1 | 73966 | 73988 | - | 0 | AACTATAACAACAACAACAC | CGG | ||||||||[||||||||||||]PAM | NA | Intergenic | NA | NA |

T28631 AACAACAACACCGGCGCTGCCGG 1000
Oligo fwd CACCAACAACAACACCGGCGCTGC
Oligo rev AAACGCAGCGCCGGTGTTGTTGTT

| Chromosome | start | end | strand | MM | target_seq | PAM | alignment | distance | position | gene name | gene id |
|---|---|---|---|---|---|---|---|---|---|---|---|
| GU305914.1 | 73957 | 73979 | - | 0 | AACAACAACACCGGCGCTGC | CGG | ||||||||[||||||||||||]PAM | NA | Intergenic | NA | NA |

T28632 GTCGACCAACGTTAGTAGCGCGG 1000
Oligo fwd CACCGTCGACCAACGTTAGTAGCG
Oligo rev AAACCGCTACTAACGTTGGTCGAC

| Chromosome | start | end | strand | MM | target_seq | PAM | alignment | distance | position | gene name | gene id |
|---|---|---|---|---|---|---|---|---|---|---|---|
| GU305914.1 | 73847 | 73869 | - | 0 | GTCGACCAACGTTAGTAGCG | CGG | ||||||||[||||||||||||]PAM | NA | Intergenic | NA | NA |

T28633 CAACGTTAGTAGCGCGGCCTCGG 1000
Oligo fwd CACCCAACGTTAGTAGCGCGGCCT
Oligo rev AAACAGGCCGCGCTACTAACGTTG

| Chromosome | start | end | strand | MM | target_seq | PAM | alignment | distance | position | gene name | gene id |
|---|---|---|---|---|---|---|---|---|---|---|---|
| GU305914.1 | 73841 | 73863 | - | 0 | CAACGTTAGTAGCGCGGCCT | CGG | ||||||||[||||||||||||]PAM | NA | Intergenic | NA | NA |

T28634 AACGTTAGTAGCGCGGCCTCGGG 1000
Oligo fwd CACCAACGTTAGTAGCGCGGCCTC

<table>
<tr><td>Oligo rev</td><td>AAACGAGGCCGCGCTACTAACGTT</td><td></td><td></td><td></td><td></td><td></td><td></td><td></td><td></td><td></td><td></td></tr>
<tr><td>Chromosome</td><td>start</td><td>end</td><td>strand</td><td>MM</td><td>target_seq</td><td>PAM</td><td>alignment</td><td>distance</td><td>position</td><td>gene name</td><td>gene id</td></tr>
<tr><td>GU305914.1</td><td>73840</td><td>73862</td><td>-</td><td>0</td><td>AACGTTAGTAGCGCGGCCTC</td><td>GGG</td><td>||||||||[||||||||||||]PAM</td><td>NA</td><td>Intergenic</td><td>NA</td><td>NA</td></tr>
<tr><td>T28635</td><td>GTTAGTAGCGCGGCCTCGGGAGG</td><td>1000</td><td></td><td></td><td></td><td></td><td></td><td></td><td></td><td></td><td></td></tr>
<tr><td>Oligo fwd</td><td>CACCGTTAGTAGCGCGGCCTCGGG</td><td></td><td></td><td></td><td></td><td></td><td></td><td></td><td></td><td></td><td></td></tr>
<tr><td>Oligo rev</td><td>AAACCCCGAGGCCGCGCTACTAAC</td><td></td><td></td><td></td><td></td><td></td><td></td><td></td><td></td><td></td><td></td></tr>
<tr><td>Chromosome</td><td>start</td><td>end</td><td>strand</td><td>MM</td><td>target_seq</td><td>PAM</td><td>alignment</td><td>distance</td><td>position</td><td>gene name</td><td>gene id</td></tr>
<tr><td>GU305914.1</td><td>73837</td><td>73859</td><td>-</td><td>0</td><td>GTTAGTAGCGCGGCCTCGGG</td><td>AGG</td><td>||||||||[||||||||||||]PAM</td><td>NA</td><td>Intergenic</td><td>NA</td><td>NA</td></tr>
<tr><td>T28636</td><td>AGTAGCGCGGCCTCGGGAGGCGG</td><td>1000</td><td></td><td></td><td></td><td></td><td></td><td></td><td></td><td></td><td></td></tr>
<tr><td>Oligo fwd</td><td>CACCAGTAGCGCGGCCTCGGGAGG</td><td></td><td></td><td></td><td></td><td></td><td></td><td></td><td></td><td></td><td></td></tr>
<tr><td>Oligo rev</td><td>AAACCCTCCCGAGGCCGCGCTACT</td><td></td><td></td><td></td><td></td><td></td><td></td><td></td><td></td><td></td><td></td></tr>
<tr><td>Chromosome</td><td>start</td><td>end</td><td>strand</td><td>MM</td><td>target_seq</td><td>PAM</td><td>alignment</td><td>distance</td><td>position</td><td>gene name</td><td>gene id</td></tr>
<tr><td>GU305914.1</td><td>73834</td><td>73856</td><td>-</td><td>0</td><td>AGTAGCGCGGCCTCGGGAGG</td><td>CGG</td><td>||||||||[||||||||||||]PAM</td><td>NA</td><td>Intergenic</td><td>NA</td><td>NA</td></tr>
<tr><td>T28637</td><td>GTAGCGCGGCCTCGGGAGGCGGG</td><td>1000</td><td></td><td></td><td></td><td></td><td></td><td></td><td></td><td></td><td></td></tr>
<tr><td>Oligo fwd</td><td>CACCGTAGCGCGGCCTCGGGAGGC</td><td></td><td></td><td></td><td></td><td></td><td></td><td></td><td></td><td></td><td></td></tr>
<tr><td>Oligo rev</td><td>AAACGCCTCCCGAGGCCGCGCTAC</td><td></td><td></td><td></td><td></td><td></td><td></td><td></td><td></td><td></td><td></td></tr>
<tr><td>Chromosome</td><td>start</td><td>end</td><td>strand</td><td>MM</td><td>target_seq</td><td>PAM</td><td>alignment</td><td>distance</td><td>position</td><td>gene name</td><td>gene id</td></tr>
<tr><td>GU305914.1</td><td>73833</td><td>73855</td><td>-</td><td>0</td><td>GTAGCGCGGCCTCGGGAGGC</td><td>GGG</td><td>||||||||[||||||||||||]PAM</td><td>NA</td><td>Intergenic</td><td>NA</td><td>NA</td></tr>
<tr><td>T28638</td><td>GCCTCGGGAGGCGGGAGTAGCGG</td><td>1000</td><td></td><td></td><td></td><td></td><td></td><td></td><td></td><td></td><td></td></tr>
<tr><td>Oligo fwd</td><td>CACCGCCTCGGGAGGCGGGAGTAG</td><td></td><td></td><td></td><td></td><td></td><td></td><td></td><td></td><td></td><td></td></tr>
<tr><td>Oligo rev</td><td>AAACCTACTCCCGCCTCCCGAGGC</td><td></td><td></td><td></td><td></td><td></td><td></td><td></td><td></td><td></td><td></td></tr>
<tr><td>Chromosome</td><td>start</td><td>end</td><td>strand</td><td>MM</td><td>target_seq</td><td>PAM</td><td>alignment</td><td>distance</td><td>position</td><td>gene name</td><td>gene id</td></tr>
<tr><td>GU305914.1</td><td>73825</td><td>73847</td><td>-</td><td>0</td><td>GCCTCGGGAGGCGGGAGTAG</td><td>CGG</td><td>||||||||[||||||||||||]PAM</td><td>NA</td><td>Intergenic</td><td>NA</td><td>NA</td></tr>
<tr><td>T28639</td><td>TCGGGAGGCGGGAGTAGCGGCGG</td><td>1000</td><td></td><td></td><td></td><td></td><td></td><td></td><td></td><td></td><td></td></tr>
<tr><td>Oligo fwd</td><td>CACCTCGGGAGGCGGGAGTAGCGG</td><td></td><td></td><td></td><td></td><td></td><td></td><td></td><td></td><td></td><td></td></tr>
<tr><td>Oligo rev</td><td>AAACCCGCTACTCCCGCCTCCCGA</td><td></td><td></td><td></td><td></td><td></td><td></td><td></td><td></td><td></td><td></td></tr>
<tr><td>Chromosome</td><td>start</td><td>end</td><td>strand</td><td>MM</td><td>target_seq</td><td>PAM</td><td>alignment</td><td>distance</td><td>position</td><td>gene name</td><td>gene id</td></tr>
<tr><td>GU305914.1</td><td>73822</td><td>73844</td><td>-</td><td>0</td><td>TCGGGAGGCGGGAGTAGCGG</td><td>CGG</td><td>||||||||[||||||||||||]PAM</td><td>NA</td><td>Intergenic</td><td>NA</td><td>NA</td></tr>
<tr><td>T28640</td><td>GGGAGTAGCGGCGGCAGCAGTGG</td><td>1000</td><td></td><td></td><td></td><td></td><td></td><td></td><td></td><td></td><td></td></tr>
<tr><td>Oligo fwd</td><td>CACCGGGAGTAGCGGCGGCAGCAG</td><td></td><td></td><td></td><td></td><td></td><td></td><td></td><td></td><td></td><td></td></tr>
<tr><td>Oligo rev</td><td>AAACCTGCTGCCGCCGCTACTCCC</td><td></td><td></td><td></td><td></td><td></td><td></td><td></td><td></td><td></td><td></td></tr>
<tr><td>Chromosome</td><td>start</td><td>end</td><td>strand</td><td>MM</td><td>target_seq</td><td>PAM</td><td>alignment</td><td>distance</td><td>position</td><td>gene name</td><td>gene id</td></tr>
<tr><td>GU305914.1</td><td>73813</td><td>73835</td><td>-</td><td>0</td><td>GGGAGTAGCGGCGGCAGCAG</td><td>TGG</td><td>||||||||[||||||||||||]PAM</td><td>NA</td><td>Intergenic</td><td>NA</td><td>NA</td></tr>
<tr><td>GU305914.1</td><td>27408</td><td>27430</td><td>+</td><td>4</td><td>GGCAGCAGCAGCAGCAGCAG</td><td>CGG</td><td>||-||-||[|-||-|||||||]PAM</td><td>NA</td><td>Intergenic</td><td>NA</td><td>NA</td></tr>
<tr><td>GU305914.1</td><td>73801</td><td>73823</td><td>-</td><td>4</td><td>GGCAGCAGTGGCGCCAGCAG</td><td>CGG</td><td>||-||-||[-||||-||||||]PAM</td><td>NA</td><td>Intergenic</td><td>NA</td><td>NA</td></tr>
<tr><td>GU305914.1</td><td>67742</td><td>67764</td><td>+</td><td>4</td><td>GGGAGCGGCGGCGGCGGCGG</td><td>CGG</td><td>|||||--|[|||||||-||-|]PAM</td><td>NA</td><td>Intergenic</td><td>NA</td><td>NA</td></tr>
<tr><td>T28641</td><td>GGCAGCAGTGGCGCCAGCAGCGG</td><td>1000</td><td></td><td></td><td></td><td></td><td></td><td></td><td></td><td></td><td></td></tr>
<tr><td>Oligo fwd</td><td>CACCGGCAGCAGTGGCGCCAGCAG</td><td></td><td></td><td></td><td></td><td></td><td></td><td></td><td></td><td></td><td></td></tr>
<tr><td>Oligo rev</td><td>AAACCTGCTGGCGCCACTGCTGCC</td><td></td><td></td><td></td><td></td><td></td><td></td><td></td><td></td><td></td><td></td></tr>
<tr><td>Chromosome</td><td>start</td><td>end</td><td>strand</td><td>MM</td><td>target_seq</td><td>PAM</td><td>alignment</td><td>distance</td><td>position</td><td>gene name</td><td>gene id</td></tr>
<tr><td>GU305914.1</td><td>73801</td><td>73823</td><td>-</td><td>0</td><td>GGCAGCAGTGGCGCCAGCAG</td><td>CGG</td><td>||||||||[||||||||||||]PAM</td><td>NA</td><td>Intergenic</td><td>NA</td><td>NA</td></tr>
<tr><td>GU305914.1</td><td>73813</td><td>73835</td><td>-</td><td>4</td><td>GGGAGTAGCGGCGGCAGCAG</td><td>TGG</td><td>||-||-||[-||||-||||||]PAM</td><td>NA</td><td>Intergenic</td><td>NA</td><td>NA</td></tr>
<tr><td>T28642</td><td>AGCAGTGGCGCCAGCAGCGGCGG</td><td>1000</td><td></td><td></td><td></td><td></td><td></td><td></td><td></td><td></td><td></td></tr>
<tr><td>Oligo fwd</td><td>CACCAGCAGTGGCGCCAGCAGCGG</td><td></td><td></td><td></td><td></td><td></td><td></td><td></td><td></td><td></td><td></td></tr>
<tr><td>Oligo rev</td><td>AAACCCGCTGCTGGCGCCACTGCT</td><td></td><td></td><td></td><td></td><td></td><td></td><td></td><td></td><td></td><td></td></tr>
<tr><td>Chromosome</td><td>start</td><td>end</td><td>strand</td><td>MM</td><td>target_seq</td><td>PAM</td><td>alignment</td><td>distance</td><td>position</td><td>gene name</td><td>gene id</td></tr>
<tr><td>GU305914.1</td><td>73798</td><td>73820</td><td>-</td><td>0</td><td>AGCAGTGGCGCCAGCAGCGG</td><td>CGG</td><td>||||||||[||||||||||||]PAM</td><td>NA</td><td>Intergenic</td><td>NA</td><td>NA</td></tr>
<tr><td>GU305914.1</td><td>27411</td><td>27433</td><td>+</td><td>4</td><td>AGCAGCAGCAGCAGCAGCGG</td><td>CGG</td><td>|||||--|[|--|||||||||]PAM</td><td>NA</td><td>Intergenic</td><td>NA</td><td>NA</td></tr>
<tr><td>T28643</td><td>AGTGGCGCCAGCAGCGGCGGCGG</td><td>1000</td><td></td><td></td><td></td><td></td><td></td><td></td><td></td><td></td><td></td></tr>
<tr><td>Oligo fwd</td><td>CACCAGTGGCGCCAGCAGCGGCGG</td><td></td><td></td><td></td><td></td><td></td><td></td><td></td><td></td><td></td><td></td></tr>
<tr><td>Oligo rev</td><td>AAACCCGCCGCTGCTGGCGCCACT</td><td></td><td></td><td></td><td></td><td></td><td></td><td></td><td></td><td></td><td></td></tr>
<tr><td>Chromosome</td><td>start</td><td>end</td><td>strand</td><td>MM</td><td>target_seq</td><td>PAM</td><td>alignment</td><td>distance</td><td>position</td><td>gene name</td><td>gene id</td></tr>
<tr><td>GU305914.1</td><td>73795</td><td>73817</td><td>-</td><td>0</td><td>AGTGGCGCCAGCAGCGGCGG</td><td>CGG</td><td>||||||||[||||||||||||]PAM</td><td>NA</td><td>Intergenic</td><td>NA</td><td>NA</td></tr>
<tr><td>GU305914.1</td><td>168820</td><td>168842</td><td>+</td><td>4</td><td>GGTGGTGGCAGCGGCGGCGG</td><td>CGG</td><td>-||||-|-[||||-|||||||]PAM</td><td>NA</td><td>Intergenic</td><td>NA</td><td>NA</td></tr>
<tr><td>GU305914.1</td><td>154407</td><td>154429</td><td>+</td><td>4</td><td>GGTGGCGGCGGCGGCGGCGG</td><td>CGG</td><td>-||||||-[|-||-|||||||]PAM</td><td>NA</td><td>Intergenic</td><td>NA</td><td>NA</td></tr>
<tr><td>GU305914.1</td><td>58123</td><td>58145</td><td>-</td><td>4</td><td>GGTGGCGGCGGCGGCGGCGG</td><td>AGG</td><td>-||||||-[|-||-|||||||]PAM</td><td>NA</td><td>Intergenic</td><td>NA</td><td>NA</td></tr>
<tr><td>GU305914.1</td><td>62152</td><td>62174</td><td>+</td><td>4</td><td>AGGGGCGACGGCAGCGGCAG</td><td>CGG</td><td>||-||||-[|-||||||||-|]PAM</td><td>NA</td><td>Intergenic</td><td>NA</td><td>NA</td></tr>
<tr><td>T28644</td><td>GGCGCCAGCAGCGGCGGCGGAGG</td><td>1000</td><td></td><td></td><td></td><td></td><td></td><td></td><td></td><td></td><td></td></tr>
<tr><td>Oligo fwd</td><td>CACCGGCGCCAGCAGCGGCGGCGG</td><td></td><td></td><td></td><td></td><td></td><td></td><td></td><td></td><td></td><td></td></tr>
<tr><td>Oligo rev</td><td>AAACCCGCCGCCGCTGCTGGCGCC</td><td></td><td></td><td></td><td></td><td></td><td></td><td></td><td></td><td></td><td></td></tr>
<tr><td>Chromosome</td><td>start</td><td>end</td><td>strand</td><td>MM</td><td>target_seq</td><td>PAM</td><td>alignment</td><td>distance</td><td>position</td><td>gene name</td><td>gene id</td></tr>
<tr><td>GU305914.1</td><td>73792</td><td>73814</td><td>-</td><td>0</td><td>GGCGCCAGCAGCGGCGGCGG</td><td>AGG</td><td>||||||||[||||||||||||]PAM</td><td>NA</td><td>Intergenic</td><td>NA</td><td>NA</td></tr>
<tr><td>GU305914.1</td><td>168820</td><td>168842</td><td>+</td><td>4</td><td>GGTGGTGGCAGCGGCGGCGG</td><td>CGG</td><td>||-|---|[||||||||||||]PAM</td><td>NA</td><td>Intergenic</td><td>NA</td><td>NA</td></tr>
<tr><td>GU305914.1</td><td>56773</td><td>56795</td><td>+</td><td>4</td><td>CGCCGCAGCGGCGGCGGCGG</td><td>CGG</td><td>-||--|||[|-||||||||||]PAM</td><td>NA</td><td>Intergenic</td><td>NA</td><td>NA</td></tr>
<tr><td>GU305914.1</td><td>149482</td><td>149504</td><td>-</td><td>3</td><td>GGCGACGGCGGCGGCGGCGG</td><td>CGG</td><td>||||-|-|[|-||||||||||]PAM</td><td>NA</td><td>Intergenic</td><td>NA</td><td>NA</td></tr>
<tr><td>GU305914.1</td><td>163896</td><td>163918</td><td>+</td><td>4</td><td>GGATCGAGCGGCGGCGGCGG</td><td>CGG</td><td>||--|-||[|-||||||||||]PAM</td><td>NA</td><td>Intergenic</td><td>NA</td><td>NA</td></tr>
<tr><td>GU305914.1</td><td>154407</td><td>154429</td><td>+</td><td>4</td><td>GGTGGCGGCGGCGGCGGCGG</td><td>CGG</td><td>||-|-|-|[|-||||||||||]PAM</td><td>NA</td><td>Intergenic</td><td>NA</td><td>NA</td></tr>
<tr><td>GU305914.1</td><td>58123</td><td>58145</td><td>-</td><td>4</td><td>GGTGGCGGCGGCGGCGGCGG</td><td>AGG</td><td>||-|-|-|[|-||||||||||]PAM</td><td>NA</td><td>Intergenic</td><td>NA</td><td>NA</td></tr>
<tr><td>GU305914.1</td><td>30000</td><td>30022</td><td>+</td><td>4</td><td>GGTGCTGGCGGCGGCGGCGG</td><td>CGG</td><td>||-||--|[|-||||||||||]PAM</td><td>NA</td><td>Intergenic</td><td>NA</td><td>NA</td></tr>
<tr><td>GU305914.1</td><td>154404</td><td>154426</td><td>+</td><td>4</td><td>GGCGGTGGCGGCGGCGGCGG</td><td>CGG</td><td>||||---|[|-||||||||||]PAM</td><td>NA</td><td>Intergenic</td><td>NA</td><td>NA</td></tr>
<tr><td>GU305914.1</td><td>58126</td><td>58148</td><td>-</td><td>4</td><td>GGCGGTGGCGGCGGCGGCGG</td><td>CGG</td><td>||||---|[|-||||||||||]PAM</td><td>NA</td><td>Intergenic</td><td>NA</td><td>NA</td></tr>
<tr><td>GU305914.1</td><td>48159</td><td>48181</td><td>+</td><td>4</td><td>GGCGGCGGTGGCGGCGGCGG</td><td>TGG</td><td>||||-|-|[--||||||||||]PAM</td><td>NA</td><td>Intergenic</td><td>NA</td><td>NA</td></tr>
<tr><td>GU305914.1</td><td>70184</td><td>70206</td><td>+</td><td>4</td><td>AGCACCGGCAGCGCCGGCGG</td><td>CGG</td><td>-||-||-|[|||||-||||||]PAM</td><td>NA</td><td>Intergenic</td><td>NA</td><td>NA</td></tr>
<tr><td>GU305914.1</td><td>48156</td><td>48178</td><td>+</td><td>4</td><td>GGCGGCGGCGGTGGCGGCGG</td><td>CGG</td><td>||||-|-|[|-|-||||||||]PAM</td><td>NA</td><td>Intergenic</td><td>NA</td><td>NA</td></tr>
<tr><td>GU305914.1</td><td>62155</td><td>62177</td><td>+</td><td>3</td><td>GGCGACGGCAGCGGCAGCGG</td><td>CGG</td><td>||||-|-|[|||||||-|||||]PAM</td><td>NA</td><td>Intergenic</td><td>NA</td><td>NA</td></tr>
<tr><td>GU305914.1</td><td>163911</td><td>163933</td><td>+</td><td>3</td><td>GGCGGCGGCAGCGGCAGCGG</td><td>AGG</td><td>||||-|-|[|||||||-|||||]PAM</td><td>NA</td><td>Intergenic</td><td>NA</td><td>NA</td></tr>
<tr><td>GU305914.1</td><td>62542</td><td>62564</td><td>+</td><td>4</td><td>GACGGCAGGAGCGGCCGCGG</td><td>CGG</td><td>|-||-|||[-||||||-||||]PAM</td><td>NA</td><td>Intergenic</td><td>NA</td><td>NA</td></tr>
<tr><td>GU305914.1</td><td>151368</td><td>151390</td><td>+</td><td>4</td><td>GACGGCAGCAGCGAAGGCGG</td><td>CGG</td><td>|-||-|||[|||||--|||||]PAM</td><td>NA</td><td>Intergenic</td><td>NA</td><td>NA</td></tr>
<tr><td>GU305914.1</td><td>30006</td><td>30028</td><td>+</td><td>4</td><td>GGCGGCGGCGGCGGCGGCAG</td><td>TGG</td><td>||||-|-|[|-||||||||-|]PAM</td><td>NA</td><td>Intergenic</td><td>NA</td><td>NA</td></tr>
</table>

| | | | | | | | | | | | |
|---|---|---|---|---|---|---|---|---|---|---|---|
| GU305914.1 | 149476 | 149498 | - | 4 | GGCGGCGGCGGCGGCGGCAG | CGG | \|\|\|\|-\|-\|[\|-\|\|\|\|\|\|\|\|-\|]PAM | NA | Intergenic | NA | NA |
| GU305914.1 | 128721 | 128743 | - | 3 | GGCGGCAGCAGCAGCGGCGC | TGG | \|\|\|\|-\|\|\|[\|\|\|\|-\|\|\|\|\|\|-]PAM | NA | Intergenic | NA | NA |
| T28645 | GCCAGCAGCGGCGGCGGAGGAGG | 1000 | | | | | | | | | |
| Oligo fwd | CACCGCCAGCAGCGGCGGCGGAGG | | | | | | | | | | |
| Oligo rev | AAACCCTCCGCCGCCGCTGCTGGC | | | | | | | | | | |
| Chromosome | start | end | strand | MM | target_seq | PAM | alignment | distance | position | gene name | gene id |
| GU305914.1 | 73789 | 73811 | - | 0 | GCCAGCAGCGGCGGCGGAGG | AGG | \|\|\|\|\|\|\|\|[\|\|\|\|\|\|\|\|\|\|\|\|]PAM | NA | Intergenic | NA | NA |
| GU305914.1 | 18656 | 18678 | - | 4 | GACGGCAGAGGTGGCGGAGG | CGG | \|-\|-\|\|\|\|[-\|\|-\|\|\|\|\|\|\|\|]PAM | NA | Intergenic | NA | NA |
| GU305914.1 | 73786 | 73808 | - | 4 | AGCAGCGGCGGCGGAGGAGG | AGG | --\|\|\|\|-\|[\|\|\|\|\|\|-\|\|\|\|\|]PAM | NA | Intergenic | NA | NA |
| GU305914.1 | 56773 | 56795 | + | 4 | CGCCGCAGCGGCGGCGGCGG | CGG | --\|-\|\|\|\|[\|\|\|\|\|\|\|\|\|-\|\|]PAM | NA | Intergenic | NA | NA |
| GU305914.1 | 67742 | 67764 | + | 4 | GGGAGCGGCGGCGGCGGCGG | CGG | \|--\|\|\|-\|[\|\|\|\|\|\|\|\|\|-\|\|]PAM | NA | Intergenic | NA | NA |
| T28646 | AGCAGCGGCGGCGGAGGAGGAGG | 1000 | | | | | | | | | |
| Oligo fwd | CACCAGCAGCGGCGGCGGAGGAGG | | | | | | | | | | |
| Oligo rev | AAACCCTCCTCCGCCGCCGCTGCT | | | | | | | | | | |
| Chromosome | start | end | strand | MM | target_seq | PAM | alignment | distance | position | gene name | gene id |
| GU305914.1 | 73786 | 73808 | - | 0 | AGCAGCGGCGGCGGAGGAGG | AGG | \|\|\|\|\|\|\|\|[\|\|\|\|\|\|\|\|\|\|\|\|]PAM | NA | Intergenic | NA | NA |
| GU305914.1 | 73789 | 73811 | - | 4 | GCCAGCAGCGGCGGCGGAGG | AGG | --\|\|\|\|-\|[\|\|\|\|\|\|-\|\|\|\|\|]PAM | NA | Intergenic | NA | NA |
| GU305914.1 | 102673 | 102695 | - | 4 | AGTAGTGGCCGCGTAGGAGG | AGG | \|\|-\|\|-\|\|[\|-\|\|\|-\|\|\|\|\|\|]PAM | NA | Intergenic | NA | NA |
| GU305914.1 | 18653 | 18675 | - | 4 | GGCAGAGGTGGCGGAGGCGG | GGG | -\|\|\|\|-\|\|[-\|\|\|\|\|\|\|\|-\|\|]PAM | NA | Intergenic | NA | NA |
| GU305914.1 | 67742 | 67764 | + | 4 | GGGAGCGGCGGCGGCGGCGG | CGG | -\|-\|\|\|\|\|[\|\|\|\|\|\|-\|\|-\|\|]PAM | NA | Intergenic | NA | NA |
| GU305914.1 | 24624 | 24646 | - | 4 | AACACCGGCGGCGGCGGTGG | CGG | \|-\|\|-\|\|\|[\|\|\|\|\|\|-\|\|-\|\|]PAM | NA | Intergenic | NA | NA |
| T28647 | GGCGGCGGAGGAGGAGGAAGCGG | 1000 | | | | | | | | | |
| Oligo fwd | CACCGGCGGCGGAGGAGGAGGAAG | | | | | | | | | | |
| Oligo rev | AAACCTTCCTCCTCCTCCGCCGCC | | | | | | | | | | |
| Chromosome | start | end | strand | MM | target_seq | PAM | alignment | distance | position | gene name | gene id |
| GU305914.1 | 73780 | 73802 | - | 0 | GGCGGCGGAGGAGGAGGAAG | CGG | \|\|\|\|\|\|\|\|[\|\|\|\|\|\|\|\|\|\|\|\|]PAM | NA | Intergenic | NA | NA |
| GU305914.1 | 75721 | 75743 | + | 4 | CGCCGGGGAGGAGGTGGAAG | TGG | -\|\|-\|-\|\|[\|\|\|\|\|\|-\|\|\|\|\|]PAM | NA | Intergenic | NA | NA |
| GU305914.1 | 145969 | 145991 | + | 4 | GTCGGTGGAGGAGGCGGGAG | CGG | \|-\|\|\|-\|\|[\|\|\|\|\|\|-\|\|-\|\|]PAM | NA | Intergenic | NA | NA |
| T28648 | GGCGGAGGAGGAGGAAGCGGCGG | 1000 | | | | | | | | | |
| Oligo fwd | CACCGGCGGAGGAGGAGGAAGCGG | | | | | | | | | | |
| Oligo rev | AAACCCGCTTCCTCCTCCTCCGCC | | | | | | | | | | |
| Chromosome | start | end | strand | MM | target_seq | PAM | alignment | distance | position | gene name | gene id |
| GU305914.1 | 73777 | 73799 | - | 0 | GGCGGAGGAGGAGGAAGCGG | CGG | \|\|\|\|\|\|\|\|[\|\|\|\|\|\|\|\|\|\|\|\|]PAM | NA | Intergenic | NA | NA |
| GU305914.1 | 75724 | 75746 | + | 4 | CGGGGAGGAGGTGGAAGTGG | AGG | -\|-\|\|\|\|\|[\|\|\|-\|\|\|\|\|-\|\|]PAM | NA | Intergenic | NA | NA |
| T28649 | CGGCAGCAGCAAGAAAGAAGAGG | 1000 | | | | | | | | | |
| Oligo fwd | CACCCGGCAGCAGCAAGAAAGAAG | | | | | | | | | | |
| Oligo rev | AAACCTTCTTTCTTGCTGCTGCCG | | | | | | | | | | |
| Chromosome | start | end | strand | MM | target_seq | PAM | alignment | distance | position | gene name | gene id |
| GU305914.1 | 73757 | 73779 | - | 0 | CGGCAGCAGCAAGAAAGAAG | AGG | \|\|\|\|\|\|\|\|[\|\|\|\|\|\|\|\|\|\|\|\|]PAM | NA | Intergenic | NA | NA |
| T28650 | GCAGCAAGAAAGAAGAGGAAAGG | 1000 | | | | | | | | | |
| Oligo fwd | CACCGCAGCAAGAAAGAAGAGGAA | | | | | | | | | | |
| Oligo rev | AAACTTCCTCTTCTTTCTTGCTGC | | | | | | | | | | |
| Chromosome | start | end | strand | MM | target_seq | PAM | alignment | distance | position | gene name | gene id |
| GU305914.1 | 73752 | 73774 | - | 0 | GCAGCAAGAAAGAAGAGGAA | AGG | \|\|\|\|\|\|\|\|[\|\|\|\|\|\|\|\|\|\|\|\|]PAM | NA | Intergenic | NA | NA |
| T28651 | GCAAGAAAGAAGAGGAAAGGAGG | 1000 | | | | | | | | | |
| Oligo fwd | CACCGCAAGAAAGAAGAGGAAAGG | | | | | | | | | | |
| Oligo rev | AAACCCTTTCCTCTTCTTTCTTGC | | | | | | | | | | |
| Chromosome | start | end | strand | MM | target_seq | PAM | alignment | distance | position | gene name | gene id |
| GU305914.1 | 73749 | 73771 | - | 0 | GCAAGAAAGAAGAGGAAAGG | AGG | \|\|\|\|\|\|\|\|[\|\|\|\|\|\|\|\|\|\|\|\|]PAM | NA | Intergenic | NA | NA |
| T28652 | GAGGAAAGGAGGAAGCAATTCGG | 1000 | | | | | | | | | |
| Oligo fwd | CACCGAGGAAAGGAGGAAGCAATT | | | | | | | | | | |
| Oligo rev | AAACAATTGCTTCCTCCTTTCCTC | | | | | | | | | | |
| Chromosome | start | end | strand | MM | target_seq | PAM | alignment | distance | position | gene name | gene id |
| GU305914.1 | 73738 | 73760 | - | 0 | GAGGAAAGGAGGAAGCAATT | CGG | \|\|\|\|\|\|\|\|[\|\|\|\|\|\|\|\|\|\|\|\|]PAM | NA | Intergenic | NA | NA |
| T28653 | CGGCGCCAACGTACTCAACCTGG | 1000 | | | | | | | | | |
| Oligo fwd | CACCCGGCGCCAACGTACTCAACC | | | | | | | | | | |
| Oligo rev | AAACGGTTGAGTACGTTGGCGCCG | | | | | | | | | | |
| Chromosome | start | end | strand | MM | target_seq | PAM | alignment | distance | position | gene name | gene id |
| GU305914.1 | 73718 | 73740 | - | 0 | CGGCGCCAACGTACTCAACC | TGG | \|\|\|\|\|\|\|\|[\|\|\|\|\|\|\|\|\|\|\|\|]PAM | NA | Intergenic | NA | NA |
| T28654 | ACTCAACCTGGCGCCTGCGATGG | 1000 | | | | | | | | | |
| Oligo fwd | CACCACTCAACCTGGCGCCTGCGA | | | | | | | | | | |
| Oligo rev | AAACTCGCAGGCGCCAGGTTGAGT | | | | | | | | | | |
| Chromosome | start | end | strand | MM | target_seq | PAM | alignment | distance | position | gene name | gene id |
| GU305914.1 | 73706 | 73728 | - | 0 | ACTCAACCTGGCGCCTGCGA | TGG | \|\|\|\|\|\|\|\|[\|\|\|\|\|\|\|\|\|\|\|\|]PAM | NA | Intergenic | NA | NA |
| T28655 | GATTCCGAAATATATGAAGATGG | 1000 | | | | | | | | | |
| Oligo fwd | CACCGATTCCGAAATATATGAAGA | | | | | | | | | | |
| Oligo rev | AAACTCTTCATATATTTCGGAATC | | | | | | | | | | |
| Chromosome | start | end | strand | MM | target_seq | PAM | alignment | distance | position | gene name | gene id |
| GU305914.1 | 73658 | 73680 | - | 0 | GATTCCGAAATATATGAAGA | TGG | \|\|\|\|\|\|\|\|[\|\|\|\|\|\|\|\|\|\|\|\|]PAM | NA | Intergenic | NA | NA |
| T28656 | ATTCCGAAATATATGAAGATGGG | 1000 | | | | | | | | | |
| Oligo fwd | CACCATTCCGAAATATATGAAGAT | | | | | | | | | | |
| Oligo rev | AAACATCTTCATATATTTCGGAAT | | | | | | | | | | |
| Chromosome | start | end | strand | MM | target_seq | PAM | alignment | distance | position | gene name | gene id |
| GU305914.1 | 73657 | 73679 | - | 0 | ATTCCGAAATATATGAAGAT | GGG | \|\|\|\|\|\|\|\|[\|\|\|\|\|\|\|\|\|\|\|\|]PAM | NA | Intergenic | NA | NA |
| T28657 | CCGAAATATATGAAGATGGGAGG | 1000 | | | | | | | | | |
| Oligo fwd | CACCCCGAAATATATGAAGATGGG | | | | | | | | | | |

Oligo rev AAACCCCATCTTCATATATTTCGG
Chromosome start end strand MM target_seq PAM alignment distance position gene name gene id
GU305914.1 73654 73676 - 0 CCGAAATATATGAAGATGGG AGG ||||||||[||||||||||||]PAM NA Intergenic NA NA

T28658 TATGAAGATGGGAGGACACGAGG 1000
Oligo fwd CACCTATGAAGATGGGAGGACACG
Oligo rev AAACCGTGTCCTCCCATCTTCATA
Chromosome start end strand MM target_seq PAM alignment distance position gene name gene id
GU305914.1 73646 73668 - 0 TATGAAGATGGGAGGACACG AGG ||||||||[||||||||||||]PAM NA Intergenic NA NA

T28659 GGACAAACTCGCCTACCAGCTGG 1000
Oligo fwd CACCGGACAAACTCGCCTACCAGC
Oligo rev AAACGCTGGTAGGCGAGTTTGTCC
Chromosome start end strand MM target_seq PAM alignment distance position gene name gene id
GU305914.1 73625 73647 - 0 GGACAAACTCGCCTACCAGC TGG ||||||||[||||||||||||]PAM NA Intergenic NA NA

T28660 CCTACCAGCTGGACCTTCTGCGG 1000
Oligo fwd CACCCCTACCAGCTGGACCTTCTG
Oligo rev AAACCAGAAGGTCCAGCTGGTAGG
Chromosome start end strand MM target_seq PAM alignment distance position gene name gene id
GU305914.1 73614 73636 - 0 CCTACCAGCTGGACCTTCTG CGG ||||||||[||||||||||||]PAM NA Intergenic NA NA

T28661 AAAAAATAAACCTCTGTGTGTGG 1000
Oligo fwd CACCAAAAAATAAACCTCTGTGTG
Oligo rev AAACCACACAGAGGTTTATTTTTT
Chromosome start end strand MM target_seq PAM alignment distance position gene name gene id
GU305914.1 73517 73539 - 0 AAAAAATAAACCTCTGTGTG TGG ||||||||[||||||||||||]PAM NA Intergenic NA NA

T28662 ATAAACCTCTGTGTGTGGTGTGG 1000
Oligo fwd CACCATAAACCTCTGTGTGTGGTG
Oligo rev AAACCACCACACACAGAGGTTTAT
Chromosome start end strand MM target_seq PAM alignment distance position gene name gene id
GU305914.1 73512 73534 - 0 ATAAACCTCTGTGTGTGGTG TGG ||||||||[||||||||||||]PAM NA Intergenic NA NA

T28663 TAAACCTCTGTGTGTGGTGTGGG 1000
Oligo fwd CACCTAAACCTCTGTGTGTGGTGT
Oligo rev AAACACACCACACACAGAGGTTTA
Chromosome start end strand MM target_seq PAM alignment distance position gene name gene id
GU305914.1 73511 73533 - 0 TAAACCTCTGTGTGTGGTGT GGG ||||||||[||||||||||||]PAM NA Intergenic NA NA

T28664 TTTTGTTTGTGTGTCAGTGCAGG 1000
Oligo fwd CACCTTTTGTTTGTGTGTCAGTGC
Oligo rev AAACGCACTGACACACAAACAAAA
Chromosome start end strand MM target_seq PAM alignment distance position gene name gene id
GU305914.1 73486 73508 - 0 TTTTGTTTGTGTGTCAGTGC AGG ||||||||[||||||||||||]PAM NA Intergenic NA NA

T28665 TTGTGTGTCAGTGCAGGTAGAGG 1000
Oligo fwd CACCTTGTGTGTCAGTGCAGGTAG
Oligo rev AAACCTACCTGCACTGACACACAA
Chromosome start end strand MM target_seq PAM alignment distance position gene name gene id
GU305914.1 73480 73502 - 0 TTGTGTGTCAGTGCAGGTAG AGG ||||||||[||||||||||||]PAM NA Intergenic NA NA

T28666 GAGGATCTTGATCCTCCTGATGG 1000
Oligo fwd CACCGAGGATCTTGATCCTCCTGA
Oligo rev AAACTCAGGAGGATCAAGATCCTC
Chromosome start end strand MM target_seq PAM alignment distance position gene name gene id
GU305914.1 73461 73483 - 0 GAGGATCTTGATCCTCCTGA TGG ||||||||[||||||||||||]PAM NA Intergenic NA NA

T28667 CGAAAGTGTCCAGCTGCAGTCGG 1000
Oligo fwd CACCCGAAAGTGTCCAGCTGCAGT
Oligo rev AAACACTGCAGCTGGACACTTTCG
Chromosome start end strand MM target_seq PAM alignment distance position gene name gene id
GU305914.1 73420 73442 - 0 CGAAAGTGTCCAGCTGCAGT CGG ||||||||[||||||||||||]PAM NA Intergenic NA NA

T28668 GAAAGTGTCCAGCTGCAGTCGGG 1000
Oligo fwd CACCGAAAGTGTCCAGCTGCAGTC
Oligo rev AAACGACTGCAGCTGGACACTTTC
Chromosome start end strand MM target_seq PAM alignment distance position gene name gene id
GU305914.1 73419 73441 - 0 GAAAGTGTCCAGCTGCAGTC GGG ||||||||[||||||||||||]PAM NA Intergenic NA NA

T28669 GCTGCAGTCGGGTCTCTTGCAGG 1000
Oligo fwd CACCGCTGCAGTCGGGTCTCTTGC
Oligo rev AAACGCAAGAGACCCGACTGCAGC
Chromosome start end strand MM target_seq PAM alignment distance position gene name gene id
GU305914.1 73408 73430 - 0 GCTGCAGTCGGGTCTCTTGC AGG ||||||||[||||||||||||]PAM NA Intergenic NA NA

T28670 CCGCCATGATCGCGCTAGACAGG 1000
Oligo fwd CACCCCGCCATGATCGCGCTAGAC
Oligo rev AAACGTCTAGCGCGATCATGGCGG
Chromosome start end strand MM target_seq PAM alignment distance position gene name gene id
GU305914.1 73375 73397 - 0 CCGCCATGATCGCGCTAGAC AGG ||||||||[||||||||||||]PAM NA Intergenic NA NA

T28671 CGCCATGATCGCGCTAGACAGGG 1000
Oligo fwd CACCCGCCATGATCGCGCTAGACA
Oligo rev AAACTGTCTAGCGCGATCATGGCG
Chromosome start end strand MM target_seq PAM alignment distance position gene name gene id
GU305914.1 73374 73396 - 0 CGCCATGATCGCGCTAGACA GGG ||||||||[||||||||||||]PAM NA Intergenic NA NA

T28672 TGATCGCGCTAGACAGGGCTCGG 1000

Oligo fwd CACCTGATCGCGCTAGACAGGGCT
Oligo rev AAACAGCCCTGTCTAGCGCGATCA
Chromosome start end strand MM target_seq PAM alignment distance position gene name gene id
GU305914.1 73369 73391 - 0 TGATCGCGCTAGACAGGGCT CGG ||||||||[||||||||||||]PAM NA Intergenic NA NA

T28673 GCGCTAGACAGGGCTCGGAACGG 1000
Oligo fwd CACCGCGCTAGACAGGGCTCGGAA
Oligo rev AAACTTCCGAGCCCTGTCTAGCGC
Chromosome start end strand MM target_seq PAM alignment distance position gene name gene id
GU305914.1 73364 73386 - 0 GCGCTAGACAGGGCTCGGAA CGG ||||||||[||||||||||||]PAM NA Intergenic NA NA

T28674 CGCTAGACAGGGCTCGGAACGGG 1000
Oligo fwd CACCCGCTAGACAGGGCTCGGAAC
Oligo rev AAACGTTCCGAGCCCTGTCTAGCG
Chromosome start end strand MM target_seq PAM alignment distance position gene name gene id
GU305914.1 73363 73385 - 0 CGCTAGACAGGGCTCGGAAC GGG ||||||||[||||||||||||]PAM NA Intergenic NA NA

T28675 TAGACAGGGCTCGGAACGGGTGG 1000
Oligo fwd CACCTAGACAGGGCTCGGAACGGG
Oligo rev AAACCCCGTTCCGAGCCCTGTCTA
Chromosome start end strand MM target_seq PAM alignment distance position gene name gene id
GU305914.1 73360 73382 - 0 TAGACAGGGCTCGGAACGGG TGG ||||||||[||||||||||||]PAM NA Intergenic NA NA

T28676 GGGCTCGGAACGGGTGGATCTGG 1000
Oligo fwd CACCGGGCTCGGAACGGGTGGATC
Oligo rev AAACGATCCACCCGTTCCGAGCCC
Chromosome start end strand MM target_seq PAM alignment distance position gene name gene id
GU305914.1 73354 73376 - 0 GGGCTCGGAACGGGTGGATC TGG ||||||||[||||||||||||]PAM NA Intergenic NA NA

T28677 GCGAGATAGATTCGTTAGACAGG 1000
Oligo fwd CACCGCGAGATAGATTCGTTAGAC
Oligo rev AAACGTCTAACGAATCTATCTCGC
Chromosome start end strand MM target_seq PAM alignment distance position gene name gene id
GU305914.1 73324 73346 - 0 GCGAGATAGATTCGTTAGAC AGG ||||||||[||||||||||||]PAM NA Intergenic NA NA

T28678 CGAGATAGATTCGTTAGACAGGG 1000
Oligo fwd CACCCGAGATAGATTCGTTAGACA
Oligo rev AAACTGTCTAACGAATCTATCTCG
Chromosome start end strand MM target_seq PAM alignment distance position gene name gene id
GU305914.1 73323 73345 - 0 CGAGATAGATTCGTTAGACA GGG ||||||||[||||||||||||]PAM NA Intergenic NA NA

T28679 AGGGTCGTCACCGCGTCCTTCGG 1000
Oligo fwd CACCAGGGTCGTCACCGCGTCCTT
Oligo rev AAACAAGGACGCGGTGACGACCCT
Chromosome start end strand MM target_seq PAM alignment distance position gene name gene id
GU305914.1 73304 73326 - 0 AGGGTCGTCACCGCGTCCTT CGG ||||||||[||||||||||||]PAM NA Intergenic NA NA

T28680 CGCGTCCTTCGGCACGATGCTGG 1000
Oligo fwd CACCCGCGTCCTTCGGCACGATGC
Oligo rev AAACGCATCGTGCCGAAGGACGCG
Chromosome start end strand MM target_seq PAM alignment distance position gene name gene id
GU305914.1 73293 73315 - 0 CGCGTCCTTCGGCACGATGC TGG ||||||||[||||||||||||]PAM NA Intergenic NA NA

T28681 CGGCACGATGCTGGACACCTTGG 1000
Oligo fwd CACCCGGCACGATGCTGGACACCT
Oligo rev AAACAGGTGTCCAGCATCGTGCCG
Chromosome start end strand MM target_seq PAM alignment distance position gene name gene id
GU305914.1 73284 73306 - 0 CGGCACGATGCTGGACACCT TGG ||||||||[||||||||||||]PAM NA Intergenic NA NA

T28682 ACACCTTGGTCTGCGAGAACAGG 1000
Oligo fwd CACCACACCTTGGTCTGCGAGAAC
Oligo rev AAACGTTCTCGCAGACCAAGGTGT
Chromosome start end strand MM target_seq PAM alignment distance position gene name gene id
GU305914.1 73270 73292 - 0 ACACCTTGGTCTGCGAGAAC AGG ||||||||[||||||||||||]PAM NA Intergenic NA NA

T28683 CACCTTGGTCTGCGAGAACAGGG 1000
Oligo fwd CACCCACCTTGGTCTGCGAGAACA
Oligo rev AAACTGTTCTCGCAGACCAAGGTG
Chromosome start end strand MM target_seq PAM alignment distance position gene name gene id
GU305914.1 73269 73291 - 0 CACCTTGGTCTGCGAGAACA GGG ||||||||[||||||||||||]PAM NA Intergenic NA NA

T28684 ACCTTGGTCTGCGAGAACAGGGG 1000
Oligo fwd CACCACCTTGGTCTGCGAGAACAG
Oligo rev AAACCTGTTCTCGCAGACCAAGGT
Chromosome start end strand MM target_seq PAM alignment distance position gene name gene id
GU305914.1 73268 73290 - 0 ACCTTGGTCTGCGAGAACAG GGG ||||||||[||||||||||||]PAM NA Intergenic NA NA

T28685 GCGAGAACAGGGGTTCGCTGAGG 1000
Oligo fwd CACCGCGAGAACAGGGGTTCGCTG
Oligo rev AAACCAGCGAACCCCTGTTCTCGC
Chromosome start end strand MM target_seq PAM alignment distance position gene name gene id
GU305914.1 73258 73280 - 0 GCGAGAACAGGGGTTCGCTG AGG ||||||||[||||||||||||]PAM NA Intergenic NA NA

T28686 CGAGAACAGGGGTTCGCTGAGGG 1000
Oligo fwd CACCCGAGAACAGGGGTTCGCTGA
Oligo rev AAACTCAGCGAACCCCTGTTCTCG
Chromosome start end strand MM target_seq PAM alignment distance position gene name gene id
GU305914.1 73257 73279 - 0 CGAGAACAGGGGTTCGCTGA GGG ||||||||[||||||||||||]PAM NA Intergenic NA NA

<table>
<tr><td>T28687</td><td>ACAGGGGTTCGCTGAGGGACTGG</td><td>1000</td><td></td><td></td><td></td><td></td><td></td><td></td><td></td><td></td><td></td></tr>
<tr><td>Oligo fwd</td><td>CACCACAGGGGTTCGCTGAGGGAC</td><td></td><td></td><td></td><td></td><td></td><td></td><td></td><td></td><td></td><td></td></tr>
<tr><td>Oligo rev</td><td>AAACGTCCCTCAGCGAACCCCTGT</td><td></td><td></td><td></td><td></td><td></td><td></td><td></td><td></td><td></td><td></td></tr>
<tr><td>Chromosome</td><td>start</td><td>end</td><td>strand</td><td>MM</td><td>target_seq</td><td>PAM</td><td>alignment</td><td>distance</td><td>position</td><td>gene name</td><td>gene id</td></tr>
<tr><td>GU305914.1</td><td>73252</td><td>73274</td><td>-</td><td>0</td><td>ACAGGGGTTCGCTGAGGGAC</td><td>TGG</td><td>||||||||[||||||||||||]PAM</td><td>NA</td><td>Intergenic</td><td>NA</td><td>NA</td></tr>
<tr><td>T28688</td><td>GGGTTCGCTGAGGGACTGGTCGG</td><td>1000</td><td></td><td></td><td></td><td></td><td></td><td></td><td></td><td></td><td></td></tr>
<tr><td>Oligo fwd</td><td>CACCGGGTTCGCTGAGGGACTGGT</td><td></td><td></td><td></td><td></td><td></td><td></td><td></td><td></td><td></td><td></td></tr>
<tr><td>Oligo rev</td><td>AAACACCAGTCCCTCAGCGAACCC</td><td></td><td></td><td></td><td></td><td></td><td></td><td></td><td></td><td></td><td></td></tr>
<tr><td>Chromosome</td><td>start</td><td>end</td><td>strand</td><td>MM</td><td>target_seq</td><td>PAM</td><td>alignment</td><td>distance</td><td>position</td><td>gene name</td><td>gene id</td></tr>
<tr><td>GU305914.1</td><td>73248</td><td>73270</td><td>-</td><td>0</td><td>GGGTTCGCTGAGGGACTGGT</td><td>CGG</td><td>||||||||[||||||||||||]PAM</td><td>NA</td><td>Intergenic</td><td>NA</td><td>NA</td></tr>
<tr><td>T28689</td><td>GGTTCGCTGAGGGACTGGTCGGG</td><td>1000</td><td></td><td></td><td></td><td></td><td></td><td></td><td></td><td></td><td></td></tr>
<tr><td>Oligo fwd</td><td>CACCGGTTCGCTGAGGGACTGGTC</td><td></td><td></td><td></td><td></td><td></td><td></td><td></td><td></td><td></td><td></td></tr>
<tr><td>Oligo rev</td><td>AAACGACCAGTCCCTCAGCGAACC</td><td></td><td></td><td></td><td></td><td></td><td></td><td></td><td></td><td></td><td></td></tr>
<tr><td>Chromosome</td><td>start</td><td>end</td><td>strand</td><td>MM</td><td>target_seq</td><td>PAM</td><td>alignment</td><td>distance</td><td>position</td><td>gene name</td><td>gene id</td></tr>
<tr><td>GU305914.1</td><td>73247</td><td>73269</td><td>-</td><td>0</td><td>GGTTCGCTGAGGGACTGGTC</td><td>GGG</td><td>||||||||[||||||||||||]PAM</td><td>NA</td><td>Intergenic</td><td>NA</td><td>NA</td></tr>
<tr><td>T28690</td><td>GTTCGCTGAGGGACTGGTCGGGG</td><td>1000</td><td></td><td></td><td></td><td></td><td></td><td></td><td></td><td></td><td></td></tr>
<tr><td>Oligo fwd</td><td>CACCGTTCGCTGAGGGACTGGTCG</td><td></td><td></td><td></td><td></td><td></td><td></td><td></td><td></td><td></td><td></td></tr>
<tr><td>Oligo rev</td><td>AAACCGACCAGTCCCTCAGCGAAC</td><td></td><td></td><td></td><td></td><td></td><td></td><td></td><td></td><td></td><td></td></tr>
<tr><td>Chromosome</td><td>start</td><td>end</td><td>strand</td><td>MM</td><td>target_seq</td><td>PAM</td><td>alignment</td><td>distance</td><td>position</td><td>gene name</td><td>gene id</td></tr>
<tr><td>GU305914.1</td><td>73246</td><td>73268</td><td>-</td><td>0</td><td>GTTCGCTGAGGGACTGGTCG</td><td>GGG</td><td>||||||||[||||||||||||]PAM</td><td>NA</td><td>Intergenic</td><td>NA</td><td>NA</td></tr>
<tr><td>T28691</td><td>ACTGGTCGGGGACCGAATCCTGG</td><td>1000</td><td></td><td></td><td></td><td></td><td></td><td></td><td></td><td></td><td></td></tr>
<tr><td>Oligo fwd</td><td>CACCACTGGTCGGGGACCGAATCC</td><td></td><td></td><td></td><td></td><td></td><td></td><td></td><td></td><td></td><td></td></tr>
<tr><td>Oligo rev</td><td>AAACGGATTCGGTCCCCGACCAGT</td><td></td><td></td><td></td><td></td><td></td><td></td><td></td><td></td><td></td><td></td></tr>
<tr><td>Chromosome</td><td>start</td><td>end</td><td>strand</td><td>MM</td><td>target_seq</td><td>PAM</td><td>alignment</td><td>distance</td><td>position</td><td>gene name</td><td>gene id</td></tr>
<tr><td>GU305914.1</td><td>73234</td><td>73256</td><td>-</td><td>0</td><td>ACTGGTCGGGGACCGAATCC</td><td>TGG</td><td>||||||||[||||||||||||]PAM</td><td>NA</td><td>Intergenic</td><td>NA</td><td>NA</td></tr>
<tr><td>T28692</td><td>CTGGTCGGGGACCGAATCCTGGG</td><td>1000</td><td></td><td></td><td></td><td></td><td></td><td></td><td></td><td></td><td></td></tr>
<tr><td>Oligo fwd</td><td>CACCCTGGTCGGGGACCGAATCCT</td><td></td><td></td><td></td><td></td><td></td><td></td><td></td><td></td><td></td><td></td></tr>
<tr><td>Oligo rev</td><td>AAACAGGATTCGGTCCCCGACCAG</td><td></td><td></td><td></td><td></td><td></td><td></td><td></td><td></td><td></td><td></td></tr>
<tr><td>Chromosome</td><td>start</td><td>end</td><td>strand</td><td>MM</td><td>target_seq</td><td>PAM</td><td>alignment</td><td>distance</td><td>position</td><td>gene name</td><td>gene id</td></tr>
<tr><td>GU305914.1</td><td>73233</td><td>73255</td><td>-</td><td>0</td><td>CTGGTCGGGGACCGAATCCT</td><td>GGG</td><td>||||||||[||||||||||||]PAM</td><td>NA</td><td>Intergenic</td><td>NA</td><td>NA</td></tr>
<tr><td>T28693</td><td>TGGTCGGGGACCGAATCCTGGGG</td><td>1000</td><td></td><td></td><td></td><td></td><td></td><td></td><td></td><td></td><td></td></tr>
<tr><td>Oligo fwd</td><td>CACCTGGTCGGGGACCGAATCCTG</td><td></td><td></td><td></td><td></td><td></td><td></td><td></td><td></td><td></td><td></td></tr>
<tr><td>Oligo rev</td><td>AAACCAGGATTCGGTCCCCGACCA</td><td></td><td></td><td></td><td></td><td></td><td></td><td></td><td></td><td></td><td></td></tr>
<tr><td>Chromosome</td><td>start</td><td>end</td><td>strand</td><td>MM</td><td>target_seq</td><td>PAM</td><td>alignment</td><td>distance</td><td>position</td><td>gene name</td><td>gene id</td></tr>
<tr><td>GU305914.1</td><td>73232</td><td>73254</td><td>-</td><td>0</td><td>TGGTCGGGGACCGAATCCTG</td><td>GGG</td><td>||||||||[||||||||||||]PAM</td><td>NA</td><td>Intergenic</td><td>NA</td><td>NA</td></tr>
<tr><td>T28694</td><td>CGAATCCTGGGGTCGCGCGTCGG</td><td>1000</td><td></td><td></td><td></td><td></td><td></td><td></td><td></td><td></td><td></td></tr>
<tr><td>Oligo fwd</td><td>CACCCGAATCCTGGGGTCGCGCGT</td><td></td><td></td><td></td><td></td><td></td><td></td><td></td><td></td><td></td><td></td></tr>
<tr><td>Oligo rev</td><td>AAACACGCGCGACCCCAGGATTCG</td><td></td><td></td><td></td><td></td><td></td><td></td><td></td><td></td><td></td><td></td></tr>
<tr><td>Chromosome</td><td>start</td><td>end</td><td>strand</td><td>MM</td><td>target_seq</td><td>PAM</td><td>alignment</td><td>distance</td><td>position</td><td>gene name</td><td>gene id</td></tr>
<tr><td>GU305914.1</td><td>73221</td><td>73243</td><td>-</td><td>0</td><td>CGAATCCTGGGGTCGCGCGT</td><td>CGG</td><td>||||||||[||||||||||||]PAM</td><td>NA</td><td>Intergenic</td><td>NA</td><td>NA</td></tr>
<tr><td>T28695</td><td>GAATCCTGGGGTCGCGCGTCGGG</td><td>1000</td><td></td><td></td><td></td><td></td><td></td><td></td><td></td><td></td><td></td></tr>
<tr><td>Oligo fwd</td><td>CACCGAATCCTGGGGTCGCGCGTC</td><td></td><td></td><td></td><td></td><td></td><td></td><td></td><td></td><td></td><td></td></tr>
<tr><td>Oligo rev</td><td>AAACGACGCGCGACCCCAGGATTC</td><td></td><td></td><td></td><td></td><td></td><td></td><td></td><td></td><td></td><td></td></tr>
<tr><td>Chromosome</td><td>start</td><td>end</td><td>strand</td><td>MM</td><td>target_seq</td><td>PAM</td><td>alignment</td><td>distance</td><td>position</td><td>gene name</td><td>gene id</td></tr>
<tr><td>GU305914.1</td><td>73220</td><td>73242</td><td>-</td><td>0</td><td>GAATCCTGGGGTCGCGCGTC</td><td>GGG</td><td>||||||||[||||||||||||]PAM</td><td>NA</td><td>Intergenic</td><td>NA</td><td>NA</td></tr>
<tr><td>T28696</td><td>AATCCTGGGGTCGCGCGTCGGGG</td><td>1000</td><td></td><td></td><td></td><td></td><td></td><td></td><td></td><td></td><td></td></tr>
<tr><td>Oligo fwd</td><td>CACCAATCCTGGGGTCGCGCGTCG</td><td></td><td></td><td></td><td></td><td></td><td></td><td></td><td></td><td></td><td></td></tr>
<tr><td>Oligo rev</td><td>AAACCGACGCGCGACCCCAGGATT</td><td></td><td></td><td></td><td></td><td></td><td></td><td></td><td></td><td></td><td></td></tr>
<tr><td>Chromosome</td><td>start</td><td>end</td><td>strand</td><td>MM</td><td>target_seq</td><td>PAM</td><td>alignment</td><td>distance</td><td>position</td><td>gene name</td><td>gene id</td></tr>
<tr><td>GU305914.1</td><td>73219</td><td>73241</td><td>-</td><td>0</td><td>AATCCTGGGGTCGCGCGTCG</td><td>GGG</td><td>||||||||[||||||||||||]PAM</td><td>NA</td><td>Intergenic</td><td>NA</td><td>NA</td></tr>
<tr><td>T28697</td><td>TCGGGGTCCCGATCCTCCGCCGG</td><td>1000</td><td></td><td></td><td></td><td></td><td></td><td></td><td></td><td></td><td></td></tr>
<tr><td>Oligo fwd</td><td>CACCTCGGGGTCCCGATCCTCCGC</td><td></td><td></td><td></td><td></td><td></td><td></td><td></td><td></td><td></td><td></td></tr>
<tr><td>Oligo rev</td><td>AAACGCGGAGGATCGGGACCCCGA</td><td></td><td></td><td></td><td></td><td></td><td></td><td></td><td></td><td></td><td></td></tr>
<tr><td>Chromosome</td><td>start</td><td>end</td><td>strand</td><td>MM</td><td>target_seq</td><td>PAM</td><td>alignment</td><td>distance</td><td>position</td><td>gene name</td><td>gene id</td></tr>
<tr><td>GU305914.1</td><td>73202</td><td>73224</td><td>-</td><td>0</td><td>TCGGGGTCCCGATCCTCCGC</td><td>CGG</td><td>||||||||[||||||||||||]PAM</td><td>NA</td><td>Intergenic</td><td>NA</td><td>NA</td></tr>
<tr><td>T28698</td><td>ATCCTCCGCCGGCACCGACGAGG</td><td>1000</td><td></td><td></td><td></td><td></td><td></td><td></td><td></td><td></td><td></td></tr>
<tr><td>Oligo fwd</td><td>CACCATCCTCCGCCGGCACCGACG</td><td></td><td></td><td></td><td></td><td></td><td></td><td></td><td></td><td></td><td></td></tr>
<tr><td>Oligo rev</td><td>AAACCGTCGGTGCCGGCGGAGGAT</td><td></td><td></td><td></td><td></td><td></td><td></td><td></td><td></td><td></td><td></td></tr>
<tr><td>Chromosome</td><td>start</td><td>end</td><td>strand</td><td>MM</td><td>target_seq</td><td>PAM</td><td>alignment</td><td>distance</td><td>position</td><td>gene name</td><td>gene id</td></tr>
<tr><td>GU305914.1</td><td>73191</td><td>73213</td><td>-</td><td>0</td><td>ATCCTCCGCCGGCACCGACG</td><td>AGG</td><td>||||||||[||||||||||||]PAM</td><td>NA</td><td>Intergenic</td><td>NA</td><td>NA</td></tr>
<tr><td>T28699</td><td>CTCCCGACGCTCTCTTCAGTCGG</td><td>1000</td><td></td><td></td><td></td><td></td><td></td><td></td><td></td><td></td><td></td></tr>
<tr><td>Oligo fwd</td><td>CACCCTCCCGACGCTCTCTTCAGT</td><td></td><td></td><td></td><td></td><td></td><td></td><td></td><td></td><td></td><td></td></tr>
<tr><td>Oligo rev</td><td>AAACACTGAAGAGAGCGTCGGGAG</td><td></td><td></td><td></td><td></td><td></td><td></td><td></td><td></td><td></td><td></td></tr>
<tr><td>Chromosome</td><td>start</td><td>end</td><td>strand</td><td>MM</td><td>target_seq</td><td>PAM</td><td>alignment</td><td>distance</td><td>position</td><td>gene name</td><td>gene id</td></tr>
<tr><td>GU305914.1</td><td>73162</td><td>73184</td><td>-</td><td>0</td><td>CTCCCGACGCTCTCTTCAGT</td><td>CGG</td><td>||||||||[||||||||||||]PAM</td><td>NA</td><td>Intergenic</td><td>NA</td><td>NA</td></tr>
<tr><td>T28700</td><td>TCCCGACGCTCTCTTCAGTCGGG</td><td>1000</td><td></td><td></td><td></td><td></td><td></td><td></td><td></td><td></td><td></td></tr>
<tr><td>Oligo fwd</td><td>CACCTCCCGACGCTCTCTTCAGTC</td><td></td><td></td><td></td><td></td><td></td><td></td><td></td><td></td><td></td><td></td></tr>
<tr><td>Oligo rev</td><td>AAACGACTGAAGAGAGCGTCGGGA</td><td></td><td></td><td></td><td></td><td></td><td></td><td></td><td></td><td></td><td></td></tr>
<tr><td>Chromosome</td><td>start</td><td>end</td><td>strand</td><td>MM</td><td>target_seq</td><td>PAM</td><td>alignment</td><td>distance</td><td>position</td><td>gene name</td><td>gene id</td></tr>
<tr><td>GU305914.1</td><td>73161</td><td>73183</td><td>-</td><td>0</td><td>TCCCGACGCTCTCTTCAGTC</td><td>GGG</td><td>||||||||[||||||||||||]PAM</td><td>NA</td><td>Intergenic</td><td>NA</td><td>NA</td></tr>
<tr><td>T28701</td><td>CCCGACGCTCTCTTCAGTCGGGG</td><td>1000</td><td></td><td></td><td></td><td></td><td></td><td></td><td></td><td></td><td></td></tr>
<tr><td>Oligo fwd</td><td>CACCCCCGACGCTCTCTTCAGTCG</td><td></td><td></td><td></td><td></td><td></td><td></td><td></td><td></td><td></td><td></td></tr>
<tr><td>Oligo rev</td><td>AAACCGACTGAAGAGAGCGTCGGG</td><td></td><td></td><td></td><td></td><td></td><td></td><td></td><td></td><td></td><td></td></tr>
<tr><td>Chromosome</td><td>start</td><td>end</td><td>strand</td><td>MM</td><td>target_seq</td><td>PAM</td><td>alignment</td><td>distance</td><td>position</td><td>gene name</td><td>gene id</td></tr>
<tr><td>GU305914.1</td><td>73160</td><td>73182</td><td>-</td><td>0</td><td>CCCGACGCTCTCTTCAGTCG</td><td>GGG</td><td>||||||||[||||||||||||]PAM</td><td>NA</td><td>Intergenic</td><td>NA</td><td>NA</td></tr>
</table>

| | | | | | | | | | | | |
|---|---|---|---|---|---|---|---|---|---|---|---|
| T28702 | GACGCTCTCTTCAGTCGGGGAGG | 1000 | | | | | | | | | |
| Oligo fwd | CACCGACGCTCTCTTCAGTCGGGG | | | | | | | | | | |
| Oligo rev | AAACCCCCGACTGAAGAGAGCGTC | | | | | | | | | | |
| Chromosome | start | end | strand | MM | target_seq | PAM | alignment | distance | position | gene name | gene id |
| GU305914.1 | 73157 | 73179 | - | 0 | GACGCTCTCTTCAGTCGGGG | AGG | \|\|\|\|\|\|\|\|[\|\|\|\|\|\|\|\|\|\|\|\|]PAM | NA | Intergenic | NA | NA |
| GU305914.1 | 18867 | 18889 | - | 4 | GTCGCACTCGTCAGTCGGTG | GGG | \|-\|\|\|-\|\|[\|-\|\|\|\|\|\|\|\|-\|]PAM | NA | Intergenic | NA | NA |
| T28703 | ACGCTCTCTTCAGTCGGGGAGGG | 1000 | | | | | | | | | |
| Oligo fwd | CACCACGCTCTCTTCAGTCGGGGA | | | | | | | | | | |
| Oligo rev | AAACTCCCCGACTGAAGAGAGCGT | | | | | | | | | | |
| Chromosome | start | end | strand | MM | target_seq | PAM | alignment | distance | position | gene name | gene id |
| GU305914.1 | 73156 | 73178 | - | 0 | ACGCTCTCTTCAGTCGGGGA | GGG | \|\|\|\|\|\|\|\|[\|\|\|\|\|\|\|\|\|\|\|\|]PAM | NA | Intergenic | NA | NA |
| T28704 | CGCAGAAAGTCGATGTTGATCGG | 1000 | | | | | | | | | |
| Oligo fwd | CACCCGCAGAAAGTCGATGTTGAT | | | | | | | | | | |
| Oligo rev | AAACATCAACATCGACTTTCTGCG | | | | | | | | | | |
| Chromosome | start | end | strand | MM | target_seq | PAM | alignment | distance | position | gene name | gene id |
| GU305914.1 | 73127 | 73149 | - | 0 | CGCAGAAAGTCGATGTTGAT | CGG | \|\|\|\|\|\|\|\|[\|\|\|\|\|\|\|\|\|\|\|\|]PAM | NA | Intergenic | NA | NA |
| T28705 | GTCGATGTTGATCGGCAACGAGG | 1000 | | | | | | | | | |
| Oligo fwd | CACCGTCGATGTTGATCGGCAACG | | | | | | | | | | |
| Oligo rev | AAACCGTTGCCGATCAACATCGAC | | | | | | | | | | |
| Chromosome | start | end | strand | MM | target_seq | PAM | alignment | distance | position | gene name | gene id |
| GU305914.1 | 73119 | 73141 | - | 0 | GTCGATGTTGATCGGCAACG | AGG | \|\|\|\|\|\|\|\|[\|\|\|\|\|\|\|\|\|\|\|\|]PAM | NA | Intergenic | NA | NA |
| T28706 | GTTGATCGGCAACGAGGTGAAGG | 1000 | | | | | | | | | |
| Oligo fwd | CACCGTTGATCGGCAACGAGGTGA | | | | | | | | | | |
| Oligo rev | AAACTCACCTCGTTGCCGATCAAC | | | | | | | | | | |
| Chromosome | start | end | strand | MM | target_seq | PAM | alignment | distance | position | gene name | gene id |
| GU305914.1 | 73113 | 73135 | - | 0 | GTTGATCGGCAACGAGGTGA | AGG | \|\|\|\|\|\|\|\|[\|\|\|\|\|\|\|\|\|\|\|\|]PAM | NA | Intergenic | NA | NA |
| T28707 | TTGATCGGCAACGAGGTGAAGGG | 1000 | | | | | | | | | |
| Oligo fwd | CACCTTGATCGGCAACGAGGTGAA | | | | | | | | | | |
| Oligo rev | AAACTTCACCTCGTTGCCGATCAA | | | | | | | | | | |
| Chromosome | start | end | strand | MM | target_seq | PAM | alignment | distance | position | gene name | gene id |
| GU305914.1 | 73112 | 73134 | - | 0 | TTGATCGGCAACGAGGTGAA | GGG | \|\|\|\|\|\|\|\|[\|\|\|\|\|\|\|\|\|\|\|\|]PAM | NA | Intergenic | NA | NA |
| T28708 | GGTGAAGGGCGTCTCGCTCGTGG | 1000 | | | | | | | | | |
| Oligo fwd | CACCGGTGAAGGGCGTCTCGCTCG | | | | | | | | | | |
| Oligo rev | AAACCGAGCGAGACGCCCTTCACC | | | | | | | | | | |
| Chromosome | start | end | strand | MM | target_seq | PAM | alignment | distance | position | gene name | gene id |
| GU305914.1 | 73098 | 73120 | - | 0 | GGTGAAGGGCGTCTCGCTCG | TGG | \|\|\|\|\|\|\|\|[\|\|\|\|\|\|\|\|\|\|\|\|]PAM | NA | Intergenic | NA | NA |
| T28709 | GAAGGGCGTCTCGCTCGTGGCGG | 1000 | | | | | | | | | |
| Oligo fwd | CACCGAAGGGCGTCTCGCTCGTGG | | | | | | | | | | |
| Oligo rev | AAACCCACGAGCGAGACGCCCTTC | | | | | | | | | | |
| Chromosome | start | end | strand | MM | target_seq | PAM | alignment | distance | position | gene name | gene id |
| GU305914.1 | 73095 | 73117 | - | 0 | GAAGGGCGTCTCGCTCGTGG | CGG | \|\|\|\|\|\|\|\|[\|\|\|\|\|\|\|\|\|\|\|\|]PAM | NA | Intergenic | NA | NA |
| GU305914.1 | 224303 | 224325 | + | 4 | GATGGCCGTCCCGCTCGTGT | AGG | \|\|-\|\|-\|\|[\|\|-\|\|\|\|\|\|\|\|-]PAM | NA | Intergenic | NA | NA |
| T28710 | CATCGTCGCCCTCGTCCTCGTGG | 1000 | | | | | | | | | |
| Oligo fwd | CACCCATCGTCGCCCTCGTCCTCG | | | | | | | | | | |
| Oligo rev | AAACCGAGGACGAGGGCGACGATG | | | | | | | | | | |
| Chromosome | start | end | strand | MM | target_seq | PAM | alignment | distance | position | gene name | gene id |
| GU305914.1 | 73072 | 73094 | - | 0 | CATCGTCGCCCTCGTCCTCG | TGG | \|\|\|\|\|\|\|\|[\|\|\|\|\|\|\|\|\|\|\|\|]PAM | NA | Intergenic | NA | NA |
| T28711 | ATCGTCGCCCTCGTCCTCGTGGG | 1000 | | | | | | | | | |
| Oligo fwd | CACCATCGTCGCCCTCGTCCTCGT | | | | | | | | | | |
| Oligo rev | AAACACGAGGACGAGGGCGACGAT | | | | | | | | | | |
| Chromosome | start | end | strand | MM | target_seq | PAM | alignment | distance | position | gene name | gene id |
| GU305914.1 | 73071 | 73093 | - | 0 | ATCGTCGCCCTCGTCCTCGT | GGG | \|\|\|\|\|\|\|\|[\|\|\|\|\|\|\|\|\|\|\|\|]PAM | NA | Intergenic | NA | NA |
| GU305914.1 | 112237 | 112259 | - | 4 | CTTGTCGTCGTCGTCCTCGT | CGG | -\|-\|\|\|\|-[\|-\|\|\|\|\|\|\|\|\|\|]PAM | NA | Intergenic | NA | NA |
| GU305914.1 | 27140 | 27162 | - | 3 | ATCGTCGTCATCGTCGTCGT | TGG | \|\|\|\|\|\|\|-[\|-\|\|\|\|\|-\|\|\|\|]PAM | NA | Intergenic | NA | NA |
| T28712 | TCGTCGCCCTCGTCCTCGTGGGG | 1000 | | | | | | | | | |
| Oligo fwd | CACCTCGTCGCCCTCGTCCTCGTG | | | | | | | | | | |
| Oligo rev | AAACCACGAGGACGAGGGCGACGA | | | | | | | | | | |
| Chromosome | start | end | strand | MM | target_seq | PAM | alignment | distance | position | gene name | gene id |
| GU305914.1 | 73070 | 73092 | - | 0 | TCGTCGCCCTCGTCCTCGTG | GGG | \|\|\|\|\|\|\|\|[\|\|\|\|\|\|\|\|\|\|\|\|]PAM | NA | Intergenic | NA | NA |
| GU305914.1 | 112236 | 112258 | - | 4 | TTGTCGTCGTCGTCCTCGTC | GGG | \|-\|\|\|\|-\|[-\|\|\|\|\|\|\|\|\|\|-]PAM | NA | Intergenic | NA | NA |
| GU305914.1 | 45183 | 45205 | + | 4 | TCGTCGGACTCGTCCACGGG | TGG | \|\|\|\|\|\|--[\|\|\|\|\|\|\|-\|\|-\|]PAM | NA | Intergenic | NA | NA |
| T28713 | CGTCGCCCTCGTCCTCGTGGGGG | 1000 | | | | | | | | | |
| Oligo fwd | CACCCGTCGCCCTCGTCCTCGTGG | | | | | | | | | | |
| Oligo rev | AAACCCACGAGGACGAGGGCGACG | | | | | | | | | | |
| Chromosome | start | end | strand | MM | target_seq | PAM | alignment | distance | position | gene name | gene id |
| GU305914.1 | 73069 | 73091 | - | 0 | CGTCGCCCTCGTCCTCGTGG | GGG | \|\|\|\|\|\|\|\|[\|\|\|\|\|\|\|\|\|\|\|\|]PAM | NA | Intergenic | NA | NA |
| T28714 | ATCATCCGCATCGTCGAGAACGG | 1000 | | | | | | | | | |
| Oligo fwd | CACCATCATCCGCATCGTCGAGAA | | | | | | | | | | |
| Oligo rev | AAACTTCTCGACGATGCGGATGAT | | | | | | | | | | |
| Chromosome | start | end | strand | MM | target_seq | PAM | alignment | distance | position | gene name | gene id |
| GU305914.1 | 73046 | 73068 | - | 0 | ATCATCCGCATCGTCGAGAA | CGG | \|\|\|\|\|\|\|\|[\|\|\|\|\|\|\|\|\|\|\|\|]PAM | NA | Intergenic | NA | NA |
| T28715 | TCATCCGCATCGTCGAGAACGGG | 1000 | | | | | | | | | |
| Oligo fwd | CACCTCATCCGCATCGTCGAGAAC | | | | | | | | | | |
| Oligo rev | AAACGTTCTCGACGATGCGGATGA | | | | | | | | | | |
| Chromosome | start | end | strand | MM | target_seq | PAM | alignment | distance | position | gene name | gene id |

| | | | | | | | | | | | |
|---|---|---|---|---|---|---|---|---|---|---|---|
| GU305914.1 | 73045 | 73067 | - | 0 | TCATCCGCATCGTCGAGAAC | GGG | \|\|\|\|\|\|\|\|[\|\|\|\|\|\|\|\|\|\|\|\|]PAM | NA | Intergenic | NA | NA |
| T28716 | AGAACGGGTTCTTGACCGTTAGG | 1000 | | | | | | | | | |
| Oligo fwd | CACCAGAACGGGTTCTTGACCGTT | | | | | | | | | | |
| Oligo rev | AAACAACGGTCAAGAACCCGTTCT | | | | | | | | | | |
| Chromosome | start | end | strand | MM | target_seq | PAM | alignment | distance | position | gene name | gene id |
| GU305914.1 | 73030 | 73052 | - | 0 | AGAACGGGTTCTTGACCGTT | AGG | \|\|\|\|\|\|\|\|[\|\|\|\|\|\|\|\|\|\|\|\|]PAM | NA | Intergenic | NA | NA |
| GU305914.1 | 58946 | 58968 | + | 4 | TGAACTGGTTCTTGACGGTC | TGG | -\|\|\|\|-\|\|[\|\|\|\|\|\|\|\|-\|\|-]PAM | NA | Intergenic | NA | NA |
| T28717 | ACGGGTTCTTGACCGTTAGGTGG | 1000 | | | | | | | | | |
| Oligo fwd | CACCACGGGTTCTTGACCGTTAGG | | | | | | | | | | |
| Oligo rev | AAACCCTAACGGTCAAGAACCCGT | | | | | | | | | | |
| Chromosome | start | end | strand | MM | target_seq | PAM | alignment | distance | position | gene name | gene id |
| GU305914.1 | 73027 | 73049 | - | 0 | ACGGGTTCTTGACCGTTAGG | TGG | \|\|\|\|\|\|\|\|[\|\|\|\|\|\|\|\|\|\|\|\|]PAM | NA | Intergenic | NA | NA |
| T28718 | AGGTGGACCCGACTCGTCGCCGG | 1000 | | | | | | | | | |
| Oligo fwd | CACCAGGTGGACCCGACTCGTCGC | | | | | | | | | | |
| Oligo rev | AAACGCGACGAGTCGGGTCCACCT | | | | | | | | | | |
| Chromosome | start | end | strand | MM | target_seq | PAM | alignment | distance | position | gene name | gene id |
| GU305914.1 | 73010 | 73032 | - | 0 | AGGTGGACCCGACTCGTCGC | CGG | \|\|\|\|\|\|\|\|[\|\|\|\|\|\|\|\|\|\|\|\|]PAM | NA | Intergenic | NA | NA |
| T28719 | GGTGGACCCGACTCGTCGCCGGG | 1000 | | | | | | | | | |
| Oligo fwd | CACCGGTGGACCCGACTCGTCGCC | | | | | | | | | | |
| Oligo rev | AAACGGCGACGAGTCGGGTCCACC | | | | | | | | | | |
| Chromosome | start | end | strand | MM | target_seq | PAM | alignment | distance | position | gene name | gene id |
| GU305914.1 | 73009 | 73031 | - | 0 | GGTGGACCCGACTCGTCGCC | GGG | \|\|\|\|\|\|\|\|[\|\|\|\|\|\|\|\|\|\|\|\|]PAM | NA | Intergenic | NA | NA |
| T28720 | GTGGACCCGACTCGTCGCCGGGG | 1000 | | | | | | | | | |
| Oligo fwd | CACCGTGGACCCGACTCGTCGCCG | | | | | | | | | | |
| Oligo rev | AAACCGGCGACGAGTCGGGTCCAC | | | | | | | | | | |
| Chromosome | start | end | strand | MM | target_seq | PAM | alignment | distance | position | gene name | gene id |
| GU305914.1 | 73008 | 73030 | - | 0 | GTGGACCCGACTCGTCGCCG | GGG | \|\|\|\|\|\|\|\|[\|\|\|\|\|\|\|\|\|\|\|\|]PAM | NA | Intergenic | NA | NA |
| T28721 | TGGACCCGACTCGTCGCCGGGGG | 1000 | | | | | | | | | |
| Oligo fwd | CACCTGGACCCGACTCGTCGCCGG | | | | | | | | | | |
| Oligo rev | AAACCCGGCGACGAGTCGGGTCCA | | | | | | | | | | |
| Chromosome | start | end | strand | MM | target_seq | PAM | alignment | distance | position | gene name | gene id |
| GU305914.1 | 73007 | 73029 | - | 0 | TGGACCCGACTCGTCGCCGG | GGG | \|\|\|\|\|\|\|\|[\|\|\|\|\|\|\|\|\|\|\|\|]PAM | NA | Intergenic | NA | NA |
| T28722 | GTCGCCGGGGGCGCCTTCAGCGG | 1000 | | | | | | | | | |
| Oligo fwd | CACCGTCGCCGGGGGCGCCTTCAG | | | | | | | | | | |
| Oligo rev | AAACCTGAAGGCGCCCCCGGCGAC | | | | | | | | | | |
| Chromosome | start | end | strand | MM | target_seq | PAM | alignment | distance | position | gene name | gene id |
| GU305914.1 | 72995 | 73017 | - | 0 | GTCGCCGGGGGCGCCTTCAG | CGG | \|\|\|\|\|\|\|\|[\|\|\|\|\|\|\|\|\|\|\|\|]PAM | NA | Intergenic | NA | NA |
| GU305914.1 | 169712 | 169734 | + | 4 | TTCGTCCGGGGCGCCTTCCG | CGG | -\|\|\|-\|-\|[\|\|\|\|\|\|\|\|\|\|-\|]PAM | NA | Intergenic | NA | NA |
| T28723 | TCGCCGGGGGCGCCTTCAGCGGG | 1000 | | | | | | | | | |
| Oligo fwd | CACCTCGCCGGGGGCGCCTTCAGC | | | | | | | | | | |
| Oligo rev | AAACGCTGAAGGCGCCCCCGGCGA | | | | | | | | | | |
| Chromosome | start | end | strand | MM | target_seq | PAM | alignment | distance | position | gene name | gene id |
| GU305914.1 | 72994 | 73016 | - | 0 | TCGCCGGGGGCGCCTTCAGC | GGG | \|\|\|\|\|\|\|\|[\|\|\|\|\|\|\|\|\|\|\|\|]PAM | NA | Intergenic | NA | NA |
| T28724 | CGCCGGGGGCGCCTTCAGCGGGG | 1000 | | | | | | | | | |
| Oligo fwd | CACCCGCCGGGGGCGCCTTCAGCG | | | | | | | | | | |
| Oligo rev | AAACCGCTGAAGGCGCCCCCGGCG | | | | | | | | | | |
| Chromosome | start | end | strand | MM | target_seq | PAM | alignment | distance | position | gene name | gene id |
| GU305914.1 | 72993 | 73015 | - | 0 | CGCCGGGGGCGCCTTCAGCG | GGG | \|\|\|\|\|\|\|\|[\|\|\|\|\|\|\|\|\|\|\|\|]PAM | NA | Intergenic | NA | NA |
| T28725 | GCCGGGGGCGCCTTCAGCGGGGG | 1000 | | | | | | | | | |
| Oligo fwd | CACCGCCGGGGGCGCCTTCAGCGG | | | | | | | | | | |
| Oligo rev | AAACCCGCTGAAGGCGCCCCCGGC | | | | | | | | | | |
| Chromosome | start | end | strand | MM | target_seq | PAM | alignment | distance | position | gene name | gene id |
| GU305914.1 | 72992 | 73014 | - | 0 | GCCGGGGGCGCCTTCAGCGG | GGG | \|\|\|\|\|\|\|\|[\|\|\|\|\|\|\|\|\|\|\|\|]PAM | NA | Intergenic | NA | NA |
| T28726 | CATCGCGTTCACCGCGTAGTTGG | 1000 | | | | | | | | | |
| Oligo fwd | CACCCATCGCGTTCACCGCGTAGT | | | | | | | | | | |
| Oligo rev | AAACACTACGCGGTGAACGCGATG | | | | | | | | | | |
| Chromosome | start | end | strand | MM | target_seq | PAM | alignment | distance | position | gene name | gene id |
| GU305914.1 | 72966 | 72988 | - | 0 | CATCGCGTTCACCGCGTAGT | TGG | \|\|\|\|\|\|\|\|[\|\|\|\|\|\|\|\|\|\|\|\|]PAM | NA | Intergenic | NA | NA |
| T28727 | GTTCACCGCGTAGTTGGTGTTGG | 1000 | | | | | | | | | |
| Oligo fwd | CACCGTTCACCGCGTAGTTGGTGT | | | | | | | | | | |
| Oligo rev | AAACACACCAACTACGCGGTGAAC | | | | | | | | | | |
| Chromosome | start | end | strand | MM | target_seq | PAM | alignment | distance | position | gene name | gene id |
| GU305914.1 | 72960 | 72982 | - | 0 | GTTCACCGCGTAGTTGGTGT | TGG | \|\|\|\|\|\|\|\|[\|\|\|\|\|\|\|\|\|\|\|\|]PAM | NA | Intergenic | NA | NA |
| T28728 | TTGGAGTCGTACACCACCAACGG | 1000 | | | | | | | | | |
| Oligo fwd | CACCTTGGAGTCGTACACCACCAA | | | | | | | | | | |
| Oligo rev | AAACTTGGTGGTGTACGACTCCAA | | | | | | | | | | |
| Chromosome | start | end | strand | MM | target_seq | PAM | alignment | distance | position | gene name | gene id |
| GU305914.1 | 72941 | 72963 | - | 0 | TTGGAGTCGTACACCACCAA | CGG | \|\|\|\|\|\|\|\|[\|\|\|\|\|\|\|\|\|\|\|\|]PAM | NA | Intergenic | NA | NA |
| T28729 | AACGGCGTTCTGTCCTGCTGTGG | 1000 | | | | | | | | | |
| Oligo fwd | CACCAACGGCGTTCTGTCCTGCTG | | | | | | | | | | |
| Oligo rev | AAACCAGCAGGACAGAACGCCGTT | | | | | | | | | | |
| Chromosome | start | end | strand | MM | target_seq | PAM | alignment | distance | position | gene name | gene id |
| GU305914.1 | 72923 | 72945 | - | 0 | AACGGCGTTCTGTCCTGCTG | TGG | \|\|\|\|\|\|\|\|[\|\|\|\|\|\|\|\|\|\|\|\|]PAM | NA | Intergenic | NA | NA |
| T28730 | ACGGCGTTCTGTCCTGCTGTGGG | 1000 | | | | | | | | | |

<table>
<tr><td>Oligo fwd</td><td>CACCACGGCGTTCTGTCCTGCTGT</td></tr>
<tr><td>Oligo rev</td><td>AAACACAGCAGGACAGAACGCCGT</td></tr>
<tr><td>Chromosome</td><td>start</td><td></td><td>end</td><td>strand</td><td>MM</td><td>target_seq</td><td>PAM</td><td>alignment</td><td>distance</td><td>position</td><td>gene name</td><td>gene id</td></tr>
<tr><td>GU305914.1</td><td></td><td>72922</td><td>72944</td><td>-</td><td>0</td><td>ACGGCGTTCTGTCCTGCTGT</td><td>GGG</td><td>||||||||[||||||||||||]PAM</td><td>NA</td><td>Intergenic</td><td>NA</td><td>NA</td></tr>
<tr><td></td></tr>
<tr><td>T28731</td><td>CGGCGTTCTGTCCTGCTGTGGGG</td><td></td><td>1000</td></tr>
<tr><td>Oligo fwd</td><td>CACCCGGCGTTCTGTCCTGCTGTG</td></tr>
<tr><td>Oligo rev</td><td>AAACCACAGCAGGACAGAACGCCG</td></tr>
<tr><td>Chromosome</td><td>start</td><td></td><td>end</td><td>strand</td><td>MM</td><td>target_seq</td><td>PAM</td><td>alignment</td><td>distance</td><td>position</td><td>gene name</td><td>gene id</td></tr>
<tr><td>GU305914.1</td><td></td><td>72921</td><td>72943</td><td>-</td><td>0</td><td>CGGCGTTCTGTCCTGCTGTG</td><td>GGG</td><td>||||||||[||||||||||||]PAM</td><td>NA</td><td>Intergenic</td><td>NA</td><td>NA</td></tr>
<tr><td></td></tr>
<tr><td>T28732</td><td>GGCGTTCTGTCCTGCTGTGGGGG</td><td></td><td>1000</td></tr>
<tr><td>Oligo fwd</td><td>CACCGGCGTTCTGTCCTGCTGTGG</td></tr>
<tr><td>Oligo rev</td><td>AAACCCACAGCAGGACAGAACGCC</td></tr>
<tr><td>Chromosome</td><td>start</td><td></td><td>end</td><td>strand</td><td>MM</td><td>target_seq</td><td>PAM</td><td>alignment</td><td>distance</td><td>position</td><td>gene name</td><td>gene id</td></tr>
<tr><td>GU305914.1</td><td></td><td>72920</td><td>72942</td><td>-</td><td>0</td><td>GGCGTTCTGTCCTGCTGTGG</td><td>GGG</td><td>||||||||[||||||||||||]PAM</td><td>NA</td><td>Intergenic</td><td>NA</td><td>NA</td></tr>
<tr><td></td></tr>
<tr><td>T28733</td><td>GCGTTCTGTCCTGCTGTGGGGGG</td><td></td><td>1000</td></tr>
<tr><td>Oligo fwd</td><td>CACCGCGTTCTGTCCTGCTGTGGG</td></tr>
<tr><td>Oligo rev</td><td>AAACCCCACAGCAGGACAGAACGC</td></tr>
<tr><td>Chromosome</td><td>start</td><td></td><td>end</td><td>strand</td><td>MM</td><td>target_seq</td><td>PAM</td><td>alignment</td><td>distance</td><td>position</td><td>gene name</td><td>gene id</td></tr>
<tr><td>GU305914.1</td><td></td><td>72919</td><td>72941</td><td>-</td><td>0</td><td>GCGTTCTGTCCTGCTGTGGG</td><td>GGG</td><td>||||||||[||||||||||||]PAM</td><td>NA</td><td>Intergenic</td><td>NA</td><td>NA</td></tr>
<tr><td></td></tr>
<tr><td>T28734</td><td>GCTGTGGGGGGTTAGTCAGCAGG</td><td></td><td>1000</td></tr>
<tr><td>Oligo fwd</td><td>CACCGCTGTGGGGGGTTAGTCAGC</td></tr>
<tr><td>Oligo rev</td><td>AAACGCTGACTAACCCCCCACAGC</td></tr>
<tr><td>Chromosome</td><td>start</td><td></td><td>end</td><td>strand</td><td>MM</td><td>target_seq</td><td>PAM</td><td>alignment</td><td>distance</td><td>position</td><td>gene name</td><td>gene id</td></tr>
<tr><td>GU305914.1</td><td></td><td>72907</td><td>72929</td><td>-</td><td>0</td><td>GCTGTGGGGGGTTAGTCAGC</td><td>AGG</td><td>||||||||[||||||||||||]PAM</td><td>NA</td><td>Intergenic</td><td>NA</td><td>NA</td></tr>
<tr><td></td></tr>
<tr><td>T28735</td><td>GTGGGGGGTTAGTCAGCAGGTGG</td><td></td><td>1000</td></tr>
<tr><td>Oligo fwd</td><td>CACCGTGGGGGGTTAGTCAGCAGG</td></tr>
<tr><td>Oligo rev</td><td>AAACCCTGCTGACTAACCCCCCAC</td></tr>
<tr><td>Chromosome</td><td>start</td><td></td><td>end</td><td>strand</td><td>MM</td><td>target_seq</td><td>PAM</td><td>alignment</td><td>distance</td><td>position</td><td>gene name</td><td>gene id</td></tr>
<tr><td>GU305914.1</td><td></td><td>72904</td><td>72926</td><td>-</td><td>0</td><td>GTGGGGGGTTAGTCAGCAGG</td><td>TGG</td><td>||||||||[||||||||||||]PAM</td><td>NA</td><td>Intergenic</td><td>NA</td><td>NA</td></tr>
<tr><td></td></tr>
<tr><td>T28736</td><td>ATCCTCTTGCAGCGAGAACCAGG</td><td></td><td>1000</td></tr>
<tr><td>Oligo fwd</td><td>CACCATCCTCTTGCAGCGAGAACC</td></tr>
<tr><td>Oligo rev</td><td>AAACGGTTCTCGCTGCAAGAGGAT</td></tr>
<tr><td>Chromosome</td><td>start</td><td></td><td>end</td><td>strand</td><td>MM</td><td>target_seq</td><td>PAM</td><td>alignment</td><td>distance</td><td>position</td><td>gene name</td><td>gene id</td></tr>
<tr><td>GU305914.1</td><td></td><td>72873</td><td>72895</td><td>-</td><td>0</td><td>ATCCTCTTGCAGCGAGAACC</td><td>AGG</td><td>||||||||[||||||||||||]PAM</td><td>NA</td><td>Intergenic</td><td>NA</td><td>NA</td></tr>
<tr><td></td></tr>
<tr><td>T28737</td><td>CGAGAACCAGGAGCGTTCGACGG</td><td></td><td>1000</td></tr>
<tr><td>Oligo fwd</td><td>CACCCGAGAACCAGGAGCGTTCGA</td></tr>
<tr><td>Oligo rev</td><td>AAACTCGAACGCTCCTGGTTCTCG</td></tr>
<tr><td>Chromosome</td><td>start</td><td></td><td>end</td><td>strand</td><td>MM</td><td>target_seq</td><td>PAM</td><td>alignment</td><td>distance</td><td>position</td><td>gene name</td><td>gene id</td></tr>
<tr><td>GU305914.1</td><td></td><td>72861</td><td>72883</td><td>-</td><td>0</td><td>CGAGAACCAGGAGCGTTCGA</td><td>CGG</td><td>||||||||[||||||||||||]PAM</td><td>NA</td><td>Intergenic</td><td>NA</td><td>NA</td></tr>
<tr><td></td></tr>
<tr><td>T28738</td><td>GAGAACCAGGAGCGTTCGACGGG</td><td></td><td>1000</td></tr>
<tr><td>Oligo fwd</td><td>CACCGAGAACCAGGAGCGTTCGAC</td></tr>
<tr><td>Oligo rev</td><td>AAACGTCGAACGCTCCTGGTTCTC</td></tr>
<tr><td>Chromosome</td><td>start</td><td></td><td>end</td><td>strand</td><td>MM</td><td>target_seq</td><td>PAM</td><td>alignment</td><td>distance</td><td>position</td><td>gene name</td><td>gene id</td></tr>
<tr><td>GU305914.1</td><td></td><td>72860</td><td>72882</td><td>-</td><td>0</td><td>GAGAACCAGGAGCGTTCGAC</td><td>GGG</td><td>||||||||[||||||||||||]PAM</td><td>NA</td><td>Intergenic</td><td>NA</td><td>NA</td></tr>
<tr><td></td></tr>
<tr><td>T28739</td><td>AGAACCAGGAGCGTTCGACGGGG</td><td></td><td>1000</td></tr>
<tr><td>Oligo fwd</td><td>CACCAGAACCAGGAGCGTTCGACG</td></tr>
<tr><td>Oligo rev</td><td>AAACCGTCGAACGCTCCTGGTTCT</td></tr>
<tr><td>Chromosome</td><td>start</td><td></td><td>end</td><td>strand</td><td>MM</td><td>target_seq</td><td>PAM</td><td>alignment</td><td>distance</td><td>position</td><td>gene name</td><td>gene id</td></tr>
<tr><td>GU305914.1</td><td></td><td>72859</td><td>72881</td><td>-</td><td>0</td><td>AGAACCAGGAGCGTTCGACG</td><td>GGG</td><td>||||||||[||||||||||||]PAM</td><td>NA</td><td>Intergenic</td><td>NA</td><td>NA</td></tr>
<tr><td></td></tr>
<tr><td>T28740</td><td>AGGAGCGTTCGACGGGGTTTTGG</td><td></td><td>1000</td></tr>
<tr><td>Oligo fwd</td><td>CACCAGGAGCGTTCGACGGGGTTT</td></tr>
<tr><td>Oligo rev</td><td>AAACAAACCCCGTCGAACGCTCCT</td></tr>
<tr><td>Chromosome</td><td>start</td><td></td><td>end</td><td>strand</td><td>MM</td><td>target_seq</td><td>PAM</td><td>alignment</td><td>distance</td><td>position</td><td>gene name</td><td>gene id</td></tr>
<tr><td>GU305914.1</td><td></td><td>72853</td><td>72875</td><td>-</td><td>0</td><td>AGGAGCGTTCGACGGGGTTT</td><td>TGG</td><td>||||||||[||||||||||||]PAM</td><td>NA</td><td>Intergenic</td><td>NA</td><td>NA</td></tr>
<tr><td></td></tr>
<tr><td>T28741</td><td>GCGTTCGACGGGGTTTTGGACGG</td><td></td><td>1000</td></tr>
<tr><td>Oligo fwd</td><td>CACCGCGTTCGACGGGGTTTTGGA</td></tr>
<tr><td>Oligo rev</td><td>AAACTCCAAAACCCCGTCGAACGC</td></tr>
<tr><td>Chromosome</td><td>start</td><td></td><td>end</td><td>strand</td><td>MM</td><td>target_seq</td><td>PAM</td><td>alignment</td><td>distance</td><td>position</td><td>gene name</td><td>gene id</td></tr>
<tr><td>GU305914.1</td><td></td><td>72849</td><td>72871</td><td>-</td><td>0</td><td>GCGTTCGACGGGGTTTTGGA</td><td>CGG</td><td>||||||||[||||||||||||]PAM</td><td>NA</td><td>Intergenic</td><td>NA</td><td>NA</td></tr>
<tr><td></td></tr>
<tr><td>T28742</td><td>CGGTGTATGACCATCCGAAGCGG</td><td></td><td>1000</td></tr>
<tr><td>Oligo fwd</td><td>CACCCGGTGTATGACCATCCGAAG</td></tr>
<tr><td>Oligo rev</td><td>AAACCTTCGGATGGTCATACACCG</td></tr>
<tr><td>Chromosome</td><td>start</td><td></td><td>end</td><td>strand</td><td>MM</td><td>target_seq</td><td>PAM</td><td>alignment</td><td>distance</td><td>position</td><td>gene name</td><td>gene id</td></tr>
<tr><td>GU305914.1</td><td></td><td>72829</td><td>72851</td><td>-</td><td>0</td><td>CGGTGTATGACCATCCGAAG</td><td>CGG</td><td>||||||||[||||||||||||]PAM</td><td>NA</td><td>Intergenic</td><td>NA</td><td>NA</td></tr>
<tr><td></td></tr>
<tr><td>T28743</td><td>ATCCGAAGCGGTCGAAGCTCAGG</td><td></td><td>1000</td></tr>
<tr><td>Oligo fwd</td><td>CACCATCCGAAGCGGTCGAAGCTC</td></tr>
<tr><td>Oligo rev</td><td>AAACGAGCTTCGACCGCTTCGGAT</td></tr>
<tr><td>Chromosome</td><td>start</td><td></td><td>end</td><td>strand</td><td>MM</td><td>target_seq</td><td>PAM</td><td>alignment</td><td>distance</td><td>position</td><td>gene name</td><td>gene id</td></tr>
<tr><td>GU305914.1</td><td></td><td>72817</td><td>72839</td><td>-</td><td>0</td><td>ATCCGAAGCGGTCGAAGCTC</td><td>AGG</td><td>||||||||[||||||||||||]PAM</td><td>NA</td><td>Intergenic</td><td>NA</td><td>NA</td></tr>
<tr><td></td></tr>
<tr><td>T28744</td><td>CGCACCACCTTGACGAAGTCCGG</td><td></td><td>1000</td></tr>
<tr><td>Oligo fwd</td><td>CACCCGCACCACCTTGACGAAGTC</td></tr>
<tr><td>Oligo rev</td><td>AAACGACTTCGTCAAGGTGGTGCG</td></tr>
<tr><td>Chromosome</td><td>start</td><td></td><td>end</td><td>strand</td><td>MM</td><td>target_seq</td><td>PAM</td><td>alignment</td><td>distance</td><td>position</td><td>gene name</td><td>gene id</td></tr>
<tr><td>GU305914.1</td><td></td><td>72785</td><td>72807</td><td>-</td><td>0</td><td>CGCACCACCTTGACGAAGTC</td><td>CGG</td><td>||||||||[||||||||||||]PAM</td><td>NA</td><td>Intergenic</td><td>NA</td><td>NA</td></tr>
</table>

<table>
<tr><td>T28745</td><td>GCACCACCTTGACGAAGTCCGGG</td><td>1000</td><td></td><td></td><td></td><td></td><td></td><td></td><td></td><td></td><td></td></tr>
<tr><td>Oligo fwd</td><td>CACCGCACCACCTTGACGAAGTCC</td><td></td><td></td><td></td><td></td><td></td><td></td><td></td><td></td><td></td><td></td></tr>
<tr><td>Oligo rev</td><td>AAACGGACTTCGTCAAGGTGGTGC</td><td></td><td></td><td></td><td></td><td></td><td></td><td></td><td></td><td></td><td></td></tr>
<tr><td>Chromosome</td><td>start</td><td>end</td><td>strand</td><td>MM</td><td>target_seq</td><td>PAM</td><td>alignment</td><td>distance</td><td>position</td><td>gene name</td><td>gene id</td></tr>
<tr><td>GU305914.1</td><td>72784</td><td>72806</td><td>-</td><td>0</td><td>GCACCACCTTGACGAAGTCC</td><td>GGG</td><td>||||||||[||||||||||||]PAM</td><td>NA</td><td>Intergenic</td><td>NA</td><td>NA</td></tr>
<tr><td>T28746</td><td>TTGACGAAGTCCGGGTCGCTCGG</td><td>1000</td><td></td><td></td><td></td><td></td><td></td><td></td><td></td><td></td><td></td></tr>
<tr><td>Oligo fwd</td><td>CACCTTGACGAAGTCCGGGTCGCT</td><td></td><td></td><td></td><td></td><td></td><td></td><td></td><td></td><td></td><td></td></tr>
<tr><td>Oligo rev</td><td>AAACAGCGACCCGGACTTCGTCAA</td><td></td><td></td><td></td><td></td><td></td><td></td><td></td><td></td><td></td><td></td></tr>
<tr><td>Chromosome</td><td>start</td><td>end</td><td>strand</td><td>MM</td><td>target_seq</td><td>PAM</td><td>alignment</td><td>distance</td><td>position</td><td>gene name</td><td>gene id</td></tr>
<tr><td>GU305914.1</td><td>72776</td><td>72798</td><td>-</td><td>0</td><td>TTGACGAAGTCCGGGTCGCT</td><td>CGG</td><td>||||||||[||||||||||||]PAM</td><td>NA</td><td>Intergenic</td><td>NA</td><td>NA</td></tr>
<tr><td>T28747</td><td>CGAAGTCCGGGTCGCTCGGCAGG</td><td>1000</td><td></td><td></td><td></td><td></td><td></td><td></td><td></td><td></td><td></td></tr>
<tr><td>Oligo fwd</td><td>CACCCGAAGTCCGGGTCGCTCGGC</td><td></td><td></td><td></td><td></td><td></td><td></td><td></td><td></td><td></td><td></td></tr>
<tr><td>Oligo rev</td><td>AAACGCCGAGCGACCCGGACTTCG</td><td></td><td></td><td></td><td></td><td></td><td></td><td></td><td></td><td></td><td></td></tr>
<tr><td>Chromosome</td><td>start</td><td>end</td><td>strand</td><td>MM</td><td>target_seq</td><td>PAM</td><td>alignment</td><td>distance</td><td>position</td><td>gene name</td><td>gene id</td></tr>
<tr><td>GU305914.1</td><td>72772</td><td>72794</td><td>-</td><td>0</td><td>CGAAGTCCGGGTCGCTCGGC</td><td>AGG</td><td>||||||||[||||||||||||]PAM</td><td>NA</td><td>Intergenic</td><td>NA</td><td>NA</td></tr>
<tr><td>T28748</td><td>GGTCGCTCGGCAGGTTCTCCAGG</td><td>1000</td><td></td><td></td><td></td><td></td><td></td><td></td><td></td><td></td><td></td></tr>
<tr><td>Oligo fwd</td><td>CACCGGTCGCTCGGCAGGTTCTCC</td><td></td><td></td><td></td><td></td><td></td><td></td><td></td><td></td><td></td><td></td></tr>
<tr><td>Oligo rev</td><td>AAACGGAGAACCTGCCGAGCGACC</td><td></td><td></td><td></td><td></td><td></td><td></td><td></td><td></td><td></td><td></td></tr>
<tr><td>Chromosome</td><td>start</td><td>end</td><td>strand</td><td>MM</td><td>target_seq</td><td>PAM</td><td>alignment</td><td>distance</td><td>position</td><td>gene name</td><td>gene id</td></tr>
<tr><td>GU305914.1</td><td>72763</td><td>72785</td><td>-</td><td>0</td><td>GGTCGCTCGGCAGGTTCTCC</td><td>AGG</td><td>||||||||[||||||||||||]PAM</td><td>NA</td><td>Intergenic</td><td>NA</td><td>NA</td></tr>
<tr><td>T28749</td><td>CCAGGTGCCGCACGATCAGCCGG</td><td>1000</td><td></td><td></td><td></td><td></td><td></td><td></td><td></td><td></td><td></td></tr>
<tr><td>Oligo fwd</td><td>CACCCCAGGTGCCGCACGATCAGC</td><td></td><td></td><td></td><td></td><td></td><td></td><td></td><td></td><td></td><td></td></tr>
<tr><td>Oligo rev</td><td>AAACGCTGATCGTGCGGCACCTGG</td><td></td><td></td><td></td><td></td><td></td><td></td><td></td><td></td><td></td><td></td></tr>
<tr><td>Chromosome</td><td>start</td><td>end</td><td>strand</td><td>MM</td><td>target_seq</td><td>PAM</td><td>alignment</td><td>distance</td><td>position</td><td>gene name</td><td>gene id</td></tr>
<tr><td>GU305914.1</td><td>72745</td><td>72767</td><td>-</td><td>0</td><td>CCAGGTGCCGCACGATCAGC</td><td>CGG</td><td>||||||||[||||||||||||]PAM</td><td>NA</td><td>Intergenic</td><td>NA</td><td>NA</td></tr>
<tr><td>T28750</td><td>GCCGCACGATCAGCCGGTATCGG</td><td>1000</td><td></td><td></td><td></td><td></td><td></td><td></td><td></td><td></td><td></td></tr>
<tr><td>Oligo fwd</td><td>CACCGCCGCACGATCAGCCGGTAT</td><td></td><td></td><td></td><td></td><td></td><td></td><td></td><td></td><td></td><td></td></tr>
<tr><td>Oligo rev</td><td>AAACATACCGGCTGATCGTGCGGC</td><td></td><td></td><td></td><td></td><td></td><td></td><td></td><td></td><td></td><td></td></tr>
<tr><td>Chromosome</td><td>start</td><td>end</td><td>strand</td><td>MM</td><td>target_seq</td><td>PAM</td><td>alignment</td><td>distance</td><td>position</td><td>gene name</td><td>gene id</td></tr>
<tr><td>GU305914.1</td><td>72739</td><td>72761</td><td>-</td><td>0</td><td>GCCGCACGATCAGCCGGTAT</td><td>CGG</td><td>||||||||[||||||||||||]PAM</td><td>NA</td><td>Intergenic</td><td>NA</td><td>NA</td></tr>
<tr><td>T28751</td><td>CGTTTACGATGTTCGTGACATGG</td><td>1000</td><td></td><td></td><td></td><td></td><td></td><td></td><td></td><td></td><td></td></tr>
<tr><td>Oligo fwd</td><td>CACCCGTTTACGATGTTCGTGACA</td><td></td><td></td><td></td><td></td><td></td><td></td><td></td><td></td><td></td><td></td></tr>
<tr><td>Oligo rev</td><td>AAACTGTCACGAACATCGTAAACG</td><td></td><td></td><td></td><td></td><td></td><td></td><td></td><td></td><td></td><td></td></tr>
<tr><td>Chromosome</td><td>start</td><td>end</td><td>strand</td><td>MM</td><td>target_seq</td><td>PAM</td><td>alignment</td><td>distance</td><td>position</td><td>gene name</td><td>gene id</td></tr>
<tr><td>GU305914.1</td><td>72715</td><td>72737</td><td>-</td><td>0</td><td>CGTTTACGATGTTCGTGACA</td><td>TGG</td><td>||||||||[||||||||||||]PAM</td><td>NA</td><td>Intergenic</td><td>NA</td><td>NA</td></tr>
<tr><td>T28752</td><td>ACATGGATGCCCACCGTCATCGG</td><td>1000</td><td></td><td></td><td></td><td></td><td></td><td></td><td></td><td></td><td></td></tr>
<tr><td>Oligo fwd</td><td>CACCACATGGATGCCCACCGTCAT</td><td></td><td></td><td></td><td></td><td></td><td></td><td></td><td></td><td></td><td></td></tr>
<tr><td>Oligo rev</td><td>AAACATGACGGTGGGCATCCATGT</td><td></td><td></td><td></td><td></td><td></td><td></td><td></td><td></td><td></td><td></td></tr>
<tr><td>Chromosome</td><td>start</td><td>end</td><td>strand</td><td>MM</td><td>target_seq</td><td>PAM</td><td>alignment</td><td>distance</td><td>position</td><td>gene name</td><td>gene id</td></tr>
<tr><td>GU305914.1</td><td>72698</td><td>72720</td><td>-</td><td>0</td><td>ACATGGATGCCCACCGTCAT</td><td>CGG</td><td>||||||||[||||||||||||]PAM</td><td>NA</td><td>Intergenic</td><td>NA</td><td>NA</td></tr>
<tr><td>T28753</td><td>CATGGATGCCCACCGTCATCGGG</td><td>1000</td><td></td><td></td><td></td><td></td><td></td><td></td><td></td><td></td><td></td></tr>
<tr><td>Oligo fwd</td><td>CACCCATGGATGCCCACCGTCATC</td><td></td><td></td><td></td><td></td><td></td><td></td><td></td><td></td><td></td><td></td></tr>
<tr><td>Oligo rev</td><td>AAACGATGACGGTGGGCATCCATG</td><td></td><td></td><td></td><td></td><td></td><td></td><td></td><td></td><td></td><td></td></tr>
<tr><td>Chromosome</td><td>start</td><td>end</td><td>strand</td><td>MM</td><td>target_seq</td><td>PAM</td><td>alignment</td><td>distance</td><td>position</td><td>gene name</td><td>gene id</td></tr>
<tr><td>GU305914.1</td><td>72697</td><td>72719</td><td>-</td><td>0</td><td>CATGGATGCCCACCGTCATC</td><td>GGG</td><td>||||||||[||||||||||||]PAM</td><td>NA</td><td>Intergenic</td><td>NA</td><td>NA</td></tr>
<tr><td>T28754</td><td>TCGCGAACACGTACGACCCCAGG</td><td>1000</td><td></td><td></td><td></td><td></td><td></td><td></td><td></td><td></td><td></td></tr>
<tr><td>Oligo fwd</td><td>CACCTCGCGAACACGTACGACCCC</td><td></td><td></td><td></td><td></td><td></td><td></td><td></td><td></td><td></td><td></td></tr>
<tr><td>Oligo rev</td><td>AAACGGGGTCGTACGTGTTCGCGA</td><td></td><td></td><td></td><td></td><td></td><td></td><td></td><td></td><td></td><td></td></tr>
<tr><td>Chromosome</td><td>start</td><td>end</td><td>strand</td><td>MM</td><td>target_seq</td><td>PAM</td><td>alignment</td><td>distance</td><td>position</td><td>gene name</td><td>gene id</td></tr>
<tr><td>GU305914.1</td><td>72673</td><td>72695</td><td>-</td><td>0</td><td>TCGCGAACACGTACGACCCC</td><td>AGG</td><td>||||||||[||||||||||||]PAM</td><td>NA</td><td>Intergenic</td><td>NA</td><td>NA</td></tr>
<tr><td>T28755</td><td>TCGTCCAGCAGCGACATCGTCGG</td><td>1000</td><td></td><td></td><td></td><td></td><td></td><td></td><td></td><td></td><td></td></tr>
<tr><td>Oligo fwd</td><td>CACCTCGTCCAGCAGCGACATCGT</td><td></td><td></td><td></td><td></td><td></td><td></td><td></td><td></td><td></td><td></td></tr>
<tr><td>Oligo rev</td><td>AAACACGATGTCGCTGCTGGACGA</td><td></td><td></td><td></td><td></td><td></td><td></td><td></td><td></td><td></td><td></td></tr>
<tr><td>Chromosome</td><td>start</td><td>end</td><td>strand</td><td>MM</td><td>target_seq</td><td>PAM</td><td>alignment</td><td>distance</td><td>position</td><td>gene name</td><td>gene id</td></tr>
<tr><td>GU305914.1</td><td>72617</td><td>72639</td><td>-</td><td>0</td><td>TCGTCCAGCAGCGACATCGT</td><td>CGG</td><td>||||||||[||||||||||||]PAM</td><td>NA</td><td>Intergenic</td><td>NA</td><td>NA</td></tr>
<tr><td>T28756</td><td>CTGCCGTTCCACCTTCTCCCCGG</td><td>1000</td><td></td><td></td><td></td><td></td><td></td><td></td><td></td><td></td><td></td></tr>
<tr><td>Oligo fwd</td><td>CACCCTGCCGTTCCACCTTCTCCC</td><td></td><td></td><td></td><td></td><td></td><td></td><td></td><td></td><td></td><td></td></tr>
<tr><td>Oligo rev</td><td>AAACGGGAGAAGGTGGAACGGCAG</td><td></td><td></td><td></td><td></td><td></td><td></td><td></td><td></td><td></td><td></td></tr>
<tr><td>Chromosome</td><td>start</td><td>end</td><td>strand</td><td>MM</td><td>target_seq</td><td>PAM</td><td>alignment</td><td>distance</td><td>position</td><td>gene name</td><td>gene id</td></tr>
<tr><td>GU305914.1</td><td>72576</td><td>72598</td><td>-</td><td>0</td><td>CTGCCGTTCCACCTTCTCCC</td><td>CGG</td><td>||||||||[||||||||||||]PAM</td><td>NA</td><td>Intergenic</td><td>NA</td><td>NA</td></tr>
<tr><td>GU305914.1</td><td>75723</td><td>75745</td><td>-</td><td>4</td><td>CTCCACTTCCACCTCCTCCC</td><td>CGG</td><td>||-|--||[||||||-|||||]PAM</td><td>NA</td><td>Intergenic</td><td>NA</td><td>NA</td></tr>
<tr><td>T28757</td><td>CGTACCGTAGATCTTGAAGAAGG</td><td>1000</td><td></td><td></td><td></td><td></td><td></td><td></td><td></td><td></td><td></td></tr>
<tr><td>Oligo fwd</td><td>CACCCGTACCGTAGATCTTGAAGA</td><td></td><td></td><td></td><td></td><td></td><td></td><td></td><td></td><td></td><td></td></tr>
<tr><td>Oligo rev</td><td>AAACTCTTCAAGATCTACGGTACG</td><td></td><td></td><td></td><td></td><td></td><td></td><td></td><td></td><td></td><td></td></tr>
<tr><td>Chromosome</td><td>start</td><td>end</td><td>strand</td><td>MM</td><td>target_seq</td><td>PAM</td><td>alignment</td><td>distance</td><td>position</td><td>gene name</td><td>gene id</td></tr>
<tr><td>GU305914.1</td><td>72522</td><td>72544</td><td>-</td><td>0</td><td>CGTACCGTAGATCTTGAAGA</td><td>AGG</td><td>||||||||[||||||||||||]PAM</td><td>NA</td><td>Intergenic</td><td>NA</td><td>NA</td></tr>
<tr><td>T28758</td><td>AGCGCTCACCAGCTCTTCTAAGG</td><td>1000</td><td></td><td></td><td></td><td></td><td></td><td></td><td></td><td></td><td></td></tr>
<tr><td>Oligo fwd</td><td>CACCAGCGCTCACCAGCTCTTCTA</td><td></td><td></td><td></td><td></td><td></td><td></td><td></td><td></td><td></td><td></td></tr>
<tr><td>Oligo rev</td><td>AAACTAGAAGAGCTGGTGAGCGCT</td><td></td><td></td><td></td><td></td><td></td><td></td><td></td><td></td><td></td><td></td></tr>
<tr><td>Chromosome</td><td>start</td><td>end</td><td>strand</td><td>MM</td><td>target_seq</td><td>PAM</td><td>alignment</td><td>distance</td><td>position</td><td>gene name</td><td>gene id</td></tr>
<tr><td>GU305914.1</td><td>72489</td><td>72511</td><td>-</td><td>0</td><td>AGCGCTCACCAGCTCTTCTA</td><td>AGG</td><td>||||||||[||||||||||||]PAM</td><td>NA</td><td>Intergenic</td><td>NA</td><td>NA</td></tr>
<tr><td>T28759</td><td>CTCTTCTAAGGACGCTATCGAGG</td><td>1000</td><td></td><td></td><td></td><td></td><td></td><td></td><td></td><td></td><td></td></tr>
<tr><td>Oligo fwd</td><td>CACCCTCTTCTAAGGACGCTATCG</td><td></td><td></td><td></td><td></td><td></td><td></td><td></td><td></td><td></td><td></td></tr>
<tr><td>Oligo rev</td><td>AAACCGATAGCGTCCTTAGAAGAG</td><td></td><td></td><td></td><td></td><td></td><td></td><td></td><td></td><td></td><td></td></tr>
<tr><td>Chromosome</td><td>start</td><td>end</td><td>strand</td><td>MM</td><td>target_seq</td><td>PAM</td><td>alignment</td><td>distance</td><td>position</td><td>gene name</td><td>gene id</td></tr>
</table>

| | | | | | | | | | | | |
|---|---|---|---|---|---|---|---|---|---|---|---|
| GU305914.1 | 72477 | 72499 | - | 0 | CTCTTCTAAGGACGCTATCG | AGG | \|\|\|\|\|\|\|\|[\|\|\|\|\|\|\|\|\|\|\|\|]PAM | NA | Intergenic | NA | NA |
| T28760 | TTCTAAGGACGCTATCGAGGAGG | 1000 | | | | | | | | | |
| Oligo fwd | CACCTTCTAAGGACGCTATCGAGG | | | | | | | | | | |
| Oligo rev | AAACCCTCGATAGCGTCCTTAGAA | | | | | | | | | | |
| Chromosome | start | end | strand | MM | target_seq | PAM | alignment | distance | position | gene name | gene id |
| GU305914.1 | 72474 | 72496 | - | 0 | TTCTAAGGACGCTATCGAGG | AGG | \|\|\|\|\|\|\|\|[\|\|\|\|\|\|\|\|\|\|\|\|]PAM | NA | Intergenic | NA | NA |
| T28761 | GGACGCTATCGAGGAGGCGTAGG | 1000 | | | | | | | | | |
| Oligo fwd | CACCGGACGCTATCGAGGAGGCGT | | | | | | | | | | |
| Oligo rev | AAACACGCCTCCTCGATAGCGTCC | | | | | | | | | | |
| Chromosome | start | end | strand | MM | target_seq | PAM | alignment | distance | position | gene name | gene id |
| GU305914.1 | 72468 | 72490 | - | 0 | GGACGCTATCGAGGAGGCGT | AGG | \|\|\|\|\|\|\|\|[\|\|\|\|\|\|\|\|\|\|\|\|]PAM | NA | Intergenic | NA | NA |
| GU305914.1 | 81252 | 81274 | - | 4 | CGCCGCGATCGAGGAGGCGG | CGG | -\|-\|\|\|-\|[\|\|\|\|\|\|\|\|\|\|\|-]PAM | NA | Intergenic | NA | NA |
| T28762 | GCTATCGAGGAGGCGTAGGCCGG | 1000 | | | | | | | | | |
| Oligo fwd | CACCGCTATCGAGGAGGCGTAGGC | | | | | | | | | | |
| Oligo rev | AAACGCCTACGCCTCCTCGATAGC | | | | | | | | | | |
| Chromosome | start | end | strand | MM | target_seq | PAM | alignment | distance | position | gene name | gene id |
| GU305914.1 | 72464 | 72486 | - | 0 | GCTATCGAGGAGGCGTAGGC | CGG | \|\|\|\|\|\|\|\|[\|\|\|\|\|\|\|\|\|\|\|\|]PAM | NA | Intergenic | NA | NA |
| GU305914.1 | 81248 | 81270 | - | 3 | GCGATCGAGGAGGCGGCGGC | GGG | \|\|-\|\|\|\|\|[\|\|\|\|\|\|\|--\|\|\|]PAM | NA | Intergenic | NA | NA |
| T28763 | ACTGCACCCAGAGACACCGAAGG | 1000 | | | | | | | | | |
| Oligo fwd | CACCACTGCACCCAGAGACACCGA | | | | | | | | | | |
| Oligo rev | AAACTCGGTGTCTCTGGGTGCAGT | | | | | | | | | | |
| Chromosome | start | end | strand | MM | target_seq | PAM | alignment | distance | position | gene name | gene id |
| GU305914.1 | 72436 | 72458 | - | 0 | ACTGCACCCAGAGACACCGA | AGG | \|\|\|\|\|\|\|\|[\|\|\|\|\|\|\|\|\|\|\|\|]PAM | NA | Intergenic | NA | NA |
| T28764 | GAGACACCGAAGGATGTCCCTGG | 1000 | | | | | | | | | |
| Oligo fwd | CACCGAGACACCGAAGGATGTCCC | | | | | | | | | | |
| Oligo rev | AAACGGGACATCCTTCGGTGTCTC | | | | | | | | | | |
| Chromosome | start | end | strand | MM | target_seq | PAM | alignment | distance | position | gene name | gene id |
| GU305914.1 | 72426 | 72448 | - | 0 | GAGACACCGAAGGATGTCCC | TGG | \|\|\|\|\|\|\|\|[\|\|\|\|\|\|\|\|\|\|\|\|]PAM | NA | Intergenic | NA | NA |
| T28765 | AGACACCGAAGGATGTCCCTGGG | 1000 | | | | | | | | | |
| Oligo fwd | CACCAGACACCGAAGGATGTCCCT | | | | | | | | | | |
| Oligo rev | AAACAGGGACATCCTTCGGTGTCT | | | | | | | | | | |
| Chromosome | start | end | strand | MM | target_seq | PAM | alignment | distance | position | gene name | gene id |
| GU305914.1 | 72425 | 72447 | - | 0 | AGACACCGAAGGATGTCCCT | GGG | \|\|\|\|\|\|\|\|[\|\|\|\|\|\|\|\|\|\|\|\|]PAM | NA | Intergenic | NA | NA |
| T28766 | GTCCCTGGGCAGCACCTTTGTGG | 1000 | | | | | | | | | |
| Oligo fwd | CACCGTCCCTGGGCAGCACCTTTG | | | | | | | | | | |
| Oligo rev | AAACCAAAGGTGCTGCCCAGGGAC | | | | | | | | | | |
| Chromosome | start | end | strand | MM | target_seq | PAM | alignment | distance | position | gene name | gene id |
| GU305914.1 | 72411 | 72433 | - | 0 | GTCCCTGGGCAGCACCTTTG | TGG | \|\|\|\|\|\|\|\|[\|\|\|\|\|\|\|\|\|\|\|\|]PAM | NA | Intergenic | NA | NA |
| T28767 | TAACATGTATAGAAACAGCTTGG | 1000 | | | | | | | | | |
| Oligo fwd | CACCTAACATGTATAGAAACAGCT | | | | | | | | | | |
| Oligo rev | AAACAGCTGTTTCTATACATGTTA | | | | | | | | | | |
| Chromosome | start | end | strand | MM | target_seq | PAM | alignment | distance | position | gene name | gene id |
| GU305914.1 | 72387 | 72409 | - | 0 | TAACATGTATAGAAACAGCT | TGG | \|\|\|\|\|\|\|\|[\|\|\|\|\|\|\|\|\|\|\|\|]PAM | NA | Intergenic | NA | NA |
| T28768 | TGGTCGAAGTGCTCGCGTCGCGG | 1000 | | | | | | | | | |
| Oligo fwd | CACCTGGTCGAAGTGCTCGCGTCG | | | | | | | | | | |
| Oligo rev | AAACCGACGCGAGCACTTCGACCA | | | | | | | | | | |
| Chromosome | start | end | strand | MM | target_seq | PAM | alignment | distance | position | gene name | gene id |
| GU305914.1 | 72367 | 72389 | - | 0 | TGGTCGAAGTGCTCGCGTCG | CGG | \|\|\|\|\|\|\|\|[\|\|\|\|\|\|\|\|\|\|\|\|]PAM | NA | Intergenic | NA | NA |
| T28769 | GTGCTCGCGTCGCGGTTTCTCGG | 1000 | | | | | | | | | |
| Oligo fwd | CACCGTGCTCGCGTCGCGGTTTCT | | | | | | | | | | |
| Oligo rev | AAACAGAAACCGCGACGCGAGCAC | | | | | | | | | | |
| Chromosome | start | end | strand | MM | target_seq | PAM | alignment | distance | position | gene name | gene id |
| GU305914.1 | 72359 | 72381 | - | 0 | GTGCTCGCGTCGCGGTTTCT | CGG | \|\|\|\|\|\|\|\|[\|\|\|\|\|\|\|\|\|\|\|\|]PAM | NA | Intergenic | NA | NA |
| T28770 | TGCTCGCGTCGCGGTTTCTCGGG | 1000 | | | | | | | | | |
| Oligo fwd | CACCTGCTCGCGTCGCGGTTTCTC | | | | | | | | | | |
| Oligo rev | AAACGAGAAACCGCGACGCGAGCA | | | | | | | | | | |
| Chromosome | start | end | strand | MM | target_seq | PAM | alignment | distance | position | gene name | gene id |
| GU305914.1 | 72358 | 72380 | - | 0 | TGCTCGCGTCGCGGTTTCTC | GGG | \|\|\|\|\|\|\|\|[\|\|\|\|\|\|\|\|\|\|\|\|]PAM | NA | Intergenic | NA | NA |
| T28771 | ACGTGCAGATCTGCTTCACCAGG | 1000 | | | | | | | | | |
| Oligo fwd | CACCACGTGCAGATCTGCTTCACC | | | | | | | | | | |
| Oligo rev | AAACGGTGAAGCAGATCTGCACGT | | | | | | | | | | |
| Chromosome | start | end | strand | MM | target_seq | PAM | alignment | distance | position | gene name | gene id |
| GU305914.1 | 72331 | 72353 | - | 0 | ACGTGCAGATCTGCTTCACC | AGG | \|\|\|\|\|\|\|\|[\|\|\|\|\|\|\|\|\|\|\|\|]PAM | NA | Intergenic | NA | NA |
| T28772 | AGGTCCGACATCCACATCTGCGG | 1000 | | | | | | | | | |
| Oligo fwd | CACCAGGTCCGACATCCACATCTG | | | | | | | | | | |
| Oligo rev | AAACCAGATGTGGATGTCGGACCT | | | | | | | | | | |
| Chromosome | start | end | strand | MM | target_seq | PAM | alignment | distance | position | gene name | gene id |
| GU305914.1 | 72311 | 72333 | - | 0 | AGGTCCGACATCCACATCTG | CGG | \|\|\|\|\|\|\|\|[\|\|\|\|\|\|\|\|\|\|\|\|]PAM | NA | Intergenic | NA | NA |
| T28773 | GGTCCGACATCCACATCTGCGGG | 1000 | | | | | | | | | |
| Oligo fwd | CACCGGTCCGACATCCACATCTGC | | | | | | | | | | |
| Oligo rev | AAACGCAGATGTGGATGTCGGACC | | | | | | | | | | |
| Chromosome | start | end | strand | MM | target_seq | PAM | alignment | distance | position | gene name | gene id |
| GU305914.1 | 72310 | 72332 | - | 0 | GGTCCGACATCCACATCTGC | GGG | \|\|\|\|\|\|\|\|[\|\|\|\|\|\|\|\|\|\|\|\|]PAM | NA | Intergenic | NA | NA |
| T28774 | CCGACATCCACATCTGCGGGCGG | 1000 | | | | | | | | | |

<table>
<tr><td>Oligo fwd</td><td>CACCCCGACATCCACATCTGCGGG</td><td></td><td></td><td></td><td></td><td></td><td></td><td></td><td></td><td></td><td></td></tr>
<tr><td>Oligo rev</td><td>AAACCCCGCAGATGTGGATGTCGG</td><td></td><td></td><td></td><td></td><td></td><td></td><td></td><td></td><td></td><td></td></tr>
<tr><td>Chromosome</td><td>start</td><td>end</td><td>strand</td><td>MM</td><td>target_seq</td><td>PAM</td><td>alignment</td><td>distance</td><td>position</td><td>gene name</td><td>gene id</td></tr>
<tr><td>GU305914.1</td><td>72307</td><td>72329</td><td>-</td><td>0</td><td>CCGACATCCACATCTGCGGG</td><td>CGG</td><td>||||||||[||||||||||||]PAM</td><td>NA</td><td>Intergenic</td><td>NA</td><td>NA</td></tr>
<tr><td>T28775</td><td>TTTCAATTCGCGCATATCGTAGG</td><td>1000</td><td></td><td></td><td></td><td></td><td></td><td></td><td></td><td></td><td></td></tr>
<tr><td>Oligo fwd</td><td>CACCTTTCAATTCGCGCATATCGT</td><td></td><td></td><td></td><td></td><td></td><td></td><td></td><td></td><td></td><td></td></tr>
<tr><td>Oligo rev</td><td>AAACACGATATGCGCGAATTGAAA</td><td></td><td></td><td></td><td></td><td></td><td></td><td></td><td></td><td></td><td></td></tr>
<tr><td>Chromosome</td><td>start</td><td>end</td><td>strand</td><td>MM</td><td>target_seq</td><td>PAM</td><td>alignment</td><td>distance</td><td>position</td><td>gene name</td><td>gene id</td></tr>
<tr><td>GU305914.1</td><td>72243</td><td>72265</td><td>-</td><td>0</td><td>TTTCAATTCGCGCATATCGT</td><td>AGG</td><td>||||||||[||||||||||||]PAM</td><td>NA</td><td>Intergenic</td><td>NA</td><td>NA</td></tr>
<tr><td>T28776</td><td>TTCAATTCGCGCATATCGTAGGG</td><td>1000</td><td></td><td></td><td></td><td></td><td></td><td></td><td></td><td></td><td></td></tr>
<tr><td>Oligo fwd</td><td>CACCTTCAATTCGCGCATATCGTA</td><td></td><td></td><td></td><td></td><td></td><td></td><td></td><td></td><td></td><td></td></tr>
<tr><td>Oligo rev</td><td>AAACTACGATATGCGCGAATTGAA</td><td></td><td></td><td></td><td></td><td></td><td></td><td></td><td></td><td></td><td></td></tr>
<tr><td>Chromosome</td><td>start</td><td>end</td><td>strand</td><td>MM</td><td>target_seq</td><td>PAM</td><td>alignment</td><td>distance</td><td>position</td><td>gene name</td><td>gene id</td></tr>
<tr><td>GU305914.1</td><td>72242</td><td>72264</td><td>-</td><td>0</td><td>TTCAATTCGCGCATATCGTA</td><td>GGG</td><td>||||||||[||||||||||||]PAM</td><td>NA</td><td>Intergenic</td><td>NA</td><td>NA</td></tr>
<tr><td>T28777</td><td>AATTCGCGCATATCGTAGGGCGG</td><td>1000</td><td></td><td></td><td></td><td></td><td></td><td></td><td></td><td></td><td></td></tr>
<tr><td>Oligo fwd</td><td>CACCAATTCGCGCATATCGTAGGG</td><td></td><td></td><td></td><td></td><td></td><td></td><td></td><td></td><td></td><td></td></tr>
<tr><td>Oligo rev</td><td>AAACCCCTACGATATGCGCGAATT</td><td></td><td></td><td></td><td></td><td></td><td></td><td></td><td></td><td></td><td></td></tr>
<tr><td>Chromosome</td><td>start</td><td>end</td><td>strand</td><td>MM</td><td>target_seq</td><td>PAM</td><td>alignment</td><td>distance</td><td>position</td><td>gene name</td><td>gene id</td></tr>
<tr><td>GU305914.1</td><td>72239</td><td>72261</td><td>-</td><td>0</td><td>AATTCGCGCATATCGTAGGG</td><td>CGG</td><td>||||||||[||||||||||||]PAM</td><td>NA</td><td>Intergenic</td><td>NA</td><td>NA</td></tr>
<tr><td>T28778</td><td>TCGCGCATATCGTAGGGCGGCGG</td><td>1000</td><td></td><td></td><td></td><td></td><td></td><td></td><td></td><td></td><td></td></tr>
<tr><td>Oligo fwd</td><td>CACCTCGCGCATATCGTAGGGCGG</td><td></td><td></td><td></td><td></td><td></td><td></td><td></td><td></td><td></td><td></td></tr>
<tr><td>Oligo rev</td><td>AAACCCGCCCTACGATATGCGCGA</td><td></td><td></td><td></td><td></td><td></td><td></td><td></td><td></td><td></td><td></td></tr>
<tr><td>Chromosome</td><td>start</td><td>end</td><td>strand</td><td>MM</td><td>target_seq</td><td>PAM</td><td>alignment</td><td>distance</td><td>position</td><td>gene name</td><td>gene id</td></tr>
<tr><td>GU305914.1</td><td>72236</td><td>72258</td><td>-</td><td>0</td><td>TCGCGCATATCGTAGGGCGG</td><td>CGG</td><td>||||||||[||||||||||||]PAM</td><td>NA</td><td>Intergenic</td><td>NA</td><td>NA</td></tr>
<tr><td>T28779</td><td>CGCGCATATCGTAGGGCGGCGGG</td><td>1000</td><td></td><td></td><td></td><td></td><td></td><td></td><td></td><td></td><td></td></tr>
<tr><td>Oligo fwd</td><td>CACCCGCGCATATCGTAGGGCGGC</td><td></td><td></td><td></td><td></td><td></td><td></td><td></td><td></td><td></td><td></td></tr>
<tr><td>Oligo rev</td><td>AAACGCCGCCCTACGATATGCGCG</td><td></td><td></td><td></td><td></td><td></td><td></td><td></td><td></td><td></td><td></td></tr>
<tr><td>Chromosome</td><td>start</td><td>end</td><td>strand</td><td>MM</td><td>target_seq</td><td>PAM</td><td>alignment</td><td>distance</td><td>position</td><td>gene name</td><td>gene id</td></tr>
<tr><td>GU305914.1</td><td>72235</td><td>72257</td><td>-</td><td>0</td><td>CGCGCATATCGTAGGGCGGC</td><td>GGG</td><td>||||||||[||||||||||||]PAM</td><td>NA</td><td>Intergenic</td><td>NA</td><td>NA</td></tr>
<tr><td>T28780</td><td>CACCGTCCTCCTGTCCAGTCTGG</td><td>1000</td><td></td><td></td><td></td><td></td><td></td><td></td><td></td><td></td><td></td></tr>
<tr><td>Oligo fwd</td><td>CACCCACCGTCCTCCTGTCCAGTC</td><td></td><td></td><td></td><td></td><td></td><td></td><td></td><td></td><td></td><td></td></tr>
<tr><td>Oligo rev</td><td>AAACGACTGGACAGGAGGACGGTG</td><td></td><td></td><td></td><td></td><td></td><td></td><td></td><td></td><td></td><td></td></tr>
<tr><td>Chromosome</td><td>start</td><td>end</td><td>strand</td><td>MM</td><td>target_seq</td><td>PAM</td><td>alignment</td><td>distance</td><td>position</td><td>gene name</td><td>gene id</td></tr>
<tr><td>GU305914.1</td><td>72186</td><td>72208</td><td>-</td><td>0</td><td>CACCGTCCTCCTGTCCAGTC</td><td>TGG</td><td>||||||||[||||||||||||]PAM</td><td>NA</td><td>Intergenic</td><td>NA</td><td>NA</td></tr>
<tr><td>T28781</td><td>GTCTGGCGATCAAGCCGCGCAGG</td><td>1000</td><td></td><td></td><td></td><td></td><td></td><td></td><td></td><td></td><td></td></tr>
<tr><td>Oligo fwd</td><td>CACCGTCTGGCGATCAAGCCGCGC</td><td></td><td></td><td></td><td></td><td></td><td></td><td></td><td></td><td></td><td></td></tr>
<tr><td>Oligo rev</td><td>AAACGCGCGGCTTGATCGCCAGAC</td><td></td><td></td><td></td><td></td><td></td><td></td><td></td><td></td><td></td><td></td></tr>
<tr><td>Chromosome</td><td>start</td><td>end</td><td>strand</td><td>MM</td><td>target_seq</td><td>PAM</td><td>alignment</td><td>distance</td><td>position</td><td>gene name</td><td>gene id</td></tr>
<tr><td>GU305914.1</td><td>72169</td><td>72191</td><td>-</td><td>0</td><td>GTCTGGCGATCAAGCCGCGC</td><td>AGG</td><td>||||||||[||||||||||||]PAM</td><td>NA</td><td>Intergenic</td><td>NA</td><td>NA</td></tr>
<tr><td>T28782</td><td>TCTGGCGATCAAGCCGCGCAGGG</td><td>1000</td><td></td><td></td><td></td><td></td><td></td><td></td><td></td><td></td><td></td></tr>
<tr><td>Oligo fwd</td><td>CACCTCTGGCGATCAAGCCGCGCA</td><td></td><td></td><td></td><td></td><td></td><td></td><td></td><td></td><td></td><td></td></tr>
<tr><td>Oligo rev</td><td>AAACTGCGCGGCTTGATCGCCAGA</td><td></td><td></td><td></td><td></td><td></td><td></td><td></td><td></td><td></td><td></td></tr>
<tr><td>Chromosome</td><td>start</td><td>end</td><td>strand</td><td>MM</td><td>target_seq</td><td>PAM</td><td>alignment</td><td>distance</td><td>position</td><td>gene name</td><td>gene id</td></tr>
<tr><td>GU305914.1</td><td>72168</td><td>72190</td><td>-</td><td>0</td><td>TCTGGCGATCAAGCCGCGCA</td><td>GGG</td><td>||||||||[||||||||||||]PAM</td><td>NA</td><td>Intergenic</td><td>NA</td><td>NA</td></tr>
<tr><td>T28783</td><td>GGCGATCAAGCCGCGCAGGGTGG</td><td>1000</td><td></td><td></td><td></td><td></td><td></td><td></td><td></td><td></td><td></td></tr>
<tr><td>Oligo fwd</td><td>CACCGGCGATCAAGCCGCGCAGGG</td><td></td><td></td><td></td><td></td><td></td><td></td><td></td><td></td><td></td><td></td></tr>
<tr><td>Oligo rev</td><td>AAACCCCTGCGCGGCTTGATCGCC</td><td></td><td></td><td></td><td></td><td></td><td></td><td></td><td></td><td></td><td></td></tr>
<tr><td>Chromosome</td><td>start</td><td>end</td><td>strand</td><td>MM</td><td>target_seq</td><td>PAM</td><td>alignment</td><td>distance</td><td>position</td><td>gene name</td><td>gene id</td></tr>
<tr><td>GU305914.1</td><td>72165</td><td>72187</td><td>-</td><td>0</td><td>GGCGATCAAGCCGCGCAGGG</td><td>TGG</td><td>||||||||[||||||||||||]PAM</td><td>NA</td><td>Intergenic</td><td>NA</td><td>NA</td></tr>
<tr><td>T28784</td><td>CCGCGCAGGGTGGCGTCCACCGG</td><td>1000</td><td></td><td></td><td></td><td></td><td></td><td></td><td></td><td></td><td></td></tr>
<tr><td>Oligo fwd</td><td>CACCCCGCGCAGGGTGGCGTCCAC</td><td></td><td></td><td></td><td></td><td></td><td></td><td></td><td></td><td></td><td></td></tr>
<tr><td>Oligo rev</td><td>AAACGTGGACGCCACCCTGCGCGG</td><td></td><td></td><td></td><td></td><td></td><td></td><td></td><td></td><td></td><td></td></tr>
<tr><td>Chromosome</td><td>start</td><td>end</td><td>strand</td><td>MM</td><td>target_seq</td><td>PAM</td><td>alignment</td><td>distance</td><td>position</td><td>gene name</td><td>gene id</td></tr>
<tr><td>GU305914.1</td><td>72155</td><td>72177</td><td>-</td><td>0</td><td>CCGCGCAGGGTGGCGTCCAC</td><td>CGG</td><td>||||||||[||||||||||||]PAM</td><td>NA</td><td>Intergenic</td><td>NA</td><td>NA</td></tr>
<tr><td>T28785</td><td>GCAGGGTGGCGTCCACCGGCTGG</td><td>1000</td><td></td><td></td><td></td><td></td><td></td><td></td><td></td><td></td><td></td></tr>
<tr><td>Oligo fwd</td><td>CACCGCAGGGTGGCGTCCACCGGC</td><td></td><td></td><td></td><td></td><td></td><td></td><td></td><td></td><td></td><td></td></tr>
<tr><td>Oligo rev</td><td>AAACGCCGGTGGACGCCACCCTGC</td><td></td><td></td><td></td><td></td><td></td><td></td><td></td><td></td><td></td><td></td></tr>
<tr><td>Chromosome</td><td>start</td><td>end</td><td>strand</td><td>MM</td><td>target_seq</td><td>PAM</td><td>alignment</td><td>distance</td><td>position</td><td>gene name</td><td>gene id</td></tr>
<tr><td>GU305914.1</td><td>72151</td><td>72173</td><td>-</td><td>0</td><td>GCAGGGTGGCGTCCACCGGC</td><td>TGG</td><td>||||||||[||||||||||||]PAM</td><td>NA</td><td>Intergenic</td><td>NA</td><td>NA</td></tr>
<tr><td>T28786</td><td>CAGGGTGGCGTCCACCGGCTGGG</td><td>1000</td><td></td><td></td><td></td><td></td><td></td><td></td><td></td><td></td><td></td></tr>
<tr><td>Oligo fwd</td><td>CACCCAGGGTGGCGTCCACCGGCT</td><td></td><td></td><td></td><td></td><td></td><td></td><td></td><td></td><td></td><td></td></tr>
<tr><td>Oligo rev</td><td>AAACAGCCGGTGGACGCCACCCTG</td><td></td><td></td><td></td><td></td><td></td><td></td><td></td><td></td><td></td><td></td></tr>
<tr><td>Chromosome</td><td>start</td><td>end</td><td>strand</td><td>MM</td><td>target_seq</td><td>PAM</td><td>alignment</td><td>distance</td><td>position</td><td>gene name</td><td>gene id</td></tr>
<tr><td>GU305914.1</td><td>72150</td><td>72172</td><td>-</td><td>0</td><td>CAGGGTGGCGTCCACCGGCT</td><td>GGG</td><td>||||||||[||||||||||||]PAM</td><td>NA</td><td>Intergenic</td><td>NA</td><td>NA</td></tr>
<tr><td>T28787</td><td>GTCCACCGGCTGGGTCGTGATGG</td><td>1000</td><td></td><td></td><td></td><td></td><td></td><td></td><td></td><td></td><td></td></tr>
<tr><td>Oligo fwd</td><td>CACCGTCCACCGGCTGGGTCGTGA</td><td></td><td></td><td></td><td></td><td></td><td></td><td></td><td></td><td></td><td></td></tr>
<tr><td>Oligo rev</td><td>AAACTCACGACCCAGCCGGTGGAC</td><td></td><td></td><td></td><td></td><td></td><td></td><td></td><td></td><td></td><td></td></tr>
<tr><td>Chromosome</td><td>start</td><td>end</td><td>strand</td><td>MM</td><td>target_seq</td><td>PAM</td><td>alignment</td><td>distance</td><td>position</td><td>gene name</td><td>gene id</td></tr>
<tr><td>GU305914.1</td><td>72141</td><td>72163</td><td>-</td><td>0</td><td>GTCCACCGGCTGGGTCGTGA</td><td>TGG</td><td>||||||||[||||||||||||]PAM</td><td>NA</td><td>Intergenic</td><td>NA</td><td>NA</td></tr>
<tr><td>T28788</td><td>GTGATGGCGTACACGTACTCCGG</td><td>1000</td><td></td><td></td><td></td><td></td><td></td><td></td><td></td><td></td><td></td></tr>
<tr><td>Oligo fwd</td><td>CACCGTGATGGCGTACACGTACTC</td><td></td><td></td><td></td><td></td><td></td><td></td><td></td><td></td><td></td><td></td></tr>
<tr><td>Oligo rev</td><td>AAACGAGTACGTGTACGCCATCAC</td><td></td><td></td><td></td><td></td><td></td><td></td><td></td><td></td><td></td><td></td></tr>
<tr><td>Chromosome</td><td>start</td><td>end</td><td>strand</td><td>MM</td><td>target_seq</td><td>PAM</td><td>alignment</td><td>distance</td><td>position</td><td>gene name</td><td>gene id</td></tr>
<tr><td>GU305914.1</td><td>72125</td><td>72147</td><td>-</td><td>0</td><td>GTGATGGCGTACACGTACTC</td><td>CGG</td><td>||||||||[||||||||||||]PAM</td><td>NA</td><td>Intergenic</td><td>NA</td><td>NA</td></tr>
</table>

<table>
<tr><td>T28789</td><td>TGATGGCGTACACGTACTCCGGG</td><td>1000</td><td></td><td></td><td></td><td></td><td></td><td></td><td></td><td></td><td></td></tr>
<tr><td>Oligo fwd</td><td>CACCTGATGGCGTACACGTACTCC</td><td></td><td></td><td></td><td></td><td></td><td></td><td></td><td></td><td></td><td></td></tr>
<tr><td>Oligo rev</td><td>AAACGGAGTACGTGTACGCCATCA</td><td></td><td></td><td></td><td></td><td></td><td></td><td></td><td></td><td></td><td></td></tr>
<tr><td>Chromosome</td><td>start</td><td>end</td><td>strand</td><td>MM</td><td>target_seq</td><td>PAM</td><td>alignment</td><td>distance</td><td>position</td><td>gene name</td><td>gene id</td></tr>
<tr><td>GU305914.1</td><td>72124</td><td>72146</td><td>-</td><td>0</td><td>TGATGGCGTACACGTACTCC</td><td>GGG</td><td>||||||||[||||||||||||]PAM</td><td>NA</td><td>Intergenic</td><td>NA</td><td>NA</td></tr>
<tr><td>T28790</td><td>TCAGCGTCGAGAAGTCCAGCAGG</td><td>1000</td><td></td><td></td><td></td><td></td><td></td><td></td><td></td><td></td><td></td></tr>
<tr><td>Oligo fwd</td><td>CACCTCAGCGTCGAGAAGTCCAGC</td><td></td><td></td><td></td><td></td><td></td><td></td><td></td><td></td><td></td><td></td></tr>
<tr><td>Oligo rev</td><td>AAACGCTGGACTTCTCGACGCTGA</td><td></td><td></td><td></td><td></td><td></td><td></td><td></td><td></td><td></td><td></td></tr>
<tr><td>Chromosome</td><td>start</td><td>end</td><td>strand</td><td>MM</td><td>target_seq</td><td>PAM</td><td>alignment</td><td>distance</td><td>position</td><td>gene name</td><td>gene id</td></tr>
<tr><td>GU305914.1</td><td>72088</td><td>72110</td><td>-</td><td>0</td><td>TCAGCGTCGAGAAGTCCAGC</td><td>AGG</td><td>||||||||[||||||||||||]PAM</td><td>NA</td><td>Intergenic</td><td>NA</td><td>NA</td></tr>
<tr><td>T28791</td><td>AGTCCAGCAGGCTCACCTCCAGG</td><td>1000</td><td></td><td></td><td></td><td></td><td></td><td></td><td></td><td></td><td></td></tr>
<tr><td>Oligo fwd</td><td>CACCAGTCCAGCAGGCTCACCTCC</td><td></td><td></td><td></td><td></td><td></td><td></td><td></td><td></td><td></td><td></td></tr>
<tr><td>Oligo rev</td><td>AAACGGAGGTGAGCCTGCTGGACT</td><td></td><td></td><td></td><td></td><td></td><td></td><td></td><td></td><td></td><td></td></tr>
<tr><td>Chromosome</td><td>start</td><td>end</td><td>strand</td><td>MM</td><td>target_seq</td><td>PAM</td><td>alignment</td><td>distance</td><td>position</td><td>gene name</td><td>gene id</td></tr>
<tr><td>GU305914.1</td><td>72076</td><td>72098</td><td>-</td><td>0</td><td>AGTCCAGCAGGCTCACCTCC</td><td>AGG</td><td>||||||||[||||||||||||]PAM</td><td>NA</td><td>Intergenic</td><td>NA</td><td>NA</td></tr>
<tr><td>T28792</td><td>GTCCAGCAGGCTCACCTCCAGGG</td><td>1000</td><td></td><td></td><td></td><td></td><td></td><td></td><td></td><td></td><td></td></tr>
<tr><td>Oligo fwd</td><td>CACCGTCCAGCAGGCTCACCTCCA</td><td></td><td></td><td></td><td></td><td></td><td></td><td></td><td></td><td></td><td></td></tr>
<tr><td>Oligo rev</td><td>AAACTGGAGGTGAGCCTGCTGGAC</td><td></td><td></td><td></td><td></td><td></td><td></td><td></td><td></td><td></td><td></td></tr>
<tr><td>Chromosome</td><td>start</td><td>end</td><td>strand</td><td>MM</td><td>target_seq</td><td>PAM</td><td>alignment</td><td>distance</td><td>position</td><td>gene name</td><td>gene id</td></tr>
<tr><td>GU305914.1</td><td>72075</td><td>72097</td><td>-</td><td>0</td><td>GTCCAGCAGGCTCACCTCCA</td><td>GGG</td><td>||||||||[||||||||||||]PAM</td><td>NA</td><td>Intergenic</td><td>NA</td><td>NA</td></tr>
<tr><td>T28793</td><td>TCCAGGGCATGCGTCGCCTCCGG</td><td>1000</td><td></td><td></td><td></td><td></td><td></td><td></td><td></td><td></td><td></td></tr>
<tr><td>Oligo fwd</td><td>CACCTCCAGGGCATGCGTCGCCTC</td><td></td><td></td><td></td><td></td><td></td><td></td><td></td><td></td><td></td><td></td></tr>
<tr><td>Oligo rev</td><td>AAACGAGGCGACGCATGCCCTGGA</td><td></td><td></td><td></td><td></td><td></td><td></td><td></td><td></td><td></td><td></td></tr>
<tr><td>Chromosome</td><td>start</td><td>end</td><td>strand</td><td>MM</td><td>target_seq</td><td>PAM</td><td>alignment</td><td>distance</td><td>position</td><td>gene name</td><td>gene id</td></tr>
<tr><td>GU305914.1</td><td>72059</td><td>72081</td><td>-</td><td>0</td><td>TCCAGGGCATGCGTCGCCTC</td><td>CGG</td><td>||||||||[||||||||||||]PAM</td><td>NA</td><td>Intergenic</td><td>NA</td><td>NA</td></tr>
<tr><td>T28794</td><td>CCAGGGCATGCGTCGCCTCCGGG</td><td>1000</td><td></td><td></td><td></td><td></td><td></td><td></td><td></td><td></td><td></td></tr>
<tr><td>Oligo fwd</td><td>CACCCCAGGGCATGCGTCGCCTCC</td><td></td><td></td><td></td><td></td><td></td><td></td><td></td><td></td><td></td><td></td></tr>
<tr><td>Oligo rev</td><td>AAACGGAGGCGACGCATGCCCTGG</td><td></td><td></td><td></td><td></td><td></td><td></td><td></td><td></td><td></td><td></td></tr>
<tr><td>Chromosome</td><td>start</td><td>end</td><td>strand</td><td>MM</td><td>target_seq</td><td>PAM</td><td>alignment</td><td>distance</td><td>position</td><td>gene name</td><td>gene id</td></tr>
<tr><td>GU305914.1</td><td>72058</td><td>72080</td><td>-</td><td>0</td><td>CCAGGGCATGCGTCGCCTCC</td><td>GGG</td><td>||||||||[||||||||||||]PAM</td><td>NA</td><td>Intergenic</td><td>NA</td><td>NA</td></tr>
<tr><td>T28795</td><td>TGCGTCGCCTCCGGGCTGCCCGG</td><td>1000</td><td></td><td></td><td></td><td></td><td></td><td></td><td></td><td></td><td></td></tr>
<tr><td>Oligo fwd</td><td>CACCTGCGTCGCCTCCGGGCTGCC</td><td></td><td></td><td></td><td></td><td></td><td></td><td></td><td></td><td></td><td></td></tr>
<tr><td>Oligo rev</td><td>AAACGGCAGCCCGGAGGCGACGCA</td><td></td><td></td><td></td><td></td><td></td><td></td><td></td><td></td><td></td><td></td></tr>
<tr><td>Chromosome</td><td>start</td><td>end</td><td>strand</td><td>MM</td><td>target_seq</td><td>PAM</td><td>alignment</td><td>distance</td><td>position</td><td>gene name</td><td>gene id</td></tr>
<tr><td>GU305914.1</td><td>72050</td><td>72072</td><td>-</td><td>0</td><td>TGCGTCGCCTCCGGGCTGCC</td><td>CGG</td><td>||||||||[||||||||||||]PAM</td><td>NA</td><td>Intergenic</td><td>NA</td><td>NA</td></tr>
<tr><td>T28796</td><td>TCCGGGCTGCCCGGCTCCACCGG</td><td>1000</td><td></td><td></td><td></td><td></td><td></td><td></td><td></td><td></td><td></td></tr>
<tr><td>Oligo fwd</td><td>CACCTCCGGGCTGCCCGGCTCCAC</td><td></td><td></td><td></td><td></td><td></td><td></td><td></td><td></td><td></td><td></td></tr>
<tr><td>Oligo rev</td><td>AAACGTGGAGCCGGGCAGCCCGGA</td><td></td><td></td><td></td><td></td><td></td><td></td><td></td><td></td><td></td><td></td></tr>
<tr><td>Chromosome</td><td>start</td><td>end</td><td>strand</td><td>MM</td><td>target_seq</td><td>PAM</td><td>alignment</td><td>distance</td><td>position</td><td>gene name</td><td>gene id</td></tr>
<tr><td>GU305914.1</td><td>72041</td><td>72063</td><td>-</td><td>0</td><td>TCCGGGCTGCCCGGCTCCAC</td><td>CGG</td><td>||||||||[||||||||||||]PAM</td><td>NA</td><td>Intergenic</td><td>NA</td><td>NA</td></tr>
<tr><td>T28797</td><td>GGGCTGCCCGGCTCCACCGGCGG</td><td>1000</td><td></td><td></td><td></td><td></td><td></td><td></td><td></td><td></td><td></td></tr>
<tr><td>Oligo fwd</td><td>CACCGGGCTGCCCGGCTCCACCGG</td><td></td><td></td><td></td><td></td><td></td><td></td><td></td><td></td><td></td><td></td></tr>
<tr><td>Oligo rev</td><td>AAACCCGGTGGAGCCGGGCAGCCC</td><td></td><td></td><td></td><td></td><td></td><td></td><td></td><td></td><td></td><td></td></tr>
<tr><td>Chromosome</td><td>start</td><td>end</td><td>strand</td><td>MM</td><td>target_seq</td><td>PAM</td><td>alignment</td><td>distance</td><td>position</td><td>gene name</td><td>gene id</td></tr>
<tr><td>GU305914.1</td><td>72038</td><td>72060</td><td>-</td><td>0</td><td>GGGCTGCCCGGCTCCACCGG</td><td>CGG</td><td>||||||||[||||||||||||]PAM</td><td>NA</td><td>Intergenic</td><td>NA</td><td>NA</td></tr>
<tr><td>T28798</td><td>CGGCGGCGTCGCCTCCGACCAGG</td><td>1000</td><td></td><td></td><td></td><td></td><td></td><td></td><td></td><td></td><td></td></tr>
<tr><td>Oligo fwd</td><td>CACCCGGCGGCGTCGCCTCCGACC</td><td></td><td></td><td></td><td></td><td></td><td></td><td></td><td></td><td></td><td></td></tr>
<tr><td>Oligo rev</td><td>AAACGGTCGGAGGCGACGCCGCCG</td><td></td><td></td><td></td><td></td><td></td><td></td><td></td><td></td><td></td><td></td></tr>
<tr><td>Chromosome</td><td>start</td><td>end</td><td>strand</td><td>MM</td><td>target_seq</td><td>PAM</td><td>alignment</td><td>distance</td><td>position</td><td>gene name</td><td>gene id</td></tr>
<tr><td>GU305914.1</td><td>72021</td><td>72043</td><td>-</td><td>0</td><td>CGGCGGCGTCGCCTCCGACC</td><td>AGG</td><td>||||||||[||||||||||||]PAM</td><td>NA</td><td>Intergenic</td><td>NA</td><td>NA</td></tr>
<tr><td>T28799</td><td>TCACATACACGAACAGTTTCAGG</td><td>1000</td><td></td><td></td><td></td><td></td><td></td><td></td><td></td><td></td><td></td></tr>
<tr><td>Oligo fwd</td><td>CACCTCACATACACGAACAGTTTC</td><td></td><td></td><td></td><td></td><td></td><td></td><td></td><td></td><td></td><td></td></tr>
<tr><td>Oligo rev</td><td>AAACGAAACTGTTCGTGTATGTGA</td><td></td><td></td><td></td><td></td><td></td><td></td><td></td><td></td><td></td><td></td></tr>
<tr><td>Chromosome</td><td>start</td><td>end</td><td>strand</td><td>MM</td><td>target_seq</td><td>PAM</td><td>alignment</td><td>distance</td><td>position</td><td>gene name</td><td>gene id</td></tr>
<tr><td>GU305914.1</td><td>71989</td><td>72011</td><td>-</td><td>0</td><td>TCACATACACGAACAGTTTC</td><td>AGG</td><td>||||||||[||||||||||||]PAM</td><td>NA</td><td>Intergenic</td><td>NA</td><td>NA</td></tr>
<tr><td>T28800</td><td>TGTTCAGCCGTCTGTCGCGCTGG</td><td>1000</td><td></td><td></td><td></td><td></td><td></td><td></td><td></td><td></td><td></td></tr>
<tr><td>Oligo fwd</td><td>CACCTGTTCAGCCGTCTGTCGCGC</td><td></td><td></td><td></td><td></td><td></td><td></td><td></td><td></td><td></td><td></td></tr>
<tr><td>Oligo rev</td><td>AAACGCGCGACAGACGGCTGAACA</td><td></td><td></td><td></td><td></td><td></td><td></td><td></td><td></td><td></td><td></td></tr>
<tr><td>Chromosome</td><td>start</td><td>end</td><td>strand</td><td>MM</td><td>target_seq</td><td>PAM</td><td>alignment</td><td>distance</td><td>position</td><td>gene name</td><td>gene id</td></tr>
<tr><td>GU305914.1</td><td>71956</td><td>71978</td><td>-</td><td>0</td><td>TGTTCAGCCGTCTGTCGCGC</td><td>TGG</td><td>||||||||[||||||||||||]PAM</td><td>NA</td><td>Intergenic</td><td>NA</td><td>NA</td></tr>
<tr><td>T28801</td><td>CAGCCGTCTGTCGCGCTGGTCGG</td><td>1000</td><td></td><td></td><td></td><td></td><td></td><td></td><td></td><td></td><td></td></tr>
<tr><td>Oligo fwd</td><td>CACCCAGCCGTCTGTCGCGCTGGT</td><td></td><td></td><td></td><td></td><td></td><td></td><td></td><td></td><td></td><td></td></tr>
<tr><td>Oligo rev</td><td>AAACACCAGCGCGACAGACGGCTG</td><td></td><td></td><td></td><td></td><td></td><td></td><td></td><td></td><td></td><td></td></tr>
<tr><td>Chromosome</td><td>start</td><td>end</td><td>strand</td><td>MM</td><td>target_seq</td><td>PAM</td><td>alignment</td><td>distance</td><td>position</td><td>gene name</td><td>gene id</td></tr>
<tr><td>GU305914.1</td><td>71952</td><td>71974</td><td>-</td><td>0</td><td>CAGCCGTCTGTCGCGCTGGT</td><td>CGG</td><td>||||||||[||||||||||||]PAM</td><td>NA</td><td>Intergenic</td><td>NA</td><td>NA</td></tr>
<tr><td>T28802</td><td>CGCTGGTCGGCCGCGCTTTCCGG</td><td>1000</td><td></td><td></td><td></td><td></td><td></td><td></td><td></td><td></td><td></td></tr>
<tr><td>Oligo fwd</td><td>CACCCGCTGGTCGGCCGCGCTTTC</td><td></td><td></td><td></td><td></td><td></td><td></td><td></td><td></td><td></td><td></td></tr>
<tr><td>Oligo rev</td><td>AAACGAAAGCGCGGCCGACCAGCG</td><td></td><td></td><td></td><td></td><td></td><td></td><td></td><td></td><td></td><td></td></tr>
<tr><td>Chromosome</td><td>start</td><td>end</td><td>strand</td><td>MM</td><td>target_seq</td><td>PAM</td><td>alignment</td><td>distance</td><td>position</td><td>gene name</td><td>gene id</td></tr>
<tr><td>GU305914.1</td><td>71939</td><td>71961</td><td>-</td><td>0</td><td>CGCTGGTCGGCCGCGCTTTC</td><td>CGG</td><td>||||||||[||||||||||||]PAM</td><td>NA</td><td>Intergenic</td><td>NA</td><td>NA</td></tr>
<tr><td>T28803</td><td>CGACCAGAACGTCTGCAGAGTGG</td><td>1000</td><td></td><td></td><td></td><td></td><td></td><td></td><td></td><td></td><td></td></tr>
<tr><td>Oligo fwd</td><td>CACCCGACCAGAACGTCTGCAGAG</td><td></td><td></td><td></td><td></td><td></td><td></td><td></td><td></td><td></td><td></td></tr>
<tr><td>Oligo rev</td><td>AAACCTCTGCAGACGTTCTGGTCG</td><td></td><td></td><td></td><td></td><td></td><td></td><td></td><td></td><td></td><td></td></tr>
<tr><td>Chromosome</td><td>start</td><td>end</td><td>strand</td><td>MM</td><td>target_seq</td><td>PAM</td><td>alignment</td><td>distance</td><td>position</td><td>gene name</td><td>gene id</td></tr>
<tr><td>GU305914.1</td><td>71865</td><td>71887</td><td>-</td><td>0</td><td>CGACCAGAACGTCTGCAGAG</td><td>TGG</td><td>||||||||[||||||||||||]PAM</td><td>NA</td><td>Intergenic</td><td>NA</td><td>NA</td></tr>
</table>

<table>
<tr><td>T28804</td><td>CCAGAACGTCTGCAGAGTGGCGG</td><td>1000</td><td></td><td></td><td></td><td></td><td></td><td></td><td></td><td></td><td></td></tr>
<tr><td>Oligo fwd</td><td>CACCCCAGAACGTCTGCAGAGTGG</td><td></td><td></td><td></td><td></td><td></td><td></td><td></td><td></td><td></td><td></td></tr>
<tr><td>Oligo rev</td><td>AAACCCACTCTGCAGACGTTCTGG</td><td></td><td></td><td></td><td></td><td></td><td></td><td></td><td></td><td></td><td></td></tr>
<tr><td>Chromosome</td><td>start</td><td>end</td><td>strand</td><td>MM</td><td>target_seq</td><td>PAM</td><td>alignment</td><td>distance</td><td>position</td><td>gene name</td><td>gene id</td></tr>
<tr><td>GU305914.1</td><td>71862</td><td>71884</td><td>-</td><td>0</td><td>CCAGAACGTCTGCAGAGTGG</td><td>CGG</td><td>||||||||[||||||||||||]PAM</td><td>NA</td><td>Intergenic</td><td>NA</td><td>NA</td></tr>
<tr><td>T28805</td><td>GGCGGCGATCGACGCCAACACGG</td><td>1000</td><td></td><td></td><td></td><td></td><td></td><td></td><td></td><td></td><td></td></tr>
<tr><td>Oligo fwd</td><td>CACCGGCGGCGATCGACGCCAACA</td><td></td><td></td><td></td><td></td><td></td><td></td><td></td><td></td><td></td><td></td></tr>
<tr><td>Oligo rev</td><td>AAACTGTTGGCGTCGATCGCCGCC</td><td></td><td></td><td></td><td></td><td></td><td></td><td></td><td></td><td></td><td></td></tr>
<tr><td>Chromosome</td><td>start</td><td>end</td><td>strand</td><td>MM</td><td>target_seq</td><td>PAM</td><td>alignment</td><td>distance</td><td>position</td><td>gene name</td><td>gene id</td></tr>
<tr><td>GU305914.1</td><td>71844</td><td>71866</td><td>-</td><td>0</td><td>GGCGGCGATCGACGCCAACA</td><td>CGG</td><td>||||||||[||||||||||||]PAM</td><td>NA</td><td>Intergenic</td><td>NA</td><td>NA</td></tr>
<tr><td>T28806</td><td>ACGCCAACACGGTCACCGACCGG</td><td>1000</td><td></td><td></td><td></td><td></td><td></td><td></td><td></td><td></td><td></td></tr>
<tr><td>Oligo fwd</td><td>CACCACGCCAACACGGTCACCGAC</td><td></td><td></td><td></td><td></td><td></td><td></td><td></td><td></td><td></td><td></td></tr>
<tr><td>Oligo rev</td><td>AAACGTCGGTGACCGTGTTGGCGT</td><td></td><td></td><td></td><td></td><td></td><td></td><td></td><td></td><td></td><td></td></tr>
<tr><td>Chromosome</td><td>start</td><td>end</td><td>strand</td><td>MM</td><td>target_seq</td><td>PAM</td><td>alignment</td><td>distance</td><td>position</td><td>gene name</td><td>gene id</td></tr>
<tr><td>GU305914.1</td><td>71833</td><td>71855</td><td>-</td><td>0</td><td>ACGCCAACACGGTCACCGAC</td><td>CGG</td><td>||||||||[||||||||||||]PAM</td><td>NA</td><td>Intergenic</td><td>NA</td><td>NA</td></tr>
<tr><td>T28807</td><td>GCGTCACCGTCGCATTCTGCTGG</td><td>1000</td><td></td><td></td><td></td><td></td><td></td><td></td><td></td><td></td><td></td></tr>
<tr><td>Oligo fwd</td><td>CACCGCGTCACCGTCGCATTCTGC</td><td></td><td></td><td></td><td></td><td></td><td></td><td></td><td></td><td></td><td></td></tr>
<tr><td>Oligo rev</td><td>AAACGCAGAATGCGACGGTGACGC</td><td></td><td></td><td></td><td></td><td></td><td></td><td></td><td></td><td></td><td></td></tr>
<tr><td>Chromosome</td><td>start</td><td>end</td><td>strand</td><td>MM</td><td>target_seq</td><td>PAM</td><td>alignment</td><td>distance</td><td>position</td><td>gene name</td><td>gene id</td></tr>
<tr><td>GU305914.1</td><td>71803</td><td>71825</td><td>-</td><td>0</td><td>GCGTCACCGTCGCATTCTGC</td><td>TGG</td><td>||||||||[||||||||||||]PAM</td><td>NA</td><td>Intergenic</td><td>NA</td><td>NA</td></tr>
<tr><td>T28808</td><td>CTGCTGGTCTTTCTGTCGAGAGG</td><td>1000</td><td></td><td></td><td></td><td></td><td></td><td></td><td></td><td></td><td></td></tr>
<tr><td>Oligo fwd</td><td>CACCCTGCTGGTCTTTCTGTCGAG</td><td></td><td></td><td></td><td></td><td></td><td></td><td></td><td></td><td></td><td></td></tr>
<tr><td>Oligo rev</td><td>AAACCTCGACAGAAAGACCAGCAG</td><td></td><td></td><td></td><td></td><td></td><td></td><td></td><td></td><td></td><td></td></tr>
<tr><td>Chromosome</td><td>start</td><td>end</td><td>strand</td><td>MM</td><td>target_seq</td><td>PAM</td><td>alignment</td><td>distance</td><td>position</td><td>gene name</td><td>gene id</td></tr>
<tr><td>GU305914.1</td><td>71787</td><td>71809</td><td>-</td><td>0</td><td>CTGCTGGTCTTTCTGTCGAG</td><td>AGG</td><td>||||||||[||||||||||||]PAM</td><td>NA</td><td>Intergenic</td><td>NA</td><td>NA</td></tr>
<tr><td>T28809</td><td>CGCCGCCGCTGCTGCTGCTGAGG</td><td>1000</td><td></td><td></td><td></td><td></td><td></td><td></td><td></td><td></td><td></td></tr>
<tr><td>Oligo fwd</td><td>CACCCGCCGCCGCTGCTGCTGCTG</td><td></td><td></td><td></td><td></td><td></td><td></td><td></td><td></td><td></td><td></td></tr>
<tr><td>Oligo rev</td><td>AAACCAGCAGCAGCAGCGGCGGCG</td><td></td><td></td><td></td><td></td><td></td><td></td><td></td><td></td><td></td><td></td></tr>
<tr><td>Chromosome</td><td>start</td><td>end</td><td>strand</td><td>MM</td><td>target_seq</td><td>PAM</td><td>alignment</td><td>distance</td><td>position</td><td>gene name</td><td>gene id</td></tr>
<tr><td>GU305914.1</td><td>71746</td><td>71768</td><td>-</td><td>0</td><td>CGCCGCCGCTGCTGCTGCTG</td><td>AGG</td><td>||||||||[||||||||||||]PAM</td><td>NA</td><td>Intergenic</td><td>NA</td><td>NA</td></tr>
<tr><td>GU305914.1</td><td>71743</td><td>71765</td><td>-</td><td>3</td><td>CGCCGCTGCTGCTGCTGAGG</td><td>AGG</td><td>||||||-|[|||||||||--|]PAM</td><td>NA</td><td>Intergenic</td><td>NA</td><td>NA</td></tr>
<tr><td>T28810</td><td>CGCCGCTGCTGCTGCTGAGGAGG</td><td>1000</td><td></td><td></td><td></td><td></td><td></td><td></td><td></td><td></td><td></td></tr>
<tr><td>Oligo fwd</td><td>CACCCGCCGCTGCTGCTGCTGAGG</td><td></td><td></td><td></td><td></td><td></td><td></td><td></td><td></td><td></td><td></td></tr>
<tr><td>Oligo rev</td><td>AAACCCTCAGCAGCAGCAGCGGCG</td><td></td><td></td><td></td><td></td><td></td><td></td><td></td><td></td><td></td><td></td></tr>
<tr><td>Chromosome</td><td>start</td><td>end</td><td>strand</td><td>MM</td><td>target_seq</td><td>PAM</td><td>alignment</td><td>distance</td><td>position</td><td>gene name</td><td>gene id</td></tr>
<tr><td>GU305914.1</td><td>71743</td><td>71765</td><td>-</td><td>0</td><td>CGCCGCTGCTGCTGCTGAGG</td><td>AGG</td><td>||||||||[||||||||||||]PAM</td><td>NA</td><td>Intergenic</td><td>NA</td><td>NA</td></tr>
<tr><td>GU305914.1</td><td>71746</td><td>71768</td><td>-</td><td>3</td><td>CGCCGCCGCTGCTGCTGCTG</td><td>AGG</td><td>||||||-|[|||||||||--|]PAM</td><td>NA</td><td>Intergenic</td><td>NA</td><td>NA</td></tr>
<tr><td>T28811</td><td>TGAGGAGGCCCTTCGTCCCGCGG</td><td>1000</td><td></td><td></td><td></td><td></td><td></td><td></td><td></td><td></td><td></td></tr>
<tr><td>Oligo fwd</td><td>CACCTGAGGAGGCCCTTCGTCCCG</td><td></td><td></td><td></td><td></td><td></td><td></td><td></td><td></td><td></td><td></td></tr>
<tr><td>Oligo rev</td><td>AAACCGGGACGAAGGGCCTCCTCA</td><td></td><td></td><td></td><td></td><td></td><td></td><td></td><td></td><td></td><td></td></tr>
<tr><td>Chromosome</td><td>start</td><td>end</td><td>strand</td><td>MM</td><td>target_seq</td><td>PAM</td><td>alignment</td><td>distance</td><td>position</td><td>gene name</td><td>gene id</td></tr>
<tr><td>GU305914.1</td><td>71728</td><td>71750</td><td>-</td><td>0</td><td>TGAGGAGGCCCTTCGTCCCG</td><td>CGG</td><td>||||||||[||||||||||||]PAM</td><td>NA</td><td>Intergenic</td><td>NA</td><td>NA</td></tr>
<tr><td>T28812</td><td>AGTTCCCGTGCGTCGCGTACAGG</td><td>1000</td><td></td><td></td><td></td><td></td><td></td><td></td><td></td><td></td><td></td></tr>
<tr><td>Oligo fwd</td><td>CACCAGTTCCCGTGCGTCGCGTAC</td><td></td><td></td><td></td><td></td><td></td><td></td><td></td><td></td><td></td><td></td></tr>
<tr><td>Oligo rev</td><td>AAACGTACGCGACGCACGGGAACT</td><td></td><td></td><td></td><td></td><td></td><td></td><td></td><td></td><td></td><td></td></tr>
<tr><td>Chromosome</td><td>start</td><td>end</td><td>strand</td><td>MM</td><td>target_seq</td><td>PAM</td><td>alignment</td><td>distance</td><td>position</td><td>gene name</td><td>gene id</td></tr>
<tr><td>GU305914.1</td><td>71680</td><td>71702</td><td>-</td><td>0</td><td>AGTTCCCGTGCGTCGCGTAC</td><td>AGG</td><td>||||||||[||||||||||||]PAM</td><td>NA</td><td>Intergenic</td><td>NA</td><td>NA</td></tr>
<tr><td>T28813</td><td>GCGTCGCGTACAGGTACAACTGG</td><td>1000</td><td></td><td></td><td></td><td></td><td></td><td></td><td></td><td></td><td></td></tr>
<tr><td>Oligo fwd</td><td>CACCGCGTCGCGTACAGGTACAAC</td><td></td><td></td><td></td><td></td><td></td><td></td><td></td><td></td><td></td><td></td></tr>
<tr><td>Oligo rev</td><td>AAACGTTGTACCTGTACGCGACGC</td><td></td><td></td><td></td><td></td><td></td><td></td><td></td><td></td><td></td><td></td></tr>
<tr><td>Chromosome</td><td>start</td><td>end</td><td>strand</td><td>MM</td><td>target_seq</td><td>PAM</td><td>alignment</td><td>distance</td><td>position</td><td>gene name</td><td>gene id</td></tr>
<tr><td>GU305914.1</td><td>71671</td><td>71693</td><td>-</td><td>0</td><td>GCGTCGCGTACAGGTACAAC</td><td>TGG</td><td>||||||||[||||||||||||]PAM</td><td>NA</td><td>Intergenic</td><td>NA</td><td>NA</td></tr>
<tr><td>T28814</td><td>CACGTCCGTCCCGCGATATAAGG</td><td>1000</td><td></td><td></td><td></td><td></td><td></td><td></td><td></td><td></td><td></td></tr>
<tr><td>Oligo fwd</td><td>CACCCACGTCCGTCCCGCGATATA</td><td></td><td></td><td></td><td></td><td></td><td></td><td></td><td></td><td></td><td></td></tr>
<tr><td>Oligo rev</td><td>AAACTATATCGCGGGACGGACGTG</td><td></td><td></td><td></td><td></td><td></td><td></td><td></td><td></td><td></td><td></td></tr>
<tr><td>Chromosome</td><td>start</td><td>end</td><td>strand</td><td>MM</td><td>target_seq</td><td>PAM</td><td>alignment</td><td>distance</td><td>position</td><td>gene name</td><td>gene id</td></tr>
<tr><td>GU305914.1</td><td>71613</td><td>71635</td><td>-</td><td>0</td><td>CACGTCCGTCCCGCGATATA</td><td>AGG</td><td>||||||||[||||||||||||]PAM</td><td>NA</td><td>Intergenic</td><td>NA</td><td>NA</td></tr>
<tr><td>T28815</td><td>GTCCGTCCCGCGATATAAGGCGG</td><td>1000</td><td></td><td></td><td></td><td></td><td></td><td></td><td></td><td></td><td></td></tr>
<tr><td>Oligo fwd</td><td>CACCGTCCGTCCCGCGATATAAGG</td><td></td><td></td><td></td><td></td><td></td><td></td><td></td><td></td><td></td><td></td></tr>
<tr><td>Oligo rev</td><td>AAACCCTTATATCGCGGGACGGAC</td><td></td><td></td><td></td><td></td><td></td><td></td><td></td><td></td><td></td><td></td></tr>
<tr><td>Chromosome</td><td>start</td><td>end</td><td>strand</td><td>MM</td><td>target_seq</td><td>PAM</td><td>alignment</td><td>distance</td><td>position</td><td>gene name</td><td>gene id</td></tr>
<tr><td>GU305914.1</td><td>71610</td><td>71632</td><td>-</td><td>0</td><td>GTCCGTCCCGCGATATAAGG</td><td>CGG</td><td>||||||||[||||||||||||]PAM</td><td>NA</td><td>Intergenic</td><td>NA</td><td>NA</td></tr>
<tr><td>T28816</td><td>TGACGTCGCATGCTGACAGTCGG</td><td>1000</td><td></td><td></td><td></td><td></td><td></td><td></td><td></td><td></td><td></td></tr>
<tr><td>Oligo fwd</td><td>CACCTGACGTCGCATGCTGACAGT</td><td></td><td></td><td></td><td></td><td></td><td></td><td></td><td></td><td></td><td></td></tr>
<tr><td>Oligo rev</td><td>AAACACTGTCAGCATGCGACGTCA</td><td></td><td></td><td></td><td></td><td></td><td></td><td></td><td></td><td></td><td></td></tr>
<tr><td>Chromosome</td><td>start</td><td>end</td><td>strand</td><td>MM</td><td>target_seq</td><td>PAM</td><td>alignment</td><td>distance</td><td>position</td><td>gene name</td><td>gene id</td></tr>
<tr><td>GU305914.1</td><td>71584</td><td>71606</td><td>-</td><td>0</td><td>TGACGTCGCATGCTGACAGT</td><td>CGG</td><td>||||||||[||||||||||||]PAM</td><td>NA</td><td>Intergenic</td><td>NA</td><td>NA</td></tr>
<tr><td>T28817</td><td>GACGTCGCATGCTGACAGTCGGG</td><td>1000</td><td></td><td></td><td></td><td></td><td></td><td></td><td></td><td></td><td></td></tr>
<tr><td>Oligo fwd</td><td>CACCGACGTCGCATGCTGACAGTC</td><td></td><td></td><td></td><td></td><td></td><td></td><td></td><td></td><td></td><td></td></tr>
<tr><td>Oligo rev</td><td>AAACGACTGTCAGCATGCGACGTC</td><td></td><td></td><td></td><td></td><td></td><td></td><td></td><td></td><td></td><td></td></tr>
<tr><td>Chromosome</td><td>start</td><td>end</td><td>strand</td><td>MM</td><td>target_seq</td><td>PAM</td><td>alignment</td><td>distance</td><td>position</td><td>gene name</td><td>gene id</td></tr>
<tr><td>GU305914.1</td><td>71583</td><td>71605</td><td>-</td><td>0</td><td>GACGTCGCATGCTGACAGTC</td><td>GGG</td><td>||||||||[||||||||||||]PAM</td><td>NA</td><td>Intergenic</td><td>NA</td><td>NA</td></tr>
<tr><td>T28818</td><td>CGCATGCTGACAGTCGGGACAGG</td><td>1000</td><td></td><td></td><td></td><td></td><td></td><td></td><td></td><td></td><td></td></tr>
<tr><td>Oligo fwd</td><td>CACCCGCATGCTGACAGTCGGGAC</td><td></td><td></td><td></td><td></td><td></td><td></td><td></td><td></td><td></td><td></td></tr>
</table>

Oligo rev AAACGTCCCGACTGTCAGCATGCG
Chromosome start end strand MM target_seq PAM alignment distance position gene name gene id
GU305914.1 71578 71600 - 0 CGCATGCTGACAGTCGGGAC AGG ||||||||[||||||||||||]PAM NA Intergenic NA NA

T28819 ACAGTCGGGACAGGTCGATACGG 1000
Oligo fwd CACCACAGTCGGGACAGGTCGATA
Oligo rev AAACTATCGACCTGTCCCGACTGT
Chromosome start end strand MM target_seq PAM alignment distance position gene name gene id
GU305914.1 71569 71591 - 0 ACAGTCGGGACAGGTCGATA CGG ||||||||[||||||||||||]PAM NA Intergenic NA NA

T28820 CAGTCGGGACAGGTCGATACGGG 1000
Oligo fwd CACCCAGTCGGGACAGGTCGATAC
Oligo rev AAACGTATCGACCTGTCCCGACTG
Chromosome start end strand MM target_seq PAM alignment distance position gene name gene id
GU305914.1 71568 71590 - 0 CAGTCGGGACAGGTCGATAC GGG ||||||||[||||||||||||]PAM NA Intergenic NA NA

T28821 TCGATACGGGTCCAGTCGAACGG 1000
Oligo fwd CACCTCGATACGGGTCCAGTCGAA
Oligo rev AAACTTCGACTGGACCCGTATCGA
Chromosome start end strand MM target_seq PAM alignment distance position gene name gene id
GU305914.1 71555 71577 - 0 TCGATACGGGTCCAGTCGAA CGG ||||||||[||||||||||||]PAM NA Intergenic NA NA

T28822 CAGTCGAACGGCTCCTGCCCTGG 1000
Oligo fwd CACCCAGTCGAACGGCTCCTGCCC
Oligo rev AAACGGGCAGGAGCCGTTCGACTG
Chromosome start end strand MM target_seq PAM alignment distance position gene name gene id
GU305914.1 71543 71565 - 0 CAGTCGAACGGCTCCTGCCC TGG ||||||||[||||||||||||]PAM NA Intergenic NA NA

T28823 CCCTGGCTTATCGTCCTCACCGG 1000
Oligo fwd CACCCCCTGGCTTATCGTCCTCAC
Oligo rev AAACGTGAGGACGATAAGCCAGGG
Chromosome start end strand MM target_seq PAM alignment distance position gene name gene id
GU305914.1 71526 71548 - 0 CCCTGGCTTATCGTCCTCAC CGG ||||||||[||||||||||||]PAM NA Intergenic NA NA

T28824 CCTGGCTTATCGTCCTCACCGGG 1000
Oligo fwd CACCCCTGGCTTATCGTCCTCACC
Oligo rev AAACGGTGAGGACGATAAGCCAGG
Chromosome start end strand MM target_seq PAM alignment distance position gene name gene id
GU305914.1 71525 71547 - 0 CCTGGCTTATCGTCCTCACC GGG ||||||||[||||||||||||]PAM NA Intergenic NA NA

T28825 CTGGCTTATCGTCCTCACCGGGG 1000
Oligo fwd CACCCTGGCTTATCGTCCTCACCG
Oligo rev AAACCGGTGAGGACGATAAGCCAG
Chromosome start end strand MM target_seq PAM alignment distance position gene name gene id
GU305914.1 71524 71546 - 0 CTGGCTTATCGTCCTCACCG GGG ||||||||[||||||||||||]PAM NA Intergenic NA NA

T28826 CCGGGGCCACCCACCAGAGCCGG 1000
Oligo fwd CACCCCGGGGCCACCCACCAGAGC
Oligo rev AAACGCTCTGGTGGGTGGCCCCGG
Chromosome start end strand MM target_seq PAM alignment distance position gene name gene id
GU305914.1 71507 71529 - 0 CCGGGGCCACCCACCAGAGC CGG ||||||||[||||||||||||]PAM NA Intergenic NA NA

T28827 CGGGGCCACCCACCAGAGCCGGG 1000
Oligo fwd CACCCGGGGCCACCCACCAGAGCC
Oligo rev AAACGGCTCTGGTGGGTGGCCCCG
Chromosome start end strand MM target_seq PAM alignment distance position gene name gene id
GU305914.1 71506 71528 - 0 CGGGGCCACCCACCAGAGCC GGG ||||||||[||||||||||||]PAM NA Intergenic NA NA

T28828 TTTTCCGCCTGACTGATCAACGG 1000
Oligo fwd CACCTTTTCCGCCTGACTGATCAA
Oligo rev AAACTTGATCAGTCAGGCGGAAAA
Chromosome start end strand MM target_seq PAM alignment distance position gene name gene id
GU305914.1 71480 71502 - 0 TTTTCCGCCTGACTGATCAA CGG ||||||||[||||||||||||]PAM NA Intergenic NA NA

T28829 CTGACTGATCAACGGCTGCGAGG 1000
Oligo fwd CACCCTGACTGATCAACGGCTGCG
Oligo rev AAACCGCAGCCGTTGATCAGTCAG
Chromosome start end strand MM target_seq PAM alignment distance position gene name gene id
GU305914.1 71472 71494 - 0 CTGACTGATCAACGGCTGCG AGG ||||||||[||||||||||||]PAM NA Intergenic NA NA

T28830 AGGACACGAAGTTATCCAGCAGG 1000
Oligo fwd CACCAGGACACGAAGTTATCCAGC
Oligo rev AAACGCTGGATAACTTCGTGTCCT
Chromosome start end strand MM target_seq PAM alignment distance position gene name gene id
GU305914.1 71452 71474 - 0 AGGACACGAAGTTATCCAGC AGG ||||||||[||||||||||||]PAM NA Intergenic NA NA

T28831 GGACACGAAGTTATCCAGCAGGG 1000
Oligo fwd CACCGGACACGAAGTTATCCAGCA
Oligo rev AAACTGCTGGATAACTTCGTGTCC
Chromosome start end strand MM target_seq PAM alignment distance position gene name gene id
GU305914.1 71451 71473 - 0 GGACACGAAGTTATCCAGCA GGG ||||||||[||||||||||||]PAM NA Intergenic NA NA

T28832 CGAAGTTATCCAGCAGGGTGCGG 1000
Oligo fwd CACCCGAAGTTATCCAGCAGGGTG
Oligo rev AAACCACCCTGCTGGATAACTTCG
Chromosome start end strand MM target_seq PAM alignment distance position gene name gene id
GU305914.1 71446 71468 - 0 CGAAGTTATCCAGCAGGGTG CGG ||||||||[||||||||||||]PAM NA Intergenic NA NA

T28833 CCAGCGCCGCGATGCCCAACAGG 1000

<table>
<tr><td>Oligo fwd</td><td>CACCCCAGCGCCGCGATGCCCAAC</td><td></td><td></td><td></td><td></td><td></td><td></td><td></td><td></td><td></td><td></td></tr>
<tr><td>Oligo rev</td><td>AAACGTTGGGCATCGCGGCGCTGG</td><td></td><td></td><td></td><td></td><td></td><td></td><td></td><td></td><td></td><td></td></tr>
<tr><td>Chromosome</td><td>start</td><td>end</td><td>strand</td><td>MM</td><td>target_seq</td><td>PAM</td><td>alignment</td><td>distance</td><td>position</td><td>gene name</td><td>gene id</td></tr>
<tr><td>GU305914.1</td><td>71404</td><td>71426</td><td>-</td><td>0</td><td>CCAGCGCCGCGATGCCCAAC</td><td>AGG</td><td>||||||||[||||||||||||]PAM</td><td>NA</td><td>Intergenic</td><td>NA</td><td>NA</td></tr>
<tr><td></td><td></td><td></td><td></td><td></td><td></td><td></td><td></td><td></td><td></td><td></td><td></td></tr>
<tr><td>T28834</td><td>CGCGATGCCCAACAGGAGCGAGG</td><td>1000</td><td></td><td></td><td></td><td></td><td></td><td></td><td></td><td></td><td></td></tr>
<tr><td>Oligo fwd</td><td>CACCCGCGATGCCCAACAGGAGCG</td><td></td><td></td><td></td><td></td><td></td><td></td><td></td><td></td><td></td><td></td></tr>
<tr><td>Oligo rev</td><td>AAACCGCTCCTGTTGGGCATCGCG</td><td></td><td></td><td></td><td></td><td></td><td></td><td></td><td></td><td></td><td></td></tr>
<tr><td>Chromosome</td><td>start</td><td>end</td><td>strand</td><td>MM</td><td>target_seq</td><td>PAM</td><td>alignment</td><td>distance</td><td>position</td><td>gene name</td><td>gene id</td></tr>
<tr><td>GU305914.1</td><td>71397</td><td>71419</td><td>-</td><td>0</td><td>CGCGATGCCCAACAGGAGCG</td><td>AGG</td><td>||||||||[||||||||||||]PAM</td><td>NA</td><td>Intergenic</td><td>NA</td><td>NA</td></tr>
<tr><td></td><td></td><td></td><td></td><td></td><td></td><td></td><td></td><td></td><td></td><td></td><td></td></tr>
<tr><td>T28835</td><td>GAGCGAGGAGATCGAGATCTTGG</td><td>1000</td><td></td><td></td><td></td><td></td><td></td><td></td><td></td><td></td><td></td></tr>
<tr><td>Oligo fwd</td><td>CACCGAGCGAGGAGATCGAGATCT</td><td></td><td></td><td></td><td></td><td></td><td></td><td></td><td></td><td></td><td></td></tr>
<tr><td>Oligo rev</td><td>AAACAGATCTCGATCTCCTCGCTC</td><td></td><td></td><td></td><td></td><td></td><td></td><td></td><td></td><td></td><td></td></tr>
<tr><td>Chromosome</td><td>start</td><td>end</td><td>strand</td><td>MM</td><td>target_seq</td><td>PAM</td><td>alignment</td><td>distance</td><td>position</td><td>gene name</td><td>gene id</td></tr>
<tr><td>GU305914.1</td><td>71382</td><td>71404</td><td>-</td><td>0</td><td>GAGCGAGGAGATCGAGATCT</td><td>TGG</td><td>||||||||[||||||||||||]PAM</td><td>NA</td><td>Intergenic</td><td>NA</td><td>NA</td></tr>
<tr><td></td><td></td><td></td><td></td><td></td><td></td><td></td><td></td><td></td><td></td><td></td><td></td></tr>
<tr><td>T28836</td><td>GCCGCCGTCGTGCTCTCTTCAGG</td><td>1000</td><td></td><td></td><td></td><td></td><td></td><td></td><td></td><td></td><td></td></tr>
<tr><td>Oligo fwd</td><td>CACCGCCGCCGTCGTGCTCTCTTC</td><td></td><td></td><td></td><td></td><td></td><td></td><td></td><td></td><td></td><td></td></tr>
<tr><td>Oligo rev</td><td>AAACGAAGAGAGCACGACGGCGGC</td><td></td><td></td><td></td><td></td><td></td><td></td><td></td><td></td><td></td><td></td></tr>
<tr><td>Chromosome</td><td>start</td><td>end</td><td>strand</td><td>MM</td><td>target_seq</td><td>PAM</td><td>alignment</td><td>distance</td><td>position</td><td>gene name</td><td>gene id</td></tr>
<tr><td>GU305914.1</td><td>71360</td><td>71382</td><td>-</td><td>0</td><td>GCCGCCGTCGTGCTCTCTTC</td><td>AGG</td><td>||||||||[||||||||||||]PAM</td><td>NA</td><td>Intergenic</td><td>NA</td><td>NA</td></tr>
<tr><td>GU305914.1</td><td>168983</td><td>169005</td><td>+</td><td>4</td><td>GACGACATCGTGCTCTTTTC</td><td>GGG</td><td>|-||-|-|[||||||||-|||]PAM</td><td>NA</td><td>Intergenic</td><td>NA</td><td>NA</td></tr>
<tr><td></td><td></td><td></td><td></td><td></td><td></td><td></td><td></td><td></td><td></td><td></td><td></td></tr>
<tr><td>T28837</td><td>GCCCACCTCCTCCCCCGTCGAGG</td><td>1000</td><td></td><td></td><td></td><td></td><td></td><td></td><td></td><td></td><td></td></tr>
<tr><td>Oligo fwd</td><td>CACCGCCCACCTCCTCCCCCGTCG</td><td></td><td></td><td></td><td></td><td></td><td></td><td></td><td></td><td></td><td></td></tr>
<tr><td>Oligo rev</td><td>AAACCGACGGGGGAGGAGGTGGGC</td><td></td><td></td><td></td><td></td><td></td><td></td><td></td><td></td><td></td><td></td></tr>
<tr><td>Chromosome</td><td>start</td><td>end</td><td>strand</td><td>MM</td><td>target_seq</td><td>PAM</td><td>alignment</td><td>distance</td><td>position</td><td>gene name</td><td>gene id</td></tr>
<tr><td>GU305914.1</td><td>71334</td><td>71356</td><td>-</td><td>0</td><td>GCCCACCTCCTCCCCCGTCG</td><td>AGG</td><td>||||||||[||||||||||||]PAM</td><td>NA</td><td>Intergenic</td><td>NA</td><td>NA</td></tr>
<tr><td></td><td></td><td></td><td></td><td></td><td></td><td></td><td></td><td></td><td></td><td></td><td></td></tr>
<tr><td>T28838</td><td>CCCACCTCCTCCCCCGTCGAGGG</td><td>1000</td><td></td><td></td><td></td><td></td><td></td><td></td><td></td><td></td><td></td></tr>
<tr><td>Oligo fwd</td><td>CACCCCCACCTCCTCCCCCGTCGA</td><td></td><td></td><td></td><td></td><td></td><td></td><td></td><td></td><td></td><td></td></tr>
<tr><td>Oligo rev</td><td>AAACTCGACGGGGGAGGAGGTGGG</td><td></td><td></td><td></td><td></td><td></td><td></td><td></td><td></td><td></td><td></td></tr>
<tr><td>Chromosome</td><td>start</td><td>end</td><td>strand</td><td>MM</td><td>target_seq</td><td>PAM</td><td>alignment</td><td>distance</td><td>position</td><td>gene name</td><td>gene id</td></tr>
<tr><td>GU305914.1</td><td>71333</td><td>71355</td><td>-</td><td>0</td><td>CCCACCTCCTCCCCCGTCGA</td><td>GGG</td><td>||||||||[||||||||||||]PAM</td><td>NA</td><td>Intergenic</td><td>NA</td><td>NA</td></tr>
<tr><td></td><td></td><td></td><td></td><td></td><td></td><td></td><td></td><td></td><td></td><td></td><td></td></tr>
<tr><td>T28839</td><td>CCACCTCCTCCCCCGTCGAGGGG</td><td>1000</td><td></td><td></td><td></td><td></td><td></td><td></td><td></td><td></td><td></td></tr>
<tr><td>Oligo fwd</td><td>CACCCCACCTCCTCCCCCGTCGAG</td><td></td><td></td><td></td><td></td><td></td><td></td><td></td><td></td><td></td><td></td></tr>
<tr><td>Oligo rev</td><td>AAACCTCGACGGGGGAGGAGGTGG</td><td></td><td></td><td></td><td></td><td></td><td></td><td></td><td></td><td></td><td></td></tr>
<tr><td>Chromosome</td><td>start</td><td>end</td><td>strand</td><td>MM</td><td>target_seq</td><td>PAM</td><td>alignment</td><td>distance</td><td>position</td><td>gene name</td><td>gene id</td></tr>
<tr><td>GU305914.1</td><td>71332</td><td>71354</td><td>-</td><td>0</td><td>CCACCTCCTCCCCCGTCGAG</td><td>GGG</td><td>||||||||[||||||||||||]PAM</td><td>NA</td><td>Intergenic</td><td>NA</td><td>NA</td></tr>
<tr><td></td><td></td><td></td><td></td><td></td><td></td><td></td><td></td><td></td><td></td><td></td><td></td></tr>
<tr><td>T28840</td><td>CACCTCCTCCCCCGTCGAGGGGG</td><td>1000</td><td></td><td></td><td></td><td></td><td></td><td></td><td></td><td></td><td></td></tr>
<tr><td>Oligo fwd</td><td>CACCCACCTCCTCCCCCGTCGAGG</td><td></td><td></td><td></td><td></td><td></td><td></td><td></td><td></td><td></td><td></td></tr>
<tr><td>Oligo rev</td><td>AAACCCTCGACGGGGGAGGAGGTG</td><td></td><td></td><td></td><td></td><td></td><td></td><td></td><td></td><td></td><td></td></tr>
<tr><td>Chromosome</td><td>start</td><td>end</td><td>strand</td><td>MM</td><td>target_seq</td><td>PAM</td><td>alignment</td><td>distance</td><td>position</td><td>gene name</td><td>gene id</td></tr>
<tr><td>GU305914.1</td><td>71331</td><td>71353</td><td>-</td><td>0</td><td>CACCTCCTCCCCCGTCGAGG</td><td>GGG</td><td>||||||||[||||||||||||]PAM</td><td>NA</td><td>Intergenic</td><td>NA</td><td>NA</td></tr>
<tr><td></td><td></td><td></td><td></td><td></td><td></td><td></td><td></td><td></td><td></td><td></td><td></td></tr>
<tr><td>T28841</td><td>ACCTCCTCCCCCGTCGAGGGGGG</td><td>1000</td><td></td><td></td><td></td><td></td><td></td><td></td><td></td><td></td><td></td></tr>
<tr><td>Oligo fwd</td><td>CACCACCTCCTCCCCCGTCGAGGG</td><td></td><td></td><td></td><td></td><td></td><td></td><td></td><td></td><td></td><td></td></tr>
<tr><td>Oligo rev</td><td>AAACCCCTCGACGGGGGAGGAGGT</td><td></td><td></td><td></td><td></td><td></td><td></td><td></td><td></td><td></td><td></td></tr>
<tr><td>Chromosome</td><td>start</td><td>end</td><td>strand</td><td>MM</td><td>target_seq</td><td>PAM</td><td>alignment</td><td>distance</td><td>position</td><td>gene name</td><td>gene id</td></tr>
<tr><td>GU305914.1</td><td>71330</td><td>71352</td><td>-</td><td>0</td><td>ACCTCCTCCCCCGTCGAGGG</td><td>GGG</td><td>||||||||[||||||||||||]PAM</td><td>NA</td><td>Intergenic</td><td>NA</td><td>NA</td></tr>
<tr><td></td><td></td><td></td><td></td><td></td><td></td><td></td><td></td><td></td><td></td><td></td><td></td></tr>
<tr><td>T28842</td><td>GCTGCCCGATCTCCGACTTCAGG</td><td>1000</td><td></td><td></td><td></td><td></td><td></td><td></td><td></td><td></td><td></td></tr>
<tr><td>Oligo fwd</td><td>CACCGCTGCCCGATCTCCGACTTC</td><td></td><td></td><td></td><td></td><td></td><td></td><td></td><td></td><td></td><td></td></tr>
<tr><td>Oligo rev</td><td>AAACGAAGTCGGAGATCGGGCAGC</td><td></td><td></td><td></td><td></td><td></td><td></td><td></td><td></td><td></td><td></td></tr>
<tr><td>Chromosome</td><td>start</td><td>end</td><td>strand</td><td>MM</td><td>target_seq</td><td>PAM</td><td>alignment</td><td>distance</td><td>position</td><td>gene name</td><td>gene id</td></tr>
<tr><td>GU305914.1</td><td>71296</td><td>71318</td><td>-</td><td>0</td><td>GCTGCCCGATCTCCGACTTC</td><td>AGG</td><td>||||||||[||||||||||||]PAM</td><td>NA</td><td>Intergenic</td><td>NA</td><td>NA</td></tr>
<tr><td></td><td></td><td></td><td></td><td></td><td></td><td></td><td></td><td></td><td></td><td></td><td></td></tr>
<tr><td>T28843</td><td>GCCCGATCTCCGACTTCAGGAGG</td><td>1000</td><td></td><td></td><td></td><td></td><td></td><td></td><td></td><td></td><td></td></tr>
<tr><td>Oligo fwd</td><td>CACCGCCCGATCTCCGACTTCAGG</td><td></td><td></td><td></td><td></td><td></td><td></td><td></td><td></td><td></td><td></td></tr>
<tr><td>Oligo rev</td><td>AAACCCTGAAGTCGGAGATCGGGC</td><td></td><td></td><td></td><td></td><td></td><td></td><td></td><td></td><td></td><td></td></tr>
<tr><td>Chromosome</td><td>start</td><td>end</td><td>strand</td><td>MM</td><td>target_seq</td><td>PAM</td><td>alignment</td><td>distance</td><td>position</td><td>gene name</td><td>gene id</td></tr>
<tr><td>GU305914.1</td><td>71293</td><td>71315</td><td>-</td><td>0</td><td>GCCCGATCTCCGACTTCAGG</td><td>AGG</td><td>||||||||[||||||||||||]PAM</td><td>NA</td><td>Intergenic</td><td>NA</td><td>NA</td></tr>
<tr><td></td><td></td><td></td><td></td><td></td><td></td><td></td><td></td><td></td><td></td><td></td><td></td></tr>
<tr><td>T28844</td><td>CCAGCCCGTACTTCGAGCTGCGG</td><td>1000</td><td></td><td></td><td></td><td></td><td></td><td></td><td></td><td></td><td></td></tr>
<tr><td>Oligo fwd</td><td>CACCCCAGCCCGTACTTCGAGCTG</td><td></td><td></td><td></td><td></td><td></td><td></td><td></td><td></td><td></td><td></td></tr>
<tr><td>Oligo rev</td><td>AAACCAGCTCGAAGTACGGGCTGG</td><td></td><td></td><td></td><td></td><td></td><td></td><td></td><td></td><td></td><td></td></tr>
<tr><td>Chromosome</td><td>start</td><td>end</td><td>strand</td><td>MM</td><td>target_seq</td><td>PAM</td><td>alignment</td><td>distance</td><td>position</td><td>gene name</td><td>gene id</td></tr>
<tr><td>GU305914.1</td><td>71242</td><td>71264</td><td>-</td><td>0</td><td>CCAGCCCGTACTTCGAGCTG</td><td>CGG</td><td>||||||||[||||||||||||]PAM</td><td>NA</td><td>Intergenic</td><td>NA</td><td>NA</td></tr>
<tr><td></td><td></td><td></td><td></td><td></td><td></td><td></td><td></td><td></td><td></td><td></td><td></td></tr>
<tr><td>T28845</td><td>CGACAGCTTCTCCAGATCTTCGG</td><td>1000</td><td></td><td></td><td></td><td></td><td></td><td></td><td></td><td></td><td></td></tr>
<tr><td>Oligo fwd</td><td>CACCCGACAGCTTCTCCAGATCTT</td><td></td><td></td><td></td><td></td><td></td><td></td><td></td><td></td><td></td><td></td></tr>
<tr><td>Oligo rev</td><td>AAACAAGATCTGGAGAAGCTGTCG</td><td></td><td></td><td></td><td></td><td></td><td></td><td></td><td></td><td></td><td></td></tr>
<tr><td>Chromosome</td><td>start</td><td>end</td><td>strand</td><td>MM</td><td>target_seq</td><td>PAM</td><td>alignment</td><td>distance</td><td>position</td><td>gene name</td><td>gene id</td></tr>
<tr><td>GU305914.1</td><td>71190</td><td>71212</td><td>-</td><td>0</td><td>CGACAGCTTCTCCAGATCTT</td><td>CGG</td><td>||||||||[||||||||||||]PAM</td><td>NA</td><td>Intergenic</td><td>NA</td><td>NA</td></tr>
<tr><td></td><td></td><td></td><td></td><td></td><td></td><td></td><td></td><td></td><td></td><td></td><td></td></tr>
<tr><td>T28846</td><td>CTTCTCCAGATCTTCGGCGAAGG</td><td>1000</td><td></td><td></td><td></td><td></td><td></td><td></td><td></td><td></td><td></td></tr>
<tr><td>Oligo fwd</td><td>CACCCTTCTCCAGATCTTCGGCGA</td><td></td><td></td><td></td><td></td><td></td><td></td><td></td><td></td><td></td><td></td></tr>
<tr><td>Oligo rev</td><td>AAACTCGCCGAAGATCTGGAGAAG</td><td></td><td></td><td></td><td></td><td></td><td></td><td></td><td></td><td></td><td></td></tr>
<tr><td>Chromosome</td><td>start</td><td>end</td><td>strand</td><td>MM</td><td>target_seq</td><td>PAM</td><td>alignment</td><td>distance</td><td>position</td><td>gene name</td><td>gene id</td></tr>
<tr><td>GU305914.1</td><td>71184</td><td>71206</td><td>-</td><td>0</td><td>CTTCTCCAGATCTTCGGCGA</td><td>AGG</td><td>||||||||[||||||||||||]PAM</td><td>NA</td><td>Intergenic</td><td>NA</td><td>NA</td></tr>
<tr><td>GU305914.1</td><td>52637</td><td>52659</td><td>+</td><td>3</td><td>CATCTCCAGAGCATCGGCGA</td><td>TGG</td><td>|-||||||[||-|-|||||||]PAM</td><td>NA</td><td>Intergenic</td><td>NA</td><td>NA</td></tr>
<tr><td></td><td></td><td></td><td></td><td></td><td></td><td></td><td></td><td></td><td></td><td></td><td></td></tr>
<tr><td>T28847</td><td>GAAGGCCAGTTTGTTCTGTTTGG</td><td>1000</td><td></td><td></td><td></td><td></td><td></td><td></td><td></td><td></td><td></td></tr>
<tr><td>Oligo fwd</td><td>CACCGAAGGCCAGTTTGTTCTGTT</td><td></td><td></td><td></td><td></td><td></td><td></td><td></td><td></td><td></td><td></td></tr>
<tr><td>Oligo rev</td><td>AAACAACAGAACAAACTGGCCTTC</td><td></td><td></td><td></td><td></td><td></td><td></td><td></td><td></td><td></td><td></td></tr>
<tr><td>Chromosome</td><td>start</td><td>end</td><td>strand</td><td>MM</td><td>target_seq</td><td>PAM</td><td>alignment</td><td>distance</td><td>position</td><td>gene name</td><td>gene id</td></tr>
</table>

| | | | | | | | | | | | | |
|---|---|---|---|---|---|---|---|---|---|---|---|---|
| GU305914.1 | | 71166 | 71188 | - | 0 | GAAGGCCAGTTTGTTCTGTT | TGG | \|\|\|\|\|\|\|\|[\|\|\|\|\|\|\|\|\|\|\|\|]PAM | NA | Intergenic | NA | NA |
| T28848 | CCAGTTTGTTCTGTTTGGTCAGG | | 1000 | | | | | | | | | |
| Oligo fwd | CACCCCAGTTTGTTCTGTTTGGTC | | | | | | | | | | | |
| Oligo rev | AAACGACCAAACAGAACAAACTGG | | | | | | | | | | | |
| Chromosome | start | | end | strand | MM | target_seq | PAM | alignment | distance | position | gene name | gene id |
| GU305914.1 | | 71161 | 71183 | - | 0 | CCAGTTTGTTCTGTTTGGTC | AGG | \|\|\|\|\|\|\|\|[\|\|\|\|\|\|\|\|\|\|\|\|]PAM | NA | Intergenic | NA | NA |
| T28849 | TGGTCAGGATCGCGTCGCTCAGG | | 1000 | | | | | | | | | |
| Oligo fwd | CACCTGGTCAGGATCGCGTCGCTC | | | | | | | | | | | |
| Oligo rev | AAACGAGCGACGCGATCCTGACCA | | | | | | | | | | | |
| Chromosome | start | | end | strand | MM | target_seq | PAM | alignment | distance | position | gene name | gene id |
| GU305914.1 | | 71146 | 71168 | - | 0 | TGGTCAGGATCGCGTCGCTC | AGG | \|\|\|\|\|\|\|\|[\|\|\|\|\|\|\|\|\|\|\|\|]PAM | NA | Intergenic | NA | NA |
| T28850 | CTCCCCGCCGACCACTCTCTTGG | | 1000 | | | | | | | | | |
| Oligo fwd | CACCCTCCCCGCCGACCACTCTCT | | | | | | | | | | | |
| Oligo rev | AAACAGAGAGTGGTCGGCGGGGAG | | | | | | | | | | | |
| Chromosome | start | | end | strand | MM | target_seq | PAM | alignment | distance | position | gene name | gene id |
| GU305914.1 | | 71109 | 71131 | - | 0 | CTCCCCGCCGACCACTCTCT | TGG | \|\|\|\|\|\|\|\|[\|\|\|\|\|\|\|\|\|\|\|\|]PAM | NA | Intergenic | NA | NA |
| T28851 | GCTGACCCAGCATCAGCTGCAGG | | 1000 | | | | | | | | | |
| Oligo fwd | CACCGCTGACCCAGCATCAGCTGC | | | | | | | | | | | |
| Oligo rev | AAACGCAGCTGATGCTGGGTCAGC | | | | | | | | | | | |
| Chromosome | start | | end | strand | MM | target_seq | PAM | alignment | distance | position | gene name | gene id |
| GU305914.1 | | 71080 | 71102 | - | 0 | GCTGACCCAGCATCAGCTGC | AGG | \|\|\|\|\|\|\|\|[\|\|\|\|\|\|\|\|\|\|\|\|]PAM | NA | Intergenic | NA | NA |
| T28852 | AGCGATTCCCTCGTGTGCTCCGG | | 1000 | | | | | | | | | |
| Oligo fwd | CACCAGCGATTCCCTCGTGTGCTC | | | | | | | | | | | |
| Oligo rev | AAACGAGCACACGAGGGAATCGCT | | | | | | | | | | | |
| Chromosome | start | | end | strand | MM | target_seq | PAM | alignment | distance | position | gene name | gene id |
| GU305914.1 | | 71051 | 71073 | - | 0 | AGCGATTCCCTCGTGTGCTC | CGG | \|\|\|\|\|\|\|\|[\|\|\|\|\|\|\|\|\|\|\|\|]PAM | NA | Intergenic | NA | NA |
| T28853 | GATCTTCTCCTCGATCTTCCTGG | | 1000 | | | | | | | | | |
| Oligo fwd | CACCGATCTTCTCCTCGATCTTCC | | | | | | | | | | | |
| Oligo rev | AAACGGAAGATCGAGGAGAAGATC | | | | | | | | | | | |
| Chromosome | start | | end | strand | MM | target_seq | PAM | alignment | distance | position | gene name | gene id |
| GU305914.1 | | 70986 | 71008 | - | 0 | GATCTTCTCCTCGATCTTCC | TGG | \|\|\|\|\|\|\|\|[\|\|\|\|\|\|\|\|\|\|\|\|]PAM | NA | Intergenic | NA | NA |
| T28854 | TCGCCTGCGAGTTGTCCAGCAGG | | 1000 | | | | | | | | | |
| Oligo fwd | CACCTCGCCTGCGAGTTGTCCAGC | | | | | | | | | | | |
| Oligo rev | AAACGCTGGACAACTCGCAGGCGA | | | | | | | | | | | |
| Chromosome | start | | end | strand | MM | target_seq | PAM | alignment | distance | position | gene name | gene id |
| GU305914.1 | | 70960 | 70982 | - | 0 | TCGCCTGCGAGTTGTCCAGC | AGG | \|\|\|\|\|\|\|\|[\|\|\|\|\|\|\|\|\|\|\|\|]PAM | NA | Intergenic | NA | NA |
| GU305914.1 | | 132240 | 132262 | + | 4 | ACGTCTCCGAGCTGTCCAGC | AGG | -\|\|-\|\|-\|[\|\|\|-\|\|\|\|\|\|\|\|]PAM | NA | Intergenic | NA | NA |
| T28855 | CGCCTGCGAGTTGTCCAGCAGGG | | 1000 | | | | | | | | | |
| Oligo fwd | CACCCGCCTGCGAGTTGTCCAGCA | | | | | | | | | | | |
| Oligo rev | AAACTGCTGGACAACTCGCAGGCG | | | | | | | | | | | |
| Chromosome | start | | end | strand | MM | target_seq | PAM | alignment | distance | position | gene name | gene id |
| GU305914.1 | | 70959 | 70981 | - | 0 | CGCCTGCGAGTTGTCCAGCA | GGG | \|\|\|\|\|\|\|\|[\|\|\|\|\|\|\|\|\|\|\|\|]PAM | NA | Intergenic | NA | NA |
| T28856 | TGTCCAGCAGGGTCGCCAGCTGG | | 1000 | | | | | | | | | |
| Oligo fwd | CACCTGTCCAGCAGGGTCGCCAGC | | | | | | | | | | | |
| Oligo rev | AAACGCTGGCGACCCTGCTGGACA | | | | | | | | | | | |
| Chromosome | start | | end | strand | MM | target_seq | PAM | alignment | distance | position | gene name | gene id |
| GU305914.1 | | 70948 | 70970 | - | 0 | TGTCCAGCAGGGTCGCCAGC | TGG | \|\|\|\|\|\|\|\|[\|\|\|\|\|\|\|\|\|\|\|\|]PAM | NA | Intergenic | NA | NA |
| T28857 | GTCCAGCAGGGTCGCCAGCTGGG | | 1000 | | | | | | | | | |
| Oligo fwd | CACCGTCCAGCAGGGTCGCCAGCT | | | | | | | | | | | |
| Oligo rev | AAACAGCTGGCGACCCTGCTGGAC | | | | | | | | | | | |
| Chromosome | start | | end | strand | MM | target_seq | PAM | alignment | distance | position | gene name | gene id |
| GU305914.1 | | 70947 | 70969 | - | 0 | GTCCAGCAGGGTCGCCAGCT | GGG | \|\|\|\|\|\|\|\|[\|\|\|\|\|\|\|\|\|\|\|\|]PAM | NA | Intergenic | NA | NA |
| T28858 | CGCCAGCTGGGCGAACATCCCGG | | 1000 | | | | | | | | | |
| Oligo fwd | CACCCGCCAGCTGGGCGAACATCC | | | | | | | | | | | |
| Oligo rev | AAACGGATGTTCGCCCAGCTGGCG | | | | | | | | | | | |
| Chromosome | start | | end | strand | MM | target_seq | PAM | alignment | distance | position | gene name | gene id |
| GU305914.1 | | 70935 | 70957 | - | 0 | CGCCAGCTGGGCGAACATCC | CGG | \|\|\|\|\|\|\|\|[\|\|\|\|\|\|\|\|\|\|\|\|]PAM | NA | Intergenic | NA | NA |
| T28859 | GCCAGCTGGGCGAACATCCCGGG | | 1000 | | | | | | | | | |
| Oligo fwd | CACCGCCAGCTGGGCGAACATCCC | | | | | | | | | | | |
| Oligo rev | AAACGGGATGTTCGCCCAGCTGGC | | | | | | | | | | | |
| Chromosome | start | | end | strand | MM | target_seq | PAM | alignment | distance | position | gene name | gene id |
| GU305914.1 | | 70934 | 70956 | - | 0 | GCCAGCTGGGCGAACATCCC | GGG | \|\|\|\|\|\|\|\|[\|\|\|\|\|\|\|\|\|\|\|\|]PAM | NA | Intergenic | NA | NA |
| T28860 | CCAGCTGGGCGAACATCCCGGGG | | 1000 | | | | | | | | | |
| Oligo fwd | CACCCCAGCTGGGCGAACATCCCG | | | | | | | | | | | |
| Oligo rev | AAACCGGGATGTTCGCCCAGCTGG | | | | | | | | | | | |
| Chromosome | start | | end | strand | MM | target_seq | PAM | alignment | distance | position | gene name | gene id |
| GU305914.1 | | 70933 | 70955 | - | 0 | CCAGCTGGGCGAACATCCCG | GGG | \|\|\|\|\|\|\|\|[\|\|\|\|\|\|\|\|\|\|\|\|]PAM | NA | Intergenic | NA | NA |
| T28861 | TCTTCTCTTCGATCGCCGCCAGG | | 1000 | | | | | | | | | |
| Oligo fwd | CACCTCTTCTCTTCGATCGCCGCC | | | | | | | | | | | |
| Oligo rev | AAACGGCGGCGATCGAAGAGAAGA | | | | | | | | | | | |
| Chromosome | start | | end | strand | MM | target_seq | PAM | alignment | distance | position | gene name | gene id |
| GU305914.1 | | 70900 | 70922 | - | 0 | TCTTCTCTTCGATCGCCGCC | AGG | \|\|\|\|\|\|\|\|[\|\|\|\|\|\|\|\|\|\|\|\|]PAM | NA | Intergenic | NA | NA |
| T28862 | TGAATTCCGTCGTCCACTCCAGG | | 1000 | | | | | | | | | |
| Oligo fwd | CACCTGAATTCCGTCGTCCACTCC | | | | | | | | | | | |

<table>
<tr><td>Oligo rev</td><td>AAACGGAGTGGACGACGGAATTCA</td><td></td><td></td><td></td><td></td><td></td><td></td><td></td><td></td><td></td><td></td></tr>
<tr><td>Chromosome</td><td>start</td><td>end</td><td>strand</td><td>MM</td><td>target_seq</td><td>PAM</td><td>alignment</td><td>distance</td><td>position</td><td>gene name</td><td>gene id</td></tr>
<tr><td>GU305914.1</td><td>70876</td><td>70898</td><td>-</td><td>0</td><td>TGAATTCCGTCGTCCACTCC</td><td>AGG</td><td>||||||||[||||||||||||]PAM</td><td>NA</td><td>Intergenic</td><td>NA</td><td>NA</td></tr>
<tr><td></td><td></td><td></td><td></td><td></td><td></td><td></td><td></td><td></td><td></td><td></td><td></td></tr>
<tr><td>T28863</td><td>CGTCCACTCCAGGCCCCTCTTGG</td><td>1000</td><td></td><td></td><td></td><td></td><td></td><td></td><td></td><td></td><td></td></tr>
<tr><td>Oligo fwd</td><td>CACCCGTCCACTCCAGGCCCCTCT</td><td></td><td></td><td></td><td></td><td></td><td></td><td></td><td></td><td></td><td></td></tr>
<tr><td>Oligo rev</td><td>AAACAGAGGGGCCTGGAGTGGACG</td><td></td><td></td><td></td><td></td><td></td><td></td><td></td><td></td><td></td><td></td></tr>
<tr><td>Chromosome</td><td>start</td><td>end</td><td>strand</td><td>MM</td><td>target_seq</td><td>PAM</td><td>alignment</td><td>distance</td><td>position</td><td>gene name</td><td>gene id</td></tr>
<tr><td>GU305914.1</td><td>70866</td><td>70888</td><td>-</td><td>0</td><td>CGTCCACTCCAGGCCCCTCT</td><td>TGG</td><td>||||||||[||||||||||||]PAM</td><td>NA</td><td>Intergenic</td><td>NA</td><td>NA</td></tr>
<tr><td></td><td></td><td></td><td></td><td></td><td></td><td></td><td></td><td></td><td></td><td></td><td></td></tr>
<tr><td>T28864</td><td>GCCCCTCTTGGCCGTGATGTAGG</td><td>1000</td><td></td><td></td><td></td><td></td><td></td><td></td><td></td><td></td><td></td></tr>
<tr><td>Oligo fwd</td><td>CACCGCCCCTCTTGGCCGTGATGT</td><td></td><td></td><td></td><td></td><td></td><td></td><td></td><td></td><td></td><td></td></tr>
<tr><td>Oligo rev</td><td>AAACACATCACGGCCAAGAGGGGC</td><td></td><td></td><td></td><td></td><td></td><td></td><td></td><td></td><td></td><td></td></tr>
<tr><td>Chromosome</td><td>start</td><td>end</td><td>strand</td><td>MM</td><td>target_seq</td><td>PAM</td><td>alignment</td><td>distance</td><td>position</td><td>gene name</td><td>gene id</td></tr>
<tr><td>GU305914.1</td><td>70854</td><td>70876</td><td>-</td><td>0</td><td>GCCCCTCTTGGCCGTGATGT</td><td>AGG</td><td>||||||||[||||||||||||]PAM</td><td>NA</td><td>Intergenic</td><td>NA</td><td>NA</td></tr>
<tr><td></td><td></td><td></td><td></td><td></td><td></td><td></td><td></td><td></td><td></td><td></td><td></td></tr>
<tr><td>T28865</td><td>CCCCTCTTGGCCGTGATGTAGGG</td><td>1000</td><td></td><td></td><td></td><td></td><td></td><td></td><td></td><td></td><td></td></tr>
<tr><td>Oligo fwd</td><td>CACCCCCCTCTTGGCCGTGATGTA</td><td></td><td></td><td></td><td></td><td></td><td></td><td></td><td></td><td></td><td></td></tr>
<tr><td>Oligo rev</td><td>AAACTACATCACGGCCAAGAGGGG</td><td></td><td></td><td></td><td></td><td></td><td></td><td></td><td></td><td></td><td></td></tr>
<tr><td>Chromosome</td><td>start</td><td>end</td><td>strand</td><td>MM</td><td>target_seq</td><td>PAM</td><td>alignment</td><td>distance</td><td>position</td><td>gene name</td><td>gene id</td></tr>
<tr><td>GU305914.1</td><td>70853</td><td>70875</td><td>-</td><td>0</td><td>CCCCTCTTGGCCGTGATGTA</td><td>GGG</td><td>||||||||[||||||||||||]PAM</td><td>NA</td><td>Intergenic</td><td>NA</td><td>NA</td></tr>
<tr><td></td><td></td><td></td><td></td><td></td><td></td><td></td><td></td><td></td><td></td><td></td><td></td></tr>
<tr><td>T28866</td><td>TCTTGGCCGTGATGTAGGGCAGG</td><td>1000</td><td></td><td></td><td></td><td></td><td></td><td></td><td></td><td></td><td></td></tr>
<tr><td>Oligo fwd</td><td>CACCTCTTGGCCGTGATGTAGGGC</td><td></td><td></td><td></td><td></td><td></td><td></td><td></td><td></td><td></td><td></td></tr>
<tr><td>Oligo rev</td><td>AAACGCCCTACATCACGGCCAAGA</td><td></td><td></td><td></td><td></td><td></td><td></td><td></td><td></td><td></td><td></td></tr>
<tr><td>Chromosome</td><td>start</td><td>end</td><td>strand</td><td>MM</td><td>target_seq</td><td>PAM</td><td>alignment</td><td>distance</td><td>position</td><td>gene name</td><td>gene id</td></tr>
<tr><td>GU305914.1</td><td>70849</td><td>70871</td><td>-</td><td>0</td><td>TCTTGGCCGTGATGTAGGGC</td><td>AGG</td><td>||||||||[||||||||||||]PAM</td><td>NA</td><td>Intergenic</td><td>NA</td><td>NA</td></tr>
<tr><td>GU305914.1</td><td>126128</td><td>126150</td><td>+</td><td>4</td><td>TCTCCGCCGTGCTGTAGGCC</td><td>CGG</td><td>|||--|||[|||-||||||-|]PAM</td><td>NA</td><td>Intergenic</td><td>NA</td><td>NA</td></tr>
<tr><td></td><td></td><td></td><td></td><td></td><td></td><td></td><td></td><td></td><td></td><td></td><td></td></tr>
<tr><td>T28867</td><td>CTTGGCCGTGATGTAGGGCAGGG</td><td>1000</td><td></td><td></td><td></td><td></td><td></td><td></td><td></td><td></td><td></td></tr>
<tr><td>Oligo fwd</td><td>CACCCTTGGCCGTGATGTAGGGCA</td><td></td><td></td><td></td><td></td><td></td><td></td><td></td><td></td><td></td><td></td></tr>
<tr><td>Oligo rev</td><td>AAACTGCCCTACATCACGGCCAAG</td><td></td><td></td><td></td><td></td><td></td><td></td><td></td><td></td><td></td><td></td></tr>
<tr><td>Chromosome</td><td>start</td><td>end</td><td>strand</td><td>MM</td><td>target_seq</td><td>PAM</td><td>alignment</td><td>distance</td><td>position</td><td>gene name</td><td>gene id</td></tr>
<tr><td>GU305914.1</td><td>70848</td><td>70870</td><td>-</td><td>0</td><td>CTTGGCCGTGATGTAGGGCA</td><td>GGG</td><td>||||||||[||||||||||||]PAM</td><td>NA</td><td>Intergenic</td><td>NA</td><td>NA</td></tr>
<tr><td></td><td></td><td></td><td></td><td></td><td></td><td></td><td></td><td></td><td></td><td></td><td></td></tr>
<tr><td>T28868</td><td>GGCCGTGATGTAGGGCAGGGTGG</td><td>1000</td><td></td><td></td><td></td><td></td><td></td><td></td><td></td><td></td><td></td></tr>
<tr><td>Oligo fwd</td><td>CACCGGCCGTGATGTAGGGCAGGG</td><td></td><td></td><td></td><td></td><td></td><td></td><td></td><td></td><td></td><td></td></tr>
<tr><td>Oligo rev</td><td>AAACCCCTGCCCTACATCACGGCC</td><td></td><td></td><td></td><td></td><td></td><td></td><td></td><td></td><td></td><td></td></tr>
<tr><td>Chromosome</td><td>start</td><td>end</td><td>strand</td><td>MM</td><td>target_seq</td><td>PAM</td><td>alignment</td><td>distance</td><td>position</td><td>gene name</td><td>gene id</td></tr>
<tr><td>GU305914.1</td><td>70845</td><td>70867</td><td>-</td><td>0</td><td>GGCCGTGATGTAGGGCAGGG</td><td>TGG</td><td>||||||||[||||||||||||]PAM</td><td>NA</td><td>Intergenic</td><td>NA</td><td>NA</td></tr>
<tr><td></td><td></td><td></td><td></td><td></td><td></td><td></td><td></td><td></td><td></td><td></td><td></td></tr>
<tr><td>T28869</td><td>CGTGATGTAGGGCAGGGTGGTGG</td><td>1000</td><td></td><td></td><td></td><td></td><td></td><td></td><td></td><td></td><td></td></tr>
<tr><td>Oligo fwd</td><td>CACCCGTGATGTAGGGCAGGGTGG</td><td></td><td></td><td></td><td></td><td></td><td></td><td></td><td></td><td></td><td></td></tr>
<tr><td>Oligo rev</td><td>AAACCCACCCTGCCCTACATCACG</td><td></td><td></td><td></td><td></td><td></td><td></td><td></td><td></td><td></td><td></td></tr>
<tr><td>Chromosome</td><td>start</td><td>end</td><td>strand</td><td>MM</td><td>target_seq</td><td>PAM</td><td>alignment</td><td>distance</td><td>position</td><td>gene name</td><td>gene id</td></tr>
<tr><td>GU305914.1</td><td>70842</td><td>70864</td><td>-</td><td>0</td><td>CGTGATGTAGGGCAGGGTGG</td><td>TGG</td><td>||||||||[||||||||||||]PAM</td><td>NA</td><td>Intergenic</td><td>NA</td><td>NA</td></tr>
<tr><td></td><td></td><td></td><td></td><td></td><td></td><td></td><td></td><td></td><td></td><td></td><td></td></tr>
<tr><td>T28870</td><td>GCAGGGTGGTGGCCGCCGAGCGG</td><td>1000</td><td></td><td></td><td></td><td></td><td></td><td></td><td></td><td></td><td></td></tr>
<tr><td>Oligo fwd</td><td>CACCGCAGGGTGGTGGCCGCCGAG</td><td></td><td></td><td></td><td></td><td></td><td></td><td></td><td></td><td></td><td></td></tr>
<tr><td>Oligo rev</td><td>AAACCTCGGCGGCCACCACCCTGC</td><td></td><td></td><td></td><td></td><td></td><td></td><td></td><td></td><td></td><td></td></tr>
<tr><td>Chromosome</td><td>start</td><td>end</td><td>strand</td><td>MM</td><td>target_seq</td><td>PAM</td><td>alignment</td><td>distance</td><td>position</td><td>gene name</td><td>gene id</td></tr>
<tr><td>GU305914.1</td><td>70831</td><td>70853</td><td>-</td><td>0</td><td>GCAGGGTGGTGGCCGCCGAG</td><td>CGG</td><td>||||||||[||||||||||||]PAM</td><td>NA</td><td>Intergenic</td><td>NA</td><td>NA</td></tr>
<tr><td></td><td></td><td></td><td></td><td></td><td></td><td></td><td></td><td></td><td></td><td></td><td></td></tr>
<tr><td>T28871</td><td>TCACCACCGTGTCCGTGCGCAGG</td><td>1000</td><td></td><td></td><td></td><td></td><td></td><td></td><td></td><td></td><td></td></tr>
<tr><td>Oligo fwd</td><td>CACCTCACCACCGTGTCCGTGCGC</td><td></td><td></td><td></td><td></td><td></td><td></td><td></td><td></td><td></td><td></td></tr>
<tr><td>Oligo rev</td><td>AAACGCGCACGGACACGGTGGTGA</td><td></td><td></td><td></td><td></td><td></td><td></td><td></td><td></td><td></td><td></td></tr>
<tr><td>Chromosome</td><td>start</td><td>end</td><td>strand</td><td>MM</td><td>target_seq</td><td>PAM</td><td>alignment</td><td>distance</td><td>position</td><td>gene name</td><td>gene id</td></tr>
<tr><td>GU305914.1</td><td>70792</td><td>70814</td><td>-</td><td>0</td><td>TCACCACCGTGTCCGTGCGC</td><td>AGG</td><td>||||||||[||||||||||||]PAM</td><td>NA</td><td>Intergenic</td><td>NA</td><td>NA</td></tr>
<tr><td></td><td></td><td></td><td></td><td></td><td></td><td></td><td></td><td></td><td></td><td></td><td></td></tr>
<tr><td>T28872</td><td>CCACCGTGTCCGTGCGCAGGCGG</td><td>1000</td><td></td><td></td><td></td><td></td><td></td><td></td><td></td><td></td><td></td></tr>
<tr><td>Oligo fwd</td><td>CACCCCACCGTGTCCGTGCGCAGG</td><td></td><td></td><td></td><td></td><td></td><td></td><td></td><td></td><td></td><td></td></tr>
<tr><td>Oligo rev</td><td>AAACCCTGCGCACGGACACGGTGG</td><td></td><td></td><td></td><td></td><td></td><td></td><td></td><td></td><td></td><td></td></tr>
<tr><td>Chromosome</td><td>start</td><td>end</td><td>strand</td><td>MM</td><td>target_seq</td><td>PAM</td><td>alignment</td><td>distance</td><td>position</td><td>gene name</td><td>gene id</td></tr>
<tr><td>GU305914.1</td><td>70789</td><td>70811</td><td>-</td><td>0</td><td>CCACCGTGTCCGTGCGCAGG</td><td>CGG</td><td>||||||||[||||||||||||]PAM</td><td>NA</td><td>Intergenic</td><td>NA</td><td>NA</td></tr>
<tr><td></td><td></td><td></td><td></td><td></td><td></td><td></td><td></td><td></td><td></td><td></td><td></td></tr>
<tr><td>T28873</td><td>CGTGTCCGTGCGCAGGCGGTTGG</td><td>1000</td><td></td><td></td><td></td><td></td><td></td><td></td><td></td><td></td><td></td></tr>
<tr><td>Oligo fwd</td><td>CACCCGTGTCCGTGCGCAGGCGGT</td><td></td><td></td><td></td><td></td><td></td><td></td><td></td><td></td><td></td><td></td></tr>
<tr><td>Oligo rev</td><td>AAACACCGCCTGCGCACGGACACG</td><td></td><td></td><td></td><td></td><td></td><td></td><td></td><td></td><td></td><td></td></tr>
<tr><td>Chromosome</td><td>start</td><td>end</td><td>strand</td><td>MM</td><td>target_seq</td><td>PAM</td><td>alignment</td><td>distance</td><td>position</td><td>gene name</td><td>gene id</td></tr>
<tr><td>GU305914.1</td><td>70785</td><td>70807</td><td>-</td><td>0</td><td>CGTGTCCGTGCGCAGGCGGT</td><td>TGG</td><td>||||||||[||||||||||||]PAM</td><td>NA</td><td>Intergenic</td><td>NA</td><td>NA</td></tr>
<tr><td></td><td></td><td></td><td></td><td></td><td></td><td></td><td></td><td></td><td></td><td></td><td></td></tr>
<tr><td>T28874</td><td>GGCGGTTGGTGTCCACGTCCAGG</td><td>1000</td><td></td><td></td><td></td><td></td><td></td><td></td><td></td><td></td><td></td></tr>
<tr><td>Oligo fwd</td><td>CACCGGCGGTTGGTGTCCACGTCC</td><td></td><td></td><td></td><td></td><td></td><td></td><td></td><td></td><td></td><td></td></tr>
<tr><td>Oligo rev</td><td>AAACGGACGTGGACACCAACCGCC</td><td></td><td></td><td></td><td></td><td></td><td></td><td></td><td></td><td></td><td></td></tr>
<tr><td>Chromosome</td><td>start</td><td>end</td><td>strand</td><td>MM</td><td>target_seq</td><td>PAM</td><td>alignment</td><td>distance</td><td>position</td><td>gene name</td><td>gene id</td></tr>
<tr><td>GU305914.1</td><td>70771</td><td>70793</td><td>-</td><td>0</td><td>GGCGGTTGGTGTCCACGTCC</td><td>AGG</td><td>||||||||[||||||||||||]PAM</td><td>NA</td><td>Intergenic</td><td>NA</td><td>NA</td></tr>
<tr><td></td><td></td><td></td><td></td><td></td><td></td><td></td><td></td><td></td><td></td><td></td><td></td></tr>
<tr><td>T28875</td><td>GGTGTCCACGTCCAGGCCCTCGG</td><td>1000</td><td></td><td></td><td></td><td></td><td></td><td></td><td></td><td></td><td></td></tr>
<tr><td>Oligo fwd</td><td>CACCGGTGTCCACGTCCAGGCCCT</td><td></td><td></td><td></td><td></td><td></td><td></td><td></td><td></td><td></td><td></td></tr>
<tr><td>Oligo rev</td><td>AAACAGGGCCTGGACGTGGACACC</td><td></td><td></td><td></td><td></td><td></td><td></td><td></td><td></td><td></td><td></td></tr>
<tr><td>Chromosome</td><td>start</td><td>end</td><td>strand</td><td>MM</td><td>target_seq</td><td>PAM</td><td>alignment</td><td>distance</td><td>position</td><td>gene name</td><td>gene id</td></tr>
<tr><td>GU305914.1</td><td>70764</td><td>70786</td><td>-</td><td>0</td><td>GGTGTCCACGTCCAGGCCCT</td><td>CGG</td><td>||||||||[||||||||||||]PAM</td><td>NA</td><td>Intergenic</td><td>NA</td><td>NA</td></tr>
<tr><td>GU305914.1</td><td>113834</td><td>113856</td><td>-</td><td>4</td><td>GGGGTCCGCTTCCAGCCCCT</td><td>TGG</td><td>||-||||-[|-|||||-||||]PAM</td><td>NA</td><td>Intergenic</td><td>NA</td><td>NA</td></tr>
<tr><td></td><td></td><td></td><td></td><td></td><td></td><td></td><td></td><td></td><td></td><td></td><td></td></tr>
<tr><td>T28876</td><td>GTGTCCACGTCCAGGCCCTCGGG</td><td>1000</td><td></td><td></td><td></td><td></td><td></td><td></td><td></td><td></td><td></td></tr>
<tr><td>Oligo fwd</td><td>CACCGTGTCCACGTCCAGGCCCTC</td><td></td><td></td><td></td><td></td><td></td><td></td><td></td><td></td><td></td><td></td></tr>
<tr><td>Oligo rev</td><td>AAACGAGGGCCTGGACGTGGACAC</td><td></td><td></td><td></td><td></td><td></td><td></td><td></td><td></td><td></td><td></td></tr>
<tr><td>Chromosome</td><td>start</td><td>end</td><td>strand</td><td>MM</td><td>target_seq</td><td>PAM</td><td>alignment</td><td>distance</td><td>position</td><td>gene name</td><td>gene id</td></tr>
<tr><td>GU305914.1</td><td>70763</td><td>70785</td><td>-</td><td>0</td><td>GTGTCCACGTCCAGGCCCTC</td><td>GGG</td><td>||||||||[||||||||||||]PAM</td><td>NA</td><td>Intergenic</td><td>NA</td><td>NA</td></tr>
</table>

<table>
<tr><td>T28877</td><td>GGCCCTCGGGCAGCCTGATGCGG</td><td>1000</td><td></td><td></td><td></td><td></td><td></td><td></td><td></td><td></td><td></td></tr>
<tr><td>Oligo fwd</td><td>CACCGGCCCTCGGGCAGCCTGATG</td><td></td><td></td><td></td><td></td><td></td><td></td><td></td><td></td><td></td><td></td></tr>
<tr><td>Oligo rev</td><td>AAACCATCAGGCTGCCCGAGGGCC</td><td></td><td></td><td></td><td></td><td></td><td></td><td></td><td></td><td></td><td></td></tr>
<tr><td>Chromosome</td><td>start</td><td>end</td><td>strand</td><td>MM</td><td>target_seq</td><td>PAM</td><td>alignment</td><td>distance</td><td>position</td><td>gene name</td><td>gene id</td></tr>
<tr><td>GU305914.1</td><td>70750</td><td>70772</td><td>-</td><td>0</td><td>GGCCCTCGGGCAGCCTGATG</td><td>CGG</td><td>||||||||[||||||||||||]PAM</td><td>NA</td><td>Intergenic</td><td>NA</td><td>NA</td></tr>
<tr><td>T28878</td><td>CGGGCAGCCTGATGCGGTGCTGG</td><td>1000</td><td></td><td></td><td></td><td></td><td></td><td></td><td></td><td></td><td></td></tr>
<tr><td>Oligo fwd</td><td>CACCCGGGCAGCCTGATGCGGTGC</td><td></td><td></td><td></td><td></td><td></td><td></td><td></td><td></td><td></td><td></td></tr>
<tr><td>Oligo rev</td><td>AAACGCACCGCATCAGGCTGCCCG</td><td></td><td></td><td></td><td></td><td></td><td></td><td></td><td></td><td></td><td></td></tr>
<tr><td>Chromosome</td><td>start</td><td>end</td><td>strand</td><td>MM</td><td>target_seq</td><td>PAM</td><td>alignment</td><td>distance</td><td>position</td><td>gene name</td><td>gene id</td></tr>
<tr><td>GU305914.1</td><td>70744</td><td>70766</td><td>-</td><td>0</td><td>CGGGCAGCCTGATGCGGTGC</td><td>TGG</td><td>||||||||[||||||||||||]PAM</td><td>NA</td><td>Intergenic</td><td>NA</td><td>NA</td></tr>
<tr><td>T28879</td><td>GGGCAGCCTGATGCGGTGCTGGG</td><td>1000</td><td></td><td></td><td></td><td></td><td></td><td></td><td></td><td></td><td></td></tr>
<tr><td>Oligo fwd</td><td>CACCGGGCAGCCTGATGCGGTGCT</td><td></td><td></td><td></td><td></td><td></td><td></td><td></td><td></td><td></td><td></td></tr>
<tr><td>Oligo rev</td><td>AAACAGCACCGCATCAGGCTGCCC</td><td></td><td></td><td></td><td></td><td></td><td></td><td></td><td></td><td></td><td></td></tr>
<tr><td>Chromosome</td><td>start</td><td>end</td><td>strand</td><td>MM</td><td>target_seq</td><td>PAM</td><td>alignment</td><td>distance</td><td>position</td><td>gene name</td><td>gene id</td></tr>
<tr><td>GU305914.1</td><td>70743</td><td>70765</td><td>-</td><td>0</td><td>GGGCAGCCTGATGCGGTGCT</td><td>GGG</td><td>||||||||[||||||||||||]PAM</td><td>NA</td><td>Intergenic</td><td>NA</td><td>NA</td></tr>
<tr><td>T28880</td><td>CCTGAGAGCTGCGCTTAGCCAGG</td><td>1000</td><td></td><td></td><td></td><td></td><td></td><td></td><td></td><td></td><td></td></tr>
<tr><td>Oligo fwd</td><td>CACCCCTGAGAGCTGCGCTTAGCC</td><td></td><td></td><td></td><td></td><td></td><td></td><td></td><td></td><td></td><td></td></tr>
<tr><td>Oligo rev</td><td>AAACGGCTAAGCGCAGCTCTCAGG</td><td></td><td></td><td></td><td></td><td></td><td></td><td></td><td></td><td></td><td></td></tr>
<tr><td>Chromosome</td><td>start</td><td>end</td><td>strand</td><td>MM</td><td>target_seq</td><td>PAM</td><td>alignment</td><td>distance</td><td>position</td><td>gene name</td><td>gene id</td></tr>
<tr><td>GU305914.1</td><td>70696</td><td>70718</td><td>-</td><td>0</td><td>CCTGAGAGCTGCGCTTAGCC</td><td>AGG</td><td>||||||||[||||||||||||]PAM</td><td>NA</td><td>Intergenic</td><td>NA</td><td>NA</td></tr>
<tr><td>T28881</td><td>TCGTCTTGAGCTTCACGTTCAGG</td><td>1000</td><td></td><td></td><td></td><td></td><td></td><td></td><td></td><td></td><td></td></tr>
<tr><td>Oligo fwd</td><td>CACCTCGTCTTGAGCTTCACGTTC</td><td></td><td></td><td></td><td></td><td></td><td></td><td></td><td></td><td></td><td></td></tr>
<tr><td>Oligo rev</td><td>AAACGAACGTGAAGCTCAAGACGA</td><td></td><td></td><td></td><td></td><td></td><td></td><td></td><td></td><td></td><td></td></tr>
<tr><td>Chromosome</td><td>start</td><td>end</td><td>strand</td><td>MM</td><td>target_seq</td><td>PAM</td><td>alignment</td><td>distance</td><td>position</td><td>gene name</td><td>gene id</td></tr>
<tr><td>GU305914.1</td><td>70660</td><td>70682</td><td>-</td><td>0</td><td>TCGTCTTGAGCTTCACGTTC</td><td>AGG</td><td>||||||||[||||||||||||]PAM</td><td>NA</td><td>Intergenic</td><td>NA</td><td>NA</td></tr>
<tr><td>T28882</td><td>GCTCGTTCTCTCGCGCCTGCAGG</td><td>1000</td><td></td><td></td><td></td><td></td><td></td><td></td><td></td><td></td><td></td></tr>
<tr><td>Oligo fwd</td><td>CACCGCTCGTTCTCTCGCGCCTGC</td><td></td><td></td><td></td><td></td><td></td><td></td><td></td><td></td><td></td><td></td></tr>
<tr><td>Oligo rev</td><td>AAACGCAGGCGCGAGAGAACGAGC</td><td></td><td></td><td></td><td></td><td></td><td></td><td></td><td></td><td></td><td></td></tr>
<tr><td>Chromosome</td><td>start</td><td>end</td><td>strand</td><td>MM</td><td>target_seq</td><td>PAM</td><td>alignment</td><td>distance</td><td>position</td><td>gene name</td><td>gene id</td></tr>
<tr><td>GU305914.1</td><td>70618</td><td>70640</td><td>-</td><td>0</td><td>GCTCGTTCTCTCGCGCCTGC</td><td>AGG</td><td>||||||||[||||||||||||]PAM</td><td>NA</td><td>Intergenic</td><td>NA</td><td>NA</td></tr>
<tr><td>T28883</td><td>CGCCTGCAGCTCCGTCGAGAAGG</td><td>1000</td><td></td><td></td><td></td><td></td><td></td><td></td><td></td><td></td><td></td></tr>
<tr><td>Oligo fwd</td><td>CACCCGCCTGCAGCTCCGTCGAGA</td><td></td><td></td><td></td><td></td><td></td><td></td><td></td><td></td><td></td><td></td></tr>
<tr><td>Oligo rev</td><td>AAACTCTCGACGGAGCTGCAGGCG</td><td></td><td></td><td></td><td></td><td></td><td></td><td></td><td></td><td></td><td></td></tr>
<tr><td>Chromosome</td><td>start</td><td>end</td><td>strand</td><td>MM</td><td>target_seq</td><td>PAM</td><td>alignment</td><td>distance</td><td>position</td><td>gene name</td><td>gene id</td></tr>
<tr><td>GU305914.1</td><td>70593</td><td>70615</td><td>-</td><td>0</td><td>CGCCTGCAGCTCCGTCGAGA</td><td>AGG</td><td>||||||||[||||||||||||]PAM</td><td>NA</td><td>Intergenic</td><td>NA</td><td>NA</td></tr>
<tr><td>T28884</td><td>GCCTGCAGCTCCGTCGAGAAGGG</td><td>1000</td><td></td><td></td><td></td><td></td><td></td><td></td><td></td><td></td><td></td></tr>
<tr><td>Oligo fwd</td><td>CACCGCCTGCAGCTCCGTCGAGAA</td><td></td><td></td><td></td><td></td><td></td><td></td><td></td><td></td><td></td><td></td></tr>
<tr><td>Oligo rev</td><td>AAACTTCTCGACGGAGCTGCAGGC</td><td></td><td></td><td></td><td></td><td></td><td></td><td></td><td></td><td></td><td></td></tr>
<tr><td>Chromosome</td><td>start</td><td>end</td><td>strand</td><td>MM</td><td>target_seq</td><td>PAM</td><td>alignment</td><td>distance</td><td>position</td><td>gene name</td><td>gene id</td></tr>
<tr><td>GU305914.1</td><td>70592</td><td>70614</td><td>-</td><td>0</td><td>GCCTGCAGCTCCGTCGAGAA</td><td>GGG</td><td>||||||||[||||||||||||]PAM</td><td>NA</td><td>Intergenic</td><td>NA</td><td>NA</td></tr>
<tr><td>T28885</td><td>GGGCGCCGCCATCGCCTTCACGG</td><td>1000</td><td></td><td></td><td></td><td></td><td></td><td></td><td></td><td></td><td></td></tr>
<tr><td>Oligo fwd</td><td>CACCGGGCGCCGCCATCGCCTTCA</td><td></td><td></td><td></td><td></td><td></td><td></td><td></td><td></td><td></td><td></td></tr>
<tr><td>Oligo rev</td><td>AAACTGAAGGCGATGGCGGCGCCC</td><td></td><td></td><td></td><td></td><td></td><td></td><td></td><td></td><td></td><td></td></tr>
<tr><td>Chromosome</td><td>start</td><td>end</td><td>strand</td><td>MM</td><td>target_seq</td><td>PAM</td><td>alignment</td><td>distance</td><td>position</td><td>gene name</td><td>gene id</td></tr>
<tr><td>GU305914.1</td><td>70572</td><td>70594</td><td>-</td><td>0</td><td>GGGCGCCGCCATCGCCTTCA</td><td>CGG</td><td>||||||||[||||||||||||]PAM</td><td>NA</td><td>Intergenic</td><td>NA</td><td>NA</td></tr>
<tr><td>T28886</td><td>GTCCCTGAACGCCGCCGTGAAGG</td><td>1000</td><td></td><td></td><td></td><td></td><td></td><td></td><td></td><td></td><td></td></tr>
<tr><td>Oligo fwd</td><td>CACCGTCCCTGAACGCCGCCGTGA</td><td></td><td></td><td></td><td></td><td></td><td></td><td></td><td></td><td></td><td></td></tr>
<tr><td>Oligo rev</td><td>AAACTCACGGCGGCGTTCAGGGAC</td><td></td><td></td><td></td><td></td><td></td><td></td><td></td><td></td><td></td><td></td></tr>
<tr><td>Chromosome</td><td>start</td><td>end</td><td>strand</td><td>MM</td><td>target_seq</td><td>PAM</td><td>alignment</td><td>distance</td><td>position</td><td>gene name</td><td>gene id</td></tr>
<tr><td>GU305914.1</td><td>70548</td><td>70570</td><td>-</td><td>0</td><td>GTCCCTGAACGCCGCCGTGA</td><td>AGG</td><td>||||||||[||||||||||||]PAM</td><td>NA</td><td>Intergenic</td><td>NA</td><td>NA</td></tr>
<tr><td>GU305914.1</td><td>70389</td><td>70411</td><td>-</td><td>4</td><td>GTTCCTGTACGCGGCCGTGT</td><td>GGG</td><td>||-||||-[||||-||||||-]PAM</td><td>NA</td><td>Intergenic</td><td>NA</td><td>NA</td></tr>
<tr><td>T28887</td><td>CGCCGCCGTGAAGGTGTCCTCGG</td><td>1000</td><td></td><td></td><td></td><td></td><td></td><td></td><td></td><td></td><td></td></tr>
<tr><td>Oligo fwd</td><td>CACCCGCCGCCGTGAAGGTGTCCT</td><td></td><td></td><td></td><td></td><td></td><td></td><td></td><td></td><td></td><td></td></tr>
<tr><td>Oligo rev</td><td>AAACAGGACACCTTCACGGCGGCG</td><td></td><td></td><td></td><td></td><td></td><td></td><td></td><td></td><td></td><td></td></tr>
<tr><td>Chromosome</td><td>start</td><td>end</td><td>strand</td><td>MM</td><td>target_seq</td><td>PAM</td><td>alignment</td><td>distance</td><td>position</td><td>gene name</td><td>gene id</td></tr>
<tr><td>GU305914.1</td><td>70539</td><td>70561</td><td>-</td><td>0</td><td>CGCCGCCGTGAAGGTGTCCT</td><td>CGG</td><td>||||||||[||||||||||||]PAM</td><td>NA</td><td>Intergenic</td><td>NA</td><td>NA</td></tr>
<tr><td>T28888</td><td>GCCGCCGTGAAGGTGTCCTCGGG</td><td>1000</td><td></td><td></td><td></td><td></td><td></td><td></td><td></td><td></td><td></td></tr>
<tr><td>Oligo fwd</td><td>CACCGCCGCCGTGAAGGTGTCCTC</td><td></td><td></td><td></td><td></td><td></td><td></td><td></td><td></td><td></td><td></td></tr>
<tr><td>Oligo rev</td><td>AAACGAGGACACCTTCACGGCGGC</td><td></td><td></td><td></td><td></td><td></td><td></td><td></td><td></td><td></td><td></td></tr>
<tr><td>Chromosome</td><td>start</td><td>end</td><td>strand</td><td>MM</td><td>target_seq</td><td>PAM</td><td>alignment</td><td>distance</td><td>position</td><td>gene name</td><td>gene id</td></tr>
<tr><td>GU305914.1</td><td>70538</td><td>70560</td><td>-</td><td>0</td><td>GCCGCCGTGAAGGTGTCCTC</td><td>GGG</td><td>||||||||[||||||||||||]PAM</td><td>NA</td><td>Intergenic</td><td>NA</td><td>NA</td></tr>
<tr><td>T28889</td><td>CGTTCGCTTTCGTGTTGTCGTGG</td><td>1000</td><td></td><td></td><td></td><td></td><td></td><td></td><td></td><td></td><td></td></tr>
<tr><td>Oligo fwd</td><td>CACCCGTTCGCTTTCGTGTTGTCG</td><td></td><td></td><td></td><td></td><td></td><td></td><td></td><td></td><td></td><td></td></tr>
<tr><td>Oligo rev</td><td>AAACCGACAACACGAAAGCGAACG</td><td></td><td></td><td></td><td></td><td></td><td></td><td></td><td></td><td></td><td></td></tr>
<tr><td>Chromosome</td><td>start</td><td>end</td><td>strand</td><td>MM</td><td>target_seq</td><td>PAM</td><td>alignment</td><td>distance</td><td>position</td><td>gene name</td><td>gene id</td></tr>
<tr><td>GU305914.1</td><td>70501</td><td>70523</td><td>-</td><td>0</td><td>CGTTCGCTTTCGTGTTGTCG</td><td>TGG</td><td>||||||||[||||||||||||]PAM</td><td>NA</td><td>Intergenic</td><td>NA</td><td>NA</td></tr>
<tr><td>T28890</td><td>GCGCCAGCGTCTCCGTCCTGCGG</td><td>1000</td><td></td><td></td><td></td><td></td><td></td><td></td><td></td><td></td><td></td></tr>
<tr><td>Oligo fwd</td><td>CACCGCGCCAGCGTCTCCGTCCTG</td><td></td><td></td><td></td><td></td><td></td><td></td><td></td><td></td><td></td><td></td></tr>
<tr><td>Oligo rev</td><td>AAACCAGGACGGAGACGCTGGCGC</td><td></td><td></td><td></td><td></td><td></td><td></td><td></td><td></td><td></td><td></td></tr>
<tr><td>Chromosome</td><td>start</td><td>end</td><td>strand</td><td>MM</td><td>target_seq</td><td>PAM</td><td>alignment</td><td>distance</td><td>position</td><td>gene name</td><td>gene id</td></tr>
<tr><td>GU305914.1</td><td>70441</td><td>70463</td><td>-</td><td>0</td><td>GCGCCAGCGTCTCCGTCCTG</td><td>CGG</td><td>||||||||[||||||||||||]PAM</td><td>NA</td><td>Intergenic</td><td>NA</td><td>NA</td></tr>
<tr><td>T28891</td><td>CTCGATCTCGTTCGCCAGTTCGG</td><td>1000</td><td></td><td></td><td></td><td></td><td></td><td></td><td></td><td></td><td></td></tr>
<tr><td>Oligo fwd</td><td>CACCCTCGATCTCGTTCGCCAGTT</td><td></td><td></td><td></td><td></td><td></td><td></td><td></td><td></td><td></td><td></td></tr>
<tr><td>Oligo rev</td><td>AAACAACTGGCGAACGAGATCGAG</td><td></td><td></td><td></td><td></td><td></td><td></td><td></td><td></td><td></td><td></td></tr>
</table>

<table>
<tr><td>Chromosome</td><td>start</td><td>end</td><td>strand</td><td>MM</td><td>target_seq</td><td>PAM</td><td>alignment</td><td>distance</td><td>position</td><td>gene name</td><td>gene id</td></tr>
<tr><td>GU305914.1</td><td>70416</td><td>70438</td><td>-</td><td>0</td><td>CTCGATCTCGTTCGCCAGTT</td><td>CGG</td><td>||||||||[||||||||||||]PAM</td><td>NA</td><td>Intergenic</td><td>NA</td><td>NA</td></tr>
<tr><td>GU305914.1</td><td>28322</td><td>28344</td><td>-</td><td>4</td><td>CGTGATCTCGTTCGCCACCT</td><td>CGG</td><td>|--|||||[|||||||||--|]PAM</td><td>NA</td><td>Intergenic</td><td>NA</td><td>NA</td></tr>
<tr><td>T28892</td><td>TTCGGCGAAGTTCCTGTACGCGG</td><td>1000</td><td></td><td></td><td></td><td></td><td></td><td></td><td></td><td></td><td></td></tr>
<tr><td>Oligo fwd</td><td>CACCTTCGGCGAAGTTCCTGTACG</td><td></td><td></td><td></td><td></td><td></td><td></td><td></td><td></td><td></td><td></td></tr>
<tr><td>Oligo rev</td><td>AAACCGTACAGGAACTTCGCCGAA</td><td></td><td></td><td></td><td></td><td></td><td></td><td></td><td></td><td></td><td></td></tr>
<tr><td>Chromosome</td><td>start</td><td>end</td><td>strand</td><td>MM</td><td>target_seq</td><td>PAM</td><td>alignment</td><td>distance</td><td>position</td><td>gene name</td><td>gene id</td></tr>
<tr><td>GU305914.1</td><td>70398</td><td>70420</td><td>-</td><td>0</td><td>TTCGGCGAAGTTCCTGTACG</td><td>CGG</td><td>||||||||[||||||||||||]PAM</td><td>NA</td><td>Intergenic</td><td>NA</td><td>NA</td></tr>
<tr><td>T28893</td><td>AGTTCCTGTACGCGGCCGTGTGG</td><td>1000</td><td></td><td></td><td></td><td></td><td></td><td></td><td></td><td></td><td></td></tr>
<tr><td>Oligo fwd</td><td>CACCAGTTCCTGTACGCGGCCGTG</td><td></td><td></td><td></td><td></td><td></td><td></td><td></td><td></td><td></td><td></td></tr>
<tr><td>Oligo rev</td><td>AAACCACGGCCGCGTACAGGAACT</td><td></td><td></td><td></td><td></td><td></td><td></td><td></td><td></td><td></td><td></td></tr>
<tr><td>Chromosome</td><td>start</td><td>end</td><td>strand</td><td>MM</td><td>target_seq</td><td>PAM</td><td>alignment</td><td>distance</td><td>position</td><td>gene name</td><td>gene id</td></tr>
<tr><td>GU305914.1</td><td>70390</td><td>70412</td><td>-</td><td>0</td><td>AGTTCCTGTACGCGGCCGTG</td><td>TGG</td><td>||||||||[||||||||||||]PAM</td><td>NA</td><td>Intergenic</td><td>NA</td><td>NA</td></tr>
<tr><td>T28894</td><td>GTTCCTGTACGCGGCCGTGTGGG</td><td>1000</td><td></td><td></td><td></td><td></td><td></td><td></td><td></td><td></td><td></td></tr>
<tr><td>Oligo fwd</td><td>CACCGTTCCTGTACGCGGCCGTGT</td><td></td><td></td><td></td><td></td><td></td><td></td><td></td><td></td><td></td><td></td></tr>
<tr><td>Oligo rev</td><td>AAACACACGGCCGCGTACAGGAAC</td><td></td><td></td><td></td><td></td><td></td><td></td><td></td><td></td><td></td><td></td></tr>
<tr><td>Chromosome</td><td>start</td><td>end</td><td>strand</td><td>MM</td><td>target_seq</td><td>PAM</td><td>alignment</td><td>distance</td><td>position</td><td>gene name</td><td>gene id</td></tr>
<tr><td>GU305914.1</td><td>70389</td><td>70411</td><td>-</td><td>0</td><td>GTTCCTGTACGCGGCCGTGT</td><td>GGG</td><td>||||||||[||||||||||||]PAM</td><td>NA</td><td>Intergenic</td><td>NA</td><td>NA</td></tr>
<tr><td>GU305914.1</td><td>70548</td><td>70570</td><td>-</td><td>4</td><td>GTCCCTGAACGCCGCCGTGA</td><td>AGG</td><td>||-||||-[||||-||||||-]PAM</td><td>NA</td><td>Intergenic</td><td>NA</td><td>NA</td></tr>
<tr><td>T28895</td><td>TTCCTGTACGCGGCCGTGTGGGG</td><td>1000</td><td></td><td></td><td></td><td></td><td></td><td></td><td></td><td></td><td></td></tr>
<tr><td>Oligo fwd</td><td>CACCTTCCTGTACGCGGCCGTGTG</td><td></td><td></td><td></td><td></td><td></td><td></td><td></td><td></td><td></td><td></td></tr>
<tr><td>Oligo rev</td><td>AAACCACACGGCCGCGTACAGGAA</td><td></td><td></td><td></td><td></td><td></td><td></td><td></td><td></td><td></td><td></td></tr>
<tr><td>Chromosome</td><td>start</td><td>end</td><td>strand</td><td>MM</td><td>target_seq</td><td>PAM</td><td>alignment</td><td>distance</td><td>position</td><td>gene name</td><td>gene id</td></tr>
<tr><td>GU305914.1</td><td>70388</td><td>70410</td><td>-</td><td>0</td><td>TTCCTGTACGCGGCCGTGTG</td><td>GGG</td><td>||||||||[||||||||||||]PAM</td><td>NA</td><td>Intergenic</td><td>NA</td><td>NA</td></tr>
<tr><td>T28896</td><td>TCCTGTACGCGGCCGTGTGGGGG</td><td>1000</td><td></td><td></td><td></td><td></td><td></td><td></td><td></td><td></td><td></td></tr>
<tr><td>Oligo fwd</td><td>CACCTCCTGTACGCGGCCGTGTGG</td><td></td><td></td><td></td><td></td><td></td><td></td><td></td><td></td><td></td><td></td></tr>
<tr><td>Oligo rev</td><td>AAACCCACACGGCCGCGTACAGGA</td><td></td><td></td><td></td><td></td><td></td><td></td><td></td><td></td><td></td><td></td></tr>
<tr><td>Chromosome</td><td>start</td><td>end</td><td>strand</td><td>MM</td><td>target_seq</td><td>PAM</td><td>alignment</td><td>distance</td><td>position</td><td>gene name</td><td>gene id</td></tr>
<tr><td>GU305914.1</td><td>70387</td><td>70409</td><td>-</td><td>0</td><td>TCCTGTACGCGGCCGTGTGG</td><td>GGG</td><td>||||||||[||||||||||||]PAM</td><td>NA</td><td>Intergenic</td><td>NA</td><td>NA</td></tr>
<tr><td>T28897</td><td>CCTGTACGCGGCCGTGTGGGGGG</td><td>1000</td><td></td><td></td><td></td><td></td><td></td><td></td><td></td><td></td><td></td></tr>
<tr><td>Oligo fwd</td><td>CACCCCTGTACGCGGCCGTGTGGG</td><td></td><td></td><td></td><td></td><td></td><td></td><td></td><td></td><td></td><td></td></tr>
<tr><td>Oligo rev</td><td>AAACCCCACACGGCCGCGTACAGG</td><td></td><td></td><td></td><td></td><td></td><td></td><td></td><td></td><td></td><td></td></tr>
<tr><td>Chromosome</td><td>start</td><td>end</td><td>strand</td><td>MM</td><td>target_seq</td><td>PAM</td><td>alignment</td><td>distance</td><td>position</td><td>gene name</td><td>gene id</td></tr>
<tr><td>GU305914.1</td><td>70386</td><td>70408</td><td>-</td><td>0</td><td>CCTGTACGCGGCCGTGTGGG</td><td>GGG</td><td>||||||||[||||||||||||]PAM</td><td>NA</td><td>Intergenic</td><td>NA</td><td>NA</td></tr>
<tr><td>T28898</td><td>CGTGTGGGGGGTGCTGCTGAAGG</td><td>1000</td><td></td><td></td><td></td><td></td><td></td><td></td><td></td><td></td><td></td></tr>
<tr><td>Oligo fwd</td><td>CACCCGTGTGGGGGGTGCTGCTGA</td><td></td><td></td><td></td><td></td><td></td><td></td><td></td><td></td><td></td><td></td></tr>
<tr><td>Oligo rev</td><td>AAACTCAGCAGCACCCCCCACACG</td><td></td><td></td><td></td><td></td><td></td><td></td><td></td><td></td><td></td><td></td></tr>
<tr><td>Chromosome</td><td>start</td><td>end</td><td>strand</td><td>MM</td><td>target_seq</td><td>PAM</td><td>alignment</td><td>distance</td><td>position</td><td>gene name</td><td>gene id</td></tr>
<tr><td>GU305914.1</td><td>70374</td><td>70396</td><td>-</td><td>0</td><td>CGTGTGGGGGGTGCTGCTGA</td><td>AGG</td><td>||||||||[||||||||||||]PAM</td><td>NA</td><td>Intergenic</td><td>NA</td><td>NA</td></tr>
<tr><td>T28899</td><td>GGTGCTGCTGAAGGCCGCCGCGG</td><td>1000</td><td></td><td></td><td></td><td></td><td></td><td></td><td></td><td></td><td></td></tr>
<tr><td>Oligo fwd</td><td>CACCGGTGCTGCTGAAGGCCGCCG</td><td></td><td></td><td></td><td></td><td></td><td></td><td></td><td></td><td></td><td></td></tr>
<tr><td>Oligo rev</td><td>AAACCGGCGGCCTTCAGCAGCACC</td><td></td><td></td><td></td><td></td><td></td><td></td><td></td><td></td><td></td><td></td></tr>
<tr><td>Chromosome</td><td>start</td><td>end</td><td>strand</td><td>MM</td><td>target_seq</td><td>PAM</td><td>alignment</td><td>distance</td><td>position</td><td>gene name</td><td>gene id</td></tr>
<tr><td>GU305914.1</td><td>70365</td><td>70387</td><td>-</td><td>0</td><td>GGTGCTGCTGAAGGCCGCCG</td><td>CGG</td><td>||||||||[||||||||||||]PAM</td><td>NA</td><td>Intergenic</td><td>NA</td><td>NA</td></tr>
<tr><td>T28900</td><td>GTGCTGCTGAAGGCCGCCGCGGG</td><td>1000</td><td></td><td></td><td></td><td></td><td></td><td></td><td></td><td></td><td></td></tr>
<tr><td>Oligo fwd</td><td>CACCGTGCTGCTGAAGGCCGCCGC</td><td></td><td></td><td></td><td></td><td></td><td></td><td></td><td></td><td></td><td></td></tr>
<tr><td>Oligo rev</td><td>AAACGCGGCGGCCTTCAGCAGCAC</td><td></td><td></td><td></td><td></td><td></td><td></td><td></td><td></td><td></td><td></td></tr>
<tr><td>Chromosome</td><td>start</td><td>end</td><td>strand</td><td>MM</td><td>target_seq</td><td>PAM</td><td>alignment</td><td>distance</td><td>position</td><td>gene name</td><td>gene id</td></tr>
<tr><td>GU305914.1</td><td>70364</td><td>70386</td><td>-</td><td>0</td><td>GTGCTGCTGAAGGCCGCCGC</td><td>GGG</td><td>||||||||[||||||||||||]PAM</td><td>NA</td><td>Intergenic</td><td>NA</td><td>NA</td></tr>
<tr><td>T28901</td><td>TGCTGCTGAAGGCCGCCGCGGGG</td><td>1000</td><td></td><td></td><td></td><td></td><td></td><td></td><td></td><td></td><td></td></tr>
<tr><td>Oligo fwd</td><td>CACCTGCTGCTGAAGGCCGCCGCG</td><td></td><td></td><td></td><td></td><td></td><td></td><td></td><td></td><td></td><td></td></tr>
<tr><td>Oligo rev</td><td>AAACCGCGGCGGCCTTCAGCAGCA</td><td></td><td></td><td></td><td></td><td></td><td></td><td></td><td></td><td></td><td></td></tr>
<tr><td>Chromosome</td><td>start</td><td>end</td><td>strand</td><td>MM</td><td>target_seq</td><td>PAM</td><td>alignment</td><td>distance</td><td>position</td><td>gene name</td><td>gene id</td></tr>
<tr><td>GU305914.1</td><td>70363</td><td>70385</td><td>-</td><td>0</td><td>TGCTGCTGAAGGCCGCCGCG</td><td>GGG</td><td>||||||||[||||||||||||]PAM</td><td>NA</td><td>Intergenic</td><td>NA</td><td>NA</td></tr>
<tr><td>T28902</td><td>AGTTCTTTCGCATGTGCGTCAGG</td><td>1000</td><td></td><td></td><td></td><td></td><td></td><td></td><td></td><td></td><td></td></tr>
<tr><td>Oligo fwd</td><td>CACCAGTTCTTTCGCATGTGCGTC</td><td></td><td></td><td></td><td></td><td></td><td></td><td></td><td></td><td></td><td></td></tr>
<tr><td>Oligo rev</td><td>AAACGACGCACATGCGAAAGAACT</td><td></td><td></td><td></td><td></td><td></td><td></td><td></td><td></td><td></td><td></td></tr>
<tr><td>Chromosome</td><td>start</td><td>end</td><td>strand</td><td>MM</td><td>target_seq</td><td>PAM</td><td>alignment</td><td>distance</td><td>position</td><td>gene name</td><td>gene id</td></tr>
<tr><td>GU305914.1</td><td>70333</td><td>70355</td><td>-</td><td>0</td><td>AGTTCTTTCGCATGTGCGTC</td><td>AGG</td><td>||||||||[||||||||||||]PAM</td><td>NA</td><td>Intergenic</td><td>NA</td><td>NA</td></tr>
<tr><td>T28903</td><td>GTTCTTTCGCATGTGCGTCAGGG</td><td>1000</td><td></td><td></td><td></td><td></td><td></td><td></td><td></td><td></td><td></td></tr>
<tr><td>Oligo fwd</td><td>CACCGTTCTTTCGCATGTGCGTCA</td><td></td><td></td><td></td><td></td><td></td><td></td><td></td><td></td><td></td><td></td></tr>
<tr><td>Oligo rev</td><td>AAACTGACGCACATGCGAAAGAAC</td><td></td><td></td><td></td><td></td><td></td><td></td><td></td><td></td><td></td><td></td></tr>
<tr><td>Chromosome</td><td>start</td><td>end</td><td>strand</td><td>MM</td><td>target_seq</td><td>PAM</td><td>alignment</td><td>distance</td><td>position</td><td>gene name</td><td>gene id</td></tr>
<tr><td>GU305914.1</td><td>70332</td><td>70354</td><td>-</td><td>0</td><td>GTTCTTTCGCATGTGCGTCA</td><td>GGG</td><td>||||||||[||||||||||||]PAM</td><td>NA</td><td>Intergenic</td><td>NA</td><td>NA</td></tr>
<tr><td>T28904</td><td>CTTTCGCATGTGCGTCAGGGCGG</td><td>1000</td><td></td><td></td><td></td><td></td><td></td><td></td><td></td><td></td><td></td></tr>
<tr><td>Oligo fwd</td><td>CACCCTTTCGCATGTGCGTCAGGG</td><td></td><td></td><td></td><td></td><td></td><td></td><td></td><td></td><td></td><td></td></tr>
<tr><td>Oligo rev</td><td>AAACCCCTGACGCACATGCGAAAG</td><td></td><td></td><td></td><td></td><td></td><td></td><td></td><td></td><td></td><td></td></tr>
<tr><td>Chromosome</td><td>start</td><td>end</td><td>strand</td><td>MM</td><td>target_seq</td><td>PAM</td><td>alignment</td><td>distance</td><td>position</td><td>gene name</td><td>gene id</td></tr>
<tr><td>GU305914.1</td><td>70329</td><td>70351</td><td>-</td><td>0</td><td>CTTTCGCATGTGCGTCAGGG</td><td>CGG</td><td>||||||||[||||||||||||]PAM</td><td>NA</td><td>Intergenic</td><td>NA</td><td>NA</td></tr>
<tr><td>T28905</td><td>GCATGTGCGTCAGGGCGGTCAGG</td><td>1000</td><td></td><td></td><td></td><td></td><td></td><td></td><td></td><td></td><td></td></tr>
<tr><td>Oligo fwd</td><td>CACCGCATGTGCGTCAGGGCGGTC</td><td></td><td></td><td></td><td></td><td></td><td></td><td></td><td></td><td></td><td></td></tr>
<tr><td>Oligo rev</td><td>AAACGACCGCCCTGACGCACATGC</td><td></td><td></td><td></td><td></td><td></td><td></td><td></td><td></td><td></td><td></td></tr>
<tr><td>Chromosome</td><td>start</td><td>end</td><td>strand</td><td>MM</td><td>target_seq</td><td>PAM</td><td>alignment</td><td>distance</td><td>position</td><td>gene name</td><td>gene id</td></tr>
<tr><td>GU305914.1</td><td>70324</td><td>70346</td><td>-</td><td>0</td><td>GCATGTGCGTCAGGGCGGTC</td><td>AGG</td><td>||||||||[||||||||||||]PAM</td><td>NA</td><td>Intergenic</td><td>NA</td><td>NA</td></tr>
</table>

| | | | | | | | | | | | |
|---|---|---|---|---|---|---|---|---|---|---|---|
| T28906 | TGTGCGTCAGGGCGGTCAGGAGG | 1000 | | | | | | | | | |
| Oligo fwd | CACCTGTGCGTCAGGGCGGTCAGG | | | | | | | | | | |
| Oligo rev | AAACCCTGACCGCCCTGACGCACA | | | | | | | | | | |
| Chromosome | start | end | strand | MM | target_seq | PAM | alignment | distance | position | gene name | gene id |
| GU305914.1 | 70321 | 70343 | - | 0 | TGTGCGTCAGGGCGGTCAGG | AGG | \|\|\|\|\|\|\|\|[\|\|\|\|\|\|\|\|\|\|\|\|]PAM | NA | Intergenic | NA | NA |
| T28907 | GGAGGTGCGCGATCGTCAGCAGG | 1000 | | | | | | | | | |
| Oligo fwd | CACCGGAGGTGCGCGATCGTCAGC | | | | | | | | | | |
| Oligo rev | AAACGCTGACGATCGCGCACCTCC | | | | | | | | | | |
| Chromosome | start | end | strand | MM | target_seq | PAM | alignment | distance | position | gene name | gene id |
| GU305914.1 | 70303 | 70325 | - | 0 | GGAGGTGCGCGATCGTCAGC | AGG | \|\|\|\|\|\|\|\|[\|\|\|\|\|\|\|\|\|\|\|\|]PAM | NA | Intergenic | NA | NA |
| T28908 | AGCAGGTCTCGTTCCGTCTCCGG | 1000 | | | | | | | | | |
| Oligo fwd | CACCAGCAGGTCTCGTTCCGTCTC | | | | | | | | | | |
| Oligo rev | AAACGAGACGGAACGAGACCTGCT | | | | | | | | | | |
| Chromosome | start | end | strand | MM | target_seq | PAM | alignment | distance | position | gene name | gene id |
| GU305914.1 | 70286 | 70308 | - | 0 | AGCAGGTCTCGTTCCGTCTC | CGG | \|\|\|\|\|\|\|\|[\|\|\|\|\|\|\|\|\|\|\|\|]PAM | NA | Intergenic | NA | NA |
| T28909 | GCTCGTGTGTTTCTGTCGATCGG | 1000 | | | | | | | | | |
| Oligo fwd | CACCGCTCGTGTGTTTCTGTCGAT | | | | | | | | | | |
| Oligo rev | AAACATCGACAGAAACACACGAGC | | | | | | | | | | |
| Chromosome | start | end | strand | MM | target_seq | PAM | alignment | distance | position | gene name | gene id |
| GU305914.1 | 70209 | 70231 | - | 0 | GCTCGTGTGTTTCTGTCGAT | CGG | \|\|\|\|\|\|\|\|[\|\|\|\|\|\|\|\|\|\|\|\|]PAM | NA | Intergenic | NA | NA |
| T28910 | CTGTCGATCGGCGCCGCCGCCGG | 1000 | | | | | | | | | |
| Oligo fwd | CACCCTGTCGATCGGCGCCGCCGC | | | | | | | | | | |
| Oligo rev | AAACGCGGCGGCGCCGATCGACAG | | | | | | | | | | |
| Chromosome | start | end | strand | MM | target_seq | PAM | alignment | distance | position | gene name | gene id |
| GU305914.1 | 70197 | 70219 | - | 0 | CTGTCGATCGGCGCCGCCGC | CGG | \|\|\|\|\|\|\|\|[\|\|\|\|\|\|\|\|\|\|\|\|]PAM | NA | Intergenic | NA | NA |
| T28911 | GGCGCCGCCGCCGGCGCTGCCGG | 1000 | | | | | | | | | |
| Oligo fwd | CACCGGCGCCGCCGCCGGCGCTGC | | | | | | | | | | |
| Oligo rev | AAACGCAGCGCCGGCGGCGGCGCC | | | | | | | | | | |
| Chromosome | start | end | strand | MM | target_seq | PAM | alignment | distance | position | gene name | gene id |
| GU305914.1 | 70188 | 70210 | - | 0 | GGCGCCGCCGCCGGCGCTGC | CGG | \|\|\|\|\|\|\|\|[\|\|\|\|\|\|\|\|\|\|\|\|]PAM | NA | Intergenic | NA | NA |
| GU305914.1 | 69121 | 69143 | + | 4 | CCCGCCGCCGCCGGAGATGC | TGG | --\|\|\|\|\|\|[\|\|\|\|\|\|-\|-\|\|\|]PAM | NA | Intergenic | NA | NA |
| GU305914.1 | 100822 | 100844 | - | 4 | GCCGCTGCCGCCGACGCTGA | TGG | \|-\|\|\|-\|\|[\|\|\|\|\|-\|\|\|\|\|-]PAM | NA | Intergenic | NA | NA |
| T28912 | GCCGGTGCTGCTCGTCGCCTCGG | 1000 | | | | | | | | | |
| Oligo fwd | CACCGCCGGTGCTGCTCGTCGCCT | | | | | | | | | | |
| Oligo rev | AAACAGGCGACGAGCAGCACCGGC | | | | | | | | | | |
| Chromosome | start | end | strand | MM | target_seq | PAM | alignment | distance | position | gene name | gene id |
| GU305914.1 | 70170 | 70192 | - | 0 | GCCGGTGCTGCTCGTCGCCT | CGG | \|\|\|\|\|\|\|\|[\|\|\|\|\|\|\|\|\|\|\|\|]PAM | NA | Intergenic | NA | NA |
| T28913 | CCGGTGCTGCTCGTCGCCTCGGG | 1000 | | | | | | | | | |
| Oligo fwd | CACCCCGGTGCTGCTCGTCGCCTC | | | | | | | | | | |
| Oligo rev | AAACGAGGCGACGAGCAGCACCGG | | | | | | | | | | |
| Chromosome | start | end | strand | MM | target_seq | PAM | alignment | distance | position | gene name | gene id |
| GU305914.1 | 70169 | 70191 | - | 0 | CCGGTGCTGCTCGTCGCCTC | GGG | \|\|\|\|\|\|\|\|[\|\|\|\|\|\|\|\|\|\|\|\|]PAM | NA | Intergenic | NA | NA |
| GU305914.1 | 52303 | 52325 | - | 3 | CCAGCGCTGCTCTTCGCCTC | TGG | \|\|-\|-\|\|\|[\|\|\|\|-\|\|\|\|\|\|\|]PAM | NA | Intergenic | NA | NA |
| T28914 | CGGTGCTGCTCGTCGCCTCGGGG | 1000 | | | | | | | | | |
| Oligo fwd | CACCCGGTGCTGCTCGTCGCCTCG | | | | | | | | | | |
| Oligo rev | AAACCGAGGCGACGAGCAGCACCG | | | | | | | | | | |
| Chromosome | start | end | strand | MM | target_seq | PAM | alignment | distance | position | gene name | gene id |
| GU305914.1 | 70168 | 70190 | - | 0 | CGGTGCTGCTCGTCGCCTCG | GGG | \|\|\|\|\|\|\|\|[\|\|\|\|\|\|\|\|\|\|\|\|]PAM | NA | Intergenic | NA | NA |
| GU305914.1 | 190140 | 190162 | + | 4 | GGGACCTGCTCGTCGCCGCG | AGG | -\|\|--\|\|\|[\|\|\|\|\|\|\|\|\|-\|\|]PAM | NA | Intergenic | NA | NA |
| GU305914.1 | 52302 | 52324 | - | 4 | CAGCGCTGCTCTTCGCCTCT | GGG | \|-\|-\|\|\|\|[\|\|\|-\|\|\|\|\|\|\|-]PAM | NA | Intergenic | NA | NA |
| T28915 | CCTCGGGGTCCGCGCCCTTGAGG | 1000 | | | | | | | | | |
| Oligo fwd | CACCCCTCGGGGTCCGCGCCCTTG | | | | | | | | | | |
| Oligo rev | AAACCAAGGGCGCGGACCCCGAGG | | | | | | | | | | |
| Chromosome | start | end | strand | MM | target_seq | PAM | alignment | distance | position | gene name | gene id |
| GU305914.1 | 70153 | 70175 | - | 0 | CCTCGGGGTCCGCGCCCTTG | AGG | \|\|\|\|\|\|\|\|[\|\|\|\|\|\|\|\|\|\|\|\|]PAM | NA | Intergenic | NA | NA |
| T28916 | GTCCGCGCCCTTGAGGATCGCGG | 1000 | | | | | | | | | |
| Oligo fwd | CACCGTCCGCGCCCTTGAGGATCG | | | | | | | | | | |
| Oligo rev | AAACCGATCCTCAAGGGCGCGGAC | | | | | | | | | | |
| Chromosome | start | end | strand | MM | target_seq | PAM | alignment | distance | position | gene name | gene id |
| GU305914.1 | 70146 | 70168 | - | 0 | GTCCGCGCCCTTGAGGATCG | CGG | \|\|\|\|\|\|\|\|[\|\|\|\|\|\|\|\|\|\|\|\|]PAM | NA | Intergenic | NA | NA |
| T28917 | CTTGAGGATCGCGGCGATCGTGG | 1000 | | | | | | | | | |
| Oligo fwd | CACCCTTGAGGATCGCGGCGATCG | | | | | | | | | | |
| Oligo rev | AAACCGATCGCCGCGATCCTCAAG | | | | | | | | | | |
| Chromosome | start | end | strand | MM | target_seq | PAM | alignment | distance | position | gene name | gene id |
| GU305914.1 | 70137 | 70159 | - | 0 | CTTGAGGATCGCGGCGATCG | TGG | \|\|\|\|\|\|\|\|[\|\|\|\|\|\|\|\|\|\|\|\|]PAM | NA | Intergenic | NA | NA |
| T28918 | CCGTGACGAGCGACGCCAGCAGG | 1000 | | | | | | | | | |
| Oligo fwd | CACCCCGTGACGAGCGACGCCAGC | | | | | | | | | | |
| Oligo rev | AAACGCTGGCGTCGCTCGTCACGG | | | | | | | | | | |
| Chromosome | start | end | strand | MM | target_seq | PAM | alignment | distance | position | gene name | gene id |
| GU305914.1 | 70105 | 70127 | - | 0 | CCGTGACGAGCGACGCCAGC | AGG | \|\|\|\|\|\|\|\|[\|\|\|\|\|\|\|\|\|\|\|\|]PAM | NA | Intergenic | NA | NA |
| T28919 | CGTGACGAGCGACGCCAGCAGGG | 1000 | | | | | | | | | |
| Oligo fwd | CACCCGTGACGAGCGACGCCAGCA | | | | | | | | | | |
| Oligo rev | AAACTGCTGGCGTCGCTCGTCACG | | | | | | | | | | |
| Chromosome | start | end | strand | MM | target_seq | PAM | alignment | distance | position | gene name | gene id |
| GU305914.1 | 70104 | 70126 | - | 0 | CGTGACGAGCGACGCCAGCA | GGG | \|\|\|\|\|\|\|\|[\|\|\|\|\|\|\|\|\|\|\|\|]PAM | NA | Intergenic | NA | NA |

<table>
<tr><td>T28920</td><td>GCGACGCCAGCAGGGAGTTCAGG</td><td></td><td>1000</td><td></td><td></td><td></td><td></td><td></td><td></td><td></td><td></td><td></td></tr>
<tr><td>Oligo fwd</td><td>CACCGCGACGCCAGCAGGGAGTTC</td><td></td><td></td><td></td><td></td><td></td><td></td><td></td><td></td><td></td><td></td><td></td></tr>
<tr><td>Oligo rev</td><td>AAACGAACTCCCTGCTGGCGTCGC</td><td></td><td></td><td></td><td></td><td></td><td></td><td></td><td></td><td></td><td></td><td></td></tr>
<tr><td>Chromosome</td><td>start</td><td></td><td>end</td><td>strand</td><td>MM</td><td>target_seq</td><td>PAM</td><td>alignment</td><td>distance</td><td>position</td><td>gene name</td><td>gene id</td></tr>
<tr><td>GU305914.1</td><td></td><td>70096</td><td>70118</td><td>-</td><td>0</td><td>GCGACGCCAGCAGGGAGTTC</td><td>AGG</td><td>||||||||[||||||||||||]PAM</td><td>NA</td><td>Intergenic</td><td>NA</td><td>NA</td></tr>
<tr><td>T28921</td><td>CGACGCCAGCAGGGAGTTCAGGG</td><td></td><td>1000</td><td></td><td></td><td></td><td></td><td></td><td></td><td></td><td></td><td></td></tr>
<tr><td>Oligo fwd</td><td>CACCCGACGCCAGCAGGGAGTTCA</td><td></td><td></td><td></td><td></td><td></td><td></td><td></td><td></td><td></td><td></td><td></td></tr>
<tr><td>Oligo rev</td><td>AAACTGAACTCCCTGCTGGCGTCG</td><td></td><td></td><td></td><td></td><td></td><td></td><td></td><td></td><td></td><td></td><td></td></tr>
<tr><td>Chromosome</td><td>start</td><td></td><td>end</td><td>strand</td><td>MM</td><td>target_seq</td><td>PAM</td><td>alignment</td><td>distance</td><td>position</td><td>gene name</td><td>gene id</td></tr>
<tr><td>GU305914.1</td><td></td><td>70095</td><td>70117</td><td>-</td><td>0</td><td>CGACGCCAGCAGGGAGTTCA</td><td>GGG</td><td>||||||||[||||||||||||]PAM</td><td>NA</td><td>Intergenic</td><td>NA</td><td>NA</td></tr>
<tr><td>T28922</td><td>CAGCAGGGAGTTCAGGGATTTGG</td><td></td><td>1000</td><td></td><td></td><td></td><td></td><td></td><td></td><td></td><td></td><td></td></tr>
<tr><td>Oligo fwd</td><td>CACCCAGCAGGGAGTTCAGGGATT</td><td></td><td></td><td></td><td></td><td></td><td></td><td></td><td></td><td></td><td></td><td></td></tr>
<tr><td>Oligo rev</td><td>AAACAATCCCTGAACTCCCTGCTG</td><td></td><td></td><td></td><td></td><td></td><td></td><td></td><td></td><td></td><td></td><td></td></tr>
<tr><td>Chromosome</td><td>start</td><td></td><td>end</td><td>strand</td><td>MM</td><td>target_seq</td><td>PAM</td><td>alignment</td><td>distance</td><td>position</td><td>gene name</td><td>gene id</td></tr>
<tr><td>GU305914.1</td><td></td><td>70089</td><td>70111</td><td>-</td><td>0</td><td>CAGCAGGGAGTTCAGGGATT</td><td>TGG</td><td>||||||||[||||||||||||]PAM</td><td>NA</td><td>Intergenic</td><td>NA</td><td>NA</td></tr>
<tr><td>T28923</td><td>GGGAGTTCAGGGATTTGGTCAGG</td><td></td><td>1000</td><td></td><td></td><td></td><td></td><td></td><td></td><td></td><td></td><td></td></tr>
<tr><td>Oligo fwd</td><td>CACCGGGAGTTCAGGGATTTGGTC</td><td></td><td></td><td></td><td></td><td></td><td></td><td></td><td></td><td></td><td></td><td></td></tr>
<tr><td>Oligo rev</td><td>AAACGACCAAATCCCTGAACTCCC</td><td></td><td></td><td></td><td></td><td></td><td></td><td></td><td></td><td></td><td></td><td></td></tr>
<tr><td>Chromosome</td><td>start</td><td></td><td>end</td><td>strand</td><td>MM</td><td>target_seq</td><td>PAM</td><td>alignment</td><td>distance</td><td>position</td><td>gene name</td><td>gene id</td></tr>
<tr><td>GU305914.1</td><td></td><td>70084</td><td>70106</td><td>-</td><td>0</td><td>GGGAGTTCAGGGATTTGGTC</td><td>AGG</td><td>||||||||[||||||||||||]PAM</td><td>NA</td><td>Intergenic</td><td>NA</td><td>NA</td></tr>
<tr><td>T28924</td><td>GTTCAGGGATTTGGTCAGGTTGG</td><td></td><td>1000</td><td></td><td></td><td></td><td></td><td></td><td></td><td></td><td></td><td></td></tr>
<tr><td>Oligo fwd</td><td>CACCGTTCAGGGATTTGGTCAGGT</td><td></td><td></td><td></td><td></td><td></td><td></td><td></td><td></td><td></td><td></td><td></td></tr>
<tr><td>Oligo rev</td><td>AAACACCTGACCAAATCCCTGAAC</td><td></td><td></td><td></td><td></td><td></td><td></td><td></td><td></td><td></td><td></td><td></td></tr>
<tr><td>Chromosome</td><td>start</td><td></td><td>end</td><td>strand</td><td>MM</td><td>target_seq</td><td>PAM</td><td>alignment</td><td>distance</td><td>position</td><td>gene name</td><td>gene id</td></tr>
<tr><td>GU305914.1</td><td></td><td>70080</td><td>70102</td><td>-</td><td>0</td><td>GTTCAGGGATTTGGTCAGGT</td><td>TGG</td><td>||||||||[||||||||||||]PAM</td><td>NA</td><td>Intergenic</td><td>NA</td><td>NA</td></tr>
<tr><td>T28925</td><td>TTTGGTCAGGTTGGCGTTGAAGG</td><td></td><td>1000</td><td></td><td></td><td></td><td></td><td></td><td></td><td></td><td></td><td></td></tr>
<tr><td>Oligo fwd</td><td>CACCTTTGGTCAGGTTGGCGTTGA</td><td></td><td></td><td></td><td></td><td></td><td></td><td></td><td></td><td></td><td></td><td></td></tr>
<tr><td>Oligo rev</td><td>AAACTCAACGCCAACCTGACCAAA</td><td></td><td></td><td></td><td></td><td></td><td></td><td></td><td></td><td></td><td></td><td></td></tr>
<tr><td>Chromosome</td><td>start</td><td></td><td>end</td><td>strand</td><td>MM</td><td>target_seq</td><td>PAM</td><td>alignment</td><td>distance</td><td>position</td><td>gene name</td><td>gene id</td></tr>
<tr><td>GU305914.1</td><td></td><td>70071</td><td>70093</td><td>-</td><td>0</td><td>TTTGGTCAGGTTGGCGTTGA</td><td>AGG</td><td>||||||||[||||||||||||]PAM</td><td>NA</td><td>Intergenic</td><td>NA</td><td>NA</td></tr>
<tr><td>T28926</td><td>GGCGTTGAAGGTCGCGATCGCGG</td><td></td><td>1000</td><td></td><td></td><td></td><td></td><td></td><td></td><td></td><td></td><td></td></tr>
<tr><td>Oligo fwd</td><td>CACCGGCGTTGAAGGTCGCGATCG</td><td></td><td></td><td></td><td></td><td></td><td></td><td></td><td></td><td></td><td></td><td></td></tr>
<tr><td>Oligo rev</td><td>AAACCGATCGCGACCTTCAACGCC</td><td></td><td></td><td></td><td></td><td></td><td></td><td></td><td></td><td></td><td></td><td></td></tr>
<tr><td>Chromosome</td><td>start</td><td></td><td>end</td><td>strand</td><td>MM</td><td>target_seq</td><td>PAM</td><td>alignment</td><td>distance</td><td>position</td><td>gene name</td><td>gene id</td></tr>
<tr><td>GU305914.1</td><td></td><td>70059</td><td>70081</td><td>-</td><td>0</td><td>GGCGTTGAAGGTCGCGATCG</td><td>CGG</td><td>||||||||[||||||||||||]PAM</td><td>NA</td><td>Intergenic</td><td>NA</td><td>NA</td></tr>
<tr><td>T28927</td><td>CGCGGCTTTCCTGTCCTCCTTGG</td><td></td><td>1000</td><td></td><td></td><td></td><td></td><td></td><td></td><td></td><td></td><td></td></tr>
<tr><td>Oligo fwd</td><td>CACCCGCGGCTTTCCTGTCCTCCT</td><td></td><td></td><td></td><td></td><td></td><td></td><td></td><td></td><td></td><td></td><td></td></tr>
<tr><td>Oligo rev</td><td>AAACAGGAGGACAGGAAAGCCGCG</td><td></td><td></td><td></td><td></td><td></td><td></td><td></td><td></td><td></td><td></td><td></td></tr>
<tr><td>Chromosome</td><td>start</td><td></td><td>end</td><td>strand</td><td>MM</td><td>target_seq</td><td>PAM</td><td>alignment</td><td>distance</td><td>position</td><td>gene name</td><td>gene id</td></tr>
<tr><td>GU305914.1</td><td></td><td>70041</td><td>70063</td><td>-</td><td>0</td><td>CGCGGCTTTCCTGTCCTCCT</td><td>TGG</td><td>||||||||[||||||||||||]PAM</td><td>NA</td><td>Intergenic</td><td>NA</td><td>NA</td></tr>
<tr><td>T28928</td><td>CTTGGTCAGCCTCACGAGCGAGG</td><td></td><td>1000</td><td></td><td></td><td></td><td></td><td></td><td></td><td></td><td></td><td></td></tr>
<tr><td>Oligo fwd</td><td>CACCCTTGGTCAGCCTCACGAGCG</td><td></td><td></td><td></td><td></td><td></td><td></td><td></td><td></td><td></td><td></td><td></td></tr>
<tr><td>Oligo rev</td><td>AAACCGCTCGTGAGGCTGACCAAG</td><td></td><td></td><td></td><td></td><td></td><td></td><td></td><td></td><td></td><td></td><td></td></tr>
<tr><td>Chromosome</td><td>start</td><td></td><td>end</td><td>strand</td><td>MM</td><td>target_seq</td><td>PAM</td><td>alignment</td><td>distance</td><td>position</td><td>gene name</td><td>gene id</td></tr>
<tr><td>GU305914.1</td><td></td><td>70023</td><td>70045</td><td>-</td><td>0</td><td>CTTGGTCAGCCTCACGAGCG</td><td>AGG</td><td>||||||||[||||||||||||]PAM</td><td>NA</td><td>Intergenic</td><td>NA</td><td>NA</td></tr>
<tr><td>T28929</td><td>GCGAGGTGCCGCAGCTTTTCAGG</td><td></td><td>1000</td><td></td><td></td><td></td><td></td><td></td><td></td><td></td><td></td><td></td></tr>
<tr><td>Oligo fwd</td><td>CACCGCGAGGTGCCGCAGCTTTTC</td><td></td><td></td><td></td><td></td><td></td><td></td><td></td><td></td><td></td><td></td><td></td></tr>
<tr><td>Oligo rev</td><td>AAACGAAAAGCTGCGGCACCTCGC</td><td></td><td></td><td></td><td></td><td></td><td></td><td></td><td></td><td></td><td></td><td></td></tr>
<tr><td>Chromosome</td><td>start</td><td></td><td>end</td><td>strand</td><td>MM</td><td>target_seq</td><td>PAM</td><td>alignment</td><td>distance</td><td>position</td><td>gene name</td><td>gene id</td></tr>
<tr><td>GU305914.1</td><td></td><td>70006</td><td>70028</td><td>-</td><td>0</td><td>GCGAGGTGCCGCAGCTTTTC</td><td>AGG</td><td>||||||||[||||||||||||]PAM</td><td>NA</td><td>Intergenic</td><td>NA</td><td>NA</td></tr>
<tr><td>T28930</td><td>TCGATCGAAGAGAACATCGACGG</td><td></td><td>1000</td><td></td><td></td><td></td><td></td><td></td><td></td><td></td><td></td><td></td></tr>
<tr><td>Oligo fwd</td><td>CACCTCGATCGAAGAGAACATCGA</td><td></td><td></td><td></td><td></td><td></td><td></td><td></td><td></td><td></td><td></td><td></td></tr>
<tr><td>Oligo rev</td><td>AAACTCGATGTTCTCTTCGATCGA</td><td></td><td></td><td></td><td></td><td></td><td></td><td></td><td></td><td></td><td></td><td></td></tr>
<tr><td>Chromosome</td><td>start</td><td></td><td>end</td><td>strand</td><td>MM</td><td>target_seq</td><td>PAM</td><td>alignment</td><td>distance</td><td>position</td><td>gene name</td><td>gene id</td></tr>
<tr><td>GU305914.1</td><td></td><td>69983</td><td>70005</td><td>-</td><td>0</td><td>TCGATCGAAGAGAACATCGA</td><td>CGG</td><td>||||||||[||||||||||||]PAM</td><td>NA</td><td>Intergenic</td><td>NA</td><td>NA</td></tr>
<tr><td>T28931</td><td>GAACATCGACGGCGTCTGCGAGG</td><td></td><td>1000</td><td></td><td></td><td></td><td></td><td></td><td></td><td></td><td></td><td></td></tr>
<tr><td>Oligo fwd</td><td>CACCGAACATCGACGGCGTCTGCG</td><td></td><td></td><td></td><td></td><td></td><td></td><td></td><td></td><td></td><td></td><td></td></tr>
<tr><td>Oligo rev</td><td>AAACCGCAGACGCCGTCGATGTTC</td><td></td><td></td><td></td><td></td><td></td><td></td><td></td><td></td><td></td><td></td><td></td></tr>
<tr><td>Chromosome</td><td>start</td><td></td><td>end</td><td>strand</td><td>MM</td><td>target_seq</td><td>PAM</td><td>alignment</td><td>distance</td><td>position</td><td>gene name</td><td>gene id</td></tr>
<tr><td>GU305914.1</td><td></td><td>69972</td><td>69994</td><td>-</td><td>0</td><td>GAACATCGACGGCGTCTGCG</td><td>AGG</td><td>||||||||[||||||||||||]PAM</td><td>NA</td><td>Intergenic</td><td>NA</td><td>NA</td></tr>
<tr><td>T28932</td><td>ACGAGTCCGACACCACCTTCAGG</td><td></td><td>1000</td><td></td><td></td><td></td><td></td><td></td><td></td><td></td><td></td><td></td></tr>
<tr><td>Oligo fwd</td><td>CACCACGAGTCCGACACCACCTTC</td><td></td><td></td><td></td><td></td><td></td><td></td><td></td><td></td><td></td><td></td><td></td></tr>
<tr><td>Oligo rev</td><td>AAACGAAGGTGGTGTCGGACTCGT</td><td></td><td></td><td></td><td></td><td></td><td></td><td></td><td></td><td></td><td></td><td></td></tr>
<tr><td>Chromosome</td><td>start</td><td></td><td>end</td><td>strand</td><td>MM</td><td>target_seq</td><td>PAM</td><td>alignment</td><td>distance</td><td>position</td><td>gene name</td><td>gene id</td></tr>
<tr><td>GU305914.1</td><td></td><td>69946</td><td>69968</td><td>-</td><td>0</td><td>ACGAGTCCGACACCACCTTC</td><td>AGG</td><td>||||||||[||||||||||||]PAM</td><td>NA</td><td>Intergenic</td><td>NA</td><td>NA</td></tr>
<tr><td>T28933</td><td>GTCCGACACCACCTTCAGGTCGG</td><td></td><td>1000</td><td></td><td></td><td></td><td></td><td></td><td></td><td></td><td></td><td></td></tr>
<tr><td>Oligo fwd</td><td>CACCGTCCGACACCACCTTCAGGT</td><td></td><td></td><td></td><td></td><td></td><td></td><td></td><td></td><td></td><td></td><td></td></tr>
<tr><td>Oligo rev</td><td>AAACACCTGAAGGTGGTGTCGGAC</td><td></td><td></td><td></td><td></td><td></td><td></td><td></td><td></td><td></td><td></td><td></td></tr>
<tr><td>Chromosome</td><td>start</td><td></td><td>end</td><td>strand</td><td>MM</td><td>target_seq</td><td>PAM</td><td>alignment</td><td>distance</td><td>position</td><td>gene name</td><td>gene id</td></tr>
<tr><td>GU305914.1</td><td></td><td>69942</td><td>69964</td><td>-</td><td>0</td><td>GTCCGACACCACCTTCAGGT</td><td>CGG</td><td>||||||||[||||||||||||]PAM</td><td>NA</td><td>Intergenic</td><td>NA</td><td>NA</td></tr>
<tr><td>T28934</td><td>TTCAGTCGCGCGTGCATGTCCGG</td><td></td><td>1000</td><td></td><td></td><td></td><td></td><td></td><td></td><td></td><td></td><td></td></tr>
<tr><td>Oligo fwd</td><td>CACCTTCAGTCGCGCGTGCATGTC</td><td></td><td></td><td></td><td></td><td></td><td></td><td></td><td></td><td></td><td></td><td></td></tr>
<tr><td>Oligo rev</td><td>AAACGACATGCACGCGCGACTGAA</td><td></td><td></td><td></td><td></td><td></td><td></td><td></td><td></td><td></td><td></td><td></td></tr>
<tr><td>Chromosome</td><td>start</td><td></td><td>end</td><td>strand</td><td>MM</td><td>target_seq</td><td>PAM</td><td>alignment</td><td>distance</td><td>position</td><td>gene name</td><td>gene id</td></tr>
<tr><td>GU305914.1</td><td></td><td>69839</td><td>69861</td><td>-</td><td>0</td><td>TTCAGTCGCGCGTGCATGTC</td><td>CGG</td><td>||||||||[||||||||||||]PAM</td><td>NA</td><td>Intergenic</td><td>NA</td><td>NA</td></tr>
</table>

T28935 TGTCCGGCGCGTGCTTCGCCAGG 1000
Oligo fwd CACCTGTCCGGCGCGTGCTTCGCC
Oligo rev AAACGGCGAAGCACGCGCCGGACA
Chromosome start end strand MM target_seq PAM alignment distance position gene name gene id
GU305914.1 69823 69845 - 0 TGTCCGGCGCGTGCTTCGCC AGG ||||||||[||||||||||||]PAM NA Intergenic NA NA

T28936 TTCGCCAGGATCGTTTCGTTCGG 1000
Oligo fwd CACCTTCGCCAGGATCGTTTCGTT
Oligo rev AAACAACGAAACGATCCTGGCGAA
Chromosome start end strand MM target_seq PAM alignment distance position gene name gene id
GU305914.1 69809 69831 - 0 TTCGCCAGGATCGTTTCGTT CGG ||||||||[||||||||||||]PAM NA Intergenic NA NA

T28937 AGCAGCCTCTCCAGCGTCTGCGG 1000
Oligo fwd CACCAGCAGCCTCTCCAGCGTCTG
Oligo rev AAACCAGACGCTGGAGAGGCTGCT
Chromosome start end strand MM target_seq PAM alignment distance position gene name gene id
GU305914.1 69776 69798 - 0 AGCAGCCTCTCCAGCGTCTG CGG ||||||||[||||||||||||]PAM NA Intergenic NA NA

T28938 GCAGCCTCTCCAGCGTCTGCGGG 1000
Oligo fwd CACCGCAGCCTCTCCAGCGTCTGC
Oligo rev AAACGCAGACGCTGGAGAGGCTGC
Chromosome start end strand MM target_seq PAM alignment distance position gene name gene id
GU305914.1 69775 69797 - 0 GCAGCCTCTCCAGCGTCTGC GGG ||||||||[||||||||||||]PAM NA Intergenic NA NA

T28939 CAGCCTCTCCAGCGTCTGCGGGG 1000
Oligo fwd CACCCAGCCTCTCCAGCGTCTGCG
Oligo rev AAACCGCAGACGCTGGAGAGGCTG
Chromosome start end strand MM target_seq PAM alignment distance position gene name gene id
GU305914.1 69774 69796 - 0 CAGCCTCTCCAGCGTCTGCG GGG ||||||||[||||||||||||]PAM NA Intergenic NA NA

T28940 CTTCACGCGCTTCAGCCACTCGG 1000
Oligo fwd CACCCTTCACGCGCTTCAGCCACT
Oligo rev AAACAGTGGCTGAAGCGCGTGAAG
Chromosome start end strand MM target_seq PAM alignment distance position gene name gene id
GU305914.1 69735 69757 - 0 CTTCACGCGCTTCAGCCACT CGG ||||||||[||||||||||||]PAM NA Intergenic NA NA

T28941 CGCGCAGCTCCTCCATCTTGCGG 1000
Oligo fwd CACCCGCGCAGCTCCTCCATCTTG
Oligo rev AAACCAAGATGGAGGAGCTGCGCG
Chromosome start end strand MM target_seq PAM alignment distance position gene name gene id
GU305914.1 69709 69731 - 0 CGCGCAGCTCCTCCATCTTG CGG ||||||||[||||||||||||]PAM NA Intergenic NA NA

T28942 GGTCGTATTCCACCTGCACCCGG 1000
Oligo fwd CACCGGTCGTATTCCACCTGCACC
Oligo rev AAACGGTGCAGGTGGAATACGACC
Chromosome start end strand MM target_seq PAM alignment distance position gene name gene id
GU305914.1 69688 69710 - 0 GGTCGTATTCCACCTGCACC CGG ||||||||[||||||||||||]PAM NA Intergenic NA NA

T28943 CGTATTCCACCTGCACCCGGCGG 1000
Oligo fwd CACCCGTATTCCACCTGCACCCGG
Oligo rev AAACCCGGGTGCAGGTGGAATACG
Chromosome start end strand MM target_seq PAM alignment distance position gene name gene id
GU305914.1 69685 69707 - 0 CGTATTCCACCTGCACCCGG CGG ||||||||[||||||||||||]PAM NA Intergenic NA NA

T28944 CCGGCGGTACACCGTCGCCACGG 1000
Oligo fwd CACCCCGGCGGTACACCGTCGCCA
Oligo rev AAACTGGCGACGGTGTACCGCCGG
Chromosome start end strand MM target_seq PAM alignment distance position gene name gene id
GU305914.1 69669 69691 - 0 CCGGCGGTACACCGTCGCCA CGG ||||||||[||||||||||||]PAM NA Intergenic NA NA

T28945 CGGCGGTACACCGTCGCCACGGG 1000
Oligo fwd CACCCGGCGGTACACCGTCGCCAC
Oligo rev AAACGTGGCGACGGTGTACCGCCG
Chromosome start end strand MM target_seq PAM alignment distance position gene name gene id
GU305914.1 69668 69690 - 0 CGGCGGTACACCGTCGCCAC GGG ||||||||[||||||||||||]PAM NA Intergenic NA NA

T28946 CCTGATCATGTCGTCGTTCTCGG 1000
Oligo fwd CACCCCTGATCATGTCGTCGTTCT
Oligo rev AAACAGAACGACGACATGATCAGG
Chromosome start end strand MM target_seq PAM alignment distance position gene name gene id
GU305914.1 69618 69640 - 0 CCTGATCATGTCGTCGTTCT CGG ||||||||[||||||||||||]PAM NA Intergenic NA NA

T28947 GCGACTGCGTCGAGTTCTTCTGG 1000
Oligo fwd CACCGCGACTGCGTCGAGTTCTTC
Oligo rev AAACGAAGAACTCGACGCAGTCGC
Chromosome start end strand MM target_seq PAM alignment distance position gene name gene id
GU305914.1 69571 69593 - 0 GCGACTGCGTCGAGTTCTTC TGG ||||||||[||||||||||||]PAM NA Intergenic NA NA

T28948 CTGCGTCGAGTTCTTCTGGTTGG 1000
Oligo fwd CACCCTGCGTCGAGTTCTTCTGGT
Oligo rev AAACACCAGAAGAACTCGACGCAG
Chromosome start end strand MM target_seq PAM alignment distance position gene name gene id
GU305914.1 69567 69589 - 0 CTGCGTCGAGTTCTTCTGGT TGG ||||||||[||||||||||||]PAM NA Intergenic NA NA

T28949 GGTTGGCCGCGTACCGCAGCCGG 1000
Oligo fwd CACCGGTTGGCCGCGTACCGCAGC
Oligo rev AAACGCTGCGGTACGCGGCCAACC
Chromosome start end strand MM target_seq PAM alignment distance position gene name gene id

<table>
<tr><td>GU305914.1</td><td>69550</td><td>69572</td><td>-</td><td>0</td><td>GGTTGGCCGCGTACCGCAGC</td><td>CGG</td><td>||||||||[||||||||||||]PAM</td><td>NA</td><td>Intergenic</td><td>NA</td><td>NA</td></tr>
<tr><td>T28950</td><td>CGATGTCCGTCGAGTCCAGCAGG</td><td>1000</td><td></td><td></td><td></td><td></td><td></td><td></td><td></td><td></td><td></td></tr>
<tr><td>Oligo fwd</td><td>CACCCGATGTCCGTCGAGTCCAGC</td><td></td><td></td><td></td><td></td><td></td><td></td><td></td><td></td><td></td><td></td></tr>
<tr><td>Oligo rev</td><td>AAACGCTGGACTCGACGGACATCG</td><td></td><td></td><td></td><td></td><td></td><td></td><td></td><td></td><td></td><td></td></tr>
<tr><td>Chromosome</td><td>start</td><td>end</td><td>strand</td><td>MM</td><td>target_seq</td><td>PAM</td><td>alignment</td><td>distance</td><td>position</td><td>gene name</td><td>gene id</td></tr>
<tr><td>GU305914.1</td><td>69511</td><td>69533</td><td>-</td><td>0</td><td>CGATGTCCGTCGAGTCCAGC</td><td>AGG</td><td>||||||||[||||||||||||]PAM</td><td>NA</td><td>Intergenic</td><td>NA</td><td>NA</td></tr>
<tr><td>T28951</td><td>GTCCGTCGAGTCCAGCAGGTCGG</td><td>1000</td><td></td><td></td><td></td><td></td><td></td><td></td><td></td><td></td><td></td></tr>
<tr><td>Oligo fwd</td><td>CACCGTCCGTCGAGTCCAGCAGGT</td><td></td><td></td><td></td><td></td><td></td><td></td><td></td><td></td><td></td><td></td></tr>
<tr><td>Oligo rev</td><td>AAACACCTGCTGGACTCGACGGAC</td><td></td><td></td><td></td><td></td><td></td><td></td><td></td><td></td><td></td><td></td></tr>
<tr><td>Chromosome</td><td>start</td><td>end</td><td>strand</td><td>MM</td><td>target_seq</td><td>PAM</td><td>alignment</td><td>distance</td><td>position</td><td>gene name</td><td>gene id</td></tr>
<tr><td>GU305914.1</td><td>69507</td><td>69529</td><td>-</td><td>0</td><td>GTCCGTCGAGTCCAGCAGGT</td><td>CGG</td><td>||||||||[||||||||||||]PAM</td><td>NA</td><td>Intergenic</td><td>NA</td><td>NA</td></tr>
<tr><td>T28952</td><td>CCAGCAGGTCGGCGCGCACCTGG</td><td>1000</td><td></td><td></td><td></td><td></td><td></td><td></td><td></td><td></td><td></td></tr>
<tr><td>Oligo fwd</td><td>CACCCCAGCAGGTCGGCGCGCACC</td><td></td><td></td><td></td><td></td><td></td><td></td><td></td><td></td><td></td><td></td></tr>
<tr><td>Oligo rev</td><td>AAACGGTGCGCGCCGACCTGCTGG</td><td></td><td></td><td></td><td></td><td></td><td></td><td></td><td></td><td></td><td></td></tr>
<tr><td>Chromosome</td><td>start</td><td>end</td><td>strand</td><td>MM</td><td>target_seq</td><td>PAM</td><td>alignment</td><td>distance</td><td>position</td><td>gene name</td><td>gene id</td></tr>
<tr><td>GU305914.1</td><td>69496</td><td>69518</td><td>-</td><td>0</td><td>CCAGCAGGTCGGCGCGCACC</td><td>TGG</td><td>||||||||[||||||||||||]PAM</td><td>NA</td><td>Intergenic</td><td>NA</td><td>NA</td></tr>
<tr><td>T28953</td><td>GCGCGCACCTGGCGCCCCGTCGG</td><td>1000</td><td></td><td></td><td></td><td></td><td></td><td></td><td></td><td></td><td></td></tr>
<tr><td>Oligo fwd</td><td>CACCGCGCGCACCTGGCGCCCCGT</td><td></td><td></td><td></td><td></td><td></td><td></td><td></td><td></td><td></td><td></td></tr>
<tr><td>Oligo rev</td><td>AAACACGGGGCGCCAGGTGCGCGC</td><td></td><td></td><td></td><td></td><td></td><td></td><td></td><td></td><td></td><td></td></tr>
<tr><td>Chromosome</td><td>start</td><td>end</td><td>strand</td><td>MM</td><td>target_seq</td><td>PAM</td><td>alignment</td><td>distance</td><td>position</td><td>gene name</td><td>gene id</td></tr>
<tr><td>GU305914.1</td><td>69485</td><td>69507</td><td>-</td><td>0</td><td>GCGCGCACCTGGCGCCCCGT</td><td>CGG</td><td>||||||||[||||||||||||]PAM</td><td>NA</td><td>Intergenic</td><td>NA</td><td>NA</td></tr>
<tr><td>T28954</td><td>CCTCGACATCATGTCGTCCGAGG</td><td>1000</td><td></td><td></td><td></td><td></td><td></td><td></td><td></td><td></td><td></td></tr>
<tr><td>Oligo fwd</td><td>CACCCCTCGACATCATGTCGTCCG</td><td></td><td></td><td></td><td></td><td></td><td></td><td></td><td></td><td></td><td></td></tr>
<tr><td>Oligo rev</td><td>AAACCGGACGACATGATGTCGAGG</td><td></td><td></td><td></td><td></td><td></td><td></td><td></td><td></td><td></td><td></td></tr>
<tr><td>Chromosome</td><td>start</td><td>end</td><td>strand</td><td>MM</td><td>target_seq</td><td>PAM</td><td>alignment</td><td>distance</td><td>position</td><td>gene name</td><td>gene id</td></tr>
<tr><td>GU305914.1</td><td>69438</td><td>69460</td><td>-</td><td>0</td><td>CCTCGACATCATGTCGTCCG</td><td>AGG</td><td>||||||||[||||||||||||]PAM</td><td>NA</td><td>Intergenic</td><td>NA</td><td>NA</td></tr>
<tr><td>T28955</td><td>CTCGACATCATGTCGTCCGAGGG</td><td>1000</td><td></td><td></td><td></td><td></td><td></td><td></td><td></td><td></td><td></td></tr>
<tr><td>Oligo fwd</td><td>CACCCTCGACATCATGTCGTCCGA</td><td></td><td></td><td></td><td></td><td></td><td></td><td></td><td></td><td></td><td></td></tr>
<tr><td>Oligo rev</td><td>AAACTCGGACGACATGATGTCGAG</td><td></td><td></td><td></td><td></td><td></td><td></td><td></td><td></td><td></td><td></td></tr>
<tr><td>Chromosome</td><td>start</td><td>end</td><td>strand</td><td>MM</td><td>target_seq</td><td>PAM</td><td>alignment</td><td>distance</td><td>position</td><td>gene name</td><td>gene id</td></tr>
<tr><td>GU305914.1</td><td>69437</td><td>69459</td><td>-</td><td>0</td><td>CTCGACATCATGTCGTCCGA</td><td>GGG</td><td>||||||||[||||||||||||]PAM</td><td>NA</td><td>Intergenic</td><td>NA</td><td>NA</td></tr>
<tr><td>T28956</td><td>ACATCATGTCGTCCGAGGGCAGG</td><td>1000</td><td></td><td></td><td></td><td></td><td></td><td></td><td></td><td></td><td></td></tr>
<tr><td>Oligo fwd</td><td>CACCACATCATGTCGTCCGAGGGC</td><td></td><td></td><td></td><td></td><td></td><td></td><td></td><td></td><td></td><td></td></tr>
<tr><td>Oligo rev</td><td>AAACGCCCTCGGACGACATGATGT</td><td></td><td></td><td></td><td></td><td></td><td></td><td></td><td></td><td></td><td></td></tr>
<tr><td>Chromosome</td><td>start</td><td>end</td><td>strand</td><td>MM</td><td>target_seq</td><td>PAM</td><td>alignment</td><td>distance</td><td>position</td><td>gene name</td><td>gene id</td></tr>
<tr><td>GU305914.1</td><td>69433</td><td>69455</td><td>-</td><td>0</td><td>ACATCATGTCGTCCGAGGGC</td><td>AGG</td><td>||||||||[||||||||||||]PAM</td><td>NA</td><td>Intergenic</td><td>NA</td><td>NA</td></tr>
<tr><td>T28957</td><td>CCCTTGATCTCGTTCAGCGACGG</td><td>1000</td><td></td><td></td><td></td><td></td><td></td><td></td><td></td><td></td><td></td></tr>
<tr><td>Oligo fwd</td><td>CACCCCCTTGATCTCGTTCAGCGA</td><td></td><td></td><td></td><td></td><td></td><td></td><td></td><td></td><td></td><td></td></tr>
<tr><td>Oligo rev</td><td>AAACTCGCTGAACGAGATCAAGGG</td><td></td><td></td><td></td><td></td><td></td><td></td><td></td><td></td><td></td><td></td></tr>
<tr><td>Chromosome</td><td>start</td><td>end</td><td>strand</td><td>MM</td><td>target_seq</td><td>PAM</td><td>alignment</td><td>distance</td><td>position</td><td>gene name</td><td>gene id</td></tr>
<tr><td>GU305914.1</td><td>69299</td><td>69321</td><td>-</td><td>0</td><td>CCCTTGATCTCGTTCAGCGA</td><td>CGG</td><td>||||||||[||||||||||||]PAM</td><td>NA</td><td>Intergenic</td><td>NA</td><td>NA</td></tr>
<tr><td>T28958</td><td>CCTTGATCTCGTTCAGCGACGGG</td><td>1000</td><td></td><td></td><td></td><td></td><td></td><td></td><td></td><td></td><td></td></tr>
<tr><td>Oligo fwd</td><td>CACCCCTTGATCTCGTTCAGCGAC</td><td></td><td></td><td></td><td></td><td></td><td></td><td></td><td></td><td></td><td></td></tr>
<tr><td>Oligo rev</td><td>AAACGTCGCTGAACGAGATCAAGG</td><td></td><td></td><td></td><td></td><td></td><td></td><td></td><td></td><td></td><td></td></tr>
<tr><td>Chromosome</td><td>start</td><td>end</td><td>strand</td><td>MM</td><td>target_seq</td><td>PAM</td><td>alignment</td><td>distance</td><td>position</td><td>gene name</td><td>gene id</td></tr>
<tr><td>GU305914.1</td><td>69298</td><td>69320</td><td>-</td><td>0</td><td>CCTTGATCTCGTTCAGCGAC</td><td>GGG</td><td>||||||||[||||||||||||]PAM</td><td>NA</td><td>Intergenic</td><td>NA</td><td>NA</td></tr>
<tr><td>T28959</td><td>CAGCTTCCCCAGCACCTTGTCGG</td><td>1000</td><td></td><td></td><td></td><td></td><td></td><td></td><td></td><td></td><td></td></tr>
<tr><td>Oligo fwd</td><td>CACCCAGCTTCCCCAGCACCTTGT</td><td></td><td></td><td></td><td></td><td></td><td></td><td></td><td></td><td></td><td></td></tr>
<tr><td>Oligo rev</td><td>AAACACAAGGTGCTGGGGAAGCTG</td><td></td><td></td><td></td><td></td><td></td><td></td><td></td><td></td><td></td><td></td></tr>
<tr><td>Chromosome</td><td>start</td><td>end</td><td>strand</td><td>MM</td><td>target_seq</td><td>PAM</td><td>alignment</td><td>distance</td><td>position</td><td>gene name</td><td>gene id</td></tr>
<tr><td>GU305914.1</td><td>69210</td><td>69232</td><td>-</td><td>0</td><td>CAGCTTCCCCAGCACCTTGT</td><td>CGG</td><td>||||||||[||||||||||||]PAM</td><td>NA</td><td>Intergenic</td><td>NA</td><td>NA</td></tr>
<tr><td>T28960</td><td>CGTAACATATCCAGCATCTCCGG</td><td>1000</td><td></td><td></td><td></td><td></td><td></td><td></td><td></td><td></td><td></td></tr>
<tr><td>Oligo fwd</td><td>CACCCGTAACATATCCAGCATCTC</td><td></td><td></td><td></td><td></td><td></td><td></td><td></td><td></td><td></td><td></td></tr>
<tr><td>Oligo rev</td><td>AAACGAGATGCTGGATATGTTACG</td><td></td><td></td><td></td><td></td><td></td><td></td><td></td><td></td><td></td><td></td></tr>
<tr><td>Chromosome</td><td>start</td><td>end</td><td>strand</td><td>MM</td><td>target_seq</td><td>PAM</td><td>alignment</td><td>distance</td><td>position</td><td>gene name</td><td>gene id</td></tr>
<tr><td>GU305914.1</td><td>69131</td><td>69153</td><td>-</td><td>0</td><td>CGTAACATATCCAGCATCTC</td><td>CGG</td><td>||||||||[||||||||||||]PAM</td><td>NA</td><td>Intergenic</td><td>NA</td><td>NA</td></tr>
<tr><td>T28961</td><td>AACATATCCAGCATCTCCGGCGG</td><td>1000</td><td></td><td></td><td></td><td></td><td></td><td></td><td></td><td></td><td></td></tr>
<tr><td>Oligo fwd</td><td>CACCAACATATCCAGCATCTCCGG</td><td></td><td></td><td></td><td></td><td></td><td></td><td></td><td></td><td></td><td></td></tr>
<tr><td>Oligo rev</td><td>AAACCCGGAGATGCTGGATATGTT</td><td></td><td></td><td></td><td></td><td></td><td></td><td></td><td></td><td></td><td></td></tr>
<tr><td>Chromosome</td><td>start</td><td>end</td><td>strand</td><td>MM</td><td>target_seq</td><td>PAM</td><td>alignment</td><td>distance</td><td>position</td><td>gene name</td><td>gene id</td></tr>
<tr><td>GU305914.1</td><td>69128</td><td>69150</td><td>-</td><td>0</td><td>AACATATCCAGCATCTCCGG</td><td>CGG</td><td>||||||||[||||||||||||]PAM</td><td>NA</td><td>Intergenic</td><td>NA</td><td>NA</td></tr>
<tr><td>T28962</td><td>ATATCCAGCATCTCCGGCGGCGG</td><td>1000</td><td></td><td></td><td></td><td></td><td></td><td></td><td></td><td></td><td></td></tr>
<tr><td>Oligo fwd</td><td>CACCATATCCAGCATCTCCGGCGG</td><td></td><td></td><td></td><td></td><td></td><td></td><td></td><td></td><td></td><td></td></tr>
<tr><td>Oligo rev</td><td>AAACCCGCCGGAGATGCTGGATAT</td><td></td><td></td><td></td><td></td><td></td><td></td><td></td><td></td><td></td><td></td></tr>
<tr><td>Chromosome</td><td>start</td><td>end</td><td>strand</td><td>MM</td><td>target_seq</td><td>PAM</td><td>alignment</td><td>distance</td><td>position</td><td>gene name</td><td>gene id</td></tr>
<tr><td>GU305914.1</td><td>69125</td><td>69147</td><td>-</td><td>0</td><td>ATATCCAGCATCTCCGGCGG</td><td>CGG</td><td>||||||||[||||||||||||]PAM</td><td>NA</td><td>Intergenic</td><td>NA</td><td>NA</td></tr>
<tr><td>T28963</td><td>TCCAGCATCTCCGGCGGCGGCGG</td><td>1000</td><td></td><td></td><td></td><td></td><td></td><td></td><td></td><td></td><td></td></tr>
<tr><td>Oligo fwd</td><td>CACCTCCAGCATCTCCGGCGGCGG</td><td></td><td></td><td></td><td></td><td></td><td></td><td></td><td></td><td></td><td></td></tr>
<tr><td>Oligo rev</td><td>AAACCCGCCGCCGGAGATGCTGGA</td><td></td><td></td><td></td><td></td><td></td><td></td><td></td><td></td><td></td><td></td></tr>
<tr><td>Chromosome</td><td>start</td><td>end</td><td>strand</td><td>MM</td><td>target_seq</td><td>PAM</td><td>alignment</td><td>distance</td><td>position</td><td>gene name</td><td>gene id</td></tr>
<tr><td>GU305914.1</td><td>69122</td><td>69144</td><td>-</td><td>0</td><td>TCCAGCATCTCCGGCGGCGG</td><td>CGG</td><td>||||||||[||||||||||||]PAM</td><td>NA</td><td>Intergenic</td><td>NA</td><td>NA</td></tr>
<tr><td>T28964</td><td>CCAGCATCTCCGGCGGCGGCGGG</td><td>1000</td><td></td><td></td><td></td><td></td><td></td><td></td><td></td><td></td><td></td></tr>
<tr><td>Oligo fwd</td><td>CACCCCAGCATCTCCGGCGGCGGC</td><td></td><td></td><td></td><td></td><td></td><td></td><td></td><td></td><td></td><td></td></tr>
<tr><td>Oligo rev</td><td>AAACGCCGCCGCCGGAGATGCTGG</td><td></td><td></td><td></td><td></td><td></td><td></td><td></td><td></td><td></td><td></td></tr>
</table>

<table>
<tr><td>Chromosome</td><td>start</td><td>end</td><td>strand</td><td>MM</td><td>target_seq</td><td>PAM</td><td>alignment</td><td>distance</td><td>position</td><td>gene name</td><td>gene id</td></tr>
<tr><td>GU305914.1</td><td>69121</td><td>69143</td><td>-</td><td>0</td><td>CCAGCATCTCCGGCGGCGGC</td><td>GGG</td><td>||||||||[||||||||||||]PAM</td><td>NA</td><td>Intergenic</td><td>NA</td><td>NA</td></tr>
<tr><td>T28965</td><td>AACTTCTTGATCTCCGCGCTCGG</td><td>1000</td><td></td><td></td><td></td><td></td><td></td><td></td><td></td><td></td><td></td></tr>
<tr><td>Oligo fwd</td><td>CACCAACTTCTTGATCTCCGCGCT</td><td></td><td></td><td></td><td></td><td></td><td></td><td></td><td></td><td></td><td></td></tr>
<tr><td>Oligo rev</td><td>AAACAGCGCGGAGATCAAGAAGTT</td><td></td><td></td><td></td><td></td><td></td><td></td><td></td><td></td><td></td><td></td></tr>
<tr><td>Chromosome</td><td>start</td><td>end</td><td>strand</td><td>MM</td><td>target_seq</td><td>PAM</td><td>alignment</td><td>distance</td><td>position</td><td>gene name</td><td>gene id</td></tr>
<tr><td>GU305914.1</td><td>68993</td><td>69015</td><td>-</td><td>0</td><td>AACTTCTTGATCTCCGCGCT</td><td>CGG</td><td>||||||||[||||||||||||]PAM</td><td>NA</td><td>Intergenic</td><td>NA</td><td>NA</td></tr>
<tr><td>T28966</td><td>CGTCACCTCCAGCTTCACCTCGG</td><td>1000</td><td></td><td></td><td></td><td></td><td></td><td></td><td></td><td></td><td></td></tr>
<tr><td>Oligo fwd</td><td>CACCCGTCACCTCCAGCTTCACCT</td><td></td><td></td><td></td><td></td><td></td><td></td><td></td><td></td><td></td><td></td></tr>
<tr><td>Oligo rev</td><td>AAACAGGTGAAGCTGGAGGTGACG</td><td></td><td></td><td></td><td></td><td></td><td></td><td></td><td></td><td></td><td></td></tr>
<tr><td>Chromosome</td><td>start</td><td>end</td><td>strand</td><td>MM</td><td>target_seq</td><td>PAM</td><td>alignment</td><td>distance</td><td>position</td><td>gene name</td><td>gene id</td></tr>
<tr><td>GU305914.1</td><td>68970</td><td>68992</td><td>-</td><td>0</td><td>CGTCACCTCCAGCTTCACCT</td><td>CGG</td><td>||||||||[||||||||||||]PAM</td><td>NA</td><td>Intergenic</td><td>NA</td><td>NA</td></tr>
<tr><td>T28967</td><td>GCTTCACCTCGGCCGTCTGCAGG</td><td>1000</td><td></td><td></td><td></td><td></td><td></td><td></td><td></td><td></td><td></td></tr>
<tr><td>Oligo fwd</td><td>CACCGCTTCACCTCGGCCGTCTGC</td><td></td><td></td><td></td><td></td><td></td><td></td><td></td><td></td><td></td><td></td></tr>
<tr><td>Oligo rev</td><td>AAACGCAGACGGCCGAGGTGAAGC</td><td></td><td></td><td></td><td></td><td></td><td></td><td></td><td></td><td></td><td></td></tr>
<tr><td>Chromosome</td><td>start</td><td>end</td><td>strand</td><td>MM</td><td>target_seq</td><td>PAM</td><td>alignment</td><td>distance</td><td>position</td><td>gene name</td><td>gene id</td></tr>
<tr><td>GU305914.1</td><td>68959</td><td>68981</td><td>-</td><td>0</td><td>GCTTCACCTCGGCCGTCTGC</td><td>AGG</td><td>||||||||[||||||||||||]PAM</td><td>NA</td><td>Intergenic</td><td>NA</td><td>NA</td></tr>
<tr><td>T28968</td><td>CGGCCGTCTGCAGGTACAGCAGG</td><td>1000</td><td></td><td></td><td></td><td></td><td></td><td></td><td></td><td></td><td></td></tr>
<tr><td>Oligo fwd</td><td>CACCCGGCCGTCTGCAGGTACAGC</td><td></td><td></td><td></td><td></td><td></td><td></td><td></td><td></td><td></td><td></td></tr>
<tr><td>Oligo rev</td><td>AAACGCTGTACCTGCAGACGGCCG</td><td></td><td></td><td></td><td></td><td></td><td></td><td></td><td></td><td></td><td></td></tr>
<tr><td>Chromosome</td><td>start</td><td>end</td><td>strand</td><td>MM</td><td>target_seq</td><td>PAM</td><td>alignment</td><td>distance</td><td>position</td><td>gene name</td><td>gene id</td></tr>
<tr><td>GU305914.1</td><td>68950</td><td>68972</td><td>-</td><td>0</td><td>CGGCCGTCTGCAGGTACAGC</td><td>AGG</td><td>||||||||[||||||||||||]PAM</td><td>NA</td><td>Intergenic</td><td>NA</td><td>NA</td></tr>
<tr><td>T28969</td><td>CTGCAGGTACAGCAGGTCCTCGG</td><td>1000</td><td></td><td></td><td></td><td></td><td></td><td></td><td></td><td></td><td></td></tr>
<tr><td>Oligo fwd</td><td>CACCCTGCAGGTACAGCAGGTCCT</td><td></td><td></td><td></td><td></td><td></td><td></td><td></td><td></td><td></td><td></td></tr>
<tr><td>Oligo rev</td><td>AAACAGGACCTGCTGTACCTGCAG</td><td></td><td></td><td></td><td></td><td></td><td></td><td></td><td></td><td></td><td></td></tr>
<tr><td>Chromosome</td><td>start</td><td>end</td><td>strand</td><td>MM</td><td>target_seq</td><td>PAM</td><td>alignment</td><td>distance</td><td>position</td><td>gene name</td><td>gene id</td></tr>
<tr><td>GU305914.1</td><td>68943</td><td>68965</td><td>-</td><td>0</td><td>CTGCAGGTACAGCAGGTCCT</td><td>CGG</td><td>||||||||[||||||||||||]PAM</td><td>NA</td><td>Intergenic</td><td>NA</td><td>NA</td></tr>
<tr><td>T28970</td><td>CGCCCGCACGTTCTCCGACACGG</td><td>1000</td><td></td><td></td><td></td><td></td><td></td><td></td><td></td><td></td><td></td></tr>
<tr><td>Oligo fwd</td><td>CACCCGCCCGCACGTTCTCCGACA</td><td></td><td></td><td></td><td></td><td></td><td></td><td></td><td></td><td></td><td></td></tr>
<tr><td>Oligo rev</td><td>AAACTGTCGGAGAACGTGCGGGCG</td><td></td><td></td><td></td><td></td><td></td><td></td><td></td><td></td><td></td><td></td></tr>
<tr><td>Chromosome</td><td>start</td><td>end</td><td>strand</td><td>MM</td><td>target_seq</td><td>PAM</td><td>alignment</td><td>distance</td><td>position</td><td>gene name</td><td>gene id</td></tr>
<tr><td>GU305914.1</td><td>68865</td><td>68887</td><td>-</td><td>0</td><td>CGCCCGCACGTTCTCCGACA</td><td>CGG</td><td>||||||||[||||||||||||]PAM</td><td>NA</td><td>Intergenic</td><td>NA</td><td>NA</td></tr>
<tr><td>T28971</td><td>GTTCTCCGACACGGCCTTGAAGG</td><td>1000</td><td></td><td></td><td></td><td></td><td></td><td></td><td></td><td></td><td></td></tr>
<tr><td>Oligo fwd</td><td>CACCGTTCTCCGACACGGCCTTGA</td><td></td><td></td><td></td><td></td><td></td><td></td><td></td><td></td><td></td><td></td></tr>
<tr><td>Oligo rev</td><td>AAACTCAAGGCCGTGTCGGAGAAC</td><td></td><td></td><td></td><td></td><td></td><td></td><td></td><td></td><td></td><td></td></tr>
<tr><td>Chromosome</td><td>start</td><td>end</td><td>strand</td><td>MM</td><td>target_seq</td><td>PAM</td><td>alignment</td><td>distance</td><td>position</td><td>gene name</td><td>gene id</td></tr>
<tr><td>GU305914.1</td><td>68856</td><td>68878</td><td>-</td><td>0</td><td>GTTCTCCGACACGGCCTTGA</td><td>AGG</td><td>||||||||[||||||||||||]PAM</td><td>NA</td><td>Intergenic</td><td>NA</td><td>NA</td></tr>
<tr><td>T28972</td><td>ACCGCCGCGATGTCCGAGTCCGG</td><td>1000</td><td></td><td></td><td></td><td></td><td></td><td></td><td></td><td></td><td></td></tr>
<tr><td>Oligo fwd</td><td>CACCACCGCCGCGATGTCCGAGTC</td><td></td><td></td><td></td><td></td><td></td><td></td><td></td><td></td><td></td><td></td></tr>
<tr><td>Oligo rev</td><td>AAACGACTCGGACATCGCGGCGGT</td><td></td><td></td><td></td><td></td><td></td><td></td><td></td><td></td><td></td><td></td></tr>
<tr><td>Chromosome</td><td>start</td><td>end</td><td>strand</td><td>MM</td><td>target_seq</td><td>PAM</td><td>alignment</td><td>distance</td><td>position</td><td>gene name</td><td>gene id</td></tr>
<tr><td>GU305914.1</td><td>68825</td><td>68847</td><td>-</td><td>0</td><td>ACCGCCGCGATGTCCGAGTC</td><td>CGG</td><td>||||||||[||||||||||||]PAM</td><td>NA</td><td>Intergenic</td><td>NA</td><td>NA</td></tr>
<tr><td>T28973</td><td>AGTCCGGCAGCGACCCCGTCAGG</td><td>1000</td><td></td><td></td><td></td><td></td><td></td><td></td><td></td><td></td><td></td></tr>
<tr><td>Oligo fwd</td><td>CACCAGTCCGGCAGCGACCCCGTC</td><td></td><td></td><td></td><td></td><td></td><td></td><td></td><td></td><td></td><td></td></tr>
<tr><td>Oligo rev</td><td>AAACGACGGGGTCGCTGCCGGACT</td><td></td><td></td><td></td><td></td><td></td><td></td><td></td><td></td><td></td><td></td></tr>
<tr><td>Chromosome</td><td>start</td><td>end</td><td>strand</td><td>MM</td><td>target_seq</td><td>PAM</td><td>alignment</td><td>distance</td><td>position</td><td>gene name</td><td>gene id</td></tr>
<tr><td>GU305914.1</td><td>68809</td><td>68831</td><td>-</td><td>0</td><td>AGTCCGGCAGCGACCCCGTC</td><td>AGG</td><td>||||||||[||||||||||||]PAM</td><td>NA</td><td>Intergenic</td><td>NA</td><td>NA</td></tr>
<tr><td>T28974</td><td>GCAGCGACCCCGTCAGGATGCGG</td><td>1000</td><td></td><td></td><td></td><td></td><td></td><td></td><td></td><td></td><td></td></tr>
<tr><td>Oligo fwd</td><td>CACCGCAGCGACCCCGTCAGGATG</td><td></td><td></td><td></td><td></td><td></td><td></td><td></td><td></td><td></td><td></td></tr>
<tr><td>Oligo rev</td><td>AAACCATCCTGACGGGGTCGCTGC</td><td></td><td></td><td></td><td></td><td></td><td></td><td></td><td></td><td></td><td></td></tr>
<tr><td>Chromosome</td><td>start</td><td>end</td><td>strand</td><td>MM</td><td>target_seq</td><td>PAM</td><td>alignment</td><td>distance</td><td>position</td><td>gene name</td><td>gene id</td></tr>
<tr><td>GU305914.1</td><td>68803</td><td>68825</td><td>-</td><td>0</td><td>GCAGCGACCCCGTCAGGATG</td><td>CGG</td><td>||||||||[||||||||||||]PAM</td><td>NA</td><td>Intergenic</td><td>NA</td><td>NA</td></tr>
<tr><td>T28975</td><td>ACCCCGTCAGGATGCGGTTGTGG</td><td>1000</td><td></td><td></td><td></td><td></td><td></td><td></td><td></td><td></td><td></td></tr>
<tr><td>Oligo fwd</td><td>CACCACCCCGTCAGGATGCGGTTG</td><td></td><td></td><td></td><td></td><td></td><td></td><td></td><td></td><td></td><td></td></tr>
<tr><td>Oligo rev</td><td>AAACCAACCGCATCCTGACGGGGT</td><td></td><td></td><td></td><td></td><td></td><td></td><td></td><td></td><td></td><td></td></tr>
<tr><td>Chromosome</td><td>start</td><td>end</td><td>strand</td><td>MM</td><td>target_seq</td><td>PAM</td><td>alignment</td><td>distance</td><td>position</td><td>gene name</td><td>gene id</td></tr>
<tr><td>GU305914.1</td><td>68797</td><td>68819</td><td>-</td><td>0</td><td>ACCCCGTCAGGATGCGGTTG</td><td>TGG</td><td>||||||||[||||||||||||]PAM</td><td>NA</td><td>Intergenic</td><td>NA</td><td>NA</td></tr>
<tr><td>T28976</td><td>GCAGCTCCGTCATCTTCGCCTGG</td><td>1000</td><td></td><td></td><td></td><td></td><td></td><td></td><td></td><td></td><td></td></tr>
<tr><td>Oligo fwd</td><td>CACCGCAGCTCCGTCATCTTCGCC</td><td></td><td></td><td></td><td></td><td></td><td></td><td></td><td></td><td></td><td></td></tr>
<tr><td>Oligo rev</td><td>AAACGGCGAAGATGACGGAGCTGC</td><td></td><td></td><td></td><td></td><td></td><td></td><td></td><td></td><td></td><td></td></tr>
<tr><td>Chromosome</td><td>start</td><td>end</td><td>strand</td><td>MM</td><td>target_seq</td><td>PAM</td><td>alignment</td><td>distance</td><td>position</td><td>gene name</td><td>gene id</td></tr>
<tr><td>GU305914.1</td><td>68764</td><td>68786</td><td>-</td><td>0</td><td>GCAGCTCCGTCATCTTCGCC</td><td>TGG</td><td>||||||||[||||||||||||]PAM</td><td>NA</td><td>Intergenic</td><td>NA</td><td>NA</td></tr>
<tr><td>T28977</td><td>CGCCGACTGCGCGTACGCCTCGG</td><td>1000</td><td></td><td></td><td></td><td></td><td></td><td></td><td></td><td></td><td></td></tr>
<tr><td>Oligo fwd</td><td>CACCCGCCGACTGCGCGTACGCCT</td><td></td><td></td><td></td><td></td><td></td><td></td><td></td><td></td><td></td><td></td></tr>
<tr><td>Oligo rev</td><td>AAACAGGCGTACGCGCAGTCGGCG</td><td></td><td></td><td></td><td></td><td></td><td></td><td></td><td></td><td></td><td></td></tr>
<tr><td>Chromosome</td><td>start</td><td>end</td><td>strand</td><td>MM</td><td>target_seq</td><td>PAM</td><td>alignment</td><td>distance</td><td>position</td><td>gene name</td><td>gene id</td></tr>
<tr><td>GU305914.1</td><td>68730</td><td>68752</td><td>-</td><td>0</td><td>CGCCGACTGCGCGTACGCCT</td><td>CGG</td><td>||||||||[||||||||||||]PAM</td><td>NA</td><td>Intergenic</td><td>NA</td><td>NA</td></tr>
<tr><td>T28978</td><td>ACGCCAGCTGCTCGCTGTCCAGG</td><td>1000</td><td></td><td></td><td></td><td></td><td></td><td></td><td></td><td></td><td></td></tr>
<tr><td>Oligo fwd</td><td>CACCACGCCAGCTGCTCGCTGTCC</td><td></td><td></td><td></td><td></td><td></td><td></td><td></td><td></td><td></td><td></td></tr>
<tr><td>Oligo rev</td><td>AAACGGACAGCGAGCAGCTGGCGT</td><td></td><td></td><td></td><td></td><td></td><td></td><td></td><td></td><td></td><td></td></tr>
<tr><td>Chromosome</td><td>start</td><td>end</td><td>strand</td><td>MM</td><td>target_seq</td><td>PAM</td><td>alignment</td><td>distance</td><td>position</td><td>gene name</td><td>gene id</td></tr>
<tr><td>GU305914.1</td><td>68692</td><td>68714</td><td>-</td><td>0</td><td>ACGCCAGCTGCTCGCTGTCC</td><td>AGG</td><td>||||||||[||||||||||||]PAM</td><td>NA</td><td>Intergenic</td><td>NA</td><td>NA</td></tr>
<tr><td>GU305914.1</td><td>126641</td><td>126663</td><td>+</td><td>4</td><td>GCGTCAGCTGCCCGCGGTCC</td><td>CGG</td><td>-||-||||[|||-|||-||||]PAM</td><td>NA</td><td>Intergenic</td><td>NA</td><td>NA</td></tr>
<tr><td>T28979</td><td>GCTGTCCAGGTGCGACGTGTCGG</td><td>1000</td><td></td><td></td><td></td><td></td><td></td><td></td><td></td><td></td><td></td></tr>
</table>

<table>
<tr><td>Oligo fwd</td><td>CACCGCTGTCCAGGTGCGACGTGT</td><td></td><td></td><td></td><td></td><td></td><td></td><td></td><td></td><td></td><td></td></tr>
<tr><td>Oligo rev</td><td>AAACACACGTCGCACCTGGACAGC</td><td></td><td></td><td></td><td></td><td></td><td></td><td></td><td></td><td></td><td></td></tr>
<tr><td>Chromosome</td><td>start</td><td>end</td><td>strand</td><td>MM</td><td>target_seq</td><td>PAM</td><td>alignment</td><td>distance</td><td>position</td><td>gene name</td><td>gene id</td></tr>
<tr><td>GU305914.1</td><td>68679</td><td>68701</td><td>-</td><td>0</td><td>GCTGTCCAGGTGCGACGTGT</td><td>CGG</td><td>||||||||[||||||||||||]PAM</td><td>NA</td><td>Intergenic</td><td>NA</td><td>NA</td></tr>
<tr><td></td><td></td><td></td><td></td><td></td><td></td><td></td><td></td><td></td><td></td><td></td><td></td></tr>
<tr><td>T28980</td><td>GGTGCGACGTGTCGGCGCTCTGG</td><td>1000</td><td></td><td></td><td></td><td></td><td></td><td></td><td></td><td></td><td></td></tr>
<tr><td>Oligo fwd</td><td>CACCGGTGCGACGTGTCGGCGCTC</td><td></td><td></td><td></td><td></td><td></td><td></td><td></td><td></td><td></td><td></td></tr>
<tr><td>Oligo rev</td><td>AAACGAGCGCCGACACGTCGCACC</td><td></td><td></td><td></td><td></td><td></td><td></td><td></td><td></td><td></td><td></td></tr>
<tr><td>Chromosome</td><td>start</td><td>end</td><td>strand</td><td>MM</td><td>target_seq</td><td>PAM</td><td>alignment</td><td>distance</td><td>position</td><td>gene name</td><td>gene id</td></tr>
<tr><td>GU305914.1</td><td>68671</td><td>68693</td><td>-</td><td>0</td><td>GGTGCGACGTGTCGGCGCTC</td><td>TGG</td><td>||||||||[||||||||||||]PAM</td><td>NA</td><td>Intergenic</td><td>NA</td><td>NA</td></tr>
<tr><td></td><td></td><td></td><td></td><td></td><td></td><td></td><td></td><td></td><td></td><td></td><td></td></tr>
<tr><td>T28981</td><td>CCGCCGCCGCGTTGCTCGCCAGG</td><td>1000</td><td></td><td></td><td></td><td></td><td></td><td></td><td></td><td></td><td></td></tr>
<tr><td>Oligo fwd</td><td>CACCCCGCCGCCGCGTTGCTCGCC</td><td></td><td></td><td></td><td></td><td></td><td></td><td></td><td></td><td></td><td></td></tr>
<tr><td>Oligo rev</td><td>AAACGGCGAGCAACGCGGCGGCGG</td><td></td><td></td><td></td><td></td><td></td><td></td><td></td><td></td><td></td><td></td></tr>
<tr><td>Chromosome</td><td>start</td><td>end</td><td>strand</td><td>MM</td><td>target_seq</td><td>PAM</td><td>alignment</td><td>distance</td><td>position</td><td>gene name</td><td>gene id</td></tr>
<tr><td>GU305914.1</td><td>68644</td><td>68666</td><td>-</td><td>0</td><td>CCGCCGCCGCGTTGCTCGCC</td><td>AGG</td><td>||||||||[||||||||||||]PAM</td><td>NA</td><td>Intergenic</td><td>NA</td><td>NA</td></tr>
<tr><td></td><td></td><td></td><td></td><td></td><td></td><td></td><td></td><td></td><td></td><td></td><td></td></tr>
<tr><td>T28982</td><td>CCAGCTTCGAGATCGCGTCCAGG</td><td>1000</td><td></td><td></td><td></td><td></td><td></td><td></td><td></td><td></td><td></td></tr>
<tr><td>Oligo fwd</td><td>CACCCCAGCTTCGAGATCGCGTCC</td><td></td><td></td><td></td><td></td><td></td><td></td><td></td><td></td><td></td><td></td></tr>
<tr><td>Oligo rev</td><td>AAACGGACGCGATCTCGAAGCTGG</td><td></td><td></td><td></td><td></td><td></td><td></td><td></td><td></td><td></td><td></td></tr>
<tr><td>Chromosome</td><td>start</td><td>end</td><td>strand</td><td>MM</td><td>target_seq</td><td>PAM</td><td>alignment</td><td>distance</td><td>position</td><td>gene name</td><td>gene id</td></tr>
<tr><td>GU305914.1</td><td>68620</td><td>68642</td><td>-</td><td>0</td><td>CCAGCTTCGAGATCGCGTCC</td><td>AGG</td><td>||||||||[||||||||||||]PAM</td><td>NA</td><td>Intergenic</td><td>NA</td><td>NA</td></tr>
<tr><td></td><td></td><td></td><td></td><td></td><td></td><td></td><td></td><td></td><td></td><td></td><td></td></tr>
<tr><td>T28983</td><td>AGATCGCGTCCAGGATCTTGCGG</td><td>1000</td><td></td><td></td><td></td><td></td><td></td><td></td><td></td><td></td><td></td></tr>
<tr><td>Oligo fwd</td><td>CACCAGATCGCGTCCAGGATCTTG</td><td></td><td></td><td></td><td></td><td></td><td></td><td></td><td></td><td></td><td></td></tr>
<tr><td>Oligo rev</td><td>AAACCAAGATCCTGGACGCGATCT</td><td></td><td></td><td></td><td></td><td></td><td></td><td></td><td></td><td></td><td></td></tr>
<tr><td>Chromosome</td><td>start</td><td>end</td><td>strand</td><td>MM</td><td>target_seq</td><td>PAM</td><td>alignment</td><td>distance</td><td>position</td><td>gene name</td><td>gene id</td></tr>
<tr><td>GU305914.1</td><td>68611</td><td>68633</td><td>-</td><td>0</td><td>AGATCGCGTCCAGGATCTTG</td><td>CGG</td><td>||||||||[||||||||||||]PAM</td><td>NA</td><td>Intergenic</td><td>NA</td><td>NA</td></tr>
<tr><td></td><td></td><td></td><td></td><td></td><td></td><td></td><td></td><td></td><td></td><td></td><td></td></tr>
<tr><td>T28984</td><td>TCTCCATGTATTTCACCTTCAGG</td><td>1000</td><td></td><td></td><td></td><td></td><td></td><td></td><td></td><td></td><td></td></tr>
<tr><td>Oligo fwd</td><td>CACCTCTCCATGTATTTCACCTTC</td><td></td><td></td><td></td><td></td><td></td><td></td><td></td><td></td><td></td><td></td></tr>
<tr><td>Oligo rev</td><td>AAACGAAGGTGAAATACATGGAGA</td><td></td><td></td><td></td><td></td><td></td><td></td><td></td><td></td><td></td><td></td></tr>
<tr><td>Chromosome</td><td>start</td><td>end</td><td>strand</td><td>MM</td><td>target_seq</td><td>PAM</td><td>alignment</td><td>distance</td><td>position</td><td>gene name</td><td>gene id</td></tr>
<tr><td>GU305914.1</td><td>68575</td><td>68597</td><td>-</td><td>0</td><td>TCTCCATGTATTTCACCTTC</td><td>AGG</td><td>||||||||[||||||||||||]PAM</td><td>NA</td><td>Intergenic</td><td>NA</td><td>NA</td></tr>
<tr><td></td><td></td><td></td><td></td><td></td><td></td><td></td><td></td><td></td><td></td><td></td><td></td></tr>
<tr><td>T28985</td><td>CTCCATGTATTTCACCTTCAGGG</td><td>1000</td><td></td><td></td><td></td><td></td><td></td><td></td><td></td><td></td><td></td></tr>
<tr><td>Oligo fwd</td><td>CACCCTCCATGTATTTCACCTTCA</td><td></td><td></td><td></td><td></td><td></td><td></td><td></td><td></td><td></td><td></td></tr>
<tr><td>Oligo rev</td><td>AAACTGAAGGTGAAATACATGGAG</td><td></td><td></td><td></td><td></td><td></td><td></td><td></td><td></td><td></td><td></td></tr>
<tr><td>Chromosome</td><td>start</td><td>end</td><td>strand</td><td>MM</td><td>target_seq</td><td>PAM</td><td>alignment</td><td>distance</td><td>position</td><td>gene name</td><td>gene id</td></tr>
<tr><td>GU305914.1</td><td>68574</td><td>68596</td><td>-</td><td>0</td><td>CTCCATGTATTTCACCTTCA</td><td>GGG</td><td>||||||||[||||||||||||]PAM</td><td>NA</td><td>Intergenic</td><td>NA</td><td>NA</td></tr>
<tr><td></td><td></td><td></td><td></td><td></td><td></td><td></td><td></td><td></td><td></td><td></td><td></td></tr>
<tr><td>T28986</td><td>TCACCGACAGCAGCTCGCCCAGG</td><td>1000</td><td></td><td></td><td></td><td></td><td></td><td></td><td></td><td></td><td></td></tr>
<tr><td>Oligo fwd</td><td>CACCTCACCGACAGCAGCTCGCCC</td><td></td><td></td><td></td><td></td><td></td><td></td><td></td><td></td><td></td><td></td></tr>
<tr><td>Oligo rev</td><td>AAACGGGCGAGCTGCTGTCGGTGA</td><td></td><td></td><td></td><td></td><td></td><td></td><td></td><td></td><td></td><td></td></tr>
<tr><td>Chromosome</td><td>start</td><td>end</td><td>strand</td><td>MM</td><td>target_seq</td><td>PAM</td><td>alignment</td><td>distance</td><td>position</td><td>gene name</td><td>gene id</td></tr>
<tr><td>GU305914.1</td><td>68542</td><td>68564</td><td>-</td><td>0</td><td>TCACCGACAGCAGCTCGCCC</td><td>AGG</td><td>||||||||[||||||||||||]PAM</td><td>NA</td><td>Intergenic</td><td>NA</td><td>NA</td></tr>
<tr><td></td><td></td><td></td><td></td><td></td><td></td><td></td><td></td><td></td><td></td><td></td><td></td></tr>
<tr><td>T28987</td><td>TCTTCACCTGCGAGTTGCGCAGG</td><td>1000</td><td></td><td></td><td></td><td></td><td></td><td></td><td></td><td></td><td></td></tr>
<tr><td>Oligo fwd</td><td>CACCTCTTCACCTGCGAGTTGCGC</td><td></td><td></td><td></td><td></td><td></td><td></td><td></td><td></td><td></td><td></td></tr>
<tr><td>Oligo rev</td><td>AAACGCGCAACTCGCAGGTGAAGA</td><td></td><td></td><td></td><td></td><td></td><td></td><td></td><td></td><td></td><td></td></tr>
<tr><td>Chromosome</td><td>start</td><td>end</td><td>strand</td><td>MM</td><td>target_seq</td><td>PAM</td><td>alignment</td><td>distance</td><td>position</td><td>gene name</td><td>gene id</td></tr>
<tr><td>GU305914.1</td><td>68518</td><td>68540</td><td>-</td><td>0</td><td>TCTTCACCTGCGAGTTGCGC</td><td>AGG</td><td>||||||||[||||||||||||]PAM</td><td>NA</td><td>Intergenic</td><td>NA</td><td>NA</td></tr>
<tr><td></td><td></td><td></td><td></td><td></td><td></td><td></td><td></td><td></td><td></td><td></td><td></td></tr>
<tr><td>T28988</td><td>GCACACGAACGCGCGCTCCCCGG</td><td>1000</td><td></td><td></td><td></td><td></td><td></td><td></td><td></td><td></td><td></td></tr>
<tr><td>Oligo fwd</td><td>CACCGCACACGAACGCGCGCTCCC</td><td></td><td></td><td></td><td></td><td></td><td></td><td></td><td></td><td></td><td></td></tr>
<tr><td>Oligo rev</td><td>AAACGGGAGCGCGCGTTCGTGTGC</td><td></td><td></td><td></td><td></td><td></td><td></td><td></td><td></td><td></td><td></td></tr>
<tr><td>Chromosome</td><td>start</td><td>end</td><td>strand</td><td>MM</td><td>target_seq</td><td>PAM</td><td>alignment</td><td>distance</td><td>position</td><td>gene name</td><td>gene id</td></tr>
<tr><td>GU305914.1</td><td>68496</td><td>68518</td><td>-</td><td>0</td><td>GCACACGAACGCGCGCTCCC</td><td>CGG</td><td>||||||||[||||||||||||]PAM</td><td>NA</td><td>Intergenic</td><td>NA</td><td>NA</td></tr>
<tr><td></td><td></td><td></td><td></td><td></td><td></td><td></td><td></td><td></td><td></td><td></td><td></td></tr>
<tr><td>T28989</td><td>CACACGAACGCGCGCTCCCCGGG</td><td>1000</td><td></td><td></td><td></td><td></td><td></td><td></td><td></td><td></td><td></td></tr>
<tr><td>Oligo fwd</td><td>CACCCACACGAACGCGCGCTCCCC</td><td></td><td></td><td></td><td></td><td></td><td></td><td></td><td></td><td></td><td></td></tr>
<tr><td>Oligo rev</td><td>AAACGGGGAGCGCGCGTTCGTGTG</td><td></td><td></td><td></td><td></td><td></td><td></td><td></td><td></td><td></td><td></td></tr>
<tr><td>Chromosome</td><td>start</td><td>end</td><td>strand</td><td>MM</td><td>target_seq</td><td>PAM</td><td>alignment</td><td>distance</td><td>position</td><td>gene name</td><td>gene id</td></tr>
<tr><td>GU305914.1</td><td>68495</td><td>68517</td><td>-</td><td>0</td><td>CACACGAACGCGCGCTCCCC</td><td>GGG</td><td>||||||||[||||||||||||]PAM</td><td>NA</td><td>Intergenic</td><td>NA</td><td>NA</td></tr>
<tr><td></td><td></td><td></td><td></td><td></td><td></td><td></td><td></td><td></td><td></td><td></td><td></td></tr>
<tr><td>T28990</td><td>ACACGAACGCGCGCTCCCCGGGG</td><td>1000</td><td></td><td></td><td></td><td></td><td></td><td></td><td></td><td></td><td></td></tr>
<tr><td>Oligo fwd</td><td>CACCACACGAACGCGCGCTCCCCG</td><td></td><td></td><td></td><td></td><td></td><td></td><td></td><td></td><td></td><td></td></tr>
<tr><td>Oligo rev</td><td>AAACCGGGGAGCGCGCGTTCGTGT</td><td></td><td></td><td></td><td></td><td></td><td></td><td></td><td></td><td></td><td></td></tr>
<tr><td>Chromosome</td><td>start</td><td>end</td><td>strand</td><td>MM</td><td>target_seq</td><td>PAM</td><td>alignment</td><td>distance</td><td>position</td><td>gene name</td><td>gene id</td></tr>
<tr><td>GU305914.1</td><td>68494</td><td>68516</td><td>-</td><td>0</td><td>ACACGAACGCGCGCTCCCCG</td><td>GGG</td><td>||||||||[||||||||||||]PAM</td><td>NA</td><td>Intergenic</td><td>NA</td><td>NA</td></tr>
<tr><td></td><td></td><td></td><td></td><td></td><td></td><td></td><td></td><td></td><td></td><td></td><td></td></tr>
<tr><td>T28991</td><td>GAACGCGCGCTCCCCGGGGATGG</td><td>1000</td><td></td><td></td><td></td><td></td><td></td><td></td><td></td><td></td><td></td></tr>
<tr><td>Oligo fwd</td><td>CACCGAACGCGCGCTCCCCGGGGA</td><td></td><td></td><td></td><td></td><td></td><td></td><td></td><td></td><td></td><td></td></tr>
<tr><td>Oligo rev</td><td>AAACTCCCCGGGGAGCGCGCGTTC</td><td></td><td></td><td></td><td></td><td></td><td></td><td></td><td></td><td></td><td></td></tr>
<tr><td>Chromosome</td><td>start</td><td>end</td><td>strand</td><td>MM</td><td>target_seq</td><td>PAM</td><td>alignment</td><td>distance</td><td>position</td><td>gene name</td><td>gene id</td></tr>
<tr><td>GU305914.1</td><td>68490</td><td>68512</td><td>-</td><td>0</td><td>GAACGCGCGCTCCCCGGGGA</td><td>TGG</td><td>||||||||[||||||||||||]PAM</td><td>NA</td><td>Intergenic</td><td>NA</td><td>NA</td></tr>
<tr><td></td><td></td><td></td><td></td><td></td><td></td><td></td><td></td><td></td><td></td><td></td><td></td></tr>
<tr><td>T28992</td><td>AACGCGCGCTCCCCGGGGATGGG</td><td>1000</td><td></td><td></td><td></td><td></td><td></td><td></td><td></td><td></td><td></td></tr>
<tr><td>Oligo fwd</td><td>CACCAACGCGCGCTCCCCGGGGAT</td><td></td><td></td><td></td><td></td><td></td><td></td><td></td><td></td><td></td><td></td></tr>
<tr><td>Oligo rev</td><td>AAACATCCCCGGGGAGCGCGCGTT</td><td></td><td></td><td></td><td></td><td></td><td></td><td></td><td></td><td></td><td></td></tr>
<tr><td>Chromosome</td><td>start</td><td>end</td><td>strand</td><td>MM</td><td>target_seq</td><td>PAM</td><td>alignment</td><td>distance</td><td>position</td><td>gene name</td><td>gene id</td></tr>
<tr><td>GU305914.1</td><td>68489</td><td>68511</td><td>-</td><td>0</td><td>AACGCGCGCTCCCCGGGGAT</td><td>GGG</td><td>||||||||[||||||||||||]PAM</td><td>NA</td><td>Intergenic</td><td>NA</td><td>NA</td></tr>
<tr><td></td><td></td><td></td><td></td><td></td><td></td><td></td><td></td><td></td><td></td><td></td><td></td></tr>
<tr><td>T28993</td><td>ACGCGCGCTCCCCGGGGATGGGG</td><td>1000</td><td></td><td></td><td></td><td></td><td></td><td></td><td></td><td></td><td></td></tr>
<tr><td>Oligo fwd</td><td>CACCACGCGCGCTCCCCGGGGATG</td><td></td><td></td><td></td><td></td><td></td><td></td><td></td><td></td><td></td><td></td></tr>
<tr><td>Oligo rev</td><td>AAACCATCCCCGGGGAGCGCGCGT</td><td></td><td></td><td></td><td></td><td></td><td></td><td></td><td></td><td></td><td></td></tr>
<tr><td>Chromosome</td><td>start</td><td>end</td><td>strand</td><td>MM</td><td>target_seq</td><td>PAM</td><td>alignment</td><td>distance</td><td>position</td><td>gene name</td><td>gene id</td></tr>
<tr><td>GU305914.1</td><td>68488</td><td>68510</td><td>-</td><td>0</td><td>ACGCGCGCTCCCCGGGGATG</td><td>GGG</td><td>||||||||[||||||||||||]PAM</td><td>NA</td><td>Intergenic</td><td>NA</td><td>NA</td></tr>
</table>

<table>
<tr><td>T28994</td><td>GGGGTGCTGCGCCACGAACGCGG</td><td></td><td>1000</td><td></td><td></td><td></td><td></td><td></td><td></td><td></td><td></td><td></td></tr>
<tr><td>Oligo fwd</td><td>CACCGGGGTGCTGCGCCACGAACG</td><td></td><td></td><td></td><td></td><td></td><td></td><td></td><td></td><td></td><td></td><td></td></tr>
<tr><td>Oligo rev</td><td>AAACCGTTCGTGGCGCAGCACCCC</td><td></td><td></td><td></td><td></td><td></td><td></td><td></td><td></td><td></td><td></td><td></td></tr>
<tr><td>Chromosome</td><td>start</td><td></td><td>end</td><td>strand</td><td>MM</td><td>target_seq</td><td>PAM</td><td>alignment</td><td>distance</td><td>position</td><td>gene name</td><td>gene id</td></tr>
<tr><td>GU305914.1</td><td></td><td>68469</td><td>68491</td><td>-</td><td>0</td><td>GGGGTGCTGCGCCACGAACG</td><td>CGG</td><td>||||||||[||||||||||||]PAM</td><td>NA</td><td>Intergenic</td><td>NA</td><td>NA</td></tr>
<tr><td>T28995</td><td>GAACGCGGCGATCTTCGCCTCGG</td><td></td><td>1000</td><td></td><td></td><td></td><td></td><td></td><td></td><td></td><td></td><td></td></tr>
<tr><td>Oligo fwd</td><td>CACCGAACGCGGCGATCTTCGCCT</td><td></td><td></td><td></td><td></td><td></td><td></td><td></td><td></td><td></td><td></td><td></td></tr>
<tr><td>Oligo rev</td><td>AAACAGGCGAAGATCGCCGCGTTC</td><td></td><td></td><td></td><td></td><td></td><td></td><td></td><td></td><td></td><td></td><td></td></tr>
<tr><td>Chromosome</td><td>start</td><td></td><td>end</td><td>strand</td><td>MM</td><td>target_seq</td><td>PAM</td><td>alignment</td><td>distance</td><td>position</td><td>gene name</td><td>gene id</td></tr>
<tr><td>GU305914.1</td><td></td><td>68454</td><td>68476</td><td>-</td><td>0</td><td>GAACGCGGCGATCTTCGCCT</td><td>CGG</td><td>||||||||[||||||||||||]PAM</td><td>NA</td><td>Intergenic</td><td>NA</td><td>NA</td></tr>
<tr><td>T28996</td><td>GGTCGTCGTCGTGTTCTCCACGG</td><td></td><td>1000</td><td></td><td></td><td></td><td></td><td></td><td></td><td></td><td></td><td></td></tr>
<tr><td>Oligo fwd</td><td>CACCGGTCGTCGTCGTGTTCTCCA</td><td></td><td></td><td></td><td></td><td></td><td></td><td></td><td></td><td></td><td></td><td></td></tr>
<tr><td>Oligo rev</td><td>AAACTGGAGAACACGACGACGACC</td><td></td><td></td><td></td><td></td><td></td><td></td><td></td><td></td><td></td><td></td><td></td></tr>
<tr><td>Chromosome</td><td>start</td><td></td><td>end</td><td>strand</td><td>MM</td><td>target_seq</td><td>PAM</td><td>alignment</td><td>distance</td><td>position</td><td>gene name</td><td>gene id</td></tr>
<tr><td>GU305914.1</td><td></td><td>68433</td><td>68455</td><td>-</td><td>0</td><td>GGTCGTCGTCGTGTTCTCCA</td><td>CGG</td><td>||||||||[||||||||||||]PAM</td><td>NA</td><td>Intergenic</td><td>NA</td><td>NA</td></tr>
<tr><td>T28997</td><td>GTCGTCGTCGTGTTCTCCACGGG</td><td></td><td>1000</td><td></td><td></td><td></td><td></td><td></td><td></td><td></td><td></td><td></td></tr>
<tr><td>Oligo fwd</td><td>CACCGTCGTCGTCGTGTTCTCCAC</td><td></td><td></td><td></td><td></td><td></td><td></td><td></td><td></td><td></td><td></td><td></td></tr>
<tr><td>Oligo rev</td><td>AAACGTGGAGAACACGACGACGAC</td><td></td><td></td><td></td><td></td><td></td><td></td><td></td><td></td><td></td><td></td><td></td></tr>
<tr><td>Chromosome</td><td>start</td><td></td><td>end</td><td>strand</td><td>MM</td><td>target_seq</td><td>PAM</td><td>alignment</td><td>distance</td><td>position</td><td>gene name</td><td>gene id</td></tr>
<tr><td>GU305914.1</td><td></td><td>68432</td><td>68454</td><td>-</td><td>0</td><td>GTCGTCGTCGTGTTCTCCAC</td><td>GGG</td><td>||||||||[||||||||||||]PAM</td><td>NA</td><td>Intergenic</td><td>NA</td><td>NA</td></tr>
<tr><td>T28998</td><td>AGCCCTCCTCGAACAGCGCCAGG</td><td></td><td>1000</td><td></td><td></td><td></td><td></td><td></td><td></td><td></td><td></td><td></td></tr>
<tr><td>Oligo fwd</td><td>CACCAGCCCTCCTCGAACAGCGCC</td><td></td><td></td><td></td><td></td><td></td><td></td><td></td><td></td><td></td><td></td><td></td></tr>
<tr><td>Oligo rev</td><td>AAACGGCGCTGTTCGAGGAGGGCT</td><td></td><td></td><td></td><td></td><td></td><td></td><td></td><td></td><td></td><td></td><td></td></tr>
<tr><td>Chromosome</td><td>start</td><td></td><td>end</td><td>strand</td><td>MM</td><td>target_seq</td><td>PAM</td><td>alignment</td><td>distance</td><td>position</td><td>gene name</td><td>gene id</td></tr>
<tr><td>GU305914.1</td><td></td><td>68353</td><td>68375</td><td>-</td><td>0</td><td>AGCCCTCCTCGAACAGCGCC</td><td>AGG</td><td>||||||||[||||||||||||]PAM</td><td>NA</td><td>Intergenic</td><td>NA</td><td>NA</td></tr>
<tr><td>T28999</td><td>GGATCTTCGACGCCAGCAGCCGG</td><td></td><td>1000</td><td></td><td></td><td></td><td></td><td></td><td></td><td></td><td></td><td></td></tr>
<tr><td>Oligo fwd</td><td>CACCGGATCTTCGACGCCAGCAGC</td><td></td><td></td><td></td><td></td><td></td><td></td><td></td><td></td><td></td><td></td><td></td></tr>
<tr><td>Oligo rev</td><td>AAACGCTGCTGGCGTCGAAGATCC</td><td></td><td></td><td></td><td></td><td></td><td></td><td></td><td></td><td></td><td></td><td></td></tr>
<tr><td>Chromosome</td><td>start</td><td></td><td>end</td><td>strand</td><td>MM</td><td>target_seq</td><td>PAM</td><td>alignment</td><td>distance</td><td>position</td><td>gene name</td><td>gene id</td></tr>
<tr><td>GU305914.1</td><td></td><td>68332</td><td>68354</td><td>-</td><td>0</td><td>GGATCTTCGACGCCAGCAGC</td><td>CGG</td><td>||||||||[||||||||||||]PAM</td><td>NA</td><td>Intergenic</td><td>NA</td><td>NA</td></tr>
<tr><td>T29000</td><td>ACGCGCTCACCCGCTGAGACAGG</td><td></td><td>1000</td><td></td><td></td><td></td><td></td><td></td><td></td><td></td><td></td><td></td></tr>
<tr><td>Oligo fwd</td><td>CACCACGCGCTCACCCGCTGAGAC</td><td></td><td></td><td></td><td></td><td></td><td></td><td></td><td></td><td></td><td></td><td></td></tr>
<tr><td>Oligo rev</td><td>AAACGTCTCAGCGGGTGAGCGCGT</td><td></td><td></td><td></td><td></td><td></td><td></td><td></td><td></td><td></td><td></td><td></td></tr>
<tr><td>Chromosome</td><td>start</td><td></td><td>end</td><td>strand</td><td>MM</td><td>target_seq</td><td>PAM</td><td>alignment</td><td>distance</td><td>position</td><td>gene name</td><td>gene id</td></tr>
<tr><td>GU305914.1</td><td></td><td>68290</td><td>68312</td><td>-</td><td>0</td><td>ACGCGCTCACCCGCTGAGAC</td><td>AGG</td><td>||||||||[||||||||||||]PAM</td><td>NA</td><td>Intergenic</td><td>NA</td><td>NA</td></tr>
<tr><td>T29001</td><td>GACAGGTAAGTCTTGTACAGCGG</td><td></td><td>1000</td><td></td><td></td><td></td><td></td><td></td><td></td><td></td><td></td><td></td></tr>
<tr><td>Oligo fwd</td><td>CACCGACAGGTAAGTCTTGTACAG</td><td></td><td></td><td></td><td></td><td></td><td></td><td></td><td></td><td></td><td></td><td></td></tr>
<tr><td>Oligo rev</td><td>AAACCTGTACAAGACTTACCTGTC</td><td></td><td></td><td></td><td></td><td></td><td></td><td></td><td></td><td></td><td></td><td></td></tr>
<tr><td>Chromosome</td><td>start</td><td></td><td>end</td><td>strand</td><td>MM</td><td>target_seq</td><td>PAM</td><td>alignment</td><td>distance</td><td>position</td><td>gene name</td><td>gene id</td></tr>
<tr><td>GU305914.1</td><td></td><td>68273</td><td>68295</td><td>-</td><td>0</td><td>GACAGGTAAGTCTTGTACAG</td><td>CGG</td><td>||||||||[||||||||||||]PAM</td><td>NA</td><td>Intergenic</td><td>NA</td><td>NA</td></tr>
<tr><td>T29002</td><td>AGGTAAGTCTTGTACAGCGGCGG</td><td></td><td>1000</td><td></td><td></td><td></td><td></td><td></td><td></td><td></td><td></td><td></td></tr>
<tr><td>Oligo fwd</td><td>CACCAGGTAAGTCTTGTACAGCGG</td><td></td><td></td><td></td><td></td><td></td><td></td><td></td><td></td><td></td><td></td><td></td></tr>
<tr><td>Oligo rev</td><td>AAACCCGCTGTACAAGACTTACCT</td><td></td><td></td><td></td><td></td><td></td><td></td><td></td><td></td><td></td><td></td><td></td></tr>
<tr><td>Chromosome</td><td>start</td><td></td><td>end</td><td>strand</td><td>MM</td><td>target_seq</td><td>PAM</td><td>alignment</td><td>distance</td><td>position</td><td>gene name</td><td>gene id</td></tr>
<tr><td>GU305914.1</td><td></td><td>68270</td><td>68292</td><td>-</td><td>0</td><td>AGGTAAGTCTTGTACAGCGG</td><td>CGG</td><td>||||||||[||||||||||||]PAM</td><td>NA</td><td>Intergenic</td><td>NA</td><td>NA</td></tr>
<tr><td>T29003</td><td>GGTAAGTCTTGTACAGCGGCGGG</td><td></td><td>1000</td><td></td><td></td><td></td><td></td><td></td><td></td><td></td><td></td><td></td></tr>
<tr><td>Oligo fwd</td><td>CACCGGTAAGTCTTGTACAGCGGC</td><td></td><td></td><td></td><td></td><td></td><td></td><td></td><td></td><td></td><td></td><td></td></tr>
<tr><td>Oligo rev</td><td>AAACGCCGCTGTACAAGACTTACC</td><td></td><td></td><td></td><td></td><td></td><td></td><td></td><td></td><td></td><td></td><td></td></tr>
<tr><td>Chromosome</td><td>start</td><td></td><td>end</td><td>strand</td><td>MM</td><td>target_seq</td><td>PAM</td><td>alignment</td><td>distance</td><td>position</td><td>gene name</td><td>gene id</td></tr>
<tr><td>GU305914.1</td><td></td><td>68269</td><td>68291</td><td>-</td><td>0</td><td>GGTAAGTCTTGTACAGCGGC</td><td>GGG</td><td>||||||||[||||||||||||]PAM</td><td>NA</td><td>Intergenic</td><td>NA</td><td>NA</td></tr>
<tr><td>T29004</td><td>GTAAGTCTTGTACAGCGGCGGGG</td><td></td><td>1000</td><td></td><td></td><td></td><td></td><td></td><td></td><td></td><td></td><td></td></tr>
<tr><td>Oligo fwd</td><td>CACCGTAAGTCTTGTACAGCGGCG</td><td></td><td></td><td></td><td></td><td></td><td></td><td></td><td></td><td></td><td></td><td></td></tr>
<tr><td>Oligo rev</td><td>AAACCGCCGCTGTACAAGACTTAC</td><td></td><td></td><td></td><td></td><td></td><td></td><td></td><td></td><td></td><td></td><td></td></tr>
<tr><td>Chromosome</td><td>start</td><td></td><td>end</td><td>strand</td><td>MM</td><td>target_seq</td><td>PAM</td><td>alignment</td><td>distance</td><td>position</td><td>gene name</td><td>gene id</td></tr>
<tr><td>GU305914.1</td><td></td><td>68268</td><td>68290</td><td>-</td><td>0</td><td>GTAAGTCTTGTACAGCGGCG</td><td>GGG</td><td>||||||||[||||||||||||]PAM</td><td>NA</td><td>Intergenic</td><td>NA</td><td>NA</td></tr>
<tr><td>T29005</td><td>GCTCGTCCGACAGACCCATCAGG</td><td></td><td>1000</td><td></td><td></td><td></td><td></td><td></td><td></td><td></td><td></td><td></td></tr>
<tr><td>Oligo fwd</td><td>CACCGCTCGTCCGACAGACCCATC</td><td></td><td></td><td></td><td></td><td></td><td></td><td></td><td></td><td></td><td></td><td></td></tr>
<tr><td>Oligo rev</td><td>AAACGATGGGTCTGTCGGACGAGC</td><td></td><td></td><td></td><td></td><td></td><td></td><td></td><td></td><td></td><td></td><td></td></tr>
<tr><td>Chromosome</td><td>start</td><td></td><td>end</td><td>strand</td><td>MM</td><td>target_seq</td><td>PAM</td><td>alignment</td><td>distance</td><td>position</td><td>gene name</td><td>gene id</td></tr>
<tr><td>GU305914.1</td><td></td><td>68203</td><td>68225</td><td>-</td><td>0</td><td>GCTCGTCCGACAGACCCATC</td><td>AGG</td><td>||||||||[||||||||||||]PAM</td><td>NA</td><td>Intergenic</td><td>NA</td><td>NA</td></tr>
<tr><td>T29006</td><td>CGATCGCGTGCGCCAGCAGCTGG</td><td></td><td>1000</td><td></td><td></td><td></td><td></td><td></td><td></td><td></td><td></td><td></td></tr>
<tr><td>Oligo fwd</td><td>CACCCGATCGCGTGCGCCAGCAGC</td><td></td><td></td><td></td><td></td><td></td><td></td><td></td><td></td><td></td><td></td><td></td></tr>
<tr><td>Oligo rev</td><td>AAACGCTGCTGGCGCACGCGATCG</td><td></td><td></td><td></td><td></td><td></td><td></td><td></td><td></td><td></td><td></td><td></td></tr>
<tr><td>Chromosome</td><td>start</td><td></td><td>end</td><td>strand</td><td>MM</td><td>target_seq</td><td>PAM</td><td>alignment</td><td>distance</td><td>position</td><td>gene name</td><td>gene id</td></tr>
<tr><td>GU305914.1</td><td></td><td>68122</td><td>68144</td><td>-</td><td>0</td><td>CGATCGCGTGCGCCAGCAGC</td><td>TGG</td><td>||||||||[||||||||||||]PAM</td><td>NA</td><td>Intergenic</td><td>NA</td><td>NA</td></tr>
<tr><td>T29007</td><td>TCGCGTGCGCCAGCAGCTGGCGG</td><td></td><td>1000</td><td></td><td></td><td></td><td></td><td></td><td></td><td></td><td></td><td></td></tr>
<tr><td>Oligo fwd</td><td>CACCTCGCGTGCGCCAGCAGCTGG</td><td></td><td></td><td></td><td></td><td></td><td></td><td></td><td></td><td></td><td></td><td></td></tr>
<tr><td>Oligo rev</td><td>AAACCCAGCTGCTGGCGCACGCGA</td><td></td><td></td><td></td><td></td><td></td><td></td><td></td><td></td><td></td><td></td><td></td></tr>
<tr><td>Chromosome</td><td>start</td><td></td><td>end</td><td>strand</td><td>MM</td><td>target_seq</td><td>PAM</td><td>alignment</td><td>distance</td><td>position</td><td>gene name</td><td>gene id</td></tr>
<tr><td>GU305914.1</td><td></td><td>68119</td><td>68141</td><td>-</td><td>0</td><td>TCGCGTGCGCCAGCAGCTGG</td><td>CGG</td><td>||||||||[||||||||||||]PAM</td><td>NA</td><td>Intergenic</td><td>NA</td><td>NA</td></tr>
<tr><td>T29008</td><td>CCACGTCCACGAAGTACTGCCGG</td><td></td><td>1000</td><td></td><td></td><td></td><td></td><td></td><td></td><td></td><td></td><td></td></tr>
<tr><td>Oligo fwd</td><td>CACCCCACGTCCACGAAGTACTGC</td><td></td><td></td><td></td><td></td><td></td><td></td><td></td><td></td><td></td><td></td><td></td></tr>
<tr><td>Oligo rev</td><td>AAACGCAGTACTTCGTGGACGTGG</td><td></td><td></td><td></td><td></td><td></td><td></td><td></td><td></td><td></td><td></td><td></td></tr>
<tr><td>Chromosome</td><td>start</td><td></td><td>end</td><td>strand</td><td>MM</td><td>target_seq</td><td>PAM</td><td>alignment</td><td>distance</td><td>position</td><td>gene name</td><td>gene id</td></tr>
<tr><td>GU305914.1</td><td></td><td>68095</td><td>68117</td><td>-</td><td>0</td><td>CCACGTCCACGAAGTACTGC</td><td>CGG</td><td>||||||||[||||||||||||]PAM</td><td>NA</td><td>Intergenic</td><td>NA</td><td>NA</td></tr>
</table>

<table>
<tr><td>T29009</td><td>ACGAAGTACTGCCGGTAGAACGG</td><td>1000</td><td></td><td></td><td></td><td></td><td></td><td></td><td></td><td></td><td></td></tr>
<tr><td>Oligo fwd</td><td>CACCACGAAGTACTGCCGGTAGAA</td><td></td><td></td><td></td><td></td><td></td><td></td><td></td><td></td><td></td><td></td></tr>
<tr><td>Oligo rev</td><td>AAACTTCTACCGGCAGTACTTCGT</td><td></td><td></td><td></td><td></td><td></td><td></td><td></td><td></td><td></td><td></td></tr>
<tr><td>Chromosome</td><td>start</td><td>end</td><td>strand</td><td>MM</td><td>target_seq</td><td>PAM</td><td>alignment</td><td>distance</td><td>position</td><td>gene name</td><td>gene id</td></tr>
<tr><td>GU305914.1</td><td>68087</td><td>68109</td><td>-</td><td>0</td><td>ACGAAGTACTGCCGGTAGAA</td><td>CGG</td><td>||||||||[||||||||||||]PAM</td><td>NA</td><td>Intergenic</td><td>NA</td><td>NA</td></tr>
<tr><td>T29010</td><td>CGGCTCCTCCCCGCAGTACATGG</td><td>1000</td><td></td><td></td><td></td><td></td><td></td><td></td><td></td><td></td><td></td></tr>
<tr><td>Oligo fwd</td><td>CACCCGGCTCCTCCCCGCAGTACA</td><td></td><td></td><td></td><td></td><td></td><td></td><td></td><td></td><td></td><td></td></tr>
<tr><td>Oligo rev</td><td>AAACTGTACTGCGGGGAGGAGCCG</td><td></td><td></td><td></td><td></td><td></td><td></td><td></td><td></td><td></td><td></td></tr>
<tr><td>Chromosome</td><td>start</td><td>end</td><td>strand</td><td>MM</td><td>target_seq</td><td>PAM</td><td>alignment</td><td>distance</td><td>position</td><td>gene name</td><td>gene id</td></tr>
<tr><td>GU305914.1</td><td>68067</td><td>68089</td><td>-</td><td>0</td><td>CGGCTCCTCCCCGCAGTACA</td><td>TGG</td><td>||||||||[||||||||||||]PAM</td><td>NA</td><td>Intergenic</td><td>NA</td><td>NA</td></tr>
<tr><td>T29011</td><td>CAGTACATGGTCCATTCCTGCGG</td><td>1000</td><td></td><td></td><td></td><td></td><td></td><td></td><td></td><td></td><td></td></tr>
<tr><td>Oligo fwd</td><td>CACCCAGTACATGGTCCATTCCTG</td><td></td><td></td><td></td><td></td><td></td><td></td><td></td><td></td><td></td><td></td></tr>
<tr><td>Oligo rev</td><td>AAACCAGGAATGGACCATGTACTG</td><td></td><td></td><td></td><td></td><td></td><td></td><td></td><td></td><td></td><td></td></tr>
<tr><td>Chromosome</td><td>start</td><td>end</td><td>strand</td><td>MM</td><td>target_seq</td><td>PAM</td><td>alignment</td><td>distance</td><td>position</td><td>gene name</td><td>gene id</td></tr>
<tr><td>GU305914.1</td><td>68054</td><td>68076</td><td>-</td><td>0</td><td>CAGTACATGGTCCATTCCTG</td><td>CGG</td><td>||||||||[||||||||||||]PAM</td><td>NA</td><td>Intergenic</td><td>NA</td><td>NA</td></tr>
<tr><td>T29012</td><td>AGTACATGGTCCATTCCTGCGGG</td><td>1000</td><td></td><td></td><td></td><td></td><td></td><td></td><td></td><td></td><td></td></tr>
<tr><td>Oligo fwd</td><td>CACCAGTACATGGTCCATTCCTGC</td><td></td><td></td><td></td><td></td><td></td><td></td><td></td><td></td><td></td><td></td></tr>
<tr><td>Oligo rev</td><td>AAACGCAGGAATGGACCATGTACT</td><td></td><td></td><td></td><td></td><td></td><td></td><td></td><td></td><td></td><td></td></tr>
<tr><td>Chromosome</td><td>start</td><td>end</td><td>strand</td><td>MM</td><td>target_seq</td><td>PAM</td><td>alignment</td><td>distance</td><td>position</td><td>gene name</td><td>gene id</td></tr>
<tr><td>GU305914.1</td><td>68053</td><td>68075</td><td>-</td><td>0</td><td>AGTACATGGTCCATTCCTGC</td><td>GGG</td><td>||||||||[||||||||||||]PAM</td><td>NA</td><td>Intergenic</td><td>NA</td><td>NA</td></tr>
<tr><td>T29013</td><td>GTACATGGTCCATTCCTGCGGGG</td><td>1000</td><td></td><td></td><td></td><td></td><td></td><td></td><td></td><td></td><td></td></tr>
<tr><td>Oligo fwd</td><td>CACCGTACATGGTCCATTCCTGCG</td><td></td><td></td><td></td><td></td><td></td><td></td><td></td><td></td><td></td><td></td></tr>
<tr><td>Oligo rev</td><td>AAACCGCAGGAATGGACCATGTAC</td><td></td><td></td><td></td><td></td><td></td><td></td><td></td><td></td><td></td><td></td></tr>
<tr><td>Chromosome</td><td>start</td><td>end</td><td>strand</td><td>MM</td><td>target_seq</td><td>PAM</td><td>alignment</td><td>distance</td><td>position</td><td>gene name</td><td>gene id</td></tr>
<tr><td>GU305914.1</td><td>68052</td><td>68074</td><td>-</td><td>0</td><td>GTACATGGTCCATTCCTGCG</td><td>GGG</td><td>||||||||[||||||||||||]PAM</td><td>NA</td><td>Intergenic</td><td>NA</td><td>NA</td></tr>
<tr><td>T29014</td><td>TACATGGTCCATTCCTGCGGGGG</td><td>1000</td><td></td><td></td><td></td><td></td><td></td><td></td><td></td><td></td><td></td></tr>
<tr><td>Oligo fwd</td><td>CACCTACATGGTCCATTCCTGCGG</td><td></td><td></td><td></td><td></td><td></td><td></td><td></td><td></td><td></td><td></td></tr>
<tr><td>Oligo rev</td><td>AAACCCGCAGGAATGGACCATGTA</td><td></td><td></td><td></td><td></td><td></td><td></td><td></td><td></td><td></td><td></td></tr>
<tr><td>Chromosome</td><td>start</td><td>end</td><td>strand</td><td>MM</td><td>target_seq</td><td>PAM</td><td>alignment</td><td>distance</td><td>position</td><td>gene name</td><td>gene id</td></tr>
<tr><td>GU305914.1</td><td>68051</td><td>68073</td><td>-</td><td>0</td><td>TACATGGTCCATTCCTGCGG</td><td>GGG</td><td>||||||||[||||||||||||]PAM</td><td>NA</td><td>Intergenic</td><td>NA</td><td>NA</td></tr>
<tr><td>T29015</td><td>CATTCCTGCGGGGGTTTCACCGG</td><td>1000</td><td></td><td></td><td></td><td></td><td></td><td></td><td></td><td></td><td></td></tr>
<tr><td>Oligo fwd</td><td>CACCCATTCCTGCGGGGGTTTCAC</td><td></td><td></td><td></td><td></td><td></td><td></td><td></td><td></td><td></td><td></td></tr>
<tr><td>Oligo rev</td><td>AAACGTGAAACCCCCGCAGGAATG</td><td></td><td></td><td></td><td></td><td></td><td></td><td></td><td></td><td></td><td></td></tr>
<tr><td>Chromosome</td><td>start</td><td>end</td><td>strand</td><td>MM</td><td>target_seq</td><td>PAM</td><td>alignment</td><td>distance</td><td>position</td><td>gene name</td><td>gene id</td></tr>
<tr><td>GU305914.1</td><td>68042</td><td>68064</td><td>-</td><td>0</td><td>CATTCCTGCGGGGGTTTCAC</td><td>CGG</td><td>||||||||[||||||||||||]PAM</td><td>NA</td><td>Intergenic</td><td>NA</td><td>NA</td></tr>
<tr><td>T29016</td><td>TTTCACCGGCAGCGACTGCGTGG</td><td>1000</td><td></td><td></td><td></td><td></td><td></td><td></td><td></td><td></td><td></td></tr>
<tr><td>Oligo fwd</td><td>CACCTTTCACCGGCAGCGACTGCG</td><td></td><td></td><td></td><td></td><td></td><td></td><td></td><td></td><td></td><td></td></tr>
<tr><td>Oligo rev</td><td>AAACCGCAGTCGCTGCCGGTGAAA</td><td></td><td></td><td></td><td></td><td></td><td></td><td></td><td></td><td></td><td></td></tr>
<tr><td>Chromosome</td><td>start</td><td>end</td><td>strand</td><td>MM</td><td>target_seq</td><td>PAM</td><td>alignment</td><td>distance</td><td>position</td><td>gene name</td><td>gene id</td></tr>
<tr><td>GU305914.1</td><td>68028</td><td>68050</td><td>-</td><td>0</td><td>TTTCACCGGCAGCGACTGCG</td><td>TGG</td><td>||||||||[||||||||||||]PAM</td><td>NA</td><td>Intergenic</td><td>NA</td><td>NA</td></tr>
<tr><td>T29017</td><td>CACCGGCAGCGACTGCGTGGCGG</td><td>1000</td><td></td><td></td><td></td><td></td><td></td><td></td><td></td><td></td><td></td></tr>
<tr><td>Oligo fwd</td><td>CACCCACCGGCAGCGACTGCGTGG</td><td></td><td></td><td></td><td></td><td></td><td></td><td></td><td></td><td></td><td></td></tr>
<tr><td>Oligo rev</td><td>AAACCCACGCAGTCGCTGCCGGTG</td><td></td><td></td><td></td><td></td><td></td><td></td><td></td><td></td><td></td><td></td></tr>
<tr><td>Chromosome</td><td>start</td><td>end</td><td>strand</td><td>MM</td><td>target_seq</td><td>PAM</td><td>alignment</td><td>distance</td><td>position</td><td>gene name</td><td>gene id</td></tr>
<tr><td>GU305914.1</td><td>68025</td><td>68047</td><td>-</td><td>0</td><td>CACCGGCAGCGACTGCGTGG</td><td>CGG</td><td>||||||||[||||||||||||]PAM</td><td>NA</td><td>Intergenic</td><td>NA</td><td>NA</td></tr>
<tr><td>T29018</td><td>GCAGCGACTGCGTGGCGGTGCGG</td><td>1000</td><td></td><td></td><td></td><td></td><td></td><td></td><td></td><td></td><td></td></tr>
<tr><td>Oligo fwd</td><td>CACCGCAGCGACTGCGTGGCGGTG</td><td></td><td></td><td></td><td></td><td></td><td></td><td></td><td></td><td></td><td></td></tr>
<tr><td>Oligo rev</td><td>AAACCACCGCCACGCAGTCGCTGC</td><td></td><td></td><td></td><td></td><td></td><td></td><td></td><td></td><td></td><td></td></tr>
<tr><td>Chromosome</td><td>start</td><td>end</td><td>strand</td><td>MM</td><td>target_seq</td><td>PAM</td><td>alignment</td><td>distance</td><td>position</td><td>gene name</td><td>gene id</td></tr>
<tr><td>GU305914.1</td><td>68020</td><td>68042</td><td>-</td><td>0</td><td>GCAGCGACTGCGTGGCGGTG</td><td>CGG</td><td>||||||||[||||||||||||]PAM</td><td>NA</td><td>Intergenic</td><td>NA</td><td>NA</td></tr>
<tr><td>T29019</td><td>CACGTCTTCCACGATCACCTCGG</td><td>1000</td><td></td><td></td><td></td><td></td><td></td><td></td><td></td><td></td><td></td></tr>
<tr><td>Oligo fwd</td><td>CACCCACGTCTTCCACGATCACCT</td><td></td><td></td><td></td><td></td><td></td><td></td><td></td><td></td><td></td><td></td></tr>
<tr><td>Oligo rev</td><td>AAACAGGTGATCGTGGAAGACGTG</td><td></td><td></td><td></td><td></td><td></td><td></td><td></td><td></td><td></td><td></td></tr>
<tr><td>Chromosome</td><td>start</td><td>end</td><td>strand</td><td>MM</td><td>target_seq</td><td>PAM</td><td>alignment</td><td>distance</td><td>position</td><td>gene name</td><td>gene id</td></tr>
<tr><td>GU305914.1</td><td>67980</td><td>68002</td><td>-</td><td>0</td><td>CACGTCTTCCACGATCACCT</td><td>CGG</td><td>||||||||[||||||||||||]PAM</td><td>NA</td><td>Intergenic</td><td>NA</td><td>NA</td></tr>
<tr><td>T29020</td><td>GATCACCTCGGACAGACGCTCGG</td><td>1000</td><td></td><td></td><td></td><td></td><td></td><td></td><td></td><td></td><td></td></tr>
<tr><td>Oligo fwd</td><td>CACCGATCACCTCGGACAGACGCT</td><td></td><td></td><td></td><td></td><td></td><td></td><td></td><td></td><td></td><td></td></tr>
<tr><td>Oligo rev</td><td>AAACAGCGTCTGTCCGAGGTGATC</td><td></td><td></td><td></td><td></td><td></td><td></td><td></td><td></td><td></td><td></td></tr>
<tr><td>Chromosome</td><td>start</td><td>end</td><td>strand</td><td>MM</td><td>target_seq</td><td>PAM</td><td>alignment</td><td>distance</td><td>position</td><td>gene name</td><td>gene id</td></tr>
<tr><td>GU305914.1</td><td>67968</td><td>67990</td><td>-</td><td>0</td><td>GATCACCTCGGACAGACGCT</td><td>CGG</td><td>||||||||[||||||||||||]PAM</td><td>NA</td><td>Intergenic</td><td>NA</td><td>NA</td></tr>
<tr><td>T29021</td><td>ATCACCTCGGACAGACGCTCGGG</td><td>1000</td><td></td><td></td><td></td><td></td><td></td><td></td><td></td><td></td><td></td></tr>
<tr><td>Oligo fwd</td><td>CACCATCACCTCGGACAGACGCTC</td><td></td><td></td><td></td><td></td><td></td><td></td><td></td><td></td><td></td><td></td></tr>
<tr><td>Oligo rev</td><td>AAACGAGCGTCTGTCCGAGGTGAT</td><td></td><td></td><td></td><td></td><td></td><td></td><td></td><td></td><td></td><td></td></tr>
<tr><td>Chromosome</td><td>start</td><td>end</td><td>strand</td><td>MM</td><td>target_seq</td><td>PAM</td><td>alignment</td><td>distance</td><td>position</td><td>gene name</td><td>gene id</td></tr>
<tr><td>GU305914.1</td><td>67967</td><td>67989</td><td>-</td><td>0</td><td>ATCACCTCGGACAGACGCTC</td><td>GGG</td><td>||||||||[||||||||||||]PAM</td><td>NA</td><td>Intergenic</td><td>NA</td><td>NA</td></tr>
<tr><td>T29022</td><td>AGACGCTCGGGCAACCTGATCGG</td><td>1000</td><td></td><td></td><td></td><td></td><td></td><td></td><td></td><td></td><td></td></tr>
<tr><td>Oligo fwd</td><td>CACCAGACGCTCGGGCAACCTGAT</td><td></td><td></td><td></td><td></td><td></td><td></td><td></td><td></td><td></td><td></td></tr>
<tr><td>Oligo rev</td><td>AAACATCAGGTTGCCCGAGCGTCT</td><td></td><td></td><td></td><td></td><td></td><td></td><td></td><td></td><td></td><td></td></tr>
<tr><td>Chromosome</td><td>start</td><td>end</td><td>strand</td><td>MM</td><td>target_seq</td><td>PAM</td><td>alignment</td><td>distance</td><td>position</td><td>gene name</td><td>gene id</td></tr>
<tr><td>GU305914.1</td><td>67955</td><td>67977</td><td>-</td><td>0</td><td>AGACGCTCGGGCAACCTGAT</td><td>CGG</td><td>||||||||[||||||||||||]PAM</td><td>NA</td><td>Intergenic</td><td>NA</td><td>NA</td></tr>
<tr><td>T29023</td><td>CGGGCAACCTGATCGGCCGCCGG</td><td>1000</td><td></td><td></td><td></td><td></td><td></td><td></td><td></td><td></td><td></td></tr>
<tr><td>Oligo fwd</td><td>CACCCGGGCAACCTGATCGGCCGC</td><td></td><td></td><td></td><td></td><td></td><td></td><td></td><td></td><td></td><td></td></tr>
<tr><td>Oligo rev</td><td>AAACGCGGCCGATCAGGTTGCCCG</td><td></td><td></td><td></td><td></td><td></td><td></td><td></td><td></td><td></td><td></td></tr>
<tr><td>Chromosome</td><td>start</td><td>end</td><td>strand</td><td>MM</td><td>target_seq</td><td>PAM</td><td>alignment</td><td>distance</td><td>position</td><td>gene name</td><td>gene id</td></tr>
</table>

GU305914.1 67948 67970 - 0 CGGGCAACCTGATCGGCCGC CGG ||||||||[||||||||||||]PAM NA Intergenic NA NA

T29024 ACCTGATCGGCCGCCGGCTGCGG 1000
Oligo fwd CACCACCTGATCGGCCGCCGGCTG
Oligo rev AAACCAGCCGGCGGCCGATCAGGT
Chromosome start end strand MM target_seq PAM alignment distance position gene name gene id
GU305914.1 67942 67964 - 0 ACCTGATCGGCCGCCGGCTG CGG ||||||||[||||||||||||]PAM NA Intergenic NA NA

T29025 TTCGCCGTGCCCTTCTGCTGCGG 1000
Oligo fwd CACCTTCGCCGTGCCCTTCTGCTG
Oligo rev AAACCAGCAGAAGGGCACGGCGAA
Chromosome start end strand MM target_seq PAM alignment distance position gene name gene id
GU305914.1 67910 67932 - 0 TTCGCCGTGCCCTTCTGCTG CGG ||||||||[||||||||||||]PAM NA Intergenic NA NA

T29026 GCCGTGCCCTTCTGCTGCGGCGG 1000
Oligo fwd CACCGCCGTGCCCTTCTGCTGCGG
Oligo rev AAACCCGCAGCAGAAGGGCACGGC
Chromosome start end strand MM target_seq PAM alignment distance position gene name gene id
GU305914.1 67907 67929 - 0 GCCGTGCCCTTCTGCTGCGG CGG ||||||||[||||||||||||]PAM NA Intergenic NA NA

T29027 TTCTGCTGCGGCGGCACGTCCGG 1000
Oligo fwd CACCTTCTGCTGCGGCGGCACGTC
Oligo rev AAACGACGTGCCGCCGCAGCAGAA
Chromosome start end strand MM target_seq PAM alignment distance position gene name gene id
GU305914.1 67898 67920 - 0 TTCTGCTGCGGCGGCACGTC CGG ||||||||[||||||||||||]PAM NA Intergenic NA NA

T29028 CCGTCTCTTCTTGCCCGACTTGG 1000
Oligo fwd CACCCCGTCTCTTCTTGCCCGACT
Oligo rev AAACAGTCGGGCAAGAAGAGACGG
Chromosome start end strand MM target_seq PAM alignment distance position gene name gene id
GU305914.1 67872 67894 - 0 CCGTCTCTTCTTGCCCGACT TGG ||||||||[||||||||||||]PAM NA Intergenic NA NA

T29029 TGCCCGACTTGGACGAAGTCTGG 1000
Oligo fwd CACCTGCCCGACTTGGACGAAGTC
Oligo rev AAACGACTTCGTCCAAGTCGGGCA
Chromosome start end strand MM target_seq PAM alignment distance position gene name gene id
GU305914.1 67861 67883 - 0 TGCCCGACTTGGACGAAGTC TGG ||||||||[||||||||||||]PAM NA Intergenic NA NA

T29030 GCCCGACTTGGACGAAGTCTGGG 1000
Oligo fwd CACCGCCCGACTTGGACGAAGTCT
Oligo rev AAACAGACTTCGTCCAAGTCGGGC
Chromosome start end strand MM target_seq PAM alignment distance position gene name gene id
GU305914.1 67860 67882 - 0 GCCCGACTTGGACGAAGTCT GGG ||||||||[||||||||||||]PAM NA Intergenic NA NA

T29031 CGCGCCGTTCGTCACGCCGTCGG 1000
Oligo fwd CACCCGCGCCGTTCGTCACGCCGT
Oligo rev AAACACGGCGTGACGAACGGCGCG
Chromosome start end strand MM target_seq PAM alignment distance position gene name gene id
GU305914.1 67833 67855 - 0 CGCGCCGTTCGTCACGCCGT CGG ||||||||[||||||||||||]PAM NA Intergenic NA NA

T29032 GTTCGTCACGCCGTCGGACGCGG 1000
Oligo fwd CACCGTTCGTCACGCCGTCGGACG
Oligo rev AAACCGTCCGACGGCGTGACGAAC
Chromosome start end strand MM target_seq PAM alignment distance position gene name gene id
GU305914.1 67827 67849 - 0 GTTCGTCACGCCGTCGGACG CGG ||||||||[||||||||||||]PAM NA Intergenic NA NA

T29033 CCGTCGGACGCGGCGCCGCTCGG 1000
Oligo fwd CACCCCGTCGGACGCGGCGCCGCT
Oligo rev AAACAGCGGCGCCGCGTCCGACGG
Chromosome start end strand MM target_seq PAM alignment distance position gene name gene id
GU305914.1 67817 67839 - 0 CCGTCGGACGCGGCGCCGCT CGG ||||||||[||||||||||||]PAM NA Intergenic NA NA

T29034 GACGCGGCGCCGCTCGGCAAAGG 1000
Oligo fwd CACCGACGCGGCGCCGCTCGGCAA
Oligo rev AAACTTGCCGAGCGGCGCCGCGTC
Chromosome start end strand MM target_seq PAM alignment distance position gene name gene id
GU305914.1 67811 67833 - 0 GACGCGGCGCCGCTCGGCAA AGG ||||||||[||||||||||||]PAM NA Intergenic NA NA

T29035 CAAAGGAGAAACAGAAGCAGAGG 1000
Oligo fwd CACCCAAAGGAGAAACAGAAGCAG
Oligo rev AAACCTGCTTCTGTTTCTCCTTTG
Chromosome start end strand MM target_seq PAM alignment distance position gene name gene id
GU305914.1 67794 67816 - 0 CAAAGGAGAAACAGAAGCAG AGG ||||||||[||||||||||||]PAM NA Intergenic NA NA

T29036 AGAAACAGAAGCAGAGGCAGAGG 1000
Oligo fwd CACCAGAAACAGAAGCAGAGGCAG
Oligo rev AAACCTGCCTCTGCTTCTGTTTCT
Chromosome start end strand MM target_seq PAM alignment distance position gene name gene id
GU305914.1 67788 67810 - 0 AGAAACAGAAGCAGAGGCAG AGG ||||||||[||||||||||||]PAM NA Intergenic NA NA
GU305914.1 67782 67804 - 3 AGAAGCAGAGGCAGAGGCGG CGG ||||-|||[|-||||||||-|]PAM NA Intergenic NA NA

T29037 AACAGAAGCAGAGGCAGAGGCGG 1000
Oligo fwd CACCAACAGAAGCAGAGGCAGAGG
Oligo rev AAACCCTCTGCCTCTGCTTCTGTT
Chromosome start end strand MM target_seq PAM alignment distance position gene name gene id
GU305914.1 67785 67807 - 0 AACAGAAGCAGAGGCAGAGG CGG ||||||||[||||||||||||]PAM NA Intergenic NA NA

T29038 AGAAGCAGAGGCAGAGGCGGCGG 1000
Oligo fwd CACCAGAAGCAGAGGCAGAGGCGG

<table>
<tr><td>Oligo rev</td><td>AAACCCGCCTCTGCCTCTGCTTCT</td><td></td><td></td><td></td><td></td><td></td><td></td><td></td><td></td><td></td><td></td></tr>
<tr><td>Chromosome</td><td>start</td><td>end</td><td>strand</td><td>MM</td><td>target_seq</td><td>PAM</td><td>alignment</td><td>distance</td><td>position</td><td>gene name</td><td>gene id</td></tr>
<tr><td>GU305914.1</td><td>67782</td><td>67804</td><td>-</td><td>0</td><td>AGAAGCAGAGGCAGAGGCGG</td><td>CGG</td><td>||||||||[||||||||||||]PAM</td><td>NA</td><td>Intergenic</td><td>NA</td><td>NA</td></tr>
<tr><td>GU305914.1</td><td>67788</td><td>67810</td><td>-</td><td>3</td><td>AGAAACAGAAGCAGAGGCAG</td><td>AGG</td><td>||||-|||[|-||||||||-|]PAM</td><td>NA</td><td>Intergenic</td><td>NA</td><td>NA</td></tr>
<tr><td>T29039</td><td>AGAGGCGGCGGAAGAAGAGATGG</td><td>1000</td><td></td><td></td><td></td><td></td><td></td><td></td><td></td><td></td><td></td></tr>
<tr><td>Oligo fwd</td><td>CACCAGAGGCGGCGGAAGAAGAGA</td><td></td><td></td><td></td><td></td><td></td><td></td><td></td><td></td><td></td><td></td></tr>
<tr><td>Oligo rev</td><td>AAACTCTCTTCTTCCGCCGCCTCT</td><td></td><td></td><td></td><td></td><td></td><td></td><td></td><td></td><td></td><td></td></tr>
<tr><td>Chromosome</td><td>start</td><td>end</td><td>strand</td><td>MM</td><td>target_seq</td><td>PAM</td><td>alignment</td><td>distance</td><td>position</td><td>gene name</td><td>gene id</td></tr>
<tr><td>GU305914.1</td><td>67770</td><td>67792</td><td>-</td><td>0</td><td>AGAGGCGGCGGAAGAAGAGA</td><td>TGG</td><td>||||||||[||||||||||||]PAM</td><td>NA</td><td>Intergenic</td><td>NA</td><td>NA</td></tr>
<tr><td>T29040</td><td>GGCGGCGGAAGAAGAGATGGAGG</td><td>1000</td><td></td><td></td><td></td><td></td><td></td><td></td><td></td><td></td><td></td></tr>
<tr><td>Oligo fwd</td><td>CACCGGCGGCGGAAGAAGAGATGG</td><td></td><td></td><td></td><td></td><td></td><td></td><td></td><td></td><td></td><td></td></tr>
<tr><td>Oligo rev</td><td>AAACCCATCTCTTCTTCCGCCGCC</td><td></td><td></td><td></td><td></td><td></td><td></td><td></td><td></td><td></td><td></td></tr>
<tr><td>Chromosome</td><td>start</td><td>end</td><td>strand</td><td>MM</td><td>target_seq</td><td>PAM</td><td>alignment</td><td>distance</td><td>position</td><td>gene name</td><td>gene id</td></tr>
<tr><td>GU305914.1</td><td>67767</td><td>67789</td><td>-</td><td>0</td><td>GGCGGCGGAAGAAGAGATGG</td><td>AGG</td><td>||||||||[||||||||||||]PAM</td><td>NA</td><td>Intergenic</td><td>NA</td><td>NA</td></tr>
<tr><td>GU305914.1</td><td>125024</td><td>125046</td><td>-</td><td>4</td><td>GACGTCGGACGAGGAGATGG</td><td>TGG</td><td>|-||-|||[|-||-|||||||]PAM</td><td>NA</td><td>Intergenic</td><td>NA</td><td>NA</td></tr>
<tr><td>T29041</td><td>GCGGCGGAAGAAGAGATGGAGGG</td><td>1000</td><td></td><td></td><td></td><td></td><td></td><td></td><td></td><td></td><td></td></tr>
<tr><td>Oligo fwd</td><td>CACCGCGGCGGAAGAAGAGATGGA</td><td></td><td></td><td></td><td></td><td></td><td></td><td></td><td></td><td></td><td></td></tr>
<tr><td>Oligo rev</td><td>AAACTCCATCTCTTCTTCCGCCGC</td><td></td><td></td><td></td><td></td><td></td><td></td><td></td><td></td><td></td><td></td></tr>
<tr><td>Chromosome</td><td>start</td><td>end</td><td>strand</td><td>MM</td><td>target_seq</td><td>PAM</td><td>alignment</td><td>distance</td><td>position</td><td>gene name</td><td>gene id</td></tr>
<tr><td>GU305914.1</td><td>67766</td><td>67788</td><td>-</td><td>0</td><td>GCGGCGGAAGAAGAGATGGA</td><td>GGG</td><td>||||||||[||||||||||||]PAM</td><td>NA</td><td>Intergenic</td><td>NA</td><td>NA</td></tr>
<tr><td>T29042</td><td>ATCACCGACGCGTCGATGTCCGG</td><td>1000</td><td></td><td></td><td></td><td></td><td></td><td></td><td></td><td></td><td></td></tr>
<tr><td>Oligo fwd</td><td>CACCATCACCGACGCGTCGATGTC</td><td></td><td></td><td></td><td></td><td></td><td></td><td></td><td></td><td></td><td></td></tr>
<tr><td>Oligo rev</td><td>AAACGACATCGACGCGTCGGTGAT</td><td></td><td></td><td></td><td></td><td></td><td></td><td></td><td></td><td></td><td></td></tr>
<tr><td>Chromosome</td><td>start</td><td>end</td><td>strand</td><td>MM</td><td>target_seq</td><td>PAM</td><td>alignment</td><td>distance</td><td>position</td><td>gene name</td><td>gene id</td></tr>
<tr><td>GU305914.1</td><td>67715</td><td>67737</td><td>-</td><td>0</td><td>ATCACCGACGCGTCGATGTC</td><td>CGG</td><td>||||||||[||||||||||||]PAM</td><td>NA</td><td>Intergenic</td><td>NA</td><td>NA</td></tr>
<tr><td>T29043</td><td>TCACCGACGCGTCGATGTCCGGG</td><td>1000</td><td></td><td></td><td></td><td></td><td></td><td></td><td></td><td></td><td></td></tr>
<tr><td>Oligo fwd</td><td>CACCTCACCGACGCGTCGATGTCC</td><td></td><td></td><td></td><td></td><td></td><td></td><td></td><td></td><td></td><td></td></tr>
<tr><td>Oligo rev</td><td>AAACGGACATCGACGCGTCGGTGA</td><td></td><td></td><td></td><td></td><td></td><td></td><td></td><td></td><td></td><td></td></tr>
<tr><td>Chromosome</td><td>start</td><td>end</td><td>strand</td><td>MM</td><td>target_seq</td><td>PAM</td><td>alignment</td><td>distance</td><td>position</td><td>gene name</td><td>gene id</td></tr>
<tr><td>GU305914.1</td><td>67714</td><td>67736</td><td>-</td><td>0</td><td>TCACCGACGCGTCGATGTCC</td><td>GGG</td><td>||||||||[||||||||||||]PAM</td><td>NA</td><td>Intergenic</td><td>NA</td><td>NA</td></tr>
<tr><td>T29044</td><td>CGATGTCCGGGTCCTTGTACTGG</td><td>1000</td><td></td><td></td><td></td><td></td><td></td><td></td><td></td><td></td><td></td></tr>
<tr><td>Oligo fwd</td><td>CACCCGATGTCCGGGTCCTTGTAC</td><td></td><td></td><td></td><td></td><td></td><td></td><td></td><td></td><td></td><td></td></tr>
<tr><td>Oligo rev</td><td>AAACGTACAAGGACCCGGACATCG</td><td></td><td></td><td></td><td></td><td></td><td></td><td></td><td></td><td></td><td></td></tr>
<tr><td>Chromosome</td><td>start</td><td>end</td><td>strand</td><td>MM</td><td>target_seq</td><td>PAM</td><td>alignment</td><td>distance</td><td>position</td><td>gene name</td><td>gene id</td></tr>
<tr><td>GU305914.1</td><td>67702</td><td>67724</td><td>-</td><td>0</td><td>CGATGTCCGGGTCCTTGTAC</td><td>TGG</td><td>||||||||[||||||||||||]PAM</td><td>NA</td><td>Intergenic</td><td>NA</td><td>NA</td></tr>
<tr><td>GU305914.1</td><td>47045</td><td>47067</td><td>-</td><td>4</td><td>AGATGTTCTGGTCGTTGTAC</td><td>GGG</td><td>-|||||-|[-||||-||||||]PAM</td><td>NA</td><td>Intergenic</td><td>NA</td><td>NA</td></tr>
<tr><td>T29045</td><td>ATCTGCACCAGCAGCTCCGACGG</td><td>1000</td><td></td><td></td><td></td><td></td><td></td><td></td><td></td><td></td><td></td></tr>
<tr><td>Oligo fwd</td><td>CACCATCTGCACCAGCAGCTCCGA</td><td></td><td></td><td></td><td></td><td></td><td></td><td></td><td></td><td></td><td></td></tr>
<tr><td>Oligo rev</td><td>AAACTCGGAGCTGCTGGTGCAGAT</td><td></td><td></td><td></td><td></td><td></td><td></td><td></td><td></td><td></td><td></td></tr>
<tr><td>Chromosome</td><td>start</td><td>end</td><td>strand</td><td>MM</td><td>target_seq</td><td>PAM</td><td>alignment</td><td>distance</td><td>position</td><td>gene name</td><td>gene id</td></tr>
<tr><td>GU305914.1</td><td>67673</td><td>67695</td><td>-</td><td>0</td><td>ATCTGCACCAGCAGCTCCGA</td><td>CGG</td><td>||||||||[||||||||||||]PAM</td><td>NA</td><td>Intergenic</td><td>NA</td><td>NA</td></tr>
<tr><td>T29046</td><td>ACGGCGAGATCAGCGTCGTGCGG</td><td>1000</td><td></td><td></td><td></td><td></td><td></td><td></td><td></td><td></td><td></td></tr>
<tr><td>Oligo fwd</td><td>CACCACGGCGAGATCAGCGTCGTG</td><td></td><td></td><td></td><td></td><td></td><td></td><td></td><td></td><td></td><td></td></tr>
<tr><td>Oligo rev</td><td>AAACCACGACGCTGATCTCGCCGT</td><td></td><td></td><td></td><td></td><td></td><td></td><td></td><td></td><td></td><td></td></tr>
<tr><td>Chromosome</td><td>start</td><td>end</td><td>strand</td><td>MM</td><td>target_seq</td><td>PAM</td><td>alignment</td><td>distance</td><td>position</td><td>gene name</td><td>gene id</td></tr>
<tr><td>GU305914.1</td><td>67654</td><td>67676</td><td>-</td><td>0</td><td>ACGGCGAGATCAGCGTCGTG</td><td>CGG</td><td>||||||||[||||||||||||]PAM</td><td>NA</td><td>Intergenic</td><td>NA</td><td>NA</td></tr>
<tr><td>T29047</td><td>GGCAGCGCGTGAAGTACACCAGG</td><td>1000</td><td></td><td></td><td></td><td></td><td></td><td></td><td></td><td></td><td></td></tr>
<tr><td>Oligo fwd</td><td>CACCGGCAGCGCGTGAAGTACACC</td><td></td><td></td><td></td><td></td><td></td><td></td><td></td><td></td><td></td><td></td></tr>
<tr><td>Oligo rev</td><td>AAACGGTGTACTTCACGCGCTGCC</td><td></td><td></td><td></td><td></td><td></td><td></td><td></td><td></td><td></td><td></td></tr>
<tr><td>Chromosome</td><td>start</td><td>end</td><td>strand</td><td>MM</td><td>target_seq</td><td>PAM</td><td>alignment</td><td>distance</td><td>position</td><td>gene name</td><td>gene id</td></tr>
<tr><td>GU305914.1</td><td>67633</td><td>67655</td><td>-</td><td>0</td><td>GGCAGCGCGTGAAGTACACC</td><td>AGG</td><td>||||||||[||||||||||||]PAM</td><td>NA</td><td>Intergenic</td><td>NA</td><td>NA</td></tr>
<tr><td>T29048</td><td>GTAGTAGAAGTCTCCGATCATGG</td><td>1000</td><td></td><td></td><td></td><td></td><td></td><td></td><td></td><td></td><td></td></tr>
<tr><td>Oligo fwd</td><td>CACCGTAGTAGAAGTCTCCGATCA</td><td></td><td></td><td></td><td></td><td></td><td></td><td></td><td></td><td></td><td></td></tr>
<tr><td>Oligo rev</td><td>AAACTGATCGGAGACTTCTACTAC</td><td></td><td></td><td></td><td></td><td></td><td></td><td></td><td></td><td></td><td></td></tr>
<tr><td>Chromosome</td><td>start</td><td>end</td><td>strand</td><td>MM</td><td>target_seq</td><td>PAM</td><td>alignment</td><td>distance</td><td>position</td><td>gene name</td><td>gene id</td></tr>
<tr><td>GU305914.1</td><td>67602</td><td>67624</td><td>-</td><td>0</td><td>GTAGTAGAAGTCTCCGATCA</td><td>TGG</td><td>||||||||[||||||||||||]PAM</td><td>NA</td><td>Intergenic</td><td>NA</td><td>NA</td></tr>
<tr><td>T29049</td><td>CGACAGCGCCGAGATCGCCTCGG</td><td>1000</td><td></td><td></td><td></td><td></td><td></td><td></td><td></td><td></td><td></td></tr>
<tr><td>Oligo fwd</td><td>CACCCGACAGCGCCGAGATCGCCT</td><td></td><td></td><td></td><td></td><td></td><td></td><td></td><td></td><td></td><td></td></tr>
<tr><td>Oligo rev</td><td>AAACAGGCGATCTCGGCGCTGTCG</td><td></td><td></td><td></td><td></td><td></td><td></td><td></td><td></td><td></td><td></td></tr>
<tr><td>Chromosome</td><td>start</td><td>end</td><td>strand</td><td>MM</td><td>target_seq</td><td>PAM</td><td>alignment</td><td>distance</td><td>position</td><td>gene name</td><td>gene id</td></tr>
<tr><td>GU305914.1</td><td>67569</td><td>67591</td><td>-</td><td>0</td><td>CGACAGCGCCGAGATCGCCT</td><td>CGG</td><td>||||||||[||||||||||||]PAM</td><td>NA</td><td>Intergenic</td><td>NA</td><td>NA</td></tr>
<tr><td>T29050</td><td>GCACACGTACACCGCCGCCTCGG</td><td>1000</td><td></td><td></td><td></td><td></td><td></td><td></td><td></td><td></td><td></td></tr>
<tr><td>Oligo fwd</td><td>CACCGCACACGTACACCGCCGCCT</td><td></td><td></td><td></td><td></td><td></td><td></td><td></td><td></td><td></td><td></td></tr>
<tr><td>Oligo rev</td><td>AAACAGGCGGCGGTGTACGTGTGC</td><td></td><td></td><td></td><td></td><td></td><td></td><td></td><td></td><td></td><td></td></tr>
<tr><td>Chromosome</td><td>start</td><td>end</td><td>strand</td><td>MM</td><td>target_seq</td><td>PAM</td><td>alignment</td><td>distance</td><td>position</td><td>gene name</td><td>gene id</td></tr>
<tr><td>GU305914.1</td><td>67536</td><td>67558</td><td>-</td><td>0</td><td>GCACACGTACACCGCCGCCT</td><td>CGG</td><td>||||||||[||||||||||||]PAM</td><td>NA</td><td>Intergenic</td><td>NA</td><td>NA</td></tr>
<tr><td>T29051</td><td>ACACCGCCGCCTCGGCCGACAGG</td><td>1000</td><td></td><td></td><td></td><td></td><td></td><td></td><td></td><td></td><td></td></tr>
<tr><td>Oligo fwd</td><td>CACCACACCGCCGCCTCGGCCGAC</td><td></td><td></td><td></td><td></td><td></td><td></td><td></td><td></td><td></td><td></td></tr>
<tr><td>Oligo rev</td><td>AAACGTCGGCCGAGGCGGCGGTGT</td><td></td><td></td><td></td><td></td><td></td><td></td><td></td><td></td><td></td><td></td></tr>
<tr><td>Chromosome</td><td>start</td><td>end</td><td>strand</td><td>MM</td><td>target_seq</td><td>PAM</td><td>alignment</td><td>distance</td><td>position</td><td>gene name</td><td>gene id</td></tr>
<tr><td>GU305914.1</td><td>67528</td><td>67550</td><td>-</td><td>0</td><td>ACACCGCCGCCTCGGCCGAC</td><td>AGG</td><td>||||||||[||||||||||||]PAM</td><td>NA</td><td>Intergenic</td><td>NA</td><td>NA</td></tr>
<tr><td>T29052</td><td>GCCGACAGGTCCGTCGTGTGCGG</td><td>1000</td><td></td><td></td><td></td><td></td><td></td><td></td><td></td><td></td><td></td></tr>
<tr><td>Oligo fwd</td><td>CACCGCCGACAGGTCCGTCGTGTG</td><td></td><td></td><td></td><td></td><td></td><td></td><td></td><td></td><td></td><td></td></tr>
<tr><td>Oligo rev</td><td>AAACCACACGACGGACCTGTCGGC</td><td></td><td></td><td></td><td></td><td></td><td></td><td></td><td></td><td></td><td></td></tr>
<tr><td>Chromosome</td><td>start</td><td>end</td><td>strand</td><td>MM</td><td>target_seq</td><td>PAM</td><td>alignment</td><td>distance</td><td>position</td><td>gene name</td><td>gene id</td></tr>
</table>

<table>
<tr><td>GU305914.1</td><td></td><td>67514</td><td>67536</td><td>-</td><td>0</td><td>GCCGACAGGTCCGTCGTGTG</td><td>CGG</td><td>||||||||[||||||||||||]PAM</td><td>NA</td><td>Intergenic</td><td>NA</td><td>NA</td></tr>
<tr><td>T29053</td><td>CCGACAGGTCCGTCGTGTGCGGG</td><td></td><td>1000</td><td></td><td></td><td></td><td></td><td></td><td></td><td></td><td></td><td></td></tr>
<tr><td>Oligo fwd</td><td>CACCCCGACAGGTCCGTCGTGTGC</td><td></td><td></td><td></td><td></td><td></td><td></td><td></td><td></td><td></td><td></td><td></td></tr>
<tr><td>Oligo rev</td><td>AAACGCACACGACGGACCTGTCGG</td><td></td><td></td><td></td><td></td><td></td><td></td><td></td><td></td><td></td><td></td><td></td></tr>
<tr><td>Chromosome</td><td>start</td><td></td><td>end</td><td>strand</td><td>MM</td><td>target_seq</td><td>PAM</td><td>alignment</td><td>distance</td><td>position</td><td>gene name</td><td>gene id</td></tr>
<tr><td>GU305914.1</td><td></td><td>67513</td><td>67535</td><td>-</td><td>0</td><td>CCGACAGGTCCGTCGTGTGC</td><td>GGG</td><td>||||||||[||||||||||||]PAM</td><td>NA</td><td>Intergenic</td><td>NA</td><td>NA</td></tr>
<tr><td>T29054</td><td>CGTGTGCGGGTCGAACACGTAGG</td><td></td><td>1000</td><td></td><td></td><td></td><td></td><td></td><td></td><td></td><td></td><td></td></tr>
<tr><td>Oligo fwd</td><td>CACCCGTGTGCGGGTCGAACACGT</td><td></td><td></td><td></td><td></td><td></td><td></td><td></td><td></td><td></td><td></td><td></td></tr>
<tr><td>Oligo rev</td><td>AAACACGTGTTCGACCCGCACACG</td><td></td><td></td><td></td><td></td><td></td><td></td><td></td><td></td><td></td><td></td><td></td></tr>
<tr><td>Chromosome</td><td>start</td><td></td><td>end</td><td>strand</td><td>MM</td><td>target_seq</td><td>PAM</td><td>alignment</td><td>distance</td><td>position</td><td>gene name</td><td>gene id</td></tr>
<tr><td>GU305914.1</td><td></td><td>67500</td><td>67522</td><td>-</td><td>0</td><td>CGTGTGCGGGTCGAACACGT</td><td>AGG</td><td>||||||||[||||||||||||]PAM</td><td>NA</td><td>Intergenic</td><td>NA</td><td>NA</td></tr>
<tr><td>T29055</td><td>ACCGTCACGATGATGTACACCGG</td><td></td><td>1000</td><td></td><td></td><td></td><td></td><td></td><td></td><td></td><td></td><td></td></tr>
<tr><td>Oligo fwd</td><td>CACCACCGTCACGATGATGTACAC</td><td></td><td></td><td></td><td></td><td></td><td></td><td></td><td></td><td></td><td></td><td></td></tr>
<tr><td>Oligo rev</td><td>AAACGTGTACATCATCGTGACGGT</td><td></td><td></td><td></td><td></td><td></td><td></td><td></td><td></td><td></td><td></td><td></td></tr>
<tr><td>Chromosome</td><td>start</td><td></td><td>end</td><td>strand</td><td>MM</td><td>target_seq</td><td>PAM</td><td>alignment</td><td>distance</td><td>position</td><td>gene name</td><td>gene id</td></tr>
<tr><td>GU305914.1</td><td></td><td>67436</td><td>67458</td><td>-</td><td>0</td><td>ACCGTCACGATGATGTACAC</td><td>CGG</td><td>||||||||[||||||||||||]PAM</td><td>NA</td><td>Intergenic</td><td>NA</td><td>NA</td></tr>
<tr><td>T29056</td><td>ACGATGATGTACACCGGCAGCGG</td><td></td><td>1000</td><td></td><td></td><td></td><td></td><td></td><td></td><td></td><td></td><td></td></tr>
<tr><td>Oligo fwd</td><td>CACCACGATGATGTACACCGGCAG</td><td></td><td></td><td></td><td></td><td></td><td></td><td></td><td></td><td></td><td></td><td></td></tr>
<tr><td>Oligo rev</td><td>AAACCTGCCGGTGTACATCATCGT</td><td></td><td></td><td></td><td></td><td></td><td></td><td></td><td></td><td></td><td></td><td></td></tr>
<tr><td>Chromosome</td><td>start</td><td></td><td>end</td><td>strand</td><td>MM</td><td>target_seq</td><td>PAM</td><td>alignment</td><td>distance</td><td>position</td><td>gene name</td><td>gene id</td></tr>
<tr><td>GU305914.1</td><td></td><td>67430</td><td>67452</td><td>-</td><td>0</td><td>ACGATGATGTACACCGGCAG</td><td>CGG</td><td>||||||||[||||||||||||]PAM</td><td>NA</td><td>Intergenic</td><td>NA</td><td>NA</td></tr>
<tr><td>T29057</td><td>GCTTCCCTCTCGCGTACGTCAGG</td><td></td><td>1000</td><td></td><td></td><td></td><td></td><td></td><td></td><td></td><td></td><td></td></tr>
<tr><td>Oligo fwd</td><td>CACCGCTTCCCTCTCGCGTACGTC</td><td></td><td></td><td></td><td></td><td></td><td></td><td></td><td></td><td></td><td></td><td></td></tr>
<tr><td>Oligo rev</td><td>AAACGACGTACGCGAGAGGGAAGC</td><td></td><td></td><td></td><td></td><td></td><td></td><td></td><td></td><td></td><td></td><td></td></tr>
<tr><td>Chromosome</td><td>start</td><td></td><td>end</td><td>strand</td><td>MM</td><td>target_seq</td><td>PAM</td><td>alignment</td><td>distance</td><td>position</td><td>gene name</td><td>gene id</td></tr>
<tr><td>GU305914.1</td><td></td><td>67408</td><td>67430</td><td>-</td><td>0</td><td>GCTTCCCTCTCGCGTACGTC</td><td>AGG</td><td>||||||||[||||||||||||]PAM</td><td>NA</td><td>Intergenic</td><td>NA</td><td>NA</td></tr>
<tr><td>T29058</td><td>TCGCGTACGTCAGGAAGTCCAGG</td><td></td><td>1000</td><td></td><td></td><td></td><td></td><td></td><td></td><td></td><td></td><td></td></tr>
<tr><td>Oligo fwd</td><td>CACCTCGCGTACGTCAGGAAGTCC</td><td></td><td></td><td></td><td></td><td></td><td></td><td></td><td></td><td></td><td></td><td></td></tr>
<tr><td>Oligo rev</td><td>AAACGGACTTCCTGACGTACGCGA</td><td></td><td></td><td></td><td></td><td></td><td></td><td></td><td></td><td></td><td></td><td></td></tr>
<tr><td>Chromosome</td><td>start</td><td></td><td>end</td><td>strand</td><td>MM</td><td>target_seq</td><td>PAM</td><td>alignment</td><td>distance</td><td>position</td><td>gene name</td><td>gene id</td></tr>
<tr><td>GU305914.1</td><td></td><td>67399</td><td>67421</td><td>-</td><td>0</td><td>TCGCGTACGTCAGGAAGTCC</td><td>AGG</td><td>||||||||[||||||||||||]PAM</td><td>NA</td><td>Intergenic</td><td>NA</td><td>NA</td></tr>
<tr><td>GU305914.1</td><td></td><td>229096</td><td>229118</td><td>-</td><td>4</td><td>CCGCGTCCGTCATGAGGTCC</td><td>AGG</td><td>-|||||-|[||||-||-||||]PAM</td><td>NA</td><td>Intergenic</td><td>NA</td><td>NA</td></tr>
<tr><td>T29059</td><td>TCAGGAAGTCCAGGATGCCCAGG</td><td></td><td>1000</td><td></td><td></td><td></td><td></td><td></td><td></td><td></td><td></td><td></td></tr>
<tr><td>Oligo fwd</td><td>CACCTCAGGAAGTCCAGGATGCCC</td><td></td><td></td><td></td><td></td><td></td><td></td><td></td><td></td><td></td><td></td><td></td></tr>
<tr><td>Oligo rev</td><td>AAACGGGCATCCTGGACTTCCTGA</td><td></td><td></td><td></td><td></td><td></td><td></td><td></td><td></td><td></td><td></td><td></td></tr>
<tr><td>Chromosome</td><td>start</td><td></td><td>end</td><td>strand</td><td>MM</td><td>target_seq</td><td>PAM</td><td>alignment</td><td>distance</td><td>position</td><td>gene name</td><td>gene id</td></tr>
<tr><td>GU305914.1</td><td></td><td>67390</td><td>67412</td><td>-</td><td>0</td><td>TCAGGAAGTCCAGGATGCCC</td><td>AGG</td><td>||||||||[||||||||||||]PAM</td><td>NA</td><td>Intergenic</td><td>NA</td><td>NA</td></tr>
<tr><td>T29060</td><td>CCAGGCACTTGTAGCCCCCCAGG</td><td></td><td>1000</td><td></td><td></td><td></td><td></td><td></td><td></td><td></td><td></td><td></td></tr>
<tr><td>Oligo fwd</td><td>CACCCCAGGCACTTGTAGCCCCCC</td><td></td><td></td><td></td><td></td><td></td><td></td><td></td><td></td><td></td><td></td><td></td></tr>
<tr><td>Oligo rev</td><td>AAACGGGGGGCTACAAGTGCCTGG</td><td></td><td></td><td></td><td></td><td></td><td></td><td></td><td></td><td></td><td></td><td></td></tr>
<tr><td>Chromosome</td><td>start</td><td></td><td>end</td><td>strand</td><td>MM</td><td>target_seq</td><td>PAM</td><td>alignment</td><td>distance</td><td>position</td><td>gene name</td><td>gene id</td></tr>
<tr><td>GU305914.1</td><td></td><td>67372</td><td>67394</td><td>-</td><td>0</td><td>CCAGGCACTTGTAGCCCCCC</td><td>AGG</td><td>||||||||[||||||||||||]PAM</td><td>NA</td><td>Intergenic</td><td>NA</td><td>NA</td></tr>
<tr><td>T29061</td><td>GTCTCCGCCGTGCCGTTGAACGG</td><td></td><td>1000</td><td></td><td></td><td></td><td></td><td></td><td></td><td></td><td></td><td></td></tr>
<tr><td>Oligo fwd</td><td>CACCGTCTCCGCCGTGCCGTTGAA</td><td></td><td></td><td></td><td></td><td></td><td></td><td></td><td></td><td></td><td></td><td></td></tr>
<tr><td>Oligo rev</td><td>AAACTTCAACGGCACGGCGGAGAC</td><td></td><td></td><td></td><td></td><td></td><td></td><td></td><td></td><td></td><td></td><td></td></tr>
<tr><td>Chromosome</td><td>start</td><td></td><td>end</td><td>strand</td><td>MM</td><td>target_seq</td><td>PAM</td><td>alignment</td><td>distance</td><td>position</td><td>gene name</td><td>gene id</td></tr>
<tr><td>GU305914.1</td><td></td><td>67343</td><td>67365</td><td>-</td><td>0</td><td>GTCTCCGCCGTGCCGTTGAA</td><td>CGG</td><td>||||||||[||||||||||||]PAM</td><td>NA</td><td>Intergenic</td><td>NA</td><td>NA</td></tr>
<tr><td>T29062</td><td>CGCGTGCCCCGTGATACCCGCGG</td><td></td><td>1000</td><td></td><td></td><td></td><td></td><td></td><td></td><td></td><td></td><td></td></tr>
<tr><td>Oligo fwd</td><td>CACCCGCGTGCCCCGTGATACCCG</td><td></td><td></td><td></td><td></td><td></td><td></td><td></td><td></td><td></td><td></td><td></td></tr>
<tr><td>Oligo rev</td><td>AAACCGGGTATCACGGGGCACGCG</td><td></td><td></td><td></td><td></td><td></td><td></td><td></td><td></td><td></td><td></td><td></td></tr>
<tr><td>Chromosome</td><td>start</td><td></td><td>end</td><td>strand</td><td>MM</td><td>target_seq</td><td>PAM</td><td>alignment</td><td>distance</td><td>position</td><td>gene name</td><td>gene id</td></tr>
<tr><td>GU305914.1</td><td></td><td>67311</td><td>67333</td><td>-</td><td>0</td><td>CGCGTGCCCCGTGATACCCG</td><td>CGG</td><td>||||||||[||||||||||||]PAM</td><td>NA</td><td>Intergenic</td><td>NA</td><td>NA</td></tr>
<tr><td>T29063</td><td>GCGTGCCCCGTGATACCCGCGGG</td><td></td><td>1000</td><td></td><td></td><td></td><td></td><td></td><td></td><td></td><td></td><td></td></tr>
<tr><td>Oligo fwd</td><td>CACCGCGTGCCCCGTGATACCCGC</td><td></td><td></td><td></td><td></td><td></td><td></td><td></td><td></td><td></td><td></td><td></td></tr>
<tr><td>Oligo rev</td><td>AAACGCGGGTATCACGGGGCACGC</td><td></td><td></td><td></td><td></td><td></td><td></td><td></td><td></td><td></td><td></td><td></td></tr>
<tr><td>Chromosome</td><td>start</td><td></td><td>end</td><td>strand</td><td>MM</td><td>target_seq</td><td>PAM</td><td>alignment</td><td>distance</td><td>position</td><td>gene name</td><td>gene id</td></tr>
<tr><td>GU305914.1</td><td></td><td>67310</td><td>67332</td><td>-</td><td>0</td><td>GCGTGCCCCGTGATACCCGC</td><td>GGG</td><td>||||||||[||||||||||||]PAM</td><td>NA</td><td>Intergenic</td><td>NA</td><td>NA</td></tr>
<tr><td>T29064</td><td>CGCGGGCGTCTCGATCACCGTGG</td><td></td><td>1000</td><td></td><td></td><td></td><td></td><td></td><td></td><td></td><td></td><td></td></tr>
<tr><td>Oligo fwd</td><td>CACCCGCGGGCGTCTCGATCACCG</td><td></td><td></td><td></td><td></td><td></td><td></td><td></td><td></td><td></td><td></td><td></td></tr>
<tr><td>Oligo rev</td><td>AAACCGGTGATCGAGACGCCCGCG</td><td></td><td></td><td></td><td></td><td></td><td></td><td></td><td></td><td></td><td></td><td></td></tr>
<tr><td>Chromosome</td><td>start</td><td></td><td>end</td><td>strand</td><td>MM</td><td>target_seq</td><td>PAM</td><td>alignment</td><td>distance</td><td>position</td><td>gene name</td><td>gene id</td></tr>
<tr><td>GU305914.1</td><td></td><td>67293</td><td>67315</td><td>-</td><td>0</td><td>CGCGGGCGTCTCGATCACCG</td><td>TGG</td><td>||||||||[||||||||||||]PAM</td><td>NA</td><td>Intergenic</td><td>NA</td><td>NA</td></tr>
<tr><td>T29065</td><td>GCGGGCGTCTCGATCACCGTGGG</td><td></td><td>1000</td><td></td><td></td><td></td><td></td><td></td><td></td><td></td><td></td><td></td></tr>
<tr><td>Oligo fwd</td><td>CACCGCGGGCGTCTCGATCACCGT</td><td></td><td></td><td></td><td></td><td></td><td></td><td></td><td></td><td></td><td></td><td></td></tr>
<tr><td>Oligo rev</td><td>AAACACGGTGATCGAGACGCCCGC</td><td></td><td></td><td></td><td></td><td></td><td></td><td></td><td></td><td></td><td></td><td></td></tr>
<tr><td>Chromosome</td><td>start</td><td></td><td>end</td><td>strand</td><td>MM</td><td>target_seq</td><td>PAM</td><td>alignment</td><td>distance</td><td>position</td><td>gene name</td><td>gene id</td></tr>
<tr><td>GU305914.1</td><td></td><td>67292</td><td>67314</td><td>-</td><td>0</td><td>GCGGGCGTCTCGATCACCGT</td><td>GGG</td><td>||||||||[||||||||||||]PAM</td><td>NA</td><td>Intergenic</td><td>NA</td><td>NA</td></tr>
<tr><td>T29066</td><td>CGGGCGTCTCGATCACCGTGGGG</td><td></td><td>1000</td><td></td><td></td><td></td><td></td><td></td><td></td><td></td><td></td><td></td></tr>
<tr><td>Oligo fwd</td><td>CACCCGGGCGTCTCGATCACCGTG</td><td></td><td></td><td></td><td></td><td></td><td></td><td></td><td></td><td></td><td></td><td></td></tr>
<tr><td>Oligo rev</td><td>AAACCACGGTGATCGAGACGCCCG</td><td></td><td></td><td></td><td></td><td></td><td></td><td></td><td></td><td></td><td></td><td></td></tr>
<tr><td>Chromosome</td><td>start</td><td></td><td>end</td><td>strand</td><td>MM</td><td>target_seq</td><td>PAM</td><td>alignment</td><td>distance</td><td>position</td><td>gene name</td><td>gene id</td></tr>
<tr><td>GU305914.1</td><td></td><td>67291</td><td>67313</td><td>-</td><td>0</td><td>CGGGCGTCTCGATCACCGTG</td><td>GGG</td><td>||||||||[||||||||||||]PAM</td><td>NA</td><td>Intergenic</td><td>NA</td><td>NA</td></tr>
<tr><td>T29067</td><td>TGGGGATCTCCGTCCCCAGACGG</td><td></td><td>1000</td><td></td><td></td><td></td><td></td><td></td><td></td><td></td><td></td><td></td></tr>
<tr><td>Oligo fwd</td><td>CACCTGGGGATCTCCGTCCCCAGA</td><td></td><td></td><td></td><td></td><td></td><td></td><td></td><td></td><td></td><td></td><td></td></tr>
</table>

<table>
<tr><td>Oligo rev</td><td>AAACTCTGGGGACGGAGATCCCCA</td><td></td><td></td><td></td><td></td><td></td><td></td><td></td><td></td><td></td><td></td></tr>
<tr><td>Chromosome</td><td>start</td><td>end</td><td>strand</td><td>MM</td><td>target_seq</td><td>PAM</td><td>alignment</td><td>distance</td><td>position</td><td>gene name</td><td>gene id</td></tr>
<tr><td>GU305914.1</td><td>67273</td><td>67295</td><td>-</td><td>0</td><td>TGGGGATCTCCGTCCCCAGA</td><td>CGG</td><td>||||||||[||||||||||||]PAM</td><td>NA</td><td>Intergenic</td><td>NA</td><td>NA</td></tr>
<tr><td>T29068</td><td>ATCTCCGTCCCCAGACGGTACGG</td><td>1000</td><td></td><td></td><td></td><td></td><td></td><td></td><td></td><td></td><td></td></tr>
<tr><td>Oligo fwd</td><td>CACCATCTCCGTCCCCAGACGGTA</td><td></td><td></td><td></td><td></td><td></td><td></td><td></td><td></td><td></td><td></td></tr>
<tr><td>Oligo rev</td><td>AAACTACCGTCTGGGGACGGAGAT</td><td></td><td></td><td></td><td></td><td></td><td></td><td></td><td></td><td></td><td></td></tr>
<tr><td>Chromosome</td><td>start</td><td>end</td><td>strand</td><td>MM</td><td>target_seq</td><td>PAM</td><td>alignment</td><td>distance</td><td>position</td><td>gene name</td><td>gene id</td></tr>
<tr><td>GU305914.1</td><td>67268</td><td>67290</td><td>-</td><td>0</td><td>ATCTCCGTCCCCAGACGGTA</td><td>CGG</td><td>||||||||[||||||||||||]PAM</td><td>NA</td><td>Intergenic</td><td>NA</td><td>NA</td></tr>
<tr><td>T29069</td><td>TCTCCGTCCCCAGACGGTACGGG</td><td>1000</td><td></td><td></td><td></td><td></td><td></td><td></td><td></td><td></td><td></td></tr>
<tr><td>Oligo fwd</td><td>CACCTCTCCGTCCCCAGACGGTAC</td><td></td><td></td><td></td><td></td><td></td><td></td><td></td><td></td><td></td><td></td></tr>
<tr><td>Oligo rev</td><td>AAACGTACCGTCTGGGGACGGAGA</td><td></td><td></td><td></td><td></td><td></td><td></td><td></td><td></td><td></td><td></td></tr>
<tr><td>Chromosome</td><td>start</td><td>end</td><td>strand</td><td>MM</td><td>target_seq</td><td>PAM</td><td>alignment</td><td>distance</td><td>position</td><td>gene name</td><td>gene id</td></tr>
<tr><td>GU305914.1</td><td>67267</td><td>67289</td><td>-</td><td>0</td><td>TCTCCGTCCCCAGACGGTAC</td><td>GGG</td><td>||||||||[||||||||||||]PAM</td><td>NA</td><td>Intergenic</td><td>NA</td><td>NA</td></tr>
<tr><td>T29070</td><td>CCGTCCCCAGACGGTACGGGTGG</td><td>1000</td><td></td><td></td><td></td><td></td><td></td><td></td><td></td><td></td><td></td></tr>
<tr><td>Oligo fwd</td><td>CACCCCGTCCCCAGACGGTACGGG</td><td></td><td></td><td></td><td></td><td></td><td></td><td></td><td></td><td></td><td></td></tr>
<tr><td>Oligo rev</td><td>AAACCCCGTACCGTCTGGGGACGG</td><td></td><td></td><td></td><td></td><td></td><td></td><td></td><td></td><td></td><td></td></tr>
<tr><td>Chromosome</td><td>start</td><td>end</td><td>strand</td><td>MM</td><td>target_seq</td><td>PAM</td><td>alignment</td><td>distance</td><td>position</td><td>gene name</td><td>gene id</td></tr>
<tr><td>GU305914.1</td><td>67264</td><td>67286</td><td>-</td><td>0</td><td>CCGTCCCCAGACGGTACGGG</td><td>TGG</td><td>||||||||[||||||||||||]PAM</td><td>NA</td><td>Intergenic</td><td>NA</td><td>NA</td></tr>
<tr><td>T29071</td><td>TCTCGTCCGCGATCGCGTCCAGG</td><td>1000</td><td></td><td></td><td></td><td></td><td></td><td></td><td></td><td></td><td></td></tr>
<tr><td>Oligo fwd</td><td>CACCTCTCGTCCGCGATCGCGTCC</td><td></td><td></td><td></td><td></td><td></td><td></td><td></td><td></td><td></td><td></td></tr>
<tr><td>Oligo rev</td><td>AAACGGACGCGATCGCGGACGAGA</td><td></td><td></td><td></td><td></td><td></td><td></td><td></td><td></td><td></td><td></td></tr>
<tr><td>Chromosome</td><td>start</td><td>end</td><td>strand</td><td>MM</td><td>target_seq</td><td>PAM</td><td>alignment</td><td>distance</td><td>position</td><td>gene name</td><td>gene id</td></tr>
<tr><td>GU305914.1</td><td>67216</td><td>67238</td><td>-</td><td>0</td><td>TCTCGTCCGCGATCGCGTCC</td><td>AGG</td><td>||||||||[||||||||||||]PAM</td><td>NA</td><td>Intergenic</td><td>NA</td><td>NA</td></tr>
<tr><td>T29072</td><td>GCGCGCGCCGCTGTCCAGCACGG</td><td>1000</td><td></td><td></td><td></td><td></td><td></td><td></td><td></td><td></td><td></td></tr>
<tr><td>Oligo fwd</td><td>CACCGCGCGCGCCGCTGTCCAGCA</td><td></td><td></td><td></td><td></td><td></td><td></td><td></td><td></td><td></td><td></td></tr>
<tr><td>Oligo rev</td><td>AAACTGCTGGACAGCGGCGCGCGC</td><td></td><td></td><td></td><td></td><td></td><td></td><td></td><td></td><td></td><td></td></tr>
<tr><td>Chromosome</td><td>start</td><td>end</td><td>strand</td><td>MM</td><td>target_seq</td><td>PAM</td><td>alignment</td><td>distance</td><td>position</td><td>gene name</td><td>gene id</td></tr>
<tr><td>GU305914.1</td><td>67194</td><td>67216</td><td>-</td><td>0</td><td>GCGCGCGCCGCTGTCCAGCA</td><td>CGG</td><td>||||||||[||||||||||||]PAM</td><td>NA</td><td>Intergenic</td><td>NA</td><td>NA</td></tr>
<tr><td>T29073</td><td>AGCGACGTCGTGTCCAGCACCGG</td><td>1000</td><td></td><td></td><td></td><td></td><td></td><td></td><td></td><td></td><td></td></tr>
<tr><td>Oligo fwd</td><td>CACCAGCGACGTCGTGTCCAGCAC</td><td></td><td></td><td></td><td></td><td></td><td></td><td></td><td></td><td></td><td></td></tr>
<tr><td>Oligo rev</td><td>AAACGTGCTGGACACGACGTCGCT</td><td></td><td></td><td></td><td></td><td></td><td></td><td></td><td></td><td></td><td></td></tr>
<tr><td>Chromosome</td><td>start</td><td>end</td><td>strand</td><td>MM</td><td>target_seq</td><td>PAM</td><td>alignment</td><td>distance</td><td>position</td><td>gene name</td><td>gene id</td></tr>
<tr><td>GU305914.1</td><td>67166</td><td>67188</td><td>-</td><td>0</td><td>AGCGACGTCGTGTCCAGCAC</td><td>CGG</td><td>||||||||[||||||||||||]PAM</td><td>NA</td><td>Intergenic</td><td>NA</td><td>NA</td></tr>
<tr><td>T29074</td><td>GCACCGGATCGATGCCCATCAGG</td><td>1000</td><td></td><td></td><td></td><td></td><td></td><td></td><td></td><td></td><td></td></tr>
<tr><td>Oligo fwd</td><td>CACCGCACCGGATCGATGCCCATC</td><td></td><td></td><td></td><td></td><td></td><td></td><td></td><td></td><td></td><td></td></tr>
<tr><td>Oligo rev</td><td>AAACGATGGGCATCGATCCGGTGC</td><td></td><td></td><td></td><td></td><td></td><td></td><td></td><td></td><td></td><td></td></tr>
<tr><td>Chromosome</td><td>start</td><td>end</td><td>strand</td><td>MM</td><td>target_seq</td><td>PAM</td><td>alignment</td><td>distance</td><td>position</td><td>gene name</td><td>gene id</td></tr>
<tr><td>GU305914.1</td><td>67150</td><td>67172</td><td>-</td><td>0</td><td>GCACCGGATCGATGCCCATC</td><td>AGG</td><td>||||||||[||||||||||||]PAM</td><td>NA</td><td>Intergenic</td><td>NA</td><td>NA</td></tr>
<tr><td>T29075</td><td>ATCGATGCCCATCAGGTAACAGG</td><td>1000</td><td></td><td></td><td></td><td></td><td></td><td></td><td></td><td></td><td></td></tr>
<tr><td>Oligo fwd</td><td>CACCATCGATGCCCATCAGGTAAC</td><td></td><td></td><td></td><td></td><td></td><td></td><td></td><td></td><td></td><td></td></tr>
<tr><td>Oligo rev</td><td>AAACGTTACCTGATGGGCATCGAT</td><td></td><td></td><td></td><td></td><td></td><td></td><td></td><td></td><td></td><td></td></tr>
<tr><td>Chromosome</td><td>start</td><td>end</td><td>strand</td><td>MM</td><td>target_seq</td><td>PAM</td><td>alignment</td><td>distance</td><td>position</td><td>gene name</td><td>gene id</td></tr>
<tr><td>GU305914.1</td><td>67143</td><td>67165</td><td>-</td><td>0</td><td>ATCGATGCCCATCAGGTAAC</td><td>AGG</td><td>||||||||[||||||||||||]PAM</td><td>NA</td><td>Intergenic</td><td>NA</td><td>NA</td></tr>
<tr><td>T29076</td><td>CCATCAGGTAACAGGTGTGTAGG</td><td>1000</td><td></td><td></td><td></td><td></td><td></td><td></td><td></td><td></td><td></td></tr>
<tr><td>Oligo fwd</td><td>CACCCCATCAGGTAACAGGTGTGT</td><td></td><td></td><td></td><td></td><td></td><td></td><td></td><td></td><td></td><td></td></tr>
<tr><td>Oligo rev</td><td>AAACACACACCTGTTACCTGATGG</td><td></td><td></td><td></td><td></td><td></td><td></td><td></td><td></td><td></td><td></td></tr>
<tr><td>Chromosome</td><td>start</td><td>end</td><td>strand</td><td>MM</td><td>target_seq</td><td>PAM</td><td>alignment</td><td>distance</td><td>position</td><td>gene name</td><td>gene id</td></tr>
<tr><td>GU305914.1</td><td>67135</td><td>67157</td><td>-</td><td>0</td><td>CCATCAGGTAACAGGTGTGT</td><td>AGG</td><td>||||||||[||||||||||||]PAM</td><td>NA</td><td>Intergenic</td><td>NA</td><td>NA</td></tr>
<tr><td>T29077</td><td>CAGGTAACAGGTGTGTAGGAAGG</td><td>1000</td><td></td><td></td><td></td><td></td><td></td><td></td><td></td><td></td><td></td></tr>
<tr><td>Oligo fwd</td><td>CACCCAGGTAACAGGTGTGTAGGA</td><td></td><td></td><td></td><td></td><td></td><td></td><td></td><td></td><td></td><td></td></tr>
<tr><td>Oligo rev</td><td>AAACTCCTACACACCTGTTACCTG</td><td></td><td></td><td></td><td></td><td></td><td></td><td></td><td></td><td></td><td></td></tr>
<tr><td>Chromosome</td><td>start</td><td>end</td><td>strand</td><td>MM</td><td>target_seq</td><td>PAM</td><td>alignment</td><td>distance</td><td>position</td><td>gene name</td><td>gene id</td></tr>
<tr><td>GU305914.1</td><td>67131</td><td>67153</td><td>-</td><td>0</td><td>CAGGTAACAGGTGTGTAGGA</td><td>AGG</td><td>||||||||[||||||||||||]PAM</td><td>NA</td><td>Intergenic</td><td>NA</td><td>NA</td></tr>
<tr><td>T29078</td><td>TAAAGCAGTTCGACATGCACTGG</td><td>1000</td><td></td><td></td><td></td><td></td><td></td><td></td><td></td><td></td><td></td></tr>
<tr><td>Oligo fwd</td><td>CACCTAAAGCAGTTCGACATGCAC</td><td></td><td></td><td></td><td></td><td></td><td></td><td></td><td></td><td></td><td></td></tr>
<tr><td>Oligo rev</td><td>AAACGTGCATGTCGAACTGCTTTA</td><td></td><td></td><td></td><td></td><td></td><td></td><td></td><td></td><td></td><td></td></tr>
<tr><td>Chromosome</td><td>start</td><td>end</td><td>strand</td><td>MM</td><td>target_seq</td><td>PAM</td><td>alignment</td><td>distance</td><td>position</td><td>gene name</td><td>gene id</td></tr>
<tr><td>GU305914.1</td><td>67108</td><td>67130</td><td>-</td><td>0</td><td>TAAAGCAGTTCGACATGCAC</td><td>TGG</td><td>||||||||[||||||||||||]PAM</td><td>NA</td><td>Intergenic</td><td>NA</td><td>NA</td></tr>
<tr><td>T29079</td><td>ACATGCACTGGCTGCCCGCGCGG</td><td>1000</td><td></td><td></td><td></td><td></td><td></td><td></td><td></td><td></td><td></td></tr>
<tr><td>Oligo fwd</td><td>CACCACATGCACTGGCTGCCCGCG</td><td></td><td></td><td></td><td></td><td></td><td></td><td></td><td></td><td></td><td></td></tr>
<tr><td>Oligo rev</td><td>AAACCGCGGGCAGCCAGTGCATGT</td><td></td><td></td><td></td><td></td><td></td><td></td><td></td><td></td><td></td><td></td></tr>
<tr><td>Chromosome</td><td>start</td><td>end</td><td>strand</td><td>MM</td><td>target_seq</td><td>PAM</td><td>alignment</td><td>distance</td><td>position</td><td>gene name</td><td>gene id</td></tr>
<tr><td>GU305914.1</td><td>67096</td><td>67118</td><td>-</td><td>0</td><td>ACATGCACTGGCTGCCCGCG</td><td>CGG</td><td>||||||||[||||||||||||]PAM</td><td>NA</td><td>Intergenic</td><td>NA</td><td>NA</td></tr>
<tr><td>T29080</td><td>CATGCACTGGCTGCCCGCGCGGG</td><td>1000</td><td></td><td></td><td></td><td></td><td></td><td></td><td></td><td></td><td></td></tr>
<tr><td>Oligo fwd</td><td>CACCCATGCACTGGCTGCCCGCGC</td><td></td><td></td><td></td><td></td><td></td><td></td><td></td><td></td><td></td><td></td></tr>
<tr><td>Oligo rev</td><td>AAACGCGCGGGCAGCCAGTGCATG</td><td></td><td></td><td></td><td></td><td></td><td></td><td></td><td></td><td></td><td></td></tr>
<tr><td>Chromosome</td><td>start</td><td>end</td><td>strand</td><td>MM</td><td>target_seq</td><td>PAM</td><td>alignment</td><td>distance</td><td>position</td><td>gene name</td><td>gene id</td></tr>
<tr><td>GU305914.1</td><td>67095</td><td>67117</td><td>-</td><td>0</td><td>CATGCACTGGCTGCCCGCGC</td><td>GGG</td><td>||||||||[||||||||||||]PAM</td><td>NA</td><td>Intergenic</td><td>NA</td><td>NA</td></tr>
<tr><td>T29081</td><td>ATGCACTGGCTGCCCGCGCGGGG</td><td>1000</td><td></td><td></td><td></td><td></td><td></td><td></td><td></td><td></td><td></td></tr>
<tr><td>Oligo fwd</td><td>CACCATGCACTGGCTGCCCGCGCG</td><td></td><td></td><td></td><td></td><td></td><td></td><td></td><td></td><td></td><td></td></tr>
<tr><td>Oligo rev</td><td>AAACCGCGCGGGCAGCCAGTGCAT</td><td></td><td></td><td></td><td></td><td></td><td></td><td></td><td></td><td></td><td></td></tr>
<tr><td>Chromosome</td><td>start</td><td>end</td><td>strand</td><td>MM</td><td>target_seq</td><td>PAM</td><td>alignment</td><td>distance</td><td>position</td><td>gene name</td><td>gene id</td></tr>
<tr><td>GU305914.1</td><td>67094</td><td>67116</td><td>-</td><td>0</td><td>ATGCACTGGCTGCCCGCGCG</td><td>GGG</td><td>||||||||[||||||||||||]PAM</td><td>NA</td><td>Intergenic</td><td>NA</td><td>NA</td></tr>
<tr><td>T29082</td><td>GCCCGCGCGGGGCCCGTACACGG</td><td>1000</td><td></td><td></td><td></td><td></td><td></td><td></td><td></td><td></td><td></td></tr>
</table>

<table>
<tr><td>Oligo fwd</td><td>CACCGCCCGCGCGGGGCCCGTACA</td><td></td><td></td><td></td><td></td><td></td><td></td><td></td><td></td><td></td><td></td></tr>
<tr><td>Oligo rev</td><td>AAACTGTACGGGCCCCGCGCGGGC</td><td></td><td></td><td></td><td></td><td></td><td></td><td></td><td></td><td></td><td></td></tr>
<tr><td>Chromosome</td><td>start</td><td>end</td><td>strand</td><td>MM</td><td>target_seq</td><td>PAM</td><td>alignment</td><td>distance</td><td>position</td><td>gene name</td><td>gene id</td></tr>
<tr><td>GU305914.1</td><td>67083</td><td>67105</td><td>-</td><td>0</td><td>GCCCGCGCGGGGCCCGTACA</td><td>CGG</td><td>||||||||[||||||||||||]PAM</td><td>NA</td><td>Intergenic</td><td>NA</td><td>NA</td></tr>
<tr><td>T29083</td><td>CCCGCGCGGGGCCCGTACACGGG</td><td>1000</td><td></td><td></td><td></td><td></td><td></td><td></td><td></td><td></td><td></td></tr>
<tr><td>Oligo fwd</td><td>CACCCCCGCGCGGGGCCCGTACAC</td><td></td><td></td><td></td><td></td><td></td><td></td><td></td><td></td><td></td><td></td></tr>
<tr><td>Oligo rev</td><td>AAACGTGTACGGGCCCCGCGCGGG</td><td></td><td></td><td></td><td></td><td></td><td></td><td></td><td></td><td></td><td></td></tr>
<tr><td>Chromosome</td><td>start</td><td>end</td><td>strand</td><td>MM</td><td>target_seq</td><td>PAM</td><td>alignment</td><td>distance</td><td>position</td><td>gene name</td><td>gene id</td></tr>
<tr><td>GU305914.1</td><td>67082</td><td>67104</td><td>-</td><td>0</td><td>CCCGCGCGGGGCCCGTACAC</td><td>GGG</td><td>||||||||[||||||||||||]PAM</td><td>NA</td><td>Intergenic</td><td>NA</td><td>NA</td></tr>
<tr><td>T29084</td><td>GCCCGTACACGGGCGCCGACTGG</td><td>1000</td><td></td><td></td><td></td><td></td><td></td><td></td><td></td><td></td><td></td></tr>
<tr><td>Oligo fwd</td><td>CACCGCCCGTACACGGGCGCCGAC</td><td></td><td></td><td></td><td></td><td></td><td></td><td></td><td></td><td></td><td></td></tr>
<tr><td>Oligo rev</td><td>AAACGTCGGCGCCCGTGTACGGGC</td><td></td><td></td><td></td><td></td><td></td><td></td><td></td><td></td><td></td><td></td></tr>
<tr><td>Chromosome</td><td>start</td><td>end</td><td>strand</td><td>MM</td><td>target_seq</td><td>PAM</td><td>alignment</td><td>distance</td><td>position</td><td>gene name</td><td>gene id</td></tr>
<tr><td>GU305914.1</td><td>67072</td><td>67094</td><td>-</td><td>0</td><td>GCCCGTACACGGGCGCCGAC</td><td>TGG</td><td>||||||||[||||||||||||]PAM</td><td>NA</td><td>Intergenic</td><td>NA</td><td>NA</td></tr>
<tr><td>T29085</td><td>ACACGGGCGCCGACTGGTCACGG</td><td>1000</td><td></td><td></td><td></td><td></td><td></td><td></td><td></td><td></td><td></td></tr>
<tr><td>Oligo fwd</td><td>CACCACACGGGCGCCGACTGGTCA</td><td></td><td></td><td></td><td></td><td></td><td></td><td></td><td></td><td></td><td></td></tr>
<tr><td>Oligo rev</td><td>AAACTGACCAGTCGGCGCCCGTGT</td><td></td><td></td><td></td><td></td><td></td><td></td><td></td><td></td><td></td><td></td></tr>
<tr><td>Chromosome</td><td>start</td><td>end</td><td>strand</td><td>MM</td><td>target_seq</td><td>PAM</td><td>alignment</td><td>distance</td><td>position</td><td>gene name</td><td>gene id</td></tr>
<tr><td>GU305914.1</td><td>67066</td><td>67088</td><td>-</td><td>0</td><td>ACACGGGCGCCGACTGGTCA</td><td>CGG</td><td>||||||||[||||||||||||]PAM</td><td>NA</td><td>Intergenic</td><td>NA</td><td>NA</td></tr>
<tr><td>T29086</td><td>CCGACTGGTCACGGCTTGCGCGG</td><td>1000</td><td></td><td></td><td></td><td></td><td></td><td></td><td></td><td></td><td></td></tr>
<tr><td>Oligo fwd</td><td>CACCCCGACTGGTCACGGCTTGCG</td><td></td><td></td><td></td><td></td><td></td><td></td><td></td><td></td><td></td><td></td></tr>
<tr><td>Oligo rev</td><td>AAACCGCAAGCCGTGACCAGTCGG</td><td></td><td></td><td></td><td></td><td></td><td></td><td></td><td></td><td></td><td></td></tr>
<tr><td>Chromosome</td><td>start</td><td>end</td><td>strand</td><td>MM</td><td>target_seq</td><td>PAM</td><td>alignment</td><td>distance</td><td>position</td><td>gene name</td><td>gene id</td></tr>
<tr><td>GU305914.1</td><td>67057</td><td>67079</td><td>-</td><td>0</td><td>CCGACTGGTCACGGCTTGCG</td><td>CGG</td><td>||||||||[||||||||||||]PAM</td><td>NA</td><td>Intergenic</td><td>NA</td><td>NA</td></tr>
<tr><td>T29087</td><td>TGTCGCCTTCTGTACGTTTCTGG</td><td>1000</td><td></td><td></td><td></td><td></td><td></td><td></td><td></td><td></td><td></td></tr>
<tr><td>Oligo fwd</td><td>CACCTGTCGCCTTCTGTACGTTTC</td><td></td><td></td><td></td><td></td><td></td><td></td><td></td><td></td><td></td><td></td></tr>
<tr><td>Oligo rev</td><td>AAACGAAACGTACAGAAGGCGACA</td><td></td><td></td><td></td><td></td><td></td><td></td><td></td><td></td><td></td><td></td></tr>
<tr><td>Chromosome</td><td>start</td><td>end</td><td>strand</td><td>MM</td><td>target_seq</td><td>PAM</td><td>alignment</td><td>distance</td><td>position</td><td>gene name</td><td>gene id</td></tr>
<tr><td>GU305914.1</td><td>67021</td><td>67043</td><td>-</td><td>0</td><td>TGTCGCCTTCTGTACGTTTC</td><td>TGG</td><td>||||||||[||||||||||||]PAM</td><td>NA</td><td>Intergenic</td><td>NA</td><td>NA</td></tr>
<tr><td>T29088</td><td>CCTTCTGTACGTTTCTGGCCAGG</td><td>1000</td><td></td><td></td><td></td><td></td><td></td><td></td><td></td><td></td><td></td></tr>
<tr><td>Oligo fwd</td><td>CACCCCTTCTGTACGTTTCTGGCC</td><td></td><td></td><td></td><td></td><td></td><td></td><td></td><td></td><td></td><td></td></tr>
<tr><td>Oligo rev</td><td>AAACGGCCAGAAACGTACAGAAGG</td><td></td><td></td><td></td><td></td><td></td><td></td><td></td><td></td><td></td><td></td></tr>
<tr><td>Chromosome</td><td>start</td><td>end</td><td>strand</td><td>MM</td><td>target_seq</td><td>PAM</td><td>alignment</td><td>distance</td><td>position</td><td>gene name</td><td>gene id</td></tr>
<tr><td>GU305914.1</td><td>67016</td><td>67038</td><td>-</td><td>0</td><td>CCTTCTGTACGTTTCTGGCC</td><td>AGG</td><td>||||||||[||||||||||||]PAM</td><td>NA</td><td>Intergenic</td><td>NA</td><td>NA</td></tr>
<tr><td>T29089</td><td>ACGTTTCTGGCCAGGCGTAGCGG</td><td>1000</td><td></td><td></td><td></td><td></td><td></td><td></td><td></td><td></td><td></td></tr>
<tr><td>Oligo fwd</td><td>CACCACGTTTCTGGCCAGGCGTAG</td><td></td><td></td><td></td><td></td><td></td><td></td><td></td><td></td><td></td><td></td></tr>
<tr><td>Oligo rev</td><td>AAACCTACGCCTGGCCAGAAACGT</td><td></td><td></td><td></td><td></td><td></td><td></td><td></td><td></td><td></td><td></td></tr>
<tr><td>Chromosome</td><td>start</td><td>end</td><td>strand</td><td>MM</td><td>target_seq</td><td>PAM</td><td>alignment</td><td>distance</td><td>position</td><td>gene name</td><td>gene id</td></tr>
<tr><td>GU305914.1</td><td>67008</td><td>67030</td><td>-</td><td>0</td><td>ACGTTTCTGGCCAGGCGTAG</td><td>CGG</td><td>||||||||[||||||||||||]PAM</td><td>NA</td><td>Intergenic</td><td>NA</td><td>NA</td></tr>
<tr><td>T29090</td><td>CGTTTCTGGCCAGGCGTAGCGGG</td><td>1000</td><td></td><td></td><td></td><td></td><td></td><td></td><td></td><td></td><td></td></tr>
<tr><td>Oligo fwd</td><td>CACCCGTTTCTGGCCAGGCGTAGC</td><td></td><td></td><td></td><td></td><td></td><td></td><td></td><td></td><td></td><td></td></tr>
<tr><td>Oligo rev</td><td>AAACGCTACGCCTGGCCAGAAACG</td><td></td><td></td><td></td><td></td><td></td><td></td><td></td><td></td><td></td><td></td></tr>
<tr><td>Chromosome</td><td>start</td><td>end</td><td>strand</td><td>MM</td><td>target_seq</td><td>PAM</td><td>alignment</td><td>distance</td><td>position</td><td>gene name</td><td>gene id</td></tr>
<tr><td>GU305914.1</td><td>67007</td><td>67029</td><td>-</td><td>0</td><td>CGTTTCTGGCCAGGCGTAGC</td><td>GGG</td><td>||||||||[||||||||||||]PAM</td><td>NA</td><td>Intergenic</td><td>NA</td><td>NA</td></tr>
<tr><td>T29091</td><td>CCAGGCGTAGCGGGTTCGTCAGG</td><td>1000</td><td></td><td></td><td></td><td></td><td></td><td></td><td></td><td></td><td></td></tr>
<tr><td>Oligo fwd</td><td>CACCCCAGGCGTAGCGGGTTCGTC</td><td></td><td></td><td></td><td></td><td></td><td></td><td></td><td></td><td></td><td></td></tr>
<tr><td>Oligo rev</td><td>AAACGACGAACCCGCTACGCCTGG</td><td></td><td></td><td></td><td></td><td></td><td></td><td></td><td></td><td></td><td></td></tr>
<tr><td>Chromosome</td><td>start</td><td>end</td><td>strand</td><td>MM</td><td>target_seq</td><td>PAM</td><td>alignment</td><td>distance</td><td>position</td><td>gene name</td><td>gene id</td></tr>
<tr><td>GU305914.1</td><td>66998</td><td>67020</td><td>-</td><td>0</td><td>CCAGGCGTAGCGGGTTCGTC</td><td>AGG</td><td>||||||||[||||||||||||]PAM</td><td>NA</td><td>Intergenic</td><td>NA</td><td>NA</td></tr>
<tr><td>T29092</td><td>TCTCCACGTACCACTCGCGCAGG</td><td>1000</td><td></td><td></td><td></td><td></td><td></td><td></td><td></td><td></td><td></td></tr>
<tr><td>Oligo fwd</td><td>CACCTCTCCACGTACCACTCGCGC</td><td></td><td></td><td></td><td></td><td></td><td></td><td></td><td></td><td></td><td></td></tr>
<tr><td>Oligo rev</td><td>AAACGCGCGAGTGGTACGTGGAGA</td><td></td><td></td><td></td><td></td><td></td><td></td><td></td><td></td><td></td><td></td></tr>
<tr><td>Chromosome</td><td>start</td><td>end</td><td>strand</td><td>MM</td><td>target_seq</td><td>PAM</td><td>alignment</td><td>distance</td><td>position</td><td>gene name</td><td>gene id</td></tr>
<tr><td>GU305914.1</td><td>66953</td><td>66975</td><td>-</td><td>0</td><td>TCTCCACGTACCACTCGCGC</td><td>AGG</td><td>||||||||[||||||||||||]PAM</td><td>NA</td><td>Intergenic</td><td>NA</td><td>NA</td></tr>
<tr><td>T29093</td><td>TACCACTCGCGCAGGTCGTTCGG</td><td>1000</td><td></td><td></td><td></td><td></td><td></td><td></td><td></td><td></td><td></td></tr>
<tr><td>Oligo fwd</td><td>CACCTACCACTCGCGCAGGTCGTT</td><td></td><td></td><td></td><td></td><td></td><td></td><td></td><td></td><td></td><td></td></tr>
<tr><td>Oligo rev</td><td>AAACAACGACCTGCGCGAGTGGTA</td><td></td><td></td><td></td><td></td><td></td><td></td><td></td><td></td><td></td><td></td></tr>
<tr><td>Chromosome</td><td>start</td><td>end</td><td>strand</td><td>MM</td><td>target_seq</td><td>PAM</td><td>alignment</td><td>distance</td><td>position</td><td>gene name</td><td>gene id</td></tr>
<tr><td>GU305914.1</td><td>66945</td><td>66967</td><td>-</td><td>0</td><td>TACCACTCGCGCAGGTCGTT</td><td>CGG</td><td>||||||||[||||||||||||]PAM</td><td>NA</td><td>Intergenic</td><td>NA</td><td>NA</td></tr>
<tr><td>T29094</td><td>AGGTCGTTCGGCGATTTCACCGG</td><td>1000</td><td></td><td></td><td></td><td></td><td></td><td></td><td></td><td></td><td></td></tr>
<tr><td>Oligo fwd</td><td>CACCAGGTCGTTCGGCGATTTCAC</td><td></td><td></td><td></td><td></td><td></td><td></td><td></td><td></td><td></td><td></td></tr>
<tr><td>Oligo rev</td><td>AAACGTGAAATCGCCGAACGACCT</td><td></td><td></td><td></td><td></td><td></td><td></td><td></td><td></td><td></td><td></td></tr>
<tr><td>Chromosome</td><td>start</td><td>end</td><td>strand</td><td>MM</td><td>target_seq</td><td>PAM</td><td>alignment</td><td>distance</td><td>position</td><td>gene name</td><td>gene id</td></tr>
<tr><td>GU305914.1</td><td>66933</td><td>66955</td><td>-</td><td>0</td><td>AGGTCGTTCGGCGATTTCAC</td><td>CGG</td><td>||||||||[||||||||||||]PAM</td><td>NA</td><td>Intergenic</td><td>NA</td><td>NA</td></tr>
<tr><td>T29095</td><td>CGTACCTCTCCGCCTCGTACCGG</td><td>1000</td><td></td><td></td><td></td><td></td><td></td><td></td><td></td><td></td><td></td></tr>
<tr><td>Oligo fwd</td><td>CACCCGTACCTCTCCGCCTCGTAC</td><td></td><td></td><td></td><td></td><td></td><td></td><td></td><td></td><td></td><td></td></tr>
<tr><td>Oligo rev</td><td>AAACGTACGAGGCGGAGAGGTACG</td><td></td><td></td><td></td><td></td><td></td><td></td><td></td><td></td><td></td><td></td></tr>
<tr><td>Chromosome</td><td>start</td><td>end</td><td>strand</td><td>MM</td><td>target_seq</td><td>PAM</td><td>alignment</td><td>distance</td><td>position</td><td>gene name</td><td>gene id</td></tr>
<tr><td>GU305914.1</td><td>66908</td><td>66930</td><td>-</td><td>0</td><td>CGTACCTCTCCGCCTCGTAC</td><td>CGG</td><td>||||||||[||||||||||||]PAM</td><td>NA</td><td>Intergenic</td><td>NA</td><td>NA</td></tr>
<tr><td>T29096</td><td>CGCCTCGTACCGGAAATCAACGG</td><td>1000</td><td></td><td></td><td></td><td></td><td></td><td></td><td></td><td></td><td></td></tr>
<tr><td>Oligo fwd</td><td>CACCCGCCTCGTACCGGAAATCAA</td><td></td><td></td><td></td><td></td><td></td><td></td><td></td><td></td><td></td><td></td></tr>
<tr><td>Oligo rev</td><td>AAACTTGATTTCCGGTACGAGGCG</td><td></td><td></td><td></td><td></td><td></td><td></td><td></td><td></td><td></td><td></td></tr>
<tr><td>Chromosome</td><td>start</td><td>end</td><td>strand</td><td>MM</td><td>target_seq</td><td>PAM</td><td>alignment</td><td>distance</td><td>position</td><td>gene name</td><td>gene id</td></tr>
<tr><td>GU305914.1</td><td>66898</td><td>66920</td><td>-</td><td>0</td><td>CGCCTCGTACCGGAAATCAA</td><td>CGG</td><td>||||||||[||||||||||||]PAM</td><td>NA</td><td>Intergenic</td><td>NA</td><td>NA</td></tr>
</table>

<table>
<tr><td>T29097</td><td>AAATCAACGGACGCGTCCCTCGG</td><td>1000</td><td></td><td></td><td></td><td></td><td></td><td></td><td></td><td></td><td></td></tr>
<tr><td>Oligo fwd</td><td>CACCAAATCAACGGACGCGTCCCT</td><td></td><td></td><td></td><td></td><td></td><td></td><td></td><td></td><td></td><td></td></tr>
<tr><td>Oligo rev</td><td>AAACAGGGACGCGTCCGTTGATTT</td><td></td><td></td><td></td><td></td><td></td><td></td><td></td><td></td><td></td><td></td></tr>
<tr><td>Chromosome</td><td>start</td><td>end</td><td>strand</td><td>MM</td><td>target_seq</td><td>PAM</td><td>alignment</td><td>distance</td><td>position</td><td>gene name</td><td>gene id</td></tr>
<tr><td>GU305914.1</td><td>66885</td><td>66907</td><td>-</td><td>0</td><td>AAATCAACGGACGCGTCCCT</td><td>CGG</td><td>||||||||[||||||||||||]PAM</td><td>NA</td><td>Intergenic</td><td>NA</td><td>NA</td></tr>
<tr><td>T29098</td><td>GACGTCCGCTCATGTTTCGTCGG</td><td>1000</td><td></td><td></td><td></td><td></td><td></td><td></td><td></td><td></td><td></td></tr>
<tr><td>Oligo fwd</td><td>CACCGACGTCCGCTCATGTTTCGT</td><td></td><td></td><td></td><td></td><td></td><td></td><td></td><td></td><td></td><td></td></tr>
<tr><td>Oligo rev</td><td>AAACACGAAACATGAGCGGACGTC</td><td></td><td></td><td></td><td></td><td></td><td></td><td></td><td></td><td></td><td></td></tr>
<tr><td>Chromosome</td><td>start</td><td>end</td><td>strand</td><td>MM</td><td>target_seq</td><td>PAM</td><td>alignment</td><td>distance</td><td>position</td><td>gene name</td><td>gene id</td></tr>
<tr><td>GU305914.1</td><td>66863</td><td>66885</td><td>-</td><td>0</td><td>GACGTCCGCTCATGTTTCGT</td><td>CGG</td><td>||||||||[||||||||||||]PAM</td><td>NA</td><td>Intergenic</td><td>NA</td><td>NA</td></tr>
<tr><td>T29099</td><td>CTCATGTTTCGTCGGCGCAGCGG</td><td>1000</td><td></td><td></td><td></td><td></td><td></td><td></td><td></td><td></td><td></td></tr>
<tr><td>Oligo fwd</td><td>CACCCTCATGTTTCGTCGGCGCAG</td><td></td><td></td><td></td><td></td><td></td><td></td><td></td><td></td><td></td><td></td></tr>
<tr><td>Oligo rev</td><td>AAACCTGCGCCGACGAAACATGAG</td><td></td><td></td><td></td><td></td><td></td><td></td><td></td><td></td><td></td><td></td></tr>
<tr><td>Chromosome</td><td>start</td><td>end</td><td>strand</td><td>MM</td><td>target_seq</td><td>PAM</td><td>alignment</td><td>distance</td><td>position</td><td>gene name</td><td>gene id</td></tr>
<tr><td>GU305914.1</td><td>66855</td><td>66877</td><td>-</td><td>0</td><td>CTCATGTTTCGTCGGCGCAG</td><td>CGG</td><td>||||||||[||||||||||||]PAM</td><td>NA</td><td>Intergenic</td><td>NA</td><td>NA</td></tr>
<tr><td>T29100</td><td>ATGTTTCGTCGGCGCAGCGGCGG</td><td>1000</td><td></td><td></td><td></td><td></td><td></td><td></td><td></td><td></td><td></td></tr>
<tr><td>Oligo fwd</td><td>CACCATGTTTCGTCGGCGCAGCGG</td><td></td><td></td><td></td><td></td><td></td><td></td><td></td><td></td><td></td><td></td></tr>
<tr><td>Oligo rev</td><td>AAACCCGCTGCGCCGACGAAACAT</td><td></td><td></td><td></td><td></td><td></td><td></td><td></td><td></td><td></td><td></td></tr>
<tr><td>Chromosome</td><td>start</td><td>end</td><td>strand</td><td>MM</td><td>target_seq</td><td>PAM</td><td>alignment</td><td>distance</td><td>position</td><td>gene name</td><td>gene id</td></tr>
<tr><td>GU305914.1</td><td>66852</td><td>66874</td><td>-</td><td>0</td><td>ATGTTTCGTCGGCGCAGCGG</td><td>CGG</td><td>||||||||[||||||||||||]PAM</td><td>NA</td><td>Intergenic</td><td>NA</td><td>NA</td></tr>
<tr><td>T29101</td><td>CGTCGGCGCAGCGGCGGCGCCGG</td><td>1000</td><td></td><td></td><td></td><td></td><td></td><td></td><td></td><td></td><td></td></tr>
<tr><td>Oligo fwd</td><td>CACCCGTCGGCGCAGCGGCGGCGC</td><td></td><td></td><td></td><td></td><td></td><td></td><td></td><td></td><td></td><td></td></tr>
<tr><td>Oligo rev</td><td>AAACGCGCCGCCGCTGCGCCGACG</td><td></td><td></td><td></td><td></td><td></td><td></td><td></td><td></td><td></td><td></td></tr>
<tr><td>Chromosome</td><td>start</td><td>end</td><td>strand</td><td>MM</td><td>target_seq</td><td>PAM</td><td>alignment</td><td>distance</td><td>position</td><td>gene name</td><td>gene id</td></tr>
<tr><td>GU305914.1</td><td>66846</td><td>66868</td><td>-</td><td>0</td><td>CGTCGGCGCAGCGGCGGCGC</td><td>CGG</td><td>||||||||[||||||||||||]PAM</td><td>NA</td><td>Intergenic</td><td>NA</td><td>NA</td></tr>
<tr><td>GU305914.1</td><td>56770</td><td>56792</td><td>+</td><td>4</td><td>CTCCGCCGCAGCGGCGGCGG</td><td>CGG</td><td>|--||-||[|||||||||||-]PAM</td><td>NA</td><td>Intergenic</td><td>NA</td><td>NA</td></tr>
<tr><td>T29102</td><td>CCGGCGCCTCCTCGCGATTCCGG</td><td>1000</td><td></td><td></td><td></td><td></td><td></td><td></td><td></td><td></td><td></td></tr>
<tr><td>Oligo fwd</td><td>CACCCCGGCGCCTCCTCGCGATTC</td><td></td><td></td><td></td><td></td><td></td><td></td><td></td><td></td><td></td><td></td></tr>
<tr><td>Oligo rev</td><td>AAACGAATCGCGAGGAGGCGCCGG</td><td></td><td></td><td></td><td></td><td></td><td></td><td></td><td></td><td></td><td></td></tr>
<tr><td>Chromosome</td><td>start</td><td>end</td><td>strand</td><td>MM</td><td>target_seq</td><td>PAM</td><td>alignment</td><td>distance</td><td>position</td><td>gene name</td><td>gene id</td></tr>
<tr><td>GU305914.1</td><td>66827</td><td>66849</td><td>-</td><td>0</td><td>CCGGCGCCTCCTCGCGATTC</td><td>CGG</td><td>||||||||[||||||||||||]PAM</td><td>NA</td><td>Intergenic</td><td>NA</td><td>NA</td></tr>
<tr><td>T29103</td><td>GCCTCCTCGCGATTCCGGTACGG</td><td>1000</td><td></td><td></td><td></td><td></td><td></td><td></td><td></td><td></td><td></td></tr>
<tr><td>Oligo fwd</td><td>CACCGCCTCCTCGCGATTCCGGTA</td><td></td><td></td><td></td><td></td><td></td><td></td><td></td><td></td><td></td><td></td></tr>
<tr><td>Oligo rev</td><td>AAACTACCGGAATCGCGAGGAGGC</td><td></td><td></td><td></td><td></td><td></td><td></td><td></td><td></td><td></td><td></td></tr>
<tr><td>Chromosome</td><td>start</td><td>end</td><td>strand</td><td>MM</td><td>target_seq</td><td>PAM</td><td>alignment</td><td>distance</td><td>position</td><td>gene name</td><td>gene id</td></tr>
<tr><td>GU305914.1</td><td>66822</td><td>66844</td><td>-</td><td>0</td><td>GCCTCCTCGCGATTCCGGTA</td><td>CGG</td><td>||||||||[||||||||||||]PAM</td><td>NA</td><td>Intergenic</td><td>NA</td><td>NA</td></tr>
<tr><td>T29104</td><td>TTAATACGCCTATCGATTCCCGG</td><td>1000</td><td></td><td></td><td></td><td></td><td></td><td></td><td></td><td></td><td></td></tr>
<tr><td>Oligo fwd</td><td>CACCTTAATACGCCTATCGATTCC</td><td></td><td></td><td></td><td></td><td></td><td></td><td></td><td></td><td></td><td></td></tr>
<tr><td>Oligo rev</td><td>AAACGGAATCGATAGGCGTATTAA</td><td></td><td></td><td></td><td></td><td></td><td></td><td></td><td></td><td></td><td></td></tr>
<tr><td>Chromosome</td><td>start</td><td>end</td><td>strand</td><td>MM</td><td>target_seq</td><td>PAM</td><td>alignment</td><td>distance</td><td>position</td><td>gene name</td><td>gene id</td></tr>
<tr><td>GU305914.1</td><td>66785</td><td>66807</td><td>-</td><td>0</td><td>TTAATACGCCTATCGATTCC</td><td>CGG</td><td>||||||||[||||||||||||]PAM</td><td>NA</td><td>Intergenic</td><td>NA</td><td>NA</td></tr>
<tr><td>T29105</td><td>CCGGTCGAGCTCCTGTCTCTTGG</td><td>1000</td><td></td><td></td><td></td><td></td><td></td><td></td><td></td><td></td><td></td></tr>
<tr><td>Oligo fwd</td><td>CACCCCGGTCGAGCTCCTGTCTCT</td><td></td><td></td><td></td><td></td><td></td><td></td><td></td><td></td><td></td><td></td></tr>
<tr><td>Oligo rev</td><td>AAACAGAGACAGGAGCTCGACCGG</td><td></td><td></td><td></td><td></td><td></td><td></td><td></td><td></td><td></td><td></td></tr>
<tr><td>Chromosome</td><td>start</td><td>end</td><td>strand</td><td>MM</td><td>target_seq</td><td>PAM</td><td>alignment</td><td>distance</td><td>position</td><td>gene name</td><td>gene id</td></tr>
<tr><td>GU305914.1</td><td>66766</td><td>66788</td><td>-</td><td>0</td><td>CCGGTCGAGCTCCTGTCTCT</td><td>TGG</td><td>||||||||[||||||||||||]PAM</td><td>NA</td><td>Intergenic</td><td>NA</td><td>NA</td></tr>
<tr><td>T29106</td><td>CGGTCGAGCTCCTGTCTCTTGGG</td><td>1000</td><td></td><td></td><td></td><td></td><td></td><td></td><td></td><td></td><td></td></tr>
<tr><td>Oligo fwd</td><td>CACCCGGTCGAGCTCCTGTCTCTT</td><td></td><td></td><td></td><td></td><td></td><td></td><td></td><td></td><td></td><td></td></tr>
<tr><td>Oligo rev</td><td>AAACAAGAGACAGGAGCTCGACCG</td><td></td><td></td><td></td><td></td><td></td><td></td><td></td><td></td><td></td><td></td></tr>
<tr><td>Chromosome</td><td>start</td><td>end</td><td>strand</td><td>MM</td><td>target_seq</td><td>PAM</td><td>alignment</td><td>distance</td><td>position</td><td>gene name</td><td>gene id</td></tr>
<tr><td>GU305914.1</td><td>66765</td><td>66787</td><td>-</td><td>0</td><td>CGGTCGAGCTCCTGTCTCTT</td><td>GGG</td><td>||||||||[||||||||||||]PAM</td><td>NA</td><td>Intergenic</td><td>NA</td><td>NA</td></tr>
<tr><td>T29107</td><td>GGTCGAGCTCCTGTCTCTTGGGG</td><td>1000</td><td></td><td></td><td></td><td></td><td></td><td></td><td></td><td></td><td></td></tr>
<tr><td>Oligo fwd</td><td>CACCGGTCGAGCTCCTGTCTCTTG</td><td></td><td></td><td></td><td></td><td></td><td></td><td></td><td></td><td></td><td></td></tr>
<tr><td>Oligo rev</td><td>AAACCAAGAGACAGGAGCTCGACC</td><td></td><td></td><td></td><td></td><td></td><td></td><td></td><td></td><td></td><td></td></tr>
<tr><td>Chromosome</td><td>start</td><td>end</td><td>strand</td><td>MM</td><td>target_seq</td><td>PAM</td><td>alignment</td><td>distance</td><td>position</td><td>gene name</td><td>gene id</td></tr>
<tr><td>GU305914.1</td><td>66764</td><td>66786</td><td>-</td><td>0</td><td>GGTCGAGCTCCTGTCTCTTG</td><td>GGG</td><td>||||||||[||||||||||||]PAM</td><td>NA</td><td>Intergenic</td><td>NA</td><td>NA</td></tr>
<tr><td>T29108</td><td>GTCGAGCTCCTGTCTCTTGGGGG</td><td>1000</td><td></td><td></td><td></td><td></td><td></td><td></td><td></td><td></td><td></td></tr>
<tr><td>Oligo fwd</td><td>CACCGTCGAGCTCCTGTCTCTTGG</td><td></td><td></td><td></td><td></td><td></td><td></td><td></td><td></td><td></td><td></td></tr>
<tr><td>Oligo rev</td><td>AAACCCAAGAGACAGGAGCTCGAC</td><td></td><td></td><td></td><td></td><td></td><td></td><td></td><td></td><td></td><td></td></tr>
<tr><td>Chromosome</td><td>start</td><td>end</td><td>strand</td><td>MM</td><td>target_seq</td><td>PAM</td><td>alignment</td><td>distance</td><td>position</td><td>gene name</td><td>gene id</td></tr>
<tr><td>GU305914.1</td><td>66763</td><td>66785</td><td>-</td><td>0</td><td>GTCGAGCTCCTGTCTCTTGG</td><td>GGG</td><td>||||||||[||||||||||||]PAM</td><td>NA</td><td>Intergenic</td><td>NA</td><td>NA</td></tr>
<tr><td>T29109</td><td>TCGAGCTCCTGTCTCTTGGGGGG</td><td>1000</td><td></td><td></td><td></td><td></td><td></td><td></td><td></td><td></td><td></td></tr>
<tr><td>Oligo fwd</td><td>CACCTCGAGCTCCTGTCTCTTGGG</td><td></td><td></td><td></td><td></td><td></td><td></td><td></td><td></td><td></td><td></td></tr>
<tr><td>Oligo rev</td><td>AAACCCCAAGAGACAGGAGCTCGA</td><td></td><td></td><td></td><td></td><td></td><td></td><td></td><td></td><td></td><td></td></tr>
<tr><td>Chromosome</td><td>start</td><td>end</td><td>strand</td><td>MM</td><td>target_seq</td><td>PAM</td><td>alignment</td><td>distance</td><td>position</td><td>gene name</td><td>gene id</td></tr>
<tr><td>GU305914.1</td><td>66762</td><td>66784</td><td>-</td><td>0</td><td>TCGAGCTCCTGTCTCTTGGG</td><td>GGG</td><td>||||||||[||||||||||||]PAM</td><td>NA</td><td>Intergenic</td><td>NA</td><td>NA</td></tr>
<tr><td>T29110</td><td>CGAGCTCCTGTCTCTTGGGGGGG</td><td>1000</td><td></td><td></td><td></td><td></td><td></td><td></td><td></td><td></td><td></td></tr>
<tr><td>Oligo fwd</td><td>CACCCGAGCTCCTGTCTCTTGGGG</td><td></td><td></td><td></td><td></td><td></td><td></td><td></td><td></td><td></td><td></td></tr>
<tr><td>Oligo rev</td><td>AAACCCCCAAGAGACAGGAGCTCG</td><td></td><td></td><td></td><td></td><td></td><td></td><td></td><td></td><td></td><td></td></tr>
<tr><td>Chromosome</td><td>start</td><td>end</td><td>strand</td><td>MM</td><td>target_seq</td><td>PAM</td><td>alignment</td><td>distance</td><td>position</td><td>gene name</td><td>gene id</td></tr>
<tr><td>GU305914.1</td><td>66761</td><td>66783</td><td>-</td><td>0</td><td>CGAGCTCCTGTCTCTTGGGG</td><td>GGG</td><td>||||||||[||||||||||||]PAM</td><td>NA</td><td>Intergenic</td><td>NA</td><td>NA</td></tr>
<tr><td>T29111</td><td>CCTGTCTCTTGGGGGGGCTGTGG</td><td>1000</td><td></td><td></td><td></td><td></td><td></td><td></td><td></td><td></td><td></td></tr>
<tr><td>Oligo fwd</td><td>CACCCCTGTCTCTTGGGGGGGCTG</td><td></td><td></td><td></td><td></td><td></td><td></td><td></td><td></td><td></td><td></td></tr>
<tr><td>Oligo rev</td><td>AAACCAGCCCCCCCAAGAGACAGG</td><td></td><td></td><td></td><td></td><td></td><td></td><td></td><td></td><td></td><td></td></tr>
<tr><td>Chromosome</td><td>start</td><td>end</td><td>strand</td><td>MM</td><td>target_seq</td><td>PAM</td><td>alignment</td><td>distance</td><td>position</td><td>gene name</td><td>gene id</td></tr>
</table>

GU305914.1 66755 66777 - 0 CCTGTCTCTTGGGGGGGCTG TGG ||||||||[||||||||||||]PAM NA Intergenic NA NA

T29112 CTGTCTCTTGGGGGGGCTGTGGG 1000
Oligo fwd CACCCTGTCTCTTGGGGGGGCTGT
Oligo rev AAACACAGCCCCCCCAAGAGACAG
Chromosome start end strand MM target_seq PAM alignment distance position gene name gene id
GU305914.1 66754 66776 - 0 CTGTCTCTTGGGGGGGCTGT GGG ||||||||[||||||||||||]PAM NA Intergenic NA NA

T29113 TGTCTCTTGGGGGGGCTGTGGGG 1000
Oligo fwd CACCTGTCTCTTGGGGGGGCTGTG
Oligo rev AAACCACAGCCCCCCCAAGAGACA
Chromosome start end strand MM target_seq PAM alignment distance position gene name gene id
GU305914.1 66753 66775 - 0 TGTCTCTTGGGGGGGCTGTG GGG ||||||||[||||||||||||]PAM NA Intergenic NA NA

T29114 GTCTCTTGGGGGGGCTGTGGGGG 1000
Oligo fwd CACCGTCTCTTGGGGGGGCTGTGG
Oligo rev AAACCCACAGCCCCCCCAAGAGAC
Chromosome start end strand MM target_seq PAM alignment distance position gene name gene id
GU305914.1 66752 66774 - 0 GTCTCTTGGGGGGGCTGTGG GGG ||||||||[||||||||||||]PAM NA Intergenic NA NA

T29115 TCTCTTGGGGGGGCTGTGGGGGG 1000
Oligo fwd CACCTCTCTTGGGGGGGCTGTGGG
Oligo rev AAACCCCACAGCCCCCCCAAGAGA
Chromosome start end strand MM target_seq PAM alignment distance position gene name gene id
GU305914.1 66751 66773 - 0 TCTCTTGGGGGGGCTGTGGG GGG ||||||||[||||||||||||]PAM NA Intergenic NA NA

T29116 TGGGGGGGCTGTGGGGGGCCCGG 1000
Oligo fwd CACCTGGGGGGGCTGTGGGGGGCC
Oligo rev AAACGGCCCCCCACAGCCCCCCCA
Chromosome start end strand MM target_seq PAM alignment distance position gene name gene id
GU305914.1 66746 66768 - 0 TGGGGGGGCTGTGGGGGGCC CGG ||||||||[||||||||||||]PAM NA Intergenic NA NA
GU305914.1 53364 53386 - 4 TGGTGCTGCTGTGTGGGGCC TGG |||-|--|[|||||-||||||]PAM NA Intergenic NA NA

T29117 GGGGGCTGTGGGGGGCCCGGTGG 1000
Oligo fwd CACCGGGGGCTGTGGGGGGCCCGG
Oligo rev AAACCCGGGCCCCCCACAGCCCCC
Chromosome start end strand MM target_seq PAM alignment distance position gene name gene id
GU305914.1 66743 66765 - 0 GGGGGCTGTGGGGGGCCCGG TGG ||||||||[||||||||||||]PAM NA Intergenic NA NA

T29118 CCACGCAGCGCCGCAGCCCGCGG 1000
Oligo fwd CACCCCACGCAGCGCCGCAGCCCG
Oligo rev AAACCGGGCTGCGGCGCTGCGTGG
Chromosome start end strand MM target_seq PAM alignment distance position gene name gene id
GU305914.1 66695 66717 - 0 CCACGCAGCGCCGCAGCCCG CGG ||||||||[||||||||||||]PAM NA Intergenic NA NA
GU305914.1 67932 67954 + 4 AGACGCTGCGCCGCAGCCGG CGG --||||-|[||||||||||-|]PAM NA Intergenic NA NA

T29119 AGCGCCGCAGCCCGCGGCTCCGG 1000
Oligo fwd CACCAGCGCCGCAGCCCGCGGCTC
Oligo rev AAACGAGCCGCGGGCTGCGGCGCT
Chromosome start end strand MM target_seq PAM alignment distance position gene name gene id
GU305914.1 66689 66711 - 0 AGCGCCGCAGCCCGCGGCTC CGG ||||||||[||||||||||||]PAM NA Intergenic NA NA
GU305914.1 100336 100358 + 4 TCCGCCGCAGCGCGCGGCAC AGG --||||||[|||-||||||-|]PAM NA Intergenic NA NA

T29120 TCCTGTCCGCGTCCATCGTGCGG 1000
Oligo fwd CACCTCCTGTCCGCGTCCATCGTG
Oligo rev AAACCACGATGGACGCGGACAGGA
Chromosome start end strand MM target_seq PAM alignment distance position gene name gene id
GU305914.1 66665 66687 - 0 TCCTGTCCGCGTCCATCGTG CGG ||||||||[||||||||||||]PAM NA Intergenic NA NA

T29121 GTCCGCGTCCATCGTGCGGCAGG 1000
Oligo fwd CACCGTCCGCGTCCATCGTGCGGC
Oligo rev AAACGCCGCACGATGGACGCGGAC
Chromosome start end strand MM target_seq PAM alignment distance position gene name gene id
GU305914.1 66661 66683 - 0 GTCCGCGTCCATCGTGCGGC AGG ||||||||[||||||||||||]PAM NA Intergenic NA NA

T29122 AGAACGACTCGCACGCCGCGCGG 1000
Oligo fwd CACCAGAACGACTCGCACGCCGCG
Oligo rev AAACCGCGGCGTGCGAGTCGTTCT
Chromosome start end strand MM target_seq PAM alignment distance position gene name gene id
GU305914.1 66623 66645 - 0 AGAACGACTCGCACGCCGCG CGG ||||||||[||||||||||||]PAM NA Intergenic NA NA

T29123 ACGCCGCGCGGTTGAAGCGCAGG 1000
Oligo fwd CACCACGCCGCGCGGTTGAAGCGC
Oligo rev AAACGCGCTTCAACCGCGCGGCGT
Chromosome start end strand MM target_seq PAM alignment distance position gene name gene id
GU305914.1 66611 66633 - 0 ACGCCGCGCGGTTGAAGCGC AGG ||||||||[||||||||||||]PAM NA Intergenic NA NA

T29124 ACAGCGATTCCAGCGTCGCCTGG 1000
Oligo fwd CACCACAGCGATTCCAGCGTCGCC
Oligo rev AAACGGCGACGCTGGAATCGCTGT
Chromosome start end strand MM target_seq PAM alignment distance position gene name gene id
GU305914.1 66578 66600 - 0 ACAGCGATTCCAGCGTCGCC TGG ||||||||[||||||||||||]PAM NA Intergenic NA NA

T29125 TCGACCGCGTCGAGAAAGTCAGG 1000
Oligo fwd CACCTCGACCGCGTCGAGAAAGTC
Oligo rev AAACGACTTTCTCGACGCGGTCGA
Chromosome start end strand MM target_seq PAM alignment distance position gene name gene id
GU305914.1 66512 66534 - 0 TCGACCGCGTCGAGAAAGTC AGG ||||||||[||||||||||||]PAM NA Intergenic NA NA

<table>
<tr><td>T29126</td><td>CCGCGTCGAGAAAGTCAGGTCGG</td><td>1000</td></tr>
<tr><td>Oligo fwd</td><td>CACCCCGCGTCGAGAAAGTCAGGT</td></tr>
<tr><td>Oligo rev</td><td>AAACACCTGACTTTCTCGACGCGG</td></tr>
<tr><td>Chromosome</td><td>start</td><td>end</td><td>strand</td><td>MM</td><td>target_seq</td><td>PAM</td><td>alignment</td><td>distance</td><td>position</td><td>gene name</td><td>gene id</td></tr>
<tr><td>GU305914.1</td><td>66508</td><td>66530</td><td>-</td><td>0</td><td>CCGCGTCGAGAAAGTCAGGT</td><td>CGG</td><td>||||||||[||||||||||||]PAM</td><td>NA</td><td>Intergenic</td><td>NA</td><td>NA</td></tr>
<tr><td>T29127</td><td>CGAGAAAGTCAGGTCGGTGAAGG</td><td>1000</td></tr>
<tr><td>Oligo fwd</td><td>CACCCGAGAAAGTCAGGTCGGTGA</td></tr>
<tr><td>Oligo rev</td><td>AAACTCACCGACCTGACTTTCTCG</td></tr>
<tr><td>Chromosome</td><td>start</td><td>end</td><td>strand</td><td>MM</td><td>target_seq</td><td>PAM</td><td>alignment</td><td>distance</td><td>position</td><td>gene name</td><td>gene id</td></tr>
<tr><td>GU305914.1</td><td>66502</td><td>66524</td><td>-</td><td>0</td><td>CGAGAAAGTCAGGTCGGTGA</td><td>AGG</td><td>||||||||[||||||||||||]PAM</td><td>NA</td><td>Intergenic</td><td>NA</td><td>NA</td></tr>
<tr><td>T29128</td><td>AAAGTCAGGTCGGTGAAGGTCGG</td><td>1000</td></tr>
<tr><td>Oligo fwd</td><td>CACCAAAGTCAGGTCGGTGAAGGT</td></tr>
<tr><td>Oligo rev</td><td>AAACACCTTCACCGACCTGACTTT</td></tr>
<tr><td>Chromosome</td><td>start</td><td>end</td><td>strand</td><td>MM</td><td>target_seq</td><td>PAM</td><td>alignment</td><td>distance</td><td>position</td><td>gene name</td><td>gene id</td></tr>
<tr><td>GU305914.1</td><td>66498</td><td>66520</td><td>-</td><td>0</td><td>AAAGTCAGGTCGGTGAAGGT</td><td>CGG</td><td>||||||||[||||||||||||]PAM</td><td>NA</td><td>Intergenic</td><td>NA</td><td>NA</td></tr>
<tr><td>T29129</td><td>AAGTCAGGTCGGTGAAGGTCGGG</td><td>1000</td></tr>
<tr><td>Oligo fwd</td><td>CACCAAGTCAGGTCGGTGAAGGTC</td></tr>
<tr><td>Oligo rev</td><td>AAACGACCTTCACCGACCTGACTT</td></tr>
<tr><td>Chromosome</td><td>start</td><td>end</td><td>strand</td><td>MM</td><td>target_seq</td><td>PAM</td><td>alignment</td><td>distance</td><td>position</td><td>gene name</td><td>gene id</td></tr>
<tr><td>GU305914.1</td><td>66497</td><td>66519</td><td>-</td><td>0</td><td>AAGTCAGGTCGGTGAAGGTC</td><td>GGG</td><td>||||||||[||||||||||||]PAM</td><td>NA</td><td>Intergenic</td><td>NA</td><td>NA</td></tr>
<tr><td>T29130</td><td>CGGGATGCTGCGCGTCTCCTCGG</td><td>1000</td></tr>
<tr><td>Oligo fwd</td><td>CACCCGGGATGCTGCGCGTCTCCT</td></tr>
<tr><td>Oligo rev</td><td>AAACAGGAGACGCGCAGCATCCCG</td></tr>
<tr><td>Chromosome</td><td>start</td><td>end</td><td>strand</td><td>MM</td><td>target_seq</td><td>PAM</td><td>alignment</td><td>distance</td><td>position</td><td>gene name</td><td>gene id</td></tr>
<tr><td>GU305914.1</td><td>66478</td><td>66500</td><td>-</td><td>0</td><td>CGGGATGCTGCGCGTCTCCT</td><td>CGG</td><td>||||||||[||||||||||||]PAM</td><td>NA</td><td>Intergenic</td><td>NA</td><td>NA</td></tr>
<tr><td>T29131</td><td>CCGCTCGACGATCAGCGCGCCGG</td><td>1000</td></tr>
<tr><td>Oligo fwd</td><td>CACCCCGCTCGACGATCAGCGCGC</td></tr>
<tr><td>Oligo rev</td><td>AAACGCGCGCTGATCGTCGAGCGG</td></tr>
<tr><td>Chromosome</td><td>start</td><td>end</td><td>strand</td><td>MM</td><td>target_seq</td><td>PAM</td><td>alignment</td><td>distance</td><td>position</td><td>gene name</td><td>gene id</td></tr>
<tr><td>GU305914.1</td><td>66442</td><td>66464</td><td>-</td><td>0</td><td>CCGCTCGACGATCAGCGCGC</td><td>CGG</td><td>||||||||[||||||||||||]PAM</td><td>NA</td><td>Intergenic</td><td>NA</td><td>NA</td></tr>
<tr><td>T29132</td><td>TCAGCGCGCCGGTGTTGCTCAGG</td><td>1000</td></tr>
<tr><td>Oligo fwd</td><td>CACCTCAGCGCGCCGGTGTTGCTC</td></tr>
<tr><td>Oligo rev</td><td>AAACGAGCAACACCGGCGCGCTGA</td></tr>
<tr><td>Chromosome</td><td>start</td><td>end</td><td>strand</td><td>MM</td><td>target_seq</td><td>PAM</td><td>alignment</td><td>distance</td><td>position</td><td>gene name</td><td>gene id</td></tr>
<tr><td>GU305914.1</td><td>66431</td><td>66453</td><td>-</td><td>0</td><td>TCAGCGCGCCGGTGTTGCTC</td><td>AGG</td><td>||||||||[||||||||||||]PAM</td><td>NA</td><td>Intergenic</td><td>NA</td><td>NA</td></tr>
<tr><td>T29133</td><td>CCGCAGCGTGTCCGTGAACTCGG</td><td>1000</td></tr>
<tr><td>Oligo fwd</td><td>CACCCCGCAGCGTGTCCGTGAACT</td></tr>
<tr><td>Oligo rev</td><td>AAACAGTTCACGGACACGCTGCGG</td></tr>
<tr><td>Chromosome</td><td>start</td><td>end</td><td>strand</td><td>MM</td><td>target_seq</td><td>PAM</td><td>alignment</td><td>distance</td><td>position</td><td>gene name</td><td>gene id</td></tr>
<tr><td>GU305914.1</td><td>66364</td><td>66386</td><td>-</td><td>0</td><td>CCGCAGCGTGTCCGTGAACT</td><td>CGG</td><td>||||||||[||||||||||||]PAM</td><td>NA</td><td>Intergenic</td><td>NA</td><td>NA</td></tr>
<tr><td>T29134</td><td>CGCAGCGTGTCCGTGAACTCGGG</td><td>1000</td></tr>
<tr><td>Oligo fwd</td><td>CACCCGCAGCGTGTCCGTGAACTC</td></tr>
<tr><td>Oligo rev</td><td>AAACGAGTTCACGGACACGCTGCG</td></tr>
<tr><td>Chromosome</td><td>start</td><td>end</td><td>strand</td><td>MM</td><td>target_seq</td><td>PAM</td><td>alignment</td><td>distance</td><td>position</td><td>gene name</td><td>gene id</td></tr>
<tr><td>GU305914.1</td><td>66363</td><td>66385</td><td>-</td><td>0</td><td>CGCAGCGTGTCCGTGAACTC</td><td>GGG</td><td>||||||||[||||||||||||]PAM</td><td>NA</td><td>Intergenic</td><td>NA</td><td>NA</td></tr>
<tr><td>T29135</td><td>GCGTGTCCGTGAACTCGGGCAGG</td><td>1000</td></tr>
<tr><td>Oligo fwd</td><td>CACCGCGTGTCCGTGAACTCGGGC</td></tr>
<tr><td>Oligo rev</td><td>AAACGCCCGAGTTCACGGACACGC</td></tr>
<tr><td>Chromosome</td><td>start</td><td>end</td><td>strand</td><td>MM</td><td>target_seq</td><td>PAM</td><td>alignment</td><td>distance</td><td>position</td><td>gene name</td><td>gene id</td></tr>
<tr><td>GU305914.1</td><td>66359</td><td>66381</td><td>-</td><td>0</td><td>GCGTGTCCGTGAACTCGGGC</td><td>AGG</td><td>||||||||[||||||||||||]PAM</td><td>NA</td><td>Intergenic</td><td>NA</td><td>NA</td></tr>
<tr><td>T29136</td><td>CGTGTCCGTGAACTCGGGCAGGG</td><td>1000</td></tr>
<tr><td>Oligo fwd</td><td>CACCCGTGTCCGTGAACTCGGGCA</td></tr>
<tr><td>Oligo rev</td><td>AAACTGCCCGAGTTCACGGACACG</td></tr>
<tr><td>Chromosome</td><td>start</td><td>end</td><td>strand</td><td>MM</td><td>target_seq</td><td>PAM</td><td>alignment</td><td>distance</td><td>position</td><td>gene name</td><td>gene id</td></tr>
<tr><td>GU305914.1</td><td>66358</td><td>66380</td><td>-</td><td>0</td><td>CGTGTCCGTGAACTCGGGCA</td><td>GGG</td><td>||||||||[||||||||||||]PAM</td><td>NA</td><td>Intergenic</td><td>NA</td><td>NA</td></tr>
<tr><td>T29137</td><td>GCAGTGTTCCGCGAACCGCTCGG</td><td>1000</td></tr>
<tr><td>Oligo fwd</td><td>CACCGCAGTGTTCCGCGAACCGCT</td></tr>
<tr><td>Oligo rev</td><td>AAACAGCGGTTCGCGGAACACTGC</td></tr>
<tr><td>Chromosome</td><td>start</td><td>end</td><td>strand</td><td>MM</td><td>target_seq</td><td>PAM</td><td>alignment</td><td>distance</td><td>position</td><td>gene name</td><td>gene id</td></tr>
<tr><td>GU305914.1</td><td>66238</td><td>66260</td><td>-</td><td>0</td><td>GCAGTGTTCCGCGAACCGCT</td><td>CGG</td><td>||||||||[||||||||||||]PAM</td><td>NA</td><td>Intergenic</td><td>NA</td><td>NA</td></tr>
<tr><td>T29138</td><td>CCGCGAACCGCTCGGCCAGCTGG</td><td>1000</td></tr>
<tr><td>Oligo fwd</td><td>CACCCCGCGAACCGCTCGGCCAGC</td></tr>
<tr><td>Oligo rev</td><td>AAACGCTGGCCGAGCGGTTCGCGG</td></tr>
<tr><td>Chromosome</td><td>start</td><td>end</td><td>strand</td><td>MM</td><td>target_seq</td><td>PAM</td><td>alignment</td><td>distance</td><td>position</td><td>gene name</td><td>gene id</td></tr>
<tr><td>GU305914.1</td><td>66230</td><td>66252</td><td>-</td><td>0</td><td>CCGCGAACCGCTCGGCCAGC</td><td>TGG</td><td>||||||||[||||||||||||]PAM</td><td>NA</td><td>Intergenic</td><td>NA</td><td>NA</td></tr>
<tr><td>T29139</td><td>ACCGCTCGGCCAGCTGGCGCCGG</td><td>1000</td></tr>
<tr><td>Oligo fwd</td><td>CACCACCGCTCGGCCAGCTGGCGC</td></tr>
<tr><td>Oligo rev</td><td>AAACGCGCCAGCTGGCCGAGCGGT</td></tr>
<tr><td>Chromosome</td><td>start</td><td>end</td><td>strand</td><td>MM</td><td>target_seq</td><td>PAM</td><td>alignment</td><td>distance</td><td>position</td><td>gene name</td><td>gene id</td></tr>
<tr><td>GU305914.1</td><td>66224</td><td>66246</td><td>-</td><td>0</td><td>ACCGCTCGGCCAGCTGGCGC</td><td>CGG</td><td>||||||||[||||||||||||]PAM</td><td>NA</td><td>Intergenic</td><td>NA</td><td>NA</td></tr>
<tr><td>T29140</td><td>GGCCAGCTGGCGCCGGTCCGCGG</td><td>1000</td></tr>
<tr><td>Oligo fwd</td><td>CACCGGCCAGCTGGCGCCGGTCCG</td></tr>
<tr><td>Oligo rev</td><td>AAACCGGACCGGCGCCAGCTGGCC</td></tr>
<tr><td>Chromosome</td><td>start</td><td>end</td><td>strand</td><td>MM</td><td>target_seq</td><td>PAM</td><td>alignment</td><td>distance</td><td>position</td><td>gene name</td><td>gene id</td></tr>
<tr><td>GU305914.1</td><td>66217</td><td>66239</td><td>-</td><td>0</td><td>GGCCAGCTGGCGCCGGTCCG</td><td>CGG</td><td>||||||||[||||||||||||]PAM</td><td>NA</td><td>Intergenic</td><td>NA</td><td>NA</td></tr>
</table>

<table>
<tr><td>T29141</td><td>GGCCCAGCGCGCCAGCGCCTCGG</td><td>1000</td><td></td><td></td><td></td><td></td><td></td><td></td><td></td><td></td><td></td></tr>
<tr><td>Oligo fwd</td><td>CACCGGCCCAGCGCGCCAGCGCCT</td><td></td><td></td><td></td><td></td><td></td><td></td><td></td><td></td><td></td><td></td></tr>
<tr><td>Oligo rev</td><td>AAACAGGCGCTGGCGCGCTGGGCC</td><td></td><td></td><td></td><td></td><td></td><td></td><td></td><td></td><td></td><td></td></tr>
<tr><td>Chromosome</td><td>start</td><td>end</td><td>strand</td><td>MM</td><td>target_seq</td><td>PAM</td><td>alignment</td><td>distance</td><td>position</td><td>gene name</td><td>gene id</td></tr>
<tr><td>GU305914.1</td><td>66196</td><td>66218</td><td>-</td><td>0</td><td>GGCCCAGCGCGCCAGCGCCT</td><td>CGG</td><td>||||||||[||||||||||||]PAM</td><td>NA</td><td>Intergenic</td><td>NA</td><td>NA</td></tr>
<tr><td>T29142</td><td>TGTCGCCCCCCAGCGCGTGCAGG</td><td>1000</td><td></td><td></td><td></td><td></td><td></td><td></td><td></td><td></td><td></td></tr>
<tr><td>Oligo fwd</td><td>CACCTGTCGCCCCCCAGCGCGTGC</td><td></td><td></td><td></td><td></td><td></td><td></td><td></td><td></td><td></td><td></td></tr>
<tr><td>Oligo rev</td><td>AAACGCACGCGCTGGGGGGCGACA</td><td></td><td></td><td></td><td></td><td></td><td></td><td></td><td></td><td></td><td></td></tr>
<tr><td>Chromosome</td><td>start</td><td>end</td><td>strand</td><td>MM</td><td>target_seq</td><td>PAM</td><td>alignment</td><td>distance</td><td>position</td><td>gene name</td><td>gene id</td></tr>
<tr><td>GU305914.1</td><td>66170</td><td>66192</td><td>-</td><td>0</td><td>TGTCGCCCCCCAGCGCGTGC</td><td>AGG</td><td>||||||||[||||||||||||]PAM</td><td>NA</td><td>Intergenic</td><td>NA</td><td>NA</td></tr>
<tr><td>T29143</td><td>GCAGGAACGACTCGATCGCCAGG</td><td>1000</td><td></td><td></td><td></td><td></td><td></td><td></td><td></td><td></td><td></td></tr>
<tr><td>Oligo fwd</td><td>CACCGCAGGAACGACTCGATCGCC</td><td></td><td></td><td></td><td></td><td></td><td></td><td></td><td></td><td></td><td></td></tr>
<tr><td>Oligo rev</td><td>AAACGGCGATCGAGTCGTTCCTGC</td><td></td><td></td><td></td><td></td><td></td><td></td><td></td><td></td><td></td><td></td></tr>
<tr><td>Chromosome</td><td>start</td><td>end</td><td>strand</td><td>MM</td><td>target_seq</td><td>PAM</td><td>alignment</td><td>distance</td><td>position</td><td>gene name</td><td>gene id</td></tr>
<tr><td>GU305914.1</td><td>66152</td><td>66174</td><td>-</td><td>0</td><td>GCAGGAACGACTCGATCGCC</td><td>AGG</td><td>||||||||[||||||||||||]PAM</td><td>NA</td><td>Intergenic</td><td>NA</td><td>NA</td></tr>
<tr><td>T29144</td><td>GTTGAACGAGTACGAGTCCCCGG</td><td>1000</td><td></td><td></td><td></td><td></td><td></td><td></td><td></td><td></td><td></td></tr>
<tr><td>Oligo fwd</td><td>CACCGTTGAACGAGTACGAGTCCC</td><td></td><td></td><td></td><td></td><td></td><td></td><td></td><td></td><td></td><td></td></tr>
<tr><td>Oligo rev</td><td>AAACGGGACTCGTACTCGTTCAAC</td><td></td><td></td><td></td><td></td><td></td><td></td><td></td><td></td><td></td><td></td></tr>
<tr><td>Chromosome</td><td>start</td><td>end</td><td>strand</td><td>MM</td><td>target_seq</td><td>PAM</td><td>alignment</td><td>distance</td><td>position</td><td>gene name</td><td>gene id</td></tr>
<tr><td>GU305914.1</td><td>66112</td><td>66134</td><td>-</td><td>0</td><td>GTTGAACGAGTACGAGTCCC</td><td>CGG</td><td>||||||||[||||||||||||]PAM</td><td>NA</td><td>Intergenic</td><td>NA</td><td>NA</td></tr>
<tr><td>T29145</td><td>AGTCCCCGGTGATGTCGCCGAGG</td><td>1000</td><td></td><td></td><td></td><td></td><td></td><td></td><td></td><td></td><td></td></tr>
<tr><td>Oligo fwd</td><td>CACCAGTCCCCGGTGATGTCGCCG</td><td></td><td></td><td></td><td></td><td></td><td></td><td></td><td></td><td></td><td></td></tr>
<tr><td>Oligo rev</td><td>AAACCGGCGACATCACCGGGGACT</td><td></td><td></td><td></td><td></td><td></td><td></td><td></td><td></td><td></td><td></td></tr>
<tr><td>Chromosome</td><td>start</td><td>end</td><td>strand</td><td>MM</td><td>target_seq</td><td>PAM</td><td>alignment</td><td>distance</td><td>position</td><td>gene name</td><td>gene id</td></tr>
<tr><td>GU305914.1</td><td>66098</td><td>66120</td><td>-</td><td>0</td><td>AGTCCCCGGTGATGTCGCCG</td><td>AGG</td><td>||||||||[||||||||||||]PAM</td><td>NA</td><td>Intergenic</td><td>NA</td><td>NA</td></tr>
<tr><td>T29146</td><td>CGGTGATGTCGCCGAGGTGCAGG</td><td>1000</td><td></td><td></td><td></td><td></td><td></td><td></td><td></td><td></td><td></td></tr>
<tr><td>Oligo fwd</td><td>CACCCGGTGATGTCGCCGAGGTGC</td><td></td><td></td><td></td><td></td><td></td><td></td><td></td><td></td><td></td><td></td></tr>
<tr><td>Oligo rev</td><td>AAACGCACCTCGGCGACATCACCG</td><td></td><td></td><td></td><td></td><td></td><td></td><td></td><td></td><td></td><td></td></tr>
<tr><td>Chromosome</td><td>start</td><td>end</td><td>strand</td><td>MM</td><td>target_seq</td><td>PAM</td><td>alignment</td><td>distance</td><td>position</td><td>gene name</td><td>gene id</td></tr>
<tr><td>GU305914.1</td><td>66092</td><td>66114</td><td>-</td><td>0</td><td>CGGTGATGTCGCCGAGGTGC</td><td>AGG</td><td>||||||||[||||||||||||]PAM</td><td>NA</td><td>Intergenic</td><td>NA</td><td>NA</td></tr>
<tr><td>T29147</td><td>CGAGGTGCAGGACCACGTTGCGG</td><td>1000</td><td></td><td></td><td></td><td></td><td></td><td></td><td></td><td></td><td></td></tr>
<tr><td>Oligo fwd</td><td>CACCCGAGGTGCAGGACCACGTTG</td><td></td><td></td><td></td><td></td><td></td><td></td><td></td><td></td><td></td><td></td></tr>
<tr><td>Oligo rev</td><td>AAACCAACGTGGTCCTGCACCTCG</td><td></td><td></td><td></td><td></td><td></td><td></td><td></td><td></td><td></td><td></td></tr>
<tr><td>Chromosome</td><td>start</td><td>end</td><td>strand</td><td>MM</td><td>target_seq</td><td>PAM</td><td>alignment</td><td>distance</td><td>position</td><td>gene name</td><td>gene id</td></tr>
<tr><td>GU305914.1</td><td>66080</td><td>66102</td><td>-</td><td>0</td><td>CGAGGTGCAGGACCACGTTG</td><td>CGG</td><td>||||||||[||||||||||||]PAM</td><td>NA</td><td>Intergenic</td><td>NA</td><td>NA</td></tr>
<tr><td>T29148</td><td>ACAGCACCTTCACCAGCCCGAGG</td><td>1000</td><td></td><td></td><td></td><td></td><td></td><td></td><td></td><td></td><td></td></tr>
<tr><td>Oligo fwd</td><td>CACCACAGCACCTTCACCAGCCCG</td><td></td><td></td><td></td><td></td><td></td><td></td><td></td><td></td><td></td><td></td></tr>
<tr><td>Oligo rev</td><td>AAACCGGGCTGGTGAAGGTGCTGT</td><td></td><td></td><td></td><td></td><td></td><td></td><td></td><td></td><td></td><td></td></tr>
<tr><td>Chromosome</td><td>start</td><td>end</td><td>strand</td><td>MM</td><td>target_seq</td><td>PAM</td><td>alignment</td><td>distance</td><td>position</td><td>gene name</td><td>gene id</td></tr>
<tr><td>GU305914.1</td><td>66047</td><td>66069</td><td>-</td><td>0</td><td>ACAGCACCTTCACCAGCCCG</td><td>AGG</td><td>||||||||[||||||||||||]PAM</td><td>NA</td><td>Intergenic</td><td>NA</td><td>NA</td></tr>
<tr><td>T29149</td><td>TCACCAGCCCGAGGTCCGTCTGG</td><td>1000</td><td></td><td></td><td></td><td></td><td></td><td></td><td></td><td></td><td></td></tr>
<tr><td>Oligo fwd</td><td>CACCTCACCAGCCCGAGGTCCGTC</td><td></td><td></td><td></td><td></td><td></td><td></td><td></td><td></td><td></td><td></td></tr>
<tr><td>Oligo rev</td><td>AAACGACGGACCTCGGGCTGGTGA</td><td></td><td></td><td></td><td></td><td></td><td></td><td></td><td></td><td></td><td></td></tr>
<tr><td>Chromosome</td><td>start</td><td>end</td><td>strand</td><td>MM</td><td>target_seq</td><td>PAM</td><td>alignment</td><td>distance</td><td>position</td><td>gene name</td><td>gene id</td></tr>
<tr><td>GU305914.1</td><td>66038</td><td>66060</td><td>-</td><td>0</td><td>TCACCAGCCCGAGGTCCGTC</td><td>TGG</td><td>||||||||[||||||||||||]PAM</td><td>NA</td><td>Intergenic</td><td>NA</td><td>NA</td></tr>
<tr><td>T29150</td><td>TCTGGCTCACGTAGTGCAGCAGG</td><td>1000</td><td></td><td></td><td></td><td></td><td></td><td></td><td></td><td></td><td></td></tr>
<tr><td>Oligo fwd</td><td>CACCTCTGGCTCACGTAGTGCAGC</td><td></td><td></td><td></td><td></td><td></td><td></td><td></td><td></td><td></td><td></td></tr>
<tr><td>Oligo rev</td><td>AAACGCTGCACTACGTGAGCCAGA</td><td></td><td></td><td></td><td></td><td></td><td></td><td></td><td></td><td></td><td></td></tr>
<tr><td>Chromosome</td><td>start</td><td>end</td><td>strand</td><td>MM</td><td>target_seq</td><td>PAM</td><td>alignment</td><td>distance</td><td>position</td><td>gene name</td><td>gene id</td></tr>
<tr><td>GU305914.1</td><td>66020</td><td>66042</td><td>-</td><td>0</td><td>TCTGGCTCACGTAGTGCAGC</td><td>AGG</td><td>||||||||[||||||||||||]PAM</td><td>NA</td><td>Intergenic</td><td>NA</td><td>NA</td></tr>
<tr><td>T29151</td><td>GCAGGTTGTACAGCGACACCAGG</td><td>1000</td><td></td><td></td><td></td><td></td><td></td><td></td><td></td><td></td><td></td></tr>
<tr><td>Oligo fwd</td><td>CACCGCAGGTTGTACAGCGACACC</td><td></td><td></td><td></td><td></td><td></td><td></td><td></td><td></td><td></td><td></td></tr>
<tr><td>Oligo rev</td><td>AAACGGTGTCGCTGTACAACCTGC</td><td></td><td></td><td></td><td></td><td></td><td></td><td></td><td></td><td></td><td></td></tr>
<tr><td>Chromosome</td><td>start</td><td>end</td><td>strand</td><td>MM</td><td>target_seq</td><td>PAM</td><td>alignment</td><td>distance</td><td>position</td><td>gene name</td><td>gene id</td></tr>
<tr><td>GU305914.1</td><td>66002</td><td>66024</td><td>-</td><td>0</td><td>GCAGGTTGTACAGCGACACC</td><td>AGG</td><td>||||||||[||||||||||||]PAM</td><td>NA</td><td>Intergenic</td><td>NA</td><td>NA</td></tr>
<tr><td>T29152</td><td>TGTACAGCGACACCAGGTTGCGG</td><td>1000</td><td></td><td></td><td></td><td></td><td></td><td></td><td></td><td></td><td></td></tr>
<tr><td>Oligo fwd</td><td>CACCTGTACAGCGACACCAGGTTG</td><td></td><td></td><td></td><td></td><td></td><td></td><td></td><td></td><td></td><td></td></tr>
<tr><td>Oligo rev</td><td>AAACCAACCTGGTGTCGCTGTACA</td><td></td><td></td><td></td><td></td><td></td><td></td><td></td><td></td><td></td><td></td></tr>
<tr><td>Chromosome</td><td>start</td><td>end</td><td>strand</td><td>MM</td><td>target_seq</td><td>PAM</td><td>alignment</td><td>distance</td><td>position</td><td>gene name</td><td>gene id</td></tr>
<tr><td>GU305914.1</td><td>65996</td><td>66018</td><td>-</td><td>0</td><td>TGTACAGCGACACCAGGTTG</td><td>CGG</td><td>||||||||[||||||||||||]PAM</td><td>NA</td><td>Intergenic</td><td>NA</td><td>NA</td></tr>
<tr><td>T29153</td><td>CCAGGTTGCGGTACAGTTCCAGG</td><td>1000</td><td></td><td></td><td></td><td></td><td></td><td></td><td></td><td></td><td></td></tr>
<tr><td>Oligo fwd</td><td>CACCCCAGGTTGCGGTACAGTTCC</td><td></td><td></td><td></td><td></td><td></td><td></td><td></td><td></td><td></td><td></td></tr>
<tr><td>Oligo rev</td><td>AAACGGAACTGTACCGCAACCTGG</td><td></td><td></td><td></td><td></td><td></td><td></td><td></td><td></td><td></td><td></td></tr>
<tr><td>Chromosome</td><td>start</td><td>end</td><td>strand</td><td>MM</td><td>target_seq</td><td>PAM</td><td>alignment</td><td>distance</td><td>position</td><td>gene name</td><td>gene id</td></tr>
<tr><td>GU305914.1</td><td>65984</td><td>66006</td><td>-</td><td>0</td><td>CCAGGTTGCGGTACAGTTCC</td><td>AGG</td><td>||||||||[||||||||||||]PAM</td><td>NA</td><td>Intergenic</td><td>NA</td><td>NA</td></tr>
<tr><td>T29154</td><td>CGGTACAGTTCCAGGCGCCGCGG</td><td>1000</td><td></td><td></td><td></td><td></td><td></td><td></td><td></td><td></td><td></td></tr>
<tr><td>Oligo fwd</td><td>CACCCGGTACAGTTCCAGGCGCCG</td><td></td><td></td><td></td><td></td><td></td><td></td><td></td><td></td><td></td><td></td></tr>
<tr><td>Oligo rev</td><td>AAACCGGCGCCTGGAACTGTACCG</td><td></td><td></td><td></td><td></td><td></td><td></td><td></td><td></td><td></td><td></td></tr>
<tr><td>Chromosome</td><td>start</td><td>end</td><td>strand</td><td>MM</td><td>target_seq</td><td>PAM</td><td>alignment</td><td>distance</td><td>position</td><td>gene name</td><td>gene id</td></tr>
<tr><td>GU305914.1</td><td>65976</td><td>65998</td><td>-</td><td>0</td><td>CGGTACAGTTCCAGGCGCCG</td><td>CGG</td><td>||||||||[||||||||||||]PAM</td><td>NA</td><td>Intergenic</td><td>NA</td><td>NA</td></tr>
<tr><td>T29155</td><td>CCAGGCGCCGCGGCATCAGCAGG</td><td>1000</td><td></td><td></td><td></td><td></td><td></td><td></td><td></td><td></td><td></td></tr>
<tr><td>Oligo fwd</td><td>CACCCCAGGCGCCGCGGCATCAGC</td><td></td><td></td><td></td><td></td><td></td><td></td><td></td><td></td><td></td><td></td></tr>
<tr><td>Oligo rev</td><td>AAACGCTGATGCCGCGGCGCCTGG</td><td></td><td></td><td></td><td></td><td></td><td></td><td></td><td></td><td></td><td></td></tr>
<tr><td>Chromosome</td><td>start</td><td>end</td><td>strand</td><td>MM</td><td>target_seq</td><td>PAM</td><td>alignment</td><td>distance</td><td>position</td><td>gene name</td><td>gene id</td></tr>
</table>

GU305914.1 65966 65988 - 0 CCAGGCGCCGCGGCATCAGC AGG ||||||||[||||||||||||]PAM NA Intergenic NA NA

T29156 GGCGCCGCGGCATCAGCAGGCGG 1000
Oligo fwd CACCGGCGCCGCGGCATCAGCAGG
Oligo rev AAACCCTGCTGATGCCGCGGCGCC
Chromosome start end strand MM target_seq PAM alignment distance position gene name gene id
GU305914.1 65963 65985 - 0 GGCGCCGCGGCATCAGCAGG CGG ||||||||[||||||||||||]PAM NA Intergenic NA NA

T29157 GCATCAGCAGGCGGCAGATGTGG 1000
Oligo fwd CACCGCATCAGCAGGCGGCAGATG
Oligo rev AAACCATCTGCCGCCTGCTGATGC
Chromosome start end strand MM target_seq PAM alignment distance position gene name gene id
GU305914.1 65954 65976 - 0 GCATCAGCAGGCGGCAGATG TGG ||||||||[||||||||||||]PAM NA Intergenic NA NA

T29158 GCCGCATGTCGTCGTGCGCGCGG 1000
Oligo fwd CACCGCCGCATGTCGTCGTGCGCG
Oligo rev AAACCGCGCACGACGACATGCGGC
Chromosome start end strand MM target_seq PAM alignment distance position gene name gene id
GU305914.1 65918 65940 - 0 GCCGCATGTCGTCGTGCGCG CGG ||||||||[||||||||||||]PAM NA Intergenic NA NA

T29159 TCGTGCGCGCGGATCTCCGCCGG 1000
Oligo fwd CACCTCGTGCGCGCGGATCTCCGC
Oligo rev AAACGCGGAGATCCGCGCGCACGA
Chromosome start end strand MM target_seq PAM alignment distance position gene name gene id
GU305914.1 65907 65929 - 0 TCGTGCGCGCGGATCTCCGC CGG ||||||||[||||||||||||]PAM NA Intergenic NA NA
GU305914.1 43159 43181 - 4 TCGCGAGCGCGGATCGGCGC CGG |||-|-||[|||||||--|||]PAM NA Intergenic NA NA

T29160 TTCACGATCGCCTCGATCCGCGG 1000
Oligo fwd CACCTTCACGATCGCCTCGATCCG
Oligo rev AAACCGGATCGAGGCGATCGTGAA
Chromosome start end strand MM target_seq PAM alignment distance position gene name gene id
GU305914.1 65877 65899 - 0 TTCACGATCGCCTCGATCCG CGG ||||||||[||||||||||||]PAM NA Intergenic NA NA

T29161 TCACGATCGCCTCGATCCGCGGG 1000
Oligo fwd CACCTCACGATCGCCTCGATCCGC
Oligo rev AAACGCGGATCGAGGCGATCGTGA
Chromosome start end strand MM target_seq PAM alignment distance position gene name gene id
GU305914.1 65876 65898 - 0 TCACGATCGCCTCGATCCGC GGG ||||||||[||||||||||||]PAM NA Intergenic NA NA

T29162 CGTCAGCGTCTTGTCGAAGTCGG 1000
Oligo fwd CACCCGTCAGCGTCTTGTCGAAGT
Oligo rev AAACACTTCGACAAGACGCTGACG
Chromosome start end strand MM target_seq PAM alignment distance position gene name gene id
GU305914.1 65839 65861 - 0 CGTCAGCGTCTTGTCGAAGT CGG ||||||||[||||||||||||]PAM NA Intergenic NA NA

T29163 GTCAGCGTCTTGTCGAAGTCGGG 1000
Oligo fwd CACCGTCAGCGTCTTGTCGAAGTC
Oligo rev AAACGACTTCGACAAGACGCTGAC
Chromosome start end strand MM target_seq PAM alignment distance position gene name gene id
GU305914.1 65838 65860 - 0 GTCAGCGTCTTGTCGAAGTC GGG ||||||||[||||||||||||]PAM NA Intergenic NA NA

T29164 TCAGCGTCTTGTCGAAGTCGGGG 1000
Oligo fwd CACCTCAGCGTCTTGTCGAAGTCG
Oligo rev AAACCGACTTCGACAAGACGCTGA
Chromosome start end strand MM target_seq PAM alignment distance position gene name gene id
GU305914.1 65837 65859 - 0 TCAGCGTCTTGTCGAAGTCG GGG ||||||||[||||||||||||]PAM NA Intergenic NA NA

T29165 CGTCTTGTCGAAGTCGGGGTAGG 1000
Oligo fwd CACCCGTCTTGTCGAAGTCGGGGT
Oligo rev AAACACCCCGACTTCGACAAGACG
Chromosome start end strand MM target_seq PAM alignment distance position gene name gene id
GU305914.1 65833 65855 - 0 CGTCTTGTCGAAGTCGGGGT AGG ||||||||[||||||||||||]PAM NA Intergenic NA NA

T29166 AAGTCGGGGTAGGTGCTCAGCGG 1000
Oligo fwd CACCAAGTCGGGGTAGGTGCTCAG
Oligo rev AAACCTGAGCACCTACCCCGACTT
Chromosome start end strand MM target_seq PAM alignment distance position gene name gene id
GU305914.1 65823 65845 - 0 AAGTCGGGGTAGGTGCTCAG CGG ||||||||[||||||||||||]PAM NA Intergenic NA NA

T29167 GCACGCCCGCGCGATGTCCTCGG 1000
Oligo fwd CACCGCACGCCCGCGCGATGTCCT
Oligo rev AAACAGGACATCGCGCGGGCGTGC
Chromosome start end strand MM target_seq PAM alignment distance position gene name gene id
GU305914.1 65755 65777 - 0 GCACGCCCGCGCGATGTCCT CGG ||||||||[||||||||||||]PAM NA Intergenic NA NA
GU305914.1 29939 29961 + 4 GACGGCCCGCGCGAAGTCCT CGG |---||||[||||||-|||||]PAM NA Intergenic NA NA

T29168 CACGCCCGCGCGATGTCCTCGGG 1000
Oligo fwd CACCCACGCCCGCGCGATGTCCTC
Oligo rev AAACGAGGACATCGCGCGGGCGTG
Chromosome start end strand MM target_seq PAM alignment distance position gene name gene id
GU305914.1 65754 65776 - 0 CACGCCCGCGCGATGTCCTC GGG ||||||||[||||||||||||]PAM NA Intergenic NA NA
GU305914.1 29940 29962 + 4 ACGGCCCGCGCGAAGTCCTC GGG ---|||||[|||||-||||||]PAM NA Intergenic NA NA

T29169 CGCGCGATGTCCTCGGGCGTCGG 1000
Oligo fwd CACCCGCGCGATGTCCTCGGGCGT
Oligo rev AAACACGCCCGAGGACATCGCGCG
Chromosome start end strand MM target_seq PAM alignment distance position gene name gene id
GU305914.1 65748 65770 - 0 CGCGCGATGTCCTCGGGCGT CGG ||||||||[||||||||||||]PAM NA Intergenic NA NA

<table>
<tr><td>T29170</td><td>TCGGGCGTCGGATCTTCCTGCGG</td><td>1000</td><td></td><td></td><td></td><td></td><td></td><td></td><td></td><td></td><td></td></tr>
<tr><td>Oligo fwd</td><td>CACCTCGGGCGTCGGATCTTCCTG</td><td></td><td></td><td></td><td></td><td></td><td></td><td></td><td></td><td></td><td></td></tr>
<tr><td>Oligo rev</td><td>AAACCAGGAAGATCCGACGCCCGA</td><td></td><td></td><td></td><td></td><td></td><td></td><td></td><td></td><td></td><td></td></tr>
<tr><td>Chromosome</td><td>start</td><td>end</td><td>strand</td><td>MM</td><td>target_seq</td><td>PAM</td><td>alignment</td><td>distance</td><td>position</td><td>gene name</td><td>gene id</td></tr>
<tr><td>GU305914.1</td><td>65736</td><td>65758</td><td>-</td><td>0</td><td>TCGGGCGTCGGATCTTCCTG</td><td>CGG</td><td>||||||||[||||||||||||]PAM</td><td>NA</td><td>Intergenic</td><td>NA</td><td>NA</td></tr>
<tr><td>T29171</td><td>GGCGTCGGATCTTCCTGCGGCGG</td><td>1000</td><td></td><td></td><td></td><td></td><td></td><td></td><td></td><td></td><td></td></tr>
<tr><td>Oligo fwd</td><td>CACCGGCGTCGGATCTTCCTGCGG</td><td></td><td></td><td></td><td></td><td></td><td></td><td></td><td></td><td></td><td></td></tr>
<tr><td>Oligo rev</td><td>AAACCCGCAGGAAGATCCGACGCC</td><td></td><td></td><td></td><td></td><td></td><td></td><td></td><td></td><td></td><td></td></tr>
<tr><td>Chromosome</td><td>start</td><td>end</td><td>strand</td><td>MM</td><td>target_seq</td><td>PAM</td><td>alignment</td><td>distance</td><td>position</td><td>gene name</td><td>gene id</td></tr>
<tr><td>GU305914.1</td><td>65733</td><td>65755</td><td>-</td><td>0</td><td>GGCGTCGGATCTTCCTGCGG</td><td>CGG</td><td>||||||||[||||||||||||]PAM</td><td>NA</td><td>Intergenic</td><td>NA</td><td>NA</td></tr>
<tr><td>T29172</td><td>GGATCTTCCTGCGGCGGCAGCGG</td><td>1000</td><td></td><td></td><td></td><td></td><td></td><td></td><td></td><td></td><td></td></tr>
<tr><td>Oligo fwd</td><td>CACCGGATCTTCCTGCGGCGGCAG</td><td></td><td></td><td></td><td></td><td></td><td></td><td></td><td></td><td></td><td></td></tr>
<tr><td>Oligo rev</td><td>AAACCTGCCGCCGCAGGAAGATCC</td><td></td><td></td><td></td><td></td><td></td><td></td><td></td><td></td><td></td><td></td></tr>
<tr><td>Chromosome</td><td>start</td><td>end</td><td>strand</td><td>MM</td><td>target_seq</td><td>PAM</td><td>alignment</td><td>distance</td><td>position</td><td>gene name</td><td>gene id</td></tr>
<tr><td>GU305914.1</td><td>65727</td><td>65749</td><td>-</td><td>0</td><td>GGATCTTCCTGCGGCGGCAG</td><td>CGG</td><td>||||||||[||||||||||||]PAM</td><td>NA</td><td>Intergenic</td><td>NA</td><td>NA</td></tr>
<tr><td>T29173</td><td>TCCTGCGGCGGCAGCGGCAGCGG</td><td>1000</td><td></td><td></td><td></td><td></td><td></td><td></td><td></td><td></td><td></td></tr>
<tr><td>Oligo fwd</td><td>CACCTCCTGCGGCGGCAGCGGCAG</td><td></td><td></td><td></td><td></td><td></td><td></td><td></td><td></td><td></td><td></td></tr>
<tr><td>Oligo rev</td><td>AAACCTGCCGCTGCCGCCGCAGGA</td><td></td><td></td><td></td><td></td><td></td><td></td><td></td><td></td><td></td><td></td></tr>
<tr><td>Chromosome</td><td>start</td><td>end</td><td>strand</td><td>MM</td><td>target_seq</td><td>PAM</td><td>alignment</td><td>distance</td><td>position</td><td>gene name</td><td>gene id</td></tr>
<tr><td>GU305914.1</td><td>65721</td><td>65743</td><td>-</td><td>0</td><td>TCCTGCGGCGGCAGCGGCAG</td><td>CGG</td><td>||||||||[||||||||||||]PAM</td><td>NA</td><td>Intergenic</td><td>NA</td><td>NA</td></tr>
<tr><td>GU305914.1</td><td>163908</td><td>163930</td><td>+</td><td>3</td><td>GGCGGCGGCGGCAGCGGCAG</td><td>CGG</td><td>--|-||||[||||||||||||]PAM</td><td>NA</td><td>Intergenic</td><td>NA</td><td>NA</td></tr>
<tr><td>GU305914.1</td><td>163902</td><td>163924</td><td>+</td><td>4</td><td>AGCGGCGGCGGCGGCGGCAG</td><td>CGG</td><td>--|-||||[||||-|||||||]PAM</td><td>NA</td><td>Intergenic</td><td>NA</td><td>NA</td></tr>
<tr><td>GU305914.1</td><td>30006</td><td>30028</td><td>+</td><td>4</td><td>GGCGGCGGCGGCGGCGGCAG</td><td>TGG</td><td>--|-||||[||||-|||||||]PAM</td><td>NA</td><td>Intergenic</td><td>NA</td><td>NA</td></tr>
<tr><td>GU305914.1</td><td>149476</td><td>149498</td><td>-</td><td>4</td><td>GGCGGCGGCGGCGGCGGCAG</td><td>CGG</td><td>--|-||||[||||-|||||||]PAM</td><td>NA</td><td>Intergenic</td><td>NA</td><td>NA</td></tr>
<tr><td>T29174</td><td>CCTGCGGCGGCAGCGGCAGCGGG</td><td>1000</td><td></td><td></td><td></td><td></td><td></td><td></td><td></td><td></td><td></td></tr>
<tr><td>Oligo fwd</td><td>CACCCCTGCGGCGGCAGCGGCAGC</td><td></td><td></td><td></td><td></td><td></td><td></td><td></td><td></td><td></td><td></td></tr>
<tr><td>Oligo rev</td><td>AAACGCTGCCGCTGCCGCCGCAGG</td><td></td><td></td><td></td><td></td><td></td><td></td><td></td><td></td><td></td><td></td></tr>
<tr><td>Chromosome</td><td>start</td><td>end</td><td>strand</td><td>MM</td><td>target_seq</td><td>PAM</td><td>alignment</td><td>distance</td><td>position</td><td>gene name</td><td>gene id</td></tr>
<tr><td>GU305914.1</td><td>65720</td><td>65742</td><td>-</td><td>0</td><td>CCTGCGGCGGCAGCGGCAGC</td><td>GGG</td><td>||||||||[||||||||||||]PAM</td><td>NA</td><td>Intergenic</td><td>NA</td><td>NA</td></tr>
<tr><td>GU305914.1</td><td>149475</td><td>149497</td><td>-</td><td>3</td><td>GCGGCGGCGGCGGCGGCAGC</td><td>GGG</td><td>-|-|||||[|||-||||||||]PAM</td><td>NA</td><td>Intergenic</td><td>NA</td><td>NA</td></tr>
<tr><td>GU305914.1</td><td>168830</td><td>168852</td><td>+</td><td>4</td><td>GCGGCGGCGGCGGCGACAGC</td><td>GGG</td><td>-|-|||||[|||-|||-||||]PAM</td><td>NA</td><td>Intergenic</td><td>NA</td><td>NA</td></tr>
<tr><td>GU305914.1</td><td>164958</td><td>164980</td><td>-</td><td>4</td><td>GCGGCGGCGGCAGCAGCTGC</td><td>GGG</td><td>-|-|||||[||||||-||-||]PAM</td><td>NA</td><td>Intergenic</td><td>NA</td><td>NA</td></tr>
<tr><td>T29175</td><td>GCAGCGGCAGCGGGTCCGTCAGG</td><td>1000</td><td></td><td></td><td></td><td></td><td></td><td></td><td></td><td></td><td></td></tr>
<tr><td>Oligo fwd</td><td>CACCGCAGCGGCAGCGGGTCCGTC</td><td></td><td></td><td></td><td></td><td></td><td></td><td></td><td></td><td></td><td></td></tr>
<tr><td>Oligo rev</td><td>AAACGACGGACCCGCTGCCGCTGC</td><td></td><td></td><td></td><td></td><td></td><td></td><td></td><td></td><td></td><td></td></tr>
<tr><td>Chromosome</td><td>start</td><td>end</td><td>strand</td><td>MM</td><td>target_seq</td><td>PAM</td><td>alignment</td><td>distance</td><td>position</td><td>gene name</td><td>gene id</td></tr>
<tr><td>GU305914.1</td><td>65711</td><td>65733</td><td>-</td><td>0</td><td>GCAGCGGCAGCGGGTCCGTC</td><td>AGG</td><td>||||||||[||||||||||||]PAM</td><td>NA</td><td>Intergenic</td><td>NA</td><td>NA</td></tr>
<tr><td>T29176</td><td>GCGGGTCCGTCAGGTCCTCCTGG</td><td>1000</td><td></td><td></td><td></td><td></td><td></td><td></td><td></td><td></td><td></td></tr>
<tr><td>Oligo fwd</td><td>CACCGCGGGTCCGTCAGGTCCTCC</td><td></td><td></td><td></td><td></td><td></td><td></td><td></td><td></td><td></td><td></td></tr>
<tr><td>Oligo rev</td><td>AAACGGAGGACCTGACGGACCCGC</td><td></td><td></td><td></td><td></td><td></td><td></td><td></td><td></td><td></td><td></td></tr>
<tr><td>Chromosome</td><td>start</td><td>end</td><td>strand</td><td>MM</td><td>target_seq</td><td>PAM</td><td>alignment</td><td>distance</td><td>position</td><td>gene name</td><td>gene id</td></tr>
<tr><td>GU305914.1</td><td>65702</td><td>65724</td><td>-</td><td>0</td><td>GCGGGTCCGTCAGGTCCTCC</td><td>TGG</td><td>||||||||[||||||||||||]PAM</td><td>NA</td><td>Intergenic</td><td>NA</td><td>NA</td></tr>
<tr><td>T29177</td><td>CCCACTTGTTCGCCGCGTCGCGG</td><td>1000</td><td></td><td></td><td></td><td></td><td></td><td></td><td></td><td></td><td></td></tr>
<tr><td>Oligo fwd</td><td>CACCCCCACTTGTTCGCCGCGTCG</td><td></td><td></td><td></td><td></td><td></td><td></td><td></td><td></td><td></td><td></td></tr>
<tr><td>Oligo rev</td><td>AAACCGACGCGGCGAACAAGTGGG</td><td></td><td></td><td></td><td></td><td></td><td></td><td></td><td></td><td></td><td></td></tr>
<tr><td>Chromosome</td><td>start</td><td>end</td><td>strand</td><td>MM</td><td>target_seq</td><td>PAM</td><td>alignment</td><td>distance</td><td>position</td><td>gene name</td><td>gene id</td></tr>
<tr><td>GU305914.1</td><td>65657</td><td>65679</td><td>-</td><td>0</td><td>CCCACTTGTTCGCCGCGTCG</td><td>CGG</td><td>||||||||[||||||||||||]PAM</td><td>NA</td><td>Intergenic</td><td>NA</td><td>NA</td></tr>
<tr><td>T29178</td><td>TGTTCGCCGCGTCGCGGTACAGG</td><td>1000</td><td></td><td></td><td></td><td></td><td></td><td></td><td></td><td></td><td></td></tr>
<tr><td>Oligo fwd</td><td>CACCTGTTCGCCGCGTCGCGGTAC</td><td></td><td></td><td></td><td></td><td></td><td></td><td></td><td></td><td></td><td></td></tr>
<tr><td>Oligo rev</td><td>AAACGTACCGCGACGCGGCGAACA</td><td></td><td></td><td></td><td></td><td></td><td></td><td></td><td></td><td></td><td></td></tr>
<tr><td>Chromosome</td><td>start</td><td>end</td><td>strand</td><td>MM</td><td>target_seq</td><td>PAM</td><td>alignment</td><td>distance</td><td>position</td><td>gene name</td><td>gene id</td></tr>
<tr><td>GU305914.1</td><td>65651</td><td>65673</td><td>-</td><td>0</td><td>TGTTCGCCGCGTCGCGGTAC</td><td>AGG</td><td>||||||||[||||||||||||]PAM</td><td>NA</td><td>Intergenic</td><td>NA</td><td>NA</td></tr>
<tr><td>T29179</td><td>TACAGGTACTCGCGACCCGCCGG</td><td>1000</td><td></td><td></td><td></td><td></td><td></td><td></td><td></td><td></td><td></td></tr>
<tr><td>Oligo fwd</td><td>CACCTACAGGTACTCGCGACCCGC</td><td></td><td></td><td></td><td></td><td></td><td></td><td></td><td></td><td></td><td></td></tr>
<tr><td>Oligo rev</td><td>AAACGCGGGTCGCGAGTACCTGTA</td><td></td><td></td><td></td><td></td><td></td><td></td><td></td><td></td><td></td><td></td></tr>
<tr><td>Chromosome</td><td>start</td><td>end</td><td>strand</td><td>MM</td><td>target_seq</td><td>PAM</td><td>alignment</td><td>distance</td><td>position</td><td>gene name</td><td>gene id</td></tr>
<tr><td>GU305914.1</td><td>65634</td><td>65656</td><td>-</td><td>0</td><td>TACAGGTACTCGCGACCCGC</td><td>CGG</td><td>||||||||[||||||||||||]PAM</td><td>NA</td><td>Intergenic</td><td>NA</td><td>NA</td></tr>
<tr><td>T29180</td><td>CGTAGCGCAGCGCGAACAGCCGG</td><td>1000</td><td></td><td></td><td></td><td></td><td></td><td></td><td></td><td></td><td></td></tr>
<tr><td>Oligo fwd</td><td>CACCCGTAGCGCAGCGCGAACAGC</td><td></td><td></td><td></td><td></td><td></td><td></td><td></td><td></td><td></td><td></td></tr>
<tr><td>Oligo rev</td><td>AAACGCTGTTCGCGCTGCGCTACG</td><td></td><td></td><td></td><td></td><td></td><td></td><td></td><td></td><td></td><td></td></tr>
<tr><td>Chromosome</td><td>start</td><td>end</td><td>strand</td><td>MM</td><td>target_seq</td><td>PAM</td><td>alignment</td><td>distance</td><td>position</td><td>gene name</td><td>gene id</td></tr>
<tr><td>GU305914.1</td><td>65573</td><td>65595</td><td>-</td><td>0</td><td>CGTAGCGCAGCGCGAACAGC</td><td>CGG</td><td>||||||||[||||||||||||]PAM</td><td>NA</td><td>Intergenic</td><td>NA</td><td>NA</td></tr>
<tr><td>GU305914.1</td><td>221430</td><td>221452</td><td>+</td><td>4</td><td>CGTAGATAAGCGCGAACAGT</td><td>AGG</td><td>|||||---[|||||||||||-]PAM</td><td>NA</td><td>Intergenic</td><td>NA</td><td>NA</td></tr>
<tr><td>T29181</td><td>GTTGATGTTGAACGCCACGTTGG</td><td>1000</td><td></td><td></td><td></td><td></td><td></td><td></td><td></td><td></td><td></td></tr>
<tr><td>Oligo fwd</td><td>CACCGTTGATGTTGAACGCCACGT</td><td></td><td></td><td></td><td></td><td></td><td></td><td></td><td></td><td></td><td></td></tr>
<tr><td>Oligo rev</td><td>AAACACGTGGCGTTCAACATCAAC</td><td></td><td></td><td></td><td></td><td></td><td></td><td></td><td></td><td></td><td></td></tr>
<tr><td>Chromosome</td><td>start</td><td>end</td><td>strand</td><td>MM</td><td>target_seq</td><td>PAM</td><td>alignment</td><td>distance</td><td>position</td><td>gene name</td><td>gene id</td></tr>
<tr><td>GU305914.1</td><td>65551</td><td>65573</td><td>-</td><td>0</td><td>GTTGATGTTGAACGCCACGT</td><td>TGG</td><td>||||||||[||||||||||||]PAM</td><td>NA</td><td>Intergenic</td><td>NA</td><td>NA</td></tr>
<tr><td>T29182</td><td>CGTTGGCCCACACGTCCTGCAGG</td><td>1000</td><td></td><td></td><td></td><td></td><td></td><td></td><td></td><td></td><td></td></tr>
<tr><td>Oligo fwd</td><td>CACCCGTTGGCCCACACGTCCTGC</td><td></td><td></td><td></td><td></td><td></td><td></td><td></td><td></td><td></td><td></td></tr>
<tr><td>Oligo rev</td><td>AAACGCAGGACGTGTGGGCCAACG</td><td></td><td></td><td></td><td></td><td></td><td></td><td></td><td></td><td></td><td></td></tr>
<tr><td>Chromosome</td><td>start</td><td>end</td><td>strand</td><td>MM</td><td>target_seq</td><td>PAM</td><td>alignment</td><td>distance</td><td>position</td><td>gene name</td><td>gene id</td></tr>
<tr><td>GU305914.1</td><td>65534</td><td>65556</td><td>-</td><td>0</td><td>CGTTGGCCCACACGTCCTGC</td><td>AGG</td><td>||||||||[||||||||||||]PAM</td><td>NA</td><td>Intergenic</td><td>NA</td><td>NA</td></tr>
<tr><td>T29183</td><td>GTTGGCCCACACGTCCTGCAGGG</td><td>1000</td><td></td><td></td><td></td><td></td><td></td><td></td><td></td><td></td><td></td></tr>
<tr><td>Oligo fwd</td><td>CACCGTTGGCCCACACGTCCTGCA</td><td></td><td></td><td></td><td></td><td></td><td></td><td></td><td></td><td></td><td></td></tr>
<tr><td>Oligo rev</td><td>AAACTGCAGGACGTGTGGGCCAAC</td><td></td><td></td><td></td><td></td><td></td><td></td><td></td><td></td><td></td><td></td></tr>
</table>

| | | | | | | | | | | | | |
|---|---|---|---|---|---|---|---|---|---|---|---|---|
| Chromosome | start | | end | strand | MM | target_seq | PAM | alignment | distance | position | gene name | gene id |
| GU305914.1 | | 65533 | 65555 | - | 0 | GTTGGCCCACACGTCCTGCA | GGG | \|\|\|\|\|\|\|\|[\|\|\|\|\|\|\|\|\|\|\|\|]PAM | NA | Intergenic | NA | NA |
| T29184 | CCCACACGTCCTGCAGGGACCGG | | 1000 | | | | | | | | | |
| Oligo fwd | CACCCCCACACGTCCTGCAGGGAC | | | | | | | | | | | |
| Oligo rev | AAACGTCCCTGCAGGACGTGTGGG | | | | | | | | | | | |
| Chromosome | start | | end | strand | MM | target_seq | PAM | alignment | distance | position | gene name | gene id |
| GU305914.1 | | 65528 | 65550 | - | 0 | CCCACACGTCCTGCAGGGAC | CGG | \|\|\|\|\|\|\|\|[\|\|\|\|\|\|\|\|\|\|\|\|]PAM | NA | Intergenic | NA | NA |
| T29185 | GGGACCGGTTGCCCGTCAACAGG | | 1000 | | | | | | | | | |
| Oligo fwd | CACCGGGACCGGTTGCCCGTCAAC | | | | | | | | | | | |
| Oligo rev | AAACGTTGACGGGCAACCGGTCCC | | | | | | | | | | | |
| Chromosome | start | | end | strand | MM | target_seq | PAM | alignment | distance | position | gene name | gene id |
| GU305914.1 | | 65513 | 65535 | - | 0 | GGGACCGGTTGCCCGTCAAC | AGG | \|\|\|\|\|\|\|\|[\|\|\|\|\|\|\|\|\|\|\|\|]PAM | NA | Intergenic | NA | NA |
| T29186 | GGACCGGTTGCCCGTCAACAGGG | | 1000 | | | | | | | | | |
| Oligo fwd | CACCGGACCGGTTGCCCGTCAACA | | | | | | | | | | | |
| Oligo rev | AAACTGTTGACGGGCAACCGGTCC | | | | | | | | | | | |
| Chromosome | start | | end | strand | MM | target_seq | PAM | alignment | distance | position | gene name | gene id |
| GU305914.1 | | 65512 | 65534 | - | 0 | GGACCGGTTGCCCGTCAACA | GGG | \|\|\|\|\|\|\|\|[\|\|\|\|\|\|\|\|\|\|\|\|]PAM | NA | Intergenic | NA | NA |
| T29187 | ACAGGGAGCGCTGTTGTTCCAGG | | 1000 | | | | | | | | | |
| Oligo fwd | CACCACAGGGAGCGCTGTTGTTCC | | | | | | | | | | | |
| Oligo rev | AAACGGAACAACAGCGCTCCCTGT | | | | | | | | | | | |
| Chromosome | start | | end | strand | MM | target_seq | PAM | alignment | distance | position | gene name | gene id |
| GU305914.1 | | 65495 | 65517 | - | 0 | ACAGGGAGCGCTGTTGTTCC | AGG | \|\|\|\|\|\|\|\|[\|\|\|\|\|\|\|\|\|\|\|\|]PAM | NA | Intergenic | NA | NA |
| T29188 | CCAGGATCAGCCGCTTCGCGTGG | | 1000 | | | | | | | | | |
| Oligo fwd | CACCCCAGGATCAGCCGCTTCGCG | | | | | | | | | | | |
| Oligo rev | AAACCGCGAAGCGGCTGATCCTGG | | | | | | | | | | | |
| Chromosome | start | | end | strand | MM | target_seq | PAM | alignment | distance | position | gene name | gene id |
| GU305914.1 | | 65477 | 65499 | - | 0 | CCAGGATCAGCCGCTTCGCG | TGG | \|\|\|\|\|\|\|\|[\|\|\|\|\|\|\|\|\|\|\|\|]PAM | NA | Intergenic | NA | NA |
| T29189 | GCCGCTTCGCGTGGTAAAACAGG | | 1000 | | | | | | | | | |
| Oligo fwd | CACCGCCGCTTCGCGTGGTAAAAC | | | | | | | | | | | |
| Oligo rev | AAACGTTTTACCACGCGAAGCGGC | | | | | | | | | | | |
| Chromosome | start | | end | strand | MM | target_seq | PAM | alignment | distance | position | gene name | gene id |
| GU305914.1 | | 65468 | 65490 | - | 0 | GCCGCTTCGCGTGGTAAAAC | AGG | \|\|\|\|\|\|\|\|[\|\|\|\|\|\|\|\|\|\|\|\|]PAM | NA | Intergenic | NA | NA |
| T29190 | CAGGAACGTGTTGCTGTATCTGG | | 1000 | | | | | | | | | |
| Oligo fwd | CACCCAGGAACGTGTTGCTGTATC | | | | | | | | | | | |
| Oligo rev | AAACGATACAGCAACACGTTCCTG | | | | | | | | | | | |
| Chromosome | start | | end | strand | MM | target_seq | PAM | alignment | distance | position | gene name | gene id |
| GU305914.1 | | 65449 | 65471 | - | 0 | CAGGAACGTGTTGCTGTATC | TGG | \|\|\|\|\|\|\|\|[\|\|\|\|\|\|\|\|\|\|\|\|]PAM | NA | Intergenic | NA | NA |
| T29191 | TGGACAGACAGAGAAAGTGCAGG | | 1000 | | | | | | | | | |
| Oligo fwd | CACCTGGACAGACAGAGAAAGTGC | | | | | | | | | | | |
| Oligo rev | AAACGCACTTTCTCTGTCTGTCCA | | | | | | | | | | | |
| Chromosome | start | | end | strand | MM | target_seq | PAM | alignment | distance | position | gene name | gene id |
| GU305914.1 | | 65429 | 65451 | - | 0 | TGGACAGACAGAGAAAGTGC | AGG | \|\|\|\|\|\|\|\|[\|\|\|\|\|\|\|\|\|\|\|\|]PAM | NA | Intergenic | NA | NA |
| T29192 | GTGCGCGTTGTACACCAGAAAGG | | 1000 | | | | | | | | | |
| Oligo fwd | CACCGTGCGCGTTGTACACCAGAA | | | | | | | | | | | |
| Oligo rev | AAACTTCTGGTGTACAACGCGCAC | | | | | | | | | | | |
| Chromosome | start | | end | strand | MM | target_seq | PAM | alignment | distance | position | gene name | gene id |
| GU305914.1 | | 65407 | 65429 | - | 0 | GTGCGCGTTGTACACCAGAA | AGG | \|\|\|\|\|\|\|\|[\|\|\|\|\|\|\|\|\|\|\|\|]PAM | NA | Intergenic | NA | NA |
| T29193 | TGTACACCAGAAAGGTGATGTGG | | 1000 | | | | | | | | | |
| Oligo fwd | CACCTGTACACCAGAAAGGTGATG | | | | | | | | | | | |
| Oligo rev | AAACCATCACCTTTCTGGTGTACA | | | | | | | | | | | |
| Chromosome | start | | end | strand | MM | target_seq | PAM | alignment | distance | position | gene name | gene id |
| GU305914.1 | | 65399 | 65421 | - | 0 | TGTACACCAGAAAGGTGATG | TGG | \|\|\|\|\|\|\|\|[\|\|\|\|\|\|\|\|\|\|\|\|]PAM | NA | Intergenic | NA | NA |
| T29194 | GCTGAACTCGTCGATGCTTCTGG | | 1000 | | | | | | | | | |
| Oligo fwd | CACCGCTGAACTCGTCGATGCTTC | | | | | | | | | | | |
| Oligo rev | AAACGAAGCATCGACGAGTTCAGC | | | | | | | | | | | |
| Chromosome | start | | end | strand | MM | target_seq | PAM | alignment | distance | position | gene name | gene id |
| GU305914.1 | | 65365 | 65387 | - | 0 | GCTGAACTCGTCGATGCTTC | TGG | \|\|\|\|\|\|\|\|[\|\|\|\|\|\|\|\|\|\|\|\|]PAM | NA | Intergenic | NA | NA |
| T29195 | GTCGATGCTTCTGGACGCCGCGG | | 1000 | | | | | | | | | |
| Oligo fwd | CACCGTCGATGCTTCTGGACGCCG | | | | | | | | | | | |
| Oligo rev | AAACCGGCGTCCAGAAGCATCGAC | | | | | | | | | | | |
| Chromosome | start | | end | strand | MM | target_seq | PAM | alignment | distance | position | gene name | gene id |
| GU305914.1 | | 65356 | 65378 | - | 0 | GTCGATGCTTCTGGACGCCG | CGG | \|\|\|\|\|\|\|\|[\|\|\|\|\|\|\|\|\|\|\|\|]PAM | NA | Intergenic | NA | NA |
| T29196 | TTCTGGACGCCGCGGCCGTGCGG | | 1000 | | | | | | | | | |
| Oligo fwd | CACCTTCTGGACGCCGCGGCCGTG | | | | | | | | | | | |
| Oligo rev | AAACCACGGCCGCGGCGTCCAGAA | | | | | | | | | | | |
| Chromosome | start | | end | strand | MM | target_seq | PAM | alignment | distance | position | gene name | gene id |
| GU305914.1 | | 65348 | 65370 | - | 0 | TTCTGGACGCCGCGGCCGTG | CGG | \|\|\|\|\|\|\|\|[\|\|\|\|\|\|\|\|\|\|\|\|]PAM | NA | Intergenic | NA | NA |
| T29197 | CGCCGCGGCCGTGCGGTACGTGG | | 1000 | | | | | | | | | |
| Oligo fwd | CACCCGCCGCGGCCGTGCGGTACG | | | | | | | | | | | |
| Oligo rev | AAACCGTACCGCACGGCCGCGGCG | | | | | | | | | | | |
| Chromosome | start | | end | strand | MM | target_seq | PAM | alignment | distance | position | gene name | gene id |
| GU305914.1 | | 65341 | 65363 | - | 0 | CGCCGCGGCCGTGCGGTACG | TGG | \|\|\|\|\|\|\|\|[\|\|\|\|\|\|\|\|\|\|\|\|]PAM | NA | Intergenic | NA | NA |
| T29198 | CCGTGCGGTACGTGGCGTGCAGG | | 1000 | | | | | | | | | |
| Oligo fwd | CACCCCGTGCGGTACGTGGCGTGC | | | | | | | | | | | |

Oligo rev AAACGCACGCCACGTACCGCACGG
Chromosome start end strand MM target_seq PAM alignment distance position gene name gene id
GU305914.1 65333 65355 - 0 CCGTGCGGTACGTGGCGTGC AGG ||||||||[||||||||||||]PAM NA Intergenic NA NA

T29199 CGTGCGGTACGTGGCGTGCAGGG 1000
Oligo fwd CACCCGTGCGGTACGTGGCGTGCA
Oligo rev AAACTGCACGCCACGTACCGCACG
Chromosome start end strand MM target_seq PAM alignment distance position gene name gene id
GU305914.1 65332 65354 - 0 CGTGCGGTACGTGGCGTGCA GGG ||||||||[||||||||||||]PAM NA Intergenic NA NA

T29200 TGGCGTGCAGGGAGATCATGCGG 1000
Oligo fwd CACCTGGCGTGCAGGGAGATCATG
Oligo rev AAACCATGATCTCCCTGCACGCCA
Chromosome start end strand MM target_seq PAM alignment distance position gene name gene id
GU305914.1 65321 65343 - 0 TGGCGTGCAGGGAGATCATG CGG ||||||||[||||||||||||]PAM NA Intergenic NA NA

T29201 CAGGGAGATCATGCGGATGTAGG 1000
Oligo fwd CACCCAGGGAGATCATGCGGATGT
Oligo rev AAACACATCCGCATGATCTCCCTG
Chromosome start end strand MM target_seq PAM alignment distance position gene name gene id
GU305914.1 65314 65336 - 0 CAGGGAGATCATGCGGATGT AGG ||||||||[||||||||||||]PAM NA Intergenic NA NA

T29202 ATGTAGGTCGCCGTGCTCTTCGG 1000
Oligo fwd CACCATGTAGGTCGCCGTGCTCTT
Oligo rev AAACAAGAGCACGGCGACCTACAT
Chromosome start end strand MM target_seq PAM alignment distance position gene name gene id
GU305914.1 65298 65320 - 0 ATGTAGGTCGCCGTGCTCTT CGG ||||||||[||||||||||||]PAM NA Intergenic NA NA

T29203 TGTAGGTCGCCGTGCTCTTCGGG 1000
Oligo fwd CACCTGTAGGTCGCCGTGCTCTTC
Oligo rev AAACGAAGAGCACGGCGACCTACA
Chromosome start end strand MM target_seq PAM alignment distance position gene name gene id
GU305914.1 65297 65319 - 0 TGTAGGTCGCCGTGCTCTTC GGG ||||||||[||||||||||||]PAM NA Intergenic NA NA

T29204 GTAGGTCGCCGTGCTCTTCGGGG 1000
Oligo fwd CACCGTAGGTCGCCGTGCTCTTCG
Oligo rev AAACCGAAGAGCACGGCGACCTAC
Chromosome start end strand MM target_seq PAM alignment distance position gene name gene id
GU305914.1 65296 65318 - 0 GTAGGTCGCCGTGCTCTTCG GGG ||||||||[||||||||||||]PAM NA Intergenic NA NA

T29205 TGCTCTTCGGGGTCAACCCGAGG 1000
Oligo fwd CACCTGCTCTTCGGGGTCAACCCG
Oligo rev AAACCGGGTTGACCCCGAAGAGCA
Chromosome start end strand MM target_seq PAM alignment distance position gene name gene id
GU305914.1 65285 65307 - 0 TGCTCTTCGGGGTCAACCCG AGG ||||||||[||||||||||||]PAM NA Intergenic NA NA

T29206 TCTTCGGGGTCAACCCGAGGCGG 1000
Oligo fwd CACCTCTTCGGGGTCAACCCGAGG
Oligo rev AAACCCTCGGGTTGACCCCGAAGA
Chromosome start end strand MM target_seq PAM alignment distance position gene name gene id
GU305914.1 65282 65304 - 0 TCTTCGGGGTCAACCCGAGG CGG ||||||||[||||||||||||]PAM NA Intergenic NA NA

T29207 TGACGCTGACGAGCCGCGAGCGG 1000
Oligo fwd CACCTGACGCTGACGAGCCGCGAG
Oligo rev AAACCTCGCGGCTCGTCAGCGTCA
Chromosome start end strand MM target_seq PAM alignment distance position gene name gene id
GU305914.1 65228 65250 - 0 TGACGCTGACGAGCCGCGAG CGG ||||||||[||||||||||||]PAM NA Intergenic NA NA

T29208 GACGCTGACGAGCCGCGAGCGGG 1000
Oligo fwd CACCGACGCTGACGAGCCGCGAGC
Oligo rev AAACGCTCGCGGCTCGTCAGCGTC
Chromosome start end strand MM target_seq PAM alignment distance position gene name gene id
GU305914.1 65227 65249 - 0 GACGCTGACGAGCCGCGAGC GGG ||||||||[||||||||||||]PAM NA Intergenic NA NA

T29209 GCCGCGAGCGGGACGATTCCTGG 1000
Oligo fwd CACCGCCGCGAGCGGGACGATTCC
Oligo rev AAACGGAATCGTCCCGCTCGCGGC
Chromosome start end strand MM target_seq PAM alignment distance position gene name gene id
GU305914.1 65216 65238 - 0 GCCGCGAGCGGGACGATTCC TGG ||||||||[||||||||||||]PAM NA Intergenic NA NA

T29210 CCGCGAGCGGGACGATTCCTGGG 1000
Oligo fwd CACCCCGCGAGCGGGACGATTCCT
Oligo rev AAACAGGAATCGTCCCGCTCGCGG
Chromosome start end strand MM target_seq PAM alignment distance position gene name gene id
GU305914.1 65215 65237 - 0 CCGCGAGCGGGACGATTCCT GGG ||||||||[||||||||||||]PAM NA Intergenic NA NA

T29211 AGCGGGACGATTCCTGGGTCTGG 1000
Oligo fwd CACCAGCGGGACGATTCCTGGGTC
Oligo rev AAACGACCCAGGAATCGTCCCGCT
Chromosome start end strand MM target_seq PAM alignment distance position gene name gene id
GU305914.1 65210 65232 - 0 AGCGGGACGATTCCTGGGTC TGG ||||||||[||||||||||||]PAM NA Intergenic NA NA

T29212 GCGGGACGATTCCTGGGTCTGGG 1000
Oligo fwd CACCGCGGGACGATTCCTGGGTCT
Oligo rev AAACAGACCCAGGAATCGTCCCGC
Chromosome start end strand MM target_seq PAM alignment distance position gene name gene id
GU305914.1 65209 65231 - 0 GCGGGACGATTCCTGGGTCT GGG ||||||||[||||||||||||]PAM NA Intergenic NA NA

T29213 GGTGAATAAAATGTCCGCGATGG 1000

<table>
<tr><td>Oligo fwd</td><td>CACCGGTGAATAAAATGTCCGCGA</td><td></td><td></td><td></td><td></td><td></td><td></td><td></td><td></td><td></td><td></td></tr>
<tr><td>Oligo rev</td><td>AAACTCGCGGACATTTTATTCACC</td><td></td><td></td><td></td><td></td><td></td><td></td><td></td><td></td><td></td><td></td></tr>
<tr><td>Chromosome</td><td>start</td><td>end</td><td>strand</td><td>MM</td><td>target_seq</td><td>PAM</td><td>alignment</td><td>distance</td><td>position</td><td>gene name</td><td>gene id</td></tr>
<tr><td>GU305914.1</td><td>65188</td><td>65210</td><td>-</td><td>0</td><td>GGTGAATAAAATGTCCGCGA</td><td>TGG</td><td>||||||||[||||||||||||]PAM</td><td>NA</td><td>Intergenic</td><td>NA</td><td>NA</td></tr>
<tr><td></td><td></td><td></td><td></td><td></td><td></td><td></td><td></td><td></td><td></td><td></td><td></td></tr>
<tr><td>T29214</td><td>CTCTTGCGCGTCGAAAAAAGAGG</td><td>1000</td><td></td><td></td><td></td><td></td><td></td><td></td><td></td><td></td><td></td></tr>
<tr><td>Oligo fwd</td><td>CACCCTCTTGCGCGTCGAAAAAAG</td><td></td><td></td><td></td><td></td><td></td><td></td><td></td><td></td><td></td><td></td></tr>
<tr><td>Oligo rev</td><td>AAACCTTTTTTCGACGCGCAAGAG</td><td></td><td></td><td></td><td></td><td></td><td></td><td></td><td></td><td></td><td></td></tr>
<tr><td>Chromosome</td><td>start</td><td>end</td><td>strand</td><td>MM</td><td>target_seq</td><td>PAM</td><td>alignment</td><td>distance</td><td>position</td><td>gene name</td><td>gene id</td></tr>
<tr><td>GU305914.1</td><td>65038</td><td>65060</td><td>-</td><td>0</td><td>CTCTTGCGCGTCGAAAAAAG</td><td>AGG</td><td>||||||||[||||||||||||]PAM</td><td>NA</td><td>Intergenic</td><td>NA</td><td>NA</td></tr>
<tr><td></td><td></td><td></td><td></td><td></td><td></td><td></td><td></td><td></td><td></td><td></td><td></td></tr>
<tr><td>T29215</td><td>GCGCGTCGAAAAAAGAGGACAGG</td><td>1000</td><td></td><td></td><td></td><td></td><td></td><td></td><td></td><td></td><td></td></tr>
<tr><td>Oligo fwd</td><td>CACCGCGCGTCGAAAAAAGAGGAC</td><td></td><td></td><td></td><td></td><td></td><td></td><td></td><td></td><td></td><td></td></tr>
<tr><td>Oligo rev</td><td>AAACGTCCTCTTTTTTCGACGCGC</td><td></td><td></td><td></td><td></td><td></td><td></td><td></td><td></td><td></td><td></td></tr>
<tr><td>Chromosome</td><td>start</td><td>end</td><td>strand</td><td>MM</td><td>target_seq</td><td>PAM</td><td>alignment</td><td>distance</td><td>position</td><td>gene name</td><td>gene id</td></tr>
<tr><td>GU305914.1</td><td>65033</td><td>65055</td><td>-</td><td>0</td><td>GCGCGTCGAAAAAAGAGGAC</td><td>AGG</td><td>||||||||[||||||||||||]PAM</td><td>NA</td><td>Intergenic</td><td>NA</td><td>NA</td></tr>
<tr><td></td><td></td><td></td><td></td><td></td><td></td><td></td><td></td><td></td><td></td><td></td><td></td></tr>
<tr><td>T29216</td><td>CGTCGAAAAAAGAGGACAGGCGG</td><td>1000</td><td></td><td></td><td></td><td></td><td></td><td></td><td></td><td></td><td></td></tr>
<tr><td>Oligo fwd</td><td>CACCCGTCGAAAAAAGAGGACAGG</td><td></td><td></td><td></td><td></td><td></td><td></td><td></td><td></td><td></td><td></td></tr>
<tr><td>Oligo rev</td><td>AAACCCTGTCCTCTTTTTTCGACG</td><td></td><td></td><td></td><td></td><td></td><td></td><td></td><td></td><td></td><td></td></tr>
<tr><td>Chromosome</td><td>start</td><td>end</td><td>strand</td><td>MM</td><td>target_seq</td><td>PAM</td><td>alignment</td><td>distance</td><td>position</td><td>gene name</td><td>gene id</td></tr>
<tr><td>GU305914.1</td><td>65030</td><td>65052</td><td>-</td><td>0</td><td>CGTCGAAAAAAGAGGACAGG</td><td>CGG</td><td>||||||||[||||||||||||]PAM</td><td>NA</td><td>Intergenic</td><td>NA</td><td>NA</td></tr>
<tr><td></td><td></td><td></td><td></td><td></td><td></td><td></td><td></td><td></td><td></td><td></td><td></td></tr>
<tr><td>T29217</td><td>CGAAAAAAGAGGACAGGCGGTGG</td><td>1000</td><td></td><td></td><td></td><td></td><td></td><td></td><td></td><td></td><td></td></tr>
<tr><td>Oligo fwd</td><td>CACCCGAAAAAAGAGGACAGGCGG</td><td></td><td></td><td></td><td></td><td></td><td></td><td></td><td></td><td></td><td></td></tr>
<tr><td>Oligo rev</td><td>AAACCCGCCTGTCCTCTTTTTTCG</td><td></td><td></td><td></td><td></td><td></td><td></td><td></td><td></td><td></td><td></td></tr>
<tr><td>Chromosome</td><td>start</td><td>end</td><td>strand</td><td>MM</td><td>target_seq</td><td>PAM</td><td>alignment</td><td>distance</td><td>position</td><td>gene name</td><td>gene id</td></tr>
<tr><td>GU305914.1</td><td>65027</td><td>65049</td><td>-</td><td>0</td><td>CGAAAAAAGAGGACAGGCGG</td><td>TGG</td><td>||||||||[||||||||||||]PAM</td><td>NA</td><td>Intergenic</td><td>NA</td><td>NA</td></tr>
<tr><td></td><td></td><td></td><td></td><td></td><td></td><td></td><td></td><td></td><td></td><td></td><td></td></tr>
<tr><td>T29218</td><td>GAAAAAAGAGGACAGGCGGTGGG</td><td>1000</td><td></td><td></td><td></td><td></td><td></td><td></td><td></td><td></td><td></td></tr>
<tr><td>Oligo fwd</td><td>CACCGAAAAAAGAGGACAGGCGGT</td><td></td><td></td><td></td><td></td><td></td><td></td><td></td><td></td><td></td><td></td></tr>
<tr><td>Oligo rev</td><td>AAACACCGCCTGTCCTCTTTTTTC</td><td></td><td></td><td></td><td></td><td></td><td></td><td></td><td></td><td></td><td></td></tr>
<tr><td>Chromosome</td><td>start</td><td>end</td><td>strand</td><td>MM</td><td>target_seq</td><td>PAM</td><td>alignment</td><td>distance</td><td>position</td><td>gene name</td><td>gene id</td></tr>
<tr><td>GU305914.1</td><td>65026</td><td>65048</td><td>-</td><td>0</td><td>GAAAAAAGAGGACAGGCGGT</td><td>GGG</td><td>||||||||[||||||||||||]PAM</td><td>NA</td><td>Intergenic</td><td>NA</td><td>NA</td></tr>
<tr><td></td><td></td><td></td><td></td><td></td><td></td><td></td><td></td><td></td><td></td><td></td><td></td></tr>
<tr><td>T29219</td><td>AGGACAGGCGGTGGGTCAGTAGG</td><td>1000</td><td></td><td></td><td></td><td></td><td></td><td></td><td></td><td></td><td></td></tr>
<tr><td>Oligo fwd</td><td>CACCAGGACAGGCGGTGGGTCAGT</td><td></td><td></td><td></td><td></td><td></td><td></td><td></td><td></td><td></td><td></td></tr>
<tr><td>Oligo rev</td><td>AAACACTGACCCACCGCCTGTCCT</td><td></td><td></td><td></td><td></td><td></td><td></td><td></td><td></td><td></td><td></td></tr>
<tr><td>Chromosome</td><td>start</td><td>end</td><td>strand</td><td>MM</td><td>target_seq</td><td>PAM</td><td>alignment</td><td>distance</td><td>position</td><td>gene name</td><td>gene id</td></tr>
<tr><td>GU305914.1</td><td>65018</td><td>65040</td><td>-</td><td>0</td><td>AGGACAGGCGGTGGGTCAGT</td><td>AGG</td><td>||||||||[||||||||||||]PAM</td><td>NA</td><td>Intergenic</td><td>NA</td><td>NA</td></tr>
<tr><td></td><td></td><td></td><td></td><td></td><td></td><td></td><td></td><td></td><td></td><td></td><td></td></tr>
<tr><td>T29220</td><td>AGTAGGCGCCGTCGTCTCTGAGG</td><td>1000</td><td></td><td></td><td></td><td></td><td></td><td></td><td></td><td></td><td></td></tr>
<tr><td>Oligo fwd</td><td>CACCAGTAGGCGCCGTCGTCTCTG</td><td></td><td></td><td></td><td></td><td></td><td></td><td></td><td></td><td></td><td></td></tr>
<tr><td>Oligo rev</td><td>AAACCAGAGACGACGGCGCCTACT</td><td></td><td></td><td></td><td></td><td></td><td></td><td></td><td></td><td></td><td></td></tr>
<tr><td>Chromosome</td><td>start</td><td>end</td><td>strand</td><td>MM</td><td>target_seq</td><td>PAM</td><td>alignment</td><td>distance</td><td>position</td><td>gene name</td><td>gene id</td></tr>
<tr><td>GU305914.1</td><td>65001</td><td>65023</td><td>-</td><td>0</td><td>AGTAGGCGCCGTCGTCTCTG</td><td>AGG</td><td>||||||||[||||||||||||]PAM</td><td>NA</td><td>Intergenic</td><td>NA</td><td>NA</td></tr>
<tr><td></td><td></td><td></td><td></td><td></td><td></td><td></td><td></td><td></td><td></td><td></td><td></td></tr>
<tr><td>T29221</td><td>AGGCGCCGTCGTCTCTGAGGCGG</td><td>1000</td><td></td><td></td><td></td><td></td><td></td><td></td><td></td><td></td><td></td></tr>
<tr><td>Oligo fwd</td><td>CACCAGGCGCCGTCGTCTCTGAGG</td><td></td><td></td><td></td><td></td><td></td><td></td><td></td><td></td><td></td><td></td></tr>
<tr><td>Oligo rev</td><td>AAACCCTCAGAGACGACGGCGCCT</td><td></td><td></td><td></td><td></td><td></td><td></td><td></td><td></td><td></td><td></td></tr>
<tr><td>Chromosome</td><td>start</td><td>end</td><td>strand</td><td>MM</td><td>target_seq</td><td>PAM</td><td>alignment</td><td>distance</td><td>position</td><td>gene name</td><td>gene id</td></tr>
<tr><td>GU305914.1</td><td>64998</td><td>65020</td><td>-</td><td>0</td><td>AGGCGCCGTCGTCTCTGAGG</td><td>CGG</td><td>||||||||[||||||||||||]PAM</td><td>NA</td><td>Intergenic</td><td>NA</td><td>NA</td></tr>
<tr><td>GU305914.1</td><td>226851</td><td>226873</td><td>+</td><td>4</td><td>AGGCCCCTCCGTCTCTGAGC</td><td>AGG</td><td>||||-||-[-||||||||||-]PAM</td><td>NA</td><td>Intergenic</td><td>NA</td><td>NA</td></tr>
<tr><td></td><td></td><td></td><td></td><td></td><td></td><td></td><td></td><td></td><td></td><td></td><td></td></tr>
<tr><td>T29222</td><td>CTCTGAGGCGGCGTCCCGCCCGG</td><td>1000</td><td></td><td></td><td></td><td></td><td></td><td></td><td></td><td></td><td></td></tr>
<tr><td>Oligo fwd</td><td>CACCCTCTGAGGCGGCGTCCCGCC</td><td></td><td></td><td></td><td></td><td></td><td></td><td></td><td></td><td></td><td></td></tr>
<tr><td>Oligo rev</td><td>AAACGGCGGGACGCCGCCTCAGAG</td><td></td><td></td><td></td><td></td><td></td><td></td><td></td><td></td><td></td><td></td></tr>
<tr><td>Chromosome</td><td>start</td><td>end</td><td>strand</td><td>MM</td><td>target_seq</td><td>PAM</td><td>alignment</td><td>distance</td><td>position</td><td>gene name</td><td>gene id</td></tr>
<tr><td>GU305914.1</td><td>64986</td><td>65008</td><td>-</td><td>0</td><td>CTCTGAGGCGGCGTCCCGCC</td><td>CGG</td><td>||||||||[||||||||||||]PAM</td><td>NA</td><td>Intergenic</td><td>NA</td><td>NA</td></tr>
<tr><td></td><td></td><td></td><td></td><td></td><td></td><td></td><td></td><td></td><td></td><td></td><td></td></tr>
<tr><td>T29223</td><td>GCGGCGTCCCGCCCGGCGACTGG</td><td>1000</td><td></td><td></td><td></td><td></td><td></td><td></td><td></td><td></td><td></td></tr>
<tr><td>Oligo fwd</td><td>CACCGCGGCGTCCCGCCCGGCGAC</td><td></td><td></td><td></td><td></td><td></td><td></td><td></td><td></td><td></td><td></td></tr>
<tr><td>Oligo rev</td><td>AAACGTCGCCGGGCGGGACGCCGC</td><td></td><td></td><td></td><td></td><td></td><td></td><td></td><td></td><td></td><td></td></tr>
<tr><td>Chromosome</td><td>start</td><td>end</td><td>strand</td><td>MM</td><td>target_seq</td><td>PAM</td><td>alignment</td><td>distance</td><td>position</td><td>gene name</td><td>gene id</td></tr>
<tr><td>GU305914.1</td><td>64979</td><td>65001</td><td>-</td><td>0</td><td>GCGGCGTCCCGCCCGGCGAC</td><td>TGG</td><td>||||||||[||||||||||||]PAM</td><td>NA</td><td>Intergenic</td><td>NA</td><td>NA</td></tr>
<tr><td></td><td></td><td></td><td></td><td></td><td></td><td></td><td></td><td></td><td></td><td></td><td></td></tr>
<tr><td>T29224</td><td>CTCTTCTCCGACGCGCGCCGCGG</td><td>1000</td><td></td><td></td><td></td><td></td><td></td><td></td><td></td><td></td><td></td></tr>
<tr><td>Oligo fwd</td><td>CACCCTCTTCTCCGACGCGCGCCG</td><td></td><td></td><td></td><td></td><td></td><td></td><td></td><td></td><td></td><td></td></tr>
<tr><td>Oligo rev</td><td>AAACCGGCGCGCGTCGGAGAAGAG</td><td></td><td></td><td></td><td></td><td></td><td></td><td></td><td></td><td></td><td></td></tr>
<tr><td>Chromosome</td><td>start</td><td>end</td><td>strand</td><td>MM</td><td>target_seq</td><td>PAM</td><td>alignment</td><td>distance</td><td>position</td><td>gene name</td><td>gene id</td></tr>
<tr><td>GU305914.1</td><td>64951</td><td>64973</td><td>-</td><td>0</td><td>CTCTTCTCCGACGCGCGCCG</td><td>CGG</td><td>||||||||[||||||||||||]PAM</td><td>NA</td><td>Intergenic</td><td>NA</td><td>NA</td></tr>
<tr><td>GU305914.1</td><td>197663</td><td>197685</td><td>+</td><td>4</td><td>CGATTCTCCGCCGCGGGCCG</td><td>TGG</td><td>|--|||||[||-||||-||||]PAM</td><td>NA</td><td>Intergenic</td><td>NA</td><td>NA</td></tr>
<tr><td></td><td></td><td></td><td></td><td></td><td></td><td></td><td></td><td></td><td></td><td></td><td></td></tr>
<tr><td>T29225</td><td>CGACGCGCGCCGCGGTCCGCAGG</td><td>1000</td><td></td><td></td><td></td><td></td><td></td><td></td><td></td><td></td><td></td></tr>
<tr><td>Oligo fwd</td><td>CACCCGACGCGCGCCGCGGTCCGC</td><td></td><td></td><td></td><td></td><td></td><td></td><td></td><td></td><td></td><td></td></tr>
<tr><td>Oligo rev</td><td>AAACGCGGACCGCGGCGCGCGTCG</td><td></td><td></td><td></td><td></td><td></td><td></td><td></td><td></td><td></td><td></td></tr>
<tr><td>Chromosome</td><td>start</td><td>end</td><td>strand</td><td>MM</td><td>target_seq</td><td>PAM</td><td>alignment</td><td>distance</td><td>position</td><td>gene name</td><td>gene id</td></tr>
<tr><td>GU305914.1</td><td>64943</td><td>64965</td><td>-</td><td>0</td><td>CGACGCGCGCCGCGGTCCGC</td><td>AGG</td><td>||||||||[||||||||||||]PAM</td><td>NA</td><td>Intergenic</td><td>NA</td><td>NA</td></tr>
<tr><td></td><td></td><td></td><td></td><td></td><td></td><td></td><td></td><td></td><td></td><td></td><td></td></tr>
<tr><td>T29226</td><td>GCGCGCCGCGGTCCGCAGGATGG</td><td>1000</td><td></td><td></td><td></td><td></td><td></td><td></td><td></td><td></td><td></td></tr>
<tr><td>Oligo fwd</td><td>CACCGCGCGCCGCGGTCCGCAGGA</td><td></td><td></td><td></td><td></td><td></td><td></td><td></td><td></td><td></td><td></td></tr>
<tr><td>Oligo rev</td><td>AAACTCCTGCGGACCGCGGCGCGC</td><td></td><td></td><td></td><td></td><td></td><td></td><td></td><td></td><td></td><td></td></tr>
<tr><td>Chromosome</td><td>start</td><td>end</td><td>strand</td><td>MM</td><td>target_seq</td><td>PAM</td><td>alignment</td><td>distance</td><td>position</td><td>gene name</td><td>gene id</td></tr>
<tr><td>GU305914.1</td><td>64939</td><td>64961</td><td>-</td><td>0</td><td>GCGCGCCGCGGTCCGCAGGA</td><td>TGG</td><td>||||||||[||||||||||||]PAM</td><td>NA</td><td>Intergenic</td><td>NA</td><td>NA</td></tr>
<tr><td></td><td></td><td></td><td></td><td></td><td></td><td></td><td></td><td></td><td></td><td></td><td></td></tr>
<tr><td>T29227</td><td>CAGGATGGCGAACACCACGCCGG</td><td>1000</td><td></td><td></td><td></td><td></td><td></td><td></td><td></td><td></td><td></td></tr>
<tr><td>Oligo fwd</td><td>CACCCAGGATGGCGAACACCACGC</td><td></td><td></td><td></td><td></td><td></td><td></td><td></td><td></td><td></td><td></td></tr>
<tr><td>Oligo rev</td><td>AAACGCGTGGTGTTCGCCATCCTG</td><td></td><td></td><td></td><td></td><td></td><td></td><td></td><td></td><td></td><td></td></tr>
<tr><td>Chromosome</td><td>start</td><td>end</td><td>strand</td><td>MM</td><td>target_seq</td><td>PAM</td><td>alignment</td><td>distance</td><td>position</td><td>gene name</td><td>gene id</td></tr>
</table>

| | | | | | | | | | | | |
|---|---|---|---|---|---|---|---|---|---|---|---|
| GU305914.1 | 64924 | 64946 | - | 0 | CAGGATGGCGAACACCACGC | CGG | \|\|\|\|\|\|\|\|[\|\|\|\|\|\|\|\|\|\|\|\|]PAM | NA | Intergenic | NA | NA |
| T29228 | AGGATGGCGAACACCACGCCGGG | 1000 | | | | | | | | | |
| Oligo fwd | CACCAGGATGGCGAACACCACGCC | | | | | | | | | | |
| Oligo rev | AAACGGCGTGGTGTTCGCCATCCT | | | | | | | | | | |
| Chromosome | start | end | strand | MM | target_seq | PAM | alignment | distance | position | gene name | gene id |
| GU305914.1 | 64923 | 64945 | - | 0 | AGGATGGCGAACACCACGCC | GGG | \|\|\|\|\|\|\|\|[\|\|\|\|\|\|\|\|\|\|\|\|]PAM | NA | Intergenic | NA | NA |
| T29229 | GGATGGCGAACACCACGCCGGGG | 1000 | | | | | | | | | |
| Oligo fwd | CACCGGATGGCGAACACCACGCCG | | | | | | | | | | |
| Oligo rev | AAACCGGCGTGGTGTTCGCCATCC | | | | | | | | | | |
| Chromosome | start | end | strand | MM | target_seq | PAM | alignment | distance | position | gene name | gene id |
| GU305914.1 | 64922 | 64944 | - | 0 | GGATGGCGAACACCACGCCG | GGG | \|\|\|\|\|\|\|\|[\|\|\|\|\|\|\|\|\|\|\|\|]PAM | NA | Intergenic | NA | NA |
| T29230 | GCCGGGGTAGAAGAGCACGTTGG | 1000 | | | | | | | | | |
| Oligo fwd | CACCGCCGGGGTAGAAGAGCACGT | | | | | | | | | | |
| Oligo rev | AAACACGTGCTCTTCTACCCCGGC | | | | | | | | | | |
| Chromosome | start | end | strand | MM | target_seq | PAM | alignment | distance | position | gene name | gene id |
| GU305914.1 | 64906 | 64928 | - | 0 | GCCGGGGTAGAAGAGCACGT | TGG | \|\|\|\|\|\|\|\|[\|\|\|\|\|\|\|\|\|\|\|\|]PAM | NA | Intergenic | NA | NA |
| GU305914.1 | 137865 | 137887 | + | 3 | GGCGAGGAAGAAGAGCACGT | GGG | \|-\|\|-\|\|-[\|\|\|\|\|\|\|\|\|\|\|\|]PAM | NA | Intergenic | NA | NA |
| T29231 | GGGGTAGAAGAGCACGTTGGCGG | 1000 | | | | | | | | | |
| Oligo fwd | CACCGGGGTAGAAGAGCACGTTGG | | | | | | | | | | |
| Oligo rev | AAACCCAACGTGCTCTTCTACCCC | | | | | | | | | | |
| Chromosome | start | end | strand | MM | target_seq | PAM | alignment | distance | position | gene name | gene id |
| GU305914.1 | 64903 | 64925 | - | 0 | GGGGTAGAAGAGCACGTTGG | CGG | \|\|\|\|\|\|\|\|[\|\|\|\|\|\|\|\|\|\|\|\|]PAM | NA | Intergenic | NA | NA |
| T29232 | GGGTAGAAGAGCACGTTGGCGGG | 1000 | | | | | | | | | |
| Oligo fwd | CACCGGGTAGAAGAGCACGTTGGC | | | | | | | | | | |
| Oligo rev | AAACGCCAACGTGCTCTTCTACCC | | | | | | | | | | |
| Chromosome | start | end | strand | MM | target_seq | PAM | alignment | distance | position | gene name | gene id |
| GU305914.1 | 64902 | 64924 | - | 0 | GGGTAGAAGAGCACGTTGGC | GGG | \|\|\|\|\|\|\|\|[\|\|\|\|\|\|\|\|\|\|\|\|]PAM | NA | Intergenic | NA | NA |
| T29233 | GCACGTTGGCGGGCGAGATGTGG | 1000 | | | | | | | | | |
| Oligo fwd | CACCGCACGTTGGCGGGCGAGATG | | | | | | | | | | |
| Oligo rev | AAACCATCTCGCCCGCCAACGTGC | | | | | | | | | | |
| Chromosome | start | end | strand | MM | target_seq | PAM | alignment | distance | position | gene name | gene id |
| GU305914.1 | 64892 | 64914 | - | 0 | GCACGTTGGCGGGCGAGATG | TGG | \|\|\|\|\|\|\|\|[\|\|\|\|\|\|\|\|\|\|\|\|]PAM | NA | Intergenic | NA | NA |
| T29234 | GATGTGGCGCAGCACGCTCACGG | 1000 | | | | | | | | | |
| Oligo fwd | CACCGATGTGGCGCAGCACGCTCA | | | | | | | | | | |
| Oligo rev | AAACTGAGCGTGCTGCGCCACATC | | | | | | | | | | |
| Chromosome | start | end | strand | MM | target_seq | PAM | alignment | distance | position | gene name | gene id |
| GU305914.1 | 64876 | 64898 | - | 0 | GATGTGGCGCAGCACGCTCA | CGG | \|\|\|\|\|\|\|\|[\|\|\|\|\|\|\|\|\|\|\|\|]PAM | NA | Intergenic | NA | NA |
| T29235 | TGGCGCAGCACGCTCACGGATGG | 1000 | | | | | | | | | |
| Oligo fwd | CACCTGGCGCAGCACGCTCACGGA | | | | | | | | | | |
| Oligo rev | AAACTCCGTGAGCGTGCTGCGCCA | | | | | | | | | | |
| Chromosome | start | end | strand | MM | target_seq | PAM | alignment | distance | position | gene name | gene id |
| GU305914.1 | 64872 | 64894 | - | 0 | TGGCGCAGCACGCTCACGGA | TGG | \|\|\|\|\|\|\|\|[\|\|\|\|\|\|\|\|\|\|\|\|]PAM | NA | Intergenic | NA | NA |
| T29236 | GGCGCAGCACGCTCACGGATGGG | 1000 | | | | | | | | | |
| Oligo fwd | CACCGGCGCAGCACGCTCACGGAT | | | | | | | | | | |
| Oligo rev | AAACATCCGTGAGCGTGCTGCGCC | | | | | | | | | | |
| Chromosome | start | end | strand | MM | target_seq | PAM | alignment | distance | position | gene name | gene id |
| GU305914.1 | 64871 | 64893 | - | 0 | GGCGCAGCACGCTCACGGAT | GGG | \|\|\|\|\|\|\|\|[\|\|\|\|\|\|\|\|\|\|\|\|]PAM | NA | Intergenic | NA | NA |
| T29237 | GCGCAGCACGCTCACGGATGGGG | 1000 | | | | | | | | | |
| Oligo fwd | CACCGCGCAGCACGCTCACGGATG | | | | | | | | | | |
| Oligo rev | AAACCATCCGTGAGCGTGCTGCGC | | | | | | | | | | |
| Chromosome | start | end | strand | MM | target_seq | PAM | alignment | distance | position | gene name | gene id |
| GU305914.1 | 64870 | 64892 | - | 0 | GCGCAGCACGCTCACGGATG | GGG | \|\|\|\|\|\|\|\|[\|\|\|\|\|\|\|\|\|\|\|\|]PAM | NA | Intergenic | NA | NA |
| T29238 | GCACGCTCACGGATGGGGTCAGG | 1000 | | | | | | | | | |
| Oligo fwd | CACCGCACGCTCACGGATGGGGTC | | | | | | | | | | |
| Oligo rev | AAACGACCCCATCCGTGAGCGTGC | | | | | | | | | | |
| Chromosome | start | end | strand | MM | target_seq | PAM | alignment | distance | position | gene name | gene id |
| GU305914.1 | 64865 | 64887 | - | 0 | GCACGCTCACGGATGGGGTC | AGG | \|\|\|\|\|\|\|\|[\|\|\|\|\|\|\|\|\|\|\|\|]PAM | NA | Intergenic | NA | NA |
| T29239 | GCTCACGGATGGGGTCAGGATGG | 1000 | | | | | | | | | |
| Oligo fwd | CACCGCTCACGGATGGGGTCAGGA | | | | | | | | | | |
| Oligo rev | AAACTCCTGACCCCATCCGTGAGC | | | | | | | | | | |
| Chromosome | start | end | strand | MM | target_seq | PAM | alignment | distance | position | gene name | gene id |
| GU305914.1 | 64861 | 64883 | - | 0 | GCTCACGGATGGGGTCAGGA | TGG | \|\|\|\|\|\|\|\|[\|\|\|\|\|\|\|\|\|\|\|\|]PAM | NA | Intergenic | NA | NA |
| T29240 | CTCACGGATGGGGTCAGGATGGG | 1000 | | | | | | | | | |
| Oligo fwd | CACCCTCACGGATGGGGTCAGGAT | | | | | | | | | | |
| Oligo rev | AAACATCCTGACCCCATCCGTGAG | | | | | | | | | | |
| Chromosome | start | end | strand | MM | target_seq | PAM | alignment | distance | position | gene name | gene id |
| GU305914.1 | 64860 | 64882 | - | 0 | CTCACGGATGGGGTCAGGAT | GGG | \|\|\|\|\|\|\|\|[\|\|\|\|\|\|\|\|\|\|\|\|]PAM | NA | Intergenic | NA | NA |
| T29241 | TCACGGATGGGGTCAGGATGGGG | 1000 | | | | | | | | | |
| Oligo fwd | CACCTCACGGATGGGGTCAGGATG | | | | | | | | | | |
| Oligo rev | AAACCATCCTGACCCCATCCGTGA | | | | | | | | | | |
| Chromosome | start | end | strand | MM | target_seq | PAM | alignment | distance | position | gene name | gene id |
| GU305914.1 | 64859 | 64881 | - | 0 | TCACGGATGGGGTCAGGATG | GGG | \|\|\|\|\|\|\|\|[\|\|\|\|\|\|\|\|\|\|\|\|]PAM | NA | Intergenic | NA | NA |
| T29242 | GGGTCAGGATGGGGAAACGCAGG | 1000 | | | | | | | | | |
| Oligo fwd | CACCGGGTCAGGATGGGGAAACGC | | | | | | | | | | |

<table>
<tr><td>Oligo rev</td><td>AAACGCGTTTCCCCATCCTGACCC</td><td></td><td></td><td></td><td></td><td></td><td></td><td></td><td></td><td></td><td></td></tr>
<tr><td>Chromosome</td><td>start</td><td>end</td><td>strand</td><td>MM</td><td>target_seq</td><td>PAM</td><td>alignment</td><td>distance</td><td>position</td><td>gene name</td><td>gene id</td></tr>
<tr><td>GU305914.1</td><td>64850</td><td>64872</td><td>-</td><td>0</td><td>GGGTCAGGATGGGGAAACGC</td><td>AGG</td><td>||||||||[||||||||||||]PAM</td><td>NA</td><td>Intergenic</td><td>NA</td><td>NA</td></tr>
<tr><td></td><td></td><td></td><td></td><td></td><td></td><td></td><td></td><td></td><td></td><td></td><td></td></tr>
<tr><td>T29243</td><td>CAGGATGGGGAAACGCAGGTCGG</td><td>1000</td><td></td><td></td><td></td><td></td><td></td><td></td><td></td><td></td><td></td></tr>
<tr><td>Oligo fwd</td><td>CACCCAGGATGGGGAAACGCAGGT</td><td></td><td></td><td></td><td></td><td></td><td></td><td></td><td></td><td></td><td></td></tr>
<tr><td>Oligo rev</td><td>AAACACCTGCGTTTCCCCATCCTG</td><td></td><td></td><td></td><td></td><td></td><td></td><td></td><td></td><td></td><td></td></tr>
<tr><td>Chromosome</td><td>start</td><td>end</td><td>strand</td><td>MM</td><td>target_seq</td><td>PAM</td><td>alignment</td><td>distance</td><td>position</td><td>gene name</td><td>gene id</td></tr>
<tr><td>GU305914.1</td><td>64846</td><td>64868</td><td>-</td><td>0</td><td>CAGGATGGGGAAACGCAGGT</td><td>CGG</td><td>||||||||[||||||||||||]PAM</td><td>NA</td><td>Intergenic</td><td>NA</td><td>NA</td></tr>
<tr><td></td><td></td><td></td><td></td><td></td><td></td><td></td><td></td><td></td><td></td><td></td><td></td></tr>
<tr><td>T29244</td><td>ACGCAGGTCGGAGCGCTCCTTGG</td><td>1000</td><td></td><td></td><td></td><td></td><td></td><td></td><td></td><td></td><td></td></tr>
<tr><td>Oligo fwd</td><td>CACCACGCAGGTCGGAGCGCTCCT</td><td></td><td></td><td></td><td></td><td></td><td></td><td></td><td></td><td></td><td></td></tr>
<tr><td>Oligo rev</td><td>AAACAGGAGCGCTCCGACCTGCGT</td><td></td><td></td><td></td><td></td><td></td><td></td><td></td><td></td><td></td><td></td></tr>
<tr><td>Chromosome</td><td>start</td><td>end</td><td>strand</td><td>MM</td><td>target_seq</td><td>PAM</td><td>alignment</td><td>distance</td><td>position</td><td>gene name</td><td>gene id</td></tr>
<tr><td>GU305914.1</td><td>64834</td><td>64856</td><td>-</td><td>0</td><td>ACGCAGGTCGGAGCGCTCCT</td><td>TGG</td><td>||||||||[||||||||||||]PAM</td><td>NA</td><td>Intergenic</td><td>NA</td><td>NA</td></tr>
<tr><td></td><td></td><td></td><td></td><td></td><td></td><td></td><td></td><td></td><td></td><td></td><td></td></tr>
<tr><td>T29245</td><td>CTTGGCCGACAGCGAGCGCGAGG</td><td>1000</td><td></td><td></td><td></td><td></td><td></td><td></td><td></td><td></td><td></td></tr>
<tr><td>Oligo fwd</td><td>CACCCTTGGCCGACAGCGAGCGCG</td><td></td><td></td><td></td><td></td><td></td><td></td><td></td><td></td><td></td><td></td></tr>
<tr><td>Oligo rev</td><td>AAACCGCGCTCGCTGTCGGCCAAG</td><td></td><td></td><td></td><td></td><td></td><td></td><td></td><td></td><td></td><td></td></tr>
<tr><td>Chromosome</td><td>start</td><td>end</td><td>strand</td><td>MM</td><td>target_seq</td><td>PAM</td><td>alignment</td><td>distance</td><td>position</td><td>gene name</td><td>gene id</td></tr>
<tr><td>GU305914.1</td><td>64816</td><td>64838</td><td>-</td><td>0</td><td>CTTGGCCGACAGCGAGCGCG</td><td>AGG</td><td>||||||||[||||||||||||]PAM</td><td>NA</td><td>Intergenic</td><td>NA</td><td>NA</td></tr>
<tr><td></td><td></td><td></td><td></td><td></td><td></td><td></td><td></td><td></td><td></td><td></td><td></td></tr>
<tr><td>T29246</td><td>TGTACTCCGAGCCGAGCTGCTGG</td><td>1000</td><td></td><td></td><td></td><td></td><td></td><td></td><td></td><td></td><td></td></tr>
<tr><td>Oligo fwd</td><td>CACCTGTACTCCGAGCCGAGCTGC</td><td></td><td></td><td></td><td></td><td></td><td></td><td></td><td></td><td></td><td></td></tr>
<tr><td>Oligo rev</td><td>AAACGCAGCTCGGCTCGGAGTACA</td><td></td><td></td><td></td><td></td><td></td><td></td><td></td><td></td><td></td><td></td></tr>
<tr><td>Chromosome</td><td>start</td><td>end</td><td>strand</td><td>MM</td><td>target_seq</td><td>PAM</td><td>alignment</td><td>distance</td><td>position</td><td>gene name</td><td>gene id</td></tr>
<tr><td>GU305914.1</td><td>64769</td><td>64791</td><td>-</td><td>0</td><td>TGTACTCCGAGCCGAGCTGC</td><td>TGG</td><td>||||||||[||||||||||||]PAM</td><td>NA</td><td>Intergenic</td><td>NA</td><td>NA</td></tr>
<tr><td></td><td></td><td></td><td></td><td></td><td></td><td></td><td></td><td></td><td></td><td></td><td></td></tr>
<tr><td>T29247</td><td>CCGAGCCGAGCTGCTGGATCTGG</td><td>1000</td><td></td><td></td><td></td><td></td><td></td><td></td><td></td><td></td><td></td></tr>
<tr><td>Oligo fwd</td><td>CACCCCGAGCCGAGCTGCTGGATC</td><td></td><td></td><td></td><td></td><td></td><td></td><td></td><td></td><td></td><td></td></tr>
<tr><td>Oligo rev</td><td>AAACGATCCAGCAGCTCGGCTCGG</td><td></td><td></td><td></td><td></td><td></td><td></td><td></td><td></td><td></td><td></td></tr>
<tr><td>Chromosome</td><td>start</td><td>end</td><td>strand</td><td>MM</td><td>target_seq</td><td>PAM</td><td>alignment</td><td>distance</td><td>position</td><td>gene name</td><td>gene id</td></tr>
<tr><td>GU305914.1</td><td>64763</td><td>64785</td><td>-</td><td>0</td><td>CCGAGCCGAGCTGCTGGATC</td><td>TGG</td><td>||||||||[||||||||||||]PAM</td><td>NA</td><td>Intergenic</td><td>NA</td><td>NA</td></tr>
<tr><td></td><td></td><td></td><td></td><td></td><td></td><td></td><td></td><td></td><td></td><td></td><td></td></tr>
<tr><td>T29248</td><td>AGCCGAGCTGCTGGATCTGGTGG</td><td>1000</td><td></td><td></td><td></td><td></td><td></td><td></td><td></td><td></td><td></td></tr>
<tr><td>Oligo fwd</td><td>CACCAGCCGAGCTGCTGGATCTGG</td><td></td><td></td><td></td><td></td><td></td><td></td><td></td><td></td><td></td><td></td></tr>
<tr><td>Oligo rev</td><td>AAACCCAGATCCAGCAGCTCGGCT</td><td></td><td></td><td></td><td></td><td></td><td></td><td></td><td></td><td></td><td></td></tr>
<tr><td>Chromosome</td><td>start</td><td>end</td><td>strand</td><td>MM</td><td>target_seq</td><td>PAM</td><td>alignment</td><td>distance</td><td>position</td><td>gene name</td><td>gene id</td></tr>
<tr><td>GU305914.1</td><td>64760</td><td>64782</td><td>-</td><td>0</td><td>AGCCGAGCTGCTGGATCTGG</td><td>TGG</td><td>||||||||[||||||||||||]PAM</td><td>NA</td><td>Intergenic</td><td>NA</td><td>NA</td></tr>
<tr><td></td><td></td><td></td><td></td><td></td><td></td><td></td><td></td><td></td><td></td><td></td><td></td></tr>
<tr><td>T29249</td><td>CTGCTGGATCTGGTGGATGTCGG</td><td>1000</td><td></td><td></td><td></td><td></td><td></td><td></td><td></td><td></td><td></td></tr>
<tr><td>Oligo fwd</td><td>CACCCTGCTGGATCTGGTGGATGT</td><td></td><td></td><td></td><td></td><td></td><td></td><td></td><td></td><td></td><td></td></tr>
<tr><td>Oligo rev</td><td>AAACACATCCACCAGATCCAGCAG</td><td></td><td></td><td></td><td></td><td></td><td></td><td></td><td></td><td></td><td></td></tr>
<tr><td>Chromosome</td><td>start</td><td>end</td><td>strand</td><td>MM</td><td>target_seq</td><td>PAM</td><td>alignment</td><td>distance</td><td>position</td><td>gene name</td><td>gene id</td></tr>
<tr><td>GU305914.1</td><td>64753</td><td>64775</td><td>-</td><td>0</td><td>CTGCTGGATCTGGTGGATGT</td><td>CGG</td><td>||||||||[||||||||||||]PAM</td><td>NA</td><td>Intergenic</td><td>NA</td><td>NA</td></tr>
<tr><td></td><td></td><td></td><td></td><td></td><td></td><td></td><td></td><td></td><td></td><td></td><td></td></tr>
<tr><td>T29250</td><td>CTGGATCTGGTGGATGTCGGTGG</td><td>1000</td><td></td><td></td><td></td><td></td><td></td><td></td><td></td><td></td><td></td></tr>
<tr><td>Oligo fwd</td><td>CACCCTGGATCTGGTGGATGTCGG</td><td></td><td></td><td></td><td></td><td></td><td></td><td></td><td></td><td></td><td></td></tr>
<tr><td>Oligo rev</td><td>AAACCCGACATCCACCAGATCCAG</td><td></td><td></td><td></td><td></td><td></td><td></td><td></td><td></td><td></td><td></td></tr>
<tr><td>Chromosome</td><td>start</td><td>end</td><td>strand</td><td>MM</td><td>target_seq</td><td>PAM</td><td>alignment</td><td>distance</td><td>position</td><td>gene name</td><td>gene id</td></tr>
<tr><td>GU305914.1</td><td>64750</td><td>64772</td><td>-</td><td>0</td><td>CTGGATCTGGTGGATGTCGG</td><td>TGG</td><td>||||||||[||||||||||||]PAM</td><td>NA</td><td>Intergenic</td><td>NA</td><td>NA</td></tr>
<tr><td>GU305914.1</td><td>162825</td><td>162847</td><td>+</td><td>3</td><td>CTCGATCTCGTGCATGTCGG</td><td>CGG</td><td>||-|||||[-|||-|||||||]PAM</td><td>NA</td><td>Intergenic</td><td>NA</td><td>NA</td></tr>
<tr><td></td><td></td><td></td><td></td><td></td><td></td><td></td><td></td><td></td><td></td><td></td><td></td></tr>
<tr><td>T29251</td><td>GATGTCGGTGGTGCAGATCTCGG</td><td>1000</td><td></td><td></td><td></td><td></td><td></td><td></td><td></td><td></td><td></td></tr>
<tr><td>Oligo fwd</td><td>CACCGATGTCGGTGGTGCAGATCT</td><td></td><td></td><td></td><td></td><td></td><td></td><td></td><td></td><td></td><td></td></tr>
<tr><td>Oligo rev</td><td>AAACAGATCTGCACCACCGACATC</td><td></td><td></td><td></td><td></td><td></td><td></td><td></td><td></td><td></td><td></td></tr>
<tr><td>Chromosome</td><td>start</td><td>end</td><td>strand</td><td>MM</td><td>target_seq</td><td>PAM</td><td>alignment</td><td>distance</td><td>position</td><td>gene name</td><td>gene id</td></tr>
<tr><td>GU305914.1</td><td>64738</td><td>64760</td><td>-</td><td>0</td><td>GATGTCGGTGGTGCAGATCT</td><td>CGG</td><td>||||||||[||||||||||||]PAM</td><td>NA</td><td>Intergenic</td><td>NA</td><td>NA</td></tr>
<tr><td>GU305914.1</td><td>121316</td><td>121338</td><td>-</td><td>4</td><td>GGTGACGGTGGTCCAGAGCT</td><td>CGG</td><td>|-||-|||[||||-||||-||]PAM</td><td>NA</td><td>Intergenic</td><td>NA</td><td>NA</td></tr>
<tr><td></td><td></td><td></td><td></td><td></td><td></td><td></td><td></td><td></td><td></td><td></td><td></td></tr>
<tr><td>T29252</td><td>GGTGCAGATCTCGGCCAGCACGG</td><td>1000</td><td></td><td></td><td></td><td></td><td></td><td></td><td></td><td></td><td></td></tr>
<tr><td>Oligo fwd</td><td>CACCGGTGCAGATCTCGGCCAGCA</td><td></td><td></td><td></td><td></td><td></td><td></td><td></td><td></td><td></td><td></td></tr>
<tr><td>Oligo rev</td><td>AAACTGCTGGCCGAGATCTGCACC</td><td></td><td></td><td></td><td></td><td></td><td></td><td></td><td></td><td></td><td></td></tr>
<tr><td>Chromosome</td><td>start</td><td>end</td><td>strand</td><td>MM</td><td>target_seq</td><td>PAM</td><td>alignment</td><td>distance</td><td>position</td><td>gene name</td><td>gene id</td></tr>
<tr><td>GU305914.1</td><td>64729</td><td>64751</td><td>-</td><td>0</td><td>GGTGCAGATCTCGGCCAGCA</td><td>CGG</td><td>||||||||[||||||||||||]PAM</td><td>NA</td><td>Intergenic</td><td>NA</td><td>NA</td></tr>
<tr><td></td><td></td><td></td><td></td><td></td><td></td><td></td><td></td><td></td><td></td><td></td><td></td></tr>
<tr><td>T29253</td><td>GCAGATCTCGGCCAGCACGGCGG</td><td>1000</td><td></td><td></td><td></td><td></td><td></td><td></td><td></td><td></td><td></td></tr>
<tr><td>Oligo fwd</td><td>CACCGCAGATCTCGGCCAGCACGG</td><td></td><td></td><td></td><td></td><td></td><td></td><td></td><td></td><td></td><td></td></tr>
<tr><td>Oligo rev</td><td>AAACCCGTGCTGGCCGAGATCTGC</td><td></td><td></td><td></td><td></td><td></td><td></td><td></td><td></td><td></td><td></td></tr>
<tr><td>Chromosome</td><td>start</td><td>end</td><td>strand</td><td>MM</td><td>target_seq</td><td>PAM</td><td>alignment</td><td>distance</td><td>position</td><td>gene name</td><td>gene id</td></tr>
<tr><td>GU305914.1</td><td>64726</td><td>64748</td><td>-</td><td>0</td><td>GCAGATCTCGGCCAGCACGG</td><td>CGG</td><td>||||||||[||||||||||||]PAM</td><td>NA</td><td>Intergenic</td><td>NA</td><td>NA</td></tr>
<tr><td></td><td></td><td></td><td></td><td></td><td></td><td></td><td></td><td></td><td></td><td></td><td></td></tr>
<tr><td>T29254</td><td>CGGCCAGCACGGCGGACGCGCGG</td><td>1000</td><td></td><td></td><td></td><td></td><td></td><td></td><td></td><td></td><td></td></tr>
<tr><td>Oligo fwd</td><td>CACCCGGCCAGCACGGCGGACGCG</td><td></td><td></td><td></td><td></td><td></td><td></td><td></td><td></td><td></td><td></td></tr>
<tr><td>Oligo rev</td><td>AAACCGCGTCCGCCGTGCTGGCCG</td><td></td><td></td><td></td><td></td><td></td><td></td><td></td><td></td><td></td><td></td></tr>
<tr><td>Chromosome</td><td>start</td><td>end</td><td>strand</td><td>MM</td><td>target_seq</td><td>PAM</td><td>alignment</td><td>distance</td><td>position</td><td>gene name</td><td>gene id</td></tr>
<tr><td>GU305914.1</td><td>64718</td><td>64740</td><td>-</td><td>0</td><td>CGGCCAGCACGGCGGACGCG</td><td>CGG</td><td>||||||||[||||||||||||]PAM</td><td>NA</td><td>Intergenic</td><td>NA</td><td>NA</td></tr>
<tr><td></td><td></td><td></td><td></td><td></td><td></td><td></td><td></td><td></td><td></td><td></td><td></td></tr>
<tr><td>T29255</td><td>CACGGCGGACGCGCGGTCGATGG</td><td>1000</td><td></td><td></td><td></td><td></td><td></td><td></td><td></td><td></td><td></td></tr>
<tr><td>Oligo fwd</td><td>CACCCACGGCGGACGCGCGGTCGA</td><td></td><td></td><td></td><td></td><td></td><td></td><td></td><td></td><td></td><td></td></tr>
<tr><td>Oligo rev</td><td>AAACTCGACCGCGCGTCCGCCGTG</td><td></td><td></td><td></td><td></td><td></td><td></td><td></td><td></td><td></td><td></td></tr>
<tr><td>Chromosome</td><td>start</td><td>end</td><td>strand</td><td>MM</td><td>target_seq</td><td>PAM</td><td>alignment</td><td>distance</td><td>position</td><td>gene name</td><td>gene id</td></tr>
<tr><td>GU305914.1</td><td>64711</td><td>64733</td><td>-</td><td>0</td><td>CACGGCGGACGCGCGGTCGA</td><td>TGG</td><td>||||||||[||||||||||||]PAM</td><td>NA</td><td>Intergenic</td><td>NA</td><td>NA</td></tr>
<tr><td>GU305914.1</td><td>135784</td><td>135806</td><td>+</td><td>4</td><td>GACGTCGGACGGGCGGACGA</td><td>CGG</td><td>-|||-|||[|||-||||-|||]PAM</td><td>NA</td><td>Intergenic</td><td>NA</td><td>NA</td></tr>
<tr><td></td><td></td><td></td><td></td><td></td><td></td><td></td><td></td><td></td><td></td><td></td><td></td></tr>
<tr><td>T29256</td><td>ACGCGCGGTCGATGGCCTTCTGG</td><td>1000</td><td></td><td></td><td></td><td></td><td></td><td></td><td></td><td></td><td></td></tr>
<tr><td>Oligo fwd</td><td>CACCACGCGCGGTCGATGGCCTTC</td><td></td><td></td><td></td><td></td><td></td><td></td><td></td><td></td><td></td><td></td></tr>
<tr><td>Oligo rev</td><td>AAACGAAGGCCATCGACCGCGCGT</td><td></td><td></td><td></td><td></td><td></td><td></td><td></td><td></td><td></td><td></td></tr>
<tr><td>Chromosome</td><td>start</td><td>end</td><td>strand</td><td>MM</td><td>target_seq</td><td>PAM</td><td>alignment</td><td>distance</td><td>position</td><td>gene name</td><td>gene id</td></tr>
</table>

<table>
<tr><td>GU305914.1</td><td></td><td>64703</td><td>64725</td><td>-</td><td>0</td><td>ACGCGCGGTCGATGGCCTTC</td><td>TGG</td><td>||||||||[||||||||||||]PAM</td><td>NA</td><td>Intergenic</td><td>NA</td><td>NA</td></tr>
<tr><td>T29257</td><td>CCTTCTGGAAGTCCGTCTCTAGG</td><td></td><td>1000</td><td></td><td></td><td></td><td></td><td></td><td></td><td></td><td></td><td></td></tr>
<tr><td>Oligo fwd</td><td>CACCCCTTCTGGAAGTCCGTCTCT</td><td></td><td></td><td></td><td></td><td></td><td></td><td></td><td></td><td></td><td></td><td></td></tr>
<tr><td>Oligo rev</td><td>AAACAGAGACGGACTTCCAGAAGG</td><td></td><td></td><td></td><td></td><td></td><td></td><td></td><td></td><td></td><td></td><td></td></tr>
<tr><td>Chromosome</td><td>start</td><td></td><td>end</td><td>strand</td><td>MM</td><td>target_seq</td><td>PAM</td><td>alignment</td><td>distance</td><td>position</td><td>gene name</td><td>gene id</td></tr>
<tr><td>GU305914.1</td><td></td><td>64688</td><td>64710</td><td>-</td><td>0</td><td>CCTTCTGGAAGTCCGTCTCT</td><td>AGG</td><td>||||||||[||||||||||||]PAM</td><td>NA</td><td>Intergenic</td><td>NA</td><td>NA</td></tr>
<tr><td>T29258</td><td>CTTCTGGAAGTCCGTCTCTAGGG</td><td></td><td>1000</td><td></td><td></td><td></td><td></td><td></td><td></td><td></td><td></td><td></td></tr>
<tr><td>Oligo fwd</td><td>CACCCTTCTGGAAGTCCGTCTCTA</td><td></td><td></td><td></td><td></td><td></td><td></td><td></td><td></td><td></td><td></td><td></td></tr>
<tr><td>Oligo rev</td><td>AAACTAGAGACGGACTTCCAGAAG</td><td></td><td></td><td></td><td></td><td></td><td></td><td></td><td></td><td></td><td></td><td></td></tr>
<tr><td>Chromosome</td><td>start</td><td></td><td>end</td><td>strand</td><td>MM</td><td>target_seq</td><td>PAM</td><td>alignment</td><td>distance</td><td>position</td><td>gene name</td><td>gene id</td></tr>
<tr><td>GU305914.1</td><td></td><td>64687</td><td>64709</td><td>-</td><td>0</td><td>CTTCTGGAAGTCCGTCTCTA</td><td>GGG</td><td>||||||||[||||||||||||]PAM</td><td>NA</td><td>Intergenic</td><td>NA</td><td>NA</td></tr>
<tr><td>T29259</td><td>CTGGAAGTCCGTCTCTAGGGCGG</td><td></td><td>1000</td><td></td><td></td><td></td><td></td><td></td><td></td><td></td><td></td><td></td></tr>
<tr><td>Oligo fwd</td><td>CACCCTGGAAGTCCGTCTCTAGGG</td><td></td><td></td><td></td><td></td><td></td><td></td><td></td><td></td><td></td><td></td><td></td></tr>
<tr><td>Oligo rev</td><td>AAACCCCTAGAGACGGACTTCCAG</td><td></td><td></td><td></td><td></td><td></td><td></td><td></td><td></td><td></td><td></td><td></td></tr>
<tr><td>Chromosome</td><td>start</td><td></td><td>end</td><td>strand</td><td>MM</td><td>target_seq</td><td>PAM</td><td>alignment</td><td>distance</td><td>position</td><td>gene name</td><td>gene id</td></tr>
<tr><td>GU305914.1</td><td></td><td>64684</td><td>64706</td><td>-</td><td>0</td><td>CTGGAAGTCCGTCTCTAGGG</td><td>CGG</td><td>||||||||[||||||||||||]PAM</td><td>NA</td><td>Intergenic</td><td>NA</td><td>NA</td></tr>
<tr><td>T29260</td><td>TGGAAGTCCGTCTCTAGGGCGGG</td><td></td><td>1000</td><td></td><td></td><td></td><td></td><td></td><td></td><td></td><td></td><td></td></tr>
<tr><td>Oligo fwd</td><td>CACCTGGAAGTCCGTCTCTAGGGC</td><td></td><td></td><td></td><td></td><td></td><td></td><td></td><td></td><td></td><td></td><td></td></tr>
<tr><td>Oligo rev</td><td>AAACGCCCTAGAGACGGACTTCCA</td><td></td><td></td><td></td><td></td><td></td><td></td><td></td><td></td><td></td><td></td><td></td></tr>
<tr><td>Chromosome</td><td>start</td><td></td><td>end</td><td>strand</td><td>MM</td><td>target_seq</td><td>PAM</td><td>alignment</td><td>distance</td><td>position</td><td>gene name</td><td>gene id</td></tr>
<tr><td>GU305914.1</td><td></td><td>64683</td><td>64705</td><td>-</td><td>0</td><td>TGGAAGTCCGTCTCTAGGGC</td><td>GGG</td><td>||||||||[||||||||||||]PAM</td><td>NA</td><td>Intergenic</td><td>NA</td><td>NA</td></tr>
<tr><td>GU305914.1</td><td></td><td>166099</td><td>166121</td><td>+</td><td>4</td><td>TTGAAGACCGTCACTAGGGA</td><td>CGG</td><td>|-||||-|[||||-||||||-]PAM</td><td>NA</td><td>Intergenic</td><td>NA</td><td>NA</td></tr>
<tr><td>T29261</td><td>CTAGGGCGGGCGTCGCCGCGTGG</td><td></td><td>1000</td><td></td><td></td><td></td><td></td><td></td><td></td><td></td><td></td><td></td></tr>
<tr><td>Oligo fwd</td><td>CACCCTAGGGCGGGCGTCGCCGCG</td><td></td><td></td><td></td><td></td><td></td><td></td><td></td><td></td><td></td><td></td><td></td></tr>
<tr><td>Oligo rev</td><td>AAACCGCGGCGACGCCCGCCCTAG</td><td></td><td></td><td></td><td></td><td></td><td></td><td></td><td></td><td></td><td></td><td></td></tr>
<tr><td>Chromosome</td><td>start</td><td></td><td>end</td><td>strand</td><td>MM</td><td>target_seq</td><td>PAM</td><td>alignment</td><td>distance</td><td>position</td><td>gene name</td><td>gene id</td></tr>
<tr><td>GU305914.1</td><td></td><td>64670</td><td>64692</td><td>-</td><td>0</td><td>CTAGGGCGGGCGTCGCCGCG</td><td>TGG</td><td>||||||||[||||||||||||]PAM</td><td>NA</td><td>Intergenic</td><td>NA</td><td>NA</td></tr>
<tr><td>T29262</td><td>GCGTCGCCGCGTGGTTATCGTGG</td><td></td><td>1000</td><td></td><td></td><td></td><td></td><td></td><td></td><td></td><td></td><td></td></tr>
<tr><td>Oligo fwd</td><td>CACCGCGTCGCCGCGTGGTTATCG</td><td></td><td></td><td></td><td></td><td></td><td></td><td></td><td></td><td></td><td></td><td></td></tr>
<tr><td>Oligo rev</td><td>AAACCGATAACCACGCGGCGACGC</td><td></td><td></td><td></td><td></td><td></td><td></td><td></td><td></td><td></td><td></td><td></td></tr>
<tr><td>Chromosome</td><td>start</td><td></td><td>end</td><td>strand</td><td>MM</td><td>target_seq</td><td>PAM</td><td>alignment</td><td>distance</td><td>position</td><td>gene name</td><td>gene id</td></tr>
<tr><td>GU305914.1</td><td></td><td>64661</td><td>64683</td><td>-</td><td>0</td><td>GCGTCGCCGCGTGGTTATCG</td><td>TGG</td><td>||||||||[||||||||||||]PAM</td><td>NA</td><td>Intergenic</td><td>NA</td><td>NA</td></tr>
<tr><td>T29263</td><td>CCGCGTGGTTATCGTGGCAGTGG</td><td></td><td>1000</td><td></td><td></td><td></td><td></td><td></td><td></td><td></td><td></td><td></td></tr>
<tr><td>Oligo fwd</td><td>CACCCCGCGTGGTTATCGTGGCAG</td><td></td><td></td><td></td><td></td><td></td><td></td><td></td><td></td><td></td><td></td><td></td></tr>
<tr><td>Oligo rev</td><td>AAACCTGCCACGATAACCACGCGG</td><td></td><td></td><td></td><td></td><td></td><td></td><td></td><td></td><td></td><td></td><td></td></tr>
<tr><td>Chromosome</td><td>start</td><td></td><td>end</td><td>strand</td><td>MM</td><td>target_seq</td><td>PAM</td><td>alignment</td><td>distance</td><td>position</td><td>gene name</td><td>gene id</td></tr>
<tr><td>GU305914.1</td><td></td><td>64655</td><td>64677</td><td>-</td><td>0</td><td>CCGCGTGGTTATCGTGGCAG</td><td>TGG</td><td>||||||||[||||||||||||]PAM</td><td>NA</td><td>Intergenic</td><td>NA</td><td>NA</td></tr>
<tr><td>T29264</td><td>GTCGCGCCGCAGCTGCTGCGTGG</td><td></td><td>1000</td><td></td><td></td><td></td><td></td><td></td><td></td><td></td><td></td><td></td></tr>
<tr><td>Oligo fwd</td><td>CACCGTCGCGCCGCAGCTGCTGCG</td><td></td><td></td><td></td><td></td><td></td><td></td><td></td><td></td><td></td><td></td><td></td></tr>
<tr><td>Oligo rev</td><td>AAACCGCAGCAGCTGCGGCGCGAC</td><td></td><td></td><td></td><td></td><td></td><td></td><td></td><td></td><td></td><td></td><td></td></tr>
<tr><td>Chromosome</td><td>start</td><td></td><td>end</td><td>strand</td><td>MM</td><td>target_seq</td><td>PAM</td><td>alignment</td><td>distance</td><td>position</td><td>gene name</td><td>gene id</td></tr>
<tr><td>GU305914.1</td><td></td><td>64597</td><td>64619</td><td>-</td><td>0</td><td>GTCGCGCCGCAGCTGCTGCG</td><td>TGG</td><td>||||||||[||||||||||||]PAM</td><td>NA</td><td>Intergenic</td><td>NA</td><td>NA</td></tr>
<tr><td>T29265</td><td>GAAAGAATTCGAAGAGCGTGTGG</td><td></td><td>1000</td><td></td><td></td><td></td><td></td><td></td><td></td><td></td><td></td><td></td></tr>
<tr><td>Oligo fwd</td><td>CACCGAAAGAATTCGAAGAGCGTG</td><td></td><td></td><td></td><td></td><td></td><td></td><td></td><td></td><td></td><td></td><td></td></tr>
<tr><td>Oligo rev</td><td>AAACCACGCTCTTCGAATTCTTTC</td><td></td><td></td><td></td><td></td><td></td><td></td><td></td><td></td><td></td><td></td><td></td></tr>
<tr><td>Chromosome</td><td>start</td><td></td><td>end</td><td>strand</td><td>MM</td><td>target_seq</td><td>PAM</td><td>alignment</td><td>distance</td><td>position</td><td>gene name</td><td>gene id</td></tr>
<tr><td>GU305914.1</td><td></td><td>64562</td><td>64584</td><td>-</td><td>0</td><td>GAAAGAATTCGAAGAGCGTG</td><td>TGG</td><td>||||||||[||||||||||||]PAM</td><td>NA</td><td>Intergenic</td><td>NA</td><td>NA</td></tr>
<tr><td>T29266</td><td>GTGCAGCTCCTCCAGCAGCTCGG</td><td></td><td>1000</td><td></td><td></td><td></td><td></td><td></td><td></td><td></td><td></td><td></td></tr>
<tr><td>Oligo fwd</td><td>CACCGTGCAGCTCCTCCAGCAGCT</td><td></td><td></td><td></td><td></td><td></td><td></td><td></td><td></td><td></td><td></td><td></td></tr>
<tr><td>Oligo rev</td><td>AAACAGCTGCTGGAGGAGCTGCAC</td><td></td><td></td><td></td><td></td><td></td><td></td><td></td><td></td><td></td><td></td><td></td></tr>
<tr><td>Chromosome</td><td>start</td><td></td><td>end</td><td>strand</td><td>MM</td><td>target_seq</td><td>PAM</td><td>alignment</td><td>distance</td><td>position</td><td>gene name</td><td>gene id</td></tr>
<tr><td>GU305914.1</td><td></td><td>64540</td><td>64562</td><td>-</td><td>0</td><td>GTGCAGCTCCTCCAGCAGCT</td><td>CGG</td><td>||||||||[||||||||||||]PAM</td><td>NA</td><td>Intergenic</td><td>NA</td><td>NA</td></tr>
<tr><td>T29267</td><td>CTCCTCCAGCAGCTCGGCCACGG</td><td></td><td>1000</td><td></td><td></td><td></td><td></td><td></td><td></td><td></td><td></td><td></td></tr>
<tr><td>Oligo fwd</td><td>CACCCTCCTCCAGCAGCTCGGCCA</td><td></td><td></td><td></td><td></td><td></td><td></td><td></td><td></td><td></td><td></td><td></td></tr>
<tr><td>Oligo rev</td><td>AAACTGGCCGAGCTGCTGGAGGAG</td><td></td><td></td><td></td><td></td><td></td><td></td><td></td><td></td><td></td><td></td><td></td></tr>
<tr><td>Chromosome</td><td>start</td><td></td><td>end</td><td>strand</td><td>MM</td><td>target_seq</td><td>PAM</td><td>alignment</td><td>distance</td><td>position</td><td>gene name</td><td>gene id</td></tr>
<tr><td>GU305914.1</td><td></td><td>64534</td><td>64556</td><td>-</td><td>0</td><td>CTCCTCCAGCAGCTCGGCCA</td><td>CGG</td><td>||||||||[||||||||||||]PAM</td><td>NA</td><td>Intergenic</td><td>NA</td><td>NA</td></tr>
<tr><td>T29268</td><td>ACAGTTCCTCAGCGTCTGCGTGG</td><td></td><td>1000</td><td></td><td></td><td></td><td></td><td></td><td></td><td></td><td></td><td></td></tr>
<tr><td>Oligo fwd</td><td>CACCACAGTTCCTCAGCGTCTGCG</td><td></td><td></td><td></td><td></td><td></td><td></td><td></td><td></td><td></td><td></td><td></td></tr>
<tr><td>Oligo rev</td><td>AAACCGCAGACGCTGAGGAACTGT</td><td></td><td></td><td></td><td></td><td></td><td></td><td></td><td></td><td></td><td></td><td></td></tr>
<tr><td>Chromosome</td><td>start</td><td></td><td>end</td><td>strand</td><td>MM</td><td>target_seq</td><td>PAM</td><td>alignment</td><td>distance</td><td>position</td><td>gene name</td><td>gene id</td></tr>
<tr><td>GU305914.1</td><td></td><td>64507</td><td>64529</td><td>-</td><td>0</td><td>ACAGTTCCTCAGCGTCTGCG</td><td>TGG</td><td>||||||||[||||||||||||]PAM</td><td>NA</td><td>Intergenic</td><td>NA</td><td>NA</td></tr>
<tr><td>T29269</td><td>CCAGCTTCTCGACCGTCTTTAGG</td><td></td><td>1000</td><td></td><td></td><td></td><td></td><td></td><td></td><td></td><td></td><td></td></tr>
<tr><td>Oligo fwd</td><td>CACCCCAGCTTCTCGACCGTCTTT</td><td></td><td></td><td></td><td></td><td></td><td></td><td></td><td></td><td></td><td></td><td></td></tr>
<tr><td>Oligo rev</td><td>AAACAAAGACGGTCGAGAAGCTGG</td><td></td><td></td><td></td><td></td><td></td><td></td><td></td><td></td><td></td><td></td><td></td></tr>
<tr><td>Chromosome</td><td>start</td><td></td><td>end</td><td>strand</td><td>MM</td><td>target_seq</td><td>PAM</td><td>alignment</td><td>distance</td><td>position</td><td>gene name</td><td>gene id</td></tr>
<tr><td>GU305914.1</td><td></td><td>64478</td><td>64500</td><td>-</td><td>0</td><td>CCAGCTTCTCGACCGTCTTT</td><td>AGG</td><td>||||||||[||||||||||||]PAM</td><td>NA</td><td>Intergenic</td><td>NA</td><td>NA</td></tr>
<tr><td>T29270</td><td>CTTCTCGACCGTCTTTAGGTCGG</td><td></td><td>1000</td><td></td><td></td><td></td><td></td><td></td><td></td><td></td><td></td><td></td></tr>
<tr><td>Oligo fwd</td><td>CACCCTTCTCGACCGTCTTTAGGT</td><td></td><td></td><td></td><td></td><td></td><td></td><td></td><td></td><td></td><td></td><td></td></tr>
<tr><td>Oligo rev</td><td>AAACACCTAAAGACGGTCGAGAAG</td><td></td><td></td><td></td><td></td><td></td><td></td><td></td><td></td><td></td><td></td><td></td></tr>
<tr><td>Chromosome</td><td>start</td><td></td><td>end</td><td>strand</td><td>MM</td><td>target_seq</td><td>PAM</td><td>alignment</td><td>distance</td><td>position</td><td>gene name</td><td>gene id</td></tr>
<tr><td>GU305914.1</td><td></td><td>64474</td><td>64496</td><td>-</td><td>0</td><td>CTTCTCGACCGTCTTTAGGT</td><td>CGG</td><td>||||||||[||||||||||||]PAM</td><td>NA</td><td>Intergenic</td><td>NA</td><td>NA</td></tr>
<tr><td>T29271</td><td>GACCGTCTTTAGGTCGGCGTCGG</td><td></td><td>1000</td><td></td><td></td><td></td><td></td><td></td><td></td><td></td><td></td><td></td></tr>
<tr><td>Oligo fwd</td><td>CACCGACCGTCTTTAGGTCGGCGT</td><td></td><td></td><td></td><td></td><td></td><td></td><td></td><td></td><td></td><td></td><td></td></tr>
</table>

<table>
<tr><td>Oligo rev</td><td>AAACACGCCGACCTAAAGACGGTC</td></tr>
<tr><td>Chromosome</td><td>start</td><td></td><td>end</td><td>strand</td><td>MM</td><td>target_seq</td><td>PAM</td><td>alignment</td><td>distance</td><td>position</td><td>gene name</td><td>gene id</td></tr>
<tr><td>GU305914.1</td><td></td><td>64468</td><td>64490</td><td>-</td><td>0</td><td>GACCGTCTTTAGGTCGGCGT</td><td>CGG</td><td>||||||||[||||||||||||]PAM</td><td>NA</td><td>Intergenic</td><td>NA</td><td>NA</td></tr>
<tr><td>T29272</td><td>CGGCGTCGGTGAATTCCCTCCGG</td><td></td><td>1000</td></tr>
<tr><td>Oligo fwd</td><td>CACCCGGCGTCGGTGAATTCCCTC</td></tr>
<tr><td>Oligo rev</td><td>AAACGAGGGAATTCACCGACGCCG</td></tr>
<tr><td>Chromosome</td><td>start</td><td></td><td>end</td><td>strand</td><td>MM</td><td>target_seq</td><td>PAM</td><td>alignment</td><td>distance</td><td>position</td><td>gene name</td><td>gene id</td></tr>
<tr><td>GU305914.1</td><td></td><td>64454</td><td>64476</td><td>-</td><td>0</td><td>CGGCGTCGGTGAATTCCCTC</td><td>CGG</td><td>||||||||[||||||||||||]PAM</td><td>NA</td><td>Intergenic</td><td>NA</td><td>NA</td></tr>
<tr><td>T29273</td><td>CGGTGAATTCCCTCCGGCCGCGG</td><td></td><td>1000</td></tr>
<tr><td>Oligo fwd</td><td>CACCCGGTGAATTCCCTCCGGCCG</td></tr>
<tr><td>Oligo rev</td><td>AAACCGGCCGGAGGGAATTCACCG</td></tr>
<tr><td>Chromosome</td><td>start</td><td></td><td>end</td><td>strand</td><td>MM</td><td>target_seq</td><td>PAM</td><td>alignment</td><td>distance</td><td>position</td><td>gene name</td><td>gene id</td></tr>
<tr><td>GU305914.1</td><td></td><td>64448</td><td>64470</td><td>-</td><td>0</td><td>CGGTGAATTCCCTCCGGCCG</td><td>CGG</td><td>||||||||[||||||||||||]PAM</td><td>NA</td><td>Intergenic</td><td>NA</td><td>NA</td></tr>
<tr><td>T29274</td><td>TCCCTCCGGCCGCGGCCCTGAGG</td><td></td><td>1000</td></tr>
<tr><td>Oligo fwd</td><td>CACCTCCCTCCGGCCGCGGCCCTG</td></tr>
<tr><td>Oligo rev</td><td>AAACCAGGGCCGCGGCCGGAGGGA</td></tr>
<tr><td>Chromosome</td><td>start</td><td></td><td>end</td><td>strand</td><td>MM</td><td>target_seq</td><td>PAM</td><td>alignment</td><td>distance</td><td>position</td><td>gene name</td><td>gene id</td></tr>
<tr><td>GU305914.1</td><td></td><td>64440</td><td>64462</td><td>-</td><td>0</td><td>TCCCTCCGGCCGCGGCCCTG</td><td>AGG</td><td>||||||||[||||||||||||]PAM</td><td>NA</td><td>Intergenic</td><td>NA</td><td>NA</td></tr>
<tr><td>T29275</td><td>TCCGGCCGCGGCCCTGAGGACGG</td><td></td><td>1000</td></tr>
<tr><td>Oligo fwd</td><td>CACCTCCGGCCGCGGCCCTGAGGA</td></tr>
<tr><td>Oligo rev</td><td>AAACTCCTCAGGGCCGCGGCCGGA</td></tr>
<tr><td>Chromosome</td><td>start</td><td></td><td>end</td><td>strand</td><td>MM</td><td>target_seq</td><td>PAM</td><td>alignment</td><td>distance</td><td>position</td><td>gene name</td><td>gene id</td></tr>
<tr><td>GU305914.1</td><td></td><td>64436</td><td>64458</td><td>-</td><td>0</td><td>TCCGGCCGCGGCCCTGAGGA</td><td>CGG</td><td>||||||||[||||||||||||]PAM</td><td>NA</td><td>Intergenic</td><td>NA</td><td>NA</td></tr>
<tr><td>T29276</td><td>GCCGCGGCCCTGAGGACGGTCGG</td><td></td><td>1000</td></tr>
<tr><td>Oligo fwd</td><td>CACCGCCGCGGCCCTGAGGACGGT</td></tr>
<tr><td>Oligo rev</td><td>AAACACCGTCCTCAGGGCCGCGGC</td></tr>
<tr><td>Chromosome</td><td>start</td><td></td><td>end</td><td>strand</td><td>MM</td><td>target_seq</td><td>PAM</td><td>alignment</td><td>distance</td><td>position</td><td>gene name</td><td>gene id</td></tr>
<tr><td>GU305914.1</td><td></td><td>64432</td><td>64454</td><td>-</td><td>0</td><td>GCCGCGGCCCTGAGGACGGT</td><td>CGG</td><td>||||||||[||||||||||||]PAM</td><td>NA</td><td>Intergenic</td><td>NA</td><td>NA</td></tr>
<tr><td>T29277</td><td>TTTCTTCGTCTATTAGACACTGG</td><td></td><td>1000</td></tr>
<tr><td>Oligo fwd</td><td>CACCTTTCTTCGTCTATTAGACAC</td></tr>
<tr><td>Oligo rev</td><td>AAACGTGTCTAATAGACGAAGAAA</td></tr>
<tr><td>Chromosome</td><td>start</td><td></td><td>end</td><td>strand</td><td>MM</td><td>target_seq</td><td>PAM</td><td>alignment</td><td>distance</td><td>position</td><td>gene name</td><td>gene id</td></tr>
<tr><td>GU305914.1</td><td></td><td>64409</td><td>64431</td><td>-</td><td>0</td><td>TTTCTTCGTCTATTAGACAC</td><td>TGG</td><td>||||||||[||||||||||||]PAM</td><td>NA</td><td>Intergenic</td><td>NA</td><td>NA</td></tr>
<tr><td>T29278</td><td>CCGAGTCCAGCAGACCCTGTTGG</td><td></td><td>1000</td></tr>
<tr><td>Oligo fwd</td><td>CACCCCGAGTCCAGCAGACCCTGT</td></tr>
<tr><td>Oligo rev</td><td>AAACACAGGGTCTGCTGGACTCGG</td></tr>
<tr><td>Chromosome</td><td>start</td><td></td><td>end</td><td>strand</td><td>MM</td><td>target_seq</td><td>PAM</td><td>alignment</td><td>distance</td><td>position</td><td>gene name</td><td>gene id</td></tr>
<tr><td>GU305914.1</td><td></td><td>64385</td><td>64407</td><td>-</td><td>0</td><td>CCGAGTCCAGCAGACCCTGT</td><td>TGG</td><td>||||||||[||||||||||||]PAM</td><td>NA</td><td>Intergenic</td><td>NA</td><td>NA</td></tr>
<tr><td>T29279</td><td>AGACGCGCGCCGCGCCTTCCAGG</td><td></td><td>1000</td></tr>
<tr><td>Oligo fwd</td><td>CACCAGACGCGCGCCGCGCCTTCC</td></tr>
<tr><td>Oligo rev</td><td>AAACGGAAGGCGCGGCGCGCGTCT</td></tr>
<tr><td>Chromosome</td><td>start</td><td></td><td>end</td><td>strand</td><td>MM</td><td>target_seq</td><td>PAM</td><td>alignment</td><td>distance</td><td>position</td><td>gene name</td><td>gene id</td></tr>
<tr><td>GU305914.1</td><td></td><td>64355</td><td>64377</td><td>-</td><td>0</td><td>AGACGCGCGCCGCGCCTTCC</td><td>AGG</td><td>||||||||[||||||||||||]PAM</td><td>NA</td><td>Intergenic</td><td>NA</td><td>NA</td></tr>
<tr><td>T29280</td><td>CGCCGCGCCTTCCAGGAACTCGG</td><td></td><td>1000</td></tr>
<tr><td>Oligo fwd</td><td>CACCCGCCGCGCCTTCCAGGAACT</td></tr>
<tr><td>Oligo rev</td><td>AAACAGTTCCTGGAAGGCGCGGCG</td></tr>
<tr><td>Chromosome</td><td>start</td><td></td><td>end</td><td>strand</td><td>MM</td><td>target_seq</td><td>PAM</td><td>alignment</td><td>distance</td><td>position</td><td>gene name</td><td>gene id</td></tr>
<tr><td>GU305914.1</td><td></td><td>64348</td><td>64370</td><td>-</td><td>0</td><td>CGCCGCGCCTTCCAGGAACT</td><td>CGG</td><td>||||||||[||||||||||||]PAM</td><td>NA</td><td>Intergenic</td><td>NA</td><td>NA</td></tr>
<tr><td>T29281</td><td>CGTCTGTAGCACCCGCTCGTTGG</td><td></td><td>1000</td></tr>
<tr><td>Oligo fwd</td><td>CACCCGTCTGTAGCACCCGCTCGT</td></tr>
<tr><td>Oligo rev</td><td>AAACACGAGCGGGTGCTACAGACG</td></tr>
<tr><td>Chromosome</td><td>start</td><td></td><td>end</td><td>strand</td><td>MM</td><td>target_seq</td><td>PAM</td><td>alignment</td><td>distance</td><td>position</td><td>gene name</td><td>gene id</td></tr>
<tr><td>GU305914.1</td><td></td><td>64321</td><td>64343</td><td>-</td><td>0</td><td>CGTCTGTAGCACCCGCTCGT</td><td>TGG</td><td>||||||||[||||||||||||]PAM</td><td>NA</td><td>Intergenic</td><td>NA</td><td>NA</td></tr>
<tr><td>T29282</td><td>CCGCTCGTTGGAGAGTTTCGAGG</td><td></td><td>1000</td></tr>
<tr><td>Oligo fwd</td><td>CACCCCGCTCGTTGGAGAGTTTCG</td></tr>
<tr><td>Oligo rev</td><td>AAACCGAAACTCTCCAACGAGCGG</td></tr>
<tr><td>Chromosome</td><td>start</td><td></td><td>end</td><td>strand</td><td>MM</td><td>target_seq</td><td>PAM</td><td>alignment</td><td>distance</td><td>position</td><td>gene name</td><td>gene id</td></tr>
<tr><td>GU305914.1</td><td></td><td>64309</td><td>64331</td><td>-</td><td>0</td><td>CCGCTCGTTGGAGAGTTTCG</td><td>AGG</td><td>||||||||[||||||||||||]PAM</td><td>NA</td><td>Intergenic</td><td>NA</td><td>NA</td></tr>
<tr><td>T29283</td><td>GCCCGCCGCCTGCGACTTCTCGG</td><td></td><td>1000</td></tr>
<tr><td>Oligo fwd</td><td>CACCGCCCGCCGCCTGCGACTTCT</td></tr>
<tr><td>Oligo rev</td><td>AAACAGAAGTCGCAGGCGGCGGGC</td></tr>
<tr><td>Chromosome</td><td>start</td><td></td><td>end</td><td>strand</td><td>MM</td><td>target_seq</td><td>PAM</td><td>alignment</td><td>distance</td><td>position</td><td>gene name</td><td>gene id</td></tr>
<tr><td>GU305914.1</td><td></td><td>64258</td><td>64280</td><td>-</td><td>0</td><td>GCCCGCCGCCTGCGACTTCT</td><td>CGG</td><td>||||||||[||||||||||||]PAM</td><td>NA</td><td>Intergenic</td><td>NA</td><td>NA</td></tr>
<tr><td>T29284</td><td>ACTTGCGTAACTCCGAGACCAGG</td><td></td><td>1000</td></tr>
<tr><td>Oligo fwd</td><td>CACCACTTGCGTAACTCCGAGACC</td></tr>
<tr><td>Oligo rev</td><td>AAACGGTCTCGGAGTTACGCAAGT</td></tr>
<tr><td>Chromosome</td><td>start</td><td></td><td>end</td><td>strand</td><td>MM</td><td>target_seq</td><td>PAM</td><td>alignment</td><td>distance</td><td>position</td><td>gene name</td><td>gene id</td></tr>
<tr><td>GU305914.1</td><td></td><td>64232</td><td>64254</td><td>-</td><td>0</td><td>ACTTGCGTAACTCCGAGACC</td><td>AGG</td><td>||||||||[||||||||||||]PAM</td><td>NA</td><td>Intergenic</td><td>NA</td><td>NA</td></tr>
<tr><td>T29285</td><td>GCGTAACTCCGAGACCAGGTCGG</td><td></td><td>1000</td></tr>
<tr><td>Oligo fwd</td><td>CACCGCGTAACTCCGAGACCAGGT</td></tr>
<tr><td>Oligo rev</td><td>AAACACCTGGTCTCGGAGTTACGC</td></tr>
<tr><td>Chromosome</td><td>start</td><td></td><td>end</td><td>strand</td><td>MM</td><td>target_seq</td><td>PAM</td><td>alignment</td><td>distance</td><td>position</td><td>gene name</td><td>gene id</td></tr>
<tr><td>GU305914.1</td><td></td><td>64228</td><td>64250</td><td>-</td><td>0</td><td>GCGTAACTCCGAGACCAGGT</td><td>CGG</td><td>||||||||[||||||||||||]PAM</td><td>NA</td><td>Intergenic</td><td>NA</td><td>NA</td></tr>
<tr><td>T29286</td><td>TAACTCCGAGACCAGGTCGGCGG</td><td></td><td>1000</td></tr>
</table>

<table>
<tr><td>Oligo fwd</td><td>CACCTAACTCCGAGACCAGGTCGG</td><td></td><td></td><td></td><td></td><td></td><td></td><td></td><td></td><td></td><td></td></tr>
<tr><td>Oligo rev</td><td>AAACCCGACCTGGTCTCGGAGTTA</td><td></td><td></td><td></td><td></td><td></td><td></td><td></td><td></td><td></td><td></td></tr>
<tr><td>Chromosome</td><td>start</td><td>end</td><td>strand</td><td>MM</td><td>target_seq</td><td>PAM</td><td>alignment</td><td>distance</td><td>position</td><td>gene name</td><td>gene id</td></tr>
<tr><td>GU305914.1</td><td>64225</td><td>64247</td><td>-</td><td>0</td><td>TAACTCCGAGACCAGGTCGG</td><td>CGG</td><td>||||||||[||||||||||||]PAM</td><td>NA</td><td>Intergenic</td><td>NA</td><td>NA</td></tr>
<tr><td></td><td></td><td></td><td></td><td></td><td></td><td></td><td></td><td></td><td></td><td></td><td></td></tr>
<tr><td>T29287</td><td>GAGACCAGGTCGGCGGCCAGCGG</td><td>1000</td><td></td><td></td><td></td><td></td><td></td><td></td><td></td><td></td><td></td></tr>
<tr><td>Oligo fwd</td><td>CACCGAGACCAGGTCGGCGGCCAG</td><td></td><td></td><td></td><td></td><td></td><td></td><td></td><td></td><td></td><td></td></tr>
<tr><td>Oligo rev</td><td>AAACCTGGCCGCCGACCTGGTCTC</td><td></td><td></td><td></td><td></td><td></td><td></td><td></td><td></td><td></td><td></td></tr>
<tr><td>Chromosome</td><td>start</td><td>end</td><td>strand</td><td>MM</td><td>target_seq</td><td>PAM</td><td>alignment</td><td>distance</td><td>position</td><td>gene name</td><td>gene id</td></tr>
<tr><td>GU305914.1</td><td>64218</td><td>64240</td><td>-</td><td>0</td><td>GAGACCAGGTCGGCGGCCAG</td><td>CGG</td><td>||||||||[||||||||||||]PAM</td><td>NA</td><td>Intergenic</td><td>NA</td><td>NA</td></tr>
<tr><td>GU305914.1</td><td>56933</td><td>56955</td><td>-</td><td>3</td><td>GATACCAGGTCGGAGGCCCG</td><td>TGG</td><td>||-|||||[|||||-||||-|]PAM</td><td>NA</td><td>Intergenic</td><td>NA</td><td>NA</td></tr>
<tr><td></td><td></td><td></td><td></td><td></td><td></td><td></td><td></td><td></td><td></td><td></td><td></td></tr>
<tr><td>T29288</td><td>AGACCAGGTCGGCGGCCAGCGGG</td><td>1000</td><td></td><td></td><td></td><td></td><td></td><td></td><td></td><td></td><td></td></tr>
<tr><td>Oligo fwd</td><td>CACCAGACCAGGTCGGCGGCCAGC</td><td></td><td></td><td></td><td></td><td></td><td></td><td></td><td></td><td></td><td></td></tr>
<tr><td>Oligo rev</td><td>AAACGCTGGCCGCCGACCTGGTCT</td><td></td><td></td><td></td><td></td><td></td><td></td><td></td><td></td><td></td><td></td></tr>
<tr><td>Chromosome</td><td>start</td><td>end</td><td>strand</td><td>MM</td><td>target_seq</td><td>PAM</td><td>alignment</td><td>distance</td><td>position</td><td>gene name</td><td>gene id</td></tr>
<tr><td>GU305914.1</td><td>64217</td><td>64239</td><td>-</td><td>0</td><td>AGACCAGGTCGGCGGCCAGC</td><td>GGG</td><td>||||||||[||||||||||||]PAM</td><td>NA</td><td>Intergenic</td><td>NA</td><td>NA</td></tr>
<tr><td></td><td></td><td></td><td></td><td></td><td></td><td></td><td></td><td></td><td></td><td></td><td></td></tr>
<tr><td>T29289</td><td>CCAGGTCGGCGGCCAGCGGGAGG</td><td>1000</td><td></td><td></td><td></td><td></td><td></td><td></td><td></td><td></td><td></td></tr>
<tr><td>Oligo fwd</td><td>CACCCCAGGTCGGCGGCCAGCGGG</td><td></td><td></td><td></td><td></td><td></td><td></td><td></td><td></td><td></td><td></td></tr>
<tr><td>Oligo rev</td><td>AAACCCCGCTGGCCGCCGACCTGG</td><td></td><td></td><td></td><td></td><td></td><td></td><td></td><td></td><td></td><td></td></tr>
<tr><td>Chromosome</td><td>start</td><td>end</td><td>strand</td><td>MM</td><td>target_seq</td><td>PAM</td><td>alignment</td><td>distance</td><td>position</td><td>gene name</td><td>gene id</td></tr>
<tr><td>GU305914.1</td><td>64214</td><td>64236</td><td>-</td><td>0</td><td>CCAGGTCGGCGGCCAGCGGG</td><td>AGG</td><td>||||||||[||||||||||||]PAM</td><td>NA</td><td>Intergenic</td><td>NA</td><td>NA</td></tr>
<tr><td></td><td></td><td></td><td></td><td></td><td></td><td></td><td></td><td></td><td></td><td></td><td></td></tr>
<tr><td>T29290</td><td>AGGAGAAGCCCTTTCTGCGCCGG</td><td>1000</td><td></td><td></td><td></td><td></td><td></td><td></td><td></td><td></td><td></td></tr>
<tr><td>Oligo fwd</td><td>CACCAGGAGAAGCCCTTTCTGCGC</td><td></td><td></td><td></td><td></td><td></td><td></td><td></td><td></td><td></td><td></td></tr>
<tr><td>Oligo rev</td><td>AAACGCGCAGAAAGGGCTTCTCCT</td><td></td><td></td><td></td><td></td><td></td><td></td><td></td><td></td><td></td><td></td></tr>
<tr><td>Chromosome</td><td>start</td><td>end</td><td>strand</td><td>MM</td><td>target_seq</td><td>PAM</td><td>alignment</td><td>distance</td><td>position</td><td>gene name</td><td>gene id</td></tr>
<tr><td>GU305914.1</td><td>64194</td><td>64216</td><td>-</td><td>0</td><td>AGGAGAAGCCCTTTCTGCGC</td><td>CGG</td><td>||||||||[||||||||||||]PAM</td><td>NA</td><td>Intergenic</td><td>NA</td><td>NA</td></tr>
<tr><td></td><td></td><td></td><td></td><td></td><td></td><td></td><td></td><td></td><td></td><td></td><td></td></tr>
<tr><td>T29291</td><td>CCCTTTCTGCGCCGGCCCGAAGG</td><td>1000</td><td></td><td></td><td></td><td></td><td></td><td></td><td></td><td></td><td></td></tr>
<tr><td>Oligo fwd</td><td>CACCCCCTTTCTGCGCCGGCCCGA</td><td></td><td></td><td></td><td></td><td></td><td></td><td></td><td></td><td></td><td></td></tr>
<tr><td>Oligo rev</td><td>AAACTCGGGCCGGCGCAGAAAGGG</td><td></td><td></td><td></td><td></td><td></td><td></td><td></td><td></td><td></td><td></td></tr>
<tr><td>Chromosome</td><td>start</td><td>end</td><td>strand</td><td>MM</td><td>target_seq</td><td>PAM</td><td>alignment</td><td>distance</td><td>position</td><td>gene name</td><td>gene id</td></tr>
<tr><td>GU305914.1</td><td>64186</td><td>64208</td><td>-</td><td>0</td><td>CCCTTTCTGCGCCGGCCCGA</td><td>AGG</td><td>||||||||[||||||||||||]PAM</td><td>NA</td><td>Intergenic</td><td>NA</td><td>NA</td></tr>
<tr><td></td><td></td><td></td><td></td><td></td><td></td><td></td><td></td><td></td><td></td><td></td><td></td></tr>
<tr><td>T29292</td><td>GCGCCGGCCCGAAGGTGCCGTGG</td><td>1000</td><td></td><td></td><td></td><td></td><td></td><td></td><td></td><td></td><td></td></tr>
<tr><td>Oligo fwd</td><td>CACCGCGCCGGCCCGAAGGTGCCG</td><td></td><td></td><td></td><td></td><td></td><td></td><td></td><td></td><td></td><td></td></tr>
<tr><td>Oligo rev</td><td>AAACCGGCACCTTCGGGCCGGCGC</td><td></td><td></td><td></td><td></td><td></td><td></td><td></td><td></td><td></td><td></td></tr>
<tr><td>Chromosome</td><td>start</td><td>end</td><td>strand</td><td>MM</td><td>target_seq</td><td>PAM</td><td>alignment</td><td>distance</td><td>position</td><td>gene name</td><td>gene id</td></tr>
<tr><td>GU305914.1</td><td>64178</td><td>64200</td><td>-</td><td>0</td><td>GCGCCGGCCCGAAGGTGCCG</td><td>TGG</td><td>||||||||[||||||||||||]PAM</td><td>NA</td><td>Intergenic</td><td>NA</td><td>NA</td></tr>
<tr><td></td><td></td><td></td><td></td><td></td><td></td><td></td><td></td><td></td><td></td><td></td><td></td></tr>
<tr><td>T29293</td><td>TGGTTCAGAAAGTAGAACACCGG</td><td>1000</td><td></td><td></td><td></td><td></td><td></td><td></td><td></td><td></td><td></td></tr>
<tr><td>Oligo fwd</td><td>CACCTGGTTCAGAAAGTAGAACAC</td><td></td><td></td><td></td><td></td><td></td><td></td><td></td><td></td><td></td><td></td></tr>
<tr><td>Oligo rev</td><td>AAACGTGTTCTACTTTCTGAACCA</td><td></td><td></td><td></td><td></td><td></td><td></td><td></td><td></td><td></td><td></td></tr>
<tr><td>Chromosome</td><td>start</td><td>end</td><td>strand</td><td>MM</td><td>target_seq</td><td>PAM</td><td>alignment</td><td>distance</td><td>position</td><td>gene name</td><td>gene id</td></tr>
<tr><td>GU305914.1</td><td>64158</td><td>64180</td><td>-</td><td>0</td><td>TGGTTCAGAAAGTAGAACAC</td><td>CGG</td><td>||||||||[||||||||||||]PAM</td><td>NA</td><td>Intergenic</td><td>NA</td><td>NA</td></tr>
<tr><td></td><td></td><td></td><td></td><td></td><td></td><td></td><td></td><td></td><td></td><td></td><td></td></tr>
<tr><td>T29294</td><td>GGTTCAGAAAGTAGAACACCGGG</td><td>1000</td><td></td><td></td><td></td><td></td><td></td><td></td><td></td><td></td><td></td></tr>
<tr><td>Oligo fwd</td><td>CACCGGTTCAGAAAGTAGAACACC</td><td></td><td></td><td></td><td></td><td></td><td></td><td></td><td></td><td></td><td></td></tr>
<tr><td>Oligo rev</td><td>AAACGGTGTTCTACTTTCTGAACC</td><td></td><td></td><td></td><td></td><td></td><td></td><td></td><td></td><td></td><td></td></tr>
<tr><td>Chromosome</td><td>start</td><td>end</td><td>strand</td><td>MM</td><td>target_seq</td><td>PAM</td><td>alignment</td><td>distance</td><td>position</td><td>gene name</td><td>gene id</td></tr>
<tr><td>GU305914.1</td><td>64157</td><td>64179</td><td>-</td><td>0</td><td>GGTTCAGAAAGTAGAACACC</td><td>GGG</td><td>||||||||[||||||||||||]PAM</td><td>NA</td><td>Intergenic</td><td>NA</td><td>NA</td></tr>
<tr><td></td><td></td><td></td><td></td><td></td><td></td><td></td><td></td><td></td><td></td><td></td><td></td></tr>
<tr><td>T29295</td><td>AGTAGAACACCGGGTGTCTGCGG</td><td>1000</td><td></td><td></td><td></td><td></td><td></td><td></td><td></td><td></td><td></td></tr>
<tr><td>Oligo fwd</td><td>CACCAGTAGAACACCGGGTGTCTG</td><td></td><td></td><td></td><td></td><td></td><td></td><td></td><td></td><td></td><td></td></tr>
<tr><td>Oligo rev</td><td>AAACCAGACACCCGGTGTTCTACT</td><td></td><td></td><td></td><td></td><td></td><td></td><td></td><td></td><td></td><td></td></tr>
<tr><td>Chromosome</td><td>start</td><td>end</td><td>strand</td><td>MM</td><td>target_seq</td><td>PAM</td><td>alignment</td><td>distance</td><td>position</td><td>gene name</td><td>gene id</td></tr>
<tr><td>GU305914.1</td><td>64148</td><td>64170</td><td>-</td><td>0</td><td>AGTAGAACACCGGGTGTCTG</td><td>CGG</td><td>||||||||[||||||||||||]PAM</td><td>NA</td><td>Intergenic</td><td>NA</td><td>NA</td></tr>
<tr><td></td><td></td><td></td><td></td><td></td><td></td><td></td><td></td><td></td><td></td><td></td><td></td></tr>
<tr><td>T29296</td><td>GTCTGCGGATGAACTCGAAGTGG</td><td>1000</td><td></td><td></td><td></td><td></td><td></td><td></td><td></td><td></td><td></td></tr>
<tr><td>Oligo fwd</td><td>CACCGTCTGCGGATGAACTCGAAG</td><td></td><td></td><td></td><td></td><td></td><td></td><td></td><td></td><td></td><td></td></tr>
<tr><td>Oligo rev</td><td>AAACCTTCGAGTTCATCCGCAGAC</td><td></td><td></td><td></td><td></td><td></td><td></td><td></td><td></td><td></td><td></td></tr>
<tr><td>Chromosome</td><td>start</td><td>end</td><td>strand</td><td>MM</td><td>target_seq</td><td>PAM</td><td>alignment</td><td>distance</td><td>position</td><td>gene name</td><td>gene id</td></tr>
<tr><td>GU305914.1</td><td>64133</td><td>64155</td><td>-</td><td>0</td><td>GTCTGCGGATGAACTCGAAG</td><td>TGG</td><td>||||||||[||||||||||||]PAM</td><td>NA</td><td>Intergenic</td><td>NA</td><td>NA</td></tr>
<tr><td></td><td></td><td></td><td></td><td></td><td></td><td></td><td></td><td></td><td></td><td></td><td></td></tr>
<tr><td>T29297</td><td>CTCGAAGTGGTAGCCCGTCTCGG</td><td>1000</td><td></td><td></td><td></td><td></td><td></td><td></td><td></td><td></td><td></td></tr>
<tr><td>Oligo fwd</td><td>CACCCTCGAAGTGGTAGCCCGTCT</td><td></td><td></td><td></td><td></td><td></td><td></td><td></td><td></td><td></td><td></td></tr>
<tr><td>Oligo rev</td><td>AAACAGACGGGCTACCACTTCGAG</td><td></td><td></td><td></td><td></td><td></td><td></td><td></td><td></td><td></td><td></td></tr>
<tr><td>Chromosome</td><td>start</td><td>end</td><td>strand</td><td>MM</td><td>target_seq</td><td>PAM</td><td>alignment</td><td>distance</td><td>position</td><td>gene name</td><td>gene id</td></tr>
<tr><td>GU305914.1</td><td>64120</td><td>64142</td><td>-</td><td>0</td><td>CTCGAAGTGGTAGCCCGTCT</td><td>CGG</td><td>||||||||[||||||||||||]PAM</td><td>NA</td><td>Intergenic</td><td>NA</td><td>NA</td></tr>
<tr><td></td><td></td><td></td><td></td><td></td><td></td><td></td><td></td><td></td><td></td><td></td><td></td></tr>
<tr><td>T29298</td><td>TCGAAGTGGTAGCCCGTCTCGGG</td><td>1000</td><td></td><td></td><td></td><td></td><td></td><td></td><td></td><td></td><td></td></tr>
<tr><td>Oligo fwd</td><td>CACCTCGAAGTGGTAGCCCGTCTC</td><td></td><td></td><td></td><td></td><td></td><td></td><td></td><td></td><td></td><td></td></tr>
<tr><td>Oligo rev</td><td>AAACGAGACGGGCTACCACTTCGA</td><td></td><td></td><td></td><td></td><td></td><td></td><td></td><td></td><td></td><td></td></tr>
<tr><td>Chromosome</td><td>start</td><td>end</td><td>strand</td><td>MM</td><td>target_seq</td><td>PAM</td><td>alignment</td><td>distance</td><td>position</td><td>gene name</td><td>gene id</td></tr>
<tr><td>GU305914.1</td><td>64119</td><td>64141</td><td>-</td><td>0</td><td>TCGAAGTGGTAGCCCGTCTC</td><td>GGG</td><td>||||||||[||||||||||||]PAM</td><td>NA</td><td>Intergenic</td><td>NA</td><td>NA</td></tr>
<tr><td></td><td></td><td></td><td></td><td></td><td></td><td></td><td></td><td></td><td></td><td></td><td></td></tr>
<tr><td>T29299</td><td>CGAAGTGGTAGCCCGTCTCGGGG</td><td>1000</td><td></td><td></td><td></td><td></td><td></td><td></td><td></td><td></td><td></td></tr>
<tr><td>Oligo fwd</td><td>CACCCGAAGTGGTAGCCCGTCTCG</td><td></td><td></td><td></td><td></td><td></td><td></td><td></td><td></td><td></td><td></td></tr>
<tr><td>Oligo rev</td><td>AAACCGAGACGGGCTACCACTTCG</td><td></td><td></td><td></td><td></td><td></td><td></td><td></td><td></td><td></td><td></td></tr>
<tr><td>Chromosome</td><td>start</td><td>end</td><td>strand</td><td>MM</td><td>target_seq</td><td>PAM</td><td>alignment</td><td>distance</td><td>position</td><td>gene name</td><td>gene id</td></tr>
<tr><td>GU305914.1</td><td>64118</td><td>64140</td><td>-</td><td>0</td><td>CGAAGTGGTAGCCCGTCTCG</td><td>GGG</td><td>||||||||[||||||||||||]PAM</td><td>NA</td><td>Intergenic</td><td>NA</td><td>NA</td></tr>
<tr><td></td><td></td><td></td><td></td><td></td><td></td><td></td><td></td><td></td><td></td><td></td><td></td></tr>
<tr><td>T29300</td><td>CCGTCTCGGGGACGTGCTGCAGG</td><td>1000</td><td></td><td></td><td></td><td></td><td></td><td></td><td></td><td></td><td></td></tr>
<tr><td>Oligo fwd</td><td>CACCCCGTCTCGGGGACGTGCTGC</td><td></td><td></td><td></td><td></td><td></td><td></td><td></td><td></td><td></td><td></td></tr>
<tr><td>Oligo rev</td><td>AAACGCAGCACGTCCCCGAGACGG</td><td></td><td></td><td></td><td></td><td></td><td></td><td></td><td></td><td></td><td></td></tr>
<tr><td>Chromosome</td><td>start</td><td>end</td><td>strand</td><td>MM</td><td>target_seq</td><td>PAM</td><td>alignment</td><td>distance</td><td>position</td><td>gene name</td><td>gene id</td></tr>
<tr><td>GU305914.1</td><td>64106</td><td>64128</td><td>-</td><td>0</td><td>CCGTCTCGGGGACGTGCTGC</td><td>AGG</td><td>||||||||[||||||||||||]PAM</td><td>NA</td><td>Intergenic</td><td>NA</td><td>NA</td></tr>
</table>

| | | | | | | | | | | | |
|---|---|---|---|---|---|---|---|---|---|---|---|
| T29301 | GGACGTGCTGCAGGAAACGACGG | 1000 | | | | | | | | | |
| Oligo fwd | CACCGGACGTGCTGCAGGAAACGA | | | | | | | | | | |
| Oligo rev | AAACTCGTTTCCTGCAGCACGTCC | | | | | | | | | | |
| Chromosome | start | end | strand | MM | target_seq | PAM | alignment | distance | position | gene name | gene id |
| GU305914.1 | 64097 | 64119 | - | 0 | GGACGTGCTGCAGGAAACGA | CGG | \|\|\|\|\|\|\|\|[\|\|\|\|\|\|\|\|\|\|\|\|]PAM | NA | Intergenic | NA | NA |
| T29302 | GCTGCAGGAAACGACGGACGCGG | 1000 | | | | | | | | | |
| Oligo fwd | CACCGCTGCAGGAAACGACGGACG | | | | | | | | | | |
| Oligo rev | AAACCGTCCGTCGTTTCCTGCAGC | | | | | | | | | | |
| Chromosome | start | end | strand | MM | target_seq | PAM | alignment | distance | position | gene name | gene id |
| GU305914.1 | 64091 | 64113 | - | 0 | GCTGCAGGAAACGACGGACG | CGG | \|\|\|\|\|\|\|\|[\|\|\|\|\|\|\|\|\|\|\|\|]PAM | NA | Intergenic | NA | NA |
| T29303 | CTGCAGGAAACGACGGACGCGGG | 1000 | | | | | | | | | |
| Oligo fwd | CACCCTGCAGGAAACGACGGACGC | | | | | | | | | | |
| Oligo rev | AAACGCGTCCGTCGTTTCCTGCAG | | | | | | | | | | |
| Chromosome | start | end | strand | MM | target_seq | PAM | alignment | distance | position | gene name | gene id |
| GU305914.1 | 64090 | 64112 | - | 0 | CTGCAGGAAACGACGGACGC | GGG | \|\|\|\|\|\|\|\|[\|\|\|\|\|\|\|\|\|\|\|\|]PAM | NA | Intergenic | NA | NA |
| T29304 | CAGGAAACGACGGACGCGGGAGG | 1000 | | | | | | | | | |
| Oligo fwd | CACCCAGGAAACGACGGACGCGGG | | | | | | | | | | |
| Oligo rev | AAACCCCGCGTCCGTCGTTTCCTG | | | | | | | | | | |
| Chromosome | start | end | strand | MM | target_seq | PAM | alignment | distance | position | gene name | gene id |
| GU305914.1 | 64087 | 64109 | - | 0 | CAGGAAACGACGGACGCGGG | AGG | \|\|\|\|\|\|\|\|[\|\|\|\|\|\|\|\|\|\|\|\|]PAM | NA | Intergenic | NA | NA |
| T29305 | CGTCACCGTCTGCAGTTGCACGG | 1000 | | | | | | | | | |
| Oligo fwd | CACCCGTCACCGTCTGCAGTTGCA | | | | | | | | | | |
| Oligo rev | AAACTGCAACTGCAGACGGTGACG | | | | | | | | | | |
| Chromosome | start | end | strand | MM | target_seq | PAM | alignment | distance | position | gene name | gene id |
| GU305914.1 | 64063 | 64085 | - | 0 | CGTCACCGTCTGCAGTTGCA | CGG | \|\|\|\|\|\|\|\|[\|\|\|\|\|\|\|\|\|\|\|\|]PAM | NA | Intergenic | NA | NA |
| T29306 | CACCGTCTGCAGTTGCACGGCGG | 1000 | | | | | | | | | |
| Oligo fwd | CACCCACCGTCTGCAGTTGCACGG | | | | | | | | | | |
| Oligo rev | AAACCCGTGCAACTGCAGACGGTG | | | | | | | | | | |
| Chromosome | start | end | strand | MM | target_seq | PAM | alignment | distance | position | gene name | gene id |
| GU305914.1 | 64060 | 64082 | - | 0 | CACCGTCTGCAGTTGCACGG | CGG | \|\|\|\|\|\|\|\|[\|\|\|\|\|\|\|\|\|\|\|\|]PAM | NA | Intergenic | NA | NA |
| GU305914.1 | 221332 | 221354 | + | 4 | GACCGCCCGCAGTTGCAAGG | GGG | -\|\|\|\|-\|-[\|\|\|\|\|\|\|\|\|-\|\|]PAM | NA | Intergenic | NA | NA |
| T29307 | CACGGCGGAGATTTCTATCTTGG | 1000 | | | | | | | | | |
| Oligo fwd | CACCCACGGCGGAGATTTCTATCT | | | | | | | | | | |
| Oligo rev | AAACAGATAGAAATCTCCGCCGTG | | | | | | | | | | |
| Chromosome | start | end | strand | MM | target_seq | PAM | alignment | distance | position | gene name | gene id |
| GU305914.1 | 64045 | 64067 | - | 0 | CACGGCGGAGATTTCTATCT | TGG | \|\|\|\|\|\|\|\|[\|\|\|\|\|\|\|\|\|\|\|\|]PAM | NA | Intergenic | NA | NA |
| T29308 | GATCTTCCAGCGTCTGTTCCAGG | 1000 | | | | | | | | | |
| Oligo fwd | CACCGATCTTCCAGCGTCTGTTCC | | | | | | | | | | |
| Oligo rev | AAACGGAACAGACGCTGGAAGATC | | | | | | | | | | |
| Chromosome | start | end | strand | MM | target_seq | PAM | alignment | distance | position | gene name | gene id |
| GU305914.1 | 63977 | 63999 | - | 0 | GATCTTCCAGCGTCTGTTCC | AGG | \|\|\|\|\|\|\|\|[\|\|\|\|\|\|\|\|\|\|\|\|]PAM | NA | Intergenic | NA | NA |
| T29309 | CCAGCGTCTGTTCCAGGTTGAGG | 1000 | | | | | | | | | |
| Oligo fwd | CACCCCAGCGTCTGTTCCAGGTTG | | | | | | | | | | |
| Oligo rev | AAACCAACCTGGAACAGACGCTGG | | | | | | | | | | |
| Chromosome | start | end | strand | MM | target_seq | PAM | alignment | distance | position | gene name | gene id |
| GU305914.1 | 63971 | 63993 | - | 0 | CCAGCGTCTGTTCCAGGTTG | AGG | \|\|\|\|\|\|\|\|[\|\|\|\|\|\|\|\|\|\|\|\|]PAM | NA | Intergenic | NA | NA |
| T29310 | CAGCGTCTGTTCCAGGTTGAGGG | 1000 | | | | | | | | | |
| Oligo fwd | CACCCAGCGTCTGTTCCAGGTTGA | | | | | | | | | | |
| Oligo rev | AAACTCAACCTGGAACAGACGCTG | | | | | | | | | | |
| Chromosome | start | end | strand | MM | target_seq | PAM | alignment | distance | position | gene name | gene id |
| GU305914.1 | 63970 | 63992 | - | 0 | CAGCGTCTGTTCCAGGTTGA | GGG | \|\|\|\|\|\|\|\|[\|\|\|\|\|\|\|\|\|\|\|\|]PAM | NA | Intergenic | NA | NA |
| T29311 | TCTGTTCCAGGTTGAGGGCTCGG | 1000 | | | | | | | | | |
| Oligo fwd | CACCTCTGTTCCAGGTTGAGGGCT | | | | | | | | | | |
| Oligo rev | AAACAGCCCTCAACCTGGAACAGA | | | | | | | | | | |
| Chromosome | start | end | strand | MM | target_seq | PAM | alignment | distance | position | gene name | gene id |
| GU305914.1 | 63965 | 63987 | - | 0 | TCTGTTCCAGGTTGAGGGCT | CGG | \|\|\|\|\|\|\|\|[\|\|\|\|\|\|\|\|\|\|\|\|]PAM | NA | Intergenic | NA | NA |
| GU305914.1 | 46295 | 46317 | - | 4 | TATCTTCCAGGTTCAGGGCG | CGG | \|-\|-\|\|\|\|[\|\|\|\|\|-\|\|\|\|\|-]PAM | NA | Intergenic | NA | NA |
| T29312 | CTGTTCCAGGTTGAGGGCTCGGG | 1000 | | | | | | | | | |
| Oligo fwd | CACCCTGTTCCAGGTTGAGGGCTC | | | | | | | | | | |
| Oligo rev | AAACGAGCCCTCAACCTGGAACAG | | | | | | | | | | |
| Chromosome | start | end | strand | MM | target_seq | PAM | alignment | distance | position | gene name | gene id |
| GU305914.1 | 63964 | 63986 | - | 0 | CTGTTCCAGGTTGAGGGCTC | GGG | \|\|\|\|\|\|\|\|[\|\|\|\|\|\|\|\|\|\|\|\|]PAM | NA | Intergenic | NA | NA |
| T29313 | TTGAGGGCTCGGGTGACGTGCGG | 1000 | | | | | | | | | |
| Oligo fwd | CACCTTGAGGGCTCGGGTGACGTG | | | | | | | | | | |
| Oligo rev | AAACCACGTCACCCGAGCCCTCAA | | | | | | | | | | |
| Chromosome | start | end | strand | MM | target_seq | PAM | alignment | distance | position | gene name | gene id |
| GU305914.1 | 63954 | 63976 | - | 0 | TTGAGGGCTCGGGTGACGTG | CGG | \|\|\|\|\|\|\|\|[\|\|\|\|\|\|\|\|\|\|\|\|]PAM | NA | Intergenic | NA | NA |
| T29314 | TGAGGGCTCGGGTGACGTGCGGG | 1000 | | | | | | | | | |
| Oligo fwd | CACCTGAGGGCTCGGGTGACGTGC | | | | | | | | | | |
| Oligo rev | AAACGCACGTCACCCGAGCCCTCA | | | | | | | | | | |
| Chromosome | start | end | strand | MM | target_seq | PAM | alignment | distance | position | gene name | gene id |
| GU305914.1 | 63953 | 63975 | - | 0 | TGAGGGCTCGGGTGACGTGC | GGG | \|\|\|\|\|\|\|\|[\|\|\|\|\|\|\|\|\|\|\|\|]PAM | NA | Intergenic | NA | NA |
| T29315 | GGGCTCGGGTGACGTGCGGGTGG | 1000 | | | | | | | | | |
| Oligo fwd | CACCGGGCTCGGGTGACGTGCGGG | | | | | | | | | | |

| | | | | | | | | | | | |
|---|---|---|---|---|---|---|---|---|---|---|---|
| Oligo rev | AAACCCCGCACGTCACCCGAGCCC | | | | | | | | | | |
| Chromosome | start | end | strand | MM | target_seq | PAM | alignment | distance | position | gene name | gene id |
| GU305914.1 | 63950 | 63972 | - | 0 | GGGCTCGGGTGACGTGCGGG | TGG | \|\|\|\|\|\|\|\|[\|\|\|\|\|\|\|\|\|\|\|\|]PAM | NA | Intergenic | NA | NA |
| T29316 | CTCGGGTGACGTGCGGGTGGTGG | 1000 | | | | | | | | | |
| Oligo fwd | CACCCTCGGGTGACGTGCGGGTGG | | | | | | | | | | |
| Oligo rev | AAACCCACCCGCACGTCACCCGAG | | | | | | | | | | |
| Chromosome | start | end | strand | MM | target_seq | PAM | alignment | distance | position | gene name | gene id |
| GU305914.1 | 63947 | 63969 | - | 0 | CTCGGGTGACGTGCGGGTGG | TGG | \|\|\|\|\|\|\|\|[\|\|\|\|\|\|\|\|\|\|\|\|]PAM | NA | Intergenic | NA | NA |
| T29317 | GTGGTGGTCCTTCGCGAACGCGG | 1000 | | | | | | | | | |
| Oligo fwd | CACCGTGGTGGTCCTTCGCGAACG | | | | | | | | | | |
| Oligo rev | AAACCGTTCGCGAAGGACCACCAC | | | | | | | | | | |
| Chromosome | start | end | strand | MM | target_seq | PAM | alignment | distance | position | gene name | gene id |
| GU305914.1 | 63931 | 63953 | - | 0 | GTGGTGGTCCTTCGCGAACG | CGG | \|\|\|\|\|\|\|\|[\|\|\|\|\|\|\|\|\|\|\|\|]PAM | NA | Intergenic | NA | NA |
| T29318 | GTCCTTCGCGAACGCGGCCATGG | 1000 | | | | | | | | | |
| Oligo fwd | CACCGTCCTTCGCGAACGCGGCCA | | | | | | | | | | |
| Oligo rev | AAACTGGCCGCGTTCGCGAAGGAC | | | | | | | | | | |
| Chromosome | start | end | strand | MM | target_seq | PAM | alignment | distance | position | gene name | gene id |
| GU305914.1 | 63925 | 63947 | - | 0 | GTCCTTCGCGAACGCGGCCA | TGG | \|\|\|\|\|\|\|\|[\|\|\|\|\|\|\|\|\|\|\|\|]PAM | NA | Intergenic | NA | NA |
| T29319 | ATGGCGAACGTCAGCAGCTTCGG | 1000 | | | | | | | | | |
| Oligo fwd | CACCATGGCGAACGTCAGCAGCTT | | | | | | | | | | |
| Oligo rev | AAACAAGCTGCTGACGTTCGCCAT | | | | | | | | | | |
| Chromosome | start | end | strand | MM | target_seq | PAM | alignment | distance | position | gene name | gene id |
| GU305914.1 | 63906 | 63928 | - | 0 | ATGGCGAACGTCAGCAGCTT | CGG | \|\|\|\|\|\|\|\|[\|\|\|\|\|\|\|\|\|\|\|\|]PAM | NA | Intergenic | NA | NA |
| T29320 | GCAGCTTCGGCCCGATGAAGCGG | 1000 | | | | | | | | | |
| Oligo fwd | CACCGCAGCTTCGGCCCGATGAAG | | | | | | | | | | |
| Oligo rev | AAACCTTCATCGGGCCGAAGCTGC | | | | | | | | | | |
| Chromosome | start | end | strand | MM | target_seq | PAM | alignment | distance | position | gene name | gene id |
| GU305914.1 | 63893 | 63915 | - | 0 | GCAGCTTCGGCCCGATGAAG | CGG | \|\|\|\|\|\|\|\|[\|\|\|\|\|\|\|\|\|\|\|\|]PAM | NA | Intergenic | NA | NA |
| T29321 | CAGCTTCGGCCCGATGAAGCGGG | 1000 | | | | | | | | | |
| Oligo fwd | CACCCAGCTTCGGCCCGATGAAGC | | | | | | | | | | |
| Oligo rev | AAACGCTTCATCGGGCCGAAGCTG | | | | | | | | | | |
| Chromosome | start | end | strand | MM | target_seq | PAM | alignment | distance | position | gene name | gene id |
| GU305914.1 | 63892 | 63914 | - | 0 | CAGCTTCGGCCCGATGAAGC | GGG | \|\|\|\|\|\|\|\|[\|\|\|\|\|\|\|\|\|\|\|\|]PAM | NA | Intergenic | NA | NA |
| T29322 | CTTCGGCCCGATGAAGCGGGAGG | 1000 | | | | | | | | | |
| Oligo fwd | CACCCTTCGGCCCGATGAAGCGGG | | | | | | | | | | |
| Oligo rev | AAACCCCGCTTCATCGGGCCGAAG | | | | | | | | | | |
| Chromosome | start | end | strand | MM | target_seq | PAM | alignment | distance | position | gene name | gene id |
| GU305914.1 | 63889 | 63911 | - | 0 | CTTCGGCCCGATGAAGCGGG | AGG | \|\|\|\|\|\|\|\|[\|\|\|\|\|\|\|\|\|\|\|\|]PAM | NA | Intergenic | NA | NA |
| T29323 | CCCGATGAAGCGGGAGGTGATGG | 1000 | | | | | | | | | |
| Oligo fwd | CACCCCCGATGAAGCGGGAGGTGA | | | | | | | | | | |
| Oligo rev | AAACTCACCTCCCGCTTCATCGGG | | | | | | | | | | |
| Chromosome | start | end | strand | MM | target_seq | PAM | alignment | distance | position | gene name | gene id |
| GU305914.1 | 63883 | 63905 | - | 0 | CCCGATGAAGCGGGAGGTGA | TGG | \|\|\|\|\|\|\|\|[\|\|\|\|\|\|\|\|\|\|\|\|]PAM | NA | Intergenic | NA | NA |
| T29324 | GGTGATGGAGTTCGACCCCGAGG | 1000 | | | | | | | | | |
| Oligo fwd | CACCGGTGATGGAGTTCGACCCCG | | | | | | | | | | |
| Oligo rev | AAACCGGGGTCGAACTCCATCACC | | | | | | | | | | |
| Chromosome | start | end | strand | MM | target_seq | PAM | alignment | distance | position | gene name | gene id |
| GU305914.1 | 63868 | 63890 | - | 0 | GGTGATGGAGTTCGACCCCG | AGG | \|\|\|\|\|\|\|\|[\|\|\|\|\|\|\|\|\|\|\|\|]PAM | NA | Intergenic | NA | NA |
| T29325 | CGACCCCGAGGACCCCTACAAGG | 1000 | | | | | | | | | |
| Oligo fwd | CACCCGACCCCGAGGACCCCTACA | | | | | | | | | | |
| Oligo rev | AAACTGTAGGGGTCCTCGGGGTCG | | | | | | | | | | |
| Chromosome | start | end | strand | MM | target_seq | PAM | alignment | distance | position | gene name | gene id |
| GU305914.1 | 63856 | 63878 | - | 0 | CGACCCCGAGGACCCCTACA | AGG | \|\|\|\|\|\|\|\|[\|\|\|\|\|\|\|\|\|\|\|\|]PAM | NA | Intergenic | NA | NA |
| GU305914.1 | 70534 | 70556 | + | 4 | CAAACCCGAGGACACCTTCA | CGG | \|-\|-\|\|\|\|[\|\|\|\|\|-\|\|\|-\|\|]PAM | NA | Intergenic | NA | NA |
| T29326 | CCTACAAGGTGTCTAAGATGAGG | 1000 | | | | | | | | | |
| Oligo fwd | CACCCCTACAAGGTGTCTAAGATG | | | | | | | | | | |
| Oligo rev | AAACCATCTTAGACACCTTGTAGG | | | | | | | | | | |
| Chromosome | start | end | strand | MM | target_seq | PAM | alignment | distance | position | gene name | gene id |
| GU305914.1 | 63842 | 63864 | - | 0 | CCTACAAGGTGTCTAAGATG | AGG | \|\|\|\|\|\|\|\|[\|\|\|\|\|\|\|\|\|\|\|\|]PAM | NA | Intergenic | NA | NA |
| T29327 | CTAAGATGAGGAAACTTGAGAGG | 1000 | | | | | | | | | |
| Oligo fwd | CACCCTAAGATGAGGAAACTTGAG | | | | | | | | | | |
| Oligo rev | AAACCTCAAGTTTCCTCATCTTAG | | | | | | | | | | |
| Chromosome | start | end | strand | MM | target_seq | PAM | alignment | distance | position | gene name | gene id |
| GU305914.1 | 63830 | 63852 | - | 0 | CTAAGATGAGGAAACTTGAG | AGG | \|\|\|\|\|\|\|\|[\|\|\|\|\|\|\|\|\|\|\|\|]PAM | NA | Intergenic | NA | NA |
| T29328 | ACTTGAGAGGTCGATAGCCAAGG | 1000 | | | | | | | | | |
| Oligo fwd | CACCACTTGAGAGGTCGATAGCCA | | | | | | | | | | |
| Oligo rev | AAACTGGCTATCGACCTCTCAAGT | | | | | | | | | | |
| Chromosome | start | end | strand | MM | target_seq | PAM | alignment | distance | position | gene name | gene id |
| GU305914.1 | 63817 | 63839 | - | 0 | ACTTGAGAGGTCGATAGCCA | AGG | \|\|\|\|\|\|\|\|[\|\|\|\|\|\|\|\|\|\|\|\|]PAM | NA | Intergenic | NA | NA |
| T29329 | CTTGAGAGGTCGATAGCCAAGGG | 1000 | | | | | | | | | |
| Oligo fwd | CACCCTTGAGAGGTCGATAGCCAA | | | | | | | | | | |
| Oligo rev | AAACTTGGCTATCGACCTCTCAAG | | | | | | | | | | |
| Chromosome | start | end | strand | MM | target_seq | PAM | alignment | distance | position | gene name | gene id |
| GU305914.1 | 63816 | 63838 | - | 0 | CTTGAGAGGTCGATAGCCAA | GGG | \|\|\|\|\|\|\|\|[\|\|\|\|\|\|\|\|\|\|\|\|]PAM | NA | Intergenic | NA | NA |

<table>
<tr><td>T29330</td><td>ATAGCCAAGGGCTATGTCTACGG</td><td></td><td>1000</td><td></td><td></td><td></td><td></td><td></td><td></td><td></td><td></td><td></td></tr>
<tr><td>Oligo fwd</td><td>CACCATAGCCAAGGGCTATGTCTA</td><td></td><td></td><td></td><td></td><td></td><td></td><td></td><td></td><td></td><td></td><td></td></tr>
<tr><td>Oligo rev</td><td>AAACTAGACATAGCCCTTGGCTAT</td><td></td><td></td><td></td><td></td><td></td><td></td><td></td><td></td><td></td><td></td><td></td></tr>
<tr><td>Chromosome</td><td>start</td><td></td><td>end</td><td>strand</td><td>MM</td><td>target_seq</td><td>PAM</td><td>alignment</td><td>distance</td><td>position</td><td>gene name</td><td>gene id</td></tr>
<tr><td>GU305914.1</td><td></td><td>63804</td><td>63826</td><td>-</td><td>0</td><td>ATAGCCAAGGGCTATGTCTA</td><td>CGG</td><td>||||||||[||||||||||||]PAM</td><td>NA</td><td>Intergenic</td><td>NA</td><td>NA</td></tr>
<tr><td></td><td></td><td></td><td></td><td></td><td></td><td></td><td></td><td></td><td></td><td></td><td></td><td></td></tr>
<tr><td>T29331</td><td>TGTCTACGGCGCCGACCACCAGG</td><td></td><td>1000</td><td></td><td></td><td></td><td></td><td></td><td></td><td></td><td></td><td></td></tr>
<tr><td>Oligo fwd</td><td>CACCTGTCTACGGCGCCGACCACC</td><td></td><td></td><td></td><td></td><td></td><td></td><td></td><td></td><td></td><td></td><td></td></tr>
<tr><td>Oligo rev</td><td>AAACGGTGGTCGGCGCCGTAGACA</td><td></td><td></td><td></td><td></td><td></td><td></td><td></td><td></td><td></td><td></td><td></td></tr>
<tr><td>Chromosome</td><td>start</td><td></td><td>end</td><td>strand</td><td>MM</td><td>target_seq</td><td>PAM</td><td>alignment</td><td>distance</td><td>position</td><td>gene name</td><td>gene id</td></tr>
<tr><td>GU305914.1</td><td></td><td>63790</td><td>63812</td><td>-</td><td>0</td><td>TGTCTACGGCGCCGACCACC</td><td>AGG</td><td>||||||||[||||||||||||]PAM</td><td>NA</td><td>Intergenic</td><td>NA</td><td>NA</td></tr>
<tr><td></td><td></td><td></td><td></td><td></td><td></td><td></td><td></td><td></td><td></td><td></td><td></td><td></td></tr>
<tr><td>T29332</td><td>CGCCGACCACCAGGCGATCACGG</td><td></td><td>1000</td><td></td><td></td><td></td><td></td><td></td><td></td><td></td><td></td><td></td></tr>
<tr><td>Oligo fwd</td><td>CACCCGCCGACCACCAGGCGATCA</td><td></td><td></td><td></td><td></td><td></td><td></td><td></td><td></td><td></td><td></td><td></td></tr>
<tr><td>Oligo rev</td><td>AAACTGATCGCCTGGTGGTCGGCG</td><td></td><td></td><td></td><td></td><td></td><td></td><td></td><td></td><td></td><td></td><td></td></tr>
<tr><td>Chromosome</td><td>start</td><td></td><td>end</td><td>strand</td><td>MM</td><td>target_seq</td><td>PAM</td><td>alignment</td><td>distance</td><td>position</td><td>gene name</td><td>gene id</td></tr>
<tr><td>GU305914.1</td><td></td><td>63781</td><td>63803</td><td>-</td><td>0</td><td>CGCCGACCACCAGGCGATCA</td><td>CGG</td><td>||||||||[||||||||||||]PAM</td><td>NA</td><td>Intergenic</td><td>NA</td><td>NA</td></tr>
<tr><td></td><td></td><td></td><td></td><td></td><td></td><td></td><td></td><td></td><td></td><td></td><td></td><td></td></tr>
<tr><td>T29333</td><td>TCACGGCCCGCTTCTTCGTGCGG</td><td></td><td>1000</td><td></td><td></td><td></td><td></td><td></td><td></td><td></td><td></td><td></td></tr>
<tr><td>Oligo fwd</td><td>CACCTCACGGCCCGCTTCTTCGTG</td><td></td><td></td><td></td><td></td><td></td><td></td><td></td><td></td><td></td><td></td><td></td></tr>
<tr><td>Oligo rev</td><td>AAACCACGAAGAAGCGGGCCGTGA</td><td></td><td></td><td></td><td></td><td></td><td></td><td></td><td></td><td></td><td></td><td></td></tr>
<tr><td>Chromosome</td><td>start</td><td></td><td>end</td><td>strand</td><td>MM</td><td>target_seq</td><td>PAM</td><td>alignment</td><td>distance</td><td>position</td><td>gene name</td><td>gene id</td></tr>
<tr><td>GU305914.1</td><td></td><td>63764</td><td>63786</td><td>-</td><td>0</td><td>TCACGGCCCGCTTCTTCGTG</td><td>CGG</td><td>||||||||[||||||||||||]PAM</td><td>NA</td><td>Intergenic</td><td>NA</td><td>NA</td></tr>
<tr><td>GU305914.1</td><td></td><td>152012</td><td>152034</td><td>-</td><td>4</td><td>GCACGTCCCGATTCTGCGTG</td><td>CGG</td><td>-||||-||[||-||||-||||]PAM</td><td>NA</td><td>Intergenic</td><td>NA</td><td>NA</td></tr>
<tr><td></td><td></td><td></td><td></td><td></td><td></td><td></td><td></td><td></td><td></td><td></td><td></td><td></td></tr>
<tr><td>T29334</td><td>CACGGCCCGCTTCTTCGTGCGGG</td><td></td><td>1000</td><td></td><td></td><td></td><td></td><td></td><td></td><td></td><td></td><td></td></tr>
<tr><td>Oligo fwd</td><td>CACCCACGGCCCGCTTCTTCGTGC</td><td></td><td></td><td></td><td></td><td></td><td></td><td></td><td></td><td></td><td></td><td></td></tr>
<tr><td>Oligo rev</td><td>AAACGCACGAAGAAGCGGGCCGTG</td><td></td><td></td><td></td><td></td><td></td><td></td><td></td><td></td><td></td><td></td><td></td></tr>
<tr><td>Chromosome</td><td>start</td><td></td><td>end</td><td>strand</td><td>MM</td><td>target_seq</td><td>PAM</td><td>alignment</td><td>distance</td><td>position</td><td>gene name</td><td>gene id</td></tr>
<tr><td>GU305914.1</td><td></td><td>63763</td><td>63785</td><td>-</td><td>0</td><td>CACGGCCCGCTTCTTCGTGC</td><td>GGG</td><td>||||||||[||||||||||||]PAM</td><td>NA</td><td>Intergenic</td><td>NA</td><td>NA</td></tr>
<tr><td></td><td></td><td></td><td></td><td></td><td></td><td></td><td></td><td></td><td></td><td></td><td></td><td></td></tr>
<tr><td>T29335</td><td>TTCTTCGTGCGGGAGAGTCTCGG</td><td></td><td>1000</td><td></td><td></td><td></td><td></td><td></td><td></td><td></td><td></td><td></td></tr>
<tr><td>Oligo fwd</td><td>CACCTTCTTCGTGCGGGAGAGTCT</td><td></td><td></td><td></td><td></td><td></td><td></td><td></td><td></td><td></td><td></td><td></td></tr>
<tr><td>Oligo rev</td><td>AAACAGACTCTCCCGCACGAAGAA</td><td></td><td></td><td></td><td></td><td></td><td></td><td></td><td></td><td></td><td></td><td></td></tr>
<tr><td>Chromosome</td><td>start</td><td></td><td>end</td><td>strand</td><td>MM</td><td>target_seq</td><td>PAM</td><td>alignment</td><td>distance</td><td>position</td><td>gene name</td><td>gene id</td></tr>
<tr><td>GU305914.1</td><td></td><td>63753</td><td>63775</td><td>-</td><td>0</td><td>TTCTTCGTGCGGGAGAGTCT</td><td>CGG</td><td>||||||||[||||||||||||]PAM</td><td>NA</td><td>Intergenic</td><td>NA</td><td>NA</td></tr>
<tr><td></td><td></td><td></td><td></td><td></td><td></td><td></td><td></td><td></td><td></td><td></td><td></td><td></td></tr>
<tr><td>T29336</td><td>CGTGCGGGAGAGTCTCGGCGAGG</td><td></td><td>1000</td><td></td><td></td><td></td><td></td><td></td><td></td><td></td><td></td><td></td></tr>
<tr><td>Oligo fwd</td><td>CACCCGTGCGGGAGAGTCTCGGCG</td><td></td><td></td><td></td><td></td><td></td><td></td><td></td><td></td><td></td><td></td><td></td></tr>
<tr><td>Oligo rev</td><td>AAACCGCCGAGACTCTCCCGCACG</td><td></td><td></td><td></td><td></td><td></td><td></td><td></td><td></td><td></td><td></td><td></td></tr>
<tr><td>Chromosome</td><td>start</td><td></td><td>end</td><td>strand</td><td>MM</td><td>target_seq</td><td>PAM</td><td>alignment</td><td>distance</td><td>position</td><td>gene name</td><td>gene id</td></tr>
<tr><td>GU305914.1</td><td></td><td>63748</td><td>63770</td><td>-</td><td>0</td><td>CGTGCGGGAGAGTCTCGGCG</td><td>AGG</td><td>||||||||[||||||||||||]PAM</td><td>NA</td><td>Intergenic</td><td>NA</td><td>NA</td></tr>
<tr><td></td><td></td><td></td><td></td><td></td><td></td><td></td><td></td><td></td><td></td><td></td><td></td><td></td></tr>
<tr><td>T29337</td><td>CGAGGTCGAGCAGAAGAACCTGG</td><td></td><td>1000</td><td></td><td></td><td></td><td></td><td></td><td></td><td></td><td></td><td></td></tr>
<tr><td>Oligo fwd</td><td>CACCCGAGGTCGAGCAGAAGAACC</td><td></td><td></td><td></td><td></td><td></td><td></td><td></td><td></td><td></td><td></td><td></td></tr>
<tr><td>Oligo rev</td><td>AAACGGTTCTTCTGCTCGACCTCG</td><td></td><td></td><td></td><td></td><td></td><td></td><td></td><td></td><td></td><td></td><td></td></tr>
<tr><td>Chromosome</td><td>start</td><td></td><td>end</td><td>strand</td><td>MM</td><td>target_seq</td><td>PAM</td><td>alignment</td><td>distance</td><td>position</td><td>gene name</td><td>gene id</td></tr>
<tr><td>GU305914.1</td><td></td><td>63730</td><td>63752</td><td>-</td><td>0</td><td>CGAGGTCGAGCAGAAGAACC</td><td>TGG</td><td>||||||||[||||||||||||]PAM</td><td>NA</td><td>Intergenic</td><td>NA</td><td>NA</td></tr>
<tr><td></td><td></td><td></td><td></td><td></td><td></td><td></td><td></td><td></td><td></td><td></td><td></td><td></td></tr>
<tr><td>T29338</td><td>GAGGTCGAGCAGAAGAACCTGGG</td><td></td><td>1000</td><td></td><td></td><td></td><td></td><td></td><td></td><td></td><td></td><td></td></tr>
<tr><td>Oligo fwd</td><td>CACCGAGGTCGAGCAGAAGAACCT</td><td></td><td></td><td></td><td></td><td></td><td></td><td></td><td></td><td></td><td></td><td></td></tr>
<tr><td>Oligo rev</td><td>AAACAGGTTCTTCTGCTCGACCTC</td><td></td><td></td><td></td><td></td><td></td><td></td><td></td><td></td><td></td><td></td><td></td></tr>
<tr><td>Chromosome</td><td>start</td><td></td><td>end</td><td>strand</td><td>MM</td><td>target_seq</td><td>PAM</td><td>alignment</td><td>distance</td><td>position</td><td>gene name</td><td>gene id</td></tr>
<tr><td>GU305914.1</td><td></td><td>63729</td><td>63751</td><td>-</td><td>0</td><td>GAGGTCGAGCAGAAGAACCT</td><td>GGG</td><td>||||||||[||||||||||||]PAM</td><td>NA</td><td>Intergenic</td><td>NA</td><td>NA</td></tr>
<tr><td></td><td></td><td></td><td></td><td></td><td></td><td></td><td></td><td></td><td></td><td></td><td></td><td></td></tr>
<tr><td>T29339</td><td>AGGTCGAGCAGAAGAACCTGGGG</td><td></td><td>1000</td><td></td><td></td><td></td><td></td><td></td><td></td><td></td><td></td><td></td></tr>
<tr><td>Oligo fwd</td><td>CACCAGGTCGAGCAGAAGAACCTG</td><td></td><td></td><td></td><td></td><td></td><td></td><td></td><td></td><td></td><td></td><td></td></tr>
<tr><td>Oligo rev</td><td>AAACCAGGTTCTTCTGCTCGACCT</td><td></td><td></td><td></td><td></td><td></td><td></td><td></td><td></td><td></td><td></td><td></td></tr>
<tr><td>Chromosome</td><td>start</td><td></td><td>end</td><td>strand</td><td>MM</td><td>target_seq</td><td>PAM</td><td>alignment</td><td>distance</td><td>position</td><td>gene name</td><td>gene id</td></tr>
<tr><td>GU305914.1</td><td></td><td>63728</td><td>63750</td><td>-</td><td>0</td><td>AGGTCGAGCAGAAGAACCTG</td><td>GGG</td><td>||||||||[||||||||||||]PAM</td><td>NA</td><td>Intergenic</td><td>NA</td><td>NA</td></tr>
<tr><td></td><td></td><td></td><td></td><td></td><td></td><td></td><td></td><td></td><td></td><td></td><td></td><td></td></tr>
<tr><td>T29340</td><td>GGTCGAGCAGAAGAACCTGGGGG</td><td></td><td>1000</td><td></td><td></td><td></td><td></td><td></td><td></td><td></td><td></td><td></td></tr>
<tr><td>Oligo fwd</td><td>CACCGGTCGAGCAGAAGAACCTGG</td><td></td><td></td><td></td><td></td><td></td><td></td><td></td><td></td><td></td><td></td><td></td></tr>
<tr><td>Oligo rev</td><td>AAACCCAGGTTCTTCTGCTCGACC</td><td></td><td></td><td></td><td></td><td></td><td></td><td></td><td></td><td></td><td></td><td></td></tr>
<tr><td>Chromosome</td><td>start</td><td></td><td>end</td><td>strand</td><td>MM</td><td>target_seq</td><td>PAM</td><td>alignment</td><td>distance</td><td>position</td><td>gene name</td><td>gene id</td></tr>
<tr><td>GU305914.1</td><td></td><td>63727</td><td>63749</td><td>-</td><td>0</td><td>GGTCGAGCAGAAGAACCTGG</td><td>GGG</td><td>||||||||[||||||||||||]PAM</td><td>NA</td><td>Intergenic</td><td>NA</td><td>NA</td></tr>
<tr><td></td><td></td><td></td><td></td><td></td><td></td><td></td><td></td><td></td><td></td><td></td><td></td><td></td></tr>
<tr><td>T29341</td><td>ACCTGGGGGTCCTGATGTTCAGG</td><td></td><td>1000</td><td></td><td></td><td></td><td></td><td></td><td></td><td></td><td></td><td></td></tr>
<tr><td>Oligo fwd</td><td>CACCACCTGGGGGTCCTGATGTTC</td><td></td><td></td><td></td><td></td><td></td><td></td><td></td><td></td><td></td><td></td><td></td></tr>
<tr><td>Oligo rev</td><td>AAACGAACATCAGGACCCCCAGGT</td><td></td><td></td><td></td><td></td><td></td><td></td><td></td><td></td><td></td><td></td><td></td></tr>
<tr><td>Chromosome</td><td>start</td><td></td><td>end</td><td>strand</td><td>MM</td><td>target_seq</td><td>PAM</td><td>alignment</td><td>distance</td><td>position</td><td>gene name</td><td>gene id</td></tr>
<tr><td>GU305914.1</td><td></td><td>63713</td><td>63735</td><td>-</td><td>0</td><td>ACCTGGGGGTCCTGATGTTC</td><td>AGG</td><td>||||||||[||||||||||||]PAM</td><td>NA</td><td>Intergenic</td><td>NA</td><td>NA</td></tr>
<tr><td></td><td></td><td></td><td></td><td></td><td></td><td></td><td></td><td></td><td></td><td></td><td></td><td></td></tr>
<tr><td>T29342</td><td>CCTGATGTTCAGGCTCGACACGG</td><td></td><td>1000</td><td></td><td></td><td></td><td></td><td></td><td></td><td></td><td></td><td></td></tr>
<tr><td>Oligo fwd</td><td>CACCCCTGATGTTCAGGCTCGACA</td><td></td><td></td><td></td><td></td><td></td><td></td><td></td><td></td><td></td><td></td><td></td></tr>
<tr><td>Oligo rev</td><td>AAACTGTCGAGCCTGAACATCAGG</td><td></td><td></td><td></td><td></td><td></td><td></td><td></td><td></td><td></td><td></td><td></td></tr>
<tr><td>Chromosome</td><td>start</td><td></td><td>end</td><td>strand</td><td>MM</td><td>target_seq</td><td>PAM</td><td>alignment</td><td>distance</td><td>position</td><td>gene name</td><td>gene id</td></tr>
<tr><td>GU305914.1</td><td></td><td>63703</td><td>63725</td><td>-</td><td>0</td><td>CCTGATGTTCAGGCTCGACA</td><td>CGG</td><td>||||||||[||||||||||||]PAM</td><td>NA</td><td>Intergenic</td><td>NA</td><td>NA</td></tr>
<tr><td></td><td></td><td></td><td></td><td></td><td></td><td></td><td></td><td></td><td></td><td></td><td></td><td></td></tr>
<tr><td>T29343</td><td>CTGATGTTCAGGCTCGACACGGG</td><td></td><td>1000</td><td></td><td></td><td></td><td></td><td></td><td></td><td></td><td></td><td></td></tr>
<tr><td>Oligo fwd</td><td>CACCCTGATGTTCAGGCTCGACAC</td><td></td><td></td><td></td><td></td><td></td><td></td><td></td><td></td><td></td><td></td><td></td></tr>
<tr><td>Oligo rev</td><td>AAACGTGTCGAGCCTGAACATCAG</td><td></td><td></td><td></td><td></td><td></td><td></td><td></td><td></td><td></td><td></td><td></td></tr>
<tr><td>Chromosome</td><td>start</td><td></td><td>end</td><td>strand</td><td>MM</td><td>target_seq</td><td>PAM</td><td>alignment</td><td>distance</td><td>position</td><td>gene name</td><td>gene id</td></tr>
<tr><td>GU305914.1</td><td></td><td>63702</td><td>63724</td><td>-</td><td>0</td><td>CTGATGTTCAGGCTCGACAC</td><td>GGG</td><td>||||||||[||||||||||||]PAM</td><td>NA</td><td>Intergenic</td><td>NA</td><td>NA</td></tr>
<tr><td></td><td></td><td></td><td></td><td></td><td></td><td></td><td></td><td></td><td></td><td></td><td></td><td></td></tr>
<tr><td>T29344</td><td>TGATGTTCAGGCTCGACACGGGG</td><td></td><td>1000</td><td></td><td></td><td></td><td></td><td></td><td></td><td></td><td></td><td></td></tr>
<tr><td>Oligo fwd</td><td>CACCTGATGTTCAGGCTCGACACG</td><td></td><td></td><td></td><td></td><td></td><td></td><td></td><td></td><td></td><td></td><td></td></tr>
<tr><td>Oligo rev</td><td>AAACCGTGTCGAGCCTGAACATCA</td><td></td><td></td><td></td><td></td><td></td><td></td><td></td><td></td><td></td><td></td><td></td></tr>
<tr><td>Chromosome</td><td>start</td><td></td><td>end</td><td>strand</td><td>MM</td><td>target_seq</td><td>PAM</td><td>alignment</td><td>distance</td><td>position</td><td>gene name</td><td>gene id</td></tr>
</table>

GU305914.1 63701 63723 - 0 TGATGTTCAGGCTCGACACG GGG ||||||||[||||||||||||]PAM NA Intergenic NA NA

T29345 CAGCCTCTTCTTCCTCTCGATGG 1000
Oligo fwd CACCCAGCCTCTTCTTCCTCTCGA
Oligo rev AAACTCGAGAGGAAGAAGAGGCTG
Chromosome start end strand MM target_seq PAM alignment distance position gene name gene id
GU305914.1 63652 63674 - 0 CAGCCTCTTCTTCCTCTCGA TGG ||||||||[||||||||||||]PAM NA Intergenic NA NA

T29346 CCTCTTCTTCCTCTCGATGGTGG 1000
Oligo fwd CACCCCTCTTCTTCCTCTCGATGG
Oligo rev AAACCCATCGAGAGGAAGAAGAGG
Chromosome start end strand MM target_seq PAM alignment distance position gene name gene id
GU305914.1 63649 63671 - 0 CCTCTTCTTCCTCTCGATGG TGG ||||||||[||||||||||||]PAM NA Intergenic NA NA
GU305914.1 118239 118261 + 4 CCTCGTCCTCCTCCCGACGG AGG ||||-||-[|||||-|||-||]PAM NA Intergenic NA NA

T29347 CGCCACCAAGAACACGCTCGCGG 1000
Oligo fwd CACCCGCCACCAAGAACACGCTCG
Oligo rev AAACCGAGCGTGTTCTTGGTGGCG
Chromosome start end strand MM target_seq PAM alignment distance position gene name gene id
GU305914.1 63610 63632 - 0 CGCCACCAAGAACACGCTCG CGG ||||||||[||||||||||||]PAM NA Intergenic NA NA

T29348 AACACGCTCGCGGCGATCTACGG 1000
Oligo fwd CACCAACACGCTCGCGGCGATCTA
Oligo rev AAACTAGATCGCCGCGAGCGTGTT
Chromosome start end strand MM target_seq PAM alignment distance position gene name gene id
GU305914.1 63600 63622 - 0 AACACGCTCGCGGCGATCTA CGG ||||||||[||||||||||||]PAM NA Intergenic NA NA

T29349 CGCGGCGATCTACGGCCGCGAGG 1000
Oligo fwd CACCCGCGGCGATCTACGGCCGCG
Oligo rev AAACCGCGGCCGTAGATCGCCGCG
Chromosome start end strand MM target_seq PAM alignment distance position gene name gene id
GU305914.1 63592 63614 - 0 CGCGGCGATCTACGGCCGCG AGG ||||||||[||||||||||||]PAM NA Intergenic NA NA

T29350 GCGGCGATCTACGGCCGCGAGGG 1000
Oligo fwd CACCGCGGCGATCTACGGCCGCGA
Oligo rev AAACTCGCGGCCGTAGATCGCCGC
Chromosome start end strand MM target_seq PAM alignment distance position gene name gene id
GU305914.1 63591 63613 - 0 GCGGCGATCTACGGCCGCGA GGG ||||||||[||||||||||||]PAM NA Intergenic NA NA

T29351 GATCTACGGCCGCGAGGGCGAGG 1000
Oligo fwd CACCGATCTACGGCCGCGAGGGCG
Oligo rev AAACCGCCCTCGCGGCCGTAGATC
Chromosome start end strand MM target_seq PAM alignment distance position gene name gene id
GU305914.1 63586 63608 - 0 GATCTACGGCCGCGAGGGCG AGG ||||||||[||||||||||||]PAM NA Intergenic NA NA

T29352 AGGGCGAGGCCATCCGCACCTGG 1000
Oligo fwd CACCAGGGCGAGGCCATCCGCACC
Oligo rev AAACGGTGCGGATGGCCTCGCCCT
Chromosome start end strand MM target_seq PAM alignment distance position gene name gene id
GU305914.1 63572 63594 - 0 AGGGCGAGGCCATCCGCACC TGG ||||||||[||||||||||||]PAM NA Intergenic NA NA

T29353 ATCCGCACCTGGCTGCGCGACGG 1000
Oligo fwd CACCATCCGCACCTGGCTGCGCGA
Oligo rev AAACTCGCGCAGCCAGGTGCGGAT
Chromosome start end strand MM target_seq PAM alignment distance position gene name gene id
GU305914.1 63561 63583 - 0 ATCCGCACCTGGCTGCGCGA CGG ||||||||[||||||||||||]PAM NA Intergenic NA NA

T29354 CCTGGCTGCGCGACGGCGCCTGG 1000
Oligo fwd CACCCCTGGCTGCGCGACGGCGCC
Oligo rev AAACGGCGCCGTCGCGCAGCCAGG
Chromosome start end strand MM target_seq PAM alignment distance position gene name gene id
GU305914.1 63554 63576 - 0 CCTGGCTGCGCGACGGCGCC TGG ||||||||[||||||||||||]PAM NA Intergenic NA NA

T29355 GGCTGCGCGACGGCGCCTGGCGG 1000
Oligo fwd CACCGGCTGCGCGACGGCGCCTGG
Oligo rev AAACCCAGGCGCCGTCGCGCAGCC
Chromosome start end strand MM target_seq PAM alignment distance position gene name gene id
GU305914.1 63551 63573 - 0 GGCTGCGCGACGGCGCCTGG CGG ||||||||[||||||||||||]PAM NA Intergenic NA NA

T29356 ACGGCGCCTGGCGGCTGCACCGG 1000
Oligo fwd CACCACGGCGCCTGGCGGCTGCAC
Oligo rev AAACGTGCAGCCGCCAGGCGCCGT
Chromosome start end strand MM target_seq PAM alignment distance position gene name gene id
GU305914.1 63542 63564 - 0 ACGGCGCCTGGCGGCTGCAC CGG ||||||||[||||||||||||]PAM NA Intergenic NA NA

T29357 CGGCGCCTGGCGGCTGCACCGGG 1000
Oligo fwd CACCCGGCGCCTGGCGGCTGCACC
Oligo rev AAACGGTGCAGCCGCCAGGCGCCG
Chromosome start end strand MM target_seq PAM alignment distance position gene name gene id
GU305914.1 63541 63563 - 0 CGGCGCCTGGCGGCTGCACC GGG ||||||||[||||||||||||]PAM NA Intergenic NA NA

T29358 CACCGGGTCGTCCACCCGCTCGG 1000
Oligo fwd CACCCACCGGGTCGTCCACCCGCT
Oligo rev AAACAGCGGGTGGACGACCCGGTG
Chromosome start end strand MM target_seq PAM alignment distance position gene name gene id
GU305914.1 63525 63547 - 0 CACCGGGTCGTCCACCCGCT CGG ||||||||[||||||||||||]PAM NA Intergenic NA NA

T29359 TGCACCAACTCGATCACGCCCGG 1000
Oligo fwd CACCTGCACCAACTCGATCACGCC

<table>
<tr><td>Oligo rev</td><td>AAACGGCGTGATCGAGTTGGTGCA</td><td></td><td></td><td></td><td></td><td></td><td></td><td></td><td></td><td></td><td></td><td></td></tr>
<tr><td>Chromosome</td><td>start</td><td></td><td>end</td><td>strand</td><td>MM</td><td>target_seq</td><td>PAM</td><td>alignment</td><td>distance</td><td>position</td><td>gene name</td><td>gene id</td></tr>
<tr><td>GU305914.1</td><td></td><td>63501</td><td>63523</td><td>-</td><td>0</td><td>TGCACCAACTCGATCACGCC</td><td>CGG</td><td>||||||||[||||||||||||]PAM</td><td>NA</td><td>Intergenic</td><td>NA</td><td>NA</td></tr>
<tr><td>T29360</td><td>CTGATCACCTGCTCGATGCGCGG</td><td></td><td>1000</td><td></td><td></td><td></td><td></td><td></td><td></td><td></td><td></td><td></td></tr>
<tr><td>Oligo fwd</td><td>CACCCTGATCACCTGCTCGATGCG</td><td></td><td></td><td></td><td></td><td></td><td></td><td></td><td></td><td></td><td></td><td></td></tr>
<tr><td>Oligo rev</td><td>AAACCGCATCGAGCAGGTGATCAG</td><td></td><td></td><td></td><td></td><td></td><td></td><td></td><td></td><td></td><td></td><td></td></tr>
<tr><td>Chromosome</td><td>start</td><td></td><td>end</td><td>strand</td><td>MM</td><td>target_seq</td><td>PAM</td><td>alignment</td><td>distance</td><td>position</td><td>gene name</td><td>gene id</td></tr>
<tr><td>GU305914.1</td><td></td><td>63468</td><td>63490</td><td>-</td><td>0</td><td>CTGATCACCTGCTCGATGCG</td><td>CGG</td><td>||||||||[||||||||||||]PAM</td><td>NA</td><td>Intergenic</td><td>NA</td><td>NA</td></tr>
<tr><td>T29361</td><td>CTATCCCCTGCTCGTTCCCAAGG</td><td></td><td>1000</td><td></td><td></td><td></td><td></td><td></td><td></td><td></td><td></td><td></td></tr>
<tr><td>Oligo fwd</td><td>CACCCTATCCCCTGCTCGTTCCCA</td><td></td><td></td><td></td><td></td><td></td><td></td><td></td><td></td><td></td><td></td><td></td></tr>
<tr><td>Oligo rev</td><td>AAACTGGGAACGAGCAGGGGATAG</td><td></td><td></td><td></td><td></td><td></td><td></td><td></td><td></td><td></td><td></td><td></td></tr>
<tr><td>Chromosome</td><td>start</td><td></td><td>end</td><td>strand</td><td>MM</td><td>target_seq</td><td>PAM</td><td>alignment</td><td>distance</td><td>position</td><td>gene name</td><td>gene id</td></tr>
<tr><td>GU305914.1</td><td></td><td>63415</td><td>63437</td><td>-</td><td>0</td><td>CTATCCCCTGCTCGTTCCCA</td><td>AGG</td><td>||||||||[||||||||||||]PAM</td><td>NA</td><td>Intergenic</td><td>NA</td><td>NA</td></tr>
<tr><td>T29362</td><td>GAGATCTACCTCGACCTCGATGG</td><td></td><td>1000</td><td></td><td></td><td></td><td></td><td></td><td></td><td></td><td></td><td></td></tr>
<tr><td>Oligo fwd</td><td>CACCGAGATCTACCTCGACCTCGA</td><td></td><td></td><td></td><td></td><td></td><td></td><td></td><td></td><td></td><td></td><td></td></tr>
<tr><td>Oligo rev</td><td>AAACTCGAGGTCGAGGTAGATCTC</td><td></td><td></td><td></td><td></td><td></td><td></td><td></td><td></td><td></td><td></td><td></td></tr>
<tr><td>Chromosome</td><td>start</td><td></td><td>end</td><td>strand</td><td>MM</td><td>target_seq</td><td>PAM</td><td>alignment</td><td>distance</td><td>position</td><td>gene name</td><td>gene id</td></tr>
<tr><td>GU305914.1</td><td></td><td>63393</td><td>63415</td><td>-</td><td>0</td><td>GAGATCTACCTCGACCTCGA</td><td>TGG</td><td>||||||||[||||||||||||]PAM</td><td>NA</td><td>Intergenic</td><td>NA</td><td>NA</td></tr>
<tr><td>T29363</td><td>AGATCTACCTCGACCTCGATGGG</td><td></td><td>1000</td><td></td><td></td><td></td><td></td><td></td><td></td><td></td><td></td><td></td></tr>
<tr><td>Oligo fwd</td><td>CACCAGATCTACCTCGACCTCGAT</td><td></td><td></td><td></td><td></td><td></td><td></td><td></td><td></td><td></td><td></td><td></td></tr>
<tr><td>Oligo rev</td><td>AAACATCGAGGTCGAGGTAGATCT</td><td></td><td></td><td></td><td></td><td></td><td></td><td></td><td></td><td></td><td></td><td></td></tr>
<tr><td>Chromosome</td><td>start</td><td></td><td>end</td><td>strand</td><td>MM</td><td>target_seq</td><td>PAM</td><td>alignment</td><td>distance</td><td>position</td><td>gene name</td><td>gene id</td></tr>
<tr><td>GU305914.1</td><td></td><td>63392</td><td>63414</td><td>-</td><td>0</td><td>AGATCTACCTCGACCTCGAT</td><td>GGG</td><td>||||||||[||||||||||||]PAM</td><td>NA</td><td>Intergenic</td><td>NA</td><td>NA</td></tr>
<tr><td>T29364</td><td>GATCTACCTCGACCTCGATGGGG</td><td></td><td>1000</td><td></td><td></td><td></td><td></td><td></td><td></td><td></td><td></td><td></td></tr>
<tr><td>Oligo fwd</td><td>CACCGATCTACCTCGACCTCGATG</td><td></td><td></td><td></td><td></td><td></td><td></td><td></td><td></td><td></td><td></td><td></td></tr>
<tr><td>Oligo rev</td><td>AAACCATCGAGGTCGAGGTAGATC</td><td></td><td></td><td></td><td></td><td></td><td></td><td></td><td></td><td></td><td></td><td></td></tr>
<tr><td>Chromosome</td><td>start</td><td></td><td>end</td><td>strand</td><td>MM</td><td>target_seq</td><td>PAM</td><td>alignment</td><td>distance</td><td>position</td><td>gene name</td><td>gene id</td></tr>
<tr><td>GU305914.1</td><td></td><td>63391</td><td>63413</td><td>-</td><td>0</td><td>GATCTACCTCGACCTCGATG</td><td>GGG</td><td>||||||||[||||||||||||]PAM</td><td>NA</td><td>Intergenic</td><td>NA</td><td>NA</td></tr>
<tr><td>T29365</td><td>CGTATGACTATAATAGCGATCGG</td><td></td><td>1000</td><td></td><td></td><td></td><td></td><td></td><td></td><td></td><td></td><td></td></tr>
<tr><td>Oligo fwd</td><td>CACCCGTATGACTATAATAGCGAT</td><td></td><td></td><td></td><td></td><td></td><td></td><td></td><td></td><td></td><td></td><td></td></tr>
<tr><td>Oligo rev</td><td>AAACATCGCTATTATAGTCATACG</td><td></td><td></td><td></td><td></td><td></td><td></td><td></td><td></td><td></td><td></td><td></td></tr>
<tr><td>Chromosome</td><td>start</td><td></td><td>end</td><td>strand</td><td>MM</td><td>target_seq</td><td>PAM</td><td>alignment</td><td>distance</td><td>position</td><td>gene name</td><td>gene id</td></tr>
<tr><td>GU305914.1</td><td></td><td>63329</td><td>63351</td><td>-</td><td>0</td><td>CGTATGACTATAATAGCGAT</td><td>CGG</td><td>||||||||[||||||||||||]PAM</td><td>NA</td><td>Intergenic</td><td>NA</td><td>NA</td></tr>
<tr><td>T29366</td><td>TGACTATAATAGCGATCGGCAGG</td><td></td><td>1000</td><td></td><td></td><td></td><td></td><td></td><td></td><td></td><td></td><td></td></tr>
<tr><td>Oligo fwd</td><td>CACCTGACTATAATAGCGATCGGC</td><td></td><td></td><td></td><td></td><td></td><td></td><td></td><td></td><td></td><td></td><td></td></tr>
<tr><td>Oligo rev</td><td>AAACGCCGATCGCTATTATAGTCA</td><td></td><td></td><td></td><td></td><td></td><td></td><td></td><td></td><td></td><td></td><td></td></tr>
<tr><td>Chromosome</td><td>start</td><td></td><td>end</td><td>strand</td><td>MM</td><td>target_seq</td><td>PAM</td><td>alignment</td><td>distance</td><td>position</td><td>gene name</td><td>gene id</td></tr>
<tr><td>GU305914.1</td><td></td><td>63325</td><td>63347</td><td>-</td><td>0</td><td>TGACTATAATAGCGATCGGC</td><td>AGG</td><td>||||||||[||||||||||||]PAM</td><td>NA</td><td>Intergenic</td><td>NA</td><td>NA</td></tr>
<tr><td>T29367</td><td>GACTATAATAGCGATCGGCAGGG</td><td></td><td>1000</td><td></td><td></td><td></td><td></td><td></td><td></td><td></td><td></td><td></td></tr>
<tr><td>Oligo fwd</td><td>CACCGACTATAATAGCGATCGGCA</td><td></td><td></td><td></td><td></td><td></td><td></td><td></td><td></td><td></td><td></td><td></td></tr>
<tr><td>Oligo rev</td><td>AAACTGCCGATCGCTATTATAGTC</td><td></td><td></td><td></td><td></td><td></td><td></td><td></td><td></td><td></td><td></td><td></td></tr>
<tr><td>Chromosome</td><td>start</td><td></td><td>end</td><td>strand</td><td>MM</td><td>target_seq</td><td>PAM</td><td>alignment</td><td>distance</td><td>position</td><td>gene name</td><td>gene id</td></tr>
<tr><td>GU305914.1</td><td></td><td>63324</td><td>63346</td><td>-</td><td>0</td><td>GACTATAATAGCGATCGGCA</td><td>GGG</td><td>||||||||[||||||||||||]PAM</td><td>NA</td><td>Intergenic</td><td>NA</td><td>NA</td></tr>
<tr><td>T29368</td><td>GGGTCGACCGTCCGCCTTTGTGG</td><td></td><td>1000</td><td></td><td></td><td></td><td></td><td></td><td></td><td></td><td></td><td></td></tr>
<tr><td>Oligo fwd</td><td>CACCGGGTCGACCGTCCGCCTTTG</td><td></td><td></td><td></td><td></td><td></td><td></td><td></td><td></td><td></td><td></td><td></td></tr>
<tr><td>Oligo rev</td><td>AAACCAAAGGCGGACGGTCGACCC</td><td></td><td></td><td></td><td></td><td></td><td></td><td></td><td></td><td></td><td></td><td></td></tr>
<tr><td>Chromosome</td><td>start</td><td></td><td>end</td><td>strand</td><td>MM</td><td>target_seq</td><td>PAM</td><td>alignment</td><td>distance</td><td>position</td><td>gene name</td><td>gene id</td></tr>
<tr><td>GU305914.1</td><td></td><td>63304</td><td>63326</td><td>-</td><td>0</td><td>GGGTCGACCGTCCGCCTTTG</td><td>TGG</td><td>||||||||[||||||||||||]PAM</td><td>NA</td><td>Intergenic</td><td>NA</td><td>NA</td></tr>
<tr><td>T29369</td><td>TCGACCGTCCGCCTTTGTGGTGG</td><td></td><td>1000</td><td></td><td></td><td></td><td></td><td></td><td></td><td></td><td></td><td></td></tr>
<tr><td>Oligo fwd</td><td>CACCTCGACCGTCCGCCTTTGTGG</td><td></td><td></td><td></td><td></td><td></td><td></td><td></td><td></td><td></td><td></td><td></td></tr>
<tr><td>Oligo rev</td><td>AAACCCACAAAGGCGGACGGTCGA</td><td></td><td></td><td></td><td></td><td></td><td></td><td></td><td></td><td></td><td></td><td></td></tr>
<tr><td>Chromosome</td><td>start</td><td></td><td>end</td><td>strand</td><td>MM</td><td>target_seq</td><td>PAM</td><td>alignment</td><td>distance</td><td>position</td><td>gene name</td><td>gene id</td></tr>
<tr><td>GU305914.1</td><td></td><td>63301</td><td>63323</td><td>-</td><td>0</td><td>TCGACCGTCCGCCTTTGTGG</td><td>TGG</td><td>||||||||[||||||||||||]PAM</td><td>NA</td><td>Intergenic</td><td>NA</td><td>NA</td></tr>
<tr><td>T29370</td><td>CCGCCTTTGTGGTGGTCTCCAGG</td><td></td><td>1000</td><td></td><td></td><td></td><td></td><td></td><td></td><td></td><td></td><td></td></tr>
<tr><td>Oligo fwd</td><td>CACCCCGCCTTTGTGGTGGTCTCC</td><td></td><td></td><td></td><td></td><td></td><td></td><td></td><td></td><td></td><td></td><td></td></tr>
<tr><td>Oligo rev</td><td>AAACGGAGACCACCACAAAGGCGG</td><td></td><td></td><td></td><td></td><td></td><td></td><td></td><td></td><td></td><td></td><td></td></tr>
<tr><td>Chromosome</td><td>start</td><td></td><td>end</td><td>strand</td><td>MM</td><td>target_seq</td><td>PAM</td><td>alignment</td><td>distance</td><td>position</td><td>gene name</td><td>gene id</td></tr>
<tr><td>GU305914.1</td><td></td><td>63293</td><td>63315</td><td>-</td><td>0</td><td>CCGCCTTTGTGGTGGTCTCC</td><td>AGG</td><td>||||||||[||||||||||||]PAM</td><td>NA</td><td>Intergenic</td><td>NA</td><td>NA</td></tr>
<tr><td>T29371</td><td>TGGTCTCCAGGATCACACACAGG</td><td></td><td>1000</td><td></td><td></td><td></td><td></td><td></td><td></td><td></td><td></td><td></td></tr>
<tr><td>Oligo fwd</td><td>CACCTGGTCTCCAGGATCACACAC</td><td></td><td></td><td></td><td></td><td></td><td></td><td></td><td></td><td></td><td></td><td></td></tr>
<tr><td>Oligo rev</td><td>AAACGTGTGTGATCCTGGAGACCA</td><td></td><td></td><td></td><td></td><td></td><td></td><td></td><td></td><td></td><td></td><td></td></tr>
<tr><td>Chromosome</td><td>start</td><td></td><td>end</td><td>strand</td><td>MM</td><td>target_seq</td><td>PAM</td><td>alignment</td><td>distance</td><td>position</td><td>gene name</td><td>gene id</td></tr>
<tr><td>GU305914.1</td><td></td><td>63281</td><td>63303</td><td>-</td><td>0</td><td>TGGTCTCCAGGATCACACAC</td><td>AGG</td><td>||||||||[||||||||||||]PAM</td><td>NA</td><td>Intergenic</td><td>NA</td><td>NA</td></tr>
<tr><td>T29372</td><td>TGATCAACGTCCTGCGCTATAGG</td><td></td><td>1000</td><td></td><td></td><td></td><td></td><td></td><td></td><td></td><td></td><td></td></tr>
<tr><td>Oligo fwd</td><td>CACCTGATCAACGTCCTGCGCTAT</td><td></td><td></td><td></td><td></td><td></td><td></td><td></td><td></td><td></td><td></td><td></td></tr>
<tr><td>Oligo rev</td><td>AAACATAGCGCAGGACGTTGATCA</td><td></td><td></td><td></td><td></td><td></td><td></td><td></td><td></td><td></td><td></td><td></td></tr>
<tr><td>Chromosome</td><td>start</td><td></td><td>end</td><td>strand</td><td>MM</td><td>target_seq</td><td>PAM</td><td>alignment</td><td>distance</td><td>position</td><td>gene name</td><td>gene id</td></tr>
<tr><td>GU305914.1</td><td></td><td>63251</td><td>63273</td><td>-</td><td>0</td><td>TGATCAACGTCCTGCGCTAT</td><td>AGG</td><td>||||||||[||||||||||||]PAM</td><td>NA</td><td>Intergenic</td><td>NA</td><td>NA</td></tr>
<tr><td>T29373</td><td>ACGTCCTGCGCTATAGGTTCAGG</td><td></td><td>1000</td><td></td><td></td><td></td><td></td><td></td><td></td><td></td><td></td><td></td></tr>
<tr><td>Oligo fwd</td><td>CACCACGTCCTGCGCTATAGGTTC</td><td></td><td></td><td></td><td></td><td></td><td></td><td></td><td></td><td></td><td></td><td></td></tr>
<tr><td>Oligo rev</td><td>AAACGAACCTATAGCGCAGGACGT</td><td></td><td></td><td></td><td></td><td></td><td></td><td></td><td></td><td></td><td></td><td></td></tr>
<tr><td>Chromosome</td><td>start</td><td></td><td>end</td><td>strand</td><td>MM</td><td>target_seq</td><td>PAM</td><td>alignment</td><td>distance</td><td>position</td><td>gene name</td><td>gene id</td></tr>
<tr><td>GU305914.1</td><td></td><td>63245</td><td>63267</td><td>-</td><td>0</td><td>ACGTCCTGCGCTATAGGTTC</td><td>AGG</td><td>||||||||[||||||||||||]PAM</td><td>NA</td><td>Intergenic</td><td>NA</td><td>NA</td></tr>
<tr><td>T29374</td><td>CGTCCTGCGCTATAGGTTCAGGG</td><td></td><td>1000</td><td></td><td></td><td></td><td></td><td></td><td></td><td></td><td></td><td></td></tr>
</table>

<table>
<tr><td>Oligo fwd</td><td>CACCCGTCCTGCGCTATAGGTTCA</td><td></td><td></td><td></td><td></td><td></td><td></td><td></td><td></td><td></td><td></td></tr>
<tr><td>Oligo rev</td><td>AAACTGAACCTATAGCGCAGGACG</td><td></td><td></td><td></td><td></td><td></td><td></td><td></td><td></td><td></td><td></td></tr>
<tr><td>Chromosome</td><td>start</td><td>end</td><td>strand</td><td>MM</td><td>target_seq</td><td>PAM</td><td>alignment</td><td>distance</td><td>position</td><td>gene name</td><td>gene id</td></tr>
<tr><td>GU305914.1</td><td>63244</td><td>63266</td><td>-</td><td>0</td><td>CGTCCTGCGCTATAGGTTCA</td><td>GGG</td><td>||||||||[||||||||||||]PAM</td><td>NA</td><td>Intergenic</td><td>NA</td><td>NA</td></tr>
<tr><td></td><td></td><td></td><td></td><td></td><td></td><td></td><td></td><td></td><td></td><td></td><td></td></tr>
<tr><td>T29375</td><td>CTTCTTAAATAACAGTATTTCGG</td><td>1000</td><td></td><td></td><td></td><td></td><td></td><td></td><td></td><td></td><td></td></tr>
<tr><td>Oligo fwd</td><td>CACCCTTCTTAAATAACAGTATTT</td><td></td><td></td><td></td><td></td><td></td><td></td><td></td><td></td><td></td><td></td></tr>
<tr><td>Oligo rev</td><td>AAACAAATACTGTTATTTAAGAAG</td><td></td><td></td><td></td><td></td><td></td><td></td><td></td><td></td><td></td><td></td></tr>
<tr><td>Chromosome</td><td>start</td><td>end</td><td>strand</td><td>MM</td><td>target_seq</td><td>PAM</td><td>alignment</td><td>distance</td><td>position</td><td>gene name</td><td>gene id</td></tr>
<tr><td>GU305914.1</td><td>63208</td><td>63230</td><td>-</td><td>0</td><td>CTTCTTAAATAACAGTATTT</td><td>CGG</td><td>||||||||[||||||||||||]PAM</td><td>NA</td><td>Intergenic</td><td>NA</td><td>NA</td></tr>
<tr><td></td><td></td><td></td><td></td><td></td><td></td><td></td><td></td><td></td><td></td><td></td><td></td></tr>
<tr><td>T29376</td><td>TTCTTAAATAACAGTATTTCGGG</td><td>1000</td><td></td><td></td><td></td><td></td><td></td><td></td><td></td><td></td><td></td></tr>
<tr><td>Oligo fwd</td><td>CACCTTCTTAAATAACAGTATTTC</td><td></td><td></td><td></td><td></td><td></td><td></td><td></td><td></td><td></td><td></td></tr>
<tr><td>Oligo rev</td><td>AAACGAAATACTGTTATTTAAGAA</td><td></td><td></td><td></td><td></td><td></td><td></td><td></td><td></td><td></td><td></td></tr>
<tr><td>Chromosome</td><td>start</td><td>end</td><td>strand</td><td>MM</td><td>target_seq</td><td>PAM</td><td>alignment</td><td>distance</td><td>position</td><td>gene name</td><td>gene id</td></tr>
<tr><td>GU305914.1</td><td>63207</td><td>63229</td><td>-</td><td>0</td><td>TTCTTAAATAACAGTATTTC</td><td>GGG</td><td>||||||||[||||||||||||]PAM</td><td>NA</td><td>Intergenic</td><td>NA</td><td>NA</td></tr>
<tr><td></td><td></td><td></td><td></td><td></td><td></td><td></td><td></td><td></td><td></td><td></td><td></td></tr>
<tr><td>T29377</td><td>AATAACAGTATTTCGGGCTACGG</td><td>1000</td><td></td><td></td><td></td><td></td><td></td><td></td><td></td><td></td><td></td></tr>
<tr><td>Oligo fwd</td><td>CACCAATAACAGTATTTCGGGCTA</td><td></td><td></td><td></td><td></td><td></td><td></td><td></td><td></td><td></td><td></td></tr>
<tr><td>Oligo rev</td><td>AAACTAGCCCGAAATACTGTTATT</td><td></td><td></td><td></td><td></td><td></td><td></td><td></td><td></td><td></td><td></td></tr>
<tr><td>Chromosome</td><td>start</td><td>end</td><td>strand</td><td>MM</td><td>target_seq</td><td>PAM</td><td>alignment</td><td>distance</td><td>position</td><td>gene name</td><td>gene id</td></tr>
<tr><td>GU305914.1</td><td>63201</td><td>63223</td><td>-</td><td>0</td><td>AATAACAGTATTTCGGGCTA</td><td>CGG</td><td>||||||||[||||||||||||]PAM</td><td>NA</td><td>Intergenic</td><td>NA</td><td>NA</td></tr>
<tr><td></td><td></td><td></td><td></td><td></td><td></td><td></td><td></td><td></td><td></td><td></td><td></td></tr>
<tr><td>T29378</td><td>ATAACAGTATTTCGGGCTACGGG</td><td>1000</td><td></td><td></td><td></td><td></td><td></td><td></td><td></td><td></td><td></td></tr>
<tr><td>Oligo fwd</td><td>CACCATAACAGTATTTCGGGCTAC</td><td></td><td></td><td></td><td></td><td></td><td></td><td></td><td></td><td></td><td></td></tr>
<tr><td>Oligo rev</td><td>AAACGTAGCCCGAAATACTGTTAT</td><td></td><td></td><td></td><td></td><td></td><td></td><td></td><td></td><td></td><td></td></tr>
<tr><td>Chromosome</td><td>start</td><td>end</td><td>strand</td><td>MM</td><td>target_seq</td><td>PAM</td><td>alignment</td><td>distance</td><td>position</td><td>gene name</td><td>gene id</td></tr>
<tr><td>GU305914.1</td><td>63200</td><td>63222</td><td>-</td><td>0</td><td>ATAACAGTATTTCGGGCTAC</td><td>GGG</td><td>||||||||[||||||||||||]PAM</td><td>NA</td><td>Intergenic</td><td>NA</td><td>NA</td></tr>
<tr><td></td><td></td><td></td><td></td><td></td><td></td><td></td><td></td><td></td><td></td><td></td><td></td></tr>
<tr><td>T29379</td><td>TCGGGCTACGGGCCGTCCACCGG</td><td>1000</td><td></td><td></td><td></td><td></td><td></td><td></td><td></td><td></td><td></td></tr>
<tr><td>Oligo fwd</td><td>CACCTCGGGCTACGGGCCGTCCAC</td><td></td><td></td><td></td><td></td><td></td><td></td><td></td><td></td><td></td><td></td></tr>
<tr><td>Oligo rev</td><td>AAACGTGGACGGCCCGTAGCCCGA</td><td></td><td></td><td></td><td></td><td></td><td></td><td></td><td></td><td></td><td></td></tr>
<tr><td>Chromosome</td><td>start</td><td>end</td><td>strand</td><td>MM</td><td>target_seq</td><td>PAM</td><td>alignment</td><td>distance</td><td>position</td><td>gene name</td><td>gene id</td></tr>
<tr><td>GU305914.1</td><td>63189</td><td>63211</td><td>-</td><td>0</td><td>TCGGGCTACGGGCCGTCCAC</td><td>CGG</td><td>||||||||[||||||||||||]PAM</td><td>NA</td><td>Intergenic</td><td>NA</td><td>NA</td></tr>
<tr><td></td><td></td><td></td><td></td><td></td><td></td><td></td><td></td><td></td><td></td><td></td><td></td></tr>
<tr><td>T29380</td><td>GGGCCGTCCACCGGCTGTCTCGG</td><td>1000</td><td></td><td></td><td></td><td></td><td></td><td></td><td></td><td></td><td></td></tr>
<tr><td>Oligo fwd</td><td>CACCGGGCCGTCCACCGGCTGTCT</td><td></td><td></td><td></td><td></td><td></td><td></td><td></td><td></td><td></td><td></td></tr>
<tr><td>Oligo rev</td><td>AAACAGACAGCCGGTGGACGGCCC</td><td></td><td></td><td></td><td></td><td></td><td></td><td></td><td></td><td></td><td></td></tr>
<tr><td>Chromosome</td><td>start</td><td>end</td><td>strand</td><td>MM</td><td>target_seq</td><td>PAM</td><td>alignment</td><td>distance</td><td>position</td><td>gene name</td><td>gene id</td></tr>
<tr><td>GU305914.1</td><td>63180</td><td>63202</td><td>-</td><td>0</td><td>GGGCCGTCCACCGGCTGTCT</td><td>CGG</td><td>||||||||[||||||||||||]PAM</td><td>NA</td><td>Intergenic</td><td>NA</td><td>NA</td></tr>
<tr><td></td><td></td><td></td><td></td><td></td><td></td><td></td><td></td><td></td><td></td><td></td><td></td></tr>
<tr><td>T29381</td><td>TCTCGGCACGCTACAGCGACTGG</td><td>1000</td><td></td><td></td><td></td><td></td><td></td><td></td><td></td><td></td><td></td></tr>
<tr><td>Oligo fwd</td><td>CACCTCTCGGCACGCTACAGCGAC</td><td></td><td></td><td></td><td></td><td></td><td></td><td></td><td></td><td></td><td></td></tr>
<tr><td>Oligo rev</td><td>AAACGTCGCTGTAGCGTGCCGAGA</td><td></td><td></td><td></td><td></td><td></td><td></td><td></td><td></td><td></td><td></td></tr>
<tr><td>Chromosome</td><td>start</td><td>end</td><td>strand</td><td>MM</td><td>target_seq</td><td>PAM</td><td>alignment</td><td>distance</td><td>position</td><td>gene name</td><td>gene id</td></tr>
<tr><td>GU305914.1</td><td>63163</td><td>63185</td><td>-</td><td>0</td><td>TCTCGGCACGCTACAGCGAC</td><td>TGG</td><td>||||||||[||||||||||||]PAM</td><td>NA</td><td>Intergenic</td><td>NA</td><td>NA</td></tr>
<tr><td></td><td></td><td></td><td></td><td></td><td></td><td></td><td></td><td></td><td></td><td></td><td></td></tr>
<tr><td>T29382</td><td>CTCGGCACGCTACAGCGACTGGG</td><td>1000</td><td></td><td></td><td></td><td></td><td></td><td></td><td></td><td></td><td></td></tr>
<tr><td>Oligo fwd</td><td>CACCCTCGGCACGCTACAGCGACT</td><td></td><td></td><td></td><td></td><td></td><td></td><td></td><td></td><td></td><td></td></tr>
<tr><td>Oligo rev</td><td>AAACAGTCGCTGTAGCGTGCCGAG</td><td></td><td></td><td></td><td></td><td></td><td></td><td></td><td></td><td></td><td></td></tr>
<tr><td>Chromosome</td><td>start</td><td>end</td><td>strand</td><td>MM</td><td>target_seq</td><td>PAM</td><td>alignment</td><td>distance</td><td>position</td><td>gene name</td><td>gene id</td></tr>
<tr><td>GU305914.1</td><td>63162</td><td>63184</td><td>-</td><td>0</td><td>CTCGGCACGCTACAGCGACT</td><td>GGG</td><td>||||||||[||||||||||||]PAM</td><td>NA</td><td>Intergenic</td><td>NA</td><td>NA</td></tr>
<tr><td></td><td></td><td></td><td></td><td></td><td></td><td></td><td></td><td></td><td></td><td></td><td></td></tr>
<tr><td>T29383</td><td>GCACGCTACAGCGACTGGGCTGG</td><td>1000</td><td></td><td></td><td></td><td></td><td></td><td></td><td></td><td></td><td></td></tr>
<tr><td>Oligo fwd</td><td>CACCGCACGCTACAGCGACTGGGC</td><td></td><td></td><td></td><td></td><td></td><td></td><td></td><td></td><td></td><td></td></tr>
<tr><td>Oligo rev</td><td>AAACGCCCAGTCGCTGTAGCGTGC</td><td></td><td></td><td></td><td></td><td></td><td></td><td></td><td></td><td></td><td></td></tr>
<tr><td>Chromosome</td><td>start</td><td>end</td><td>strand</td><td>MM</td><td>target_seq</td><td>PAM</td><td>alignment</td><td>distance</td><td>position</td><td>gene name</td><td>gene id</td></tr>
<tr><td>GU305914.1</td><td>63158</td><td>63180</td><td>-</td><td>0</td><td>GCACGCTACAGCGACTGGGC</td><td>TGG</td><td>||||||||[||||||||||||]PAM</td><td>NA</td><td>Intergenic</td><td>NA</td><td>NA</td></tr>
<tr><td></td><td></td><td></td><td></td><td></td><td></td><td></td><td></td><td></td><td></td><td></td><td></td></tr>
<tr><td>T29384</td><td>GGTTCTGCTCGCGCGACAGCAGG</td><td>1000</td><td></td><td></td><td></td><td></td><td></td><td></td><td></td><td></td><td></td></tr>
<tr><td>Oligo fwd</td><td>CACCGGTTCTGCTCGCGCGACAGC</td><td></td><td></td><td></td><td></td><td></td><td></td><td></td><td></td><td></td><td></td></tr>
<tr><td>Oligo rev</td><td>AAACGCTGTCGCGCGAGCAGAACC</td><td></td><td></td><td></td><td></td><td></td><td></td><td></td><td></td><td></td><td></td></tr>
<tr><td>Chromosome</td><td>start</td><td>end</td><td>strand</td><td>MM</td><td>target_seq</td><td>PAM</td><td>alignment</td><td>distance</td><td>position</td><td>gene name</td><td>gene id</td></tr>
<tr><td>GU305914.1</td><td>63137</td><td>63159</td><td>-</td><td>0</td><td>GGTTCTGCTCGCGCGACAGC</td><td>AGG</td><td>||||||||[||||||||||||]PAM</td><td>NA</td><td>Intergenic</td><td>NA</td><td>NA</td></tr>
<tr><td></td><td></td><td></td><td></td><td></td><td></td><td></td><td></td><td></td><td></td><td></td><td></td></tr>
<tr><td>T29385</td><td>TGCTCGCGCGACAGCAGGTCCGG</td><td>1000</td><td></td><td></td><td></td><td></td><td></td><td></td><td></td><td></td><td></td></tr>
<tr><td>Oligo fwd</td><td>CACCTGCTCGCGCGACAGCAGGTC</td><td></td><td></td><td></td><td></td><td></td><td></td><td></td><td></td><td></td><td></td></tr>
<tr><td>Oligo rev</td><td>AAACGACCTGCTGTCGCGCGAGCA</td><td></td><td></td><td></td><td></td><td></td><td></td><td></td><td></td><td></td><td></td></tr>
<tr><td>Chromosome</td><td>start</td><td>end</td><td>strand</td><td>MM</td><td>target_seq</td><td>PAM</td><td>alignment</td><td>distance</td><td>position</td><td>gene name</td><td>gene id</td></tr>
<tr><td>GU305914.1</td><td>63132</td><td>63154</td><td>-</td><td>0</td><td>TGCTCGCGCGACAGCAGGTC</td><td>CGG</td><td>||||||||[||||||||||||]PAM</td><td>NA</td><td>Intergenic</td><td>NA</td><td>NA</td></tr>
<tr><td></td><td></td><td></td><td></td><td></td><td></td><td></td><td></td><td></td><td></td><td></td><td></td></tr>
<tr><td>T29386</td><td>GCTCGCGCGACAGCAGGTCCGGG</td><td>1000</td><td></td><td></td><td></td><td></td><td></td><td></td><td></td><td></td><td></td></tr>
<tr><td>Oligo fwd</td><td>CACCGCTCGCGCGACAGCAGGTCC</td><td></td><td></td><td></td><td></td><td></td><td></td><td></td><td></td><td></td><td></td></tr>
<tr><td>Oligo rev</td><td>AAACGGACCTGCTGTCGCGCGAGC</td><td></td><td></td><td></td><td></td><td></td><td></td><td></td><td></td><td></td><td></td></tr>
<tr><td>Chromosome</td><td>start</td><td>end</td><td>strand</td><td>MM</td><td>target_seq</td><td>PAM</td><td>alignment</td><td>distance</td><td>position</td><td>gene name</td><td>gene id</td></tr>
<tr><td>GU305914.1</td><td>63131</td><td>63153</td><td>-</td><td>0</td><td>GCTCGCGCGACAGCAGGTCC</td><td>GGG</td><td>||||||||[||||||||||||]PAM</td><td>NA</td><td>Intergenic</td><td>NA</td><td>NA</td></tr>
<tr><td></td><td></td><td></td><td></td><td></td><td></td><td></td><td></td><td></td><td></td><td></td><td></td></tr>
<tr><td>T29387</td><td>GGGATCGTCGCCTCGCGCGCCGG</td><td>1000</td><td></td><td></td><td></td><td></td><td></td><td></td><td></td><td></td><td></td></tr>
<tr><td>Oligo fwd</td><td>CACCGGGATCGTCGCCTCGCGCGC</td><td></td><td></td><td></td><td></td><td></td><td></td><td></td><td></td><td></td><td></td></tr>
<tr><td>Oligo rev</td><td>AAACGCGCGCGAGGCGACGATCCC</td><td></td><td></td><td></td><td></td><td></td><td></td><td></td><td></td><td></td><td></td></tr>
<tr><td>Chromosome</td><td>start</td><td>end</td><td>strand</td><td>MM</td><td>target_seq</td><td>PAM</td><td>alignment</td><td>distance</td><td>position</td><td>gene name</td><td>gene id</td></tr>
<tr><td>GU305914.1</td><td>63111</td><td>63133</td><td>-</td><td>0</td><td>GGGATCGTCGCCTCGCGCGC</td><td>CGG</td><td>||||||||[||||||||||||]PAM</td><td>NA</td><td>Intergenic</td><td>NA</td><td>NA</td></tr>
<tr><td></td><td></td><td></td><td></td><td></td><td></td><td></td><td></td><td></td><td></td><td></td><td></td></tr>
<tr><td>T29388</td><td>GCGCGCCGGACAGCTCTCCGTGG</td><td>1000</td><td></td><td></td><td></td><td></td><td></td><td></td><td></td><td></td><td></td></tr>
<tr><td>Oligo fwd</td><td>CACCGCGCGCCGGACAGCTCTCCG</td><td></td><td></td><td></td><td></td><td></td><td></td><td></td><td></td><td></td><td></td></tr>
<tr><td>Oligo rev</td><td>AAACCGGAGAGCTGTCCGGCGCGC</td><td></td><td></td><td></td><td></td><td></td><td></td><td></td><td></td><td></td><td></td></tr>
<tr><td>Chromosome</td><td>start</td><td>end</td><td>strand</td><td>MM</td><td>target_seq</td><td>PAM</td><td>alignment</td><td>distance</td><td>position</td><td>gene name</td><td>gene id</td></tr>
<tr><td>GU305914.1</td><td>63097</td><td>63119</td><td>-</td><td>0</td><td>GCGCGCCGGACAGCTCTCCG</td><td>TGG</td><td>||||||||[||||||||||||]PAM</td><td>NA</td><td>Intergenic</td><td>NA</td><td>NA</td></tr>
</table>

| | | | | | | | | | | | | |
|---|---|---|---|---|---|---|---|---|---|---|---|---|
| T29389 | GAGAAGTTCTACGTCGACGTCGG | | 1000 | | | | | | | | | |
| Oligo fwd | CACCGAGAAGTTCTACGTCGACGT | | | | | | | | | | | |
| Oligo rev | AAACACGTCGACGTAGAACTTCTC | | | | | | | | | | | |
| Chromosome | start | | end | strand | MM | target_seq | PAM | alignment | distance | position | gene name | gene id |
| GU305914.1 | | 63066 | 63088 | - | 0 | GAGAAGTTCTACGTCGACGT | CGG | \|\|\|\|\|\|\|\|[\|\|\|\|\|\|\|\|\|\|\|\|]PAM | NA | Intergenic | NA | NA |
| T29390 | AGAAGTTCTACGTCGACGTCGGG | | 1000 | | | | | | | | | |
| Oligo fwd | CACCAGAAGTTCTACGTCGACGTC | | | | | | | | | | | |
| Oligo rev | AAACGACGTCGACGTAGAACTTCT | | | | | | | | | | | |
| Chromosome | start | | end | strand | MM | target_seq | PAM | alignment | distance | position | gene name | gene id |
| GU305914.1 | | 63065 | 63087 | - | 0 | AGAAGTTCTACGTCGACGTC | GGG | \|\|\|\|\|\|\|\|[\|\|\|\|\|\|\|\|\|\|\|\|]PAM | NA | Intergenic | NA | NA |
| T29391 | CTCGTCGAGTTCGCGTGACATGG | | 1000 | | | | | | | | | |
| Oligo fwd | CACCCTCGTCGAGTTCGCGTGACA | | | | | | | | | | | |
| Oligo rev | AAACTGTCACGCGAACTCGACGAG | | | | | | | | | | | |
| Chromosome | start | | end | strand | MM | target_seq | PAM | alignment | distance | position | gene name | gene id |
| GU305914.1 | | 63039 | 63061 | - | 0 | CTCGTCGAGTTCGCGTGACA | TGG | \|\|\|\|\|\|\|\|[\|\|\|\|\|\|\|\|\|\|\|\|]PAM | NA | Intergenic | NA | NA |
| T29392 | GCCCAAAGTCTACTCCGACCCGG | | 1000 | | | | | | | | | |
| Oligo fwd | CACCGCCCAAAGTCTACTCCGACC | | | | | | | | | | | |
| Oligo rev | AAACGGTCGGAGTAGACTTTGGGC | | | | | | | | | | | |
| Chromosome | start | | end | strand | MM | target_seq | PAM | alignment | distance | position | gene name | gene id |
| GU305914.1 | | 63009 | 63031 | - | 0 | GCCCAAAGTCTACTCCGACC | CGG | \|\|\|\|\|\|\|\|[\|\|\|\|\|\|\|\|\|\|\|\|]PAM | NA | Intergenic | NA | NA |
| T29393 | GTCTACTCCGACCCGGACAACGG | | 1000 | | | | | | | | | |
| Oligo fwd | CACCGTCTACTCCGACCCGGACAA | | | | | | | | | | | |
| Oligo rev | AAACTTGTCCGGGTCGGAGTAGAC | | | | | | | | | | | |
| Chromosome | start | | end | strand | MM | target_seq | PAM | alignment | distance | position | gene name | gene id |
| GU305914.1 | | 63002 | 63024 | - | 0 | GTCTACTCCGACCCGGACAA | CGG | \|\|\|\|\|\|\|\|[\|\|\|\|\|\|\|\|\|\|\|\|]PAM | NA | Intergenic | NA | NA |
| T29394 | GGACAACGGATTCTTCTTTTTGG | | 1000 | | | | | | | | | |
| Oligo fwd | CACCGGACAACGGATTCTTCTTTT | | | | | | | | | | | |
| Oligo rev | AAACAAAAGAAGAATCCGTTGTCC | | | | | | | | | | | |
| Chromosome | start | | end | strand | MM | target_seq | PAM | alignment | distance | position | gene name | gene id |
| GU305914.1 | | 62988 | 63010 | - | 0 | GGACAACGGATTCTTCTTTT | TGG | \|\|\|\|\|\|\|\|[\|\|\|\|\|\|\|\|\|\|\|\|]PAM | NA | Intergenic | NA | NA |
| T29395 | GATGTCCCCATGCCTGACGACGG | | 1000 | | | | | | | | | |
| Oligo fwd | CACCGATGTCCCCATGCCTGACGA | | | | | | | | | | | |
| Oligo rev | AAACTCGTCAGGCATGGGGACATC | | | | | | | | | | | |
| Chromosome | start | | end | strand | MM | target_seq | PAM | alignment | distance | position | gene name | gene id |
| GU305914.1 | | 62966 | 62988 | - | 0 | GATGTCCCCATGCCTGACGA | CGG | \|\|\|\|\|\|\|\|[\|\|\|\|\|\|\|\|\|\|\|\|]PAM | NA | Intergenic | NA | NA |
| T29396 | ATGTCCCCATGCCTGACGACGGG | | 1000 | | | | | | | | | |
| Oligo fwd | CACCATGTCCCCATGCCTGACGAC | | | | | | | | | | | |
| Oligo rev | AAACGTCGTCAGGCATGGGGACAT | | | | | | | | | | | |
| Chromosome | start | | end | strand | MM | target_seq | PAM | alignment | distance | position | gene name | gene id |
| GU305914.1 | | 62965 | 62987 | - | 0 | ATGTCCCCATGCCTGACGAC | GGG | \|\|\|\|\|\|\|\|[\|\|\|\|\|\|\|\|\|\|\|\|]PAM | NA | Intergenic | NA | NA |
| T29397 | CCCCATGCCTGACGACGGGCAGG | | 1000 | | | | | | | | | |
| Oligo fwd | CACCCCCCATGCCTGACGACGGGC | | | | | | | | | | | |
| Oligo rev | AAACGCCCGTCGTCAGGCATGGGG | | | | | | | | | | | |
| Chromosome | start | | end | strand | MM | target_seq | PAM | alignment | distance | position | gene name | gene id |
| GU305914.1 | | 62961 | 62983 | - | 0 | CCCCATGCCTGACGACGGGC | AGG | \|\|\|\|\|\|\|\|[\|\|\|\|\|\|\|\|\|\|\|\|]PAM | NA | Intergenic | NA | NA |
| T29398 | CCCATGCCTGACGACGGGCAGGG | | 1000 | | | | | | | | | |
| Oligo fwd | CACCCCCATGCCTGACGACGGGCA | | | | | | | | | | | |
| Oligo rev | AAACTGCCCGTCGTCAGGCATGGG | | | | | | | | | | | |
| Chromosome | start | | end | strand | MM | target_seq | PAM | alignment | distance | position | gene name | gene id |
| GU305914.1 | | 62960 | 62982 | - | 0 | CCCATGCCTGACGACGGGCA | GGG | \|\|\|\|\|\|\|\|[\|\|\|\|\|\|\|\|\|\|\|\|]PAM | NA | Intergenic | NA | NA |
| T29399 | ATGCCTGACGACGGGCAGGGAGG | | 1000 | | | | | | | | | |
| Oligo fwd | CACCATGCCTGACGACGGGCAGGG | | | | | | | | | | | |
| Oligo rev | AAACCCCTGCCCGTCGTCAGGCAT | | | | | | | | | | | |
| Chromosome | start | | end | strand | MM | target_seq | PAM | alignment | distance | position | gene name | gene id |
| GU305914.1 | | 62957 | 62979 | - | 0 | ATGCCTGACGACGGGCAGGG | AGG | \|\|\|\|\|\|\|\|[\|\|\|\|\|\|\|\|\|\|\|\|]PAM | NA | Intergenic | NA | NA |
| T29400 | CAGACCGCCACCACCGCTGCCGG | | 1000 | | | | | | | | | |
| Oligo fwd | CACCCAGACCGCCACCACCGCTGC | | | | | | | | | | | |
| Oligo rev | AAACGCAGCGGTGGTGGCGGTCTG | | | | | | | | | | | |
| Chromosome | start | | end | strand | MM | target_seq | PAM | alignment | distance | position | gene name | gene id |
| GU305914.1 | | 62930 | 62952 | - | 0 | CAGACCGCCACCACCGCTGC | CGG | \|\|\|\|\|\|\|\|[\|\|\|\|\|\|\|\|\|\|\|\|]PAM | NA | Intergenic | NA | NA |
| T29401 | AGACCGCCACCACCGCTGCCGGG | | 1000 | | | | | | | | | |
| Oligo fwd | CACCAGACCGCCACCACCGCTGCC | | | | | | | | | | | |
| Oligo rev | AAACGGCAGCGGTGGTGGCGGTCT | | | | | | | | | | | |
| Chromosome | start | | end | strand | MM | target_seq | PAM | alignment | distance | position | gene name | gene id |
| GU305914.1 | | 62929 | 62951 | - | 0 | AGACCGCCACCACCGCTGCC | GGG | \|\|\|\|\|\|\|\|[\|\|\|\|\|\|\|\|\|\|\|\|]PAM | NA | Intergenic | NA | NA |
| T29402 | GACCGCCACCACCGCTGCCGGGG | | 1000 | | | | | | | | | |
| Oligo fwd | CACCGACCGCCACCACCGCTGCCG | | | | | | | | | | | |
| Oligo rev | AAACCGGCAGCGGTGGTGGCGGTC | | | | | | | | | | | |
| Chromosome | start | | end | strand | MM | target_seq | PAM | alignment | distance | position | gene name | gene id |
| GU305914.1 | | 62928 | 62950 | - | 0 | GACCGCCACCACCGCTGCCG | GGG | \|\|\|\|\|\|\|\|[\|\|\|\|\|\|\|\|\|\|\|\|]PAM | NA | Intergenic | NA | NA |
| T29403 | ACCGCCACCACCGCTGCCGGGGG | | 1000 | | | | | | | | | |
| Oligo fwd | CACCACCGCCACCACCGCTGCCGG | | | | | | | | | | | |
| Oligo rev | AAACCCGGCAGCGGTGGTGGCGGT | | | | | | | | | | | |
| Chromosome | start | | end | strand | MM | target_seq | PAM | alignment | distance | position | gene name | gene id |
| GU305914.1 | | 62927 | 62949 | - | 0 | ACCGCCACCACCGCTGCCGG | GGG | \|\|\|\|\|\|\|\|[\|\|\|\|\|\|\|\|\|\|\|\|]PAM | NA | Intergenic | NA | NA |

<table>
<tr><td>T29404</td><td>ACCGCTGCCGGGGGAGCCTTCGG</td><td>1000</td><td></td><td></td><td></td><td></td><td></td><td></td><td></td><td></td><td></td></tr>
<tr><td>Oligo fwd</td><td>CACCACCGCTGCCGGGGGAGCCTT</td><td></td><td></td><td></td><td></td><td></td><td></td><td></td><td></td><td></td><td></td></tr>
<tr><td>Oligo rev</td><td>AAACAAGGCTCCCCCGGCAGCGGT</td><td></td><td></td><td></td><td></td><td></td><td></td><td></td><td></td><td></td><td></td></tr>
<tr><td>Chromosome</td><td>start</td><td>end</td><td>strand</td><td>MM</td><td>target_seq</td><td>PAM</td><td>alignment</td><td>distance</td><td>position</td><td>gene name</td><td>gene id</td></tr>
<tr><td>GU305914.1</td><td>62918</td><td>62940</td><td>-</td><td>0</td><td>ACCGCTGCCGGGGGAGCCTT</td><td>CGG</td><td>||||||||[||||||||||||]PAM</td><td>NA</td><td>Intergenic</td><td>NA</td><td>NA</td></tr>
<tr><td>T29405</td><td>CCGCTGCCGGGGGAGCCTTCGGG</td><td>1000</td><td></td><td></td><td></td><td></td><td></td><td></td><td></td><td></td><td></td></tr>
<tr><td>Oligo fwd</td><td>CACCCCGCTGCCGGGGGAGCCTTC</td><td></td><td></td><td></td><td></td><td></td><td></td><td></td><td></td><td></td><td></td></tr>
<tr><td>Oligo rev</td><td>AAACGAAGGCTCCCCCGGCAGCGG</td><td></td><td></td><td></td><td></td><td></td><td></td><td></td><td></td><td></td><td></td></tr>
<tr><td>Chromosome</td><td>start</td><td>end</td><td>strand</td><td>MM</td><td>target_seq</td><td>PAM</td><td>alignment</td><td>distance</td><td>position</td><td>gene name</td><td>gene id</td></tr>
<tr><td>GU305914.1</td><td>62917</td><td>62939</td><td>-</td><td>0</td><td>CCGCTGCCGGGGGAGCCTTC</td><td>GGG</td><td>||||||||[||||||||||||]PAM</td><td>NA</td><td>Intergenic</td><td>NA</td><td>NA</td></tr>
<tr><td>T29406</td><td>CGCTGCCGGGGGAGCCTTCGGGG</td><td>1000</td><td></td><td></td><td></td><td></td><td></td><td></td><td></td><td></td><td></td></tr>
<tr><td>Oligo fwd</td><td>CACCCGCTGCCGGGGGAGCCTTCG</td><td></td><td></td><td></td><td></td><td></td><td></td><td></td><td></td><td></td><td></td></tr>
<tr><td>Oligo rev</td><td>AAACCGAAGGCTCCCCCGGCAGCG</td><td></td><td></td><td></td><td></td><td></td><td></td><td></td><td></td><td></td><td></td></tr>
<tr><td>Chromosome</td><td>start</td><td>end</td><td>strand</td><td>MM</td><td>target_seq</td><td>PAM</td><td>alignment</td><td>distance</td><td>position</td><td>gene name</td><td>gene id</td></tr>
<tr><td>GU305914.1</td><td>62916</td><td>62938</td><td>-</td><td>0</td><td>CGCTGCCGGGGGAGCCTTCG</td><td>GGG</td><td>||||||||[||||||||||||]PAM</td><td>NA</td><td>Intergenic</td><td>NA</td><td>NA</td></tr>
<tr><td>T29407</td><td>TGCCGGGGGAGCCTTCGGGGTGG</td><td>1000</td><td></td><td></td><td></td><td></td><td></td><td></td><td></td><td></td><td></td></tr>
<tr><td>Oligo fwd</td><td>CACCTGCCGGGGGAGCCTTCGGGG</td><td></td><td></td><td></td><td></td><td></td><td></td><td></td><td></td><td></td><td></td></tr>
<tr><td>Oligo rev</td><td>AAACCCCCGAAGGCTCCCCCGGCA</td><td></td><td></td><td></td><td></td><td></td><td></td><td></td><td></td><td></td><td></td></tr>
<tr><td>Chromosome</td><td>start</td><td>end</td><td>strand</td><td>MM</td><td>target_seq</td><td>PAM</td><td>alignment</td><td>distance</td><td>position</td><td>gene name</td><td>gene id</td></tr>
<tr><td>GU305914.1</td><td>62913</td><td>62935</td><td>-</td><td>0</td><td>TGCCGGGGGAGCCTTCGGGG</td><td>TGG</td><td>||||||||[||||||||||||]PAM</td><td>NA</td><td>Intergenic</td><td>NA</td><td>NA</td></tr>
<tr><td>T29408</td><td>GCCGGGGGAGCCTTCGGGGTGGG</td><td>1000</td><td></td><td></td><td></td><td></td><td></td><td></td><td></td><td></td><td></td></tr>
<tr><td>Oligo fwd</td><td>CACCGCCGGGGGAGCCTTCGGGGT</td><td></td><td></td><td></td><td></td><td></td><td></td><td></td><td></td><td></td><td></td></tr>
<tr><td>Oligo rev</td><td>AAACACCCCGAAGGCTCCCCCGGC</td><td></td><td></td><td></td><td></td><td></td><td></td><td></td><td></td><td></td><td></td></tr>
<tr><td>Chromosome</td><td>start</td><td>end</td><td>strand</td><td>MM</td><td>target_seq</td><td>PAM</td><td>alignment</td><td>distance</td><td>position</td><td>gene name</td><td>gene id</td></tr>
<tr><td>GU305914.1</td><td>62912</td><td>62934</td><td>-</td><td>0</td><td>GCCGGGGGAGCCTTCGGGGT</td><td>GGG</td><td>||||||||[||||||||||||]PAM</td><td>NA</td><td>Intergenic</td><td>NA</td><td>NA</td></tr>
<tr><td>T29409</td><td>GGGGGAGCCTTCGGGGTGGGTGG</td><td>1000</td><td></td><td></td><td></td><td></td><td></td><td></td><td></td><td></td><td></td></tr>
<tr><td>Oligo fwd</td><td>CACCGGGGGAGCCTTCGGGGTGGG</td><td></td><td></td><td></td><td></td><td></td><td></td><td></td><td></td><td></td><td></td></tr>
<tr><td>Oligo rev</td><td>AAACCCCACCCCGAAGGCTCCCCC</td><td></td><td></td><td></td><td></td><td></td><td></td><td></td><td></td><td></td><td></td></tr>
<tr><td>Chromosome</td><td>start</td><td>end</td><td>strand</td><td>MM</td><td>target_seq</td><td>PAM</td><td>alignment</td><td>distance</td><td>position</td><td>gene name</td><td>gene id</td></tr>
<tr><td>GU305914.1</td><td>62909</td><td>62931</td><td>-</td><td>0</td><td>GGGGGAGCCTTCGGGGTGGG</td><td>TGG</td><td>||||||||[||||||||||||]PAM</td><td>NA</td><td>Intergenic</td><td>NA</td><td>NA</td></tr>
<tr><td>T29410</td><td>GGGGAGCCTTCGGGGTGGGTGGG</td><td>1000</td><td></td><td></td><td></td><td></td><td></td><td></td><td></td><td></td><td></td></tr>
<tr><td>Oligo fwd</td><td>CACCGGGGAGCCTTCGGGGTGGGT</td><td></td><td></td><td></td><td></td><td></td><td></td><td></td><td></td><td></td><td></td></tr>
<tr><td>Oligo rev</td><td>AAACACCCACCCCGAAGGCTCCCC</td><td></td><td></td><td></td><td></td><td></td><td></td><td></td><td></td><td></td><td></td></tr>
<tr><td>Chromosome</td><td>start</td><td>end</td><td>strand</td><td>MM</td><td>target_seq</td><td>PAM</td><td>alignment</td><td>distance</td><td>position</td><td>gene name</td><td>gene id</td></tr>
<tr><td>GU305914.1</td><td>62908</td><td>62930</td><td>-</td><td>0</td><td>GGGGAGCCTTCGGGGTGGGT</td><td>GGG</td><td>||||||||[||||||||||||]PAM</td><td>NA</td><td>Intergenic</td><td>NA</td><td>NA</td></tr>
<tr><td>T29411</td><td>GGGAGCCTTCGGGGTGGGTGGGG</td><td>1000</td><td></td><td></td><td></td><td></td><td></td><td></td><td></td><td></td><td></td></tr>
<tr><td>Oligo fwd</td><td>CACCGGGAGCCTTCGGGGTGGGTG</td><td></td><td></td><td></td><td></td><td></td><td></td><td></td><td></td><td></td><td></td></tr>
<tr><td>Oligo rev</td><td>AAACCACCCACCCCGAAGGCTCCC</td><td></td><td></td><td></td><td></td><td></td><td></td><td></td><td></td><td></td><td></td></tr>
<tr><td>Chromosome</td><td>start</td><td>end</td><td>strand</td><td>MM</td><td>target_seq</td><td>PAM</td><td>alignment</td><td>distance</td><td>position</td><td>gene name</td><td>gene id</td></tr>
<tr><td>GU305914.1</td><td>62907</td><td>62929</td><td>-</td><td>0</td><td>GGGAGCCTTCGGGGTGGGTG</td><td>GGG</td><td>||||||||[||||||||||||]PAM</td><td>NA</td><td>Intergenic</td><td>NA</td><td>NA</td></tr>
<tr><td>T29412</td><td>GGAGCCTTCGGGGTGGGTGGGGG</td><td>1000</td><td></td><td></td><td></td><td></td><td></td><td></td><td></td><td></td><td></td></tr>
<tr><td>Oligo fwd</td><td>CACCGGAGCCTTCGGGGTGGGTGG</td><td></td><td></td><td></td><td></td><td></td><td></td><td></td><td></td><td></td><td></td></tr>
<tr><td>Oligo rev</td><td>AAACCCACCCACCCCGAAGGCTCC</td><td></td><td></td><td></td><td></td><td></td><td></td><td></td><td></td><td></td><td></td></tr>
<tr><td>Chromosome</td><td>start</td><td>end</td><td>strand</td><td>MM</td><td>target_seq</td><td>PAM</td><td>alignment</td><td>distance</td><td>position</td><td>gene name</td><td>gene id</td></tr>
<tr><td>GU305914.1</td><td>62906</td><td>62928</td><td>-</td><td>0</td><td>GGAGCCTTCGGGGTGGGTGG</td><td>GGG</td><td>||||||||[||||||||||||]PAM</td><td>NA</td><td>Intergenic</td><td>NA</td><td>NA</td></tr>
<tr><td>T29413</td><td>GAGCCTTCGGGGTGGGTGGGGGG</td><td>1000</td><td></td><td></td><td></td><td></td><td></td><td></td><td></td><td></td><td></td></tr>
<tr><td>Oligo fwd</td><td>CACCGAGCCTTCGGGGTGGGTGGG</td><td></td><td></td><td></td><td></td><td></td><td></td><td></td><td></td><td></td><td></td></tr>
<tr><td>Oligo rev</td><td>AAACCCCACCCACCCCGAAGGCTC</td><td></td><td></td><td></td><td></td><td></td><td></td><td></td><td></td><td></td><td></td></tr>
<tr><td>Chromosome</td><td>start</td><td>end</td><td>strand</td><td>MM</td><td>target_seq</td><td>PAM</td><td>alignment</td><td>distance</td><td>position</td><td>gene name</td><td>gene id</td></tr>
<tr><td>GU305914.1</td><td>62905</td><td>62927</td><td>-</td><td>0</td><td>GAGCCTTCGGGGTGGGTGGG</td><td>GGG</td><td>||||||||[||||||||||||]PAM</td><td>NA</td><td>Intergenic</td><td>NA</td><td>NA</td></tr>
<tr><td>T29414</td><td>GGCACAGCGTGCCCTACGTCAGG</td><td>1000</td><td></td><td></td><td></td><td></td><td></td><td></td><td></td><td></td><td></td></tr>
<tr><td>Oligo fwd</td><td>CACCGGCACAGCGTGCCCTACGTC</td><td></td><td></td><td></td><td></td><td></td><td></td><td></td><td></td><td></td><td></td></tr>
<tr><td>Oligo rev</td><td>AAACGACGTAGGGCACGCTGTGCC</td><td></td><td></td><td></td><td></td><td></td><td></td><td></td><td></td><td></td><td></td></tr>
<tr><td>Chromosome</td><td>start</td><td>end</td><td>strand</td><td>MM</td><td>target_seq</td><td>PAM</td><td>alignment</td><td>distance</td><td>position</td><td>gene name</td><td>gene id</td></tr>
<tr><td>GU305914.1</td><td>62884</td><td>62906</td><td>-</td><td>0</td><td>GGCACAGCGTGCCCTACGTC</td><td>AGG</td><td>||||||||[||||||||||||]PAM</td><td>NA</td><td>Intergenic</td><td>NA</td><td>NA</td></tr>
<tr><td>T29415</td><td>CCCTACGTCAGGATCATGAATGG</td><td>1000</td><td></td><td></td><td></td><td></td><td></td><td></td><td></td><td></td><td></td></tr>
<tr><td>Oligo fwd</td><td>CACCCCCTACGTCAGGATCATGAA</td><td></td><td></td><td></td><td></td><td></td><td></td><td></td><td></td><td></td><td></td></tr>
<tr><td>Oligo rev</td><td>AAACTTCATGATCCTGACGTAGGG</td><td></td><td></td><td></td><td></td><td></td><td></td><td></td><td></td><td></td><td></td></tr>
<tr><td>Chromosome</td><td>start</td><td>end</td><td>strand</td><td>MM</td><td>target_seq</td><td>PAM</td><td>alignment</td><td>distance</td><td>position</td><td>gene name</td><td>gene id</td></tr>
<tr><td>GU305914.1</td><td>62873</td><td>62895</td><td>-</td><td>0</td><td>CCCTACGTCAGGATCATGAA</td><td>TGG</td><td>||||||||[||||||||||||]PAM</td><td>NA</td><td>Intergenic</td><td>NA</td><td>NA</td></tr>
<tr><td>T29416</td><td>AGGATCATGAATGGAGTCTCTGG</td><td>1000</td><td></td><td></td><td></td><td></td><td></td><td></td><td></td><td></td><td></td></tr>
<tr><td>Oligo fwd</td><td>CACCAGGATCATGAATGGAGTCTC</td><td></td><td></td><td></td><td></td><td></td><td></td><td></td><td></td><td></td><td></td></tr>
<tr><td>Oligo rev</td><td>AAACGAGACTCCATTCATGATCCT</td><td></td><td></td><td></td><td></td><td></td><td></td><td></td><td></td><td></td><td></td></tr>
<tr><td>Chromosome</td><td>start</td><td>end</td><td>strand</td><td>MM</td><td>target_seq</td><td>PAM</td><td>alignment</td><td>distance</td><td>position</td><td>gene name</td><td>gene id</td></tr>
<tr><td>GU305914.1</td><td>62864</td><td>62886</td><td>-</td><td>0</td><td>AGGATCATGAATGGAGTCTC</td><td>TGG</td><td>||||||||[||||||||||||]PAM</td><td>NA</td><td>Intergenic</td><td>NA</td><td>NA</td></tr>
<tr><td>T29417</td><td>GGAGTCTCTGGAATACAGATCGG</td><td>1000</td><td></td><td></td><td></td><td></td><td></td><td></td><td></td><td></td><td></td></tr>
<tr><td>Oligo fwd</td><td>CACCGGAGTCTCTGGAATACAGAT</td><td></td><td></td><td></td><td></td><td></td><td></td><td></td><td></td><td></td><td></td></tr>
<tr><td>Oligo rev</td><td>AAACATCTGTATTCCAGAGACTCC</td><td></td><td></td><td></td><td></td><td></td><td></td><td></td><td></td><td></td><td></td></tr>
<tr><td>Chromosome</td><td>start</td><td>end</td><td>strand</td><td>MM</td><td>target_seq</td><td>PAM</td><td>alignment</td><td>distance</td><td>position</td><td>gene name</td><td>gene id</td></tr>
<tr><td>GU305914.1</td><td>62852</td><td>62874</td><td>-</td><td>0</td><td>GGAGTCTCTGGAATACAGAT</td><td>CGG</td><td>||||||||[||||||||||||]PAM</td><td>NA</td><td>Intergenic</td><td>NA</td><td>NA</td></tr>
<tr><td>T29418</td><td>CTATGAGCATCGCCTCTTGCTGG</td><td>1000</td><td></td><td></td><td></td><td></td><td></td><td></td><td></td><td></td><td></td></tr>
<tr><td>Oligo fwd</td><td>CACCCTATGAGCATCGCCTCTTGC</td><td></td><td></td><td></td><td></td><td></td><td></td><td></td><td></td><td></td><td></td></tr>
<tr><td>Oligo rev</td><td>AAACGCAAGAGGCGATGCTCATAG</td><td></td><td></td><td></td><td></td><td></td><td></td><td></td><td></td><td></td><td></td></tr>
<tr><td>Chromosome</td><td>start</td><td>end</td><td>strand</td><td>MM</td><td>target_seq</td><td>PAM</td><td>alignment</td><td>distance</td><td>position</td><td>gene name</td><td>gene id</td></tr>
</table>

GU305914.1 62818 62840 - 0 CTATGAGCATCGCCTCTTGC TGG ||||||||[||||||||||||]PAM NA Intergenic NA NA

T29419 CCTACACTGACCGACGCCGCAGG 1000
Oligo fwd CACCCCTACACTGACCGACGCCGC
Oligo rev AAACGCGGCGTCGGTCAGTGTAGG
Chromosome start end strand MM target_seq PAM alignment distance position gene name gene id
GU305914.1 62788 62810 - 0 CCTACACTGACCGACGCCGC AGG ||||||||[||||||||||||]PAM NA Intergenic NA NA

T29420 CCCCAAGACCGCGACCAACGCGG 1000
Oligo fwd CACCCCCCAAGACCGCGACCAACG
Oligo rev AAACCGTTGGTCGCGGTCTTGGGG
Chromosome start end strand MM target_seq PAM alignment distance position gene name gene id
GU305914.1 62760 62782 - 0 CCCCAAGACCGCGACCAACG CGG ||||||||[||||||||||||]PAM NA Intergenic NA NA
GU305914.1 218352 218374 + 4 CACTAAGACCGCGACGATCG CGG |-|-||||[|||||||-|-||]PAM NA Intergenic NA NA

T29421 CAAGACCGCGACCAACGCGGCGG 1000
Oligo fwd CACCCAAGACCGCGACCAACGCGG
Oligo rev AAACCCGCGTTGGTCGCGGTCTTG
Chromosome start end strand MM target_seq PAM alignment distance position gene name gene id
GU305914.1 62757 62779 - 0 CAAGACCGCGACCAACGCGG CGG ||||||||[||||||||||||]PAM NA Intergenic NA NA

T29422 CGACCAACGCGGCGGCAGACAGG 1000
Oligo fwd CACCCGACCAACGCGGCGGCAGAC
Oligo rev AAACGTCTGCCGCCGCGTTGGTCG
Chromosome start end strand MM target_seq PAM alignment distance position gene name gene id
GU305914.1 62749 62771 - 0 CGACCAACGCGGCGGCAGAC AGG ||||||||[||||||||||||]PAM NA Intergenic NA NA

T29423 GACCAACGCGGCGGCAGACAGGG 1000
Oligo fwd CACCGACCAACGCGGCGGCAGACA
Oligo rev AAACTGTCTGCCGCCGCGTTGGTC
Chromosome start end strand MM target_seq PAM alignment distance position gene name gene id
GU305914.1 62748 62770 - 0 GACCAACGCGGCGGCAGACA GGG ||||||||[||||||||||||]PAM NA Intergenic NA NA

T29424 TGCTGTCAATGCTGCTGCCGCGG 1000
Oligo fwd CACCTGCTGTCAATGCTGCTGCCG
Oligo rev AAACCGGCAGCAGCATTGACAGCA
Chromosome start end strand MM target_seq PAM alignment distance position gene name gene id
GU305914.1 62697 62719 - 0 TGCTGTCAATGCTGCTGCCG CGG ||||||||[||||||||||||]PAM NA Intergenic NA NA

T29425 GCTGCCGCGGCTGCTGCCGCCGG 1000
Oligo fwd CACCGCTGCCGCGGCTGCTGCCGC
Oligo rev AAACGCGGCAGCAGCCGCGGCAGC
Chromosome start end strand MM target_seq PAM alignment distance position gene name gene id
GU305914.1 62684 62706 - 0 GCTGCCGCGGCTGCTGCCGC CGG ||||||||[||||||||||||]PAM NA Intergenic NA NA
GU305914.1 38224 38246 - 4 GATGCCGAAGCTGATGCCGC TGG |-|||||-[-||||-||||||]PAM NA Intergenic NA NA
GU305914.1 168890 168912 + 4 GCTGCTGGCGCTGGTGCCGC GGG |||||-|-[-||||-||||||]PAM NA Intergenic NA NA
GU305914.1 168881 168903 + 4 GCAGAAGCGGCTGCTGGCGC TGG ||-|--||[||||||||-|||]PAM NA Intergenic NA NA
GU305914.1 27406 27428 - 3 GCTGCTGCTGCTGCTGCTGC CGG |||||-||[-||||||||-||]PAM NA Intergenic NA NA
GU305914.1 113357 113379 - 4 GCTGGCGTGCCTGCTGACGC TGG ||||-||-[|-||||||-|||]PAM NA Intergenic NA NA

T29426 GCCGCGGCTGCTGCCGCCGGCGG 1000
Oligo fwd CACCGCCGCGGCTGCTGCCGCCGG
Oligo rev AAACCCGGCGGCAGCAGCCGCGGC
Chromosome start end strand MM target_seq PAM alignment distance position gene name gene id
GU305914.1 62681 62703 - 0 GCCGCGGCTGCTGCCGCCGG CGG ||||||||[||||||||||||]PAM NA Intergenic NA NA
GU305914.1 62678 62700 - 4 GCGGCTGCTGCCGCCGGCGG GGG ||-||-||[|||-||||-|||]PAM NA Intergenic NA NA
GU305914.1 65720 65742 + 4 CCCGCTGCCGCTGCCGCCGC AGG -||||-||[-||||||||||-]PAM NA Intergenic NA NA

T29427 CCGCGGCTGCTGCCGCCGGCGGG 1000
Oligo fwd CACCCCGCGGCTGCTGCCGCCGGC
Oligo rev AAACGCCGGCGGCAGCAGCCGCGG
Chromosome start end strand MM target_seq PAM alignment distance position gene name gene id
GU305914.1 62680 62702 - 0 CCGCGGCTGCTGCCGCCGGC GGG ||||||||[||||||||||||]PAM NA Intergenic NA NA

T29428 CGCGGCTGCTGCCGCCGGCGGGG 1000
Oligo fwd CACCCGCGGCTGCTGCCGCCGGCG
Oligo rev AAACCGCCGGCGGCAGCAGCCGCG
Chromosome start end strand MM target_seq PAM alignment distance position gene name gene id
GU305914.1 62679 62701 - 0 CGCGGCTGCTGCCGCCGGCG GGG ||||||||[||||||||||||]PAM NA Intergenic NA NA
GU305914.1 38219 38241 - 4 CGAAGCTGATGCCGCTGGCG CGG ||--||||[-||||||-||||]PAM NA Intergenic NA NA
GU305914.1 45592 45614 - 3 GGCGGCTGCCGCCGCCGCCG CGG -|||||||[|-|||||||-||]PAM NA Intergenic NA NA
GU305914.1 130312 130334 + 4 CGCAGCAGCAGCCGCCGCCG CGG |||-||-|[|-|||||||-||]PAM NA Intergenic NA NA

T29429 GCGGCTGCTGCCGCCGGCGGGGG 1000
Oligo fwd CACCGCGGCTGCTGCCGCCGGCGG
Oligo rev AAACCCGCCGGCGGCAGCAGCCGC
Chromosome start end strand MM target_seq PAM alignment distance position gene name gene id
GU305914.1 62678 62700 - 0 GCGGCTGCTGCCGCCGGCGG GGG ||||||||[||||||||||||]PAM NA Intergenic NA NA
GU305914.1 62681 62703 - 4 GCCGCGGCTGCTGCCGCCGG CGG ||-||-||[|||-||||-|||]PAM NA Intergenic NA NA

T29430 GCTGCTGCCGCCGGCGGGGGCGG 1000
Oligo fwd CACCGCTGCTGCCGCCGGCGGGGG
Oligo rev AAACCCCCCGCCGGCGGCAGCAGC
Chromosome start end strand MM target_seq PAM alignment distance position gene name gene id
GU305914.1 62675 62697 - 0 GCTGCTGCCGCCGGCGGGGG CGG ||||||||[||||||||||||]PAM NA Intergenic NA NA
GU305914.1 30000 30022 + 4 GGTGCTGGCGGCGGCGGCGG CGG |-|||||-[||-||||||-||]PAM NA Intergenic NA NA
GU305914.1 229979 230001 - 4 GCCACTGCCGCCGGCCGGGA TGG ||--||||[|||||||-|||-]PAM NA Intergenic NA NA

T29431 GCTGCCGCCGGCGGGGGCGGCGG 1000

<table>
<tr><td>Oligo fwd</td><td>CACCGCTGCCGCCGGCGGGGGCGG</td><td></td><td></td><td></td><td></td><td></td><td></td><td></td><td></td><td></td><td></td></tr>
<tr><td>Oligo rev</td><td>AAACCCGCCCCCGCCGGCGGCAGC</td><td></td><td></td><td></td><td></td><td></td><td></td><td></td><td></td><td></td><td></td></tr>
<tr><td>Chromosome</td><td>start</td><td>end</td><td>strand</td><td>MM</td><td>target_seq</td><td>PAM</td><td>alignment</td><td>distance</td><td>position</td><td>gene name</td><td>gene id</td></tr>
<tr><td>GU305914.1</td><td>62672</td><td>62694</td><td>-</td><td>0</td><td>GCTGCCGCCGGCGGGGGCGG</td><td>CGG</td><td>||||||||[||||||||||||]PAM</td><td>NA</td><td>Intergenic</td><td>NA</td><td>NA</td></tr>
<tr><td>GU305914.1</td><td>52716</td><td>52738</td><td>+</td><td>3</td><td>GCCGCCGCTGGCGGTGGCGG</td><td>TGG</td><td>||-|||||[-|||||-|||||]PAM</td><td>NA</td><td>Intergenic</td><td>NA</td><td>NA</td></tr>
<tr><td>GU305914.1</td><td>154407</td><td>154429</td><td>+</td><td>4</td><td>GGTGGCGGCGGCGGCGGCGG</td><td>CGG</td><td>|-||-||-[||||||-|||||]PAM</td><td>NA</td><td>Intergenic</td><td>NA</td><td>NA</td></tr>
<tr><td>GU305914.1</td><td>58123</td><td>58145</td><td>-</td><td>4</td><td>GGTGGCGGCGGCGGCGGCGG</td><td>AGG</td><td>|-||-||-[||||||-|||||]PAM</td><td>NA</td><td>Intergenic</td><td>NA</td><td>NA</td></tr>
<tr><td>GU305914.1</td><td>149485</td><td>149507</td><td>-</td><td>4</td><td>GCAGGCGACGGCGGCGGCGG</td><td>CGG</td><td>||-|-||-[||||||-|||||]PAM</td><td>NA</td><td>Intergenic</td><td>NA</td><td>NA</td></tr>
<tr><td>GU305914.1</td><td>90259</td><td>90281</td><td>-</td><td>4</td><td>GCGGGCGACGGCGGAGGCGG</td><td>CGG</td><td>||-|-||-[||||||-|||||]PAM</td><td>NA</td><td>Intergenic</td><td>NA</td><td>NA</td></tr>
<tr><td>GU305914.1</td><td>30000</td><td>30022</td><td>+</td><td>4</td><td>GGTGCTGGCGGCGGCGGCGG</td><td>CGG</td><td>|-|||-|-[||||||-|||||]PAM</td><td>NA</td><td>Intergenic</td><td>NA</td><td>NA</td></tr>
<tr><td></td><td></td><td></td><td></td><td></td><td></td><td></td><td></td><td></td><td></td><td></td><td></td></tr>
<tr><td>T29432</td><td>GGGCGGCGGCGCTAACCTACTGG</td><td>1000</td><td></td><td></td><td></td><td></td><td></td><td></td><td></td><td></td><td></td></tr>
<tr><td>Oligo fwd</td><td>CACCGGGCGGCGGCGCTAACCTAC</td><td></td><td></td><td></td><td></td><td></td><td></td><td></td><td></td><td></td><td></td></tr>
<tr><td>Oligo rev</td><td>AAACGTAGGTTAGCGCCGCCGCCC</td><td></td><td></td><td></td><td></td><td></td><td></td><td></td><td></td><td></td><td></td></tr>
<tr><td>Chromosome</td><td>start</td><td>end</td><td>strand</td><td>MM</td><td>target_seq</td><td>PAM</td><td>alignment</td><td>distance</td><td>position</td><td>gene name</td><td>gene id</td></tr>
<tr><td>GU305914.1</td><td>62658</td><td>62680</td><td>-</td><td>0</td><td>GGGCGGCGGCGCTAACCTAC</td><td>TGG</td><td>||||||||[||||||||||||]PAM</td><td>NA</td><td>Intergenic</td><td>NA</td><td>NA</td></tr>
<tr><td></td><td></td><td></td><td></td><td></td><td></td><td></td><td></td><td></td><td></td><td></td><td></td></tr>
<tr><td>T29433</td><td>GTCACTTGTGCAAATCAGCGAGG</td><td>1000</td><td></td><td></td><td></td><td></td><td></td><td></td><td></td><td></td><td></td></tr>
<tr><td>Oligo fwd</td><td>CACCGTCACTTGTGCAAATCAGCG</td><td></td><td></td><td></td><td></td><td></td><td></td><td></td><td></td><td></td><td></td></tr>
<tr><td>Oligo rev</td><td>AAACCGCTGATTTGCACAAGTGAC</td><td></td><td></td><td></td><td></td><td></td><td></td><td></td><td></td><td></td><td></td></tr>
<tr><td>Chromosome</td><td>start</td><td>end</td><td>strand</td><td>MM</td><td>target_seq</td><td>PAM</td><td>alignment</td><td>distance</td><td>position</td><td>gene name</td><td>gene id</td></tr>
<tr><td>GU305914.1</td><td>62627</td><td>62649</td><td>-</td><td>0</td><td>GTCACTTGTGCAAATCAGCG</td><td>AGG</td><td>||||||||[||||||||||||]PAM</td><td>NA</td><td>Intergenic</td><td>NA</td><td>NA</td></tr>
<tr><td></td><td></td><td></td><td></td><td></td><td></td><td></td><td></td><td></td><td></td><td></td><td></td></tr>
<tr><td>T29434</td><td>GCAAATCAGCGAGGTTGCTGCGG</td><td>1000</td><td></td><td></td><td></td><td></td><td></td><td></td><td></td><td></td><td></td></tr>
<tr><td>Oligo fwd</td><td>CACCGCAAATCAGCGAGGTTGCTG</td><td></td><td></td><td></td><td></td><td></td><td></td><td></td><td></td><td></td><td></td></tr>
<tr><td>Oligo rev</td><td>AAACCAGCAACCTCGCTGATTTGC</td><td></td><td></td><td></td><td></td><td></td><td></td><td></td><td></td><td></td><td></td></tr>
<tr><td>Chromosome</td><td>start</td><td>end</td><td>strand</td><td>MM</td><td>target_seq</td><td>PAM</td><td>alignment</td><td>distance</td><td>position</td><td>gene name</td><td>gene id</td></tr>
<tr><td>GU305914.1</td><td>62618</td><td>62640</td><td>-</td><td>0</td><td>GCAAATCAGCGAGGTTGCTG</td><td>CGG</td><td>||||||||[||||||||||||]PAM</td><td>NA</td><td>Intergenic</td><td>NA</td><td>NA</td></tr>
<tr><td></td><td></td><td></td><td></td><td></td><td></td><td></td><td></td><td></td><td></td><td></td><td></td></tr>
<tr><td>T29435</td><td>AATCAGCGAGGTTGCTGCGGAGG</td><td>1000</td><td></td><td></td><td></td><td></td><td></td><td></td><td></td><td></td><td></td></tr>
<tr><td>Oligo fwd</td><td>CACCAATCAGCGAGGTTGCTGCGG</td><td></td><td></td><td></td><td></td><td></td><td></td><td></td><td></td><td></td><td></td></tr>
<tr><td>Oligo rev</td><td>AAACCCGCAGCAACCTCGCTGATT</td><td></td><td></td><td></td><td></td><td></td><td></td><td></td><td></td><td></td><td></td></tr>
<tr><td>Chromosome</td><td>start</td><td>end</td><td>strand</td><td>MM</td><td>target_seq</td><td>PAM</td><td>alignment</td><td>distance</td><td>position</td><td>gene name</td><td>gene id</td></tr>
<tr><td>GU305914.1</td><td>62615</td><td>62637</td><td>-</td><td>0</td><td>AATCAGCGAGGTTGCTGCGG</td><td>AGG</td><td>||||||||[||||||||||||]PAM</td><td>NA</td><td>Intergenic</td><td>NA</td><td>NA</td></tr>
<tr><td></td><td></td><td></td><td></td><td></td><td></td><td></td><td></td><td></td><td></td><td></td><td></td></tr>
<tr><td>T29436</td><td>CGAGGTTGCTGCGGAGGAAATGG</td><td>1000</td><td></td><td></td><td></td><td></td><td></td><td></td><td></td><td></td><td></td></tr>
<tr><td>Oligo fwd</td><td>CACCCGAGGTTGCTGCGGAGGAAA</td><td></td><td></td><td></td><td></td><td></td><td></td><td></td><td></td><td></td><td></td></tr>
<tr><td>Oligo rev</td><td>AAACTTTCCTCCGCAGCAACCTCG</td><td></td><td></td><td></td><td></td><td></td><td></td><td></td><td></td><td></td><td></td></tr>
<tr><td>Chromosome</td><td>start</td><td>end</td><td>strand</td><td>MM</td><td>target_seq</td><td>PAM</td><td>alignment</td><td>distance</td><td>position</td><td>gene name</td><td>gene id</td></tr>
<tr><td>GU305914.1</td><td>62609</td><td>62631</td><td>-</td><td>0</td><td>CGAGGTTGCTGCGGAGGAAA</td><td>TGG</td><td>||||||||[||||||||||||]PAM</td><td>NA</td><td>Intergenic</td><td>NA</td><td>NA</td></tr>
<tr><td></td><td></td><td></td><td></td><td></td><td></td><td></td><td></td><td></td><td></td><td></td><td></td></tr>
<tr><td>T29437</td><td>GAGGTTGCTGCGGAGGAAATGGG</td><td>1000</td><td></td><td></td><td></td><td></td><td></td><td></td><td></td><td></td><td></td></tr>
<tr><td>Oligo fwd</td><td>CACCGAGGTTGCTGCGGAGGAAAT</td><td></td><td></td><td></td><td></td><td></td><td></td><td></td><td></td><td></td><td></td></tr>
<tr><td>Oligo rev</td><td>AAACATTTCCTCCGCAGCAACCTC</td><td></td><td></td><td></td><td></td><td></td><td></td><td></td><td></td><td></td><td></td></tr>
<tr><td>Chromosome</td><td>start</td><td>end</td><td>strand</td><td>MM</td><td>target_seq</td><td>PAM</td><td>alignment</td><td>distance</td><td>position</td><td>gene name</td><td>gene id</td></tr>
<tr><td>GU305914.1</td><td>62608</td><td>62630</td><td>-</td><td>0</td><td>GAGGTTGCTGCGGAGGAAAT</td><td>GGG</td><td>||||||||[||||||||||||]PAM</td><td>NA</td><td>Intergenic</td><td>NA</td><td>NA</td></tr>
<tr><td></td><td></td><td></td><td></td><td></td><td></td><td></td><td></td><td></td><td></td><td></td><td></td></tr>
<tr><td>T29438</td><td>AGGTTGCTGCGGAGGAAATGGGG</td><td>1000</td><td></td><td></td><td></td><td></td><td></td><td></td><td></td><td></td><td></td></tr>
<tr><td>Oligo fwd</td><td>CACCAGGTTGCTGCGGAGGAAATG</td><td></td><td></td><td></td><td></td><td></td><td></td><td></td><td></td><td></td><td></td></tr>
<tr><td>Oligo rev</td><td>AAACCATTTCCTCCGCAGCAACCT</td><td></td><td></td><td></td><td></td><td></td><td></td><td></td><td></td><td></td><td></td></tr>
<tr><td>Chromosome</td><td>start</td><td>end</td><td>strand</td><td>MM</td><td>target_seq</td><td>PAM</td><td>alignment</td><td>distance</td><td>position</td><td>gene name</td><td>gene id</td></tr>
<tr><td>GU305914.1</td><td>62607</td><td>62629</td><td>-</td><td>0</td><td>AGGTTGCTGCGGAGGAAATG</td><td>GGG</td><td>||||||||[||||||||||||]PAM</td><td>NA</td><td>Intergenic</td><td>NA</td><td>NA</td></tr>
<tr><td></td><td></td><td></td><td></td><td></td><td></td><td></td><td></td><td></td><td></td><td></td><td></td></tr>
<tr><td>T29439</td><td>GGTTGCTGCGGAGGAAATGGGGG</td><td>1000</td><td></td><td></td><td></td><td></td><td></td><td></td><td></td><td></td><td></td></tr>
<tr><td>Oligo fwd</td><td>CACCGGTTGCTGCGGAGGAAATGG</td><td></td><td></td><td></td><td></td><td></td><td></td><td></td><td></td><td></td><td></td></tr>
<tr><td>Oligo rev</td><td>AAACCCATTTCCTCCGCAGCAACC</td><td></td><td></td><td></td><td></td><td></td><td></td><td></td><td></td><td></td><td></td></tr>
<tr><td>Chromosome</td><td>start</td><td>end</td><td>strand</td><td>MM</td><td>target_seq</td><td>PAM</td><td>alignment</td><td>distance</td><td>position</td><td>gene name</td><td>gene id</td></tr>
<tr><td>GU305914.1</td><td>62606</td><td>62628</td><td>-</td><td>0</td><td>GGTTGCTGCGGAGGAAATGG</td><td>GGG</td><td>||||||||[||||||||||||]PAM</td><td>NA</td><td>Intergenic</td><td>NA</td><td>NA</td></tr>
<tr><td></td><td></td><td></td><td></td><td></td><td></td><td></td><td></td><td></td><td></td><td></td><td></td></tr>
<tr><td>T29440</td><td>GTTGCTGCGGAGGAAATGGGGGG</td><td>1000</td><td></td><td></td><td></td><td></td><td></td><td></td><td></td><td></td><td></td></tr>
<tr><td>Oligo fwd</td><td>CACCGTTGCTGCGGAGGAAATGGG</td><td></td><td></td><td></td><td></td><td></td><td></td><td></td><td></td><td></td><td></td></tr>
<tr><td>Oligo rev</td><td>AAACCCCATTTCCTCCGCAGCAAC</td><td></td><td></td><td></td><td></td><td></td><td></td><td></td><td></td><td></td><td></td></tr>
<tr><td>Chromosome</td><td>start</td><td>end</td><td>strand</td><td>MM</td><td>target_seq</td><td>PAM</td><td>alignment</td><td>distance</td><td>position</td><td>gene name</td><td>gene id</td></tr>
<tr><td>GU305914.1</td><td>62605</td><td>62627</td><td>-</td><td>0</td><td>GTTGCTGCGGAGGAAATGGG</td><td>GGG</td><td>||||||||[||||||||||||]PAM</td><td>NA</td><td>Intergenic</td><td>NA</td><td>NA</td></tr>
<tr><td></td><td></td><td></td><td></td><td></td><td></td><td></td><td></td><td></td><td></td><td></td><td></td></tr>
<tr><td>T29441</td><td>GAAGACCAACGCTACCGCCGCGG</td><td>1000</td><td></td><td></td><td></td><td></td><td></td><td></td><td></td><td></td><td></td></tr>
<tr><td>Oligo fwd</td><td>CACCGAAGACCAACGCTACCGCCG</td><td></td><td></td><td></td><td></td><td></td><td></td><td></td><td></td><td></td><td></td></tr>
<tr><td>Oligo rev</td><td>AAACCGGCGGTAGCGTTGGTCTTC</td><td></td><td></td><td></td><td></td><td></td><td></td><td></td><td></td><td></td><td></td></tr>
<tr><td>Chromosome</td><td>start</td><td>end</td><td>strand</td><td>MM</td><td>target_seq</td><td>PAM</td><td>alignment</td><td>distance</td><td>position</td><td>gene name</td><td>gene id</td></tr>
<tr><td>GU305914.1</td><td>62556</td><td>62578</td><td>-</td><td>0</td><td>GAAGACCAACGCTACCGCCG</td><td>CGG</td><td>||||||||[||||||||||||]PAM</td><td>NA</td><td>Intergenic</td><td>NA</td><td>NA</td></tr>
<tr><td></td><td></td><td></td><td></td><td></td><td></td><td></td><td></td><td></td><td></td><td></td><td></td></tr>
<tr><td>T29442</td><td>GCGGCCGCTCCTGCCGTCGCCGG</td><td>1000</td><td></td><td></td><td></td><td></td><td></td><td></td><td></td><td></td><td></td></tr>
<tr><td>Oligo fwd</td><td>CACCGCGGCCGCTCCTGCCGTCGC</td><td></td><td></td><td></td><td></td><td></td><td></td><td></td><td></td><td></td><td></td></tr>
<tr><td>Oligo rev</td><td>AAACGCGACGGCAGGAGCGGCCGC</td><td></td><td></td><td></td><td></td><td></td><td></td><td></td><td></td><td></td><td></td></tr>
<tr><td>Chromosome</td><td>start</td><td>end</td><td>strand</td><td>MM</td><td>target_seq</td><td>PAM</td><td>alignment</td><td>distance</td><td>position</td><td>gene name</td><td>gene id</td></tr>
<tr><td>GU305914.1</td><td>62537</td><td>62559</td><td>-</td><td>0</td><td>GCGGCCGCTCCTGCCGTCGC</td><td>CGG</td><td>||||||||[||||||||||||]PAM</td><td>NA</td><td>Intergenic</td><td>NA</td><td>NA</td></tr>
<tr><td></td><td></td><td></td><td></td><td></td><td></td><td></td><td></td><td></td><td></td><td></td><td></td></tr>
<tr><td>T29443</td><td>AATCACGCTAACGCCACCTCTGG</td><td>1000</td><td></td><td></td><td></td><td></td><td></td><td></td><td></td><td></td><td></td></tr>
<tr><td>Oligo fwd</td><td>CACCAATCACGCTAACGCCACCTC</td><td></td><td></td><td></td><td></td><td></td><td></td><td></td><td></td><td></td><td></td></tr>
<tr><td>Oligo rev</td><td>AAACGAGGTGGCGTTAGCGTGATT</td><td></td><td></td><td></td><td></td><td></td><td></td><td></td><td></td><td></td><td></td></tr>
<tr><td>Chromosome</td><td>start</td><td>end</td><td>strand</td><td>MM</td><td>target_seq</td><td>PAM</td><td>alignment</td><td>distance</td><td>position</td><td>gene name</td><td>gene id</td></tr>
<tr><td>GU305914.1</td><td>62489</td><td>62511</td><td>-</td><td>0</td><td>AATCACGCTAACGCCACCTC</td><td>TGG</td><td>||||||||[||||||||||||]PAM</td><td>NA</td><td>Intergenic</td><td>NA</td><td>NA</td></tr>
<tr><td></td><td></td><td></td><td></td><td></td><td></td><td></td><td></td><td></td><td></td><td></td><td></td></tr>
<tr><td>T29444</td><td>GCTAACGCCACCTCTGGTGCTGG</td><td>1000</td><td></td><td></td><td></td><td></td><td></td><td></td><td></td><td></td><td></td></tr>
<tr><td>Oligo fwd</td><td>CACCGCTAACGCCACCTCTGGTGC</td><td></td><td></td><td></td><td></td><td></td><td></td><td></td><td></td><td></td><td></td></tr>
<tr><td>Oligo rev</td><td>AAACGCACCAGAGGTGGCGTTAGC</td><td></td><td></td><td></td><td></td><td></td><td></td><td></td><td></td><td></td><td></td></tr>
<tr><td>Chromosome</td><td>start</td><td>end</td><td>strand</td><td>MM</td><td>target_seq</td><td>PAM</td><td>alignment</td><td>distance</td><td>position</td><td>gene name</td><td>gene id</td></tr>
<tr><td>GU305914.1</td><td>62483</td><td>62505</td><td>-</td><td>0</td><td>GCTAACGCCACCTCTGGTGC</td><td>TGG</td><td>||||||||[||||||||||||]PAM</td><td>NA</td><td>Intergenic</td><td>NA</td><td>NA</td></tr>
</table>

<table>
<tr><td>T29445</td><td>CACCTCTGGTGCTGGATCCGCGG</td><td>1000</td><td></td><td></td><td></td><td></td><td></td><td></td><td></td><td></td><td></td></tr>
<tr><td>Oligo fwd</td><td>CACCCACCTCTGGTGCTGGATCCG</td><td></td><td></td><td></td><td></td><td></td><td></td><td></td><td></td><td></td><td></td></tr>
<tr><td>Oligo rev</td><td>AAACCGGATCCAGCACCAGAGGTG</td><td></td><td></td><td></td><td></td><td></td><td></td><td></td><td></td><td></td><td></td></tr>
<tr><td>Chromosome</td><td>start</td><td>end</td><td>strand</td><td>MM</td><td>target_seq</td><td>PAM</td><td>alignment</td><td>distance</td><td>position</td><td>gene name</td><td>gene id</td></tr>
<tr><td>GU305914.1</td><td>62475</td><td>62497</td><td>-</td><td>0</td><td>CACCTCTGGTGCTGGATCCG</td><td>CGG</td><td>||||||||[||||||||||||]PAM</td><td>NA</td><td>Intergenic</td><td>NA</td><td>NA</td></tr>
<tr><td>T29446</td><td>CTCGACTGCGCCCTGTGACGAGG</td><td>1000</td><td></td><td></td><td></td><td></td><td></td><td></td><td></td><td></td><td></td></tr>
<tr><td>Oligo fwd</td><td>CACCCTCGACTGCGCCCTGTGACG</td><td></td><td></td><td></td><td></td><td></td><td></td><td></td><td></td><td></td><td></td></tr>
<tr><td>Oligo rev</td><td>AAACCGTCACAGGGCGCAGTCGAG</td><td></td><td></td><td></td><td></td><td></td><td></td><td></td><td></td><td></td><td></td></tr>
<tr><td>Chromosome</td><td>start</td><td>end</td><td>strand</td><td>MM</td><td>target_seq</td><td>PAM</td><td>alignment</td><td>distance</td><td>position</td><td>gene name</td><td>gene id</td></tr>
<tr><td>GU305914.1</td><td>62376</td><td>62398</td><td>-</td><td>0</td><td>CTCGACTGCGCCCTGTGACG</td><td>AGG</td><td>||||||||[||||||||||||]PAM</td><td>NA</td><td>Intergenic</td><td>NA</td><td>NA</td></tr>
<tr><td>T29447</td><td>TCGACTGCGCCCTGTGACGAGGG</td><td>1000</td><td></td><td></td><td></td><td></td><td></td><td></td><td></td><td></td><td></td></tr>
<tr><td>Oligo fwd</td><td>CACCTCGACTGCGCCCTGTGACGA</td><td></td><td></td><td></td><td></td><td></td><td></td><td></td><td></td><td></td><td></td></tr>
<tr><td>Oligo rev</td><td>AAACTCGTCACAGGGCGCAGTCGA</td><td></td><td></td><td></td><td></td><td></td><td></td><td></td><td></td><td></td><td></td></tr>
<tr><td>Chromosome</td><td>start</td><td>end</td><td>strand</td><td>MM</td><td>target_seq</td><td>PAM</td><td>alignment</td><td>distance</td><td>position</td><td>gene name</td><td>gene id</td></tr>
<tr><td>GU305914.1</td><td>62375</td><td>62397</td><td>-</td><td>0</td><td>TCGACTGCGCCCTGTGACGA</td><td>GGG</td><td>||||||||[||||||||||||]PAM</td><td>NA</td><td>Intergenic</td><td>NA</td><td>NA</td></tr>
<tr><td>T29448</td><td>CGTCCTAGTCACGTTAGTGTCGG</td><td>1000</td><td></td><td></td><td></td><td></td><td></td><td></td><td></td><td></td><td></td></tr>
<tr><td>Oligo fwd</td><td>CACCCGTCCTAGTCACGTTAGTGT</td><td></td><td></td><td></td><td></td><td></td><td></td><td></td><td></td><td></td><td></td></tr>
<tr><td>Oligo rev</td><td>AAACACACTAACGTGACTAGGACG</td><td></td><td></td><td></td><td></td><td></td><td></td><td></td><td></td><td></td><td></td></tr>
<tr><td>Chromosome</td><td>start</td><td>end</td><td>strand</td><td>MM</td><td>target_seq</td><td>PAM</td><td>alignment</td><td>distance</td><td>position</td><td>gene name</td><td>gene id</td></tr>
<tr><td>GU305914.1</td><td>62333</td><td>62355</td><td>-</td><td>0</td><td>CGTCCTAGTCACGTTAGTGT</td><td>CGG</td><td>||||||||[||||||||||||]PAM</td><td>NA</td><td>Intergenic</td><td>NA</td><td>NA</td></tr>
<tr><td>T29449</td><td>TCACGTTAGTGTCGGCACTCAGG</td><td>1000</td><td></td><td></td><td></td><td></td><td></td><td></td><td></td><td></td><td></td></tr>
<tr><td>Oligo fwd</td><td>CACCTCACGTTAGTGTCGGCACTC</td><td></td><td></td><td></td><td></td><td></td><td></td><td></td><td></td><td></td><td></td></tr>
<tr><td>Oligo rev</td><td>AAACGAGTGCCGACACTAACGTGA</td><td></td><td></td><td></td><td></td><td></td><td></td><td></td><td></td><td></td><td></td></tr>
<tr><td>Chromosome</td><td>start</td><td>end</td><td>strand</td><td>MM</td><td>target_seq</td><td>PAM</td><td>alignment</td><td>distance</td><td>position</td><td>gene name</td><td>gene id</td></tr>
<tr><td>GU305914.1</td><td>62325</td><td>62347</td><td>-</td><td>0</td><td>TCACGTTAGTGTCGGCACTC</td><td>AGG</td><td>||||||||[||||||||||||]PAM</td><td>NA</td><td>Intergenic</td><td>NA</td><td>NA</td></tr>
<tr><td>T29450</td><td>CTATCCCGATTCCCGCTCCCCGG</td><td>1000</td><td></td><td></td><td></td><td></td><td></td><td></td><td></td><td></td><td></td></tr>
<tr><td>Oligo fwd</td><td>CACCCTATCCCGATTCCCGCTCCC</td><td></td><td></td><td></td><td></td><td></td><td></td><td></td><td></td><td></td><td></td></tr>
<tr><td>Oligo rev</td><td>AAACGGGAGCGGGAATCGGGATAG</td><td></td><td></td><td></td><td></td><td></td><td></td><td></td><td></td><td></td><td></td></tr>
<tr><td>Chromosome</td><td>start</td><td>end</td><td>strand</td><td>MM</td><td>target_seq</td><td>PAM</td><td>alignment</td><td>distance</td><td>position</td><td>gene name</td><td>gene id</td></tr>
<tr><td>GU305914.1</td><td>62287</td><td>62309</td><td>-</td><td>0</td><td>CTATCCCGATTCCCGCTCCC</td><td>CGG</td><td>||||||||[||||||||||||]PAM</td><td>NA</td><td>Intergenic</td><td>NA</td><td>NA</td></tr>
<tr><td>T29451</td><td>CCCGCTCCCCGGTGCAGCACAGG</td><td>1000</td><td></td><td></td><td></td><td></td><td></td><td></td><td></td><td></td><td></td></tr>
<tr><td>Oligo fwd</td><td>CACCCCCGCTCCCCGGTGCAGCAC</td><td></td><td></td><td></td><td></td><td></td><td></td><td></td><td></td><td></td><td></td></tr>
<tr><td>Oligo rev</td><td>AAACGTGCTGCACCGGGGAGCGGG</td><td></td><td></td><td></td><td></td><td></td><td></td><td></td><td></td><td></td><td></td></tr>
<tr><td>Chromosome</td><td>start</td><td>end</td><td>strand</td><td>MM</td><td>target_seq</td><td>PAM</td><td>alignment</td><td>distance</td><td>position</td><td>gene name</td><td>gene id</td></tr>
<tr><td>GU305914.1</td><td>62276</td><td>62298</td><td>-</td><td>0</td><td>CCCGCTCCCCGGTGCAGCAC</td><td>AGG</td><td>||||||||[||||||||||||]PAM</td><td>NA</td><td>Intergenic</td><td>NA</td><td>NA</td></tr>
<tr><td>T29452</td><td>AGCTGATCCCCTCCTGTACGCGG</td><td>1000</td><td></td><td></td><td></td><td></td><td></td><td></td><td></td><td></td><td></td></tr>
<tr><td>Oligo fwd</td><td>CACCAGCTGATCCCCTCCTGTACG</td><td></td><td></td><td></td><td></td><td></td><td></td><td></td><td></td><td></td><td></td></tr>
<tr><td>Oligo rev</td><td>AAACCGTACAGGAGGGGATCAGCT</td><td></td><td></td><td></td><td></td><td></td><td></td><td></td><td></td><td></td><td></td></tr>
<tr><td>Chromosome</td><td>start</td><td>end</td><td>strand</td><td>MM</td><td>target_seq</td><td>PAM</td><td>alignment</td><td>distance</td><td>position</td><td>gene name</td><td>gene id</td></tr>
<tr><td>GU305914.1</td><td>62208</td><td>62230</td><td>-</td><td>0</td><td>AGCTGATCCCCTCCTGTACG</td><td>CGG</td><td>||||||||[||||||||||||]PAM</td><td>NA</td><td>Intergenic</td><td>NA</td><td>NA</td></tr>
<tr><td>T29453</td><td>CGAATCCTCACCTGCCGCTTCGG</td><td>1000</td><td></td><td></td><td></td><td></td><td></td><td></td><td></td><td></td><td></td></tr>
<tr><td>Oligo fwd</td><td>CACCCGAATCCTCACCTGCCGCTT</td><td></td><td></td><td></td><td></td><td></td><td></td><td></td><td></td><td></td><td></td></tr>
<tr><td>Oligo rev</td><td>AAACAAGCGGCAGGTGAGGATTCG</td><td></td><td></td><td></td><td></td><td></td><td></td><td></td><td></td><td></td><td></td></tr>
<tr><td>Chromosome</td><td>start</td><td>end</td><td>strand</td><td>MM</td><td>target_seq</td><td>PAM</td><td>alignment</td><td>distance</td><td>position</td><td>gene name</td><td>gene id</td></tr>
<tr><td>GU305914.1</td><td>62124</td><td>62146</td><td>-</td><td>0</td><td>CGAATCCTCACCTGCCGCTT</td><td>CGG</td><td>||||||||[||||||||||||]PAM</td><td>NA</td><td>Intergenic</td><td>NA</td><td>NA</td></tr>
<tr><td>T29454</td><td>ACCACCAGCAGCAGCAGCGATGG</td><td>1000</td><td></td><td></td><td></td><td></td><td></td><td></td><td></td><td></td><td></td></tr>
<tr><td>Oligo fwd</td><td>CACCACCACCAGCAGCAGCAGCGA</td><td></td><td></td><td></td><td></td><td></td><td></td><td></td><td></td><td></td><td></td></tr>
<tr><td>Oligo rev</td><td>AAACTCGCTGCTGCTGCTGGTGGT</td><td></td><td></td><td></td><td></td><td></td><td></td><td></td><td></td><td></td><td></td></tr>
<tr><td>Chromosome</td><td>start</td><td>end</td><td>strand</td><td>MM</td><td>target_seq</td><td>PAM</td><td>alignment</td><td>distance</td><td>position</td><td>gene name</td><td>gene id</td></tr>
<tr><td>GU305914.1</td><td>62100</td><td>62122</td><td>-</td><td>0</td><td>ACCACCAGCAGCAGCAGCGA</td><td>TGG</td><td>||||||||[||||||||||||]PAM</td><td>NA</td><td>Intergenic</td><td>NA</td><td>NA</td></tr>
<tr><td>GU305914.1</td><td>27411</td><td>27433</td><td>+</td><td>3</td><td>AGCAGCAGCAGCAGCAGCGG</td><td>CGG</td><td>|-||-|||[|||||||||||-]PAM</td><td>NA</td><td>Intergenic</td><td>NA</td><td>NA</td></tr>
<tr><td>T29455</td><td>GCAGCAGCAGCGATGGCGACCGG</td><td>1000</td><td></td><td></td><td></td><td></td><td></td><td></td><td></td><td></td><td></td></tr>
<tr><td>Oligo fwd</td><td>CACCGCAGCAGCAGCGATGGCGAC</td><td></td><td></td><td></td><td></td><td></td><td></td><td></td><td></td><td></td><td></td></tr>
<tr><td>Oligo rev</td><td>AAACGTCGCCATCGCTGCTGCTGC</td><td></td><td></td><td></td><td></td><td></td><td></td><td></td><td></td><td></td><td></td></tr>
<tr><td>Chromosome</td><td>start</td><td>end</td><td>strand</td><td>MM</td><td>target_seq</td><td>PAM</td><td>alignment</td><td>distance</td><td>position</td><td>gene name</td><td>gene id</td></tr>
<tr><td>GU305914.1</td><td>62093</td><td>62115</td><td>-</td><td>0</td><td>GCAGCAGCAGCGATGGCGAC</td><td>CGG</td><td>||||||||[||||||||||||]PAM</td><td>NA</td><td>Intergenic</td><td>NA</td><td>NA</td></tr>
<tr><td>T29456</td><td>CAGCAGCAGCGATGGCGACCGGG</td><td>1000</td><td></td><td></td><td></td><td></td><td></td><td></td><td></td><td></td><td></td></tr>
<tr><td>Oligo fwd</td><td>CACCCAGCAGCAGCGATGGCGACC</td><td></td><td></td><td></td><td></td><td></td><td></td><td></td><td></td><td></td><td></td></tr>
<tr><td>Oligo rev</td><td>AAACGGTCGCCATCGCTGCTGCTG</td><td></td><td></td><td></td><td></td><td></td><td></td><td></td><td></td><td></td><td></td></tr>
<tr><td>Chromosome</td><td>start</td><td>end</td><td>strand</td><td>MM</td><td>target_seq</td><td>PAM</td><td>alignment</td><td>distance</td><td>position</td><td>gene name</td><td>gene id</td></tr>
<tr><td>GU305914.1</td><td>62092</td><td>62114</td><td>-</td><td>0</td><td>CAGCAGCAGCGATGGCGACC</td><td>GGG</td><td>||||||||[||||||||||||]PAM</td><td>NA</td><td>Intergenic</td><td>NA</td><td>NA</td></tr>
<tr><td>T29457</td><td>AGCAGCAGCGATGGCGACCGGGG</td><td>1000</td><td></td><td></td><td></td><td></td><td></td><td></td><td></td><td></td><td></td></tr>
<tr><td>Oligo fwd</td><td>CACCAGCAGCAGCGATGGCGACCG</td><td></td><td></td><td></td><td></td><td></td><td></td><td></td><td></td><td></td><td></td></tr>
<tr><td>Oligo rev</td><td>AAACCGGTCGCCATCGCTGCTGCT</td><td></td><td></td><td></td><td></td><td></td><td></td><td></td><td></td><td></td><td></td></tr>
<tr><td>Chromosome</td><td>start</td><td>end</td><td>strand</td><td>MM</td><td>target_seq</td><td>PAM</td><td>alignment</td><td>distance</td><td>position</td><td>gene name</td><td>gene id</td></tr>
<tr><td>GU305914.1</td><td>62091</td><td>62113</td><td>-</td><td>0</td><td>AGCAGCAGCGATGGCGACCG</td><td>GGG</td><td>||||||||[||||||||||||]PAM</td><td>NA</td><td>Intergenic</td><td>NA</td><td>NA</td></tr>
<tr><td>T29458</td><td>GCAGCAGCGATGGCGACCGGGGG</td><td>1000</td><td></td><td></td><td></td><td></td><td></td><td></td><td></td><td></td><td></td></tr>
<tr><td>Oligo fwd</td><td>CACCGCAGCAGCGATGGCGACCGG</td><td></td><td></td><td></td><td></td><td></td><td></td><td></td><td></td><td></td><td></td></tr>
<tr><td>Oligo rev</td><td>AAACCCGGTCGCCATCGCTGCTGC</td><td></td><td></td><td></td><td></td><td></td><td></td><td></td><td></td><td></td><td></td></tr>
<tr><td>Chromosome</td><td>start</td><td>end</td><td>strand</td><td>MM</td><td>target_seq</td><td>PAM</td><td>alignment</td><td>distance</td><td>position</td><td>gene name</td><td>gene id</td></tr>
<tr><td>GU305914.1</td><td>62090</td><td>62112</td><td>-</td><td>0</td><td>GCAGCAGCGATGGCGACCGG</td><td>GGG</td><td>||||||||[||||||||||||]PAM</td><td>NA</td><td>Intergenic</td><td>NA</td><td>NA</td></tr>
<tr><td>T29459</td><td>TTCGTGAGCGACGACGTCCTCGG</td><td>1000</td><td></td><td></td><td></td><td></td><td></td><td></td><td></td><td></td><td></td></tr>
<tr><td>Oligo fwd</td><td>CACCTTCGTGAGCGACGACGTCCT</td><td></td><td></td><td></td><td></td><td></td><td></td><td></td><td></td><td></td><td></td></tr>
<tr><td>Oligo rev</td><td>AAACAGGACGTCGTCGCTCACGAA</td><td></td><td></td><td></td><td></td><td></td><td></td><td></td><td></td><td></td><td></td></tr>
<tr><td>Chromosome</td><td>start</td><td>end</td><td>strand</td><td>MM</td><td>target_seq</td><td>PAM</td><td>alignment</td><td>distance</td><td>position</td><td>gene name</td><td>gene id</td></tr>
</table>

| | | | | | | | | | | | |
|---|---|---|---|---|---|---|---|---|---|---|---|
| GU305914.1 | 62039 | 62061 | - | 0 | TTCGTGAGCGACGACGTCCT | CGG | \|\|\|\|\|\|\|\|[\|\|\|\|\|\|\|\|\|\|\|\|]PAM | NA | Intergenic | NA | NA |
| T29460 | TTCGAAGATCTGCGCATCGCCGG | 1000 | | | | | | | | | |
| Oligo fwd | CACCTTCGAAGATCTGCGCATCGC | | | | | | | | | | |
| Oligo rev | AAACGCGATGCGCAGATCTTCGAA | | | | | | | | | | |
| Chromosome | start | end | strand | MM | target_seq | PAM | alignment | distance | position | gene name | gene id |
| GU305914.1 | 62012 | 62034 | - | 0 | TTCGAAGATCTGCGCATCGC | CGG | \|\|\|\|\|\|\|\|[\|\|\|\|\|\|\|\|\|\|\|\|]PAM | NA | Intergenic | NA | NA |
| T29461 | TCGAAGATCTGCGCATCGCCGGG | 1000 | | | | | | | | | |
| Oligo fwd | CACCTCGAAGATCTGCGCATCGCC | | | | | | | | | | |
| Oligo rev | AAACGGCGATGCGCAGATCTTCGA | | | | | | | | | | |
| Chromosome | start | end | strand | MM | target_seq | PAM | alignment | distance | position | gene name | gene id |
| GU305914.1 | 62011 | 62033 | - | 0 | TCGAAGATCTGCGCATCGCC | GGG | \|\|\|\|\|\|\|\|[\|\|\|\|\|\|\|\|\|\|\|\|]PAM | NA | Intergenic | NA | NA |
| T29462 | GCATCGCCGGGCTCCCCGTCAGG | 1000 | | | | | | | | | |
| Oligo fwd | CACCGCATCGCCGGGCTCCCCGTC | | | | | | | | | | |
| Oligo rev | AAACGACGGGGAGCCCGGCGATGC | | | | | | | | | | |
| Chromosome | start | end | strand | MM | target_seq | PAM | alignment | distance | position | gene name | gene id |
| GU305914.1 | 61999 | 62021 | - | 0 | GCATCGCCGGGCTCCCCGTC | AGG | \|\|\|\|\|\|\|\|[\|\|\|\|\|\|\|\|\|\|\|\|]PAM | NA | Intergenic | NA | NA |
| T29463 | CATCGCCGGGCTCCCCGTCAGGG | 1000 | | | | | | | | | |
| Oligo fwd | CACCCATCGCCGGGCTCCCCGTCA | | | | | | | | | | |
| Oligo rev | AAACTGACGGGGAGCCCGGCGATG | | | | | | | | | | |
| Chromosome | start | end | strand | MM | target_seq | PAM | alignment | distance | position | gene name | gene id |
| GU305914.1 | 61998 | 62020 | - | 0 | CATCGCCGGGCTCCCCGTCA | GGG | \|\|\|\|\|\|\|\|[\|\|\|\|\|\|\|\|\|\|\|\|]PAM | NA | Intergenic | NA | NA |
| T29464 | CCCCCCGACCCCGACGATGACGG | 1000 | | | | | | | | | |
| Oligo fwd | CACCCCCCCCGACCCCGACGATGA | | | | | | | | | | |
| Oligo rev | AAACTCATCGTCGGGGTCGGGGGG | | | | | | | | | | |
| Chromosome | start | end | strand | MM | target_seq | PAM | alignment | distance | position | gene name | gene id |
| GU305914.1 | 61965 | 61987 | - | 0 | CCCCCCGACCCCGACGATGA | CGG | \|\|\|\|\|\|\|\|[\|\|\|\|\|\|\|\|\|\|\|\|]PAM | NA | Intergenic | NA | NA |
| GU305914.1 | 146798 | 146820 | - | 4 | ACCCGCGACGCCGACGATGT | CGG | -\|\|\|-\|\|\|[\|-\|\|\|\|\|\|\|\|\|-]PAM | NA | Intergenic | NA | NA |
| T29465 | TGACGGTCATTCCCGTCAGCAGG | 1000 | | | | | | | | | |
| Oligo fwd | CACCTGACGGTCATTCCCGTCAGC | | | | | | | | | | |
| Oligo rev | AAACGCTGACGGGAATGACCGTCA | | | | | | | | | | |
| Chromosome | start | end | strand | MM | target_seq | PAM | alignment | distance | position | gene name | gene id |
| GU305914.1 | 61948 | 61970 | - | 0 | TGACGGTCATTCCCGTCAGC | AGG | \|\|\|\|\|\|\|\|[\|\|\|\|\|\|\|\|\|\|\|\|]PAM | NA | Intergenic | NA | NA |
| T29466 | CGTCAGCAGGCCCTTCCGCGCGG | 1000 | | | | | | | | | |
| Oligo fwd | CACCCGTCAGCAGGCCCTTCCGCG | | | | | | | | | | |
| Oligo rev | AAACCGCGGAAGGGCCTGCTGACG | | | | | | | | | | |
| Chromosome | start | end | strand | MM | target_seq | PAM | alignment | distance | position | gene name | gene id |
| GU305914.1 | 61935 | 61957 | - | 0 | CGTCAGCAGGCCCTTCCGCG | CGG | \|\|\|\|\|\|\|\|[\|\|\|\|\|\|\|\|\|\|\|\|]PAM | NA | Intergenic | NA | NA |
| T29467 | GTCAGCAGGCCCTTCCGCGCGGG | 1000 | | | | | | | | | |
| Oligo fwd | CACCGTCAGCAGGCCCTTCCGCGC | | | | | | | | | | |
| Oligo rev | AAACGCGCGGAAGGGCCTGCTGAC | | | | | | | | | | |
| Chromosome | start | end | strand | MM | target_seq | PAM | alignment | distance | position | gene name | gene id |
| GU305914.1 | 61934 | 61956 | - | 0 | GTCAGCAGGCCCTTCCGCGC | GGG | \|\|\|\|\|\|\|\|[\|\|\|\|\|\|\|\|\|\|\|\|]PAM | NA | Intergenic | NA | NA |
| T29468 | TCAGCAGGCCCTTCCGCGCGGGG | 1000 | | | | | | | | | |
| Oligo fwd | CACCTCAGCAGGCCCTTCCGCGCG | | | | | | | | | | |
| Oligo rev | AAACCGCGCGGAAGGGCCTGCTGA | | | | | | | | | | |
| Chromosome | start | end | strand | MM | target_seq | PAM | alignment | distance | position | gene name | gene id |
| GU305914.1 | 61933 | 61955 | - | 0 | TCAGCAGGCCCTTCCGCGCG | GGG | \|\|\|\|\|\|\|\|[\|\|\|\|\|\|\|\|\|\|\|\|]PAM | NA | Intergenic | NA | NA |
| T29469 | TTCCGCGCGGGGCGCGACAGCGG | 1000 | | | | | | | | | |
| Oligo fwd | CACCTTCCGCGCGGGGCGCGACAG | | | | | | | | | | |
| Oligo rev | AAACCTGTCGCGCCCCGCGCGGAA | | | | | | | | | | |
| Chromosome | start | end | strand | MM | target_seq | PAM | alignment | distance | position | gene name | gene id |
| GU305914.1 | 61922 | 61944 | - | 0 | TTCCGCGCGGGGCGCGACAG | CGG | \|\|\|\|\|\|\|\|[\|\|\|\|\|\|\|\|\|\|\|\|]PAM | NA | Intergenic | NA | NA |
| GU305914.1 | 188079 | 188101 | - | 4 | ATCGGCGGGGGGCGCGATAG | GGG | -\|\|-\|\|\|-[\|\|\|\|\|\|\|\|\|-\|\|]PAM | NA | Intergenic | NA | NA |
| T29470 | TCCGCGCGGGGCGCGACAGCGGG | 1000 | | | | | | | | | |
| Oligo fwd | CACCTCCGCGCGGGGCGCGACAGC | | | | | | | | | | |
| Oligo rev | AAACGCTGTCGCGCCCCGCGCGGA | | | | | | | | | | |
| Chromosome | start | end | strand | MM | target_seq | PAM | alignment | distance | position | gene name | gene id |
| GU305914.1 | 61921 | 61943 | - | 0 | TCCGCGCGGGGCGCGACAGC | GGG | \|\|\|\|\|\|\|\|[\|\|\|\|\|\|\|\|\|\|\|\|]PAM | NA | Intergenic | NA | NA |
| GU305914.1 | 188078 | 188100 | - | 4 | TCGGCGGGGGGCGCGATAGG | GGG | \|\|-\|\|\|-\|[\|\|\|\|\|\|\|\|-\|\|-]PAM | NA | Intergenic | NA | NA |
| T29471 | TCCGTGCGCTGCTACTGCCACGG | 1000 | | | | | | | | | |
| Oligo fwd | CACCTCCGTGCGCTGCTACTGCCA | | | | | | | | | | |
| Oligo rev | AAACTGGCAGTAGCAGCGCACGGA | | | | | | | | | | |
| Chromosome | start | end | strand | MM | target_seq | PAM | alignment | distance | position | gene name | gene id |
| GU305914.1 | 61868 | 61890 | - | 0 | TCCGTGCGCTGCTACTGCCA | CGG | \|\|\|\|\|\|\|\|[\|\|\|\|\|\|\|\|\|\|\|\|]PAM | NA | Intergenic | NA | NA |
| T29472 | TGCTACTGCCACGGCATCCTCGG | 1000 | | | | | | | | | |
| Oligo fwd | CACCTGCTACTGCCACGGCATCCT | | | | | | | | | | |
| Oligo rev | AAACAGGATGCCGTGGCAGTAGCA | | | | | | | | | | |
| Chromosome | start | end | strand | MM | target_seq | PAM | alignment | distance | position | gene name | gene id |
| GU305914.1 | 61859 | 61881 | - | 0 | TGCTACTGCCACGGCATCCT | CGG | \|\|\|\|\|\|\|\|[\|\|\|\|\|\|\|\|\|\|\|\|]PAM | NA | Intergenic | NA | NA |
| T29473 | GAAGTGCTCCGAACCCGCCAAGG | 1000 | | | | | | | | | |
| Oligo fwd | CACCGAAGTGCTCCGAACCCGCCA | | | | | | | | | | |
| Oligo rev | AAACTGGCGGGTTCGGAGCACTTC | | | | | | | | | | |
| Chromosome | start | end | strand | MM | target_seq | PAM | alignment | distance | position | gene name | gene id |
| GU305914.1 | 61806 | 61828 | - | 0 | GAAGTGCTCCGAACCCGCCA | AGG | \|\|\|\|\|\|\|\|[\|\|\|\|\|\|\|\|\|\|\|\|]PAM | NA | Intergenic | NA | NA |

| | | | | | | | | | | | | |
|---|---|---|---|---|---|---|---|---|---|---|---|---|
| T29474 | CGAACCCGCCAAGGAGCGCATGG | | 1000 | | | | | | | | | |
| Oligo fwd | CACCCGAACCCGCCAAGGAGCGCA | | | | | | | | | | | |
| Oligo rev | AAACTGCGCTCCTTGGCGGGTTCG | | | | | | | | | | | |
| Chromosome | start | | end | strand | MM | target_seq | PAM | alignment | distance | position | gene name | gene id |
| GU305914.1 | | 61797 | 61819 | - | 0 | CGAACCCGCCAAGGAGCGCA | TGG | \|\|\|\|\|\|\|\|[\|\|\|\|\|\|\|\|\|\|\|\|]PAM | NA | Intergenic | NA | NA |
| T29475 | TCGCATCCGCCGCTACGCCGCGG | | 1000 | | | | | | | | | |
| Oligo fwd | CACCTCGCATCCGCCGCTACGCCG | | | | | | | | | | | |
| Oligo rev | AAACCGGCGTAGCGGCGGATGCGA | | | | | | | | | | | |
| Chromosome | start | | end | strand | MM | target_seq | PAM | alignment | distance | position | gene name | gene id |
| GU305914.1 | | 61773 | 61795 | - | 0 | TCGCATCCGCCGCTACGCCG | CGG | \|\|\|\|\|\|\|\|[\|\|\|\|\|\|\|\|\|\|\|\|]PAM | NA | Intergenic | NA | NA |
| GU305914.1 | | 52115 | 52137 | - | 4 | TCGCGAACGCCGCTTCGCCG | AGG | \|\|\|\|---\|[\|\|\|\|\|\|-\|\|\|\|\|]PAM | NA | Intergenic | NA | NA |
| T29476 | GCCGCGGACGTGACGCGCTGCGG | | 1000 | | | | | | | | | |
| Oligo fwd | CACCGCCGCGGACGTGACGCGCTG | | | | | | | | | | | |
| Oligo rev | AAACCAGCGCGTCACGTCCGCGGC | | | | | | | | | | | |
| Chromosome | start | | end | strand | MM | target_seq | PAM | alignment | distance | position | gene name | gene id |
| GU305914.1 | | 61757 | 61779 | - | 0 | GCCGCGGACGTGACGCGCTG | CGG | \|\|\|\|\|\|\|\|[\|\|\|\|\|\|\|\|\|\|\|\|]PAM | NA | Intergenic | NA | NA |
| T29477 | GACCGACCCGTCGTGCGACCTGG | | 1000 | | | | | | | | | |
| Oligo fwd | CACCGACCGACCCGTCGTGCGACC | | | | | | | | | | | |
| Oligo rev | AAACGGTCGCACGACGGGTCGGTC | | | | | | | | | | | |
| Chromosome | start | | end | strand | MM | target_seq | PAM | alignment | distance | position | gene name | gene id |
| GU305914.1 | | 61683 | 61705 | - | 0 | GACCGACCCGTCGTGCGACC | TGG | \|\|\|\|\|\|\|\|[\|\|\|\|\|\|\|\|\|\|\|\|]PAM | NA | Intergenic | NA | NA |
| T29478 | CGCCGAATGTTACGTCGAGACGG | | 1000 | | | | | | | | | |
| Oligo fwd | CACCCGCCGAATGTTACGTCGAGA | | | | | | | | | | | |
| Oligo rev | AAACTCTCGACGTAACATTCGGCG | | | | | | | | | | | |
| Chromosome | start | | end | strand | MM | target_seq | PAM | alignment | distance | position | gene name | gene id |
| GU305914.1 | | 61656 | 61678 | - | 0 | CGCCGAATGTTACGTCGAGA | CGG | \|\|\|\|\|\|\|\|[\|\|\|\|\|\|\|\|\|\|\|\|]PAM | NA | Intergenic | NA | NA |
| T29479 | ACGTCGAGACGGCGTCCAAGAGG | | 1000 | | | | | | | | | |
| Oligo fwd | CACCACGTCGAGACGGCGTCCAAG | | | | | | | | | | | |
| Oligo rev | AAACCTTGGACGCCGTCTCGACGT | | | | | | | | | | | |
| Chromosome | start | | end | strand | MM | target_seq | PAM | alignment | distance | position | gene name | gene id |
| GU305914.1 | | 61645 | 61667 | - | 0 | ACGTCGAGACGGCGTCCAAG | AGG | \|\|\|\|\|\|\|\|[\|\|\|\|\|\|\|\|\|\|\|\|]PAM | NA | Intergenic | NA | NA |
| T29480 | GAGGAACGCCGTCCCCCTCTCGG | | 1000 | | | | | | | | | |
| Oligo fwd | CACCGAGGAACGCCGTCCCCCTCT | | | | | | | | | | | |
| Oligo rev | AAACAGAGGGGGACGGCGTTCCTC | | | | | | | | | | | |
| Chromosome | start | | end | strand | MM | target_seq | PAM | alignment | distance | position | gene name | gene id |
| GU305914.1 | | 61626 | 61648 | - | 0 | GAGGAACGCCGTCCCCCTCT | CGG | \|\|\|\|\|\|\|\|[\|\|\|\|\|\|\|\|\|\|\|\|]PAM | NA | Intergenic | NA | NA |
| T29481 | ACCGCGACTGCGATCGCCTGCGG | | 1000 | | | | | | | | | |
| Oligo fwd | CACCACCGCGACTGCGATCGCCTG | | | | | | | | | | | |
| Oligo rev | AAACCAGGCGATCGCAGTCGCGGT | | | | | | | | | | | |
| Chromosome | start | | end | strand | MM | target_seq | PAM | alignment | distance | position | gene name | gene id |
| GU305914.1 | | 61597 | 61619 | - | 0 | ACCGCGACTGCGATCGCCTG | CGG | \|\|\|\|\|\|\|\|[\|\|\|\|\|\|\|\|\|\|\|\|]PAM | NA | Intergenic | NA | NA |
| T29482 | CCGCGACTGCGATCGCCTGCGGG | | 1000 | | | | | | | | | |
| Oligo fwd | CACCCCGCGACTGCGATCGCCTGC | | | | | | | | | | | |
| Oligo rev | AAACGCAGGCGATCGCAGTCGCGG | | | | | | | | | | | |
| Chromosome | start | | end | strand | MM | target_seq | PAM | alignment | distance | position | gene name | gene id |
| GU305914.1 | | 61596 | 61618 | - | 0 | CCGCGACTGCGATCGCCTGC | GGG | \|\|\|\|\|\|\|\|[\|\|\|\|\|\|\|\|\|\|\|\|]PAM | NA | Intergenic | NA | NA |
| T29483 | GCCGCGTTCTTCAAGACCTACGG | | 1000 | | | | | | | | | |
| Oligo fwd | CACCGCCGCGTTCTTCAAGACCTA | | | | | | | | | | | |
| Oligo rev | AAACTAGGTCTTGAAGAACGCGGC | | | | | | | | | | | |
| Chromosome | start | | end | strand | MM | target_seq | PAM | alignment | distance | position | gene name | gene id |
| GU305914.1 | | 61565 | 61587 | - | 0 | GCCGCGTTCTTCAAGACCTA | CGG | \|\|\|\|\|\|\|\|[\|\|\|\|\|\|\|\|\|\|\|\|]PAM | NA | Intergenic | NA | NA |
| GU305914.1 | | 72518 | 72540 | + | 4 | GACTCCTTCTTCAAGATCTA | CGG | \|-\|-\|-\|\|[\|\|\|\|\|\|\|\|-\|\|\|]PAM | NA | Intergenic | NA | NA |
| T29484 | GTTCTTCAAGACCTACGGCATGG | | 1000 | | | | | | | | | |
| Oligo fwd | CACCGTTCTTCAAGACCTACGGCA | | | | | | | | | | | |
| Oligo rev | AAACTGCCGTAGGTCTTGAAGAAC | | | | | | | | | | | |
| Chromosome | start | | end | strand | MM | target_seq | PAM | alignment | distance | position | gene name | gene id |
| GU305914.1 | | 61560 | 61582 | - | 0 | GTTCTTCAAGACCTACGGCA | TGG | \|\|\|\|\|\|\|\|[\|\|\|\|\|\|\|\|\|\|\|\|]PAM | NA | Intergenic | NA | NA |
| T29485 | CTTCAAGACCTACGGCATGGTGG | | 1000 | | | | | | | | | |
| Oligo fwd | CACCCTTCAAGACCTACGGCATGG | | | | | | | | | | | |
| Oligo rev | AAACCCATGCCGTAGGTCTTGAAG | | | | | | | | | | | |
| Chromosome | start | | end | strand | MM | target_seq | PAM | alignment | distance | position | gene name | gene id |
| GU305914.1 | | 61557 | 61579 | - | 0 | CTTCAAGACCTACGGCATGG | TGG | \|\|\|\|\|\|\|\|[\|\|\|\|\|\|\|\|\|\|\|\|]PAM | NA | Intergenic | NA | NA |
| T29486 | CAAGACCTACGGCATGGTGGTGG | | 1000 | | | | | | | | | |
| Oligo fwd | CACCCAAGACCTACGGCATGGTGG | | | | | | | | | | | |
| Oligo rev | AAACCCACCATGCCGTAGGTCTTG | | | | | | | | | | | |
| Chromosome | start | | end | strand | MM | target_seq | PAM | alignment | distance | position | gene name | gene id |
| GU305914.1 | | 61554 | 61576 | - | 0 | CAAGACCTACGGCATGGTGG | TGG | \|\|\|\|\|\|\|\|[\|\|\|\|\|\|\|\|\|\|\|\|]PAM | NA | Intergenic | NA | NA |
| GU305914.1 | | 148005 | 148027 | + | 3 | CAACACCGTCGGCATGGTGG | TGG | \|\|\|-\|\|\|-[-\|\|\|\|\|\|\|\|\|\|\|]PAM | NA | Intergenic | NA | NA |
| T29487 | CGGCATGGTGGTGGACGCCATGG | | 1000 | | | | | | | | | |
| Oligo fwd | CACCCGGCATGGTGGTGGACGCCA | | | | | | | | | | | |
| Oligo rev | AAACTGGCGTCCACCACCATGCCG | | | | | | | | | | | |
| Chromosome | start | | end | strand | MM | target_seq | PAM | alignment | distance | position | gene name | gene id |
| GU305914.1 | | 61545 | 61567 | - | 0 | CGGCATGGTGGTGGACGCCA | TGG | \|\|\|\|\|\|\|\|[\|\|\|\|\|\|\|\|\|\|\|\|]PAM | NA | Intergenic | NA | NA |
| T29488 | CGCCATGGCGCAGCGCATCACGG | | 1000 | | | | | | | | | |
| Oligo fwd | CACCCGCCATGGCGCAGCGCATCA | | | | | | | | | | | |

| | | | | | | | | | | | |
|---|---|---|---|---|---|---|---|---|---|---|---|
| Oligo rev | AAACTGATGCGCTGCGCCATGGCG | | | | | | | | | | |
| Chromosome | start | end | strand | MM | target_seq | PAM | alignment | distance | position | gene name | gene id |
| GU305914.1 | 61530 | 61552 | - | 0 | CGCCATGGCGCAGCGCATCA | CGG | \|\|\|\|\|\|\|\|[\|\|\|\|\|\|\|\|\|\|\|\|]PAM | NA | Intergenic | NA | NA |
| T29489 | TGGCGCAGCGCATCACGGAGCGG | 1000 | | | | | | | | | |
| Oligo fwd | CACCTGGCGCAGCGCATCACGGAG | | | | | | | | | | |
| Oligo rev | AAACCTCCGTGATGCGCTGCGCCA | | | | | | | | | | |
| Chromosome | start | end | strand | MM | target_seq | PAM | alignment | distance | position | gene name | gene id |
| GU305914.1 | 61525 | 61547 | - | 0 | TGGCGCAGCGCATCACGGAG | CGG | \|\|\|\|\|\|\|\|[\|\|\|\|\|\|\|\|\|\|\|\|]PAM | NA | Intergenic | NA | NA |
| T29490 | GGCGCAGCGCATCACGGAGCGGG | 1000 | | | | | | | | | |
| Oligo fwd | CACCGGCGCAGCGCATCACGGAGC | | | | | | | | | | |
| Oligo rev | AAACGCTCCGTGATGCGCTGCGCC | | | | | | | | | | |
| Chromosome | start | end | strand | MM | target_seq | PAM | alignment | distance | position | gene name | gene id |
| GU305914.1 | 61524 | 61546 | - | 0 | GGCGCAGCGCATCACGGAGC | GGG | \|\|\|\|\|\|\|\|[\|\|\|\|\|\|\|\|\|\|\|\|]PAM | NA | Intergenic | NA | NA |
| T29491 | CAGCGCATCACGGAGCGGGTCGG | 1000 | | | | | | | | | |
| Oligo fwd | CACCCAGCGCATCACGGAGCGGGT | | | | | | | | | | |
| Oligo rev | AAACACCCGCTCCGTGATGCGCTG | | | | | | | | | | |
| Chromosome | start | end | strand | MM | target_seq | PAM | alignment | distance | position | gene name | gene id |
| GU305914.1 | 61520 | 61542 | - | 0 | CAGCGCATCACGGAGCGGGT | CGG | \|\|\|\|\|\|\|\|[\|\|\|\|\|\|\|\|\|\|\|\|]PAM | NA | Intergenic | NA | NA |
| T29492 | AGCGCATCACGGAGCGGGTCGGG | 1000 | | | | | | | | | |
| Oligo fwd | CACCAGCGCATCACGGAGCGGGTC | | | | | | | | | | |
| Oligo rev | AAACGACCCGCTCCGTGATGCGCT | | | | | | | | | | |
| Chromosome | start | end | strand | MM | target_seq | PAM | alignment | distance | position | gene name | gene id |
| GU305914.1 | 61519 | 61541 | - | 0 | AGCGCATCACGGAGCGGGTC | GGG | \|\|\|\|\|\|\|\|[\|\|\|\|\|\|\|\|\|\|\|\|]PAM | NA | Intergenic | NA | NA |
| T29493 | CATCACGGAGCGGGTCGGGCCGG | 1000 | | | | | | | | | |
| Oligo fwd | CACCCATCACGGAGCGGGTCGGGC | | | | | | | | | | |
| Oligo rev | AAACGCCCGACCCGCTCCGTGATG | | | | | | | | | | |
| Chromosome | start | end | strand | MM | target_seq | PAM | alignment | distance | position | gene name | gene id |
| GU305914.1 | 61515 | 61537 | - | 0 | CATCACGGAGCGGGTCGGGC | CGG | \|\|\|\|\|\|\|\|[\|\|\|\|\|\|\|\|\|\|\|\|]PAM | NA | Intergenic | NA | NA |
| T29494 | GGAGCGGGTCGGGCCGGCCCTGG | 1000 | | | | | | | | | |
| Oligo fwd | CACCGGAGCGGGTCGGGCCGGCCC | | | | | | | | | | |
| Oligo rev | AAACGGGCCGGCCCGACCCGCTCC | | | | | | | | | | |
| Chromosome | start | end | strand | MM | target_seq | PAM | alignment | distance | position | gene name | gene id |
| GU305914.1 | 61509 | 61531 | - | 0 | GGAGCGGGTCGGGCCGGCCC | TGG | \|\|\|\|\|\|\|\|[\|\|\|\|\|\|\|\|\|\|\|\|]PAM | NA | Intergenic | NA | NA |
| T29495 | GAGCGGGTCGGGCCGGCCCTGGG | 1000 | | | | | | | | | |
| Oligo fwd | CACCGAGCGGGTCGGGCCGGCCCT | | | | | | | | | | |
| Oligo rev | AAACAGGGCCGGCCCGACCCGCTC | | | | | | | | | | |
| Chromosome | start | end | strand | MM | target_seq | PAM | alignment | distance | position | gene name | gene id |
| GU305914.1 | 61508 | 61530 | - | 0 | GAGCGGGTCGGGCCGGCCCT | GGG | \|\|\|\|\|\|\|\|[\|\|\|\|\|\|\|\|\|\|\|\|]PAM | NA | Intergenic | NA | NA |
| T29496 | GGGTCGGGCCGGCCCTGGGTAGG | 1000 | | | | | | | | | |
| Oligo fwd | CACCGGGTCGGGCCGGCCCTGGGT | | | | | | | | | | |
| Oligo rev | AAACACCCAGGGCCGGCCCGACCC | | | | | | | | | | |
| Chromosome | start | end | strand | MM | target_seq | PAM | alignment | distance | position | gene name | gene id |
| GU305914.1 | 61504 | 61526 | - | 0 | GGGTCGGGCCGGCCCTGGGT | AGG | \|\|\|\|\|\|\|\|[\|\|\|\|\|\|\|\|\|\|\|\|]PAM | NA | Intergenic | NA | NA |
| T29497 | GGTCGGGCCGGCCCTGGGTAGGG | 1000 | | | | | | | | | |
| Oligo fwd | CACCGGTCGGGCCGGCCCTGGGTA | | | | | | | | | | |
| Oligo rev | AAACTACCCAGGGCCGGCCCGACC | | | | | | | | | | |
| Chromosome | start | end | strand | MM | target_seq | PAM | alignment | distance | position | gene name | gene id |
| GU305914.1 | 61503 | 61525 | - | 0 | GGTCGGGCCGGCCCTGGGTA | GGG | \|\|\|\|\|\|\|\|[\|\|\|\|\|\|\|\|\|\|\|\|]PAM | NA | Intergenic | NA | NA |
| T29498 | GTCGGGCCGGCCCTGGGTAGGGG | 1000 | | | | | | | | | |
| Oligo fwd | CACCGTCGGGCCGGCCCTGGGTAG | | | | | | | | | | |
| Oligo rev | AAACCTACCCAGGGCCGGCCCGAC | | | | | | | | | | |
| Chromosome | start | end | strand | MM | target_seq | PAM | alignment | distance | position | gene name | gene id |
| GU305914.1 | 61502 | 61524 | - | 0 | GTCGGGCCGGCCCTGGGTAG | GGG | \|\|\|\|\|\|\|\|[\|\|\|\|\|\|\|\|\|\|\|\|]PAM | NA | Intergenic | NA | NA |
| T29499 | CTACTCGACCGTCGTCATGATGG | 1000 | | | | | | | | | |
| Oligo fwd | CACCCTACTCGACCGTCGTCATGA | | | | | | | | | | |
| Oligo rev | AAACTCATGACGACGGTCGAGTAG | | | | | | | | | | |
| Chromosome | start | end | strand | MM | target_seq | PAM | alignment | distance | position | gene name | gene id |
| GU305914.1 | 61476 | 61498 | - | 0 | CTACTCGACCGTCGTCATGA | TGG | \|\|\|\|\|\|\|\|[\|\|\|\|\|\|\|\|\|\|\|\|]PAM | NA | Intergenic | NA | NA |
| T29500 | GTCGTCATGATGGATCGCTGCGG | 1000 | | | | | | | | | |
| Oligo fwd | CACCGTCGTCATGATGGATCGCTG | | | | | | | | | | |
| Oligo rev | AAACCAGCGATCCATCATGACGAC | | | | | | | | | | |
| Chromosome | start | end | strand | MM | target_seq | PAM | alignment | distance | position | gene name | gene id |
| GU305914.1 | 61466 | 61488 | - | 0 | GTCGTCATGATGGATCGCTG | CGG | \|\|\|\|\|\|\|\|[\|\|\|\|\|\|\|\|\|\|\|\|]PAM | NA | Intergenic | NA | NA |
| T29501 | TCGCTGCGGAAACAGCTTCCAGG | 1000 | | | | | | | | | |
| Oligo fwd | CACCTCGCTGCGGAAACAGCTTCC | | | | | | | | | | |
| Oligo rev | AAACGGAAGCTGTTTCCGCAGCGA | | | | | | | | | | |
| Chromosome | start | end | strand | MM | target_seq | PAM | alignment | distance | position | gene name | gene id |
| GU305914.1 | 61452 | 61474 | - | 0 | TCGCTGCGGAAACAGCTTCC | AGG | \|\|\|\|\|\|\|\|[\|\|\|\|\|\|\|\|\|\|\|\|]PAM | NA | Intergenic | NA | NA |
| T29502 | CGCTGCGGAAACAGCTTCCAGGG | 1000 | | | | | | | | | |
| Oligo fwd | CACCCGCTGCGGAAACAGCTTCCA | | | | | | | | | | |
| Oligo rev | AAACTGGAAGCTGTTTCCGCAGCG | | | | | | | | | | |
| Chromosome | start | end | strand | MM | target_seq | PAM | alignment | distance | position | gene name | gene id |
| GU305914.1 | 61451 | 61473 | - | 0 | CGCTGCGGAAACAGCTTCCA | GGG | \|\|\|\|\|\|\|\|[\|\|\|\|\|\|\|\|\|\|\|\|]PAM | NA | Intergenic | NA | NA |
| T29503 | AAACAGCTTCCAGGGACGCGAGG | 1000 | | | | | | | | | |

Oligo fwd CACCAAACAGCTTCCAGGGACGCG
Oligo rev AAACCGCGTCCCTGGAAGCTGTTT
Chromosome start end strand MM target_seq PAM alignment distance position gene name gene id
GU305914.1 61443 61465 - 0 AAACAGCTTCCAGGGACGCG AGG ||||||||[||||||||||||]PAM NA Intergenic NA NA

T29504 CCCCCATCTCCGTCTTCGCCCGG 1000
Oligo fwd CACCCCCCCATCTCCGTCTTCGCC
Oligo rev AAACGGCGAAGACGGAGATGGGGG
Chromosome start end strand MM target_seq PAM alignment distance position gene name gene id
GU305914.1 61417 61439 - 0 CCCCCATCTCCGTCTTCGCC CGG ||||||||[||||||||||||]PAM NA Intergenic NA NA

T29505 CCCCATCTCCGTCTTCGCCCGGG 1000
Oligo fwd CACCCCCCATCTCCGTCTTCGCCC
Oligo rev AAACGGGCGAAGACGGAGATGGGG
Chromosome start end strand MM target_seq PAM alignment distance position gene name gene id
GU305914.1 61416 61438 - 0 CCCCATCTCCGTCTTCGCCC GGG ||||||||[||||||||||||]PAM NA Intergenic NA NA

T29506 CGTCTTCGCCCGGGTCGCCGCGG 1000
Oligo fwd CACCCGTCTTCGCCCGGGTCGCCG
Oligo rev AAACCGGCGACCCGGGCGAAGACG
Chromosome start end strand MM target_seq PAM alignment distance position gene name gene id
GU305914.1 61407 61429 - 0 CGTCTTCGCCCGGGTCGCCG CGG ||||||||[||||||||||||]PAM NA Intergenic NA NA

T29507 GGCTCTCGCCGTCGAGTGCGAGG 1000
Oligo fwd CACCGGCTCTCGCCGTCGAGTGCG
Oligo rev AAACCGCACTCGACGGCGAGAGCC
Chromosome start end strand MM target_seq PAM alignment distance position gene name gene id
GU305914.1 61386 61408 - 0 GGCTCTCGCCGTCGAGTGCG AGG ||||||||[||||||||||||]PAM NA Intergenic NA NA
GU305914.1 206825 206847 - 4 CGCTATCGCCGTCTGGTGCG TGG -|||-|||[|||||--|||||]PAM NA Intergenic NA NA

T29508 GCCGTCGAGTGCGAGGTCGACGG 1000
Oligo fwd CACCGCCGTCGAGTGCGAGGTCGA
Oligo rev AAACTCGACCTCGCACTCGACGGC
Chromosome start end strand MM target_seq PAM alignment distance position gene name gene id
GU305914.1 61379 61401 - 0 GCCGTCGAGTGCGAGGTCGA CGG ||||||||[||||||||||||]PAM NA Intergenic NA NA

T29509 CCGTCGAGTGCGAGGTCGACGGG 1000
Oligo fwd CACCCCGTCGAGTGCGAGGTCGAC
Oligo rev AAACGTCGACCTCGCACTCGACGG
Chromosome start end strand MM target_seq PAM alignment distance position gene name gene id
GU305914.1 61378 61400 - 0 CCGTCGAGTGCGAGGTCGAC GGG ||||||||[||||||||||||]PAM NA Intergenic NA NA

T29510 CGTCGAGTGCGAGGTCGACGGGG 1000
Oligo fwd CACCCGTCGAGTGCGAGGTCGACG
Oligo rev AAACCGTCGACCTCGCACTCGACG
Chromosome start end strand MM target_seq PAM alignment distance position gene name gene id
GU305914.1 61377 61399 - 0 CGTCGAGTGCGAGGTCGACG GGG ||||||||[||||||||||||]PAM NA Intergenic NA NA
GU305914.1 74048 74070 + 4 CGGCGAGGTCGAGGTCGTCG CGG ||-||||-[-||||||||-||]PAM NA Intergenic NA NA

T29511 GTCGAGTGCGAGGTCGACGGGGG 1000
Oligo fwd CACCGTCGAGTGCGAGGTCGACGG
Oligo rev AAACCCGTCGACCTCGCACTCGAC
Chromosome start end strand MM target_seq PAM alignment distance position gene name gene id
GU305914.1 61376 61398 - 0 GTCGAGTGCGAGGTCGACGG GGG ||||||||[||||||||||||]PAM NA Intergenic NA NA
GU305914.1 201151 201173 + 4 GTGGAGTTCGGCGTCGACGG GGG ||-||||-[||--||||||||]PAM NA Intergenic NA NA

T29512 CAAGCCCGTCGACGCCGCGCAGG 1000
Oligo fwd CACCCAAGCCCGTCGACGCCGCGC
Oligo rev AAACGCGCGGCGTCGACGGGCTTG
Chromosome start end strand MM target_seq PAM alignment distance position gene name gene id
GU305914.1 61329 61351 - 0 CAAGCCCGTCGACGCCGCGC AGG ||||||||[||||||||||||]PAM NA Intergenic NA NA

T29513 TCGCCATCATCCCCTCGCCGCGG 1000
Oligo fwd CACCTCGCCATCATCCCCTCGCCG
Oligo rev AAACCGGCGAGGGGATGATGGCGA
Chromosome start end strand MM target_seq PAM alignment distance position gene name gene id
GU305914.1 61273 61295 - 0 TCGCCATCATCCCCTCGCCG CGG ||||||||[||||||||||||]PAM NA Intergenic NA NA

T29514 CGCCATCATCCCCTCGCCGCGGG 1000
Oligo fwd CACCCGCCATCATCCCCTCGCCGC
Oligo rev AAACGCGGCGAGGGGATGATGGCG
Chromosome start end strand MM target_seq PAM alignment distance position gene name gene id
GU305914.1 61272 61294 - 0 CGCCATCATCCCCTCGCCGC GGG ||||||||[||||||||||||]PAM NA Intergenic NA NA

T29515 CATCCCCTCGCCGCGGGTCCTGG 1000
Oligo fwd CACCCATCCCCTCGCCGCGGGTCC
Oligo rev AAACGGACCCGCGGCGAGGGGATG
Chromosome start end strand MM target_seq PAM alignment distance position gene name gene id
GU305914.1 61266 61288 - 0 CATCCCCTCGCCGCGGGTCC TGG ||||||||[||||||||||||]PAM NA Intergenic NA NA

T29516 CCGCGGGTCCTGGCCTACGCCGG 1000
Oligo fwd CACCCCGCGGGTCCTGGCCTACGC
Oligo rev AAACGCGTAGGCCAGGACCCGCGG
Chromosome start end strand MM target_seq PAM alignment distance position gene name gene id
GU305914.1 61256 61278 - 0 CCGCGGGTCCTGGCCTACGC CGG ||||||||[||||||||||||]PAM NA Intergenic NA NA

T29517 CGCGGGTCCTGGCCTACGCCGGG 1000
Oligo fwd CACCCGCGGGTCCTGGCCTACGCC
Oligo rev AAACGGCGTAGGCCAGGACCCGCG

Chromosome start end strand MM target_seq PAM alignment distance position gene name gene id
GU305914.1 61255 61277 - 0 CGCGGGTCCTGGCCTACGCC GGG ||||||||[||||||||||||]PAM NA Intergenic NA NA

T29518 GTCCTGGCCTACGCCGGGTTCGG 1000
Oligo fwd CACCGTCCTGGCCTACGCCGGGTT
Oligo rev AAACAACCCGGCGTAGGCCAGGAC
Chromosome start end strand MM target_seq PAM alignment distance position gene name gene id
GU305914.1 61250 61272 - 0 GTCCTGGCCTACGCCGGGTT CGG ||||||||[||||||||||||]PAM NA Intergenic NA NA

T29519 CTGGCCTACGCCGGGTTCGGCGG 1000
Oligo fwd CACCCTGGCCTACGCCGGGTTCGG
Oligo rev AAACCCGAACCCGGCGTAGGCCAG
Chromosome start end strand MM target_seq PAM alignment distance position gene name gene id
GU305914.1 61247 61269 - 0 CTGGCCTACGCCGGGTTCGG CGG ||||||||[||||||||||||]PAM NA Intergenic NA NA

T29520 TTACCGCGCTCAGTTCTACTCGG 1000
Oligo fwd CACCTTACCGCGCTCAGTTCTACT
Oligo rev AAACAGTAGAACTGAGCGCGGTAA
Chromosome start end strand MM target_seq PAM alignment distance position gene name gene id
GU305914.1 61197 61219 - 0 TTACCGCGCTCAGTTCTACT CGG ||||||||[||||||||||||]PAM NA Intergenic NA NA

T29521 TCGGCCGAGAGCACGATCAACGG 1000
Oligo fwd CACCTCGGCCGAGAGCACGATCAA
Oligo rev AAACTTGATCGTGCTCTCGGCCGA
Chromosome start end strand MM target_seq PAM alignment distance position gene name gene id
GU305914.1 61178 61200 - 0 TCGGCCGAGAGCACGATCAA CGG ||||||||[||||||||||||]PAM NA Intergenic NA NA

T29522 CCTGCACGACATCTGCGACATGG 1000
Oligo fwd CACCCCTGCACGACATCTGCGACA
Oligo rev AAACTGTCGCAGATGTCGTGCAGG
Chromosome start end strand MM target_seq PAM alignment distance position gene name gene id
GU305914.1 61152 61174 - 0 CCTGCACGACATCTGCGACA TGG ||||||||[||||||||||||]PAM NA Intergenic NA NA

T29523 ATCTGCGACATGGTGACCAACGG 1000
Oligo fwd CACCATCTGCGACATGGTGACCAA
Oligo rev AAACTTGGTCACCATGTCGCAGAT
Chromosome start end strand MM target_seq PAM alignment distance position gene name gene id
GU305914.1 61142 61164 - 0 ATCTGCGACATGGTGACCAA CGG ||||||||[||||||||||||]PAM NA Intergenic NA NA

T29524 CATGGTGACCAACGGCCTCTCGG 1000
Oligo fwd CACCCATGGTGACCAACGGCCTCT
Oligo rev AAACAGAGGCCGTTGGTCACCATG
Chromosome start end strand MM target_seq PAM alignment distance position gene name gene id
GU305914.1 61134 61156 - 0 CATGGTGACCAACGGCCTCT CGG ||||||||[||||||||||||]PAM NA Intergenic NA NA

T29525 CCTCTCGGTGTCCGTCAGCGCGG 1000
Oligo fwd CACCCCTCTCGGTGTCCGTCAGCG
Oligo rev AAACCGCTGACGGACACCGAGAGG
Chromosome start end strand MM target_seq PAM alignment distance position gene name gene id
GU305914.1 61119 61141 - 0 CCTCTCGGTGTCCGTCAGCG CGG ||||||||[||||||||||||]PAM NA Intergenic NA NA

T29526 CTCGGTGTCCGTCAGCGCGGCGG 1000
Oligo fwd CACCCTCGGTGTCCGTCAGCGCGG
Oligo rev AAACCCGCGCTGACGGACACCGAG
Chromosome start end strand MM target_seq PAM alignment distance position gene name gene id
GU305914.1 61116 61138 - 0 CTCGGTGTCCGTCAGCGCGG CGG ||||||||[||||||||||||]PAM NA Intergenic NA NA

T29527 TCCGTCAGCGCGGCGGACCTCGG 1000
Oligo fwd CACCTCCGTCAGCGCGGCGGACCT
Oligo rev AAACAGGTCCGCCGCGCTGACGGA
Chromosome start end strand MM target_seq PAM alignment distance position gene name gene id
GU305914.1 61109 61131 - 0 TCCGTCAGCGCGGCGGACCT CGG ||||||||[||||||||||||]PAM NA Intergenic NA NA

T29528 GTCAGCGCGGCGGACCTCGGAGG 1000
Oligo fwd CACCGTCAGCGCGGCGGACCTCGG
Oligo rev AAACCCGAGGTCCGCCGCGCTGAC
Chromosome start end strand MM target_seq PAM alignment distance position gene name gene id
GU305914.1 61106 61128 - 0 GTCAGCGCGGCGGACCTCGG AGG ||||||||[||||||||||||]PAM NA Intergenic NA NA

T29529 GGACCTCGGAGGCGACATCGTGG 1000
Oligo fwd CACCGGACCTCGGAGGCGACATCG
Oligo rev AAACCGATGTCGCCTCCGAGGTCC
Chromosome start end strand MM target_seq PAM alignment distance position gene name gene id
GU305914.1 61095 61117 - 0 GGACCTCGGAGGCGACATCG TGG ||||||||[||||||||||||]PAM NA Intergenic NA NA
GU305914.1 96953 96975 + 4 CGACGGCGGAGGCGGCATCG CGG -|||--||[||||||-|||||]PAM NA Intergenic NA NA

T29530 GTGGCCTCTCTGCACATCCTCGG 1000
Oligo fwd CACCGTGGCCTCTCTGCACATCCT
Oligo rev AAACAGGATGTGCAGAGAGGCCAC
Chromosome start end strand MM target_seq PAM alignment distance position gene name gene id
GU305914.1 61076 61098 - 0 GTGGCCTCTCTGCACATCCT CGG ||||||||[||||||||||||]PAM NA Intergenic NA NA

T29531 CATCCTCGGACAGCAGTGCAAGG 1000
Oligo fwd CACCCATCCTCGGACAGCAGTGCA
Oligo rev AAACTGCACTGCTGTCCGAGGATG
Chromosome start end strand MM target_seq PAM alignment distance position gene name gene id
GU305914.1 61062 61084 - 0 CATCCTCGGACAGCAGTGCA AGG ||||||||[||||||||||||]PAM NA Intergenic NA NA

T29532 GACAGCAGTGCAAGGCGCTGCGG 1000

| | | | | | | | | | | | |
|---|---|---|---|---|---|---|---|---|---|---|---|
| Oligo fwd | CACCGACAGCAGTGCAAGGCGCTG | | | | | | | | | | |
| Oligo rev | AAACCAGCGCCTTGCACTGCTGTC | | | | | | | | | | |
| Chromosome | start | end | strand | MM | target_seq | PAM | alignment | distance | position | gene name | gene id |
| GU305914.1 | 61054 | 61076 | - | 0 | GACAGCAGTGCAAGGCGCTG | CGG | \|\|\|\|\|\|\|\|[\|\|\|\|\|\|\|\|\|\|\|\|]PAM | NA | Intergenic | NA | NA |
| T29533 | CGCGATTCAAGACCGTCTTGAGG | 1000 | | | | | | | | | |
| Oligo fwd | CACCCGCGATTCAAGACCGTCTTG | | | | | | | | | | |
| Oligo rev | AAACCAAGACGGTCTTGAATCGCG | | | | | | | | | | |
| Chromosome | start | end | strand | MM | target_seq | PAM | alignment | distance | position | gene name | gene id |
| GU305914.1 | 61024 | 61046 | - | 0 | CGCGATTCAAGACCGTCTTG | AGG | \|\|\|\|\|\|\|\|[\|\|\|\|\|\|\|\|\|\|\|\|]PAM | NA | Intergenic | NA | NA |
| T29534 | TGAGGATCTACTTCGACATCTGG | 1000 | | | | | | | | | |
| Oligo fwd | CACCTGAGGATCTACTTCGACATC | | | | | | | | | | |
| Oligo rev | AAACGATGTCGAAGTAGATCCTCA | | | | | | | | | | |
| Chromosome | start | end | strand | MM | target_seq | PAM | alignment | distance | position | gene name | gene id |
| GU305914.1 | 61006 | 61028 | - | 0 | TGAGGATCTACTTCGACATC | TGG | \|\|\|\|\|\|\|\|[\|\|\|\|\|\|\|\|\|\|\|\|]PAM | NA | Intergenic | NA | NA |
| T29535 | TTCTCTTTCATCCTCGACGTCGG | 1000 | | | | | | | | | |
| Oligo fwd | CACCTTCTCTTTCATCCTCGACGT | | | | | | | | | | |
| Oligo rev | AAACACGTCGAGGATGAAAGAGAA | | | | | | | | | | |
| Chromosome | start | end | strand | MM | target_seq | PAM | alignment | distance | position | gene name | gene id |
| GU305914.1 | 60962 | 60984 | - | 0 | TTCTCTTTCATCCTCGACGT | CGG | \|\|\|\|\|\|\|\|[\|\|\|\|\|\|\|\|\|\|\|\|]PAM | NA | Intergenic | NA | NA |
| T29536 | TCTCTTTCATCCTCGACGTCGGG | 1000 | | | | | | | | | |
| Oligo fwd | CACCTCTCTTTCATCCTCGACGTC | | | | | | | | | | |
| Oligo rev | AAACGACGTCGAGGATGAAAGAGA | | | | | | | | | | |
| Chromosome | start | end | strand | MM | target_seq | PAM | alignment | distance | position | gene name | gene id |
| GU305914.1 | 60961 | 60983 | - | 0 | TCTCTTTCATCCTCGACGTC | GGG | \|\|\|\|\|\|\|\|[\|\|\|\|\|\|\|\|\|\|\|\|]PAM | NA | Intergenic | NA | NA |
| T29537 | CTTTCATCCTCGACGTCGGGCGG | 1000 | | | | | | | | | |
| Oligo fwd | CACCCTTTCATCCTCGACGTCGGG | | | | | | | | | | |
| Oligo rev | AAACCCCGACGTCGAGGATGAAAG | | | | | | | | | | |
| Chromosome | start | end | strand | MM | target_seq | PAM | alignment | distance | position | gene name | gene id |
| GU305914.1 | 60958 | 60980 | - | 0 | CTTTCATCCTCGACGTCGGG | CGG | \|\|\|\|\|\|\|\|[\|\|\|\|\|\|\|\|\|\|\|\|]PAM | NA | Intergenic | NA | NA |
| T29538 | TTTCATCCTCGACGTCGGGCGGG | 1000 | | | | | | | | | |
| Oligo fwd | CACCTTTCATCCTCGACGTCGGGC | | | | | | | | | | |
| Oligo rev | AAACGCCCGACGTCGAGGATGAAA | | | | | | | | | | |
| Chromosome | start | end | strand | MM | target_seq | PAM | alignment | distance | position | gene name | gene id |
| GU305914.1 | 60957 | 60979 | - | 0 | TTTCATCCTCGACGTCGGGC | GGG | \|\|\|\|\|\|\|\|[\|\|\|\|\|\|\|\|\|\|\|\|]PAM | NA | Intergenic | NA | NA |
| T29539 | CGACGTCGGGCGGGAATACGAGG | 1000 | | | | | | | | | |
| Oligo fwd | CACCCGACGTCGGGCGGGAATACG | | | | | | | | | | |
| Oligo rev | AAACCGTATTCCCGCCCGACGTCG | | | | | | | | | | |
| Chromosome | start | end | strand | MM | target_seq | PAM | alignment | distance | position | gene name | gene id |
| GU305914.1 | 60948 | 60970 | - | 0 | CGACGTCGGGCGGGAATACG | AGG | \|\|\|\|\|\|\|\|[\|\|\|\|\|\|\|\|\|\|\|\|]PAM | NA | Intergenic | NA | NA |
| T29540 | GACGTCGGGCGGGAATACGAGGG | 1000 | | | | | | | | | |
| Oligo fwd | CACCGACGTCGGGCGGGAATACGA | | | | | | | | | | |
| Oligo rev | AAACTCGTATTCCCGCCCGACGTC | | | | | | | | | | |
| Chromosome | start | end | strand | MM | target_seq | PAM | alignment | distance | position | gene name | gene id |
| GU305914.1 | 60947 | 60969 | - | 0 | GACGTCGGGCGGGAATACGA | GGG | \|\|\|\|\|\|\|\|[\|\|\|\|\|\|\|\|\|\|\|\|]PAM | NA | Intergenic | NA | NA |
| T29541 | GCGGGAATACGAGGGCCTGATGG | 1000 | | | | | | | | | |
| Oligo fwd | CACCGCGGGAATACGAGGGCCTGA | | | | | | | | | | |
| Oligo rev | AAACTCAGGCCCTCGTATTCCCGC | | | | | | | | | | |
| Chromosome | start | end | strand | MM | target_seq | PAM | alignment | distance | position | gene name | gene id |
| GU305914.1 | 60939 | 60961 | - | 0 | GCGGGAATACGAGGGCCTGA | TGG | \|\|\|\|\|\|\|\|[\|\|\|\|\|\|\|\|\|\|\|\|]PAM | NA | Intergenic | NA | NA |
| T29542 | CGAGGGCCTGATGGCCTTCGCGG | 1000 | | | | | | | | | |
| Oligo fwd | CACCCGAGGGCCTGATGGCCTTCG | | | | | | | | | | |
| Oligo rev | AAACCGAAGGCCATCAGGCCCTCG | | | | | | | | | | |
| Chromosome | start | end | strand | MM | target_seq | PAM | alignment | distance | position | gene name | gene id |
| GU305914.1 | 60930 | 60952 | - | 0 | CGAGGGCCTGATGGCCTTCG | CGG | \|\|\|\|\|\|\|\|[\|\|\|\|\|\|\|\|\|\|\|\|]PAM | NA | Intergenic | NA | NA |
| T29543 | CCTTCGCGGTCAACACGCCGAGG | 1000 | | | | | | | | | |
| Oligo fwd | CACCCCTTCGCGGTCAACACGCCG | | | | | | | | | | |
| Oligo rev | AAACCGGCGTGTTGACCGCGAAGG | | | | | | | | | | |
| Chromosome | start | end | strand | MM | target_seq | PAM | alignment | distance | position | gene name | gene id |
| GU305914.1 | 60916 | 60938 | - | 0 | CCTTCGCGGTCAACACGCCG | AGG | \|\|\|\|\|\|\|\|[\|\|\|\|\|\|\|\|\|\|\|\|]PAM | NA | Intergenic | NA | NA |
| T29544 | TCAACACGCCGAGGATCTTCTGG | 1000 | | | | | | | | | |
| Oligo fwd | CACCTCAACACGCCGAGGATCTTC | | | | | | | | | | |
| Oligo rev | AAACGAAGATCCTCGGCGTGTTGA | | | | | | | | | | |
| Chromosome | start | end | strand | MM | target_seq | PAM | alignment | distance | position | gene name | gene id |
| GU305914.1 | 60907 | 60929 | - | 0 | TCAACACGCCGAGGATCTTC | TGG | \|\|\|\|\|\|\|\|[\|\|\|\|\|\|\|\|\|\|\|\|]PAM | NA | Intergenic | NA | NA |
| T29545 | CAACACGCCGAGGATCTTCTGGG | 1000 | | | | | | | | | |
| Oligo fwd | CACCCAACACGCCGAGGATCTTCT | | | | | | | | | | |
| Oligo rev | AAACAGAAGATCCTCGGCGTGTTG | | | | | | | | | | |
| Chromosome | start | end | strand | MM | target_seq | PAM | alignment | distance | position | gene name | gene id |
| GU305914.1 | 60906 | 60928 | - | 0 | CAACACGCCGAGGATCTTCT | GGG | \|\|\|\|\|\|\|\|[\|\|\|\|\|\|\|\|\|\|\|\|]PAM | NA | Intergenic | NA | NA |
| T29546 | GATCGCTACCTAGACAGCTCCGG | 1000 | | | | | | | | | |
| Oligo fwd | CACCGATCGCTACCTAGACAGCTC | | | | | | | | | | |
| Oligo rev | AAACGAGCTGTCTAGGTAGCGATC | | | | | | | | | | |
| Chromosome | start | end | strand | MM | target_seq | PAM | alignment | distance | position | gene name | gene id |
| GU305914.1 | 60884 | 60906 | - | 0 | GATCGCTACCTAGACAGCTC | CGG | \|\|\|\|\|\|\|\|[\|\|\|\|\|\|\|\|\|\|\|\|]PAM | NA | Intergenic | NA | NA |

<table>
<tr><td>T29547</td><td>ACAGCTCCGGCGACAAGATGTGG</td><td></td><td>1000</td><td></td><td></td><td></td><td></td><td></td><td></td><td></td><td></td><td></td></tr>
<tr><td>Oligo fwd</td><td>CACCACAGCTCCGGCGACAAGATG</td><td></td><td></td><td></td><td></td><td></td><td></td><td></td><td></td><td></td><td></td><td></td></tr>
<tr><td>Oligo rev</td><td>AAACCATCTTGTCGCCGGAGCTGT</td><td></td><td></td><td></td><td></td><td></td><td></td><td></td><td></td><td></td><td></td><td></td></tr>
<tr><td>Chromosome</td><td>start</td><td></td><td>end</td><td>strand</td><td>MM</td><td>target_seq</td><td>PAM</td><td>alignment</td><td>distance</td><td>position</td><td>gene name</td><td>gene id</td></tr>
<tr><td>GU305914.1</td><td></td><td>60871</td><td>60893</td><td>-</td><td>0</td><td>ACAGCTCCGGCGACAAGATG</td><td>TGG</td><td>||||||||[||||||||||||]PAM</td><td>NA</td><td>Intergenic</td><td>NA</td><td>NA</td></tr>
<tr><td>T29548</td><td>AGATGTGGCTCATGTTCGCGAGG</td><td></td><td>1000</td><td></td><td></td><td></td><td></td><td></td><td></td><td></td><td></td><td></td></tr>
<tr><td>Oligo fwd</td><td>CACCAGATGTGGCTCATGTTCGCG</td><td></td><td></td><td></td><td></td><td></td><td></td><td></td><td></td><td></td><td></td><td></td></tr>
<tr><td>Oligo rev</td><td>AAACCGCGAACATGAGCCACATCT</td><td></td><td></td><td></td><td></td><td></td><td></td><td></td><td></td><td></td><td></td><td></td></tr>
<tr><td>Chromosome</td><td>start</td><td></td><td>end</td><td>strand</td><td>MM</td><td>target_seq</td><td>PAM</td><td>alignment</td><td>distance</td><td>position</td><td>gene name</td><td>gene id</td></tr>
<tr><td>GU305914.1</td><td></td><td>60856</td><td>60878</td><td>-</td><td>0</td><td>AGATGTGGCTCATGTTCGCG</td><td>AGG</td><td>||||||||[||||||||||||]PAM</td><td>NA</td><td>Intergenic</td><td>NA</td><td>NA</td></tr>
<tr><td>T29549</td><td>TGTGGCTCATGTTCGCGAGGCGG</td><td></td><td>1000</td><td></td><td></td><td></td><td></td><td></td><td></td><td></td><td></td><td></td></tr>
<tr><td>Oligo fwd</td><td>CACCTGTGGCTCATGTTCGCGAGG</td><td></td><td></td><td></td><td></td><td></td><td></td><td></td><td></td><td></td><td></td><td></td></tr>
<tr><td>Oligo rev</td><td>AAACCCTCGCGAACATGAGCCACA</td><td></td><td></td><td></td><td></td><td></td><td></td><td></td><td></td><td></td><td></td><td></td></tr>
<tr><td>Chromosome</td><td>start</td><td></td><td>end</td><td>strand</td><td>MM</td><td>target_seq</td><td>PAM</td><td>alignment</td><td>distance</td><td>position</td><td>gene name</td><td>gene id</td></tr>
<tr><td>GU305914.1</td><td></td><td>60853</td><td>60875</td><td>-</td><td>0</td><td>TGTGGCTCATGTTCGCGAGG</td><td>CGG</td><td>||||||||[||||||||||||]PAM</td><td>NA</td><td>Intergenic</td><td>NA</td><td>NA</td></tr>
<tr><td>T29550</td><td>GTGGCTCATGTTCGCGAGGCGGG</td><td></td><td>1000</td><td></td><td></td><td></td><td></td><td></td><td></td><td></td><td></td><td></td></tr>
<tr><td>Oligo fwd</td><td>CACCGTGGCTCATGTTCGCGAGGC</td><td></td><td></td><td></td><td></td><td></td><td></td><td></td><td></td><td></td><td></td><td></td></tr>
<tr><td>Oligo rev</td><td>AAACGCCTCGCGAACATGAGCCAC</td><td></td><td></td><td></td><td></td><td></td><td></td><td></td><td></td><td></td><td></td><td></td></tr>
<tr><td>Chromosome</td><td>start</td><td></td><td>end</td><td>strand</td><td>MM</td><td>target_seq</td><td>PAM</td><td>alignment</td><td>distance</td><td>position</td><td>gene name</td><td>gene id</td></tr>
<tr><td>GU305914.1</td><td></td><td>60852</td><td>60874</td><td>-</td><td>0</td><td>GTGGCTCATGTTCGCGAGGC</td><td>GGG</td><td>||||||||[||||||||||||]PAM</td><td>NA</td><td>Intergenic</td><td>NA</td><td>NA</td></tr>
<tr><td>T29551</td><td>GCTCATGTTCGCGAGGCGGGAGG</td><td></td><td>1000</td><td></td><td></td><td></td><td></td><td></td><td></td><td></td><td></td><td></td></tr>
<tr><td>Oligo fwd</td><td>CACCGCTCATGTTCGCGAGGCGGG</td><td></td><td></td><td></td><td></td><td></td><td></td><td></td><td></td><td></td><td></td><td></td></tr>
<tr><td>Oligo rev</td><td>AAACCCCGCCTCGCGAACATGAGC</td><td></td><td></td><td></td><td></td><td></td><td></td><td></td><td></td><td></td><td></td><td></td></tr>
<tr><td>Chromosome</td><td>start</td><td></td><td>end</td><td>strand</td><td>MM</td><td>target_seq</td><td>PAM</td><td>alignment</td><td>distance</td><td>position</td><td>gene name</td><td>gene id</td></tr>
<tr><td>GU305914.1</td><td></td><td>60849</td><td>60871</td><td>-</td><td>0</td><td>GCTCATGTTCGCGAGGCGGG</td><td>AGG</td><td>||||||||[||||||||||||]PAM</td><td>NA</td><td>Intergenic</td><td>NA</td><td>NA</td></tr>
<tr><td>T29552</td><td>GTTCGCGAGGCGGGAGGCCGCGG</td><td></td><td>1000</td><td></td><td></td><td></td><td></td><td></td><td></td><td></td><td></td><td></td></tr>
<tr><td>Oligo fwd</td><td>CACCGTTCGCGAGGCGGGAGGCCG</td><td></td><td></td><td></td><td></td><td></td><td></td><td></td><td></td><td></td><td></td><td></td></tr>
<tr><td>Oligo rev</td><td>AAACCGGCCTCCCGCCTCGCGAAC</td><td></td><td></td><td></td><td></td><td></td><td></td><td></td><td></td><td></td><td></td><td></td></tr>
<tr><td>Chromosome</td><td>start</td><td></td><td>end</td><td>strand</td><td>MM</td><td>target_seq</td><td>PAM</td><td>alignment</td><td>distance</td><td>position</td><td>gene name</td><td>gene id</td></tr>
<tr><td>GU305914.1</td><td></td><td>60843</td><td>60865</td><td>-</td><td>0</td><td>GTTCGCGAGGCGGGAGGCCG</td><td>CGG</td><td>||||||||[||||||||||||]PAM</td><td>NA</td><td>Intergenic</td><td>NA</td><td>NA</td></tr>
<tr><td>T29553</td><td>CGGGAGGCCGCGGCCCTGTGCGG</td><td></td><td>1000</td><td></td><td></td><td></td><td></td><td></td><td></td><td></td><td></td><td></td></tr>
<tr><td>Oligo fwd</td><td>CACCCGGGAGGCCGCGGCCCTGTG</td><td></td><td></td><td></td><td></td><td></td><td></td><td></td><td></td><td></td><td></td><td></td></tr>
<tr><td>Oligo rev</td><td>AAACCACAGGGCCGCGGCCTCCCG</td><td></td><td></td><td></td><td></td><td></td><td></td><td></td><td></td><td></td><td></td><td></td></tr>
<tr><td>Chromosome</td><td>start</td><td></td><td>end</td><td>strand</td><td>MM</td><td>target_seq</td><td>PAM</td><td>alignment</td><td>distance</td><td>position</td><td>gene name</td><td>gene id</td></tr>
<tr><td>GU305914.1</td><td></td><td>60833</td><td>60855</td><td>-</td><td>0</td><td>CGGGAGGCCGCGGCCCTGTG</td><td>CGG</td><td>||||||||[||||||||||||]PAM</td><td>NA</td><td>Intergenic</td><td>NA</td><td>NA</td></tr>
<tr><td>T29554</td><td>CCGTAACGTCTACGAGAAGATGG</td><td></td><td>1000</td><td></td><td></td><td></td><td></td><td></td><td></td><td></td><td></td><td></td></tr>
<tr><td>Oligo fwd</td><td>CACCCCGTAACGTCTACGAGAAGA</td><td></td><td></td><td></td><td></td><td></td><td></td><td></td><td></td><td></td><td></td><td></td></tr>
<tr><td>Oligo rev</td><td>AAACTCTTCTCGTAGACGTTACGG</td><td></td><td></td><td></td><td></td><td></td><td></td><td></td><td></td><td></td><td></td><td></td></tr>
<tr><td>Chromosome</td><td>start</td><td></td><td>end</td><td>strand</td><td>MM</td><td>target_seq</td><td>PAM</td><td>alignment</td><td>distance</td><td>position</td><td>gene name</td><td>gene id</td></tr>
<tr><td>GU305914.1</td><td></td><td>60792</td><td>60814</td><td>-</td><td>0</td><td>CCGTAACGTCTACGAGAAGA</td><td>TGG</td><td>||||||||[||||||||||||]PAM</td><td>NA</td><td>Intergenic</td><td>NA</td><td>NA</td></tr>
<tr><td>T29555</td><td>TACGAGAAGATGGAGCGTGACGG</td><td></td><td>1000</td><td></td><td></td><td></td><td></td><td></td><td></td><td></td><td></td><td></td></tr>
<tr><td>Oligo fwd</td><td>CACCTACGAGAAGATGGAGCGTGA</td><td></td><td></td><td></td><td></td><td></td><td></td><td></td><td></td><td></td><td></td><td></td></tr>
<tr><td>Oligo rev</td><td>AAACTCACGCTCCATCTTCTCGTA</td><td></td><td></td><td></td><td></td><td></td><td></td><td></td><td></td><td></td><td></td><td></td></tr>
<tr><td>Chromosome</td><td>start</td><td></td><td>end</td><td>strand</td><td>MM</td><td>target_seq</td><td>PAM</td><td>alignment</td><td>distance</td><td>position</td><td>gene name</td><td>gene id</td></tr>
<tr><td>GU305914.1</td><td></td><td>60782</td><td>60804</td><td>-</td><td>0</td><td>TACGAGAAGATGGAGCGTGA</td><td>CGG</td><td>||||||||[||||||||||||]PAM</td><td>NA</td><td>Intergenic</td><td>NA</td><td>NA</td></tr>
<tr><td>T29556</td><td>ACGAGAAGATGGAGCGTGACGGG</td><td></td><td>1000</td><td></td><td></td><td></td><td></td><td></td><td></td><td></td><td></td><td></td></tr>
<tr><td>Oligo fwd</td><td>CACCACGAGAAGATGGAGCGTGAC</td><td></td><td></td><td></td><td></td><td></td><td></td><td></td><td></td><td></td><td></td><td></td></tr>
<tr><td>Oligo rev</td><td>AAACGTCACGCTCCATCTTCTCGT</td><td></td><td></td><td></td><td></td><td></td><td></td><td></td><td></td><td></td><td></td><td></td></tr>
<tr><td>Chromosome</td><td>start</td><td></td><td>end</td><td>strand</td><td>MM</td><td>target_seq</td><td>PAM</td><td>alignment</td><td>distance</td><td>position</td><td>gene name</td><td>gene id</td></tr>
<tr><td>GU305914.1</td><td></td><td>60781</td><td>60803</td><td>-</td><td>0</td><td>ACGAGAAGATGGAGCGTGAC</td><td>GGG</td><td>||||||||[||||||||||||]PAM</td><td>NA</td><td>Intergenic</td><td>NA</td><td>NA</td></tr>
<tr><td>T29557</td><td>GCGCCATCACCGTCTCGCCCTGG</td><td></td><td>1000</td><td></td><td></td><td></td><td></td><td></td><td></td><td></td><td></td><td></td></tr>
<tr><td>Oligo fwd</td><td>CACCGCGCCATCACCGTCTCGCCC</td><td></td><td></td><td></td><td></td><td></td><td></td><td></td><td></td><td></td><td></td><td></td></tr>
<tr><td>Oligo rev</td><td>AAACGGGCGAGACGGTGATGGCGC</td><td></td><td></td><td></td><td></td><td></td><td></td><td></td><td></td><td></td><td></td><td></td></tr>
<tr><td>Chromosome</td><td>start</td><td></td><td>end</td><td>strand</td><td>MM</td><td>target_seq</td><td>PAM</td><td>alignment</td><td>distance</td><td>position</td><td>gene name</td><td>gene id</td></tr>
<tr><td>GU305914.1</td><td></td><td>60754</td><td>60776</td><td>-</td><td>0</td><td>GCGCCATCACCGTCTCGCCC</td><td>TGG</td><td>||||||||[||||||||||||]PAM</td><td>NA</td><td>Intergenic</td><td>NA</td><td>NA</td></tr>
<tr><td>T29558</td><td>CCATCACCGTCTCGCCCTGGTGG</td><td></td><td>1000</td><td></td><td></td><td></td><td></td><td></td><td></td><td></td><td></td><td></td></tr>
<tr><td>Oligo fwd</td><td>CACCCCATCACCGTCTCGCCCTGG</td><td></td><td></td><td></td><td></td><td></td><td></td><td></td><td></td><td></td><td></td><td></td></tr>
<tr><td>Oligo rev</td><td>AAACCCAGGGCGAGACGGTGATGG</td><td></td><td></td><td></td><td></td><td></td><td></td><td></td><td></td><td></td><td></td><td></td></tr>
<tr><td>Chromosome</td><td>start</td><td></td><td>end</td><td>strand</td><td>MM</td><td>target_seq</td><td>PAM</td><td>alignment</td><td>distance</td><td>position</td><td>gene name</td><td>gene id</td></tr>
<tr><td>GU305914.1</td><td></td><td>60751</td><td>60773</td><td>-</td><td>0</td><td>CCATCACCGTCTCGCCCTGG</td><td>TGG</td><td>||||||||[||||||||||||]PAM</td><td>NA</td><td>Intergenic</td><td>NA</td><td>NA</td></tr>
<tr><td>T29559</td><td>CATCACCGTCTCGCCCTGGTGGG</td><td></td><td>1000</td><td></td><td></td><td></td><td></td><td></td><td></td><td></td><td></td><td></td></tr>
<tr><td>Oligo fwd</td><td>CACCCATCACCGTCTCGCCCTGGT</td><td></td><td></td><td></td><td></td><td></td><td></td><td></td><td></td><td></td><td></td><td></td></tr>
<tr><td>Oligo rev</td><td>AAACACCAGGGCGAGACGGTGATG</td><td></td><td></td><td></td><td></td><td></td><td></td><td></td><td></td><td></td><td></td><td></td></tr>
<tr><td>Chromosome</td><td>start</td><td></td><td>end</td><td>strand</td><td>MM</td><td>target_seq</td><td>PAM</td><td>alignment</td><td>distance</td><td>position</td><td>gene name</td><td>gene id</td></tr>
<tr><td>GU305914.1</td><td></td><td>60750</td><td>60772</td><td>-</td><td>0</td><td>CATCACCGTCTCGCCCTGGT</td><td>GGG</td><td>||||||||[||||||||||||]PAM</td><td>NA</td><td>Intergenic</td><td>NA</td><td>NA</td></tr>
<tr><td>T29560</td><td>CTGTCAGCTCGACGCGTGCGTGG</td><td></td><td>1000</td><td></td><td></td><td></td><td></td><td></td><td></td><td></td><td></td><td></td></tr>
<tr><td>Oligo fwd</td><td>CACCCTGTCAGCTCGACGCGTGCG</td><td></td><td></td><td></td><td></td><td></td><td></td><td></td><td></td><td></td><td></td><td></td></tr>
<tr><td>Oligo rev</td><td>AAACCGCACGCGTCGAGCTGACAG</td><td></td><td></td><td></td><td></td><td></td><td></td><td></td><td></td><td></td><td></td><td></td></tr>
<tr><td>Chromosome</td><td>start</td><td></td><td>end</td><td>strand</td><td>MM</td><td>target_seq</td><td>PAM</td><td>alignment</td><td>distance</td><td>position</td><td>gene name</td><td>gene id</td></tr>
<tr><td>GU305914.1</td><td></td><td>60723</td><td>60745</td><td>-</td><td>0</td><td>CTGTCAGCTCGACGCGTGCG</td><td>TGG</td><td>||||||||[||||||||||||]PAM</td><td>NA</td><td>Intergenic</td><td>NA</td><td>NA</td></tr>
<tr><td>T29561</td><td>AGCTCGACGCGTGCGTGGCGCGG</td><td></td><td>1000</td><td></td><td></td><td></td><td></td><td></td><td></td><td></td><td></td><td></td></tr>
<tr><td>Oligo fwd</td><td>CACCAGCTCGACGCGTGCGTGGCG</td><td></td><td></td><td></td><td></td><td></td><td></td><td></td><td></td><td></td><td></td><td></td></tr>
<tr><td>Oligo rev</td><td>AAACCGCCACGCACGCGTCGAGCT</td><td></td><td></td><td></td><td></td><td></td><td></td><td></td><td></td><td></td><td></td><td></td></tr>
<tr><td>Chromosome</td><td>start</td><td></td><td>end</td><td>strand</td><td>MM</td><td>target_seq</td><td>PAM</td><td>alignment</td><td>distance</td><td>position</td><td>gene name</td><td>gene id</td></tr>
<tr><td>GU305914.1</td><td></td><td>60718</td><td>60740</td><td>-</td><td>0</td><td>AGCTCGACGCGTGCGTGGCG</td><td>CGG</td><td>||||||||[||||||||||||]PAM</td><td>NA</td><td>Intergenic</td><td>NA</td><td>NA</td></tr>
</table>

T29562 GCTCGACGCGTGCGTGGCGCGGG 1000
Oligo fwd CACCGCTCGACGCGTGCGTGGCGC
Oligo rev AAACGCGCCACGCACGCGTCGAGC
Chromosome start end strand MM target_seq PAM alignment distance position gene name gene id
GU305914.1 60717 60739 - 0 GCTCGACGCGTGCGTGGCGC GGG ||||||||[||||||||||||]PAM NA Intergenic NA NA

T29563 CTCGACGCGTGCGTGGCGCGGGG 1000
Oligo fwd CACCCTCGACGCGTGCGTGGCGCG
Oligo rev AAACCGCGCCACGCACGCGTCGAG
Chromosome start end strand MM target_seq PAM alignment distance position gene name gene id
GU305914.1 60716 60738 - 0 CTCGACGCGTGCGTGGCGCG GGG ||||||||[||||||||||||]PAM NA Intergenic NA NA

T29564 GTGCGTGGCGCGGGGCAACACGG 1000
Oligo fwd CACCGTGCGTGGCGCGGGGCAACA
Oligo rev AAACTGTTGCCCCGCGCCACGCAC
Chromosome start end strand MM target_seq PAM alignment distance position gene name gene id
GU305914.1 60708 60730 - 0 GTGCGTGGCGCGGGGCAACA CGG ||||||||[||||||||||||]PAM NA Intergenic NA NA

T29565 GGCGCGGGGCAACACGGCCGTGG 1000
Oligo fwd CACCGGCGCGGGGCAACACGGCCG
Oligo rev AAACCGGCCGTGTTGCCCCGCGCC
Chromosome start end strand MM target_seq PAM alignment distance position gene name gene id
GU305914.1 60702 60724 - 0 GGCGCGGGGCAACACGGCCG TGG ||||||||[||||||||||||]PAM NA Intergenic NA NA

T29566 AGTATGATCCCCGAGAACATCGG 1000
Oligo fwd CACCAGTATGATCCCCGAGAACAT
Oligo rev AAACATGTTCTCGGGGATCATACT
Chromosome start end strand MM target_seq PAM alignment distance position gene name gene id
GU305914.1 60659 60681 - 0 AGTATGATCCCCGAGAACAT CGG ||||||||[||||||||||||]PAM NA Intergenic NA NA

T29567 GTATGATCCCCGAGAACATCGGG 1000
Oligo fwd CACCGTATGATCCCCGAGAACATC
Oligo rev AAACGATGTTCTCGGGGATCATAC
Chromosome start end strand MM target_seq PAM alignment distance position gene name gene id
GU305914.1 60658 60680 - 0 GTATGATCCCCGAGAACATC GGG ||||||||[||||||||||||]PAM NA Intergenic NA NA

T29568 ATCGGGCGCCCCGCCGTGTGCGG 1000
Oligo fwd CACCATCGGGCGCCCCGCCGTGTG
Oligo rev AAACCACACGGCGGGGCGCCCGAT
Chromosome start end strand MM target_seq PAM alignment distance position gene name gene id
GU305914.1 60641 60663 - 0 ATCGGGCGCCCCGCCGTGTG CGG ||||||||[||||||||||||]PAM NA Intergenic NA NA

T29569 CGCCCCGCCGTGTGCGGACCCGG 1000
Oligo fwd CACCCGCCCCGCCGTGTGCGGACC
Oligo rev AAACGGTCCGCACACGGCGGGGCG
Chromosome start end strand MM target_seq PAM alignment distance position gene name gene id
GU305914.1 60635 60657 - 0 CGCCCCGCCGTGTGCGGACC CGG ||||||||[||||||||||||]PAM NA Intergenic NA NA

T29570 CCCGGCGTCTCCGTCGTCTCCGG 1000
Oligo fwd CACCCCCGGCGTCTCCGTCGTCTC
Oligo rev AAACGAGACGACGGAGACGCCGGG
Chromosome start end strand MM target_seq PAM alignment distance position gene name gene id
GU305914.1 60617 60639 - 0 CCCGGCGTCTCCGTCGTCTC CGG ||||||||[||||||||||||]PAM NA Intergenic NA NA

T29571 GGCGTCTCCGTCGTCTCCGGCGG 1000
Oligo fwd CACCGGCGTCTCCGTCGTCTCCGG
Oligo rev AAACCCGGAGACGACGGAGACGCC
Chromosome start end strand MM target_seq PAM alignment distance position gene name gene id
GU305914.1 60614 60636 - 0 GGCGTCTCCGTCGTCTCCGG CGG ||||||||[||||||||||||]PAM NA Intergenic NA NA
GU305914.1 26356 26378 - 4 GACGACGACGTCGTCTCCGG GGG |-||-|--[||||||||||||]PAM NA Intergenic NA NA
GU305914.1 142216 142238 - 4 GTCATCGCCGTCGCCTCCGG AGG |-|-||-|[|||||-||||||]PAM NA Intergenic NA NA

T29572 GTCGTCTCCGGCGGCTTCGTCGG 1000
Oligo fwd CACCGTCGTCTCCGGCGGCTTCGT
Oligo rev AAACACGAAGCCGCCGGAGACGAC
Chromosome start end strand MM target_seq PAM alignment distance position gene name gene id
GU305914.1 60605 60627 - 0 GTCGTCTCCGGCGGCTTCGT CGG ||||||||[||||||||||||]PAM NA Intergenic NA NA

T29573 CCACGAGCTGTGCATCAACCTGG 1000
Oligo fwd CACCCCACGAGCTGTGCATCAACC
Oligo rev AAACGGTTGATGCACAGCTCGTGG
Chromosome start end strand MM target_seq PAM alignment distance position gene name gene id
GU305914.1 60570 60592 - 0 CCACGAGCTGTGCATCAACC TGG ||||||||[||||||||||||]PAM NA Intergenic NA NA

T29574 CAACCTGGAGAACTGCGTCCTGG 1000
Oligo fwd CACCCAACCTGGAGAACTGCGTCC
Oligo rev AAACGGACGCAGTTCTCCAGGTTG
Chromosome start end strand MM target_seq PAM alignment distance position gene name gene id
GU305914.1 60555 60577 - 0 CAACCTGGAGAACTGCGTCC TGG ||||||||[||||||||||||]PAM NA Intergenic NA NA

T29575 CCTGGAGAACTGCGTCCTGGAGG 1000
Oligo fwd CACCCCTGGAGAACTGCGTCCTGG
Oligo rev AAACCCAGGACGCAGTTCTCCAGG
Chromosome start end strand MM target_seq PAM alignment distance position gene name gene id
GU305914.1 60552 60574 - 0 CCTGGAGAACTGCGTCCTGG AGG ||||||||[||||||||||||]PAM NA Intergenic NA NA

T29576 CTGGAGAACTGCGTCCTGGAGGG 1000
Oligo fwd CACCCTGGAGAACTGCGTCCTGGA

<table>
<tr><td>Oligo rev</td><td>AAACTCCAGGACGCAGTTCTCCAG</td><td></td><td></td><td></td><td></td><td></td><td></td><td></td><td></td><td></td><td></td></tr>
<tr><td>Chromosome</td><td>start</td><td>end</td><td>strand</td><td>MM</td><td>target_seq</td><td>PAM</td><td>alignment</td><td>distance</td><td>position</td><td>gene name</td><td>gene id</td></tr>
<tr><td>GU305914.1</td><td>60551</td><td>60573</td><td>-</td><td>0</td><td>CTGGAGAACTGCGTCCTGGA</td><td>GGG</td><td>||||||||[||||||||||||]PAM</td><td>NA</td><td>Intergenic</td><td>NA</td><td>NA</td></tr>
<tr><td>T29577</td><td>GAACTGCGTCCTGGAGGGCGCGG</td><td>1000</td><td></td><td></td><td></td><td></td><td></td><td></td><td></td><td></td><td></td></tr>
<tr><td>Oligo fwd</td><td>CACCGAACTGCGTCCTGGAGGGCG</td><td></td><td></td><td></td><td></td><td></td><td></td><td></td><td></td><td></td><td></td></tr>
<tr><td>Oligo rev</td><td>AAACCGCCCTCCAGGACGCAGTTC</td><td></td><td></td><td></td><td></td><td></td><td></td><td></td><td></td><td></td><td></td></tr>
<tr><td>Chromosome</td><td>start</td><td>end</td><td>strand</td><td>MM</td><td>target_seq</td><td>PAM</td><td>alignment</td><td>distance</td><td>position</td><td>gene name</td><td>gene id</td></tr>
<tr><td>GU305914.1</td><td>60546</td><td>60568</td><td>-</td><td>0</td><td>GAACTGCGTCCTGGAGGGCG</td><td>CGG</td><td>||||||||[||||||||||||]PAM</td><td>NA</td><td>Intergenic</td><td>NA</td><td>NA</td></tr>
<tr><td>T29578</td><td>AGTCGAGAGCTCCGTCGACGTGG</td><td>1000</td><td></td><td></td><td></td><td></td><td></td><td></td><td></td><td></td><td></td></tr>
<tr><td>Oligo fwd</td><td>CACCAGTCGAGAGCTCCGTCGACG</td><td></td><td></td><td></td><td></td><td></td><td></td><td></td><td></td><td></td><td></td></tr>
<tr><td>Oligo rev</td><td>AAACCGTCGACGGAGCTCTCGACT</td><td></td><td></td><td></td><td></td><td></td><td></td><td></td><td></td><td></td><td></td></tr>
<tr><td>Chromosome</td><td>start</td><td>end</td><td>strand</td><td>MM</td><td>target_seq</td><td>PAM</td><td>alignment</td><td>distance</td><td>position</td><td>gene name</td><td>gene id</td></tr>
<tr><td>GU305914.1</td><td>60522</td><td>60544</td><td>-</td><td>0</td><td>AGTCGAGAGCTCCGTCGACG</td><td>TGG</td><td>||||||||[||||||||||||]PAM</td><td>NA</td><td>Intergenic</td><td>NA</td><td>NA</td></tr>
<tr><td>T29579</td><td>AGCTCCGTCGACGTGGTCCTCGG</td><td>1000</td><td></td><td></td><td></td><td></td><td></td><td></td><td></td><td></td><td></td></tr>
<tr><td>Oligo fwd</td><td>CACCAGCTCCGTCGACGTGGTCCT</td><td></td><td></td><td></td><td></td><td></td><td></td><td></td><td></td><td></td><td></td></tr>
<tr><td>Oligo rev</td><td>AAACAGGACCACGTCGACGGAGCT</td><td></td><td></td><td></td><td></td><td></td><td></td><td></td><td></td><td></td><td></td></tr>
<tr><td>Chromosome</td><td>start</td><td>end</td><td>strand</td><td>MM</td><td>target_seq</td><td>PAM</td><td>alignment</td><td>distance</td><td>position</td><td>gene name</td><td>gene id</td></tr>
<tr><td>GU305914.1</td><td>60515</td><td>60537</td><td>-</td><td>0</td><td>AGCTCCGTCGACGTGGTCCT</td><td>CGG</td><td>||||||||[||||||||||||]PAM</td><td>NA</td><td>Intergenic</td><td>NA</td><td>NA</td></tr>
<tr><td>T29580</td><td>GTCGACGTGGTCCTCGGTCTCGG</td><td>1000</td><td></td><td></td><td></td><td></td><td></td><td></td><td></td><td></td><td></td></tr>
<tr><td>Oligo fwd</td><td>CACCGTCGACGTGGTCCTCGGTCT</td><td></td><td></td><td></td><td></td><td></td><td></td><td></td><td></td><td></td><td></td></tr>
<tr><td>Oligo rev</td><td>AAACAGACCGAGGACCACGTCGAC</td><td></td><td></td><td></td><td></td><td></td><td></td><td></td><td></td><td></td><td></td></tr>
<tr><td>Chromosome</td><td>start</td><td>end</td><td>strand</td><td>MM</td><td>target_seq</td><td>PAM</td><td>alignment</td><td>distance</td><td>position</td><td>gene name</td><td>gene id</td></tr>
<tr><td>GU305914.1</td><td>60509</td><td>60531</td><td>-</td><td>0</td><td>GTCGACGTGGTCCTCGGTCT</td><td>CGG</td><td>||||||||[||||||||||||]PAM</td><td>NA</td><td>Intergenic</td><td>NA</td><td>NA</td></tr>
<tr><td>T29581</td><td>CGGTTGCCGCTTCAGCTTCAAGG</td><td>1000</td><td></td><td></td><td></td><td></td><td></td><td></td><td></td><td></td><td></td></tr>
<tr><td>Oligo fwd</td><td>CACCCGGTTGCCGCTTCAGCTTCA</td><td></td><td></td><td></td><td></td><td></td><td></td><td></td><td></td><td></td><td></td></tr>
<tr><td>Oligo rev</td><td>AAACTGAAGCTGAAGCGGCAACCG</td><td></td><td></td><td></td><td></td><td></td><td></td><td></td><td></td><td></td><td></td></tr>
<tr><td>Chromosome</td><td>start</td><td>end</td><td>strand</td><td>MM</td><td>target_seq</td><td>PAM</td><td>alignment</td><td>distance</td><td>position</td><td>gene name</td><td>gene id</td></tr>
<tr><td>GU305914.1</td><td>60489</td><td>60511</td><td>-</td><td>0</td><td>CGGTTGCCGCTTCAGCTTCA</td><td>AGG</td><td>||||||||[||||||||||||]PAM</td><td>NA</td><td>Intergenic</td><td>NA</td><td>NA</td></tr>
<tr><td>T29582</td><td>CCGCTTCAGCTTCAAGGCCCTGG</td><td>1000</td><td></td><td></td><td></td><td></td><td></td><td></td><td></td><td></td><td></td></tr>
<tr><td>Oligo fwd</td><td>CACCCCGCTTCAGCTTCAAGGCCC</td><td></td><td></td><td></td><td></td><td></td><td></td><td></td><td></td><td></td><td></td></tr>
<tr><td>Oligo rev</td><td>AAACGGGCCTTGAAGCTGAAGCGG</td><td></td><td></td><td></td><td></td><td></td><td></td><td></td><td></td><td></td><td></td></tr>
<tr><td>Chromosome</td><td>start</td><td>end</td><td>strand</td><td>MM</td><td>target_seq</td><td>PAM</td><td>alignment</td><td>distance</td><td>position</td><td>gene name</td><td>gene id</td></tr>
<tr><td>GU305914.1</td><td>60483</td><td>60505</td><td>-</td><td>0</td><td>CCGCTTCAGCTTCAAGGCCC</td><td>TGG</td><td>||||||||[||||||||||||]PAM</td><td>NA</td><td>Intergenic</td><td>NA</td><td>NA</td></tr>
<tr><td>T29583</td><td>CTTCAAGGCCCTGGAGTCCCTGG</td><td>1000</td><td></td><td></td><td></td><td></td><td></td><td></td><td></td><td></td><td></td></tr>
<tr><td>Oligo fwd</td><td>CACCCTTCAAGGCCCTGGAGTCCC</td><td></td><td></td><td></td><td></td><td></td><td></td><td></td><td></td><td></td><td></td></tr>
<tr><td>Oligo rev</td><td>AAACGGGACTCCAGGGCCTTGAAG</td><td></td><td></td><td></td><td></td><td></td><td></td><td></td><td></td><td></td><td></td></tr>
<tr><td>Chromosome</td><td>start</td><td>end</td><td>strand</td><td>MM</td><td>target_seq</td><td>PAM</td><td>alignment</td><td>distance</td><td>position</td><td>gene name</td><td>gene id</td></tr>
<tr><td>GU305914.1</td><td>60474</td><td>60496</td><td>-</td><td>0</td><td>CTTCAAGGCCCTGGAGTCCC</td><td>TGG</td><td>||||||||[||||||||||||]PAM</td><td>NA</td><td>Intergenic</td><td>NA</td><td>NA</td></tr>
<tr><td>T29584</td><td>GGAGTCCCTGGTCCGCGACGCGG</td><td>1000</td><td></td><td></td><td></td><td></td><td></td><td></td><td></td><td></td><td></td></tr>
<tr><td>Oligo fwd</td><td>CACCGGAGTCCCTGGTCCGCGACG</td><td></td><td></td><td></td><td></td><td></td><td></td><td></td><td></td><td></td><td></td></tr>
<tr><td>Oligo rev</td><td>AAACCGTCGCGGACCAGGGACTCC</td><td></td><td></td><td></td><td></td><td></td><td></td><td></td><td></td><td></td><td></td></tr>
<tr><td>Chromosome</td><td>start</td><td>end</td><td>strand</td><td>MM</td><td>target_seq</td><td>PAM</td><td>alignment</td><td>distance</td><td>position</td><td>gene name</td><td>gene id</td></tr>
<tr><td>GU305914.1</td><td>60462</td><td>60484</td><td>-</td><td>0</td><td>GGAGTCCCTGGTCCGCGACG</td><td>CGG</td><td>||||||||[||||||||||||]PAM</td><td>NA</td><td>Intergenic</td><td>NA</td><td>NA</td></tr>
<tr><td>GU305914.1</td><td>65645</td><td>65667</td><td>+</td><td>4</td><td>CGAGTACCTGTACCGCGACG</td><td>CGG</td><td>-||||-||[||--||||||||]PAM</td><td>NA</td><td>Intergenic</td><td>NA</td><td>NA</td></tr>
<tr><td>T29585</td><td>GTCCCTGGTCCGCGACGCGGTGG</td><td>1000</td><td></td><td></td><td></td><td></td><td></td><td></td><td></td><td></td><td></td></tr>
<tr><td>Oligo fwd</td><td>CACCGTCCCTGGTCCGCGACGCGG</td><td></td><td></td><td></td><td></td><td></td><td></td><td></td><td></td><td></td><td></td></tr>
<tr><td>Oligo rev</td><td>AAACCCGCGTCGCGGACCAGGGAC</td><td></td><td></td><td></td><td></td><td></td><td></td><td></td><td></td><td></td><td></td></tr>
<tr><td>Chromosome</td><td>start</td><td>end</td><td>strand</td><td>MM</td><td>target_seq</td><td>PAM</td><td>alignment</td><td>distance</td><td>position</td><td>gene name</td><td>gene id</td></tr>
<tr><td>GU305914.1</td><td>60459</td><td>60481</td><td>-</td><td>0</td><td>GTCCCTGGTCCGCGACGCGG</td><td>TGG</td><td>||||||||[||||||||||||]PAM</td><td>NA</td><td>Intergenic</td><td>NA</td><td>NA</td></tr>
<tr><td>T29586</td><td>GGTCCGCGACGCGGTGGTGCTGG</td><td>1000</td><td></td><td></td><td></td><td></td><td></td><td></td><td></td><td></td><td></td></tr>
<tr><td>Oligo fwd</td><td>CACCGGTCCGCGACGCGGTGGTGC</td><td></td><td></td><td></td><td></td><td></td><td></td><td></td><td></td><td></td><td></td></tr>
<tr><td>Oligo rev</td><td>AAACGCACCACCGCGTCGCGGACC</td><td></td><td></td><td></td><td></td><td></td><td></td><td></td><td></td><td></td><td></td></tr>
<tr><td>Chromosome</td><td>start</td><td>end</td><td>strand</td><td>MM</td><td>target_seq</td><td>PAM</td><td>alignment</td><td>distance</td><td>position</td><td>gene name</td><td>gene id</td></tr>
<tr><td>GU305914.1</td><td>60453</td><td>60475</td><td>-</td><td>0</td><td>GGTCCGCGACGCGGTGGTGC</td><td>TGG</td><td>||||||||[||||||||||||]PAM</td><td>NA</td><td>Intergenic</td><td>NA</td><td>NA</td></tr>
<tr><td>GU305914.1</td><td>125354</td><td>125376</td><td>-</td><td>4</td><td>GGGCGACGGCGCGGTGGTGC</td><td>CGG</td><td>||-|--||[-|||||||||||]PAM</td><td>NA</td><td>Intergenic</td><td>NA</td><td>NA</td></tr>
<tr><td>T29587</td><td>GTCCGCGACGCGGTGGTGCTGGG</td><td>1000</td><td></td><td></td><td></td><td></td><td></td><td></td><td></td><td></td><td></td></tr>
<tr><td>Oligo fwd</td><td>CACCGTCCGCGACGCGGTGGTGCT</td><td></td><td></td><td></td><td></td><td></td><td></td><td></td><td></td><td></td><td></td></tr>
<tr><td>Oligo rev</td><td>AAACAGCACCACCGCGTCGCGGAC</td><td></td><td></td><td></td><td></td><td></td><td></td><td></td><td></td><td></td><td></td></tr>
<tr><td>Chromosome</td><td>start</td><td>end</td><td>strand</td><td>MM</td><td>target_seq</td><td>PAM</td><td>alignment</td><td>distance</td><td>position</td><td>gene name</td><td>gene id</td></tr>
<tr><td>GU305914.1</td><td>60452</td><td>60474</td><td>-</td><td>0</td><td>GTCCGCGACGCGGTGGTGCT</td><td>GGG</td><td>||||||||[||||||||||||]PAM</td><td>NA</td><td>Intergenic</td><td>NA</td><td>NA</td></tr>
<tr><td>T29588</td><td>ACCGTGCGCACCAACGCGTACGG</td><td>1000</td><td></td><td></td><td></td><td></td><td></td><td></td><td></td><td></td><td></td></tr>
<tr><td>Oligo fwd</td><td>CACCACCGTGCGCACCAACGCGTA</td><td></td><td></td><td></td><td></td><td></td><td></td><td></td><td></td><td></td><td></td></tr>
<tr><td>Oligo rev</td><td>AAACTACGCGTTGGTGCGCACGGT</td><td></td><td></td><td></td><td></td><td></td><td></td><td></td><td></td><td></td><td></td></tr>
<tr><td>Chromosome</td><td>start</td><td>end</td><td>strand</td><td>MM</td><td>target_seq</td><td>PAM</td><td>alignment</td><td>distance</td><td>position</td><td>gene name</td><td>gene id</td></tr>
<tr><td>GU305914.1</td><td>60410</td><td>60432</td><td>-</td><td>0</td><td>ACCGTGCGCACCAACGCGTA</td><td>CGG</td><td>||||||||[||||||||||||]PAM</td><td>NA</td><td>Intergenic</td><td>NA</td><td>NA</td></tr>
<tr><td>T29589</td><td>CGCACCAACGCGTACGGCGCCGG</td><td>1000</td><td></td><td></td><td></td><td></td><td></td><td></td><td></td><td></td><td></td></tr>
<tr><td>Oligo fwd</td><td>CACCCGCACCAACGCGTACGGCGC</td><td></td><td></td><td></td><td></td><td></td><td></td><td></td><td></td><td></td><td></td></tr>
<tr><td>Oligo rev</td><td>AAACGCGCCGTACGCGTTGGTGCG</td><td></td><td></td><td></td><td></td><td></td><td></td><td></td><td></td><td></td><td></td></tr>
<tr><td>Chromosome</td><td>start</td><td>end</td><td>strand</td><td>MM</td><td>target_seq</td><td>PAM</td><td>alignment</td><td>distance</td><td>position</td><td>gene name</td><td>gene id</td></tr>
<tr><td>GU305914.1</td><td>60404</td><td>60426</td><td>-</td><td>0</td><td>CGCACCAACGCGTACGGCGC</td><td>CGG</td><td>||||||||[||||||||||||]PAM</td><td>NA</td><td>Intergenic</td><td>NA</td><td>NA</td></tr>
<tr><td>T29590</td><td>CTCTATCGCGACCTGCACATCGG</td><td>1000</td><td></td><td></td><td></td><td></td><td></td><td></td><td></td><td></td><td></td></tr>
<tr><td>Oligo fwd</td><td>CACCCTCTATCGCGACCTGCACAT</td><td></td><td></td><td></td><td></td><td></td><td></td><td></td><td></td><td></td><td></td></tr>
<tr><td>Oligo rev</td><td>AAACATGTGCAGGTCGCGATAGAG</td><td></td><td></td><td></td><td></td><td></td><td></td><td></td><td></td><td></td><td></td></tr>
<tr><td>Chromosome</td><td>start</td><td>end</td><td>strand</td><td>MM</td><td>target_seq</td><td>PAM</td><td>alignment</td><td>distance</td><td>position</td><td>gene name</td><td>gene id</td></tr>
<tr><td>GU305914.1</td><td>60368</td><td>60390</td><td>-</td><td>0</td><td>CTCTATCGCGACCTGCACAT</td><td>CGG</td><td>||||||||[||||||||||||]PAM</td><td>NA</td><td>Intergenic</td><td>NA</td><td>NA</td></tr>
</table>

<table>
<tr><td>T29591</td><td>TCTATCGCGACCTGCACATCGGG</td><td>1000</td><td></td><td></td><td></td><td></td><td></td><td></td><td></td><td></td><td></td></tr>
<tr><td>Oligo fwd</td><td>CACCTCTATCGCGACCTGCACATC</td><td></td><td></td><td></td><td></td><td></td><td></td><td></td><td></td><td></td><td></td></tr>
<tr><td>Oligo rev</td><td>AAACGATGTGCAGGTCGCGATAGA</td><td></td><td></td><td></td><td></td><td></td><td></td><td></td><td></td><td></td><td></td></tr>
<tr><td>Chromosome</td><td>start</td><td>end</td><td>strand</td><td>MM</td><td>target_seq</td><td>PAM</td><td>alignment</td><td>distance</td><td>position</td><td>gene name</td><td>gene id</td></tr>
<tr><td>GU305914.1</td><td>60367</td><td>60389</td><td>-</td><td>0</td><td>TCTATCGCGACCTGCACATC</td><td>GGG</td><td>||||||||[||||||||||||]PAM</td><td>NA</td><td>Intergenic</td><td>NA</td><td>NA</td></tr>
<tr><td>T29592</td><td>CTATCGCGACCTGCACATCGGGG</td><td>1000</td><td></td><td></td><td></td><td></td><td></td><td></td><td></td><td></td><td></td></tr>
<tr><td>Oligo fwd</td><td>CACCCTATCGCGACCTGCACATCG</td><td></td><td></td><td></td><td></td><td></td><td></td><td></td><td></td><td></td><td></td></tr>
<tr><td>Oligo rev</td><td>AAACCGATGTGCAGGTCGCGATAG</td><td></td><td></td><td></td><td></td><td></td><td></td><td></td><td></td><td></td><td></td></tr>
<tr><td>Chromosome</td><td>start</td><td>end</td><td>strand</td><td>MM</td><td>target_seq</td><td>PAM</td><td>alignment</td><td>distance</td><td>position</td><td>gene name</td><td>gene id</td></tr>
<tr><td>GU305914.1</td><td>60366</td><td>60388</td><td>-</td><td>0</td><td>CTATCGCGACCTGCACATCG</td><td>GGG</td><td>||||||||[||||||||||||]PAM</td><td>NA</td><td>Intergenic</td><td>NA</td><td>NA</td></tr>
<tr><td>T29593</td><td>GACCTGCACATCGGGGTCGTCGG</td><td>1000</td><td></td><td></td><td></td><td></td><td></td><td></td><td></td><td></td><td></td></tr>
<tr><td>Oligo fwd</td><td>CACCGACCTGCACATCGGGGTCGT</td><td></td><td></td><td></td><td></td><td></td><td></td><td></td><td></td><td></td><td></td></tr>
<tr><td>Oligo rev</td><td>AAACACGACCCCGATGTGCAGGTC</td><td></td><td></td><td></td><td></td><td></td><td></td><td></td><td></td><td></td><td></td></tr>
<tr><td>Chromosome</td><td>start</td><td>end</td><td>strand</td><td>MM</td><td>target_seq</td><td>PAM</td><td>alignment</td><td>distance</td><td>position</td><td>gene name</td><td>gene id</td></tr>
<tr><td>GU305914.1</td><td>60359</td><td>60381</td><td>-</td><td>0</td><td>GACCTGCACATCGGGGTCGT</td><td>CGG</td><td>||||||||[||||||||||||]PAM</td><td>NA</td><td>Intergenic</td><td>NA</td><td>NA</td></tr>
<tr><td>T29594</td><td>CACGCTGTGATGAATCGCCTCGG</td><td>1000</td><td></td><td></td><td></td><td></td><td></td><td></td><td></td><td></td><td></td></tr>
<tr><td>Oligo fwd</td><td>CACCCACGCTGTGATGAATCGCCT</td><td></td><td></td><td></td><td></td><td></td><td></td><td></td><td></td><td></td><td></td></tr>
<tr><td>Oligo rev</td><td>AAACAGGCGATTCATCACAGCGTG</td><td></td><td></td><td></td><td></td><td></td><td></td><td></td><td></td><td></td><td></td></tr>
<tr><td>Chromosome</td><td>start</td><td>end</td><td>strand</td><td>MM</td><td>target_seq</td><td>PAM</td><td>alignment</td><td>distance</td><td>position</td><td>gene name</td><td>gene id</td></tr>
<tr><td>GU305914.1</td><td>60332</td><td>60354</td><td>-</td><td>0</td><td>CACGCTGTGATGAATCGCCT</td><td>CGG</td><td>||||||||[||||||||||||]PAM</td><td>NA</td><td>Intergenic</td><td>NA</td><td>NA</td></tr>
<tr><td>T29595</td><td>ACGCTGTGATGAATCGCCTCGGG</td><td>1000</td><td></td><td></td><td></td><td></td><td></td><td></td><td></td><td></td><td></td></tr>
<tr><td>Oligo fwd</td><td>CACCACGCTGTGATGAATCGCCTC</td><td></td><td></td><td></td><td></td><td></td><td></td><td></td><td></td><td></td><td></td></tr>
<tr><td>Oligo rev</td><td>AAACGAGGCGATTCATCACAGCGT</td><td></td><td></td><td></td><td></td><td></td><td></td><td></td><td></td><td></td><td></td></tr>
<tr><td>Chromosome</td><td>start</td><td>end</td><td>strand</td><td>MM</td><td>target_seq</td><td>PAM</td><td>alignment</td><td>distance</td><td>position</td><td>gene name</td><td>gene id</td></tr>
<tr><td>GU305914.1</td><td>60331</td><td>60353</td><td>-</td><td>0</td><td>ACGCTGTGATGAATCGCCTC</td><td>GGG</td><td>||||||||[||||||||||||]PAM</td><td>NA</td><td>Intergenic</td><td>NA</td><td>NA</td></tr>
<tr><td>T29596</td><td>CGGGCAGAAGTTCGCCGACATGG</td><td>1000</td><td></td><td></td><td></td><td></td><td></td><td></td><td></td><td></td><td></td></tr>
<tr><td>Oligo fwd</td><td>CACCCGGGCAGAAGTTCGCCGACA</td><td></td><td></td><td></td><td></td><td></td><td></td><td></td><td></td><td></td><td></td></tr>
<tr><td>Oligo rev</td><td>AAACTGTCGGCGAACTTCTGCCCG</td><td></td><td></td><td></td><td></td><td></td><td></td><td></td><td></td><td></td><td></td></tr>
<tr><td>Chromosome</td><td>start</td><td>end</td><td>strand</td><td>MM</td><td>target_seq</td><td>PAM</td><td>alignment</td><td>distance</td><td>position</td><td>gene name</td><td>gene id</td></tr>
<tr><td>GU305914.1</td><td>60312</td><td>60334</td><td>-</td><td>0</td><td>CGGGCAGAAGTTCGCCGACA</td><td>TGG</td><td>||||||||[||||||||||||]PAM</td><td>NA</td><td>Intergenic</td><td>NA</td><td>NA</td></tr>
<tr><td>T29597</td><td>CGACCTCAACCAGCGTATCGCGG</td><td>1000</td><td></td><td></td><td></td><td></td><td></td><td></td><td></td><td></td><td></td></tr>
<tr><td>Oligo fwd</td><td>CACCCGACCTCAACCAGCGTATCG</td><td></td><td></td><td></td><td></td><td></td><td></td><td></td><td></td><td></td><td></td></tr>
<tr><td>Oligo rev</td><td>AAACCGATACGCTGGTTGAGGTCG</td><td></td><td></td><td></td><td></td><td></td><td></td><td></td><td></td><td></td><td></td></tr>
<tr><td>Chromosome</td><td>start</td><td>end</td><td>strand</td><td>MM</td><td>target_seq</td><td>PAM</td><td>alignment</td><td>distance</td><td>position</td><td>gene name</td><td>gene id</td></tr>
<tr><td>GU305914.1</td><td>60282</td><td>60304</td><td>-</td><td>0</td><td>CGACCTCAACCAGCGTATCG</td><td>CGG</td><td>||||||||[||||||||||||]PAM</td><td>NA</td><td>Intergenic</td><td>NA</td><td>NA</td></tr>
<tr><td>T29598</td><td>TTATCTACTACACCGCCGTGCGG</td><td>1000</td><td></td><td></td><td></td><td></td><td></td><td></td><td></td><td></td><td></td></tr>
<tr><td>Oligo fwd</td><td>CACCTTATCTACTACACCGCCGTG</td><td></td><td></td><td></td><td></td><td></td><td></td><td></td><td></td><td></td><td></td></tr>
<tr><td>Oligo rev</td><td>AAACCACGGCGGTGTAGTAGATAA</td><td></td><td></td><td></td><td></td><td></td><td></td><td></td><td></td><td></td><td></td></tr>
<tr><td>Chromosome</td><td>start</td><td>end</td><td>strand</td><td>MM</td><td>target_seq</td><td>PAM</td><td>alignment</td><td>distance</td><td>position</td><td>gene name</td><td>gene id</td></tr>
<tr><td>GU305914.1</td><td>60256</td><td>60278</td><td>-</td><td>0</td><td>TTATCTACTACACCGCCGTG</td><td>CGG</td><td>||||||||[||||||||||||]PAM</td><td>NA</td><td>Intergenic</td><td>NA</td><td>NA</td></tr>
<tr><td>T29599</td><td>TATCTACTACACCGCCGTGCGGG</td><td>1000</td><td></td><td></td><td></td><td></td><td></td><td></td><td></td><td></td><td></td></tr>
<tr><td>Oligo fwd</td><td>CACCTATCTACTACACCGCCGTGC</td><td></td><td></td><td></td><td></td><td></td><td></td><td></td><td></td><td></td><td></td></tr>
<tr><td>Oligo rev</td><td>AAACGCACGGCGGTGTAGTAGATA</td><td></td><td></td><td></td><td></td><td></td><td></td><td></td><td></td><td></td><td></td></tr>
<tr><td>Chromosome</td><td>start</td><td>end</td><td>strand</td><td>MM</td><td>target_seq</td><td>PAM</td><td>alignment</td><td>distance</td><td>position</td><td>gene name</td><td>gene id</td></tr>
<tr><td>GU305914.1</td><td>60255</td><td>60277</td><td>-</td><td>0</td><td>TATCTACTACACCGCCGTGC</td><td>GGG</td><td>||||||||[||||||||||||]PAM</td><td>NA</td><td>Intergenic</td><td>NA</td><td>NA</td></tr>
<tr><td>T29600</td><td>GGCCAGCGTCGACCTGTGCATGG</td><td>1000</td><td></td><td></td><td></td><td></td><td></td><td></td><td></td><td></td><td></td></tr>
<tr><td>Oligo fwd</td><td>CACCGGCCAGCGTCGACCTGTGCA</td><td></td><td></td><td></td><td></td><td></td><td></td><td></td><td></td><td></td><td></td></tr>
<tr><td>Oligo rev</td><td>AAACTGCACAGGTCGACGCTGGCC</td><td></td><td></td><td></td><td></td><td></td><td></td><td></td><td></td><td></td><td></td></tr>
<tr><td>Chromosome</td><td>start</td><td>end</td><td>strand</td><td>MM</td><td>target_seq</td><td>PAM</td><td>alignment</td><td>distance</td><td>position</td><td>gene name</td><td>gene id</td></tr>
<tr><td>GU305914.1</td><td>60234</td><td>60256</td><td>-</td><td>0</td><td>GGCCAGCGTCGACCTGTGCA</td><td>TGG</td><td>||||||||[||||||||||||]PAM</td><td>NA</td><td>Intergenic</td><td>NA</td><td>NA</td></tr>
<tr><td>T29601</td><td>CAGCGTCGACCTGTGCATGGCGG</td><td>1000</td><td></td><td></td><td></td><td></td><td></td><td></td><td></td><td></td><td></td></tr>
<tr><td>Oligo fwd</td><td>CACCCAGCGTCGACCTGTGCATGG</td><td></td><td></td><td></td><td></td><td></td><td></td><td></td><td></td><td></td><td></td></tr>
<tr><td>Oligo rev</td><td>AAACCCATGCACAGGTCGACGCTG</td><td></td><td></td><td></td><td></td><td></td><td></td><td></td><td></td><td></td><td></td></tr>
<tr><td>Chromosome</td><td>start</td><td>end</td><td>strand</td><td>MM</td><td>target_seq</td><td>PAM</td><td>alignment</td><td>distance</td><td>position</td><td>gene name</td><td>gene id</td></tr>
<tr><td>GU305914.1</td><td>60231</td><td>60253</td><td>-</td><td>0</td><td>CAGCGTCGACCTGTGCATGG</td><td>CGG</td><td>||||||||[||||||||||||]PAM</td><td>NA</td><td>Intergenic</td><td>NA</td><td>NA</td></tr>
<tr><td>T29602</td><td>AGCGTCGACCTGTGCATGGCGGG</td><td>1000</td><td></td><td></td><td></td><td></td><td></td><td></td><td></td><td></td><td></td></tr>
<tr><td>Oligo fwd</td><td>CACCAGCGTCGACCTGTGCATGGC</td><td></td><td></td><td></td><td></td><td></td><td></td><td></td><td></td><td></td><td></td></tr>
<tr><td>Oligo rev</td><td>AAACGCCATGCACAGGTCGACGCT</td><td></td><td></td><td></td><td></td><td></td><td></td><td></td><td></td><td></td><td></td></tr>
<tr><td>Chromosome</td><td>start</td><td>end</td><td>strand</td><td>MM</td><td>target_seq</td><td>PAM</td><td>alignment</td><td>distance</td><td>position</td><td>gene name</td><td>gene id</td></tr>
<tr><td>GU305914.1</td><td>60230</td><td>60252</td><td>-</td><td>0</td><td>AGCGTCGACCTGTGCATGGC</td><td>GGG</td><td>||||||||[||||||||||||]PAM</td><td>NA</td><td>Intergenic</td><td>NA</td><td>NA</td></tr>
<tr><td>T29603</td><td>GTTCCCTAAGAGCCTGTACGCGG</td><td>1000</td><td></td><td></td><td></td><td></td><td></td><td></td><td></td><td></td><td></td></tr>
<tr><td>Oligo fwd</td><td>CACCGTTCCCTAAGAGCCTGTACG</td><td></td><td></td><td></td><td></td><td></td><td></td><td></td><td></td><td></td><td></td></tr>
<tr><td>Oligo rev</td><td>AAACCGTACAGGCTCTTAGGGAAC</td><td></td><td></td><td></td><td></td><td></td><td></td><td></td><td></td><td></td><td></td></tr>
<tr><td>Chromosome</td><td>start</td><td>end</td><td>strand</td><td>MM</td><td>target_seq</td><td>PAM</td><td>alignment</td><td>distance</td><td>position</td><td>gene name</td><td>gene id</td></tr>
<tr><td>GU305914.1</td><td>60189</td><td>60211</td><td>-</td><td>0</td><td>GTTCCCTAAGAGCCTGTACG</td><td>CGG</td><td>||||||||[||||||||||||]PAM</td><td>NA</td><td>Intergenic</td><td>NA</td><td>NA</td></tr>
<tr><td>T29604</td><td>CCTAAGAGCCTGTACGCGGCCGG</td><td>1000</td><td></td><td></td><td></td><td></td><td></td><td></td><td></td><td></td><td></td></tr>
<tr><td>Oligo fwd</td><td>CACCCCTAAGAGCCTGTACGCGGC</td><td></td><td></td><td></td><td></td><td></td><td></td><td></td><td></td><td></td><td></td></tr>
<tr><td>Oligo rev</td><td>AAACGCCGCGTACAGGCTCTTAGG</td><td></td><td></td><td></td><td></td><td></td><td></td><td></td><td></td><td></td><td></td></tr>
<tr><td>Chromosome</td><td>start</td><td>end</td><td>strand</td><td>MM</td><td>target_seq</td><td>PAM</td><td>alignment</td><td>distance</td><td>position</td><td>gene name</td><td>gene id</td></tr>
<tr><td>GU305914.1</td><td>60185</td><td>60207</td><td>-</td><td>0</td><td>CCTAAGAGCCTGTACGCGGC</td><td>CGG</td><td>||||||||[||||||||||||]PAM</td><td>NA</td><td>Intergenic</td><td>NA</td><td>NA</td></tr>
<tr><td>T29605</td><td>CTCTTCGACGACGACGAGCGCGG</td><td>1000</td><td></td><td></td><td></td><td></td><td></td><td></td><td></td><td></td><td></td></tr>
<tr><td>Oligo fwd</td><td>CACCCTCTTCGACGACGACGAGCG</td><td></td><td></td><td></td><td></td><td></td><td></td><td></td><td></td><td></td><td></td></tr>
<tr><td>Oligo rev</td><td>AAACCGCTCGTCGTCGTCGAAGAG</td><td></td><td></td><td></td><td></td><td></td><td></td><td></td><td></td><td></td><td></td></tr>
<tr><td>Chromosome</td><td>start</td><td>end</td><td>strand</td><td>MM</td><td>target_seq</td><td>PAM</td><td>alignment</td><td>distance</td><td>position</td><td>gene name</td><td>gene id</td></tr>
</table>

| | | | | | | | | | | | |
|---|---|---|---|---|---|---|---|---|---|---|---|
| GU305914.1 | 60146 | 60168 | - | 0 | CTCTTCGACGACGACGAGCG | CGG | \|\|\|\|\|\|\|\|[\|\|\|\|\|\|\|\|\|\|\|\|]PAM | NA | Intergenic | NA | NA |
| T29606 | CGGCCCGCGTCGCATGACCAAGG | 1000 | | | | | | | | | |
| Oligo fwd | CACCCGGCCCGCGTCGCATGACCA | | | | | | | | | | |
| Oligo rev | AAACTGGTCATGCGACGCGGGCCG | | | | | | | | | | |
| Chromosome | start | end | strand | MM | target_seq | PAM | alignment | distance | position | gene name | gene id |
| GU305914.1 | 60126 | 60148 | - | 0 | CGGCCCGCGTCGCATGACCA | AGG | \|\|\|\|\|\|\|\|[\|\|\|\|\|\|\|\|\|\|\|\|]PAM | NA | Intergenic | NA | NA |
| T29607 | CGAGAAACTACGTGAAGACGTGG | 1000 | | | | | | | | | |
| Oligo fwd | CACCCGAGAAACTACGTGAAGACG | | | | | | | | | | |
| Oligo rev | AAACCGTCTTCACGTAGTTTCTCG | | | | | | | | | | |
| Chromosome | start | end | strand | MM | target_seq | PAM | alignment | distance | position | gene name | gene id |
| GU305914.1 | 60096 | 60118 | - | 0 | CGAGAAACTACGTGAAGACG | TGG | \|\|\|\|\|\|\|\|[\|\|\|\|\|\|\|\|\|\|\|\|]PAM | NA | Intergenic | NA | NA |
| T29608 | CGTGAAGACGTGGTGAAACACGG | 1000 | | | | | | | | | |
| Oligo fwd | CACCCGTGAAGACGTGGTGAAACA | | | | | | | | | | |
| Oligo rev | AAACTGTTTCACCACGTCTTCACG | | | | | | | | | | |
| Chromosome | start | end | strand | MM | target_seq | PAM | alignment | distance | position | gene name | gene id |
| GU305914.1 | 60086 | 60108 | - | 0 | CGTGAAGACGTGGTGAAACA | CGG | \|\|\|\|\|\|\|\|[\|\|\|\|\|\|\|\|\|\|\|\|]PAM | NA | Intergenic | NA | NA |
| T29609 | ACGTGGTGAAACACGGCATCAGG | 1000 | | | | | | | | | |
| Oligo fwd | CACCACGTGGTGAAACACGGCATC | | | | | | | | | | |
| Oligo rev | AAACGATGCCGTGTTTCACCACGT | | | | | | | | | | |
| Chromosome | start | end | strand | MM | target_seq | PAM | alignment | distance | position | gene name | gene id |
| GU305914.1 | 60079 | 60101 | - | 0 | ACGTGGTGAAACACGGCATC | AGG | \|\|\|\|\|\|\|\|[\|\|\|\|\|\|\|\|\|\|\|\|]PAM | NA | Intergenic | NA | NA |
| T29610 | AGGAACGCCTCCTTCATCACCGG | 1000 | | | | | | | | | |
| Oligo fwd | CACCAGGAACGCCTCCTTCATCAC | | | | | | | | | | |
| Oligo rev | AAACGTGATGAAGGAGGCGTTCCT | | | | | | | | | | |
| Chromosome | start | end | strand | MM | target_seq | PAM | alignment | distance | position | gene name | gene id |
| GU305914.1 | 60059 | 60081 | - | 0 | AGGAACGCCTCCTTCATCAC | CGG | \|\|\|\|\|\|\|\|[\|\|\|\|\|\|\|\|\|\|\|\|]PAM | NA | Intergenic | NA | NA |
| T29611 | CTCCTTCATCACCGGCTGCTCGG | 1000 | | | | | | | | | |
| Oligo fwd | CACCCTCCTTCATCACCGGCTGCT | | | | | | | | | | |
| Oligo rev | AAACAGCAGCCGGTGATGAAGGAG | | | | | | | | | | |
| Chromosome | start | end | strand | MM | target_seq | PAM | alignment | distance | position | gene name | gene id |
| GU305914.1 | 60051 | 60073 | - | 0 | CTCCTTCATCACCGGCTGCT | CGG | \|\|\|\|\|\|\|\|[\|\|\|\|\|\|\|\|\|\|\|\|]PAM | NA | Intergenic | NA | NA |
| T29612 | GGCCGACGAAGCCGCCAACCTGG | 1000 | | | | | | | | | |
| Oligo fwd | CACCGGCCGACGAAGCCGCCAACC | | | | | | | | | | |
| Oligo rev | AAACGGTTGGCGGCTTCGTCGGCC | | | | | | | | | | |
| Chromosome | start | end | strand | MM | target_seq | PAM | alignment | distance | position | gene name | gene id |
| GU305914.1 | 60030 | 60052 | - | 0 | GGCCGACGAAGCCGCCAACC | TGG | \|\|\|\|\|\|\|\|[\|\|\|\|\|\|\|\|\|\|\|\|]PAM | NA | Intergenic | NA | NA |
| T29613 | CGACGAAGCCGCCAACCTGGCGG | 1000 | | | | | | | | | |
| Oligo fwd | CACCCGACGAAGCCGCCAACCTGG | | | | | | | | | | |
| Oligo rev | AAACCCAGGTTGGCGGCTTCGTCG | | | | | | | | | | |
| Chromosome | start | end | strand | MM | target_seq | PAM | alignment | distance | position | gene name | gene id |
| GU305914.1 | 60027 | 60049 | - | 0 | CGACGAAGCCGCCAACCTGG | CGG | \|\|\|\|\|\|\|\|[\|\|\|\|\|\|\|\|\|\|\|\|]PAM | NA | Intergenic | NA | NA |
| T29614 | GACGAAGCCGCCAACCTGGCGGG | 1000 | | | | | | | | | |
| Oligo fwd | CACCGACGAAGCCGCCAACCTGGC | | | | | | | | | | |
| Oligo rev | AAACGCCAGGTTGGCGGCTTCGTC | | | | | | | | | | |
| Chromosome | start | end | strand | MM | target_seq | PAM | alignment | distance | position | gene name | gene id |
| GU305914.1 | 60026 | 60048 | - | 0 | GACGAAGCCGCCAACCTGGC | GGG | \|\|\|\|\|\|\|\|[\|\|\|\|\|\|\|\|\|\|\|\|]PAM | NA | Intergenic | NA | NA |
| T29615 | CAACCTGGCGGGCACCACTCCGG | 1000 | | | | | | | | | |
| Oligo fwd | CACCCAACCTGGCGGGCACCACTC | | | | | | | | | | |
| Oligo rev | AAACGAGTGGTGCCCGCCAGGTTG | | | | | | | | | | |
| Chromosome | start | end | strand | MM | target_seq | PAM | alignment | distance | position | gene name | gene id |
| GU305914.1 | 60015 | 60037 | - | 0 | CAACCTGGCGGGCACCACTC | CGG | \|\|\|\|\|\|\|\|[\|\|\|\|\|\|\|\|\|\|\|\|]PAM | NA | Intergenic | NA | NA |
| T29616 | AACCTGGCGGGCACCACTCCGGG | 1000 | | | | | | | | | |
| Oligo fwd | CACCAACCTGGCGGGCACCACTCC | | | | | | | | | | |
| Oligo rev | AAACGGAGTGGTGCCCGCCAGGTT | | | | | | | | | | |
| Chromosome | start | end | strand | MM | target_seq | PAM | alignment | distance | position | gene name | gene id |
| GU305914.1 | 60014 | 60036 | - | 0 | AACCTGGCGGGCACCACTCC | GGG | \|\|\|\|\|\|\|\|[\|\|\|\|\|\|\|\|\|\|\|\|]PAM | NA | Intergenic | NA | NA |
| T29617 | CGGGCACCACTCCGGGCTTCTGG | 1000 | | | | | | | | | |
| Oligo fwd | CACCCGGGCACCACTCCGGGCTTC | | | | | | | | | | |
| Oligo rev | AAACGAAGCCCGGAGTGGTGCCCG | | | | | | | | | | |
| Chromosome | start | end | strand | MM | target_seq | PAM | alignment | distance | position | gene name | gene id |
| GU305914.1 | 60007 | 60029 | - | 0 | CGGGCACCACTCCGGGCTTC | TGG | \|\|\|\|\|\|\|\|[\|\|\|\|\|\|\|\|\|\|\|\|]PAM | NA | Intergenic | NA | NA |
| T29618 | GCTCATGATGACCCCGACGAAGG | 1000 | | | | | | | | | |
| Oligo fwd | CACCGCTCATGATGACCCCGACGA | | | | | | | | | | |
| Oligo rev | AAACTCGTCGGGGTCATCATGAGC | | | | | | | | | | |
| Chromosome | start | end | strand | MM | target_seq | PAM | alignment | distance | position | gene name | gene id |
| GU305914.1 | 59949 | 59971 | - | 0 | GCTCATGATGACCCCGACGA | AGG | \|\|\|\|\|\|\|\|[\|\|\|\|\|\|\|\|\|\|\|\|]PAM | NA | Intergenic | NA | NA |
| T29619 | CGAGCCCCATCGCCTGCACGAGG | 1000 | | | | | | | | | |
| Oligo fwd | CACCCGAGCCCCATCGCCTGCACG | | | | | | | | | | |
| Oligo rev | AAACCGTGCAGGCGATGGGGCTCG | | | | | | | | | | |
| Chromosome | start | end | strand | MM | target_seq | PAM | alignment | distance | position | gene name | gene id |
| GU305914.1 | 59889 | 59911 | - | 0 | CGAGCCCCATCGCCTGCACG | AGG | \|\|\|\|\|\|\|\|[\|\|\|\|\|\|\|\|\|\|\|\|]PAM | NA | Intergenic | NA | NA |
| T29620 | GCCCCATCGCCTGCACGAGGAGG | 1000 | | | | | | | | | |
| Oligo fwd | CACCGCCCCATCGCCTGCACGAGG | | | | | | | | | | |
| Oligo rev | AAACCCTCGTGCAGGCGATGGGGC | | | | | | | | | | |

Chromosome start end strand MM target_seq PAM alignment distance position gene name gene id
GU305914.1 59886 59908 - 0 GCCCCATCGCCTGCACGAGG AGG ||||||||[||||||||||||]PAM NA Intergenic NA NA

T29621 GAGGAGGATCTCTCCTGTCTCGG 1000
Oligo fwd CACCGAGGAGGATCTCTCCTGTCT
Oligo rev AAACAGACAGGAGAGATCCTCCTC
Chromosome start end strand MM target_seq PAM alignment distance position gene name gene id
GU305914.1 59870 59892 - 0 GAGGAGGATCTCTCCTGTCT CGG ||||||||[||||||||||||]PAM NA Intergenic NA NA

T29622 TCTCCTGTCTCGGCGAGAACCGG 1000
Oligo fwd CACCTCTCCTGTCTCGGCGAGAAC
Oligo rev AAACGTTCTCGCCGAGACAGGAGA
Chromosome start end strand MM target_seq PAM alignment distance position gene name gene id
GU305914.1 59860 59882 - 0 TCTCCTGTCTCGGCGAGAAC CGG ||||||||[||||||||||||]PAM NA Intergenic NA NA

T29623 CCGTGCTCAACAGCCGCCTCAGG 1000
Oligo fwd CACCCCGTGCTCAACAGCCGCCTC
Oligo rev AAACGAGGCGGCTGTTGAGCACGG
Chromosome start end strand MM target_seq PAM alignment distance position gene name gene id
GU305914.1 59824 59846 - 0 CCGTGCTCAACAGCCGCCTC AGG ||||||||[||||||||||||]PAM NA Intergenic NA NA

T29624 GATCTCGAAAGAGTCCGCGACGG 1000
Oligo fwd CACCGATCTCGAAAGAGTCCGCGA
Oligo rev AAACTCGCGGACTCTTTCGAGATC
Chromosome start end strand MM target_seq PAM alignment distance position gene name gene id
GU305914.1 59799 59821 - 0 GATCTCGAAAGAGTCCGCGA CGG ||||||||[||||||||||||]PAM NA Intergenic NA NA

T29625 GCGACGGTCGCCGTGCGCCGCGG 1000
Oligo fwd CACCGCGACGGTCGCCGTGCGCCG
Oligo rev AAACCGGCGCACGGCGACCGTCGC
Chromosome start end strand MM target_seq PAM alignment distance position gene name gene id
GU305914.1 59783 59805 - 0 GCGACGGTCGCCGTGCGCCG CGG ||||||||[||||||||||||]PAM NA Intergenic NA NA

T29626 CGCCGTGCGCCGCGGCCGCTCGG 1000
Oligo fwd CACCCGCCGTGCGCCGCGGCCGCT
Oligo rev AAACAGCGGCCGCGGCGCACGGCG
Chromosome start end strand MM target_seq PAM alignment distance position gene name gene id
GU305914.1 59775 59797 - 0 CGCCGTGCGCCGCGGCCGCT CGG ||||||||[||||||||||||]PAM NA Intergenic NA NA

T29627 GGCGCCCTTCTACGACGACTCGG 1000
Oligo fwd CACCGGCGCCCTTCTACGACGACT
Oligo rev AAACAGTCGTCGTAGAAGGGCGCC
Chromosome start end strand MM target_seq PAM alignment distance position gene name gene id
GU305914.1 59754 59776 - 0 GGCGCCCTTCTACGACGACT CGG ||||||||[||||||||||||]PAM NA Intergenic NA NA

T29628 CTACGACGACTCGGACGACGAGG 1000
Oligo fwd CACCCTACGACGACTCGGACGACG
Oligo rev AAACCGTCGTCCGAGTCGTCGTAG
Chromosome start end strand MM target_seq PAM alignment distance position gene name gene id
GU305914.1 59745 59767 - 0 CTACGACGACTCGGACGACG AGG ||||||||[||||||||||||]PAM NA Intergenic NA NA

T29629 CGACTCGGACGACGAGGACGAGG 1000
Oligo fwd CACCCGACTCGGACGACGAGGACG
Oligo rev AAACCGTCCTCGTCGTCCGAGTCG
Chromosome start end strand MM target_seq PAM alignment distance position gene name gene id
GU305914.1 59739 59761 - 0 CGACTCGGACGACGAGGACG AGG ||||||||[||||||||||||]PAM NA Intergenic NA NA
GU305914.1 123785 123807 - 2 CGACCCGTACGACGAGGACG CGG ||||-||-[||||||||||||]PAM NA Intergenic NA NA
GU305914.1 77043 77065 - 4 GGAGTCGGTCGACGAGGATG AGG -||-||||[-|||||||||-|]PAM NA Intergenic NA NA
GU305914.1 116060 116082 - 3 CGACTCGTACGAGGAGGAGG CGG |||||||-[||||-|||||-|]PAM NA Intergenic NA NA

T29630 CTCGGACGACGAGGACGAGGTGG 1000
Oligo fwd CACCCTCGGACGACGAGGACGAGG
Oligo rev AAACCCTCGTCCTCGTCGTCCGAG
Chromosome start end strand MM target_seq PAM alignment distance position gene name gene id
GU305914.1 59736 59758 - 0 CTCGGACGACGAGGACGAGG TGG ||||||||[||||||||||||]PAM NA Intergenic NA NA
GU305914.1 77040 77062 - 3 GTCGGTCGACGAGGATGAGG AGG -||||-||[|||||||-||||]PAM NA Intergenic NA NA
GU305914.1 96811 96833 - 4 CGCCGGCGACGAGGACGAAG GGG |-|-|-||[||||||||||-|]PAM NA Intergenic NA NA

T29631 GAGGTGGCGTGCTCCGAGACCGG 1000
Oligo fwd CACCGAGGTGGCGTGCTCCGAGAC
Oligo rev AAACGTCTCGGAGCACGCCACCTC
Chromosome start end strand MM target_seq PAM alignment distance position gene name gene id
GU305914.1 59720 59742 - 0 GAGGTGGCGTGCTCCGAGAC CGG ||||||||[||||||||||||]PAM NA Intergenic NA NA

T29632 TGGCGTGCTCCGAGACCGGCTGG 1000
Oligo fwd CACCTGGCGTGCTCCGAGACCGGC
Oligo rev AAACGCCGGTCTCGGAGCACGCCA
Chromosome start end strand MM target_seq PAM alignment distance position gene name gene id
GU305914.1 59716 59738 - 0 TGGCGTGCTCCGAGACCGGC TGG ||||||||[||||||||||||]PAM NA Intergenic NA NA

T29633 CTGGACCGTGTCGACCGACGCGG 1000
Oligo fwd CACCCTGGACCGTGTCGACCGACG
Oligo rev AAACCGTCGGTCGACACGGTCCAG
Chromosome start end strand MM target_seq PAM alignment distance position gene name gene id
GU305914.1 59697 59719 - 0 CTGGACCGTGTCGACCGACG CGG ||||||||[||||||||||||]PAM NA Intergenic NA NA

T29634 CAGTCGCTGCCCGTCGCCATCGG 1000
Oligo fwd CACCCAGTCGCTGCCCGTCGCCAT
Oligo rev AAACATGGCGACGGGCAGCGACTG

<table>
<tr><td>Chromosome</td><td>start</td><td>end</td><td>strand</td><td>MM</td><td>target_seq</td><td>PAM</td><td>alignment</td><td>distance</td><td>position</td><td>gene name</td><td>gene id</td></tr>
<tr><td>GU305914.1</td><td>59630</td><td>59652</td><td>-</td><td>0</td><td>CAGTCGCTGCCCGTCGCCAT</td><td>CGG</td><td>||||||||[||||||||||||]PAM</td><td>NA</td><td>Intergenic</td><td>NA</td><td>NA</td></tr>
<tr><td></td><td></td><td></td><td></td><td></td><td></td><td></td><td></td><td></td><td></td><td></td><td></td></tr>
<tr><td>T29635</td><td>CTGCCCGTCGCCATCGGCTTCGG</td><td>1000</td><td></td><td></td><td></td><td></td><td></td><td></td><td></td><td></td><td></td></tr>
<tr><td>Oligo fwd</td><td>CACCCTGCCCGTCGCCATCGGCTT</td><td></td><td></td><td></td><td></td><td></td><td></td><td></td><td></td><td></td><td></td></tr>
<tr><td>Oligo rev</td><td>AAACAAGCCGATGGCGACGGGCAG</td><td></td><td></td><td></td><td></td><td></td><td></td><td></td><td></td><td></td><td></td></tr>
<tr><td>Chromosome</td><td>start</td><td>end</td><td>strand</td><td>MM</td><td>target_seq</td><td>PAM</td><td>alignment</td><td>distance</td><td>position</td><td>gene name</td><td>gene id</td></tr>
<tr><td>GU305914.1</td><td>59624</td><td>59646</td><td>-</td><td>0</td><td>CTGCCCGTCGCCATCGGCTT</td><td>CGG</td><td>||||||||[||||||||||||]PAM</td><td>NA</td><td>Intergenic</td><td>NA</td><td>NA</td></tr>
<tr><td></td><td></td><td></td><td></td><td></td><td></td><td></td><td></td><td></td><td></td><td></td><td></td></tr>
<tr><td>T29636</td><td>TGCCCGTCGCCATCGGCTTCGGG</td><td>1000</td><td></td><td></td><td></td><td></td><td></td><td></td><td></td><td></td><td></td></tr>
<tr><td>Oligo fwd</td><td>CACCTGCCCGTCGCCATCGGCTTC</td><td></td><td></td><td></td><td></td><td></td><td></td><td></td><td></td><td></td><td></td></tr>
<tr><td>Oligo rev</td><td>AAACGAAGCCGATGGCGACGGGCA</td><td></td><td></td><td></td><td></td><td></td><td></td><td></td><td></td><td></td><td></td></tr>
<tr><td>Chromosome</td><td>start</td><td>end</td><td>strand</td><td>MM</td><td>target_seq</td><td>PAM</td><td>alignment</td><td>distance</td><td>position</td><td>gene name</td><td>gene id</td></tr>
<tr><td>GU305914.1</td><td>59623</td><td>59645</td><td>-</td><td>0</td><td>TGCCCGTCGCCATCGGCTTC</td><td>GGG</td><td>||||||||[||||||||||||]PAM</td><td>NA</td><td>Intergenic</td><td>NA</td><td>NA</td></tr>
<tr><td></td><td></td><td></td><td></td><td></td><td></td><td></td><td></td><td></td><td></td><td></td><td></td></tr>
<tr><td>T29637</td><td>GCCCGTCGCCATCGGCTTCGGGG</td><td>1000</td><td></td><td></td><td></td><td></td><td></td><td></td><td></td><td></td><td></td></tr>
<tr><td>Oligo fwd</td><td>CACCGCCCGTCGCCATCGGCTTCG</td><td></td><td></td><td></td><td></td><td></td><td></td><td></td><td></td><td></td><td></td></tr>
<tr><td>Oligo rev</td><td>AAACCGAAGCCGATGGCGACGGGC</td><td></td><td></td><td></td><td></td><td></td><td></td><td></td><td></td><td></td><td></td></tr>
<tr><td>Chromosome</td><td>start</td><td>end</td><td>strand</td><td>MM</td><td>target_seq</td><td>PAM</td><td>alignment</td><td>distance</td><td>position</td><td>gene name</td><td>gene id</td></tr>
<tr><td>GU305914.1</td><td>59622</td><td>59644</td><td>-</td><td>0</td><td>GCCCGTCGCCATCGGCTTCG</td><td>GGG</td><td>||||||||[||||||||||||]PAM</td><td>NA</td><td>Intergenic</td><td>NA</td><td>NA</td></tr>
<tr><td></td><td></td><td></td><td></td><td></td><td></td><td></td><td></td><td></td><td></td><td></td><td></td></tr>
<tr><td>T29638</td><td>CCCGTCGCCATCGGCTTCGGGGG</td><td>1000</td><td></td><td></td><td></td><td></td><td></td><td></td><td></td><td></td><td></td></tr>
<tr><td>Oligo fwd</td><td>CACCCCCGTCGCCATCGGCTTCGG</td><td></td><td></td><td></td><td></td><td></td><td></td><td></td><td></td><td></td><td></td></tr>
<tr><td>Oligo rev</td><td>AAACCCGAAGCCGATGGCGACGGG</td><td></td><td></td><td></td><td></td><td></td><td></td><td></td><td></td><td></td><td></td></tr>
<tr><td>Chromosome</td><td>start</td><td>end</td><td>strand</td><td>MM</td><td>target_seq</td><td>PAM</td><td>alignment</td><td>distance</td><td>position</td><td>gene name</td><td>gene id</td></tr>
<tr><td>GU305914.1</td><td>59621</td><td>59643</td><td>-</td><td>0</td><td>CCCGTCGCCATCGGCTTCGG</td><td>GGG</td><td>||||||||[||||||||||||]PAM</td><td>NA</td><td>Intergenic</td><td>NA</td><td>NA</td></tr>
<tr><td></td><td></td><td></td><td></td><td></td><td></td><td></td><td></td><td></td><td></td><td></td><td></td></tr>
<tr><td>T29639</td><td>CCGTCGCCATCGGCTTCGGGGGG</td><td>1000</td><td></td><td></td><td></td><td></td><td></td><td></td><td></td><td></td><td></td></tr>
<tr><td>Oligo fwd</td><td>CACCCCGTCGCCATCGGCTTCGGG</td><td></td><td></td><td></td><td></td><td></td><td></td><td></td><td></td><td></td><td></td></tr>
<tr><td>Oligo rev</td><td>AAACCCCGAAGCCGATGGCGACGG</td><td></td><td></td><td></td><td></td><td></td><td></td><td></td><td></td><td></td><td></td></tr>
<tr><td>Chromosome</td><td>start</td><td>end</td><td>strand</td><td>MM</td><td>target_seq</td><td>PAM</td><td>alignment</td><td>distance</td><td>position</td><td>gene name</td><td>gene id</td></tr>
<tr><td>GU305914.1</td><td>59620</td><td>59642</td><td>-</td><td>0</td><td>CCGTCGCCATCGGCTTCGGG</td><td>GGG</td><td>||||||||[||||||||||||]PAM</td><td>NA</td><td>Intergenic</td><td>NA</td><td>NA</td></tr>
<tr><td></td><td></td><td></td><td></td><td></td><td></td><td></td><td></td><td></td><td></td><td></td><td></td></tr>
<tr><td>T29640</td><td>CATCGGCTTCGGGGGGTCTTCGG</td><td>1000</td><td></td><td></td><td></td><td></td><td></td><td></td><td></td><td></td><td></td></tr>
<tr><td>Oligo fwd</td><td>CACCCATCGGCTTCGGGGGGTCTT</td><td></td><td></td><td></td><td></td><td></td><td></td><td></td><td></td><td></td><td></td></tr>
<tr><td>Oligo rev</td><td>AAACAAGACCCCCCGAAGCCGATG</td><td></td><td></td><td></td><td></td><td></td><td></td><td></td><td></td><td></td><td></td></tr>
<tr><td>Chromosome</td><td>start</td><td>end</td><td>strand</td><td>MM</td><td>target_seq</td><td>PAM</td><td>alignment</td><td>distance</td><td>position</td><td>gene name</td><td>gene id</td></tr>
<tr><td>GU305914.1</td><td>59613</td><td>59635</td><td>-</td><td>0</td><td>CATCGGCTTCGGGGGGTCTT</td><td>CGG</td><td>||||||||[||||||||||||]PAM</td><td>NA</td><td>Intergenic</td><td>NA</td><td>NA</td></tr>
<tr><td></td><td></td><td></td><td></td><td></td><td></td><td></td><td></td><td></td><td></td><td></td><td></td></tr>
<tr><td>T29641</td><td>CGGCTTCGGGGGGTCTTCGGTGG</td><td>1000</td><td></td><td></td><td></td><td></td><td></td><td></td><td></td><td></td><td></td></tr>
<tr><td>Oligo fwd</td><td>CACCCGGCTTCGGGGGGTCTTCGG</td><td></td><td></td><td></td><td></td><td></td><td></td><td></td><td></td><td></td><td></td></tr>
<tr><td>Oligo rev</td><td>AAACCCGAAGACCCCCCGAAGCCG</td><td></td><td></td><td></td><td></td><td></td><td></td><td></td><td></td><td></td><td></td></tr>
<tr><td>Chromosome</td><td>start</td><td>end</td><td>strand</td><td>MM</td><td>target_seq</td><td>PAM</td><td>alignment</td><td>distance</td><td>position</td><td>gene name</td><td>gene id</td></tr>
<tr><td>GU305914.1</td><td>59610</td><td>59632</td><td>-</td><td>0</td><td>CGGCTTCGGGGGGTCTTCGG</td><td>TGG</td><td>||||||||[||||||||||||]PAM</td><td>NA</td><td>Intergenic</td><td>NA</td><td>NA</td></tr>
<tr><td></td><td></td><td></td><td></td><td></td><td></td><td></td><td></td><td></td><td></td><td></td><td></td></tr>
<tr><td>T29642</td><td>CGGGGGGTCTTCGGTGGAATTGG</td><td>1000</td><td></td><td></td><td></td><td></td><td></td><td></td><td></td><td></td><td></td></tr>
<tr><td>Oligo fwd</td><td>CACCCGGGGGGTCTTCGGTGGAAT</td><td></td><td></td><td></td><td></td><td></td><td></td><td></td><td></td><td></td><td></td></tr>
<tr><td>Oligo rev</td><td>AAACATTCCACCGAAGACCCCCCG</td><td></td><td></td><td></td><td></td><td></td><td></td><td></td><td></td><td></td><td></td></tr>
<tr><td>Chromosome</td><td>start</td><td>end</td><td>strand</td><td>MM</td><td>target_seq</td><td>PAM</td><td>alignment</td><td>distance</td><td>position</td><td>gene name</td><td>gene id</td></tr>
<tr><td>GU305914.1</td><td>59604</td><td>59626</td><td>-</td><td>0</td><td>CGGGGGGTCTTCGGTGGAAT</td><td>TGG</td><td>||||||||[||||||||||||]PAM</td><td>NA</td><td>Intergenic</td><td>NA</td><td>NA</td></tr>
<tr><td></td><td></td><td></td><td></td><td></td><td></td><td></td><td></td><td></td><td></td><td></td><td></td></tr>
<tr><td>T29643</td><td>TTGGCGCGTCATCTGAGACGAGG</td><td>1000</td><td></td><td></td><td></td><td></td><td></td><td></td><td></td><td></td><td></td></tr>
<tr><td>Oligo fwd</td><td>CACCTTGGCGCGTCATCTGAGACG</td><td></td><td></td><td></td><td></td><td></td><td></td><td></td><td></td><td></td><td></td></tr>
<tr><td>Oligo rev</td><td>AAACCGTCTCAGATGACGCGCCAA</td><td></td><td></td><td></td><td></td><td></td><td></td><td></td><td></td><td></td><td></td></tr>
<tr><td>Chromosome</td><td>start</td><td>end</td><td>strand</td><td>MM</td><td>target_seq</td><td>PAM</td><td>alignment</td><td>distance</td><td>position</td><td>gene name</td><td>gene id</td></tr>
<tr><td>GU305914.1</td><td>59585</td><td>59607</td><td>-</td><td>0</td><td>TTGGCGCGTCATCTGAGACG</td><td>AGG</td><td>||||||||[||||||||||||]PAM</td><td>NA</td><td>Intergenic</td><td>NA</td><td>NA</td></tr>
<tr><td></td><td></td><td></td><td></td><td></td><td></td><td></td><td></td><td></td><td></td><td></td><td></td></tr>
<tr><td>T29644</td><td>TGGCGCGTCATCTGAGACGAGGG</td><td>1000</td><td></td><td></td><td></td><td></td><td></td><td></td><td></td><td></td><td></td></tr>
<tr><td>Oligo fwd</td><td>CACCTGGCGCGTCATCTGAGACGA</td><td></td><td></td><td></td><td></td><td></td><td></td><td></td><td></td><td></td><td></td></tr>
<tr><td>Oligo rev</td><td>AAACTCGTCTCAGATGACGCGCCA</td><td></td><td></td><td></td><td></td><td></td><td></td><td></td><td></td><td></td><td></td></tr>
<tr><td>Chromosome</td><td>start</td><td>end</td><td>strand</td><td>MM</td><td>target_seq</td><td>PAM</td><td>alignment</td><td>distance</td><td>position</td><td>gene name</td><td>gene id</td></tr>
<tr><td>GU305914.1</td><td>59584</td><td>59606</td><td>-</td><td>0</td><td>TGGCGCGTCATCTGAGACGA</td><td>GGG</td><td>||||||||[||||||||||||]PAM</td><td>NA</td><td>Intergenic</td><td>NA</td><td>NA</td></tr>
<tr><td></td><td></td><td></td><td></td><td></td><td></td><td></td><td></td><td></td><td></td><td></td><td></td></tr>
<tr><td>T29645</td><td>TCTGAGACGAGGGAACGCTCTGG</td><td>1000</td><td></td><td></td><td></td><td></td><td></td><td></td><td></td><td></td><td></td></tr>
<tr><td>Oligo fwd</td><td>CACCTCTGAGACGAGGGAACGCTC</td><td></td><td></td><td></td><td></td><td></td><td></td><td></td><td></td><td></td><td></td></tr>
<tr><td>Oligo rev</td><td>AAACGAGCGTTCCCTCGTCTCAGA</td><td></td><td></td><td></td><td></td><td></td><td></td><td></td><td></td><td></td><td></td></tr>
<tr><td>Chromosome</td><td>start</td><td>end</td><td>strand</td><td>MM</td><td>target_seq</td><td>PAM</td><td>alignment</td><td>distance</td><td>position</td><td>gene name</td><td>gene id</td></tr>
<tr><td>GU305914.1</td><td>59574</td><td>59596</td><td>-</td><td>0</td><td>TCTGAGACGAGGGAACGCTC</td><td>TGG</td><td>||||||||[||||||||||||]PAM</td><td>NA</td><td>Intergenic</td><td>NA</td><td>NA</td></tr>
<tr><td></td><td></td><td></td><td></td><td></td><td></td><td></td><td></td><td></td><td></td><td></td><td></td></tr>
<tr><td>T29646</td><td>CTGAGACGAGGGAACGCTCTGGG</td><td>1000</td><td></td><td></td><td></td><td></td><td></td><td></td><td></td><td></td><td></td></tr>
<tr><td>Oligo fwd</td><td>CACCCTGAGACGAGGGAACGCTCT</td><td></td><td></td><td></td><td></td><td></td><td></td><td></td><td></td><td></td><td></td></tr>
<tr><td>Oligo rev</td><td>AAACAGAGCGTTCCCTCGTCTCAG</td><td></td><td></td><td></td><td></td><td></td><td></td><td></td><td></td><td></td><td></td></tr>
<tr><td>Chromosome</td><td>start</td><td>end</td><td>strand</td><td>MM</td><td>target_seq</td><td>PAM</td><td>alignment</td><td>distance</td><td>position</td><td>gene name</td><td>gene id</td></tr>
<tr><td>GU305914.1</td><td>59573</td><td>59595</td><td>-</td><td>0</td><td>CTGAGACGAGGGAACGCTCT</td><td>GGG</td><td>||||||||[||||||||||||]PAM</td><td>NA</td><td>Intergenic</td><td>NA</td><td>NA</td></tr>
<tr><td></td><td></td><td></td><td></td><td></td><td></td><td></td><td></td><td></td><td></td><td></td><td></td></tr>
<tr><td>T29647</td><td>AACGCTCTGGGACTGTCCGTCGG</td><td>1000</td><td></td><td></td><td></td><td></td><td></td><td></td><td></td><td></td><td></td></tr>
<tr><td>Oligo fwd</td><td>CACCAACGCTCTGGGACTGTCCGT</td><td></td><td></td><td></td><td></td><td></td><td></td><td></td><td></td><td></td><td></td></tr>
<tr><td>Oligo rev</td><td>AAACACGGACAGTCCCAGAGCGTT</td><td></td><td></td><td></td><td></td><td></td><td></td><td></td><td></td><td></td><td></td></tr>
<tr><td>Chromosome</td><td>start</td><td>end</td><td>strand</td><td>MM</td><td>target_seq</td><td>PAM</td><td>alignment</td><td>distance</td><td>position</td><td>gene name</td><td>gene id</td></tr>
<tr><td>GU305914.1</td><td>59561</td><td>59583</td><td>-</td><td>0</td><td>AACGCTCTGGGACTGTCCGT</td><td>CGG</td><td>||||||||[||||||||||||]PAM</td><td>NA</td><td>Intergenic</td><td>NA</td><td>NA</td></tr>
<tr><td></td><td></td><td></td><td></td><td></td><td></td><td></td><td></td><td></td><td></td><td></td><td></td></tr>
<tr><td>T29648</td><td>CTTCCGTGAATTATCGCTGACGG</td><td>1000</td><td></td><td></td><td></td><td></td><td></td><td></td><td></td><td></td><td></td></tr>
<tr><td>Oligo fwd</td><td>CACCCTTCCGTGAATTATCGCTGA</td><td></td><td></td><td></td><td></td><td></td><td></td><td></td><td></td><td></td><td></td></tr>
<tr><td>Oligo rev</td><td>AAACTCAGCGATAATTCACGGAAG</td><td></td><td></td><td></td><td></td><td></td><td></td><td></td><td></td><td></td><td></td></tr>
<tr><td>Chromosome</td><td>start</td><td>end</td><td>strand</td><td>MM</td><td>target_seq</td><td>PAM</td><td>alignment</td><td>distance</td><td>position</td><td>gene name</td><td>gene id</td></tr>
<tr><td>GU305914.1</td><td>59515</td><td>59537</td><td>-</td><td>0</td><td>CTTCCGTGAATTATCGCTGA</td><td>CGG</td><td>||||||||[||||||||||||]PAM</td><td>NA</td><td>Intergenic</td><td>NA</td><td>NA</td></tr>
<tr><td></td><td></td><td></td><td></td><td></td><td></td><td></td><td></td><td></td><td></td><td></td><td></td></tr>
<tr><td>T29649</td><td>AAAAACGAGTTCTATTGCTCTGG</td><td>1000</td><td></td><td></td><td></td><td></td><td></td><td></td><td></td><td></td><td></td></tr>
<tr><td>Oligo fwd</td><td>CACCAAAAACGAGTTCTATTGCTC</td><td></td><td></td><td></td><td></td><td></td><td></td><td></td><td></td><td></td><td></td></tr>
</table>

<table>
<tr><td>Oligo rev</td><td>AAACGAGCAATAGAACTCGTTTTT</td><td></td><td></td><td></td><td></td><td></td><td></td><td></td><td></td><td></td><td></td><td></td></tr>
<tr><td>Chromosome</td><td>start</td><td></td><td>end</td><td>strand</td><td>MM</td><td>target_seq</td><td>PAM</td><td>alignment</td><td>distance</td><td>position</td><td>gene name</td><td>gene id</td></tr>
<tr><td>GU305914.1</td><td></td><td>59478</td><td>59500</td><td>-</td><td>0</td><td>AAAAACGAGTTCTATTGCTC</td><td>TGG</td><td>||||||||[||||||||||||]PAM</td><td>NA</td><td>Intergenic</td><td>NA</td><td>NA</td></tr>
<tr><td>T29650</td><td>CTATTGCTCTGGCGCTCGAGTGG</td><td></td><td>1000</td><td></td><td></td><td></td><td></td><td></td><td></td><td></td><td></td><td></td></tr>
<tr><td>Oligo fwd</td><td>CACCCTATTGCTCTGGCGCTCGAG</td><td></td><td></td><td></td><td></td><td></td><td></td><td></td><td></td><td></td><td></td><td></td></tr>
<tr><td>Oligo rev</td><td>AAACCTCGAGCGCCAGAGCAATAG</td><td></td><td></td><td></td><td></td><td></td><td></td><td></td><td></td><td></td><td></td><td></td></tr>
<tr><td>Chromosome</td><td>start</td><td></td><td>end</td><td>strand</td><td>MM</td><td>target_seq</td><td>PAM</td><td>alignment</td><td>distance</td><td>position</td><td>gene name</td><td>gene id</td></tr>
<tr><td>GU305914.1</td><td></td><td>59467</td><td>59489</td><td>-</td><td>0</td><td>CTATTGCTCTGGCGCTCGAG</td><td>TGG</td><td>||||||||[||||||||||||]PAM</td><td>NA</td><td>Intergenic</td><td>NA</td><td>NA</td></tr>
<tr><td>T29651</td><td>TATTGCTCTGGCGCTCGAGTGGG</td><td></td><td>1000</td><td></td><td></td><td></td><td></td><td></td><td></td><td></td><td></td><td></td></tr>
<tr><td>Oligo fwd</td><td>CACCTATTGCTCTGGCGCTCGAGT</td><td></td><td></td><td></td><td></td><td></td><td></td><td></td><td></td><td></td><td></td><td></td></tr>
<tr><td>Oligo rev</td><td>AAACACTCGAGCGCCAGAGCAATA</td><td></td><td></td><td></td><td></td><td></td><td></td><td></td><td></td><td></td><td></td><td></td></tr>
<tr><td>Chromosome</td><td>start</td><td></td><td>end</td><td>strand</td><td>MM</td><td>target_seq</td><td>PAM</td><td>alignment</td><td>distance</td><td>position</td><td>gene name</td><td>gene id</td></tr>
<tr><td>GU305914.1</td><td></td><td>59466</td><td>59488</td><td>-</td><td>0</td><td>TATTGCTCTGGCGCTCGAGT</td><td>GGG</td><td>||||||||[||||||||||||]PAM</td><td>NA</td><td>Intergenic</td><td>NA</td><td>NA</td></tr>
<tr><td>T29652</td><td>TTGTTTTTGTGCTTAGCGTGCGG</td><td></td><td>1000</td><td></td><td></td><td></td><td></td><td></td><td></td><td></td><td></td><td></td></tr>
<tr><td>Oligo fwd</td><td>CACCTTGTTTTTGTGCTTAGCGTG</td><td></td><td></td><td></td><td></td><td></td><td></td><td></td><td></td><td></td><td></td><td></td></tr>
<tr><td>Oligo rev</td><td>AAACCACGCTAAGCACAAAAACAA</td><td></td><td></td><td></td><td></td><td></td><td></td><td></td><td></td><td></td><td></td><td></td></tr>
<tr><td>Chromosome</td><td>start</td><td></td><td>end</td><td>strand</td><td>MM</td><td>target_seq</td><td>PAM</td><td>alignment</td><td>distance</td><td>position</td><td>gene name</td><td>gene id</td></tr>
<tr><td>GU305914.1</td><td></td><td>59421</td><td>59443</td><td>-</td><td>0</td><td>TTGTTTTTGTGCTTAGCGTG</td><td>CGG</td><td>||||||||[||||||||||||]PAM</td><td>NA</td><td>Intergenic</td><td>NA</td><td>NA</td></tr>
<tr><td>T29653</td><td>TTAGCGTGCGGCCAGACGATTGG</td><td></td><td>1000</td><td></td><td></td><td></td><td></td><td></td><td></td><td></td><td></td><td></td></tr>
<tr><td>Oligo fwd</td><td>CACCTTAGCGTGCGGCCAGACGAT</td><td></td><td></td><td></td><td></td><td></td><td></td><td></td><td></td><td></td><td></td><td></td></tr>
<tr><td>Oligo rev</td><td>AAACATCGTCTGGCCGCACGCTAA</td><td></td><td></td><td></td><td></td><td></td><td></td><td></td><td></td><td></td><td></td><td></td></tr>
<tr><td>Chromosome</td><td>start</td><td></td><td>end</td><td>strand</td><td>MM</td><td>target_seq</td><td>PAM</td><td>alignment</td><td>distance</td><td>position</td><td>gene name</td><td>gene id</td></tr>
<tr><td>GU305914.1</td><td></td><td>59409</td><td>59431</td><td>-</td><td>0</td><td>TTAGCGTGCGGCCAGACGAT</td><td>TGG</td><td>||||||||[||||||||||||]PAM</td><td>NA</td><td>Intergenic</td><td>NA</td><td>NA</td></tr>
<tr><td>T29654</td><td>ACGATTGGTTTTCGTGACAGCGG</td><td></td><td>1000</td><td></td><td></td><td></td><td></td><td></td><td></td><td></td><td></td><td></td></tr>
<tr><td>Oligo fwd</td><td>CACCACGATTGGTTTTCGTGACAG</td><td></td><td></td><td></td><td></td><td></td><td></td><td></td><td></td><td></td><td></td><td></td></tr>
<tr><td>Oligo rev</td><td>AAACCTGTCACGAAAACCAATCGT</td><td></td><td></td><td></td><td></td><td></td><td></td><td></td><td></td><td></td><td></td><td></td></tr>
<tr><td>Chromosome</td><td>start</td><td></td><td>end</td><td>strand</td><td>MM</td><td>target_seq</td><td>PAM</td><td>alignment</td><td>distance</td><td>position</td><td>gene name</td><td>gene id</td></tr>
<tr><td>GU305914.1</td><td></td><td>59394</td><td>59416</td><td>-</td><td>0</td><td>ACGATTGGTTTTCGTGACAG</td><td>CGG</td><td>||||||||[||||||||||||]PAM</td><td>NA</td><td>Intergenic</td><td>NA</td><td>NA</td></tr>
<tr><td>T29655</td><td>GAGCGCTTCATTCCCCTTCGAGG</td><td></td><td>1000</td><td></td><td></td><td></td><td></td><td></td><td></td><td></td><td></td><td></td></tr>
<tr><td>Oligo fwd</td><td>CACCGAGCGCTTCATTCCCCTTCG</td><td></td><td></td><td></td><td></td><td></td><td></td><td></td><td></td><td></td><td></td><td></td></tr>
<tr><td>Oligo rev</td><td>AAACCGAAGGGGAATGAAGCGCTC</td><td></td><td></td><td></td><td></td><td></td><td></td><td></td><td></td><td></td><td></td><td></td></tr>
<tr><td>Chromosome</td><td>start</td><td></td><td>end</td><td>strand</td><td>MM</td><td>target_seq</td><td>PAM</td><td>alignment</td><td>distance</td><td>position</td><td>gene name</td><td>gene id</td></tr>
<tr><td>GU305914.1</td><td></td><td>59339</td><td>59361</td><td>-</td><td>0</td><td>GAGCGCTTCATTCCCCTTCG</td><td>AGG</td><td>||||||||[||||||||||||]PAM</td><td>NA</td><td>Intergenic</td><td>NA</td><td>NA</td></tr>
<tr><td>T29656</td><td>TTCCCCTTCGAGGAGATCCGCGG</td><td></td><td>1000</td><td></td><td></td><td></td><td></td><td></td><td></td><td></td><td></td><td></td></tr>
<tr><td>Oligo fwd</td><td>CACCTTCCCCTTCGAGGAGATCCG</td><td></td><td></td><td></td><td></td><td></td><td></td><td></td><td></td><td></td><td></td><td></td></tr>
<tr><td>Oligo rev</td><td>AAACCGGATCTCCTCGAAGGGGAA</td><td></td><td></td><td></td><td></td><td></td><td></td><td></td><td></td><td></td><td></td><td></td></tr>
<tr><td>Chromosome</td><td>start</td><td></td><td>end</td><td>strand</td><td>MM</td><td>target_seq</td><td>PAM</td><td>alignment</td><td>distance</td><td>position</td><td>gene name</td><td>gene id</td></tr>
<tr><td>GU305914.1</td><td></td><td>59329</td><td>59351</td><td>-</td><td>0</td><td>TTCCCCTTCGAGGAGATCCG</td><td>CGG</td><td>||||||||[||||||||||||]PAM</td><td>NA</td><td>Intergenic</td><td>NA</td><td>NA</td></tr>
<tr><td>T29657</td><td>TTCGAGGAGATCCGCGGTGCAGG</td><td></td><td>1000</td><td></td><td></td><td></td><td></td><td></td><td></td><td></td><td></td><td></td></tr>
<tr><td>Oligo fwd</td><td>CACCTTCGAGGAGATCCGCGGTGC</td><td></td><td></td><td></td><td></td><td></td><td></td><td></td><td></td><td></td><td></td><td></td></tr>
<tr><td>Oligo rev</td><td>AAACGCACCGCGGATCTCCTCGAA</td><td></td><td></td><td></td><td></td><td></td><td></td><td></td><td></td><td></td><td></td><td></td></tr>
<tr><td>Chromosome</td><td>start</td><td></td><td>end</td><td>strand</td><td>MM</td><td>target_seq</td><td>PAM</td><td>alignment</td><td>distance</td><td>position</td><td>gene name</td><td>gene id</td></tr>
<tr><td>GU305914.1</td><td></td><td>59323</td><td>59345</td><td>-</td><td>0</td><td>TTCGAGGAGATCCGCGGTGC</td><td>AGG</td><td>||||||||[||||||||||||]PAM</td><td>NA</td><td>Intergenic</td><td>NA</td><td>NA</td></tr>
<tr><td>T29658</td><td>TCGAGGAGATCCGCGGTGCAGGG</td><td></td><td>1000</td><td></td><td></td><td></td><td></td><td></td><td></td><td></td><td></td><td></td></tr>
<tr><td>Oligo fwd</td><td>CACCTCGAGGAGATCCGCGGTGCA</td><td></td><td></td><td></td><td></td><td></td><td></td><td></td><td></td><td></td><td></td><td></td></tr>
<tr><td>Oligo rev</td><td>AAACTGCACCGCGGATCTCCTCGA</td><td></td><td></td><td></td><td></td><td></td><td></td><td></td><td></td><td></td><td></td><td></td></tr>
<tr><td>Chromosome</td><td>start</td><td></td><td>end</td><td>strand</td><td>MM</td><td>target_seq</td><td>PAM</td><td>alignment</td><td>distance</td><td>position</td><td>gene name</td><td>gene id</td></tr>
<tr><td>GU305914.1</td><td></td><td>59322</td><td>59344</td><td>-</td><td>0</td><td>TCGAGGAGATCCGCGGTGCA</td><td>GGG</td><td>||||||||[||||||||||||]PAM</td><td>NA</td><td>Intergenic</td><td>NA</td><td>NA</td></tr>
<tr><td>T29659</td><td>TTTTTTTCTGACGCATCCCGTGG</td><td></td><td>1000</td><td></td><td></td><td></td><td></td><td></td><td></td><td></td><td></td><td></td></tr>
<tr><td>Oligo fwd</td><td>CACCTTTTTTTCTGACGCATCCCG</td><td></td><td></td><td></td><td></td><td></td><td></td><td></td><td></td><td></td><td></td><td></td></tr>
<tr><td>Oligo rev</td><td>AAACCGGGATGCGTCAGAAAAAAA</td><td></td><td></td><td></td><td></td><td></td><td></td><td></td><td></td><td></td><td></td><td></td></tr>
<tr><td>Chromosome</td><td>start</td><td></td><td>end</td><td>strand</td><td>MM</td><td>target_seq</td><td>PAM</td><td>alignment</td><td>distance</td><td>position</td><td>gene name</td><td>gene id</td></tr>
<tr><td>GU305914.1</td><td></td><td>59265</td><td>59287</td><td>-</td><td>0</td><td>TTTTTTTCTGACGCATCCCG</td><td>TGG</td><td>||||||||[||||||||||||]PAM</td><td>NA</td><td>Intergenic</td><td>NA</td><td>NA</td></tr>
<tr><td>T29660</td><td>CTGTTCGCCGCGCGCCCGATCGG</td><td></td><td>1000</td><td></td><td></td><td></td><td></td><td></td><td></td><td></td><td></td><td></td></tr>
<tr><td>Oligo fwd</td><td>CACCCTGTTCGCCGCGCGCCCGAT</td><td></td><td></td><td></td><td></td><td></td><td></td><td></td><td></td><td></td><td></td><td></td></tr>
<tr><td>Oligo rev</td><td>AAACATCGGGCGCGCGGCGAACAG</td><td></td><td></td><td></td><td></td><td></td><td></td><td></td><td></td><td></td><td></td><td></td></tr>
<tr><td>Chromosome</td><td>start</td><td></td><td>end</td><td>strand</td><td>MM</td><td>target_seq</td><td>PAM</td><td>alignment</td><td>distance</td><td>position</td><td>gene name</td><td>gene id</td></tr>
<tr><td>GU305914.1</td><td></td><td>59230</td><td>59252</td><td>-</td><td>0</td><td>CTGTTCGCCGCGCGCCCGAT</td><td>CGG</td><td>||||||||[||||||||||||]PAM</td><td>NA</td><td>Intergenic</td><td>NA</td><td>NA</td></tr>
<tr><td>T29661</td><td>TCGCCTCGCGCTCGCTCGCTCGG</td><td></td><td>1000</td><td></td><td></td><td></td><td></td><td></td><td></td><td></td><td></td><td></td></tr>
<tr><td>Oligo fwd</td><td>CACCTCGCCTCGCGCTCGCTCGCT</td><td></td><td></td><td></td><td></td><td></td><td></td><td></td><td></td><td></td><td></td><td></td></tr>
<tr><td>Oligo rev</td><td>AAACAGCGAGCGAGCGCGAGGCGA</td><td></td><td></td><td></td><td></td><td></td><td></td><td></td><td></td><td></td><td></td><td></td></tr>
<tr><td>Chromosome</td><td>start</td><td></td><td>end</td><td>strand</td><td>MM</td><td>target_seq</td><td>PAM</td><td>alignment</td><td>distance</td><td>position</td><td>gene name</td><td>gene id</td></tr>
<tr><td>GU305914.1</td><td></td><td>59194</td><td>59216</td><td>-</td><td>0</td><td>TCGCCTCGCGCTCGCTCGCT</td><td>CGG</td><td>||||||||[||||||||||||]PAM</td><td>NA</td><td>Intergenic</td><td>NA</td><td>NA</td></tr>
<tr><td>T29662</td><td>CTCGCGCTCGCTCGCTCGGTCGG</td><td></td><td>1000</td><td></td><td></td><td></td><td></td><td></td><td></td><td></td><td></td><td></td></tr>
<tr><td>Oligo fwd</td><td>CACCCTCGCGCTCGCTCGCTCGGT</td><td></td><td></td><td></td><td></td><td></td><td></td><td></td><td></td><td></td><td></td><td></td></tr>
<tr><td>Oligo rev</td><td>AAACACCGAGCGAGCGAGCGCGAG</td><td></td><td></td><td></td><td></td><td></td><td></td><td></td><td></td><td></td><td></td><td></td></tr>
<tr><td>Chromosome</td><td>start</td><td></td><td>end</td><td>strand</td><td>MM</td><td>target_seq</td><td>PAM</td><td>alignment</td><td>distance</td><td>position</td><td>gene name</td><td>gene id</td></tr>
<tr><td>GU305914.1</td><td></td><td>59190</td><td>59212</td><td>-</td><td>0</td><td>CTCGCGCTCGCTCGCTCGGT</td><td>CGG</td><td>||||||||[||||||||||||]PAM</td><td>NA</td><td>Intergenic</td><td>NA</td><td>NA</td></tr>
<tr><td>GU305914.1</td><td></td><td>59186</td><td>59208</td><td>-</td><td>3</td><td>CGCTCGCTCGCTCGGTCGGT</td><td>CGG</td><td>|-|-||||[||||||-|||||]PAM</td><td>NA</td><td>Intergenic</td><td>NA</td><td>NA</td></tr>
<tr><td>T29663</td><td>CGCTCGCTCGCTCGGTCGGTCGG</td><td></td><td>1000</td><td></td><td></td><td></td><td></td><td></td><td></td><td></td><td></td><td></td></tr>
<tr><td>Oligo fwd</td><td>CACCCGCTCGCTCGCTCGGTCGGT</td><td></td><td></td><td></td><td></td><td></td><td></td><td></td><td></td><td></td><td></td><td></td></tr>
<tr><td>Oligo rev</td><td>AAACACCGACCGAGCGAGCGAGCG</td><td></td><td></td><td></td><td></td><td></td><td></td><td></td><td></td><td></td><td></td><td></td></tr>
<tr><td>Chromosome</td><td>start</td><td></td><td>end</td><td>strand</td><td>MM</td><td>target_seq</td><td>PAM</td><td>alignment</td><td>distance</td><td>position</td><td>gene name</td><td>gene id</td></tr>
<tr><td>GU305914.1</td><td></td><td>59186</td><td>59208</td><td>-</td><td>0</td><td>CGCTCGCTCGCTCGGTCGGT</td><td>CGG</td><td>||||||||[||||||||||||]PAM</td><td>NA</td><td>Intergenic</td><td>NA</td><td>NA</td></tr>
<tr><td>GU305914.1</td><td></td><td>59190</td><td>59212</td><td>-</td><td>3</td><td>CTCGCGCTCGCTCGCTCGGT</td><td>CGG</td><td>|-|-||||[||||||-|||||]PAM</td><td>NA</td><td>Intergenic</td><td>NA</td><td>NA</td></tr>
</table>

T29664 GTCTCGTGTCCTTCTTTCGCCGG 1000
Oligo fwd CACCGTCTCGTGTCCTTCTTTCGC
Oligo rev AAACGCGAAAGAAGGACACGAGAC
Chromosome start end strand MM target_seq PAM alignment distance position gene name gene id
GU305914.1 59160 59182 - 0 GTCTCGTGTCCTTCTTTCGC CGG ||||||||[||||||||||||]PAM NA Intergenic NA NA

T29665 CTCGCTCACATACATCTCCGCGG 1000
Oligo fwd CACCCTCGCTCACATACATCTCCG
Oligo rev AAACCGGAGATGTATGTGAGCGAG
Chromosome start end strand MM target_seq PAM alignment distance position gene name gene id
GU305914.1 59126 59148 - 0 CTCGCTCACATACATCTCCG CGG ||||||||[||||||||||||]PAM NA Intergenic NA NA

T29666 TCGCTCACATACATCTCCGCGGG 1000
Oligo fwd CACCTCGCTCACATACATCTCCGC
Oligo rev AAACGCGGAGATGTATGTGAGCGA
Chromosome start end strand MM target_seq PAM alignment distance position gene name gene id
GU305914.1 59125 59147 - 0 TCGCTCACATACATCTCCGC GGG ||||||||[||||||||||||]PAM NA Intergenic NA NA

T29667 CACATACATCTCCGCGGGAATGG 1000
Oligo fwd CACCCACATACATCTCCGCGGGAA
Oligo rev AAACTTCCCGCGGAGATGTATGTG
Chromosome start end strand MM target_seq PAM alignment distance position gene name gene id
GU305914.1 59120 59142 - 0 CACATACATCTCCGCGGGAA TGG ||||||||[||||||||||||]PAM NA Intergenic NA NA

T29668 ATACATCTCCGCGGGAATGGAGG 1000
Oligo fwd CACCATACATCTCCGCGGGAATGG
Oligo rev AAACCCATTCCCGCGGAGATGTAT
Chromosome start end strand MM target_seq PAM alignment distance position gene name gene id
GU305914.1 59117 59139 - 0 ATACATCTCCGCGGGAATGG AGG ||||||||[||||||||||||]PAM NA Intergenic NA NA

T29669 TACATCTCCGCGGGAATGGAGGG 1000
Oligo fwd CACCTACATCTCCGCGGGAATGGA
Oligo rev AAACTCCATTCCCGCGGAGATGTA
Chromosome start end strand MM target_seq PAM alignment distance position gene name gene id
GU305914.1 59116 59138 - 0 TACATCTCCGCGGGAATGGA GGG ||||||||[||||||||||||]PAM NA Intergenic NA NA

T29670 ATCTCCGCGGGAATGGAGGGTGG 1000
Oligo fwd CACCATCTCCGCGGGAATGGAGGG
Oligo rev AAACCCCTCCATTCCCGCGGAGAT
Chromosome start end strand MM target_seq PAM alignment distance position gene name gene id
GU305914.1 59113 59135 - 0 ATCTCCGCGGGAATGGAGGG TGG ||||||||[||||||||||||]PAM NA Intergenic NA NA

T29671 CCGCGGGAATGGAGGGTGGTAGG 1000
Oligo fwd CACCCCGCGGGAATGGAGGGTGGT
Oligo rev AAACACCACCCTCCATTCCCGCGG
Chromosome start end strand MM target_seq PAM alignment distance position gene name gene id
GU305914.1 59109 59131 - 0 CCGCGGGAATGGAGGGTGGT AGG ||||||||[||||||||||||]PAM NA Intergenic NA NA

T29672 CGAGCACGAACCGCCGACCCTGG 1000
Oligo fwd CACCCGAGCACGAACCGCCGACCC
Oligo rev AAACGGGTCGGCGGTTCGTGCTCG
Chromosome start end strand MM target_seq PAM alignment distance position gene name gene id
GU305914.1 59078 59100 - 0 CGAGCACGAACCGCCGACCC TGG ||||||||[||||||||||||]PAM NA Intergenic NA NA

T29673 TTGCGTCGTACGCTCGTTGAAGG 1000
Oligo fwd CACCTTGCGTCGTACGCTCGTTGA
Oligo rev AAACTCAACGAGCGTACGACGCAA
Chromosome start end strand MM target_seq PAM alignment distance position gene name gene id
GU305914.1 59012 59034 - 0 TTGCGTCGTACGCTCGTTGA AGG ||||||||[||||||||||||]PAM NA Intergenic NA NA

T29674 CTCGTTGAAGGAGAATACCACGG 1000
Oligo fwd CACCCTCGTTGAAGGAGAATACCA
Oligo rev AAACTGGTATTCTCCTTCAACGAG
Chromosome start end strand MM target_seq PAM alignment distance position gene name gene id
GU305914.1 59000 59022 - 0 CTCGTTGAAGGAGAATACCA CGG ||||||||[||||||||||||]PAM NA Intergenic NA NA

T29675 CATCACCGACAAGTCTTTCTCGG 1000
Oligo fwd CACCCATCACCGACAAGTCTTTCT
Oligo rev AAACAGAAAGACTTGTCGGTGATG
Chromosome start end strand MM target_seq PAM alignment distance position gene name gene id
GU305914.1 58889 58911 - 0 CATCACCGACAAGTCTTTCT CGG ||||||||[||||||||||||]PAM NA Intergenic NA NA

T29676 CAACAACTCCATCCCCCTGCTGG 1000
Oligo fwd CACCCAACAACTCCATCCCCCTGC
Oligo rev AAACGCAGGGGGATGGAGTTGTTG
Chromosome start end strand MM target_seq PAM alignment distance position gene name gene id
GU305914.1 58856 58878 - 0 CAACAACTCCATCCCCCTGC TGG ||||||||[||||||||||||]PAM NA Intergenic NA NA

T29677 AACAACTCCATCCCCCTGCTGGG 1000
Oligo fwd CACCAACAACTCCATCCCCCTGCT
Oligo rev AAACAGCAGGGGGATGGAGTTGTT
Chromosome start end strand MM target_seq PAM alignment distance position gene name gene id
GU305914.1 58855 58877 - 0 AACAACTCCATCCCCCTGCT GGG ||||||||[||||||||||||]PAM NA Intergenic NA NA

T29678 TCTGACCAAATTCACCGTGCAGG 1000
Oligo fwd CACCTCTGACCAAATTCACCGTGC
Oligo rev AAACGCACGGTGAATTTGGTCAGA
Chromosome start end strand MM target_seq PAM alignment distance position gene name gene id

<table>
<tr><td>GU305914.1</td><td>58802</td><td>58824</td><td>-</td><td>0</td><td>TCTGACCAAATTCACCGTGC</td><td>AGG</td><td>||||||||[||||||||||||]PAM</td><td>NA</td><td>Intergenic</td><td>NA</td><td>NA</td></tr>
<tr><td>T29679</td><td>ATTCACCGTGCAGGACACCTCGG</td><td>1000</td><td></td><td></td><td></td><td></td><td></td><td></td><td></td><td></td><td></td></tr>
<tr><td>Oligo fwd</td><td>CACCATTCACCGTGCAGGACACCT</td><td></td><td></td><td></td><td></td><td></td><td></td><td></td><td></td><td></td><td></td></tr>
<tr><td>Oligo rev</td><td>AAACAGGTGTCCTGCACGGTGAAT</td><td></td><td></td><td></td><td></td><td></td><td></td><td></td><td></td><td></td><td></td></tr>
<tr><td>Chromosome</td><td>start</td><td>end</td><td>strand</td><td>MM</td><td>target_seq</td><td>PAM</td><td>alignment</td><td>distance</td><td>position</td><td>gene name</td><td>gene id</td></tr>
<tr><td>GU305914.1</td><td>58793</td><td>58815</td><td>-</td><td>0</td><td>ATTCACCGTGCAGGACACCT</td><td>CGG</td><td>||||||||[||||||||||||]PAM</td><td>NA</td><td>Intergenic</td><td>NA</td><td>NA</td></tr>
<tr><td>T29680</td><td>CACCTCGGACCTCAGCGCCAAGG</td><td>1000</td><td></td><td></td><td></td><td></td><td></td><td></td><td></td><td></td><td></td></tr>
<tr><td>Oligo fwd</td><td>CACCCACCTCGGACCTCAGCGCCA</td><td></td><td></td><td></td><td></td><td></td><td></td><td></td><td></td><td></td><td></td></tr>
<tr><td>Oligo rev</td><td>AAACTGGCGCTGAGGTCCGAGGTG</td><td></td><td></td><td></td><td></td><td></td><td></td><td></td><td></td><td></td><td></td></tr>
<tr><td>Chromosome</td><td>start</td><td>end</td><td>strand</td><td>MM</td><td>target_seq</td><td>PAM</td><td>alignment</td><td>distance</td><td>position</td><td>gene name</td><td>gene id</td></tr>
<tr><td>GU305914.1</td><td>58778</td><td>58800</td><td>-</td><td>0</td><td>CACCTCGGACCTCAGCGCCA</td><td>AGG</td><td>||||||||[||||||||||||]PAM</td><td>NA</td><td>Intergenic</td><td>NA</td><td>NA</td></tr>
<tr><td>GU305914.1</td><td>141692</td><td>141714</td><td>+</td><td>4</td><td>CGTCTCGGACCTCTGCGCCC</td><td>TGG</td><td>|--|||||[|||||-|||||-]PAM</td><td>NA</td><td>Intergenic</td><td>NA</td><td>NA</td></tr>
<tr><td>T29681</td><td>CAGCGCCAAGGTCTGCATGTCGG</td><td>1000</td><td></td><td></td><td></td><td></td><td></td><td></td><td></td><td></td><td></td></tr>
<tr><td>Oligo fwd</td><td>CACCCAGCGCCAAGGTCTGCATGT</td><td></td><td></td><td></td><td></td><td></td><td></td><td></td><td></td><td></td><td></td></tr>
<tr><td>Oligo rev</td><td>AAACACATGCAGACCTTGGCGCTG</td><td></td><td></td><td></td><td></td><td></td><td></td><td></td><td></td><td></td><td></td></tr>
<tr><td>Chromosome</td><td>start</td><td>end</td><td>strand</td><td>MM</td><td>target_seq</td><td>PAM</td><td>alignment</td><td>distance</td><td>position</td><td>gene name</td><td>gene id</td></tr>
<tr><td>GU305914.1</td><td>58766</td><td>58788</td><td>-</td><td>0</td><td>CAGCGCCAAGGTCTGCATGT</td><td>CGG</td><td>||||||||[||||||||||||]PAM</td><td>NA</td><td>Intergenic</td><td>NA</td><td>NA</td></tr>
<tr><td>T29682</td><td>GTCGGCGCCCGACTACAACATGG</td><td>1000</td><td></td><td></td><td></td><td></td><td></td><td></td><td></td><td></td><td></td></tr>
<tr><td>Oligo fwd</td><td>CACCGTCGGCGCCCGACTACAACA</td><td></td><td></td><td></td><td></td><td></td><td></td><td></td><td></td><td></td><td></td></tr>
<tr><td>Oligo rev</td><td>AAACTGTTGTAGTCGGGCGCCGAC</td><td></td><td></td><td></td><td></td><td></td><td></td><td></td><td></td><td></td><td></td></tr>
<tr><td>Chromosome</td><td>start</td><td>end</td><td>strand</td><td>MM</td><td>target_seq</td><td>PAM</td><td>alignment</td><td>distance</td><td>position</td><td>gene name</td><td>gene id</td></tr>
<tr><td>GU305914.1</td><td>58748</td><td>58770</td><td>-</td><td>0</td><td>GTCGGCGCCCGACTACAACA</td><td>TGG</td><td>||||||||[||||||||||||]PAM</td><td>NA</td><td>Intergenic</td><td>NA</td><td>NA</td></tr>
<tr><td>T29683</td><td>CTCCGCCTGCGTGCACAACCAGG</td><td>1000</td><td></td><td></td><td></td><td></td><td></td><td></td><td></td><td></td><td></td></tr>
<tr><td>Oligo fwd</td><td>CACCCTCCGCCTGCGTGCACAACC</td><td></td><td></td><td></td><td></td><td></td><td></td><td></td><td></td><td></td><td></td></tr>
<tr><td>Oligo rev</td><td>AAACGGTTGTGCACGCAGGCGGAG</td><td></td><td></td><td></td><td></td><td></td><td></td><td></td><td></td><td></td><td></td></tr>
<tr><td>Chromosome</td><td>start</td><td>end</td><td>strand</td><td>MM</td><td>target_seq</td><td>PAM</td><td>alignment</td><td>distance</td><td>position</td><td>gene name</td><td>gene id</td></tr>
<tr><td>GU305914.1</td><td>58718</td><td>58740</td><td>-</td><td>0</td><td>CTCCGCCTGCGTGCACAACC</td><td>AGG</td><td>||||||||[||||||||||||]PAM</td><td>NA</td><td>Intergenic</td><td>NA</td><td>NA</td></tr>
<tr><td>T29684</td><td>CAGGACATCATCCGCGAGACCGG</td><td>1000</td><td></td><td></td><td></td><td></td><td></td><td></td><td></td><td></td><td></td></tr>
<tr><td>Oligo fwd</td><td>CACCCAGGACATCATCCGCGAGAC</td><td></td><td></td><td></td><td></td><td></td><td></td><td></td><td></td><td></td><td></td></tr>
<tr><td>Oligo rev</td><td>AAACGTCTCGCGGATGATGTCCTG</td><td></td><td></td><td></td><td></td><td></td><td></td><td></td><td></td><td></td><td></td></tr>
<tr><td>Chromosome</td><td>start</td><td>end</td><td>strand</td><td>MM</td><td>target_seq</td><td>PAM</td><td>alignment</td><td>distance</td><td>position</td><td>gene name</td><td>gene id</td></tr>
<tr><td>GU305914.1</td><td>58699</td><td>58721</td><td>-</td><td>0</td><td>CAGGACATCATCCGCGAGAC</td><td>CGG</td><td>||||||||[||||||||||||]PAM</td><td>NA</td><td>Intergenic</td><td>NA</td><td>NA</td></tr>
<tr><td>T29685</td><td>GATCTCGACTCCGCCGTCGTCGG</td><td>1000</td><td></td><td></td><td></td><td></td><td></td><td></td><td></td><td></td><td></td></tr>
<tr><td>Oligo fwd</td><td>CACCGATCTCGACTCCGCCGTCGT</td><td></td><td></td><td></td><td></td><td></td><td></td><td></td><td></td><td></td><td></td></tr>
<tr><td>Oligo rev</td><td>AAACACGACGGCGGAGTCGAGATC</td><td></td><td></td><td></td><td></td><td></td><td></td><td></td><td></td><td></td><td></td></tr>
<tr><td>Chromosome</td><td>start</td><td>end</td><td>strand</td><td>MM</td><td>target_seq</td><td>PAM</td><td>alignment</td><td>distance</td><td>position</td><td>gene name</td><td>gene id</td></tr>
<tr><td>GU305914.1</td><td>58657</td><td>58679</td><td>-</td><td>0</td><td>GATCTCGACTCCGCCGTCGT</td><td>CGG</td><td>||||||||[||||||||||||]PAM</td><td>NA</td><td>Intergenic</td><td>NA</td><td>NA</td></tr>
<tr><td>T29686</td><td>TCGTCGGCGAGCTCATCCGCTGG</td><td>1000</td><td></td><td></td><td></td><td></td><td></td><td></td><td></td><td></td><td></td></tr>
<tr><td>Oligo fwd</td><td>CACCTCGTCGGCGAGCTCATCCGC</td><td></td><td></td><td></td><td></td><td></td><td></td><td></td><td></td><td></td><td></td></tr>
<tr><td>Oligo rev</td><td>AAACGCGGATGAGCTCGCCGACGA</td><td></td><td></td><td></td><td></td><td></td><td></td><td></td><td></td><td></td><td></td></tr>
<tr><td>Chromosome</td><td>start</td><td>end</td><td>strand</td><td>MM</td><td>target_seq</td><td>PAM</td><td>alignment</td><td>distance</td><td>position</td><td>gene name</td><td>gene id</td></tr>
<tr><td>GU305914.1</td><td>58641</td><td>58663</td><td>-</td><td>0</td><td>TCGTCGGCGAGCTCATCCGC</td><td>TGG</td><td>||||||||[||||||||||||]PAM</td><td>NA</td><td>Intergenic</td><td>NA</td><td>NA</td></tr>
<tr><td>T29687</td><td>GCAATCCACCTCCAGCAGCACGG</td><td>1000</td><td></td><td></td><td></td><td></td><td></td><td></td><td></td><td></td><td></td></tr>
<tr><td>Oligo fwd</td><td>CACCGCAATCCACCTCCAGCAGCA</td><td></td><td></td><td></td><td></td><td></td><td></td><td></td><td></td><td></td><td></td></tr>
<tr><td>Oligo rev</td><td>AAACTGCTGCTGGAGGTGGATTGC</td><td></td><td></td><td></td><td></td><td></td><td></td><td></td><td></td><td></td><td></td></tr>
<tr><td>Chromosome</td><td>start</td><td>end</td><td>strand</td><td>MM</td><td>target_seq</td><td>PAM</td><td>alignment</td><td>distance</td><td>position</td><td>gene name</td><td>gene id</td></tr>
<tr><td>GU305914.1</td><td>58580</td><td>58602</td><td>-</td><td>0</td><td>GCAATCCACCTCCAGCAGCA</td><td>CGG</td><td>||||||||[||||||||||||]PAM</td><td>NA</td><td>Intergenic</td><td>NA</td><td>NA</td></tr>
<tr><td>T29688</td><td>GCTGCACGCTAATCCGCCCACGG</td><td>1000</td><td></td><td></td><td></td><td></td><td></td><td></td><td></td><td></td><td></td></tr>
<tr><td>Oligo fwd</td><td>CACCGCTGCACGCTAATCCGCCCA</td><td></td><td></td><td></td><td></td><td></td><td></td><td></td><td></td><td></td><td></td></tr>
<tr><td>Oligo rev</td><td>AAACTGGGCGGATTAGCGTGCAGC</td><td></td><td></td><td></td><td></td><td></td><td></td><td></td><td></td><td></td><td></td></tr>
<tr><td>Chromosome</td><td>start</td><td>end</td><td>strand</td><td>MM</td><td>target_seq</td><td>PAM</td><td>alignment</td><td>distance</td><td>position</td><td>gene name</td><td>gene id</td></tr>
<tr><td>GU305914.1</td><td>58547</td><td>58569</td><td>-</td><td>0</td><td>GCTGCACGCTAATCCGCCCA</td><td>CGG</td><td>||||||||[||||||||||||]PAM</td><td>NA</td><td>Intergenic</td><td>NA</td><td>NA</td></tr>
<tr><td>T29689</td><td>CCCACGGTCAAGTTCTCGCTCGG</td><td>1000</td><td></td><td></td><td></td><td></td><td></td><td></td><td></td><td></td><td></td></tr>
<tr><td>Oligo fwd</td><td>CACCCCCACGGTCAAGTTCTCGCT</td><td></td><td></td><td></td><td></td><td></td><td></td><td></td><td></td><td></td><td></td></tr>
<tr><td>Oligo rev</td><td>AAACAGCGAGAACTTGACCGTGGG</td><td></td><td></td><td></td><td></td><td></td><td></td><td></td><td></td><td></td><td></td></tr>
<tr><td>Chromosome</td><td>start</td><td>end</td><td>strand</td><td>MM</td><td>target_seq</td><td>PAM</td><td>alignment</td><td>distance</td><td>position</td><td>gene name</td><td>gene id</td></tr>
<tr><td>GU305914.1</td><td>58531</td><td>58553</td><td>-</td><td>0</td><td>CCCACGGTCAAGTTCTCGCT</td><td>CGG</td><td>||||||||[||||||||||||]PAM</td><td>NA</td><td>Intergenic</td><td>NA</td><td>NA</td></tr>
<tr><td>T29690</td><td>CAACCGCATCGCCTTTCACGAGG</td><td>1000</td><td></td><td></td><td></td><td></td><td></td><td></td><td></td><td></td><td></td></tr>
<tr><td>Oligo fwd</td><td>CACCCAACCGCATCGCCTTTCACG</td><td></td><td></td><td></td><td></td><td></td><td></td><td></td><td></td><td></td><td></td></tr>
<tr><td>Oligo rev</td><td>AAACCGTGAAAGGCGATGCGGTTG</td><td></td><td></td><td></td><td></td><td></td><td></td><td></td><td></td><td></td><td></td></tr>
<tr><td>Chromosome</td><td>start</td><td>end</td><td>strand</td><td>MM</td><td>target_seq</td><td>PAM</td><td>alignment</td><td>distance</td><td>position</td><td>gene name</td><td>gene id</td></tr>
<tr><td>GU305914.1</td><td>58478</td><td>58500</td><td>-</td><td>0</td><td>CAACCGCATCGCCTTTCACG</td><td>AGG</td><td>||||||||[||||||||||||]PAM</td><td>NA</td><td>Intergenic</td><td>NA</td><td>NA</td></tr>
<tr><td>T29691</td><td>GGTCAAGAACCTGCGCATCACGG</td><td>1000</td><td></td><td></td><td></td><td></td><td></td><td></td><td></td><td></td><td></td></tr>
<tr><td>Oligo fwd</td><td>CACCGGTCAAGAACCTGCGCATCA</td><td></td><td></td><td></td><td></td><td></td><td></td><td></td><td></td><td></td><td></td></tr>
<tr><td>Oligo rev</td><td>AAACTGATGCGCAGGTTCTTGACC</td><td></td><td></td><td></td><td></td><td></td><td></td><td></td><td></td><td></td><td></td></tr>
<tr><td>Chromosome</td><td>start</td><td>end</td><td>strand</td><td>MM</td><td>target_seq</td><td>PAM</td><td>alignment</td><td>distance</td><td>position</td><td>gene name</td><td>gene id</td></tr>
<tr><td>GU305914.1</td><td>58457</td><td>58479</td><td>-</td><td>0</td><td>GGTCAAGAACCTGCGCATCA</td><td>CGG</td><td>||||||||[||||||||||||]PAM</td><td>NA</td><td>Intergenic</td><td>NA</td><td>NA</td></tr>
<tr><td>T29692</td><td>GAACCTGCGCATCACGGTGCAGG</td><td>1000</td><td></td><td></td><td></td><td></td><td></td><td></td><td></td><td></td><td></td></tr>
<tr><td>Oligo fwd</td><td>CACCGAACCTGCGCATCACGGTGC</td><td></td><td></td><td></td><td></td><td></td><td></td><td></td><td></td><td></td><td></td></tr>
<tr><td>Oligo rev</td><td>AAACGCACCGTGATGCGCAGGTTC</td><td></td><td></td><td></td><td></td><td></td><td></td><td></td><td></td><td></td><td></td></tr>
<tr><td>Chromosome</td><td>start</td><td>end</td><td>strand</td><td>MM</td><td>target_seq</td><td>PAM</td><td>alignment</td><td>distance</td><td>position</td><td>gene name</td><td>gene id</td></tr>
<tr><td>GU305914.1</td><td>58451</td><td>58473</td><td>-</td><td>0</td><td>GAACCTGCGCATCACGGTGC</td><td>AGG</td><td>||||||||[||||||||||||]PAM</td><td>NA</td><td>Intergenic</td><td>NA</td><td>NA</td></tr>
<tr><td>GU305914.1</td><td>149133</td><td>149155</td><td>-</td><td>4</td><td>GATCGTGAGCATCACGGTGT</td><td>CGG</td><td>||-|-||-[|||||||||||-]PAM</td><td>NA</td><td>Intergenic</td><td>NA</td><td>NA</td></tr>
<tr><td>T29693</td><td>GCAGGCCAAGAACCTGCACCAGG</td><td>1000</td><td></td><td></td><td></td><td></td><td></td><td></td><td></td><td></td><td></td></tr>
</table>

<table>
<tr><td>Oligo fwd</td><td>CACCGCAGGCCAAGAACCTGCACC</td><td></td><td></td><td></td><td></td><td></td><td></td><td></td><td></td><td></td><td></td></tr>
<tr><td>Oligo rev</td><td>AAACGGTGCAGGTTCTTGGCCTGC</td><td></td><td></td><td></td><td></td><td></td><td></td><td></td><td></td><td></td><td></td></tr>
<tr><td>Chromosome</td><td>start</td><td>end</td><td>strand</td><td>MM</td><td>target_seq</td><td>PAM</td><td>alignment</td><td>distance</td><td>position</td><td>gene name</td><td>gene id</td></tr>
<tr><td>GU305914.1</td><td>58433</td><td>58455</td><td>-</td><td>0</td><td>GCAGGCCAAGAACCTGCACC</td><td>AGG</td><td>||||||||[||||||||||||]PAM</td><td>NA</td><td>Intergenic</td><td>NA</td><td>NA</td></tr>
<tr><td>T29694</td><td>CCAGGCGCTCTGTAACTGTGTGG</td><td>1000</td><td></td><td></td><td></td><td></td><td></td><td></td><td></td><td></td><td></td></tr>
<tr><td>Oligo fwd</td><td>CACCCCAGGCGCTCTGTAACTGTG</td><td></td><td></td><td></td><td></td><td></td><td></td><td></td><td></td><td></td><td></td></tr>
<tr><td>Oligo rev</td><td>AAACCACAGTTACAGAGCGCCTGG</td><td></td><td></td><td></td><td></td><td></td><td></td><td></td><td></td><td></td><td></td></tr>
<tr><td>Chromosome</td><td>start</td><td>end</td><td>strand</td><td>MM</td><td>target_seq</td><td>PAM</td><td>alignment</td><td>distance</td><td>position</td><td>gene name</td><td>gene id</td></tr>
<tr><td>GU305914.1</td><td>58415</td><td>58437</td><td>-</td><td>0</td><td>CCAGGCGCTCTGTAACTGTG</td><td>TGG</td><td>||||||||[||||||||||||]PAM</td><td>NA</td><td>Intergenic</td><td>NA</td><td>NA</td></tr>
<tr><td>T29695</td><td>TAACTGTGTGGTCACTAAGCTGG</td><td>1000</td><td></td><td></td><td></td><td></td><td></td><td></td><td></td><td></td><td></td></tr>
<tr><td>Oligo fwd</td><td>CACCTAACTGTGTGGTCACTAAGC</td><td></td><td></td><td></td><td></td><td></td><td></td><td></td><td></td><td></td><td></td></tr>
<tr><td>Oligo rev</td><td>AAACGCTTAGTGACCACACAGTTA</td><td></td><td></td><td></td><td></td><td></td><td></td><td></td><td></td><td></td><td></td></tr>
<tr><td>Chromosome</td><td>start</td><td>end</td><td>strand</td><td>MM</td><td>target_seq</td><td>PAM</td><td>alignment</td><td>distance</td><td>position</td><td>gene name</td><td>gene id</td></tr>
<tr><td>GU305914.1</td><td>58403</td><td>58425</td><td>-</td><td>0</td><td>TAACTGTGTGGTCACTAAGC</td><td>TGG</td><td>||||||||[||||||||||||]PAM</td><td>NA</td><td>Intergenic</td><td>NA</td><td>NA</td></tr>
<tr><td>T29696</td><td>CTGCACGCTGCGCGTCATGACGG</td><td>1000</td><td></td><td></td><td></td><td></td><td></td><td></td><td></td><td></td><td></td></tr>
<tr><td>Oligo fwd</td><td>CACCCTGCACGCTGCGCGTCATGA</td><td></td><td></td><td></td><td></td><td></td><td></td><td></td><td></td><td></td><td></td></tr>
<tr><td>Oligo rev</td><td>AAACTCATGACGCGCAGCGTGCAG</td><td></td><td></td><td></td><td></td><td></td><td></td><td></td><td></td><td></td><td></td></tr>
<tr><td>Chromosome</td><td>start</td><td>end</td><td>strand</td><td>MM</td><td>target_seq</td><td>PAM</td><td>alignment</td><td>distance</td><td>position</td><td>gene name</td><td>gene id</td></tr>
<tr><td>GU305914.1</td><td>58379</td><td>58401</td><td>-</td><td>0</td><td>CTGCACGCTGCGCGTCATGA</td><td>CGG</td><td>||||||||[||||||||||||]PAM</td><td>NA</td><td>Intergenic</td><td>NA</td><td>NA</td></tr>
<tr><td>T29697</td><td>CATCGAGAACTTCCTCAGCGAGG</td><td>1000</td><td></td><td></td><td></td><td></td><td></td><td></td><td></td><td></td><td></td></tr>
<tr><td>Oligo fwd</td><td>CACCCATCGAGAACTTCCTCAGCG</td><td></td><td></td><td></td><td></td><td></td><td></td><td></td><td></td><td></td><td></td></tr>
<tr><td>Oligo rev</td><td>AAACCGCTGAGGAAGTTCTCGATG</td><td></td><td></td><td></td><td></td><td></td><td></td><td></td><td></td><td></td><td></td></tr>
<tr><td>Chromosome</td><td>start</td><td>end</td><td>strand</td><td>MM</td><td>target_seq</td><td>PAM</td><td>alignment</td><td>distance</td><td>position</td><td>gene name</td><td>gene id</td></tr>
<tr><td>GU305914.1</td><td>58310</td><td>58332</td><td>-</td><td>0</td><td>CATCGAGAACTTCCTCAGCG</td><td>AGG</td><td>||||||||[||||||||||||]PAM</td><td>NA</td><td>Intergenic</td><td>NA</td><td>NA</td></tr>
<tr><td>T29698</td><td>AGCGAGGAGCCCTTCGTGCGCGG</td><td>1000</td><td></td><td></td><td></td><td></td><td></td><td></td><td></td><td></td><td></td></tr>
<tr><td>Oligo fwd</td><td>CACCAGCGAGGAGCCCTTCGTGCG</td><td></td><td></td><td></td><td></td><td></td><td></td><td></td><td></td><td></td><td></td></tr>
<tr><td>Oligo rev</td><td>AAACCGCACGAAGGGCTCCTCGCT</td><td></td><td></td><td></td><td></td><td></td><td></td><td></td><td></td><td></td><td></td></tr>
<tr><td>Chromosome</td><td>start</td><td>end</td><td>strand</td><td>MM</td><td>target_seq</td><td>PAM</td><td>alignment</td><td>distance</td><td>position</td><td>gene name</td><td>gene id</td></tr>
<tr><td>GU305914.1</td><td>58294</td><td>58316</td><td>-</td><td>0</td><td>AGCGAGGAGCCCTTCGTGCG</td><td>CGG</td><td>||||||||[||||||||||||]PAM</td><td>NA</td><td>Intergenic</td><td>NA</td><td>NA</td></tr>
<tr><td>T29699</td><td>GCCCTTCGTGCGCGGCGACGTGG</td><td>1000</td><td></td><td></td><td></td><td></td><td></td><td></td><td></td><td></td><td></td></tr>
<tr><td>Oligo fwd</td><td>CACCGCCCTTCGTGCGCGGCGACG</td><td></td><td></td><td></td><td></td><td></td><td></td><td></td><td></td><td></td><td></td></tr>
<tr><td>Oligo rev</td><td>AAACCGTCGCCGCGCACGAAGGGC</td><td></td><td></td><td></td><td></td><td></td><td></td><td></td><td></td><td></td><td></td></tr>
<tr><td>Chromosome</td><td>start</td><td>end</td><td>strand</td><td>MM</td><td>target_seq</td><td>PAM</td><td>alignment</td><td>distance</td><td>position</td><td>gene name</td><td>gene id</td></tr>
<tr><td>GU305914.1</td><td>58286</td><td>58308</td><td>-</td><td>0</td><td>GCCCTTCGTGCGCGGCGACG</td><td>TGG</td><td>||||||||[||||||||||||]PAM</td><td>NA</td><td>Intergenic</td><td>NA</td><td>NA</td></tr>
<tr><td>T29700</td><td>CCCTTCGTGCGCGGCGACGTGGG</td><td>1000</td><td></td><td></td><td></td><td></td><td></td><td></td><td></td><td></td><td></td></tr>
<tr><td>Oligo fwd</td><td>CACCCCCTTCGTGCGCGGCGACGT</td><td></td><td></td><td></td><td></td><td></td><td></td><td></td><td></td><td></td><td></td></tr>
<tr><td>Oligo rev</td><td>AAACACGTCGCCGCGCACGAAGGG</td><td></td><td></td><td></td><td></td><td></td><td></td><td></td><td></td><td></td><td></td></tr>
<tr><td>Chromosome</td><td>start</td><td>end</td><td>strand</td><td>MM</td><td>target_seq</td><td>PAM</td><td>alignment</td><td>distance</td><td>position</td><td>gene name</td><td>gene id</td></tr>
<tr><td>GU305914.1</td><td>58285</td><td>58307</td><td>-</td><td>0</td><td>CCCTTCGTGCGCGGCGACGT</td><td>GGG</td><td>||||||||[||||||||||||]PAM</td><td>NA</td><td>Intergenic</td><td>NA</td><td>NA</td></tr>
<tr><td>T29701</td><td>CAGAACTCCAGCAGCTCCGCCGG</td><td>1000</td><td></td><td></td><td></td><td></td><td></td><td></td><td></td><td></td><td></td></tr>
<tr><td>Oligo fwd</td><td>CACCCAGAACTCCAGCAGCTCCGC</td><td></td><td></td><td></td><td></td><td></td><td></td><td></td><td></td><td></td><td></td></tr>
<tr><td>Oligo rev</td><td>AAACGCGGAGCTGCTGGAGTTCTG</td><td></td><td></td><td></td><td></td><td></td><td></td><td></td><td></td><td></td><td></td></tr>
<tr><td>Chromosome</td><td>start</td><td>end</td><td>strand</td><td>MM</td><td>target_seq</td><td>PAM</td><td>alignment</td><td>distance</td><td>position</td><td>gene name</td><td>gene id</td></tr>
<tr><td>GU305914.1</td><td>58225</td><td>58247</td><td>-</td><td>0</td><td>CAGAACTCCAGCAGCTCCGC</td><td>CGG</td><td>||||||||[||||||||||||]PAM</td><td>NA</td><td>Intergenic</td><td>NA</td><td>NA</td></tr>
<tr><td>T29702</td><td>ACTGTACCAAGAAACACGAGAGG</td><td>1000</td><td></td><td></td><td></td><td></td><td></td><td></td><td></td><td></td><td></td></tr>
<tr><td>Oligo fwd</td><td>CACCACTGTACCAAGAAACACGAG</td><td></td><td></td><td></td><td></td><td></td><td></td><td></td><td></td><td></td><td></td></tr>
<tr><td>Oligo rev</td><td>AAACCTCGTGTTTCTTGGTACAGT</td><td></td><td></td><td></td><td></td><td></td><td></td><td></td><td></td><td></td><td></td></tr>
<tr><td>Chromosome</td><td>start</td><td>end</td><td>strand</td><td>MM</td><td>target_seq</td><td>PAM</td><td>alignment</td><td>distance</td><td>position</td><td>gene name</td><td>gene id</td></tr>
<tr><td>GU305914.1</td><td>58164</td><td>58186</td><td>-</td><td>0</td><td>ACTGTACCAAGAAACACGAG</td><td>AGG</td><td>||||||||[||||||||||||]PAM</td><td>NA</td><td>Intergenic</td><td>NA</td><td>NA</td></tr>
<tr><td>T29703</td><td>CTGTACCAAGAAACACGAGAGGG</td><td>1000</td><td></td><td></td><td></td><td></td><td></td><td></td><td></td><td></td><td></td></tr>
<tr><td>Oligo fwd</td><td>CACCCTGTACCAAGAAACACGAGA</td><td></td><td></td><td></td><td></td><td></td><td></td><td></td><td></td><td></td><td></td></tr>
<tr><td>Oligo rev</td><td>AAACTCTCGTGTTTCTTGGTACAG</td><td></td><td></td><td></td><td></td><td></td><td></td><td></td><td></td><td></td><td></td></tr>
<tr><td>Chromosome</td><td>start</td><td>end</td><td>strand</td><td>MM</td><td>target_seq</td><td>PAM</td><td>alignment</td><td>distance</td><td>position</td><td>gene name</td><td>gene id</td></tr>
<tr><td>GU305914.1</td><td>58163</td><td>58185</td><td>-</td><td>0</td><td>CTGTACCAAGAAACACGAGA</td><td>GGG</td><td>||||||||[||||||||||||]PAM</td><td>NA</td><td>Intergenic</td><td>NA</td><td>NA</td></tr>
<tr><td>T29704</td><td>ACACGAGAGGGTCTCGCGCAAGG</td><td>1000</td><td></td><td></td><td></td><td></td><td></td><td></td><td></td><td></td><td></td></tr>
<tr><td>Oligo fwd</td><td>CACCACACGAGAGGGTCTCGCGCA</td><td></td><td></td><td></td><td></td><td></td><td></td><td></td><td></td><td></td><td></td></tr>
<tr><td>Oligo rev</td><td>AAACTGCGCGAGACCCTCTCGTGT</td><td></td><td></td><td></td><td></td><td></td><td></td><td></td><td></td><td></td><td></td></tr>
<tr><td>Chromosome</td><td>start</td><td>end</td><td>strand</td><td>MM</td><td>target_seq</td><td>PAM</td><td>alignment</td><td>distance</td><td>position</td><td>gene name</td><td>gene id</td></tr>
<tr><td>GU305914.1</td><td>58151</td><td>58173</td><td>-</td><td>0</td><td>ACACGAGAGGGTCTCGCGCA</td><td>AGG</td><td>||||||||[||||||||||||]PAM</td><td>NA</td><td>Intergenic</td><td>NA</td><td>NA</td></tr>
<tr><td>T29705</td><td>CGAGAGGGTCTCGCGCAAGGCGG</td><td>1000</td><td></td><td></td><td></td><td></td><td></td><td></td><td></td><td></td><td></td></tr>
<tr><td>Oligo fwd</td><td>CACCCGAGAGGGTCTCGCGCAAGG</td><td></td><td></td><td></td><td></td><td></td><td></td><td></td><td></td><td></td><td></td></tr>
<tr><td>Oligo rev</td><td>AAACCCTTGCGCGAGACCCTCTCG</td><td></td><td></td><td></td><td></td><td></td><td></td><td></td><td></td><td></td><td></td></tr>
<tr><td>Chromosome</td><td>start</td><td>end</td><td>strand</td><td>MM</td><td>target_seq</td><td>PAM</td><td>alignment</td><td>distance</td><td>position</td><td>gene name</td><td>gene id</td></tr>
<tr><td>GU305914.1</td><td>58148</td><td>58170</td><td>-</td><td>0</td><td>CGAGAGGGTCTCGCGCAAGG</td><td>CGG</td><td>||||||||[||||||||||||]PAM</td><td>NA</td><td>Intergenic</td><td>NA</td><td>NA</td></tr>
<tr><td>T29706</td><td>GAGAGGGTCTCGCGCAAGGCGGG</td><td>1000</td><td></td><td></td><td></td><td></td><td></td><td></td><td></td><td></td><td></td></tr>
<tr><td>Oligo fwd</td><td>CACCGAGAGGGTCTCGCGCAAGGC</td><td></td><td></td><td></td><td></td><td></td><td></td><td></td><td></td><td></td><td></td></tr>
<tr><td>Oligo rev</td><td>AAACGCCTTGCGCGAGACCCTCTC</td><td></td><td></td><td></td><td></td><td></td><td></td><td></td><td></td><td></td><td></td></tr>
<tr><td>Chromosome</td><td>start</td><td>end</td><td>strand</td><td>MM</td><td>target_seq</td><td>PAM</td><td>alignment</td><td>distance</td><td>position</td><td>gene name</td><td>gene id</td></tr>
<tr><td>GU305914.1</td><td>58147</td><td>58169</td><td>-</td><td>0</td><td>GAGAGGGTCTCGCGCAAGGC</td><td>GGG</td><td>||||||||[||||||||||||]PAM</td><td>NA</td><td>Intergenic</td><td>NA</td><td>NA</td></tr>
<tr><td>T29707</td><td>AGGGTCTCGCGCAAGGCGGGCGG</td><td>1000</td><td></td><td></td><td></td><td></td><td></td><td></td><td></td><td></td><td></td></tr>
<tr><td>Oligo fwd</td><td>CACCAGGGTCTCGCGCAAGGCGGG</td><td></td><td></td><td></td><td></td><td></td><td></td><td></td><td></td><td></td><td></td></tr>
<tr><td>Oligo rev</td><td>AAACCCCGCCTTGCGCGAGACCCT</td><td></td><td></td><td></td><td></td><td></td><td></td><td></td><td></td><td></td><td></td></tr>
<tr><td>Chromosome</td><td>start</td><td>end</td><td>strand</td><td>MM</td><td>target_seq</td><td>PAM</td><td>alignment</td><td>distance</td><td>position</td><td>gene name</td><td>gene id</td></tr>
<tr><td>GU305914.1</td><td>58144</td><td>58166</td><td>-</td><td>0</td><td>AGGGTCTCGCGCAAGGCGGG</td><td>CGG</td><td>||||||||[||||||||||||]PAM</td><td>NA</td><td>Intergenic</td><td>NA</td><td>NA</td></tr>
</table>

<table>
<tr><td>T29708</td><td>GTCTCGCGCAAGGCGGGCGGTGG</td><td>1000</td><td></td><td></td><td></td><td></td><td></td><td></td><td></td><td></td><td></td></tr>
<tr><td>Oligo fwd</td><td>CACCGTCTCGCGCAAGGCGGGCGG</td><td></td><td></td><td></td><td></td><td></td><td></td><td></td><td></td><td></td><td></td></tr>
<tr><td>Oligo rev</td><td>AAACCCGCCCGCCTTGCGCGAGAC</td><td></td><td></td><td></td><td></td><td></td><td></td><td></td><td></td><td></td><td></td></tr>
<tr><td>Chromosome</td><td>start</td><td>end</td><td>strand</td><td>MM</td><td>target_seq</td><td>PAM</td><td>alignment</td><td>distance</td><td>position</td><td>gene name</td><td>gene id</td></tr>
<tr><td>GU305914.1</td><td>58141</td><td>58163</td><td>-</td><td>0</td><td>GTCTCGCGCAAGGCGGGCGG</td><td>TGG</td><td>||||||||[||||||||||||]PAM</td><td>NA</td><td>Intergenic</td><td>NA</td><td>NA</td></tr>
<tr><td>T29709</td><td>TCGCGCAAGGCGGGCGGTGGCGG</td><td>1000</td><td></td><td></td><td></td><td></td><td></td><td></td><td></td><td></td><td></td></tr>
<tr><td>Oligo fwd</td><td>CACCTCGCGCAAGGCGGGCGGTGG</td><td></td><td></td><td></td><td></td><td></td><td></td><td></td><td></td><td></td><td></td></tr>
<tr><td>Oligo rev</td><td>AAACCCACCGCCCGCCTTGCGCGA</td><td></td><td></td><td></td><td></td><td></td><td></td><td></td><td></td><td></td><td></td></tr>
<tr><td>Chromosome</td><td>start</td><td>end</td><td>strand</td><td>MM</td><td>target_seq</td><td>PAM</td><td>alignment</td><td>distance</td><td>position</td><td>gene name</td><td>gene id</td></tr>
<tr><td>GU305914.1</td><td>58138</td><td>58160</td><td>-</td><td>0</td><td>TCGCGCAAGGCGGGCGGTGG</td><td>CGG</td><td>||||||||[||||||||||||]PAM</td><td>NA</td><td>Intergenic</td><td>NA</td><td>NA</td></tr>
<tr><td>T29710</td><td>CGCAAGGCGGGCGGTGGCGGCGG</td><td>1000</td><td></td><td></td><td></td><td></td><td></td><td></td><td></td><td></td><td></td></tr>
<tr><td>Oligo fwd</td><td>CACCCGCAAGGCGGGCGGTGGCGG</td><td></td><td></td><td></td><td></td><td></td><td></td><td></td><td></td><td></td><td></td></tr>
<tr><td>Oligo rev</td><td>AAACCCGCCACCGCCCGCCTTGCG</td><td></td><td></td><td></td><td></td><td></td><td></td><td></td><td></td><td></td><td></td></tr>
<tr><td>Chromosome</td><td>start</td><td>end</td><td>strand</td><td>MM</td><td>target_seq</td><td>PAM</td><td>alignment</td><td>distance</td><td>position</td><td>gene name</td><td>gene id</td></tr>
<tr><td>GU305914.1</td><td>58135</td><td>58157</td><td>-</td><td>0</td><td>CGCAAGGCGGGCGGTGGCGG</td><td>CGG</td><td>||||||||[||||||||||||]PAM</td><td>NA</td><td>Intergenic</td><td>NA</td><td>NA</td></tr>
<tr><td>GU305914.1</td><td>22926</td><td>22948</td><td>+</td><td>4</td><td>CCCAACGCGGTCAGTGGCGG</td><td>CGG</td><td>|-|||-||[||-|-|||||||]PAM</td><td>NA</td><td>Intergenic</td><td>NA</td><td>NA</td></tr>
<tr><td>GU305914.1</td><td>24627</td><td>24649</td><td>-</td><td>4</td><td>CGCAACACCGGCGGCGGCGG</td><td>TGG</td><td>|||||--|[-|||||-|||||]PAM</td><td>NA</td><td>Intergenic</td><td>NA</td><td>NA</td></tr>
<tr><td>GU305914.1</td><td>90265</td><td>90287</td><td>-</td><td>4</td><td>CGGGAGGCGGGCGACGGCGG</td><td>AGG</td><td>||--||||[|||||--|||||]PAM</td><td>NA</td><td>Intergenic</td><td>NA</td><td>NA</td></tr>
<tr><td>T29711</td><td>AAGGCGGGCGGTGGCGGCGGCGG</td><td>1000</td><td></td><td></td><td></td><td></td><td></td><td></td><td></td><td></td><td></td></tr>
<tr><td>Oligo fwd</td><td>CACCAAGGCGGGCGGTGGCGGCGG</td><td></td><td></td><td></td><td></td><td></td><td></td><td></td><td></td><td></td><td></td></tr>
<tr><td>Oligo rev</td><td>AAACCCGCCGCCACCGCCCGCCTT</td><td></td><td></td><td></td><td></td><td></td><td></td><td></td><td></td><td></td><td></td></tr>
<tr><td>Chromosome</td><td>start</td><td>end</td><td>strand</td><td>MM</td><td>target_seq</td><td>PAM</td><td>alignment</td><td>distance</td><td>position</td><td>gene name</td><td>gene id</td></tr>
<tr><td>GU305914.1</td><td>58132</td><td>58154</td><td>-</td><td>0</td><td>AAGGCGGGCGGTGGCGGCGG</td><td>CGG</td><td>||||||||[||||||||||||]PAM</td><td>NA</td><td>Intergenic</td><td>NA</td><td>NA</td></tr>
<tr><td>GU305914.1</td><td>154398</td><td>154420</td><td>+</td><td>4</td><td>GCCCCGGGCGGTGGCGGCGG</td><td>CGG</td><td>----||||[||||||||||||]PAM</td><td>NA</td><td>Intergenic</td><td>NA</td><td>NA</td></tr>
<tr><td>GU305914.1</td><td>45258</td><td>45280</td><td>-</td><td>4</td><td>AACGGTGGCGGTGGCGGCAG</td><td>CGG</td><td>||-|--||[||||||||||-|]PAM</td><td>NA</td><td>Intergenic</td><td>NA</td><td>NA</td></tr>
<tr><td>T29712</td><td>GCGGGCGGTGGCGGCGGCGGCGG</td><td>1000</td><td></td><td></td><td></td><td></td><td></td><td></td><td></td><td></td><td></td></tr>
<tr><td>Oligo fwd</td><td>CACCGCGGGCGGTGGCGGCGGCGG</td><td></td><td></td><td></td><td></td><td></td><td></td><td></td><td></td><td></td><td></td></tr>
<tr><td>Oligo rev</td><td>AAACCCGCCGCCGCCACCGCCCGC</td><td></td><td></td><td></td><td></td><td></td><td></td><td></td><td></td><td></td><td></td></tr>
<tr><td>Chromosome</td><td>start</td><td>end</td><td>strand</td><td>MM</td><td>target_seq</td><td>PAM</td><td>alignment</td><td>distance</td><td>position</td><td>gene name</td><td>gene id</td></tr>
<tr><td>GU305914.1</td><td>58129</td><td>58151</td><td>-</td><td>0</td><td>GCGGGCGGTGGCGGCGGCGG</td><td>CGG</td><td>||||||||[||||||||||||]PAM</td><td>NA</td><td>Intergenic</td><td>NA</td><td>NA</td></tr>
<tr><td>GU305914.1</td><td>154401</td><td>154423</td><td>+</td><td>1</td><td>CCGGGCGGTGGCGGCGGCGG</td><td>CGG</td><td>-|||||||[||||||||||||]PAM</td><td>NA</td><td>Intergenic</td><td>NA</td><td>NA</td></tr>
<tr><td>GU305914.1</td><td>48159</td><td>48181</td><td>+</td><td>2</td><td>GGCGGCGGTGGCGGCGGCGG</td><td>TGG</td><td>|--|||||[||||||||||||]PAM</td><td>NA</td><td>Intergenic</td><td>NA</td><td>NA</td></tr>
<tr><td>GU305914.1</td><td>154407</td><td>154429</td><td>+</td><td>3</td><td>GGTGGCGGCGGCGGCGGCGG</td><td>CGG</td><td>|--|||||[-|||||||||||]PAM</td><td>NA</td><td>Intergenic</td><td>NA</td><td>NA</td></tr>
<tr><td>GU305914.1</td><td>58123</td><td>58145</td><td>-</td><td>3</td><td>GGTGGCGGCGGCGGCGGCGG</td><td>AGG</td><td>|--|||||[-|||||||||||]PAM</td><td>NA</td><td>Intergenic</td><td>NA</td><td>NA</td></tr>
<tr><td>GU305914.1</td><td>67742</td><td>67764</td><td>+</td><td>3</td><td>GGGAGCGGCGGCGGCGGCGG</td><td>CGG</td><td>|-|-||||[-|||||||||||]PAM</td><td>NA</td><td>Intergenic</td><td>NA</td><td>NA</td></tr>
<tr><td>GU305914.1</td><td>29997</td><td>30019</td><td>+</td><td>4</td><td>ACCGGTGCTGGCGGCGGCGG</td><td>CGG</td><td>-|-||-|-[||||||||||||]PAM</td><td>NA</td><td>Intergenic</td><td>NA</td><td>NA</td></tr>
<tr><td>GU305914.1</td><td>149482</td><td>149504</td><td>-</td><td>4</td><td>GGCGACGGCGGCGGCGGCGG</td><td>CGG</td><td>|--|-|||[-|||||||||||]PAM</td><td>NA</td><td>Intergenic</td><td>NA</td><td>NA</td></tr>
<tr><td>GU305914.1</td><td>149485</td><td>149507</td><td>-</td><td>3</td><td>GCAGGCGACGGCGGCGGCGG</td><td>CGG</td><td>||-||||-[-|||||||||||]PAM</td><td>NA</td><td>Intergenic</td><td>NA</td><td>NA</td></tr>
<tr><td>GU305914.1</td><td>154404</td><td>154426</td><td>+</td><td>4</td><td>GGCGGTGGCGGCGGCGGCGG</td><td>CGG</td><td>|--||-||[-|||||||||||]PAM</td><td>NA</td><td>Intergenic</td><td>NA</td><td>NA</td></tr>
<tr><td>GU305914.1</td><td>58126</td><td>58148</td><td>-</td><td>4</td><td>GGCGGTGGCGGCGGCGGCGG</td><td>CGG</td><td>|--||-||[-|||||||||||]PAM</td><td>NA</td><td>Intergenic</td><td>NA</td><td>NA</td></tr>
<tr><td>GU305914.1</td><td>67739</td><td>67761</td><td>+</td><td>4</td><td>TCGGGGAGCGGCGGCGGCGG</td><td>CGG</td><td>-||||--|[-|||||||||||]PAM</td><td>NA</td><td>Intergenic</td><td>NA</td><td>NA</td></tr>
<tr><td>GU305914.1</td><td>62446</td><td>62468</td><td>+</td><td>4</td><td>GGCGGGGGTGGTGGCGGCGG</td><td>GGG</td><td>|--||-||[|||-||||||||]PAM</td><td>NA</td><td>Intergenic</td><td>NA</td><td>NA</td></tr>
<tr><td>GU305914.1</td><td>114658</td><td>114680</td><td>-</td><td>4</td><td>GGCGGTGGTGGTGGCGGCGG</td><td>CGG</td><td>|--||-||[|||-||||||||]PAM</td><td>NA</td><td>Intergenic</td><td>NA</td><td>NA</td></tr>
<tr><td>GU305914.1</td><td>168817</td><td>168839</td><td>+</td><td>4</td><td>GGCGGTGGTGGCAGCGGCGG</td><td>CGG</td><td>|--||-||[||||-|||||||]PAM</td><td>NA</td><td>Intergenic</td><td>NA</td><td>NA</td></tr>
<tr><td>GU305914.1</td><td>170536</td><td>170558</td><td>-</td><td>4</td><td>GTGGCCGATGGTGGCGGCGG</td><td>CGG</td><td>|-||-||-[|||-||||||||]PAM</td><td>NA</td><td>Intergenic</td><td>NA</td><td>NA</td></tr>
<tr><td>GU305914.1</td><td>48156</td><td>48178</td><td>+</td><td>4</td><td>GGCGGCGGCGGTGGCGGCGG</td><td>CGG</td><td>|--|||||[-||-||||||||]PAM</td><td>NA</td><td>Intergenic</td><td>NA</td><td>NA</td></tr>
<tr><td>GU305914.1</td><td>45255</td><td>45277</td><td>-</td><td>3</td><td>GGTGGCGGTGGCGGCAGCGG</td><td>CGG</td><td>|--|||||[|||||||-||||]PAM</td><td>NA</td><td>Intergenic</td><td>NA</td><td>NA</td></tr>
<tr><td>GU305914.1</td><td>52716</td><td>52738</td><td>+</td><td>4</td><td>GCCGCCGCTGGCGGTGGCGG</td><td>TGG</td><td>||-|-||-[||||||-|||||]PAM</td><td>NA</td><td>Intergenic</td><td>NA</td><td>NA</td></tr>
<tr><td>GU305914.1</td><td>90259</td><td>90281</td><td>-</td><td>3</td><td>GCGGGCGACGGCGGAGGCGG</td><td>CGG</td><td>|||||||-[-|||||-|||||]PAM</td><td>NA</td><td>Intergenic</td><td>NA</td><td>NA</td></tr>
<tr><td>GU305914.1</td><td>114661</td><td>114683</td><td>-</td><td>4</td><td>GACGGCGGTGGTGGTGGCGG</td><td>CGG</td><td>|--|||||[|||-||-|||||]PAM</td><td>NA</td><td>Intergenic</td><td>NA</td><td>NA</td></tr>
<tr><td>GU305914.1</td><td>62434</td><td>62456</td><td>+</td><td>3</td><td>GACGGCGGTGGCGGCGGGGG</td><td>TGG</td><td>|--|||||[|||||||||-||]PAM</td><td>NA</td><td>Intergenic</td><td>NA</td><td>NA</td></tr>
<tr><td>GU305914.1</td><td>62449</td><td>62471</td><td>+</td><td>3</td><td>GGGGGTGGTGGCGGCGGGGG</td><td>TGG</td><td>|-|||-||[|||||||||-||]PAM</td><td>NA</td><td>Intergenic</td><td>NA</td><td>NA</td></tr>
<tr><td>GU305914.1</td><td>30006</td><td>30028</td><td>+</td><td>4</td><td>GGCGGCGGCGGCGGCGGCAG</td><td>TGG</td><td>|--|||||[-|||||||||-|]PAM</td><td>NA</td><td>Intergenic</td><td>NA</td><td>NA</td></tr>
<tr><td>GU305914.1</td><td>149476</td><td>149498</td><td>-</td><td>4</td><td>GGCGGCGGCGGCGGCGGCAG</td><td>CGG</td><td>|--|||||[-|||||||||-|]PAM</td><td>NA</td><td>Intergenic</td><td>NA</td><td>NA</td></tr>
<tr><td>GU305914.1</td><td>143022</td><td>143044</td><td>+</td><td>4</td><td>GCTGGTGGTGGCGCCGGGGG</td><td>CGG</td><td>||-||-||[|||||-|||-||]PAM</td><td>NA</td><td>Intergenic</td><td>NA</td><td>NA</td></tr>
<tr><td>T29713</td><td>GGCGGTGGCGGCGGCGGCGGCGG</td><td>1000</td><td></td><td></td><td></td><td></td><td></td><td></td><td></td><td></td><td></td></tr>
<tr><td>Oligo fwd</td><td>CACCGGCGGTGGCGGCGGCGGCGG</td><td></td><td></td><td></td><td></td><td></td><td></td><td></td><td></td><td></td><td></td></tr>
<tr><td>Oligo rev</td><td>AAACCCGCCGCCGCCGCCACCGCC</td><td></td><td></td><td></td><td></td><td></td><td></td><td></td><td></td><td></td><td></td></tr>
<tr><td>Chromosome</td><td>start</td><td>end</td><td>strand</td><td>MM</td><td>target_seq</td><td>PAM</td><td>alignment</td><td>distance</td><td>position</td><td>gene name</td><td>gene id</td></tr>
<tr><td>GU305914.1</td><td>154404</td><td>154426</td><td>+</td><td>0</td><td>GGCGGTGGCGGCGGCGGCGG</td><td>CGG</td><td>||||||||[||||||||||||]PAM</td><td>NA</td><td>Intergenic</td><td>NA</td><td>NA</td></tr>
<tr><td>GU305914.1</td><td>58126</td><td>58148</td><td>-</td><td>0</td><td>GGCGGTGGCGGCGGCGGCGG</td><td>CGG</td><td>||||||||[||||||||||||]PAM</td><td>NA</td><td>Intergenic</td><td>NA</td><td>NA</td></tr>
<tr><td>GU305914.1</td><td>30000</td><td>30022</td><td>+</td><td>2</td><td>GGTGCTGGCGGCGGCGGCGG</td><td>CGG</td><td>||-|-|||[||||||||||||]PAM</td><td>NA</td><td>Intergenic</td><td>NA</td><td>NA</td></tr>
<tr><td>GU305914.1</td><td>154407</td><td>154429</td><td>+</td><td>2</td><td>GGTGGCGGCGGCGGCGGCGG</td><td>CGG</td><td>||-||-||[||||||||||||]PAM</td><td>NA</td><td>Intergenic</td><td>NA</td><td>NA</td></tr>
<tr><td>GU305914.1</td><td>58123</td><td>58145</td><td>-</td><td>2</td><td>GGTGGCGGCGGCGGCGGCGG</td><td>AGG</td><td>||-||-||[||||||||||||]PAM</td><td>NA</td><td>Intergenic</td><td>NA</td><td>NA</td></tr>
<tr><td>GU305914.1</td><td>149482</td><td>149504</td><td>-</td><td>2</td><td>GGCGACGGCGGCGGCGGCGG</td><td>CGG</td><td>||||--||[||||||||||||]PAM</td><td>NA</td><td>Intergenic</td><td>NA</td><td>NA</td></tr>
<tr><td>GU305914.1</td><td>67742</td><td>67764</td><td>+</td><td>3</td><td>GGGAGCGGCGGCGGCGGCGG</td><td>CGG</td><td>||--|-||[||||||||||||]PAM</td><td>NA</td><td>Intergenic</td><td>NA</td><td>NA</td></tr>
<tr><td>GU305914.1</td><td>168820</td><td>168842</td><td>+</td><td>2</td><td>GGTGGTGGCAGCGGCGGCGG</td><td>CGG</td><td>||-|||||[|-||||||||||]PAM</td><td>NA</td><td>Intergenic</td><td>NA</td><td>NA</td></tr>
<tr><td>GU305914.1</td><td>48159</td><td>48181</td><td>+</td><td>2</td><td>GGCGGCGGTGGCGGCGGCGG</td><td>TGG</td><td>|||||-||[-|||||||||||]PAM</td><td>NA</td><td>Intergenic</td><td>NA</td><td>NA</td></tr>
<tr><td>GU305914.1</td><td>56773</td><td>56795</td><td>+</td><td>4</td><td>CGCCGCAGCGGCGGCGGCGG</td><td>CGG</td><td>-||-|--|[||||||||||||]PAM</td><td>NA</td><td>Intergenic</td><td>NA</td><td>NA</td></tr>
<tr><td>GU305914.1</td><td>161754</td><td>161776</td><td>-</td><td>3</td><td>GGCCGAGGAGGCGGCGGCGG</td><td>AGG</td><td>|||-|-||[-|||||||||||]PAM</td><td>NA</td><td>Intergenic</td><td>NA</td><td>NA</td></tr>
<tr><td>GU305914.1</td><td>149485</td><td>149507</td><td>-</td><td>4</td><td>GCAGGCGACGGCGGCGGCGG</td><td>CGG</td><td>|--||-|-[||||||||||||]PAM</td><td>NA</td><td>Intergenic</td><td>NA</td><td>NA</td></tr>
<tr><td>GU305914.1</td><td>58129</td><td>58151</td><td>-</td><td>4</td><td>GCGGGCGGTGGCGGCGGCGG</td><td>CGG</td><td>|--||-||[-|||||||||||]PAM</td><td>NA</td><td>Intergenic</td><td>NA</td><td>NA</td></tr>
<tr><td>GU305914.1</td><td>176454</td><td>176476</td><td>+</td><td>3</td><td>GACGATGGCGACGGCGGCGG</td><td>AGG</td><td>|-||-|||[||-|||||||||]PAM</td><td>NA</td><td>Intergenic</td><td>NA</td><td>NA</td></tr>
<tr><td>GU305914.1</td><td>48156</td><td>48178</td><td>+</td><td>2</td><td>GGCGGCGGCGGTGGCGGCGG</td><td>CGG</td><td>|||||-||[|||-||||||||]PAM</td><td>NA</td><td>Intergenic</td><td>NA</td><td>NA</td></tr>
<tr><td>GU305914.1</td><td>29997</td><td>30019</td><td>+</td><td>4</td><td>ACCGGTGCTGGCGGCGGCGG</td><td>CGG</td><td>--|||||-[-|||||||||||]PAM</td><td>NA</td><td>Intergenic</td><td>NA</td><td>NA</td></tr>
<tr><td>GU305914.1</td><td>164971</td><td>164993</td><td>-</td><td>4</td><td>GGTGGGACCGGCGGCGGCGG</td><td>CGG</td><td>||-||---[||||||||||||]PAM</td><td>NA</td><td>Intergenic</td><td>NA</td><td>NA</td></tr>
<tr><td>GU305914.1</td><td>114658</td><td>114680</td><td>-</td><td>2</td><td>GGCGGTGGTGGTGGCGGCGG</td><td>CGG</td><td>||||||||[-||-||||||||]PAM</td><td>NA</td><td>Intergenic</td><td>NA</td><td>NA</td></tr>
<tr><td>GU305914.1</td><td>149488</td><td>149510</td><td>-</td><td>4</td><td>GGAGCAGGCGACGGCGGCGG</td><td>CGG</td><td>||-|--||[||-|||||||||]PAM</td><td>NA</td><td>Intergenic</td><td>NA</td><td>NA</td></tr>
<tr><td>GU305914.1</td><td>73792</td><td>73814</td><td>-</td><td>4</td><td>GGCGCCAGCAGCGGCGGCGG</td><td>AGG</td><td>||||---|[|-||||||||||]PAM</td><td>NA</td><td>Intergenic</td><td>NA</td><td>NA</td></tr>
<tr><td>GU305914.1</td><td>168817</td><td>168839</td><td>+</td><td>2</td><td>GGCGGTGGTGGCAGCGGCGG</td><td>CGG</td><td>||||||||[-|||-|||||||]PAM</td><td>NA</td><td>Intergenic</td><td>NA</td><td>NA</td></tr>
<tr><td>GU305914.1</td><td>62446</td><td>62468</td><td>+</td><td>3</td><td>GGCGGGGGTGGTGGCGGCGG</td><td>GGG</td><td>|||||-||[-||-||||||||]PAM</td><td>NA</td><td>Intergenic</td><td>NA</td><td>NA</td></tr>
<tr><td>GU305914.1</td><td>113930</td><td>113952</td><td>-</td><td>4</td><td>CGAGGTGTCGGCGTCGGCGG</td><td>CGG</td><td>-|-||||-[|||||-||||||]PAM</td><td>NA</td><td>Intergenic</td><td>NA</td><td>NA</td></tr>
<tr><td>GU305914.1</td><td>48153</td><td>48175</td><td>+</td><td>4</td><td>TGTGGCGGCGGCGGTGGCGG</td><td>CGG</td><td>-|-||-||[||||||-|||||]PAM</td><td>NA</td><td>Intergenic</td><td>NA</td><td>NA</td></tr>
<tr><td>GU305914.1</td><td>18653</td><td>18675</td><td>-</td><td>4</td><td>GGCAGAGGTGGCGGAGGCGG</td><td>GGG</td><td>|||-|-||[-|||||-|||||]PAM</td><td>NA</td><td>Intergenic</td><td>NA</td><td>NA</td></tr>
</table>

<table>
<tr><td>GU305914.1</td><td>163749</td><td>163771</td><td>+</td><td>4</td><td>GTCGGCGGCGCCGGTGGCGG</td><td>CGG</td><td>|-|||-||[||-|||-|||||]PAM</td><td>NA</td><td>Intergenic</td><td>NA</td><td>NA</td></tr>
<tr><td>GU305914.1</td><td>163911</td><td>163933</td><td>+</td><td>3</td><td>GGCGGCGGCAGCGGCAGCGG</td><td>AGG</td><td>|||||-||[|-|||||-||||]PAM</td><td>NA</td><td>Intergenic</td><td>NA</td><td>NA</td></tr>
<tr><td>GU305914.1</td><td>45255</td><td>45277</td><td>-</td><td>4</td><td>GGTGGCGGTGGCGGCAGCGG</td><td>CGG</td><td>||-||-||[-||||||-||||]PAM</td><td>NA</td><td>Intergenic</td><td>NA</td><td>NA</td></tr>
<tr><td>GU305914.1</td><td>19936</td><td>19958</td><td>+</td><td>4</td><td>GACGGTAGCGGTGGTGGCGG</td><td>TGG</td><td>|-||||-|[|||-||-|||||]PAM</td><td>NA</td><td>Intergenic</td><td>NA</td><td>NA</td></tr>
<tr><td>GU305914.1</td><td>62155</td><td>62177</td><td>+</td><td>4</td><td>GGCGACGGCAGCGGCAGCGG</td><td>CGG</td><td>||||--||[|-|||||-||||]PAM</td><td>NA</td><td>Intergenic</td><td>NA</td><td>NA</td></tr>
<tr><td>GU305914.1</td><td>48150</td><td>48172</td><td>+</td><td>4</td><td>ACCTGTGGCGGCGGCGGTGG</td><td>CGG</td><td>--|-||||[|||||||||-||]PAM</td><td>NA</td><td>Intergenic</td><td>NA</td><td>NA</td></tr>
<tr><td>GU305914.1</td><td>62449</td><td>62471</td><td>+</td><td>3</td><td>GGGGGTGGTGGCGGCGGGGG</td><td>TGG</td><td>||-|||||[-||||||||-||]PAM</td><td>NA</td><td>Intergenic</td><td>NA</td><td>NA</td></tr>
<tr><td>GU305914.1</td><td>30006</td><td>30028</td><td>+</td><td>2</td><td>GGCGGCGGCGGCGGCGGCAG</td><td>TGG</td><td>|||||-||[||||||||||-|]PAM</td><td>NA</td><td>Intergenic</td><td>NA</td><td>NA</td></tr>
<tr><td>GU305914.1</td><td>149476</td><td>149498</td><td>-</td><td>2</td><td>GGCGGCGGCGGCGGCGGCAG</td><td>CGG</td><td>|||||-||[||||||||||-|]PAM</td><td>NA</td><td>Intergenic</td><td>NA</td><td>NA</td></tr>
<tr><td>GU305914.1</td><td>163902</td><td>163924</td><td>+</td><td>3</td><td>AGCGGCGGCGGCGGCGGCAG</td><td>CGG</td><td>-||||-||[||||||||||-|]PAM</td><td>NA</td><td>Intergenic</td><td>NA</td><td>NA</td></tr>
<tr><td>GU305914.1</td><td>62434</td><td>62456</td><td>+</td><td>4</td><td>GACGGCGGTGGCGGCGGGGG</td><td>TGG</td><td>|-|||-||[-||||||||-||]PAM</td><td>NA</td><td>Intergenic</td><td>NA</td><td>NA</td></tr>
<tr><td>GU305914.1</td><td>52719</td><td>52741</td><td>+</td><td>4</td><td>GCCGCTGGCGGTGGCGGTGG</td><td>CGG</td><td>|-||-|||[|||-|||||-||]PAM</td><td>NA</td><td>Intergenic</td><td>NA</td><td>NA</td></tr>
<tr><td>GU305914.1</td><td>90250</td><td>90272</td><td>-</td><td>3</td><td>GGCGGAGGCGGCGGAGCCGG</td><td>CGG</td><td>|||||-||[||||||-|-|||]PAM</td><td>NA</td><td>Intergenic</td><td>NA</td><td>NA</td></tr>
<tr><td>GU305914.1</td><td>113924</td><td>113946</td><td>-</td><td>4</td><td>GTCGGCGTCGGCGGCGGCAG</td><td>CGG</td><td>|-|||-|-[||||||||||-|]PAM</td><td>NA</td><td>Intergenic</td><td>NA</td><td>NA</td></tr>
<tr><td>GU305914.1</td><td>152553</td><td>152575</td><td>+</td><td>4</td><td>TGCGGCGGCGGCGGTGCCGG</td><td>CGG</td><td>-||||-||[||||||-|-|||]PAM</td><td>NA</td><td>Intergenic</td><td>NA</td><td>NA</td></tr>
<tr><td>GU305914.1</td><td>62437</td><td>62459</td><td>+</td><td>2</td><td>GGCGGTGGCGGCGGGGGTGG</td><td>TGG</td><td>||||||||[||||||-||-||]PAM</td><td>NA</td><td>Intergenic</td><td>NA</td><td>NA</td></tr>
<tr><td>GU305914.1</td><td>45258</td><td>45280</td><td>-</td><td>4</td><td>AACGGTGGCGGTGGCGGCAG</td><td>CGG</td><td>--||||||[|||-||||||-|]PAM</td><td>NA</td><td>Intergenic</td><td>NA</td><td>NA</td></tr>
<tr><td>GU305914.1</td><td>62452</td><td>62474</td><td>+</td><td>3</td><td>GGTGGTGGCGGCGGGGGTGG</td><td>CGG</td><td>||-|||||[||||||-||-||]PAM</td><td>NA</td><td>Intergenic</td><td>NA</td><td>NA</td></tr>
<tr><td>GU305914.1</td><td>163908</td><td>163930</td><td>+</td><td>3</td><td>GGCGGCGGCGGCAGCGGCAG</td><td>CGG</td><td>|||||-||[||||-|||||-|]PAM</td><td>NA</td><td>Intergenic</td><td>NA</td><td>NA</td></tr>
<tr><td>GU305914.1</td><td>113918</td><td>113940</td><td>-</td><td>4</td><td>GTCGGCGGCGGCAGCGGCGA</td><td>CGG</td><td>|-|||-||[||||-||||||-]PAM</td><td>NA</td><td>Intergenic</td><td>NA</td><td>NA</td></tr>
<tr><td>GU305914.1</td><td>113912</td><td>113934</td><td>-</td><td>4</td><td>GGCGGCAGCGGCGACGGCGT</td><td>CGG</td><td>|||||--|[|||||-|||||-]PAM</td><td>NA</td><td>Intergenic</td><td>NA</td><td>NA</td></tr>
<tr><td>GU305914.1</td><td>168829</td><td>168851</td><td>+</td><td>4</td><td>AGCGGCGGCGGCGGCGACAG</td><td>CGG</td><td>-||||-||[||||||||-|-|]PAM</td><td>NA</td><td>Intergenic</td><td>NA</td><td>NA</td></tr>
<tr><td>GU305914.1</td><td>170530</td><td>170552</td><td>-</td><td>4</td><td>GATGGTGGCGGCGGCGGCCA</td><td>CGG</td><td>|--|||||[||||||||||--]PAM</td><td>NA</td><td>Intergenic</td><td>NA</td><td>NA</td></tr>
</table>

T29714 GGTGGCGGCGGCGGCGGCGGAGG 1000
Oligo fwd CACCGGTGGCGGCGGCGGCGGCGG
Oligo rev AAACCCGCCGCCGCCGCCGCCACC

<table>
<tr><th>Chromosome</th><th>start</th><th>end</th><th>strand</th><th>MM</th><th>target_seq</th><th>PAM</th><th>alignment</th><th>distance</th><th>position</th><th>gene name</th><th>gene id</th></tr>
<tr><td>GU305914.1</td><td>154407</td><td>154429</td><td>+</td><td>0</td><td>GGTGGCGGCGGCGGCGGCGG</td><td>CGG</td><td>||||||||[||||||||||||]PAM</td><td>NA</td><td>Intergenic</td><td>NA</td><td>NA</td></tr>
<tr><td>GU305914.1</td><td>58123</td><td>58145</td><td>-</td><td>0</td><td>GGTGGCGGCGGCGGCGGCGG</td><td>AGG</td><td>||||||||[||||||||||||]PAM</td><td>NA</td><td>Intergenic</td><td>NA</td><td>NA</td></tr>
<tr><td>GU305914.1</td><td>67742</td><td>67764</td><td>+</td><td>2</td><td>GGGAGCGGCGGCGGCGGCGG</td><td>CGG</td><td>||--||||[||||||||||||]PAM</td><td>NA</td><td>Intergenic</td><td>NA</td><td>NA</td></tr>
<tr><td>GU305914.1</td><td>149482</td><td>149504</td><td>-</td><td>2</td><td>GGCGACGGCGGCGGCGGCGG</td><td>CGG</td><td>||-|-|||[||||||||||||]PAM</td><td>NA</td><td>Intergenic</td><td>NA</td><td>NA</td></tr>
<tr><td>GU305914.1</td><td>154404</td><td>154426</td><td>+</td><td>2</td><td>GGCGGTGGCGGCGGCGGCGG</td><td>CGG</td><td>||-||-||[||||||||||||]PAM</td><td>NA</td><td>Intergenic</td><td>NA</td><td>NA</td></tr>
<tr><td>GU305914.1</td><td>58126</td><td>58148</td><td>-</td><td>2</td><td>GGCGGTGGCGGCGGCGGCGG</td><td>CGG</td><td>||-||-||[||||||||||||]PAM</td><td>NA</td><td>Intergenic</td><td>NA</td><td>NA</td></tr>
<tr><td>GU305914.1</td><td>30000</td><td>30022</td><td>+</td><td>2</td><td>GGTGCTGGCGGCGGCGGCGG</td><td>CGG</td><td>||||--||[||||||||||||]PAM</td><td>NA</td><td>Intergenic</td><td>NA</td><td>NA</td></tr>
<tr><td>GU305914.1</td><td>48159</td><td>48181</td><td>+</td><td>2</td><td>GGCGGCGGTGGCGGCGGCGG</td><td>TGG</td><td>||-|||||[-|||||||||||]PAM</td><td>NA</td><td>Intergenic</td><td>NA</td><td>NA</td></tr>
<tr><td>GU305914.1</td><td>149485</td><td>149507</td><td>-</td><td>3</td><td>GCAGGCGACGGCGGCGGCGG</td><td>CGG</td><td>|--||||-[||||||||||||]PAM</td><td>NA</td><td>Intergenic</td><td>NA</td><td>NA</td></tr>
<tr><td>GU305914.1</td><td>58129</td><td>58151</td><td>-</td><td>3</td><td>GCGGGCGGTGGCGGCGGCGG</td><td>CGG</td><td>|--|||||[-|||||||||||]PAM</td><td>NA</td><td>Intergenic</td><td>NA</td><td>NA</td></tr>
<tr><td>GU305914.1</td><td>56773</td><td>56795</td><td>+</td><td>4</td><td>CGCCGCAGCGGCGGCGGCGG</td><td>CGG</td><td>-|--||-|[||||||||||||]PAM</td><td>NA</td><td>Intergenic</td><td>NA</td><td>NA</td></tr>
<tr><td>GU305914.1</td><td>168820</td><td>168842</td><td>+</td><td>2</td><td>GGTGGTGGCAGCGGCGGCGG</td><td>CGG</td><td>|||||-||[|-||||||||||]PAM</td><td>NA</td><td>Intergenic</td><td>NA</td><td>NA</td></tr>
<tr><td>GU305914.1</td><td>154401</td><td>154423</td><td>+</td><td>4</td><td>CCGGGCGGTGGCGGCGGCGG</td><td>CGG</td><td>---|||||[-|||||||||||]PAM</td><td>NA</td><td>Intergenic</td><td>NA</td><td>NA</td></tr>
<tr><td>GU305914.1</td><td>48156</td><td>48178</td><td>+</td><td>2</td><td>GGCGGCGGCGGTGGCGGCGG</td><td>CGG</td><td>||-|||||[|||-||||||||]PAM</td><td>NA</td><td>Intergenic</td><td>NA</td><td>NA</td></tr>
<tr><td>GU305914.1</td><td>164971</td><td>164993</td><td>-</td><td>3</td><td>GGTGGGACCGGCGGCGGCGG</td><td>CGG</td><td>|||||---[||||||||||||]PAM</td><td>NA</td><td>Intergenic</td><td>NA</td><td>NA</td></tr>
<tr><td>GU305914.1</td><td>81246</td><td>81268</td><td>-</td><td>4</td><td>GATCGAGGAGGCGGCGGCGG</td><td>GGG</td><td>|-|-|-||[-|||||||||||]PAM</td><td>NA</td><td>Intergenic</td><td>NA</td><td>NA</td></tr>
<tr><td>GU305914.1</td><td>161754</td><td>161776</td><td>-</td><td>4</td><td>GGCCGAGGAGGCGGCGGCGG</td><td>AGG</td><td>||--|-||[-|||||||||||]PAM</td><td>NA</td><td>Intergenic</td><td>NA</td><td>NA</td></tr>
<tr><td>GU305914.1</td><td>73792</td><td>73814</td><td>-</td><td>4</td><td>GGCGCCAGCAGCGGCGGCGG</td><td>AGG</td><td>||-|-|-|[|-||||||||||]PAM</td><td>NA</td><td>Intergenic</td><td>NA</td><td>NA</td></tr>
<tr><td>GU305914.1</td><td>149488</td><td>149510</td><td>-</td><td>4</td><td>GGAGCAGGCGACGGCGGCGG</td><td>CGG</td><td>||-|--||[||-|||||||||]PAM</td><td>NA</td><td>Intergenic</td><td>NA</td><td>NA</td></tr>
<tr><td>GU305914.1</td><td>48153</td><td>48175</td><td>+</td><td>2</td><td>TGTGGCGGCGGCGGTGGCGG</td><td>CGG</td><td>-|||||||[||||||-|||||]PAM</td><td>NA</td><td>Intergenic</td><td>NA</td><td>NA</td></tr>
<tr><td>GU305914.1</td><td>62446</td><td>62468</td><td>+</td><td>4</td><td>GGCGGGGGTGGTGGCGGCGG</td><td>GGG</td><td>||-||-||[-||-||||||||]PAM</td><td>NA</td><td>Intergenic</td><td>NA</td><td>NA</td></tr>
<tr><td>GU305914.1</td><td>114658</td><td>114680</td><td>-</td><td>4</td><td>GGCGGTGGTGGTGGCGGCGG</td><td>CGG</td><td>||-||-||[-||-||||||||]PAM</td><td>NA</td><td>Intergenic</td><td>NA</td><td>NA</td></tr>
<tr><td>GU305914.1</td><td>161757</td><td>161779</td><td>-</td><td>4</td><td>GGAGGCCGAGGAGGCGGCGG</td><td>CGG</td><td>||-|||-|[-||-||||||||]PAM</td><td>NA</td><td>Intergenic</td><td>NA</td><td>NA</td></tr>
<tr><td>GU305914.1</td><td>168817</td><td>168839</td><td>+</td><td>4</td><td>GGCGGTGGTGGCAGCGGCGG</td><td>CGG</td><td>||-||-||[-|||-|||||||]PAM</td><td>NA</td><td>Intergenic</td><td>NA</td><td>NA</td></tr>
<tr><td>GU305914.1</td><td>73795</td><td>73817</td><td>-</td><td>4</td><td>AGTGGCGCCAGCAGCGGCGG</td><td>CGG</td><td>-||||||-[|-||-|||||||]PAM</td><td>NA</td><td>Intergenic</td><td>NA</td><td>NA</td></tr>
<tr><td>GU305914.1</td><td>90259</td><td>90281</td><td>-</td><td>4</td><td>GCGGGCGACGGCGGAGGCGG</td><td>CGG</td><td>|--||||-[||||||-|||||]PAM</td><td>NA</td><td>Intergenic</td><td>NA</td><td>NA</td></tr>
<tr><td>GU305914.1</td><td>62672</td><td>62694</td><td>-</td><td>4</td><td>GCTGCCGCCGGCGGGGGCGG</td><td>CGG</td><td>|-||-||-[||||||-|||||]PAM</td><td>NA</td><td>Intergenic</td><td>NA</td><td>NA</td></tr>
<tr><td>GU305914.1</td><td>45255</td><td>45277</td><td>-</td><td>2</td><td>GGTGGCGGTGGCGGCAGCGG</td><td>CGG</td><td>||||||||[-||||||-||||]PAM</td><td>NA</td><td>Intergenic</td><td>NA</td><td>NA</td></tr>
<tr><td>GU305914.1</td><td>163749</td><td>163771</td><td>+</td><td>4</td><td>GTCGGCGGCGCCGGTGGCGG</td><td>CGG</td><td>|--|||||[||-|||-|||||]PAM</td><td>NA</td><td>Intergenic</td><td>NA</td><td>NA</td></tr>
<tr><td>GU305914.1</td><td>163911</td><td>163933</td><td>+</td><td>3</td><td>GGCGGCGGCAGCGGCAGCGG</td><td>AGG</td><td>||-|||||[|-|||||-||||]PAM</td><td>NA</td><td>Intergenic</td><td>NA</td><td>NA</td></tr>
<tr><td>GU305914.1</td><td>110125</td><td>110147</td><td>-</td><td>4</td><td>GGACGCGACGGCGGCCGCGG</td><td>CGG</td><td>||--|||-[|||||||-||||]PAM</td><td>NA</td><td>Intergenic</td><td>NA</td><td>NA</td></tr>
<tr><td>GU305914.1</td><td>62155</td><td>62177</td><td>+</td><td>4</td><td>GGCGACGGCAGCGGCAGCGG</td><td>CGG</td><td>||-|-|||[|-|||||-||||]PAM</td><td>NA</td><td>Intergenic</td><td>NA</td><td>NA</td></tr>
<tr><td>GU305914.1</td><td>30006</td><td>30028</td><td>+</td><td>2</td><td>GGCGGCGGCGGCGGCGGCAG</td><td>TGG</td><td>||-|||||[||||||||||-|]PAM</td><td>NA</td><td>Intergenic</td><td>NA</td><td>NA</td></tr>
<tr><td>GU305914.1</td><td>149476</td><td>149498</td><td>-</td><td>2</td><td>GGCGGCGGCGGCGGCGGCAG</td><td>CGG</td><td>||-|||||[||||||||||-|]PAM</td><td>NA</td><td>Intergenic</td><td>NA</td><td>NA</td></tr>
<tr><td>GU305914.1</td><td>163902</td><td>163924</td><td>+</td><td>3</td><td>AGCGGCGGCGGCGGCGGCAG</td><td>CGG</td><td>-|-|||||[||||||||||-|]PAM</td><td>NA</td><td>Intergenic</td><td>NA</td><td>NA</td></tr>
<tr><td>GU305914.1</td><td>62434</td><td>62456</td><td>+</td><td>4</td><td>GACGGCGGTGGCGGCGGGGG</td><td>TGG</td><td>|--|||||[-||||||||-||]PAM</td><td>NA</td><td>Intergenic</td><td>NA</td><td>NA</td></tr>
<tr><td>GU305914.1</td><td>62449</td><td>62471</td><td>+</td><td>4</td><td>GGGGGTGGTGGCGGCGGGGG</td><td>TGG</td><td>||-||-||[-||||||||-||]PAM</td><td>NA</td><td>Intergenic</td><td>NA</td><td>NA</td></tr>
<tr><td>GU305914.1</td><td>113924</td><td>113946</td><td>-</td><td>4</td><td>GTCGGCGTCGGCGGCGGCAG</td><td>CGG</td><td>|--||||-[||||||||||-|]PAM</td><td>NA</td><td>Intergenic</td><td>NA</td><td>NA</td></tr>
<tr><td>GU305914.1</td><td>152553</td><td>152575</td><td>+</td><td>4</td><td>TGCGGCGGCGGCGGTGCCGG</td><td>CGG</td><td>-|-|||||[||||||-|-|||]PAM</td><td>NA</td><td>Intergenic</td><td>NA</td><td>NA</td></tr>
<tr><td>GU305914.1</td><td>90250</td><td>90272</td><td>-</td><td>4</td><td>GGCGGAGGCGGCGGAGCCGG</td><td>CGG</td><td>||-||-||[||||||-|-|||]PAM</td><td>NA</td><td>Intergenic</td><td>NA</td><td>NA</td></tr>
<tr><td>GU305914.1</td><td>74673</td><td>74695</td><td>+</td><td>4</td><td>GGTGCCGTCGGTGGCGGTGG</td><td>CGG</td><td>||||-||-[|||-|||||-||]PAM</td><td>NA</td><td>Intergenic</td><td>NA</td><td>NA</td></tr>
<tr><td>GU305914.1</td><td>163908</td><td>163930</td><td>+</td><td>3</td><td>GGCGGCGGCGGCAGCGGCAG</td><td>CGG</td><td>||-|||||[||||-|||||-|]PAM</td><td>NA</td><td>Intergenic</td><td>NA</td><td>NA</td></tr>
<tr><td>GU305914.1</td><td>62452</td><td>62474</td><td>+</td><td>3</td><td>GGTGGTGGCGGCGGGGGTGG</td><td>CGG</td><td>|||||-||[||||||-||-||]PAM</td><td>NA</td><td>Intergenic</td><td>NA</td><td>NA</td></tr>
<tr><td>GU305914.1</td><td>62437</td><td>62459</td><td>+</td><td>4</td><td>GGCGGTGGCGGCGGGGGTGG</td><td>TGG</td><td>||-||-||[||||||-||-||]PAM</td><td>NA</td><td>Intergenic</td><td>NA</td><td>NA</td></tr>
<tr><td>GU305914.1</td><td>113918</td><td>113940</td><td>-</td><td>4</td><td>GTCGGCGGCGGCAGCGGCGA</td><td>CGG</td><td>|--|||||[||||-||||||-]PAM</td><td>NA</td><td>Intergenic</td><td>NA</td><td>NA</td></tr>
<tr><td>GU305914.1</td><td>113912</td><td>113934</td><td>-</td><td>4</td><td>GGCGGCAGCGGCGACGGCGT</td><td>CGG</td><td>||-|||-|[|||||-|||||-]PAM</td><td>NA</td><td>Intergenic</td><td>NA</td><td>NA</td></tr>
<tr><td>GU305914.1</td><td>168829</td><td>168851</td><td>+</td><td>4</td><td>AGCGGCGGCGGCGGCGACAG</td><td>CGG</td><td>-|-|||||[||||||||-|-|]PAM</td><td>NA</td><td>Intergenic</td><td>NA</td><td>NA</td></tr>
<tr><td>GU305914.1</td><td>170530</td><td>170552</td><td>-</td><td>4</td><td>GATGGTGGCGGCGGCGGCCA</td><td>CGG</td><td>|-|||-||[||||||||||--]PAM</td><td>NA</td><td>Intergenic</td><td>NA</td><td>NA</td></tr>
</table>

T29715 GTGGCGGCGGCGGCGGCGGAGGG 1000
Oligo fwd CACCGTGGCGGCGGCGGCGGCGGA
Oligo rev AAACTCCGCCGCCGCCGCCGCCAC

<table>
<tr><th>Chromosome</th><th>start</th><th>end</th><th>strand</th><th>MM</th><th>target_seq</th><th>PAM</th><th>alignment</th><th>distance</th><th>position</th><th>gene name</th><th>gene id</th></tr>
<tr><td>GU305914.1</td><td>58122</td><td>58144</td><td>-</td><td>0</td><td>GTGGCGGCGGCGGCGGCGGA</td><td>GGG</td><td>||||||||[||||||||||||]PAM</td><td>NA</td><td>Intergenic</td><td>NA</td><td>NA</td></tr>
<tr><td>GU305914.1</td><td>67743</td><td>67765</td><td>+</td><td>3</td><td>GGAGCGGCGGCGGCGGCGGC</td><td>GGG</td><td>|--|||||[|||||||||||-]PAM</td><td>NA</td><td>Intergenic</td><td>NA</td><td>NA</td></tr>
<tr><td>GU305914.1</td><td>48160</td><td>48182</td><td>+</td><td>3</td><td>GCGGCGGTGGCGGCGGCGGT</td><td>GGG</td><td>|-|||||-[|||||||||||-]PAM</td><td>NA</td><td>Intergenic</td><td>NA</td><td>NA</td></tr>
<tr><td>GU305914.1</td><td>75852</td><td>75874</td><td>+</td><td>4</td><td>GGGGAGGCGGAGGCGGCGGT</td><td>CGG</td><td>|-||-|||[||-||||||||-]PAM</td><td>NA</td><td>Intergenic</td><td>NA</td><td>NA</td></tr>
<tr><td>GU305914.1</td><td>86780</td><td>86802</td><td>-</td><td>4</td><td>GAGGGGGCGGGGGCGGCGGC</td><td>AGG</td><td>|-||-|||[||-||||||||-]PAM</td><td>NA</td><td>Intergenic</td><td>NA</td><td>NA</td></tr>
<tr><td>GU305914.1</td><td>149475</td><td>149497</td><td>-</td><td>3</td><td>GCGGCGGCGGCGGCGGCAGC</td><td>GGG</td><td>|-||||||[|||||||||-|-]PAM</td><td>NA</td><td>Intergenic</td><td>NA</td><td>NA</td></tr>
</table>

T29716 TGGCGGCGGCGGCGGCGGAGGGG 1000

| | | | | | | | | | | | |
|---|---|---|---|---|---|---|---|---|---|---|---|
| Oligo fwd | CACCTGGCGGCGGCGGCGGCGGAG | | | | | | | | | | |
| Oligo rev | AAACCTCCGCCGCCGCCGCCGCCA | | | | | | | | | | |
| Chromosome | start | end | strand | MM | target_seq | PAM | alignment | distance | position | gene name | gene id |
| GU305914.1 | 58121 | 58143 | - | 0 | TGGCGGCGGCGGCGGCGGAG | GGG | \|\|\|\|\|\|\|\|[\|\|\|\|\|\|\|\|\|\|\|\|]PAM | NA | Intergenic | NA | NA |
| GU305914.1 | 35655 | 35677 | + | 4 | CCACGGCGGCGGCGGCGGCG | GGG | ---\|\|\|\|\|[\|\|\|\|\|\|\|\|\|\|-\|]PAM | NA | Intergenic | NA | NA |
| GU305914.1 | 159954 | 159976 | - | 4 | TATCGGCGGCGACGGCGGCG | AGG | \|--\|\|\|\|\|[\|\|\|-\|\|\|\|\|\|-\|]PAM | NA | Intergenic | NA | NA |
| T29717 | GTCGTGAACGACGACCATCAAGG | 1000 | | | | | | | | | |
| Oligo fwd | CACCGTCGTGAACGACGACCATCA | | | | | | | | | | |
| Oligo rev | AAACTGATGGTCGTCGTTCACGAC | | | | | | | | | | |
| Chromosome | start | end | strand | MM | target_seq | PAM | alignment | distance | position | gene name | gene id |
| GU305914.1 | 58096 | 58118 | - | 0 | GTCGTGAACGACGACCATCA | AGG | \|\|\|\|\|\|\|\|[\|\|\|\|\|\|\|\|\|\|\|\|]PAM | NA | Intergenic | NA | NA |
| T29718 | GTGAACGACGACCATCAAGGTGG | 1000 | | | | | | | | | |
| Oligo fwd | CACCGTGAACGACGACCATCAAGG | | | | | | | | | | |
| Oligo rev | AAACCCTTGATGGTCGTCGTTCAC | | | | | | | | | | |
| Chromosome | start | end | strand | MM | target_seq | PAM | alignment | distance | position | gene name | gene id |
| GU305914.1 | 58093 | 58115 | - | 0 | GTGAACGACGACCATCAAGG | TGG | \|\|\|\|\|\|\|\|[\|\|\|\|\|\|\|\|\|\|\|\|]PAM | NA | Intergenic | NA | NA |
| T29719 | AACGACGACCATCAAGGTGGCGG | 1000 | | | | | | | | | |
| Oligo fwd | CACCAACGACGACCATCAAGGTGG | | | | | | | | | | |
| Oligo rev | AAACCCACCTTGATGGTCGTCGTT | | | | | | | | | | |
| Chromosome | start | end | strand | MM | target_seq | PAM | alignment | distance | position | gene name | gene id |
| GU305914.1 | 58090 | 58112 | - | 0 | AACGACGACCATCAAGGTGG | CGG | \|\|\|\|\|\|\|\|[\|\|\|\|\|\|\|\|\|\|\|\|]PAM | NA | Intergenic | NA | NA |
| T29720 | GACGACCATCAAGGTGGCGGCGG | 1000 | | | | | | | | | |
| Oligo fwd | CACCGACGACCATCAAGGTGGCGG | | | | | | | | | | |
| Oligo rev | AAACCCGCCACCTTGATGGTCGTC | | | | | | | | | | |
| Chromosome | start | end | strand | MM | target_seq | PAM | alignment | distance | position | gene name | gene id |
| GU305914.1 | 58087 | 58109 | - | 0 | GACGACCATCAAGGTGGCGG | CGG | \|\|\|\|\|\|\|\|[\|\|\|\|\|\|\|\|\|\|\|\|]PAM | NA | Intergenic | NA | NA |
| T29721 | CATCAAGGTGGCGGCGGAAGCGG | 1000 | | | | | | | | | |
| Oligo fwd | CACCCATCAAGGTGGCGGCGGAAG | | | | | | | | | | |
| Oligo rev | AAACCTTCCGCCGCCACCTTGATG | | | | | | | | | | |
| Chromosome | start | end | strand | MM | target_seq | PAM | alignment | distance | position | gene name | gene id |
| GU305914.1 | 58081 | 58103 | - | 0 | CATCAAGGTGGCGGCGGAAG | CGG | \|\|\|\|\|\|\|\|[\|\|\|\|\|\|\|\|\|\|\|\|]PAM | NA | Intergenic | NA | NA |
| T29722 | GCACAAGATCACCAGTTTCATGG | 1000 | | | | | | | | | |
| Oligo fwd | CACCGCACAAGATCACCAGTTTCA | | | | | | | | | | |
| Oligo rev | AAACTGAAACTGGTGATCTTGTGC | | | | | | | | | | |
| Chromosome | start | end | strand | MM | target_seq | PAM | alignment | distance | position | gene name | gene id |
| GU305914.1 | 58037 | 58059 | - | 0 | GCACAAGATCACCAGTTTCA | TGG | \|\|\|\|\|\|\|\|[\|\|\|\|\|\|\|\|\|\|\|\|]PAM | NA | Intergenic | NA | NA |
| T29723 | CACCAGTTTCATGGTCTCCAAGG | 1000 | | | | | | | | | |
| Oligo fwd | CACCCACCAGTTTCATGGTCTCCA | | | | | | | | | | |
| Oligo rev | AAACTGGAGACCATGAAACTGGTG | | | | | | | | | | |
| Chromosome | start | end | strand | MM | target_seq | PAM | alignment | distance | position | gene name | gene id |
| GU305914.1 | 58028 | 58050 | - | 0 | CACCAGTTTCATGGTCTCCA | AGG | \|\|\|\|\|\|\|\|[\|\|\|\|\|\|\|\|\|\|\|\|]PAM | NA | Intergenic | NA | NA |
| T29724 | ACCAGTTTCATGGTCTCCAAGGG | 1000 | | | | | | | | | |
| Oligo fwd | CACCACCAGTTTCATGGTCTCCAA | | | | | | | | | | |
| Oligo rev | AAACTTGGAGACCATGAAACTGGT | | | | | | | | | | |
| Chromosome | start | end | strand | MM | target_seq | PAM | alignment | distance | position | gene name | gene id |
| GU305914.1 | 58027 | 58049 | - | 0 | ACCAGTTTCATGGTCTCCAA | GGG | \|\|\|\|\|\|\|\|[\|\|\|\|\|\|\|\|\|\|\|\|]PAM | NA | Intergenic | NA | NA |
| T29725 | CCAGTTTCATGGTCTCCAAGGGG | 1000 | | | | | | | | | |
| Oligo fwd | CACCCCAGTTTCATGGTCTCCAAG | | | | | | | | | | |
| Oligo rev | AAACCTTGGAGACCATGAAACTGG | | | | | | | | | | |
| Chromosome | start | end | strand | MM | target_seq | PAM | alignment | distance | position | gene name | gene id |
| GU305914.1 | 58026 | 58048 | - | 0 | CCAGTTTCATGGTCTCCAAG | GGG | \|\|\|\|\|\|\|\|[\|\|\|\|\|\|\|\|\|\|\|\|]PAM | NA | Intergenic | NA | NA |
| T29726 | CAGTTTCATGGTCTCCAAGGGGG | 1000 | | | | | | | | | |
| Oligo fwd | CACCCAGTTTCATGGTCTCCAAGG | | | | | | | | | | |
| Oligo rev | AAACCCTTGGAGACCATGAAACTG | | | | | | | | | | |
| Chromosome | start | end | strand | MM | target_seq | PAM | alignment | distance | position | gene name | gene id |
| GU305914.1 | 58025 | 58047 | - | 0 | CAGTTTCATGGTCTCCAAGG | GGG | \|\|\|\|\|\|\|\|[\|\|\|\|\|\|\|\|\|\|\|\|]PAM | NA | Intergenic | NA | NA |
| T29727 | CATGGTCTCCAAGGGGGCTGTGG | 1000 | | | | | | | | | |
| Oligo fwd | CACCCATGGTCTCCAAGGGGGCTG | | | | | | | | | | |
| Oligo rev | AAACCAGCCCCCTTGGAGACCATG | | | | | | | | | | |
| Chromosome | start | end | strand | MM | target_seq | PAM | alignment | distance | position | gene name | gene id |
| GU305914.1 | 58019 | 58041 | - | 0 | CATGGTCTCCAAGGGGGCTG | TGG | \|\|\|\|\|\|\|\|[\|\|\|\|\|\|\|\|\|\|\|\|]PAM | NA | Intergenic | NA | NA |
| T29728 | ATGGTCTCCAAGGGGGCTGTGGG | 1000 | | | | | | | | | |
| Oligo fwd | CACCATGGTCTCCAAGGGGGCTGT | | | | | | | | | | |
| Oligo rev | AAACACAGCCCCCTTGGAGACCAT | | | | | | | | | | |
| Chromosome | start | end | strand | MM | target_seq | PAM | alignment | distance | position | gene name | gene id |
| GU305914.1 | 58018 | 58040 | - | 0 | ATGGTCTCCAAGGGGGCTGT | GGG | \|\|\|\|\|\|\|\|[\|\|\|\|\|\|\|\|\|\|\|\|]PAM | NA | Intergenic | NA | NA |
| T29729 | GTCTCCAAGGGGGCTGTGGGTGG | 1000 | | | | | | | | | |
| Oligo fwd | CACCGTCTCCAAGGGGGCTGTGGG | | | | | | | | | | |
| Oligo rev | AAACCCCACAGCCCCCTTGGAGAC | | | | | | | | | | |
| Chromosome | start | end | strand | MM | target_seq | PAM | alignment | distance | position | gene name | gene id |
| GU305914.1 | 58015 | 58037 | - | 0 | GTCTCCAAGGGGGCTGTGGG | TGG | \|\|\|\|\|\|\|\|[\|\|\|\|\|\|\|\|\|\|\|\|]PAM | NA | Intergenic | NA | NA |
| T29730 | TCCAAGGGGGCTGTGGGTGGAGG | 1000 | | | | | | | | | |
| Oligo fwd | CACCTCCAAGGGGGCTGTGGGTGG | | | | | | | | | | |
| Oligo rev | AAACCCACCCACAGCCCCCTTGGA | | | | | | | | | | |
| Chromosome | start | end | strand | MM | target_seq | PAM | alignment | distance | position | gene name | gene id |

<table>
<tr><td>GU305914.1</td><td>58012</td><td>58034</td><td>-</td><td>0</td><td>TCCAAGGGGGCTGTGGGTGG</td><td>AGG</td><td>||||||||[||||||||||||]PAM</td><td>NA</td><td>Intergenic</td><td>NA</td><td>NA</td></tr>
<tr><td></td><td></td><td></td><td></td><td></td><td></td><td></td><td></td><td></td><td></td><td></td><td></td></tr>
<tr><td>T29731</td><td>CCAAGGGGGCTGTGGGTGGAGGG</td><td>1000</td><td></td><td></td><td></td><td></td><td></td><td></td><td></td><td></td><td></td></tr>
<tr><td>Oligo fwd</td><td>CACCCCAAGGGGGCTGTGGGTGGA</td><td></td><td></td><td></td><td></td><td></td><td></td><td></td><td></td><td></td><td></td></tr>
<tr><td>Oligo rev</td><td>AAACTCCACCCACAGCCCCCTTGG</td><td></td><td></td><td></td><td></td><td></td><td></td><td></td><td></td><td></td><td></td></tr>
<tr><td>Chromosome</td><td>start</td><td>end</td><td>strand</td><td>MM</td><td>target_seq</td><td>PAM</td><td>alignment</td><td>distance</td><td>position</td><td>gene name</td><td>gene id</td></tr>
<tr><td>GU305914.1</td><td>58011</td><td>58033</td><td>-</td><td>0</td><td>CCAAGGGGGCTGTGGGTGGA</td><td>GGG</td><td>||||||||[||||||||||||]PAM</td><td>NA</td><td>Intergenic</td><td>NA</td><td>NA</td></tr>
<tr><td></td><td></td><td></td><td></td><td></td><td></td><td></td><td></td><td></td><td></td><td></td><td></td></tr>
<tr><td>T29732</td><td>CAAGGGGGCTGTGGGTGGAGGGG</td><td>1000</td><td></td><td></td><td></td><td></td><td></td><td></td><td></td><td></td><td></td></tr>
<tr><td>Oligo fwd</td><td>CACCCAAGGGGGCTGTGGGTGGAG</td><td></td><td></td><td></td><td></td><td></td><td></td><td></td><td></td><td></td><td></td></tr>
<tr><td>Oligo rev</td><td>AAACCTCCACCCACAGCCCCCTTG</td><td></td><td></td><td></td><td></td><td></td><td></td><td></td><td></td><td></td><td></td></tr>
<tr><td>Chromosome</td><td>start</td><td>end</td><td>strand</td><td>MM</td><td>target_seq</td><td>PAM</td><td>alignment</td><td>distance</td><td>position</td><td>gene name</td><td>gene id</td></tr>
<tr><td>GU305914.1</td><td>58010</td><td>58032</td><td>-</td><td>0</td><td>CAAGGGGGCTGTGGGTGGAG</td><td>GGG</td><td>||||||||[||||||||||||]PAM</td><td>NA</td><td>Intergenic</td><td>NA</td><td>NA</td></tr>
<tr><td></td><td></td><td></td><td></td><td></td><td></td><td></td><td></td><td></td><td></td><td></td><td></td></tr>
<tr><td>T29733</td><td>GGGGCTGTGGGTGGAGGGGCCGG</td><td>1000</td><td></td><td></td><td></td><td></td><td></td><td></td><td></td><td></td><td></td></tr>
<tr><td>Oligo fwd</td><td>CACCGGGGCTGTGGGTGGAGGGGC</td><td></td><td></td><td></td><td></td><td></td><td></td><td></td><td></td><td></td><td></td></tr>
<tr><td>Oligo rev</td><td>AAACGCCCCTCCACCCACAGCCCC</td><td></td><td></td><td></td><td></td><td></td><td></td><td></td><td></td><td></td><td></td></tr>
<tr><td>Chromosome</td><td>start</td><td>end</td><td>strand</td><td>MM</td><td>target_seq</td><td>PAM</td><td>alignment</td><td>distance</td><td>position</td><td>gene name</td><td>gene id</td></tr>
<tr><td>GU305914.1</td><td>58006</td><td>58028</td><td>-</td><td>0</td><td>GGGGCTGTGGGTGGAGGGGC</td><td>CGG</td><td>||||||||[||||||||||||]PAM</td><td>NA</td><td>Intergenic</td><td>NA</td><td>NA</td></tr>
<tr><td></td><td></td><td></td><td></td><td></td><td></td><td></td><td></td><td></td><td></td><td></td><td></td></tr>
<tr><td>T29734</td><td>GGGCTGTGGGTGGAGGGGCCGGG</td><td>1000</td><td></td><td></td><td></td><td></td><td></td><td></td><td></td><td></td><td></td></tr>
<tr><td>Oligo fwd</td><td>CACCGGGCTGTGGGTGGAGGGGCC</td><td></td><td></td><td></td><td></td><td></td><td></td><td></td><td></td><td></td><td></td></tr>
<tr><td>Oligo rev</td><td>AAACGGCCCCTCCACCCACAGCCC</td><td></td><td></td><td></td><td></td><td></td><td></td><td></td><td></td><td></td><td></td></tr>
<tr><td>Chromosome</td><td>start</td><td>end</td><td>strand</td><td>MM</td><td>target_seq</td><td>PAM</td><td>alignment</td><td>distance</td><td>position</td><td>gene name</td><td>gene id</td></tr>
<tr><td>GU305914.1</td><td>58005</td><td>58027</td><td>-</td><td>0</td><td>GGGCTGTGGGTGGAGGGGCC</td><td>GGG</td><td>||||||||[||||||||||||]PAM</td><td>NA</td><td>Intergenic</td><td>NA</td><td>NA</td></tr>
<tr><td></td><td></td><td></td><td></td><td></td><td></td><td></td><td></td><td></td><td></td><td></td><td></td></tr>
<tr><td>T29735</td><td>GGCTGTGGGTGGAGGGGCCGGGG</td><td>1000</td><td></td><td></td><td></td><td></td><td></td><td></td><td></td><td></td><td></td></tr>
<tr><td>Oligo fwd</td><td>CACCGGCTGTGGGTGGAGGGGCCG</td><td></td><td></td><td></td><td></td><td></td><td></td><td></td><td></td><td></td><td></td></tr>
<tr><td>Oligo rev</td><td>AAACCGGCCCCTCCACCCACAGCC</td><td></td><td></td><td></td><td></td><td></td><td></td><td></td><td></td><td></td><td></td></tr>
<tr><td>Chromosome</td><td>start</td><td>end</td><td>strand</td><td>MM</td><td>target_seq</td><td>PAM</td><td>alignment</td><td>distance</td><td>position</td><td>gene name</td><td>gene id</td></tr>
<tr><td>GU305914.1</td><td>58004</td><td>58026</td><td>-</td><td>0</td><td>GGCTGTGGGTGGAGGGGCCG</td><td>GGG</td><td>||||||||[||||||||||||]PAM</td><td>NA</td><td>Intergenic</td><td>NA</td><td>NA</td></tr>
<tr><td></td><td></td><td></td><td></td><td></td><td></td><td></td><td></td><td></td><td></td><td></td><td></td></tr>
<tr><td>T29736</td><td>GCTGTGGGTGGAGGGGCCGGGGG</td><td>1000</td><td></td><td></td><td></td><td></td><td></td><td></td><td></td><td></td><td></td></tr>
<tr><td>Oligo fwd</td><td>CACCGCTGTGGGTGGAGGGGCCGG</td><td></td><td></td><td></td><td></td><td></td><td></td><td></td><td></td><td></td><td></td></tr>
<tr><td>Oligo rev</td><td>AAACCCGGCCCCTCCACCCACAGC</td><td></td><td></td><td></td><td></td><td></td><td></td><td></td><td></td><td></td><td></td></tr>
<tr><td>Chromosome</td><td>start</td><td>end</td><td>strand</td><td>MM</td><td>target_seq</td><td>PAM</td><td>alignment</td><td>distance</td><td>position</td><td>gene name</td><td>gene id</td></tr>
<tr><td>GU305914.1</td><td>58003</td><td>58025</td><td>-</td><td>0</td><td>GCTGTGGGTGGAGGGGCCGG</td><td>GGG</td><td>||||||||[||||||||||||]PAM</td><td>NA</td><td>Intergenic</td><td>NA</td><td>NA</td></tr>
<tr><td></td><td></td><td></td><td></td><td></td><td></td><td></td><td></td><td></td><td></td><td></td><td></td></tr>
<tr><td>T29737</td><td>GTGGGTGGAGGGGCCGGGGGAGG</td><td>1000</td><td></td><td></td><td></td><td></td><td></td><td></td><td></td><td></td><td></td></tr>
<tr><td>Oligo fwd</td><td>CACCGTGGGTGGAGGGGCCGGGGG</td><td></td><td></td><td></td><td></td><td></td><td></td><td></td><td></td><td></td><td></td></tr>
<tr><td>Oligo rev</td><td>AAACCCCCCGGCCCCTCCACCCAC</td><td></td><td></td><td></td><td></td><td></td><td></td><td></td><td></td><td></td><td></td></tr>
<tr><td>Chromosome</td><td>start</td><td>end</td><td>strand</td><td>MM</td><td>target_seq</td><td>PAM</td><td>alignment</td><td>distance</td><td>position</td><td>gene name</td><td>gene id</td></tr>
<tr><td>GU305914.1</td><td>58000</td><td>58022</td><td>-</td><td>0</td><td>GTGGGTGGAGGGGCCGGGGG</td><td>AGG</td><td>||||||||[||||||||||||]PAM</td><td>NA</td><td>Intergenic</td><td>NA</td><td>NA</td></tr>
<tr><td>GU305914.1</td><td>143022</td><td>143044</td><td>+</td><td>4</td><td>GCTGGTGGTGGCGCCGGGGG</td><td>CGG</td><td>|--|||||[-||-||||||||]PAM</td><td>NA</td><td>Intergenic</td><td>NA</td><td>NA</td></tr>
<tr><td></td><td></td><td></td><td></td><td></td><td></td><td></td><td></td><td></td><td></td><td></td><td></td></tr>
<tr><td>T29738</td><td>GGTGGAGGGGCCGGGGGAGGCGG</td><td>1000</td><td></td><td></td><td></td><td></td><td></td><td></td><td></td><td></td><td></td></tr>
<tr><td>Oligo fwd</td><td>CACCGGTGGAGGGGCCGGGGGAGG</td><td></td><td></td><td></td><td></td><td></td><td></td><td></td><td></td><td></td><td></td></tr>
<tr><td>Oligo rev</td><td>AAACCCTCCCCCGGCCCCTCCACC</td><td></td><td></td><td></td><td></td><td></td><td></td><td></td><td></td><td></td><td></td></tr>
<tr><td>Chromosome</td><td>start</td><td>end</td><td>strand</td><td>MM</td><td>target_seq</td><td>PAM</td><td>alignment</td><td>distance</td><td>position</td><td>gene name</td><td>gene id</td></tr>
<tr><td>GU305914.1</td><td>57997</td><td>58019</td><td>-</td><td>0</td><td>GGTGGAGGGGCCGGGGGAGG</td><td>CGG</td><td>||||||||[||||||||||||]PAM</td><td>NA</td><td>Intergenic</td><td>NA</td><td>NA</td></tr>
<tr><td>GU305914.1</td><td>145960</td><td>145982</td><td>+</td><td>4</td><td>GGAGGAGAGGTCGGTGGAGG</td><td>AGG</td><td>||-||||-[||-|||-|||||]PAM</td><td>NA</td><td>Intergenic</td><td>NA</td><td>NA</td></tr>
<tr><td>GU305914.1</td><td>86784</td><td>86806</td><td>-</td><td>4</td><td>GGAAGAGGGGGCGGGGGCGG</td><td>CGG</td><td>||--||||[||-||||||-||]PAM</td><td>NA</td><td>Intergenic</td><td>NA</td><td>NA</td></tr>
<tr><td></td><td></td><td></td><td></td><td></td><td></td><td></td><td></td><td></td><td></td><td></td><td></td></tr>
<tr><td>T29739</td><td>GGGGGAGGCGGCTCTGACCGCGG</td><td>1000</td><td></td><td></td><td></td><td></td><td></td><td></td><td></td><td></td><td></td></tr>
<tr><td>Oligo fwd</td><td>CACCGGGGGAGGCGGCTCTGACCG</td><td></td><td></td><td></td><td></td><td></td><td></td><td></td><td></td><td></td><td></td></tr>
<tr><td>Oligo rev</td><td>AAACCGGTCAGAGCCGCCTCCCCC</td><td></td><td></td><td></td><td></td><td></td><td></td><td></td><td></td><td></td><td></td></tr>
<tr><td>Chromosome</td><td>start</td><td>end</td><td>strand</td><td>MM</td><td>target_seq</td><td>PAM</td><td>alignment</td><td>distance</td><td>position</td><td>gene name</td><td>gene id</td></tr>
<tr><td>GU305914.1</td><td>57985</td><td>58007</td><td>-</td><td>0</td><td>GGGGGAGGCGGCTCTGACCG</td><td>CGG</td><td>||||||||[||||||||||||]PAM</td><td>NA</td><td>Intergenic</td><td>NA</td><td>NA</td></tr>
<tr><td></td><td></td><td></td><td></td><td></td><td></td><td></td><td></td><td></td><td></td><td></td><td></td></tr>
<tr><td>T29740</td><td>GGAGGCGGCTCTGACCGCGGTGG</td><td>1000</td><td></td><td></td><td></td><td></td><td></td><td></td><td></td><td></td><td></td></tr>
<tr><td>Oligo fwd</td><td>CACCGGAGGCGGCTCTGACCGCGG</td><td></td><td></td><td></td><td></td><td></td><td></td><td></td><td></td><td></td><td></td></tr>
<tr><td>Oligo rev</td><td>AAACCCGCGGTCAGAGCCGCCTCC</td><td></td><td></td><td></td><td></td><td></td><td></td><td></td><td></td><td></td><td></td></tr>
<tr><td>Chromosome</td><td>start</td><td>end</td><td>strand</td><td>MM</td><td>target_seq</td><td>PAM</td><td>alignment</td><td>distance</td><td>position</td><td>gene name</td><td>gene id</td></tr>
<tr><td>GU305914.1</td><td>57982</td><td>58004</td><td>-</td><td>0</td><td>GGAGGCGGCTCTGACCGCGG</td><td>TGG</td><td>||||||||[||||||||||||]PAM</td><td>NA</td><td>Intergenic</td><td>NA</td><td>NA</td></tr>
<tr><td></td><td></td><td></td><td></td><td></td><td></td><td></td><td></td><td></td><td></td><td></td><td></td></tr>
<tr><td>T29741</td><td>TGGCTACTTCAACGACACCAAGG</td><td>1000</td><td></td><td></td><td></td><td></td><td></td><td></td><td></td><td></td><td></td></tr>
<tr><td>Oligo fwd</td><td>CACCTGGCTACTTCAACGACACCA</td><td></td><td></td><td></td><td></td><td></td><td></td><td></td><td></td><td></td><td></td></tr>
<tr><td>Oligo rev</td><td>AAACTGGTGTCGTTGAAGTAGCCA</td><td></td><td></td><td></td><td></td><td></td><td></td><td></td><td></td><td></td><td></td></tr>
<tr><td>Chromosome</td><td>start</td><td>end</td><td>strand</td><td>MM</td><td>target_seq</td><td>PAM</td><td>alignment</td><td>distance</td><td>position</td><td>gene name</td><td>gene id</td></tr>
<tr><td>GU305914.1</td><td>57962</td><td>57984</td><td>-</td><td>0</td><td>TGGCTACTTCAACGACACCA</td><td>AGG</td><td>||||||||[||||||||||||]PAM</td><td>NA</td><td>Intergenic</td><td>NA</td><td>NA</td></tr>
<tr><td></td><td></td><td></td><td></td><td></td><td></td><td></td><td></td><td></td><td></td><td></td><td></td></tr>
<tr><td>T29742</td><td>AGAGAGCGACAGTGAAGACTCGG</td><td>1000</td><td></td><td></td><td></td><td></td><td></td><td></td><td></td><td></td><td></td></tr>
<tr><td>Oligo fwd</td><td>CACCAGAGAGCGACAGTGAAGACT</td><td></td><td></td><td></td><td></td><td></td><td></td><td></td><td></td><td></td><td></td></tr>
<tr><td>Oligo rev</td><td>AAACAGTCTTCACTGTCGCTCTCT</td><td></td><td></td><td></td><td></td><td></td><td></td><td></td><td></td><td></td><td></td></tr>
<tr><td>Chromosome</td><td>start</td><td>end</td><td>strand</td><td>MM</td><td>target_seq</td><td>PAM</td><td>alignment</td><td>distance</td><td>position</td><td>gene name</td><td>gene id</td></tr>
<tr><td>GU305914.1</td><td>57938</td><td>57960</td><td>-</td><td>0</td><td>AGAGAGCGACAGTGAAGACT</td><td>CGG</td><td>||||||||[||||||||||||]PAM</td><td>NA</td><td>Intergenic</td><td>NA</td><td>NA</td></tr>
<tr><td></td><td></td><td></td><td></td><td></td><td></td><td></td><td></td><td></td><td></td><td></td><td></td></tr>
<tr><td>T29743</td><td>CACTAAAAAACAAAAGTGCGCGG</td><td>1000</td><td></td><td></td><td></td><td></td><td></td><td></td><td></td><td></td><td></td></tr>
<tr><td>Oligo fwd</td><td>CACCCACTAAAAAACAAAAGTGCG</td><td></td><td></td><td></td><td></td><td></td><td></td><td></td><td></td><td></td><td></td></tr>
<tr><td>Oligo rev</td><td>AAACCGCACTTTTGTTTTTTAGTG</td><td></td><td></td><td></td><td></td><td></td><td></td><td></td><td></td><td></td><td></td></tr>
<tr><td>Chromosome</td><td>start</td><td>end</td><td>strand</td><td>MM</td><td>target_seq</td><td>PAM</td><td>alignment</td><td>distance</td><td>position</td><td>gene name</td><td>gene id</td></tr>
<tr><td>GU305914.1</td><td>57893</td><td>57915</td><td>-</td><td>0</td><td>CACTAAAAAACAAAAGTGCG</td><td>CGG</td><td>||||||||[||||||||||||]PAM</td><td>NA</td><td>Intergenic</td><td>NA</td><td>NA</td></tr>
<tr><td></td><td></td><td></td><td></td><td></td><td></td><td></td><td></td><td></td><td></td><td></td><td></td></tr>
<tr><td>T29744</td><td>GCGGCCTGAATGCCTTATTCTGG</td><td>1000</td><td></td><td></td><td></td><td></td><td></td><td></td><td></td><td></td><td></td></tr>
<tr><td>Oligo fwd</td><td>CACCGCGGCCTGAATGCCTTATTC</td><td></td><td></td><td></td><td></td><td></td><td></td><td></td><td></td><td></td><td></td></tr>
<tr><td>Oligo rev</td><td>AAACGAATAAGGCATTCAGGCCGC</td><td></td><td></td><td></td><td></td><td></td><td></td><td></td><td></td><td></td><td></td></tr>
<tr><td>Chromosome</td><td>start</td><td>end</td><td>strand</td><td>MM</td><td>target_seq</td><td>PAM</td><td>alignment</td><td>distance</td><td>position</td><td>gene name</td><td>gene id</td></tr>
<tr><td>GU305914.1</td><td>57874</td><td>57896</td><td>-</td><td>0</td><td>GCGGCCTGAATGCCTTATTC</td><td>TGG</td><td>||||||||[||||||||||||]PAM</td><td>NA</td><td>Intergenic</td><td>NA</td><td>NA</td></tr>
</table>

<table>
<tr><td>T29745</td><td>CTGAATGCCTTATTCTGGCCCGG</td><td>1000</td><td></td><td></td><td></td><td></td><td></td><td></td><td></td><td></td><td></td></tr>
<tr><td>Oligo fwd</td><td>CACCCTGAATGCCTTATTCTGGCC</td><td></td><td></td><td></td><td></td><td></td><td></td><td></td><td></td><td></td><td></td></tr>
<tr><td>Oligo rev</td><td>AAACGGCCAGAATAAGGCATTCAG</td><td></td><td></td><td></td><td></td><td></td><td></td><td></td><td></td><td></td><td></td></tr>
<tr><td>Chromosome</td><td>start</td><td>end</td><td>strand</td><td>MM</td><td>target_seq</td><td>PAM</td><td>alignment</td><td>distance</td><td>position</td><td>gene name</td><td>gene id</td></tr>
<tr><td>GU305914.1</td><td>57869</td><td>57891</td><td>-</td><td>0</td><td>CTGAATGCCTTATTCTGGCC</td><td>CGG</td><td>||||||||[||||||||||||]PAM</td><td>NA</td><td>Intergenic</td><td>NA</td><td>NA</td></tr>
<tr><td>T29746</td><td>TGAATGCCTTATTCTGGCCCGGG</td><td>1000</td><td></td><td></td><td></td><td></td><td></td><td></td><td></td><td></td><td></td></tr>
<tr><td>Oligo fwd</td><td>CACCTGAATGCCTTATTCTGGCCC</td><td></td><td></td><td></td><td></td><td></td><td></td><td></td><td></td><td></td><td></td></tr>
<tr><td>Oligo rev</td><td>AAACGGGCCAGAATAAGGCATTCA</td><td></td><td></td><td></td><td></td><td></td><td></td><td></td><td></td><td></td><td></td></tr>
<tr><td>Chromosome</td><td>start</td><td>end</td><td>strand</td><td>MM</td><td>target_seq</td><td>PAM</td><td>alignment</td><td>distance</td><td>position</td><td>gene name</td><td>gene id</td></tr>
<tr><td>GU305914.1</td><td>57868</td><td>57890</td><td>-</td><td>0</td><td>TGAATGCCTTATTCTGGCCC</td><td>GGG</td><td>||||||||[||||||||||||]PAM</td><td>NA</td><td>Intergenic</td><td>NA</td><td>NA</td></tr>
<tr><td>T29747</td><td>GAATGCCTTATTCTGGCCCGGGG</td><td>1000</td><td></td><td></td><td></td><td></td><td></td><td></td><td></td><td></td><td></td></tr>
<tr><td>Oligo fwd</td><td>CACCGAATGCCTTATTCTGGCCCG</td><td></td><td></td><td></td><td></td><td></td><td></td><td></td><td></td><td></td><td></td></tr>
<tr><td>Oligo rev</td><td>AAACCGGGCCAGAATAAGGCATTC</td><td></td><td></td><td></td><td></td><td></td><td></td><td></td><td></td><td></td><td></td></tr>
<tr><td>Chromosome</td><td>start</td><td>end</td><td>strand</td><td>MM</td><td>target_seq</td><td>PAM</td><td>alignment</td><td>distance</td><td>position</td><td>gene name</td><td>gene id</td></tr>
<tr><td>GU305914.1</td><td>57867</td><td>57889</td><td>-</td><td>0</td><td>GAATGCCTTATTCTGGCCCG</td><td>GGG</td><td>||||||||[||||||||||||]PAM</td><td>NA</td><td>Intergenic</td><td>NA</td><td>NA</td></tr>
<tr><td>T29748</td><td>CGACTGACCACCGAGAGCGACGG</td><td>1000</td><td></td><td></td><td></td><td></td><td></td><td></td><td></td><td></td><td></td></tr>
<tr><td>Oligo fwd</td><td>CACCCGACTGACCACCGAGAGCGA</td><td></td><td></td><td></td><td></td><td></td><td></td><td></td><td></td><td></td><td></td></tr>
<tr><td>Oligo rev</td><td>AAACTCGCTCTCGGTGGTCAGTCG</td><td></td><td></td><td></td><td></td><td></td><td></td><td></td><td></td><td></td><td></td></tr>
<tr><td>Chromosome</td><td>start</td><td>end</td><td>strand</td><td>MM</td><td>target_seq</td><td>PAM</td><td>alignment</td><td>distance</td><td>position</td><td>gene name</td><td>gene id</td></tr>
<tr><td>GU305914.1</td><td>57838</td><td>57860</td><td>-</td><td>0</td><td>CGACTGACCACCGAGAGCGA</td><td>CGG</td><td>||||||||[||||||||||||]PAM</td><td>NA</td><td>Intergenic</td><td>NA</td><td>NA</td></tr>
<tr><td>T29749</td><td>CTGACCACCGAGAGCGACGGCGG</td><td>1000</td><td></td><td></td><td></td><td></td><td></td><td></td><td></td><td></td><td></td></tr>
<tr><td>Oligo fwd</td><td>CACCCTGACCACCGAGAGCGACGG</td><td></td><td></td><td></td><td></td><td></td><td></td><td></td><td></td><td></td><td></td></tr>
<tr><td>Oligo rev</td><td>AAACCCGTCGCTCTCGGTGGTCAG</td><td></td><td></td><td></td><td></td><td></td><td></td><td></td><td></td><td></td><td></td></tr>
<tr><td>Chromosome</td><td>start</td><td>end</td><td>strand</td><td>MM</td><td>target_seq</td><td>PAM</td><td>alignment</td><td>distance</td><td>position</td><td>gene name</td><td>gene id</td></tr>
<tr><td>GU305914.1</td><td>57835</td><td>57857</td><td>-</td><td>0</td><td>CTGACCACCGAGAGCGACGG</td><td>CGG</td><td>||||||||[||||||||||||]PAM</td><td>NA</td><td>Intergenic</td><td>NA</td><td>NA</td></tr>
<tr><td>T29750</td><td>ACCACCGAGAGCGACGGCGGCGG</td><td>1000</td><td></td><td></td><td></td><td></td><td></td><td></td><td></td><td></td><td></td></tr>
<tr><td>Oligo fwd</td><td>CACCACCACCGAGAGCGACGGCGG</td><td></td><td></td><td></td><td></td><td></td><td></td><td></td><td></td><td></td><td></td></tr>
<tr><td>Oligo rev</td><td>AAACCCGCCGTCGCTCTCGGTGGT</td><td></td><td></td><td></td><td></td><td></td><td></td><td></td><td></td><td></td><td></td></tr>
<tr><td>Chromosome</td><td>start</td><td>end</td><td>strand</td><td>MM</td><td>target_seq</td><td>PAM</td><td>alignment</td><td>distance</td><td>position</td><td>gene name</td><td>gene id</td></tr>
<tr><td>GU305914.1</td><td>57832</td><td>57854</td><td>-</td><td>0</td><td>ACCACCGAGAGCGACGGCGG</td><td>CGG</td><td>||||||||[||||||||||||]PAM</td><td>NA</td><td>Intergenic</td><td>NA</td><td>NA</td></tr>
<tr><td>GU305914.1</td><td>70184</td><td>70206</td><td>+</td><td>4</td><td>AGCACCGGCAGCGCCGGCGG</td><td>CGG</td><td>|-|||||-[-||||-||||||]PAM</td><td>NA</td><td>Intergenic</td><td>NA</td><td>NA</td></tr>
<tr><td>T29751</td><td>TCCTCCCCTCCTCTCGCCCGCGG</td><td>1000</td><td></td><td></td><td></td><td></td><td></td><td></td><td></td><td></td><td></td></tr>
<tr><td>Oligo fwd</td><td>CACCTCCTCCCCTCCTCTCGCCCG</td><td></td><td></td><td></td><td></td><td></td><td></td><td></td><td></td><td></td><td></td></tr>
<tr><td>Oligo rev</td><td>AAACCGGGCGAGAGGAGGGGAGGA</td><td></td><td></td><td></td><td></td><td></td><td></td><td></td><td></td><td></td><td></td></tr>
<tr><td>Chromosome</td><td>start</td><td>end</td><td>strand</td><td>MM</td><td>target_seq</td><td>PAM</td><td>alignment</td><td>distance</td><td>position</td><td>gene name</td><td>gene id</td></tr>
<tr><td>GU305914.1</td><td>57794</td><td>57816</td><td>-</td><td>0</td><td>TCCTCCCCTCCTCTCGCCCG</td><td>CGG</td><td>||||||||[||||||||||||]PAM</td><td>NA</td><td>Intergenic</td><td>NA</td><td>NA</td></tr>
<tr><td>T29752</td><td>GAACGTCTCGACGCATGACTTGG</td><td>1000</td><td></td><td></td><td></td><td></td><td></td><td></td><td></td><td></td><td></td></tr>
<tr><td>Oligo fwd</td><td>CACCGAACGTCTCGACGCATGACT</td><td></td><td></td><td></td><td></td><td></td><td></td><td></td><td></td><td></td><td></td></tr>
<tr><td>Oligo rev</td><td>AAACAGTCATGCGTCGAGACGTTC</td><td></td><td></td><td></td><td></td><td></td><td></td><td></td><td></td><td></td><td></td></tr>
<tr><td>Chromosome</td><td>start</td><td>end</td><td>strand</td><td>MM</td><td>target_seq</td><td>PAM</td><td>alignment</td><td>distance</td><td>position</td><td>gene name</td><td>gene id</td></tr>
<tr><td>GU305914.1</td><td>57753</td><td>57775</td><td>-</td><td>0</td><td>GAACGTCTCGACGCATGACT</td><td>TGG</td><td>||||||||[||||||||||||]PAM</td><td>NA</td><td>Intergenic</td><td>NA</td><td>NA</td></tr>
<tr><td>T29753</td><td>AGATAAATTCCCCGAGCGTGCGG</td><td>1000</td><td></td><td></td><td></td><td></td><td></td><td></td><td></td><td></td><td></td></tr>
<tr><td>Oligo fwd</td><td>CACCAGATAAATTCCCCGAGCGTG</td><td></td><td></td><td></td><td></td><td></td><td></td><td></td><td></td><td></td><td></td></tr>
<tr><td>Oligo rev</td><td>AAACCACGCTCGGGGAATTTATCT</td><td></td><td></td><td></td><td></td><td></td><td></td><td></td><td></td><td></td><td></td></tr>
<tr><td>Chromosome</td><td>start</td><td>end</td><td>strand</td><td>MM</td><td>target_seq</td><td>PAM</td><td>alignment</td><td>distance</td><td>position</td><td>gene name</td><td>gene id</td></tr>
<tr><td>GU305914.1</td><td>57704</td><td>57726</td><td>-</td><td>0</td><td>AGATAAATTCCCCGAGCGTG</td><td>CGG</td><td>||||||||[||||||||||||]PAM</td><td>NA</td><td>Intergenic</td><td>NA</td><td>NA</td></tr>
<tr><td>T29754</td><td>GCGGTCTTTCTTTCAGTCCGAGG</td><td>1000</td><td></td><td></td><td></td><td></td><td></td><td></td><td></td><td></td><td></td></tr>
<tr><td>Oligo fwd</td><td>CACCGCGGTCTTTCTTTCAGTCCG</td><td></td><td></td><td></td><td></td><td></td><td></td><td></td><td></td><td></td><td></td></tr>
<tr><td>Oligo rev</td><td>AAACCGGACTGAAAGAAAGACCGC</td><td></td><td></td><td></td><td></td><td></td><td></td><td></td><td></td><td></td><td></td></tr>
<tr><td>Chromosome</td><td>start</td><td>end</td><td>strand</td><td>MM</td><td>target_seq</td><td>PAM</td><td>alignment</td><td>distance</td><td>position</td><td>gene name</td><td>gene id</td></tr>
<tr><td>GU305914.1</td><td>57685</td><td>57707</td><td>-</td><td>0</td><td>GCGGTCTTTCTTTCAGTCCG</td><td>AGG</td><td>||||||||[||||||||||||]PAM</td><td>NA</td><td>Intergenic</td><td>NA</td><td>NA</td></tr>
<tr><td>T29755</td><td>GTCTTTCTTTCAGTCCGAGGAGG</td><td>1000</td><td></td><td></td><td></td><td></td><td></td><td></td><td></td><td></td><td></td></tr>
<tr><td>Oligo fwd</td><td>CACCGTCTTTCTTTCAGTCCGAGG</td><td></td><td></td><td></td><td></td><td></td><td></td><td></td><td></td><td></td><td></td></tr>
<tr><td>Oligo rev</td><td>AAACCCTCGGACTGAAAGAAAGAC</td><td></td><td></td><td></td><td></td><td></td><td></td><td></td><td></td><td></td><td></td></tr>
<tr><td>Chromosome</td><td>start</td><td>end</td><td>strand</td><td>MM</td><td>target_seq</td><td>PAM</td><td>alignment</td><td>distance</td><td>position</td><td>gene name</td><td>gene id</td></tr>
<tr><td>GU305914.1</td><td>57682</td><td>57704</td><td>-</td><td>0</td><td>GTCTTTCTTTCAGTCCGAGG</td><td>AGG</td><td>||||||||[||||||||||||]PAM</td><td>NA</td><td>Intergenic</td><td>NA</td><td>NA</td></tr>
<tr><td>T29756</td><td>GGAGGATCTGAACCCCCCTCAGG</td><td>1000</td><td></td><td></td><td></td><td></td><td></td><td></td><td></td><td></td><td></td></tr>
<tr><td>Oligo fwd</td><td>CACCGGAGGATCTGAACCCCCCTC</td><td></td><td></td><td></td><td></td><td></td><td></td><td></td><td></td><td></td><td></td></tr>
<tr><td>Oligo rev</td><td>AAACGAGGGGGGTTCAGATCCTCC</td><td></td><td></td><td></td><td></td><td></td><td></td><td></td><td></td><td></td><td></td></tr>
<tr><td>Chromosome</td><td>start</td><td>end</td><td>strand</td><td>MM</td><td>target_seq</td><td>PAM</td><td>alignment</td><td>distance</td><td>position</td><td>gene name</td><td>gene id</td></tr>
<tr><td>GU305914.1</td><td>57664</td><td>57686</td><td>-</td><td>0</td><td>GGAGGATCTGAACCCCCCTC</td><td>AGG</td><td>||||||||[||||||||||||]PAM</td><td>NA</td><td>Intergenic</td><td>NA</td><td>NA</td></tr>
<tr><td>GU305914.1</td><td>123542</td><td>123564</td><td>-</td><td>4</td><td>GGAGTACCTGAACCGCTCTC</td><td>AGG</td><td>||||-|-|[||||||-|-|||]PAM</td><td>NA</td><td>Intergenic</td><td>NA</td><td>NA</td></tr>
<tr><td>T29757</td><td>TCTGAACCCCCCTCAGGTGCCGG</td><td>1000</td><td></td><td></td><td></td><td></td><td></td><td></td><td></td><td></td><td></td></tr>
<tr><td>Oligo fwd</td><td>CACCTCTGAACCCCCCTCAGGTGC</td><td></td><td></td><td></td><td></td><td></td><td></td><td></td><td></td><td></td><td></td></tr>
<tr><td>Oligo rev</td><td>AAACGCACCTGAGGGGGGTTCAGA</td><td></td><td></td><td></td><td></td><td></td><td></td><td></td><td></td><td></td><td></td></tr>
<tr><td>Chromosome</td><td>start</td><td>end</td><td>strand</td><td>MM</td><td>target_seq</td><td>PAM</td><td>alignment</td><td>distance</td><td>position</td><td>gene name</td><td>gene id</td></tr>
<tr><td>GU305914.1</td><td>57658</td><td>57680</td><td>-</td><td>0</td><td>TCTGAACCCCCCTCAGGTGC</td><td>CGG</td><td>||||||||[||||||||||||]PAM</td><td>NA</td><td>Intergenic</td><td>NA</td><td>NA</td></tr>
<tr><td>T29758</td><td>CCGCCGCGCCATCGCCACGACGG</td><td>1000</td><td></td><td></td><td></td><td></td><td></td><td></td><td></td><td></td><td></td></tr>
<tr><td>Oligo fwd</td><td>CACCCCGCCGCGCCATCGCCACGA</td><td></td><td></td><td></td><td></td><td></td><td></td><td></td><td></td><td></td><td></td></tr>
<tr><td>Oligo rev</td><td>AAACTCGTGGCGATGGCGCGGCGG</td><td></td><td></td><td></td><td></td><td></td><td></td><td></td><td></td><td></td><td></td></tr>
<tr><td>Chromosome</td><td>start</td><td>end</td><td>strand</td><td>MM</td><td>target_seq</td><td>PAM</td><td>alignment</td><td>distance</td><td>position</td><td>gene name</td><td>gene id</td></tr>
<tr><td>GU305914.1</td><td>57522</td><td>57544</td><td>-</td><td>0</td><td>CCGCCGCGCCATCGCCACGA</td><td>CGG</td><td>||||||||[||||||||||||]PAM</td><td>NA</td><td>Intergenic</td><td>NA</td><td>NA</td></tr>
<tr><td>T29759</td><td>GCCGTGACGACCGCGAGCTCCGG</td><td>1000</td><td></td><td></td><td></td><td></td><td></td><td></td><td></td><td></td><td></td></tr>
<tr><td>Oligo fwd</td><td>CACCGCCGTGACGACCGCGAGCTC</td><td></td><td></td><td></td><td></td><td></td><td></td><td></td><td></td><td></td><td></td></tr>
<tr><td>Oligo rev</td><td>AAACGAGCTCGCGGTCGTCACGGC</td><td></td><td></td><td></td><td></td><td></td><td></td><td></td><td></td><td></td><td></td></tr>
</table>

<table>
<tr><td>Chromosome</td><td>start</td><td></td><td>end</td><td>strand</td><td>MM</td><td>target_seq</td><td>PAM</td><td>alignment</td><td>distance</td><td>position</td><td>gene name</td><td>gene id</td></tr>
<tr><td>GU305914.1</td><td></td><td>57451</td><td>57473</td><td>-</td><td>0</td><td>GCCGTGACGACCGCGAGCTC</td><td>CGG</td><td>||||||||[||||||||||||]PAM</td><td>NA</td><td>Intergenic</td><td>NA</td><td>NA</td></tr>
<tr><td>T29760</td><td>ACGACCGCGAGCTCCGGAGAAGG</td><td></td><td>1000</td><td></td><td></td><td></td><td></td><td></td><td></td><td></td><td></td><td></td></tr>
<tr><td>Oligo fwd</td><td>CACCACGACCGCGAGCTCCGGAGA</td><td></td><td></td><td></td><td></td><td></td><td></td><td></td><td></td><td></td><td></td><td></td></tr>
<tr><td>Oligo rev</td><td>AAACTCTCCGGAGCTCGCGGTCGT</td><td></td><td></td><td></td><td></td><td></td><td></td><td></td><td></td><td></td><td></td><td></td></tr>
<tr><td>Chromosome</td><td>start</td><td></td><td>end</td><td>strand</td><td>MM</td><td>target_seq</td><td>PAM</td><td>alignment</td><td>distance</td><td>position</td><td>gene name</td><td>gene id</td></tr>
<tr><td>GU305914.1</td><td></td><td>57445</td><td>57467</td><td>-</td><td>0</td><td>ACGACCGCGAGCTCCGGAGA</td><td>AGG</td><td>||||||||[||||||||||||]PAM</td><td>NA</td><td>Intergenic</td><td>NA</td><td>NA</td></tr>
<tr><td>T29761</td><td>CGTATCTTTTTCCAGAGCCGCGG</td><td></td><td>1000</td><td></td><td></td><td></td><td></td><td></td><td></td><td></td><td></td><td></td></tr>
<tr><td>Oligo fwd</td><td>CACCCGTATCTTTTTCCAGAGCCG</td><td></td><td></td><td></td><td></td><td></td><td></td><td></td><td></td><td></td><td></td><td></td></tr>
<tr><td>Oligo rev</td><td>AAACCGGCTCTGGAAAAAGATACG</td><td></td><td></td><td></td><td></td><td></td><td></td><td></td><td></td><td></td><td></td><td></td></tr>
<tr><td>Chromosome</td><td>start</td><td></td><td>end</td><td>strand</td><td>MM</td><td>target_seq</td><td>PAM</td><td>alignment</td><td>distance</td><td>position</td><td>gene name</td><td>gene id</td></tr>
<tr><td>GU305914.1</td><td></td><td>57406</td><td>57428</td><td>-</td><td>0</td><td>CGTATCTTTTTCCAGAGCCG</td><td>CGG</td><td>||||||||[||||||||||||]PAM</td><td>NA</td><td>Intergenic</td><td>NA</td><td>NA</td></tr>
<tr><td>T29762</td><td>TCCAGAGCCGCGGTCGTCGCCGG</td><td></td><td>1000</td><td></td><td></td><td></td><td></td><td></td><td></td><td></td><td></td><td></td></tr>
<tr><td>Oligo fwd</td><td>CACCTCCAGAGCCGCGGTCGTCGC</td><td></td><td></td><td></td><td></td><td></td><td></td><td></td><td></td><td></td><td></td><td></td></tr>
<tr><td>Oligo rev</td><td>AAACGCGACGACCGCGGCTCTGGA</td><td></td><td></td><td></td><td></td><td></td><td></td><td></td><td></td><td></td><td></td><td></td></tr>
<tr><td>Chromosome</td><td>start</td><td></td><td>end</td><td>strand</td><td>MM</td><td>target_seq</td><td>PAM</td><td>alignment</td><td>distance</td><td>position</td><td>gene name</td><td>gene id</td></tr>
<tr><td>GU305914.1</td><td></td><td>57396</td><td>57418</td><td>-</td><td>0</td><td>TCCAGAGCCGCGGTCGTCGC</td><td>CGG</td><td>||||||||[||||||||||||]PAM</td><td>NA</td><td>Intergenic</td><td>NA</td><td>NA</td></tr>
<tr><td>T29763</td><td>GCATCGAGAATCCGTCTTCACGG</td><td></td><td>1000</td><td></td><td></td><td></td><td></td><td></td><td></td><td></td><td></td><td></td></tr>
<tr><td>Oligo fwd</td><td>CACCGCATCGAGAATCCGTCTTCA</td><td></td><td></td><td></td><td></td><td></td><td></td><td></td><td></td><td></td><td></td><td></td></tr>
<tr><td>Oligo rev</td><td>AAACTGAAGACGGATTCTCGATGC</td><td></td><td></td><td></td><td></td><td></td><td></td><td></td><td></td><td></td><td></td><td></td></tr>
<tr><td>Chromosome</td><td>start</td><td></td><td>end</td><td>strand</td><td>MM</td><td>target_seq</td><td>PAM</td><td>alignment</td><td>distance</td><td>position</td><td>gene name</td><td>gene id</td></tr>
<tr><td>GU305914.1</td><td></td><td>57347</td><td>57369</td><td>-</td><td>0</td><td>GCATCGAGAATCCGTCTTCA</td><td>CGG</td><td>||||||||[||||||||||||]PAM</td><td>NA</td><td>Intergenic</td><td>NA</td><td>NA</td></tr>
<tr><td>T29764</td><td>CCGATCCGTTAGCTAGAGACCGG</td><td></td><td>1000</td><td></td><td></td><td></td><td></td><td></td><td></td><td></td><td></td><td></td></tr>
<tr><td>Oligo fwd</td><td>CACCCCGATCCGTTAGCTAGAGAC</td><td></td><td></td><td></td><td></td><td></td><td></td><td></td><td></td><td></td><td></td><td></td></tr>
<tr><td>Oligo rev</td><td>AAACGTCTCTAGCTAACGGATCGG</td><td></td><td></td><td></td><td></td><td></td><td></td><td></td><td></td><td></td><td></td><td></td></tr>
<tr><td>Chromosome</td><td>start</td><td></td><td>end</td><td>strand</td><td>MM</td><td>target_seq</td><td>PAM</td><td>alignment</td><td>distance</td><td>position</td><td>gene name</td><td>gene id</td></tr>
<tr><td>GU305914.1</td><td></td><td>57321</td><td>57343</td><td>-</td><td>0</td><td>CCGATCCGTTAGCTAGAGAC</td><td>CGG</td><td>||||||||[||||||||||||]PAM</td><td>NA</td><td>Intergenic</td><td>NA</td><td>NA</td></tr>
<tr><td>T29765</td><td>GCTAGAGACCGGCGTCTTCGAGG</td><td></td><td>1000</td><td></td><td></td><td></td><td></td><td></td><td></td><td></td><td></td><td></td></tr>
<tr><td>Oligo fwd</td><td>CACCGCTAGAGACCGGCGTCTTCG</td><td></td><td></td><td></td><td></td><td></td><td></td><td></td><td></td><td></td><td></td><td></td></tr>
<tr><td>Oligo rev</td><td>AAACCGAAGACGCCGGTCTCTAGC</td><td></td><td></td><td></td><td></td><td></td><td></td><td></td><td></td><td></td><td></td><td></td></tr>
<tr><td>Chromosome</td><td>start</td><td></td><td>end</td><td>strand</td><td>MM</td><td>target_seq</td><td>PAM</td><td>alignment</td><td>distance</td><td>position</td><td>gene name</td><td>gene id</td></tr>
<tr><td>GU305914.1</td><td></td><td>57310</td><td>57332</td><td>-</td><td>0</td><td>GCTAGAGACCGGCGTCTTCG</td><td>AGG</td><td>||||||||[||||||||||||]PAM</td><td>NA</td><td>Intergenic</td><td>NA</td><td>NA</td></tr>
<tr><td>T29766</td><td>CTAGAGACCGGCGTCTTCGAGGG</td><td></td><td>1000</td><td></td><td></td><td></td><td></td><td></td><td></td><td></td><td></td><td></td></tr>
<tr><td>Oligo fwd</td><td>CACCCTAGAGACCGGCGTCTTCGA</td><td></td><td></td><td></td><td></td><td></td><td></td><td></td><td></td><td></td><td></td><td></td></tr>
<tr><td>Oligo rev</td><td>AAACTCGAAGACGCCGGTCTCTAG</td><td></td><td></td><td></td><td></td><td></td><td></td><td></td><td></td><td></td><td></td><td></td></tr>
<tr><td>Chromosome</td><td>start</td><td></td><td>end</td><td>strand</td><td>MM</td><td>target_seq</td><td>PAM</td><td>alignment</td><td>distance</td><td>position</td><td>gene name</td><td>gene id</td></tr>
<tr><td>GU305914.1</td><td></td><td>57309</td><td>57331</td><td>-</td><td>0</td><td>CTAGAGACCGGCGTCTTCGA</td><td>GGG</td><td>||||||||[||||||||||||]PAM</td><td>NA</td><td>Intergenic</td><td>NA</td><td>NA</td></tr>
<tr><td>T29767</td><td>TAGAGACCGGCGTCTTCGAGGGG</td><td></td><td>1000</td><td></td><td></td><td></td><td></td><td></td><td></td><td></td><td></td><td></td></tr>
<tr><td>Oligo fwd</td><td>CACCTAGAGACCGGCGTCTTCGAG</td><td></td><td></td><td></td><td></td><td></td><td></td><td></td><td></td><td></td><td></td><td></td></tr>
<tr><td>Oligo rev</td><td>AAACCTCGAAGACGCCGGTCTCTA</td><td></td><td></td><td></td><td></td><td></td><td></td><td></td><td></td><td></td><td></td><td></td></tr>
<tr><td>Chromosome</td><td>start</td><td></td><td>end</td><td>strand</td><td>MM</td><td>target_seq</td><td>PAM</td><td>alignment</td><td>distance</td><td>position</td><td>gene name</td><td>gene id</td></tr>
<tr><td>GU305914.1</td><td></td><td>57308</td><td>57330</td><td>-</td><td>0</td><td>TAGAGACCGGCGTCTTCGAG</td><td>GGG</td><td>||||||||[||||||||||||]PAM</td><td>NA</td><td>Intergenic</td><td>NA</td><td>NA</td></tr>
<tr><td>T29768</td><td>AGAGACCGGCGTCTTCGAGGGGG</td><td></td><td>1000</td><td></td><td></td><td></td><td></td><td></td><td></td><td></td><td></td><td></td></tr>
<tr><td>Oligo fwd</td><td>CACCAGAGACCGGCGTCTTCGAGG</td><td></td><td></td><td></td><td></td><td></td><td></td><td></td><td></td><td></td><td></td><td></td></tr>
<tr><td>Oligo rev</td><td>AAACCCTCGAAGACGCCGGTCTCT</td><td></td><td></td><td></td><td></td><td></td><td></td><td></td><td></td><td></td><td></td><td></td></tr>
<tr><td>Chromosome</td><td>start</td><td></td><td>end</td><td>strand</td><td>MM</td><td>target_seq</td><td>PAM</td><td>alignment</td><td>distance</td><td>position</td><td>gene name</td><td>gene id</td></tr>
<tr><td>GU305914.1</td><td></td><td>57307</td><td>57329</td><td>-</td><td>0</td><td>AGAGACCGGCGTCTTCGAGG</td><td>GGG</td><td>||||||||[||||||||||||]PAM</td><td>NA</td><td>Intergenic</td><td>NA</td><td>NA</td></tr>
<tr><td>T29769</td><td>TTGTCGTCTACGCTCGTTTCCGG</td><td></td><td>1000</td><td></td><td></td><td></td><td></td><td></td><td></td><td></td><td></td><td></td></tr>
<tr><td>Oligo fwd</td><td>CACCTTGTCGTCTACGCTCGTTTC</td><td></td><td></td><td></td><td></td><td></td><td></td><td></td><td></td><td></td><td></td><td></td></tr>
<tr><td>Oligo rev</td><td>AAACGAAACGAGCGTAGACGACAA</td><td></td><td></td><td></td><td></td><td></td><td></td><td></td><td></td><td></td><td></td><td></td></tr>
<tr><td>Chromosome</td><td>start</td><td></td><td>end</td><td>strand</td><td>MM</td><td>target_seq</td><td>PAM</td><td>alignment</td><td>distance</td><td>position</td><td>gene name</td><td>gene id</td></tr>
<tr><td>GU305914.1</td><td></td><td>57226</td><td>57248</td><td>-</td><td>0</td><td>TTGTCGTCTACGCTCGTTTC</td><td>CGG</td><td>||||||||[||||||||||||]PAM</td><td>NA</td><td>Intergenic</td><td>NA</td><td>NA</td></tr>
<tr><td>T29770</td><td>TACGCTCGTTTCCGGCGTGACGG</td><td></td><td>1000</td><td></td><td></td><td></td><td></td><td></td><td></td><td></td><td></td><td></td></tr>
<tr><td>Oligo fwd</td><td>CACCTACGCTCGTTTCCGGCGTGA</td><td></td><td></td><td></td><td></td><td></td><td></td><td></td><td></td><td></td><td></td><td></td></tr>
<tr><td>Oligo rev</td><td>AAACTCACGCCGGAAACGAGCGTA</td><td></td><td></td><td></td><td></td><td></td><td></td><td></td><td></td><td></td><td></td><td></td></tr>
<tr><td>Chromosome</td><td>start</td><td></td><td>end</td><td>strand</td><td>MM</td><td>target_seq</td><td>PAM</td><td>alignment</td><td>distance</td><td>position</td><td>gene name</td><td>gene id</td></tr>
<tr><td>GU305914.1</td><td></td><td>57218</td><td>57240</td><td>-</td><td>0</td><td>TACGCTCGTTTCCGGCGTGA</td><td>CGG</td><td>||||||||[||||||||||||]PAM</td><td>NA</td><td>Intergenic</td><td>NA</td><td>NA</td></tr>
<tr><td>T29771</td><td>GTAGACCTCGCCACGTTCGTTGG</td><td></td><td>1000</td><td></td><td></td><td></td><td></td><td></td><td></td><td></td><td></td><td></td></tr>
<tr><td>Oligo fwd</td><td>CACCGTAGACCTCGCCACGTTCGT</td><td></td><td></td><td></td><td></td><td></td><td></td><td></td><td></td><td></td><td></td><td></td></tr>
<tr><td>Oligo rev</td><td>AAACACGAACGTGGCGAGGTCTAC</td><td></td><td></td><td></td><td></td><td></td><td></td><td></td><td></td><td></td><td></td><td></td></tr>
<tr><td>Chromosome</td><td>start</td><td></td><td>end</td><td>strand</td><td>MM</td><td>target_seq</td><td>PAM</td><td>alignment</td><td>distance</td><td>position</td><td>gene name</td><td>gene id</td></tr>
<tr><td>GU305914.1</td><td></td><td>57151</td><td>57173</td><td>-</td><td>0</td><td>GTAGACCTCGCCACGTTCGT</td><td>TGG</td><td>||||||||[||||||||||||]PAM</td><td>NA</td><td>Intergenic</td><td>NA</td><td>NA</td></tr>
<tr><td>T29772</td><td>GTCTATGACAACGACGTTGACGG</td><td></td><td>1000</td><td></td><td></td><td></td><td></td><td></td><td></td><td></td><td></td><td></td></tr>
<tr><td>Oligo fwd</td><td>CACCGTCTATGACAACGACGTTGA</td><td></td><td></td><td></td><td></td><td></td><td></td><td></td><td></td><td></td><td></td><td></td></tr>
<tr><td>Oligo rev</td><td>AAACTCAACGTCGTTGTCATAGAC</td><td></td><td></td><td></td><td></td><td></td><td></td><td></td><td></td><td></td><td></td><td></td></tr>
<tr><td>Chromosome</td><td>start</td><td></td><td>end</td><td>strand</td><td>MM</td><td>target_seq</td><td>PAM</td><td>alignment</td><td>distance</td><td>position</td><td>gene name</td><td>gene id</td></tr>
<tr><td>GU305914.1</td><td></td><td>57129</td><td>57151</td><td>-</td><td>0</td><td>GTCTATGACAACGACGTTGA</td><td>CGG</td><td>||||||||[||||||||||||]PAM</td><td>NA</td><td>Intergenic</td><td>NA</td><td>NA</td></tr>
<tr><td>T29773</td><td>TCTATGACAACGACGTTGACGGG</td><td></td><td>1000</td><td></td><td></td><td></td><td></td><td></td><td></td><td></td><td></td><td></td></tr>
<tr><td>Oligo fwd</td><td>CACCTCTATGACAACGACGTTGAC</td><td></td><td></td><td></td><td></td><td></td><td></td><td></td><td></td><td></td><td></td><td></td></tr>
<tr><td>Oligo rev</td><td>AAACGTCAACGTCGTTGTCATAGA</td><td></td><td></td><td></td><td></td><td></td><td></td><td></td><td></td><td></td><td></td><td></td></tr>
<tr><td>Chromosome</td><td>start</td><td></td><td>end</td><td>strand</td><td>MM</td><td>target_seq</td><td>PAM</td><td>alignment</td><td>distance</td><td>position</td><td>gene name</td><td>gene id</td></tr>
<tr><td>GU305914.1</td><td></td><td>57128</td><td>57150</td><td>-</td><td>0</td><td>TCTATGACAACGACGTTGAC</td><td>GGG</td><td>||||||||[||||||||||||]PAM</td><td>NA</td><td>Intergenic</td><td>NA</td><td>NA</td></tr>
<tr><td>T29774</td><td>CTATGACAACGACGTTGACGGGG</td><td></td><td>1000</td><td></td><td></td><td></td><td></td><td></td><td></td><td></td><td></td><td></td></tr>
<tr><td>Oligo fwd</td><td>CACCCTATGACAACGACGTTGACG</td><td></td><td></td><td></td><td></td><td></td><td></td><td></td><td></td><td></td><td></td><td></td></tr>
</table>

<table>
<tr><td>Oligo rev</td><td>AAACCGTCAACGTCGTTGTCATAG</td><td></td><td></td><td></td><td></td><td></td><td></td><td></td><td></td><td></td><td></td><td></td></tr>
<tr><td>Chromosome</td><td>start</td><td></td><td>end</td><td>strand</td><td>MM</td><td>target_seq</td><td>PAM</td><td>alignment</td><td>distance</td><td>position</td><td>gene name</td><td>gene id</td></tr>
<tr><td>GU305914.1</td><td></td><td>57127</td><td>57149</td><td>-</td><td>0</td><td>CTATGACAACGACGTTGACG</td><td>GGG</td><td>||||||||[||||||||||||]PAM</td><td>NA</td><td>Intergenic</td><td>NA</td><td>NA</td></tr>
<tr><td></td><td></td><td></td><td></td><td></td><td></td><td></td><td></td><td></td><td></td><td></td><td></td><td></td></tr>
<tr><td>T29775</td><td>GTCGTGCCGACGACGAGCCATGG</td><td></td><td>1000</td><td></td><td></td><td></td><td></td><td></td><td></td><td></td><td></td><td></td></tr>
<tr><td>Oligo fwd</td><td>CACCGTCGTGCCGACGACGAGCCA</td><td></td><td></td><td></td><td></td><td></td><td></td><td></td><td></td><td></td><td></td><td></td></tr>
<tr><td>Oligo rev</td><td>AAACTGGCTCGTCGTCGGCACGAC</td><td></td><td></td><td></td><td></td><td></td><td></td><td></td><td></td><td></td><td></td><td></td></tr>
<tr><td>Chromosome</td><td>start</td><td></td><td>end</td><td>strand</td><td>MM</td><td>target_seq</td><td>PAM</td><td>alignment</td><td>distance</td><td>position</td><td>gene name</td><td>gene id</td></tr>
<tr><td>GU305914.1</td><td></td><td>57096</td><td>57118</td><td>-</td><td>0</td><td>GTCGTGCCGACGACGAGCCA</td><td>TGG</td><td>||||||||[||||||||||||]PAM</td><td>NA</td><td>Intergenic</td><td>NA</td><td>NA</td></tr>
<tr><td></td><td></td><td></td><td></td><td></td><td></td><td></td><td></td><td></td><td></td><td></td><td></td><td></td></tr>
<tr><td>T29776</td><td>CGAGCCATGGCTCGAGAGCTCGG</td><td></td><td>1000</td><td></td><td></td><td></td><td></td><td></td><td></td><td></td><td></td><td></td></tr>
<tr><td>Oligo fwd</td><td>CACCCGAGCCATGGCTCGAGAGCT</td><td></td><td></td><td></td><td></td><td></td><td></td><td></td><td></td><td></td><td></td><td></td></tr>
<tr><td>Oligo rev</td><td>AAACAGCTCTCGAGCCATGGCTCG</td><td></td><td></td><td></td><td></td><td></td><td></td><td></td><td></td><td></td><td></td><td></td></tr>
<tr><td>Chromosome</td><td>start</td><td></td><td>end</td><td>strand</td><td>MM</td><td>target_seq</td><td>PAM</td><td>alignment</td><td>distance</td><td>position</td><td>gene name</td><td>gene id</td></tr>
<tr><td>GU305914.1</td><td></td><td>57083</td><td>57105</td><td>-</td><td>0</td><td>CGAGCCATGGCTCGAGAGCT</td><td>CGG</td><td>||||||||[||||||||||||]PAM</td><td>NA</td><td>Intergenic</td><td>NA</td><td>NA</td></tr>
<tr><td></td><td></td><td></td><td></td><td></td><td></td><td></td><td></td><td></td><td></td><td></td><td></td><td></td></tr>
<tr><td>T29777</td><td>GAGCCATGGCTCGAGAGCTCGGG</td><td></td><td>1000</td><td></td><td></td><td></td><td></td><td></td><td></td><td></td><td></td><td></td></tr>
<tr><td>Oligo fwd</td><td>CACCGAGCCATGGCTCGAGAGCTC</td><td></td><td></td><td></td><td></td><td></td><td></td><td></td><td></td><td></td><td></td><td></td></tr>
<tr><td>Oligo rev</td><td>AAACGAGCTCTCGAGCCATGGCTC</td><td></td><td></td><td></td><td></td><td></td><td></td><td></td><td></td><td></td><td></td><td></td></tr>
<tr><td>Chromosome</td><td>start</td><td></td><td>end</td><td>strand</td><td>MM</td><td>target_seq</td><td>PAM</td><td>alignment</td><td>distance</td><td>position</td><td>gene name</td><td>gene id</td></tr>
<tr><td>GU305914.1</td><td></td><td>57082</td><td>57104</td><td>-</td><td>0</td><td>GAGCCATGGCTCGAGAGCTC</td><td>GGG</td><td>||||||||[||||||||||||]PAM</td><td>NA</td><td>Intergenic</td><td>NA</td><td>NA</td></tr>
<tr><td></td><td></td><td></td><td></td><td></td><td></td><td></td><td></td><td></td><td></td><td></td><td></td><td></td></tr>
<tr><td>T29778</td><td>TCGAGAGCTCGGGTACACGTTGG</td><td></td><td>1000</td><td></td><td></td><td></td><td></td><td></td><td></td><td></td><td></td><td></td></tr>
<tr><td>Oligo fwd</td><td>CACCTCGAGAGCTCGGGTACACGT</td><td></td><td></td><td></td><td></td><td></td><td></td><td></td><td></td><td></td><td></td><td></td></tr>
<tr><td>Oligo rev</td><td>AAACACGTGTACCCGAGCTCTCGA</td><td></td><td></td><td></td><td></td><td></td><td></td><td></td><td></td><td></td><td></td><td></td></tr>
<tr><td>Chromosome</td><td>start</td><td></td><td>end</td><td>strand</td><td>MM</td><td>target_seq</td><td>PAM</td><td>alignment</td><td>distance</td><td>position</td><td>gene name</td><td>gene id</td></tr>
<tr><td>GU305914.1</td><td></td><td>57072</td><td>57094</td><td>-</td><td>0</td><td>TCGAGAGCTCGGGTACACGT</td><td>TGG</td><td>||||||||[||||||||||||]PAM</td><td>NA</td><td>Intergenic</td><td>NA</td><td>NA</td></tr>
<tr><td></td><td></td><td></td><td></td><td></td><td></td><td></td><td></td><td></td><td></td><td></td><td></td><td></td></tr>
<tr><td>T29779</td><td>ACACGTTGGCTGTTACCGCTAGG</td><td></td><td>1000</td><td></td><td></td><td></td><td></td><td></td><td></td><td></td><td></td><td></td></tr>
<tr><td>Oligo fwd</td><td>CACCACACGTTGGCTGTTACCGCT</td><td></td><td></td><td></td><td></td><td></td><td></td><td></td><td></td><td></td><td></td><td></td></tr>
<tr><td>Oligo rev</td><td>AAACAGCGGTAACAGCCAACGTGT</td><td></td><td></td><td></td><td></td><td></td><td></td><td></td><td></td><td></td><td></td><td></td></tr>
<tr><td>Chromosome</td><td>start</td><td></td><td>end</td><td>strand</td><td>MM</td><td>target_seq</td><td>PAM</td><td>alignment</td><td>distance</td><td>position</td><td>gene name</td><td>gene id</td></tr>
<tr><td>GU305914.1</td><td></td><td>57058</td><td>57080</td><td>-</td><td>0</td><td>ACACGTTGGCTGTTACCGCT</td><td>AGG</td><td>||||||||[||||||||||||]PAM</td><td>NA</td><td>Intergenic</td><td>NA</td><td>NA</td></tr>
<tr><td></td><td></td><td></td><td></td><td></td><td></td><td></td><td></td><td></td><td></td><td></td><td></td><td></td></tr>
<tr><td>T29780</td><td>CGTTGGCTGTTACCGCTAGGTGG</td><td></td><td>1000</td><td></td><td></td><td></td><td></td><td></td><td></td><td></td><td></td><td></td></tr>
<tr><td>Oligo fwd</td><td>CACCCGTTGGCTGTTACCGCTAGG</td><td></td><td></td><td></td><td></td><td></td><td></td><td></td><td></td><td></td><td></td><td></td></tr>
<tr><td>Oligo rev</td><td>AAACCCTAGCGGTAACAGCCAACG</td><td></td><td></td><td></td><td></td><td></td><td></td><td></td><td></td><td></td><td></td><td></td></tr>
<tr><td>Chromosome</td><td>start</td><td></td><td>end</td><td>strand</td><td>MM</td><td>target_seq</td><td>PAM</td><td>alignment</td><td>distance</td><td>position</td><td>gene name</td><td>gene id</td></tr>
<tr><td>GU305914.1</td><td></td><td>57055</td><td>57077</td><td>-</td><td>0</td><td>CGTTGGCTGTTACCGCTAGG</td><td>TGG</td><td>||||||||[||||||||||||]PAM</td><td>NA</td><td>Intergenic</td><td>NA</td><td>NA</td></tr>
<tr><td></td><td></td><td></td><td></td><td></td><td></td><td></td><td></td><td></td><td></td><td></td><td></td><td></td></tr>
<tr><td>T29781</td><td>CTGTTACCGCTAGGTGGCGACGG</td><td></td><td>1000</td><td></td><td></td><td></td><td></td><td></td><td></td><td></td><td></td><td></td></tr>
<tr><td>Oligo fwd</td><td>CACCCTGTTACCGCTAGGTGGCGA</td><td></td><td></td><td></td><td></td><td></td><td></td><td></td><td></td><td></td><td></td><td></td></tr>
<tr><td>Oligo rev</td><td>AAACTCGCCACCTAGCGGTAACAG</td><td></td><td></td><td></td><td></td><td></td><td></td><td></td><td></td><td></td><td></td><td></td></tr>
<tr><td>Chromosome</td><td>start</td><td></td><td>end</td><td>strand</td><td>MM</td><td>target_seq</td><td>PAM</td><td>alignment</td><td>distance</td><td>position</td><td>gene name</td><td>gene id</td></tr>
<tr><td>GU305914.1</td><td></td><td>57049</td><td>57071</td><td>-</td><td>0</td><td>CTGTTACCGCTAGGTGGCGA</td><td>CGG</td><td>||||||||[||||||||||||]PAM</td><td>NA</td><td>Intergenic</td><td>NA</td><td>NA</td></tr>
<tr><td></td><td></td><td></td><td></td><td></td><td></td><td></td><td></td><td></td><td></td><td></td><td></td><td></td></tr>
<tr><td>T29782</td><td>CTCTATGTATCCCGAACGACTGG</td><td></td><td>1000</td><td></td><td></td><td></td><td></td><td></td><td></td><td></td><td></td><td></td></tr>
<tr><td>Oligo fwd</td><td>CACCCTCTATGTATCCCGAACGAC</td><td></td><td></td><td></td><td></td><td></td><td></td><td></td><td></td><td></td><td></td><td></td></tr>
<tr><td>Oligo rev</td><td>AAACGTCGTTCGGGATACATAGAG</td><td></td><td></td><td></td><td></td><td></td><td></td><td></td><td></td><td></td><td></td><td></td></tr>
<tr><td>Chromosome</td><td>start</td><td></td><td>end</td><td>strand</td><td>MM</td><td>target_seq</td><td>PAM</td><td>alignment</td><td>distance</td><td>position</td><td>gene name</td><td>gene id</td></tr>
<tr><td>GU305914.1</td><td></td><td>57000</td><td>57022</td><td>-</td><td>0</td><td>CTCTATGTATCCCGAACGAC</td><td>TGG</td><td>||||||||[||||||||||||]PAM</td><td>NA</td><td>Intergenic</td><td>NA</td><td>NA</td></tr>
<tr><td></td><td></td><td></td><td></td><td></td><td></td><td></td><td></td><td></td><td></td><td></td><td></td><td></td></tr>
<tr><td>T29783</td><td>CCGAACGACTGGCCGACGCGCGG</td><td></td><td>1000</td><td></td><td></td><td></td><td></td><td></td><td></td><td></td><td></td><td></td></tr>
<tr><td>Oligo fwd</td><td>CACCCCGAACGACTGGCCGACGCG</td><td></td><td></td><td></td><td></td><td></td><td></td><td></td><td></td><td></td><td></td><td></td></tr>
<tr><td>Oligo rev</td><td>AAACCGCGTCGGCCAGTCGTTCGG</td><td></td><td></td><td></td><td></td><td></td><td></td><td></td><td></td><td></td><td></td><td></td></tr>
<tr><td>Chromosome</td><td>start</td><td></td><td>end</td><td>strand</td><td>MM</td><td>target_seq</td><td>PAM</td><td>alignment</td><td>distance</td><td>position</td><td>gene name</td><td>gene id</td></tr>
<tr><td>GU305914.1</td><td></td><td>56989</td><td>57011</td><td>-</td><td>0</td><td>CCGAACGACTGGCCGACGCG</td><td>CGG</td><td>||||||||[||||||||||||]PAM</td><td>NA</td><td>Intergenic</td><td>NA</td><td>NA</td></tr>
<tr><td></td><td></td><td></td><td></td><td></td><td></td><td></td><td></td><td></td><td></td><td></td><td></td><td></td></tr>
<tr><td>T29784</td><td>CGCGGTTCCGACCGCAGCCGAGG</td><td></td><td>1000</td><td></td><td></td><td></td><td></td><td></td><td></td><td></td><td></td><td></td></tr>
<tr><td>Oligo fwd</td><td>CACCCGCGGTTCCGACCGCAGCCG</td><td></td><td></td><td></td><td></td><td></td><td></td><td></td><td></td><td></td><td></td><td></td></tr>
<tr><td>Oligo rev</td><td>AAACCGGCTGCGGTCGGAACCGCG</td><td></td><td></td><td></td><td></td><td></td><td></td><td></td><td></td><td></td><td></td><td></td></tr>
<tr><td>Chromosome</td><td>start</td><td></td><td>end</td><td>strand</td><td>MM</td><td>target_seq</td><td>PAM</td><td>alignment</td><td>distance</td><td>position</td><td>gene name</td><td>gene id</td></tr>
<tr><td>GU305914.1</td><td></td><td>56971</td><td>56993</td><td>-</td><td>0</td><td>CGCGGTTCCGACCGCAGCCG</td><td>AGG</td><td>||||||||[||||||||||||]PAM</td><td>NA</td><td>Intergenic</td><td>NA</td><td>NA</td></tr>
<tr><td></td><td></td><td></td><td></td><td></td><td></td><td></td><td></td><td></td><td></td><td></td><td></td><td></td></tr>
<tr><td>T29785</td><td>GCCGAGGATCATCCGCACGCTGG</td><td></td><td>1000</td><td></td><td></td><td></td><td></td><td></td><td></td><td></td><td></td><td></td></tr>
<tr><td>Oligo fwd</td><td>CACCGCCGAGGATCATCCGCACGC</td><td></td><td></td><td></td><td></td><td></td><td></td><td></td><td></td><td></td><td></td><td></td></tr>
<tr><td>Oligo rev</td><td>AAACGCGTGCGGATGATCCTCGGC</td><td></td><td></td><td></td><td></td><td></td><td></td><td></td><td></td><td></td><td></td><td></td></tr>
<tr><td>Chromosome</td><td>start</td><td></td><td>end</td><td>strand</td><td>MM</td><td>target_seq</td><td>PAM</td><td>alignment</td><td>distance</td><td>position</td><td>gene name</td><td>gene id</td></tr>
<tr><td>GU305914.1</td><td></td><td>56955</td><td>56977</td><td>-</td><td>0</td><td>GCCGAGGATCATCCGCACGC</td><td>TGG</td><td>||||||||[||||||||||||]PAM</td><td>NA</td><td>Intergenic</td><td>NA</td><td>NA</td></tr>
<tr><td></td><td></td><td></td><td></td><td></td><td></td><td></td><td></td><td></td><td></td><td></td><td></td><td></td></tr>
<tr><td>T29786</td><td>TCATCCGCACGCTGGATACCAGG</td><td></td><td>1000</td><td></td><td></td><td></td><td></td><td></td><td></td><td></td><td></td><td></td></tr>
<tr><td>Oligo fwd</td><td>CACCTCATCCGCACGCTGGATACC</td><td></td><td></td><td></td><td></td><td></td><td></td><td></td><td></td><td></td><td></td><td></td></tr>
<tr><td>Oligo rev</td><td>AAACGGTATCCAGCGTGCGGATGA</td><td></td><td></td><td></td><td></td><td></td><td></td><td></td><td></td><td></td><td></td><td></td></tr>
<tr><td>Chromosome</td><td>start</td><td></td><td>end</td><td>strand</td><td>MM</td><td>target_seq</td><td>PAM</td><td>alignment</td><td>distance</td><td>position</td><td>gene name</td><td>gene id</td></tr>
<tr><td>GU305914.1</td><td></td><td>56947</td><td>56969</td><td>-</td><td>0</td><td>TCATCCGCACGCTGGATACC</td><td>AGG</td><td>||||||||[||||||||||||]PAM</td><td>NA</td><td>Intergenic</td><td>NA</td><td>NA</td></tr>
<tr><td></td><td></td><td></td><td></td><td></td><td></td><td></td><td></td><td></td><td></td><td></td><td></td><td></td></tr>
<tr><td>T29787</td><td>CCGCACGCTGGATACCAGGTCGG</td><td></td><td>1000</td><td></td><td></td><td></td><td></td><td></td><td></td><td></td><td></td><td></td></tr>
<tr><td>Oligo fwd</td><td>CACCCCGCACGCTGGATACCAGGT</td><td></td><td></td><td></td><td></td><td></td><td></td><td></td><td></td><td></td><td></td><td></td></tr>
<tr><td>Oligo rev</td><td>AAACACCTGGTATCCAGCGTGCGG</td><td></td><td></td><td></td><td></td><td></td><td></td><td></td><td></td><td></td><td></td><td></td></tr>
<tr><td>Chromosome</td><td>start</td><td></td><td>end</td><td>strand</td><td>MM</td><td>target_seq</td><td>PAM</td><td>alignment</td><td>distance</td><td>position</td><td>gene name</td><td>gene id</td></tr>
<tr><td>GU305914.1</td><td></td><td>56943</td><td>56965</td><td>-</td><td>0</td><td>CCGCACGCTGGATACCAGGT</td><td>CGG</td><td>||||||||[||||||||||||]PAM</td><td>NA</td><td>Intergenic</td><td>NA</td><td>NA</td></tr>
<tr><td></td><td></td><td></td><td></td><td></td><td></td><td></td><td></td><td></td><td></td><td></td><td></td><td></td></tr>
<tr><td>T29788</td><td>CACGCTGGATACCAGGTCGGAGG</td><td></td><td>1000</td><td></td><td></td><td></td><td></td><td></td><td></td><td></td><td></td><td></td></tr>
<tr><td>Oligo fwd</td><td>CACCCACGCTGGATACCAGGTCGG</td><td></td><td></td><td></td><td></td><td></td><td></td><td></td><td></td><td></td><td></td><td></td></tr>
<tr><td>Oligo rev</td><td>AAACCCGACCTGGTATCCAGCGTG</td><td></td><td></td><td></td><td></td><td></td><td></td><td></td><td></td><td></td><td></td><td></td></tr>
<tr><td>Chromosome</td><td>start</td><td></td><td>end</td><td>strand</td><td>MM</td><td>target_seq</td><td>PAM</td><td>alignment</td><td>distance</td><td>position</td><td>gene name</td><td>gene id</td></tr>
<tr><td>GU305914.1</td><td></td><td>56940</td><td>56962</td><td>-</td><td>0</td><td>CACGCTGGATACCAGGTCGG</td><td>AGG</td><td>||||||||[||||||||||||]PAM</td><td>NA</td><td>Intergenic</td><td>NA</td><td>NA</td></tr>
<tr><td></td><td></td><td></td><td></td><td></td><td></td><td></td><td></td><td></td><td></td><td></td><td></td><td></td></tr>
<tr><td>T29789</td><td>GATACCAGGTCGGAGGCCCGTGG</td><td></td><td>1000</td><td></td><td></td><td></td><td></td><td></td><td></td><td></td><td></td><td></td></tr>
</table>

<table>
<tr><td>Oligo fwd</td><td>CACCGATACCAGGTCGGAGGCCCG</td><td></td><td></td><td></td><td></td><td></td><td></td><td></td><td></td><td></td><td></td></tr>
<tr><td>Oligo rev</td><td>AAACCGGGCCTCCGACCTGGTATC</td><td></td><td></td><td></td><td></td><td></td><td></td><td></td><td></td><td></td><td></td></tr>
<tr><td>Chromosome</td><td>start</td><td>end</td><td>strand</td><td>MM</td><td>target_seq</td><td>PAM</td><td>alignment</td><td>distance</td><td>position</td><td>gene name</td><td>gene id</td></tr>
<tr><td>GU305914.1</td><td>56933</td><td>56955</td><td>-</td><td>0</td><td>GATACCAGGTCGGAGGCCCG</td><td>TGG</td><td>||||||||[||||||||||||]PAM</td><td>NA</td><td>Intergenic</td><td>NA</td><td>NA</td></tr>
<tr><td>GU305914.1</td><td>64218</td><td>64240</td><td>-</td><td>3</td><td>GAGACCAGGTCGGCGGCCAG</td><td>CGG</td><td>||-|||||[|||||-||||-|]PAM</td><td>NA</td><td>Intergenic</td><td>NA</td><td>NA</td></tr>
<tr><td></td><td></td><td></td><td></td><td></td><td></td><td></td><td></td><td></td><td></td><td></td><td></td></tr>
<tr><td>T29790</td><td>ATACCAGGTCGGAGGCCCGTGGG</td><td>1000</td><td></td><td></td><td></td><td></td><td></td><td></td><td></td><td></td><td></td></tr>
<tr><td>Oligo fwd</td><td>CACCATACCAGGTCGGAGGCCCGT</td><td></td><td></td><td></td><td></td><td></td><td></td><td></td><td></td><td></td><td></td></tr>
<tr><td>Oligo rev</td><td>AAACACGGGCCTCCGACCTGGTAT</td><td></td><td></td><td></td><td></td><td></td><td></td><td></td><td></td><td></td><td></td></tr>
<tr><td>Chromosome</td><td>start</td><td>end</td><td>strand</td><td>MM</td><td>target_seq</td><td>PAM</td><td>alignment</td><td>distance</td><td>position</td><td>gene name</td><td>gene id</td></tr>
<tr><td>GU305914.1</td><td>56932</td><td>56954</td><td>-</td><td>0</td><td>ATACCAGGTCGGAGGCCCGT</td><td>GGG</td><td>||||||||[||||||||||||]PAM</td><td>NA</td><td>Intergenic</td><td>NA</td><td>NA</td></tr>
<tr><td></td><td></td><td></td><td></td><td></td><td></td><td></td><td></td><td></td><td></td><td></td><td></td></tr>
<tr><td>T29791</td><td>CCAGGTCGGAGGCCCGTGGGAGG</td><td>1000</td><td></td><td></td><td></td><td></td><td></td><td></td><td></td><td></td><td></td></tr>
<tr><td>Oligo fwd</td><td>CACCCCAGGTCGGAGGCCCGTGGG</td><td></td><td></td><td></td><td></td><td></td><td></td><td></td><td></td><td></td><td></td></tr>
<tr><td>Oligo rev</td><td>AAACCCCACGGGCCTCCGACCTGG</td><td></td><td></td><td></td><td></td><td></td><td></td><td></td><td></td><td></td><td></td></tr>
<tr><td>Chromosome</td><td>start</td><td>end</td><td>strand</td><td>MM</td><td>target_seq</td><td>PAM</td><td>alignment</td><td>distance</td><td>position</td><td>gene name</td><td>gene id</td></tr>
<tr><td>GU305914.1</td><td>56929</td><td>56951</td><td>-</td><td>0</td><td>CCAGGTCGGAGGCCCGTGGG</td><td>AGG</td><td>||||||||[||||||||||||]PAM</td><td>NA</td><td>Intergenic</td><td>NA</td><td>NA</td></tr>
<tr><td></td><td></td><td></td><td></td><td></td><td></td><td></td><td></td><td></td><td></td><td></td><td></td></tr>
<tr><td>T29792</td><td>CAGGTCGGAGGCCCGTGGGAGGG</td><td>1000</td><td></td><td></td><td></td><td></td><td></td><td></td><td></td><td></td><td></td></tr>
<tr><td>Oligo fwd</td><td>CACCCAGGTCGGAGGCCCGTGGGA</td><td></td><td></td><td></td><td></td><td></td><td></td><td></td><td></td><td></td><td></td></tr>
<tr><td>Oligo rev</td><td>AAACTCCCACGGGCCTCCGACCTG</td><td></td><td></td><td></td><td></td><td></td><td></td><td></td><td></td><td></td><td></td></tr>
<tr><td>Chromosome</td><td>start</td><td>end</td><td>strand</td><td>MM</td><td>target_seq</td><td>PAM</td><td>alignment</td><td>distance</td><td>position</td><td>gene name</td><td>gene id</td></tr>
<tr><td>GU305914.1</td><td>56928</td><td>56950</td><td>-</td><td>0</td><td>CAGGTCGGAGGCCCGTGGGA</td><td>GGG</td><td>||||||||[||||||||||||]PAM</td><td>NA</td><td>Intergenic</td><td>NA</td><td>NA</td></tr>
<tr><td></td><td></td><td></td><td></td><td></td><td></td><td></td><td></td><td></td><td></td><td></td><td></td></tr>
<tr><td>T29793</td><td>GAGGGCCGTCGTGCGATTTTCGG</td><td>1000</td><td></td><td></td><td></td><td></td><td></td><td></td><td></td><td></td><td></td></tr>
<tr><td>Oligo fwd</td><td>CACCGAGGGCCGTCGTGCGATTTT</td><td></td><td></td><td></td><td></td><td></td><td></td><td></td><td></td><td></td><td></td></tr>
<tr><td>Oligo rev</td><td>AAACAAAATCGCACGACGGCCCTC</td><td></td><td></td><td></td><td></td><td></td><td></td><td></td><td></td><td></td><td></td></tr>
<tr><td>Chromosome</td><td>start</td><td>end</td><td>strand</td><td>MM</td><td>target_seq</td><td>PAM</td><td>alignment</td><td>distance</td><td>position</td><td>gene name</td><td>gene id</td></tr>
<tr><td>GU305914.1</td><td>56910</td><td>56932</td><td>-</td><td>0</td><td>GAGGGCCGTCGTGCGATTTT</td><td>CGG</td><td>||||||||[||||||||||||]PAM</td><td>NA</td><td>Intergenic</td><td>NA</td><td>NA</td></tr>
<tr><td></td><td></td><td></td><td></td><td></td><td></td><td></td><td></td><td></td><td></td><td></td><td></td></tr>
<tr><td>T29794</td><td>GTGCGATTTTCGGCACCTCGTGG</td><td>1000</td><td></td><td></td><td></td><td></td><td></td><td></td><td></td><td></td><td></td></tr>
<tr><td>Oligo fwd</td><td>CACCGTGCGATTTTCGGCACCTCG</td><td></td><td></td><td></td><td></td><td></td><td></td><td></td><td></td><td></td><td></td></tr>
<tr><td>Oligo rev</td><td>AAACCGAGGTGCCGAAAATCGCAC</td><td></td><td></td><td></td><td></td><td></td><td></td><td></td><td></td><td></td><td></td></tr>
<tr><td>Chromosome</td><td>start</td><td>end</td><td>strand</td><td>MM</td><td>target_seq</td><td>PAM</td><td>alignment</td><td>distance</td><td>position</td><td>gene name</td><td>gene id</td></tr>
<tr><td>GU305914.1</td><td>56900</td><td>56922</td><td>-</td><td>0</td><td>GTGCGATTTTCGGCACCTCG</td><td>TGG</td><td>||||||||[||||||||||||]PAM</td><td>NA</td><td>Intergenic</td><td>NA</td><td>NA</td></tr>
<tr><td></td><td></td><td></td><td></td><td></td><td></td><td></td><td></td><td></td><td></td><td></td><td></td></tr>
<tr><td>T29795</td><td>CGGCACCTCGTGGACGAGATTGG</td><td>1000</td><td></td><td></td><td></td><td></td><td></td><td></td><td></td><td></td><td></td></tr>
<tr><td>Oligo fwd</td><td>CACCCGGCACCTCGTGGACGAGAT</td><td></td><td></td><td></td><td></td><td></td><td></td><td></td><td></td><td></td><td></td></tr>
<tr><td>Oligo rev</td><td>AAACATCTCGTCCACGAGGTGCCG</td><td></td><td></td><td></td><td></td><td></td><td></td><td></td><td></td><td></td><td></td></tr>
<tr><td>Chromosome</td><td>start</td><td>end</td><td>strand</td><td>MM</td><td>target_seq</td><td>PAM</td><td>alignment</td><td>distance</td><td>position</td><td>gene name</td><td>gene id</td></tr>
<tr><td>GU305914.1</td><td>56890</td><td>56912</td><td>-</td><td>0</td><td>CGGCACCTCGTGGACGAGAT</td><td>TGG</td><td>||||||||[||||||||||||]PAM</td><td>NA</td><td>Intergenic</td><td>NA</td><td>NA</td></tr>
<tr><td></td><td></td><td></td><td></td><td></td><td></td><td></td><td></td><td></td><td></td><td></td><td></td></tr>
<tr><td>T29796</td><td>GTCTCCGTCCGACGATCTCACGG</td><td>1000</td><td></td><td></td><td></td><td></td><td></td><td></td><td></td><td></td><td></td></tr>
<tr><td>Oligo fwd</td><td>CACCGTCTCCGTCCGACGATCTCA</td><td></td><td></td><td></td><td></td><td></td><td></td><td></td><td></td><td></td><td></td></tr>
<tr><td>Oligo rev</td><td>AAACTGAGATCGTCGGACGGAGAC</td><td></td><td></td><td></td><td></td><td></td><td></td><td></td><td></td><td></td><td></td></tr>
<tr><td>Chromosome</td><td>start</td><td>end</td><td>strand</td><td>MM</td><td>target_seq</td><td>PAM</td><td>alignment</td><td>distance</td><td>position</td><td>gene name</td><td>gene id</td></tr>
<tr><td>GU305914.1</td><td>56868</td><td>56890</td><td>-</td><td>0</td><td>GTCTCCGTCCGACGATCTCA</td><td>CGG</td><td>||||||||[||||||||||||]PAM</td><td>NA</td><td>Intergenic</td><td>NA</td><td>NA</td></tr>
<tr><td></td><td></td><td></td><td></td><td></td><td></td><td></td><td></td><td></td><td></td><td></td><td></td></tr>
<tr><td>T29797</td><td>AGAGAGATCTTTTCGAAAGAAGG</td><td>1000</td><td></td><td></td><td></td><td></td><td></td><td></td><td></td><td></td><td></td></tr>
<tr><td>Oligo fwd</td><td>CACCAGAGAGATCTTTTCGAAAGA</td><td></td><td></td><td></td><td></td><td></td><td></td><td></td><td></td><td></td><td></td></tr>
<tr><td>Oligo rev</td><td>AAACTCTTTCGAAAAGATCTCTCT</td><td></td><td></td><td></td><td></td><td></td><td></td><td></td><td></td><td></td><td></td></tr>
<tr><td>Chromosome</td><td>start</td><td>end</td><td>strand</td><td>MM</td><td>target_seq</td><td>PAM</td><td>alignment</td><td>distance</td><td>position</td><td>gene name</td><td>gene id</td></tr>
<tr><td>GU305914.1</td><td>56820</td><td>56842</td><td>-</td><td>0</td><td>AGAGAGATCTTTTCGAAAGA</td><td>AGG</td><td>||||||||[||||||||||||]PAM</td><td>NA</td><td>Intergenic</td><td>NA</td><td>NA</td></tr>
<tr><td></td><td></td><td></td><td></td><td></td><td></td><td></td><td></td><td></td><td></td><td></td><td></td></tr>
<tr><td>T29798</td><td>GAGAGATCTTTTCGAAAGAAGGG</td><td>1000</td><td></td><td></td><td></td><td></td><td></td><td></td><td></td><td></td><td></td></tr>
<tr><td>Oligo fwd</td><td>CACCGAGAGATCTTTTCGAAAGAA</td><td></td><td></td><td></td><td></td><td></td><td></td><td></td><td></td><td></td><td></td></tr>
<tr><td>Oligo rev</td><td>AAACTTCTTTCGAAAAGATCTCTC</td><td></td><td></td><td></td><td></td><td></td><td></td><td></td><td></td><td></td><td></td></tr>
<tr><td>Chromosome</td><td>start</td><td>end</td><td>strand</td><td>MM</td><td>target_seq</td><td>PAM</td><td>alignment</td><td>distance</td><td>position</td><td>gene name</td><td>gene id</td></tr>
<tr><td>GU305914.1</td><td>56819</td><td>56841</td><td>-</td><td>0</td><td>GAGAGATCTTTTCGAAAGAA</td><td>GGG</td><td>||||||||[||||||||||||]PAM</td><td>NA</td><td>Intergenic</td><td>NA</td><td>NA</td></tr>
<tr><td></td><td></td><td></td><td></td><td></td><td></td><td></td><td></td><td></td><td></td><td></td><td></td></tr>
<tr><td>T29799</td><td>AGAGATCTTTTCGAAAGAAGGGG</td><td>1000</td><td></td><td></td><td></td><td></td><td></td><td></td><td></td><td></td><td></td></tr>
<tr><td>Oligo fwd</td><td>CACCAGAGATCTTTTCGAAAGAAG</td><td></td><td></td><td></td><td></td><td></td><td></td><td></td><td></td><td></td><td></td></tr>
<tr><td>Oligo rev</td><td>AAACCTTCTTTCGAAAAGATCTCT</td><td></td><td></td><td></td><td></td><td></td><td></td><td></td><td></td><td></td><td></td></tr>
<tr><td>Chromosome</td><td>start</td><td>end</td><td>strand</td><td>MM</td><td>target_seq</td><td>PAM</td><td>alignment</td><td>distance</td><td>position</td><td>gene name</td><td>gene id</td></tr>
<tr><td>GU305914.1</td><td>56818</td><td>56840</td><td>-</td><td>0</td><td>AGAGATCTTTTCGAAAGAAG</td><td>GGG</td><td>||||||||[||||||||||||]PAM</td><td>NA</td><td>Intergenic</td><td>NA</td><td>NA</td></tr>
<tr><td></td><td></td><td></td><td></td><td></td><td></td><td></td><td></td><td></td><td></td><td></td><td></td></tr>
<tr><td>T29800</td><td>TCTTTTCGAAAGAAGGGGAGAGG</td><td>1000</td><td></td><td></td><td></td><td></td><td></td><td></td><td></td><td></td><td></td></tr>
<tr><td>Oligo fwd</td><td>CACCTCTTTTCGAAAGAAGGGGAG</td><td></td><td></td><td></td><td></td><td></td><td></td><td></td><td></td><td></td><td></td></tr>
<tr><td>Oligo rev</td><td>AAACCTCCCCTTCTTTCGAAAAGA</td><td></td><td></td><td></td><td></td><td></td><td></td><td></td><td></td><td></td><td></td></tr>
<tr><td>Chromosome</td><td>start</td><td>end</td><td>strand</td><td>MM</td><td>target_seq</td><td>PAM</td><td>alignment</td><td>distance</td><td>position</td><td>gene name</td><td>gene id</td></tr>
<tr><td>GU305914.1</td><td>56813</td><td>56835</td><td>-</td><td>0</td><td>TCTTTTCGAAAGAAGGGGAG</td><td>AGG</td><td>||||||||[||||||||||||]PAM</td><td>NA</td><td>Intergenic</td><td>NA</td><td>NA</td></tr>
<tr><td></td><td></td><td></td><td></td><td></td><td></td><td></td><td></td><td></td><td></td><td></td><td></td></tr>
<tr><td>T29801</td><td>TTTCGAAAGAAGGGGAGAGGAGG</td><td>1000</td><td></td><td></td><td></td><td></td><td></td><td></td><td></td><td></td><td></td></tr>
<tr><td>Oligo fwd</td><td>CACCTTTCGAAAGAAGGGGAGAGG</td><td></td><td></td><td></td><td></td><td></td><td></td><td></td><td></td><td></td><td></td></tr>
<tr><td>Oligo rev</td><td>AAACCCTCTCCCCTTCTTTCGAAA</td><td></td><td></td><td></td><td></td><td></td><td></td><td></td><td></td><td></td><td></td></tr>
<tr><td>Chromosome</td><td>start</td><td>end</td><td>strand</td><td>MM</td><td>target_seq</td><td>PAM</td><td>alignment</td><td>distance</td><td>position</td><td>gene name</td><td>gene id</td></tr>
<tr><td>GU305914.1</td><td>56810</td><td>56832</td><td>-</td><td>0</td><td>TTTCGAAAGAAGGGGAGAGG</td><td>AGG</td><td>||||||||[||||||||||||]PAM</td><td>NA</td><td>Intergenic</td><td>NA</td><td>NA</td></tr>
<tr><td></td><td></td><td></td><td></td><td></td><td></td><td></td><td></td><td></td><td></td><td></td><td></td></tr>
<tr><td>T29802</td><td>TGCCGCCGCCGCCGCCGCTGCGG</td><td>1000</td><td></td><td></td><td></td><td></td><td></td><td></td><td></td><td></td><td></td></tr>
<tr><td>Oligo fwd</td><td>CACCTGCCGCCGCCGCCGCCGCTG</td><td></td><td></td><td></td><td></td><td></td><td></td><td></td><td></td><td></td><td></td></tr>
<tr><td>Oligo rev</td><td>AAACCAGCGGCGGCGGCGGCGGCA</td><td></td><td></td><td></td><td></td><td></td><td></td><td></td><td></td><td></td><td></td></tr>
<tr><td>Chromosome</td><td>start</td><td>end</td><td>strand</td><td>MM</td><td>target_seq</td><td>PAM</td><td>alignment</td><td>distance</td><td>position</td><td>gene name</td><td>gene id</td></tr>
<tr><td>GU305914.1</td><td>56775</td><td>56797</td><td>-</td><td>0</td><td>TGCCGCCGCCGCCGCCGCTG</td><td>CGG</td><td>||||||||[||||||||||||]PAM</td><td>NA</td><td>Intergenic</td><td>NA</td><td>NA</td></tr>
<tr><td>GU305914.1</td><td>128802</td><td>128824</td><td>-</td><td>4</td><td>CGCCGACGTCGCTGCCGCTG</td><td>CGG</td><td>-||||-||[-|||-|||||||]PAM</td><td>NA</td><td>Intergenic</td><td>NA</td><td>NA</td></tr>
<tr><td>GU305914.1</td><td>35655</td><td>35677</td><td>-</td><td>3</td><td>CCCCGCCGCCGCCGCCGCCG</td><td>TGG</td><td>--||||||[||||||||||-|]PAM</td><td>NA</td><td>Intergenic</td><td>NA</td><td>NA</td></tr>
<tr><td>GU305914.1</td><td>130315</td><td>130337</td><td>+</td><td>4</td><td>AGCAGCAGCCGCCGCCGCGG</td><td>CGG</td><td>-||-||-|[||||||||||-|]PAM</td><td>NA</td><td>Intergenic</td><td>NA</td><td>NA</td></tr>
<tr><td>GU305914.1</td><td>45592</td><td>45614</td><td>-</td><td>4</td><td>GGCGGCTGCCGCCGCCGCCG</td><td>CGG</td><td>-||-||-|[||||||||||-|]PAM</td><td>NA</td><td>Intergenic</td><td>NA</td><td>NA</td></tr>
<tr><td>GU305914.1</td><td>56772</td><td>56794</td><td>-</td><td>3</td><td>CGCCGCCGCCGCCGCTGCGG</td><td>CGG</td><td>-|||||||[|||||||-||-|]PAM</td><td>NA</td><td>Intergenic</td><td>NA</td><td>NA</td></tr>
</table>

| | | | | | | | | | | | |
|---|---|---|---|---|---|---|---|---|---|---|---|
| T29803 | CGCCGCCGCCGCCGCTGCGGCGG | 1000 | | | | | | | | | |
| Oligo fwd | CACCCGCCGCCGCCGCCGCTGCGG | | | | | | | | | | |
| Oligo rev | AAACCCGCAGCGGCGGCGGCGGCG | | | | | | | | | | |
| Chromosome | start | end | strand | MM | target_seq | PAM | alignment | distance | position | gene name | gene id |
| GU305914.1 | 56772 | 56794 | - | 0 | CGCCGCCGCCGCCGCTGCGG | CGG | \|\|\|\|\|\|\|\|[\|\|\|\|\|\|\|\|\|\|\|\|]PAM | NA | Intergenic | NA | NA |
| GU305914.1 | 130315 | 130337 | + | 4 | AGCAGCAGCCGCCGCCGCGG | CGG | -\|\|-\|\|-\|[\|\|\|\|\|\|\|-\|\|\|\|]PAM | NA | Intergenic | NA | NA |
| GU305914.1 | 56775 | 56797 | - | 3 | TGCCGCCGCCGCCGCCGCTG | CGG | -\|\|\|\|\|\|\|[\|\|\|\|\|\|\|-\|\|-\|]PAM | NA | Intergenic | NA | NA |
| GU305914.1 | 35655 | 35677 | - | 3 | CCCCGCCGCCGCCGCCGCCG | TGG | \|-\|\|\|\|\|\|[\|\|\|\|\|\|\|-\|\|-\|]PAM | NA | Intergenic | NA | NA |
| GU305914.1 | 78745 | 78767 | - | 4 | CGCCGCGCCGGCCGCTGCGC | TGG | \|\|\|\|\|\|--[\|-\|\|\|\|\|\|\|\|\|-]PAM | NA | Intergenic | NA | NA |
| GU305914.1 | 45586 | 45608 | - | 3 | TGCCGCCGCCGCCGCGGCGC | CGG | -\|\|\|\|\|\|\|[\|\|\|\|\|\|\|-\|\|\|-]PAM | NA | Intergenic | NA | NA |
| T29804 | TGCGGCGGAGAAGAGTCGTGAGG | 1000 | | | | | | | | | |
| Oligo fwd | CACCTGCGGCGGAGAAGAGTCGTG | | | | | | | | | | |
| Oligo rev | AAACCACGACTCTTCTCCGCCGCA | | | | | | | | | | |
| Chromosome | start | end | strand | MM | target_seq | PAM | alignment | distance | position | gene name | gene id |
| GU305914.1 | 56757 | 56779 | - | 0 | TGCGGCGGAGAAGAGTCGTG | AGG | \|\|\|\|\|\|\|\|[\|\|\|\|\|\|\|\|\|\|\|\|]PAM | NA | Intergenic | NA | NA |
| T29805 | TTACGAGAACGTCGAACCGACGG | 1000 | | | | | | | | | |
| Oligo fwd | CACCTTACGAGAACGTCGAACCGA | | | | | | | | | | |
| Oligo rev | AAACTCGGTTCGACGTTCTCGTAA | | | | | | | | | | |
| Chromosome | start | end | strand | MM | target_seq | PAM | alignment | distance | position | gene name | gene id |
| GU305914.1 | 56724 | 56746 | - | 0 | TTACGAGAACGTCGAACCGA | CGG | \|\|\|\|\|\|\|\|[\|\|\|\|\|\|\|\|\|\|\|\|]PAM | NA | Intergenic | NA | NA |
| T29806 | GAACGTCGAACCGACGGTGAAGG | 1000 | | | | | | | | | |
| Oligo fwd | CACCGAACGTCGAACCGACGGTGA | | | | | | | | | | |
| Oligo rev | AAACTCACCGTCGGTTCGACGTTC | | | | | | | | | | |
| Chromosome | start | end | strand | MM | target_seq | PAM | alignment | distance | position | gene name | gene id |
| GU305914.1 | 56718 | 56740 | - | 0 | GAACGTCGAACCGACGGTGA | AGG | \|\|\|\|\|\|\|\|[\|\|\|\|\|\|\|\|\|\|\|\|]PAM | NA | Intergenic | NA | NA |
| T29807 | AACGTCGAACCGACGGTGAAGGG | 1000 | | | | | | | | | |
| Oligo fwd | CACCAACGTCGAACCGACGGTGAA | | | | | | | | | | |
| Oligo rev | AAACTTCACCGTCGGTTCGACGTT | | | | | | | | | | |
| Chromosome | start | end | strand | MM | target_seq | PAM | alignment | distance | position | gene name | gene id |
| GU305914.1 | 56717 | 56739 | - | 0 | AACGTCGAACCGACGGTGAA | GGG | \|\|\|\|\|\|\|\|[\|\|\|\|\|\|\|\|\|\|\|\|]PAM | NA | Intergenic | NA | NA |
| T29808 | ACCGACGGTGAAGGGTGCCGCGG | 1000 | | | | | | | | | |
| Oligo fwd | CACCACCGACGGTGAAGGGTGCCG | | | | | | | | | | |
| Oligo rev | AAACCGGCACCCTTCACCGTCGGT | | | | | | | | | | |
| Chromosome | start | end | strand | MM | target_seq | PAM | alignment | distance | position | gene name | gene id |
| GU305914.1 | 56709 | 56731 | - | 0 | ACCGACGGTGAAGGGTGCCG | CGG | \|\|\|\|\|\|\|\|[\|\|\|\|\|\|\|\|\|\|\|\|]PAM | NA | Intergenic | NA | NA |
| T29809 | TGCCACCGCATCCACATCCGCGG | 1000 | | | | | | | | | |
| Oligo fwd | CACCTGCCACCGCATCCACATCCG | | | | | | | | | | |
| Oligo rev | AAACCGGATGTGGATGCGGTGGCA | | | | | | | | | | |
| Chromosome | start | end | strand | MM | target_seq | PAM | alignment | distance | position | gene name | gene id |
| GU305914.1 | 56682 | 56704 | - | 0 | TGCCACCGCATCCACATCCG | CGG | \|\|\|\|\|\|\|\|[\|\|\|\|\|\|\|\|\|\|\|\|]PAM | NA | Intergenic | NA | NA |
| T29810 | GCCGCCTGCTCGCTCGACGACGG | 1000 | | | | | | | | | |
| Oligo fwd | CACCGCCGCCTGCTCGCTCGACGA | | | | | | | | | | |
| Oligo rev | AAACTCGTCGAGCGAGCAGGCGGC | | | | | | | | | | |
| Chromosome | start | end | strand | MM | target_seq | PAM | alignment | distance | position | gene name | gene id |
| GU305914.1 | 56660 | 56682 | - | 0 | GCCGCCTGCTCGCTCGACGA | CGG | \|\|\|\|\|\|\|\|[\|\|\|\|\|\|\|\|\|\|\|\|]PAM | NA | Intergenic | NA | NA |
| T29811 | GAAGTAGTAACAAAGACAAGAGG | 1000 | | | | | | | | | |
| Oligo fwd | CACCGAAGTAGTAACAAAGACAAG | | | | | | | | | | |
| Oligo rev | AAACCTTGTCTTTGTTACTACTTC | | | | | | | | | | |
| Chromosome | start | end | strand | MM | target_seq | PAM | alignment | distance | position | gene name | gene id |
| GU305914.1 | 56635 | 56657 | - | 0 | GAAGTAGTAACAAAGACAAG | AGG | \|\|\|\|\|\|\|\|[\|\|\|\|\|\|\|\|\|\|\|\|]PAM | NA | Intergenic | NA | NA |
| T29812 | AAAAGTATCATCAGCGCCTGCGG | 1000 | | | | | | | | | |
| Oligo fwd | CACCAAAAGTATCATCAGCGCCTG | | | | | | | | | | |
| Oligo rev | AAACCAGGCGCTGATGATACTTTT | | | | | | | | | | |
| Chromosome | start | end | strand | MM | target_seq | PAM | alignment | distance | position | gene name | gene id |
| GU305914.1 | 56612 | 56634 | - | 0 | AAAAGTATCATCAGCGCCTG | CGG | \|\|\|\|\|\|\|\|[\|\|\|\|\|\|\|\|\|\|\|\|]PAM | NA | Intergenic | NA | NA |
| T29813 | TCCGTCGCCGCAGCAGTACTAGG | 1000 | | | | | | | | | |
| Oligo fwd | CACCTCCGTCGCCGCAGCAGTACT | | | | | | | | | | |
| Oligo rev | AAACAGTACTGCTGCGGCGACGGA | | | | | | | | | | |
| Chromosome | start | end | strand | MM | target_seq | PAM | alignment | distance | position | gene name | gene id |
| GU305914.1 | 56560 | 56582 | - | 0 | TCCGTCGCCGCAGCAGTACT | AGG | \|\|\|\|\|\|\|\|[\|\|\|\|\|\|\|\|\|\|\|\|]PAM | NA | Intergenic | NA | NA |
| GU305914.1 | 152545 | 152567 | - | 4 | ACCGCCGCCGCCGCAGGACT | GGG | -\|\|\|-\|\|\|[\|\|\|-\|\|\|\|-\|\|\|]PAM | NA | Intergenic | NA | NA |
| T29814 | ATCACACTACTAGCAACAAGAGG | 1000 | | | | | | | | | |
| Oligo fwd | CACCATCACACTACTAGCAACAAG | | | | | | | | | | |
| Oligo rev | AAACCTTGTTGCTAGTAGTGTGAT | | | | | | | | | | |
| Chromosome | start | end | strand | MM | target_seq | PAM | alignment | distance | position | gene name | gene id |
| GU305914.1 | 56512 | 56534 | - | 0 | ATCACACTACTAGCAACAAG | AGG | \|\|\|\|\|\|\|\|[\|\|\|\|\|\|\|\|\|\|\|\|]PAM | NA | Intergenic | NA | NA |
| T29815 | TAGCAACAAGAGGAAGAGTACGG | 1000 | | | | | | | | | |
| Oligo fwd | CACCTAGCAACAAGAGGAAGAGTA | | | | | | | | | | |
| Oligo rev | AAACTACTCTTCCTCTTGTTGCTA | | | | | | | | | | |
| Chromosome | start | end | strand | MM | target_seq | PAM | alignment | distance | position | gene name | gene id |
| GU305914.1 | 56502 | 56524 | - | 0 | TAGCAACAAGAGGAAGAGTA | CGG | \|\|\|\|\|\|\|\|[\|\|\|\|\|\|\|\|\|\|\|\|]PAM | NA | Intergenic | NA | NA |
| T29816 | CTCGATCGCCACTCGCTGATCGG | 1000 | | | | | | | | | |
| Oligo fwd | CACCCTCGATCGCCACTCGCTGAT | | | | | | | | | | |
| Oligo rev | AAACATCAGCGAGTGGCGATCGAG | | | | | | | | | | |
| Chromosome | start | end | strand | MM | target_seq | PAM | alignment | distance | position | gene name | gene id |
| GU305914.1 | 56462 | 56484 | - | 0 | CTCGATCGCCACTCGCTGAT | CGG | \|\|\|\|\|\|\|\|[\|\|\|\|\|\|\|\|\|\|\|\|]PAM | NA | Intergenic | NA | NA |

| | | | | | | | | | | | |
|---|---|---|---|---|---|---|---|---|---|---|---|
| T29817 | CGGAGCGCCTCCGACCCTGCCGG | 1000 | | | | | | | | | |
| Oligo fwd | CACCCGGAGCGCCTCCGACCCTGC | | | | | | | | | | |
| Oligo rev | AAACGCAGGGTCGGAGGCGCTCCG | | | | | | | | | | |
| Chromosome | start | end | strand | MM | target_seq | PAM | alignment | distance | position | gene name | gene id |
| GU305914.1 | 56442 | 56464 | - | 0 | CGGAGCGCCTCCGACCCTGC | CGG | \|\|\|\|\|\|\|\|[\|\|\|\|\|\|\|\|\|\|\|\|]PAM | NA | Intergenic | NA | NA |
| T29818 | TAAAGATTTCGTCGAGAAGTTGG | 1000 | | | | | | | | | |
| Oligo fwd | CACCTAAAGATTTCGTCGAGAAGT | | | | | | | | | | |
| Oligo rev | AAACACTTCTCGACGAAATCTTTA | | | | | | | | | | |
| Chromosome | start | end | strand | MM | target_seq | PAM | alignment | distance | position | gene name | gene id |
| GU305914.1 | 56403 | 56425 | - | 0 | TAAAGATTTCGTCGAGAAGT | TGG | \|\|\|\|\|\|\|\|[\|\|\|\|\|\|\|\|\|\|\|\|]PAM | NA | Intergenic | NA | NA |
| T29819 | GTTGGCTGCCAGACTCTCTCAGG | 1000 | | | | | | | | | |
| Oligo fwd | CACCGTTGGCTGCCAGACTCTCTC | | | | | | | | | | |
| Oligo rev | AAACGAGAGAGTCTGGCAGCCAAC | | | | | | | | | | |
| Chromosome | start | end | strand | MM | target_seq | PAM | alignment | distance | position | gene name | gene id |
| GU305914.1 | 56385 | 56407 | - | 0 | GTTGGCTGCCAGACTCTCTC | AGG | \|\|\|\|\|\|\|\|[\|\|\|\|\|\|\|\|\|\|\|\|]PAM | NA | Intergenic | NA | NA |
| T29820 | CTCTAAGATGATTCTCTGCACGG | 1000 | | | | | | | | | |
| Oligo fwd | CACCCTCTAAGATGATTCTCTGCA | | | | | | | | | | |
| Oligo rev | AAACTGCAGAGAATCATCTTAGAG | | | | | | | | | | |
| Chromosome | start | end | strand | MM | target_seq | PAM | alignment | distance | position | gene name | gene id |
| GU305914.1 | 56358 | 56380 | - | 0 | CTCTAAGATGATTCTCTGCA | CGG | \|\|\|\|\|\|\|\|[\|\|\|\|\|\|\|\|\|\|\|\|]PAM | NA | Intergenic | NA | NA |
| T29821 | TCCACGAATTGATCGCCGCCAGG | 1000 | | | | | | | | | |
| Oligo fwd | CACCTCCACGAATTGATCGCCGCC | | | | | | | | | | |
| Oligo rev | AAACGGCGGCGATCAATTCGTGGA | | | | | | | | | | |
| Chromosome | start | end | strand | MM | target_seq | PAM | alignment | distance | position | gene name | gene id |
| GU305914.1 | 56320 | 56342 | - | 0 | TCCACGAATTGATCGCCGCC | AGG | \|\|\|\|\|\|\|\|[\|\|\|\|\|\|\|\|\|\|\|\|]PAM | NA | Intergenic | NA | NA |
| T29822 | CCACGAATTGATCGCCGCCAGGG | 1000 | | | | | | | | | |
| Oligo fwd | CACCCCACGAATTGATCGCCGCCA | | | | | | | | | | |
| Oligo rev | AAACTGGCGGCGATCAATTCGTGG | | | | | | | | | | |
| Chromosome | start | end | strand | MM | target_seq | PAM | alignment | distance | position | gene name | gene id |
| GU305914.1 | 56319 | 56341 | - | 0 | CCACGAATTGATCGCCGCCA | GGG | \|\|\|\|\|\|\|\|[\|\|\|\|\|\|\|\|\|\|\|\|]PAM | NA | Intergenic | NA | NA |
| T29823 | TGATCGCCGCCAGGGACATCCGG | 1000 | | | | | | | | | |
| Oligo fwd | CACCTGATCGCCGCCAGGGACATC | | | | | | | | | | |
| Oligo rev | AAACGATGTCCCTGGCGGCGATCA | | | | | | | | | | |
| Chromosome | start | end | strand | MM | target_seq | PAM | alignment | distance | position | gene name | gene id |
| GU305914.1 | 56311 | 56333 | - | 0 | TGATCGCCGCCAGGGACATC | CGG | \|\|\|\|\|\|\|\|[\|\|\|\|\|\|\|\|\|\|\|\|]PAM | NA | Intergenic | NA | NA |
| T29824 | GACTCAACTGCCTCATCTACAGG | 1000 | | | | | | | | | |
| Oligo fwd | CACCGACTCAACTGCCTCATCTAC | | | | | | | | | | |
| Oligo rev | AAACGTAGATGAGGCAGTTGAGTC | | | | | | | | | | |
| Chromosome | start | end | strand | MM | target_seq | PAM | alignment | distance | position | gene name | gene id |
| GU305914.1 | 56287 | 56309 | - | 0 | GACTCAACTGCCTCATCTAC | AGG | \|\|\|\|\|\|\|\|[\|\|\|\|\|\|\|\|\|\|\|\|]PAM | NA | Intergenic | NA | NA |
| T29825 | TCCATCACGTGCTGTTGACCTGG | 1000 | | | | | | | | | |
| Oligo fwd | CACCTCCATCACGTGCTGTTGACC | | | | | | | | | | |
| Oligo rev | AAACGGTCAACAGCACGTGATGGA | | | | | | | | | | |
| Chromosome | start | end | strand | MM | target_seq | PAM | alignment | distance | position | gene name | gene id |
| GU305914.1 | 56257 | 56279 | - | 0 | TCCATCACGTGCTGTTGACC | TGG | \|\|\|\|\|\|\|\|[\|\|\|\|\|\|\|\|\|\|\|\|]PAM | NA | Intergenic | NA | NA |
| T29826 | GTGCTGTTGACCTGGCCCCCCGG | 1000 | | | | | | | | | |
| Oligo fwd | CACCGTGCTGTTGACCTGGCCCCC | | | | | | | | | | |
| Oligo rev | AAACGGGGGCCAGGTCAACAGCAC | | | | | | | | | | |
| Chromosome | start | end | strand | MM | target_seq | PAM | alignment | distance | position | gene name | gene id |
| GU305914.1 | 56249 | 56271 | - | 0 | GTGCTGTTGACCTGGCCCCC | CGG | \|\|\|\|\|\|\|\|[\|\|\|\|\|\|\|\|\|\|\|\|]PAM | NA | Intergenic | NA | NA |
| T29827 | TGTTGACCTGGCCCCCCGGCTGG | 1000 | | | | | | | | | |
| Oligo fwd | CACCTGTTGACCTGGCCCCCCGGC | | | | | | | | | | |
| Oligo rev | AAACGCCGGGGGGCCAGGTCAACA | | | | | | | | | | |
| Chromosome | start | end | strand | MM | target_seq | PAM | alignment | distance | position | gene name | gene id |
| GU305914.1 | 56245 | 56267 | - | 0 | TGTTGACCTGGCCCCCCGGC | TGG | \|\|\|\|\|\|\|\|[\|\|\|\|\|\|\|\|\|\|\|\|]PAM | NA | Intergenic | NA | NA |
| T29828 | GTTGACCTGGCCCCCCGGCTGGG | 1000 | | | | | | | | | |
| Oligo fwd | CACCGTTGACCTGGCCCCCCGGCT | | | | | | | | | | |
| Oligo rev | AAACAGCCGGGGGGCCAGGTCAAC | | | | | | | | | | |
| Chromosome | start | end | strand | MM | target_seq | PAM | alignment | distance | position | gene name | gene id |
| GU305914.1 | 56244 | 56266 | - | 0 | GTTGACCTGGCCCCCCGGCT | GGG | \|\|\|\|\|\|\|\|[\|\|\|\|\|\|\|\|\|\|\|\|]PAM | NA | Intergenic | NA | NA |
| T29829 | TCGATGCGACTCGCTCGCCGAGG | 1000 | | | | | | | | | |
| Oligo fwd | CACCTCGATGCGACTCGCTCGCCG | | | | | | | | | | |
| Oligo rev | AAACCGGCGAGCGAGTCGCATCGA | | | | | | | | | | |
| Chromosome | start | end | strand | MM | target_seq | PAM | alignment | distance | position | gene name | gene id |
| GU305914.1 | 56209 | 56231 | - | 0 | TCGATGCGACTCGCTCGCCG | AGG | \|\|\|\|\|\|\|\|[\|\|\|\|\|\|\|\|\|\|\|\|]PAM | NA | Intergenic | NA | NA |
| T29830 | TCGCTCGCCGAGGCTGCAGAAGG | 1000 | | | | | | | | | |
| Oligo fwd | CACCTCGCTCGCCGAGGCTGCAGA | | | | | | | | | | |
| Oligo rev | AAACTCTGCAGCCTCGGCGAGCGA | | | | | | | | | | |
| Chromosome | start | end | strand | MM | target_seq | PAM | alignment | distance | position | gene name | gene id |
| GU305914.1 | 56199 | 56221 | - | 0 | TCGCTCGCCGAGGCTGCAGA | AGG | \|\|\|\|\|\|\|\|[\|\|\|\|\|\|\|\|\|\|\|\|]PAM | NA | Intergenic | NA | NA |
| T29831 | AGGCGAGCGCCGCCGTCGTCCGG | 1000 | | | | | | | | | |
| Oligo fwd | CACCAGGCGAGCGCCGCCGTCGTC | | | | | | | | | | |
| Oligo rev | AAACGACGACGGCGGCGCTCGCCT | | | | | | | | | | |
| Chromosome | start | end | strand | MM | target_seq | PAM | alignment | distance | position | gene name | gene id |

<table>
<tr><td>GU305914.1</td><td>56179</td><td>56201</td><td>-</td><td>0</td><td>AGGCGAGCGCCGCCGTCGTC</td><td>CGG</td><td>||||||||[||||||||||||]PAM</td><td>NA</td><td>Intergenic</td><td>NA</td><td>NA</td></tr>
<tr><td>T29832</td><td>GGCGAGCGCCGCCGTCGTCCGGG</td><td>1000</td><td></td><td></td><td></td><td></td><td></td><td></td><td></td><td></td><td></td></tr>
<tr><td>Oligo fwd</td><td>CACCGGCGAGCGCCGCCGTCGTCC</td><td></td><td></td><td></td><td></td><td></td><td></td><td></td><td></td><td></td><td></td></tr>
<tr><td>Oligo rev</td><td>AAACGGACGACGGCGGCGCTCGCC</td><td></td><td></td><td></td><td></td><td></td><td></td><td></td><td></td><td></td><td></td></tr>
<tr><td>Chromosome</td><td>start</td><td>end</td><td>strand</td><td>MM</td><td>target_seq</td><td>PAM</td><td>alignment</td><td>distance</td><td>position</td><td>gene name</td><td>gene id</td></tr>
<tr><td>GU305914.1</td><td>56178</td><td>56200</td><td>-</td><td>0</td><td>GGCGAGCGCCGCCGTCGTCC</td><td>GGG</td><td>||||||||[||||||||||||]PAM</td><td>NA</td><td>Intergenic</td><td>NA</td><td>NA</td></tr>
<tr><td>T29833</td><td>AGCGCCGCCGTCGTCCGGGACGG</td><td>1000</td><td></td><td></td><td></td><td></td><td></td><td></td><td></td><td></td><td></td></tr>
<tr><td>Oligo fwd</td><td>CACCAGCGCCGCCGTCGTCCGGGA</td><td></td><td></td><td></td><td></td><td></td><td></td><td></td><td></td><td></td><td></td></tr>
<tr><td>Oligo rev</td><td>AAACTCCCGGACGACGGCGGCGCT</td><td></td><td></td><td></td><td></td><td></td><td></td><td></td><td></td><td></td><td></td></tr>
<tr><td>Chromosome</td><td>start</td><td>end</td><td>strand</td><td>MM</td><td>target_seq</td><td>PAM</td><td>alignment</td><td>distance</td><td>position</td><td>gene name</td><td>gene id</td></tr>
<tr><td>GU305914.1</td><td>56174</td><td>56196</td><td>-</td><td>0</td><td>AGCGCCGCCGTCGTCCGGGA</td><td>CGG</td><td>||||||||[||||||||||||]PAM</td><td>NA</td><td>Intergenic</td><td>NA</td><td>NA</td></tr>
<tr><td>T29834</td><td>GAGCAGAACCTCATCGCCATCGG</td><td>1000</td><td></td><td></td><td></td><td></td><td></td><td></td><td></td><td></td><td></td></tr>
<tr><td>Oligo fwd</td><td>CACCGAGCAGAACCTCATCGCCAT</td><td></td><td></td><td></td><td></td><td></td><td></td><td></td><td></td><td></td><td></td></tr>
<tr><td>Oligo rev</td><td>AAACATGGCGATGAGGTTCTGCTC</td><td></td><td></td><td></td><td></td><td></td><td></td><td></td><td></td><td></td><td></td></tr>
<tr><td>Chromosome</td><td>start</td><td>end</td><td>strand</td><td>MM</td><td>target_seq</td><td>PAM</td><td>alignment</td><td>distance</td><td>position</td><td>gene name</td><td>gene id</td></tr>
<tr><td>GU305914.1</td><td>56138</td><td>56160</td><td>-</td><td>0</td><td>GAGCAGAACCTCATCGCCAT</td><td>CGG</td><td>||||||||[||||||||||||]PAM</td><td>NA</td><td>Intergenic</td><td>NA</td><td>NA</td></tr>
<tr><td>T29835</td><td>AGCAGAACCTCATCGCCATCGGG</td><td>1000</td><td></td><td></td><td></td><td></td><td></td><td></td><td></td><td></td><td></td></tr>
<tr><td>Oligo fwd</td><td>CACCAGCAGAACCTCATCGCCATC</td><td></td><td></td><td></td><td></td><td></td><td></td><td></td><td></td><td></td><td></td></tr>
<tr><td>Oligo rev</td><td>AAACGATGGCGATGAGGTTCTGCT</td><td></td><td></td><td></td><td></td><td></td><td></td><td></td><td></td><td></td><td></td></tr>
<tr><td>Chromosome</td><td>start</td><td>end</td><td>strand</td><td>MM</td><td>target_seq</td><td>PAM</td><td>alignment</td><td>distance</td><td>position</td><td>gene name</td><td>gene id</td></tr>
<tr><td>GU305914.1</td><td>56137</td><td>56159</td><td>-</td><td>0</td><td>AGCAGAACCTCATCGCCATC</td><td>GGG</td><td>||||||||[||||||||||||]PAM</td><td>NA</td><td>Intergenic</td><td>NA</td><td>NA</td></tr>
<tr><td>T29836</td><td>GCAGAACCTCATCGCCATCGGGG</td><td>1000</td><td></td><td></td><td></td><td></td><td></td><td></td><td></td><td></td><td></td></tr>
<tr><td>Oligo fwd</td><td>CACCGCAGAACCTCATCGCCATCG</td><td></td><td></td><td></td><td></td><td></td><td></td><td></td><td></td><td></td><td></td></tr>
<tr><td>Oligo rev</td><td>AAACCGATGGCGATGAGGTTCTGC</td><td></td><td></td><td></td><td></td><td></td><td></td><td></td><td></td><td></td><td></td></tr>
<tr><td>Chromosome</td><td>start</td><td>end</td><td>strand</td><td>MM</td><td>target_seq</td><td>PAM</td><td>alignment</td><td>distance</td><td>position</td><td>gene name</td><td>gene id</td></tr>
<tr><td>GU305914.1</td><td>56136</td><td>56158</td><td>-</td><td>0</td><td>GCAGAACCTCATCGCCATCG</td><td>GGG</td><td>||||||||[||||||||||||]PAM</td><td>NA</td><td>Intergenic</td><td>NA</td><td>NA</td></tr>
<tr><td>T29837</td><td>GAACCTCATCGCCATCGGGGTGG</td><td>1000</td><td></td><td></td><td></td><td></td><td></td><td></td><td></td><td></td><td></td></tr>
<tr><td>Oligo fwd</td><td>CACCGAACCTCATCGCCATCGGGG</td><td></td><td></td><td></td><td></td><td></td><td></td><td></td><td></td><td></td><td></td></tr>
<tr><td>Oligo rev</td><td>AAACCCCCGATGGCGATGAGGTTC</td><td></td><td></td><td></td><td></td><td></td><td></td><td></td><td></td><td></td><td></td></tr>
<tr><td>Chromosome</td><td>start</td><td>end</td><td>strand</td><td>MM</td><td>target_seq</td><td>PAM</td><td>alignment</td><td>distance</td><td>position</td><td>gene name</td><td>gene id</td></tr>
<tr><td>GU305914.1</td><td>56133</td><td>56155</td><td>-</td><td>0</td><td>GAACCTCATCGCCATCGGGG</td><td>TGG</td><td>||||||||[||||||||||||]PAM</td><td>NA</td><td>Intergenic</td><td>NA</td><td>NA</td></tr>
<tr><td>T29838</td><td>CATCGGGGTGGTCCACGACCTGG</td><td>1000</td><td></td><td></td><td></td><td></td><td></td><td></td><td></td><td></td><td></td></tr>
<tr><td>Oligo fwd</td><td>CACCCATCGGGGTGGTCCACGACC</td><td></td><td></td><td></td><td></td><td></td><td></td><td></td><td></td><td></td><td></td></tr>
<tr><td>Oligo rev</td><td>AAACGGTCGTGGACCACCCCGATG</td><td></td><td></td><td></td><td></td><td></td><td></td><td></td><td></td><td></td><td></td></tr>
<tr><td>Chromosome</td><td>start</td><td>end</td><td>strand</td><td>MM</td><td>target_seq</td><td>PAM</td><td>alignment</td><td>distance</td><td>position</td><td>gene name</td><td>gene id</td></tr>
<tr><td>GU305914.1</td><td>56121</td><td>56143</td><td>-</td><td>0</td><td>CATCGGGGTGGTCCACGACC</td><td>TGG</td><td>||||||||[||||||||||||]PAM</td><td>NA</td><td>Intergenic</td><td>NA</td><td>NA</td></tr>
<tr><td>T29839</td><td>ATCGGGGTGGTCCACGACCTGGG</td><td>1000</td><td></td><td></td><td></td><td></td><td></td><td></td><td></td><td></td><td></td></tr>
<tr><td>Oligo fwd</td><td>CACCATCGGGGTGGTCCACGACCT</td><td></td><td></td><td></td><td></td><td></td><td></td><td></td><td></td><td></td><td></td></tr>
<tr><td>Oligo rev</td><td>AAACAGGTCGTGGACCACCCCGAT</td><td></td><td></td><td></td><td></td><td></td><td></td><td></td><td></td><td></td><td></td></tr>
<tr><td>Chromosome</td><td>start</td><td>end</td><td>strand</td><td>MM</td><td>target_seq</td><td>PAM</td><td>alignment</td><td>distance</td><td>position</td><td>gene name</td><td>gene id</td></tr>
<tr><td>GU305914.1</td><td>56120</td><td>56142</td><td>-</td><td>0</td><td>ATCGGGGTGGTCCACGACCT</td><td>GGG</td><td>||||||||[||||||||||||]PAM</td><td>NA</td><td>Intergenic</td><td>NA</td><td>NA</td></tr>
<tr><td>T29840</td><td>TCGGGGTGGTCCACGACCTGGGG</td><td>1000</td><td></td><td></td><td></td><td></td><td></td><td></td><td></td><td></td><td></td></tr>
<tr><td>Oligo fwd</td><td>CACCTCGGGGTGGTCCACGACCTG</td><td></td><td></td><td></td><td></td><td></td><td></td><td></td><td></td><td></td><td></td></tr>
<tr><td>Oligo rev</td><td>AAACCAGGTCGTGGACCACCCCGA</td><td></td><td></td><td></td><td></td><td></td><td></td><td></td><td></td><td></td><td></td></tr>
<tr><td>Chromosome</td><td>start</td><td>end</td><td>strand</td><td>MM</td><td>target_seq</td><td>PAM</td><td>alignment</td><td>distance</td><td>position</td><td>gene name</td><td>gene id</td></tr>
<tr><td>GU305914.1</td><td>56119</td><td>56141</td><td>-</td><td>0</td><td>TCGGGGTGGTCCACGACCTG</td><td>GGG</td><td>||||||||[||||||||||||]PAM</td><td>NA</td><td>Intergenic</td><td>NA</td><td>NA</td></tr>
<tr><td>T29841</td><td>CCTGGGGCGCGAACCCTCCGAGG</td><td>1000</td><td></td><td></td><td></td><td></td><td></td><td></td><td></td><td></td><td></td></tr>
<tr><td>Oligo fwd</td><td>CACCCCTGGGGCGCGAACCCTCCG</td><td></td><td></td><td></td><td></td><td></td><td></td><td></td><td></td><td></td><td></td></tr>
<tr><td>Oligo rev</td><td>AAACCGGAGGGTTCGCGCCCCAGG</td><td></td><td></td><td></td><td></td><td></td><td></td><td></td><td></td><td></td><td></td></tr>
<tr><td>Chromosome</td><td>start</td><td>end</td><td>strand</td><td>MM</td><td>target_seq</td><td>PAM</td><td>alignment</td><td>distance</td><td>position</td><td>gene name</td><td>gene id</td></tr>
<tr><td>GU305914.1</td><td>56103</td><td>56125</td><td>-</td><td>0</td><td>CCTGGGGCGCGAACCCTCCG</td><td>AGG</td><td>||||||||[||||||||||||]PAM</td><td>NA</td><td>Intergenic</td><td>NA</td><td>NA</td></tr>
<tr><td>T29842</td><td>ACCCTCCGAGGCCGAGTACCCGG</td><td>1000</td><td></td><td></td><td></td><td></td><td></td><td></td><td></td><td></td><td></td></tr>
<tr><td>Oligo fwd</td><td>CACCACCCTCCGAGGCCGAGTACC</td><td></td><td></td><td></td><td></td><td></td><td></td><td></td><td></td><td></td><td></td></tr>
<tr><td>Oligo rev</td><td>AAACGGTACTCGGCCTCGGAGGGT</td><td></td><td></td><td></td><td></td><td></td><td></td><td></td><td></td><td></td><td></td></tr>
<tr><td>Chromosome</td><td>start</td><td>end</td><td>strand</td><td>MM</td><td>target_seq</td><td>PAM</td><td>alignment</td><td>distance</td><td>position</td><td>gene name</td><td>gene id</td></tr>
<tr><td>GU305914.1</td><td>56091</td><td>56113</td><td>-</td><td>0</td><td>ACCCTCCGAGGCCGAGTACC</td><td>CGG</td><td>||||||||[||||||||||||]PAM</td><td>NA</td><td>Intergenic</td><td>NA</td><td>NA</td></tr>
<tr><td>T29843</td><td>GTCATCCTCGTCAACCTCGATGG</td><td>1000</td><td></td><td></td><td></td><td></td><td></td><td></td><td></td><td></td><td></td></tr>
<tr><td>Oligo fwd</td><td>CACCGTCATCCTCGTCAACCTCGA</td><td></td><td></td><td></td><td></td><td></td><td></td><td></td><td></td><td></td><td></td></tr>
<tr><td>Oligo rev</td><td>AAACTCGAGGTTGACGAGGATGAC</td><td></td><td></td><td></td><td></td><td></td><td></td><td></td><td></td><td></td><td></td></tr>
<tr><td>Chromosome</td><td>start</td><td>end</td><td>strand</td><td>MM</td><td>target_seq</td><td>PAM</td><td>alignment</td><td>distance</td><td>position</td><td>gene name</td><td>gene id</td></tr>
<tr><td>GU305914.1</td><td>56069</td><td>56091</td><td>-</td><td>0</td><td>GTCATCCTCGTCAACCTCGA</td><td>TGG</td><td>||||||||[||||||||||||]PAM</td><td>NA</td><td>Intergenic</td><td>NA</td><td>NA</td></tr>
<tr><td>T29844</td><td>TTCGTGTACATGTACGACGATGG</td><td>1000</td><td></td><td></td><td></td><td></td><td></td><td></td><td></td><td></td><td></td></tr>
<tr><td>Oligo fwd</td><td>CACCTTCGTGTACATGTACGACGA</td><td></td><td></td><td></td><td></td><td></td><td></td><td></td><td></td><td></td><td></td></tr>
<tr><td>Oligo rev</td><td>AAACTCGTCGTACATGTACACGAA</td><td></td><td></td><td></td><td></td><td></td><td></td><td></td><td></td><td></td><td></td></tr>
<tr><td>Chromosome</td><td>start</td><td>end</td><td>strand</td><td>MM</td><td>target_seq</td><td>PAM</td><td>alignment</td><td>distance</td><td>position</td><td>gene name</td><td>gene id</td></tr>
<tr><td>GU305914.1</td><td>56045</td><td>56067</td><td>-</td><td>0</td><td>TTCGTGTACATGTACGACGA</td><td>TGG</td><td>||||||||[||||||||||||]PAM</td><td>NA</td><td>Intergenic</td><td>NA</td><td>NA</td></tr>
<tr><td>T29845</td><td>GTGTACATGTACGACGATGGCGG</td><td>1000</td><td></td><td></td><td></td><td></td><td></td><td></td><td></td><td></td><td></td></tr>
<tr><td>Oligo fwd</td><td>CACCGTGTACATGTACGACGATGG</td><td></td><td></td><td></td><td></td><td></td><td></td><td></td><td></td><td></td><td></td></tr>
<tr><td>Oligo rev</td><td>AAACCCATCGTCGTACATGTACAC</td><td></td><td></td><td></td><td></td><td></td><td></td><td></td><td></td><td></td><td></td></tr>
<tr><td>Chromosome</td><td>start</td><td>end</td><td>strand</td><td>MM</td><td>target_seq</td><td>PAM</td><td>alignment</td><td>distance</td><td>position</td><td>gene name</td><td>gene id</td></tr>
<tr><td>GU305914.1</td><td>56042</td><td>56064</td><td>-</td><td>0</td><td>GTGTACATGTACGACGATGG</td><td>CGG</td><td>||||||||[||||||||||||]PAM</td><td>NA</td><td>Intergenic</td><td>NA</td><td>NA</td></tr>
<tr><td>T29846</td><td>CTGCACCTGCTCACCAAGTGCGG</td><td>1000</td><td></td><td></td><td></td><td></td><td></td><td></td><td></td><td></td><td></td></tr>
<tr><td>Oligo fwd</td><td>CACCCTGCACCTGCTCACCAAGTG</td><td></td><td></td><td></td><td></td><td></td><td></td><td></td><td></td><td></td><td></td></tr>
<tr><td>Oligo rev</td><td>AAACCACTTGGTGAGCAGGTGCAG</td><td></td><td></td><td></td><td></td><td></td><td></td><td></td><td></td><td></td><td></td></tr>
</table>

<table>
<tr><td>Chromosome</td><td>start</td><td>end</td><td>strand</td><td>MM</td><td>target_seq</td><td>PAM</td><td>alignment</td><td>distance</td><td>position</td><td>gene name</td><td>gene id</td></tr>
<tr><td>GU305914.1</td><td>56006</td><td>56028</td><td>-</td><td>0</td><td>CTGCACCTGCTCACCAAGTG</td><td>CGG</td><td>||||||||[||||||||||||]PAM</td><td>NA</td><td>Intergenic</td><td>NA</td><td>NA</td></tr>
<tr><td></td><td></td><td></td><td></td><td></td><td></td><td></td><td></td><td></td><td></td><td></td><td></td></tr>
<tr><td>T29847</td><td>GCGGCTTCTCCGACTTCTGCCGG</td><td>1000</td><td></td><td></td><td></td><td></td><td></td><td></td><td></td><td></td><td></td></tr>
<tr><td>Oligo fwd</td><td>CACCGCGGCTTCTCCGACTTCTGC</td><td></td><td></td><td></td><td></td><td></td><td></td><td></td><td></td><td></td><td></td></tr>
<tr><td>Oligo rev</td><td>AAACGCAGAAGTCGGAGAAGCCGC</td><td></td><td></td><td></td><td></td><td></td><td></td><td></td><td></td><td></td><td></td></tr>
<tr><td>Chromosome</td><td>start</td><td>end</td><td>strand</td><td>MM</td><td>target_seq</td><td>PAM</td><td>alignment</td><td>distance</td><td>position</td><td>gene name</td><td>gene id</td></tr>
<tr><td>GU305914.1</td><td>55987</td><td>56009</td><td>-</td><td>0</td><td>GCGGCTTCTCCGACTTCTGC</td><td>CGG</td><td>||||||||[||||||||||||]PAM</td><td>NA</td><td>Intergenic</td><td>NA</td><td>NA</td></tr>
<tr><td></td><td></td><td></td><td></td><td></td><td></td><td></td><td></td><td></td><td></td><td></td><td></td></tr>
<tr><td>T29848</td><td>TTCTCCGACTTCTGCCGGCGCGG</td><td>1000</td><td></td><td></td><td></td><td></td><td></td><td></td><td></td><td></td><td></td></tr>
<tr><td>Oligo fwd</td><td>CACCTTCTCCGACTTCTGCCGGCG</td><td></td><td></td><td></td><td></td><td></td><td></td><td></td><td></td><td></td><td></td></tr>
<tr><td>Oligo rev</td><td>AAACCGCCGGCAGAAGTCGGAGAA</td><td></td><td></td><td></td><td></td><td></td><td></td><td></td><td></td><td></td><td></td></tr>
<tr><td>Chromosome</td><td>start</td><td>end</td><td>strand</td><td>MM</td><td>target_seq</td><td>PAM</td><td>alignment</td><td>distance</td><td>position</td><td>gene name</td><td>gene id</td></tr>
<tr><td>GU305914.1</td><td>55982</td><td>56004</td><td>-</td><td>0</td><td>TTCTCCGACTTCTGCCGGCG</td><td>CGG</td><td>||||||||[||||||||||||]PAM</td><td>NA</td><td>Intergenic</td><td>NA</td><td>NA</td></tr>
<tr><td></td><td></td><td></td><td></td><td></td><td></td><td></td><td></td><td></td><td></td><td></td><td></td></tr>
<tr><td>T29849</td><td>GCCGGCGCGGACTCCGTGAGAGG</td><td>1000</td><td></td><td></td><td></td><td></td><td></td><td></td><td></td><td></td><td></td></tr>
<tr><td>Oligo fwd</td><td>CACCGCCGGCGCGGACTCCGTGAG</td><td></td><td></td><td></td><td></td><td></td><td></td><td></td><td></td><td></td><td></td></tr>
<tr><td>Oligo rev</td><td>AAACCTCACGGAGTCCGCGCCGGC</td><td></td><td></td><td></td><td></td><td></td><td></td><td></td><td></td><td></td><td></td></tr>
<tr><td>Chromosome</td><td>start</td><td>end</td><td>strand</td><td>MM</td><td>target_seq</td><td>PAM</td><td>alignment</td><td>distance</td><td>position</td><td>gene name</td><td>gene id</td></tr>
<tr><td>GU305914.1</td><td>55969</td><td>55991</td><td>-</td><td>0</td><td>GCCGGCGCGGACTCCGTGAG</td><td>AGG</td><td>||||||||[||||||||||||]PAM</td><td>NA</td><td>Intergenic</td><td>NA</td><td>NA</td></tr>
<tr><td></td><td></td><td></td><td></td><td></td><td></td><td></td><td></td><td></td><td></td><td></td><td></td></tr>
<tr><td>T29850</td><td>CGCGGACTCCGTGAGAGGTGCGG</td><td>1000</td><td></td><td></td><td></td><td></td><td></td><td></td><td></td><td></td><td></td></tr>
<tr><td>Oligo fwd</td><td>CACCCGCGGACTCCGTGAGAGGTG</td><td></td><td></td><td></td><td></td><td></td><td></td><td></td><td></td><td></td><td></td></tr>
<tr><td>Oligo rev</td><td>AAACCACCTCTCACGGAGTCCGCG</td><td></td><td></td><td></td><td></td><td></td><td></td><td></td><td></td><td></td><td></td></tr>
<tr><td>Chromosome</td><td>start</td><td>end</td><td>strand</td><td>MM</td><td>target_seq</td><td>PAM</td><td>alignment</td><td>distance</td><td>position</td><td>gene name</td><td>gene id</td></tr>
<tr><td>GU305914.1</td><td>55964</td><td>55986</td><td>-</td><td>0</td><td>CGCGGACTCCGTGAGAGGTG</td><td>CGG</td><td>||||||||[||||||||||||]PAM</td><td>NA</td><td>Intergenic</td><td>NA</td><td>NA</td></tr>
<tr><td></td><td></td><td></td><td></td><td></td><td></td><td></td><td></td><td></td><td></td><td></td><td></td></tr>
<tr><td>T29851</td><td>CTCCGTGAGAGGTGCGGCATCGG</td><td>1000</td><td></td><td></td><td></td><td></td><td></td><td></td><td></td><td></td><td></td></tr>
<tr><td>Oligo fwd</td><td>CACCCTCCGTGAGAGGTGCGGCAT</td><td></td><td></td><td></td><td></td><td></td><td></td><td></td><td></td><td></td><td></td></tr>
<tr><td>Oligo rev</td><td>AAACATGCCGCACCTCTCACGGAG</td><td></td><td></td><td></td><td></td><td></td><td></td><td></td><td></td><td></td><td></td></tr>
<tr><td>Chromosome</td><td>start</td><td>end</td><td>strand</td><td>MM</td><td>target_seq</td><td>PAM</td><td>alignment</td><td>distance</td><td>position</td><td>gene name</td><td>gene id</td></tr>
<tr><td>GU305914.1</td><td>55958</td><td>55980</td><td>-</td><td>0</td><td>CTCCGTGAGAGGTGCGGCAT</td><td>CGG</td><td>||||||||[||||||||||||]PAM</td><td>NA</td><td>Intergenic</td><td>NA</td><td>NA</td></tr>
<tr><td></td><td></td><td></td><td></td><td></td><td></td><td></td><td></td><td></td><td></td><td></td><td></td></tr>
<tr><td>T29852</td><td>TGAGAGGTGCGGCATCGGCCTGG</td><td>1000</td><td></td><td></td><td></td><td></td><td></td><td></td><td></td><td></td><td></td></tr>
<tr><td>Oligo fwd</td><td>CACCTGAGAGGTGCGGCATCGGCC</td><td></td><td></td><td></td><td></td><td></td><td></td><td></td><td></td><td></td><td></td></tr>
<tr><td>Oligo rev</td><td>AAACGGCCGATGCCGCACCTCTCA</td><td></td><td></td><td></td><td></td><td></td><td></td><td></td><td></td><td></td><td></td></tr>
<tr><td>Chromosome</td><td>start</td><td>end</td><td>strand</td><td>MM</td><td>target_seq</td><td>PAM</td><td>alignment</td><td>distance</td><td>position</td><td>gene name</td><td>gene id</td></tr>
<tr><td>GU305914.1</td><td>55953</td><td>55975</td><td>-</td><td>0</td><td>TGAGAGGTGCGGCATCGGCC</td><td>TGG</td><td>||||||||[||||||||||||]PAM</td><td>NA</td><td>Intergenic</td><td>NA</td><td>NA</td></tr>
<tr><td></td><td></td><td></td><td></td><td></td><td></td><td></td><td></td><td></td><td></td><td></td><td></td></tr>
<tr><td>T29853</td><td>TGCGGCATCGGCCTGGACGCCGG</td><td>1000</td><td></td><td></td><td></td><td></td><td></td><td></td><td></td><td></td><td></td></tr>
<tr><td>Oligo fwd</td><td>CACCTGCGGCATCGGCCTGGACGC</td><td></td><td></td><td></td><td></td><td></td><td></td><td></td><td></td><td></td><td></td></tr>
<tr><td>Oligo rev</td><td>AAACGCGTCCAGGCCGATGCCGCA</td><td></td><td></td><td></td><td></td><td></td><td></td><td></td><td></td><td></td><td></td></tr>
<tr><td>Chromosome</td><td>start</td><td>end</td><td>strand</td><td>MM</td><td>target_seq</td><td>PAM</td><td>alignment</td><td>distance</td><td>position</td><td>gene name</td><td>gene id</td></tr>
<tr><td>GU305914.1</td><td>55946</td><td>55968</td><td>-</td><td>0</td><td>TGCGGCATCGGCCTGGACGC</td><td>CGG</td><td>||||||||[||||||||||||]PAM</td><td>NA</td><td>Intergenic</td><td>NA</td><td>NA</td></tr>
<tr><td></td><td></td><td></td><td></td><td></td><td></td><td></td><td></td><td></td><td></td><td></td><td></td></tr>
<tr><td>T29854</td><td>ATCGACGCCGATCGACTATACGG</td><td>1000</td><td></td><td></td><td></td><td></td><td></td><td></td><td></td><td></td><td></td></tr>
<tr><td>Oligo fwd</td><td>CACCATCGACGCCGATCGACTATA</td><td></td><td></td><td></td><td></td><td></td><td></td><td></td><td></td><td></td><td></td></tr>
<tr><td>Oligo rev</td><td>AAACTATAGTCGATCGGCGTCGAT</td><td></td><td></td><td></td><td></td><td></td><td></td><td></td><td></td><td></td><td></td></tr>
<tr><td>Chromosome</td><td>start</td><td>end</td><td>strand</td><td>MM</td><td>target_seq</td><td>PAM</td><td>alignment</td><td>distance</td><td>position</td><td>gene name</td><td>gene id</td></tr>
<tr><td>GU305914.1</td><td>55923</td><td>55945</td><td>-</td><td>0</td><td>ATCGACGCCGATCGACTATA</td><td>CGG</td><td>||||||||[||||||||||||]PAM</td><td>NA</td><td>Intergenic</td><td>NA</td><td>NA</td></tr>
<tr><td></td><td></td><td></td><td></td><td></td><td></td><td></td><td></td><td></td><td></td><td></td><td></td></tr>
<tr><td>T29855</td><td>GACGCCGATCGACTATACGGAGG</td><td>1000</td><td></td><td></td><td></td><td></td><td></td><td></td><td></td><td></td><td></td></tr>
<tr><td>Oligo fwd</td><td>CACCGACGCCGATCGACTATACGG</td><td></td><td></td><td></td><td></td><td></td><td></td><td></td><td></td><td></td><td></td></tr>
<tr><td>Oligo rev</td><td>AAACCCGTATAGTCGATCGGCGTC</td><td></td><td></td><td></td><td></td><td></td><td></td><td></td><td></td><td></td><td></td></tr>
<tr><td>Chromosome</td><td>start</td><td>end</td><td>strand</td><td>MM</td><td>target_seq</td><td>PAM</td><td>alignment</td><td>distance</td><td>position</td><td>gene name</td><td>gene id</td></tr>
<tr><td>GU305914.1</td><td>55920</td><td>55942</td><td>-</td><td>0</td><td>GACGCCGATCGACTATACGG</td><td>AGG</td><td>||||||||[||||||||||||]PAM</td><td>NA</td><td>Intergenic</td><td>NA</td><td>NA</td></tr>
<tr><td></td><td></td><td></td><td></td><td></td><td></td><td></td><td></td><td></td><td></td><td></td><td></td></tr>
<tr><td>T29856</td><td>TCAAGTCCATCCTCGAATCCAGG</td><td>1000</td><td></td><td></td><td></td><td></td><td></td><td></td><td></td><td></td><td></td></tr>
<tr><td>Oligo fwd</td><td>CACCTCAAGTCCATCCTCGAATCC</td><td></td><td></td><td></td><td></td><td></td><td></td><td></td><td></td><td></td><td></td></tr>
<tr><td>Oligo rev</td><td>AAACGGATTCGAGGATGGACTTGA</td><td></td><td></td><td></td><td></td><td></td><td></td><td></td><td></td><td></td><td></td></tr>
<tr><td>Chromosome</td><td>start</td><td>end</td><td>strand</td><td>MM</td><td>target_seq</td><td>PAM</td><td>alignment</td><td>distance</td><td>position</td><td>gene name</td><td>gene id</td></tr>
<tr><td>GU305914.1</td><td>55891</td><td>55913</td><td>-</td><td>0</td><td>TCAAGTCCATCCTCGAATCC</td><td>AGG</td><td>||||||||[||||||||||||]PAM</td><td>NA</td><td>Intergenic</td><td>NA</td><td>NA</td></tr>
<tr><td></td><td></td><td></td><td></td><td></td><td></td><td></td><td></td><td></td><td></td><td></td><td></td></tr>
<tr><td>T29857</td><td>CAGGAGCTCGATAGACGACCTGG</td><td>1000</td><td></td><td></td><td></td><td></td><td></td><td></td><td></td><td></td><td></td></tr>
<tr><td>Oligo fwd</td><td>CACCCAGGAGCTCGATAGACGACC</td><td></td><td></td><td></td><td></td><td></td><td></td><td></td><td></td><td></td><td></td></tr>
<tr><td>Oligo rev</td><td>AAACGGTCGTCTATCGAGCTCCTG</td><td></td><td></td><td></td><td></td><td></td><td></td><td></td><td></td><td></td><td></td></tr>
<tr><td>Chromosome</td><td>start</td><td>end</td><td>strand</td><td>MM</td><td>target_seq</td><td>PAM</td><td>alignment</td><td>distance</td><td>position</td><td>gene name</td><td>gene id</td></tr>
<tr><td>GU305914.1</td><td>55872</td><td>55894</td><td>-</td><td>0</td><td>CAGGAGCTCGATAGACGACC</td><td>TGG</td><td>||||||||[||||||||||||]PAM</td><td>NA</td><td>Intergenic</td><td>NA</td><td>NA</td></tr>
<tr><td></td><td></td><td></td><td></td><td></td><td></td><td></td><td></td><td></td><td></td><td></td><td></td></tr>
<tr><td>T29858</td><td>GTCAGCCGCGATCGCCTGCTCGG</td><td>1000</td><td></td><td></td><td></td><td></td><td></td><td></td><td></td><td></td><td></td></tr>
<tr><td>Oligo fwd</td><td>CACCGTCAGCCGCGATCGCCTGCT</td><td></td><td></td><td></td><td></td><td></td><td></td><td></td><td></td><td></td><td></td></tr>
<tr><td>Oligo rev</td><td>AAACAGCAGGCGATCGCGGCTGAC</td><td></td><td></td><td></td><td></td><td></td><td></td><td></td><td></td><td></td><td></td></tr>
<tr><td>Chromosome</td><td>start</td><td>end</td><td>strand</td><td>MM</td><td>target_seq</td><td>PAM</td><td>alignment</td><td>distance</td><td>position</td><td>gene name</td><td>gene id</td></tr>
<tr><td>GU305914.1</td><td>55847</td><td>55869</td><td>-</td><td>0</td><td>GTCAGCCGCGATCGCCTGCT</td><td>CGG</td><td>||||||||[||||||||||||]PAM</td><td>NA</td><td>Intergenic</td><td>NA</td><td>NA</td></tr>
<tr><td></td><td></td><td></td><td></td><td></td><td></td><td></td><td></td><td></td><td></td><td></td><td></td></tr>
<tr><td>T29859</td><td>CGAGACCGCCATCTTCCACTCGG</td><td>1000</td><td></td><td></td><td></td><td></td><td></td><td></td><td></td><td></td><td></td></tr>
<tr><td>Oligo fwd</td><td>CACCCGAGACCGCCATCTTCCACT</td><td></td><td></td><td></td><td></td><td></td><td></td><td></td><td></td><td></td><td></td></tr>
<tr><td>Oligo rev</td><td>AAACAGTGGAAGATGGCGGTCTCG</td><td></td><td></td><td></td><td></td><td></td><td></td><td></td><td></td><td></td><td></td></tr>
<tr><td>Chromosome</td><td>start</td><td>end</td><td>strand</td><td>MM</td><td>target_seq</td><td>PAM</td><td>alignment</td><td>distance</td><td>position</td><td>gene name</td><td>gene id</td></tr>
<tr><td>GU305914.1</td><td>55821</td><td>55843</td><td>-</td><td>0</td><td>CGAGACCGCCATCTTCCACT</td><td>CGG</td><td>||||||||[||||||||||||]PAM</td><td>NA</td><td>Intergenic</td><td>NA</td><td>NA</td></tr>
<tr><td></td><td></td><td></td><td></td><td></td><td></td><td></td><td></td><td></td><td></td><td></td><td></td></tr>
<tr><td>T29860</td><td>TCTTCCACTCGGATAGCATGTGG</td><td>1000</td><td></td><td></td><td></td><td></td><td></td><td></td><td></td><td></td><td></td></tr>
<tr><td>Oligo fwd</td><td>CACCTCTTCCACTCGGATAGCATG</td><td></td><td></td><td></td><td></td><td></td><td></td><td></td><td></td><td></td><td></td></tr>
<tr><td>Oligo rev</td><td>AAACCATGCTATCCGAGTGGAAGA</td><td></td><td></td><td></td><td></td><td></td><td></td><td></td><td></td><td></td><td></td></tr>
<tr><td>Chromosome</td><td>start</td><td>end</td><td>strand</td><td>MM</td><td>target_seq</td><td>PAM</td><td>alignment</td><td>distance</td><td>position</td><td>gene name</td><td>gene id</td></tr>
<tr><td>GU305914.1</td><td>55810</td><td>55832</td><td>-</td><td>0</td><td>TCTTCCACTCGGATAGCATG</td><td>TGG</td><td>||||||||[||||||||||||]PAM</td><td>NA</td><td>Intergenic</td><td>NA</td><td>NA</td></tr>
<tr><td></td><td></td><td></td><td></td><td></td><td></td><td></td><td></td><td></td><td></td><td></td><td></td></tr>
<tr><td>T29861</td><td>GTCAGCAACCTGTGCGACACCGG</td><td>1000</td><td></td><td></td><td></td><td></td><td></td><td></td><td></td><td></td><td></td></tr>
<tr><td>Oligo fwd</td><td>CACCGTCAGCAACCTGTGCGACAC</td><td></td><td></td><td></td><td></td><td></td><td></td><td></td><td></td><td></td><td></td></tr>
</table>

Oligo rev AAACGTGTCGCACAGGTTGCTGAC
Chromosome start end strand MM target_seq PAM alignment distance position gene name gene id
GU305914.1 55775 55797 - 0 GTCAGCAACCTGTGCGACAC CGG ||||||||[||||||||||||]PAM NA Intergenic NA NA

T29862 ACAGCTTCGACTACGAGTGCTGG 1000
Oligo fwd CACCACAGCTTCGACTACGAGTGC
Oligo rev AAACGCACTCGTAGTCGAAGCTGT
Chromosome start end strand MM target_seq PAM alignment distance position gene name gene id
GU305914.1 55747 55769 - 0 ACAGCTTCGACTACGAGTGC TGG ||||||||[||||||||||||]PAM NA Intergenic NA NA

T29863 GAGTGCTGGAAGCACGAGACCGG 1000
Oligo fwd CACCGAGTGCTGGAAGCACGAGAC
Oligo rev AAACGTCTCGTGCTTCCAGCACTC
Chromosome start end strand MM target_seq PAM alignment distance position gene name gene id
GU305914.1 55733 55755 - 0 GAGTGCTGGAAGCACGAGAC CGG ||||||||[||||||||||||]PAM NA Intergenic NA NA

T29864 CCGACTCGAGCCCGTCTTCATGG 1000
Oligo fwd CACCCCGACTCGAGCCCGTCTTCA
Oligo rev AAACTGAAGACGGGCTCGAGTCGG
Chromosome start end strand MM target_seq PAM alignment distance position gene name gene id
GU305914.1 55704 55726 - 0 CCGACTCGAGCCCGTCTTCA TGG ||||||||[||||||||||||]PAM NA Intergenic NA NA

T29865 CGAGCCCGTCTTCATGGTCAAGG 1000
Oligo fwd CACCCGAGCCCGTCTTCATGGTCA
Oligo rev AAACTGACCATGAAGACGGGCTCG
Chromosome start end strand MM target_seq PAM alignment distance position gene name gene id
GU305914.1 55698 55720 - 0 CGAGCCCGTCTTCATGGTCA AGG ||||||||[||||||||||||]PAM NA Intergenic NA NA

T29866 ATGGTCAAGGCGTGCGTGTCCGG 1000
Oligo fwd CACCATGGTCAAGGCGTGCGTGTC
Oligo rev AAACGACACGCACGCCTTGACCAT
Chromosome start end strand MM target_seq PAM alignment distance position gene name gene id
GU305914.1 55685 55707 - 0 ATGGTCAAGGCGTGCGTGTC CGG ||||||||[||||||||||||]PAM NA Intergenic NA NA

T29867 TGGTCAAGGCGTGCGTGTCCGGG 1000
Oligo fwd CACCTGGTCAAGGCGTGCGTGTCC
Oligo rev AAACGGACACGCACGCCTTGACCA
Chromosome start end strand MM target_seq PAM alignment distance position gene name gene id
GU305914.1 55684 55706 - 0 TGGTCAAGGCGTGCGTGTCC GGG ||||||||[||||||||||||]PAM NA Intergenic NA NA

T29868 GGTCAAGGCGTGCGTGTCCGGGG 1000
Oligo fwd CACCGGTCAAGGCGTGCGTGTCCG
Oligo rev AAACCGGACACGCACGCCTTGACC
Chromosome start end strand MM target_seq PAM alignment distance position gene name gene id
GU305914.1 55683 55705 - 0 GGTCAAGGCGTGCGTGTCCG GGG ||||||||[||||||||||||]PAM NA Intergenic NA NA

T29869 AGGCGTGCGTGTCCGGGGTCTGG 1000
Oligo fwd CACCAGGCGTGCGTGTCCGGGGTC
Oligo rev AAACGACCCCGGACACGCACGCCT
Chromosome start end strand MM target_seq PAM alignment distance position gene name gene id
GU305914.1 55678 55700 - 0 AGGCGTGCGTGTCCGGGGTC TGG ||||||||[||||||||||||]PAM NA Intergenic NA NA

T29870 GGCGTGCGTGTCCGGGGTCTGGG 1000
Oligo fwd CACCGGCGTGCGTGTCCGGGGTCT
Oligo rev AAACAGACCCCGGACACGCACGCC
Chromosome start end strand MM target_seq PAM alignment distance position gene name gene id
GU305914.1 55677 55699 - 0 GGCGTGCGTGTCCGGGGTCT GGG ||||||||[||||||||||||]PAM NA Intergenic NA NA

T29871 GCGCCCATCCTCGTGTCCGACGG 1000
Oligo fwd CACCGCGCCCATCCTCGTGTCCGA
Oligo rev AAACTCGGACACGAGGATGGGCGC
Chromosome start end strand MM target_seq PAM alignment distance position gene name gene id
GU305914.1 55649 55671 - 0 GCGCCCATCCTCGTGTCCGA CGG ||||||||[||||||||||||]PAM NA Intergenic NA NA

T29872 CCCATCCTCGTGTCCGACGGCGG 1000
Oligo fwd CACCCCCATCCTCGTGTCCGACGG
Oligo rev AAACCCGTCGGACACGAGGATGGG
Chromosome start end strand MM target_seq PAM alignment distance position gene name gene id
GU305914.1 55646 55668 - 0 CCCATCCTCGTGTCCGACGG CGG ||||||||[||||||||||||]PAM NA Intergenic NA NA

T29873 CCATCCTCGTGTCCGACGGCGGG 1000
Oligo fwd CACCCCATCCTCGTGTCCGACGGC
Oligo rev AAACGCCGTCGGACACGAGGATGG
Chromosome start end strand MM target_seq PAM alignment distance position gene name gene id
GU305914.1 55645 55667 - 0 CCATCCTCGTGTCCGACGGC GGG ||||||||[||||||||||||]PAM NA Intergenic NA NA

T29874 GACCGTCTACTACGTCGACCCGG 1000
Oligo fwd CACCGACCGTCTACTACGTCGACC
Oligo rev AAACGGTCGACGTAGTAGACGGTC
Chromosome start end strand MM target_seq PAM alignment distance position gene name gene id
GU305914.1 55623 55645 - 0 GACCGTCTACTACGTCGACC CGG ||||||||[||||||||||||]PAM NA Intergenic NA NA

T29875 ACCGTCTACTACGTCGACCCGGG 1000
Oligo fwd CACCACCGTCTACTACGTCGACCC
Oligo rev AAACGGGTCGACGTAGTAGACGGT
Chromosome start end strand MM target_seq PAM alignment distance position gene name gene id
GU305914.1 55622 55644 - 0 ACCGTCTACTACGTCGACCC GGG ||||||||[||||||||||||]PAM NA Intergenic NA NA

T29876 CCGTCTACTACGTCGACCCGGGG 1000

<table>
<tr><td>Oligo fwd</td><td>CACCCCGTCTACTACGTCGACCCG</td><td></td><td></td><td></td><td></td><td></td><td></td><td></td><td></td><td></td><td></td></tr>
<tr><td>Oligo rev</td><td>AAACCGGGTCGACGTAGTAGACGG</td><td></td><td></td><td></td><td></td><td></td><td></td><td></td><td></td><td></td><td></td></tr>
<tr><td>Chromosome</td><td>start</td><td>end</td><td>strand</td><td>MM</td><td>target_seq</td><td>PAM</td><td>alignment</td><td>distance</td><td>position</td><td>gene name</td><td>gene id</td></tr>
<tr><td>GU305914.1</td><td>55621</td><td>55643</td><td>-</td><td>0</td><td>CCGTCTACTACGTCGACCCG</td><td>GGG</td><td>||||||||[||||||||||||]PAM</td><td>NA</td><td>Intergenic</td><td>NA</td><td>NA</td></tr>
<tr><td></td><td></td><td></td><td></td><td></td><td></td><td></td><td></td><td></td><td></td><td></td><td></td></tr>
<tr><td>T29877</td><td>TACTACGTCGACCCGGGGCACGG</td><td>1000</td><td></td><td></td><td></td><td></td><td></td><td></td><td></td><td></td><td></td></tr>
<tr><td>Oligo fwd</td><td>CACCTACTACGTCGACCCGGGGCA</td><td></td><td></td><td></td><td></td><td></td><td></td><td></td><td></td><td></td><td></td></tr>
<tr><td>Oligo rev</td><td>AAACTGCCCCGGGTCGACGTAGTA</td><td></td><td></td><td></td><td></td><td></td><td></td><td></td><td></td><td></td><td></td></tr>
<tr><td>Chromosome</td><td>start</td><td>end</td><td>strand</td><td>MM</td><td>target_seq</td><td>PAM</td><td>alignment</td><td>distance</td><td>position</td><td>gene name</td><td>gene id</td></tr>
<tr><td>GU305914.1</td><td>55616</td><td>55638</td><td>-</td><td>0</td><td>TACTACGTCGACCCGGGGCA</td><td>CGG</td><td>||||||||[||||||||||||]PAM</td><td>NA</td><td>Intergenic</td><td>NA</td><td>NA</td></tr>
<tr><td></td><td></td><td></td><td></td><td></td><td></td><td></td><td></td><td></td><td></td><td></td><td></td></tr>
<tr><td>T29878</td><td>CTCTTCACCTTCCTCGTGCTCGG</td><td>1000</td><td></td><td></td><td></td><td></td><td></td><td></td><td></td><td></td><td></td></tr>
<tr><td>Oligo fwd</td><td>CACCCTCTTCACCTTCCTCGTGCT</td><td></td><td></td><td></td><td></td><td></td><td></td><td></td><td></td><td></td><td></td></tr>
<tr><td>Oligo rev</td><td>AAACAGCACGAGGAAGGTGAAGAG</td><td></td><td></td><td></td><td></td><td></td><td></td><td></td><td></td><td></td><td></td></tr>
<tr><td>Chromosome</td><td>start</td><td>end</td><td>strand</td><td>MM</td><td>target_seq</td><td>PAM</td><td>alignment</td><td>distance</td><td>position</td><td>gene name</td><td>gene id</td></tr>
<tr><td>GU305914.1</td><td>55568</td><td>55590</td><td>-</td><td>0</td><td>CTCTTCACCTTCCTCGTGCT</td><td>CGG</td><td>||||||||[||||||||||||]PAM</td><td>NA</td><td>Intergenic</td><td>NA</td><td>NA</td></tr>
<tr><td></td><td></td><td></td><td></td><td></td><td></td><td></td><td></td><td></td><td></td><td></td><td></td></tr>
<tr><td>T29879</td><td>CGACTCGAAGACCCGCGCCCCGG</td><td>1000</td><td></td><td></td><td></td><td></td><td></td><td></td><td></td><td></td><td></td></tr>
<tr><td>Oligo fwd</td><td>CACCCGACTCGAAGACCCGCGCCC</td><td></td><td></td><td></td><td></td><td></td><td></td><td></td><td></td><td></td><td></td></tr>
<tr><td>Oligo rev</td><td>AAACGGGCGCGGGTCTTCGAGTCG</td><td></td><td></td><td></td><td></td><td></td><td></td><td></td><td></td><td></td><td></td></tr>
<tr><td>Chromosome</td><td>start</td><td>end</td><td>strand</td><td>MM</td><td>target_seq</td><td>PAM</td><td>alignment</td><td>distance</td><td>position</td><td>gene name</td><td>gene id</td></tr>
<tr><td>GU305914.1</td><td>55490</td><td>55512</td><td>-</td><td>0</td><td>CGACTCGAAGACCCGCGCCC</td><td>CGG</td><td>||||||||[||||||||||||]PAM</td><td>NA</td><td>Intergenic</td><td>NA</td><td>NA</td></tr>
<tr><td></td><td></td><td></td><td></td><td></td><td></td><td></td><td></td><td></td><td></td><td></td><td></td></tr>
<tr><td>T29880</td><td>CGCGCCCCGGACAGCCCATCAGG</td><td>1000</td><td></td><td></td><td></td><td></td><td></td><td></td><td></td><td></td><td></td></tr>
<tr><td>Oligo fwd</td><td>CACCCGCGCCCCGGACAGCCCATC</td><td></td><td></td><td></td><td></td><td></td><td></td><td></td><td></td><td></td><td></td></tr>
<tr><td>Oligo rev</td><td>AAACGATGGGCTGTCCGGGGCGCG</td><td></td><td></td><td></td><td></td><td></td><td></td><td></td><td></td><td></td><td></td></tr>
<tr><td>Chromosome</td><td>start</td><td>end</td><td>strand</td><td>MM</td><td>target_seq</td><td>PAM</td><td>alignment</td><td>distance</td><td>position</td><td>gene name</td><td>gene id</td></tr>
<tr><td>GU305914.1</td><td>55477</td><td>55499</td><td>-</td><td>0</td><td>CGCGCCCCGGACAGCCCATC</td><td>AGG</td><td>||||||||[||||||||||||]PAM</td><td>NA</td><td>Intergenic</td><td>NA</td><td>NA</td></tr>
<tr><td></td><td></td><td></td><td></td><td></td><td></td><td></td><td></td><td></td><td></td><td></td><td></td></tr>
<tr><td>T29881</td><td>CCCCGGACAGCCCATCAGGCCGG</td><td>1000</td><td></td><td></td><td></td><td></td><td></td><td></td><td></td><td></td><td></td></tr>
<tr><td>Oligo fwd</td><td>CACCCCCCGGACAGCCCATCAGGC</td><td></td><td></td><td></td><td></td><td></td><td></td><td></td><td></td><td></td><td></td></tr>
<tr><td>Oligo rev</td><td>AAACGCCTGATGGGCTGTCCGGGG</td><td></td><td></td><td></td><td></td><td></td><td></td><td></td><td></td><td></td><td></td></tr>
<tr><td>Chromosome</td><td>start</td><td>end</td><td>strand</td><td>MM</td><td>target_seq</td><td>PAM</td><td>alignment</td><td>distance</td><td>position</td><td>gene name</td><td>gene id</td></tr>
<tr><td>GU305914.1</td><td>55473</td><td>55495</td><td>-</td><td>0</td><td>CCCCGGACAGCCCATCAGGC</td><td>CGG</td><td>||||||||[||||||||||||]PAM</td><td>NA</td><td>Intergenic</td><td>NA</td><td>NA</td></tr>
<tr><td></td><td></td><td></td><td></td><td></td><td></td><td></td><td></td><td></td><td></td><td></td><td></td></tr>
<tr><td>T29882</td><td>AGGCCGGTCTTCTGTCCGCTCGG</td><td>1000</td><td></td><td></td><td></td><td></td><td></td><td></td><td></td><td></td><td></td></tr>
<tr><td>Oligo fwd</td><td>CACCAGGCCGGTCTTCTGTCCGCT</td><td></td><td></td><td></td><td></td><td></td><td></td><td></td><td></td><td></td><td></td></tr>
<tr><td>Oligo rev</td><td>AAACAGCGGACAGAAGACCGGCCT</td><td></td><td></td><td></td><td></td><td></td><td></td><td></td><td></td><td></td><td></td></tr>
<tr><td>Chromosome</td><td>start</td><td>end</td><td>strand</td><td>MM</td><td>target_seq</td><td>PAM</td><td>alignment</td><td>distance</td><td>position</td><td>gene name</td><td>gene id</td></tr>
<tr><td>GU305914.1</td><td>55457</td><td>55479</td><td>-</td><td>0</td><td>AGGCCGGTCTTCTGTCCGCT</td><td>CGG</td><td>||||||||[||||||||||||]PAM</td><td>NA</td><td>Intergenic</td><td>NA</td><td>NA</td></tr>
<tr><td></td><td></td><td></td><td></td><td></td><td></td><td></td><td></td><td></td><td></td><td></td><td></td></tr>
<tr><td>T29883</td><td>GGCCGGTCTTCTGTCCGCTCGGG</td><td>1000</td><td></td><td></td><td></td><td></td><td></td><td></td><td></td><td></td><td></td></tr>
<tr><td>Oligo fwd</td><td>CACCGGCCGGTCTTCTGTCCGCTC</td><td></td><td></td><td></td><td></td><td></td><td></td><td></td><td></td><td></td><td></td></tr>
<tr><td>Oligo rev</td><td>AAACGAGCGGACAGAAGACCGGCC</td><td></td><td></td><td></td><td></td><td></td><td></td><td></td><td></td><td></td><td></td></tr>
<tr><td>Chromosome</td><td>start</td><td>end</td><td>strand</td><td>MM</td><td>target_seq</td><td>PAM</td><td>alignment</td><td>distance</td><td>position</td><td>gene name</td><td>gene id</td></tr>
<tr><td>GU305914.1</td><td>55456</td><td>55478</td><td>-</td><td>0</td><td>GGCCGGTCTTCTGTCCGCTC</td><td>GGG</td><td>||||||||[||||||||||||]PAM</td><td>NA</td><td>Intergenic</td><td>NA</td><td>NA</td></tr>
<tr><td></td><td></td><td></td><td></td><td></td><td></td><td></td><td></td><td></td><td></td><td></td><td></td></tr>
<tr><td>T29884</td><td>TCTTCTGTCCGCTCGGGCAGAGG</td><td>1000</td><td></td><td></td><td></td><td></td><td></td><td></td><td></td><td></td><td></td></tr>
<tr><td>Oligo fwd</td><td>CACCTCTTCTGTCCGCTCGGGCAG</td><td></td><td></td><td></td><td></td><td></td><td></td><td></td><td></td><td></td><td></td></tr>
<tr><td>Oligo rev</td><td>AAACCTGCCCGAGCGGACAGAAGA</td><td></td><td></td><td></td><td></td><td></td><td></td><td></td><td></td><td></td><td></td></tr>
<tr><td>Chromosome</td><td>start</td><td>end</td><td>strand</td><td>MM</td><td>target_seq</td><td>PAM</td><td>alignment</td><td>distance</td><td>position</td><td>gene name</td><td>gene id</td></tr>
<tr><td>GU305914.1</td><td>55450</td><td>55472</td><td>-</td><td>0</td><td>TCTTCTGTCCGCTCGGGCAG</td><td>AGG</td><td>||||||||[||||||||||||]PAM</td><td>NA</td><td>Intergenic</td><td>NA</td><td>NA</td></tr>
<tr><td></td><td></td><td></td><td></td><td></td><td></td><td></td><td></td><td></td><td></td><td></td><td></td></tr>
<tr><td>T29885</td><td>AGAGGTGCCGACGCAAGAGAAGG</td><td>1000</td><td></td><td></td><td></td><td></td><td></td><td></td><td></td><td></td><td></td></tr>
<tr><td>Oligo fwd</td><td>CACCAGAGGTGCCGACGCAAGAGA</td><td></td><td></td><td></td><td></td><td></td><td></td><td></td><td></td><td></td><td></td></tr>
<tr><td>Oligo rev</td><td>AAACTCTCTTGCGTCGGCACCTCT</td><td></td><td></td><td></td><td></td><td></td><td></td><td></td><td></td><td></td><td></td></tr>
<tr><td>Chromosome</td><td>start</td><td>end</td><td>strand</td><td>MM</td><td>target_seq</td><td>PAM</td><td>alignment</td><td>distance</td><td>position</td><td>gene name</td><td>gene id</td></tr>
<tr><td>GU305914.1</td><td>55432</td><td>55454</td><td>-</td><td>0</td><td>AGAGGTGCCGACGCAAGAGA</td><td>AGG</td><td>||||||||[||||||||||||]PAM</td><td>NA</td><td>Intergenic</td><td>NA</td><td>NA</td></tr>
<tr><td></td><td></td><td></td><td></td><td></td><td></td><td></td><td></td><td></td><td></td><td></td><td></td></tr>
<tr><td>T29886</td><td>GAGGTGCCGACGCAAGAGAAGGG</td><td>1000</td><td></td><td></td><td></td><td></td><td></td><td></td><td></td><td></td><td></td></tr>
<tr><td>Oligo fwd</td><td>CACCGAGGTGCCGACGCAAGAGAA</td><td></td><td></td><td></td><td></td><td></td><td></td><td></td><td></td><td></td><td></td></tr>
<tr><td>Oligo rev</td><td>AAACTTCTCTTGCGTCGGCACCTC</td><td></td><td></td><td></td><td></td><td></td><td></td><td></td><td></td><td></td><td></td></tr>
<tr><td>Chromosome</td><td>start</td><td>end</td><td>strand</td><td>MM</td><td>target_seq</td><td>PAM</td><td>alignment</td><td>distance</td><td>position</td><td>gene name</td><td>gene id</td></tr>
<tr><td>GU305914.1</td><td>55431</td><td>55453</td><td>-</td><td>0</td><td>GAGGTGCCGACGCAAGAGAA</td><td>GGG</td><td>||||||||[||||||||||||]PAM</td><td>NA</td><td>Intergenic</td><td>NA</td><td>NA</td></tr>
<tr><td></td><td></td><td></td><td></td><td></td><td></td><td></td><td></td><td></td><td></td><td></td><td></td></tr>
<tr><td>T29887</td><td>TCTCGATCATCAGTCGTTTCTGG</td><td>1000</td><td></td><td></td><td></td><td></td><td></td><td></td><td></td><td></td><td></td></tr>
<tr><td>Oligo fwd</td><td>CACCTCTCGATCATCAGTCGTTTC</td><td></td><td></td><td></td><td></td><td></td><td></td><td></td><td></td><td></td><td></td></tr>
<tr><td>Oligo rev</td><td>AAACGAAACGACTGATGATCGAGA</td><td></td><td></td><td></td><td></td><td></td><td></td><td></td><td></td><td></td><td></td></tr>
<tr><td>Chromosome</td><td>start</td><td>end</td><td>strand</td><td>MM</td><td>target_seq</td><td>PAM</td><td>alignment</td><td>distance</td><td>position</td><td>gene name</td><td>gene id</td></tr>
<tr><td>GU305914.1</td><td>55408</td><td>55430</td><td>-</td><td>0</td><td>TCTCGATCATCAGTCGTTTC</td><td>TGG</td><td>||||||||[||||||||||||]PAM</td><td>NA</td><td>Intergenic</td><td>NA</td><td>NA</td></tr>
<tr><td></td><td></td><td></td><td></td><td></td><td></td><td></td><td></td><td></td><td></td><td></td><td></td></tr>
<tr><td>T29888</td><td>TCATCAGTCGTTTCTGGCACTGG</td><td>1000</td><td></td><td></td><td></td><td></td><td></td><td></td><td></td><td></td><td></td></tr>
<tr><td>Oligo fwd</td><td>CACCTCATCAGTCGTTTCTGGCAC</td><td></td><td></td><td></td><td></td><td></td><td></td><td></td><td></td><td></td><td></td></tr>
<tr><td>Oligo rev</td><td>AAACGTGCCAGAAACGACTGATGA</td><td></td><td></td><td></td><td></td><td></td><td></td><td></td><td></td><td></td><td></td></tr>
<tr><td>Chromosome</td><td>start</td><td>end</td><td>strand</td><td>MM</td><td>target_seq</td><td>PAM</td><td>alignment</td><td>distance</td><td>position</td><td>gene name</td><td>gene id</td></tr>
<tr><td>GU305914.1</td><td>55402</td><td>55424</td><td>-</td><td>0</td><td>TCATCAGTCGTTTCTGGCAC</td><td>TGG</td><td>||||||||[||||||||||||]PAM</td><td>NA</td><td>Intergenic</td><td>NA</td><td>NA</td></tr>
<tr><td></td><td></td><td></td><td></td><td></td><td></td><td></td><td></td><td></td><td></td><td></td><td></td></tr>
<tr><td>T29889</td><td>GTCGTTTCTGGCACTGGCTCTGG</td><td>1000</td><td></td><td></td><td></td><td></td><td></td><td></td><td></td><td></td><td></td></tr>
<tr><td>Oligo fwd</td><td>CACCGTCGTTTCTGGCACTGGCTC</td><td></td><td></td><td></td><td></td><td></td><td></td><td></td><td></td><td></td><td></td></tr>
<tr><td>Oligo rev</td><td>AAACGAGCCAGTGCCAGAAACGAC</td><td></td><td></td><td></td><td></td><td></td><td></td><td></td><td></td><td></td><td></td></tr>
<tr><td>Chromosome</td><td>start</td><td>end</td><td>strand</td><td>MM</td><td>target_seq</td><td>PAM</td><td>alignment</td><td>distance</td><td>position</td><td>gene name</td><td>gene id</td></tr>
<tr><td>GU305914.1</td><td>55396</td><td>55418</td><td>-</td><td>0</td><td>GTCGTTTCTGGCACTGGCTC</td><td>TGG</td><td>||||||||[||||||||||||]PAM</td><td>NA</td><td>Intergenic</td><td>NA</td><td>NA</td></tr>
<tr><td></td><td></td><td></td><td></td><td></td><td></td><td></td><td></td><td></td><td></td><td></td><td></td></tr>
<tr><td>T29890</td><td>TCGTTTCTGGCACTGGCTCTGGG</td><td>1000</td><td></td><td></td><td></td><td></td><td></td><td></td><td></td><td></td><td></td></tr>
<tr><td>Oligo fwd</td><td>CACCTCGTTTCTGGCACTGGCTCT</td><td></td><td></td><td></td><td></td><td></td><td></td><td></td><td></td><td></td><td></td></tr>
<tr><td>Oligo rev</td><td>AAACAGAGCCAGTGCCAGAAACGA</td><td></td><td></td><td></td><td></td><td></td><td></td><td></td><td></td><td></td><td></td></tr>
<tr><td>Chromosome</td><td>start</td><td>end</td><td>strand</td><td>MM</td><td>target_seq</td><td>PAM</td><td>alignment</td><td>distance</td><td>position</td><td>gene name</td><td>gene id</td></tr>
<tr><td>GU305914.1</td><td>55395</td><td>55417</td><td>-</td><td>0</td><td>TCGTTTCTGGCACTGGCTCT</td><td>GGG</td><td>||||||||[||||||||||||]PAM</td><td>NA</td><td>Intergenic</td><td>NA</td><td>NA</td></tr>
</table>

<table>
<tr><td>T29891</td><td>CGTTTCTGGCACTGGCTCTGGGG</td><td>1000</td><td></td><td></td><td></td><td></td><td></td><td></td><td></td><td></td><td></td></tr>
<tr><td>Oligo fwd</td><td>CACCCGTTTCTGGCACTGGCTCTG</td><td></td><td></td><td></td><td></td><td></td><td></td><td></td><td></td><td></td><td></td></tr>
<tr><td>Oligo rev</td><td>AAACCAGAGCCAGTGCCAGAAACG</td><td></td><td></td><td></td><td></td><td></td><td></td><td></td><td></td><td></td><td></td></tr>
<tr><td>Chromosome</td><td>start</td><td>end</td><td>strand</td><td>MM</td><td>target_seq</td><td>PAM</td><td>alignment</td><td>distance</td><td>position</td><td>gene name</td><td>gene id</td></tr>
<tr><td>GU305914.1</td><td>55394</td><td>55416</td><td>-</td><td>0</td><td>CGTTTCTGGCACTGGCTCTG</td><td>GGG</td><td>||||||||[||||||||||||]PAM</td><td>NA</td><td>Intergenic</td><td>NA</td><td>NA</td></tr>
<tr><td>T29892</td><td>GGCACTGGCTCTGGGGCACGTGG</td><td>1000</td><td></td><td></td><td></td><td></td><td></td><td></td><td></td><td></td><td></td></tr>
<tr><td>Oligo fwd</td><td>CACCGGCACTGGCTCTGGGGCACG</td><td></td><td></td><td></td><td></td><td></td><td></td><td></td><td></td><td></td><td></td></tr>
<tr><td>Oligo rev</td><td>AAACCGTGCCCCAGAGCCAGTGCC</td><td></td><td></td><td></td><td></td><td></td><td></td><td></td><td></td><td></td><td></td></tr>
<tr><td>Chromosome</td><td>start</td><td>end</td><td>strand</td><td>MM</td><td>target_seq</td><td>PAM</td><td>alignment</td><td>distance</td><td>position</td><td>gene name</td><td>gene id</td></tr>
<tr><td>GU305914.1</td><td>55387</td><td>55409</td><td>-</td><td>0</td><td>GGCACTGGCTCTGGGGCACG</td><td>TGG</td><td>||||||||[||||||||||||]PAM</td><td>NA</td><td>Intergenic</td><td>NA</td><td>NA</td></tr>
<tr><td>T29893</td><td>GCACTGGCTCTGGGGCACGTGGG</td><td>1000</td><td></td><td></td><td></td><td></td><td></td><td></td><td></td><td></td><td></td></tr>
<tr><td>Oligo fwd</td><td>CACCGCACTGGCTCTGGGGCACGT</td><td></td><td></td><td></td><td></td><td></td><td></td><td></td><td></td><td></td><td></td></tr>
<tr><td>Oligo rev</td><td>AAACACGTGCCCCAGAGCCAGTGC</td><td></td><td></td><td></td><td></td><td></td><td></td><td></td><td></td><td></td><td></td></tr>
<tr><td>Chromosome</td><td>start</td><td>end</td><td>strand</td><td>MM</td><td>target_seq</td><td>PAM</td><td>alignment</td><td>distance</td><td>position</td><td>gene name</td><td>gene id</td></tr>
<tr><td>GU305914.1</td><td>55386</td><td>55408</td><td>-</td><td>0</td><td>GCACTGGCTCTGGGGCACGT</td><td>GGG</td><td>||||||||[||||||||||||]PAM</td><td>NA</td><td>Intergenic</td><td>NA</td><td>NA</td></tr>
<tr><td>T29894</td><td>CACTGGCTCTGGGGCACGTGGGG</td><td>1000</td><td></td><td></td><td></td><td></td><td></td><td></td><td></td><td></td><td></td></tr>
<tr><td>Oligo fwd</td><td>CACCCACTGGCTCTGGGGCACGTG</td><td></td><td></td><td></td><td></td><td></td><td></td><td></td><td></td><td></td><td></td></tr>
<tr><td>Oligo rev</td><td>AAACCACGTGCCCCAGAGCCAGTG</td><td></td><td></td><td></td><td></td><td></td><td></td><td></td><td></td><td></td><td></td></tr>
<tr><td>Chromosome</td><td>start</td><td>end</td><td>strand</td><td>MM</td><td>target_seq</td><td>PAM</td><td>alignment</td><td>distance</td><td>position</td><td>gene name</td><td>gene id</td></tr>
<tr><td>GU305914.1</td><td>55385</td><td>55407</td><td>-</td><td>0</td><td>CACTGGCTCTGGGGCACGTG</td><td>GGG</td><td>||||||||[||||||||||||]PAM</td><td>NA</td><td>Intergenic</td><td>NA</td><td>NA</td></tr>
<tr><td>T29895</td><td>GGGCACGTGGGGCTATGTGACGG</td><td>1000</td><td></td><td></td><td></td><td></td><td></td><td></td><td></td><td></td><td></td></tr>
<tr><td>Oligo fwd</td><td>CACCGGGCACGTGGGGCTATGTGA</td><td></td><td></td><td></td><td></td><td></td><td></td><td></td><td></td><td></td><td></td></tr>
<tr><td>Oligo rev</td><td>AAACTCACATAGCCCCACGTGCCC</td><td></td><td></td><td></td><td></td><td></td><td></td><td></td><td></td><td></td><td></td></tr>
<tr><td>Chromosome</td><td>start</td><td>end</td><td>strand</td><td>MM</td><td>target_seq</td><td>PAM</td><td>alignment</td><td>distance</td><td>position</td><td>gene name</td><td>gene id</td></tr>
<tr><td>GU305914.1</td><td>55374</td><td>55396</td><td>-</td><td>0</td><td>GGGCACGTGGGGCTATGTGA</td><td>CGG</td><td>||||||||[||||||||||||]PAM</td><td>NA</td><td>Intergenic</td><td>NA</td><td>NA</td></tr>
<tr><td>T29896</td><td>GTTGCCGACATCAATAAAAACGG</td><td>1000</td><td></td><td></td><td></td><td></td><td></td><td></td><td></td><td></td><td></td></tr>
<tr><td>Oligo fwd</td><td>CACCGTTGCCGACATCAATAAAAA</td><td></td><td></td><td></td><td></td><td></td><td></td><td></td><td></td><td></td><td></td></tr>
<tr><td>Oligo rev</td><td>AAACTTTTTATTGATGTCGGCAAC</td><td></td><td></td><td></td><td></td><td></td><td></td><td></td><td></td><td></td><td></td></tr>
<tr><td>Chromosome</td><td>start</td><td>end</td><td>strand</td><td>MM</td><td>target_seq</td><td>PAM</td><td>alignment</td><td>distance</td><td>position</td><td>gene name</td><td>gene id</td></tr>
<tr><td>GU305914.1</td><td>55348</td><td>55370</td><td>-</td><td>0</td><td>GTTGCCGACATCAATAAAAA</td><td>CGG</td><td>||||||||[||||||||||||]PAM</td><td>NA</td><td>Intergenic</td><td>NA</td><td>NA</td></tr>
<tr><td>T29897</td><td>TTGCCGACATCAATAAAAACGGG</td><td>1000</td><td></td><td></td><td></td><td></td><td></td><td></td><td></td><td></td><td></td></tr>
<tr><td>Oligo fwd</td><td>CACCTTGCCGACATCAATAAAAAC</td><td></td><td></td><td></td><td></td><td></td><td></td><td></td><td></td><td></td><td></td></tr>
<tr><td>Oligo rev</td><td>AAACGTTTTTATTGATGTCGGCAA</td><td></td><td></td><td></td><td></td><td></td><td></td><td></td><td></td><td></td><td></td></tr>
<tr><td>Chromosome</td><td>start</td><td>end</td><td>strand</td><td>MM</td><td>target_seq</td><td>PAM</td><td>alignment</td><td>distance</td><td>position</td><td>gene name</td><td>gene id</td></tr>
<tr><td>GU305914.1</td><td>55347</td><td>55369</td><td>-</td><td>0</td><td>TTGCCGACATCAATAAAAAC</td><td>GGG</td><td>||||||||[||||||||||||]PAM</td><td>NA</td><td>Intergenic</td><td>NA</td><td>NA</td></tr>
<tr><td>T29898</td><td>TCAATAAAAACGGGACGCTGCGG</td><td>1000</td><td></td><td></td><td></td><td></td><td></td><td></td><td></td><td></td><td></td></tr>
<tr><td>Oligo fwd</td><td>CACCTCAATAAAAACGGGACGCTG</td><td></td><td></td><td></td><td></td><td></td><td></td><td></td><td></td><td></td><td></td></tr>
<tr><td>Oligo rev</td><td>AAACCAGCGTCCCGTTTTTATTGA</td><td></td><td></td><td></td><td></td><td></td><td></td><td></td><td></td><td></td><td></td></tr>
<tr><td>Chromosome</td><td>start</td><td>end</td><td>strand</td><td>MM</td><td>target_seq</td><td>PAM</td><td>alignment</td><td>distance</td><td>position</td><td>gene name</td><td>gene id</td></tr>
<tr><td>GU305914.1</td><td>55338</td><td>55360</td><td>-</td><td>0</td><td>TCAATAAAAACGGGACGCTG</td><td>CGG</td><td>||||||||[||||||||||||]PAM</td><td>NA</td><td>Intergenic</td><td>NA</td><td>NA</td></tr>
<tr><td>T29899</td><td>GCGCTCTCCTTCCTCTCTCCCGG</td><td>1000</td><td></td><td></td><td></td><td></td><td></td><td></td><td></td><td></td><td></td></tr>
<tr><td>Oligo fwd</td><td>CACCGCGCTCTCCTTCCTCTCTCC</td><td></td><td></td><td></td><td></td><td></td><td></td><td></td><td></td><td></td><td></td></tr>
<tr><td>Oligo rev</td><td>AAACGGAGAGAGGAAGGAGAGCGC</td><td></td><td></td><td></td><td></td><td></td><td></td><td></td><td></td><td></td><td></td></tr>
<tr><td>Chromosome</td><td>start</td><td>end</td><td>strand</td><td>MM</td><td>target_seq</td><td>PAM</td><td>alignment</td><td>distance</td><td>position</td><td>gene name</td><td>gene id</td></tr>
<tr><td>GU305914.1</td><td>55236</td><td>55258</td><td>-</td><td>0</td><td>GCGCTCTCCTTCCTCTCTCC</td><td>CGG</td><td>||||||||[||||||||||||]PAM</td><td>NA</td><td>Intergenic</td><td>NA</td><td>NA</td></tr>
<tr><td>T29900</td><td>CGCTCTCCTTCCTCTCTCCCGGG</td><td>1000</td><td></td><td></td><td></td><td></td><td></td><td></td><td></td><td></td><td></td></tr>
<tr><td>Oligo fwd</td><td>CACCCGCTCTCCTTCCTCTCTCCC</td><td></td><td></td><td></td><td></td><td></td><td></td><td></td><td></td><td></td><td></td></tr>
<tr><td>Oligo rev</td><td>AAACGGGAGAGAGGAAGGAGAGCG</td><td></td><td></td><td></td><td></td><td></td><td></td><td></td><td></td><td></td><td></td></tr>
<tr><td>Chromosome</td><td>start</td><td>end</td><td>strand</td><td>MM</td><td>target_seq</td><td>PAM</td><td>alignment</td><td>distance</td><td>position</td><td>gene name</td><td>gene id</td></tr>
<tr><td>GU305914.1</td><td>55235</td><td>55257</td><td>-</td><td>0</td><td>CGCTCTCCTTCCTCTCTCCC</td><td>GGG</td><td>||||||||[||||||||||||]PAM</td><td>NA</td><td>Intergenic</td><td>NA</td><td>NA</td></tr>
<tr><td>T29901</td><td>GCTCTCCTTCCTCTCTCCCGGGG</td><td>1000</td><td></td><td></td><td></td><td></td><td></td><td></td><td></td><td></td><td></td></tr>
<tr><td>Oligo fwd</td><td>CACCGCTCTCCTTCCTCTCTCCCG</td><td></td><td></td><td></td><td></td><td></td><td></td><td></td><td></td><td></td><td></td></tr>
<tr><td>Oligo rev</td><td>AAACCGGGAGAGAGGAAGGAGAGC</td><td></td><td></td><td></td><td></td><td></td><td></td><td></td><td></td><td></td><td></td></tr>
<tr><td>Chromosome</td><td>start</td><td>end</td><td>strand</td><td>MM</td><td>target_seq</td><td>PAM</td><td>alignment</td><td>distance</td><td>position</td><td>gene name</td><td>gene id</td></tr>
<tr><td>GU305914.1</td><td>55234</td><td>55256</td><td>-</td><td>0</td><td>GCTCTCCTTCCTCTCTCCCG</td><td>GGG</td><td>||||||||[||||||||||||]PAM</td><td>NA</td><td>Intergenic</td><td>NA</td><td>NA</td></tr>
<tr><td>T29902</td><td>TCCTCTCTCCCGGGGAGAAGAGG</td><td>1000</td><td></td><td></td><td></td><td></td><td></td><td></td><td></td><td></td><td></td></tr>
<tr><td>Oligo fwd</td><td>CACCTCCTCTCTCCCGGGGAGAAG</td><td></td><td></td><td></td><td></td><td></td><td></td><td></td><td></td><td></td><td></td></tr>
<tr><td>Oligo rev</td><td>AAACCTTCTCCCCGGGAGAGAGGA</td><td></td><td></td><td></td><td></td><td></td><td></td><td></td><td></td><td></td><td></td></tr>
<tr><td>Chromosome</td><td>start</td><td>end</td><td>strand</td><td>MM</td><td>target_seq</td><td>PAM</td><td>alignment</td><td>distance</td><td>position</td><td>gene name</td><td>gene id</td></tr>
<tr><td>GU305914.1</td><td>55226</td><td>55248</td><td>-</td><td>0</td><td>TCCTCTCTCCCGGGGAGAAG</td><td>AGG</td><td>||||||||[||||||||||||]PAM</td><td>NA</td><td>Intergenic</td><td>NA</td><td>NA</td></tr>
<tr><td>T29903</td><td>CCTCTCTCCCGGGGAGAAGAGGG</td><td>1000</td><td></td><td></td><td></td><td></td><td></td><td></td><td></td><td></td><td></td></tr>
<tr><td>Oligo fwd</td><td>CACCCCTCTCTCCCGGGGAGAAGA</td><td></td><td></td><td></td><td></td><td></td><td></td><td></td><td></td><td></td><td></td></tr>
<tr><td>Oligo rev</td><td>AAACTCTTCTCCCCGGGAGAGAGG</td><td></td><td></td><td></td><td></td><td></td><td></td><td></td><td></td><td></td><td></td></tr>
<tr><td>Chromosome</td><td>start</td><td>end</td><td>strand</td><td>MM</td><td>target_seq</td><td>PAM</td><td>alignment</td><td>distance</td><td>position</td><td>gene name</td><td>gene id</td></tr>
<tr><td>GU305914.1</td><td>55225</td><td>55247</td><td>-</td><td>0</td><td>CCTCTCTCCCGGGGAGAAGA</td><td>GGG</td><td>||||||||[||||||||||||]PAM</td><td>NA</td><td>Intergenic</td><td>NA</td><td>NA</td></tr>
<tr><td>T29904</td><td>TCTCCCGGGGAGAAGAGGGCCGG</td><td>1000</td><td></td><td></td><td></td><td></td><td></td><td></td><td></td><td></td><td></td></tr>
<tr><td>Oligo fwd</td><td>CACCTCTCCCGGGGAGAAGAGGGC</td><td></td><td></td><td></td><td></td><td></td><td></td><td></td><td></td><td></td><td></td></tr>
<tr><td>Oligo rev</td><td>AAACGCCCTCTTCTCCCCGGGAGA</td><td></td><td></td><td></td><td></td><td></td><td></td><td></td><td></td><td></td><td></td></tr>
<tr><td>Chromosome</td><td>start</td><td>end</td><td>strand</td><td>MM</td><td>target_seq</td><td>PAM</td><td>alignment</td><td>distance</td><td>position</td><td>gene name</td><td>gene id</td></tr>
<tr><td>GU305914.1</td><td>55221</td><td>55243</td><td>-</td><td>0</td><td>TCTCCCGGGGAGAAGAGGGC</td><td>CGG</td><td>||||||||[||||||||||||]PAM</td><td>NA</td><td>Intergenic</td><td>NA</td><td>NA</td></tr>
<tr><td>T29905</td><td>CTCCCGGGGAGAAGAGGGCCGGG</td><td>1000</td><td></td><td></td><td></td><td></td><td></td><td></td><td></td><td></td><td></td></tr>
<tr><td>Oligo fwd</td><td>CACCCTCCCGGGGAGAAGAGGGCC</td><td></td><td></td><td></td><td></td><td></td><td></td><td></td><td></td><td></td><td></td></tr>
<tr><td>Oligo rev</td><td>AAACGGCCCTCTTCTCCCCGGGAG</td><td></td><td></td><td></td><td></td><td></td><td></td><td></td><td></td><td></td><td></td></tr>
<tr><td>Chromosome</td><td>start</td><td>end</td><td>strand</td><td>MM</td><td>target_seq</td><td>PAM</td><td>alignment</td><td>distance</td><td>position</td><td>gene name</td><td>gene id</td></tr>
<tr><td>GU305914.1</td><td>55220</td><td>55242</td><td>-</td><td>0</td><td>CTCCCGGGGAGAAGAGGGCC</td><td>GGG</td><td>||||||||[||||||||||||]PAM</td><td>NA</td><td>Intergenic</td><td>NA</td><td>NA</td></tr>
</table>

T29906 GGGGAGAAGAGGGCCGGGAACGG 1000
Oligo fwd CACCGGGGAGAAGAGGGCCGGGAA
Oligo rev AAACTTCCCGGCCCTCTTCTCCCC
Chromosome start end strand MM target_seq PAM alignment distance position gene name gene id
GU305914.1 55215 55237 - 0 GGGGAGAAGAGGGCCGGGAA CGG ||||||||[||||||||||||]PAM NA Intergenic NA NA

T29907 GAGAAGAGGGCCGGGAACGGCGG 1000
Oligo fwd CACCGAGAAGAGGGCCGGGAACGG
Oligo rev AAACCCGTTCCCGGCCCTCTTCTC
Chromosome start end strand MM target_seq PAM alignment distance position gene name gene id
GU305914.1 55212 55234 - 0 GAGAAGAGGGCCGGGAACGG CGG ||||||||[||||||||||||]PAM NA Intergenic NA NA

T29908 CCTTCCGTAGATAACTGTCTCGG 1000
Oligo fwd CACCCCTTCCGTAGATAACTGTCT
Oligo rev AAACAGACAGTTATCTACGGAAGG
Chromosome start end strand MM target_seq PAM alignment distance position gene name gene id
GU305914.1 55186 55208 - 0 CCTTCCGTAGATAACTGTCT CGG ||||||||[||||||||||||]PAM NA Intergenic NA NA

T29909 TAACTGTCTCGGCGCCGCCGCGG 1000
Oligo fwd CACCTAACTGTCTCGGCGCCGCCG
Oligo rev AAACCGGCGGCGCCGAGACAGTTA
Chromosome start end strand MM target_seq PAM alignment distance position gene name gene id
GU305914.1 55175 55197 - 0 TAACTGTCTCGGCGCCGCCG CGG ||||||||[||||||||||||]PAM NA Intergenic NA NA

T29910 AACTGTCTCGGCGCCGCCGCGGG 1000
Oligo fwd CACCAACTGTCTCGGCGCCGCCGC
Oligo rev AAACGCGGCGGCGCCGAGACAGTT
Chromosome start end strand MM target_seq PAM alignment distance position gene name gene id
GU305914.1 55174 55196 - 0 AACTGTCTCGGCGCCGCCGC GGG ||||||||[||||||||||||]PAM NA Intergenic NA NA

T29911 TCGGCGCCGCCGCGGGCGCCAGG 1000
Oligo fwd CACCTCGGCGCCGCCGCGGGCGCC
Oligo rev AAACGGCGCCCGCGGCGGCGCCGA
Chromosome start end strand MM target_seq PAM alignment distance position gene name gene id
GU305914.1 55167 55189 - 0 TCGGCGCCGCCGCGGGCGCC AGG ||||||||[||||||||||||]PAM NA Intergenic NA NA
GU305914.1 149953 149975 - 4 CCGGCGCGGCCGCGGCCCCC CGG -||||||-[|||||||-|-||]PAM NA Intergenic NA NA

T29912 TCGCGAAGCACAACTCCGATAGG 1000
Oligo fwd CACCTCGCGAAGCACAACTCCGAT
Oligo rev AAACATCGGAGTTGTGCTTCGCGA
Chromosome start end strand MM target_seq PAM alignment distance position gene name gene id
GU305914.1 55127 55149 - 0 TCGCGAAGCACAACTCCGAT AGG ||||||||[||||||||||||]PAM NA Intergenic NA NA

T29913 AAGCACAACTCCGATAGGTACGG 1000
Oligo fwd CACCAAGCACAACTCCGATAGGTA
Oligo rev AAACTACCTATCGGAGTTGTGCTT
Chromosome start end strand MM target_seq PAM alignment distance position gene name gene id
GU305914.1 55122 55144 - 0 AAGCACAACTCCGATAGGTA CGG ||||||||[||||||||||||]PAM NA Intergenic NA NA

T29914 AGCACAACTCCGATAGGTACGGG 1000
Oligo fwd CACCAGCACAACTCCGATAGGTAC
Oligo rev AAACGTACCTATCGGAGTTGTGCT
Chromosome start end strand MM target_seq PAM alignment distance position gene name gene id
GU305914.1 55121 55143 - 0 AGCACAACTCCGATAGGTAC GGG ||||||||[||||||||||||]PAM NA Intergenic NA NA

T29915 AGGTACGGGAACCGAGACTCTGG 1000
Oligo fwd CACCAGGTACGGGAACCGAGACTC
Oligo rev AAACGAGTCTCGGTTCCCGTACCT
Chromosome start end strand MM target_seq PAM alignment distance position gene name gene id
GU305914.1 55107 55129 - 0 AGGTACGGGAACCGAGACTC TGG ||||||||[||||||||||||]PAM NA Intergenic NA NA

T29916 TTCTTCCTCTCTGAACGCTTCGG 1000
Oligo fwd CACCTTCTTCCTCTCTGAACGCTT
Oligo rev AAACAAGCGTTCAGAGAGGAAGAA
Chromosome start end strand MM target_seq PAM alignment distance position gene name gene id
GU305914.1 55072 55094 - 0 TTCTTCCTCTCTGAACGCTT CGG ||||||||[||||||||||||]PAM NA Intergenic NA NA

T29917 TCCTCTCTGAACGCTTCGGAAGG 1000
Oligo fwd CACCTCCTCTCTGAACGCTTCGGA
Oligo rev AAACTCCGAAGCGTTCAGAGAGGA
Chromosome start end strand MM target_seq PAM alignment distance position gene name gene id
GU305914.1 55068 55090 - 0 TCCTCTCTGAACGCTTCGGA AGG ||||||||[||||||||||||]PAM NA Intergenic NA NA

T29918 GAACGCTTCGGAAGGTCAAGAGG 1000
Oligo fwd CACCGAACGCTTCGGAAGGTCAAG
Oligo rev AAACCTTGACCTTCCGAAGCGTTC
Chromosome start end strand MM target_seq PAM alignment distance position gene name gene id
GU305914.1 55060 55082 - 0 GAACGCTTCGGAAGGTCAAG AGG ||||||||[||||||||||||]PAM NA Intergenic NA NA

T29919 GCTTCGGAAGGTCAAGAGGTAGG 1000
Oligo fwd CACCGCTTCGGAAGGTCAAGAGGT
Oligo rev AAACACCTCTTGACCTTCCGAAGC
Chromosome start end strand MM target_seq PAM alignment distance position gene name gene id
GU305914.1 55056 55078 - 0 GCTTCGGAAGGTCAAGAGGT AGG ||||||||[||||||||||||]PAM NA Intergenic NA NA
GU305914.1 130077 130099 - 4 GCGACAGAGGGTCAAGAGGT TGG ||--|-||[-|||||||||||]PAM NA Intergenic NA NA

T29920 CTTCGGAAGGTCAAGAGGTAGGG 1000
Oligo fwd CACCCTTCGGAAGGTCAAGAGGTA

<table>
<tr><td>Oligo rev</td><td>AAACTACCTCTTGACCTTCCGAAG</td></tr>
<tr><td>Chromosome</td><td>start</td><td></td><td>end</td><td>strand</td><td>MM</td><td>target_seq</td><td>PAM</td><td>alignment</td><td>distance</td><td>position</td><td>gene name</td><td>gene id</td></tr>
<tr><td>GU305914.1</td><td></td><td>55055</td><td>55077</td><td>-</td><td>0</td><td>CTTCGGAAGGTCAAGAGGTA</td><td>GGG</td><td>||||||||[||||||||||||]PAM</td><td>NA</td><td>Intergenic</td><td>NA</td><td>NA</td></tr>
<tr><td></td></tr>
<tr><td>T29921</td><td>CGGAAGGTCAAGAGGTAGGGTGG</td><td></td><td>1000</td></tr>
<tr><td>Oligo fwd</td><td>CACCCGGAAGGTCAAGAGGTAGGG</td></tr>
<tr><td>Oligo rev</td><td>AAACCCCTACCTCTTGACCTTCCG</td></tr>
<tr><td>Chromosome</td><td>start</td><td></td><td>end</td><td>strand</td><td>MM</td><td>target_seq</td><td>PAM</td><td>alignment</td><td>distance</td><td>position</td><td>gene name</td><td>gene id</td></tr>
<tr><td>GU305914.1</td><td></td><td>55052</td><td>55074</td><td>-</td><td>0</td><td>CGGAAGGTCAAGAGGTAGGG</td><td>TGG</td><td>||||||||[||||||||||||]PAM</td><td>NA</td><td>Intergenic</td><td>NA</td><td>NA</td></tr>
<tr><td></td></tr>
<tr><td>T29922</td><td>GTCAAGAGGTAGGGTGGTTGTGG</td><td></td><td>1000</td></tr>
<tr><td>Oligo fwd</td><td>CACCGTCAAGAGGTAGGGTGGTTG</td></tr>
<tr><td>Oligo rev</td><td>AAACCAACCACCCTACCTCTTGAC</td></tr>
<tr><td>Chromosome</td><td>start</td><td></td><td>end</td><td>strand</td><td>MM</td><td>target_seq</td><td>PAM</td><td>alignment</td><td>distance</td><td>position</td><td>gene name</td><td>gene id</td></tr>
<tr><td>GU305914.1</td><td></td><td>55046</td><td>55068</td><td>-</td><td>0</td><td>GTCAAGAGGTAGGGTGGTTG</td><td>TGG</td><td>||||||||[||||||||||||]PAM</td><td>NA</td><td>Intergenic</td><td>NA</td><td>NA</td></tr>
<tr><td></td></tr>
<tr><td>T29923</td><td>TCAAGAGGTAGGGTGGTTGTGGG</td><td></td><td>1000</td></tr>
<tr><td>Oligo fwd</td><td>CACCTCAAGAGGTAGGGTGGTTGT</td></tr>
<tr><td>Oligo rev</td><td>AAACACAACCACCCTACCTCTTGA</td></tr>
<tr><td>Chromosome</td><td>start</td><td></td><td>end</td><td>strand</td><td>MM</td><td>target_seq</td><td>PAM</td><td>alignment</td><td>distance</td><td>position</td><td>gene name</td><td>gene id</td></tr>
<tr><td>GU305914.1</td><td></td><td>55045</td><td>55067</td><td>-</td><td>0</td><td>TCAAGAGGTAGGGTGGTTGT</td><td>GGG</td><td>||||||||[||||||||||||]PAM</td><td>NA</td><td>Intergenic</td><td>NA</td><td>NA</td></tr>
<tr><td></td></tr>
<tr><td>T29924</td><td>GAGGTAGGGTGGTTGTGGGTCGG</td><td></td><td>1000</td></tr>
<tr><td>Oligo fwd</td><td>CACCGAGGTAGGGTGGTTGTGGGT</td></tr>
<tr><td>Oligo rev</td><td>AAACACCCACAACCACCCTACCTC</td></tr>
<tr><td>Chromosome</td><td>start</td><td></td><td>end</td><td>strand</td><td>MM</td><td>target_seq</td><td>PAM</td><td>alignment</td><td>distance</td><td>position</td><td>gene name</td><td>gene id</td></tr>
<tr><td>GU305914.1</td><td></td><td>55041</td><td>55063</td><td>-</td><td>0</td><td>GAGGTAGGGTGGTTGTGGGT</td><td>CGG</td><td>||||||||[||||||||||||]PAM</td><td>NA</td><td>Intergenic</td><td>NA</td><td>NA</td></tr>
<tr><td></td></tr>
<tr><td>T29925</td><td>AGGGTGGTTGTGGGTCGGAACGG</td><td></td><td>1000</td></tr>
<tr><td>Oligo fwd</td><td>CACCAGGGTGGTTGTGGGTCGGAA</td></tr>
<tr><td>Oligo rev</td><td>AAACTTCCGACCCACAACCACCCT</td></tr>
<tr><td>Chromosome</td><td>start</td><td></td><td>end</td><td>strand</td><td>MM</td><td>target_seq</td><td>PAM</td><td>alignment</td><td>distance</td><td>position</td><td>gene name</td><td>gene id</td></tr>
<tr><td>GU305914.1</td><td></td><td>55036</td><td>55058</td><td>-</td><td>0</td><td>AGGGTGGTTGTGGGTCGGAA</td><td>CGG</td><td>||||||||[||||||||||||]PAM</td><td>NA</td><td>Intergenic</td><td>NA</td><td>NA</td></tr>
<tr><td></td></tr>
<tr><td>T29926</td><td>GGGTGGTTGTGGGTCGGAACGGG</td><td></td><td>1000</td></tr>
<tr><td>Oligo fwd</td><td>CACCGGGTGGTTGTGGGTCGGAAC</td></tr>
<tr><td>Oligo rev</td><td>AAACGTTCCGACCCACAACCACCC</td></tr>
<tr><td>Chromosome</td><td>start</td><td></td><td>end</td><td>strand</td><td>MM</td><td>target_seq</td><td>PAM</td><td>alignment</td><td>distance</td><td>position</td><td>gene name</td><td>gene id</td></tr>
<tr><td>GU305914.1</td><td></td><td>55035</td><td>55057</td><td>-</td><td>0</td><td>GGGTGGTTGTGGGTCGGAAC</td><td>GGG</td><td>||||||||[||||||||||||]PAM</td><td>NA</td><td>Intergenic</td><td>NA</td><td>NA</td></tr>
<tr><td></td></tr>
<tr><td>T29927</td><td>TGGTTGTGGGTCGGAACGGGTGG</td><td></td><td>1000</td></tr>
<tr><td>Oligo fwd</td><td>CACCTGGTTGTGGGTCGGAACGGG</td></tr>
<tr><td>Oligo rev</td><td>AAACCCCGTTCCGACCCACAACCA</td></tr>
<tr><td>Chromosome</td><td>start</td><td></td><td>end</td><td>strand</td><td>MM</td><td>target_seq</td><td>PAM</td><td>alignment</td><td>distance</td><td>position</td><td>gene name</td><td>gene id</td></tr>
<tr><td>GU305914.1</td><td></td><td>55032</td><td>55054</td><td>-</td><td>0</td><td>TGGTTGTGGGTCGGAACGGG</td><td>TGG</td><td>||||||||[||||||||||||]PAM</td><td>NA</td><td>Intergenic</td><td>NA</td><td>NA</td></tr>
<tr><td></td></tr>
<tr><td>T29928</td><td>TGTGGGTCGGAACGGGTGGTTGG</td><td></td><td>1000</td></tr>
<tr><td>Oligo fwd</td><td>CACCTGTGGGTCGGAACGGGTGGT</td></tr>
<tr><td>Oligo rev</td><td>AAACACCACCCGTTCCGACCCACA</td></tr>
<tr><td>Chromosome</td><td>start</td><td></td><td>end</td><td>strand</td><td>MM</td><td>target_seq</td><td>PAM</td><td>alignment</td><td>distance</td><td>position</td><td>gene name</td><td>gene id</td></tr>
<tr><td>GU305914.1</td><td></td><td>55028</td><td>55050</td><td>-</td><td>0</td><td>TGTGGGTCGGAACGGGTGGT</td><td>TGG</td><td>||||||||[||||||||||||]PAM</td><td>NA</td><td>Intergenic</td><td>NA</td><td>NA</td></tr>
<tr><td></td></tr>
<tr><td>T29929</td><td>TTGCAGATCCGATTGCCTTTCGG</td><td></td><td>1000</td></tr>
<tr><td>Oligo fwd</td><td>CACCTTGCAGATCCGATTGCCTTT</td></tr>
<tr><td>Oligo rev</td><td>AAACAAAGGCAATCGGATCTGCAA</td></tr>
<tr><td>Chromosome</td><td>start</td><td></td><td>end</td><td>strand</td><td>MM</td><td>target_seq</td><td>PAM</td><td>alignment</td><td>distance</td><td>position</td><td>gene name</td><td>gene id</td></tr>
<tr><td>GU305914.1</td><td></td><td>55005</td><td>55027</td><td>-</td><td>0</td><td>TTGCAGATCCGATTGCCTTT</td><td>CGG</td><td>||||||||[||||||||||||]PAM</td><td>NA</td><td>Intergenic</td><td>NA</td><td>NA</td></tr>
<tr><td></td></tr>
<tr><td>T29930</td><td>TGCAGATCCGATTGCCTTTCGGG</td><td></td><td>1000</td></tr>
<tr><td>Oligo fwd</td><td>CACCTGCAGATCCGATTGCCTTTC</td></tr>
<tr><td>Oligo rev</td><td>AAACGAAAGGCAATCGGATCTGCA</td></tr>
<tr><td>Chromosome</td><td>start</td><td></td><td>end</td><td>strand</td><td>MM</td><td>target_seq</td><td>PAM</td><td>alignment</td><td>distance</td><td>position</td><td>gene name</td><td>gene id</td></tr>
<tr><td>GU305914.1</td><td></td><td>55004</td><td>55026</td><td>-</td><td>0</td><td>TGCAGATCCGATTGCCTTTC</td><td>GGG</td><td>||||||||[||||||||||||]PAM</td><td>NA</td><td>Intergenic</td><td>NA</td><td>NA</td></tr>
<tr><td></td></tr>
<tr><td>T29931</td><td>CGATTGCCTTTCGGGATAGACGG</td><td></td><td>1000</td></tr>
<tr><td>Oligo fwd</td><td>CACCCGATTGCCTTTCGGGATAGA</td></tr>
<tr><td>Oligo rev</td><td>AAACTCTATCCCGAAAGGCAATCG</td></tr>
<tr><td>Chromosome</td><td>start</td><td></td><td>end</td><td>strand</td><td>MM</td><td>target_seq</td><td>PAM</td><td>alignment</td><td>distance</td><td>position</td><td>gene name</td><td>gene id</td></tr>
<tr><td>GU305914.1</td><td></td><td>54996</td><td>55018</td><td>-</td><td>0</td><td>CGATTGCCTTTCGGGATAGA</td><td>CGG</td><td>||||||||[||||||||||||]PAM</td><td>NA</td><td>Intergenic</td><td>NA</td><td>NA</td></tr>
<tr><td></td></tr>
<tr><td>T29932</td><td>GGGATAGACGGAGTATGCCTAGG</td><td></td><td>1000</td></tr>
<tr><td>Oligo fwd</td><td>CACCGGGATAGACGGAGTATGCCT</td></tr>
<tr><td>Oligo rev</td><td>AAACAGGCATACTCCGTCTATCCC</td></tr>
<tr><td>Chromosome</td><td>start</td><td></td><td>end</td><td>strand</td><td>MM</td><td>target_seq</td><td>PAM</td><td>alignment</td><td>distance</td><td>position</td><td>gene name</td><td>gene id</td></tr>
<tr><td>GU305914.1</td><td></td><td>54984</td><td>55006</td><td>-</td><td>0</td><td>GGGATAGACGGAGTATGCCT</td><td>AGG</td><td>||||||||[||||||||||||]PAM</td><td>NA</td><td>Intergenic</td><td>NA</td><td>NA</td></tr>
<tr><td></td></tr>
<tr><td>T29933</td><td>ATAGACGGAGTATGCCTAGGCGG</td><td></td><td>1000</td></tr>
<tr><td>Oligo fwd</td><td>CACCATAGACGGAGTATGCCTAGG</td></tr>
<tr><td>Oligo rev</td><td>AAACCCTAGGCATACTCCGTCTAT</td></tr>
<tr><td>Chromosome</td><td>start</td><td></td><td>end</td><td>strand</td><td>MM</td><td>target_seq</td><td>PAM</td><td>alignment</td><td>distance</td><td>position</td><td>gene name</td><td>gene id</td></tr>
<tr><td>GU305914.1</td><td></td><td>54981</td><td>55003</td><td>-</td><td>0</td><td>ATAGACGGAGTATGCCTAGG</td><td>CGG</td><td>||||||||[||||||||||||]PAM</td><td>NA</td><td>Intergenic</td><td>NA</td><td>NA</td></tr>
<tr><td></td></tr>
<tr><td>T29934</td><td>TAGACGGAGTATGCCTAGGCGGG</td><td></td><td>1000</td></tr>
<tr><td>Oligo fwd</td><td>CACCTAGACGGAGTATGCCTAGGC</td></tr>
<tr><td>Oligo rev</td><td>AAACGCCTAGGCATACTCCGTCTA</td></tr>
<tr><td>Chromosome</td><td>start</td><td></td><td>end</td><td>strand</td><td>MM</td><td>target_seq</td><td>PAM</td><td>alignment</td><td>distance</td><td>position</td><td>gene name</td><td>gene id</td></tr>
<tr><td>GU305914.1</td><td></td><td>54980</td><td>55002</td><td>-</td><td>0</td><td>TAGACGGAGTATGCCTAGGC</td><td>GGG</td><td>||||||||[||||||||||||]PAM</td><td>NA</td><td>Intergenic</td><td>NA</td><td>NA</td></tr>
<tr><td></td></tr>
<tr><td>T29935</td><td>GGAGTATGCCTAGGCGGGTCAGG</td><td></td><td>1000</td></tr>
</table>

<table>
<tr><td>Oligo fwd</td><td>CACCGGAGTATGCCTAGGCGGGTC</td><td></td><td></td><td></td><td></td><td></td><td></td><td></td><td></td><td></td><td></td></tr>
<tr><td>Oligo rev</td><td>AAACGACCCGCCTAGGCATACTCC</td><td></td><td></td><td></td><td></td><td></td><td></td><td></td><td></td><td></td><td></td></tr>
<tr><td>Chromosome</td><td>start</td><td>end</td><td>strand</td><td>MM</td><td>target_seq</td><td>PAM</td><td>alignment</td><td>distance</td><td>position</td><td>gene name</td><td>gene id</td></tr>
<tr><td>GU305914.1</td><td>54975</td><td>54997</td><td>-</td><td>0</td><td>GGAGTATGCCTAGGCGGGTC</td><td>AGG</td><td>||||||||[||||||||||||]PAM</td><td>NA</td><td>Intergenic</td><td>NA</td><td>NA</td></tr>
<tr><td>T29936</td><td>TATGCCTAGGCGGGTCAGGCAGG</td><td>1000</td><td></td><td></td><td></td><td></td><td></td><td></td><td></td><td></td><td></td></tr>
<tr><td>Oligo fwd</td><td>CACCTATGCCTAGGCGGGTCAGGC</td><td></td><td></td><td></td><td></td><td></td><td></td><td></td><td></td><td></td><td></td></tr>
<tr><td>Oligo rev</td><td>AAACGCCTGACCCGCCTAGGCATA</td><td></td><td></td><td></td><td></td><td></td><td></td><td></td><td></td><td></td><td></td></tr>
<tr><td>Chromosome</td><td>start</td><td>end</td><td>strand</td><td>MM</td><td>target_seq</td><td>PAM</td><td>alignment</td><td>distance</td><td>position</td><td>gene name</td><td>gene id</td></tr>
<tr><td>GU305914.1</td><td>54971</td><td>54993</td><td>-</td><td>0</td><td>TATGCCTAGGCGGGTCAGGC</td><td>AGG</td><td>||||||||[||||||||||||]PAM</td><td>NA</td><td>Intergenic</td><td>NA</td><td>NA</td></tr>
<tr><td>T29937</td><td>ATGCCTAGGCGGGTCAGGCAGGG</td><td>1000</td><td></td><td></td><td></td><td></td><td></td><td></td><td></td><td></td><td></td></tr>
<tr><td>Oligo fwd</td><td>CACCATGCCTAGGCGGGTCAGGCA</td><td></td><td></td><td></td><td></td><td></td><td></td><td></td><td></td><td></td><td></td></tr>
<tr><td>Oligo rev</td><td>AAACTGCCTGACCCGCCTAGGCAT</td><td></td><td></td><td></td><td></td><td></td><td></td><td></td><td></td><td></td><td></td></tr>
<tr><td>Chromosome</td><td>start</td><td>end</td><td>strand</td><td>MM</td><td>target_seq</td><td>PAM</td><td>alignment</td><td>distance</td><td>position</td><td>gene name</td><td>gene id</td></tr>
<tr><td>GU305914.1</td><td>54970</td><td>54992</td><td>-</td><td>0</td><td>ATGCCTAGGCGGGTCAGGCA</td><td>GGG</td><td>||||||||[||||||||||||]PAM</td><td>NA</td><td>Intergenic</td><td>NA</td><td>NA</td></tr>
<tr><td>T29938</td><td>TCAGGCAGGGCCACAACCCGTGG</td><td>1000</td><td></td><td></td><td></td><td></td><td></td><td></td><td></td><td></td><td></td></tr>
<tr><td>Oligo fwd</td><td>CACCTCAGGCAGGGCCACAACCCG</td><td></td><td></td><td></td><td></td><td></td><td></td><td></td><td></td><td></td><td></td></tr>
<tr><td>Oligo rev</td><td>AAACCGGGTTGTGGCCCTGCCTGA</td><td></td><td></td><td></td><td></td><td></td><td></td><td></td><td></td><td></td><td></td></tr>
<tr><td>Chromosome</td><td>start</td><td>end</td><td>strand</td><td>MM</td><td>target_seq</td><td>PAM</td><td>alignment</td><td>distance</td><td>position</td><td>gene name</td><td>gene id</td></tr>
<tr><td>GU305914.1</td><td>54957</td><td>54979</td><td>-</td><td>0</td><td>TCAGGCAGGGCCACAACCCG</td><td>TGG</td><td>||||||||[||||||||||||]PAM</td><td>NA</td><td>Intergenic</td><td>NA</td><td>NA</td></tr>
<tr><td>T29939</td><td>GTACGATCTCCTGAAGATCGTGG</td><td>1000</td><td></td><td></td><td></td><td></td><td></td><td></td><td></td><td></td><td></td></tr>
<tr><td>Oligo fwd</td><td>CACCGTACGATCTCCTGAAGATCG</td><td></td><td></td><td></td><td></td><td></td><td></td><td></td><td></td><td></td><td></td></tr>
<tr><td>Oligo rev</td><td>AAACCGATCTTCAGGAGATCGTAC</td><td></td><td></td><td></td><td></td><td></td><td></td><td></td><td></td><td></td><td></td></tr>
<tr><td>Chromosome</td><td>start</td><td>end</td><td>strand</td><td>MM</td><td>target_seq</td><td>PAM</td><td>alignment</td><td>distance</td><td>position</td><td>gene name</td><td>gene id</td></tr>
<tr><td>GU305914.1</td><td>54935</td><td>54957</td><td>-</td><td>0</td><td>GTACGATCTCCTGAAGATCG</td><td>TGG</td><td>||||||||[||||||||||||]PAM</td><td>NA</td><td>Intergenic</td><td>NA</td><td>NA</td></tr>
<tr><td>T29940</td><td>AAGATCGTGGCTCTCATTGTAGG</td><td>1000</td><td></td><td></td><td></td><td></td><td></td><td></td><td></td><td></td><td></td></tr>
<tr><td>Oligo fwd</td><td>CACCAAGATCGTGGCTCTCATTGT</td><td></td><td></td><td></td><td></td><td></td><td></td><td></td><td></td><td></td><td></td></tr>
<tr><td>Oligo rev</td><td>AAACACAATGAGAGCCACGATCTT</td><td></td><td></td><td></td><td></td><td></td><td></td><td></td><td></td><td></td><td></td></tr>
<tr><td>Chromosome</td><td>start</td><td>end</td><td>strand</td><td>MM</td><td>target_seq</td><td>PAM</td><td>alignment</td><td>distance</td><td>position</td><td>gene name</td><td>gene id</td></tr>
<tr><td>GU305914.1</td><td>54922</td><td>54944</td><td>-</td><td>0</td><td>AAGATCGTGGCTCTCATTGT</td><td>AGG</td><td>||||||||[||||||||||||]PAM</td><td>NA</td><td>Intergenic</td><td>NA</td><td>NA</td></tr>
<tr><td>T29941</td><td>CGTGGCTCTCATTGTAGGCTTGG</td><td>1000</td><td></td><td></td><td></td><td></td><td></td><td></td><td></td><td></td><td></td></tr>
<tr><td>Oligo fwd</td><td>CACCCGTGGCTCTCATTGTAGGCT</td><td></td><td></td><td></td><td></td><td></td><td></td><td></td><td></td><td></td><td></td></tr>
<tr><td>Oligo rev</td><td>AAACAGCCTACAATGAGAGCCACG</td><td></td><td></td><td></td><td></td><td></td><td></td><td></td><td></td><td></td><td></td></tr>
<tr><td>Chromosome</td><td>start</td><td>end</td><td>strand</td><td>MM</td><td>target_seq</td><td>PAM</td><td>alignment</td><td>distance</td><td>position</td><td>gene name</td><td>gene id</td></tr>
<tr><td>GU305914.1</td><td>54917</td><td>54939</td><td>-</td><td>0</td><td>CGTGGCTCTCATTGTAGGCT</td><td>TGG</td><td>||||||||[||||||||||||]PAM</td><td>NA</td><td>Intergenic</td><td>NA</td><td>NA</td></tr>
<tr><td>T29942</td><td>GATACTATATATGCTGATAGAGG</td><td>1000</td><td></td><td></td><td></td><td></td><td></td><td></td><td></td><td></td><td></td></tr>
<tr><td>Oligo fwd</td><td>CACCGATACTATATATGCTGATAG</td><td></td><td></td><td></td><td></td><td></td><td></td><td></td><td></td><td></td><td></td></tr>
<tr><td>Oligo rev</td><td>AAACCTATCAGCATATATAGTATC</td><td></td><td></td><td></td><td></td><td></td><td></td><td></td><td></td><td></td><td></td></tr>
<tr><td>Chromosome</td><td>start</td><td>end</td><td>strand</td><td>MM</td><td>target_seq</td><td>PAM</td><td>alignment</td><td>distance</td><td>position</td><td>gene name</td><td>gene id</td></tr>
<tr><td>GU305914.1</td><td>54890</td><td>54912</td><td>-</td><td>0</td><td>GATACTATATATGCTGATAG</td><td>AGG</td><td>||||||||[||||||||||||]PAM</td><td>NA</td><td>Intergenic</td><td>NA</td><td>NA</td></tr>
<tr><td>T29943</td><td>ATATATGCTGATAGAGGACGTGG</td><td>1000</td><td></td><td></td><td></td><td></td><td></td><td></td><td></td><td></td><td></td></tr>
<tr><td>Oligo fwd</td><td>CACCATATATGCTGATAGAGGACG</td><td></td><td></td><td></td><td></td><td></td><td></td><td></td><td></td><td></td><td></td></tr>
<tr><td>Oligo rev</td><td>AAACCGTCCTCTATCAGCATATAT</td><td></td><td></td><td></td><td></td><td></td><td></td><td></td><td></td><td></td><td></td></tr>
<tr><td>Chromosome</td><td>start</td><td>end</td><td>strand</td><td>MM</td><td>target_seq</td><td>PAM</td><td>alignment</td><td>distance</td><td>position</td><td>gene name</td><td>gene id</td></tr>
<tr><td>GU305914.1</td><td>54884</td><td>54906</td><td>-</td><td>0</td><td>ATATATGCTGATAGAGGACG</td><td>TGG</td><td>||||||||[||||||||||||]PAM</td><td>NA</td><td>Intergenic</td><td>NA</td><td>NA</td></tr>
<tr><td>T29944</td><td>GGTTATGCTACTTTTGTGAACGG</td><td>1000</td><td></td><td></td><td></td><td></td><td></td><td></td><td></td><td></td><td></td></tr>
<tr><td>Oligo fwd</td><td>CACCGGTTATGCTACTTTTGTGAA</td><td></td><td></td><td></td><td></td><td></td><td></td><td></td><td></td><td></td><td></td></tr>
<tr><td>Oligo rev</td><td>AAACTTCACAAAAGTAGCATAACC</td><td></td><td></td><td></td><td></td><td></td><td></td><td></td><td></td><td></td><td></td></tr>
<tr><td>Chromosome</td><td>start</td><td>end</td><td>strand</td><td>MM</td><td>target_seq</td><td>PAM</td><td>alignment</td><td>distance</td><td>position</td><td>gene name</td><td>gene id</td></tr>
<tr><td>GU305914.1</td><td>54863</td><td>54885</td><td>-</td><td>0</td><td>GGTTATGCTACTTTTGTGAA</td><td>CGG</td><td>||||||||[||||||||||||]PAM</td><td>NA</td><td>Intergenic</td><td>NA</td><td>NA</td></tr>
<tr><td>T29945</td><td>GCTACTTTTGTGAACGGACGCGG</td><td>1000</td><td></td><td></td><td></td><td></td><td></td><td></td><td></td><td></td><td></td></tr>
<tr><td>Oligo fwd</td><td>CACCGCTACTTTTGTGAACGGACG</td><td></td><td></td><td></td><td></td><td></td><td></td><td></td><td></td><td></td><td></td></tr>
<tr><td>Oligo rev</td><td>AAACCGTCCGTTCACAAAAGTAGC</td><td></td><td></td><td></td><td></td><td></td><td></td><td></td><td></td><td></td><td></td></tr>
<tr><td>Chromosome</td><td>start</td><td>end</td><td>strand</td><td>MM</td><td>target_seq</td><td>PAM</td><td>alignment</td><td>distance</td><td>position</td><td>gene name</td><td>gene id</td></tr>
<tr><td>GU305914.1</td><td>54857</td><td>54879</td><td>-</td><td>0</td><td>GCTACTTTTGTGAACGGACG</td><td>CGG</td><td>||||||||[||||||||||||]PAM</td><td>NA</td><td>Intergenic</td><td>NA</td><td>NA</td></tr>
<tr><td>T29946</td><td>CTTTTGTGAACGGACGCGGTTGG</td><td>1000</td><td></td><td></td><td></td><td></td><td></td><td></td><td></td><td></td><td></td></tr>
<tr><td>Oligo fwd</td><td>CACCCTTTTGTGAACGGACGCGGT</td><td></td><td></td><td></td><td></td><td></td><td></td><td></td><td></td><td></td><td></td></tr>
<tr><td>Oligo rev</td><td>AAACACCGCGTCCGTTCACAAAAG</td><td></td><td></td><td></td><td></td><td></td><td></td><td></td><td></td><td></td><td></td></tr>
<tr><td>Chromosome</td><td>start</td><td>end</td><td>strand</td><td>MM</td><td>target_seq</td><td>PAM</td><td>alignment</td><td>distance</td><td>position</td><td>gene name</td><td>gene id</td></tr>
<tr><td>GU305914.1</td><td>54853</td><td>54875</td><td>-</td><td>0</td><td>CTTTTGTGAACGGACGCGGT</td><td>TGG</td><td>||||||||[||||||||||||]PAM</td><td>NA</td><td>Intergenic</td><td>NA</td><td>NA</td></tr>
<tr><td>T29947</td><td>TTTTGTGAACGGACGCGGTTGGG</td><td>1000</td><td></td><td></td><td></td><td></td><td></td><td></td><td></td><td></td><td></td></tr>
<tr><td>Oligo fwd</td><td>CACCTTTTGTGAACGGACGCGGTT</td><td></td><td></td><td></td><td></td><td></td><td></td><td></td><td></td><td></td><td></td></tr>
<tr><td>Oligo rev</td><td>AAACAACCGCGTCCGTTCACAAAA</td><td></td><td></td><td></td><td></td><td></td><td></td><td></td><td></td><td></td><td></td></tr>
<tr><td>Chromosome</td><td>start</td><td>end</td><td>strand</td><td>MM</td><td>target_seq</td><td>PAM</td><td>alignment</td><td>distance</td><td>position</td><td>gene name</td><td>gene id</td></tr>
<tr><td>GU305914.1</td><td>54852</td><td>54874</td><td>-</td><td>0</td><td>TTTTGTGAACGGACGCGGTT</td><td>GGG</td><td>||||||||[||||||||||||]PAM</td><td>NA</td><td>Intergenic</td><td>NA</td><td>NA</td></tr>
<tr><td>T29948</td><td>TGTGAACGGACGCGGTTGGGAGG</td><td>1000</td><td></td><td></td><td></td><td></td><td></td><td></td><td></td><td></td><td></td></tr>
<tr><td>Oligo fwd</td><td>CACCTGTGAACGGACGCGGTTGGG</td><td></td><td></td><td></td><td></td><td></td><td></td><td></td><td></td><td></td><td></td></tr>
<tr><td>Oligo rev</td><td>AAACCCCAACCGCGTCCGTTCACA</td><td></td><td></td><td></td><td></td><td></td><td></td><td></td><td></td><td></td><td></td></tr>
<tr><td>Chromosome</td><td>start</td><td>end</td><td>strand</td><td>MM</td><td>target_seq</td><td>PAM</td><td>alignment</td><td>distance</td><td>position</td><td>gene name</td><td>gene id</td></tr>
<tr><td>GU305914.1</td><td>54849</td><td>54871</td><td>-</td><td>0</td><td>TGTGAACGGACGCGGTTGGG</td><td>AGG</td><td>||||||||[||||||||||||]PAM</td><td>NA</td><td>Intergenic</td><td>NA</td><td>NA</td></tr>
<tr><td>T29949</td><td>GGACATGACTTCTTCCGAACCGG</td><td>1000</td><td></td><td></td><td></td><td></td><td></td><td></td><td></td><td></td><td></td></tr>
<tr><td>Oligo fwd</td><td>CACCGGACATGACTTCTTCCGAAC</td><td></td><td></td><td></td><td></td><td></td><td></td><td></td><td></td><td></td><td></td></tr>
<tr><td>Oligo rev</td><td>AAACGTTCGGAAGAAGTCATGTCC</td><td></td><td></td><td></td><td></td><td></td><td></td><td></td><td></td><td></td><td></td></tr>
<tr><td>Chromosome</td><td>start</td><td>end</td><td>strand</td><td>MM</td><td>target_seq</td><td>PAM</td><td>alignment</td><td>distance</td><td>position</td><td>gene name</td><td>gene id</td></tr>
<tr><td>GU305914.1</td><td>54828</td><td>54850</td><td>-</td><td>0</td><td>GGACATGACTTCTTCCGAAC</td><td>CGG</td><td>||||||||[||||||||||||]PAM</td><td>NA</td><td>Intergenic</td><td>NA</td><td>NA</td></tr>
</table>

T29950 GCACCGTTTGCCTACTTATCTGG 1000
Oligo fwd CACCGCACCGTTTGCCTACTTATC
Oligo rev AAACGATAAGTAGGCAAACGGTGC
Chromosome start end strand MM target_seq PAM alignment distance position gene name gene id
GU305914.1 54804 54826 - 0 GCACCGTTTGCCTACTTATC TGG ||||||||[||||||||||||]PAM NA Intergenic NA NA

T29951 TTTGCCTACTTATCTGGAAGCGG 1000
Oligo fwd CACCTTTGCCTACTTATCTGGAAG
Oligo rev AAACCTTCCAGATAAGTAGGCAAA
Chromosome start end strand MM target_seq PAM alignment distance position gene name gene id
GU305914.1 54798 54820 - 0 TTTGCCTACTTATCTGGAAG CGG ||||||||[||||||||||||]PAM NA Intergenic NA NA

T29952 CCTACTTATCTGGAAGCGGTCGG 1000
Oligo fwd CACCCCTACTTATCTGGAAGCGGT
Oligo rev AAACACCGCTTCCAGATAAGTAGG
Chromosome start end strand MM target_seq PAM alignment distance position gene name gene id
GU305914.1 54794 54816 - 0 CCTACTTATCTGGAAGCGGT CGG ||||||||[||||||||||||]PAM NA Intergenic NA NA

T29953 CGACACTGCCACCACGCGACTGG 1000
Oligo fwd CACCCGACACTGCCACCACGCGAC
Oligo rev AAACGTCGCGTGGTGGCAGTGTCG
Chromosome start end strand MM target_seq PAM alignment distance position gene name gene id
GU305914.1 54738 54760 - 0 CGACACTGCCACCACGCGAC TGG ||||||||[||||||||||||]PAM NA Intergenic NA NA

T29954 CACTGCCACCACGCGACTGGTGG 1000
Oligo fwd CACCCACTGCCACCACGCGACTGG
Oligo rev AAACCCAGTCGCGTGGTGGCAGTG
Chromosome start end strand MM target_seq PAM alignment distance position gene name gene id
GU305914.1 54735 54757 - 0 CACTGCCACCACGCGACTGG TGG ||||||||[||||||||||||]PAM NA Intergenic NA NA

T29955 GACTGGTGGTCACGAATCTGCGG 1000
Oligo fwd CACCGACTGGTGGTCACGAATCTG
Oligo rev AAACCAGATTCGTGACCACCAGTC
Chromosome start end strand MM target_seq PAM alignment distance position gene name gene id
GU305914.1 54721 54743 - 0 GACTGGTGGTCACGAATCTG CGG ||||||||[||||||||||||]PAM NA Intergenic NA NA

T29956 GGTCACGAATCTGCGGCAGCAGG 1000
Oligo fwd CACCGGTCACGAATCTGCGGCAGC
Oligo rev AAACGCTGCCGCAGATTCGTGACC
Chromosome start end strand MM target_seq PAM alignment distance position gene name gene id
GU305914.1 54714 54736 - 0 GGTCACGAATCTGCGGCAGC AGG ||||||||[||||||||||||]PAM NA Intergenic NA NA

T29957 CACGAATCTGCGGCAGCAGGCGG 1000
Oligo fwd CACCCACGAATCTGCGGCAGCAGG
Oligo rev AAACCCTGCTGCCGCAGATTCGTG
Chromosome start end strand MM target_seq PAM alignment distance position gene name gene id
GU305914.1 54711 54733 - 0 CACGAATCTGCGGCAGCAGG CGG ||||||||[||||||||||||]PAM NA Intergenic NA NA

T29958 CCGTCCTACGAGAGTCTCTTGG 1000
Oligo fwd CACCCCGTCCTACGAGAGTCTCTT
Oligo rev AAACAAGAGACTCTCGTAGGACGG
Chromosome start end strand MM target_seq PAM alignment distance position gene name gene id
GU305914.1 54674 54696 - 0 CCGTCCTACGAGAGTCTCTT TGG ||||||||[||||||||||||]PAM NA Intergenic NA NA

T29959 TCTTTGGTAACAACGACGAGCGG 1000
Oligo fwd CACCTCTTTGGTAACAACGACGAG
Oligo rev AAACCTCGTCGTTGTTACCAAAGA
Chromosome start end strand MM target_seq PAM alignment distance position gene name gene id
GU305914.1 54658 54680 - 0 TCTTTGGTAACAACGACGAG CGG ||||||||[||||||||||||]PAM NA Intergenic NA NA

T29960 AGTCTCCGCGCCGCGTTACGCGG 1000
Oligo fwd CACCAGTCTCCGCGCCGCGTTACG
Oligo rev AAACCGTAACGCGGCGCGGAGACT
Chromosome start end strand MM target_seq PAM alignment distance position gene name gene id
GU305914.1 54617 54639 - 0 AGTCTCCGCGCCGCGTTACG CGG ||||||||[||||||||||||]PAM NA Intergenic NA NA

T29961 CTCCACCACCTCGTCCTCGACGG 1000
Oligo fwd CACCCTCCACCACCTCGTCCTCGA
Oligo rev AAACTCGAGGACGAGGTGGTGGAG
Chromosome start end strand MM target_seq PAM alignment distance position gene name gene id
GU305914.1 54555 54577 - 0 CTCCACCACCTCGTCCTCGA CGG ||||||||[||||||||||||]PAM NA Intergenic NA NA

T29962 CACCACCTCGTCCTCGACGGAGG 1000
Oligo fwd CACCCACCACCTCGTCCTCGACGG
Oligo rev AAACCCGTCGAGGACGAGGTGGTG
Chromosome start end strand MM target_seq PAM alignment distance position gene name gene id
GU305914.1 54552 54574 - 0 CACCACCTCGTCCTCGACGG AGG ||||||||[||||||||||||]PAM NA Intergenic NA NA

T29963 CACCTCGTCCTCGACGGAGGTGG 1000
Oligo fwd CACCCACCTCGTCCTCGACGGAGG
Oligo rev AAACCCTCCGTCGAGGACGAGGTG
Chromosome start end strand MM target_seq PAM alignment distance position gene name gene id
GU305914.1 54549 54571 - 0 CACCTCGTCCTCGACGGAGG TGG ||||||||[||||||||||||]PAM NA Intergenic NA NA

T29964 GGTGGCTGCGCGTGCGCTCGCGG 1000
Oligo fwd CACCGGTGGCTGCGCGTGCGCTCG
Oligo rev AAACCGAGCGCACGCGCAGCCACC
Chromosome start end strand MM target_seq PAM alignment distance position gene name gene id
GU305914.1 54531 54553 - 0 GGTGGCTGCGCGTGCGCTCG CGG ||||||||[||||||||||||]PAM NA Intergenic NA NA

| | | | | | | | | | | | | |
|---|---|---|---|---|---|---|---|---|---|---|---|---|
| T29965 | GCGCGCGAACAGCGAGCGAGAGG | | 1000 | | | | | | | | | |
| Oligo fwd | CACCGCGCGCGAACAGCGAGCGAG | | | | | | | | | | | |
| Oligo rev | AAACCTCGCTCGCTGTTCGCGCGC | | | | | | | | | | | |
| Chromosome | start | | end | strand | MM | target_seq | PAM | alignment | distance | position | gene name | gene id |
| GU305914.1 | | 54502 | 54524 | - | 0 | GCGCGCGAACAGCGAGCGAG | AGG | ||||||||[||||||||||||]PAM | NA | Intergenic | NA | NA |
| T29966 | GAACAGCGAGCGAGAGGCTGAGG | | 1000 | | | | | | | | | |
| Oligo fwd | CACCGAACAGCGAGCGAGAGGCTG | | | | | | | | | | | |
| Oligo rev | AAACCAGCCTCTCGCTCGCTGTTC | | | | | | | | | | | |
| Chromosome | start | | end | strand | MM | target_seq | PAM | alignment | distance | position | gene name | gene id |
| GU305914.1 | | 54496 | 54518 | - | 0 | GAACAGCGAGCGAGAGGCTG | AGG | ||||||||[||||||||||||]PAM | NA | Intergenic | NA | NA |
| T29967 | AACCCTGTCTCTCGAACGCATGG | | 1000 | | | | | | | | | |
| Oligo fwd | CACCAACCCTGTCTCTCGAACGCA | | | | | | | | | | | |
| Oligo rev | AAACTGCGTTCGAGAGACAGGGTT | | | | | | | | | | | |
| Chromosome | start | | end | strand | MM | target_seq | PAM | alignment | distance | position | gene name | gene id |
| GU305914.1 | | 54471 | 54493 | - | 0 | AACCCTGTCTCTCGAACGCA | TGG | ||||||||[||||||||||||]PAM | NA | Intergenic | NA | NA |
| T29968 | GCATGGATAAAGAGTTTCAGTGG | | 1000 | | | | | | | | | |
| Oligo fwd | CACCGCATGGATAAAGAGTTTCAG | | | | | | | | | | | |
| Oligo rev | AAACCTGAAACTCTTTATCCATGC | | | | | | | | | | | |
| Chromosome | start | | end | strand | MM | target_seq | PAM | alignment | distance | position | gene name | gene id |
| GU305914.1 | | 54454 | 54476 | - | 0 | GCATGGATAAAGAGTTTCAG | TGG | ||||||||[||||||||||||]PAM | NA | Intergenic | NA | NA |
| T29969 | GTTTCAGTGGACCGCCATCACGG | | 1000 | | | | | | | | | |
| Oligo fwd | CACCGTTTCAGTGGACCGCCATCA | | | | | | | | | | | |
| Oligo rev | AAACTGATGGCGGTCCACTGAAAC | | | | | | | | | | | |
| Chromosome | start | | end | strand | MM | target_seq | PAM | alignment | distance | position | gene name | gene id |
| GU305914.1 | | 54441 | 54463 | - | 0 | GTTTCAGTGGACCGCCATCA | CGG | ||||||||[||||||||||||]PAM | NA | Intergenic | NA | NA |
| T29970 | CAGTGGACCGCCATCACGGTCGG | | 1000 | | | | | | | | | |
| Oligo fwd | CACCCAGTGGACCGCCATCACGGT | | | | | | | | | | | |
| Oligo rev | AAACACCGTGATGGCGGTCCACTG | | | | | | | | | | | |
| Chromosome | start | | end | strand | MM | target_seq | PAM | alignment | distance | position | gene name | gene id |
| GU305914.1 | | 54437 | 54459 | - | 0 | CAGTGGACCGCCATCACGGT | CGG | ||||||||[||||||||||||]PAM | NA | Intergenic | NA | NA |
| T29971 | CGGAGTCTGTGTGTCCCTCTTGG | | 1000 | | | | | | | | | |
| Oligo fwd | CACCCGGAGTCTGTGTGTCCCTCT | | | | | | | | | | | |
| Oligo rev | AAACAGAGGGACACACAGACTCCG | | | | | | | | | | | |
| Chromosome | start | | end | strand | MM | target_seq | PAM | alignment | distance | position | gene name | gene id |
| GU305914.1 | | 54417 | 54439 | - | 0 | CGGAGTCTGTGTGTCCCTCT | TGG | ||||||||[||||||||||||]PAM | NA | Intergenic | NA | NA |
| T29972 | CATACTCGTTCTCGTCATCACGG | | 1000 | | | | | | | | | |
| Oligo fwd | CACCCATACTCGTTCTCGTCATCA | | | | | | | | | | | |
| Oligo rev | AAACTGATGACGAGAACGAGTATG | | | | | | | | | | | |
| Chromosome | start | | end | strand | MM | target_seq | PAM | alignment | distance | position | gene name | gene id |
| GU305914.1 | | 54393 | 54415 | - | 0 | CATACTCGTTCTCGTCATCA | CGG | ||||||||[||||||||||||]PAM | NA | Intergenic | NA | NA |
| T29973 | CGTTCTCGTCATCACGGCCGTGG | | 1000 | | | | | | | | | |
| Oligo fwd | CACCCGTTCTCGTCATCACGGCCG | | | | | | | | | | | |
| Oligo rev | AAACCGGCCGTGATGACGAGAACG | | | | | | | | | | | |
| Chromosome | start | | end | strand | MM | target_seq | PAM | alignment | distance | position | gene name | gene id |
| GU305914.1 | | 54387 | 54409 | - | 0 | CGTTCTCGTCATCACGGCCG | TGG | ||||||||[||||||||||||]PAM | NA | Intergenic | NA | NA |
| T29974 | GTCATCACGGCCGTGGTCACTGG | | 1000 | | | | | | | | | |
| Oligo fwd | CACCGTCATCACGGCCGTGGTCAC | | | | | | | | | | | |
| Oligo rev | AAACGTGACCACGGCCGTGATGAC | | | | | | | | | | | |
| Chromosome | start | | end | strand | MM | target_seq | PAM | alignment | distance | position | gene name | gene id |
| GU305914.1 | | 54380 | 54402 | - | 0 | GTCATCACGGCCGTGGTCAC | TGG | ||||||||[||||||||||||]PAM | NA | Intergenic | NA | NA |
| T29975 | TCACGGCCGTGGTCACTGGTCGG | | 1000 | | | | | | | | | |
| Oligo fwd | CACCTCACGGCCGTGGTCACTGGT | | | | | | | | | | | |
| Oligo rev | AAACACCAGTGACCACGGCCGTGA | | | | | | | | | | | |
| Chromosome | start | | end | strand | MM | target_seq | PAM | alignment | distance | position | gene name | gene id |
| GU305914.1 | | 54376 | 54398 | - | 0 | TCACGGCCGTGGTCACTGGT | CGG | ||||||||[||||||||||||]PAM | NA | Intergenic | NA | NA |
| T29976 | CACGGCCGTGGTCACTGGTCGGG | | 1000 | | | | | | | | | |
| Oligo fwd | CACCCACGGCCGTGGTCACTGGTC | | | | | | | | | | | |
| Oligo rev | AAACGACCAGTGACCACGGCCGTG | | | | | | | | | | | |
| Chromosome | start | | end | strand | MM | target_seq | PAM | alignment | distance | position | gene name | gene id |
| GU305914.1 | | 54375 | 54397 | - | 0 | CACGGCCGTGGTCACTGGTC | GGG | ||||||||[||||||||||||]PAM | NA | Intergenic | NA | NA |
| T29977 | ACGGCCGTGGTCACTGGTCGGGG | | 1000 | | | | | | | | | |
| Oligo fwd | CACCACGGCCGTGGTCACTGGTCG | | | | | | | | | | | |
| Oligo rev | AAACCGACCAGTGACCACGGCCGT | | | | | | | | | | | |
| Chromosome | start | | end | strand | MM | target_seq | PAM | alignment | distance | position | gene name | gene id |
| GU305914.1 | | 54374 | 54396 | - | 0 | ACGGCCGTGGTCACTGGTCG | GGG | ||||||||[||||||||||||]PAM | NA | Intergenic | NA | NA |
| T29978 | GCCGTGGTCACTGGTCGGGGAGG | | 1000 | | | | | | | | | |
| Oligo fwd | CACCGCCGTGGTCACTGGTCGGGG | | | | | | | | | | | |
| Oligo rev | AAACCCCCGACCAGTGACCACGGC | | | | | | | | | | | |
| Chromosome | start | | end | strand | MM | target_seq | PAM | alignment | distance | position | gene name | gene id |
| GU305914.1 | | 54371 | 54393 | - | 0 | GCCGTGGTCACTGGTCGGGG | AGG | ||||||||[||||||||||||]PAM | NA | Intergenic | NA | NA |
| T29979 | GTGGTCACTGGTCGGGGAGGTGG | | 1000 | | | | | | | | | |
| Oligo fwd | CACCGTGGTCACTGGTCGGGGAGG | | | | | | | | | | | |
| Oligo rev | AAACCCTCCCCGACCAGTGACCAC | | | | | | | | | | | |
| Chromosome | start | | end | strand | MM | target_seq | PAM | alignment | distance | position | gene name | gene id |

| | | | | | | | | | | | |
|---|---|---|---|---|---|---|---|---|---|---|---|
| GU305914.1 | | 54368 | 54390 | - | 0 | GTGGTCACTGGTCGGGGAGG | TGG | \|\|\|\|\|\|\|\|[\|\|\|\|\|\|\|\|\|\|\|\|]PAM | NA | Intergenic | NA | NA |
| T29980 | GTCACTGGTCGGGGAGGTGGCGG | 1000 | | | | | | | | | |
| Oligo fwd | CACCGTCACTGGTCGGGGAGGTGG | | | | | | | | | | |
| Oligo rev | AAACCCACCTCCCCGACCAGTGAC | | | | | | | | | | |
| Chromosome | start | end | strand | MM | target_seq | PAM | alignment | distance | position | gene name | gene id |
| GU305914.1 | 54365 | 54387 | - | 0 | GTCACTGGTCGGGGAGGTGG | CGG | \|\|\|\|\|\|\|\|[\|\|\|\|\|\|\|\|\|\|\|\|]PAM | NA | Intergenic | NA | NA |
| T29981 | GGCGGAGACGATTAGACTTGAGG | 1000 | | | | | | | | | |
| Oligo fwd | CACCGGCGGAGACGATTAGACTTG | | | | | | | | | | |
| Oligo rev | AAACCAAGTCTAATCGTCTCCGCC | | | | | | | | | | |
| Chromosome | start | end | strand | MM | target_seq | PAM | alignment | distance | position | gene name | gene id |
| GU305914.1 | 54347 | 54369 | - | 0 | GGCGGAGACGATTAGACTTG | AGG | \|\|\|\|\|\|\|\|[\|\|\|\|\|\|\|\|\|\|\|\|]PAM | NA | Intergenic | NA | NA |
| T29982 | GGAGACGATTAGACTTGAGGAGG | 1000 | | | | | | | | | |
| Oligo fwd | CACCGGAGACGATTAGACTTGAGG | | | | | | | | | | |
| Oligo rev | AAACCCTCAAGTCTAATCGTCTCC | | | | | | | | | | |
| Chromosome | start | end | strand | MM | target_seq | PAM | alignment | distance | position | gene name | gene id |
| GU305914.1 | 54344 | 54366 | - | 0 | GGAGACGATTAGACTTGAGG | AGG | \|\|\|\|\|\|\|\|[\|\|\|\|\|\|\|\|\|\|\|\|]PAM | NA | Intergenic | NA | NA |
| T29983 | TAGACTTGAGGAGGAACGACCGG | 1000 | | | | | | | | | |
| Oligo fwd | CACCTAGACTTGAGGAGGAACGAC | | | | | | | | | | |
| Oligo rev | AAACGTCGTTCCTCCTCAAGTCTA | | | | | | | | | | |
| Chromosome | start | end | strand | MM | target_seq | PAM | alignment | distance | position | gene name | gene id |
| GU305914.1 | 54335 | 54357 | - | 0 | TAGACTTGAGGAGGAACGAC | CGG | \|\|\|\|\|\|\|\|[\|\|\|\|\|\|\|\|\|\|\|\|]PAM | NA | Intergenic | NA | NA |
| T29984 | GAGGAGGAACGACCGGTCGCCGG | 1000 | | | | | | | | | |
| Oligo fwd | CACCGAGGAGGAACGACCGGTCGC | | | | | | | | | | |
| Oligo rev | AAACGCGACCGGTCGTTCCTCCTC | | | | | | | | | | |
| Chromosome | start | end | strand | MM | target_seq | PAM | alignment | distance | position | gene name | gene id |
| GU305914.1 | 54328 | 54350 | - | 0 | GAGGAGGAACGACCGGTCGC | CGG | \|\|\|\|\|\|\|\|[\|\|\|\|\|\|\|\|\|\|\|\|]PAM | NA | Intergenic | NA | NA |
| T29985 | AGGAACGACCGGTCGCCGGACGG | 1000 | | | | | | | | | |
| Oligo fwd | CACCAGGAACGACCGGTCGCCGGA | | | | | | | | | | |
| Oligo rev | AAACTCCGGCGACCGGTCGTTCCT | | | | | | | | | | |
| Chromosome | start | end | strand | MM | target_seq | PAM | alignment | distance | position | gene name | gene id |
| GU305914.1 | 54324 | 54346 | - | 0 | AGGAACGACCGGTCGCCGGA | CGG | \|\|\|\|\|\|\|\|[\|\|\|\|\|\|\|\|\|\|\|\|]PAM | NA | Intergenic | NA | NA |
| T29986 | GGAACGACCGGTCGCCGGACGGG | 1000 | | | | | | | | | |
| Oligo fwd | CACCGGAACGACCGGTCGCCGGAC | | | | | | | | | | |
| Oligo rev | AAACGTCCGGCGACCGGTCGTTCC | | | | | | | | | | |
| Chromosome | start | end | strand | MM | target_seq | PAM | alignment | distance | position | gene name | gene id |
| GU305914.1 | 54323 | 54345 | - | 0 | GGAACGACCGGTCGCCGGAC | GGG | \|\|\|\|\|\|\|\|[\|\|\|\|\|\|\|\|\|\|\|\|]PAM | NA | Intergenic | NA | NA |
| T29987 | CACATTCCGTTGTCGTGCGCAGG | 1000 | | | | | | | | | |
| Oligo fwd | CACCCACATTCCGTTGTCGTGCGC | | | | | | | | | | |
| Oligo rev | AAACGCGCACGACAACGGAATGTG | | | | | | | | | | |
| Chromosome | start | end | strand | MM | target_seq | PAM | alignment | distance | position | gene name | gene id |
| GU305914.1 | 54219 | 54241 | - | 0 | CACATTCCGTTGTCGTGCGC | AGG | \|\|\|\|\|\|\|\|[\|\|\|\|\|\|\|\|\|\|\|\|]PAM | NA | Intergenic | NA | NA |
| T29988 | AGGTTCCTCCGAACCTTTGATGG | 1000 | | | | | | | | | |
| Oligo fwd | CACCAGGTTCCTCCGAACCTTTGA | | | | | | | | | | |
| Oligo rev | AAACTCAAAGGTTCGGAGGAACCT | | | | | | | | | | |
| Chromosome | start | end | strand | MM | target_seq | PAM | alignment | distance | position | gene name | gene id |
| GU305914.1 | 54199 | 54221 | - | 0 | AGGTTCCTCCGAACCTTTGA | TGG | \|\|\|\|\|\|\|\|[\|\|\|\|\|\|\|\|\|\|\|\|]PAM | NA | Intergenic | NA | NA |
| T29989 | GGTTCCTCCGAACCTTTGATGGG | 1000 | | | | | | | | | |
| Oligo fwd | CACCGGTTCCTCCGAACCTTTGAT | | | | | | | | | | |
| Oligo rev | AAACATCAAAGGTTCGGAGGAACC | | | | | | | | | | |
| Chromosome | start | end | strand | MM | target_seq | PAM | alignment | distance | position | gene name | gene id |
| GU305914.1 | 54198 | 54220 | - | 0 | GGTTCCTCCGAACCTTTGAT | GGG | \|\|\|\|\|\|\|\|[\|\|\|\|\|\|\|\|\|\|\|\|]PAM | NA | Intergenic | NA | NA |
| T29990 | ATGGGAGACGATGATCGTCGCGG | 1000 | | | | | | | | | |
| Oligo fwd | CACCATGGGAGACGATGATCGTCG | | | | | | | | | | |
| Oligo rev | AAACCGACGATCATCGTCTCCCAT | | | | | | | | | | |
| Chromosome | start | end | strand | MM | target_seq | PAM | alignment | distance | position | gene name | gene id |
| GU305914.1 | 54180 | 54202 | - | 0 | ATGGGAGACGATGATCGTCG | CGG | \|\|\|\|\|\|\|\|[\|\|\|\|\|\|\|\|\|\|\|\|]PAM | NA | Intergenic | NA | NA |
| T29991 | GATGATCGTCGCGGCGATGACGG | 1000 | | | | | | | | | |
| Oligo fwd | CACCGATGATCGTCGCGGCGATGA | | | | | | | | | | |
| Oligo rev | AAACTCATCGCCGCGACGATCATC | | | | | | | | | | |
| Chromosome | start | end | strand | MM | target_seq | PAM | alignment | distance | position | gene name | gene id |
| GU305914.1 | 54171 | 54193 | - | 0 | GATGATCGTCGCGGCGATGA | CGG | \|\|\|\|\|\|\|\|[\|\|\|\|\|\|\|\|\|\|\|\|]PAM | NA | Intergenic | NA | NA |
| T29992 | GATCGTCGCGGCGATGACGGCGG | 1000 | | | | | | | | | |
| Oligo fwd | CACCGATCGTCGCGGCGATGACGG | | | | | | | | | | |
| Oligo rev | AAACCCGTCATCGCCGCGACGATC | | | | | | | | | | |
| Chromosome | start | end | strand | MM | target_seq | PAM | alignment | distance | position | gene name | gene id |
| GU305914.1 | 54168 | 54190 | - | 0 | GATCGTCGCGGCGATGACGG | CGG | \|\|\|\|\|\|\|\|[\|\|\|\|\|\|\|\|\|\|\|\|]PAM | NA | Intergenic | NA | NA |
| T29993 | GGCGATGACGGCGGCGTATATGG | 1000 | | | | | | | | | |
| Oligo fwd | CACCGGCGATGACGGCGGCGTATA | | | | | | | | | | |
| Oligo rev | AAACTATACGCCGCCGTCATCGCC | | | | | | | | | | |
| Chromosome | start | end | strand | MM | target_seq | PAM | alignment | distance | position | gene name | gene id |
| GU305914.1 | 54159 | 54181 | - | 0 | GGCGATGACGGCGGCGTATA | TGG | \|\|\|\|\|\|\|\|[\|\|\|\|\|\|\|\|\|\|\|\|]PAM | NA | Intergenic | NA | NA |
| T29994 | GACGGCGGCGTATATGGCTCTGG | 1000 | | | | | | | | | |
| Oligo fwd | CACCGACGGCGGCGTATATGGCTC | | | | | | | | | | |
| Oligo rev | AAACGAGCCATATACGCCGCCGTC | | | | | | | | | | |

| | | | | | | | | | | | |
|---|---|---|---|---|---|---|---|---|---|---|---|
| Chromosome | start | end | strand | MM | target_seq | PAM | alignment | distance | position | gene name | gene id |
| GU305914.1 | 54153 | 54175 | - | 0 | GACGGCGGCGTATATGGCTC | TGG | \|\|\|\|\|\|\|\|[\|\|\|\|\|\|\|\|\|\|\|\|]PAM | NA | Intergenic | NA | NA |
| T29995 | TATGGCTCTGGCTCTGCCGACGG | 1000 | | | | | | | | | |
| Oligo fwd | CACCTATGGCTCTGGCTCTGCCGA | | | | | | | | | | |
| Oligo rev | AAACTCGGCAGAGCCAGAGCCATA | | | | | | | | | | |
| Chromosome | start | end | strand | MM | target_seq | PAM | alignment | distance | position | gene name | gene id |
| GU305914.1 | 54141 | 54163 | - | 0 | TATGGCTCTGGCTCTGCCGA | CGG | \|\|\|\|\|\|\|\|[\|\|\|\|\|\|\|\|\|\|\|\|]PAM | NA | Intergenic | NA | NA |
| T29996 | AGACCTAATCTCCGTCGCCTTGG | 1000 | | | | | | | | | |
| Oligo fwd | CACCAGACCTAATCTCCGTCGCCT | | | | | | | | | | |
| Oligo rev | AAACAGGCGACGGAGATTAGGTCT | | | | | | | | | | |
| Chromosome | start | end | strand | MM | target_seq | PAM | alignment | distance | position | gene name | gene id |
| GU305914.1 | 54072 | 54094 | - | 0 | AGACCTAATCTCCGTCGCCT | TGG | \|\|\|\|\|\|\|\|[\|\|\|\|\|\|\|\|\|\|\|\|]PAM | NA | Intergenic | NA | NA |
| T29997 | CCTAATCTCCGTCGCCTTGGCGG | 1000 | | | | | | | | | |
| Oligo fwd | CACCCCTAATCTCCGTCGCCTTGG | | | | | | | | | | |
| Oligo rev | AAACCCAAGGCGACGGAGATTAGG | | | | | | | | | | |
| Chromosome | start | end | strand | MM | target_seq | PAM | alignment | distance | position | gene name | gene id |
| GU305914.1 | 54069 | 54091 | - | 0 | CCTAATCTCCGTCGCCTTGG | CGG | \|\|\|\|\|\|\|\|[\|\|\|\|\|\|\|\|\|\|\|\|]PAM | NA | Intergenic | NA | NA |
| T29998 | ACTATGCCTTACCGTGAGCTCGG | 1000 | | | | | | | | | |
| Oligo fwd | CACCACTATGCCTTACCGTGAGCT | | | | | | | | | | |
| Oligo rev | AAACAGCTCACGGTAAGGCATAGT | | | | | | | | | | |
| Chromosome | start | end | strand | MM | target_seq | PAM | alignment | distance | position | gene name | gene id |
| GU305914.1 | 54039 | 54061 | - | 0 | ACTATGCCTTACCGTGAGCT | CGG | \|\|\|\|\|\|\|\|[\|\|\|\|\|\|\|\|\|\|\|\|]PAM | NA | Intergenic | NA | NA |
| T29999 | CTATGCCTTACCGTGAGCTCGGG | 1000 | | | | | | | | | |
| Oligo fwd | CACCCTATGCCTTACCGTGAGCTC | | | | | | | | | | |
| Oligo rev | AAACGAGCTCACGGTAAGGCATAG | | | | | | | | | | |
| Chromosome | start | end | strand | MM | target_seq | PAM | alignment | distance | position | gene name | gene id |
| GU305914.1 | 54038 | 54060 | - | 0 | CTATGCCTTACCGTGAGCTC | GGG | \|\|\|\|\|\|\|\|[\|\|\|\|\|\|\|\|\|\|\|\|]PAM | NA | Intergenic | NA | NA |
| T30000 | TATGCCTTACCGTGAGCTCGGGG | 1000 | | | | | | | | | |
| Oligo fwd | CACCTATGCCTTACCGTGAGCTCG | | | | | | | | | | |
| Oligo rev | AAACCGAGCTCACGGTAAGGCATA | | | | | | | | | | |
| Chromosome | start | end | strand | MM | target_seq | PAM | alignment | distance | position | gene name | gene id |
| GU305914.1 | 54037 | 54059 | - | 0 | TATGCCTTACCGTGAGCTCG | GGG | \|\|\|\|\|\|\|\|[\|\|\|\|\|\|\|\|\|\|\|\|]PAM | NA | Intergenic | NA | NA |
| T30001 | ATGCCTTACCGTGAGCTCGGGGG | 1000 | | | | | | | | | |
| Oligo fwd | CACCATGCCTTACCGTGAGCTCGG | | | | | | | | | | |
| Oligo rev | AAACCCGAGCTCACGGTAAGGCAT | | | | | | | | | | |
| Chromosome | start | end | strand | MM | target_seq | PAM | alignment | distance | position | gene name | gene id |
| GU305914.1 | 54036 | 54058 | - | 0 | ATGCCTTACCGTGAGCTCGG | GGG | \|\|\|\|\|\|\|\|[\|\|\|\|\|\|\|\|\|\|\|\|]PAM | NA | Intergenic | NA | NA |
| T30002 | GGGGGCTCTACGCCGTCGTTAGG | 1000 | | | | | | | | | |
| Oligo fwd | CACCGGGGGCTCTACGCCGTCGTT | | | | | | | | | | |
| Oligo rev | AAACAACGACGGCGTAGAGCCCCC | | | | | | | | | | |
| Chromosome | start | end | strand | MM | target_seq | PAM | alignment | distance | position | gene name | gene id |
| GU305914.1 | 54018 | 54040 | - | 0 | GGGGGCTCTACGCCGTCGTT | AGG | \|\|\|\|\|\|\|\|[\|\|\|\|\|\|\|\|\|\|\|\|]PAM | NA | Intergenic | NA | NA |
| T30003 | GTTAGGAGACAACGTTTCCTCGG | 1000 | | | | | | | | | |
| Oligo fwd | CACCGTTAGGAGACAACGTTTCCT | | | | | | | | | | |
| Oligo rev | AAACAGGAAACGTTGTCTCCTAAC | | | | | | | | | | |
| Chromosome | start | end | strand | MM | target_seq | PAM | alignment | distance | position | gene name | gene id |
| GU305914.1 | 54001 | 54023 | - | 0 | GTTAGGAGACAACGTTTCCT | CGG | \|\|\|\|\|\|\|\|[\|\|\|\|\|\|\|\|\|\|\|\|]PAM | NA | Intergenic | NA | NA |
| T30004 | GGAGACAACGTTTCCTCGGAAGG | 1000 | | | | | | | | | |
| Oligo fwd | CACCGGAGACAACGTTTCCTCGGA | | | | | | | | | | |
| Oligo rev | AAACTCCGAGGAAACGTTGTCTCC | | | | | | | | | | |
| Chromosome | start | end | strand | MM | target_seq | PAM | alignment | distance | position | gene name | gene id |
| GU305914.1 | 53997 | 54019 | - | 0 | GGAGACAACGTTTCCTCGGA | AGG | \|\|\|\|\|\|\|\|[\|\|\|\|\|\|\|\|\|\|\|\|]PAM | NA | Intergenic | NA | NA |
| T30005 | TCGGAAGGCGTCATCTTCGTCGG | 1000 | | | | | | | | | |
| Oligo fwd | CACCTCGGAAGGCGTCATCTTCGT | | | | | | | | | | |
| Oligo rev | AAACACGAAGATGACGCCTTCCGA | | | | | | | | | | |
| Chromosome | start | end | strand | MM | target_seq | PAM | alignment | distance | position | gene name | gene id |
| GU305914.1 | 53982 | 54004 | - | 0 | TCGGAAGGCGTCATCTTCGT | CGG | \|\|\|\|\|\|\|\|[\|\|\|\|\|\|\|\|\|\|\|\|]PAM | NA | Intergenic | NA | NA |
| T30006 | CTGTTTCCCGTCGACGCCAACGG | 1000 | | | | | | | | | |
| Oligo fwd | CACCCTGTTTCCCGTCGACGCCAA | | | | | | | | | | |
| Oligo rev | AAACTTGGCGTCGACGGGAAACAG | | | | | | | | | | |
| Chromosome | start | end | strand | MM | target_seq | PAM | alignment | distance | position | gene name | gene id |
| GU305914.1 | 53940 | 53962 | - | 0 | CTGTTTCCCGTCGACGCCAA | CGG | \|\|\|\|\|\|\|\|[\|\|\|\|\|\|\|\|\|\|\|\|]PAM | NA | Intergenic | NA | NA |
| T30007 | TCCCGTCGACGCCAACGGAGTGG | 1000 | | | | | | | | | |
| Oligo fwd | CACCTCCCGTCGACGCCAACGGAG | | | | | | | | | | |
| Oligo rev | AAACCTCCGTTGGCGTCGACGGGA | | | | | | | | | | |
| Chromosome | start | end | strand | MM | target_seq | PAM | alignment | distance | position | gene name | gene id |
| GU305914.1 | 53935 | 53957 | - | 0 | TCCCGTCGACGCCAACGGAG | TGG | \|\|\|\|\|\|\|\|[\|\|\|\|\|\|\|\|\|\|\|\|]PAM | NA | Intergenic | NA | NA |
| T30008 | CGCCAACGGAGTGGCCATCCCGG | 1000 | | | | | | | | | |
| Oligo fwd | CACCCGCCAACGGAGTGGCCATCC | | | | | | | | | | |
| Oligo rev | AAACGGATGGCCACTCCGTTGGCG | | | | | | | | | | |
| Chromosome | start | end | strand | MM | target_seq | PAM | alignment | distance | position | gene name | gene id |
| GU305914.1 | 53926 | 53948 | - | 0 | CGCCAACGGAGTGGCCATCC | CGG | \|\|\|\|\|\|\|\|[\|\|\|\|\|\|\|\|\|\|\|\|]PAM | NA | Intergenic | NA | NA |
| T30009 | GCGCGACCGCCGCGAAGAGTTGG | 1000 | | | | | | | | | |
| Oligo fwd | CACCGCGCGACCGCCGCGAAGAGT | | | | | | | | | | |

<table>
<tr><td>Oligo rev</td><td>AAACACTCTTCGCGGCGGTCGCGC</td><td></td><td></td><td></td><td></td><td></td><td></td><td></td><td></td><td></td><td></td></tr>
<tr><td>Chromosome</td><td>start</td><td>end</td><td>strand</td><td>MM</td><td>target_seq</td><td>PAM</td><td>alignment</td><td>distance</td><td>position</td><td>gene name</td><td>gene id</td></tr>
<tr><td>GU305914.1</td><td>53904</td><td>53926</td><td>-</td><td>0</td><td>GCGCGACCGCCGCGAAGAGT</td><td>TGG</td><td>||||||||[||||||||||||]PAM</td><td>NA</td><td>Intergenic</td><td>NA</td><td>NA</td></tr>
<tr><td>T30010</td><td>ACCGCCGCGAAGAGTTGGCACGG</td><td>1000</td><td></td><td></td><td></td><td></td><td></td><td></td><td></td><td></td><td></td></tr>
<tr><td>Oligo fwd</td><td>CACCACCGCCGCGAAGAGTTGGCA</td><td></td><td></td><td></td><td></td><td></td><td></td><td></td><td></td><td></td><td></td></tr>
<tr><td>Oligo rev</td><td>AAACTGCCAACTCTTCGCGGCGGT</td><td></td><td></td><td></td><td></td><td></td><td></td><td></td><td></td><td></td><td></td></tr>
<tr><td>Chromosome</td><td>start</td><td>end</td><td>strand</td><td>MM</td><td>target_seq</td><td>PAM</td><td>alignment</td><td>distance</td><td>position</td><td>gene name</td><td>gene id</td></tr>
<tr><td>GU305914.1</td><td>53899</td><td>53921</td><td>-</td><td>0</td><td>ACCGCCGCGAAGAGTTGGCA</td><td>CGG</td><td>||||||||[||||||||||||]PAM</td><td>NA</td><td>Intergenic</td><td>NA</td><td>NA</td></tr>
<tr><td>T30011</td><td>ACGGTCGCCAAACTCTGTAACGG</td><td>1000</td><td></td><td></td><td></td><td></td><td></td><td></td><td></td><td></td><td></td></tr>
<tr><td>Oligo fwd</td><td>CACCACGGTCGCCAAACTCTGTAA</td><td></td><td></td><td></td><td></td><td></td><td></td><td></td><td></td><td></td><td></td></tr>
<tr><td>Oligo rev</td><td>AAACTTACAGAGTTTGGCGACCGT</td><td></td><td></td><td></td><td></td><td></td><td></td><td></td><td></td><td></td><td></td></tr>
<tr><td>Chromosome</td><td>start</td><td>end</td><td>strand</td><td>MM</td><td>target_seq</td><td>PAM</td><td>alignment</td><td>distance</td><td>position</td><td>gene name</td><td>gene id</td></tr>
<tr><td>GU305914.1</td><td>53880</td><td>53902</td><td>-</td><td>0</td><td>ACGGTCGCCAAACTCTGTAA</td><td>CGG</td><td>||||||||[||||||||||||]PAM</td><td>NA</td><td>Intergenic</td><td>NA</td><td>NA</td></tr>
<tr><td>T30012</td><td>TAAAAAAATCACTCCCGACGTGG</td><td>1000</td><td></td><td></td><td></td><td></td><td></td><td></td><td></td><td></td><td></td></tr>
<tr><td>Oligo fwd</td><td>CACCTAAAAAAATCACTCCCGACG</td><td></td><td></td><td></td><td></td><td></td><td></td><td></td><td></td><td></td><td></td></tr>
<tr><td>Oligo rev</td><td>AAACCGTCGGGAGTGATTTTTTTA</td><td></td><td></td><td></td><td></td><td></td><td></td><td></td><td></td><td></td><td></td></tr>
<tr><td>Chromosome</td><td>start</td><td>end</td><td>strand</td><td>MM</td><td>target_seq</td><td>PAM</td><td>alignment</td><td>distance</td><td>position</td><td>gene name</td><td>gene id</td></tr>
<tr><td>GU305914.1</td><td>53851</td><td>53873</td><td>-</td><td>0</td><td>TAAAAAAATCACTCCCGACG</td><td>TGG</td><td>||||||||[||||||||||||]PAM</td><td>NA</td><td>Intergenic</td><td>NA</td><td>NA</td></tr>
<tr><td>T30013</td><td>GTGATCATTGACAGATGAATTGG</td><td>1000</td><td></td><td></td><td></td><td></td><td></td><td></td><td></td><td></td><td></td></tr>
<tr><td>Oligo fwd</td><td>CACCGTGATCATTGACAGATGAAT</td><td></td><td></td><td></td><td></td><td></td><td></td><td></td><td></td><td></td><td></td></tr>
<tr><td>Oligo rev</td><td>AAACATTCATCTGTCAATGATCAC</td><td></td><td></td><td></td><td></td><td></td><td></td><td></td><td></td><td></td><td></td></tr>
<tr><td>Chromosome</td><td>start</td><td>end</td><td>strand</td><td>MM</td><td>target_seq</td><td>PAM</td><td>alignment</td><td>distance</td><td>position</td><td>gene name</td><td>gene id</td></tr>
<tr><td>GU305914.1</td><td>53781</td><td>53803</td><td>-</td><td>0</td><td>GTGATCATTGACAGATGAAT</td><td>TGG</td><td>||||||||[||||||||||||]PAM</td><td>NA</td><td>Intergenic</td><td>NA</td><td>NA</td></tr>
<tr><td>T30014</td><td>TGATCATTGACAGATGAATTGGG</td><td>1000</td><td></td><td></td><td></td><td></td><td></td><td></td><td></td><td></td><td></td></tr>
<tr><td>Oligo fwd</td><td>CACCTGATCATTGACAGATGAATT</td><td></td><td></td><td></td><td></td><td></td><td></td><td></td><td></td><td></td><td></td></tr>
<tr><td>Oligo rev</td><td>AAACAATTCATCTGTCAATGATCA</td><td></td><td></td><td></td><td></td><td></td><td></td><td></td><td></td><td></td><td></td></tr>
<tr><td>Chromosome</td><td>start</td><td>end</td><td>strand</td><td>MM</td><td>target_seq</td><td>PAM</td><td>alignment</td><td>distance</td><td>position</td><td>gene name</td><td>gene id</td></tr>
<tr><td>GU305914.1</td><td>53780</td><td>53802</td><td>-</td><td>0</td><td>TGATCATTGACAGATGAATT</td><td>GGG</td><td>||||||||[||||||||||||]PAM</td><td>NA</td><td>Intergenic</td><td>NA</td><td>NA</td></tr>
<tr><td>T30015</td><td>GATCATTGACAGATGAATTGGGG</td><td>1000</td><td></td><td></td><td></td><td></td><td></td><td></td><td></td><td></td><td></td></tr>
<tr><td>Oligo fwd</td><td>CACCGATCATTGACAGATGAATTG</td><td></td><td></td><td></td><td></td><td></td><td></td><td></td><td></td><td></td><td></td></tr>
<tr><td>Oligo rev</td><td>AAACCAATTCATCTGTCAATGATC</td><td></td><td></td><td></td><td></td><td></td><td></td><td></td><td></td><td></td><td></td></tr>
<tr><td>Chromosome</td><td>start</td><td>end</td><td>strand</td><td>MM</td><td>target_seq</td><td>PAM</td><td>alignment</td><td>distance</td><td>position</td><td>gene name</td><td>gene id</td></tr>
<tr><td>GU305914.1</td><td>53779</td><td>53801</td><td>-</td><td>0</td><td>GATCATTGACAGATGAATTG</td><td>GGG</td><td>||||||||[||||||||||||]PAM</td><td>NA</td><td>Intergenic</td><td>NA</td><td>NA</td></tr>
<tr><td>T30016</td><td>ATCATTGACAGATGAATTGGGGG</td><td>1000</td><td></td><td></td><td></td><td></td><td></td><td></td><td></td><td></td><td></td></tr>
<tr><td>Oligo fwd</td><td>CACCATCATTGACAGATGAATTGG</td><td></td><td></td><td></td><td></td><td></td><td></td><td></td><td></td><td></td><td></td></tr>
<tr><td>Oligo rev</td><td>AAACCCAATTCATCTGTCAATGAT</td><td></td><td></td><td></td><td></td><td></td><td></td><td></td><td></td><td></td><td></td></tr>
<tr><td>Chromosome</td><td>start</td><td>end</td><td>strand</td><td>MM</td><td>target_seq</td><td>PAM</td><td>alignment</td><td>distance</td><td>position</td><td>gene name</td><td>gene id</td></tr>
<tr><td>GU305914.1</td><td>53778</td><td>53800</td><td>-</td><td>0</td><td>ATCATTGACAGATGAATTGG</td><td>GGG</td><td>||||||||[||||||||||||]PAM</td><td>NA</td><td>Intergenic</td><td>NA</td><td>NA</td></tr>
<tr><td>T30017</td><td>ATTGACAGATGAATTGGGGGAGG</td><td>1000</td><td></td><td></td><td></td><td></td><td></td><td></td><td></td><td></td><td></td></tr>
<tr><td>Oligo fwd</td><td>CACCATTGACAGATGAATTGGGGG</td><td></td><td></td><td></td><td></td><td></td><td></td><td></td><td></td><td></td><td></td></tr>
<tr><td>Oligo rev</td><td>AAACCCCCCAATTCATCTGTCAAT</td><td></td><td></td><td></td><td></td><td></td><td></td><td></td><td></td><td></td><td></td></tr>
<tr><td>Chromosome</td><td>start</td><td>end</td><td>strand</td><td>MM</td><td>target_seq</td><td>PAM</td><td>alignment</td><td>distance</td><td>position</td><td>gene name</td><td>gene id</td></tr>
<tr><td>GU305914.1</td><td>53775</td><td>53797</td><td>-</td><td>0</td><td>ATTGACAGATGAATTGGGGG</td><td>AGG</td><td>||||||||[||||||||||||]PAM</td><td>NA</td><td>Intergenic</td><td>NA</td><td>NA</td></tr>
<tr><td>T30018</td><td>GACAGATGAATTGGGGGAGGAGG</td><td>1000</td><td></td><td></td><td></td><td></td><td></td><td></td><td></td><td></td><td></td></tr>
<tr><td>Oligo fwd</td><td>CACCGACAGATGAATTGGGGGAGG</td><td></td><td></td><td></td><td></td><td></td><td></td><td></td><td></td><td></td><td></td></tr>
<tr><td>Oligo rev</td><td>AAACCCTCCCCCAATTCATCTGTC</td><td></td><td></td><td></td><td></td><td></td><td></td><td></td><td></td><td></td><td></td></tr>
<tr><td>Chromosome</td><td>start</td><td>end</td><td>strand</td><td>MM</td><td>target_seq</td><td>PAM</td><td>alignment</td><td>distance</td><td>position</td><td>gene name</td><td>gene id</td></tr>
<tr><td>GU305914.1</td><td>53772</td><td>53794</td><td>-</td><td>0</td><td>GACAGATGAATTGGGGGAGG</td><td>AGG</td><td>||||||||[||||||||||||]PAM</td><td>NA</td><td>Intergenic</td><td>NA</td><td>NA</td></tr>
<tr><td>T30019</td><td>ACAGATGAATTGGGGGAGGAGGG</td><td>1000</td><td></td><td></td><td></td><td></td><td></td><td></td><td></td><td></td><td></td></tr>
<tr><td>Oligo fwd</td><td>CACCACAGATGAATTGGGGGAGGA</td><td></td><td></td><td></td><td></td><td></td><td></td><td></td><td></td><td></td><td></td></tr>
<tr><td>Oligo rev</td><td>AAACTCCTCCCCCAATTCATCTGT</td><td></td><td></td><td></td><td></td><td></td><td></td><td></td><td></td><td></td><td></td></tr>
<tr><td>Chromosome</td><td>start</td><td>end</td><td>strand</td><td>MM</td><td>target_seq</td><td>PAM</td><td>alignment</td><td>distance</td><td>position</td><td>gene name</td><td>gene id</td></tr>
<tr><td>GU305914.1</td><td>53771</td><td>53793</td><td>-</td><td>0</td><td>ACAGATGAATTGGGGGAGGA</td><td>GGG</td><td>||||||||[||||||||||||]PAM</td><td>NA</td><td>Intergenic</td><td>NA</td><td>NA</td></tr>
<tr><td>T30020</td><td>GATGAATTGGGGGAGGAGGGCGG</td><td>1000</td><td></td><td></td><td></td><td></td><td></td><td></td><td></td><td></td><td></td></tr>
<tr><td>Oligo fwd</td><td>CACCGATGAATTGGGGGAGGAGGG</td><td></td><td></td><td></td><td></td><td></td><td></td><td></td><td></td><td></td><td></td></tr>
<tr><td>Oligo rev</td><td>AAACCCCTCCTCCCCCAATTCATC</td><td></td><td></td><td></td><td></td><td></td><td></td><td></td><td></td><td></td><td></td></tr>
<tr><td>Chromosome</td><td>start</td><td>end</td><td>strand</td><td>MM</td><td>target_seq</td><td>PAM</td><td>alignment</td><td>distance</td><td>position</td><td>gene name</td><td>gene id</td></tr>
<tr><td>GU305914.1</td><td>53768</td><td>53790</td><td>-</td><td>0</td><td>GATGAATTGGGGGAGGAGGG</td><td>CGG</td><td>||||||||[||||||||||||]PAM</td><td>NA</td><td>Intergenic</td><td>NA</td><td>NA</td></tr>
<tr><td>T30021</td><td>AATTGGGGGAGGAGGGCGGTCGG</td><td>1000</td><td></td><td></td><td></td><td></td><td></td><td></td><td></td><td></td><td></td></tr>
<tr><td>Oligo fwd</td><td>CACCAATTGGGGGAGGAGGGCGGT</td><td></td><td></td><td></td><td></td><td></td><td></td><td></td><td></td><td></td><td></td></tr>
<tr><td>Oligo rev</td><td>AAACACCGCCCTCCTCCCCCAATT</td><td></td><td></td><td></td><td></td><td></td><td></td><td></td><td></td><td></td><td></td></tr>
<tr><td>Chromosome</td><td>start</td><td>end</td><td>strand</td><td>MM</td><td>target_seq</td><td>PAM</td><td>alignment</td><td>distance</td><td>position</td><td>gene name</td><td>gene id</td></tr>
<tr><td>GU305914.1</td><td>53764</td><td>53786</td><td>-</td><td>0</td><td>AATTGGGGGAGGAGGGCGGT</td><td>CGG</td><td>||||||||[||||||||||||]PAM</td><td>NA</td><td>Intergenic</td><td>NA</td><td>NA</td></tr>
<tr><td>T30022</td><td>ATTGGGGGAGGAGGGCGGTCGGG</td><td>1000</td><td></td><td></td><td></td><td></td><td></td><td></td><td></td><td></td><td></td></tr>
<tr><td>Oligo fwd</td><td>CACCATTGGGGGAGGAGGGCGGTC</td><td></td><td></td><td></td><td></td><td></td><td></td><td></td><td></td><td></td><td></td></tr>
<tr><td>Oligo rev</td><td>AAACGACCGCCCTCCTCCCCCAAT</td><td></td><td></td><td></td><td></td><td></td><td></td><td></td><td></td><td></td><td></td></tr>
<tr><td>Chromosome</td><td>start</td><td>end</td><td>strand</td><td>MM</td><td>target_seq</td><td>PAM</td><td>alignment</td><td>distance</td><td>position</td><td>gene name</td><td>gene id</td></tr>
<tr><td>GU305914.1</td><td>53763</td><td>53785</td><td>-</td><td>0</td><td>ATTGGGGGAGGAGGGCGGTC</td><td>GGG</td><td>||||||||[||||||||||||]PAM</td><td>NA</td><td>Intergenic</td><td>NA</td><td>NA</td></tr>
<tr><td>T30023</td><td>GGGGGAGGAGGGCGGTCGGGCGG</td><td>1000</td><td></td><td></td><td></td><td></td><td></td><td></td><td></td><td></td><td></td></tr>
<tr><td>Oligo fwd</td><td>CACCGGGGGAGGAGGGCGGTCGGG</td><td></td><td></td><td></td><td></td><td></td><td></td><td></td><td></td><td></td><td></td></tr>
<tr><td>Oligo rev</td><td>AAACCCCGACCGCCCTCCTCCCCC</td><td></td><td></td><td></td><td></td><td></td><td></td><td></td><td></td><td></td><td></td></tr>
<tr><td>Chromosome</td><td>start</td><td>end</td><td>strand</td><td>MM</td><td>target_seq</td><td>PAM</td><td>alignment</td><td>distance</td><td>position</td><td>gene name</td><td>gene id</td></tr>
<tr><td>GU305914.1</td><td>53760</td><td>53782</td><td>-</td><td>0</td><td>GGGGGAGGAGGGCGGTCGGG</td><td>CGG</td><td>||||||||[||||||||||||]PAM</td><td>NA</td><td>Intergenic</td><td>NA</td><td>NA</td></tr>
<tr><td>T30024</td><td>TCGGGCGGCGACACGATCATCGG</td><td>1000</td><td></td><td></td><td></td><td></td><td></td><td></td><td></td><td></td><td></td></tr>
</table>

Oligo fwd CACCTCGGGCGGCGACACGATCAT
Oligo rev AAACATGATCGTGTCGCCGCCCGA
Chromosome start end strand MM target_seq PAM alignment distance position gene name gene id
GU305914.1 53745 53767 - 0 TCGGGCGGCGACACGATCAT CGG ||||||||[||||||||||||]PAM NA Intergenic NA NA

T30025 GATCTTCAATAAAAATCAATCGG 1000
Oligo fwd CACCGATCTTCAATAAAAATCAAT
Oligo rev AAACATTGATTTTTATTGAAGATC
Chromosome start end strand MM target_seq PAM alignment distance position gene name gene id
GU305914.1 53694 53716 - 0 GATCTTCAATAAAAATCAAT CGG ||||||||[||||||||||||]PAM NA Intergenic NA NA

T30026 ATCTTCAATAAAAATCAATCGGG 1000
Oligo fwd CACCATCTTCAATAAAAATCAATC
Oligo rev AAACGATTGATTTTTATTGAAGAT
Chromosome start end strand MM target_seq PAM alignment distance position gene name gene id
GU305914.1 53693 53715 - 0 ATCTTCAATAAAAATCAATC GGG ||||||||[||||||||||||]PAM NA Intergenic NA NA

T30027 TCGCGTCTTCTCTGTCGATCTGG 1000
Oligo fwd CACCTCGCGTCTTCTCTGTCGATC
Oligo rev AAACGATCGACAGAGAAGACGCGA
Chromosome start end strand MM target_seq PAM alignment distance position gene name gene id
GU305914.1 53647 53669 - 0 TCGCGTCTTCTCTGTCGATC TGG ||||||||[||||||||||||]PAM NA Intergenic NA NA

T30028 CGCGTCTTCTCTGTCGATCTGGG 1000
Oligo fwd CACCCGCGTCTTCTCTGTCGATCT
Oligo rev AAACAGATCGACAGAGAAGACGCG
Chromosome start end strand MM target_seq PAM alignment distance position gene name gene id
GU305914.1 53646 53668 - 0 CGCGTCTTCTCTGTCGATCT GGG ||||||||[||||||||||||]PAM NA Intergenic NA NA

T30029 TTCTCTGTCGATCTGGGTGTCGG 1000
Oligo fwd CACCTTCTCTGTCGATCTGGGTGT
Oligo rev AAACACACCCAGATCGACAGAGAA
Chromosome start end strand MM target_seq PAM alignment distance position gene name gene id
GU305914.1 53640 53662 - 0 TTCTCTGTCGATCTGGGTGT CGG ||||||||[||||||||||||]PAM NA Intergenic NA NA

T30030 TCTCTGTCGATCTGGGTGTCGGG 1000
Oligo fwd CACCTCTCTGTCGATCTGGGTGTC
Oligo rev AAACGACACCCAGATCGACAGAGA
Chromosome start end strand MM target_seq PAM alignment distance position gene name gene id
GU305914.1 53639 53661 - 0 TCTCTGTCGATCTGGGTGTC GGG ||||||||[||||||||||||]PAM NA Intergenic NA NA

T30031 CTCTGTCGATCTGGGTGTCGGGG 1000
Oligo fwd CACCCTCTGTCGATCTGGGTGTCG
Oligo rev AAACCGACACCCAGATCGACAGAG
Chromosome start end strand MM target_seq PAM alignment distance position gene name gene id
GU305914.1 53638 53660 - 0 CTCTGTCGATCTGGGTGTCG GGG ||||||||[||||||||||||]PAM NA Intergenic NA NA

T30032 TGTCGATCTGGGTGTCGGGGAGG 1000
Oligo fwd CACCTGTCGATCTGGGTGTCGGGG
Oligo rev AAACCCCCGACACCCAGATCGACA
Chromosome start end strand MM target_seq PAM alignment distance position gene name gene id
GU305914.1 53635 53657 - 0 TGTCGATCTGGGTGTCGGGG AGG ||||||||[||||||||||||]PAM NA Intergenic NA NA

T30033 GTCGATCTGGGTGTCGGGGAGGG 1000
Oligo fwd CACCGTCGATCTGGGTGTCGGGGA
Oligo rev AAACTCCCCGACACCCAGATCGAC
Chromosome start end strand MM target_seq PAM alignment distance position gene name gene id
GU305914.1 53634 53656 - 0 GTCGATCTGGGTGTCGGGGA GGG ||||||||[||||||||||||]PAM NA Intergenic NA NA

T30034 TGGGTGTCGGGGAGGGATGTCGG 1000
Oligo fwd CACCTGGGTGTCGGGGAGGGATGT
Oligo rev AAACACATCCCTCCCCGACACCCA
Chromosome start end strand MM target_seq PAM alignment distance position gene name gene id
GU305914.1 53627 53649 - 0 TGGGTGTCGGGGAGGGATGT CGG ||||||||[||||||||||||]PAM NA Intergenic NA NA

T30035 GGGTGTCGGGGAGGGATGTCGGG 1000
Oligo fwd CACCGGGTGTCGGGGAGGGATGTC
Oligo rev AAACGACATCCCTCCCCGACACCC
Chromosome start end strand MM target_seq PAM alignment distance position gene name gene id
GU305914.1 53626 53648 - 0 GGGTGTCGGGGAGGGATGTC GGG ||||||||[||||||||||||]PAM NA Intergenic NA NA

T30036 CGGGGAGGGATGTCGGGCAGAGG 1000
Oligo fwd CACCCGGGGAGGGATGTCGGGCAG
Oligo rev AAACCTGCCCGACATCCCTCCCCG
Chromosome start end strand MM target_seq PAM alignment distance position gene name gene id
GU305914.1 53620 53642 - 0 CGGGGAGGGATGTCGGGCAG AGG ||||||||[||||||||||||]PAM NA Intergenic NA NA

T30037 GGAGGGATGTCGGGCAGAGGCGG 1000
Oligo fwd CACCGGAGGGATGTCGGGCAGAGG
Oligo rev AAACCCTCTGCCCGACATCCCTCC
Chromosome start end strand MM target_seq PAM alignment distance position gene name gene id
GU305914.1 53617 53639 - 0 GGAGGGATGTCGGGCAGAGG CGG ||||||||[||||||||||||]PAM NA Intergenic NA NA

T30038 GGGATGTCGGGCAGAGGCGGAGG 1000
Oligo fwd CACCGGGATGTCGGGCAGAGGCGG
Oligo rev AAACCCGCCTCTGCCCGACATCCC
Chromosome start end strand MM target_seq PAM alignment distance position gene name gene id
GU305914.1 53614 53636 - 0 GGGATGTCGGGCAGAGGCGG AGG ||||||||[||||||||||||]PAM NA Intergenic NA NA
GU305914.1 67733 67755 + 3 GTGATGTCGGGGAGCGGCGG CGG |-||||||[|||-||-|||||]PAM NA Intergenic NA NA

| | | | | | | | | | | | |
|---|---|---|---|---|---|---|---|---|---|---|---|
| T30039 | TCGGGCAGAGGCGGAGGCACCGG | 1000 | | | | | | | | | |
| Oligo fwd | CACCTCGGGCAGAGGCGGAGGCAC | | | | | | | | | | |
| Oligo rev | AAACGTGCCTCCGCCTCTGCCCGA | | | | | | | | | | |
| Chromosome | start | end | strand | MM | target_seq | PAM | alignment | distance | position | gene name | gene id |
| GU305914.1 | 53608 | 53630 | - | 0 | TCGGGCAGAGGCGGAGGCAC | CGG | \|\|\|\|\|\|\|\|[\|\|\|\|\|\|\|\|\|\|\|\|]PAM | NA | Intergenic | NA | NA |

| | | | | | | | | | | | |
|---|---|---|---|---|---|---|---|---|---|---|---|
| T30040 | CGGGCAGAGGCGGAGGCACCGGG | 1000 | | | | | | | | | |
| Oligo fwd | CACCCGGGCAGAGGCGGAGGCACC | | | | | | | | | | |
| Oligo rev | AAACGGTGCCTCCGCCTCTGCCCG | | | | | | | | | | |
| Chromosome | start | end | strand | MM | target_seq | PAM | alignment | distance | position | gene name | gene id |
| GU305914.1 | 53607 | 53629 | - | 0 | CGGGCAGAGGCGGAGGCACC | GGG | \|\|\|\|\|\|\|\|[\|\|\|\|\|\|\|\|\|\|\|\|]PAM | NA | Intergenic | NA | NA |

| | | | | | | | | | | | |
|---|---|---|---|---|---|---|---|---|---|---|---|
| T30041 | CAGAGGCGGAGGCACCGGGACGG | 1000 | | | | | | | | | |
| Oligo fwd | CACCCAGAGGCGGAGGCACCGGGA | | | | | | | | | | |
| Oligo rev | AAACTCCCGGTGCCTCCGCCTCTG | | | | | | | | | | |
| Chromosome | start | end | strand | MM | target_seq | PAM | alignment | distance | position | gene name | gene id |
| GU305914.1 | 53603 | 53625 | - | 0 | CAGAGGCGGAGGCACCGGGA | CGG | \|\|\|\|\|\|\|\|[\|\|\|\|\|\|\|\|\|\|\|\|]PAM | NA | Intergenic | NA | NA |

| | | | | | | | | | | | |
|---|---|---|---|---|---|---|---|---|---|---|---|
| T30042 | AGGCGGAGGCACCGGGACGGTGG | 1000 | | | | | | | | | |
| Oligo fwd | CACCAGGCGGAGGCACCGGGACGG | | | | | | | | | | |
| Oligo rev | AAACCCGTCCCGGTGCCTCCGCCT | | | | | | | | | | |
| Chromosome | start | end | strand | MM | target_seq | PAM | alignment | distance | position | gene name | gene id |
| GU305914.1 | 53600 | 53622 | - | 0 | AGGCGGAGGCACCGGGACGG | TGG | \|\|\|\|\|\|\|\|[\|\|\|\|\|\|\|\|\|\|\|\|]PAM | NA | Intergenic | NA | NA |

| | | | | | | | | | | | |
|---|---|---|---|---|---|---|---|---|---|---|---|
| T30043 | GGCGGAGGCACCGGGACGGTGGG | 1000 | | | | | | | | | |
| Oligo fwd | CACCGGCGGAGGCACCGGGACGGT | | | | | | | | | | |
| Oligo rev | AAACACCGTCCCGGTGCCTCCGCC | | | | | | | | | | |
| Chromosome | start | end | strand | MM | target_seq | PAM | alignment | distance | position | gene name | gene id |
| GU305914.1 | 53599 | 53621 | - | 0 | GGCGGAGGCACCGGGACGGT | GGG | \|\|\|\|\|\|\|\|[\|\|\|\|\|\|\|\|\|\|\|\|]PAM | NA | Intergenic | NA | NA |

| | | | | | | | | | | | |
|---|---|---|---|---|---|---|---|---|---|---|---|
| T30044 | GGACGGTGGGTGTCCTGTCTCGG | 1000 | | | | | | | | | |
| Oligo fwd | CACCGGACGGTGGGTGTCCTGTCT | | | | | | | | | | |
| Oligo rev | AAACAGACAGGACACCCACCGTCC | | | | | | | | | | |
| Chromosome | start | end | strand | MM | target_seq | PAM | alignment | distance | position | gene name | gene id |
| GU305914.1 | 53586 | 53608 | - | 0 | GGACGGTGGGTGTCCTGTCT | CGG | \|\|\|\|\|\|\|\|[\|\|\|\|\|\|\|\|\|\|\|\|]PAM | NA | Intergenic | NA | NA |

| | | | | | | | | | | | |
|---|---|---|---|---|---|---|---|---|---|---|---|
| T30045 | GACGATCCTCTCCGATGAAGAGG | 1000 | | | | | | | | | |
| Oligo fwd | CACCGACGATCCTCTCCGATGAAG | | | | | | | | | | |
| Oligo rev | AAACCTTCATCGGAGAGGATCGTC | | | | | | | | | | |
| Chromosome | start | end | strand | MM | target_seq | PAM | alignment | distance | position | gene name | gene id |
| GU305914.1 | 53564 | 53586 | - | 0 | GACGATCCTCTCCGATGAAG | AGG | \|\|\|\|\|\|\|\|[\|\|\|\|\|\|\|\|\|\|\|\|]PAM | NA | Intergenic | NA | NA |

| | | | | | | | | | | | |
|---|---|---|---|---|---|---|---|---|---|---|---|
| T30046 | TGAAGAGGTCACCATCTTCCCGG | 1000 | | | | | | | | | |
| Oligo fwd | CACCTGAAGAGGTCACCATCTTCC | | | | | | | | | | |
| Oligo rev | AAACGGAAGATGGTGACCTCTTCA | | | | | | | | | | |
| Chromosome | start | end | strand | MM | target_seq | PAM | alignment | distance | position | gene name | gene id |
| GU305914.1 | 53549 | 53571 | - | 0 | TGAAGAGGTCACCATCTTCC | CGG | \|\|\|\|\|\|\|\|[\|\|\|\|\|\|\|\|\|\|\|\|]PAM | NA | Intergenic | NA | NA |

| | | | | | | | | | | | |
|---|---|---|---|---|---|---|---|---|---|---|---|
| T30047 | CACCATCTTCCCGGAGAGCTCGG | 1000 | | | | | | | | | |
| Oligo fwd | CACCCACCATCTTCCCGGAGAGCT | | | | | | | | | | |
| Oligo rev | AAACAGCTCTCCGGGAAGATGGTG | | | | | | | | | | |
| Chromosome | start | end | strand | MM | target_seq | PAM | alignment | distance | position | gene name | gene id |
| GU305914.1 | 53540 | 53562 | - | 0 | CACCATCTTCCCGGAGAGCT | CGG | \|\|\|\|\|\|\|\|[\|\|\|\|\|\|\|\|\|\|\|\|]PAM | NA | Intergenic | NA | NA |

| | | | | | | | | | | | |
|---|---|---|---|---|---|---|---|---|---|---|---|
| T30048 | CATCTTCCCGGAGAGCTCGGCGG | 1000 | | | | | | | | | |
| Oligo fwd | CACCCATCTTCCCGGAGAGCTCGG | | | | | | | | | | |
| Oligo rev | AAACCCGAGCTCTCCGGGAAGATG | | | | | | | | | | |
| Chromosome | start | end | strand | MM | target_seq | PAM | alignment | distance | position | gene name | gene id |
| GU305914.1 | 53537 | 53559 | - | 0 | CATCTTCCCGGAGAGCTCGG | CGG | \|\|\|\|\|\|\|\|[\|\|\|\|\|\|\|\|\|\|\|\|]PAM | NA | Intergenic | NA | NA |

| | | | | | | | | | | | |
|---|---|---|---|---|---|---|---|---|---|---|---|
| T30049 | CTCGGCGGTCTTGAACCTCTCGG | 1000 | | | | | | | | | |
| Oligo fwd | CACCCTCGGCGGTCTTGAACCTCT | | | | | | | | | | |
| Oligo rev | AAACAGAGGTTCAAGACCGCCGAG | | | | | | | | | | |
| Chromosome | start | end | strand | MM | target_seq | PAM | alignment | distance | position | gene name | gene id |
| GU305914.1 | 53522 | 53544 | - | 0 | CTCGGCGGTCTTGAACCTCT | CGG | \|\|\|\|\|\|\|\|[\|\|\|\|\|\|\|\|\|\|\|\|]PAM | NA | Intergenic | NA | NA |

| | | | | | | | | | | | |
|---|---|---|---|---|---|---|---|---|---|---|---|
| T30050 | TCGGCGGTCTTGAACCTCTCGGG | 1000 | | | | | | | | | |
| Oligo fwd | CACCTCGGCGGTCTTGAACCTCTC | | | | | | | | | | |
| Oligo rev | AAACGAGAGGTTCAAGACCGCCGA | | | | | | | | | | |
| Chromosome | start | end | strand | MM | target_seq | PAM | alignment | distance | position | gene name | gene id |
| GU305914.1 | 53521 | 53543 | - | 0 | TCGGCGGTCTTGAACCTCTC | GGG | \|\|\|\|\|\|\|\|[\|\|\|\|\|\|\|\|\|\|\|\|]PAM | NA | Intergenic | NA | NA |

| | | | | | | | | | | | |
|---|---|---|---|---|---|---|---|---|---|---|---|
| T30051 | CTCTCGGGCACGACCACCGTCGG | 1000 | | | | | | | | | |
| Oligo fwd | CACCCTCTCGGGCACGACCACCGT | | | | | | | | | | |
| Oligo rev | AAACACGGTGGTCGTGCCCGAGAG | | | | | | | | | | |
| Chromosome | start | end | strand | MM | target_seq | PAM | alignment | distance | position | gene name | gene id |
| GU305914.1 | 53506 | 53528 | - | 0 | CTCTCGGGCACGACCACCGT | CGG | \|\|\|\|\|\|\|\|[\|\|\|\|\|\|\|\|\|\|\|\|]PAM | NA | Intergenic | NA | NA |
| GU305914.1 | 230092 | 230114 | - | 4 | CTGTCCGGCACGACTACCGC | GGG | \|\|-\|\|-\|\|[\|\|\|\|\|\|-\|\|\|\|-]PAM | NA | Intergenic | NA | NA |

| | | | | | | | | | | | |
|---|---|---|---|---|---|---|---|---|---|---|---|
| T30052 | CGGGCACGACCACCGTCGGTCGG | 1000 | | | | | | | | | |
| Oligo fwd | CACCCGGGCACGACCACCGTCGGT | | | | | | | | | | |
| Oligo rev | AAACACCGACGGTGGTCGTGCCCG | | | | | | | | | | |
| Chromosome | start | end | strand | MM | target_seq | PAM | alignment | distance | position | gene name | gene id |
| GU305914.1 | 53502 | 53524 | - | 0 | CGGGCACGACCACCGTCGGT | CGG | \|\|\|\|\|\|\|\|[\|\|\|\|\|\|\|\|\|\|\|\|]PAM | NA | Intergenic | NA | NA |

| | | |
|---|---|---|
| T30053 | GGGCACGACCACCGTCGGTCGGG | 1000 |
| Oligo fwd | CACCGGGCACGACCACCGTCGGTC | |
| Oligo rev | AAACGACCGACGGTGGTCGTGCCC | |

<table>
<tr><td>Chromosome</td><td>start</td><td>end</td><td>strand</td><td>MM</td><td>target_seq</td><td>PAM</td><td>alignment</td><td>distance</td><td>position</td><td>gene name</td><td>gene id</td></tr>
<tr><td>GU305914.1</td><td>53501</td><td>53523</td><td>-</td><td>0</td><td>GGGCACGACCACCGTCGGTC</td><td>GGG</td><td>||||||||[||||||||||||]PAM</td><td>NA</td><td>Intergenic</td><td>NA</td><td>NA</td></tr>
<tr><td>T30054</td><td>GGCACGACCACCGTCGGTCGGGG</td><td>1000</td><td></td><td></td><td></td><td></td><td></td><td></td><td></td><td></td><td></td></tr>
<tr><td>Oligo fwd</td><td>CACCGGCACGACCACCGTCGGTCG</td><td></td><td></td><td></td><td></td><td></td><td></td><td></td><td></td><td></td><td></td></tr>
<tr><td>Oligo rev</td><td>AAACCGACCGACGGTGGTCGTGCC</td><td></td><td></td><td></td><td></td><td></td><td></td><td></td><td></td><td></td><td></td></tr>
<tr><td>Chromosome</td><td>start</td><td>end</td><td>strand</td><td>MM</td><td>target_seq</td><td>PAM</td><td>alignment</td><td>distance</td><td>position</td><td>gene name</td><td>gene id</td></tr>
<tr><td>GU305914.1</td><td>53500</td><td>53522</td><td>-</td><td>0</td><td>GGCACGACCACCGTCGGTCG</td><td>GGG</td><td>||||||||[||||||||||||]PAM</td><td>NA</td><td>Intergenic</td><td>NA</td><td>NA</td></tr>
<tr><td>T30055</td><td>CGGTCGGGGCGAGAACGACGTGG</td><td>1000</td><td></td><td></td><td></td><td></td><td></td><td></td><td></td><td></td><td></td></tr>
<tr><td>Oligo fwd</td><td>CACCCGGTCGGGGCGAGAACGACG</td><td></td><td></td><td></td><td></td><td></td><td></td><td></td><td></td><td></td><td></td></tr>
<tr><td>Oligo rev</td><td>AAACCGTCGTTCTCGCCCCGACCG</td><td></td><td></td><td></td><td></td><td></td><td></td><td></td><td></td><td></td><td></td></tr>
<tr><td>Chromosome</td><td>start</td><td>end</td><td>strand</td><td>MM</td><td>target_seq</td><td>PAM</td><td>alignment</td><td>distance</td><td>position</td><td>gene name</td><td>gene id</td></tr>
<tr><td>GU305914.1</td><td>53486</td><td>53508</td><td>-</td><td>0</td><td>CGGTCGGGGCGAGAACGACG</td><td>TGG</td><td>||||||||[||||||||||||]PAM</td><td>NA</td><td>Intergenic</td><td>NA</td><td>NA</td></tr>
<tr><td>T30056</td><td>GGTCGGGGCGAGAACGACGTGGG</td><td>1000</td><td></td><td></td><td></td><td></td><td></td><td></td><td></td><td></td><td></td></tr>
<tr><td>Oligo fwd</td><td>CACCGGTCGGGGCGAGAACGACGT</td><td></td><td></td><td></td><td></td><td></td><td></td><td></td><td></td><td></td><td></td></tr>
<tr><td>Oligo rev</td><td>AAACACGTCGTTCTCGCCCCGACC</td><td></td><td></td><td></td><td></td><td></td><td></td><td></td><td></td><td></td><td></td></tr>
<tr><td>Chromosome</td><td>start</td><td>end</td><td>strand</td><td>MM</td><td>target_seq</td><td>PAM</td><td>alignment</td><td>distance</td><td>position</td><td>gene name</td><td>gene id</td></tr>
<tr><td>GU305914.1</td><td>53485</td><td>53507</td><td>-</td><td>0</td><td>GGTCGGGGCGAGAACGACGT</td><td>GGG</td><td>||||||||[||||||||||||]PAM</td><td>NA</td><td>Intergenic</td><td>NA</td><td>NA</td></tr>
<tr><td>T30057</td><td>GTCGGGGCGAGAACGACGTGGGG</td><td>1000</td><td></td><td></td><td></td><td></td><td></td><td></td><td></td><td></td><td></td></tr>
<tr><td>Oligo fwd</td><td>CACCGTCGGGGCGAGAACGACGTG</td><td></td><td></td><td></td><td></td><td></td><td></td><td></td><td></td><td></td><td></td></tr>
<tr><td>Oligo rev</td><td>AAACCACGTCGTTCTCGCCCCGAC</td><td></td><td></td><td></td><td></td><td></td><td></td><td></td><td></td><td></td><td></td></tr>
<tr><td>Chromosome</td><td>start</td><td>end</td><td>strand</td><td>MM</td><td>target_seq</td><td>PAM</td><td>alignment</td><td>distance</td><td>position</td><td>gene name</td><td>gene id</td></tr>
<tr><td>GU305914.1</td><td>53484</td><td>53506</td><td>-</td><td>0</td><td>GTCGGGGCGAGAACGACGTG</td><td>GGG</td><td>||||||||[||||||||||||]PAM</td><td>NA</td><td>Intergenic</td><td>NA</td><td>NA</td></tr>
<tr><td>T30058</td><td>GACGTGGGGACTATCCGTGACGG</td><td>1000</td><td></td><td></td><td></td><td></td><td></td><td></td><td></td><td></td><td></td></tr>
<tr><td>Oligo fwd</td><td>CACCGACGTGGGGACTATCCGTGA</td><td></td><td></td><td></td><td></td><td></td><td></td><td></td><td></td><td></td><td></td></tr>
<tr><td>Oligo rev</td><td>AAACTCACGGATAGTCCCCACGTC</td><td></td><td></td><td></td><td></td><td></td><td></td><td></td><td></td><td></td><td></td></tr>
<tr><td>Chromosome</td><td>start</td><td>end</td><td>strand</td><td>MM</td><td>target_seq</td><td>PAM</td><td>alignment</td><td>distance</td><td>position</td><td>gene name</td><td>gene id</td></tr>
<tr><td>GU305914.1</td><td>53470</td><td>53492</td><td>-</td><td>0</td><td>GACGTGGGGACTATCCGTGA</td><td>CGG</td><td>||||||||[||||||||||||]PAM</td><td>NA</td><td>Intergenic</td><td>NA</td><td>NA</td></tr>
<tr><td>T30059</td><td>ACGTGGGGACTATCCGTGACGGG</td><td>1000</td><td></td><td></td><td></td><td></td><td></td><td></td><td></td><td></td><td></td></tr>
<tr><td>Oligo fwd</td><td>CACCACGTGGGGACTATCCGTGAC</td><td></td><td></td><td></td><td></td><td></td><td></td><td></td><td></td><td></td><td></td></tr>
<tr><td>Oligo rev</td><td>AAACGTCACGGATAGTCCCCACGT</td><td></td><td></td><td></td><td></td><td></td><td></td><td></td><td></td><td></td><td></td></tr>
<tr><td>Chromosome</td><td>start</td><td>end</td><td>strand</td><td>MM</td><td>target_seq</td><td>PAM</td><td>alignment</td><td>distance</td><td>position</td><td>gene name</td><td>gene id</td></tr>
<tr><td>GU305914.1</td><td>53469</td><td>53491</td><td>-</td><td>0</td><td>ACGTGGGGACTATCCGTGAC</td><td>GGG</td><td>||||||||[||||||||||||]PAM</td><td>NA</td><td>Intergenic</td><td>NA</td><td>NA</td></tr>
<tr><td>T30060</td><td>GGGACTATCCGTGACGGGTCCGG</td><td>1000</td><td></td><td></td><td></td><td></td><td></td><td></td><td></td><td></td><td></td></tr>
<tr><td>Oligo fwd</td><td>CACCGGGACTATCCGTGACGGGTC</td><td></td><td></td><td></td><td></td><td></td><td></td><td></td><td></td><td></td><td></td></tr>
<tr><td>Oligo rev</td><td>AAACGACCCGTCACGGATAGTCCC</td><td></td><td></td><td></td><td></td><td></td><td></td><td></td><td></td><td></td><td></td></tr>
<tr><td>Chromosome</td><td>start</td><td>end</td><td>strand</td><td>MM</td><td>target_seq</td><td>PAM</td><td>alignment</td><td>distance</td><td>position</td><td>gene name</td><td>gene id</td></tr>
<tr><td>GU305914.1</td><td>53464</td><td>53486</td><td>-</td><td>0</td><td>GGGACTATCCGTGACGGGTC</td><td>CGG</td><td>||||||||[||||||||||||]PAM</td><td>NA</td><td>Intergenic</td><td>NA</td><td>NA</td></tr>
<tr><td>T30061</td><td>GGACTATCCGTGACGGGTCCGGG</td><td>1000</td><td></td><td></td><td></td><td></td><td></td><td></td><td></td><td></td><td></td></tr>
<tr><td>Oligo fwd</td><td>CACCGGACTATCCGTGACGGGTCC</td><td></td><td></td><td></td><td></td><td></td><td></td><td></td><td></td><td></td><td></td></tr>
<tr><td>Oligo rev</td><td>AAACGGACCCGTCACGGATAGTCC</td><td></td><td></td><td></td><td></td><td></td><td></td><td></td><td></td><td></td><td></td></tr>
<tr><td>Chromosome</td><td>start</td><td>end</td><td>strand</td><td>MM</td><td>target_seq</td><td>PAM</td><td>alignment</td><td>distance</td><td>position</td><td>gene name</td><td>gene id</td></tr>
<tr><td>GU305914.1</td><td>53463</td><td>53485</td><td>-</td><td>0</td><td>GGACTATCCGTGACGGGTCC</td><td>GGG</td><td>||||||||[||||||||||||]PAM</td><td>NA</td><td>Intergenic</td><td>NA</td><td>NA</td></tr>
<tr><td>T30062</td><td>GACTATCCGTGACGGGTCCGGGG</td><td>1000</td><td></td><td></td><td></td><td></td><td></td><td></td><td></td><td></td><td></td></tr>
<tr><td>Oligo fwd</td><td>CACCGACTATCCGTGACGGGTCCG</td><td></td><td></td><td></td><td></td><td></td><td></td><td></td><td></td><td></td><td></td></tr>
<tr><td>Oligo rev</td><td>AAACCGGACCCGTCACGGATAGTC</td><td></td><td></td><td></td><td></td><td></td><td></td><td></td><td></td><td></td><td></td></tr>
<tr><td>Chromosome</td><td>start</td><td>end</td><td>strand</td><td>MM</td><td>target_seq</td><td>PAM</td><td>alignment</td><td>distance</td><td>position</td><td>gene name</td><td>gene id</td></tr>
<tr><td>GU305914.1</td><td>53462</td><td>53484</td><td>-</td><td>0</td><td>GACTATCCGTGACGGGTCCG</td><td>GGG</td><td>||||||||[||||||||||||]PAM</td><td>NA</td><td>Intergenic</td><td>NA</td><td>NA</td></tr>
<tr><td>T30063</td><td>GGACGCCTCACTCTTGCATCTGG</td><td>1000</td><td></td><td></td><td></td><td></td><td></td><td></td><td></td><td></td><td></td></tr>
<tr><td>Oligo fwd</td><td>CACCGGACGCCTCACTCTTGCATC</td><td></td><td></td><td></td><td></td><td></td><td></td><td></td><td></td><td></td><td></td></tr>
<tr><td>Oligo rev</td><td>AAACGATGCAAGAGTGAGGCGTCC</td><td></td><td></td><td></td><td></td><td></td><td></td><td></td><td></td><td></td><td></td></tr>
<tr><td>Chromosome</td><td>start</td><td>end</td><td>strand</td><td>MM</td><td>target_seq</td><td>PAM</td><td>alignment</td><td>distance</td><td>position</td><td>gene name</td><td>gene id</td></tr>
<tr><td>GU305914.1</td><td>53441</td><td>53463</td><td>-</td><td>0</td><td>GGACGCCTCACTCTTGCATC</td><td>TGG</td><td>||||||||[||||||||||||]PAM</td><td>NA</td><td>Intergenic</td><td>NA</td><td>NA</td></tr>
<tr><td>T30064</td><td>CCTCACTCTTGCATCTGGCGTGG</td><td>1000</td><td></td><td></td><td></td><td></td><td></td><td></td><td></td><td></td><td></td></tr>
<tr><td>Oligo fwd</td><td>CACCCCTCACTCTTGCATCTGGCG</td><td></td><td></td><td></td><td></td><td></td><td></td><td></td><td></td><td></td><td></td></tr>
<tr><td>Oligo rev</td><td>AAACCGCCAGATGCAAGAGTGAGG</td><td></td><td></td><td></td><td></td><td></td><td></td><td></td><td></td><td></td><td></td></tr>
<tr><td>Chromosome</td><td>start</td><td>end</td><td>strand</td><td>MM</td><td>target_seq</td><td>PAM</td><td>alignment</td><td>distance</td><td>position</td><td>gene name</td><td>gene id</td></tr>
<tr><td>GU305914.1</td><td>53436</td><td>53458</td><td>-</td><td>0</td><td>CCTCACTCTTGCATCTGGCG</td><td>TGG</td><td>||||||||[||||||||||||]PAM</td><td>NA</td><td>Intergenic</td><td>NA</td><td>NA</td></tr>
<tr><td>T30065</td><td>CACTCTTGCATCTGGCGTGGTGG</td><td>1000</td><td></td><td></td><td></td><td></td><td></td><td></td><td></td><td></td><td></td></tr>
<tr><td>Oligo fwd</td><td>CACCCACTCTTGCATCTGGCGTGG</td><td></td><td></td><td></td><td></td><td></td><td></td><td></td><td></td><td></td><td></td></tr>
<tr><td>Oligo rev</td><td>AAACCCACGCCAGATGCAAGAGTG</td><td></td><td></td><td></td><td></td><td></td><td></td><td></td><td></td><td></td><td></td></tr>
<tr><td>Chromosome</td><td>start</td><td>end</td><td>strand</td><td>MM</td><td>target_seq</td><td>PAM</td><td>alignment</td><td>distance</td><td>position</td><td>gene name</td><td>gene id</td></tr>
<tr><td>GU305914.1</td><td>53433</td><td>53455</td><td>-</td><td>0</td><td>CACTCTTGCATCTGGCGTGG</td><td>TGG</td><td>||||||||[||||||||||||]PAM</td><td>NA</td><td>Intergenic</td><td>NA</td><td>NA</td></tr>
<tr><td>T30066</td><td>TCTTGCATCTGGCGTGGTGGAGG</td><td>1000</td><td></td><td></td><td></td><td></td><td></td><td></td><td></td><td></td><td></td></tr>
<tr><td>Oligo fwd</td><td>CACCTCTTGCATCTGGCGTGGTGG</td><td></td><td></td><td></td><td></td><td></td><td></td><td></td><td></td><td></td><td></td></tr>
<tr><td>Oligo rev</td><td>AAACCCACCACGCCAGATGCAAGA</td><td></td><td></td><td></td><td></td><td></td><td></td><td></td><td></td><td></td><td></td></tr>
<tr><td>Chromosome</td><td>start</td><td>end</td><td>strand</td><td>MM</td><td>target_seq</td><td>PAM</td><td>alignment</td><td>distance</td><td>position</td><td>gene name</td><td>gene id</td></tr>
<tr><td>GU305914.1</td><td>53430</td><td>53452</td><td>-</td><td>0</td><td>TCTTGCATCTGGCGTGGTGG</td><td>AGG</td><td>||||||||[||||||||||||]PAM</td><td>NA</td><td>Intergenic</td><td>NA</td><td>NA</td></tr>
<tr><td>T30067</td><td>GCATCTGGCGTGGTGGAGGACGG</td><td>1000</td><td></td><td></td><td></td><td></td><td></td><td></td><td></td><td></td><td></td></tr>
<tr><td>Oligo fwd</td><td>CACCGCATCTGGCGTGGTGGAGGA</td><td></td><td></td><td></td><td></td><td></td><td></td><td></td><td></td><td></td><td></td></tr>
<tr><td>Oligo rev</td><td>AAACTCCTCCACCACGCCAGATGC</td><td></td><td></td><td></td><td></td><td></td><td></td><td></td><td></td><td></td><td></td></tr>
<tr><td>Chromosome</td><td>start</td><td>end</td><td>strand</td><td>MM</td><td>target_seq</td><td>PAM</td><td>alignment</td><td>distance</td><td>position</td><td>gene name</td><td>gene id</td></tr>
<tr><td>GU305914.1</td><td>53426</td><td>53448</td><td>-</td><td>0</td><td>GCATCTGGCGTGGTGGAGGA</td><td>CGG</td><td>||||||||[||||||||||||]PAM</td><td>NA</td><td>Intergenic</td><td>NA</td><td>NA</td></tr>
<tr><td>GU305914.1</td><td>186729</td><td>186751</td><td>-</td><td>3</td><td>GCCTCTGGCGTGGAGGAAGA</td><td>TGG</td><td>||-|||||[|||||-|||-||]PAM</td><td>NA</td><td>Intergenic</td><td>NA</td><td>NA</td></tr>
<tr><td>T30068</td><td>CTTCTACTTCATGATCGCCGTGG</td><td>1000</td><td></td><td></td><td></td><td></td><td></td><td></td><td></td><td></td><td></td></tr>
</table>

Oligo fwd CACCCTTCTACTTCATGATCGCCG
Oligo rev AAACCGGCGATCATGAAGTAGAAG
Chromosome start end strand MM target_seq PAM alignment distance position gene name gene id
GU305914.1 53396 53418 - 0 CTTCTACTTCATGATCGCCG TGG ||||||||[||||||||||||]PAM NA Intergenic NA NA

T30069 GATCGCCGTGGTTTTGTCCGTGG 1000
Oligo fwd CACCGATCGCCGTGGTTTTGTCCG
Oligo rev AAACCGGACAAAACCACGGCGATC
Chromosome start end strand MM target_seq PAM alignment distance position gene name gene id
GU305914.1 53384 53406 - 0 GATCGCCGTGGTTTTGTCCG TGG ||||||||[||||||||||||]PAM NA Intergenic NA NA

T30070 TTGTCCGTGGTGCTGCTGTGTGG 1000
Oligo fwd CACCTTGTCCGTGGTGCTGCTGTG
Oligo rev AAACCACAGCAGCACCACGGACAA
Chromosome start end strand MM target_seq PAM alignment distance position gene name gene id
GU305914.1 53371 53393 - 0 TTGTCCGTGGTGCTGCTGTG TGG ||||||||[||||||||||||]PAM NA Intergenic NA NA

T30071 TGTCCGTGGTGCTGCTGTGTGGG 1000
Oligo fwd CACCTGTCCGTGGTGCTGCTGTGT
Oligo rev AAACACACAGCAGCACCACGGACA
Chromosome start end strand MM target_seq PAM alignment distance position gene name gene id
GU305914.1 53370 53392 - 0 TGTCCGTGGTGCTGCTGTGT GGG ||||||||[||||||||||||]PAM NA Intergenic NA NA

T30072 GTCCGTGGTGCTGCTGTGTGGGG 1000
Oligo fwd CACCGTCCGTGGTGCTGCTGTGTG
Oligo rev AAACCACACAGCAGCACCACGGAC
Chromosome start end strand MM target_seq PAM alignment distance position gene name gene id
GU305914.1 53369 53391 - 0 GTCCGTGGTGCTGCTGTGTG GGG ||||||||[||||||||||||]PAM NA Intergenic NA NA

T30073 TGGTGCTGCTGTGTGGGGCCTGG 1000
Oligo fwd CACCTGGTGCTGCTGTGTGGGGCC
Oligo rev AAACGGCCCCACACAGCAGCACCA
Chromosome start end strand MM target_seq PAM alignment distance position gene name gene id
GU305914.1 53364 53386 - 0 TGGTGCTGCTGTGTGGGGCC TGG ||||||||[||||||||||||]PAM NA Intergenic NA NA
GU305914.1 66746 66768 - 4 TGGGGGGGCTGTGGGGGGCC CGG |||-|--|[|||||-||||||]PAM NA Intergenic NA NA

T30074 TTCGTTCGACACGATCGCGCCGG 1000
Oligo fwd CACCTTCGTTCGACACGATCGCGC
Oligo rev AAACGCGCGATCGTGTCGAACGAA
Chromosome start end strand MM target_seq PAM alignment distance position gene name gene id
GU305914.1 53308 53330 - 0 TTCGTTCGACACGATCGCGC CGG ||||||||[||||||||||||]PAM NA Intergenic NA NA

T30075 GTTCGACACGATCGCGCCGGCGG 1000
Oligo fwd CACCGTTCGACACGATCGCGCCGG
Oligo rev AAACCCGGCGCGATCGTGTCGAAC
Chromosome start end strand MM target_seq PAM alignment distance position gene name gene id
GU305914.1 53305 53327 - 0 GTTCGACACGATCGCGCCGG CGG ||||||||[||||||||||||]PAM NA Intergenic NA NA

T30076 ATCGCGCCGGCGGAGCGCGTTGG 1000
Oligo fwd CACCATCGCGCCGGCGGAGCGCGT
Oligo rev AAACACGCGCTCCGCCGGCGCGAT
Chromosome start end strand MM target_seq PAM alignment distance position gene name gene id
GU305914.1 53295 53317 - 0 ATCGCGCCGGCGGAGCGCGT TGG ||||||||[||||||||||||]PAM NA Intergenic NA NA

T30077 CCGGCGGAGCGCGTTGGACCAGG 1000
Oligo fwd CACCCCGGCGGAGCGCGTTGGACC
Oligo rev AAACGGTCCAACGCGCTCCGCCGG
Chromosome start end strand MM target_seq PAM alignment distance position gene name gene id
GU305914.1 53289 53311 - 0 CCGGCGGAGCGCGTTGGACC AGG ||||||||[||||||||||||]PAM NA Intergenic NA NA

T30078 GAGCGCGTTGGACCAGGTTGAGG 1000
Oligo fwd CACCGAGCGCGTTGGACCAGGTTG
Oligo rev AAACCAACCTGGTCCAACGCGCTC
Chromosome start end strand MM target_seq PAM alignment distance position gene name gene id
GU305914.1 53283 53305 - 0 GAGCGCGTTGGACCAGGTTG AGG ||||||||[||||||||||||]PAM NA Intergenic NA NA

T30079 AGCGCGTTGGACCAGGTTGAGGG 1000
Oligo fwd CACCAGCGCGTTGGACCAGGTTGA
Oligo rev AAACTCAACCTGGTCCAACGCGCT
Chromosome start end strand MM target_seq PAM alignment distance position gene name gene id
GU305914.1 53282 53304 - 0 AGCGCGTTGGACCAGGTTGA GGG ||||||||[||||||||||||]PAM NA Intergenic NA NA

T30080 CCAGGTTGAGGGAAGAGAGTCGG 1000
Oligo fwd CACCCCAGGTTGAGGGAAGAGAGT
Oligo rev AAACACTCTCTTCCCTCAACCTGG
Chromosome start end strand MM target_seq PAM alignment distance position gene name gene id
GU305914.1 53271 53293 - 0 CCAGGTTGAGGGAAGAGAGT CGG ||||||||[||||||||||||]PAM NA Intergenic NA NA

T30081 AAGAGAGTCGGTGAGATGAGTGG 1000
Oligo fwd CACCAAGAGAGTCGGTGAGATGAG
Oligo rev AAACCTCATCTCACCGACTCTCTT
Chromosome start end strand MM target_seq PAM alignment distance position gene name gene id
GU305914.1 53259 53281 - 0 AAGAGAGTCGGTGAGATGAG TGG ||||||||[||||||||||||]PAM NA Intergenic NA NA

T30082 AGAGAGTCGGTGAGATGAGTGGG 1000
Oligo fwd CACCAGAGAGTCGGTGAGATGAGT
Oligo rev AAACACTCATCTCACCGACTCTCT
Chromosome start end strand MM target_seq PAM alignment distance position gene name gene id
GU305914.1 53258 53280 - 0 AGAGAGTCGGTGAGATGAGT GGG ||||||||[||||||||||||]PAM NA Intergenic NA NA

T30083 GATGCTGTGTACTCAGTTTTTGG 1000
Oligo fwd CACCGATGCTGTGTACTCAGTTTT
Oligo rev AAACAAAACTGAGTACACAGCATC
Chromosome start end strand MM target_seq PAM alignment distance position gene name gene id
GU305914.1 53182 53204 - 0 GATGCTGTGTACTCAGTTTT TGG ||||||||[||||||||||||]PAM NA Intergenic NA NA

T30084 ATGCTGTGTACTCAGTTTTTGGG 1000
Oligo fwd CACCATGCTGTGTACTCAGTTTTT
Oligo rev AAACAAAAACTGAGTACACAGCAT
Chromosome start end strand MM target_seq PAM alignment distance position gene name gene id
GU305914.1 53181 53203 - 0 ATGCTGTGTACTCAGTTTTT GGG ||||||||[||||||||||||]PAM NA Intergenic NA NA

T30085 TGCTGTGTACTCAGTTTTTGGGG 1000
Oligo fwd CACCTGCTGTGTACTCAGTTTTTG
Oligo rev AAACCAAAAACTGAGTACACAGCA
Chromosome start end strand MM target_seq PAM alignment distance position gene name gene id
GU305914.1 53180 53202 - 0 TGCTGTGTACTCAGTTTTTG GGG ||||||||[||||||||||||]PAM NA Intergenic NA NA

T30086 CAGTTTTTGGGGATATTGTTCGG 1000
Oligo fwd CACCCAGTTTTTGGGGATATTGTT
Oligo rev AAACAACAATATCCCCAAAAACTG
Chromosome start end strand MM target_seq PAM alignment distance position gene name gene id
GU305914.1 53169 53191 - 0 CAGTTTTTGGGGATATTGTT CGG ||||||||[||||||||||||]PAM NA Intergenic NA NA

T30087 GCTACTCATCTTTTAGTCCGCGG 1000
Oligo fwd CACCGCTACTCATCTTTTAGTCCG
Oligo rev AAACCGGACTAAAAGATGAGTAGC
Chromosome start end strand MM target_seq PAM alignment distance position gene name gene id
GU305914.1 53145 53167 - 0 GCTACTCATCTTTTAGTCCG CGG ||||||||[||||||||||||]PAM NA Intergenic NA NA

T30088 GCCGTTCGTATCTTCGTCCGCGG 1000
Oligo fwd CACCGCCGTTCGTATCTTCGTCCG
Oligo rev AAACCGGACGAAGATACGAACGGC
Chromosome start end strand MM target_seq PAM alignment distance position gene name gene id
GU305914.1 53115 53137 - 0 GCCGTTCGTATCTTCGTCCG CGG ||||||||[||||||||||||]PAM NA Intergenic NA NA

T30089 TCGTATCTTCGTCCGCGGCCAGG 1000
Oligo fwd CACCTCGTATCTTCGTCCGCGGCC
Oligo rev AAACGGCCGCGGACGAAGATACGA
Chromosome start end strand MM target_seq PAM alignment distance position gene name gene id
GU305914.1 53110 53132 - 0 TCGTATCTTCGTCCGCGGCC AGG ||||||||[||||||||||||]PAM NA Intergenic NA NA

T30090 AACGAGTGATCAAAACTTATCGG 1000
Oligo fwd CACCAACGAGTGATCAAAACTTAT
Oligo rev AAACATAAGTTTTGATCACTCGTT
Chromosome start end strand MM target_seq PAM alignment distance position gene name gene id
GU305914.1 53087 53109 - 0 AACGAGTGATCAAAACTTAT CGG ||||||||[||||||||||||]PAM NA Intergenic NA NA

T30091 GTGATCAAAACTTATCGGTACGG 1000
Oligo fwd CACCGTGATCAAAACTTATCGGTA
Oligo rev AAACTACCGATAAGTTTTGATCAC
Chromosome start end strand MM target_seq PAM alignment distance position gene name gene id
GU305914.1 53082 53104 - 0 GTGATCAAAACTTATCGGTA CGG ||||||||[||||||||||||]PAM NA Intergenic NA NA

T30092 AAGATGCCCGATTTCCCTAACGG 1000
Oligo fwd CACCAAGATGCCCGATTTCCCTAA
Oligo rev AAACTTAGGGAAATCGGGCATCTT
Chromosome start end strand MM target_seq PAM alignment distance position gene name gene id
GU305914.1 53052 53074 - 0 AAGATGCCCGATTTCCCTAA CGG ||||||||[||||||||||||]PAM NA Intergenic NA NA

T30093 AGATGCCCGATTTCCCTAACGGG 1000
Oligo fwd CACCAGATGCCCGATTTCCCTAAC
Oligo rev AAACGTTAGGGAAATCGGGCATCT
Chromosome start end strand MM target_seq PAM alignment distance position gene name gene id
GU305914.1 53051 53073 - 0 AGATGCCCGATTTCCCTAAC GGG ||||||||[||||||||||||]PAM NA Intergenic NA NA

T30094 AACGGGTCATCCATCTGTTGTGG 1000
Oligo fwd CACCAACGGGTCATCCATCTGTTG
Oligo rev AAACCAACAGATGGATGACCCGTT
Chromosome start end strand MM target_seq PAM alignment distance position gene name gene id
GU305914.1 53034 53056 - 0 AACGGGTCATCCATCTGTTG TGG ||||||||[||||||||||||]PAM NA Intergenic NA NA

T30095 TGTGGAGCGTGTGACCTTATCGG 1000
Oligo fwd CACCTGTGGAGCGTGTGACCTTAT
Oligo rev AAACATAAGGTCACACGCTCCACA
Chromosome start end strand MM target_seq PAM alignment distance position gene name gene id
GU305914.1 53016 53038 - 0 TGTGGAGCGTGTGACCTTAT CGG ||||||||[||||||||||||]PAM NA Intergenic NA NA

T30096 CACAATACCATTGTGAGACAAGG 1000
Oligo fwd CACCCACAATACCATTGTGAGACA
Oligo rev AAACTGTCTCACAATGGTATTGTG
Chromosome start end strand MM target_seq PAM alignment distance position gene name gene id
GU305914.1 52992 53014 - 0 CACAATACCATTGTGAGACA AGG ||||||||[||||||||||||]PAM NA Intergenic NA NA

T30097 TACCATTGTGAGACAAGGATTGG 1000
Oligo fwd CACCTACCATTGTGAGACAAGGAT
Oligo rev AAACATCCTTGTCTCACAATGGTA
Chromosome start end strand MM target_seq PAM alignment distance position gene name gene id

GU305914.1 52987 53009 - 0 TACCATTGTGAGACAAGGAT TGG ||||||||[||||||||||||]PAM NA Intergenic NA NA

T30098 AGGATTGGCCCCAGAGTATGCGG 1000
Oligo fwd CACCAGGATTGGCCCCAGAGTATG
Oligo rev AAACCATACTCTGGGGCCAATCCT
Chromosome start end strand MM target_seq PAM alignment distance position gene name gene id
GU305914.1 52972 52994 - 0 AGGATTGGCCCCAGAGTATG CGG ||||||||[||||||||||||]PAM NA Intergenic NA NA

T30099 TATGCGGTGTTGTGTGTCATCGG 1000
Oligo fwd CACCTATGCGGTGTTGTGTGTCAT
Oligo rev AAACATGACACACAACACCGCATA
Chromosome start end strand MM target_seq PAM alignment distance position gene name gene id
GU305914.1 52956 52978 - 0 TATGCGGTGTTGTGTGTCAT CGG ||||||||[||||||||||||]PAM NA Intergenic NA NA

T30100 GCGGTGTTGTGTGTCATCGGAGG 1000
Oligo fwd CACCGCGGTGTTGTGTGTCATCGG
Oligo rev AAACCCGATGACACACAACACCGC
Chromosome start end strand MM target_seq PAM alignment distance position gene name gene id
GU305914.1 52953 52975 - 0 GCGGTGTTGTGTGTCATCGG AGG ||||||||[||||||||||||]PAM NA Intergenic NA NA

T30101 TGTCATCGGAGGCTTTACCGCGG 1000
Oligo fwd CACCTGTCATCGGAGGCTTTACCG
Oligo rev AAACCGGTAAAGCCTCCGATGACA
Chromosome start end strand MM target_seq PAM alignment distance position gene name gene id
GU305914.1 52942 52964 - 0 TGTCATCGGAGGCTTTACCG CGG ||||||||[||||||||||||]PAM NA Intergenic NA NA

T30102 GTCATCGGAGGCTTTACCGCGGG 1000
Oligo fwd CACCGTCATCGGAGGCTTTACCGC
Oligo rev AAACGCGGTAAAGCCTCCGATGAC
Chromosome start end strand MM target_seq PAM alignment distance position gene name gene id
GU305914.1 52941 52963 - 0 GTCATCGGAGGCTTTACCGC GGG ||||||||[||||||||||||]PAM NA Intergenic NA NA

T30103 TCTTCGCCTTTCTCGTCGACAGG 1000
Oligo fwd CACCTCTTCGCCTTTCTCGTCGAC
Oligo rev AAACGTCGACGAGAAAGGCGAAGA
Chromosome start end strand MM target_seq PAM alignment distance position gene name gene id
GU305914.1 52913 52935 - 0 TCTTCGCCTTTCTCGTCGAC AGG ||||||||[||||||||||||]PAM NA Intergenic NA NA

T30104 TTTCTCGTCGACAGGATCGTCGG 1000
Oligo fwd CACCTTTCTCGTCGACAGGATCGT
Oligo rev AAACACGATCCTGTCGACGAGAAA
Chromosome start end strand MM target_seq PAM alignment distance position gene name gene id
GU305914.1 52905 52927 - 0 TTTCTCGTCGACAGGATCGT CGG ||||||||[||||||||||||]PAM NA Intergenic NA NA

T30105 GATCGTCGGCTCTCCCCTACCGG 1000
Oligo fwd CACCGATCGTCGGCTCTCCCCTAC
Oligo rev AAACGTAGGGGAGAGCCGACGATC
Chromosome start end strand MM target_seq PAM alignment distance position gene name gene id
GU305914.1 52891 52913 - 0 GATCGTCGGCTCTCCCCTAC CGG ||||||||[||||||||||||]PAM NA Intergenic NA NA

T30106 CCTACCGGTCACTTTTGCCGAGG 1000
Oligo fwd CACCCCTACCGGTCACTTTTGCCG
Oligo rev AAACCGGCAAAAGTGACCGGTAGG
Chromosome start end strand MM target_seq PAM alignment distance position gene name gene id
GU305914.1 52876 52898 - 0 CCTACCGGTCACTTTTGCCG AGG ||||||||[||||||||||||]PAM NA Intergenic NA NA

T30107 CTACCGGTCACTTTTGCCGAGGG 1000
Oligo fwd CACCCTACCGGTCACTTTTGCCGA
Oligo rev AAACTCGGCAAAAGTGACCGGTAG
Chromosome start end strand MM target_seq PAM alignment distance position gene name gene id
GU305914.1 52875 52897 - 0 CTACCGGTCACTTTTGCCGA GGG ||||||||[||||||||||||]PAM NA Intergenic NA NA

T30108 CCGGTCACTTTTGCCGAGGGCGG 1000
Oligo fwd CACCCCGGTCACTTTTGCCGAGGG
Oligo rev AAACCCCTCGGCAAAAGTGACCGG
Chromosome start end strand MM target_seq PAM alignment distance position gene name gene id
GU305914.1 52872 52894 - 0 CCGGTCACTTTTGCCGAGGG CGG ||||||||[||||||||||||]PAM NA Intergenic NA NA

T30109 CGGTCACTTTTGCCGAGGGCGGG 1000
Oligo fwd CACCCGGTCACTTTTGCCGAGGGC
Oligo rev AAACGCCCTCGGCAAAAGTGACCG
Chromosome start end strand MM target_seq PAM alignment distance position gene name gene id
GU305914.1 52871 52893 - 0 CGGTCACTTTTGCCGAGGGC GGG ||||||||[||||||||||||]PAM NA Intergenic NA NA

T30110 CCTCTGCTTTCCCCTTTCTCAGG 1000
Oligo fwd CACCCCTCTGCTTTCCCCTTTCTC
Oligo rev AAACGAGAAAGGGGAAAGCAGAGG
Chromosome start end strand MM target_seq PAM alignment distance position gene name gene id
GU305914.1 52835 52857 - 0 CCTCTGCTTTCCCCTTTCTC AGG ||||||||[||||||||||||]PAM NA Intergenic NA NA

T30111 TCCCCTTTCTCAGGAAGAAATGG 1000
Oligo fwd CACCTCCCCTTTCTCAGGAAGAAA
Oligo rev AAACTTTCTTCCTGAGAAAGGGGA
Chromosome start end strand MM target_seq PAM alignment distance position gene name gene id
GU305914.1 52826 52848 - 0 TCCCCTTTCTCAGGAAGAAA TGG ||||||||[||||||||||||]PAM NA Intergenic NA NA

T30112 TCTCAGGAAGAAATGGTACTCGG 1000
Oligo fwd CACCTCTCAGGAAGAAATGGTACT
Oligo rev AAACAGTACCATTTCTTCCTGAGA

<table>
<tr><td>Chromosome</td><td>start</td><td>end</td><td>strand</td><td>MM</td><td>target_seq</td><td>PAM</td><td>alignment</td><td>distance</td><td>position</td><td>gene name</td><td>gene id</td></tr>
<tr><td>GU305914.1</td><td>52819</td><td>52841</td><td>-</td><td>0</td><td>TCTCAGGAAGAAATGGTACT</td><td>CGG</td><td>||||||||[||||||||||||]PAM</td><td>NA</td><td>Intergenic</td><td>NA</td><td>NA</td></tr>
<tr><td>T30113</td><td>GAAGAAATGGTACTCGGACGAGG</td><td>1000</td><td></td><td></td><td></td><td></td><td></td><td></td><td></td><td></td><td></td></tr>
<tr><td>Oligo fwd</td><td>CACCGAAGAAATGGTACTCGGACG</td><td></td><td></td><td></td><td></td><td></td><td></td><td></td><td></td><td></td><td></td></tr>
<tr><td>Oligo rev</td><td>AAACCGTCCGAGTACCATTTCTTC</td><td></td><td></td><td></td><td></td><td></td><td></td><td></td><td></td><td></td><td></td></tr>
<tr><td>Chromosome</td><td>start</td><td>end</td><td>strand</td><td>MM</td><td>target_seq</td><td>PAM</td><td>alignment</td><td>distance</td><td>position</td><td>gene name</td><td>gene id</td></tr>
<tr><td>GU305914.1</td><td>52813</td><td>52835</td><td>-</td><td>0</td><td>GAAGAAATGGTACTCGGACG</td><td>AGG</td><td>||||||||[||||||||||||]PAM</td><td>NA</td><td>Intergenic</td><td>NA</td><td>NA</td></tr>
<tr><td>T30114</td><td>ACTTTCTCCCACCGTGCAAGTGG</td><td>1000</td><td></td><td></td><td></td><td></td><td></td><td></td><td></td><td></td><td></td></tr>
<tr><td>Oligo fwd</td><td>CACCACTTTCTCCCACCGTGCAAG</td><td></td><td></td><td></td><td></td><td></td><td></td><td></td><td></td><td></td><td></td></tr>
<tr><td>Oligo rev</td><td>AAACCTTGCACGGTGGGAGAAAGT</td><td></td><td></td><td></td><td></td><td></td><td></td><td></td><td></td><td></td><td></td></tr>
<tr><td>Chromosome</td><td>start</td><td>end</td><td>strand</td><td>MM</td><td>target_seq</td><td>PAM</td><td>alignment</td><td>distance</td><td>position</td><td>gene name</td><td>gene id</td></tr>
<tr><td>GU305914.1</td><td>52789</td><td>52811</td><td>-</td><td>0</td><td>ACTTTCTCCCACCGTGCAAG</td><td>TGG</td><td>||||||||[||||||||||||]PAM</td><td>NA</td><td>Intergenic</td><td>NA</td><td>NA</td></tr>
<tr><td>T30115</td><td>CACCGTGCAAGTGGTCTGCACGG</td><td>1000</td><td></td><td></td><td></td><td></td><td></td><td></td><td></td><td></td><td></td></tr>
<tr><td>Oligo fwd</td><td>CACCCACCGTGCAAGTGGTCTGCA</td><td></td><td></td><td></td><td></td><td></td><td></td><td></td><td></td><td></td><td></td></tr>
<tr><td>Oligo rev</td><td>AAACTGCAGACCACTTGCACGGTG</td><td></td><td></td><td></td><td></td><td></td><td></td><td></td><td></td><td></td><td></td></tr>
<tr><td>Chromosome</td><td>start</td><td>end</td><td>strand</td><td>MM</td><td>target_seq</td><td>PAM</td><td>alignment</td><td>distance</td><td>position</td><td>gene name</td><td>gene id</td></tr>
<tr><td>GU305914.1</td><td>52780</td><td>52802</td><td>-</td><td>0</td><td>CACCGTGCAAGTGGTCTGCA</td><td>CGG</td><td>||||||||[||||||||||||]PAM</td><td>NA</td><td>Intergenic</td><td>NA</td><td>NA</td></tr>
<tr><td>T30116</td><td>GCAAGTGGTCTGCACGGATCTGG</td><td>1000</td><td></td><td></td><td></td><td></td><td></td><td></td><td></td><td></td><td></td></tr>
<tr><td>Oligo fwd</td><td>CACCGCAAGTGGTCTGCACGGATC</td><td></td><td></td><td></td><td></td><td></td><td></td><td></td><td></td><td></td><td></td></tr>
<tr><td>Oligo rev</td><td>AAACGATCCGTGCAGACCACTTGC</td><td></td><td></td><td></td><td></td><td></td><td></td><td></td><td></td><td></td><td></td></tr>
<tr><td>Chromosome</td><td>start</td><td>end</td><td>strand</td><td>MM</td><td>target_seq</td><td>PAM</td><td>alignment</td><td>distance</td><td>position</td><td>gene name</td><td>gene id</td></tr>
<tr><td>GU305914.1</td><td>52774</td><td>52796</td><td>-</td><td>0</td><td>GCAAGTGGTCTGCACGGATC</td><td>TGG</td><td>||||||||[||||||||||||]PAM</td><td>NA</td><td>Intergenic</td><td>NA</td><td>NA</td></tr>
<tr><td>T30117</td><td>CACGGATCTGGATCCAGAGATGG</td><td>1000</td><td></td><td></td><td></td><td></td><td></td><td></td><td></td><td></td><td></td></tr>
<tr><td>Oligo fwd</td><td>CACCCACGGATCTGGATCCAGAGA</td><td></td><td></td><td></td><td></td><td></td><td></td><td></td><td></td><td></td><td></td></tr>
<tr><td>Oligo rev</td><td>AAACTCTCTGGATCCAGATCCGTG</td><td></td><td></td><td></td><td></td><td></td><td></td><td></td><td></td><td></td><td></td></tr>
<tr><td>Chromosome</td><td>start</td><td>end</td><td>strand</td><td>MM</td><td>target_seq</td><td>PAM</td><td>alignment</td><td>distance</td><td>position</td><td>gene name</td><td>gene id</td></tr>
<tr><td>GU305914.1</td><td>52762</td><td>52784</td><td>-</td><td>0</td><td>CACGGATCTGGATCCAGAGA</td><td>TGG</td><td>||||||||[||||||||||||]PAM</td><td>NA</td><td>Intergenic</td><td>NA</td><td>NA</td></tr>
<tr><td>T30118</td><td>ACGGATCTGGATCCAGAGATGGG</td><td>1000</td><td></td><td></td><td></td><td></td><td></td><td></td><td></td><td></td><td></td></tr>
<tr><td>Oligo fwd</td><td>CACCACGGATCTGGATCCAGAGAT</td><td></td><td></td><td></td><td></td><td></td><td></td><td></td><td></td><td></td><td></td></tr>
<tr><td>Oligo rev</td><td>AAACATCTCTGGATCCAGATCCGT</td><td></td><td></td><td></td><td></td><td></td><td></td><td></td><td></td><td></td><td></td></tr>
<tr><td>Chromosome</td><td>start</td><td>end</td><td>strand</td><td>MM</td><td>target_seq</td><td>PAM</td><td>alignment</td><td>distance</td><td>position</td><td>gene name</td><td>gene id</td></tr>
<tr><td>GU305914.1</td><td>52761</td><td>52783</td><td>-</td><td>0</td><td>ACGGATCTGGATCCAGAGAT</td><td>GGG</td><td>||||||||[||||||||||||]PAM</td><td>NA</td><td>Intergenic</td><td>NA</td><td>NA</td></tr>
<tr><td>T30119</td><td>TCCGCCACCGCCACCGCCAGCGG</td><td>1000</td><td></td><td></td><td></td><td></td><td></td><td></td><td></td><td></td><td></td></tr>
<tr><td>Oligo fwd</td><td>CACCTCCGCCACCGCCACCGCCAG</td><td></td><td></td><td></td><td></td><td></td><td></td><td></td><td></td><td></td><td></td></tr>
<tr><td>Oligo rev</td><td>AAACCTGGCGGTGGCGGTGGCGGA</td><td></td><td></td><td></td><td></td><td></td><td></td><td></td><td></td><td></td><td></td></tr>
<tr><td>Chromosome</td><td>start</td><td>end</td><td>strand</td><td>MM</td><td>target_seq</td><td>PAM</td><td>alignment</td><td>distance</td><td>position</td><td>gene name</td><td>gene id</td></tr>
<tr><td>GU305914.1</td><td>52720</td><td>52742</td><td>-</td><td>0</td><td>TCCGCCACCGCCACCGCCAG</td><td>CGG</td><td>||||||||[||||||||||||]PAM</td><td>NA</td><td>Intergenic</td><td>NA</td><td>NA</td></tr>
<tr><td>GU305914.1</td><td>154399</td><td>154421</td><td>-</td><td>3</td><td>GCCGCCGCCGCCACCGCCCG</td><td>GGG</td><td>-|||||-|[||||||||||-|]PAM</td><td>NA</td><td>Intergenic</td><td>NA</td><td>NA</td></tr>
<tr><td>GU305914.1</td><td>48151</td><td>48173</td><td>-</td><td>3</td><td>GCCGCCACCGCCGCCGCCAC</td><td>AGG</td><td>-|||||||[||||-||||||-]PAM</td><td>NA</td><td>Intergenic</td><td>NA</td><td>NA</td></tr>
<tr><td>T30120</td><td>GCCACCGCCACCGCCAGCGGCGG</td><td>1000</td><td></td><td></td><td></td><td></td><td></td><td></td><td></td><td></td><td></td></tr>
<tr><td>Oligo fwd</td><td>CACCGCCACCGCCACCGCCAGCGG</td><td></td><td></td><td></td><td></td><td></td><td></td><td></td><td></td><td></td><td></td></tr>
<tr><td>Oligo rev</td><td>AAACCCGCTGGCGGTGGCGGTGGC</td><td></td><td></td><td></td><td></td><td></td><td></td><td></td><td></td><td></td><td></td></tr>
<tr><td>Chromosome</td><td>start</td><td>end</td><td>strand</td><td>MM</td><td>target_seq</td><td>PAM</td><td>alignment</td><td>distance</td><td>position</td><td>gene name</td><td>gene id</td></tr>
<tr><td>GU305914.1</td><td>52717</td><td>52739</td><td>-</td><td>0</td><td>GCCACCGCCACCGCCAGCGG</td><td>CGG</td><td>||||||||[||||||||||||]PAM</td><td>NA</td><td>Intergenic</td><td>NA</td><td>NA</td></tr>
<tr><td>T30121</td><td>GCCACCGCCAGCGGCGGCCAAGG</td><td>1000</td><td></td><td></td><td></td><td></td><td></td><td></td><td></td><td></td><td></td></tr>
<tr><td>Oligo fwd</td><td>CACCGCCACCGCCAGCGGCGGCCA</td><td></td><td></td><td></td><td></td><td></td><td></td><td></td><td></td><td></td><td></td></tr>
<tr><td>Oligo rev</td><td>AAACTGGCCGCCGCTGGCGGTGGC</td><td></td><td></td><td></td><td></td><td></td><td></td><td></td><td></td><td></td><td></td></tr>
<tr><td>Chromosome</td><td>start</td><td>end</td><td>strand</td><td>MM</td><td>target_seq</td><td>PAM</td><td>alignment</td><td>distance</td><td>position</td><td>gene name</td><td>gene id</td></tr>
<tr><td>GU305914.1</td><td>52711</td><td>52733</td><td>-</td><td>0</td><td>GCCACCGCCAGCGGCGGCCA</td><td>AGG</td><td>||||||||[||||||||||||]PAM</td><td>NA</td><td>Intergenic</td><td>NA</td><td>NA</td></tr>
<tr><td>T30122</td><td>GCGGCGGCCAAGGTGTTCGTCGG</td><td>1000</td><td></td><td></td><td></td><td></td><td></td><td></td><td></td><td></td><td></td></tr>
<tr><td>Oligo fwd</td><td>CACCGCGGCGGCCAAGGTGTTCGT</td><td></td><td></td><td></td><td></td><td></td><td></td><td></td><td></td><td></td><td></td></tr>
<tr><td>Oligo rev</td><td>AAACACGAACACCTTGGCCGCCGC</td><td></td><td></td><td></td><td></td><td></td><td></td><td></td><td></td><td></td><td></td></tr>
<tr><td>Chromosome</td><td>start</td><td>end</td><td>strand</td><td>MM</td><td>target_seq</td><td>PAM</td><td>alignment</td><td>distance</td><td>position</td><td>gene name</td><td>gene id</td></tr>
<tr><td>GU305914.1</td><td>52701</td><td>52723</td><td>-</td><td>0</td><td>GCGGCGGCCAAGGTGTTCGT</td><td>CGG</td><td>||||||||[||||||||||||]PAM</td><td>NA</td><td>Intergenic</td><td>NA</td><td>NA</td></tr>
<tr><td>T30123</td><td>GCGGCCAAGGTGTTCGTCGGTGG</td><td>1000</td><td></td><td></td><td></td><td></td><td></td><td></td><td></td><td></td><td></td></tr>
<tr><td>Oligo fwd</td><td>CACCGCGGCCAAGGTGTTCGTCGG</td><td></td><td></td><td></td><td></td><td></td><td></td><td></td><td></td><td></td><td></td></tr>
<tr><td>Oligo rev</td><td>AAACCCGACGAACACCTTGGCCGC</td><td></td><td></td><td></td><td></td><td></td><td></td><td></td><td></td><td></td><td></td></tr>
<tr><td>Chromosome</td><td>start</td><td>end</td><td>strand</td><td>MM</td><td>target_seq</td><td>PAM</td><td>alignment</td><td>distance</td><td>position</td><td>gene name</td><td>gene id</td></tr>
<tr><td>GU305914.1</td><td>52698</td><td>52720</td><td>-</td><td>0</td><td>GCGGCCAAGGTGTTCGTCGG</td><td>TGG</td><td>||||||||[||||||||||||]PAM</td><td>NA</td><td>Intergenic</td><td>NA</td><td>NA</td></tr>
<tr><td>T30124</td><td>CGACCTGAGTAAAAAATCGAAGG</td><td>1000</td><td></td><td></td><td></td><td></td><td></td><td></td><td></td><td></td><td></td></tr>
<tr><td>Oligo fwd</td><td>CACCCGACCTGAGTAAAAAATCGA</td><td></td><td></td><td></td><td></td><td></td><td></td><td></td><td></td><td></td><td></td></tr>
<tr><td>Oligo rev</td><td>AAACTCGATTTTTTACTCAGGTCG</td><td></td><td></td><td></td><td></td><td></td><td></td><td></td><td></td><td></td><td></td></tr>
<tr><td>Chromosome</td><td>start</td><td>end</td><td>strand</td><td>MM</td><td>target_seq</td><td>PAM</td><td>alignment</td><td>distance</td><td>position</td><td>gene name</td><td>gene id</td></tr>
<tr><td>GU305914.1</td><td>52675</td><td>52697</td><td>-</td><td>0</td><td>CGACCTGAGTAAAAAATCGA</td><td>AGG</td><td>||||||||[||||||||||||]PAM</td><td>NA</td><td>Intergenic</td><td>NA</td><td>NA</td></tr>
<tr><td>T30125</td><td>TGTCGCCATCGCCGATGCTCTGG</td><td>1000</td><td></td><td></td><td></td><td></td><td></td><td></td><td></td><td></td><td></td></tr>
<tr><td>Oligo fwd</td><td>CACCTGTCGCCATCGCCGATGCTC</td><td></td><td></td><td></td><td></td><td></td><td></td><td></td><td></td><td></td><td></td></tr>
<tr><td>Oligo rev</td><td>AAACGAGCATCGGCGATGGCGACA</td><td></td><td></td><td></td><td></td><td></td><td></td><td></td><td></td><td></td><td></td></tr>
<tr><td>Chromosome</td><td>start</td><td>end</td><td>strand</td><td>MM</td><td>target_seq</td><td>PAM</td><td>alignment</td><td>distance</td><td>position</td><td>gene name</td><td>gene id</td></tr>
<tr><td>GU305914.1</td><td>52642</td><td>52664</td><td>-</td><td>0</td><td>TGTCGCCATCGCCGATGCTC</td><td>TGG</td><td>||||||||[||||||||||||]PAM</td><td>NA</td><td>Intergenic</td><td>NA</td><td>NA</td></tr>
<tr><td>T30126</td><td>GGAGATGCAGATGATGCGCTCGG</td><td>1000</td><td></td><td></td><td></td><td></td><td></td><td></td><td></td><td></td><td></td></tr>
<tr><td>Oligo fwd</td><td>CACCGGAGATGCAGATGATGCGCT</td><td></td><td></td><td></td><td></td><td></td><td></td><td></td><td></td><td></td><td></td></tr>
<tr><td>Oligo rev</td><td>AAACAGCGCATCATCTGCATCTCC</td><td></td><td></td><td></td><td></td><td></td><td></td><td></td><td></td><td></td><td></td></tr>
<tr><td>Chromosome</td><td>start</td><td>end</td><td>strand</td><td>MM</td><td>target_seq</td><td>PAM</td><td>alignment</td><td>distance</td><td>position</td><td>gene name</td><td>gene id</td></tr>
<tr><td>GU305914.1</td><td>52621</td><td>52643</td><td>-</td><td>0</td><td>GGAGATGCAGATGATGCGCT</td><td>CGG</td><td>||||||||[||||||||||||]PAM</td><td>NA</td><td>Intergenic</td><td>NA</td><td>NA</td></tr>
</table>

| | | | | | | | | | | | |
|---|---|---|---|---|---|---|---|---|---|---|---|
| T30127 | GATGCAGATGATGCGCTCGGCGG | 1000 | | | | | | | | | |
| Oligo fwd | CACCGATGCAGATGATGCGCTCGG | | | | | | | | | | |
| Oligo rev | AAACCCGAGCGCATCATCTGCATC | | | | | | | | | | |
| Chromosome | start | end | strand | MM | target_seq | PAM | alignment | distance | position | gene name | gene id |
| GU305914.1 | 52618 | 52640 | - | 0 | GATGCAGATGATGCGCTCGG | CGG | \|\|\|\|\|\|\|\|[\|\|\|\|\|\|\|\|\|\|\|\|]PAM | NA | Intergenic | NA | NA |
| T30128 | ATGCAGATGATGCGCTCGGCGGG | 1000 | | | | | | | | | |
| Oligo fwd | CACCATGCAGATGATGCGCTCGGC | | | | | | | | | | |
| Oligo rev | AAACGCCGAGCGCATCATCTGCAT | | | | | | | | | | |
| Chromosome | start | end | strand | MM | target_seq | PAM | alignment | distance | position | gene name | gene id |
| GU305914.1 | 52617 | 52639 | - | 0 | ATGCAGATGATGCGCTCGGC | GGG | \|\|\|\|\|\|\|\|[\|\|\|\|\|\|\|\|\|\|\|\|]PAM | NA | Intergenic | NA | NA |
| T30129 | TGCAGATGATGCGCTCGGCGGGG | 1000 | | | | | | | | | |
| Oligo fwd | CACCTGCAGATGATGCGCTCGGCG | | | | | | | | | | |
| Oligo rev | AAACCGCCGAGCGCATCATCTGCA | | | | | | | | | | |
| Chromosome | start | end | strand | MM | target_seq | PAM | alignment | distance | position | gene name | gene id |
| GU305914.1 | 52616 | 52638 | - | 0 | TGCAGATGATGCGCTCGGCG | GGG | \|\|\|\|\|\|\|\|[\|\|\|\|\|\|\|\|\|\|\|\|]PAM | NA | Intergenic | NA | NA |
| T30130 | GCAGATGATGCGCTCGGCGGGGG | 1000 | | | | | | | | | |
| Oligo fwd | CACCGCAGATGATGCGCTCGGCGG | | | | | | | | | | |
| Oligo rev | AAACCCGCCGAGCGCATCATCTGC | | | | | | | | | | |
| Chromosome | start | end | strand | MM | target_seq | PAM | alignment | distance | position | gene name | gene id |
| GU305914.1 | 52615 | 52637 | - | 0 | GCAGATGATGCGCTCGGCGG | GGG | \|\|\|\|\|\|\|\|[\|\|\|\|\|\|\|\|\|\|\|\|]PAM | NA | Intergenic | NA | NA |
| T30131 | ATGCGCTCGGCGGGGGCGCACGG | 1000 | | | | | | | | | |
| Oligo fwd | CACCATGCGCTCGGCGGGGGCGCA | | | | | | | | | | |
| Oligo rev | AAACTGCGCCCCCGCCGAGCGCAT | | | | | | | | | | |
| Chromosome | start | end | strand | MM | target_seq | PAM | alignment | distance | position | gene name | gene id |
| GU305914.1 | 52608 | 52630 | - | 0 | ATGCGCTCGGCGGGGGCGCA | CGG | \|\|\|\|\|\|\|\|[\|\|\|\|\|\|\|\|\|\|\|\|]PAM | NA | Intergenic | NA | NA |
| T30132 | CGCTCGGCGGGGGCGCACGGCGG | 1000 | | | | | | | | | |
| Oligo fwd | CACCCGCTCGGCGGGGGCGCACGG | | | | | | | | | | |
| Oligo rev | AAACCCGTGCGCCCCCGCCGAGCG | | | | | | | | | | |
| Chromosome | start | end | strand | MM | target_seq | PAM | alignment | distance | position | gene name | gene id |
| GU305914.1 | 52605 | 52627 | - | 0 | CGCTCGGCGGGGGCGCACGG | CGG | \|\|\|\|\|\|\|\|[\|\|\|\|\|\|\|\|\|\|\|\|]PAM | NA | Intergenic | NA | NA |
| T30133 | GCTCGGCGGGGGCGCACGGCGGG | 1000 | | | | | | | | | |
| Oligo fwd | CACCGCTCGGCGGGGGCGCACGGC | | | | | | | | | | |
| Oligo rev | AAACGCCGTGCGCCCCCGCCGAGC | | | | | | | | | | |
| Chromosome | start | end | strand | MM | target_seq | PAM | alignment | distance | position | gene name | gene id |
| GU305914.1 | 52604 | 52626 | - | 0 | GCTCGGCGGGGGCGCACGGC | GGG | \|\|\|\|\|\|\|\|[\|\|\|\|\|\|\|\|\|\|\|\|]PAM | NA | Intergenic | NA | NA |
| T30134 | CTCGGCGGGGGCGCACGGCGGGG | 1000 | | | | | | | | | |
| Oligo fwd | CACCCTCGGCGGGGGCGCACGGCG | | | | | | | | | | |
| Oligo rev | AAACCGCCGTGCGCCCCCGCCGAG | | | | | | | | | | |
| Chromosome | start | end | strand | MM | target_seq | PAM | alignment | distance | position | gene name | gene id |
| GU305914.1 | 52603 | 52625 | - | 0 | CTCGGCGGGGGCGCACGGCG | GGG | \|\|\|\|\|\|\|\|[\|\|\|\|\|\|\|\|\|\|\|\|]PAM | NA | Intergenic | NA | NA |
| T30135 | GGGGCGCACGGCGGGGCCGCCGG | 1000 | | | | | | | | | |
| Oligo fwd | CACCGGGGCGCACGGCGGGGCCGC | | | | | | | | | | |
| Oligo rev | AAACGCGGCCCCGCCGTGCGCCCC | | | | | | | | | | |
| Chromosome | start | end | strand | MM | target_seq | PAM | alignment | distance | position | gene name | gene id |
| GU305914.1 | 52596 | 52618 | - | 0 | GGGGCGCACGGCGGGGCCGC | CGG | \|\|\|\|\|\|\|\|[\|\|\|\|\|\|\|\|\|\|\|\|]PAM | NA | Intergenic | NA | NA |
| GU305914.1 | 103945 | 103967 | + | 4 | GGTTCGCGCGGCGGGGCCTC | CGG | \|\|--\|\|\|-[\|\|\|\|\|\|\|\|\|\|-\|]PAM | NA | Intergenic | NA | NA |
| GU305914.1 | 113657 | 113679 | - | 4 | GAGGGGCACGGCGGGCCCCC | CGG | \|-\|\|-\|\|\|[\|\|\|\|\|\|\|-\|\|-\|]PAM | NA | Intergenic | NA | NA |
| T30136 | GGGCGCACGGCGGGGCCGCCGGG | 1000 | | | | | | | | | |
| Oligo fwd | CACCGGGCGCACGGCGGGGCCGCC | | | | | | | | | | |
| Oligo rev | AAACGGCGGCCCCGCCGTGCGCCC | | | | | | | | | | |
| Chromosome | start | end | strand | MM | target_seq | PAM | alignment | distance | position | gene name | gene id |
| GU305914.1 | 52595 | 52617 | - | 0 | GGGCGCACGGCGGGGCCGCC | GGG | \|\|\|\|\|\|\|\|[\|\|\|\|\|\|\|\|\|\|\|\|]PAM | NA | Intergenic | NA | NA |
| GU305914.1 | 96457 | 96479 | + | 4 | GGATGGACGGCGGGGCGGCC | AGG | \|\|--\|-\|\|[\|\|\|\|\|\|\|\|-\|\|\|]PAM | NA | Intergenic | NA | NA |
| T30137 | GGCGCACGGCGGGGCCGCCGGGG | 1000 | | | | | | | | | |
| Oligo fwd | CACCGGCGCACGGCGGGGCCGCCG | | | | | | | | | | |
| Oligo rev | AAACCGGCGGCCCCGCCGTGCGCC | | | | | | | | | | |
| Chromosome | start | end | strand | MM | target_seq | PAM | alignment | distance | position | gene name | gene id |
| GU305914.1 | 52594 | 52616 | - | 0 | GGCGCACGGCGGGGCCGCCG | GGG | \|\|\|\|\|\|\|\|[\|\|\|\|\|\|\|\|\|\|\|\|]PAM | NA | Intergenic | NA | NA |
| T30138 | GCGCACGGCGGGGCCGCCGGGGG | 1000 | | | | | | | | | |
| Oligo fwd | CACCGCGCACGGCGGGGCCGCCGG | | | | | | | | | | |
| Oligo rev | AAACCCGGCGGCCCCGCCGTGCGC | | | | | | | | | | |
| Chromosome | start | end | strand | MM | target_seq | PAM | alignment | distance | position | gene name | gene id |
| GU305914.1 | 52593 | 52615 | - | 0 | GCGCACGGCGGGGCCGCCGG | GGG | \|\|\|\|\|\|\|\|[\|\|\|\|\|\|\|\|\|\|\|\|]PAM | NA | Intergenic | NA | NA |
| T30139 | GGGGTCAGTATCCCTTGCCTCGG | 1000 | | | | | | | | | |
| Oligo fwd | CACCGGGGTCAGTATCCCTTGCCT | | | | | | | | | | |
| Oligo rev | AAACAGGCAAGGGATACTGACCCC | | | | | | | | | | |
| Chromosome | start | end | strand | MM | target_seq | PAM | alignment | distance | position | gene name | gene id |
| GU305914.1 | 52574 | 52596 | - | 0 | GGGGTCAGTATCCCTTGCCT | CGG | \|\|\|\|\|\|\|\|[\|\|\|\|\|\|\|\|\|\|\|\|]PAM | NA | Intergenic | NA | NA |
| T30140 | GATGAAGATGACGACTACGATGG | 1000 | | | | | | | | | |
| Oligo fwd | CACCGATGAAGATGACGACTACGA | | | | | | | | | | |
| Oligo rev | AAACTCGTAGTCGTCATCTTCATC | | | | | | | | | | |
| Chromosome | start | end | strand | MM | target_seq | PAM | alignment | distance | position | gene name | gene id |
| GU305914.1 | 52533 | 52555 | - | 0 | GATGAAGATGACGACTACGA | TGG | \|\|\|\|\|\|\|\|[\|\|\|\|\|\|\|\|\|\|\|\|]PAM | NA | Intergenic | NA | NA |
| GU305914.1 | 118472 | 118494 | - | 3 | GATGATGATGACGACGAAGA | CGG | \|\|\|\|\|-\|\|[\|\|\|\|\|\|\|-\|-\|\|]PAM | NA | Intergenic | NA | NA |
| T30141 | ATGAAGATGACGACTACGATGGG | 1000 | | | | | | | | | |

<table>
<tr><td>Oligo fwd</td><td>CACCATGAAGATGACGACTACGAT</td><td></td><td></td><td></td><td></td><td></td><td></td><td></td><td></td><td></td><td></td></tr>
<tr><td>Oligo rev</td><td>AAACATCGTAGTCGTCATCTTCAT</td><td></td><td></td><td></td><td></td><td></td><td></td><td></td><td></td><td></td><td></td></tr>
<tr><td>Chromosome</td><td>start</td><td>end</td><td>strand</td><td>MM</td><td>target_seq</td><td>PAM</td><td>alignment</td><td>distance</td><td>position</td><td>gene name</td><td>gene id</td></tr>
<tr><td>GU305914.1</td><td>52532</td><td>52554</td><td>-</td><td>0</td><td>ATGAAGATGACGACTACGAT</td><td>GGG</td><td>||||||||[||||||||||||]PAM</td><td>NA</td><td>Intergenic</td><td>NA</td><td>NA</td></tr>
<tr><td>T30142</td><td>TGAAGATGACGACTACGATGGGG</td><td>1000</td><td></td><td></td><td></td><td></td><td></td><td></td><td></td><td></td><td></td></tr>
<tr><td>Oligo fwd</td><td>CACCTGAAGATGACGACTACGATG</td><td></td><td></td><td></td><td></td><td></td><td></td><td></td><td></td><td></td><td></td></tr>
<tr><td>Oligo rev</td><td>AAACCATCGTAGTCGTCATCTTCA</td><td></td><td></td><td></td><td></td><td></td><td></td><td></td><td></td><td></td><td></td></tr>
<tr><td>Chromosome</td><td>start</td><td>end</td><td>strand</td><td>MM</td><td>target_seq</td><td>PAM</td><td>alignment</td><td>distance</td><td>position</td><td>gene name</td><td>gene id</td></tr>
<tr><td>GU305914.1</td><td>52531</td><td>52553</td><td>-</td><td>0</td><td>TGAAGATGACGACTACGATG</td><td>GGG</td><td>||||||||[||||||||||||]PAM</td><td>NA</td><td>Intergenic</td><td>NA</td><td>NA</td></tr>
<tr><td>GU305914.1</td><td>65111</td><td>65133</td><td>+</td><td>4</td><td>GGAAGAAGACGACGACGAGG</td><td>AGG</td><td>-|||||-|[|||||-||||-|]PAM</td><td>NA</td><td>Intergenic</td><td>NA</td><td>NA</td></tr>
<tr><td>T30143</td><td>GACTACGATGGGGTCGATGATGG</td><td>1000</td><td></td><td></td><td></td><td></td><td></td><td></td><td></td><td></td><td></td></tr>
<tr><td>Oligo fwd</td><td>CACCGACTACGATGGGGTCGATGA</td><td></td><td></td><td></td><td></td><td></td><td></td><td></td><td></td><td></td><td></td></tr>
<tr><td>Oligo rev</td><td>AAACTCATCGACCCCATCGTAGTC</td><td></td><td></td><td></td><td></td><td></td><td></td><td></td><td></td><td></td><td></td></tr>
<tr><td>Chromosome</td><td>start</td><td>end</td><td>strand</td><td>MM</td><td>target_seq</td><td>PAM</td><td>alignment</td><td>distance</td><td>position</td><td>gene name</td><td>gene id</td></tr>
<tr><td>GU305914.1</td><td>52521</td><td>52543</td><td>-</td><td>0</td><td>GACTACGATGGGGTCGATGA</td><td>TGG</td><td>||||||||[||||||||||||]PAM</td><td>NA</td><td>Intergenic</td><td>NA</td><td>NA</td></tr>
<tr><td>T30144</td><td>ACAACTTCTGACCCTGACCGTGG</td><td>1000</td><td></td><td></td><td></td><td></td><td></td><td></td><td></td><td></td><td></td></tr>
<tr><td>Oligo fwd</td><td>CACCACAACTTCTGACCCTGACCG</td><td></td><td></td><td></td><td></td><td></td><td></td><td></td><td></td><td></td><td></td></tr>
<tr><td>Oligo rev</td><td>AAACCGGTCAGGGTCAGAAGTTGT</td><td></td><td></td><td></td><td></td><td></td><td></td><td></td><td></td><td></td><td></td></tr>
<tr><td>Chromosome</td><td>start</td><td>end</td><td>strand</td><td>MM</td><td>target_seq</td><td>PAM</td><td>alignment</td><td>distance</td><td>position</td><td>gene name</td><td>gene id</td></tr>
<tr><td>GU305914.1</td><td>52475</td><td>52497</td><td>-</td><td>0</td><td>ACAACTTCTGACCCTGACCG</td><td>TGG</td><td>||||||||[||||||||||||]PAM</td><td>NA</td><td>Intergenic</td><td>NA</td><td>NA</td></tr>
<tr><td>GU305914.1</td><td>137109</td><td>137131</td><td>-</td><td>4</td><td>CCAGCTTCTGACGCTGACCA</td><td>CGG</td><td>-||-||||[||||-||||||-]PAM</td><td>NA</td><td>Intergenic</td><td>NA</td><td>NA</td></tr>
<tr><td>T30145</td><td>ACTTCTGACCCTGACCGTGGCGG</td><td>1000</td><td></td><td></td><td></td><td></td><td></td><td></td><td></td><td></td><td></td></tr>
<tr><td>Oligo fwd</td><td>CACCACTTCTGACCCTGACCGTGG</td><td></td><td></td><td></td><td></td><td></td><td></td><td></td><td></td><td></td><td></td></tr>
<tr><td>Oligo rev</td><td>AAACCCACGGTCAGGGTCAGAAGT</td><td></td><td></td><td></td><td></td><td></td><td></td><td></td><td></td><td></td><td></td></tr>
<tr><td>Chromosome</td><td>start</td><td>end</td><td>strand</td><td>MM</td><td>target_seq</td><td>PAM</td><td>alignment</td><td>distance</td><td>position</td><td>gene name</td><td>gene id</td></tr>
<tr><td>GU305914.1</td><td>52472</td><td>52494</td><td>-</td><td>0</td><td>ACTTCTGACCCTGACCGTGG</td><td>CGG</td><td>||||||||[||||||||||||]PAM</td><td>NA</td><td>Intergenic</td><td>NA</td><td>NA</td></tr>
<tr><td>T30146</td><td>TGACCCTGACCGTGGCGGCTCGG</td><td>1000</td><td></td><td></td><td></td><td></td><td></td><td></td><td></td><td></td><td></td></tr>
<tr><td>Oligo fwd</td><td>CACCTGACCCTGACCGTGGCGGCT</td><td></td><td></td><td></td><td></td><td></td><td></td><td></td><td></td><td></td><td></td></tr>
<tr><td>Oligo rev</td><td>AAACAGCCGCCACGGTCAGGGTCA</td><td></td><td></td><td></td><td></td><td></td><td></td><td></td><td></td><td></td><td></td></tr>
<tr><td>Chromosome</td><td>start</td><td>end</td><td>strand</td><td>MM</td><td>target_seq</td><td>PAM</td><td>alignment</td><td>distance</td><td>position</td><td>gene name</td><td>gene id</td></tr>
<tr><td>GU305914.1</td><td>52467</td><td>52489</td><td>-</td><td>0</td><td>TGACCCTGACCGTGGCGGCT</td><td>CGG</td><td>||||||||[||||||||||||]PAM</td><td>NA</td><td>Intergenic</td><td>NA</td><td>NA</td></tr>
<tr><td>T30147</td><td>CCGTGGCGGCTCGGATTGATTGG</td><td>1000</td><td></td><td></td><td></td><td></td><td></td><td></td><td></td><td></td><td></td></tr>
<tr><td>Oligo fwd</td><td>CACCCCGTGGCGGCTCGGATTGAT</td><td></td><td></td><td></td><td></td><td></td><td></td><td></td><td></td><td></td><td></td></tr>
<tr><td>Oligo rev</td><td>AAACATCAATCCGAGCCGCCACGG</td><td></td><td></td><td></td><td></td><td></td><td></td><td></td><td></td><td></td><td></td></tr>
<tr><td>Chromosome</td><td>start</td><td>end</td><td>strand</td><td>MM</td><td>target_seq</td><td>PAM</td><td>alignment</td><td>distance</td><td>position</td><td>gene name</td><td>gene id</td></tr>
<tr><td>GU305914.1</td><td>52458</td><td>52480</td><td>-</td><td>0</td><td>CCGTGGCGGCTCGGATTGAT</td><td>TGG</td><td>||||||||[||||||||||||]PAM</td><td>NA</td><td>Intergenic</td><td>NA</td><td>NA</td></tr>
<tr><td>T30148</td><td>GCGGCTCGGATTGATTGGACAGG</td><td>1000</td><td></td><td></td><td></td><td></td><td></td><td></td><td></td><td></td><td></td></tr>
<tr><td>Oligo fwd</td><td>CACCGCGGCTCGGATTGATTGGAC</td><td></td><td></td><td></td><td></td><td></td><td></td><td></td><td></td><td></td><td></td></tr>
<tr><td>Oligo rev</td><td>AAACGTCCAATCAATCCGAGCCGC</td><td></td><td></td><td></td><td></td><td></td><td></td><td></td><td></td><td></td><td></td></tr>
<tr><td>Chromosome</td><td>start</td><td>end</td><td>strand</td><td>MM</td><td>target_seq</td><td>PAM</td><td>alignment</td><td>distance</td><td>position</td><td>gene name</td><td>gene id</td></tr>
<tr><td>GU305914.1</td><td>52453</td><td>52475</td><td>-</td><td>0</td><td>GCGGCTCGGATTGATTGGAC</td><td>AGG</td><td>||||||||[||||||||||||]PAM</td><td>NA</td><td>Intergenic</td><td>NA</td><td>NA</td></tr>
<tr><td>T30149</td><td>CGGATTGATTGGACAGGAAGTGG</td><td>1000</td><td></td><td></td><td></td><td></td><td></td><td></td><td></td><td></td><td></td></tr>
<tr><td>Oligo fwd</td><td>CACCCGGATTGATTGGACAGGAAG</td><td></td><td></td><td></td><td></td><td></td><td></td><td></td><td></td><td></td><td></td></tr>
<tr><td>Oligo rev</td><td>AAACCTTCCTGTCCAATCAATCCG</td><td></td><td></td><td></td><td></td><td></td><td></td><td></td><td></td><td></td><td></td></tr>
<tr><td>Chromosome</td><td>start</td><td>end</td><td>strand</td><td>MM</td><td>target_seq</td><td>PAM</td><td>alignment</td><td>distance</td><td>position</td><td>gene name</td><td>gene id</td></tr>
<tr><td>GU305914.1</td><td>52447</td><td>52469</td><td>-</td><td>0</td><td>CGGATTGATTGGACAGGAAG</td><td>TGG</td><td>||||||||[||||||||||||]PAM</td><td>NA</td><td>Intergenic</td><td>NA</td><td>NA</td></tr>
<tr><td>T30150</td><td>GATTGGACAGGAAGTGGAACCGG</td><td>1000</td><td></td><td></td><td></td><td></td><td></td><td></td><td></td><td></td><td></td></tr>
<tr><td>Oligo fwd</td><td>CACCGATTGGACAGGAAGTGGAAC</td><td></td><td></td><td></td><td></td><td></td><td></td><td></td><td></td><td></td><td></td></tr>
<tr><td>Oligo rev</td><td>AAACGTTCCACTTCCTGTCCAATC</td><td></td><td></td><td></td><td></td><td></td><td></td><td></td><td></td><td></td><td></td></tr>
<tr><td>Chromosome</td><td>start</td><td>end</td><td>strand</td><td>MM</td><td>target_seq</td><td>PAM</td><td>alignment</td><td>distance</td><td>position</td><td>gene name</td><td>gene id</td></tr>
<tr><td>GU305914.1</td><td>52441</td><td>52463</td><td>-</td><td>0</td><td>GATTGGACAGGAAGTGGAAC</td><td>CGG</td><td>||||||||[||||||||||||]PAM</td><td>NA</td><td>Intergenic</td><td>NA</td><td>NA</td></tr>
<tr><td>T30151</td><td>ACAGGAAGTGGAACCGGAGACGG</td><td>1000</td><td></td><td></td><td></td><td></td><td></td><td></td><td></td><td></td><td></td></tr>
<tr><td>Oligo fwd</td><td>CACCACAGGAAGTGGAACCGGAGA</td><td></td><td></td><td></td><td></td><td></td><td></td><td></td><td></td><td></td><td></td></tr>
<tr><td>Oligo rev</td><td>AAACTCTCCGGTTCCACTTCCTGT</td><td></td><td></td><td></td><td></td><td></td><td></td><td></td><td></td><td></td><td></td></tr>
<tr><td>Chromosome</td><td>start</td><td>end</td><td>strand</td><td>MM</td><td>target_seq</td><td>PAM</td><td>alignment</td><td>distance</td><td>position</td><td>gene name</td><td>gene id</td></tr>
<tr><td>GU305914.1</td><td>52435</td><td>52457</td><td>-</td><td>0</td><td>ACAGGAAGTGGAACCGGAGA</td><td>CGG</td><td>||||||||[||||||||||||]PAM</td><td>NA</td><td>Intergenic</td><td>NA</td><td>NA</td></tr>
<tr><td>T30152</td><td>GAAGTGGAACCGGAGACGGACGG</td><td>1000</td><td></td><td></td><td></td><td></td><td></td><td></td><td></td><td></td><td></td></tr>
<tr><td>Oligo fwd</td><td>CACCGAAGTGGAACCGGAGACGGA</td><td></td><td></td><td></td><td></td><td></td><td></td><td></td><td></td><td></td><td></td></tr>
<tr><td>Oligo rev</td><td>AAACTCCGTCTCCGGTTCCACTTC</td><td></td><td></td><td></td><td></td><td></td><td></td><td></td><td></td><td></td><td></td></tr>
<tr><td>Chromosome</td><td>start</td><td>end</td><td>strand</td><td>MM</td><td>target_seq</td><td>PAM</td><td>alignment</td><td>distance</td><td>position</td><td>gene name</td><td>gene id</td></tr>
<tr><td>GU305914.1</td><td>52431</td><td>52453</td><td>-</td><td>0</td><td>GAAGTGGAACCGGAGACGGA</td><td>CGG</td><td>||||||||[||||||||||||]PAM</td><td>NA</td><td>Intergenic</td><td>NA</td><td>NA</td></tr>
<tr><td>T30153</td><td>AAGTGGAACCGGAGACGGACGGG</td><td>1000</td><td></td><td></td><td></td><td></td><td></td><td></td><td></td><td></td><td></td></tr>
<tr><td>Oligo fwd</td><td>CACCAAGTGGAACCGGAGACGGAC</td><td></td><td></td><td></td><td></td><td></td><td></td><td></td><td></td><td></td><td></td></tr>
<tr><td>Oligo rev</td><td>AAACGTCCGTCTCCGGTTCCACTT</td><td></td><td></td><td></td><td></td><td></td><td></td><td></td><td></td><td></td><td></td></tr>
<tr><td>Chromosome</td><td>start</td><td>end</td><td>strand</td><td>MM</td><td>target_seq</td><td>PAM</td><td>alignment</td><td>distance</td><td>position</td><td>gene name</td><td>gene id</td></tr>
<tr><td>GU305914.1</td><td>52430</td><td>52452</td><td>-</td><td>0</td><td>AAGTGGAACCGGAGACGGAC</td><td>GGG</td><td>||||||||[||||||||||||]PAM</td><td>NA</td><td>Intergenic</td><td>NA</td><td>NA</td></tr>
<tr><td>T30154</td><td>AGTGGAACCGGAGACGGACGGGG</td><td>1000</td><td></td><td></td><td></td><td></td><td></td><td></td><td></td><td></td><td></td></tr>
<tr><td>Oligo fwd</td><td>CACCAGTGGAACCGGAGACGGACG</td><td></td><td></td><td></td><td></td><td></td><td></td><td></td><td></td><td></td><td></td></tr>
<tr><td>Oligo rev</td><td>AAACCGTCCGTCTCCGGTTCCACT</td><td></td><td></td><td></td><td></td><td></td><td></td><td></td><td></td><td></td><td></td></tr>
<tr><td>Chromosome</td><td>start</td><td>end</td><td>strand</td><td>MM</td><td>target_seq</td><td>PAM</td><td>alignment</td><td>distance</td><td>position</td><td>gene name</td><td>gene id</td></tr>
<tr><td>GU305914.1</td><td>52429</td><td>52451</td><td>-</td><td>0</td><td>AGTGGAACCGGAGACGGACG</td><td>GGG</td><td>||||||||[||||||||||||]PAM</td><td>NA</td><td>Intergenic</td><td>NA</td><td>NA</td></tr>
<tr><td>T30155</td><td>GTGGAACCGGAGACGGACGGGGG</td><td>1000</td><td></td><td></td><td></td><td></td><td></td><td></td><td></td><td></td><td></td></tr>
<tr><td>Oligo fwd</td><td>CACCGTGGAACCGGAGACGGACGG</td><td></td><td></td><td></td><td></td><td></td><td></td><td></td><td></td><td></td><td></td></tr>
<tr><td>Oligo rev</td><td>AAACCCGTCCGTCTCCGGTTCCAC</td><td></td><td></td><td></td><td></td><td></td><td></td><td></td><td></td><td></td><td></td></tr>
<tr><td>Chromosome</td><td>start</td><td>end</td><td>strand</td><td>MM</td><td>target_seq</td><td>PAM</td><td>alignment</td><td>distance</td><td>position</td><td>gene name</td><td>gene id</td></tr>
</table>

GU305914.1 52428 52450 - 0 GTGGAACCGGAGACGGACGG GGG ||||||||[||||||||||||]PAM NA Intergenic NA NA

T30156 GGACACATATAAAAGTTGCGCGG 1000
Oligo fwd CACCGGACACATATAAAAGTTGCG
Oligo rev AAACCGCAACTTTTATATGTGTCC
Chromosome start end strand MM target_seq PAM alignment distance position gene name gene id
GU305914.1 52407 52429 - 0 GGACACATATAAAAGTTGCG CGG ||||||||[||||||||||||]PAM NA Intergenic NA NA

T30157 ACATATAAAAGTTGCGCGGACGG 1000
Oligo fwd CACCACATATAAAAGTTGCGCGGA
Oligo rev AAACTCCGCGCAACTTTTATATGT
Chromosome start end strand MM target_seq PAM alignment distance position gene name gene id
GU305914.1 52403 52425 - 0 ACATATAAAAGTTGCGCGGA CGG ||||||||[||||||||||||]PAM NA Intergenic NA NA

T30158 AAGTTGCGCGGACGGTGCCGCGG 1000
Oligo fwd CACCAAGTTGCGCGGACGGTGCCG
Oligo rev AAACCGGCACCGTCCGCGCAACTT
Chromosome start end strand MM target_seq PAM alignment distance position gene name gene id
GU305914.1 52395 52417 - 0 AAGTTGCGCGGACGGTGCCG CGG ||||||||[||||||||||||]PAM NA Intergenic NA NA

T30159 AGTTGCGCGGACGGTGCCGCGGG 1000
Oligo fwd CACCAGTTGCGCGGACGGTGCCGC
Oligo rev AAACGCGGCACCGTCCGCGCAACT
Chromosome start end strand MM target_seq PAM alignment distance position gene name gene id
GU305914.1 52394 52416 - 0 AGTTGCGCGGACGGTGCCGC GGG ||||||||[||||||||||||]PAM NA Intergenic NA NA

T30160 GGTGCCGCGGGTTGTAGTTGTGG 1000
Oligo fwd CACCGGTGCCGCGGGTTGTAGTTG
Oligo rev AAACCAACTACAACCCGCGGCACC
Chromosome start end strand MM target_seq PAM alignment distance position gene name gene id
GU305914.1 52382 52404 - 0 GGTGCCGCGGGTTGTAGTTG TGG ||||||||[||||||||||||]PAM NA Intergenic NA NA

T30161 GCCGCGGGTTGTAGTTGTGGAGG 1000
Oligo fwd CACCGCCGCGGGTTGTAGTTGTGG
Oligo rev AAACCCACAACTACAACCCGCGGC
Chromosome start end strand MM target_seq PAM alignment distance position gene name gene id
GU305914.1 52379 52401 - 0 GCCGCGGGTTGTAGTTGTGG AGG ||||||||[||||||||||||]PAM NA Intergenic NA NA

T30162 CCGCGGGTTGTAGTTGTGGAGGG 1000
Oligo fwd CACCCCGCGGGTTGTAGTTGTGGA
Oligo rev AAACTCCACAACTACAACCCGCGG
Chromosome start end strand MM target_seq PAM alignment distance position gene name gene id
GU305914.1 52378 52400 - 0 CCGCGGGTTGTAGTTGTGGA GGG ||||||||[||||||||||||]PAM NA Intergenic NA NA

T30163 CGCGGGTTGTAGTTGTGGAGGGG 1000
Oligo fwd CACCCGCGGGTTGTAGTTGTGGAG
Oligo rev AAACCTCCACAACTACAACCCGCG
Chromosome start end strand MM target_seq PAM alignment distance position gene name gene id
GU305914.1 52377 52399 - 0 CGCGGGTTGTAGTTGTGGAG GGG ||||||||[||||||||||||]PAM NA Intergenic NA NA

T30164 AGTTGTGGAGGGGACAGCGATGG 1000
Oligo fwd CACCAGTTGTGGAGGGGACAGCGA
Oligo rev AAACTCGCTGTCCCCTCCACAACT
Chromosome start end strand MM target_seq PAM alignment distance position gene name gene id
GU305914.1 52367 52389 - 0 AGTTGTGGAGGGGACAGCGA TGG ||||||||[||||||||||||]PAM NA Intergenic NA NA

T30165 AGTGTGCGCCGACCCTTCTTTGG 1000
Oligo fwd CACCAGTGTGCGCCGACCCTTCTT
Oligo rev AAACAAGAAGGGTCGGCGCACACT
Chromosome start end strand MM target_seq PAM alignment distance position gene name gene id
GU305914.1 52342 52364 - 0 AGTGTGCGCCGACCCTTCTT TGG ||||||||[||||||||||||]PAM NA Intergenic NA NA

T30166 GTGTGCGCCGACCCTTCTTTGGG 1000
Oligo fwd CACCGTGTGCGCCGACCCTTCTTT
Oligo rev AAACAAAGAAGGGTCGGCGCACAC
Chromosome start end strand MM target_seq PAM alignment distance position gene name gene id
GU305914.1 52341 52363 - 0 GTGTGCGCCGACCCTTCTTT GGG ||||||||[||||||||||||]PAM NA Intergenic NA NA

T30167 CCAGCGCTGCTCTTCGCCTCTGG 1000
Oligo fwd CACCCCAGCGCTGCTCTTCGCCTC
Oligo rev AAACGAGGCGAAGAGCAGCGCTGG
Chromosome start end strand MM target_seq PAM alignment distance position gene name gene id
GU305914.1 52303 52325 - 0 CCAGCGCTGCTCTTCGCCTC TGG ||||||||[||||||||||||]PAM NA Intergenic NA NA
GU305914.1 70169 70191 - 3 CCGGTGCTGCTCGTCGCCTC GGG ||-|-|||[||||-||||||||]PAM NA Intergenic NA NA

T30168 CAGCGCTGCTCTTCGCCTCTGGG 1000
Oligo fwd CACCCAGCGCTGCTCTTCGCCTCT
Oligo rev AAACAGAGGCGAAGAGCAGCGCTG
Chromosome start end strand MM target_seq PAM alignment distance position gene name gene id
GU305914.1 52302 52324 - 0 CAGCGCTGCTCTTCGCCTCT GGG ||||||||[||||||||||||]PAM NA Intergenic NA NA
GU305914.1 70168 70190 - 4 CGGTGCTGCTCGTCGCCTCG GGG |-|-||||[|||-|||||||-]PAM NA Intergenic NA NA

T30169 AGCGCTGCTCTTCGCCTCTGGGG 1000
Oligo fwd CACCAGCGCTGCTCTTCGCCTCTG
Oligo rev AAACCAGAGGCGAAGAGCAGCGCT
Chromosome start end strand MM target_seq PAM alignment distance position gene name gene id
GU305914.1 52301 52323 - 0 AGCGCTGCTCTTCGCCTCTG GGG ||||||||[||||||||||||]PAM NA Intergenic NA NA

T30170 CTCTTCGCCTCTGGGGCGCTTGG 1000

Oligo fwd CACCCTCTTCGCCTCTGGGGCGCT
Oligo rev AAACAGCGCCCCAGAGGCGAAGAG
Chromosome start end strand MM target_seq PAM alignment distance position gene name gene id
GU305914.1 52294 52316 - 0 CTCTTCGCCTCTGGGGCGCT TGG ||||||||[||||||||||||]PAM NA Intergenic NA NA

T30171 GGGGCGCTTGGTTTCGTAGCCGG 1000
Oligo fwd CACCGGGGCGCTTGGTTTCGTAGC
Oligo rev AAACGCTACGAAACCAAGCGCCCC
Chromosome start end strand MM target_seq PAM alignment distance position gene name gene id
GU305914.1 52282 52304 - 0 GGGGCGCTTGGTTTCGTAGC CGG ||||||||[||||||||||||]PAM NA Intergenic NA NA

T30172 CGCTTGGTTTCGTAGCCGGTTGG 1000
Oligo fwd CACCCGCTTGGTTTCGTAGCCGGT
Oligo rev AAACACCGGCTACGAAACCAAGCG
Chromosome start end strand MM target_seq PAM alignment distance position gene name gene id
GU305914.1 52278 52300 - 0 CGCTTGGTTTCGTAGCCGGT TGG ||||||||[||||||||||||]PAM NA Intergenic NA NA

T30173 CCGGTTGGATTCTCTCTAATTGG 1000
Oligo fwd CACCCCGGTTGGATTCTCTCTAAT
Oligo rev AAACATTAGAGAGAATCCAACCGG
Chromosome start end strand MM target_seq PAM alignment distance position gene name gene id
GU305914.1 52263 52285 - 0 CCGGTTGGATTCTCTCTAAT TGG ||||||||[||||||||||||]PAM NA Intergenic NA NA

T30174 TTCTCTCTAATTGGTTAACCAGG 1000
Oligo fwd CACCTTCTCTCTAATTGGTTAACC
Oligo rev AAACGGTTAACCAATTAGAGAGAA
Chromosome start end strand MM target_seq PAM alignment distance position gene name gene id
GU305914.1 52254 52276 - 0 TTCTCTCTAATTGGTTAACC AGG ||||||||[||||||||||||]PAM NA Intergenic NA NA

T30175 TCTCTAATTGGTTAACCAGGCGG 1000
Oligo fwd CACCTCTCTAATTGGTTAACCAGG
Oligo rev AAACCCTGGTTAACCAATTAGAGA
Chromosome start end strand MM target_seq PAM alignment distance position gene name gene id
GU305914.1 52251 52273 - 0 TCTCTAATTGGTTAACCAGG CGG ||||||||[||||||||||||]PAM NA Intergenic NA NA

T30176 CTAATTGGTTAACCAGGCGGCGG 1000
Oligo fwd CACCCTAATTGGTTAACCAGGCGG
Oligo rev AAACCCGCCTGGTTAACCAATTAG
Chromosome start end strand MM target_seq PAM alignment distance position gene name gene id
GU305914.1 52248 52270 - 0 CTAATTGGTTAACCAGGCGG CGG ||||||||[||||||||||||]PAM NA Intergenic NA NA

T30177 TAATTGGTTAACCAGGCGGCGGG 1000
Oligo fwd CACCTAATTGGTTAACCAGGCGGC
Oligo rev AAACGCCGCCTGGTTAACCAATTA
Chromosome start end strand MM target_seq PAM alignment distance position gene name gene id
GU305914.1 52247 52269 - 0 TAATTGGTTAACCAGGCGGC GGG ||||||||[||||||||||||]PAM NA Intergenic NA NA

T30178 GCGGGACAGTCAGAACTGCTCGG 1000
Oligo fwd CACCGCGGGACAGTCAGAACTGCT
Oligo rev AAACAGCAGTTCTGACTGTCCCGC
Chromosome start end strand MM target_seq PAM alignment distance position gene name gene id
GU305914.1 52229 52251 - 0 GCGGGACAGTCAGAACTGCT CGG ||||||||[||||||||||||]PAM NA Intergenic NA NA

T30179 CGGGACAGTCAGAACTGCTCGGG 1000
Oligo fwd CACCCGGGACAGTCAGAACTGCTC
Oligo rev AAACGAGCAGTTCTGACTGTCCCG
Chromosome start end strand MM target_seq PAM alignment distance position gene name gene id
GU305914.1 52228 52250 - 0 CGGGACAGTCAGAACTGCTC GGG ||||||||[||||||||||||]PAM NA Intergenic NA NA

T30180 GCTCGGGAAGCAGCTGTGATCGG 1000
Oligo fwd CACCGCTCGGGAAGCAGCTGTGAT
Oligo rev AAACATCACAGCTGCTTCCCGAGC
Chromosome start end strand MM target_seq PAM alignment distance position gene name gene id
GU305914.1 52212 52234 - 0 GCTCGGGAAGCAGCTGTGAT CGG ||||||||[||||||||||||]PAM NA Intergenic NA NA

T30181 AGCAGCTGTGATCGGCCCGCTGG 1000
Oligo fwd CACCAGCAGCTGTGATCGGCCCGC
Oligo rev AAACGCGGGCCGATCACAGCTGCT
Chromosome start end strand MM target_seq PAM alignment distance position gene name gene id
GU305914.1 52204 52226 - 0 AGCAGCTGTGATCGGCCCGC TGG ||||||||[||||||||||||]PAM NA Intergenic NA NA

T30182 TCGGCCCGCTGGAGACGTTACGG 1000
Oligo fwd CACCTCGGCCCGCTGGAGACGTTA
Oligo rev AAACTAACGTCTCCAGCGGGCCGA
Chromosome start end strand MM target_seq PAM alignment distance position gene name gene id
GU305914.1 52193 52215 - 0 TCGGCCCGCTGGAGACGTTA CGG ||||||||[||||||||||||]PAM NA Intergenic NA NA

T30183 GCTGGAGACGTTACGGACGACGG 1000
Oligo fwd CACCGCTGGAGACGTTACGGACGA
Oligo rev AAACTCGTCCGTAACGTCTCCAGC
Chromosome start end strand MM target_seq PAM alignment distance position gene name gene id
GU305914.1 52186 52208 - 0 GCTGGAGACGTTACGGACGA CGG ||||||||[||||||||||||]PAM NA Intergenic NA NA

T30184 CGTTACGGACGACGGTTCAGCGG 1000
Oligo fwd CACCCGTTACGGACGACGGTTCAG
Oligo rev AAACCTGAACCGTCGTCCGTAACG
Chromosome start end strand MM target_seq PAM alignment distance position gene name gene id
GU305914.1 52178 52200 - 0 CGTTACGGACGACGGTTCAG CGG ||||||||[||||||||||||]PAM NA Intergenic NA NA

<table>
<tr><td>T30185</td><td>GACGGTTCAGCGGTCTACGTCGG</td><td>1000</td><td></td><td></td><td></td><td></td><td></td><td></td><td></td><td></td><td></td></tr>
<tr><td>Oligo fwd</td><td>CACCGACGGTTCAGCGGTCTACGT</td><td></td><td></td><td></td><td></td><td></td><td></td><td></td><td></td><td></td><td></td></tr>
<tr><td>Oligo rev</td><td>AAACACGTAGACCGCTGAACCGTC</td><td></td><td></td><td></td><td></td><td></td><td></td><td></td><td></td><td></td><td></td></tr>
<tr><td>Chromosome</td><td>start</td><td>end</td><td>strand</td><td>MM</td><td>target_seq</td><td>PAM</td><td>alignment</td><td>distance</td><td>position</td><td>gene name</td><td>gene id</td></tr>
<tr><td>GU305914.1</td><td>52168</td><td>52190</td><td>-</td><td>0</td><td>GACGGTTCAGCGGTCTACGT</td><td>CGG</td><td>||||||||[||||||||||||]PAM</td><td>NA</td><td>Intergenic</td><td>NA</td><td>NA</td></tr>
<tr><td>T30186</td><td>CAGCGGTCTACGTCGGTCTTCGG</td><td>1000</td><td></td><td></td><td></td><td></td><td></td><td></td><td></td><td></td><td></td></tr>
<tr><td>Oligo fwd</td><td>CACCCAGCGGTCTACGTCGGTCTT</td><td></td><td></td><td></td><td></td><td></td><td></td><td></td><td></td><td></td><td></td></tr>
<tr><td>Oligo rev</td><td>AAACAAGACCGACGTAGACCGCTG</td><td></td><td></td><td></td><td></td><td></td><td></td><td></td><td></td><td></td><td></td></tr>
<tr><td>Chromosome</td><td>start</td><td>end</td><td>strand</td><td>MM</td><td>target_seq</td><td>PAM</td><td>alignment</td><td>distance</td><td>position</td><td>gene name</td><td>gene id</td></tr>
<tr><td>GU305914.1</td><td>52161</td><td>52183</td><td>-</td><td>0</td><td>CAGCGGTCTACGTCGGTCTT</td><td>CGG</td><td>||||||||[||||||||||||]PAM</td><td>NA</td><td>Intergenic</td><td>NA</td><td>NA</td></tr>
<tr><td>T30187</td><td>TTCGGCGCCGCCGCTTCTCCTGG</td><td>1000</td><td></td><td></td><td></td><td></td><td></td><td></td><td></td><td></td><td></td></tr>
<tr><td>Oligo fwd</td><td>CACCTTCGGCGCCGCCGCTTCTCC</td><td></td><td></td><td></td><td></td><td></td><td></td><td></td><td></td><td></td><td></td></tr>
<tr><td>Oligo rev</td><td>AAACGGAGAAGCGGCGGCGCCGAA</td><td></td><td></td><td></td><td></td><td></td><td></td><td></td><td></td><td></td><td></td></tr>
<tr><td>Chromosome</td><td>start</td><td>end</td><td>strand</td><td>MM</td><td>target_seq</td><td>PAM</td><td>alignment</td><td>distance</td><td>position</td><td>gene name</td><td>gene id</td></tr>
<tr><td>GU305914.1</td><td>52143</td><td>52165</td><td>-</td><td>0</td><td>TTCGGCGCCGCCGCTTCTCC</td><td>TGG</td><td>||||||||[||||||||||||]PAM</td><td>NA</td><td>Intergenic</td><td>NA</td><td>NA</td></tr>
<tr><td>T30188</td><td>TCGGCGCCGCCGCTTCTCCTGGG</td><td>1000</td><td></td><td></td><td></td><td></td><td></td><td></td><td></td><td></td><td></td></tr>
<tr><td>Oligo fwd</td><td>CACCTCGGCGCCGCCGCTTCTCCT</td><td></td><td></td><td></td><td></td><td></td><td></td><td></td><td></td><td></td><td></td></tr>
<tr><td>Oligo rev</td><td>AAACAGGAGAAGCGGCGGCGCCGA</td><td></td><td></td><td></td><td></td><td></td><td></td><td></td><td></td><td></td><td></td></tr>
<tr><td>Chromosome</td><td>start</td><td>end</td><td>strand</td><td>MM</td><td>target_seq</td><td>PAM</td><td>alignment</td><td>distance</td><td>position</td><td>gene name</td><td>gene id</td></tr>
<tr><td>GU305914.1</td><td>52142</td><td>52164</td><td>-</td><td>0</td><td>TCGGCGCCGCCGCTTCTCCT</td><td>GGG</td><td>||||||||[||||||||||||]PAM</td><td>NA</td><td>Intergenic</td><td>NA</td><td>NA</td></tr>
<tr><td>T30189</td><td>CGGCGCCGCCGCTTCTCCTGGGG</td><td>1000</td><td></td><td></td><td></td><td></td><td></td><td></td><td></td><td></td><td></td></tr>
<tr><td>Oligo fwd</td><td>CACCCGGCGCCGCCGCTTCTCCTG</td><td></td><td></td><td></td><td></td><td></td><td></td><td></td><td></td><td></td><td></td></tr>
<tr><td>Oligo rev</td><td>AAACCAGGAGAAGCGGCGGCGCCG</td><td></td><td></td><td></td><td></td><td></td><td></td><td></td><td></td><td></td><td></td></tr>
<tr><td>Chromosome</td><td>start</td><td>end</td><td>strand</td><td>MM</td><td>target_seq</td><td>PAM</td><td>alignment</td><td>distance</td><td>position</td><td>gene name</td><td>gene id</td></tr>
<tr><td>GU305914.1</td><td>52141</td><td>52163</td><td>-</td><td>0</td><td>CGGCGCCGCCGCTTCTCCTG</td><td>GGG</td><td>||||||||[||||||||||||]PAM</td><td>NA</td><td>Intergenic</td><td>NA</td><td>NA</td></tr>
<tr><td>T30190</td><td>TCGCGAACGCCGCTTCGCCGAGG</td><td>1000</td><td></td><td></td><td></td><td></td><td></td><td></td><td></td><td></td><td></td></tr>
<tr><td>Oligo fwd</td><td>CACCTCGCGAACGCCGCTTCGCCG</td><td></td><td></td><td></td><td></td><td></td><td></td><td></td><td></td><td></td><td></td></tr>
<tr><td>Oligo rev</td><td>AAACCGGCGAAGCGGCGTTCGCGA</td><td></td><td></td><td></td><td></td><td></td><td></td><td></td><td></td><td></td><td></td></tr>
<tr><td>Chromosome</td><td>start</td><td>end</td><td>strand</td><td>MM</td><td>target_seq</td><td>PAM</td><td>alignment</td><td>distance</td><td>position</td><td>gene name</td><td>gene id</td></tr>
<tr><td>GU305914.1</td><td>52115</td><td>52137</td><td>-</td><td>0</td><td>TCGCGAACGCCGCTTCGCCG</td><td>AGG</td><td>||||||||[||||||||||||]PAM</td><td>NA</td><td>Intergenic</td><td>NA</td><td>NA</td></tr>
<tr><td>GU305914.1</td><td>61773</td><td>61795</td><td>-</td><td>4</td><td>TCGCATCCGCCGCTACGCCG</td><td>CGG</td><td>||||---|[||||||-|||||]PAM</td><td>NA</td><td>Intergenic</td><td>NA</td><td>NA</td></tr>
<tr><td>T30191</td><td>ACGAGCTCATACGCGTCTTGCGG</td><td>1000</td><td></td><td></td><td></td><td></td><td></td><td></td><td></td><td></td><td></td></tr>
<tr><td>Oligo fwd</td><td>CACCACGAGCTCATACGCGTCTTG</td><td></td><td></td><td></td><td></td><td></td><td></td><td></td><td></td><td></td><td></td></tr>
<tr><td>Oligo rev</td><td>AAACCAAGACGCGTATGAGCTCGT</td><td></td><td></td><td></td><td></td><td></td><td></td><td></td><td></td><td></td><td></td></tr>
<tr><td>Chromosome</td><td>start</td><td>end</td><td>strand</td><td>MM</td><td>target_seq</td><td>PAM</td><td>alignment</td><td>distance</td><td>position</td><td>gene name</td><td>gene id</td></tr>
<tr><td>GU305914.1</td><td>52083</td><td>52105</td><td>-</td><td>0</td><td>ACGAGCTCATACGCGTCTTG</td><td>CGG</td><td>||||||||[||||||||||||]PAM</td><td>NA</td><td>Intergenic</td><td>NA</td><td>NA</td></tr>
<tr><td>T30192</td><td>CGAGCTCATACGCGTCTTGCGGG</td><td>1000</td><td></td><td></td><td></td><td></td><td></td><td></td><td></td><td></td><td></td></tr>
<tr><td>Oligo fwd</td><td>CACCCGAGCTCATACGCGTCTTGC</td><td></td><td></td><td></td><td></td><td></td><td></td><td></td><td></td><td></td><td></td></tr>
<tr><td>Oligo rev</td><td>AAACGCAAGACGCGTATGAGCTCG</td><td></td><td></td><td></td><td></td><td></td><td></td><td></td><td></td><td></td><td></td></tr>
<tr><td>Chromosome</td><td>start</td><td>end</td><td>strand</td><td>MM</td><td>target_seq</td><td>PAM</td><td>alignment</td><td>distance</td><td>position</td><td>gene name</td><td>gene id</td></tr>
<tr><td>GU305914.1</td><td>52082</td><td>52104</td><td>-</td><td>0</td><td>CGAGCTCATACGCGTCTTGC</td><td>GGG</td><td>||||||||[||||||||||||]PAM</td><td>NA</td><td>Intergenic</td><td>NA</td><td>NA</td></tr>
<tr><td>T30193</td><td>TACGCGTCTTGCGGGACATCAGG</td><td>1000</td><td></td><td></td><td></td><td></td><td></td><td></td><td></td><td></td><td></td></tr>
<tr><td>Oligo fwd</td><td>CACCTACGCGTCTTGCGGGACATC</td><td></td><td></td><td></td><td></td><td></td><td></td><td></td><td></td><td></td><td></td></tr>
<tr><td>Oligo rev</td><td>AAACGATGTCCCGCAAGACGCGTA</td><td></td><td></td><td></td><td></td><td></td><td></td><td></td><td></td><td></td><td></td></tr>
<tr><td>Chromosome</td><td>start</td><td>end</td><td>strand</td><td>MM</td><td>target_seq</td><td>PAM</td><td>alignment</td><td>distance</td><td>position</td><td>gene name</td><td>gene id</td></tr>
<tr><td>GU305914.1</td><td>52074</td><td>52096</td><td>-</td><td>0</td><td>TACGCGTCTTGCGGGACATC</td><td>AGG</td><td>||||||||[||||||||||||]PAM</td><td>NA</td><td>Intergenic</td><td>NA</td><td>NA</td></tr>
<tr><td>T30194</td><td>GCAGCAGCTACAACAACAGCAGG</td><td>1000</td><td></td><td></td><td></td><td></td><td></td><td></td><td></td><td></td><td></td></tr>
<tr><td>Oligo fwd</td><td>CACCGCAGCAGCTACAACAACAGC</td><td></td><td></td><td></td><td></td><td></td><td></td><td></td><td></td><td></td><td></td></tr>
<tr><td>Oligo rev</td><td>AAACGCTGTTGTTGTAGCTGCTGC</td><td></td><td></td><td></td><td></td><td></td><td></td><td></td><td></td><td></td><td></td></tr>
<tr><td>Chromosome</td><td>start</td><td>end</td><td>strand</td><td>MM</td><td>target_seq</td><td>PAM</td><td>alignment</td><td>distance</td><td>position</td><td>gene name</td><td>gene id</td></tr>
<tr><td>GU305914.1</td><td>52040</td><td>52062</td><td>-</td><td>0</td><td>GCAGCAGCTACAACAACAGC</td><td>AGG</td><td>||||||||[||||||||||||]PAM</td><td>NA</td><td>Intergenic</td><td>NA</td><td>NA</td></tr>
<tr><td>GU305914.1</td><td>52025</td><td>52047</td><td>-</td><td>4</td><td>ACAGCAGGAACAACAGCAGC</td><td>AGG</td><td>-||||||-[-||||||-||||]PAM</td><td>NA</td><td>Intergenic</td><td>NA</td><td>NA</td></tr>
<tr><td>T30195</td><td>ACAGCAGGAACAACAGCAGCAGG</td><td>1000</td><td></td><td></td><td></td><td></td><td></td><td></td><td></td><td></td><td></td></tr>
<tr><td>Oligo fwd</td><td>CACCACAGCAGGAACAACAGCAGC</td><td></td><td></td><td></td><td></td><td></td><td></td><td></td><td></td><td></td><td></td></tr>
<tr><td>Oligo rev</td><td>AAACGCTGCTGTTGTTCCTGCTGT</td><td></td><td></td><td></td><td></td><td></td><td></td><td></td><td></td><td></td><td></td></tr>
<tr><td>Chromosome</td><td>start</td><td>end</td><td>strand</td><td>MM</td><td>target_seq</td><td>PAM</td><td>alignment</td><td>distance</td><td>position</td><td>gene name</td><td>gene id</td></tr>
<tr><td>GU305914.1</td><td>52025</td><td>52047</td><td>-</td><td>0</td><td>ACAGCAGGAACAACAGCAGC</td><td>AGG</td><td>||||||||[||||||||||||]PAM</td><td>NA</td><td>Intergenic</td><td>NA</td><td>NA</td></tr>
<tr><td>GU305914.1</td><td>52040</td><td>52062</td><td>-</td><td>4</td><td>GCAGCAGCTACAACAACAGC</td><td>AGG</td><td>-||||||-[-||||||-||||]PAM</td><td>NA</td><td>Intergenic</td><td>NA</td><td>NA</td></tr>
<tr><td>T30196</td><td>TCGTCCGCAGTCTCAGCGTCAGG</td><td>1000</td><td></td><td></td><td></td><td></td><td></td><td></td><td></td><td></td><td></td></tr>
<tr><td>Oligo fwd</td><td>CACCTCGTCCGCAGTCTCAGCGTC</td><td></td><td></td><td></td><td></td><td></td><td></td><td></td><td></td><td></td><td></td></tr>
<tr><td>Oligo rev</td><td>AAACGACGCTGAGACTGCGGACGA</td><td></td><td></td><td></td><td></td><td></td><td></td><td></td><td></td><td></td><td></td></tr>
<tr><td>Chromosome</td><td>start</td><td>end</td><td>strand</td><td>MM</td><td>target_seq</td><td>PAM</td><td>alignment</td><td>distance</td><td>position</td><td>gene name</td><td>gene id</td></tr>
<tr><td>GU305914.1</td><td>51968</td><td>51990</td><td>-</td><td>0</td><td>TCGTCCGCAGTCTCAGCGTC</td><td>AGG</td><td>||||||||[||||||||||||]PAM</td><td>NA</td><td>Intergenic</td><td>NA</td><td>NA</td></tr>
<tr><td>T30197</td><td>CAGTCTCAGCGTCAGGAAGCCGG</td><td>1000</td><td></td><td></td><td></td><td></td><td></td><td></td><td></td><td></td><td></td></tr>
<tr><td>Oligo fwd</td><td>CACCCAGTCTCAGCGTCAGGAAGC</td><td></td><td></td><td></td><td></td><td></td><td></td><td></td><td></td><td></td><td></td></tr>
<tr><td>Oligo rev</td><td>AAACGCTTCCTGACGCTGAGACTG</td><td></td><td></td><td></td><td></td><td></td><td></td><td></td><td></td><td></td><td></td></tr>
<tr><td>Chromosome</td><td>start</td><td>end</td><td>strand</td><td>MM</td><td>target_seq</td><td>PAM</td><td>alignment</td><td>distance</td><td>position</td><td>gene name</td><td>gene id</td></tr>
<tr><td>GU305914.1</td><td>51961</td><td>51983</td><td>-</td><td>0</td><td>CAGTCTCAGCGTCAGGAAGC</td><td>CGG</td><td>||||||||[||||||||||||]PAM</td><td>NA</td><td>Intergenic</td><td>NA</td><td>NA</td></tr>
<tr><td>T30198</td><td>CAGCGTCAGGAAGCCGGCGATGG</td><td>1000</td><td></td><td></td><td></td><td></td><td></td><td></td><td></td><td></td><td></td></tr>
<tr><td>Oligo fwd</td><td>CACCCAGCGTCAGGAAGCCGGCGA</td><td></td><td></td><td></td><td></td><td></td><td></td><td></td><td></td><td></td><td></td></tr>
<tr><td>Oligo rev</td><td>AAACTCGCCGGCTTCCTGACGCTG</td><td></td><td></td><td></td><td></td><td></td><td></td><td></td><td></td><td></td><td></td></tr>
<tr><td>Chromosome</td><td>start</td><td>end</td><td>strand</td><td>MM</td><td>target_seq</td><td>PAM</td><td>alignment</td><td>distance</td><td>position</td><td>gene name</td><td>gene id</td></tr>
<tr><td>GU305914.1</td><td>51955</td><td>51977</td><td>-</td><td>0</td><td>CAGCGTCAGGAAGCCGGCGA</td><td>TGG</td><td>||||||||[||||||||||||]PAM</td><td>NA</td><td>Intergenic</td><td>NA</td><td>NA</td></tr>
<tr><td>T30199</td><td>CGTCAGGAAGCCGGCGATGGCGG</td><td>1000</td><td></td><td></td><td></td><td></td><td></td><td></td><td></td><td></td><td></td></tr>
<tr><td>Oligo fwd</td><td>CACCCGTCAGGAAGCCGGCGATGG</td><td></td><td></td><td></td><td></td><td></td><td></td><td></td><td></td><td></td><td></td></tr>
</table>

<table>
<tr><td>Oligo rev</td><td>AAACCCATCGCCGGCTTCCTGACG</td><td></td><td></td><td></td><td></td><td></td><td></td><td></td><td></td><td></td><td></td></tr>
<tr><td>Chromosome</td><td>start</td><td>end</td><td>strand</td><td>MM</td><td>target_seq</td><td>PAM</td><td>alignment</td><td>distance</td><td>position</td><td>gene name</td><td>gene id</td></tr>
<tr><td>GU305914.1</td><td>51952</td><td>51974</td><td>-</td><td>0</td><td>CGTCAGGAAGCCGGCGATGG</td><td>CGG</td><td>||||||||[||||||||||||]PAM</td><td>NA</td><td>Intergenic</td><td>NA</td><td>NA</td></tr>
<tr><td></td><td></td><td></td><td></td><td></td><td></td><td></td><td></td><td></td><td></td><td></td><td></td></tr>
<tr><td>T30200</td><td>CAATCGTGCCGAACACCGAGAGG</td><td>1000</td><td></td><td></td><td></td><td></td><td></td><td></td><td></td><td></td><td></td></tr>
<tr><td>Oligo fwd</td><td>CACCCAATCGTGCCGAACACCGAG</td><td></td><td></td><td></td><td></td><td></td><td></td><td></td><td></td><td></td><td></td></tr>
<tr><td>Oligo rev</td><td>AAACCTCGGTGTTCGGCACGATTG</td><td></td><td></td><td></td><td></td><td></td><td></td><td></td><td></td><td></td><td></td></tr>
<tr><td>Chromosome</td><td>start</td><td>end</td><td>strand</td><td>MM</td><td>target_seq</td><td>PAM</td><td>alignment</td><td>distance</td><td>position</td><td>gene name</td><td>gene id</td></tr>
<tr><td>GU305914.1</td><td>51917</td><td>51939</td><td>-</td><td>0</td><td>CAATCGTGCCGAACACCGAG</td><td>AGG</td><td>||||||||[||||||||||||]PAM</td><td>NA</td><td>Intergenic</td><td>NA</td><td>NA</td></tr>
<tr><td></td><td></td><td></td><td></td><td></td><td></td><td></td><td></td><td></td><td></td><td></td><td></td></tr>
<tr><td>T30201</td><td>GAGGATGTTCGCCAACGCGATGG</td><td>1000</td><td></td><td></td><td></td><td></td><td></td><td></td><td></td><td></td><td></td></tr>
<tr><td>Oligo fwd</td><td>CACCGAGGATGTTCGCCAACGCGA</td><td></td><td></td><td></td><td></td><td></td><td></td><td></td><td></td><td></td><td></td></tr>
<tr><td>Oligo rev</td><td>AAACTCGCGTTGGCGAACATCCTC</td><td></td><td></td><td></td><td></td><td></td><td></td><td></td><td></td><td></td><td></td></tr>
<tr><td>Chromosome</td><td>start</td><td>end</td><td>strand</td><td>MM</td><td>target_seq</td><td>PAM</td><td>alignment</td><td>distance</td><td>position</td><td>gene name</td><td>gene id</td></tr>
<tr><td>GU305914.1</td><td>51898</td><td>51920</td><td>-</td><td>0</td><td>GAGGATGTTCGCCAACGCGA</td><td>TGG</td><td>||||||||[||||||||||||]PAM</td><td>NA</td><td>Intergenic</td><td>NA</td><td>NA</td></tr>
<tr><td></td><td></td><td></td><td></td><td></td><td></td><td></td><td></td><td></td><td></td><td></td><td></td></tr>
<tr><td>T30202</td><td>TGTTCGCCAACGCGATGGATCGG</td><td>1000</td><td></td><td></td><td></td><td></td><td></td><td></td><td></td><td></td><td></td></tr>
<tr><td>Oligo fwd</td><td>CACCTGTTCGCCAACGCGATGGAT</td><td></td><td></td><td></td><td></td><td></td><td></td><td></td><td></td><td></td><td></td></tr>
<tr><td>Oligo rev</td><td>AAACATCCATCGCGTTGGCGAACA</td><td></td><td></td><td></td><td></td><td></td><td></td><td></td><td></td><td></td><td></td></tr>
<tr><td>Chromosome</td><td>start</td><td>end</td><td>strand</td><td>MM</td><td>target_seq</td><td>PAM</td><td>alignment</td><td>distance</td><td>position</td><td>gene name</td><td>gene id</td></tr>
<tr><td>GU305914.1</td><td>51893</td><td>51915</td><td>-</td><td>0</td><td>TGTTCGCCAACGCGATGGAT</td><td>CGG</td><td>||||||||[||||||||||||]PAM</td><td>NA</td><td>Intergenic</td><td>NA</td><td>NA</td></tr>
<tr><td></td><td></td><td></td><td></td><td></td><td></td><td></td><td></td><td></td><td></td><td></td><td></td></tr>
<tr><td>T30203</td><td>GTTCGCCAACGCGATGGATCGGG</td><td>1000</td><td></td><td></td><td></td><td></td><td></td><td></td><td></td><td></td><td></td></tr>
<tr><td>Oligo fwd</td><td>CACCGTTCGCCAACGCGATGGATC</td><td></td><td></td><td></td><td></td><td></td><td></td><td></td><td></td><td></td><td></td></tr>
<tr><td>Oligo rev</td><td>AAACGATCCATCGCGTTGGCGAAC</td><td></td><td></td><td></td><td></td><td></td><td></td><td></td><td></td><td></td><td></td></tr>
<tr><td>Chromosome</td><td>start</td><td>end</td><td>strand</td><td>MM</td><td>target_seq</td><td>PAM</td><td>alignment</td><td>distance</td><td>position</td><td>gene name</td><td>gene id</td></tr>
<tr><td>GU305914.1</td><td>51892</td><td>51914</td><td>-</td><td>0</td><td>GTTCGCCAACGCGATGGATC</td><td>GGG</td><td>||||||||[||||||||||||]PAM</td><td>NA</td><td>Intergenic</td><td>NA</td><td>NA</td></tr>
<tr><td></td><td></td><td></td><td></td><td></td><td></td><td></td><td></td><td></td><td></td><td></td><td></td></tr>
<tr><td>T30204</td><td>TGGATCGGGAGTCCGCCATCCGG</td><td>1000</td><td></td><td></td><td></td><td></td><td></td><td></td><td></td><td></td><td></td></tr>
<tr><td>Oligo fwd</td><td>CACCTGGATCGGGAGTCCGCCATC</td><td></td><td></td><td></td><td></td><td></td><td></td><td></td><td></td><td></td><td></td></tr>
<tr><td>Oligo rev</td><td>AAACGATGGCGGACTCCCGATCCA</td><td></td><td></td><td></td><td></td><td></td><td></td><td></td><td></td><td></td><td></td></tr>
<tr><td>Chromosome</td><td>start</td><td>end</td><td>strand</td><td>MM</td><td>target_seq</td><td>PAM</td><td>alignment</td><td>distance</td><td>position</td><td>gene name</td><td>gene id</td></tr>
<tr><td>GU305914.1</td><td>51878</td><td>51900</td><td>-</td><td>0</td><td>TGGATCGGGAGTCCGCCATC</td><td>CGG</td><td>||||||||[||||||||||||]PAM</td><td>NA</td><td>Intergenic</td><td>NA</td><td>NA</td></tr>
<tr><td></td><td></td><td></td><td></td><td></td><td></td><td></td><td></td><td></td><td></td><td></td><td></td></tr>
<tr><td>T30205</td><td>GGATCGGGAGTCCGCCATCCGGG</td><td>1000</td><td></td><td></td><td></td><td></td><td></td><td></td><td></td><td></td><td></td></tr>
<tr><td>Oligo fwd</td><td>CACCGGATCGGGAGTCCGCCATCC</td><td></td><td></td><td></td><td></td><td></td><td></td><td></td><td></td><td></td><td></td></tr>
<tr><td>Oligo rev</td><td>AAACGGATGGCGGACTCCCGATCC</td><td></td><td></td><td></td><td></td><td></td><td></td><td></td><td></td><td></td><td></td></tr>
<tr><td>Chromosome</td><td>start</td><td>end</td><td>strand</td><td>MM</td><td>target_seq</td><td>PAM</td><td>alignment</td><td>distance</td><td>position</td><td>gene name</td><td>gene id</td></tr>
<tr><td>GU305914.1</td><td>51877</td><td>51899</td><td>-</td><td>0</td><td>GGATCGGGAGTCCGCCATCC</td><td>GGG</td><td>||||||||[||||||||||||]PAM</td><td>NA</td><td>Intergenic</td><td>NA</td><td>NA</td></tr>
<tr><td></td><td></td><td></td><td></td><td></td><td></td><td></td><td></td><td></td><td></td><td></td><td></td></tr>
<tr><td>T30206</td><td>TCGGGAGTCCGCCATCCGGGCGG</td><td>1000</td><td></td><td></td><td></td><td></td><td></td><td></td><td></td><td></td><td></td></tr>
<tr><td>Oligo fwd</td><td>CACCTCGGGAGTCCGCCATCCGGG</td><td></td><td></td><td></td><td></td><td></td><td></td><td></td><td></td><td></td><td></td></tr>
<tr><td>Oligo rev</td><td>AAACCCCGGATGGCGGACTCCCGA</td><td></td><td></td><td></td><td></td><td></td><td></td><td></td><td></td><td></td><td></td></tr>
<tr><td>Chromosome</td><td>start</td><td>end</td><td>strand</td><td>MM</td><td>target_seq</td><td>PAM</td><td>alignment</td><td>distance</td><td>position</td><td>gene name</td><td>gene id</td></tr>
<tr><td>GU305914.1</td><td>51874</td><td>51896</td><td>-</td><td>0</td><td>TCGGGAGTCCGCCATCCGGG</td><td>CGG</td><td>||||||||[||||||||||||]PAM</td><td>NA</td><td>Intergenic</td><td>NA</td><td>NA</td></tr>
<tr><td></td><td></td><td></td><td></td><td></td><td></td><td></td><td></td><td></td><td></td><td></td><td></td></tr>
<tr><td>T30207</td><td>CGCCATCCGGGCGGCGAAACTGG</td><td>1000</td><td></td><td></td><td></td><td></td><td></td><td></td><td></td><td></td><td></td></tr>
<tr><td>Oligo fwd</td><td>CACCCGCCATCCGGGCGGCGAAAC</td><td></td><td></td><td></td><td></td><td></td><td></td><td></td><td></td><td></td><td></td></tr>
<tr><td>Oligo rev</td><td>AAACGTTTCGCCGCCCGGATGGCG</td><td></td><td></td><td></td><td></td><td></td><td></td><td></td><td></td><td></td><td></td></tr>
<tr><td>Chromosome</td><td>start</td><td>end</td><td>strand</td><td>MM</td><td>target_seq</td><td>PAM</td><td>alignment</td><td>distance</td><td>position</td><td>gene name</td><td>gene id</td></tr>
<tr><td>GU305914.1</td><td>51865</td><td>51887</td><td>-</td><td>0</td><td>CGCCATCCGGGCGGCGAAAC</td><td>TGG</td><td>||||||||[||||||||||||]PAM</td><td>NA</td><td>Intergenic</td><td>NA</td><td>NA</td></tr>
<tr><td></td><td></td><td></td><td></td><td></td><td></td><td></td><td></td><td></td><td></td><td></td><td></td></tr>
<tr><td>T30208</td><td>CCGGGCGGCGAAACTGGTACAGG</td><td>1000</td><td></td><td></td><td></td><td></td><td></td><td></td><td></td><td></td><td></td></tr>
<tr><td>Oligo fwd</td><td>CACCCCGGGCGGCGAAACTGGTAC</td><td></td><td></td><td></td><td></td><td></td><td></td><td></td><td></td><td></td><td></td></tr>
<tr><td>Oligo rev</td><td>AAACGTACCAGTTTCGCCGCCCGG</td><td></td><td></td><td></td><td></td><td></td><td></td><td></td><td></td><td></td><td></td></tr>
<tr><td>Chromosome</td><td>start</td><td>end</td><td>strand</td><td>MM</td><td>target_seq</td><td>PAM</td><td>alignment</td><td>distance</td><td>position</td><td>gene name</td><td>gene id</td></tr>
<tr><td>GU305914.1</td><td>51859</td><td>51881</td><td>-</td><td>0</td><td>CCGGGCGGCGAAACTGGTAC</td><td>AGG</td><td>||||||||[||||||||||||]PAM</td><td>NA</td><td>Intergenic</td><td>NA</td><td>NA</td></tr>
<tr><td></td><td></td><td></td><td></td><td></td><td></td><td></td><td></td><td></td><td></td><td></td><td></td></tr>
<tr><td>T30209</td><td>GGCGAAACTGGTACAGGCTTTGG</td><td>1000</td><td></td><td></td><td></td><td></td><td></td><td></td><td></td><td></td><td></td></tr>
<tr><td>Oligo fwd</td><td>CACCGGCGAAACTGGTACAGGCTT</td><td></td><td></td><td></td><td></td><td></td><td></td><td></td><td></td><td></td><td></td></tr>
<tr><td>Oligo rev</td><td>AAACAAGCCTGTACCAGTTTCGCC</td><td></td><td></td><td></td><td></td><td></td><td></td><td></td><td></td><td></td><td></td></tr>
<tr><td>Chromosome</td><td>start</td><td>end</td><td>strand</td><td>MM</td><td>target_seq</td><td>PAM</td><td>alignment</td><td>distance</td><td>position</td><td>gene name</td><td>gene id</td></tr>
<tr><td>GU305914.1</td><td>51853</td><td>51875</td><td>-</td><td>0</td><td>GGCGAAACTGGTACAGGCTT</td><td>TGG</td><td>||||||||[||||||||||||]PAM</td><td>NA</td><td>Intergenic</td><td>NA</td><td>NA</td></tr>
<tr><td></td><td></td><td></td><td></td><td></td><td></td><td></td><td></td><td></td><td></td><td></td><td></td></tr>
<tr><td>T30210</td><td>CGAGCACCTATACGTTTGTGAGG</td><td>1000</td><td></td><td></td><td></td><td></td><td></td><td></td><td></td><td></td><td></td></tr>
<tr><td>Oligo fwd</td><td>CACCCGAGCACCTATACGTTTGTG</td><td></td><td></td><td></td><td></td><td></td><td></td><td></td><td></td><td></td><td></td></tr>
<tr><td>Oligo rev</td><td>AAACCACAAACGTATAGGTGCTCG</td><td></td><td></td><td></td><td></td><td></td><td></td><td></td><td></td><td></td><td></td></tr>
<tr><td>Chromosome</td><td>start</td><td>end</td><td>strand</td><td>MM</td><td>target_seq</td><td>PAM</td><td>alignment</td><td>distance</td><td>position</td><td>gene name</td><td>gene id</td></tr>
<tr><td>GU305914.1</td><td>51827</td><td>51849</td><td>-</td><td>0</td><td>CGAGCACCTATACGTTTGTG</td><td>AGG</td><td>||||||||[||||||||||||]PAM</td><td>NA</td><td>Intergenic</td><td>NA</td><td>NA</td></tr>
<tr><td></td><td></td><td></td><td></td><td></td><td></td><td></td><td></td><td></td><td></td><td></td><td></td></tr>
<tr><td>T30211</td><td>ATACGTTTGTGAGGACTTTGAGG</td><td>1000</td><td></td><td></td><td></td><td></td><td></td><td></td><td></td><td></td><td></td></tr>
<tr><td>Oligo fwd</td><td>CACCATACGTTTGTGAGGACTTTG</td><td></td><td></td><td></td><td></td><td></td><td></td><td></td><td></td><td></td><td></td></tr>
<tr><td>Oligo rev</td><td>AAACCAAAGTCCTCACAAACGTAT</td><td></td><td></td><td></td><td></td><td></td><td></td><td></td><td></td><td></td><td></td></tr>
<tr><td>Chromosome</td><td>start</td><td>end</td><td>strand</td><td>MM</td><td>target_seq</td><td>PAM</td><td>alignment</td><td>distance</td><td>position</td><td>gene name</td><td>gene id</td></tr>
<tr><td>GU305914.1</td><td>51818</td><td>51840</td><td>-</td><td>0</td><td>ATACGTTTGTGAGGACTTTG</td><td>AGG</td><td>||||||||[||||||||||||]PAM</td><td>NA</td><td>Intergenic</td><td>NA</td><td>NA</td></tr>
<tr><td></td><td></td><td></td><td></td><td></td><td></td><td></td><td></td><td></td><td></td><td></td><td></td></tr>
<tr><td>T30212</td><td>TACGTTTGTGAGGACTTTGAGGG</td><td>1000</td><td></td><td></td><td></td><td></td><td></td><td></td><td></td><td></td><td></td></tr>
<tr><td>Oligo fwd</td><td>CACCTACGTTTGTGAGGACTTTGA</td><td></td><td></td><td></td><td></td><td></td><td></td><td></td><td></td><td></td><td></td></tr>
<tr><td>Oligo rev</td><td>AAACTCAAAGTCCTCACAAACGTA</td><td></td><td></td><td></td><td></td><td></td><td></td><td></td><td></td><td></td><td></td></tr>
<tr><td>Chromosome</td><td>start</td><td>end</td><td>strand</td><td>MM</td><td>target_seq</td><td>PAM</td><td>alignment</td><td>distance</td><td>position</td><td>gene name</td><td>gene id</td></tr>
<tr><td>GU305914.1</td><td>51817</td><td>51839</td><td>-</td><td>0</td><td>TACGTTTGTGAGGACTTTGA</td><td>GGG</td><td>||||||||[||||||||||||]PAM</td><td>NA</td><td>Intergenic</td><td>NA</td><td>NA</td></tr>
<tr><td></td><td></td><td></td><td></td><td></td><td></td><td></td><td></td><td></td><td></td><td></td><td></td></tr>
<tr><td>T30213</td><td>GGAGTGTACGATTCAGAAGATGG</td><td>1000</td><td></td><td></td><td></td><td></td><td></td><td></td><td></td><td></td><td></td></tr>
<tr><td>Oligo fwd</td><td>CACCGGAGTGTACGATTCAGAAGA</td><td></td><td></td><td></td><td></td><td></td><td></td><td></td><td></td><td></td><td></td></tr>
<tr><td>Oligo rev</td><td>AAACTCTTCTGAATCGTACACTCC</td><td></td><td></td><td></td><td></td><td></td><td></td><td></td><td></td><td></td><td></td></tr>
<tr><td>Chromosome</td><td>start</td><td>end</td><td>strand</td><td>MM</td><td>target_seq</td><td>PAM</td><td>alignment</td><td>distance</td><td>position</td><td>gene name</td><td>gene id</td></tr>
<tr><td>GU305914.1</td><td>51796</td><td>51818</td><td>-</td><td>0</td><td>GGAGTGTACGATTCAGAAGA</td><td>TGG</td><td>||||||||[||||||||||||]PAM</td><td>NA</td><td>Intergenic</td><td>NA</td><td>NA</td></tr>
<tr><td></td><td></td><td></td><td></td><td></td><td></td><td></td><td></td><td></td><td></td><td></td><td></td></tr>
<tr><td>T30214</td><td>ATGGAGATTACACATTCTAGCGG</td><td>1000</td><td></td><td></td><td></td><td></td><td></td><td></td><td></td><td></td><td></td></tr>
</table>

<table>
<tr><td>Oligo fwd</td><td>CACCATGGAGATTACACATTCTAG</td><td></td><td></td><td></td><td></td><td></td><td></td><td></td><td></td><td></td><td></td></tr>
<tr><td>Oligo rev</td><td>AAACCTAGAATGTGTAATCTCCAT</td><td></td><td></td><td></td><td></td><td></td><td></td><td></td><td></td><td></td><td></td></tr>
<tr><td>Chromosome</td><td>start</td><td>end</td><td>strand</td><td>MM</td><td>target_seq</td><td>PAM</td><td>alignment</td><td>distance</td><td>position</td><td>gene name</td><td>gene id</td></tr>
<tr><td>GU305914.1</td><td>51777</td><td>51799</td><td>-</td><td>0</td><td>ATGGAGATTACACATTCTAG</td><td>CGG</td><td>||||||||[||||||||||||]PAM</td><td>NA</td><td>Intergenic</td><td>NA</td><td>NA</td></tr>
<tr><td>T30215</td><td>CTACGAACACCTGACCATCAAGG</td><td>1000</td><td></td><td></td><td></td><td></td><td></td><td></td><td></td><td></td><td></td></tr>
<tr><td>Oligo fwd</td><td>CACCCTACGAACACCTGACCATCA</td><td></td><td></td><td></td><td></td><td></td><td></td><td></td><td></td><td></td><td></td></tr>
<tr><td>Oligo rev</td><td>AAACTGATGGTCAGGTGTTCGTAG</td><td></td><td></td><td></td><td></td><td></td><td></td><td></td><td></td><td></td><td></td></tr>
<tr><td>Chromosome</td><td>start</td><td>end</td><td>strand</td><td>MM</td><td>target_seq</td><td>PAM</td><td>alignment</td><td>distance</td><td>position</td><td>gene name</td><td>gene id</td></tr>
<tr><td>GU305914.1</td><td>51727</td><td>51749</td><td>-</td><td>0</td><td>CTACGAACACCTGACCATCA</td><td>AGG</td><td>||||||||[||||||||||||]PAM</td><td>NA</td><td>Intergenic</td><td>NA</td><td>NA</td></tr>
<tr><td>T30216</td><td>TCAAGGAACCCGCGCTCTCCTGG</td><td>1000</td><td></td><td></td><td></td><td></td><td></td><td></td><td></td><td></td><td></td></tr>
<tr><td>Oligo fwd</td><td>CACCTCAAGGAACCCGCGCTCTCC</td><td></td><td></td><td></td><td></td><td></td><td></td><td></td><td></td><td></td><td></td></tr>
<tr><td>Oligo rev</td><td>AAACGGAGAGCGCGGGTTCCTTGA</td><td></td><td></td><td></td><td></td><td></td><td></td><td></td><td></td><td></td><td></td></tr>
<tr><td>Chromosome</td><td>start</td><td>end</td><td>strand</td><td>MM</td><td>target_seq</td><td>PAM</td><td>alignment</td><td>distance</td><td>position</td><td>gene name</td><td>gene id</td></tr>
<tr><td>GU305914.1</td><td>51710</td><td>51732</td><td>-</td><td>0</td><td>TCAAGGAACCCGCGCTCTCC</td><td>TGG</td><td>||||||||[||||||||||||]PAM</td><td>NA</td><td>Intergenic</td><td>NA</td><td>NA</td></tr>
<tr><td>T30217</td><td>CGCGACTTCTGCCCCCTGTCCGG</td><td>1000</td><td></td><td></td><td></td><td></td><td></td><td></td><td></td><td></td><td></td></tr>
<tr><td>Oligo fwd</td><td>CACCCGCGACTTCTGCCCCCTGTC</td><td></td><td></td><td></td><td></td><td></td><td></td><td></td><td></td><td></td><td></td></tr>
<tr><td>Oligo rev</td><td>AAACGACAGGGGGCAGAAGTCGCG</td><td></td><td></td><td></td><td></td><td></td><td></td><td></td><td></td><td></td><td></td></tr>
<tr><td>Chromosome</td><td>start</td><td>end</td><td>strand</td><td>MM</td><td>target_seq</td><td>PAM</td><td>alignment</td><td>distance</td><td>position</td><td>gene name</td><td>gene id</td></tr>
<tr><td>GU305914.1</td><td>51681</td><td>51703</td><td>-</td><td>0</td><td>CGCGACTTCTGCCCCCTGTC</td><td>CGG</td><td>||||||||[||||||||||||]PAM</td><td>NA</td><td>Intergenic</td><td>NA</td><td>NA</td></tr>
<tr><td>T30218</td><td>GGACCGCTCATGATCATCTTCGG</td><td>1000</td><td></td><td></td><td></td><td></td><td></td><td></td><td></td><td></td><td></td></tr>
<tr><td>Oligo fwd</td><td>CACCGGACCGCTCATGATCATCTT</td><td></td><td></td><td></td><td></td><td></td><td></td><td></td><td></td><td></td><td></td></tr>
<tr><td>Oligo rev</td><td>AAACAAGATGATCATGAGCGGTCC</td><td></td><td></td><td></td><td></td><td></td><td></td><td></td><td></td><td></td><td></td></tr>
<tr><td>Chromosome</td><td>start</td><td>end</td><td>strand</td><td>MM</td><td>target_seq</td><td>PAM</td><td>alignment</td><td>distance</td><td>position</td><td>gene name</td><td>gene id</td></tr>
<tr><td>GU305914.1</td><td>51660</td><td>51682</td><td>-</td><td>0</td><td>GGACCGCTCATGATCATCTT</td><td>CGG</td><td>||||||||[||||||||||||]PAM</td><td>NA</td><td>Intergenic</td><td>NA</td><td>NA</td></tr>
<tr><td>T30219</td><td>GACCGCTCATGATCATCTTCGGG</td><td>1000</td><td></td><td></td><td></td><td></td><td></td><td></td><td></td><td></td><td></td></tr>
<tr><td>Oligo fwd</td><td>CACCGACCGCTCATGATCATCTTC</td><td></td><td></td><td></td><td></td><td></td><td></td><td></td><td></td><td></td><td></td></tr>
<tr><td>Oligo rev</td><td>AAACGAAGATGATCATGAGCGGTC</td><td></td><td></td><td></td><td></td><td></td><td></td><td></td><td></td><td></td><td></td></tr>
<tr><td>Chromosome</td><td>start</td><td>end</td><td>strand</td><td>MM</td><td>target_seq</td><td>PAM</td><td>alignment</td><td>distance</td><td>position</td><td>gene name</td><td>gene id</td></tr>
<tr><td>GU305914.1</td><td>51659</td><td>51681</td><td>-</td><td>0</td><td>GACCGCTCATGATCATCTTC</td><td>GGG</td><td>||||||||[||||||||||||]PAM</td><td>NA</td><td>Intergenic</td><td>NA</td><td>NA</td></tr>
<tr><td>T30220</td><td>ACCGCTCATGATCATCTTCGGGG</td><td>1000</td><td></td><td></td><td></td><td></td><td></td><td></td><td></td><td></td><td></td></tr>
<tr><td>Oligo fwd</td><td>CACCACCGCTCATGATCATCTTCG</td><td></td><td></td><td></td><td></td><td></td><td></td><td></td><td></td><td></td><td></td></tr>
<tr><td>Oligo rev</td><td>AAACCGAAGATGATCATGAGCGGT</td><td></td><td></td><td></td><td></td><td></td><td></td><td></td><td></td><td></td><td></td></tr>
<tr><td>Chromosome</td><td>start</td><td>end</td><td>strand</td><td>MM</td><td>target_seq</td><td>PAM</td><td>alignment</td><td>distance</td><td>position</td><td>gene name</td><td>gene id</td></tr>
<tr><td>GU305914.1</td><td>51658</td><td>51680</td><td>-</td><td>0</td><td>ACCGCTCATGATCATCTTCG</td><td>GGG</td><td>||||||||[||||||||||||]PAM</td><td>NA</td><td>Intergenic</td><td>NA</td><td>NA</td></tr>
<tr><td>T30221</td><td>TCTTCGGGGTCGCCGAAGAGTGG</td><td>1000</td><td></td><td></td><td></td><td></td><td></td><td></td><td></td><td></td><td></td></tr>
<tr><td>Oligo fwd</td><td>CACCTCTTCGGGGTCGCCGAAGAG</td><td></td><td></td><td></td><td></td><td></td><td></td><td></td><td></td><td></td><td></td></tr>
<tr><td>Oligo rev</td><td>AAACCTCTTCGGCGACCCCGAAGA</td><td></td><td></td><td></td><td></td><td></td><td></td><td></td><td></td><td></td><td></td></tr>
<tr><td>Chromosome</td><td>start</td><td>end</td><td>strand</td><td>MM</td><td>target_seq</td><td>PAM</td><td>alignment</td><td>distance</td><td>position</td><td>gene name</td><td>gene id</td></tr>
<tr><td>GU305914.1</td><td>51644</td><td>51666</td><td>-</td><td>0</td><td>TCTTCGGGGTCGCCGAAGAG</td><td>TGG</td><td>||||||||[||||||||||||]PAM</td><td>NA</td><td>Intergenic</td><td>NA</td><td>NA</td></tr>
<tr><td>T30222</td><td>TCGGGGTCGCCGAAGAGTGGCGG</td><td>1000</td><td></td><td></td><td></td><td></td><td></td><td></td><td></td><td></td><td></td></tr>
<tr><td>Oligo fwd</td><td>CACCTCGGGGTCGCCGAAGAGTGG</td><td></td><td></td><td></td><td></td><td></td><td></td><td></td><td></td><td></td><td></td></tr>
<tr><td>Oligo rev</td><td>AAACCCACTCTTCGGCGACCCCGA</td><td></td><td></td><td></td><td></td><td></td><td></td><td></td><td></td><td></td><td></td></tr>
<tr><td>Chromosome</td><td>start</td><td>end</td><td>strand</td><td>MM</td><td>target_seq</td><td>PAM</td><td>alignment</td><td>distance</td><td>position</td><td>gene name</td><td>gene id</td></tr>
<tr><td>GU305914.1</td><td>51641</td><td>51663</td><td>-</td><td>0</td><td>TCGGGGTCGCCGAAGAGTGG</td><td>CGG</td><td>||||||||[||||||||||||]PAM</td><td>NA</td><td>Intergenic</td><td>NA</td><td>NA</td></tr>
<tr><td>T30223</td><td>CTCCAATCGACCCTCGAGACAGG</td><td>1000</td><td></td><td></td><td></td><td></td><td></td><td></td><td></td><td></td><td></td></tr>
<tr><td>Oligo fwd</td><td>CACCCTCCAATCGACCCTCGAGAC</td><td></td><td></td><td></td><td></td><td></td><td></td><td></td><td></td><td></td><td></td></tr>
<tr><td>Oligo rev</td><td>AAACGTCTCGAGGGTCGATTGGAG</td><td></td><td></td><td></td><td></td><td></td><td></td><td></td><td></td><td></td><td></td></tr>
<tr><td>Chromosome</td><td>start</td><td>end</td><td>strand</td><td>MM</td><td>target_seq</td><td>PAM</td><td>alignment</td><td>distance</td><td>position</td><td>gene name</td><td>gene id</td></tr>
<tr><td>GU305914.1</td><td>51613</td><td>51635</td><td>-</td><td>0</td><td>CTCCAATCGACCCTCGAGAC</td><td>AGG</td><td>||||||||[||||||||||||]PAM</td><td>NA</td><td>Intergenic</td><td>NA</td><td>NA</td></tr>
<tr><td>T30224</td><td>AGACAGGTCATCTTCATCATCGG</td><td>1000</td><td></td><td></td><td></td><td></td><td></td><td></td><td></td><td></td><td></td></tr>
<tr><td>Oligo fwd</td><td>CACCAGACAGGTCATCTTCATCAT</td><td></td><td></td><td></td><td></td><td></td><td></td><td></td><td></td><td></td><td></td></tr>
<tr><td>Oligo rev</td><td>AAACATGATGAAGATGACCTGTCT</td><td></td><td></td><td></td><td></td><td></td><td></td><td></td><td></td><td></td><td></td></tr>
<tr><td>Chromosome</td><td>start</td><td>end</td><td>strand</td><td>MM</td><td>target_seq</td><td>PAM</td><td>alignment</td><td>distance</td><td>position</td><td>gene name</td><td>gene id</td></tr>
<tr><td>GU305914.1</td><td>51597</td><td>51619</td><td>-</td><td>0</td><td>AGACAGGTCATCTTCATCAT</td><td>CGG</td><td>||||||||[||||||||||||]PAM</td><td>NA</td><td>Intergenic</td><td>NA</td><td>NA</td></tr>
<tr><td>T30225</td><td>ATCTTCATCATCGGACGCGACGG</td><td>1000</td><td></td><td></td><td></td><td></td><td></td><td></td><td></td><td></td><td></td></tr>
<tr><td>Oligo fwd</td><td>CACCATCTTCATCATCGGACGCGA</td><td></td><td></td><td></td><td></td><td></td><td></td><td></td><td></td><td></td><td></td></tr>
<tr><td>Oligo rev</td><td>AAACTCGCGTCCGATGATGAAGAT</td><td></td><td></td><td></td><td></td><td></td><td></td><td></td><td></td><td></td><td></td></tr>
<tr><td>Chromosome</td><td>start</td><td>end</td><td>strand</td><td>MM</td><td>target_seq</td><td>PAM</td><td>alignment</td><td>distance</td><td>position</td><td>gene name</td><td>gene id</td></tr>
<tr><td>GU305914.1</td><td>51588</td><td>51610</td><td>-</td><td>0</td><td>ATCTTCATCATCGGACGCGA</td><td>CGG</td><td>||||||||[||||||||||||]PAM</td><td>NA</td><td>Intergenic</td><td>NA</td><td>NA</td></tr>
<tr><td>T30226</td><td>CGGACGCGACGGCGTCGTCATGG</td><td>1000</td><td></td><td></td><td></td><td></td><td></td><td></td><td></td><td></td><td></td></tr>
<tr><td>Oligo fwd</td><td>CACCCGGACGCGACGGCGTCGTCA</td><td></td><td></td><td></td><td></td><td></td><td></td><td></td><td></td><td></td><td></td></tr>
<tr><td>Oligo rev</td><td>AAACTGACGACGCCGTCGCGTCCG</td><td></td><td></td><td></td><td></td><td></td><td></td><td></td><td></td><td></td><td></td></tr>
<tr><td>Chromosome</td><td>start</td><td>end</td><td>strand</td><td>MM</td><td>target_seq</td><td>PAM</td><td>alignment</td><td>distance</td><td>position</td><td>gene name</td><td>gene id</td></tr>
<tr><td>GU305914.1</td><td>51577</td><td>51599</td><td>-</td><td>0</td><td>CGGACGCGACGGCGTCGTCA</td><td>TGG</td><td>||||||||[||||||||||||]PAM</td><td>NA</td><td>Intergenic</td><td>NA</td><td>NA</td></tr>
<tr><td>T30227</td><td>GCGTCGTCATGGCCTACGACAGG</td><td>1000</td><td></td><td></td><td></td><td></td><td></td><td></td><td></td><td></td><td></td></tr>
<tr><td>Oligo fwd</td><td>CACCGCGTCGTCATGGCCTACGAC</td><td></td><td></td><td></td><td></td><td></td><td></td><td></td><td></td><td></td><td></td></tr>
<tr><td>Oligo rev</td><td>AAACGTCGTAGGCCATGACGACGC</td><td></td><td></td><td></td><td></td><td></td><td></td><td></td><td></td><td></td><td></td></tr>
<tr><td>Chromosome</td><td>start</td><td>end</td><td>strand</td><td>MM</td><td>target_seq</td><td>PAM</td><td>alignment</td><td>distance</td><td>position</td><td>gene name</td><td>gene id</td></tr>
<tr><td>GU305914.1</td><td>51566</td><td>51588</td><td>-</td><td>0</td><td>GCGTCGTCATGGCCTACGAC</td><td>AGG</td><td>||||||||[||||||||||||]PAM</td><td>NA</td><td>Intergenic</td><td>NA</td><td>NA</td></tr>
<tr><td>T30228</td><td>CGTCGTCATGGCCTACGACAGGG</td><td>1000</td><td></td><td></td><td></td><td></td><td></td><td></td><td></td><td></td><td></td></tr>
<tr><td>Oligo fwd</td><td>CACCCGTCGTCATGGCCTACGACA</td><td></td><td></td><td></td><td></td><td></td><td></td><td></td><td></td><td></td><td></td></tr>
<tr><td>Oligo rev</td><td>AAACTGTCGTAGGCCATGACGACG</td><td></td><td></td><td></td><td></td><td></td><td></td><td></td><td></td><td></td><td></td></tr>
<tr><td>Chromosome</td><td>start</td><td>end</td><td>strand</td><td>MM</td><td>target_seq</td><td>PAM</td><td>alignment</td><td>distance</td><td>position</td><td>gene name</td><td>gene id</td></tr>
<tr><td>GU305914.1</td><td>51565</td><td>51587</td><td>-</td><td>0</td><td>CGTCGTCATGGCCTACGACA</td><td>GGG</td><td>||||||||[||||||||||||]PAM</td><td>NA</td><td>Intergenic</td><td>NA</td><td>NA</td></tr>
</table>

<table>
<tr><td>T30229</td><td>GTCGTCATGGCCTACGACAGGGG</td><td></td><td>1000</td><td></td><td></td><td></td><td></td><td></td><td></td><td></td><td></td><td></td></tr>
<tr><td>Oligo fwd</td><td>CACCGTCGTCATGGCCTACGACAG</td><td></td><td></td><td></td><td></td><td></td><td></td><td></td><td></td><td></td><td></td><td></td></tr>
<tr><td>Oligo rev</td><td>AAACCTGTCGTAGGCCATGACGAC</td><td></td><td></td><td></td><td></td><td></td><td></td><td></td><td></td><td></td><td></td><td></td></tr>
<tr><td>Chromosome</td><td>start</td><td></td><td>end</td><td>strand</td><td>MM</td><td>target_seq</td><td>PAM</td><td>alignment</td><td>distance</td><td>position</td><td>gene name</td><td>gene id</td></tr>
<tr><td>GU305914.1</td><td></td><td>51564</td><td>51586</td><td>-</td><td>0</td><td>GTCGTCATGGCCTACGACAG</td><td>GGG</td><td>||||||||[||||||||||||]PAM</td><td>NA</td><td>Intergenic</td><td>NA</td><td>NA</td></tr>
<tr><td>T30230</td><td>GTCCCAGCCTGCAAGACTTCTGG</td><td></td><td>1000</td><td></td><td></td><td></td><td></td><td></td><td></td><td></td><td></td><td></td></tr>
<tr><td>Oligo fwd</td><td>CACCGTCCCAGCCTGCAAGACTTC</td><td></td><td></td><td></td><td></td><td></td><td></td><td></td><td></td><td></td><td></td><td></td></tr>
<tr><td>Oligo rev</td><td>AAACGAAGTCTTGCAGGCTGGGAC</td><td></td><td></td><td></td><td></td><td></td><td></td><td></td><td></td><td></td><td></td><td></td></tr>
<tr><td>Chromosome</td><td>start</td><td></td><td>end</td><td>strand</td><td>MM</td><td>target_seq</td><td>PAM</td><td>alignment</td><td>distance</td><td>position</td><td>gene name</td><td>gene id</td></tr>
<tr><td>GU305914.1</td><td></td><td>51524</td><td>51546</td><td>-</td><td>0</td><td>GTCCCAGCCTGCAAGACTTC</td><td>TGG</td><td>||||||||[||||||||||||]PAM</td><td>NA</td><td>Intergenic</td><td>NA</td><td>NA</td></tr>
<tr><td>T30231</td><td>CCATCTTTCCTTCGCCCCTGCGG</td><td></td><td>1000</td><td></td><td></td><td></td><td></td><td></td><td></td><td></td><td></td><td></td></tr>
<tr><td>Oligo fwd</td><td>CACCCCATCTTTCCTTCGCCCCTG</td><td></td><td></td><td></td><td></td><td></td><td></td><td></td><td></td><td></td><td></td><td></td></tr>
<tr><td>Oligo rev</td><td>AAACCAGGGGCGAAGGAAAGATGG</td><td></td><td></td><td></td><td></td><td></td><td></td><td></td><td></td><td></td><td></td><td></td></tr>
<tr><td>Chromosome</td><td>start</td><td></td><td>end</td><td>strand</td><td>MM</td><td>target_seq</td><td>PAM</td><td>alignment</td><td>distance</td><td>position</td><td>gene name</td><td>gene id</td></tr>
<tr><td>GU305914.1</td><td></td><td>51470</td><td>51492</td><td>-</td><td>0</td><td>CCATCTTTCCTTCGCCCCTG</td><td>CGG</td><td>||||||||[||||||||||||]PAM</td><td>NA</td><td>Intergenic</td><td>NA</td><td>NA</td></tr>
<tr><td>T30232</td><td>TCTTTCCTTCGCCCCTGCGGAGG</td><td></td><td>1000</td><td></td><td></td><td></td><td></td><td></td><td></td><td></td><td></td><td></td></tr>
<tr><td>Oligo fwd</td><td>CACCTCTTTCCTTCGCCCCTGCGG</td><td></td><td></td><td></td><td></td><td></td><td></td><td></td><td></td><td></td><td></td><td></td></tr>
<tr><td>Oligo rev</td><td>AAACCCGCAGGGGCGAAGGAAAGA</td><td></td><td></td><td></td><td></td><td></td><td></td><td></td><td></td><td></td><td></td><td></td></tr>
<tr><td>Chromosome</td><td>start</td><td></td><td>end</td><td>strand</td><td>MM</td><td>target_seq</td><td>PAM</td><td>alignment</td><td>distance</td><td>position</td><td>gene name</td><td>gene id</td></tr>
<tr><td>GU305914.1</td><td></td><td>51467</td><td>51489</td><td>-</td><td>0</td><td>TCTTTCCTTCGCCCCTGCGG</td><td>AGG</td><td>||||||||[||||||||||||]PAM</td><td>NA</td><td>Intergenic</td><td>NA</td><td>NA</td></tr>
<tr><td>T30233</td><td>GCAGCTCAATATCGACCTCAAGG</td><td></td><td>1000</td><td></td><td></td><td></td><td></td><td></td><td></td><td></td><td></td><td></td></tr>
<tr><td>Oligo fwd</td><td>CACCGCAGCTCAATATCGACCTCA</td><td></td><td></td><td></td><td></td><td></td><td></td><td></td><td></td><td></td><td></td><td></td></tr>
<tr><td>Oligo rev</td><td>AAACTGAGGTCGATATTGAGCTGC</td><td></td><td></td><td></td><td></td><td></td><td></td><td></td><td></td><td></td><td></td><td></td></tr>
<tr><td>Chromosome</td><td>start</td><td></td><td>end</td><td>strand</td><td>MM</td><td>target_seq</td><td>PAM</td><td>alignment</td><td>distance</td><td>position</td><td>gene name</td><td>gene id</td></tr>
<tr><td>GU305914.1</td><td></td><td>51436</td><td>51458</td><td>-</td><td>0</td><td>GCAGCTCAATATCGACCTCA</td><td>AGG</td><td>||||||||[||||||||||||]PAM</td><td>NA</td><td>Intergenic</td><td>NA</td><td>NA</td></tr>
<tr><td>T30234</td><td>TATCGACCTCAAGGATCTCATGG</td><td></td><td>1000</td><td></td><td></td><td></td><td></td><td></td><td></td><td></td><td></td><td></td></tr>
<tr><td>Oligo fwd</td><td>CACCTATCGACCTCAAGGATCTCA</td><td></td><td></td><td></td><td></td><td></td><td></td><td></td><td></td><td></td><td></td><td></td></tr>
<tr><td>Oligo rev</td><td>AAACTGAGATCCTTGAGGTCGATA</td><td></td><td></td><td></td><td></td><td></td><td></td><td></td><td></td><td></td><td></td><td></td></tr>
<tr><td>Chromosome</td><td>start</td><td></td><td>end</td><td>strand</td><td>MM</td><td>target_seq</td><td>PAM</td><td>alignment</td><td>distance</td><td>position</td><td>gene name</td><td>gene id</td></tr>
<tr><td>GU305914.1</td><td></td><td>51427</td><td>51449</td><td>-</td><td>0</td><td>TATCGACCTCAAGGATCTCA</td><td>TGG</td><td>||||||||[||||||||||||]PAM</td><td>NA</td><td>Intergenic</td><td>NA</td><td>NA</td></tr>
<tr><td>T30235</td><td>ATCGACCTCAAGGATCTCATGGG</td><td></td><td>1000</td><td></td><td></td><td></td><td></td><td></td><td></td><td></td><td></td><td></td></tr>
<tr><td>Oligo fwd</td><td>CACCATCGACCTCAAGGATCTCAT</td><td></td><td></td><td></td><td></td><td></td><td></td><td></td><td></td><td></td><td></td><td></td></tr>
<tr><td>Oligo rev</td><td>AAACATGAGATCCTTGAGGTCGAT</td><td></td><td></td><td></td><td></td><td></td><td></td><td></td><td></td><td></td><td></td><td></td></tr>
<tr><td>Chromosome</td><td>start</td><td></td><td>end</td><td>strand</td><td>MM</td><td>target_seq</td><td>PAM</td><td>alignment</td><td>distance</td><td>position</td><td>gene name</td><td>gene id</td></tr>
<tr><td>GU305914.1</td><td></td><td>51426</td><td>51448</td><td>-</td><td>0</td><td>ATCGACCTCAAGGATCTCAT</td><td>GGG</td><td>||||||||[||||||||||||]PAM</td><td>NA</td><td>Intergenic</td><td>NA</td><td>NA</td></tr>
<tr><td>T30236</td><td>GCGCGTCATCATCGAATCCAAGG</td><td></td><td>1000</td><td></td><td></td><td></td><td></td><td></td><td></td><td></td><td></td><td></td></tr>
<tr><td>Oligo fwd</td><td>CACCGCGCGTCATCATCGAATCCA</td><td></td><td></td><td></td><td></td><td></td><td></td><td></td><td></td><td></td><td></td><td></td></tr>
<tr><td>Oligo rev</td><td>AAACTGGATTCGATGATGACGCGC</td><td></td><td></td><td></td><td></td><td></td><td></td><td></td><td></td><td></td><td></td><td></td></tr>
<tr><td>Chromosome</td><td>start</td><td></td><td>end</td><td>strand</td><td>MM</td><td>target_seq</td><td>PAM</td><td>alignment</td><td>distance</td><td>position</td><td>gene name</td><td>gene id</td></tr>
<tr><td>GU305914.1</td><td></td><td>51379</td><td>51401</td><td>-</td><td>0</td><td>GCGCGTCATCATCGAATCCA</td><td>AGG</td><td>||||||||[||||||||||||]PAM</td><td>NA</td><td>Intergenic</td><td>NA</td><td>NA</td></tr>
<tr><td>T30237</td><td>TCATCGAATCCAAGGAGAAGAGG</td><td></td><td>1000</td><td></td><td></td><td></td><td></td><td></td><td></td><td></td><td></td><td></td></tr>
<tr><td>Oligo fwd</td><td>CACCTCATCGAATCCAAGGAGAAG</td><td></td><td></td><td></td><td></td><td></td><td></td><td></td><td></td><td></td><td></td><td></td></tr>
<tr><td>Oligo rev</td><td>AAACCTTCTCCTTGGATTCGATGA</td><td></td><td></td><td></td><td></td><td></td><td></td><td></td><td></td><td></td><td></td><td></td></tr>
<tr><td>Chromosome</td><td>start</td><td></td><td>end</td><td>strand</td><td>MM</td><td>target_seq</td><td>PAM</td><td>alignment</td><td>distance</td><td>position</td><td>gene name</td><td>gene id</td></tr>
<tr><td>GU305914.1</td><td></td><td>51371</td><td>51393</td><td>-</td><td>0</td><td>TCATCGAATCCAAGGAGAAG</td><td>AGG</td><td>||||||||[||||||||||||]PAM</td><td>NA</td><td>Intergenic</td><td>NA</td><td>NA</td></tr>
<tr><td>T30238</td><td>AAGGAGAAGAGGCAGAAGTCTGG</td><td></td><td>1000</td><td></td><td></td><td></td><td></td><td></td><td></td><td></td><td></td><td></td></tr>
<tr><td>Oligo fwd</td><td>CACCAAGGAGAAGAGGCAGAAGTC</td><td></td><td></td><td></td><td></td><td></td><td></td><td></td><td></td><td></td><td></td><td></td></tr>
<tr><td>Oligo rev</td><td>AAACGACTTCTGCCTCTTCTCCTT</td><td></td><td></td><td></td><td></td><td></td><td></td><td></td><td></td><td></td><td></td><td></td></tr>
<tr><td>Chromosome</td><td>start</td><td></td><td>end</td><td>strand</td><td>MM</td><td>target_seq</td><td>PAM</td><td>alignment</td><td>distance</td><td>position</td><td>gene name</td><td>gene id</td></tr>
<tr><td>GU305914.1</td><td></td><td>51360</td><td>51382</td><td>-</td><td>0</td><td>AAGGAGAAGAGGCAGAAGTC</td><td>TGG</td><td>||||||||[||||||||||||]PAM</td><td>NA</td><td>Intergenic</td><td>NA</td><td>NA</td></tr>
<tr><td>T30239</td><td>AGGAGAAGAGGCAGAAGTCTGGG</td><td></td><td>1000</td><td></td><td></td><td></td><td></td><td></td><td></td><td></td><td></td><td></td></tr>
<tr><td>Oligo fwd</td><td>CACCAGGAGAAGAGGCAGAAGTCT</td><td></td><td></td><td></td><td></td><td></td><td></td><td></td><td></td><td></td><td></td><td></td></tr>
<tr><td>Oligo rev</td><td>AAACAGACTTCTGCCTCTTCTCCT</td><td></td><td></td><td></td><td></td><td></td><td></td><td></td><td></td><td></td><td></td><td></td></tr>
<tr><td>Chromosome</td><td>start</td><td></td><td>end</td><td>strand</td><td>MM</td><td>target_seq</td><td>PAM</td><td>alignment</td><td>distance</td><td>position</td><td>gene name</td><td>gene id</td></tr>
<tr><td>GU305914.1</td><td></td><td>51359</td><td>51381</td><td>-</td><td>0</td><td>AGGAGAAGAGGCAGAAGTCT</td><td>GGG</td><td>||||||||[||||||||||||]PAM</td><td>NA</td><td>Intergenic</td><td>NA</td><td>NA</td></tr>
<tr><td>T30240</td><td>CATGCGCATCCTCATTTCCGCGG</td><td></td><td>1000</td><td></td><td></td><td></td><td></td><td></td><td></td><td></td><td></td><td></td></tr>
<tr><td>Oligo fwd</td><td>CACCCATGCGCATCCTCATTTCCG</td><td></td><td></td><td></td><td></td><td></td><td></td><td></td><td></td><td></td><td></td><td></td></tr>
<tr><td>Oligo rev</td><td>AAACCGGAAATGAGGATGCGCATG</td><td></td><td></td><td></td><td></td><td></td><td></td><td></td><td></td><td></td><td></td><td></td></tr>
<tr><td>Chromosome</td><td>start</td><td></td><td>end</td><td>strand</td><td>MM</td><td>target_seq</td><td>PAM</td><td>alignment</td><td>distance</td><td>position</td><td>gene name</td><td>gene id</td></tr>
<tr><td>GU305914.1</td><td></td><td>51307</td><td>51329</td><td>-</td><td>0</td><td>CATGCGCATCCTCATTTCCG</td><td>CGG</td><td>||||||||[||||||||||||]PAM</td><td>NA</td><td>Intergenic</td><td>NA</td><td>NA</td></tr>
<tr><td>T30241</td><td>CGCGGCCAGCGCCATCCAGCAGG</td><td></td><td>1000</td><td></td><td></td><td></td><td></td><td></td><td></td><td></td><td></td><td></td></tr>
<tr><td>Oligo fwd</td><td>CACCCGCGGCCAGCGCCATCCAGC</td><td></td><td></td><td></td><td></td><td></td><td></td><td></td><td></td><td></td><td></td><td></td></tr>
<tr><td>Oligo rev</td><td>AAACGCTGGATGGCGCTGGCCGCG</td><td></td><td></td><td></td><td></td><td></td><td></td><td></td><td></td><td></td><td></td><td></td></tr>
<tr><td>Chromosome</td><td>start</td><td></td><td>end</td><td>strand</td><td>MM</td><td>target_seq</td><td>PAM</td><td>alignment</td><td>distance</td><td>position</td><td>gene name</td><td>gene id</td></tr>
<tr><td>GU305914.1</td><td></td><td>51289</td><td>51311</td><td>-</td><td>0</td><td>CGCGGCCAGCGCCATCCAGC</td><td>AGG</td><td>||||||||[||||||||||||]PAM</td><td>NA</td><td>Intergenic</td><td>NA</td><td>NA</td></tr>
<tr><td>T30242</td><td>GCGGCCAGCGCCATCCAGCAGGG</td><td></td><td>1000</td><td></td><td></td><td></td><td></td><td></td><td></td><td></td><td></td><td></td></tr>
<tr><td>Oligo fwd</td><td>CACCGCGGCCAGCGCCATCCAGCA</td><td></td><td></td><td></td><td></td><td></td><td></td><td></td><td></td><td></td><td></td><td></td></tr>
<tr><td>Oligo rev</td><td>AAACTGCTGGATGGCGCTGGCCGC</td><td></td><td></td><td></td><td></td><td></td><td></td><td></td><td></td><td></td><td></td><td></td></tr>
<tr><td>Chromosome</td><td>start</td><td></td><td>end</td><td>strand</td><td>MM</td><td>target_seq</td><td>PAM</td><td>alignment</td><td>distance</td><td>position</td><td>gene name</td><td>gene id</td></tr>
<tr><td>GU305914.1</td><td></td><td>51288</td><td>51310</td><td>-</td><td>0</td><td>GCGGCCAGCGCCATCCAGCA</td><td>GGG</td><td>||||||||[||||||||||||]PAM</td><td>NA</td><td>Intergenic</td><td>NA</td><td>NA</td></tr>
<tr><td>T30243</td><td>CGCCACCCTCCACCAAAACGTGG</td><td></td><td>1000</td><td></td><td></td><td></td><td></td><td></td><td></td><td></td><td></td><td></td></tr>
<tr><td>Oligo fwd</td><td>CACCCGCCACCCTCCACCAAAACG</td><td></td><td></td><td></td><td></td><td></td><td></td><td></td><td></td><td></td><td></td><td></td></tr>
<tr><td>Oligo rev</td><td>AAACCGTTTTGGTGGAGGGTGGCG</td><td></td><td></td><td></td><td></td><td></td><td></td><td></td><td></td><td></td><td></td><td></td></tr>
<tr><td>Chromosome</td><td>start</td><td></td><td>end</td><td>strand</td><td>MM</td><td>target_seq</td><td>PAM</td><td>alignment</td><td>distance</td><td>position</td><td>gene name</td><td>gene id</td></tr>
<tr><td>GU305914.1</td><td></td><td>51238</td><td>51260</td><td>-</td><td>0</td><td>CGCCACCCTCCACCAAAACG</td><td>TGG</td><td>||||||||[||||||||||||]PAM</td><td>NA</td><td>Intergenic</td><td>NA</td><td>NA</td></tr>
</table>

| | | | | | | | | | | | |
|---|---|---|---|---|---|---|---|---|---|---|---|
| T30244 | CCTCCACCAAAACGTGGACATGG | 1000 | | | | | | | | | |
| Oligo fwd | CACCCCTCCACCAAAACGTGGACA | | | | | | | | | | |
| Oligo rev | AAACTGTCCACGTTTTGGTGGAGG | | | | | | | | | | |
| Chromosome | start | end | strand | MM | target_seq | PAM | alignment | distance | position | gene name | gene id |
| GU305914.1 | 51232 | 51254 | - | 0 | CCTCCACCAAAACGTGGACA | TGG | \|\|\|\|\|\|\|\|[\|\|\|\|\|\|\|\|\|\|\|\|]PAM | NA | Intergenic | NA | NA |
| T30245 | CTCCACCAAAACGTGGACATGGG | 1000 | | | | | | | | | |
| Oligo fwd | CACCCTCCACCAAAACGTGGACAT | | | | | | | | | | |
| Oligo rev | AAACATGTCCACGTTTTGGTGGAG | | | | | | | | | | |
| Chromosome | start | end | strand | MM | target_seq | PAM | alignment | distance | position | gene name | gene id |
| GU305914.1 | 51231 | 51253 | - | 0 | CTCCACCAAAACGTGGACAT | GGG | \|\|\|\|\|\|\|\|[\|\|\|\|\|\|\|\|\|\|\|\|]PAM | NA | Intergenic | NA | NA |
| T30246 | TGGGCTTCCTCGAGATGTACTGG | 1000 | | | | | | | | | |
| Oligo fwd | CACCTGGGCTTCCTCGAGATGTAC | | | | | | | | | | |
| Oligo rev | AAACGTACATCTCGAGGAAGCCCA | | | | | | | | | | |
| Chromosome | start | end | strand | MM | target_seq | PAM | alignment | distance | position | gene name | gene id |
| GU305914.1 | 51212 | 51234 | - | 0 | TGGGCTTCCTCGAGATGTAC | TGG | \|\|\|\|\|\|\|\|[\|\|\|\|\|\|\|\|\|\|\|\|]PAM | NA | Intergenic | NA | NA |
| T30247 | CCGCTACGTGCCGCAGCAATCGG | 1000 | | | | | | | | | |
| Oligo fwd | CACCCCGCTACGTGCCGCAGCAAT | | | | | | | | | | |
| Oligo rev | AAACATTGCTGCGGCACGTAGCGG | | | | | | | | | | |
| Chromosome | start | end | strand | MM | target_seq | PAM | alignment | distance | position | gene name | gene id |
| GU305914.1 | 51181 | 51203 | - | 0 | CCGCTACGTGCCGCAGCAAT | CGG | \|\|\|\|\|\|\|\|[\|\|\|\|\|\|\|\|\|\|\|\|]PAM | NA | Intergenic | NA | NA |
| T30248 | CCGCAGCAATCGGCGCAGAACGG | 1000 | | | | | | | | | |
| Oligo fwd | CACCCCGCAGCAATCGGCGCAGAA | | | | | | | | | | |
| Oligo rev | AAACTTCTGCGCCGATTGCTGCGG | | | | | | | | | | |
| Chromosome | start | end | strand | MM | target_seq | PAM | alignment | distance | position | gene name | gene id |
| GU305914.1 | 51171 | 51193 | - | 0 | CCGCAGCAATCGGCGCAGAA | CGG | \|\|\|\|\|\|\|\|[\|\|\|\|\|\|\|\|\|\|\|\|]PAM | NA | Intergenic | NA | NA |
| T30249 | CGCAGAACGGCGTCGTGCGTAGG | 1000 | | | | | | | | | |
| Oligo fwd | CACCCGCAGAACGGCGTCGTGCGT | | | | | | | | | | |
| Oligo rev | AAACACGCACGACGCCGTTCTGCG | | | | | | | | | | |
| Chromosome | start | end | strand | MM | target_seq | PAM | alignment | distance | position | gene name | gene id |
| GU305914.1 | 51158 | 51180 | - | 0 | CGCAGAACGGCGTCGTGCGT | AGG | \|\|\|\|\|\|\|\|[\|\|\|\|\|\|\|\|\|\|\|\|]PAM | NA | Intergenic | NA | NA |
| T30250 | AACGGCGTCGTGCGTAGGAACGG | 1000 | | | | | | | | | |
| Oligo fwd | CACCAACGGCGTCGTGCGTAGGAA | | | | | | | | | | |
| Oligo rev | AAACTTCCTACGCACGACGCCGTT | | | | | | | | | | |
| Chromosome | start | end | strand | MM | target_seq | PAM | alignment | distance | position | gene name | gene id |
| GU305914.1 | 51153 | 51175 | - | 0 | AACGGCGTCGTGCGTAGGAA | CGG | \|\|\|\|\|\|\|\|[\|\|\|\|\|\|\|\|\|\|\|\|]PAM | NA | Intergenic | NA | NA |
| T30251 | TCATCCTCGAATCTACTCCTCGG | 1000 | | | | | | | | | |
| Oligo fwd | CACCTCATCCTCGAATCTACTCCT | | | | | | | | | | |
| Oligo rev | AAACAGGAGTAGATTCGAGGATGA | | | | | | | | | | |
| Chromosome | start | end | strand | MM | target_seq | PAM | alignment | distance | position | gene name | gene id |
| GU305914.1 | 51125 | 51147 | - | 0 | TCATCCTCGAATCTACTCCT | CGG | \|\|\|\|\|\|\|\|[\|\|\|\|\|\|\|\|\|\|\|\|]PAM | NA | Intergenic | NA | NA |
| T30252 | CATCCTCGAATCTACTCCTCGGG | 1000 | | | | | | | | | |
| Oligo fwd | CACCCATCCTCGAATCTACTCCTC | | | | | | | | | | |
| Oligo rev | AAACGAGGAGTAGATTCGAGGATG | | | | | | | | | | |
| Chromosome | start | end | strand | MM | target_seq | PAM | alignment | distance | position | gene name | gene id |
| GU305914.1 | 51124 | 51146 | - | 0 | CATCCTCGAATCTACTCCTC | GGG | \|\|\|\|\|\|\|\|[\|\|\|\|\|\|\|\|\|\|\|\|]PAM | NA | Intergenic | NA | NA |
| T30253 | ATCCTCGAATCTACTCCTCGGGG | 1000 | | | | | | | | | |
| Oligo fwd | CACCATCCTCGAATCTACTCCTCG | | | | | | | | | | |
| Oligo rev | AAACCGAGGAGTAGATTCGAGGAT | | | | | | | | | | |
| Chromosome | start | end | strand | MM | target_seq | PAM | alignment | distance | position | gene name | gene id |
| GU305914.1 | 51123 | 51145 | - | 0 | ATCCTCGAATCTACTCCTCG | GGG | \|\|\|\|\|\|\|\|[\|\|\|\|\|\|\|\|\|\|\|\|]PAM | NA | Intergenic | NA | NA |
| T30254 | CCCGTGCATCGATCTCGCCGCGG | 1000 | | | | | | | | | |
| Oligo fwd | CACCCCCGTGCATCGATCTCGCCG | | | | | | | | | | |
| Oligo rev | AAACCGGCGAGATCGATGCACGGG | | | | | | | | | | |
| Chromosome | start | end | strand | MM | target_seq | PAM | alignment | distance | position | gene name | gene id |
| GU305914.1 | 51079 | 51101 | - | 0 | CCCGTGCATCGATCTCGCCG | CGG | \|\|\|\|\|\|\|\|[\|\|\|\|\|\|\|\|\|\|\|\|]PAM | NA | Intergenic | NA | NA |
| T30255 | CGGCGTCTCCTCCCATCGCTCGG | 1000 | | | | | | | | | |
| Oligo fwd | CACCCGGCGTCTCCTCCCATCGCT | | | | | | | | | | |
| Oligo rev | AAACAGCGATGGGAGGAGACGCCG | | | | | | | | | | |
| Chromosome | start | end | strand | MM | target_seq | PAM | alignment | distance | position | gene name | gene id |
| GU305914.1 | 51059 | 51081 | - | 0 | CGGCGTCTCCTCCCATCGCT | CGG | \|\|\|\|\|\|\|\|[\|\|\|\|\|\|\|\|\|\|\|\|]PAM | NA | Intergenic | NA | NA |
| T30256 | CGTCTCCTCCCATCGCTCGGAGG | 1000 | | | | | | | | | |
| Oligo fwd | CACCCGTCTCCTCCCATCGCTCGG | | | | | | | | | | |
| Oligo rev | AAACCCGAGCGATGGGAGGAGACG | | | | | | | | | | |
| Chromosome | start | end | strand | MM | target_seq | PAM | alignment | distance | position | gene name | gene id |
| GU305914.1 | 51056 | 51078 | - | 0 | CGTCTCCTCCCATCGCTCGG | AGG | \|\|\|\|\|\|\|\|[\|\|\|\|\|\|\|\|\|\|\|\|]PAM | NA | Intergenic | NA | NA |
| T30257 | GTCTCCTCCCATCGCTCGGAGGG | 1000 | | | | | | | | | |
| Oligo fwd | CACCGTCTCCTCCCATCGCTCGGA | | | | | | | | | | |
| Oligo rev | AAACTCCGAGCGATGGGAGGAGAC | | | | | | | | | | |
| Chromosome | start | end | strand | MM | target_seq | PAM | alignment | distance | position | gene name | gene id |
| GU305914.1 | 51055 | 51077 | - | 0 | GTCTCCTCCCATCGCTCGGA | GGG | \|\|\|\|\|\|\|\|[\|\|\|\|\|\|\|\|\|\|\|\|]PAM | NA | Intergenic | NA | NA |
| T30258 | CATCGCTCGGAGGGCTATCGAGG | 1000 | | | | | | | | | |
| Oligo fwd | CACCCATCGCTCGGAGGGCTATCG | | | | | | | | | | |
| Oligo rev | AAACCGATAGCCCTCCGAGCGATG | | | | | | | | | | |
| Chromosome | start | end | strand | MM | target_seq | PAM | alignment | distance | position | gene name | gene id |

<table>
<tr><td>GU305914.1</td><td>51046</td><td>51068</td><td>-</td><td>0</td><td>CATCGCTCGGAGGGCTATCG</td><td>AGG</td><td>||||||||[||||||||||||]PAM</td><td>NA</td><td>Intergenic</td><td>NA</td><td>NA</td></tr>
<tr><td>GU305914.1</td><td>114751</td><td>114773</td><td>+</td><td>4</td><td>CATCACCCGGACGCCTATCG</td><td>GGG</td><td>||||-|-|[|||-|-||||||]PAM</td><td>NA</td><td>Intergenic</td><td>NA</td><td>NA</td></tr>
<tr><td></td><td></td><td></td><td></td><td></td><td></td><td></td><td></td><td></td><td></td><td></td><td></td></tr>
<tr><td>T30259</td><td>ATCGCTCGGAGGGCTATCGAGGG</td><td>1000</td><td></td><td></td><td></td><td></td><td></td><td></td><td></td><td></td><td></td></tr>
<tr><td>Oligo fwd</td><td>CACCATCGCTCGGAGGGCTATCGA</td><td></td><td></td><td></td><td></td><td></td><td></td><td></td><td></td><td></td><td></td></tr>
<tr><td>Oligo rev</td><td>AAACTCGATAGCCCTCCGAGCGAT</td><td></td><td></td><td></td><td></td><td></td><td></td><td></td><td></td><td></td><td></td></tr>
<tr><td>Chromosome</td><td>start</td><td>end</td><td>strand</td><td>MM</td><td>target_seq</td><td>PAM</td><td>alignment</td><td>distance</td><td>position</td><td>gene name</td><td>gene id</td></tr>
<tr><td>GU305914.1</td><td>51045</td><td>51067</td><td>-</td><td>0</td><td>ATCGCTCGGAGGGCTATCGA</td><td>GGG</td><td>||||||||[||||||||||||]PAM</td><td>NA</td><td>Intergenic</td><td>NA</td><td>NA</td></tr>
<tr><td></td><td></td><td></td><td></td><td></td><td></td><td></td><td></td><td></td><td></td><td></td><td></td></tr>
<tr><td>T30260</td><td>GTGCCCGTGAACCCCGTGCCTGG</td><td>1000</td><td></td><td></td><td></td><td></td><td></td><td></td><td></td><td></td><td></td></tr>
<tr><td>Oligo fwd</td><td>CACCGTGCCCGTGAACCCCGTGCC</td><td></td><td></td><td></td><td></td><td></td><td></td><td></td><td></td><td></td><td></td></tr>
<tr><td>Oligo rev</td><td>AAACGGCACGGGGTTCACGGGCAC</td><td></td><td></td><td></td><td></td><td></td><td></td><td></td><td></td><td></td><td></td></tr>
<tr><td>Chromosome</td><td>start</td><td>end</td><td>strand</td><td>MM</td><td>target_seq</td><td>PAM</td><td>alignment</td><td>distance</td><td>position</td><td>gene name</td><td>gene id</td></tr>
<tr><td>GU305914.1</td><td>50970</td><td>50992</td><td>-</td><td>0</td><td>GTGCCCGTGAACCCCGTGCC</td><td>TGG</td><td>||||||||[||||||||||||]PAM</td><td>NA</td><td>Intergenic</td><td>NA</td><td>NA</td></tr>
<tr><td></td><td></td><td></td><td></td><td></td><td></td><td></td><td></td><td></td><td></td><td></td><td></td></tr>
<tr><td>T30261</td><td>CCCGTGAACCCCGTGCCTGGTGG</td><td>1000</td><td></td><td></td><td></td><td></td><td></td><td></td><td></td><td></td><td></td></tr>
<tr><td>Oligo fwd</td><td>CACCCCCGTGAACCCCGTGCCTGG</td><td></td><td></td><td></td><td></td><td></td><td></td><td></td><td></td><td></td><td></td></tr>
<tr><td>Oligo rev</td><td>AAACCCAGGCACGGGGTTCACGGG</td><td></td><td></td><td></td><td></td><td></td><td></td><td></td><td></td><td></td><td></td></tr>
<tr><td>Chromosome</td><td>start</td><td>end</td><td>strand</td><td>MM</td><td>target_seq</td><td>PAM</td><td>alignment</td><td>distance</td><td>position</td><td>gene name</td><td>gene id</td></tr>
<tr><td>GU305914.1</td><td>50967</td><td>50989</td><td>-</td><td>0</td><td>CCCGTGAACCCCGTGCCTGG</td><td>TGG</td><td>||||||||[||||||||||||]PAM</td><td>NA</td><td>Intergenic</td><td>NA</td><td>NA</td></tr>
<tr><td></td><td></td><td></td><td></td><td></td><td></td><td></td><td></td><td></td><td></td><td></td><td></td></tr>
<tr><td>T30262</td><td>GTGAACCCCGTGCCTGGTGGCGG</td><td>1000</td><td></td><td></td><td></td><td></td><td></td><td></td><td></td><td></td><td></td></tr>
<tr><td>Oligo fwd</td><td>CACCGTGAACCCCGTGCCTGGTGG</td><td></td><td></td><td></td><td></td><td></td><td></td><td></td><td></td><td></td><td></td></tr>
<tr><td>Oligo rev</td><td>AAACCCACCAGGCACGGGGTTCAC</td><td></td><td></td><td></td><td></td><td></td><td></td><td></td><td></td><td></td><td></td></tr>
<tr><td>Chromosome</td><td>start</td><td>end</td><td>strand</td><td>MM</td><td>target_seq</td><td>PAM</td><td>alignment</td><td>distance</td><td>position</td><td>gene name</td><td>gene id</td></tr>
<tr><td>GU305914.1</td><td>50964</td><td>50986</td><td>-</td><td>0</td><td>GTGAACCCCGTGCCTGGTGG</td><td>CGG</td><td>||||||||[||||||||||||]PAM</td><td>NA</td><td>Intergenic</td><td>NA</td><td>NA</td></tr>
<tr><td></td><td></td><td></td><td></td><td></td><td></td><td></td><td></td><td></td><td></td><td></td><td></td></tr>
<tr><td>T30263</td><td>AACCCCGTGCCTGGTGGCGGTGG</td><td>1000</td><td></td><td></td><td></td><td></td><td></td><td></td><td></td><td></td><td></td></tr>
<tr><td>Oligo fwd</td><td>CACCAACCCCGTGCCTGGTGGCGG</td><td></td><td></td><td></td><td></td><td></td><td></td><td></td><td></td><td></td><td></td></tr>
<tr><td>Oligo rev</td><td>AAACCCGCCACCAGGCACGGGGTT</td><td></td><td></td><td></td><td></td><td></td><td></td><td></td><td></td><td></td><td></td></tr>
<tr><td>Chromosome</td><td>start</td><td>end</td><td>strand</td><td>MM</td><td>target_seq</td><td>PAM</td><td>alignment</td><td>distance</td><td>position</td><td>gene name</td><td>gene id</td></tr>
<tr><td>GU305914.1</td><td>50961</td><td>50983</td><td>-</td><td>0</td><td>AACCCCGTGCCTGGTGGCGG</td><td>TGG</td><td>||||||||[||||||||||||]PAM</td><td>NA</td><td>Intergenic</td><td>NA</td><td>NA</td></tr>
<tr><td></td><td></td><td></td><td></td><td></td><td></td><td></td><td></td><td></td><td></td><td></td><td></td></tr>
<tr><td>T30264</td><td>CCCGTCGATCTCTCCGTCAGAGG</td><td>1000</td><td></td><td></td><td></td><td></td><td></td><td></td><td></td><td></td><td></td></tr>
<tr><td>Oligo fwd</td><td>CACCCCCGTCGATCTCTCCGTCAG</td><td></td><td></td><td></td><td></td><td></td><td></td><td></td><td></td><td></td><td></td></tr>
<tr><td>Oligo rev</td><td>AAACCTGACGGAGAGATCGACGGG</td><td></td><td></td><td></td><td></td><td></td><td></td><td></td><td></td><td></td><td></td></tr>
<tr><td>Chromosome</td><td>start</td><td>end</td><td>strand</td><td>MM</td><td>target_seq</td><td>PAM</td><td>alignment</td><td>distance</td><td>position</td><td>gene name</td><td>gene id</td></tr>
<tr><td>GU305914.1</td><td>50919</td><td>50941</td><td>-</td><td>0</td><td>CCCGTCGATCTCTCCGTCAG</td><td>AGG</td><td>||||||||[||||||||||||]PAM</td><td>NA</td><td>Intergenic</td><td>NA</td><td>NA</td></tr>
<tr><td></td><td></td><td></td><td></td><td></td><td></td><td></td><td></td><td></td><td></td><td></td><td></td></tr>
<tr><td>T30265</td><td>TCCGTCAGAGGCCCCGCGCCCGG</td><td>1000</td><td></td><td></td><td></td><td></td><td></td><td></td><td></td><td></td><td></td></tr>
<tr><td>Oligo fwd</td><td>CACCTCCGTCAGAGGCCCCGCGCC</td><td></td><td></td><td></td><td></td><td></td><td></td><td></td><td></td><td></td><td></td></tr>
<tr><td>Oligo rev</td><td>AAACGGCGCGGGGCCTCTGACGGA</td><td></td><td></td><td></td><td></td><td></td><td></td><td></td><td></td><td></td><td></td></tr>
<tr><td>Chromosome</td><td>start</td><td>end</td><td>strand</td><td>MM</td><td>target_seq</td><td>PAM</td><td>alignment</td><td>distance</td><td>position</td><td>gene name</td><td>gene id</td></tr>
<tr><td>GU305914.1</td><td>50907</td><td>50929</td><td>-</td><td>0</td><td>TCCGTCAGAGGCCCCGCGCC</td><td>CGG</td><td>||||||||[||||||||||||]PAM</td><td>NA</td><td>Intergenic</td><td>NA</td><td>NA</td></tr>
<tr><td></td><td></td><td></td><td></td><td></td><td></td><td></td><td></td><td></td><td></td><td></td><td></td></tr>
<tr><td>T30266</td><td>CCCGGCGTTCACGTTCTCTTCGG</td><td>1000</td><td></td><td></td><td></td><td></td><td></td><td></td><td></td><td></td><td></td></tr>
<tr><td>Oligo fwd</td><td>CACCCCCGGCGTTCACGTTCTCTT</td><td></td><td></td><td></td><td></td><td></td><td></td><td></td><td></td><td></td><td></td></tr>
<tr><td>Oligo rev</td><td>AAACAAGAGAACGTGAACGCCGGG</td><td></td><td></td><td></td><td></td><td></td><td></td><td></td><td></td><td></td><td></td></tr>
<tr><td>Chromosome</td><td>start</td><td>end</td><td>strand</td><td>MM</td><td>target_seq</td><td>PAM</td><td>alignment</td><td>distance</td><td>position</td><td>gene name</td><td>gene id</td></tr>
<tr><td>GU305914.1</td><td>50889</td><td>50911</td><td>-</td><td>0</td><td>CCCGGCGTTCACGTTCTCTT</td><td>CGG</td><td>||||||||[||||||||||||]PAM</td><td>NA</td><td>Intergenic</td><td>NA</td><td>NA</td></tr>
<tr><td></td><td></td><td></td><td></td><td></td><td></td><td></td><td></td><td></td><td></td><td></td><td></td></tr>
<tr><td>T30267</td><td>CCGGCGTTCACGTTCTCTTCGGG</td><td>1000</td><td></td><td></td><td></td><td></td><td></td><td></td><td></td><td></td><td></td></tr>
<tr><td>Oligo fwd</td><td>CACCCCGGCGTTCACGTTCTCTTC</td><td></td><td></td><td></td><td></td><td></td><td></td><td></td><td></td><td></td><td></td></tr>
<tr><td>Oligo rev</td><td>AAACGAAGAGAACGTGAACGCCGG</td><td></td><td></td><td></td><td></td><td></td><td></td><td></td><td></td><td></td><td></td></tr>
<tr><td>Chromosome</td><td>start</td><td>end</td><td>strand</td><td>MM</td><td>target_seq</td><td>PAM</td><td>alignment</td><td>distance</td><td>position</td><td>gene name</td><td>gene id</td></tr>
<tr><td>GU305914.1</td><td>50888</td><td>50910</td><td>-</td><td>0</td><td>CCGGCGTTCACGTTCTCTTC</td><td>GGG</td><td>||||||||[||||||||||||]PAM</td><td>NA</td><td>Intergenic</td><td>NA</td><td>NA</td></tr>
<tr><td></td><td></td><td></td><td></td><td></td><td></td><td></td><td></td><td></td><td></td><td></td><td></td></tr>
<tr><td>T30268</td><td>TCACGTTCTCTTCGGGCCCGTGG</td><td>1000</td><td></td><td></td><td></td><td></td><td></td><td></td><td></td><td></td><td></td></tr>
<tr><td>Oligo fwd</td><td>CACCTCACGTTCTCTTCGGGCCCG</td><td></td><td></td><td></td><td></td><td></td><td></td><td></td><td></td><td></td><td></td></tr>
<tr><td>Oligo rev</td><td>AAACCGGGCCCGAAGAGAACGTGA</td><td></td><td></td><td></td><td></td><td></td><td></td><td></td><td></td><td></td><td></td></tr>
<tr><td>Chromosome</td><td>start</td><td>end</td><td>strand</td><td>MM</td><td>target_seq</td><td>PAM</td><td>alignment</td><td>distance</td><td>position</td><td>gene name</td><td>gene id</td></tr>
<tr><td>GU305914.1</td><td>50881</td><td>50903</td><td>-</td><td>0</td><td>TCACGTTCTCTTCGGGCCCG</td><td>TGG</td><td>||||||||[||||||||||||]PAM</td><td>NA</td><td>Intergenic</td><td>NA</td><td>NA</td></tr>
<tr><td>GU305914.1</td><td>91376</td><td>91398</td><td>+</td><td>4</td><td>TCACGGCCTCTTCGGGACAG</td><td>GGG</td><td>|||||--|[||||||||-|-|]PAM</td><td>NA</td><td>Intergenic</td><td>NA</td><td>NA</td></tr>
<tr><td></td><td></td><td></td><td></td><td></td><td></td><td></td><td></td><td></td><td></td><td></td><td></td></tr>
<tr><td>T30269</td><td>TCTCTTCGGGCCCGTGGACGAGG</td><td>1000</td><td></td><td></td><td></td><td></td><td></td><td></td><td></td><td></td><td></td></tr>
<tr><td>Oligo fwd</td><td>CACCTCTCTTCGGGCCCGTGGACG</td><td></td><td></td><td></td><td></td><td></td><td></td><td></td><td></td><td></td><td></td></tr>
<tr><td>Oligo rev</td><td>AAACCGTCCACGGGCCCGAAGAGA</td><td></td><td></td><td></td><td></td><td></td><td></td><td></td><td></td><td></td><td></td></tr>
<tr><td>Chromosome</td><td>start</td><td>end</td><td>strand</td><td>MM</td><td>target_seq</td><td>PAM</td><td>alignment</td><td>distance</td><td>position</td><td>gene name</td><td>gene id</td></tr>
<tr><td>GU305914.1</td><td>50875</td><td>50897</td><td>-</td><td>0</td><td>TCTCTTCGGGCCCGTGGACG</td><td>AGG</td><td>||||||||[||||||||||||]PAM</td><td>NA</td><td>Intergenic</td><td>NA</td><td>NA</td></tr>
<tr><td></td><td></td><td></td><td></td><td></td><td></td><td></td><td></td><td></td><td></td><td></td><td></td></tr>
<tr><td>T30270</td><td>CGATGACAACGCCTTCAACTCGG</td><td>1000</td><td></td><td></td><td></td><td></td><td></td><td></td><td></td><td></td><td></td></tr>
<tr><td>Oligo fwd</td><td>CACCCGATGACAACGCCTTCAACT</td><td></td><td></td><td></td><td></td><td></td><td></td><td></td><td></td><td></td><td></td></tr>
<tr><td>Oligo rev</td><td>AAACAGTTGAAGGCGTTGTCATCG</td><td></td><td></td><td></td><td></td><td></td><td></td><td></td><td></td><td></td><td></td></tr>
<tr><td>Chromosome</td><td>start</td><td>end</td><td>strand</td><td>MM</td><td>target_seq</td><td>PAM</td><td>alignment</td><td>distance</td><td>position</td><td>gene name</td><td>gene id</td></tr>
<tr><td>GU305914.1</td><td>50842</td><td>50864</td><td>-</td><td>0</td><td>CGATGACAACGCCTTCAACT</td><td>CGG</td><td>||||||||[||||||||||||]PAM</td><td>NA</td><td>Intergenic</td><td>NA</td><td>NA</td></tr>
<tr><td></td><td></td><td></td><td></td><td></td><td></td><td></td><td></td><td></td><td></td><td></td><td></td></tr>
<tr><td>T30271</td><td>GTCTCCTCCACGCGACCACACGG</td><td>1000</td><td></td><td></td><td></td><td></td><td></td><td></td><td></td><td></td><td></td></tr>
<tr><td>Oligo fwd</td><td>CACCGTCTCCTCCACGCGACCACA</td><td></td><td></td><td></td><td></td><td></td><td></td><td></td><td></td><td></td><td></td></tr>
<tr><td>Oligo rev</td><td>AAACTGTGGTCGCGTGGAGGAGAC</td><td></td><td></td><td></td><td></td><td></td><td></td><td></td><td></td><td></td><td></td></tr>
<tr><td>Chromosome</td><td>start</td><td>end</td><td>strand</td><td>MM</td><td>target_seq</td><td>PAM</td><td>alignment</td><td>distance</td><td>position</td><td>gene name</td><td>gene id</td></tr>
<tr><td>GU305914.1</td><td>50806</td><td>50828</td><td>-</td><td>0</td><td>GTCTCCTCCACGCGACCACA</td><td>CGG</td><td>||||||||[||||||||||||]PAM</td><td>NA</td><td>Intergenic</td><td>NA</td><td>NA</td></tr>
<tr><td></td><td></td><td></td><td></td><td></td><td></td><td></td><td></td><td></td><td></td><td></td><td></td></tr>
<tr><td>T30272</td><td>GCCGTGCATAGACCTCACCCTGG</td><td>1000</td><td></td><td></td><td></td><td></td><td></td><td></td><td></td><td></td><td></td></tr>
<tr><td>Oligo fwd</td><td>CACCGCCGTGCATAGACCTCACCC</td><td></td><td></td><td></td><td></td><td></td><td></td><td></td><td></td><td></td><td></td></tr>
<tr><td>Oligo rev</td><td>AAACGGGTGAGGTCTATGCACGGC</td><td></td><td></td><td></td><td></td><td></td><td></td><td></td><td></td><td></td><td></td></tr>
<tr><td>Chromosome</td><td>start</td><td>end</td><td>strand</td><td>MM</td><td>target_seq</td><td>PAM</td><td>alignment</td><td>distance</td><td>position</td><td>gene name</td><td>gene id</td></tr>
<tr><td>GU305914.1</td><td>50779</td><td>50801</td><td>-</td><td>0</td><td>GCCGTGCATAGACCTCACCC</td><td>TGG</td><td>||||||||[||||||||||||]PAM</td><td>NA</td><td>Intergenic</td><td>NA</td><td>NA</td></tr>
<tr><td></td><td></td><td></td><td></td><td></td><td></td><td></td><td></td><td></td><td></td><td></td><td></td></tr>
<tr><td>T30273</td><td>AGACCTCACCCTGGACTCTTCGG</td><td>1000</td><td></td><td></td><td></td><td></td><td></td><td></td><td></td><td></td><td></td></tr>
</table>

<table>
<tr><td>Oligo fwd</td><td>CACCAGACCTCACCCTGGACTCTT</td><td></td><td></td><td></td><td></td><td></td><td></td><td></td><td></td><td></td><td></td></tr>
<tr><td>Oligo rev</td><td>AAACAAGAGTCCAGGGTGAGGTCT</td><td></td><td></td><td></td><td></td><td></td><td></td><td></td><td></td><td></td><td></td></tr>
<tr><td>Chromosome</td><td>start</td><td>end</td><td>strand</td><td>MM</td><td>target_seq</td><td>PAM</td><td>alignment</td><td>distance</td><td>position</td><td>gene name</td><td>gene id</td></tr>
<tr><td>GU305914.1</td><td>50770</td><td>50792</td><td>-</td><td>0</td><td>AGACCTCACCCTGGACTCTT</td><td>CGG</td><td>||||||||[||||||||||||]PAM</td><td>NA</td><td>Intergenic</td><td>NA</td><td>NA</td></tr>
<tr><td></td><td></td><td></td><td></td><td></td><td></td><td></td><td></td><td></td><td></td><td></td><td></td></tr>
<tr><td>T30274</td><td>GGACTCTTCGGAAGATCAGCAGG</td><td>1000</td><td></td><td></td><td></td><td></td><td></td><td></td><td></td><td></td><td></td></tr>
<tr><td>Oligo fwd</td><td>CACCGGACTCTTCGGAAGATCAGC</td><td></td><td></td><td></td><td></td><td></td><td></td><td></td><td></td><td></td><td></td></tr>
<tr><td>Oligo rev</td><td>AAACGCTGATCTTCCGAAGAGTCC</td><td></td><td></td><td></td><td></td><td></td><td></td><td></td><td></td><td></td><td></td></tr>
<tr><td>Chromosome</td><td>start</td><td>end</td><td>strand</td><td>MM</td><td>target_seq</td><td>PAM</td><td>alignment</td><td>distance</td><td>position</td><td>gene name</td><td>gene id</td></tr>
<tr><td>GU305914.1</td><td>50758</td><td>50780</td><td>-</td><td>0</td><td>GGACTCTTCGGAAGATCAGC</td><td>AGG</td><td>||||||||[||||||||||||]PAM</td><td>NA</td><td>Intergenic</td><td>NA</td><td>NA</td></tr>
<tr><td></td><td></td><td></td><td></td><td></td><td></td><td></td><td></td><td></td><td></td><td></td><td></td></tr>
<tr><td>T30275</td><td>AGATCAGCAGGAGCAGCCCATGG</td><td>1000</td><td></td><td></td><td></td><td></td><td></td><td></td><td></td><td></td><td></td></tr>
<tr><td>Oligo fwd</td><td>CACCAGATCAGCAGGAGCAGCCCA</td><td></td><td></td><td></td><td></td><td></td><td></td><td></td><td></td><td></td><td></td></tr>
<tr><td>Oligo rev</td><td>AAACTGGGCTGCTCCTGCTGATCT</td><td></td><td></td><td></td><td></td><td></td><td></td><td></td><td></td><td></td><td></td></tr>
<tr><td>Chromosome</td><td>start</td><td>end</td><td>strand</td><td>MM</td><td>target_seq</td><td>PAM</td><td>alignment</td><td>distance</td><td>position</td><td>gene name</td><td>gene id</td></tr>
<tr><td>GU305914.1</td><td>50746</td><td>50768</td><td>-</td><td>0</td><td>AGATCAGCAGGAGCAGCCCA</td><td>TGG</td><td>||||||||[||||||||||||]PAM</td><td>NA</td><td>Intergenic</td><td>NA</td><td>NA</td></tr>
<tr><td></td><td></td><td></td><td></td><td></td><td></td><td></td><td></td><td></td><td></td><td></td><td></td></tr>
<tr><td>T30276</td><td>CCCATGGACCTCACCGTGAACGG</td><td>1000</td><td></td><td></td><td></td><td></td><td></td><td></td><td></td><td></td><td></td></tr>
<tr><td>Oligo fwd</td><td>CACCCCCATGGACCTCACCGTGAA</td><td></td><td></td><td></td><td></td><td></td><td></td><td></td><td></td><td></td><td></td></tr>
<tr><td>Oligo rev</td><td>AAACTTCACGGTGAGGTCCATGGG</td><td></td><td></td><td></td><td></td><td></td><td></td><td></td><td></td><td></td><td></td></tr>
<tr><td>Chromosome</td><td>start</td><td>end</td><td>strand</td><td>MM</td><td>target_seq</td><td>PAM</td><td>alignment</td><td>distance</td><td>position</td><td>gene name</td><td>gene id</td></tr>
<tr><td>GU305914.1</td><td>50730</td><td>50752</td><td>-</td><td>0</td><td>CCCATGGACCTCACCGTGAA</td><td>CGG</td><td>||||||||[||||||||||||]PAM</td><td>NA</td><td>Intergenic</td><td>NA</td><td>NA</td></tr>
<tr><td></td><td></td><td></td><td></td><td></td><td></td><td></td><td></td><td></td><td></td><td></td><td></td></tr>
<tr><td>T30277</td><td>CGTGAACGGCTCTCCTCCTGTGG</td><td>1000</td><td></td><td></td><td></td><td></td><td></td><td></td><td></td><td></td><td></td></tr>
<tr><td>Oligo fwd</td><td>CACCCGTGAACGGCTCTCCTCCTG</td><td></td><td></td><td></td><td></td><td></td><td></td><td></td><td></td><td></td><td></td></tr>
<tr><td>Oligo rev</td><td>AAACCAGGAGGAGAGCCGTTCACG</td><td></td><td></td><td></td><td></td><td></td><td></td><td></td><td></td><td></td><td></td></tr>
<tr><td>Chromosome</td><td>start</td><td>end</td><td>strand</td><td>MM</td><td>target_seq</td><td>PAM</td><td>alignment</td><td>distance</td><td>position</td><td>gene name</td><td>gene id</td></tr>
<tr><td>GU305914.1</td><td>50716</td><td>50738</td><td>-</td><td>0</td><td>CGTGAACGGCTCTCCTCCTG</td><td>TGG</td><td>||||||||[||||||||||||]PAM</td><td>NA</td><td>Intergenic</td><td>NA</td><td>NA</td></tr>
<tr><td></td><td></td><td></td><td></td><td></td><td></td><td></td><td></td><td></td><td></td><td></td><td></td></tr>
<tr><td>T30278</td><td>CCTGTGGCCCTGTCGTACAACGG</td><td>1000</td><td></td><td></td><td></td><td></td><td></td><td></td><td></td><td></td><td></td></tr>
<tr><td>Oligo fwd</td><td>CACCCCTGTGGCCCTGTCGTACAA</td><td></td><td></td><td></td><td></td><td></td><td></td><td></td><td></td><td></td><td></td></tr>
<tr><td>Oligo rev</td><td>AAACTTGTACGACAGGGCCACAGG</td><td></td><td></td><td></td><td></td><td></td><td></td><td></td><td></td><td></td><td></td></tr>
<tr><td>Chromosome</td><td>start</td><td>end</td><td>strand</td><td>MM</td><td>target_seq</td><td>PAM</td><td>alignment</td><td>distance</td><td>position</td><td>gene name</td><td>gene id</td></tr>
<tr><td>GU305914.1</td><td>50700</td><td>50722</td><td>-</td><td>0</td><td>CCTGTGGCCCTGTCGTACAA</td><td>CGG</td><td>||||||||[||||||||||||]PAM</td><td>NA</td><td>Intergenic</td><td>NA</td><td>NA</td></tr>
<tr><td></td><td></td><td></td><td></td><td></td><td></td><td></td><td></td><td></td><td></td><td></td><td></td></tr>
<tr><td>T30279</td><td>GTCGTACAACGGCTGTATCCAGG</td><td>1000</td><td></td><td></td><td></td><td></td><td></td><td></td><td></td><td></td><td></td></tr>
<tr><td>Oligo fwd</td><td>CACCGTCGTACAACGGCTGTATCC</td><td></td><td></td><td></td><td></td><td></td><td></td><td></td><td></td><td></td><td></td></tr>
<tr><td>Oligo rev</td><td>AAACGGATACAGCCGTTGTACGAC</td><td></td><td></td><td></td><td></td><td></td><td></td><td></td><td></td><td></td><td></td></tr>
<tr><td>Chromosome</td><td>start</td><td>end</td><td>strand</td><td>MM</td><td>target_seq</td><td>PAM</td><td>alignment</td><td>distance</td><td>position</td><td>gene name</td><td>gene id</td></tr>
<tr><td>GU305914.1</td><td>50689</td><td>50711</td><td>-</td><td>0</td><td>GTCGTACAACGGCTGTATCC</td><td>AGG</td><td>||||||||[||||||||||||]PAM</td><td>NA</td><td>Intergenic</td><td>NA</td><td>NA</td></tr>
<tr><td></td><td></td><td></td><td></td><td></td><td></td><td></td><td></td><td></td><td></td><td></td><td></td></tr>
<tr><td>T30280</td><td>GTACAACGGCTGTATCCAGGTGG</td><td>1000</td><td></td><td></td><td></td><td></td><td></td><td></td><td></td><td></td><td></td></tr>
<tr><td>Oligo fwd</td><td>CACCGTACAACGGCTGTATCCAGG</td><td></td><td></td><td></td><td></td><td></td><td></td><td></td><td></td><td></td><td></td></tr>
<tr><td>Oligo rev</td><td>AAACCCTGGATACAGCCGTTGTAC</td><td></td><td></td><td></td><td></td><td></td><td></td><td></td><td></td><td></td><td></td></tr>
<tr><td>Chromosome</td><td>start</td><td>end</td><td>strand</td><td>MM</td><td>target_seq</td><td>PAM</td><td>alignment</td><td>distance</td><td>position</td><td>gene name</td><td>gene id</td></tr>
<tr><td>GU305914.1</td><td>50686</td><td>50708</td><td>-</td><td>0</td><td>GTACAACGGCTGTATCCAGG</td><td>TGG</td><td>||||||||[||||||||||||]PAM</td><td>NA</td><td>Intergenic</td><td>NA</td><td>NA</td></tr>
<tr><td></td><td></td><td></td><td></td><td></td><td></td><td></td><td></td><td></td><td></td><td></td><td></td></tr>
<tr><td>T30281</td><td>TCCAGGTGGCCTCCGTCCCGAGG</td><td>1000</td><td></td><td></td><td></td><td></td><td></td><td></td><td></td><td></td><td></td></tr>
<tr><td>Oligo fwd</td><td>CACCTCCAGGTGGCCTCCGTCCCG</td><td></td><td></td><td></td><td></td><td></td><td></td><td></td><td></td><td></td><td></td></tr>
<tr><td>Oligo rev</td><td>AAACCGGGACGGAGGCCACCTGGA</td><td></td><td></td><td></td><td></td><td></td><td></td><td></td><td></td><td></td><td></td></tr>
<tr><td>Chromosome</td><td>start</td><td>end</td><td>strand</td><td>MM</td><td>target_seq</td><td>PAM</td><td>alignment</td><td>distance</td><td>position</td><td>gene name</td><td>gene id</td></tr>
<tr><td>GU305914.1</td><td>50672</td><td>50694</td><td>-</td><td>0</td><td>TCCAGGTGGCCTCCGTCCCG</td><td>AGG</td><td>||||||||[||||||||||||]PAM</td><td>NA</td><td>Intergenic</td><td>NA</td><td>NA</td></tr>
<tr><td></td><td></td><td></td><td></td><td></td><td></td><td></td><td></td><td></td><td></td><td></td><td></td></tr>
<tr><td>T30282</td><td>TGGCCTCCGTCCCGAGGAAGAGG</td><td>1000</td><td></td><td></td><td></td><td></td><td></td><td></td><td></td><td></td><td></td></tr>
<tr><td>Oligo fwd</td><td>CACCTGGCCTCCGTCCCGAGGAAG</td><td></td><td></td><td></td><td></td><td></td><td></td><td></td><td></td><td></td><td></td></tr>
<tr><td>Oligo rev</td><td>AAACCTTCCTCGGGACGGAGGCCA</td><td></td><td></td><td></td><td></td><td></td><td></td><td></td><td></td><td></td><td></td></tr>
<tr><td>Chromosome</td><td>start</td><td>end</td><td>strand</td><td>MM</td><td>target_seq</td><td>PAM</td><td>alignment</td><td>distance</td><td>position</td><td>gene name</td><td>gene id</td></tr>
<tr><td>GU305914.1</td><td>50666</td><td>50688</td><td>-</td><td>0</td><td>TGGCCTCCGTCCCGAGGAAG</td><td>AGG</td><td>||||||||[||||||||||||]PAM</td><td>NA</td><td>Intergenic</td><td>NA</td><td>NA</td></tr>
<tr><td></td><td></td><td></td><td></td><td></td><td></td><td></td><td></td><td></td><td></td><td></td><td></td></tr>
<tr><td>T30283</td><td>CTCCGTCCCGAGGAAGAGGCCGG</td><td>1000</td><td></td><td></td><td></td><td></td><td></td><td></td><td></td><td></td><td></td></tr>
<tr><td>Oligo fwd</td><td>CACCCTCCGTCCCGAGGAAGAGGC</td><td></td><td></td><td></td><td></td><td></td><td></td><td></td><td></td><td></td><td></td></tr>
<tr><td>Oligo rev</td><td>AAACGCCTCTTCCTCGGGACGGAG</td><td></td><td></td><td></td><td></td><td></td><td></td><td></td><td></td><td></td><td></td></tr>
<tr><td>Chromosome</td><td>start</td><td>end</td><td>strand</td><td>MM</td><td>target_seq</td><td>PAM</td><td>alignment</td><td>distance</td><td>position</td><td>gene name</td><td>gene id</td></tr>
<tr><td>GU305914.1</td><td>50662</td><td>50684</td><td>-</td><td>0</td><td>CTCCGTCCCGAGGAAGAGGC</td><td>CGG</td><td>||||||||[||||||||||||]PAM</td><td>NA</td><td>Intergenic</td><td>NA</td><td>NA</td></tr>
<tr><td></td><td></td><td></td><td></td><td></td><td></td><td></td><td></td><td></td><td></td><td></td><td></td></tr>
<tr><td>T30284</td><td>GGCCGGCGCCCGAAGACAGTAGG</td><td>1000</td><td></td><td></td><td></td><td></td><td></td><td></td><td></td><td></td><td></td></tr>
<tr><td>Oligo fwd</td><td>CACCGGCCGGCGCCCGAAGACAGT</td><td></td><td></td><td></td><td></td><td></td><td></td><td></td><td></td><td></td><td></td></tr>
<tr><td>Oligo rev</td><td>AAACACTGTCTTCGGGCGCCGGCC</td><td></td><td></td><td></td><td></td><td></td><td></td><td></td><td></td><td></td><td></td></tr>
<tr><td>Chromosome</td><td>start</td><td>end</td><td>strand</td><td>MM</td><td>target_seq</td><td>PAM</td><td>alignment</td><td>distance</td><td>position</td><td>gene name</td><td>gene id</td></tr>
<tr><td>GU305914.1</td><td>50645</td><td>50667</td><td>-</td><td>0</td><td>GGCCGGCGCCCGAAGACAGT</td><td>AGG</td><td>||||||||[||||||||||||]PAM</td><td>NA</td><td>Intergenic</td><td>NA</td><td>NA</td></tr>
<tr><td></td><td></td><td></td><td></td><td></td><td></td><td></td><td></td><td></td><td></td><td></td><td></td></tr>
<tr><td>T30285</td><td>GCCGGCGCCCGAAGACAGTAGGG</td><td>1000</td><td></td><td></td><td></td><td></td><td></td><td></td><td></td><td></td><td></td></tr>
<tr><td>Oligo fwd</td><td>CACCGCCGGCGCCCGAAGACAGTA</td><td></td><td></td><td></td><td></td><td></td><td></td><td></td><td></td><td></td><td></td></tr>
<tr><td>Oligo rev</td><td>AAACTACTGTCTTCGGGCGCCGGC</td><td></td><td></td><td></td><td></td><td></td><td></td><td></td><td></td><td></td><td></td></tr>
<tr><td>Chromosome</td><td>start</td><td>end</td><td>strand</td><td>MM</td><td>target_seq</td><td>PAM</td><td>alignment</td><td>distance</td><td>position</td><td>gene name</td><td>gene id</td></tr>
<tr><td>GU305914.1</td><td>50644</td><td>50666</td><td>-</td><td>0</td><td>GCCGGCGCCCGAAGACAGTA</td><td>GGG</td><td>||||||||[||||||||||||]PAM</td><td>NA</td><td>Intergenic</td><td>NA</td><td>NA</td></tr>
<tr><td></td><td></td><td></td><td></td><td></td><td></td><td></td><td></td><td></td><td></td><td></td><td></td></tr>
<tr><td>T30286</td><td>CGAAGACAGTAGGGTGCTCGAGG</td><td>1000</td><td></td><td></td><td></td><td></td><td></td><td></td><td></td><td></td><td></td></tr>
<tr><td>Oligo fwd</td><td>CACCCGAAGACAGTAGGGTGCTCG</td><td></td><td></td><td></td><td></td><td></td><td></td><td></td><td></td><td></td><td></td></tr>
<tr><td>Oligo rev</td><td>AAACCGAGCACCCTACTGTCTTCG</td><td></td><td></td><td></td><td></td><td></td><td></td><td></td><td></td><td></td><td></td></tr>
<tr><td>Chromosome</td><td>start</td><td>end</td><td>strand</td><td>MM</td><td>target_seq</td><td>PAM</td><td>alignment</td><td>distance</td><td>position</td><td>gene name</td><td>gene id</td></tr>
<tr><td>GU305914.1</td><td>50635</td><td>50657</td><td>-</td><td>0</td><td>CGAAGACAGTAGGGTGCTCG</td><td>AGG</td><td>||||||||[||||||||||||]PAM</td><td>NA</td><td>Intergenic</td><td>NA</td><td>NA</td></tr>
<tr><td></td><td></td><td></td><td></td><td></td><td></td><td></td><td></td><td></td><td></td><td></td><td></td></tr>
<tr><td>T30287</td><td>GACAGTAGGGTGCTCGAGGATGG</td><td>1000</td><td></td><td></td><td></td><td></td><td></td><td></td><td></td><td></td><td></td></tr>
<tr><td>Oligo fwd</td><td>CACCGACAGTAGGGTGCTCGAGGA</td><td></td><td></td><td></td><td></td><td></td><td></td><td></td><td></td><td></td><td></td></tr>
<tr><td>Oligo rev</td><td>AAACTCCTCGAGCACCCTACTGTC</td><td></td><td></td><td></td><td></td><td></td><td></td><td></td><td></td><td></td><td></td></tr>
<tr><td>Chromosome</td><td>start</td><td>end</td><td>strand</td><td>MM</td><td>target_seq</td><td>PAM</td><td>alignment</td><td>distance</td><td>position</td><td>gene name</td><td>gene id</td></tr>
<tr><td>GU305914.1</td><td>50631</td><td>50653</td><td>-</td><td>0</td><td>GACAGTAGGGTGCTCGAGGA</td><td>TGG</td><td>||||||||[||||||||||||]PAM</td><td>NA</td><td>Intergenic</td><td>NA</td><td>NA</td></tr>
</table>

<table>
<tr><td>T30288</td><td>ACAGTAGGGTGCTCGAGGATGGG</td><td></td><td>1000</td></tr>
<tr><td>Oligo fwd</td><td>CACCACAGTAGGGTGCTCGAGGAT</td></tr>
<tr><td>Oligo rev</td><td>AAACATCCTCGAGCACCCTACTGT</td></tr>
<tr><td>Chromosome</td><td>start</td><td></td><td>end</td><td>strand</td><td>MM</td><td>target_seq</td><td>PAM</td><td>alignment</td><td>distance</td><td>position</td><td>gene name</td><td>gene id</td></tr>
<tr><td>GU305914.1</td><td></td><td>50630</td><td>50652</td><td>-</td><td>0</td><td>ACAGTAGGGTGCTCGAGGAT</td><td>GGG</td><td>||||||||[||||||||||||]PAM</td><td>NA</td><td>Intergenic</td><td>NA</td><td>NA</td></tr>
<tr><td>T30289</td><td>GGTGCTCGAGGATGGGAGCGTGG</td><td></td><td>1000</td></tr>
<tr><td>Oligo fwd</td><td>CACCGGTGCTCGAGGATGGGAGCG</td></tr>
<tr><td>Oligo rev</td><td>AAACCGCTCCCATCCTCGAGCACC</td></tr>
<tr><td>Chromosome</td><td>start</td><td></td><td>end</td><td>strand</td><td>MM</td><td>target_seq</td><td>PAM</td><td>alignment</td><td>distance</td><td>position</td><td>gene name</td><td>gene id</td></tr>
<tr><td>GU305914.1</td><td></td><td>50623</td><td>50645</td><td>-</td><td>0</td><td>GGTGCTCGAGGATGGGAGCG</td><td>TGG</td><td>||||||||[||||||||||||]PAM</td><td>NA</td><td>Intergenic</td><td>NA</td><td>NA</td></tr>
<tr><td>T30290</td><td>GTGCTCGAGGATGGGAGCGTGGG</td><td></td><td>1000</td></tr>
<tr><td>Oligo fwd</td><td>CACCGTGCTCGAGGATGGGAGCGT</td></tr>
<tr><td>Oligo rev</td><td>AAACACGCTCCCATCCTCGAGCAC</td></tr>
<tr><td>Chromosome</td><td>start</td><td></td><td>end</td><td>strand</td><td>MM</td><td>target_seq</td><td>PAM</td><td>alignment</td><td>distance</td><td>position</td><td>gene name</td><td>gene id</td></tr>
<tr><td>GU305914.1</td><td></td><td>50622</td><td>50644</td><td>-</td><td>0</td><td>GTGCTCGAGGATGGGAGCGT</td><td>GGG</td><td>||||||||[||||||||||||]PAM</td><td>NA</td><td>Intergenic</td><td>NA</td><td>NA</td></tr>
<tr><td>T30291</td><td>TGCTCGAGGATGGGAGCGTGGGG</td><td></td><td>1000</td></tr>
<tr><td>Oligo fwd</td><td>CACCTGCTCGAGGATGGGAGCGTG</td></tr>
<tr><td>Oligo rev</td><td>AAACCACGCTCCCATCCTCGAGCA</td></tr>
<tr><td>Chromosome</td><td>start</td><td></td><td>end</td><td>strand</td><td>MM</td><td>target_seq</td><td>PAM</td><td>alignment</td><td>distance</td><td>position</td><td>gene name</td><td>gene id</td></tr>
<tr><td>GU305914.1</td><td></td><td>50621</td><td>50643</td><td>-</td><td>0</td><td>TGCTCGAGGATGGGAGCGTG</td><td>GGG</td><td>||||||||[||||||||||||]PAM</td><td>NA</td><td>Intergenic</td><td>NA</td><td>NA</td></tr>
<tr><td>T30292</td><td>GCTCGAGGATGGGAGCGTGGGGG</td><td></td><td>1000</td></tr>
<tr><td>Oligo fwd</td><td>CACCGCTCGAGGATGGGAGCGTGG</td></tr>
<tr><td>Oligo rev</td><td>AAACCCACGCTCCCATCCTCGAGC</td></tr>
<tr><td>Chromosome</td><td>start</td><td></td><td>end</td><td>strand</td><td>MM</td><td>target_seq</td><td>PAM</td><td>alignment</td><td>distance</td><td>position</td><td>gene name</td><td>gene id</td></tr>
<tr><td>GU305914.1</td><td></td><td>50620</td><td>50642</td><td>-</td><td>0</td><td>GCTCGAGGATGGGAGCGTGG</td><td>GGG</td><td>||||||||[||||||||||||]PAM</td><td>NA</td><td>Intergenic</td><td>NA</td><td>NA</td></tr>
<tr><td>T30293</td><td>GGGGGCCAAGATCACCAAGACGG</td><td></td><td>1000</td></tr>
<tr><td>Oligo fwd</td><td>CACCGGGGGCCAAGATCACCAAGA</td></tr>
<tr><td>Oligo rev</td><td>AAACTCTTGGTGATCTTGGCCCCC</td></tr>
<tr><td>Chromosome</td><td>start</td><td></td><td>end</td><td>strand</td><td>MM</td><td>target_seq</td><td>PAM</td><td>alignment</td><td>distance</td><td>position</td><td>gene name</td><td>gene id</td></tr>
<tr><td>GU305914.1</td><td></td><td>50602</td><td>50624</td><td>-</td><td>0</td><td>GGGGGCCAAGATCACCAAGA</td><td>CGG</td><td>||||||||[||||||||||||]PAM</td><td>NA</td><td>Intergenic</td><td>NA</td><td>NA</td></tr>
<tr><td>T30294</td><td>GGAAGAATGCTTCGAGCAGAAGG</td><td></td><td>1000</td></tr>
<tr><td>Oligo fwd</td><td>CACCGGAAGAATGCTTCGAGCAGA</td></tr>
<tr><td>Oligo rev</td><td>AAACTCTGCTCGAAGCATTCTTCC</td></tr>
<tr><td>Chromosome</td><td>start</td><td></td><td>end</td><td>strand</td><td>MM</td><td>target_seq</td><td>PAM</td><td>alignment</td><td>distance</td><td>position</td><td>gene name</td><td>gene id</td></tr>
<tr><td>GU305914.1</td><td></td><td>50581</td><td>50603</td><td>-</td><td>0</td><td>GGAAGAATGCTTCGAGCAGA</td><td>AGG</td><td>||||||||[||||||||||||]PAM</td><td>NA</td><td>Intergenic</td><td>NA</td><td>NA</td></tr>
<tr><td>T30295</td><td>TGCTTCGAGCAGAAGGAACGAGG</td><td></td><td>1000</td></tr>
<tr><td>Oligo fwd</td><td>CACCTGCTTCGAGCAGAAGGAACG</td></tr>
<tr><td>Oligo rev</td><td>AAACCGTTCCTTCTGCTCGAAGCA</td></tr>
<tr><td>Chromosome</td><td>start</td><td></td><td>end</td><td>strand</td><td>MM</td><td>target_seq</td><td>PAM</td><td>alignment</td><td>distance</td><td>position</td><td>gene name</td><td>gene id</td></tr>
<tr><td>GU305914.1</td><td></td><td>50574</td><td>50596</td><td>-</td><td>0</td><td>TGCTTCGAGCAGAAGGAACG</td><td>AGG</td><td>||||||||[||||||||||||]PAM</td><td>NA</td><td>Intergenic</td><td>NA</td><td>NA</td></tr>
<tr><td>T30296</td><td>GCTTCGAGCAGAAGGAACGAGGG</td><td></td><td>1000</td></tr>
<tr><td>Oligo fwd</td><td>CACCGCTTCGAGCAGAAGGAACGA</td></tr>
<tr><td>Oligo rev</td><td>AAACTCGTTCCTTCTGCTCGAAGC</td></tr>
<tr><td>Chromosome</td><td>start</td><td></td><td>end</td><td>strand</td><td>MM</td><td>target_seq</td><td>PAM</td><td>alignment</td><td>distance</td><td>position</td><td>gene name</td><td>gene id</td></tr>
<tr><td>GU305914.1</td><td></td><td>50573</td><td>50595</td><td>-</td><td>0</td><td>GCTTCGAGCAGAAGGAACGA</td><td>GGG</td><td>||||||||[||||||||||||]PAM</td><td>NA</td><td>Intergenic</td><td>NA</td><td>NA</td></tr>
<tr><td>T30297</td><td>CTTCGAGCAGAAGGAACGAGGGG</td><td></td><td>1000</td></tr>
<tr><td>Oligo fwd</td><td>CACCCTTCGAGCAGAAGGAACGAG</td></tr>
<tr><td>Oligo rev</td><td>AAACCTCGTTCCTTCTGCTCGAAG</td></tr>
<tr><td>Chromosome</td><td>start</td><td></td><td>end</td><td>strand</td><td>MM</td><td>target_seq</td><td>PAM</td><td>alignment</td><td>distance</td><td>position</td><td>gene name</td><td>gene id</td></tr>
<tr><td>GU305914.1</td><td></td><td>50572</td><td>50594</td><td>-</td><td>0</td><td>CTTCGAGCAGAAGGAACGAG</td><td>GGG</td><td>||||||||[||||||||||||]PAM</td><td>NA</td><td>Intergenic</td><td>NA</td><td>NA</td></tr>
<tr><td>T30298</td><td>AACGAGGGGACGCCCAACTGTGG</td><td></td><td>1000</td></tr>
<tr><td>Oligo fwd</td><td>CACCAACGAGGGGACGCCCAACTG</td></tr>
<tr><td>Oligo rev</td><td>AAACCAGTTGGGCGTCCCCTCGTT</td></tr>
<tr><td>Chromosome</td><td>start</td><td></td><td>end</td><td>strand</td><td>MM</td><td>target_seq</td><td>PAM</td><td>alignment</td><td>distance</td><td>position</td><td>gene name</td><td>gene id</td></tr>
<tr><td>GU305914.1</td><td></td><td>50558</td><td>50580</td><td>-</td><td>0</td><td>AACGAGGGGACGCCCAACTG</td><td>TGG</td><td>||||||||[||||||||||||]PAM</td><td>NA</td><td>Intergenic</td><td>NA</td><td>NA</td></tr>
<tr><td>T30299</td><td>ACGAGGGGACGCCCAACTGTGGG</td><td></td><td>1000</td></tr>
<tr><td>Oligo fwd</td><td>CACCACGAGGGGACGCCCAACTGT</td></tr>
<tr><td>Oligo rev</td><td>AAACACAGTTGGGCGTCCCCTCGT</td></tr>
<tr><td>Chromosome</td><td>start</td><td></td><td>end</td><td>strand</td><td>MM</td><td>target_seq</td><td>PAM</td><td>alignment</td><td>distance</td><td>position</td><td>gene name</td><td>gene id</td></tr>
<tr><td>GU305914.1</td><td></td><td>50557</td><td>50579</td><td>-</td><td>0</td><td>ACGAGGGGACGCCCAACTGT</td><td>GGG</td><td>||||||||[||||||||||||]PAM</td><td>NA</td><td>Intergenic</td><td>NA</td><td>NA</td></tr>
<tr><td>T30300</td><td>CGAGGGGACGCCCAACTGTGGGG</td><td></td><td>1000</td></tr>
<tr><td>Oligo fwd</td><td>CACCCGAGGGGACGCCCAACTGTG</td></tr>
<tr><td>Oligo rev</td><td>AAACCACAGTTGGGCGTCCCCTCG</td></tr>
<tr><td>Chromosome</td><td>start</td><td></td><td>end</td><td>strand</td><td>MM</td><td>target_seq</td><td>PAM</td><td>alignment</td><td>distance</td><td>position</td><td>gene name</td><td>gene id</td></tr>
<tr><td>GU305914.1</td><td></td><td>50556</td><td>50578</td><td>-</td><td>0</td><td>CGAGGGGACGCCCAACTGTG</td><td>GGG</td><td>||||||||[||||||||||||]PAM</td><td>NA</td><td>Intergenic</td><td>NA</td><td>NA</td></tr>
<tr><td>T30301</td><td>CGCCCAACTGTGGGGACCTGCGG</td><td></td><td>1000</td></tr>
<tr><td>Oligo fwd</td><td>CACCCGCCCAACTGTGGGGACCTG</td></tr>
<tr><td>Oligo rev</td><td>AAACCAGGTCCCCACAGTTGGGCG</td></tr>
<tr><td>Chromosome</td><td>start</td><td></td><td>end</td><td>strand</td><td>MM</td><td>target_seq</td><td>PAM</td><td>alignment</td><td>distance</td><td>position</td><td>gene name</td><td>gene id</td></tr>
<tr><td>GU305914.1</td><td></td><td>50548</td><td>50570</td><td>-</td><td>0</td><td>CGCCCAACTGTGGGGACCTG</td><td>CGG</td><td>||||||||[||||||||||||]PAM</td><td>NA</td><td>Intergenic</td><td>NA</td><td>NA</td></tr>
<tr><td>T30302</td><td>GATGACGACGAGCATCGTGCTGG</td><td></td><td>1000</td></tr>
<tr><td>Oligo fwd</td><td>CACCGATGACGACGAGCATCGTGC</td></tr>
<tr><td>Oligo rev</td><td>AAACGCACGATGCTCGTCGTCATC</td></tr>
<tr><td>Chromosome</td><td>start</td><td></td><td>end</td><td>strand</td><td>MM</td><td>target_seq</td><td>PAM</td><td>alignment</td><td>distance</td><td>position</td><td>gene name</td><td>gene id</td></tr>
<tr><td>GU305914.1</td><td></td><td>50524</td><td>50546</td><td>-</td><td>0</td><td>GATGACGACGAGCATCGTGC</td><td>TGG</td><td>||||||||[||||||||||||]PAM</td><td>NA</td><td>Intergenic</td><td>NA</td><td>NA</td></tr>
</table>

| | | | | | | | | | | | |
|---|---|---|---|---|---|---|---|---|---|---|---|
| T30303 | ATGACGACGAGCATCGTGCTGGG | 1000 | | | | | | | | | |
| Oligo fwd | CACCATGACGACGAGCATCGTGCT | | | | | | | | | | |
| Oligo rev | AAACAGCACGATGCTCGTCGTCAT | | | | | | | | | | |
| Chromosome | start | end | strand | MM | target_seq | PAM | alignment | distance | position | gene name | gene id |
| GU305914.1 | 50523 | 50545 | - | 0 | ATGACGACGAGCATCGTGCT | GGG | ||||||||[||||||||||||]PAM | NA | Intergenic | NA | NA |

| | | | | | | | | | | | |
|---|---|---|---|---|---|---|---|---|---|---|---|
| T30304 | AGCATCGTGCTGGGCGCTGAAGG | 1000 | | | | | | | | | |
| Oligo fwd | CACCAGCATCGTGCTGGGCGCTGA | | | | | | | | | | |
| Oligo rev | AAACTCAGCGCCCAGCACGATGCT | | | | | | | | | | |
| Chromosome | start | end | strand | MM | target_seq | PAM | alignment | distance | position | gene name | gene id |
| GU305914.1 | 50514 | 50536 | - | 0 | AGCATCGTGCTGGGCGCTGA | AGG | ||||||||[||||||||||||]PAM | NA | Intergenic | NA | NA |

| | | | | | | | | | | | |
|---|---|---|---|---|---|---|---|---|---|---|---|
| T30305 | GCATCGTGCTGGGCGCTGAAGGG | 1000 | | | | | | | | | |
| Oligo fwd | CACCGCATCGTGCTGGGCGCTGAA | | | | | | | | | | |
| Oligo rev | AAACTTCAGCGCCCAGCACGATGC | | | | | | | | | | |
| Chromosome | start | end | strand | MM | target_seq | PAM | alignment | distance | position | gene name | gene id |
| GU305914.1 | 50513 | 50535 | - | 0 | GCATCGTGCTGGGCGCTGAA | GGG | ||||||||[||||||||||||]PAM | NA | Intergenic | NA | NA |

| | | | | | | | | | | | |
|---|---|---|---|---|---|---|---|---|---|---|---|
| T30306 | CATCGTGCTGGGCGCTGAAGGGG | 1000 | | | | | | | | | |
| Oligo fwd | CACCCATCGTGCTGGGCGCTGAAG | | | | | | | | | | |
| Oligo rev | AAACCTTCAGCGCCCAGCACGATG | | | | | | | | | | |
| Chromosome | start | end | strand | MM | target_seq | PAM | alignment | distance | position | gene name | gene id |
| GU305914.1 | 50512 | 50534 | - | 0 | CATCGTGCTGGGCGCTGAAG | GGG | ||||||||[||||||||||||]PAM | NA | Intergenic | NA | NA |

| | | | | | | | | | | | |
|---|---|---|---|---|---|---|---|---|---|---|---|
| T30307 | TGCGACCGCGCCTCGAGCGATGG | 1000 | | | | | | | | | |
| Oligo fwd | CACCTGCGACCGCGCCTCGAGCGA | | | | | | | | | | |
| Oligo rev | AAACTCGCTCGAGGCGCGGTCGCA | | | | | | | | | | |
| Chromosome | start | end | strand | MM | target_seq | PAM | alignment | distance | position | gene name | gene id |
| GU305914.1 | 50478 | 50500 | - | 0 | TGCGACCGCGCCTCGAGCGA | TGG | ||||||||[||||||||||||]PAM | NA | Intergenic | NA | NA |

| | | | | | | | | | | | |
|---|---|---|---|---|---|---|---|---|---|---|---|
| T30308 | TGACACATAAGTCCGAATCTCGG | 1000 | | | | | | | | | |
| Oligo fwd | CACCTGACACATAAGTCCGAATCT | | | | | | | | | | |
| Oligo rev | AAACAGATTCGGACTTATGTGTCA | | | | | | | | | | |
| Chromosome | start | end | strand | MM | target_seq | PAM | alignment | distance | position | gene name | gene id |
| GU305914.1 | 50452 | 50474 | - | 0 | TGACACATAAGTCCGAATCT | CGG | ||||||||[||||||||||||]PAM | NA | Intergenic | NA | NA |

| | | | | | | | | | | | |
|---|---|---|---|---|---|---|---|---|---|---|---|
| T30309 | GACACATAAGTCCGAATCTCGGG | 1000 | | | | | | | | | |
| Oligo fwd | CACCGACACATAAGTCCGAATCTC | | | | | | | | | | |
| Oligo rev | AAACGAGATTCGGACTTATGTGTC | | | | | | | | | | |
| Chromosome | start | end | strand | MM | target_seq | PAM | alignment | distance | position | gene name | gene id |
| GU305914.1 | 50451 | 50473 | - | 0 | GACACATAAGTCCGAATCTC | GGG | ||||||||[||||||||||||]PAM | NA | Intergenic | NA | NA |

| | | | | | | | | | | | |
|---|---|---|---|---|---|---|---|---|---|---|---|
| T30310 | ACATAAGTCCGAATCTCGGGCGG | 1000 | | | | | | | | | |
| Oligo fwd | CACCACATAAGTCCGAATCTCGGG | | | | | | | | | | |
| Oligo rev | AAACCCCGAGATTCGGACTTATGT | | | | | | | | | | |
| Chromosome | start | end | strand | MM | target_seq | PAM | alignment | distance | position | gene name | gene id |
| GU305914.1 | 50448 | 50470 | - | 0 | ACATAAGTCCGAATCTCGGG | CGG | ||||||||[||||||||||||]PAM | NA | Intergenic | NA | NA |

| | | | | | | | | | | | |
|---|---|---|---|---|---|---|---|---|---|---|---|
| T30311 | CTCGGGCGGATTCTCTCGTCAGG | 1000 | | | | | | | | | |
| Oligo fwd | CACCCTCGGGCGGATTCTCTCGTC | | | | | | | | | | |
| Oligo rev | AAACGACGAGAGAATCCGCCCGAG | | | | | | | | | | |
| Chromosome | start | end | strand | MM | target_seq | PAM | alignment | distance | position | gene name | gene id |
| GU305914.1 | 50434 | 50456 | - | 0 | CTCGGGCGGATTCTCTCGTC | AGG | ||||||||[||||||||||||]PAM | NA | Intergenic | NA | NA |

| | | | | | | | | | | | |
|---|---|---|---|---|---|---|---|---|---|---|---|
| T30312 | TTCTCTCGTCAGGATGACAGCGG | 1000 | | | | | | | | | |
| Oligo fwd | CACCTTCTCTCGTCAGGATGACAG | | | | | | | | | | |
| Oligo rev | AAACCTGTCATCCTGACGAGAGAA | | | | | | | | | | |
| Chromosome | start | end | strand | MM | target_seq | PAM | alignment | distance | position | gene name | gene id |
| GU305914.1 | 50424 | 50446 | - | 0 | TTCTCTCGTCAGGATGACAG | CGG | ||||||||[||||||||||||]PAM | NA | Intergenic | NA | NA |

| | | | | | | | | | | | |
|---|---|---|---|---|---|---|---|---|---|---|---|
| T30313 | TCTCTCGTCAGGATGACAGCGGG | 1000 | | | | | | | | | |
| Oligo fwd | CACCTCTCTCGTCAGGATGACAGC | | | | | | | | | | |
| Oligo rev | AAACGCTGTCATCCTGACGAGAGA | | | | | | | | | | |
| Chromosome | start | end | strand | MM | target_seq | PAM | alignment | distance | position | gene name | gene id |
| GU305914.1 | 50423 | 50445 | - | 0 | TCTCTCGTCAGGATGACAGC | GGG | ||||||||[||||||||||||]PAM | NA | Intergenic | NA | NA |

| | | | | | | | | | | | |
|---|---|---|---|---|---|---|---|---|---|---|---|
| T30314 | CTCTCGTCAGGATGACAGCGGGG | 1000 | | | | | | | | | |
| Oligo fwd | CACCCTCTCGTCAGGATGACAGCG | | | | | | | | | | |
| Oligo rev | AAACCGCTGTCATCCTGACGAGAG | | | | | | | | | | |
| Chromosome | start | end | strand | MM | target_seq | PAM | alignment | distance | position | gene name | gene id |
| GU305914.1 | 50422 | 50444 | - | 0 | CTCTCGTCAGGATGACAGCG | GGG | ||||||||[||||||||||||]PAM | NA | Intergenic | NA | NA |

| | | | | | | | | | | | |
|---|---|---|---|---|---|---|---|---|---|---|---|
| T30315 | TCTCGTCAGGATGACAGCGGGGG | 1000 | | | | | | | | | |
| Oligo fwd | CACCTCTCGTCAGGATGACAGCGG | | | | | | | | | | |
| Oligo rev | AAACCCGCTGTCATCCTGACGAGA | | | | | | | | | | |
| Chromosome | start | end | strand | MM | target_seq | PAM | alignment | distance | position | gene name | gene id |
| GU305914.1 | 50421 | 50443 | - | 0 | TCTCGTCAGGATGACAGCGG | GGG | ||||||||[||||||||||||]PAM | NA | Intergenic | NA | NA |

| | | | | | | | | | | | |
|---|---|---|---|---|---|---|---|---|---|---|---|
| T30316 | CTCGTCAGGATGACAGCGGGGGG | 1000 | | | | | | | | | |
| Oligo fwd | CACCCTCGTCAGGATGACAGCGGG | | | | | | | | | | |
| Oligo rev | AAACCCCGCTGTCATCCTGACGAG | | | | | | | | | | |
| Chromosome | start | end | strand | MM | target_seq | PAM | alignment | distance | position | gene name | gene id |
| GU305914.1 | 50420 | 50442 | - | 0 | CTCGTCAGGATGACAGCGGG | GGG | ||||||||[||||||||||||]PAM | NA | Intergenic | NA | NA |

| | | | | | | | | | | | |
|---|---|---|---|---|---|---|---|---|---|---|---|
| T30317 | TCGTCAGGATGACAGCGGGGGGG | 1000 | | | | | | | | | |
| Oligo fwd | CACCTCGTCAGGATGACAGCGGGG | | | | | | | | | | |
| Oligo rev | AAACCCCCGCTGTCATCCTGACGA | | | | | | | | | | |
| Chromosome | start | end | strand | MM | target_seq | PAM | alignment | distance | position | gene name | gene id |

<table>
<tr><td>GU305914.1</td><td>50419</td><td>50441</td><td>-</td><td>0</td><td>TCGTCAGGATGACAGCGGGG</td><td>GGG</td><td>||||||||[||||||||||||]PAM</td><td>NA</td><td>Intergenic</td><td>NA</td><td>NA</td></tr>
<tr><td>T30318</td><td>AGGATGACAGCGGGGGGGCTCGG</td><td>1000</td><td></td><td></td><td></td><td></td><td></td><td></td><td></td><td></td><td></td></tr>
<tr><td>Oligo fwd</td><td>CACCAGGATGACAGCGGGGGGGCT</td><td></td><td></td><td></td><td></td><td></td><td></td><td></td><td></td><td></td><td></td></tr>
<tr><td>Oligo rev</td><td>AAACAGCCCCCCCGCTGTCATCCT</td><td></td><td></td><td></td><td></td><td></td><td></td><td></td><td></td><td></td><td></td></tr>
<tr><td>Chromosome</td><td>start</td><td>end</td><td>strand</td><td>MM</td><td>target_seq</td><td>PAM</td><td>alignment</td><td>distance</td><td>position</td><td>gene name</td><td>gene id</td></tr>
<tr><td>GU305914.1</td><td>50414</td><td>50436</td><td>-</td><td>0</td><td>AGGATGACAGCGGGGGGGCT</td><td>CGG</td><td>||||||||[||||||||||||]PAM</td><td>NA</td><td>Intergenic</td><td>NA</td><td>NA</td></tr>
<tr><td>T30319</td><td>GGATGACAGCGGGGGGGCTCGGG</td><td>1000</td><td></td><td></td><td></td><td></td><td></td><td></td><td></td><td></td><td></td></tr>
<tr><td>Oligo fwd</td><td>CACCGGATGACAGCGGGGGGGCTC</td><td></td><td></td><td></td><td></td><td></td><td></td><td></td><td></td><td></td><td></td></tr>
<tr><td>Oligo rev</td><td>AAACGAGCCCCCCCGCTGTCATCC</td><td></td><td></td><td></td><td></td><td></td><td></td><td></td><td></td><td></td><td></td></tr>
<tr><td>Chromosome</td><td>start</td><td>end</td><td>strand</td><td>MM</td><td>target_seq</td><td>PAM</td><td>alignment</td><td>distance</td><td>position</td><td>gene name</td><td>gene id</td></tr>
<tr><td>GU305914.1</td><td>50413</td><td>50435</td><td>-</td><td>0</td><td>GGATGACAGCGGGGGGGCTC</td><td>GGG</td><td>||||||||[||||||||||||]PAM</td><td>NA</td><td>Intergenic</td><td>NA</td><td>NA</td></tr>
<tr><td>T30320</td><td>AGCGGGGGGGCTCGGGATCGCGG</td><td>1000</td><td></td><td></td><td></td><td></td><td></td><td></td><td></td><td></td><td></td></tr>
<tr><td>Oligo fwd</td><td>CACCAGCGGGGGGGCTCGGGATCG</td><td></td><td></td><td></td><td></td><td></td><td></td><td></td><td></td><td></td><td></td></tr>
<tr><td>Oligo rev</td><td>AAACCGATCCCGAGCCCCCCCGCT</td><td></td><td></td><td></td><td></td><td></td><td></td><td></td><td></td><td></td><td></td></tr>
<tr><td>Chromosome</td><td>start</td><td>end</td><td>strand</td><td>MM</td><td>target_seq</td><td>PAM</td><td>alignment</td><td>distance</td><td>position</td><td>gene name</td><td>gene id</td></tr>
<tr><td>GU305914.1</td><td>50406</td><td>50428</td><td>-</td><td>0</td><td>AGCGGGGGGGCTCGGGATCG</td><td>CGG</td><td>||||||||[||||||||||||]PAM</td><td>NA</td><td>Intergenic</td><td>NA</td><td>NA</td></tr>
<tr><td>T30321</td><td>GGGGGGGCTCGGGATCGCGGCGG</td><td>1000</td><td></td><td></td><td></td><td></td><td></td><td></td><td></td><td></td><td></td></tr>
<tr><td>Oligo fwd</td><td>CACCGGGGGGGCTCGGGATCGCGG</td><td></td><td></td><td></td><td></td><td></td><td></td><td></td><td></td><td></td><td></td></tr>
<tr><td>Oligo rev</td><td>AAACCCGCGATCCCGAGCCCCCCC</td><td></td><td></td><td></td><td></td><td></td><td></td><td></td><td></td><td></td><td></td></tr>
<tr><td>Chromosome</td><td>start</td><td>end</td><td>strand</td><td>MM</td><td>target_seq</td><td>PAM</td><td>alignment</td><td>distance</td><td>position</td><td>gene name</td><td>gene id</td></tr>
<tr><td>GU305914.1</td><td>50403</td><td>50425</td><td>-</td><td>0</td><td>GGGGGGGCTCGGGATCGCGG</td><td>CGG</td><td>||||||||[||||||||||||]PAM</td><td>NA</td><td>Intergenic</td><td>NA</td><td>NA</td></tr>
<tr><td>T30322</td><td>GGGGCTCGGGATCGCGGCGGTGG</td><td>1000</td><td></td><td></td><td></td><td></td><td></td><td></td><td></td><td></td><td></td></tr>
<tr><td>Oligo fwd</td><td>CACCGGGGCTCGGGATCGCGGCGG</td><td></td><td></td><td></td><td></td><td></td><td></td><td></td><td></td><td></td><td></td></tr>
<tr><td>Oligo rev</td><td>AAACCCGCCGCGATCCCGAGCCCC</td><td></td><td></td><td></td><td></td><td></td><td></td><td></td><td></td><td></td><td></td></tr>
<tr><td>Chromosome</td><td>start</td><td>end</td><td>strand</td><td>MM</td><td>target_seq</td><td>PAM</td><td>alignment</td><td>distance</td><td>position</td><td>gene name</td><td>gene id</td></tr>
<tr><td>GU305914.1</td><td>50400</td><td>50422</td><td>-</td><td>0</td><td>GGGGCTCGGGATCGCGGCGG</td><td>TGG</td><td>||||||||[||||||||||||]PAM</td><td>NA</td><td>Intergenic</td><td>NA</td><td>NA</td></tr>
<tr><td>T30323</td><td>GTTGCCTCGCGATCGCCGCGCGG</td><td>1000</td><td></td><td></td><td></td><td></td><td></td><td></td><td></td><td></td><td></td></tr>
<tr><td>Oligo fwd</td><td>CACCGTTGCCTCGCGATCGCCGCG</td><td></td><td></td><td></td><td></td><td></td><td></td><td></td><td></td><td></td><td></td></tr>
<tr><td>Oligo rev</td><td>AAACCGCGGCGATCGCGAGGCAAC</td><td></td><td></td><td></td><td></td><td></td><td></td><td></td><td></td><td></td><td></td></tr>
<tr><td>Chromosome</td><td>start</td><td>end</td><td>strand</td><td>MM</td><td>target_seq</td><td>PAM</td><td>alignment</td><td>distance</td><td>position</td><td>gene name</td><td>gene id</td></tr>
<tr><td>GU305914.1</td><td>50353</td><td>50375</td><td>-</td><td>0</td><td>GTTGCCTCGCGATCGCCGCG</td><td>CGG</td><td>||||||||[||||||||||||]PAM</td><td>NA</td><td>Intergenic</td><td>NA</td><td>NA</td></tr>
<tr><td>T30324</td><td>CCTCGCGATCGCCGCGCGGACGG</td><td>1000</td><td></td><td></td><td></td><td></td><td></td><td></td><td></td><td></td><td></td></tr>
<tr><td>Oligo fwd</td><td>CACCCCTCGCGATCGCCGCGCGGA</td><td></td><td></td><td></td><td></td><td></td><td></td><td></td><td></td><td></td><td></td></tr>
<tr><td>Oligo rev</td><td>AAACTCCGCGCGGCGATCGCGAGG</td><td></td><td></td><td></td><td></td><td></td><td></td><td></td><td></td><td></td><td></td></tr>
<tr><td>Chromosome</td><td>start</td><td>end</td><td>strand</td><td>MM</td><td>target_seq</td><td>PAM</td><td>alignment</td><td>distance</td><td>position</td><td>gene name</td><td>gene id</td></tr>
<tr><td>GU305914.1</td><td>50349</td><td>50371</td><td>-</td><td>0</td><td>CCTCGCGATCGCCGCGCGGA</td><td>CGG</td><td>||||||||[||||||||||||]PAM</td><td>NA</td><td>Intergenic</td><td>NA</td><td>NA</td></tr>
<tr><td>T30325</td><td>CGCGATCGCCGCGCGGACGGCGG</td><td>1000</td><td></td><td></td><td></td><td></td><td></td><td></td><td></td><td></td><td></td></tr>
<tr><td>Oligo fwd</td><td>CACCCGCGATCGCCGCGCGGACGG</td><td></td><td></td><td></td><td></td><td></td><td></td><td></td><td></td><td></td><td></td></tr>
<tr><td>Oligo rev</td><td>AAACCCGTCCGCGCGGCGATCGCG</td><td></td><td></td><td></td><td></td><td></td><td></td><td></td><td></td><td></td><td></td></tr>
<tr><td>Chromosome</td><td>start</td><td>end</td><td>strand</td><td>MM</td><td>target_seq</td><td>PAM</td><td>alignment</td><td>distance</td><td>position</td><td>gene name</td><td>gene id</td></tr>
<tr><td>GU305914.1</td><td>50346</td><td>50368</td><td>-</td><td>0</td><td>CGCGATCGCCGCGCGGACGG</td><td>CGG</td><td>||||||||[||||||||||||]PAM</td><td>NA</td><td>Intergenic</td><td>NA</td><td>NA</td></tr>
<tr><td>GU305914.1</td><td>45908</td><td>45930</td><td>+</td><td>3</td><td>CGCGATGGCCGCTCCGACGG</td><td>AGG</td><td>||||||-|[||||-|-|||||]PAM</td><td>NA</td><td>Intergenic</td><td>NA</td><td>NA</td></tr>
<tr><td>T30326</td><td>CGCCGCGCGGACGGCGGTCTCGG</td><td>1000</td><td></td><td></td><td></td><td></td><td></td><td></td><td></td><td></td><td></td></tr>
<tr><td>Oligo fwd</td><td>CACCCGCCGCGCGGACGGCGGTCT</td><td></td><td></td><td></td><td></td><td></td><td></td><td></td><td></td><td></td><td></td></tr>
<tr><td>Oligo rev</td><td>AAACAGACCGCCGTCCGCGCGGCG</td><td></td><td></td><td></td><td></td><td></td><td></td><td></td><td></td><td></td><td></td></tr>
<tr><td>Chromosome</td><td>start</td><td>end</td><td>strand</td><td>MM</td><td>target_seq</td><td>PAM</td><td>alignment</td><td>distance</td><td>position</td><td>gene name</td><td>gene id</td></tr>
<tr><td>GU305914.1</td><td>50340</td><td>50362</td><td>-</td><td>0</td><td>CGCCGCGCGGACGGCGGTCT</td><td>CGG</td><td>||||||||[||||||||||||]PAM</td><td>NA</td><td>Intergenic</td><td>NA</td><td>NA</td></tr>
<tr><td>T30327</td><td>GCCGCGCGGACGGCGGTCTCGGG</td><td>1000</td><td></td><td></td><td></td><td></td><td></td><td></td><td></td><td></td><td></td></tr>
<tr><td>Oligo fwd</td><td>CACCGCCGCGCGGACGGCGGTCTC</td><td></td><td></td><td></td><td></td><td></td><td></td><td></td><td></td><td></td><td></td></tr>
<tr><td>Oligo rev</td><td>AAACGAGACCGCCGTCCGCGCGGC</td><td></td><td></td><td></td><td></td><td></td><td></td><td></td><td></td><td></td><td></td></tr>
<tr><td>Chromosome</td><td>start</td><td>end</td><td>strand</td><td>MM</td><td>target_seq</td><td>PAM</td><td>alignment</td><td>distance</td><td>position</td><td>gene name</td><td>gene id</td></tr>
<tr><td>GU305914.1</td><td>50339</td><td>50361</td><td>-</td><td>0</td><td>GCCGCGCGGACGGCGGTCTC</td><td>GGG</td><td>||||||||[||||||||||||]PAM</td><td>NA</td><td>Intergenic</td><td>NA</td><td>NA</td></tr>
<tr><td>T30328</td><td>CCGCGCGGACGGCGGTCTCGGGG</td><td>1000</td><td></td><td></td><td></td><td></td><td></td><td></td><td></td><td></td><td></td></tr>
<tr><td>Oligo fwd</td><td>CACCCCGCGCGGACGGCGGTCTCG</td><td></td><td></td><td></td><td></td><td></td><td></td><td></td><td></td><td></td><td></td></tr>
<tr><td>Oligo rev</td><td>AAACCGAGACCGCCGTCCGCGCGG</td><td></td><td></td><td></td><td></td><td></td><td></td><td></td><td></td><td></td><td></td></tr>
<tr><td>Chromosome</td><td>start</td><td>end</td><td>strand</td><td>MM</td><td>target_seq</td><td>PAM</td><td>alignment</td><td>distance</td><td>position</td><td>gene name</td><td>gene id</td></tr>
<tr><td>GU305914.1</td><td>50338</td><td>50360</td><td>-</td><td>0</td><td>CCGCGCGGACGGCGGTCTCG</td><td>GGG</td><td>||||||||[||||||||||||]PAM</td><td>NA</td><td>Intergenic</td><td>NA</td><td>NA</td></tr>
<tr><td>T30329</td><td>GGCGGTCTCGGGGCCCGAATCGG</td><td>1000</td><td></td><td></td><td></td><td></td><td></td><td></td><td></td><td></td><td></td></tr>
<tr><td>Oligo fwd</td><td>CACCGGCGGTCTCGGGGCCCGAAT</td><td></td><td></td><td></td><td></td><td></td><td></td><td></td><td></td><td></td><td></td></tr>
<tr><td>Oligo rev</td><td>AAACATTCGGGCCCCGAGACCGCC</td><td></td><td></td><td></td><td></td><td></td><td></td><td></td><td></td><td></td><td></td></tr>
<tr><td>Chromosome</td><td>start</td><td>end</td><td>strand</td><td>MM</td><td>target_seq</td><td>PAM</td><td>alignment</td><td>distance</td><td>position</td><td>gene name</td><td>gene id</td></tr>
<tr><td>GU305914.1</td><td>50328</td><td>50350</td><td>-</td><td>0</td><td>GGCGGTCTCGGGGCCCGAAT</td><td>CGG</td><td>||||||||[||||||||||||]PAM</td><td>NA</td><td>Intergenic</td><td>NA</td><td>NA</td></tr>
<tr><td>T30330</td><td>GCGGTCTCGGGGCCCGAATCGGG</td><td>1000</td><td></td><td></td><td></td><td></td><td></td><td></td><td></td><td></td><td></td></tr>
<tr><td>Oligo fwd</td><td>CACCGCGGTCTCGGGGCCCGAATC</td><td></td><td></td><td></td><td></td><td></td><td></td><td></td><td></td><td></td><td></td></tr>
<tr><td>Oligo rev</td><td>AAACGATTCGGGCCCCGAGACCGC</td><td></td><td></td><td></td><td></td><td></td><td></td><td></td><td></td><td></td><td></td></tr>
<tr><td>Chromosome</td><td>start</td><td>end</td><td>strand</td><td>MM</td><td>target_seq</td><td>PAM</td><td>alignment</td><td>distance</td><td>position</td><td>gene name</td><td>gene id</td></tr>
<tr><td>GU305914.1</td><td>50327</td><td>50349</td><td>-</td><td>0</td><td>GCGGTCTCGGGGCCCGAATC</td><td>GGG</td><td>||||||||[||||||||||||]PAM</td><td>NA</td><td>Intergenic</td><td>NA</td><td>NA</td></tr>
<tr><td>T30331</td><td>CGTCTGTCTCGTGCAGCGCAAGG</td><td>1000</td><td></td><td></td><td></td><td></td><td></td><td></td><td></td><td></td><td></td></tr>
<tr><td>Oligo fwd</td><td>CACCCGTCTGTCTCGTGCAGCGCA</td><td></td><td></td><td></td><td></td><td></td><td></td><td></td><td></td><td></td><td></td></tr>
<tr><td>Oligo rev</td><td>AAACTGCGCTGCACGAGACAGACG</td><td></td><td></td><td></td><td></td><td></td><td></td><td></td><td></td><td></td><td></td></tr>
<tr><td>Chromosome</td><td>start</td><td>end</td><td>strand</td><td>MM</td><td>target_seq</td><td>PAM</td><td>alignment</td><td>distance</td><td>position</td><td>gene name</td><td>gene id</td></tr>
<tr><td>GU305914.1</td><td>50283</td><td>50305</td><td>-</td><td>0</td><td>CGTCTGTCTCGTGCAGCGCA</td><td>AGG</td><td>||||||||[||||||||||||]PAM</td><td>NA</td><td>Intergenic</td><td>NA</td><td>NA</td></tr>
<tr><td>T30332</td><td>CAAGGACAACCGCACCAGTGTGG</td><td>1000</td><td></td><td></td><td></td><td></td><td></td><td></td><td></td><td></td><td></td></tr>
<tr><td>Oligo fwd</td><td>CACCCAAGGACAACCGCACCAGTG</td><td></td><td></td><td></td><td></td><td></td><td></td><td></td><td></td><td></td><td></td></tr>
</table>

<table>
<tr><td>Oligo rev</td><td>AAACCACTGGTGCGGTTGTCCTTG</td><td></td><td></td><td></td><td></td><td></td><td></td><td></td><td></td><td></td><td></td></tr>
<tr><td>Chromosome</td><td>start</td><td>end</td><td>strand</td><td>MM</td><td>target_seq</td><td>PAM</td><td>alignment</td><td>distance</td><td>position</td><td>gene name</td><td>gene id</td></tr>
<tr><td>GU305914.1</td><td>50265</td><td>50287</td><td>-</td><td>0</td><td>CAAGGACAACCGCACCAGTG</td><td>TGG</td><td>||||||||[||||||||||||]PAM</td><td>NA</td><td>Intergenic</td><td>NA</td><td>NA</td></tr>
<tr><td></td><td></td><td></td><td></td><td></td><td></td><td></td><td></td><td></td><td></td><td></td><td></td></tr>
<tr><td>T30333</td><td>AAGGACAACCGCACCAGTGTGGG</td><td>1000</td><td></td><td></td><td></td><td></td><td></td><td></td><td></td><td></td><td></td></tr>
<tr><td>Oligo fwd</td><td>CACCAAGGACAACCGCACCAGTGT</td><td></td><td></td><td></td><td></td><td></td><td></td><td></td><td></td><td></td><td></td></tr>
<tr><td>Oligo rev</td><td>AAACACACTGGTGCGGTTGTCCTT</td><td></td><td></td><td></td><td></td><td></td><td></td><td></td><td></td><td></td><td></td></tr>
<tr><td>Chromosome</td><td>start</td><td>end</td><td>strand</td><td>MM</td><td>target_seq</td><td>PAM</td><td>alignment</td><td>distance</td><td>position</td><td>gene name</td><td>gene id</td></tr>
<tr><td>GU305914.1</td><td>50264</td><td>50286</td><td>-</td><td>0</td><td>AAGGACAACCGCACCAGTGT</td><td>GGG</td><td>||||||||[||||||||||||]PAM</td><td>NA</td><td>Intergenic</td><td>NA</td><td>NA</td></tr>
<tr><td></td><td></td><td></td><td></td><td></td><td></td><td></td><td></td><td></td><td></td><td></td><td></td></tr>
<tr><td>T30334</td><td>TGTATCGTCGACTGCAAGTACGG</td><td>1000</td><td></td><td></td><td></td><td></td><td></td><td></td><td></td><td></td><td></td></tr>
<tr><td>Oligo fwd</td><td>CACCTGTATCGTCGACTGCAAGTA</td><td></td><td></td><td></td><td></td><td></td><td></td><td></td><td></td><td></td><td></td></tr>
<tr><td>Oligo rev</td><td>AAACTACTTGCAGTCGACGATACA</td><td></td><td></td><td></td><td></td><td></td><td></td><td></td><td></td><td></td><td></td></tr>
<tr><td>Chromosome</td><td>start</td><td>end</td><td>strand</td><td>MM</td><td>target_seq</td><td>PAM</td><td>alignment</td><td>distance</td><td>position</td><td>gene name</td><td>gene id</td></tr>
<tr><td>GU305914.1</td><td>50240</td><td>50262</td><td>-</td><td>0</td><td>TGTATCGTCGACTGCAAGTA</td><td>CGG</td><td>||||||||[||||||||||||]PAM</td><td>NA</td><td>Intergenic</td><td>NA</td><td>NA</td></tr>
<tr><td>GU305914.1</td><td>220343</td><td>220365</td><td>-</td><td>4</td><td>TGGATCATCGACTACAACTA</td><td>TGG</td><td>||-|||-|[|||||-|||-||]PAM</td><td>NA</td><td>Intergenic</td><td>NA</td><td>NA</td></tr>
<tr><td></td><td></td><td></td><td></td><td></td><td></td><td></td><td></td><td></td><td></td><td></td><td></td></tr>
<tr><td>T30335</td><td>GGATCCGAACTCGTCTTCTCCGG</td><td>1000</td><td></td><td></td><td></td><td></td><td></td><td></td><td></td><td></td><td></td></tr>
<tr><td>Oligo fwd</td><td>CACCGGATCCGAACTCGTCTTCTC</td><td></td><td></td><td></td><td></td><td></td><td></td><td></td><td></td><td></td><td></td></tr>
<tr><td>Oligo rev</td><td>AAACGAGAAGACGAGTTCGGATCC</td><td></td><td></td><td></td><td></td><td></td><td></td><td></td><td></td><td></td><td></td></tr>
<tr><td>Chromosome</td><td>start</td><td>end</td><td>strand</td><td>MM</td><td>target_seq</td><td>PAM</td><td>alignment</td><td>distance</td><td>position</td><td>gene name</td><td>gene id</td></tr>
<tr><td>GU305914.1</td><td>50219</td><td>50241</td><td>-</td><td>0</td><td>GGATCCGAACTCGTCTTCTC</td><td>CGG</td><td>||||||||[||||||||||||]PAM</td><td>NA</td><td>Intergenic</td><td>NA</td><td>NA</td></tr>
<tr><td></td><td></td><td></td><td></td><td></td><td></td><td></td><td></td><td></td><td></td><td></td><td></td></tr>
<tr><td>T30336</td><td>CTGTCAAGATGCCTTCACGTTGG</td><td>1000</td><td></td><td></td><td></td><td></td><td></td><td></td><td></td><td></td><td></td></tr>
<tr><td>Oligo fwd</td><td>CACCCTGTCAAGATGCCTTCACGT</td><td></td><td></td><td></td><td></td><td></td><td></td><td></td><td></td><td></td><td></td></tr>
<tr><td>Oligo rev</td><td>AAACACGTGAAGGCATCTTGACAG</td><td></td><td></td><td></td><td></td><td></td><td></td><td></td><td></td><td></td><td></td></tr>
<tr><td>Chromosome</td><td>start</td><td>end</td><td>strand</td><td>MM</td><td>target_seq</td><td>PAM</td><td>alignment</td><td>distance</td><td>position</td><td>gene name</td><td>gene id</td></tr>
<tr><td>GU305914.1</td><td>50193</td><td>50215</td><td>-</td><td>0</td><td>CTGTCAAGATGCCTTCACGT</td><td>TGG</td><td>||||||||[||||||||||||]PAM</td><td>NA</td><td>Intergenic</td><td>NA</td><td>NA</td></tr>
<tr><td></td><td></td><td></td><td></td><td></td><td></td><td></td><td></td><td></td><td></td><td></td><td></td></tr>
<tr><td>T30337</td><td>CCTTCACGTTGGTCAGTACCTGG</td><td>1000</td><td></td><td></td><td></td><td></td><td></td><td></td><td></td><td></td><td></td></tr>
<tr><td>Oligo fwd</td><td>CACCCCTTCACGTTGGTCAGTACC</td><td></td><td></td><td></td><td></td><td></td><td></td><td></td><td></td><td></td><td></td></tr>
<tr><td>Oligo rev</td><td>AAACGGTACTGACCAACGTGAAGG</td><td></td><td></td><td></td><td></td><td></td><td></td><td></td><td></td><td></td><td></td></tr>
<tr><td>Chromosome</td><td>start</td><td>end</td><td>strand</td><td>MM</td><td>target_seq</td><td>PAM</td><td>alignment</td><td>distance</td><td>position</td><td>gene name</td><td>gene id</td></tr>
<tr><td>GU305914.1</td><td>50182</td><td>50204</td><td>-</td><td>0</td><td>CCTTCACGTTGGTCAGTACC</td><td>TGG</td><td>||||||||[||||||||||||]PAM</td><td>NA</td><td>Intergenic</td><td>NA</td><td>NA</td></tr>
<tr><td></td><td></td><td></td><td></td><td></td><td></td><td></td><td></td><td></td><td></td><td></td><td></td></tr>
<tr><td>T30338</td><td>ACGTTGGTCAGTACCTGGTTCGG</td><td>1000</td><td></td><td></td><td></td><td></td><td></td><td></td><td></td><td></td><td></td></tr>
<tr><td>Oligo fwd</td><td>CACCACGTTGGTCAGTACCTGGTT</td><td></td><td></td><td></td><td></td><td></td><td></td><td></td><td></td><td></td><td></td></tr>
<tr><td>Oligo rev</td><td>AAACAACCAGGTACTGACCAACGT</td><td></td><td></td><td></td><td></td><td></td><td></td><td></td><td></td><td></td><td></td></tr>
<tr><td>Chromosome</td><td>start</td><td>end</td><td>strand</td><td>MM</td><td>target_seq</td><td>PAM</td><td>alignment</td><td>distance</td><td>position</td><td>gene name</td><td>gene id</td></tr>
<tr><td>GU305914.1</td><td>50177</td><td>50199</td><td>-</td><td>0</td><td>ACGTTGGTCAGTACCTGGTT</td><td>CGG</td><td>||||||||[||||||||||||]PAM</td><td>NA</td><td>Intergenic</td><td>NA</td><td>NA</td></tr>
<tr><td></td><td></td><td></td><td></td><td></td><td></td><td></td><td></td><td></td><td></td><td></td><td></td></tr>
<tr><td>T30339</td><td>GGTCAGTACCTGGTTCGGCAAGG</td><td>1000</td><td></td><td></td><td></td><td></td><td></td><td></td><td></td><td></td><td></td></tr>
<tr><td>Oligo fwd</td><td>CACCGGTCAGTACCTGGTTCGGCA</td><td></td><td></td><td></td><td></td><td></td><td></td><td></td><td></td><td></td><td></td></tr>
<tr><td>Oligo rev</td><td>AAACTGCCGAACCAGGTACTGACC</td><td></td><td></td><td></td><td></td><td></td><td></td><td></td><td></td><td></td><td></td></tr>
<tr><td>Chromosome</td><td>start</td><td>end</td><td>strand</td><td>MM</td><td>target_seq</td><td>PAM</td><td>alignment</td><td>distance</td><td>position</td><td>gene name</td><td>gene id</td></tr>
<tr><td>GU305914.1</td><td>50172</td><td>50194</td><td>-</td><td>0</td><td>GGTCAGTACCTGGTTCGGCA</td><td>AGG</td><td>||||||||[||||||||||||]PAM</td><td>NA</td><td>Intergenic</td><td>NA</td><td>NA</td></tr>
<tr><td></td><td></td><td></td><td></td><td></td><td></td><td></td><td></td><td></td><td></td><td></td><td></td></tr>
<tr><td>T30340</td><td>CCTGGTTCGGCAAGGTTAACCGG</td><td>1000</td><td></td><td></td><td></td><td></td><td></td><td></td><td></td><td></td><td></td></tr>
<tr><td>Oligo fwd</td><td>CACCCCTGGTTCGGCAAGGTTAAC</td><td></td><td></td><td></td><td></td><td></td><td></td><td></td><td></td><td></td><td></td></tr>
<tr><td>Oligo rev</td><td>AAACGTTAACCTTGCCGAACCAGG</td><td></td><td></td><td></td><td></td><td></td><td></td><td></td><td></td><td></td><td></td></tr>
<tr><td>Chromosome</td><td>start</td><td>end</td><td>strand</td><td>MM</td><td>target_seq</td><td>PAM</td><td>alignment</td><td>distance</td><td>position</td><td>gene name</td><td>gene id</td></tr>
<tr><td>GU305914.1</td><td>50164</td><td>50186</td><td>-</td><td>0</td><td>CCTGGTTCGGCAAGGTTAAC</td><td>CGG</td><td>||||||||[||||||||||||]PAM</td><td>NA</td><td>Intergenic</td><td>NA</td><td>NA</td></tr>
<tr><td></td><td></td><td></td><td></td><td></td><td></td><td></td><td></td><td></td><td></td><td></td><td></td></tr>
<tr><td>T30341</td><td>TCGGCAAGGTTAACCGGCGCTGG</td><td>1000</td><td></td><td></td><td></td><td></td><td></td><td></td><td></td><td></td><td></td></tr>
<tr><td>Oligo fwd</td><td>CACCTCGGCAAGGTTAACCGGCGC</td><td></td><td></td><td></td><td></td><td></td><td></td><td></td><td></td><td></td><td></td></tr>
<tr><td>Oligo rev</td><td>AAACGCGCCGGTTAACCTTGCCGA</td><td></td><td></td><td></td><td></td><td></td><td></td><td></td><td></td><td></td><td></td></tr>
<tr><td>Chromosome</td><td>start</td><td>end</td><td>strand</td><td>MM</td><td>target_seq</td><td>PAM</td><td>alignment</td><td>distance</td><td>position</td><td>gene name</td><td>gene id</td></tr>
<tr><td>GU305914.1</td><td>50158</td><td>50180</td><td>-</td><td>0</td><td>TCGGCAAGGTTAACCGGCGC</td><td>TGG</td><td>||||||||[||||||||||||]PAM</td><td>NA</td><td>Intergenic</td><td>NA</td><td>NA</td></tr>
<tr><td></td><td></td><td></td><td></td><td></td><td></td><td></td><td></td><td></td><td></td><td></td><td></td></tr>
<tr><td>T30342</td><td>TTAACCGGCGCTGGTCGTTTCGG</td><td>1000</td><td></td><td></td><td></td><td></td><td></td><td></td><td></td><td></td><td></td></tr>
<tr><td>Oligo fwd</td><td>CACCTTAACCGGCGCTGGTCGTTT</td><td></td><td></td><td></td><td></td><td></td><td></td><td></td><td></td><td></td><td></td></tr>
<tr><td>Oligo rev</td><td>AAACAAACGACCAGCGCCGGTTAA</td><td></td><td></td><td></td><td></td><td></td><td></td><td></td><td></td><td></td><td></td></tr>
<tr><td>Chromosome</td><td>start</td><td>end</td><td>strand</td><td>MM</td><td>target_seq</td><td>PAM</td><td>alignment</td><td>distance</td><td>position</td><td>gene name</td><td>gene id</td></tr>
<tr><td>GU305914.1</td><td>50149</td><td>50171</td><td>-</td><td>0</td><td>TTAACCGGCGCTGGTCGTTT</td><td>CGG</td><td>||||||||[||||||||||||]PAM</td><td>NA</td><td>Intergenic</td><td>NA</td><td>NA</td></tr>
<tr><td></td><td></td><td></td><td></td><td></td><td></td><td></td><td></td><td></td><td></td><td></td><td></td></tr>
<tr><td>T30343</td><td>TAACCGGCGCTGGTCGTTTCGGG</td><td>1000</td><td></td><td></td><td></td><td></td><td></td><td></td><td></td><td></td><td></td></tr>
<tr><td>Oligo fwd</td><td>CACCTAACCGGCGCTGGTCGTTTC</td><td></td><td></td><td></td><td></td><td></td><td></td><td></td><td></td><td></td><td></td></tr>
<tr><td>Oligo rev</td><td>AAACGAAACGACCAGCGCCGGTTA</td><td></td><td></td><td></td><td></td><td></td><td></td><td></td><td></td><td></td><td></td></tr>
<tr><td>Chromosome</td><td>start</td><td>end</td><td>strand</td><td>MM</td><td>target_seq</td><td>PAM</td><td>alignment</td><td>distance</td><td>position</td><td>gene name</td><td>gene id</td></tr>
<tr><td>GU305914.1</td><td>50148</td><td>50170</td><td>-</td><td>0</td><td>TAACCGGCGCTGGTCGTTTC</td><td>GGG</td><td>||||||||[||||||||||||]PAM</td><td>NA</td><td>Intergenic</td><td>NA</td><td>NA</td></tr>
<tr><td></td><td></td><td></td><td></td><td></td><td></td><td></td><td></td><td></td><td></td><td></td><td></td></tr>
<tr><td>T30344</td><td>GTCGTTTCGGGATATCCAGCTGG</td><td>1000</td><td></td><td></td><td></td><td></td><td></td><td></td><td></td><td></td><td></td></tr>
<tr><td>Oligo fwd</td><td>CACCGTCGTTTCGGGATATCCAGC</td><td></td><td></td><td></td><td></td><td></td><td></td><td></td><td></td><td></td><td></td></tr>
<tr><td>Oligo rev</td><td>AAACGCTGGATATCCCGAAACGAC</td><td></td><td></td><td></td><td></td><td></td><td></td><td></td><td></td><td></td><td></td></tr>
<tr><td>Chromosome</td><td>start</td><td>end</td><td>strand</td><td>MM</td><td>target_seq</td><td>PAM</td><td>alignment</td><td>distance</td><td>position</td><td>gene name</td><td>gene id</td></tr>
<tr><td>GU305914.1</td><td>50136</td><td>50158</td><td>-</td><td>0</td><td>GTCGTTTCGGGATATCCAGC</td><td>TGG</td><td>||||||||[||||||||||||]PAM</td><td>NA</td><td>Intergenic</td><td>NA</td><td>NA</td></tr>
<tr><td></td><td></td><td></td><td></td><td></td><td></td><td></td><td></td><td></td><td></td><td></td><td></td></tr>
<tr><td>T30345</td><td>GATATCCAGCTGGTTGACATCGG</td><td>1000</td><td></td><td></td><td></td><td></td><td></td><td></td><td></td><td></td><td></td></tr>
<tr><td>Oligo fwd</td><td>CACCGATATCCAGCTGGTTGACAT</td><td></td><td></td><td></td><td></td><td></td><td></td><td></td><td></td><td></td><td></td></tr>
<tr><td>Oligo rev</td><td>AAACATGTCAACCAGCTGGATATC</td><td></td><td></td><td></td><td></td><td></td><td></td><td></td><td></td><td></td><td></td></tr>
<tr><td>Chromosome</td><td>start</td><td>end</td><td>strand</td><td>MM</td><td>target_seq</td><td>PAM</td><td>alignment</td><td>distance</td><td>position</td><td>gene name</td><td>gene id</td></tr>
<tr><td>GU305914.1</td><td>50126</td><td>50148</td><td>-</td><td>0</td><td>GATATCCAGCTGGTTGACAT</td><td>CGG</td><td>||||||||[||||||||||||]PAM</td><td>NA</td><td>Intergenic</td><td>NA</td><td>NA</td></tr>
<tr><td></td><td></td><td></td><td></td><td></td><td></td><td></td><td></td><td></td><td></td><td></td><td></td></tr>
<tr><td>T30346</td><td>GGCATCGCTTACTACTTCACCGG</td><td>1000</td><td></td><td></td><td></td><td></td><td></td><td></td><td></td><td></td><td></td></tr>
<tr><td>Oligo fwd</td><td>CACCGGCATCGCTTACTACTTCAC</td><td></td><td></td><td></td><td></td><td></td><td></td><td></td><td></td><td></td><td></td></tr>
<tr><td>Oligo rev</td><td>AAACGTGAAGTAGTAAGCGATGCC</td><td></td><td></td><td></td><td></td><td></td><td></td><td></td><td></td><td></td><td></td></tr>
<tr><td>Chromosome</td><td>start</td><td>end</td><td>strand</td><td>MM</td><td>target_seq</td><td>PAM</td><td>alignment</td><td>distance</td><td>position</td><td>gene name</td><td>gene id</td></tr>
<tr><td>GU305914.1</td><td>50105</td><td>50127</td><td>-</td><td>0</td><td>GGCATCGCTTACTACTTCAC</td><td>CGG</td><td>||||||||[||||||||||||]PAM</td><td>NA</td><td>Intergenic</td><td>NA</td><td>NA</td></tr>
</table>

<table>
<tr><td>T30347</td><td>ACTTCACCGGTTACATCCTCAGG</td><td>1000</td><td></td><td></td><td></td><td></td><td></td><td></td><td></td><td></td><td></td></tr>
<tr><td>Oligo fwd</td><td>CACCACTTCACCGGTTACATCCTC</td><td></td><td></td><td></td><td></td><td></td><td></td><td></td><td></td><td></td><td></td></tr>
<tr><td>Oligo rev</td><td>AAACGAGGATGTAACCGGTGAAGT</td><td></td><td></td><td></td><td></td><td></td><td></td><td></td><td></td><td></td><td></td></tr>
<tr><td>Chromosome</td><td>start</td><td>end</td><td>strand</td><td>MM</td><td>target_seq</td><td>PAM</td><td>alignment</td><td>distance</td><td>position</td><td>gene name</td><td>gene id</td></tr>
<tr><td>GU305914.1</td><td>50092</td><td>50114</td><td>-</td><td>0</td><td>ACTTCACCGGTTACATCCTC</td><td>AGG</td><td>||||||||[||||||||||||]PAM</td><td>NA</td><td>Intergenic</td><td>NA</td><td>NA</td></tr>
<tr><td>T30348</td><td>AGGCCCCTGAGCTCGTGTCCCGG</td><td>1000</td><td></td><td></td><td></td><td></td><td></td><td></td><td></td><td></td><td></td></tr>
<tr><td>Oligo fwd</td><td>CACCAGGCCCCTGAGCTCGTGTCC</td><td></td><td></td><td></td><td></td><td></td><td></td><td></td><td></td><td></td><td></td></tr>
<tr><td>Oligo rev</td><td>AAACGGACACGAGCTCAGGGGCCT</td><td></td><td></td><td></td><td></td><td></td><td></td><td></td><td></td><td></td><td></td></tr>
<tr><td>Chromosome</td><td>start</td><td>end</td><td>strand</td><td>MM</td><td>target_seq</td><td>PAM</td><td>alignment</td><td>distance</td><td>position</td><td>gene name</td><td>gene id</td></tr>
<tr><td>GU305914.1</td><td>50072</td><td>50094</td><td>-</td><td>0</td><td>AGGCCCCTGAGCTCGTGTCC</td><td>CGG</td><td>||||||||[||||||||||||]PAM</td><td>NA</td><td>Intergenic</td><td>NA</td><td>NA</td></tr>
<tr><td>T30349</td><td>CGGACTCGTCTCGCACCGTGCGG</td><td>1000</td><td></td><td></td><td></td><td></td><td></td><td></td><td></td><td></td><td></td></tr>
<tr><td>Oligo fwd</td><td>CACCCGGACTCGTCTCGCACCGTG</td><td></td><td></td><td></td><td></td><td></td><td></td><td></td><td></td><td></td><td></td></tr>
<tr><td>Oligo rev</td><td>AAACCACGGTGCGAGACGAGTCCG</td><td></td><td></td><td></td><td></td><td></td><td></td><td></td><td></td><td></td><td></td></tr>
<tr><td>Chromosome</td><td>start</td><td>end</td><td>strand</td><td>MM</td><td>target_seq</td><td>PAM</td><td>alignment</td><td>distance</td><td>position</td><td>gene name</td><td>gene id</td></tr>
<tr><td>GU305914.1</td><td>50052</td><td>50074</td><td>-</td><td>0</td><td>CGGACTCGTCTCGCACCGTG</td><td>CGG</td><td>||||||||[||||||||||||]PAM</td><td>NA</td><td>Intergenic</td><td>NA</td><td>NA</td></tr>
<tr><td>T30350</td><td>CTCGTCTCGCACCGTGCGGATGG</td><td>1000</td><td></td><td></td><td></td><td></td><td></td><td></td><td></td><td></td><td></td></tr>
<tr><td>Oligo fwd</td><td>CACCCTCGTCTCGCACCGTGCGGA</td><td></td><td></td><td></td><td></td><td></td><td></td><td></td><td></td><td></td><td></td></tr>
<tr><td>Oligo rev</td><td>AAACTCCGCACGGTGCGAGACGAG</td><td></td><td></td><td></td><td></td><td></td><td></td><td></td><td></td><td></td><td></td></tr>
<tr><td>Chromosome</td><td>start</td><td>end</td><td>strand</td><td>MM</td><td>target_seq</td><td>PAM</td><td>alignment</td><td>distance</td><td>position</td><td>gene name</td><td>gene id</td></tr>
<tr><td>GU305914.1</td><td>50048</td><td>50070</td><td>-</td><td>0</td><td>CTCGTCTCGCACCGTGCGGA</td><td>TGG</td><td>||||||||[||||||||||||]PAM</td><td>NA</td><td>Intergenic</td><td>NA</td><td>NA</td></tr>
<tr><td>T30351</td><td>TCTCGCACCGTGCGGATGGAAGG</td><td>1000</td><td></td><td></td><td></td><td></td><td></td><td></td><td></td><td></td><td></td></tr>
<tr><td>Oligo fwd</td><td>CACCTCTCGCACCGTGCGGATGGA</td><td></td><td></td><td></td><td></td><td></td><td></td><td></td><td></td><td></td><td></td></tr>
<tr><td>Oligo rev</td><td>AAACTCCATCCGCACGGTGCGAGA</td><td></td><td></td><td></td><td></td><td></td><td></td><td></td><td></td><td></td><td></td></tr>
<tr><td>Chromosome</td><td>start</td><td>end</td><td>strand</td><td>MM</td><td>target_seq</td><td>PAM</td><td>alignment</td><td>distance</td><td>position</td><td>gene name</td><td>gene id</td></tr>
<tr><td>GU305914.1</td><td>50044</td><td>50066</td><td>-</td><td>0</td><td>TCTCGCACCGTGCGGATGGA</td><td>AGG</td><td>||||||||[||||||||||||]PAM</td><td>NA</td><td>Intergenic</td><td>NA</td><td>NA</td></tr>
<tr><td>T30352</td><td>GGAAGGACTTCGTCGTTCTCCGG</td><td>1000</td><td></td><td></td><td></td><td></td><td></td><td></td><td></td><td></td><td></td></tr>
<tr><td>Oligo fwd</td><td>CACCGGAAGGACTTCGTCGTTCTC</td><td></td><td></td><td></td><td></td><td></td><td></td><td></td><td></td><td></td><td></td></tr>
<tr><td>Oligo rev</td><td>AAACGAGAACGACGAAGTCCTTCC</td><td></td><td></td><td></td><td></td><td></td><td></td><td></td><td></td><td></td><td></td></tr>
<tr><td>Chromosome</td><td>start</td><td>end</td><td>strand</td><td>MM</td><td>target_seq</td><td>PAM</td><td>alignment</td><td>distance</td><td>position</td><td>gene name</td><td>gene id</td></tr>
<tr><td>GU305914.1</td><td>50027</td><td>50049</td><td>-</td><td>0</td><td>GGAAGGACTTCGTCGTTCTC</td><td>CGG</td><td>||||||||[||||||||||||]PAM</td><td>NA</td><td>Intergenic</td><td>NA</td><td>NA</td></tr>
<tr><td>T30353</td><td>GACTTCGTCGTTCTCCGGCAAGG</td><td>1000</td><td></td><td></td><td></td><td></td><td></td><td></td><td></td><td></td><td></td></tr>
<tr><td>Oligo fwd</td><td>CACCGACTTCGTCGTTCTCCGGCA</td><td></td><td></td><td></td><td></td><td></td><td></td><td></td><td></td><td></td><td></td></tr>
<tr><td>Oligo rev</td><td>AAACTGCCGGAGAACGACGAAGTC</td><td></td><td></td><td></td><td></td><td></td><td></td><td></td><td></td><td></td><td></td></tr>
<tr><td>Chromosome</td><td>start</td><td>end</td><td>strand</td><td>MM</td><td>target_seq</td><td>PAM</td><td>alignment</td><td>distance</td><td>position</td><td>gene name</td><td>gene id</td></tr>
<tr><td>GU305914.1</td><td>50022</td><td>50044</td><td>-</td><td>0</td><td>GACTTCGTCGTTCTCCGGCA</td><td>AGG</td><td>||||||||[||||||||||||]PAM</td><td>NA</td><td>Intergenic</td><td>NA</td><td>NA</td></tr>
<tr><td>T30354</td><td>ACTTCGTCGTTCTCCGGCAAGGG</td><td>1000</td><td></td><td></td><td></td><td></td><td></td><td></td><td></td><td></td><td></td></tr>
<tr><td>Oligo fwd</td><td>CACCACTTCGTCGTTCTCCGGCAA</td><td></td><td></td><td></td><td></td><td></td><td></td><td></td><td></td><td></td><td></td></tr>
<tr><td>Oligo rev</td><td>AAACTTGCCGGAGAACGACGAAGT</td><td></td><td></td><td></td><td></td><td></td><td></td><td></td><td></td><td></td><td></td></tr>
<tr><td>Chromosome</td><td>start</td><td>end</td><td>strand</td><td>MM</td><td>target_seq</td><td>PAM</td><td>alignment</td><td>distance</td><td>position</td><td>gene name</td><td>gene id</td></tr>
<tr><td>GU305914.1</td><td>50021</td><td>50043</td><td>-</td><td>0</td><td>ACTTCGTCGTTCTCCGGCAA</td><td>GGG</td><td>||||||||[||||||||||||]PAM</td><td>NA</td><td>Intergenic</td><td>NA</td><td>NA</td></tr>
<tr><td>T30355</td><td>CGGCAAGGGCAGATGTCATCCGG</td><td>1000</td><td></td><td></td><td></td><td></td><td></td><td></td><td></td><td></td><td></td></tr>
<tr><td>Oligo fwd</td><td>CACCCGGCAAGGGCAGATGTCATC</td><td></td><td></td><td></td><td></td><td></td><td></td><td></td><td></td><td></td><td></td></tr>
<tr><td>Oligo rev</td><td>AAACGATGACATCTGCCCTTGCCG</td><td></td><td></td><td></td><td></td><td></td><td></td><td></td><td></td><td></td><td></td></tr>
<tr><td>Chromosome</td><td>start</td><td>end</td><td>strand</td><td>MM</td><td>target_seq</td><td>PAM</td><td>alignment</td><td>distance</td><td>position</td><td>gene name</td><td>gene id</td></tr>
<tr><td>GU305914.1</td><td>50007</td><td>50029</td><td>-</td><td>0</td><td>CGGCAAGGGCAGATGTCATC</td><td>CGG</td><td>||||||||[||||||||||||]PAM</td><td>NA</td><td>Intergenic</td><td>NA</td><td>NA</td></tr>
<tr><td>T30356</td><td>AAGGGCAGATGTCATCCGGACGG</td><td>1000</td><td></td><td></td><td></td><td></td><td></td><td></td><td></td><td></td><td></td></tr>
<tr><td>Oligo fwd</td><td>CACCAAGGGCAGATGTCATCCGGA</td><td></td><td></td><td></td><td></td><td></td><td></td><td></td><td></td><td></td><td></td></tr>
<tr><td>Oligo rev</td><td>AAACTCCGGATGACATCTGCCCTT</td><td></td><td></td><td></td><td></td><td></td><td></td><td></td><td></td><td></td><td></td></tr>
<tr><td>Chromosome</td><td>start</td><td>end</td><td>strand</td><td>MM</td><td>target_seq</td><td>PAM</td><td>alignment</td><td>distance</td><td>position</td><td>gene name</td><td>gene id</td></tr>
<tr><td>GU305914.1</td><td>50003</td><td>50025</td><td>-</td><td>0</td><td>AAGGGCAGATGTCATCCGGA</td><td>CGG</td><td>||||||||[||||||||||||]PAM</td><td>NA</td><td>Intergenic</td><td>NA</td><td>NA</td></tr>
<tr><td>T30357</td><td>AGATGTCATCCGGACGGTCGCGG</td><td>1000</td><td></td><td></td><td></td><td></td><td></td><td></td><td></td><td></td><td></td></tr>
<tr><td>Oligo fwd</td><td>CACCAGATGTCATCCGGACGGTCG</td><td></td><td></td><td></td><td></td><td></td><td></td><td></td><td></td><td></td><td></td></tr>
<tr><td>Oligo rev</td><td>AAACCGACCGTCCGGATGACATCT</td><td></td><td></td><td></td><td></td><td></td><td></td><td></td><td></td><td></td><td></td></tr>
<tr><td>Chromosome</td><td>start</td><td>end</td><td>strand</td><td>MM</td><td>target_seq</td><td>PAM</td><td>alignment</td><td>distance</td><td>position</td><td>gene name</td><td>gene id</td></tr>
<tr><td>GU305914.1</td><td>49997</td><td>50019</td><td>-</td><td>0</td><td>AGATGTCATCCGGACGGTCG</td><td>CGG</td><td>||||||||[||||||||||||]PAM</td><td>NA</td><td>Intergenic</td><td>NA</td><td>NA</td></tr>
<tr><td>T30358</td><td>TCATCCGGACGGTCGCGGTTCGG</td><td>1000</td><td></td><td></td><td></td><td></td><td></td><td></td><td></td><td></td><td></td></tr>
<tr><td>Oligo fwd</td><td>CACCTCATCCGGACGGTCGCGGTT</td><td></td><td></td><td></td><td></td><td></td><td></td><td></td><td></td><td></td><td></td></tr>
<tr><td>Oligo rev</td><td>AAACAACCGCGACCGTCCGGATGA</td><td></td><td></td><td></td><td></td><td></td><td></td><td></td><td></td><td></td><td></td></tr>
<tr><td>Chromosome</td><td>start</td><td>end</td><td>strand</td><td>MM</td><td>target_seq</td><td>PAM</td><td>alignment</td><td>distance</td><td>position</td><td>gene name</td><td>gene id</td></tr>
<tr><td>GU305914.1</td><td>49992</td><td>50014</td><td>-</td><td>0</td><td>TCATCCGGACGGTCGCGGTT</td><td>CGG</td><td>||||||||[||||||||||||]PAM</td><td>NA</td><td>Intergenic</td><td>NA</td><td>NA</td></tr>
<tr><td>T30359</td><td>GGTAGAGCTCAAGACGACGATGG</td><td>1000</td><td></td><td></td><td></td><td></td><td></td><td></td><td></td><td></td><td></td></tr>
<tr><td>Oligo fwd</td><td>CACCGGTAGAGCTCAAGACGACGA</td><td></td><td></td><td></td><td></td><td></td><td></td><td></td><td></td><td></td><td></td></tr>
<tr><td>Oligo rev</td><td>AAACTCGTCGTCTTGAGCTCTACC</td><td></td><td></td><td></td><td></td><td></td><td></td><td></td><td></td><td></td><td></td></tr>
<tr><td>Chromosome</td><td>start</td><td>end</td><td>strand</td><td>MM</td><td>target_seq</td><td>PAM</td><td>alignment</td><td>distance</td><td>position</td><td>gene name</td><td>gene id</td></tr>
<tr><td>GU305914.1</td><td>49971</td><td>49993</td><td>-</td><td>0</td><td>GGTAGAGCTCAAGACGACGA</td><td>TGG</td><td>||||||||[||||||||||||]PAM</td><td>NA</td><td>Intergenic</td><td>NA</td><td>NA</td></tr>
<tr><td>T30360</td><td>GAGCTCAAGACGACGATGGACGG</td><td>1000</td><td></td><td></td><td></td><td></td><td></td><td></td><td></td><td></td><td></td></tr>
<tr><td>Oligo fwd</td><td>CACCGAGCTCAAGACGACGATGGA</td><td></td><td></td><td></td><td></td><td></td><td></td><td></td><td></td><td></td><td></td></tr>
<tr><td>Oligo rev</td><td>AAACTCCATCGTCGTCTTGAGCTC</td><td></td><td></td><td></td><td></td><td></td><td></td><td></td><td></td><td></td><td></td></tr>
<tr><td>Chromosome</td><td>start</td><td>end</td><td>strand</td><td>MM</td><td>target_seq</td><td>PAM</td><td>alignment</td><td>distance</td><td>position</td><td>gene name</td><td>gene id</td></tr>
<tr><td>GU305914.1</td><td>49967</td><td>49989</td><td>-</td><td>0</td><td>GAGCTCAAGACGACGATGGA</td><td>CGG</td><td>||||||||[||||||||||||]PAM</td><td>NA</td><td>Intergenic</td><td>NA</td><td>NA</td></tr>
<tr><td>T30361</td><td>CTCAAGACGACGATGGACGGCGG</td><td>1000</td><td></td><td></td><td></td><td></td><td></td><td></td><td></td><td></td><td></td></tr>
<tr><td>Oligo fwd</td><td>CACCCTCAAGACGACGATGGACGG</td><td></td><td></td><td></td><td></td><td></td><td></td><td></td><td></td><td></td><td></td></tr>
<tr><td>Oligo rev</td><td>AAACCCGTCCATCGTCGTCTTGAG</td><td></td><td></td><td></td><td></td><td></td><td></td><td></td><td></td><td></td><td></td></tr>
<tr><td>Chromosome</td><td>start</td><td>end</td><td>strand</td><td>MM</td><td>target_seq</td><td>PAM</td><td>alignment</td><td>distance</td><td>position</td><td>gene name</td><td>gene id</td></tr>
<tr><td>GU305914.1</td><td>49964</td><td>49986</td><td>-</td><td>0</td><td>CTCAAGACGACGATGGACGG</td><td>CGG</td><td>||||||||[||||||||||||]PAM</td><td>NA</td><td>Intergenic</td><td>NA</td><td>NA</td></tr>
</table>

| | | | | | | | | | | | |
|---|---|---|---|---|---|---|---|---|---|---|---|
| T30362 | TCAAGACGACGATGGACGGCGGG | 1000 | | | | | | | | | |
| Oligo fwd | CACCTCAAGACGACGATGGACGGC | | | | | | | | | | |
| Oligo rev | AAACGCCGTCCATCGTCGTCTTGA | | | | | | | | | | |
| Chromosome | start | end | strand | MM | target_seq | PAM | alignment | distance | position | gene name | gene id |
| GU305914.1 | 49963 | 49985 | - | 0 | TCAAGACGACGATGGACGGC | GGG | \|\|\|\|\|\|\|\|[\|\|\|\|\|\|\|\|\|\|\|\|]PAM | NA | Intergenic | NA | NA |
| T30363 | GATGGACGGCGGGTTCCGTCTGG | 1000 | | | | | | | | | |
| Oligo fwd | CACCGATGGACGGCGGGTTCCGTC | | | | | | | | | | |
| Oligo rev | AAACGACGGAACCCGCCGTCCATC | | | | | | | | | | |
| Chromosome | start | end | strand | MM | target_seq | PAM | alignment | distance | position | gene name | gene id |
| GU305914.1 | 49953 | 49975 | - | 0 | GATGGACGGCGGGTTCCGTC | TGG | \|\|\|\|\|\|\|\|[\|\|\|\|\|\|\|\|\|\|\|\|]PAM | NA | Intergenic | NA | NA |
| T30364 | GGACGGCGGGTTCCGTCTGGAGG | 1000 | | | | | | | | | |
| Oligo fwd | CACCGGACGGCGGGTTCCGTCTGG | | | | | | | | | | |
| Oligo rev | AAACCCAGACGGAACCCGCCGTCC | | | | | | | | | | |
| Chromosome | start | end | strand | MM | target_seq | PAM | alignment | distance | position | gene name | gene id |
| GU305914.1 | 49950 | 49972 | - | 0 | GGACGGCGGGTTCCGTCTGG | AGG | \|\|\|\|\|\|\|\|[\|\|\|\|\|\|\|\|\|\|\|\|]PAM | NA | Intergenic | NA | NA |
| T30365 | GCAAGAGCAGCGACACGAACTGG | 1000 | | | | | | | | | |
| Oligo fwd | CACCGCAAGAGCAGCGACACGAAC | | | | | | | | | | |
| Oligo rev | AAACGTTCGTGTCGCTGCTCTTGC | | | | | | | | | | |
| Chromosome | start | end | strand | MM | target_seq | PAM | alignment | distance | position | gene name | gene id |
| GU305914.1 | 49918 | 49940 | - | 0 | GCAAGAGCAGCGACACGAAC | TGG | \|\|\|\|\|\|\|\|[\|\|\|\|\|\|\|\|\|\|\|\|]PAM | NA | Intergenic | NA | NA |
| T30366 | CAAGAGCAGCGACACGAACTGGG | 1000 | | | | | | | | | |
| Oligo fwd | CACCCAAGAGCAGCGACACGAACT | | | | | | | | | | |
| Oligo rev | AAACAGTTCGTGTCGCTGCTCTTG | | | | | | | | | | |
| Chromosome | start | end | strand | MM | target_seq | PAM | alignment | distance | position | gene name | gene id |
| GU305914.1 | 49917 | 49939 | - | 0 | CAAGAGCAGCGACACGAACT | GGG | \|\|\|\|\|\|\|\|[\|\|\|\|\|\|\|\|\|\|\|\|]PAM | NA | Intergenic | NA | NA |
| T30367 | CAGCGACACGAACTGGGTTTCGG | 1000 | | | | | | | | | |
| Oligo fwd | CACCCAGCGACACGAACTGGGTTT | | | | | | | | | | |
| Oligo rev | AAACAAACCCAGTTCGTGTCGCTG | | | | | | | | | | |
| Chromosome | start | end | strand | MM | target_seq | PAM | alignment | distance | position | gene name | gene id |
| GU305914.1 | 49911 | 49933 | - | 0 | CAGCGACACGAACTGGGTTT | CGG | \|\|\|\|\|\|\|\|[\|\|\|\|\|\|\|\|\|\|\|\|]PAM | NA | Intergenic | NA | NA |
| T30368 | ACGAACTGGGTTTCGGACGACGG | 1000 | | | | | | | | | |
| Oligo fwd | CACCACGAACTGGGTTTCGGACGA | | | | | | | | | | |
| Oligo rev | AAACTCGTCCGAAACCCAGTTCGT | | | | | | | | | | |
| Chromosome | start | end | strand | MM | target_seq | PAM | alignment | distance | position | gene name | gene id |
| GU305914.1 | 49904 | 49926 | - | 0 | ACGAACTGGGTTTCGGACGA | CGG | \|\|\|\|\|\|\|\|[\|\|\|\|\|\|\|\|\|\|\|\|]PAM | NA | Intergenic | NA | NA |
| T30369 | CGACGGAAATCACACCGAGACGG | 1000 | | | | | | | | | |
| Oligo fwd | CACCCGACGGAAATCACACCGAGA | | | | | | | | | | |
| Oligo rev | AAACTCTCGGTGTGATTTCCGTCG | | | | | | | | | | |
| Chromosome | start | end | strand | MM | target_seq | PAM | alignment | distance | position | gene name | gene id |
| GU305914.1 | 49887 | 49909 | - | 0 | CGACGGAAATCACACCGAGA | CGG | \|\|\|\|\|\|\|\|[\|\|\|\|\|\|\|\|\|\|\|\|]PAM | NA | Intergenic | NA | NA |
| T30370 | CCTGATGAAAAACATTAGTCTGG | 1000 | | | | | | | | | |
| Oligo fwd | CACCCCTGATGAAAAACATTAGTC | | | | | | | | | | |
| Oligo rev | AAACGACTAATGTTTTTCATCAGG | | | | | | | | | | |
| Chromosome | start | end | strand | MM | target_seq | PAM | alignment | distance | position | gene name | gene id |
| GU305914.1 | 49857 | 49879 | - | 0 | CCTGATGAAAAACATTAGTC | TGG | \|\|\|\|\|\|\|\|[\|\|\|\|\|\|\|\|\|\|\|\|]PAM | NA | Intergenic | NA | NA |
| T30371 | CTGCTGCGCGCCGTCAGTGTCGG | 1000 | | | | | | | | | |
| Oligo fwd | CACCCTGCTGCGCGCCGTCAGTGT | | | | | | | | | | |
| Oligo rev | AAACACACTGACGGCGCGCAGCAG | | | | | | | | | | |
| Chromosome | start | end | strand | MM | target_seq | PAM | alignment | distance | position | gene name | gene id |
| GU305914.1 | 49805 | 49827 | - | 0 | CTGCTGCGCGCCGTCAGTGT | CGG | \|\|\|\|\|\|\|\|[\|\|\|\|\|\|\|\|\|\|\|\|]PAM | NA | Intergenic | NA | NA |
| T30372 | GCGCCGTCAGTGTCGGACACCGG | 1000 | | | | | | | | | |
| Oligo fwd | CACCGCGCCGTCAGTGTCGGACAC | | | | | | | | | | |
| Oligo rev | AAACGTGTCCGACACTGACGGCGC | | | | | | | | | | |
| Chromosome | start | end | strand | MM | target_seq | PAM | alignment | distance | position | gene name | gene id |
| GU305914.1 | 49798 | 49820 | - | 0 | GCGCCGTCAGTGTCGGACAC | CGG | \|\|\|\|\|\|\|\|[\|\|\|\|\|\|\|\|\|\|\|\|]PAM | NA | Intergenic | NA | NA |
| T30373 | ACACCGGTCCGAACTCTCGCCGG | 1000 | | | | | | | | | |
| Oligo fwd | CACCACACCGGTCCGAACTCTCGC | | | | | | | | | | |
| Oligo rev | AAACGCGAGAGTTCGGACCGGTGT | | | | | | | | | | |
| Chromosome | start | end | strand | MM | target_seq | PAM | alignment | distance | position | gene name | gene id |
| GU305914.1 | 49782 | 49804 | - | 0 | ACACCGGTCCGAACTCTCGC | CGG | \|\|\|\|\|\|\|\|[\|\|\|\|\|\|\|\|\|\|\|\|]PAM | NA | Intergenic | NA | NA |
| T30374 | CACCGGTCCGAACTCTCGCCGGG | 1000 | | | | | | | | | |
| Oligo fwd | CACCCACCGGTCCGAACTCTCGCC | | | | | | | | | | |
| Oligo rev | AAACGGCGAGAGTTCGGACCGGTG | | | | | | | | | | |
| Chromosome | start | end | strand | MM | target_seq | PAM | alignment | distance | position | gene name | gene id |
| GU305914.1 | 49781 | 49803 | - | 0 | CACCGGTCCGAACTCTCGCC | GGG | \|\|\|\|\|\|\|\|[\|\|\|\|\|\|\|\|\|\|\|\|]PAM | NA | Intergenic | NA | NA |
| T30375 | CGAACTCTCGCCGGGCATCATGG | 1000 | | | | | | | | | |
| Oligo fwd | CACCCGAACTCTCGCCGGGCATCA | | | | | | | | | | |
| Oligo rev | AAACTGATGCCCGGCGAGAGTTCG | | | | | | | | | | |
| Chromosome | start | end | strand | MM | target_seq | PAM | alignment | distance | position | gene name | gene id |
| GU305914.1 | 49773 | 49795 | - | 0 | CGAACTCTCGCCGGGCATCA | TGG | \|\|\|\|\|\|\|\|[\|\|\|\|\|\|\|\|\|\|\|\|]PAM | NA | Intergenic | NA | NA |
| T30376 | TCTCGCCGGGCATCATGGAGAGG | 1000 | | | | | | | | | |
| Oligo fwd | CACCTCTCGCCGGGCATCATGGAG | | | | | | | | | | |
| Oligo rev | AAACCTCCATGATGCCCGGCGAGA | | | | | | | | | | |
| Chromosome | start | end | strand | MM | target_seq | PAM | alignment | distance | position | gene name | gene id |

| | | | | | | | | | | | |
|---|---|---|---|---|---|---|---|---|---|---|---|
| GU305914.1 | 49768 | 49790 | - | 0 | TCTCGCCGGGCATCATGGAG | AGG | \|\|\|\|\|\|\|\|[\|\|\|\|\|\|\|\|\|\|\|\|]PAM | NA | Intergenic | NA | NA |
| T30377 | TGGAGAGGACCGCCGTCATCAGG | 1000 | | | | | | | | | |
| Oligo fwd | CACCTGGAGAGGACCGCCGTCATC | | | | | | | | | | |
| Oligo rev | AAACGATGACGGCGGTCCTCTCCA | | | | | | | | | | |
| Chromosome | start | end | strand | MM | target_seq | PAM | alignment | distance | position | gene name | gene id |
| GU305914.1 | 49753 | 49775 | - | 0 | TGGAGAGGACCGCCGTCATC | AGG | \|\|\|\|\|\|\|\|[\|\|\|\|\|\|\|\|\|\|\|\|]PAM | NA | Intergenic | NA | NA |
| T30378 | AGAGGACCGCCGTCATCAGGAGG | 1000 | | | | | | | | | |
| Oligo fwd | CACCAGAGGACCGCCGTCATCAGG | | | | | | | | | | |
| Oligo rev | AAACCCTGATGACGGCGGTCCTCT | | | | | | | | | | |
| Chromosome | start | end | strand | MM | target_seq | PAM | alignment | distance | position | gene name | gene id |
| GU305914.1 | 49750 | 49772 | - | 0 | AGAGGACCGCCGTCATCAGG | AGG | \|\|\|\|\|\|\|\|[\|\|\|\|\|\|\|\|\|\|\|\|]PAM | NA | Intergenic | NA | NA |
| T30379 | GAGGACCGCCGTCATCAGGAGGG | 1000 | | | | | | | | | |
| Oligo fwd | CACCGAGGACCGCCGTCATCAGGA | | | | | | | | | | |
| Oligo rev | AAACTCCTGATGACGGCGGTCCTC | | | | | | | | | | |
| Chromosome | start | end | strand | MM | target_seq | PAM | alignment | distance | position | gene name | gene id |
| GU305914.1 | 49749 | 49771 | - | 0 | GAGGACCGCCGTCATCAGGA | GGG | \|\|\|\|\|\|\|\|[\|\|\|\|\|\|\|\|\|\|\|\|]PAM | NA | Intergenic | NA | NA |
| T30380 | ATCAGGAGGGACGCCTGCTCCGG | 1000 | | | | | | | | | |
| Oligo fwd | CACCATCAGGAGGGACGCCTGCTC | | | | | | | | | | |
| Oligo rev | AAACGAGCAGGCGTCCCTCCTGAT | | | | | | | | | | |
| Chromosome | start | end | strand | MM | target_seq | PAM | alignment | distance | position | gene name | gene id |
| GU305914.1 | 49736 | 49758 | - | 0 | ATCAGGAGGGACGCCTGCTC | CGG | \|\|\|\|\|\|\|\|[\|\|\|\|\|\|\|\|\|\|\|\|]PAM | NA | Intergenic | NA | NA |
| T30381 | AGGAGGGACGCCTGCTCCGGCGG | 1000 | | | | | | | | | |
| Oligo fwd | CACCAGGAGGGACGCCTGCTCCGG | | | | | | | | | | |
| Oligo rev | AAACCCGGAGCAGGCGTCCCTCCT | | | | | | | | | | |
| Chromosome | start | end | strand | MM | target_seq | PAM | alignment | distance | position | gene name | gene id |
| GU305914.1 | 49733 | 49755 | - | 0 | AGGAGGGACGCCTGCTCCGG | CGG | \|\|\|\|\|\|\|\|[\|\|\|\|\|\|\|\|\|\|\|\|]PAM | NA | Intergenic | NA | NA |
| T30382 | CTGCTCCGGCGGCACCGCCACGG | 1000 | | | | | | | | | |
| Oligo fwd | CACCCTGCTCCGGCGGCACCGCCA | | | | | | | | | | |
| Oligo rev | AAACTGGCGGTGCCGCCGGAGCAG | | | | | | | | | | |
| Chromosome | start | end | strand | MM | target_seq | PAM | alignment | distance | position | gene name | gene id |
| GU305914.1 | 49722 | 49744 | - | 0 | CTGCTCCGGCGGCACCGCCA | CGG | \|\|\|\|\|\|\|\|[\|\|\|\|\|\|\|\|\|\|\|\|]PAM | NA | Intergenic | NA | NA |
| T30383 | TGCTCCGGCGGCACCGCCACGGG | 1000 | | | | | | | | | |
| Oligo fwd | CACCTGCTCCGGCGGCACCGCCAC | | | | | | | | | | |
| Oligo rev | AAACGTGGCGGTGCCGCCGGAGCA | | | | | | | | | | |
| Chromosome | start | end | strand | MM | target_seq | PAM | alignment | distance | position | gene name | gene id |
| GU305914.1 | 49721 | 49743 | - | 0 | TGCTCCGGCGGCACCGCCAC | GGG | \|\|\|\|\|\|\|\|[\|\|\|\|\|\|\|\|\|\|\|\|]PAM | NA | Intergenic | NA | NA |
| T30384 | GCGGCACCGCCACGGGAAACTGG | 1000 | | | | | | | | | |
| Oligo fwd | CACCGCGGCACCGCCACGGGAAAC | | | | | | | | | | |
| Oligo rev | AAACGTTTCCCGTGGCGGTGCCGC | | | | | | | | | | |
| Chromosome | start | end | strand | MM | target_seq | PAM | alignment | distance | position | gene name | gene id |
| GU305914.1 | 49714 | 49736 | - | 0 | GCGGCACCGCCACGGGAAAC | TGG | \|\|\|\|\|\|\|\|[\|\|\|\|\|\|\|\|\|\|\|\|]PAM | NA | Intergenic | NA | NA |
| T30385 | CGGGAAACTGGTCCACCGTCTGG | 1000 | | | | | | | | | |
| Oligo fwd | CACCCGGGAAACTGGTCCACCGTC | | | | | | | | | | |
| Oligo rev | AAACGACGGTGGACCAGTTTCCCG | | | | | | | | | | |
| Chromosome | start | end | strand | MM | target_seq | PAM | alignment | distance | position | gene name | gene id |
| GU305914.1 | 49702 | 49724 | - | 0 | CGGGAAACTGGTCCACCGTC | TGG | \|\|\|\|\|\|\|\|[\|\|\|\|\|\|\|\|\|\|\|\|]PAM | NA | Intergenic | NA | NA |
| T30386 | GGGAAACTGGTCCACCGTCTGGG | 1000 | | | | | | | | | |
| Oligo fwd | CACCGGGAAACTGGTCCACCGTCT | | | | | | | | | | |
| Oligo rev | AAACAGACGGTGGACCAGTTTCCC | | | | | | | | | | |
| Chromosome | start | end | strand | MM | target_seq | PAM | alignment | distance | position | gene name | gene id |
| GU305914.1 | 49701 | 49723 | - | 0 | GGGAAACTGGTCCACCGTCT | GGG | \|\|\|\|\|\|\|\|[\|\|\|\|\|\|\|\|\|\|\|\|]PAM | NA | Intergenic | NA | NA |
| T30387 | GGAAACTGGTCCACCGTCTGGGG | 1000 | | | | | | | | | |
| Oligo fwd | CACCGGAAACTGGTCCACCGTCTG | | | | | | | | | | |
| Oligo rev | AAACCAGACGGTGGACCAGTTTCC | | | | | | | | | | |
| Chromosome | start | end | strand | MM | target_seq | PAM | alignment | distance | position | gene name | gene id |
| GU305914.1 | 49700 | 49722 | - | 0 | GGAAACTGGTCCACCGTCTG | GGG | \|\|\|\|\|\|\|\|[\|\|\|\|\|\|\|\|\|\|\|\|]PAM | NA | Intergenic | NA | NA |
| T30388 | GGTCCACCGTCTGGGGCAACTGG | 1000 | | | | | | | | | |
| Oligo fwd | CACCGGTCCACCGTCTGGGGCAAC | | | | | | | | | | |
| Oligo rev | AAACGTTGCCCCAGACGGTGGACC | | | | | | | | | | |
| Chromosome | start | end | strand | MM | target_seq | PAM | alignment | distance | position | gene name | gene id |
| GU305914.1 | 49693 | 49715 | - | 0 | GGTCCACCGTCTGGGGCAAC | TGG | \|\|\|\|\|\|\|\|[\|\|\|\|\|\|\|\|\|\|\|\|]PAM | NA | Intergenic | NA | NA |
| T30389 | AGTACTGCGAGATGAACGAGCGG | 1000 | | | | | | | | | |
| Oligo fwd | CACCAGTACTGCGAGATGAACGAG | | | | | | | | | | |
| Oligo rev | AAACCTCGTTCATCTCGCAGTACT | | | | | | | | | | |
| Chromosome | start | end | strand | MM | target_seq | PAM | alignment | distance | position | gene name | gene id |
| GU305914.1 | 49666 | 49688 | - | 0 | AGTACTGCGAGATGAACGAG | CGG | \|\|\|\|\|\|\|\|[\|\|\|\|\|\|\|\|\|\|\|\|]PAM | NA | Intergenic | NA | NA |
| T30390 | AGATGAACGAGCGGTTCGTCAGG | 1000 | | | | | | | | | |
| Oligo fwd | CACCAGATGAACGAGCGGTTCGTC | | | | | | | | | | |
| Oligo rev | AAACGACGAACCGCTCGTTCATCT | | | | | | | | | | |
| Chromosome | start | end | strand | MM | target_seq | PAM | alignment | distance | position | gene name | gene id |
| GU305914.1 | 49657 | 49679 | - | 0 | AGATGAACGAGCGGTTCGTC | AGG | \|\|\|\|\|\|\|\|[\|\|\|\|\|\|\|\|\|\|\|\|]PAM | NA | Intergenic | NA | NA |
| T30391 | GATGAACGAGCGGTTCGTCAGGG | 1000 | | | | | | | | | |
| Oligo fwd | CACCGATGAACGAGCGGTTCGTCA | | | | | | | | | | |
| Oligo rev | AAACTGACGAACCGCTCGTTCATC | | | | | | | | | | |

<table>
<tr><td>Chromosome</td><td>start</td><td>end</td><td>strand</td><td>MM</td><td>target_seq</td><td>PAM</td><td>alignment</td><td>distance</td><td>position</td><td>gene name</td><td>gene id</td></tr>
<tr><td>GU305914.1</td><td>49656</td><td>49678</td><td>-</td><td>0</td><td>GATGAACGAGCGGTTCGTCA</td><td>GGG</td><td>||||||||[||||||||||||]PAM</td><td>NA</td><td>Intergenic</td><td>NA</td><td>NA</td></tr>
<tr><td>T30392</td><td>CGAGCGGTTCGTCAGGGACCCGG</td><td>1000</td><td></td><td></td><td></td><td></td><td></td><td></td><td></td><td></td><td></td></tr>
<tr><td>Oligo fwd</td><td>CACCCGAGCGGTTCGTCAGGGACC</td><td></td><td></td><td></td><td></td><td></td><td></td><td></td><td></td><td></td><td></td></tr>
<tr><td>Oligo rev</td><td>AAACGGTCCCTGACGAACCGCTCG</td><td></td><td></td><td></td><td></td><td></td><td></td><td></td><td></td><td></td><td></td></tr>
<tr><td>Chromosome</td><td>start</td><td>end</td><td>strand</td><td>MM</td><td>target_seq</td><td>PAM</td><td>alignment</td><td>distance</td><td>position</td><td>gene name</td><td>gene id</td></tr>
<tr><td>GU305914.1</td><td>49650</td><td>49672</td><td>-</td><td>0</td><td>CGAGCGGTTCGTCAGGGACC</td><td>CGG</td><td>||||||||[||||||||||||]PAM</td><td>NA</td><td>Intergenic</td><td>NA</td><td>NA</td></tr>
<tr><td>T30393</td><td>GAGCGGTTCGTCAGGGACCCGGG</td><td>1000</td><td></td><td></td><td></td><td></td><td></td><td></td><td></td><td></td><td></td></tr>
<tr><td>Oligo fwd</td><td>CACCGAGCGGTTCGTCAGGGACCC</td><td></td><td></td><td></td><td></td><td></td><td></td><td></td><td></td><td></td><td></td></tr>
<tr><td>Oligo rev</td><td>AAACGGGTCCCTGACGAACCGCTC</td><td></td><td></td><td></td><td></td><td></td><td></td><td></td><td></td><td></td><td></td></tr>
<tr><td>Chromosome</td><td>start</td><td>end</td><td>strand</td><td>MM</td><td>target_seq</td><td>PAM</td><td>alignment</td><td>distance</td><td>position</td><td>gene name</td><td>gene id</td></tr>
<tr><td>GU305914.1</td><td>49649</td><td>49671</td><td>-</td><td>0</td><td>GAGCGGTTCGTCAGGGACCC</td><td>GGG</td><td>||||||||[||||||||||||]PAM</td><td>NA</td><td>Intergenic</td><td>NA</td><td>NA</td></tr>
<tr><td>T30394</td><td>GGCTTTCTCGTCGCGATGCACGG</td><td>1000</td><td></td><td></td><td></td><td></td><td></td><td></td><td></td><td></td><td></td></tr>
<tr><td>Oligo fwd</td><td>CACCGGCTTTCTCGTCGCGATGCA</td><td></td><td></td><td></td><td></td><td></td><td></td><td></td><td></td><td></td><td></td></tr>
<tr><td>Oligo rev</td><td>AAACTGCATCGCGACGAGAAAGCC</td><td></td><td></td><td></td><td></td><td></td><td></td><td></td><td></td><td></td><td></td></tr>
<tr><td>Chromosome</td><td>start</td><td>end</td><td>strand</td><td>MM</td><td>target_seq</td><td>PAM</td><td>alignment</td><td>distance</td><td>position</td><td>gene name</td><td>gene id</td></tr>
<tr><td>GU305914.1</td><td>49628</td><td>49650</td><td>-</td><td>0</td><td>GGCTTTCTCGTCGCGATGCA</td><td>CGG</td><td>||||||||[||||||||||||]PAM</td><td>NA</td><td>Intergenic</td><td>NA</td><td>NA</td></tr>
<tr><td>T30395</td><td>CGTCGCGATGCACGGCTACGTGG</td><td>1000</td><td></td><td></td><td></td><td></td><td></td><td></td><td></td><td></td><td></td></tr>
<tr><td>Oligo fwd</td><td>CACCCGTCGCGATGCACGGCTACG</td><td></td><td></td><td></td><td></td><td></td><td></td><td></td><td></td><td></td><td></td></tr>
<tr><td>Oligo rev</td><td>AAACCGTAGCCGTGCATCGCGACG</td><td></td><td></td><td></td><td></td><td></td><td></td><td></td><td></td><td></td><td></td></tr>
<tr><td>Chromosome</td><td>start</td><td>end</td><td>strand</td><td>MM</td><td>target_seq</td><td>PAM</td><td>alignment</td><td>distance</td><td>position</td><td>gene name</td><td>gene id</td></tr>
<tr><td>GU305914.1</td><td>49620</td><td>49642</td><td>-</td><td>0</td><td>CGTCGCGATGCACGGCTACG</td><td>TGG</td><td>||||||||[||||||||||||]PAM</td><td>NA</td><td>Intergenic</td><td>NA</td><td>NA</td></tr>
<tr><td>T30396</td><td>GCGATGCACGGCTACGTGGCCGG</td><td>1000</td><td></td><td></td><td></td><td></td><td></td><td></td><td></td><td></td><td></td></tr>
<tr><td>Oligo fwd</td><td>CACCGCGATGCACGGCTACGTGGC</td><td></td><td></td><td></td><td></td><td></td><td></td><td></td><td></td><td></td><td></td></tr>
<tr><td>Oligo rev</td><td>AAACGCCACGTAGCCGTGCATCGC</td><td></td><td></td><td></td><td></td><td></td><td></td><td></td><td></td><td></td><td></td></tr>
<tr><td>Chromosome</td><td>start</td><td>end</td><td>strand</td><td>MM</td><td>target_seq</td><td>PAM</td><td>alignment</td><td>distance</td><td>position</td><td>gene name</td><td>gene id</td></tr>
<tr><td>GU305914.1</td><td>49616</td><td>49638</td><td>-</td><td>0</td><td>GCGATGCACGGCTACGTGGC</td><td>CGG</td><td>||||||||[||||||||||||]PAM</td><td>NA</td><td>Intergenic</td><td>NA</td><td>NA</td></tr>
<tr><td>T30397</td><td>CGGCGATCCCCTGCCCAAGATGG</td><td>1000</td><td></td><td></td><td></td><td></td><td></td><td></td><td></td><td></td><td></td></tr>
<tr><td>Oligo fwd</td><td>CACCCGGCGATCCCCTGCCCAAGA</td><td></td><td></td><td></td><td></td><td></td><td></td><td></td><td></td><td></td><td></td></tr>
<tr><td>Oligo rev</td><td>AAACTCTTGGGCAGGGGATCGCCG</td><td></td><td></td><td></td><td></td><td></td><td></td><td></td><td></td><td></td><td></td></tr>
<tr><td>Chromosome</td><td>start</td><td>end</td><td>strand</td><td>MM</td><td>target_seq</td><td>PAM</td><td>alignment</td><td>distance</td><td>position</td><td>gene name</td><td>gene id</td></tr>
<tr><td>GU305914.1</td><td>49596</td><td>49618</td><td>-</td><td>0</td><td>CGGCGATCCCCTGCCCAAGA</td><td>TGG</td><td>||||||||[||||||||||||]PAM</td><td>NA</td><td>Intergenic</td><td>NA</td><td>NA</td></tr>
<tr><td>T30398</td><td>GATCCCCTGCCCAAGATGGTCGG</td><td>1000</td><td></td><td></td><td></td><td></td><td></td><td></td><td></td><td></td><td></td></tr>
<tr><td>Oligo fwd</td><td>CACCGATCCCCTGCCCAAGATGGT</td><td></td><td></td><td></td><td></td><td></td><td></td><td></td><td></td><td></td><td></td></tr>
<tr><td>Oligo rev</td><td>AAACACCATCTTGGGCAGGGGATC</td><td></td><td></td><td></td><td></td><td></td><td></td><td></td><td></td><td></td><td></td></tr>
<tr><td>Chromosome</td><td>start</td><td>end</td><td>strand</td><td>MM</td><td>target_seq</td><td>PAM</td><td>alignment</td><td>distance</td><td>position</td><td>gene name</td><td>gene id</td></tr>
<tr><td>GU305914.1</td><td>49592</td><td>49614</td><td>-</td><td>0</td><td>GATCCCCTGCCCAAGATGGT</td><td>CGG</td><td>||||||||[||||||||||||]PAM</td><td>NA</td><td>Intergenic</td><td>NA</td><td>NA</td></tr>
<tr><td>T30399</td><td>GGCATGTTCTTCTTCGTGTTCGG</td><td>1000</td><td></td><td></td><td></td><td></td><td></td><td></td><td></td><td></td><td></td></tr>
<tr><td>Oligo fwd</td><td>CACCGGCATGTTCTTCTTCGTGTT</td><td></td><td></td><td></td><td></td><td></td><td></td><td></td><td></td><td></td><td></td></tr>
<tr><td>Oligo rev</td><td>AAACAACACGAAGAAGAACATGCC</td><td></td><td></td><td></td><td></td><td></td><td></td><td></td><td></td><td></td><td></td></tr>
<tr><td>Chromosome</td><td>start</td><td>end</td><td>strand</td><td>MM</td><td>target_seq</td><td>PAM</td><td>alignment</td><td>distance</td><td>position</td><td>gene name</td><td>gene id</td></tr>
<tr><td>GU305914.1</td><td>49571</td><td>49593</td><td>-</td><td>0</td><td>GGCATGTTCTTCTTCGTGTT</td><td>CGG</td><td>||||||||[||||||||||||]PAM</td><td>NA</td><td>Intergenic</td><td>NA</td><td>NA</td></tr>
<tr><td>GU305914.1</td><td>148394</td><td>148416</td><td>+</td><td>4</td><td>GTCATATTCTTCTTAGTGGT</td><td>GGG</td><td>|-|||-||[||||||-|||-|]PAM</td><td>NA</td><td>Intergenic</td><td>NA</td><td>NA</td></tr>
<tr><td>T30400</td><td>ATGTTCTTCTTCGTGTTCGGCGG</td><td>1000</td><td></td><td></td><td></td><td></td><td></td><td></td><td></td><td></td><td></td></tr>
<tr><td>Oligo fwd</td><td>CACCATGTTCTTCTTCGTGTTCGG</td><td></td><td></td><td></td><td></td><td></td><td></td><td></td><td></td><td></td><td></td></tr>
<tr><td>Oligo rev</td><td>AAACCCGAACACGAAGAAGAACAT</td><td></td><td></td><td></td><td></td><td></td><td></td><td></td><td></td><td></td><td></td></tr>
<tr><td>Chromosome</td><td>start</td><td>end</td><td>strand</td><td>MM</td><td>target_seq</td><td>PAM</td><td>alignment</td><td>distance</td><td>position</td><td>gene name</td><td>gene id</td></tr>
<tr><td>GU305914.1</td><td>49568</td><td>49590</td><td>-</td><td>0</td><td>ATGTTCTTCTTCGTGTTCGG</td><td>CGG</td><td>||||||||[||||||||||||]PAM</td><td>NA</td><td>Intergenic</td><td>NA</td><td>NA</td></tr>
<tr><td>T30401</td><td>CTTCTTCGTGTTCGGCGGCCTGG</td><td>1000</td><td></td><td></td><td></td><td></td><td></td><td></td><td></td><td></td><td></td></tr>
<tr><td>Oligo fwd</td><td>CACCCTTCTTCGTGTTCGGCGGCC</td><td></td><td></td><td></td><td></td><td></td><td></td><td></td><td></td><td></td><td></td></tr>
<tr><td>Oligo rev</td><td>AAACGGCCGCCGAACACGAAGAAG</td><td></td><td></td><td></td><td></td><td></td><td></td><td></td><td></td><td></td><td></td></tr>
<tr><td>Chromosome</td><td>start</td><td>end</td><td>strand</td><td>MM</td><td>target_seq</td><td>PAM</td><td>alignment</td><td>distance</td><td>position</td><td>gene name</td><td>gene id</td></tr>
<tr><td>GU305914.1</td><td>49563</td><td>49585</td><td>-</td><td>0</td><td>CTTCTTCGTGTTCGGCGGCC</td><td>TGG</td><td>||||||||[||||||||||||]PAM</td><td>NA</td><td>Intergenic</td><td>NA</td><td>NA</td></tr>
<tr><td>T30402</td><td>AGTCGCTGTTCAAAGATATGAGG</td><td>1000</td><td></td><td></td><td></td><td></td><td></td><td></td><td></td><td></td><td></td></tr>
<tr><td>Oligo fwd</td><td>CACCAGTCGCTGTTCAAAGATATG</td><td></td><td></td><td></td><td></td><td></td><td></td><td></td><td></td><td></td><td></td></tr>
<tr><td>Oligo rev</td><td>AAACCATATCTTTGAACAGCGACT</td><td></td><td></td><td></td><td></td><td></td><td></td><td></td><td></td><td></td><td></td></tr>
<tr><td>Chromosome</td><td>start</td><td>end</td><td>strand</td><td>MM</td><td>target_seq</td><td>PAM</td><td>alignment</td><td>distance</td><td>position</td><td>gene name</td><td>gene id</td></tr>
<tr><td>GU305914.1</td><td>49498</td><td>49520</td><td>-</td><td>0</td><td>AGTCGCTGTTCAAAGATATG</td><td>AGG</td><td>||||||||[||||||||||||]PAM</td><td>NA</td><td>Intergenic</td><td>NA</td><td>NA</td></tr>
<tr><td>T30403</td><td>GTCGCTGTTCAAAGATATGAGGG</td><td>1000</td><td></td><td></td><td></td><td></td><td></td><td></td><td></td><td></td><td></td></tr>
<tr><td>Oligo fwd</td><td>CACCGTCGCTGTTCAAAGATATGA</td><td></td><td></td><td></td><td></td><td></td><td></td><td></td><td></td><td></td><td></td></tr>
<tr><td>Oligo rev</td><td>AAACTCATATCTTTGAACAGCGAC</td><td></td><td></td><td></td><td></td><td></td><td></td><td></td><td></td><td></td><td></td></tr>
<tr><td>Chromosome</td><td>start</td><td>end</td><td>strand</td><td>MM</td><td>target_seq</td><td>PAM</td><td>alignment</td><td>distance</td><td>position</td><td>gene name</td><td>gene id</td></tr>
<tr><td>GU305914.1</td><td>49497</td><td>49519</td><td>-</td><td>0</td><td>GTCGCTGTTCAAAGATATGA</td><td>GGG</td><td>||||||||[||||||||||||]PAM</td><td>NA</td><td>Intergenic</td><td>NA</td><td>NA</td></tr>
<tr><td>T30404</td><td>TCGCTGTTCAAAGATATGAGGGG</td><td>1000</td><td></td><td></td><td></td><td></td><td></td><td></td><td></td><td></td><td></td></tr>
<tr><td>Oligo fwd</td><td>CACCTCGCTGTTCAAAGATATGAG</td><td></td><td></td><td></td><td></td><td></td><td></td><td></td><td></td><td></td><td></td></tr>
<tr><td>Oligo rev</td><td>AAACCTCATATCTTTGAACAGCGA</td><td></td><td></td><td></td><td></td><td></td><td></td><td></td><td></td><td></td><td></td></tr>
<tr><td>Chromosome</td><td>start</td><td>end</td><td>strand</td><td>MM</td><td>target_seq</td><td>PAM</td><td>alignment</td><td>distance</td><td>position</td><td>gene name</td><td>gene id</td></tr>
<tr><td>GU305914.1</td><td>49496</td><td>49518</td><td>-</td><td>0</td><td>TCGCTGTTCAAAGATATGAG</td><td>GGG</td><td>||||||||[||||||||||||]PAM</td><td>NA</td><td>Intergenic</td><td>NA</td><td>NA</td></tr>
<tr><td>T30405</td><td>GACAGTCGTCCCTCCGTCTCCGG</td><td>1000</td><td></td><td></td><td></td><td></td><td></td><td></td><td></td><td></td><td></td></tr>
<tr><td>Oligo fwd</td><td>CACCGACAGTCGTCCCTCCGTCTC</td><td></td><td></td><td></td><td></td><td></td><td></td><td></td><td></td><td></td><td></td></tr>
<tr><td>Oligo rev</td><td>AAACGAGACGGAGGGACGACTGTC</td><td></td><td></td><td></td><td></td><td></td><td></td><td></td><td></td><td></td><td></td></tr>
<tr><td>Chromosome</td><td>start</td><td>end</td><td>strand</td><td>MM</td><td>target_seq</td><td>PAM</td><td>alignment</td><td>distance</td><td>position</td><td>gene name</td><td>gene id</td></tr>
<tr><td>GU305914.1</td><td>49474</td><td>49496</td><td>-</td><td>0</td><td>GACAGTCGTCCCTCCGTCTC</td><td>CGG</td><td>||||||||[||||||||||||]PAM</td><td>NA</td><td>Intergenic</td><td>NA</td><td>NA</td></tr>
<tr><td>T30406</td><td>GTCCCTCCGTCTCCGGAACGAGG</td><td>1000</td><td></td><td></td><td></td><td></td><td></td><td></td><td></td><td></td><td></td></tr>
</table>

Oligo fwd CACCGTCCCTCCGTCTCCGGAACG
Oligo rev AAACCGTTCCGGAGACGGAGGGAC
Chromosome start end strand MM target_seq PAM alignment distance position gene name gene id
GU305914.1 49467 49489 - 0 GTCCCTCCGTCTCCGGAACG AGG ||||||||[||||||||||||]PAM NA Intergenic NA NA

T30407 CCGGAACGAGGATTTCGCCGCGG 1000
Oligo fwd CACCCCGGAACGAGGATTTCGCCG
Oligo rev AAACCGGCGAAATCCTCGTTCCGG
Chromosome start end strand MM target_seq PAM alignment distance position gene name gene id
GU305914.1 49455 49477 - 0 CCGGAACGAGGATTTCGCCG CGG ||||||||[||||||||||||]PAM NA Intergenic NA NA

T30408 TGCGTCAAAATGCTCGAATGCGG 1000
Oligo fwd CACCTGCGTCAAAATGCTCGAATG
Oligo rev AAACCATTCGAGCATTTTGACGCA
Chromosome start end strand MM target_seq PAM alignment distance position gene name gene id
GU305914.1 49377 49399 - 0 TGCGTCAAAATGCTCGAATG CGG ||||||||[||||||||||||]PAM NA Intergenic NA NA

T30409 CGTGCGTCATACACGCTATAAGG 1000
Oligo fwd CACCCGTGCGTCATACACGCTATA
Oligo rev AAACTATAGCGTGTATGACGCACG
Chromosome start end strand MM target_seq PAM alignment distance position gene name gene id
GU305914.1 49342 49364 - 0 CGTGCGTCATACACGCTATA AGG ||||||||[||||||||||||]PAM NA Intergenic NA NA

T30410 GCGTCATACACGCTATAAGGAGG 1000
Oligo fwd CACCGCGTCATACACGCTATAAGG
Oligo rev AAACCCTTATAGCGTGTATGACGC
Chromosome start end strand MM target_seq PAM alignment distance position gene name gene id
GU305914.1 49339 49361 - 0 GCGTCATACACGCTATAAGG AGG ||||||||[||||||||||||]PAM NA Intergenic NA NA

T30411 TATAAGGAGGTCACATTCTTTGG 1000
Oligo fwd CACCTATAAGGAGGTCACATTCTT
Oligo rev AAACAAGAATGTGACCTCCTTATA
Chromosome start end strand MM target_seq PAM alignment distance position gene name gene id
GU305914.1 49326 49348 - 0 TATAAGGAGGTCACATTCTT TGG ||||||||[||||||||||||]PAM NA Intergenic NA NA
GU305914.1 9894 9916 - 4 TATAAAGCGGTCGCATTCAT TGG |||||-|-[||||-|||||-|]PAM NA Intergenic NA NA

T30412 AGGAGGTCACATTCTTTGGACGG 1000
Oligo fwd CACCAGGAGGTCACATTCTTTGGA
Oligo rev AAACTCCAAAGAATGTGACCTCCT
Chromosome start end strand MM target_seq PAM alignment distance position gene name gene id
GU305914.1 49322 49344 - 0 AGGAGGTCACATTCTTTGGA CGG ||||||||[||||||||||||]PAM NA Intergenic NA NA

T30413 TTGGACGGAGCTCATTCTTTCGG 1000
Oligo fwd CACCTTGGACGGAGCTCATTCTTT
Oligo rev AAACAAAGAATGAGCTCCGTCCAA
Chromosome start end strand MM target_seq PAM alignment distance position gene name gene id
GU305914.1 49307 49329 - 0 TTGGACGGAGCTCATTCTTT CGG ||||||||[||||||||||||]PAM NA Intergenic NA NA

T30414 TGGACGGAGCTCATTCTTTCGGG 1000
Oligo fwd CACCTGGACGGAGCTCATTCTTTC
Oligo rev AAACGAAAGAATGAGCTCCGTCCA
Chromosome start end strand MM target_seq PAM alignment distance position gene name gene id
GU305914.1 49306 49328 - 0 TGGACGGAGCTCATTCTTTC GGG ||||||||[||||||||||||]PAM NA Intergenic NA NA

T30415 GGAGCTCATTCTTTCGGGAAAGG 1000
Oligo fwd CACCGGAGCTCATTCTTTCGGGAA
Oligo rev AAACTTCCCGAAAGAATGAGCTCC
Chromosome start end strand MM target_seq PAM alignment distance position gene name gene id
GU305914.1 49301 49323 - 0 GGAGCTCATTCTTTCGGGAA AGG ||||||||[||||||||||||]PAM NA Intergenic NA NA

T30416 GAGCTCATTCTTTCGGGAAAGGG 1000
Oligo fwd CACCGAGCTCATTCTTTCGGGAAA
Oligo rev AAACTTTCCCGAAAGAATGAGCTC
Chromosome start end strand MM target_seq PAM alignment distance position gene name gene id
GU305914.1 49300 49322 - 0 GAGCTCATTCTTTCGGGAAA GGG ||||||||[||||||||||||]PAM NA Intergenic NA NA

T30417 AGCTCATTCTTTCGGGAAAGGGG 1000
Oligo fwd CACCAGCTCATTCTTTCGGGAAAG
Oligo rev AAACCTTTCCCGAAAGAATGAGCT
Chromosome start end strand MM target_seq PAM alignment distance position gene name gene id
GU305914.1 49299 49321 - 0 AGCTCATTCTTTCGGGAAAG GGG ||||||||[||||||||||||]PAM NA Intergenic NA NA

T30418 TCATTCTTTCGGGAAAGGGGTGG 1000
Oligo fwd CACCTCATTCTTTCGGGAAAGGGG
Oligo rev AAACCCCCTTTCCCGAAAGAATGA
Chromosome start end strand MM target_seq PAM alignment distance position gene name gene id
GU305914.1 49296 49318 - 0 TCATTCTTTCGGGAAAGGGG TGG ||||||||[||||||||||||]PAM NA Intergenic NA NA

T30419 TTCTTTCGGGAAAGGGGTGGAGG 1000
Oligo fwd CACCTTCTTTCGGGAAAGGGGTGG
Oligo rev AAACCCACCCCTTTCCCGAAAGAA
Chromosome start end strand MM target_seq PAM alignment distance position gene name gene id
GU305914.1 49293 49315 - 0 TTCTTTCGGGAAAGGGGTGG AGG ||||||||[||||||||||||]PAM NA Intergenic NA NA

T30420 TTTCGGGAAAGGGGTGGAGGAGG 1000
Oligo fwd CACCTTTCGGGAAAGGGGTGGAGG
Oligo rev AAACCCTCCACCCCTTTCCCGAAA
Chromosome start end strand MM target_seq PAM alignment distance position gene name gene id
GU305914.1 49290 49312 - 0 TTTCGGGAAAGGGGTGGAGG AGG ||||||||[||||||||||||]PAM NA Intergenic NA NA

<table>
<tr><td>T30421</td><td>TTCGGGAAAGGGGTGGAGGAGGG</td><td>1000</td><td></td><td></td><td></td><td></td><td></td><td></td><td></td><td></td><td></td></tr>
<tr><td>Oligo fwd</td><td>CACCTTCGGGAAAGGGGTGGAGGA</td><td></td><td></td><td></td><td></td><td></td><td></td><td></td><td></td><td></td><td></td></tr>
<tr><td>Oligo rev</td><td>AAACTCCTCCACCCCTTTCCCGAA</td><td></td><td></td><td></td><td></td><td></td><td></td><td></td><td></td><td></td><td></td></tr>
<tr><td>Chromosome</td><td>start</td><td>end</td><td>strand</td><td>MM</td><td>target_seq</td><td>PAM</td><td>alignment</td><td>distance</td><td>position</td><td>gene name</td><td>gene id</td></tr>
<tr><td>GU305914.1</td><td>49289</td><td>49311</td><td>-</td><td>0</td><td>TTCGGGAAAGGGGTGGAGGA</td><td>GGG</td><td>||||||||[||||||||||||]PAM</td><td>NA</td><td>Intergenic</td><td>NA</td><td>NA</td></tr>
<tr><td>T30422</td><td>GAGGGTCGTTTGACAGTGAAAGG</td><td>1000</td><td></td><td></td><td></td><td></td><td></td><td></td><td></td><td></td><td></td></tr>
<tr><td>Oligo fwd</td><td>CACCGAGGGTCGTTTGACAGTGAA</td><td></td><td></td><td></td><td></td><td></td><td></td><td></td><td></td><td></td><td></td></tr>
<tr><td>Oligo rev</td><td>AAACTTCACTGTCAAACGACCCTC</td><td></td><td></td><td></td><td></td><td></td><td></td><td></td><td></td><td></td><td></td></tr>
<tr><td>Chromosome</td><td>start</td><td>end</td><td>strand</td><td>MM</td><td>target_seq</td><td>PAM</td><td>alignment</td><td>distance</td><td>position</td><td>gene name</td><td>gene id</td></tr>
<tr><td>GU305914.1</td><td>49271</td><td>49293</td><td>-</td><td>0</td><td>GAGGGTCGTTTGACAGTGAA</td><td>AGG</td><td>||||||||[||||||||||||]PAM</td><td>NA</td><td>Intergenic</td><td>NA</td><td>NA</td></tr>
<tr><td>T30423</td><td>TGAAAGGATGTATGAGCAAGAGG</td><td>1000</td><td></td><td></td><td></td><td></td><td></td><td></td><td></td><td></td><td></td></tr>
<tr><td>Oligo fwd</td><td>CACCTGAAAGGATGTATGAGCAAG</td><td></td><td></td><td></td><td></td><td></td><td></td><td></td><td></td><td></td><td></td></tr>
<tr><td>Oligo rev</td><td>AAACCTTGCTCATACATCCTTTCA</td><td></td><td></td><td></td><td></td><td></td><td></td><td></td><td></td><td></td><td></td></tr>
<tr><td>Chromosome</td><td>start</td><td>end</td><td>strand</td><td>MM</td><td>target_seq</td><td>PAM</td><td>alignment</td><td>distance</td><td>position</td><td>gene name</td><td>gene id</td></tr>
<tr><td>GU305914.1</td><td>49255</td><td>49277</td><td>-</td><td>0</td><td>TGAAAGGATGTATGAGCAAG</td><td>AGG</td><td>||||||||[||||||||||||]PAM</td><td>NA</td><td>Intergenic</td><td>NA</td><td>NA</td></tr>
<tr><td>T30424</td><td>ACCCGAGATCGTCCCTTACCAGG</td><td>1000</td><td></td><td></td><td></td><td></td><td></td><td></td><td></td><td></td><td></td></tr>
<tr><td>Oligo fwd</td><td>CACCACCCGAGATCGTCCCTTACC</td><td></td><td></td><td></td><td></td><td></td><td></td><td></td><td></td><td></td><td></td></tr>
<tr><td>Oligo rev</td><td>AAACGGTAAGGGACGATCTCGGGT</td><td></td><td></td><td></td><td></td><td></td><td></td><td></td><td></td><td></td><td></td></tr>
<tr><td>Chromosome</td><td>start</td><td>end</td><td>strand</td><td>MM</td><td>target_seq</td><td>PAM</td><td>alignment</td><td>distance</td><td>position</td><td>gene name</td><td>gene id</td></tr>
<tr><td>GU305914.1</td><td>49225</td><td>49247</td><td>-</td><td>0</td><td>ACCCGAGATCGTCCCTTACC</td><td>AGG</td><td>||||||||[||||||||||||]PAM</td><td>NA</td><td>Intergenic</td><td>NA</td><td>NA</td></tr>
<tr><td>T30425</td><td>ATCGTCCCTTACCAGGACATCGG</td><td>1000</td><td></td><td></td><td></td><td></td><td></td><td></td><td></td><td></td><td></td></tr>
<tr><td>Oligo fwd</td><td>CACCATCGTCCCTTACCAGGACAT</td><td></td><td></td><td></td><td></td><td></td><td></td><td></td><td></td><td></td><td></td></tr>
<tr><td>Oligo rev</td><td>AAACATGTCCTGGTAAGGGACGAT</td><td></td><td></td><td></td><td></td><td></td><td></td><td></td><td></td><td></td><td></td></tr>
<tr><td>Chromosome</td><td>start</td><td>end</td><td>strand</td><td>MM</td><td>target_seq</td><td>PAM</td><td>alignment</td><td>distance</td><td>position</td><td>gene name</td><td>gene id</td></tr>
<tr><td>GU305914.1</td><td>49218</td><td>49240</td><td>-</td><td>0</td><td>ATCGTCCCTTACCAGGACAT</td><td>CGG</td><td>||||||||[||||||||||||]PAM</td><td>NA</td><td>Intergenic</td><td>NA</td><td>NA</td></tr>
<tr><td>T30426</td><td>TCGTCCCTTACCAGGACATCGGG</td><td>1000</td><td></td><td></td><td></td><td></td><td></td><td></td><td></td><td></td><td></td></tr>
<tr><td>Oligo fwd</td><td>CACCTCGTCCCTTACCAGGACATC</td><td></td><td></td><td></td><td></td><td></td><td></td><td></td><td></td><td></td><td></td></tr>
<tr><td>Oligo rev</td><td>AAACGATGTCCTGGTAAGGGACGA</td><td></td><td></td><td></td><td></td><td></td><td></td><td></td><td></td><td></td><td></td></tr>
<tr><td>Chromosome</td><td>start</td><td>end</td><td>strand</td><td>MM</td><td>target_seq</td><td>PAM</td><td>alignment</td><td>distance</td><td>position</td><td>gene name</td><td>gene id</td></tr>
<tr><td>GU305914.1</td><td>49217</td><td>49239</td><td>-</td><td>0</td><td>TCGTCCCTTACCAGGACATC</td><td>GGG</td><td>||||||||[||||||||||||]PAM</td><td>NA</td><td>Intergenic</td><td>NA</td><td>NA</td></tr>
<tr><td>T30427</td><td>CGTCCCTTACCAGGACATCGGGG</td><td>1000</td><td></td><td></td><td></td><td></td><td></td><td></td><td></td><td></td><td></td></tr>
<tr><td>Oligo fwd</td><td>CACCCGTCCCTTACCAGGACATCG</td><td></td><td></td><td></td><td></td><td></td><td></td><td></td><td></td><td></td><td></td></tr>
<tr><td>Oligo rev</td><td>AAACCGATGTCCTGGTAAGGGACG</td><td></td><td></td><td></td><td></td><td></td><td></td><td></td><td></td><td></td><td></td></tr>
<tr><td>Chromosome</td><td>start</td><td>end</td><td>strand</td><td>MM</td><td>target_seq</td><td>PAM</td><td>alignment</td><td>distance</td><td>position</td><td>gene name</td><td>gene id</td></tr>
<tr><td>GU305914.1</td><td>49216</td><td>49238</td><td>-</td><td>0</td><td>CGTCCCTTACCAGGACATCG</td><td>GGG</td><td>||||||||[||||||||||||]PAM</td><td>NA</td><td>Intergenic</td><td>NA</td><td>NA</td></tr>
<tr><td>T30428</td><td>CCCTTACCAGGACATCGGGGAGG</td><td>1000</td><td></td><td></td><td></td><td></td><td></td><td></td><td></td><td></td><td></td></tr>
<tr><td>Oligo fwd</td><td>CACCCCCTTACCAGGACATCGGGG</td><td></td><td></td><td></td><td></td><td></td><td></td><td></td><td></td><td></td><td></td></tr>
<tr><td>Oligo rev</td><td>AAACCCCCGATGTCCTGGTAAGGG</td><td></td><td></td><td></td><td></td><td></td><td></td><td></td><td></td><td></td><td></td></tr>
<tr><td>Chromosome</td><td>start</td><td>end</td><td>strand</td><td>MM</td><td>target_seq</td><td>PAM</td><td>alignment</td><td>distance</td><td>position</td><td>gene name</td><td>gene id</td></tr>
<tr><td>GU305914.1</td><td>49213</td><td>49235</td><td>-</td><td>0</td><td>CCCTTACCAGGACATCGGGG</td><td>AGG</td><td>||||||||[||||||||||||]PAM</td><td>NA</td><td>Intergenic</td><td>NA</td><td>NA</td></tr>
<tr><td>T30429</td><td>ACATCGGGGAGGCTTATCGCTGG</td><td>1000</td><td></td><td></td><td></td><td></td><td></td><td></td><td></td><td></td><td></td></tr>
<tr><td>Oligo fwd</td><td>CACCACATCGGGGAGGCTTATCGC</td><td></td><td></td><td></td><td></td><td></td><td></td><td></td><td></td><td></td><td></td></tr>
<tr><td>Oligo rev</td><td>AAACGCGATAAGCCTCCCCGATGT</td><td></td><td></td><td></td><td></td><td></td><td></td><td></td><td></td><td></td><td></td></tr>
<tr><td>Chromosome</td><td>start</td><td>end</td><td>strand</td><td>MM</td><td>target_seq</td><td>PAM</td><td>alignment</td><td>distance</td><td>position</td><td>gene name</td><td>gene id</td></tr>
<tr><td>GU305914.1</td><td>49202</td><td>49224</td><td>-</td><td>0</td><td>ACATCGGGGAGGCTTATCGC</td><td>TGG</td><td>||||||||[||||||||||||]PAM</td><td>NA</td><td>Intergenic</td><td>NA</td><td>NA</td></tr>
<tr><td>T30430</td><td>TATCGCTGGCTCGCCAACATCGG</td><td>1000</td><td></td><td></td><td></td><td></td><td></td><td></td><td></td><td></td><td></td></tr>
<tr><td>Oligo fwd</td><td>CACCTATCGCTGGCTCGCCAACAT</td><td></td><td></td><td></td><td></td><td></td><td></td><td></td><td></td><td></td><td></td></tr>
<tr><td>Oligo rev</td><td>AAACATGTTGGCGAGCCAGCGATA</td><td></td><td></td><td></td><td></td><td></td><td></td><td></td><td></td><td></td><td></td></tr>
<tr><td>Chromosome</td><td>start</td><td>end</td><td>strand</td><td>MM</td><td>target_seq</td><td>PAM</td><td>alignment</td><td>distance</td><td>position</td><td>gene name</td><td>gene id</td></tr>
<tr><td>GU305914.1</td><td>49188</td><td>49210</td><td>-</td><td>0</td><td>TATCGCTGGCTCGCCAACAT</td><td>CGG</td><td>||||||||[||||||||||||]PAM</td><td>NA</td><td>Intergenic</td><td>NA</td><td>NA</td></tr>
<tr><td>T30431</td><td>CGCCAACATCGGCTTCAATCCGG</td><td>1000</td><td></td><td></td><td></td><td></td><td></td><td></td><td></td><td></td><td></td></tr>
<tr><td>Oligo fwd</td><td>CACCCGCCAACATCGGCTTCAATC</td><td></td><td></td><td></td><td></td><td></td><td></td><td></td><td></td><td></td><td></td></tr>
<tr><td>Oligo rev</td><td>AAACGATTGAAGCCGATGTTGGCG</td><td></td><td></td><td></td><td></td><td></td><td></td><td></td><td></td><td></td><td></td></tr>
<tr><td>Chromosome</td><td>start</td><td>end</td><td>strand</td><td>MM</td><td>target_seq</td><td>PAM</td><td>alignment</td><td>distance</td><td>position</td><td>gene name</td><td>gene id</td></tr>
<tr><td>GU305914.1</td><td>49177</td><td>49199</td><td>-</td><td>0</td><td>CGCCAACATCGGCTTCAATC</td><td>CGG</td><td>||||||||[||||||||||||]PAM</td><td>NA</td><td>Intergenic</td><td>NA</td><td>NA</td></tr>
<tr><td>T30432</td><td>CAACATCGGCTTCAATCCGGAGG</td><td>1000</td><td></td><td></td><td></td><td></td><td></td><td></td><td></td><td></td><td></td></tr>
<tr><td>Oligo fwd</td><td>CACCCAACATCGGCTTCAATCCGG</td><td></td><td></td><td></td><td></td><td></td><td></td><td></td><td></td><td></td><td></td></tr>
<tr><td>Oligo rev</td><td>AAACCCGGATTGAAGCCGATGTTG</td><td></td><td></td><td></td><td></td><td></td><td></td><td></td><td></td><td></td><td></td></tr>
<tr><td>Chromosome</td><td>start</td><td>end</td><td>strand</td><td>MM</td><td>target_seq</td><td>PAM</td><td>alignment</td><td>distance</td><td>position</td><td>gene name</td><td>gene id</td></tr>
<tr><td>GU305914.1</td><td>49174</td><td>49196</td><td>-</td><td>0</td><td>CAACATCGGCTTCAATCCGG</td><td>AGG</td><td>||||||||[||||||||||||]PAM</td><td>NA</td><td>Intergenic</td><td>NA</td><td>NA</td></tr>
<tr><td>T30433</td><td>AGCTATGTCTCCGAGCGCGCCGG</td><td>1000</td><td></td><td></td><td></td><td></td><td></td><td></td><td></td><td></td><td></td></tr>
<tr><td>Oligo fwd</td><td>CACCAGCTATGTCTCCGAGCGCGC</td><td></td><td></td><td></td><td></td><td></td><td></td><td></td><td></td><td></td><td></td></tr>
<tr><td>Oligo rev</td><td>AAACGCGCGCTCGGAGACATAGCT</td><td></td><td></td><td></td><td></td><td></td><td></td><td></td><td></td><td></td><td></td></tr>
<tr><td>Chromosome</td><td>start</td><td>end</td><td>strand</td><td>MM</td><td>target_seq</td><td>PAM</td><td>alignment</td><td>distance</td><td>position</td><td>gene name</td><td>gene id</td></tr>
<tr><td>GU305914.1</td><td>49143</td><td>49165</td><td>-</td><td>0</td><td>AGCTATGTCTCCGAGCGCGC</td><td>CGG</td><td>||||||||[||||||||||||]PAM</td><td>NA</td><td>Intergenic</td><td>NA</td><td>NA</td></tr>
<tr><td>T30434</td><td>GCGCGCCGGTACCTCCTTGCCGG</td><td>1000</td><td></td><td></td><td></td><td></td><td></td><td></td><td></td><td></td><td></td></tr>
<tr><td>Oligo fwd</td><td>CACCGCGCGCCGGTACCTCCTTGC</td><td></td><td></td><td></td><td></td><td></td><td></td><td></td><td></td><td></td><td></td></tr>
<tr><td>Oligo rev</td><td>AAACGCAAGGAGGTACCGGCGCGC</td><td></td><td></td><td></td><td></td><td></td><td></td><td></td><td></td><td></td><td></td></tr>
<tr><td>Chromosome</td><td>start</td><td>end</td><td>strand</td><td>MM</td><td>target_seq</td><td>PAM</td><td>alignment</td><td>distance</td><td>position</td><td>gene name</td><td>gene id</td></tr>
<tr><td>GU305914.1</td><td>49129</td><td>49151</td><td>-</td><td>0</td><td>GCGCGCCGGTACCTCCTTGC</td><td>CGG</td><td>||||||||[||||||||||||]PAM</td><td>NA</td><td>Intergenic</td><td>NA</td><td>NA</td></tr>
<tr><td>T30435</td><td>GTACCTCCTTGCCGGTCAGCTGG</td><td>1000</td><td></td><td></td><td></td><td></td><td></td><td></td><td></td><td></td><td></td></tr>
<tr><td>Oligo fwd</td><td>CACCGTACCTCCTTGCCGGTCAGC</td><td></td><td></td><td></td><td></td><td></td><td></td><td></td><td></td><td></td><td></td></tr>
<tr><td>Oligo rev</td><td>AAACGCTGACCGGCAAGGAGGTAC</td><td></td><td></td><td></td><td></td><td></td><td></td><td></td><td></td><td></td><td></td></tr>
<tr><td>Chromosome</td><td>start</td><td>end</td><td>strand</td><td>MM</td><td>target_seq</td><td>PAM</td><td>alignment</td><td>distance</td><td>position</td><td>gene name</td><td>gene id</td></tr>
</table>

| | | | | | | | | | | | |
|---|---|---|---|---|---|---|---|---|---|---|---|
| GU305914.1 | 49121 | 49143 | - | 0 | GTACCTCCTTGCCGGTCAGC | TGG | \|\|\|\|\|\|\|\|[\|\|\|\|\|\|\|\|\|\|\|\|]PAM | NA | Intergenic | NA | NA |
| T30436 | CTCCTTGCCGGTCAGCTGGCCGG | 1000 | | | | | | | | | |
| Oligo fwd | CACCCTCCTTGCCGGTCAGCTGGC | | | | | | | | | | |
| Oligo rev | AAACGCCAGCTGACCGGCAAGGAG | | | | | | | | | | |
| Chromosome | start | end | strand | MM | target_seq | PAM | alignment | distance | position | gene name | gene id |
| GU305914.1 | 49117 | 49139 | - | 0 | CTCCTTGCCGGTCAGCTGGC | CGG | \|\|\|\|\|\|\|\|[\|\|\|\|\|\|\|\|\|\|\|\|]PAM | NA | Intergenic | NA | NA |
| T30437 | TTGCCGGTCAGCTGGCCGGACGG | 1000 | | | | | | | | | |
| Oligo fwd | CACCTTGCCGGTCAGCTGGCCGGA | | | | | | | | | | |
| Oligo rev | AAACTCCGGCCAGCTGACCGGCAA | | | | | | | | | | |
| Chromosome | start | end | strand | MM | target_seq | PAM | alignment | distance | position | gene name | gene id |
| GU305914.1 | 49113 | 49135 | - | 0 | TTGCCGGTCAGCTGGCCGGA | CGG | \|\|\|\|\|\|\|\|[\|\|\|\|\|\|\|\|\|\|\|\|]PAM | NA | Intergenic | NA | NA |
| T30438 | TGCCGGTCAGCTGGCCGGACGGG | 1000 | | | | | | | | | |
| Oligo fwd | CACCTGCCGGTCAGCTGGCCGGAC | | | | | | | | | | |
| Oligo rev | AAACGTCCGGCCAGCTGACCGGCA | | | | | | | | | | |
| Chromosome | start | end | strand | MM | target_seq | PAM | alignment | distance | position | gene name | gene id |
| GU305914.1 | 49112 | 49134 | - | 0 | TGCCGGTCAGCTGGCCGGAC | GGG | \|\|\|\|\|\|\|\|[\|\|\|\|\|\|\|\|\|\|\|\|]PAM | NA | Intergenic | NA | NA |
| T30439 | GCCGGTCAGCTGGCCGGACGGGG | 1000 | | | | | | | | | |
| Oligo fwd | CACCGCCGGTCAGCTGGCCGGACG | | | | | | | | | | |
| Oligo rev | AAACCGTCCGGCCAGCTGACCGGC | | | | | | | | | | |
| Chromosome | start | end | strand | MM | target_seq | PAM | alignment | distance | position | gene name | gene id |
| GU305914.1 | 49111 | 49133 | - | 0 | GCCGGTCAGCTGGCCGGACG | GGG | \|\|\|\|\|\|\|\|[\|\|\|\|\|\|\|\|\|\|\|\|]PAM | NA | Intergenic | NA | NA |
| T30440 | CAGCTGGCCGGACGGGGTTTCGG | 1000 | | | | | | | | | |
| Oligo fwd | CACCCAGCTGGCCGGACGGGGTTT | | | | | | | | | | |
| Oligo rev | AAACAAACCCCGTCCGGCCAGCTG | | | | | | | | | | |
| Chromosome | start | end | strand | MM | target_seq | PAM | alignment | distance | position | gene name | gene id |
| GU305914.1 | 49105 | 49127 | - | 0 | CAGCTGGCCGGACGGGGTTT | CGG | \|\|\|\|\|\|\|\|[\|\|\|\|\|\|\|\|\|\|\|\|]PAM | NA | Intergenic | NA | NA |
| T30441 | GGCCGGACGGGGTTTCGGTCCGG | 1000 | | | | | | | | | |
| Oligo fwd | CACCGGCCGGACGGGGTTTCGGTC | | | | | | | | | | |
| Oligo rev | AAACGACCGAAACCCCGTCCGGCC | | | | | | | | | | |
| Chromosome | start | end | strand | MM | target_seq | PAM | alignment | distance | position | gene name | gene id |
| GU305914.1 | 49100 | 49122 | - | 0 | GGCCGGACGGGGTTTCGGTC | CGG | \|\|\|\|\|\|\|\|[\|\|\|\|\|\|\|\|\|\|\|\|]PAM | NA | Intergenic | NA | NA |
| GU305914.1 | 35355 | 35377 | - | 4 | CACCGGACGGCGTTTCGGGC | TGG | --\|\|\|\|\|\|[\|\|-\|\|\|\|\|\|\|-\|]PAM | NA | Intergenic | NA | NA |
| T30442 | ACGTATTTCTTGAAGAGATACGG | 1000 | | | | | | | | | |
| Oligo fwd | CACCACGTATTTCTTGAAGAGATA | | | | | | | | | | |
| Oligo rev | AAACTATCTCTTCAAGAAATACGT | | | | | | | | | | |
| Chromosome | start | end | strand | MM | target_seq | PAM | alignment | distance | position | gene name | gene id |
| GU305914.1 | 49056 | 49078 | - | 0 | ACGTATTTCTTGAAGAGATA | CGG | \|\|\|\|\|\|\|\|[\|\|\|\|\|\|\|\|\|\|\|\|]PAM | NA | Intergenic | NA | NA |
| T30443 | TATTTCTTGAAGAGATACGGTGG | 1000 | | | | | | | | | |
| Oligo fwd | CACCTATTTCTTGAAGAGATACGG | | | | | | | | | | |
| Oligo rev | AAACCCGTATCTCTTCAAGAAATA | | | | | | | | | | |
| Chromosome | start | end | strand | MM | target_seq | PAM | alignment | distance | position | gene name | gene id |
| GU305914.1 | 49053 | 49075 | - | 0 | TATTTCTTGAAGAGATACGG | TGG | \|\|\|\|\|\|\|\|[\|\|\|\|\|\|\|\|\|\|\|\|]PAM | NA | Intergenic | NA | NA |
| T30444 | TACGGTGGTCTGTGTGATGAAGG | 1000 | | | | | | | | | |
| Oligo fwd | CACCTACGGTGGTCTGTGTGATGA | | | | | | | | | | |
| Oligo rev | AAACTCATCACACAGACCACCGTA | | | | | | | | | | |
| Chromosome | start | end | strand | MM | target_seq | PAM | alignment | distance | position | gene name | gene id |
| GU305914.1 | 49038 | 49060 | - | 0 | TACGGTGGTCTGTGTGATGA | AGG | \|\|\|\|\|\|\|\|[\|\|\|\|\|\|\|\|\|\|\|\|]PAM | NA | Intergenic | NA | NA |
| T30445 | ATGAAGGTGAGTGCTACGTGCGG | 1000 | | | | | | | | | |
| Oligo fwd | CACCATGAAGGTGAGTGCTACGTG | | | | | | | | | | |
| Oligo rev | AAACCACGTAGCACTCACCTTCAT | | | | | | | | | | |
| Chromosome | start | end | strand | MM | target_seq | PAM | alignment | distance | position | gene name | gene id |
| GU305914.1 | 49022 | 49044 | - | 0 | ATGAAGGTGAGTGCTACGTG | CGG | \|\|\|\|\|\|\|\|[\|\|\|\|\|\|\|\|\|\|\|\|]PAM | NA | Intergenic | NA | NA |
| T30446 | AGGTGAGTGCTACGTGCGGACGG | 1000 | | | | | | | | | |
| Oligo fwd | CACCAGGTGAGTGCTACGTGCGGA | | | | | | | | | | |
| Oligo rev | AAACTCCGCACGTAGCACTCACCT | | | | | | | | | | |
| Chromosome | start | end | strand | MM | target_seq | PAM | alignment | distance | position | gene name | gene id |
| GU305914.1 | 49018 | 49040 | - | 0 | AGGTGAGTGCTACGTGCGGA | CGG | \|\|\|\|\|\|\|\|[\|\|\|\|\|\|\|\|\|\|\|\|]PAM | NA | Intergenic | NA | NA |
| T30447 | GCTACGTGCGGACGGACCCACGG | 1000 | | | | | | | | | |
| Oligo fwd | CACCGCTACGTGCGGACGGACCCA | | | | | | | | | | |
| Oligo rev | AAACTGGGTCCGTCCGCACGTAGC | | | | | | | | | | |
| Chromosome | start | end | strand | MM | target_seq | PAM | alignment | distance | position | gene name | gene id |
| GU305914.1 | 49010 | 49032 | - | 0 | GCTACGTGCGGACGGACCCA | CGG | \|\|\|\|\|\|\|\|[\|\|\|\|\|\|\|\|\|\|\|\|]PAM | NA | Intergenic | NA | NA |
| T30448 | ACGTGCGGACGGACCCACGGAGG | 1000 | | | | | | | | | |
| Oligo fwd | CACCACGTGCGGACGGACCCACGG | | | | | | | | | | |
| Oligo rev | AAACCCGTGGGTCCGTCCGCACGT | | | | | | | | | | |
| Chromosome | start | end | strand | MM | target_seq | PAM | alignment | distance | position | gene name | gene id |
| GU305914.1 | 49007 | 49029 | - | 0 | ACGTGCGGACGGACCCACGG | AGG | \|\|\|\|\|\|\|\|[\|\|\|\|\|\|\|\|\|\|\|\|]PAM | NA | Intergenic | NA | NA |
| T30449 | CGTGCGGACGGACCCACGGAGGG | 1000 | | | | | | | | | |
| Oligo fwd | CACCCGTGCGGACGGACCCACGGA | | | | | | | | | | |
| Oligo rev | AAACTCCGTGGGTCCGTCCGCACG | | | | | | | | | | |
| Chromosome | start | end | strand | MM | target_seq | PAM | alignment | distance | position | gene name | gene id |
| GU305914.1 | 49006 | 49028 | - | 0 | CGTGCGGACGGACCCACGGA | GGG | \|\|\|\|\|\|\|\|[\|\|\|\|\|\|\|\|\|\|\|\|]PAM | NA | Intergenic | NA | NA |
| T30450 | GTGCGGACGGACCCACGGAGGGG | 1000 | | | | | | | | | |
| Oligo fwd | CACCGTGCGGACGGACCCACGGAG | | | | | | | | | | |

| | | | | | | | | | | | |
|---|---|---|---|---|---|---|---|---|---|---|---|
| Oligo rev | AAACCTCCGTGGGTCCGTCCGCAC | | | | | | | | | | |
| Chromosome | start | end | strand | MM | target_seq | PAM | alignment | distance | position | gene name | gene id |
| GU305914.1 | 49005 | 49027 | - | 0 | GTGCGGACGGACCCACGGAG | GGG | \|\|\|\|\|\|\|\|[\|\|\|\|\|\|\|\|\|\|\|\|]PAM | NA | Intergenic | NA | NA |
| T30451 | CGGACGGACCCACGGAGGGGAGG | 1000 | | | | | | | | | |
| Oligo fwd | CACCCGGACGGACCCACGGAGGGG | | | | | | | | | | |
| Oligo rev | AAACCCCCTCCGTGGGTCCGTCCG | | | | | | | | | | |
| Chromosome | start | end | strand | MM | target_seq | PAM | alignment | distance | position | gene name | gene id |
| GU305914.1 | 49002 | 49024 | - | 0 | CGGACGGACCCACGGAGGGG | AGG | \|\|\|\|\|\|\|\|[\|\|\|\|\|\|\|\|\|\|\|\|]PAM | NA | Intergenic | NA | NA |
| T30452 | CGGACCCACGGAGGGGAGGATGG | 1000 | | | | | | | | | |
| Oligo fwd | CACCCGGACCCACGGAGGGGAGGA | | | | | | | | | | |
| Oligo rev | AAACTCCTCCCCTCCGTGGGTCCG | | | | | | | | | | |
| Chromosome | start | end | strand | MM | target_seq | PAM | alignment | distance | position | gene name | gene id |
| GU305914.1 | 48998 | 49020 | - | 0 | CGGACCCACGGAGGGGAGGA | TGG | \|\|\|\|\|\|\|\|[\|\|\|\|\|\|\|\|\|\|\|\|]PAM | NA | Intergenic | NA | NA |
| T30453 | CCCACGGAGGGGAGGATGGATGG | 1000 | | | | | | | | | |
| Oligo fwd | CACCCCCACGGAGGGGAGGATGGA | | | | | | | | | | |
| Oligo rev | AAACTCCATCCTCCCCTCCGTGGG | | | | | | | | | | |
| Chromosome | start | end | strand | MM | target_seq | PAM | alignment | distance | position | gene name | gene id |
| GU305914.1 | 48994 | 49016 | - | 0 | CCCACGGAGGGGAGGATGGA | TGG | \|\|\|\|\|\|\|\|[\|\|\|\|\|\|\|\|\|\|\|\|]PAM | NA | Intergenic | NA | NA |
| T30454 | CGGAGGGGAGGATGGATGGATGG | 1000 | | | | | | | | | |
| Oligo fwd | CACCCGGAGGGGAGGATGGATGGA | | | | | | | | | | |
| Oligo rev | AAACTCCATCCATCCTCCCCTCCG | | | | | | | | | | |
| Chromosome | start | end | strand | MM | target_seq | PAM | alignment | distance | position | gene name | gene id |
| GU305914.1 | 48990 | 49012 | - | 0 | CGGAGGGGAGGATGGATGGA | TGG | \|\|\|\|\|\|\|\|[\|\|\|\|\|\|\|\|\|\|\|\|]PAM | NA | Intergenic | NA | NA |
| T30455 | GGGGAGGATGGATGGATGGATGG | 1000 | | | | | | | | | |
| Oligo fwd | CACCGGGGAGGATGGATGGATGGA | | | | | | | | | | |
| Oligo rev | AAACTCCATCCATCCATCCTCCCC | | | | | | | | | | |
| Chromosome | start | end | strand | MM | target_seq | PAM | alignment | distance | position | gene name | gene id |
| GU305914.1 | 48986 | 49008 | - | 0 | GGGGAGGATGGATGGATGGA | TGG | \|\|\|\|\|\|\|\|[\|\|\|\|\|\|\|\|\|\|\|\|]PAM | NA | Intergenic | NA | NA |
| T30456 | ATGGATGGATGGATGCATGACGG | 1000 | | | | | | | | | |
| Oligo fwd | CACCATGGATGGATGGATGCATGA | | | | | | | | | | |
| Oligo rev | AAACTCATGCATCCATCCATCCAT | | | | | | | | | | |
| Chromosome | start | end | strand | MM | target_seq | PAM | alignment | distance | position | gene name | gene id |
| GU305914.1 | 48975 | 48997 | - | 0 | ATGGATGGATGGATGCATGA | CGG | \|\|\|\|\|\|\|\|[\|\|\|\|\|\|\|\|\|\|\|\|]PAM | NA | Intergenic | NA | NA |
| T30457 | TGGATGGATGGATGCATGACGGG | 1000 | | | | | | | | | |
| Oligo fwd | CACCTGGATGGATGGATGCATGAC | | | | | | | | | | |
| Oligo rev | AAACGTCATGCATCCATCCATCCA | | | | | | | | | | |
| Chromosome | start | end | strand | MM | target_seq | PAM | alignment | distance | position | gene name | gene id |
| GU305914.1 | 48974 | 48996 | - | 0 | TGGATGGATGGATGCATGAC | GGG | \|\|\|\|\|\|\|\|[\|\|\|\|\|\|\|\|\|\|\|\|]PAM | NA | Intergenic | NA | NA |
| T30458 | GGATGGATGGATGCATGACGGGG | 1000 | | | | | | | | | |
| Oligo fwd | CACCGGATGGATGGATGCATGACG | | | | | | | | | | |
| Oligo rev | AAACCGTCATGCATCCATCCATCC | | | | | | | | | | |
| Chromosome | start | end | strand | MM | target_seq | PAM | alignment | distance | position | gene name | gene id |
| GU305914.1 | 48973 | 48995 | - | 0 | GGATGGATGGATGCATGACG | GGG | \|\|\|\|\|\|\|\|[\|\|\|\|\|\|\|\|\|\|\|\|]PAM | NA | Intergenic | NA | NA |
| T30459 | TGGATGGATGCATGACGGGGTGG | 1000 | | | | | | | | | |
| Oligo fwd | CACCTGGATGGATGCATGACGGGG | | | | | | | | | | |
| Oligo rev | AAACCCCCGTCATGCATCCATCCA | | | | | | | | | | |
| Chromosome | start | end | strand | MM | target_seq | PAM | alignment | distance | position | gene name | gene id |
| GU305914.1 | 48970 | 48992 | - | 0 | TGGATGGATGCATGACGGGG | TGG | \|\|\|\|\|\|\|\|[\|\|\|\|\|\|\|\|\|\|\|\|]PAM | NA | Intergenic | NA | NA |
| T30460 | ATGGATGCATGACGGGGTGGCGG | 1000 | | | | | | | | | |
| Oligo fwd | CACCATGGATGCATGACGGGGTGG | | | | | | | | | | |
| Oligo rev | AAACCCACCCCGTCATGCATCCAT | | | | | | | | | | |
| Chromosome | start | end | strand | MM | target_seq | PAM | alignment | distance | position | gene name | gene id |
| GU305914.1 | 48967 | 48989 | - | 0 | ATGGATGCATGACGGGGTGG | CGG | \|\|\|\|\|\|\|\|[\|\|\|\|\|\|\|\|\|\|\|\|]PAM | NA | Intergenic | NA | NA |
| T30461 | GATGCATGACGGGGTGGCGGCGG | 1000 | | | | | | | | | |
| Oligo fwd | CACCGATGCATGACGGGGTGGCGG | | | | | | | | | | |
| Oligo rev | AAACCCGCCACCCCGTCATGCATC | | | | | | | | | | |
| Chromosome | start | end | strand | MM | target_seq | PAM | alignment | distance | position | gene name | gene id |
| GU305914.1 | 48964 | 48986 | - | 0 | GATGCATGACGGGGTGGCGG | CGG | \|\|\|\|\|\|\|\|[\|\|\|\|\|\|\|\|\|\|\|\|]PAM | NA | Intergenic | NA | NA |
| T30462 | GCATGACGGGGTGGCGGCGGCGG | 1000 | | | | | | | | | |
| Oligo fwd | CACCGCATGACGGGGTGGCGGCGG | | | | | | | | | | |
| Oligo rev | AAACCCGCCGCCACCCCGTCATGC | | | | | | | | | | |
| Chromosome | start | end | strand | MM | target_seq | PAM | alignment | distance | position | gene name | gene id |
| GU305914.1 | 48961 | 48983 | - | 0 | GCATGACGGGGTGGCGGCGG | CGG | \|\|\|\|\|\|\|\|[\|\|\|\|\|\|\|\|\|\|\|\|]PAM | NA | Intergenic | NA | NA |
| T30463 | CATGACGGGGTGGCGGCGGCGGG | 1000 | | | | | | | | | |
| Oligo fwd | CACCCATGACGGGGTGGCGGCGGC | | | | | | | | | | |
| Oligo rev | AAACGCCGCCGCCACCCCGTCATG | | | | | | | | | | |
| Chromosome | start | end | strand | MM | target_seq | PAM | alignment | distance | position | gene name | gene id |
| GU305914.1 | 48960 | 48982 | - | 0 | CATGACGGGGTGGCGGCGGC | GGG | \|\|\|\|\|\|\|\|[\|\|\|\|\|\|\|\|\|\|\|\|]PAM | NA | Intergenic | NA | NA |
| GU305914.1 | 221599 | 221621 | - | 4 | GAGGCCGTGGTGGCGGCGGC | AGG | -\|-\|-\|\|-[\|\|\|\|\|\|\|\|\|\|\|\|]PAM | NA | Intergenic | NA | NA |
| GU305914.1 | 132935 | 132957 | + | 4 | CCCGACGGGGTCGGGGCGGC | TGG | \|--\|\|\|\|\|[\|\|\|-\|-\|\|\|\|\|\|]PAM | NA | Intergenic | NA | NA |
| T30464 | ATGACGGGGTGGCGGCGGCGGGG | 1000 | | | | | | | | | |
| Oligo fwd | CACCATGACGGGGTGGCGGCGGCG | | | | | | | | | | |
| Oligo rev | AAACCGCCGCCGCCACCCCGTCAT | | | | | | | | | | |
| Chromosome | start | end | strand | MM | target_seq | PAM | alignment | distance | position | gene name | gene id |
| GU305914.1 | 48959 | 48981 | - | 0 | ATGACGGGGTGGCGGCGGCG | GGG | \|\|\|\|\|\|\|\|[\|\|\|\|\|\|\|\|\|\|\|\|]PAM | NA | Intergenic | NA | NA |

<table>
<tr><td>GU305914.1</td><td></td><td>221598</td><td>221620</td><td>-</td><td>4</td><td>AGGCCGTGGTGGCGGCGGCA</td><td>GGG</td><td>|-|-||-|[||||||||||||-]PAM</td><td>NA</td><td>Intergenic</td><td>NA</td><td>NA</td></tr>
<tr><td>T30465</td><td>GCGGCGGGGAGAGAAAACGTAGG</td><td></td><td>1000</td><td></td><td></td><td></td><td></td><td></td><td></td><td></td><td></td><td></td></tr>
<tr><td>Oligo fwd</td><td>CACCGCGGCGGGGAGAGAAAACGT</td><td></td><td></td><td></td><td></td><td></td><td></td><td></td><td></td><td></td><td></td><td></td></tr>
<tr><td>Oligo rev</td><td>AAACACGTTTTCTCTCCCCGCCGC</td><td></td><td></td><td></td><td></td><td></td><td></td><td></td><td></td><td></td><td></td><td></td></tr>
<tr><td>Chromosome</td><td>start</td><td></td><td>end</td><td>strand</td><td>MM</td><td>target_seq</td><td>PAM</td><td>alignment</td><td>distance</td><td>position</td><td>gene name</td><td>gene id</td></tr>
<tr><td>GU305914.1</td><td></td><td>48945</td><td>48967</td><td>-</td><td>0</td><td>GCGGCGGGGAGAGAAAACGT</td><td>AGG</td><td>||||||||[||||||||||||]PAM</td><td>NA</td><td>Intergenic</td><td>NA</td><td>NA</td></tr>
<tr><td>T30466</td><td>CGGCGGGGAGAGAAAACGTAGGG</td><td></td><td>1000</td><td></td><td></td><td></td><td></td><td></td><td></td><td></td><td></td><td></td></tr>
<tr><td>Oligo fwd</td><td>CACCCGGCGGGGAGAGAAAACGTA</td><td></td><td></td><td></td><td></td><td></td><td></td><td></td><td></td><td></td><td></td><td></td></tr>
<tr><td>Oligo rev</td><td>AAACTACGTTTTCTCTCCCCGCCG</td><td></td><td></td><td></td><td></td><td></td><td></td><td></td><td></td><td></td><td></td><td></td></tr>
<tr><td>Chromosome</td><td>start</td><td></td><td>end</td><td>strand</td><td>MM</td><td>target_seq</td><td>PAM</td><td>alignment</td><td>distance</td><td>position</td><td>gene name</td><td>gene id</td></tr>
<tr><td>GU305914.1</td><td></td><td>48944</td><td>48966</td><td>-</td><td>0</td><td>CGGCGGGGAGAGAAAACGTA</td><td>GGG</td><td>||||||||[||||||||||||]PAM</td><td>NA</td><td>Intergenic</td><td>NA</td><td>NA</td></tr>
<tr><td>T30467</td><td>CGCACGACGTCCTCCCTAAAAGG</td><td></td><td>1000</td><td></td><td></td><td></td><td></td><td></td><td></td><td></td><td></td><td></td></tr>
<tr><td>Oligo fwd</td><td>CACCCGCACGACGTCCTCCCTAAA</td><td></td><td></td><td></td><td></td><td></td><td></td><td></td><td></td><td></td><td></td><td></td></tr>
<tr><td>Oligo rev</td><td>AAACTTTAGGGAGGACGTCGTGCG</td><td></td><td></td><td></td><td></td><td></td><td></td><td></td><td></td><td></td><td></td><td></td></tr>
<tr><td>Chromosome</td><td>start</td><td></td><td>end</td><td>strand</td><td>MM</td><td>target_seq</td><td>PAM</td><td>alignment</td><td>distance</td><td>position</td><td>gene name</td><td>gene id</td></tr>
<tr><td>GU305914.1</td><td></td><td>48912</td><td>48934</td><td>-</td><td>0</td><td>CGCACGACGTCCTCCCTAAA</td><td>AGG</td><td>||||||||[||||||||||||]PAM</td><td>NA</td><td>Intergenic</td><td>NA</td><td>NA</td></tr>
<tr><td>T30468</td><td>AAAGGAAAGTACGTCTGCTGCGG</td><td></td><td>1000</td><td></td><td></td><td></td><td></td><td></td><td></td><td></td><td></td><td></td></tr>
<tr><td>Oligo fwd</td><td>CACCAAAGGAAAGTACGTCTGCTG</td><td></td><td></td><td></td><td></td><td></td><td></td><td></td><td></td><td></td><td></td><td></td></tr>
<tr><td>Oligo rev</td><td>AAACCAGCAGACGTACTTTCCTTT</td><td></td><td></td><td></td><td></td><td></td><td></td><td></td><td></td><td></td><td></td><td></td></tr>
<tr><td>Chromosome</td><td>start</td><td></td><td>end</td><td>strand</td><td>MM</td><td>target_seq</td><td>PAM</td><td>alignment</td><td>distance</td><td>position</td><td>gene name</td><td>gene id</td></tr>
<tr><td>GU305914.1</td><td></td><td>48894</td><td>48916</td><td>-</td><td>0</td><td>AAAGGAAAGTACGTCTGCTG</td><td>CGG</td><td>||||||||[||||||||||||]PAM</td><td>NA</td><td>Intergenic</td><td>NA</td><td>NA</td></tr>
<tr><td>T30469</td><td>AGTACGTCTGCTGCGGTGAGAGG</td><td></td><td>1000</td><td></td><td></td><td></td><td></td><td></td><td></td><td></td><td></td><td></td></tr>
<tr><td>Oligo fwd</td><td>CACCAGTACGTCTGCTGCGGTGAG</td><td></td><td></td><td></td><td></td><td></td><td></td><td></td><td></td><td></td><td></td><td></td></tr>
<tr><td>Oligo rev</td><td>AAACCTCACCGCAGCAGACGTACT</td><td></td><td></td><td></td><td></td><td></td><td></td><td></td><td></td><td></td><td></td><td></td></tr>
<tr><td>Chromosome</td><td>start</td><td></td><td>end</td><td>strand</td><td>MM</td><td>target_seq</td><td>PAM</td><td>alignment</td><td>distance</td><td>position</td><td>gene name</td><td>gene id</td></tr>
<tr><td>GU305914.1</td><td></td><td>48887</td><td>48909</td><td>-</td><td>0</td><td>AGTACGTCTGCTGCGGTGAG</td><td>AGG</td><td>||||||||[||||||||||||]PAM</td><td>NA</td><td>Intergenic</td><td>NA</td><td>NA</td></tr>
<tr><td>T30470</td><td>CGTCTGCTGCGGTGAGAGGATGG</td><td></td><td>1000</td><td></td><td></td><td></td><td></td><td></td><td></td><td></td><td></td><td></td></tr>
<tr><td>Oligo fwd</td><td>CACCCGTCTGCTGCGGTGAGAGGA</td><td></td><td></td><td></td><td></td><td></td><td></td><td></td><td></td><td></td><td></td><td></td></tr>
<tr><td>Oligo rev</td><td>AAACTCCTCTCACCGCAGCAGACG</td><td></td><td></td><td></td><td></td><td></td><td></td><td></td><td></td><td></td><td></td><td></td></tr>
<tr><td>Chromosome</td><td>start</td><td></td><td>end</td><td>strand</td><td>MM</td><td>target_seq</td><td>PAM</td><td>alignment</td><td>distance</td><td>position</td><td>gene name</td><td>gene id</td></tr>
<tr><td>GU305914.1</td><td></td><td>48883</td><td>48905</td><td>-</td><td>0</td><td>CGTCTGCTGCGGTGAGAGGA</td><td>TGG</td><td>||||||||[||||||||||||]PAM</td><td>NA</td><td>Intergenic</td><td>NA</td><td>NA</td></tr>
<tr><td>T30471</td><td>GGTGAGAGGATGGTGCCGTTCGG</td><td></td><td>1000</td><td></td><td></td><td></td><td></td><td></td><td></td><td></td><td></td><td></td></tr>
<tr><td>Oligo fwd</td><td>CACCGGTGAGAGGATGGTGCCGTT</td><td></td><td></td><td></td><td></td><td></td><td></td><td></td><td></td><td></td><td></td><td></td></tr>
<tr><td>Oligo rev</td><td>AAACAACGGCACCATCCTCTCACC</td><td></td><td></td><td></td><td></td><td></td><td></td><td></td><td></td><td></td><td></td><td></td></tr>
<tr><td>Chromosome</td><td>start</td><td></td><td>end</td><td>strand</td><td>MM</td><td>target_seq</td><td>PAM</td><td>alignment</td><td>distance</td><td>position</td><td>gene name</td><td>gene id</td></tr>
<tr><td>GU305914.1</td><td></td><td>48873</td><td>48895</td><td>-</td><td>0</td><td>GGTGAGAGGATGGTGCCGTT</td><td>CGG</td><td>||||||||[||||||||||||]PAM</td><td>NA</td><td>Intergenic</td><td>NA</td><td>NA</td></tr>
<tr><td>T30472</td><td>AGAGGATGGTGCCGTTCGGCCGG</td><td></td><td>1000</td><td></td><td></td><td></td><td></td><td></td><td></td><td></td><td></td><td></td></tr>
<tr><td>Oligo fwd</td><td>CACCAGAGGATGGTGCCGTTCGGC</td><td></td><td></td><td></td><td></td><td></td><td></td><td></td><td></td><td></td><td></td><td></td></tr>
<tr><td>Oligo rev</td><td>AAACGCCGAACGGCACCATCCTCT</td><td></td><td></td><td></td><td></td><td></td><td></td><td></td><td></td><td></td><td></td><td></td></tr>
<tr><td>Chromosome</td><td>start</td><td></td><td>end</td><td>strand</td><td>MM</td><td>target_seq</td><td>PAM</td><td>alignment</td><td>distance</td><td>position</td><td>gene name</td><td>gene id</td></tr>
<tr><td>GU305914.1</td><td></td><td>48869</td><td>48891</td><td>-</td><td>0</td><td>AGAGGATGGTGCCGTTCGGC</td><td>CGG</td><td>||||||||[||||||||||||]PAM</td><td>NA</td><td>Intergenic</td><td>NA</td><td>NA</td></tr>
<tr><td>T30473</td><td>GTGCCGTTCGGCCGGCTGCGTGG</td><td></td><td>1000</td><td></td><td></td><td></td><td></td><td></td><td></td><td></td><td></td><td></td></tr>
<tr><td>Oligo fwd</td><td>CACCGTGCCGTTCGGCCGGCTGCG</td><td></td><td></td><td></td><td></td><td></td><td></td><td></td><td></td><td></td><td></td><td></td></tr>
<tr><td>Oligo rev</td><td>AAACCGCAGCCGGCCGAACGGCAC</td><td></td><td></td><td></td><td></td><td></td><td></td><td></td><td></td><td></td><td></td><td></td></tr>
<tr><td>Chromosome</td><td>start</td><td></td><td>end</td><td>strand</td><td>MM</td><td>target_seq</td><td>PAM</td><td>alignment</td><td>distance</td><td>position</td><td>gene name</td><td>gene id</td></tr>
<tr><td>GU305914.1</td><td></td><td>48861</td><td>48883</td><td>-</td><td>0</td><td>GTGCCGTTCGGCCGGCTGCG</td><td>TGG</td><td>||||||||[||||||||||||]PAM</td><td>NA</td><td>Intergenic</td><td>NA</td><td>NA</td></tr>
<tr><td>T30474</td><td>CGTTCGGCCGGCTGCGTGGACGG</td><td></td><td>1000</td><td></td><td></td><td></td><td></td><td></td><td></td><td></td><td></td><td></td></tr>
<tr><td>Oligo fwd</td><td>CACCCGTTCGGCCGGCTGCGTGGA</td><td></td><td></td><td></td><td></td><td></td><td></td><td></td><td></td><td></td><td></td><td></td></tr>
<tr><td>Oligo rev</td><td>AAACTCCACGCAGCCGGCCGAACG</td><td></td><td></td><td></td><td></td><td></td><td></td><td></td><td></td><td></td><td></td><td></td></tr>
<tr><td>Chromosome</td><td>start</td><td></td><td>end</td><td>strand</td><td>MM</td><td>target_seq</td><td>PAM</td><td>alignment</td><td>distance</td><td>position</td><td>gene name</td><td>gene id</td></tr>
<tr><td>GU305914.1</td><td></td><td>48857</td><td>48879</td><td>-</td><td>0</td><td>CGTTCGGCCGGCTGCGTGGA</td><td>CGG</td><td>||||||||[||||||||||||]PAM</td><td>NA</td><td>Intergenic</td><td>NA</td><td>NA</td></tr>
<tr><td>T30475</td><td>GTTCGGCCGGCTGCGTGGACGGG</td><td></td><td>1000</td><td></td><td></td><td></td><td></td><td></td><td></td><td></td><td></td><td></td></tr>
<tr><td>Oligo fwd</td><td>CACCGTTCGGCCGGCTGCGTGGAC</td><td></td><td></td><td></td><td></td><td></td><td></td><td></td><td></td><td></td><td></td><td></td></tr>
<tr><td>Oligo rev</td><td>AAACGTCCACGCAGCCGGCCGAAC</td><td></td><td></td><td></td><td></td><td></td><td></td><td></td><td></td><td></td><td></td><td></td></tr>
<tr><td>Chromosome</td><td>start</td><td></td><td>end</td><td>strand</td><td>MM</td><td>target_seq</td><td>PAM</td><td>alignment</td><td>distance</td><td>position</td><td>gene name</td><td>gene id</td></tr>
<tr><td>GU305914.1</td><td></td><td>48856</td><td>48878</td><td>-</td><td>0</td><td>GTTCGGCCGGCTGCGTGGAC</td><td>GGG</td><td>||||||||[||||||||||||]PAM</td><td>NA</td><td>Intergenic</td><td>NA</td><td>NA</td></tr>
<tr><td>T30476</td><td>CCGGCTGCGTGGACGGGACCTGG</td><td></td><td>1000</td><td></td><td></td><td></td><td></td><td></td><td></td><td></td><td></td><td></td></tr>
<tr><td>Oligo fwd</td><td>CACCCCGGCTGCGTGGACGGGACC</td><td></td><td></td><td></td><td></td><td></td><td></td><td></td><td></td><td></td><td></td><td></td></tr>
<tr><td>Oligo rev</td><td>AAACGGTCCCGTCCACGCAGCCGG</td><td></td><td></td><td></td><td></td><td></td><td></td><td></td><td></td><td></td><td></td><td></td></tr>
<tr><td>Chromosome</td><td>start</td><td></td><td>end</td><td>strand</td><td>MM</td><td>target_seq</td><td>PAM</td><td>alignment</td><td>distance</td><td>position</td><td>gene name</td><td>gene id</td></tr>
<tr><td>GU305914.1</td><td></td><td>48850</td><td>48872</td><td>-</td><td>0</td><td>CCGGCTGCGTGGACGGGACC</td><td>TGG</td><td>||||||||[||||||||||||]PAM</td><td>NA</td><td>Intergenic</td><td>NA</td><td>NA</td></tr>
<tr><td>T30477</td><td>GACGGGACCTGGACACCACGAGG</td><td></td><td>1000</td><td></td><td></td><td></td><td></td><td></td><td></td><td></td><td></td><td></td></tr>
<tr><td>Oligo fwd</td><td>CACCGACGGGACCTGGACACCACG</td><td></td><td></td><td></td><td></td><td></td><td></td><td></td><td></td><td></td><td></td><td></td></tr>
<tr><td>Oligo rev</td><td>AAACCGTGGTGTCCAGGTCCCGTC</td><td></td><td></td><td></td><td></td><td></td><td></td><td></td><td></td><td></td><td></td><td></td></tr>
<tr><td>Chromosome</td><td>start</td><td></td><td>end</td><td>strand</td><td>MM</td><td>target_seq</td><td>PAM</td><td>alignment</td><td>distance</td><td>position</td><td>gene name</td><td>gene id</td></tr>
<tr><td>GU305914.1</td><td></td><td>48839</td><td>48861</td><td>-</td><td>0</td><td>GACGGGACCTGGACACCACG</td><td>AGG</td><td>||||||||[||||||||||||]PAM</td><td>NA</td><td>Intergenic</td><td>NA</td><td>NA</td></tr>
<tr><td>T30478</td><td>ACGAGGTTCGTGATGCTCTTCGG</td><td></td><td>1000</td><td></td><td></td><td></td><td></td><td></td><td></td><td></td><td></td><td></td></tr>
<tr><td>Oligo fwd</td><td>CACCACGAGGTTCGTGATGCTCTT</td><td></td><td></td><td></td><td></td><td></td><td></td><td></td><td></td><td></td><td></td><td></td></tr>
<tr><td>Oligo rev</td><td>AAACAAGAGCATCACGAACCTCGT</td><td></td><td></td><td></td><td></td><td></td><td></td><td></td><td></td><td></td><td></td><td></td></tr>
<tr><td>Chromosome</td><td>start</td><td></td><td>end</td><td>strand</td><td>MM</td><td>target_seq</td><td>PAM</td><td>alignment</td><td>distance</td><td>position</td><td>gene name</td><td>gene id</td></tr>
<tr><td>GU305914.1</td><td></td><td>48822</td><td>48844</td><td>-</td><td>0</td><td>ACGAGGTTCGTGATGCTCTT</td><td>CGG</td><td>||||||||[||||||||||||]PAM</td><td>NA</td><td>Intergenic</td><td>NA</td><td>NA</td></tr>
<tr><td>T30479</td><td>GTGATGCTCTTCGGCGCCACCGG</td><td></td><td>1000</td><td></td><td></td><td></td><td></td><td></td><td></td><td></td><td></td><td></td></tr>
<tr><td>Oligo fwd</td><td>CACCGTGATGCTCTTCGGCGCCAC</td><td></td><td></td><td></td><td></td><td></td><td></td><td></td><td></td><td></td><td></td><td></td></tr>
<tr><td>Oligo rev</td><td>AAACGTGGCGCCGAAGAGCATCAC</td><td></td><td></td><td></td><td></td><td></td><td></td><td></td><td></td><td></td><td></td><td></td></tr>
</table>

Chromosome start end strand MM target_seq PAM alignment distance position gene name gene id
GU305914.1 48813 48835 - 0 GTGATGCTCTTCGGCGCCAC CGG ||||||||[||||||||||||]PAM NA Intergenic NA NA

T30480 ATGCTCTTCGGCGCCACCGGCGG 1000
Oligo fwd CACCATGCTCTTCGGCGCCACCGG
Oligo rev AAACCCGGTGGCGCCGAAGAGCAT
Chromosome start end strand MM target_seq PAM alignment distance position gene name gene id
GU305914.1 48810 48832 - 0 ATGCTCTTCGGCGCCACCGG CGG ||||||||[||||||||||||]PAM NA Intergenic NA NA

T30481 GGCGGCATCTACTGCTACGACGG 1000
Oligo fwd CACCGGCGGCATCTACTGCTACGA
Oligo rev AAACTCGTAGCAGTAGATGCCGCC
Chromosome start end strand MM target_seq PAM alignment distance position gene name gene id
GU305914.1 48792 48814 - 0 GGCGGCATCTACTGCTACGA CGG ||||||||[||||||||||||]PAM NA Intergenic NA NA

T30482 CTACTGCTACGACGGACACGAGG 1000
Oligo fwd CACCCTACTGCTACGACGGACACG
Oligo rev AAACCGTGTCCGTCGTAGCAGTAG
Chromosome start end strand MM target_seq PAM alignment distance position gene name gene id
GU305914.1 48784 48806 - 0 CTACTGCTACGACGGACACG AGG ||||||||[||||||||||||]PAM NA Intergenic NA NA

T30483 CGAGGACGTCGTCTTCCATCTGG 1000
Oligo fwd CACCCGAGGACGTCGTCTTCCATC
Oligo rev AAACGATGGAAGACGACGTCCTCG
Chromosome start end strand MM target_seq PAM alignment distance position gene name gene id
GU305914.1 48766 48788 - 0 CGAGGACGTCGTCTTCCATC TGG ||||||||[||||||||||||]PAM NA Intergenic NA NA

T30484 GAGGACGTCGTCTTCCATCTGGG 1000
Oligo fwd CACCGAGGACGTCGTCTTCCATCT
Oligo rev AAACAGATGGAAGACGACGTCCTC
Chromosome start end strand MM target_seq PAM alignment distance position gene name gene id
GU305914.1 48765 48787 - 0 GAGGACGTCGTCTTCCATCT GGG ||||||||[||||||||||||]PAM NA Intergenic NA NA

T30485 AGGACGTCGTCTTCCATCTGGGG 1000
Oligo fwd CACCAGGACGTCGTCTTCCATCTG
Oligo rev AAACCAGATGGAAGACGACGTCCT
Chromosome start end strand MM target_seq PAM alignment distance position gene name gene id
GU305914.1 48764 48786 - 0 AGGACGTCGTCTTCCATCTG GGG ||||||||[||||||||||||]PAM NA Intergenic NA NA

T30486 GGACGTCGTCTTCCATCTGGGGG 1000
Oligo fwd CACCGGACGTCGTCTTCCATCTGG
Oligo rev AAACCCAGATGGAAGACGACGTCC
Chromosome start end strand MM target_seq PAM alignment distance position gene name gene id
GU305914.1 48763 48785 - 0 GGACGTCGTCTTCCATCTGG GGG ||||||||[||||||||||||]PAM NA Intergenic NA NA

T30487 CAGTCTAGTGTCCTTCTTCCAGG 1000
Oligo fwd CACCCAGTCTAGTGTCCTTCTTCC
Oligo rev AAACGGAAGAAGGACACTAGACTG
Chromosome start end strand MM target_seq PAM alignment distance position gene name gene id
GU305914.1 48739 48761 - 0 CAGTCTAGTGTCCTTCTTCC AGG ||||||||[||||||||||||]PAM NA Intergenic NA NA

T30488 CTAGTGTCCTTCTTCCAGGACGG 1000
Oligo fwd CACCCTAGTGTCCTTCTTCCAGGA
Oligo rev AAACTCCTGGAAGAAGGACACTAG
Chromosome start end strand MM target_seq PAM alignment distance position gene name gene id
GU305914.1 48735 48757 - 0 CTAGTGTCCTTCTTCCAGGA CGG ||||||||[||||||||||||]PAM NA Intergenic NA NA

T30489 TCTTCCAGGACGGCCTGCGTCGG 1000
Oligo fwd CACCTCTTCCAGGACGGCCTGCGT
Oligo rev AAACACGCAGGCCGTCCTGGAAGA
Chromosome start end strand MM target_seq PAM alignment distance position gene name gene id
GU305914.1 48725 48747 - 0 TCTTCCAGGACGGCCTGCGT CGG ||||||||[||||||||||||]PAM NA Intergenic NA NA

T30490 TCCCATCTTCGCCACGTCGCTGG 1000
Oligo fwd CACCTCCCATCTTCGCCACGTCGC
Oligo rev AAACGCGACGTGGCGAAGATGGGA
Chromosome start end strand MM target_seq PAM alignment distance position gene name gene id
GU305914.1 48697 48719 - 0 TCCCATCTTCGCCACGTCGC TGG ||||||||[||||||||||||]PAM NA Intergenic NA NA

T30491 GCGCCCATACTTCTCGTCGACGG 1000
Oligo fwd CACCGCGCCCATACTTCTCGTCGA
Oligo rev AAACTCGACGAGAAGTATGGGCGC
Chromosome start end strand MM target_seq PAM alignment distance position gene name gene id
GU305914.1 48672 48694 - 0 GCGCCCATACTTCTCGTCGA CGG ||||||||[||||||||||||]PAM NA Intergenic NA NA

T30492 CGGTGTCATCGCCAGCCTCCTGG 1000
Oligo fwd CACCCGGTGTCATCGCCAGCCTCC
Oligo rev AAACGGAGGCTGGCGATGACACCG
Chromosome start end strand MM target_seq PAM alignment distance position gene name gene id
GU305914.1 48652 48674 - 0 CGGTGTCATCGCCAGCCTCC TGG ||||||||[||||||||||||]PAM NA Intergenic NA NA

T30493 TGTCATCGCCAGCCTCCTGGAGG 1000
Oligo fwd CACCTGTCATCGCCAGCCTCCTGG
Oligo rev AAACCCAGGAGGCTGGCGATGACA
Chromosome start end strand MM target_seq PAM alignment distance position gene name gene id
GU305914.1 48649 48671 - 0 TGTCATCGCCAGCCTCCTGG AGG ||||||||[||||||||||||]PAM NA Intergenic NA NA

T30494 CGCTTTCATCTCCATCGTCGCGG 1000
Oligo fwd CACCCGCTTTCATCTCCATCGTCG

Oligo rev AAACCGACGATGGAGATGAAAGCG
Chromosome start end strand MM target_seq PAM alignment distance position gene name gene id
GU305914.1 48613 48635 - 0 CGCTTTCATCTCCATCGTCG CGG ||||||||[||||||||||||]PAM NA Intergenic NA NA

T30495 CTCCATCGTCGCGGAGAATCAGG 1000
Oligo fwd CACCCTCCATCGTCGCGGAGAATC
Oligo rev AAACGATTCTCCGCGACGATGGAG
Chromosome start end strand MM target_seq PAM alignment distance position gene name gene id
GU305914.1 48604 48626 - 0 CTCCATCGTCGCGGAGAATC AGG ||||||||[||||||||||||]PAM NA Intergenic NA NA

T30496 TCCATCGTCGCGGAGAATCAGGG 1000
Oligo fwd CACCTCCATCGTCGCGGAGAATCA
Oligo rev AAACTGATTCTCCGCGACGATGGA
Chromosome start end strand MM target_seq PAM alignment distance position gene name gene id
GU305914.1 48603 48625 - 0 TCCATCGTCGCGGAGAATCA GGG ||||||||[||||||||||||]PAM NA Intergenic NA NA

T30497 TCAGGGTGCCGTCTACCAGATGG 1000
Oligo fwd CACCTCAGGGTGCCGTCTACCAGA
Oligo rev AAACTCTGGTAGACGGCACCCTGA
Chromosome start end strand MM target_seq PAM alignment distance position gene name gene id
GU305914.1 48586 48608 - 0 TCAGGGTGCCGTCTACCAGA TGG ||||||||[||||||||||||]PAM NA Intergenic NA NA

T30498 GGTGCCGTCTACCAGATGGTCGG 1000
Oligo fwd CACCGGTGCCGTCTACCAGATGGT
Oligo rev AAACACCATCTGGTAGACGGCACC
Chromosome start end strand MM target_seq PAM alignment distance position gene name gene id
GU305914.1 48582 48604 - 0 GGTGCCGTCTACCAGATGGT CGG ||||||||[||||||||||||]PAM NA Intergenic NA NA

T30499 CAGATGGTCGGCAACATCGCCGG 1000
Oligo fwd CACCCAGATGGTCGGCAACATCGC
Oligo rev AAACGCGATGTTGCCGACCATCTG
Chromosome start end strand MM target_seq PAM alignment distance position gene name gene id
GU305914.1 48570 48592 - 0 CAGATGGTCGGCAACATCGC CGG ||||||||[||||||||||||]PAM NA Intergenic NA NA

T30500 GCCCAGATGATGCTCTATCGCGG 1000
Oligo fwd CACCGCCCAGATGATGCTCTATCG
Oligo rev AAACCGATAGAGCATCATCTGGGC
Chromosome start end strand MM target_seq PAM alignment distance position gene name gene id
GU305914.1 48540 48562 - 0 GCCCAGATGATGCTCTATCG CGG ||||||||[||||||||||||]PAM NA Intergenic NA NA

T30501 ATGCTCTATCGCGGCGACTACGG 1000
Oligo fwd CACCATGCTCTATCGCGGCGACTA
Oligo rev AAACTAGTCGCCGCGATAGAGCAT
Chromosome start end strand MM target_seq PAM alignment distance position gene name gene id
GU305914.1 48531 48553 - 0 ATGCTCTATCGCGGCGACTA CGG ||||||||[||||||||||||]PAM NA Intergenic NA NA

T30502 CACCGTCACCTCCTGCAGCCTGG 1000
Oligo fwd CACCCACCGTCACCTCCTGCAGCC
Oligo rev AAACGGCTGCAGGAGGTGACGGTG
Chromosome start end strand MM target_seq PAM alignment distance position gene name gene id
GU305914.1 48508 48530 - 0 CACCGTCACCTCCTGCAGCC TGG ||||||||[||||||||||||]PAM NA Intergenic NA NA

T30503 ACCGTCACCTCCTGCAGCCTGGG 1000
Oligo fwd CACCACCGTCACCTCCTGCAGCCT
Oligo rev AAACAGGCTGCAGGAGGTGACGGT
Chromosome start end strand MM target_seq PAM alignment distance position gene name gene id
GU305914.1 48507 48529 - 0 ACCGTCACCTCCTGCAGCCT GGG ||||||||[||||||||||||]PAM NA Intergenic NA NA

T30504 CACCTCCTGCAGCCTGGGTACGG 1000
Oligo fwd CACCCACCTCCTGCAGCCTGGGTA
Oligo rev AAACTACCCAGGCTGCAGGAGGTG
Chromosome start end strand MM target_seq PAM alignment distance position gene name gene id
GU305914.1 48502 48524 - 0 CACCTCCTGCAGCCTGGGTA CGG ||||||||[||||||||||||]PAM NA Intergenic NA NA

T30505 CCGTCATGTGCTCCGTCGTCCGG 1000
Oligo fwd CACCCCGTCATGTGCTCCGTCGTC
Oligo rev AAACGACGACGGAGCACATGACGG
Chromosome start end strand MM target_seq PAM alignment distance position gene name gene id
GU305914.1 48464 48486 - 0 CCGTCATGTGCTCCGTCGTC CGG ||||||||[||||||||||||]PAM NA Intergenic NA NA

T30506 TCATGTGCTCCGTCGTCCGGAGG 1000
Oligo fwd CACCTCATGTGCTCCGTCGTCCGG
Oligo rev AAACCCGGACGACGGAGCACATGA
Chromosome start end strand MM target_seq PAM alignment distance position gene name gene id
GU305914.1 48461 48483 - 0 TCATGTGCTCCGTCGTCCGG AGG ||||||||[||||||||||||]PAM NA Intergenic NA NA

T30507 CACCTTTGACATCTTCGCCGTGG 1000
Oligo fwd CACCCACCTTTGACATCTTCGCCG
Oligo rev AAACCGGCGAAGATGTCAAAGGTG
Chromosome start end strand MM target_seq PAM alignment distance position gene name gene id
GU305914.1 48430 48452 - 0 CACCTTTGACATCTTCGCCG TGG ||||||||[||||||||||||]PAM NA Intergenic NA NA

T30508 TTTGACATCTTCGCCGTGGTCGG 1000
Oligo fwd CACCTTTGACATCTTCGCCGTGGT
Oligo rev AAACACCACGGCGAAGATGTCAAA
Chromosome start end strand MM target_seq PAM alignment distance position gene name gene id
GU305914.1 48426 48448 - 0 TTTGACATCTTCGCCGTGGT CGG ||||||||[||||||||||||]PAM NA Intergenic NA NA

T30509 CTTCGCCGTGGTCGGATACAAGG 1000

Oligo fwd CACCCTTCGCCGTGGTCGGATACA
Oligo rev AAACTGTATCCGACCACGGCGAAG
Chromosome start end strand MM target_seq PAM alignment distance position gene name gene id
GU305914.1 48418 48440 - 0 CTTCGCCGTGGTCGGATACA AGG ||||||||[||||||||||||]PAM NA Intergenic NA NA

T30510 CGGATACAAGGCCGCGACCTCGG 1000
Oligo fwd CACCCGGATACAAGGCCGCGACCT
Oligo rev AAACAGGTCGCGGCCTTGTATCCG
Chromosome start end strand MM target_seq PAM alignment distance position gene name gene id
GU305914.1 48406 48428 - 0 CGGATACAAGGCCGCGACCT CGG ||||||||[||||||||||||]PAM NA Intergenic NA NA

T30511 CGACCTCGGTCTTCCGACCCCGG 1000
Oligo fwd CACCCGACCTCGGTCTTCCGACCC
Oligo rev AAACGGGTCGGAAGACCGAGGTCG
Chromosome start end strand MM target_seq PAM alignment distance position gene name gene id
GU305914.1 48392 48414 - 0 CGACCTCGGTCTTCCGACCC CGG ||||||||[||||||||||||]PAM NA Intergenic NA NA

T30512 ATCCTCATCATCGATCCCTTCGG 1000
Oligo fwd CACCATCCTCATCATCGATCCCTT
Oligo rev AAACAAGGGATCGATGATGAGGAT
Chromosome start end strand MM target_seq PAM alignment distance position gene name gene id
GU305914.1 48366 48388 - 0 ATCCTCATCATCGATCCCTT CGG ||||||||[||||||||||||]PAM NA Intergenic NA NA

T30513 GATCCCTTCGGCGCCATCTTCGG 1000
Oligo fwd CACCGATCCCTTCGGCGCCATCTT
Oligo rev AAACAAGATGGCGCCGAAGGGATC
Chromosome start end strand MM target_seq PAM alignment distance position gene name gene id
GU305914.1 48354 48376 - 0 GATCCCTTCGGCGCCATCTT CGG ||||||||[||||||||||||]PAM NA Intergenic NA NA

T30514 TCTCAACAGAGTCACTCGCCTGG 1000
Oligo fwd CACCTCTCAACAGAGTCACTCGCC
Oligo rev AAACGGCGAGTGACTCTGTTGAGA
Chromosome start end strand MM target_seq PAM alignment distance position gene name gene id
GU305914.1 48319 48341 - 0 TCTCAACAGAGTCACTCGCC TGG ||||||||[||||||||||||]PAM NA Intergenic NA NA

T30515 TTTAATATGTTCATCCGCATCGG 1000
Oligo fwd CACCTTTAATATGTTCATCCGCAT
Oligo rev AAACATGCGGATGAACATATTAAA
Chromosome start end strand MM target_seq PAM alignment distance position gene name gene id
GU305914.1 48288 48310 - 0 TTTAATATGTTCATCCGCAT CGG ||||||||[||||||||||||]PAM NA Intergenic NA NA

T30516 CCTCAACTTCAGACACGACAAGG 1000
Oligo fwd CACCCCTCAACTTCAGACACGACA
Oligo rev AAACTGTCGTGTCTGAAGTTGAGG
Chromosome start end strand MM target_seq PAM alignment distance position gene name gene id
GU305914.1 48250 48272 - 0 CCTCAACTTCAGACACGACA AGG ||||||||[||||||||||||]PAM NA Intergenic NA NA

T30517 CTCAACTTCAGACACGACAAGGG 1000
Oligo fwd CACCCTCAACTTCAGACACGACAA
Oligo rev AAACTTGTCGTGTCTGAAGTTGAG
Chromosome start end strand MM target_seq PAM alignment distance position gene name gene id
GU305914.1 48249 48271 - 0 CTCAACTTCAGACACGACAA GGG ||||||||[||||||||||||]PAM NA Intergenic NA NA

T30518 AGACACGACAAGGGCTTCCGCGG 1000
Oligo fwd CACCAGACACGACAAGGGCTTCCG
Oligo rev AAACCGGAAGCCCTTGTCGTGTCT
Chromosome start end strand MM target_seq PAM alignment distance position gene name gene id
GU305914.1 48240 48262 - 0 AGACACGACAAGGGCTTCCG CGG ||||||||[||||||||||||]PAM NA Intergenic NA NA

T30519 GACAAGGGCTTCCGCGGTATCGG 1000
Oligo fwd CACCGACAAGGGCTTCCGCGGTAT
Oligo rev AAACATACCGCGGAAGCCCTTGTC
Chromosome start end strand MM target_seq PAM alignment distance position gene name gene id
GU305914.1 48234 48256 - 0 GACAAGGGCTTCCGCGGTAT CGG ||||||||[||||||||||||]PAM NA Intergenic NA NA

T30520 CGGTATCGGACGCCTCGAGAAGG 1000
Oligo fwd CACCCGGTATCGGACGCCTCGAGA
Oligo rev AAACTCTCGAGGCGTCCGATACCG
Chromosome start end strand MM target_seq PAM alignment distance position gene name gene id
GU305914.1 48220 48242 - 0 CGGTATCGGACGCCTCGAGA AGG ||||||||[||||||||||||]PAM NA Intergenic NA NA

T30521 GTTCCCTACTGTCCGCACGTCGG 1000
Oligo fwd CACCGTTCCCTACTGTCCGCACGT
Oligo rev AAACACGTGCGGACAGTAGGGAAC
Chromosome start end strand MM target_seq PAM alignment distance position gene name gene id
GU305914.1 48198 48220 - 0 GTTCCCTACTGTCCGCACGT CGG ||||||||[||||||||||||]PAM NA Intergenic NA NA

T30522 GCACGTCGGCAACGTGCGCGAGG 1000
Oligo fwd CACCGCACGTCGGCAACGTGCGCG
Oligo rev AAACCGCGCACGTTGCCGACGTGC
Chromosome start end strand MM target_seq PAM alignment distance position gene name gene id
GU305914.1 48184 48206 - 0 GCACGTCGGCAACGTGCGCG AGG ||||||||[||||||||||||]PAM NA Intergenic NA NA

T30523 GCCGCCACCGCCGCCGCCACAGG 1000
Oligo fwd CACCGCCGCCACCGCCGCCGCCAC
Oligo rev AAACGTGGCGGCGGCGGTGGCGGC
Chromosome start end strand MM target_seq PAM alignment distance position gene name gene id
GU305914.1 48151 48173 - 0 GCCGCCACCGCCGCCGCCAC AGG ||||||||[||||||||||||]PAM NA Intergenic NA NA
GU305914.1 152551 152573 - 2 GCCGGCACCGCCGCCGCCGC AGG ||||-|||[||||||||||-|]PAM NA Intergenic NA NA

| | | | | | | | | | | | |
|---|---|---|---|---|---|---|---|---|---|---|---|
| GU305914.1 | 164964 | 164986 | + | 4 | GCTGCTGCCGCCGCCGCCGC | CGG | \|\|-\|\|--\|[\|\|\|\|\|\|\|\|\|\|-\|]PAM | NA | Intergenic | NA | NA |
| GU305914.1 | 24620 | 24642 | + | 4 | GTCGCCGCCACCGCCGCCGC | CGG | \|-\|\|\|\|-\|[\|-\|\|\|\|\|\|\|\|-\|]PAM | NA | Intergenic | NA | NA |
| GU305914.1 | 29998 | 30020 | - | 3 | GCCGCCGCCGCCGCCAGCAC | CGG | \|\|\|\|\|\|-\|[\|\|\|\|\|\|\|--\|\|\|]PAM | NA | Intergenic | NA | NA |
| GU305914.1 | 52720 | 52742 | - | 3 | TCCGCCACCGCCACCGCCAG | CGG | -\|\|\|\|\|\|\|[\|\|\|\|-\|\|\|\|\|\|-]PAM | NA | Intergenic | NA | NA |
| T30524 | GCCGCCGCCACAGGTCCGCGAGG | 1000 | | | | | | | | | |
| Oligo fwd | CACCGCCGCCGCCACAGGTCCGCG | | | | | | | | | | |
| Oligo rev | AAACCGCGGACCTGTGGCGGCGGC | | | | | | | | | | |
| Chromosome | start | end | strand | MM | target_seq | PAM | alignment | distance | position | gene name | gene id |
| GU305914.1 | 48142 | 48164 | - | 0 | GCCGCCGCCACAGGTCCGCG | AGG | \|\|\|\|\|\|\|\|[\|\|\|\|\|\|\|\|\|\|\|\|]PAM | NA | Intergenic | NA | NA |
| T30525 | CCGCCGCCACAGGTCCGCGAGGG | 1000 | | | | | | | | | |
| Oligo fwd | CACCCCGCCGCCACAGGTCCGCGA | | | | | | | | | | |
| Oligo rev | AAACTCGCGGACCTGTGGCGGCGG | | | | | | | | | | |
| Chromosome | start | end | strand | MM | target_seq | PAM | alignment | distance | position | gene name | gene id |
| GU305914.1 | 48141 | 48163 | - | 0 | CCGCCGCCACAGGTCCGCGA | GGG | \|\|\|\|\|\|\|\|[\|\|\|\|\|\|\|\|\|\|\|\|]PAM | NA | Intergenic | NA | NA |
| T30526 | ACAGGTCCGCGAGGGTGAAGTGG | 1000 | | | | | | | | | |
| Oligo fwd | CACCACAGGTCCGCGAGGGTGAAG | | | | | | | | | | |
| Oligo rev | AAACCTTCACCCTCGCGGACCTGT | | | | | | | | | | |
| Chromosome | start | end | strand | MM | target_seq | PAM | alignment | distance | position | gene name | gene id |
| GU305914.1 | 48133 | 48155 | - | 0 | ACAGGTCCGCGAGGGTGAAG | TGG | \|\|\|\|\|\|\|\|[\|\|\|\|\|\|\|\|\|\|\|\|]PAM | NA | Intergenic | NA | NA |
| T30527 | GTGAAGTGGTTCTTCAGCGCAGG | 1000 | | | | | | | | | |
| Oligo fwd | CACCGTGAAGTGGTTCTTCAGCGC | | | | | | | | | | |
| Oligo rev | AAACGCGCTGAAGAACCACTTCAC | | | | | | | | | | |
| Chromosome | start | end | strand | MM | target_seq | PAM | alignment | distance | position | gene name | gene id |
| GU305914.1 | 48119 | 48141 | - | 0 | GTGAAGTGGTTCTTCAGCGC | AGG | \|\|\|\|\|\|\|\|[\|\|\|\|\|\|\|\|\|\|\|\|]PAM | NA | Intergenic | NA | NA |
| T30528 | TGAAGTGGTTCTTCAGCGCAGGG | 1000 | | | | | | | | | |
| Oligo fwd | CACCTGAAGTGGTTCTTCAGCGCA | | | | | | | | | | |
| Oligo rev | AAACTGCGCTGAAGAACCACTTCA | | | | | | | | | | |
| Chromosome | start | end | strand | MM | target_seq | PAM | alignment | distance | position | gene name | gene id |
| GU305914.1 | 48118 | 48140 | - | 0 | TGAAGTGGTTCTTCAGCGCA | GGG | \|\|\|\|\|\|\|\|[\|\|\|\|\|\|\|\|\|\|\|\|]PAM | NA | Intergenic | NA | NA |
| T30529 | TCTTCAGCGCAGGGTGTCCGTGG | 1000 | | | | | | | | | |
| Oligo fwd | CACCTCTTCAGCGCAGGGTGTCCG | | | | | | | | | | |
| Oligo rev | AAACCGGACACCCTGCGCTGAAGA | | | | | | | | | | |
| Chromosome | start | end | strand | MM | target_seq | PAM | alignment | distance | position | gene name | gene id |
| GU305914.1 | 48109 | 48131 | - | 0 | TCTTCAGCGCAGGGTGTCCG | TGG | \|\|\|\|\|\|\|\|[\|\|\|\|\|\|\|\|\|\|\|\|]PAM | NA | Intergenic | NA | NA |
| T30530 | CTTCAGCGCAGGGTGTCCGTGGG | 1000 | | | | | | | | | |
| Oligo fwd | CACCCTTCAGCGCAGGGTGTCCGT | | | | | | | | | | |
| Oligo rev | AAACACGGACACCCTGCGCTGAAG | | | | | | | | | | |
| Chromosome | start | end | strand | MM | target_seq | PAM | alignment | distance | position | gene name | gene id |
| GU305914.1 | 48108 | 48130 | - | 0 | CTTCAGCGCAGGGTGTCCGT | GGG | \|\|\|\|\|\|\|\|[\|\|\|\|\|\|\|\|\|\|\|\|]PAM | NA | Intergenic | NA | NA |
| T30531 | TTCAGCGCAGGGTGTCCGTGGGG | 1000 | | | | | | | | | |
| Oligo fwd | CACCTTCAGCGCAGGGTGTCCGTG | | | | | | | | | | |
| Oligo rev | AAACCACGGACACCCTGCGCTGAA | | | | | | | | | | |
| Chromosome | start | end | strand | MM | target_seq | PAM | alignment | distance | position | gene name | gene id |
| GU305914.1 | 48107 | 48129 | - | 0 | TTCAGCGCAGGGTGTCCGTG | GGG | \|\|\|\|\|\|\|\|[\|\|\|\|\|\|\|\|\|\|\|\|]PAM | NA | Intergenic | NA | NA |
| T30532 | TCAGCGCAGGGTGTCCGTGGGGG | 1000 | | | | | | | | | |
| Oligo fwd | CACCTCAGCGCAGGGTGTCCGTGG | | | | | | | | | | |
| Oligo rev | AAACCCACGGACACCCTGCGCTGA | | | | | | | | | | |
| Chromosome | start | end | strand | MM | target_seq | PAM | alignment | distance | position | gene name | gene id |
| GU305914.1 | 48106 | 48128 | - | 0 | TCAGCGCAGGGTGTCCGTGG | GGG | \|\|\|\|\|\|\|\|[\|\|\|\|\|\|\|\|\|\|\|\|]PAM | NA | Intergenic | NA | NA |
| T30533 | CAGGGTGTCCGTGGGGGATGAGG | 1000 | | | | | | | | | |
| Oligo fwd | CACCCAGGGTGTCCGTGGGGGATG | | | | | | | | | | |
| Oligo rev | AAACCATCCCCCACGGACACCCTG | | | | | | | | | | |
| Chromosome | start | end | strand | MM | target_seq | PAM | alignment | distance | position | gene name | gene id |
| GU305914.1 | 48100 | 48122 | - | 0 | CAGGGTGTCCGTGGGGGATG | AGG | \|\|\|\|\|\|\|\|[\|\|\|\|\|\|\|\|\|\|\|\|]PAM | NA | Intergenic | NA | NA |
| T30534 | AGGACCCGCAGTACGTCGTCAGG | 1000 | | | | | | | | | |
| Oligo fwd | CACCAGGACCCGCAGTACGTCGTC | | | | | | | | | | |
| Oligo rev | AAACGACGACGTACTGCGGGTCCT | | | | | | | | | | |
| Chromosome | start | end | strand | MM | target_seq | PAM | alignment | distance | position | gene name | gene id |
| GU305914.1 | 48080 | 48102 | - | 0 | AGGACCCGCAGTACGTCGTC | AGG | \|\|\|\|\|\|\|\|[\|\|\|\|\|\|\|\|\|\|\|\|]PAM | NA | Intergenic | NA | NA |
| T30535 | GGACCCGCAGTACGTCGTCAGGG | 1000 | | | | | | | | | |
| Oligo fwd | CACCGGACCCGCAGTACGTCGTCA | | | | | | | | | | |
| Oligo rev | AAACTGACGACGTACTGCGGGTCC | | | | | | | | | | |
| Chromosome | start | end | strand | MM | target_seq | PAM | alignment | distance | position | gene name | gene id |
| GU305914.1 | 48079 | 48101 | - | 0 | GGACCCGCAGTACGTCGTCA | GGG | \|\|\|\|\|\|\|\|[\|\|\|\|\|\|\|\|\|\|\|\|]PAM | NA | Intergenic | NA | NA |
| GU305914.1 | 21293 | 21315 | + | 4 | GGACTCGGTGTACGGCGTCA | TGG | \|\|\|\|-\|\|-[-\|\|\|\|\|-\|\|\|\|\|]PAM | NA | Intergenic | NA | NA |
| T30536 | ACAGCTCCTTCTGCATGTCGCGG | 1000 | | | | | | | | | |
| Oligo fwd | CACCACAGCTCCTTCTGCATGTCG | | | | | | | | | | |
| Oligo rev | AAACCGACATGCAGAAGGAGCTGT | | | | | | | | | | |
| Chromosome | start | end | strand | MM | target_seq | PAM | alignment | distance | position | gene name | gene id |
| GU305914.1 | 48050 | 48072 | - | 0 | ACAGCTCCTTCTGCATGTCG | CGG | \|\|\|\|\|\|\|\|[\|\|\|\|\|\|\|\|\|\|\|\|]PAM | NA | Intergenic | NA | NA |
| T30537 | ATGTCGCGGATGACCAGTTTCGG | 1000 | | | | | | | | | |
| Oligo fwd | CACCATGTCGCGGATGACCAGTTT | | | | | | | | | | |
| Oligo rev | AAACAAACTGGTCATCCGCGACAT | | | | | | | | | | |
| Chromosome | start | end | strand | MM | target_seq | PAM | alignment | distance | position | gene name | gene id |
| GU305914.1 | 48036 | 48058 | - | 0 | ATGTCGCGGATGACCAGTTT | CGG | \|\|\|\|\|\|\|\|[\|\|\|\|\|\|\|\|\|\|\|\|]PAM | NA | Intergenic | NA | NA |

T30538 TGTCGCGGATGACCAGTTTCGGG 1000
Oligo fwd CACCTGTCGCGGATGACCAGTTTC
Oligo rev AAACGAAACTGGTCATCCGCGACA
Chromosome start end strand MM target_seq PAM alignment distance position gene name gene id
GU305914.1 48035 48057 - 0 TGTCGCGGATGACCAGTTTC GGG ||||||||[|||||||||||||]PAM NA Intergenic NA NA

T30539 GTCGCGGATGACCAGTTTCGGGG 1000
Oligo fwd CACCGTCGCGGATGACCAGTTTCG
Oligo rev AAACCGAAACTGGTCATCCGCGAC
Chromosome start end strand MM target_seq PAM alignment distance position gene name gene id
GU305914.1 48034 48056 - 0 GTCGCGGATGACCAGTTTCG GGG ||||||||[|||||||||||||]PAM NA Intergenic NA NA

T30540 ACCAGTTTCGGGGAGCAGATCGG 1000
Oligo fwd CACCACCAGTTTCGGGGAGCAGAT
Oligo rev AAACATCTGCTCCCCGAAACTGGT
Chromosome start end strand MM target_seq PAM alignment distance position gene name gene id
GU305914.1 48024 48046 - 0 ACCAGTTTCGGGGAGCAGAT CGG ||||||||[|||||||||||||]PAM NA Intergenic NA NA

T30541 ATCGGCCACAATCTCTTCCTCGG 1000
Oligo fwd CACCATCGGCCACAATCTCTTCCT
Oligo rev AAACAGGAAGAGATTGTGGCCGAT
Chromosome start end strand MM target_seq PAM alignment distance position gene name gene id
GU305914.1 48006 48028 - 0 ATCGGCCACAATCTCTTCCT CGG ||||||||[|||||||||||||]PAM NA Intergenic NA NA

T30542 TCGGCCACAATCTCTTCCTCGGG 1000
Oligo fwd CACCTCGGCCACAATCTCTTCCTC
Oligo rev AAACGAGGAAGAGATTGTGGCCGA
Chromosome start end strand MM target_seq PAM alignment distance position gene name gene id
GU305914.1 48005 48027 - 0 TCGGCCACAATCTCTTCCTC GGG ||||||||[|||||||||||||]PAM NA Intergenic NA NA

T30543 GCCACAATCTCTTCCTCGGGAGG 1000
Oligo fwd CACCGCCACAATCTCTTCCTCGGG
Oligo rev AAACCCCGAGGAAGAGATTGTGGC
Chromosome start end strand MM target_seq PAM alignment distance position gene name gene id
GU305914.1 48002 48024 - 0 GCCACAATCTCTTCCTCGGG AGG ||||||||[|||||||||||||]PAM NA Intergenic NA NA

T30544 CAACAGAAATCAGTGCGTCGAGG 1000
Oligo fwd CACCCAACAGAAATCAGTGCGTCG
Oligo rev AAACCGACGCACTGATTTCTGTTG
Chromosome start end strand MM target_seq PAM alignment distance position gene name gene id
GU305914.1 47950 47972 - 0 CAACAGAAATCAGTGCGTCG AGG ||||||||[|||||||||||||]PAM NA Intergenic NA NA

T30545 AACAGAAATCAGTGCGTCGAGGG 1000
Oligo fwd CACCAACAGAAATCAGTGCGTCGA
Oligo rev AAACTCGACGCACTGATTTCTGTT
Chromosome start end strand MM target_seq PAM alignment distance position gene name gene id
GU305914.1 47949 47971 - 0 AACAGAAATCAGTGCGTCGA GGG ||||||||[|||||||||||||]PAM NA Intergenic NA NA

T30546 TCAGTGCGTCGAGGGCTTCGTGG 1000
Oligo fwd CACCTCAGTGCGTCGAGGGCTTCG
Oligo rev AAACCGAAGCCCTCGACGCACTGA
Chromosome start end strand MM target_seq PAM alignment distance position gene name gene id
GU305914.1 47941 47963 - 0 TCAGTGCGTCGAGGGCTTCG TGG ||||||||[|||||||||||||]PAM NA Intergenic NA NA

T30547 ACGACGCCGCCAACATCATGAGG 1000
Oligo fwd CACCACGACGCCGCCAACATCATG
Oligo rev AAACCATGATGTTGGCGGCGTCGT
Chromosome start end strand MM target_seq PAM alignment distance position gene name gene id
GU305914.1 47900 47922 - 0 ACGACGCCGCCAACATCATG AGG ||||||||[|||||||||||||]PAM NA Intergenic NA NA

T30548 CGACGCCGCCAACATCATGAGGG 1000
Oligo fwd CACCCGACGCCGCCAACATCATGA
Oligo rev AAACTCATGATGTTGGCGGCGTCG
Chromosome start end strand MM target_seq PAM alignment distance position gene name gene id
GU305914.1 47899 47921 - 0 CGACGCCGCCAACATCATGA GGG ||||||||[|||||||||||||]PAM NA Intergenic NA NA

T30549 ATCATGAGGGCCGCTTACTTCGG 1000
Oligo fwd CACCATCATGAGGGCCGCTTACTT
Oligo rev AAACAAGTAAGCGGCCCTCATGAT
Chromosome start end strand MM target_seq PAM alignment distance position gene name gene id
GU305914.1 47886 47908 - 0 ATCATGAGGGCCGCTTACTT CGG ||||||||[|||||||||||||]PAM NA Intergenic NA NA

T30550 AGCAGATCACCGATCTCCTCTGG 1000
Oligo fwd CACCAGCAGATCACCGATCTCCTC
Oligo rev AAACGAGGAGATCGGTGATCTGCT
Chromosome start end strand MM target_seq PAM alignment distance position gene name gene id
GU305914.1 47858 47880 - 0 AGCAGATCACCGATCTCCTC TGG ||||||||[|||||||||||||]PAM NA Intergenic NA NA

T30551 CACCGATCTCCTCTGGCCCGAGG 1000
Oligo fwd CACCCACCGATCTCCTCTGGCCCG
Oligo rev AAACCGGGCCAGAGGAGATCGGTG
Chromosome start end strand MM target_seq PAM alignment distance position gene name gene id
GU305914.1 47851 47873 - 0 CACCGATCTCCTCTGGCCCG AGG ||||||||[|||||||||||||]PAM NA Intergenic NA NA
GU305914.1 47842 47864 + 4 CCTCGATCTCCTCGGGCCAG AGG |--|||||[|||||-||||-|]PAM NA Intergenic NA NA

T30552 CCTCTGGCCCGAGGAGATCGAGG 1000
Oligo fwd CACCCCTCTGGCCCGAGGAGATCG
Oligo rev AAACCGATCTCCTCGGGCCAGAGG

| Chromosome | start | end | strand | MM | target_seq | PAM | alignment | distance | position | gene name | gene id |
|---|---|---|---|---|---|---|---|---|---|---|---|
| GU305914.1 | 47842 | 47864 | - | 0 | CCTCTGGCCCGAGGAGATCG | AGG | \|\|\|\|\|\|\|[\|\|\|\|\|\|\|\|\|\|\|\|\|]PAM | NA | Intergenic | NA | NA |
| T30553 | CTCTGGCCCGAGGAGATCGAGGG | 1000 | | | | | | | | | |
| Oligo fwd | CACCCTCTGGCCCGAGGAGATCGA | | | | | | | | | | |
| Oligo rev | AAACTCGATCTCCTCGGGCCAGAG | | | | | | | | | | |
| Chromosome | start | end | strand | MM | target_seq | PAM | alignment | distance | position | gene name | gene id |
| GU305914.1 | 47841 | 47863 | - | 0 | CTCTGGCCCGAGGAGATCGA | GGG | \|\|\|\|\|\|\|[\|\|\|\|\|\|\|\|\|\|\|\|\|]PAM | NA | Intergenic | NA | NA |
| T30554 | TCTGGCCCGAGGAGATCGAGGGG | 1000 | | | | | | | | | |
| Oligo fwd | CACCTCTGGCCCGAGGAGATCGAG | | | | | | | | | | |
| Oligo rev | AAACCTCGATCTCCTCGGGCCAGA | | | | | | | | | | |
| Chromosome | start | end | strand | MM | target_seq | PAM | alignment | distance | position | gene name | gene id |
| GU305914.1 | 47840 | 47862 | - | 0 | TCTGGCCCGAGGAGATCGAG | GGG | \|\|\|\|\|\|\|[\|\|\|\|\|\|\|\|\|\|\|\|\|]PAM | NA | Intergenic | NA | NA |
| T30555 | GAGATCGAGGGGCTCCTCGTCGG | 1000 | | | | | | | | | |
| Oligo fwd | CACCGAGATCGAGGGGCTCCTCGT | | | | | | | | | | |
| Oligo rev | AAACACGAGGAGCCCCTCGATCTC | | | | | | | | | | |
| Chromosome | start | end | strand | MM | target_seq | PAM | alignment | distance | position | gene name | gene id |
| GU305914.1 | 47829 | 47851 | - | 0 | GAGATCGAGGGGCTCCTCGT | CGG | \|\|\|\|\|\|\|[\|\|\|\|\|\|\|\|\|\|\|\|\|]PAM | NA | Intergenic | NA | NA |
| T30556 | CGTCGGCCACCCCTCCTGCCCGG | 1000 | | | | | | | | | |
| Oligo fwd | CACCCGTCGGCCACCCCTCCTGCC | | | | | | | | | | |
| Oligo rev | AAACGGCAGGAGGGGTGGCCGACG | | | | | | | | | | |
| Chromosome | start | end | strand | MM | target_seq | PAM | alignment | distance | position | gene name | gene id |
| GU305914.1 | 47812 | 47834 | - | 0 | CGTCGGCCACCCCTCCTGCC | CGG | \|\|\|\|\|\|\|[\|\|\|\|\|\|\|\|\|\|\|\|\|]PAM | NA | Intergenic | NA | NA |
| T30557 | CCACCCCTCCTGCCCGGCCATGG | 1000 | | | | | | | | | |
| Oligo fwd | CACCCCACCCCTCCTGCCCGGCCA | | | | | | | | | | |
| Oligo rev | AAACTGGCCGGGCAGGAGGGGTGG | | | | | | | | | | |
| Chromosome | start | end | strand | MM | target_seq | PAM | alignment | distance | position | gene name | gene id |
| GU305914.1 | 47806 | 47828 | - | 0 | CCACCCCTCCTGCCCGGCCA | TGG | \|\|\|\|\|\|\|[\|\|\|\|\|\|\|\|\|\|\|\|\|]PAM | NA | Intergenic | NA | NA |
| T30558 | GGCCTCTGACCCCGTGATCAAGG | 1000 | | | | | | | | | |
| Oligo fwd | CACCGGCCTCTGACCCCGTGATCA | | | | | | | | | | |
| Oligo rev | AAACTGATCACGGGGTCAGAGGCC | | | | | | | | | | |
| Chromosome | start | end | strand | MM | target_seq | PAM | alignment | distance | position | gene name | gene id |
| GU305914.1 | 47785 | 47807 | - | 0 | GGCCTCTGACCCCGTGATCA | AGG | \|\|\|\|\|\|\|[\|\|\|\|\|\|\|\|\|\|\|\|\|]PAM | NA | Intergenic | NA | NA |
| T30559 | CGTGATCAAGGCCACGCTCGAGG | 1000 | | | | | | | | | |
| Oligo fwd | CACCCGTGATCAAGGCCACGCTCG | | | | | | | | | | |
| Oligo rev | AAACCGAGCGTGGCCTTGATCACG | | | | | | | | | | |
| Chromosome | start | end | strand | MM | target_seq | PAM | alignment | distance | position | gene name | gene id |
| GU305914.1 | 47773 | 47795 | - | 0 | CGTGATCAAGGCCACGCTCG | AGG | \|\|\|\|\|\|\|[\|\|\|\|\|\|\|\|\|\|\|\|\|]PAM | NA | Intergenic | NA | NA |
| T30560 | GTGATCAAGGCCACGCTCGAGGG | 1000 | | | | | | | | | |
| Oligo fwd | CACCGTGATCAAGGCCACGCTCGA | | | | | | | | | | |
| Oligo rev | AAACTCGAGCGTGGCCTTGATCAC | | | | | | | | | | |
| Chromosome | start | end | strand | MM | target_seq | PAM | alignment | distance | position | gene name | gene id |
| GU305914.1 | 47772 | 47794 | - | 0 | GTGATCAAGGCCACGCTCGA | GGG | \|\|\|\|\|\|\|[\|\|\|\|\|\|\|\|\|\|\|\|\|]PAM | NA | Intergenic | NA | NA |
| T30561 | TGATCAAGGCCACGCTCGAGGGG | 1000 | | | | | | | | | |
| Oligo fwd | CACCTGATCAAGGCCACGCTCGAG | | | | | | | | | | |
| Oligo rev | AAACCTCGAGCGTGGCCTTGATCA | | | | | | | | | | |
| Chromosome | start | end | strand | MM | target_seq | PAM | alignment | distance | position | gene name | gene id |
| GU305914.1 | 47771 | 47793 | - | 0 | TGATCAAGGCCACGCTCGAG | GGG | \|\|\|\|\|\|\|[\|\|\|\|\|\|\|\|\|\|\|\|\|]PAM | NA | Intergenic | NA | NA |
| T30562 | AGGGGATGCGCAGCGTCATGAGG | 1000 | | | | | | | | | |
| Oligo fwd | CACCAGGGGATGCGCAGCGTCATG | | | | | | | | | | |
| Oligo rev | AAACCATGACGCTGCGCATCCCCT | | | | | | | | | | |
| Chromosome | start | end | strand | MM | target_seq | PAM | alignment | distance | position | gene name | gene id |
| GU305914.1 | 47753 | 47775 | - | 0 | AGGGGATGCGCAGCGTCATG | AGG | \|\|\|\|\|\|\|[\|\|\|\|\|\|\|\|\|\|\|\|\|]PAM | NA | Intergenic | NA | NA |
| T30563 | GATGCGCAGCGTCATGAGGTCGG | 1000 | | | | | | | | | |
| Oligo fwd | CACCGATGCGCAGCGTCATGAGGT | | | | | | | | | | |
| Oligo rev | AAACACCTCATGACGCTGCGCATC | | | | | | | | | | |
| Chromosome | start | end | strand | MM | target_seq | PAM | alignment | distance | position | gene name | gene id |
| GU305914.1 | 47749 | 47771 | - | 0 | GATGCGCAGCGTCATGAGGT | CGG | \|\|\|\|\|\|\|[\|\|\|\|\|\|\|\|\|\|\|\|\|]PAM | NA | Intergenic | NA | NA |
| T30564 | GCGCAGCGTCATGAGGTCGGTGG | 1000 | | | | | | | | | |
| Oligo fwd | CACCGCGCAGCGTCATGAGGTCGG | | | | | | | | | | |
| Oligo rev | AAACCCGACCTCATGACGCTGCGC | | | | | | | | | | |
| Chromosome | start | end | strand | MM | target_seq | PAM | alignment | distance | position | gene name | gene id |
| GU305914.1 | 47746 | 47768 | - | 0 | GCGCAGCGTCATGAGGTCGG | TGG | \|\|\|\|\|\|\|[\|\|\|\|\|\|\|\|\|\|\|\|\|]PAM | NA | Intergenic | NA | NA |
| T30565 | GACACCAGCCTGACCACCATCGG | 1000 | | | | | | | | | |
| Oligo fwd | CACCGACACCAGCCTGACCACCAT | | | | | | | | | | |
| Oligo rev | AAACATGGTGGTCAGGCTGGTGTC | | | | | | | | | | |
| Chromosome | start | end | strand | MM | target_seq | PAM | alignment | distance | position | gene name | gene id |
| GU305914.1 | 47700 | 47722 | - | 0 | GACACCAGCCTGACCACCAT | CGG | \|\|\|\|\|\|\|[\|\|\|\|\|\|\|\|\|\|\|\|\|]PAM | NA | Intergenic | NA | NA |
| T30566 | ACACCAGCCTGACCACCATCGGG | 1000 | | | | | | | | | |
| Oligo fwd | CACCACACCAGCCTGACCACCATC | | | | | | | | | | |
| Oligo rev | AAACGATGGTGGTCAGGCTGGTGT | | | | | | | | | | |
| Chromosome | start | end | strand | MM | target_seq | PAM | alignment | distance | position | gene name | gene id |
| GU305914.1 | 47699 | 47721 | - | 0 | ACACCAGCCTGACCACCATC | GGG | \|\|\|\|\|\|\|[\|\|\|\|\|\|\|\|\|\|\|\|\|]PAM | NA | Intergenic | NA | NA |
| T30567 | GGATCAACGATCCACCGTGTCGG | 1000 | | | | | | | | | |
| Oligo fwd | CACCGGATCAACGATCCACCGTGT | | | | | | | | | | |

Oligo rev AAACACACGGTGGATCGTTGATCC
Chromosome start end strand MM target_seq PAM alignment distance position gene name gene id
GU305914.1 47678 47700 - 0 GGATCAACGATCCACCGTGT CGG ||||||||[||||||||||||]PAM NA Intergenic NA NA

T30568 CGTGTCGGCGATGTATCGAGAGG 1000
Oligo fwd CACCCGTGTCGGCGATGTATCGAG
Oligo rev AAACCTCGATACATCGCCGACACG
Chromosome start end strand MM target_seq PAM alignment distance position gene name gene id
GU305914.1 47663 47685 - 0 CGTGTCGGCGATGTATCGAG AGG ||||||||[||||||||||||]PAM NA Intergenic NA NA

T30569 GTCGGCGATGTATCGAGAGGAGG 1000
Oligo fwd CACCGTCGGCGATGTATCGAGAGG
Oligo rev AAACCCTCTCGATACATCGCCGAC
Chromosome start end strand MM target_seq PAM alignment distance position gene name gene id
GU305914.1 47660 47682 - 0 GTCGGCGATGTATCGAGAGG AGG ||||||||[||||||||||||]PAM NA Intergenic NA NA

T30570 GGCGATGTATCGAGAGGAGGAGG 1000
Oligo fwd CACCGGCGATGTATCGAGAGGAGG
Oligo rev AAACCCTCCTCTCGATACATCGCC
Chromosome start end strand MM target_seq PAM alignment distance position gene name gene id
GU305914.1 47657 47679 - 0 GGCGATGTATCGAGAGGAGG AGG ||||||||[||||||||||||]PAM NA Intergenic NA NA

T30571 GCGATGTATCGAGAGGAGGAGGG 1000
Oligo fwd CACCGCGATGTATCGAGAGGAGGA
Oligo rev AAACTCCTCCTCTCGATACATCGC
Chromosome start end strand MM target_seq PAM alignment distance position gene name gene id
GU305914.1 47656 47678 - 0 GCGATGTATCGAGAGGAGGA GGG ||||||||[||||||||||||]PAM NA Intergenic NA NA

T30572 TCAAGCTCTTTAAGATGACACGG 1000
Oligo fwd CACCTCAAGCTCTTTAAGATGACA
Oligo rev AAACTGTCATCTTAAAGAGCTTGA
Chromosome start end strand MM target_seq PAM alignment distance position gene name gene id
GU305914.1 47633 47655 - 0 TCAAGCTCTTTAAGATGACA CGG ||||||||[||||||||||||]PAM NA Intergenic NA NA

T30573 CAAGCTCTTTAAGATGACACGGG 1000
Oligo fwd CACCCAAGCTCTTTAAGATGACAC
Oligo rev AAACGTGTCATCTTAAAGAGCTTG
Chromosome start end strand MM target_seq PAM alignment distance position gene name gene id
GU305914.1 47632 47654 - 0 CAAGCTCTTTAAGATGACAC GGG ||||||||[||||||||||||]PAM NA Intergenic NA NA

T30574 AAGCTCTTTAAGATGACACGGGG 1000
Oligo fwd CACCAAGCTCTTTAAGATGACACG
Oligo rev AAACCGTGTCATCTTAAAGAGCTT
Chromosome start end strand MM target_seq PAM alignment distance position gene name gene id
GU305914.1 47631 47653 - 0 AAGCTCTTTAAGATGACACG GGG ||||||||[||||||||||||]PAM NA Intergenic NA NA

T30575 TGACACGGGGATATCGATAGAGG 1000
Oligo fwd CACCTGACACGGGGATATCGATAG
Oligo rev AAACCTATCGATATCCCCGTGTCA
Chromosome start end strand MM target_seq PAM alignment distance position gene name gene id
GU305914.1 47618 47640 - 0 TGACACGGGGATATCGATAG AGG ||||||||[||||||||||||]PAM NA Intergenic NA NA

T30576 GACACGGGGATATCGATAGAGGG 1000
Oligo fwd CACCGACACGGGGATATCGATAGA
Oligo rev AAACTCTATCGATATCCCCGTGTC
Chromosome start end strand MM target_seq PAM alignment distance position gene name gene id
GU305914.1 47617 47639 - 0 GACACGGGGATATCGATAGA GGG ||||||||[||||||||||||]PAM NA Intergenic NA NA

T30577 GGATATCGATAGAGGGAAAACGG 1000
Oligo fwd CACCGGATATCGATAGAGGGAAAA
Oligo rev AAACTTTTCCCTCTATCGATATCC
Chromosome start end strand MM target_seq PAM alignment distance position gene name gene id
GU305914.1 47610 47632 - 0 GGATATCGATAGAGGGAAAA CGG ||||||||[||||||||||||]PAM NA Intergenic NA NA

T30578 AGAGGGAAAACGGATAAGAAAGG 1000
Oligo fwd CACCAGAGGGAAAACGGATAAGAA
Oligo rev AAACTTCTTATCCGTTTTCCCTCT
Chromosome start end strand MM target_seq PAM alignment distance position gene name gene id
GU305914.1 47600 47622 - 0 AGAGGGAAAACGGATAAGAA AGG ||||||||[||||||||||||]PAM NA Intergenic NA NA

T30579 GAGGGAAAACGGATAAGAAAGGG 1000
Oligo fwd CACCGAGGGAAAACGGATAAGAAA
Oligo rev AAACTTTCTTATCCGTTTTCCCTC
Chromosome start end strand MM target_seq PAM alignment distance position gene name gene id
GU305914.1 47599 47621 - 0 GAGGGAAAACGGATAAGAAA GGG ||||||||[||||||||||||]PAM NA Intergenic NA NA

T30580 GAAAACGGATAAGAAAGGGACGG 1000
Oligo fwd CACCGAAAACGGATAAGAAAGGGA
Oligo rev AAACTCCCTTTCTTATCCGTTTTC
Chromosome start end strand MM target_seq PAM alignment distance position gene name gene id
GU305914.1 47595 47617 - 0 GAAAACGGATAAGAAAGGGA CGG ||||||||[||||||||||||]PAM NA Intergenic NA NA

T30581 CGGATAAGAAAGGGACGGAGAGG 1000
Oligo fwd CACCCGGATAAGAAAGGGACGGAG
Oligo rev AAACCTCCGTCCCTTTCTTATCCG
Chromosome start end strand MM target_seq PAM alignment distance position gene name gene id
GU305914.1 47590 47612 - 0 CGGATAAGAAAGGGACGGAG AGG ||||||||[||||||||||||]PAM NA Intergenic NA NA

T30582 GGATAAGAAAGGGACGGAGAGGG 1000

<table>
<tr><td>Oligo fwd</td><td>CACCGGATAAGAAAGGGACGGAGA</td><td></td><td></td><td></td><td></td><td></td><td></td><td></td><td></td><td></td><td></td></tr>
<tr><td>Oligo rev</td><td>AAACTCTCCGTCCCTTTCTTATCC</td><td></td><td></td><td></td><td></td><td></td><td></td><td></td><td></td><td></td><td></td></tr>
<tr><td>Chromosome</td><td>start</td><td>end</td><td>strand</td><td>MM</td><td>target_seq</td><td>PAM</td><td>alignment</td><td>distance</td><td>position</td><td>gene name</td><td>gene id</td></tr>
<tr><td>GU305914.1</td><td>47589</td><td>47611</td><td>-</td><td>0</td><td>GGATAAGAAAGGGACGGAGA</td><td>GGG</td><td>||||||||[||||||||||||]PAM</td><td>NA</td><td>Intergenic</td><td>NA</td><td>NA</td></tr>
<tr><td></td><td></td><td></td><td></td><td></td><td></td><td></td><td></td><td></td><td></td><td></td><td></td></tr>
<tr><td>T30583</td><td>AGAAAGGGACGGAGAGGGTGAGG</td><td>1000</td><td></td><td></td><td></td><td></td><td></td><td></td><td></td><td></td><td></td></tr>
<tr><td>Oligo fwd</td><td>CACCAGAAAGGGACGGAGAGGGTG</td><td></td><td></td><td></td><td></td><td></td><td></td><td></td><td></td><td></td><td></td></tr>
<tr><td>Oligo rev</td><td>AAACCACCCTCTCCGTCCCTTTCT</td><td></td><td></td><td></td><td></td><td></td><td></td><td></td><td></td><td></td><td></td></tr>
<tr><td>Chromosome</td><td>start</td><td>end</td><td>strand</td><td>MM</td><td>target_seq</td><td>PAM</td><td>alignment</td><td>distance</td><td>position</td><td>gene name</td><td>gene id</td></tr>
<tr><td>GU305914.1</td><td>47584</td><td>47606</td><td>-</td><td>0</td><td>AGAAAGGGACGGAGAGGGTG</td><td>AGG</td><td>||||||||[||||||||||||]PAM</td><td>NA</td><td>Intergenic</td><td>NA</td><td>NA</td></tr>
<tr><td></td><td></td><td></td><td></td><td></td><td></td><td></td><td></td><td></td><td></td><td></td><td></td></tr>
<tr><td>T30584</td><td>GAAAGGGACGGAGAGGGTGAGGG</td><td>1000</td><td></td><td></td><td></td><td></td><td></td><td></td><td></td><td></td><td></td></tr>
<tr><td>Oligo fwd</td><td>CACCGAAAGGGACGGAGAGGGTGA</td><td></td><td></td><td></td><td></td><td></td><td></td><td></td><td></td><td></td><td></td></tr>
<tr><td>Oligo rev</td><td>AAACTCACCCTCTCCGTCCCTTTC</td><td></td><td></td><td></td><td></td><td></td><td></td><td></td><td></td><td></td><td></td></tr>
<tr><td>Chromosome</td><td>start</td><td>end</td><td>strand</td><td>MM</td><td>target_seq</td><td>PAM</td><td>alignment</td><td>distance</td><td>position</td><td>gene name</td><td>gene id</td></tr>
<tr><td>GU305914.1</td><td>47583</td><td>47605</td><td>-</td><td>0</td><td>GAAAGGGACGGAGAGGGTGA</td><td>GGG</td><td>||||||||[||||||||||||]PAM</td><td>NA</td><td>Intergenic</td><td>NA</td><td>NA</td></tr>
<tr><td></td><td></td><td></td><td></td><td></td><td></td><td></td><td></td><td></td><td></td><td></td><td></td></tr>
<tr><td>T30585</td><td>AAAGGGACGGAGAGGGTGAGGGG</td><td>1000</td><td></td><td></td><td></td><td></td><td></td><td></td><td></td><td></td><td></td></tr>
<tr><td>Oligo fwd</td><td>CACCAAAGGGACGGAGAGGGTGAG</td><td></td><td></td><td></td><td></td><td></td><td></td><td></td><td></td><td></td><td></td></tr>
<tr><td>Oligo rev</td><td>AAACCTCACCCTCTCCGTCCCTTT</td><td></td><td></td><td></td><td></td><td></td><td></td><td></td><td></td><td></td><td></td></tr>
<tr><td>Chromosome</td><td>start</td><td>end</td><td>strand</td><td>MM</td><td>target_seq</td><td>PAM</td><td>alignment</td><td>distance</td><td>position</td><td>gene name</td><td>gene id</td></tr>
<tr><td>GU305914.1</td><td>47582</td><td>47604</td><td>-</td><td>0</td><td>AAAGGGACGGAGAGGGTGAG</td><td>GGG</td><td>||||||||[||||||||||||]PAM</td><td>NA</td><td>Intergenic</td><td>NA</td><td>NA</td></tr>
<tr><td></td><td></td><td></td><td></td><td></td><td></td><td></td><td></td><td></td><td></td><td></td><td></td></tr>
<tr><td>T30586</td><td>AGAAAAAAAAATAAACAAAACGG</td><td>1000</td><td></td><td></td><td></td><td></td><td></td><td></td><td></td><td></td><td></td></tr>
<tr><td>Oligo fwd</td><td>CACCAGAAAAAAAAATAAACAAAA</td><td></td><td></td><td></td><td></td><td></td><td></td><td></td><td></td><td></td><td></td></tr>
<tr><td>Oligo rev</td><td>AAACTTTTGTTTATTTTTTTTTCT</td><td></td><td></td><td></td><td></td><td></td><td></td><td></td><td></td><td></td><td></td></tr>
<tr><td>Chromosome</td><td>start</td><td>end</td><td>strand</td><td>MM</td><td>target_seq</td><td>PAM</td><td>alignment</td><td>distance</td><td>position</td><td>gene name</td><td>gene id</td></tr>
<tr><td>GU305914.1</td><td>47559</td><td>47581</td><td>-</td><td>0</td><td>AGAAAAAAAAATAAACAAAA</td><td>CGG</td><td>||||||||[||||||||||||]PAM</td><td>NA</td><td>Intergenic</td><td>NA</td><td>NA</td></tr>
<tr><td></td><td></td><td></td><td></td><td></td><td></td><td></td><td></td><td></td><td></td><td></td><td></td></tr>
<tr><td>T30587</td><td>AAAAAAAATAAACAAAACGGAGG</td><td>1000</td><td></td><td></td><td></td><td></td><td></td><td></td><td></td><td></td><td></td></tr>
<tr><td>Oligo fwd</td><td>CACCAAAAAAAATAAACAAAACGG</td><td></td><td></td><td></td><td></td><td></td><td></td><td></td><td></td><td></td><td></td></tr>
<tr><td>Oligo rev</td><td>AAACCCGTTTTGTTTATTTTTTTT</td><td></td><td></td><td></td><td></td><td></td><td></td><td></td><td></td><td></td><td></td></tr>
<tr><td>Chromosome</td><td>start</td><td>end</td><td>strand</td><td>MM</td><td>target_seq</td><td>PAM</td><td>alignment</td><td>distance</td><td>position</td><td>gene name</td><td>gene id</td></tr>
<tr><td>GU305914.1</td><td>47556</td><td>47578</td><td>-</td><td>0</td><td>AAAAAAAATAAACAAAACGG</td><td>AGG</td><td>||||||||[||||||||||||]PAM</td><td>NA</td><td>Intergenic</td><td>NA</td><td>NA</td></tr>
<tr><td></td><td></td><td></td><td></td><td></td><td></td><td></td><td></td><td></td><td></td><td></td><td></td></tr>
<tr><td>T30588</td><td>AAAAAGCGACACATAACACACGG</td><td>1000</td><td></td><td></td><td></td><td></td><td></td><td></td><td></td><td></td><td></td></tr>
<tr><td>Oligo fwd</td><td>CACCAAAAAGCGACACATAACACA</td><td></td><td></td><td></td><td></td><td></td><td></td><td></td><td></td><td></td><td></td></tr>
<tr><td>Oligo rev</td><td>AAACTGTGTTATGTGTCGCTTTTT</td><td></td><td></td><td></td><td></td><td></td><td></td><td></td><td></td><td></td><td></td></tr>
<tr><td>Chromosome</td><td>start</td><td>end</td><td>strand</td><td>MM</td><td>target_seq</td><td>PAM</td><td>alignment</td><td>distance</td><td>position</td><td>gene name</td><td>gene id</td></tr>
<tr><td>GU305914.1</td><td>47528</td><td>47550</td><td>-</td><td>0</td><td>AAAAAGCGACACATAACACA</td><td>CGG</td><td>||||||||[||||||||||||]PAM</td><td>NA</td><td>Intergenic</td><td>NA</td><td>NA</td></tr>
<tr><td></td><td></td><td></td><td></td><td></td><td></td><td></td><td></td><td></td><td></td><td></td><td></td></tr>
<tr><td>T30589</td><td>ACACTTCGCTCTCGTGCCGACGG</td><td>1000</td><td></td><td></td><td></td><td></td><td></td><td></td><td></td><td></td><td></td></tr>
<tr><td>Oligo fwd</td><td>CACCACACTTCGCTCTCGTGCCGA</td><td></td><td></td><td></td><td></td><td></td><td></td><td></td><td></td><td></td><td></td></tr>
<tr><td>Oligo rev</td><td>AAACTCGGCACGAGAGCGAAGTGT</td><td></td><td></td><td></td><td></td><td></td><td></td><td></td><td></td><td></td><td></td></tr>
<tr><td>Chromosome</td><td>start</td><td>end</td><td>strand</td><td>MM</td><td>target_seq</td><td>PAM</td><td>alignment</td><td>distance</td><td>position</td><td>gene name</td><td>gene id</td></tr>
<tr><td>GU305914.1</td><td>47447</td><td>47469</td><td>-</td><td>0</td><td>ACACTTCGCTCTCGTGCCGA</td><td>CGG</td><td>||||||||[||||||||||||]PAM</td><td>NA</td><td>Intergenic</td><td>NA</td><td>NA</td></tr>
<tr><td></td><td></td><td></td><td></td><td></td><td></td><td></td><td></td><td></td><td></td><td></td><td></td></tr>
<tr><td>T30590</td><td>CTTCGCTCTCGTGCCGACGGCGG</td><td>1000</td><td></td><td></td><td></td><td></td><td></td><td></td><td></td><td></td><td></td></tr>
<tr><td>Oligo fwd</td><td>CACCCTTCGCTCTCGTGCCGACGG</td><td></td><td></td><td></td><td></td><td></td><td></td><td></td><td></td><td></td><td></td></tr>
<tr><td>Oligo rev</td><td>AAACCCGTCGGCACGAGAGCGAAG</td><td></td><td></td><td></td><td></td><td></td><td></td><td></td><td></td><td></td><td></td></tr>
<tr><td>Chromosome</td><td>start</td><td>end</td><td>strand</td><td>MM</td><td>target_seq</td><td>PAM</td><td>alignment</td><td>distance</td><td>position</td><td>gene name</td><td>gene id</td></tr>
<tr><td>GU305914.1</td><td>47444</td><td>47466</td><td>-</td><td>0</td><td>CTTCGCTCTCGTGCCGACGG</td><td>CGG</td><td>||||||||[||||||||||||]PAM</td><td>NA</td><td>Intergenic</td><td>NA</td><td>NA</td></tr>
<tr><td></td><td></td><td></td><td></td><td></td><td></td><td></td><td></td><td></td><td></td><td></td><td></td></tr>
<tr><td>T30591</td><td>CGTGCCGACGGCGGCCGCGCTGG</td><td>1000</td><td></td><td></td><td></td><td></td><td></td><td></td><td></td><td></td><td></td></tr>
<tr><td>Oligo fwd</td><td>CACCCGTGCCGACGGCGGCCGCGC</td><td></td><td></td><td></td><td></td><td></td><td></td><td></td><td></td><td></td><td></td></tr>
<tr><td>Oligo rev</td><td>AAACGCGCGGCCGCCGTCGGCACG</td><td></td><td></td><td></td><td></td><td></td><td></td><td></td><td></td><td></td><td></td></tr>
<tr><td>Chromosome</td><td>start</td><td>end</td><td>strand</td><td>MM</td><td>target_seq</td><td>PAM</td><td>alignment</td><td>distance</td><td>position</td><td>gene name</td><td>gene id</td></tr>
<tr><td>GU305914.1</td><td>47435</td><td>47457</td><td>-</td><td>0</td><td>CGTGCCGACGGCGGCCGCGC</td><td>TGG</td><td>||||||||[||||||||||||]PAM</td><td>NA</td><td>Intergenic</td><td>NA</td><td>NA</td></tr>
<tr><td>GU305914.1</td><td>168918</td><td>168940</td><td>-</td><td>4</td><td>CGAGACGACGGTCGCCGCGC</td><td>CGG</td><td>||-|-|||[|||--|||||||]PAM</td><td>NA</td><td>Intergenic</td><td>NA</td><td>NA</td></tr>
<tr><td></td><td></td><td></td><td></td><td></td><td></td><td></td><td></td><td></td><td></td><td></td><td></td></tr>
<tr><td>T30592</td><td>ACGGCGGCCGCGCTGGACTGTGG</td><td>1000</td><td></td><td></td><td></td><td></td><td></td><td></td><td></td><td></td><td></td></tr>
<tr><td>Oligo fwd</td><td>CACCACGGCGGCCGCGCTGGACTG</td><td></td><td></td><td></td><td></td><td></td><td></td><td></td><td></td><td></td><td></td></tr>
<tr><td>Oligo rev</td><td>AAACCAGTCCAGCGCGGCCGCCGT</td><td></td><td></td><td></td><td></td><td></td><td></td><td></td><td></td><td></td><td></td></tr>
<tr><td>Chromosome</td><td>start</td><td>end</td><td>strand</td><td>MM</td><td>target_seq</td><td>PAM</td><td>alignment</td><td>distance</td><td>position</td><td>gene name</td><td>gene id</td></tr>
<tr><td>GU305914.1</td><td>47428</td><td>47450</td><td>-</td><td>0</td><td>ACGGCGGCCGCGCTGGACTG</td><td>TGG</td><td>||||||||[||||||||||||]PAM</td><td>NA</td><td>Intergenic</td><td>NA</td><td>NA</td></tr>
<tr><td></td><td></td><td></td><td></td><td></td><td></td><td></td><td></td><td></td><td></td><td></td><td></td></tr>
<tr><td>T30593</td><td>CCGCGCTGGACTGTGGAGTACGG</td><td>1000</td><td></td><td></td><td></td><td></td><td></td><td></td><td></td><td></td><td></td></tr>
<tr><td>Oligo fwd</td><td>CACCCCGCGCTGGACTGTGGAGTA</td><td></td><td></td><td></td><td></td><td></td><td></td><td></td><td></td><td></td><td></td></tr>
<tr><td>Oligo rev</td><td>AAACTACTCCACAGTCCAGCGCGG</td><td></td><td></td><td></td><td></td><td></td><td></td><td></td><td></td><td></td><td></td></tr>
<tr><td>Chromosome</td><td>start</td><td>end</td><td>strand</td><td>MM</td><td>target_seq</td><td>PAM</td><td>alignment</td><td>distance</td><td>position</td><td>gene name</td><td>gene id</td></tr>
<tr><td>GU305914.1</td><td>47421</td><td>47443</td><td>-</td><td>0</td><td>CCGCGCTGGACTGTGGAGTA</td><td>CGG</td><td>||||||||[||||||||||||]PAM</td><td>NA</td><td>Intergenic</td><td>NA</td><td>NA</td></tr>
<tr><td></td><td></td><td></td><td></td><td></td><td></td><td></td><td></td><td></td><td></td><td></td><td></td></tr>
<tr><td>T30594</td><td>CTGGACTGTGGAGTACGGTCTGG</td><td>1000</td><td></td><td></td><td></td><td></td><td></td><td></td><td></td><td></td><td></td></tr>
<tr><td>Oligo fwd</td><td>CACCCTGGACTGTGGAGTACGGTC</td><td></td><td></td><td></td><td></td><td></td><td></td><td></td><td></td><td></td><td></td></tr>
<tr><td>Oligo rev</td><td>AAACGACCGTACTCCACAGTCCAG</td><td></td><td></td><td></td><td></td><td></td><td></td><td></td><td></td><td></td><td></td></tr>
<tr><td>Chromosome</td><td>start</td><td>end</td><td>strand</td><td>MM</td><td>target_seq</td><td>PAM</td><td>alignment</td><td>distance</td><td>position</td><td>gene name</td><td>gene id</td></tr>
<tr><td>GU305914.1</td><td>47416</td><td>47438</td><td>-</td><td>0</td><td>CTGGACTGTGGAGTACGGTC</td><td>TGG</td><td>||||||||[||||||||||||]PAM</td><td>NA</td><td>Intergenic</td><td>NA</td><td>NA</td></tr>
<tr><td></td><td></td><td></td><td></td><td></td><td></td><td></td><td></td><td></td><td></td><td></td><td></td></tr>
<tr><td>T30595</td><td>CGGTCTGGCACCTCTCATGTTGG</td><td>1000</td><td></td><td></td><td></td><td></td><td></td><td></td><td></td><td></td><td></td></tr>
<tr><td>Oligo fwd</td><td>CACCCGGTCTGGCACCTCTCATGT</td><td></td><td></td><td></td><td></td><td></td><td></td><td></td><td></td><td></td><td></td></tr>
<tr><td>Oligo rev</td><td>AAACACATGAGAGGTGCCAGACCG</td><td></td><td></td><td></td><td></td><td></td><td></td><td></td><td></td><td></td><td></td></tr>
<tr><td>Chromosome</td><td>start</td><td>end</td><td>strand</td><td>MM</td><td>target_seq</td><td>PAM</td><td>alignment</td><td>distance</td><td>position</td><td>gene name</td><td>gene id</td></tr>
<tr><td>GU305914.1</td><td>47401</td><td>47423</td><td>-</td><td>0</td><td>CGGTCTGGCACCTCTCATGT</td><td>TGG</td><td>||||||||[||||||||||||]PAM</td><td>NA</td><td>Intergenic</td><td>NA</td><td>NA</td></tr>
<tr><td></td><td></td><td></td><td></td><td></td><td></td><td></td><td></td><td></td><td></td><td></td><td></td></tr>
<tr><td>T30596</td><td>CCATCCGTTGCCCCCCGCCGCGG</td><td>1000</td><td></td><td></td><td></td><td></td><td></td><td></td><td></td><td></td><td></td></tr>
<tr><td>Oligo fwd</td><td>CACCCCATCCGTTGCCCCCCGCCG</td><td></td><td></td><td></td><td></td><td></td><td></td><td></td><td></td><td></td><td></td></tr>
<tr><td>Oligo rev</td><td>AAACCGGCGGGGGGCAACGGATGG</td><td></td><td></td><td></td><td></td><td></td><td></td><td></td><td></td><td></td><td></td></tr>
<tr><td>Chromosome</td><td>start</td><td>end</td><td>strand</td><td>MM</td><td>target_seq</td><td>PAM</td><td>alignment</td><td>distance</td><td>position</td><td>gene name</td><td>gene id</td></tr>
<tr><td>GU305914.1</td><td>47378</td><td>47400</td><td>-</td><td>0</td><td>CCATCCGTTGCCCCCCGCCG</td><td>CGG</td><td>||||||||[||||||||||||]PAM</td><td>NA</td><td>Intergenic</td><td>NA</td><td>NA</td></tr>
</table>

| | | | | | | | | | | | |
|---|---|---|---|---|---|---|---|---|---|---|---|
| T30597 | CGTTGCCCCCCGCCGCGGATCGG | 1000 | | | | | | | | | |
| Oligo fwd | CACCCGTTGCCCCCCGCCGCGGAT | | | | | | | | | | |
| Oligo rev | AAACATCCGCGGCGGGGGGCAACG | | | | | | | | | | |
| Chromosome | start | end | strand | MM | target_seq | PAM | alignment | distance | position | gene name | gene id |
| GU305914.1 | 47373 | 47395 | - | 0 | CGTTGCCCCCCGCCGCGGAT | CGG | \|\|\|\|\|\|\|\|[\|\|\|\|\|\|\|\|\|\|\|\|]PAM | NA | Intergenic | NA | NA |
| T30598 | CCCGCCGCGGATCGGCCTCACGG | 1000 | | | | | | | | | |
| Oligo fwd | CACCCCCGCCGCGGATCGGCCTCA | | | | | | | | | | |
| Oligo rev | AAACTGAGGCCGATCCGCGGCGGG | | | | | | | | | | |
| Chromosome | start | end | strand | MM | target_seq | PAM | alignment | distance | position | gene name | gene id |
| GU305914.1 | 47365 | 47387 | - | 0 | CCCGCCGCGGATCGGCCTCA | CGG | \|\|\|\|\|\|\|\|[\|\|\|\|\|\|\|\|\|\|\|\|]PAM | NA | Intergenic | NA | NA |
| T30599 | CCGCCGCGGATCGGCCTCACGGG | 1000 | | | | | | | | | |
| Oligo fwd | CACCCCGCCGCGGATCGGCCTCAC | | | | | | | | | | |
| Oligo rev | AAACGTGAGGCCGATCCGCGGCGG | | | | | | | | | | |
| Chromosome | start | end | strand | MM | target_seq | PAM | alignment | distance | position | gene name | gene id |
| GU305914.1 | 47364 | 47386 | - | 0 | CCGCCGCGGATCGGCCTCAC | GGG | \|\|\|\|\|\|\|\|[\|\|\|\|\|\|\|\|\|\|\|\|]PAM | NA | Intergenic | NA | NA |
| T30600 | ATCGGCCTCACGGGCGACGCCGG | 1000 | | | | | | | | | |
| Oligo fwd | CACCATCGGCCTCACGGGCGACGC | | | | | | | | | | |
| Oligo rev | AAACGCGTCGCCCGTGAGGCCGAT | | | | | | | | | | |
| Chromosome | start | end | strand | MM | target_seq | PAM | alignment | distance | position | gene name | gene id |
| GU305914.1 | 47355 | 47377 | - | 0 | ATCGGCCTCACGGGCGACGC | CGG | \|\|\|\|\|\|\|\|[\|\|\|\|\|\|\|\|\|\|\|\|]PAM | NA | Intergenic | NA | NA |
| T30601 | TCGGCCTCACGGGCGACGCCGGG | 1000 | | | | | | | | | |
| Oligo fwd | CACCTCGGCCTCACGGGCGACGCC | | | | | | | | | | |
| Oligo rev | AAACGGCGTCGCCCGTGAGGCCGA | | | | | | | | | | |
| Chromosome | start | end | strand | MM | target_seq | PAM | alignment | distance | position | gene name | gene id |
| GU305914.1 | 47354 | 47376 | - | 0 | TCGGCCTCACGGGCGACGCC | GGG | \|\|\|\|\|\|\|\|[\|\|\|\|\|\|\|\|\|\|\|\|]PAM | NA | Intergenic | NA | NA |
| T30602 | CGGCCTCACGGGCGACGCCGGGG | 1000 | | | | | | | | | |
| Oligo fwd | CACCCGGCCTCACGGGCGACGCCG | | | | | | | | | | |
| Oligo rev | AAACCGGCGTCGCCCGTGAGGCCG | | | | | | | | | | |
| Chromosome | start | end | strand | MM | target_seq | PAM | alignment | distance | position | gene name | gene id |
| GU305914.1 | 47353 | 47375 | - | 0 | CGGCCTCACGGGCGACGCCG | GGG | \|\|\|\|\|\|\|\|[\|\|\|\|\|\|\|\|\|\|\|\|]PAM | NA | Intergenic | NA | NA |
| T30603 | CGCCGGGGCCACCGACACGTCGG | 1000 | | | | | | | | | |
| Oligo fwd | CACCCGCCGGGGCCACCGACACGT | | | | | | | | | | |
| Oligo rev | AAACACGTGTCGGTGGCCCCGGCG | | | | | | | | | | |
| Chromosome | start | end | strand | MM | target_seq | PAM | alignment | distance | position | gene name | gene id |
| GU305914.1 | 47338 | 47360 | - | 0 | CGCCGGGGCCACCGACACGT | CGG | \|\|\|\|\|\|\|\|[\|\|\|\|\|\|\|\|\|\|\|\|]PAM | NA | Intergenic | NA | NA |
| T30604 | TCGTCTGTGACGACTTCGACCGG | 1000 | | | | | | | | | |
| Oligo fwd | CACCTCGTCTGTGACGACTTCGAC | | | | | | | | | | |
| Oligo rev | AAACGTCGAAGTCGTCACAGACGA | | | | | | | | | | |
| Chromosome | start | end | strand | MM | target_seq | PAM | alignment | distance | position | gene name | gene id |
| GU305914.1 | 47280 | 47302 | - | 0 | TCGTCTGTGACGACTTCGAC | CGG | \|\|\|\|\|\|\|\|[\|\|\|\|\|\|\|\|\|\|\|\|]PAM | NA | Intergenic | NA | NA |
| T30605 | CGTCTGTGACGACTTCGACCGGG | 1000 | | | | | | | | | |
| Oligo fwd | CACCCGTCTGTGACGACTTCGACC | | | | | | | | | | |
| Oligo rev | AAACGGTCGAAGTCGTCACAGACG | | | | | | | | | | |
| Chromosome | start | end | strand | MM | target_seq | PAM | alignment | distance | position | gene name | gene id |
| GU305914.1 | 47279 | 47301 | - | 0 | CGTCTGTGACGACTTCGACC | GGG | \|\|\|\|\|\|\|\|[\|\|\|\|\|\|\|\|\|\|\|\|]PAM | NA | Intergenic | NA | NA |
| T30606 | GTCTGTGACGACTTCGACCGGGG | 1000 | | | | | | | | | |
| Oligo fwd | CACCGTCTGTGACGACTTCGACCG | | | | | | | | | | |
| Oligo rev | AAACCGGTCGAAGTCGTCACAGAC | | | | | | | | | | |
| Chromosome | start | end | strand | MM | target_seq | PAM | alignment | distance | position | gene name | gene id |
| GU305914.1 | 47278 | 47300 | - | 0 | GTCTGTGACGACTTCGACCG | GGG | \|\|\|\|\|\|\|\|[\|\|\|\|\|\|\|\|\|\|\|\|]PAM | NA | Intergenic | NA | NA |
| T30607 | TTCGACCGGGGTGCTTACCACGG | 1000 | | | | | | | | | |
| Oligo fwd | CACCTTCGACCGGGGTGCTTACCA | | | | | | | | | | |
| Oligo rev | AAACTGGTAAGCACCCCGGTCGAA | | | | | | | | | | |
| Chromosome | start | end | strand | MM | target_seq | PAM | alignment | distance | position | gene name | gene id |
| GU305914.1 | 47266 | 47288 | - | 0 | TTCGACCGGGGTGCTTACCA | CGG | \|\|\|\|\|\|\|\|[\|\|\|\|\|\|\|\|\|\|\|\|]PAM | NA | Intergenic | NA | NA |
| T30608 | CCGGGGTGCTTACCACGGACAGG | 1000 | | | | | | | | | |
| Oligo fwd | CACCCCGGGGTGCTTACCACGGAC | | | | | | | | | | |
| Oligo rev | AAACGTCCGTGGTAAGCACCCCGG | | | | | | | | | | |
| Chromosome | start | end | strand | MM | target_seq | PAM | alignment | distance | position | gene name | gene id |
| GU305914.1 | 47261 | 47283 | - | 0 | CCGGGGTGCTTACCACGGAC | AGG | \|\|\|\|\|\|\|\|[\|\|\|\|\|\|\|\|\|\|\|\|]PAM | NA | Intergenic | NA | NA |
| T30609 | CGGACAGGTAGTGCGACAGCTGG | 1000 | | | | | | | | | |
| Oligo fwd | CACCCGGACAGGTAGTGCGACAGC | | | | | | | | | | |
| Oligo rev | AAACGCTGTCGCACTACCTGTCCG | | | | | | | | | | |
| Chromosome | start | end | strand | MM | target_seq | PAM | alignment | distance | position | gene name | gene id |
| GU305914.1 | 47246 | 47268 | - | 0 | CGGACAGGTAGTGCGACAGC | TGG | \|\|\|\|\|\|\|\|[\|\|\|\|\|\|\|\|\|\|\|\|]PAM | NA | Intergenic | NA | NA |
| T30610 | AGGTAGTGCGACAGCTGGTCTGG | 1000 | | | | | | | | | |
| Oligo fwd | CACCAGGTAGTGCGACAGCTGGTC | | | | | | | | | | |
| Oligo rev | AAACGACCAGCTGTCGCACTACCT | | | | | | | | | | |
| Chromosome | start | end | strand | MM | target_seq | PAM | alignment | distance | position | gene name | gene id |
| GU305914.1 | 47241 | 47263 | - | 0 | AGGTAGTGCGACAGCTGGTC | TGG | \|\|\|\|\|\|\|\|[\|\|\|\|\|\|\|\|\|\|\|\|]PAM | NA | Intergenic | NA | NA |
| T30611 | GGTAGTGCGACAGCTGGTCTGGG | 1000 | | | | | | | | | |
| Oligo fwd | CACCGGTAGTGCGACAGCTGGTCT | | | | | | | | | | |
| Oligo rev | AAACAGACCAGCTGTCGCACTACC | | | | | | | | | | |
| Chromosome | start | end | strand | MM | target_seq | PAM | alignment | distance | position | gene name | gene id |

GU305914.1 47240 47262 - 0 GGTAGTGCGACAGCTGGTCT GGG ||||||||[||||||||||||]PAM NA Intergenic NA NA

T30612 GTAGTGCGACAGCTGGTCTGGGG 1000
Oligo fwd CACCGTAGTGCGACAGCTGGTCTG
Oligo rev AAACCAGACCAGCTGTCGCACTAC
Chromosome start end strand MM target_seq PAM alignment distance position gene name gene id
GU305914.1 47239 47261 - 0 GTAGTGCGACAGCTGGTCTG GGG ||||||||[||||||||||||]PAM NA Intergenic NA NA

T30613 GCGACAGCTGGTCTGGGGTGAGG 1000
Oligo fwd CACCGCGACAGCTGGTCTGGGGTG
Oligo rev AAACCACCCCAGACCAGCTGTCGC
Chromosome start end strand MM target_seq PAM alignment distance position gene name gene id
GU305914.1 47234 47256 - 0 GCGACAGCTGGTCTGGGGTG AGG ||||||||[||||||||||||]PAM NA Intergenic NA NA

T30614 CACGCCTTCGCCCACGAGCTCGG 1000
Oligo fwd CACCCACGCCTTCGCCCACGAGCT
Oligo rev AAACAGCTCGTGGGCGAAGGCGTG
Chromosome start end strand MM target_seq PAM alignment distance position gene name gene id
GU305914.1 47197 47219 - 0 CACGCCTTCGCCCACGAGCT CGG ||||||||[||||||||||||]PAM NA Intergenic NA NA

T30615 CCCACGAGCTCGGAGACCGACGG 1000
Oligo fwd CACCCCCACGAGCTCGGAGACCGA
Oligo rev AAACTCGGTCTCCGAGCTCGTGGG
Chromosome start end strand MM target_seq PAM alignment distance position gene name gene id
GU305914.1 47187 47209 - 0 CCCACGAGCTCGGAGACCGA CGG ||||||||[||||||||||||]PAM NA Intergenic NA NA

T30616 GGAGACCGACGGCAGCTCGTAGG 1000
Oligo fwd CACCGGAGACCGACGGCAGCTCGT
Oligo rev AAACACGAGCTGCCGTCGGTCTCC
Chromosome start end strand MM target_seq PAM alignment distance position gene name gene id
GU305914.1 47176 47198 - 0 GGAGACCGACGGCAGCTCGT AGG ||||||||[||||||||||||]PAM NA Intergenic NA NA

T30617 CCGACGGCAGCTCGTAGGTCTGG 1000
Oligo fwd CACCCCGACGGCAGCTCGTAGGTC
Oligo rev AAACGACCTACGAGCTGCCGTCGG
Chromosome start end strand MM target_seq PAM alignment distance position gene name gene id
GU305914.1 47171 47193 - 0 CCGACGGCAGCTCGTAGGTC TGG ||||||||[||||||||||||]PAM NA Intergenic NA NA

T30618 AGGTCTGGTACGTCGCCTCTTGG 1000
Oligo fwd CACCAGGTCTGGTACGTCGCCTCT
Oligo rev AAACAGAGGCGACGTACCAGACCT
Chromosome start end strand MM target_seq PAM alignment distance position gene name gene id
GU305914.1 47156 47178 - 0 AGGTCTGGTACGTCGCCTCT TGG ||||||||[||||||||||||]PAM NA Intergenic NA NA

T30619 TGCAGCTCACTAGCAGCTCCTGG 1000
Oligo fwd CACCTGCAGCTCACTAGCAGCTCC
Oligo rev AAACGGAGCTGCTAGTGAGCTGCA
Chromosome start end strand MM target_seq PAM alignment distance position gene name gene id
GU305914.1 47129 47151 - 0 TGCAGCTCACTAGCAGCTCC TGG ||||||||[||||||||||||]PAM NA Intergenic NA NA

T30620 GCAGCTCACTAGCAGCTCCTGGG 1000
Oligo fwd CACCGCAGCTCACTAGCAGCTCCT
Oligo rev AAACAGGAGCTGCTAGTGAGCTGC
Chromosome start end strand MM target_seq PAM alignment distance position gene name gene id
GU305914.1 47128 47150 - 0 GCAGCTCACTAGCAGCTCCT GGG ||||||||[||||||||||||]PAM NA Intergenic NA NA

T30621 CACTAGCAGCTCCTGGGTCATGG 1000
Oligo fwd CACCCACTAGCAGCTCCTGGGTCA
Oligo rev AAACTGACCCAGGAGCTGCTAGTG
Chromosome start end strand MM target_seq PAM alignment distance position gene name gene id
GU305914.1 47122 47144 - 0 CACTAGCAGCTCCTGGGTCA TGG ||||||||[||||||||||||]PAM NA Intergenic NA NA

T30622 CATCGCCCACGTAATGCACCAGG 1000
Oligo fwd CACCCATCGCCCACGTAATGCACC
Oligo rev AAACGGTGCATTACGTGGGCGATG
Chromosome start end strand MM target_seq PAM alignment distance position gene name gene id
GU305914.1 47084 47106 - 0 CATCGCCCACGTAATGCACC AGG ||||||||[||||||||||||]PAM NA Intergenic NA NA

T30623 CACGTAATGCACCAGGCTCCTGG 1000
Oligo fwd CACCCACGTAATGCACCAGGCTCC
Oligo rev AAACGGAGCCTGGTGCATTACGTG
Chromosome start end strand MM target_seq PAM alignment distance position gene name gene id
GU305914.1 47077 47099 - 0 CACGTAATGCACCAGGCTCC TGG ||||||||[||||||||||||]PAM NA Intergenic NA NA

T30624 ACGTAATGCACCAGGCTCCTGGG 1000
Oligo fwd CACCACGTAATGCACCAGGCTCCT
Oligo rev AAACAGGAGCCTGGTGCATTACGT
Chromosome start end strand MM target_seq PAM alignment distance position gene name gene id
GU305914.1 47076 47098 - 0 ACGTAATGCACCAGGCTCCT GGG ||||||||[||||||||||||]PAM NA Intergenic NA NA

T30625 CGTAATGCACCAGGCTCCTGGGG 1000
Oligo fwd CACCCGTAATGCACCAGGCTCCTG
Oligo rev AAACCAGGAGCCTGGTGCATTACG
Chromosome start end strand MM target_seq PAM alignment distance position gene name gene id
GU305914.1 47075 47097 - 0 CGTAATGCACCAGGCTCCTG GGG ||||||||[||||||||||||]PAM NA Intergenic NA NA

T30626 TGGGGCACTTGAAGATGTTCTGG 1000
Oligo fwd CACCTGGGGCACTTGAAGATGTTC
Oligo rev AAACGAACATCTTCAAGTGCCCCA

Chromosome start end strand MM target_seq PAM alignment distance position gene name gene id
GU305914.1 47057 47079 - 0 TGGGGCACTTGAAGATGTTC TGG ||||||||[||||||||||||]PAM NA Intergenic NA NA

T30627 AAGATGTTCTGGTCGTTGTACGG 1000
Oligo fwd CACCAAGATGTTCTGGTCGTTGTA
Oligo rev AAACTACAACGACCAGAACATCTT
Chromosome start end strand MM target_seq PAM alignment distance position gene name gene id
GU305914.1 47046 47068 - 0 AAGATGTTCTGGTCGTTGTA CGG ||||||||[||||||||||||]PAM NA Intergenic NA NA

T30628 AGATGTTCTGGTCGTTGTACGGG 1000
Oligo fwd CACCAGATGTTCTGGTCGTTGTAC
Oligo rev AAACGTACAACGACCAGAACATCT
Chromosome start end strand MM target_seq PAM alignment distance position gene name gene id
GU305914.1 47045 47067 - 0 AGATGTTCTGGTCGTTGTAC GGG ||||||||[||||||||||||]PAM NA Intergenic NA NA
GU305914.1 67702 67724 - 4 CGATGTCCGGGTCCTTGTAC TGG -|||||-|[-||||-||||||]PAM NA Intergenic NA NA

T30629 CGTTGTACGGGTTGTTCATGCGG 1000
Oligo fwd CACCCGTTGTACGGGTTGTTCATG
Oligo rev AAACCATGAACAACCCGTACAACG
Chromosome start end strand MM target_seq PAM alignment distance position gene name gene id
GU305914.1 47033 47055 - 0 CGTTGTACGGGTTGTTCATG CGG ||||||||[||||||||||||]PAM NA Intergenic NA NA

T30630 CGGGTTGTTCATGCGGTACTTGG 1000
Oligo fwd CACCCGGGTTGTTCATGCGGTACT
Oligo rev AAACAGTACCGCATGAACAACCCG
Chromosome start end strand MM target_seq PAM alignment distance position gene name gene id
GU305914.1 47026 47048 - 0 CGGGTTGTTCATGCGGTACT TGG ||||||||[||||||||||||]PAM NA Intergenic NA NA

T30631 ATGCGGTACTTGGCGCTGTAAGG 1000
Oligo fwd CACCATGCGGTACTTGGCGCTGTA
Oligo rev AAACTACAGCGCCAAGTACCGCAT
Chromosome start end strand MM target_seq PAM alignment distance position gene name gene id
GU305914.1 47016 47038 - 0 ATGCGGTACTTGGCGCTGTA AGG ||||||||[||||||||||||]PAM NA Intergenic NA NA

T30632 TGCGGTACTTGGCGCTGTAAGGG 1000
Oligo fwd CACCTGCGGTACTTGGCGCTGTAA
Oligo rev AAACTTACAGCGCCAAGTACCGCA
Chromosome start end strand MM target_seq PAM alignment distance position gene name gene id
GU305914.1 47015 47037 - 0 TGCGGTACTTGGCGCTGTAA GGG ||||||||[||||||||||||]PAM NA Intergenic NA NA

T30633 TTGGCGCTGTAAGGGTTTCCCGG 1000
Oligo fwd CACCTTGGCGCTGTAAGGGTTTCC
Oligo rev AAACGGAAACCCTTACAGCGCCAA
Chromosome start end strand MM target_seq PAM alignment distance position gene name gene id
GU305914.1 47007 47029 - 0 TTGGCGCTGTAAGGGTTTCC CGG ||||||||[||||||||||||]PAM NA Intergenic NA NA

T30634 TGGCGCTGTAAGGGTTTCCCGGG 1000
Oligo fwd CACCTGGCGCTGTAAGGGTTTCCC
Oligo rev AAACGGGAAACCCTTACAGCGCCA
Chromosome start end strand MM target_seq PAM alignment distance position gene name gene id
GU305914.1 47006 47028 - 0 TGGCGCTGTAAGGGTTTCCC GGG ||||||||[||||||||||||]PAM NA Intergenic NA NA

T30635 CCGGGTCCACCTCGATGTCCAGG 1000
Oligo fwd CACCCCGGGTCCACCTCGATGTCC
Oligo rev AAACGGACATCGAGGTGGACCCGG
Chromosome start end strand MM target_seq PAM alignment distance position gene name gene id
GU305914.1 46988 47010 - 0 CCGGGTCCACCTCGATGTCC AGG ||||||||[||||||||||||]PAM NA Intergenic NA NA

T30636 GTCGTCGTCGTGACCCATCGCGG 1000
Oligo fwd CACCGTCGTCGTCGTGACCCATCG
Oligo rev AAACCGATGGGTCACGACGACGAC
Chromosome start end strand MM target_seq PAM alignment distance position gene name gene id
GU305914.1 46966 46988 - 0 GTCGTCGTCGTGACCCATCG CGG ||||||||[||||||||||||]PAM NA Intergenic NA NA

T30637 GACCCATCGCGGCCCCCTCCAGG 1000
Oligo fwd CACCGACCCATCGCGGCCCCCTCC
Oligo rev AAACGGAGGGGGCCGCGATGGGTC
Chromosome start end strand MM target_seq PAM alignment distance position gene name gene id
GU305914.1 46955 46977 - 0 GACCCATCGCGGCCCCCTCC AGG ||||||||[||||||||||||]PAM NA Intergenic NA NA

T30638 GCCGTGCATCCTCGCCGTCACGG 1000
Oligo fwd CACCGCCGTGCATCCTCGCCGTCA
Oligo rev AAACTGACGGCGAGGATGCACGGC
Chromosome start end strand MM target_seq PAM alignment distance position gene name gene id
GU305914.1 46933 46955 - 0 GCCGTGCATCCTCGCCGTCA CGG ||||||||[||||||||||||]PAM NA Intergenic NA NA

T30639 GCATCCTCGCCGTCACGGCCAGG 1000
Oligo fwd CACCGCATCCTCGCCGTCACGGCC
Oligo rev AAACGGCCGTGACGGCGAGGATGC
Chromosome start end strand MM target_seq PAM alignment distance position gene name gene id
GU305914.1 46928 46950 - 0 GCATCCTCGCCGTCACGGCC AGG ||||||||[||||||||||||]PAM NA Intergenic NA NA

T30640 TCGCCGTCACGGCCAGGATCTGG 1000
Oligo fwd CACCTCGCCGTCACGGCCAGGATC
Oligo rev AAACGATCCTGGCCGTGACGGCGA
Chromosome start end strand MM target_seq PAM alignment distance position gene name gene id
GU305914.1 46922 46944 - 0 TCGCCGTCACGGCCAGGATC TGG ||||||||[||||||||||||]PAM NA Intergenic NA NA

T30641 CGTCACGGCCAGGATCTGGTTGG 1000

<table>
<tr><td>Oligo fwd</td><td>CACCCGTCACGGCCAGGATCTGGT</td><td></td><td></td><td></td><td></td><td></td><td></td><td></td><td></td><td></td><td></td></tr>
<tr><td>Oligo rev</td><td>AAACACCAGATCCTGGCCGTGACG</td><td></td><td></td><td></td><td></td><td></td><td></td><td></td><td></td><td></td><td></td></tr>
<tr><td>Chromosome</td><td>start</td><td>end</td><td>strand</td><td>MM</td><td>target_seq</td><td>PAM</td><td>alignment</td><td>distance</td><td>position</td><td>gene name</td><td>gene id</td></tr>
<tr><td>GU305914.1</td><td>46918</td><td>46940</td><td>-</td><td>0</td><td>CGTCACGGCCAGGATCTGGT</td><td>TGG</td><td>||||||||[||||||||||||]PAM</td><td>NA</td><td>Intergenic</td><td>NA</td><td>NA</td></tr>
<tr><td></td><td></td><td></td><td></td><td></td><td></td><td></td><td></td><td></td><td></td><td></td><td></td></tr>
<tr><td>T30642</td><td>CGGCCAGGATCTGGTTGGTCAGG</td><td>1000</td><td></td><td></td><td></td><td></td><td></td><td></td><td></td><td></td><td></td></tr>
<tr><td>Oligo fwd</td><td>CACCCGGCCAGGATCTGGTTGGTC</td><td></td><td></td><td></td><td></td><td></td><td></td><td></td><td></td><td></td><td></td></tr>
<tr><td>Oligo rev</td><td>AAACGACCAACCAGATCCTGGCCG</td><td></td><td></td><td></td><td></td><td></td><td></td><td></td><td></td><td></td><td></td></tr>
<tr><td>Chromosome</td><td>start</td><td>end</td><td>strand</td><td>MM</td><td>target_seq</td><td>PAM</td><td>alignment</td><td>distance</td><td>position</td><td>gene name</td><td>gene id</td></tr>
<tr><td>GU305914.1</td><td>46913</td><td>46935</td><td>-</td><td>0</td><td>CGGCCAGGATCTGGTTGGTC</td><td>AGG</td><td>||||||||[||||||||||||]PAM</td><td>NA</td><td>Intergenic</td><td>NA</td><td>NA</td></tr>
<tr><td></td><td></td><td></td><td></td><td></td><td></td><td></td><td></td><td></td><td></td><td></td><td></td></tr>
<tr><td>T30643</td><td>GATCTGGTTGGTCAGGTACCTGG</td><td>1000</td><td></td><td></td><td></td><td></td><td></td><td></td><td></td><td></td><td></td></tr>
<tr><td>Oligo fwd</td><td>CACCGATCTGGTTGGTCAGGTACC</td><td></td><td></td><td></td><td></td><td></td><td></td><td></td><td></td><td></td><td></td></tr>
<tr><td>Oligo rev</td><td>AAACGGTACCTGACCAACCAGATC</td><td></td><td></td><td></td><td></td><td></td><td></td><td></td><td></td><td></td><td></td></tr>
<tr><td>Chromosome</td><td>start</td><td>end</td><td>strand</td><td>MM</td><td>target_seq</td><td>PAM</td><td>alignment</td><td>distance</td><td>position</td><td>gene name</td><td>gene id</td></tr>
<tr><td>GU305914.1</td><td>46906</td><td>46928</td><td>-</td><td>0</td><td>GATCTGGTTGGTCAGGTACC</td><td>TGG</td><td>||||||||[||||||||||||]PAM</td><td>NA</td><td>Intergenic</td><td>NA</td><td>NA</td></tr>
<tr><td></td><td></td><td></td><td></td><td></td><td></td><td></td><td></td><td></td><td></td><td></td><td></td></tr>
<tr><td>T30644</td><td>ATCTGGTTGGTCAGGTACCTGGG</td><td>1000</td><td></td><td></td><td></td><td></td><td></td><td></td><td></td><td></td><td></td></tr>
<tr><td>Oligo fwd</td><td>CACCATCTGGTTGGTCAGGTACCT</td><td></td><td></td><td></td><td></td><td></td><td></td><td></td><td></td><td></td><td></td></tr>
<tr><td>Oligo rev</td><td>AAACAGGTACCTGACCAACCAGAT</td><td></td><td></td><td></td><td></td><td></td><td></td><td></td><td></td><td></td><td></td></tr>
<tr><td>Chromosome</td><td>start</td><td>end</td><td>strand</td><td>MM</td><td>target_seq</td><td>PAM</td><td>alignment</td><td>distance</td><td>position</td><td>gene name</td><td>gene id</td></tr>
<tr><td>GU305914.1</td><td>46905</td><td>46927</td><td>-</td><td>0</td><td>ATCTGGTTGGTCAGGTACCT</td><td>GGG</td><td>||||||||[||||||||||||]PAM</td><td>NA</td><td>Intergenic</td><td>NA</td><td>NA</td></tr>
<tr><td></td><td></td><td></td><td></td><td></td><td></td><td></td><td></td><td></td><td></td><td></td><td></td></tr>
<tr><td>T30645</td><td>TCTGGTTGGTCAGGTACCTGGGG</td><td>1000</td><td></td><td></td><td></td><td></td><td></td><td></td><td></td><td></td><td></td></tr>
<tr><td>Oligo fwd</td><td>CACCTCTGGTTGGTCAGGTACCTG</td><td></td><td></td><td></td><td></td><td></td><td></td><td></td><td></td><td></td><td></td></tr>
<tr><td>Oligo rev</td><td>AAACCAGGTACCTGACCAACCAGA</td><td></td><td></td><td></td><td></td><td></td><td></td><td></td><td></td><td></td><td></td></tr>
<tr><td>Chromosome</td><td>start</td><td>end</td><td>strand</td><td>MM</td><td>target_seq</td><td>PAM</td><td>alignment</td><td>distance</td><td>position</td><td>gene name</td><td>gene id</td></tr>
<tr><td>GU305914.1</td><td>46904</td><td>46926</td><td>-</td><td>0</td><td>TCTGGTTGGTCAGGTACCTG</td><td>GGG</td><td>||||||||[||||||||||||]PAM</td><td>NA</td><td>Intergenic</td><td>NA</td><td>NA</td></tr>
<tr><td></td><td></td><td></td><td></td><td></td><td></td><td></td><td></td><td></td><td></td><td></td><td></td></tr>
<tr><td>T30646</td><td>GTTGGTCAGGTACCTGGGGCAGG</td><td>1000</td><td></td><td></td><td></td><td></td><td></td><td></td><td></td><td></td><td></td></tr>
<tr><td>Oligo fwd</td><td>CACCGTTGGTCAGGTACCTGGGGC</td><td></td><td></td><td></td><td></td><td></td><td></td><td></td><td></td><td></td><td></td></tr>
<tr><td>Oligo rev</td><td>AAACGCCCCAGGTACCTGACCAAC</td><td></td><td></td><td></td><td></td><td></td><td></td><td></td><td></td><td></td><td></td></tr>
<tr><td>Chromosome</td><td>start</td><td>end</td><td>strand</td><td>MM</td><td>target_seq</td><td>PAM</td><td>alignment</td><td>distance</td><td>position</td><td>gene name</td><td>gene id</td></tr>
<tr><td>GU305914.1</td><td>46900</td><td>46922</td><td>-</td><td>0</td><td>GTTGGTCAGGTACCTGGGGC</td><td>AGG</td><td>||||||||[||||||||||||]PAM</td><td>NA</td><td>Intergenic</td><td>NA</td><td>NA</td></tr>
<tr><td></td><td></td><td></td><td></td><td></td><td></td><td></td><td></td><td></td><td></td><td></td><td></td></tr>
<tr><td>T30647</td><td>CCTGGGGCAGGTGTACACGTCGG</td><td>1000</td><td></td><td></td><td></td><td></td><td></td><td></td><td></td><td></td><td></td></tr>
<tr><td>Oligo fwd</td><td>CACCCCTGGGGCAGGTGTACACGT</td><td></td><td></td><td></td><td></td><td></td><td></td><td></td><td></td><td></td><td></td></tr>
<tr><td>Oligo rev</td><td>AAACACGTGTACACCTGCCCCAGG</td><td></td><td></td><td></td><td></td><td></td><td></td><td></td><td></td><td></td><td></td></tr>
<tr><td>Chromosome</td><td>start</td><td>end</td><td>strand</td><td>MM</td><td>target_seq</td><td>PAM</td><td>alignment</td><td>distance</td><td>position</td><td>gene name</td><td>gene id</td></tr>
<tr><td>GU305914.1</td><td>46888</td><td>46910</td><td>-</td><td>0</td><td>CCTGGGGCAGGTGTACACGT</td><td>CGG</td><td>||||||||[||||||||||||]PAM</td><td>NA</td><td>Intergenic</td><td>NA</td><td>NA</td></tr>
<tr><td></td><td></td><td></td><td></td><td></td><td></td><td></td><td></td><td></td><td></td><td></td><td></td></tr>
<tr><td>T30648</td><td>CTTGATGAACTGCAACATGCAGG</td><td>1000</td><td></td><td></td><td></td><td></td><td></td><td></td><td></td><td></td><td></td></tr>
<tr><td>Oligo fwd</td><td>CACCCTTGATGAACTGCAACATGC</td><td></td><td></td><td></td><td></td><td></td><td></td><td></td><td></td><td></td><td></td></tr>
<tr><td>Oligo rev</td><td>AAACGCATGTTGCAGTTCATCAAG</td><td></td><td></td><td></td><td></td><td></td><td></td><td></td><td></td><td></td><td></td></tr>
<tr><td>Chromosome</td><td>start</td><td>end</td><td>strand</td><td>MM</td><td>target_seq</td><td>PAM</td><td>alignment</td><td>distance</td><td>position</td><td>gene name</td><td>gene id</td></tr>
<tr><td>GU305914.1</td><td>46852</td><td>46874</td><td>-</td><td>0</td><td>CTTGATGAACTGCAACATGC</td><td>AGG</td><td>||||||||[||||||||||||]PAM</td><td>NA</td><td>Intergenic</td><td>NA</td><td>NA</td></tr>
<tr><td></td><td></td><td></td><td></td><td></td><td></td><td></td><td></td><td></td><td></td><td></td><td></td></tr>
<tr><td>T30649</td><td>TGCAACATGCAGGCGACGAACGG</td><td>1000</td><td></td><td></td><td></td><td></td><td></td><td></td><td></td><td></td><td></td></tr>
<tr><td>Oligo fwd</td><td>CACCTGCAACATGCAGGCGACGAA</td><td></td><td></td><td></td><td></td><td></td><td></td><td></td><td></td><td></td><td></td></tr>
<tr><td>Oligo rev</td><td>AAACTTCGTCGCCTGCATGTTGCA</td><td></td><td></td><td></td><td></td><td></td><td></td><td></td><td></td><td></td><td></td></tr>
<tr><td>Chromosome</td><td>start</td><td>end</td><td>strand</td><td>MM</td><td>target_seq</td><td>PAM</td><td>alignment</td><td>distance</td><td>position</td><td>gene name</td><td>gene id</td></tr>
<tr><td>GU305914.1</td><td>46842</td><td>46864</td><td>-</td><td>0</td><td>TGCAACATGCAGGCGACGAA</td><td>CGG</td><td>||||||||[||||||||||||]PAM</td><td>NA</td><td>Intergenic</td><td>NA</td><td>NA</td></tr>
<tr><td></td><td></td><td></td><td></td><td></td><td></td><td></td><td></td><td></td><td></td><td></td><td></td></tr>
<tr><td>T30650</td><td>GCGACGAACGGTCTGAAGTTCGG</td><td>1000</td><td></td><td></td><td></td><td></td><td></td><td></td><td></td><td></td><td></td></tr>
<tr><td>Oligo fwd</td><td>CACCGCGACGAACGGTCTGAAGTT</td><td></td><td></td><td></td><td></td><td></td><td></td><td></td><td></td><td></td><td></td></tr>
<tr><td>Oligo rev</td><td>AAACAACTTCAGACCGTTCGTCGC</td><td></td><td></td><td></td><td></td><td></td><td></td><td></td><td></td><td></td><td></td></tr>
<tr><td>Chromosome</td><td>start</td><td>end</td><td>strand</td><td>MM</td><td>target_seq</td><td>PAM</td><td>alignment</td><td>distance</td><td>position</td><td>gene name</td><td>gene id</td></tr>
<tr><td>GU305914.1</td><td>46830</td><td>46852</td><td>-</td><td>0</td><td>GCGACGAACGGTCTGAAGTT</td><td>CGG</td><td>||||||||[||||||||||||]PAM</td><td>NA</td><td>Intergenic</td><td>NA</td><td>NA</td></tr>
<tr><td></td><td></td><td></td><td></td><td></td><td></td><td></td><td></td><td></td><td></td><td></td><td></td></tr>
<tr><td>T30651</td><td>CGACGAACGGTCTGAAGTTCGGG</td><td>1000</td><td></td><td></td><td></td><td></td><td></td><td></td><td></td><td></td><td></td></tr>
<tr><td>Oligo fwd</td><td>CACCCGACGAACGGTCTGAAGTTC</td><td></td><td></td><td></td><td></td><td></td><td></td><td></td><td></td><td></td><td></td></tr>
<tr><td>Oligo rev</td><td>AAACGAACTTCAGACCGTTCGTCG</td><td></td><td></td><td></td><td></td><td></td><td></td><td></td><td></td><td></td><td></td></tr>
<tr><td>Chromosome</td><td>start</td><td>end</td><td>strand</td><td>MM</td><td>target_seq</td><td>PAM</td><td>alignment</td><td>distance</td><td>position</td><td>gene name</td><td>gene id</td></tr>
<tr><td>GU305914.1</td><td>46829</td><td>46851</td><td>-</td><td>0</td><td>CGACGAACGGTCTGAAGTTC</td><td>GGG</td><td>||||||||[||||||||||||]PAM</td><td>NA</td><td>Intergenic</td><td>NA</td><td>NA</td></tr>
<tr><td></td><td></td><td></td><td></td><td></td><td></td><td></td><td></td><td></td><td></td><td></td><td></td></tr>
<tr><td>T30652</td><td>GACGAACGGTCTGAAGTTCGGGG</td><td>1000</td><td></td><td></td><td></td><td></td><td></td><td></td><td></td><td></td><td></td></tr>
<tr><td>Oligo fwd</td><td>CACCGACGAACGGTCTGAAGTTCG</td><td></td><td></td><td></td><td></td><td></td><td></td><td></td><td></td><td></td><td></td></tr>
<tr><td>Oligo rev</td><td>AAACCGAACTTCAGACCGTTCGTC</td><td></td><td></td><td></td><td></td><td></td><td></td><td></td><td></td><td></td><td></td></tr>
<tr><td>Chromosome</td><td>start</td><td>end</td><td>strand</td><td>MM</td><td>target_seq</td><td>PAM</td><td>alignment</td><td>distance</td><td>position</td><td>gene name</td><td>gene id</td></tr>
<tr><td>GU305914.1</td><td>46828</td><td>46850</td><td>-</td><td>0</td><td>GACGAACGGTCTGAAGTTCG</td><td>GGG</td><td>||||||||[||||||||||||]PAM</td><td>NA</td><td>Intergenic</td><td>NA</td><td>NA</td></tr>
<tr><td></td><td></td><td></td><td></td><td></td><td></td><td></td><td></td><td></td><td></td><td></td><td></td></tr>
<tr><td>T30653</td><td>CGGTCTGAAGTTCGGGGTGTCGG</td><td>1000</td><td></td><td></td><td></td><td></td><td></td><td></td><td></td><td></td><td></td></tr>
<tr><td>Oligo fwd</td><td>CACCCGGTCTGAAGTTCGGGGTGT</td><td></td><td></td><td></td><td></td><td></td><td></td><td></td><td></td><td></td><td></td></tr>
<tr><td>Oligo rev</td><td>AAACACACCCCGAACTTCAGACCG</td><td></td><td></td><td></td><td></td><td></td><td></td><td></td><td></td><td></td><td></td></tr>
<tr><td>Chromosome</td><td>start</td><td>end</td><td>strand</td><td>MM</td><td>target_seq</td><td>PAM</td><td>alignment</td><td>distance</td><td>position</td><td>gene name</td><td>gene id</td></tr>
<tr><td>GU305914.1</td><td>46822</td><td>46844</td><td>-</td><td>0</td><td>CGGTCTGAAGTTCGGGGTGT</td><td>CGG</td><td>||||||||[||||||||||||]PAM</td><td>NA</td><td>Intergenic</td><td>NA</td><td>NA</td></tr>
<tr><td></td><td></td><td></td><td></td><td></td><td></td><td></td><td></td><td></td><td></td><td></td><td></td></tr>
<tr><td>T30654</td><td>GGTCTGAAGTTCGGGGTGTCGGG</td><td>1000</td><td></td><td></td><td></td><td></td><td></td><td></td><td></td><td></td><td></td></tr>
<tr><td>Oligo fwd</td><td>CACCGGTCTGAAGTTCGGGGTGTC</td><td></td><td></td><td></td><td></td><td></td><td></td><td></td><td></td><td></td><td></td></tr>
<tr><td>Oligo rev</td><td>AAACGACACCCCGAACTTCAGACC</td><td></td><td></td><td></td><td></td><td></td><td></td><td></td><td></td><td></td><td></td></tr>
<tr><td>Chromosome</td><td>start</td><td>end</td><td>strand</td><td>MM</td><td>target_seq</td><td>PAM</td><td>alignment</td><td>distance</td><td>position</td><td>gene name</td><td>gene id</td></tr>
<tr><td>GU305914.1</td><td>46821</td><td>46843</td><td>-</td><td>0</td><td>GGTCTGAAGTTCGGGGTGTC</td><td>GGG</td><td>||||||||[||||||||||||]PAM</td><td>NA</td><td>Intergenic</td><td>NA</td><td>NA</td></tr>
<tr><td></td><td></td><td></td><td></td><td></td><td></td><td></td><td></td><td></td><td></td><td></td><td></td></tr>
<tr><td>T30655</td><td>GTCTGAAGTTCGGGGTGTCGGGG</td><td>1000</td><td></td><td></td><td></td><td></td><td></td><td></td><td></td><td></td><td></td></tr>
<tr><td>Oligo fwd</td><td>CACCGTCTGAAGTTCGGGGTGTCG</td><td></td><td></td><td></td><td></td><td></td><td></td><td></td><td></td><td></td><td></td></tr>
<tr><td>Oligo rev</td><td>AAACCGACACCCCGAACTTCAGAC</td><td></td><td></td><td></td><td></td><td></td><td></td><td></td><td></td><td></td><td></td></tr>
<tr><td>Chromosome</td><td>start</td><td>end</td><td>strand</td><td>MM</td><td>target_seq</td><td>PAM</td><td>alignment</td><td>distance</td><td>position</td><td>gene name</td><td>gene id</td></tr>
<tr><td>GU305914.1</td><td>46820</td><td>46842</td><td>-</td><td>0</td><td>GTCTGAAGTTCGGGGTGTCG</td><td>GGG</td><td>||||||||[||||||||||||]PAM</td><td>NA</td><td>Intergenic</td><td>NA</td><td>NA</td></tr>
</table>

| | | | | | | | | | | | |
|---|---|---|---|---|---|---|---|---|---|---|---|
| T30656 | TCTGAAGTTCGGGGTGTCGGGGG | 1000 | | | | | | | | | |
| Oligo fwd | CACCTCTGAAGTTCGGGGTGTCGG | | | | | | | | | | |
| Oligo rev | AAACCCGACACCCCGAACTTCAGA | | | | | | | | | | |
| Chromosome | start | end | strand | MM | target_seq | PAM | alignment | distance | position | gene name | gene id |
| GU305914.1 | 46819 | 46841 | - | 0 | TCTGAAGTTCGGGGTGTCGG | GGG | ||||||||[||||||||||||]PAM | NA | Intergenic | NA | NA |
| T30657 | GTTCGGGGTGTCGGGGGACCTGG | 1000 | | | | | | | | | |
| Oligo fwd | CACCGTTCGGGGTGTCGGGGGACC | | | | | | | | | | |
| Oligo rev | AAACGGTCCCCCGACACCCCGAAC | | | | | | | | | | |
| Chromosome | start | end | strand | MM | target_seq | PAM | alignment | distance | position | gene name | gene id |
| GU305914.1 | 46813 | 46835 | - | 0 | GTTCGGGGTGTCGGGGGACC | TGG | ||||||||[||||||||||||]PAM | NA | Intergenic | NA | NA |
| T30658 | CGGGGTGTCGGGGGACCTGGAGG | 1000 | | | | | | | | | |
| Oligo fwd | CACCCGGGGTGTCGGGGGACCTGG | | | | | | | | | | |
| Oligo rev | AAACCCAGGTCCCCCGACACCCCG | | | | | | | | | | |
| Chromosome | start | end | strand | MM | target_seq | PAM | alignment | distance | position | gene name | gene id |
| GU305914.1 | 46810 | 46832 | - | 0 | CGGGGTGTCGGGGGACCTGG | AGG | ||||||||[||||||||||||]PAM | NA | Intergenic | NA | NA |
| T30659 | GCGCCTCGTGCGCGACCTCGCGG | 1000 | | | | | | | | | |
| Oligo fwd | CACCGCGCCTCGTGCGCGACCTCG | | | | | | | | | | |
| Oligo rev | AAACCGAGGTCGCGCACGAGGCGC | | | | | | | | | | |
| Chromosome | start | end | strand | MM | target_seq | PAM | alignment | distance | position | gene name | gene id |
| GU305914.1 | 46775 | 46797 | - | 0 | GCGCCTCGTGCGCGACCTCG | CGG | ||||||||[||||||||||||]PAM | NA | Intergenic | NA | NA |
| T30660 | CGTGCGCGACCTCGCGGAACAGG | 1000 | | | | | | | | | |
| Oligo fwd | CACCCGTGCGCGACCTCGCGGAAC | | | | | | | | | | |
| Oligo rev | AAACGTTCCGCGAGGTCGCGCACG | | | | | | | | | | |
| Chromosome | start | end | strand | MM | target_seq | PAM | alignment | distance | position | gene name | gene id |
| GU305914.1 | 46769 | 46791 | - | 0 | CGTGCGCGACCTCGCGGAAC | AGG | ||||||||[||||||||||||]PAM | NA | Intergenic | NA | NA |
| T30661 | GTGCGCGACCTCGCGGAACAGGG | 1000 | | | | | | | | | |
| Oligo fwd | CACCGTGCGCGACCTCGCGGAACA | | | | | | | | | | |
| Oligo rev | AAACTGTTCCGCGAGGTCGCGCAC | | | | | | | | | | |
| Chromosome | start | end | strand | MM | target_seq | PAM | alignment | distance | position | gene name | gene id |
| GU305914.1 | 46768 | 46790 | - | 0 | GTGCGCGACCTCGCGGAACA | GGG | ||||||||[||||||||||||]PAM | NA | Intergenic | NA | NA |
| T30662 | CGGAACAGGGATTTCGTCTCCGG | 1000 | | | | | | | | | |
| Oligo fwd | CACCCGGAACAGGGATTTCGTCTC | | | | | | | | | | |
| Oligo rev | AAACGAGACGAAATCCCTGTTCCG | | | | | | | | | | |
| Chromosome | start | end | strand | MM | target_seq | PAM | alignment | distance | position | gene name | gene id |
| GU305914.1 | 46755 | 46777 | - | 0 | CGGAACAGGGATTTCGTCTC | CGG | ||||||||[||||||||||||]PAM | NA | Intergenic | NA | NA |
| T30663 | CGTCTCCGGTACCTGAGTGTAGG | 1000 | | | | | | | | | |
| Oligo fwd | CACCCGTCTCCGGTACCTGAGTGT | | | | | | | | | | |
| Oligo rev | AAACACACTCAGGTACCGGAGACG | | | | | | | | | | |
| Chromosome | start | end | strand | MM | target_seq | PAM | alignment | distance | position | gene name | gene id |
| GU305914.1 | 46741 | 46763 | - | 0 | CGTCTCCGGTACCTGAGTGT | AGG | ||||||||[||||||||||||]PAM | NA | Intergenic | NA | NA |
| T30664 | CTCCGGTACCTGAGTGTAGGCGG | 1000 | | | | | | | | | |
| Oligo fwd | CACCCTCCGGTACCTGAGTGTAGG | | | | | | | | | | |
| Oligo rev | AAACCCTACACTCAGGTACCGGAG | | | | | | | | | | |
| Chromosome | start | end | strand | MM | target_seq | PAM | alignment | distance | position | gene name | gene id |
| GU305914.1 | 46738 | 46760 | - | 0 | CTCCGGTACCTGAGTGTAGG | CGG | ||||||||[||||||||||||]PAM | NA | Intergenic | NA | NA |
| T30665 | GCGCGCTCGCCGTCAAGATCAGG | 1000 | | | | | | | | | |
| Oligo fwd | CACCGCGCGCTCGCCGTCAAGATC | | | | | | | | | | |
| Oligo rev | AAACGATCTTGACGGCGAGCGCGC | | | | | | | | | | |
| Chromosome | start | end | strand | MM | target_seq | PAM | alignment | distance | position | gene name | gene id |
| GU305914.1 | 46697 | 46719 | - | 0 | GCGCGCTCGCCGTCAAGATC | AGG | ||||||||[||||||||||||]PAM | NA | Intergenic | NA | NA |
| T30666 | TCAAGATCAGGTTCTTGCGCAGG | 1000 | | | | | | | | | |
| Oligo fwd | CACCTCAAGATCAGGTTCTTGCGC | | | | | | | | | | |
| Oligo rev | AAACGCGCAAGAACCTGATCTTGA | | | | | | | | | | |
| Chromosome | start | end | strand | MM | target_seq | PAM | alignment | distance | position | gene name | gene id |
| GU305914.1 | 46685 | 46707 | - | 0 | TCAAGATCAGGTTCTTGCGC | AGG | ||||||||[||||||||||||]PAM | NA | Intergenic | NA | NA |
| T30667 | CAAGATCAGGTTCTTGCGCAGGG | 1000 | | | | | | | | | |
| Oligo fwd | CACCCAAGATCAGGTTCTTGCGCA | | | | | | | | | | |
| Oligo rev | AAACTGCGCAAGAACCTGATCTTG | | | | | | | | | | |
| Chromosome | start | end | strand | MM | target_seq | PAM | alignment | distance | position | gene name | gene id |
| GU305914.1 | 46684 | 46706 | - | 0 | CAAGATCAGGTTCTTGCGCA | GGG | ||||||||[||||||||||||]PAM | NA | Intergenic | NA | NA |
| T30668 | GCAGGGCTCTCAGCGATTCCTGG | 1000 | | | | | | | | | |
| Oligo fwd | CACCGCAGGGCTCTCAGCGATTCC | | | | | | | | | | |
| Oligo rev | AAACGGAATCGCTGAGAGCCCTGC | | | | | | | | | | |
| Chromosome | start | end | strand | MM | target_seq | PAM | alignment | distance | position | gene name | gene id |
| GU305914.1 | 46667 | 46689 | - | 0 | GCAGGGCTCTCAGCGATTCC | TGG | ||||||||[||||||||||||]PAM | NA | Intergenic | NA | NA |
| T30669 | GGGCTCTCAGCGATTCCTGGTGG | 1000 | | | | | | | | | |
| Oligo fwd | CACCGGGCTCTCAGCGATTCCTGG | | | | | | | | | | |
| Oligo rev | AAACCCAGGAATCGCTGAGAGCCC | | | | | | | | | | |
| Chromosome | start | end | strand | MM | target_seq | PAM | alignment | distance | position | gene name | gene id |
| GU305914.1 | 46664 | 46686 | - | 0 | GGGCTCTCAGCGATTCCTGG | TGG | ||||||||[||||||||||||]PAM | NA | Intergenic | NA | NA |
| T30670 | CCTGGTGGAGCACGATCATGCGG | 1000 | | | | | | | | | |
| Oligo fwd | CACCCCTGGTGGAGCACGATCATG | | | | | | | | | | |
| Oligo rev | AAACCATGATCGTGCTCCACCAGG | | | | | | | | | | |
| Chromosome | start | end | strand | MM | target_seq | PAM | alignment | distance | position | gene name | gene id |
| GU305914.1 | 46649 | 46671 | - | 0 | CCTGGTGGAGCACGATCATG | CGG | ||||||||[||||||||||||]PAM | NA | Intergenic | NA | NA |

T30671 CGATCATGCGGCCGACCGACAGG 1000
Oligo fwd CACCCGATCATGCGGCCGACCGAC
Oligo rev AAACGTCGGTCGGCCGCATGATCG

| Chromosome | start | end | strand | MM | target_seq | PAM | alignment | distance | position | gene name | gene id |
|---|---|---|---|---|---|---|---|---|---|---|---|
| GU305914.1 | 46637 | 46659 | - | 0 | CGATCATGCGGCCGACCGAC | AGG | \|\|\|\|\|\|\|\|[\|\|\|\|\|\|\|\|\|\|\|\|]PAM | NA | Intergenic | NA | NA |

T30672 TCATGCGGCCGACCGACAGGAGG 1000
Oligo fwd CACCTCATGCGGCCGACCGACAGG
Oligo rev AAACCCTGTCGGTCGGCCGCATGA

| Chromosome | start | end | strand | MM | target_seq | PAM | alignment | distance | position | gene name | gene id |
|---|---|---|---|---|---|---|---|---|---|---|---|
| GU305914.1 | 46634 | 46656 | - | 0 | TCATGCGGCCGACCGACAGG | AGG | \|\|\|\|\|\|\|\|[\|\|\|\|\|\|\|\|\|\|\|\|]PAM | NA | Intergenic | NA | NA |

T30673 GGAGGAACAGCATGTCGTTCAGG 1000
Oligo fwd CACCGGAGGAACAGCATGTCGTTC
Oligo rev AAACGAACGACATGCTGTTCCTCC

| Chromosome | start | end | strand | MM | target_seq | PAM | alignment | distance | position | gene name | gene id |
|---|---|---|---|---|---|---|---|---|---|---|---|
| GU305914.1 | 46616 | 46638 | - | 0 | GGAGGAACAGCATGTCGTTC | AGG | \|\|\|\|\|\|\|\|[\|\|\|\|\|\|\|\|\|\|\|\|]PAM | NA | Intergenic | NA | NA |
| GU305914.1 | 84017 | 84039 | + | 4 | GGAAGATCCGCATGTCCTTC | AGG | \|\|\|-\|\|-\|[-\|\|\|\|\|\|\|-\|\|\|]PAM | NA | Intergenic | NA | NA |

T30674 CAGCATGTCGTTCAGGATCTCGG 1000
Oligo fwd CACCCAGCATGTCGTTCAGGATCT
Oligo rev AAACAGATCCTGAACGACATGCTG

| Chromosome | start | end | strand | MM | target_seq | PAM | alignment | distance | position | gene name | gene id |
|---|---|---|---|---|---|---|---|---|---|---|---|
| GU305914.1 | 46609 | 46631 | - | 0 | CAGCATGTCGTTCAGGATCT | CGG | \|\|\|\|\|\|\|\|[\|\|\|\|\|\|\|\|\|\|\|\|]PAM | NA | Intergenic | NA | NA |

T30675 AGCATGTCGTTCAGGATCTCGGG 1000
Oligo fwd CACCAGCATGTCGTTCAGGATCTC
Oligo rev AAACGAGATCCTGAACGACATGCT

| Chromosome | start | end | strand | MM | target_seq | PAM | alignment | distance | position | gene name | gene id |
|---|---|---|---|---|---|---|---|---|---|---|---|
| GU305914.1 | 46608 | 46630 | - | 0 | AGCATGTCGTTCAGGATCTC | GGG | \|\|\|\|\|\|\|\|[\|\|\|\|\|\|\|\|\|\|\|\|]PAM | NA | Intergenic | NA | NA |
| GU305914.1 | 119147 | 119169 | - | 4 | AACCTGTCGTTCACGATCAC | CGG | \|-\|-\|\|\|\|[\|\|\|\|\|-\|\|\|\|-\|]PAM | NA | Intergenic | NA | NA |

T30676 ATGTCGTTCAGGATCTCGGGAGG 1000
Oligo fwd CACCATGTCGTTCAGGATCTCGGG
Oligo rev AAACCCCGAGATCCTGAACGACAT

| Chromosome | start | end | strand | MM | target_seq | PAM | alignment | distance | position | gene name | gene id |
|---|---|---|---|---|---|---|---|---|---|---|---|
| GU305914.1 | 46605 | 46627 | - | 0 | ATGTCGTTCAGGATCTCGGG | AGG | \|\|\|\|\|\|\|\|[\|\|\|\|\|\|\|\|\|\|\|\|]PAM | NA | Intergenic | NA | NA |

T30677 TCTCGGGAGGTTCTTTGCTCCGG 1000
Oligo fwd CACCTCTCGGGAGGTTCTTTGCTC
Oligo rev AAACGAGCAAAGAACCTCCCGAGA

| Chromosome | start | end | strand | MM | target_seq | PAM | alignment | distance | position | gene name | gene id |
|---|---|---|---|---|---|---|---|---|---|---|---|
| GU305914.1 | 46592 | 46614 | - | 0 | TCTCGGGAGGTTCTTTGCTC | CGG | \|\|\|\|\|\|\|\|[\|\|\|\|\|\|\|\|\|\|\|\|]PAM | NA | Intergenic | NA | NA |

T30678 TGCTCCGGTGCACGACCGACAGG 1000
Oligo fwd CACCTGCTCCGGTGCACGACCGAC
Oligo rev AAACGTCGGTCGTGCACCGGAGCA

| Chromosome | start | end | strand | MM | target_seq | PAM | alignment | distance | position | gene name | gene id |
|---|---|---|---|---|---|---|---|---|---|---|---|
| GU305914.1 | 46577 | 46599 | - | 0 | TGCTCCGGTGCACGACCGAC | AGG | \|\|\|\|\|\|\|\|[\|\|\|\|\|\|\|\|\|\|\|\|]PAM | NA | Intergenic | NA | NA |

T30679 GCTCCGGTGCACGACCGACAGGG 1000
Oligo fwd CACCGCTCCGGTGCACGACCGACA
Oligo rev AAACTGTCGGTCGTGCACCGGAGC

| Chromosome | start | end | strand | MM | target_seq | PAM | alignment | distance | position | gene name | gene id |
|---|---|---|---|---|---|---|---|---|---|---|---|
| GU305914.1 | 46576 | 46598 | - | 0 | GCTCCGGTGCACGACCGACA | GGG | \|\|\|\|\|\|\|\|[\|\|\|\|\|\|\|\|\|\|\|\|]PAM | NA | Intergenic | NA | NA |

T30680 ACGACCGACAGGGTCTCGAACGG 1000
Oligo fwd CACCACGACCGACAGGGTCTCGAA
Oligo rev AAACTTCGAGACCCTGTCGGTCGT

| Chromosome | start | end | strand | MM | target_seq | PAM | alignment | distance | position | gene name | gene id |
|---|---|---|---|---|---|---|---|---|---|---|---|
| GU305914.1 | 46566 | 46588 | - | 0 | ACGACCGACAGGGTCTCGAA | CGG | \|\|\|\|\|\|\|\|[\|\|\|\|\|\|\|\|\|\|\|\|]PAM | NA | Intergenic | NA | NA |

T30681 GAACGGCTCGCGCGACATCTTGG 1000
Oligo fwd CACCGAACGGCTCGCGCGACATCT
Oligo rev AAACAGATGTCGCGCGAGCCGTTC

| Chromosome | start | end | strand | MM | target_seq | PAM | alignment | distance | position | gene name | gene id |
|---|---|---|---|---|---|---|---|---|---|---|---|
| GU305914.1 | 46549 | 46571 | - | 0 | GAACGGCTCGCGCGACATCT | TGG | \|\|\|\|\|\|\|\|[\|\|\|\|\|\|\|\|\|\|\|\|]PAM | NA | Intergenic | NA | NA |

T30682 GCGCGACATCTTGGTGATGCCGG 1000
Oligo fwd CACCGCGCGACATCTTGGTGATGC
Oligo rev AAACGCATCACCAAGATGTCGCGC

| Chromosome | start | end | strand | MM | target_seq | PAM | alignment | distance | position | gene name | gene id |
|---|---|---|---|---|---|---|---|---|---|---|---|
| GU305914.1 | 46540 | 46562 | - | 0 | GCGCGACATCTTGGTGATGC | CGG | \|\|\|\|\|\|\|\|[\|\|\|\|\|\|\|\|\|\|\|\|]PAM | NA | Intergenic | NA | NA |

T30683 ACATCTTGGTGATGCCGGCGAGG 1000
Oligo fwd CACCACATCTTGGTGATGCCGGCG
Oligo rev AAACCGCCGGCATCACCAAGATGT

| Chromosome | start | end | strand | MM | target_seq | PAM | alignment | distance | position | gene name | gene id |
|---|---|---|---|---|---|---|---|---|---|---|---|
| GU305914.1 | 46535 | 46557 | - | 0 | ACATCTTGGTGATGCCGGCG | AGG | \|\|\|\|\|\|\|\|[\|\|\|\|\|\|\|\|\|\|\|\|]PAM | NA | Intergenic | NA | NA |

T30684 GGTGATGCCGGCGAGGTTCCTGG 1000
Oligo fwd CACCGGTGATGCCGGCGAGGTTCC
Oligo rev AAACGGAACCTCGCCGGCATCACC

| Chromosome | start | end | strand | MM | target_seq | PAM | alignment | distance | position | gene name | gene id |
|---|---|---|---|---|---|---|---|---|---|---|---|
| GU305914.1 | 46528 | 46550 | - | 0 | GGTGATGCCGGCGAGGTTCC | TGG | \|\|\|\|\|\|\|\|[\|\|\|\|\|\|\|\|\|\|\|\|]PAM | NA | Intergenic | NA | NA |

T30685 GTTCCTGGAGTACATCGTCACGG 1000
Oligo fwd CACCGTTCCTGGAGTACATCGTCA

Oligo rev AAACTGACGATGTACTCCAGGAAC
Chromosome start end strand MM target_seq PAM alignment distance position gene name gene id
GU305914.1 46513 46535 - 0 GTTCCTGGAGTACATCGTCA CGG ||||||||[||||||||||||]PAM NA Intergenic NA NA

T30686 AGTACATCGTCACGGCCTCCTGG 1000
Oligo fwd CACCAGTACATCGTCACGGCCTCC
Oligo rev AAACGGAGGCCGTGACGATGTACT
Chromosome start end strand MM target_seq PAM alignment distance position gene name gene id
GU305914.1 46505 46527 - 0 AGTACATCGTCACGGCCTCC TGG ||||||||[||||||||||||]PAM NA Intergenic NA NA

T30687 CGGCCTCCTGGCTCGTGAACAGG 1000
Oligo fwd CACCCGGCCTCCTGGCTCGTGAAC
Oligo rev AAACGTTCACGAGCCAGGAGGCCG
Chromosome start end strand MM target_seq PAM alignment distance position gene name gene id
GU305914.1 46493 46515 - 0 CGGCCTCCTGGCTCGTGAAC AGG ||||||||[||||||||||||]PAM NA Intergenic NA NA

T30688 GGCCTCCTGGCTCGTGAACAGGG 1000
Oligo fwd CACCGGCCTCCTGGCTCGTGAACA
Oligo rev AAACTGTTCACGAGCCAGGAGGCC
Chromosome start end strand MM target_seq PAM alignment distance position gene name gene id
GU305914.1 46492 46514 - 0 GGCCTCCTGGCTCGTGAACA GGG ||||||||[||||||||||||]PAM NA Intergenic NA NA

T30689 GCCTCCTGGCTCGTGAACAGGGG 1000
Oligo fwd CACCGCCTCCTGGCTCGTGAACAG
Oligo rev AAACCTGTTCACGAGCCAGGAGGC
Chromosome start end strand MM target_seq PAM alignment distance position gene name gene id
GU305914.1 46491 46513 - 0 GCCTCCTGGCTCGTGAACAG GGG ||||||||[||||||||||||]PAM NA Intergenic NA NA

T30690 CCTCCTGGCTCGTGAACAGGGGG 1000
Oligo fwd CACCCCTCCTGGCTCGTGAACAGG
Oligo rev AAACCCTGTTCACGAGCCAGGAGG
Chromosome start end strand MM target_seq PAM alignment distance position gene name gene id
GU305914.1 46490 46512 - 0 CCTCCTGGCTCGTGAACAGG GGG ||||||||[||||||||||||]PAM NA Intergenic NA NA

T30691 CTCCTGGCTCGTGAACAGGGGGG 1000
Oligo fwd CACCCTCCTGGCTCGTGAACAGGG
Oligo rev AAACCCCTGTTCACGAGCCAGGAG
Chromosome start end strand MM target_seq PAM alignment distance position gene name gene id
GU305914.1 46489 46511 - 0 CTCCTGGCTCGTGAACAGGG GGG ||||||||[||||||||||||]PAM NA Intergenic NA NA

T30692 TGAACAGGGGGGATTCGTAGAGG 1000
Oligo fwd CACCTGAACAGGGGGGATTCGTAG
Oligo rev AAACCTACGAATCCCCCCTGTTCA
Chromosome start end strand MM target_seq PAM alignment distance position gene name gene id
GU305914.1 46478 46500 - 0 TGAACAGGGGGGATTCGTAG AGG ||||||||[||||||||||||]PAM NA Intergenic NA NA

T30693 GTAGTTGAACACCGTCATCGTGG 1000
Oligo fwd CACCGTAGTTGAACACCGTCATCG
Oligo rev AAACCGATGACGGTGTTCAACTAC
Chromosome start end strand MM target_seq PAM alignment distance position gene name gene id
GU305914.1 46456 46478 - 0 GTAGTTGAACACCGTCATCG TGG ||||||||[||||||||||||]PAM NA Intergenic NA NA

T30694 CCGTCATCGTGGAGTAGTCCAGG 1000
Oligo fwd CACCCCGTCATCGTGGAGTAGTCC
Oligo rev AAACGGACTACTCCACGATGACGG
Chromosome start end strand MM target_seq PAM alignment distance position gene name gene id
GU305914.1 46445 46467 - 0 CCGTCATCGTGGAGTAGTCC AGG ||||||||[||||||||||||]PAM NA Intergenic NA NA

T30695 CGTCATCGTGGAGTAGTCCAGGG 1000
Oligo fwd CACCCGTCATCGTGGAGTAGTCCA
Oligo rev AAACTGGACTACTCCACGATGACG
Chromosome start end strand MM target_seq PAM alignment distance position gene name gene id
GU305914.1 46444 46466 - 0 CGTCATCGTGGAGTAGTCCA GGG ||||||||[||||||||||||]PAM NA Intergenic NA NA

T30696 TCGTGGAGTAGTCCAGGGACCGG 1000
Oligo fwd CACCTCGTGGAGTAGTCCAGGGAC
Oligo rev AAACGTCCCTGGACTACTCCACGA
Chromosome start end strand MM target_seq PAM alignment distance position gene name gene id
GU305914.1 46439 46461 - 0 TCGTGGAGTAGTCCAGGGAC CGG ||||||||[||||||||||||]PAM NA Intergenic NA NA

T30697 AGTAGTCCAGGGACCGGCTGAGG 1000
Oligo fwd CACCAGTAGTCCAGGGACCGGCTG
Oligo rev AAACCAGCCGGTCCCTGGACTACT
Chromosome start end strand MM target_seq PAM alignment distance position gene name gene id
GU305914.1 46433 46455 - 0 AGTAGTCCAGGGACCGGCTG AGG ||||||||[||||||||||||]PAM NA Intergenic NA NA

T30698 AGTCCAGGGACCGGCTGAGGTGG 1000
Oligo fwd CACCAGTCCAGGGACCGGCTGAGG
Oligo rev AAACCCTCAGCCGGTCCCTGGACT
Chromosome start end strand MM target_seq PAM alignment distance position gene name gene id
GU305914.1 46430 46452 - 0 AGTCCAGGGACCGGCTGAGG TGG ||||||||[||||||||||||]PAM NA Intergenic NA NA

T30699 GGTGGCCCACATCGTGTCCCAGG 1000
Oligo fwd CACCGGTGGCCCACATCGTGTCCC
Oligo rev AAACGGGACACGATGTGGGCCACC
Chromosome start end strand MM target_seq PAM alignment distance position gene name gene id
GU305914.1 46412 46434 - 0 GGTGGCCCACATCGTGTCCC AGG ||||||||[||||||||||||]PAM NA Intergenic NA NA

T30700 GTGGCCCACATCGTGTCCCAGGG 1000

Oligo fwd CACCGTGGCCCACATCGTGTCCCA
Oligo rev AAACTGGGACACGATGTGGGCCAC
Chromosome start end strand MM target_seq PAM alignment distance position gene name gene id
GU305914.1 46411 46433 - 0 GTGGCCCACATCGTGTCCCA GGG ||||||||[||||||||||||]PAM NA Intergenic NA NA

T30701 ACTTCATGAGCGAGTAGATGTGG 1000
Oligo fwd CACCACTTCATGAGCGAGTAGATG
Oligo rev AAACCATCTACTCGCTCATGAAGT
Chromosome start end strand MM target_seq PAM alignment distance position gene name gene id
GU305914.1 46388 46410 - 0 ACTTCATGAGCGAGTAGATG TGG ||||||||[||||||||||||]PAM NA Intergenic NA NA

T30702 GTGGCGCACGTAGCCCTCGTAGG 1000
Oligo fwd CACCGTGGCGCACGTAGCCCTCGT
Oligo rev AAACACGAGGGCTACGTGCGCCAC
Chromosome start end strand MM target_seq PAM alignment distance position gene name gene id
GU305914.1 46369 46391 - 0 GTGGCGCACGTAGCCCTCGT AGG ||||||||[||||||||||||]PAM NA Intergenic NA NA

T30703 CCTCGTAGGCGCCCTGAGACAGG 1000
Oligo fwd CACCCCTCGTAGGCGCCCTGAGAC
Oligo rev AAACGTCTCAGGGCGCCTACGAGG
Chromosome start end strand MM target_seq PAM alignment distance position gene name gene id
GU305914.1 46355 46377 - 0 CCTCGTAGGCGCCCTGAGAC AGG ||||||||[||||||||||||]PAM NA Intergenic NA NA

T30704 AGGCGCCCTGAGACAGGTCCAGG 1000
Oligo fwd CACCAGGCGCCCTGAGACAGGTCC
Oligo rev AAACGGACCTGTCTCAGGGCGCCT
Chromosome start end strand MM target_seq PAM alignment distance position gene name gene id
GU305914.1 46349 46371 - 0 AGGCGCCCTGAGACAGGTCC AGG ||||||||[||||||||||||]PAM NA Intergenic NA NA

T30705 AGACAGGTCCAGGTCGCCCACGG 1000
Oligo fwd CACCAGACAGGTCCAGGTCGCCCA
Oligo rev AAACTGGGCGACCTGGACCTGTCT
Chromosome start end strand MM target_seq PAM alignment distance position gene name gene id
GU305914.1 46339 46361 - 0 AGACAGGTCCAGGTCGCCCA CGG ||||||||[||||||||||||]PAM NA Intergenic NA NA

T30706 GGTCGCCCACGGCCTCGACCAGG 1000
Oligo fwd CACCGGTCGCCCACGGCCTCGACC
Oligo rev AAACGGTCGAGGCCGTGGGCGACC
Chromosome start end strand MM target_seq PAM alignment distance position gene name gene id
GU305914.1 46328 46350 - 0 GGTCGCCCACGGCCTCGACC AGG ||||||||[||||||||||||]PAM NA Intergenic NA NA

T30707 ACGGCCTCGACCAGGTCGAACGG 1000
Oligo fwd CACCACGGCCTCGACCAGGTCGAA
Oligo rev AAACTTCGACCTGGTCGAGGCCGT
Chromosome start end strand MM target_seq PAM alignment distance position gene name gene id
GU305914.1 46320 46342 - 0 ACGGCCTCGACCAGGTCGAA CGG ||||||||[||||||||||||]PAM NA Intergenic NA NA

T30708 CGGCCTCGACCAGGTCGAACGGG 1000
Oligo fwd CACCCGGCCTCGACCAGGTCGAAC
Oligo rev AAACGTTCGACCTGGTCGAGGCCG
Chromosome start end strand MM target_seq PAM alignment distance position gene name gene id
GU305914.1 46319 46341 - 0 CGGCCTCGACCAGGTCGAAC GGG ||||||||[||||||||||||]PAM NA Intergenic NA NA

T30709 GGTCGAACGGGTTATCTTCCAGG 1000
Oligo fwd CACCGGTCGAACGGGTTATCTTCC
Oligo rev AAACGGAAGATAACCCGTTCGACC
Chromosome start end strand MM target_seq PAM alignment distance position gene name gene id
GU305914.1 46307 46329 - 0 GGTCGAACGGGTTATCTTCC AGG ||||||||[||||||||||||]PAM NA Intergenic NA NA

T30710 ACGGGTTATCTTCCAGGTTCAGG 1000
Oligo fwd CACCACGGGTTATCTTCCAGGTTC
Oligo rev AAACGAACCTGGAAGATAACCCGT
Chromosome start end strand MM target_seq PAM alignment distance position gene name gene id
GU305914.1 46301 46323 - 0 ACGGGTTATCTTCCAGGTTC AGG ||||||||[||||||||||||]PAM NA Intergenic NA NA

T30711 CGGGTTATCTTCCAGGTTCAGGG 1000
Oligo fwd CACCCGGGTTATCTTCCAGGTTCA
Oligo rev AAACTGAACCTGGAAGATAACCCG
Chromosome start end strand MM target_seq PAM alignment distance position gene name gene id
GU305914.1 46300 46322 - 0 CGGGTTATCTTCCAGGTTCA GGG ||||||||[||||||||||||]PAM NA Intergenic NA NA

T30712 TATCTTCCAGGTTCAGGGCGCGG 1000
Oligo fwd CACCTATCTTCCAGGTTCAGGGCG
Oligo rev AAACCGCCCTGAACCTGGAAGATA
Chromosome start end strand MM target_seq PAM alignment distance position gene name gene id
GU305914.1 46295 46317 - 0 TATCTTCCAGGTTCAGGGCG CGG ||||||||[||||||||||||]PAM NA Intergenic NA NA
GU305914.1 63965 63987 - 4 TCTGTTCCAGGTTGAGGGCT CGG |-|-||||[|||||-|||||-]PAM NA Intergenic NA NA

T30713 TCAGGGCGCGGTAGATCCTTCGG 1000
Oligo fwd CACCTCAGGGCGCGGTAGATCCTT
Oligo rev AAACAAGGATCTACCGCGCCCTGA
Chromosome start end strand MM target_seq PAM alignment distance position gene name gene id
GU305914.1 46283 46305 - 0 TCAGGGCGCGGTAGATCCTT CGG ||||||||[||||||||||||]PAM NA Intergenic NA NA

T30714 GCGGTAGATCCTTCGGAGCCCGG 1000
Oligo fwd CACCGCGGTAGATCCTTCGGAGCC
Oligo rev AAACGGCTCCGAAGGATCTACCGC
Chromosome start end strand MM target_seq PAM alignment distance position gene name gene id
GU305914.1 46276 46298 - 0 GCGGTAGATCCTTCGGAGCC CGG ||||||||[||||||||||||]PAM NA Intergenic NA NA

| | | | | | | | | | | | |
|---|---|---|---|---|---|---|---|---|---|---|---|
| T30715 | TTCGGAGCCCGGACTCCGCGCGG | 1000 | | | | | | | | | |
| Oligo fwd | CACCTTCGGAGCCCGGACTCCGCG | | | | | | | | | | |
| Oligo rev | AAACCGCGGAGTCCGGGCTCCGAA | | | | | | | | | | |
| Chromosome | start | end | strand | MM | target_seq | PAM | alignment | distance | position | gene name | gene id |
| GU305914.1 | 46265 | 46287 | - | 0 | TTCGGAGCCCGGACTCCGCG | CGG | ||||||||[||||||||||||]PAM | NA | Intergenic | NA | NA |
| T30716 | TCCGCGCGGTGTTTGTCGTTCGG | 1000 | | | | | | | | | |
| Oligo fwd | CACCTCCGCGCGGTGTTTGTCGTT | | | | | | | | | | |
| Oligo rev | AAACAACGACAAACACCGCGCGGA | | | | | | | | | | |
| Chromosome | start | end | strand | MM | target_seq | PAM | alignment | distance | position | gene name | gene id |
| GU305914.1 | 46251 | 46273 | - | 0 | TCCGCGCGGTGTTTGTCGTT | CGG | ||||||||[||||||||||||]PAM | NA | Intergenic | NA | NA |
| T30717 | CCCCCACGCCGTCGCGTGTTTGG | 1000 | | | | | | | | | |
| Oligo fwd | CACCCCCCCACGCCGTCGCGTGTT | | | | | | | | | | |
| Oligo rev | AAACAACACGCGACGGCGTGGGGG | | | | | | | | | | |
| Chromosome | start | end | strand | MM | target_seq | PAM | alignment | distance | position | gene name | gene id |
| GU305914.1 | 46228 | 46250 | - | 0 | CCCCCACGCCGTCGCGTGTT | TGG | ||||||||[||||||||||||]PAM | NA | Intergenic | NA | NA |
| T30718 | CGCCGTCGCGTGTTTGGCCGTGG | 1000 | | | | | | | | | |
| Oligo fwd | CACCCGCCGTCGCGTGTTTGGCCG | | | | | | | | | | |
| Oligo rev | AAACCGGCCAAACACGCGACGGCG | | | | | | | | | | |
| Chromosome | start | end | strand | MM | target_seq | PAM | alignment | distance | position | gene name | gene id |
| GU305914.1 | 46222 | 46244 | - | 0 | CGCCGTCGCGTGTTTGGCCG | TGG | ||||||||[||||||||||||]PAM | NA | Intergenic | NA | NA |
| T30719 | AGATGCGCGTACCGTTGATGAGG | 1000 | | | | | | | | | |
| Oligo fwd | CACCAGATGCGCGTACCGTTGATG | | | | | | | | | | |
| Oligo rev | AAACCATCAACGGTACGCGCATCT | | | | | | | | | | |
| Chromosome | start | end | strand | MM | target_seq | PAM | alignment | distance | position | gene name | gene id |
| GU305914.1 | 46190 | 46212 | - | 0 | AGATGCGCGTACCGTTGATG | AGG | ||||||||[||||||||||||]PAM | NA | Intergenic | NA | NA |
| T30720 | GATGAGGTGCGCCACGCTGATGG | 1000 | | | | | | | | | |
| Oligo fwd | CACCGATGAGGTGCGCCACGCTGA | | | | | | | | | | |
| Oligo rev | AAACTCAGCGTGGCGCACCTCATC | | | | | | | | | | |
| Chromosome | start | end | strand | MM | target_seq | PAM | alignment | distance | position | gene name | gene id |
| GU305914.1 | 46174 | 46196 | - | 0 | GATGAGGTGCGCCACGCTGA | TGG | ||||||||[||||||||||||]PAM | NA | Intergenic | NA | NA |
| T30721 | GCTGATGGTGTAACAGATGACGG | 1000 | | | | | | | | | |
| Oligo fwd | CACCGCTGATGGTGTAACAGATGA | | | | | | | | | | |
| Oligo rev | AAACTCATCTGTTACACCATCAGC | | | | | | | | | | |
| Chromosome | start | end | strand | MM | target_seq | PAM | alignment | distance | position | gene name | gene id |
| GU305914.1 | 46159 | 46181 | - | 0 | GCTGATGGTGTAACAGATGA | CGG | ||||||||[||||||||||||]PAM | NA | Intergenic | NA | NA |
| T30722 | CTGATGGTGTAACAGATGACGGG | 1000 | | | | | | | | | |
| Oligo fwd | CACCCTGATGGTGTAACAGATGAC | | | | | | | | | | |
| Oligo rev | AAACGTCATCTGTTACACCATCAG | | | | | | | | | | |
| Chromosome | start | end | strand | MM | target_seq | PAM | alignment | distance | position | gene name | gene id |
| GU305914.1 | 46158 | 46180 | - | 0 | CTGATGGTGTAACAGATGAC | GGG | ||||||||[||||||||||||]PAM | NA | Intergenic | NA | NA |
| T30723 | TCGTCAGCCTGTGTTTGACGCGG | 1000 | | | | | | | | | |
| Oligo fwd | CACCTCGTCAGCCTGTGTTTGACG | | | | | | | | | | |
| Oligo rev | AAACCGTCAAACACAGGCTGACGA | | | | | | | | | | |
| Chromosome | start | end | strand | MM | target_seq | PAM | alignment | distance | position | gene name | gene id |
| GU305914.1 | 46115 | 46137 | - | 0 | TCGTCAGCCTGTGTTTGACG | CGG | ||||||||[||||||||||||]PAM | NA | Intergenic | NA | NA |
| T30724 | TGTGTTTGACGCGGACCGCCAGG | 1000 | | | | | | | | | |
| Oligo fwd | CACCTGTGTTTGACGCGGACCGCC | | | | | | | | | | |
| Oligo rev | AAACGGCGGTCCGCGTCAAACACA | | | | | | | | | | |
| Chromosome | start | end | strand | MM | target_seq | PAM | alignment | distance | position | gene name | gene id |
| GU305914.1 | 46106 | 46128 | - | 0 | TGTGTTTGACGCGGACCGCC | AGG | ||||||||[||||||||||||]PAM | NA | Intergenic | NA | NA |
| T30725 | GTTTGACGCGGACCGCCAGGAGG | 1000 | | | | | | | | | |
| Oligo fwd | CACCGTTTGACGCGGACCGCCAGG | | | | | | | | | | |
| Oligo rev | AAACCCTGGCGGTCCGCGTCAAAC | | | | | | | | | | |
| Chromosome | start | end | strand | MM | target_seq | PAM | alignment | distance | position | gene name | gene id |
| GU305914.1 | 46103 | 46125 | - | 0 | GTTTGACGCGGACCGCCAGG | AGG | ||||||||[||||||||||||]PAM | NA | Intergenic | NA | NA |
| T30726 | TTTGACGCGGACCGCCAGGAGGG | 1000 | | | | | | | | | |
| Oligo fwd | CACCTTTGACGCGGACCGCCAGGA | | | | | | | | | | |
| Oligo rev | AAACTCCTGGCGGTCCGCGTCAAA | | | | | | | | | | |
| Chromosome | start | end | strand | MM | target_seq | PAM | alignment | distance | position | gene name | gene id |
| GU305914.1 | 46102 | 46124 | - | 0 | TTTGACGCGGACCGCCAGGA | GGG | ||||||||[||||||||||||]PAM | NA | Intergenic | NA | NA |
| T30727 | AGGAGGGCCTCTAGCATGTACGG | 1000 | | | | | | | | | |
| Oligo fwd | CACCAGGAGGGCCTCTAGCATGTA | | | | | | | | | | |
| Oligo rev | AAACTACATGCTAGAGGCCCTCCT | | | | | | | | | | |
| Chromosome | start | end | strand | MM | target_seq | PAM | alignment | distance | position | gene name | gene id |
| GU305914.1 | 46086 | 46108 | - | 0 | AGGAGGGCCTCTAGCATGTA | CGG | ||||||||[||||||||||||]PAM | NA | Intergenic | NA | NA |
| T30728 | GGAGGGCCTCTAGCATGTACGGG | 1000 | | | | | | | | | |
| Oligo fwd | CACCGGAGGGCCTCTAGCATGTAC | | | | | | | | | | |
| Oligo rev | AAACGTACATGCTAGAGGCCCTCC | | | | | | | | | | |
| Chromosome | start | end | strand | MM | target_seq | PAM | alignment | distance | position | gene name | gene id |
| GU305914.1 | 46085 | 46107 | - | 0 | GGAGGGCCTCTAGCATGTAC | GGG | ||||||||[||||||||||||]PAM | NA | Intergenic | NA | NA |
| T30729 | GAGGGCCTCTAGCATGTACGGGG | 1000 | | | | | | | | | |
| Oligo fwd | CACCGAGGGCCTCTAGCATGTACG | | | | | | | | | | |
| Oligo rev | AAACCGTACATGCTAGAGGCCCTC | | | | | | | | | | |
| Chromosome | start | end | strand | MM | target_seq | PAM | alignment | distance | position | gene name | gene id |

<table>
<tr><td>GU305914.1</td><td></td><td>46084</td><td>46106</td><td>-</td><td>0</td><td>GAGGGCCTCTAGCATGTACG</td><td>GGG</td><td>||||||||[||||||||||||]PAM</td><td>NA</td><td>Intergenic</td><td>NA</td><td>NA</td></tr>
<tr><td>T30730</td><td>TCTAGCATGTACGGGGCCATCGG</td><td></td><td>1000</td><td></td><td></td><td></td><td></td><td></td><td></td><td></td><td></td><td></td></tr>
<tr><td>Oligo fwd</td><td>CACCTCTAGCATGTACGGGGCCAT</td><td></td><td></td><td></td><td></td><td></td><td></td><td></td><td></td><td></td><td></td><td></td></tr>
<tr><td>Oligo rev</td><td>AAACATGGCCCCGTACATGCTAGA</td><td></td><td></td><td></td><td></td><td></td><td></td><td></td><td></td><td></td><td></td><td></td></tr>
<tr><td>Chromosome</td><td>start</td><td></td><td>end</td><td>strand</td><td>MM</td><td>target_seq</td><td>PAM</td><td>alignment</td><td>distance</td><td>position</td><td>gene name</td><td>gene id</td></tr>
<tr><td>GU305914.1</td><td></td><td>46077</td><td>46099</td><td>-</td><td>0</td><td>TCTAGCATGTACGGGGCCAT</td><td>CGG</td><td>||||||||[||||||||||||]PAM</td><td>NA</td><td>Intergenic</td><td>NA</td><td>NA</td></tr>
<tr><td>GU305914.1</td><td></td><td>27610</td><td>27632</td><td>-</td><td>4</td><td>TCTAGGGTGTACGGCGGCAT</td><td>CGG</td><td>|||||--|[||||||-|-|||]PAM</td><td>NA</td><td>Intergenic</td><td>NA</td><td>NA</td></tr>
<tr><td>T30731</td><td>ACCCGAGTTCAACGCCGCCACGG</td><td></td><td>1000</td><td></td><td></td><td></td><td></td><td></td><td></td><td></td><td></td><td></td></tr>
<tr><td>Oligo fwd</td><td>CACCACCCGAGTTCAACGCCGCCA</td><td></td><td></td><td></td><td></td><td></td><td></td><td></td><td></td><td></td><td></td><td></td></tr>
<tr><td>Oligo rev</td><td>AAACTGGCGGCGTTGAACTCGGGT</td><td></td><td></td><td></td><td></td><td></td><td></td><td></td><td></td><td></td><td></td><td></td></tr>
<tr><td>Chromosome</td><td>start</td><td></td><td>end</td><td>strand</td><td>MM</td><td>target_seq</td><td>PAM</td><td>alignment</td><td>distance</td><td>position</td><td>gene name</td><td>gene id</td></tr>
<tr><td>GU305914.1</td><td></td><td>46051</td><td>46073</td><td>-</td><td>0</td><td>ACCCGAGTTCAACGCCGCCA</td><td>CGG</td><td>||||||||[||||||||||||]PAM</td><td>NA</td><td>Intergenic</td><td>NA</td><td>NA</td></tr>
<tr><td>T30732</td><td>TCAACGCCGCCACGGCGCCCAGG</td><td></td><td>1000</td><td></td><td></td><td></td><td></td><td></td><td></td><td></td><td></td><td></td></tr>
<tr><td>Oligo fwd</td><td>CACCTCAACGCCGCCACGGCGCCC</td><td></td><td></td><td></td><td></td><td></td><td></td><td></td><td></td><td></td><td></td><td></td></tr>
<tr><td>Oligo rev</td><td>AAACGGGCGCCGTGGCGGCGTTGA</td><td></td><td></td><td></td><td></td><td></td><td></td><td></td><td></td><td></td><td></td><td></td></tr>
<tr><td>Chromosome</td><td>start</td><td></td><td>end</td><td>strand</td><td>MM</td><td>target_seq</td><td>PAM</td><td>alignment</td><td>distance</td><td>position</td><td>gene name</td><td>gene id</td></tr>
<tr><td>GU305914.1</td><td></td><td>46043</td><td>46065</td><td>-</td><td>0</td><td>TCAACGCCGCCACGGCGCCC</td><td>AGG</td><td>||||||||[||||||||||||]PAM</td><td>NA</td><td>Intergenic</td><td>NA</td><td>NA</td></tr>
<tr><td>T30733</td><td>GCCCAGGTCGCCGTTACACACGG</td><td></td><td>1000</td><td></td><td></td><td></td><td></td><td></td><td></td><td></td><td></td><td></td></tr>
<tr><td>Oligo fwd</td><td>CACCGCCCAGGTCGCCGTTACACA</td><td></td><td></td><td></td><td></td><td></td><td></td><td></td><td></td><td></td><td></td><td></td></tr>
<tr><td>Oligo rev</td><td>AAACTGTGTAACGGCGACCTGGGC</td><td></td><td></td><td></td><td></td><td></td><td></td><td></td><td></td><td></td><td></td><td></td></tr>
<tr><td>Chromosome</td><td>start</td><td></td><td>end</td><td>strand</td><td>MM</td><td>target_seq</td><td>PAM</td><td>alignment</td><td>distance</td><td>position</td><td>gene name</td><td>gene id</td></tr>
<tr><td>GU305914.1</td><td></td><td>46027</td><td>46049</td><td>-</td><td>0</td><td>GCCCAGGTCGCCGTTACACA</td><td>CGG</td><td>||||||||[||||||||||||]PAM</td><td>NA</td><td>Intergenic</td><td>NA</td><td>NA</td></tr>
<tr><td>T30734</td><td>TTTCCTCCGACAGCATATTGTGG</td><td></td><td>1000</td><td></td><td></td><td></td><td></td><td></td><td></td><td></td><td></td><td></td></tr>
<tr><td>Oligo fwd</td><td>CACCTTTCCTCCGACAGCATATTG</td><td></td><td></td><td></td><td></td><td></td><td></td><td></td><td></td><td></td><td></td><td></td></tr>
<tr><td>Oligo rev</td><td>AAACCAATATGCTGTCGGAGGAAA</td><td></td><td></td><td></td><td></td><td></td><td></td><td></td><td></td><td></td><td></td><td></td></tr>
<tr><td>Chromosome</td><td>start</td><td></td><td>end</td><td>strand</td><td>MM</td><td>target_seq</td><td>PAM</td><td>alignment</td><td>distance</td><td>position</td><td>gene name</td><td>gene id</td></tr>
<tr><td>GU305914.1</td><td></td><td>45983</td><td>46005</td><td>-</td><td>0</td><td>TTTCCTCCGACAGCATATTG</td><td>TGG</td><td>||||||||[||||||||||||]PAM</td><td>NA</td><td>Intergenic</td><td>NA</td><td>NA</td></tr>
<tr><td>T30735</td><td>TTCCTCCGACAGCATATTGTGGG</td><td></td><td>1000</td><td></td><td></td><td></td><td></td><td></td><td></td><td></td><td></td><td></td></tr>
<tr><td>Oligo fwd</td><td>CACCTTCCTCCGACAGCATATTGT</td><td></td><td></td><td></td><td></td><td></td><td></td><td></td><td></td><td></td><td></td><td></td></tr>
<tr><td>Oligo rev</td><td>AAACACAATATGCTGTCGGAGGAA</td><td></td><td></td><td></td><td></td><td></td><td></td><td></td><td></td><td></td><td></td><td></td></tr>
<tr><td>Chromosome</td><td>start</td><td></td><td>end</td><td>strand</td><td>MM</td><td>target_seq</td><td>PAM</td><td>alignment</td><td>distance</td><td>position</td><td>gene name</td><td>gene id</td></tr>
<tr><td>GU305914.1</td><td></td><td>45982</td><td>46004</td><td>-</td><td>0</td><td>TTCCTCCGACAGCATATTGT</td><td>GGG</td><td>||||||||[||||||||||||]PAM</td><td>NA</td><td>Intergenic</td><td>NA</td><td>NA</td></tr>
<tr><td>T30736</td><td>TCCTCCGACAGCATATTGTGGGG</td><td></td><td>1000</td><td></td><td></td><td></td><td></td><td></td><td></td><td></td><td></td><td></td></tr>
<tr><td>Oligo fwd</td><td>CACCTCCTCCGACAGCATATTGTG</td><td></td><td></td><td></td><td></td><td></td><td></td><td></td><td></td><td></td><td></td><td></td></tr>
<tr><td>Oligo rev</td><td>AAACCACAATATGCTGTCGGAGGA</td><td></td><td></td><td></td><td></td><td></td><td></td><td></td><td></td><td></td><td></td><td></td></tr>
<tr><td>Chromosome</td><td>start</td><td></td><td>end</td><td>strand</td><td>MM</td><td>target_seq</td><td>PAM</td><td>alignment</td><td>distance</td><td>position</td><td>gene name</td><td>gene id</td></tr>
<tr><td>GU305914.1</td><td></td><td>45981</td><td>46003</td><td>-</td><td>0</td><td>TCCTCCGACAGCATATTGTG</td><td>GGG</td><td>||||||||[||||||||||||]PAM</td><td>NA</td><td>Intergenic</td><td>NA</td><td>NA</td></tr>
<tr><td>T30737</td><td>CCTCCGACAGCATATTGTGGGGG</td><td></td><td>1000</td><td></td><td></td><td></td><td></td><td></td><td></td><td></td><td></td><td></td></tr>
<tr><td>Oligo fwd</td><td>CACCCCTCCGACAGCATATTGTGG</td><td></td><td></td><td></td><td></td><td></td><td></td><td></td><td></td><td></td><td></td><td></td></tr>
<tr><td>Oligo rev</td><td>AAACCCACAATATGCTGTCGGAGG</td><td></td><td></td><td></td><td></td><td></td><td></td><td></td><td></td><td></td><td></td><td></td></tr>
<tr><td>Chromosome</td><td>start</td><td></td><td>end</td><td>strand</td><td>MM</td><td>target_seq</td><td>PAM</td><td>alignment</td><td>distance</td><td>position</td><td>gene name</td><td>gene id</td></tr>
<tr><td>GU305914.1</td><td></td><td>45980</td><td>46002</td><td>-</td><td>0</td><td>CCTCCGACAGCATATTGTGG</td><td>GGG</td><td>||||||||[||||||||||||]PAM</td><td>NA</td><td>Intergenic</td><td>NA</td><td>NA</td></tr>
<tr><td>T30738</td><td>CATCAGACTCGTGTCGCGTCTGG</td><td></td><td>1000</td><td></td><td></td><td></td><td></td><td></td><td></td><td></td><td></td><td></td></tr>
<tr><td>Oligo fwd</td><td>CACCCATCAGACTCGTGTCGCGTC</td><td></td><td></td><td></td><td></td><td></td><td></td><td></td><td></td><td></td><td></td><td></td></tr>
<tr><td>Oligo rev</td><td>AAACGACGCGACACGAGTCTGATG</td><td></td><td></td><td></td><td></td><td></td><td></td><td></td><td></td><td></td><td></td><td></td></tr>
<tr><td>Chromosome</td><td>start</td><td></td><td>end</td><td>strand</td><td>MM</td><td>target_seq</td><td>PAM</td><td>alignment</td><td>distance</td><td>position</td><td>gene name</td><td>gene id</td></tr>
<tr><td>GU305914.1</td><td></td><td>45937</td><td>45959</td><td>-</td><td>0</td><td>CATCAGACTCGTGTCGCGTC</td><td>TGG</td><td>||||||||[||||||||||||]PAM</td><td>NA</td><td>Intergenic</td><td>NA</td><td>NA</td></tr>
<tr><td>T30739</td><td>ATCAGACTCGTGTCGCGTCTGGG</td><td></td><td>1000</td><td></td><td></td><td></td><td></td><td></td><td></td><td></td><td></td><td></td></tr>
<tr><td>Oligo fwd</td><td>CACCATCAGACTCGTGTCGCGTCT</td><td></td><td></td><td></td><td></td><td></td><td></td><td></td><td></td><td></td><td></td><td></td></tr>
<tr><td>Oligo rev</td><td>AAACAGACGCGACACGAGTCTGAT</td><td></td><td></td><td></td><td></td><td></td><td></td><td></td><td></td><td></td><td></td><td></td></tr>
<tr><td>Chromosome</td><td>start</td><td></td><td>end</td><td>strand</td><td>MM</td><td>target_seq</td><td>PAM</td><td>alignment</td><td>distance</td><td>position</td><td>gene name</td><td>gene id</td></tr>
<tr><td>GU305914.1</td><td></td><td>45936</td><td>45958</td><td>-</td><td>0</td><td>ATCAGACTCGTGTCGCGTCT</td><td>GGG</td><td>||||||||[||||||||||||]PAM</td><td>NA</td><td>Intergenic</td><td>NA</td><td>NA</td></tr>
<tr><td>T30740</td><td>TCAGACTCGTGTCGCGTCTGGGG</td><td></td><td>1000</td><td></td><td></td><td></td><td></td><td></td><td></td><td></td><td></td><td></td></tr>
<tr><td>Oligo fwd</td><td>CACCTCAGACTCGTGTCGCGTCTG</td><td></td><td></td><td></td><td></td><td></td><td></td><td></td><td></td><td></td><td></td><td></td></tr>
<tr><td>Oligo rev</td><td>AAACCAGACGCGACACGAGTCTGA</td><td></td><td></td><td></td><td></td><td></td><td></td><td></td><td></td><td></td><td></td><td></td></tr>
<tr><td>Chromosome</td><td>start</td><td></td><td>end</td><td>strand</td><td>MM</td><td>target_seq</td><td>PAM</td><td>alignment</td><td>distance</td><td>position</td><td>gene name</td><td>gene id</td></tr>
<tr><td>GU305914.1</td><td></td><td>45935</td><td>45957</td><td>-</td><td>0</td><td>TCAGACTCGTGTCGCGTCTG</td><td>GGG</td><td>||||||||[||||||||||||]PAM</td><td>NA</td><td>Intergenic</td><td>NA</td><td>NA</td></tr>
<tr><td>T30741</td><td>CGTCTGGGGTCTTCCTCCGTCGG</td><td></td><td>1000</td><td></td><td></td><td></td><td></td><td></td><td></td><td></td><td></td><td></td></tr>
<tr><td>Oligo fwd</td><td>CACCCGTCTGGGGTCTTCCTCCGT</td><td></td><td></td><td></td><td></td><td></td><td></td><td></td><td></td><td></td><td></td><td></td></tr>
<tr><td>Oligo rev</td><td>AAACACGGAGGAAGACCCCAGACG</td><td></td><td></td><td></td><td></td><td></td><td></td><td></td><td></td><td></td><td></td><td></td></tr>
<tr><td>Chromosome</td><td>start</td><td></td><td>end</td><td>strand</td><td>MM</td><td>target_seq</td><td>PAM</td><td>alignment</td><td>distance</td><td>position</td><td>gene name</td><td>gene id</td></tr>
<tr><td>GU305914.1</td><td></td><td>45921</td><td>45943</td><td>-</td><td>0</td><td>CGTCTGGGGTCTTCCTCCGT</td><td>CGG</td><td>||||||||[||||||||||||]PAM</td><td>NA</td><td>Intergenic</td><td>NA</td><td>NA</td></tr>
<tr><td>T30742</td><td>GGGGTCTTCCTCCGTCGGAGCGG</td><td></td><td>1000</td><td></td><td></td><td></td><td></td><td></td><td></td><td></td><td></td><td></td></tr>
<tr><td>Oligo fwd</td><td>CACCGGGGTCTTCCTCCGTCGGAG</td><td></td><td></td><td></td><td></td><td></td><td></td><td></td><td></td><td></td><td></td><td></td></tr>
<tr><td>Oligo rev</td><td>AAACCTCCGACGGAGGAAGACCCC</td><td></td><td></td><td></td><td></td><td></td><td></td><td></td><td></td><td></td><td></td><td></td></tr>
<tr><td>Chromosome</td><td>start</td><td></td><td>end</td><td>strand</td><td>MM</td><td>target_seq</td><td>PAM</td><td>alignment</td><td>distance</td><td>position</td><td>gene name</td><td>gene id</td></tr>
<tr><td>GU305914.1</td><td></td><td>45916</td><td>45938</td><td>-</td><td>0</td><td>GGGGTCTTCCTCCGTCGGAG</td><td>CGG</td><td>||||||||[||||||||||||]PAM</td><td>NA</td><td>Intergenic</td><td>NA</td><td>NA</td></tr>
<tr><td>T30743</td><td>AGCGGCCATCGCGTCCCGTGCGG</td><td></td><td>1000</td><td></td><td></td><td></td><td></td><td></td><td></td><td></td><td></td><td></td></tr>
<tr><td>Oligo fwd</td><td>CACCAGCGGCCATCGCGTCCCGTG</td><td></td><td></td><td></td><td></td><td></td><td></td><td></td><td></td><td></td><td></td><td></td></tr>
<tr><td>Oligo rev</td><td>AAACCACGGGACGCGATGGCCGCT</td><td></td><td></td><td></td><td></td><td></td><td></td><td></td><td></td><td></td><td></td><td></td></tr>
<tr><td>Chromosome</td><td>start</td><td></td><td>end</td><td>strand</td><td>MM</td><td>target_seq</td><td>PAM</td><td>alignment</td><td>distance</td><td>position</td><td>gene name</td><td>gene id</td></tr>
<tr><td>GU305914.1</td><td></td><td>45898</td><td>45920</td><td>-</td><td>0</td><td>AGCGGCCATCGCGTCCCGTG</td><td>CGG</td><td>||||||||[||||||||||||]PAM</td><td>NA</td><td>Intergenic</td><td>NA</td><td>NA</td></tr>
<tr><td>GU305914.1</td><td></td><td>110132</td><td>110154</td><td>+</td><td>4</td><td>GGCCGCCGTCGCGTCCCGGG</td><td>AGG</td><td>-||-|||-[||||||||||-|]PAM</td><td>NA</td><td>Intergenic</td><td>NA</td><td>NA</td></tr>
<tr><td>T30744</td><td>GCCATCGCGTCCCGTGCGGCCGG</td><td></td><td>1000</td><td></td><td></td><td></td><td></td><td></td><td></td><td></td><td></td><td></td></tr>
</table>

Oligo fwd CACCGCCATCGCGTCCCGTGCGGC
Oligo rev AAACGCCGCACGGGACGCGATGGC
Chromosome start end strand MM target_seq PAM alignment distance position gene name gene id
GU305914.1 45894 45916 - 0 GCCATCGCGTCCCGTGCGGC CGG ||||||||[||||||||||||]PAM NA Intergenic NA NA

T30745 GATACATGCCCACCCGCACGCGG 1000
Oligo fwd CACCGATACATGCCCACCCGCACG
Oligo rev AAACCGTGCGGGTGGGCATGTATC
Chromosome start end strand MM target_seq PAM alignment distance position gene name gene id
GU305914.1 45868 45890 - 0 GATACATGCCCACCCGCACG CGG ||||||||[||||||||||||]PAM NA Intergenic NA NA

T30746 GCCCACCCGCACGCGGCGTGTGG 1000
Oligo fwd CACCGCCCACCCGCACGCGGCGTG
Oligo rev AAACCACGCCGCGTGCGGGTGGGC
Chromosome start end strand MM target_seq PAM alignment distance position gene name gene id
GU305914.1 45861 45883 - 0 GCCCACCCGCACGCGGCGTG TGG ||||||||[||||||||||||]PAM NA Intergenic NA NA

T30747 CGTGTGGCGCCAGCCCCCGATGG 1000
Oligo fwd CACCCGTGTGGCGCCAGCCCCCGA
Oligo rev AAACTCGGGGGCTGGCGCCACACG
Chromosome start end strand MM target_seq PAM alignment distance position gene name gene id
GU305914.1 45845 45867 - 0 CGTGTGGCGCCAGCCCCCGA TGG ||||||||[||||||||||||]PAM NA Intergenic NA NA

T30748 TGGCGCCAGCCCCCGATGGATGG 1000
Oligo fwd CACCTGGCGCCAGCCCCCGATGGA
Oligo rev AAACTCCATCGGGGGCTGGCGCCA
Chromosome start end strand MM target_seq PAM alignment distance position gene name gene id
GU305914.1 45841 45863 - 0 TGGCGCCAGCCCCCGATGGA TGG ||||||||[||||||||||||]PAM NA Intergenic NA NA

T30749 GGCGCCAGCCCCCGATGGATGGG 1000
Oligo fwd CACCGGCGCCAGCCCCCGATGGAT
Oligo rev AAACATCCATCGGGGGCTGGCGCC
Chromosome start end strand MM target_seq PAM alignment distance position gene name gene id
GU305914.1 45840 45862 - 0 GGCGCCAGCCCCCGATGGAT GGG ||||||||[||||||||||||]PAM NA Intergenic NA NA

T30750 ACACGATTCCAATTAGCTTTTGG 1000
Oligo fwd CACCACACGATTCCAATTAGCTTT
Oligo rev AAACAAAGCTAATTGGAATCGTGT
Chromosome start end strand MM target_seq PAM alignment distance position gene name gene id
GU305914.1 45804 45826 - 0 ACACGATTCCAATTAGCTTT TGG ||||||||[||||||||||||]PAM NA Intergenic NA NA

T30751 CACGATTCCAATTAGCTTTTGGG 1000
Oligo fwd CACCCACGATTCCAATTAGCTTTT
Oligo rev AAACAAAAGCTAATTGGAATCGTG
Chromosome start end strand MM target_seq PAM alignment distance position gene name gene id
GU305914.1 45803 45825 - 0 CACGATTCCAATTAGCTTTT GGG ||||||||[||||||||||||]PAM NA Intergenic NA NA

T30752 CAATTAGCTTTTGGGCCCAGAGG 1000
Oligo fwd CACCCAATTAGCTTTTGGGCCCAG
Oligo rev AAACCTGGGCCCAAAAGCTAATTG
Chromosome start end strand MM target_seq PAM alignment distance position gene name gene id
GU305914.1 45795 45817 - 0 CAATTAGCTTTTGGGCCCAG AGG ||||||||[||||||||||||]PAM NA Intergenic NA NA

T30753 CCCTATGCTACAAAACGACACGG 1000
Oligo fwd CACCCCCTATGCTACAAAACGACA
Oligo rev AAACTGTCGTTTTGTAGCATAGGG
Chromosome start end strand MM target_seq PAM alignment distance position gene name gene id
GU305914.1 45716 45738 - 0 CCCTATGCTACAAAACGACA CGG ||||||||[||||||||||||]PAM NA Intergenic NA NA

T30754 GCTACAAAACGACACGGTACTGG 1000
Oligo fwd CACCGCTACAAAACGACACGGTAC
Oligo rev AAACGTACCGTGTCGTTTTGTAGC
Chromosome start end strand MM target_seq PAM alignment distance position gene name gene id
GU305914.1 45710 45732 - 0 GCTACAAAACGACACGGTAC TGG ||||||||[||||||||||||]PAM NA Intergenic NA NA

T30755 AAAACGACACGGTACTGGTGAGG 1000
Oligo fwd CACCAAAACGACACGGTACTGGTG
Oligo rev AAACCACCAGTACCGTGTCGTTTT
Chromosome start end strand MM target_seq PAM alignment distance position gene name gene id
GU305914.1 45705 45727 - 0 AAAACGACACGGTACTGGTG AGG ||||||||[||||||||||||]PAM NA Intergenic NA NA

T30756 AAACGACACGGTACTGGTGAGGG 1000
Oligo fwd CACCAAACGACACGGTACTGGTGA
Oligo rev AAACTCACCAGTACCGTGTCGTTT
Chromosome start end strand MM target_seq PAM alignment distance position gene name gene id
GU305914.1 45704 45726 - 0 AAACGACACGGTACTGGTGA GGG ||||||||[||||||||||||]PAM NA Intergenic NA NA

T30757 CACGGTACTGGTGAGGGAGACGG 1000
Oligo fwd CACCCACGGTACTGGTGAGGGAGA
Oligo rev AAACTCTCCCTCACCAGTACCGTG
Chromosome start end strand MM target_seq PAM alignment distance position gene name gene id
GU305914.1 45698 45720 - 0 CACGGTACTGGTGAGGGAGA CGG ||||||||[||||||||||||]PAM NA Intergenic NA NA

T30758 TGGTGAGGGAGACGGTGTCGCGG 1000
Oligo fwd CACCTGGTGAGGGAGACGGTGTCG
Oligo rev AAACCGACACCGTCTCCCTCACCA
Chromosome start end strand MM target_seq PAM alignment distance position gene name gene id
GU305914.1 45690 45712 - 0 TGGTGAGGGAGACGGTGTCG CGG ||||||||[||||||||||||]PAM NA Intergenic NA NA

| | | | | | | | | | | | |
|---|---|---|---|---|---|---|---|---|---|---|---|
| T30759 | TCCCGCGCCCCCCTTAGAGCAGG | 1000 | | | | | | | | | |
| Oligo fwd | CACCTCCCGCGCCCCCCTTAGAGC | | | | | | | | | | |
| Oligo rev | AAACGCTCTAAGGGGGGCGCGGGA | | | | | | | | | | |
| Chromosome | start | end | strand | MM | target_seq | PAM | alignment | distance | position | gene name | gene id |
| GU305914.1 | 45641 | 45663 | - | 0 | TCCCGCGCCCCCCTTAGAGC | AGG | \|\|\|\|\|\|\|\|[\|\|\|\|\|\|\|\|\|\|\|\|]PAM | NA | Intergenic | NA | NA |
| T30760 | TGATGCACTCAGAATGTTTCTGG | 1000 | | | | | | | | | |
| Oligo fwd | CACCTGATGCACTCAGAATGTTTC | | | | | | | | | | |
| Oligo rev | AAACGAAACATTCTGAGTGCATCA | | | | | | | | | | |
| Chromosome | start | end | strand | MM | target_seq | PAM | alignment | distance | position | gene name | gene id |
| GU305914.1 | 45614 | 45636 | - | 0 | TGATGCACTCAGAATGTTTC | TGG | \|\|\|\|\|\|\|\|[\|\|\|\|\|\|\|\|\|\|\|\|]PAM | NA | Intergenic | NA | NA |
| T30761 | GATGCACTCAGAATGTTTCTGGG | 1000 | | | | | | | | | |
| Oligo fwd | CACCGATGCACTCAGAATGTTTCT | | | | | | | | | | |
| Oligo rev | AAACAGAAACATTCTGAGTGCATC | | | | | | | | | | |
| Chromosome | start | end | strand | MM | target_seq | PAM | alignment | distance | position | gene name | gene id |
| GU305914.1 | 45613 | 45635 | - | 0 | GATGCACTCAGAATGTTTCT | GGG | \|\|\|\|\|\|\|\|[\|\|\|\|\|\|\|\|\|\|\|\|]PAM | NA | Intergenic | NA | NA |
| T30762 | GCACTCAGAATGTTTCTGGGCGG | 1000 | | | | | | | | | |
| Oligo fwd | CACCGCACTCAGAATGTTTCTGGG | | | | | | | | | | |
| Oligo rev | AAACCCCAGAAACATTCTGAGTGC | | | | | | | | | | |
| Chromosome | start | end | strand | MM | target_seq | PAM | alignment | distance | position | gene name | gene id |
| GU305914.1 | 45610 | 45632 | - | 0 | GCACTCAGAATGTTTCTGGG | CGG | \|\|\|\|\|\|\|\|[\|\|\|\|\|\|\|\|\|\|\|\|]PAM | NA | Intergenic | NA | NA |
| T30763 | GGCGGCTGCCGCCGCCGCCGCGG | 1000 | | | | | | | | | |
| Oligo fwd | CACCGGCGGCTGCCGCCGCCGCCG | | | | | | | | | | |
| Oligo rev | AAACCGGCGGCGGCGGCAGCCGCC | | | | | | | | | | |
| Chromosome | start | end | strand | MM | target_seq | PAM | alignment | distance | position | gene name | gene id |
| GU305914.1 | 45592 | 45614 | - | 0 | GGCGGCTGCCGCCGCCGCCG | CGG | \|\|\|\|\|\|\|\|[\|\|\|\|\|\|\|\|\|\|\|\|]PAM | NA | Intergenic | NA | NA |
| GU305914.1 | 35655 | 35677 | - | 4 | CCCCGCCGCCGCCGCCGCCG | TGG | --\|-\|\|-\|[\|\|\|\|\|\|\|\|\|\|\|\|]PAM | NA | Intergenic | NA | NA |
| GU305914.1 | 26444 | 26466 | - | 4 | CTCGGAGGCCGCCGCCGCCG | TGG | --\|\|\|--\|[\|\|\|\|\|\|\|\|\|\|\|\|]PAM | NA | Intergenic | NA | NA |
| GU305914.1 | 130312 | 130334 | + | 4 | CGCAGCAGCAGCCGCCGCCG | CGG | -\|\|-\|\|-\|[\|-\|\|\|\|\|\|\|\|\|\|]PAM | NA | Intergenic | NA | NA |
| GU305914.1 | 86842 | 86864 | + | 4 | TGAAGCTGCCGCCCCCGCCG | CGG | -\|--\|\|\|\|[\|\|\|\|\|-\|\|\|\|\|\|]PAM | NA | Intergenic | NA | NA |
| GU305914.1 | 62679 | 62701 | - | 3 | CGCGGCTGCTGCCGCCGGCG | GGG | -\|\|\|\|\|\|\|[\|-\|\|\|\|\|\|\|-\|\|]PAM | NA | Intergenic | NA | NA |
| GU305914.1 | 130315 | 130337 | + | 4 | AGCAGCAGCCGCCGCCGCGG | CGG | -\|\|-\|\|-\|[\|\|\|\|\|\|\|\|\|\|-\|]PAM | NA | Intergenic | NA | NA |
| GU305914.1 | 56775 | 56797 | - | 4 | TGCCGCCGCCGCCGCCGCTG | CGG | -\|\|-\|\|-\|[\|\|\|\|\|\|\|\|\|\|-\|]PAM | NA | Intergenic | NA | NA |
| GU305914.1 | 97885 | 97907 | + | 3 | GGTCGCTGCCGCCGCCGCCC | GGG | \|\|--\|\|\|\|[\|\|\|\|\|\|\|\|\|\|\|-]PAM | NA | Intergenic | NA | NA |
| GU305914.1 | 26441 | 26463 | - | 4 | GGAGGCCGCCGCCGCCGTGG | CGG | \|\|-\|\|\|-\|[\|\|\|\|\|\|\|\|\|--\|]PAM | NA | Intergenic | NA | NA |
| T30764 | TGCCGCCGCCGCCGCGGCGCCGG | 1000 | | | | | | | | | |
| Oligo fwd | CACCTGCCGCCGCCGCCGCGGCGC | | | | | | | | | | |
| Oligo rev | AAACGCGCCGCGGCGGCGGCGGCA | | | | | | | | | | |
| Chromosome | start | end | strand | MM | target_seq | PAM | alignment | distance | position | gene name | gene id |
| GU305914.1 | 45586 | 45608 | - | 0 | TGCCGCCGCCGCCGCGGCGC | CGG | \|\|\|\|\|\|\|\|[\|\|\|\|\|\|\|\|\|\|\|\|]PAM | NA | Intergenic | NA | NA |
| GU305914.1 | 56772 | 56794 | - | 3 | CGCCGCCGCCGCCGCTGCGG | CGG | -\|\|\|\|\|\|\|[\|\|\|\|\|\|\|-\|\|\|-]PAM | NA | Intergenic | NA | NA |
| T30765 | GCCGCCGCCGCCGCGGCGCCGGG | 1000 | | | | | | | | | |
| Oligo fwd | CACCGCCGCCGCCGCCGCGGCGCC | | | | | | | | | | |
| Oligo rev | AAACGGCGCCGCGGCGGCGGCGGC | | | | | | | | | | |
| Chromosome | start | end | strand | MM | target_seq | PAM | alignment | distance | position | gene name | gene id |
| GU305914.1 | 45585 | 45607 | - | 0 | GCCGCCGCCGCCGCGGCGCC | GGG | \|\|\|\|\|\|\|\|[\|\|\|\|\|\|\|\|\|\|\|\|]PAM | NA | Intergenic | NA | NA |
| T30766 | GCCGCCGCCGCGGCGCCGGGTGG | 1000 | | | | | | | | | |
| Oligo fwd | CACCGCCGCCGCCGCGGCGCCGGG | | | | | | | | | | |
| Oligo rev | AAACCCCGGCGCCGCGGCGGCGGC | | | | | | | | | | |
| Chromosome | start | end | strand | MM | target_seq | PAM | alignment | distance | position | gene name | gene id |
| GU305914.1 | 45582 | 45604 | - | 0 | GCCGCCGCCGCGGCGCCGGG | TGG | \|\|\|\|\|\|\|\|[\|\|\|\|\|\|\|\|\|\|\|\|]PAM | NA | Intergenic | NA | NA |
| GU305914.1 | 169849 | 169871 | + | 4 | GCCGCTGTCGCGGCTCCGGC | CGG | \|\|\|\|\|-\|-[\|\|\|\|\|\|-\|\|\|\|-]PAM | NA | Intergenic | NA | NA |
| T30767 | GTGGCGTAGACCGTCCGACGTGG | 1000 | | | | | | | | | |
| Oligo fwd | CACCGTGGCGTAGACCGTCCGACG | | | | | | | | | | |
| Oligo rev | AAACCGTCGGACGGTCTACGCCAC | | | | | | | | | | |
| Chromosome | start | end | strand | MM | target_seq | PAM | alignment | distance | position | gene name | gene id |
| GU305914.1 | 45563 | 45585 | - | 0 | GTGGCGTAGACCGTCCGACG | TGG | \|\|\|\|\|\|\|\|[\|\|\|\|\|\|\|\|\|\|\|\|]PAM | NA | Intergenic | NA | NA |
| T30768 | CGTAGACCGTCCGACGTGGACGG | 1000 | | | | | | | | | |
| Oligo fwd | CACCCGTAGACCGTCCGACGTGGA | | | | | | | | | | |
| Oligo rev | AAACTCCACGTCGGACGGTCTACG | | | | | | | | | | |
| Chromosome | start | end | strand | MM | target_seq | PAM | alignment | distance | position | gene name | gene id |
| GU305914.1 | 45559 | 45581 | - | 0 | CGTAGACCGTCCGACGTGGA | CGG | \|\|\|\|\|\|\|\|[\|\|\|\|\|\|\|\|\|\|\|\|]PAM | NA | Intergenic | NA | NA |
| T30769 | TGGACGGAGATGAAGTTTTTCGG | 1000 | | | | | | | | | |
| Oligo fwd | CACCTGGACGGAGATGAAGTTTTT | | | | | | | | | | |
| Oligo rev | AAACAAAAACTTCATCTCCGTCCA | | | | | | | | | | |
| Chromosome | start | end | strand | MM | target_seq | PAM | alignment | distance | position | gene name | gene id |
| GU305914.1 | 45543 | 45565 | - | 0 | TGGACGGAGATGAAGTTTTT | CGG | \|\|\|\|\|\|\|\|[\|\|\|\|\|\|\|\|\|\|\|\|]PAM | NA | Intergenic | NA | NA |
| T30770 | ACGGAGATGAAGTTTTTCGGAGG | 1000 | | | | | | | | | |
| Oligo fwd | CACCACGGAGATGAAGTTTTTCGG | | | | | | | | | | |
| Oligo rev | AAACCCGAAAAACTTCATCTCCGT | | | | | | | | | | |
| Chromosome | start | end | strand | MM | target_seq | PAM | alignment | distance | position | gene name | gene id |
| GU305914.1 | 45540 | 45562 | - | 0 | ACGGAGATGAAGTTTTTCGG | AGG | \|\|\|\|\|\|\|\|[\|\|\|\|\|\|\|\|\|\|\|\|]PAM | NA | Intergenic | NA | NA |
| T30771 | CGGAGATGAAGTTTTTCGGAGGG | 1000 | | | | | | | | | |
| Oligo fwd | CACCCGGAGATGAAGTTTTTCGGA | | | | | | | | | | |
| Oligo rev | AAACTCCGAAAAACTTCATCTCCG | | | | | | | | | | |
| Chromosome | start | end | strand | MM | target_seq | PAM | alignment | distance | position | gene name | gene id |
| GU305914.1 | 45539 | 45561 | - | 0 | CGGAGATGAAGTTTTTCGGA | GGG | \|\|\|\|\|\|\|\|[\|\|\|\|\|\|\|\|\|\|\|\|]PAM | NA | Intergenic | NA | NA |

T30772 GGGCTTCCGTTTATCTCGTTCGG 1000
Oligo fwd CACCGGGCTTCCGTTTATCTCGTT
Oligo rev AAACAACGAGATAAACGGAAGCCC
Chromosome start end strand MM target_seq PAM alignment distance position gene name gene id
GU305914.1 45519 45541 - 0 GGGCTTCCGTTTATCTCGTT CGG ||||||||[||||||||||||]PAM NA Intergenic NA NA

T30773 GAAAAGTACCCGAGTTCGTGTGG 1000
Oligo fwd CACCGAAAAGTACCCGAGTTCGTG
Oligo rev AAACCACGAACTCGGGTACTTTTC
Chromosome start end strand MM target_seq PAM alignment distance position gene name gene id
GU305914.1 45495 45517 - 0 GAAAAGTACCCGAGTTCGTG TGG ||||||||[||||||||||||]PAM NA Intergenic NA NA

T30774 AAAAGTACCCGAGTTCGTGTGGG 1000
Oligo fwd CACCAAAAGTACCCGAGTTCGTGT
Oligo rev AAACACACGAACTCGGGTACTTTT
Chromosome start end strand MM target_seq PAM alignment distance position gene name gene id
GU305914.1 45494 45516 - 0 AAAAGTACCCGAGTTCGTGT GGG ||||||||[||||||||||||]PAM NA Intergenic NA NA

T30775 TACCCGAGTTCGTGTGGGCTTGG 1000
Oligo fwd CACCTACCCGAGTTCGTGTGGGCT
Oligo rev AAACAGCCCACACGAACTCGGGTA
Chromosome start end strand MM target_seq PAM alignment distance position gene name gene id
GU305914.1 45489 45511 - 0 TACCCGAGTTCGTGTGGGCT TGG ||||||||[||||||||||||]PAM NA Intergenic NA NA

T30776 CGTGTGGGCTTGGCGAGTAATGG 1000
Oligo fwd CACCCGTGTGGGCTTGGCGAGTAA
Oligo rev AAACTTACTCGCCAAGCCCACACG
Chromosome start end strand MM target_seq PAM alignment distance position gene name gene id
GU305914.1 45479 45501 - 0 CGTGTGGGCTTGGCGAGTAA TGG ||||||||[||||||||||||]PAM NA Intergenic NA NA

T30777 GCTTGGCGAGTAATGGTGATAGG 1000
Oligo fwd CACCGCTTGGCGAGTAATGGTGAT
Oligo rev AAACATCACCATTACTCGCCAAGC
Chromosome start end strand MM target_seq PAM alignment distance position gene name gene id
GU305914.1 45472 45494 - 0 GCTTGGCGAGTAATGGTGAT AGG ||||||||[||||||||||||]PAM NA Intergenic NA NA

T30778 CTTGGCGAGTAATGGTGATAGGG 1000
Oligo fwd CACCCTTGGCGAGTAATGGTGATA
Oligo rev AAACTATCACCATTACTCGCCAAG
Chromosome start end strand MM target_seq PAM alignment distance position gene name gene id
GU305914.1 45471 45493 - 0 CTTGGCGAGTAATGGTGATA GGG ||||||||[||||||||||||]PAM NA Intergenic NA NA

T30779 TTGGCGAGTAATGGTGATAGGGG 1000
Oligo fwd CACCTTGGCGAGTAATGGTGATAG
Oligo rev AAACCTATCACCATTACTCGCCAA
Chromosome start end strand MM target_seq PAM alignment distance position gene name gene id
GU305914.1 45470 45492 - 0 TTGGCGAGTAATGGTGATAG GGG ||||||||[||||||||||||]PAM NA Intergenic NA NA

T30780 GGTGATGAAACTGCTGCGCTTGG 1000
Oligo fwd CACCGGTGATGAAACTGCTGCGCT
Oligo rev AAACAGCGCAGCAGTTTCATCACC
Chromosome start end strand MM target_seq PAM alignment distance position gene name gene id
GU305914.1 45449 45471 - 0 GGTGATGAAACTGCTGCGCT TGG ||||||||[||||||||||||]PAM NA Intergenic NA NA

T30781 GATGAAACTGCTGCGCTTGGTGG 1000
Oligo fwd CACCGATGAAACTGCTGCGCTTGG
Oligo rev AAACCCAAGCGCAGCAGTTTCATC
Chromosome start end strand MM target_seq PAM alignment distance position gene name gene id
GU305914.1 45446 45468 - 0 GATGAAACTGCTGCGCTTGG TGG ||||||||[||||||||||||]PAM NA Intergenic NA NA

T30782 CCCCCATTGTCATCATGTCGTGG 1000
Oligo fwd CACCCCCCCATTGTCATCATGTCG
Oligo rev AAACCGACATGATGACAATGGGGG
Chromosome start end strand MM target_seq PAM alignment distance position gene name gene id
GU305914.1 45374 45396 - 0 CCCCCATTGTCATCATGTCG TGG ||||||||[||||||||||||]PAM NA Intergenic NA NA

T30783 TGGCTGCCGATGCTGTCGATCGG 1000
Oligo fwd CACCTGGCTGCCGATGCTGTCGAT
Oligo rev AAACATCGACAGCATCGGCAGCCA
Chromosome start end strand MM target_seq PAM alignment distance position gene name gene id
GU305914.1 45354 45376 - 0 TGGCTGCCGATGCTGTCGAT CGG ||||||||[||||||||||||]PAM NA Intergenic NA NA

T30784 CTGCCGATGCTGTCGATCGGAGG 1000
Oligo fwd CACCCTGCCGATGCTGTCGATCGG
Oligo rev AAACCCGATCGACAGCATCGGCAG
Chromosome start end strand MM target_seq PAM alignment distance position gene name gene id
GU305914.1 45351 45373 - 0 CTGCCGATGCTGTCGATCGG AGG ||||||||[||||||||||||]PAM NA Intergenic NA NA

T30785 CCGATGCTGTCGATCGGAGGAGG 1000
Oligo fwd CACCCCGATGCTGTCGATCGGAGG
Oligo rev AAACCCTCCGATCGACAGCATCGG
Chromosome start end strand MM target_seq PAM alignment distance position gene name gene id
GU305914.1 45348 45370 - 0 CCGATGCTGTCGATCGGAGG AGG ||||||||[||||||||||||]PAM NA Intergenic NA NA

T30786 GGAGGAGGAAGATGATGAAGTGG 1000
Oligo fwd CACCGGAGGAGGAAGATGATGAAG
Oligo rev AAACCTTCATCATCTTCCTCCTCC
Chromosome start end strand MM target_seq PAM alignment distance position gene name gene id
GU305914.1 45333 45355 - 0 GGAGGAGGAAGATGATGAAG TGG ||||||||[||||||||||||]PAM NA Intergenic NA NA

| | | | | | | | | | | | |
|---|---|---|---|---|---|---|---|---|---|---|---|
| GU305914.1 | 118515 | 118537 | - | 3 | CGATGAGGACGATGATGAAG | AGG | -\|\|-\|\|\|\|[\|-\|\|\|\|\|\|\|\|\|\|]PAM | NA | Intergenic | NA | NA |
| GU305914.1 | 118851 | 118873 | - | 2 | GGAGGAGGAAGAAGACGAAG | AGG | \|\|\|\|\|\|\|\|[\|\|\|\|-\|\|-\|\|\|\|]PAM | NA | Intergenic | NA | NA |
| T30787 | GGAGGAAGATGATGAAGTGGAGG | 1000 | | | | | | | | | |
| Oligo fwd | CACCGGAGGAAGATGATGAAGTGG | | | | | | | | | | |
| Oligo rev | AAACCCACTTCATCATCTTCCTCC | | | | | | | | | | |
| Chromosome | start | end | strand | MM | target_seq | PAM | alignment | distance | position | gene name | gene id |
| GU305914.1 | 45330 | 45352 | - | 0 | GGAGGAAGATGATGAAGTGG | AGG | \|\|\|\|\|\|\|\|[\|\|\|\|\|\|\|\|\|\|\|\|]PAM | NA | Intergenic | NA | NA |
| GU305914.1 | 118512 | 118534 | - | 3 | TGAGGACGATGATGAAGAGG | AGG | -\|\|\|\|\|-\|[\|\|\|\|\|\|\|\|\|-\|\|]PAM | NA | Intergenic | NA | NA |
| T30788 | GGAAGATGATGAAGTGGAGGAGG | 1000 | | | | | | | | | |
| Oligo fwd | CACCGGAAGATGATGAAGTGGAGG | | | | | | | | | | |
| Oligo rev | AAACCCTCCACTTCATCATCTTCC | | | | | | | | | | |
| Chromosome | start | end | strand | MM | target_seq | PAM | alignment | distance | position | gene name | gene id |
| GU305914.1 | 45327 | 45349 | - | 0 | GGAAGATGATGAAGTGGAGG | AGG | \|\|\|\|\|\|\|\|[\|\|\|\|\|\|\|\|\|\|\|\|]PAM | NA | Intergenic | NA | NA |
| T30789 | AGATGATGAAGTGGAGGAGGAGG | 1000 | | | | | | | | | |
| Oligo fwd | CACCAGATGATGAAGTGGAGGAGG | | | | | | | | | | |
| Oligo rev | AAACCCTCCTCCACTTCATCATCT | | | | | | | | | | |
| Chromosome | start | end | strand | MM | target_seq | PAM | alignment | distance | position | gene name | gene id |
| GU305914.1 | 45324 | 45346 | - | 0 | AGATGATGAAGTGGAGGAGG | AGG | \|\|\|\|\|\|\|\|[\|\|\|\|\|\|\|\|\|\|\|\|]PAM | NA | Intergenic | NA | NA |
| GU305914.1 | 118506 | 118528 | - | 3 | CGATGATGAAGAGGAGGACG | AGG | -\|\|\|\|\|\|\|[\|\|\|-\|\|\|\|\|\|-\|]PAM | NA | Intergenic | NA | NA |
| T30790 | TGATGAAGTGGAGGAGGAGGAGG | 1000 | | | | | | | | | |
| Oligo fwd | CACCTGATGAAGTGGAGGAGGAGG | | | | | | | | | | |
| Oligo rev | AAACCCTCCTCCTCCACTTCATCA | | | | | | | | | | |
| Chromosome | start | end | strand | MM | target_seq | PAM | alignment | distance | position | gene name | gene id |
| GU305914.1 | 45321 | 45343 | - | 0 | TGATGAAGTGGAGGAGGAGG | AGG | \|\|\|\|\|\|\|\|[\|\|\|\|\|\|\|\|\|\|\|\|]PAM | NA | Intergenic | NA | NA |
| T30791 | AGTGGAGGAGGAGGAGGAGAAGG | 1000 | | | | | | | | | |
| Oligo fwd | CACCAGTGGAGGAGGAGGAGGAGA | | | | | | | | | | |
| Oligo rev | AAACTCTCCTCCTCCTCCTCCACT | | | | | | | | | | |
| Chromosome | start | end | strand | MM | target_seq | PAM | alignment | distance | position | gene name | gene id |
| GU305914.1 | 45315 | 45337 | - | 0 | AGTGGAGGAGGAGGAGGAGA | AGG | \|\|\|\|\|\|\|\|[\|\|\|\|\|\|\|\|\|\|\|\|]PAM | NA | Intergenic | NA | NA |
| GU305914.1 | 99528 | 99550 | - | 4 | GGGCGAGGAGGCGGAGGAGA | AGG | -\|--\|\|\|\|[\|\|\|-\|\|\|\|\|\|\|\|]PAM | NA | Intergenic | NA | NA |
| GU305914.1 | 42858 | 42880 | - | 4 | GGGGGAGGATGAGGAAGAGA | AGG | -\|-\|\|\|\|\|[\|-\|\|\|\|\|-\|\|\|\|]PAM | NA | Intergenic | NA | NA |
| GU305914.1 | 18851 | 18873 | - | 4 | GGTGGGGGAGGAGGGGAAGA | CGG | -\|\|\|\|-\|\|[\|\|\|\|\|\|-\|-\|\|\|]PAM | NA | Intergenic | NA | NA |
| T30792 | AGGAGAAGGAGACGACGCGACGG | 1000 | | | | | | | | | |
| Oligo fwd | CACCAGGAGAAGGAGACGACGCGA | | | | | | | | | | |
| Oligo rev | AAACTCGCGTCGTCTCCTTCTCCT | | | | | | | | | | |
| Chromosome | start | end | strand | MM | target_seq | PAM | alignment | distance | position | gene name | gene id |
| GU305914.1 | 45301 | 45323 | - | 0 | AGGAGAAGGAGACGACGCGA | CGG | \|\|\|\|\|\|\|\|[\|\|\|\|\|\|\|\|\|\|\|\|]PAM | NA | Intergenic | NA | NA |
| T30793 | GGAGAAGGAGACGACGCGACGGG | 1000 | | | | | | | | | |
| Oligo fwd | CACCGGAGAAGGAGACGACGCGAC | | | | | | | | | | |
| Oligo rev | AAACGTCGCGTCGTCTCCTTCTCC | | | | | | | | | | |
| Chromosome | start | end | strand | MM | target_seq | PAM | alignment | distance | position | gene name | gene id |
| GU305914.1 | 45300 | 45322 | - | 0 | GGAGAAGGAGACGACGCGAC | GGG | \|\|\|\|\|\|\|\|[\|\|\|\|\|\|\|\|\|\|\|\|]PAM | NA | Intergenic | NA | NA |
| T30794 | GAAGGAGACGACGCGACGGGCGG | 1000 | | | | | | | | | |
| Oligo fwd | CACCGAAGGAGACGACGCGACGGG | | | | | | | | | | |
| Oligo rev | AAACCCCGTCGCGTCGTCTCCTTC | | | | | | | | | | |
| Chromosome | start | end | strand | MM | target_seq | PAM | alignment | distance | position | gene name | gene id |
| GU305914.1 | 45297 | 45319 | - | 0 | GAAGGAGACGACGCGACGGG | CGG | \|\|\|\|\|\|\|\|[\|\|\|\|\|\|\|\|\|\|\|\|]PAM | NA | Intergenic | NA | NA |
| T30795 | GGAGACGACGCGACGGGCGGAGG | 1000 | | | | | | | | | |
| Oligo fwd | CACCGGAGACGACGCGACGGGCGG | | | | | | | | | | |
| Oligo rev | AAACCCGCCCGTCGCGTCGTCTCC | | | | | | | | | | |
| Chromosome | start | end | strand | MM | target_seq | PAM | alignment | distance | position | gene name | gene id |
| GU305914.1 | 45294 | 45316 | - | 0 | GGAGACGACGCGACGGGCGG | AGG | \|\|\|\|\|\|\|\|[\|\|\|\|\|\|\|\|\|\|\|\|]PAM | NA | Intergenic | NA | NA |
| T30796 | GACGACGCGACGGGCGGAGGAGG | 1000 | | | | | | | | | |
| Oligo fwd | CACCGACGACGCGACGGGCGGAGG | | | | | | | | | | |
| Oligo rev | AAACCCTCCGCCCGTCGCGTCGTC | | | | | | | | | | |
| Chromosome | start | end | strand | MM | target_seq | PAM | alignment | distance | position | gene name | gene id |
| GU305914.1 | 45291 | 45313 | - | 0 | GACGACGCGACGGGCGGAGG | AGG | \|\|\|\|\|\|\|\|[\|\|\|\|\|\|\|\|\|\|\|\|]PAM | NA | Intergenic | NA | NA |
| T30797 | ACGCGACGGGCGGAGGAGGACGG | 1000 | | | | | | | | | |
| Oligo fwd | CACCACGCGACGGGCGGAGGAGGA | | | | | | | | | | |
| Oligo rev | AAACTCCTCCTCCGCCCGTCGCGT | | | | | | | | | | |
| Chromosome | start | end | strand | MM | target_seq | PAM | alignment | distance | position | gene name | gene id |
| GU305914.1 | 45287 | 45309 | - | 0 | ACGCGACGGGCGGAGGAGGA | CGG | \|\|\|\|\|\|\|\|[\|\|\|\|\|\|\|\|\|\|\|\|]PAM | NA | Intergenic | NA | NA |
| T30798 | GGAGGAGGACGGTGACGCAACGG | 1000 | | | | | | | | | |
| Oligo fwd | CACCGGAGGAGGACGGTGACGCAA | | | | | | | | | | |
| Oligo rev | AAACTTGCGTCACCGTCCTCCTCC | | | | | | | | | | |
| Chromosome | start | end | strand | MM | target_seq | PAM | alignment | distance | position | gene name | gene id |
| GU305914.1 | 45276 | 45298 | - | 0 | GGAGGAGGACGGTGACGCAA | CGG | \|\|\|\|\|\|\|\|[\|\|\|\|\|\|\|\|\|\|\|\|]PAM | NA | Intergenic | NA | NA |
| T30799 | GGAGGACGGTGACGCAACGGTGG | 1000 | | | | | | | | | |
| Oligo fwd | CACCGGAGGACGGTGACGCAACGG | | | | | | | | | | |
| Oligo rev | AAACCCGTTGCGTCACCGTCCTCC | | | | | | | | | | |
| Chromosome | start | end | strand | MM | target_seq | PAM | alignment | distance | position | gene name | gene id |
| GU305914.1 | 45273 | 45295 | - | 0 | GGAGGACGGTGACGCAACGG | TGG | \|\|\|\|\|\|\|\|[\|\|\|\|\|\|\|\|\|\|\|\|]PAM | NA | Intergenic | NA | NA |
| T30800 | GGACGGTGACGCAACGGTGGCGG | 1000 | | | | | | | | | |
| Oligo fwd | CACCGGACGGTGACGCAACGGTGG | | | | | | | | | | |
| Oligo rev | AAACCCACCGTTGCGTCACCGTCC | | | | | | | | | | |

| Chromosome | start | end | strand | MM | target_seq | PAM | alignment | distance | position | gene name | gene id |
|---|---|---|---|---|---|---|---|---|---|---|---|
| GU305914.1 | 45270 | 45292 | - | 0 | GGACGGTGACGCAACGGTGG | CGG | \|\|\|\|\|\|\|\|[\|\|\|\|\|\|\|\|\|\|\|\|]PAM | NA | Intergenic | NA | NA |
| T30801 | CGGTGACGCAACGGTGGCGGTGG | 1000 | | | | | | | | | |
| Oligo fwd | CACCCGGTGACGCAACGGTGGCGG | | | | | | | | | | |
| Oligo rev | AAACCCGCCACCGTTGCGTCACCG | | | | | | | | | | |
| Chromosome | start | end | strand | MM | target_seq | PAM | alignment | distance | position | gene name | gene id |
| GU305914.1 | 45267 | 45289 | - | 0 | CGGTGACGCAACGGTGGCGG | TGG | \|\|\|\|\|\|\|\|[\|\|\|\|\|\|\|\|\|\|\|\|]PAM | NA | Intergenic | NA | NA |
| T30802 | TGACGCAACGGTGGCGGTGGCGG | 1000 | | | | | | | | | |
| Oligo fwd | CACCTGACGCAACGGTGGCGGTGG | | | | | | | | | | |
| Oligo rev | AAACCCACCGCCACCGTTGCGTCA | | | | | | | | | | |
| Chromosome | start | end | strand | MM | target_seq | PAM | alignment | distance | position | gene name | gene id |
| GU305914.1 | 45264 | 45286 | - | 0 | TGACGCAACGGTGGCGGTGG | CGG | \|\|\|\|\|\|\|\|[\|\|\|\|\|\|\|\|\|\|\|\|]PAM | NA | Intergenic | NA | NA |
| T30803 | AACGGTGGCGGTGGCGGCAGCGG | 1000 | | | | | | | | | |
| Oligo fwd | CACCAACGGTGGCGGTGGCGGCAG | | | | | | | | | | |
| Oligo rev | AAACCTGCCGCCACCGCCACCGTT | | | | | | | | | | |
| Chromosome | start | end | strand | MM | target_seq | PAM | alignment | distance | position | gene name | gene id |
| GU305914.1 | 45258 | 45280 | - | 0 | AACGGTGGCGGTGGCGGCAG | CGG | \|\|\|\|\|\|\|\|[\|\|\|\|\|\|\|\|\|\|\|\|]PAM | NA | Intergenic | NA | NA |
| GU305914.1 | 40310 | 40332 | - | 4 | GAGGATTGCGGTGGCGGCAG | TGG | -\|-\|-\|-\|[\|\|\|\|\|\|\|\|\|\|\|\|]PAM | NA | Intergenic | NA | NA |
| GU305914.1 | 163902 | 163924 | + | 3 | AGCGGCGGCGGCGGCGGCAG | CGG | \|-\|\|\|-\|\|[\|\|\|-\|\|\|\|\|\|\|\|]PAM | NA | Intergenic | NA | NA |
| GU305914.1 | 30006 | 30028 | + | 4 | GGCGGCGGCGGCGGCGGCAG | TGG | --\|\|\|-\|\|[\|\|\|-\|\|\|\|\|\|\|\|]PAM | NA | Intergenic | NA | NA |
| GU305914.1 | 149476 | 149498 | - | 4 | GGCGGCGGCGGCGGCGGCAG | CGG | --\|\|\|-\|\|[\|\|\|-\|\|\|\|\|\|\|\|]PAM | NA | Intergenic | NA | NA |
| GU305914.1 | 168829 | 168851 | + | 4 | AGCGGCGGCGGCGGCGACAG | CGG | \|-\|\|\|-\|\|[\|\|\|-\|\|\|\|-\|\|\|]PAM | NA | Intergenic | NA | NA |
| GU305914.1 | 48156 | 48178 | + | 4 | GGCGGCGGCGGTGGCGGCGG | CGG | --\|\|\|-\|\|[\|\|\|\|\|\|\|\|\|\|-\|]PAM | NA | Intergenic | NA | NA |
| GU305914.1 | 58132 | 58154 | - | 4 | AAGGCGGGCGGTGGCGGCGG | CGG | \|\|-\|--\|\|[\|\|\|\|\|\|\|\|\|\|-\|]PAM | NA | Intergenic | NA | NA |
| GU305914.1 | 114658 | 114680 | - | 4 | GGCGGTGGTGGTGGCGGCGG | CGG | --\|\|\|\|\|\|[-\|\|\|\|\|\|\|\|\|-\|]PAM | NA | Intergenic | NA | NA |
| GU305914.1 | 154404 | 154426 | + | 4 | GGCGGTGGCGGCGGCGGCGG | CGG | --\|\|\|\|\|\|[\|\|\|-\|\|\|\|\|\|-\|]PAM | NA | Intergenic | NA | NA |
| GU305914.1 | 58126 | 58148 | - | 4 | GGCGGTGGCGGCGGCGGCGG | CGG | --\|\|\|\|\|\|[\|\|\|-\|\|\|\|\|\|-\|]PAM | NA | Intergenic | NA | NA |
| GU305914.1 | 19936 | 19958 | + | 4 | GACGGTAGCGGTGGTGGCGG | TGG | -\|\|\|\|\|-\|[\|\|\|\|\|\|-\|\|\|-\|]PAM | NA | Intergenic | NA | NA |
| T30804 | GGTGGCGGTGGCGGCAGCGGCGG | 1000 | | | | | | | | | |
| Oligo fwd | CACCGGTGGCGGTGGCGGCAGCGG | | | | | | | | | | |
| Oligo rev | AAACCCGCTGCCGCCACCGCCACC | | | | | | | | | | |
| Chromosome | start | end | strand | MM | target_seq | PAM | alignment | distance | position | gene name | gene id |
| GU305914.1 | 45255 | 45277 | - | 0 | GGTGGCGGTGGCGGCAGCGG | CGG | \|\|\|\|\|\|\|\|[\|\|\|\|\|\|\|\|\|\|\|\|]PAM | NA | Intergenic | NA | NA |
| GU305914.1 | 168814 | 168836 | + | 3 | AGCGGCGGTGGTGGCAGCGG | CGG | -\|-\|\|\|\|\|[\|\|\|-\|\|\|\|\|\|\|\|]PAM | NA | Intergenic | NA | NA |
| GU305914.1 | 163911 | 163933 | + | 3 | GGCGGCGGCAGCGGCAGCGG | AGG | \|\|-\|\|\|\|\|[--\|\|\|\|\|\|\|\|\|\|]PAM | NA | Intergenic | NA | NA |
| GU305914.1 | 62155 | 62177 | + | 4 | GGCGACGGCAGCGGCAGCGG | CGG | \|\|-\|-\|\|\|[--\|\|\|\|\|\|\|\|\|\|]PAM | NA | Intergenic | NA | NA |
| GU305914.1 | 48159 | 48181 | + | 2 | GGCGGCGGTGGCGGCGGCGG | TGG | \|\|-\|\|\|\|\|[\|\|\|\|\|\|\|-\|\|\|\|]PAM | NA | Intergenic | NA | NA |
| GU305914.1 | 58129 | 58151 | - | 3 | GCGGGCGGTGGCGGCGGCGG | CGG | \|--\|\|\|\|\|[\|\|\|\|\|\|\|-\|\|\|\|]PAM | NA | Intergenic | NA | NA |
| GU305914.1 | 154401 | 154423 | + | 4 | CCGGGCGGTGGCGGCGGCGG | CGG | ---\|\|\|\|\|[\|\|\|\|\|\|\|-\|\|\|\|]PAM | NA | Intergenic | NA | NA |
| GU305914.1 | 154407 | 154429 | + | 2 | GGTGGCGGCGGCGGCGGCGG | CGG | \|\|\|\|\|\|\|\|[-\|\|\|\|\|\|-\|\|\|\|]PAM | NA | Intergenic | NA | NA |
| GU305914.1 | 58123 | 58145 | - | 2 | GGTGGCGGCGGCGGCGGCGG | AGG | \|\|\|\|\|\|\|\|[-\|\|\|\|\|\|-\|\|\|\|]PAM | NA | Intergenic | NA | NA |
| GU305914.1 | 67742 | 67764 | + | 4 | GGGAGCGGCGGCGGCGGCGG | CGG | \|\|--\|\|\|\|[-\|\|\|\|\|\|-\|\|\|\|]PAM | NA | Intergenic | NA | NA |
| GU305914.1 | 149482 | 149504 | - | 4 | GGCGACGGCGGCGGCGGCGG | CGG | \|\|-\|-\|\|\|[-\|\|\|\|\|\|-\|\|\|\|]PAM | NA | Intergenic | NA | NA |
| GU305914.1 | 154404 | 154426 | + | 4 | GGCGGTGGCGGCGGCGGCGG | CGG | \|\|-\|\|-\|\|[-\|\|\|\|\|\|-\|\|\|\|]PAM | NA | Intergenic | NA | NA |
| GU305914.1 | 58126 | 58148 | - | 4 | GGCGGTGGCGGCGGCGGCGG | CGG | \|\|-\|\|-\|\|[-\|\|\|\|\|\|-\|\|\|\|]PAM | NA | Intergenic | NA | NA |
| GU305914.1 | 30000 | 30022 | + | 4 | GGTGCTGGCGGCGGCGGCGG | CGG | \|\|\|\|--\|\|[-\|\|\|\|\|\|-\|\|\|\|]PAM | NA | Intergenic | NA | NA |
| GU305914.1 | 62446 | 62468 | + | 4 | GGCGGGGGTGGTGGCGGCGG | GGG | \|\|-\|\|-\|\|[\|\|\|-\|\|\|-\|\|\|\|]PAM | NA | Intergenic | NA | NA |
| GU305914.1 | 114658 | 114680 | - | 4 | GGCGGTGGTGGTGGCGGCGG | CGG | \|\|-\|\|-\|\|[\|\|\|-\|\|\|-\|\|\|\|]PAM | NA | Intergenic | NA | NA |
| GU305914.1 | 168817 | 168839 | + | 4 | GGCGGTGGTGGCAGCGGCGG | CGG | \|\|-\|\|-\|\|[\|\|\|\|-\|\|-\|\|\|\|]PAM | NA | Intergenic | NA | NA |
| GU305914.1 | 62434 | 62456 | + | 4 | GACGGCGGTGGCGGCGGGGG | TGG | \|--\|\|\|\|\|[\|\|\|\|\|\|\|-\|-\|\|]PAM | NA | Intergenic | NA | NA |
| GU305914.1 | 62449 | 62471 | + | 4 | GGGGGTGGTGGCGGCGGGGG | TGG | \|\|-\|\|-\|\|[\|\|\|\|\|\|\|-\|-\|\|]PAM | NA | Intergenic | NA | NA |
| T30805 | CGGCGGCGTGTTGCCCCTGTCGG | 1000 | | | | | | | | | |
| Oligo fwd | CACCCGGCGGCGTGTTGCCCCTGT | | | | | | | | | | |
| Oligo rev | AAACACAGGGGCAACACGCCGCCG | | | | | | | | | | |
| Chromosome | start | end | strand | MM | target_seq | PAM | alignment | distance | position | gene name | gene id |
| GU305914.1 | 45238 | 45260 | - | 0 | CGGCGGCGTGTTGCCCCTGT | CGG | \|\|\|\|\|\|\|\|[\|\|\|\|\|\|\|\|\|\|\|\|]PAM | NA | Intergenic | NA | NA |
| T30806 | TTGCCCCTGTCGGATAACGTCGG | 1000 | | | | | | | | | |
| Oligo fwd | CACCTTGCCCCTGTCGGATAACGT | | | | | | | | | | |
| Oligo rev | AAACACGTTATCCGACAGGGGCAA | | | | | | | | | | |
| Chromosome | start | end | strand | MM | target_seq | PAM | alignment | distance | position | gene name | gene id |
| GU305914.1 | 45228 | 45250 | - | 0 | TTGCCCCTGTCGGATAACGT | CGG | \|\|\|\|\|\|\|\|[\|\|\|\|\|\|\|\|\|\|\|\|]PAM | NA | Intergenic | NA | NA |
| T30807 | GTCGGATAACGTCGGTGAACCGG | 1000 | | | | | | | | | |
| Oligo fwd | CACCGTCGGATAACGTCGGTGAAC | | | | | | | | | | |
| Oligo rev | AAACGTTCACCGACGTTATCCGAC | | | | | | | | | | |
| Chromosome | start | end | strand | MM | target_seq | PAM | alignment | distance | position | gene name | gene id |
| GU305914.1 | 45220 | 45242 | - | 0 | GTCGGATAACGTCGGTGAAC | CGG | \|\|\|\|\|\|\|\|[\|\|\|\|\|\|\|\|\|\|\|\|]PAM | NA | Intergenic | NA | NA |
| T30808 | ATAACGTCGGTGAACCGGTGTGG | 1000 | | | | | | | | | |
| Oligo fwd | CACCATAACGTCGGTGAACCGGTG | | | | | | | | | | |
| Oligo rev | AAACCACCGGTTCACCGACGTTAT | | | | | | | | | | |
| Chromosome | start | end | strand | MM | target_seq | PAM | alignment | distance | position | gene name | gene id |
| GU305914.1 | 45215 | 45237 | - | 0 | ATAACGTCGGTGAACCGGTG | TGG | \|\|\|\|\|\|\|\|[\|\|\|\|\|\|\|\|\|\|\|\|]PAM | NA | Intergenic | NA | NA |
| T30809 | TAACGTCGGTGAACCGGTGTGGG | 1000 | | | | | | | | | |
| Oligo fwd | CACCTAACGTCGGTGAACCGGTGT | | | | | | | | | | |
| Oligo rev | AAACACACCGGTTCACCGACGTTA | | | | | | | | | | |
| Chromosome | start | end | strand | MM | target_seq | PAM | alignment | distance | position | gene name | gene id |
| GU305914.1 | 45214 | 45236 | - | 0 | TAACGTCGGTGAACCGGTGT | GGG | \|\|\|\|\|\|\|\|[\|\|\|\|\|\|\|\|\|\|\|\|]PAM | NA | Intergenic | NA | NA |
| T30810 | GTGGGACCGAGAACCACCCGTGG | 1000 | | | | | | | | | |
| Oligo fwd | CACCGTGGGACCGAGAACCACCCG | | | | | | | | | | |
| Oligo rev | AAACCGGGTGGTTCTCGGTCCCAC | | | | | | | | | | |

| | | | | | | | | | | | |
|---|---|---|---|---|---|---|---|---|---|---|---|
| Chromosome | start | end | strand | MM | target_seq | PAM | alignment | distance | position | gene name | gene id |
| GU305914.1 | 45196 | 45218 | - | 0 | GTGGGACCGAGAACCACCCG | TGG | \|\|\|\|\|\|\|\|[\|\|\|\|\|\|\|\|\|\|\|\|]PAM | NA | Intergenic | NA | NA |
| T30811 | CGAGTCGCCTTCGCCGCCGCTGG | 1000 | | | | | | | | | |
| Oligo fwd | CACCCGAGTCGCCTTCGCCGCCGC | | | | | | | | | | |
| Oligo rev | AAACGCGGCGGCGAAGGCGACTCG | | | | | | | | | | |
| Chromosome | start | end | strand | MM | target_seq | PAM | alignment | distance | position | gene name | gene id |
| GU305914.1 | 45163 | 45185 | - | 0 | CGAGTCGCCTTCGCCGCCGC | TGG | \|\|\|\|\|\|\|\|[\|\|\|\|\|\|\|\|\|\|\|\|]PAM | NA | Intergenic | NA | NA |
| T30812 | GTCGCCTTCGCCGCCGCTGGAGG | 1000 | | | | | | | | | |
| Oligo fwd | CACCGTCGCCTTCGCCGCCGCTGG | | | | | | | | | | |
| Oligo rev | AAACCCAGCGGCGGCGAAGGCGAC | | | | | | | | | | |
| Chromosome | start | end | strand | MM | target_seq | PAM | alignment | distance | position | gene name | gene id |
| GU305914.1 | 45160 | 45182 | - | 0 | GTCGCCTTCGCCGCCGCTGG | AGG | \|\|\|\|\|\|\|\|[\|\|\|\|\|\|\|\|\|\|\|\|]PAM | NA | Intergenic | NA | NA |
| GU305914.1 | 52707 | 52729 | + | 4 | AACACCTTGGCCGCCGCTGG | CGG | --\|-\|\|\|\|[-\|\|\|\|\|\|\|\|\|\|\|]PAM | NA | Intergenic | NA | NA |
| T30813 | ACGCGAACTCGATTTCGACGAGG | 1000 | | | | | | | | | |
| Oligo fwd | CACCACGCGAACTCGATTTCGACG | | | | | | | | | | |
| Oligo rev | AAACCGTCGAAATCGAGTTCGCGT | | | | | | | | | | |
| Chromosome | start | end | strand | MM | target_seq | PAM | alignment | distance | position | gene name | gene id |
| GU305914.1 | 45109 | 45131 | - | 0 | ACGCGAACTCGATTTCGACG | AGG | \|\|\|\|\|\|\|\|[\|\|\|\|\|\|\|\|\|\|\|\|]PAM | NA | Intergenic | NA | NA |
| T30814 | CTCCTCCTACTACTCCTCCTCGG | 1000 | | | | | | | | | |
| Oligo fwd | CACCCTCCTCCTACTACTCCTCCT | | | | | | | | | | |
| Oligo rev | AAACAGGAGGAGTAGTAGGAGGAG | | | | | | | | | | |
| Chromosome | start | end | strand | MM | target_seq | PAM | alignment | distance | position | gene name | gene id |
| GU305914.1 | 45047 | 45069 | - | 0 | CTCCTCCTACTACTCCTCCT | CGG | \|\|\|\|\|\|\|\|[\|\|\|\|\|\|\|\|\|\|\|\|]PAM | NA | Intergenic | NA | NA |
| T30815 | TCCTACTACTCCTCCTCGGCCGG | 1000 | | | | | | | | | |
| Oligo fwd | CACCTCCTACTACTCCTCCTCGGC | | | | | | | | | | |
| Oligo rev | AAACGCCGAGGAGGAGTAGTAGGA | | | | | | | | | | |
| Chromosome | start | end | strand | MM | target_seq | PAM | alignment | distance | position | gene name | gene id |
| GU305914.1 | 45043 | 45065 | - | 0 | TCCTACTACTCCTCCTCGGC | CGG | \|\|\|\|\|\|\|\|[\|\|\|\|\|\|\|\|\|\|\|\|]PAM | NA | Intergenic | NA | NA |
| T30816 | CCTACTACTCCTCCTCGGCCGGG | 1000 | | | | | | | | | |
| Oligo fwd | CACCCCTACTACTCCTCCTCGGCC | | | | | | | | | | |
| Oligo rev | AAACGGCCGAGGAGGAGTAGTAGG | | | | | | | | | | |
| Chromosome | start | end | strand | MM | target_seq | PAM | alignment | distance | position | gene name | gene id |
| GU305914.1 | 45042 | 45064 | - | 0 | CCTACTACTCCTCCTCGGCC | GGG | \|\|\|\|\|\|\|\|[\|\|\|\|\|\|\|\|\|\|\|\|]PAM | NA | Intergenic | NA | NA |
| T30817 | CTACTACTCCTCCTCGGCCGGGG | 1000 | | | | | | | | | |
| Oligo fwd | CACCCTACTACTCCTCCTCGGCCG | | | | | | | | | | |
| Oligo rev | AAACCGGCCGAGGAGGAGTAGTAG | | | | | | | | | | |
| Chromosome | start | end | strand | MM | target_seq | PAM | alignment | distance | position | gene name | gene id |
| GU305914.1 | 45041 | 45063 | - | 0 | CTACTACTCCTCCTCGGCCG | GGG | \|\|\|\|\|\|\|\|[\|\|\|\|\|\|\|\|\|\|\|\|]PAM | NA | Intergenic | NA | NA |
| T30818 | TACTACTCCTCCTCGGCCGGGGG | 1000 | | | | | | | | | |
| Oligo fwd | CACCTACTACTCCTCCTCGGCCGG | | | | | | | | | | |
| Oligo rev | AAACCCGGCCGAGGAGGAGTAGTA | | | | | | | | | | |
| Chromosome | start | end | strand | MM | target_seq | PAM | alignment | distance | position | gene name | gene id |
| GU305914.1 | 45040 | 45062 | - | 0 | TACTACTCCTCCTCGGCCGG | GGG | \|\|\|\|\|\|\|\|[\|\|\|\|\|\|\|\|\|\|\|\|]PAM | NA | Intergenic | NA | NA |
| T30819 | CCTCCTCGGCCGGGGGAGAACGG | 1000 | | | | | | | | | |
| Oligo fwd | CACCCCTCCTCGGCCGGGGGAGAA | | | | | | | | | | |
| Oligo rev | AAACTTCTCCCCCGGCCGAGGAGG | | | | | | | | | | |
| Chromosome | start | end | strand | MM | target_seq | PAM | alignment | distance | position | gene name | gene id |
| GU305914.1 | 45033 | 45055 | - | 0 | CCTCCTCGGCCGGGGGAGAA | CGG | \|\|\|\|\|\|\|\|[\|\|\|\|\|\|\|\|\|\|\|\|]PAM | NA | Intergenic | NA | NA |
| T30820 | CTCCTCGGCCGGGGGAGAACGGG | 1000 | | | | | | | | | |
| Oligo fwd | CACCCTCCTCGGCCGGGGGAGAAC | | | | | | | | | | |
| Oligo rev | AAACGTTCTCCCCCGGCCGAGGAG | | | | | | | | | | |
| Chromosome | start | end | strand | MM | target_seq | PAM | alignment | distance | position | gene name | gene id |
| GU305914.1 | 45032 | 45054 | - | 0 | CTCCTCGGCCGGGGGAGAAC | GGG | \|\|\|\|\|\|\|\|[\|\|\|\|\|\|\|\|\|\|\|\|]PAM | NA | Intergenic | NA | NA |
| T30821 | GGAGAACGGGTCGCTGTCGAAGG | 1000 | | | | | | | | | |
| Oligo fwd | CACCGGAGAACGGGTCGCTGTCGA | | | | | | | | | | |
| Oligo rev | AAACTCGACAGCGACCCGTTCTCC | | | | | | | | | | |
| Chromosome | start | end | strand | MM | target_seq | PAM | alignment | distance | position | gene name | gene id |
| GU305914.1 | 45019 | 45041 | - | 0 | GGAGAACGGGTCGCTGTCGA | AGG | \|\|\|\|\|\|\|\|[\|\|\|\|\|\|\|\|\|\|\|\|]PAM | NA | Intergenic | NA | NA |
| T30822 | AACGGGTCGCTGTCGAAGGTCGG | 1000 | | | | | | | | | |
| Oligo fwd | CACCAACGGGTCGCTGTCGAAGGT | | | | | | | | | | |
| Oligo rev | AAACACCTTCGACAGCGACCCGTT | | | | | | | | | | |
| Chromosome | start | end | strand | MM | target_seq | PAM | alignment | distance | position | gene name | gene id |
| GU305914.1 | 45015 | 45037 | - | 0 | AACGGGTCGCTGTCGAAGGT | CGG | \|\|\|\|\|\|\|\|[\|\|\|\|\|\|\|\|\|\|\|\|]PAM | NA | Intergenic | NA | NA |
| T30823 | GTCTTCCTCCTCTAGATCCGAGG | 1000 | | | | | | | | | |
| Oligo fwd | CACCGTCTTCCTCCTCTAGATCCG | | | | | | | | | | |
| Oligo rev | AAACCGGATCTAGAGGAGGAAGAC | | | | | | | | | | |
| Chromosome | start | end | strand | MM | target_seq | PAM | alignment | distance | position | gene name | gene id |
| GU305914.1 | 44968 | 44990 | - | 0 | GTCTTCCTCCTCTAGATCCG | AGG | \|\|\|\|\|\|\|\|[\|\|\|\|\|\|\|\|\|\|\|\|]PAM | NA | Intergenic | NA | NA |
| T30824 | TCTTCTGCTGCCTCCGCGAACGG | 1000 | | | | | | | | | |
| Oligo fwd | CACCTCTTCTGCTGCCTCCGCGAA | | | | | | | | | | |
| Oligo rev | AAACTTCGCGGAGGCAGCAGAAGA | | | | | | | | | | |
| Chromosome | start | end | strand | MM | target_seq | PAM | alignment | distance | position | gene name | gene id |
| GU305914.1 | 44943 | 44965 | - | 0 | TCTTCTGCTGCCTCCGCGAA | CGG | \|\|\|\|\|\|\|\|[\|\|\|\|\|\|\|\|\|\|\|\|]PAM | NA | Intergenic | NA | NA |
| T30825 | TCTGCTGCCTCCGCGAACGGAGG | 1000 | | | | | | | | | |

<table>
<tr><td>Oligo fwd</td><td>CACCTCTGCTGCCTCCGCGAACGG</td><td></td><td></td><td></td><td></td><td></td><td></td><td></td><td></td><td></td><td></td></tr>
<tr><td>Oligo rev</td><td>AAACCCGTTCGCGGAGGCAGCAGA</td><td></td><td></td><td></td><td></td><td></td><td></td><td></td><td></td><td></td><td></td></tr>
<tr><td>Chromosome</td><td>start</td><td>end</td><td>strand</td><td>MM</td><td>target_seq</td><td>PAM</td><td>alignment</td><td>distance</td><td>position</td><td>gene name</td><td>gene id</td></tr>
<tr><td>GU305914.1</td><td>44940</td><td>44962</td><td>-</td><td>0</td><td>TCTGCTGCCTCCGCGAACGG</td><td>AGG</td><td>||||||||[||||||||||||]PAM</td><td>NA</td><td>Intergenic</td><td>NA</td><td>NA</td></tr>
<tr><td>T30826</td><td>AGGCGTCCTGCTCGAGCGAGAGG</td><td>1000</td><td></td><td></td><td></td><td></td><td></td><td></td><td></td><td></td><td></td></tr>
<tr><td>Oligo fwd</td><td>CACCAGGCGTCCTGCTCGAGCGAG</td><td></td><td></td><td></td><td></td><td></td><td></td><td></td><td></td><td></td><td></td></tr>
<tr><td>Oligo rev</td><td>AAACCTCGCTCGAGCAGGACGCCT</td><td></td><td></td><td></td><td></td><td></td><td></td><td></td><td></td><td></td><td></td></tr>
<tr><td>Chromosome</td><td>start</td><td>end</td><td>strand</td><td>MM</td><td>target_seq</td><td>PAM</td><td>alignment</td><td>distance</td><td>position</td><td>gene name</td><td>gene id</td></tr>
<tr><td>GU305914.1</td><td>44920</td><td>44942</td><td>-</td><td>0</td><td>AGGCGTCCTGCTCGAGCGAG</td><td>AGG</td><td>||||||||[||||||||||||]PAM</td><td>NA</td><td>Intergenic</td><td>NA</td><td>NA</td></tr>
<tr><td>T30827</td><td>AGCGAGAGGCGACGCTGCTCCGG</td><td>1000</td><td></td><td></td><td></td><td></td><td></td><td></td><td></td><td></td><td></td></tr>
<tr><td>Oligo fwd</td><td>CACCAGCGAGAGGCGACGCTGCTC</td><td></td><td></td><td></td><td></td><td></td><td></td><td></td><td></td><td></td><td></td></tr>
<tr><td>Oligo rev</td><td>AAACGAGCAGCGTCGCCTCTCGCT</td><td></td><td></td><td></td><td></td><td></td><td></td><td></td><td></td><td></td><td></td></tr>
<tr><td>Chromosome</td><td>start</td><td>end</td><td>strand</td><td>MM</td><td>target_seq</td><td>PAM</td><td>alignment</td><td>distance</td><td>position</td><td>gene name</td><td>gene id</td></tr>
<tr><td>GU305914.1</td><td>44906</td><td>44928</td><td>-</td><td>0</td><td>AGCGAGAGGCGACGCTGCTC</td><td>CGG</td><td>||||||||[||||||||||||]PAM</td><td>NA</td><td>Intergenic</td><td>NA</td><td>NA</td></tr>
<tr><td>GU305914.1</td><td>141031</td><td>141053</td><td>+</td><td>4</td><td>AGAGAGACGCGACGCTCATC</td><td>GGG</td><td>||-||||-[||||||||--||]PAM</td><td>NA</td><td>Intergenic</td><td>NA</td><td>NA</td></tr>
<tr><td>T30828</td><td>CCGTGTCGCTGTCCGTGTCGCGG</td><td>1000</td><td></td><td></td><td></td><td></td><td></td><td></td><td></td><td></td><td></td></tr>
<tr><td>Oligo fwd</td><td>CACCCCGTGTCGCTGTCCGTGTCG</td><td></td><td></td><td></td><td></td><td></td><td></td><td></td><td></td><td></td><td></td></tr>
<tr><td>Oligo rev</td><td>AAACCGACACGGACAGCGACACGG</td><td></td><td></td><td></td><td></td><td></td><td></td><td></td><td></td><td></td><td></td></tr>
<tr><td>Chromosome</td><td>start</td><td>end</td><td>strand</td><td>MM</td><td>target_seq</td><td>PAM</td><td>alignment</td><td>distance</td><td>position</td><td>gene name</td><td>gene id</td></tr>
<tr><td>GU305914.1</td><td>44882</td><td>44904</td><td>-</td><td>0</td><td>CCGTGTCGCTGTCCGTGTCG</td><td>CGG</td><td>||||||||[||||||||||||]PAM</td><td>NA</td><td>Intergenic</td><td>NA</td><td>NA</td></tr>
<tr><td>T30829</td><td>TGTCGCTGTCCGTGTCGCGGTGG</td><td>1000</td><td></td><td></td><td></td><td></td><td></td><td></td><td></td><td></td><td></td></tr>
<tr><td>Oligo fwd</td><td>CACCTGTCGCTGTCCGTGTCGCGG</td><td></td><td></td><td></td><td></td><td></td><td></td><td></td><td></td><td></td><td></td></tr>
<tr><td>Oligo rev</td><td>AAACCCGCGACACGGACAGCGACA</td><td></td><td></td><td></td><td></td><td></td><td></td><td></td><td></td><td></td><td></td></tr>
<tr><td>Chromosome</td><td>start</td><td>end</td><td>strand</td><td>MM</td><td>target_seq</td><td>PAM</td><td>alignment</td><td>distance</td><td>position</td><td>gene name</td><td>gene id</td></tr>
<tr><td>GU305914.1</td><td>44879</td><td>44901</td><td>-</td><td>0</td><td>TGTCGCTGTCCGTGTCGCGG</td><td>TGG</td><td>||||||||[||||||||||||]PAM</td><td>NA</td><td>Intergenic</td><td>NA</td><td>NA</td></tr>
<tr><td>T30830</td><td>CGCTGTCCGTGTCGCGGTGGTGG</td><td>1000</td><td></td><td></td><td></td><td></td><td></td><td></td><td></td><td></td><td></td></tr>
<tr><td>Oligo fwd</td><td>CACCCGCTGTCCGTGTCGCGGTGG</td><td></td><td></td><td></td><td></td><td></td><td></td><td></td><td></td><td></td><td></td></tr>
<tr><td>Oligo rev</td><td>AAACCCACCGCGACACGGACAGCG</td><td></td><td></td><td></td><td></td><td></td><td></td><td></td><td></td><td></td><td></td></tr>
<tr><td>Chromosome</td><td>start</td><td>end</td><td>strand</td><td>MM</td><td>target_seq</td><td>PAM</td><td>alignment</td><td>distance</td><td>position</td><td>gene name</td><td>gene id</td></tr>
<tr><td>GU305914.1</td><td>44876</td><td>44898</td><td>-</td><td>0</td><td>CGCTGTCCGTGTCGCGGTGG</td><td>TGG</td><td>||||||||[||||||||||||]PAM</td><td>NA</td><td>Intergenic</td><td>NA</td><td>NA</td></tr>
<tr><td>T30831</td><td>TCCGTGTCGCGGTGGTGGCGCGG</td><td>1000</td><td></td><td></td><td></td><td></td><td></td><td></td><td></td><td></td><td></td></tr>
<tr><td>Oligo fwd</td><td>CACCTCCGTGTCGCGGTGGTGGCG</td><td></td><td></td><td></td><td></td><td></td><td></td><td></td><td></td><td></td><td></td></tr>
<tr><td>Oligo rev</td><td>AAACCGCCACCACCGCGACACGGA</td><td></td><td></td><td></td><td></td><td></td><td></td><td></td><td></td><td></td><td></td></tr>
<tr><td>Chromosome</td><td>start</td><td>end</td><td>strand</td><td>MM</td><td>target_seq</td><td>PAM</td><td>alignment</td><td>distance</td><td>position</td><td>gene name</td><td>gene id</td></tr>
<tr><td>GU305914.1</td><td>44871</td><td>44893</td><td>-</td><td>0</td><td>TCCGTGTCGCGGTGGTGGCG</td><td>CGG</td><td>||||||||[||||||||||||]PAM</td><td>NA</td><td>Intergenic</td><td>NA</td><td>NA</td></tr>
<tr><td>T30832</td><td>GTGGCGCGGAGAATCCGTATCGG</td><td>1000</td><td></td><td></td><td></td><td></td><td></td><td></td><td></td><td></td><td></td></tr>
<tr><td>Oligo fwd</td><td>CACCGTGGCGCGGAGAATCCGTAT</td><td></td><td></td><td></td><td></td><td></td><td></td><td></td><td></td><td></td><td></td></tr>
<tr><td>Oligo rev</td><td>AAACATACGGATTCTCCGCGCCAC</td><td></td><td></td><td></td><td></td><td></td><td></td><td></td><td></td><td></td><td></td></tr>
<tr><td>Chromosome</td><td>start</td><td>end</td><td>strand</td><td>MM</td><td>target_seq</td><td>PAM</td><td>alignment</td><td>distance</td><td>position</td><td>gene name</td><td>gene id</td></tr>
<tr><td>GU305914.1</td><td>44857</td><td>44879</td><td>-</td><td>0</td><td>GTGGCGCGGAGAATCCGTAT</td><td>CGG</td><td>||||||||[||||||||||||]PAM</td><td>NA</td><td>Intergenic</td><td>NA</td><td>NA</td></tr>
<tr><td>T30833</td><td>GCGCGGAGAATCCGTATCGGAGG</td><td>1000</td><td></td><td></td><td></td><td></td><td></td><td></td><td></td><td></td><td></td></tr>
<tr><td>Oligo fwd</td><td>CACCGCGCGGAGAATCCGTATCGG</td><td></td><td></td><td></td><td></td><td></td><td></td><td></td><td></td><td></td><td></td></tr>
<tr><td>Oligo rev</td><td>AAACCCGATACGGATTCTCCGCGC</td><td></td><td></td><td></td><td></td><td></td><td></td><td></td><td></td><td></td><td></td></tr>
<tr><td>Chromosome</td><td>start</td><td>end</td><td>strand</td><td>MM</td><td>target_seq</td><td>PAM</td><td>alignment</td><td>distance</td><td>position</td><td>gene name</td><td>gene id</td></tr>
<tr><td>GU305914.1</td><td>44854</td><td>44876</td><td>-</td><td>0</td><td>GCGCGGAGAATCCGTATCGG</td><td>AGG</td><td>||||||||[||||||||||||]PAM</td><td>NA</td><td>Intergenic</td><td>NA</td><td>NA</td></tr>
<tr><td>T30834</td><td>ATCCGTATCGGAGGACACGTCGG</td><td>1000</td><td></td><td></td><td></td><td></td><td></td><td></td><td></td><td></td><td></td></tr>
<tr><td>Oligo fwd</td><td>CACCATCCGTATCGGAGGACACGT</td><td></td><td></td><td></td><td></td><td></td><td></td><td></td><td></td><td></td><td></td></tr>
<tr><td>Oligo rev</td><td>AAACACGTGTCCTCCGATACGGAT</td><td></td><td></td><td></td><td></td><td></td><td></td><td></td><td></td><td></td><td></td></tr>
<tr><td>Chromosome</td><td>start</td><td>end</td><td>strand</td><td>MM</td><td>target_seq</td><td>PAM</td><td>alignment</td><td>distance</td><td>position</td><td>gene name</td><td>gene id</td></tr>
<tr><td>GU305914.1</td><td>44845</td><td>44867</td><td>-</td><td>0</td><td>ATCCGTATCGGAGGACACGT</td><td>CGG</td><td>||||||||[||||||||||||]PAM</td><td>NA</td><td>Intergenic</td><td>NA</td><td>NA</td></tr>
<tr><td>T30835</td><td>GACACGTCGGACAACGTTCCCGG</td><td>1000</td><td></td><td></td><td></td><td></td><td></td><td></td><td></td><td></td><td></td></tr>
<tr><td>Oligo fwd</td><td>CACCGACACGTCGGACAACGTTCC</td><td></td><td></td><td></td><td></td><td></td><td></td><td></td><td></td><td></td><td></td></tr>
<tr><td>Oligo rev</td><td>AAACGGAACGTTGTCCGACGTGTC</td><td></td><td></td><td></td><td></td><td></td><td></td><td></td><td></td><td></td><td></td></tr>
<tr><td>Chromosome</td><td>start</td><td>end</td><td>strand</td><td>MM</td><td>target_seq</td><td>PAM</td><td>alignment</td><td>distance</td><td>position</td><td>gene name</td><td>gene id</td></tr>
<tr><td>GU305914.1</td><td>44832</td><td>44854</td><td>-</td><td>0</td><td>GACACGTCGGACAACGTTCC</td><td>CGG</td><td>||||||||[||||||||||||]PAM</td><td>NA</td><td>Intergenic</td><td>NA</td><td>NA</td></tr>
<tr><td>T30836</td><td>ACACGTCGGACAACGTTCCCGGG</td><td>1000</td><td></td><td></td><td></td><td></td><td></td><td></td><td></td><td></td><td></td></tr>
<tr><td>Oligo fwd</td><td>CACCACACGTCGGACAACGTTCCC</td><td></td><td></td><td></td><td></td><td></td><td></td><td></td><td></td><td></td><td></td></tr>
<tr><td>Oligo rev</td><td>AAACGGGAACGTTGTCCGACGTGT</td><td></td><td></td><td></td><td></td><td></td><td></td><td></td><td></td><td></td><td></td></tr>
<tr><td>Chromosome</td><td>start</td><td>end</td><td>strand</td><td>MM</td><td>target_seq</td><td>PAM</td><td>alignment</td><td>distance</td><td>position</td><td>gene name</td><td>gene id</td></tr>
<tr><td>GU305914.1</td><td>44831</td><td>44853</td><td>-</td><td>0</td><td>ACACGTCGGACAACGTTCCC</td><td>GGG</td><td>||||||||[||||||||||||]PAM</td><td>NA</td><td>Intergenic</td><td>NA</td><td>NA</td></tr>
<tr><td>T30837</td><td>TCGGACAACGTTCCCGGGCGCGG</td><td>1000</td><td></td><td></td><td></td><td></td><td></td><td></td><td></td><td></td><td></td></tr>
<tr><td>Oligo fwd</td><td>CACCTCGGACAACGTTCCCGGGCG</td><td></td><td></td><td></td><td></td><td></td><td></td><td></td><td></td><td></td><td></td></tr>
<tr><td>Oligo rev</td><td>AAACCGCCCGGGAACGTTGTCCGA</td><td></td><td></td><td></td><td></td><td></td><td></td><td></td><td></td><td></td><td></td></tr>
<tr><td>Chromosome</td><td>start</td><td>end</td><td>strand</td><td>MM</td><td>target_seq</td><td>PAM</td><td>alignment</td><td>distance</td><td>position</td><td>gene name</td><td>gene id</td></tr>
<tr><td>GU305914.1</td><td>44826</td><td>44848</td><td>-</td><td>0</td><td>TCGGACAACGTTCCCGGGCG</td><td>CGG</td><td>||||||||[||||||||||||]PAM</td><td>NA</td><td>Intergenic</td><td>NA</td><td>NA</td></tr>
<tr><td>T30838</td><td>CGGACAACGTTCCCGGGCGCGGG</td><td>1000</td><td></td><td></td><td></td><td></td><td></td><td></td><td></td><td></td><td></td></tr>
<tr><td>Oligo fwd</td><td>CACCCGGACAACGTTCCCGGGCGC</td><td></td><td></td><td></td><td></td><td></td><td></td><td></td><td></td><td></td><td></td></tr>
<tr><td>Oligo rev</td><td>AAACGCGCCCGGGAACGTTGTCCG</td><td></td><td></td><td></td><td></td><td></td><td></td><td></td><td></td><td></td><td></td></tr>
<tr><td>Chromosome</td><td>start</td><td>end</td><td>strand</td><td>MM</td><td>target_seq</td><td>PAM</td><td>alignment</td><td>distance</td><td>position</td><td>gene name</td><td>gene id</td></tr>
<tr><td>GU305914.1</td><td>44825</td><td>44847</td><td>-</td><td>0</td><td>CGGACAACGTTCCCGGGCGC</td><td>GGG</td><td>||||||||[||||||||||||]PAM</td><td>NA</td><td>Intergenic</td><td>NA</td><td>NA</td></tr>
<tr><td>T30839</td><td>GGTAGAAGATGTACTCGTCGAGG</td><td>1000</td><td></td><td></td><td></td><td></td><td></td><td></td><td></td><td></td><td></td></tr>
<tr><td>Oligo fwd</td><td>CACCGGTAGAAGATGTACTCGTCG</td><td></td><td></td><td></td><td></td><td></td><td></td><td></td><td></td><td></td><td></td></tr>
<tr><td>Oligo rev</td><td>AAACCGACGAGTACATCTTCTACC</td><td></td><td></td><td></td><td></td><td></td><td></td><td></td><td></td><td></td><td></td></tr>
<tr><td>Chromosome</td><td>start</td><td>end</td><td>strand</td><td>MM</td><td>target_seq</td><td>PAM</td><td>alignment</td><td>distance</td><td>position</td><td>gene name</td><td>gene id</td></tr>
<tr><td>GU305914.1</td><td>44804</td><td>44826</td><td>-</td><td>0</td><td>GGTAGAAGATGTACTCGTCG</td><td>AGG</td><td>||||||||[||||||||||||]PAM</td><td>NA</td><td>Intergenic</td><td>NA</td><td>NA</td></tr>
</table>

| | | | | | | | | | | | |
|---|---|---|---|---|---|---|---|---|---|---|---|
| T30840 | ACTCGTCGAGGATGTCTTCTCGG | 1000 | | | | | | | | | |
| Oligo fwd | CACCACTCGTCGAGGATGTCTTCT | | | | | | | | | | |
| Oligo rev | AAACAGAAGACATCCTCGACGAGT | | | | | | | | | | |
| Chromosome | start | end | strand | MM | target_seq | PAM | alignment | distance | position | gene name | gene id |
| GU305914.1 | 44792 | 44814 | - | 0 | ACTCGTCGAGGATGTCTTCT | CGG | \|\|\|\|\|\|\|\|[\|\|\|\|\|\|\|\|\|\|\|\|]PAM | NA | Intergenic | NA | NA |
| T30841 | AGGATGTCTTCTCGGATGATCGG | 1000 | | | | | | | | | |
| Oligo fwd | CACCAGGATGTCTTCTCGGATGAT | | | | | | | | | | |
| Oligo rev | AAACATCATCCGAGAAGACATCCT | | | | | | | | | | |
| Chromosome | start | end | strand | MM | target_seq | PAM | alignment | distance | position | gene name | gene id |
| GU305914.1 | 44784 | 44806 | - | 0 | AGGATGTCTTCTCGGATGAT | CGG | \|\|\|\|\|\|\|\|[\|\|\|\|\|\|\|\|\|\|\|\|]PAM | NA | Intergenic | NA | NA |
| T30842 | TTCTCGGATGATCGGAGTCTTGG | 1000 | | | | | | | | | |
| Oligo fwd | CACCTTCTCGGATGATCGGAGTCT | | | | | | | | | | |
| Oligo rev | AAACAGACTCCGATCATCCGAGAA | | | | | | | | | | |
| Chromosome | start | end | strand | MM | target_seq | PAM | alignment | distance | position | gene name | gene id |
| GU305914.1 | 44776 | 44798 | - | 0 | TTCTCGGATGATCGGAGTCT | TGG | \|\|\|\|\|\|\|\|[\|\|\|\|\|\|\|\|\|\|\|\|]PAM | NA | Intergenic | NA | NA |
| T30843 | TCTCGGATGATCGGAGTCTTGGG | 1000 | | | | | | | | | |
| Oligo fwd | CACCTCTCGGATGATCGGAGTCTT | | | | | | | | | | |
| Oligo rev | AAACAAGACTCCGATCATCCGAGA | | | | | | | | | | |
| Chromosome | start | end | strand | MM | target_seq | PAM | alignment | distance | position | gene name | gene id |
| GU305914.1 | 44775 | 44797 | - | 0 | TCTCGGATGATCGGAGTCTT | GGG | \|\|\|\|\|\|\|\|[\|\|\|\|\|\|\|\|\|\|\|\|]PAM | NA | Intergenic | NA | NA |
| T30844 | CGGATGATCGGAGTCTTGGGCGG | 1000 | | | | | | | | | |
| Oligo fwd | CACCCGGATGATCGGAGTCTTGGG | | | | | | | | | | |
| Oligo rev | AAACCCCAAGACTCCGATCATCCG | | | | | | | | | | |
| Chromosome | start | end | strand | MM | target_seq | PAM | alignment | distance | position | gene name | gene id |
| GU305914.1 | 44772 | 44794 | - | 0 | CGGATGATCGGAGTCTTGGG | CGG | \|\|\|\|\|\|\|\|[\|\|\|\|\|\|\|\|\|\|\|\|]PAM | NA | Intergenic | NA | NA |
| T30845 | AGTCTTGGGCGGAGAGAACGAGG | 1000 | | | | | | | | | |
| Oligo fwd | CACCAGTCTTGGGCGGAGAGAACG | | | | | | | | | | |
| Oligo rev | AAACCGTTCTCTCCGCCCAAGACT | | | | | | | | | | |
| Chromosome | start | end | strand | MM | target_seq | PAM | alignment | distance | position | gene name | gene id |
| GU305914.1 | 44761 | 44783 | - | 0 | AGTCTTGGGCGGAGAGAACG | AGG | \|\|\|\|\|\|\|\|[\|\|\|\|\|\|\|\|\|\|\|\|]PAM | NA | Intergenic | NA | NA |
| T30846 | GTCTTGGGCGGAGAGAACGAGGG | 1000 | | | | | | | | | |
| Oligo fwd | CACCGTCTTGGGCGGAGAGAACGA | | | | | | | | | | |
| Oligo rev | AAACTCGTTCTCTCCGCCCAAGAC | | | | | | | | | | |
| Chromosome | start | end | strand | MM | target_seq | PAM | alignment | distance | position | gene name | gene id |
| GU305914.1 | 44760 | 44782 | - | 0 | GTCTTGGGCGGAGAGAACGA | GGG | \|\|\|\|\|\|\|\|[\|\|\|\|\|\|\|\|\|\|\|\|]PAM | NA | Intergenic | NA | NA |
| T30847 | TCTTGGGCGGAGAGAACGAGGGG | 1000 | | | | | | | | | |
| Oligo fwd | CACCTCTTGGGCGGAGAGAACGAG | | | | | | | | | | |
| Oligo rev | AAACCTCGTTCTCTCCGCCCAAGA | | | | | | | | | | |
| Chromosome | start | end | strand | MM | target_seq | PAM | alignment | distance | position | gene name | gene id |
| GU305914.1 | 44759 | 44781 | - | 0 | TCTTGGGCGGAGAGAACGAG | GGG | \|\|\|\|\|\|\|\|[\|\|\|\|\|\|\|\|\|\|\|\|]PAM | NA | Intergenic | NA | NA |
| T30848 | AGAACGAGGGGAAACCGCTCTGG | 1000 | | | | | | | | | |
| Oligo fwd | CACCAGAACGAGGGGAAACCGCTC | | | | | | | | | | |
| Oligo rev | AAACGAGCGGTTTCCCCTCGTTCT | | | | | | | | | | |
| Chromosome | start | end | strand | MM | target_seq | PAM | alignment | distance | position | gene name | gene id |
| GU305914.1 | 44747 | 44769 | - | 0 | AGAACGAGGGGAAACCGCTC | TGG | \|\|\|\|\|\|\|\|[\|\|\|\|\|\|\|\|\|\|\|\|]PAM | NA | Intergenic | NA | NA |
| T30849 | CCGTCGTCTTCGAAGCGATGAGG | 1000 | | | | | | | | | |
| Oligo fwd | CACCCCGTCGTCTTCGAAGCGATG | | | | | | | | | | |
| Oligo rev | AAACCATCGCTTCGAAGACGACGG | | | | | | | | | | |
| Chromosome | start | end | strand | MM | target_seq | PAM | alignment | distance | position | gene name | gene id |
| GU305914.1 | 44694 | 44716 | - | 0 | CCGTCGTCTTCGAAGCGATG | AGG | \|\|\|\|\|\|\|\|[\|\|\|\|\|\|\|\|\|\|\|\|]PAM | NA | Intergenic | NA | NA |
| T30850 | CGAAGCGATGAGGCGAGTGATGG | 1000 | | | | | | | | | |
| Oligo fwd | CACCCGAAGCGATGAGGCGAGTGA | | | | | | | | | | |
| Oligo rev | AAACTCACTCGCCTCATCGCTTCG | | | | | | | | | | |
| Chromosome | start | end | strand | MM | target_seq | PAM | alignment | distance | position | gene name | gene id |
| GU305914.1 | 44684 | 44706 | - | 0 | CGAAGCGATGAGGCGAGTGA | TGG | \|\|\|\|\|\|\|\|[\|\|\|\|\|\|\|\|\|\|\|\|]PAM | NA | Intergenic | NA | NA |
| T30851 | AGCGATGAGGCGAGTGATGGTGG | 1000 | | | | | | | | | |
| Oligo fwd | CACCAGCGATGAGGCGAGTGATGG | | | | | | | | | | |
| Oligo rev | AAACCCATCACTCGCCTCATCGCT | | | | | | | | | | |
| Chromosome | start | end | strand | MM | target_seq | PAM | alignment | distance | position | gene name | gene id |
| GU305914.1 | 44681 | 44703 | - | 0 | AGCGATGAGGCGAGTGATGG | TGG | \|\|\|\|\|\|\|\|[\|\|\|\|\|\|\|\|\|\|\|\|]PAM | NA | Intergenic | NA | NA |
| T30852 | GATGAGGCGAGTGATGGTGGTGG | 1000 | | | | | | | | | |
| Oligo fwd | CACCGATGAGGCGAGTGATGGTGG | | | | | | | | | | |
| Oligo rev | AAACCCACCATCACTCGCCTCATC | | | | | | | | | | |
| Chromosome | start | end | strand | MM | target_seq | PAM | alignment | distance | position | gene name | gene id |
| GU305914.1 | 44678 | 44700 | - | 0 | GATGAGGCGAGTGATGGTGG | TGG | \|\|\|\|\|\|\|\|[\|\|\|\|\|\|\|\|\|\|\|\|]PAM | NA | Intergenic | NA | NA |
| T30853 | ATGAGGCGAGTGATGGTGGTGGG | 1000 | | | | | | | | | |
| Oligo fwd | CACCATGAGGCGAGTGATGGTGGT | | | | | | | | | | |
| Oligo rev | AAACACCACCATCACTCGCCTCAT | | | | | | | | | | |
| Chromosome | start | end | strand | MM | target_seq | PAM | alignment | distance | position | gene name | gene id |
| GU305914.1 | 44677 | 44699 | - | 0 | ATGAGGCGAGTGATGGTGGT | GGG | \|\|\|\|\|\|\|\|[\|\|\|\|\|\|\|\|\|\|\|\|]PAM | NA | Intergenic | NA | NA |
| T30854 | TGAGGCGAGTGATGGTGGTGGGG | 1000 | | | | | | | | | |
| Oligo fwd | CACCTGAGGCGAGTGATGGTGGTG | | | | | | | | | | |
| Oligo rev | AAACCACCACCATCACTCGCCTCA | | | | | | | | | | |
| Chromosome | start | end | strand | MM | target_seq | PAM | alignment | distance | position | gene name | gene id |

<table>
<tr><td>GU305914.1</td><td></td><td>44676</td><td>44698</td><td>-</td><td>0</td><td>TGAGGCGAGTGATGGTGGTG</td><td>GGG</td><td>||||||||[||||||||||||]PAM</td><td>NA</td><td>Intergenic</td><td>NA</td><td>NA</td></tr>
<tr><td>GU305914.1</td><td></td><td>192579</td><td>192601</td><td>+</td><td>3</td><td>TGAGGGGGGTGATGGTGTTG</td><td>TGG</td><td>|||||-|-[|||||||||-||]PAM</td><td>NA</td><td>Intergenic</td><td>NA</td><td>NA</td></tr>
<tr><td>T30855</td><td>AGTGATGGTGGTGGGGATGATGG</td><td></td><td>1000</td><td></td><td></td><td></td><td></td><td></td><td></td><td></td><td></td><td></td></tr>
<tr><td>Oligo fwd</td><td>CACCAGTGATGGTGGTGGGGATGA</td><td></td><td></td><td></td><td></td><td></td><td></td><td></td><td></td><td></td><td></td><td></td></tr>
<tr><td>Oligo rev</td><td>AAACTCATCCCCACCACCATCACT</td><td></td><td></td><td></td><td></td><td></td><td></td><td></td><td></td><td></td><td></td><td></td></tr>
<tr><td>Chromosome</td><td>start</td><td></td><td>end</td><td>strand</td><td>MM</td><td>target_seq</td><td>PAM</td><td>alignment</td><td>distance</td><td>position</td><td>gene name</td><td>gene id</td></tr>
<tr><td>GU305914.1</td><td></td><td>44669</td><td>44691</td><td>-</td><td>0</td><td>AGTGATGGTGGTGGGGATGA</td><td>TGG</td><td>||||||||[||||||||||||]PAM</td><td>NA</td><td>Intergenic</td><td>NA</td><td>NA</td></tr>
<tr><td>T30856</td><td>GATGGTGGTGGGGATGATGGTGG</td><td></td><td>1000</td><td></td><td></td><td></td><td></td><td></td><td></td><td></td><td></td><td></td></tr>
<tr><td>Oligo fwd</td><td>CACCGATGGTGGTGGGGATGATGG</td><td></td><td></td><td></td><td></td><td></td><td></td><td></td><td></td><td></td><td></td><td></td></tr>
<tr><td>Oligo rev</td><td>AAACCCATCATCCCCACCACCATC</td><td></td><td></td><td></td><td></td><td></td><td></td><td></td><td></td><td></td><td></td><td></td></tr>
<tr><td>Chromosome</td><td>start</td><td></td><td>end</td><td>strand</td><td>MM</td><td>target_seq</td><td>PAM</td><td>alignment</td><td>distance</td><td>position</td><td>gene name</td><td>gene id</td></tr>
<tr><td>GU305914.1</td><td></td><td>44666</td><td>44688</td><td>-</td><td>0</td><td>GATGGTGGTGGGGATGATGG</td><td>TGG</td><td>||||||||[||||||||||||]PAM</td><td>NA</td><td>Intergenic</td><td>NA</td><td>NA</td></tr>
<tr><td>GU305914.1</td><td></td><td>106019</td><td>106041</td><td>+</td><td>4</td><td>GATCATGGTGGCGATGATGT</td><td>TGG</td><td>|||--|||[|||-|||||||-]PAM</td><td>NA</td><td>Intergenic</td><td>NA</td><td>NA</td></tr>
<tr><td>T30857</td><td>GGTGGGGATGATGGTGGTGATGG</td><td></td><td>1000</td><td></td><td></td><td></td><td></td><td></td><td></td><td></td><td></td><td></td></tr>
<tr><td>Oligo fwd</td><td>CACCGGTGGGGATGATGGTGGTGA</td><td></td><td></td><td></td><td></td><td></td><td></td><td></td><td></td><td></td><td></td><td></td></tr>
<tr><td>Oligo rev</td><td>AAACTCACCACCATCATCCCCACC</td><td></td><td></td><td></td><td></td><td></td><td></td><td></td><td></td><td></td><td></td><td></td></tr>
<tr><td>Chromosome</td><td>start</td><td></td><td>end</td><td>strand</td><td>MM</td><td>target_seq</td><td>PAM</td><td>alignment</td><td>distance</td><td>position</td><td>gene name</td><td>gene id</td></tr>
<tr><td>GU305914.1</td><td></td><td>44660</td><td>44682</td><td>-</td><td>0</td><td>GGTGGGGATGATGGTGGTGA</td><td>TGG</td><td>||||||||[||||||||||||]PAM</td><td>NA</td><td>Intergenic</td><td>NA</td><td>NA</td></tr>
<tr><td>T30858</td><td>GGGGATGATGGTGGTGATGGTGG</td><td></td><td>1000</td><td></td><td></td><td></td><td></td><td></td><td></td><td></td><td></td><td></td></tr>
<tr><td>Oligo fwd</td><td>CACCGGGGATGATGGTGGTGATGG</td><td></td><td></td><td></td><td></td><td></td><td></td><td></td><td></td><td></td><td></td><td></td></tr>
<tr><td>Oligo rev</td><td>AAACCCATCACCACCATCATCCCC</td><td></td><td></td><td></td><td></td><td></td><td></td><td></td><td></td><td></td><td></td><td></td></tr>
<tr><td>Chromosome</td><td>start</td><td></td><td>end</td><td>strand</td><td>MM</td><td>target_seq</td><td>PAM</td><td>alignment</td><td>distance</td><td>position</td><td>gene name</td><td>gene id</td></tr>
<tr><td>GU305914.1</td><td></td><td>44657</td><td>44679</td><td>-</td><td>0</td><td>GGGGATGATGGTGGTGATGG</td><td>TGG</td><td>||||||||[||||||||||||]PAM</td><td>NA</td><td>Intergenic</td><td>NA</td><td>NA</td></tr>
<tr><td>GU305914.1</td><td></td><td>167809</td><td>167831</td><td>+</td><td>4</td><td>GGGGTCCTTGGTGGTGATGG</td><td>CGG</td><td>||||----[||||||||||||]PAM</td><td>NA</td><td>Intergenic</td><td>NA</td><td>NA</td></tr>
<tr><td>T30859</td><td>GGGATGATGGTGGTGATGGTGGG</td><td></td><td>1000</td><td></td><td></td><td></td><td></td><td></td><td></td><td></td><td></td><td></td></tr>
<tr><td>Oligo fwd</td><td>CACCGGGATGATGGTGGTGATGGT</td><td></td><td></td><td></td><td></td><td></td><td></td><td></td><td></td><td></td><td></td><td></td></tr>
<tr><td>Oligo rev</td><td>AAACACCATCACCACCATCATCCC</td><td></td><td></td><td></td><td></td><td></td><td></td><td></td><td></td><td></td><td></td><td></td></tr>
<tr><td>Chromosome</td><td>start</td><td></td><td>end</td><td>strand</td><td>MM</td><td>target_seq</td><td>PAM</td><td>alignment</td><td>distance</td><td>position</td><td>gene name</td><td>gene id</td></tr>
<tr><td>GU305914.1</td><td></td><td>44656</td><td>44678</td><td>-</td><td>0</td><td>GGGATGATGGTGGTGATGGT</td><td>GGG</td><td>||||||||[||||||||||||]PAM</td><td>NA</td><td>Intergenic</td><td>NA</td><td>NA</td></tr>
<tr><td>T30860</td><td>ATGATGGTGGTGATGGTGGGAGG</td><td></td><td>1000</td><td></td><td></td><td></td><td></td><td></td><td></td><td></td><td></td><td></td></tr>
<tr><td>Oligo fwd</td><td>CACCATGATGGTGGTGATGGTGGG</td><td></td><td></td><td></td><td></td><td></td><td></td><td></td><td></td><td></td><td></td><td></td></tr>
<tr><td>Oligo rev</td><td>AAACCCCACCATCACCACCATCAT</td><td></td><td></td><td></td><td></td><td></td><td></td><td></td><td></td><td></td><td></td><td></td></tr>
<tr><td>Chromosome</td><td>start</td><td></td><td>end</td><td>strand</td><td>MM</td><td>target_seq</td><td>PAM</td><td>alignment</td><td>distance</td><td>position</td><td>gene name</td><td>gene id</td></tr>
<tr><td>GU305914.1</td><td></td><td>44653</td><td>44675</td><td>-</td><td>0</td><td>ATGATGGTGGTGATGGTGGG</td><td>AGG</td><td>||||||||[||||||||||||]PAM</td><td>NA</td><td>Intergenic</td><td>NA</td><td>NA</td></tr>
<tr><td>T30861</td><td>ATGGTGGTGATGGTGGGAGGAGG</td><td></td><td>1000</td><td></td><td></td><td></td><td></td><td></td><td></td><td></td><td></td><td></td></tr>
<tr><td>Oligo fwd</td><td>CACCATGGTGGTGATGGTGGGAGG</td><td></td><td></td><td></td><td></td><td></td><td></td><td></td><td></td><td></td><td></td><td></td></tr>
<tr><td>Oligo rev</td><td>AAACCCTCCCACCATCACCACCAT</td><td></td><td></td><td></td><td></td><td></td><td></td><td></td><td></td><td></td><td></td><td></td></tr>
<tr><td>Chromosome</td><td>start</td><td></td><td>end</td><td>strand</td><td>MM</td><td>target_seq</td><td>PAM</td><td>alignment</td><td>distance</td><td>position</td><td>gene name</td><td>gene id</td></tr>
<tr><td>GU305914.1</td><td></td><td>44650</td><td>44672</td><td>-</td><td>0</td><td>ATGGTGGTGATGGTGGGAGG</td><td>AGG</td><td>||||||||[||||||||||||]PAM</td><td>NA</td><td>Intergenic</td><td>NA</td><td>NA</td></tr>
<tr><td>T30862</td><td>GTGGTGATGGTGGGAGGAGGAGG</td><td></td><td>1000</td><td></td><td></td><td></td><td></td><td></td><td></td><td></td><td></td><td></td></tr>
<tr><td>Oligo fwd</td><td>CACCGTGGTGATGGTGGGAGGAGG</td><td></td><td></td><td></td><td></td><td></td><td></td><td></td><td></td><td></td><td></td><td></td></tr>
<tr><td>Oligo rev</td><td>AAACCCTCCTCCCACCATCACCAC</td><td></td><td></td><td></td><td></td><td></td><td></td><td></td><td></td><td></td><td></td><td></td></tr>
<tr><td>Chromosome</td><td>start</td><td></td><td>end</td><td>strand</td><td>MM</td><td>target_seq</td><td>PAM</td><td>alignment</td><td>distance</td><td>position</td><td>gene name</td><td>gene id</td></tr>
<tr><td>GU305914.1</td><td></td><td>44647</td><td>44669</td><td>-</td><td>0</td><td>GTGGTGATGGTGGGAGGAGG</td><td>AGG</td><td>||||||||[||||||||||||]PAM</td><td>NA</td><td>Intergenic</td><td>NA</td><td>NA</td></tr>
<tr><td>T30863</td><td>ATGGTGGGAGGAGGAGGACGAGG</td><td></td><td>1000</td><td></td><td></td><td></td><td></td><td></td><td></td><td></td><td></td><td></td></tr>
<tr><td>Oligo fwd</td><td>CACCATGGTGGGAGGAGGAGGACG</td><td></td><td></td><td></td><td></td><td></td><td></td><td></td><td></td><td></td><td></td><td></td></tr>
<tr><td>Oligo rev</td><td>AAACCGTCCTCCTCCTCCCACCAT</td><td></td><td></td><td></td><td></td><td></td><td></td><td></td><td></td><td></td><td></td><td></td></tr>
<tr><td>Chromosome</td><td>start</td><td></td><td>end</td><td>strand</td><td>MM</td><td>target_seq</td><td>PAM</td><td>alignment</td><td>distance</td><td>position</td><td>gene name</td><td>gene id</td></tr>
<tr><td>GU305914.1</td><td></td><td>44641</td><td>44663</td><td>-</td><td>0</td><td>ATGGTGGGAGGAGGAGGACG</td><td>AGG</td><td>||||||||[||||||||||||]PAM</td><td>NA</td><td>Intergenic</td><td>NA</td><td>NA</td></tr>
<tr><td>T30864</td><td>AGGACGAGGAGTGATGATGAAGG</td><td></td><td>1000</td><td></td><td></td><td></td><td></td><td></td><td></td><td></td><td></td><td></td></tr>
<tr><td>Oligo fwd</td><td>CACCAGGACGAGGAGTGATGATGA</td><td></td><td></td><td></td><td></td><td></td><td></td><td></td><td></td><td></td><td></td><td></td></tr>
<tr><td>Oligo rev</td><td>AAACTCATCATCACTCCTCGTCCT</td><td></td><td></td><td></td><td></td><td></td><td></td><td></td><td></td><td></td><td></td><td></td></tr>
<tr><td>Chromosome</td><td>start</td><td></td><td>end</td><td>strand</td><td>MM</td><td>target_seq</td><td>PAM</td><td>alignment</td><td>distance</td><td>position</td><td>gene name</td><td>gene id</td></tr>
<tr><td>GU305914.1</td><td></td><td>44627</td><td>44649</td><td>-</td><td>0</td><td>AGGACGAGGAGTGATGATGA</td><td>AGG</td><td>||||||||[||||||||||||]PAM</td><td>NA</td><td>Intergenic</td><td>NA</td><td>NA</td></tr>
<tr><td>GU305914.1</td><td></td><td>188470</td><td>188492</td><td>-</td><td>4</td><td>AGGATGACGTGTGACGATGA</td><td>AGG</td><td>||||-||-[|-||||-|||||]PAM</td><td>NA</td><td>Intergenic</td><td>NA</td><td>NA</td></tr>
<tr><td>T30865</td><td>ACGAGGAGTGATGATGAAGGTGG</td><td></td><td>1000</td><td></td><td></td><td></td><td></td><td></td><td></td><td></td><td></td><td></td></tr>
<tr><td>Oligo fwd</td><td>CACCACGAGGAGTGATGATGAAGG</td><td></td><td></td><td></td><td></td><td></td><td></td><td></td><td></td><td></td><td></td><td></td></tr>
<tr><td>Oligo rev</td><td>AAACCCTTCATCATCACTCCTCGT</td><td></td><td></td><td></td><td></td><td></td><td></td><td></td><td></td><td></td><td></td><td></td></tr>
<tr><td>Chromosome</td><td>start</td><td></td><td>end</td><td>strand</td><td>MM</td><td>target_seq</td><td>PAM</td><td>alignment</td><td>distance</td><td>position</td><td>gene name</td><td>gene id</td></tr>
<tr><td>GU305914.1</td><td></td><td>44624</td><td>44646</td><td>-</td><td>0</td><td>ACGAGGAGTGATGATGAAGG</td><td>TGG</td><td>||||||||[||||||||||||]PAM</td><td>NA</td><td>Intergenic</td><td>NA</td><td>NA</td></tr>
<tr><td>T30866</td><td>AGGAGTGATGATGAAGGTGGTGG</td><td></td><td>1000</td><td></td><td></td><td></td><td></td><td></td><td></td><td></td><td></td><td></td></tr>
<tr><td>Oligo fwd</td><td>CACCAGGAGTGATGATGAAGGTGG</td><td></td><td></td><td></td><td></td><td></td><td></td><td></td><td></td><td></td><td></td><td></td></tr>
<tr><td>Oligo rev</td><td>AAACCCACCTTCATCATCACTCCT</td><td></td><td></td><td></td><td></td><td></td><td></td><td></td><td></td><td></td><td></td><td></td></tr>
<tr><td>Chromosome</td><td>start</td><td></td><td>end</td><td>strand</td><td>MM</td><td>target_seq</td><td>PAM</td><td>alignment</td><td>distance</td><td>position</td><td>gene name</td><td>gene id</td></tr>
<tr><td>GU305914.1</td><td></td><td>44621</td><td>44643</td><td>-</td><td>0</td><td>AGGAGTGATGATGAAGGTGG</td><td>TGG</td><td>||||||||[||||||||||||]PAM</td><td>NA</td><td>Intergenic</td><td>NA</td><td>NA</td></tr>
<tr><td>T30867</td><td>AGTGATGATGAAGGTGGTGGTGG</td><td></td><td>1000</td><td></td><td></td><td></td><td></td><td></td><td></td><td></td><td></td><td></td></tr>
<tr><td>Oligo fwd</td><td>CACCAGTGATGATGAAGGTGGTGG</td><td></td><td></td><td></td><td></td><td></td><td></td><td></td><td></td><td></td><td></td><td></td></tr>
<tr><td>Oligo rev</td><td>AAACCCACCACCTTCATCATCACT</td><td></td><td></td><td></td><td></td><td></td><td></td><td></td><td></td><td></td><td></td><td></td></tr>
<tr><td>Chromosome</td><td>start</td><td></td><td>end</td><td>strand</td><td>MM</td><td>target_seq</td><td>PAM</td><td>alignment</td><td>distance</td><td>position</td><td>gene name</td><td>gene id</td></tr>
<tr><td>GU305914.1</td><td></td><td>44618</td><td>44640</td><td>-</td><td>0</td><td>AGTGATGATGAAGGTGGTGG</td><td>TGG</td><td>||||||||[||||||||||||]PAM</td><td>NA</td><td>Intergenic</td><td>NA</td><td>NA</td></tr>
<tr><td>T30868</td><td>TGATGAAGGTGGTGGTGGTGCGG</td><td></td><td>1000</td><td></td><td></td><td></td><td></td><td></td><td></td><td></td><td></td><td></td></tr>
<tr><td>Oligo fwd</td><td>CACCTGATGAAGGTGGTGGTGGTG</td><td></td><td></td><td></td><td></td><td></td><td></td><td></td><td></td><td></td><td></td><td></td></tr>
<tr><td>Oligo rev</td><td>AAACCACCACCACCACCTTCATCA</td><td></td><td></td><td></td><td></td><td></td><td></td><td></td><td></td><td></td><td></td><td></td></tr>
<tr><td>Chromosome</td><td>start</td><td></td><td>end</td><td>strand</td><td>MM</td><td>target_seq</td><td>PAM</td><td>alignment</td><td>distance</td><td>position</td><td>gene name</td><td>gene id</td></tr>
<tr><td>GU305914.1</td><td></td><td>44613</td><td>44635</td><td>-</td><td>0</td><td>TGATGAAGGTGGTGGTGGTG</td><td>CGG</td><td>||||||||[||||||||||||]PAM</td><td>NA</td><td>Intergenic</td><td>NA</td><td>NA</td></tr>
</table>

T30869 GATGAAGGTGGTGGTGGTGCGGG 1000
Oligo fwd CACCGATGAAGGTGGTGGTGGTGC
Oligo rev AAACGCACCACCACCACCTTCATC
Chromosome start end strand MM target_seq PAM alignment distance position gene name gene id
GU305914.1 44612 44634 - 0 GATGAAGGTGGTGGTGGTGC GGG ||||||||[||||||||||||]PAM NA Intergenic NA NA

T30870 AAGGTGGTGGTGGTGCGGGTCGG 1000
Oligo fwd CACCAAGGTGGTGGTGGTGCGGGT
Oligo rev AAACACCCGCACCACCACCACCTT
Chromosome start end strand MM target_seq PAM alignment distance position gene name gene id
GU305914.1 44608 44630 - 0 AAGGTGGTGGTGGTGCGGGT CGG ||||||||[||||||||||||]PAM NA Intergenic NA NA

T30871 AGGTGGTGGTGGTGCGGGTCGGG 1000
Oligo fwd CACCAGGTGGTGGTGGTGCGGGTC
Oligo rev AAACGACCCGCACCACCACCACCT
Chromosome start end strand MM target_seq PAM alignment distance position gene name gene id
GU305914.1 44607 44629 - 0 AGGTGGTGGTGGTGCGGGTC GGG ||||||||[||||||||||||]PAM NA Intergenic NA NA

T30872 TGGTGGTGGTGCGGGTCGGGAGG 1000
Oligo fwd CACCTGGTGGTGGTGCGGGTCGGG
Oligo rev AAACCCCGACCCGCACCACCACCA
Chromosome start end strand MM target_seq PAM alignment distance position gene name gene id
GU305914.1 44604 44626 - 0 TGGTGGTGGTGCGGGTCGGG AGG ||||||||[||||||||||||]PAM NA Intergenic NA NA

T30873 GGCAGCGCCGCCACCGTCCCCGG 1000
Oligo fwd CACCGGCAGCGCCGCCACCGTCCC
Oligo rev AAACGGGACGGTGGCGGCGCTGCC
Chromosome start end strand MM target_seq PAM alignment distance position gene name gene id
GU305914.1 44583 44605 - 0 GGCAGCGCCGCCACCGTCCC CGG ||||||||[||||||||||||]PAM NA Intergenic NA NA

T30874 CCCCGGTAACATCGCGCCGCCGG 1000
Oligo fwd CACCCCCCGGTAACATCGCGCCGC
Oligo rev AAACGCGGCGCGATGTTACCGGGG
Chromosome start end strand MM target_seq PAM alignment distance position gene name gene id
GU305914.1 44566 44588 - 0 CCCCGGTAACATCGCGCCGC CGG ||||||||[||||||||||||]PAM NA Intergenic NA NA

T30875 CGCGTCGTTCAAGACGAACGCGG 1000
Oligo fwd CACCCGCGTCGTTCAAGACGAACG
Oligo rev AAACCGTTCGTCTTGAACGACGCG
Chromosome start end strand MM target_seq PAM alignment distance position gene name gene id
GU305914.1 44497 44519 - 0 CGCGTCGTTCAAGACGAACG CGG ||||||||[||||||||||||]PAM NA Intergenic NA NA

T30876 GTTCAAGACGAACGCGGAGCTGG 1000
Oligo fwd CACCGTTCAAGACGAACGCGGAGC
Oligo rev AAACGCTCCGCGTTCGTCTTGAAC
Chromosome start end strand MM target_seq PAM alignment distance position gene name gene id
GU305914.1 44491 44513 - 0 GTTCAAGACGAACGCGGAGC TGG ||||||||[||||||||||||]PAM NA Intergenic NA NA

T30877 CGAACGCGGAGCTGGTGTACTGG 1000
Oligo fwd CACCCGAACGCGGAGCTGGTGTAC
Oligo rev AAACGTACACCAGCTCCGCGTTCG
Chromosome start end strand MM target_seq PAM alignment distance position gene name gene id
GU305914.1 44483 44505 - 0 CGAACGCGGAGCTGGTGTAC TGG ||||||||[||||||||||||]PAM NA Intergenic NA NA

T30878 ACGCGGAGCTGGTGTACTGGCGG 1000
Oligo fwd CACCACGCGGAGCTGGTGTACTGG
Oligo rev AAACCCAGTACACCAGCTCCGCGT
Chromosome start end strand MM target_seq PAM alignment distance position gene name gene id
GU305914.1 44480 44502 - 0 ACGCGGAGCTGGTGTACTGG CGG ||||||||[||||||||||||]PAM NA Intergenic NA NA

T30879 TGTACTGGCGGATCTGCGACTGG 1000
Oligo fwd CACCTGTACTGGCGGATCTGCGAC
Oligo rev AAACGTCGCAGATCCGCCAGTACA
Chromosome start end strand MM target_seq PAM alignment distance position gene name gene id
GU305914.1 44468 44490 - 0 TGTACTGGCGGATCTGCGAC TGG ||||||||[||||||||||||]PAM NA Intergenic NA NA

T30880 GGATCTTGTATCCCGTCGAGTGG 1000
Oligo fwd CACCGGATCTTGTATCCCGTCGAG
Oligo rev AAACCTCGACGGGATACAAGATCC
Chromosome start end strand MM target_seq PAM alignment distance position gene name gene id
GU305914.1 44447 44469 - 0 GGATCTTGTATCCCGTCGAG TGG ||||||||[||||||||||||]PAM NA Intergenic NA NA

T30881 GCAGCTCGAAGACGTCCCTGAGG 1000
Oligo fwd CACCGCAGCTCGAAGACGTCCCTG
Oligo rev AAACCAGGGACGTCTTCGAGCTGC
Chromosome start end strand MM target_seq PAM alignment distance position gene name gene id
GU305914.1 44420 44442 - 0 GCAGCTCGAAGACGTCCCTG AGG ||||||||[||||||||||||]PAM NA Intergenic NA NA

T30882 CCTGAGGTCGATCTTCTCGTCGG 1000
Oligo fwd CACCCCTGAGGTCGATCTTCTCGT
Oligo rev AAACACGAGAAGATCGACCTCAGG
Chromosome start end strand MM target_seq PAM alignment distance position gene name gene id
GU305914.1 44404 44426 - 0 CCTGAGGTCGATCTTCTCGT CGG ||||||||[||||||||||||]PAM NA Intergenic NA NA

T30883 CGTCGGCGAACATCTTGATCAGG 1000
Oligo fwd CACCCGTCGGCGAACATCTTGATC
Oligo rev AAACGATCAAGATGTTCGCCGACG
Chromosome start end strand MM target_seq PAM alignment distance position gene name gene id

<table>
<tr><td>GU305914.1</td><td>44387</td><td>44409</td><td>-</td><td>0</td><td>CGTCGGCGAACATCTTGATC</td><td>AGG</td><td>||||||||[||||||||||||]PAM</td><td>NA</td><td>Intergenic</td><td>NA</td><td>NA</td></tr>
<tr><td></td><td></td><td></td><td></td><td></td><td></td><td></td><td></td><td></td><td></td><td></td><td></td></tr>
<tr><td>T30884</td><td>ACATCTTGATCAGGACGTTCTGG</td><td>1000</td><td></td><td></td><td></td><td></td><td></td><td></td><td></td><td></td><td></td></tr>
<tr><td>Oligo fwd</td><td>CACCACATCTTGATCAGGACGTTC</td><td></td><td></td><td></td><td></td><td></td><td></td><td></td><td></td><td></td><td></td></tr>
<tr><td>Oligo rev</td><td>AAACGAACGTCCTGATCAAGATGT</td><td></td><td></td><td></td><td></td><td></td><td></td><td></td><td></td><td></td><td></td></tr>
<tr><td>Chromosome</td><td>start</td><td>end</td><td>strand</td><td>MM</td><td>target_seq</td><td>PAM</td><td>alignment</td><td>distance</td><td>position</td><td>gene name</td><td>gene id</td></tr>
<tr><td>GU305914.1</td><td>44378</td><td>44400</td><td>-</td><td>0</td><td>ACATCTTGATCAGGACGTTC</td><td>TGG</td><td>||||||||[||||||||||||]PAM</td><td>NA</td><td>Intergenic</td><td>NA</td><td>NA</td></tr>
<tr><td></td><td></td><td></td><td></td><td></td><td></td><td></td><td></td><td></td><td></td><td></td><td></td></tr>
<tr><td>T30885</td><td>CGTTCTGGTAGAAGATCTTGAGG</td><td>1000</td><td></td><td></td><td></td><td></td><td></td><td></td><td></td><td></td><td></td></tr>
<tr><td>Oligo fwd</td><td>CACCCGTTCTGGTAGAAGATCTTG</td><td></td><td></td><td></td><td></td><td></td><td></td><td></td><td></td><td></td><td></td></tr>
<tr><td>Oligo rev</td><td>AAACCAAGATCTTCTACCAGAACG</td><td></td><td></td><td></td><td></td><td></td><td></td><td></td><td></td><td></td><td></td></tr>
<tr><td>Chromosome</td><td>start</td><td>end</td><td>strand</td><td>MM</td><td>target_seq</td><td>PAM</td><td>alignment</td><td>distance</td><td>position</td><td>gene name</td><td>gene id</td></tr>
<tr><td>GU305914.1</td><td>44363</td><td>44385</td><td>-</td><td>0</td><td>CGTTCTGGTAGAAGATCTTG</td><td>AGG</td><td>||||||||[||||||||||||]PAM</td><td>NA</td><td>Intergenic</td><td>NA</td><td>NA</td></tr>
<tr><td></td><td></td><td></td><td></td><td></td><td></td><td></td><td></td><td></td><td></td><td></td><td></td></tr>
<tr><td>T30886</td><td>TCTGGTAGAAGATCTTGAGGCGG</td><td>1000</td><td></td><td></td><td></td><td></td><td></td><td></td><td></td><td></td><td></td></tr>
<tr><td>Oligo fwd</td><td>CACCTCTGGTAGAAGATCTTGAGG</td><td></td><td></td><td></td><td></td><td></td><td></td><td></td><td></td><td></td><td></td></tr>
<tr><td>Oligo rev</td><td>AAACCCTCAAGATCTTCTACCAGA</td><td></td><td></td><td></td><td></td><td></td><td></td><td></td><td></td><td></td><td></td></tr>
<tr><td>Chromosome</td><td>start</td><td>end</td><td>strand</td><td>MM</td><td>target_seq</td><td>PAM</td><td>alignment</td><td>distance</td><td>position</td><td>gene name</td><td>gene id</td></tr>
<tr><td>GU305914.1</td><td>44360</td><td>44382</td><td>-</td><td>0</td><td>TCTGGTAGAAGATCTTGAGG</td><td>CGG</td><td>||||||||[||||||||||||]PAM</td><td>NA</td><td>Intergenic</td><td>NA</td><td>NA</td></tr>
<tr><td></td><td></td><td></td><td></td><td></td><td></td><td></td><td></td><td></td><td></td><td></td><td></td></tr>
<tr><td>T30887</td><td>AGATCTTGAGGCGGCACAGCAGG</td><td>1000</td><td></td><td></td><td></td><td></td><td></td><td></td><td></td><td></td><td></td></tr>
<tr><td>Oligo fwd</td><td>CACCAGATCTTGAGGCGGCACAGC</td><td></td><td></td><td></td><td></td><td></td><td></td><td></td><td></td><td></td><td></td></tr>
<tr><td>Oligo rev</td><td>AAACGCTGTGCCGCCTCAAGATCT</td><td></td><td></td><td></td><td></td><td></td><td></td><td></td><td></td><td></td><td></td></tr>
<tr><td>Chromosome</td><td>start</td><td>end</td><td>strand</td><td>MM</td><td>target_seq</td><td>PAM</td><td>alignment</td><td>distance</td><td>position</td><td>gene name</td><td>gene id</td></tr>
<tr><td>GU305914.1</td><td>44351</td><td>44373</td><td>-</td><td>0</td><td>AGATCTTGAGGCGGCACAGC</td><td>AGG</td><td>||||||||[||||||||||||]PAM</td><td>NA</td><td>Intergenic</td><td>NA</td><td>NA</td></tr>
<tr><td></td><td></td><td></td><td></td><td></td><td></td><td></td><td></td><td></td><td></td><td></td><td></td></tr>
<tr><td>T30888</td><td>TCTTGAGGCGGCACAGCAGGCGG</td><td>1000</td><td></td><td></td><td></td><td></td><td></td><td></td><td></td><td></td><td></td></tr>
<tr><td>Oligo fwd</td><td>CACCTCTTGAGGCGGCACAGCAGG</td><td></td><td></td><td></td><td></td><td></td><td></td><td></td><td></td><td></td><td></td></tr>
<tr><td>Oligo rev</td><td>AAACCCTGCTGTGCCGCCTCAAGA</td><td></td><td></td><td></td><td></td><td></td><td></td><td></td><td></td><td></td><td></td></tr>
<tr><td>Chromosome</td><td>start</td><td>end</td><td>strand</td><td>MM</td><td>target_seq</td><td>PAM</td><td>alignment</td><td>distance</td><td>position</td><td>gene name</td><td>gene id</td></tr>
<tr><td>GU305914.1</td><td>44348</td><td>44370</td><td>-</td><td>0</td><td>TCTTGAGGCGGCACAGCAGG</td><td>CGG</td><td>||||||||[||||||||||||]PAM</td><td>NA</td><td>Intergenic</td><td>NA</td><td>NA</td></tr>
<tr><td></td><td></td><td></td><td></td><td></td><td></td><td></td><td></td><td></td><td></td><td></td><td></td></tr>
<tr><td>T30889</td><td>GGCACAGCAGGCGGATCGACCGG</td><td>1000</td><td></td><td></td><td></td><td></td><td></td><td></td><td></td><td></td><td></td></tr>
<tr><td>Oligo fwd</td><td>CACCGGCACAGCAGGCGGATCGAC</td><td></td><td></td><td></td><td></td><td></td><td></td><td></td><td></td><td></td><td></td></tr>
<tr><td>Oligo rev</td><td>AAACGTCGATCCGCCTGCTGTGCC</td><td></td><td></td><td></td><td></td><td></td><td></td><td></td><td></td><td></td><td></td></tr>
<tr><td>Chromosome</td><td>start</td><td>end</td><td>strand</td><td>MM</td><td>target_seq</td><td>PAM</td><td>alignment</td><td>distance</td><td>position</td><td>gene name</td><td>gene id</td></tr>
<tr><td>GU305914.1</td><td>44339</td><td>44361</td><td>-</td><td>0</td><td>GGCACAGCAGGCGGATCGAC</td><td>CGG</td><td>||||||||[||||||||||||]PAM</td><td>NA</td><td>Intergenic</td><td>NA</td><td>NA</td></tr>
<tr><td></td><td></td><td></td><td></td><td></td><td></td><td></td><td></td><td></td><td></td><td></td><td></td></tr>
<tr><td>T30890</td><td>GCACAGCAGGCGGATCGACCGGG</td><td>1000</td><td></td><td></td><td></td><td></td><td></td><td></td><td></td><td></td><td></td></tr>
<tr><td>Oligo fwd</td><td>CACCGCACAGCAGGCGGATCGACC</td><td></td><td></td><td></td><td></td><td></td><td></td><td></td><td></td><td></td><td></td></tr>
<tr><td>Oligo rev</td><td>AAACGGTCGATCCGCCTGCTGTGC</td><td></td><td></td><td></td><td></td><td></td><td></td><td></td><td></td><td></td><td></td></tr>
<tr><td>Chromosome</td><td>start</td><td>end</td><td>strand</td><td>MM</td><td>target_seq</td><td>PAM</td><td>alignment</td><td>distance</td><td>position</td><td>gene name</td><td>gene id</td></tr>
<tr><td>GU305914.1</td><td>44338</td><td>44360</td><td>-</td><td>0</td><td>GCACAGCAGGCGGATCGACC</td><td>GGG</td><td>||||||||[||||||||||||]PAM</td><td>NA</td><td>Intergenic</td><td>NA</td><td>NA</td></tr>
<tr><td></td><td></td><td></td><td></td><td></td><td></td><td></td><td></td><td></td><td></td><td></td><td></td></tr>
<tr><td>T30891</td><td>GGATCGACCGGGCCCACATGTGG</td><td>1000</td><td></td><td></td><td></td><td></td><td></td><td></td><td></td><td></td><td></td></tr>
<tr><td>Oligo fwd</td><td>CACCGGATCGACCGGGCCCACATG</td><td></td><td></td><td></td><td></td><td></td><td></td><td></td><td></td><td></td><td></td></tr>
<tr><td>Oligo rev</td><td>AAACCATGTGGGCCCGGTCGATCC</td><td></td><td></td><td></td><td></td><td></td><td></td><td></td><td></td><td></td><td></td></tr>
<tr><td>Chromosome</td><td>start</td><td>end</td><td>strand</td><td>MM</td><td>target_seq</td><td>PAM</td><td>alignment</td><td>distance</td><td>position</td><td>gene name</td><td>gene id</td></tr>
<tr><td>GU305914.1</td><td>44327</td><td>44349</td><td>-</td><td>0</td><td>GGATCGACCGGGCCCACATG</td><td>TGG</td><td>||||||||[||||||||||||]PAM</td><td>NA</td><td>Intergenic</td><td>NA</td><td>NA</td></tr>
<tr><td></td><td></td><td></td><td></td><td></td><td></td><td></td><td></td><td></td><td></td><td></td><td></td></tr>
<tr><td>T30892</td><td>GATCGACCGGGCCCACATGTGGG</td><td>1000</td><td></td><td></td><td></td><td></td><td></td><td></td><td></td><td></td><td></td></tr>
<tr><td>Oligo fwd</td><td>CACCGATCGACCGGGCCCACATGT</td><td></td><td></td><td></td><td></td><td></td><td></td><td></td><td></td><td></td><td></td></tr>
<tr><td>Oligo rev</td><td>AAACACATGTGGGCCCGGTCGATC</td><td></td><td></td><td></td><td></td><td></td><td></td><td></td><td></td><td></td><td></td></tr>
<tr><td>Chromosome</td><td>start</td><td>end</td><td>strand</td><td>MM</td><td>target_seq</td><td>PAM</td><td>alignment</td><td>distance</td><td>position</td><td>gene name</td><td>gene id</td></tr>
<tr><td>GU305914.1</td><td>44326</td><td>44348</td><td>-</td><td>0</td><td>GATCGACCGGGCCCACATGT</td><td>GGG</td><td>||||||||[||||||||||||]PAM</td><td>NA</td><td>Intergenic</td><td>NA</td><td>NA</td></tr>
<tr><td></td><td></td><td></td><td></td><td></td><td></td><td></td><td></td><td></td><td></td><td></td><td></td></tr>
<tr><td>T30893</td><td>ATCGACCGGGCCCACATGTGGGG</td><td>1000</td><td></td><td></td><td></td><td></td><td></td><td></td><td></td><td></td><td></td></tr>
<tr><td>Oligo fwd</td><td>CACCATCGACCGGGCCCACATGTG</td><td></td><td></td><td></td><td></td><td></td><td></td><td></td><td></td><td></td><td></td></tr>
<tr><td>Oligo rev</td><td>AAACCACATGTGGGCCCGGTCGAT</td><td></td><td></td><td></td><td></td><td></td><td></td><td></td><td></td><td></td><td></td></tr>
<tr><td>Chromosome</td><td>start</td><td>end</td><td>strand</td><td>MM</td><td>target_seq</td><td>PAM</td><td>alignment</td><td>distance</td><td>position</td><td>gene name</td><td>gene id</td></tr>
<tr><td>GU305914.1</td><td>44325</td><td>44347</td><td>-</td><td>0</td><td>ATCGACCGGGCCCACATGTG</td><td>GGG</td><td>||||||||[||||||||||||]PAM</td><td>NA</td><td>Intergenic</td><td>NA</td><td>NA</td></tr>
<tr><td></td><td></td><td></td><td></td><td></td><td></td><td></td><td></td><td></td><td></td><td></td><td></td></tr>
<tr><td>T30894</td><td>GGGCCCACATGTGGGGCGTCTGG</td><td>1000</td><td></td><td></td><td></td><td></td><td></td><td></td><td></td><td></td><td></td></tr>
<tr><td>Oligo fwd</td><td>CACCGGGCCCACATGTGGGGCGTC</td><td></td><td></td><td></td><td></td><td></td><td></td><td></td><td></td><td></td><td></td></tr>
<tr><td>Oligo rev</td><td>AAACGACGCCCCACATGTGGGCCC</td><td></td><td></td><td></td><td></td><td></td><td></td><td></td><td></td><td></td><td></td></tr>
<tr><td>Chromosome</td><td>start</td><td>end</td><td>strand</td><td>MM</td><td>target_seq</td><td>PAM</td><td>alignment</td><td>distance</td><td>position</td><td>gene name</td><td>gene id</td></tr>
<tr><td>GU305914.1</td><td>44318</td><td>44340</td><td>-</td><td>0</td><td>GGGCCCACATGTGGGGCGTC</td><td>TGG</td><td>||||||||[||||||||||||]PAM</td><td>NA</td><td>Intergenic</td><td>NA</td><td>NA</td></tr>
<tr><td></td><td></td><td></td><td></td><td></td><td></td><td></td><td></td><td></td><td></td><td></td><td></td></tr>
<tr><td>T30895</td><td>TCTGGCACAGCATCCCCACCAGG</td><td>1000</td><td></td><td></td><td></td><td></td><td></td><td></td><td></td><td></td><td></td></tr>
<tr><td>Oligo fwd</td><td>CACCTCTGGCACAGCATCCCCACC</td><td></td><td></td><td></td><td></td><td></td><td></td><td></td><td></td><td></td><td></td></tr>
<tr><td>Oligo rev</td><td>AAACGGTGGGGATGCTGTGCCAGA</td><td></td><td></td><td></td><td></td><td></td><td></td><td></td><td></td><td></td><td></td></tr>
<tr><td>Chromosome</td><td>start</td><td>end</td><td>strand</td><td>MM</td><td>target_seq</td><td>PAM</td><td>alignment</td><td>distance</td><td>position</td><td>gene name</td><td>gene id</td></tr>
<tr><td>GU305914.1</td><td>44300</td><td>44322</td><td>-</td><td>0</td><td>TCTGGCACAGCATCCCCACC</td><td>AGG</td><td>||||||||[||||||||||||]PAM</td><td>NA</td><td>Intergenic</td><td>NA</td><td>NA</td></tr>
<tr><td></td><td></td><td></td><td></td><td></td><td></td><td></td><td></td><td></td><td></td><td></td><td></td></tr>
<tr><td>T30896</td><td>TCCCCACCAGGATCATGCAGAGG</td><td>1000</td><td></td><td></td><td></td><td></td><td></td><td></td><td></td><td></td><td></td></tr>
<tr><td>Oligo fwd</td><td>CACCTCCCCACCAGGATCATGCAG</td><td></td><td></td><td></td><td></td><td></td><td></td><td></td><td></td><td></td><td></td></tr>
<tr><td>Oligo rev</td><td>AAACCTGCATGATCCTGGTGGGGA</td><td></td><td></td><td></td><td></td><td></td><td></td><td></td><td></td><td></td><td></td></tr>
<tr><td>Chromosome</td><td>start</td><td>end</td><td>strand</td><td>MM</td><td>target_seq</td><td>PAM</td><td>alignment</td><td>distance</td><td>position</td><td>gene name</td><td>gene id</td></tr>
<tr><td>GU305914.1</td><td>44288</td><td>44310</td><td>-</td><td>0</td><td>TCCCCACCAGGATCATGCAG</td><td>AGG</td><td>||||||||[||||||||||||]PAM</td><td>NA</td><td>Intergenic</td><td>NA</td><td>NA</td></tr>
<tr><td></td><td></td><td></td><td></td><td></td><td></td><td></td><td></td><td></td><td></td><td></td><td></td></tr>
<tr><td>T30897</td><td>CCAGGATCATGCAGAGGTCTCGG</td><td>1000</td><td></td><td></td><td></td><td></td><td></td><td></td><td></td><td></td><td></td></tr>
<tr><td>Oligo fwd</td><td>CACCCCAGGATCATGCAGAGGTCT</td><td></td><td></td><td></td><td></td><td></td><td></td><td></td><td></td><td></td><td></td></tr>
<tr><td>Oligo rev</td><td>AAACAGACCTCTGCATGATCCTGG</td><td></td><td></td><td></td><td></td><td></td><td></td><td></td><td></td><td></td><td></td></tr>
<tr><td>Chromosome</td><td>start</td><td>end</td><td>strand</td><td>MM</td><td>target_seq</td><td>PAM</td><td>alignment</td><td>distance</td><td>position</td><td>gene name</td><td>gene id</td></tr>
<tr><td>GU305914.1</td><td>44282</td><td>44304</td><td>-</td><td>0</td><td>CCAGGATCATGCAGAGGTCT</td><td>CGG</td><td>||||||||[||||||||||||]PAM</td><td>NA</td><td>Intergenic</td><td>NA</td><td>NA</td></tr>
<tr><td></td><td></td><td></td><td></td><td></td><td></td><td></td><td></td><td></td><td></td><td></td><td></td></tr>
<tr><td>T30898</td><td>TGATGAAACGACGTATCCCGAGG</td><td>1000</td><td></td><td></td><td></td><td></td><td></td><td></td><td></td><td></td><td></td></tr>
<tr><td>Oligo fwd</td><td>CACCTGATGAAACGACGTATCCCG</td><td></td><td></td><td></td><td></td><td></td><td></td><td></td><td></td><td></td><td></td></tr>
<tr><td>Oligo rev</td><td>AAACCGGGATACGTCGTTTCATCA</td><td></td><td></td><td></td><td></td><td></td><td></td><td></td><td></td><td></td><td></td></tr>
</table>

<table>
<tr><td>Chromosome</td><td>start</td><td>end</td><td>strand</td><td>MM</td><td>target_seq</td><td>PAM</td><td>alignment</td><td>distance</td><td>position</td><td>gene name</td><td>gene id</td></tr>
<tr><td>GU305914.1</td><td>44237</td><td>44259</td><td>-</td><td>0</td><td>TGATGAAACGACGTATCCCG</td><td>AGG</td><td>||||||||[||||||||||||]PAM</td><td>NA</td><td>Intergenic</td><td>NA</td><td>NA</td></tr>
<tr><td>T30899</td><td>AACGACGTATCCCGAGGTTGAGG</td><td>1000</td><td></td><td></td><td></td><td></td><td></td><td></td><td></td><td></td><td></td></tr>
<tr><td>Oligo fwd</td><td>CACCAACGACGTATCCCGAGGTTG</td><td></td><td></td><td></td><td></td><td></td><td></td><td></td><td></td><td></td><td></td></tr>
<tr><td>Oligo rev</td><td>AAACCAACCTCGGGATACGTCGTT</td><td></td><td></td><td></td><td></td><td></td><td></td><td></td><td></td><td></td><td></td></tr>
<tr><td>Chromosome</td><td>start</td><td>end</td><td>strand</td><td>MM</td><td>target_seq</td><td>PAM</td><td>alignment</td><td>distance</td><td>position</td><td>gene name</td><td>gene id</td></tr>
<tr><td>GU305914.1</td><td>44231</td><td>44253</td><td>-</td><td>0</td><td>AACGACGTATCCCGAGGTTG</td><td>AGG</td><td>||||||||[||||||||||||]PAM</td><td>NA</td><td>Intergenic</td><td>NA</td><td>NA</td></tr>
<tr><td>T30900</td><td>CGAGCCTTCGCTGCACCGTTTGG</td><td>1000</td><td></td><td></td><td></td><td></td><td></td><td></td><td></td><td></td><td></td></tr>
<tr><td>Oligo fwd</td><td>CACCCGAGCCTTCGCTGCACCGTT</td><td></td><td></td><td></td><td></td><td></td><td></td><td></td><td></td><td></td><td></td></tr>
<tr><td>Oligo rev</td><td>AAACAACGGTGCAGCGAAGGCTCG</td><td></td><td></td><td></td><td></td><td></td><td></td><td></td><td></td><td></td><td></td></tr>
<tr><td>Chromosome</td><td>start</td><td>end</td><td>strand</td><td>MM</td><td>target_seq</td><td>PAM</td><td>alignment</td><td>distance</td><td>position</td><td>gene name</td><td>gene id</td></tr>
<tr><td>GU305914.1</td><td>44206</td><td>44228</td><td>-</td><td>0</td><td>CGAGCCTTCGCTGCACCGTT</td><td>TGG</td><td>||||||||[||||||||||||]PAM</td><td>NA</td><td>Intergenic</td><td>NA</td><td>NA</td></tr>
<tr><td>T30901</td><td>CGCTGCACCGTTTGGTGAGCTGG</td><td>1000</td><td></td><td></td><td></td><td></td><td></td><td></td><td></td><td></td><td></td></tr>
<tr><td>Oligo fwd</td><td>CACCCGCTGCACCGTTTGGTGAGC</td><td></td><td></td><td></td><td></td><td></td><td></td><td></td><td></td><td></td><td></td></tr>
<tr><td>Oligo rev</td><td>AAACGCTCACCAAACGGTGCAGCG</td><td></td><td></td><td></td><td></td><td></td><td></td><td></td><td></td><td></td><td></td></tr>
<tr><td>Chromosome</td><td>start</td><td>end</td><td>strand</td><td>MM</td><td>target_seq</td><td>PAM</td><td>alignment</td><td>distance</td><td>position</td><td>gene name</td><td>gene id</td></tr>
<tr><td>GU305914.1</td><td>44198</td><td>44220</td><td>-</td><td>0</td><td>CGCTGCACCGTTTGGTGAGC</td><td>TGG</td><td>||||||||[||||||||||||]PAM</td><td>NA</td><td>Intergenic</td><td>NA</td><td>NA</td></tr>
<tr><td>T30902</td><td>ACCGTTTGGTGAGCTGGTCGCGG</td><td>1000</td><td></td><td></td><td></td><td></td><td></td><td></td><td></td><td></td><td></td></tr>
<tr><td>Oligo fwd</td><td>CACCACCGTTTGGTGAGCTGGTCG</td><td></td><td></td><td></td><td></td><td></td><td></td><td></td><td></td><td></td><td></td></tr>
<tr><td>Oligo rev</td><td>AAACCGACCAGCTCACCAAACGGT</td><td></td><td></td><td></td><td></td><td></td><td></td><td></td><td></td><td></td><td></td></tr>
<tr><td>Chromosome</td><td>start</td><td>end</td><td>strand</td><td>MM</td><td>target_seq</td><td>PAM</td><td>alignment</td><td>distance</td><td>position</td><td>gene name</td><td>gene id</td></tr>
<tr><td>GU305914.1</td><td>44192</td><td>44214</td><td>-</td><td>0</td><td>ACCGTTTGGTGAGCTGGTCG</td><td>CGG</td><td>||||||||[||||||||||||]PAM</td><td>NA</td><td>Intergenic</td><td>NA</td><td>NA</td></tr>
<tr><td>T30903</td><td>GTTTGGTGAGCTGGTCGCGGAGG</td><td>1000</td><td></td><td></td><td></td><td></td><td></td><td></td><td></td><td></td><td></td></tr>
<tr><td>Oligo fwd</td><td>CACCGTTTGGTGAGCTGGTCGCGG</td><td></td><td></td><td></td><td></td><td></td><td></td><td></td><td></td><td></td><td></td></tr>
<tr><td>Oligo rev</td><td>AAACCCGCGACCAGCTCACCAAAC</td><td></td><td></td><td></td><td></td><td></td><td></td><td></td><td></td><td></td><td></td></tr>
<tr><td>Chromosome</td><td>start</td><td>end</td><td>strand</td><td>MM</td><td>target_seq</td><td>PAM</td><td>alignment</td><td>distance</td><td>position</td><td>gene name</td><td>gene id</td></tr>
<tr><td>GU305914.1</td><td>44189</td><td>44211</td><td>-</td><td>0</td><td>GTTTGGTGAGCTGGTCGCGG</td><td>AGG</td><td>||||||||[||||||||||||]PAM</td><td>NA</td><td>Intergenic</td><td>NA</td><td>NA</td></tr>
<tr><td>T30904</td><td>GAGCTGGTCGCGGAGGTAGTTGG</td><td>1000</td><td></td><td></td><td></td><td></td><td></td><td></td><td></td><td></td><td></td></tr>
<tr><td>Oligo fwd</td><td>CACCGAGCTGGTCGCGGAGGTAGT</td><td></td><td></td><td></td><td></td><td></td><td></td><td></td><td></td><td></td><td></td></tr>
<tr><td>Oligo rev</td><td>AAACACTACCTCCGCGACCAGCTC</td><td></td><td></td><td></td><td></td><td></td><td></td><td></td><td></td><td></td><td></td></tr>
<tr><td>Chromosome</td><td>start</td><td>end</td><td>strand</td><td>MM</td><td>target_seq</td><td>PAM</td><td>alignment</td><td>distance</td><td>position</td><td>gene name</td><td>gene id</td></tr>
<tr><td>GU305914.1</td><td>44182</td><td>44204</td><td>-</td><td>0</td><td>GAGCTGGTCGCGGAGGTAGT</td><td>TGG</td><td>||||||||[||||||||||||]PAM</td><td>NA</td><td>Intergenic</td><td>NA</td><td>NA</td></tr>
<tr><td>T30905</td><td>GCGGAGGTAGTTGGAGAACCTGG</td><td>1000</td><td></td><td></td><td></td><td></td><td></td><td></td><td></td><td></td><td></td></tr>
<tr><td>Oligo fwd</td><td>CACCGCGGAGGTAGTTGGAGAACC</td><td></td><td></td><td></td><td></td><td></td><td></td><td></td><td></td><td></td><td></td></tr>
<tr><td>Oligo rev</td><td>AAACGGTTCTCCAACTACCTCCGC</td><td></td><td></td><td></td><td></td><td></td><td></td><td></td><td></td><td></td><td></td></tr>
<tr><td>Chromosome</td><td>start</td><td>end</td><td>strand</td><td>MM</td><td>target_seq</td><td>PAM</td><td>alignment</td><td>distance</td><td>position</td><td>gene name</td><td>gene id</td></tr>
<tr><td>GU305914.1</td><td>44173</td><td>44195</td><td>-</td><td>0</td><td>GCGGAGGTAGTTGGAGAACC</td><td>TGG</td><td>||||||||[||||||||||||]PAM</td><td>NA</td><td>Intergenic</td><td>NA</td><td>NA</td></tr>
<tr><td>T30906</td><td>CGCGCGCCACGTACGTGATGCGG</td><td>1000</td><td></td><td></td><td></td><td></td><td></td><td></td><td></td><td></td><td></td></tr>
<tr><td>Oligo fwd</td><td>CACCCGCGCGCCACGTACGTGATG</td><td></td><td></td><td></td><td></td><td></td><td></td><td></td><td></td><td></td><td></td></tr>
<tr><td>Oligo rev</td><td>AAACCATCACGTACGTGGCGCGCG</td><td></td><td></td><td></td><td></td><td></td><td></td><td></td><td></td><td></td><td></td></tr>
<tr><td>Chromosome</td><td>start</td><td>end</td><td>strand</td><td>MM</td><td>target_seq</td><td>PAM</td><td>alignment</td><td>distance</td><td>position</td><td>gene name</td><td>gene id</td></tr>
<tr><td>GU305914.1</td><td>44126</td><td>44148</td><td>-</td><td>0</td><td>CGCGCGCCACGTACGTGATG</td><td>CGG</td><td>||||||||[||||||||||||]PAM</td><td>NA</td><td>Intergenic</td><td>NA</td><td>NA</td></tr>
<tr><td>T30907</td><td>CACGTACGTGATGCGGTTGAAGG</td><td>1000</td><td></td><td></td><td></td><td></td><td></td><td></td><td></td><td></td><td></td></tr>
<tr><td>Oligo fwd</td><td>CACCCACGTACGTGATGCGGTTGA</td><td></td><td></td><td></td><td></td><td></td><td></td><td></td><td></td><td></td><td></td></tr>
<tr><td>Oligo rev</td><td>AAACTCAACCGCATCACGTACGTG</td><td></td><td></td><td></td><td></td><td></td><td></td><td></td><td></td><td></td><td></td></tr>
<tr><td>Chromosome</td><td>start</td><td>end</td><td>strand</td><td>MM</td><td>target_seq</td><td>PAM</td><td>alignment</td><td>distance</td><td>position</td><td>gene name</td><td>gene id</td></tr>
<tr><td>GU305914.1</td><td>44119</td><td>44141</td><td>-</td><td>0</td><td>CACGTACGTGATGCGGTTGA</td><td>AGG</td><td>||||||||[||||||||||||]PAM</td><td>NA</td><td>Intergenic</td><td>NA</td><td>NA</td></tr>
<tr><td>T30908</td><td>CGTGATGCGGTTGAAGGAGATGG</td><td>1000</td><td></td><td></td><td></td><td></td><td></td><td></td><td></td><td></td><td></td></tr>
<tr><td>Oligo fwd</td><td>CACCCGTGATGCGGTTGAAGGAGA</td><td></td><td></td><td></td><td></td><td></td><td></td><td></td><td></td><td></td><td></td></tr>
<tr><td>Oligo rev</td><td>AAACTCTCCTTCAACCGCATCACG</td><td></td><td></td><td></td><td></td><td></td><td></td><td></td><td></td><td></td><td></td></tr>
<tr><td>Chromosome</td><td>start</td><td>end</td><td>strand</td><td>MM</td><td>target_seq</td><td>PAM</td><td>alignment</td><td>distance</td><td>position</td><td>gene name</td><td>gene id</td></tr>
<tr><td>GU305914.1</td><td>44113</td><td>44135</td><td>-</td><td>0</td><td>CGTGATGCGGTTGAAGGAGA</td><td>TGG</td><td>||||||||[||||||||||||]PAM</td><td>NA</td><td>Intergenic</td><td>NA</td><td>NA</td></tr>
<tr><td>T30909</td><td>TGAAGGAGATGGCGATGATGCGG</td><td>1000</td><td></td><td></td><td></td><td></td><td></td><td></td><td></td><td></td><td></td></tr>
<tr><td>Oligo fwd</td><td>CACCTGAAGGAGATGGCGATGATG</td><td></td><td></td><td></td><td></td><td></td><td></td><td></td><td></td><td></td><td></td></tr>
<tr><td>Oligo rev</td><td>AAACCATCATCGCCATCTCCTTCA</td><td></td><td></td><td></td><td></td><td></td><td></td><td></td><td></td><td></td><td></td></tr>
<tr><td>Chromosome</td><td>start</td><td>end</td><td>strand</td><td>MM</td><td>target_seq</td><td>PAM</td><td>alignment</td><td>distance</td><td>position</td><td>gene name</td><td>gene id</td></tr>
<tr><td>GU305914.1</td><td>44102</td><td>44124</td><td>-</td><td>0</td><td>TGAAGGAGATGGCGATGATG</td><td>CGG</td><td>||||||||[||||||||||||]PAM</td><td>NA</td><td>Intergenic</td><td>NA</td><td>NA</td></tr>
<tr><td>T30910</td><td>GGAGATGGCGATGATGCGGATGG</td><td>1000</td><td></td><td></td><td></td><td></td><td></td><td></td><td></td><td></td><td></td></tr>
<tr><td>Oligo fwd</td><td>CACCGGAGATGGCGATGATGCGGA</td><td></td><td></td><td></td><td></td><td></td><td></td><td></td><td></td><td></td><td></td></tr>
<tr><td>Oligo rev</td><td>AAACTCCGCATCATCGCCATCTCC</td><td></td><td></td><td></td><td></td><td></td><td></td><td></td><td></td><td></td><td></td></tr>
<tr><td>Chromosome</td><td>start</td><td>end</td><td>strand</td><td>MM</td><td>target_seq</td><td>PAM</td><td>alignment</td><td>distance</td><td>position</td><td>gene name</td><td>gene id</td></tr>
<tr><td>GU305914.1</td><td>44098</td><td>44120</td><td>-</td><td>0</td><td>GGAGATGGCGATGATGCGGA</td><td>TGG</td><td>||||||||[||||||||||||]PAM</td><td>NA</td><td>Intergenic</td><td>NA</td><td>NA</td></tr>
<tr><td>T30911</td><td>TGGCGATGATGCGGATGGCGCGG</td><td>1000</td><td></td><td></td><td></td><td></td><td></td><td></td><td></td><td></td><td></td></tr>
<tr><td>Oligo fwd</td><td>CACCTGGCGATGATGCGGATGGCG</td><td></td><td></td><td></td><td></td><td></td><td></td><td></td><td></td><td></td><td></td></tr>
<tr><td>Oligo rev</td><td>AAACCGCCATCCGCATCATCGCCA</td><td></td><td></td><td></td><td></td><td></td><td></td><td></td><td></td><td></td><td></td></tr>
<tr><td>Chromosome</td><td>start</td><td>end</td><td>strand</td><td>MM</td><td>target_seq</td><td>PAM</td><td>alignment</td><td>distance</td><td>position</td><td>gene name</td><td>gene id</td></tr>
<tr><td>GU305914.1</td><td>44093</td><td>44115</td><td>-</td><td>0</td><td>TGGCGATGATGCGGATGGCG</td><td>CGG</td><td>||||||||[||||||||||||]PAM</td><td>NA</td><td>Intergenic</td><td>NA</td><td>NA</td></tr>
<tr><td>T30912</td><td>TCGCCACCTGCTGTTCGCTCAGG</td><td>1000</td><td></td><td></td><td></td><td></td><td></td><td></td><td></td><td></td><td></td></tr>
<tr><td>Oligo fwd</td><td>CACCTCGCCACCTGCTGTTCGCTC</td><td></td><td></td><td></td><td></td><td></td><td></td><td></td><td></td><td></td><td></td></tr>
<tr><td>Oligo rev</td><td>AAACGAGCGAACAGCAGGTGGCGA</td><td></td><td></td><td></td><td></td><td></td><td></td><td></td><td></td><td></td><td></td></tr>
<tr><td>Chromosome</td><td>start</td><td>end</td><td>strand</td><td>MM</td><td>target_seq</td><td>PAM</td><td>alignment</td><td>distance</td><td>position</td><td>gene name</td><td>gene id</td></tr>
<tr><td>GU305914.1</td><td>44066</td><td>44088</td><td>-</td><td>0</td><td>TCGCCACCTGCTGTTCGCTC</td><td>AGG</td><td>||||||||[||||||||||||]PAM</td><td>NA</td><td>Intergenic</td><td>NA</td><td>NA</td></tr>
<tr><td>T30913</td><td>ACCTGCTGTTCGCTCAGGTTCGG</td><td>1000</td><td></td><td></td><td></td><td></td><td></td><td></td><td></td><td></td><td></td></tr>
<tr><td>Oligo fwd</td><td>CACCACCTGCTGTTCGCTCAGGTT</td><td></td><td></td><td></td><td></td><td></td><td></td><td></td><td></td><td></td><td></td></tr>
</table>

Oligo rev AAACAACCTGAGCGAACAGCAGGT
Chromosome start end strand MM target_seq PAM alignment distance position gene name gene id
GU305914.1 44061 44083 - 0 ACCTGCTGTTCGCTCAGGTT CGG ||||||||[||||||||||||]PAM NA Intergenic NA NA

T30914 CCTGCTGTTCGCTCAGGTTCGGG 1000
Oligo fwd CACCCCTGCTGTTCGCTCAGGTTC
Oligo rev AAACGAACCTGAGCGAACAGCAGG
Chromosome start end strand MM target_seq PAM alignment distance position gene name gene id
GU305914.1 44060 44082 - 0 CCTGCTGTTCGCTCAGGTTC GGG ||||||||[||||||||||||]PAM NA Intergenic NA NA

T30915 CTGCTGTTCGCTCAGGTTCGGGG 1000
Oligo fwd CACCCTGCTGTTCGCTCAGGTTCG
Oligo rev AAACCGAACCTGAGCGAACAGCAG
Chromosome start end strand MM target_seq PAM alignment distance position gene name gene id
GU305914.1 44059 44081 - 0 CTGCTGTTCGCTCAGGTTCG GGG ||||||||[||||||||||||]PAM NA Intergenic NA NA

T30916 TCGCTCAGGTTCGGGGAGATCGG 1000
Oligo fwd CACCTCGCTCAGGTTCGGGGAGAT
Oligo rev AAACATCTCCCCGAACCTGAGCGA
Chromosome start end strand MM target_seq PAM alignment distance position gene name gene id
GU305914.1 44052 44074 - 0 TCGCTCAGGTTCGGGGAGAT CGG ||||||||[||||||||||||]PAM NA Intergenic NA NA

T30917 CGCTCAGGTTCGGGGAGATCGGG 1000
Oligo fwd CACCCGCTCAGGTTCGGGGAGATC
Oligo rev AAACGATCTCCCCGAACCTGAGCG
Chromosome start end strand MM target_seq PAM alignment distance position gene name gene id
GU305914.1 44051 44073 - 0 CGCTCAGGTTCGGGGAGATC GGG ||||||||[||||||||||||]PAM NA Intergenic NA NA

T30918 TGATCCTGCAGCTCACCCCGTGG 1000
Oligo fwd CACCTGATCCTGCAGCTCACCCCG
Oligo rev AAACCGGGGTGAGCTGCAGGATCA
Chromosome start end strand MM target_seq PAM alignment distance position gene name gene id
GU305914.1 44021 44043 - 0 TGATCCTGCAGCTCACCCCG TGG ||||||||[||||||||||||]PAM NA Intergenic NA NA

T30919 AGCTCACCCCGTGGATCGCGCGG 1000
Oligo fwd CACCAGCTCACCCCGTGGATCGCG
Oligo rev AAACCGCGATCCACGGGGTGAGCT
Chromosome start end strand MM target_seq PAM alignment distance position gene name gene id
GU305914.1 44012 44034 - 0 AGCTCACCCCGTGGATCGCG CGG ||||||||[||||||||||||]PAM NA Intergenic NA NA

T30920 TGAAATCCGTGATGTGTTTCTGG 1000
Oligo fwd CACCTGAAATCCGTGATGTGTTTC
Oligo rev AAACGAAACACATCACGGATTTCA
Chromosome start end strand MM target_seq PAM alignment distance position gene name gene id
GU305914.1 43979 44001 - 0 TGAAATCCGTGATGTGTTTC TGG ||||||||[||||||||||||]PAM NA Intergenic NA NA

T30921 AATCCGTGATGTGTTTCTGGTGG 1000
Oligo fwd CACCAATCCGTGATGTGTTTCTGG
Oligo rev AAACCCAGAAACACATCACGGATT
Chromosome start end strand MM target_seq PAM alignment distance position gene name gene id
GU305914.1 43976 43998 - 0 AATCCGTGATGTGTTTCTGG TGG ||||||||[||||||||||||]PAM NA Intergenic NA NA

T30922 CCGTGATGTGTTTCTGGTGGCGG 1000
Oligo fwd CACCCCGTGATGTGTTTCTGGTGG
Oligo rev AAACCCACCAGAAACACATCACGG
Chromosome start end strand MM target_seq PAM alignment distance position gene name gene id
GU305914.1 43973 43995 - 0 CCGTGATGTGTTTCTGGTGG CGG ||||||||[||||||||||||]PAM NA Intergenic NA NA

T30923 TCGACGCGATCTCCGACTCCAGG 1000
Oligo fwd CACCTCGACGCGATCTCCGACTCC
Oligo rev AAACGGAGTCGGAGATCGCGTCGA
Chromosome start end strand MM target_seq PAM alignment distance position gene name gene id
GU305914.1 43946 43968 - 0 TCGACGCGATCTCCGACTCC AGG ||||||||[||||||||||||]PAM NA Intergenic NA NA

T30924 CGTGCAGAAGATGAACGAGTTGG 1000
Oligo fwd CACCCGTGCAGAAGATGAACGAGT
Oligo rev AAACACTCGTTCATCTTCTGCACG
Chromosome start end strand MM target_seq PAM alignment distance position gene name gene id
GU305914.1 43906 43928 - 0 CGTGCAGAAGATGAACGAGT TGG ||||||||[||||||||||||]PAM NA Intergenic NA NA

T30925 GAACGAGTTGGCGATCGACTTGG 1000
Oligo fwd CACCGAACGAGTTGGCGATCGACT
Oligo rev AAACAGTCGATCGCCAACTCGTTC
Chromosome start end strand MM target_seq PAM alignment distance position gene name gene id
GU305914.1 43894 43916 - 0 GAACGAGTTGGCGATCGACT TGG ||||||||[||||||||||||]PAM NA Intergenic NA NA

T30926 GACTTGGACTTGATGCACGAAGG 1000
Oligo fwd CACCGACTTGGACTTGATGCACGA
Oligo rev AAACTCGTGCATCAAGTCCAAGTC
Chromosome start end strand MM target_seq PAM alignment distance position gene name gene id
GU305914.1 43878 43900 - 0 GACTTGGACTTGATGCACGA AGG ||||||||[||||||||||||]PAM NA Intergenic NA NA

T30927 ACTTGATGCACGAAGGCAGCTGG 1000
Oligo fwd CACCACTTGATGCACGAAGGCAGC
Oligo rev AAACGCTGCCTTCGTGCATCAAGT
Chromosome start end strand MM target_seq PAM alignment distance position gene name gene id
GU305914.1 43871 43893 - 0 ACTTGATGCACGAAGGCAGC TGG ||||||||[||||||||||||]PAM NA Intergenic NA NA

T30928 GATGCACGAAGGCAGCTGGTTGG 1000

<table>
<tr><td>Oligo fwd</td><td>CACCGATGCACGAAGGCAGCTGGT</td><td></td><td></td><td></td><td></td><td></td><td></td><td></td><td></td><td></td><td></td></tr>
<tr><td>Oligo rev</td><td>AAACACCAGCTGCCTTCGTGCATC</td><td></td><td></td><td></td><td></td><td></td><td></td><td></td><td></td><td></td><td></td></tr>
<tr><td>Chromosome</td><td>start</td><td>end</td><td>strand</td><td>MM</td><td>target_seq</td><td>PAM</td><td>alignment</td><td>distance</td><td>position</td><td>gene name</td><td>gene id</td></tr>
<tr><td>GU305914.1</td><td>43867</td><td>43889</td><td>-</td><td>0</td><td>GATGCACGAAGGCAGCTGGT</td><td>TGG</td><td>||||||||[||||||||||||]PAM</td><td>NA</td><td>Intergenic</td><td>NA</td><td>NA</td></tr>
<tr><td>T30929</td><td>CGCTCGATTCATCATCTTGTAGG</td><td>1000</td><td></td><td></td><td></td><td></td><td></td><td></td><td></td><td></td><td></td></tr>
<tr><td>Oligo fwd</td><td>CACCCGCTCGATTCATCATCTTGT</td><td></td><td></td><td></td><td></td><td></td><td></td><td></td><td></td><td></td><td></td></tr>
<tr><td>Oligo rev</td><td>AAACACAAGATGATGAATCGAGCG</td><td></td><td></td><td></td><td></td><td></td><td></td><td></td><td></td><td></td><td></td></tr>
<tr><td>Chromosome</td><td>start</td><td>end</td><td>strand</td><td>MM</td><td>target_seq</td><td>PAM</td><td>alignment</td><td>distance</td><td>position</td><td>gene name</td><td>gene id</td></tr>
<tr><td>GU305914.1</td><td>43822</td><td>43844</td><td>-</td><td>0</td><td>CGCTCGATTCATCATCTTGT</td><td>AGG</td><td>||||||||[||||||||||||]PAM</td><td>NA</td><td>Intergenic</td><td>NA</td><td>NA</td></tr>
<tr><td>T30930</td><td>ATTCATCATCTTGTAGGTCCCGG</td><td>1000</td><td></td><td></td><td></td><td></td><td></td><td></td><td></td><td></td><td></td></tr>
<tr><td>Oligo fwd</td><td>CACCATTCATCATCTTGTAGGTCC</td><td></td><td></td><td></td><td></td><td></td><td></td><td></td><td></td><td></td><td></td></tr>
<tr><td>Oligo rev</td><td>AAACGGACCTACAAGATGATGAAT</td><td></td><td></td><td></td><td></td><td></td><td></td><td></td><td></td><td></td><td></td></tr>
<tr><td>Chromosome</td><td>start</td><td>end</td><td>strand</td><td>MM</td><td>target_seq</td><td>PAM</td><td>alignment</td><td>distance</td><td>position</td><td>gene name</td><td>gene id</td></tr>
<tr><td>GU305914.1</td><td>43816</td><td>43838</td><td>-</td><td>0</td><td>ATTCATCATCTTGTAGGTCC</td><td>CGG</td><td>||||||||[||||||||||||]PAM</td><td>NA</td><td>Intergenic</td><td>NA</td><td>NA</td></tr>
<tr><td>GU305914.1</td><td>40461</td><td>40483</td><td>+</td><td>4</td><td>CTTCTTCATCTTCGAGGTCC</td><td>AGG</td><td>-|||-|||[||||--||||||]PAM</td><td>NA</td><td>Intergenic</td><td>NA</td><td>NA</td></tr>
<tr><td>T30931</td><td>CATCTTGTAGGTCCCGGACACGG</td><td>1000</td><td></td><td></td><td></td><td></td><td></td><td></td><td></td><td></td><td></td></tr>
<tr><td>Oligo fwd</td><td>CACCCATCTTGTAGGTCCCGGACA</td><td></td><td></td><td></td><td></td><td></td><td></td><td></td><td></td><td></td><td></td></tr>
<tr><td>Oligo rev</td><td>AAACTGTCCGGGACCTACAAGATG</td><td></td><td></td><td></td><td></td><td></td><td></td><td></td><td></td><td></td><td></td></tr>
<tr><td>Chromosome</td><td>start</td><td>end</td><td>strand</td><td>MM</td><td>target_seq</td><td>PAM</td><td>alignment</td><td>distance</td><td>position</td><td>gene name</td><td>gene id</td></tr>
<tr><td>GU305914.1</td><td>43810</td><td>43832</td><td>-</td><td>0</td><td>CATCTTGTAGGTCCCGGACA</td><td>CGG</td><td>||||||||[||||||||||||]PAM</td><td>NA</td><td>Intergenic</td><td>NA</td><td>NA</td></tr>
<tr><td>T30932</td><td>TGTAGGTCCCGGACACGGAGCGG</td><td>1000</td><td></td><td></td><td></td><td></td><td></td><td></td><td></td><td></td><td></td></tr>
<tr><td>Oligo fwd</td><td>CACCTGTAGGTCCCGGACACGGAG</td><td></td><td></td><td></td><td></td><td></td><td></td><td></td><td></td><td></td><td></td></tr>
<tr><td>Oligo rev</td><td>AAACCTCCGTGTCCGGGACCTACA</td><td></td><td></td><td></td><td></td><td></td><td></td><td></td><td></td><td></td><td></td></tr>
<tr><td>Chromosome</td><td>start</td><td>end</td><td>strand</td><td>MM</td><td>target_seq</td><td>PAM</td><td>alignment</td><td>distance</td><td>position</td><td>gene name</td><td>gene id</td></tr>
<tr><td>GU305914.1</td><td>43805</td><td>43827</td><td>-</td><td>0</td><td>TGTAGGTCCCGGACACGGAG</td><td>CGG</td><td>||||||||[||||||||||||]PAM</td><td>NA</td><td>Intergenic</td><td>NA</td><td>NA</td></tr>
<tr><td>T30933</td><td>CACGGAGCGGATGAAACCCTCGG</td><td>1000</td><td></td><td></td><td></td><td></td><td></td><td></td><td></td><td></td><td></td></tr>
<tr><td>Oligo fwd</td><td>CACCCACGGAGCGGATGAAACCCT</td><td></td><td></td><td></td><td></td><td></td><td></td><td></td><td></td><td></td><td></td></tr>
<tr><td>Oligo rev</td><td>AAACAGGGTTTCATCCGCTCCGTG</td><td></td><td></td><td></td><td></td><td></td><td></td><td></td><td></td><td></td><td></td></tr>
<tr><td>Chromosome</td><td>start</td><td>end</td><td>strand</td><td>MM</td><td>target_seq</td><td>PAM</td><td>alignment</td><td>distance</td><td>position</td><td>gene name</td><td>gene id</td></tr>
<tr><td>GU305914.1</td><td>43792</td><td>43814</td><td>-</td><td>0</td><td>CACGGAGCGGATGAAACCCT</td><td>CGG</td><td>||||||||[||||||||||||]PAM</td><td>NA</td><td>Intergenic</td><td>NA</td><td>NA</td></tr>
<tr><td>T30934</td><td>TCGTACCTCACGATCTTCGACGG</td><td>1000</td><td></td><td></td><td></td><td></td><td></td><td></td><td></td><td></td><td></td></tr>
<tr><td>Oligo fwd</td><td>CACCTCGTACCTCACGATCTTCGA</td><td></td><td></td><td></td><td></td><td></td><td></td><td></td><td></td><td></td><td></td></tr>
<tr><td>Oligo rev</td><td>AAACTCGAAGATCGTGAGGTACGA</td><td></td><td></td><td></td><td></td><td></td><td></td><td></td><td></td><td></td><td></td></tr>
<tr><td>Chromosome</td><td>start</td><td>end</td><td>strand</td><td>MM</td><td>target_seq</td><td>PAM</td><td>alignment</td><td>distance</td><td>position</td><td>gene name</td><td>gene id</td></tr>
<tr><td>GU305914.1</td><td>43746</td><td>43768</td><td>-</td><td>0</td><td>TCGTACCTCACGATCTTCGA</td><td>CGG</td><td>||||||||[||||||||||||]PAM</td><td>NA</td><td>Intergenic</td><td>NA</td><td>NA</td></tr>
<tr><td>T30935</td><td>CCTCACGATCTTCGACGGCGCGG</td><td>1000</td><td></td><td></td><td></td><td></td><td></td><td></td><td></td><td></td><td></td></tr>
<tr><td>Oligo fwd</td><td>CACCCCTCACGATCTTCGACGGCG</td><td></td><td></td><td></td><td></td><td></td><td></td><td></td><td></td><td></td><td></td></tr>
<tr><td>Oligo rev</td><td>AAACCGCCGTCGAAGATCGTGAGG</td><td></td><td></td><td></td><td></td><td></td><td></td><td></td><td></td><td></td><td></td></tr>
<tr><td>Chromosome</td><td>start</td><td>end</td><td>strand</td><td>MM</td><td>target_seq</td><td>PAM</td><td>alignment</td><td>distance</td><td>position</td><td>gene name</td><td>gene id</td></tr>
<tr><td>GU305914.1</td><td>43741</td><td>43763</td><td>-</td><td>0</td><td>CCTCACGATCTTCGACGGCG</td><td>CGG</td><td>||||||||[||||||||||||]PAM</td><td>NA</td><td>Intergenic</td><td>NA</td><td>NA</td></tr>
<tr><td>T30936</td><td>ATCTTCGACGGCGCGGACGACGG</td><td>1000</td><td></td><td></td><td></td><td></td><td></td><td></td><td></td><td></td><td></td></tr>
<tr><td>Oligo fwd</td><td>CACCATCTTCGACGGCGCGGACGA</td><td></td><td></td><td></td><td></td><td></td><td></td><td></td><td></td><td></td><td></td></tr>
<tr><td>Oligo rev</td><td>AAACTCGTCCGCGCCGTCGAAGAT</td><td></td><td></td><td></td><td></td><td></td><td></td><td></td><td></td><td></td><td></td></tr>
<tr><td>Chromosome</td><td>start</td><td>end</td><td>strand</td><td>MM</td><td>target_seq</td><td>PAM</td><td>alignment</td><td>distance</td><td>position</td><td>gene name</td><td>gene id</td></tr>
<tr><td>GU305914.1</td><td>43734</td><td>43756</td><td>-</td><td>0</td><td>ATCTTCGACGGCGCGGACGA</td><td>CGG</td><td>||||||||[||||||||||||]PAM</td><td>NA</td><td>Intergenic</td><td>NA</td><td>NA</td></tr>
<tr><td>T30937</td><td>ACGACGGCATTGTCGTTTCGCGG</td><td>1000</td><td></td><td></td><td></td><td></td><td></td><td></td><td></td><td></td><td></td></tr>
<tr><td>Oligo fwd</td><td>CACCACGACGGCATTGTCGTTTCG</td><td></td><td></td><td></td><td></td><td></td><td></td><td></td><td></td><td></td><td></td></tr>
<tr><td>Oligo rev</td><td>AAACCGAAACGACAATGCCGTCGT</td><td></td><td></td><td></td><td></td><td></td><td></td><td></td><td></td><td></td><td></td></tr>
<tr><td>Chromosome</td><td>start</td><td>end</td><td>strand</td><td>MM</td><td>target_seq</td><td>PAM</td><td>alignment</td><td>distance</td><td>position</td><td>gene name</td><td>gene id</td></tr>
<tr><td>GU305914.1</td><td>43718</td><td>43740</td><td>-</td><td>0</td><td>ACGACGGCATTGTCGTTTCG</td><td>CGG</td><td>||||||||[||||||||||||]PAM</td><td>NA</td><td>Intergenic</td><td>NA</td><td>NA</td></tr>
<tr><td>T30938</td><td>GTCGTTTCGCGGTGATGCGACGG</td><td>1000</td><td></td><td></td><td></td><td></td><td></td><td></td><td></td><td></td><td></td></tr>
<tr><td>Oligo fwd</td><td>CACCGTCGTTTCGCGGTGATGCGA</td><td></td><td></td><td></td><td></td><td></td><td></td><td></td><td></td><td></td><td></td></tr>
<tr><td>Oligo rev</td><td>AAACTCGCATCACCGCGAAACGAC</td><td></td><td></td><td></td><td></td><td></td><td></td><td></td><td></td><td></td><td></td></tr>
<tr><td>Chromosome</td><td>start</td><td>end</td><td>strand</td><td>MM</td><td>target_seq</td><td>PAM</td><td>alignment</td><td>distance</td><td>position</td><td>gene name</td><td>gene id</td></tr>
<tr><td>GU305914.1</td><td>43707</td><td>43729</td><td>-</td><td>0</td><td>GTCGTTTCGCGGTGATGCGA</td><td>CGG</td><td>||||||||[||||||||||||]PAM</td><td>NA</td><td>Intergenic</td><td>NA</td><td>NA</td></tr>
<tr><td>T30939</td><td>GTTTCGCGGTGATGCGACGGCGG</td><td>1000</td><td></td><td></td><td></td><td></td><td></td><td></td><td></td><td></td><td></td></tr>
<tr><td>Oligo fwd</td><td>CACCGTTTCGCGGTGATGCGACGG</td><td></td><td></td><td></td><td></td><td></td><td></td><td></td><td></td><td></td><td></td></tr>
<tr><td>Oligo rev</td><td>AAACCCGTCGCATCACCGCGAAAC</td><td></td><td></td><td></td><td></td><td></td><td></td><td></td><td></td><td></td><td></td></tr>
<tr><td>Chromosome</td><td>start</td><td>end</td><td>strand</td><td>MM</td><td>target_seq</td><td>PAM</td><td>alignment</td><td>distance</td><td>position</td><td>gene name</td><td>gene id</td></tr>
<tr><td>GU305914.1</td><td>43704</td><td>43726</td><td>-</td><td>0</td><td>GTTTCGCGGTGATGCGACGG</td><td>CGG</td><td>||||||||[||||||||||||]PAM</td><td>NA</td><td>Intergenic</td><td>NA</td><td>NA</td></tr>
<tr><td>T30940</td><td>TCGCGGTGATGCGACGGCGGAGG</td><td>1000</td><td></td><td></td><td></td><td></td><td></td><td></td><td></td><td></td><td></td></tr>
<tr><td>Oligo fwd</td><td>CACCTCGCGGTGATGCGACGGCGG</td><td></td><td></td><td></td><td></td><td></td><td></td><td></td><td></td><td></td><td></td></tr>
<tr><td>Oligo rev</td><td>AAACCCGCCGTCGCATCACCGCGA</td><td></td><td></td><td></td><td></td><td></td><td></td><td></td><td></td><td></td><td></td></tr>
<tr><td>Chromosome</td><td>start</td><td>end</td><td>strand</td><td>MM</td><td>target_seq</td><td>PAM</td><td>alignment</td><td>distance</td><td>position</td><td>gene name</td><td>gene id</td></tr>
<tr><td>GU305914.1</td><td>43701</td><td>43723</td><td>-</td><td>0</td><td>TCGCGGTGATGCGACGGCGG</td><td>AGG</td><td>||||||||[||||||||||||]PAM</td><td>NA</td><td>Intergenic</td><td>NA</td><td>NA</td></tr>
<tr><td>GU305914.1</td><td>114673</td><td>114695</td><td>-</td><td>4</td><td>CCGCGATGGTGAGACGGCGG</td><td>TGG</td><td>-||||-||[-||-||||||||]PAM</td><td>NA</td><td>Intergenic</td><td>NA</td><td>NA</td></tr>
<tr><td>T30941</td><td>TGCGACGGCGGAGGCTGCGTTGG</td><td>1000</td><td></td><td></td><td></td><td></td><td></td><td></td><td></td><td></td><td></td></tr>
<tr><td>Oligo fwd</td><td>CACCTGCGACGGCGGAGGCTGCGT</td><td></td><td></td><td></td><td></td><td></td><td></td><td></td><td></td><td></td><td></td></tr>
<tr><td>Oligo rev</td><td>AAACACGCAGCCTCCGCCGTCGCA</td><td></td><td></td><td></td><td></td><td></td><td></td><td></td><td></td><td></td><td></td></tr>
<tr><td>Chromosome</td><td>start</td><td>end</td><td>strand</td><td>MM</td><td>target_seq</td><td>PAM</td><td>alignment</td><td>distance</td><td>position</td><td>gene name</td><td>gene id</td></tr>
<tr><td>GU305914.1</td><td>43692</td><td>43714</td><td>-</td><td>0</td><td>TGCGACGGCGGAGGCTGCGT</td><td>TGG</td><td>||||||||[||||||||||||]PAM</td><td>NA</td><td>Intergenic</td><td>NA</td><td>NA</td></tr>
<tr><td>T30942</td><td>GACGGCGGAGGCTGCGTTGGTGG</td><td>1000</td><td></td><td></td><td></td><td></td><td></td><td></td><td></td><td></td><td></td></tr>
<tr><td>Oligo fwd</td><td>CACCGACGGCGGAGGCTGCGTTGG</td><td></td><td></td><td></td><td></td><td></td><td></td><td></td><td></td><td></td><td></td></tr>
<tr><td>Oligo rev</td><td>AAACCCAACGCAGCCTCCGCCGTC</td><td></td><td></td><td></td><td></td><td></td><td></td><td></td><td></td><td></td><td></td></tr>
<tr><td>Chromosome</td><td>start</td><td>end</td><td>strand</td><td>MM</td><td>target_seq</td><td>PAM</td><td>alignment</td><td>distance</td><td>position</td><td>gene name</td><td>gene id</td></tr>
</table>

GU305914.1 43689 43711 - 0 GACGGCGGAGGCTGCGTTGG TGG ||||||||[||||||||||||]PAM NA Intergenic NA NA

T30943 CTGCGTTGGTGGTTCGTGTCTGG 1000
Oligo fwd CACCCTGCGTTGGTGGTTCGTGTC
Oligo rev AAACGACACGAACCACCAACGCAG
Chromosome start end strand MM target_seq PAM alignment distance position gene name gene id
GU305914.1 43678 43700 - 0 CTGCGTTGGTGGTTCGTGTC TGG ||||||||[||||||||||||]PAM NA Intergenic NA NA

T30944 CGCCGCCCTGTTGTTGTTGCTGG 1000
Oligo fwd CACCCGCCGCCCTGTTGTTGTTGC
Oligo rev AAACGCAACAACAACAGGGCGGCG
Chromosome start end strand MM target_seq PAM alignment distance position gene name gene id
GU305914.1 43622 43644 - 0 CGCCGCCCTGTTGTTGTTGC TGG ||||||||[||||||||||||]PAM NA Intergenic NA NA

T30945 CAGAGTAGCGTCTCTGCGCGCGG 1000
Oligo fwd CACCCAGAGTAGCGTCTCTGCGCG
Oligo rev AAACCGCGCAGAGACGCTACTCTG
Chromosome start end strand MM target_seq PAM alignment distance position gene name gene id
GU305914.1 43583 43605 - 0 CAGAGTAGCGTCTCTGCGCG CGG ||||||||[||||||||||||]PAM NA Intergenic NA NA

T30946 TAGCGTCTCTGCGCGCGGACCGG 1000
Oligo fwd CACCTAGCGTCTCTGCGCGCGGAC
Oligo rev AAACGTCCGCGCGCAGAGACGCTA
Chromosome start end strand MM target_seq PAM alignment distance position gene name gene id
GU305914.1 43578 43600 - 0 TAGCGTCTCTGCGCGCGGAC CGG ||||||||[||||||||||||]PAM NA Intergenic NA NA

T30947 AGCGTCTCTGCGCGCGGACCGGG 1000
Oligo fwd CACCAGCGTCTCTGCGCGCGGACC
Oligo rev AAACGGTCCGCGCGCAGAGACGCT
Chromosome start end strand MM target_seq PAM alignment distance position gene name gene id
GU305914.1 43577 43599 - 0 AGCGTCTCTGCGCGCGGACC GGG ||||||||[||||||||||||]PAM NA Intergenic NA NA

T30948 TCTCTGCGCGCGGACCGGGTCGG 1000
Oligo fwd CACCTCTCTGCGCGCGGACCGGGT
Oligo rev AAACACCCGGTCCGCGCGCAGAGA
Chromosome start end strand MM target_seq PAM alignment distance position gene name gene id
GU305914.1 43573 43595 - 0 TCTCTGCGCGCGGACCGGGT CGG ||||||||[||||||||||||]PAM NA Intergenic NA NA

T30949 CGTCTCTCCAAGTGATGTATCGG 1000
Oligo fwd CACCCGTCTCTCCAAGTGATGTAT
Oligo rev AAACATACATCACTTGGAGAGACG
Chromosome start end strand MM target_seq PAM alignment distance position gene name gene id
GU305914.1 43545 43567 - 0 CGTCTCTCCAAGTGATGTAT CGG ||||||||[||||||||||||]PAM NA Intergenic NA NA

T30950 CTCTCCAAGTGATGTATCGGCGG 1000
Oligo fwd CACCCTCTCCAAGTGATGTATCGG
Oligo rev AAACCCGATACATCACTTGGAGAG
Chromosome start end strand MM target_seq PAM alignment distance position gene name gene id
GU305914.1 43542 43564 - 0 CTCTCCAAGTGATGTATCGG CGG ||||||||[||||||||||||]PAM NA Intergenic NA NA

T30951 GTGATGTATCGGCGGCGCCGAGG 1000
Oligo fwd CACCGTGATGTATCGGCGGCGCCG
Oligo rev AAACCGGCGCCGCCGATACATCAC
Chromosome start end strand MM target_seq PAM alignment distance position gene name gene id
GU305914.1 43534 43556 - 0 GTGATGTATCGGCGGCGCCG AGG ||||||||[||||||||||||]PAM NA Intergenic NA NA

T30952 GGCGCCGAGGACACGTGCTGCGG 1000
Oligo fwd CACCGGCGCCGAGGACACGTGCTG
Oligo rev AAACCAGCACGTGTCCTCGGCGCC
Chromosome start end strand MM target_seq PAM alignment distance position gene name gene id
GU305914.1 43521 43543 - 0 GGCGCCGAGGACACGTGCTG CGG ||||||||[||||||||||||]PAM NA Intergenic NA NA

T30953 GCCGAGGACACGTGCTGCGGCGG 1000
Oligo fwd CACCGCCGAGGACACGTGCTGCGG
Oligo rev AAACCCGCAGCACGTGTCCTCGGC
Chromosome start end strand MM target_seq PAM alignment distance position gene name gene id
GU305914.1 43518 43540 - 0 GCCGAGGACACGTGCTGCGG CGG ||||||||[||||||||||||]PAM NA Intergenic NA NA

T30954 ACGTGCTGCGGCGGATGATGTGG 1000
Oligo fwd CACCACGTGCTGCGGCGGATGATG
Oligo rev AAACCATCATCCGCCGCAGCACGT
Chromosome start end strand MM target_seq PAM alignment distance position gene name gene id
GU305914.1 43509 43531 - 0 ACGTGCTGCGGCGGATGATG TGG ||||||||[||||||||||||]PAM NA Intergenic NA NA

T30955 TGGCTGCGACGACGACGATGAGG 1000
Oligo fwd CACCTGGCTGCGACGACGACGATG
Oligo rev AAACCATCGTCGTCGTCGCAGCCA
Chromosome start end strand MM target_seq PAM alignment distance position gene name gene id
GU305914.1 43489 43511 - 0 TGGCTGCGACGACGACGATG AGG ||||||||[||||||||||||]PAM NA Intergenic NA NA

T30956 ACGACGATGAGGAGTGTCGATGG 1000
Oligo fwd CACCACGACGATGAGGAGTGTCGA
Oligo rev AAACTCGACACTCCTCATCGTCGT
Chromosome start end strand MM target_seq PAM alignment distance position gene name gene id
GU305914.1 43478 43500 - 0 ACGACGATGAGGAGTGTCGA TGG ||||||||[||||||||||||]PAM NA Intergenic NA NA

T30957 GATGAGGAGTGTCGATGGTGAGG 1000
Oligo fwd CACCGATGAGGAGTGTCGATGGTG
Oligo rev AAACCACCATCGACACTCCTCATC

<table>
<tr><td>Chromosome</td><td>start</td><td>end</td><td>strand</td><td>MM</td><td>target_seq</td><td>PAM</td><td>alignment</td><td>distance</td><td>position</td><td>gene name</td><td>gene id</td></tr>
<tr><td>GU305914.1</td><td>43473</td><td>43495</td><td>-</td><td>0</td><td>GATGAGGAGTGTCGATGGTG</td><td>AGG</td><td>||||||||[||||||||||||]PAM</td><td>NA</td><td>Intergenic</td><td>NA</td><td>NA</td></tr>
<tr><td></td><td></td><td></td><td></td><td></td><td></td><td></td><td></td><td></td><td></td><td></td><td></td></tr>
<tr><td>T30958</td><td>GAGGAGTGTCGATGGTGAGGAGG</td><td>1000</td><td></td><td></td><td></td><td></td><td></td><td></td><td></td><td></td><td></td></tr>
<tr><td>Oligo fwd</td><td>CACCGAGGAGTGTCGATGGTGAGG</td><td></td><td></td><td></td><td></td><td></td><td></td><td></td><td></td><td></td><td></td></tr>
<tr><td>Oligo rev</td><td>AAACCCTCACCATCGACACTCCTC</td><td></td><td></td><td></td><td></td><td></td><td></td><td></td><td></td><td></td><td></td></tr>
<tr><td>Chromosome</td><td>start</td><td>end</td><td>strand</td><td>MM</td><td>target_seq</td><td>PAM</td><td>alignment</td><td>distance</td><td>position</td><td>gene name</td><td>gene id</td></tr>
<tr><td>GU305914.1</td><td>43470</td><td>43492</td><td>-</td><td>0</td><td>GAGGAGTGTCGATGGTGAGG</td><td>AGG</td><td>||||||||[||||||||||||]PAM</td><td>NA</td><td>Intergenic</td><td>NA</td><td>NA</td></tr>
<tr><td></td><td></td><td></td><td></td><td></td><td></td><td></td><td></td><td></td><td></td><td></td><td></td></tr>
<tr><td>T30959</td><td>AGTGTCGATGGTGAGGAGGATGG</td><td>1000</td><td></td><td></td><td></td><td></td><td></td><td></td><td></td><td></td><td></td></tr>
<tr><td>Oligo fwd</td><td>CACCAGTGTCGATGGTGAGGAGGA</td><td></td><td></td><td></td><td></td><td></td><td></td><td></td><td></td><td></td><td></td></tr>
<tr><td>Oligo rev</td><td>AAACTCCTCCTCACCATCGACACT</td><td></td><td></td><td></td><td></td><td></td><td></td><td></td><td></td><td></td><td></td></tr>
<tr><td>Chromosome</td><td>start</td><td>end</td><td>strand</td><td>MM</td><td>target_seq</td><td>PAM</td><td>alignment</td><td>distance</td><td>position</td><td>gene name</td><td>gene id</td></tr>
<tr><td>GU305914.1</td><td>43466</td><td>43488</td><td>-</td><td>0</td><td>AGTGTCGATGGTGAGGAGGA</td><td>TGG</td><td>||||||||[||||||||||||]PAM</td><td>NA</td><td>Intergenic</td><td>NA</td><td>NA</td></tr>
<tr><td></td><td></td><td></td><td></td><td></td><td></td><td></td><td></td><td></td><td></td><td></td><td></td></tr>
<tr><td>T30960</td><td>GGAGGATGGTGACGATCGAACGG</td><td>1000</td><td></td><td></td><td></td><td></td><td></td><td></td><td></td><td></td><td></td></tr>
<tr><td>Oligo fwd</td><td>CACCGGAGGATGGTGACGATCGAA</td><td></td><td></td><td></td><td></td><td></td><td></td><td></td><td></td><td></td><td></td></tr>
<tr><td>Oligo rev</td><td>AAACTTCGATCGTCACCATCCTCC</td><td></td><td></td><td></td><td></td><td></td><td></td><td></td><td></td><td></td><td></td></tr>
<tr><td>Chromosome</td><td>start</td><td>end</td><td>strand</td><td>MM</td><td>target_seq</td><td>PAM</td><td>alignment</td><td>distance</td><td>position</td><td>gene name</td><td>gene id</td></tr>
<tr><td>GU305914.1</td><td>43452</td><td>43474</td><td>-</td><td>0</td><td>GGAGGATGGTGACGATCGAA</td><td>CGG</td><td>||||||||[||||||||||||]PAM</td><td>NA</td><td>Intergenic</td><td>NA</td><td>NA</td></tr>
<tr><td></td><td></td><td></td><td></td><td></td><td></td><td></td><td></td><td></td><td></td><td></td><td></td></tr>
<tr><td>T30961</td><td>GAGGATGGTGACGATCGAACGGG</td><td>1000</td><td></td><td></td><td></td><td></td><td></td><td></td><td></td><td></td><td></td></tr>
<tr><td>Oligo fwd</td><td>CACCGAGGATGGTGACGATCGAAC</td><td></td><td></td><td></td><td></td><td></td><td></td><td></td><td></td><td></td><td></td></tr>
<tr><td>Oligo rev</td><td>AAACGTTCGATCGTCACCATCCTC</td><td></td><td></td><td></td><td></td><td></td><td></td><td></td><td></td><td></td><td></td></tr>
<tr><td>Chromosome</td><td>start</td><td>end</td><td>strand</td><td>MM</td><td>target_seq</td><td>PAM</td><td>alignment</td><td>distance</td><td>position</td><td>gene name</td><td>gene id</td></tr>
<tr><td>GU305914.1</td><td>43451</td><td>43473</td><td>-</td><td>0</td><td>GAGGATGGTGACGATCGAAC</td><td>GGG</td><td>||||||||[||||||||||||]PAM</td><td>NA</td><td>Intergenic</td><td>NA</td><td>NA</td></tr>
<tr><td></td><td></td><td></td><td></td><td></td><td></td><td></td><td></td><td></td><td></td><td></td><td></td></tr>
<tr><td>T30962</td><td>GATGGTGACGATCGAACGGGTGG</td><td>1000</td><td></td><td></td><td></td><td></td><td></td><td></td><td></td><td></td><td></td></tr>
<tr><td>Oligo fwd</td><td>CACCGATGGTGACGATCGAACGGG</td><td></td><td></td><td></td><td></td><td></td><td></td><td></td><td></td><td></td><td></td></tr>
<tr><td>Oligo rev</td><td>AAACCCCGTTCGATCGTCACCATC</td><td></td><td></td><td></td><td></td><td></td><td></td><td></td><td></td><td></td><td></td></tr>
<tr><td>Chromosome</td><td>start</td><td>end</td><td>strand</td><td>MM</td><td>target_seq</td><td>PAM</td><td>alignment</td><td>distance</td><td>position</td><td>gene name</td><td>gene id</td></tr>
<tr><td>GU305914.1</td><td>43448</td><td>43470</td><td>-</td><td>0</td><td>GATGGTGACGATCGAACGGG</td><td>TGG</td><td>||||||||[||||||||||||]PAM</td><td>NA</td><td>Intergenic</td><td>NA</td><td>NA</td></tr>
<tr><td></td><td></td><td></td><td></td><td></td><td></td><td></td><td></td><td></td><td></td><td></td><td></td></tr>
<tr><td>T30963</td><td>GTGACGATCGAACGGGTGGTCGG</td><td>1000</td><td></td><td></td><td></td><td></td><td></td><td></td><td></td><td></td><td></td></tr>
<tr><td>Oligo fwd</td><td>CACCGTGACGATCGAACGGGTGGT</td><td></td><td></td><td></td><td></td><td></td><td></td><td></td><td></td><td></td><td></td></tr>
<tr><td>Oligo rev</td><td>AAACACCACCCGTTCGATCGTCAC</td><td></td><td></td><td></td><td></td><td></td><td></td><td></td><td></td><td></td><td></td></tr>
<tr><td>Chromosome</td><td>start</td><td>end</td><td>strand</td><td>MM</td><td>target_seq</td><td>PAM</td><td>alignment</td><td>distance</td><td>position</td><td>gene name</td><td>gene id</td></tr>
<tr><td>GU305914.1</td><td>43444</td><td>43466</td><td>-</td><td>0</td><td>GTGACGATCGAACGGGTGGT</td><td>CGG</td><td>||||||||[||||||||||||]PAM</td><td>NA</td><td>Intergenic</td><td>NA</td><td>NA</td></tr>
<tr><td></td><td></td><td></td><td></td><td></td><td></td><td></td><td></td><td></td><td></td><td></td><td></td></tr>
<tr><td>T30964</td><td>TGACGATCGAACGGGTGGTCGGG</td><td>1000</td><td></td><td></td><td></td><td></td><td></td><td></td><td></td><td></td><td></td></tr>
<tr><td>Oligo fwd</td><td>CACCTGACGATCGAACGGGTGGTC</td><td></td><td></td><td></td><td></td><td></td><td></td><td></td><td></td><td></td><td></td></tr>
<tr><td>Oligo rev</td><td>AAACGACCACCCGTTCGATCGTCA</td><td></td><td></td><td></td><td></td><td></td><td></td><td></td><td></td><td></td><td></td></tr>
<tr><td>Chromosome</td><td>start</td><td>end</td><td>strand</td><td>MM</td><td>target_seq</td><td>PAM</td><td>alignment</td><td>distance</td><td>position</td><td>gene name</td><td>gene id</td></tr>
<tr><td>GU305914.1</td><td>43443</td><td>43465</td><td>-</td><td>0</td><td>TGACGATCGAACGGGTGGTC</td><td>GGG</td><td>||||||||[||||||||||||]PAM</td><td>NA</td><td>Intergenic</td><td>NA</td><td>NA</td></tr>
<tr><td></td><td></td><td></td><td></td><td></td><td></td><td></td><td></td><td></td><td></td><td></td><td></td></tr>
<tr><td>T30965</td><td>CCGCCGATACGTCGTCGACTCGG</td><td>1000</td><td></td><td></td><td></td><td></td><td></td><td></td><td></td><td></td><td></td></tr>
<tr><td>Oligo fwd</td><td>CACCCCGCCGATACGTCGTCGACT</td><td></td><td></td><td></td><td></td><td></td><td></td><td></td><td></td><td></td><td></td></tr>
<tr><td>Oligo rev</td><td>AAACAGTCGACGACGTATCGGCGG</td><td></td><td></td><td></td><td></td><td></td><td></td><td></td><td></td><td></td><td></td></tr>
<tr><td>Chromosome</td><td>start</td><td>end</td><td>strand</td><td>MM</td><td>target_seq</td><td>PAM</td><td>alignment</td><td>distance</td><td>position</td><td>gene name</td><td>gene id</td></tr>
<tr><td>GU305914.1</td><td>43420</td><td>43442</td><td>-</td><td>0</td><td>CCGCCGATACGTCGTCGACT</td><td>CGG</td><td>||||||||[||||||||||||]PAM</td><td>NA</td><td>Intergenic</td><td>NA</td><td>NA</td></tr>
<tr><td></td><td></td><td></td><td></td><td></td><td></td><td></td><td></td><td></td><td></td><td></td><td></td></tr>
<tr><td>T30966</td><td>TCGTCGACTCGGACGAGTGTCGG</td><td>1000</td><td></td><td></td><td></td><td></td><td></td><td></td><td></td><td></td><td></td></tr>
<tr><td>Oligo fwd</td><td>CACCTCGTCGACTCGGACGAGTGT</td><td></td><td></td><td></td><td></td><td></td><td></td><td></td><td></td><td></td><td></td></tr>
<tr><td>Oligo rev</td><td>AAACACACTCGTCCGAGTCGACGA</td><td></td><td></td><td></td><td></td><td></td><td></td><td></td><td></td><td></td><td></td></tr>
<tr><td>Chromosome</td><td>start</td><td>end</td><td>strand</td><td>MM</td><td>target_seq</td><td>PAM</td><td>alignment</td><td>distance</td><td>position</td><td>gene name</td><td>gene id</td></tr>
<tr><td>GU305914.1</td><td>43409</td><td>43431</td><td>-</td><td>0</td><td>TCGTCGACTCGGACGAGTGT</td><td>CGG</td><td>||||||||[||||||||||||]PAM</td><td>NA</td><td>Intergenic</td><td>NA</td><td>NA</td></tr>
<tr><td></td><td></td><td></td><td></td><td></td><td></td><td></td><td></td><td></td><td></td><td></td><td></td></tr>
<tr><td>T30967</td><td>CGTCGACTCGGACGAGTGTCGGG</td><td>1000</td><td></td><td></td><td></td><td></td><td></td><td></td><td></td><td></td><td></td></tr>
<tr><td>Oligo fwd</td><td>CACCCGTCGACTCGGACGAGTGTC</td><td></td><td></td><td></td><td></td><td></td><td></td><td></td><td></td><td></td><td></td></tr>
<tr><td>Oligo rev</td><td>AAACGACACTCGTCCGAGTCGACG</td><td></td><td></td><td></td><td></td><td></td><td></td><td></td><td></td><td></td><td></td></tr>
<tr><td>Chromosome</td><td>start</td><td>end</td><td>strand</td><td>MM</td><td>target_seq</td><td>PAM</td><td>alignment</td><td>distance</td><td>position</td><td>gene name</td><td>gene id</td></tr>
<tr><td>GU305914.1</td><td>43408</td><td>43430</td><td>-</td><td>0</td><td>CGTCGACTCGGACGAGTGTC</td><td>GGG</td><td>||||||||[||||||||||||]PAM</td><td>NA</td><td>Intergenic</td><td>NA</td><td>NA</td></tr>
<tr><td></td><td></td><td></td><td></td><td></td><td></td><td></td><td></td><td></td><td></td><td></td><td></td></tr>
<tr><td>T30968</td><td>GTCGACTCGGACGAGTGTCGGGG</td><td>1000</td><td></td><td></td><td></td><td></td><td></td><td></td><td></td><td></td><td></td></tr>
<tr><td>Oligo fwd</td><td>CACCGTCGACTCGGACGAGTGTCG</td><td></td><td></td><td></td><td></td><td></td><td></td><td></td><td></td><td></td><td></td></tr>
<tr><td>Oligo rev</td><td>AAACCGACACTCGTCCGAGTCGAC</td><td></td><td></td><td></td><td></td><td></td><td></td><td></td><td></td><td></td><td></td></tr>
<tr><td>Chromosome</td><td>start</td><td>end</td><td>strand</td><td>MM</td><td>target_seq</td><td>PAM</td><td>alignment</td><td>distance</td><td>position</td><td>gene name</td><td>gene id</td></tr>
<tr><td>GU305914.1</td><td>43407</td><td>43429</td><td>-</td><td>0</td><td>GTCGACTCGGACGAGTGTCG</td><td>GGG</td><td>||||||||[||||||||||||]PAM</td><td>NA</td><td>Intergenic</td><td>NA</td><td>NA</td></tr>
<tr><td></td><td></td><td></td><td></td><td></td><td></td><td></td><td></td><td></td><td></td><td></td><td></td></tr>
<tr><td>T30969</td><td>TCGGACGAGTGTCGGGGACGCGG</td><td>1000</td><td></td><td></td><td></td><td></td><td></td><td></td><td></td><td></td><td></td></tr>
<tr><td>Oligo fwd</td><td>CACCTCGGACGAGTGTCGGGGACG</td><td></td><td></td><td></td><td></td><td></td><td></td><td></td><td></td><td></td><td></td></tr>
<tr><td>Oligo rev</td><td>AAACCGTCCCCGACACTCGTCCGA</td><td></td><td></td><td></td><td></td><td></td><td></td><td></td><td></td><td></td><td></td></tr>
<tr><td>Chromosome</td><td>start</td><td>end</td><td>strand</td><td>MM</td><td>target_seq</td><td>PAM</td><td>alignment</td><td>distance</td><td>position</td><td>gene name</td><td>gene id</td></tr>
<tr><td>GU305914.1</td><td>43401</td><td>43423</td><td>-</td><td>0</td><td>TCGGACGAGTGTCGGGGACG</td><td>CGG</td><td>||||||||[||||||||||||]PAM</td><td>NA</td><td>Intergenic</td><td>NA</td><td>NA</td></tr>
<tr><td></td><td></td><td></td><td></td><td></td><td></td><td></td><td></td><td></td><td></td><td></td><td></td></tr>
<tr><td>T30970</td><td>AGTGTCGGGGACGCGGCGCGTGG</td><td>1000</td><td></td><td></td><td></td><td></td><td></td><td></td><td></td><td></td><td></td></tr>
<tr><td>Oligo fwd</td><td>CACCAGTGTCGGGGACGCGGCGCG</td><td></td><td></td><td></td><td></td><td></td><td></td><td></td><td></td><td></td><td></td></tr>
<tr><td>Oligo rev</td><td>AAACCGCGCCGCGTCCCCGACACT</td><td></td><td></td><td></td><td></td><td></td><td></td><td></td><td></td><td></td><td></td></tr>
<tr><td>Chromosome</td><td>start</td><td>end</td><td>strand</td><td>MM</td><td>target_seq</td><td>PAM</td><td>alignment</td><td>distance</td><td>position</td><td>gene name</td><td>gene id</td></tr>
<tr><td>GU305914.1</td><td>43394</td><td>43416</td><td>-</td><td>0</td><td>AGTGTCGGGGACGCGGCGCG</td><td>TGG</td><td>||||||||[||||||||||||]PAM</td><td>NA</td><td>Intergenic</td><td>NA</td><td>NA</td></tr>
<tr><td>GU305914.1</td><td>114142</td><td>114164</td><td>+</td><td>4</td><td>CGTGTCGACGACGCGGCGCT</td><td>CGG</td><td>-||||||-[-||||||||||-]PAM</td><td>NA</td><td>Intergenic</td><td>NA</td><td>NA</td></tr>
<tr><td></td><td></td><td></td><td></td><td></td><td></td><td></td><td></td><td></td><td></td><td></td><td></td></tr>
<tr><td>T30971</td><td>GTGTCGGGGACGCGGCGCGTGGG</td><td>1000</td><td></td><td></td><td></td><td></td><td></td><td></td><td></td><td></td><td></td></tr>
<tr><td>Oligo fwd</td><td>CACCGTGTCGGGGACGCGGCGCGT</td><td></td><td></td><td></td><td></td><td></td><td></td><td></td><td></td><td></td><td></td></tr>
<tr><td>Oligo rev</td><td>AAACACGCGCCGCGTCCCCGACAC</td><td></td><td></td><td></td><td></td><td></td><td></td><td></td><td></td><td></td><td></td></tr>
<tr><td>Chromosome</td><td>start</td><td>end</td><td>strand</td><td>MM</td><td>target_seq</td><td>PAM</td><td>alignment</td><td>distance</td><td>position</td><td>gene name</td><td>gene id</td></tr>
<tr><td>GU305914.1</td><td>43393</td><td>43415</td><td>-</td><td>0</td><td>GTGTCGGGGACGCGGCGCGT</td><td>GGG</td><td>||||||||[||||||||||||]PAM</td><td>NA</td><td>Intergenic</td><td>NA</td><td>NA</td></tr>
<tr><td>GU305914.1</td><td>114143</td><td>114165</td><td>+</td><td>4</td><td>GTGTCGACGACGCGGCGCTC</td><td>GGG</td><td>||||||--[||||||||||--]PAM</td><td>NA</td><td>Intergenic</td><td>NA</td><td>NA</td></tr>
</table>

T30972 TGTCGGGGACGCGGCGCGTGGGG 1000
Oligo fwd CACCTGTCGGGGACGCGGCGCGTG
Oligo rev AAACCACGCGCCGCGTCCCCGACA

| Chromosome | start | end | strand | MM | target_seq | PAM | alignment | distance | position | gene name | gene id |
|---|---|---|---|---|---|---|---|---|---|---|---|
| GU305914.1 | 43392 | 43414 | - | 0 | TGTCGGGGACGCGGCGCGTG | GGG | ||||||||[||||||||||||]PAM | NA | Intergenic | NA | NA |
| GU305914.1 | 198416 | 198438 | + | 3 | TGTCGGCGACGCGCCGCGTC | AGG | ||||||-|[|||||-|||||-]PAM | NA | Intergenic | NA | NA |
| GU305914.1 | 114144 | 114166 | + | 4 | TGTCGACGACGCGGCGCTCG | GGG | |||||--|[|||||||||--|]PAM | NA | Intergenic | NA | NA |

T30973 GGCGCGTGGGGCGTCCCCGCCGG 1000
Oligo fwd CACCGGCGCGTGGGGCGTCCCCGC
Oligo rev AAACGCGGGGACGCCCCACGCGCC

| Chromosome | start | end | strand | MM | target_seq | PAM | alignment | distance | position | gene name | gene id |
|---|---|---|---|---|---|---|---|---|---|---|---|
| GU305914.1 | 43380 | 43402 | - | 0 | GGCGCGTGGGGCGTCCCCGC | CGG | ||||||||[||||||||||||]PAM | NA | Intergenic | NA | NA |

T30974 GCGCGTGGGGCGTCCCCGCCGGG 1000
Oligo fwd CACCGCGCGTGGGGCGTCCCCGCC
Oligo rev AAACGGCGGGGACGCCCCACGCGC

| Chromosome | start | end | strand | MM | target_seq | PAM | alignment | distance | position | gene name | gene id |
|---|---|---|---|---|---|---|---|---|---|---|---|
| GU305914.1 | 43379 | 43401 | - | 0 | GCGCGTGGGGCGTCCCCGCC | GGG | ||||||||[||||||||||||]PAM | NA | Intergenic | NA | NA |

T30975 ACGTCGTTTTTGTCGTCGTCTGG 1000
Oligo fwd CACCACGTCGTTTTTGTCGTCGTC
Oligo rev AAACGACGACGACAAAAACGACGT

| Chromosome | start | end | strand | MM | target_seq | PAM | alignment | distance | position | gene name | gene id |
|---|---|---|---|---|---|---|---|---|---|---|---|
| GU305914.1 | 43346 | 43368 | - | 0 | ACGTCGTTTTTGTCGTCGTC | TGG | ||||||||[||||||||||||]PAM | NA | Intergenic | NA | NA |

T30976 TTTTTGTCGTCGTCTGGTCGCGG 1000
Oligo fwd CACCTTTTTGTCGTCGTCTGGTCG
Oligo rev AAACCGACCAGACGACGACAAAAA

| Chromosome | start | end | strand | MM | target_seq | PAM | alignment | distance | position | gene name | gene id |
|---|---|---|---|---|---|---|---|---|---|---|---|
| GU305914.1 | 43340 | 43362 | - | 0 | TTTTTGTCGTCGTCTGGTCG | CGG | ||||||||[||||||||||||]PAM | NA | Intergenic | NA | NA |

T30977 TTTTGTCGTCGTCTGGTCGCGGG 1000
Oligo fwd CACCTTTTGTCGTCGTCTGGTCGC
Oligo rev AAACGCGACCAGACGACGACAAAA

| Chromosome | start | end | strand | MM | target_seq | PAM | alignment | distance | position | gene name | gene id |
|---|---|---|---|---|---|---|---|---|---|---|---|
| GU305914.1 | 43339 | 43361 | - | 0 | TTTTGTCGTCGTCTGGTCGC | GGG | ||||||||[||||||||||||]PAM | NA | Intergenic | NA | NA |

T30978 GCGGGTCGACGACGACGACTTGG 1000
Oligo fwd CACCGCGGGTCGACGACGACGACT
Oligo rev AAACAGTCGTCGTCGTCGACCCGC

| Chromosome | start | end | strand | MM | target_seq | PAM | alignment | distance | position | gene name | gene id |
|---|---|---|---|---|---|---|---|---|---|---|---|
| GU305914.1 | 43321 | 43343 | - | 0 | GCGGGTCGACGACGACGACT | TGG | ||||||||[||||||||||||]PAM | NA | Intergenic | NA | NA |

T30979 GACGACTTGGAGAAGCTGCGCGG 1000
Oligo fwd CACCGACGACTTGGAGAAGCTGCG
Oligo rev AAACCGCAGCTTCTCCAAGTCGTC

| Chromosome | start | end | strand | MM | target_seq | PAM | alignment | distance | position | gene name | gene id |
|---|---|---|---|---|---|---|---|---|---|---|---|
| GU305914.1 | 43308 | 43330 | - | 0 | GACGACTTGGAGAAGCTGCG | CGG | ||||||||[||||||||||||]PAM | NA | Intergenic | NA | NA |

T30980 ACGACTTGGAGAAGCTGCGCGGG 1000
Oligo fwd CACCACGACTTGGAGAAGCTGCGC
Oligo rev AAACGCGCAGCTTCTCCAAGTCGT

| Chromosome | start | end | strand | MM | target_seq | PAM | alignment | distance | position | gene name | gene id |
|---|---|---|---|---|---|---|---|---|---|---|---|
| GU305914.1 | 43307 | 43329 | - | 0 | ACGACTTGGAGAAGCTGCGC | GGG | ||||||||[||||||||||||]PAM | NA | Intergenic | NA | NA |

T30981 GGCGCTTGATGAAAGCTCCCCGG 1000
Oligo fwd CACCGGCGCTTGATGAAAGCTCCC
Oligo rev AAACGGGAGCTTTCATCAAGCGCC

| Chromosome | start | end | strand | MM | target_seq | PAM | alignment | distance | position | gene name | gene id |
|---|---|---|---|---|---|---|---|---|---|---|---|
| GU305914.1 | 43286 | 43308 | - | 0 | GGCGCTTGATGAAAGCTCCC | CGG | ||||||||[||||||||||||]PAM | NA | Intergenic | NA | NA |

T30982 CGTTGTTATTGTTGTCAGAACGG 1000
Oligo fwd CACCCGTTGTTATTGTTGTCAGAA
Oligo rev AAACTTCTGACAACAATAACAACG

| Chromosome | start | end | strand | MM | target_seq | PAM | alignment | distance | position | gene name | gene id |
|---|---|---|---|---|---|---|---|---|---|---|---|
| GU305914.1 | 43256 | 43278 | - | 0 | CGTTGTTATTGTTGTCAGAA | CGG | ||||||||[||||||||||||]PAM | NA | Intergenic | NA | NA |

T30983 TCCTCTTGCGTCGTCTGTGTCGG 1000
Oligo fwd CACCTCCTCTTGCGTCGTCTGTGT
Oligo rev AAACACACAGACGACGCAAGAGGA

| Chromosome | start | end | strand | MM | target_seq | PAM | alignment | distance | position | gene name | gene id |
|---|---|---|---|---|---|---|---|---|---|---|---|
| GU305914.1 | 43232 | 43254 | - | 0 | TCCTCTTGCGTCGTCTGTGT | CGG | ||||||||[||||||||||||]PAM | NA | Intergenic | NA | NA |

T30984 GTCCGTGCGCGTGTCTTCCGCGG 1000
Oligo fwd CACCGTCCGTGCGCGTGTCTTCCG
Oligo rev AAACCGGAAGACACGCGCACGGAC

| Chromosome | start | end | strand | MM | target_seq | PAM | alignment | distance | position | gene name | gene id |
|---|---|---|---|---|---|---|---|---|---|---|---|
| GU305914.1 | 43192 | 43214 | - | 0 | GTCCGTGCGCGTGTCTTCCG | CGG | ||||||||[||||||||||||]PAM | NA | Intergenic | NA | NA |

T30985 GTGCGCGTGTCTTCCGCGGCCGG 1000
Oligo fwd CACCGTGCGCGTGTCTTCCGCGGC
Oligo rev AAACGCCGCGGAAGACACGCGCAC

| Chromosome | start | end | strand | MM | target_seq | PAM | alignment | distance | position | gene name | gene id |
|---|---|---|---|---|---|---|---|---|---|---|---|
| GU305914.1 | 43188 | 43210 | - | 0 | GTGCGCGTGTCTTCCGCGGC | CGG | ||||||||[||||||||||||]PAM | NA | Intergenic | NA | NA |

T30986 CGCGTGTCTTCCGCGGCCGGCGG 1000
Oligo fwd CACCCGCGTGTCTTCCGCGGCCGG
Oligo rev AAACCCGGCCGCGGAAGACACGCG

```
Chromosome    start                      end    strand  MM  target_seq            PAM  alignment                       distance  position    gene name  gene id
GU305914.1                      43185    43207  -       0   CGCGTGTCTTCCGCGGCCGG  CGG  ||||||||[||||||||||||]PAM       NA        Intergenic  NA         NA

T30987        GCCGGCGGTCTTCGCGAGCGCGG    1000
Oligo fwd     CACCGCCGGCGGTCTTCGCGAGCG
Oligo rev     AAACCGCTCGCGAAGACCGCCGGC
Chromosome    start                      end    strand  MM  target_seq            PAM  alignment                       distance  position    gene name  gene id
GU305914.1                      43170    43192  -       0   GCCGGCGGTCTTCGCGAGCG  CGG  ||||||||[||||||||||||]PAM       NA        Intergenic  NA         NA

T30988        CGGTCTTCGCGAGCGCGGATCGG    1000
Oligo fwd     CACCCGGTCTTCGCGAGCGCGGAT
Oligo rev     AAACATCCGCGCTCGCGAAGACCG
Chromosome    start                      end    strand  MM  target_seq            PAM  alignment                       distance  position    gene name  gene id
GU305914.1                      43165    43187  -       0   CGGTCTTCGCGAGCGCGGAT  CGG  ||||||||[||||||||||||]PAM       NA        Intergenic  NA         NA

T30989        TCGCGAGCGCGGATCGGCGCCGG    1000
Oligo fwd     CACCTCGCGAGCGCGGATCGGCGC
Oligo rev     AAACGCGCCGATCCGCGCTCGCGA
Chromosome    start                      end    strand  MM  target_seq            PAM  alignment                       distance  position    gene name  gene id
GU305914.1                      43159    43181  -       0   TCGCGAGCGCGGATCGGCGC  CGG  ||||||||[||||||||||||]PAM       NA        Intergenic  NA         NA
GU305914.1                      65907    65929  -       4   TCGTGCGCGCGGATCTCCGC  CGG  |||-|-||[|||||||--|||]PAM       NA        Intergenic  NA         NA

T30990        CGCGAGCGCGGATCGGCGCCGGG    1000
Oligo fwd     CACCCGCGAGCGCGGATCGGCGCC
Oligo rev     AAACGGCGCCGATCCGCGCTCGCG
Chromosome    start                      end    strand  MM  target_seq            PAM  alignment                       distance  position    gene name  gene id
GU305914.1                      43158    43180  -       0   CGCGAGCGCGGATCGGCGCC  GGG  ||||||||[||||||||||||]PAM       NA        Intergenic  NA         NA

T30991        CGCGGATCGGCGCCGGGTCGCGG    1000
Oligo fwd     CACCCGCGGATCGGCGCCGGGTCG
Oligo rev     AAACCGACCCGGCGCCGATCCGCG
Chromosome    start                      end    strand  MM  target_seq            PAM  alignment                       distance  position    gene name  gene id
GU305914.1                      43152    43174  -       0   CGCGGATCGGCGCCGGGTCG  CGG  ||||||||[||||||||||||]PAM       NA        Intergenic  NA         NA

T30992        GGCGCCGGGTCGCGGATGCCCGG    1000
Oligo fwd     CACCGGCGCCGGGTCGCGGATGCC
Oligo rev     AAACGGCATCCGCGACCCGGCGCC
Chromosome    start                      end    strand  MM  target_seq            PAM  alignment                       distance  position    gene name  gene id
GU305914.1                      43144    43166  -       0   GGCGCCGGGTCGCGGATGCC  CGG  ||||||||[||||||||||||]PAM       NA        Intergenic  NA         NA

T30993        GCCCGGACCGTCCGAACGTGCGG    1000
Oligo fwd     CACCGCCCGGACCGTCCGAACGTG
Oligo rev     AAACCACGTTCGGACGGTCCGGGC
Chromosome    start                      end    strand  MM  target_seq            PAM  alignment                       distance  position    gene name  gene id
GU305914.1                      43127    43149  -       0   GCCCGGACCGTCCGAACGTG  CGG  ||||||||[||||||||||||]PAM       NA        Intergenic  NA         NA

T30994        GACGCCTCGAGAAGCCGACGCGG    1000
Oligo fwd     CACCGACGCCTCGAGAAGCCGACG
Oligo rev     AAACCGTCGGCTTCTCGAGGCGTC
Chromosome    start                      end    strand  MM  target_seq            PAM  alignment                       distance  position    gene name  gene id
GU305914.1                      43093    43115  -       0   GACGCCTCGAGAAGCCGACG  CGG  ||||||||[||||||||||||]PAM       NA        Intergenic  NA         NA

T30995        TCGAGCGAGCAAACTGTTAGTGG    1000
Oligo fwd     CACCTCGAGCGAGCAAACTGTTAG
Oligo rev     AAACCTAACAGTTTGCTCGCTCGA
Chromosome    start                      end    strand  MM  target_seq            PAM  alignment                       distance  position    gene name  gene id
GU305914.1                      43064    43086  -       0   TCGAGCGAGCAAACTGTTAG  TGG  ||||||||[||||||||||||]PAM       NA        Intergenic  NA         NA

T30996        AGCGAGCAAACTGTTAGTGGCGG    1000
Oligo fwd     CACCAGCGAGCAAACTGTTAGTGG
Oligo rev     AAACCCACTAACAGTTTGCTCGCT
Chromosome    start                      end    strand  MM  target_seq            PAM  alignment                       distance  position    gene name  gene id
GU305914.1                      43061    43083  -       0   AGCGAGCAAACTGTTAGTGG  CGG  ||||||||[||||||||||||]PAM       NA        Intergenic  NA         NA

T30997        GAGCAAACTGTTAGTGGCGGCGG    1000
Oligo fwd     CACCGAGCAAACTGTTAGTGGCGG
Oligo rev     AAACCCGCCACTAACAGTTTGCTC
Chromosome    start                      end    strand  MM  target_seq            PAM  alignment                       distance  position    gene name  gene id
GU305914.1                      43058    43080  -       0   GAGCAAACTGTTAGTGGCGG  CGG  ||||||||[||||||||||||]PAM       NA        Intergenic  NA         NA

T30998        AACTGTTAGTGGCGGCGGCGCGG    1000
Oligo fwd     CACCAACTGTTAGTGGCGGCGGCG
Oligo rev     AAACCGCCGCCGCCACTAACAGTT
Chromosome    start                      end    strand  MM  target_seq            PAM  alignment                       distance  position    gene name  gene id
GU305914.1                      43053    43075  -       0   AACTGTTAGTGGCGGCGGCG  CGG  ||||||||[||||||||||||]PAM       NA        Intergenic  NA         NA

T30999        TGGCGGCGGCGCGGCGTCTGAGG    1000
Oligo fwd     CACCTGGCGGCGGCGCGGCGTCTG
Oligo rev     AAACCAGACGCCGCGCCGCCGCCA
Chromosome    start                      end    strand  MM  target_seq            PAM  alignment                       distance  position    gene name  gene id
GU305914.1                      43044    43066  -       0   TGGCGGCGGCGCGGCGTCTG  AGG  ||||||||[||||||||||||]PAM       NA        Intergenic  NA         NA

T31000        AGCACCGCTGCGCTTTTAGCCGG    1000
Oligo fwd     CACCAGCACCGCTGCGCTTTTAGC
Oligo rev     AAACGCTAAAAGCGCAGCGGTGCT
Chromosome    start                      end    strand  MM  target_seq            PAM  alignment                       distance  position    gene name  gene id
GU305914.1                      43013    43035  -       0   AGCACCGCTGCGCTTTTAGC  CGG  ||||||||[||||||||||||]PAM       NA        Intergenic  NA         NA

T31001        GCACCGCTGCGCTTTTAGCCGGG    1000
```

Oligo fwd CACCGCACCGCTGCGCTTTTAGCC
Oligo rev AAACGGCTAAAAGCGCAGCGGTGC
Chromosome start end strand MM target_seq PAM alignment distance position gene name gene id
GU305914.1 43012 43034 - 0 GCACCGCTGCGCTTTTAGCC GGG ||||||||[||||||||||||]PAM NA Intergenic NA NA

T31002 GCCGGGCCGCCACTCCTACACGG 1000
Oligo fwd CACCGCCGGGCCGCCACTCCTACA
Oligo rev AAACTGTAGGAGTGGCGGCCCGGC
Chromosome start end strand MM target_seq PAM alignment distance position gene name gene id
GU305914.1 42995 43017 - 0 GCCGGGCCGCCACTCCTACA CGG ||||||||[||||||||||||]PAM NA Intergenic NA NA

T31003 GTATTCTGCGAAAGATTGATTGG 1000
Oligo fwd CACCGTATTCTGCGAAAGATTGAT
Oligo rev AAACATCAATCTTTCGCAGAATAC
Chromosome start end strand MM target_seq PAM alignment distance position gene name gene id
GU305914.1 42973 42995 - 0 GTATTCTGCGAAAGATTGAT TGG ||||||||[||||||||||||]PAM NA Intergenic NA NA

T31004 GGTCTAGATGATGTTGTGCGCGG 1000
Oligo fwd CACCGGTCTAGATGATGTTGTGCG
Oligo rev AAACCGCACAACATCATCTAGACC
Chromosome start end strand MM target_seq PAM alignment distance position gene name gene id
GU305914.1 42952 42974 - 0 GGTCTAGATGATGTTGTGCG CGG ||||||||[||||||||||||]PAM NA Intergenic NA NA

T31005 GGTGATGTGTCTCTTTCGACAGG 1000
Oligo fwd CACCGGTGATGTGTCTCTTTCGAC
Oligo rev AAACGTCGAAAGAGACACATCACC
Chromosome start end strand MM target_seq PAM alignment distance position gene name gene id
GU305914.1 42931 42953 - 0 GGTGATGTGTCTCTTTCGAC AGG ||||||||[||||||||||||]PAM NA Intergenic NA NA

T31006 TGTCTCTTTCGACAGGTGCGTGG 1000
Oligo fwd CACCTGTCTCTTTCGACAGGTGCG
Oligo rev AAACCGCACCTGTCGAAAGAGACA
Chromosome start end strand MM target_seq PAM alignment distance position gene name gene id
GU305914.1 42924 42946 - 0 TGTCTCTTTCGACAGGTGCG TGG ||||||||[||||||||||||]PAM NA Intergenic NA NA

T31007 GGCGTCCCTCGCGCTCTCCGAGG 1000
Oligo fwd CACCGGCGTCCCTCGCGCTCTCCG
Oligo rev AAACCGGAGAGCGCGAGGGACGCC
Chromosome start end strand MM target_seq PAM alignment distance position gene name gene id
GU305914.1 42903 42925 - 0 GGCGTCCCTCGCGCTCTCCG AGG ||||||||[||||||||||||]PAM NA Intergenic NA NA

T31008 GCGTCCCTCGCGCTCTCCGAGGG 1000
Oligo fwd CACCGCGTCCCTCGCGCTCTCCGA
Oligo rev AAACTCGGAGAGCGCGAGGGACGC
Chromosome start end strand MM target_seq PAM alignment distance position gene name gene id
GU305914.1 42902 42924 - 0 GCGTCCCTCGCGCTCTCCGA GGG ||||||||[||||||||||||]PAM NA Intergenic NA NA

T31009 CGTCCCTCGCGCTCTCCGAGGGG 1000
Oligo fwd CACCCGTCCCTCGCGCTCTCCGAG
Oligo rev AAACCTCGGAGAGCGCGAGGGACG
Chromosome start end strand MM target_seq PAM alignment distance position gene name gene id
GU305914.1 42901 42923 - 0 CGTCCCTCGCGCTCTCCGAG GGG ||||||||[||||||||||||]PAM NA Intergenic NA NA
GU305914.1 226718 226740 - 3 CGACCTCCGCGCTCTCCGAG TGG ||-||--|[||||||||||||]PAM NA Intergenic NA NA

T31010 GTCTCGACCGACCCTCCGCACGG 1000
Oligo fwd CACCGTCTCGACCGACCCTCCGCA
Oligo rev AAACTGCGGAGGGTCGGTCGAGAC
Chromosome start end strand MM target_seq PAM alignment distance position gene name gene id
GU305914.1 42879 42901 - 0 GTCTCGACCGACCCTCCGCA CGG ||||||||[||||||||||||]PAM NA Intergenic NA NA

T31011 TCTCGACCGACCCTCCGCACGGG 1000
Oligo fwd CACCTCTCGACCGACCCTCCGCAC
Oligo rev AAACGTGCGGAGGGTCGGTCGAGA
Chromosome start end strand MM target_seq PAM alignment distance position gene name gene id
GU305914.1 42878 42900 - 0 TCTCGACCGACCCTCCGCAC GGG ||||||||[||||||||||||]PAM NA Intergenic NA NA

T31012 CTCGACCGACCCTCCGCACGGGG 1000
Oligo fwd CACCCTCGACCGACCCTCCGCACG
Oligo rev AAACCGTGCGGAGGGTCGGTCGAG
Chromosome start end strand MM target_seq PAM alignment distance position gene name gene id
GU305914.1 42877 42899 - 0 CTCGACCGACCCTCCGCACG GGG ||||||||[||||||||||||]PAM NA Intergenic NA NA

T31013 TCGACCGACCCTCCGCACGGGGG 1000
Oligo fwd CACCTCGACCGACCCTCCGCACGG
Oligo rev AAACCCGTGCGGAGGGTCGGTCGA
Chromosome start end strand MM target_seq PAM alignment distance position gene name gene id
GU305914.1 42876 42898 - 0 TCGACCGACCCTCCGCACGG GGG ||||||||[||||||||||||]PAM NA Intergenic NA NA

T31014 ACCGACCCTCCGCACGGGGGAGG 1000
Oligo fwd CACCACCGACCCTCCGCACGGGGG
Oligo rev AAACCCCCCGTGCGGAGGGTCGGT
Chromosome start end strand MM target_seq PAM alignment distance position gene name gene id
GU305914.1 42873 42895 - 0 ACCGACCCTCCGCACGGGGG AGG ||||||||[||||||||||||]PAM NA Intergenic NA NA

T31015 CCTCCGCACGGGGGAGGATGAGG 1000
Oligo fwd CACCCCTCCGCACGGGGGAGGATG
Oligo rev AAACCATCCTCCCCCGTGCGGAGG
Chromosome start end strand MM target_seq PAM alignment distance position gene name gene id
GU305914.1 42867 42889 - 0 CCTCCGCACGGGGGAGGATG AGG ||||||||[||||||||||||]PAM NA Intergenic NA NA

<table>
<tr><td>T31016</td><td>GGGGGAGGATGAGGAAGAGAAGG</td><td>1000</td><td></td><td></td><td></td><td></td><td></td><td></td><td></td><td></td><td></td></tr>
<tr><td>Oligo fwd</td><td>CACCGGGGGAGGATGAGGAAGAGA</td><td></td><td></td><td></td><td></td><td></td><td></td><td></td><td></td><td></td><td></td></tr>
<tr><td>Oligo rev</td><td>AAACTCTCTTCCTCATCCTCCCCC</td><td></td><td></td><td></td><td></td><td></td><td></td><td></td><td></td><td></td><td></td></tr>
<tr><td>Chromosome</td><td>start</td><td>end</td><td>strand</td><td>MM</td><td>target_seq</td><td>PAM</td><td>alignment</td><td>distance</td><td>position</td><td>gene name</td><td>gene id</td></tr>
<tr><td>GU305914.1</td><td>42858</td><td>42880</td><td>-</td><td>0</td><td>GGGGGAGGATGAGGAAGAGA</td><td>AGG</td><td>||||||||[||||||||||||]PAM</td><td>NA</td><td>Intergenic</td><td>NA</td><td>NA</td></tr>
<tr><td>GU305914.1</td><td>44981</td><td>45003</td><td>+</td><td>4</td><td>GGAGGAAGACGGGGAAGAGA</td><td>CGG</td><td>||-|||-|[|-|-||||||||]PAM</td><td>NA</td><td>Intergenic</td><td>NA</td><td>NA</td></tr>
<tr><td>GU305914.1</td><td>45315</td><td>45337</td><td>-</td><td>4</td><td>AGTGGAGGAGGAGGAGGAGA</td><td>AGG</td><td>-|-|||||[|-|||||-||||]PAM</td><td>NA</td><td>Intergenic</td><td>NA</td><td>NA</td></tr>
<tr><td>GU305914.1</td><td>140726</td><td>140748</td><td>+</td><td>4</td><td>CGAGGAGGATGAGGATGAGG</td><td>AGG</td><td>-|-|||||[|||||||-|||-]PAM</td><td>NA</td><td>Intergenic</td><td>NA</td><td>NA</td></tr>
<tr><td>GU305914.1</td><td>140711</td><td>140733</td><td>+</td><td>4</td><td>GGGGGGAGATGAGGACGAGG</td><td>AGG</td><td>|||||--|[|||||||-|||-]PAM</td><td>NA</td><td>Intergenic</td><td>NA</td><td>NA</td></tr>
<tr><td>T31017</td><td>ATGAGGAAGAGAAGGAGAAGAGG</td><td>1000</td><td></td><td></td><td></td><td></td><td></td><td></td><td></td><td></td><td></td></tr>
<tr><td>Oligo fwd</td><td>CACCATGAGGAAGAGAAGGAGAAG</td><td></td><td></td><td></td><td></td><td></td><td></td><td></td><td></td><td></td><td></td></tr>
<tr><td>Oligo rev</td><td>AAACCTTCTCCTTCTCTTCCTCAT</td><td></td><td></td><td></td><td></td><td></td><td></td><td></td><td></td><td></td><td></td></tr>
<tr><td>Chromosome</td><td>start</td><td>end</td><td>strand</td><td>MM</td><td>target_seq</td><td>PAM</td><td>alignment</td><td>distance</td><td>position</td><td>gene name</td><td>gene id</td></tr>
<tr><td>GU305914.1</td><td>42850</td><td>42872</td><td>-</td><td>0</td><td>ATGAGGAAGAGAAGGAGAAG</td><td>AGG</td><td>||||||||[||||||||||||]PAM</td><td>NA</td><td>Intergenic</td><td>NA</td><td>NA</td></tr>
<tr><td>T31018</td><td>GAAGGAGAAGAGGATGACGAAGG</td><td>1000</td><td></td><td></td><td></td><td></td><td></td><td></td><td></td><td></td><td></td></tr>
<tr><td>Oligo fwd</td><td>CACCGAAGGAGAAGAGGATGACGA</td><td></td><td></td><td></td><td></td><td></td><td></td><td></td><td></td><td></td><td></td></tr>
<tr><td>Oligo rev</td><td>AAACTCGTCATCCTCTTCTCCTTC</td><td></td><td></td><td></td><td></td><td></td><td></td><td></td><td></td><td></td><td></td></tr>
<tr><td>Chromosome</td><td>start</td><td>end</td><td>strand</td><td>MM</td><td>target_seq</td><td>PAM</td><td>alignment</td><td>distance</td><td>position</td><td>gene name</td><td>gene id</td></tr>
<tr><td>GU305914.1</td><td>42840</td><td>42862</td><td>-</td><td>0</td><td>GAAGGAGAAGAGGATGACGA</td><td>AGG</td><td>||||||||[||||||||||||]PAM</td><td>NA</td><td>Intergenic</td><td>NA</td><td>NA</td></tr>
<tr><td>GU305914.1</td><td>28744</td><td>28766</td><td>-</td><td>4</td><td>GGAGGAGGAGAGGATGATGG</td><td>TGG</td><td>|-|||||-[|||||||||-|-]PAM</td><td>NA</td><td>Intergenic</td><td>NA</td><td>NA</td></tr>
<tr><td>T31019</td><td>AAGGAGAAGAGGATGACGAAGGG</td><td>1000</td><td></td><td></td><td></td><td></td><td></td><td></td><td></td><td></td><td></td></tr>
<tr><td>Oligo fwd</td><td>CACCAAGGAGAAGAGGATGACGAA</td><td></td><td></td><td></td><td></td><td></td><td></td><td></td><td></td><td></td><td></td></tr>
<tr><td>Oligo rev</td><td>AAACTTCGTCATCCTCTTCTCCTT</td><td></td><td></td><td></td><td></td><td></td><td></td><td></td><td></td><td></td><td></td></tr>
<tr><td>Chromosome</td><td>start</td><td>end</td><td>strand</td><td>MM</td><td>target_seq</td><td>PAM</td><td>alignment</td><td>distance</td><td>position</td><td>gene name</td><td>gene id</td></tr>
<tr><td>GU305914.1</td><td>42839</td><td>42861</td><td>-</td><td>0</td><td>AAGGAGAAGAGGATGACGAA</td><td>GGG</td><td>||||||||[||||||||||||]PAM</td><td>NA</td><td>Intergenic</td><td>NA</td><td>NA</td></tr>
<tr><td>T31020</td><td>GACGAAGGGACGACGACGAGAGG</td><td>1000</td><td></td><td></td><td></td><td></td><td></td><td></td><td></td><td></td><td></td></tr>
<tr><td>Oligo fwd</td><td>CACCGACGAAGGGACGACGACGAG</td><td></td><td></td><td></td><td></td><td></td><td></td><td></td><td></td><td></td><td></td></tr>
<tr><td>Oligo rev</td><td>AAACCTCGTCGTCGTCCCTTCGTC</td><td></td><td></td><td></td><td></td><td></td><td></td><td></td><td></td><td></td><td></td></tr>
<tr><td>Chromosome</td><td>start</td><td>end</td><td>strand</td><td>MM</td><td>target_seq</td><td>PAM</td><td>alignment</td><td>distance</td><td>position</td><td>gene name</td><td>gene id</td></tr>
<tr><td>GU305914.1</td><td>42825</td><td>42847</td><td>-</td><td>0</td><td>GACGAAGGGACGACGACGAG</td><td>AGG</td><td>||||||||[||||||||||||]PAM</td><td>NA</td><td>Intergenic</td><td>NA</td><td>NA</td></tr>
<tr><td>T31021</td><td>AAGCGAAAGAGAAGAGAGAGCGG</td><td>1000</td><td></td><td></td><td></td><td></td><td></td><td></td><td></td><td></td><td></td></tr>
<tr><td>Oligo fwd</td><td>CACCAAGCGAAAGAGAAGAGAGAG</td><td></td><td></td><td></td><td></td><td></td><td></td><td></td><td></td><td></td><td></td></tr>
<tr><td>Oligo rev</td><td>AAACCTCTCTCTTCTCTTTCGCTT</td><td></td><td></td><td></td><td></td><td></td><td></td><td></td><td></td><td></td><td></td></tr>
<tr><td>Chromosome</td><td>start</td><td>end</td><td>strand</td><td>MM</td><td>target_seq</td><td>PAM</td><td>alignment</td><td>distance</td><td>position</td><td>gene name</td><td>gene id</td></tr>
<tr><td>GU305914.1</td><td>42798</td><td>42820</td><td>-</td><td>0</td><td>AAGCGAAAGAGAAGAGAGAG</td><td>CGG</td><td>||||||||[||||||||||||]PAM</td><td>NA</td><td>Intergenic</td><td>NA</td><td>NA</td></tr>
<tr><td>T31022</td><td>GGCCGCCGCGCTGCGATCACTGG</td><td>1000</td><td></td><td></td><td></td><td></td><td></td><td></td><td></td><td></td><td></td></tr>
<tr><td>Oligo fwd</td><td>CACCGGCCGCCGCGCTGCGATCAC</td><td></td><td></td><td></td><td></td><td></td><td></td><td></td><td></td><td></td><td></td></tr>
<tr><td>Oligo rev</td><td>AAACGTGATCGCAGCGCGGCGGCC</td><td></td><td></td><td></td><td></td><td></td><td></td><td></td><td></td><td></td><td></td></tr>
<tr><td>Chromosome</td><td>start</td><td>end</td><td>strand</td><td>MM</td><td>target_seq</td><td>PAM</td><td>alignment</td><td>distance</td><td>position</td><td>gene name</td><td>gene id</td></tr>
<tr><td>GU305914.1</td><td>42777</td><td>42799</td><td>-</td><td>0</td><td>GGCCGCCGCGCTGCGATCAC</td><td>TGG</td><td>||||||||[||||||||||||]PAM</td><td>NA</td><td>Intergenic</td><td>NA</td><td>NA</td></tr>
<tr><td>T31023</td><td>GCCGCCGCGCTGCGATCACTGGG</td><td>1000</td><td></td><td></td><td></td><td></td><td></td><td></td><td></td><td></td><td></td></tr>
<tr><td>Oligo fwd</td><td>CACCGCCGCCGCGCTGCGATCACT</td><td></td><td></td><td></td><td></td><td></td><td></td><td></td><td></td><td></td><td></td></tr>
<tr><td>Oligo rev</td><td>AAACAGTGATCGCAGCGCGGCGGC</td><td></td><td></td><td></td><td></td><td></td><td></td><td></td><td></td><td></td><td></td></tr>
<tr><td>Chromosome</td><td>start</td><td>end</td><td>strand</td><td>MM</td><td>target_seq</td><td>PAM</td><td>alignment</td><td>distance</td><td>position</td><td>gene name</td><td>gene id</td></tr>
<tr><td>GU305914.1</td><td>42776</td><td>42798</td><td>-</td><td>0</td><td>GCCGCCGCGCTGCGATCACT</td><td>GGG</td><td>||||||||[||||||||||||]PAM</td><td>NA</td><td>Intergenic</td><td>NA</td><td>NA</td></tr>
<tr><td>T31024</td><td>CCGCCGCGCTGCGATCACTGGGG</td><td>1000</td><td></td><td></td><td></td><td></td><td></td><td></td><td></td><td></td><td></td></tr>
<tr><td>Oligo fwd</td><td>CACCCCGCCGCGCTGCGATCACTG</td><td></td><td></td><td></td><td></td><td></td><td></td><td></td><td></td><td></td><td></td></tr>
<tr><td>Oligo rev</td><td>AAACCAGTGATCGCAGCGCGGCGG</td><td></td><td></td><td></td><td></td><td></td><td></td><td></td><td></td><td></td><td></td></tr>
<tr><td>Chromosome</td><td>start</td><td>end</td><td>strand</td><td>MM</td><td>target_seq</td><td>PAM</td><td>alignment</td><td>distance</td><td>position</td><td>gene name</td><td>gene id</td></tr>
<tr><td>GU305914.1</td><td>42775</td><td>42797</td><td>-</td><td>0</td><td>CCGCCGCGCTGCGATCACTG</td><td>GGG</td><td>||||||||[||||||||||||]PAM</td><td>NA</td><td>Intergenic</td><td>NA</td><td>NA</td></tr>
<tr><td>T31025</td><td>CCGCGCTGCGATCACTGGGGCGG</td><td>1000</td><td></td><td></td><td></td><td></td><td></td><td></td><td></td><td></td><td></td></tr>
<tr><td>Oligo fwd</td><td>CACCCCGCGCTGCGATCACTGGGG</td><td></td><td></td><td></td><td></td><td></td><td></td><td></td><td></td><td></td><td></td></tr>
<tr><td>Oligo rev</td><td>AAACCCCCAGTGATCGCAGCGCGG</td><td></td><td></td><td></td><td></td><td></td><td></td><td></td><td></td><td></td><td></td></tr>
<tr><td>Chromosome</td><td>start</td><td>end</td><td>strand</td><td>MM</td><td>target_seq</td><td>PAM</td><td>alignment</td><td>distance</td><td>position</td><td>gene name</td><td>gene id</td></tr>
<tr><td>GU305914.1</td><td>42772</td><td>42794</td><td>-</td><td>0</td><td>CCGCGCTGCGATCACTGGGG</td><td>CGG</td><td>||||||||[||||||||||||]PAM</td><td>NA</td><td>Intergenic</td><td>NA</td><td>NA</td></tr>
<tr><td>T31026</td><td>CGCTGCGATCACTGGGGCGGAGG</td><td>1000</td><td></td><td></td><td></td><td></td><td></td><td></td><td></td><td></td><td></td></tr>
<tr><td>Oligo fwd</td><td>CACCCGCTGCGATCACTGGGGCGG</td><td></td><td></td><td></td><td></td><td></td><td></td><td></td><td></td><td></td><td></td></tr>
<tr><td>Oligo rev</td><td>AAACCCGCCCCAGTGATCGCAGCG</td><td></td><td></td><td></td><td></td><td></td><td></td><td></td><td></td><td></td><td></td></tr>
<tr><td>Chromosome</td><td>start</td><td>end</td><td>strand</td><td>MM</td><td>target_seq</td><td>PAM</td><td>alignment</td><td>distance</td><td>position</td><td>gene name</td><td>gene id</td></tr>
<tr><td>GU305914.1</td><td>42769</td><td>42791</td><td>-</td><td>0</td><td>CGCTGCGATCACTGGGGCGG</td><td>AGG</td><td>||||||||[||||||||||||]PAM</td><td>NA</td><td>Intergenic</td><td>NA</td><td>NA</td></tr>
<tr><td>T31027</td><td>GCGTCCCGATCTTTGCCAGCTGG</td><td>1000</td><td></td><td></td><td></td><td></td><td></td><td></td><td></td><td></td><td></td></tr>
<tr><td>Oligo fwd</td><td>CACCGCGTCCCGATCTTTGCCAGC</td><td></td><td></td><td></td><td></td><td></td><td></td><td></td><td></td><td></td><td></td></tr>
<tr><td>Oligo rev</td><td>AAACGCTGGCAAAGATCGGGACGC</td><td></td><td></td><td></td><td></td><td></td><td></td><td></td><td></td><td></td><td></td></tr>
<tr><td>Chromosome</td><td>start</td><td>end</td><td>strand</td><td>MM</td><td>target_seq</td><td>PAM</td><td>alignment</td><td>distance</td><td>position</td><td>gene name</td><td>gene id</td></tr>
<tr><td>GU305914.1</td><td>42699</td><td>42721</td><td>-</td><td>0</td><td>GCGTCCCGATCTTTGCCAGC</td><td>TGG</td><td>||||||||[||||||||||||]PAM</td><td>NA</td><td>Intergenic</td><td>NA</td><td>NA</td></tr>
<tr><td>T31028</td><td>GATCTTTGCCAGCTGGCGCGCGG</td><td>1000</td><td></td><td></td><td></td><td></td><td></td><td></td><td></td><td></td><td></td></tr>
<tr><td>Oligo fwd</td><td>CACCGATCTTTGCCAGCTGGCGCG</td><td></td><td></td><td></td><td></td><td></td><td></td><td></td><td></td><td></td><td></td></tr>
<tr><td>Oligo rev</td><td>AAACCGCGCCAGCTGGCAAAGATC</td><td></td><td></td><td></td><td></td><td></td><td></td><td></td><td></td><td></td><td></td></tr>
<tr><td>Chromosome</td><td>start</td><td>end</td><td>strand</td><td>MM</td><td>target_seq</td><td>PAM</td><td>alignment</td><td>distance</td><td>position</td><td>gene name</td><td>gene id</td></tr>
<tr><td>GU305914.1</td><td>42692</td><td>42714</td><td>-</td><td>0</td><td>GATCTTTGCCAGCTGGCGCG</td><td>CGG</td><td>||||||||[||||||||||||]PAM</td><td>NA</td><td>Intergenic</td><td>NA</td><td>NA</td></tr>
<tr><td>T31029</td><td>TTGCCAGCTGGCGCGCGGCGCGG</td><td>1000</td><td></td><td></td><td></td><td></td><td></td><td></td><td></td><td></td><td></td></tr>
<tr><td>Oligo fwd</td><td>CACCTTGCCAGCTGGCGCGCGGCG</td><td></td><td></td><td></td><td></td><td></td><td></td><td></td><td></td><td></td><td></td></tr>
<tr><td>Oligo rev</td><td>AAACCGCCGCGCGCCAGCTGGCAA</td><td></td><td></td><td></td><td></td><td></td><td></td><td></td><td></td><td></td><td></td></tr>
<tr><td>Chromosome</td><td>start</td><td>end</td><td>strand</td><td>MM</td><td>target_seq</td><td>PAM</td><td>alignment</td><td>distance</td><td>position</td><td>gene name</td><td>gene id</td></tr>
<tr><td>GU305914.1</td><td>42687</td><td>42709</td><td>-</td><td>0</td><td>TTGCCAGCTGGCGCGCGGCG</td><td>CGG</td><td>||||||||[||||||||||||]PAM</td><td>NA</td><td>Intergenic</td><td>NA</td><td>NA</td></tr>
</table>

<table>
<tr><td>T31030</td><td>AGCTGGCGCGCGGCGCGGTGTGG</td><td>1000</td><td></td><td></td><td></td><td></td><td></td><td></td><td></td><td></td><td></td></tr>
<tr><td>Oligo fwd</td><td>CACCAGCTGGCGCGCGGCGCGGTG</td><td></td><td></td><td></td><td></td><td></td><td></td><td></td><td></td><td></td><td></td></tr>
<tr><td>Oligo rev</td><td>AAACCACCGCGCCGCGCGCCAGCT</td><td></td><td></td><td></td><td></td><td></td><td></td><td></td><td></td><td></td><td></td></tr>
<tr><td>Chromosome</td><td>start</td><td>end</td><td>strand</td><td>MM</td><td>target_seq</td><td>PAM</td><td>alignment</td><td>distance</td><td>position</td><td>gene name</td><td>gene id</td></tr>
<tr><td>GU305914.1</td><td>42682</td><td>42704</td><td>-</td><td>0</td><td>AGCTGGCGCGCGGCGCGGTG</td><td>TGG</td><td>||||||||[||||||||||||]PAM</td><td>NA</td><td>Intergenic</td><td>NA</td><td>NA</td></tr>
<tr><td></td><td></td><td></td><td></td><td></td><td></td><td></td><td></td><td></td><td></td><td></td><td></td></tr>
<tr><td>T31031</td><td>CGCGGCGCGGTGTGGCGTCGAGG</td><td>1000</td><td></td><td></td><td></td><td></td><td></td><td></td><td></td><td></td><td></td></tr>
<tr><td>Oligo fwd</td><td>CACCCGCGGCGCGGTGTGGCGTCG</td><td></td><td></td><td></td><td></td><td></td><td></td><td></td><td></td><td></td><td></td></tr>
<tr><td>Oligo rev</td><td>AAACCGACGCCACACCGCGCCGCG</td><td></td><td></td><td></td><td></td><td></td><td></td><td></td><td></td><td></td><td></td></tr>
<tr><td>Chromosome</td><td>start</td><td>end</td><td>strand</td><td>MM</td><td>target_seq</td><td>PAM</td><td>alignment</td><td>distance</td><td>position</td><td>gene name</td><td>gene id</td></tr>
<tr><td>GU305914.1</td><td>42674</td><td>42696</td><td>-</td><td>0</td><td>CGCGGCGCGGTGTGGCGTCG</td><td>AGG</td><td>||||||||[||||||||||||]PAM</td><td>NA</td><td>Intergenic</td><td>NA</td><td>NA</td></tr>
<tr><td></td><td></td><td></td><td></td><td></td><td></td><td></td><td></td><td></td><td></td><td></td><td></td></tr>
<tr><td>T31032</td><td>GCGGCGCGGTGTGGCGTCGAGGG</td><td>1000</td><td></td><td></td><td></td><td></td><td></td><td></td><td></td><td></td><td></td></tr>
<tr><td>Oligo fwd</td><td>CACCGCGGCGCGGTGTGGCGTCGA</td><td></td><td></td><td></td><td></td><td></td><td></td><td></td><td></td><td></td><td></td></tr>
<tr><td>Oligo rev</td><td>AAACTCGACGCCACACCGCGCCGC</td><td></td><td></td><td></td><td></td><td></td><td></td><td></td><td></td><td></td><td></td></tr>
<tr><td>Chromosome</td><td>start</td><td>end</td><td>strand</td><td>MM</td><td>target_seq</td><td>PAM</td><td>alignment</td><td>distance</td><td>position</td><td>gene name</td><td>gene id</td></tr>
<tr><td>GU305914.1</td><td>42673</td><td>42695</td><td>-</td><td>0</td><td>GCGGCGCGGTGTGGCGTCGA</td><td>GGG</td><td>||||||||[||||||||||||]PAM</td><td>NA</td><td>Intergenic</td><td>NA</td><td>NA</td></tr>
<tr><td></td><td></td><td></td><td></td><td></td><td></td><td></td><td></td><td></td><td></td><td></td><td></td></tr>
<tr><td>T31033</td><td>CAGCGCGCTCCTGAAGCATGCGG</td><td>1000</td><td></td><td></td><td></td><td></td><td></td><td></td><td></td><td></td><td></td></tr>
<tr><td>Oligo fwd</td><td>CACCCAGCGCGCTCCTGAAGCATG</td><td></td><td></td><td></td><td></td><td></td><td></td><td></td><td></td><td></td><td></td></tr>
<tr><td>Oligo rev</td><td>AAACCATGCTTCAGGAGCGCGCTG</td><td></td><td></td><td></td><td></td><td></td><td></td><td></td><td></td><td></td><td></td></tr>
<tr><td>Chromosome</td><td>start</td><td>end</td><td>strand</td><td>MM</td><td>target_seq</td><td>PAM</td><td>alignment</td><td>distance</td><td>position</td><td>gene name</td><td>gene id</td></tr>
<tr><td>GU305914.1</td><td>42639</td><td>42661</td><td>-</td><td>0</td><td>CAGCGCGCTCCTGAAGCATG</td><td>CGG</td><td>||||||||[||||||||||||]PAM</td><td>NA</td><td>Intergenic</td><td>NA</td><td>NA</td></tr>
<tr><td></td><td></td><td></td><td></td><td></td><td></td><td></td><td></td><td></td><td></td><td></td><td></td></tr>
<tr><td>T31034</td><td>GTTTGTTGAAGTCCCGCCCGAGG</td><td>1000</td><td></td><td></td><td></td><td></td><td></td><td></td><td></td><td></td><td></td></tr>
<tr><td>Oligo fwd</td><td>CACCGTTTGTTGAAGTCCCGCCCG</td><td></td><td></td><td></td><td></td><td></td><td></td><td></td><td></td><td></td><td></td></tr>
<tr><td>Oligo rev</td><td>AAACCGGGCGGGACTTCAACAAAC</td><td></td><td></td><td></td><td></td><td></td><td></td><td></td><td></td><td></td><td></td></tr>
<tr><td>Chromosome</td><td>start</td><td>end</td><td>strand</td><td>MM</td><td>target_seq</td><td>PAM</td><td>alignment</td><td>distance</td><td>position</td><td>gene name</td><td>gene id</td></tr>
<tr><td>GU305914.1</td><td>42576</td><td>42598</td><td>-</td><td>0</td><td>GTTTGTTGAAGTCCCGCCCG</td><td>AGG</td><td>||||||||[||||||||||||]PAM</td><td>NA</td><td>Intergenic</td><td>NA</td><td>NA</td></tr>
<tr><td></td><td></td><td></td><td></td><td></td><td></td><td></td><td></td><td></td><td></td><td></td><td></td></tr>
<tr><td>T31035</td><td>GTTGAAGTCCCGCCCGAGGAAGG</td><td>1000</td><td></td><td></td><td></td><td></td><td></td><td></td><td></td><td></td><td></td></tr>
<tr><td>Oligo fwd</td><td>CACCGTTGAAGTCCCGCCCGAGGA</td><td></td><td></td><td></td><td></td><td></td><td></td><td></td><td></td><td></td><td></td></tr>
<tr><td>Oligo rev</td><td>AAACTCCTCGGGCGGGACTTCAAC</td><td></td><td></td><td></td><td></td><td></td><td></td><td></td><td></td><td></td><td></td></tr>
<tr><td>Chromosome</td><td>start</td><td>end</td><td>strand</td><td>MM</td><td>target_seq</td><td>PAM</td><td>alignment</td><td>distance</td><td>position</td><td>gene name</td><td>gene id</td></tr>
<tr><td>GU305914.1</td><td>42572</td><td>42594</td><td>-</td><td>0</td><td>GTTGAAGTCCCGCCCGAGGA</td><td>AGG</td><td>||||||||[||||||||||||]PAM</td><td>NA</td><td>Intergenic</td><td>NA</td><td>NA</td></tr>
<tr><td></td><td></td><td></td><td></td><td></td><td></td><td></td><td></td><td></td><td></td><td></td><td></td></tr>
<tr><td>T31036</td><td>CCCGAGGAAGGCGTACAGCACGG</td><td>1000</td><td></td><td></td><td></td><td></td><td></td><td></td><td></td><td></td><td></td></tr>
<tr><td>Oligo fwd</td><td>CACCCCCGAGGAAGGCGTACAGCA</td><td></td><td></td><td></td><td></td><td></td><td></td><td></td><td></td><td></td><td></td></tr>
<tr><td>Oligo rev</td><td>AAACTGCTGTACGCCTTCCTCGGG</td><td></td><td></td><td></td><td></td><td></td><td></td><td></td><td></td><td></td><td></td></tr>
<tr><td>Chromosome</td><td>start</td><td>end</td><td>strand</td><td>MM</td><td>target_seq</td><td>PAM</td><td>alignment</td><td>distance</td><td>position</td><td>gene name</td><td>gene id</td></tr>
<tr><td>GU305914.1</td><td>42560</td><td>42582</td><td>-</td><td>0</td><td>CCCGAGGAAGGCGTACAGCA</td><td>CGG</td><td>||||||||[||||||||||||]PAM</td><td>NA</td><td>Intergenic</td><td>NA</td><td>NA</td></tr>
<tr><td></td><td></td><td></td><td></td><td></td><td></td><td></td><td></td><td></td><td></td><td></td><td></td></tr>
<tr><td>T31037</td><td>CCGAGGAAGGCGTACAGCACGGG</td><td>1000</td><td></td><td></td><td></td><td></td><td></td><td></td><td></td><td></td><td></td></tr>
<tr><td>Oligo fwd</td><td>CACCCCGAGGAAGGCGTACAGCAC</td><td></td><td></td><td></td><td></td><td></td><td></td><td></td><td></td><td></td><td></td></tr>
<tr><td>Oligo rev</td><td>AAACGTGCTGTACGCCTTCCTCGG</td><td></td><td></td><td></td><td></td><td></td><td></td><td></td><td></td><td></td><td></td></tr>
<tr><td>Chromosome</td><td>start</td><td>end</td><td>strand</td><td>MM</td><td>target_seq</td><td>PAM</td><td>alignment</td><td>distance</td><td>position</td><td>gene name</td><td>gene id</td></tr>
<tr><td>GU305914.1</td><td>42559</td><td>42581</td><td>-</td><td>0</td><td>CCGAGGAAGGCGTACAGCAC</td><td>GGG</td><td>||||||||[||||||||||||]PAM</td><td>NA</td><td>Intergenic</td><td>NA</td><td>NA</td></tr>
<tr><td></td><td></td><td></td><td></td><td></td><td></td><td></td><td></td><td></td><td></td><td></td><td></td></tr>
<tr><td>T31038</td><td>GCACGGGATTCAGCATGCAGTGG</td><td>1000</td><td></td><td></td><td></td><td></td><td></td><td></td><td></td><td></td><td></td></tr>
<tr><td>Oligo fwd</td><td>CACCGCACGGGATTCAGCATGCAG</td><td></td><td></td><td></td><td></td><td></td><td></td><td></td><td></td><td></td><td></td></tr>
<tr><td>Oligo rev</td><td>AAACCTGCATGCTGAATCCCGTGC</td><td></td><td></td><td></td><td></td><td></td><td></td><td></td><td></td><td></td><td></td></tr>
<tr><td>Chromosome</td><td>start</td><td>end</td><td>strand</td><td>MM</td><td>target_seq</td><td>PAM</td><td>alignment</td><td>distance</td><td>position</td><td>gene name</td><td>gene id</td></tr>
<tr><td>GU305914.1</td><td>42543</td><td>42565</td><td>-</td><td>0</td><td>GCACGGGATTCAGCATGCAG</td><td>TGG</td><td>||||||||[||||||||||||]PAM</td><td>NA</td><td>Intergenic</td><td>NA</td><td>NA</td></tr>
<tr><td></td><td></td><td></td><td></td><td></td><td></td><td></td><td></td><td></td><td></td><td></td><td></td></tr>
<tr><td>T31039</td><td>ATTCAGCATGCAGTGGAAGTTGG</td><td>1000</td><td></td><td></td><td></td><td></td><td></td><td></td><td></td><td></td><td></td></tr>
<tr><td>Oligo fwd</td><td>CACCATTCAGCATGCAGTGGAAGT</td><td></td><td></td><td></td><td></td><td></td><td></td><td></td><td></td><td></td><td></td></tr>
<tr><td>Oligo rev</td><td>AAACACTTCCACTGCATGCTGAAT</td><td></td><td></td><td></td><td></td><td></td><td></td><td></td><td></td><td></td><td></td></tr>
<tr><td>Chromosome</td><td>start</td><td>end</td><td>strand</td><td>MM</td><td>target_seq</td><td>PAM</td><td>alignment</td><td>distance</td><td>position</td><td>gene name</td><td>gene id</td></tr>
<tr><td>GU305914.1</td><td>42536</td><td>42558</td><td>-</td><td>0</td><td>ATTCAGCATGCAGTGGAAGT</td><td>TGG</td><td>||||||||[||||||||||||]PAM</td><td>NA</td><td>Intergenic</td><td>NA</td><td>NA</td></tr>
<tr><td></td><td></td><td></td><td></td><td></td><td></td><td></td><td></td><td></td><td></td><td></td><td></td></tr>
<tr><td>T31040</td><td>TTCAGCATGCAGTGGAAGTTGGG</td><td>1000</td><td></td><td></td><td></td><td></td><td></td><td></td><td></td><td></td><td></td></tr>
<tr><td>Oligo fwd</td><td>CACCTTCAGCATGCAGTGGAAGTT</td><td></td><td></td><td></td><td></td><td></td><td></td><td></td><td></td><td></td><td></td></tr>
<tr><td>Oligo rev</td><td>AAACAACTTCCACTGCATGCTGAA</td><td></td><td></td><td></td><td></td><td></td><td></td><td></td><td></td><td></td><td></td></tr>
<tr><td>Chromosome</td><td>start</td><td>end</td><td>strand</td><td>MM</td><td>target_seq</td><td>PAM</td><td>alignment</td><td>distance</td><td>position</td><td>gene name</td><td>gene id</td></tr>
<tr><td>GU305914.1</td><td>42535</td><td>42557</td><td>-</td><td>0</td><td>TTCAGCATGCAGTGGAAGTT</td><td>GGG</td><td>||||||||[||||||||||||]PAM</td><td>NA</td><td>Intergenic</td><td>NA</td><td>NA</td></tr>
<tr><td></td><td></td><td></td><td></td><td></td><td></td><td></td><td></td><td></td><td></td><td></td><td></td></tr>
<tr><td>T31041</td><td>GGGAACCACTCGCGCCAGCATGG</td><td>1000</td><td></td><td></td><td></td><td></td><td></td><td></td><td></td><td></td><td></td></tr>
<tr><td>Oligo fwd</td><td>CACCGGGAACCACTCGCGCCAGCA</td><td></td><td></td><td></td><td></td><td></td><td></td><td></td><td></td><td></td><td></td></tr>
<tr><td>Oligo rev</td><td>AAACTGCTGGCGCGAGTGGTTCCC</td><td></td><td></td><td></td><td></td><td></td><td></td><td></td><td></td><td></td><td></td></tr>
<tr><td>Chromosome</td><td>start</td><td>end</td><td>strand</td><td>MM</td><td>target_seq</td><td>PAM</td><td>alignment</td><td>distance</td><td>position</td><td>gene name</td><td>gene id</td></tr>
<tr><td>GU305914.1</td><td>42515</td><td>42537</td><td>-</td><td>0</td><td>GGGAACCACTCGCGCCAGCA</td><td>TGG</td><td>||||||||[||||||||||||]PAM</td><td>NA</td><td>Intergenic</td><td>NA</td><td>NA</td></tr>
<tr><td></td><td></td><td></td><td></td><td></td><td></td><td></td><td></td><td></td><td></td><td></td><td></td></tr>
<tr><td>T31042</td><td>CCAGCATGGCGACCGCGTCTCGG</td><td>1000</td><td></td><td></td><td></td><td></td><td></td><td></td><td></td><td></td><td></td></tr>
<tr><td>Oligo fwd</td><td>CACCCCAGCATGGCGACCGCGTCT</td><td></td><td></td><td></td><td></td><td></td><td></td><td></td><td></td><td></td><td></td></tr>
<tr><td>Oligo rev</td><td>AAACAGACGCGGTCGCCATGCTGG</td><td></td><td></td><td></td><td></td><td></td><td></td><td></td><td></td><td></td><td></td></tr>
<tr><td>Chromosome</td><td>start</td><td>end</td><td>strand</td><td>MM</td><td>target_seq</td><td>PAM</td><td>alignment</td><td>distance</td><td>position</td><td>gene name</td><td>gene id</td></tr>
<tr><td>GU305914.1</td><td>42501</td><td>42523</td><td>-</td><td>0</td><td>CCAGCATGGCGACCGCGTCT</td><td>CGG</td><td>||||||||[||||||||||||]PAM</td><td>NA</td><td>Intergenic</td><td>NA</td><td>NA</td></tr>
<tr><td></td><td></td><td></td><td></td><td></td><td></td><td></td><td></td><td></td><td></td><td></td><td></td></tr>
<tr><td>T31043</td><td>CGTCACGTCCCACTCGATCACGG</td><td>1000</td><td></td><td></td><td></td><td></td><td></td><td></td><td></td><td></td><td></td></tr>
<tr><td>Oligo fwd</td><td>CACCCGTCACGTCCCACTCGATCA</td><td></td><td></td><td></td><td></td><td></td><td></td><td></td><td></td><td></td><td></td></tr>
<tr><td>Oligo rev</td><td>AAACTGATCGAGTGGGACGTGACG</td><td></td><td></td><td></td><td></td><td></td><td></td><td></td><td></td><td></td><td></td></tr>
<tr><td>Chromosome</td><td>start</td><td>end</td><td>strand</td><td>MM</td><td>target_seq</td><td>PAM</td><td>alignment</td><td>distance</td><td>position</td><td>gene name</td><td>gene id</td></tr>
<tr><td>GU305914.1</td><td>42461</td><td>42483</td><td>-</td><td>0</td><td>CGTCACGTCCCACTCGATCA</td><td>CGG</td><td>||||||||[||||||||||||]PAM</td><td>NA</td><td>Intergenic</td><td>NA</td><td>NA</td></tr>
<tr><td></td><td></td><td></td><td></td><td></td><td></td><td></td><td></td><td></td><td></td><td></td><td></td></tr>
<tr><td>T31044</td><td>AAGATCATGATGCCCACGAACGG</td><td>1000</td><td></td><td></td><td></td><td></td><td></td><td></td><td></td><td></td><td></td></tr>
<tr><td>Oligo fwd</td><td>CACCAAGATCATGATGCCCACGAA</td><td></td><td></td><td></td><td></td><td></td><td></td><td></td><td></td><td></td><td></td></tr>
<tr><td>Oligo rev</td><td>AAACTTCGTGGGCATCATGATCTT</td><td></td><td></td><td></td><td></td><td></td><td></td><td></td><td></td><td></td><td></td></tr>
<tr><td>Chromosome</td><td>start</td><td>end</td><td>strand</td><td>MM</td><td>target_seq</td><td>PAM</td><td>alignment</td><td>distance</td><td>position</td><td>gene name</td><td>gene id</td></tr>
</table>

GU305914.1 42427 42449 - 0 AAGATCATGATGCCCACGAA CGG ||||||||[||||||||||||]PAM NA Intergenic NA NA

T31045 AGATCATGATGCCCACGAACGGG 1000
Oligo fwd CACCAGATCATGATGCCCACGAAC
Oligo rev AAACGTTCGTGGGCATCATGATCT
Chromosome start end strand MM target_seq PAM alignment distance position gene name gene id
GU305914.1 42426 42448 - 0 AGATCATGATGCCCACGAAC GGG ||||||||[||||||||||||]PAM NA Intergenic NA NA

T31046 GATCATGATGCCCACGAACGGGG 1000
Oligo fwd CACCGATCATGATGCCCACGAACG
Oligo rev AAACCGTTCGTGGGCATCATGATC
Chromosome start end strand MM target_seq PAM alignment distance position gene name gene id
GU305914.1 42425 42447 - 0 GATCATGATGCCCACGAACG GGG ||||||||[||||||||||||]PAM NA Intergenic NA NA

T31047 TCTGCAGCACTAAGAATGACAGG 1000
Oligo fwd CACCTCTGCAGCACTAAGAATGAC
Oligo rev AAACGTCATTCTTAGTGCTGCAGA
Chromosome start end strand MM target_seq PAM alignment distance position gene name gene id
GU305914.1 42402 42424 - 0 TCTGCAGCACTAAGAATGAC AGG ||||||||[||||||||||||]PAM NA Intergenic NA NA

T31048 GCACTAAGAATGACAGGATGAGG 1000
Oligo fwd CACCGCACTAAGAATGACAGGATG
Oligo rev AAACCATCCTGTCATTCTTAGTGC
Chromosome start end strand MM target_seq PAM alignment distance position gene name gene id
GU305914.1 42396 42418 - 0 GCACTAAGAATGACAGGATG AGG ||||||||[||||||||||||]PAM NA Intergenic NA NA

T31049 CACTAAGAATGACAGGATGAGGG 1000
Oligo fwd CACCCACTAAGAATGACAGGATGA
Oligo rev AAACTCATCCTGTCATTCTTAGTG
Chromosome start end strand MM target_seq PAM alignment distance position gene name gene id
GU305914.1 42395 42417 - 0 CACTAAGAATGACAGGATGA GGG ||||||||[||||||||||||]PAM NA Intergenic NA NA

T31050 CAGGATGAGGGTCGACACGAAGG 1000
Oligo fwd CACCCAGGATGAGGGTCGACACGA
Oligo rev AAACTCGTGTCGACCCTCATCCTG
Chromosome start end strand MM target_seq PAM alignment distance position gene name gene id
GU305914.1 42383 42405 - 0 CAGGATGAGGGTCGACACGA AGG ||||||||[||||||||||||]PAM NA Intergenic NA NA

T31051 TCGACACGAAGGCCAGCGTGTGG 1000
Oligo fwd CACCTCGACACGAAGGCCAGCGTG
Oligo rev AAACCACGCTGGCCTTCGTGTCGA
Chromosome start end strand MM target_seq PAM alignment distance position gene name gene id
GU305914.1 42372 42394 - 0 TCGACACGAAGGCCAGCGTG TGG ||||||||[||||||||||||]PAM NA Intergenic NA NA

T31052 CCAGCGTGTGGCTGCGCCTGCGG 1000
Oligo fwd CACCCCAGCGTGTGGCTGCGCCTG
Oligo rev AAACCAGGCGCAGCCACACGCTGG
Chromosome start end strand MM target_seq PAM alignment distance position gene name gene id
GU305914.1 42360 42382 - 0 CCAGCGTGTGGCTGCGCCTG CGG ||||||||[||||||||||||]PAM NA Intergenic NA NA

T31053 GTGGCTGCGCCTGCGGTACGAGG 1000
Oligo fwd CACCGTGGCTGCGCCTGCGGTACG
Oligo rev AAACCGTACCGCAGGCGCAGCCAC
Chromosome start end strand MM target_seq PAM alignment distance position gene name gene id
GU305914.1 42353 42375 - 0 GTGGCTGCGCCTGCGGTACG AGG ||||||||[||||||||||||]PAM NA Intergenic NA NA
GU305914.1 69544 69566 + 4 GTCGCGCCGGCTGCGGTACG CGG ||-||--|[|-||||||||||]PAM NA Intergenic NA NA

T31054 TGCGGTACGAGGTCAGTTTCAGG 1000
Oligo fwd CACCTGCGGTACGAGGTCAGTTTC
Oligo rev AAACGAAACTGACCTCGTACCGCA
Chromosome start end strand MM target_seq PAM alignment distance position gene name gene id
GU305914.1 42342 42364 - 0 TGCGGTACGAGGTCAGTTTC AGG ||||||||[||||||||||||]PAM NA Intergenic NA NA

T31055 TCAGGCGTTTGTAGAAAAAAAGG 1000
Oligo fwd CACCTCAGGCGTTTGTAGAAAAAA
Oligo rev AAACTTTTTTCTACAAACGCCTGA
Chromosome start end strand MM target_seq PAM alignment distance position gene name gene id
GU305914.1 42324 42346 - 0 TCAGGCGTTTGTAGAAAAAA AGG ||||||||[||||||||||||]PAM NA Intergenic NA NA

T31056 AACCATGTCATCATCACCACTGG 1000
Oligo fwd CACCAACCATGTCATCATCACCAC
Oligo rev AAACGTGGTGATGATGACATGGTT
Chromosome start end strand MM target_seq PAM alignment distance position gene name gene id
GU305914.1 42298 42320 - 0 AACCATGTCATCATCACCAC TGG ||||||||[||||||||||||]PAM NA Intergenic NA NA

T31057 CGCCCCATACTATGCAAATGAGG 1000
Oligo fwd CACCCGCCCCATACTATGCAAATG
Oligo rev AAACCATTTGCATAGTATGGGGCG
Chromosome start end strand MM target_seq PAM alignment distance position gene name gene id
GU305914.1 42270 42292 - 0 CGCCCCATACTATGCAAATG AGG ||||||||[||||||||||||]PAM NA Intergenic NA NA

T31058 TATGCAAATGAGGATCTTAAAGG 1000
Oligo fwd CACCTATGCAAATGAGGATCTTAA
Oligo rev AAACTTAAGATCCTCATTTGCATA
Chromosome start end strand MM target_seq PAM alignment distance position gene name gene id
GU305914.1 42260 42282 - 0 TATGCAAATGAGGATCTTAA AGG ||||||||[||||||||||||]PAM NA Intergenic NA NA

T31059 GATCTTAAAGGTCGCCAGCACGG 1000
Oligo fwd CACCGATCTTAAAGGTCGCCAGCA

| | | | | | | | | | | | |
|---|---|---|---|---|---|---|---|---|---|---|---|
| Oligo rev | AAACTGCTGGCGACCTTTAAGATC | | | | | | | | | | |
| Chromosome | start | end | strand | MM | target_seq | PAM | alignment | distance | position | gene name | gene id |
| GU305914.1 | 42248 | 42270 | - | 0 | GATCTTAAAGGTCGCCAGCA | CGG | \|\|\|\|\|\|\|\|[\|\|\|\|\|\|\|\|\|\|\|\|]PAM | NA | Intergenic | NA | NA |
| T31060 | GTAGTTGAAGAAGATGATGCAGG | 1000 | | | | | | | | | |
| Oligo fwd | CACCGTAGTTGAAGAAGATGATGC | | | | | | | | | | |
| Oligo rev | AAACGCATCATCTTCTTCAACTAC | | | | | | | | | | |
| Chromosome | start | end | strand | MM | target_seq | PAM | alignment | distance | position | gene name | gene id |
| GU305914.1 | 42212 | 42234 | - | 0 | GTAGTTGAAGAAGATGATGC | AGG | \|\|\|\|\|\|\|\|[\|\|\|\|\|\|\|\|\|\|\|\|]PAM | NA | Intergenic | NA | NA |
| GU305914.1 | 89090 | 89112 | - | 4 | GCACTTCAAGAAGATGAAGC | TGG | \|-\|-\|\|-\|[\|\|\|\|\|\|\|\|\|-\|\|]PAM | NA | Intergenic | NA | NA |
| T31061 | CAGGTCTCGTGCACGCTGTCCGG | 1000 | | | | | | | | | |
| Oligo fwd | CACCCAGGTCTCGTGCACGCTGTC | | | | | | | | | | |
| Oligo rev | AAACGACAGCGTGCACGAGACCTG | | | | | | | | | | |
| Chromosome | start | end | strand | MM | target_seq | PAM | alignment | distance | position | gene name | gene id |
| GU305914.1 | 42193 | 42215 | - | 0 | CAGGTCTCGTGCACGCTGTC | CGG | \|\|\|\|\|\|\|\|[\|\|\|\|\|\|\|\|\|\|\|\|]PAM | NA | Intergenic | NA | NA |
| T31062 | GTCCCCGTCGTGCGCCAAAACGG | 1000 | | | | | | | | | |
| Oligo fwd | CACCGTCCCCGTCGTGCGCCAAAA | | | | | | | | | | |
| Oligo rev | AAACTTTTGGCGCACGACGGGGAC | | | | | | | | | | |
| Chromosome | start | end | strand | MM | target_seq | PAM | alignment | distance | position | gene name | gene id |
| GU305914.1 | 42167 | 42189 | - | 0 | GTCCCCGTCGTGCGCCAAAA | CGG | \|\|\|\|\|\|\|\|[\|\|\|\|\|\|\|\|\|\|\|\|]PAM | NA | Intergenic | NA | NA |
| T31063 | GCCAAAACGGTCACGTACGTCGG | 1000 | | | | | | | | | |
| Oligo fwd | CACCGCCAAAACGGTCACGTACGT | | | | | | | | | | |
| Oligo rev | AAACACGTACGTGACCGTTTTGGC | | | | | | | | | | |
| Chromosome | start | end | strand | MM | target_seq | PAM | alignment | distance | position | gene name | gene id |
| GU305914.1 | 42154 | 42176 | - | 0 | GCCAAAACGGTCACGTACGT | CGG | \|\|\|\|\|\|\|\|[\|\|\|\|\|\|\|\|\|\|\|\|]PAM | NA | Intergenic | NA | NA |
| T31064 | ACGGTCACGTACGTCGGCGCCGG | 1000 | | | | | | | | | |
| Oligo fwd | CACCACGGTCACGTACGTCGGCGC | | | | | | | | | | |
| Oligo rev | AAACGCGCCGACGTACGTGACCGT | | | | | | | | | | |
| Chromosome | start | end | strand | MM | target_seq | PAM | alignment | distance | position | gene name | gene id |
| GU305914.1 | 42148 | 42170 | - | 0 | ACGGTCACGTACGTCGGCGC | CGG | \|\|\|\|\|\|\|\|[\|\|\|\|\|\|\|\|\|\|\|\|]PAM | NA | Intergenic | NA | NA |
| T31065 | CGGCGACGCGCACATCAGCGAGG | 1000 | | | | | | | | | |
| Oligo fwd | CACCCGGCGACGCGCACATCAGCG | | | | | | | | | | |
| Oligo rev | AAACCGCTGATGTGCGCGTCGCCG | | | | | | | | | | |
| Chromosome | start | end | strand | MM | target_seq | PAM | alignment | distance | position | gene name | gene id |
| GU305914.1 | 42128 | 42150 | - | 0 | CGGCGACGCGCACATCAGCG | AGG | \|\|\|\|\|\|\|\|[\|\|\|\|\|\|\|\|\|\|\|\|]PAM | NA | Intergenic | NA | NA |
| T31066 | CCGCGTTCTTGTTCGTGCTCTGG | 1000 | | | | | | | | | |
| Oligo fwd | CACCCCGCGTTCTTGTTCGTGCTC | | | | | | | | | | |
| Oligo rev | AAACGAGCACGAACAAGAACGCGG | | | | | | | | | | |
| Chromosome | start | end | strand | MM | target_seq | PAM | alignment | distance | position | gene name | gene id |
| GU305914.1 | 42057 | 42079 | - | 0 | CCGCGTTCTTGTTCGTGCTC | TGG | \|\|\|\|\|\|\|\|[\|\|\|\|\|\|\|\|\|\|\|\|]PAM | NA | Intergenic | NA | NA |
| T31067 | CGTTCTTGTTCGTGCTCTGGTGG | 1000 | | | | | | | | | |
| Oligo fwd | CACCCGTTCTTGTTCGTGCTCTGG | | | | | | | | | | |
| Oligo rev | AAACCCAGAGCACGAACAAGAACG | | | | | | | | | | |
| Chromosome | start | end | strand | MM | target_seq | PAM | alignment | distance | position | gene name | gene id |
| GU305914.1 | 42054 | 42076 | - | 0 | CGTTCTTGTTCGTGCTCTGG | TGG | \|\|\|\|\|\|\|\|[\|\|\|\|\|\|\|\|\|\|\|\|]PAM | NA | Intergenic | NA | NA |
| T31068 | CAGCGCCAGCGTCGAGAACCCGG | 1000 | | | | | | | | | |
| Oligo fwd | CACCCAGCGCCAGCGTCGAGAACC | | | | | | | | | | |
| Oligo rev | AAACGGTTCTCGACGCTGGCGCTG | | | | | | | | | | |
| Chromosome | start | end | strand | MM | target_seq | PAM | alignment | distance | position | gene name | gene id |
| GU305914.1 | 42005 | 42027 | - | 0 | CAGCGCCAGCGTCGAGAACC | CGG | \|\|\|\|\|\|\|\|[\|\|\|\|\|\|\|\|\|\|\|\|]PAM | NA | Intergenic | NA | NA |
| GU305914.1 | 100832 | 100854 | + | 4 | CGGCGGCAGCGGCGAGAACA | GGG | \|-\|\|\|-\|\|[\|\|\|-\|\|\|\|\|\|\|-]PAM | NA | Intergenic | NA | NA |
| T31069 | GGCCGTGCACGACGCGTAGTAGG | 1000 | | | | | | | | | |
| Oligo fwd | CACCGGCCGTGCACGACGCGTAGT | | | | | | | | | | |
| Oligo rev | AAACACTACGCGTCGTGCACGGCC | | | | | | | | | | |
| Chromosome | start | end | strand | MM | target_seq | PAM | alignment | distance | position | gene name | gene id |
| GU305914.1 | 41984 | 42006 | - | 0 | GGCCGTGCACGACGCGTAGT | AGG | \|\|\|\|\|\|\|\|[\|\|\|\|\|\|\|\|\|\|\|\|]PAM | NA | Intergenic | NA | NA |
| T31070 | GCCACCAGCTTGCACCCCTCCGG | 1000 | | | | | | | | | |
| Oligo fwd | CACCGCCACCAGCTTGCACCCCTC | | | | | | | | | | |
| Oligo rev | AAACGAGGGGTGCAAGCTGGTGGC | | | | | | | | | | |
| Chromosome | start | end | strand | MM | target_seq | PAM | alignment | distance | position | gene name | gene id |
| GU305914.1 | 41956 | 41978 | - | 0 | GCCACCAGCTTGCACCCCTC | CGG | \|\|\|\|\|\|\|\|[\|\|\|\|\|\|\|\|\|\|\|\|]PAM | NA | Intergenic | NA | NA |
| T31071 | CCACCAGCTTGCACCCCTCCGGG | 1000 | | | | | | | | | |
| Oligo fwd | CACCCCACCAGCTTGCACCCCTCC | | | | | | | | | | |
| Oligo rev | AAACGGAGGGGTGCAAGCTGGTGG | | | | | | | | | | |
| Chromosome | start | end | strand | MM | target_seq | PAM | alignment | distance | position | gene name | gene id |
| GU305914.1 | 41955 | 41977 | - | 0 | CCACCAGCTTGCACCCCTCC | GGG | \|\|\|\|\|\|\|\|[\|\|\|\|\|\|\|\|\|\|\|\|]PAM | NA | Intergenic | NA | NA |
| T31072 | CACCAGCTTGCACCCCTCCGGGG | 1000 | | | | | | | | | |
| Oligo fwd | CACCCACCAGCTTGCACCCCTCCG | | | | | | | | | | |
| Oligo rev | AAACCGGAGGGGTGCAAGCTGGTG | | | | | | | | | | |
| Chromosome | start | end | strand | MM | target_seq | PAM | alignment | distance | position | gene name | gene id |
| GU305914.1 | 41954 | 41976 | - | 0 | CACCAGCTTGCACCCCTCCG | GGG | \|\|\|\|\|\|\|\|[\|\|\|\|\|\|\|\|\|\|\|\|]PAM | NA | Intergenic | NA | NA |
| T31073 | TGATCTGACCCCGATTGCTCAGG | 1000 | | | | | | | | | |
| Oligo fwd | CACCTGATCTGACCCCGATTGCTC | | | | | | | | | | |
| Oligo rev | AAACGAGCAATCGGGGTCAGATCA | | | | | | | | | | |
| Chromosome | start | end | strand | MM | target_seq | PAM | alignment | distance | position | gene name | gene id |
| GU305914.1 | 41925 | 41947 | - | 0 | TGATCTGACCCCGATTGCTC | AGG | \|\|\|\|\|\|\|\|[\|\|\|\|\|\|\|\|\|\|\|\|]PAM | NA | Intergenic | NA | NA |

| | | | | | | | | | | | | |
|---|---|---|---|---|---|---|---|---|---|---|---|---|
| T31074 | GACCCCGATTGCTCAGGATCAGG | | 1000 | | | | | | | | | |
| Oligo fwd | CACCGACCCCGATTGCTCAGGATC | | | | | | | | | | | |
| Oligo rev | AAACGATCCTGAGCAATCGGGGTC | | | | | | | | | | | |
| Chromosome | start | | end | strand | MM | target_seq | PAM | alignment | distance | position | gene name | gene id |
| GU305914.1 | | 41919 | 41941 | - | 0 | GACCCCGATTGCTCAGGATC | AGG | \|\|\|\|\|\|\|\|[\|\|\|\|\|\|\|\|\|\|\|\|]PAM | NA | Intergenic | NA | NA |
| T31075 | CGATTGCTCAGGATCAGGAACGG | | 1000 | | | | | | | | | |
| Oligo fwd | CACCCGATTGCTCAGGATCAGGAA | | | | | | | | | | | |
| Oligo rev | AAACTTCCTGATCCTGAGCAATCG | | | | | | | | | | | |
| Chromosome | start | | end | strand | MM | target_seq | PAM | alignment | distance | position | gene name | gene id |
| GU305914.1 | | 41914 | 41936 | - | 0 | CGATTGCTCAGGATCAGGAA | CGG | \|\|\|\|\|\|\|\|[\|\|\|\|\|\|\|\|\|\|\|\|]PAM | NA | Intergenic | NA | NA |
| T31076 | GCAGCATCGTCACCGTCAACAGG | | 1000 | | | | | | | | | |
| Oligo fwd | CACCGCAGCATCGTCACCGTCAAC | | | | | | | | | | | |
| Oligo rev | AAACGTTGACGGTGACGATGCTGC | | | | | | | | | | | |
| Chromosome | start | | end | strand | MM | target_seq | PAM | alignment | distance | position | gene name | gene id |
| GU305914.1 | | 41892 | 41914 | - | 0 | GCAGCATCGTCACCGTCAAC | AGG | \|\|\|\|\|\|\|\|[\|\|\|\|\|\|\|\|\|\|\|\|]PAM | NA | Intergenic | NA | NA |
| T31077 | TCAACAGGTTCGCAAAGTACAGG | | 1000 | | | | | | | | | |
| Oligo fwd | CACCTCAACAGGTTCGCAAAGTAC | | | | | | | | | | | |
| Oligo rev | AAACGTACTTTGCGAACCTGTTGA | | | | | | | | | | | |
| Chromosome | start | | end | strand | MM | target_seq | PAM | alignment | distance | position | gene name | gene id |
| GU305914.1 | | 41877 | 41899 | - | 0 | TCAACAGGTTCGCAAAGTAC | AGG | \|\|\|\|\|\|\|\|[\|\|\|\|\|\|\|\|\|\|\|\|]PAM | NA | Intergenic | NA | NA |
| T31078 | CAGGTTCGCAAAGTACAGGTTGG | | 1000 | | | | | | | | | |
| Oligo fwd | CACCCAGGTTCGCAAAGTACAGGT | | | | | | | | | | | |
| Oligo rev | AAACACCTGTACTTTGCGAACCTG | | | | | | | | | | | |
| Chromosome | start | | end | strand | MM | target_seq | PAM | alignment | distance | position | gene name | gene id |
| GU305914.1 | | 41873 | 41895 | - | 0 | CAGGTTCGCAAAGTACAGGT | TGG | \|\|\|\|\|\|\|\|[\|\|\|\|\|\|\|\|\|\|\|\|]PAM | NA | Intergenic | NA | NA |
| T31079 | AGTACAGGTTGGTCATGTACAGG | | 1000 | | | | | | | | | |
| Oligo fwd | CACCAGTACAGGTTGGTCATGTAC | | | | | | | | | | | |
| Oligo rev | AAACGTACATGACCAACCTGTACT | | | | | | | | | | | |
| Chromosome | start | | end | strand | MM | target_seq | PAM | alignment | distance | position | gene name | gene id |
| GU305914.1 | | 41862 | 41884 | - | 0 | AGTACAGGTTGGTCATGTAC | AGG | \|\|\|\|\|\|\|\|[\|\|\|\|\|\|\|\|\|\|\|\|]PAM | NA | Intergenic | NA | NA |
| T31080 | GTACAGGTTGGTCATGTACAGGG | | 1000 | | | | | | | | | |
| Oligo fwd | CACCGTACAGGTTGGTCATGTACA | | | | | | | | | | | |
| Oligo rev | AAACTGTACATGACCAACCTGTAC | | | | | | | | | | | |
| Chromosome | start | | end | strand | MM | target_seq | PAM | alignment | distance | position | gene name | gene id |
| GU305914.1 | | 41861 | 41883 | - | 0 | GTACAGGTTGGTCATGTACA | GGG | \|\|\|\|\|\|\|\|[\|\|\|\|\|\|\|\|\|\|\|\|]PAM | NA | Intergenic | NA | NA |
| T31081 | AGGTTGGTCATGTACAGGGTCGG | | 1000 | | | | | | | | | |
| Oligo fwd | CACCAGGTTGGTCATGTACAGGGT | | | | | | | | | | | |
| Oligo rev | AAACACCCTGTACATGACCAACCT | | | | | | | | | | | |
| Chromosome | start | | end | strand | MM | target_seq | PAM | alignment | distance | position | gene name | gene id |
| GU305914.1 | | 41857 | 41879 | - | 0 | AGGTTGGTCATGTACAGGGT | CGG | \|\|\|\|\|\|\|\|[\|\|\|\|\|\|\|\|\|\|\|\|]PAM | NA | Intergenic | NA | NA |
| T31082 | GGTTGGTCATGTACAGGGTCGGG | | 1000 | | | | | | | | | |
| Oligo fwd | CACCGGTTGGTCATGTACAGGGTC | | | | | | | | | | | |
| Oligo rev | AAACGACCCTGTACATGACCAACC | | | | | | | | | | | |
| Chromosome | start | | end | strand | MM | target_seq | PAM | alignment | distance | position | gene name | gene id |
| GU305914.1 | | 41856 | 41878 | - | 0 | GGTTGGTCATGTACAGGGTC | GGG | \|\|\|\|\|\|\|\|[\|\|\|\|\|\|\|\|\|\|\|\|]PAM | NA | Intergenic | NA | NA |
| T31083 | GTTGGTCATGTACAGGGTCGGGG | | 1000 | | | | | | | | | |
| Oligo fwd | CACCGTTGGTCATGTACAGGGTCG | | | | | | | | | | | |
| Oligo rev | AAACCGACCCTGTACATGACCAAC | | | | | | | | | | | |
| Chromosome | start | | end | strand | MM | target_seq | PAM | alignment | distance | position | gene name | gene id |
| GU305914.1 | | 41855 | 41877 | - | 0 | GTTGGTCATGTACAGGGTCG | GGG | \|\|\|\|\|\|\|\|[\|\|\|\|\|\|\|\|\|\|\|\|]PAM | NA | Intergenic | NA | NA |
| T31084 | GGGCCGAGCTGCGATAGATGCGG | | 1000 | | | | | | | | | |
| Oligo fwd | CACCGGGCCGAGCTGCGATAGATG | | | | | | | | | | | |
| Oligo rev | AAACCATCTATCGCAGCTCGGCCC | | | | | | | | | | | |
| Chromosome | start | | end | strand | MM | target_seq | PAM | alignment | distance | position | gene name | gene id |
| GU305914.1 | | 41835 | 41857 | - | 0 | GGGCCGAGCTGCGATAGATG | CGG | \|\|\|\|\|\|\|\|[\|\|\|\|\|\|\|\|\|\|\|\|]PAM | NA | Intergenic | NA | NA |
| T31085 | GTCAGCACCACGAAGTTCATCGG | | 1000 | | | | | | | | | |
| Oligo fwd | CACCGTCAGCACCACGAAGTTCAT | | | | | | | | | | | |
| Oligo rev | AAACATGAACTTCGTGGTGCTGAC | | | | | | | | | | | |
| Chromosome | start | | end | strand | MM | target_seq | PAM | alignment | distance | position | gene name | gene id |
| GU305914.1 | | 41794 | 41816 | - | 0 | GTCAGCACCACGAAGTTCAT | CGG | \|\|\|\|\|\|\|\|[\|\|\|\|\|\|\|\|\|\|\|\|]PAM | NA | Intergenic | NA | NA |
| T31086 | AGTTCATCGGCCCGCCGATCAGG | | 1000 | | | | | | | | | |
| Oligo fwd | CACCAGTTCATCGGCCCGCCGATC | | | | | | | | | | | |
| Oligo rev | AAACGATCGGCGGGCCGATGAACT | | | | | | | | | | | |
| Chromosome | start | | end | strand | MM | target_seq | PAM | alignment | distance | position | gene name | gene id |
| GU305914.1 | | 41781 | 41803 | - | 0 | AGTTCATCGGCCCGCCGATC | AGG | \|\|\|\|\|\|\|\|[\|\|\|\|\|\|\|\|\|\|\|\|]PAM | NA | Intergenic | NA | NA |
| T31087 | CAGGATGATCACCGTGTTGATGG | | 1000 | | | | | | | | | |
| Oligo fwd | CACCCAGGATGATCACCGTGTTGA | | | | | | | | | | | |
| Oligo rev | AAACTCAACACGGTGATCATCCTG | | | | | | | | | | | |
| Chromosome | start | | end | strand | MM | target_seq | PAM | alignment | distance | position | gene name | gene id |
| GU305914.1 | | 41762 | 41784 | - | 0 | CAGGATGATCACCGTGTTGA | TGG | \|\|\|\|\|\|\|\|[\|\|\|\|\|\|\|\|\|\|\|\|]PAM | NA | Intergenic | NA | NA |
| T31088 | CGCGATGCTCAGCCCCAGCGAGG | | 1000 | | | | | | | | | |
| Oligo fwd | CACCCGCGATGCTCAGCCCCAGCG | | | | | | | | | | | |
| Oligo rev | AAACCGCTGGGGCTGAGCATCGCG | | | | | | | | | | | |
| Chromosome | start | | end | strand | MM | target_seq | PAM | alignment | distance | position | gene name | gene id |

<table>
<tr><td>GU305914.1</td><td>41723</td><td>41745</td><td>-</td><td>0</td><td>CGCGATGCTCAGCCCCAGCG</td><td>AGG</td><td>||||||||[||||||||||||]PAM</td><td>NA</td><td>Intergenic</td><td>NA</td><td>NA</td></tr>
<tr><td></td><td></td><td></td><td></td><td></td><td></td><td></td><td></td><td></td><td></td><td></td><td></td></tr>
<tr><td>T31089</td><td>GCGATGCTCAGCCCCAGCGAGGG</td><td>1000</td><td></td><td></td><td></td><td></td><td></td><td></td><td></td><td></td><td></td></tr>
<tr><td>Oligo fwd</td><td>CACCGCGATGCTCAGCCCCAGCGA</td><td></td><td></td><td></td><td></td><td></td><td></td><td></td><td></td><td></td><td></td></tr>
<tr><td>Oligo rev</td><td>AAACTCGCTGGGGCTGAGCATCGC</td><td></td><td></td><td></td><td></td><td></td><td></td><td></td><td></td><td></td><td></td></tr>
<tr><td>Chromosome</td><td>start</td><td>end</td><td>strand</td><td>MM</td><td>target_seq</td><td>PAM</td><td>alignment</td><td>distance</td><td>position</td><td>gene name</td><td>gene id</td></tr>
<tr><td>GU305914.1</td><td>41722</td><td>41744</td><td>-</td><td>0</td><td>GCGATGCTCAGCCCCAGCGA</td><td>GGG</td><td>||||||||[||||||||||||]PAM</td><td>NA</td><td>Intergenic</td><td>NA</td><td>NA</td></tr>
<tr><td></td><td></td><td></td><td></td><td></td><td></td><td></td><td></td><td></td><td></td><td></td><td></td></tr>
<tr><td>T31090</td><td>CGATGCTCAGCCCCAGCGAGGGG</td><td>1000</td><td></td><td></td><td></td><td></td><td></td><td></td><td></td><td></td><td></td></tr>
<tr><td>Oligo fwd</td><td>CACCCGATGCTCAGCCCCAGCGAG</td><td></td><td></td><td></td><td></td><td></td><td></td><td></td><td></td><td></td><td></td></tr>
<tr><td>Oligo rev</td><td>AAACCTCGCTGGGGCTGAGCATCG</td><td></td><td></td><td></td><td></td><td></td><td></td><td></td><td></td><td></td><td></td></tr>
<tr><td>Chromosome</td><td>start</td><td>end</td><td>strand</td><td>MM</td><td>target_seq</td><td>PAM</td><td>alignment</td><td>distance</td><td>position</td><td>gene name</td><td>gene id</td></tr>
<tr><td>GU305914.1</td><td>41721</td><td>41743</td><td>-</td><td>0</td><td>CGATGCTCAGCCCCAGCGAG</td><td>GGG</td><td>||||||||[||||||||||||]PAM</td><td>NA</td><td>Intergenic</td><td>NA</td><td>NA</td></tr>
<tr><td></td><td></td><td></td><td></td><td></td><td></td><td></td><td></td><td></td><td></td><td></td><td></td></tr>
<tr><td>T31091</td><td>GATGCTCAGCCCCAGCGAGGGGG</td><td>1000</td><td></td><td></td><td></td><td></td><td></td><td></td><td></td><td></td><td></td></tr>
<tr><td>Oligo fwd</td><td>CACCGATGCTCAGCCCCAGCGAGG</td><td></td><td></td><td></td><td></td><td></td><td></td><td></td><td></td><td></td><td></td></tr>
<tr><td>Oligo rev</td><td>AAACCCTCGCTGGGGCTGAGCATC</td><td></td><td></td><td></td><td></td><td></td><td></td><td></td><td></td><td></td><td></td></tr>
<tr><td>Chromosome</td><td>start</td><td>end</td><td>strand</td><td>MM</td><td>target_seq</td><td>PAM</td><td>alignment</td><td>distance</td><td>position</td><td>gene name</td><td>gene id</td></tr>
<tr><td>GU305914.1</td><td>41720</td><td>41742</td><td>-</td><td>0</td><td>GATGCTCAGCCCCAGCGAGG</td><td>GGG</td><td>||||||||[||||||||||||]PAM</td><td>NA</td><td>Intergenic</td><td>NA</td><td>NA</td></tr>
<tr><td></td><td></td><td></td><td></td><td></td><td></td><td></td><td></td><td></td><td></td><td></td><td></td></tr>
<tr><td>T31092</td><td>CGCACGTGTTGTTCACGTGCAGG</td><td>1000</td><td></td><td></td><td></td><td></td><td></td><td></td><td></td><td></td><td></td></tr>
<tr><td>Oligo fwd</td><td>CACCCGCACGTGTTGTTCACGTGC</td><td></td><td></td><td></td><td></td><td></td><td></td><td></td><td></td><td></td><td></td></tr>
<tr><td>Oligo rev</td><td>AAACGCACGTGAACAACACGTGCG</td><td></td><td></td><td></td><td></td><td></td><td></td><td></td><td></td><td></td><td></td></tr>
<tr><td>Chromosome</td><td>start</td><td>end</td><td>strand</td><td>MM</td><td>target_seq</td><td>PAM</td><td>alignment</td><td>distance</td><td>position</td><td>gene name</td><td>gene id</td></tr>
<tr><td>GU305914.1</td><td>41697</td><td>41719</td><td>-</td><td>0</td><td>CGCACGTGTTGTTCACGTGC</td><td>AGG</td><td>||||||||[||||||||||||]PAM</td><td>NA</td><td>Intergenic</td><td>NA</td><td>NA</td></tr>
<tr><td></td><td></td><td></td><td></td><td></td><td></td><td></td><td></td><td></td><td></td><td></td><td></td></tr>
<tr><td>T31093</td><td>AGGTAGTCGCTCTCGTCCTGCGG</td><td>1000</td><td></td><td></td><td></td><td></td><td></td><td></td><td></td><td></td><td></td></tr>
<tr><td>Oligo fwd</td><td>CACCAGGTAGTCGCTCTCGTCCTG</td><td></td><td></td><td></td><td></td><td></td><td></td><td></td><td></td><td></td><td></td></tr>
<tr><td>Oligo rev</td><td>AAACCAGGACGAGAGCGACTACCT</td><td></td><td></td><td></td><td></td><td></td><td></td><td></td><td></td><td></td><td></td></tr>
<tr><td>Chromosome</td><td>start</td><td>end</td><td>strand</td><td>MM</td><td>target_seq</td><td>PAM</td><td>alignment</td><td>distance</td><td>position</td><td>gene name</td><td>gene id</td></tr>
<tr><td>GU305914.1</td><td>41677</td><td>41699</td><td>-</td><td>0</td><td>AGGTAGTCGCTCTCGTCCTG</td><td>CGG</td><td>||||||||[||||||||||||]PAM</td><td>NA</td><td>Intergenic</td><td>NA</td><td>NA</td></tr>
<tr><td></td><td></td><td></td><td></td><td></td><td></td><td></td><td></td><td></td><td></td><td></td><td></td></tr>
<tr><td>T31094</td><td>GGTAGTCGCTCTCGTCCTGCGGG</td><td>1000</td><td></td><td></td><td></td><td></td><td></td><td></td><td></td><td></td><td></td></tr>
<tr><td>Oligo fwd</td><td>CACCGGTAGTCGCTCTCGTCCTGC</td><td></td><td></td><td></td><td></td><td></td><td></td><td></td><td></td><td></td><td></td></tr>
<tr><td>Oligo rev</td><td>AAACGCAGGACGAGAGCGACTACC</td><td></td><td></td><td></td><td></td><td></td><td></td><td></td><td></td><td></td><td></td></tr>
<tr><td>Chromosome</td><td>start</td><td>end</td><td>strand</td><td>MM</td><td>target_seq</td><td>PAM</td><td>alignment</td><td>distance</td><td>position</td><td>gene name</td><td>gene id</td></tr>
<tr><td>GU305914.1</td><td>41676</td><td>41698</td><td>-</td><td>0</td><td>GGTAGTCGCTCTCGTCCTGC</td><td>GGG</td><td>||||||||[||||||||||||]PAM</td><td>NA</td><td>Intergenic</td><td>NA</td><td>NA</td></tr>
<tr><td></td><td></td><td></td><td></td><td></td><td></td><td></td><td></td><td></td><td></td><td></td><td></td></tr>
<tr><td>T31095</td><td>GCTCTCGTCCTGCGGGCGAGAGG</td><td>1000</td><td></td><td></td><td></td><td></td><td></td><td></td><td></td><td></td><td></td></tr>
<tr><td>Oligo fwd</td><td>CACCGCTCTCGTCCTGCGGGCGAG</td><td></td><td></td><td></td><td></td><td></td><td></td><td></td><td></td><td></td><td></td></tr>
<tr><td>Oligo rev</td><td>AAACCTCGCCCGCAGGACGAGAGC</td><td></td><td></td><td></td><td></td><td></td><td></td><td></td><td></td><td></td><td></td></tr>
<tr><td>Chromosome</td><td>start</td><td>end</td><td>strand</td><td>MM</td><td>target_seq</td><td>PAM</td><td>alignment</td><td>distance</td><td>position</td><td>gene name</td><td>gene id</td></tr>
<tr><td>GU305914.1</td><td>41669</td><td>41691</td><td>-</td><td>0</td><td>GCTCTCGTCCTGCGGGCGAG</td><td>AGG</td><td>||||||||[||||||||||||]PAM</td><td>NA</td><td>Intergenic</td><td>NA</td><td>NA</td></tr>
<tr><td></td><td></td><td></td><td></td><td></td><td></td><td></td><td></td><td></td><td></td><td></td><td></td></tr>
<tr><td>T31096</td><td>CTCTCGTCCTGCGGGCGAGAGGG</td><td>1000</td><td></td><td></td><td></td><td></td><td></td><td></td><td></td><td></td><td></td></tr>
<tr><td>Oligo fwd</td><td>CACCCTCTCGTCCTGCGGGCGAGA</td><td></td><td></td><td></td><td></td><td></td><td></td><td></td><td></td><td></td><td></td></tr>
<tr><td>Oligo rev</td><td>AAACTCTCGCCCGCAGGACGAGAG</td><td></td><td></td><td></td><td></td><td></td><td></td><td></td><td></td><td></td><td></td></tr>
<tr><td>Chromosome</td><td>start</td><td>end</td><td>strand</td><td>MM</td><td>target_seq</td><td>PAM</td><td>alignment</td><td>distance</td><td>position</td><td>gene name</td><td>gene id</td></tr>
<tr><td>GU305914.1</td><td>41668</td><td>41690</td><td>-</td><td>0</td><td>CTCTCGTCCTGCGGGCGAGA</td><td>GGG</td><td>||||||||[||||||||||||]PAM</td><td>NA</td><td>Intergenic</td><td>NA</td><td>NA</td></tr>
<tr><td></td><td></td><td></td><td></td><td></td><td></td><td></td><td></td><td></td><td></td><td></td><td></td></tr>
<tr><td>T31097</td><td>CGAGAGGGCGAGATATGTTGTGG</td><td>1000</td><td></td><td></td><td></td><td></td><td></td><td></td><td></td><td></td><td></td></tr>
<tr><td>Oligo fwd</td><td>CACCCGAGAGGGCGAGATATGTTG</td><td></td><td></td><td></td><td></td><td></td><td></td><td></td><td></td><td></td><td></td></tr>
<tr><td>Oligo rev</td><td>AAACCAACATATCTCGCCCTCTCG</td><td></td><td></td><td></td><td></td><td></td><td></td><td></td><td></td><td></td><td></td></tr>
<tr><td>Chromosome</td><td>start</td><td>end</td><td>strand</td><td>MM</td><td>target_seq</td><td>PAM</td><td>alignment</td><td>distance</td><td>position</td><td>gene name</td><td>gene id</td></tr>
<tr><td>GU305914.1</td><td>41653</td><td>41675</td><td>-</td><td>0</td><td>CGAGAGGGCGAGATATGTTG</td><td>TGG</td><td>||||||||[||||||||||||]PAM</td><td>NA</td><td>Intergenic</td><td>NA</td><td>NA</td></tr>
<tr><td></td><td></td><td></td><td></td><td></td><td></td><td></td><td></td><td></td><td></td><td></td><td></td></tr>
<tr><td>T31098</td><td>GAGAGGGCGAGATATGTTGTGGG</td><td>1000</td><td></td><td></td><td></td><td></td><td></td><td></td><td></td><td></td><td></td></tr>
<tr><td>Oligo fwd</td><td>CACCGAGAGGGCGAGATATGTTGT</td><td></td><td></td><td></td><td></td><td></td><td></td><td></td><td></td><td></td><td></td></tr>
<tr><td>Oligo rev</td><td>AAACACAACATATCTCGCCCTCTC</td><td></td><td></td><td></td><td></td><td></td><td></td><td></td><td></td><td></td><td></td></tr>
<tr><td>Chromosome</td><td>start</td><td>end</td><td>strand</td><td>MM</td><td>target_seq</td><td>PAM</td><td>alignment</td><td>distance</td><td>position</td><td>gene name</td><td>gene id</td></tr>
<tr><td>GU305914.1</td><td>41652</td><td>41674</td><td>-</td><td>0</td><td>GAGAGGGCGAGATATGTTGT</td><td>GGG</td><td>||||||||[||||||||||||]PAM</td><td>NA</td><td>Intergenic</td><td>NA</td><td>NA</td></tr>
<tr><td></td><td></td><td></td><td></td><td></td><td></td><td></td><td></td><td></td><td></td><td></td><td></td></tr>
<tr><td>T31099</td><td>AGAGGGCGAGATATGTTGTGGGG</td><td>1000</td><td></td><td></td><td></td><td></td><td></td><td></td><td></td><td></td><td></td></tr>
<tr><td>Oligo fwd</td><td>CACCAGAGGGCGAGATATGTTGTG</td><td></td><td></td><td></td><td></td><td></td><td></td><td></td><td></td><td></td><td></td></tr>
<tr><td>Oligo rev</td><td>AAACCACAACATATCTCGCCCTCT</td><td></td><td></td><td></td><td></td><td></td><td></td><td></td><td></td><td></td><td></td></tr>
<tr><td>Chromosome</td><td>start</td><td>end</td><td>strand</td><td>MM</td><td>target_seq</td><td>PAM</td><td>alignment</td><td>distance</td><td>position</td><td>gene name</td><td>gene id</td></tr>
<tr><td>GU305914.1</td><td>41651</td><td>41673</td><td>-</td><td>0</td><td>AGAGGGCGAGATATGTTGTG</td><td>GGG</td><td>||||||||[||||||||||||]PAM</td><td>NA</td><td>Intergenic</td><td>NA</td><td>NA</td></tr>
<tr><td></td><td></td><td></td><td></td><td></td><td></td><td></td><td></td><td></td><td></td><td></td><td></td></tr>
<tr><td>T31100</td><td>TATGTTGTGGGGATAGACACCGG</td><td>1000</td><td></td><td></td><td></td><td></td><td></td><td></td><td></td><td></td><td></td></tr>
<tr><td>Oligo fwd</td><td>CACCTATGTTGTGGGGATAGACAC</td><td></td><td></td><td></td><td></td><td></td><td></td><td></td><td></td><td></td><td></td></tr>
<tr><td>Oligo rev</td><td>AAACGTGTCTATCCCCACAACATA</td><td></td><td></td><td></td><td></td><td></td><td></td><td></td><td></td><td></td><td></td></tr>
<tr><td>Chromosome</td><td>start</td><td>end</td><td>strand</td><td>MM</td><td>target_seq</td><td>PAM</td><td>alignment</td><td>distance</td><td>position</td><td>gene name</td><td>gene id</td></tr>
<tr><td>GU305914.1</td><td>41640</td><td>41662</td><td>-</td><td>0</td><td>TATGTTGTGGGGATAGACAC</td><td>CGG</td><td>||||||||[||||||||||||]PAM</td><td>NA</td><td>Intergenic</td><td>NA</td><td>NA</td></tr>
<tr><td></td><td></td><td></td><td></td><td></td><td></td><td></td><td></td><td></td><td></td><td></td><td></td></tr>
<tr><td>T31101</td><td>CACCGGACACCGAAGATAGACGG</td><td>1000</td><td></td><td></td><td></td><td></td><td></td><td></td><td></td><td></td><td></td></tr>
<tr><td>Oligo fwd</td><td>CACCCACCGGACACCGAAGATAGA</td><td></td><td></td><td></td><td></td><td></td><td></td><td></td><td></td><td></td><td></td></tr>
<tr><td>Oligo rev</td><td>AAACTCTATCTTCGGTGTCCGGTG</td><td></td><td></td><td></td><td></td><td></td><td></td><td></td><td></td><td></td><td></td></tr>
<tr><td>Chromosome</td><td>start</td><td>end</td><td>strand</td><td>MM</td><td>target_seq</td><td>PAM</td><td>alignment</td><td>distance</td><td>position</td><td>gene name</td><td>gene id</td></tr>
<tr><td>GU305914.1</td><td>41623</td><td>41645</td><td>-</td><td>0</td><td>CACCGGACACCGAAGATAGA</td><td>CGG</td><td>||||||||[||||||||||||]PAM</td><td>NA</td><td>Intergenic</td><td>NA</td><td>NA</td></tr>
<tr><td></td><td></td><td></td><td></td><td></td><td></td><td></td><td></td><td></td><td></td><td></td><td></td></tr>
<tr><td>T31102</td><td>CTGCCTTTCTCTTGACAGATAGG</td><td>1000</td><td></td><td></td><td></td><td></td><td></td><td></td><td></td><td></td><td></td></tr>
<tr><td>Oligo fwd</td><td>CACCCTGCCTTTCTCTTGACAGAT</td><td></td><td></td><td></td><td></td><td></td><td></td><td></td><td></td><td></td><td></td></tr>
<tr><td>Oligo rev</td><td>AAACATCTGTCAAGAGAAAGGCAG</td><td></td><td></td><td></td><td></td><td></td><td></td><td></td><td></td><td></td><td></td></tr>
<tr><td>Chromosome</td><td>start</td><td>end</td><td>strand</td><td>MM</td><td>target_seq</td><td>PAM</td><td>alignment</td><td>distance</td><td>position</td><td>gene name</td><td>gene id</td></tr>
<tr><td>GU305914.1</td><td>41585</td><td>41607</td><td>-</td><td>0</td><td>CTGCCTTTCTCTTGACAGAT</td><td>AGG</td><td>||||||||[||||||||||||]PAM</td><td>NA</td><td>Intergenic</td><td>NA</td><td>NA</td></tr>
<tr><td></td><td></td><td></td><td></td><td></td><td></td><td></td><td></td><td></td><td></td><td></td><td></td></tr>
<tr><td>T31103</td><td>TGACAGATAGGCCGCGACTCTGG</td><td>1000</td><td></td><td></td><td></td><td></td><td></td><td></td><td></td><td></td><td></td></tr>
<tr><td>Oligo fwd</td><td>CACCTGACAGATAGGCCGCGACTC</td><td></td><td></td><td></td><td></td><td></td><td></td><td></td><td></td><td></td><td></td></tr>
<tr><td>Oligo rev</td><td>AAACGAGTCGCGGCCTATCTGTCA</td><td></td><td></td><td></td><td></td><td></td><td></td><td></td><td></td><td></td><td></td></tr>
</table>

```
Chromosome  start  end  strand  MM  target_seq  PAM  alignment  distance  position  gene name  gene id
GU305914.1  41573  41595  -  0  TGACAGATAGGCCGCGACTC  TGG  ||||||||[||||||||||||]PAM  NA  Intergenic  NA  NA

T31104  CAGATAGGCCGCGACTCTGGCGG  1000
Oligo fwd  CACCCAGATAGGCCGCGACTCTGG
Oligo rev  AAACCCAGAGTCGCGGCCTATCTG
Chromosome  start  end  strand  MM  target_seq  PAM  alignment  distance  position  gene name  gene id
GU305914.1  41570  41592  -  0  CAGATAGGCCGCGACTCTGG  CGG  ||||||||[||||||||||||]PAM  NA  Intergenic  NA  NA

T31105  AGATAGGCCGCGACTCTGGCGGG  1000
Oligo fwd  CACCAGATAGGCCGCGACTCTGGC
Oligo rev  AAACGCCAGAGTCGCGGCCTATCT
Chromosome  start  end  strand  MM  target_seq  PAM  alignment  distance  position  gene name  gene id
GU305914.1  41569  41591  -  0  AGATAGGCCGCGACTCTGGC  GGG  ||||||||[||||||||||||]PAM  NA  Intergenic  NA  NA

T31106  CGCTCCCCGTCCCGACGCCCCGG  1000
Oligo fwd  CACCCGCTCCCCGTCCCGACGCCC
Oligo rev  AAACGGGCGTCGGGACGGGGAGCG
Chromosome  start  end  strand  MM  target_seq  PAM  alignment  distance  position  gene name  gene id
GU305914.1  41536  41558  -  0  CGCTCCCCGTCCCGACGCCC  CGG  ||||||||[||||||||||||]PAM  NA  Intergenic  NA  NA

T31107  TGCTTACCATAGTCTCGTCCCGG  1000
Oligo fwd  CACCTGCTTACCATAGTCTCGTCC
Oligo rev  AAACGGACGAGACTATGGTAAGCA
Chromosome  start  end  strand  MM  target_seq  PAM  alignment  distance  position  gene name  gene id
GU305914.1  41506  41528  -  0  TGCTTACCATAGTCTCGTCC  CGG  ||||||||[||||||||||||]PAM  NA  Intergenic  NA  NA

T31108  TAGTCTCGTCCCGGCCCAAAAGG  1000
Oligo fwd  CACCTAGTCTCGTCCCGGCCCAAA
Oligo rev  AAACTTTGGGCCGGGACGAGACTA
Chromosome  start  end  strand  MM  target_seq  PAM  alignment  distance  position  gene name  gene id
GU305914.1  41497  41519  -  0  TAGTCTCGTCCCGGCCCAAA  AGG  ||||||||[||||||||||||]PAM  NA  Intergenic  NA  NA

T31109  AAAGGACGTCCATCTCGAGCCGG  1000
Oligo fwd  CACCAAAGGACGTCCATCTCGAGC
Oligo rev  AAACGCTCGAGATGGACGTCCTTT
Chromosome  start  end  strand  MM  target_seq  PAM  alignment  distance  position  gene name  gene id
GU305914.1  41479  41501  -  0  AAAGGACGTCCATCTCGAGC  CGG  ||||||||[||||||||||||]PAM  NA  Intergenic  NA  NA

T31110  CATCTCGAGCCGGTCGTCGCCGG  1000
Oligo fwd  CACCCATCTCGAGCCGGTCGTCGC
Oligo rev  AAACGCGACGACCGGCTCGAGATG
Chromosome  start  end  strand  MM  target_seq  PAM  alignment  distance  position  gene name  gene id
GU305914.1  41469  41491  -  0  CATCTCGAGCCGGTCGTCGC  CGG  ||||||||[||||||||||||]PAM  NA  Intergenic  NA  NA

T31111  TATTTAACGATGTCCGCTCGAGG  1000
Oligo fwd  CACCTATTTAACGATGTCCGCTCG
Oligo rev  AAACCGAGCGGACATCGTTAAATA
Chromosome  start  end  strand  MM  target_seq  PAM  alignment  distance  position  gene name  gene id
GU305914.1  41426  41448  -  0  TATTTAACGATGTCCGCTCG  AGG  ||||||||[||||||||||||]PAM  NA  Intergenic  NA  NA

T31112  ATTTAACGATGTCCGCTCGAGGG  1000
Oligo fwd  CACCATTTAACGATGTCCGCTCGA
Oligo rev  AAACTCGAGCGGACATCGTTAAAT
Chromosome  start  end  strand  MM  target_seq  PAM  alignment  distance  position  gene name  gene id
GU305914.1  41425  41447  -  0  ATTTAACGATGTCCGCTCGA  GGG  ||||||||[||||||||||||]PAM  NA  Intergenic  NA  NA

T31113  GATGTCCGCTCGAGGGCGCGCGG  1000
Oligo fwd  CACCGATGTCCGCTCGAGGGCGCG
Oligo rev  AAACCGCGCCCTCGAGCGGACATC
Chromosome  start  end  strand  MM  target_seq  PAM  alignment  distance  position  gene name  gene id
GU305914.1  41418  41440  -  0  GATGTCCGCTCGAGGGCGCG  CGG  ||||||||[||||||||||||]PAM  NA  Intergenic  NA  NA

T31114  GTCCGCTCGAGGGCGCGCGGCGG  1000
Oligo fwd  CACCGTCCGCTCGAGGGCGCGCGG
Oligo rev  AAACCCGCGCGCCCTCGAGCGGAC
Chromosome  start  end  strand  MM  target_seq  PAM  alignment  distance  position  gene name  gene id
GU305914.1  41415  41437  -  0  GTCCGCTCGAGGGCGCGCGG  CGG  ||||||||[||||||||||||]PAM  NA  Intergenic  NA  NA
GU305914.1  138667  138689  -  4  GACCCCTCGAGGACGCACGG  CGG  |-||-|||[||||-|||-|||]PAM  NA  Intergenic  NA  NA

T31115  TCCGCTCGAGGGCGCGCGGCGGG  1000
Oligo fwd  CACCTCCGCTCGAGGGCGCGCGGC
Oligo rev  AAACGCCGCGCGCCCTCGAGCGGA
Chromosome  start  end  strand  MM  target_seq  PAM  alignment  distance  position  gene name  gene id
GU305914.1  41414  41436  -  0  TCCGCTCGAGGGCGCGCGGC  GGG  ||||||||[||||||||||||]PAM  NA  Intergenic  NA  NA

T31116  GGGCGCGCGGCGGGAGACGACGG  1000
Oligo fwd  CACCGGGCGCGCGGCGGGAGACGA
Oligo rev  AAACTCGTCTCCCGCCGCGCGCCC
Chromosome  start  end  strand  MM  target_seq  PAM  alignment  distance  position  gene name  gene id
GU305914.1  41405  41427  -  0  GGGCGCGCGGCGGGAGACGA  CGG  ||||||||[||||||||||||]PAM  NA  Intergenic  NA  NA

T31117  GGCGGGAGACGACGGTCGTCAGG  1000
Oligo fwd  CACCGGCGGGAGACGACGGTCGTC
Oligo rev  AAACGACGACCGTCGTCTCCCGCC
Chromosome  start  end  strand  MM  target_seq  PAM  alignment  distance  position  gene name  gene id
GU305914.1  41397  41419  -  0  GGCGGGAGACGACGGTCGTC  AGG  ||||||||[||||||||||||]PAM  NA  Intergenic  NA  NA

T31118  CGGTCGTCAGGCCGAACTGATGG  1000
```

Oligo fwd CACCCGGTCGTCAGGCCGAACTGA
Oligo rev AAACTCAGTTCGGCCTGACGACCG
Chromosome start end strand MM target_seq PAM alignment distance position gene name gene id
GU305914.1 41385 41407 - 0 CGGTCGTCAGGCCGAACTGA TGG ||||||||[||||||||||||]PAM NA Intergenic NA NA

T31119 GCCGAACTGATGGCCACCCTCGG 1000
Oligo fwd CACCGCCGAACTGATGGCCACCCT
Oligo rev AAACAGGGTGGCCATCAGTTCGGC
Chromosome start end strand MM target_seq PAM alignment distance position gene name gene id
GU305914.1 41375 41397 - 0 GCCGAACTGATGGCCACCCT CGG ||||||||[||||||||||||]PAM NA Intergenic NA NA

T31120 CCGAACTGATGGCCACCCTCGGG 1000
Oligo fwd CACCCCGAACTGATGGCCACCCTC
Oligo rev AAACGAGGGTGGCCATCAGTTCGG
Chromosome start end strand MM target_seq PAM alignment distance position gene name gene id
GU305914.1 41374 41396 - 0 CCGAACTGATGGCCACCCTC GGG ||||||||[||||||||||||]PAM NA Intergenic NA NA

T31121 TGGCCACCCTCGGGTTCGTCAGG 1000
Oligo fwd CACCTGGCCACCCTCGGGTTCGTC
Oligo rev AAACGACGAACCCGAGGGTGGCCA
Chromosome start end strand MM target_seq PAM alignment distance position gene name gene id
GU305914.1 41365 41387 - 0 TGGCCACCCTCGGGTTCGTC AGG ||||||||[||||||||||||]PAM NA Intergenic NA NA

T31122 AGGCTCTCCAAGAGCTCCGTTGG 1000
Oligo fwd CACCAGGCTCTCCAAGAGCTCCGT
Oligo rev AAACACGGAGCTCTTGGAGAGCCT
Chromosome start end strand MM target_seq PAM alignment distance position gene name gene id
GU305914.1 41345 41367 - 0 AGGCTCTCCAAGAGCTCCGT TGG ||||||||[||||||||||||]PAM NA Intergenic NA NA

T31123 CTCCAAGAGCTCCGTTGGAAAGG 1000
Oligo fwd CACCCTCCAAGAGCTCCGTTGGAA
Oligo rev AAACTTCCAACGGAGCTCTTGGAG
Chromosome start end strand MM target_seq PAM alignment distance position gene name gene id
GU305914.1 41340 41362 - 0 CTCCAAGAGCTCCGTTGGAA AGG ||||||||[||||||||||||]PAM NA Intergenic NA NA

T31124 TCGCCACCCACGTGTGATTGCGG 1000
Oligo fwd CACCTCGCCACCCACGTGTGATTG
Oligo rev AAACCAATCACACGTGGGTGGCGA
Chromosome start end strand MM target_seq PAM alignment distance position gene name gene id
GU305914.1 41268 41290 - 0 TCGCCACCCACGTGTGATTG CGG ||||||||[||||||||||||]PAM NA Intergenic NA NA

T31125 CGTGTGATTGCGGAATGTCGCGG 1000
Oligo fwd CACCCGTGTGATTGCGGAATGTCG
Oligo rev AAACCGACATTCCGCAATCACACG
Chromosome start end strand MM target_seq PAM alignment distance position gene name gene id
GU305914.1 41258 41280 - 0 CGTGTGATTGCGGAATGTCG CGG ||||||||[||||||||||||]PAM NA Intergenic NA NA

T31126 GTCGCGGCACCGACCTTTCCAGG 1000
Oligo fwd CACCGTCGCGGCACCGACCTTTCC
Oligo rev AAACGGAAAGGTCGGTGCCGCGAC
Chromosome start end strand MM target_seq PAM alignment distance position gene name gene id
GU305914.1 41242 41264 - 0 GTCGCGGCACCGACCTTTCC AGG ||||||||[||||||||||||]PAM NA Intergenic NA NA

T31127 TCGCGGCACCGACCTTTCCAGGG 1000
Oligo fwd CACCTCGCGGCACCGACCTTTCCA
Oligo rev AAACTGGAAAGGTCGGTGCCGCGA
Chromosome start end strand MM target_seq PAM alignment distance position gene name gene id
GU305914.1 41241 41263 - 0 TCGCGGCACCGACCTTTCCA GGG ||||||||[||||||||||||]PAM NA Intergenic NA NA

T31128 GACGCAGCTCTACAACGAGATGG 1000
Oligo fwd CACCGACGCAGCTCTACAACGAGA
Oligo rev AAACTCTCGTTGTAGAGCTGCGTC
Chromosome start end strand MM target_seq PAM alignment distance position gene name gene id
GU305914.1 41217 41239 - 0 GACGCAGCTCTACAACGAGA TGG ||||||||[||||||||||||]PAM NA Intergenic NA NA

T31129 TCTACAACGAGATGGTCCTCTGG 1000
Oligo fwd CACCTCTACAACGAGATGGTCCTC
Oligo rev AAACGAGGACCATCTCGTTGTAGA
Chromosome start end strand MM target_seq PAM alignment distance position gene name gene id
GU305914.1 41209 41231 - 0 TCTACAACGAGATGGTCCTC TGG ||||||||[||||||||||||]PAM NA Intergenic NA NA

T31130 ACGAGATGGTCCTCTGGCTCAGG 1000
Oligo fwd CACCACGAGATGGTCCTCTGGCTC
Oligo rev AAACGAGCCAGAGGACCATCTCGT
Chromosome start end strand MM target_seq PAM alignment distance position gene name gene id
GU305914.1 41203 41225 - 0 ACGAGATGGTCCTCTGGCTC AGG ||||||||[||||||||||||]PAM NA Intergenic NA NA

T31131 CAGGTACCACGAGAAGCTCACGG 1000
Oligo fwd CACCCAGGTACCACGAGAAGCTCA
Oligo rev AAACTGAGCTTCTCGTGGTACCTG
Chromosome start end strand MM target_seq PAM alignment distance position gene name gene id
GU305914.1 41184 41206 - 0 CAGGTACCACGAGAAGCTCA CGG ||||||||[||||||||||||]PAM NA Intergenic NA NA

T31132 AAGCTCACGGCCCGACGACCCGG 1000
Oligo fwd CACCAAGCTCACGGCCCGACGACC
Oligo rev AAACGGTCGTCGGGCCGTGAGCTT
Chromosome start end strand MM target_seq PAM alignment distance position gene name gene id
GU305914.1 41171 41193 - 0 AAGCTCACGGCCCGACGACC CGG ||||||||[||||||||||||]PAM NA Intergenic NA NA

T31133 GACACCTGCCGCTCCTCACTAGG 1000
Oligo fwd CACCGACACCTGCCGCTCCTCACT
Oligo rev AAACAGTGAGGAGCGGCAGGTGTC
Chromosome start end strand MM target_seq PAM alignment distance position gene name gene id
GU305914.1 41149 41171 - 0 GACACCTGCCGCTCCTCACT AGG ||||||||[||||||||||||]PAM NA Intergenic NA NA

T31134 CAAGATTACGACAAGCTCTTCGG 1000
Oligo fwd CACCCAAGATTACGACAAGCTCTT
Oligo rev AAACAAGAGCTTGTCGTAATCTTG
Chromosome start end strand MM target_seq PAM alignment distance position gene name gene id
GU305914.1 41120 41142 - 0 CAAGATTACGACAAGCTCTT CGG ||||||||[||||||||||||]PAM NA Intergenic NA NA

T31135 TCTTCGGCTTCGTCGCCGCGCGG 1000
Oligo fwd CACCTCTTCGGCTTCGTCGCCGCG
Oligo rev AAACCGCGGCGACGAAGCCGAAGA
Chromosome start end strand MM target_seq PAM alignment distance position gene name gene id
GU305914.1 41104 41126 - 0 TCTTCGGCTTCGTCGCCGCG CGG ||||||||[||||||||||||]PAM NA Intergenic NA NA

T31136 GCCGCGCGGCCCGAGCTCTGCGG 1000
Oligo fwd CACCGCCGCGCGGCCCGAGCTCTG
Oligo rev AAACCAGAGCTCGGGCCGCGCGGC
Chromosome start end strand MM target_seq PAM alignment distance position gene name gene id
GU305914.1 41090 41112 - 0 GCCGCGCGGCCCGAGCTCTG CGG ||||||||[||||||||||||]PAM NA Intergenic NA NA

T31137 CCCGAGCTCTGCGGCTTCGACGG 1000
Oligo fwd CACCCCCGAGCTCTGCGGCTTCGA
Oligo rev AAACTCGAAGCCGCAGAGCTCGGG
Chromosome start end strand MM target_seq PAM alignment distance position gene name gene id
GU305914.1 41081 41103 - 0 CCCGAGCTCTGCGGCTTCGA CGG ||||||||[||||||||||||]PAM NA Intergenic NA NA

T31138 CTGCGGCTTCGACGGACTCACGG 1000
Oligo fwd CACCCTGCGGCTTCGACGGACTCA
Oligo rev AAACTGAGTCCGTCGAAGCCGCAG
Chromosome start end strand MM target_seq PAM alignment distance position gene name gene id
GU305914.1 41073 41095 - 0 CTGCGGCTTCGACGGACTCA CGG ||||||||[||||||||||||]PAM NA Intergenic NA NA

T31139 CGGCTTCGACGGACTCACGGAGG 1000
Oligo fwd CACCCGGCTTCGACGGACTCACGG
Oligo rev AAACCCGTGAGTCCGTCGAAGCCG
Chromosome start end strand MM target_seq PAM alignment distance position gene name gene id
GU305914.1 41070 41092 - 0 CGGCTTCGACGGACTCACGG AGG ||||||||[||||||||||||]PAM NA Intergenic NA NA

T31140 GTGTTCGACGACGCCGTCTACGG 1000
Oligo fwd CACCGTGTTCGACGACGCCGTCTA
Oligo rev AAACTAGACGGCGTCGTCGAACAC
Chromosome start end strand MM target_seq PAM alignment distance position gene name gene id
GU305914.1 41042 41064 - 0 GTGTTCGACGACGCCGTCTA CGG ||||||||[||||||||||||]PAM NA Intergenic NA NA

T31141 CGACGATTATGTGCCTCGCGTGG 1000
Oligo fwd CACCCGACGATTATGTGCCTCGCG
Oligo rev AAACCGCGAGGCACATAATCGTCG
Chromosome start end strand MM target_seq PAM alignment distance position gene name gene id
GU305914.1 41019 41041 - 0 CGACGATTATGTGCCTCGCG TGG ||||||||[||||||||||||]PAM NA Intergenic NA NA

T31142 CTCGCGTGGACGTCTTCCTGCGG 1000
Oligo fwd CACCCTCGCGTGGACGTCTTCCTG
Oligo rev AAACCAGGAAGACGTCCACGCGAG
Chromosome start end strand MM target_seq PAM alignment distance position gene name gene id
GU305914.1 41005 41027 - 0 CTCGCGTGGACGTCTTCCTG CGG ||||||||[||||||||||||]PAM NA Intergenic NA NA

T31143 TCGCGTGGACGTCTTCCTGCGGG 1000
Oligo fwd CACCTCGCGTGGACGTCTTCCTGC
Oligo rev AAACGCAGGAAGACGTCCACGCGA
Chromosome start end strand MM target_seq PAM alignment distance position gene name gene id
GU305914.1 41004 41026 - 0 TCGCGTGGACGTCTTCCTGC GGG ||||||||[||||||||||||]PAM NA Intergenic NA NA

T31144 CGCGTGGACGTCTTCCTGCGGGG 1000
Oligo fwd CACCCGCGTGGACGTCTTCCTGCG
Oligo rev AAACCGCAGGAAGACGTCCACGCG
Chromosome start end strand MM target_seq PAM alignment distance position gene name gene id
GU305914.1 41003 41025 - 0 CGCGTGGACGTCTTCCTGCG GGG ||||||||[||||||||||||]PAM NA Intergenic NA NA

T31145 GCGTGGACGTCTTCCTGCGGGGG 1000
Oligo fwd CACCGCGTGGACGTCTTCCTGCGG
Oligo rev AAACCCGCAGGAAGACGTCCACGC
Chromosome start end strand MM target_seq PAM alignment distance position gene name gene id
GU305914.1 41002 41024 - 0 GCGTGGACGTCTTCCTGCGG GGG ||||||||[||||||||||||]PAM NA Intergenic NA NA

T31146 CGTCTTCCTGCGGGGGCTCGAGG 1000
Oligo fwd CACCCGTCTTCCTGCGGGGGCTCG
Oligo rev AAACCGAGCCCCCGCAGGAAGACG
Chromosome start end strand MM target_seq PAM alignment distance position gene name gene id
GU305914.1 40995 41017 - 0 CGTCTTCCTGCGGGGGCTCG AGG ||||||||[||||||||||||]PAM NA Intergenic NA NA
GU305914.1 116360 116382 + 3 CGGCGTCCTGCGGCGGCTCG GGG ||-|-|||[|||||-||||||]PAM NA Intergenic NA NA

T31147 GGGGGCTCGAGGACCTCGCCCGG 1000
Oligo fwd CACCGGGGGCTCGAGGACCTCGCC
Oligo rev AAACGGCGAGGTCCTCGAGCCCCC
Chromosome start end strand MM target_seq PAM alignment distance position gene name gene id

GU305914.1 40984 41006 - 0 GGGGGCTCGAGGACCTCGCC CGG ||||||||[||||||||||||]PAM NA Intergenic NA NA

T31148 CGCCCGGTGTCTCTGTGCTCAGG 1000
Oligo fwd CACCGCCCGGTGTCTCTGTGCTC
Oligo rev AAACGAGCACAGAGACACCGGGCG
Chromosome start end strand MM target_seq PAM alignment distance position gene name gene id
GU305914.1 40968 40990 - 0 CGCCCGGTGTCTCTGTGCTC AGG ||||||||[||||||||||||]PAM NA Intergenic NA NA

T31149 GCCCGGTGTCTCTGTGCTCAGGG 1000
Oligo fwd CACCGCCCGGTGTCTCTGTGCTCA
Oligo rev AAACTGAGCACAGAGACACCGGGC
Chromosome start end strand MM target_seq PAM alignment distance position gene name gene id
GU305914.1 40967 40989 - 0 GCCCGGTGTCTCTGTGCTCA GGG ||||||||[||||||||||||]PAM NA Intergenic NA NA

T31150 GTGTCTCTGTGCTCAGGGCCCGG 1000
Oligo fwd CACCGTGTCTCTGTGCTCAGGGCC
Oligo rev AAACGGCCCTGAGCACAGAGACAC
Chromosome start end strand MM target_seq PAM alignment distance position gene name gene id
GU305914.1 40962 40984 - 0 GTGTCTCTGTGCTCAGGGCC CGG ||||||||[||||||||||||]PAM NA Intergenic NA NA

T31151 TGCTCAGGGCCCGGACAAGCCGG 1000
Oligo fwd CACCTGCTCAGGGCCCGGACAAGC
Oligo rev AAACGCTTGTCCGGGCCCTGAGCA
Chromosome start end strand MM target_seq PAM alignment distance position gene name gene id
GU305914.1 40953 40975 - 0 TGCTCAGGGCCCGGACAAGC CGG ||||||||[||||||||||||]PAM NA Intergenic NA NA

T31152 CCCGGACAAGCCGGCGCGTGCGG 1000
Oligo fwd CACCCCCGGACAAGCCGGCGCGTG
Oligo rev AAACCACGCGCCGGCTTGTCCGGG
Chromosome start end strand MM target_seq PAM alignment distance position gene name gene id
GU305914.1 40944 40966 - 0 CCCGGACAAGCCGGCGCGTG CGG ||||||||[||||||||||||]PAM NA Intergenic NA NA

T31153 GCCGGCGCGTGCGGTGATCATGG 1000
Oligo fwd CACCGCCGGCGCGTGCGGTGATCA
Oligo rev AAACTGATCACCGCACGCGCCGGC
Chromosome start end strand MM target_seq PAM alignment distance position gene name gene id
GU305914.1 40935 40957 - 0 GCCGGCGCGTGCGGTGATCA TGG ||||||||[||||||||||||]PAM NA Intergenic NA NA

T31154 CCGGCGCGTGCGGTGATCATGGG 1000
Oligo fwd CACCCCGGCGCGTGCGGTGATCAT
Oligo rev AAACATGATCACCGCACGCGCCGG
Chromosome start end strand MM target_seq PAM alignment distance position gene name gene id
GU305914.1 40934 40956 - 0 CCGGCGCGTGCGGTGATCAT GGG ||||||||[||||||||||||]PAM NA Intergenic NA NA

T31155 GGGCTTCATCAACATGAGAGCGG 1000
Oligo fwd CACCGGGCTTCATCAACATGAGAG
Oligo rev AAACCTCTCATGTTGATGAAGCCC
Chromosome start end strand MM target_seq PAM alignment distance position gene name gene id
GU305914.1 40914 40936 - 0 GGGCTTCATCAACATGAGAG CGG ||||||||[||||||||||||]PAM NA Intergenic NA NA

T31156 CTTCATCAACATGAGAGCGGAGG 1000
Oligo fwd CACCCTTCATCAACATGAGAGCGG
Oligo rev AAACCCGCTCTCATGTTGATGAAG
Chromosome start end strand MM target_seq PAM alignment distance position gene name gene id
GU305914.1 40911 40933 - 0 CTTCATCAACATGAGAGCGG AGG ||||||||[||||||||||||]PAM NA Intergenic NA NA

T31157 CATCAACATGAGAGCGGAGGAGG 1000
Oligo fwd CACCCATCAACATGAGAGCGGAGG
Oligo rev AAACCCTCCGCTCTCATGTTGATG
Chromosome start end strand MM target_seq PAM alignment distance position gene name gene id
GU305914.1 40908 40930 - 0 CATCAACATGAGAGCGGAGG AGG ||||||||[||||||||||||]PAM NA Intergenic NA NA

T31158 GGAGGAGGTCAATCGCCTGATGG 1000
Oligo fwd CACCGGAGGAGGTCAATCGCCTGA
Oligo rev AAACTCAGGCGATTGACCTCCTCC
Chromosome start end strand MM target_seq PAM alignment distance position gene name gene id
GU305914.1 40893 40915 - 0 GGAGGAGGTCAATCGCCTGA TGG ||||||||[||||||||||||]PAM NA Intergenic NA NA

T31159 ATCGCCTGATGGACAACGTCAGG 1000
Oligo fwd CACCATCGCCTGATGGACAACGTC
Oligo rev AAACGACGTTGTCCATCAGGCGAT
Chromosome start end strand MM target_seq PAM alignment distance position gene name gene id
GU305914.1 40882 40904 - 0 ATCGCCTGATGGACAACGTC AGG ||||||||[||||||||||||]PAM NA Intergenic NA NA

T31160 TCGCCTGATGGACAACGTCAGGG 1000
Oligo fwd CACCTCGCCTGATGGACAACGTCA
Oligo rev AAACTGACGTTGTCCATCAGGCGA
Chromosome start end strand MM target_seq PAM alignment distance position gene name gene id
GU305914.1 40881 40903 - 0 TCGCCTGATGGACAACGTCA GGG ||||||||[||||||||||||]PAM NA Intergenic NA NA

T31161 GATGGACAACGTCAGGGACGCGG 1000
Oligo fwd CACCGATGGACAACGTCAGGGACG
Oligo rev AAACCGTCCCTGACGTTGTCCATC
Chromosome start end strand MM target_seq PAM alignment distance position gene name gene id
GU305914.1 40875 40897 - 0 GATGGACAACGTCAGGGACG CGG ||||||||[||||||||||||]PAM NA Intergenic NA NA

T31162 CGTCCTCGTCTACGAAGTCTTGG 1000
Oligo fwd CACCCGTCCTCGTCTACGAAGTCT
Oligo rev AAACAGACTTCGTAGACGAGGACG

<table>
<tr><td>Chromosome</td><td>start</td><td>end</td><td>strand</td><td>MM</td><td>target_seq</td><td>PAM</td><td>alignment</td><td>distance</td><td>position</td><td>gene name</td><td>gene id</td></tr>
<tr><td>GU305914.1</td><td>40845</td><td>40867</td><td>-</td><td>0</td><td>CGTCCTCGTCTACGAAGTCT</td><td>TGG</td><td>||||||||[||||||||||||]PAM</td><td>NA</td><td>Intergenic</td><td>NA</td><td>NA</td></tr>
<tr><td>T31163</td><td>TCTACGAAGTCTTGGACGTGAGG</td><td>1000</td></tr>
<tr><td>Oligo fwd</td><td>CACCTCTACGAAGTCTTGGACGTG</td></tr>
<tr><td>Oligo rev</td><td>AAACCACGTCCAAGACTTCGTAGA</td></tr>
<tr><td>Chromosome</td><td>start</td><td>end</td><td>strand</td><td>MM</td><td>target_seq</td><td>PAM</td><td>alignment</td><td>distance</td><td>position</td><td>gene name</td><td>gene id</td></tr>
<tr><td>GU305914.1</td><td>40837</td><td>40859</td><td>-</td><td>0</td><td>TCTACGAAGTCTTGGACGTG</td><td>AGG</td><td>||||||||[||||||||||||]PAM</td><td>NA</td><td>Intergenic</td><td>NA</td><td>NA</td></tr>
<tr><td>T31164</td><td>CTACGAAGTCTTGGACGTGAGGG</td><td>1000</td></tr>
<tr><td>Oligo fwd</td><td>CACCCTACGAAGTCTTGGACGTGA</td></tr>
<tr><td>Oligo rev</td><td>AAACTCACGTCCAAGACTTCGTAG</td></tr>
<tr><td>Chromosome</td><td>start</td><td>end</td><td>strand</td><td>MM</td><td>target_seq</td><td>PAM</td><td>alignment</td><td>distance</td><td>position</td><td>gene name</td><td>gene id</td></tr>
<tr><td>GU305914.1</td><td>40836</td><td>40858</td><td>-</td><td>0</td><td>CTACGAAGTCTTGGACGTGA</td><td>GGG</td><td>||||||||[||||||||||||]PAM</td><td>NA</td><td>Intergenic</td><td>NA</td><td>NA</td></tr>
<tr><td>T31165</td><td>CCTACGCCATCAGTAAGTCTTGG</td><td>1000</td></tr>
<tr><td>Oligo fwd</td><td>CACCCCTACGCCATCAGTAAGTCT</td></tr>
<tr><td>Oligo rev</td><td>AAACAGACTTACTGATGGCGTAGG</td></tr>
<tr><td>Chromosome</td><td>start</td><td>end</td><td>strand</td><td>MM</td><td>target_seq</td><td>PAM</td><td>alignment</td><td>distance</td><td>position</td><td>gene name</td><td>gene id</td></tr>
<tr><td>GU305914.1</td><td>40750</td><td>40772</td><td>-</td><td>0</td><td>CCTACGCCATCAGTAAGTCT</td><td>TGG</td><td>||||||||[||||||||||||]PAM</td><td>NA</td><td>Intergenic</td><td>NA</td><td>NA</td></tr>
<tr><td>T31166</td><td>GACGCTCTCCCACATGTGTCTGG</td><td>1000</td></tr>
<tr><td>Oligo fwd</td><td>CACCGACGCTCTCCCACATGTGTC</td></tr>
<tr><td>Oligo rev</td><td>AAACGACACATGTGGGAGAGCGTC</td></tr>
<tr><td>Chromosome</td><td>start</td><td>end</td><td>strand</td><td>MM</td><td>target_seq</td><td>PAM</td><td>alignment</td><td>distance</td><td>position</td><td>gene name</td><td>gene id</td></tr>
<tr><td>GU305914.1</td><td>40725</td><td>40747</td><td>-</td><td>0</td><td>GACGCTCTCCCACATGTGTC</td><td>TGG</td><td>||||||||[||||||||||||]PAM</td><td>NA</td><td>Intergenic</td><td>NA</td><td>NA</td></tr>
<tr><td>T31167</td><td>ATCGCATCAACTCGCTGCGACGG</td><td>1000</td></tr>
<tr><td>Oligo fwd</td><td>CACCATCGCATCAACTCGCTGCGA</td></tr>
<tr><td>Oligo rev</td><td>AAACTCGCAGCGAGTTGATGCGAT</td></tr>
<tr><td>Chromosome</td><td>start</td><td>end</td><td>strand</td><td>MM</td><td>target_seq</td><td>PAM</td><td>alignment</td><td>distance</td><td>position</td><td>gene name</td><td>gene id</td></tr>
<tr><td>GU305914.1</td><td>40702</td><td>40724</td><td>-</td><td>0</td><td>ATCGCATCAACTCGCTGCGA</td><td>CGG</td><td>||||||||[||||||||||||]PAM</td><td>NA</td><td>Intergenic</td><td>NA</td><td>NA</td></tr>
<tr><td>T31168</td><td>ACTAGCGTTTCACGATAGACCGG</td><td>1000</td></tr>
<tr><td>Oligo fwd</td><td>CACCACTAGCGTTTCACGATAGAC</td></tr>
<tr><td>Oligo rev</td><td>AAACGTCTATCGTGAAACGCTAGT</td></tr>
<tr><td>Chromosome</td><td>start</td><td>end</td><td>strand</td><td>MM</td><td>target_seq</td><td>PAM</td><td>alignment</td><td>distance</td><td>position</td><td>gene name</td><td>gene id</td></tr>
<tr><td>GU305914.1</td><td>40671</td><td>40693</td><td>-</td><td>0</td><td>ACTAGCGTTTCACGATAGAC</td><td>CGG</td><td>||||||||[||||||||||||]PAM</td><td>NA</td><td>Intergenic</td><td>NA</td><td>NA</td></tr>
<tr><td>T31169</td><td>GCGCTCGAGAGACCCGTCGACGG</td><td>1000</td></tr>
<tr><td>Oligo fwd</td><td>CACCGCGCTCGAGAGACCCGTCGA</td></tr>
<tr><td>Oligo rev</td><td>AAACTCGACGGGTCTCTCGAGCGC</td></tr>
<tr><td>Chromosome</td><td>start</td><td>end</td><td>strand</td><td>MM</td><td>target_seq</td><td>PAM</td><td>alignment</td><td>distance</td><td>position</td><td>gene name</td><td>gene id</td></tr>
<tr><td>GU305914.1</td><td>40622</td><td>40644</td><td>-</td><td>0</td><td>GCGCTCGAGAGACCCGTCGA</td><td>CGG</td><td>||||||||[||||||||||||]PAM</td><td>NA</td><td>Intergenic</td><td>NA</td><td>NA</td></tr>
<tr><td>T31170</td><td>CGACCGCGTATAATCTACTGCGG</td><td>1000</td></tr>
<tr><td>Oligo fwd</td><td>CACCCGACCGCGTATAATCTACTG</td></tr>
<tr><td>Oligo rev</td><td>AAACCAGTAGATTATACGCGGTCG</td></tr>
<tr><td>Chromosome</td><td>start</td><td>end</td><td>strand</td><td>MM</td><td>target_seq</td><td>PAM</td><td>alignment</td><td>distance</td><td>position</td><td>gene name</td><td>gene id</td></tr>
<tr><td>GU305914.1</td><td>40597</td><td>40619</td><td>-</td><td>0</td><td>CGACCGCGTATAATCTACTG</td><td>CGG</td><td>||||||||[||||||||||||]PAM</td><td>NA</td><td>Intergenic</td><td>NA</td><td>NA</td></tr>
<tr><td>T31171</td><td>CCGCGTATAATCTACTGCGGCGG</td><td>1000</td></tr>
<tr><td>Oligo fwd</td><td>CACCCCGCGTATAATCTACTGCGG</td></tr>
<tr><td>Oligo rev</td><td>AAACCCGCAGTAGATTATACGCGG</td></tr>
<tr><td>Chromosome</td><td>start</td><td>end</td><td>strand</td><td>MM</td><td>target_seq</td><td>PAM</td><td>alignment</td><td>distance</td><td>position</td><td>gene name</td><td>gene id</td></tr>
<tr><td>GU305914.1</td><td>40594</td><td>40616</td><td>-</td><td>0</td><td>CCGCGTATAATCTACTGCGG</td><td>CGG</td><td>||||||||[||||||||||||]PAM</td><td>NA</td><td>Intergenic</td><td>NA</td><td>NA</td></tr>
<tr><td>T31172</td><td>TAATCTACTGCGGCGGCTCGAGG</td><td>1000</td></tr>
<tr><td>Oligo fwd</td><td>CACCTAATCTACTGCGGCGGCTCG</td></tr>
<tr><td>Oligo rev</td><td>AAACCGAGCCGCCGCAGTAGATTA</td></tr>
<tr><td>Chromosome</td><td>start</td><td>end</td><td>strand</td><td>MM</td><td>target_seq</td><td>PAM</td><td>alignment</td><td>distance</td><td>position</td><td>gene name</td><td>gene id</td></tr>
<tr><td>GU305914.1</td><td>40587</td><td>40609</td><td>-</td><td>0</td><td>TAATCTACTGCGGCGGCTCG</td><td>AGG</td><td>||||||||[||||||||||||]PAM</td><td>NA</td><td>Intergenic</td><td>NA</td><td>NA</td></tr>
<tr><td>T31173</td><td>TCTACTGCGGCGGCTCGAGGAGG</td><td>1000</td></tr>
<tr><td>Oligo fwd</td><td>CACCTCTACTGCGGCGGCTCGAGG</td></tr>
<tr><td>Oligo rev</td><td>AAACCCTCGAGCCGCCGCAGTAGA</td></tr>
<tr><td>Chromosome</td><td>start</td><td>end</td><td>strand</td><td>MM</td><td>target_seq</td><td>PAM</td><td>alignment</td><td>distance</td><td>position</td><td>gene name</td><td>gene id</td></tr>
<tr><td>GU305914.1</td><td>40584</td><td>40606</td><td>-</td><td>0</td><td>TCTACTGCGGCGGCTCGAGG</td><td>AGG</td><td>||||||||[||||||||||||]PAM</td><td>NA</td><td>Intergenic</td><td>NA</td><td>NA</td></tr>
<tr><td>T31174</td><td>TCCTCTTCAGAAACGCGCTTAGG</td><td>1000</td></tr>
<tr><td>Oligo fwd</td><td>CACCTCCTCTTCAGAAACGCGCTT</td></tr>
<tr><td>Oligo rev</td><td>AAACAAGCGCGTTTCTGAAGAGGA</td></tr>
<tr><td>Chromosome</td><td>start</td><td>end</td><td>strand</td><td>MM</td><td>target_seq</td><td>PAM</td><td>alignment</td><td>distance</td><td>position</td><td>gene name</td><td>gene id</td></tr>
<tr><td>GU305914.1</td><td>40555</td><td>40577</td><td>-</td><td>0</td><td>TCCTCTTCAGAAACGCGCTT</td><td>AGG</td><td>||||||||[||||||||||||]PAM</td><td>NA</td><td>Intergenic</td><td>NA</td><td>NA</td></tr>
<tr><td>T31175</td><td>TCTTCAGAAACGCGCTTAGGTGG</td><td>1000</td></tr>
<tr><td>Oligo fwd</td><td>CACCTCTTCAGAAACGCGCTTAGG</td></tr>
<tr><td>Oligo rev</td><td>AAACCCTAAGCGCGTTTCTGAAGA</td></tr>
<tr><td>Chromosome</td><td>start</td><td>end</td><td>strand</td><td>MM</td><td>target_seq</td><td>PAM</td><td>alignment</td><td>distance</td><td>position</td><td>gene name</td><td>gene id</td></tr>
<tr><td>GU305914.1</td><td>40552</td><td>40574</td><td>-</td><td>0</td><td>TCTTCAGAAACGCGCTTAGG</td><td>TGG</td><td>||||||||[||||||||||||]PAM</td><td>NA</td><td>Intergenic</td><td>NA</td><td>NA</td></tr>
<tr><td>T31176</td><td>CTTCAGAAACGCGCTTAGGTGGG</td><td>1000</td></tr>
<tr><td>Oligo fwd</td><td>CACCCTTCAGAAACGCGCTTAGGT</td></tr>
<tr><td>Oligo rev</td><td>AAACACCTAAGCGCGTTTCTGAAG</td></tr>
<tr><td>Chromosome</td><td>start</td><td>end</td><td>strand</td><td>MM</td><td>target_seq</td><td>PAM</td><td>alignment</td><td>distance</td><td>position</td><td>gene name</td><td>gene id</td></tr>
<tr><td>GU305914.1</td><td>40551</td><td>40573</td><td>-</td><td>0</td><td>CTTCAGAAACGCGCTTAGGT</td><td>GGG</td><td>||||||||[||||||||||||]PAM</td><td>NA</td><td>Intergenic</td><td>NA</td><td>NA</td></tr>
<tr><td>T31177</td><td>TTCAGAAACGCGCTTAGGTGGGG</td><td>1000</td></tr>
<tr><td>Oligo fwd</td><td>CACCTTCAGAAACGCGCTTAGGTG</td></tr>
</table>

Oligo rev AAACCACCTAAGCGCGTTTCTGAA
Chromosome start end strand MM target_seq PAM alignment distance position gene name gene id
GU305914.1 40550 40572 - 0 TTCAGAAACGCGCTTAGGTG GGG ||||||||[||||||||||||]PAM NA Intergenic NA NA

T31178 TTAGGTGGGGTGATCCCGACTGG 1000
Oligo fwd CACCTTAGGTGGGGTGATCCCGAC
Oligo rev AAACGTCGGGATCACCCCACCTAA
Chromosome start end strand MM target_seq PAM alignment distance position gene name gene id
GU305914.1 40537 40559 - 0 TTAGGTGGGGTGATCCCGAC TGG ||||||||[||||||||||||]PAM NA Intergenic NA NA

T31179 TAGGTGGGGTGATCCCGACTGGG 1000
Oligo fwd CACCTAGGTGGGGTGATCCCGACT
Oligo rev AAACAGTCGGGATCACCCCACCTA
Chromosome start end strand MM target_seq PAM alignment distance position gene name gene id
GU305914.1 40536 40558 - 0 TAGGTGGGGTGATCCCGACT GGG ||||||||[||||||||||||]PAM NA Intergenic NA NA

T31180 AGGTGGGGTGATCCCGACTGGGG 1000
Oligo fwd CACCAGGTGGGGTGATCCCGACTG
Oligo rev AAACCAGTCGGGATCACCCCACCT
Chromosome start end strand MM target_seq PAM alignment distance position gene name gene id
GU305914.1 40535 40557 - 0 AGGTGGGGTGATCCCGACTG GGG ||||||||[||||||||||||]PAM NA Intergenic NA NA

T31181 GGTGGGGTGATCCCGACTGGGGG 1000
Oligo fwd CACCGGTGGGGTGATCCCGACTGG
Oligo rev AAACCCAGTCGGGATCACCCCACC
Chromosome start end strand MM target_seq PAM alignment distance position gene name gene id
GU305914.1 40534 40556 - 0 GGTGGGGTGATCCCGACTGG GGG ||||||||[||||||||||||]PAM NA Intergenic NA NA

T31182 AGAGTCCGAGCTCGAGAGCGAGG 1000
Oligo fwd CACCAGAGTCCGAGCTCGAGAGCG
Oligo rev AAACCGCTCTCGAGCTCGGACTCT
Chromosome start end strand MM target_seq PAM alignment distance position gene name gene id
GU305914.1 40509 40531 - 0 AGAGTCCGAGCTCGAGAGCG AGG ||||||||[||||||||||||]PAM NA Intergenic NA NA

T31183 GAGTCCGAGCTCGAGAGCGAGGG 1000
Oligo fwd CACCGAGTCCGAGCTCGAGAGCGA
Oligo rev AAACTCGCTCTCGAGCTCGGACTC
Chromosome start end strand MM target_seq PAM alignment distance position gene name gene id
GU305914.1 40508 40530 - 0 GAGTCCGAGCTCGAGAGCGA GGG ||||||||[||||||||||||]PAM NA Intergenic NA NA
GU305914.1 47193 47215 + 4 GTCTCCGAGCTCGTGGGCGA AGG |--|||||[|||||-|-||||]PAM NA Intergenic NA NA

T31184 GAGGGAGATAATAGTGACGCCGG 1000
Oligo fwd CACCGAGGGAGATAATAGTGACGC
Oligo rev AAACGCGTCACTATTATCTCCCTC
Chromosome start end strand MM target_seq PAM alignment distance position gene name gene id
GU305914.1 40490 40512 - 0 GAGGGAGATAATAGTGACGC CGG ||||||||[||||||||||||]PAM NA Intergenic NA NA

T31185 AGATAATAGTGACGCCGGCTCGG 1000
Oligo fwd CACCAGATAATAGTGACGCCGGCT
Oligo rev AAACAGCCGGCGTCACTATTATCT
Chromosome start end strand MM target_seq PAM alignment distance position gene name gene id
GU305914.1 40485 40507 - 0 AGATAATAGTGACGCCGGCT CGG ||||||||[||||||||||||]PAM NA Intergenic NA NA

T31186 TAGTGACGCCGGCTCGGACCTGG 1000
Oligo fwd CACCTAGTGACGCCGGCTCGGACC
Oligo rev AAACGGTCCGAGCCGGCGTCACTA
Chromosome start end strand MM target_seq PAM alignment distance position gene name gene id
GU305914.1 40479 40501 - 0 TAGTGACGCCGGCTCGGACC TGG ||||||||[||||||||||||]PAM NA Intergenic NA NA

T31187 GATGAAGAAGACGACGATGACGG 1000
Oligo fwd CACCGATGAAGAAGACGACGATGA
Oligo rev AAACTCATCGTCGTCTTCTTCATC
Chromosome start end strand MM target_seq PAM alignment distance position gene name gene id
GU305914.1 40448 40470 - 0 GATGAAGAAGACGACGATGA CGG ||||||||[||||||||||||]PAM NA Intergenic NA NA
GU305914.1 118472 118494 - 3 GATGATGATGACGACGAAGA CGG |||||-||[-||||||||-||]PAM NA Intergenic NA NA
GU305914.1 102657 102679 - 4 GAGGAGGAAGAAGACGACGA CGG ||-||-||[|||-|||||-||]PAM NA Intergenic NA NA

T31188 GAAGAAGACGACGATGACGGCGG 1000
Oligo fwd CACCGAAGAAGACGACGATGACGG
Oligo rev AAACCCGTCATCGTCGTCTTCTTC
Chromosome start end strand MM target_seq PAM alignment distance position gene name gene id
GU305914.1 40445 40467 - 0 GAAGAAGACGACGATGACGG CGG ||||||||[||||||||||||]PAM NA Intergenic NA NA
GU305914.1 68437 68459 + 4 GGAGAACACGACGACGACCG AGG |-||||-|[||||||-|||-|]PAM NA Intergenic NA NA

T31189 GAAGACGACGATGACGGCGGAGG 1000
Oligo fwd CACCGAAGACGACGATGACGGCGG
Oligo rev AAACCCGCCGTCATCGTCGTCTTC
Chromosome start end strand MM target_seq PAM alignment distance position gene name gene id
GU305914.1 40442 40464 - 0 GAAGACGACGATGACGGCGG AGG ||||||||[||||||||||||]PAM NA Intergenic NA NA
GU305914.1 20756 20778 + 4 GACGACGGCGATCACGGCGA GGG ||-||||-[||||-||||||-]PAM NA Intergenic NA NA

T31190 GACGATGACGGCGGAGGACCCGG 1000
Oligo fwd CACCGACGATGACGGCGGAGGACC
Oligo rev AAACGGTCCTCCGCCGTCATCGTC
Chromosome start end strand MM target_seq PAM alignment distance position gene name gene id
GU305914.1 40436 40458 - 0 GACGATGACGGCGGAGGACC CGG ||||||||[||||||||||||]PAM NA Intergenic NA NA
GU305914.1 73195 73217 + 4 GTCGGTGCCGGCGGAGGATC GGG |-||-||-[||||||||||-|]PAM NA Intergenic NA NA

T31191 GATGACGGCGGAGGACCCGGAGG 1000

<table>
<tr><td>Oligo fwd</td><td>CACCGATGACGGCGGAGGACCCGG</td><td></td><td></td><td></td><td></td><td></td><td></td><td></td><td></td><td></td><td></td></tr>
<tr><td>Oligo rev</td><td>AAACCCGGGTCCTCCGCCGTCATC</td><td></td><td></td><td></td><td></td><td></td><td></td><td></td><td></td><td></td><td></td></tr>
<tr><td>Chromosome</td><td>start</td><td>end</td><td>strand</td><td>MM</td><td>target_seq</td><td>PAM</td><td>alignment</td><td>distance</td><td>position</td><td>gene name</td><td>gene id</td></tr>
<tr><td>GU305914.1</td><td>40433</td><td>40455</td><td>-</td><td>0</td><td>GATGACGGCGGAGGACCCGG</td><td>AGG</td><td>||||||||[||||||||||||]PAM</td><td>NA</td><td>Intergenic</td><td>NA</td><td>NA</td></tr>
<tr><td></td><td></td><td></td><td></td><td></td><td></td><td></td><td></td><td></td><td></td><td></td><td></td></tr>
<tr><td>T31192</td><td>CGGAGGACATGACGACGAGTCGG</td><td>1000</td><td></td><td></td><td></td><td></td><td></td><td></td><td></td><td></td><td></td></tr>
<tr><td>Oligo fwd</td><td>CACCCGGAGGACATGACGACGAGT</td><td></td><td></td><td></td><td></td><td></td><td></td><td></td><td></td><td></td><td></td></tr>
<tr><td>Oligo rev</td><td>AAACACTCGTCGTCATGTCCTCCG</td><td></td><td></td><td></td><td></td><td></td><td></td><td></td><td></td><td></td><td></td></tr>
<tr><td>Chromosome</td><td>start</td><td>end</td><td>strand</td><td>MM</td><td>target_seq</td><td>PAM</td><td>alignment</td><td>distance</td><td>position</td><td>gene name</td><td>gene id</td></tr>
<tr><td>GU305914.1</td><td>40416</td><td>40438</td><td>-</td><td>0</td><td>CGGAGGACATGACGACGAGT</td><td>CGG</td><td>||||||||[||||||||||||]PAM</td><td>NA</td><td>Intergenic</td><td>NA</td><td>NA</td></tr>
<tr><td></td><td></td><td></td><td></td><td></td><td></td><td></td><td></td><td></td><td></td><td></td><td></td></tr>
<tr><td>T31193</td><td>GGAGGACATGACGACGAGTCGGG</td><td>1000</td><td></td><td></td><td></td><td></td><td></td><td></td><td></td><td></td><td></td></tr>
<tr><td>Oligo fwd</td><td>CACCGGAGGACATGACGACGAGTC</td><td></td><td></td><td></td><td></td><td></td><td></td><td></td><td></td><td></td><td></td></tr>
<tr><td>Oligo rev</td><td>AAACGACTCGTCGTCATGTCCTCC</td><td></td><td></td><td></td><td></td><td></td><td></td><td></td><td></td><td></td><td></td></tr>
<tr><td>Chromosome</td><td>start</td><td>end</td><td>strand</td><td>MM</td><td>target_seq</td><td>PAM</td><td>alignment</td><td>distance</td><td>position</td><td>gene name</td><td>gene id</td></tr>
<tr><td>GU305914.1</td><td>40415</td><td>40437</td><td>-</td><td>0</td><td>GGAGGACATGACGACGAGTC</td><td>GGG</td><td>||||||||[||||||||||||]PAM</td><td>NA</td><td>Intergenic</td><td>NA</td><td>NA</td></tr>
<tr><td>GU305914.1</td><td>38917</td><td>38939</td><td>-</td><td>4</td><td>GGAGTGCATGAAGAAGAGTC</td><td>TGG</td><td>||||--||[|||-||-|||||]PAM</td><td>NA</td><td>Intergenic</td><td>NA</td><td>NA</td></tr>
<tr><td></td><td></td><td></td><td></td><td></td><td></td><td></td><td></td><td></td><td></td><td></td><td></td></tr>
<tr><td>T31194</td><td>GGACATGACGACGAGTCGGGAGG</td><td>1000</td><td></td><td></td><td></td><td></td><td></td><td></td><td></td><td></td><td></td></tr>
<tr><td>Oligo fwd</td><td>CACCGGACATGACGACGAGTCGGG</td><td></td><td></td><td></td><td></td><td></td><td></td><td></td><td></td><td></td><td></td></tr>
<tr><td>Oligo rev</td><td>AAACCCCGACTCGTCGTCATGTCC</td><td></td><td></td><td></td><td></td><td></td><td></td><td></td><td></td><td></td><td></td></tr>
<tr><td>Chromosome</td><td>start</td><td>end</td><td>strand</td><td>MM</td><td>target_seq</td><td>PAM</td><td>alignment</td><td>distance</td><td>position</td><td>gene name</td><td>gene id</td></tr>
<tr><td>GU305914.1</td><td>40412</td><td>40434</td><td>-</td><td>0</td><td>GGACATGACGACGAGTCGGG</td><td>AGG</td><td>||||||||[||||||||||||]PAM</td><td>NA</td><td>Intergenic</td><td>NA</td><td>NA</td></tr>
<tr><td></td><td></td><td></td><td></td><td></td><td></td><td></td><td></td><td></td><td></td><td></td><td></td></tr>
<tr><td>T31195</td><td>GACATGACGACGAGTCGGGAGGG</td><td>1000</td><td></td><td></td><td></td><td></td><td></td><td></td><td></td><td></td><td></td></tr>
<tr><td>Oligo fwd</td><td>CACCGACATGACGACGAGTCGGGA</td><td></td><td></td><td></td><td></td><td></td><td></td><td></td><td></td><td></td><td></td></tr>
<tr><td>Oligo rev</td><td>AAACTCCCGACTCGTCGTCATGTC</td><td></td><td></td><td></td><td></td><td></td><td></td><td></td><td></td><td></td><td></td></tr>
<tr><td>Chromosome</td><td>start</td><td>end</td><td>strand</td><td>MM</td><td>target_seq</td><td>PAM</td><td>alignment</td><td>distance</td><td>position</td><td>gene name</td><td>gene id</td></tr>
<tr><td>GU305914.1</td><td>40411</td><td>40433</td><td>-</td><td>0</td><td>GACATGACGACGAGTCGGGA</td><td>GGG</td><td>||||||||[||||||||||||]PAM</td><td>NA</td><td>Intergenic</td><td>NA</td><td>NA</td></tr>
<tr><td></td><td></td><td></td><td></td><td></td><td></td><td></td><td></td><td></td><td></td><td></td><td></td></tr>
<tr><td>T31196</td><td>ACGACGAGTCGGGAGGGAATCGG</td><td>1000</td><td></td><td></td><td></td><td></td><td></td><td></td><td></td><td></td><td></td></tr>
<tr><td>Oligo fwd</td><td>CACCACGACGAGTCGGGAGGGAAT</td><td></td><td></td><td></td><td></td><td></td><td></td><td></td><td></td><td></td><td></td></tr>
<tr><td>Oligo rev</td><td>AAACATTCCCTCCCGACTCGTCGT</td><td></td><td></td><td></td><td></td><td></td><td></td><td></td><td></td><td></td><td></td></tr>
<tr><td>Chromosome</td><td>start</td><td>end</td><td>strand</td><td>MM</td><td>target_seq</td><td>PAM</td><td>alignment</td><td>distance</td><td>position</td><td>gene name</td><td>gene id</td></tr>
<tr><td>GU305914.1</td><td>40405</td><td>40427</td><td>-</td><td>0</td><td>ACGACGAGTCGGGAGGGAAT</td><td>CGG</td><td>||||||||[||||||||||||]PAM</td><td>NA</td><td>Intergenic</td><td>NA</td><td>NA</td></tr>
<tr><td></td><td></td><td></td><td></td><td></td><td></td><td></td><td></td><td></td><td></td><td></td><td></td></tr>
<tr><td>T31197</td><td>AGGGAATCGGACGCCCGATCCGG</td><td>1000</td><td></td><td></td><td></td><td></td><td></td><td></td><td></td><td></td><td></td></tr>
<tr><td>Oligo fwd</td><td>CACCAGGGAATCGGACGCCCGATC</td><td></td><td></td><td></td><td></td><td></td><td></td><td></td><td></td><td></td><td></td></tr>
<tr><td>Oligo rev</td><td>AAACGATCGGGCGTCCGATTCCCT</td><td></td><td></td><td></td><td></td><td></td><td></td><td></td><td></td><td></td><td></td></tr>
<tr><td>Chromosome</td><td>start</td><td>end</td><td>strand</td><td>MM</td><td>target_seq</td><td>PAM</td><td>alignment</td><td>distance</td><td>position</td><td>gene name</td><td>gene id</td></tr>
<tr><td>GU305914.1</td><td>40392</td><td>40414</td><td>-</td><td>0</td><td>AGGGAATCGGACGCCCGATC</td><td>CGG</td><td>||||||||[||||||||||||]PAM</td><td>NA</td><td>Intergenic</td><td>NA</td><td>NA</td></tr>
<tr><td></td><td></td><td></td><td></td><td></td><td></td><td></td><td></td><td></td><td></td><td></td><td></td></tr>
<tr><td>T31198</td><td>GGGAATCGGACGCCCGATCCGGG</td><td>1000</td><td></td><td></td><td></td><td></td><td></td><td></td><td></td><td></td><td></td></tr>
<tr><td>Oligo fwd</td><td>CACCGGGAATCGGACGCCCGATCC</td><td></td><td></td><td></td><td></td><td></td><td></td><td></td><td></td><td></td><td></td></tr>
<tr><td>Oligo rev</td><td>AAACGGATCGGGCGTCCGATTCCC</td><td></td><td></td><td></td><td></td><td></td><td></td><td></td><td></td><td></td><td></td></tr>
<tr><td>Chromosome</td><td>start</td><td>end</td><td>strand</td><td>MM</td><td>target_seq</td><td>PAM</td><td>alignment</td><td>distance</td><td>position</td><td>gene name</td><td>gene id</td></tr>
<tr><td>GU305914.1</td><td>40391</td><td>40413</td><td>-</td><td>0</td><td>GGGAATCGGACGCCCGATCC</td><td>GGG</td><td>||||||||[||||||||||||]PAM</td><td>NA</td><td>Intergenic</td><td>NA</td><td>NA</td></tr>
<tr><td></td><td></td><td></td><td></td><td></td><td></td><td></td><td></td><td></td><td></td><td></td><td></td></tr>
<tr><td>T31199</td><td>GGAATCGGACGCCCGATCCGGGG</td><td>1000</td><td></td><td></td><td></td><td></td><td></td><td></td><td></td><td></td><td></td></tr>
<tr><td>Oligo fwd</td><td>CACCGGAATCGGACGCCCGATCCG</td><td></td><td></td><td></td><td></td><td></td><td></td><td></td><td></td><td></td><td></td></tr>
<tr><td>Oligo rev</td><td>AAACCGGATCGGGCGTCCGATTCC</td><td></td><td></td><td></td><td></td><td></td><td></td><td></td><td></td><td></td><td></td></tr>
<tr><td>Chromosome</td><td>start</td><td>end</td><td>strand</td><td>MM</td><td>target_seq</td><td>PAM</td><td>alignment</td><td>distance</td><td>position</td><td>gene name</td><td>gene id</td></tr>
<tr><td>GU305914.1</td><td>40390</td><td>40412</td><td>-</td><td>0</td><td>GGAATCGGACGCCCGATCCG</td><td>GGG</td><td>||||||||[||||||||||||]PAM</td><td>NA</td><td>Intergenic</td><td>NA</td><td>NA</td></tr>
<tr><td></td><td></td><td></td><td></td><td></td><td></td><td></td><td></td><td></td><td></td><td></td><td></td></tr>
<tr><td>T31200</td><td>ATGTCCCTCCACGACGACACCGG</td><td>1000</td><td></td><td></td><td></td><td></td><td></td><td></td><td></td><td></td><td></td></tr>
<tr><td>Oligo fwd</td><td>CACCATGTCCCTCCACGACGACAC</td><td></td><td></td><td></td><td></td><td></td><td></td><td></td><td></td><td></td><td></td></tr>
<tr><td>Oligo rev</td><td>AAACGTGTCGTCGTGGAGGGACAT</td><td></td><td></td><td></td><td></td><td></td><td></td><td></td><td></td><td></td><td></td></tr>
<tr><td>Chromosome</td><td>start</td><td>end</td><td>strand</td><td>MM</td><td>target_seq</td><td>PAM</td><td>alignment</td><td>distance</td><td>position</td><td>gene name</td><td>gene id</td></tr>
<tr><td>GU305914.1</td><td>40367</td><td>40389</td><td>-</td><td>0</td><td>ATGTCCCTCCACGACGACAC</td><td>CGG</td><td>||||||||[||||||||||||]PAM</td><td>NA</td><td>Intergenic</td><td>NA</td><td>NA</td></tr>
<tr><td></td><td></td><td></td><td></td><td></td><td></td><td></td><td></td><td></td><td></td><td></td><td></td></tr>
<tr><td>T31201</td><td>ATCGCAAACACGTGTCTCATAGG</td><td>1000</td><td></td><td></td><td></td><td></td><td></td><td></td><td></td><td></td><td></td></tr>
<tr><td>Oligo fwd</td><td>CACCATCGCAAACACGTGTCTCAT</td><td></td><td></td><td></td><td></td><td></td><td></td><td></td><td></td><td></td><td></td></tr>
<tr><td>Oligo rev</td><td>AAACATGAGACACGTGTTTGCGAT</td><td></td><td></td><td></td><td></td><td></td><td></td><td></td><td></td><td></td><td></td></tr>
<tr><td>Chromosome</td><td>start</td><td>end</td><td>strand</td><td>MM</td><td>target_seq</td><td>PAM</td><td>alignment</td><td>distance</td><td>position</td><td>gene name</td><td>gene id</td></tr>
<tr><td>GU305914.1</td><td>40343</td><td>40365</td><td>-</td><td>0</td><td>ATCGCAAACACGTGTCTCAT</td><td>AGG</td><td>||||||||[||||||||||||]PAM</td><td>NA</td><td>Intergenic</td><td>NA</td><td>NA</td></tr>
<tr><td></td><td></td><td></td><td></td><td></td><td></td><td></td><td></td><td></td><td></td><td></td><td></td></tr>
<tr><td>T31202</td><td>TCGCAAACACGTGTCTCATAGGG</td><td>1000</td><td></td><td></td><td></td><td></td><td></td><td></td><td></td><td></td><td></td></tr>
<tr><td>Oligo fwd</td><td>CACCTCGCAAACACGTGTCTCATA</td><td></td><td></td><td></td><td></td><td></td><td></td><td></td><td></td><td></td><td></td></tr>
<tr><td>Oligo rev</td><td>AAACTATGAGACACGTGTTTGCGA</td><td></td><td></td><td></td><td></td><td></td><td></td><td></td><td></td><td></td><td></td></tr>
<tr><td>Chromosome</td><td>start</td><td>end</td><td>strand</td><td>MM</td><td>target_seq</td><td>PAM</td><td>alignment</td><td>distance</td><td>position</td><td>gene name</td><td>gene id</td></tr>
<tr><td>GU305914.1</td><td>40342</td><td>40364</td><td>-</td><td>0</td><td>TCGCAAACACGTGTCTCATA</td><td>GGG</td><td>||||||||[||||||||||||]PAM</td><td>NA</td><td>Intergenic</td><td>NA</td><td>NA</td></tr>
<tr><td></td><td></td><td></td><td></td><td></td><td></td><td></td><td></td><td></td><td></td><td></td><td></td></tr>
<tr><td>T31203</td><td>CGCAAACACGTGTCTCATAGGGG</td><td>1000</td><td></td><td></td><td></td><td></td><td></td><td></td><td></td><td></td><td></td></tr>
<tr><td>Oligo fwd</td><td>CACCCGCAAACACGTGTCTCATAG</td><td></td><td></td><td></td><td></td><td></td><td></td><td></td><td></td><td></td><td></td></tr>
<tr><td>Oligo rev</td><td>AAACCTATGAGACACGTGTTTGCG</td><td></td><td></td><td></td><td></td><td></td><td></td><td></td><td></td><td></td><td></td></tr>
<tr><td>Chromosome</td><td>start</td><td>end</td><td>strand</td><td>MM</td><td>target_seq</td><td>PAM</td><td>alignment</td><td>distance</td><td>position</td><td>gene name</td><td>gene id</td></tr>
<tr><td>GU305914.1</td><td>40341</td><td>40363</td><td>-</td><td>0</td><td>CGCAAACACGTGTCTCATAG</td><td>GGG</td><td>||||||||[||||||||||||]PAM</td><td>NA</td><td>Intergenic</td><td>NA</td><td>NA</td></tr>
<tr><td></td><td></td><td></td><td></td><td></td><td></td><td></td><td></td><td></td><td></td><td></td><td></td></tr>
<tr><td>T31204</td><td>GCAAACACGTGTCTCATAGGGGG</td><td>1000</td><td></td><td></td><td></td><td></td><td></td><td></td><td></td><td></td><td></td></tr>
<tr><td>Oligo fwd</td><td>CACCGCAAACACGTGTCTCATAGG</td><td></td><td></td><td></td><td></td><td></td><td></td><td></td><td></td><td></td><td></td></tr>
<tr><td>Oligo rev</td><td>AAACCCTATGAGACACGTGTTTGC</td><td></td><td></td><td></td><td></td><td></td><td></td><td></td><td></td><td></td><td></td></tr>
<tr><td>Chromosome</td><td>start</td><td>end</td><td>strand</td><td>MM</td><td>target_seq</td><td>PAM</td><td>alignment</td><td>distance</td><td>position</td><td>gene name</td><td>gene id</td></tr>
<tr><td>GU305914.1</td><td>40340</td><td>40362</td><td>-</td><td>0</td><td>GCAAACACGTGTCTCATAGG</td><td>GGG</td><td>||||||||[||||||||||||]PAM</td><td>NA</td><td>Intergenic</td><td>NA</td><td>NA</td></tr>
<tr><td></td><td></td><td></td><td></td><td></td><td></td><td></td><td></td><td></td><td></td><td></td><td></td></tr>
<tr><td>T31205</td><td>TCTCATAGGGGGAGACGATGAGG</td><td>1000</td><td></td><td></td><td></td><td></td><td></td><td></td><td></td><td></td><td></td></tr>
<tr><td>Oligo fwd</td><td>CACCTCTCATAGGGGGAGACGATG</td><td></td><td></td><td></td><td></td><td></td><td></td><td></td><td></td><td></td><td></td></tr>
<tr><td>Oligo rev</td><td>AAACCATCGTCTCCCCCTATGAGA</td><td></td><td></td><td></td><td></td><td></td><td></td><td></td><td></td><td></td><td></td></tr>
<tr><td>Chromosome</td><td>start</td><td>end</td><td>strand</td><td>MM</td><td>target_seq</td><td>PAM</td><td>alignment</td><td>distance</td><td>position</td><td>gene name</td><td>gene id</td></tr>
<tr><td>GU305914.1</td><td>40329</td><td>40351</td><td>-</td><td>0</td><td>TCTCATAGGGGGAGACGATG</td><td>AGG</td><td>||||||||[||||||||||||]PAM</td><td>NA</td><td>Intergenic</td><td>NA</td><td>NA</td></tr>
</table>

T31206 GGGGGAGACGATGAGGATTGCGG 1000
Oligo fwd CACCGGGGGAGACGATGAGGATTG
Oligo rev AAACCAATCCTCATCGTCTCCCCC

| Chromosome | start | end | strand | MM | target_seq | PAM | alignment | distance | position | gene name | gene id |
|---|---|---|---|---|---|---|---|---|---|---|---|
| GU305914.1 | 40322 | 40344 | - | 0 | GGGGGAGACGATGAGGATTG | CGG | \|\|\|\|\|\|\|\|[\|\|\|\|\|\|\|\|\|\|\|\|]PAM | NA | Intergenic | NA | NA |

T31207 GGAGACGATGAGGATTGCGGTGG 1000
Oligo fwd CACCGGAGACGATGAGGATTGCGG
Oligo rev AAACCCGCAATCCTCATCGTCTCC

| Chromosome | start | end | strand | MM | target_seq | PAM | alignment | distance | position | gene name | gene id |
|---|---|---|---|---|---|---|---|---|---|---|---|
| GU305914.1 | 40319 | 40341 | - | 0 | GGAGACGATGAGGATTGCGG | TGG | \|\|\|\|\|\|\|\|[\|\|\|\|\|\|\|\|\|\|\|\|]PAM | NA | Intergenic | NA | NA |

T31208 GACGATGAGGATTGCGGTGGCGG 1000
Oligo fwd CACCGACGATGAGGATTGCGGTGG
Oligo rev AAACCCACCGCAATCCTCATCGTC

| Chromosome | start | end | strand | MM | target_seq | PAM | alignment | distance | position | gene name | gene id |
|---|---|---|---|---|---|---|---|---|---|---|---|
| GU305914.1 | 40316 | 40338 | - | 0 | GACGATGAGGATTGCGGTGG | CGG | \|\|\|\|\|\|\|\|[\|\|\|\|\|\|\|\|\|\|\|\|]PAM | NA | Intergenic | NA | NA |

T31209 GAGGATTGCGGTGGCGGCAGTGG 1000
Oligo fwd CACCGAGGATTGCGGTGGCGGCAG
Oligo rev AAACCTGCCGCCACCGCAATCCTC

| Chromosome | start | end | strand | MM | target_seq | PAM | alignment | distance | position | gene name | gene id |
|---|---|---|---|---|---|---|---|---|---|---|---|
| GU305914.1 | 40310 | 40332 | - | 0 | GAGGATTGCGGTGGCGGCAG | TGG | \|\|\|\|\|\|\|\|[\|\|\|\|\|\|\|\|\|\|\|\|]PAM | NA | Intergenic | NA | NA |
| GU305914.1 | 45258 | 45280 | - | 4 | AACGGTGGCGGTGGCGGCAG | CGG | -\|-\|-\|-\|[\|\|\|\|\|\|\|\|\|\|\|\|]PAM | NA | Intergenic | NA | NA |

T31210 TGGCGGCAGTGGACGCTGTTTGG 1000
Oligo fwd CACCTGGCGGCAGTGGACGCTGTT
Oligo rev AAACAACAGCGTCCACTGCCGCCA

| Chromosome | start | end | strand | MM | target_seq | PAM | alignment | distance | position | gene name | gene id |
|---|---|---|---|---|---|---|---|---|---|---|---|
| GU305914.1 | 40299 | 40321 | - | 0 | TGGCGGCAGTGGACGCTGTT | TGG | \|\|\|\|\|\|\|\|[\|\|\|\|\|\|\|\|\|\|\|\|]PAM | NA | Intergenic | NA | NA |

T31211 AATTCGATCCCGATAGCGAGCGG 1000
Oligo fwd CACCAATTCGATCCCGATAGCGAG
Oligo rev AAACCTCGCTATCGGGATCGAATT

| Chromosome | start | end | strand | MM | target_seq | PAM | alignment | distance | position | gene name | gene id |
|---|---|---|---|---|---|---|---|---|---|---|---|
| GU305914.1 | 40276 | 40298 | - | 0 | AATTCGATCCCGATAGCGAG | CGG | \|\|\|\|\|\|\|\|[\|\|\|\|\|\|\|\|\|\|\|\|]PAM | NA | Intergenic | NA | NA |

T31212 CCCGATAGCGAGCGGTGTCTTGG 1000
Oligo fwd CACCCCCGATAGCGAGCGGTGTCT
Oligo rev AAACAGACACCGCTCGCTATCGGG

| Chromosome | start | end | strand | MM | target_seq | PAM | alignment | distance | position | gene name | gene id |
|---|---|---|---|---|---|---|---|---|---|---|---|
| GU305914.1 | 40268 | 40290 | - | 0 | CCCGATAGCGAGCGGTGTCT | TGG | \|\|\|\|\|\|\|\|[\|\|\|\|\|\|\|\|\|\|\|\|]PAM | NA | Intergenic | NA | NA |

T31213 CCGATAGCGAGCGGTGTCTTGGG 1000
Oligo fwd CACCCCGATAGCGAGCGGTGTCTT
Oligo rev AAACAAGACACCGCTCGCTATCGG

| Chromosome | start | end | strand | MM | target_seq | PAM | alignment | distance | position | gene name | gene id |
|---|---|---|---|---|---|---|---|---|---|---|---|
| GU305914.1 | 40267 | 40289 | - | 0 | CCGATAGCGAGCGGTGTCTT | GGG | \|\|\|\|\|\|\|\|[\|\|\|\|\|\|\|\|\|\|\|\|]PAM | NA | Intergenic | NA | NA |

T31214 CGATAGCGAGCGGTGTCTTGGGG 1000
Oligo fwd CACCCGATAGCGAGCGGTGTCTTG
Oligo rev AAACCAAGACACCGCTCGCTATCG

| Chromosome | start | end | strand | MM | target_seq | PAM | alignment | distance | position | gene name | gene id |
|---|---|---|---|---|---|---|---|---|---|---|---|
| GU305914.1 | 40266 | 40288 | - | 0 | CGATAGCGAGCGGTGTCTTG | GGG | \|\|\|\|\|\|\|\|[\|\|\|\|\|\|\|\|\|\|\|\|]PAM | NA | Intergenic | NA | NA |

T31215 GCGGTGTCTTGGGGTCAAGATGG 1000
Oligo fwd CACCGCGGTGTCTTGGGGTCAAGA
Oligo rev AAACTCTTGACCCCAAGACACCGC

| Chromosome | start | end | strand | MM | target_seq | PAM | alignment | distance | position | gene name | gene id |
|---|---|---|---|---|---|---|---|---|---|---|---|
| GU305914.1 | 40257 | 40279 | - | 0 | GCGGTGTCTTGGGGTCAAGA | TGG | \|\|\|\|\|\|\|\|[\|\|\|\|\|\|\|\|\|\|\|\|]PAM | NA | Intergenic | NA | NA |

T31216 CTTGGGGTCAAGATGGTGAACGG 1000
Oligo fwd CACCCTTGGGGTCAAGATGGTGAA
Oligo rev AAACTTCACCATCTTGACCCCAAG

| Chromosome | start | end | strand | MM | target_seq | PAM | alignment | distance | position | gene name | gene id |
|---|---|---|---|---|---|---|---|---|---|---|---|
| GU305914.1 | 40250 | 40272 | - | 0 | CTTGGGGTCAAGATGGTGAA | CGG | \|\|\|\|\|\|\|\|[\|\|\|\|\|\|\|\|\|\|\|\|]PAM | NA | Intergenic | NA | NA |

T31217 GGGTCAAGATGGTGAACGGCAGG 1000
Oligo fwd CACCGGGTCAAGATGGTGAACGGC
Oligo rev AAACGCCGTTCACCATCTTGACCC

| Chromosome | start | end | strand | MM | target_seq | PAM | alignment | distance | position | gene name | gene id |
|---|---|---|---|---|---|---|---|---|---|---|---|
| GU305914.1 | 40246 | 40268 | - | 0 | GGGTCAAGATGGTGAACGGC | AGG | \|\|\|\|\|\|\|\|[\|\|\|\|\|\|\|\|\|\|\|\|]PAM | NA | Intergenic | NA | NA |

T31218 GGTCAAGATGGTGAACGGCAGGG 1000
Oligo fwd CACCGGTCAAGATGGTGAACGGCA
Oligo rev AAACTGCCGTTCACCATCTTGACC

| Chromosome | start | end | strand | MM | target_seq | PAM | alignment | distance | position | gene name | gene id |
|---|---|---|---|---|---|---|---|---|---|---|---|
| GU305914.1 | 40245 | 40267 | - | 0 | GGTCAAGATGGTGAACGGCA | GGG | \|\|\|\|\|\|\|\|[\|\|\|\|\|\|\|\|\|\|\|\|]PAM | NA | Intergenic | NA | NA |

T31219 TGGTGAACGGCAGGGCTCTGAGG 1000
Oligo fwd CACCTGGTGAACGGCAGGGCTCTG
Oligo rev AAACCAGAGCCCTGCCGTTCACCA

| Chromosome | start | end | strand | MM | target_seq | PAM | alignment | distance | position | gene name | gene id |
|---|---|---|---|---|---|---|---|---|---|---|---|
| GU305914.1 | 40237 | 40259 | - | 0 | TGGTGAACGGCAGGGCTCTG | AGG | \|\|\|\|\|\|\|\|[\|\|\|\|\|\|\|\|\|\|\|\|]PAM | NA | Intergenic | NA | NA |

T31220 TGAACGGCAGGGCTCTGAGGTGG 1000
Oligo fwd CACCTGAACGGCAGGGCTCTGAGG
Oligo rev AAACCCTCAGAGCCCTGCCGTTCA

Chromosome start end strand MM target_seq PAM alignment distance position gene name gene id
GU305914.1 40234 40256 - 0 TGAACGGCAGGGCTCTGAGG TGG ||||||||[||||||||||||]PAM NA Intergenic NA NA

T31221 ACGGCAGGGCTCTGAGGTGGTGG 1000
Oligo fwd CACCACGGCAGGGCTCTGAGGTGG
Oligo rev AAACCCACCTCAGAGCCCTGCCGT
Chromosome start end strand MM target_seq PAM alignment distance position gene name gene id
GU305914.1 40231 40253 - 0 ACGGCAGGGCTCTGAGGTGG TGG ||||||||[||||||||||||]PAM NA Intergenic NA NA

T31222 CTGAGGTGGTGGAATCCGACCGG 1000
Oligo fwd CACCCTGAGGTGGTGGAATCCGAC
Oligo rev AAACGTCGGATTCCACCACCTCAG
Chromosome start end strand MM target_seq PAM alignment distance position gene name gene id
GU305914.1 40220 40242 - 0 CTGAGGTGGTGGAATCCGAC CGG ||||||||[||||||||||||]PAM NA Intergenic NA NA

T31223 TGAGGTGGTGGAATCCGACCGGG 1000
Oligo fwd CACCTGAGGTGGTGGAATCCGACC
Oligo rev AAACGGTCGGATTCCACCACCTCA
Chromosome start end strand MM target_seq PAM alignment distance position gene name gene id
GU305914.1 40219 40241 - 0 TGAGGTGGTGGAATCCGACC GGG ||||||||[||||||||||||]PAM NA Intergenic NA NA

T31224 GTGGAATCCGACCGGGATAATGG 1000
Oligo fwd CACCGTGGAATCCGACCGGGATAA
Oligo rev AAACTTATCCCGGTCGGATTCCAC
Chromosome start end strand MM target_seq PAM alignment distance position gene name gene id
GU305914.1 40212 40234 - 0 GTGGAATCCGACCGGGATAA TGG ||||||||[||||||||||||]PAM NA Intergenic NA NA

T31225 GAATCCGACCGGGATAATGGTGG 1000
Oligo fwd CACCGAATCCGACCGGGATAATGG
Oligo rev AAACCCATTATCCCGGTCGGATTC
Chromosome start end strand MM target_seq PAM alignment distance position gene name gene id
GU305914.1 40209 40231 - 0 GAATCCGACCGGGATAATGG TGG ||||||||[||||||||||||]PAM NA Intergenic NA NA

T31226 CGGGATAATGGTGGACAACGAGG 1000
Oligo fwd CACCCGGGATAATGGTGGACAACG
Oligo rev AAACCGTTGTCCACCATTATCCCG
Chromosome start end strand MM target_seq PAM alignment distance position gene name gene id
GU305914.1 40200 40222 - 0 CGGGATAATGGTGGACAACG AGG ||||||||[||||||||||||]PAM NA Intergenic NA NA

T31227 TGGACAACGAGGCCGCCGTCTGG 1000
Oligo fwd CACCTGGACAACGAGGCCGCCGTC
Oligo rev AAACGACGGCGGCCTCGTTGTCCA
Chromosome start end strand MM target_seq PAM alignment distance position gene name gene id
GU305914.1 40189 40211 - 0 TGGACAACGAGGCCGCCGTC TGG ||||||||[||||||||||||]PAM NA Intergenic NA NA

T31228 GCCGTCTGGATCGACGAACACGG 1000
Oligo fwd CACCGCCGTCTGGATCGACGAACA
Oligo rev AAACTGTTCGTCGATCCAGACGGC
Chromosome start end strand MM target_seq PAM alignment distance position gene name gene id
GU305914.1 40175 40197 - 0 GCCGTCTGGATCGACGAACA CGG ||||||||[||||||||||||]PAM NA Intergenic NA NA

T31229 TCTGGATCGACGAACACGGCAGG 1000
Oligo fwd CACCTCTGGATCGACGAACACGGC
Oligo rev AAACGCCGTGTTCGTCGATCCAGA
Chromosome start end strand MM target_seq PAM alignment distance position gene name gene id
GU305914.1 40171 40193 - 0 TCTGGATCGACGAACACGGC AGG ||||||||[||||||||||||]PAM NA Intergenic NA NA

T31230 CTGGATCGACGAACACGGCAGGG 1000
Oligo fwd CACCCTGGATCGACGAACACGGCA
Oligo rev AAACTGCCGTGTTCGTCGATCCAG
Chromosome start end strand MM target_seq PAM alignment distance position gene name gene id
GU305914.1 40170 40192 - 0 CTGGATCGACGAACACGGCA GGG ||||||||[||||||||||||]PAM NA Intergenic NA NA

T31231 CGACGAACACGGCAGGGTGATGG 1000
Oligo fwd CACCCGACGAACACGGCAGGGTGA
Oligo rev AAACTCACCCTGCCGTGTTCGTCG
Chromosome start end strand MM target_seq PAM alignment distance position gene name gene id
GU305914.1 40164 40186 - 0 CGACGAACACGGCAGGGTGA TGG ||||||||[||||||||||||]PAM NA Intergenic NA NA

T31232 GATGGACAAACCACCGCCCAAGG 1000
Oligo fwd CACCGATGGACAAACCACCGCCCA
Oligo rev AAACTGGGCGGTGGTTTGTCCATC
Chromosome start end strand MM target_seq PAM alignment distance position gene name gene id
GU305914.1 40146 40168 - 0 GATGGACAAACCACCGCCCA AGG ||||||||[||||||||||||]PAM NA Intergenic NA NA

T31233 ACCGCCCAAGGAACTCCGAAAGG 1000
Oligo fwd CACCACCGCCCAAGGAACTCCGAA
Oligo rev AAACTTCGGAGTTCCTTGGGCGGT
Chromosome start end strand MM target_seq PAM alignment distance position gene name gene id
GU305914.1 40134 40156 - 0 ACCGCCCAAGGAACTCCGAA AGG ||||||||[||||||||||||]PAM NA Intergenic NA NA

T31234 CGAAAGGCAAGCTCCGACGACGG 1000
Oligo fwd CACCCGAAAGGCAAGCTCCGACGA
Oligo rev AAACTCGTCGGAGCTTGCCTTTCG
Chromosome start end strand MM target_seq PAM alignment distance position gene name gene id
GU305914.1 40118 40140 - 0 CGAAAGGCAAGCTCCGACGA CGG ||||||||[||||||||||||]PAM NA Intergenic NA NA

T31235 AAGGCAAGCTCCGACGACGGTGG 1000
Oligo fwd CACCAAGGCAAGCTCCGACGACGG

| | | | | | | | | | | | |
|---|---|---|---|---|---|---|---|---|---|---|---|
| Oligo rev | AAACCCGTCGTCGGAGCTTGCCTT | | | | | | | | | | |
| Chromosome | start | end | strand | MM | target_seq | PAM | alignment | distance | position | gene name | gene id |
| GU305914.1 | 40115 | 40137 | - | 0 | AAGGCAAGCTCCGACGACGG | TGG | \|\|\|\|\|\|\|\|[\|\|\|\|\|\|\|\|\|\|\|\|]PAM | NA | Intergenic | NA | NA |
| T31236 | GAATGTCACGCCACCCGTCTCGG | 1000 | | | | | | | | | |
| Oligo fwd | CACCGAATGTCACGCCACCCGTCT | | | | | | | | | | |
| Oligo rev | AAACAGACGGGTGGCGTGACATTC | | | | | | | | | | |
| Chromosome | start | end | strand | MM | target_seq | PAM | alignment | distance | position | gene name | gene id |
| GU305914.1 | 40065 | 40087 | - | 0 | GAATGTCACGCCACCCGTCT | CGG | \|\|\|\|\|\|\|\|[\|\|\|\|\|\|\|\|\|\|\|\|]PAM | NA | Intergenic | NA | NA |
| T31237 | AATGTCACGCCACCCGTCTCGGG | 1000 | | | | | | | | | |
| Oligo fwd | CACCAATGTCACGCCACCCGTCTC | | | | | | | | | | |
| Oligo rev | AAACGAGACGGGTGGCGTGACATT | | | | | | | | | | |
| Chromosome | start | end | strand | MM | target_seq | PAM | alignment | distance | position | gene name | gene id |
| GU305914.1 | 40064 | 40086 | - | 0 | AATGTCACGCCACCCGTCTC | GGG | \|\|\|\|\|\|\|\|[\|\|\|\|\|\|\|\|\|\|\|\|]PAM | NA | Intergenic | NA | NA |
| T31238 | GTCTCGGGAAGTAATAGCGTTGG | 1000 | | | | | | | | | |
| Oligo fwd | CACCGTCTCGGGAAGTAATAGCGT | | | | | | | | | | |
| Oligo rev | AAACACGCTATTACTTCCCGAGAC | | | | | | | | | | |
| Chromosome | start | end | strand | MM | target_seq | PAM | alignment | distance | position | gene name | gene id |
| GU305914.1 | 40049 | 40071 | - | 0 | GTCTCGGGAAGTAATAGCGT | TGG | \|\|\|\|\|\|\|\|[\|\|\|\|\|\|\|\|\|\|\|\|]PAM | NA | Intergenic | NA | NA |
| T31239 | TCGGGAAGTAATAGCGTTGGCGG | 1000 | | | | | | | | | |
| Oligo fwd | CACCTCGGGAAGTAATAGCGTTGG | | | | | | | | | | |
| Oligo rev | AAACCCAACGCTATTACTTCCCGA | | | | | | | | | | |
| Chromosome | start | end | strand | MM | target_seq | PAM | alignment | distance | position | gene name | gene id |
| GU305914.1 | 40046 | 40068 | - | 0 | TCGGGAAGTAATAGCGTTGG | CGG | \|\|\|\|\|\|\|\|[\|\|\|\|\|\|\|\|\|\|\|\|]PAM | NA | Intergenic | NA | NA |
| T31240 | GGAAGTAATAGCGTTGGCGGCGG | 1000 | | | | | | | | | |
| Oligo fwd | CACCGGAAGTAATAGCGTTGGCGG | | | | | | | | | | |
| Oligo rev | AAACCCGCCAACGCTATTACTTCC | | | | | | | | | | |
| Chromosome | start | end | strand | MM | target_seq | PAM | alignment | distance | position | gene name | gene id |
| GU305914.1 | 40043 | 40065 | - | 0 | GGAAGTAATAGCGTTGGCGG | CGG | \|\|\|\|\|\|\|\|[\|\|\|\|\|\|\|\|\|\|\|\|]PAM | NA | Intergenic | NA | NA |
| T31241 | CGGTGTACAGACACCGTCAACGG | 1000 | | | | | | | | | |
| Oligo fwd | CACCCGGTGTACAGACACCGTCAA | | | | | | | | | | |
| Oligo rev | AAACTTGACGGTGTCTGTACACCG | | | | | | | | | | |
| Chromosome | start | end | strand | MM | target_seq | PAM | alignment | distance | position | gene name | gene id |
| GU305914.1 | 40023 | 40045 | - | 0 | CGGTGTACAGACACCGTCAA | CGG | \|\|\|\|\|\|\|\|[\|\|\|\|\|\|\|\|\|\|\|\|]PAM | NA | Intergenic | NA | NA |
| T31242 | ACAGACACCGTCAACGGCGTCGG | 1000 | | | | | | | | | |
| Oligo fwd | CACCACAGACACCGTCAACGGCGT | | | | | | | | | | |
| Oligo rev | AAACACGCCGTTGACGGTGTCTGT | | | | | | | | | | |
| Chromosome | start | end | strand | MM | target_seq | PAM | alignment | distance | position | gene name | gene id |
| GU305914.1 | 40017 | 40039 | - | 0 | ACAGACACCGTCAACGGCGT | CGG | \|\|\|\|\|\|\|\|[\|\|\|\|\|\|\|\|\|\|\|\|]PAM | NA | Intergenic | NA | NA |
| T31243 | CAGACACCGTCAACGGCGTCGGG | 1000 | | | | | | | | | |
| Oligo fwd | CACCCAGACACCGTCAACGGCGTC | | | | | | | | | | |
| Oligo rev | AAACGACGCCGTTGACGGTGTCTG | | | | | | | | | | |
| Chromosome | start | end | strand | MM | target_seq | PAM | alignment | distance | position | gene name | gene id |
| GU305914.1 | 40016 | 40038 | - | 0 | CAGACACCGTCAACGGCGTC | GGG | \|\|\|\|\|\|\|\|[\|\|\|\|\|\|\|\|\|\|\|\|]PAM | NA | Intergenic | NA | NA |
| T31244 | AGACACCGTCAACGGCGTCGGGG | 1000 | | | | | | | | | |
| Oligo fwd | CACCAGACACCGTCAACGGCGTCG | | | | | | | | | | |
| Oligo rev | AAACCGACGCCGTTGACGGTGTCT | | | | | | | | | | |
| Chromosome | start | end | strand | MM | target_seq | PAM | alignment | distance | position | gene name | gene id |
| GU305914.1 | 40015 | 40037 | - | 0 | AGACACCGTCAACGGCGTCG | GGG | \|\|\|\|\|\|\|\|[\|\|\|\|\|\|\|\|\|\|\|\|]PAM | NA | Intergenic | NA | NA |
| T31245 | ACGGCGTCGGGGAAACGTACAGG | 1000 | | | | | | | | | |
| Oligo fwd | CACCACGGCGTCGGGGAAACGTAC | | | | | | | | | | |
| Oligo rev | AAACGTACGTTTCCCCGACGCCGT | | | | | | | | | | |
| Chromosome | start | end | strand | MM | target_seq | PAM | alignment | distance | position | gene name | gene id |
| GU305914.1 | 40004 | 40026 | - | 0 | ACGGCGTCGGGGAAACGTAC | AGG | \|\|\|\|\|\|\|\|[\|\|\|\|\|\|\|\|\|\|\|\|]PAM | NA | Intergenic | NA | NA |
| T31246 | CGGCGTCGGGGAAACGTACAGGG | 1000 | | | | | | | | | |
| Oligo fwd | CACCCGGCGTCGGGGAAACGTACA | | | | | | | | | | |
| Oligo rev | AAACTGTACGTTTCCCCGACGCCG | | | | | | | | | | |
| Chromosome | start | end | strand | MM | target_seq | PAM | alignment | distance | position | gene name | gene id |
| GU305914.1 | 40003 | 40025 | - | 0 | CGGCGTCGGGGAAACGTACA | GGG | \|\|\|\|\|\|\|\|[\|\|\|\|\|\|\|\|\|\|\|\|]PAM | NA | Intergenic | NA | NA |
| T31247 | GAAACGTACAGGGAAGAAAAAGG | 1000 | | | | | | | | | |
| Oligo fwd | CACCGAAACGTACAGGGAAGAAAA | | | | | | | | | | |
| Oligo rev | AAACTTTTCTTCCCTGTACGTTTC | | | | | | | | | | |
| Chromosome | start | end | strand | MM | target_seq | PAM | alignment | distance | position | gene name | gene id |
| GU305914.1 | 39993 | 40015 | - | 0 | GAAACGTACAGGGAAGAAAA | AGG | \|\|\|\|\|\|\|\|[\|\|\|\|\|\|\|\|\|\|\|\|]PAM | NA | Intergenic | NA | NA |
| T31248 | CGTACAGGGAAGAAAAAGGAAGG | 1000 | | | | | | | | | |
| Oligo fwd | CACCCGTACAGGGAAGAAAAAGGA | | | | | | | | | | |
| Oligo rev | AAACTCCTTTTTCTTCCCTGTACG | | | | | | | | | | |
| Chromosome | start | end | strand | MM | target_seq | PAM | alignment | distance | position | gene name | gene id |
| GU305914.1 | 39989 | 40011 | - | 0 | CGTACAGGGAAGAAAAAGGA | AGG | \|\|\|\|\|\|\|\|[\|\|\|\|\|\|\|\|\|\|\|\|]PAM | NA | Intergenic | NA | NA |
| T31249 | ACAGGGAAGAAAAAGGAAGGTGG | 1000 | | | | | | | | | |
| Oligo fwd | CACCACAGGGAAGAAAAAGGAAGG | | | | | | | | | | |
| Oligo rev | AAACCCTTCCTTTTTCTTCCCTGT | | | | | | | | | | |
| Chromosome | start | end | strand | MM | target_seq | PAM | alignment | distance | position | gene name | gene id |
| GU305914.1 | 39986 | 40008 | - | 0 | ACAGGGAAGAAAAAGGAAGG | TGG | \|\|\|\|\|\|\|\|[\|\|\|\|\|\|\|\|\|\|\|\|]PAM | NA | Intergenic | NA | NA |
| T31250 | CAGGGAAGAAAAAGGAAGGTGGG | 1000 | | | | | | | | | |

<table>
<tr><td>Oligo fwd</td><td>CACCCAGGGAAGAAAAAGGAAGGT</td><td></td><td></td><td></td><td></td><td></td><td></td><td></td><td></td><td></td><td></td></tr>
<tr><td>Oligo rev</td><td>AAACACCTTCCTTTTTCTTCCCTG</td><td></td><td></td><td></td><td></td><td></td><td></td><td></td><td></td><td></td><td></td></tr>
<tr><td>Chromosome</td><td>start</td><td>end</td><td>strand</td><td>MM</td><td>target_seq</td><td>PAM</td><td>alignment</td><td>distance</td><td>position</td><td>gene name</td><td>gene id</td></tr>
<tr><td>GU305914.1</td><td>39985</td><td>40007</td><td>-</td><td>0</td><td>CAGGGAAGAAAAAGGAAGGT</td><td>GGG</td><td>||||||||[||||||||||||]PAM</td><td>NA</td><td>Intergenic</td><td>NA</td><td>NA</td></tr>
<tr><td></td><td></td><td></td><td></td><td></td><td></td><td></td><td></td><td></td><td></td><td></td><td></td></tr>
<tr><td>T31251</td><td>AGGGAAGAAAAAGGAAGGTGGGG</td><td>1000</td><td></td><td></td><td></td><td></td><td></td><td></td><td></td><td></td><td></td></tr>
<tr><td>Oligo fwd</td><td>CACCAGGGAAGAAAAAGGAAGGTG</td><td></td><td></td><td></td><td></td><td></td><td></td><td></td><td></td><td></td><td></td></tr>
<tr><td>Oligo rev</td><td>AAACCACCTTCCTTTTTCTTCCCT</td><td></td><td></td><td></td><td></td><td></td><td></td><td></td><td></td><td></td><td></td></tr>
<tr><td>Chromosome</td><td>start</td><td>end</td><td>strand</td><td>MM</td><td>target_seq</td><td>PAM</td><td>alignment</td><td>distance</td><td>position</td><td>gene name</td><td>gene id</td></tr>
<tr><td>GU305914.1</td><td>39984</td><td>40006</td><td>-</td><td>0</td><td>AGGGAAGAAAAAGGAAGGTG</td><td>GGG</td><td>||||||||[||||||||||||]PAM</td><td>NA</td><td>Intergenic</td><td>NA</td><td>NA</td></tr>
<tr><td></td><td></td><td></td><td></td><td></td><td></td><td></td><td></td><td></td><td></td><td></td><td></td></tr>
<tr><td>T31252</td><td>GGGAAGAAAAAGGAAGGTGGGGG</td><td>1000</td><td></td><td></td><td></td><td></td><td></td><td></td><td></td><td></td><td></td></tr>
<tr><td>Oligo fwd</td><td>CACCGGGAAGAAAAAGGAAGGTGG</td><td></td><td></td><td></td><td></td><td></td><td></td><td></td><td></td><td></td><td></td></tr>
<tr><td>Oligo rev</td><td>AAACCCACCTTCCTTTTTCTTCCC</td><td></td><td></td><td></td><td></td><td></td><td></td><td></td><td></td><td></td><td></td></tr>
<tr><td>Chromosome</td><td>start</td><td>end</td><td>strand</td><td>MM</td><td>target_seq</td><td>PAM</td><td>alignment</td><td>distance</td><td>position</td><td>gene name</td><td>gene id</td></tr>
<tr><td>GU305914.1</td><td>39983</td><td>40005</td><td>-</td><td>0</td><td>GGGAAGAAAAAGGAAGGTGG</td><td>GGG</td><td>||||||||[||||||||||||]PAM</td><td>NA</td><td>Intergenic</td><td>NA</td><td>NA</td></tr>
<tr><td></td><td></td><td></td><td></td><td></td><td></td><td></td><td></td><td></td><td></td><td></td><td></td></tr>
<tr><td>T31253</td><td>AAGAAAAAGGAAGGTGGGGGCGG</td><td>1000</td><td></td><td></td><td></td><td></td><td></td><td></td><td></td><td></td><td></td></tr>
<tr><td>Oligo fwd</td><td>CACCAAGAAAAAGGAAGGTGGGGG</td><td></td><td></td><td></td><td></td><td></td><td></td><td></td><td></td><td></td><td></td></tr>
<tr><td>Oligo rev</td><td>AAACCCCCCACCTTCCTTTTTCTT</td><td></td><td></td><td></td><td></td><td></td><td></td><td></td><td></td><td></td><td></td></tr>
<tr><td>Chromosome</td><td>start</td><td>end</td><td>strand</td><td>MM</td><td>target_seq</td><td>PAM</td><td>alignment</td><td>distance</td><td>position</td><td>gene name</td><td>gene id</td></tr>
<tr><td>GU305914.1</td><td>39980</td><td>40002</td><td>-</td><td>0</td><td>AAGAAAAAGGAAGGTGGGGG</td><td>CGG</td><td>||||||||[||||||||||||]PAM</td><td>NA</td><td>Intergenic</td><td>NA</td><td>NA</td></tr>
<tr><td></td><td></td><td></td><td></td><td></td><td></td><td></td><td></td><td></td><td></td><td></td><td></td></tr>
<tr><td>T31254</td><td>GCGGCTACTTGTTGAGAAGCAGG</td><td>1000</td><td></td><td></td><td></td><td></td><td></td><td></td><td></td><td></td><td></td></tr>
<tr><td>Oligo fwd</td><td>CACCGCGGCTACTTGTTGAGAAGC</td><td></td><td></td><td></td><td></td><td></td><td></td><td></td><td></td><td></td><td></td></tr>
<tr><td>Oligo rev</td><td>AAACGCTTCTCAACAAGTAGCCGC</td><td></td><td></td><td></td><td></td><td></td><td></td><td></td><td></td><td></td><td></td></tr>
<tr><td>Chromosome</td><td>start</td><td>end</td><td>strand</td><td>MM</td><td>target_seq</td><td>PAM</td><td>alignment</td><td>distance</td><td>position</td><td>gene name</td><td>gene id</td></tr>
<tr><td>GU305914.1</td><td>39961</td><td>39983</td><td>-</td><td>0</td><td>GCGGCTACTTGTTGAGAAGC</td><td>AGG</td><td>||||||||[||||||||||||]PAM</td><td>NA</td><td>Intergenic</td><td>NA</td><td>NA</td></tr>
<tr><td></td><td></td><td></td><td></td><td></td><td></td><td></td><td></td><td></td><td></td><td></td><td></td></tr>
<tr><td>T31255</td><td>CTTGTTGAGAAGCAGGAGTACGG</td><td>1000</td><td></td><td></td><td></td><td></td><td></td><td></td><td></td><td></td><td></td></tr>
<tr><td>Oligo fwd</td><td>CACCCTTGTTGAGAAGCAGGAGTA</td><td></td><td></td><td></td><td></td><td></td><td></td><td></td><td></td><td></td><td></td></tr>
<tr><td>Oligo rev</td><td>AAACTACTCCTGCTTCTCAACAAG</td><td></td><td></td><td></td><td></td><td></td><td></td><td></td><td></td><td></td><td></td></tr>
<tr><td>Chromosome</td><td>start</td><td>end</td><td>strand</td><td>MM</td><td>target_seq</td><td>PAM</td><td>alignment</td><td>distance</td><td>position</td><td>gene name</td><td>gene id</td></tr>
<tr><td>GU305914.1</td><td>39954</td><td>39976</td><td>-</td><td>0</td><td>CTTGTTGAGAAGCAGGAGTA</td><td>CGG</td><td>||||||||[||||||||||||]PAM</td><td>NA</td><td>Intergenic</td><td>NA</td><td>NA</td></tr>
<tr><td></td><td></td><td></td><td></td><td></td><td></td><td></td><td></td><td></td><td></td><td></td><td></td></tr>
<tr><td>T31256</td><td>CAGGAGTACGGACGATGATGAGG</td><td>1000</td><td></td><td></td><td></td><td></td><td></td><td></td><td></td><td></td><td></td></tr>
<tr><td>Oligo fwd</td><td>CACCCAGGAGTACGGACGATGATG</td><td></td><td></td><td></td><td></td><td></td><td></td><td></td><td></td><td></td><td></td></tr>
<tr><td>Oligo rev</td><td>AAACCATCATCGTCCGTACTCCTG</td><td></td><td></td><td></td><td></td><td></td><td></td><td></td><td></td><td></td><td></td></tr>
<tr><td>Chromosome</td><td>start</td><td>end</td><td>strand</td><td>MM</td><td>target_seq</td><td>PAM</td><td>alignment</td><td>distance</td><td>position</td><td>gene name</td><td>gene id</td></tr>
<tr><td>GU305914.1</td><td>39942</td><td>39964</td><td>-</td><td>0</td><td>CAGGAGTACGGACGATGATG</td><td>AGG</td><td>||||||||[||||||||||||]PAM</td><td>NA</td><td>Intergenic</td><td>NA</td><td>NA</td></tr>
<tr><td></td><td></td><td></td><td></td><td></td><td></td><td></td><td></td><td></td><td></td><td></td><td></td></tr>
<tr><td>T31257</td><td>GTACGGACGATGATGAGGTGCGG</td><td>1000</td><td></td><td></td><td></td><td></td><td></td><td></td><td></td><td></td><td></td></tr>
<tr><td>Oligo fwd</td><td>CACCGTACGGACGATGATGAGGTG</td><td></td><td></td><td></td><td></td><td></td><td></td><td></td><td></td><td></td><td></td></tr>
<tr><td>Oligo rev</td><td>AAACCACCTCATCATCGTCCGTAC</td><td></td><td></td><td></td><td></td><td></td><td></td><td></td><td></td><td></td><td></td></tr>
<tr><td>Chromosome</td><td>start</td><td>end</td><td>strand</td><td>MM</td><td>target_seq</td><td>PAM</td><td>alignment</td><td>distance</td><td>position</td><td>gene name</td><td>gene id</td></tr>
<tr><td>GU305914.1</td><td>39937</td><td>39959</td><td>-</td><td>0</td><td>GTACGGACGATGATGAGGTG</td><td>CGG</td><td>||||||||[||||||||||||]PAM</td><td>NA</td><td>Intergenic</td><td>NA</td><td>NA</td></tr>
<tr><td></td><td></td><td></td><td></td><td></td><td></td><td></td><td></td><td></td><td></td><td></td><td></td></tr>
<tr><td>T31258</td><td>GTGCGGAAGATGAAGAAAGATGG</td><td>1000</td><td></td><td></td><td></td><td></td><td></td><td></td><td></td><td></td><td></td></tr>
<tr><td>Oligo fwd</td><td>CACCGTGCGGAAGATGAAGAAAGA</td><td></td><td></td><td></td><td></td><td></td><td></td><td></td><td></td><td></td><td></td></tr>
<tr><td>Oligo rev</td><td>AAACTCTTTCTTCATCTTCCGCAC</td><td></td><td></td><td></td><td></td><td></td><td></td><td></td><td></td><td></td><td></td></tr>
<tr><td>Chromosome</td><td>start</td><td>end</td><td>strand</td><td>MM</td><td>target_seq</td><td>PAM</td><td>alignment</td><td>distance</td><td>position</td><td>gene name</td><td>gene id</td></tr>
<tr><td>GU305914.1</td><td>39920</td><td>39942</td><td>-</td><td>0</td><td>GTGCGGAAGATGAAGAAAGA</td><td>TGG</td><td>||||||||[||||||||||||]PAM</td><td>NA</td><td>Intergenic</td><td>NA</td><td>NA</td></tr>
<tr><td></td><td></td><td></td><td></td><td></td><td></td><td></td><td></td><td></td><td></td><td></td><td></td></tr>
<tr><td>T31259</td><td>TGGCACGATCGACGATCGCGCGG</td><td>1000</td><td></td><td></td><td></td><td></td><td></td><td></td><td></td><td></td><td></td></tr>
<tr><td>Oligo fwd</td><td>CACCTGGCACGATCGACGATCGCG</td><td></td><td></td><td></td><td></td><td></td><td></td><td></td><td></td><td></td><td></td></tr>
<tr><td>Oligo rev</td><td>AAACCGCGATCGTCGATCGTGCCA</td><td></td><td></td><td></td><td></td><td></td><td></td><td></td><td></td><td></td><td></td></tr>
<tr><td>Chromosome</td><td>start</td><td>end</td><td>strand</td><td>MM</td><td>target_seq</td><td>PAM</td><td>alignment</td><td>distance</td><td>position</td><td>gene name</td><td>gene id</td></tr>
<tr><td>GU305914.1</td><td>39900</td><td>39922</td><td>-</td><td>0</td><td>TGGCACGATCGACGATCGCG</td><td>CGG</td><td>||||||||[||||||||||||]PAM</td><td>NA</td><td>Intergenic</td><td>NA</td><td>NA</td></tr>
<tr><td></td><td></td><td></td><td></td><td></td><td></td><td></td><td></td><td></td><td></td><td></td><td></td></tr>
<tr><td>T31260</td><td>CGCGGACAGAGAATTAAAGATGG</td><td>1000</td><td></td><td></td><td></td><td></td><td></td><td></td><td></td><td></td><td></td></tr>
<tr><td>Oligo fwd</td><td>CACCCGCGGACAGAGAATTAAAGA</td><td></td><td></td><td></td><td></td><td></td><td></td><td></td><td></td><td></td><td></td></tr>
<tr><td>Oligo rev</td><td>AAACTCTTTAATTCTCTGTCCGCG</td><td></td><td></td><td></td><td></td><td></td><td></td><td></td><td></td><td></td><td></td></tr>
<tr><td>Chromosome</td><td>start</td><td>end</td><td>strand</td><td>MM</td><td>target_seq</td><td>PAM</td><td>alignment</td><td>distance</td><td>position</td><td>gene name</td><td>gene id</td></tr>
<tr><td>GU305914.1</td><td>39882</td><td>39904</td><td>-</td><td>0</td><td>CGCGGACAGAGAATTAAAGA</td><td>TGG</td><td>||||||||[||||||||||||]PAM</td><td>NA</td><td>Intergenic</td><td>NA</td><td>NA</td></tr>
<tr><td></td><td></td><td></td><td></td><td></td><td></td><td></td><td></td><td></td><td></td><td></td><td></td></tr>
<tr><td>T31261</td><td>ACAAAAAGCGAGAGAAAGCACGG</td><td>1000</td><td></td><td></td><td></td><td></td><td></td><td></td><td></td><td></td><td></td></tr>
<tr><td>Oligo fwd</td><td>CACCACAAAAAGCGAGAGAAAGCA</td><td></td><td></td><td></td><td></td><td></td><td></td><td></td><td></td><td></td><td></td></tr>
<tr><td>Oligo rev</td><td>AAACTGCTTTCTCTCGCTTTTTGT</td><td></td><td></td><td></td><td></td><td></td><td></td><td></td><td></td><td></td><td></td></tr>
<tr><td>Chromosome</td><td>start</td><td>end</td><td>strand</td><td>MM</td><td>target_seq</td><td>PAM</td><td>alignment</td><td>distance</td><td>position</td><td>gene name</td><td>gene id</td></tr>
<tr><td>GU305914.1</td><td>39855</td><td>39877</td><td>-</td><td>0</td><td>ACAAAAAGCGAGAGAAAGCA</td><td>CGG</td><td>||||||||[||||||||||||]PAM</td><td>NA</td><td>Intergenic</td><td>NA</td><td>NA</td></tr>
<tr><td></td><td></td><td></td><td></td><td></td><td></td><td></td><td></td><td></td><td></td><td></td><td></td></tr>
<tr><td>T31262</td><td>GAGAGAAAGCACGGCCGACTCGG</td><td>1000</td><td></td><td></td><td></td><td></td><td></td><td></td><td></td><td></td><td></td></tr>
<tr><td>Oligo fwd</td><td>CACCGAGAGAAAGCACGGCCGACT</td><td></td><td></td><td></td><td></td><td></td><td></td><td></td><td></td><td></td><td></td></tr>
<tr><td>Oligo rev</td><td>AAACAGTCGGCCGTGCTTTCTCTC</td><td></td><td></td><td></td><td></td><td></td><td></td><td></td><td></td><td></td><td></td></tr>
<tr><td>Chromosome</td><td>start</td><td>end</td><td>strand</td><td>MM</td><td>target_seq</td><td>PAM</td><td>alignment</td><td>distance</td><td>position</td><td>gene name</td><td>gene id</td></tr>
<tr><td>GU305914.1</td><td>39846</td><td>39868</td><td>-</td><td>0</td><td>GAGAGAAAGCACGGCCGACT</td><td>CGG</td><td>||||||||[||||||||||||]PAM</td><td>NA</td><td>Intergenic</td><td>NA</td><td>NA</td></tr>
<tr><td></td><td></td><td></td><td></td><td></td><td></td><td></td><td></td><td></td><td></td><td></td><td></td></tr>
<tr><td>T31263</td><td>ATCTGAGCACAATCCTTCCCCGG</td><td>1000</td><td></td><td></td><td></td><td></td><td></td><td></td><td></td><td></td><td></td></tr>
<tr><td>Oligo fwd</td><td>CACCATCTGAGCACAATCCTTCCC</td><td></td><td></td><td></td><td></td><td></td><td></td><td></td><td></td><td></td><td></td></tr>
<tr><td>Oligo rev</td><td>AAACGGGAAGGATTGTGCTCAGAT</td><td></td><td></td><td></td><td></td><td></td><td></td><td></td><td></td><td></td><td></td></tr>
<tr><td>Chromosome</td><td>start</td><td>end</td><td>strand</td><td>MM</td><td>target_seq</td><td>PAM</td><td>alignment</td><td>distance</td><td>position</td><td>gene name</td><td>gene id</td></tr>
<tr><td>GU305914.1</td><td>39823</td><td>39845</td><td>-</td><td>0</td><td>ATCTGAGCACAATCCTTCCC</td><td>CGG</td><td>||||||||[||||||||||||]PAM</td><td>NA</td><td>Intergenic</td><td>NA</td><td>NA</td></tr>
<tr><td></td><td></td><td></td><td></td><td></td><td></td><td></td><td></td><td></td><td></td><td></td><td></td></tr>
<tr><td>T31264</td><td>TTCCCCGGACCGAGCCGCTGCGG</td><td>1000</td><td></td><td></td><td></td><td></td><td></td><td></td><td></td><td></td><td></td></tr>
<tr><td>Oligo fwd</td><td>CACCTTCCCCGGACCGAGCCGCTG</td><td></td><td></td><td></td><td></td><td></td><td></td><td></td><td></td><td></td><td></td></tr>
<tr><td>Oligo rev</td><td>AAACCAGCGGCTCGGTCCGGGGAA</td><td></td><td></td><td></td><td></td><td></td><td></td><td></td><td></td><td></td><td></td></tr>
<tr><td>Chromosome</td><td>start</td><td>end</td><td>strand</td><td>MM</td><td>target_seq</td><td>PAM</td><td>alignment</td><td>distance</td><td>position</td><td>gene name</td><td>gene id</td></tr>
<tr><td>GU305914.1</td><td>39808</td><td>39830</td><td>-</td><td>0</td><td>TTCCCCGGACCGAGCCGCTG</td><td>CGG</td><td>||||||||[||||||||||||]PAM</td><td>NA</td><td>Intergenic</td><td>NA</td><td>NA</td></tr>
</table>

T31265 CTGCGGAAAGTCGCCTTCGTCGG 1000
Oligo fwd CACCCTGCGGAAAGTCGCCTTCGT
Oligo rev AAACACGAAGGCGACTTTCCGCAG
Chromosome start end strand MM target_seq PAM alignment distance position gene name gene id
GU305914.1 39791 39813 - 0 CTGCGGAAAGTCGCCTTCGT CGG ||||||||[||||||||||||]PAM NA Intergenic NA NA

T31266 AGTCGCCTTCGTCGGCGATCCGG 1000
Oligo fwd CACCAGTCGCCTTCGTCGGCGATC
Oligo rev AAACGATCGCCGACGAAGGCGACT
Chromosome start end strand MM target_seq PAM alignment distance position gene name gene id
GU305914.1 39783 39805 - 0 AGTCGCCTTCGTCGGCGATC CGG ||||||||[||||||||||||]PAM NA Intergenic NA NA

T31267 CTTCGTCGGCGATCCGGTCGCGG 1000
Oligo fwd CACCCTTCGTCGGCGATCCGGTCG
Oligo rev AAACCGACCGGATCGCCGACGAAG
Chromosome start end strand MM target_seq PAM alignment distance position gene name gene id
GU305914.1 39777 39799 - 0 CTTCGTCGGCGATCCGGTCG CGG ||||||||[||||||||||||]PAM NA Intergenic NA NA
GU305914.1 23938 23960 - 4 CTTCATCATCGATCCGCTCG AGG ||||-||-[-|||||||-|||]PAM NA Intergenic NA NA

T31268 GGCGATCCGGTCGCGGCGTTCGG 1000
Oligo fwd CACCGGCGATCCGGTCGCGGCGTT
Oligo rev AAACAACGCCGCGACCGGATCGCC
Chromosome start end strand MM target_seq PAM alignment distance position gene name gene id
GU305914.1 39770 39792 - 0 GGCGATCCGGTCGCGGCGTT CGG ||||||||[||||||||||||]PAM NA Intergenic NA NA

T31269 GGTCGCGGCGTTCGGAGATACGG 1000
Oligo fwd CACCGGTCGCGGCGTTCGGAGATA
Oligo rev AAACTATCTCCGAACGCCGCGACC
Chromosome start end strand MM target_seq PAM alignment distance position gene name gene id
GU305914.1 39762 39784 - 0 GGTCGCGGCGTTCGGAGATA CGG ||||||||[||||||||||||]PAM NA Intergenic NA NA

T31270 CGGCGTTCGGAGATACGGTGAGG 1000
Oligo fwd CACCCGGCGTTCGGAGATACGGTG
Oligo rev AAACCACCGTATCTCCGAACGCCG
Chromosome start end strand MM target_seq PAM alignment distance position gene name gene id
GU305914.1 39757 39779 - 0 CGGCGTTCGGAGATACGGTG AGG ||||||||[||||||||||||]PAM NA Intergenic NA NA

T31271 GAGGACCACCTCCTCGTCGAAGG 1000
Oligo fwd CACCGAGGACCACCTCCTCGTCGA
Oligo rev AAACTCGACGAGGAGGTGGTCCTC
Chromosome start end strand MM target_seq PAM alignment distance position gene name gene id
GU305914.1 39738 39760 - 0 GAGGACCACCTCCTCGTCGA AGG ||||||||[||||||||||||]PAM NA Intergenic NA NA

T31272 AGGACCACCTCCTCGTCGAAGGG 1000
Oligo fwd CACCAGGACCACCTCCTCGTCGAA
Oligo rev AAACTTCGACGAGGAGGTGGTCCT
Chromosome start end strand MM target_seq PAM alignment distance position gene name gene id
GU305914.1 39737 39759 - 0 AGGACCACCTCCTCGTCGAA GGG ||||||||[||||||||||||]PAM NA Intergenic NA NA

T31273 TCGTCGAAGGGATTCGACGACGG 1000
Oligo fwd CACCTCGTCGAAGGGATTCGACGA
Oligo rev AAACTCGTCGAATCCCTTCGACGA
Chromosome start end strand MM target_seq PAM alignment distance position gene name gene id
GU305914.1 39725 39747 - 0 TCGTCGAAGGGATTCGACGA CGG ||||||||[||||||||||||]PAM NA Intergenic NA NA

T31274 CGTCGAAGGGATTCGACGACGGG 1000
Oligo fwd CACCCGTCGAAGGGATTCGACGAC
Oligo rev AAACGTCGTCGAATCCCTTCGACG
Chromosome start end strand MM target_seq PAM alignment distance position gene name gene id
GU305914.1 39724 39746 - 0 CGTCGAAGGGATTCGACGAC GGG ||||||||[||||||||||||]PAM NA Intergenic NA NA

T31275 GACGGGCCTTTCACGACCCAAGG 1000
Oligo fwd CACCGACGGGCCTTTCACGACCCA
Oligo rev AAACTGGGTCGTGAAAGGCCCGTC
Chromosome start end strand MM target_seq PAM alignment distance position gene name gene id
GU305914.1 39707 39729 - 0 GACGGGCCTTTCACGACCCA AGG ||||||||[||||||||||||]PAM NA Intergenic NA NA

T31276 GTGCTCCTTCCTCCTCCCCTCGG 1000
Oligo fwd CACCGTGCTCCTTCCTCCTCCCCT
Oligo rev AAACAGGGGAGGAGGAAGGAGCAC
Chromosome start end strand MM target_seq PAM alignment distance position gene name gene id
GU305914.1 39677 39699 - 0 GTGCTCCTTCCTCCTCCCCT CGG ||||||||[||||||||||||]PAM NA Intergenic NA NA
GU305914.1 78506 78528 - 4 CTCCTCCTTCCTCCTCTTCT CGG -|-|||||[||||||||--||]PAM NA Intergenic NA NA

T31277 CTCCTTCCTCCTCCCCTCGGAGG 1000
Oligo fwd CACCCTCCTTCCTCCTCCCCTCGG
Oligo rev AAACCCGAGGGGAGGAGGAAGGAG
Chromosome start end strand MM target_seq PAM alignment distance position gene name gene id
GU305914.1 39674 39696 - 0 CTCCTTCCTCCTCCCCTCGG AGG ||||||||[||||||||||||]PAM NA Intergenic NA NA

T31278 TCCTTCCTCCTCCCCTCGGAGGG 1000
Oligo fwd CACCTCCTTCCTCCTCCCCTCGGA
Oligo rev AAACTCCGAGGGGAGGAGGAAGGA
Chromosome start end strand MM target_seq PAM alignment distance position gene name gene id
GU305914.1 39673 39695 - 0 TCCTTCCTCCTCCCCTCGGA GGG ||||||||[||||||||||||]PAM NA Intergenic NA NA

T31279 CCTCCTCCCCTCGGAGGGCCCGG 1000
Oligo fwd CACCCCTCCTCCCCTCGGAGGGCC
Oligo rev AAACGGCCCTCCGAGGGGAGGAGG

Chromosome start end strand MM target_seq PAM alignment distance position gene name gene id
GU305914.1 39668 39690 - 0 CCTCCTCCCCTCGGAGGGCC CGG ||||||||[||||||||||||]PAM NA Intergenic NA NA

T31280 GACTCTCCCCCCCGATCTGCCGG 1000
Oligo fwd CACCGACTCTCCCCCCCGATCTGC
Oligo rev AAACGCAGATCGGGGGGGAGAGTC
Chromosome start end strand MM target_seq PAM alignment distance position gene name gene id
GU305914.1 39636 39658 - 0 GACTCTCCCCCCCGATCTGC CGG ||||||||[||||||||||||]PAM NA Intergenic NA NA

T31281 CCCGATCTGCCGGATCTCTCCGG 1000
Oligo fwd CACCCCCGATCTGCCGGATCTCTC
Oligo rev AAACGAGAGATCCGGCAGATCGGG
Chromosome start end strand MM target_seq PAM alignment distance position gene name gene id
GU305914.1 39626 39648 - 0 CCCGATCTGCCGGATCTCTC CGG ||||||||[||||||||||||]PAM NA Intergenic NA NA

T31282 CCGATCTGCCGGATCTCTCCGGG 1000
Oligo fwd CACCCCGATCTGCCGGATCTCTCC
Oligo rev AAACGGAGAGATCCGGCAGATCGG
Chromosome start end strand MM target_seq PAM alignment distance position gene name gene id
GU305914.1 39625 39647 - 0 CCGATCTGCCGGATCTCTCC GGG ||||||||[||||||||||||]PAM NA Intergenic NA NA

T31283 CGATCTGCCGGATCTCTCCGGGG 1000
Oligo fwd CACCCGATCTGCCGGATCTCTCCG
Oligo rev AAACCGGAGAGATCCGGCAGATCG
Chromosome start end strand MM target_seq PAM alignment distance position gene name gene id
GU305914.1 39624 39646 - 0 CGATCTGCCGGATCTCTCCG GGG ||||||||[||||||||||||]PAM NA Intergenic NA NA

T31284 GGATCTCTCCGGGGTTCTCGCGG 1000
Oligo fwd CACCGGATCTCTCCGGGGTTCTCG
Oligo rev AAACCGAGAACCCCGGAGAGATCC
Chromosome start end strand MM target_seq PAM alignment distance position gene name gene id
GU305914.1 39615 39637 - 0 GGATCTCTCCGGGGTTCTCG CGG ||||||||[||||||||||||]PAM NA Intergenic NA NA

T31285 TCTCGCGGACGAAGACGTCCAGG 1000
Oligo fwd CACCTCTCGCGGACGAAGACGTCC
Oligo rev AAACGGACGTCTTCGTCCGCGAGA
Chromosome start end strand MM target_seq PAM alignment distance position gene name gene id
GU305914.1 39600 39622 - 0 TCTCGCGGACGAAGACGTCC AGG ||||||||[||||||||||||]PAM NA Intergenic NA NA

T31286 GACGTCCAGGACAGCAGATACGG 1000
Oligo fwd CACCGACGTCCAGGACAGCAGATA
Oligo rev AAACTATCTGCTGTCCTGGACGTC
Chromosome start end strand MM target_seq PAM alignment distance position gene name gene id
GU305914.1 39587 39609 - 0 GACGTCCAGGACAGCAGATA CGG ||||||||[||||||||||||]PAM NA Intergenic NA NA

T31287 ACGGCAAGATTCCAAAGAGCAGG 1000
Oligo fwd CACCACGGCAAGATTCCAAAGAGC
Oligo rev AAACGCTCTTTGGAATCTTGCCGT
Chromosome start end strand MM target_seq PAM alignment distance position gene name gene id
GU305914.1 39568 39590 - 0 ACGGCAAGATTCCAAAGAGC AGG ||||||||[||||||||||||]PAM NA Intergenic NA NA

T31288 ATAAAGACGACGAGTATTACTGG 1000
Oligo fwd CACCATAAAGACGACGAGTATTAC
Oligo rev AAACGTAATACTCGTCGTCTTTAT
Chromosome start end strand MM target_seq PAM alignment distance position gene name gene id
GU305914.1 39490 39512 - 0 ATAAAGACGACGAGTATTAC TGG ||||||||[||||||||||||]PAM NA Intergenic NA NA

T31289 TAAAGACGACGAGTATTACTGGG 1000
Oligo fwd CACCTAAAGACGACGAGTATTACT
Oligo rev AAACAGTAATACTCGTCGTCTTTA
Chromosome start end strand MM target_seq PAM alignment distance position gene name gene id
GU305914.1 39489 39511 - 0 TAAAGACGACGAGTATTACT GGG ||||||||[||||||||||||]PAM NA Intergenic NA NA

T31290 GAGTATTACTGGGACGAGACTGG 1000
Oligo fwd CACCGAGTATTACTGGGACGAGAC
Oligo rev AAACGTCTCGTCCCAGTAATACTC
Chromosome start end strand MM target_seq PAM alignment distance position gene name gene id
GU305914.1 39479 39501 - 0 GAGTATTACTGGGACGAGAC TGG ||||||||[||||||||||||]PAM NA Intergenic NA NA

T31291 AGTATTACTGGGACGAGACTGGG 1000
Oligo fwd CACCAGTATTACTGGGACGAGACT
Oligo rev AAACAGTCTCGTCCCAGTAATACT
Chromosome start end strand MM target_seq PAM alignment distance position gene name gene id
GU305914.1 39478 39500 - 0 AGTATTACTGGGACGAGACT GGG ||||||||[||||||||||||]PAM NA Intergenic NA NA

T31292 GTATTACTGGGACGAGACTGGGG 1000
Oligo fwd CACCGTATTACTGGGACGAGACTG
Oligo rev AAACCAGTCTCGTCCCAGTAATAC
Chromosome start end strand MM target_seq PAM alignment distance position gene name gene id
GU305914.1 39477 39499 - 0 GTATTACTGGGACGAGACTG GGG ||||||||[||||||||||||]PAM NA Intergenic NA NA

T31293 ACTGGGGATAATATGCCCGTCGG 1000
Oligo fwd CACCACTGGGGATAATATGCCCGT
Oligo rev AAACACGGGCATATTATCCCCAGT
Chromosome start end strand MM target_seq PAM alignment distance position gene name gene id
GU305914.1 39461 39483 - 0 ACTGGGGATAATATGCCCGT CGG ||||||||[||||||||||||]PAM NA Intergenic NA NA

T31294 CTGGGGATAATATGCCCGTCGGG 1000
Oligo fwd CACCCTGGGGATAATATGCCCGTC

<table>
<tr><td>Oligo rev</td><td>AAACGACGGGCATATTATCCCCAG</td><td></td><td></td><td></td><td></td><td></td><td></td><td></td><td></td><td></td><td></td></tr>
<tr><td>Chromosome</td><td>start</td><td>end</td><td>strand</td><td>MM</td><td>target_seq</td><td>PAM</td><td>alignment</td><td>distance</td><td>position</td><td>gene name</td><td>gene id</td></tr>
<tr><td>GU305914.1</td><td>39460</td><td>39482</td><td>-</td><td>0</td><td>CTGGGGATAATATGCCCGTC</td><td>GGG</td><td>||||||||[||||||||||||]PAM</td><td>NA</td><td>Intergenic</td><td>NA</td><td>NA</td></tr>
<tr><td></td><td></td><td></td><td></td><td></td><td></td><td></td><td></td><td></td><td></td><td></td><td></td></tr>
<tr><td>T31295</td><td>TGGGGATAATATGCCCGTCGGGG</td><td>1000</td><td></td><td></td><td></td><td></td><td></td><td></td><td></td><td></td><td></td></tr>
<tr><td>Oligo fwd</td><td>CACCTGGGGATAATATGCCCGTCG</td><td></td><td></td><td></td><td></td><td></td><td></td><td></td><td></td><td></td><td></td></tr>
<tr><td>Oligo rev</td><td>AAACCGACGGGCATATTATCCCCA</td><td></td><td></td><td></td><td></td><td></td><td></td><td></td><td></td><td></td><td></td></tr>
<tr><td>Chromosome</td><td>start</td><td>end</td><td>strand</td><td>MM</td><td>target_seq</td><td>PAM</td><td>alignment</td><td>distance</td><td>position</td><td>gene name</td><td>gene id</td></tr>
<tr><td>GU305914.1</td><td>39459</td><td>39481</td><td>-</td><td>0</td><td>TGGGGATAATATGCCCGTCG</td><td>GGG</td><td>||||||||[||||||||||||]PAM</td><td>NA</td><td>Intergenic</td><td>NA</td><td>NA</td></tr>
<tr><td></td><td></td><td></td><td></td><td></td><td></td><td></td><td></td><td></td><td></td><td></td><td></td></tr>
<tr><td>T31296</td><td>GGATAATATGCCCGTCGGGGAGG</td><td>1000</td><td></td><td></td><td></td><td></td><td></td><td></td><td></td><td></td><td></td></tr>
<tr><td>Oligo fwd</td><td>CACCGGATAATATGCCCGTCGGGG</td><td></td><td></td><td></td><td></td><td></td><td></td><td></td><td></td><td></td><td></td></tr>
<tr><td>Oligo rev</td><td>AAACCCCCGACGGGCATATTATCC</td><td></td><td></td><td></td><td></td><td></td><td></td><td></td><td></td><td></td><td></td></tr>
<tr><td>Chromosome</td><td>start</td><td>end</td><td>strand</td><td>MM</td><td>target_seq</td><td>PAM</td><td>alignment</td><td>distance</td><td>position</td><td>gene name</td><td>gene id</td></tr>
<tr><td>GU305914.1</td><td>39456</td><td>39478</td><td>-</td><td>0</td><td>GGATAATATGCCCGTCGGGG</td><td>AGG</td><td>||||||||[||||||||||||]PAM</td><td>NA</td><td>Intergenic</td><td>NA</td><td>NA</td></tr>
<tr><td></td><td></td><td></td><td></td><td></td><td></td><td></td><td></td><td></td><td></td><td></td><td></td></tr>
<tr><td>T31297</td><td>AATATGCCCGTCGGGGAGGATGG</td><td>1000</td><td></td><td></td><td></td><td></td><td></td><td></td><td></td><td></td><td></td></tr>
<tr><td>Oligo fwd</td><td>CACCAATATGCCCGTCGGGGAGGA</td><td></td><td></td><td></td><td></td><td></td><td></td><td></td><td></td><td></td><td></td></tr>
<tr><td>Oligo rev</td><td>AAACTCCTCCCCGACGGGCATATT</td><td></td><td></td><td></td><td></td><td></td><td></td><td></td><td></td><td></td><td></td></tr>
<tr><td>Chromosome</td><td>start</td><td>end</td><td>strand</td><td>MM</td><td>target_seq</td><td>PAM</td><td>alignment</td><td>distance</td><td>position</td><td>gene name</td><td>gene id</td></tr>
<tr><td>GU305914.1</td><td>39452</td><td>39474</td><td>-</td><td>0</td><td>AATATGCCCGTCGGGGAGGA</td><td>TGG</td><td>||||||||[||||||||||||]PAM</td><td>NA</td><td>Intergenic</td><td>NA</td><td>NA</td></tr>
<tr><td></td><td></td><td></td><td></td><td></td><td></td><td></td><td></td><td></td><td></td><td></td><td></td></tr>
<tr><td>T31298</td><td>ATGCCCGTCGGGGAGGATGGAGG</td><td>1000</td><td></td><td></td><td></td><td></td><td></td><td></td><td></td><td></td><td></td></tr>
<tr><td>Oligo fwd</td><td>CACCATGCCCGTCGGGGAGGATGG</td><td></td><td></td><td></td><td></td><td></td><td></td><td></td><td></td><td></td><td></td></tr>
<tr><td>Oligo rev</td><td>AAACCCATCCTCCCCGACGGGCAT</td><td></td><td></td><td></td><td></td><td></td><td></td><td></td><td></td><td></td><td></td></tr>
<tr><td>Chromosome</td><td>start</td><td>end</td><td>strand</td><td>MM</td><td>target_seq</td><td>PAM</td><td>alignment</td><td>distance</td><td>position</td><td>gene name</td><td>gene id</td></tr>
<tr><td>GU305914.1</td><td>39449</td><td>39471</td><td>-</td><td>0</td><td>ATGCCCGTCGGGGAGGATGG</td><td>AGG</td><td>||||||||[||||||||||||]PAM</td><td>NA</td><td>Intergenic</td><td>NA</td><td>NA</td></tr>
<tr><td></td><td></td><td></td><td></td><td></td><td></td><td></td><td></td><td></td><td></td><td></td><td></td></tr>
<tr><td>T31299</td><td>CCCGTCGGGGAGGATGGAGGTGG</td><td>1000</td><td></td><td></td><td></td><td></td><td></td><td></td><td></td><td></td><td></td></tr>
<tr><td>Oligo fwd</td><td>CACCCCCGTCGGGGAGGATGGAGG</td><td></td><td></td><td></td><td></td><td></td><td></td><td></td><td></td><td></td><td></td></tr>
<tr><td>Oligo rev</td><td>AAACCCTCCATCCTCCCCGACGGG</td><td></td><td></td><td></td><td></td><td></td><td></td><td></td><td></td><td></td><td></td></tr>
<tr><td>Chromosome</td><td>start</td><td>end</td><td>strand</td><td>MM</td><td>target_seq</td><td>PAM</td><td>alignment</td><td>distance</td><td>position</td><td>gene name</td><td>gene id</td></tr>
<tr><td>GU305914.1</td><td>39446</td><td>39468</td><td>-</td><td>0</td><td>CCCGTCGGGGAGGATGGAGG</td><td>TGG</td><td>||||||||[||||||||||||]PAM</td><td>NA</td><td>Intergenic</td><td>NA</td><td>NA</td></tr>
<tr><td></td><td></td><td></td><td></td><td></td><td></td><td></td><td></td><td></td><td></td><td></td><td></td></tr>
<tr><td>T31300</td><td>GTTCTAGAAGACCTCCGTAAAGG</td><td>1000</td><td></td><td></td><td></td><td></td><td></td><td></td><td></td><td></td><td></td></tr>
<tr><td>Oligo fwd</td><td>CACCGTTCTAGAAGACCTCCGTAA</td><td></td><td></td><td></td><td></td><td></td><td></td><td></td><td></td><td></td><td></td></tr>
<tr><td>Oligo rev</td><td>AAACTTACGGAGGTCTTCTAGAAC</td><td></td><td></td><td></td><td></td><td></td><td></td><td></td><td></td><td></td><td></td></tr>
<tr><td>Chromosome</td><td>start</td><td>end</td><td>strand</td><td>MM</td><td>target_seq</td><td>PAM</td><td>alignment</td><td>distance</td><td>position</td><td>gene name</td><td>gene id</td></tr>
<tr><td>GU305914.1</td><td>39422</td><td>39444</td><td>-</td><td>0</td><td>GTTCTAGAAGACCTCCGTAA</td><td>AGG</td><td>||||||||[||||||||||||]PAM</td><td>NA</td><td>Intergenic</td><td>NA</td><td>NA</td></tr>
<tr><td></td><td></td><td></td><td></td><td></td><td></td><td></td><td></td><td></td><td></td><td></td><td></td></tr>
<tr><td>T31301</td><td>AGAAGACCTCCGTAAAGGTCTGG</td><td>1000</td><td></td><td></td><td></td><td></td><td></td><td></td><td></td><td></td><td></td></tr>
<tr><td>Oligo fwd</td><td>CACCAGAAGACCTCCGTAAAGGTC</td><td></td><td></td><td></td><td></td><td></td><td></td><td></td><td></td><td></td><td></td></tr>
<tr><td>Oligo rev</td><td>AAACGACCTTTACGGAGGTCTTCT</td><td></td><td></td><td></td><td></td><td></td><td></td><td></td><td></td><td></td><td></td></tr>
<tr><td>Chromosome</td><td>start</td><td>end</td><td>strand</td><td>MM</td><td>target_seq</td><td>PAM</td><td>alignment</td><td>distance</td><td>position</td><td>gene name</td><td>gene id</td></tr>
<tr><td>GU305914.1</td><td>39417</td><td>39439</td><td>-</td><td>0</td><td>AGAAGACCTCCGTAAAGGTC</td><td>TGG</td><td>||||||||[||||||||||||]PAM</td><td>NA</td><td>Intergenic</td><td>NA</td><td>NA</td></tr>
<tr><td></td><td></td><td></td><td></td><td></td><td></td><td></td><td></td><td></td><td></td><td></td><td></td></tr>
<tr><td>T31302</td><td>AGACCTCCGTAAAGGTCTGGAGG</td><td>1000</td><td></td><td></td><td></td><td></td><td></td><td></td><td></td><td></td><td></td></tr>
<tr><td>Oligo fwd</td><td>CACCAGACCTCCGTAAAGGTCTGG</td><td></td><td></td><td></td><td></td><td></td><td></td><td></td><td></td><td></td><td></td></tr>
<tr><td>Oligo rev</td><td>AAACCCAGACCTTTACGGAGGTCT</td><td></td><td></td><td></td><td></td><td></td><td></td><td></td><td></td><td></td><td></td></tr>
<tr><td>Chromosome</td><td>start</td><td>end</td><td>strand</td><td>MM</td><td>target_seq</td><td>PAM</td><td>alignment</td><td>distance</td><td>position</td><td>gene name</td><td>gene id</td></tr>
<tr><td>GU305914.1</td><td>39414</td><td>39436</td><td>-</td><td>0</td><td>AGACCTCCGTAAAGGTCTGG</td><td>AGG</td><td>||||||||[||||||||||||]PAM</td><td>NA</td><td>Intergenic</td><td>NA</td><td>NA</td></tr>
<tr><td></td><td></td><td></td><td></td><td></td><td></td><td></td><td></td><td></td><td></td><td></td><td></td></tr>
<tr><td>T31303</td><td>GACCTCCGTAAAGGTCTGGAGGG</td><td>1000</td><td></td><td></td><td></td><td></td><td></td><td></td><td></td><td></td><td></td></tr>
<tr><td>Oligo fwd</td><td>CACCGACCTCCGTAAAGGTCTGGA</td><td></td><td></td><td></td><td></td><td></td><td></td><td></td><td></td><td></td><td></td></tr>
<tr><td>Oligo rev</td><td>AAACTCCAGACCTTTACGGAGGTC</td><td></td><td></td><td></td><td></td><td></td><td></td><td></td><td></td><td></td><td></td></tr>
<tr><td>Chromosome</td><td>start</td><td>end</td><td>strand</td><td>MM</td><td>target_seq</td><td>PAM</td><td>alignment</td><td>distance</td><td>position</td><td>gene name</td><td>gene id</td></tr>
<tr><td>GU305914.1</td><td>39413</td><td>39435</td><td>-</td><td>0</td><td>GACCTCCGTAAAGGTCTGGA</td><td>GGG</td><td>||||||||[||||||||||||]PAM</td><td>NA</td><td>Intergenic</td><td>NA</td><td>NA</td></tr>
<tr><td></td><td></td><td></td><td></td><td></td><td></td><td></td><td></td><td></td><td></td><td></td><td></td></tr>
<tr><td>T31304</td><td>GAGGGTATAGATCTCAAAACCGG</td><td>1000</td><td></td><td></td><td></td><td></td><td></td><td></td><td></td><td></td><td></td></tr>
<tr><td>Oligo fwd</td><td>CACCGAGGGTATAGATCTCAAAAC</td><td></td><td></td><td></td><td></td><td></td><td></td><td></td><td></td><td></td><td></td></tr>
<tr><td>Oligo rev</td><td>AAACGTTTTGAGATCTATACCCTC</td><td></td><td></td><td></td><td></td><td></td><td></td><td></td><td></td><td></td><td></td></tr>
<tr><td>Chromosome</td><td>start</td><td>end</td><td>strand</td><td>MM</td><td>target_seq</td><td>PAM</td><td>alignment</td><td>distance</td><td>position</td><td>gene name</td><td>gene id</td></tr>
<tr><td>GU305914.1</td><td>39395</td><td>39417</td><td>-</td><td>0</td><td>GAGGGTATAGATCTCAAAAC</td><td>CGG</td><td>||||||||[||||||||||||]PAM</td><td>NA</td><td>Intergenic</td><td>NA</td><td>NA</td></tr>
<tr><td></td><td></td><td></td><td></td><td></td><td></td><td></td><td></td><td></td><td></td><td></td><td></td></tr>
<tr><td>T31305</td><td>GGTATAGATCTCAAAACCGGTGG</td><td>1000</td><td></td><td></td><td></td><td></td><td></td><td></td><td></td><td></td><td></td></tr>
<tr><td>Oligo fwd</td><td>CACCGGTATAGATCTCAAAACCGG</td><td></td><td></td><td></td><td></td><td></td><td></td><td></td><td></td><td></td><td></td></tr>
<tr><td>Oligo rev</td><td>AAACCCGGTTTTGAGATCTATACC</td><td></td><td></td><td></td><td></td><td></td><td></td><td></td><td></td><td></td><td></td></tr>
<tr><td>Chromosome</td><td>start</td><td>end</td><td>strand</td><td>MM</td><td>target_seq</td><td>PAM</td><td>alignment</td><td>distance</td><td>position</td><td>gene name</td><td>gene id</td></tr>
<tr><td>GU305914.1</td><td>39392</td><td>39414</td><td>-</td><td>0</td><td>GGTATAGATCTCAAAACCGG</td><td>TGG</td><td>||||||||[||||||||||||]PAM</td><td>NA</td><td>Intergenic</td><td>NA</td><td>NA</td></tr>
<tr><td></td><td></td><td></td><td></td><td></td><td></td><td></td><td></td><td></td><td></td><td></td><td></td></tr>
<tr><td>T31306</td><td>ATAGATCTCAAAACCGGTGGCGG</td><td>1000</td><td></td><td></td><td></td><td></td><td></td><td></td><td></td><td></td><td></td></tr>
<tr><td>Oligo fwd</td><td>CACCATAGATCTCAAAACCGGTGG</td><td></td><td></td><td></td><td></td><td></td><td></td><td></td><td></td><td></td><td></td></tr>
<tr><td>Oligo rev</td><td>AAACCCACCGGTTTTGAGATCTAT</td><td></td><td></td><td></td><td></td><td></td><td></td><td></td><td></td><td></td><td></td></tr>
<tr><td>Chromosome</td><td>start</td><td>end</td><td>strand</td><td>MM</td><td>target_seq</td><td>PAM</td><td>alignment</td><td>distance</td><td>position</td><td>gene name</td><td>gene id</td></tr>
<tr><td>GU305914.1</td><td>39389</td><td>39411</td><td>-</td><td>0</td><td>ATAGATCTCAAAACCGGTGG</td><td>CGG</td><td>||||||||[||||||||||||]PAM</td><td>NA</td><td>Intergenic</td><td>NA</td><td>NA</td></tr>
<tr><td></td><td></td><td></td><td></td><td></td><td></td><td></td><td></td><td></td><td></td><td></td><td></td></tr>
<tr><td>T31307</td><td>GATCTCAAAACCGGTGGCGGTGG</td><td>1000</td><td></td><td></td><td></td><td></td><td></td><td></td><td></td><td></td><td></td></tr>
<tr><td>Oligo fwd</td><td>CACCGATCTCAAAACCGGTGGCGG</td><td></td><td></td><td></td><td></td><td></td><td></td><td></td><td></td><td></td><td></td></tr>
<tr><td>Oligo rev</td><td>AAACCCGCCACCGGTTTTGAGATC</td><td></td><td></td><td></td><td></td><td></td><td></td><td></td><td></td><td></td><td></td></tr>
<tr><td>Chromosome</td><td>start</td><td>end</td><td>strand</td><td>MM</td><td>target_seq</td><td>PAM</td><td>alignment</td><td>distance</td><td>position</td><td>gene name</td><td>gene id</td></tr>
<tr><td>GU305914.1</td><td>39386</td><td>39408</td><td>-</td><td>0</td><td>GATCTCAAAACCGGTGGCGG</td><td>TGG</td><td>||||||||[||||||||||||]PAM</td><td>NA</td><td>Intergenic</td><td>NA</td><td>NA</td></tr>
<tr><td></td><td></td><td></td><td></td><td></td><td></td><td></td><td></td><td></td><td></td><td></td><td></td></tr>
<tr><td>T31308</td><td>CCACTCTCCCAACAGTTCGCCGG</td><td>1000</td><td></td><td></td><td></td><td></td><td></td><td></td><td></td><td></td><td></td></tr>
<tr><td>Oligo fwd</td><td>CACCCCACTCTCCCAACAGTTCGC</td><td></td><td></td><td></td><td></td><td></td><td></td><td></td><td></td><td></td><td></td></tr>
<tr><td>Oligo rev</td><td>AAACGCGAACTGTTGGGAGAGTGG</td><td></td><td></td><td></td><td></td><td></td><td></td><td></td><td></td><td></td><td></td></tr>
<tr><td>Chromosome</td><td>start</td><td>end</td><td>strand</td><td>MM</td><td>target_seq</td><td>PAM</td><td>alignment</td><td>distance</td><td>position</td><td>gene name</td><td>gene id</td></tr>
<tr><td>GU305914.1</td><td>39350</td><td>39372</td><td>-</td><td>0</td><td>CCACTCTCCCAACAGTTCGC</td><td>CGG</td><td>||||||||[||||||||||||]PAM</td><td>NA</td><td>Intergenic</td><td>NA</td><td>NA</td></tr>
<tr><td></td><td></td><td></td><td></td><td></td><td></td><td></td><td></td><td></td><td></td><td></td><td></td></tr>
<tr><td>T31309</td><td>TTCGCCGGCTCCCCATTCGCCGG</td><td>1000</td><td></td><td></td><td></td><td></td><td></td><td></td><td></td><td></td><td></td></tr>
</table>

<table>
<tr><td>Oligo fwd</td><td>CACCTTCGCCGGCTCCCCATTCGC</td><td></td><td></td><td></td><td></td><td></td><td></td><td></td><td></td><td></td><td></td></tr>
<tr><td>Oligo rev</td><td>AAACGCGAATGGGGAGCCGGCGAA</td><td></td><td></td><td></td><td></td><td></td><td></td><td></td><td></td><td></td><td></td></tr>
<tr><td>Chromosome</td><td>start</td><td>end</td><td>strand</td><td>MM</td><td>target_seq</td><td>PAM</td><td>alignment</td><td>distance</td><td>position</td><td>gene name</td><td>gene id</td></tr>
<tr><td>GU305914.1</td><td>39335</td><td>39357</td><td>-</td><td>0</td><td>TTCGCCGGCTCCCCATTCGC</td><td>CGG</td><td>||||||||[||||||||||||]PAM</td><td>NA</td><td>Intergenic</td><td>NA</td><td>NA</td></tr>
<tr><td></td><td></td><td></td><td></td><td></td><td></td><td></td><td></td><td></td><td></td><td></td><td></td></tr>
<tr><td>T31310</td><td>TCCCCATTCGCCGGTTCTGATGG</td><td>1000</td><td></td><td></td><td></td><td></td><td></td><td></td><td></td><td></td><td></td></tr>
<tr><td>Oligo fwd</td><td>CACCTCCCCATTCGCCGGTTCTGA</td><td></td><td></td><td></td><td></td><td></td><td></td><td></td><td></td><td></td><td></td></tr>
<tr><td>Oligo rev</td><td>AAACTCAGAACCGGCGAATGGGGA</td><td></td><td></td><td></td><td></td><td></td><td></td><td></td><td></td><td></td><td></td></tr>
<tr><td>Chromosome</td><td>start</td><td>end</td><td>strand</td><td>MM</td><td>target_seq</td><td>PAM</td><td>alignment</td><td>distance</td><td>position</td><td>gene name</td><td>gene id</td></tr>
<tr><td>GU305914.1</td><td>39326</td><td>39348</td><td>-</td><td>0</td><td>TCCCCATTCGCCGGTTCTGA</td><td>TGG</td><td>||||||||[||||||||||||]PAM</td><td>NA</td><td>Intergenic</td><td>NA</td><td>NA</td></tr>
<tr><td></td><td></td><td></td><td></td><td></td><td></td><td></td><td></td><td></td><td></td><td></td><td></td></tr>
<tr><td>T31311</td><td>CCCCATTCGCCGGTTCTGATGGG</td><td>1000</td><td></td><td></td><td></td><td></td><td></td><td></td><td></td><td></td><td></td></tr>
<tr><td>Oligo fwd</td><td>CACCCCCCATTCGCCGGTTCTGAT</td><td></td><td></td><td></td><td></td><td></td><td></td><td></td><td></td><td></td><td></td></tr>
<tr><td>Oligo rev</td><td>AAACATCAGAACCGGCGAATGGGG</td><td></td><td></td><td></td><td></td><td></td><td></td><td></td><td></td><td></td><td></td></tr>
<tr><td>Chromosome</td><td>start</td><td>end</td><td>strand</td><td>MM</td><td>target_seq</td><td>PAM</td><td>alignment</td><td>distance</td><td>position</td><td>gene name</td><td>gene id</td></tr>
<tr><td>GU305914.1</td><td>39325</td><td>39347</td><td>-</td><td>0</td><td>CCCCATTCGCCGGTTCTGAT</td><td>GGG</td><td>||||||||[||||||||||||]PAM</td><td>NA</td><td>Intergenic</td><td>NA</td><td>NA</td></tr>
<tr><td></td><td></td><td></td><td></td><td></td><td></td><td></td><td></td><td></td><td></td><td></td><td></td></tr>
<tr><td>T31312</td><td>CCCATTCGCCGGTTCTGATGGGG</td><td>1000</td><td></td><td></td><td></td><td></td><td></td><td></td><td></td><td></td><td></td></tr>
<tr><td>Oligo fwd</td><td>CACCCCCATTCGCCGGTTCTGATG</td><td></td><td></td><td></td><td></td><td></td><td></td><td></td><td></td><td></td><td></td></tr>
<tr><td>Oligo rev</td><td>AAACCATCAGAACCGGCGAATGGG</td><td></td><td></td><td></td><td></td><td></td><td></td><td></td><td></td><td></td><td></td></tr>
<tr><td>Chromosome</td><td>start</td><td>end</td><td>strand</td><td>MM</td><td>target_seq</td><td>PAM</td><td>alignment</td><td>distance</td><td>position</td><td>gene name</td><td>gene id</td></tr>
<tr><td>GU305914.1</td><td>39324</td><td>39346</td><td>-</td><td>0</td><td>CCCATTCGCCGGTTCTGATG</td><td>GGG</td><td>||||||||[||||||||||||]PAM</td><td>NA</td><td>Intergenic</td><td>NA</td><td>NA</td></tr>
<tr><td></td><td></td><td></td><td></td><td></td><td></td><td></td><td></td><td></td><td></td><td></td><td></td></tr>
<tr><td>T31313</td><td>TTCGCCGGTTCTGATGGGGACGG</td><td>1000</td><td></td><td></td><td></td><td></td><td></td><td></td><td></td><td></td><td></td></tr>
<tr><td>Oligo fwd</td><td>CACCTTCGCCGGTTCTGATGGGGA</td><td></td><td></td><td></td><td></td><td></td><td></td><td></td><td></td><td></td><td></td></tr>
<tr><td>Oligo rev</td><td>AAACTCCCCATCAGAACCGGCGAA</td><td></td><td></td><td></td><td></td><td></td><td></td><td></td><td></td><td></td><td></td></tr>
<tr><td>Chromosome</td><td>start</td><td>end</td><td>strand</td><td>MM</td><td>target_seq</td><td>PAM</td><td>alignment</td><td>distance</td><td>position</td><td>gene name</td><td>gene id</td></tr>
<tr><td>GU305914.1</td><td>39320</td><td>39342</td><td>-</td><td>0</td><td>TTCGCCGGTTCTGATGGGGA</td><td>CGG</td><td>||||||||[||||||||||||]PAM</td><td>NA</td><td>Intergenic</td><td>NA</td><td>NA</td></tr>
<tr><td></td><td></td><td></td><td></td><td></td><td></td><td></td><td></td><td></td><td></td><td></td><td></td></tr>
<tr><td>T31314</td><td>GCCGGTTCTGATGGGGACGGTGG</td><td>1000</td><td></td><td></td><td></td><td></td><td></td><td></td><td></td><td></td><td></td></tr>
<tr><td>Oligo fwd</td><td>CACCGCCGGTTCTGATGGGGACGG</td><td></td><td></td><td></td><td></td><td></td><td></td><td></td><td></td><td></td><td></td></tr>
<tr><td>Oligo rev</td><td>AAACCCGTCCCCATCAGAACCGGC</td><td></td><td></td><td></td><td></td><td></td><td></td><td></td><td></td><td></td><td></td></tr>
<tr><td>Chromosome</td><td>start</td><td>end</td><td>strand</td><td>MM</td><td>target_seq</td><td>PAM</td><td>alignment</td><td>distance</td><td>position</td><td>gene name</td><td>gene id</td></tr>
<tr><td>GU305914.1</td><td>39317</td><td>39339</td><td>-</td><td>0</td><td>GCCGGTTCTGATGGGGACGG</td><td>TGG</td><td>||||||||[||||||||||||]PAM</td><td>NA</td><td>Intergenic</td><td>NA</td><td>NA</td></tr>
<tr><td></td><td></td><td></td><td></td><td></td><td></td><td></td><td></td><td></td><td></td><td></td><td></td></tr>
<tr><td>T31315</td><td>GGTTCTGATGGGGACGGTGGCGG</td><td>1000</td><td></td><td></td><td></td><td></td><td></td><td></td><td></td><td></td><td></td></tr>
<tr><td>Oligo fwd</td><td>CACCGGTTCTGATGGGGACGGTGG</td><td></td><td></td><td></td><td></td><td></td><td></td><td></td><td></td><td></td><td></td></tr>
<tr><td>Oligo rev</td><td>AAACCCACCGTCCCCATCAGAACC</td><td></td><td></td><td></td><td></td><td></td><td></td><td></td><td></td><td></td><td></td></tr>
<tr><td>Chromosome</td><td>start</td><td>end</td><td>strand</td><td>MM</td><td>target_seq</td><td>PAM</td><td>alignment</td><td>distance</td><td>position</td><td>gene name</td><td>gene id</td></tr>
<tr><td>GU305914.1</td><td>39314</td><td>39336</td><td>-</td><td>0</td><td>GGTTCTGATGGGGACGGTGG</td><td>CGG</td><td>||||||||[||||||||||||]PAM</td><td>NA</td><td>Intergenic</td><td>NA</td><td>NA</td></tr>
<tr><td></td><td></td><td></td><td></td><td></td><td></td><td></td><td></td><td></td><td></td><td></td><td></td></tr>
<tr><td>T31316</td><td>TGATGGGGACGGTGGCGGCCTGG</td><td>1000</td><td></td><td></td><td></td><td></td><td></td><td></td><td></td><td></td><td></td></tr>
<tr><td>Oligo fwd</td><td>CACCTGATGGGGACGGTGGCGGCC</td><td></td><td></td><td></td><td></td><td></td><td></td><td></td><td></td><td></td><td></td></tr>
<tr><td>Oligo rev</td><td>AAACGGCCGCCACCGTCCCCATCA</td><td></td><td></td><td></td><td></td><td></td><td></td><td></td><td></td><td></td><td></td></tr>
<tr><td>Chromosome</td><td>start</td><td>end</td><td>strand</td><td>MM</td><td>target_seq</td><td>PAM</td><td>alignment</td><td>distance</td><td>position</td><td>gene name</td><td>gene id</td></tr>
<tr><td>GU305914.1</td><td>39309</td><td>39331</td><td>-</td><td>0</td><td>TGATGGGGACGGTGGCGGCC</td><td>TGG</td><td>||||||||[||||||||||||]PAM</td><td>NA</td><td>Intergenic</td><td>NA</td><td>NA</td></tr>
<tr><td></td><td></td><td></td><td></td><td></td><td></td><td></td><td></td><td></td><td></td><td></td><td></td></tr>
<tr><td>T31317</td><td>GATCGATGATCGCTCGCTCCTGG</td><td>1000</td><td></td><td></td><td></td><td></td><td></td><td></td><td></td><td></td><td></td></tr>
<tr><td>Oligo fwd</td><td>CACCGATCGATGATCGCTCGCTCC</td><td></td><td></td><td></td><td></td><td></td><td></td><td></td><td></td><td></td><td></td></tr>
<tr><td>Oligo rev</td><td>AAACGGAGCGAGCGATCATCGATC</td><td></td><td></td><td></td><td></td><td></td><td></td><td></td><td></td><td></td><td></td></tr>
<tr><td>Chromosome</td><td>start</td><td>end</td><td>strand</td><td>MM</td><td>target_seq</td><td>PAM</td><td>alignment</td><td>distance</td><td>position</td><td>gene name</td><td>gene id</td></tr>
<tr><td>GU305914.1</td><td>39256</td><td>39278</td><td>-</td><td>0</td><td>GATCGATGATCGCTCGCTCC</td><td>TGG</td><td>||||||||[||||||||||||]PAM</td><td>NA</td><td>Intergenic</td><td>NA</td><td>NA</td></tr>
<tr><td></td><td></td><td></td><td></td><td></td><td></td><td></td><td></td><td></td><td></td><td></td><td></td></tr>
<tr><td>T31318</td><td>GATGATCGCTCGCTCCTGGTTGG</td><td>1000</td><td></td><td></td><td></td><td></td><td></td><td></td><td></td><td></td><td></td></tr>
<tr><td>Oligo fwd</td><td>CACCGATGATCGCTCGCTCCTGGT</td><td></td><td></td><td></td><td></td><td></td><td></td><td></td><td></td><td></td><td></td></tr>
<tr><td>Oligo rev</td><td>AAACACCAGGAGCGAGCGATCATC</td><td></td><td></td><td></td><td></td><td></td><td></td><td></td><td></td><td></td><td></td></tr>
<tr><td>Chromosome</td><td>start</td><td>end</td><td>strand</td><td>MM</td><td>target_seq</td><td>PAM</td><td>alignment</td><td>distance</td><td>position</td><td>gene name</td><td>gene id</td></tr>
<tr><td>GU305914.1</td><td>39252</td><td>39274</td><td>-</td><td>0</td><td>GATGATCGCTCGCTCCTGGT</td><td>TGG</td><td>||||||||[||||||||||||]PAM</td><td>NA</td><td>Intergenic</td><td>NA</td><td>NA</td></tr>
<tr><td></td><td></td><td></td><td></td><td></td><td></td><td></td><td></td><td></td><td></td><td></td><td></td></tr>
<tr><td>T31319</td><td>GTGTTTTTCTTGATTTATTGAGG</td><td>1000</td><td></td><td></td><td></td><td></td><td></td><td></td><td></td><td></td><td></td></tr>
<tr><td>Oligo fwd</td><td>CACCGTGTTTTTCTTGATTTATTG</td><td></td><td></td><td></td><td></td><td></td><td></td><td></td><td></td><td></td><td></td></tr>
<tr><td>Oligo rev</td><td>AAACCAATAAATCAAGAAAAACAC</td><td></td><td></td><td></td><td></td><td></td><td></td><td></td><td></td><td></td><td></td></tr>
<tr><td>Chromosome</td><td>start</td><td>end</td><td>strand</td><td>MM</td><td>target_seq</td><td>PAM</td><td>alignment</td><td>distance</td><td>position</td><td>gene name</td><td>gene id</td></tr>
<tr><td>GU305914.1</td><td>39163</td><td>39185</td><td>-</td><td>0</td><td>GTGTTTTTCTTGATTTATTG</td><td>AGG</td><td>||||||||[||||||||||||]PAM</td><td>NA</td><td>Intergenic</td><td>NA</td><td>NA</td></tr>
<tr><td>GU305914.1</td><td>156986</td><td>157008</td><td>+</td><td>4</td><td>TTGTCTTTCTTTATTTATCG</td><td>CGG</td><td>-|||-|||[|||-||||||-|]PAM</td><td>NA</td><td>Intergenic</td><td>NA</td><td>NA</td></tr>
<tr><td></td><td></td><td></td><td></td><td></td><td></td><td></td><td></td><td></td><td></td><td></td><td></td></tr>
<tr><td>T31320</td><td>TGTTTTTCTTGATTTATTGAGGG</td><td>1000</td><td></td><td></td><td></td><td></td><td></td><td></td><td></td><td></td><td></td></tr>
<tr><td>Oligo fwd</td><td>CACCTGTTTTTCTTGATTTATTGA</td><td></td><td></td><td></td><td></td><td></td><td></td><td></td><td></td><td></td><td></td></tr>
<tr><td>Oligo rev</td><td>AAACTCAATAAATCAAGAAAAACA</td><td></td><td></td><td></td><td></td><td></td><td></td><td></td><td></td><td></td><td></td></tr>
<tr><td>Chromosome</td><td>start</td><td>end</td><td>strand</td><td>MM</td><td>target_seq</td><td>PAM</td><td>alignment</td><td>distance</td><td>position</td><td>gene name</td><td>gene id</td></tr>
<tr><td>GU305914.1</td><td>39162</td><td>39184</td><td>-</td><td>0</td><td>TGTTTTTCTTGATTTATTGA</td><td>GGG</td><td>||||||||[||||||||||||]PAM</td><td>NA</td><td>Intergenic</td><td>NA</td><td>NA</td></tr>
<tr><td></td><td></td><td></td><td></td><td></td><td></td><td></td><td></td><td></td><td></td><td></td><td></td></tr>
<tr><td>T31321</td><td>TTCTTGATTTATTGAGGGTTTGG</td><td>1000</td><td></td><td></td><td></td><td></td><td></td><td></td><td></td><td></td><td></td></tr>
<tr><td>Oligo fwd</td><td>CACCTTCTTGATTTATTGAGGGTT</td><td></td><td></td><td></td><td></td><td></td><td></td><td></td><td></td><td></td><td></td></tr>
<tr><td>Oligo rev</td><td>AAACAACCCTCAATAAATCAAGAA</td><td></td><td></td><td></td><td></td><td></td><td></td><td></td><td></td><td></td><td></td></tr>
<tr><td>Chromosome</td><td>start</td><td>end</td><td>strand</td><td>MM</td><td>target_seq</td><td>PAM</td><td>alignment</td><td>distance</td><td>position</td><td>gene name</td><td>gene id</td></tr>
<tr><td>GU305914.1</td><td>39157</td><td>39179</td><td>-</td><td>0</td><td>TTCTTGATTTATTGAGGGTT</td><td>TGG</td><td>||||||||[||||||||||||]PAM</td><td>NA</td><td>Intergenic</td><td>NA</td><td>NA</td></tr>
<tr><td></td><td></td><td></td><td></td><td></td><td></td><td></td><td></td><td></td><td></td><td></td><td></td></tr>
<tr><td>T31322</td><td>ATTTATTGAGGGTTTGGCAGTGG</td><td>1000</td><td></td><td></td><td></td><td></td><td></td><td></td><td></td><td></td><td></td></tr>
<tr><td>Oligo fwd</td><td>CACCATTTATTGAGGGTTTGGCAG</td><td></td><td></td><td></td><td></td><td></td><td></td><td></td><td></td><td></td><td></td></tr>
<tr><td>Oligo rev</td><td>AAACCTGCCAAACCCTCAATAAAT</td><td></td><td></td><td></td><td></td><td></td><td></td><td></td><td></td><td></td><td></td></tr>
<tr><td>Chromosome</td><td>start</td><td>end</td><td>strand</td><td>MM</td><td>target_seq</td><td>PAM</td><td>alignment</td><td>distance</td><td>position</td><td>gene name</td><td>gene id</td></tr>
<tr><td>GU305914.1</td><td>39151</td><td>39173</td><td>-</td><td>0</td><td>ATTTATTGAGGGTTTGGCAG</td><td>TGG</td><td>||||||||[||||||||||||]PAM</td><td>NA</td><td>Intergenic</td><td>NA</td><td>NA</td></tr>
<tr><td></td><td></td><td></td><td></td><td></td><td></td><td></td><td></td><td></td><td></td><td></td><td></td></tr>
<tr><td>T31323</td><td>TTTATTGAGGGTTTGGCAGTGGG</td><td>1000</td><td></td><td></td><td></td><td></td><td></td><td></td><td></td><td></td><td></td></tr>
<tr><td>Oligo fwd</td><td>CACCTTTATTGAGGGTTTGGCAGT</td><td></td><td></td><td></td><td></td><td></td><td></td><td></td><td></td><td></td><td></td></tr>
<tr><td>Oligo rev</td><td>AAACACTGCCAAACCCTCAATAAA</td><td></td><td></td><td></td><td></td><td></td><td></td><td></td><td></td><td></td><td></td></tr>
<tr><td>Chromosome</td><td>start</td><td>end</td><td>strand</td><td>MM</td><td>target_seq</td><td>PAM</td><td>alignment</td><td>distance</td><td>position</td><td>gene name</td><td>gene id</td></tr>
<tr><td>GU305914.1</td><td>39150</td><td>39172</td><td>-</td><td>0</td><td>TTTATTGAGGGTTTGGCAGT</td><td>GGG</td><td>||||||||[||||||||||||]PAM</td><td>NA</td><td>Intergenic</td><td>NA</td><td>NA</td></tr>
</table>

<table>
<tr><td>T31324</td><td>GTGGGTATTAATTACACTACAGG</td><td></td><td>1000</td></tr>
<tr><td>Oligo fwd</td><td>CACCGTGGGTATTAATTACACTAC</td></tr>
<tr><td>Oligo rev</td><td>AAACGTAGTGTAATTAATACCCAC</td></tr>
<tr><td>Chromosome</td><td>start</td><td></td><td>end</td><td>strand</td><td>MM</td><td>target_seq</td><td>PAM</td><td>alignment</td><td>distance</td><td>position</td><td>gene name</td><td>gene id</td></tr>
<tr><td>GU305914.1</td><td></td><td>39132</td><td>39154</td><td>-</td><td>0</td><td>GTGGGTATTAATTACACTAC</td><td>AGG</td><td>||||||||[||||||||||||]PAM</td><td>NA</td><td>Intergenic</td><td>NA</td><td>NA</td></tr>
<tr><td>T31325</td><td>ATATAATCACAACTCGTCCGAGG</td><td></td><td>1000</td></tr>
<tr><td>Oligo fwd</td><td>CACCATATAATCACAACTCGTCCG</td></tr>
<tr><td>Oligo rev</td><td>AAACCGGACGAGTTGTGATTATAT</td></tr>
<tr><td>Chromosome</td><td>start</td><td></td><td>end</td><td>strand</td><td>MM</td><td>target_seq</td><td>PAM</td><td>alignment</td><td>distance</td><td>position</td><td>gene name</td><td>gene id</td></tr>
<tr><td>GU305914.1</td><td></td><td>39055</td><td>39077</td><td>-</td><td>0</td><td>ATATAATCACAACTCGTCCG</td><td>AGG</td><td>||||||||[||||||||||||]PAM</td><td>NA</td><td>Intergenic</td><td>NA</td><td>NA</td></tr>
<tr><td>T31326</td><td>GAGGTTGTAAAGTTCTGATAAGG</td><td></td><td>1000</td></tr>
<tr><td>Oligo fwd</td><td>CACCGAGGTTGTAAAGTTCTGATA</td></tr>
<tr><td>Oligo rev</td><td>AAACTATCAGAACTTTACAACCTC</td></tr>
<tr><td>Chromosome</td><td>start</td><td></td><td>end</td><td>strand</td><td>MM</td><td>target_seq</td><td>PAM</td><td>alignment</td><td>distance</td><td>position</td><td>gene name</td><td>gene id</td></tr>
<tr><td>GU305914.1</td><td></td><td>39036</td><td>39058</td><td>-</td><td>0</td><td>GAGGTTGTAAAGTTCTGATA</td><td>AGG</td><td>||||||||[||||||||||||]PAM</td><td>NA</td><td>Intergenic</td><td>NA</td><td>NA</td></tr>
<tr><td>T31327</td><td>AGGTTGTAAAGTTCTGATAAGGG</td><td></td><td>1000</td></tr>
<tr><td>Oligo fwd</td><td>CACCAGGTTGTAAAGTTCTGATAA</td></tr>
<tr><td>Oligo rev</td><td>AAACTTATCAGAACTTTACAACCT</td></tr>
<tr><td>Chromosome</td><td>start</td><td></td><td>end</td><td>strand</td><td>MM</td><td>target_seq</td><td>PAM</td><td>alignment</td><td>distance</td><td>position</td><td>gene name</td><td>gene id</td></tr>
<tr><td>GU305914.1</td><td></td><td>39035</td><td>39057</td><td>-</td><td>0</td><td>AGGTTGTAAAGTTCTGATAA</td><td>GGG</td><td>||||||||[||||||||||||]PAM</td><td>NA</td><td>Intergenic</td><td>NA</td><td>NA</td></tr>
<tr><td>T31328</td><td>AGTTCTGATAAGGGAGTCTCAGG</td><td></td><td>1000</td></tr>
<tr><td>Oligo fwd</td><td>CACCAGTTCTGATAAGGGAGTCTC</td></tr>
<tr><td>Oligo rev</td><td>AAACGAGACTCCCTTATCAGAACT</td></tr>
<tr><td>Chromosome</td><td>start</td><td></td><td>end</td><td>strand</td><td>MM</td><td>target_seq</td><td>PAM</td><td>alignment</td><td>distance</td><td>position</td><td>gene name</td><td>gene id</td></tr>
<tr><td>GU305914.1</td><td></td><td>39026</td><td>39048</td><td>-</td><td>0</td><td>AGTTCTGATAAGGGAGTCTC</td><td>AGG</td><td>||||||||[||||||||||||]PAM</td><td>NA</td><td>Intergenic</td><td>NA</td><td>NA</td></tr>
<tr><td>T31329</td><td>GTCTCAGGTCTCCCAAGTGAAGG</td><td></td><td>1000</td></tr>
<tr><td>Oligo fwd</td><td>CACCGTCTCAGGTCTCCCAAGTGA</td></tr>
<tr><td>Oligo rev</td><td>AAACTCACTTGGGAGACCTGAGAC</td></tr>
<tr><td>Chromosome</td><td>start</td><td></td><td>end</td><td>strand</td><td>MM</td><td>target_seq</td><td>PAM</td><td>alignment</td><td>distance</td><td>position</td><td>gene name</td><td>gene id</td></tr>
<tr><td>GU305914.1</td><td></td><td>39011</td><td>39033</td><td>-</td><td>0</td><td>GTCTCAGGTCTCCCAAGTGA</td><td>AGG</td><td>||||||||[||||||||||||]PAM</td><td>NA</td><td>Intergenic</td><td>NA</td><td>NA</td></tr>
<tr><td>T31330</td><td>CTCCCAAGTGAAGGTTGATTCGG</td><td></td><td>1000</td></tr>
<tr><td>Oligo fwd</td><td>CACCCTCCCAAGTGAAGGTTGATT</td></tr>
<tr><td>Oligo rev</td><td>AAACAATCAACCTTCACTTGGGAG</td></tr>
<tr><td>Chromosome</td><td>start</td><td></td><td>end</td><td>strand</td><td>MM</td><td>target_seq</td><td>PAM</td><td>alignment</td><td>distance</td><td>position</td><td>gene name</td><td>gene id</td></tr>
<tr><td>GU305914.1</td><td></td><td>39002</td><td>39024</td><td>-</td><td>0</td><td>CTCCCAAGTGAAGGTTGATT</td><td>CGG</td><td>||||||||[||||||||||||]PAM</td><td>NA</td><td>Intergenic</td><td>NA</td><td>NA</td></tr>
<tr><td>T31331</td><td>GGTCTCCCGTCTTGATCACTCGG</td><td></td><td>1000</td></tr>
<tr><td>Oligo fwd</td><td>CACCGGTCTCCCGTCTTGATCACT</td></tr>
<tr><td>Oligo rev</td><td>AAACAGTGATCAAGACGGGAGACC</td></tr>
<tr><td>Chromosome</td><td>start</td><td></td><td>end</td><td>strand</td><td>MM</td><td>target_seq</td><td>PAM</td><td>alignment</td><td>distance</td><td>position</td><td>gene name</td><td>gene id</td></tr>
<tr><td>GU305914.1</td><td></td><td>38981</td><td>39003</td><td>-</td><td>0</td><td>GGTCTCCCGTCTTGATCACT</td><td>CGG</td><td>||||||||[||||||||||||]PAM</td><td>NA</td><td>Intergenic</td><td>NA</td><td>NA</td></tr>
<tr><td>T31332</td><td>GTCTCCCGTCTTGATCACTCGGG</td><td></td><td>1000</td></tr>
<tr><td>Oligo fwd</td><td>CACCGTCTCCCGTCTTGATCACTC</td></tr>
<tr><td>Oligo rev</td><td>AAACGAGTGATCAAGACGGGAGAC</td></tr>
<tr><td>Chromosome</td><td>start</td><td></td><td>end</td><td>strand</td><td>MM</td><td>target_seq</td><td>PAM</td><td>alignment</td><td>distance</td><td>position</td><td>gene name</td><td>gene id</td></tr>
<tr><td>GU305914.1</td><td></td><td>38980</td><td>39002</td><td>-</td><td>0</td><td>GTCTCCCGTCTTGATCACTC</td><td>GGG</td><td>||||||||[||||||||||||]PAM</td><td>NA</td><td>Intergenic</td><td>NA</td><td>NA</td></tr>
<tr><td>T31333</td><td>TCGCCGTGCGTTCGTTGTGTTGG</td><td></td><td>1000</td></tr>
<tr><td>Oligo fwd</td><td>CACCTCGCCGTGCGTTCGTTGTGT</td></tr>
<tr><td>Oligo rev</td><td>AAACACACAACGAACGCACGGCGA</td></tr>
<tr><td>Chromosome</td><td>start</td><td></td><td>end</td><td>strand</td><td>MM</td><td>target_seq</td><td>PAM</td><td>alignment</td><td>distance</td><td>position</td><td>gene name</td><td>gene id</td></tr>
<tr><td>GU305914.1</td><td></td><td>38957</td><td>38979</td><td>-</td><td>0</td><td>TCGCCGTGCGTTCGTTGTGT</td><td>TGG</td><td>||||||||[||||||||||||]PAM</td><td>NA</td><td>Intergenic</td><td>NA</td><td>NA</td></tr>
<tr><td>T31334</td><td>TTGGTAGATGTTGCCCTCTGTGG</td><td></td><td>1000</td></tr>
<tr><td>Oligo fwd</td><td>CACCTTGGTAGATGTTGCCCTCTG</td></tr>
<tr><td>Oligo rev</td><td>AAACCAGAGGGCAACATCTACCAA</td></tr>
<tr><td>Chromosome</td><td>start</td><td></td><td>end</td><td>strand</td><td>MM</td><td>target_seq</td><td>PAM</td><td>alignment</td><td>distance</td><td>position</td><td>gene name</td><td>gene id</td></tr>
<tr><td>GU305914.1</td><td></td><td>38938</td><td>38960</td><td>-</td><td>0</td><td>TTGGTAGATGTTGCCCTCTG</td><td>TGG</td><td>||||||||[||||||||||||]PAM</td><td>NA</td><td>Intergenic</td><td>NA</td><td>NA</td></tr>
<tr><td>T31335</td><td>GGAGTGCATGAAGAAGAGTCTGG</td><td></td><td>1000</td></tr>
<tr><td>Oligo fwd</td><td>CACCGGAGTGCATGAAGAAGAGTC</td></tr>
<tr><td>Oligo rev</td><td>AAACGACTCTTCTTCATGCACTCC</td></tr>
<tr><td>Chromosome</td><td>start</td><td></td><td>end</td><td>strand</td><td>MM</td><td>target_seq</td><td>PAM</td><td>alignment</td><td>distance</td><td>position</td><td>gene name</td><td>gene id</td></tr>
<tr><td>GU305914.1</td><td></td><td>38917</td><td>38939</td><td>-</td><td>0</td><td>GGAGTGCATGAAGAAGAGTC</td><td>TGG</td><td>||||||||[||||||||||||]PAM</td><td>NA</td><td>Intergenic</td><td>NA</td><td>NA</td></tr>
<tr><td>GU305914.1</td><td></td><td>40415</td><td>40437</td><td>-</td><td>4</td><td>GGAGGACATGACGACGAGTC</td><td>GGG</td><td>||||--||[|||-||-|||||]PAM</td><td>NA</td><td>Intergenic</td><td>NA</td><td>NA</td></tr>
<tr><td>T31336</td><td>CATGAAGAAGAGTCTGGCGATGG</td><td></td><td>1000</td></tr>
<tr><td>Oligo fwd</td><td>CACCCATGAAGAAGAGTCTGGCGA</td></tr>
<tr><td>Oligo rev</td><td>AAACTCGCCAGACTCTTCTTCATG</td></tr>
<tr><td>Chromosome</td><td>start</td><td></td><td>end</td><td>strand</td><td>MM</td><td>target_seq</td><td>PAM</td><td>alignment</td><td>distance</td><td>position</td><td>gene name</td><td>gene id</td></tr>
<tr><td>GU305914.1</td><td></td><td>38911</td><td>38933</td><td>-</td><td>0</td><td>CATGAAGAAGAGTCTGGCGA</td><td>TGG</td><td>||||||||[||||||||||||]PAM</td><td>NA</td><td>Intergenic</td><td>NA</td><td>NA</td></tr>
<tr><td>T31337</td><td>ATGAAGAAGAGTCTGGCGATGGG</td><td></td><td>1000</td></tr>
<tr><td>Oligo fwd</td><td>CACCATGAAGAAGAGTCTGGCGAT</td></tr>
<tr><td>Oligo rev</td><td>AAACATCGCCAGACTCTTCTTCAT</td></tr>
<tr><td>Chromosome</td><td>start</td><td></td><td>end</td><td>strand</td><td>MM</td><td>target_seq</td><td>PAM</td><td>alignment</td><td>distance</td><td>position</td><td>gene name</td><td>gene id</td></tr>
<tr><td>GU305914.1</td><td></td><td>38910</td><td>38932</td><td>-</td><td>0</td><td>ATGAAGAAGAGTCTGGCGAT</td><td>GGG</td><td>||||||||[||||||||||||]PAM</td><td>NA</td><td>Intergenic</td><td>NA</td><td>NA</td></tr>
<tr><td>T31338</td><td>GTCTGGCGATGGGCGTCCCGCGG</td><td></td><td>1000</td></tr>
<tr><td>Oligo fwd</td><td>CACCGTCTGGCGATGGGCGTCCCG</td></tr>
<tr><td>Oligo rev</td><td>AAACCGGGACGCCCATCGCCAGAC</td></tr>
</table>

| | | | | | | | | | | | |
|---|---|---|---|---|---|---|---|---|---|---|---|
| Chromosome | start | end | strand | MM | target_seq | PAM | alignment | distance | position | gene name | gene id |
| GU305914.1 | 38900 | 38922 | - | 0 | GTCTGGCGATGGGCGTCCCG | CGG | \|\|\|\|\|\|\|\|[\|\|\|\|\|\|\|\|\|\|\|\|]PAM | NA | Intergenic | NA | NA |
| T31339 | TGGGCGTCCCGCGGCAGATGTGG | 1000 | | | | | | | | | |
| Oligo fwd | CACCTGGGCGTCCCGCGGCAGATG | | | | | | | | | | |
| Oligo rev | AAACCATCTGCCGCGGGACGCCCA | | | | | | | | | | |
| Chromosome | start | end | strand | MM | target_seq | PAM | alignment | distance | position | gene name | gene id |
| GU305914.1 | 38891 | 38913 | - | 0 | TGGGCGTCCCGCGGCAGATG | TGG | \|\|\|\|\|\|\|\|[\|\|\|\|\|\|\|\|\|\|\|\|]PAM | NA | Intergenic | NA | NA |
| T31340 | GCGTCCCGCGGCAGATGTGGAGG | 1000 | | | | | | | | | |
| Oligo fwd | CACCGCGTCCCGCGGCAGATGTGG | | | | | | | | | | |
| Oligo rev | AAACCCACATCTGCCGCGGGACGC | | | | | | | | | | |
| Chromosome | start | end | strand | MM | target_seq | PAM | alignment | distance | position | gene name | gene id |
| GU305914.1 | 38888 | 38910 | - | 0 | GCGTCCCGCGGCAGATGTGG | AGG | \|\|\|\|\|\|\|\|[\|\|\|\|\|\|\|\|\|\|\|\|]PAM | NA | Intergenic | NA | NA |
| T31341 | CCCGCGGCAGATGTGGAGGCTGG | 1000 | | | | | | | | | |
| Oligo fwd | CACCCCCGCGGCAGATGTGGAGGC | | | | | | | | | | |
| Oligo rev | AAACGCCTCCACATCTGCCGCGGG | | | | | | | | | | |
| Chromosome | start | end | strand | MM | target_seq | PAM | alignment | distance | position | gene name | gene id |
| GU305914.1 | 38884 | 38906 | - | 0 | CCCGCGGCAGATGTGGAGGC | TGG | \|\|\|\|\|\|\|\|[\|\|\|\|\|\|\|\|\|\|\|\|]PAM | NA | Intergenic | NA | NA |
| GU305914.1 | 180320 | 180342 | - | 4 | CCCCCTGCAGATGAGGAGGA | AGG | \|\|\|-\|-\|\|[\|\|\|\|\|-\|\|\|\|\|-]PAM | NA | Intergenic | NA | NA |
| T31342 | CCGCGGCAGATGTGGAGGCTGGG | 1000 | | | | | | | | | |
| Oligo fwd | CACCCCGCGGCAGATGTGGAGGCT | | | | | | | | | | |
| Oligo rev | AAACAGCCTCCACATCTGCCGCGG | | | | | | | | | | |
| Chromosome | start | end | strand | MM | target_seq | PAM | alignment | distance | position | gene name | gene id |
| GU305914.1 | 38883 | 38905 | - | 0 | CCGCGGCAGATGTGGAGGCT | GGG | \|\|\|\|\|\|\|\|[\|\|\|\|\|\|\|\|\|\|\|\|]PAM | NA | Intergenic | NA | NA |
| T31343 | CGCGGCAGATGTGGAGGCTGGGG | 1000 | | | | | | | | | |
| Oligo fwd | CACCCGCGGCAGATGTGGAGGCTG | | | | | | | | | | |
| Oligo rev | AAACCAGCCTCCACATCTGCCGCG | | | | | | | | | | |
| Chromosome | start | end | strand | MM | target_seq | PAM | alignment | distance | position | gene name | gene id |
| GU305914.1 | 38882 | 38904 | - | 0 | CGCGGCAGATGTGGAGGCTG | GGG | \|\|\|\|\|\|\|\|[\|\|\|\|\|\|\|\|\|\|\|\|]PAM | NA | Intergenic | NA | NA |
| T31344 | GCGGCAGATGTGGAGGCTGGGGG | 1000 | | | | | | | | | |
| Oligo fwd | CACCGCGGCAGATGTGGAGGCTGG | | | | | | | | | | |
| Oligo rev | AAACCCAGCCTCCACATCTGCCGC | | | | | | | | | | |
| Chromosome | start | end | strand | MM | target_seq | PAM | alignment | distance | position | gene name | gene id |
| GU305914.1 | 38881 | 38903 | - | 0 | GCGGCAGATGTGGAGGCTGG | GGG | \|\|\|\|\|\|\|\|[\|\|\|\|\|\|\|\|\|\|\|\|]PAM | NA | Intergenic | NA | NA |
| T31345 | CAGATGTGGAGGCTGGGGTCGG | 1000 | | | | | | | | | |
| Oligo fwd | CACCCAGATGTGGAGGCTGGGGGT | | | | | | | | | | |
| Oligo rev | AAACACCCCCAGCCTCCACATCTG | | | | | | | | | | |
| Chromosome | start | end | strand | MM | target_seq | PAM | alignment | distance | position | gene name | gene id |
| GU305914.1 | 38877 | 38899 | - | 0 | CAGATGTGGAGGCTGGGGGT | CGG | \|\|\|\|\|\|\|\|[\|\|\|\|\|\|\|\|\|\|\|\|]PAM | NA | Intergenic | NA | NA |
| T31346 | AGATGTGGAGGCTGGGGGTCGGG | 1000 | | | | | | | | | |
| Oligo fwd | CACCAGATGTGGAGGCTGGGGGTC | | | | | | | | | | |
| Oligo rev | AAACGACCCCCAGCCTCCACATCT | | | | | | | | | | |
| Chromosome | start | end | strand | MM | target_seq | PAM | alignment | distance | position | gene name | gene id |
| GU305914.1 | 38876 | 38898 | - | 0 | AGATGTGGAGGCTGGGGGTC | GGG | \|\|\|\|\|\|\|\|[\|\|\|\|\|\|\|\|\|\|\|\|]PAM | NA | Intergenic | NA | NA |
| T31347 | GATGTGGAGGCTGGGGGTCGGGG | 1000 | | | | | | | | | |
| Oligo fwd | CACCGATGTGGAGGCTGGGGGTCG | | | | | | | | | | |
| Oligo rev | AAACCGACCCCCAGCCTCCACATC | | | | | | | | | | |
| Chromosome | start | end | strand | MM | target_seq | PAM | alignment | distance | position | gene name | gene id |
| GU305914.1 | 38875 | 38897 | - | 0 | GATGTGGAGGCTGGGGGTCG | GGG | \|\|\|\|\|\|\|\|[\|\|\|\|\|\|\|\|\|\|\|\|]PAM | NA | Intergenic | NA | NA |
| T31348 | GGGTCGGGGAGATCAGCGAGAGG | 1000 | | | | | | | | | |
| Oligo fwd | CACCGGGTCGGGGAGATCAGCGAG | | | | | | | | | | |
| Oligo rev | AAACCTCGCTGATCTCCCCGACCC | | | | | | | | | | |
| Chromosome | start | end | strand | MM | target_seq | PAM | alignment | distance | position | gene name | gene id |
| GU305914.1 | 38861 | 38883 | - | 0 | GGGTCGGGGAGATCAGCGAG | AGG | \|\|\|\|\|\|\|\|[\|\|\|\|\|\|\|\|\|\|\|\|]PAM | NA | Intergenic | NA | NA |
| T31349 | TCGGGGAGATCAGCGAGAGGCGG | 1000 | | | | | | | | | |
| Oligo fwd | CACCTCGGGGAGATCAGCGAGAGG | | | | | | | | | | |
| Oligo rev | AAACCCTCTCGCTGATCTCCCCGA | | | | | | | | | | |
| Chromosome | start | end | strand | MM | target_seq | PAM | alignment | distance | position | gene name | gene id |
| GU305914.1 | 38858 | 38880 | - | 0 | TCGGGGAGATCAGCGAGAGG | CGG | \|\|\|\|\|\|\|\|[\|\|\|\|\|\|\|\|\|\|\|\|]PAM | NA | Intergenic | NA | NA |
| T31350 | GGAGATCAGCGAGAGGCGGATGG | 1000 | | | | | | | | | |
| Oligo fwd | CACCGGAGATCAGCGAGAGGCGGA | | | | | | | | | | |
| Oligo rev | AAACTCCGCCTCTCGCTGATCTCC | | | | | | | | | | |
| Chromosome | start | end | strand | MM | target_seq | PAM | alignment | distance | position | gene name | gene id |
| GU305914.1 | 38854 | 38876 | - | 0 | GGAGATCAGCGAGAGGCGGA | TGG | \|\|\|\|\|\|\|\|[\|\|\|\|\|\|\|\|\|\|\|\|]PAM | NA | Intergenic | NA | NA |
| T31351 | GAGATCAGCGAGAGGCGGATGGG | 1000 | | | | | | | | | |
| Oligo fwd | CACCGAGATCAGCGAGAGGCGGAT | | | | | | | | | | |
| Oligo rev | AAACATCCGCCTCTCGCTGATCTC | | | | | | | | | | |
| Chromosome | start | end | strand | MM | target_seq | PAM | alignment | distance | position | gene name | gene id |
| GU305914.1 | 38853 | 38875 | - | 0 | GAGATCAGCGAGAGGCGGAT | GGG | \|\|\|\|\|\|\|\|[\|\|\|\|\|\|\|\|\|\|\|\|]PAM | NA | Intergenic | NA | NA |
| T31352 | AGATCAGCGAGAGGCGGATGGGG | 1000 | | | | | | | | | |
| Oligo fwd | CACCAGATCAGCGAGAGGCGGATG | | | | | | | | | | |
| Oligo rev | AAACCATCCGCCTCTCGCTGATCT | | | | | | | | | | |
| Chromosome | start | end | strand | MM | target_seq | PAM | alignment | distance | position | gene name | gene id |
| GU305914.1 | 38852 | 38874 | - | 0 | AGATCAGCGAGAGGCGGATG | GGG | \|\|\|\|\|\|\|\|[\|\|\|\|\|\|\|\|\|\|\|\|]PAM | NA | Intergenic | NA | NA |
| T31353 | GAGAGGCGGATGGGGCTCTGCGG | 1000 | | | | | | | | | |

<table>
<tr><td>Oligo fwd</td><td>CACCGAGAGGCGGATGGGGCTCTG</td><td></td><td></td><td></td><td></td><td></td><td></td><td></td><td></td><td></td><td></td></tr>
<tr><td>Oligo rev</td><td>AAACCAGAGCCCCATCCGCCTCTC</td><td></td><td></td><td></td><td></td><td></td><td></td><td></td><td></td><td></td><td></td></tr>
<tr><td>Chromosome</td><td>start</td><td>end</td><td>strand</td><td>MM</td><td>target_seq</td><td>PAM</td><td>alignment</td><td>distance</td><td>position</td><td>gene name</td><td>gene id</td></tr>
<tr><td>GU305914.1</td><td>38844</td><td>38866</td><td>-</td><td>0</td><td>GAGAGGCGGATGGGGCTCTG</td><td>CGG</td><td>||||||||[||||||||||||]PAM</td><td>NA</td><td>Intergenic</td><td>NA</td><td>NA</td></tr>
<tr><td>T31354</td><td>AGAGGCGGATGGGGCTCTGCGGG</td><td>1000</td><td></td><td></td><td></td><td></td><td></td><td></td><td></td><td></td><td></td></tr>
<tr><td>Oligo fwd</td><td>CACCAGAGGCGGATGGGGCTCTGC</td><td></td><td></td><td></td><td></td><td></td><td></td><td></td><td></td><td></td><td></td></tr>
<tr><td>Oligo rev</td><td>AAACGCAGAGCCCCATCCGCCTCT</td><td></td><td></td><td></td><td></td><td></td><td></td><td></td><td></td><td></td><td></td></tr>
<tr><td>Chromosome</td><td>start</td><td>end</td><td>strand</td><td>MM</td><td>target_seq</td><td>PAM</td><td>alignment</td><td>distance</td><td>position</td><td>gene name</td><td>gene id</td></tr>
<tr><td>GU305914.1</td><td>38843</td><td>38865</td><td>-</td><td>0</td><td>AGAGGCGGATGGGGCTCTGC</td><td>GGG</td><td>||||||||[||||||||||||]PAM</td><td>NA</td><td>Intergenic</td><td>NA</td><td>NA</td></tr>
<tr><td>T31355</td><td>CGGGCGCCAGACCGTCAGCCCGG</td><td>1000</td><td></td><td></td><td></td><td></td><td></td><td></td><td></td><td></td><td></td></tr>
<tr><td>Oligo fwd</td><td>CACCCGGGCGCCAGACCGTCAGCC</td><td></td><td></td><td></td><td></td><td></td><td></td><td></td><td></td><td></td><td></td></tr>
<tr><td>Oligo rev</td><td>AAACGGCTGACGGTCTGGCGCCCG</td><td></td><td></td><td></td><td></td><td></td><td></td><td></td><td></td><td></td><td></td></tr>
<tr><td>Chromosome</td><td>start</td><td>end</td><td>strand</td><td>MM</td><td>target_seq</td><td>PAM</td><td>alignment</td><td>distance</td><td>position</td><td>gene name</td><td>gene id</td></tr>
<tr><td>GU305914.1</td><td>38824</td><td>38846</td><td>-</td><td>0</td><td>CGGGCGCCAGACCGTCAGCC</td><td>CGG</td><td>||||||||[||||||||||||]PAM</td><td>NA</td><td>Intergenic</td><td>NA</td><td>NA</td></tr>
<tr><td>T31356</td><td>GCCAGACCGTCAGCCCGGTGTGG</td><td>1000</td><td></td><td></td><td></td><td></td><td></td><td></td><td></td><td></td><td></td></tr>
<tr><td>Oligo fwd</td><td>CACCGCCAGACCGTCAGCCCGGTG</td><td></td><td></td><td></td><td></td><td></td><td></td><td></td><td></td><td></td><td></td></tr>
<tr><td>Oligo rev</td><td>AAACCACCGGGCTGACGGTCTGGC</td><td></td><td></td><td></td><td></td><td></td><td></td><td></td><td></td><td></td><td></td></tr>
<tr><td>Chromosome</td><td>start</td><td>end</td><td>strand</td><td>MM</td><td>target_seq</td><td>PAM</td><td>alignment</td><td>distance</td><td>position</td><td>gene name</td><td>gene id</td></tr>
<tr><td>GU305914.1</td><td>38819</td><td>38841</td><td>-</td><td>0</td><td>GCCAGACCGTCAGCCCGGTG</td><td>TGG</td><td>||||||||[||||||||||||]PAM</td><td>NA</td><td>Intergenic</td><td>NA</td><td>NA</td></tr>
<tr><td>T31357</td><td>CCGTCAGCCCGGTGTGGAAACGG</td><td>1000</td><td></td><td></td><td></td><td></td><td></td><td></td><td></td><td></td><td></td></tr>
<tr><td>Oligo fwd</td><td>CACCCCGTCAGCCCGGTGTGGAAA</td><td></td><td></td><td></td><td></td><td></td><td></td><td></td><td></td><td></td><td></td></tr>
<tr><td>Oligo rev</td><td>AAACTTTCCACACCGGGCTGACGG</td><td></td><td></td><td></td><td></td><td></td><td></td><td></td><td></td><td></td><td></td></tr>
<tr><td>Chromosome</td><td>start</td><td>end</td><td>strand</td><td>MM</td><td>target_seq</td><td>PAM</td><td>alignment</td><td>distance</td><td>position</td><td>gene name</td><td>gene id</td></tr>
<tr><td>GU305914.1</td><td>38813</td><td>38835</td><td>-</td><td>0</td><td>CCGTCAGCCCGGTGTGGAAA</td><td>CGG</td><td>||||||||[||||||||||||]PAM</td><td>NA</td><td>Intergenic</td><td>NA</td><td>NA</td></tr>
<tr><td>T31358</td><td>GGTGTGGAAACGGTACTCGCCGG</td><td>1000</td><td></td><td></td><td></td><td></td><td></td><td></td><td></td><td></td><td></td></tr>
<tr><td>Oligo fwd</td><td>CACCGGTGTGGAAACGGTACTCGC</td><td></td><td></td><td></td><td></td><td></td><td></td><td></td><td></td><td></td><td></td></tr>
<tr><td>Oligo rev</td><td>AAACGCGAGTACCGTTTCCACACC</td><td></td><td></td><td></td><td></td><td></td><td></td><td></td><td></td><td></td><td></td></tr>
<tr><td>Chromosome</td><td>start</td><td>end</td><td>strand</td><td>MM</td><td>target_seq</td><td>PAM</td><td>alignment</td><td>distance</td><td>position</td><td>gene name</td><td>gene id</td></tr>
<tr><td>GU305914.1</td><td>38803</td><td>38825</td><td>-</td><td>0</td><td>GGTGTGGAAACGGTACTCGC</td><td>CGG</td><td>||||||||[||||||||||||]PAM</td><td>NA</td><td>Intergenic</td><td>NA</td><td>NA</td></tr>
<tr><td>T31359</td><td>GTGTGGAAACGGTACTCGCCGGG</td><td>1000</td><td></td><td></td><td></td><td></td><td></td><td></td><td></td><td></td><td></td></tr>
<tr><td>Oligo fwd</td><td>CACCGTGTGGAAACGGTACTCGCC</td><td></td><td></td><td></td><td></td><td></td><td></td><td></td><td></td><td></td><td></td></tr>
<tr><td>Oligo rev</td><td>AAACGGCGAGTACCGTTTCCACAC</td><td></td><td></td><td></td><td></td><td></td><td></td><td></td><td></td><td></td><td></td></tr>
<tr><td>Chromosome</td><td>start</td><td>end</td><td>strand</td><td>MM</td><td>target_seq</td><td>PAM</td><td>alignment</td><td>distance</td><td>position</td><td>gene name</td><td>gene id</td></tr>
<tr><td>GU305914.1</td><td>38802</td><td>38824</td><td>-</td><td>0</td><td>GTGTGGAAACGGTACTCGCC</td><td>GGG</td><td>||||||||[||||||||||||]PAM</td><td>NA</td><td>Intergenic</td><td>NA</td><td>NA</td></tr>
<tr><td>T31360</td><td>ACGGTACTCGCCGGGCACGCCGG</td><td>1000</td><td></td><td></td><td></td><td></td><td></td><td></td><td></td><td></td><td></td></tr>
<tr><td>Oligo fwd</td><td>CACCACGGTACTCGCCGGGCACGC</td><td></td><td></td><td></td><td></td><td></td><td></td><td></td><td></td><td></td><td></td></tr>
<tr><td>Oligo rev</td><td>AAACGCGTGCCCGGCGAGTACCGT</td><td></td><td></td><td></td><td></td><td></td><td></td><td></td><td></td><td></td><td></td></tr>
<tr><td>Chromosome</td><td>start</td><td>end</td><td>strand</td><td>MM</td><td>target_seq</td><td>PAM</td><td>alignment</td><td>distance</td><td>position</td><td>gene name</td><td>gene id</td></tr>
<tr><td>GU305914.1</td><td>38794</td><td>38816</td><td>-</td><td>0</td><td>ACGGTACTCGCCGGGCACGC</td><td>CGG</td><td>||||||||[||||||||||||]PAM</td><td>NA</td><td>Intergenic</td><td>NA</td><td>NA</td></tr>
<tr><td>T31361</td><td>CGCCGGAGATGAAAAAGCAGCGG</td><td>1000</td><td></td><td></td><td></td><td></td><td></td><td></td><td></td><td></td><td></td></tr>
<tr><td>Oligo fwd</td><td>CACCCGCCGGAGATGAAAAAGCAG</td><td></td><td></td><td></td><td></td><td></td><td></td><td></td><td></td><td></td><td></td></tr>
<tr><td>Oligo rev</td><td>AAACCTGCTTTTTCATCTCCGGCG</td><td></td><td></td><td></td><td></td><td></td><td></td><td></td><td></td><td></td><td></td></tr>
<tr><td>Chromosome</td><td>start</td><td>end</td><td>strand</td><td>MM</td><td>target_seq</td><td>PAM</td><td>alignment</td><td>distance</td><td>position</td><td>gene name</td><td>gene id</td></tr>
<tr><td>GU305914.1</td><td>38777</td><td>38799</td><td>-</td><td>0</td><td>CGCCGGAGATGAAAAAGCAG</td><td>CGG</td><td>||||||||[||||||||||||]PAM</td><td>NA</td><td>Intergenic</td><td>NA</td><td>NA</td></tr>
<tr><td>T31362</td><td>GTATCTGATGTCCATGAAGACGG</td><td>1000</td><td></td><td></td><td></td><td></td><td></td><td></td><td></td><td></td><td></td></tr>
<tr><td>Oligo fwd</td><td>CACCGTATCTGATGTCCATGAAGA</td><td></td><td></td><td></td><td></td><td></td><td></td><td></td><td></td><td></td><td></td></tr>
<tr><td>Oligo rev</td><td>AAACTCTTCATGGACATCAGATAC</td><td></td><td></td><td></td><td></td><td></td><td></td><td></td><td></td><td></td><td></td></tr>
<tr><td>Chromosome</td><td>start</td><td>end</td><td>strand</td><td>MM</td><td>target_seq</td><td>PAM</td><td>alignment</td><td>distance</td><td>position</td><td>gene name</td><td>gene id</td></tr>
<tr><td>GU305914.1</td><td>38737</td><td>38759</td><td>-</td><td>0</td><td>GTATCTGATGTCCATGAAGA</td><td>CGG</td><td>||||||||[||||||||||||]PAM</td><td>NA</td><td>Intergenic</td><td>NA</td><td>NA</td></tr>
<tr><td>T31363</td><td>CTCTCTGAAGCAGAGATCGTAGG</td><td>1000</td><td></td><td></td><td></td><td></td><td></td><td></td><td></td><td></td><td></td></tr>
<tr><td>Oligo fwd</td><td>CACCCTCTCTGAAGCAGAGATCGT</td><td></td><td></td><td></td><td></td><td></td><td></td><td></td><td></td><td></td><td></td></tr>
<tr><td>Oligo rev</td><td>AAACACGATCTCTGCTTCAGAGAG</td><td></td><td></td><td></td><td></td><td></td><td></td><td></td><td></td><td></td><td></td></tr>
<tr><td>Chromosome</td><td>start</td><td>end</td><td>strand</td><td>MM</td><td>target_seq</td><td>PAM</td><td>alignment</td><td>distance</td><td>position</td><td>gene name</td><td>gene id</td></tr>
<tr><td>GU305914.1</td><td>38704</td><td>38726</td><td>-</td><td>0</td><td>CTCTCTGAAGCAGAGATCGT</td><td>AGG</td><td>||||||||[||||||||||||]PAM</td><td>NA</td><td>Intergenic</td><td>NA</td><td>NA</td></tr>
<tr><td>T31364</td><td>TCTCTGAAGCAGAGATCGTAGGG</td><td>1000</td><td></td><td></td><td></td><td></td><td></td><td></td><td></td><td></td><td></td></tr>
<tr><td>Oligo fwd</td><td>CACCTCTCTGAAGCAGAGATCGTA</td><td></td><td></td><td></td><td></td><td></td><td></td><td></td><td></td><td></td><td></td></tr>
<tr><td>Oligo rev</td><td>AAACTACGATCTCTGCTTCAGAGA</td><td></td><td></td><td></td><td></td><td></td><td></td><td></td><td></td><td></td><td></td></tr>
<tr><td>Chromosome</td><td>start</td><td>end</td><td>strand</td><td>MM</td><td>target_seq</td><td>PAM</td><td>alignment</td><td>distance</td><td>position</td><td>gene name</td><td>gene id</td></tr>
<tr><td>GU305914.1</td><td>38703</td><td>38725</td><td>-</td><td>0</td><td>TCTCTGAAGCAGAGATCGTA</td><td>GGG</td><td>||||||||[||||||||||||]PAM</td><td>NA</td><td>Intergenic</td><td>NA</td><td>NA</td></tr>
<tr><td>T31365</td><td>GTAGGGCGCGTACATCGTGATGG</td><td>1000</td><td></td><td></td><td></td><td></td><td></td><td></td><td></td><td></td><td></td></tr>
<tr><td>Oligo fwd</td><td>CACCGTAGGGCGCGTACATCGTGA</td><td></td><td></td><td></td><td></td><td></td><td></td><td></td><td></td><td></td><td></td></tr>
<tr><td>Oligo rev</td><td>AAACTCACGATGTACGCGCCCTAC</td><td></td><td></td><td></td><td></td><td></td><td></td><td></td><td></td><td></td><td></td></tr>
<tr><td>Chromosome</td><td>start</td><td>end</td><td>strand</td><td>MM</td><td>target_seq</td><td>PAM</td><td>alignment</td><td>distance</td><td>position</td><td>gene name</td><td>gene id</td></tr>
<tr><td>GU305914.1</td><td>38686</td><td>38708</td><td>-</td><td>0</td><td>GTAGGGCGCGTACATCGTGA</td><td>TGG</td><td>||||||||[||||||||||||]PAM</td><td>NA</td><td>Intergenic</td><td>NA</td><td>NA</td></tr>
<tr><td>T31366</td><td>TAGGGCGCGTACATCGTGATGGG</td><td>1000</td><td></td><td></td><td></td><td></td><td></td><td></td><td></td><td></td><td></td></tr>
<tr><td>Oligo fwd</td><td>CACCTAGGGCGCGTACATCGTGAT</td><td></td><td></td><td></td><td></td><td></td><td></td><td></td><td></td><td></td><td></td></tr>
<tr><td>Oligo rev</td><td>AAACATCACGATGTACGCGCCCTA</td><td></td><td></td><td></td><td></td><td></td><td></td><td></td><td></td><td></td><td></td></tr>
<tr><td>Chromosome</td><td>start</td><td>end</td><td>strand</td><td>MM</td><td>target_seq</td><td>PAM</td><td>alignment</td><td>distance</td><td>position</td><td>gene name</td><td>gene id</td></tr>
<tr><td>GU305914.1</td><td>38685</td><td>38707</td><td>-</td><td>0</td><td>TAGGGCGCGTACATCGTGAT</td><td>GGG</td><td>||||||||[||||||||||||]PAM</td><td>NA</td><td>Intergenic</td><td>NA</td><td>NA</td></tr>
<tr><td>T31367</td><td>GTACATCGTGATGGGATCGTCGG</td><td>1000</td><td></td><td></td><td></td><td></td><td></td><td></td><td></td><td></td><td></td></tr>
<tr><td>Oligo fwd</td><td>CACCGTACATCGTGATGGGATCGT</td><td></td><td></td><td></td><td></td><td></td><td></td><td></td><td></td><td></td><td></td></tr>
<tr><td>Oligo rev</td><td>AAACACGATCCCATCACGATGTAC</td><td></td><td></td><td></td><td></td><td></td><td></td><td></td><td></td><td></td><td></td></tr>
<tr><td>Chromosome</td><td>start</td><td>end</td><td>strand</td><td>MM</td><td>target_seq</td><td>PAM</td><td>alignment</td><td>distance</td><td>position</td><td>gene name</td><td>gene id</td></tr>
<tr><td>GU305914.1</td><td>38677</td><td>38699</td><td>-</td><td>0</td><td>GTACATCGTGATGGGATCGT</td><td>CGG</td><td>||||||||[||||||||||||]PAM</td><td>NA</td><td>Intergenic</td><td>NA</td><td>NA</td></tr>
</table>

| | | | | | | | | | | | | |
|---|---|---|---|---|---|---|---|---|---|---|---|---|
| T31368 | TACATCGTGATGGGATCGTCGGG | | 1000 | | | | | | | | | |
| Oligo fwd | CACCTACATCGTGATGGGATCGTC | | | | | | | | | | | |
| Oligo rev | AAACGACGATCCCATCACGATGTA | | | | | | | | | | | |
| Chromosome | start | | end | strand | MM | target_seq | PAM | alignment | distance | position | gene name | gene id |
| GU305914.1 | | 38676 | 38698 | - | 0 | TACATCGTGATGGGATCGTC | GGG | \|\|\|\|\|\|\|\|[\|\|\|\|\|\|\|\|\|\|\|\|]PAM | NA | Intergenic | NA | NA |
| T31369 | ACATCGTGATGGGATCGTCGGGG | | 1000 | | | | | | | | | |
| Oligo fwd | CACCACATCGTGATGGGATCGTCG | | | | | | | | | | | |
| Oligo rev | AAACCGACGATCCCATCACGATGT | | | | | | | | | | | |
| Chromosome | start | | end | strand | MM | target_seq | PAM | alignment | distance | position | gene name | gene id |
| GU305914.1 | | 38675 | 38697 | - | 0 | ACATCGTGATGGGATCGTCG | GGG | \|\|\|\|\|\|\|\|[\|\|\|\|\|\|\|\|\|\|\|\|]PAM | NA | Intergenic | NA | NA |
| T31370 | GGATCGTCGGGGCTTTCGCTCGG | | 1000 | | | | | | | | | |
| Oligo fwd | CACCGGATCGTCGGGGCTTTCGCT | | | | | | | | | | | |
| Oligo rev | AAACAGCGAAAGCCCCGACGATCC | | | | | | | | | | | |
| Chromosome | start | | end | strand | MM | target_seq | PAM | alignment | distance | position | gene name | gene id |
| GU305914.1 | | 38664 | 38686 | - | 0 | GGATCGTCGGGGCTTTCGCT | CGG | \|\|\|\|\|\|\|\|[\|\|\|\|\|\|\|\|\|\|\|\|]PAM | NA | Intergenic | NA | NA |
| T31371 | GTCGGGGCTTTCGCTCGGCCAGG | | 1000 | | | | | | | | | |
| Oligo fwd | CACCGTCGGGGCTTTCGCTCGGCC | | | | | | | | | | | |
| Oligo rev | AAACGGCCGAGCGAAAGCCCCGAC | | | | | | | | | | | |
| Chromosome | start | | end | strand | MM | target_seq | PAM | alignment | distance | position | gene name | gene id |
| GU305914.1 | | 38659 | 38681 | - | 0 | GTCGGGGCTTTCGCTCGGCC | AGG | \|\|\|\|\|\|\|\|[\|\|\|\|\|\|\|\|\|\|\|\|]PAM | NA | Intergenic | NA | NA |
| T31372 | GGCTTTCGCTCGGCCAGGTGAGG | | 1000 | | | | | | | | | |
| Oligo fwd | CACCGGCTTTCGCTCGGCCAGGTG | | | | | | | | | | | |
| Oligo rev | AAACCACCTGGCCGAGCGAAAGCC | | | | | | | | | | | |
| Chromosome | start | | end | strand | MM | target_seq | PAM | alignment | distance | position | gene name | gene id |
| GU305914.1 | | 38654 | 38676 | - | 0 | GGCTTTCGCTCGGCCAGGTG | AGG | \|\|\|\|\|\|\|\|[\|\|\|\|\|\|\|\|\|\|\|\|]PAM | NA | Intergenic | NA | NA |
| T31373 | CGCTCGGCCAGGTGAGGTGCAGG | | 1000 | | | | | | | | | |
| Oligo fwd | CACCCGCTCGGCCAGGTGAGGTGC | | | | | | | | | | | |
| Oligo rev | AAACGCACCTCACCTGGCCGAGCG | | | | | | | | | | | |
| Chromosome | start | | end | strand | MM | target_seq | PAM | alignment | distance | position | gene name | gene id |
| GU305914.1 | | 38648 | 38670 | - | 0 | CGCTCGGCCAGGTGAGGTGC | AGG | \|\|\|\|\|\|\|\|[\|\|\|\|\|\|\|\|\|\|\|\|]PAM | NA | Intergenic | NA | NA |
| T31374 | GCTCGGCCAGGTGAGGTGCAGGG | | 1000 | | | | | | | | | |
| Oligo fwd | CACCGCTCGGCCAGGTGAGGTGCA | | | | | | | | | | | |
| Oligo rev | AAACTGCACCTCACCTGGCCGAGC | | | | | | | | | | | |
| Chromosome | start | | end | strand | MM | target_seq | PAM | alignment | distance | position | gene name | gene id |
| GU305914.1 | | 38647 | 38669 | - | 0 | GCTCGGCCAGGTGAGGTGCA | GGG | \|\|\|\|\|\|\|\|[\|\|\|\|\|\|\|\|\|\|\|\|]PAM | NA | Intergenic | NA | NA |
| T31375 | CGGCCAGGTGAGGTGCAGGGTGG | | 1000 | | | | | | | | | |
| Oligo fwd | CACCCGGCCAGGTGAGGTGCAGGG | | | | | | | | | | | |
| Oligo rev | AAACCCCTGCACCTCACCTGGCCG | | | | | | | | | | | |
| Chromosome | start | | end | strand | MM | target_seq | PAM | alignment | distance | position | gene name | gene id |
| GU305914.1 | | 38644 | 38666 | - | 0 | CGGCCAGGTGAGGTGCAGGG | TGG | \|\|\|\|\|\|\|\|[\|\|\|\|\|\|\|\|\|\|\|\|]PAM | NA | Intergenic | NA | NA |
| T31376 | GGCCAGGTGAGGTGCAGGGTGGG | | 1000 | | | | | | | | | |
| Oligo fwd | CACCGGCCAGGTGAGGTGCAGGGT | | | | | | | | | | | |
| Oligo rev | AAACACCCTGCACCTCACCTGGCC | | | | | | | | | | | |
| Chromosome | start | | end | strand | MM | target_seq | PAM | alignment | distance | position | gene name | gene id |
| GU305914.1 | | 38643 | 38665 | - | 0 | GGCCAGGTGAGGTGCAGGGT | GGG | \|\|\|\|\|\|\|\|[\|\|\|\|\|\|\|\|\|\|\|\|]PAM | NA | Intergenic | NA | NA |
| T31377 | GCCAGGTGAGGTGCAGGGTGGGG | | 1000 | | | | | | | | | |
| Oligo fwd | CACCGCCAGGTGAGGTGCAGGGTG | | | | | | | | | | | |
| Oligo rev | AAACCACCCTGCACCTCACCTGGC | | | | | | | | | | | |
| Chromosome | start | | end | strand | MM | target_seq | PAM | alignment | distance | position | gene name | gene id |
| GU305914.1 | | 38642 | 38664 | - | 0 | GCCAGGTGAGGTGCAGGGTG | GGG | \|\|\|\|\|\|\|\|[\|\|\|\|\|\|\|\|\|\|\|\|]PAM | NA | Intergenic | NA | NA |
| T31378 | GGTGAGGTGCAGGGTGGGGTTGG | | 1000 | | | | | | | | | |
| Oligo fwd | CACCGGTGAGGTGCAGGGTGGGGT | | | | | | | | | | | |
| Oligo rev | AAACACCCCACCCTGCACCTCACC | | | | | | | | | | | |
| Chromosome | start | | end | strand | MM | target_seq | PAM | alignment | distance | position | gene name | gene id |
| GU305914.1 | | 38638 | 38660 | - | 0 | GGTGAGGTGCAGGGTGGGGT | TGG | \|\|\|\|\|\|\|\|[\|\|\|\|\|\|\|\|\|\|\|\|]PAM | NA | Intergenic | NA | NA |
| T31379 | GGGGTTGGTCGAGAACACGATGG | | 1000 | | | | | | | | | |
| Oligo fwd | CACCGGGGTTGGTCGAGAACACGA | | | | | | | | | | | |
| Oligo rev | AAACTCGTGTTCTCGACCAACCCC | | | | | | | | | | | |
| Chromosome | start | | end | strand | MM | target_seq | PAM | alignment | distance | position | gene name | gene id |
| GU305914.1 | | 38623 | 38645 | - | 0 | GGGGTTGGTCGAGAACACGA | TGG | \|\|\|\|\|\|\|\|[\|\|\|\|\|\|\|\|\|\|\|\|]PAM | NA | Intergenic | NA | NA |
| T31380 | TTGGTCGAGAACACGATGGTCGG | | 1000 | | | | | | | | | |
| Oligo fwd | CACCTTGGTCGAGAACACGATGGT | | | | | | | | | | | |
| Oligo rev | AAACACCATCGTGTTCTCGACCAA | | | | | | | | | | | |
| Chromosome | start | | end | strand | MM | target_seq | PAM | alignment | distance | position | gene name | gene id |
| GU305914.1 | | 38619 | 38641 | - | 0 | TTGGTCGAGAACACGATGGT | CGG | \|\|\|\|\|\|\|\|[\|\|\|\|\|\|\|\|\|\|\|\|]PAM | NA | Intergenic | NA | NA |
| T31381 | AGAACACGATGGTCGGTTCTCGG | | 1000 | | | | | | | | | |
| Oligo fwd | CACCAGAACACGATGGTCGGTTCT | | | | | | | | | | | |
| Oligo rev | AAACAGAACCGACCATCGTGTTCT | | | | | | | | | | | |
| Chromosome | start | | end | strand | MM | target_seq | PAM | alignment | distance | position | gene name | gene id |
| GU305914.1 | | 38612 | 38634 | - | 0 | AGAACACGATGGTCGGTTCT | CGG | \|\|\|\|\|\|\|\|[\|\|\|\|\|\|\|\|\|\|\|\|]PAM | NA | Intergenic | NA | NA |
| T31382 | AAAGCGCAGCACGATCATGCTGG | | 1000 | | | | | | | | | |
| Oligo fwd | CACCAAAGCGCAGCACGATCATGC | | | | | | | | | | | |
| Oligo rev | AAACGCATGATCGTGCTGCGCTTT | | | | | | | | | | | |
| Chromosome | start | | end | strand | MM | target_seq | PAM | alignment | distance | position | gene name | gene id |
| GU305914.1 | | 38569 | 38591 | - | 0 | AAAGCGCAGCACGATCATGC | TGG | \|\|\|\|\|\|\|\|[\|\|\|\|\|\|\|\|\|\|\|\|]PAM | NA | Intergenic | NA | NA |

| | | | | | | | | | | | | |
|---|---|---|---|---|---|---|---|---|---|---|---|---|
| T31383 | AAGCGCAGCACGATCATGCTGGG | | 1000 | | | | | | | | | |
| Oligo fwd | CACCAAGCGCAGCACGATCATGCT | | | | | | | | | | | |
| Oligo rev | AAACAGCATGATCGTGCTGCGCTT | | | | | | | | | | | |
| Chromosome | start | | end | strand | MM | target_seq | PAM | alignment | distance | position | gene name | gene id |
| GU305914.1 | | 38568 | 38590 | - | 0 | AAGCGCAGCACGATCATGCT | GGG | \|\|\|\|\|\|\|\|[\|\|\|\|\|\|\|\|\|\|\|\|]PAM | NA | Intergenic | NA | NA |
| T31384 | ATCATGCTGGGCATCAACTCCGG | | 1000 | | | | | | | | | |
| Oligo fwd | CACCATCATGCTGGGCATCAACTC | | | | | | | | | | | |
| Oligo rev | AAACGAGTTGATGCCCAGCATGAT | | | | | | | | | | | |
| Chromosome | start | | end | strand | MM | target_seq | PAM | alignment | distance | position | gene name | gene id |
| GU305914.1 | | 38556 | 38578 | - | 0 | ATCATGCTGGGCATCAACTC | CGG | \|\|\|\|\|\|\|\|[\|\|\|\|\|\|\|\|\|\|\|\|]PAM | NA | Intergenic | NA | NA |
| T31385 | ATCAACTCCGGTGACAGCGACGG | | 1000 | | | | | | | | | |
| Oligo fwd | CACCATCAACTCCGGTGACAGCGA | | | | | | | | | | | |
| Oligo rev | AAACTCGCTGTCACCGGAGTTGAT | | | | | | | | | | | |
| Chromosome | start | | end | strand | MM | target_seq | PAM | alignment | distance | position | gene name | gene id |
| GU305914.1 | | 38544 | 38566 | - | 0 | ATCAACTCCGGTGACAGCGA | CGG | \|\|\|\|\|\|\|\|[\|\|\|\|\|\|\|\|\|\|\|\|]PAM | NA | Intergenic | NA | NA |
| T31386 | CAGCGACGGCGTGTTCGTAGTGG | | 1000 | | | | | | | | | |
| Oligo fwd | CACCCAGCGACGGCGTGTTCGTAG | | | | | | | | | | | |
| Oligo rev | AAACCTACGAACACGCCGTCGCTG | | | | | | | | | | | |
| Chromosome | start | | end | strand | MM | target_seq | PAM | alignment | distance | position | gene name | gene id |
| GU305914.1 | | 38530 | 38552 | - | 0 | CAGCGACGGCGTGTTCGTAG | TGG | \|\|\|\|\|\|\|\|[\|\|\|\|\|\|\|\|\|\|\|\|]PAM | NA | Intergenic | NA | NA |
| T31387 | CATGCTCTCGACGTTGATACCGG | | 1000 | | | | | | | | | |
| Oligo fwd | CACCCATGCTCTCGACGTTGATAC | | | | | | | | | | | |
| Oligo rev | AAACGTATCAACGTCGAGAGCATG | | | | | | | | | | | |
| Chromosome | start | | end | strand | MM | target_seq | PAM | alignment | distance | position | gene name | gene id |
| GU305914.1 | | 38452 | 38474 | - | 0 | CATGCTCTCGACGTTGATAC | CGG | \|\|\|\|\|\|\|\|[\|\|\|\|\|\|\|\|\|\|\|\|]PAM | NA | Intergenic | NA | NA |
| T31388 | CTCGAAATCGCCGCACTCGTAGG | | 1000 | | | | | | | | | |
| Oligo fwd | CACCCTCGAAATCGCCGCACTCGT | | | | | | | | | | | |
| Oligo rev | AAACACGAGTGCGGCGATTTCGAG | | | | | | | | | | | |
| Chromosome | start | | end | strand | MM | target_seq | PAM | alignment | distance | position | gene name | gene id |
| GU305914.1 | | 38428 | 38450 | - | 0 | CTCGAAATCGCCGCACTCGT | AGG | \|\|\|\|\|\|\|\|[\|\|\|\|\|\|\|\|\|\|\|\|]PAM | NA | Intergenic | NA | NA |
| T31389 | TCGTAGGTCACCTTCCTGTACGG | | 1000 | | | | | | | | | |
| Oligo fwd | CACCTCGTAGGTCACCTTCCTGTA | | | | | | | | | | | |
| Oligo rev | AAACTACAGGAAGGTGACCTACGA | | | | | | | | | | | |
| Chromosome | start | | end | strand | MM | target_seq | PAM | alignment | distance | position | gene name | gene id |
| GU305914.1 | | 38412 | 38434 | - | 0 | TCGTAGGTCACCTTCCTGTA | CGG | \|\|\|\|\|\|\|\|[\|\|\|\|\|\|\|\|\|\|\|\|]PAM | NA | Intergenic | NA | NA |
| T31390 | ACGGTCGCCAGTACTTGCATAGG | | 1000 | | | | | | | | | |
| Oligo fwd | CACCACGGTCGCCAGTACTTGCAT | | | | | | | | | | | |
| Oligo rev | AAACATGCAAGTACTGGCGACCGT | | | | | | | | | | | |
| Chromosome | start | | end | strand | MM | target_seq | PAM | alignment | distance | position | gene name | gene id |
| GU305914.1 | | 38393 | 38415 | - | 0 | ACGGTCGCCAGTACTTGCAT | AGG | \|\|\|\|\|\|\|\|[\|\|\|\|\|\|\|\|\|\|\|\|]PAM | NA | Intergenic | NA | NA |
| T31391 | CGGTCGCCAGTACTTGCATAGGG | | 1000 | | | | | | | | | |
| Oligo fwd | CACCCGGTCGCCAGTACTTGCATA | | | | | | | | | | | |
| Oligo rev | AAACTATGCAAGTACTGGCGACCG | | | | | | | | | | | |
| Chromosome | start | | end | strand | MM | target_seq | PAM | alignment | distance | position | gene name | gene id |
| GU305914.1 | | 38392 | 38414 | - | 0 | CGGTCGCCAGTACTTGCATA | GGG | \|\|\|\|\|\|\|\|[\|\|\|\|\|\|\|\|\|\|\|\|]PAM | NA | Intergenic | NA | NA |
| T31392 | ATAGGGAGTAGCTGTTCTCCTGG | | 1000 | | | | | | | | | |
| Oligo fwd | CACCATAGGGAGTAGCTGTTCTCC | | | | | | | | | | | |
| Oligo rev | AAACGGAGAACAGCTACTCCCTAT | | | | | | | | | | | |
| Chromosome | start | | end | strand | MM | target_seq | PAM | alignment | distance | position | gene name | gene id |
| GU305914.1 | | 38375 | 38397 | - | 0 | ATAGGGAGTAGCTGTTCTCC | TGG | \|\|\|\|\|\|\|\|[\|\|\|\|\|\|\|\|\|\|\|\|]PAM | NA | Intergenic | NA | NA |
| T31393 | TAGGGAGTAGCTGTTCTCCTGGG | | 1000 | | | | | | | | | |
| Oligo fwd | CACCTAGGGAGTAGCTGTTCTCCT | | | | | | | | | | | |
| Oligo rev | AAACAGGAGAACAGCTACTCCCTA | | | | | | | | | | | |
| Chromosome | start | | end | strand | MM | target_seq | PAM | alignment | distance | position | gene name | gene id |
| GU305914.1 | | 38374 | 38396 | - | 0 | TAGGGAGTAGCTGTTCTCCT | GGG | \|\|\|\|\|\|\|\|[\|\|\|\|\|\|\|\|\|\|\|\|]PAM | NA | Intergenic | NA | NA |
| T31394 | CTCCTGGGTGCACTCGAACCAGG | | 1000 | | | | | | | | | |
| Oligo fwd | CACCCTCCTGGGTGCACTCGAACC | | | | | | | | | | | |
| Oligo rev | AAACGGTTCGAGTGCACCCAGGAG | | | | | | | | | | | |
| Chromosome | start | | end | strand | MM | target_seq | PAM | alignment | distance | position | gene name | gene id |
| GU305914.1 | | 38359 | 38381 | - | 0 | CTCCTGGGTGCACTCGAACC | AGG | \|\|\|\|\|\|\|\|[\|\|\|\|\|\|\|\|\|\|\|\|]PAM | NA | Intergenic | NA | NA |
| T31395 | GAACCAGGCTACGTAGCTCGTGG | | 1000 | | | | | | | | | |
| Oligo fwd | CACCGAACCAGGCTACGTAGCTCG | | | | | | | | | | | |
| Oligo rev | AAACCGAGCTACGTAGCCTGGTTC | | | | | | | | | | | |
| Chromosome | start | | end | strand | MM | target_seq | PAM | alignment | distance | position | gene name | gene id |
| GU305914.1 | | 38344 | 38366 | - | 0 | GAACCAGGCTACGTAGCTCG | TGG | \|\|\|\|\|\|\|\|[\|\|\|\|\|\|\|\|\|\|\|\|]PAM | NA | Intergenic | NA | NA |
| T31396 | TACGCCGTTTTCGTTCATGATGG | | 1000 | | | | | | | | | |
| Oligo fwd | CACCTACGCCGTTTTCGTTCATGA | | | | | | | | | | | |
| Oligo rev | AAACTCATGAACGAAAACGGCGTA | | | | | | | | | | | |
| Chromosome | start | | end | strand | MM | target_seq | PAM | alignment | distance | position | gene name | gene id |
| GU305914.1 | | 38317 | 38339 | - | 0 | TACGCCGTTTTCGTTCATGA | TGG | \|\|\|\|\|\|\|\|[\|\|\|\|\|\|\|\|\|\|\|\|]PAM | NA | Intergenic | NA | NA |
| T31397 | TTTTCGTTCATGATGGTGCGCGG | | 1000 | | | | | | | | | |
| Oligo fwd | CACCTTTTCGTTCATGATGGTGCG | | | | | | | | | | | |
| Oligo rev | AAACCGCACCATCATGAACGAAAA | | | | | | | | | | | |
| Chromosome | start | | end | strand | MM | target_seq | PAM | alignment | distance | position | gene name | gene id |

| | | | | | | | | | | | |
|---|---|---|---|---|---|---|---|---|---|---|---|
| GU305914.1 | 38310 | 38332 | - | 0 | TTTTCGTTCATGATGGTGCG | CGG | \|\|\|\|\|\|\|\|[\|\|\|\|\|\|\|\|\|\|\|\|]PAM | NA | Intergenic | NA | NA |
| T31398 | TGGTGCGCGGAGTCCAGCTCAGG | 1000 | | | | | | | | | |
| Oligo fwd | CACCTGGTGCGCGGAGTCCAGCTC | | | | | | | | | | |
| Oligo rev | AAACGAGCTGGACTCCGCGCACCA | | | | | | | | | | |
| Chromosome | start | end | strand | MM | target_seq | PAM | alignment | distance | position | gene name | gene id |
| GU305914.1 | 38297 | 38319 | - | 0 | TGGTGCGCGGAGTCCAGCTC | AGG | \|\|\|\|\|\|\|\|[\|\|\|\|\|\|\|\|\|\|\|\|]PAM | NA | Intergenic | NA | NA |
| T31399 | GGTGCGCGGAGTCCAGCTCAGGG | 1000 | | | | | | | | | |
| Oligo fwd | CACCGGTGCGCGGAGTCCAGCTCA | | | | | | | | | | |
| Oligo rev | AAACTGAGCTGGACTCCGCGCACC | | | | | | | | | | |
| Chromosome | start | end | strand | MM | target_seq | PAM | alignment | distance | position | gene name | gene id |
| GU305914.1 | 38296 | 38318 | - | 0 | GGTGCGCGGAGTCCAGCTCA | GGG | \|\|\|\|\|\|\|\|[\|\|\|\|\|\|\|\|\|\|\|\|]PAM | NA | Intergenic | NA | NA |
| T31400 | CACCCCGCCCGTCCTCCGCACGG | 1000 | | | | | | | | | |
| Oligo fwd | CACCCACCCCGCCCGTCCTCCGCA | | | | | | | | | | |
| Oligo rev | AAACTGCGGAGGACGGGCGGGGTG | | | | | | | | | | |
| Chromosome | start | end | strand | MM | target_seq | PAM | alignment | distance | position | gene name | gene id |
| GU305914.1 | 38254 | 38276 | - | 0 | CACCCCGCCCGTCCTCCGCA | CGG | \|\|\|\|\|\|\|\|[\|\|\|\|\|\|\|\|\|\|\|\|]PAM | NA | Intergenic | NA | NA |
| T31401 | ACCCCGCCCGTCCTCCGCACGGG | 1000 | | | | | | | | | |
| Oligo fwd | CACCACCCCGCCCGTCCTCCGCAC | | | | | | | | | | |
| Oligo rev | AAACGTGCGGAGGACGGGCGGGGT | | | | | | | | | | |
| Chromosome | start | end | strand | MM | target_seq | PAM | alignment | distance | position | gene name | gene id |
| GU305914.1 | 38253 | 38275 | - | 0 | ACCCCGCCCGTCCTCCGCAC | GGG | \|\|\|\|\|\|\|\|[\|\|\|\|\|\|\|\|\|\|\|\|]PAM | NA | Intergenic | NA | NA |
| T31402 | CCCCGCCCGTCCTCCGCACGGGG | 1000 | | | | | | | | | |
| Oligo fwd | CACCCCCCGCCCGTCCTCCGCACG | | | | | | | | | | |
| Oligo rev | AAACCGTGCGGAGGACGGGCGGGG | | | | | | | | | | |
| Chromosome | start | end | strand | MM | target_seq | PAM | alignment | distance | position | gene name | gene id |
| GU305914.1 | 38252 | 38274 | - | 0 | CCCCGCCCGTCCTCCGCACG | GGG | \|\|\|\|\|\|\|\|[\|\|\|\|\|\|\|\|\|\|\|\|]PAM | NA | Intergenic | NA | NA |
| T31403 | CCCGCCCGTCCTCCGCACGGGGG | 1000 | | | | | | | | | |
| Oligo fwd | CACCCCCGCCCGTCCTCCGCACGG | | | | | | | | | | |
| Oligo rev | AAACCCGTGCGGAGGACGGGCGGG | | | | | | | | | | |
| Chromosome | start | end | strand | MM | target_seq | PAM | alignment | distance | position | gene name | gene id |
| GU305914.1 | 38251 | 38273 | - | 0 | CCCGCCCGTCCTCCGCACGG | GGG | \|\|\|\|\|\|\|\|[\|\|\|\|\|\|\|\|\|\|\|\|]PAM | NA | Intergenic | NA | NA |
| T31404 | GATGCCGAAGCTGATGCCGCTGG | 1000 | | | | | | | | | |
| Oligo fwd | CACCGATGCCGAAGCTGATGCCGC | | | | | | | | | | |
| Oligo rev | AAACGCGGCATCAGCTTCGGCATC | | | | | | | | | | |
| Chromosome | start | end | strand | MM | target_seq | PAM | alignment | distance | position | gene name | gene id |
| GU305914.1 | 38224 | 38246 | - | 0 | GATGCCGAAGCTGATGCCGC | TGG | \|\|\|\|\|\|\|\|[\|\|\|\|\|\|\|\|\|\|\|\|]PAM | NA | Intergenic | NA | NA |
| GU305914.1 | 62684 | 62706 | - | 4 | GCTGCCGCGGCTGCTGCCGC | CGG | \|-\|\|\|\|\|-[-\|\|\|\|-\|\|\|\|\|\|]PAM | NA | Intergenic | NA | NA |
| T31405 | CGAAGCTGATGCCGCTGGCGCGG | 1000 | | | | | | | | | |
| Oligo fwd | CACCCGAAGCTGATGCCGCTGGCG | | | | | | | | | | |
| Oligo rev | AAACCGCCAGCGGCATCAGCTTCG | | | | | | | | | | |
| Chromosome | start | end | strand | MM | target_seq | PAM | alignment | distance | position | gene name | gene id |
| GU305914.1 | 38219 | 38241 | - | 0 | CGAAGCTGATGCCGCTGGCG | CGG | \|\|\|\|\|\|\|\|[\|\|\|\|\|\|\|\|\|\|\|\|]PAM | NA | Intergenic | NA | NA |
| GU305914.1 | 62679 | 62701 | - | 4 | CGCGGCTGCTGCCGCCGGCG | GGG | \|\|--\|\|\|\|[-\|\|\|\|\|\|-\|\|\|\|]PAM | NA | Intergenic | NA | NA |
| T31406 | AGCATGCGCACGCCGAAGTCCGG | 1000 | | | | | | | | | |
| Oligo fwd | CACCAGCATGCGCACGCCGAAGTC | | | | | | | | | | |
| Oligo rev | AAACGACTTCGGCGTGCGCATGCT | | | | | | | | | | |
| Chromosome | start | end | strand | MM | target_seq | PAM | alignment | distance | position | gene name | gene id |
| GU305914.1 | 38175 | 38197 | - | 0 | AGCATGCGCACGCCGAAGTC | CGG | \|\|\|\|\|\|\|\|[\|\|\|\|\|\|\|\|\|\|\|\|]PAM | NA | Intergenic | NA | NA |
| T31407 | CCGAAGTCCGGCGCGACGTACGG | 1000 | | | | | | | | | |
| Oligo fwd | CACCCCGAAGTCCGGCGCGACGTA | | | | | | | | | | |
| Oligo rev | AAACTACGTCGCGCCGGACTTCGG | | | | | | | | | | |
| Chromosome | start | end | strand | MM | target_seq | PAM | alignment | distance | position | gene name | gene id |
| GU305914.1 | 38163 | 38185 | - | 0 | CCGAAGTCCGGCGCGACGTA | CGG | \|\|\|\|\|\|\|\|[\|\|\|\|\|\|\|\|\|\|\|\|]PAM | NA | Intergenic | NA | NA |
| T31408 | CGCCGACAGCTCCAGCCTCACGG | 1000 | | | | | | | | | |
| Oligo fwd | CACCCGCCGACAGCTCCAGCCTCA | | | | | | | | | | |
| Oligo rev | AAACTGAGGCTGGAGCTGTCGGCG | | | | | | | | | | |
| Chromosome | start | end | strand | MM | target_seq | PAM | alignment | distance | position | gene name | gene id |
| GU305914.1 | 38128 | 38150 | - | 0 | CGCCGACAGCTCCAGCCTCA | CGG | \|\|\|\|\|\|\|\|[\|\|\|\|\|\|\|\|\|\|\|\|]PAM | NA | Intergenic | NA | NA |
| GU305914.1 | 140236 | 140258 | - | 4 | CGGCGTCAGCTCCACGCTCA | TGG | \|\|-\|\|-\|\|[\|\|\|\|\|\|--\|\|\|\|]PAM | NA | Intergenic | NA | NA |
| T31409 | GCCGACAGCTCCAGCCTCACGGG | 1000 | | | | | | | | | |
| Oligo fwd | CACCGCCGACAGCTCCAGCCTCAC | | | | | | | | | | |
| Oligo rev | AAACGTGAGGCTGGAGCTGTCGGC | | | | | | | | | | |
| Chromosome | start | end | strand | MM | target_seq | PAM | alignment | distance | position | gene name | gene id |
| GU305914.1 | 38127 | 38149 | - | 0 | GCCGACAGCTCCAGCCTCAC | GGG | \|\|\|\|\|\|\|\|[\|\|\|\|\|\|\|\|\|\|\|\|]PAM | NA | Intergenic | NA | NA |
| T31410 | AGCCTCACGGGCCTGTTGTCCGG | 1000 | | | | | | | | | |
| Oligo fwd | CACCAGCCTCACGGGCCTGTTGTC | | | | | | | | | | |
| Oligo rev | AAACGACAACAGGCCCGTGAGGCT | | | | | | | | | | |
| Chromosome | start | end | strand | MM | target_seq | PAM | alignment | distance | position | gene name | gene id |
| GU305914.1 | 38115 | 38137 | - | 0 | AGCCTCACGGGCCTGTTGTC | CGG | \|\|\|\|\|\|\|\|[\|\|\|\|\|\|\|\|\|\|\|\|]PAM | NA | Intergenic | NA | NA |
| T31411 | CTCACGGGCCTGTTGTCCGGCGG | 1000 | | | | | | | | | |
| Oligo fwd | CACCCTCACGGGCCTGTTGTCCGG | | | | | | | | | | |
| Oligo rev | AAACCCGGACAACAGGCCCGTGAG | | | | | | | | | | |
| Chromosome | start | end | strand | MM | target_seq | PAM | alignment | distance | position | gene name | gene id |
| GU305914.1 | 38112 | 38134 | - | 0 | CTCACGGGCCTGTTGTCCGG | CGG | \|\|\|\|\|\|\|\|[\|\|\|\|\|\|\|\|\|\|\|\|]PAM | NA | Intergenic | NA | NA |

<table>
<tr><td>T31412</td><td>CCTGTTGTCCGGCGGCACGTTGG</td><td>1000</td><td></td><td></td><td></td><td></td><td></td><td></td><td></td><td></td><td></td></tr>
<tr><td>Oligo fwd</td><td>CACCCCTGTTGTCCGGCGGCACGT</td><td></td><td></td><td></td><td></td><td></td><td></td><td></td><td></td><td></td><td></td></tr>
<tr><td>Oligo rev</td><td>AAACACGTGCCGCCGGACAACAGG</td><td></td><td></td><td></td><td></td><td></td><td></td><td></td><td></td><td></td><td></td></tr>
<tr><td>Chromosome</td><td>start</td><td>end</td><td>strand</td><td>MM</td><td>target_seq</td><td>PAM</td><td>alignment</td><td>distance</td><td>position</td><td>gene name</td><td>gene id</td></tr>
<tr><td>GU305914.1</td><td>38104</td><td>38126</td><td>-</td><td>0</td><td>CCTGTTGTCCGGCGGCACGT</td><td>TGG</td><td>||||||||[||||||||||||]PAM</td><td>NA</td><td>Intergenic</td><td>NA</td><td>NA</td></tr>
<tr><td>T31413</td><td>TTGTCCGGCGGCACGTTGGACGG</td><td>1000</td><td></td><td></td><td></td><td></td><td></td><td></td><td></td><td></td><td></td></tr>
<tr><td>Oligo fwd</td><td>CACCTTGTCCGGCGGCACGTTGGA</td><td></td><td></td><td></td><td></td><td></td><td></td><td></td><td></td><td></td><td></td></tr>
<tr><td>Oligo rev</td><td>AAACTCCAACGTGCCGCCGGACAA</td><td></td><td></td><td></td><td></td><td></td><td></td><td></td><td></td><td></td><td></td></tr>
<tr><td>Chromosome</td><td>start</td><td>end</td><td>strand</td><td>MM</td><td>target_seq</td><td>PAM</td><td>alignment</td><td>distance</td><td>position</td><td>gene name</td><td>gene id</td></tr>
<tr><td>GU305914.1</td><td>38100</td><td>38122</td><td>-</td><td>0</td><td>TTGTCCGGCGGCACGTTGGA</td><td>CGG</td><td>||||||||[||||||||||||]PAM</td><td>NA</td><td>Intergenic</td><td>NA</td><td>NA</td></tr>
<tr><td>T31414</td><td>TGTCCGGCGGCACGTTGGACGGG</td><td>1000</td><td></td><td></td><td></td><td></td><td></td><td></td><td></td><td></td><td></td></tr>
<tr><td>Oligo fwd</td><td>CACCTGTCCGGCGGCACGTTGGAC</td><td></td><td></td><td></td><td></td><td></td><td></td><td></td><td></td><td></td><td></td></tr>
<tr><td>Oligo rev</td><td>AAACGTCCAACGTGCCGCCGGACA</td><td></td><td></td><td></td><td></td><td></td><td></td><td></td><td></td><td></td><td></td></tr>
<tr><td>Chromosome</td><td>start</td><td>end</td><td>strand</td><td>MM</td><td>target_seq</td><td>PAM</td><td>alignment</td><td>distance</td><td>position</td><td>gene name</td><td>gene id</td></tr>
<tr><td>GU305914.1</td><td>38099</td><td>38121</td><td>-</td><td>0</td><td>TGTCCGGCGGCACGTTGGAC</td><td>GGG</td><td>||||||||[||||||||||||]PAM</td><td>NA</td><td>Intergenic</td><td>NA</td><td>NA</td></tr>
<tr><td>T31415</td><td>GCGGCACGTTGGACGGGTCGTGG</td><td>1000</td><td></td><td></td><td></td><td></td><td></td><td></td><td></td><td></td><td></td></tr>
<tr><td>Oligo fwd</td><td>CACCGCGGCACGTTGGACGGGTCG</td><td></td><td></td><td></td><td></td><td></td><td></td><td></td><td></td><td></td><td></td></tr>
<tr><td>Oligo rev</td><td>AAACCGACCCGTCCAACGTGCCGC</td><td></td><td></td><td></td><td></td><td></td><td></td><td></td><td></td><td></td><td></td></tr>
<tr><td>Chromosome</td><td>start</td><td>end</td><td>strand</td><td>MM</td><td>target_seq</td><td>PAM</td><td>alignment</td><td>distance</td><td>position</td><td>gene name</td><td>gene id</td></tr>
<tr><td>GU305914.1</td><td>38093</td><td>38115</td><td>-</td><td>0</td><td>GCGGCACGTTGGACGGGTCG</td><td>TGG</td><td>||||||||[||||||||||||]PAM</td><td>NA</td><td>Intergenic</td><td>NA</td><td>NA</td></tr>
<tr><td>T31416</td><td>CGGCACGTTGGACGGGTCGTGGG</td><td>1000</td><td></td><td></td><td></td><td></td><td></td><td></td><td></td><td></td><td></td></tr>
<tr><td>Oligo fwd</td><td>CACCCGGCACGTTGGACGGGTCGT</td><td></td><td></td><td></td><td></td><td></td><td></td><td></td><td></td><td></td><td></td></tr>
<tr><td>Oligo rev</td><td>AAACACGACCCGTCCAACGTGCCG</td><td></td><td></td><td></td><td></td><td></td><td></td><td></td><td></td><td></td><td></td></tr>
<tr><td>Chromosome</td><td>start</td><td>end</td><td>strand</td><td>MM</td><td>target_seq</td><td>PAM</td><td>alignment</td><td>distance</td><td>position</td><td>gene name</td><td>gene id</td></tr>
<tr><td>GU305914.1</td><td>38092</td><td>38114</td><td>-</td><td>0</td><td>CGGCACGTTGGACGGGTCGT</td><td>GGG</td><td>||||||||[||||||||||||]PAM</td><td>NA</td><td>Intergenic</td><td>NA</td><td>NA</td></tr>
<tr><td>T31417</td><td>TGGACGGGTCGTGGGCCAGCAGG</td><td>1000</td><td></td><td></td><td></td><td></td><td></td><td></td><td></td><td></td><td></td></tr>
<tr><td>Oligo fwd</td><td>CACCTGGACGGGTCGTGGGCCAGC</td><td></td><td></td><td></td><td></td><td></td><td></td><td></td><td></td><td></td><td></td></tr>
<tr><td>Oligo rev</td><td>AAACGCTGGCCCACGACCCGTCCA</td><td></td><td></td><td></td><td></td><td></td><td></td><td></td><td></td><td></td><td></td></tr>
<tr><td>Chromosome</td><td>start</td><td>end</td><td>strand</td><td>MM</td><td>target_seq</td><td>PAM</td><td>alignment</td><td>distance</td><td>position</td><td>gene name</td><td>gene id</td></tr>
<tr><td>GU305914.1</td><td>38084</td><td>38106</td><td>-</td><td>0</td><td>TGGACGGGTCGTGGGCCAGC</td><td>AGG</td><td>||||||||[||||||||||||]PAM</td><td>NA</td><td>Intergenic</td><td>NA</td><td>NA</td></tr>
<tr><td>T31418</td><td>CGGGTCGTGGGCCAGCAGGATGG</td><td>1000</td><td></td><td></td><td></td><td></td><td></td><td></td><td></td><td></td><td></td></tr>
<tr><td>Oligo fwd</td><td>CACCCGGGTCGTGGGCCAGCAGGA</td><td></td><td></td><td></td><td></td><td></td><td></td><td></td><td></td><td></td><td></td></tr>
<tr><td>Oligo rev</td><td>AAACTCCTGCTGGCCCACGACCCG</td><td></td><td></td><td></td><td></td><td></td><td></td><td></td><td></td><td></td><td></td></tr>
<tr><td>Chromosome</td><td>start</td><td>end</td><td>strand</td><td>MM</td><td>target_seq</td><td>PAM</td><td>alignment</td><td>distance</td><td>position</td><td>gene name</td><td>gene id</td></tr>
<tr><td>GU305914.1</td><td>38080</td><td>38102</td><td>-</td><td>0</td><td>CGGGTCGTGGGCCAGCAGGA</td><td>TGG</td><td>||||||||[||||||||||||]PAM</td><td>NA</td><td>Intergenic</td><td>NA</td><td>NA</td></tr>
<tr><td>T31419</td><td>TGGGCCAGCAGGATGGTGTGCGG</td><td>1000</td><td></td><td></td><td></td><td></td><td></td><td></td><td></td><td></td><td></td></tr>
<tr><td>Oligo fwd</td><td>CACCTGGGCCAGCAGGATGGTGTG</td><td></td><td></td><td></td><td></td><td></td><td></td><td></td><td></td><td></td><td></td></tr>
<tr><td>Oligo rev</td><td>AAACCACACCATCCTGCTGGCCCA</td><td></td><td></td><td></td><td></td><td></td><td></td><td></td><td></td><td></td><td></td></tr>
<tr><td>Chromosome</td><td>start</td><td>end</td><td>strand</td><td>MM</td><td>target_seq</td><td>PAM</td><td>alignment</td><td>distance</td><td>position</td><td>gene name</td><td>gene id</td></tr>
<tr><td>GU305914.1</td><td>38073</td><td>38095</td><td>-</td><td>0</td><td>TGGGCCAGCAGGATGGTGTG</td><td>CGG</td><td>||||||||[||||||||||||]PAM</td><td>NA</td><td>Intergenic</td><td>NA</td><td>NA</td></tr>
<tr><td>T31420</td><td>GGGCCAGCAGGATGGTGTGCGGG</td><td>1000</td><td></td><td></td><td></td><td></td><td></td><td></td><td></td><td></td><td></td></tr>
<tr><td>Oligo fwd</td><td>CACCGGGCCAGCAGGATGGTGTGC</td><td></td><td></td><td></td><td></td><td></td><td></td><td></td><td></td><td></td><td></td></tr>
<tr><td>Oligo rev</td><td>AAACGCACACCATCCTGCTGGCCC</td><td></td><td></td><td></td><td></td><td></td><td></td><td></td><td></td><td></td><td></td></tr>
<tr><td>Chromosome</td><td>start</td><td>end</td><td>strand</td><td>MM</td><td>target_seq</td><td>PAM</td><td>alignment</td><td>distance</td><td>position</td><td>gene name</td><td>gene id</td></tr>
<tr><td>GU305914.1</td><td>38072</td><td>38094</td><td>-</td><td>0</td><td>GGGCCAGCAGGATGGTGTGC</td><td>GGG</td><td>||||||||[||||||||||||]PAM</td><td>NA</td><td>Intergenic</td><td>NA</td><td>NA</td></tr>
<tr><td>T31421</td><td>GATGGTGTGCGGGCCACCCGTGG</td><td>1000</td><td></td><td></td><td></td><td></td><td></td><td></td><td></td><td></td><td></td></tr>
<tr><td>Oligo fwd</td><td>CACCGATGGTGTGCGGGCCACCCG</td><td></td><td></td><td></td><td></td><td></td><td></td><td></td><td></td><td></td><td></td></tr>
<tr><td>Oligo rev</td><td>AAACCGGGTGGCCCGCACACCATC</td><td></td><td></td><td></td><td></td><td></td><td></td><td></td><td></td><td></td><td></td></tr>
<tr><td>Chromosome</td><td>start</td><td>end</td><td>strand</td><td>MM</td><td>target_seq</td><td>PAM</td><td>alignment</td><td>distance</td><td>position</td><td>gene name</td><td>gene id</td></tr>
<tr><td>GU305914.1</td><td>38062</td><td>38084</td><td>-</td><td>0</td><td>GATGGTGTGCGGGCCACCCG</td><td>TGG</td><td>||||||||[||||||||||||]PAM</td><td>NA</td><td>Intergenic</td><td>NA</td><td>NA</td></tr>
<tr><td>T31422</td><td>GTGCGGGCCACCCGTGGTCACGG</td><td>1000</td><td></td><td></td><td></td><td></td><td></td><td></td><td></td><td></td><td></td></tr>
<tr><td>Oligo fwd</td><td>CACCGTGCGGGCCACCCGTGGTCA</td><td></td><td></td><td></td><td></td><td></td><td></td><td></td><td></td><td></td><td></td></tr>
<tr><td>Oligo rev</td><td>AAACTGACCACGGGTGGCCCGCAC</td><td></td><td></td><td></td><td></td><td></td><td></td><td></td><td></td><td></td><td></td></tr>
<tr><td>Chromosome</td><td>start</td><td>end</td><td>strand</td><td>MM</td><td>target_seq</td><td>PAM</td><td>alignment</td><td>distance</td><td>position</td><td>gene name</td><td>gene id</td></tr>
<tr><td>GU305914.1</td><td>38056</td><td>38078</td><td>-</td><td>0</td><td>GTGCGGGCCACCCGTGGTCA</td><td>CGG</td><td>||||||||[||||||||||||]PAM</td><td>NA</td><td>Intergenic</td><td>NA</td><td>NA</td></tr>
<tr><td>T31423</td><td>GGGCCACCCGTGGTCACGGTCGG</td><td>1000</td><td></td><td></td><td></td><td></td><td></td><td></td><td></td><td></td><td></td></tr>
<tr><td>Oligo fwd</td><td>CACCGGGCCACCCGTGGTCACGGT</td><td></td><td></td><td></td><td></td><td></td><td></td><td></td><td></td><td></td><td></td></tr>
<tr><td>Oligo rev</td><td>AAACACCGTGACCACGGGTGGCCC</td><td></td><td></td><td></td><td></td><td></td><td></td><td></td><td></td><td></td><td></td></tr>
<tr><td>Chromosome</td><td>start</td><td>end</td><td>strand</td><td>MM</td><td>target_seq</td><td>PAM</td><td>alignment</td><td>distance</td><td>position</td><td>gene name</td><td>gene id</td></tr>
<tr><td>GU305914.1</td><td>38052</td><td>38074</td><td>-</td><td>0</td><td>GGGCCACCCGTGGTCACGGT</td><td>CGG</td><td>||||||||[||||||||||||]PAM</td><td>NA</td><td>Intergenic</td><td>NA</td><td>NA</td></tr>
<tr><td>T31424</td><td>ACACCGCTGCGTTTCCCTTCAGG</td><td>1000</td><td></td><td></td><td></td><td></td><td></td><td></td><td></td><td></td><td></td></tr>
<tr><td>Oligo fwd</td><td>CACCACACCGCTGCGTTTCCCTTC</td><td></td><td></td><td></td><td></td><td></td><td></td><td></td><td></td><td></td><td></td></tr>
<tr><td>Oligo rev</td><td>AAACGAAGGGAAACGCAGCGGTGT</td><td></td><td></td><td></td><td></td><td></td><td></td><td></td><td></td><td></td><td></td></tr>
<tr><td>Chromosome</td><td>start</td><td>end</td><td>strand</td><td>MM</td><td>target_seq</td><td>PAM</td><td>alignment</td><td>distance</td><td>position</td><td>gene name</td><td>gene id</td></tr>
<tr><td>GU305914.1</td><td>38024</td><td>38046</td><td>-</td><td>0</td><td>ACACCGCTGCGTTTCCCTTC</td><td>AGG</td><td>||||||||[||||||||||||]PAM</td><td>NA</td><td>Intergenic</td><td>NA</td><td>NA</td></tr>
<tr><td>T31425</td><td>CACCGCTGCGTTTCCCTTCAGGG</td><td>1000</td><td></td><td></td><td></td><td></td><td></td><td></td><td></td><td></td><td></td></tr>
<tr><td>Oligo fwd</td><td>CACCCACCGCTGCGTTTCCCTTCA</td><td></td><td></td><td></td><td></td><td></td><td></td><td></td><td></td><td></td><td></td></tr>
<tr><td>Oligo rev</td><td>AAACTGAAGGGAAACGCAGCGGTG</td><td></td><td></td><td></td><td></td><td></td><td></td><td></td><td></td><td></td><td></td></tr>
<tr><td>Chromosome</td><td>start</td><td>end</td><td>strand</td><td>MM</td><td>target_seq</td><td>PAM</td><td>alignment</td><td>distance</td><td>position</td><td>gene name</td><td>gene id</td></tr>
<tr><td>GU305914.1</td><td>38023</td><td>38045</td><td>-</td><td>0</td><td>CACCGCTGCGTTTCCCTTCA</td><td>GGG</td><td>||||||||[||||||||||||]PAM</td><td>NA</td><td>Intergenic</td><td>NA</td><td>NA</td></tr>
<tr><td>T31426</td><td>TTCCCTTCAGGGAGCACAGCAGG</td><td>1000</td><td></td><td></td><td></td><td></td><td></td><td></td><td></td><td></td><td></td></tr>
<tr><td>Oligo fwd</td><td>CACCTTCCCTTCAGGGAGCACAGC</td><td></td><td></td><td></td><td></td><td></td><td></td><td></td><td></td><td></td><td></td></tr>
<tr><td>Oligo rev</td><td>AAACGCTGTGCTCCCTGAAGGGAA</td><td></td><td></td><td></td><td></td><td></td><td></td><td></td><td></td><td></td><td></td></tr>
<tr><td>Chromosome</td><td>start</td><td>end</td><td>strand</td><td>MM</td><td>target_seq</td><td>PAM</td><td>alignment</td><td>distance</td><td>position</td><td>gene name</td><td>gene id</td></tr>
<tr><td>GU305914.1</td><td>38012</td><td>38034</td><td>-</td><td>0</td><td>TTCCCTTCAGGGAGCACAGC</td><td>AGG</td><td>||||||||[||||||||||||]PAM</td><td>NA</td><td>Intergenic</td><td>NA</td><td>NA</td></tr>
</table>

<table>
<tr><td>T31427</td><td>CACAGCAGGAACTGTCCGCCCGG</td><td>1000</td><td></td><td></td><td></td><td></td><td></td><td></td><td></td><td></td><td></td></tr>
<tr><td>Oligo fwd</td><td>CACCCACAGCAGGAACTGTCCGCC</td><td></td><td></td><td></td><td></td><td></td><td></td><td></td><td></td><td></td><td></td></tr>
<tr><td>Oligo rev</td><td>AAACGGCGGACAGTTCCTGCTGTG</td><td></td><td></td><td></td><td></td><td></td><td></td><td></td><td></td><td></td><td></td></tr>
<tr><td>Chromosome</td><td>start</td><td>end</td><td>strand</td><td>MM</td><td>target_seq</td><td>PAM</td><td>alignment</td><td>distance</td><td>position</td><td>gene name</td><td>gene id</td></tr>
<tr><td>GU305914.1</td><td>37998</td><td>38020</td><td>-</td><td>0</td><td>CACAGCAGGAACTGTCCGCC</td><td>CGG</td><td>||||||||[||||||||||||]PAM</td><td>NA</td><td>Intergenic</td><td>NA</td><td>NA</td></tr>
<tr><td>T31428</td><td>ACAGCAGGAACTGTCCGCCCGGG</td><td>1000</td><td></td><td></td><td></td><td></td><td></td><td></td><td></td><td></td><td></td></tr>
<tr><td>Oligo fwd</td><td>CACCACAGCAGGAACTGTCCGCCC</td><td></td><td></td><td></td><td></td><td></td><td></td><td></td><td></td><td></td><td></td></tr>
<tr><td>Oligo rev</td><td>AAACGGGCGGACAGTTCCTGCTGT</td><td></td><td></td><td></td><td></td><td></td><td></td><td></td><td></td><td></td><td></td></tr>
<tr><td>Chromosome</td><td>start</td><td>end</td><td>strand</td><td>MM</td><td>target_seq</td><td>PAM</td><td>alignment</td><td>distance</td><td>position</td><td>gene name</td><td>gene id</td></tr>
<tr><td>GU305914.1</td><td>37997</td><td>38019</td><td>-</td><td>0</td><td>ACAGCAGGAACTGTCCGCCC</td><td>GGG</td><td>||||||||[||||||||||||]PAM</td><td>NA</td><td>Intergenic</td><td>NA</td><td>NA</td></tr>
<tr><td>T31429</td><td>CCCGGGATGTTCGAGACGCGAGG</td><td>1000</td><td></td><td></td><td></td><td></td><td></td><td></td><td></td><td></td><td></td></tr>
<tr><td>Oligo fwd</td><td>CACCCCCGGGATGTTCGAGACGCG</td><td></td><td></td><td></td><td></td><td></td><td></td><td></td><td></td><td></td><td></td></tr>
<tr><td>Oligo rev</td><td>AAACCGCGTCTCGAACATCCCGGG</td><td></td><td></td><td></td><td></td><td></td><td></td><td></td><td></td><td></td><td></td></tr>
<tr><td>Chromosome</td><td>start</td><td>end</td><td>strand</td><td>MM</td><td>target_seq</td><td>PAM</td><td>alignment</td><td>distance</td><td>position</td><td>gene name</td><td>gene id</td></tr>
<tr><td>GU305914.1</td><td>37980</td><td>38002</td><td>-</td><td>0</td><td>CCCGGGATGTTCGAGACGCG</td><td>AGG</td><td>||||||||[||||||||||||]PAM</td><td>NA</td><td>Intergenic</td><td>NA</td><td>NA</td></tr>
<tr><td>T31430</td><td>CCGGGATGTTCGAGACGCGAGGG</td><td>1000</td><td></td><td></td><td></td><td></td><td></td><td></td><td></td><td></td><td></td></tr>
<tr><td>Oligo fwd</td><td>CACCCCGGGATGTTCGAGACGCGA</td><td></td><td></td><td></td><td></td><td></td><td></td><td></td><td></td><td></td><td></td></tr>
<tr><td>Oligo rev</td><td>AAACTCGCGTCTCGAACATCCCGG</td><td></td><td></td><td></td><td></td><td></td><td></td><td></td><td></td><td></td><td></td></tr>
<tr><td>Chromosome</td><td>start</td><td>end</td><td>strand</td><td>MM</td><td>target_seq</td><td>PAM</td><td>alignment</td><td>distance</td><td>position</td><td>gene name</td><td>gene id</td></tr>
<tr><td>GU305914.1</td><td>37979</td><td>38001</td><td>-</td><td>0</td><td>CCGGGATGTTCGAGACGCGA</td><td>GGG</td><td>||||||||[||||||||||||]PAM</td><td>NA</td><td>Intergenic</td><td>NA</td><td>NA</td></tr>
<tr><td>T31431</td><td>GACGCGAGGGACCAGCAGCGTGG</td><td>1000</td><td></td><td></td><td></td><td></td><td></td><td></td><td></td><td></td><td></td></tr>
<tr><td>Oligo fwd</td><td>CACCGACGCGAGGGACCAGCAGCG</td><td></td><td></td><td></td><td></td><td></td><td></td><td></td><td></td><td></td><td></td></tr>
<tr><td>Oligo rev</td><td>AAACCGCTGCTGGTCCCTCGCGTC</td><td></td><td></td><td></td><td></td><td></td><td></td><td></td><td></td><td></td><td></td></tr>
<tr><td>Chromosome</td><td>start</td><td>end</td><td>strand</td><td>MM</td><td>target_seq</td><td>PAM</td><td>alignment</td><td>distance</td><td>position</td><td>gene name</td><td>gene id</td></tr>
<tr><td>GU305914.1</td><td>37966</td><td>37988</td><td>-</td><td>0</td><td>GACGCGAGGGACCAGCAGCG</td><td>TGG</td><td>||||||||[||||||||||||]PAM</td><td>NA</td><td>Intergenic</td><td>NA</td><td>NA</td></tr>
<tr><td>T31432</td><td>AGGGACCAGCAGCGTGGTCCAGG</td><td>1000</td><td></td><td></td><td></td><td></td><td></td><td></td><td></td><td></td><td></td></tr>
<tr><td>Oligo fwd</td><td>CACCAGGGACCAGCAGCGTGGTCC</td><td></td><td></td><td></td><td></td><td></td><td></td><td></td><td></td><td></td><td></td></tr>
<tr><td>Oligo rev</td><td>AAACGGACCACGCTGCTGGTCCCT</td><td></td><td></td><td></td><td></td><td></td><td></td><td></td><td></td><td></td><td></td></tr>
<tr><td>Chromosome</td><td>start</td><td>end</td><td>strand</td><td>MM</td><td>target_seq</td><td>PAM</td><td>alignment</td><td>distance</td><td>position</td><td>gene name</td><td>gene id</td></tr>
<tr><td>GU305914.1</td><td>37960</td><td>37982</td><td>-</td><td>0</td><td>AGGGACCAGCAGCGTGGTCC</td><td>AGG</td><td>||||||||[||||||||||||]PAM</td><td>NA</td><td>Intergenic</td><td>NA</td><td>NA</td></tr>
<tr><td>T31433</td><td>GGTGTACGTCAGCTCGTCCGAGG</td><td>1000</td><td></td><td></td><td></td><td></td><td></td><td></td><td></td><td></td><td></td></tr>
<tr><td>Oligo fwd</td><td>CACCGGTGTACGTCAGCTCGTCCG</td><td></td><td></td><td></td><td></td><td></td><td></td><td></td><td></td><td></td><td></td></tr>
<tr><td>Oligo rev</td><td>AAACCGGACGAGCTGACGTACACC</td><td></td><td></td><td></td><td></td><td></td><td></td><td></td><td></td><td></td><td></td></tr>
<tr><td>Chromosome</td><td>start</td><td>end</td><td>strand</td><td>MM</td><td>target_seq</td><td>PAM</td><td>alignment</td><td>distance</td><td>position</td><td>gene name</td><td>gene id</td></tr>
<tr><td>GU305914.1</td><td>37939</td><td>37961</td><td>-</td><td>0</td><td>GGTGTACGTCAGCTCGTCCG</td><td>AGG</td><td>||||||||[||||||||||||]PAM</td><td>NA</td><td>Intergenic</td><td>NA</td><td>NA</td></tr>
<tr><td>T31434</td><td>ACGTCAGCTCGTCCGAGGTGCGG</td><td>1000</td><td></td><td></td><td></td><td></td><td></td><td></td><td></td><td></td><td></td></tr>
<tr><td>Oligo fwd</td><td>CACCACGTCAGCTCGTCCGAGGTG</td><td></td><td></td><td></td><td></td><td></td><td></td><td></td><td></td><td></td><td></td></tr>
<tr><td>Oligo rev</td><td>AAACCACCTCGGACGAGCTGACGT</td><td></td><td></td><td></td><td></td><td></td><td></td><td></td><td></td><td></td><td></td></tr>
<tr><td>Chromosome</td><td>start</td><td>end</td><td>strand</td><td>MM</td><td>target_seq</td><td>PAM</td><td>alignment</td><td>distance</td><td>position</td><td>gene name</td><td>gene id</td></tr>
<tr><td>GU305914.1</td><td>37934</td><td>37956</td><td>-</td><td>0</td><td>ACGTCAGCTCGTCCGAGGTG</td><td>CGG</td><td>||||||||[||||||||||||]PAM</td><td>NA</td><td>Intergenic</td><td>NA</td><td>NA</td></tr>
<tr><td>T31435</td><td>CACCGCCGTCGCGAAGTGAATGG</td><td>1000</td><td></td><td></td><td></td><td></td><td></td><td></td><td></td><td></td><td></td></tr>
<tr><td>Oligo fwd</td><td>CACCCACCGCCGTCGCGAAGTGAA</td><td></td><td></td><td></td><td></td><td></td><td></td><td></td><td></td><td></td><td></td></tr>
<tr><td>Oligo rev</td><td>AAACTTCACTTCGCGACGGCGGTG</td><td></td><td></td><td></td><td></td><td></td><td></td><td></td><td></td><td></td><td></td></tr>
<tr><td>Chromosome</td><td>start</td><td>end</td><td>strand</td><td>MM</td><td>target_seq</td><td>PAM</td><td>alignment</td><td>distance</td><td>position</td><td>gene name</td><td>gene id</td></tr>
<tr><td>GU305914.1</td><td>37897</td><td>37919</td><td>-</td><td>0</td><td>CACCGCCGTCGCGAAGTGAA</td><td>TGG</td><td>||||||||[||||||||||||]PAM</td><td>NA</td><td>Intergenic</td><td>NA</td><td>NA</td></tr>
<tr><td>T31436</td><td>TAGCGAGCGCGCAGCTCCGACGG</td><td>1000</td><td></td><td></td><td></td><td></td><td></td><td></td><td></td><td></td><td></td></tr>
<tr><td>Oligo fwd</td><td>CACCTAGCGAGCGCGCAGCTCCGA</td><td></td><td></td><td></td><td></td><td></td><td></td><td></td><td></td><td></td><td></td></tr>
<tr><td>Oligo rev</td><td>AAACTCGGAGCTGCGCGCTCGCTA</td><td></td><td></td><td></td><td></td><td></td><td></td><td></td><td></td><td></td><td></td></tr>
<tr><td>Chromosome</td><td>start</td><td>end</td><td>strand</td><td>MM</td><td>target_seq</td><td>PAM</td><td>alignment</td><td>distance</td><td>position</td><td>gene name</td><td>gene id</td></tr>
<tr><td>GU305914.1</td><td>37863</td><td>37885</td><td>-</td><td>0</td><td>TAGCGAGCGCGCAGCTCCGA</td><td>CGG</td><td>||||||||[||||||||||||]PAM</td><td>NA</td><td>Intergenic</td><td>NA</td><td>NA</td></tr>
<tr><td>T31437</td><td>ACGGCACCACCTCGTTGTACTGG</td><td>1000</td><td></td><td></td><td></td><td></td><td></td><td></td><td></td><td></td><td></td></tr>
<tr><td>Oligo fwd</td><td>CACCACGGCACCACCTCGTTGTAC</td><td></td><td></td><td></td><td></td><td></td><td></td><td></td><td></td><td></td><td></td></tr>
<tr><td>Oligo rev</td><td>AAACGTACAACGAGGTGGTGCCGT</td><td></td><td></td><td></td><td></td><td></td><td></td><td></td><td></td><td></td><td></td></tr>
<tr><td>Chromosome</td><td>start</td><td>end</td><td>strand</td><td>MM</td><td>target_seq</td><td>PAM</td><td>alignment</td><td>distance</td><td>position</td><td>gene name</td><td>gene id</td></tr>
<tr><td>GU305914.1</td><td>37844</td><td>37866</td><td>-</td><td>0</td><td>ACGGCACCACCTCGTTGTAC</td><td>TGG</td><td>||||||||[||||||||||||]PAM</td><td>NA</td><td>Intergenic</td><td>NA</td><td>NA</td></tr>
<tr><td>T31438</td><td>CACCACCTCGTTGTACTGGCTGG</td><td>1000</td><td></td><td></td><td></td><td></td><td></td><td></td><td></td><td></td><td></td></tr>
<tr><td>Oligo fwd</td><td>CACCCACCACCTCGTTGTACTGGC</td><td></td><td></td><td></td><td></td><td></td><td></td><td></td><td></td><td></td><td></td></tr>
<tr><td>Oligo rev</td><td>AAACGCCAGTACAACGAGGTGGTG</td><td></td><td></td><td></td><td></td><td></td><td></td><td></td><td></td><td></td><td></td></tr>
<tr><td>Chromosome</td><td>start</td><td>end</td><td>strand</td><td>MM</td><td>target_seq</td><td>PAM</td><td>alignment</td><td>distance</td><td>position</td><td>gene name</td><td>gene id</td></tr>
<tr><td>GU305914.1</td><td>37840</td><td>37862</td><td>-</td><td>0</td><td>CACCACCTCGTTGTACTGGC</td><td>TGG</td><td>||||||||[||||||||||||]PAM</td><td>NA</td><td>Intergenic</td><td>NA</td><td>NA</td></tr>
<tr><td>T31439</td><td>GGCTGGCGCTGAGTGCCAGTCGG</td><td>1000</td><td></td><td></td><td></td><td></td><td></td><td></td><td></td><td></td><td></td></tr>
<tr><td>Oligo fwd</td><td>CACCGGCTGGCGCTGAGTGCCAGT</td><td></td><td></td><td></td><td></td><td></td><td></td><td></td><td></td><td></td><td></td></tr>
<tr><td>Oligo rev</td><td>AAACACTGGCACTCAGCGCCAGCC</td><td></td><td></td><td></td><td></td><td></td><td></td><td></td><td></td><td></td><td></td></tr>
<tr><td>Chromosome</td><td>start</td><td>end</td><td>strand</td><td>MM</td><td>target_seq</td><td>PAM</td><td>alignment</td><td>distance</td><td>position</td><td>gene name</td><td>gene id</td></tr>
<tr><td>GU305914.1</td><td>37823</td><td>37845</td><td>-</td><td>0</td><td>GGCTGGCGCTGAGTGCCAGT</td><td>CGG</td><td>||||||||[||||||||||||]PAM</td><td>NA</td><td>Intergenic</td><td>NA</td><td>NA</td></tr>
<tr><td>T31440</td><td>CGCTGAGTGCCAGTCGGTAGCGG</td><td>1000</td><td></td><td></td><td></td><td></td><td></td><td></td><td></td><td></td><td></td></tr>
<tr><td>Oligo fwd</td><td>CACCCGCTGAGTGCCAGTCGGTAG</td><td></td><td></td><td></td><td></td><td></td><td></td><td></td><td></td><td></td><td></td></tr>
<tr><td>Oligo rev</td><td>AAACCTACCGACTGGCACTCAGCG</td><td></td><td></td><td></td><td></td><td></td><td></td><td></td><td></td><td></td><td></td></tr>
<tr><td>Chromosome</td><td>start</td><td>end</td><td>strand</td><td>MM</td><td>target_seq</td><td>PAM</td><td>alignment</td><td>distance</td><td>position</td><td>gene name</td><td>gene id</td></tr>
<tr><td>GU305914.1</td><td>37817</td><td>37839</td><td>-</td><td>0</td><td>CGCTGAGTGCCAGTCGGTAG</td><td>CGG</td><td>||||||||[||||||||||||]PAM</td><td>NA</td><td>Intergenic</td><td>NA</td><td>NA</td></tr>
<tr><td>T31441</td><td>TGCCAGTCGGTAGCGGTAGTTGG</td><td>1000</td><td></td><td></td><td></td><td></td><td></td><td></td><td></td><td></td><td></td></tr>
<tr><td>Oligo fwd</td><td>CACCTGCCAGTCGGTAGCGGTAGT</td><td></td><td></td><td></td><td></td><td></td><td></td><td></td><td></td><td></td><td></td></tr>
<tr><td>Oligo rev</td><td>AAACACTACCGCTACCGACTGGCA</td><td></td><td></td><td></td><td></td><td></td><td></td><td></td><td></td><td></td><td></td></tr>
<tr><td>Chromosome</td><td>start</td><td>end</td><td>strand</td><td>MM</td><td>target_seq</td><td>PAM</td><td>alignment</td><td>distance</td><td>position</td><td>gene name</td><td>gene id</td></tr>
</table>

<table>
<tr><td>GU305914.1</td><td></td><td>37810</td><td>37832</td><td>-</td><td>0</td><td>TGCCAGTCGGTAGCGGTAGT</td><td>TGG</td><td>||||||||[||||||||||||]PAM</td><td>NA</td><td>Intergenic</td><td>NA</td><td>NA</td></tr>
<tr><td></td></tr>
<tr><td>T31442</td><td>GTTGGAAACGTGCCGTCTCTCGG</td><td></td><td>1000</td></tr>
<tr><td>Oligo fwd</td><td>CACCGTTGGAAACGTGCCGTCTCT</td></tr>
<tr><td>Oligo rev</td><td>AAACAGAGACGGCACGTTTCCAAC</td></tr>
<tr><td>Chromosome</td><td>start</td><td></td><td>end</td><td>strand</td><td>MM</td><td>target_seq</td><td>PAM</td><td>alignment</td><td>distance</td><td>position</td><td>gene name</td><td>gene id</td></tr>
<tr><td>GU305914.1</td><td></td><td>37792</td><td>37814</td><td>-</td><td>0</td><td>GTTGGAAACGTGCCGTCTCT</td><td>CGG</td><td>||||||||[||||||||||||]PAM</td><td>NA</td><td>Intergenic</td><td>NA</td><td>NA</td></tr>
<tr><td></td></tr>
<tr><td>T31443</td><td>TCCAGCTTCACGCGCGTGTCCGG</td><td></td><td>1000</td></tr>
<tr><td>Oligo fwd</td><td>CACCTCCAGCTTCACGCGCGTGTC</td></tr>
<tr><td>Oligo rev</td><td>AAACGACACGCGCGTGAAGCTGGA</td></tr>
<tr><td>Chromosome</td><td>start</td><td></td><td>end</td><td>strand</td><td>MM</td><td>target_seq</td><td>PAM</td><td>alignment</td><td>distance</td><td>position</td><td>gene name</td><td>gene id</td></tr>
<tr><td>GU305914.1</td><td></td><td>37764</td><td>37786</td><td>-</td><td>0</td><td>TCCAGCTTCACGCGCGTGTC</td><td>CGG</td><td>||||||||[||||||||||||]PAM</td><td>NA</td><td>Intergenic</td><td>NA</td><td>NA</td></tr>
<tr><td></td></tr>
<tr><td>T31444</td><td>CCAGCTTCACGCGCGTGTCCGGG</td><td></td><td>1000</td></tr>
<tr><td>Oligo fwd</td><td>CACCCCAGCTTCACGCGCGTGTCC</td></tr>
<tr><td>Oligo rev</td><td>AAACGGACACGCGCGTGAAGCTGG</td></tr>
<tr><td>Chromosome</td><td>start</td><td></td><td>end</td><td>strand</td><td>MM</td><td>target_seq</td><td>PAM</td><td>alignment</td><td>distance</td><td>position</td><td>gene name</td><td>gene id</td></tr>
<tr><td>GU305914.1</td><td></td><td>37763</td><td>37785</td><td>-</td><td>0</td><td>CCAGCTTCACGCGCGTGTCC</td><td>GGG</td><td>||||||||[||||||||||||]PAM</td><td>NA</td><td>Intergenic</td><td>NA</td><td>NA</td></tr>
<tr><td></td></tr>
<tr><td>T31445</td><td>GCACGACGCGTCGATCGAGTTGG</td><td></td><td>1000</td></tr>
<tr><td>Oligo fwd</td><td>CACCGCACGACGCGTCGATCGAGT</td></tr>
<tr><td>Oligo rev</td><td>AAACACTCGATCGACGCGTCGTGC</td></tr>
<tr><td>Chromosome</td><td>start</td><td></td><td>end</td><td>strand</td><td>MM</td><td>target_seq</td><td>PAM</td><td>alignment</td><td>distance</td><td>position</td><td>gene name</td><td>gene id</td></tr>
<tr><td>GU305914.1</td><td></td><td>37741</td><td>37763</td><td>-</td><td>0</td><td>GCACGACGCGTCGATCGAGT</td><td>TGG</td><td>||||||||[||||||||||||]PAM</td><td>NA</td><td>Intergenic</td><td>NA</td><td>NA</td></tr>
<tr><td></td></tr>
<tr><td>T31446</td><td>GTACGCGATCATGCTGTTGCAGG</td><td></td><td>1000</td></tr>
<tr><td>Oligo fwd</td><td>CACCGTACGCGATCATGCTGTTGC</td></tr>
<tr><td>Oligo rev</td><td>AAACGCAACAGCATGATCGCGTAC</td></tr>
<tr><td>Chromosome</td><td>start</td><td></td><td>end</td><td>strand</td><td>MM</td><td>target_seq</td><td>PAM</td><td>alignment</td><td>distance</td><td>position</td><td>gene name</td><td>gene id</td></tr>
<tr><td>GU305914.1</td><td></td><td>37717</td><td>37739</td><td>-</td><td>0</td><td>GTACGCGATCATGCTGTTGC</td><td>AGG</td><td>||||||||[||||||||||||]PAM</td><td>NA</td><td>Intergenic</td><td>NA</td><td>NA</td></tr>
<tr><td></td></tr>
<tr><td>T31447</td><td>GGTATCGCGCATAGACTCTGCGG</td><td></td><td>1000</td></tr>
<tr><td>Oligo fwd</td><td>CACCGGTATCGCGCATAGACTCTG</td></tr>
<tr><td>Oligo rev</td><td>AAACCAGAGTCTATGCGCGATACC</td></tr>
<tr><td>Chromosome</td><td>start</td><td></td><td>end</td><td>strand</td><td>MM</td><td>target_seq</td><td>PAM</td><td>alignment</td><td>distance</td><td>position</td><td>gene name</td><td>gene id</td></tr>
<tr><td>GU305914.1</td><td></td><td>37696</td><td>37718</td><td>-</td><td>0</td><td>GGTATCGCGCATAGACTCTG</td><td>CGG</td><td>||||||||[||||||||||||]PAM</td><td>NA</td><td>Intergenic</td><td>NA</td><td>NA</td></tr>
<tr><td></td></tr>
<tr><td>T31448</td><td>TAGACTCTGCGGCCGTGTCCAGG</td><td></td><td>1000</td></tr>
<tr><td>Oligo fwd</td><td>CACCTAGACTCTGCGGCCGTGTCC</td></tr>
<tr><td>Oligo rev</td><td>AAACGGACACGGCCGCAGAGTCTA</td></tr>
<tr><td>Chromosome</td><td>start</td><td></td><td>end</td><td>strand</td><td>MM</td><td>target_seq</td><td>PAM</td><td>alignment</td><td>distance</td><td>position</td><td>gene name</td><td>gene id</td></tr>
<tr><td>GU305914.1</td><td></td><td>37685</td><td>37707</td><td>-</td><td>0</td><td>TAGACTCTGCGGCCGTGTCC</td><td>AGG</td><td>||||||||[||||||||||||]PAM</td><td>NA</td><td>Intergenic</td><td>NA</td><td>NA</td></tr>
<tr><td></td></tr>
<tr><td>T31449</td><td>CTCTGCGGCCGTGTCCAGGACGG</td><td></td><td>1000</td></tr>
<tr><td>Oligo fwd</td><td>CACCCTCTGCGGCCGTGTCCAGGA</td></tr>
<tr><td>Oligo rev</td><td>AAACTCCTGGACACGGCCGCAGAG</td></tr>
<tr><td>Chromosome</td><td>start</td><td></td><td>end</td><td>strand</td><td>MM</td><td>target_seq</td><td>PAM</td><td>alignment</td><td>distance</td><td>position</td><td>gene name</td><td>gene id</td></tr>
<tr><td>GU305914.1</td><td></td><td>37681</td><td>37703</td><td>-</td><td>0</td><td>CTCTGCGGCCGTGTCCAGGA</td><td>CGG</td><td>||||||||[||||||||||||]PAM</td><td>NA</td><td>Intergenic</td><td>NA</td><td>NA</td></tr>
<tr><td></td></tr>
<tr><td>T31450</td><td>TGCGGCCGTGTCCAGGACGGTGG</td><td></td><td>1000</td></tr>
<tr><td>Oligo fwd</td><td>CACCTGCGGCCGTGTCCAGGACGG</td></tr>
<tr><td>Oligo rev</td><td>AAACCCGTCCTGGACACGGCCGCA</td></tr>
<tr><td>Chromosome</td><td>start</td><td></td><td>end</td><td>strand</td><td>MM</td><td>target_seq</td><td>PAM</td><td>alignment</td><td>distance</td><td>position</td><td>gene name</td><td>gene id</td></tr>
<tr><td>GU305914.1</td><td></td><td>37678</td><td>37700</td><td>-</td><td>0</td><td>TGCGGCCGTGTCCAGGACGG</td><td>TGG</td><td>||||||||[||||||||||||]PAM</td><td>NA</td><td>Intergenic</td><td>NA</td><td>NA</td></tr>
<tr><td></td></tr>
<tr><td>T31451</td><td>ACAGCATGACGACCTGAGCATGG</td><td></td><td>1000</td></tr>
<tr><td>Oligo fwd</td><td>CACCACAGCATGACGACCTGAGCA</td></tr>
<tr><td>Oligo rev</td><td>AAACTGCTCAGGTCGTCATGCTGT</td></tr>
<tr><td>Chromosome</td><td>start</td><td></td><td>end</td><td>strand</td><td>MM</td><td>target_seq</td><td>PAM</td><td>alignment</td><td>distance</td><td>position</td><td>gene name</td><td>gene id</td></tr>
<tr><td>GU305914.1</td><td></td><td>37643</td><td>37665</td><td>-</td><td>0</td><td>ACAGCATGACGACCTGAGCA</td><td>TGG</td><td>||||||||[||||||||||||]PAM</td><td>NA</td><td>Intergenic</td><td>NA</td><td>NA</td></tr>
<tr><td></td></tr>
<tr><td>T31452</td><td>GACCTGAGCATGGAAATCTCCGG</td><td></td><td>1000</td></tr>
<tr><td>Oligo fwd</td><td>CACCGACCTGAGCATGGAAATCTC</td></tr>
<tr><td>Oligo rev</td><td>AAACGAGATTTCCATGCTCAGGTC</td></tr>
<tr><td>Chromosome</td><td>start</td><td></td><td>end</td><td>strand</td><td>MM</td><td>target_seq</td><td>PAM</td><td>alignment</td><td>distance</td><td>position</td><td>gene name</td><td>gene id</td></tr>
<tr><td>GU305914.1</td><td></td><td>37633</td><td>37655</td><td>-</td><td>0</td><td>GACCTGAGCATGGAAATCTC</td><td>CGG</td><td>||||||||[||||||||||||]PAM</td><td>NA</td><td>Intergenic</td><td>NA</td><td>NA</td></tr>
<tr><td></td></tr>
<tr><td>T31453</td><td>ATTGAGCGCCACCGACGAACTGG</td><td></td><td>1000</td></tr>
<tr><td>Oligo fwd</td><td>CACCATTGAGCGCCACCGACGAAC</td></tr>
<tr><td>Oligo rev</td><td>AAACGTTCGTCGGTGGCGCTCAAT</td></tr>
<tr><td>Chromosome</td><td>start</td><td></td><td>end</td><td>strand</td><td>MM</td><td>target_seq</td><td>PAM</td><td>alignment</td><td>distance</td><td>position</td><td>gene name</td><td>gene id</td></tr>
<tr><td>GU305914.1</td><td></td><td>37609</td><td>37631</td><td>-</td><td>0</td><td>ATTGAGCGCCACCGACGAAC</td><td>TGG</td><td>||||||||[||||||||||||]PAM</td><td>NA</td><td>Intergenic</td><td>NA</td><td>NA</td></tr>
<tr><td></td></tr>
<tr><td>T31454</td><td>TTGAGCGCCACCGACGAACTGGG</td><td></td><td>1000</td></tr>
<tr><td>Oligo fwd</td><td>CACCTTGAGCGCCACCGACGAACT</td></tr>
<tr><td>Oligo rev</td><td>AAACAGTTCGTCGGTGGCGCTCAA</td></tr>
<tr><td>Chromosome</td><td>start</td><td></td><td>end</td><td>strand</td><td>MM</td><td>target_seq</td><td>PAM</td><td>alignment</td><td>distance</td><td>position</td><td>gene name</td><td>gene id</td></tr>
<tr><td>GU305914.1</td><td></td><td>37608</td><td>37630</td><td>-</td><td>0</td><td>TTGAGCGCCACCGACGAACT</td><td>GGG</td><td>||||||||[||||||||||||]PAM</td><td>NA</td><td>Intergenic</td><td>NA</td><td>NA</td></tr>
<tr><td></td></tr>
<tr><td>T31455</td><td>ACTGGGTGCCCCGACGATGATGG</td><td></td><td>1000</td></tr>
<tr><td>Oligo fwd</td><td>CACCACTGGGTGCCCCGACGATGA</td></tr>
<tr><td>Oligo rev</td><td>AAACTCATCGTCGGGGCACCCAGT</td></tr>
<tr><td>Chromosome</td><td>start</td><td></td><td>end</td><td>strand</td><td>MM</td><td>target_seq</td><td>PAM</td><td>alignment</td><td>distance</td><td>position</td><td>gene name</td><td>gene id</td></tr>
<tr><td>GU305914.1</td><td></td><td>37591</td><td>37613</td><td>-</td><td>0</td><td>ACTGGGTGCCCCGACGATGA</td><td>TGG</td><td>||||||||[||||||||||||]PAM</td><td>NA</td><td>Intergenic</td><td>NA</td><td>NA</td></tr>
<tr><td>GU305914.1</td><td></td><td>111571</td><td>111593</td><td>-</td><td>4</td><td>AGCGGGAGCGCCGACGATGA</td><td>AGG</td><td>|--|||-|[|-||||||||||]PAM</td><td>NA</td><td>Intergenic</td><td>NA</td><td>NA</td></tr>
<tr><td></td></tr>
<tr><td>T31456</td><td>GGGTGCCCCGACGATGATGGAGG</td><td></td><td>1000</td></tr>
<tr><td>Oligo fwd</td><td>CACCGGGTGCCCCGACGATGATGG</td></tr>
</table>

<table>
<tr><td>Oligo rev</td><td>AAACCCATCATCGTCGGGGCACCC</td><td></td><td></td><td></td><td></td><td></td><td></td><td></td><td></td><td></td><td></td></tr>
<tr><td>Chromosome</td><td>start</td><td>end</td><td>strand</td><td>MM</td><td>target_seq</td><td>PAM</td><td>alignment</td><td>distance</td><td>position</td><td>gene name</td><td>gene id</td></tr>
<tr><td>GU305914.1</td><td>37588</td><td>37610</td><td>-</td><td>0</td><td>GGGTGCCCCGACGATGATGG</td><td>AGG</td><td>||||||||[||||||||||||]PAM</td><td>NA</td><td>Intergenic</td><td>NA</td><td>NA</td></tr>
<tr><td>GU305914.1</td><td>111568</td><td>111590</td><td>-</td><td>3</td><td>GGGAGCGCCGACGATGAAGG</td><td>CGG</td><td>|||-||-|[|||||||||-||]PAM</td><td>NA</td><td>Intergenic</td><td>NA</td><td>NA</td></tr>
<tr><td></td><td></td><td></td><td></td><td></td><td></td><td></td><td></td><td></td><td></td><td></td><td></td></tr>
<tr><td>T31457</td><td>GGTGCCCCGACGATGATGGAGGG</td><td>1000</td><td></td><td></td><td></td><td></td><td></td><td></td><td></td><td></td><td></td></tr>
<tr><td>Oligo fwd</td><td>CACCGGTGCCCCGACGATGATGGA</td><td></td><td></td><td></td><td></td><td></td><td></td><td></td><td></td><td></td><td></td></tr>
<tr><td>Oligo rev</td><td>AAACTCCATCATCGTCGGGGCACC</td><td></td><td></td><td></td><td></td><td></td><td></td><td></td><td></td><td></td><td></td></tr>
<tr><td>Chromosome</td><td>start</td><td>end</td><td>strand</td><td>MM</td><td>target_seq</td><td>PAM</td><td>alignment</td><td>distance</td><td>position</td><td>gene name</td><td>gene id</td></tr>
<tr><td>GU305914.1</td><td>37587</td><td>37609</td><td>-</td><td>0</td><td>GGTGCCCCGACGATGATGGA</td><td>GGG</td><td>||||||||[||||||||||||]PAM</td><td>NA</td><td>Intergenic</td><td>NA</td><td>NA</td></tr>
<tr><td></td><td></td><td></td><td></td><td></td><td></td><td></td><td></td><td></td><td></td><td></td><td></td></tr>
<tr><td>T31458</td><td>TGGAGGGCTGTTGCACGCTGAGG</td><td>1000</td><td></td><td></td><td></td><td></td><td></td><td></td><td></td><td></td><td></td></tr>
<tr><td>Oligo fwd</td><td>CACCTGGAGGGCTGTTGCACGCTG</td><td></td><td></td><td></td><td></td><td></td><td></td><td></td><td></td><td></td><td></td></tr>
<tr><td>Oligo rev</td><td>AAACCAGCGTGCAACAGCCCTCCA</td><td></td><td></td><td></td><td></td><td></td><td></td><td></td><td></td><td></td><td></td></tr>
<tr><td>Chromosome</td><td>start</td><td>end</td><td>strand</td><td>MM</td><td>target_seq</td><td>PAM</td><td>alignment</td><td>distance</td><td>position</td><td>gene name</td><td>gene id</td></tr>
<tr><td>GU305914.1</td><td>37571</td><td>37593</td><td>-</td><td>0</td><td>TGGAGGGCTGTTGCACGCTG</td><td>AGG</td><td>||||||||[||||||||||||]PAM</td><td>NA</td><td>Intergenic</td><td>NA</td><td>NA</td></tr>
<tr><td></td><td></td><td></td><td></td><td></td><td></td><td></td><td></td><td></td><td></td><td></td><td></td></tr>
<tr><td>T31459</td><td>GCACGCTGAGGTCCACCGCTTGG</td><td>1000</td><td></td><td></td><td></td><td></td><td></td><td></td><td></td><td></td><td></td></tr>
<tr><td>Oligo fwd</td><td>CACCGCACGCTGAGGTCCACCGCT</td><td></td><td></td><td></td><td></td><td></td><td></td><td></td><td></td><td></td><td></td></tr>
<tr><td>Oligo rev</td><td>AAACAGCGGTGGACCTCAGCGTGC</td><td></td><td></td><td></td><td></td><td></td><td></td><td></td><td></td><td></td><td></td></tr>
<tr><td>Chromosome</td><td>start</td><td>end</td><td>strand</td><td>MM</td><td>target_seq</td><td>PAM</td><td>alignment</td><td>distance</td><td>position</td><td>gene name</td><td>gene id</td></tr>
<tr><td>GU305914.1</td><td>37559</td><td>37581</td><td>-</td><td>0</td><td>GCACGCTGAGGTCCACCGCT</td><td>TGG</td><td>||||||||[||||||||||||]PAM</td><td>NA</td><td>Intergenic</td><td>NA</td><td>NA</td></tr>
<tr><td></td><td></td><td></td><td></td><td></td><td></td><td></td><td></td><td></td><td></td><td></td><td></td></tr>
<tr><td>T31460</td><td>CGCTGAGGTCCACCGCTTGGCGG</td><td>1000</td><td></td><td></td><td></td><td></td><td></td><td></td><td></td><td></td><td></td></tr>
<tr><td>Oligo fwd</td><td>CACCCGCTGAGGTCCACCGCTTGG</td><td></td><td></td><td></td><td></td><td></td><td></td><td></td><td></td><td></td><td></td></tr>
<tr><td>Oligo rev</td><td>AAACCCAAGCGGTGGACCTCAGCG</td><td></td><td></td><td></td><td></td><td></td><td></td><td></td><td></td><td></td><td></td></tr>
<tr><td>Chromosome</td><td>start</td><td>end</td><td>strand</td><td>MM</td><td>target_seq</td><td>PAM</td><td>alignment</td><td>distance</td><td>position</td><td>gene name</td><td>gene id</td></tr>
<tr><td>GU305914.1</td><td>37556</td><td>37578</td><td>-</td><td>0</td><td>CGCTGAGGTCCACCGCTTGG</td><td>CGG</td><td>||||||||[||||||||||||]PAM</td><td>NA</td><td>Intergenic</td><td>NA</td><td>NA</td></tr>
<tr><td></td><td></td><td></td><td></td><td></td><td></td><td></td><td></td><td></td><td></td><td></td><td></td></tr>
<tr><td>T31461</td><td>CCGCTTGGCGGAGAGCCGCCAGG</td><td>1000</td><td></td><td></td><td></td><td></td><td></td><td></td><td></td><td></td><td></td></tr>
<tr><td>Oligo fwd</td><td>CACCCCGCTTGGCGGAGAGCCGCC</td><td></td><td></td><td></td><td></td><td></td><td></td><td></td><td></td><td></td><td></td></tr>
<tr><td>Oligo rev</td><td>AAACGGCGGCTCTCCGCCAAGCGG</td><td></td><td></td><td></td><td></td><td></td><td></td><td></td><td></td><td></td><td></td></tr>
<tr><td>Chromosome</td><td>start</td><td>end</td><td>strand</td><td>MM</td><td>target_seq</td><td>PAM</td><td>alignment</td><td>distance</td><td>position</td><td>gene name</td><td>gene id</td></tr>
<tr><td>GU305914.1</td><td>37544</td><td>37566</td><td>-</td><td>0</td><td>CCGCTTGGCGGAGAGCCGCC</td><td>AGG</td><td>||||||||[||||||||||||]PAM</td><td>NA</td><td>Intergenic</td><td>NA</td><td>NA</td></tr>
<tr><td></td><td></td><td></td><td></td><td></td><td></td><td></td><td></td><td></td><td></td><td></td><td></td></tr>
<tr><td>T31462</td><td>CGCTTGGCGGAGAGCCGCCAGGG</td><td>1000</td><td></td><td></td><td></td><td></td><td></td><td></td><td></td><td></td><td></td></tr>
<tr><td>Oligo fwd</td><td>CACCCGCTTGGCGGAGAGCCGCCA</td><td></td><td></td><td></td><td></td><td></td><td></td><td></td><td></td><td></td><td></td></tr>
<tr><td>Oligo rev</td><td>AAACTGGCGGCTCTCCGCCAAGCG</td><td></td><td></td><td></td><td></td><td></td><td></td><td></td><td></td><td></td><td></td></tr>
<tr><td>Chromosome</td><td>start</td><td>end</td><td>strand</td><td>MM</td><td>target_seq</td><td>PAM</td><td>alignment</td><td>distance</td><td>position</td><td>gene name</td><td>gene id</td></tr>
<tr><td>GU305914.1</td><td>37543</td><td>37565</td><td>-</td><td>0</td><td>CGCTTGGCGGAGAGCCGCCA</td><td>GGG</td><td>||||||||[||||||||||||]PAM</td><td>NA</td><td>Intergenic</td><td>NA</td><td>NA</td></tr>
<tr><td></td><td></td><td></td><td></td><td></td><td></td><td></td><td></td><td></td><td></td><td></td><td></td></tr>
<tr><td>T31463</td><td>GCTTGGCGGAGAGCCGCCAGGGG</td><td>1000</td><td></td><td></td><td></td><td></td><td></td><td></td><td></td><td></td><td></td></tr>
<tr><td>Oligo fwd</td><td>CACCGCTTGGCGGAGAGCCGCCAG</td><td></td><td></td><td></td><td></td><td></td><td></td><td></td><td></td><td></td><td></td></tr>
<tr><td>Oligo rev</td><td>AAACCTGGCGGCTCTCCGCCAAGC</td><td></td><td></td><td></td><td></td><td></td><td></td><td></td><td></td><td></td><td></td></tr>
<tr><td>Chromosome</td><td>start</td><td>end</td><td>strand</td><td>MM</td><td>target_seq</td><td>PAM</td><td>alignment</td><td>distance</td><td>position</td><td>gene name</td><td>gene id</td></tr>
<tr><td>GU305914.1</td><td>37542</td><td>37564</td><td>-</td><td>0</td><td>GCTTGGCGGAGAGCCGCCAG</td><td>GGG</td><td>||||||||[||||||||||||]PAM</td><td>NA</td><td>Intergenic</td><td>NA</td><td>NA</td></tr>
<tr><td></td><td></td><td></td><td></td><td></td><td></td><td></td><td></td><td></td><td></td><td></td><td></td></tr>
<tr><td>T31464</td><td>CAGGGGTCCGCACAGCCCTGCGG</td><td>1000</td><td></td><td></td><td></td><td></td><td></td><td></td><td></td><td></td><td></td></tr>
<tr><td>Oligo fwd</td><td>CACCCAGGGGTCCGCACAGCCCTG</td><td></td><td></td><td></td><td></td><td></td><td></td><td></td><td></td><td></td><td></td></tr>
<tr><td>Oligo rev</td><td>AAACCAGGGCTGTGCGGACCCCTG</td><td></td><td></td><td></td><td></td><td></td><td></td><td></td><td></td><td></td><td></td></tr>
<tr><td>Chromosome</td><td>start</td><td>end</td><td>strand</td><td>MM</td><td>target_seq</td><td>PAM</td><td>alignment</td><td>distance</td><td>position</td><td>gene name</td><td>gene id</td></tr>
<tr><td>GU305914.1</td><td>37525</td><td>37547</td><td>-</td><td>0</td><td>CAGGGGTCCGCACAGCCCTG</td><td>CGG</td><td>||||||||[||||||||||||]PAM</td><td>NA</td><td>Intergenic</td><td>NA</td><td>NA</td></tr>
<tr><td></td><td></td><td></td><td></td><td></td><td></td><td></td><td></td><td></td><td></td><td></td><td></td></tr>
<tr><td>T31465</td><td>AGGGGTCCGCACAGCCCTGCGGG</td><td>1000</td><td></td><td></td><td></td><td></td><td></td><td></td><td></td><td></td><td></td></tr>
<tr><td>Oligo fwd</td><td>CACCAGGGGTCCGCACAGCCCTGC</td><td></td><td></td><td></td><td></td><td></td><td></td><td></td><td></td><td></td><td></td></tr>
<tr><td>Oligo rev</td><td>AAACGCAGGGCTGTGCGGACCCCT</td><td></td><td></td><td></td><td></td><td></td><td></td><td></td><td></td><td></td><td></td></tr>
<tr><td>Chromosome</td><td>start</td><td>end</td><td>strand</td><td>MM</td><td>target_seq</td><td>PAM</td><td>alignment</td><td>distance</td><td>position</td><td>gene name</td><td>gene id</td></tr>
<tr><td>GU305914.1</td><td>37524</td><td>37546</td><td>-</td><td>0</td><td>AGGGGTCCGCACAGCCCTGC</td><td>GGG</td><td>||||||||[||||||||||||]PAM</td><td>NA</td><td>Intergenic</td><td>NA</td><td>NA</td></tr>
<tr><td></td><td></td><td></td><td></td><td></td><td></td><td></td><td></td><td></td><td></td><td></td><td></td></tr>
<tr><td>T31466</td><td>CAGCCCTGCGGGATGAGCTTCGG</td><td>1000</td><td></td><td></td><td></td><td></td><td></td><td></td><td></td><td></td><td></td></tr>
<tr><td>Oligo fwd</td><td>CACCCAGCCCTGCGGGATGAGCTT</td><td></td><td></td><td></td><td></td><td></td><td></td><td></td><td></td><td></td><td></td></tr>
<tr><td>Oligo rev</td><td>AAACAAGCTCATCCCGCAGGGCTG</td><td></td><td></td><td></td><td></td><td></td><td></td><td></td><td></td><td></td><td></td></tr>
<tr><td>Chromosome</td><td>start</td><td>end</td><td>strand</td><td>MM</td><td>target_seq</td><td>PAM</td><td>alignment</td><td>distance</td><td>position</td><td>gene name</td><td>gene id</td></tr>
<tr><td>GU305914.1</td><td>37513</td><td>37535</td><td>-</td><td>0</td><td>CAGCCCTGCGGGATGAGCTT</td><td>CGG</td><td>||||||||[||||||||||||]PAM</td><td>NA</td><td>Intergenic</td><td>NA</td><td>NA</td></tr>
<tr><td></td><td></td><td></td><td></td><td></td><td></td><td></td><td></td><td></td><td></td><td></td><td></td></tr>
<tr><td>T31467</td><td>TGCGGGATGAGCTTCGGAGAAGG</td><td>1000</td><td></td><td></td><td></td><td></td><td></td><td></td><td></td><td></td><td></td></tr>
<tr><td>Oligo fwd</td><td>CACCTGCGGGATGAGCTTCGGAGA</td><td></td><td></td><td></td><td></td><td></td><td></td><td></td><td></td><td></td><td></td></tr>
<tr><td>Oligo rev</td><td>AAACTCTCCGAAGCTCATCCCGCA</td><td></td><td></td><td></td><td></td><td></td><td></td><td></td><td></td><td></td><td></td></tr>
<tr><td>Chromosome</td><td>start</td><td>end</td><td>strand</td><td>MM</td><td>target_seq</td><td>PAM</td><td>alignment</td><td>distance</td><td>position</td><td>gene name</td><td>gene id</td></tr>
<tr><td>GU305914.1</td><td>37507</td><td>37529</td><td>-</td><td>0</td><td>TGCGGGATGAGCTTCGGAGA</td><td>AGG</td><td>||||||||[||||||||||||]PAM</td><td>NA</td><td>Intergenic</td><td>NA</td><td>NA</td></tr>
<tr><td></td><td></td><td></td><td></td><td></td><td></td><td></td><td></td><td></td><td></td><td></td><td></td></tr>
<tr><td>T31468</td><td>GGGATGAGCTTCGGAGAAGGCGG</td><td>1000</td><td></td><td></td><td></td><td></td><td></td><td></td><td></td><td></td><td></td></tr>
<tr><td>Oligo fwd</td><td>CACCGGGATGAGCTTCGGAGAAGG</td><td></td><td></td><td></td><td></td><td></td><td></td><td></td><td></td><td></td><td></td></tr>
<tr><td>Oligo rev</td><td>AAACCCTTCTCCGAAGCTCATCCC</td><td></td><td></td><td></td><td></td><td></td><td></td><td></td><td></td><td></td><td></td></tr>
<tr><td>Chromosome</td><td>start</td><td>end</td><td>strand</td><td>MM</td><td>target_seq</td><td>PAM</td><td>alignment</td><td>distance</td><td>position</td><td>gene name</td><td>gene id</td></tr>
<tr><td>GU305914.1</td><td>37504</td><td>37526</td><td>-</td><td>0</td><td>GGGATGAGCTTCGGAGAAGG</td><td>CGG</td><td>||||||||[||||||||||||]PAM</td><td>NA</td><td>Intergenic</td><td>NA</td><td>NA</td></tr>
<tr><td></td><td></td><td></td><td></td><td></td><td></td><td></td><td></td><td></td><td></td><td></td><td></td></tr>
<tr><td>T31469</td><td>ATGAGCTTCGGAGAAGGCGGTGG</td><td>1000</td><td></td><td></td><td></td><td></td><td></td><td></td><td></td><td></td><td></td></tr>
<tr><td>Oligo fwd</td><td>CACCATGAGCTTCGGAGAAGGCGG</td><td></td><td></td><td></td><td></td><td></td><td></td><td></td><td></td><td></td><td></td></tr>
<tr><td>Oligo rev</td><td>AAACCCGCCTTCTCCGAAGCTCAT</td><td></td><td></td><td></td><td></td><td></td><td></td><td></td><td></td><td></td><td></td></tr>
<tr><td>Chromosome</td><td>start</td><td>end</td><td>strand</td><td>MM</td><td>target_seq</td><td>PAM</td><td>alignment</td><td>distance</td><td>position</td><td>gene name</td><td>gene id</td></tr>
<tr><td>GU305914.1</td><td>37501</td><td>37523</td><td>-</td><td>0</td><td>ATGAGCTTCGGAGAAGGCGG</td><td>TGG</td><td>||||||||[||||||||||||]PAM</td><td>NA</td><td>Intergenic</td><td>NA</td><td>NA</td></tr>
<tr><td></td><td></td><td></td><td></td><td></td><td></td><td></td><td></td><td></td><td></td><td></td><td></td></tr>
<tr><td>T31470</td><td>CGCTCGTCGACCGCATGATGCGG</td><td>1000</td><td></td><td></td><td></td><td></td><td></td><td></td><td></td><td></td><td></td></tr>
<tr><td>Oligo fwd</td><td>CACCCGCTCGTCGACCGCATGATG</td><td></td><td></td><td></td><td></td><td></td><td></td><td></td><td></td><td></td><td></td></tr>
<tr><td>Oligo rev</td><td>AAACCATCATGCGGTCGACGAGCG</td><td></td><td></td><td></td><td></td><td></td><td></td><td></td><td></td><td></td><td></td></tr>
<tr><td>Chromosome</td><td>start</td><td>end</td><td>strand</td><td>MM</td><td>target_seq</td><td>PAM</td><td>alignment</td><td>distance</td><td>position</td><td>gene name</td><td>gene id</td></tr>
<tr><td>GU305914.1</td><td>37473</td><td>37495</td><td>-</td><td>0</td><td>CGCTCGTCGACCGCATGATG</td><td>CGG</td><td>||||||||[||||||||||||]PAM</td><td>NA</td><td>Intergenic</td><td>NA</td><td>NA</td></tr>
</table>

| | | | | | | | | | | | |
|---|---|---|---|---|---|---|---|---|---|---|---|
| T31471 | CGACCGCATGATGCGGATCGAGG | 1000 | | | | | | | | | |
| Oligo fwd | CACCCGACCGCATGATGCGGATCG | | | | | | | | | | |
| Oligo rev | AAACCGATCCGCATCATGCGGTCG | | | | | | | | | | |
| Chromosome | start | end | strand | MM | target_seq | PAM | alignment | distance | position | gene name | gene id |
| GU305914.1 | 37466 | 37488 | - | 0 | CGACCGCATGATGCGGATCG | AGG | \|\|\|\|\|\|\|\|[\|\|\|\|\|\|\|\|\|\|\|\|]PAM | NA | Intergenic | NA | NA |

| | | | | | | | | | | | |
|---|---|---|---|---|---|---|---|---|---|---|---|
| T31472 | GATGCGGATCGAGGATGTTCTGG | 1000 | | | | | | | | | |
| Oligo fwd | CACCGATGCGGATCGAGGATGTTC | | | | | | | | | | |
| Oligo rev | AAACGAACATCCTCGATCCGCATC | | | | | | | | | | |
| Chromosome | start | end | strand | MM | target_seq | PAM | alignment | distance | position | gene name | gene id |
| GU305914.1 | 37457 | 37479 | - | 0 | GATGCGGATCGAGGATGTTC | TGG | \|\|\|\|\|\|\|\|[\|\|\|\|\|\|\|\|\|\|\|\|]PAM | NA | Intergenic | NA | NA |

| | | | | | | | | | | | |
|---|---|---|---|---|---|---|---|---|---|---|---|
| T31473 | GGATGTTCTGGAGACGACGCAGG | 1000 | | | | | | | | | |
| Oligo fwd | CACCGGATGTTCTGGAGACGACGC | | | | | | | | | | |
| Oligo rev | AAACGCGTCGTCTCCAGAACATCC | | | | | | | | | | |
| Chromosome | start | end | strand | MM | target_seq | PAM | alignment | distance | position | gene name | gene id |
| GU305914.1 | 37445 | 37467 | - | 0 | GGATGTTCTGGAGACGACGC | AGG | \|\|\|\|\|\|\|\|[\|\|\|\|\|\|\|\|\|\|\|\|]PAM | NA | Intergenic | NA | NA |

| | | | | | | | | | | | |
|---|---|---|---|---|---|---|---|---|---|---|---|
| T31474 | ACGACGCAGGTCTTCGATAGTGG | 1000 | | | | | | | | | |
| Oligo fwd | CACCACGACGCAGGTCTTCGATAG | | | | | | | | | | |
| Oligo rev | AAACCTATCGAAGACCTGCGTCGT | | | | | | | | | | |
| Chromosome | start | end | strand | MM | target_seq | PAM | alignment | distance | position | gene name | gene id |
| GU305914.1 | 37432 | 37454 | - | 0 | ACGACGCAGGTCTTCGATAG | TGG | \|\|\|\|\|\|\|\|[\|\|\|\|\|\|\|\|\|\|\|\|]PAM | NA | Intergenic | NA | NA |

| | | | | | | | | | | | |
|---|---|---|---|---|---|---|---|---|---|---|---|
| T31475 | GCAGGTCTTCGATAGTGGCACGG | 1000 | | | | | | | | | |
| Oligo fwd | CACCGCAGGTCTTCGATAGTGGCA | | | | | | | | | | |
| Oligo rev | AAACTGCCACTATCGAAGACCTGC | | | | | | | | | | |
| Chromosome | start | end | strand | MM | target_seq | PAM | alignment | distance | position | gene name | gene id |
| GU305914.1 | 37427 | 37449 | - | 0 | GCAGGTCTTCGATAGTGGCA | CGG | \|\|\|\|\|\|\|\|[\|\|\|\|\|\|\|\|\|\|\|\|]PAM | NA | Intergenic | NA | NA |

| | | | | | | | | | | | |
|---|---|---|---|---|---|---|---|---|---|---|---|
| T31476 | GGCACGGTAATGCGTCGCGTCGG | 1000 | | | | | | | | | |
| Oligo fwd | CACCGGCACGGTAATGCGTCGCGT | | | | | | | | | | |
| Oligo rev | AAACACGCGACGCATTACCGTGCC | | | | | | | | | | |
| Chromosome | start | end | strand | MM | target_seq | PAM | alignment | distance | position | gene name | gene id |
| GU305914.1 | 37411 | 37433 | - | 0 | GGCACGGTAATGCGTCGCGT | CGG | \|\|\|\|\|\|\|\|[\|\|\|\|\|\|\|\|\|\|\|\|]PAM | NA | Intergenic | NA | NA |

| | | | | | | | | | | | |
|---|---|---|---|---|---|---|---|---|---|---|---|
| T31477 | GCACGGTAATGCGTCGCGTCGGG | 1000 | | | | | | | | | |
| Oligo fwd | CACCGCACGGTAATGCGTCGCGTC | | | | | | | | | | |
| Oligo rev | AAACGACGCGACGCATTACCGTGC | | | | | | | | | | |
| Chromosome | start | end | strand | MM | target_seq | PAM | alignment | distance | position | gene name | gene id |
| GU305914.1 | 37410 | 37432 | - | 0 | GCACGGTAATGCGTCGCGTC | GGG | \|\|\|\|\|\|\|\|[\|\|\|\|\|\|\|\|\|\|\|\|]PAM | NA | Intergenic | NA | NA |

| | | | | | | | | | | | |
|---|---|---|---|---|---|---|---|---|---|---|---|
| T31478 | CACGGTAATGCGTCGCGTCGGGG | 1000 | | | | | | | | | |
| Oligo fwd | CACCCACGGTAATGCGTCGCGTCG | | | | | | | | | | |
| Oligo rev | AAACCGACGCGACGCATTACCGTG | | | | | | | | | | |
| Chromosome | start | end | strand | MM | target_seq | PAM | alignment | distance | position | gene name | gene id |
| GU305914.1 | 37409 | 37431 | - | 0 | CACGGTAATGCGTCGCGTCG | GGG | \|\|\|\|\|\|\|\|[\|\|\|\|\|\|\|\|\|\|\|\|]PAM | NA | Intergenic | NA | NA |
| GU305914.1 | 123003 | 123025 | - | 4 | CACCATCATGCGGCGCGTCG | AGG | \|\|\|--\|-\|[\|\|\|\|-\|\|\|\|\|\|\|]PAM | NA | Intergenic | NA | NA |

| | | | | | | | | | | | |
|---|---|---|---|---|---|---|---|---|---|---|---|
| T31479 | GCGTCGGGGTTCTTCCTCTGCGG | 1000 | | | | | | | | | |
| Oligo fwd | CACCGCGTCGGGGTTCTTCCTCTG | | | | | | | | | | |
| Oligo rev | AAACCAGAGGAAGAACCCCGACGC | | | | | | | | | | |
| Chromosome | start | end | strand | MM | target_seq | PAM | alignment | distance | position | gene name | gene id |
| GU305914.1 | 37395 | 37417 | - | 0 | GCGTCGGGGTTCTTCCTCTG | CGG | \|\|\|\|\|\|\|\|[\|\|\|\|\|\|\|\|\|\|\|\|]PAM | NA | Intergenic | NA | NA |
| GU305914.1 | 131211 | 131233 | + | 4 | GCCTCGGAGGTCTGCCTCTG | TGG | \|\|-\|\|\|\|-[\|-\|\|\|-\|\|\|\|\|\|]PAM | NA | Intergenic | NA | NA |

| | | | | | | | | | | | |
|---|---|---|---|---|---|---|---|---|---|---|---|
| T31480 | TCGGGGTTCTTCCTCTGCGGTGG | 1000 | | | | | | | | | |
| Oligo fwd | CACCTCGGGGTTCTTCCTCTGCGG | | | | | | | | | | |
| Oligo rev | AAACCCGCAGAGGAAGAACCCCGA | | | | | | | | | | |
| Chromosome | start | end | strand | MM | target_seq | PAM | alignment | distance | position | gene name | gene id |
| GU305914.1 | 37392 | 37414 | - | 0 | TCGGGGTTCTTCCTCTGCGG | TGG | \|\|\|\|\|\|\|\|[\|\|\|\|\|\|\|\|\|\|\|\|]PAM | NA | Intergenic | NA | NA |

| | | | | | | | | | | | |
|---|---|---|---|---|---|---|---|---|---|---|---|
| T31481 | CGGGGTTCTTCCTCTGCGGTGGG | 1000 | | | | | | | | | |
| Oligo fwd | CACCCGGGGTTCTTCCTCTGCGGT | | | | | | | | | | |
| Oligo rev | AAACACCGCAGAGGAAGAACCCCG | | | | | | | | | | |
| Chromosome | start | end | strand | MM | target_seq | PAM | alignment | distance | position | gene name | gene id |
| GU305914.1 | 37391 | 37413 | - | 0 | CGGGGTTCTTCCTCTGCGGT | GGG | \|\|\|\|\|\|\|\|[\|\|\|\|\|\|\|\|\|\|\|\|]PAM | NA | Intergenic | NA | NA |

| | | | | | | | | | | | |
|---|---|---|---|---|---|---|---|---|---|---|---|
| T31482 | TCCTCTGCGGTGGGCTGCTCCGG | 1000 | | | | | | | | | |
| Oligo fwd | CACCTCCTCTGCGGTGGGCTGCTC | | | | | | | | | | |
| Oligo rev | AAACGAGCAGCCCACCGCAGAGGA | | | | | | | | | | |
| Chromosome | start | end | strand | MM | target_seq | PAM | alignment | distance | position | gene name | gene id |
| GU305914.1 | 37382 | 37404 | - | 0 | TCCTCTGCGGTGGGCTGCTC | CGG | \|\|\|\|\|\|\|\|[\|\|\|\|\|\|\|\|\|\|\|\|]PAM | NA | Intergenic | NA | NA |

| | | | | | | | | | | | |
|---|---|---|---|---|---|---|---|---|---|---|---|
| T31483 | CGGTGGGCTGCTCCGGATGACGG | 1000 | | | | | | | | | |
| Oligo fwd | CACCCGGTGGGCTGCTCCGGATGA | | | | | | | | | | |
| Oligo rev | AAACTCATCCGGAGCAGCCCACCG | | | | | | | | | | |
| Chromosome | start | end | strand | MM | target_seq | PAM | alignment | distance | position | gene name | gene id |
| GU305914.1 | 37375 | 37397 | - | 0 | CGGTGGGCTGCTCCGGATGA | CGG | \|\|\|\|\|\|\|\|[\|\|\|\|\|\|\|\|\|\|\|\|]PAM | NA | Intergenic | NA | NA |

| | | | | | | | | | | | |
|---|---|---|---|---|---|---|---|---|---|---|---|
| T31484 | CTGCTCCGGATGACGGAAGCTGG | 1000 | | | | | | | | | |
| Oligo fwd | CACCCTGCTCCGGATGACGGAAGC | | | | | | | | | | |
| Oligo rev | AAACGCTTCCGTCATCCGGAGCAG | | | | | | | | | | |
| Chromosome | start | end | strand | MM | target_seq | PAM | alignment | distance | position | gene name | gene id |
| GU305914.1 | 37368 | 37390 | - | 0 | CTGCTCCGGATGACGGAAGC | TGG | \|\|\|\|\|\|\|\|[\|\|\|\|\|\|\|\|\|\|\|\|]PAM | NA | Intergenic | NA | NA |

| | | |
|---|---|---|
| T31485 | TGACGGAAGCTGGTGCCGACCGG | 1000 |
| Oligo fwd | CACCTGACGGAAGCTGGTGCCGAC | |
| Oligo rev | AAACGTCGGCACCAGCTTCCGTCA | |

Chromosome start end strand MM target_seq PAM alignment distance position gene name gene id
GU305914.1 37358 37380 - 0 TGACGGAAGCTGGTGCCGAC CGG ||||||||[||||||||||||]PAM NA Intergenic NA NA

T31486 GACGGAAGCTGGTGCCGACCGGG 1000
Oligo fwd CACCGACGGAAGCTGGTGCCGACC
Oligo rev AAACGGTCGGCACCAGCTTCCGTC
Chromosome start end strand MM target_seq PAM alignment distance position gene name gene id
GU305914.1 37357 37379 - 0 GACGGAAGCTGGTGCCGACC GGG ||||||||[||||||||||||]PAM NA Intergenic NA NA

T31487 ACGGAAGCTGGTGCCGACCGGGG 1000
Oligo fwd CACCACGGAAGCTGGTGCCGACCG
Oligo rev AAACCGGTCGGCACCAGCTTCCGT
Chromosome start end strand MM target_seq PAM alignment distance position gene name gene id
GU305914.1 37356 37378 - 0 ACGGAAGCTGGTGCCGACCG GGG ||||||||[||||||||||||]PAM NA Intergenic NA NA

T31488 CGGAAGCTGGTGCCGACCGGGGG 1000
Oligo fwd CACCCGGAAGCTGGTGCCGACCGG
Oligo rev AAACCCGGTCGGCACCAGCTTCCG
Chromosome start end strand MM target_seq PAM alignment distance position gene name gene id
GU305914.1 37355 37377 - 0 CGGAAGCTGGTGCCGACCGG GGG ||||||||[||||||||||||]PAM NA Intergenic NA NA

T31489 GTGCCGACCGGGGGCTCTTGAGG 1000
Oligo fwd CACCGTGCCGACCGGGGGCTCTTG
Oligo rev AAACCAAGAGCCCCCGGTCGGCAC
Chromosome start end strand MM target_seq PAM alignment distance position gene name gene id
GU305914.1 37346 37368 - 0 GTGCCGACCGGGGGCTCTTG AGG ||||||||[||||||||||||]PAM NA Intergenic NA NA

T31490 GCTCTTGAGGCCGCCGAAACTGG 1000
Oligo fwd CACCGCTCTTGAGGCCGCCGAAAC
Oligo rev AAACGTTTCGGCGGCCTCAAGAGC
Chromosome start end strand MM target_seq PAM alignment distance position gene name gene id
GU305914.1 37333 37355 - 0 GCTCTTGAGGCCGCCGAAAC TGG ||||||||[||||||||||||]PAM NA Intergenic NA NA

T31491 CCGAAACTGGCGCTCTTCTCTGG 1000
Oligo fwd CACCCCGAAACTGGCGCTCTTCTC
Oligo rev AAACGAGAAGAGCGCCAGTTTCGG
Chromosome start end strand MM target_seq PAM alignment distance position gene name gene id
GU305914.1 37320 37342 - 0 CCGAAACTGGCGCTCTTCTC TGG ||||||||[||||||||||||]PAM NA Intergenic NA NA

T31492 CGAAACTGGCGCTCTTCTCTGGG 1000
Oligo fwd CACCCGAAACTGGCGCTCTTCTCT
Oligo rev AAACAGAGAAGAGCGCCAGTTTCG
Chromosome start end strand MM target_seq PAM alignment distance position gene name gene id
GU305914.1 37319 37341 - 0 CGAAACTGGCGCTCTTCTCT GGG ||||||||[||||||||||||]PAM NA Intergenic NA NA

T31493 GAAACTGGCGCTCTTCTCTGGGG 1000
Oligo fwd CACCGAAACTGGCGCTCTTCTCTG
Oligo rev AAACCAGAGAAGAGCGCCAGTTTC
Chromosome start end strand MM target_seq PAM alignment distance position gene name gene id
GU305914.1 37318 37340 - 0 GAAACTGGCGCTCTTCTCTG GGG ||||||||[||||||||||||]PAM NA Intergenic NA NA

T31494 ACTGGCGCTCTTCTCTGGGGTGG 1000
Oligo fwd CACCACTGGCGCTCTTCTCTGGGG
Oligo rev AAACCCCCAGAGAAGAGCGCCAGT
Chromosome start end strand MM target_seq PAM alignment distance position gene name gene id
GU305914.1 37315 37337 - 0 ACTGGCGCTCTTCTCTGGGG TGG ||||||||[||||||||||||]PAM NA Intergenic NA NA

T31495 TCTGGGGTGGTCGTCGTGTCCGG 1000
Oligo fwd CACCTCTGGGGTGGTCGTCGTGTC
Oligo rev AAACGACACGACGACCACCCCAGA
Chromosome start end strand MM target_seq PAM alignment distance position gene name gene id
GU305914.1 37302 37324 - 0 TCTGGGGTGGTCGTCGTGTC CGG ||||||||[||||||||||||]PAM NA Intergenic NA NA

T31496 TTTTCGCTGCTTCGCCTGCATGG 1000
Oligo fwd CACCTTTTCGCTGCTTCGCCTGCA
Oligo rev AAACTGCAGGCGAAGCAGCGAAAA
Chromosome start end strand MM target_seq PAM alignment distance position gene name gene id
GU305914.1 37279 37301 - 0 TTTTCGCTGCTTCGCCTGCA TGG ||||||||[||||||||||||]PAM NA Intergenic NA NA

T31497 CTGCTTCGCCTGCATGGAGACGG 1000
Oligo fwd CACCCTGCTTCGCCTGCATGGAGA
Oligo rev AAACTCTCCATGCAGGCGAAGCAG
Chromosome start end strand MM target_seq PAM alignment distance position gene name gene id
GU305914.1 37273 37295 - 0 CTGCTTCGCCTGCATGGAGA CGG ||||||||[||||||||||||]PAM NA Intergenic NA NA

T31498 CTTCGCCTGCATGGAGACGGCGG 1000
Oligo fwd CACCCTTCGCCTGCATGGAGACGG
Oligo rev AAACCCGTCTCCATGCAGGCGAAG
Chromosome start end strand MM target_seq PAM alignment distance position gene name gene id
GU305914.1 37270 37292 - 0 CTTCGCCTGCATGGAGACGG CGG ||||||||[||||||||||||]PAM NA Intergenic NA NA

T31499 TGCATGGAGACGGCGGCGTCTGG 1000
Oligo fwd CACCTGCATGGAGACGGCGGCGTC
Oligo rev AAACGACGCCGCCGTCTCCATGCA
Chromosome start end strand MM target_seq PAM alignment distance position gene name gene id
GU305914.1 37263 37285 - 0 TGCATGGAGACGGCGGCGTC TGG ||||||||[||||||||||||]PAM NA Intergenic NA NA

T31500 GCATGGAGACGGCGGCGTCTGGG 1000
Oligo fwd CACCGCATGGAGACGGCGGCGTCT

| | | | | | | | | | | | |
|---|---|---|---|---|---|---|---|---|---|---|---|
| Oligo rev | AAACAGACGCCGCCGTCTCCATGC | | | | | | | | | | |
| Chromosome | start | end | strand | MM | target_seq | PAM | alignment | distance | position | gene name | gene id |
| GU305914.1 | 37262 | 37284 | - | 0 | GCATGGAGACGGCGGCGTCT | GGG | \|\|\|\|\|\|\|\|[\|\|\|\|\|\|\|\|\|\|\|\|]PAM | NA | Intergenic | NA | NA |
| GU305914.1 | 195204 | 195226 | - | 4 | GTATCGAGACGGCGGCGCCG | TGG | \|-\|\|-\|\|\|[\|\|\|\|\|\|\|\|\|-\|-]PAM | NA | Intergenic | NA | NA |
| T31501 | CATGGAGACGGCGGCGTCTGGGG | 1000 | | | | | | | | | |
| Oligo fwd | CACCCATGGAGACGGCGGCGTCTG | | | | | | | | | | |
| Oligo rev | AAACCAGACGCCGCCGTCTCCATG | | | | | | | | | | |
| Chromosome | start | end | strand | MM | target_seq | PAM | alignment | distance | position | gene name | gene id |
| GU305914.1 | 37261 | 37283 | - | 0 | CATGGAGACGGCGGCGTCTG | GGG | \|\|\|\|\|\|\|\|[\|\|\|\|\|\|\|\|\|\|\|\|]PAM | NA | Intergenic | NA | NA |
| T31502 | CGTCTGGGGCGCTGCGCTTGAGG | 1000 | | | | | | | | | |
| Oligo fwd | CACCCGTCTGGGGCGCTGCGCTTG | | | | | | | | | | |
| Oligo rev | AAACCAAGCGCAGCGCCCCAGACG | | | | | | | | | | |
| Chromosome | start | end | strand | MM | target_seq | PAM | alignment | distance | position | gene name | gene id |
| GU305914.1 | 37247 | 37269 | - | 0 | CGTCTGGGGCGCTGCGCTTG | AGG | \|\|\|\|\|\|\|\|[\|\|\|\|\|\|\|\|\|\|\|\|]PAM | NA | Intergenic | NA | NA |
| T31503 | GCGCTTGAGGCAGCCGCTCTCGG | 1000 | | | | | | | | | |
| Oligo fwd | CACCGCGCTTGAGGCAGCCGCTCT | | | | | | | | | | |
| Oligo rev | AAACAGAGCGGCTGCCTCAAGCGC | | | | | | | | | | |
| Chromosome | start | end | strand | MM | target_seq | PAM | alignment | distance | position | gene name | gene id |
| GU305914.1 | 37234 | 37256 | - | 0 | GCGCTTGAGGCAGCCGCTCT | CGG | \|\|\|\|\|\|\|\|[\|\|\|\|\|\|\|\|\|\|\|\|]PAM | NA | Intergenic | NA | NA |
| T31504 | GAGGCAGCCGCTCTCGGTGACGG | 1000 | | | | | | | | | |
| Oligo fwd | CACCGAGGCAGCCGCTCTCGGTGA | | | | | | | | | | |
| Oligo rev | AAACTCACCGAGAGCGGCTGCCTC | | | | | | | | | | |
| Chromosome | start | end | strand | MM | target_seq | PAM | alignment | distance | position | gene name | gene id |
| GU305914.1 | 37228 | 37250 | - | 0 | GAGGCAGCCGCTCTCGGTGA | CGG | \|\|\|\|\|\|\|\|[\|\|\|\|\|\|\|\|\|\|\|\|]PAM | NA | Intergenic | NA | NA |
| T31505 | GCAGCCGCTCTCGGTGACGGCGG | 1000 | | | | | | | | | |
| Oligo fwd | CACCGCAGCCGCTCTCGGTGACGG | | | | | | | | | | |
| Oligo rev | AAACCCGTCACCGAGAGCGGCTGC | | | | | | | | | | |
| Chromosome | start | end | strand | MM | target_seq | PAM | alignment | distance | position | gene name | gene id |
| GU305914.1 | 37225 | 37247 | - | 0 | GCAGCCGCTCTCGGTGACGG | CGG | \|\|\|\|\|\|\|\|[\|\|\|\|\|\|\|\|\|\|\|\|]PAM | NA | Intergenic | NA | NA |
| T31506 | GCCGCTCTCGGTGACGGCGGCGG | 1000 | | | | | | | | | |
| Oligo fwd | CACCGCCGCTCTCGGTGACGGCGG | | | | | | | | | | |
| Oligo rev | AAACCCGCCGTCACCGAGAGCGGC | | | | | | | | | | |
| Chromosome | start | end | strand | MM | target_seq | PAM | alignment | distance | position | gene name | gene id |
| GU305914.1 | 37222 | 37244 | - | 0 | GCCGCTCTCGGTGACGGCGG | CGG | \|\|\|\|\|\|\|\|[\|\|\|\|\|\|\|\|\|\|\|\|]PAM | NA | Intergenic | NA | NA |
| GU305914.1 | 52719 | 52741 | + | 4 | GCCGCTGGCGGTGGCGGTGG | CGG | \|\|\|\|\|\|--[\|\|\|\|\|-\|\|\|-\|\|]PAM | NA | Intergenic | NA | NA |
| T31507 | CGATGTTGACACTGCAACTGCGG | 1000 | | | | | | | | | |
| Oligo fwd | CACCCGATGTTGACACTGCAACTG | | | | | | | | | | |
| Oligo rev | AAACCAGTTGCAGTGTCAACATCG | | | | | | | | | | |
| Chromosome | start | end | strand | MM | target_seq | PAM | alignment | distance | position | gene name | gene id |
| GU305914.1 | 37178 | 37200 | - | 0 | CGATGTTGACACTGCAACTG | CGG | \|\|\|\|\|\|\|\|[\|\|\|\|\|\|\|\|\|\|\|\|]PAM | NA | Intergenic | NA | NA |
| T31508 | ACACTGCAACTGCGGATCTGCGG | 1000 | | | | | | | | | |
| Oligo fwd | CACCACACTGCAACTGCGGATCTG | | | | | | | | | | |
| Oligo rev | AAACCAGATCCGCAGTTGCAGTGT | | | | | | | | | | |
| Chromosome | start | end | strand | MM | target_seq | PAM | alignment | distance | position | gene name | gene id |
| GU305914.1 | 37170 | 37192 | - | 0 | ACACTGCAACTGCGGATCTG | CGG | \|\|\|\|\|\|\|\|[\|\|\|\|\|\|\|\|\|\|\|\|]PAM | NA | Intergenic | NA | NA |
| T31509 | GCAACTGCGGATCTGCGGAGCGG | 1000 | | | | | | | | | |
| Oligo fwd | CACCGCAACTGCGGATCTGCGGAG | | | | | | | | | | |
| Oligo rev | AAACCTCCGCAGATCCGCAGTTGC | | | | | | | | | | |
| Chromosome | start | end | strand | MM | target_seq | PAM | alignment | distance | position | gene name | gene id |
| GU305914.1 | 37165 | 37187 | - | 0 | GCAACTGCGGATCTGCGGAG | CGG | \|\|\|\|\|\|\|\|[\|\|\|\|\|\|\|\|\|\|\|\|]PAM | NA | Intergenic | NA | NA |
| T31510 | CAACTGCGGATCTGCGGAGCGGG | 1000 | | | | | | | | | |
| Oligo fwd | CACCCAACTGCGGATCTGCGGAGC | | | | | | | | | | |
| Oligo rev | AAACGCTCCGCAGATCCGCAGTTG | | | | | | | | | | |
| Chromosome | start | end | strand | MM | target_seq | PAM | alignment | distance | position | gene name | gene id |
| GU305914.1 | 37164 | 37186 | - | 0 | CAACTGCGGATCTGCGGAGC | GGG | \|\|\|\|\|\|\|\|[\|\|\|\|\|\|\|\|\|\|\|\|]PAM | NA | Intergenic | NA | NA |
| T31511 | AACTGCGGATCTGCGGAGCGGGG | 1000 | | | | | | | | | |
| Oligo fwd | CACCAACTGCGGATCTGCGGAGCG | | | | | | | | | | |
| Oligo rev | AAACCGCTCCGCAGATCCGCAGTT | | | | | | | | | | |
| Chromosome | start | end | strand | MM | target_seq | PAM | alignment | distance | position | gene name | gene id |
| GU305914.1 | 37163 | 37185 | - | 0 | AACTGCGGATCTGCGGAGCG | GGG | \|\|\|\|\|\|\|\|[\|\|\|\|\|\|\|\|\|\|\|\|]PAM | NA | Intergenic | NA | NA |
| T31512 | ACTGCGGATCTGCGGAGCGGGGG | 1000 | | | | | | | | | |
| Oligo fwd | CACCACTGCGGATCTGCGGAGCGG | | | | | | | | | | |
| Oligo rev | AAACCCGCTCCGCAGATCCGCAGT | | | | | | | | | | |
| Chromosome | start | end | strand | MM | target_seq | PAM | alignment | distance | position | gene name | gene id |
| GU305914.1 | 37162 | 37184 | - | 0 | ACTGCGGATCTGCGGAGCGG | GGG | \|\|\|\|\|\|\|\|[\|\|\|\|\|\|\|\|\|\|\|\|]PAM | NA | Intergenic | NA | NA |
| T31513 | CGGATCTGCGGAGCGGGGGCCGG | 1000 | | | | | | | | | |
| Oligo fwd | CACCCGGATCTGCGGAGCGGGGGC | | | | | | | | | | |
| Oligo rev | AAACGCCCCCGCTCCGCAGATCCG | | | | | | | | | | |
| Chromosome | start | end | strand | MM | target_seq | PAM | alignment | distance | position | gene name | gene id |
| GU305914.1 | 37158 | 37180 | - | 0 | CGGATCTGCGGAGCGGGGGC | CGG | \|\|\|\|\|\|\|\|[\|\|\|\|\|\|\|\|\|\|\|\|]PAM | NA | Intergenic | NA | NA |
| T31514 | AGCGGGGGCCGGCTCTTTGATGG | 1000 | | | | | | | | | |
| Oligo fwd | CACCAGCGGGGGCCGGCTCTTTGA | | | | | | | | | | |
| Oligo rev | AAACTCAAAGAGCCGGCCCCCGCT | | | | | | | | | | |
| Chromosome | start | end | strand | MM | target_seq | PAM | alignment | distance | position | gene name | gene id |
| GU305914.1 | 37147 | 37169 | - | 0 | AGCGGGGGCCGGCTCTTTGA | TGG | \|\|\|\|\|\|\|\|[\|\|\|\|\|\|\|\|\|\|\|\|]PAM | NA | Intergenic | NA | NA |

T31515 CGCTGACCGATCTGCGCTCTCGG 1000
Oligo fwd CACCCGCTGACCGATCTGCGCTCT
Oligo rev AAACAGAGCGCAGATCGGTCAGCG
Chromosome start end strand MM target_seq PAM alignment distance position gene name gene id
GU305914.1 37124 37146 - 0 CGCTGACCGATCTGCGCTCT CGG ||||||||[||||||||||||]PAM NA Intergenic NA NA

T31516 GCTGACCGATCTGCGCTCTCGGG 1000
Oligo fwd CACCGCTGACCGATCTGCGCTCTC
Oligo rev AAACGAGAGCGCAGATCGGTCAGC
Chromosome start end strand MM target_seq PAM alignment distance position gene name gene id
GU305914.1 37123 37145 - 0 GCTGACCGATCTGCGCTCTC GGG ||||||||[||||||||||||]PAM NA Intergenic NA NA

T31517 CTGACCGATCTGCGCTCTCGGGG 1000
Oligo fwd CACCCTGACCGATCTGCGCTCTCG
Oligo rev AAACCGAGAGCGCAGATCGGTCAG
Chromosome start end strand MM target_seq PAM alignment distance position gene name gene id
GU305914.1 37122 37144 - 0 CTGACCGATCTGCGCTCTCG GGG ||||||||[||||||||||||]PAM NA Intergenic NA NA

T31518 TGACCGATCTGCGCTCTCGGGGG 1000
Oligo fwd CACCTGACCGATCTGCGCTCTCGG
Oligo rev AAACCCGAGAGCGCAGATCGGTCA
Chromosome start end strand MM target_seq PAM alignment distance position gene name gene id
GU305914.1 37121 37143 - 0 TGACCGATCTGCGCTCTCGG GGG ||||||||[||||||||||||]PAM NA Intergenic NA NA

T31519 GACCGATCTGCGCTCTCGGGGGG 1000
Oligo fwd CACCGACCGATCTGCGCTCTCGGG
Oligo rev AAACCCCGAGAGCGCAGATCGGTC
Chromosome start end strand MM target_seq PAM alignment distance position gene name gene id
GU305914.1 37120 37142 - 0 GACCGATCTGCGCTCTCGGG GGG ||||||||[||||||||||||]PAM NA Intergenic NA NA

T31520 TGCATTCTTGTGAGACGAAGAGG 1000
Oligo fwd CACCTGCATTCTTGTGAGACGAAG
Oligo rev AAACCTTCGTCTCACAAGAATGCA
Chromosome start end strand MM target_seq PAM alignment distance position gene name gene id
GU305914.1 37093 37115 - 0 TGCATTCTTGTGAGACGAAG AGG ||||||||[||||||||||||]PAM NA Intergenic NA NA

T31521 TTGTGAGACGAAGAGGACGACGG 1000
Oligo fwd CACCTTGTGAGACGAAGAGGACGA
Oligo rev AAACTCGTCCTCTTCGTCTCACAA
Chromosome start end strand MM target_seq PAM alignment distance position gene name gene id
GU305914.1 37086 37108 - 0 TTGTGAGACGAAGAGGACGA CGG ||||||||[||||||||||||]PAM NA Intergenic NA NA

T31522 GAGGACGACGGTCGACTCTCTGG 1000
Oligo fwd CACCGAGGACGACGGTCGACTCTC
Oligo rev AAACGAGAGTCGACCGTCGTCCTC
Chromosome start end strand MM target_seq PAM alignment distance position gene name gene id
GU305914.1 37074 37096 - 0 GAGGACGACGGTCGACTCTC TGG ||||||||[||||||||||||]PAM NA Intergenic NA NA

T31523 GACGACGGTCGACTCTCTGGCGG 1000
Oligo fwd CACCGACGACGGTCGACTCTCTGG
Oligo rev AAACCCAGAGAGTCGACCGTCGTC
Chromosome start end strand MM target_seq PAM alignment distance position gene name gene id
GU305914.1 37071 37093 - 0 GACGACGGTCGACTCTCTGG CGG ||||||||[||||||||||||]PAM NA Intergenic NA NA

T31524 ACGACGGTCGACTCTCTGGCGGG 1000
Oligo fwd CACCACGACGGTCGACTCTCTGGC
Oligo rev AAACGCCAGAGAGTCGACCGTCGT
Chromosome start end strand MM target_seq PAM alignment distance position gene name gene id
GU305914.1 37070 37092 - 0 ACGACGGTCGACTCTCTGGC GGG ||||||||[||||||||||||]PAM NA Intergenic NA NA

T31525 ACGGTCGACTCTCTGGCGGGCGG 1000
Oligo fwd CACCACGGTCGACTCTCTGGCGGG
Oligo rev AAACCCCGCCAGAGAGTCGACCGT
Chromosome start end strand MM target_seq PAM alignment distance position gene name gene id
GU305914.1 37067 37089 - 0 ACGGTCGACTCTCTGGCGGG CGG ||||||||[||||||||||||]PAM NA Intergenic NA NA

T31526 GACTCTCTGGCGGGCGGTAGCGG 1000
Oligo fwd CACCGACTCTCTGGCGGGCGGTAG
Oligo rev AAACCTACCGCCCGCCAGAGAGTC
Chromosome start end strand MM target_seq PAM alignment distance position gene name gene id
GU305914.1 37061 37083 - 0 GACTCTCTGGCGGGCGGTAG CGG ||||||||[||||||||||||]PAM NA Intergenic NA NA

T31527 GGGCGGTAGCGGTCTTTATAAGG 1000
Oligo fwd CACCGGGCGGTAGCGGTCTTTATA
Oligo rev AAACTATAAAGACCGCTACCGCCC
Chromosome start end strand MM target_seq PAM alignment distance position gene name gene id
GU305914.1 37050 37072 - 0 GGGCGGTAGCGGTCTTTATA AGG ||||||||[||||||||||||]PAM NA Intergenic NA NA

T31528 TAGCGGTCTTTATAAGGCTTCGG 1000
Oligo fwd CACCTAGCGGTCTTTATAAGGCTT
Oligo rev AAACAAGCCTTATAAAGACCGCTA
Chromosome start end strand MM target_seq PAM alignment distance position gene name gene id
GU305914.1 37044 37066 - 0 TAGCGGTCTTTATAAGGCTT CGG ||||||||[||||||||||||]PAM NA Intergenic NA NA

T31529 AGCGGTCTTTATAAGGCTTCGGG 1000
Oligo fwd CACCAGCGGTCTTTATAAGGCTTC
Oligo rev AAACGAAGCCTTATAAAGACCGCT
Chromosome start end strand MM target_seq PAM alignment distance position gene name gene id

GU305914.1 37043 37065 - 0 AGCGGTCTTTATAAGGCTTC GGG ||||||||[||||||||||||]PAM NA Intergenic NA NA

T31530 TTCGCGATTTGTCTCGATGATGG 1000
Oligo fwd CACCTTCGCGATTTGTCTCGATGA
Oligo rev AAACTCATCGAGACAAATCGCGAA
Chromosome start end strand MM target_seq PAM alignment distance position gene name gene id
GU305914.1 37010 37032 - 0 TTCGCGATTTGTCTCGATGA TGG ||||||||[||||||||||||]PAM NA Intergenic NA NA

T31531 TCGATGATGGTCTCGAGACACGG 1000
Oligo fwd CACCTCGATGATGGTCTCGAGACA
Oligo rev AAACTGTCTCGAGACCATCATCGA
Chromosome start end strand MM target_seq PAM alignment distance position gene name gene id
GU305914.1 36997 37019 - 0 TCGATGATGGTCTCGAGACA CGG ||||||||[||||||||||||]PAM NA Intergenic NA NA

T31532 GATGGTCTCGAGACACGGATCGG 1000
Oligo fwd CACCGATGGTCTCGAGACACGGAT
Oligo rev AAACATCCGTGTCTCGAGACCATC
Chromosome start end strand MM target_seq PAM alignment distance position gene name gene id
GU305914.1 36992 37014 - 0 GATGGTCTCGAGACACGGAT CGG ||||||||[||||||||||||]PAM NA Intergenic NA NA

T31533 ATGGTCTCGAGACACGGATCGGG 1000
Oligo fwd CACCATGGTCTCGAGACACGGATC
Oligo rev AAACGATCCGTGTCTCGAGACCAT
Chromosome start end strand MM target_seq PAM alignment distance position gene name gene id
GU305914.1 36991 37013 - 0 ATGGTCTCGAGACACGGATC GGG ||||||||[||||||||||||]PAM NA Intergenic NA NA

T31534 TCGAGACACGGATCGGGACCTGG 1000
Oligo fwd CACCTCGAGACACGGATCGGGACC
Oligo rev AAACGGTCCCGATCCGTGTCTCGA
Chromosome start end strand MM target_seq PAM alignment distance position gene name gene id
GU305914.1 36985 37007 - 0 TCGAGACACGGATCGGGACC TGG ||||||||[||||||||||||]PAM NA Intergenic NA NA

T31535 GGACCTGGATCTCTCCCTCGAGG 1000
Oligo fwd CACCGGACCTGGATCTCTCCCTCG
Oligo rev AAACCGAGGGAGAGATCCAGGTCC
Chromosome start end strand MM target_seq PAM alignment distance position gene name gene id
GU305914.1 36970 36992 - 0 GGACCTGGATCTCTCCCTCG AGG ||||||||[||||||||||||]PAM NA Intergenic NA NA

T31536 CTGCTGATGTTGTCCGCGTCAGG 1000
Oligo fwd CACCCTGCTGATGTTGTCCGCGTC
Oligo rev AAACGACGCGGACAACATCAGCAG
Chromosome start end strand MM target_seq PAM alignment distance position gene name gene id
GU305914.1 36933 36955 - 0 CTGCTGATGTTGTCCGCGTC AGG ||||||||[||||||||||||]PAM NA Intergenic NA NA

T31537 CGCGTCAGGACGCCTGCTGCTGG 1000
Oligo fwd CACCCGCGTCAGGACGCCTGCTGC
Oligo rev AAACGCAGCAGGCGTCCTGACGCG
Chromosome start end strand MM target_seq PAM alignment distance position gene name gene id
GU305914.1 36919 36941 - 0 CGCGTCAGGACGCCTGCTGC TGG ||||||||[||||||||||||]PAM NA Intergenic NA NA

T31538 GCGTCAGGACGCCTGCTGCTGGG 1000
Oligo fwd CACCGCGTCAGGACGCCTGCTGCT
Oligo rev AAACAGCAGCAGGCGTCCTGACGC
Chromosome start end strand MM target_seq PAM alignment distance position gene name gene id
GU305914.1 36918 36940 - 0 GCGTCAGGACGCCTGCTGCT GGG ||||||||[||||||||||||]PAM NA Intergenic NA NA

T31539 TCAGGACGCCTGCTGCTGGGCGG 1000
Oligo fwd CACCTCAGGACGCCTGCTGCTGGG
Oligo rev AAACCCCAGCAGCAGGCGTCCTGA
Chromosome start end strand MM target_seq PAM alignment distance position gene name gene id
GU305914.1 36915 36937 - 0 TCAGGACGCCTGCTGCTGGG CGG ||||||||[||||||||||||]PAM NA Intergenic NA NA

T31540 GGCGATGCGCGTTCTGTTTCTGG 1000
Oligo fwd CACCGGCGATGCGCGTTCTGTTTC
Oligo rev AAACGAAACAGAACGCGCATCGCC
Chromosome start end strand MM target_seq PAM alignment distance position gene name gene id
GU305914.1 36894 36916 - 0 GGCGATGCGCGTTCTGTTTC TGG ||||||||[||||||||||||]PAM NA Intergenic NA NA

T31541 TCCATCCCTCTGTTCTCGTTAGG 1000
Oligo fwd CACCTCCATCCCTCTGTTCTCGTT
Oligo rev AAACAACGAGAACAGAGGGATGGA
Chromosome start end strand MM target_seq PAM alignment distance position gene name gene id
GU305914.1 36867 36889 - 0 TCCATCCCTCTGTTCTCGTT AGG ||||||||[||||||||||||]PAM NA Intergenic NA NA

T31542 GGAGAATTCAGAGAGTCTCTCGG 1000
Oligo fwd CACCGGAGAATTCAGAGAGTCTCT
Oligo rev AAACAGAGACTCTCTGAATTCTCC
Chromosome start end strand MM target_seq PAM alignment distance position gene name gene id
GU305914.1 36846 36868 - 0 GGAGAATTCAGAGAGTCTCT CGG ||||||||[||||||||||||]PAM NA Intergenic NA NA

T31543 AGAGAGTCTCTCGGAAGTAGAGG 1000
Oligo fwd CACCAGAGAGTCTCTCGGAAGTAG
Oligo rev AAACCTACTTCCGAGAGACTCTCT
Chromosome start end strand MM target_seq PAM alignment distance position gene name gene id
GU305914.1 36837 36859 - 0 AGAGAGTCTCTCGGAAGTAG AGG ||||||||[||||||||||||]PAM NA Intergenic NA NA

T31544 TCGGAAGTAGAGGTGTGTTGTGG 1000
Oligo fwd CACCTCGGAAGTAGAGGTGTGTTG
Oligo rev AAACCAACACACCTCTACTTCCGA

<table>
<tr><td>Chromosome</td><td>start</td><td>end</td><td>strand</td><td>MM</td><td>target_seq</td><td>PAM</td><td>alignment</td><td>distance</td><td>position</td><td>gene name</td><td>gene id</td></tr>
<tr><td>GU305914.1</td><td>36827</td><td>36849</td><td>-</td><td>0</td><td>TCGGAAGTAGAGGTGTGTTG</td><td>TGG</td><td>||||||||[||||||||||||]PAM</td><td>NA</td><td>Intergenic</td><td>NA</td><td>NA</td></tr>
<tr><td></td><td></td><td></td><td></td><td></td><td></td><td></td><td></td><td></td><td></td><td></td><td></td></tr>
<tr><td>T31545</td><td>AGGTGTGTTGTGGTGATGTCAGG</td><td>1000</td><td></td><td></td><td></td><td></td><td></td><td></td><td></td><td></td><td></td></tr>
<tr><td>Oligo fwd</td><td>CACCAGGTGTGTTGTGGTGATGTC</td><td></td><td></td><td></td><td></td><td></td><td></td><td></td><td></td><td></td><td></td></tr>
<tr><td>Oligo rev</td><td>AAACGACATCACCACAACACACCT</td><td></td><td></td><td></td><td></td><td></td><td></td><td></td><td></td><td></td><td></td></tr>
<tr><td>Chromosome</td><td>start</td><td>end</td><td>strand</td><td>MM</td><td>target_seq</td><td>PAM</td><td>alignment</td><td>distance</td><td>position</td><td>gene name</td><td>gene id</td></tr>
<tr><td>GU305914.1</td><td>36817</td><td>36839</td><td>-</td><td>0</td><td>AGGTGTGTTGTGGTGATGTC</td><td>AGG</td><td>||||||||[||||||||||||]PAM</td><td>NA</td><td>Intergenic</td><td>NA</td><td>NA</td></tr>
<tr><td></td><td></td><td></td><td></td><td></td><td></td><td></td><td></td><td></td><td></td><td></td><td></td></tr>
<tr><td>T31546</td><td>TCCACGCTCGCGTATAAAATAGG</td><td>1000</td><td></td><td></td><td></td><td></td><td></td><td></td><td></td><td></td><td></td></tr>
<tr><td>Oligo fwd</td><td>CACCTCCACGCTCGCGTATAAAAT</td><td></td><td></td><td></td><td></td><td></td><td></td><td></td><td></td><td></td><td></td></tr>
<tr><td>Oligo rev</td><td>AAACATTTTATACGCGAGCGTGGA</td><td></td><td></td><td></td><td></td><td></td><td></td><td></td><td></td><td></td><td></td></tr>
<tr><td>Chromosome</td><td>start</td><td>end</td><td>strand</td><td>MM</td><td>target_seq</td><td>PAM</td><td>alignment</td><td>distance</td><td>position</td><td>gene name</td><td>gene id</td></tr>
<tr><td>GU305914.1</td><td>36775</td><td>36797</td><td>-</td><td>0</td><td>TCCACGCTCGCGTATAAAAT</td><td>AGG</td><td>||||||||[||||||||||||]PAM</td><td>NA</td><td>Intergenic</td><td>NA</td><td>NA</td></tr>
<tr><td></td><td></td><td></td><td></td><td></td><td></td><td></td><td></td><td></td><td></td><td></td><td></td></tr>
<tr><td>T31547</td><td>TGCGCTTCAGACTCTGCGACAGG</td><td>1000</td><td></td><td></td><td></td><td></td><td></td><td></td><td></td><td></td><td></td></tr>
<tr><td>Oligo fwd</td><td>CACCTGCGCTTCAGACTCTGCGAC</td><td></td><td></td><td></td><td></td><td></td><td></td><td></td><td></td><td></td><td></td></tr>
<tr><td>Oligo rev</td><td>AAACGTCGCAGAGTCTGAAGCGCA</td><td></td><td></td><td></td><td></td><td></td><td></td><td></td><td></td><td></td><td></td></tr>
<tr><td>Chromosome</td><td>start</td><td>end</td><td>strand</td><td>MM</td><td>target_seq</td><td>PAM</td><td>alignment</td><td>distance</td><td>position</td><td>gene name</td><td>gene id</td></tr>
<tr><td>GU305914.1</td><td>36739</td><td>36761</td><td>-</td><td>0</td><td>TGCGCTTCAGACTCTGCGAC</td><td>AGG</td><td>||||||||[||||||||||||]PAM</td><td>NA</td><td>Intergenic</td><td>NA</td><td>NA</td></tr>
<tr><td></td><td></td><td></td><td></td><td></td><td></td><td></td><td></td><td></td><td></td><td></td><td></td></tr>
<tr><td>T31548</td><td>TGCGACAGGTATCTTCTCACCGG</td><td>1000</td><td></td><td></td><td></td><td></td><td></td><td></td><td></td><td></td><td></td></tr>
<tr><td>Oligo fwd</td><td>CACCTGCGACAGGTATCTTCTCAC</td><td></td><td></td><td></td><td></td><td></td><td></td><td></td><td></td><td></td><td></td></tr>
<tr><td>Oligo rev</td><td>AAACGTGAGAAGATACCTGTCGCA</td><td></td><td></td><td></td><td></td><td></td><td></td><td></td><td></td><td></td><td></td></tr>
<tr><td>Chromosome</td><td>start</td><td>end</td><td>strand</td><td>MM</td><td>target_seq</td><td>PAM</td><td>alignment</td><td>distance</td><td>position</td><td>gene name</td><td>gene id</td></tr>
<tr><td>GU305914.1</td><td>36725</td><td>36747</td><td>-</td><td>0</td><td>TGCGACAGGTATCTTCTCAC</td><td>CGG</td><td>||||||||[||||||||||||]PAM</td><td>NA</td><td>Intergenic</td><td>NA</td><td>NA</td></tr>
<tr><td></td><td></td><td></td><td></td><td></td><td></td><td></td><td></td><td></td><td></td><td></td><td></td></tr>
<tr><td>T31549</td><td>GCGACAGGTATCTTCTCACCGGG</td><td>1000</td><td></td><td></td><td></td><td></td><td></td><td></td><td></td><td></td><td></td></tr>
<tr><td>Oligo fwd</td><td>CACCGCGACAGGTATCTTCTCACC</td><td></td><td></td><td></td><td></td><td></td><td></td><td></td><td></td><td></td><td></td></tr>
<tr><td>Oligo rev</td><td>AAACGGTGAGAAGATACCTGTCGC</td><td></td><td></td><td></td><td></td><td></td><td></td><td></td><td></td><td></td><td></td></tr>
<tr><td>Chromosome</td><td>start</td><td>end</td><td>strand</td><td>MM</td><td>target_seq</td><td>PAM</td><td>alignment</td><td>distance</td><td>position</td><td>gene name</td><td>gene id</td></tr>
<tr><td>GU305914.1</td><td>36724</td><td>36746</td><td>-</td><td>0</td><td>GCGACAGGTATCTTCTCACC</td><td>GGG</td><td>||||||||[||||||||||||]PAM</td><td>NA</td><td>Intergenic</td><td>NA</td><td>NA</td></tr>
<tr><td></td><td></td><td></td><td></td><td></td><td></td><td></td><td></td><td></td><td></td><td></td><td></td></tr>
<tr><td>T31550</td><td>CGACAGGTATCTTCTCACCGGGG</td><td>1000</td><td></td><td></td><td></td><td></td><td></td><td></td><td></td><td></td><td></td></tr>
<tr><td>Oligo fwd</td><td>CACCCGACAGGTATCTTCTCACCG</td><td></td><td></td><td></td><td></td><td></td><td></td><td></td><td></td><td></td><td></td></tr>
<tr><td>Oligo rev</td><td>AAACCGGTGAGAAGATACCTGTCG</td><td></td><td></td><td></td><td></td><td></td><td></td><td></td><td></td><td></td><td></td></tr>
<tr><td>Chromosome</td><td>start</td><td>end</td><td>strand</td><td>MM</td><td>target_seq</td><td>PAM</td><td>alignment</td><td>distance</td><td>position</td><td>gene name</td><td>gene id</td></tr>
<tr><td>GU305914.1</td><td>36723</td><td>36745</td><td>-</td><td>0</td><td>CGACAGGTATCTTCTCACCG</td><td>GGG</td><td>||||||||[||||||||||||]PAM</td><td>NA</td><td>Intergenic</td><td>NA</td><td>NA</td></tr>
<tr><td></td><td></td><td></td><td></td><td></td><td></td><td></td><td></td><td></td><td></td><td></td><td></td></tr>
<tr><td>T31551</td><td>CGCGATCGCCGACGTTGATCTGG</td><td>1000</td><td></td><td></td><td></td><td></td><td></td><td></td><td></td><td></td><td></td></tr>
<tr><td>Oligo fwd</td><td>CACCCGCGATCGCCGACGTTGATC</td><td></td><td></td><td></td><td></td><td></td><td></td><td></td><td></td><td></td><td></td></tr>
<tr><td>Oligo rev</td><td>AAACGATCAACGTCGGCGATCGCG</td><td></td><td></td><td></td><td></td><td></td><td></td><td></td><td></td><td></td><td></td></tr>
<tr><td>Chromosome</td><td>start</td><td>end</td><td>strand</td><td>MM</td><td>target_seq</td><td>PAM</td><td>alignment</td><td>distance</td><td>position</td><td>gene name</td><td>gene id</td></tr>
<tr><td>GU305914.1</td><td>36684</td><td>36706</td><td>-</td><td>0</td><td>CGCGATCGCCGACGTTGATC</td><td>TGG</td><td>||||||||[||||||||||||]PAM</td><td>NA</td><td>Intergenic</td><td>NA</td><td>NA</td></tr>
<tr><td></td><td></td><td></td><td></td><td></td><td></td><td></td><td></td><td></td><td></td><td></td><td></td></tr>
<tr><td>T31552</td><td>GCGATCGCCGACGTTGATCTGGG</td><td>1000</td><td></td><td></td><td></td><td></td><td></td><td></td><td></td><td></td><td></td></tr>
<tr><td>Oligo fwd</td><td>CACCGCGATCGCCGACGTTGATCT</td><td></td><td></td><td></td><td></td><td></td><td></td><td></td><td></td><td></td><td></td></tr>
<tr><td>Oligo rev</td><td>AAACAGATCAACGTCGGCGATCGC</td><td></td><td></td><td></td><td></td><td></td><td></td><td></td><td></td><td></td><td></td></tr>
<tr><td>Chromosome</td><td>start</td><td>end</td><td>strand</td><td>MM</td><td>target_seq</td><td>PAM</td><td>alignment</td><td>distance</td><td>position</td><td>gene name</td><td>gene id</td></tr>
<tr><td>GU305914.1</td><td>36683</td><td>36705</td><td>-</td><td>0</td><td>GCGATCGCCGACGTTGATCT</td><td>GGG</td><td>||||||||[||||||||||||]PAM</td><td>NA</td><td>Intergenic</td><td>NA</td><td>NA</td></tr>
<tr><td></td><td></td><td></td><td></td><td></td><td></td><td></td><td></td><td></td><td></td><td></td><td></td></tr>
<tr><td>T31553</td><td>CGATCGCCGACGTTGATCTGGGG</td><td>1000</td><td></td><td></td><td></td><td></td><td></td><td></td><td></td><td></td><td></td></tr>
<tr><td>Oligo fwd</td><td>CACCCGATCGCCGACGTTGATCTG</td><td></td><td></td><td></td><td></td><td></td><td></td><td></td><td></td><td></td><td></td></tr>
<tr><td>Oligo rev</td><td>AAACCAGATCAACGTCGGCGATCG</td><td></td><td></td><td></td><td></td><td></td><td></td><td></td><td></td><td></td><td></td></tr>
<tr><td>Chromosome</td><td>start</td><td>end</td><td>strand</td><td>MM</td><td>target_seq</td><td>PAM</td><td>alignment</td><td>distance</td><td>position</td><td>gene name</td><td>gene id</td></tr>
<tr><td>GU305914.1</td><td>36682</td><td>36704</td><td>-</td><td>0</td><td>CGATCGCCGACGTTGATCTG</td><td>GGG</td><td>||||||||[||||||||||||]PAM</td><td>NA</td><td>Intergenic</td><td>NA</td><td>NA</td></tr>
<tr><td></td><td></td><td></td><td></td><td></td><td></td><td></td><td></td><td></td><td></td><td></td><td></td></tr>
<tr><td>T31554</td><td>GATCGCCGACGTTGATCTGGGGG</td><td>1000</td><td></td><td></td><td></td><td></td><td></td><td></td><td></td><td></td><td></td></tr>
<tr><td>Oligo fwd</td><td>CACCGATCGCCGACGTTGATCTGG</td><td></td><td></td><td></td><td></td><td></td><td></td><td></td><td></td><td></td><td></td></tr>
<tr><td>Oligo rev</td><td>AAACCCAGATCAACGTCGGCGATC</td><td></td><td></td><td></td><td></td><td></td><td></td><td></td><td></td><td></td><td></td></tr>
<tr><td>Chromosome</td><td>start</td><td>end</td><td>strand</td><td>MM</td><td>target_seq</td><td>PAM</td><td>alignment</td><td>distance</td><td>position</td><td>gene name</td><td>gene id</td></tr>
<tr><td>GU305914.1</td><td>36681</td><td>36703</td><td>-</td><td>0</td><td>GATCGCCGACGTTGATCTGG</td><td>GGG</td><td>||||||||[||||||||||||]PAM</td><td>NA</td><td>Intergenic</td><td>NA</td><td>NA</td></tr>
<tr><td></td><td></td><td></td><td></td><td></td><td></td><td></td><td></td><td></td><td></td><td></td><td></td></tr>
<tr><td>T31555</td><td>ATCGCCGACGTTGATCTGGGGGG</td><td>1000</td><td></td><td></td><td></td><td></td><td></td><td></td><td></td><td></td><td></td></tr>
<tr><td>Oligo fwd</td><td>CACCATCGCCGACGTTGATCTGGG</td><td></td><td></td><td></td><td></td><td></td><td></td><td></td><td></td><td></td><td></td></tr>
<tr><td>Oligo rev</td><td>AAACCCCAGATCAACGTCGGCGAT</td><td></td><td></td><td></td><td></td><td></td><td></td><td></td><td></td><td></td><td></td></tr>
<tr><td>Chromosome</td><td>start</td><td>end</td><td>strand</td><td>MM</td><td>target_seq</td><td>PAM</td><td>alignment</td><td>distance</td><td>position</td><td>gene name</td><td>gene id</td></tr>
<tr><td>GU305914.1</td><td>36680</td><td>36702</td><td>-</td><td>0</td><td>ATCGCCGACGTTGATCTGGG</td><td>GGG</td><td>||||||||[||||||||||||]PAM</td><td>NA</td><td>Intergenic</td><td>NA</td><td>NA</td></tr>
<tr><td></td><td></td><td></td><td></td><td></td><td></td><td></td><td></td><td></td><td></td><td></td><td></td></tr>
<tr><td>T31556</td><td>TGGGGGGCGTGTTTCGCGTACGG</td><td>1000</td><td></td><td></td><td></td><td></td><td></td><td></td><td></td><td></td><td></td></tr>
<tr><td>Oligo fwd</td><td>CACCTGGGGGGCGTGTTTCGCGTA</td><td></td><td></td><td></td><td></td><td></td><td></td><td></td><td></td><td></td><td></td></tr>
<tr><td>Oligo rev</td><td>AAACTACGCGAAACACGCCCCCCA</td><td></td><td></td><td></td><td></td><td></td><td></td><td></td><td></td><td></td><td></td></tr>
<tr><td>Chromosome</td><td>start</td><td>end</td><td>strand</td><td>MM</td><td>target_seq</td><td>PAM</td><td>alignment</td><td>distance</td><td>position</td><td>gene name</td><td>gene id</td></tr>
<tr><td>GU305914.1</td><td>36664</td><td>36686</td><td>-</td><td>0</td><td>TGGGGGGCGTGTTTCGCGTA</td><td>CGG</td><td>||||||||[||||||||||||]PAM</td><td>NA</td><td>Intergenic</td><td>NA</td><td>NA</td></tr>
<tr><td></td><td></td><td></td><td></td><td></td><td></td><td></td><td></td><td></td><td></td><td></td><td></td></tr>
<tr><td>T31557</td><td>GCGTGTTTCGCGTACGGTTATGG</td><td>1000</td><td></td><td></td><td></td><td></td><td></td><td></td><td></td><td></td><td></td></tr>
<tr><td>Oligo fwd</td><td>CACCGCGTGTTTCGCGTACGGTTA</td><td></td><td></td><td></td><td></td><td></td><td></td><td></td><td></td><td></td><td></td></tr>
<tr><td>Oligo rev</td><td>AAACTAACCGTACGCGAAACACGC</td><td></td><td></td><td></td><td></td><td></td><td></td><td></td><td></td><td></td><td></td></tr>
<tr><td>Chromosome</td><td>start</td><td>end</td><td>strand</td><td>MM</td><td>target_seq</td><td>PAM</td><td>alignment</td><td>distance</td><td>position</td><td>gene name</td><td>gene id</td></tr>
<tr><td>GU305914.1</td><td>36658</td><td>36680</td><td>-</td><td>0</td><td>GCGTGTTTCGCGTACGGTTA</td><td>TGG</td><td>||||||||[||||||||||||]PAM</td><td>NA</td><td>Intergenic</td><td>NA</td><td>NA</td></tr>
<tr><td></td><td></td><td></td><td></td><td></td><td></td><td></td><td></td><td></td><td></td><td></td><td></td></tr>
<tr><td>T31558</td><td>TGGAAACTAAAGAAGAGTTGTGG</td><td>1000</td><td></td><td></td><td></td><td></td><td></td><td></td><td></td><td></td><td></td></tr>
<tr><td>Oligo fwd</td><td>CACCTGGAAACTAAAGAAGAGTTG</td><td></td><td></td><td></td><td></td><td></td><td></td><td></td><td></td><td></td><td></td></tr>
<tr><td>Oligo rev</td><td>AAACCAACTCTTCTTTAGTTTCCA</td><td></td><td></td><td></td><td></td><td></td><td></td><td></td><td></td><td></td><td></td></tr>
<tr><td>Chromosome</td><td>start</td><td>end</td><td>strand</td><td>MM</td><td>target_seq</td><td>PAM</td><td>alignment</td><td>distance</td><td>position</td><td>gene name</td><td>gene id</td></tr>
<tr><td>GU305914.1</td><td>36638</td><td>36660</td><td>-</td><td>0</td><td>TGGAAACTAAAGAAGAGTTG</td><td>TGG</td><td>||||||||[||||||||||||]PAM</td><td>NA</td><td>Intergenic</td><td>NA</td><td>NA</td></tr>
<tr><td></td><td></td><td></td><td></td><td></td><td></td><td></td><td></td><td></td><td></td><td></td><td></td></tr>
<tr><td>T31559</td><td>GGAAACTAAAGAAGAGTTGTGGG</td><td>1000</td><td></td><td></td><td></td><td></td><td></td><td></td><td></td><td></td><td></td></tr>
<tr><td>Oligo fwd</td><td>CACCGGAAACTAAAGAAGAGTTGT</td><td></td><td></td><td></td><td></td><td></td><td></td><td></td><td></td><td></td><td></td></tr>
</table>

<table>
<tr><td>Oligo rev</td><td>AAACACAACTCTTCTTTAGTTTCC</td><td></td><td></td><td></td><td></td><td></td><td></td><td></td><td></td><td></td><td></td><td></td></tr>
<tr><td>Chromosome</td><td>start</td><td></td><td>end</td><td>strand</td><td>MM</td><td>target_seq</td><td>PAM</td><td>alignment</td><td>distance</td><td>position</td><td>gene name</td><td>gene id</td></tr>
<tr><td>GU305914.1</td><td></td><td>36637</td><td>36659</td><td>-</td><td>0</td><td>GGAAACTAAAGAAGAGTTGT</td><td>GGG</td><td>||||||||[||||||||||||]PAM</td><td>NA</td><td>Intergenic</td><td>NA</td><td>NA</td></tr>
<tr><td>T31560</td><td>GAAACTAAAGAAGAGTTGTGGGG</td><td></td><td>1000</td><td></td><td></td><td></td><td></td><td></td><td></td><td></td><td></td><td></td></tr>
<tr><td>Oligo fwd</td><td>CACCGAAACTAAAGAAGAGTTGTG</td><td></td><td></td><td></td><td></td><td></td><td></td><td></td><td></td><td></td><td></td><td></td></tr>
<tr><td>Oligo rev</td><td>AAACCACAACTCTTCTTTAGTTTC</td><td></td><td></td><td></td><td></td><td></td><td></td><td></td><td></td><td></td><td></td><td></td></tr>
<tr><td>Chromosome</td><td>start</td><td></td><td>end</td><td>strand</td><td>MM</td><td>target_seq</td><td>PAM</td><td>alignment</td><td>distance</td><td>position</td><td>gene name</td><td>gene id</td></tr>
<tr><td>GU305914.1</td><td></td><td>36636</td><td>36658</td><td>-</td><td>0</td><td>GAAACTAAAGAAGAGTTGTG</td><td>GGG</td><td>||||||||[||||||||||||]PAM</td><td>NA</td><td>Intergenic</td><td>NA</td><td>NA</td></tr>
<tr><td>T31561</td><td>GGGTAAAGCCTTTGATCTCATGG</td><td></td><td>1000</td><td></td><td></td><td></td><td></td><td></td><td></td><td></td><td></td><td></td></tr>
<tr><td>Oligo fwd</td><td>CACCGGGTAAAGCCTTTGATCTCA</td><td></td><td></td><td></td><td></td><td></td><td></td><td></td><td></td><td></td><td></td><td></td></tr>
<tr><td>Oligo rev</td><td>AAACTGAGATCAAAGGCTTTACCC</td><td></td><td></td><td></td><td></td><td></td><td></td><td></td><td></td><td></td><td></td><td></td></tr>
<tr><td>Chromosome</td><td>start</td><td></td><td>end</td><td>strand</td><td>MM</td><td>target_seq</td><td>PAM</td><td>alignment</td><td>distance</td><td>position</td><td>gene name</td><td>gene id</td></tr>
<tr><td>GU305914.1</td><td></td><td>36616</td><td>36638</td><td>-</td><td>0</td><td>GGGTAAAGCCTTTGATCTCA</td><td>TGG</td><td>||||||||[||||||||||||]PAM</td><td>NA</td><td>Intergenic</td><td>NA</td><td>NA</td></tr>
<tr><td>T31562</td><td>TGATCTCATGGTCAACTTCGCGG</td><td></td><td>1000</td><td></td><td></td><td></td><td></td><td></td><td></td><td></td><td></td><td></td></tr>
<tr><td>Oligo fwd</td><td>CACCTGATCTCATGGTCAACTTCG</td><td></td><td></td><td></td><td></td><td></td><td></td><td></td><td></td><td></td><td></td><td></td></tr>
<tr><td>Oligo rev</td><td>AAACCGAAGTTGACCATGAGATCA</td><td></td><td></td><td></td><td></td><td></td><td></td><td></td><td></td><td></td><td></td><td></td></tr>
<tr><td>Chromosome</td><td>start</td><td></td><td>end</td><td>strand</td><td>MM</td><td>target_seq</td><td>PAM</td><td>alignment</td><td>distance</td><td>position</td><td>gene name</td><td>gene id</td></tr>
<tr><td>GU305914.1</td><td></td><td>36604</td><td>36626</td><td>-</td><td>0</td><td>TGATCTCATGGTCAACTTCG</td><td>CGG</td><td>||||||||[||||||||||||]PAM</td><td>NA</td><td>Intergenic</td><td>NA</td><td>NA</td></tr>
<tr><td>GU305914.1</td><td></td><td>29365</td><td>29387</td><td>-</td><td>4</td><td>TGACACCATGGTCTACTTCG</td><td>CGG</td><td>|||---||[|||||-||||||]PAM</td><td>NA</td><td>Intergenic</td><td>NA</td><td>NA</td></tr>
<tr><td>T31563</td><td>GATCTCATGGTCAACTTCGCGGG</td><td></td><td>1000</td><td></td><td></td><td></td><td></td><td></td><td></td><td></td><td></td><td></td></tr>
<tr><td>Oligo fwd</td><td>CACCGATCTCATGGTCAACTTCGC</td><td></td><td></td><td></td><td></td><td></td><td></td><td></td><td></td><td></td><td></td><td></td></tr>
<tr><td>Oligo rev</td><td>AAACGCGAAGTTGACCATGAGATC</td><td></td><td></td><td></td><td></td><td></td><td></td><td></td><td></td><td></td><td></td><td></td></tr>
<tr><td>Chromosome</td><td>start</td><td></td><td>end</td><td>strand</td><td>MM</td><td>target_seq</td><td>PAM</td><td>alignment</td><td>distance</td><td>position</td><td>gene name</td><td>gene id</td></tr>
<tr><td>GU305914.1</td><td></td><td>36603</td><td>36625</td><td>-</td><td>0</td><td>GATCTCATGGTCAACTTCGC</td><td>GGG</td><td>||||||||[||||||||||||]PAM</td><td>NA</td><td>Intergenic</td><td>NA</td><td>NA</td></tr>
<tr><td>T31564</td><td>ATGGTCAACTTCGCGGGCGCCGG</td><td></td><td>1000</td><td></td><td></td><td></td><td></td><td></td><td></td><td></td><td></td><td></td></tr>
<tr><td>Oligo fwd</td><td>CACCATGGTCAACTTCGCGGGCGC</td><td></td><td></td><td></td><td></td><td></td><td></td><td></td><td></td><td></td><td></td><td></td></tr>
<tr><td>Oligo rev</td><td>AAACGCGCCCGCGAAGTTGACCAT</td><td></td><td></td><td></td><td></td><td></td><td></td><td></td><td></td><td></td><td></td><td></td></tr>
<tr><td>Chromosome</td><td>start</td><td></td><td>end</td><td>strand</td><td>MM</td><td>target_seq</td><td>PAM</td><td>alignment</td><td>distance</td><td>position</td><td>gene name</td><td>gene id</td></tr>
<tr><td>GU305914.1</td><td></td><td>36597</td><td>36619</td><td>-</td><td>0</td><td>ATGGTCAACTTCGCGGGCGC</td><td>CGG</td><td>||||||||[||||||||||||]PAM</td><td>NA</td><td>Intergenic</td><td>NA</td><td>NA</td></tr>
<tr><td>T31565</td><td>TCAACTTCGCGGGCGCCGGACGG</td><td></td><td>1000</td><td></td><td></td><td></td><td></td><td></td><td></td><td></td><td></td><td></td></tr>
<tr><td>Oligo fwd</td><td>CACCTCAACTTCGCGGGCGCCGGA</td><td></td><td></td><td></td><td></td><td></td><td></td><td></td><td></td><td></td><td></td><td></td></tr>
<tr><td>Oligo rev</td><td>AAACTCCGGCGCCCGCGAAGTTGA</td><td></td><td></td><td></td><td></td><td></td><td></td><td></td><td></td><td></td><td></td><td></td></tr>
<tr><td>Chromosome</td><td>start</td><td></td><td>end</td><td>strand</td><td>MM</td><td>target_seq</td><td>PAM</td><td>alignment</td><td>distance</td><td>position</td><td>gene name</td><td>gene id</td></tr>
<tr><td>GU305914.1</td><td></td><td>36593</td><td>36615</td><td>-</td><td>0</td><td>TCAACTTCGCGGGCGCCGGA</td><td>CGG</td><td>||||||||[||||||||||||]PAM</td><td>NA</td><td>Intergenic</td><td>NA</td><td>NA</td></tr>
<tr><td>T31566</td><td>GATGCCCGTTAGCGACATGAAGG</td><td></td><td>1000</td><td></td><td></td><td></td><td></td><td></td><td></td><td></td><td></td><td></td></tr>
<tr><td>Oligo fwd</td><td>CACCGATGCCCGTTAGCGACATGA</td><td></td><td></td><td></td><td></td><td></td><td></td><td></td><td></td><td></td><td></td><td></td></tr>
<tr><td>Oligo rev</td><td>AAACTCATGTCGCTAACGGGCATC</td><td></td><td></td><td></td><td></td><td></td><td></td><td></td><td></td><td></td><td></td><td></td></tr>
<tr><td>Chromosome</td><td>start</td><td></td><td>end</td><td>strand</td><td>MM</td><td>target_seq</td><td>PAM</td><td>alignment</td><td>distance</td><td>position</td><td>gene name</td><td>gene id</td></tr>
<tr><td>GU305914.1</td><td></td><td>36571</td><td>36593</td><td>-</td><td>0</td><td>GATGCCCGTTAGCGACATGA</td><td>AGG</td><td>||||||||[||||||||||||]PAM</td><td>NA</td><td>Intergenic</td><td>NA</td><td>NA</td></tr>
<tr><td>T31567</td><td>GCGACATGAAGGTCGTGTCGCGG</td><td></td><td>1000</td><td></td><td></td><td></td><td></td><td></td><td></td><td></td><td></td><td></td></tr>
<tr><td>Oligo fwd</td><td>CACCGCGACATGAAGGTCGTGTCG</td><td></td><td></td><td></td><td></td><td></td><td></td><td></td><td></td><td></td><td></td><td></td></tr>
<tr><td>Oligo rev</td><td>AAACCGACACGACCTTCATGTCGC</td><td></td><td></td><td></td><td></td><td></td><td></td><td></td><td></td><td></td><td></td><td></td></tr>
<tr><td>Chromosome</td><td>start</td><td></td><td>end</td><td>strand</td><td>MM</td><td>target_seq</td><td>PAM</td><td>alignment</td><td>distance</td><td>position</td><td>gene name</td><td>gene id</td></tr>
<tr><td>GU305914.1</td><td></td><td>36560</td><td>36582</td><td>-</td><td>0</td><td>GCGACATGAAGGTCGTGTCG</td><td>CGG</td><td>||||||||[||||||||||||]PAM</td><td>NA</td><td>Intergenic</td><td>NA</td><td>NA</td></tr>
<tr><td>T31568</td><td>CATGAAGGTCGTGTCGCGGCTGG</td><td></td><td>1000</td><td></td><td></td><td></td><td></td><td></td><td></td><td></td><td></td><td></td></tr>
<tr><td>Oligo fwd</td><td>CACCCATGAAGGTCGTGTCGCGGC</td><td></td><td></td><td></td><td></td><td></td><td></td><td></td><td></td><td></td><td></td><td></td></tr>
<tr><td>Oligo rev</td><td>AAACGCCGCGACACGACCTTCATG</td><td></td><td></td><td></td><td></td><td></td><td></td><td></td><td></td><td></td><td></td><td></td></tr>
<tr><td>Chromosome</td><td>start</td><td></td><td>end</td><td>strand</td><td>MM</td><td>target_seq</td><td>PAM</td><td>alignment</td><td>distance</td><td>position</td><td>gene name</td><td>gene id</td></tr>
<tr><td>GU305914.1</td><td></td><td>36556</td><td>36578</td><td>-</td><td>0</td><td>CATGAAGGTCGTGTCGCGGC</td><td>TGG</td><td>||||||||[||||||||||||]PAM</td><td>NA</td><td>Intergenic</td><td>NA</td><td>NA</td></tr>
<tr><td>T31569</td><td>GCTGGCTACGATGTTGACTAAGG</td><td></td><td>1000</td><td></td><td></td><td></td><td></td><td></td><td></td><td></td><td></td><td></td></tr>
<tr><td>Oligo fwd</td><td>CACCGCTGGCTACGATGTTGACTA</td><td></td><td></td><td></td><td></td><td></td><td></td><td></td><td></td><td></td><td></td><td></td></tr>
<tr><td>Oligo rev</td><td>AAACTAGTCAACATCGTAGCCAGC</td><td></td><td></td><td></td><td></td><td></td><td></td><td></td><td></td><td></td><td></td><td></td></tr>
<tr><td>Chromosome</td><td>start</td><td></td><td>end</td><td>strand</td><td>MM</td><td>target_seq</td><td>PAM</td><td>alignment</td><td>distance</td><td>position</td><td>gene name</td><td>gene id</td></tr>
<tr><td>GU305914.1</td><td></td><td>36538</td><td>36560</td><td>-</td><td>0</td><td>GCTGGCTACGATGTTGACTA</td><td>AGG</td><td>||||||||[||||||||||||]PAM</td><td>NA</td><td>Intergenic</td><td>NA</td><td>NA</td></tr>
<tr><td>T31570</td><td>GATGTTGACTAAGGAAGAAGAGG</td><td></td><td>1000</td><td></td><td></td><td></td><td></td><td></td><td></td><td></td><td></td><td></td></tr>
<tr><td>Oligo fwd</td><td>CACCGATGTTGACTAAGGAAGAAG</td><td></td><td></td><td></td><td></td><td></td><td></td><td></td><td></td><td></td><td></td><td></td></tr>
<tr><td>Oligo rev</td><td>AAACCTTCTTCCTTAGTCAACATC</td><td></td><td></td><td></td><td></td><td></td><td></td><td></td><td></td><td></td><td></td><td></td></tr>
<tr><td>Chromosome</td><td>start</td><td></td><td>end</td><td>strand</td><td>MM</td><td>target_seq</td><td>PAM</td><td>alignment</td><td>distance</td><td>position</td><td>gene name</td><td>gene id</td></tr>
<tr><td>GU305914.1</td><td></td><td>36529</td><td>36551</td><td>-</td><td>0</td><td>GATGTTGACTAAGGAAGAAG</td><td>AGG</td><td>||||||||[||||||||||||]PAM</td><td>NA</td><td>Intergenic</td><td>NA</td><td>NA</td></tr>
<tr><td>T31571</td><td>AGAAGAGGCTCGCTTCGATAAGG</td><td></td><td>1000</td><td></td><td></td><td></td><td></td><td></td><td></td><td></td><td></td><td></td></tr>
<tr><td>Oligo fwd</td><td>CACCAGAAGAGGCTCGCTTCGATA</td><td></td><td></td><td></td><td></td><td></td><td></td><td></td><td></td><td></td><td></td><td></td></tr>
<tr><td>Oligo rev</td><td>AAACTATCGAAGCGAGCCTCTTCT</td><td></td><td></td><td></td><td></td><td></td><td></td><td></td><td></td><td></td><td></td><td></td></tr>
<tr><td>Chromosome</td><td>start</td><td></td><td>end</td><td>strand</td><td>MM</td><td>target_seq</td><td>PAM</td><td>alignment</td><td>distance</td><td>position</td><td>gene name</td><td>gene id</td></tr>
<tr><td>GU305914.1</td><td></td><td>36514</td><td>36536</td><td>-</td><td>0</td><td>AGAAGAGGCTCGCTTCGATA</td><td>AGG</td><td>||||||||[||||||||||||]PAM</td><td>NA</td><td>Intergenic</td><td>NA</td><td>NA</td></tr>
<tr><td>T31572</td><td>AGAGGCTCGCTTCGATAAGGAGG</td><td></td><td>1000</td><td></td><td></td><td></td><td></td><td></td><td></td><td></td><td></td><td></td></tr>
<tr><td>Oligo fwd</td><td>CACCAGAGGCTCGCTTCGATAAGG</td><td></td><td></td><td></td><td></td><td></td><td></td><td></td><td></td><td></td><td></td><td></td></tr>
<tr><td>Oligo rev</td><td>AAACCCTTATCGAAGCGAGCCTCT</td><td></td><td></td><td></td><td></td><td></td><td></td><td></td><td></td><td></td><td></td><td></td></tr>
<tr><td>Chromosome</td><td>start</td><td></td><td>end</td><td>strand</td><td>MM</td><td>target_seq</td><td>PAM</td><td>alignment</td><td>distance</td><td>position</td><td>gene name</td><td>gene id</td></tr>
<tr><td>GU305914.1</td><td></td><td>36511</td><td>36533</td><td>-</td><td>0</td><td>AGAGGCTCGCTTCGATAAGG</td><td>AGG</td><td>||||||||[||||||||||||]PAM</td><td>NA</td><td>Intergenic</td><td>NA</td><td>NA</td></tr>
<tr><td>T31573</td><td>TCGCTTCGATAAGGAGGTCGCGG</td><td></td><td>1000</td><td></td><td></td><td></td><td></td><td></td><td></td><td></td><td></td><td></td></tr>
<tr><td>Oligo fwd</td><td>CACCTCGCTTCGATAAGGAGGTCG</td><td></td><td></td><td></td><td></td><td></td><td></td><td></td><td></td><td></td><td></td><td></td></tr>
<tr><td>Oligo rev</td><td>AAACCGACCTCCTTATCGAAGCGA</td><td></td><td></td><td></td><td></td><td></td><td></td><td></td><td></td><td></td><td></td><td></td></tr>
<tr><td>Chromosome</td><td>start</td><td></td><td>end</td><td>strand</td><td>MM</td><td>target_seq</td><td>PAM</td><td>alignment</td><td>distance</td><td>position</td><td>gene name</td><td>gene id</td></tr>
<tr><td>GU305914.1</td><td></td><td>36505</td><td>36527</td><td>-</td><td>0</td><td>TCGCTTCGATAAGGAGGTCG</td><td>CGG</td><td>||||||||[||||||||||||]PAM</td><td>NA</td><td>Intergenic</td><td>NA</td><td>NA</td></tr>
</table>

| | | | | | | | | | | | |
|---|---|---|---|---|---|---|---|---|---|---|---|
| T31574 | CGATAAGGAGGTCGCGGACGAGG | 1000 | | | | | | | | | |
| Oligo fwd | CACCCGATAAGGAGGTCGCGGACG | | | | | | | | | | |
| Oligo rev | AAACCGTCCGCGACCTCCTTATCG | | | | | | | | | | |
| Chromosome | start | end | strand | MM | target_seq | PAM | alignment | distance | position | gene name | gene id |
| GU305914.1 | 36499 | 36521 | - | 0 | CGATAAGGAGGTCGCGGACG | AGG | \|\|\|\|\|\|\|\|[\|\|\|\|\|\|\|\|\|\|\|\|]PAM | NA | Intergenic | NA | NA |
| T31575 | CATACTAGCGTCTAGTGAAGAGG | 1000 | | | | | | | | | |
| Oligo fwd | CACCCATACTAGCGTCTAGTGAAG | | | | | | | | | | |
| Oligo rev | AAACCTTCACTAGACGCTAGTATG | | | | | | | | | | |
| Chromosome | start | end | strand | MM | target_seq | PAM | alignment | distance | position | gene name | gene id |
| GU305914.1 | 36457 | 36479 | - | 0 | CATACTAGCGTCTAGTGAAG | AGG | \|\|\|\|\|\|\|\|[\|\|\|\|\|\|\|\|\|\|\|\|]PAM | NA | Intergenic | NA | NA |
| T31576 | GAAGAGGATGAAAGCGAAAGTGG | 1000 | | | | | | | | | |
| Oligo fwd | CACCGAAGAGGATGAAAGCGAAAG | | | | | | | | | | |
| Oligo rev | AAACCTTTCGCTTTCATCCTCTTC | | | | | | | | | | |
| Chromosome | start | end | strand | MM | target_seq | PAM | alignment | distance | position | gene name | gene id |
| GU305914.1 | 36441 | 36463 | - | 0 | GAAGAGGATGAAAGCGAAAG | TGG | \|\|\|\|\|\|\|\|[\|\|\|\|\|\|\|\|\|\|\|\|]PAM | NA | Intergenic | NA | NA |
| T31577 | GAGGATGAAAGCGAAAGTGGCGG | 1000 | | | | | | | | | |
| Oligo fwd | CACCGAGGATGAAAGCGAAAGTGG | | | | | | | | | | |
| Oligo rev | AAACCCACTTTCGCTTTCATCCTC | | | | | | | | | | |
| Chromosome | start | end | strand | MM | target_seq | PAM | alignment | distance | position | gene name | gene id |
| GU305914.1 | 36438 | 36460 | - | 0 | GAGGATGAAAGCGAAAGTGG | CGG | \|\|\|\|\|\|\|\|[\|\|\|\|\|\|\|\|\|\|\|\|]PAM | NA | Intergenic | NA | NA |
| T31578 | GATGAAAGCGAAAGTGGCGGCGG | 1000 | | | | | | | | | |
| Oligo fwd | CACCGATGAAAGCGAAAGTGGCGG | | | | | | | | | | |
| Oligo rev | AAACCCGCCACTTTCGCTTTCATC | | | | | | | | | | |
| Chromosome | start | end | strand | MM | target_seq | PAM | alignment | distance | position | gene name | gene id |
| GU305914.1 | 36435 | 36457 | - | 0 | GATGAAAGCGAAAGTGGCGG | CGG | \|\|\|\|\|\|\|\|[\|\|\|\|\|\|\|\|\|\|\|\|]PAM | NA | Intergenic | NA | NA |
| T31579 | AGCGACGCACTCGCCTGTCGTGG | 1000 | | | | | | | | | |
| Oligo fwd | CACCAGCGACGCACTCGCCTGTCG | | | | | | | | | | |
| Oligo rev | AAACCGACAGGCGAGTGCGTCGCT | | | | | | | | | | |
| Chromosome | start | end | strand | MM | target_seq | PAM | alignment | distance | position | gene name | gene id |
| GU305914.1 | 36396 | 36418 | - | 0 | AGCGACGCACTCGCCTGTCG | TGG | \|\|\|\|\|\|\|\|[\|\|\|\|\|\|\|\|\|\|\|\|]PAM | NA | Intergenic | NA | NA |
| T31580 | CGCACTCGCCTGTCGTGGCTCGG | 1000 | | | | | | | | | |
| Oligo fwd | CACCCGCACTCGCCTGTCGTGGCT | | | | | | | | | | |
| Oligo rev | AAACAGCCACGACAGGCGAGTGCG | | | | | | | | | | |
| Chromosome | start | end | strand | MM | target_seq | PAM | alignment | distance | position | gene name | gene id |
| GU305914.1 | 36391 | 36413 | - | 0 | CGCACTCGCCTGTCGTGGCT | CGG | \|\|\|\|\|\|\|\|[\|\|\|\|\|\|\|\|\|\|\|\|]PAM | NA | Intergenic | NA | NA |
| T31581 | CAACCTTCGAAGATATCGAGAGG | 1000 | | | | | | | | | |
| Oligo fwd | CACCCAACCTTCGAAGATATCGAG | | | | | | | | | | |
| Oligo rev | AAACCTCGATATCTTCGAAGGTTG | | | | | | | | | | |
| Chromosome | start | end | strand | MM | target_seq | PAM | alignment | distance | position | gene name | gene id |
| GU305914.1 | 36350 | 36372 | - | 0 | CAACCTTCGAAGATATCGAG | AGG | \|\|\|\|\|\|\|\|[\|\|\|\|\|\|\|\|\|\|\|\|]PAM | NA | Intergenic | NA | NA |
| T31582 | GCGAAAAGCGCCGCTCCGAGCGG | 1000 | | | | | | | | | |
| Oligo fwd | CACCGCGAAAAGCGCCGCTCCGAG | | | | | | | | | | |
| Oligo rev | AAACCTCGGAGCGGCGCTTTTCGC | | | | | | | | | | |
| Chromosome | start | end | strand | MM | target_seq | PAM | alignment | distance | position | gene name | gene id |
| GU305914.1 | 36290 | 36312 | - | 0 | GCGAAAAGCGCCGCTCCGAG | CGG | \|\|\|\|\|\|\|\|[\|\|\|\|\|\|\|\|\|\|\|\|]PAM | NA | Intergenic | NA | NA |
| T31583 | AAAGCGCCGCTCCGAGCGGAAGG | 1000 | | | | | | | | | |
| Oligo fwd | CACCAAAGCGCCGCTCCGAGCGGA | | | | | | | | | | |
| Oligo rev | AAACTCCGCTCGGAGCGGCGCTTT | | | | | | | | | | |
| Chromosome | start | end | strand | MM | target_seq | PAM | alignment | distance | position | gene name | gene id |
| GU305914.1 | 36286 | 36308 | - | 0 | AAAGCGCCGCTCCGAGCGGA | AGG | \|\|\|\|\|\|\|\|[\|\|\|\|\|\|\|\|\|\|\|\|]PAM | NA | Intergenic | NA | NA |
| GU305914.1 | 94281 | 94303 | - | 4 | AAAACGCAGCGCCGAGCGAA | AGG | \|\|\|-\|\|\|-[\|\|-\|\|\|\|\|\|\|-\|]PAM | NA | Intergenic | NA | NA |
| T31584 | GAAGGCCAAGCGCAAAAGAAAGG | 1000 | | | | | | | | | |
| Oligo fwd | CACCGAAGGCCAAGCGCAAAAGAA | | | | | | | | | | |
| Oligo rev | AAACTTCTTTTGCGCTTGGCCTTC | | | | | | | | | | |
| Chromosome | start | end | strand | MM | target_seq | PAM | alignment | distance | position | gene name | gene id |
| GU305914.1 | 36268 | 36290 | - | 0 | GAAGGCCAAGCGCAAAAGAA | AGG | \|\|\|\|\|\|\|\|[\|\|\|\|\|\|\|\|\|\|\|\|]PAM | NA | Intergenic | NA | NA |
| T31585 | AAGGCCAAGCGCAAAAGAAAGGG | 1000 | | | | | | | | | |
| Oligo fwd | CACCAAGGCCAAGCGCAAAAGAAA | | | | | | | | | | |
| Oligo rev | AAACTTTCTTTTGCGCTTGGCCTT | | | | | | | | | | |
| Chromosome | start | end | strand | MM | target_seq | PAM | alignment | distance | position | gene name | gene id |
| GU305914.1 | 36267 | 36289 | - | 0 | AAGGCCAAGCGCAAAAGAAA | GGG | \|\|\|\|\|\|\|\|[\|\|\|\|\|\|\|\|\|\|\|\|]PAM | NA | Intergenic | NA | NA |
| T31586 | CGCAAAAGAAAGGGCGACCACGG | 1000 | | | | | | | | | |
| Oligo fwd | CACCCGCAAAAGAAAGGGCGACCA | | | | | | | | | | |
| Oligo rev | AAACTGGTCGCCCTTTCTTTTGCG | | | | | | | | | | |
| Chromosome | start | end | strand | MM | target_seq | PAM | alignment | distance | position | gene name | gene id |
| GU305914.1 | 36258 | 36280 | - | 0 | CGCAAAAGAAAGGGCGACCA | CGG | \|\|\|\|\|\|\|\|[\|\|\|\|\|\|\|\|\|\|\|\|]PAM | NA | Intergenic | NA | NA |
| T31587 | AAGAAAGGGCGACCACGGTGCGG | 1000 | | | | | | | | | |
| Oligo fwd | CACCAAGAAAGGGCGACCACGGTG | | | | | | | | | | |
| Oligo rev | AAACCACCGTGGTCGCCCTTTCTT | | | | | | | | | | |
| Chromosome | start | end | strand | MM | target_seq | PAM | alignment | distance | position | gene name | gene id |
| GU305914.1 | 36253 | 36275 | - | 0 | AAGAAAGGGCGACCACGGTG | CGG | \|\|\|\|\|\|\|\|[\|\|\|\|\|\|\|\|\|\|\|\|]PAM | NA | Intergenic | NA | NA |
| T31588 | TGCTGCCGTCGCTTCGTCCGCGG | 1000 | | | | | | | | | |
| Oligo fwd | CACCTGCTGCCGTCGCTTCGTCCG | | | | | | | | | | |
| Oligo rev | AAACCGGACGAAGCGACGGCAGCA | | | | | | | | | | |
| Chromosome | start | end | strand | MM | target_seq | PAM | alignment | distance | position | gene name | gene id |

| | | | | | | | | | | | |
|---|---|---|---|---|---|---|---|---|---|---|---|
| GU305914.1 | 36211 | 36233 | - | 0 | TGCTGCCGTCGCTTCGTCCG | CGG | \|\|\|\|\|\|\|\|[\|\|\|\|\|\|\|\|\|\|\|\|]PAM | NA | Intergenic | NA | NA |
| T31589 | ACAACCACCACAACCGCCGCCGG | 1000 | | | | | | | | | |
| Oligo fwd | CACCACAACCACCACAACCGCCGC | | | | | | | | | | |
| Oligo rev | AAACGCGGCGGTTGTGGTGGTTGT | | | | | | | | | | |
| Chromosome | start | end | strand | MM | target_seq | PAM | alignment | distance | position | gene name | gene id |
| GU305914.1 | 36165 | 36187 | - | 0 | ACAACCACCACAACCGCCGC | CGG | \|\|\|\|\|\|\|\|[\|\|\|\|\|\|\|\|\|\|\|\|]PAM | NA | Intergenic | NA | NA |
| GU305914.1 | 200143 | 200165 | + | 4 | AGAACGGCCACCACCGCCGC | AGG | \|-\|\|\|--\|[\|\|\|-\|\|\|\|\|\|\|\|]PAM | NA | Intergenic | NA | NA |
| T31590 | ACCACAACCGCCGCCGGCAGCGG | 1000 | | | | | | | | | |
| Oligo fwd | CACCACCACAACCGCCGCCGGCAG | | | | | | | | | | |
| Oligo rev | AAACCTGCCGGCGGCGGTTGTGGT | | | | | | | | | | |
| Chromosome | start | end | strand | MM | target_seq | PAM | alignment | distance | position | gene name | gene id |
| GU305914.1 | 36159 | 36181 | - | 0 | ACCACAACCGCCGCCGGCAG | CGG | \|\|\|\|\|\|\|\|[\|\|\|\|\|\|\|\|\|\|\|\|]PAM | NA | Intergenic | NA | NA |
| T31591 | CAGCGGCCCCGTTGCCGCCACGG | 1000 | | | | | | | | | |
| Oligo fwd | CACCCAGCGGCCCCGTTGCCGCCA | | | | | | | | | | |
| Oligo rev | AAACTGGCGGCAACGGGGCCGCTG | | | | | | | | | | |
| Chromosome | start | end | strand | MM | target_seq | PAM | alignment | distance | position | gene name | gene id |
| GU305914.1 | 36142 | 36164 | - | 0 | CAGCGGCCCCGTTGCCGCCA | CGG | \|\|\|\|\|\|\|\|[\|\|\|\|\|\|\|\|\|\|\|\|]PAM | NA | Intergenic | NA | NA |
| T31592 | ACGCCGCTCCTCTGTTCGTGAGG | 1000 | | | | | | | | | |
| Oligo fwd | CACCACGCCGCTCCTCTGTTCGTG | | | | | | | | | | |
| Oligo rev | AAACCACGAACAGAGGAGCGGCGT | | | | | | | | | | |
| Chromosome | start | end | strand | MM | target_seq | PAM | alignment | distance | position | gene name | gene id |
| GU305914.1 | 36107 | 36129 | - | 0 | ACGCCGCTCCTCTGTTCGTG | AGG | \|\|\|\|\|\|\|\|[\|\|\|\|\|\|\|\|\|\|\|\|]PAM | NA | Intergenic | NA | NA |
| T31593 | TTCGTGAGGAGTGTCCCTATCGG | 1000 | | | | | | | | | |
| Oligo fwd | CACCTTCGTGAGGAGTGTCCCTAT | | | | | | | | | | |
| Oligo rev | AAACATAGGGACACTCCTCACGAA | | | | | | | | | | |
| Chromosome | start | end | strand | MM | target_seq | PAM | alignment | distance | position | gene name | gene id |
| GU305914.1 | 36093 | 36115 | - | 0 | TTCGTGAGGAGTGTCCCTAT | CGG | \|\|\|\|\|\|\|\|[\|\|\|\|\|\|\|\|\|\|\|\|]PAM | NA | Intergenic | NA | NA |
| T31594 | TCCCGATCAGAAAATCGAGCAGG | 1000 | | | | | | | | | |
| Oligo fwd | CACCTCCCGATCAGAAAATCGAGC | | | | | | | | | | |
| Oligo rev | AAACGCTCGATTTTCTGATCGGGA | | | | | | | | | | |
| Chromosome | start | end | strand | MM | target_seq | PAM | alignment | distance | position | gene name | gene id |
| GU305914.1 | 36047 | 36069 | - | 0 | TCCCGATCAGAAAATCGAGC | AGG | \|\|\|\|\|\|\|\|[\|\|\|\|\|\|\|\|\|\|\|\|]PAM | NA | Intergenic | NA | NA |
| T31595 | ACACCACCATCTAGTAAGTGAGG | 1000 | | | | | | | | | |
| Oligo fwd | CACCACACCACCATCTAGTAAGTG | | | | | | | | | | |
| Oligo rev | AAACCACTTACTAGATGGTGGTGT | | | | | | | | | | |
| Chromosome | start | end | strand | MM | target_seq | PAM | alignment | distance | position | gene name | gene id |
| GU305914.1 | 36020 | 36042 | - | 0 | ACACCACCATCTAGTAAGTG | AGG | \|\|\|\|\|\|\|\|[\|\|\|\|\|\|\|\|\|\|\|\|]PAM | NA | Intergenic | NA | NA |
| T31596 | CACCATCTAGTAAGTGAGGCAGG | 1000 | | | | | | | | | |
| Oligo fwd | CACCCACCATCTAGTAAGTGAGGC | | | | | | | | | | |
| Oligo rev | AAACGCCTCACTTACTAGATGGTG | | | | | | | | | | |
| Chromosome | start | end | strand | MM | target_seq | PAM | alignment | distance | position | gene name | gene id |
| GU305914.1 | 36016 | 36038 | - | 0 | CACCATCTAGTAAGTGAGGC | AGG | \|\|\|\|\|\|\|\|[\|\|\|\|\|\|\|\|\|\|\|\|]PAM | NA | Intergenic | NA | NA |
| T31597 | AGTAAGTGAGGCAGGAGCAGCGG | 1000 | | | | | | | | | |
| Oligo fwd | CACCAGTAAGTGAGGCAGGAGCAG | | | | | | | | | | |
| Oligo rev | AAACCTGCTCCTGCCTCACTTACT | | | | | | | | | | |
| Chromosome | start | end | strand | MM | target_seq | PAM | alignment | distance | position | gene name | gene id |
| GU305914.1 | 36008 | 36030 | - | 0 | AGTAAGTGAGGCAGGAGCAG | CGG | \|\|\|\|\|\|\|\|[\|\|\|\|\|\|\|\|\|\|\|\|]PAM | NA | Intergenic | NA | NA |
| T31598 | GGCGCGAGTCCGACGACCTGCGG | 1000 | | | | | | | | | |
| Oligo fwd | CACCGGCGCGAGTCCGACGACCTG | | | | | | | | | | |
| Oligo rev | AAACCAGGTCGTCGGACTCGCGCC | | | | | | | | | | |
| Chromosome | start | end | strand | MM | target_seq | PAM | alignment | distance | position | gene name | gene id |
| GU305914.1 | 35987 | 36009 | - | 0 | GGCGCGAGTCCGACGACCTG | CGG | \|\|\|\|\|\|\|\|[\|\|\|\|\|\|\|\|\|\|\|\|]PAM | NA | Intergenic | NA | NA |
| T31599 | GGCCGCGACAGAGCACCGAACGG | 1000 | | | | | | | | | |
| Oligo fwd | CACCGGCCGCGACAGAGCACCGAA | | | | | | | | | | |
| Oligo rev | AAACTTCGGTGCTCTGTCGCGGCC | | | | | | | | | | |
| Chromosome | start | end | strand | MM | target_seq | PAM | alignment | distance | position | gene name | gene id |
| GU305914.1 | 35966 | 35988 | - | 0 | GGCCGCGACAGAGCACCGAA | CGG | \|\|\|\|\|\|\|\|[\|\|\|\|\|\|\|\|\|\|\|\|]PAM | NA | Intergenic | NA | NA |
| T31600 | AGCACCGAACGGTCGCCATGCGG | 1000 | | | | | | | | | |
| Oligo fwd | CACCAGCACCGAACGGTCGCCATG | | | | | | | | | | |
| Oligo rev | AAACCATGGCGACCGTTCGGTGCT | | | | | | | | | | |
| Chromosome | start | end | strand | MM | target_seq | PAM | alignment | distance | position | gene name | gene id |
| GU305914.1 | 35955 | 35977 | - | 0 | AGCACCGAACGGTCGCCATG | CGG | \|\|\|\|\|\|\|\|[\|\|\|\|\|\|\|\|\|\|\|\|]PAM | NA | Intergenic | NA | NA |
| T31601 | GCACCGAACGGTCGCCATGCGGG | 1000 | | | | | | | | | |
| Oligo fwd | CACCGCACCGAACGGTCGCCATGC | | | | | | | | | | |
| Oligo rev | AAACGCATGGCGACCGTTCGGTGC | | | | | | | | | | |
| Chromosome | start | end | strand | MM | target_seq | PAM | alignment | distance | position | gene name | gene id |
| GU305914.1 | 35954 | 35976 | - | 0 | GCACCGAACGGTCGCCATGC | GGG | \|\|\|\|\|\|\|\|[\|\|\|\|\|\|\|\|\|\|\|\|]PAM | NA | Intergenic | NA | NA |
| T31602 | CCGAACGGTCGCCATGCGGGAGG | 1000 | | | | | | | | | |
| Oligo fwd | CACCCCGAACGGTCGCCATGCGGG | | | | | | | | | | |
| Oligo rev | AAACCCCGCATGGCGACCGTTCGG | | | | | | | | | | |
| Chromosome | start | end | strand | MM | target_seq | PAM | alignment | distance | position | gene name | gene id |
| GU305914.1 | 35951 | 35973 | - | 0 | CCGAACGGTCGCCATGCGGG | AGG | \|\|\|\|\|\|\|\|[\|\|\|\|\|\|\|\|\|\|\|\|]PAM | NA | Intergenic | NA | NA |
| T31603 | TTTGCCAGCTCAGATCGTAGAGG | 1000 | | | | | | | | | |
| Oligo fwd | CACCTTTGCCAGCTCAGATCGTAG | | | | | | | | | | |

Oligo rev AAACCTACGATCTGAGCTGGCAAA
Chromosome start end strand MM target_seq PAM alignment distance position gene name gene id
GU305914.1 35920 35942 - 0 TTTGCCAGCTCAGATCGTAG AGG ||||||||[||||||||||||]PAM NA Intergenic NA NA

T31604 GCCAGCTCAGATCGTAGAGGAGG 1000
Oligo fwd CACCGCCAGCTCAGATCGTAGAGG
Oligo rev AAACCCTCTACGATCTGAGCTGGC
Chromosome start end strand MM target_seq PAM alignment distance position gene name gene id
GU305914.1 35917 35939 - 0 GCCAGCTCAGATCGTAGAGG AGG ||||||||[||||||||||||]PAM NA Intergenic NA NA

T31605 TCGTGCGATCGTATGAGTAGTGG 1000
Oligo fwd CACCTCGTGCGATCGTATGAGTAG
Oligo rev AAACCTACTCATACGATCGCACGA
Chromosome start end strand MM target_seq PAM alignment distance position gene name gene id
GU305914.1 35890 35912 - 0 TCGTGCGATCGTATGAGTAG TGG ||||||||[||||||||||||]PAM NA Intergenic NA NA

T31606 CGTGCGATCGTATGAGTAGTGGG 1000
Oligo fwd CACCCGTGCGATCGTATGAGTAGT
Oligo rev AAACACTACTCATACGATCGCACG
Chromosome start end strand MM target_seq PAM alignment distance position gene name gene id
GU305914.1 35889 35911 - 0 CGTGCGATCGTATGAGTAGT GGG ||||||||[||||||||||||]PAM NA Intergenic NA NA

T31607 AGTAGTGGGCGTTTCTATTTCGG 1000
Oligo fwd CACCAGTAGTGGGCGTTTCTATTT
Oligo rev AAACAAATAGAAACGCCCACTACT
Chromosome start end strand MM target_seq PAM alignment distance position gene name gene id
GU305914.1 35875 35897 - 0 AGTAGTGGGCGTTTCTATTT CGG ||||||||[||||||||||||]PAM NA Intergenic NA NA

T31608 GTAGTGGGCGTTTCTATTTCGGG 1000
Oligo fwd CACCGTAGTGGGCGTTTCTATTTC
Oligo rev AAACGAAATAGAAACGCCCACTAC
Chromosome start end strand MM target_seq PAM alignment distance position gene name gene id
GU305914.1 35874 35896 - 0 GTAGTGGGCGTTTCTATTTC GGG ||||||||[||||||||||||]PAM NA Intergenic NA NA

T31609 GGCGTTTCTATTTCGGGCAATGG 1000
Oligo fwd CACCGGCGTTTCTATTTCGGGCAA
Oligo rev AAACTTGCCCGAAATAGAAACGCC
Chromosome start end strand MM target_seq PAM alignment distance position gene name gene id
GU305914.1 35868 35890 - 0 GGCGTTTCTATTTCGGGCAA TGG ||||||||[||||||||||||]PAM NA Intergenic NA NA

T31610 GGCAATGGCATAAAGCCATTCGG 1000
Oligo fwd CACCGGCAATGGCATAAAGCCATT
Oligo rev AAACAATGGCTTTATGCCATTGCC
Chromosome start end strand MM target_seq PAM alignment distance position gene name gene id
GU305914.1 35853 35875 - 0 GGCAATGGCATAAAGCCATT CGG ||||||||[||||||||||||]PAM NA Intergenic NA NA

T31611 GCAATGGCATAAAGCCATTCGGG 1000
Oligo fwd CACCGCAATGGCATAAAGCCATTC
Oligo rev AAACGAATGGCTTTATGCCATTGC
Chromosome start end strand MM target_seq PAM alignment distance position gene name gene id
GU305914.1 35852 35874 - 0 GCAATGGCATAAAGCCATTC GGG ||||||||[||||||||||||]PAM NA Intergenic NA NA

T31612 AAAATCAGTCGATATGAGCCTGG 1000
Oligo fwd CACCAAAATCAGTCGATATGAGCC
Oligo rev AAACGGCTCATATCGACTGATTTT
Chromosome start end strand MM target_seq PAM alignment distance position gene name gene id
GU305914.1 35769 35791 - 0 AAAATCAGTCGATATGAGCC TGG ||||||||[||||||||||||]PAM NA Intergenic NA NA

T31613 GATATGAGCCTGGAAGCTGTCGG 1000
Oligo fwd CACCGATATGAGCCTGGAAGCTGT
Oligo rev AAACACAGCTTCCAGGCTCATATC
Chromosome start end strand MM target_seq PAM alignment distance position gene name gene id
GU305914.1 35759 35781 - 0 GATATGAGCCTGGAAGCTGT CGG ||||||||[||||||||||||]PAM NA Intergenic NA NA

T31614 GCCTGGAAGCTGTCGGTATGCGG 1000
Oligo fwd CACCGCCTGGAAGCTGTCGGTATG
Oligo rev AAACCATACCGACAGCTTCCAGGC
Chromosome start end strand MM target_seq PAM alignment distance position gene name gene id
GU305914.1 35752 35774 - 0 GCCTGGAAGCTGTCGGTATG CGG ||||||||[||||||||||||]PAM NA Intergenic NA NA

T31615 CAGCGACGACGACCTCGATGAGG 1000
Oligo fwd CACCCAGCGACGACGACCTCGATG
Oligo rev AAACCATCGAGGTCGTCGTCGCTG
Chromosome start end strand MM target_seq PAM alignment distance position gene name gene id
GU305914.1 35724 35746 - 0 CAGCGACGACGACCTCGATG AGG ||||||||[||||||||||||]PAM NA Intergenic NA NA

T31616 GTTCCTTCGCGACGATGAACAGG 1000
Oligo fwd CACCGTTCCTTCGCGACGATGAAC
Oligo rev AAACGTTCATCGTCGCGAAGGAAC
Chromosome start end strand MM target_seq PAM alignment distance position gene name gene id
GU305914.1 35700 35722 - 0 GTTCCTTCGCGACGATGAAC AGG ||||||||[||||||||||||]PAM NA Intergenic NA NA

T31617 TTCCTTCGCGACGATGAACAGGG 1000
Oligo fwd CACCTTCCTTCGCGACGATGAACA
Oligo rev AAACTGTTCATCGTCGCGAAGGAA
Chromosome start end strand MM target_seq PAM alignment distance position gene name gene id
GU305914.1 35699 35721 - 0 TTCCTTCGCGACGATGAACA GGG ||||||||[||||||||||||]PAM NA Intergenic NA NA

T31618 TCCTTCGCGACGATGAACAGGGG 1000

| | | | | | | | | | | | |
|---|---|---|---|---|---|---|---|---|---|---|---|
| Oligo fwd | CACCTCCTTCGCGACGATGAACAG | | | | | | | | | | |
| Oligo rev | AAACCTGTTCATCGTCGCGAAGGA | | | | | | | | | | |
| Chromosome | start | end | strand | MM | target_seq | PAM | alignment | distance | position | gene name | gene id |
| GU305914.1 | 35698 | 35720 | - | 0 | TCCTTCGCGACGATGAACAG | GGG | \|\|\|\|\|\|\|\|[\|\|\|\|\|\|\|\|\|\|\|\|]PAM | NA | Intergenic | NA | NA |
| T31619 | CCTTCGCGACGATGAACAGGGGG | 1000 | | | | | | | | | |
| Oligo fwd | CACCCCTTCGCGACGATGAACAGG | | | | | | | | | | |
| Oligo rev | AAACCCTGTTCATCGTCGCGAAGG | | | | | | | | | | |
| Chromosome | start | end | strand | MM | target_seq | PAM | alignment | distance | position | gene name | gene id |
| GU305914.1 | 35697 | 35719 | - | 0 | CCTTCGCGACGATGAACAGG | GGG | \|\|\|\|\|\|\|\|[\|\|\|\|\|\|\|\|\|\|\|\|]PAM | NA | Intergenic | NA | NA |
| T31620 | CCCCGCCGCCGCCGCCGCCGTGG | 1000 | | | | | | | | | |
| Oligo fwd | CACCCCCCGCCGCCGCCGCCGCCG | | | | | | | | | | |
| Oligo rev | AAACCGGCGGCGGCGGCGGCGGGG | | | | | | | | | | |
| Chromosome | start | end | strand | MM | target_seq | PAM | alignment | distance | position | gene name | gene id |
| GU305914.1 | 35655 | 35677 | - | 0 | CCCCGCCGCCGCCGCCGCCG | TGG | \|\|\|\|\|\|\|\|[\|\|\|\|\|\|\|\|\|\|\|\|]PAM | NA | Intergenic | NA | NA |
| GU305914.1 | 45592 | 45614 | - | 4 | GGCGGCTGCCGCCGCCGCCG | CGG | --\|-\|\|-\|[\|\|\|\|\|\|\|\|\|\|\|\|]PAM | NA | Intergenic | NA | NA |
| GU305914.1 | 26444 | 26466 | - | 4 | CTCGGAGGCCGCCGCCGCCG | TGG | \|-\|-\|--\|[\|\|\|\|\|\|\|\|\|\|\|\|]PAM | NA | Intergenic | NA | NA |
| GU305914.1 | 130312 | 130334 | + | 4 | CGCAGCAGCAGCCGCCGCCG | CGG | \|-\|-\|\|-\|[\|-\|\|\|\|\|\|\|\|\|\|]PAM | NA | Intergenic | NA | NA |
| GU305914.1 | 56775 | 56797 | - | 3 | TGCCGCCGCCGCCGCCGCTG | CGG | --\|\|\|\|\|\|[\|\|\|\|\|\|\|\|\|\|-\|]PAM | NA | Intergenic | NA | NA |
| GU305914.1 | 98350 | 98372 | + | 3 | CGCCGCCGCCACCGCCGCCT | AGG | \|-\|\|\|\|\|\|[\|\|-\|\|\|\|\|\|\|\|-]PAM | NA | Intergenic | NA | NA |
| GU305914.1 | 56772 | 56794 | - | 3 | CGCCGCCGCCGCCGCTGCGG | CGG | \|-\|\|\|\|\|\|[\|\|\|\|\|\|\|-\|\|-\|]PAM | NA | Intergenic | NA | NA |
| GU305914.1 | 154400 | 154422 | - | 3 | CGCCGCCGCCGCCACCGCCC | GGG | \|-\|\|\|\|\|\|[\|\|\|\|\|-\|\|\|\|\|-]PAM | NA | Intergenic | NA | NA |
| GU305914.1 | 86815 | 86837 | + | 4 | CCCGGCCCCCGACGCCGCCT | CGG | \|\|\|-\|\|\|-[\|\|\|-\|\|\|\|\|\|\|-]PAM | NA | Intergenic | NA | NA |
| GU305914.1 | 161752 | 161774 | + | 4 | CACCTCCGCCGCCGCCTCCT | CGG | \|-\|\|-\|\|\|[\|\|\|\|\|\|\|\|-\|\|-]PAM | NA | Intergenic | NA | NA |
| T31621 | GACGCCACCGATTCCGCCTGTGG | 1000 | | | | | | | | | |
| Oligo fwd | CACCGACGCCACCGATTCCGCCTG | | | | | | | | | | |
| Oligo rev | AAACCAGGCGGAATCGGTGGCGTC | | | | | | | | | | |
| Chromosome | start | end | strand | MM | target_seq | PAM | alignment | distance | position | gene name | gene id |
| GU305914.1 | 35613 | 35635 | - | 0 | GACGCCACCGATTCCGCCTG | TGG | \|\|\|\|\|\|\|\|[\|\|\|\|\|\|\|\|\|\|\|\|]PAM | NA | Intergenic | NA | NA |
| T31622 | TCCGCCTGTGGCCGCCTTCTCGG | 1000 | | | | | | | | | |
| Oligo fwd | CACCTCCGCCTGTGGCCGCCTTCT | | | | | | | | | | |
| Oligo rev | AAACAGAAGGCGGCCACAGGCGGA | | | | | | | | | | |
| Chromosome | start | end | strand | MM | target_seq | PAM | alignment | distance | position | gene name | gene id |
| GU305914.1 | 35601 | 35623 | - | 0 | TCCGCCTGTGGCCGCCTTCT | CGG | \|\|\|\|\|\|\|\|[\|\|\|\|\|\|\|\|\|\|\|\|]PAM | NA | Intergenic | NA | NA |
| T31623 | TGTGGCCGCCTTCTCGGAGATGG | 1000 | | | | | | | | | |
| Oligo fwd | CACCTGTGGCCGCCTTCTCGGAGA | | | | | | | | | | |
| Oligo rev | AAACTCTCCGAGAAGGCGGCCACA | | | | | | | | | | |
| Chromosome | start | end | strand | MM | target_seq | PAM | alignment | distance | position | gene name | gene id |
| GU305914.1 | 35595 | 35617 | - | 0 | TGTGGCCGCCTTCTCGGAGA | TGG | \|\|\|\|\|\|\|\|[\|\|\|\|\|\|\|\|\|\|\|\|]PAM | NA | Intergenic | NA | NA |
| T31624 | CTCCGTCGCCTCCAAGACGCCGG | 1000 | | | | | | | | | |
| Oligo fwd | CACCCTCCGTCGCCTCCAAGACGC | | | | | | | | | | |
| Oligo rev | AAACGCGTCTTGGAGGCGACGGAG | | | | | | | | | | |
| Chromosome | start | end | strand | MM | target_seq | PAM | alignment | distance | position | gene name | gene id |
| GU305914.1 | 35544 | 35566 | - | 0 | CTCCGTCGCCTCCAAGACGC | CGG | \|\|\|\|\|\|\|\|[\|\|\|\|\|\|\|\|\|\|\|\|]PAM | NA | Intergenic | NA | NA |
| GU305914.1 | 33779 | 33801 | - | 4 | CTCCGACTCCTGCAAGGCGC | TGG | \|\|\|\|\|-\|-[\|\|\|-\|\|\|\|-\|\|\|]PAM | NA | Intergenic | NA | NA |
| T31625 | CGTGCCCGTGAACGTCGTCACGG | 1000 | | | | | | | | | |
| Oligo fwd | CACCCGTGCCCGTGAACGTCGTCA | | | | | | | | | | |
| Oligo rev | AAACTGACGACGTTCACGGGCACG | | | | | | | | | | |
| Chromosome | start | end | strand | MM | target_seq | PAM | alignment | distance | position | gene name | gene id |
| GU305914.1 | 35520 | 35542 | - | 0 | CGTGCCCGTGAACGTCGTCA | CGG | \|\|\|\|\|\|\|\|[\|\|\|\|\|\|\|\|\|\|\|\|]PAM | NA | Intergenic | NA | NA |
| T31626 | GTGCCCGTGAACGTCGTCACGGG | 1000 | | | | | | | | | |
| Oligo fwd | CACCGTGCCCGTGAACGTCGTCAC | | | | | | | | | | |
| Oligo rev | AAACGTGACGACGTTCACGGGCAC | | | | | | | | | | |
| Chromosome | start | end | strand | MM | target_seq | PAM | alignment | distance | position | gene name | gene id |
| GU305914.1 | 35519 | 35541 | - | 0 | GTGCCCGTGAACGTCGTCAC | GGG | \|\|\|\|\|\|\|\|[\|\|\|\|\|\|\|\|\|\|\|\|]PAM | NA | Intergenic | NA | NA |
| T31627 | CACCGCGCAGCGCGTCGACTTGG | 1000 | | | | | | | | | |
| Oligo fwd | CACCCACCGCGCAGCGCGTCGACT | | | | | | | | | | |
| Oligo rev | AAACAGTCGACGCGCTGCGCGGTG | | | | | | | | | | |
| Chromosome | start | end | strand | MM | target_seq | PAM | alignment | distance | position | gene name | gene id |
| GU305914.1 | 35496 | 35518 | - | 0 | CACCGCGCAGCGCGTCGACT | TGG | \|\|\|\|\|\|\|\|[\|\|\|\|\|\|\|\|\|\|\|\|]PAM | NA | Intergenic | NA | NA |
| T31628 | CAGCGCGTCGACTTGGATTCCGG | 1000 | | | | | | | | | |
| Oligo fwd | CACCCAGCGCGTCGACTTGGATTC | | | | | | | | | | |
| Oligo rev | AAACGAATCCAAGTCGACGCGCTG | | | | | | | | | | |
| Chromosome | start | end | strand | MM | target_seq | PAM | alignment | distance | position | gene name | gene id |
| GU305914.1 | 35489 | 35511 | - | 0 | CAGCGCGTCGACTTGGATTC | CGG | \|\|\|\|\|\|\|\|[\|\|\|\|\|\|\|\|\|\|\|\|]PAM | NA | Intergenic | NA | NA |
| T31629 | GACTTGGATTCCGGCAGTCGCGG | 1000 | | | | | | | | | |
| Oligo fwd | CACCGACTTGGATTCCGGCAGTCG | | | | | | | | | | |
| Oligo rev | AAACCGACTGCCGGAATCCAAGTC | | | | | | | | | | |
| Chromosome | start | end | strand | MM | target_seq | PAM | alignment | distance | position | gene name | gene id |
| GU305914.1 | 35480 | 35502 | - | 0 | GACTTGGATTCCGGCAGTCG | CGG | \|\|\|\|\|\|\|\|[\|\|\|\|\|\|\|\|\|\|\|\|]PAM | NA | Intergenic | NA | NA |
| T31630 | GCGTCGACAACTTCCACGCGAGG | 1000 | | | | | | | | | |
| Oligo fwd | CACCGCGTCGACAACTTCCACGCG | | | | | | | | | | |
| Oligo rev | AAACCGCGTGGAAGTTGTCGACGC | | | | | | | | | | |
| Chromosome | start | end | strand | MM | target_seq | PAM | alignment | distance | position | gene name | gene id |
| GU305914.1 | 35452 | 35474 | - | 0 | GCGTCGACAACTTCCACGCG | AGG | \|\|\|\|\|\|\|\|[\|\|\|\|\|\|\|\|\|\|\|\|]PAM | NA | Intergenic | NA | NA |
| T31631 | CGTCGACAACTTCCACGCGAGGG | 1000 | | | | | | | | | |
| Oligo fwd | CACCCGTCGACAACTTCCACGCGA | | | | | | | | | | |

<table>
<tr><td>Oligo rev</td><td>AAACTCGCGTGGAAGTTGTCGACG</td><td></td><td></td><td></td><td></td><td></td><td></td><td></td><td></td><td></td><td></td><td></td></tr>
<tr><td>Chromosome</td><td>start</td><td></td><td>end</td><td>strand</td><td>MM</td><td>target_seq</td><td>PAM</td><td>alignment</td><td>distance</td><td>position</td><td>gene name</td><td>gene id</td></tr>
<tr><td>GU305914.1</td><td></td><td>35451</td><td>35473</td><td>-</td><td>0</td><td>CGTCGACAACTTCCACGCGA</td><td>GGG</td><td>||||||||[||||||||||||]PAM</td><td>NA</td><td>Intergenic</td><td>NA</td><td>NA</td></tr>
<tr><td>T31632</td><td>GTCGACAACTTCCACGCGAGGGG</td><td></td><td>1000</td><td></td><td></td><td></td><td></td><td></td><td></td><td></td><td></td><td></td></tr>
<tr><td>Oligo fwd</td><td>CACCGTCGACAACTTCCACGCGAG</td><td></td><td></td><td></td><td></td><td></td><td></td><td></td><td></td><td></td><td></td><td></td></tr>
<tr><td>Oligo rev</td><td>AAACCTCGCGTGGAAGTTGTCGAC</td><td></td><td></td><td></td><td></td><td></td><td></td><td></td><td></td><td></td><td></td><td></td></tr>
<tr><td>Chromosome</td><td>start</td><td></td><td>end</td><td>strand</td><td>MM</td><td>target_seq</td><td>PAM</td><td>alignment</td><td>distance</td><td>position</td><td>gene name</td><td>gene id</td></tr>
<tr><td>GU305914.1</td><td></td><td>35450</td><td>35472</td><td>-</td><td>0</td><td>GTCGACAACTTCCACGCGAG</td><td>GGG</td><td>||||||||[||||||||||||]PAM</td><td>NA</td><td>Intergenic</td><td>NA</td><td>NA</td></tr>
<tr><td>T31633</td><td>TCGACAACTTCCACGCGAGGGGG</td><td></td><td>1000</td><td></td><td></td><td></td><td></td><td></td><td></td><td></td><td></td><td></td></tr>
<tr><td>Oligo fwd</td><td>CACCTCGACAACTTCCACGCGAGG</td><td></td><td></td><td></td><td></td><td></td><td></td><td></td><td></td><td></td><td></td><td></td></tr>
<tr><td>Oligo rev</td><td>AAACCCTCGCGTGGAAGTTGTCGA</td><td></td><td></td><td></td><td></td><td></td><td></td><td></td><td></td><td></td><td></td><td></td></tr>
<tr><td>Chromosome</td><td>start</td><td></td><td>end</td><td>strand</td><td>MM</td><td>target_seq</td><td>PAM</td><td>alignment</td><td>distance</td><td>position</td><td>gene name</td><td>gene id</td></tr>
<tr><td>GU305914.1</td><td></td><td>35449</td><td>35471</td><td>-</td><td>0</td><td>TCGACAACTTCCACGCGAGG</td><td>GGG</td><td>||||||||[||||||||||||]PAM</td><td>NA</td><td>Intergenic</td><td>NA</td><td>NA</td></tr>
<tr><td>T31634</td><td>CGACAACTTCCACGCGAGGGGGG</td><td></td><td>1000</td><td></td><td></td><td></td><td></td><td></td><td></td><td></td><td></td><td></td></tr>
<tr><td>Oligo fwd</td><td>CACCCGACAACTTCCACGCGAGGG</td><td></td><td></td><td></td><td></td><td></td><td></td><td></td><td></td><td></td><td></td><td></td></tr>
<tr><td>Oligo rev</td><td>AAACCCCTCGCGTGGAAGTTGTCG</td><td></td><td></td><td></td><td></td><td></td><td></td><td></td><td></td><td></td><td></td><td></td></tr>
<tr><td>Chromosome</td><td>start</td><td></td><td>end</td><td>strand</td><td>MM</td><td>target_seq</td><td>PAM</td><td>alignment</td><td>distance</td><td>position</td><td>gene name</td><td>gene id</td></tr>
<tr><td>GU305914.1</td><td></td><td>35448</td><td>35470</td><td>-</td><td>0</td><td>CGACAACTTCCACGCGAGGG</td><td>GGG</td><td>||||||||[||||||||||||]PAM</td><td>NA</td><td>Intergenic</td><td>NA</td><td>NA</td></tr>
<tr><td>T31635</td><td>GGGAGCTGTTGCGCTGCGCCAGG</td><td></td><td>1000</td><td></td><td></td><td></td><td></td><td></td><td></td><td></td><td></td><td></td></tr>
<tr><td>Oligo fwd</td><td>CACCGGGAGCTGTTGCGCTGCGCC</td><td></td><td></td><td></td><td></td><td></td><td></td><td></td><td></td><td></td><td></td><td></td></tr>
<tr><td>Oligo rev</td><td>AAACGGCGCAGCGCAACAGCTCCC</td><td></td><td></td><td></td><td></td><td></td><td></td><td></td><td></td><td></td><td></td><td></td></tr>
<tr><td>Chromosome</td><td>start</td><td></td><td>end</td><td>strand</td><td>MM</td><td>target_seq</td><td>PAM</td><td>alignment</td><td>distance</td><td>position</td><td>gene name</td><td>gene id</td></tr>
<tr><td>GU305914.1</td><td></td><td>35428</td><td>35450</td><td>-</td><td>0</td><td>GGGAGCTGTTGCGCTGCGCC</td><td>AGG</td><td>||||||||[||||||||||||]PAM</td><td>NA</td><td>Intergenic</td><td>NA</td><td>NA</td></tr>
<tr><td>T31636</td><td>GGAGCTGTTGCGCTGCGCCAGGG</td><td></td><td>1000</td><td></td><td></td><td></td><td></td><td></td><td></td><td></td><td></td><td></td></tr>
<tr><td>Oligo fwd</td><td>CACCGGAGCTGTTGCGCTGCGCCA</td><td></td><td></td><td></td><td></td><td></td><td></td><td></td><td></td><td></td><td></td><td></td></tr>
<tr><td>Oligo rev</td><td>AAACTGGCGCAGCGCAACAGCTCC</td><td></td><td></td><td></td><td></td><td></td><td></td><td></td><td></td><td></td><td></td><td></td></tr>
<tr><td>Chromosome</td><td>start</td><td></td><td>end</td><td>strand</td><td>MM</td><td>target_seq</td><td>PAM</td><td>alignment</td><td>distance</td><td>position</td><td>gene name</td><td>gene id</td></tr>
<tr><td>GU305914.1</td><td></td><td>35427</td><td>35449</td><td>-</td><td>0</td><td>GGAGCTGTTGCGCTGCGCCA</td><td>GGG</td><td>||||||||[||||||||||||]PAM</td><td>NA</td><td>Intergenic</td><td>NA</td><td>NA</td></tr>
<tr><td>T31637</td><td>CGCTGCGCCAGGGCGTGCAATGG</td><td></td><td>1000</td><td></td><td></td><td></td><td></td><td></td><td></td><td></td><td></td><td></td></tr>
<tr><td>Oligo fwd</td><td>CACCCGCTGCGCCAGGGCGTGCAA</td><td></td><td></td><td></td><td></td><td></td><td></td><td></td><td></td><td></td><td></td><td></td></tr>
<tr><td>Oligo rev</td><td>AAACTTGCACGCCCTGGCGCAGCG</td><td></td><td></td><td></td><td></td><td></td><td></td><td></td><td></td><td></td><td></td><td></td></tr>
<tr><td>Chromosome</td><td>start</td><td></td><td>end</td><td>strand</td><td>MM</td><td>target_seq</td><td>PAM</td><td>alignment</td><td>distance</td><td>position</td><td>gene name</td><td>gene id</td></tr>
<tr><td>GU305914.1</td><td></td><td>35417</td><td>35439</td><td>-</td><td>0</td><td>CGCTGCGCCAGGGCGTGCAA</td><td>TGG</td><td>||||||||[||||||||||||]PAM</td><td>NA</td><td>Intergenic</td><td>NA</td><td>NA</td></tr>
<tr><td>T31638</td><td>CAATGGCAGCTCCTTCGACGAGG</td><td></td><td>1000</td><td></td><td></td><td></td><td></td><td></td><td></td><td></td><td></td><td></td></tr>
<tr><td>Oligo fwd</td><td>CACCCAATGGCAGCTCCTTCGACG</td><td></td><td></td><td></td><td></td><td></td><td></td><td></td><td></td><td></td><td></td><td></td></tr>
<tr><td>Oligo rev</td><td>AAACCGTCGAAGGAGCTGCCATTG</td><td></td><td></td><td></td><td></td><td></td><td></td><td></td><td></td><td></td><td></td><td></td></tr>
<tr><td>Chromosome</td><td>start</td><td></td><td>end</td><td>strand</td><td>MM</td><td>target_seq</td><td>PAM</td><td>alignment</td><td>distance</td><td>position</td><td>gene name</td><td>gene id</td></tr>
<tr><td>GU305914.1</td><td></td><td>35400</td><td>35422</td><td>-</td><td>0</td><td>CAATGGCAGCTCCTTCGACG</td><td>AGG</td><td>||||||||[||||||||||||]PAM</td><td>NA</td><td>Intergenic</td><td>NA</td><td>NA</td></tr>
<tr><td>T31639</td><td>CAGCTCCTTCGACGAGGTGAAGG</td><td></td><td>1000</td><td></td><td></td><td></td><td></td><td></td><td></td><td></td><td></td><td></td></tr>
<tr><td>Oligo fwd</td><td>CACCCAGCTCCTTCGACGAGGTGA</td><td></td><td></td><td></td><td></td><td></td><td></td><td></td><td></td><td></td><td></td><td></td></tr>
<tr><td>Oligo rev</td><td>AAACTCACCTCGTCGAAGGAGCTG</td><td></td><td></td><td></td><td></td><td></td><td></td><td></td><td></td><td></td><td></td><td></td></tr>
<tr><td>Chromosome</td><td>start</td><td></td><td>end</td><td>strand</td><td>MM</td><td>target_seq</td><td>PAM</td><td>alignment</td><td>distance</td><td>position</td><td>gene name</td><td>gene id</td></tr>
<tr><td>GU305914.1</td><td></td><td>35394</td><td>35416</td><td>-</td><td>0</td><td>CAGCTCCTTCGACGAGGTGA</td><td>AGG</td><td>||||||||[||||||||||||]PAM</td><td>NA</td><td>Intergenic</td><td>NA</td><td>NA</td></tr>
<tr><td>T31640</td><td>GCTTTCGTCAAGAGAAACACCGG</td><td></td><td>1000</td><td></td><td></td><td></td><td></td><td></td><td></td><td></td><td></td><td></td></tr>
<tr><td>Oligo fwd</td><td>CACCGCTTTCGTCAAGAGAAACAC</td><td></td><td></td><td></td><td></td><td></td><td></td><td></td><td></td><td></td><td></td><td></td></tr>
<tr><td>Oligo rev</td><td>AAACGTGTTTCTCTTGACGAAAGC</td><td></td><td></td><td></td><td></td><td></td><td></td><td></td><td></td><td></td><td></td><td></td></tr>
<tr><td>Chromosome</td><td>start</td><td></td><td>end</td><td>strand</td><td>MM</td><td>target_seq</td><td>PAM</td><td>alignment</td><td>distance</td><td>position</td><td>gene name</td><td>gene id</td></tr>
<tr><td>GU305914.1</td><td></td><td>35372</td><td>35394</td><td>-</td><td>0</td><td>GCTTTCGTCAAGAGAAACAC</td><td>CGG</td><td>||||||||[||||||||||||]PAM</td><td>NA</td><td>Intergenic</td><td>NA</td><td>NA</td></tr>
<tr><td>T31641</td><td>TCGTCAAGAGAAACACCGGACGG</td><td></td><td>1000</td><td></td><td></td><td></td><td></td><td></td><td></td><td></td><td></td><td></td></tr>
<tr><td>Oligo fwd</td><td>CACCTCGTCAAGAGAAACACCGGA</td><td></td><td></td><td></td><td></td><td></td><td></td><td></td><td></td><td></td><td></td><td></td></tr>
<tr><td>Oligo rev</td><td>AAACTCCGGTGTTTCTCTTGACGA</td><td></td><td></td><td></td><td></td><td></td><td></td><td></td><td></td><td></td><td></td><td></td></tr>
<tr><td>Chromosome</td><td>start</td><td></td><td>end</td><td>strand</td><td>MM</td><td>target_seq</td><td>PAM</td><td>alignment</td><td>distance</td><td>position</td><td>gene name</td><td>gene id</td></tr>
<tr><td>GU305914.1</td><td></td><td>35368</td><td>35390</td><td>-</td><td>0</td><td>TCGTCAAGAGAAACACCGGA</td><td>CGG</td><td>||||||||[||||||||||||]PAM</td><td>NA</td><td>Intergenic</td><td>NA</td><td>NA</td></tr>
<tr><td>T31642</td><td>AGAAACACCGGACGGCGTTTCGG</td><td></td><td>1000</td><td></td><td></td><td></td><td></td><td></td><td></td><td></td><td></td><td></td></tr>
<tr><td>Oligo fwd</td><td>CACCAGAAACACCGGACGGCGTTT</td><td></td><td></td><td></td><td></td><td></td><td></td><td></td><td></td><td></td><td></td><td></td></tr>
<tr><td>Oligo rev</td><td>AAACAAACGCCGTCCGGTGTTTCT</td><td></td><td></td><td></td><td></td><td></td><td></td><td></td><td></td><td></td><td></td><td></td></tr>
<tr><td>Chromosome</td><td>start</td><td></td><td>end</td><td>strand</td><td>MM</td><td>target_seq</td><td>PAM</td><td>alignment</td><td>distance</td><td>position</td><td>gene name</td><td>gene id</td></tr>
<tr><td>GU305914.1</td><td></td><td>35360</td><td>35382</td><td>-</td><td>0</td><td>AGAAACACCGGACGGCGTTT</td><td>CGG</td><td>||||||||[||||||||||||]PAM</td><td>NA</td><td>Intergenic</td><td>NA</td><td>NA</td></tr>
<tr><td>T31643</td><td>GAAACACCGGACGGCGTTTCGGG</td><td></td><td>1000</td><td></td><td></td><td></td><td></td><td></td><td></td><td></td><td></td><td></td></tr>
<tr><td>Oligo fwd</td><td>CACCGAAACACCGGACGGCGTTTC</td><td></td><td></td><td></td><td></td><td></td><td></td><td></td><td></td><td></td><td></td><td></td></tr>
<tr><td>Oligo rev</td><td>AAACGAAACGCCGTCCGGTGTTTC</td><td></td><td></td><td></td><td></td><td></td><td></td><td></td><td></td><td></td><td></td><td></td></tr>
<tr><td>Chromosome</td><td>start</td><td></td><td>end</td><td>strand</td><td>MM</td><td>target_seq</td><td>PAM</td><td>alignment</td><td>distance</td><td>position</td><td>gene name</td><td>gene id</td></tr>
<tr><td>GU305914.1</td><td></td><td>35359</td><td>35381</td><td>-</td><td>0</td><td>GAAACACCGGACGGCGTTTC</td><td>GGG</td><td>||||||||[||||||||||||]PAM</td><td>NA</td><td>Intergenic</td><td>NA</td><td>NA</td></tr>
<tr><td>T31644</td><td>CACCGGACGGCGTTTCGGGCTGG</td><td></td><td>1000</td><td></td><td></td><td></td><td></td><td></td><td></td><td></td><td></td><td></td></tr>
<tr><td>Oligo fwd</td><td>CACCCACCGGACGGCGTTTCGGGC</td><td></td><td></td><td></td><td></td><td></td><td></td><td></td><td></td><td></td><td></td><td></td></tr>
<tr><td>Oligo rev</td><td>AAACGCCCGAAACGCCGTCCGGTG</td><td></td><td></td><td></td><td></td><td></td><td></td><td></td><td></td><td></td><td></td><td></td></tr>
<tr><td>Chromosome</td><td>start</td><td></td><td>end</td><td>strand</td><td>MM</td><td>target_seq</td><td>PAM</td><td>alignment</td><td>distance</td><td>position</td><td>gene name</td><td>gene id</td></tr>
<tr><td>GU305914.1</td><td></td><td>35355</td><td>35377</td><td>-</td><td>0</td><td>CACCGGACGGCGTTTCGGGC</td><td>TGG</td><td>||||||||[||||||||||||]PAM</td><td>NA</td><td>Intergenic</td><td>NA</td><td>NA</td></tr>
<tr><td>GU305914.1</td><td></td><td>49100</td><td>49122</td><td>-</td><td>4</td><td>GGCCGGACGGGGTTTCGGTC</td><td>CGG</td><td>--||||||[||-|||||||-|]PAM</td><td>NA</td><td>Intergenic</td><td>NA</td><td>NA</td></tr>
<tr><td>T31645</td><td>GCGTTTCGGGCTGGACCACATGG</td><td></td><td>1000</td><td></td><td></td><td></td><td></td><td></td><td></td><td></td><td></td><td></td></tr>
<tr><td>Oligo fwd</td><td>CACCGCGTTTCGGGCTGGACCACA</td><td></td><td></td><td></td><td></td><td></td><td></td><td></td><td></td><td></td><td></td><td></td></tr>
<tr><td>Oligo rev</td><td>AAACTGTGGTCCAGCCCGAAACGC</td><td></td><td></td><td></td><td></td><td></td><td></td><td></td><td></td><td></td><td></td><td></td></tr>
<tr><td>Chromosome</td><td>start</td><td></td><td>end</td><td>strand</td><td>MM</td><td>target_seq</td><td>PAM</td><td>alignment</td><td>distance</td><td>position</td><td>gene name</td><td>gene id</td></tr>
<tr><td>GU305914.1</td><td></td><td>35346</td><td>35368</td><td>-</td><td>0</td><td>GCGTTTCGGGCTGGACCACA</td><td>TGG</td><td>||||||||[||||||||||||]PAM</td><td>NA</td><td>Intergenic</td><td>NA</td><td>NA</td></tr>
</table>

<table>
<tr><td>T31646</td><td>TTTCGGGCTGGACCACATGGCGG</td><td>1000</td><td></td><td></td><td></td><td></td><td></td><td></td><td></td><td></td><td></td></tr>
<tr><td>Oligo fwd</td><td>CACCTTTCGGGCTGGACCACATGG</td><td></td><td></td><td></td><td></td><td></td><td></td><td></td><td></td><td></td><td></td></tr>
<tr><td>Oligo rev</td><td>AAACCCATGTGGTCCAGCCCGAAA</td><td></td><td></td><td></td><td></td><td></td><td></td><td></td><td></td><td></td><td></td></tr>
<tr><td>Chromosome</td><td>start</td><td>end</td><td>strand</td><td>MM</td><td>target_seq</td><td>PAM</td><td>alignment</td><td>distance</td><td>position</td><td>gene name</td><td>gene id</td></tr>
<tr><td>GU305914.1</td><td>35343</td><td>35365</td><td>-</td><td>0</td><td>TTTCGGGCTGGACCACATGG</td><td>CGG</td><td>||||||||[||||||||||||]PAM</td><td>NA</td><td>Intergenic</td><td>NA</td><td>NA</td></tr>
<tr><td>T31647</td><td>ACATGGCGGATATCACGTTGCGG</td><td>1000</td><td></td><td></td><td></td><td></td><td></td><td></td><td></td><td></td><td></td></tr>
<tr><td>Oligo fwd</td><td>CACCACATGGCGGATATCACGTTG</td><td></td><td></td><td></td><td></td><td></td><td></td><td></td><td></td><td></td><td></td></tr>
<tr><td>Oligo rev</td><td>AAACCAACGTGATATCCGCCATGT</td><td></td><td></td><td></td><td></td><td></td><td></td><td></td><td></td><td></td><td></td></tr>
<tr><td>Chromosome</td><td>start</td><td>end</td><td>strand</td><td>MM</td><td>target_seq</td><td>PAM</td><td>alignment</td><td>distance</td><td>position</td><td>gene name</td><td>gene id</td></tr>
<tr><td>GU305914.1</td><td>35329</td><td>35351</td><td>-</td><td>0</td><td>ACATGGCGGATATCACGTTG</td><td>CGG</td><td>||||||||[||||||||||||]PAM</td><td>NA</td><td>Intergenic</td><td>NA</td><td>NA</td></tr>
<tr><td>T31648</td><td>GCGGATATCACGTTGCGGATCGG</td><td>1000</td><td></td><td></td><td></td><td></td><td></td><td></td><td></td><td></td><td></td></tr>
<tr><td>Oligo fwd</td><td>CACCGCGGATATCACGTTGCGGAT</td><td></td><td></td><td></td><td></td><td></td><td></td><td></td><td></td><td></td><td></td></tr>
<tr><td>Oligo rev</td><td>AAACATCCGCAACGTGATATCCGC</td><td></td><td></td><td></td><td></td><td></td><td></td><td></td><td></td><td></td><td></td></tr>
<tr><td>Chromosome</td><td>start</td><td>end</td><td>strand</td><td>MM</td><td>target_seq</td><td>PAM</td><td>alignment</td><td>distance</td><td>position</td><td>gene name</td><td>gene id</td></tr>
<tr><td>GU305914.1</td><td>35324</td><td>35346</td><td>-</td><td>0</td><td>GCGGATATCACGTTGCGGAT</td><td>CGG</td><td>||||||||[||||||||||||]PAM</td><td>NA</td><td>Intergenic</td><td>NA</td><td>NA</td></tr>
<tr><td>T31649</td><td>TTCGACGTAGAAAAGCTGCTTGG</td><td>1000</td><td></td><td></td><td></td><td></td><td></td><td></td><td></td><td></td><td></td></tr>
<tr><td>Oligo fwd</td><td>CACCTTCGACGTAGAAAAGCTGCT</td><td></td><td></td><td></td><td></td><td></td><td></td><td></td><td></td><td></td><td></td></tr>
<tr><td>Oligo rev</td><td>AAACAGCAGCTTTTCTACGTCGAA</td><td></td><td></td><td></td><td></td><td></td><td></td><td></td><td></td><td></td><td></td></tr>
<tr><td>Chromosome</td><td>start</td><td>end</td><td>strand</td><td>MM</td><td>target_seq</td><td>PAM</td><td>alignment</td><td>distance</td><td>position</td><td>gene name</td><td>gene id</td></tr>
<tr><td>GU305914.1</td><td>35276</td><td>35298</td><td>-</td><td>0</td><td>TTCGACGTAGAAAAGCTGCT</td><td>TGG</td><td>||||||||[||||||||||||]PAM</td><td>NA</td><td>Intergenic</td><td>NA</td><td>NA</td></tr>
<tr><td>T31650</td><td>TTGGCTACCGCTGTAGCGAGCGG</td><td>1000</td><td></td><td></td><td></td><td></td><td></td><td></td><td></td><td></td><td></td></tr>
<tr><td>Oligo fwd</td><td>CACCTTGGCTACCGCTGTAGCGAG</td><td></td><td></td><td></td><td></td><td></td><td></td><td></td><td></td><td></td><td></td></tr>
<tr><td>Oligo rev</td><td>AAACCTCGCTACAGCGGTAGCCAA</td><td></td><td></td><td></td><td></td><td></td><td></td><td></td><td></td><td></td><td></td></tr>
<tr><td>Chromosome</td><td>start</td><td>end</td><td>strand</td><td>MM</td><td>target_seq</td><td>PAM</td><td>alignment</td><td>distance</td><td>position</td><td>gene name</td><td>gene id</td></tr>
<tr><td>GU305914.1</td><td>35257</td><td>35279</td><td>-</td><td>0</td><td>TTGGCTACCGCTGTAGCGAG</td><td>CGG</td><td>||||||||[||||||||||||]PAM</td><td>NA</td><td>Intergenic</td><td>NA</td><td>NA</td></tr>
<tr><td>T31651</td><td>TCCTCCGAGATGACCGTCGTCGG</td><td>1000</td><td></td><td></td><td></td><td></td><td></td><td></td><td></td><td></td><td></td></tr>
<tr><td>Oligo fwd</td><td>CACCTCCTCCGAGATGACCGTCGT</td><td></td><td></td><td></td><td></td><td></td><td></td><td></td><td></td><td></td><td></td></tr>
<tr><td>Oligo rev</td><td>AAACACGACGGTCATCTCGGAGGA</td><td></td><td></td><td></td><td></td><td></td><td></td><td></td><td></td><td></td><td></td></tr>
<tr><td>Chromosome</td><td>start</td><td>end</td><td>strand</td><td>MM</td><td>target_seq</td><td>PAM</td><td>alignment</td><td>distance</td><td>position</td><td>gene name</td><td>gene id</td></tr>
<tr><td>GU305914.1</td><td>35234</td><td>35256</td><td>-</td><td>0</td><td>TCCTCCGAGATGACCGTCGT</td><td>CGG</td><td>||||||||[||||||||||||]PAM</td><td>NA</td><td>Intergenic</td><td>NA</td><td>NA</td></tr>
<tr><td>T31652</td><td>CCGAGATGACCGTCGTCGGTCGG</td><td>1000</td><td></td><td></td><td></td><td></td><td></td><td></td><td></td><td></td><td></td></tr>
<tr><td>Oligo fwd</td><td>CACCCCGAGATGACCGTCGTCGGT</td><td></td><td></td><td></td><td></td><td></td><td></td><td></td><td></td><td></td><td></td></tr>
<tr><td>Oligo rev</td><td>AAACACCGACGACGGTCATCTCGG</td><td></td><td></td><td></td><td></td><td></td><td></td><td></td><td></td><td></td><td></td></tr>
<tr><td>Chromosome</td><td>start</td><td>end</td><td>strand</td><td>MM</td><td>target_seq</td><td>PAM</td><td>alignment</td><td>distance</td><td>position</td><td>gene name</td><td>gene id</td></tr>
<tr><td>GU305914.1</td><td>35230</td><td>35252</td><td>-</td><td>0</td><td>CCGAGATGACCGTCGTCGGT</td><td>CGG</td><td>||||||||[||||||||||||]PAM</td><td>NA</td><td>Intergenic</td><td>NA</td><td>NA</td></tr>
<tr><td>T31653</td><td>GTCGTCGGTCGGATCGTTTTCGG</td><td>1000</td><td></td><td></td><td></td><td></td><td></td><td></td><td></td><td></td><td></td></tr>
<tr><td>Oligo fwd</td><td>CACCGTCGTCGGTCGGATCGTTTT</td><td></td><td></td><td></td><td></td><td></td><td></td><td></td><td></td><td></td><td></td></tr>
<tr><td>Oligo rev</td><td>AAACAAAACGATCCGACCGACGAC</td><td></td><td></td><td></td><td></td><td></td><td></td><td></td><td></td><td></td><td></td></tr>
<tr><td>Chromosome</td><td>start</td><td>end</td><td>strand</td><td>MM</td><td>target_seq</td><td>PAM</td><td>alignment</td><td>distance</td><td>position</td><td>gene name</td><td>gene id</td></tr>
<tr><td>GU305914.1</td><td>35219</td><td>35241</td><td>-</td><td>0</td><td>GTCGTCGGTCGGATCGTTTT</td><td>CGG</td><td>||||||||[||||||||||||]PAM</td><td>NA</td><td>Intergenic</td><td>NA</td><td>NA</td></tr>
<tr><td>T31654</td><td>GTCGGATCGTTTTCGGTGCCAGG</td><td>1000</td><td></td><td></td><td></td><td></td><td></td><td></td><td></td><td></td><td></td></tr>
<tr><td>Oligo fwd</td><td>CACCGTCGGATCGTTTTCGGTGCC</td><td></td><td></td><td></td><td></td><td></td><td></td><td></td><td></td><td></td><td></td></tr>
<tr><td>Oligo rev</td><td>AAACGGCACCGAAAACGATCCGAC</td><td></td><td></td><td></td><td></td><td></td><td></td><td></td><td></td><td></td><td></td></tr>
<tr><td>Chromosome</td><td>start</td><td>end</td><td>strand</td><td>MM</td><td>target_seq</td><td>PAM</td><td>alignment</td><td>distance</td><td>position</td><td>gene name</td><td>gene id</td></tr>
<tr><td>GU305914.1</td><td>35212</td><td>35234</td><td>-</td><td>0</td><td>GTCGGATCGTTTTCGGTGCC</td><td>AGG</td><td>||||||||[||||||||||||]PAM</td><td>NA</td><td>Intergenic</td><td>NA</td><td>NA</td></tr>
<tr><td>T31655</td><td>GTTTTCGGTGCCAGGAGTCACGG</td><td>1000</td><td></td><td></td><td></td><td></td><td></td><td></td><td></td><td></td><td></td></tr>
<tr><td>Oligo fwd</td><td>CACCGTTTTCGGTGCCAGGAGTCA</td><td></td><td></td><td></td><td></td><td></td><td></td><td></td><td></td><td></td><td></td></tr>
<tr><td>Oligo rev</td><td>AAACTGACTCCTGGCACCGAAAAC</td><td></td><td></td><td></td><td></td><td></td><td></td><td></td><td></td><td></td><td></td></tr>
<tr><td>Chromosome</td><td>start</td><td>end</td><td>strand</td><td>MM</td><td>target_seq</td><td>PAM</td><td>alignment</td><td>distance</td><td>position</td><td>gene name</td><td>gene id</td></tr>
<tr><td>GU305914.1</td><td>35204</td><td>35226</td><td>-</td><td>0</td><td>GTTTTCGGTGCCAGGAGTCA</td><td>CGG</td><td>||||||||[||||||||||||]PAM</td><td>NA</td><td>Intergenic</td><td>NA</td><td>NA</td></tr>
<tr><td>T31656</td><td>GAGTCACGGCGATCTGATCGCGG</td><td>1000</td><td></td><td></td><td></td><td></td><td></td><td></td><td></td><td></td><td></td></tr>
<tr><td>Oligo fwd</td><td>CACCGAGTCACGGCGATCTGATCG</td><td></td><td></td><td></td><td></td><td></td><td></td><td></td><td></td><td></td><td></td></tr>
<tr><td>Oligo rev</td><td>AAACCGATCAGATCGCCGTGACTC</td><td></td><td></td><td></td><td></td><td></td><td></td><td></td><td></td><td></td><td></td></tr>
<tr><td>Chromosome</td><td>start</td><td>end</td><td>strand</td><td>MM</td><td>target_seq</td><td>PAM</td><td>alignment</td><td>distance</td><td>position</td><td>gene name</td><td>gene id</td></tr>
<tr><td>GU305914.1</td><td>35190</td><td>35212</td><td>-</td><td>0</td><td>GAGTCACGGCGATCTGATCG</td><td>CGG</td><td>||||||||[||||||||||||]PAM</td><td>NA</td><td>Intergenic</td><td>NA</td><td>NA</td></tr>
<tr><td>T31657</td><td>GATCTGATCGCGGTGTACACCGG</td><td>1000</td><td></td><td></td><td></td><td></td><td></td><td></td><td></td><td></td><td></td></tr>
<tr><td>Oligo fwd</td><td>CACCGATCTGATCGCGGTGTACAC</td><td></td><td></td><td></td><td></td><td></td><td></td><td></td><td></td><td></td><td></td></tr>
<tr><td>Oligo rev</td><td>AAACGTGTACACCGCGATCAGATC</td><td></td><td></td><td></td><td></td><td></td><td></td><td></td><td></td><td></td><td></td></tr>
<tr><td>Chromosome</td><td>start</td><td>end</td><td>strand</td><td>MM</td><td>target_seq</td><td>PAM</td><td>alignment</td><td>distance</td><td>position</td><td>gene name</td><td>gene id</td></tr>
<tr><td>GU305914.1</td><td>35180</td><td>35202</td><td>-</td><td>0</td><td>GATCTGATCGCGGTGTACAC</td><td>CGG</td><td>||||||||[||||||||||||]PAM</td><td>NA</td><td>Intergenic</td><td>NA</td><td>NA</td></tr>
<tr><td>T31658</td><td>CTGATCGCGGTGTACACCGGCGG</td><td>1000</td><td></td><td></td><td></td><td></td><td></td><td></td><td></td><td></td><td></td></tr>
<tr><td>Oligo fwd</td><td>CACCCTGATCGCGGTGTACACCGG</td><td></td><td></td><td></td><td></td><td></td><td></td><td></td><td></td><td></td><td></td></tr>
<tr><td>Oligo rev</td><td>AAACCCGGTGTACACCGCGATCAG</td><td></td><td></td><td></td><td></td><td></td><td></td><td></td><td></td><td></td><td></td></tr>
<tr><td>Chromosome</td><td>start</td><td>end</td><td>strand</td><td>MM</td><td>target_seq</td><td>PAM</td><td>alignment</td><td>distance</td><td>position</td><td>gene name</td><td>gene id</td></tr>
<tr><td>GU305914.1</td><td>35177</td><td>35199</td><td>-</td><td>0</td><td>CTGATCGCGGTGTACACCGG</td><td>CGG</td><td>||||||||[||||||||||||]PAM</td><td>NA</td><td>Intergenic</td><td>NA</td><td>NA</td></tr>
<tr><td>T31659</td><td>TCGTTGCTACGTGCACAGCCAGG</td><td>1000</td><td></td><td></td><td></td><td></td><td></td><td></td><td></td><td></td><td></td></tr>
<tr><td>Oligo fwd</td><td>CACCTCGTTGCTACGTGCACAGCC</td><td></td><td></td><td></td><td></td><td></td><td></td><td></td><td></td><td></td><td></td></tr>
<tr><td>Oligo rev</td><td>AAACGGCTGTGCACGTAGCAACGA</td><td></td><td></td><td></td><td></td><td></td><td></td><td></td><td></td><td></td><td></td></tr>
<tr><td>Chromosome</td><td>start</td><td>end</td><td>strand</td><td>MM</td><td>target_seq</td><td>PAM</td><td>alignment</td><td>distance</td><td>position</td><td>gene name</td><td>gene id</td></tr>
<tr><td>GU305914.1</td><td>35151</td><td>35173</td><td>-</td><td>0</td><td>TCGTTGCTACGTGCACAGCC</td><td>AGG</td><td>||||||||[||||||||||||]PAM</td><td>NA</td><td>Intergenic</td><td>NA</td><td>NA</td></tr>
<tr><td>T31660</td><td>TGCTCTATATCGTCTCCGAACGG</td><td>1000</td><td></td><td></td><td></td><td></td><td></td><td></td><td></td><td></td><td></td></tr>
<tr><td>Oligo fwd</td><td>CACCTGCTCTATATCGTCTCCGAA</td><td></td><td></td><td></td><td></td><td></td><td></td><td></td><td></td><td></td><td></td></tr>
<tr><td>Oligo rev</td><td>AAACTTCGGAGACGATATAGAGCA</td><td></td><td></td><td></td><td></td><td></td><td></td><td></td><td></td><td></td><td></td></tr>
<tr><td>Chromosome</td><td>start</td><td>end</td><td>strand</td><td>MM</td><td>target_seq</td><td>PAM</td><td>alignment</td><td>distance</td><td>position</td><td>gene name</td><td>gene id</td></tr>
<tr><td>GU305914.1</td><td>35119</td><td>35141</td><td>-</td><td>0</td><td>TGCTCTATATCGTCTCCGAA</td><td>CGG</td><td>||||||||[||||||||||||]PAM</td><td>NA</td><td>Intergenic</td><td>NA</td><td>NA</td></tr>
</table>

T31661 GCTCTATATCGTCTCCGAACGGG 1000
Oligo fwd CACCGCTCTATATCGTCTCCGAAC
Oligo rev AAACGTTCGGAGACGATATAGAGC
Chromosome start end strand MM target_seq PAM alignment distance position gene name gene id
GU305914.1 35118 35140 - 0 GCTCTATATCGTCTCCGAAC GGG ||||||||[||||||||||||]PAM NA Intergenic NA NA

T31662 CTCTATATCGTCTCCGAACGGGG 1000
Oligo fwd CACCCTCTATATCGTCTCCGAACG
Oligo rev AAACCGTTCGGAGACGATATAGAG
Chromosome start end strand MM target_seq PAM alignment distance position gene name gene id
GU305914.1 35117 35139 - 0 CTCTATATCGTCTCCGAACG GGG ||||||||[||||||||||||]PAM NA Intergenic NA NA

T31663 TATCGTCTCCGAACGGGGTCTGG 1000
Oligo fwd CACCTATCGTCTCCGAACGGGGTC
Oligo rev AAACGACCCCGTTCGGAGACGATA
Chromosome start end strand MM target_seq PAM alignment distance position gene name gene id
GU305914.1 35112 35134 - 0 TATCGTCTCCGAACGGGGTC TGG ||||||||[||||||||||||]PAM NA Intergenic NA NA
GU305914.1 135643 135665 - 4 GATCGGCTCCTGACGGGGTC GGG -||||-||[||--||||||||]PAM NA Intergenic NA NA

T31664 CGTCTCCGAACGGGGTCTGGTGG 1000
Oligo fwd CACCCGTCTCCGAACGGGGTCTGG
Oligo rev AAACCCAGACCCCGTTCGGAGACG
Chromosome start end strand MM target_seq PAM alignment distance position gene name gene id
GU305914.1 35109 35131 - 0 CGTCTCCGAACGGGGTCTGG TGG ||||||||[||||||||||||]PAM NA Intergenic NA NA

T31665 GGTGGATCTTCTGTCGAGAGAGG 1000
Oligo fwd CACCGGTGGATCTTCTGTCGAGAG
Oligo rev AAACCTCTCGACAGAAGATCCACC
Chromosome start end strand MM target_seq PAM alignment distance position gene name gene id
GU305914.1 35091 35113 - 0 GGTGGATCTTCTGTCGAGAG AGG ||||||||[||||||||||||]PAM NA Intergenic NA NA

T31666 GTGGATCTTCTGTCGAGAGAGGG 1000
Oligo fwd CACCGTGGATCTTCTGTCGAGAGA
Oligo rev AAACTCTCTCGACAGAAGATCCAC
Chromosome start end strand MM target_seq PAM alignment distance position gene name gene id
GU305914.1 35090 35112 - 0 GTGGATCTTCTGTCGAGAGA GGG ||||||||[||||||||||||]PAM NA Intergenic NA NA

T31667 TTCTGTCGAGAGAGGGACACCGG 1000
Oligo fwd CACCTTCTGTCGAGAGAGGGACAC
Oligo rev AAACGTGTCCCTCTCTCGACAGAA
Chromosome start end strand MM target_seq PAM alignment distance position gene name gene id
GU305914.1 35083 35105 - 0 TTCTGTCGAGAGAGGGACAC CGG ||||||||[||||||||||||]PAM NA Intergenic NA NA

T31668 GTTCGACGCGCCGCCAACGACGG 1000
Oligo fwd CACCGTTCGACGCGCCGCCAACGA
Oligo rev AAACTCGTTGGCGGCGCGTCGAAC
Chromosome start end strand MM target_seq PAM alignment distance position gene name gene id
GU305914.1 35046 35068 - 0 GTTCGACGCGCCGCCAACGA CGG ||||||||[||||||||||||]PAM NA Intergenic NA NA

T31669 TTCGACGCGCCGCCAACGACGGG 1000
Oligo fwd CACCTTCGACGCGCCGCCAACGAC
Oligo rev AAACGTCGTTGGCGGCGCGTCGAA
Chromosome start end strand MM target_seq PAM alignment distance position gene name gene id
GU305914.1 35045 35067 - 0 TTCGACGCGCCGCCAACGAC GGG ||||||||[||||||||||||]PAM NA Intergenic NA NA

T31670 AACGACGGGCGACGACGTCGAGG 1000
Oligo fwd CACCAACGACGGGCGACGACGTCG
Oligo rev AAACCGACGTCGTCGCCCGTCGTT
Chromosome start end strand MM target_seq PAM alignment distance position gene name gene id
GU305914.1 35031 35053 - 0 AACGACGGGCGACGACGTCG AGG ||||||||[||||||||||||]PAM NA Intergenic NA NA

T31671 TACTGCCCCTGACCGAGCTCCGG 1000
Oligo fwd CACCTACTGCCCCTGACCGAGCTC
Oligo rev AAACGAGCTCGGTCAGGGGCAGTA
Chromosome start end strand MM target_seq PAM alignment distance position gene name gene id
GU305914.1 34996 35018 - 0 TACTGCCCCTGACCGAGCTC CGG ||||||||[||||||||||||]PAM NA Intergenic NA NA

T31672 ACTGCCCCTGACCGAGCTCCGGG 1000
Oligo fwd CACCACTGCCCCTGACCGAGCTCC
Oligo rev AAACGGAGCTCGGTCAGGGGCAGT
Chromosome start end strand MM target_seq PAM alignment distance position gene name gene id
GU305914.1 34995 35017 - 0 ACTGCCCCTGACCGAGCTCC GGG ||||||||[||||||||||||]PAM NA Intergenic NA NA

T31673 TGCTTCGTCCGCGACCGCACCGG 1000
Oligo fwd CACCTGCTTCGTCCGCGACCGCAC
Oligo rev AAACGTGCGGTCGCGGACGAAGCA
Chromosome start end strand MM target_seq PAM alignment distance position gene name gene id
GU305914.1 34955 34977 - 0 TGCTTCGTCCGCGACCGCAC CGG ||||||||[||||||||||||]PAM NA Intergenic NA NA

T31674 GCTTCGTCCGCGACCGCACCGGG 1000
Oligo fwd CACCGCTTCGTCCGCGACCGCACC
Oligo rev AAACGGTGCGGTCGCGGACGAAGC
Chromosome start end strand MM target_seq PAM alignment distance position gene name gene id
GU305914.1 34954 34976 - 0 GCTTCGTCCGCGACCGCACC GGG ||||||||[||||||||||||]PAM NA Intergenic NA NA

T31675 CGGGCTGTCTACTTTCCGCCAGG 1000
Oligo fwd CACCCGGGCTGTCTACTTTCCGCC
Oligo rev AAACGGCGGAAAGTAGACAGCCCG

Chromosome start end strand MM target_seq PAM alignment distance position gene name gene id
GU305914.1 34935 34957 - 0 CGGGCTGTCTACTTTCCGCC AGG ||||||||[||||||||||||]PAM NA Intergenic NA NA

T31676 TCCGCCAGGCCAGTAACCTCAGG 1000
Oligo fwd CACCTCCGCCAGGCCAGTAACCTC
Oligo rev AAACGAGGTTACTGGCCTGGCGGA
Chromosome start end strand MM target_seq PAM alignment distance position gene name gene id
GU305914.1 34921 34943 - 0 TCCGCCAGGCCAGTAACCTC AGG ||||||||[||||||||||||]PAM NA Intergenic NA NA

T31677 GGCCAGTAACCTCAGGTTTATGG 1000
Oligo fwd CACCGGCCAGTAACCTCAGGTTTA
Oligo rev AAACTAAACCTGAGGTTACTGGCC
Chromosome start end strand MM target_seq PAM alignment distance position gene name gene id
GU305914.1 34914 34936 - 0 GGCCAGTAACCTCAGGTTTA TGG ||||||||[||||||||||||]PAM NA Intergenic NA NA

T31678 GCCAGTAACCTCAGGTTTATGGG 1000
Oligo fwd CACCGCCAGTAACCTCAGGTTTAT
Oligo rev AAACATAAACCTGAGGTTACTGGC
Chromosome start end strand MM target_seq PAM alignment distance position gene name gene id
GU305914.1 34913 34935 - 0 GCCAGTAACCTCAGGTTTAT GGG ||||||||[||||||||||||]PAM NA Intergenic NA NA

T31679 AGGTTTATGGGCGAGTTCTTCGG 1000
Oligo fwd CACCAGGTTTATGGGCGAGTTCTT
Oligo rev AAACAAGAACTCGCCCATAAACCT
Chromosome start end strand MM target_seq PAM alignment distance position gene name gene id
GU305914.1 34901 34923 - 0 AGGTTTATGGGCGAGTTCTT CGG ||||||||[||||||||||||]PAM NA Intergenic NA NA

T31680 CGAGTTCTTCGGCTATTTCATGG 1000
Oligo fwd CACCCGAGTTCTTCGGCTATTTCA
Oligo rev AAACTGAAATAGCCGAAGAACTCG
Chromosome start end strand MM target_seq PAM alignment distance position gene name gene id
GU305914.1 34890 34912 - 0 CGAGTTCTTCGGCTATTTCA TGG ||||||||[||||||||||||]PAM NA Intergenic NA NA

T31681 TTCTTCGGCTATTTCATGGTCGG 1000
Oligo fwd CACCTTCTTCGGCTATTTCATGGT
Oligo rev AAACACCATGAAATAGCCGAAGAA
Chromosome start end strand MM target_seq PAM alignment distance position gene name gene id
GU305914.1 34886 34908 - 0 TTCTTCGGCTATTTCATGGT CGG ||||||||[||||||||||||]PAM NA Intergenic NA NA

T31682 TGATGAAGAGAGTTTGAGATTGG 1000
Oligo fwd CACCTGATGAAGAGAGTTTGAGAT
Oligo rev AAACATCTCAAACTCTCTTCATCA
Chromosome start end strand MM target_seq PAM alignment distance position gene name gene id
GU305914.1 34863 34885 - 0 TGATGAAGAGAGTTTGAGAT TGG ||||||||[||||||||||||]PAM NA Intergenic NA NA

T31683 GATGAAGAGAGTTTGAGATTGGG 1000
Oligo fwd CACCGATGAAGAGAGTTTGAGATT
Oligo rev AAACAATCTCAAACTCTCTTCATC
Chromosome start end strand MM target_seq PAM alignment distance position gene name gene id
GU305914.1 34862 34884 - 0 GATGAAGAGAGTTTGAGATT GGG ||||||||[||||||||||||]PAM NA Intergenic NA NA

T31684 AGAGAGTTTGAGATTGGGCCAGG 1000
Oligo fwd CACCAGAGAGTTTGAGATTGGGCC
Oligo rev AAACGGCCCAATCTCAAACTCTCT
Chromosome start end strand MM target_seq PAM alignment distance position gene name gene id
GU305914.1 34857 34879 - 0 AGAGAGTTTGAGATTGGGCC AGG ||||||||[||||||||||||]PAM NA Intergenic NA NA

T31685 GAGTTTGAGATTGGGCCAGGTGG 1000
Oligo fwd CACCGAGTTTGAGATTGGGCCAGG
Oligo rev AAACCCTGGCCCAATCTCAAACTC
Chromosome start end strand MM target_seq PAM alignment distance position gene name gene id
GU305914.1 34854 34876 - 0 GAGTTTGAGATTGGGCCAGG TGG ||||||||[||||||||||||]PAM NA Intergenic NA NA

T31686 AGATTGGGCCAGGTGGTGCCCGG 1000
Oligo fwd CACCAGATTGGGCCAGGTGGTGCC
Oligo rev AAACGGCACCACCTGGCCCAATCT
Chromosome start end strand MM target_seq PAM alignment distance position gene name gene id
GU305914.1 34847 34869 - 0 AGATTGGGCCAGGTGGTGCC CGG ||||||||[||||||||||||]PAM NA Intergenic NA NA

T31687 TGGGCCAGGTGGTGCCCGGACGG 1000
Oligo fwd CACCTGGGCCAGGTGGTGCCCGGA
Oligo rev AAACTCCGGGCACCACCTGGCCCA
Chromosome start end strand MM target_seq PAM alignment distance position gene name gene id
GU305914.1 34843 34865 - 0 TGGGCCAGGTGGTGCCCGGA CGG ||||||||[||||||||||||]PAM NA Intergenic NA NA

T31688 GGGCCAGGTGGTGCCCGGACGGG 1000
Oligo fwd CACCGGGCCAGGTGGTGCCCGGAC
Oligo rev AAACGTCCGGGCACCACCTGGCCC
Chromosome start end strand MM target_seq PAM alignment distance position gene name gene id
GU305914.1 34842 34864 - 0 GGGCCAGGTGGTGCCCGGAC GGG ||||||||[||||||||||||]PAM NA Intergenic NA NA

T31689 GGGTGTTCTTCTATCTTCGCCGG 1000
Oligo fwd CACCGGGTGTTCTTCTATCTTCGC
Oligo rev AAACGCGAAGATAGAAGAACACCC
Chromosome start end strand MM target_seq PAM alignment distance position gene name gene id
GU305914.1 34822 34844 - 0 GGGTGTTCTTCTATCTTCGC CGG ||||||||[||||||||||||]PAM NA Intergenic NA NA

T31690 GGTGTTCTTCTATCTTCGCCGGG 1000
Oligo fwd CACCGGTGTTCTTCTATCTTCGCC

<table>
<tr><td>Oligo rev</td><td>AAACGGCGAAGATAGAAGAACACC</td></tr>
<tr><td>Chromosome</td><td>start</td><td>end</td><td>strand</td><td>MM</td><td>target_seq</td><td>PAM</td><td>alignment</td><td>distance</td><td>position</td><td>gene name</td><td>gene id</td></tr>
<tr><td>GU305914.1</td><td>34821</td><td>34843</td><td>-</td><td>0</td><td>GGTGTTCTTCTATCTTCGCC</td><td>GGG</td><td>||||||||[||||||||||||]PAM</td><td>NA</td><td>Intergenic</td><td>NA</td><td>NA</td></tr>
<tr><td>T31691</td><td>GTTCTTCTATCTTCGCCGGGCGG</td><td>1000</td></tr>
<tr><td>Oligo fwd</td><td>CACCGTTCTTCTATCTTCGCCGGG</td></tr>
<tr><td>Oligo rev</td><td>AAACCCCGGCGAAGATAGAAGAAC</td></tr>
<tr><td>Chromosome</td><td>start</td><td>end</td><td>strand</td><td>MM</td><td>target_seq</td><td>PAM</td><td>alignment</td><td>distance</td><td>position</td><td>gene name</td><td>gene id</td></tr>
<tr><td>GU305914.1</td><td>34818</td><td>34840</td><td>-</td><td>0</td><td>GTTCTTCTATCTTCGCCGGG</td><td>CGG</td><td>||||||||[||||||||||||]PAM</td><td>NA</td><td>Intergenic</td><td>NA</td><td>NA</td></tr>
<tr><td>T31692</td><td>TTCTTCTATCTTCGCCGGGCGGG</td><td>1000</td></tr>
<tr><td>Oligo fwd</td><td>CACCTTCTTCTATCTTCGCCGGGC</td></tr>
<tr><td>Oligo rev</td><td>AAACGCCCGGCGAAGATAGAAGAA</td></tr>
<tr><td>Chromosome</td><td>start</td><td>end</td><td>strand</td><td>MM</td><td>target_seq</td><td>PAM</td><td>alignment</td><td>distance</td><td>position</td><td>gene name</td><td>gene id</td></tr>
<tr><td>GU305914.1</td><td>34817</td><td>34839</td><td>-</td><td>0</td><td>TTCTTCTATCTTCGCCGGGC</td><td>GGG</td><td>||||||||[||||||||||||]PAM</td><td>NA</td><td>Intergenic</td><td>NA</td><td>NA</td></tr>
<tr><td>T31693</td><td>CGGGCGGGCTATCGCGTCATCGG</td><td>1000</td></tr>
<tr><td>Oligo fwd</td><td>CACCCGGGCGGGCTATCGCGTCAT</td></tr>
<tr><td>Oligo rev</td><td>AAACATGACGCGATAGCCCGCCCG</td></tr>
<tr><td>Chromosome</td><td>start</td><td>end</td><td>strand</td><td>MM</td><td>target_seq</td><td>PAM</td><td>alignment</td><td>distance</td><td>position</td><td>gene name</td><td>gene id</td></tr>
<tr><td>GU305914.1</td><td>34802</td><td>34824</td><td>-</td><td>0</td><td>CGGGCGGGCTATCGCGTCAT</td><td>CGG</td><td>||||||||[||||||||||||]PAM</td><td>NA</td><td>Intergenic</td><td>NA</td><td>NA</td></tr>
<tr><td>T31694</td><td>GCTGTGTAACGAGAAGCTCGAGG</td><td>1000</td></tr>
<tr><td>Oligo fwd</td><td>CACCGCTGTGTAACGAGAAGCTCG</td></tr>
<tr><td>Oligo rev</td><td>AAACCGAGCTTCTCGTTACACAGC</td></tr>
<tr><td>Chromosome</td><td>start</td><td>end</td><td>strand</td><td>MM</td><td>target_seq</td><td>PAM</td><td>alignment</td><td>distance</td><td>position</td><td>gene name</td><td>gene id</td></tr>
<tr><td>GU305914.1</td><td>34755</td><td>34777</td><td>-</td><td>0</td><td>GCTGTGTAACGAGAAGCTCG</td><td>AGG</td><td>||||||||[||||||||||||]PAM</td><td>NA</td><td>Intergenic</td><td>NA</td><td>NA</td></tr>
<tr><td>T31695</td><td>GCTCGAGGTCTTCGTGCTCCTGG</td><td>1000</td></tr>
<tr><td>Oligo fwd</td><td>CACCGCTCGAGGTCTTCGTGCTCC</td></tr>
<tr><td>Oligo rev</td><td>AAACGGAGCACGAAGACCTCGAGC</td></tr>
<tr><td>Chromosome</td><td>start</td><td>end</td><td>strand</td><td>MM</td><td>target_seq</td><td>PAM</td><td>alignment</td><td>distance</td><td>position</td><td>gene name</td><td>gene id</td></tr>
<tr><td>GU305914.1</td><td>34740</td><td>34762</td><td>-</td><td>0</td><td>GCTCGAGGTCTTCGTGCTCC</td><td>TGG</td><td>||||||||[||||||||||||]PAM</td><td>NA</td><td>Intergenic</td><td>NA</td><td>NA</td></tr>
<tr><td>T31696</td><td>CTCGAGGTCTTCGTGCTCCTGGG</td><td>1000</td></tr>
<tr><td>Oligo fwd</td><td>CACCCTCGAGGTCTTCGTGCTCCT</td></tr>
<tr><td>Oligo rev</td><td>AAACAGGAGCACGAAGACCTCGAG</td></tr>
<tr><td>Chromosome</td><td>start</td><td>end</td><td>strand</td><td>MM</td><td>target_seq</td><td>PAM</td><td>alignment</td><td>distance</td><td>position</td><td>gene name</td><td>gene id</td></tr>
<tr><td>GU305914.1</td><td>34739</td><td>34761</td><td>-</td><td>0</td><td>CTCGAGGTCTTCGTGCTCCT</td><td>GGG</td><td>||||||||[||||||||||||]PAM</td><td>NA</td><td>Intergenic</td><td>NA</td><td>NA</td></tr>
<tr><td>T31697</td><td>GAGGTCTTCGTGCTCCTGGGCGG</td><td>1000</td></tr>
<tr><td>Oligo fwd</td><td>CACCGAGGTCTTCGTGCTCCTGGG</td></tr>
<tr><td>Oligo rev</td><td>AAACCCCAGGAGCACGAAGACCTC</td></tr>
<tr><td>Chromosome</td><td>start</td><td>end</td><td>strand</td><td>MM</td><td>target_seq</td><td>PAM</td><td>alignment</td><td>distance</td><td>position</td><td>gene name</td><td>gene id</td></tr>
<tr><td>GU305914.1</td><td>34736</td><td>34758</td><td>-</td><td>0</td><td>GAGGTCTTCGTGCTCCTGGG</td><td>CGG</td><td>||||||||[||||||||||||]PAM</td><td>NA</td><td>Intergenic</td><td>NA</td><td>NA</td></tr>
<tr><td>T31698</td><td>GGGCGGCGCCCGCGTCTTGAAGG</td><td>1000</td></tr>
<tr><td>Oligo fwd</td><td>CACCGGGCGGCGCCCGCGTCTTGA</td></tr>
<tr><td>Oligo rev</td><td>AAACTCAAGACGCGGGCGCCGCCC</td></tr>
<tr><td>Chromosome</td><td>start</td><td>end</td><td>strand</td><td>MM</td><td>target_seq</td><td>PAM</td><td>alignment</td><td>distance</td><td>position</td><td>gene name</td><td>gene id</td></tr>
<tr><td>GU305914.1</td><td>34719</td><td>34741</td><td>-</td><td>0</td><td>GGGCGGCGCCCGCGTCTTGA</td><td>AGG</td><td>||||||||[||||||||||||]PAM</td><td>NA</td><td>Intergenic</td><td>NA</td><td>NA</td></tr>
<tr><td>T31699</td><td>CGGCGCCCGCGTCTTGAAGGTGG</td><td>1000</td></tr>
<tr><td>Oligo fwd</td><td>CACCCGGCGCCCGCGTCTTGAAGG</td></tr>
<tr><td>Oligo rev</td><td>AAACCCTTCAAGACGCGGGCGCCG</td></tr>
<tr><td>Chromosome</td><td>start</td><td>end</td><td>strand</td><td>MM</td><td>target_seq</td><td>PAM</td><td>alignment</td><td>distance</td><td>position</td><td>gene name</td><td>gene id</td></tr>
<tr><td>GU305914.1</td><td>34716</td><td>34738</td><td>-</td><td>0</td><td>CGGCGCCCGCGTCTTGAAGG</td><td>TGG</td><td>||||||||[||||||||||||]PAM</td><td>NA</td><td>Intergenic</td><td>NA</td><td>NA</td></tr>
<tr><td>T31700</td><td>GAAGGTGGCCAACACGATAGCGG</td><td>1000</td></tr>
<tr><td>Oligo fwd</td><td>CACCGAAGGTGGCCAACACGATAG</td></tr>
<tr><td>Oligo rev</td><td>AAACCTATCGTGTTGGCCACCTTC</td></tr>
<tr><td>Chromosome</td><td>start</td><td>end</td><td>strand</td><td>MM</td><td>target_seq</td><td>PAM</td><td>alignment</td><td>distance</td><td>position</td><td>gene name</td><td>gene id</td></tr>
<tr><td>GU305914.1</td><td>34701</td><td>34723</td><td>-</td><td>0</td><td>GAAGGTGGCCAACACGATAG</td><td>CGG</td><td>||||||||[||||||||||||]PAM</td><td>NA</td><td>Intergenic</td><td>NA</td><td>NA</td></tr>
<tr><td>T31701</td><td>AAGGTGGCCAACACGATAGCGGG</td><td>1000</td></tr>
<tr><td>Oligo fwd</td><td>CACCAAGGTGGCCAACACGATAGC</td></tr>
<tr><td>Oligo rev</td><td>AAACGCTATCGTGTTGGCCACCTT</td></tr>
<tr><td>Chromosome</td><td>start</td><td>end</td><td>strand</td><td>MM</td><td>target_seq</td><td>PAM</td><td>alignment</td><td>distance</td><td>position</td><td>gene name</td><td>gene id</td></tr>
<tr><td>GU305914.1</td><td>34700</td><td>34722</td><td>-</td><td>0</td><td>AAGGTGGCCAACACGATAGC</td><td>GGG</td><td>||||||||[||||||||||||]PAM</td><td>NA</td><td>Intergenic</td><td>NA</td><td>NA</td></tr>
<tr><td>T31702</td><td>ACACGATAGCGGGATTCCTCCGG</td><td>1000</td></tr>
<tr><td>Oligo fwd</td><td>CACCACACGATAGCGGGATTCCTC</td></tr>
<tr><td>Oligo rev</td><td>AAACGAGGAATCCCGCTATCGTGT</td></tr>
<tr><td>Chromosome</td><td>start</td><td>end</td><td>strand</td><td>MM</td><td>target_seq</td><td>PAM</td><td>alignment</td><td>distance</td><td>position</td><td>gene name</td><td>gene id</td></tr>
<tr><td>GU305914.1</td><td>34690</td><td>34712</td><td>-</td><td>0</td><td>ACACGATAGCGGGATTCCTC</td><td>CGG</td><td>||||||||[||||||||||||]PAM</td><td>NA</td><td>Intergenic</td><td>NA</td><td>NA</td></tr>
<tr><td>T31703</td><td>CACGATAGCGGGATTCCTCCGGG</td><td>1000</td></tr>
<tr><td>Oligo fwd</td><td>CACCCACGATAGCGGGATTCCTCC</td></tr>
<tr><td>Oligo rev</td><td>AAACGGAGGAATCCCGCTATCGTG</td></tr>
<tr><td>Chromosome</td><td>start</td><td>end</td><td>strand</td><td>MM</td><td>target_seq</td><td>PAM</td><td>alignment</td><td>distance</td><td>position</td><td>gene name</td><td>gene id</td></tr>
<tr><td>GU305914.1</td><td>34689</td><td>34711</td><td>-</td><td>0</td><td>CACGATAGCGGGATTCCTCC</td><td>GGG</td><td>||||||||[||||||||||||]PAM</td><td>NA</td><td>Intergenic</td><td>NA</td><td>NA</td></tr>
<tr><td>T31704</td><td>TAGCGGGATTCCTCCGGGACCGG</td><td>1000</td></tr>
<tr><td>Oligo fwd</td><td>CACCTAGCGGGATTCCTCCGGGAC</td></tr>
<tr><td>Oligo rev</td><td>AAACGTCCCGGAGGAATCCCGCTA</td></tr>
<tr><td>Chromosome</td><td>start</td><td>end</td><td>strand</td><td>MM</td><td>target_seq</td><td>PAM</td><td>alignment</td><td>distance</td><td>position</td><td>gene name</td><td>gene id</td></tr>
<tr><td>GU305914.1</td><td>34684</td><td>34706</td><td>-</td><td>0</td><td>TAGCGGGATTCCTCCGGGAC</td><td>CGG</td><td>||||||||[||||||||||||]PAM</td><td>NA</td><td>Intergenic</td><td>NA</td><td>NA</td></tr>
<tr><td>T31705</td><td>GATTCCTCCGGGACCGGCTCAGG</td><td>1000</td></tr>
</table>

Oligo fwd CACCGATTCCTCCGGGACCGGCTC
Oligo rev AAACGAGCCGGTCCCGGAGGAATC
Chromosome start end strand MM target_seq PAM alignment distance position gene name gene id
GU305914.1 34678 34700 - 0 GATTCCTCCGGGACCGGCTC AGG ||||||||[||||||||||||]PAM NA Intergenic NA NA

T31706 GGATCAACCTGCAGCCCTACCGG 1000
Oligo fwd CACCGGATCAACCTGCAGCCCTAC
Oligo rev AAACGTAGGGCTGCAGGTTGATCC
Chromosome start end strand MM target_seq PAM alignment distance position gene name gene id
GU305914.1 34657 34679 - 0 GGATCAACCTGCAGCCCTAC CGG ||||||||[||||||||||||]PAM NA Intergenic NA NA

T31707 TCAACCTGCAGCCCTACCGGAGG 1000
Oligo fwd CACCTCAACCTGCAGCCCTACCGG
Oligo rev AAACCCGGTAGGGCTGCAGGTTGA
Chromosome start end strand MM target_seq PAM alignment distance position gene name gene id
GU305914.1 34654 34676 - 0 TCAACCTGCAGCCCTACCGG AGG ||||||||[||||||||||||]PAM NA Intergenic NA NA

T31708 CAACCTGCAGCCCTACCGGAGGG 1000
Oligo fwd CACCCAACCTGCAGCCCTACCGGA
Oligo rev AAACTCCGGTAGGGCTGCAGGTTG
Chromosome start end strand MM target_seq PAM alignment distance position gene name gene id
GU305914.1 34653 34675 - 0 CAACCTGCAGCCCTACCGGA GGG ||||||||[||||||||||||]PAM NA Intergenic NA NA

T31709 CCTACCGGAGGGTGTTCGCCAGG 1000
Oligo fwd CACCCCTACCGGAGGGTGTTCGCC
Oligo rev AAACGGCGAACACCCTCCGGTAGG
Chromosome start end strand MM target_seq PAM alignment distance position gene name gene id
GU305914.1 34642 34664 - 0 CCTACCGGAGGGTGTTCGCC AGG ||||||||[||||||||||||]PAM NA Intergenic NA NA

T31710 CGGAGGGTGTTCGCCAGGAACGG 1000
Oligo fwd CACCCGGAGGGTGTTCGCCAGGAA
Oligo rev AAACTTCCTGGCGAACACCCTCCG
Chromosome start end strand MM target_seq PAM alignment distance position gene name gene id
GU305914.1 34637 34659 - 0 CGGAGGGTGTTCGCCAGGAA CGG ||||||||[||||||||||||]PAM NA Intergenic NA NA

T31711 GTTCGCCAGGAACGGCGTCGCGG 1000
Oligo fwd CACCGTTCGCCAGGAACGGCGTCG
Oligo rev AAACCGACGCCGTTCCTGGCGAAC
Chromosome start end strand MM target_seq PAM alignment distance position gene name gene id
GU305914.1 34629 34651 - 0 GTTCGCCAGGAACGGCGTCG CGG ||||||||[||||||||||||]PAM NA Intergenic NA NA

T31712 GAACGGCGTCGCGGAGCGCACGG 1000
Oligo fwd CACCGAACGGCGTCGCGGAGCGCA
Oligo rev AAACTGCGCTCCGCGACGCCGTTC
Chromosome start end strand MM target_seq PAM alignment distance position gene name gene id
GU305914.1 34620 34642 - 0 GAACGGCGTCGCGGAGCGCA CGG ||||||||[||||||||||||]PAM NA Intergenic NA NA

T31713 GCGGAGCGCACGGTCTGCGTCGG 1000
Oligo fwd CACCGCGGAGCGCACGGTCTGCGT
Oligo rev AAACACGCAGACCGTGCGCTCCGC
Chromosome start end strand MM target_seq PAM alignment distance position gene name gene id
GU305914.1 34610 34632 - 0 GCGGAGCGCACGGTCTGCGT CGG ||||||||[||||||||||||]PAM NA Intergenic NA NA

T31714 GCGCACGGTCTGCGTCGGACAGG 1000
Oligo fwd CACCGCGCACGGTCTGCGTCGGAC
Oligo rev AAACGTCCGACGCAGACCGTGCGC
Chromosome start end strand MM target_seq PAM alignment distance position gene name gene id
GU305914.1 34605 34627 - 0 GCGCACGGTCTGCGTCGGAC AGG ||||||||[||||||||||||]PAM NA Intergenic NA NA

T31715 GAGATCGACTACGTCGTGCCCGG 1000
Oligo fwd CACCGAGATCGACTACGTCGTGCC
Oligo rev AAACGGCACGACGTAGTCGATCTC
Chromosome start end strand MM target_seq PAM alignment distance position gene name gene id
GU305914.1 34565 34587 - 0 GAGATCGACTACGTCGTGCC CGG ||||||||[||||||||||||]PAM NA Intergenic NA NA

T31716 ATCGACTACGTCGTGCCCGGCGG 1000
Oligo fwd CACCATCGACTACGTCGTGCCCGG
Oligo rev AAACCCGGGCACGACGTAGTCGAT
Chromosome start end strand MM target_seq PAM alignment distance position gene name gene id
GU305914.1 34562 34584 - 0 ATCGACTACGTCGTGCCCGG CGG ||||||||[||||||||||||]PAM NA Intergenic NA NA

T31717 GCGGTGCCGACTTCGCCAAGTGG 1000
Oligo fwd CACCGCGGTGCCGACTTCGCCAAG
Oligo rev AAACCTTGGCGAAGTCGGCACCGC
Chromosome start end strand MM target_seq PAM alignment distance position gene name gene id
GU305914.1 34543 34565 - 0 GCGGTGCCGACTTCGCCAAG TGG ||||||||[||||||||||||]PAM NA Intergenic NA NA

T31718 TGCCGACTTCGCCAAGTGGTTGG 1000
Oligo fwd CACCTGCCGACTTCGCCAAGTGGT
Oligo rev AAACACCACTTGGCGAAGTCGGCA
Chromosome start end strand MM target_seq PAM alignment distance position gene name gene id
GU305914.1 34539 34561 - 0 TGCCGACTTCGCCAAGTGGT TGG ||||||||[||||||||||||]PAM NA Intergenic NA NA

T31719 TCGCCAAGTGGTTGGCGTTGCGG 1000
Oligo fwd CACCTCGCCAAGTGGTTGGCGTTG
Oligo rev AAACCAACGCCAACCACTTGGCGA
Chromosome start end strand MM target_seq PAM alignment distance position gene name gene id
GU305914.1 34531 34553 - 0 TCGCCAAGTGGTTGGCGTTG CGG ||||||||[||||||||||||]PAM NA Intergenic NA NA

| | | | | | | | | | | | |
|---|---|---|---|---|---|---|---|---|---|---|---|
| T31720 | CGCCAAGTGGTTGGCGTTGCGGG | 1000 | | | | | | | | | |
| Oligo fwd | CACCCGCCAAGTGGTTGGCGTTGC | | | | | | | | | | |
| Oligo rev | AAACGCAACGCCAACCACTTGGCG | | | | | | | | | | |
| Chromosome | start | end | strand | MM | target_seq | PAM | alignment | distance | position | gene name | gene id |
| GU305914.1 | 34530 | 34552 | - | 0 | CGCCAAGTGGTTGGCGTTGC | GGG | \|\|\|\|\|\|\|\|[\|\|\|\|\|\|\|\|\|\|\|\|]PAM | NA | Intergenic | NA | NA |
| T31721 | TTGGCGTTGCGGGCTCCGTCCGG | 1000 | | | | | | | | | |
| Oligo fwd | CACCTTGGCGTTGCGGGCTCCGTC | | | | | | | | | | |
| Oligo rev | AAACGACGGAGCCCGCAACGCCAA | | | | | | | | | | |
| Chromosome | start | end | strand | MM | target_seq | PAM | alignment | distance | position | gene name | gene id |
| GU305914.1 | 34520 | 34542 | - | 0 | TTGGCGTTGCGGGCTCCGTC | CGG | \|\|\|\|\|\|\|\|[\|\|\|\|\|\|\|\|\|\|\|\|]PAM | NA | Intergenic | NA | NA |
| T31722 | TGGCGTTGCGGGCTCCGTCCGGG | 1000 | | | | | | | | | |
| Oligo fwd | CACCTGGCGTTGCGGGCTCCGTCC | | | | | | | | | | |
| Oligo rev | AAACGGACGGAGCCCGCAACGCCA | | | | | | | | | | |
| Chromosome | start | end | strand | MM | target_seq | PAM | alignment | distance | position | gene name | gene id |
| GU305914.1 | 34519 | 34541 | - | 0 | TGGCGTTGCGGGCTCCGTCC | GGG | \|\|\|\|\|\|\|\|[\|\|\|\|\|\|\|\|\|\|\|\|]PAM | NA | Intergenic | NA | NA |
| T31723 | GAGACCTGCGTGCCCACGCTCGG | 1000 | | | | | | | | | |
| Oligo fwd | CACCGAGACCTGCGTGCCCACGCT | | | | | | | | | | |
| Oligo rev | AAACAGCGTGGGCACGCAGGTCTC | | | | | | | | | | |
| Chromosome | start | end | strand | MM | target_seq | PAM | alignment | distance | position | gene name | gene id |
| GU305914.1 | 34490 | 34512 | - | 0 | GAGACCTGCGTGCCCACGCT | CGG | \|\|\|\|\|\|\|\|[\|\|\|\|\|\|\|\|\|\|\|\|]PAM | NA | Intergenic | NA | NA |
| T31724 | AGACCTGCGTGCCCACGCTCGGG | 1000 | | | | | | | | | |
| Oligo fwd | CACCAGACCTGCGTGCCCACGCTC | | | | | | | | | | |
| Oligo rev | AAACGAGCGTGGGCACGCAGGTCT | | | | | | | | | | |
| Chromosome | start | end | strand | MM | target_seq | PAM | alignment | distance | position | gene name | gene id |
| GU305914.1 | 34489 | 34511 | - | 0 | AGACCTGCGTGCCCACGCTC | GGG | \|\|\|\|\|\|\|\|[\|\|\|\|\|\|\|\|\|\|\|\|]PAM | NA | Intergenic | NA | NA |
| T31725 | GCGTGCCCACGCTCGGGTGACGG | 1000 | | | | | | | | | |
| Oligo fwd | CACCGCGTGCCCACGCTCGGGTGA | | | | | | | | | | |
| Oligo rev | AAACTCACCCGAGCGTGGGCACGC | | | | | | | | | | |
| Chromosome | start | end | strand | MM | target_seq | PAM | alignment | distance | position | gene name | gene id |
| GU305914.1 | 34483 | 34505 | - | 0 | GCGTGCCCACGCTCGGGTGA | CGG | \|\|\|\|\|\|\|\|[\|\|\|\|\|\|\|\|\|\|\|\|]PAM | NA | Intergenic | NA | NA |
| T31726 | GCCCACGCTCGGGTGACGGCTGG | 1000 | | | | | | | | | |
| Oligo fwd | CACCGCCCACGCTCGGGTGACGGC | | | | | | | | | | |
| Oligo rev | AAACGCCGTCACCCGAGCGTGGGC | | | | | | | | | | |
| Chromosome | start | end | strand | MM | target_seq | PAM | alignment | distance | position | gene name | gene id |
| GU305914.1 | 34479 | 34501 | - | 0 | GCCCACGCTCGGGTGACGGC | TGG | \|\|\|\|\|\|\|\|[\|\|\|\|\|\|\|\|\|\|\|\|]PAM | NA | Intergenic | NA | NA |
| T31727 | CCCACGCTCGGGTGACGGCTGGG | 1000 | | | | | | | | | |
| Oligo fwd | CACCCCCACGCTCGGGTGACGGCT | | | | | | | | | | |
| Oligo rev | AAACAGCCGTCACCCGAGCGTGGG | | | | | | | | | | |
| Chromosome | start | end | strand | MM | target_seq | PAM | alignment | distance | position | gene name | gene id |
| GU305914.1 | 34478 | 34500 | - | 0 | CCCACGCTCGGGTGACGGCT | GGG | \|\|\|\|\|\|\|\|[\|\|\|\|\|\|\|\|\|\|\|\|]PAM | NA | Intergenic | NA | NA |
| T31728 | GCCAGTTTCGCCTCGCCTCCCGG | 1000 | | | | | | | | | |
| Oligo fwd | CACCGCCAGTTTCGCCTCGCCTCC | | | | | | | | | | |
| Oligo rev | AAACGGAGGCGAGGCGAAACTGGC | | | | | | | | | | |
| Chromosome | start | end | strand | MM | target_seq | PAM | alignment | distance | position | gene name | gene id |
| GU305914.1 | 34421 | 34443 | - | 0 | GCCAGTTTCGCCTCGCCTCC | CGG | \|\|\|\|\|\|\|\|[\|\|\|\|\|\|\|\|\|\|\|\|]PAM | NA | Intergenic | NA | NA |
| T31729 | CTCGCCTCCCGGCAGCCGAGCGG | 1000 | | | | | | | | | |
| Oligo fwd | CACCCTCGCCTCCCGGCAGCCGAG | | | | | | | | | | |
| Oligo rev | AAACCTCGGCTGCCGGGAGGCGAG | | | | | | | | | | |
| Chromosome | start | end | strand | MM | target_seq | PAM | alignment | distance | position | gene name | gene id |
| GU305914.1 | 34410 | 34432 | - | 0 | CTCGCCTCCCGGCAGCCGAG | CGG | \|\|\|\|\|\|\|\|[\|\|\|\|\|\|\|\|\|\|\|\|]PAM | NA | Intergenic | NA | NA |
| T31730 | GCCTCCCGGCAGCCGAGCGGCGG | 1000 | | | | | | | | | |
| Oligo fwd | CACCGCCTCCCGGCAGCCGAGCGG | | | | | | | | | | |
| Oligo rev | AAACCCGCTCGGCTGCCGGGAGGC | | | | | | | | | | |
| Chromosome | start | end | strand | MM | target_seq | PAM | alignment | distance | position | gene name | gene id |
| GU305914.1 | 34407 | 34429 | - | 0 | GCCTCCCGGCAGCCGAGCGG | CGG | \|\|\|\|\|\|\|\|[\|\|\|\|\|\|\|\|\|\|\|\|]PAM | NA | Intergenic | NA | NA |
| T31731 | TCCCGGCAGCCGAGCGGCGGAGG | 1000 | | | | | | | | | |
| Oligo fwd | CACCTCCCGGCAGCCGAGCGGCGG | | | | | | | | | | |
| Oligo rev | AAACCCGCCGCTCGGCTGCCGGGA | | | | | | | | | | |
| Chromosome | start | end | strand | MM | target_seq | PAM | alignment | distance | position | gene name | gene id |
| GU305914.1 | 34404 | 34426 | - | 0 | TCCCGGCAGCCGAGCGGCGG | AGG | \|\|\|\|\|\|\|\|[\|\|\|\|\|\|\|\|\|\|\|\|]PAM | NA | Intergenic | NA | NA |
| T31732 | CCCGGCAGCCGAGCGGCGGAGGG | 1000 | | | | | | | | | |
| Oligo fwd | CACCCCCGGCAGCCGAGCGGCGGA | | | | | | | | | | |
| Oligo rev | AAACTCCGCCGCTCGGCTGCCGGG | | | | | | | | | | |
| Chromosome | start | end | strand | MM | target_seq | PAM | alignment | distance | position | gene name | gene id |
| GU305914.1 | 34403 | 34425 | - | 0 | CCCGGCAGCCGAGCGGCGGA | GGG | \|\|\|\|\|\|\|\|[\|\|\|\|\|\|\|\|\|\|\|\|]PAM | NA | Intergenic | NA | NA |
| GU305914.1 | 88250 | 88272 | + | 4 | CTCGCCGGACGAGCGGCGGA | CGG | \|-\|\|-\|-\|[-\|\|\|\|\|\|\|\|\|\|\|]PAM | NA | Intergenic | NA | NA |
| T31733 | GGCAGCCGAGCGGCGGAGGGTGG | 1000 | | | | | | | | | |
| Oligo fwd | CACCGGCAGCCGAGCGGCGGAGGG | | | | | | | | | | |
| Oligo rev | AAACCCCTCCGCCGCTCGGCTGCC | | | | | | | | | | |
| Chromosome | start | end | strand | MM | target_seq | PAM | alignment | distance | position | gene name | gene id |
| GU305914.1 | 34400 | 34422 | - | 0 | GGCAGCCGAGCGGCGGAGGG | TGG | \|\|\|\|\|\|\|\|[\|\|\|\|\|\|\|\|\|\|\|\|]PAM | NA | Intergenic | NA | NA |
| T31734 | GGGTGGCGTGCTTGCTGTGTAGG | 1000 | | | | | | | | | |
| Oligo fwd | CACCGGGTGGCGTGCTTGCTGTGT | | | | | | | | | | |
| Oligo rev | AAACACACAGCAAGCACGCCACCC | | | | | | | | | | |
| Chromosome | start | end | strand | MM | target_seq | PAM | alignment | distance | position | gene name | gene id |

GU305914.1 34383 34405 - 0 GGGTGGCGTGCTTGCTGTGT AGG ||||||||[||||||||||||]PAM NA Intergenic NA NA

T31735 GCGTGCTTGCTGTGTAGGTAAGG 1000
Oligo fwd CACCGCGTGCTTGCTGTGTAGGTA
Oligo rev AAACTACCTACACAGCAAGCACGC
Chromosome start end strand MM target_seq PAM alignment distance position gene name gene id
GU305914.1 34378 34400 - 0 GCGTGCTTGCTGTGTAGGTA AGG ||||||||[||||||||||||]PAM NA Intergenic NA NA

T31736 CGTGCTTGCTGTGTAGGTAAGGG 1000
Oligo fwd CACCCGTGCTTGCTGTGTAGGTAA
Oligo rev AAACTTACCTACACAGCAAGCACG
Chromosome start end strand MM target_seq PAM alignment distance position gene name gene id
GU305914.1 34377 34399 - 0 CGTGCTTGCTGTGTAGGTAA GGG ||||||||[||||||||||||]PAM NA Intergenic NA NA

T31737 GTGCTTGCTGTGTAGGTAAGGGG 1000
Oligo fwd CACCGTGCTTGCTGTGTAGGTAAG
Oligo rev AAACCTTACCTACACAGCAAGCAC
Chromosome start end strand MM target_seq PAM alignment distance position gene name gene id
GU305914.1 34376 34398 - 0 GTGCTTGCTGTGTAGGTAAG GGG ||||||||[||||||||||||]PAM NA Intergenic NA NA

T31738 CTTGCTGTGTAGGTAAGGGGCGG 1000
Oligo fwd CACCCTTGCTGTGTAGGTAAGGGG
Oligo rev AAACCCCCTTACCTACACAGCAAG
Chromosome start end strand MM target_seq PAM alignment distance position gene name gene id
GU305914.1 34373 34395 - 0 CTTGCTGTGTAGGTAAGGGG CGG ||||||||[||||||||||||]PAM NA Intergenic NA NA

T31739 TTGCTGTGTAGGTAAGGGGCGGG 1000
Oligo fwd CACCTTGCTGTGTAGGTAAGGGGC
Oligo rev AAACGCCCCTTACCTACACAGCAA
Chromosome start end strand MM target_seq PAM alignment distance position gene name gene id
GU305914.1 34372 34394 - 0 TTGCTGTGTAGGTAAGGGGC GGG ||||||||[||||||||||||]PAM NA Intergenic NA NA

T31740 TGCTGTGTAGGTAAGGGGCGGGG 1000
Oligo fwd CACCTGCTGTGTAGGTAAGGGGCG
Oligo rev AAACCGCCCCTTACCTACACAGCA
Chromosome start end strand MM target_seq PAM alignment distance position gene name gene id
GU305914.1 34371 34393 - 0 TGCTGTGTAGGTAAGGGGCG GGG ||||||||[||||||||||||]PAM NA Intergenic NA NA

T31741 ATTTTCAATAAAGTCGTGTGAGG 1000
Oligo fwd CACCATTTTCAATAAAGTCGTGTG
Oligo rev AAACCACACGACTTTATTGAAAAT
Chromosome start end strand MM target_seq PAM alignment distance position gene name gene id
GU305914.1 34343 34365 - 0 ATTTTCAATAAAGTCGTGTG AGG ||||||||[||||||||||||]PAM NA Intergenic NA NA

T31742 TCGTGTGAGGAAAAAGACGACGG 1000
Oligo fwd CACCTCGTGTGAGGAAAAAGACGA
Oligo rev AAACTCGTCTTTTTCCTCACACGA
Chromosome start end strand MM target_seq PAM alignment distance position gene name gene id
GU305914.1 34330 34352 - 0 TCGTGTGAGGAAAAAGACGA CGG ||||||||[||||||||||||]PAM NA Intergenic NA NA

T31743 ACGACGGTGTCATTTATCGACGG 1000
Oligo fwd CACCACGACGGTGTCATTTATCGA
Oligo rev AAACTCGATAAATGACACCGTCGT
Chromosome start end strand MM target_seq PAM alignment distance position gene name gene id
GU305914.1 34314 34336 - 0 ACGACGGTGTCATTTATCGA CGG ||||||||[||||||||||||]PAM NA Intergenic NA NA

T31744 GCGCCGTGTCGCGCTGACCATGG 1000
Oligo fwd CACCGCGCCGTGTCGCGCTGACCA
Oligo rev AAACTGGTCAGCGCGACACGGCGC
Chromosome start end strand MM target_seq PAM alignment distance position gene name gene id
GU305914.1 34292 34314 - 0 GCGCCGTGTCGCGCTGACCA TGG ||||||||[||||||||||||]PAM NA Intergenic NA NA

T31745 CCGTGTCGCGCTGACCATGGCGG 1000
Oligo fwd CACCCCGTGTCGCGCTGACCATGG
Oligo rev AAACCCATGGTCAGCGCGACACGG
Chromosome start end strand MM target_seq PAM alignment distance position gene name gene id
GU305914.1 34289 34311 - 0 CCGTGTCGCGCTGACCATGG CGG ||||||||[||||||||||||]PAM NA Intergenic NA NA

T31746 GCGCTGACCATGGCGGACCGCGG 1000
Oligo fwd CACCGCGCTGACCATGGCGGACCG
Oligo rev AAACCGGTCCGCCATGGTCAGCGC
Chromosome start end strand MM target_seq PAM alignment distance position gene name gene id
GU305914.1 34282 34304 - 0 GCGCTGACCATGGCGGACCG CGG ||||||||[||||||||||||]PAM NA Intergenic NA NA

T31747 CGCTGACCATGGCGGACCGCGGG 1000
Oligo fwd CACCCGCTGACCATGGCGGACCGC
Oligo rev AAACGCGGTCCGCCATGGTCAGCG
Chromosome start end strand MM target_seq PAM alignment distance position gene name gene id
GU305914.1 34281 34303 - 0 CGCTGACCATGGCGGACCGC GGG ||||||||[||||||||||||]PAM NA Intergenic NA NA

T31748 GACCGCGGGCCCTGCGCTGAAGG 1000
Oligo fwd CACCGACCGCGGGCCCTGCGCTGA
Oligo rev AAACTCAGCGCAGGGCCCGCGGTC
Chromosome start end strand MM target_seq PAM alignment distance position gene name gene id
GU305914.1 34267 34289 - 0 GACCGCGGGCCCTGCGCTGA AGG ||||||||[||||||||||||]PAM NA Intergenic NA NA

T31749 ACCGCGGGCCCTGCGCTGAAGGG 1000
Oligo fwd CACCACCGCGGGCCCTGCGCTGAA
Oligo rev AAACTTCAGCGCAGGGCCCGCGGT

<table>
<tr><td>Chromosome</td><td>start</td><td>end</td><td>strand</td><td>MM</td><td>target_seq</td><td>PAM</td><td>alignment</td><td>distance</td><td>position</td><td>gene name</td><td>gene id</td></tr>
<tr><td>GU305914.1</td><td>34266</td><td>34288</td><td>-</td><td>0</td><td>ACCGCGGGCCCTGCGCTGAA</td><td>GGG</td><td>||||||||[||||||||||||]PAM</td><td>NA</td><td>Intergenic</td><td>NA</td><td>NA</td></tr>
<tr><td>T31750</td><td>CGGGCCCTGCGCTGAAGGGTCGG</td><td>1000</td><td></td><td></td><td></td><td></td><td></td><td></td><td></td><td></td><td></td></tr>
<tr><td>Oligo fwd</td><td>CACCCGGGCCCTGCGCTGAAGGGT</td><td></td><td></td><td></td><td></td><td></td><td></td><td></td><td></td><td></td><td></td></tr>
<tr><td>Oligo rev</td><td>AAACACCCTTCAGCGCAGGGCCCG</td><td></td><td></td><td></td><td></td><td></td><td></td><td></td><td></td><td></td><td></td></tr>
<tr><td>Chromosome</td><td>start</td><td>end</td><td>strand</td><td>MM</td><td>target_seq</td><td>PAM</td><td>alignment</td><td>distance</td><td>position</td><td>gene name</td><td>gene id</td></tr>
<tr><td>GU305914.1</td><td>34262</td><td>34284</td><td>-</td><td>0</td><td>CGGGCCCTGCGCTGAAGGGT</td><td>CGG</td><td>||||||||[||||||||||||]PAM</td><td>NA</td><td>Intergenic</td><td>NA</td><td>NA</td></tr>
<tr><td>T31751</td><td>GGGCCCTGCGCTGAAGGGTCGGG</td><td>1000</td><td></td><td></td><td></td><td></td><td></td><td></td><td></td><td></td><td></td></tr>
<tr><td>Oligo fwd</td><td>CACCGGGCCCTGCGCTGAAGGGTC</td><td></td><td></td><td></td><td></td><td></td><td></td><td></td><td></td><td></td><td></td></tr>
<tr><td>Oligo rev</td><td>AAACGACCCTTCAGCGCAGGGCCC</td><td></td><td></td><td></td><td></td><td></td><td></td><td></td><td></td><td></td><td></td></tr>
<tr><td>Chromosome</td><td>start</td><td>end</td><td>strand</td><td>MM</td><td>target_seq</td><td>PAM</td><td>alignment</td><td>distance</td><td>position</td><td>gene name</td><td>gene id</td></tr>
<tr><td>GU305914.1</td><td>34261</td><td>34283</td><td>-</td><td>0</td><td>GGGCCCTGCGCTGAAGGGTC</td><td>GGG</td><td>||||||||[||||||||||||]PAM</td><td>NA</td><td>Intergenic</td><td>NA</td><td>NA</td></tr>
<tr><td>T31752</td><td>CCCTGCGCTGAAGGGTCGGGTGG</td><td>1000</td><td></td><td></td><td></td><td></td><td></td><td></td><td></td><td></td><td></td></tr>
<tr><td>Oligo fwd</td><td>CACCCCCTGCGCTGAAGGGTCGGG</td><td></td><td></td><td></td><td></td><td></td><td></td><td></td><td></td><td></td><td></td></tr>
<tr><td>Oligo rev</td><td>AAACCCCGACCCTTCAGCGCAGGG</td><td></td><td></td><td></td><td></td><td></td><td></td><td></td><td></td><td></td><td></td></tr>
<tr><td>Chromosome</td><td>start</td><td>end</td><td>strand</td><td>MM</td><td>target_seq</td><td>PAM</td><td>alignment</td><td>distance</td><td>position</td><td>gene name</td><td>gene id</td></tr>
<tr><td>GU305914.1</td><td>34258</td><td>34280</td><td>-</td><td>0</td><td>CCCTGCGCTGAAGGGTCGGG</td><td>TGG</td><td>||||||||[||||||||||||]PAM</td><td>NA</td><td>Intergenic</td><td>NA</td><td>NA</td></tr>
<tr><td>T31753</td><td>CCTGCGCTGAAGGGTCGGGTGGG</td><td>1000</td><td></td><td></td><td></td><td></td><td></td><td></td><td></td><td></td><td></td></tr>
<tr><td>Oligo fwd</td><td>CACCCCTGCGCTGAAGGGTCGGGT</td><td></td><td></td><td></td><td></td><td></td><td></td><td></td><td></td><td></td><td></td></tr>
<tr><td>Oligo rev</td><td>AAACACCCGACCCTTCAGCGCAGG</td><td></td><td></td><td></td><td></td><td></td><td></td><td></td><td></td><td></td><td></td></tr>
<tr><td>Chromosome</td><td>start</td><td>end</td><td>strand</td><td>MM</td><td>target_seq</td><td>PAM</td><td>alignment</td><td>distance</td><td>position</td><td>gene name</td><td>gene id</td></tr>
<tr><td>GU305914.1</td><td>34257</td><td>34279</td><td>-</td><td>0</td><td>CCTGCGCTGAAGGGTCGGGT</td><td>GGG</td><td>||||||||[||||||||||||]PAM</td><td>NA</td><td>Intergenic</td><td>NA</td><td>NA</td></tr>
<tr><td>T31754</td><td>CGCTGAAGGGTCGGGTGGGACGG</td><td>1000</td><td></td><td></td><td></td><td></td><td></td><td></td><td></td><td></td><td></td></tr>
<tr><td>Oligo fwd</td><td>CACCCGCTGAAGGGTCGGGTGGGA</td><td></td><td></td><td></td><td></td><td></td><td></td><td></td><td></td><td></td><td></td></tr>
<tr><td>Oligo rev</td><td>AAACTCCCACCCGACCCTTCAGCG</td><td></td><td></td><td></td><td></td><td></td><td></td><td></td><td></td><td></td><td></td></tr>
<tr><td>Chromosome</td><td>start</td><td>end</td><td>strand</td><td>MM</td><td>target_seq</td><td>PAM</td><td>alignment</td><td>distance</td><td>position</td><td>gene name</td><td>gene id</td></tr>
<tr><td>GU305914.1</td><td>34253</td><td>34275</td><td>-</td><td>0</td><td>CGCTGAAGGGTCGGGTGGGA</td><td>CGG</td><td>||||||||[||||||||||||]PAM</td><td>NA</td><td>Intergenic</td><td>NA</td><td>NA</td></tr>
<tr><td>T31755</td><td>GGTCGGGTGGGACGGTGAGACGG</td><td>1000</td><td></td><td></td><td></td><td></td><td></td><td></td><td></td><td></td><td></td></tr>
<tr><td>Oligo fwd</td><td>CACCGGTCGGGTGGGACGGTGAGA</td><td></td><td></td><td></td><td></td><td></td><td></td><td></td><td></td><td></td><td></td></tr>
<tr><td>Oligo rev</td><td>AAACTCTCACCGTCCCACCCGACC</td><td></td><td></td><td></td><td></td><td></td><td></td><td></td><td></td><td></td><td></td></tr>
<tr><td>Chromosome</td><td>start</td><td>end</td><td>strand</td><td>MM</td><td>target_seq</td><td>PAM</td><td>alignment</td><td>distance</td><td>position</td><td>gene name</td><td>gene id</td></tr>
<tr><td>GU305914.1</td><td>34245</td><td>34267</td><td>-</td><td>0</td><td>GGTCGGGTGGGACGGTGAGA</td><td>CGG</td><td>||||||||[||||||||||||]PAM</td><td>NA</td><td>Intergenic</td><td>NA</td><td>NA</td></tr>
<tr><td>T31756</td><td>GAGCCCGTCCGAGTCTTCCGAGG</td><td>1000</td><td></td><td></td><td></td><td></td><td></td><td></td><td></td><td></td><td></td></tr>
<tr><td>Oligo fwd</td><td>CACCGAGCCCGTCCGAGTCTTCCG</td><td></td><td></td><td></td><td></td><td></td><td></td><td></td><td></td><td></td><td></td></tr>
<tr><td>Oligo rev</td><td>AAACCGGAAGACTCGGACGGGCTC</td><td></td><td></td><td></td><td></td><td></td><td></td><td></td><td></td><td></td><td></td></tr>
<tr><td>Chromosome</td><td>start</td><td>end</td><td>strand</td><td>MM</td><td>target_seq</td><td>PAM</td><td>alignment</td><td>distance</td><td>position</td><td>gene name</td><td>gene id</td></tr>
<tr><td>GU305914.1</td><td>34217</td><td>34239</td><td>-</td><td>0</td><td>GAGCCCGTCCGAGTCTTCCG</td><td>AGG</td><td>||||||||[||||||||||||]PAM</td><td>NA</td><td>Intergenic</td><td>NA</td><td>NA</td></tr>
<tr><td>T31757</td><td>CACGCAGTTCGTCTTTGAACTGG</td><td>1000</td><td></td><td></td><td></td><td></td><td></td><td></td><td></td><td></td><td></td></tr>
<tr><td>Oligo fwd</td><td>CACCCACGCAGTTCGTCTTTGAAC</td><td></td><td></td><td></td><td></td><td></td><td></td><td></td><td></td><td></td><td></td></tr>
<tr><td>Oligo rev</td><td>AAACGTTCAAAGACGAACTGCGTG</td><td></td><td></td><td></td><td></td><td></td><td></td><td></td><td></td><td></td><td></td></tr>
<tr><td>Chromosome</td><td>start</td><td>end</td><td>strand</td><td>MM</td><td>target_seq</td><td>PAM</td><td>alignment</td><td>distance</td><td>position</td><td>gene name</td><td>gene id</td></tr>
<tr><td>GU305914.1</td><td>34187</td><td>34209</td><td>-</td><td>0</td><td>CACGCAGTTCGTCTTTGAAC</td><td>TGG</td><td>||||||||[||||||||||||]PAM</td><td>NA</td><td>Intergenic</td><td>NA</td><td>NA</td></tr>
<tr><td>T31758</td><td>TGAACTGGTCGTCGACGAGACGG</td><td>1000</td><td></td><td></td><td></td><td></td><td></td><td></td><td></td><td></td><td></td></tr>
<tr><td>Oligo fwd</td><td>CACCTGAACTGGTCGTCGACGAGA</td><td></td><td></td><td></td><td></td><td></td><td></td><td></td><td></td><td></td><td></td></tr>
<tr><td>Oligo rev</td><td>AAACTCTCGTCGACGACCAGTTCA</td><td></td><td></td><td></td><td></td><td></td><td></td><td></td><td></td><td></td><td></td></tr>
<tr><td>Chromosome</td><td>start</td><td>end</td><td>strand</td><td>MM</td><td>target_seq</td><td>PAM</td><td>alignment</td><td>distance</td><td>position</td><td>gene name</td><td>gene id</td></tr>
<tr><td>GU305914.1</td><td>34172</td><td>34194</td><td>-</td><td>0</td><td>TGAACTGGTCGTCGACGAGA</td><td>CGG</td><td>||||||||[||||||||||||]PAM</td><td>NA</td><td>Intergenic</td><td>NA</td><td>NA</td></tr>
<tr><td>T31759</td><td>ACTGGTCGTCGACGAGACGGCGG</td><td>1000</td><td></td><td></td><td></td><td></td><td></td><td></td><td></td><td></td><td></td></tr>
<tr><td>Oligo fwd</td><td>CACCACTGGTCGTCGACGAGACGG</td><td></td><td></td><td></td><td></td><td></td><td></td><td></td><td></td><td></td><td></td></tr>
<tr><td>Oligo rev</td><td>AAACCCGTCTCGTCGACGACCAGT</td><td></td><td></td><td></td><td></td><td></td><td></td><td></td><td></td><td></td><td></td></tr>
<tr><td>Chromosome</td><td>start</td><td>end</td><td>strand</td><td>MM</td><td>target_seq</td><td>PAM</td><td>alignment</td><td>distance</td><td>position</td><td>gene name</td><td>gene id</td></tr>
<tr><td>GU305914.1</td><td>34169</td><td>34191</td><td>-</td><td>0</td><td>ACTGGTCGTCGACGAGACGG</td><td>CGG</td><td>||||||||[||||||||||||]PAM</td><td>NA</td><td>Intergenic</td><td>NA</td><td>NA</td></tr>
<tr><td>T31760</td><td>AGACGGCGGACTCGTTCTGCCGG</td><td>1000</td><td></td><td></td><td></td><td></td><td></td><td></td><td></td><td></td><td></td></tr>
<tr><td>Oligo fwd</td><td>CACCAGACGGCGGACTCGTTCTGC</td><td></td><td></td><td></td><td></td><td></td><td></td><td></td><td></td><td></td><td></td></tr>
<tr><td>Oligo rev</td><td>AAACGCAGAACGAGTCCGCCGTCT</td><td></td><td></td><td></td><td></td><td></td><td></td><td></td><td></td><td></td><td></td></tr>
<tr><td>Chromosome</td><td>start</td><td>end</td><td>strand</td><td>MM</td><td>target_seq</td><td>PAM</td><td>alignment</td><td>distance</td><td>position</td><td>gene name</td><td>gene id</td></tr>
<tr><td>GU305914.1</td><td>34155</td><td>34177</td><td>-</td><td>0</td><td>AGACGGCGGACTCGTTCTGC</td><td>CGG</td><td>||||||||[||||||||||||]PAM</td><td>NA</td><td>Intergenic</td><td>NA</td><td>NA</td></tr>
<tr><td>T31761</td><td>GCGCTTTATCGCCGATCACCTGG</td><td>1000</td><td></td><td></td><td></td><td></td><td></td><td></td><td></td><td></td><td></td></tr>
<tr><td>Oligo fwd</td><td>CACCGCGCTTTATCGCCGATCACC</td><td></td><td></td><td></td><td></td><td></td><td></td><td></td><td></td><td></td><td></td></tr>
<tr><td>Oligo rev</td><td>AAACGGTGATCGGCGATAAAGCGC</td><td></td><td></td><td></td><td></td><td></td><td></td><td></td><td></td><td></td><td></td></tr>
<tr><td>Chromosome</td><td>start</td><td>end</td><td>strand</td><td>MM</td><td>target_seq</td><td>PAM</td><td>alignment</td><td>distance</td><td>position</td><td>gene name</td><td>gene id</td></tr>
<tr><td>GU305914.1</td><td>34133</td><td>34155</td><td>-</td><td>0</td><td>GCGCTTTATCGCCGATCACC</td><td>TGG</td><td>||||||||[||||||||||||]PAM</td><td>NA</td><td>Intergenic</td><td>NA</td><td>NA</td></tr>
<tr><td>T31762</td><td>CCGATCACCTGGTCCAGCTCAGG</td><td>1000</td><td></td><td></td><td></td><td></td><td></td><td></td><td></td><td></td><td></td></tr>
<tr><td>Oligo fwd</td><td>CACCCCGATCACCTGGTCCAGCTC</td><td></td><td></td><td></td><td></td><td></td><td></td><td></td><td></td><td></td><td></td></tr>
<tr><td>Oligo rev</td><td>AAACGAGCTGGACCAGGTGATCGG</td><td></td><td></td><td></td><td></td><td></td><td></td><td></td><td></td><td></td><td></td></tr>
<tr><td>Chromosome</td><td>start</td><td>end</td><td>strand</td><td>MM</td><td>target_seq</td><td>PAM</td><td>alignment</td><td>distance</td><td>position</td><td>gene name</td><td>gene id</td></tr>
<tr><td>GU305914.1</td><td>34122</td><td>34144</td><td>-</td><td>0</td><td>CCGATCACCTGGTCCAGCTC</td><td>AGG</td><td>||||||||[||||||||||||]PAM</td><td>NA</td><td>Intergenic</td><td>NA</td><td>NA</td></tr>
<tr><td>T31763</td><td>CCAGCTCAGGAACTACGAGCAGG</td><td>1000</td><td></td><td></td><td></td><td></td><td></td><td></td><td></td><td></td><td></td></tr>
<tr><td>Oligo fwd</td><td>CACCCCAGCTCAGGAACTACGAGC</td><td></td><td></td><td></td><td></td><td></td><td></td><td></td><td></td><td></td><td></td></tr>
<tr><td>Oligo rev</td><td>AAACGCTCGTAGTTCCTGAGCTGG</td><td></td><td></td><td></td><td></td><td></td><td></td><td></td><td></td><td></td><td></td></tr>
<tr><td>Chromosome</td><td>start</td><td>end</td><td>strand</td><td>MM</td><td>target_seq</td><td>PAM</td><td>alignment</td><td>distance</td><td>position</td><td>gene name</td><td>gene id</td></tr>
<tr><td>GU305914.1</td><td>34109</td><td>34131</td><td>-</td><td>0</td><td>CCAGCTCAGGAACTACGAGC</td><td>AGG</td><td>||||||||[||||||||||||]PAM</td><td>NA</td><td>Intergenic</td><td>NA</td><td>NA</td></tr>
<tr><td>T31764</td><td>TACGAGCAGGCCTCGCGCTTCGG</td><td>1000</td><td></td><td></td><td></td><td></td><td></td><td></td><td></td><td></td><td></td></tr>
<tr><td>Oligo fwd</td><td>CACCTACGAGCAGGCCTCGCGCTT</td><td></td><td></td><td></td><td></td><td></td><td></td><td></td><td></td><td></td><td></td></tr>
</table>

<table>
<tr><td>Oligo rev</td><td>AAACAAGCGCGAGGCCTGCTCGTA</td><td></td><td></td><td></td><td></td><td></td><td></td><td></td><td></td><td></td><td></td><td></td></tr>
<tr><td>Chromosome</td><td>start</td><td></td><td>end</td><td>strand</td><td>MM</td><td>target_seq</td><td>PAM</td><td>alignment</td><td>distance</td><td>position</td><td>gene name</td><td>gene id</td></tr>
<tr><td>GU305914.1</td><td></td><td>34096</td><td>34118</td><td>-</td><td>0</td><td>TACGAGCAGGCCTCGCGCTT</td><td>CGG</td><td>||||||||[||||||||||||]PAM</td><td>NA</td><td>Intergenic</td><td>NA</td><td>NA</td></tr>
<tr><td></td><td></td><td></td><td></td><td></td><td></td><td></td><td></td><td></td><td></td><td></td><td></td><td></td></tr>
<tr><td>T31765</td><td>CTCGCGCTTCGGCTTCGTCATGG</td><td></td><td>1000</td><td></td><td></td><td></td><td></td><td></td><td></td><td></td><td></td><td></td></tr>
<tr><td>Oligo fwd</td><td>CACCCTCGCGCTTCGGCTTCGTCA</td><td></td><td></td><td></td><td></td><td></td><td></td><td></td><td></td><td></td><td></td><td></td></tr>
<tr><td>Oligo rev</td><td>AAACTGACGAAGCCGAAGCGCGAG</td><td></td><td></td><td></td><td></td><td></td><td></td><td></td><td></td><td></td><td></td><td></td></tr>
<tr><td>Chromosome</td><td>start</td><td></td><td>end</td><td>strand</td><td>MM</td><td>target_seq</td><td>PAM</td><td>alignment</td><td>distance</td><td>position</td><td>gene name</td><td>gene id</td></tr>
<tr><td>GU305914.1</td><td></td><td>34085</td><td>34107</td><td>-</td><td>0</td><td>CTCGCGCTTCGGCTTCGTCA</td><td>TGG</td><td>||||||||[||||||||||||]PAM</td><td>NA</td><td>Intergenic</td><td>NA</td><td>NA</td></tr>
<tr><td></td><td></td><td></td><td></td><td></td><td></td><td></td><td></td><td></td><td></td><td></td><td></td><td></td></tr>
<tr><td>T31766</td><td>TCGGCTTCGTCATGGACGCCTGG</td><td></td><td>1000</td><td></td><td></td><td></td><td></td><td></td><td></td><td></td><td></td><td></td></tr>
<tr><td>Oligo fwd</td><td>CACCTCGGCTTCGTCATGGACGCC</td><td></td><td></td><td></td><td></td><td></td><td></td><td></td><td></td><td></td><td></td><td></td></tr>
<tr><td>Oligo rev</td><td>AAACGGCGTCCATGACGAAGCCGA</td><td></td><td></td><td></td><td></td><td></td><td></td><td></td><td></td><td></td><td></td><td></td></tr>
<tr><td>Chromosome</td><td>start</td><td></td><td>end</td><td>strand</td><td>MM</td><td>target_seq</td><td>PAM</td><td>alignment</td><td>distance</td><td>position</td><td>gene name</td><td>gene id</td></tr>
<tr><td>GU305914.1</td><td></td><td>34077</td><td>34099</td><td>-</td><td>0</td><td>TCGGCTTCGTCATGGACGCC</td><td>TGG</td><td>||||||||[||||||||||||]PAM</td><td>NA</td><td>Intergenic</td><td>NA</td><td>NA</td></tr>
<tr><td></td><td></td><td></td><td></td><td></td><td></td><td></td><td></td><td></td><td></td><td></td><td></td><td></td></tr>
<tr><td>T31767</td><td>TCGTCATGGACGCCTGGTCCAGG</td><td></td><td>1000</td><td></td><td></td><td></td><td></td><td></td><td></td><td></td><td></td><td></td></tr>
<tr><td>Oligo fwd</td><td>CACCTCGTCATGGACGCCTGGTCC</td><td></td><td></td><td></td><td></td><td></td><td></td><td></td><td></td><td></td><td></td><td></td></tr>
<tr><td>Oligo rev</td><td>AAACGGACCAGGCGTCCATGACGA</td><td></td><td></td><td></td><td></td><td></td><td></td><td></td><td></td><td></td><td></td><td></td></tr>
<tr><td>Chromosome</td><td>start</td><td></td><td>end</td><td>strand</td><td>MM</td><td>target_seq</td><td>PAM</td><td>alignment</td><td>distance</td><td>position</td><td>gene name</td><td>gene id</td></tr>
<tr><td>GU305914.1</td><td></td><td>34071</td><td>34093</td><td>-</td><td>0</td><td>TCGTCATGGACGCCTGGTCC</td><td>AGG</td><td>||||||||[||||||||||||]PAM</td><td>NA</td><td>Intergenic</td><td>NA</td><td>NA</td></tr>
<tr><td></td><td></td><td></td><td></td><td></td><td></td><td></td><td></td><td></td><td></td><td></td><td></td><td></td></tr>
<tr><td>T31768</td><td>CGTCATGGACGCCTGGTCCAGGG</td><td></td><td>1000</td><td></td><td></td><td></td><td></td><td></td><td></td><td></td><td></td><td></td></tr>
<tr><td>Oligo fwd</td><td>CACCCGTCATGGACGCCTGGTCCA</td><td></td><td></td><td></td><td></td><td></td><td></td><td></td><td></td><td></td><td></td><td></td></tr>
<tr><td>Oligo rev</td><td>AAACTGGACCAGGCGTCCATGACG</td><td></td><td></td><td></td><td></td><td></td><td></td><td></td><td></td><td></td><td></td><td></td></tr>
<tr><td>Chromosome</td><td>start</td><td></td><td>end</td><td>strand</td><td>MM</td><td>target_seq</td><td>PAM</td><td>alignment</td><td>distance</td><td>position</td><td>gene name</td><td>gene id</td></tr>
<tr><td>GU305914.1</td><td></td><td>34070</td><td>34092</td><td>-</td><td>0</td><td>CGTCATGGACGCCTGGTCCA</td><td>GGG</td><td>||||||||[||||||||||||]PAM</td><td>NA</td><td>Intergenic</td><td>NA</td><td>NA</td></tr>
<tr><td></td><td></td><td></td><td></td><td></td><td></td><td></td><td></td><td></td><td></td><td></td><td></td><td></td></tr>
<tr><td>T31769</td><td>GGACGCCTGGTCCAGGGAGTCGG</td><td></td><td>1000</td><td></td><td></td><td></td><td></td><td></td><td></td><td></td><td></td><td></td></tr>
<tr><td>Oligo fwd</td><td>CACCGGACGCCTGGTCCAGGGAGT</td><td></td><td></td><td></td><td></td><td></td><td></td><td></td><td></td><td></td><td></td><td></td></tr>
<tr><td>Oligo rev</td><td>AAACACTCCCTGGACCAGGCGTCC</td><td></td><td></td><td></td><td></td><td></td><td></td><td></td><td></td><td></td><td></td><td></td></tr>
<tr><td>Chromosome</td><td>start</td><td></td><td>end</td><td>strand</td><td>MM</td><td>target_seq</td><td>PAM</td><td>alignment</td><td>distance</td><td>position</td><td>gene name</td><td>gene id</td></tr>
<tr><td>GU305914.1</td><td></td><td>34064</td><td>34086</td><td>-</td><td>0</td><td>GGACGCCTGGTCCAGGGAGT</td><td>CGG</td><td>||||||||[||||||||||||]PAM</td><td>NA</td><td>Intergenic</td><td>NA</td><td>NA</td></tr>
<tr><td></td><td></td><td></td><td></td><td></td><td></td><td></td><td></td><td></td><td></td><td></td><td></td><td></td></tr>
<tr><td>T31770</td><td>GACGCCTGGTCCAGGGAGTCGGG</td><td></td><td>1000</td><td></td><td></td><td></td><td></td><td></td><td></td><td></td><td></td><td></td></tr>
<tr><td>Oligo fwd</td><td>CACCGACGCCTGGTCCAGGGAGTC</td><td></td><td></td><td></td><td></td><td></td><td></td><td></td><td></td><td></td><td></td><td></td></tr>
<tr><td>Oligo rev</td><td>AAACGACTCCCTGGACCAGGCGTC</td><td></td><td></td><td></td><td></td><td></td><td></td><td></td><td></td><td></td><td></td><td></td></tr>
<tr><td>Chromosome</td><td>start</td><td></td><td>end</td><td>strand</td><td>MM</td><td>target_seq</td><td>PAM</td><td>alignment</td><td>distance</td><td>position</td><td>gene name</td><td>gene id</td></tr>
<tr><td>GU305914.1</td><td></td><td>34063</td><td>34085</td><td>-</td><td>0</td><td>GACGCCTGGTCCAGGGAGTC</td><td>GGG</td><td>||||||||[||||||||||||]PAM</td><td>NA</td><td>Intergenic</td><td>NA</td><td>NA</td></tr>
<tr><td></td><td></td><td></td><td></td><td></td><td></td><td></td><td></td><td></td><td></td><td></td><td></td><td></td></tr>
<tr><td>T31771</td><td>CTGGTCCAGGGAGTCGGGCAAGG</td><td></td><td>1000</td><td></td><td></td><td></td><td></td><td></td><td></td><td></td><td></td><td></td></tr>
<tr><td>Oligo fwd</td><td>CACCCTGGTCCAGGGAGTCGGGCA</td><td></td><td></td><td></td><td></td><td></td><td></td><td></td><td></td><td></td><td></td><td></td></tr>
<tr><td>Oligo rev</td><td>AAACTGCCCGACTCCCTGGACCAG</td><td></td><td></td><td></td><td></td><td></td><td></td><td></td><td></td><td></td><td></td><td></td></tr>
<tr><td>Chromosome</td><td>start</td><td></td><td>end</td><td>strand</td><td>MM</td><td>target_seq</td><td>PAM</td><td>alignment</td><td>distance</td><td>position</td><td>gene name</td><td>gene id</td></tr>
<tr><td>GU305914.1</td><td></td><td>34058</td><td>34080</td><td>-</td><td>0</td><td>CTGGTCCAGGGAGTCGGGCA</td><td>AGG</td><td>||||||||[||||||||||||]PAM</td><td>NA</td><td>Intergenic</td><td>NA</td><td>NA</td></tr>
<tr><td></td><td></td><td></td><td></td><td></td><td></td><td></td><td></td><td></td><td></td><td></td><td></td><td></td></tr>
<tr><td>T31772</td><td>ACAGCTACCTGAAGAACGCGCGG</td><td></td><td>1000</td><td></td><td></td><td></td><td></td><td></td><td></td><td></td><td></td><td></td></tr>
<tr><td>Oligo fwd</td><td>CACCACAGCTACCTGAAGAACGCG</td><td></td><td></td><td></td><td></td><td></td><td></td><td></td><td></td><td></td><td></td><td></td></tr>
<tr><td>Oligo rev</td><td>AAACCGCGTTCTTCAGGTAGCTGT</td><td></td><td></td><td></td><td></td><td></td><td></td><td></td><td></td><td></td><td></td><td></td></tr>
<tr><td>Chromosome</td><td>start</td><td></td><td>end</td><td>strand</td><td>MM</td><td>target_seq</td><td>PAM</td><td>alignment</td><td>distance</td><td>position</td><td>gene name</td><td>gene id</td></tr>
<tr><td>GU305914.1</td><td></td><td>33975</td><td>33997</td><td>-</td><td>0</td><td>ACAGCTACCTGAAGAACGCG</td><td>CGG</td><td>||||||||[||||||||||||]PAM</td><td>NA</td><td>Intergenic</td><td>NA</td><td>NA</td></tr>
<tr><td></td><td></td><td></td><td></td><td></td><td></td><td></td><td></td><td></td><td></td><td></td><td></td><td></td></tr>
<tr><td>T31773</td><td>GCGCGGCTCGCCACCGCCAACGG</td><td></td><td>1000</td><td></td><td></td><td></td><td></td><td></td><td></td><td></td><td></td><td></td></tr>
<tr><td>Oligo fwd</td><td>CACCGCGCGGCTCGCCACCGCCAA</td><td></td><td></td><td></td><td></td><td></td><td></td><td></td><td></td><td></td><td></td><td></td></tr>
<tr><td>Oligo rev</td><td>AAACTTGGCGGTGGCGAGCCGCGC</td><td></td><td></td><td></td><td></td><td></td><td></td><td></td><td></td><td></td><td></td><td></td></tr>
<tr><td>Chromosome</td><td>start</td><td></td><td>end</td><td>strand</td><td>MM</td><td>target_seq</td><td>PAM</td><td>alignment</td><td>distance</td><td>position</td><td>gene name</td><td>gene id</td></tr>
<tr><td>GU305914.1</td><td></td><td>33958</td><td>33980</td><td>-</td><td>0</td><td>GCGCGGCTCGCCACCGCCAA</td><td>CGG</td><td>||||||||[||||||||||||]PAM</td><td>NA</td><td>Intergenic</td><td>NA</td><td>NA</td></tr>
<tr><td></td><td></td><td></td><td></td><td></td><td></td><td></td><td></td><td></td><td></td><td></td><td></td><td></td></tr>
<tr><td>T31774</td><td>CGGCTCGCCACCGCCAACGGCGG</td><td></td><td>1000</td><td></td><td></td><td></td><td></td><td></td><td></td><td></td><td></td><td></td></tr>
<tr><td>Oligo fwd</td><td>CACCCGGCTCGCCACCGCCAACGG</td><td></td><td></td><td></td><td></td><td></td><td></td><td></td><td></td><td></td><td></td><td></td></tr>
<tr><td>Oligo rev</td><td>AAACCCGTTGGCGGTGGCGAGCCG</td><td></td><td></td><td></td><td></td><td></td><td></td><td></td><td></td><td></td><td></td><td></td></tr>
<tr><td>Chromosome</td><td>start</td><td></td><td>end</td><td>strand</td><td>MM</td><td>target_seq</td><td>PAM</td><td>alignment</td><td>distance</td><td>position</td><td>gene name</td><td>gene id</td></tr>
<tr><td>GU305914.1</td><td></td><td>33955</td><td>33977</td><td>-</td><td>0</td><td>CGGCTCGCCACCGCCAACGG</td><td>CGG</td><td>||||||||[||||||||||||]PAM</td><td>NA</td><td>Intergenic</td><td>NA</td><td>NA</td></tr>
<tr><td></td><td></td><td></td><td></td><td></td><td></td><td></td><td></td><td></td><td></td><td></td><td></td><td></td></tr>
<tr><td>T31775</td><td>GGCTCGCCACCGCCAACGGCGGG</td><td></td><td>1000</td><td></td><td></td><td></td><td></td><td></td><td></td><td></td><td></td><td></td></tr>
<tr><td>Oligo fwd</td><td>CACCGGCTCGCCACCGCCAACGGC</td><td></td><td></td><td></td><td></td><td></td><td></td><td></td><td></td><td></td><td></td><td></td></tr>
<tr><td>Oligo rev</td><td>AAACGCCGTTGGCGGTGGCGAGCC</td><td></td><td></td><td></td><td></td><td></td><td></td><td></td><td></td><td></td><td></td><td></td></tr>
<tr><td>Chromosome</td><td>start</td><td></td><td>end</td><td>strand</td><td>MM</td><td>target_seq</td><td>PAM</td><td>alignment</td><td>distance</td><td>position</td><td>gene name</td><td>gene id</td></tr>
<tr><td>GU305914.1</td><td></td><td>33954</td><td>33976</td><td>-</td><td>0</td><td>GGCTCGCCACCGCCAACGGC</td><td>GGG</td><td>||||||||[||||||||||||]PAM</td><td>NA</td><td>Intergenic</td><td>NA</td><td>NA</td></tr>
<tr><td></td><td></td><td></td><td></td><td></td><td></td><td></td><td></td><td></td><td></td><td></td><td></td><td></td></tr>
<tr><td>T31776</td><td>GCCACCGCCAACGGCGGGTACGG</td><td></td><td>1000</td><td></td><td></td><td></td><td></td><td></td><td></td><td></td><td></td><td></td></tr>
<tr><td>Oligo fwd</td><td>CACCGCCACCGCCAACGGCGGGTA</td><td></td><td></td><td></td><td></td><td></td><td></td><td></td><td></td><td></td><td></td><td></td></tr>
<tr><td>Oligo rev</td><td>AAACTACCCGCCGTTGGCGGTGGC</td><td></td><td></td><td></td><td></td><td></td><td></td><td></td><td></td><td></td><td></td><td></td></tr>
<tr><td>Chromosome</td><td>start</td><td></td><td>end</td><td>strand</td><td>MM</td><td>target_seq</td><td>PAM</td><td>alignment</td><td>distance</td><td>position</td><td>gene name</td><td>gene id</td></tr>
<tr><td>GU305914.1</td><td></td><td>33949</td><td>33971</td><td>-</td><td>0</td><td>GCCACCGCCAACGGCGGGTA</td><td>CGG</td><td>||||||||[||||||||||||]PAM</td><td>NA</td><td>Intergenic</td><td>NA</td><td>NA</td></tr>
<tr><td></td><td></td><td></td><td></td><td></td><td></td><td></td><td></td><td></td><td></td><td></td><td></td><td></td></tr>
<tr><td>T31777</td><td>AACGGCGGGTACGGCATCCGCGG</td><td></td><td>1000</td><td></td><td></td><td></td><td></td><td></td><td></td><td></td><td></td><td></td></tr>
<tr><td>Oligo fwd</td><td>CACCAACGGCGGGTACGGCATCCG</td><td></td><td></td><td></td><td></td><td></td><td></td><td></td><td></td><td></td><td></td><td></td></tr>
<tr><td>Oligo rev</td><td>AAACCGGATGCCGTACCCGCCGTT</td><td></td><td></td><td></td><td></td><td></td><td></td><td></td><td></td><td></td><td></td><td></td></tr>
<tr><td>Chromosome</td><td>start</td><td></td><td>end</td><td>strand</td><td>MM</td><td>target_seq</td><td>PAM</td><td>alignment</td><td>distance</td><td>position</td><td>gene name</td><td>gene id</td></tr>
<tr><td>GU305914.1</td><td></td><td>33940</td><td>33962</td><td>-</td><td>0</td><td>AACGGCGGGTACGGCATCCG</td><td>CGG</td><td>||||||||[||||||||||||]PAM</td><td>NA</td><td>Intergenic</td><td>NA</td><td>NA</td></tr>
<tr><td></td><td></td><td></td><td></td><td></td><td></td><td></td><td></td><td></td><td></td><td></td><td></td><td></td></tr>
<tr><td>T31778</td><td>TCCGCGGTCTCACCCTCGCCTGG</td><td></td><td>1000</td><td></td><td></td><td></td><td></td><td></td><td></td><td></td><td></td><td></td></tr>
<tr><td>Oligo fwd</td><td>CACCTCCGCGGTCTCACCCTCGCC</td><td></td><td></td><td></td><td></td><td></td><td></td><td></td><td></td><td></td><td></td><td></td></tr>
<tr><td>Oligo rev</td><td>AAACGGCGAGGGTGAGACCGCGGA</td><td></td><td></td><td></td><td></td><td></td><td></td><td></td><td></td><td></td><td></td><td></td></tr>
<tr><td>Chromosome</td><td>start</td><td></td><td>end</td><td>strand</td><td>MM</td><td>target_seq</td><td>PAM</td><td>alignment</td><td>distance</td><td>position</td><td>gene name</td><td>gene id</td></tr>
<tr><td>GU305914.1</td><td></td><td>33924</td><td>33946</td><td>-</td><td>0</td><td>TCCGCGGTCTCACCCTCGCC</td><td>TGG</td><td>||||||||[||||||||||||]PAM</td><td>NA</td><td>Intergenic</td><td>NA</td><td>NA</td></tr>
<tr><td></td><td></td><td></td><td></td><td></td><td></td><td></td><td></td><td></td><td></td><td></td><td></td><td></td></tr>
<tr><td>T31779</td><td>CCGCGGTCTCACCCTCGCCTGGG</td><td></td><td>1000</td><td></td><td></td><td></td><td></td><td></td><td></td><td></td><td></td><td></td></tr>
</table>

| | | | | | | | | | | | |
|---|---|---|---|---|---|---|---|---|---|---|---|
| Oligo fwd | CACCCCGCGGTCTCACCCTCGCCT | | | | | | | | | | |
| Oligo rev | AAACAGGCGAGGGTGAGACCGCGG | | | | | | | | | | |
| Chromosome | start | end | strand | MM | target_seq | PAM | alignment | distance | position | gene name | gene id |
| GU305914.1 | 33923 | 33945 | - | 0 | CCGCGGTCTCACCCTCGCCT | GGG | \|\|\|\|\|\|\|\|[\|\|\|\|\|\|\|\|\|\|\|\|]PAM | NA | Intergenic | NA | NA |
| GU305914.1 | 132395 | 132417 | - | 4 | CCGCAGTTCCACCCTCACCT | GGG | \|\|\|\|-\|\|-[-\|\|\|\|\|\|\|-\|\|\|]PAM | NA | Intergenic | NA | NA |
| T31780 | TCGCCTGGGACTCCGCCATCTGG | 1000 | | | | | | | | | |
| Oligo fwd | CACCTCGCCTGGGACTCCGCCATC | | | | | | | | | | |
| Oligo rev | AAACGATGGCGGAGTCCCAGGCGA | | | | | | | | | | |
| Chromosome | start | end | strand | MM | target_seq | PAM | alignment | distance | position | gene name | gene id |
| GU305914.1 | 33909 | 33931 | - | 0 | TCGCCTGGGACTCCGCCATC | TGG | \|\|\|\|\|\|\|\|[\|\|\|\|\|\|\|\|\|\|\|\|]PAM | NA | Intergenic | NA | NA |
| T31781 | CGCCTGGGACTCCGCCATCTGGG | 1000 | | | | | | | | | |
| Oligo fwd | CACCCGCCTGGGACTCCGCCATCT | | | | | | | | | | |
| Oligo rev | AAACAGATGGCGGAGTCCCAGGCG | | | | | | | | | | |
| Chromosome | start | end | strand | MM | target_seq | PAM | alignment | distance | position | gene name | gene id |
| GU305914.1 | 33908 | 33930 | - | 0 | CGCCTGGGACTCCGCCATCT | GGG | \|\|\|\|\|\|\|\|[\|\|\|\|\|\|\|\|\|\|\|\|]PAM | NA | Intergenic | NA | NA |
| T31782 | GCCTGGGACTCCGCCATCTGGGG | 1000 | | | | | | | | | |
| Oligo fwd | CACCGCCTGGGACTCCGCCATCTG | | | | | | | | | | |
| Oligo rev | AAACCAGATGGCGGAGTCCCAGGC | | | | | | | | | | |
| Chromosome | start | end | strand | MM | target_seq | PAM | alignment | distance | position | gene name | gene id |
| GU305914.1 | 33907 | 33929 | - | 0 | GCCTGGGACTCCGCCATCTG | GGG | \|\|\|\|\|\|\|\|[\|\|\|\|\|\|\|\|\|\|\|\|]PAM | NA | Intergenic | NA | NA |
| T31783 | CCTGGGACTCCGCCATCTGGGGG | 1000 | | | | | | | | | |
| Oligo fwd | CACCCCTGGGACTCCGCCATCTGG | | | | | | | | | | |
| Oligo rev | AAACCCAGATGGCGGAGTCCCAGG | | | | | | | | | | |
| Chromosome | start | end | strand | MM | target_seq | PAM | alignment | distance | position | gene name | gene id |
| GU305914.1 | 33906 | 33928 | - | 0 | CCTGGGACTCCGCCATCTGG | GGG | \|\|\|\|\|\|\|\|[\|\|\|\|\|\|\|\|\|\|\|\|]PAM | NA | Intergenic | NA | NA |
| T31784 | CCGCCATCTGGGGGCTCCTCAGG | 1000 | | | | | | | | | |
| Oligo fwd | CACCCCGCCATCTGGGGGCTCCTC | | | | | | | | | | |
| Oligo rev | AAACGAGGAGCCCCCAGATGGCGG | | | | | | | | | | |
| Chromosome | start | end | strand | MM | target_seq | PAM | alignment | distance | position | gene name | gene id |
| GU305914.1 | 33897 | 33919 | - | 0 | CCGCCATCTGGGGGCTCCTC | AGG | \|\|\|\|\|\|\|\|[\|\|\|\|\|\|\|\|\|\|\|\|]PAM | NA | Intergenic | NA | NA |
| T31785 | CGCCATCTGGGGGCTCCTCAGGG | 1000 | | | | | | | | | |
| Oligo fwd | CACCCGCCATCTGGGGGCTCCTCA | | | | | | | | | | |
| Oligo rev | AAACTGAGGAGCCCCCAGATGGCG | | | | | | | | | | |
| Chromosome | start | end | strand | MM | target_seq | PAM | alignment | distance | position | gene name | gene id |
| GU305914.1 | 33896 | 33918 | - | 0 | CGCCATCTGGGGGCTCCTCA | GGG | \|\|\|\|\|\|\|\|[\|\|\|\|\|\|\|\|\|\|\|\|]PAM | NA | Intergenic | NA | NA |
| T31786 | TTCTCCTTCCCCTCGTGCGACGG | 1000 | | | | | | | | | |
| Oligo fwd | CACCTTCTCCTTCCCCTCGTGCGA | | | | | | | | | | |
| Oligo rev | AAACTCGCACGAGGGGAAGGAGAA | | | | | | | | | | |
| Chromosome | start | end | strand | MM | target_seq | PAM | alignment | distance | position | gene name | gene id |
| GU305914.1 | 33844 | 33866 | - | 0 | TTCTCCTTCCCCTCGTGCGA | CGG | \|\|\|\|\|\|\|\|[\|\|\|\|\|\|\|\|\|\|\|\|]PAM | NA | Intergenic | NA | NA |
| T31787 | TCTCCTTCCCCTCGTGCGACGGG | 1000 | | | | | | | | | |
| Oligo fwd | CACCTCTCCTTCCCCTCGTGCGAC | | | | | | | | | | |
| Oligo rev | AAACGTCGCACGAGGGGAAGGAGA | | | | | | | | | | |
| Chromosome | start | end | strand | MM | target_seq | PAM | alignment | distance | position | gene name | gene id |
| GU305914.1 | 33843 | 33865 | - | 0 | TCTCCTTCCCCTCGTGCGAC | GGG | \|\|\|\|\|\|\|\|[\|\|\|\|\|\|\|\|\|\|\|\|]PAM | NA | Intergenic | NA | NA |
| T31788 | AGATCCTGTTCGATCTCGAATGG | 1000 | | | | | | | | | |
| Oligo fwd | CACCAGATCCTGTTCGATCTCGAA | | | | | | | | | | |
| Oligo rev | AAACTTCGAGATCGAACAGGATCT | | | | | | | | | | |
| Chromosome | start | end | strand | MM | target_seq | PAM | alignment | distance | position | gene name | gene id |
| GU305914.1 | 33813 | 33835 | - | 0 | AGATCCTGTTCGATCTCGAA | TGG | \|\|\|\|\|\|\|\|[\|\|\|\|\|\|\|\|\|\|\|\|]PAM | NA | Intergenic | NA | NA |
| T31789 | TCCTGTTCGATCTCGAATGGAGG | 1000 | | | | | | | | | |
| Oligo fwd | CACCTCCTGTTCGATCTCGAATGG | | | | | | | | | | |
| Oligo rev | AAACCCATTCGAGATCGAACAGGA | | | | | | | | | | |
| Chromosome | start | end | strand | MM | target_seq | PAM | alignment | distance | position | gene name | gene id |
| GU305914.1 | 33810 | 33832 | - | 0 | TCCTGTTCGATCTCGAATGG | AGG | \|\|\|\|\|\|\|\|[\|\|\|\|\|\|\|\|\|\|\|\|]PAM | NA | Intergenic | NA | NA |
| T31790 | CGATCTCGAATGGAGGCCCGAGG | 1000 | | | | | | | | | |
| Oligo fwd | CACCCGATCTCGAATGGAGGCCCG | | | | | | | | | | |
| Oligo rev | AAACCGGGCCTCCATTCGAGATCG | | | | | | | | | | |
| Chromosome | start | end | strand | MM | target_seq | PAM | alignment | distance | position | gene name | gene id |
| GU305914.1 | 33803 | 33825 | - | 0 | CGATCTCGAATGGAGGCCCG | AGG | \|\|\|\|\|\|\|\|[\|\|\|\|\|\|\|\|\|\|\|\|]PAM | NA | Intergenic | NA | NA |
| T31791 | CGAGGCCTCCGACTCCTGCAAGG | 1000 | | | | | | | | | |
| Oligo fwd | CACCCGAGGCCTCCGACTCCTGCA | | | | | | | | | | |
| Oligo rev | AAACTGCAGGAGTCGGAGGCCTCG | | | | | | | | | | |
| Chromosome | start | end | strand | MM | target_seq | PAM | alignment | distance | position | gene name | gene id |
| GU305914.1 | 33785 | 33807 | - | 0 | CGAGGCCTCCGACTCCTGCA | AGG | \|\|\|\|\|\|\|\|[\|\|\|\|\|\|\|\|\|\|\|\|]PAM | NA | Intergenic | NA | NA |
| T31792 | CTCCGACTCCTGCAAGGCGCTGG | 1000 | | | | | | | | | |
| Oligo fwd | CACCCTCCGACTCCTGCAAGGCGC | | | | | | | | | | |
| Oligo rev | AAACGCGCCTTGCAGGAGTCGGAG | | | | | | | | | | |
| Chromosome | start | end | strand | MM | target_seq | PAM | alignment | distance | position | gene name | gene id |
| GU305914.1 | 33779 | 33801 | - | 0 | CTCCGACTCCTGCAAGGCGC | TGG | \|\|\|\|\|\|\|\|[\|\|\|\|\|\|\|\|\|\|\|\|]PAM | NA | Intergenic | NA | NA |
| GU305914.1 | 35544 | 35566 | - | 4 | CTCCGTCGCCTCCAAGACGC | CGG | \|\|\|\|\|-\|-[\|\|\|-\|\|\|\|-\|\|\|]PAM | NA | Intergenic | NA | NA |
| T31793 | TCCGACTCCTGCAAGGCGCTGGG | 1000 | | | | | | | | | |
| Oligo fwd | CACCTCCGACTCCTGCAAGGCGCT | | | | | | | | | | |
| Oligo rev | AAACAGCGCCTTGCAGGAGTCGGA | | | | | | | | | | |
| Chromosome | start | end | strand | MM | target_seq | PAM | alignment | distance | position | gene name | gene id |

<table>
<tr><td>GU305914.1</td><td>33778</td><td>33800</td><td>-</td><td>0</td><td>TCCGACTCCTGCAAGGCGCT</td><td>GGG</td><td>||||||||[||||||||||||]PAM</td><td>NA</td><td>Intergenic</td><td>NA</td><td>NA</td></tr>
<tr><td></td><td></td><td></td><td></td><td></td><td></td><td></td><td></td><td></td><td></td><td></td><td></td></tr>
<tr><td>T31794</td><td>TACGCCATCTCGCTGCTGTACGG</td><td>1000</td><td></td><td></td><td></td><td></td><td></td><td></td><td></td><td></td><td></td></tr>
<tr><td>Oligo fwd</td><td>CACCTACGCCATCTCGCTGCTGTA</td><td></td><td></td><td></td><td></td><td></td><td></td><td></td><td></td><td></td><td></td></tr>
<tr><td>Oligo rev</td><td>AAACTACAGCAGCGAGATGGCGTA</td><td></td><td></td><td></td><td></td><td></td><td></td><td></td><td></td><td></td><td></td></tr>
<tr><td>Chromosome</td><td>start</td><td>end</td><td>strand</td><td>MM</td><td>target_seq</td><td>PAM</td><td>alignment</td><td>distance</td><td>position</td><td>gene name</td><td>gene id</td></tr>
<tr><td>GU305914.1</td><td>33733</td><td>33755</td><td>-</td><td>0</td><td>TACGCCATCTCGCTGCTGTA</td><td>CGG</td><td>||||||||[||||||||||||]PAM</td><td>NA</td><td>Intergenic</td><td>NA</td><td>NA</td></tr>
<tr><td>GU305914.1</td><td>10700</td><td>10722</td><td>-</td><td>3</td><td>TGAGCCATCTCGGTGCTGTA</td><td>CGG</td><td>|--|||||[||||-|||||||]PAM</td><td>NA</td><td>Intergenic</td><td>NA</td><td>NA</td></tr>
<tr><td></td><td></td><td></td><td></td><td></td><td></td><td></td><td></td><td></td><td></td><td></td><td></td></tr>
<tr><td>T31795</td><td>GCTGTACGGCTACGATCGCTCGG</td><td>1000</td><td></td><td></td><td></td><td></td><td></td><td></td><td></td><td></td><td></td></tr>
<tr><td>Oligo fwd</td><td>CACCGCTGTACGGCTACGATCGCT</td><td></td><td></td><td></td><td></td><td></td><td></td><td></td><td></td><td></td><td></td></tr>
<tr><td>Oligo rev</td><td>AAACAGCGATCGTAGCCGTACAGC</td><td></td><td></td><td></td><td></td><td></td><td></td><td></td><td></td><td></td><td></td></tr>
<tr><td>Chromosome</td><td>start</td><td>end</td><td>strand</td><td>MM</td><td>target_seq</td><td>PAM</td><td>alignment</td><td>distance</td><td>position</td><td>gene name</td><td>gene id</td></tr>
<tr><td>GU305914.1</td><td>33719</td><td>33741</td><td>-</td><td>0</td><td>GCTGTACGGCTACGATCGCT</td><td>CGG</td><td>||||||||[||||||||||||]PAM</td><td>NA</td><td>Intergenic</td><td>NA</td><td>NA</td></tr>
<tr><td></td><td></td><td></td><td></td><td></td><td></td><td></td><td></td><td></td><td></td><td></td><td></td></tr>
<tr><td>T31796</td><td>CCATGTCCCGCTATCTCCGCCGG</td><td>1000</td><td></td><td></td><td></td><td></td><td></td><td></td><td></td><td></td><td></td></tr>
<tr><td>Oligo fwd</td><td>CACCCCATGTCCCGCTATCTCCGC</td><td></td><td></td><td></td><td></td><td></td><td></td><td></td><td></td><td></td><td></td></tr>
<tr><td>Oligo rev</td><td>AAACGCGGAGATAGCGGGACATGG</td><td></td><td></td><td></td><td></td><td></td><td></td><td></td><td></td><td></td><td></td></tr>
<tr><td>Chromosome</td><td>start</td><td>end</td><td>strand</td><td>MM</td><td>target_seq</td><td>PAM</td><td>alignment</td><td>distance</td><td>position</td><td>gene name</td><td>gene id</td></tr>
<tr><td>GU305914.1</td><td>33693</td><td>33715</td><td>-</td><td>0</td><td>CCATGTCCCGCTATCTCCGC</td><td>CGG</td><td>||||||||[||||||||||||]PAM</td><td>NA</td><td>Intergenic</td><td>NA</td><td>NA</td></tr>
<tr><td></td><td></td><td></td><td></td><td></td><td></td><td></td><td></td><td></td><td></td><td></td><td></td></tr>
<tr><td>T31797</td><td>AGCTCATCGACCTGCACGACAGG</td><td>1000</td><td></td><td></td><td></td><td></td><td></td><td></td><td></td><td></td><td></td></tr>
<tr><td>Oligo fwd</td><td>CACCAGCTCATCGACCTGCACGAC</td><td></td><td></td><td></td><td></td><td></td><td></td><td></td><td></td><td></td><td></td></tr>
<tr><td>Oligo rev</td><td>AAACGTCGTGCAGGTCGATGAGCT</td><td></td><td></td><td></td><td></td><td></td><td></td><td></td><td></td><td></td><td></td></tr>
<tr><td>Chromosome</td><td>start</td><td>end</td><td>strand</td><td>MM</td><td>target_seq</td><td>PAM</td><td>alignment</td><td>distance</td><td>position</td><td>gene name</td><td>gene id</td></tr>
<tr><td>GU305914.1</td><td>33660</td><td>33682</td><td>-</td><td>0</td><td>AGCTCATCGACCTGCACGAC</td><td>AGG</td><td>||||||||[||||||||||||]PAM</td><td>NA</td><td>Intergenic</td><td>NA</td><td>NA</td></tr>
<tr><td></td><td></td><td></td><td></td><td></td><td></td><td></td><td></td><td></td><td></td><td></td><td></td></tr>
<tr><td>T31798</td><td>CATGTCCAGCCTCCGCATCCCGG</td><td>1000</td><td></td><td></td><td></td><td></td><td></td><td></td><td></td><td></td><td></td></tr>
<tr><td>Oligo fwd</td><td>CACCCATGTCCAGCCTCCGCATCC</td><td></td><td></td><td></td><td></td><td></td><td></td><td></td><td></td><td></td><td></td></tr>
<tr><td>Oligo rev</td><td>AAACGGATGCGGAGGCTGGACATG</td><td></td><td></td><td></td><td></td><td></td><td></td><td></td><td></td><td></td><td></td></tr>
<tr><td>Chromosome</td><td>start</td><td>end</td><td>strand</td><td>MM</td><td>target_seq</td><td>PAM</td><td>alignment</td><td>distance</td><td>position</td><td>gene name</td><td>gene id</td></tr>
<tr><td>GU305914.1</td><td>33620</td><td>33642</td><td>-</td><td>0</td><td>CATGTCCAGCCTCCGCATCC</td><td>CGG</td><td>||||||||[||||||||||||]PAM</td><td>NA</td><td>Intergenic</td><td>NA</td><td>NA</td></tr>
<tr><td></td><td></td><td></td><td></td><td></td><td></td><td></td><td></td><td></td><td></td><td></td><td></td></tr>
<tr><td>T31799</td><td>ATGTCCAGCCTCCGCATCCCGGG</td><td>1000</td><td></td><td></td><td></td><td></td><td></td><td></td><td></td><td></td><td></td></tr>
<tr><td>Oligo fwd</td><td>CACCATGTCCAGCCTCCGCATCCC</td><td></td><td></td><td></td><td></td><td></td><td></td><td></td><td></td><td></td><td></td></tr>
<tr><td>Oligo rev</td><td>AAACGGGATGCGGAGGCTGGACAT</td><td></td><td></td><td></td><td></td><td></td><td></td><td></td><td></td><td></td><td></td></tr>
<tr><td>Chromosome</td><td>start</td><td>end</td><td>strand</td><td>MM</td><td>target_seq</td><td>PAM</td><td>alignment</td><td>distance</td><td>position</td><td>gene name</td><td>gene id</td></tr>
<tr><td>GU305914.1</td><td>33619</td><td>33641</td><td>-</td><td>0</td><td>ATGTCCAGCCTCCGCATCCC</td><td>GGG</td><td>||||||||[||||||||||||]PAM</td><td>NA</td><td>Intergenic</td><td>NA</td><td>NA</td></tr>
<tr><td></td><td></td><td></td><td></td><td></td><td></td><td></td><td></td><td></td><td></td><td></td><td></td></tr>
<tr><td>T31800</td><td>GAGCGCATCTGCCAGCAACTCGG</td><td>1000</td><td></td><td></td><td></td><td></td><td></td><td></td><td></td><td></td><td></td></tr>
<tr><td>Oligo fwd</td><td>CACCGAGCGCATCTGCCAGCAACT</td><td></td><td></td><td></td><td></td><td></td><td></td><td></td><td></td><td></td><td></td></tr>
<tr><td>Oligo rev</td><td>AAACAGTTGCTGGCAGATGCGCTC</td><td></td><td></td><td></td><td></td><td></td><td></td><td></td><td></td><td></td><td></td></tr>
<tr><td>Chromosome</td><td>start</td><td>end</td><td>strand</td><td>MM</td><td>target_seq</td><td>PAM</td><td>alignment</td><td>distance</td><td>position</td><td>gene name</td><td>gene id</td></tr>
<tr><td>GU305914.1</td><td>33562</td><td>33584</td><td>-</td><td>0</td><td>GAGCGCATCTGCCAGCAACT</td><td>CGG</td><td>||||||||[||||||||||||]PAM</td><td>NA</td><td>Intergenic</td><td>NA</td><td>NA</td></tr>
<tr><td></td><td></td><td></td><td></td><td></td><td></td><td></td><td></td><td></td><td></td><td></td><td></td></tr>
<tr><td>T31801</td><td>CGAGTACGTCGAGATGACCATGG</td><td>1000</td><td></td><td></td><td></td><td></td><td></td><td></td><td></td><td></td><td></td></tr>
<tr><td>Oligo fwd</td><td>CACCCGAGTACGTCGAGATGACCA</td><td></td><td></td><td></td><td></td><td></td><td></td><td></td><td></td><td></td><td></td></tr>
<tr><td>Oligo rev</td><td>AAACTGGTCATCTCGACGTACTCG</td><td></td><td></td><td></td><td></td><td></td><td></td><td></td><td></td><td></td><td></td></tr>
<tr><td>Chromosome</td><td>start</td><td>end</td><td>strand</td><td>MM</td><td>target_seq</td><td>PAM</td><td>alignment</td><td>distance</td><td>position</td><td>gene name</td><td>gene id</td></tr>
<tr><td>GU305914.1</td><td>33497</td><td>33519</td><td>-</td><td>0</td><td>CGAGTACGTCGAGATGACCA</td><td>TGG</td><td>||||||||[||||||||||||]PAM</td><td>NA</td><td>Intergenic</td><td>NA</td><td>NA</td></tr>
<tr><td></td><td></td><td></td><td></td><td></td><td></td><td></td><td></td><td></td><td></td><td></td><td></td></tr>
<tr><td>T31802</td><td>GTACGTCGAGATGACCATGGAGG</td><td>1000</td><td></td><td></td><td></td><td></td><td></td><td></td><td></td><td></td><td></td></tr>
<tr><td>Oligo fwd</td><td>CACCGTACGTCGAGATGACCATGG</td><td></td><td></td><td></td><td></td><td></td><td></td><td></td><td></td><td></td><td></td></tr>
<tr><td>Oligo rev</td><td>AAACCCATGGTCATCTCGACGTAC</td><td></td><td></td><td></td><td></td><td></td><td></td><td></td><td></td><td></td><td></td></tr>
<tr><td>Chromosome</td><td>start</td><td>end</td><td>strand</td><td>MM</td><td>target_seq</td><td>PAM</td><td>alignment</td><td>distance</td><td>position</td><td>gene name</td><td>gene id</td></tr>
<tr><td>GU305914.1</td><td>33494</td><td>33516</td><td>-</td><td>0</td><td>GTACGTCGAGATGACCATGG</td><td>AGG</td><td>||||||||[||||||||||||]PAM</td><td>NA</td><td>Intergenic</td><td>NA</td><td>NA</td></tr>
<tr><td></td><td></td><td></td><td></td><td></td><td></td><td></td><td></td><td></td><td></td><td></td><td></td></tr>
<tr><td>T31803</td><td>TCTGTTACTGTCGCGAGTGTCGG</td><td>1000</td><td></td><td></td><td></td><td></td><td></td><td></td><td></td><td></td><td></td></tr>
<tr><td>Oligo fwd</td><td>CACCTCTGTTACTGTCGCGAGTGT</td><td></td><td></td><td></td><td></td><td></td><td></td><td></td><td></td><td></td><td></td></tr>
<tr><td>Oligo rev</td><td>AAACACACTCGCGACAGTAACAGA</td><td></td><td></td><td></td><td></td><td></td><td></td><td></td><td></td><td></td><td></td></tr>
<tr><td>Chromosome</td><td>start</td><td>end</td><td>strand</td><td>MM</td><td>target_seq</td><td>PAM</td><td>alignment</td><td>distance</td><td>position</td><td>gene name</td><td>gene id</td></tr>
<tr><td>GU305914.1</td><td>33456</td><td>33478</td><td>-</td><td>0</td><td>TCTGTTACTGTCGCGAGTGT</td><td>CGG</td><td>||||||||[||||||||||||]PAM</td><td>NA</td><td>Intergenic</td><td>NA</td><td>NA</td></tr>
<tr><td></td><td></td><td></td><td></td><td></td><td></td><td></td><td></td><td></td><td></td><td></td><td></td></tr>
<tr><td>T31804</td><td>GTCGCGAGTGTCGGATGCGCCGG</td><td>1000</td><td></td><td></td><td></td><td></td><td></td><td></td><td></td><td></td><td></td></tr>
<tr><td>Oligo fwd</td><td>CACCGTCGCGAGTGTCGGATGCGC</td><td></td><td></td><td></td><td></td><td></td><td></td><td></td><td></td><td></td><td></td></tr>
<tr><td>Oligo rev</td><td>AAACGCGCATCCGACACTCGCGAC</td><td></td><td></td><td></td><td></td><td></td><td></td><td></td><td></td><td></td><td></td></tr>
<tr><td>Chromosome</td><td>start</td><td>end</td><td>strand</td><td>MM</td><td>target_seq</td><td>PAM</td><td>alignment</td><td>distance</td><td>position</td><td>gene name</td><td>gene id</td></tr>
<tr><td>GU305914.1</td><td>33447</td><td>33469</td><td>-</td><td>0</td><td>GTCGCGAGTGTCGGATGCGC</td><td>CGG</td><td>||||||||[||||||||||||]PAM</td><td>NA</td><td>Intergenic</td><td>NA</td><td>NA</td></tr>
<tr><td></td><td></td><td></td><td></td><td></td><td></td><td></td><td></td><td></td><td></td><td></td><td></td></tr>
<tr><td>T31805</td><td>TCGCGAGTGTCGGATGCGCCGGG</td><td>1000</td><td></td><td></td><td></td><td></td><td></td><td></td><td></td><td></td><td></td></tr>
<tr><td>Oligo fwd</td><td>CACCTCGCGAGTGTCGGATGCGCC</td><td></td><td></td><td></td><td></td><td></td><td></td><td></td><td></td><td></td><td></td></tr>
<tr><td>Oligo rev</td><td>AAACGGCGCATCCGACACTCGCGA</td><td></td><td></td><td></td><td></td><td></td><td></td><td></td><td></td><td></td><td></td></tr>
<tr><td>Chromosome</td><td>start</td><td>end</td><td>strand</td><td>MM</td><td>target_seq</td><td>PAM</td><td>alignment</td><td>distance</td><td>position</td><td>gene name</td><td>gene id</td></tr>
<tr><td>GU305914.1</td><td>33446</td><td>33468</td><td>-</td><td>0</td><td>TCGCGAGTGTCGGATGCGCC</td><td>GGG</td><td>||||||||[||||||||||||]PAM</td><td>NA</td><td>Intergenic</td><td>NA</td><td>NA</td></tr>
<tr><td></td><td></td><td></td><td></td><td></td><td></td><td></td><td></td><td></td><td></td><td></td><td></td></tr>
<tr><td>T31806</td><td>CGCGAGTGTCGGATGCGCCGGGG</td><td>1000</td><td></td><td></td><td></td><td></td><td></td><td></td><td></td><td></td><td></td></tr>
<tr><td>Oligo fwd</td><td>CACCCGCGAGTGTCGGATGCGCCG</td><td></td><td></td><td></td><td></td><td></td><td></td><td></td><td></td><td></td><td></td></tr>
<tr><td>Oligo rev</td><td>AAACCGGCGCATCCGACACTCGCG</td><td></td><td></td><td></td><td></td><td></td><td></td><td></td><td></td><td></td><td></td></tr>
<tr><td>Chromosome</td><td>start</td><td>end</td><td>strand</td><td>MM</td><td>target_seq</td><td>PAM</td><td>alignment</td><td>distance</td><td>position</td><td>gene name</td><td>gene id</td></tr>
<tr><td>GU305914.1</td><td>33445</td><td>33467</td><td>-</td><td>0</td><td>CGCGAGTGTCGGATGCGCCG</td><td>GGG</td><td>||||||||[||||||||||||]PAM</td><td>NA</td><td>Intergenic</td><td>NA</td><td>NA</td></tr>
<tr><td></td><td></td><td></td><td></td><td></td><td></td><td></td><td></td><td></td><td></td><td></td><td></td></tr>
<tr><td>T31807</td><td>GCCGGGGCGAGCGCCTCGAGAGG</td><td>1000</td><td></td><td></td><td></td><td></td><td></td><td></td><td></td><td></td><td></td></tr>
<tr><td>Oligo fwd</td><td>CACCGCCGGGGCGAGCGCCTCGAG</td><td></td><td></td><td></td><td></td><td></td><td></td><td></td><td></td><td></td><td></td></tr>
<tr><td>Oligo rev</td><td>AAACCTCGAGGCGCTCGCCCCGGC</td><td></td><td></td><td></td><td></td><td></td><td></td><td></td><td></td><td></td><td></td></tr>
<tr><td>Chromosome</td><td>start</td><td>end</td><td>strand</td><td>MM</td><td>target_seq</td><td>PAM</td><td>alignment</td><td>distance</td><td>position</td><td>gene name</td><td>gene id</td></tr>
<tr><td>GU305914.1</td><td>33429</td><td>33451</td><td>-</td><td>0</td><td>GCCGGGGCGAGCGCCTCGAG</td><td>AGG</td><td>||||||||[||||||||||||]PAM</td><td>NA</td><td>Intergenic</td><td>NA</td><td>NA</td></tr>
<tr><td>GU305914.1</td><td>165717</td><td>165739</td><td>-</td><td>4</td><td>GCCGCGGGGAGCGCGTCGAC</td><td>TGG</td><td>||||-||-[||||||-||||-]PAM</td><td>NA</td><td>Intergenic</td><td>NA</td><td>NA</td></tr>
<tr><td></td><td></td><td></td><td></td><td></td><td></td><td></td><td></td><td></td><td></td><td></td><td></td></tr>
<tr><td>T31808</td><td>AGGCTCCGCGCGCAGAAGATCGG</td><td>1000</td><td></td><td></td><td></td><td></td><td></td><td></td><td></td><td></td><td></td></tr>
</table>

Oligo fwd CACCAGGCTCCGCGCGCAGAAGAT
Oligo rev AAACATCTTCTGCGCGCGGAGCCT
Chromosome start end strand MM target_seq PAM alignment distance position gene name gene id
GU305914.1 33409 33431 - 0 AGGCTCCGCGCGCAGAAGAT CGG ||||||||[||||||||||||]PAM NA Intergenic NA NA

T31809 AGAAGATCGGCACGAAGTACCGG 1000
Oligo fwd CACCAGAAGATCGGCACGAAGTAC
Oligo rev AAACGTACTTCGTGCCGATCTTCT
Chromosome start end strand MM target_seq PAM alignment distance position gene name gene id
GU305914.1 33396 33418 - 0 AGAAGATCGGCACGAAGTAC CGG ||||||||[||||||||||||]PAM NA Intergenic NA NA

T31810 TCGGCACGAAGTACCGGCGTCGG 1000
Oligo fwd CACCTCGGCACGAAGTACCGGCGT
Oligo rev AAACACGCCGGTACTTCGTGCCGA
Chromosome start end strand MM target_seq PAM alignment distance position gene name gene id
GU305914.1 33390 33412 - 0 TCGGCACGAAGTACCGGCGT CGG ||||||||[||||||||||||]PAM NA Intergenic NA NA

T31811 TACCGGCGTCGGAACCCCGACGG 1000
Oligo fwd CACCTACCGGCGTCGGAACCCCGA
Oligo rev AAACTCGGGGTTCCGACGCCGGTA
Chromosome start end strand MM target_seq PAM alignment distance position gene name gene id
GU305914.1 33379 33401 - 0 TACCGGCGTCGGAACCCCGA CGG ||||||||[||||||||||||]PAM NA Intergenic NA NA

T31812 ACCGGCGTCGGAACCCCGACGGG 1000
Oligo fwd CACCACCGGCGTCGGAACCCCGAC
Oligo rev AAACGTCGGGGTTCCGACGCCGGT
Chromosome start end strand MM target_seq PAM alignment distance position gene name gene id
GU305914.1 33378 33400 - 0 ACCGGCGTCGGAACCCCGAC GGG ||||||||[||||||||||||]PAM NA Intergenic NA NA

T31813 CGACGGGATCCGCCTCTTCGAGG 1000
Oligo fwd CACCCGACGGGATCCGCCTCTTCG
Oligo rev AAACCGAAGAGGCGGATCCCGTCG
Chromosome start end strand MM target_seq PAM alignment distance position gene name gene id
GU305914.1 33362 33384 - 0 CGACGGGATCCGCCTCTTCG AGG ||||||||[||||||||||||]PAM NA Intergenic NA NA

T31814 GTCGCCGCCTACCCCAACATCGG 1000
Oligo fwd CACCGTCGCCGCCTACCCCAACAT
Oligo rev AAACATGTTGGGGTAGGCGGCGAC
Chromosome start end strand MM target_seq PAM alignment distance position gene name gene id
GU305914.1 33325 33347 - 0 GTCGCCGCCTACCCCAACAT CGG ||||||||[||||||||||||]PAM NA Intergenic NA NA

T31815 GAACACCAGACCCTCGTCTGCGG 1000
Oligo fwd CACCGAACACCAGACCCTCGTCTG
Oligo rev AAACCAGACGAGGGTCTGGTGTTC
Chromosome start end strand MM target_seq PAM alignment distance position gene name gene id
GU305914.1 33265 33287 - 0 GAACACCAGACCCTCGTCTG CGG ||||||||[||||||||||||]PAM NA Intergenic NA NA

T31816 CTCATGTACGACGTGCACTTCGG 1000
Oligo fwd CACCCTCATGTACGACGTGCACTT
Oligo rev AAACAAGTGCACGTCGTACATGAG
Chromosome start end strand MM target_seq PAM alignment distance position gene name gene id
GU305914.1 33226 33248 - 0 CTCATGTACGACGTGCACTT CGG ||||||||[||||||||||||]PAM NA Intergenic NA NA

T31817 AGTCCGCTCGCGCGCGTCAACGG 1000
Oligo fwd CACCAGTCCGCTCGCGCGCGTCAA
Oligo rev AAACTTGACGCGCGCGAGCGGACT
Chromosome start end strand MM target_seq PAM alignment distance position gene name gene id
GU305914.1 33196 33218 - 0 AGTCCGCTCGCGCGCGTCAA CGG ||||||||[||||||||||||]PAM NA Intergenic NA NA

T31818 CGGCATCCCCCTGCCCTTCACGG 1000
Oligo fwd CACCCGGCATCCCCCTGCCCTTCA
Oligo rev AAACTGAAGGGCAGGGGGATGCCG
Chromosome start end strand MM target_seq PAM alignment distance position gene name gene id
GU305914.1 33176 33198 - 0 CGGCATCCCCCTGCCCTTCA CGG ||||||||[||||||||||||]PAM NA Intergenic NA NA

T31819 CTTCACGGCCGTCGACTGCATGG 1000
Oligo fwd CACCCTTCACGGCCGTCGACTGCA
Oligo rev AAACTGCAGTCGACGGCCGTGAAG
Chromosome start end strand MM target_seq PAM alignment distance position gene name gene id
GU305914.1 33161 33183 - 0 CTTCACGGCCGTCGACTGCA TGG ||||||||[||||||||||||]PAM NA Intergenic NA NA

T31820 CGACTGCATGGACATCGATCTGG 1000
Oligo fwd CACCCGACTGCATGGACATCGATC
Oligo rev AAACGATCGATGTCCATGCAGTCG
Chromosome start end strand MM target_seq PAM alignment distance position gene name gene id
GU305914.1 33149 33171 - 0 CGACTGCATGGACATCGATC TGG ||||||||[||||||||||||]PAM NA Intergenic NA NA

T31821 GACTGCATGGACATCGATCTGGG 1000
Oligo fwd CACCGACTGCATGGACATCGATCT
Oligo rev AAACAGATCGATGTCCATGCAGTC
Chromosome start end strand MM target_seq PAM alignment distance position gene name gene id
GU305914.1 33148 33170 - 0 GACTGCATGGACATCGATCT GGG ||||||||[||||||||||||]PAM NA Intergenic NA NA

T31822 TCTGGGCTACGCGCTCTACATGG 1000
Oligo fwd CACCTCTGGGCTACGCGCTCTACA
Oligo rev AAACTGTAGAGCGCGTAGCCCAGA
Chromosome start end strand MM target_seq PAM alignment distance position gene name gene id
GU305914.1 33131 33153 - 0 TCTGGGCTACGCGCTCTACA TGG ||||||||[||||||||||||]PAM NA Intergenic NA NA

| | | | | | | | | | | | |
|---|---|---|---|---|---|---|---|---|---|---|---|
| T31823 | ACGTAGTGCGTAGCGAGCAGAGG | 1000 | | | | | | | | | |
| Oligo fwd | CACCACGTAGTGCGTAGCGAGCAG | | | | | | | | | | |
| Oligo rev | AAACCTGCTCGCTACGCACTACGT | | | | | | | | | | |
| Chromosome | start | end | strand | MM | target_seq | PAM | alignment | distance | position | gene name | gene id |
| GU305914.1 | 33063 | 33085 | - | 0 | ACGTAGTGCGTAGCGAGCAG | AGG | \|\|\|\|\|\|\|\|[\|\|\|\|\|\|\|\|\|\|\|\|]PAM | NA | Intergenic | NA | NA |
| T31824 | CGTAGTGCGTAGCGAGCAGAGGG | 1000 | | | | | | | | | |
| Oligo fwd | CACCCGTAGTGCGTAGCGAGCAGA | | | | | | | | | | |
| Oligo rev | AAACTCTGCTCGCTACGCACTACG | | | | | | | | | | |
| Chromosome | start | end | strand | MM | target_seq | PAM | alignment | distance | position | gene name | gene id |
| GU305914.1 | 33062 | 33084 | - | 0 | CGTAGTGCGTAGCGAGCAGA | GGG | \|\|\|\|\|\|\|\|[\|\|\|\|\|\|\|\|\|\|\|\|]PAM | NA | Intergenic | NA | NA |
| T31825 | GAGTACGTCGACCTCGTCACCGG | 1000 | | | | | | | | | |
| Oligo fwd | CACCGAGTACGTCGACCTCGTCAC | | | | | | | | | | |
| Oligo rev | AAACGTGACGAGGTCGACGTACTC | | | | | | | | | | |
| Chromosome | start | end | strand | MM | target_seq | PAM | alignment | distance | position | gene name | gene id |
| GU305914.1 | 33040 | 33062 | - | 0 | GAGTACGTCGACCTCGTCAC | CGG | \|\|\|\|\|\|\|\|[\|\|\|\|\|\|\|\|\|\|\|\|]PAM | NA | Intergenic | NA | NA |
| T31826 | CGTCGACCTCGTCACCGGCCTGG | 1000 | | | | | | | | | |
| Oligo fwd | CACCCGTCGACCTCGTCACCGGCC | | | | | | | | | | |
| Oligo rev | AAACGGCCGGTGACGAGGTCGACG | | | | | | | | | | |
| Chromosome | start | end | strand | MM | target_seq | PAM | alignment | distance | position | gene name | gene id |
| GU305914.1 | 33035 | 33057 | - | 0 | CGTCGACCTCGTCACCGGCC | TGG | \|\|\|\|\|\|\|\|[\|\|\|\|\|\|\|\|\|\|\|\|]PAM | NA | Intergenic | NA | NA |
| T31827 | CGTCACCGGCCTGGTCCGCGAGG | 1000 | | | | | | | | | |
| Oligo fwd | CACCCGTCACCGGCCTGGTCCGCG | | | | | | | | | | |
| Oligo rev | AAACCGCGGACCAGGCCGGTGACG | | | | | | | | | | |
| Chromosome | start | end | strand | MM | target_seq | PAM | alignment | distance | position | gene name | gene id |
| GU305914.1 | 33026 | 33048 | - | 0 | CGTCACCGGCCTGGTCCGCG | AGG | \|\|\|\|\|\|\|\|[\|\|\|\|\|\|\|\|\|\|\|\|]PAM | NA | Intergenic | NA | NA |
| T31828 | CGTCGACGCCCGCATCCGCGCGG | 1000 | | | | | | | | | |
| Oligo fwd | CACCCGTCGACGCCCGCATCCGCG | | | | | | | | | | |
| Oligo rev | AAACCGCGGATGCGGGCGTCGACG | | | | | | | | | | |
| Chromosome | start | end | strand | MM | target_seq | PAM | alignment | distance | position | gene name | gene id |
| GU305914.1 | 32981 | 33003 | - | 0 | CGTCGACGCCCGCATCCGCG | CGG | \|\|\|\|\|\|\|\|[\|\|\|\|\|\|\|\|\|\|\|\|]PAM | NA | Intergenic | NA | NA |
| T31829 | GTCGACGCCCGCATCCGCGCGGG | 1000 | | | | | | | | | |
| Oligo fwd | CACCGTCGACGCCCGCATCCGCGC | | | | | | | | | | |
| Oligo rev | AAACGCGCGGATGCGGGCGTCGAC | | | | | | | | | | |
| Chromosome | start | end | strand | MM | target_seq | PAM | alignment | distance | position | gene name | gene id |
| GU305914.1 | 32980 | 33002 | - | 0 | GTCGACGCCCGCATCCGCGC | GGG | \|\|\|\|\|\|\|\|[\|\|\|\|\|\|\|\|\|\|\|\|]PAM | NA | Intergenic | NA | NA |
| T31830 | GCATCCGCGCGGGAGCCGACCGG | 1000 | | | | | | | | | |
| Oligo fwd | CACCGCATCCGCGCGGGAGCCGAC | | | | | | | | | | |
| Oligo rev | AAACGTCGGCTCCCGCGCGGATGC | | | | | | | | | | |
| Chromosome | start | end | strand | MM | target_seq | PAM | alignment | distance | position | gene name | gene id |
| GU305914.1 | 32970 | 32992 | - | 0 | GCATCCGCGCGGGAGCCGAC | CGG | \|\|\|\|\|\|\|\|[\|\|\|\|\|\|\|\|\|\|\|\|]PAM | NA | Intergenic | NA | NA |
| T31831 | CGTCGCCTCCGACCCCCGACAGG | 1000 | | | | | | | | | |
| Oligo fwd | CACCCGTCGCCTCCGACCCCCGAC | | | | | | | | | | |
| Oligo rev | AAACGTCGGGGGTCGGAGGCGACG | | | | | | | | | | |
| Chromosome | start | end | strand | MM | target_seq | PAM | alignment | distance | position | gene name | gene id |
| GU305914.1 | 32939 | 32961 | - | 0 | CGTCGCCTCCGACCCCCGAC | AGG | \|\|\|\|\|\|\|\|[\|\|\|\|\|\|\|\|\|\|\|\|]PAM | NA | Intergenic | NA | NA |
| T31832 | GCCTCCGACCCCCGACAGGACGG | 1000 | | | | | | | | | |
| Oligo fwd | CACCGCCTCCGACCCCCGACAGGA | | | | | | | | | | |
| Oligo rev | AAACTCCTGTCGGGGGTCGGAGGC | | | | | | | | | | |
| Chromosome | start | end | strand | MM | target_seq | PAM | alignment | distance | position | gene name | gene id |
| GU305914.1 | 32935 | 32957 | - | 0 | GCCTCCGACCCCCGACAGGA | CGG | \|\|\|\|\|\|\|\|[\|\|\|\|\|\|\|\|\|\|\|\|]PAM | NA | Intergenic | NA | NA |
| T31833 | CCCCCGACAGGACGGCATCTCGG | 1000 | | | | | | | | | |
| Oligo fwd | CACCCCCCCGACAGGACGGCATCT | | | | | | | | | | |
| Oligo rev | AAACAGATGCCGTCCTGTCGGGGG | | | | | | | | | | |
| Chromosome | start | end | strand | MM | target_seq | PAM | alignment | distance | position | gene name | gene id |
| GU305914.1 | 32927 | 32949 | - | 0 | CCCCCGACAGGACGGCATCT | CGG | \|\|\|\|\|\|\|\|[\|\|\|\|\|\|\|\|\|\|\|\|]PAM | NA | Intergenic | NA | NA |
| T31834 | ACAGGACGGCATCTCGGACGAGG | 1000 | | | | | | | | | |
| Oligo fwd | CACCACAGGACGGCATCTCGGACG | | | | | | | | | | |
| Oligo rev | AAACCGTCCGAGATGCCGTCCTGT | | | | | | | | | | |
| Chromosome | start | end | strand | MM | target_seq | PAM | alignment | distance | position | gene name | gene id |
| GU305914.1 | 32921 | 32943 | - | 0 | ACAGGACGGCATCTCGGACG | AGG | \|\|\|\|\|\|\|\|[\|\|\|\|\|\|\|\|\|\|\|\|]PAM | NA | Intergenic | NA | NA |
| T31835 | CCTGTCCATCGCCAACGCGCTGG | 1000 | | | | | | | | | |
| Oligo fwd | CACCCCTGTCCATCGCCAACGCGC | | | | | | | | | | |
| Oligo rev | AAACGCGCGTTGGCGATGGACAGG | | | | | | | | | | |
| Chromosome | start | end | strand | MM | target_seq | PAM | alignment | distance | position | gene name | gene id |
| GU305914.1 | 32891 | 32913 | - | 0 | CCTGTCCATCGCCAACGCGC | TGG | \|\|\|\|\|\|\|\|[\|\|\|\|\|\|\|\|\|\|\|\|]PAM | NA | Intergenic | NA | NA |
| GU305914.1 | 76441 | 76463 | + | 4 | CTTGTCGATCTCCATCGCGC | CGG | \|-\|\|\|\|-\|[\|\|-\|\|\|-\|\|\|\|\|]PAM | NA | Intergenic | NA | NA |
| T31836 | CCATCGCCAACGCGCTGGACCGG | 1000 | | | | | | | | | |
| Oligo fwd | CACCCCATCGCCAACGCGCTGGAC | | | | | | | | | | |
| Oligo rev | AAACGTCCAGCGCGTTGGCGATGG | | | | | | | | | | |
| Chromosome | start | end | strand | MM | target_seq | PAM | alignment | distance | position | gene name | gene id |
| GU305914.1 | 32886 | 32908 | - | 0 | CCATCGCCAACGCGCTGGAC | CGG | \|\|\|\|\|\|\|\|[\|\|\|\|\|\|\|\|\|\|\|\|]PAM | NA | Intergenic | NA | NA |
| T31837 | GCTGGACCGGCTCACCTTCTCGG | 1000 | | | | | | | | | |
| Oligo fwd | CACCGCTGGACCGGCTCACCTTCT | | | | | | | | | | |
| Oligo rev | AAACAGAAGGTGAGCCGGTCCAGC | | | | | | | | | | |
| Chromosome | start | end | strand | MM | target_seq | PAM | alignment | distance | position | gene name | gene id |

| Chromosome | start | end | strand | MM | target_seq | PAM | alignment | distance | position | gene name | gene id |
|---|---|---|---|---|---|---|---|---|---|---|---|
| GU305914.1 | 32873 | 32895 | - | 0 | GCTGGACCGGCTCACCTTCT | CGG | ||||||||[||||||||||||]PAM | NA | Intergenic | NA | NA |

T31838 GGCTCACCTTCTCGGACGAGCGG 1000
Oligo fwd CACCGGCTCACCTTCTCGGACGAG
Oligo rev AAACCTCGTCCGAGAAGGTGAGCC

| Chromosome | start | end | strand | MM | target_seq | PAM | alignment | distance | position | gene name | gene id |
|---|---|---|---|---|---|---|---|---|---|---|---|
| GU305914.1 | 32865 | 32887 | - | 0 | GGCTCACCTTCTCGGACGAG | CGG | ||||||||[||||||||||||]PAM | NA | Intergenic | NA | NA |

T31839 TTCTCGGACGAGCGGCCCGACGG 1000
Oligo fwd CACCTTCTCGGACGAGCGGCCCGA
Oligo rev AAACTCGGGCCGCTCGTCCGAGAA

| Chromosome | start | end | strand | MM | target_seq | PAM | alignment | distance | position | gene name | gene id |
|---|---|---|---|---|---|---|---|---|---|---|---|
| GU305914.1 | 32857 | 32879 | - | 0 | TTCTCGGACGAGCGGCCCGA | CGG | ||||||||[||||||||||||]PAM | NA | Intergenic | NA | NA |

T31840 CGAGCGGCCCGACGGCTACGAGG 1000
Oligo fwd CACCCGAGCGGCCCGACGGCTACG
Oligo rev AAACCGTAGCCGTCGGGCCGCTCG

| Chromosome | start | end | strand | MM | target_seq | PAM | alignment | distance | position | gene name | gene id |
|---|---|---|---|---|---|---|---|---|---|---|---|
| GU305914.1 | 32849 | 32871 | - | 0 | CGAGCGGCCCGACGGCTACG | AGG | ||||||||[||||||||||||]PAM | NA | Intergenic | NA | NA |

T31841 CTACGAGGACGACGACAACGAGG 1000
Oligo fwd CACCCTACGAGGACGACGACAACG
Oligo rev AAACCGTTGTCGTCGTCCTCGTAG

| Chromosome | start | end | strand | MM | target_seq | PAM | alignment | distance | position | gene name | gene id |
|---|---|---|---|---|---|---|---|---|---|---|---|
| GU305914.1 | 32834 | 32856 | - | 0 | CTACGAGGACGACGACAACG | AGG | ||||||||[||||||||||||]PAM | NA | Intergenic | NA | NA |
| GU305914.1 | 114978 | 115000 | + | 4 | CGACGATGACGATGACGACG | AGG | |-||||-|[||||-|||-|||]PAM | NA | Intergenic | NA | NA |
| GU305914.1 | 196398 | 196420 | - | 4 | ACAGGAGGACGACGACAACA | GGG | --|-||||[|||||||||||-]PAM | NA | Intergenic | NA | NA |

T31842 CGAGCAGCGCTACGAGCGCATGG 1000
Oligo fwd CACCCGAGCAGCGCTACGAGCGCA
Oligo rev AAACTGCGCTCGTAGCGCTGCTCG

| Chromosome | start | end | strand | MM | target_seq | PAM | alignment | distance | position | gene name | gene id |
|---|---|---|---|---|---|---|---|---|---|---|---|
| GU305914.1 | 32792 | 32814 | - | 0 | CGAGCAGCGCTACGAGCGCA | TGG | ||||||||[||||||||||||]PAM | NA | Intergenic | NA | NA |

T31843 GAGCAGCGCTACGAGCGCATGGG 1000
Oligo fwd CACCGAGCAGCGCTACGAGCGCAT
Oligo rev AAACATGCGCTCGTAGCGCTGCTC

| Chromosome | start | end | strand | MM | target_seq | PAM | alignment | distance | position | gene name | gene id |
|---|---|---|---|---|---|---|---|---|---|---|---|
| GU305914.1 | 32791 | 32813 | - | 0 | GAGCAGCGCTACGAGCGCAT | GGG | ||||||||[||||||||||||]PAM | NA | Intergenic | NA | NA |

T31844 CAGCGCTACGAGCGCATGGGCGG 1000
Oligo fwd CACCCAGCGCTACGAGCGCATGGG
Oligo rev AAACCCCATGCGCTCGTAGCGCTG

| Chromosome | start | end | strand | MM | target_seq | PAM | alignment | distance | position | gene name | gene id |
|---|---|---|---|---|---|---|---|---|---|---|---|
| GU305914.1 | 32788 | 32810 | - | 0 | CAGCGCTACGAGCGCATGGG | CGG | ||||||||[||||||||||||]PAM | NA | Intergenic | NA | NA |

T31845 GCGCGACGCCGCCAAGTTCATGG 1000
Oligo fwd CACCGCGCGACGCCGCCAAGTTCA
Oligo rev AAACTGAACTTGGCGGCGTCGCGC

| Chromosome | start | end | strand | MM | target_seq | PAM | alignment | distance | position | gene name | gene id |
|---|---|---|---|---|---|---|---|---|---|---|---|
| GU305914.1 | 32759 | 32781 | - | 0 | GCGCGACGCCGCCAAGTTCA | TGG | ||||||||[||||||||||||]PAM | NA | Intergenic | NA | NA |

T31846 CGCCAAGTTCATGGACCTCTTGG 1000
Oligo fwd CACCCGCCAAGTTCATGGACCTCT
Oligo rev AAACAGAGGTCCATGAACTTGGCG

| Chromosome | start | end | strand | MM | target_seq | PAM | alignment | distance | position | gene name | gene id |
|---|---|---|---|---|---|---|---|---|---|---|---|
| GU305914.1 | 32750 | 32772 | - | 0 | CGCCAAGTTCATGGACCTCT | TGG | ||||||||[||||||||||||]PAM | NA | Intergenic | NA | NA |

T31847 GGATGCGTTCCGCGTCCCGAAGG 1000
Oligo fwd CACCGGATGCGTTCCGCGTCCCGA
Oligo rev AAACTCGGGACGCGGAACGCATCC

| Chromosome | start | end | strand | MM | target_seq | PAM | alignment | distance | position | gene name | gene id |
|---|---|---|---|---|---|---|---|---|---|---|---|
| GU305914.1 | 32729 | 32751 | - | 0 | GGATGCGTTCCGCGTCCCGA | AGG | ||||||||[||||||||||||]PAM | NA | Intergenic | NA | NA |

T31848 GCGCGAGAACCGCGCCCGCGAGG 1000
Oligo fwd CACCGCGCGAGAACCGCGCCCGCG
Oligo rev AAACCGCGGGCGCGGTTCTCGCGC

| Chromosome | start | end | strand | MM | target_seq | PAM | alignment | distance | position | gene name | gene id |
|---|---|---|---|---|---|---|---|---|---|---|---|
| GU305914.1 | 32702 | 32724 | - | 0 | GCGCGAGAACCGCGCCCGCG | AGG | ||||||||[||||||||||||]PAM | NA | Intergenic | NA | NA |

T31849 TGCACACCTTCTTCATCAAGAGG 1000
Oligo fwd CACCTGCACACCTTCTTCATCAAG
Oligo rev AAACCTTGATGAAGAAGGTGTGCA

| Chromosome | start | end | strand | MM | target_seq | PAM | alignment | distance | position | gene name | gene id |
|---|---|---|---|---|---|---|---|---|---|---|---|
| GU305914.1 | 32670 | 32692 | - | 0 | TGCACACCTTCTTCATCAAG | AGG | ||||||||[||||||||||||]PAM | NA | Intergenic | NA | NA |

T31850 GATGTACGACGAGCGCCCGCTGG 1000
Oligo fwd CACCGATGTACGACGAGCGCCCGC
Oligo rev AAACGCGGGCGCTCGTCGTACATC

| Chromosome | start | end | strand | MM | target_seq | PAM | alignment | distance | position | gene name | gene id |
|---|---|---|---|---|---|---|---|---|---|---|---|
| GU305914.1 | 32648 | 32670 | - | 0 | GATGTACGACGAGCGCCCGC | TGG | ||||||||[||||||||||||]PAM | NA | Intergenic | NA | NA |

T31851 TACGACGAGCGCCCGCTGGCCGG 1000
Oligo fwd CACCTACGACGAGCGCCCGCTGGC
Oligo rev AAACGCCAGCGGGCGCTCGTCGTA

| Chromosome | start | end | strand | MM | target_seq | PAM | alignment | distance | position | gene name | gene id |
|---|---|---|---|---|---|---|---|---|---|---|---|
| GU305914.1 | 32644 | 32666 | - | 0 | TACGACGAGCGCCCGCTGGC | CGG | ||||||||[||||||||||||]PAM | NA | Intergenic | NA | NA |

T31852 CGCCCGCTGGCCGGCTTCTACGG 1000

<table>
<tr><td>Oligo fwd</td><td>CACCCGCCCGCTGGCCGGCTTCTA</td><td></td><td></td><td></td><td></td><td></td><td></td><td></td><td></td><td></td><td></td></tr>
<tr><td>Oligo rev</td><td>AAACTAGAAGCCGGCCAGCGGGCG</td><td></td><td></td><td></td><td></td><td></td><td></td><td></td><td></td><td></td><td></td></tr>
<tr><td>Chromosome</td><td>start</td><td>end</td><td>strand</td><td>MM</td><td>target_seq</td><td>PAM</td><td>alignment</td><td>distance</td><td>position</td><td>gene name</td><td>gene id</td></tr>
<tr><td>GU305914.1</td><td>32635</td><td>32657</td><td>-</td><td>0</td><td>CGCCCGCTGGCCGGCTTCTA</td><td>CGG</td><td>||||||||[||||||||||||]PAM</td><td>NA</td><td>Intergenic</td><td>NA</td><td>NA</td></tr>
<tr><td></td><td></td><td></td><td></td><td></td><td></td><td></td><td></td><td></td><td></td><td></td><td></td></tr>
<tr><td>T31853</td><td>TGGCCGGCTTCTACGGCGACAGG</td><td>1000</td><td></td><td></td><td></td><td></td><td></td><td></td><td></td><td></td><td></td></tr>
<tr><td>Oligo fwd</td><td>CACCTGGCCGGCTTCTACGGCGAC</td><td></td><td></td><td></td><td></td><td></td><td></td><td></td><td></td><td></td><td></td></tr>
<tr><td>Oligo rev</td><td>AAACGTCGCCGTAGAAGCCGGCCA</td><td></td><td></td><td></td><td></td><td></td><td></td><td></td><td></td><td></td><td></td></tr>
<tr><td>Chromosome</td><td>start</td><td>end</td><td>strand</td><td>MM</td><td>target_seq</td><td>PAM</td><td>alignment</td><td>distance</td><td>position</td><td>gene name</td><td>gene id</td></tr>
<tr><td>GU305914.1</td><td>32628</td><td>32650</td><td>-</td><td>0</td><td>TGGCCGGCTTCTACGGCGAC</td><td>AGG</td><td>||||||||[||||||||||||]PAM</td><td>NA</td><td>Intergenic</td><td>NA</td><td>NA</td></tr>
<tr><td></td><td></td><td></td><td></td><td></td><td></td><td></td><td></td><td></td><td></td><td></td><td></td></tr>
<tr><td>T31854</td><td>ATCCCCTACTACATGTTTGTCGG</td><td>1000</td><td></td><td></td><td></td><td></td><td></td><td></td><td></td><td></td><td></td></tr>
<tr><td>Oligo fwd</td><td>CACCATCCCCTACTACATGTTTGT</td><td></td><td></td><td></td><td></td><td></td><td></td><td></td><td></td><td></td><td></td></tr>
<tr><td>Oligo rev</td><td>AAACACAAACATGTAGTAGGGGAT</td><td></td><td></td><td></td><td></td><td></td><td></td><td></td><td></td><td></td><td></td></tr>
<tr><td>Chromosome</td><td>start</td><td>end</td><td>strand</td><td>MM</td><td>target_seq</td><td>PAM</td><td>alignment</td><td>distance</td><td>position</td><td>gene name</td><td>gene id</td></tr>
<tr><td>GU305914.1</td><td>32602</td><td>32624</td><td>-</td><td>0</td><td>ATCCCCTACTACATGTTTGT</td><td>CGG</td><td>||||||||[||||||||||||]PAM</td><td>NA</td><td>Intergenic</td><td>NA</td><td>NA</td></tr>
<tr><td></td><td></td><td></td><td></td><td></td><td></td><td></td><td></td><td></td><td></td><td></td><td></td></tr>
<tr><td>T31855</td><td>CCCTACTACATGTTTGTCGGCGG</td><td>1000</td><td></td><td></td><td></td><td></td><td></td><td></td><td></td><td></td><td></td></tr>
<tr><td>Oligo fwd</td><td>CACCCCCTACTACATGTTTGTCGG</td><td></td><td></td><td></td><td></td><td></td><td></td><td></td><td></td><td></td><td></td></tr>
<tr><td>Oligo rev</td><td>AAACCCGACAAACATGTAGTAGGG</td><td></td><td></td><td></td><td></td><td></td><td></td><td></td><td></td><td></td><td></td></tr>
<tr><td>Chromosome</td><td>start</td><td>end</td><td>strand</td><td>MM</td><td>target_seq</td><td>PAM</td><td>alignment</td><td>distance</td><td>position</td><td>gene name</td><td>gene id</td></tr>
<tr><td>GU305914.1</td><td>32599</td><td>32621</td><td>-</td><td>0</td><td>CCCTACTACATGTTTGTCGG</td><td>CGG</td><td>||||||||[||||||||||||]PAM</td><td>NA</td><td>Intergenic</td><td>NA</td><td>NA</td></tr>
<tr><td></td><td></td><td></td><td></td><td></td><td></td><td></td><td></td><td></td><td></td><td></td><td></td></tr>
<tr><td>T31856</td><td>TGTTTGTCGGCGGATACCGCCGG</td><td>1000</td><td></td><td></td><td></td><td></td><td></td><td></td><td></td><td></td><td></td></tr>
<tr><td>Oligo fwd</td><td>CACCTGTTTGTCGGCGGATACCGC</td><td></td><td></td><td></td><td></td><td></td><td></td><td></td><td></td><td></td><td></td></tr>
<tr><td>Oligo rev</td><td>AAACGCGGTATCCGCCGACAAACA</td><td></td><td></td><td></td><td></td><td></td><td></td><td></td><td></td><td></td><td></td></tr>
<tr><td>Chromosome</td><td>start</td><td>end</td><td>strand</td><td>MM</td><td>target_seq</td><td>PAM</td><td>alignment</td><td>distance</td><td>position</td><td>gene name</td><td>gene id</td></tr>
<tr><td>GU305914.1</td><td>32589</td><td>32611</td><td>-</td><td>0</td><td>TGTTTGTCGGCGGATACCGC</td><td>CGG</td><td>||||||||[||||||||||||]PAM</td><td>NA</td><td>Intergenic</td><td>NA</td><td>NA</td></tr>
<tr><td></td><td></td><td></td><td></td><td></td><td></td><td></td><td></td><td></td><td></td><td></td><td></td></tr>
<tr><td>T31857</td><td>GGCGGATACCGCCGGCGCGACGG</td><td>1000</td><td></td><td></td><td></td><td></td><td></td><td></td><td></td><td></td><td></td></tr>
<tr><td>Oligo fwd</td><td>CACCGGCGGATACCGCCGGCGCGA</td><td></td><td></td><td></td><td></td><td></td><td></td><td></td><td></td><td></td><td></td></tr>
<tr><td>Oligo rev</td><td>AAACTCGCGCCGGCGGTATCCGCC</td><td></td><td></td><td></td><td></td><td></td><td></td><td></td><td></td><td></td><td></td></tr>
<tr><td>Chromosome</td><td>start</td><td>end</td><td>strand</td><td>MM</td><td>target_seq</td><td>PAM</td><td>alignment</td><td>distance</td><td>position</td><td>gene name</td><td>gene id</td></tr>
<tr><td>GU305914.1</td><td>32581</td><td>32603</td><td>-</td><td>0</td><td>GGCGGATACCGCCGGCGCGA</td><td>CGG</td><td>||||||||[||||||||||||]PAM</td><td>NA</td><td>Intergenic</td><td>NA</td><td>NA</td></tr>
<tr><td></td><td></td><td></td><td></td><td></td><td></td><td></td><td></td><td></td><td></td><td></td><td></td></tr>
<tr><td>T31858</td><td>GCGGATACCGCCGGCGCGACGGG</td><td>1000</td><td></td><td></td><td></td><td></td><td></td><td></td><td></td><td></td><td></td></tr>
<tr><td>Oligo fwd</td><td>CACCGCGGATACCGCCGGCGCGAC</td><td></td><td></td><td></td><td></td><td></td><td></td><td></td><td></td><td></td><td></td></tr>
<tr><td>Oligo rev</td><td>AAACGTCGCGCCGGCGGTATCCGC</td><td></td><td></td><td></td><td></td><td></td><td></td><td></td><td></td><td></td><td></td></tr>
<tr><td>Chromosome</td><td>start</td><td>end</td><td>strand</td><td>MM</td><td>target_seq</td><td>PAM</td><td>alignment</td><td>distance</td><td>position</td><td>gene name</td><td>gene id</td></tr>
<tr><td>GU305914.1</td><td>32580</td><td>32602</td><td>-</td><td>0</td><td>GCGGATACCGCCGGCGCGAC</td><td>GGG</td><td>||||||||[||||||||||||]PAM</td><td>NA</td><td>Intergenic</td><td>NA</td><td>NA</td></tr>
<tr><td></td><td></td><td></td><td></td><td></td><td></td><td></td><td></td><td></td><td></td><td></td><td></td></tr>
<tr><td>T31859</td><td>CGGATACCGCCGGCGCGACGGGG</td><td>1000</td><td></td><td></td><td></td><td></td><td></td><td></td><td></td><td></td><td></td></tr>
<tr><td>Oligo fwd</td><td>CACCCGGATACCGCCGGCGCGACG</td><td></td><td></td><td></td><td></td><td></td><td></td><td></td><td></td><td></td><td></td></tr>
<tr><td>Oligo rev</td><td>AAACCGTCGCGCCGGCGGTATCCG</td><td></td><td></td><td></td><td></td><td></td><td></td><td></td><td></td><td></td><td></td></tr>
<tr><td>Chromosome</td><td>start</td><td>end</td><td>strand</td><td>MM</td><td>target_seq</td><td>PAM</td><td>alignment</td><td>distance</td><td>position</td><td>gene name</td><td>gene id</td></tr>
<tr><td>GU305914.1</td><td>32579</td><td>32601</td><td>-</td><td>0</td><td>CGGATACCGCCGGCGCGACG</td><td>GGG</td><td>||||||||[||||||||||||]PAM</td><td>NA</td><td>Intergenic</td><td>NA</td><td>NA</td></tr>
<tr><td></td><td></td><td></td><td></td><td></td><td></td><td></td><td></td><td></td><td></td><td></td><td></td></tr>
<tr><td>T31860</td><td>GGCGCGACGGGGCCGTCATCAGG</td><td>1000</td><td></td><td></td><td></td><td></td><td></td><td></td><td></td><td></td><td></td></tr>
<tr><td>Oligo fwd</td><td>CACCGGCGCGACGGGGCCGTCATC</td><td></td><td></td><td></td><td></td><td></td><td></td><td></td><td></td><td></td><td></td></tr>
<tr><td>Oligo rev</td><td>AAACGATGACGGCCCCGTCGCGCC</td><td></td><td></td><td></td><td></td><td></td><td></td><td></td><td></td><td></td><td></td></tr>
<tr><td>Chromosome</td><td>start</td><td>end</td><td>strand</td><td>MM</td><td>target_seq</td><td>PAM</td><td>alignment</td><td>distance</td><td>position</td><td>gene name</td><td>gene id</td></tr>
<tr><td>GU305914.1</td><td>32568</td><td>32590</td><td>-</td><td>0</td><td>GGCGCGACGGGGCCGTCATC</td><td>AGG</td><td>||||||||[||||||||||||]PAM</td><td>NA</td><td>Intergenic</td><td>NA</td><td>NA</td></tr>
<tr><td></td><td></td><td></td><td></td><td></td><td></td><td></td><td></td><td></td><td></td><td></td><td></td></tr>
<tr><td>T31861</td><td>CGACGGGGCCGTCATCAGGATGG</td><td>1000</td><td></td><td></td><td></td><td></td><td></td><td></td><td></td><td></td><td></td></tr>
<tr><td>Oligo fwd</td><td>CACCCGACGGGGCCGTCATCAGGA</td><td></td><td></td><td></td><td></td><td></td><td></td><td></td><td></td><td></td><td></td></tr>
<tr><td>Oligo rev</td><td>AAACTCCTGATGACGGCCCCGTCG</td><td></td><td></td><td></td><td></td><td></td><td></td><td></td><td></td><td></td><td></td></tr>
<tr><td>Chromosome</td><td>start</td><td>end</td><td>strand</td><td>MM</td><td>target_seq</td><td>PAM</td><td>alignment</td><td>distance</td><td>position</td><td>gene name</td><td>gene id</td></tr>
<tr><td>GU305914.1</td><td>32564</td><td>32586</td><td>-</td><td>0</td><td>CGACGGGGCCGTCATCAGGA</td><td>TGG</td><td>||||||||[||||||||||||]PAM</td><td>NA</td><td>Intergenic</td><td>NA</td><td>NA</td></tr>
<tr><td></td><td></td><td></td><td></td><td></td><td></td><td></td><td></td><td></td><td></td><td></td><td></td></tr>
<tr><td>T31862</td><td>GACGGGGCCGTCATCAGGATGGG</td><td>1000</td><td></td><td></td><td></td><td></td><td></td><td></td><td></td><td></td><td></td></tr>
<tr><td>Oligo fwd</td><td>CACCGACGGGGCCGTCATCAGGAT</td><td></td><td></td><td></td><td></td><td></td><td></td><td></td><td></td><td></td><td></td></tr>
<tr><td>Oligo rev</td><td>AAACATCCTGATGACGGCCCCGTC</td><td></td><td></td><td></td><td></td><td></td><td></td><td></td><td></td><td></td><td></td></tr>
<tr><td>Chromosome</td><td>start</td><td>end</td><td>strand</td><td>MM</td><td>target_seq</td><td>PAM</td><td>alignment</td><td>distance</td><td>position</td><td>gene name</td><td>gene id</td></tr>
<tr><td>GU305914.1</td><td>32563</td><td>32585</td><td>-</td><td>0</td><td>GACGGGGCCGTCATCAGGAT</td><td>GGG</td><td>||||||||[||||||||||||]PAM</td><td>NA</td><td>Intergenic</td><td>NA</td><td>NA</td></tr>
<tr><td></td><td></td><td></td><td></td><td></td><td></td><td></td><td></td><td></td><td></td><td></td><td></td></tr>
<tr><td>T31863</td><td>CAGTGACACCGAGATCGTCAAGG</td><td>1000</td><td></td><td></td><td></td><td></td><td></td><td></td><td></td><td></td><td></td></tr>
<tr><td>Oligo fwd</td><td>CACCCAGTGACACCGAGATCGTCA</td><td></td><td></td><td></td><td></td><td></td><td></td><td></td><td></td><td></td><td></td></tr>
<tr><td>Oligo rev</td><td>AAACTGACGATCTCGGTGTCACTG</td><td></td><td></td><td></td><td></td><td></td><td></td><td></td><td></td><td></td><td></td></tr>
<tr><td>Chromosome</td><td>start</td><td>end</td><td>strand</td><td>MM</td><td>target_seq</td><td>PAM</td><td>alignment</td><td>distance</td><td>position</td><td>gene name</td><td>gene id</td></tr>
<tr><td>GU305914.1</td><td>32528</td><td>32550</td><td>-</td><td>0</td><td>CAGTGACACCGAGATCGTCA</td><td>AGG</td><td>||||||||[||||||||||||]PAM</td><td>NA</td><td>Intergenic</td><td>NA</td><td>NA</td></tr>
<tr><td></td><td></td><td></td><td></td><td></td><td></td><td></td><td></td><td></td><td></td><td></td><td></td></tr>
<tr><td>T31864</td><td>AGTGACACCGAGATCGTCAAGGG</td><td>1000</td><td></td><td></td><td></td><td></td><td></td><td></td><td></td><td></td><td></td></tr>
<tr><td>Oligo fwd</td><td>CACCAGTGACACCGAGATCGTCAA</td><td></td><td></td><td></td><td></td><td></td><td></td><td></td><td></td><td></td><td></td></tr>
<tr><td>Oligo rev</td><td>AAACTTGACGATCTCGGTGTCACT</td><td></td><td></td><td></td><td></td><td></td><td></td><td></td><td></td><td></td><td></td></tr>
<tr><td>Chromosome</td><td>start</td><td>end</td><td>strand</td><td>MM</td><td>target_seq</td><td>PAM</td><td>alignment</td><td>distance</td><td>position</td><td>gene name</td><td>gene id</td></tr>
<tr><td>GU305914.1</td><td>32527</td><td>32549</td><td>-</td><td>0</td><td>AGTGACACCGAGATCGTCAA</td><td>GGG</td><td>||||||||[||||||||||||]PAM</td><td>NA</td><td>Intergenic</td><td>NA</td><td>NA</td></tr>
<tr><td></td><td></td><td></td><td></td><td></td><td></td><td></td><td></td><td></td><td></td><td></td><td></td></tr>
<tr><td>T31865</td><td>CCGAGATCGTCAAGGGACACTGG</td><td>1000</td><td></td><td></td><td></td><td></td><td></td><td></td><td></td><td></td><td></td></tr>
<tr><td>Oligo fwd</td><td>CACCCCGAGATCGTCAAGGGACAC</td><td></td><td></td><td></td><td></td><td></td><td></td><td></td><td></td><td></td><td></td></tr>
<tr><td>Oligo rev</td><td>AAACGTGTCCCTTGACGATCTCGG</td><td></td><td></td><td></td><td></td><td></td><td></td><td></td><td></td><td></td><td></td></tr>
<tr><td>Chromosome</td><td>start</td><td>end</td><td>strand</td><td>MM</td><td>target_seq</td><td>PAM</td><td>alignment</td><td>distance</td><td>position</td><td>gene name</td><td>gene id</td></tr>
<tr><td>GU305914.1</td><td>32520</td><td>32542</td><td>-</td><td>0</td><td>CCGAGATCGTCAAGGGACAC</td><td>TGG</td><td>||||||||[||||||||||||]PAM</td><td>NA</td><td>Intergenic</td><td>NA</td><td>NA</td></tr>
<tr><td></td><td></td><td></td><td></td><td></td><td></td><td></td><td></td><td></td><td></td><td></td><td></td></tr>
<tr><td>T31866</td><td>ACACTGGCGCGCCGACGACCTGG</td><td>1000</td><td></td><td></td><td></td><td></td><td></td><td></td><td></td><td></td><td></td></tr>
<tr><td>Oligo fwd</td><td>CACCACACTGGCGCGCCGACGACC</td><td></td><td></td><td></td><td></td><td></td><td></td><td></td><td></td><td></td><td></td></tr>
<tr><td>Oligo rev</td><td>AAACGGTCGTCGGCGCGCCAGTGT</td><td></td><td></td><td></td><td></td><td></td><td></td><td></td><td></td><td></td><td></td></tr>
<tr><td>Chromosome</td><td>start</td><td>end</td><td>strand</td><td>MM</td><td>target_seq</td><td>PAM</td><td>alignment</td><td>distance</td><td>position</td><td>gene name</td><td>gene id</td></tr>
<tr><td>GU305914.1</td><td>32504</td><td>32526</td><td>-</td><td>0</td><td>ACACTGGCGCGCCGACGACC</td><td>TGG</td><td>||||||||[||||||||||||]PAM</td><td>NA</td><td>Intergenic</td><td>NA</td><td>NA</td></tr>
</table>

<table>
<tr><td>T31867</td><td>CTGGCGCGCCGACGACCTGGTGG</td><td>1000</td><td></td><td></td><td></td><td></td><td></td><td></td><td></td><td></td><td></td></tr>
<tr><td>Oligo fwd</td><td>CACCCTGGCGCGCCGACGACCTGG</td><td></td><td></td><td></td><td></td><td></td><td></td><td></td><td></td><td></td><td></td></tr>
<tr><td>Oligo rev</td><td>AAACCCAGGTCGTCGGCGCGCCAG</td><td></td><td></td><td></td><td></td><td></td><td></td><td></td><td></td><td></td><td></td></tr>
<tr><td>Chromosome</td><td>start</td><td>end</td><td>strand</td><td>MM</td><td>target_seq</td><td>PAM</td><td>alignment</td><td>distance</td><td>position</td><td>gene name</td><td>gene id</td></tr>
<tr><td>GU305914.1</td><td>32501</td><td>32523</td><td>-</td><td>0</td><td>CTGGCGCGCCGACGACCTGG</td><td>TGG</td><td>||||||||[||||||||||||]PAM</td><td>NA</td><td>Intergenic</td><td>NA</td><td>NA</td></tr>
<tr><td>T31868</td><td>TGGCGCGCCGACGACCTGGTGGG</td><td>1000</td><td></td><td></td><td></td><td></td><td></td><td></td><td></td><td></td><td></td></tr>
<tr><td>Oligo fwd</td><td>CACCTGGCGCGCCGACGACCTGGT</td><td></td><td></td><td></td><td></td><td></td><td></td><td></td><td></td><td></td><td></td></tr>
<tr><td>Oligo rev</td><td>AAACACCAGGTCGTCGGCGCGCCA</td><td></td><td></td><td></td><td></td><td></td><td></td><td></td><td></td><td></td><td></td></tr>
<tr><td>Chromosome</td><td>start</td><td>end</td><td>strand</td><td>MM</td><td>target_seq</td><td>PAM</td><td>alignment</td><td>distance</td><td>position</td><td>gene name</td><td>gene id</td></tr>
<tr><td>GU305914.1</td><td>32500</td><td>32522</td><td>-</td><td>0</td><td>TGGCGCGCCGACGACCTGGT</td><td>GGG</td><td>||||||||[||||||||||||]PAM</td><td>NA</td><td>Intergenic</td><td>NA</td><td>NA</td></tr>
<tr><td>T31869</td><td>CCGACGACCTGGTGGGCATGCGG</td><td>1000</td><td></td><td></td><td></td><td></td><td></td><td></td><td></td><td></td><td></td></tr>
<tr><td>Oligo fwd</td><td>CACCCCGACGACCTGGTGGGCATG</td><td></td><td></td><td></td><td></td><td></td><td></td><td></td><td></td><td></td><td></td></tr>
<tr><td>Oligo rev</td><td>AAACCATGCCCACCAGGTCGTCGG</td><td></td><td></td><td></td><td></td><td></td><td></td><td></td><td></td><td></td><td></td></tr>
<tr><td>Chromosome</td><td>start</td><td>end</td><td>strand</td><td>MM</td><td>target_seq</td><td>PAM</td><td>alignment</td><td>distance</td><td>position</td><td>gene name</td><td>gene id</td></tr>
<tr><td>GU305914.1</td><td>32493</td><td>32515</td><td>-</td><td>0</td><td>CCGACGACCTGGTGGGCATG</td><td>CGG</td><td>||||||||[||||||||||||]PAM</td><td>NA</td><td>Intergenic</td><td>NA</td><td>NA</td></tr>
<tr><td>T31870</td><td>TCGAGTACGTCGACATGCTGAGG</td><td>1000</td><td></td><td></td><td></td><td></td><td></td><td></td><td></td><td></td><td></td></tr>
<tr><td>Oligo fwd</td><td>CACCTCGAGTACGTCGACATGCTG</td><td></td><td></td><td></td><td></td><td></td><td></td><td></td><td></td><td></td><td></td></tr>
<tr><td>Oligo rev</td><td>AAACCAGCATGTCGACGTACTCGA</td><td></td><td></td><td></td><td></td><td></td><td></td><td></td><td></td><td></td><td></td></tr>
<tr><td>Chromosome</td><td>start</td><td>end</td><td>strand</td><td>MM</td><td>target_seq</td><td>PAM</td><td>alignment</td><td>distance</td><td>position</td><td>gene name</td><td>gene id</td></tr>
<tr><td>GU305914.1</td><td>32451</td><td>32473</td><td>-</td><td>0</td><td>TCGAGTACGTCGACATGCTG</td><td>AGG</td><td>||||||||[||||||||||||]PAM</td><td>NA</td><td>Intergenic</td><td>NA</td><td>NA</td></tr>
<tr><td>T31871</td><td>CAATAAACAGACGCTCATGATGG</td><td>1000</td><td></td><td></td><td></td><td></td><td></td><td></td><td></td><td></td><td></td></tr>
<tr><td>Oligo fwd</td><td>CACCCAATAAACAGACGCTCATGA</td><td></td><td></td><td></td><td></td><td></td><td></td><td></td><td></td><td></td><td></td></tr>
<tr><td>Oligo rev</td><td>AAACTCATGAGCGTCTGTTTATTG</td><td></td><td></td><td></td><td></td><td></td><td></td><td></td><td></td><td></td><td></td></tr>
<tr><td>Chromosome</td><td>start</td><td>end</td><td>strand</td><td>MM</td><td>target_seq</td><td>PAM</td><td>alignment</td><td>distance</td><td>position</td><td>gene name</td><td>gene id</td></tr>
<tr><td>GU305914.1</td><td>32417</td><td>32439</td><td>-</td><td>0</td><td>CAATAAACAGACGCTCATGA</td><td>TGG</td><td>||||||||[||||||||||||]PAM</td><td>NA</td><td>Intergenic</td><td>NA</td><td>NA</td></tr>
<tr><td>T31872</td><td>ACAGACGCTCATGATGGAGCTGG</td><td>1000</td><td></td><td></td><td></td><td></td><td></td><td></td><td></td><td></td><td></td></tr>
<tr><td>Oligo fwd</td><td>CACCACAGACGCTCATGATGGAGC</td><td></td><td></td><td></td><td></td><td></td><td></td><td></td><td></td><td></td><td></td></tr>
<tr><td>Oligo rev</td><td>AAACGCTCCATCATGAGCGTCTGT</td><td></td><td></td><td></td><td></td><td></td><td></td><td></td><td></td><td></td><td></td></tr>
<tr><td>Chromosome</td><td>start</td><td>end</td><td>strand</td><td>MM</td><td>target_seq</td><td>PAM</td><td>alignment</td><td>distance</td><td>position</td><td>gene name</td><td>gene id</td></tr>
<tr><td>GU305914.1</td><td>32411</td><td>32433</td><td>-</td><td>0</td><td>ACAGACGCTCATGATGGAGC</td><td>TGG</td><td>||||||||[||||||||||||]PAM</td><td>NA</td><td>Intergenic</td><td>NA</td><td>NA</td></tr>
<tr><td>T31873</td><td>CGCTCATGATGGAGCTGGATAGG</td><td>1000</td><td></td><td></td><td></td><td></td><td></td><td></td><td></td><td></td><td></td></tr>
<tr><td>Oligo fwd</td><td>CACCCGCTCATGATGGAGCTGGAT</td><td></td><td></td><td></td><td></td><td></td><td></td><td></td><td></td><td></td><td></td></tr>
<tr><td>Oligo rev</td><td>AAACATCCAGCTCCATCATGAGCG</td><td></td><td></td><td></td><td></td><td></td><td></td><td></td><td></td><td></td><td></td></tr>
<tr><td>Chromosome</td><td>start</td><td>end</td><td>strand</td><td>MM</td><td>target_seq</td><td>PAM</td><td>alignment</td><td>distance</td><td>position</td><td>gene name</td><td>gene id</td></tr>
<tr><td>GU305914.1</td><td>32406</td><td>32428</td><td>-</td><td>0</td><td>CGCTCATGATGGAGCTGGAT</td><td>AGG</td><td>||||||||[||||||||||||]PAM</td><td>NA</td><td>Intergenic</td><td>NA</td><td>NA</td></tr>
<tr><td>T31874</td><td>ATAGGTTATACGTCCCCAAGCGG</td><td>1000</td><td></td><td></td><td></td><td></td><td></td><td></td><td></td><td></td><td></td></tr>
<tr><td>Oligo fwd</td><td>CACCATAGGTTATACGTCCCCAAG</td><td></td><td></td><td></td><td></td><td></td><td></td><td></td><td></td><td></td><td></td></tr>
<tr><td>Oligo rev</td><td>AAACCTTGGGGACGTATAACCTAT</td><td></td><td></td><td></td><td></td><td></td><td></td><td></td><td></td><td></td><td></td></tr>
<tr><td>Chromosome</td><td>start</td><td>end</td><td>strand</td><td>MM</td><td>target_seq</td><td>PAM</td><td>alignment</td><td>distance</td><td>position</td><td>gene name</td><td>gene id</td></tr>
<tr><td>GU305914.1</td><td>32388</td><td>32410</td><td>-</td><td>0</td><td>ATAGGTTATACGTCCCCAAG</td><td>CGG</td><td>||||||||[||||||||||||]PAM</td><td>NA</td><td>Intergenic</td><td>NA</td><td>NA</td></tr>
<tr><td>T31875</td><td>TAGGTTATACGTCCCCAAGCGGG</td><td>1000</td><td></td><td></td><td></td><td></td><td></td><td></td><td></td><td></td><td></td></tr>
<tr><td>Oligo fwd</td><td>CACCTAGGTTATACGTCCCCAAGC</td><td></td><td></td><td></td><td></td><td></td><td></td><td></td><td></td><td></td><td></td></tr>
<tr><td>Oligo rev</td><td>AAACGCTTGGGGACGTATAACCTA</td><td></td><td></td><td></td><td></td><td></td><td></td><td></td><td></td><td></td><td></td></tr>
<tr><td>Chromosome</td><td>start</td><td>end</td><td>strand</td><td>MM</td><td>target_seq</td><td>PAM</td><td>alignment</td><td>distance</td><td>position</td><td>gene name</td><td>gene id</td></tr>
<tr><td>GU305914.1</td><td>32387</td><td>32409</td><td>-</td><td>0</td><td>TAGGTTATACGTCCCCAAGC</td><td>GGG</td><td>||||||||[||||||||||||]PAM</td><td>NA</td><td>Intergenic</td><td>NA</td><td>NA</td></tr>
<tr><td>T31876</td><td>GCTCGTCGAGCTCGTCATCTCGG</td><td>1000</td><td></td><td></td><td></td><td></td><td></td><td></td><td></td><td></td><td></td></tr>
<tr><td>Oligo fwd</td><td>CACCGCTCGTCGAGCTCGTCATCT</td><td></td><td></td><td></td><td></td><td></td><td></td><td></td><td></td><td></td><td></td></tr>
<tr><td>Oligo rev</td><td>AAACAGATGACGAGCTCGACGAGC</td><td></td><td></td><td></td><td></td><td></td><td></td><td></td><td></td><td></td><td></td></tr>
<tr><td>Chromosome</td><td>start</td><td>end</td><td>strand</td><td>MM</td><td>target_seq</td><td>PAM</td><td>alignment</td><td>distance</td><td>position</td><td>gene name</td><td>gene id</td></tr>
<tr><td>GU305914.1</td><td>32363</td><td>32385</td><td>-</td><td>0</td><td>GCTCGTCGAGCTCGTCATCT</td><td>CGG</td><td>||||||||[||||||||||||]PAM</td><td>NA</td><td>Intergenic</td><td>NA</td><td>NA</td></tr>
<tr><td>T31877</td><td>CGAGCTCGTCATCTCGGACAAGG</td><td>1000</td><td></td><td></td><td></td><td></td><td></td><td></td><td></td><td></td><td></td></tr>
<tr><td>Oligo fwd</td><td>CACCCGAGCTCGTCATCTCGGACA</td><td></td><td></td><td></td><td></td><td></td><td></td><td></td><td></td><td></td><td></td></tr>
<tr><td>Oligo rev</td><td>AAACTGTCCGAGATGACGAGCTCG</td><td></td><td></td><td></td><td></td><td></td><td></td><td></td><td></td><td></td><td></td></tr>
<tr><td>Chromosome</td><td>start</td><td>end</td><td>strand</td><td>MM</td><td>target_seq</td><td>PAM</td><td>alignment</td><td>distance</td><td>position</td><td>gene name</td><td>gene id</td></tr>
<tr><td>GU305914.1</td><td>32357</td><td>32379</td><td>-</td><td>0</td><td>CGAGCTCGTCATCTCGGACA</td><td>AGG</td><td>||||||||[||||||||||||]PAM</td><td>NA</td><td>Intergenic</td><td>NA</td><td>NA</td></tr>
<tr><td>T31878</td><td>CATCTCGGACAAGGATTCGTCGG</td><td>1000</td><td></td><td></td><td></td><td></td><td></td><td></td><td></td><td></td><td></td></tr>
<tr><td>Oligo fwd</td><td>CACCCATCTCGGACAAGGATTCGT</td><td></td><td></td><td></td><td></td><td></td><td></td><td></td><td></td><td></td><td></td></tr>
<tr><td>Oligo rev</td><td>AAACACGAATCCTTGTCCGAGATG</td><td></td><td></td><td></td><td></td><td></td><td></td><td></td><td></td><td></td><td></td></tr>
<tr><td>Chromosome</td><td>start</td><td>end</td><td>strand</td><td>MM</td><td>target_seq</td><td>PAM</td><td>alignment</td><td>distance</td><td>position</td><td>gene name</td><td>gene id</td></tr>
<tr><td>GU305914.1</td><td>32348</td><td>32370</td><td>-</td><td>0</td><td>CATCTCGGACAAGGATTCGT</td><td>CGG</td><td>||||||||[||||||||||||]PAM</td><td>NA</td><td>Intergenic</td><td>NA</td><td>NA</td></tr>
<tr><td>T31879</td><td>ATCTCGGACAAGGATTCGTCGGG</td><td>1000</td><td></td><td></td><td></td><td></td><td></td><td></td><td></td><td></td><td></td></tr>
<tr><td>Oligo fwd</td><td>CACCATCTCGGACAAGGATTCGTC</td><td></td><td></td><td></td><td></td><td></td><td></td><td></td><td></td><td></td><td></td></tr>
<tr><td>Oligo rev</td><td>AAACGACGAATCCTTGTCCGAGAT</td><td></td><td></td><td></td><td></td><td></td><td></td><td></td><td></td><td></td><td></td></tr>
<tr><td>Chromosome</td><td>start</td><td>end</td><td>strand</td><td>MM</td><td>target_seq</td><td>PAM</td><td>alignment</td><td>distance</td><td>position</td><td>gene name</td><td>gene id</td></tr>
<tr><td>GU305914.1</td><td>32347</td><td>32369</td><td>-</td><td>0</td><td>ATCTCGGACAAGGATTCGTC</td><td>GGG</td><td>||||||||[||||||||||||]PAM</td><td>NA</td><td>Intergenic</td><td>NA</td><td>NA</td></tr>
<tr><td>T31880</td><td>GGATTCGTCGGGCTCCGACGAGG</td><td>1000</td><td></td><td></td><td></td><td></td><td></td><td></td><td></td><td></td><td></td></tr>
<tr><td>Oligo fwd</td><td>CACCGGATTCGTCGGGCTCCGACG</td><td></td><td></td><td></td><td></td><td></td><td></td><td></td><td></td><td></td><td></td></tr>
<tr><td>Oligo rev</td><td>AAACCGTCGGAGCCCGACGAATCC</td><td></td><td></td><td></td><td></td><td></td><td></td><td></td><td></td><td></td><td></td></tr>
<tr><td>Chromosome</td><td>start</td><td>end</td><td>strand</td><td>MM</td><td>target_seq</td><td>PAM</td><td>alignment</td><td>distance</td><td>position</td><td>gene name</td><td>gene id</td></tr>
<tr><td>GU305914.1</td><td>32336</td><td>32358</td><td>-</td><td>0</td><td>GGATTCGTCGGGCTCCGACG</td><td>AGG</td><td>||||||||[||||||||||||]PAM</td><td>NA</td><td>Intergenic</td><td>NA</td><td>NA</td></tr>
<tr><td>T31881</td><td>GGGCTCCGACGAGGAGACCGAGG</td><td>1000</td><td></td><td></td><td></td><td></td><td></td><td></td><td></td><td></td><td></td></tr>
<tr><td>Oligo fwd</td><td>CACCGGGCTCCGACGAGGAGACCG</td><td></td><td></td><td></td><td></td><td></td><td></td><td></td><td></td><td></td><td></td></tr>
<tr><td>Oligo rev</td><td>AAACCGGTCTCCTCGTCGGAGCCC</td><td></td><td></td><td></td><td></td><td></td><td></td><td></td><td></td><td></td><td></td></tr>
<tr><td>Chromosome</td><td>start</td><td>end</td><td>strand</td><td>MM</td><td>target_seq</td><td>PAM</td><td>alignment</td><td>distance</td><td>position</td><td>gene name</td><td>gene id</td></tr>
<tr><td>GU305914.1</td><td>32327</td><td>32349</td><td>-</td><td>0</td><td>GGGCTCCGACGAGGAGACCG</td><td>AGG</td><td>||||||||[||||||||||||]PAM</td><td>NA</td><td>Intergenic</td><td>NA</td><td>NA</td></tr>
</table>

<table>
<tr><td>GU305914.1</td><td>113955</td><td>113977</td><td>+</td><td>4</td><td>GAACTCCGGCGAGGAGGCCG</td><td>AGG</td><td>|--|||||[-|||||||-|||]PAM</td><td>NA</td><td>Intergenic</td><td>NA</td><td>NA</td></tr>
<tr><td>T31882</td><td>CGACGAGGAGACCGAGGACGCGG</td><td>1000</td><td colspan="9"></td></tr>
<tr><td>Oligo fwd</td><td>CACCCGACGAGGAGACCGAGGACG</td><td colspan="10"></td></tr>
<tr><td>Oligo rev</td><td>AAACCGTCCTCGGTCTCCTCGTCG</td><td colspan="10"></td></tr>
<tr><td>Chromosome</td><td>start</td><td>end</td><td>strand</td><td>MM</td><td>target_seq</td><td>PAM</td><td>alignment</td><td>distance</td><td>position</td><td>gene name</td><td>gene id</td></tr>
<tr><td>GU305914.1</td><td>32321</td><td>32343</td><td>-</td><td>0</td><td>CGACGAGGAGACCGAGGACG</td><td>CGG</td><td>||||||||[||||||||||||]PAM</td><td>NA</td><td>Intergenic</td><td>NA</td><td>NA</td></tr>
<tr><td>T31883</td><td>GAGACCGAGGACGCGGACAGCGG</td><td>1000</td><td colspan="9"></td></tr>
<tr><td>Oligo fwd</td><td>CACCGAGACCGAGGACGCGGACAG</td><td colspan="10"></td></tr>
<tr><td>Oligo rev</td><td>AAACCTGTCCGCGTCCTCGGTCTC</td><td colspan="10"></td></tr>
<tr><td>Chromosome</td><td>start</td><td>end</td><td>strand</td><td>MM</td><td>target_seq</td><td>PAM</td><td>alignment</td><td>distance</td><td>position</td><td>gene name</td><td>gene id</td></tr>
<tr><td>GU305914.1</td><td>32314</td><td>32336</td><td>-</td><td>0</td><td>GAGACCGAGGACGCGGACAG</td><td>CGG</td><td>||||||||[||||||||||||]PAM</td><td>NA</td><td>Intergenic</td><td>NA</td><td>NA</td></tr>
<tr><td>T31884</td><td>AGACCGAGGACGCGGACAGCGGG</td><td>1000</td><td colspan="9"></td></tr>
<tr><td>Oligo fwd</td><td>CACCAGACCGAGGACGCGGACAGC</td><td colspan="10"></td></tr>
<tr><td>Oligo rev</td><td>AAACGCTGTCCGCGTCCTCGGTCT</td><td colspan="10"></td></tr>
<tr><td>Chromosome</td><td>start</td><td>end</td><td>strand</td><td>MM</td><td>target_seq</td><td>PAM</td><td>alignment</td><td>distance</td><td>position</td><td>gene name</td><td>gene id</td></tr>
<tr><td>GU305914.1</td><td>32313</td><td>32335</td><td>-</td><td>0</td><td>AGACCGAGGACGCGGACAGC</td><td>GGG</td><td>||||||||[||||||||||||]PAM</td><td>NA</td><td>Intergenic</td><td>NA</td><td>NA</td></tr>
<tr><td>T31885</td><td>GGACGCGGACAGCGGGCACTCGG</td><td>1000</td><td colspan="9"></td></tr>
<tr><td>Oligo fwd</td><td>CACCGGACGCGGACAGCGGGCACT</td><td colspan="10"></td></tr>
<tr><td>Oligo rev</td><td>AAACAGTGCCCGCTGTCCGCGTCC</td><td colspan="10"></td></tr>
<tr><td>Chromosome</td><td>start</td><td>end</td><td>strand</td><td>MM</td><td>target_seq</td><td>PAM</td><td>alignment</td><td>distance</td><td>position</td><td>gene name</td><td>gene id</td></tr>
<tr><td>GU305914.1</td><td>32306</td><td>32328</td><td>-</td><td>0</td><td>GGACGCGGACAGCGGGCACT</td><td>CGG</td><td>||||||||[||||||||||||]PAM</td><td>NA</td><td>Intergenic</td><td>NA</td><td>NA</td></tr>
<tr><td>T31886</td><td>CACTCGGAGCCGTCCGATGATGG</td><td>1000</td><td colspan="9"></td></tr>
<tr><td>Oligo fwd</td><td>CACCCACTCGGAGCCGTCCGATGA</td><td colspan="10"></td></tr>
<tr><td>Oligo rev</td><td>AAACTCATCGGACGGCTCCGAGTG</td><td colspan="10"></td></tr>
<tr><td>Chromosome</td><td>start</td><td>end</td><td>strand</td><td>MM</td><td>target_seq</td><td>PAM</td><td>alignment</td><td>distance</td><td>position</td><td>gene name</td><td>gene id</td></tr>
<tr><td>GU305914.1</td><td>32290</td><td>32312</td><td>-</td><td>0</td><td>CACTCGGAGCCGTCCGATGA</td><td>TGG</td><td>||||||||[||||||||||||]PAM</td><td>NA</td><td>Intergenic</td><td>NA</td><td>NA</td></tr>
<tr><td>T31887</td><td>TCGGAGCCGTCCGATGATGGCGG</td><td>1000</td><td colspan="9"></td></tr>
<tr><td>Oligo fwd</td><td>CACCTCGGAGCCGTCCGATGATGG</td><td colspan="10"></td></tr>
<tr><td>Oligo rev</td><td>AAACCCATCATCGGACGGCTCCGA</td><td colspan="10"></td></tr>
<tr><td>Chromosome</td><td>start</td><td>end</td><td>strand</td><td>MM</td><td>target_seq</td><td>PAM</td><td>alignment</td><td>distance</td><td>position</td><td>gene name</td><td>gene id</td></tr>
<tr><td>GU305914.1</td><td>32287</td><td>32309</td><td>-</td><td>0</td><td>TCGGAGCCGTCCGATGATGG</td><td>CGG</td><td>||||||||[||||||||||||]PAM</td><td>NA</td><td>Intergenic</td><td>NA</td><td>NA</td></tr>
<tr><td>T31888</td><td>GATGATGGCGGCGAGTCCGACGG</td><td>1000</td><td colspan="9"></td></tr>
<tr><td>Oligo fwd</td><td>CACCGATGATGGCGGCGAGTCCGA</td><td colspan="10"></td></tr>
<tr><td>Oligo rev</td><td>AAACTCGGACTCGCCGCCATCATC</td><td colspan="10"></td></tr>
<tr><td>Chromosome</td><td>start</td><td>end</td><td>strand</td><td>MM</td><td>target_seq</td><td>PAM</td><td>alignment</td><td>distance</td><td>position</td><td>gene name</td><td>gene id</td></tr>
<tr><td>GU305914.1</td><td>32275</td><td>32297</td><td>-</td><td>0</td><td>GATGATGGCGGCGAGTCCGA</td><td>CGG</td><td>||||||||[||||||||||||]PAM</td><td>NA</td><td>Intergenic</td><td>NA</td><td>NA</td></tr>
<tr><td>T31889</td><td>GTCCGACGGCTCCGAGTTTGTGG</td><td>1000</td><td colspan="9"></td></tr>
<tr><td>Oligo fwd</td><td>CACCGTCCGACGGCTCCGAGTTTG</td><td colspan="10"></td></tr>
<tr><td>Oligo rev</td><td>AAACCAAACTCGGAGCCGTCGGAC</td><td colspan="10"></td></tr>
<tr><td>Chromosome</td><td>start</td><td>end</td><td>strand</td><td>MM</td><td>target_seq</td><td>PAM</td><td>alignment</td><td>distance</td><td>position</td><td>gene name</td><td>gene id</td></tr>
<tr><td>GU305914.1</td><td>32261</td><td>32283</td><td>-</td><td>0</td><td>GTCCGACGGCTCCGAGTTTG</td><td>TGG</td><td>||||||||[||||||||||||]PAM</td><td>NA</td><td>Intergenic</td><td>NA</td><td>NA</td></tr>
<tr><td>T31890</td><td>CGACGGCTCCGAGTTTGTGGTGG</td><td>1000</td><td colspan="9"></td></tr>
<tr><td>Oligo fwd</td><td>CACCCGACGGCTCCGAGTTTGTGG</td><td colspan="10"></td></tr>
<tr><td>Oligo rev</td><td>AAACCCACAAACTCGGAGCCGTCG</td><td colspan="10"></td></tr>
<tr><td>Chromosome</td><td>start</td><td>end</td><td>strand</td><td>MM</td><td>target_seq</td><td>PAM</td><td>alignment</td><td>distance</td><td>position</td><td>gene name</td><td>gene id</td></tr>
<tr><td>GU305914.1</td><td>32258</td><td>32280</td><td>-</td><td>0</td><td>CGACGGCTCCGAGTTTGTGG</td><td>TGG</td><td>||||||||[||||||||||||]PAM</td><td>NA</td><td>Intergenic</td><td>NA</td><td>NA</td></tr>
<tr><td>T31891</td><td>GCTCCGAGTTTGTGGTGGAGCGG</td><td>1000</td><td colspan="9"></td></tr>
<tr><td>Oligo fwd</td><td>CACCGCTCCGAGTTTGTGGTGGAG</td><td colspan="10"></td></tr>
<tr><td>Oligo rev</td><td>AAACCTCCACCACAAACTCGGAGC</td><td colspan="10"></td></tr>
<tr><td>Chromosome</td><td>start</td><td>end</td><td>strand</td><td>MM</td><td>target_seq</td><td>PAM</td><td>alignment</td><td>distance</td><td>position</td><td>gene name</td><td>gene id</td></tr>
<tr><td>GU305914.1</td><td>32253</td><td>32275</td><td>-</td><td>0</td><td>GCTCCGAGTTTGTGGTGGAG</td><td>CGG</td><td>||||||||[||||||||||||]PAM</td><td>NA</td><td>Intergenic</td><td>NA</td><td>NA</td></tr>
<tr><td>T31892</td><td>CTCCGAGTTTGTGGTGGAGCGGG</td><td>1000</td><td colspan="9"></td></tr>
<tr><td>Oligo fwd</td><td>CACCCTCCGAGTTTGTGGTGGAGC</td><td colspan="10"></td></tr>
<tr><td>Oligo rev</td><td>AAACGCTCCACCACAAACTCGGAG</td><td colspan="10"></td></tr>
<tr><td>Chromosome</td><td>start</td><td>end</td><td>strand</td><td>MM</td><td>target_seq</td><td>PAM</td><td>alignment</td><td>distance</td><td>position</td><td>gene name</td><td>gene id</td></tr>
<tr><td>GU305914.1</td><td>32252</td><td>32274</td><td>-</td><td>0</td><td>CTCCGAGTTTGTGGTGGAGC</td><td>GGG</td><td>||||||||[||||||||||||]PAM</td><td>NA</td><td>Intergenic</td><td>NA</td><td>NA</td></tr>
<tr><td>T31893</td><td>TGTGGTGGAGCGGGTGTGACAGG</td><td>1000</td><td colspan="9"></td></tr>
<tr><td>Oligo fwd</td><td>CACCTGTGGTGGAGCGGGTGTGAC</td><td colspan="10"></td></tr>
<tr><td>Oligo rev</td><td>AAACGTCACACCCGCTCCACCACA</td><td colspan="10"></td></tr>
<tr><td>Chromosome</td><td>start</td><td>end</td><td>strand</td><td>MM</td><td>target_seq</td><td>PAM</td><td>alignment</td><td>distance</td><td>position</td><td>gene name</td><td>gene id</td></tr>
<tr><td>GU305914.1</td><td>32243</td><td>32265</td><td>-</td><td>0</td><td>TGTGGTGGAGCGGGTGTGAC</td><td>AGG</td><td>||||||||[||||||||||||]PAM</td><td>NA</td><td>Intergenic</td><td>NA</td><td>NA</td></tr>
<tr><td>T31894</td><td>GACAGGACACGAGCGACACACGG</td><td>1000</td><td colspan="9"></td></tr>
<tr><td>Oligo fwd</td><td>CACCGACAGGACACGAGCGACACA</td><td colspan="10"></td></tr>
<tr><td>Oligo rev</td><td>AAACTGTGTCGCTCGTGTCCTGTC</td><td colspan="10"></td></tr>
<tr><td>Chromosome</td><td>start</td><td>end</td><td>strand</td><td>MM</td><td>target_seq</td><td>PAM</td><td>alignment</td><td>distance</td><td>position</td><td>gene name</td><td>gene id</td></tr>
<tr><td>GU305914.1</td><td>32226</td><td>32248</td><td>-</td><td>0</td><td>GACAGGACACGAGCGACACA</td><td>CGG</td><td>||||||||[||||||||||||]PAM</td><td>NA</td><td>Intergenic</td><td>NA</td><td>NA</td></tr>
<tr><td>T31895</td><td>ACGAGCGACACACGGAAGACAGG</td><td>1000</td><td colspan="9"></td></tr>
<tr><td>Oligo fwd</td><td>CACCACGAGCGACACACGGAAGAC</td><td colspan="10"></td></tr>
<tr><td>Oligo rev</td><td>AAACGTCTTCCGTGTGTCGCTCGT</td><td colspan="10"></td></tr>
<tr><td>Chromosome</td><td>start</td><td>end</td><td>strand</td><td>MM</td><td>target_seq</td><td>PAM</td><td>alignment</td><td>distance</td><td>position</td><td>gene name</td><td>gene id</td></tr>
<tr><td>GU305914.1</td><td>32218</td><td>32240</td><td>-</td><td>0</td><td>ACGAGCGACACACGGAAGAC</td><td>AGG</td><td>||||||||[||||||||||||]PAM</td><td>NA</td><td>Intergenic</td><td>NA</td><td>NA</td></tr>
<tr><td>T31896</td><td>AGCGACACACGGAAGACAGGAGG</td><td>1000</td><td colspan="9"></td></tr>
<tr><td>Oligo fwd</td><td>CACCAGCGACACACGGAAGACAGG</td><td colspan="10"></td></tr>
<tr><td>Oligo rev</td><td>AAACCCTGTCTTCCGTGTGTCGCT</td><td colspan="10"></td></tr>
</table>

Chromosome start end strand MM target_seq PAM alignment distance position gene name gene id
GU305914.1 32215 32237 - 0 AGCGACACACGGAAGACAGG AGG ||||||||[||||||||||||]PAM NA Intergenic NA NA

T31897 CACGGAAGACAGGAGGAATATGG 1000
Oligo fwd CACCCACGGAAGACAGGAGGAATA
Oligo rev AAACTATTCCTCCTGTCTTCCGTG
Chromosome start end strand MM target_seq PAM alignment distance position gene name gene id
GU305914.1 32208 32230 - 0 CACGGAAGACAGGAGGAATA TGG ||||||||[||||||||||||]PAM NA Intergenic NA NA

T31898 GAAGACAGGAGGAATATGGCTGG 1000
Oligo fwd CACCGAAGACAGGAGGAATATGGC
Oligo rev AAACGCCATATTCCTCCTGTCTTC
Chromosome start end strand MM target_seq PAM alignment distance position gene name gene id
GU305914.1 32204 32226 - 0 GAAGACAGGAGGAATATGGC TGG ||||||||[||||||||||||]PAM NA Intergenic NA NA

T31899 CTGGTTCTGTTCGCTCCGAGTGG 1000
Oligo fwd CACCCTGGTTCTGTTCGCTCCGAG
Oligo rev AAACCTCGGAGCGAACAGAACCAG
Chromosome start end strand MM target_seq PAM alignment distance position gene name gene id
GU305914.1 32185 32207 - 0 CTGGTTCTGTTCGCTCCGAG TGG ||||||||[||||||||||||]PAM NA Intergenic NA NA

T31900 TGGTTCTGTTCGCTCCGAGTGGG 1000
Oligo fwd CACCTGGTTCTGTTCGCTCCGAGT
Oligo rev AAACACTCGGAGCGAACAGAACCA
Chromosome start end strand MM target_seq PAM alignment distance position gene name gene id
GU305914.1 32184 32206 - 0 TGGTTCTGTTCGCTCCGAGT GGG ||||||||[||||||||||||]PAM NA Intergenic NA NA

T31901 TTCTGTTCGCTCCGAGTGGGCGG 1000
Oligo fwd CACCTTCTGTTCGCTCCGAGTGGG
Oligo rev AAACCCCACTCGGAGCGAACAGAA
Chromosome start end strand MM target_seq PAM alignment distance position gene name gene id
GU305914.1 32181 32203 - 0 TTCTGTTCGCTCCGAGTGGG CGG ||||||||[||||||||||||]PAM NA Intergenic NA NA

T31902 CGGTGTTAAGTCACGTAGTCCGG 1000
Oligo fwd CACCCGGTGTTAAGTCACGTAGTC
Oligo rev AAACGACTACGTGACTTAACACCG
Chromosome start end strand MM target_seq PAM alignment distance position gene name gene id
GU305914.1 32161 32183 - 0 CGGTGTTAAGTCACGTAGTC CGG ||||||||[||||||||||||]PAM NA Intergenic NA NA

T31903 CGGTTATTATTATAAAGTTTCGG 1000
Oligo fwd CACCCGGTTATTATTATAAAGTTT
Oligo rev AAACAAACTTTATAATAATAACCG
Chromosome start end strand MM target_seq PAM alignment distance position gene name gene id
GU305914.1 32141 32163 - 0 CGGTTATTATTATAAAGTTT CGG ||||||||[||||||||||||]PAM NA Intergenic NA NA

T31904 TTATTATAAAGTTTCGGTTTTGG 1000
Oligo fwd CACCTTATTATAAAGTTTCGGTTT
Oligo rev AAACAAACCGAAACTTTATAATAA
Chromosome start end strand MM target_seq PAM alignment distance position gene name gene id
GU305914.1 32135 32157 - 0 TTATTATAAAGTTTCGGTTT TGG ||||||||[||||||||||||]PAM NA Intergenic NA NA

T31905 TTTTGGCGATAACTTCCGTGAGG 1000
Oligo fwd CACCTTTTGGCGATAACTTCCGTG
Oligo rev AAACCACGGAAGTTATCGCCAAAA
Chromosome start end strand MM target_seq PAM alignment distance position gene name gene id
GU305914.1 32118 32140 - 0 TTTTGGCGATAACTTCCGTG AGG ||||||||[||||||||||||]PAM NA Intergenic NA NA

T31906 TTTGGCGATAACTTCCGTGAGGG 1000
Oligo fwd CACCTTTGGCGATAACTTCCGTGA
Oligo rev AAACTCACGGAAGTTATCGCCAAA
Chromosome start end strand MM target_seq PAM alignment distance position gene name gene id
GU305914.1 32117 32139 - 0 TTTGGCGATAACTTCCGTGA GGG ||||||||[||||||||||||]PAM NA Intergenic NA NA

T31907 CCGTGAGGGTCTGTCTGTGTCGG 1000
Oligo fwd CACCCCGTGAGGGTCTGTCTGTGT
Oligo rev AAACACACAGACAGACCCTCACGG
Chromosome start end strand MM target_seq PAM alignment distance position gene name gene id
GU305914.1 32103 32125 - 0 CCGTGAGGGTCTGTCTGTGT CGG ||||||||[||||||||||||]PAM NA Intergenic NA NA
GU305914.1 95541 95563 + 4 CCGAGGGGGTGTGTCGGTGT AGG |||-|-||[||-||||-||||]PAM NA Intergenic NA NA

T31908 TGTCGGTCATTTTTCCAGAGTGG 1000
Oligo fwd CACCTGTCGGTCATTTTTCCAGAG
Oligo rev AAACCTCTGGAAAAATGACCGACA
Chromosome start end strand MM target_seq PAM alignment distance position gene name gene id
GU305914.1 32086 32108 - 0 TGTCGGTCATTTTTCCAGAG TGG ||||||||[||||||||||||]PAM NA Intergenic NA NA

T31909 GTCGGTCATTTTTCCAGAGTGGG 1000
Oligo fwd CACCGTCGGTCATTTTTCCAGAGT
Oligo rev AAACACTCTGGAAAAATGACCGAC
Chromosome start end strand MM target_seq PAM alignment distance position gene name gene id
GU305914.1 32085 32107 - 0 GTCGGTCATTTTTCCAGAGT GGG ||||||||[||||||||||||]PAM NA Intergenic NA NA

T31910 GTCATTTTTCCAGAGTGGGCAGG 1000
Oligo fwd CACCGTCATTTTTCCAGAGTGGGC
Oligo rev AAACGCCCACTCTGGAAAAATGAC
Chromosome start end strand MM target_seq PAM alignment distance position gene name gene id
GU305914.1 32081 32103 - 0 GTCATTTTTCCAGAGTGGGC AGG ||||||||[||||||||||||]PAM NA Intergenic NA NA

T31911 TTTTTCCAGAGTGGGCAGGACGG 1000

| | | | | | | | | | | | |
|---|---|---|---|---|---|---|---|---|---|---|---|
| Oligo fwd | CACCTTTTTCCAGAGTGGGCAGGA | | | | | | | | | | |
| Oligo rev | AAACTCCTGCCCACTCTGGAAAAA | | | | | | | | | | |
| Chromosome | start | end | strand | MM | target_seq | PAM | alignment | distance | position | gene name | gene id |
| GU305914.1 | 32077 | 32099 | - | 0 | TTTTTCCAGAGTGGGCAGGA | CGG | \|\|\|\|\|\|\|\|[\|\|\|\|\|\|\|\|\|\|\|\|]PAM | NA | Intergenic | NA | NA |
| T31912 | TTTTCCAGAGTGGGCAGGACGGG | 1000 | | | | | | | | | |
| Oligo fwd | CACCTTTTCCAGAGTGGGCAGGAC | | | | | | | | | | |
| Oligo rev | AAACGTCCTGCCCACTCTGGAAAA | | | | | | | | | | |
| Chromosome | start | end | strand | MM | target_seq | PAM | alignment | distance | position | gene name | gene id |
| GU305914.1 | 32076 | 32098 | - | 0 | TTTTCCAGAGTGGGCAGGAC | GGG | \|\|\|\|\|\|\|\|[\|\|\|\|\|\|\|\|\|\|\|\|]PAM | NA | Intergenic | NA | NA |
| T31913 | CAGGACGGGACACTGCGCGATGG | 1000 | | | | | | | | | |
| Oligo fwd | CACCCAGGACGGGACACTGCGCGA | | | | | | | | | | |
| Oligo rev | AAACTCGCGCAGTGTCCCGTCCTG | | | | | | | | | | |
| Chromosome | start | end | strand | MM | target_seq | PAM | alignment | distance | position | gene name | gene id |
| GU305914.1 | 32062 | 32084 | - | 0 | CAGGACGGGACACTGCGCGA | TGG | \|\|\|\|\|\|\|\|[\|\|\|\|\|\|\|\|\|\|\|\|]PAM | NA | Intergenic | NA | NA |
| T31914 | GGGACACTGCGCGATGGCCTCGG | 1000 | | | | | | | | | |
| Oligo fwd | CACCGGGACACTGCGCGATGGCCT | | | | | | | | | | |
| Oligo rev | AAACAGGCCATCGCGCAGTGTCCC | | | | | | | | | | |
| Chromosome | start | end | strand | MM | target_seq | PAM | alignment | distance | position | gene name | gene id |
| GU305914.1 | 32056 | 32078 | - | 0 | GGGACACTGCGCGATGGCCT | CGG | \|\|\|\|\|\|\|\|[\|\|\|\|\|\|\|\|\|\|\|\|]PAM | NA | Intergenic | NA | NA |
| T31915 | ACACTGCGCGATGGCCTCGGAGG | 1000 | | | | | | | | | |
| Oligo fwd | CACCACACTGCGCGATGGCCTCGG | | | | | | | | | | |
| Oligo rev | AAACCCGAGGCCATCGCGCAGTGT | | | | | | | | | | |
| Chromosome | start | end | strand | MM | target_seq | PAM | alignment | distance | position | gene name | gene id |
| GU305914.1 | 32053 | 32075 | - | 0 | ACACTGCGCGATGGCCTCGG | AGG | \|\|\|\|\|\|\|\|[\|\|\|\|\|\|\|\|\|\|\|\|]PAM | NA | Intergenic | NA | NA |
| T31916 | CTGCGCGATGGCCTCGGAGGCGG | 1000 | | | | | | | | | |
| Oligo fwd | CACCCTGCGCGATGGCCTCGGAGG | | | | | | | | | | |
| Oligo rev | AAACCCTCCGAGGCCATCGCGCAG | | | | | | | | | | |
| Chromosome | start | end | strand | MM | target_seq | PAM | alignment | distance | position | gene name | gene id |
| GU305914.1 | 32050 | 32072 | - | 0 | CTGCGCGATGGCCTCGGAGG | CGG | \|\|\|\|\|\|\|\|[\|\|\|\|\|\|\|\|\|\|\|\|]PAM | NA | Intergenic | NA | NA |
| T31917 | GATGGCCTCGGAGGCGGTCGCGG | 1000 | | | | | | | | | |
| Oligo fwd | CACCGATGGCCTCGGAGGCGGTCG | | | | | | | | | | |
| Oligo rev | AAACCGACCGCCTCCGAGGCCATC | | | | | | | | | | |
| Chromosome | start | end | strand | MM | target_seq | PAM | alignment | distance | position | gene name | gene id |
| GU305914.1 | 32044 | 32066 | - | 0 | GATGGCCTCGGAGGCGGTCG | CGG | \|\|\|\|\|\|\|\|[\|\|\|\|\|\|\|\|\|\|\|\|]PAM | NA | Intergenic | NA | NA |
| T31918 | TCGGAGGCGGTCGCGGTCGCCGG | 1000 | | | | | | | | | |
| Oligo fwd | CACCTCGGAGGCGGTCGCGGTCGC | | | | | | | | | | |
| Oligo rev | AAACGCGACCGCGACCGCCTCCGA | | | | | | | | | | |
| Chromosome | start | end | strand | MM | target_seq | PAM | alignment | distance | position | gene name | gene id |
| GU305914.1 | 32037 | 32059 | - | 0 | TCGGAGGCGGTCGCGGTCGC | CGG | \|\|\|\|\|\|\|\|[\|\|\|\|\|\|\|\|\|\|\|\|]PAM | NA | Intergenic | NA | NA |
| T31919 | GAGGCGGTCGCGGTCGCCGGCGG | 1000 | | | | | | | | | |
| Oligo fwd | CACCGAGGCGGTCGCGGTCGCCGG | | | | | | | | | | |
| Oligo rev | AAACCCGGCGACCGCGACCGCCTC | | | | | | | | | | |
| Chromosome | start | end | strand | MM | target_seq | PAM | alignment | distance | position | gene name | gene id |
| GU305914.1 | 32034 | 32056 | - | 0 | GAGGCGGTCGCGGTCGCCGG | CGG | \|\|\|\|\|\|\|\|[\|\|\|\|\|\|\|\|\|\|\|\|]PAM | NA | Intergenic | NA | NA |
| T31920 | CGCGGTCGCCGGCGGTTCCAAGG | 1000 | | | | | | | | | |
| Oligo fwd | CACCCGCGGTCGCCGGCGGTTCCA | | | | | | | | | | |
| Oligo rev | AAACTGGAACCGCCGGCGACCGCG | | | | | | | | | | |
| Chromosome | start | end | strand | MM | target_seq | PAM | alignment | distance | position | gene name | gene id |
| GU305914.1 | 32026 | 32048 | - | 0 | CGCGGTCGCCGGCGGTTCCA | AGG | \|\|\|\|\|\|\|\|[\|\|\|\|\|\|\|\|\|\|\|\|]PAM | NA | Intergenic | NA | NA |
| T31921 | GCGGTCGCCGGCGGTTCCAAGGG | 1000 | | | | | | | | | |
| Oligo fwd | CACCGCGGTCGCCGGCGGTTCCAA | | | | | | | | | | |
| Oligo rev | AAACTTGGAACCGCCGGCGACCGC | | | | | | | | | | |
| Chromosome | start | end | strand | MM | target_seq | PAM | alignment | distance | position | gene name | gene id |
| GU305914.1 | 32025 | 32047 | - | 0 | GCGGTCGCCGGCGGTTCCAA | GGG | \|\|\|\|\|\|\|\|[\|\|\|\|\|\|\|\|\|\|\|\|]PAM | NA | Intergenic | NA | NA |
| T31922 | CGGTCGCCGGCGGTTCCAAGGGG | 1000 | | | | | | | | | |
| Oligo fwd | CACCCGGTCGCCGGCGGTTCCAAG | | | | | | | | | | |
| Oligo rev | AAACCTTGGAACCGCCGGCGACCG | | | | | | | | | | |
| Chromosome | start | end | strand | MM | target_seq | PAM | alignment | distance | position | gene name | gene id |
| GU305914.1 | 32024 | 32046 | - | 0 | CGGTCGCCGGCGGTTCCAAG | GGG | \|\|\|\|\|\|\|\|[\|\|\|\|\|\|\|\|\|\|\|\|]PAM | NA | Intergenic | NA | NA |
| T31923 | CGCCGGCGGTTCCAAGGGGATGG | 1000 | | | | | | | | | |
| Oligo fwd | CACCCGCCGGCGGTTCCAAGGGGA | | | | | | | | | | |
| Oligo rev | AAACTCCCCTTGGAACCGCCGGCG | | | | | | | | | | |
| Chromosome | start | end | strand | MM | target_seq | PAM | alignment | distance | position | gene name | gene id |
| GU305914.1 | 32020 | 32042 | - | 0 | CGCCGGCGGTTCCAAGGGGA | TGG | \|\|\|\|\|\|\|\|[\|\|\|\|\|\|\|\|\|\|\|\|]PAM | NA | Intergenic | NA | NA |
| GU305914.1 | 223198 | 223220 | - | 4 | CGTCGCCGGTTTCACGGGGA | TGG | \|\|-\|\|-\|\|[\|\|\|-\|\|-\|\|\|\|\|]PAM | NA | Intergenic | NA | NA |
| T31924 | GTTCCAAGGGGATGGAGTTGCGG | 1000 | | | | | | | | | |
| Oligo fwd | CACCGTTCCAAGGGGATGGAGTTG | | | | | | | | | | |
| Oligo rev | AAACCAACTCCATCCCCTTGGAAC | | | | | | | | | | |
| Chromosome | start | end | strand | MM | target_seq | PAM | alignment | distance | position | gene name | gene id |
| GU305914.1 | 32012 | 32034 | - | 0 | GTTCCAAGGGGATGGAGTTG | CGG | \|\|\|\|\|\|\|\|[\|\|\|\|\|\|\|\|\|\|\|\|]PAM | NA | Intergenic | NA | NA |
| T31925 | GTTGCGGACGCAGTTGCGTCTGG | 1000 | | | | | | | | | |
| Oligo fwd | CACCGTTGCGGACGCAGTTGCGTC | | | | | | | | | | |
| Oligo rev | AAACGACGCAACTGCGTCCGCAAC | | | | | | | | | | |
| Chromosome | start | end | strand | MM | target_seq | PAM | alignment | distance | position | gene name | gene id |
| GU305914.1 | 31996 | 32018 | - | 0 | GTTGCGGACGCAGTTGCGTC | TGG | \|\|\|\|\|\|\|\|[\|\|\|\|\|\|\|\|\|\|\|\|]PAM | NA | Intergenic | NA | NA |

<table>
<tr><td>T31926</td><td>GTTGCGTCTGGATCTGATCGAGG</td><td>1000</td><td></td><td></td><td></td><td></td><td></td><td></td><td></td><td></td><td></td></tr>
<tr><td>Oligo fwd</td><td>CACCGTTGCGTCTGGATCTGATCG</td><td></td><td></td><td></td><td></td><td></td><td></td><td></td><td></td><td></td><td></td></tr>
<tr><td>Oligo rev</td><td>AAACCGATCAGATCCAGACGCAAC</td><td></td><td></td><td></td><td></td><td></td><td></td><td></td><td></td><td></td><td></td></tr>
<tr><td>Chromosome</td><td>start</td><td>end</td><td>strand</td><td>MM</td><td>target_seq</td><td>PAM</td><td>alignment</td><td>distance</td><td>position</td><td>gene name</td><td>gene id</td></tr>
<tr><td>GU305914.1</td><td>31984</td><td>32006</td><td>-</td><td>0</td><td>GTTGCGTCTGGATCTGATCG</td><td>AGG</td><td>||||||||[||||||||||||]PAM</td><td>NA</td><td>Intergenic</td><td>NA</td><td>NA</td></tr>
<tr><td>GU305914.1</td><td>172021</td><td>172043</td><td>-</td><td>4</td><td>GTTACGTTTGGTTCTGTTCG</td><td>CGG</td><td>|||-|||-[|||-||||-|||]PAM</td><td>NA</td><td>Intergenic</td><td>NA</td><td>NA</td></tr>
<tr><td></td><td></td><td></td><td></td><td></td><td></td><td></td><td></td><td></td><td></td><td></td><td></td></tr>
<tr><td>T31927</td><td>GCGTCTGGATCTGATCGAGGCGG</td><td>1000</td><td></td><td></td><td></td><td></td><td></td><td></td><td></td><td></td><td></td></tr>
<tr><td>Oligo fwd</td><td>CACCGCGTCTGGATCTGATCGAGG</td><td></td><td></td><td></td><td></td><td></td><td></td><td></td><td></td><td></td><td></td></tr>
<tr><td>Oligo rev</td><td>AAACCCTCGATCAGATCCAGACGC</td><td></td><td></td><td></td><td></td><td></td><td></td><td></td><td></td><td></td><td></td></tr>
<tr><td>Chromosome</td><td>start</td><td>end</td><td>strand</td><td>MM</td><td>target_seq</td><td>PAM</td><td>alignment</td><td>distance</td><td>position</td><td>gene name</td><td>gene id</td></tr>
<tr><td>GU305914.1</td><td>31981</td><td>32003</td><td>-</td><td>0</td><td>GCGTCTGGATCTGATCGAGG</td><td>CGG</td><td>||||||||[||||||||||||]PAM</td><td>NA</td><td>Intergenic</td><td>NA</td><td>NA</td></tr>
<tr><td></td><td></td><td></td><td></td><td></td><td></td><td></td><td></td><td></td><td></td><td></td><td></td></tr>
<tr><td>T31928</td><td>ATCGAGGCGGATTCGCCCTACGG</td><td>1000</td><td></td><td></td><td></td><td></td><td></td><td></td><td></td><td></td><td></td></tr>
<tr><td>Oligo fwd</td><td>CACCATCGAGGCGGATTCGCCCTA</td><td></td><td></td><td></td><td></td><td></td><td></td><td></td><td></td><td></td><td></td></tr>
<tr><td>Oligo rev</td><td>AAACTAGGGCGAATCCGCCTCGAT</td><td></td><td></td><td></td><td></td><td></td><td></td><td></td><td></td><td></td><td></td></tr>
<tr><td>Chromosome</td><td>start</td><td>end</td><td>strand</td><td>MM</td><td>target_seq</td><td>PAM</td><td>alignment</td><td>distance</td><td>position</td><td>gene name</td><td>gene id</td></tr>
<tr><td>GU305914.1</td><td>31968</td><td>31990</td><td>-</td><td>0</td><td>ATCGAGGCGGATTCGCCCTA</td><td>CGG</td><td>||||||||[||||||||||||]PAM</td><td>NA</td><td>Intergenic</td><td>NA</td><td>NA</td></tr>
<tr><td></td><td></td><td></td><td></td><td></td><td></td><td></td><td></td><td></td><td></td><td></td><td></td></tr>
<tr><td>T31929</td><td>CTACGGCGCGTCCCAGAACGAGG</td><td>1000</td><td></td><td></td><td></td><td></td><td></td><td></td><td></td><td></td><td></td></tr>
<tr><td>Oligo fwd</td><td>CACCCTACGGCGCGTCCCAGAACG</td><td></td><td></td><td></td><td></td><td></td><td></td><td></td><td></td><td></td><td></td></tr>
<tr><td>Oligo rev</td><td>AAACCGTTCTGGGACGCGCCGTAG</td><td></td><td></td><td></td><td></td><td></td><td></td><td></td><td></td><td></td><td></td></tr>
<tr><td>Chromosome</td><td>start</td><td>end</td><td>strand</td><td>MM</td><td>target_seq</td><td>PAM</td><td>alignment</td><td>distance</td><td>position</td><td>gene name</td><td>gene id</td></tr>
<tr><td>GU305914.1</td><td>31951</td><td>31973</td><td>-</td><td>0</td><td>CTACGGCGCGTCCCAGAACG</td><td>AGG</td><td>||||||||[||||||||||||]PAM</td><td>NA</td><td>Intergenic</td><td>NA</td><td>NA</td></tr>
<tr><td></td><td></td><td></td><td></td><td></td><td></td><td></td><td></td><td></td><td></td><td></td><td></td></tr>
<tr><td>T31930</td><td>CCAGAACGAGGTCTGCGAGCTGG</td><td>1000</td><td></td><td></td><td></td><td></td><td></td><td></td><td></td><td></td><td></td></tr>
<tr><td>Oligo fwd</td><td>CACCCCAGAACGAGGTCTGCGAGC</td><td></td><td></td><td></td><td></td><td></td><td></td><td></td><td></td><td></td><td></td></tr>
<tr><td>Oligo rev</td><td>AAACGCTCGCAGACCTCGTTCTGG</td><td></td><td></td><td></td><td></td><td></td><td></td><td></td><td></td><td></td><td></td></tr>
<tr><td>Chromosome</td><td>start</td><td>end</td><td>strand</td><td>MM</td><td>target_seq</td><td>PAM</td><td>alignment</td><td>distance</td><td>position</td><td>gene name</td><td>gene id</td></tr>
<tr><td>GU305914.1</td><td>31939</td><td>31961</td><td>-</td><td>0</td><td>CCAGAACGAGGTCTGCGAGC</td><td>TGG</td><td>||||||||[||||||||||||]PAM</td><td>NA</td><td>Intergenic</td><td>NA</td><td>NA</td></tr>
<tr><td></td><td></td><td></td><td></td><td></td><td></td><td></td><td></td><td></td><td></td><td></td><td></td></tr>
<tr><td>T31931</td><td>GCTGGTCATCGCCGCCGCCATGG</td><td>1000</td><td></td><td></td><td></td><td></td><td></td><td></td><td></td><td></td><td></td></tr>
<tr><td>Oligo fwd</td><td>CACCGCTGGTCATCGCCGCCGCCA</td><td></td><td></td><td></td><td></td><td></td><td></td><td></td><td></td><td></td><td></td></tr>
<tr><td>Oligo rev</td><td>AAACTGGCGGCGGCGATGACCAGC</td><td></td><td></td><td></td><td></td><td></td><td></td><td></td><td></td><td></td><td></td></tr>
<tr><td>Chromosome</td><td>start</td><td>end</td><td>strand</td><td>MM</td><td>target_seq</td><td>PAM</td><td>alignment</td><td>distance</td><td>position</td><td>gene name</td><td>gene id</td></tr>
<tr><td>GU305914.1</td><td>31921</td><td>31943</td><td>-</td><td>0</td><td>GCTGGTCATCGCCGCCGCCA</td><td>TGG</td><td>||||||||[||||||||||||]PAM</td><td>NA</td><td>Intergenic</td><td>NA</td><td>NA</td></tr>
<tr><td></td><td></td><td></td><td></td><td></td><td></td><td></td><td></td><td></td><td></td><td></td><td></td></tr>
<tr><td>T31932</td><td>CTGGTCATCGCCGCCGCCATGGG</td><td>1000</td><td></td><td></td><td></td><td></td><td></td><td></td><td></td><td></td><td></td></tr>
<tr><td>Oligo fwd</td><td>CACCCTGGTCATCGCCGCCGCCAT</td><td></td><td></td><td></td><td></td><td></td><td></td><td></td><td></td><td></td><td></td></tr>
<tr><td>Oligo rev</td><td>AAACATGGCGGCGGCGATGACCAG</td><td></td><td></td><td></td><td></td><td></td><td></td><td></td><td></td><td></td><td></td></tr>
<tr><td>Chromosome</td><td>start</td><td>end</td><td>strand</td><td>MM</td><td>target_seq</td><td>PAM</td><td>alignment</td><td>distance</td><td>position</td><td>gene name</td><td>gene id</td></tr>
<tr><td>GU305914.1</td><td>31920</td><td>31942</td><td>-</td><td>0</td><td>CTGGTCATCGCCGCCGCCAT</td><td>GGG</td><td>||||||||[||||||||||||]PAM</td><td>NA</td><td>Intergenic</td><td>NA</td><td>NA</td></tr>
<tr><td></td><td></td><td></td><td></td><td></td><td></td><td></td><td></td><td></td><td></td><td></td><td></td></tr>
<tr><td>T31933</td><td>GTCATCGCCGCCGCCATGGGTGG</td><td>1000</td><td></td><td></td><td></td><td></td><td></td><td></td><td></td><td></td><td></td></tr>
<tr><td>Oligo fwd</td><td>CACCGTCATCGCCGCCGCCATGGG</td><td></td><td></td><td></td><td></td><td></td><td></td><td></td><td></td><td></td><td></td></tr>
<tr><td>Oligo rev</td><td>AAACCCCATGGCGGCGGCGATGAC</td><td></td><td></td><td></td><td></td><td></td><td></td><td></td><td></td><td></td><td></td></tr>
<tr><td>Chromosome</td><td>start</td><td>end</td><td>strand</td><td>MM</td><td>target_seq</td><td>PAM</td><td>alignment</td><td>distance</td><td>position</td><td>gene name</td><td>gene id</td></tr>
<tr><td>GU305914.1</td><td>31917</td><td>31939</td><td>-</td><td>0</td><td>GTCATCGCCGCCGCCATGGG</td><td>TGG</td><td>||||||||[||||||||||||]PAM</td><td>NA</td><td>Intergenic</td><td>NA</td><td>NA</td></tr>
<tr><td></td><td></td><td></td><td></td><td></td><td></td><td></td><td></td><td></td><td></td><td></td><td></td></tr>
<tr><td>T31934</td><td>CCATGGGTGGCGAATGCCTGAGG</td><td>1000</td><td></td><td></td><td></td><td></td><td></td><td></td><td></td><td></td><td></td></tr>
<tr><td>Oligo fwd</td><td>CACCCCATGGGTGGCGAATGCCTG</td><td></td><td></td><td></td><td></td><td></td><td></td><td></td><td></td><td></td><td></td></tr>
<tr><td>Oligo rev</td><td>AAACCAGGCATTCGCCACCCATGG</td><td></td><td></td><td></td><td></td><td></td><td></td><td></td><td></td><td></td><td></td></tr>
<tr><td>Chromosome</td><td>start</td><td>end</td><td>strand</td><td>MM</td><td>target_seq</td><td>PAM</td><td>alignment</td><td>distance</td><td>position</td><td>gene name</td><td>gene id</td></tr>
<tr><td>GU305914.1</td><td>31904</td><td>31926</td><td>-</td><td>0</td><td>CCATGGGTGGCGAATGCCTG</td><td>AGG</td><td>||||||||[||||||||||||]PAM</td><td>NA</td><td>Intergenic</td><td>NA</td><td>NA</td></tr>
<tr><td></td><td></td><td></td><td></td><td></td><td></td><td></td><td></td><td></td><td></td><td></td><td></td></tr>
<tr><td>T31935</td><td>CATGGGTGGCGAATGCCTGAGGG</td><td>1000</td><td></td><td></td><td></td><td></td><td></td><td></td><td></td><td></td><td></td></tr>
<tr><td>Oligo fwd</td><td>CACCCATGGGTGGCGAATGCCTGA</td><td></td><td></td><td></td><td></td><td></td><td></td><td></td><td></td><td></td><td></td></tr>
<tr><td>Oligo rev</td><td>AAACTCAGGCATTCGCCACCCATG</td><td></td><td></td><td></td><td></td><td></td><td></td><td></td><td></td><td></td><td></td></tr>
<tr><td>Chromosome</td><td>start</td><td>end</td><td>strand</td><td>MM</td><td>target_seq</td><td>PAM</td><td>alignment</td><td>distance</td><td>position</td><td>gene name</td><td>gene id</td></tr>
<tr><td>GU305914.1</td><td>31903</td><td>31925</td><td>-</td><td>0</td><td>CATGGGTGGCGAATGCCTGA</td><td>GGG</td><td>||||||||[||||||||||||]PAM</td><td>NA</td><td>Intergenic</td><td>NA</td><td>NA</td></tr>
<tr><td></td><td></td><td></td><td></td><td></td><td></td><td></td><td></td><td></td><td></td><td></td><td></td></tr>
<tr><td>T31936</td><td>ATGGGTGGCGAATGCCTGAGGGG</td><td>1000</td><td></td><td></td><td></td><td></td><td></td><td></td><td></td><td></td><td></td></tr>
<tr><td>Oligo fwd</td><td>CACCATGGGTGGCGAATGCCTGAG</td><td></td><td></td><td></td><td></td><td></td><td></td><td></td><td></td><td></td><td></td></tr>
<tr><td>Oligo rev</td><td>AAACCTCAGGCATTCGCCACCCAT</td><td></td><td></td><td></td><td></td><td></td><td></td><td></td><td></td><td></td><td></td></tr>
<tr><td>Chromosome</td><td>start</td><td>end</td><td>strand</td><td>MM</td><td>target_seq</td><td>PAM</td><td>alignment</td><td>distance</td><td>position</td><td>gene name</td><td>gene id</td></tr>
<tr><td>GU305914.1</td><td>31902</td><td>31924</td><td>-</td><td>0</td><td>ATGGGTGGCGAATGCCTGAG</td><td>GGG</td><td>||||||||[||||||||||||]PAM</td><td>NA</td><td>Intergenic</td><td>NA</td><td>NA</td></tr>
<tr><td></td><td></td><td></td><td></td><td></td><td></td><td></td><td></td><td></td><td></td><td></td><td></td></tr>
<tr><td>T31937</td><td>GGGGATGTGTACAGAGACTCAGG</td><td>1000</td><td></td><td></td><td></td><td></td><td></td><td></td><td></td><td></td><td></td></tr>
<tr><td>Oligo fwd</td><td>CACCGGGGATGTGTACAGAGACTC</td><td></td><td></td><td></td><td></td><td></td><td></td><td></td><td></td><td></td><td></td></tr>
<tr><td>Oligo rev</td><td>AAACGAGTCTCTGTACACATCCCC</td><td></td><td></td><td></td><td></td><td></td><td></td><td></td><td></td><td></td><td></td></tr>
<tr><td>Chromosome</td><td>start</td><td>end</td><td>strand</td><td>MM</td><td>target_seq</td><td>PAM</td><td>alignment</td><td>distance</td><td>position</td><td>gene name</td><td>gene id</td></tr>
<tr><td>GU305914.1</td><td>31883</td><td>31905</td><td>-</td><td>0</td><td>GGGGATGTGTACAGAGACTC</td><td>AGG</td><td>||||||||[||||||||||||]PAM</td><td>NA</td><td>Intergenic</td><td>NA</td><td>NA</td></tr>
<tr><td></td><td></td><td></td><td></td><td></td><td></td><td></td><td></td><td></td><td></td><td></td><td></td></tr>
<tr><td>T31938</td><td>GGGATGTGTACAGAGACTCAGGG</td><td>1000</td><td></td><td></td><td></td><td></td><td></td><td></td><td></td><td></td><td></td></tr>
<tr><td>Oligo fwd</td><td>CACCGGGATGTGTACAGAGACTCA</td><td></td><td></td><td></td><td></td><td></td><td></td><td></td><td></td><td></td><td></td></tr>
<tr><td>Oligo rev</td><td>AAACTGAGTCTCTGTACACATCCC</td><td></td><td></td><td></td><td></td><td></td><td></td><td></td><td></td><td></td><td></td></tr>
<tr><td>Chromosome</td><td>start</td><td>end</td><td>strand</td><td>MM</td><td>target_seq</td><td>PAM</td><td>alignment</td><td>distance</td><td>position</td><td>gene name</td><td>gene id</td></tr>
<tr><td>GU305914.1</td><td>31882</td><td>31904</td><td>-</td><td>0</td><td>GGGATGTGTACAGAGACTCA</td><td>GGG</td><td>||||||||[||||||||||||]PAM</td><td>NA</td><td>Intergenic</td><td>NA</td><td>NA</td></tr>
<tr><td></td><td></td><td></td><td></td><td></td><td></td><td></td><td></td><td></td><td></td><td></td><td></td></tr>
<tr><td>T31939</td><td>GGATGTGTACAGAGACTCAGGGG</td><td>1000</td><td></td><td></td><td></td><td></td><td></td><td></td><td></td><td></td><td></td></tr>
<tr><td>Oligo fwd</td><td>CACCGGATGTGTACAGAGACTCAG</td><td></td><td></td><td></td><td></td><td></td><td></td><td></td><td></td><td></td><td></td></tr>
<tr><td>Oligo rev</td><td>AAACCTGAGTCTCTGTACACATCC</td><td></td><td></td><td></td><td></td><td></td><td></td><td></td><td></td><td></td><td></td></tr>
<tr><td>Chromosome</td><td>start</td><td>end</td><td>strand</td><td>MM</td><td>target_seq</td><td>PAM</td><td>alignment</td><td>distance</td><td>position</td><td>gene name</td><td>gene id</td></tr>
<tr><td>GU305914.1</td><td>31881</td><td>31903</td><td>-</td><td>0</td><td>GGATGTGTACAGAGACTCAG</td><td>GGG</td><td>||||||||[||||||||||||]PAM</td><td>NA</td><td>Intergenic</td><td>NA</td><td>NA</td></tr>
<tr><td></td><td></td><td></td><td></td><td></td><td></td><td></td><td></td><td></td><td></td><td></td><td></td></tr>
<tr><td>T31940</td><td>ACAGAGACTCAGGGGAATAGAGG</td><td>1000</td><td></td><td></td><td></td><td></td><td></td><td></td><td></td><td></td><td></td></tr>
<tr><td>Oligo fwd</td><td>CACCACAGAGACTCAGGGGAATAG</td><td></td><td></td><td></td><td></td><td></td><td></td><td></td><td></td><td></td><td></td></tr>
<tr><td>Oligo rev</td><td>AAACCTATTCCCCTGAGTCTCTGT</td><td></td><td></td><td></td><td></td><td></td><td></td><td></td><td></td><td></td><td></td></tr>
</table>

Chromosome start end strand MM target_seq PAM alignment distance position gene name gene id
GU305914.1 31873 31895 - 0 ACAGAGACTCAGGGGAATAG AGG ||||||||[||||||||||||]PAM NA Intergenic NA NA

T31941 GGTCGCGCTCCCTTTCCCCACGG 1000
Oligo fwd CACCGGTCGCGCTCCCTTTCCCCA
Oligo rev AAACTGGGGAAAGGGAGCGCGACC
Chromosome start end strand MM target_seq PAM alignment distance position gene name gene id
GU305914.1 31852 31874 - 0 GGTCGCGCTCCCTTTCCCCA CGG ||||||||[||||||||||||]PAM NA Intergenic NA NA

T31942 GTCGCGCTCCCTTTCCCCACGGG 1000
Oligo fwd CACCGTCGCGCTCCCTTTCCCCAC
Oligo rev AAACGTGGGGAAAGGGAGCGCGAC
Chromosome start end strand MM target_seq PAM alignment distance position gene name gene id
GU305914.1 31851 31873 - 0 GTCGCGCTCCCTTTCCCCAC GGG ||||||||[||||||||||||]PAM NA Intergenic NA NA

T31943 TCGCGCTCCCTTTCCCCACGGGG 1000
Oligo fwd CACCTCGCGCTCCCTTTCCCCACG
Oligo rev AAACCGTGGGGAAAGGGAGCGCGA
Chromosome start end strand MM target_seq PAM alignment distance position gene name gene id
GU305914.1 31850 31872 - 0 TCGCGCTCCCTTTCCCCACG GGG ||||||||[||||||||||||]PAM NA Intergenic NA NA

T31944 ACGGGGTACATCTTTTACATCGG 1000
Oligo fwd CACCACGGGGTACATCTTTTACAT
Oligo rev AAACATGTAAAAGATGTACCCCGT
Chromosome start end strand MM target_seq PAM alignment distance position gene name gene id
GU305914.1 31833 31855 - 0 ACGGGGTACATCTTTTACAT CGG ||||||||[||||||||||||]PAM NA Intergenic NA NA

T31945 CGGGGTACATCTTTTACATCGGG 1000
Oligo fwd CACCCGGGGTACATCTTTTACATC
Oligo rev AAACGATGTAAAAGATGTACCCCG
Chromosome start end strand MM target_seq PAM alignment distance position gene name gene id
GU305914.1 31832 31854 - 0 CGGGGTACATCTTTTACATC GGG ||||||||[||||||||||||]PAM NA Intergenic NA NA

T31946 GGGGTACATCTTTTACATCGGGG 1000
Oligo fwd CACCGGGGTACATCTTTTACATCG
Oligo rev AAACCGATGTAAAAGATGTACCCC
Chromosome start end strand MM target_seq PAM alignment distance position gene name gene id
GU305914.1 31831 31853 - 0 GGGGTACATCTTTTACATCG GGG ||||||||[||||||||||||]PAM NA Intergenic NA NA

T31947 GGGTACATCTTTTACATCGGGGG 1000
Oligo fwd CACCGGGTACATCTTTTACATCGG
Oligo rev AAACCCGATGTAAAAGATGTACCC
Chromosome start end strand MM target_seq PAM alignment distance position gene name gene id
GU305914.1 31830 31852 - 0 GGGTACATCTTTTACATCGG GGG ||||||||[||||||||||||]PAM NA Intergenic NA NA

T31948 CATCTTTTACATCGGGGGCGAGG 1000
Oligo fwd CACCCATCTTTTACATCGGGGGCG
Oligo rev AAACCGCCCCCGATGTAAAAGATG
Chromosome start end strand MM target_seq PAM alignment distance position gene name gene id
GU305914.1 31825 31847 - 0 CATCTTTTACATCGGGGGCG AGG ||||||||[||||||||||||]PAM NA Intergenic NA NA

T31949 CTTTTACATCGGGGGCGAGGAGG 1000
Oligo fwd CACCCTTTTACATCGGGGGCGAGG
Oligo rev AAACCCTCGCCCCCGATGTAAAAG
Chromosome start end strand MM target_seq PAM alignment distance position gene name gene id
GU305914.1 31822 31844 - 0 CTTTTACATCGGGGGCGAGG AGG ||||||||[||||||||||||]PAM NA Intergenic NA NA

T31950 CCTCGTGCGATCTCCATTTCTGG 1000
Oligo fwd CACCCCTCGTGCGATCTCCATTTC
Oligo rev AAACGAAATGGAGATCGCACGAGG
Chromosome start end strand MM target_seq PAM alignment distance position gene name gene id
GU305914.1 31787 31809 - 0 CCTCGTGCGATCTCCATTTC TGG ||||||||[||||||||||||]PAM NA Intergenic NA NA

T31951 CGTGCGATCTCCATTTCTGGCGG 1000
Oligo fwd CACCCGTGCGATCTCCATTTCTGG
Oligo rev AAACCCAGAAATGGAGATCGCACG
Chromosome start end strand MM target_seq PAM alignment distance position gene name gene id
GU305914.1 31784 31806 - 0 CGTGCGATCTCCATTTCTGG CGG ||||||||[||||||||||||]PAM NA Intergenic NA NA

T31952 CGGATGCACCCGCTCATGTGCGG 1000
Oligo fwd CACCCGGATGCACCCGCTCATGTG
Oligo rev AAACCACATGAGCGGGTGCATCCG
Chromosome start end strand MM target_seq PAM alignment distance position gene name gene id
GU305914.1 31764 31786 - 0 CGGATGCACCCGCTCATGTG CGG ||||||||[||||||||||||]PAM NA Intergenic NA NA

T31953 GGATGCACCCGCTCATGTGCGGG 1000
Oligo fwd CACCGGATGCACCCGCTCATGTGC
Oligo rev AAACGCACATGAGCGGGTGCATCC
Chromosome start end strand MM target_seq PAM alignment distance position gene name gene id
GU305914.1 31763 31785 - 0 GGATGCACCCGCTCATGTGC GGG ||||||||[||||||||||||]PAM NA Intergenic NA NA

T31954 GATGCACCCGCTCATGTGCGGGG 1000
Oligo fwd CACCGATGCACCCGCTCATGTGCG
Oligo rev AAACCGCACATGAGCGGGTGCATC
Chromosome start end strand MM target_seq PAM alignment distance position gene name gene id
GU305914.1 31762 31784 - 0 GATGCACCCGCTCATGTGCG GGG ||||||||[||||||||||||]PAM NA Intergenic NA NA

T31955 TGCGGGGAGCTCACCGTCATCGG 1000
Oligo fwd CACCTGCGGGGAGCTCACCGTCAT

| | | | | | | | | | | | | |
|---|---|---|---|---|---|---|---|---|---|---|---|---|
| Oligo rev | AAACATGACGGTGAGCTCCCCGCA | | | | | | | | | | | |
| Chromosome | start | | end | strand | MM | target_seq | PAM | alignment | distance | position | gene name | gene id |
| GU305914.1 | | 31746 | 31768 | - | 0 | TGCGGGGAGCTCACCGTCAT | CGG | \|\|\|\|\|\|\|\|[\|\|\|\|\|\|\|\|\|\|\|\|]PAM | NA | Intergenic | NA | NA |
| T31956 | CTCACCGTCATCGGCACGCTCGG | | 1000 | | | | | | | | | |
| Oligo fwd | CACCCTCACCGTCATCGGCACGCT | | | | | | | | | | | |
| Oligo rev | AAACAGCGTGCCGATGACGGTGAG | | | | | | | | | | | |
| Chromosome | start | | end | strand | MM | target_seq | PAM | alignment | distance | position | gene name | gene id |
| GU305914.1 | | 31737 | 31759 | - | 0 | CTCACCGTCATCGGCACGCT | CGG | \|\|\|\|\|\|\|\|[\|\|\|\|\|\|\|\|\|\|\|\|]PAM | NA | Intergenic | NA | NA |
| T31957 | TCACCGTCATCGGCACGCTCGGG | | 1000 | | | | | | | | | |
| Oligo fwd | CACCTCACCGTCATCGGCACGCTC | | | | | | | | | | | |
| Oligo rev | AAACGAGCGTGCCGATGACGGTGA | | | | | | | | | | | |
| Chromosome | start | | end | strand | MM | target_seq | PAM | alignment | distance | position | gene name | gene id |
| GU305914.1 | | 31736 | 31758 | - | 0 | TCACCGTCATCGGCACGCTC | GGG | \|\|\|\|\|\|\|\|[\|\|\|\|\|\|\|\|\|\|\|\|]PAM | NA | Intergenic | NA | NA |
| T31958 | GGCACCGACGCACCTTCGCTTGG | | 1000 | | | | | | | | | |
| Oligo fwd | CACCGGCACCGACGCACCTTCGCT | | | | | | | | | | | |
| Oligo rev | AAACAGCGAAGGTGCGTCGGTGCC | | | | | | | | | | | |
| Chromosome | start | | end | strand | MM | target_seq | PAM | alignment | distance | position | gene name | gene id |
| GU305914.1 | | 31715 | 31737 | - | 0 | GGCACCGACGCACCTTCGCT | TGG | \|\|\|\|\|\|\|\|[\|\|\|\|\|\|\|\|\|\|\|\|]PAM | NA | Intergenic | NA | NA |
| T31959 | GCACCGACGCACCTTCGCTTGGG | | 1000 | | | | | | | | | |
| Oligo fwd | CACCGCACCGACGCACCTTCGCTT | | | | | | | | | | | |
| Oligo rev | AAACAAGCGAAGGTGCGTCGGTGC | | | | | | | | | | | |
| Chromosome | start | | end | strand | MM | target_seq | PAM | alignment | distance | position | gene name | gene id |
| GU305914.1 | | 31714 | 31736 | - | 0 | GCACCGACGCACCTTCGCTT | GGG | \|\|\|\|\|\|\|\|[\|\|\|\|\|\|\|\|\|\|\|\|]PAM | NA | Intergenic | NA | NA |
| T31960 | GCACCTTCGCTTGGGACCGCCGG | | 1000 | | | | | | | | | |
| Oligo fwd | CACCGCACCTTCGCTTGGGACCGC | | | | | | | | | | | |
| Oligo rev | AAACGCGGTCCCAAGCGAAGGTGC | | | | | | | | | | | |
| Chromosome | start | | end | strand | MM | target_seq | PAM | alignment | distance | position | gene name | gene id |
| GU305914.1 | | 31706 | 31728 | - | 0 | GCACCTTCGCTTGGGACCGC | CGG | \|\|\|\|\|\|\|\|[\|\|\|\|\|\|\|\|\|\|\|\|]PAM | NA | Intergenic | NA | NA |
| T31961 | CACCTTCGCTTGGGACCGCCGGG | | 1000 | | | | | | | | | |
| Oligo fwd | CACCCACCTTCGCTTGGGACCGCC | | | | | | | | | | | |
| Oligo rev | AAACGGCGGTCCCAAGCGAAGGTG | | | | | | | | | | | |
| Chromosome | start | | end | strand | MM | target_seq | PAM | alignment | distance | position | gene name | gene id |
| GU305914.1 | | 31705 | 31727 | - | 0 | CACCTTCGCTTGGGACCGCC | GGG | \|\|\|\|\|\|\|\|[\|\|\|\|\|\|\|\|\|\|\|\|]PAM | NA | Intergenic | NA | NA |
| T31962 | CTTCGCTTGGGACCGCCGGGTGG | | 1000 | | | | | | | | | |
| Oligo fwd | CACCCTTCGCTTGGGACCGCCGGG | | | | | | | | | | | |
| Oligo rev | AAACCCCGGCGGTCCCAAGCGAAG | | | | | | | | | | | |
| Chromosome | start | | end | strand | MM | target_seq | PAM | alignment | distance | position | gene name | gene id |
| GU305914.1 | | 31702 | 31724 | - | 0 | CTTCGCTTGGGACCGCCGGG | TGG | \|\|\|\|\|\|\|\|[\|\|\|\|\|\|\|\|\|\|\|\|]PAM | NA | Intergenic | NA | NA |
| T31963 | GCTTGGGACCGCCGGGTGGTCGG | | 1000 | | | | | | | | | |
| Oligo fwd | CACCGCTTGGGACCGCCGGGTGGT | | | | | | | | | | | |
| Oligo rev | AAACACCACCCGGCGGTCCCAAGC | | | | | | | | | | | |
| Chromosome | start | | end | strand | MM | target_seq | PAM | alignment | distance | position | gene name | gene id |
| GU305914.1 | | 31698 | 31720 | - | 0 | GCTTGGGACCGCCGGGTGGT | CGG | \|\|\|\|\|\|\|\|[\|\|\|\|\|\|\|\|\|\|\|\|]PAM | NA | Intergenic | NA | NA |
| T31964 | GGTGGTCGGCGTCAACGCCGTGG | | 1000 | | | | | | | | | |
| Oligo fwd | CACCGGTGGTCGGCGTCAACGCCG | | | | | | | | | | | |
| Oligo rev | AAACCGGCGTTGACGCCGACCACC | | | | | | | | | | | |
| Chromosome | start | | end | strand | MM | target_seq | PAM | alignment | distance | position | gene name | gene id |
| GU305914.1 | | 31684 | 31706 | - | 0 | GGTGGTCGGCGTCAACGCCG | TGG | \|\|\|\|\|\|\|\|[\|\|\|\|\|\|\|\|\|\|\|\|]PAM | NA | Intergenic | NA | NA |
| T31965 | GTGGTCGGCGTCAACGCCGTGGG | | 1000 | | | | | | | | | |
| Oligo fwd | CACCGTGGTCGGCGTCAACGCCGT | | | | | | | | | | | |
| Oligo rev | AAACACGGCGTTGACGCCGACCAC | | | | | | | | | | | |
| Chromosome | start | | end | strand | MM | target_seq | PAM | alignment | distance | position | gene name | gene id |
| GU305914.1 | | 31683 | 31705 | - | 0 | GTGGTCGGCGTCAACGCCGT | GGG | \|\|\|\|\|\|\|\|[\|\|\|\|\|\|\|\|\|\|\|\|]PAM | NA | Intergenic | NA | NA |
| GU305914.1 | | 20748 | 20770 | - | 4 | GTGATCGCCGTCGTCGCCGT | CGG | \|\|\|-\|\|\|-[\|\|\|\|--\|\|\|\|\|\|]PAM | NA | Intergenic | NA | NA |
| T31966 | CGGCGTCAACGCCGTGGGCCAGG | | 1000 | | | | | | | | | |
| Oligo fwd | CACCCGGCGTCAACGCCGTGGGCC | | | | | | | | | | | |
| Oligo rev | AAACGGCCCACGGCGTTGACGCCG | | | | | | | | | | | |
| Chromosome | start | | end | strand | MM | target_seq | PAM | alignment | distance | position | gene name | gene id |
| GU305914.1 | | 31678 | 31700 | - | 0 | CGGCGTCAACGCCGTGGGCC | AGG | \|\|\|\|\|\|\|\|[\|\|\|\|\|\|\|\|\|\|\|\|]PAM | NA | Intergenic | NA | NA |
| GU305914.1 | | 109712 | 109734 | + | 4 | CGTCCTCAACGCCGTGGAGC | AGG | \|\|-\|-\|\|\|[\|\|\|\|\|\|\|\|\|--\|]PAM | NA | Intergenic | NA | NA |
| T31967 | CTGCTTCATCATGTACGTCTCGG | | 1000 | | | | | | | | | |
| Oligo fwd | CACCCTGCTTCATCATGTACGTCT | | | | | | | | | | | |
| Oligo rev | AAACAGACGTACATGATGAAGCAG | | | | | | | | | | | |
| Chromosome | start | | end | strand | MM | target_seq | PAM | alignment | distance | position | gene name | gene id |
| GU305914.1 | | 31630 | 31652 | - | 0 | CTGCTTCATCATGTACGTCT | CGG | \|\|\|\|\|\|\|\|[\|\|\|\|\|\|\|\|\|\|\|\|]PAM | NA | Intergenic | NA | NA |
| T31968 | CAACCTCACGCAGCTCATCTCGG | | 1000 | | | | | | | | | |
| Oligo fwd | CACCCAACCTCACGCAGCTCATCT | | | | | | | | | | | |
| Oligo rev | AAACAGATGAGCTGCGTGAGGTTG | | | | | | | | | | | |
| Chromosome | start | | end | strand | MM | target_seq | PAM | alignment | distance | position | gene name | gene id |
| GU305914.1 | | 31606 | 31628 | - | 0 | CAACCTCACGCAGCTCATCT | CGG | \|\|\|\|\|\|\|\|[\|\|\|\|\|\|\|\|\|\|\|\|]PAM | NA | Intergenic | NA | NA |
| GU305914.1 | | 144806 | 144828 | - | 4 | GAACCTGACCCAGCTGATCT | TGG | -\|\|\|\|\|-\|[\|-\|\|\|\|\|-\|\|\|\|]PAM | NA | Intergenic | NA | NA |
| T31969 | CTCACGCAGCTCATCTCGGACGG | | 1000 | | | | | | | | | |
| Oligo fwd | CACCCTCACGCAGCTCATCTCGGA | | | | | | | | | | | |
| Oligo rev | AAACTCCGAGATGAGCTGCGTGAG | | | | | | | | | | | |
| Chromosome | start | | end | strand | MM | target_seq | PAM | alignment | distance | position | gene name | gene id |

<table>
<tr><td>GU305914.1</td><td>31602</td><td>31624</td><td>-</td><td>0</td><td>CTCACGCAGCTCATCTCGGA</td><td>CGG</td><td>||||||||[||||||||||||]PAM</td><td>NA</td><td>Intergenic</td><td>NA</td><td>NA</td></tr>
<tr><td>T31970</td><td>TCTCGGACGGCGTCTCTAGAAGG</td><td>1000</td><td></td><td></td><td></td><td></td><td></td><td></td><td></td><td></td><td></td></tr>
<tr><td>Oligo fwd</td><td>CACCTCTCGGACGGCGTCTCTAGA</td><td></td><td></td><td></td><td></td><td></td><td></td><td></td><td></td><td></td><td></td></tr>
<tr><td>Oligo rev</td><td>AAACTCTAGAGACGCCGTCCGAGA</td><td></td><td></td><td></td><td></td><td></td><td></td><td></td><td></td><td></td><td></td></tr>
<tr><td>Chromosome</td><td>start</td><td>end</td><td>strand</td><td>MM</td><td>target_seq</td><td>PAM</td><td>alignment</td><td>distance</td><td>position</td><td>gene name</td><td>gene id</td></tr>
<tr><td>GU305914.1</td><td>31589</td><td>31611</td><td>-</td><td>0</td><td>TCTCGGACGGCGTCTCTAGA</td><td>AGG</td><td>||||||||[||||||||||||]PAM</td><td>NA</td><td>Intergenic</td><td>NA</td><td>NA</td></tr>
<tr><td>T31971</td><td>CGTCTCTAGAAGGTACATCCAGG</td><td>1000</td><td></td><td></td><td></td><td></td><td></td><td></td><td></td><td></td><td></td></tr>
<tr><td>Oligo fwd</td><td>CACCCGTCTCTAGAAGGTACATCC</td><td></td><td></td><td></td><td></td><td></td><td></td><td></td><td></td><td></td><td></td></tr>
<tr><td>Oligo rev</td><td>AAACGGATGTACCTTCTAGAGACG</td><td></td><td></td><td></td><td></td><td></td><td></td><td></td><td></td><td></td><td></td></tr>
<tr><td>Chromosome</td><td>start</td><td>end</td><td>strand</td><td>MM</td><td>target_seq</td><td>PAM</td><td>alignment</td><td>distance</td><td>position</td><td>gene name</td><td>gene id</td></tr>
<tr><td>GU305914.1</td><td>31579</td><td>31601</td><td>-</td><td>0</td><td>CGTCTCTAGAAGGTACATCC</td><td>AGG</td><td>||||||||[||||||||||||]PAM</td><td>NA</td><td>Intergenic</td><td>NA</td><td>NA</td></tr>
<tr><td>T31972</td><td>GGTACAGCGCGACCTGCGTGAGG</td><td>1000</td><td></td><td></td><td></td><td></td><td></td><td></td><td></td><td></td><td></td></tr>
<tr><td>Oligo fwd</td><td>CACCGGTACAGCGCGACCTGCGTG</td><td></td><td></td><td></td><td></td><td></td><td></td><td></td><td></td><td></td><td></td></tr>
<tr><td>Oligo rev</td><td>AAACCACGCAGGTCGCGCTGTACC</td><td></td><td></td><td></td><td></td><td></td><td></td><td></td><td></td><td></td><td></td></tr>
<tr><td>Chromosome</td><td>start</td><td>end</td><td>strand</td><td>MM</td><td>target_seq</td><td>PAM</td><td>alignment</td><td>distance</td><td>position</td><td>gene name</td><td>gene id</td></tr>
<tr><td>GU305914.1</td><td>31558</td><td>31580</td><td>-</td><td>0</td><td>GGTACAGCGCGACCTGCGTG</td><td>AGG</td><td>||||||||[||||||||||||]PAM</td><td>NA</td><td>Intergenic</td><td>NA</td><td>NA</td></tr>
<tr><td>GU305914.1</td><td>21506</td><td>21528</td><td>-</td><td>4</td><td>GCAACAGCGGCACCTGCGTG</td><td>TGG</td><td>|--|||||[|--|||||||||]PAM</td><td>NA</td><td>Intergenic</td><td>NA</td><td>NA</td></tr>
<tr><td>T31973</td><td>GTACAGCGCGACCTGCGTGAGGG</td><td>1000</td><td></td><td></td><td></td><td></td><td></td><td></td><td></td><td></td><td></td></tr>
<tr><td>Oligo fwd</td><td>CACCGTACAGCGCGACCTGCGTGA</td><td></td><td></td><td></td><td></td><td></td><td></td><td></td><td></td><td></td><td></td></tr>
<tr><td>Oligo rev</td><td>AAACTCACGCAGGTCGCGCTGTAC</td><td></td><td></td><td></td><td></td><td></td><td></td><td></td><td></td><td></td><td></td></tr>
<tr><td>Chromosome</td><td>start</td><td>end</td><td>strand</td><td>MM</td><td>target_seq</td><td>PAM</td><td>alignment</td><td>distance</td><td>position</td><td>gene name</td><td>gene id</td></tr>
<tr><td>GU305914.1</td><td>31557</td><td>31579</td><td>-</td><td>0</td><td>GTACAGCGCGACCTGCGTGA</td><td>GGG</td><td>||||||||[||||||||||||]PAM</td><td>NA</td><td>Intergenic</td><td>NA</td><td>NA</td></tr>
<tr><td>T31974</td><td>TCTGCCAGTGTAACGAGCCGCGG</td><td>1000</td><td></td><td></td><td></td><td></td><td></td><td></td><td></td><td></td><td></td></tr>
<tr><td>Oligo fwd</td><td>CACCTCTGCCAGTGTAACGAGCCG</td><td></td><td></td><td></td><td></td><td></td><td></td><td></td><td></td><td></td><td></td></tr>
<tr><td>Oligo rev</td><td>AAACCGGCTCGTTACACTGGCAGA</td><td></td><td></td><td></td><td></td><td></td><td></td><td></td><td></td><td></td><td></td></tr>
<tr><td>Chromosome</td><td>start</td><td>end</td><td>strand</td><td>MM</td><td>target_seq</td><td>PAM</td><td>alignment</td><td>distance</td><td>position</td><td>gene name</td><td>gene id</td></tr>
<tr><td>GU305914.1</td><td>31511</td><td>31533</td><td>-</td><td>0</td><td>TCTGCCAGTGTAACGAGCCG</td><td>CGG</td><td>||||||||[||||||||||||]PAM</td><td>NA</td><td>Intergenic</td><td>NA</td><td>NA</td></tr>
<tr><td>T31975</td><td>GTAACGAGCCGCGGTCTCCCTGG</td><td>1000</td><td></td><td></td><td></td><td></td><td></td><td></td><td></td><td></td><td></td></tr>
<tr><td>Oligo fwd</td><td>CACCGTAACGAGCCGCGGTCTCCC</td><td></td><td></td><td></td><td></td><td></td><td></td><td></td><td></td><td></td><td></td></tr>
<tr><td>Oligo rev</td><td>AAACGGGAGACCGCGGCTCGTTAC</td><td></td><td></td><td></td><td></td><td></td><td></td><td></td><td></td><td></td><td></td></tr>
<tr><td>Chromosome</td><td>start</td><td>end</td><td>strand</td><td>MM</td><td>target_seq</td><td>PAM</td><td>alignment</td><td>distance</td><td>position</td><td>gene name</td><td>gene id</td></tr>
<tr><td>GU305914.1</td><td>31502</td><td>31524</td><td>-</td><td>0</td><td>GTAACGAGCCGCGGTCTCCC</td><td>TGG</td><td>||||||||[||||||||||||]PAM</td><td>NA</td><td>Intergenic</td><td>NA</td><td>NA</td></tr>
<tr><td>T31976</td><td>TACCGCCGTCCTTCAGCCAGTGG</td><td>1000</td><td></td><td></td><td></td><td></td><td></td><td></td><td></td><td></td><td></td></tr>
<tr><td>Oligo fwd</td><td>CACCTACCGCCGTCCTTCAGCCAG</td><td></td><td></td><td></td><td></td><td></td><td></td><td></td><td></td><td></td><td></td></tr>
<tr><td>Oligo rev</td><td>AAACCTGGCTGAAGGACGGCGGTA</td><td></td><td></td><td></td><td></td><td></td><td></td><td></td><td></td><td></td><td></td></tr>
<tr><td>Chromosome</td><td>start</td><td>end</td><td>strand</td><td>MM</td><td>target_seq</td><td>PAM</td><td>alignment</td><td>distance</td><td>position</td><td>gene name</td><td>gene id</td></tr>
<tr><td>GU305914.1</td><td>31478</td><td>31500</td><td>-</td><td>0</td><td>TACCGCCGTCCTTCAGCCAG</td><td>TGG</td><td>||||||||[||||||||||||]PAM</td><td>NA</td><td>Intergenic</td><td>NA</td><td>NA</td></tr>
<tr><td>T31977</td><td>CAGTGGCTGCCCGTCGAGCTCGG</td><td>1000</td><td></td><td></td><td></td><td></td><td></td><td></td><td></td><td></td><td></td></tr>
<tr><td>Oligo fwd</td><td>CACCCAGTGGCTGCCCGTCGAGCT</td><td></td><td></td><td></td><td></td><td></td><td></td><td></td><td></td><td></td><td></td></tr>
<tr><td>Oligo rev</td><td>AAACAGCTCGACGGGCAGCCACTG</td><td></td><td></td><td></td><td></td><td></td><td></td><td></td><td></td><td></td><td></td></tr>
<tr><td>Chromosome</td><td>start</td><td>end</td><td>strand</td><td>MM</td><td>target_seq</td><td>PAM</td><td>alignment</td><td>distance</td><td>position</td><td>gene name</td><td>gene id</td></tr>
<tr><td>GU305914.1</td><td>31461</td><td>31483</td><td>-</td><td>0</td><td>CAGTGGCTGCCCGTCGAGCT</td><td>CGG</td><td>||||||||[||||||||||||]PAM</td><td>NA</td><td>Intergenic</td><td>NA</td><td>NA</td></tr>
<tr><td>T31978</td><td>TGGCTGCCCGTCGAGCTCGGAGG</td><td>1000</td><td></td><td></td><td></td><td></td><td></td><td></td><td></td><td></td><td></td></tr>
<tr><td>Oligo fwd</td><td>CACCTGGCTGCCCGTCGAGCTCGG</td><td></td><td></td><td></td><td></td><td></td><td></td><td></td><td></td><td></td><td></td></tr>
<tr><td>Oligo rev</td><td>AAACCCGAGCTCGACGGGCAGCCA</td><td></td><td></td><td></td><td></td><td></td><td></td><td></td><td></td><td></td><td></td></tr>
<tr><td>Chromosome</td><td>start</td><td>end</td><td>strand</td><td>MM</td><td>target_seq</td><td>PAM</td><td>alignment</td><td>distance</td><td>position</td><td>gene name</td><td>gene id</td></tr>
<tr><td>GU305914.1</td><td>31458</td><td>31480</td><td>-</td><td>0</td><td>TGGCTGCCCGTCGAGCTCGG</td><td>AGG</td><td>||||||||[||||||||||||]PAM</td><td>NA</td><td>Intergenic</td><td>NA</td><td>NA</td></tr>
<tr><td>T31979</td><td>TCGGAGGCGTACTCTGCGCCAGG</td><td>1000</td><td></td><td></td><td></td><td></td><td></td><td></td><td></td><td></td><td></td></tr>
<tr><td>Oligo fwd</td><td>CACCTCGGAGGCGTACTCTGCGCC</td><td></td><td></td><td></td><td></td><td></td><td></td><td></td><td></td><td></td><td></td></tr>
<tr><td>Oligo rev</td><td>AAACGGCGCAGAGTACGCCTCCGA</td><td></td><td></td><td></td><td></td><td></td><td></td><td></td><td></td><td></td><td></td></tr>
<tr><td>Chromosome</td><td>start</td><td>end</td><td>strand</td><td>MM</td><td>target_seq</td><td>PAM</td><td>alignment</td><td>distance</td><td>position</td><td>gene name</td><td>gene id</td></tr>
<tr><td>GU305914.1</td><td>31442</td><td>31464</td><td>-</td><td>0</td><td>TCGGAGGCGTACTCTGCGCC</td><td>AGG</td><td>||||||||[||||||||||||]PAM</td><td>NA</td><td>Intergenic</td><td>NA</td><td>NA</td></tr>
<tr><td>T31980</td><td>CGGAGGCGTACTCTGCGCCAGGG</td><td>1000</td><td></td><td></td><td></td><td></td><td></td><td></td><td></td><td></td><td></td></tr>
<tr><td>Oligo fwd</td><td>CACCCGGAGGCGTACTCTGCGCCA</td><td></td><td></td><td></td><td></td><td></td><td></td><td></td><td></td><td></td><td></td></tr>
<tr><td>Oligo rev</td><td>AAACTGGCGCAGAGTACGCCTCCG</td><td></td><td></td><td></td><td></td><td></td><td></td><td></td><td></td><td></td><td></td></tr>
<tr><td>Chromosome</td><td>start</td><td>end</td><td>strand</td><td>MM</td><td>target_seq</td><td>PAM</td><td>alignment</td><td>distance</td><td>position</td><td>gene name</td><td>gene id</td></tr>
<tr><td>GU305914.1</td><td>31441</td><td>31463</td><td>-</td><td>0</td><td>CGGAGGCGTACTCTGCGCCA</td><td>GGG</td><td>||||||||[||||||||||||]PAM</td><td>NA</td><td>Intergenic</td><td>NA</td><td>NA</td></tr>
<tr><td>T31981</td><td>CGTACTCTGCGCCAGGGTCCCGG</td><td>1000</td><td></td><td></td><td></td><td></td><td></td><td></td><td></td><td></td><td></td></tr>
<tr><td>Oligo fwd</td><td>CACCCGTACTCTGCGCCAGGGTCC</td><td></td><td></td><td></td><td></td><td></td><td></td><td></td><td></td><td></td><td></td></tr>
<tr><td>Oligo rev</td><td>AAACGGACCCTGGCGCAGAGTACG</td><td></td><td></td><td></td><td></td><td></td><td></td><td></td><td></td><td></td><td></td></tr>
<tr><td>Chromosome</td><td>start</td><td>end</td><td>strand</td><td>MM</td><td>target_seq</td><td>PAM</td><td>alignment</td><td>distance</td><td>position</td><td>gene name</td><td>gene id</td></tr>
<tr><td>GU305914.1</td><td>31435</td><td>31457</td><td>-</td><td>0</td><td>CGTACTCTGCGCCAGGGTCC</td><td>CGG</td><td>||||||||[||||||||||||]PAM</td><td>NA</td><td>Intergenic</td><td>NA</td><td>NA</td></tr>
<tr><td>T31982</td><td>GCGCCAGGGTCCCGGCCGAGAGG</td><td>1000</td><td></td><td></td><td></td><td></td><td></td><td></td><td></td><td></td><td></td></tr>
<tr><td>Oligo fwd</td><td>CACCGCGCCAGGGTCCCGGCCGAG</td><td></td><td></td><td></td><td></td><td></td><td></td><td></td><td></td><td></td><td></td></tr>
<tr><td>Oligo rev</td><td>AAACCTCGGCCGGGACCCTGGCGC</td><td></td><td></td><td></td><td></td><td></td><td></td><td></td><td></td><td></td><td></td></tr>
<tr><td>Chromosome</td><td>start</td><td>end</td><td>strand</td><td>MM</td><td>target_seq</td><td>PAM</td><td>alignment</td><td>distance</td><td>position</td><td>gene name</td><td>gene id</td></tr>
<tr><td>GU305914.1</td><td>31427</td><td>31449</td><td>-</td><td>0</td><td>GCGCCAGGGTCCCGGCCGAG</td><td>AGG</td><td>||||||||[||||||||||||]PAM</td><td>NA</td><td>Intergenic</td><td>NA</td><td>NA</td></tr>
<tr><td>T31983</td><td>TCCCGGCCGAGAGGATCCGCCGG</td><td>1000</td><td></td><td></td><td></td><td></td><td></td><td></td><td></td><td></td><td></td></tr>
<tr><td>Oligo fwd</td><td>CACCTCCCGGCCGAGAGGATCCGC</td><td></td><td></td><td></td><td></td><td></td><td></td><td></td><td></td><td></td><td></td></tr>
<tr><td>Oligo rev</td><td>AAACGCGGATCCTCTCGGCCGGGA</td><td></td><td></td><td></td><td></td><td></td><td></td><td></td><td></td><td></td><td></td></tr>
<tr><td>Chromosome</td><td>start</td><td>end</td><td>strand</td><td>MM</td><td>target_seq</td><td>PAM</td><td>alignment</td><td>distance</td><td>position</td><td>gene name</td><td>gene id</td></tr>
<tr><td>GU305914.1</td><td>31418</td><td>31440</td><td>-</td><td>0</td><td>TCCCGGCCGAGAGGATCCGC</td><td>CGG</td><td>||||||||[||||||||||||]PAM</td><td>NA</td><td>Intergenic</td><td>NA</td><td>NA</td></tr>
<tr><td>T31984</td><td>GCCGATCAGACTCCGTCCCGCGG</td><td>1000</td><td></td><td></td><td></td><td></td><td></td><td></td><td></td><td></td><td></td></tr>
<tr><td>Oligo fwd</td><td>CACCGCCGATCAGACTCCGTCCCG</td><td></td><td></td><td></td><td></td><td></td><td></td><td></td><td></td><td></td><td></td></tr>
</table>

| | | | | | | | | | | | | |
|---|---|---|---|---|---|---|---|---|---|---|---|---|
| Oligo rev | AAACCGGGACGGAGTCTGATCGGC | | | | | | | | | | | |
| Chromosome | start | | end | strand | MM | target_seq | PAM | alignment | distance | position | gene name | gene id |
| GU305914.1 | | 31396 | 31418 | - | 0 | GCCGATCAGACTCCGTCCCG | CGG | \|\|\|\|\|\|\|\|[\|\|\|\|\|\|\|\|\|\|\|\|]PAM | NA | Intergenic | NA | NA |
| T31985 | ACTCCGTCCCGCGGATTTTCCGG | | 1000 | | | | | | | | | |
| Oligo fwd | CACCACTCCGTCCCGCGGATTTTC | | | | | | | | | | | |
| Oligo rev | AAACGAAAATCCGCGGGACGGAGT | | | | | | | | | | | |
| Chromosome | start | | end | strand | MM | target_seq | PAM | alignment | distance | position | gene name | gene id |
| GU305914.1 | | 31387 | 31409 | - | 0 | ACTCCGTCCCGCGGATTTTC | CGG | \|\|\|\|\|\|\|\|[\|\|\|\|\|\|\|\|\|\|\|\|]PAM | NA | Intergenic | NA | NA |
| T31986 | CGTCCCGCGGATTTTCCGGACGG | | 1000 | | | | | | | | | |
| Oligo fwd | CACCCGTCCCGCGGATTTTCCGGA | | | | | | | | | | | |
| Oligo rev | AAACTCCGGAAAATCCGCGGGACG | | | | | | | | | | | |
| Chromosome | start | | end | strand | MM | target_seq | PAM | alignment | distance | position | gene name | gene id |
| GU305914.1 | | 31383 | 31405 | - | 0 | CGTCCCGCGGATTTTCCGGA | CGG | \|\|\|\|\|\|\|\|[\|\|\|\|\|\|\|\|\|\|\|\|]PAM | NA | Intergenic | NA | NA |
| T31987 | GTCCCGCGGATTTTCCGGACGGG | | 1000 | | | | | | | | | |
| Oligo fwd | CACCGTCCCGCGGATTTTCCGGAC | | | | | | | | | | | |
| Oligo rev | AAACGTCCGGAAAATCCGCGGGAC | | | | | | | | | | | |
| Chromosome | start | | end | strand | MM | target_seq | PAM | alignment | distance | position | gene name | gene id |
| GU305914.1 | | 31382 | 31404 | - | 0 | GTCCCGCGGATTTTCCGGAC | GGG | \|\|\|\|\|\|\|\|[\|\|\|\|\|\|\|\|\|\|\|\|]PAM | NA | Intergenic | NA | NA |
| T31988 | TCCCGCGGATTTTCCGGACGGGG | | 1000 | | | | | | | | | |
| Oligo fwd | CACCTCCCGCGGATTTTCCGGACG | | | | | | | | | | | |
| Oligo rev | AAACCGTCCGGAAAATCCGCGGGA | | | | | | | | | | | |
| Chromosome | start | | end | strand | MM | target_seq | PAM | alignment | distance | position | gene name | gene id |
| GU305914.1 | | 31381 | 31403 | - | 0 | TCCCGCGGATTTTCCGGACG | GGG | \|\|\|\|\|\|\|\|[\|\|\|\|\|\|\|\|\|\|\|\|]PAM | NA | Intergenic | NA | NA |
| T31989 | TTTTCCGGACGGGGTGTTTAAGG | | 1000 | | | | | | | | | |
| Oligo fwd | CACCTTTTCCGGACGGGGTGTTTA | | | | | | | | | | | |
| Oligo rev | AAACTAAACACCCCGTCCGGAAAA | | | | | | | | | | | |
| Chromosome | start | | end | strand | MM | target_seq | PAM | alignment | distance | position | gene name | gene id |
| GU305914.1 | | 31372 | 31394 | - | 0 | TTTTCCGGACGGGGTGTTTA | AGG | \|\|\|\|\|\|\|\|[\|\|\|\|\|\|\|\|\|\|\|\|]PAM | NA | Intergenic | NA | NA |
| T31990 | TCCGGACGGGGTGTTTAAGGCGG | | 1000 | | | | | | | | | |
| Oligo fwd | CACCTCCGGACGGGGTGTTTAAGG | | | | | | | | | | | |
| Oligo rev | AAACCCTTAAACACCCCGTCCGGA | | | | | | | | | | | |
| Chromosome | start | | end | strand | MM | target_seq | PAM | alignment | distance | position | gene name | gene id |
| GU305914.1 | | 31369 | 31391 | - | 0 | TCCGGACGGGGTGTTTAAGG | CGG | \|\|\|\|\|\|\|\|[\|\|\|\|\|\|\|\|\|\|\|\|]PAM | NA | Intergenic | NA | NA |
| T31991 | CGGGGTGTTTAAGGCGGCCCTGG | | 1000 | | | | | | | | | |
| Oligo fwd | CACCCGGGGTGTTTAAGGCGGCCC | | | | | | | | | | | |
| Oligo rev | AAACGGGCCGCCTTAAACACCCCG | | | | | | | | | | | |
| Chromosome | start | | end | strand | MM | target_seq | PAM | alignment | distance | position | gene name | gene id |
| GU305914.1 | | 31363 | 31385 | - | 0 | CGGGGTGTTTAAGGCGGCCC | TGG | \|\|\|\|\|\|\|\|[\|\|\|\|\|\|\|\|\|\|\|\|]PAM | NA | Intergenic | NA | NA |
| T31992 | GACGTCCTCCTCTCCGTCCGTGG | | 1000 | | | | | | | | | |
| Oligo fwd | CACCGACGTCCTCCTCTCCGTCCG | | | | | | | | | | | |
| Oligo rev | AAACCGGACGGAGAGGAGGACGTC | | | | | | | | | | | |
| Chromosome | start | | end | strand | MM | target_seq | PAM | alignment | distance | position | gene name | gene id |
| GU305914.1 | | 31309 | 31331 | - | 0 | GACGTCCTCCTCTCCGTCCG | TGG | \|\|\|\|\|\|\|\|[\|\|\|\|\|\|\|\|\|\|\|\|]PAM | NA | Intergenic | NA | NA |
| T31993 | GTGGCGACCGCAGAGACGATCGG | | 1000 | | | | | | | | | |
| Oligo fwd | CACCGTGGCGACCGCAGAGACGAT | | | | | | | | | | | |
| Oligo rev | AAACATCGTCTCTGCGGTCGCCAC | | | | | | | | | | | |
| Chromosome | start | | end | strand | MM | target_seq | PAM | alignment | distance | position | gene name | gene id |
| GU305914.1 | | 31290 | 31312 | - | 0 | GTGGCGACCGCAGAGACGAT | CGG | \|\|\|\|\|\|\|\|[\|\|\|\|\|\|\|\|\|\|\|\|]PAM | NA | Intergenic | NA | NA |
| T31994 | ACCGCAGAGACGATCGGACACGG | | 1000 | | | | | | | | | |
| Oligo fwd | CACCACCGCAGAGACGATCGGACA | | | | | | | | | | | |
| Oligo rev | AAACTGTCCGATCGTCTCTGCGGT | | | | | | | | | | | |
| Chromosome | start | | end | strand | MM | target_seq | PAM | alignment | distance | position | gene name | gene id |
| GU305914.1 | | 31284 | 31306 | - | 0 | ACCGCAGAGACGATCGGACA | CGG | \|\|\|\|\|\|\|\|[\|\|\|\|\|\|\|\|\|\|\|\|]PAM | NA | Intergenic | NA | NA |
| T31995 | GACGATCGGACACGGTCGCGAGG | | 1000 | | | | | | | | | |
| Oligo fwd | CACCGACGATCGGACACGGTCGCG | | | | | | | | | | | |
| Oligo rev | AAACCGCGACCGTGTCCGATCGTC | | | | | | | | | | | |
| Chromosome | start | | end | strand | MM | target_seq | PAM | alignment | distance | position | gene name | gene id |
| GU305914.1 | | 31276 | 31298 | - | 0 | GACGATCGGACACGGTCGCG | AGG | \|\|\|\|\|\|\|\|[\|\|\|\|\|\|\|\|\|\|\|\|]PAM | NA | Intergenic | NA | NA |
| T31996 | CGGACACGGTCGCGAGGAAGAGG | | 1000 | | | | | | | | | |
| Oligo fwd | CACCCGGACACGGTCGCGAGGAAG | | | | | | | | | | | |
| Oligo rev | AAACCTTCCTCGCGACCGTGTCCG | | | | | | | | | | | |
| Chromosome | start | | end | strand | MM | target_seq | PAM | alignment | distance | position | gene name | gene id |
| GU305914.1 | | 31270 | 31292 | - | 0 | CGGACACGGTCGCGAGGAAG | AGG | \|\|\|\|\|\|\|\|[\|\|\|\|\|\|\|\|\|\|\|\|]PAM | NA | Intergenic | NA | NA |
| T31997 | CGGTCGCGAGGAAGAGGACGAGG | | 1000 | | | | | | | | | |
| Oligo fwd | CACCCGGTCGCGAGGAAGAGGACG | | | | | | | | | | | |
| Oligo rev | AAACCGTCCTCTTCCTCGCGACCG | | | | | | | | | | | |
| Chromosome | start | | end | strand | MM | target_seq | PAM | alignment | distance | position | gene name | gene id |
| GU305914.1 | | 31264 | 31286 | - | 0 | CGGTCGCGAGGAAGAGGACG | AGG | \|\|\|\|\|\|\|\|[\|\|\|\|\|\|\|\|\|\|\|\|]PAM | NA | Intergenic | NA | NA |
| T31998 | CGAGGAAGAGGACGAGGAGTAGG | | 1000 | | | | | | | | | |
| Oligo fwd | CACCCGAGGAAGAGGACGAGGAGT | | | | | | | | | | | |
| Oligo rev | AAACACTCCTCGTCCTCTTCCTCG | | | | | | | | | | | |
| Chromosome | start | | end | strand | MM | target_seq | PAM | alignment | distance | position | gene name | gene id |
| GU305914.1 | | 31258 | 31280 | - | 0 | CGAGGAAGAGGACGAGGAGT | AGG | \|\|\|\|\|\|\|\|[\|\|\|\|\|\|\|\|\|\|\|\|]PAM | NA | Intergenic | NA | NA |
| T31999 | GTAGGACTAGAACTAGAATTAGG | | 1000 | | | | | | | | | |

| | | | | | | | | | | | |
|---|---|---|---|---|---|---|---|---|---|---|---|
| Oligo fwd | CACCGTAGGACTAGAACTAGAATT | | | | | | | | | | |
| Oligo rev | AAACAATTCTAGTTCTAGTCCTAC | | | | | | | | | | |
| Chromosome | start | end | strand | MM | target_seq | PAM | alignment | distance | position | gene name | gene id |
| GU305914.1 | 31240 | 31262 | - | 0 | GTAGGACTAGAACTAGAATT | AGG | \|\|\|\|\|\|\|\|[\|\|\|\|\|\|\|\|\|\|\|\|]PAM | NA | Intergenic | NA | NA |
| T32000 | ACTAGAACTAGAATTAGGAACGG | 1000 | | | | | | | | | |
| Oligo fwd | CACCACTAGAACTAGAATTAGGAA | | | | | | | | | | |
| Oligo rev | AAACTTCCTAATTCTAGTTCTAGT | | | | | | | | | | |
| Chromosome | start | end | strand | MM | target_seq | PAM | alignment | distance | position | gene name | gene id |
| GU305914.1 | 31235 | 31257 | - | 0 | ACTAGAACTAGAATTAGGAA | CGG | \|\|\|\|\|\|\|\|[\|\|\|\|\|\|\|\|\|\|\|\|]PAM | NA | Intergenic | NA | NA |
| T32001 | CTAGAACTAGAATTAGGAACGGG | 1000 | | | | | | | | | |
| Oligo fwd | CACCCTAGAACTAGAATTAGGAAC | | | | | | | | | | |
| Oligo rev | AAACGTTCCTAATTCTAGTTCTAG | | | | | | | | | | |
| Chromosome | start | end | strand | MM | target_seq | PAM | alignment | distance | position | gene name | gene id |
| GU305914.1 | 31234 | 31256 | - | 0 | CTAGAACTAGAATTAGGAAC | GGG | \|\|\|\|\|\|\|\|[\|\|\|\|\|\|\|\|\|\|\|\|]PAM | NA | Intergenic | NA | NA |
| T32002 | TAGAACTAGAATTAGGAACGGGG | 1000 | | | | | | | | | |
| Oligo fwd | CACCTAGAACTAGAATTAGGAACG | | | | | | | | | | |
| Oligo rev | AAACCGTTCCTAATTCTAGTTCTA | | | | | | | | | | |
| Chromosome | start | end | strand | MM | target_seq | PAM | alignment | distance | position | gene name | gene id |
| GU305914.1 | 31233 | 31255 | - | 0 | TAGAACTAGAATTAGGAACG | GGG | \|\|\|\|\|\|\|\|[\|\|\|\|\|\|\|\|\|\|\|\|]PAM | NA | Intergenic | NA | NA |
| T32003 | AGAATTAGGAACGGGGCGAAAGG | 1000 | | | | | | | | | |
| Oligo fwd | CACCAGAATTAGGAACGGGGCGAA | | | | | | | | | | |
| Oligo rev | AAACTTCGCCCCGTTCCTAATTCT | | | | | | | | | | |
| Chromosome | start | end | strand | MM | target_seq | PAM | alignment | distance | position | gene name | gene id |
| GU305914.1 | 31226 | 31248 | - | 0 | AGAATTAGGAACGGGGCGAA | AGG | \|\|\|\|\|\|\|\|[\|\|\|\|\|\|\|\|\|\|\|\|]PAM | NA | Intergenic | NA | NA |
| T32004 | GAATTAGGAACGGGGCGAAAGGG | 1000 | | | | | | | | | |
| Oligo fwd | CACCGAATTAGGAACGGGGCGAAA | | | | | | | | | | |
| Oligo rev | AAACTTTCGCCCCGTTCCTAATTC | | | | | | | | | | |
| Chromosome | start | end | strand | MM | target_seq | PAM | alignment | distance | position | gene name | gene id |
| GU305914.1 | 31225 | 31247 | - | 0 | GAATTAGGAACGGGGCGAAA | GGG | \|\|\|\|\|\|\|\|[\|\|\|\|\|\|\|\|\|\|\|\|]PAM | NA | Intergenic | NA | NA |
| T32005 | AATTAGGAACGGGGCGAAAGGGG | 1000 | | | | | | | | | |
| Oligo fwd | CACCAATTAGGAACGGGGCGAAAG | | | | | | | | | | |
| Oligo rev | AAACCTTTCGCCCCGTTCCTAATT | | | | | | | | | | |
| Chromosome | start | end | strand | MM | target_seq | PAM | alignment | distance | position | gene name | gene id |
| GU305914.1 | 31224 | 31246 | - | 0 | AATTAGGAACGGGGCGAAAG | GGG | \|\|\|\|\|\|\|\|[\|\|\|\|\|\|\|\|\|\|\|\|]PAM | NA | Intergenic | NA | NA |
| T32006 | GGACCAGACCATGCTAGCTCCGG | 1000 | | | | | | | | | |
| Oligo fwd | CACCGGACCAGACCATGCTAGCTC | | | | | | | | | | |
| Oligo rev | AAACGAGCTAGCATGGTCTGGTCC | | | | | | | | | | |
| Chromosome | start | end | strand | MM | target_seq | PAM | alignment | distance | position | gene name | gene id |
| GU305914.1 | 31203 | 31225 | - | 0 | GGACCAGACCATGCTAGCTC | CGG | \|\|\|\|\|\|\|\|[\|\|\|\|\|\|\|\|\|\|\|\|]PAM | NA | Intergenic | NA | NA |
| T32007 | GACCATGCTAGCTCCGGCCGCGG | 1000 | | | | | | | | | |
| Oligo fwd | CACCGACCATGCTAGCTCCGGCCG | | | | | | | | | | |
| Oligo rev | AAACCGGCCGGAGCTAGCATGGTC | | | | | | | | | | |
| Chromosome | start | end | strand | MM | target_seq | PAM | alignment | distance | position | gene name | gene id |
| GU305914.1 | 31197 | 31219 | - | 0 | GACCATGCTAGCTCCGGCCG | CGG | \|\|\|\|\|\|\|\|[\|\|\|\|\|\|\|\|\|\|\|\|]PAM | NA | Intergenic | NA | NA |
| T32008 | CATGCTAGCTCCGGCCGCGGCGG | 1000 | | | | | | | | | |
| Oligo fwd | CACCCATGCTAGCTCCGGCCGCGG | | | | | | | | | | |
| Oligo rev | AAACCCGCGGCCGGAGCTAGCATG | | | | | | | | | | |
| Chromosome | start | end | strand | MM | target_seq | PAM | alignment | distance | position | gene name | gene id |
| GU305914.1 | 31194 | 31216 | - | 0 | CATGCTAGCTCCGGCCGCGG | CGG | \|\|\|\|\|\|\|\|[\|\|\|\|\|\|\|\|\|\|\|\|]PAM | NA | Intergenic | NA | NA |
| T32009 | GGCGGTGTCGATGCCCGAGTCGG | 1000 | | | | | | | | | |
| Oligo fwd | CACCGGCGGTGTCGATGCCCGAGT | | | | | | | | | | |
| Oligo rev | AAACACTCGGGCATCGACACCGCC | | | | | | | | | | |
| Chromosome | start | end | strand | MM | target_seq | PAM | alignment | distance | position | gene name | gene id |
| GU305914.1 | 31176 | 31198 | - | 0 | GGCGGTGTCGATGCCCGAGT | CGG | \|\|\|\|\|\|\|\|[\|\|\|\|\|\|\|\|\|\|\|\|]PAM | NA | Intergenic | NA | NA |
| T32010 | GTCGATGCCCGAGTCGGAGACGG | 1000 | | | | | | | | | |
| Oligo fwd | CACCGTCGATGCCCGAGTCGGAGA | | | | | | | | | | |
| Oligo rev | AAACTCTCCGACTCGGGCATCGAC | | | | | | | | | | |
| Chromosome | start | end | strand | MM | target_seq | PAM | alignment | distance | position | gene name | gene id |
| GU305914.1 | 31170 | 31192 | - | 0 | GTCGATGCCCGAGTCGGAGA | CGG | \|\|\|\|\|\|\|\|[\|\|\|\|\|\|\|\|\|\|\|\|]PAM | NA | Intergenic | NA | NA |
| T32011 | GCCCGAGTCGGAGACGGAGACGG | 1000 | | | | | | | | | |
| Oligo fwd | CACCGCCCGAGTCGGAGACGGAGA | | | | | | | | | | |
| Oligo rev | AAACTCTCCGTCTCCGACTCGGGC | | | | | | | | | | |
| Chromosome | start | end | strand | MM | target_seq | PAM | alignment | distance | position | gene name | gene id |
| GU305914.1 | 31164 | 31186 | - | 0 | GCCCGAGTCGGAGACGGAGA | CGG | \|\|\|\|\|\|\|\|[\|\|\|\|\|\|\|\|\|\|\|\|]PAM | NA | Intergenic | NA | NA |
| GU305914.1 | 31158 | 31180 | - | 4 | GTCGGAGACGGAGACGGTGA | CGG | \|-\|-\|\|\|-[\|\|\|\|\|\|\|\|\|-\|\|]PAM | NA | Intergenic | NA | NA |
| T32012 | GTCGGAGACGGAGACGGTGACGG | 1000 | | | | | | | | | |
| Oligo fwd | CACCGTCGGAGACGGAGACGGTGA | | | | | | | | | | |
| Oligo rev | AAACTCACCGTCTCCGTCTCCGAC | | | | | | | | | | |
| Chromosome | start | end | strand | MM | target_seq | PAM | alignment | distance | position | gene name | gene id |
| GU305914.1 | 31158 | 31180 | - | 0 | GTCGGAGACGGAGACGGTGA | CGG | \|\|\|\|\|\|\|\|[\|\|\|\|\|\|\|\|\|\|\|\|]PAM | NA | Intergenic | NA | NA |
| GU305914.1 | 170252 | 170274 | + | 4 | TTCGGATTCGGGGACGGTGA | CGG | -\|\|\|\|\|--[\|\|\|-\|\|\|\|\|\|\|\|]PAM | NA | Intergenic | NA | NA |
| GU305914.1 | 31164 | 31186 | - | 4 | GCCCGAGTCGGAGACGGAGA | CGG | \|-\|-\|\|\|-[\|\|\|\|\|\|\|\|\|-\|\|]PAM | NA | Intergenic | NA | NA |
| T32013 | GGAGACGGAGACGGTGACGGTGG | 1000 | | | | | | | | | |
| Oligo fwd | CACCGGAGACGGAGACGGTGACGG | | | | | | | | | | |
| Oligo rev | AAACCCGTCACCGTCTCCGTCTCC | | | | | | | | | | |

<table>
<tr><td>Chromosome</td><td>start</td><td>end</td><td>strand</td><td>MM</td><td>target_seq</td><td>PAM</td><td>alignment</td><td>distance</td><td>position</td><td>gene name</td><td>gene id</td></tr>
<tr><td>GU305914.1</td><td>31155</td><td>31177</td><td>-</td><td>0</td><td>GGAGACGGAGACGGTGACGG</td><td>TGG</td><td>||||||||[||||||||||||]PAM</td><td>NA</td><td>Intergenic</td><td>NA</td><td>NA</td></tr>
<tr><td>GU305914.1</td><td>149875</td><td>149897</td><td>-</td><td>4</td><td>GCCGACGGCGCCGGTGACGG</td><td>TGG</td><td>|--|||||[-|-|||||||||]PAM</td><td>NA</td><td>Intergenic</td><td>NA</td><td>NA</td></tr>
<tr><td>T32014</td><td>GGTGCCTGCCCGCTCCACCCTGG</td><td>1000</td><td></td><td></td><td></td><td></td><td></td><td></td><td></td><td></td><td></td></tr>
<tr><td>Oligo fwd</td><td>CACCGGTGCCTGCCCGCTCCACCC</td><td></td><td></td><td></td><td></td><td></td><td></td><td></td><td></td><td></td><td></td></tr>
<tr><td>Oligo rev</td><td>AAACGGGTGGAGCGGGCAGGCACC</td><td></td><td></td><td></td><td></td><td></td><td></td><td></td><td></td><td></td><td></td></tr>
<tr><td>Chromosome</td><td>start</td><td>end</td><td>strand</td><td>MM</td><td>target_seq</td><td>PAM</td><td>alignment</td><td>distance</td><td>position</td><td>gene name</td><td>gene id</td></tr>
<tr><td>GU305914.1</td><td>31134</td><td>31156</td><td>-</td><td>0</td><td>GGTGCCTGCCCGCTCCACCC</td><td>TGG</td><td>||||||||[||||||||||||]PAM</td><td>NA</td><td>Intergenic</td><td>NA</td><td>NA</td></tr>
<tr><td>T32015</td><td>GCCTGCCCGCTCCACCCTGGCGG</td><td>1000</td><td></td><td></td><td></td><td></td><td></td><td></td><td></td><td></td><td></td></tr>
<tr><td>Oligo fwd</td><td>CACCGCCTGCCCGCTCCACCCTGG</td><td></td><td></td><td></td><td></td><td></td><td></td><td></td><td></td><td></td><td></td></tr>
<tr><td>Oligo rev</td><td>AAACCCAGGGTGGAGCGGGCAGGC</td><td></td><td></td><td></td><td></td><td></td><td></td><td></td><td></td><td></td><td></td></tr>
<tr><td>Chromosome</td><td>start</td><td>end</td><td>strand</td><td>MM</td><td>target_seq</td><td>PAM</td><td>alignment</td><td>distance</td><td>position</td><td>gene name</td><td>gene id</td></tr>
<tr><td>GU305914.1</td><td>31131</td><td>31153</td><td>-</td><td>0</td><td>GCCTGCCCGCTCCACCCTGG</td><td>CGG</td><td>||||||||[||||||||||||]PAM</td><td>NA</td><td>Intergenic</td><td>NA</td><td>NA</td></tr>
<tr><td>T32016</td><td>CCGCTCCACCCTGGCGGCCAAGG</td><td>1000</td><td></td><td></td><td></td><td></td><td></td><td></td><td></td><td></td><td></td></tr>
<tr><td>Oligo fwd</td><td>CACCCCGCTCCACCCTGGCGGCCA</td><td></td><td></td><td></td><td></td><td></td><td></td><td></td><td></td><td></td><td></td></tr>
<tr><td>Oligo rev</td><td>AAACTGGCCGCCAGGGTGGAGCGG</td><td></td><td></td><td></td><td></td><td></td><td></td><td></td><td></td><td></td><td></td></tr>
<tr><td>Chromosome</td><td>start</td><td>end</td><td>strand</td><td>MM</td><td>target_seq</td><td>PAM</td><td>alignment</td><td>distance</td><td>position</td><td>gene name</td><td>gene id</td></tr>
<tr><td>GU305914.1</td><td>31125</td><td>31147</td><td>-</td><td>0</td><td>CCGCTCCACCCTGGCGGCCA</td><td>AGG</td><td>||||||||[||||||||||||]PAM</td><td>NA</td><td>Intergenic</td><td>NA</td><td>NA</td></tr>
<tr><td>T32017</td><td>CGGCCAAGGCCGCCGCCATGAGG</td><td>1000</td><td></td><td></td><td></td><td></td><td></td><td></td><td></td><td></td><td></td></tr>
<tr><td>Oligo fwd</td><td>CACCCGGCCAAGGCCGCCGCCATG</td><td></td><td></td><td></td><td></td><td></td><td></td><td></td><td></td><td></td><td></td></tr>
<tr><td>Oligo rev</td><td>AAACCATGGCGGCGGCCTTGGCCG</td><td></td><td></td><td></td><td></td><td></td><td></td><td></td><td></td><td></td><td></td></tr>
<tr><td>Chromosome</td><td>start</td><td>end</td><td>strand</td><td>MM</td><td>target_seq</td><td>PAM</td><td>alignment</td><td>distance</td><td>position</td><td>gene name</td><td>gene id</td></tr>
<tr><td>GU305914.1</td><td>31111</td><td>31133</td><td>-</td><td>0</td><td>CGGCCAAGGCCGCCGCCATG</td><td>AGG</td><td>||||||||[||||||||||||]PAM</td><td>NA</td><td>Intergenic</td><td>NA</td><td>NA</td></tr>
<tr><td>T32018</td><td>CAAGGCCGCCGCCATGAGGAAGG</td><td>1000</td><td></td><td></td><td></td><td></td><td></td><td></td><td></td><td></td><td></td></tr>
<tr><td>Oligo fwd</td><td>CACCCAAGGCCGCCGCCATGAGGA</td><td></td><td></td><td></td><td></td><td></td><td></td><td></td><td></td><td></td><td></td></tr>
<tr><td>Oligo rev</td><td>AAACTCCTCATGGCGGCGGCCTTG</td><td></td><td></td><td></td><td></td><td></td><td></td><td></td><td></td><td></td><td></td></tr>
<tr><td>Chromosome</td><td>start</td><td>end</td><td>strand</td><td>MM</td><td>target_seq</td><td>PAM</td><td>alignment</td><td>distance</td><td>position</td><td>gene name</td><td>gene id</td></tr>
<tr><td>GU305914.1</td><td>31107</td><td>31129</td><td>-</td><td>0</td><td>CAAGGCCGCCGCCATGAGGA</td><td>AGG</td><td>||||||||[||||||||||||]PAM</td><td>NA</td><td>Intergenic</td><td>NA</td><td>NA</td></tr>
<tr><td>T32019</td><td>GTACGCCGTCAACAGACTCCTGG</td><td>1000</td><td></td><td></td><td></td><td></td><td></td><td></td><td></td><td></td><td></td></tr>
<tr><td>Oligo fwd</td><td>CACCGTACGCCGTCAACAGACTCC</td><td></td><td></td><td></td><td></td><td></td><td></td><td></td><td></td><td></td><td></td></tr>
<tr><td>Oligo rev</td><td>AAACGGAGTCTGTTGACGGCGTAC</td><td></td><td></td><td></td><td></td><td></td><td></td><td></td><td></td><td></td><td></td></tr>
<tr><td>Chromosome</td><td>start</td><td>end</td><td>strand</td><td>MM</td><td>target_seq</td><td>PAM</td><td>alignment</td><td>distance</td><td>position</td><td>gene name</td><td>gene id</td></tr>
<tr><td>GU305914.1</td><td>31083</td><td>31105</td><td>-</td><td>0</td><td>GTACGCCGTCAACAGACTCC</td><td>TGG</td><td>||||||||[||||||||||||]PAM</td><td>NA</td><td>Intergenic</td><td>NA</td><td>NA</td></tr>
<tr><td>T32020</td><td>GTCAACAGACTCCTGGCGCTCGG</td><td>1000</td><td></td><td></td><td></td><td></td><td></td><td></td><td></td><td></td><td></td></tr>
<tr><td>Oligo fwd</td><td>CACCGTCAACAGACTCCTGGCGCT</td><td></td><td></td><td></td><td></td><td></td><td></td><td></td><td></td><td></td><td></td></tr>
<tr><td>Oligo rev</td><td>AAACAGCGCCAGGAGTCTGTTGAC</td><td></td><td></td><td></td><td></td><td></td><td></td><td></td><td></td><td></td><td></td></tr>
<tr><td>Chromosome</td><td>start</td><td>end</td><td>strand</td><td>MM</td><td>target_seq</td><td>PAM</td><td>alignment</td><td>distance</td><td>position</td><td>gene name</td><td>gene id</td></tr>
<tr><td>GU305914.1</td><td>31076</td><td>31098</td><td>-</td><td>0</td><td>GTCAACAGACTCCTGGCGCT</td><td>CGG</td><td>||||||||[||||||||||||]PAM</td><td>NA</td><td>Intergenic</td><td>NA</td><td>NA</td></tr>
<tr><td>T32021</td><td>ACAGACTCCTGGCGCTCGGCAGG</td><td>1000</td><td></td><td></td><td></td><td></td><td></td><td></td><td></td><td></td><td></td></tr>
<tr><td>Oligo fwd</td><td>CACCACAGACTCCTGGCGCTCGGC</td><td></td><td></td><td></td><td></td><td></td><td></td><td></td><td></td><td></td><td></td></tr>
<tr><td>Oligo rev</td><td>AAACGCCGAGCGCCAGGAGTCTGT</td><td></td><td></td><td></td><td></td><td></td><td></td><td></td><td></td><td></td><td></td></tr>
<tr><td>Chromosome</td><td>start</td><td>end</td><td>strand</td><td>MM</td><td>target_seq</td><td>PAM</td><td>alignment</td><td>distance</td><td>position</td><td>gene name</td><td>gene id</td></tr>
<tr><td>GU305914.1</td><td>31072</td><td>31094</td><td>-</td><td>0</td><td>ACAGACTCCTGGCGCTCGGC</td><td>AGG</td><td>||||||||[||||||||||||]PAM</td><td>NA</td><td>Intergenic</td><td>NA</td><td>NA</td></tr>
<tr><td>T32022</td><td>CAGACTCCTGGCGCTCGGCAGGG</td><td>1000</td><td></td><td></td><td></td><td></td><td></td><td></td><td></td><td></td><td></td></tr>
<tr><td>Oligo fwd</td><td>CACCCAGACTCCTGGCGCTCGGCA</td><td></td><td></td><td></td><td></td><td></td><td></td><td></td><td></td><td></td><td></td></tr>
<tr><td>Oligo rev</td><td>AAACTGCCGAGCGCCAGGAGTCTG</td><td></td><td></td><td></td><td></td><td></td><td></td><td></td><td></td><td></td><td></td></tr>
<tr><td>Chromosome</td><td>start</td><td>end</td><td>strand</td><td>MM</td><td>target_seq</td><td>PAM</td><td>alignment</td><td>distance</td><td>position</td><td>gene name</td><td>gene id</td></tr>
<tr><td>GU305914.1</td><td>31071</td><td>31093</td><td>-</td><td>0</td><td>CAGACTCCTGGCGCTCGGCA</td><td>GGG</td><td>||||||||[||||||||||||]PAM</td><td>NA</td><td>Intergenic</td><td>NA</td><td>NA</td></tr>
<tr><td>T32023</td><td>CCTGGCGCTCGGCAGGGCCGCGG</td><td>1000</td><td></td><td></td><td></td><td></td><td></td><td></td><td></td><td></td><td></td></tr>
<tr><td>Oligo fwd</td><td>CACCCCTGGCGCTCGGCAGGGCCG</td><td></td><td></td><td></td><td></td><td></td><td></td><td></td><td></td><td></td><td></td></tr>
<tr><td>Oligo rev</td><td>AAACCGGCCCTGCCGAGCGCCAGG</td><td></td><td></td><td></td><td></td><td></td><td></td><td></td><td></td><td></td><td></td></tr>
<tr><td>Chromosome</td><td>start</td><td>end</td><td>strand</td><td>MM</td><td>target_seq</td><td>PAM</td><td>alignment</td><td>distance</td><td>position</td><td>gene name</td><td>gene id</td></tr>
<tr><td>GU305914.1</td><td>31065</td><td>31087</td><td>-</td><td>0</td><td>CCTGGCGCTCGGCAGGGCCG</td><td>CGG</td><td>||||||||[||||||||||||]PAM</td><td>NA</td><td>Intergenic</td><td>NA</td><td>NA</td></tr>
<tr><td>T32024</td><td>GCTCGGCAGGGCCGCGGCCGAGG</td><td>1000</td><td></td><td></td><td></td><td></td><td></td><td></td><td></td><td></td><td></td></tr>
<tr><td>Oligo fwd</td><td>CACCGCTCGGCAGGGCCGCGGCCG</td><td></td><td></td><td></td><td></td><td></td><td></td><td></td><td></td><td></td><td></td></tr>
<tr><td>Oligo rev</td><td>AAACCGGCCGCGGCCCTGCCGAGC</td><td></td><td></td><td></td><td></td><td></td><td></td><td></td><td></td><td></td><td></td></tr>
<tr><td>Chromosome</td><td>start</td><td>end</td><td>strand</td><td>MM</td><td>target_seq</td><td>PAM</td><td>alignment</td><td>distance</td><td>position</td><td>gene name</td><td>gene id</td></tr>
<tr><td>GU305914.1</td><td>31059</td><td>31081</td><td>-</td><td>0</td><td>GCTCGGCAGGGCCGCGGCCG</td><td>AGG</td><td>||||||||[||||||||||||]PAM</td><td>NA</td><td>Intergenic</td><td>NA</td><td>NA</td></tr>
<tr><td>GU305914.1</td><td>113640</td><td>113662</td><td>+</td><td>4</td><td>GCTGCGCAGAGCGGCGGCCG</td><td>GGG</td><td>|||--|||[|-||-|||||||]PAM</td><td>NA</td><td>Intergenic</td><td>NA</td><td>NA</td></tr>
<tr><td>T32025</td><td>CTCGGCAGGGCCGCGGCCGAGGG</td><td>1000</td><td></td><td></td><td></td><td></td><td></td><td></td><td></td><td></td><td></td></tr>
<tr><td>Oligo fwd</td><td>CACCCTCGGCAGGGCCGCGGCCGA</td><td></td><td></td><td></td><td></td><td></td><td></td><td></td><td></td><td></td><td></td></tr>
<tr><td>Oligo rev</td><td>AAACTCGGCCGCGGCCCTGCCGAG</td><td></td><td></td><td></td><td></td><td></td><td></td><td></td><td></td><td></td><td></td></tr>
<tr><td>Chromosome</td><td>start</td><td>end</td><td>strand</td><td>MM</td><td>target_seq</td><td>PAM</td><td>alignment</td><td>distance</td><td>position</td><td>gene name</td><td>gene id</td></tr>
<tr><td>GU305914.1</td><td>31058</td><td>31080</td><td>-</td><td>0</td><td>CTCGGCAGGGCCGCGGCCGA</td><td>GGG</td><td>||||||||[||||||||||||]PAM</td><td>NA</td><td>Intergenic</td><td>NA</td><td>NA</td></tr>
<tr><td>GU305914.1</td><td>64438</td><td>64460</td><td>+</td><td>4</td><td>GTCCTCAGGGCCGCGGCCGG</td><td>AGG</td><td>-||--|||[|||||||||||-]PAM</td><td>NA</td><td>Intergenic</td><td>NA</td><td>NA</td></tr>
<tr><td>T32026</td><td>CCGAGGGCAAGCTCGAACAGTGG</td><td>1000</td><td></td><td></td><td></td><td></td><td></td><td></td><td></td><td></td><td></td></tr>
<tr><td>Oligo fwd</td><td>CACCCCGAGGGCAAGCTCGAACAG</td><td></td><td></td><td></td><td></td><td></td><td></td><td></td><td></td><td></td><td></td></tr>
<tr><td>Oligo rev</td><td>AAACCTGTTCGAGCTTGCCCTCGG</td><td></td><td></td><td></td><td></td><td></td><td></td><td></td><td></td><td></td><td></td></tr>
<tr><td>Chromosome</td><td>start</td><td>end</td><td>strand</td><td>MM</td><td>target_seq</td><td>PAM</td><td>alignment</td><td>distance</td><td>position</td><td>gene name</td><td>gene id</td></tr>
<tr><td>GU305914.1</td><td>31042</td><td>31064</td><td>-</td><td>0</td><td>CCGAGGGCAAGCTCGAACAG</td><td>TGG</td><td>||||||||[||||||||||||]PAM</td><td>NA</td><td>Intergenic</td><td>NA</td><td>NA</td></tr>
<tr><td>T32027</td><td>CAGTGGCTCAGCGCCCGACGCGG</td><td>1000</td><td></td><td></td><td></td><td></td><td></td><td></td><td></td><td></td><td></td></tr>
<tr><td>Oligo fwd</td><td>CACCCAGTGGCTCAGCGCCCGACG</td><td></td><td></td><td></td><td></td><td></td><td></td><td></td><td></td><td></td><td></td></tr>
<tr><td>Oligo rev</td><td>AAACCGTCGGGCGCTGAGCCACTG</td><td></td><td></td><td></td><td></td><td></td><td></td><td></td><td></td><td></td><td></td></tr>
<tr><td>Chromosome</td><td>start</td><td>end</td><td>strand</td><td>MM</td><td>target_seq</td><td>PAM</td><td>alignment</td><td>distance</td><td>position</td><td>gene name</td><td>gene id</td></tr>
<tr><td>GU305914.1</td><td>31025</td><td>31047</td><td>-</td><td>0</td><td>CAGTGGCTCAGCGCCCGACG</td><td>CGG</td><td>||||||||[||||||||||||]PAM</td><td>NA</td><td>Intergenic</td><td>NA</td><td>NA</td></tr>
</table>

T32028 TATCTCATCCTCTGCGCTCCCGG 1000
Oligo fwd CACCTATCTCATCCTCTGCGCTCC
Oligo rev AAACGGAGCGCAGAGGATGAGATA
Chromosome start end strand MM target_seq PAM alignment distance position gene name gene id
GU305914.1 30962 30984 - 0 TATCTCATCCTCTGCGCTCC CGG ||||||||[||||||||||||]PAM NA Intergenic NA NA

T32029 TGCGCTCCCGGCGATGTCGACGG 1000
Oligo fwd CACCTGCGCTCCCGGCGATGTCGA
Oligo rev AAACTCGACATCGCCGGGAGCGCA
Chromosome start end strand MM target_seq PAM alignment distance position gene name gene id
GU305914.1 30950 30972 - 0 TGCGCTCCCGGCGATGTCGA CGG ||||||||[||||||||||||]PAM NA Intergenic NA NA

T32030 AGTGCGACCCCGACGCCCTGCGG 1000
Oligo fwd CACCAGTGCGACCCCGACGCCCTG
Oligo rev AAACCAGGGCGTCGGGGTCGCACT
Chromosome start end strand MM target_seq PAM alignment distance position gene name gene id
GU305914.1 30922 30944 - 0 AGTGCGACCCCGACGCCCTG CGG ||||||||[||||||||||||]PAM NA Intergenic NA NA

T32031 GTGCGACCCCGACGCCCTGCGGG 1000
Oligo fwd CACCGTGCGACCCCGACGCCCTGC
Oligo rev AAACGCAGGGCGTCGGGGTCGCAC
Chromosome start end strand MM target_seq PAM alignment distance position gene name gene id
GU305914.1 30921 30943 - 0 GTGCGACCCCGACGCCCTGC GGG ||||||||[||||||||||||]PAM NA Intergenic NA NA

T32032 CGCTCGAACCCGCCGTCGACAGG 1000
Oligo fwd CACCCGCTCGAACCCGCCGTCGAC
Oligo rev AAACGTCGACGGCGGGTTCGAGCG
Chromosome start end strand MM target_seq PAM alignment distance position gene name gene id
GU305914.1 30898 30920 - 0 CGCTCGAACCCGCCGTCGAC AGG ||||||||[||||||||||||]PAM NA Intergenic NA NA

T32033 CGAACCCGCCGTCGACAGGCTGG 1000
Oligo fwd CACCCGAACCCGCCGTCGACAGGC
Oligo rev AAACGCCTGTCGACGGCGGGTTCG
Chromosome start end strand MM target_seq PAM alignment distance position gene name gene id
GU305914.1 30894 30916 - 0 CGAACCCGCCGTCGACAGGC TGG ||||||||[||||||||||||]PAM NA Intergenic NA NA

T32034 GTCGACAGGCTGGTCGTCGTCGG 1000
Oligo fwd CACCGTCGACAGGCTGGTCGTCGT
Oligo rev AAACACGACGACCAGCCTGTCGAC
Chromosome start end strand MM target_seq PAM alignment distance position gene name gene id
GU305914.1 30884 30906 - 0 GTCGACAGGCTGGTCGTCGT CGG ||||||||[||||||||||||]PAM NA Intergenic NA NA

T32035 GGTCGTCGTCGGCTTCGTCGAGG 1000
Oligo fwd CACCGGTCGTCGTCGGCTTCGTCG
Oligo rev AAACCGACGAAGCCGACGACGACC
Chromosome start end strand MM target_seq PAM alignment distance position gene name gene id
GU305914.1 30873 30895 - 0 GGTCGTCGTCGGCTTCGTCG AGG ||||||||[||||||||||||]PAM NA Intergenic NA NA
GU305914.1 163655 163677 - 4 CGACATCGTCGGCTTCGGCG CGG -|-|-|||[|||||||||-||]PAM NA Intergenic NA NA

T32036 GTCGTCGTCGGCTTCGTCGAGGG 1000
Oligo fwd CACCGTCGTCGTCGGCTTCGTCGA
Oligo rev AAACTCGACGAAGCCGACGACGAC
Chromosome start end strand MM target_seq PAM alignment distance position gene name gene id
GU305914.1 30872 30894 - 0 GTCGTCGTCGGCTTCGTCGA GGG ||||||||[||||||||||||]PAM NA Intergenic NA NA

T32037 CGGCTTCGTCGAGGGTCGCGAGG 1000
Oligo fwd CACCCGGCTTCGTCGAGGGTCGCG
Oligo rev AAACCGCGACCCTCGACGAAGCCG
Chromosome start end strand MM target_seq PAM alignment distance position gene name gene id
GU305914.1 30864 30886 - 0 CGGCTTCGTCGAGGGTCGCG AGG ||||||||[||||||||||||]PAM NA Intergenic NA NA

T32038 CGCGAGGACGCGCACCGCGACGG 1000
Oligo fwd CACCCGCGAGGACGCGCACCGCGA
Oligo rev AAACTCGCGGTGCGCGTCCTCGCG
Chromosome start end strand MM target_seq PAM alignment distance position gene name gene id
GU305914.1 30848 30870 - 0 CGCGAGGACGCGCACCGCGA CGG ||||||||[||||||||||||]PAM NA Intergenic NA NA

T32039 CGCACCGCGACGGTGACGACAGG 1000
Oligo fwd CACCCGCACCGCGACGGTGACGAC
Oligo rev AAACGTCGTCACCGTCGCGGTGCG
Chromosome start end strand MM target_seq PAM alignment distance position gene name gene id
GU305914.1 30838 30860 - 0 CGCACCGCGACGGTGACGAC AGG ||||||||[||||||||||||]PAM NA Intergenic NA NA

T32040 TCTCGTCAATCCGTCGAGCGCGG 1000
Oligo fwd CACCTCTCGTCAATCCGTCGAGCG
Oligo rev AAACCGCTCGACGGATTGACGAGA
Chromosome start end strand MM target_seq PAM alignment distance position gene name gene id
GU305914.1 30807 30829 - 0 TCTCGTCAATCCGTCGAGCG CGG ||||||||[||||||||||||]PAM NA Intergenic NA NA

T32041 CGCGGTCTTCATCTACGACCCGG 1000
Oligo fwd CACCCGCGGTCTTCATCTACGACC
Oligo rev AAACGGTCGTAGATGAAGACCGCG
Chromosome start end strand MM target_seq PAM alignment distance position gene name gene id
GU305914.1 30789 30811 - 0 CGCGGTCTTCATCTACGACC CGG ||||||||[||||||||||||]PAM NA Intergenic NA NA

T32042 ATCTACGACCCGGCGCCCTACGG 1000
Oligo fwd CACCATCTACGACCCGGCGCCCTA
Oligo rev AAACTAGGGCGCCGGGTCGTAGAT

Chromosome start end strand MM target_seq PAM alignment distance position gene name gene id
GU305914.1 30779 30801 - 0 ATCTACGACCCGGCGCCCTA CGG ||||||||[||||||||||||]PAM NA Intergenic NA NA

T32043 TCTACGACCCGGCGCCCTACGGG 1000
Oligo fwd CACCTCTACGACCCGGCGCCCTAC
Oligo rev AAACGTAGGGCGCCGGGTCGTAGA
Chromosome start end strand MM target_seq PAM alignment distance position gene name gene id
GU305914.1 30778 30800 - 0 TCTACGACCCGGCGCCCTAC GGG ||||||||[||||||||||||]PAM NA Intergenic NA NA

T32044 CTACGACCCGGCGCCCTACGGGG 1000
Oligo fwd CACCCTACGACCCGGCGCCCTACG
Oligo rev AAACCGTAGGGCGCCGGGTCGTAG
Chromosome start end strand MM target_seq PAM alignment distance position gene name gene id
GU305914.1 30777 30799 - 0 CTACGACCCGGCGCCCTACG GGG ||||||||[||||||||||||]PAM NA Intergenic NA NA

T32045 TACGACCCGGCGCCCTACGGGGG 1000
Oligo fwd CACCTACGACCCGGCGCCCTACGG
Oligo rev AAACCCGTAGGGCGCCGGGTCGTA
Chromosome start end strand MM target_seq PAM alignment distance position gene name gene id
GU305914.1 30776 30798 - 0 TACGACCCGGCGCCCTACGG GGG ||||||||[||||||||||||]PAM NA Intergenic NA NA

T32046 CGCCCTACGGGGGACTATATAGG 1000
Oligo fwd CACCCGCCCTACGGGGGACTATAT
Oligo rev AAACATATAGTCCCCCGTAGGGCG
Chromosome start end strand MM target_seq PAM alignment distance position gene name gene id
GU305914.1 30766 30788 - 0 CGCCCTACGGGGGACTATAT AGG ||||||||[||||||||||||]PAM NA Intergenic NA NA
GU305914.1 215833 215855 + 4 CGTCATACCGGGGACTACAT CGG ||-|-|||[-||||||||-||]PAM NA Intergenic NA NA

T32047 AGGCTTGCCAACACTATGATCGG 1000
Oligo fwd CACCAGGCTTGCCAACACTATGAT
Oligo rev AAACATCATAGTGTTGGCAAGCCT
Chromosome start end strand MM target_seq PAM alignment distance position gene name gene id
GU305914.1 30746 30768 - 0 AGGCTTGCCAACACTATGAT CGG ||||||||[||||||||||||]PAM NA Intergenic NA NA

T32048 CTATGATCGGCTTCGTGAGACGG 1000
Oligo fwd CACCCTATGATCGGCTTCGTGAGA
Oligo rev AAACTCTCACGAAGCCGATCATAG
Chromosome start end strand MM target_seq PAM alignment distance position gene name gene id
GU305914.1 30733 30755 - 0 CTATGATCGGCTTCGTGAGA CGG ||||||||[||||||||||||]PAM NA Intergenic NA NA

T32049 TATGATCGGCTTCGTGAGACGGG 1000
Oligo fwd CACCTATGATCGGCTTCGTGAGAC
Oligo rev AAACGTCTCACGAAGCCGATCATA
Chromosome start end strand MM target_seq PAM alignment distance position gene name gene id
GU305914.1 30732 30754 - 0 TATGATCGGCTTCGTGAGAC GGG ||||||||[||||||||||||]PAM NA Intergenic NA NA

T32050 ATGATCGGCTTCGTGAGACGGGG 1000
Oligo fwd CACCATGATCGGCTTCGTGAGACG
Oligo rev AAACCGTCTCACGAAGCCGATCAT
Chromosome start end strand MM target_seq PAM alignment distance position gene name gene id
GU305914.1 30731 30753 - 0 ATGATCGGCTTCGTGAGACG GGG ||||||||[||||||||||||]PAM NA Intergenic NA NA

T32051 TCGTGAGACGGGGTCTGCGCAGG 1000
Oligo fwd CACCTCGTGAGACGGGGTCTGCGC
Oligo rev AAACGCGCAGACCCCGTCTCACGA
Chromosome start end strand MM target_seq PAM alignment distance position gene name gene id
GU305914.1 30721 30743 - 0 TCGTGAGACGGGGTCTGCGC AGG ||||||||[||||||||||||]PAM NA Intergenic NA NA

T32052 TCGTGATCCCTGCTGTGTCGAGG 1000
Oligo fwd CACCTCGTGATCCCTGCTGTGTCG
Oligo rev AAACCGACACAGCAGGGATCACGA
Chromosome start end strand MM target_seq PAM alignment distance position gene name gene id
GU305914.1 30684 30706 - 0 TCGTGATCCCTGCTGTGTCG AGG ||||||||[||||||||||||]PAM NA Intergenic NA NA

T32053 GCTGTGTCGAGGATGCAATCCGG 1000
Oligo fwd CACCGCTGTGTCGAGGATGCAATC
Oligo rev AAACGATTGCATCCTCGACACAGC
Chromosome start end strand MM target_seq PAM alignment distance position gene name gene id
GU305914.1 30673 30695 - 0 GCTGTGTCGAGGATGCAATC CGG ||||||||[||||||||||||]PAM NA Intergenic NA NA

T32054 CTGTGTCGAGGATGCAATCCGGG 1000
Oligo fwd CACCCTGTGTCGAGGATGCAATCC
Oligo rev AAACGGATTGCATCCTCGACACAG
Chromosome start end strand MM target_seq PAM alignment distance position gene name gene id
GU305914.1 30672 30694 - 0 CTGTGTCGAGGATGCAATCC GGG ||||||||[||||||||||||]PAM NA Intergenic NA NA

T32055 CGAGGATGCAATCCGGGTGAAGG 1000
Oligo fwd CACCCGAGGATGCAATCCGGGTGA
Oligo rev AAACTCACCCGGATTGCATCCTCG
Chromosome start end strand MM target_seq PAM alignment distance position gene name gene id
GU305914.1 30666 30688 - 0 CGAGGATGCAATCCGGGTGA AGG ||||||||[||||||||||||]PAM NA Intergenic NA NA

T32056 GATGCAATCCGGGTGAAGGACGG 1000
Oligo fwd CACCGATGCAATCCGGGTGAAGGA
Oligo rev AAACTCCTTCACCCGGATTGCATC
Chromosome start end strand MM target_seq PAM alignment distance position gene name gene id
GU305914.1 30662 30684 - 0 GATGCAATCCGGGTGAAGGA CGG ||||||||[||||||||||||]PAM NA Intergenic NA NA

T32057 ACGGACACGCGTCGCTGCCTCGG 1000

<table>
<tr><td>Oligo fwd</td><td>CACCACGGACACGCGTCGCTGCCT</td><td></td><td></td><td></td><td></td><td></td><td></td><td></td><td></td><td></td><td></td></tr>
<tr><td>Oligo rev</td><td>AAACAGGCAGCGACGCGTGTCCGT</td><td></td><td></td><td></td><td></td><td></td><td></td><td></td><td></td><td></td><td></td></tr>
<tr><td>Chromosome</td><td>start</td><td>end</td><td>strand</td><td>MM</td><td>target_seq</td><td>PAM</td><td>alignment</td><td>distance</td><td>position</td><td>gene name</td><td>gene id</td></tr>
<tr><td>GU305914.1</td><td>30643</td><td>30665</td><td>-</td><td>0</td><td>ACGGACACGCGTCGCTGCCT</td><td>CGG</td><td>||||||||[||||||||||||]PAM</td><td>NA</td><td>Intergenic</td><td>NA</td><td>NA</td></tr>
<tr><td>T32058</td><td>CGCGTCGCTGCCTCGGAACGCGG</td><td>1000</td><td></td><td></td><td></td><td></td><td></td><td></td><td></td><td></td><td></td></tr>
<tr><td>Oligo fwd</td><td>CACCCGCGTCGCTGCCTCGGAACG</td><td></td><td></td><td></td><td></td><td></td><td></td><td></td><td></td><td></td><td></td></tr>
<tr><td>Oligo rev</td><td>AAACCGTTCCGAGGCAGCGACGCG</td><td></td><td></td><td></td><td></td><td></td><td></td><td></td><td></td><td></td><td></td></tr>
<tr><td>Chromosome</td><td>start</td><td>end</td><td>strand</td><td>MM</td><td>target_seq</td><td>PAM</td><td>alignment</td><td>distance</td><td>position</td><td>gene name</td><td>gene id</td></tr>
<tr><td>GU305914.1</td><td>30636</td><td>30658</td><td>-</td><td>0</td><td>CGCGTCGCTGCCTCGGAACG</td><td>CGG</td><td>||||||||[||||||||||||]PAM</td><td>NA</td><td>Intergenic</td><td>NA</td><td>NA</td></tr>
<tr><td>T32059</td><td>GCTGCCTCGGAACGCGGACGAGG</td><td>1000</td><td></td><td></td><td></td><td></td><td></td><td></td><td></td><td></td><td></td></tr>
<tr><td>Oligo fwd</td><td>CACCGCTGCCTCGGAACGCGGACG</td><td></td><td></td><td></td><td></td><td></td><td></td><td></td><td></td><td></td><td></td></tr>
<tr><td>Oligo rev</td><td>AAACCGTCCGCGTTCCGAGGCAGC</td><td></td><td></td><td></td><td></td><td></td><td></td><td></td><td></td><td></td><td></td></tr>
<tr><td>Chromosome</td><td>start</td><td>end</td><td>strand</td><td>MM</td><td>target_seq</td><td>PAM</td><td>alignment</td><td>distance</td><td>position</td><td>gene name</td><td>gene id</td></tr>
<tr><td>GU305914.1</td><td>30630</td><td>30652</td><td>-</td><td>0</td><td>GCTGCCTCGGAACGCGGACG</td><td>AGG</td><td>||||||||[||||||||||||]PAM</td><td>NA</td><td>Intergenic</td><td>NA</td><td>NA</td></tr>
<tr><td>T32060</td><td>CGCGGACGAGGCGCTCGCCGCGG</td><td>1000</td><td></td><td></td><td></td><td></td><td></td><td></td><td></td><td></td><td></td></tr>
<tr><td>Oligo fwd</td><td>CACCCGCGGACGAGGCGCTCGCCG</td><td></td><td></td><td></td><td></td><td></td><td></td><td></td><td></td><td></td><td></td></tr>
<tr><td>Oligo rev</td><td>AAACCGGCGAGCGCCTCGTCCGCG</td><td></td><td></td><td></td><td></td><td></td><td></td><td></td><td></td><td></td><td></td></tr>
<tr><td>Chromosome</td><td>start</td><td>end</td><td>strand</td><td>MM</td><td>target_seq</td><td>PAM</td><td>alignment</td><td>distance</td><td>position</td><td>gene name</td><td>gene id</td></tr>
<tr><td>GU305914.1</td><td>30618</td><td>30640</td><td>-</td><td>0</td><td>CGCGGACGAGGCGCTCGCCG</td><td>CGG</td><td>||||||||[||||||||||||]PAM</td><td>NA</td><td>Intergenic</td><td>NA</td><td>NA</td></tr>
<tr><td>T32061</td><td>GGACGAGGCGCTCGCCGCGGCGG</td><td>1000</td><td></td><td></td><td></td><td></td><td></td><td></td><td></td><td></td><td></td></tr>
<tr><td>Oligo fwd</td><td>CACCGGACGAGGCGCTCGCCGCGG</td><td></td><td></td><td></td><td></td><td></td><td></td><td></td><td></td><td></td><td></td></tr>
<tr><td>Oligo rev</td><td>AAACCCGCGGCGAGCGCCTCGTCC</td><td></td><td></td><td></td><td></td><td></td><td></td><td></td><td></td><td></td><td></td></tr>
<tr><td>Chromosome</td><td>start</td><td>end</td><td>strand</td><td>MM</td><td>target_seq</td><td>PAM</td><td>alignment</td><td>distance</td><td>position</td><td>gene name</td><td>gene id</td></tr>
<tr><td>GU305914.1</td><td>30615</td><td>30637</td><td>-</td><td>0</td><td>GGACGAGGCGCTCGCCGCGG</td><td>CGG</td><td>||||||||[||||||||||||]PAM</td><td>NA</td><td>Intergenic</td><td>NA</td><td>NA</td></tr>
<tr><td>GU305914.1</td><td>165719</td><td>165741</td><td>+</td><td>4</td><td>AGTCGACGCGCTCCCCGCGG</td><td>CGG</td><td>-|-|||-|[|||||-||||||]PAM</td><td>NA</td><td>Intergenic</td><td>NA</td><td>NA</td></tr>
<tr><td>T32062</td><td>AGGCGCTCGCCGCGGCGGAACGG</td><td>1000</td><td></td><td></td><td></td><td></td><td></td><td></td><td></td><td></td><td></td></tr>
<tr><td>Oligo fwd</td><td>CACCAGGCGCTCGCCGCGGCGGAA</td><td></td><td></td><td></td><td></td><td></td><td></td><td></td><td></td><td></td><td></td></tr>
<tr><td>Oligo rev</td><td>AAACTTCCGCCGCGGCGAGCGCCT</td><td></td><td></td><td></td><td></td><td></td><td></td><td></td><td></td><td></td><td></td></tr>
<tr><td>Chromosome</td><td>start</td><td>end</td><td>strand</td><td>MM</td><td>target_seq</td><td>PAM</td><td>alignment</td><td>distance</td><td>position</td><td>gene name</td><td>gene id</td></tr>
<tr><td>GU305914.1</td><td>30610</td><td>30632</td><td>-</td><td>0</td><td>AGGCGCTCGCCGCGGCGGAA</td><td>CGG</td><td>||||||||[||||||||||||]PAM</td><td>NA</td><td>Intergenic</td><td>NA</td><td>NA</td></tr>
<tr><td>T32063</td><td>GGCGCTCGCCGCGGCGGAACGGG</td><td>1000</td><td></td><td></td><td></td><td></td><td></td><td></td><td></td><td></td><td></td></tr>
<tr><td>Oligo fwd</td><td>CACCGGCGCTCGCCGCGGCGGAAC</td><td></td><td></td><td></td><td></td><td></td><td></td><td></td><td></td><td></td><td></td></tr>
<tr><td>Oligo rev</td><td>AAACGTTCCGCCGCGGCGAGCGCC</td><td></td><td></td><td></td><td></td><td></td><td></td><td></td><td></td><td></td><td></td></tr>
<tr><td>Chromosome</td><td>start</td><td>end</td><td>strand</td><td>MM</td><td>target_seq</td><td>PAM</td><td>alignment</td><td>distance</td><td>position</td><td>gene name</td><td>gene id</td></tr>
<tr><td>GU305914.1</td><td>30609</td><td>30631</td><td>-</td><td>0</td><td>GGCGCTCGCCGCGGCGGAAC</td><td>GGG</td><td>||||||||[||||||||||||]PAM</td><td>NA</td><td>Intergenic</td><td>NA</td><td>NA</td></tr>
<tr><td>T32064</td><td>GCGCTCGCCGCGGCGGAACGGGG</td><td>1000</td><td></td><td></td><td></td><td></td><td></td><td></td><td></td><td></td><td></td></tr>
<tr><td>Oligo fwd</td><td>CACCGCGCTCGCCGCGGCGGAACG</td><td></td><td></td><td></td><td></td><td></td><td></td><td></td><td></td><td></td><td></td></tr>
<tr><td>Oligo rev</td><td>AAACCGTTCCGCCGCGGCGAGCGC</td><td></td><td></td><td></td><td></td><td></td><td></td><td></td><td></td><td></td><td></td></tr>
<tr><td>Chromosome</td><td>start</td><td>end</td><td>strand</td><td>MM</td><td>target_seq</td><td>PAM</td><td>alignment</td><td>distance</td><td>position</td><td>gene name</td><td>gene id</td></tr>
<tr><td>GU305914.1</td><td>30608</td><td>30630</td><td>-</td><td>0</td><td>GCGCTCGCCGCGGCGGAACG</td><td>GGG</td><td>||||||||[||||||||||||]PAM</td><td>NA</td><td>Intergenic</td><td>NA</td><td>NA</td></tr>
<tr><td>T32065</td><td>CGCTCGCCGCGGCGGAACGGGGG</td><td>1000</td><td></td><td></td><td></td><td></td><td></td><td></td><td></td><td></td><td></td></tr>
<tr><td>Oligo fwd</td><td>CACCCGCTCGCCGCGGCGGAACGG</td><td></td><td></td><td></td><td></td><td></td><td></td><td></td><td></td><td></td><td></td></tr>
<tr><td>Oligo rev</td><td>AAACCCGTTCCGCCGCGGCGAGCG</td><td></td><td></td><td></td><td></td><td></td><td></td><td></td><td></td><td></td><td></td></tr>
<tr><td>Chromosome</td><td>start</td><td>end</td><td>strand</td><td>MM</td><td>target_seq</td><td>PAM</td><td>alignment</td><td>distance</td><td>position</td><td>gene name</td><td>gene id</td></tr>
<tr><td>GU305914.1</td><td>30607</td><td>30629</td><td>-</td><td>0</td><td>CGCTCGCCGCGGCGGAACGG</td><td>GGG</td><td>||||||||[||||||||||||]PAM</td><td>NA</td><td>Intergenic</td><td>NA</td><td>NA</td></tr>
<tr><td>T32066</td><td>CGCCGCGGCGGAACGGGGGCTGG</td><td>1000</td><td></td><td></td><td></td><td></td><td></td><td></td><td></td><td></td><td></td></tr>
<tr><td>Oligo fwd</td><td>CACCCGCCGCGGCGGAACGGGGGC</td><td></td><td></td><td></td><td></td><td></td><td></td><td></td><td></td><td></td><td></td></tr>
<tr><td>Oligo rev</td><td>AAACGCCCCCGTTCCGCCGCGGCG</td><td></td><td></td><td></td><td></td><td></td><td></td><td></td><td></td><td></td><td></td></tr>
<tr><td>Chromosome</td><td>start</td><td>end</td><td>strand</td><td>MM</td><td>target_seq</td><td>PAM</td><td>alignment</td><td>distance</td><td>position</td><td>gene name</td><td>gene id</td></tr>
<tr><td>GU305914.1</td><td>30603</td><td>30625</td><td>-</td><td>0</td><td>CGCCGCGGCGGAACGGGGGC</td><td>TGG</td><td>||||||||[||||||||||||]PAM</td><td>NA</td><td>Intergenic</td><td>NA</td><td>NA</td></tr>
<tr><td>T32067</td><td>GGCTGGTATGTACCCTACAGTGG</td><td>1000</td><td></td><td></td><td></td><td></td><td></td><td></td><td></td><td></td><td></td></tr>
<tr><td>Oligo fwd</td><td>CACCGGCTGGTATGTACCCTACAG</td><td></td><td></td><td></td><td></td><td></td><td></td><td></td><td></td><td></td><td></td></tr>
<tr><td>Oligo rev</td><td>AAACCTGTAGGGTACATACCAGCC</td><td></td><td></td><td></td><td></td><td></td><td></td><td></td><td></td><td></td><td></td></tr>
<tr><td>Chromosome</td><td>start</td><td>end</td><td>strand</td><td>MM</td><td>target_seq</td><td>PAM</td><td>alignment</td><td>distance</td><td>position</td><td>gene name</td><td>gene id</td></tr>
<tr><td>GU305914.1</td><td>30586</td><td>30608</td><td>-</td><td>0</td><td>GGCTGGTATGTACCCTACAG</td><td>TGG</td><td>||||||||[||||||||||||]PAM</td><td>NA</td><td>Intergenic</td><td>NA</td><td>NA</td></tr>
<tr><td>T32068</td><td>GAAGATTACGACTTCGTTTTCGG</td><td>1000</td><td></td><td></td><td></td><td></td><td></td><td></td><td></td><td></td><td></td></tr>
<tr><td>Oligo fwd</td><td>CACCGAAGATTACGACTTCGTTTT</td><td></td><td></td><td></td><td></td><td></td><td></td><td></td><td></td><td></td><td></td></tr>
<tr><td>Oligo rev</td><td>AAACAAAACGAAGTCGTAATCTTC</td><td></td><td></td><td></td><td></td><td></td><td></td><td></td><td></td><td></td><td></td></tr>
<tr><td>Chromosome</td><td>start</td><td>end</td><td>strand</td><td>MM</td><td>target_seq</td><td>PAM</td><td>alignment</td><td>distance</td><td>position</td><td>gene name</td><td>gene id</td></tr>
<tr><td>GU305914.1</td><td>30560</td><td>30582</td><td>-</td><td>0</td><td>GAAGATTACGACTTCGTTTT</td><td>CGG</td><td>||||||||[||||||||||||]PAM</td><td>NA</td><td>Intergenic</td><td>NA</td><td>NA</td></tr>
<tr><td>T32069</td><td>CTTCGTTTTCGGAACGCCCCCGG</td><td>1000</td><td></td><td></td><td></td><td></td><td></td><td></td><td></td><td></td><td></td></tr>
<tr><td>Oligo fwd</td><td>CACCCTTCGTTTTCGGAACGCCCC</td><td></td><td></td><td></td><td></td><td></td><td></td><td></td><td></td><td></td><td></td></tr>
<tr><td>Oligo rev</td><td>AAACGGGGCGTTCCGAAAACGAAG</td><td></td><td></td><td></td><td></td><td></td><td></td><td></td><td></td><td></td><td></td></tr>
<tr><td>Chromosome</td><td>start</td><td>end</td><td>strand</td><td>MM</td><td>target_seq</td><td>PAM</td><td>alignment</td><td>distance</td><td>position</td><td>gene name</td><td>gene id</td></tr>
<tr><td>GU305914.1</td><td>30549</td><td>30571</td><td>-</td><td>0</td><td>CTTCGTTTTCGGAACGCCCC</td><td>CGG</td><td>||||||||[||||||||||||]PAM</td><td>NA</td><td>Intergenic</td><td>NA</td><td>NA</td></tr>
<tr><td>T32070</td><td>GCTTTACGACTGTCTACATTTGG</td><td>1000</td><td></td><td></td><td></td><td></td><td></td><td></td><td></td><td></td><td></td></tr>
<tr><td>Oligo fwd</td><td>CACCGCTTTACGACTGTCTACATT</td><td></td><td></td><td></td><td></td><td></td><td></td><td></td><td></td><td></td><td></td></tr>
<tr><td>Oligo rev</td><td>AAACAATGTAGACAGTCGTAAAGC</td><td></td><td></td><td></td><td></td><td></td><td></td><td></td><td></td><td></td><td></td></tr>
<tr><td>Chromosome</td><td>start</td><td>end</td><td>strand</td><td>MM</td><td>target_seq</td><td>PAM</td><td>alignment</td><td>distance</td><td>position</td><td>gene name</td><td>gene id</td></tr>
<tr><td>GU305914.1</td><td>30470</td><td>30492</td><td>-</td><td>0</td><td>GCTTTACGACTGTCTACATT</td><td>TGG</td><td>||||||||[||||||||||||]PAM</td><td>NA</td><td>Intergenic</td><td>NA</td><td>NA</td></tr>
<tr><td>T32071</td><td>CTGTCTACATTTGGACATTTTGG</td><td>1000</td><td></td><td></td><td></td><td></td><td></td><td></td><td></td><td></td><td></td></tr>
<tr><td>Oligo fwd</td><td>CACCCTGTCTACATTTGGACATTT</td><td></td><td></td><td></td><td></td><td></td><td></td><td></td><td></td><td></td><td></td></tr>
<tr><td>Oligo rev</td><td>AAACAAATGTCCAAATGTAGACAG</td><td></td><td></td><td></td><td></td><td></td><td></td><td></td><td></td><td></td><td></td></tr>
<tr><td>Chromosome</td><td>start</td><td>end</td><td>strand</td><td>MM</td><td>target_seq</td><td>PAM</td><td>alignment</td><td>distance</td><td>position</td><td>gene name</td><td>gene id</td></tr>
<tr><td>GU305914.1</td><td>30461</td><td>30483</td><td>-</td><td>0</td><td>CTGTCTACATTTGGACATTT</td><td>TGG</td><td>||||||||[||||||||||||]PAM</td><td>NA</td><td>Intergenic</td><td>NA</td><td>NA</td></tr>
</table>

| | | | | | | | | | | | |
|---|---|---|---|---|---|---|---|---|---|---|---|
| T32072 | TACATTTGGACATTTTGGCACGG | 1000 | | | | | | | | | |
| Oligo fwd | CACCTACATTTGGACATTTTGGCA | | | | | | | | | | |
| Oligo rev | AAACTGCCAAAATGTCCAAATGTA | | | | | | | | | | |
| Chromosome | start | end | strand | MM | target_seq | PAM | alignment | distance | position | gene name | gene id |
| GU305914.1 | 30456 | 30478 | - | 0 | TACATTTGGACATTTTGGCA | CGG | ||||||||[||||||||||||]PAM | NA | Intergenic | NA | NA |
| T32073 | ATTTGGACATTTTGGCACGGCGG | 1000 | | | | | | | | | |
| Oligo fwd | CACCATTTGGACATTTTGGCACGG | | | | | | | | | | |
| Oligo rev | AAACCCGTGCCAAAATGTCCAAAT | | | | | | | | | | |
| Chromosome | start | end | strand | MM | target_seq | PAM | alignment | distance | position | gene name | gene id |
| GU305914.1 | 30453 | 30475 | - | 0 | ATTTGGACATTTTGGCACGG | CGG | ||||||||[||||||||||||]PAM | NA | Intergenic | NA | NA |
| T32074 | GGACATTTTGGCACGGCGGCAGG | 1000 | | | | | | | | | |
| Oligo fwd | CACCGGACATTTTGGCACGGCGGC | | | | | | | | | | |
| Oligo rev | AAACGCCGCCGTGCCAAAATGTCC | | | | | | | | | | |
| Chromosome | start | end | strand | MM | target_seq | PAM | alignment | distance | position | gene name | gene id |
| GU305914.1 | 30449 | 30471 | - | 0 | GGACATTTTGGCACGGCGGC | AGG | ||||||||[||||||||||||]PAM | NA | Intergenic | NA | NA |
| T32075 | AAACTCTACCAGACCGCGTACGG | 1000 | | | | | | | | | |
| Oligo fwd | CACCAAACTCTACCAGACCGCGTA | | | | | | | | | | |
| Oligo rev | AAACTACGCGGTCTGGTAGAGTTT | | | | | | | | | | |
| Chromosome | start | end | strand | MM | target_seq | PAM | alignment | distance | position | gene name | gene id |
| GU305914.1 | 30423 | 30445 | - | 0 | AAACTCTACCAGACCGCGTA | CGG | ||||||||[||||||||||||]PAM | NA | Intergenic | NA | NA |
| T32076 | CCGCGTACGGCCGTTTTCATTGG | 1000 | | | | | | | | | |
| Oligo fwd | CACCCCGCGTACGGCCGTTTTCAT | | | | | | | | | | |
| Oligo rev | AAACATGAAAACGGCCGTACGCGG | | | | | | | | | | |
| Chromosome | start | end | strand | MM | target_seq | PAM | alignment | distance | position | gene name | gene id |
| GU305914.1 | 30410 | 30432 | - | 0 | CCGCGTACGGCCGTTTTCAT | TGG | ||||||||[||||||||||||]PAM | NA | Intergenic | NA | NA |
| T32077 | CGCGTACGGCCGTTTTCATTGGG | 1000 | | | | | | | | | |
| Oligo fwd | CACCCGCGTACGGCCGTTTTCATT | | | | | | | | | | |
| Oligo rev | AAACAATGAAAACGGCCGTACGCG | | | | | | | | | | |
| Chromosome | start | end | strand | MM | target_seq | PAM | alignment | distance | position | gene name | gene id |
| GU305914.1 | 30409 | 30431 | - | 0 | CGCGTACGGCCGTTTTCATT | GGG | ||||||||[||||||||||||]PAM | NA | Intergenic | NA | NA |
| T32078 | GCGTACGGCCGTTTTCATTGGGG | 1000 | | | | | | | | | |
| Oligo fwd | CACCGCGTACGGCCGTTTTCATTG | | | | | | | | | | |
| Oligo rev | AAACCAATGAAAACGGCCGTACGC | | | | | | | | | | |
| Chromosome | start | end | strand | MM | target_seq | PAM | alignment | distance | position | gene name | gene id |
| GU305914.1 | 30408 | 30430 | - | 0 | GCGTACGGCCGTTTTCATTG | GGG | ||||||||[||||||||||||]PAM | NA | Intergenic | NA | NA |
| T32079 | TTTCATTGGGGCCGATCACGCGG | 1000 | | | | | | | | | |
| Oligo fwd | CACCTTTCATTGGGGCCGATCACG | | | | | | | | | | |
| Oligo rev | AAACCGTGATCGGCCCCAATGAAA | | | | | | | | | | |
| Chromosome | start | end | strand | MM | target_seq | PAM | alignment | distance | position | gene name | gene id |
| GU305914.1 | 30396 | 30418 | - | 0 | TTTCATTGGGGCCGATCACG | CGG | ||||||||[||||||||||||]PAM | NA | Intergenic | NA | NA |
| T32080 | GCTCAGACTCGCAGAATCTCTGG | 1000 | | | | | | | | | |
| Oligo fwd | CACCGCTCAGACTCGCAGAATCTC | | | | | | | | | | |
| Oligo rev | AAACGAGATTCTGCGAGTCTGAGC | | | | | | | | | | |
| Chromosome | start | end | strand | MM | target_seq | PAM | alignment | distance | position | gene name | gene id |
| GU305914.1 | 30342 | 30364 | - | 0 | GCTCAGACTCGCAGAATCTC | TGG | ||||||||[||||||||||||]PAM | NA | Intergenic | NA | NA |
| T32081 | CTGGATATGTTCTACAAGATAGG | 1000 | | | | | | | | | |
| Oligo fwd | CACCCTGGATATGTTCTACAAGAT | | | | | | | | | | |
| Oligo rev | AAACATCTTGTAGAACATATCCAG | | | | | | | | | | |
| Chromosome | start | end | strand | MM | target_seq | PAM | alignment | distance | position | gene name | gene id |
| GU305914.1 | 30323 | 30345 | - | 0 | CTGGATATGTTCTACAAGAT | AGG | ||||||||[||||||||||||]PAM | NA | Intergenic | NA | NA |
| T32082 | TGTTCTACAAGATAGGCATCAGG | 1000 | | | | | | | | | |
| Oligo fwd | CACCTGTTCTACAAGATAGGCATC | | | | | | | | | | |
| Oligo rev | AAACGATGCCTATCTTGTAGAACA | | | | | | | | | | |
| Chromosome | start | end | strand | MM | target_seq | PAM | alignment | distance | position | gene name | gene id |
| GU305914.1 | 30316 | 30338 | - | 0 | TGTTCTACAAGATAGGCATC | AGG | ||||||||[||||||||||||]PAM | NA | Intergenic | NA | NA |
| T32083 | TCTACAAGATAGGCATCAGGAGG | 1000 | | | | | | | | | |
| Oligo fwd | CACCTCTACAAGATAGGCATCAGG | | | | | | | | | | |
| Oligo rev | AAACCCTGATGCCTATCTTGTAGA | | | | | | | | | | |
| Chromosome | start | end | strand | MM | target_seq | PAM | alignment | distance | position | gene name | gene id |
| GU305914.1 | 30313 | 30335 | - | 0 | TCTACAAGATAGGCATCAGG | AGG | ||||||||[||||||||||||]PAM | NA | Intergenic | NA | NA |
| T32084 | GTTTTTTCACAACTATCGCGTGG | 1000 | | | | | | | | | |
| Oligo fwd | CACCGTTTTTTCACAACTATCGCG | | | | | | | | | | |
| Oligo rev | AAACCGCGATAGTTGTGAAAAAAC | | | | | | | | | | |
| Chromosome | start | end | strand | MM | target_seq | PAM | alignment | distance | position | gene name | gene id |
| GU305914.1 | 30291 | 30313 | - | 0 | GTTTTTTCACAACTATCGCG | TGG | ||||||||[||||||||||||]PAM | NA | Intergenic | NA | NA |
| T32085 | CGCGTGGTACCGAGCAGATGTGG | 1000 | | | | | | | | | |
| Oligo fwd | CACCCGCGTGGTACCGAGCAGATG | | | | | | | | | | |
| Oligo rev | AAACCATCTGCTCGGTACCACGCG | | | | | | | | | | |
| Chromosome | start | end | strand | MM | target_seq | PAM | alignment | distance | position | gene name | gene id |
| GU305914.1 | 30275 | 30297 | - | 0 | CGCGTGGTACCGAGCAGATG | TGG | ||||||||[||||||||||||]PAM | NA | Intergenic | NA | NA |
| T32086 | GCGTGGTACCGAGCAGATGTGGG | 1000 | | | | | | | | | |
| Oligo fwd | CACCGCGTGGTACCGAGCAGATGT | | | | | | | | | | |
| Oligo rev | AAACACATCTGCTCGGTACCACGC | | | | | | | | | | |
| Chromosome | start | end | strand | MM | target_seq | PAM | alignment | distance | position | gene name | gene id |

<table>
<tr><td>GU305914.1</td><td>30274</td><td>30296</td><td>-</td><td>0</td><td>GCGTGGTACCGAGCAGATGT</td><td>GGG</td><td>||||||||[||||||||||||]PAM</td><td>NA</td><td>Intergenic</td><td>NA</td><td>NA</td></tr>
<tr><td>T32087</td><td>CGTGGTACCGAGCAGATGTGGGG</td><td>1000</td><td></td><td></td><td></td><td></td><td></td><td></td><td></td><td></td><td></td></tr>
<tr><td>Oligo fwd</td><td>CACCCGTGGTACCGAGCAGATGTG</td><td></td><td></td><td></td><td></td><td></td><td></td><td></td><td></td><td></td><td></td></tr>
<tr><td>Oligo rev</td><td>AAACCACATCTGCTCGGTACCACG</td><td></td><td></td><td></td><td></td><td></td><td></td><td></td><td></td><td></td><td></td></tr>
<tr><td>Chromosome</td><td>start</td><td>end</td><td>strand</td><td>MM</td><td>target_seq</td><td>PAM</td><td>alignment</td><td>distance</td><td>position</td><td>gene name</td><td>gene id</td></tr>
<tr><td>GU305914.1</td><td>30273</td><td>30295</td><td>-</td><td>0</td><td>CGTGGTACCGAGCAGATGTG</td><td>GGG</td><td>||||||||[||||||||||||]PAM</td><td>NA</td><td>Intergenic</td><td>NA</td><td>NA</td></tr>
<tr><td>T32088</td><td>CGAGCAGATGTGGGGACGACAGG</td><td>1000</td><td></td><td></td><td></td><td></td><td></td><td></td><td></td><td></td><td></td></tr>
<tr><td>Oligo fwd</td><td>CACCCGAGCAGATGTGGGGACGAC</td><td></td><td></td><td></td><td></td><td></td><td></td><td></td><td></td><td></td><td></td></tr>
<tr><td>Oligo rev</td><td>AAACGTCGTCCCCACATCTGCTCG</td><td></td><td></td><td></td><td></td><td></td><td></td><td></td><td></td><td></td><td></td></tr>
<tr><td>Chromosome</td><td>start</td><td>end</td><td>strand</td><td>MM</td><td>target_seq</td><td>PAM</td><td>alignment</td><td>distance</td><td>position</td><td>gene name</td><td>gene id</td></tr>
<tr><td>GU305914.1</td><td>30265</td><td>30287</td><td>-</td><td>0</td><td>CGAGCAGATGTGGGGACGAC</td><td>AGG</td><td>||||||||[||||||||||||]PAM</td><td>NA</td><td>Intergenic</td><td>NA</td><td>NA</td></tr>
<tr><td>T32089</td><td>CGACAGGTTCTTTTTGAAAGCGG</td><td>1000</td><td></td><td></td><td></td><td></td><td></td><td></td><td></td><td></td><td></td></tr>
<tr><td>Oligo fwd</td><td>CACCCGACAGGTTCTTTTTGAAAG</td><td></td><td></td><td></td><td></td><td></td><td></td><td></td><td></td><td></td><td></td></tr>
<tr><td>Oligo rev</td><td>AAACCTTTCAAAAAGAACCTGTCG</td><td></td><td></td><td></td><td></td><td></td><td></td><td></td><td></td><td></td><td></td></tr>
<tr><td>Chromosome</td><td>start</td><td>end</td><td>strand</td><td>MM</td><td>target_seq</td><td>PAM</td><td>alignment</td><td>distance</td><td>position</td><td>gene name</td><td>gene id</td></tr>
<tr><td>GU305914.1</td><td>30249</td><td>30271</td><td>-</td><td>0</td><td>CGACAGGTTCTTTTTGAAAG</td><td>CGG</td><td>||||||||[||||||||||||]PAM</td><td>NA</td><td>Intergenic</td><td>NA</td><td>NA</td></tr>
<tr><td>T32090</td><td>TTCTTTTTGAAAGCGGTCTGAGG</td><td>1000</td><td></td><td></td><td></td><td></td><td></td><td></td><td></td><td></td><td></td></tr>
<tr><td>Oligo fwd</td><td>CACCTTCTTTTTGAAAGCGGTCTG</td><td></td><td></td><td></td><td></td><td></td><td></td><td></td><td></td><td></td><td></td></tr>
<tr><td>Oligo rev</td><td>AAACCAGACCGCTTTCAAAAAGAA</td><td></td><td></td><td></td><td></td><td></td><td></td><td></td><td></td><td></td><td></td></tr>
<tr><td>Chromosome</td><td>start</td><td>end</td><td>strand</td><td>MM</td><td>target_seq</td><td>PAM</td><td>alignment</td><td>distance</td><td>position</td><td>gene name</td><td>gene id</td></tr>
<tr><td>GU305914.1</td><td>30242</td><td>30264</td><td>-</td><td>0</td><td>TTCTTTTTGAAAGCGGTCTG</td><td>AGG</td><td>||||||||[||||||||||||]PAM</td><td>NA</td><td>Intergenic</td><td>NA</td><td>NA</td></tr>
<tr><td>T32091</td><td>AGCGGTCTGAGGCGCTACGACGG</td><td>1000</td><td></td><td></td><td></td><td></td><td></td><td></td><td></td><td></td><td></td></tr>
<tr><td>Oligo fwd</td><td>CACCAGCGGTCTGAGGCGCTACGA</td><td></td><td></td><td></td><td></td><td></td><td></td><td></td><td></td><td></td><td></td></tr>
<tr><td>Oligo rev</td><td>AAACTCGTAGCGCCTCAGACCGCT</td><td></td><td></td><td></td><td></td><td></td><td></td><td></td><td></td><td></td><td></td></tr>
<tr><td>Chromosome</td><td>start</td><td>end</td><td>strand</td><td>MM</td><td>target_seq</td><td>PAM</td><td>alignment</td><td>distance</td><td>position</td><td>gene name</td><td>gene id</td></tr>
<tr><td>GU305914.1</td><td>30231</td><td>30253</td><td>-</td><td>0</td><td>AGCGGTCTGAGGCGCTACGA</td><td>CGG</td><td>||||||||[||||||||||||]PAM</td><td>NA</td><td>Intergenic</td><td>NA</td><td>NA</td></tr>
<tr><td>T32092</td><td>CTGAGGCGCTACGACGGCTTCGG</td><td>1000</td><td></td><td></td><td></td><td></td><td></td><td></td><td></td><td></td><td></td></tr>
<tr><td>Oligo fwd</td><td>CACCCTGAGGCGCTACGACGGCTT</td><td></td><td></td><td></td><td></td><td></td><td></td><td></td><td></td><td></td><td></td></tr>
<tr><td>Oligo rev</td><td>AAACAAGCCGTCGTAGCGCCTCAG</td><td></td><td></td><td></td><td></td><td></td><td></td><td></td><td></td><td></td><td></td></tr>
<tr><td>Chromosome</td><td>start</td><td>end</td><td>strand</td><td>MM</td><td>target_seq</td><td>PAM</td><td>alignment</td><td>distance</td><td>position</td><td>gene name</td><td>gene id</td></tr>
<tr><td>GU305914.1</td><td>30225</td><td>30247</td><td>-</td><td>0</td><td>CTGAGGCGCTACGACGGCTT</td><td>CGG</td><td>||||||||[||||||||||||]PAM</td><td>NA</td><td>Intergenic</td><td>NA</td><td>NA</td></tr>
<tr><td>T32093</td><td>TGAGGCGCTACGACGGCTTCGGG</td><td>1000</td><td></td><td></td><td></td><td></td><td></td><td></td><td></td><td></td><td></td></tr>
<tr><td>Oligo fwd</td><td>CACCTGAGGCGCTACGACGGCTTC</td><td></td><td></td><td></td><td></td><td></td><td></td><td></td><td></td><td></td><td></td></tr>
<tr><td>Oligo rev</td><td>AAACGAAGCCGTCGTAGCGCCTCA</td><td></td><td></td><td></td><td></td><td></td><td></td><td></td><td></td><td></td><td></td></tr>
<tr><td>Chromosome</td><td>start</td><td>end</td><td>strand</td><td>MM</td><td>target_seq</td><td>PAM</td><td>alignment</td><td>distance</td><td>position</td><td>gene name</td><td>gene id</td></tr>
<tr><td>GU305914.1</td><td>30224</td><td>30246</td><td>-</td><td>0</td><td>TGAGGCGCTACGACGGCTTC</td><td>GGG</td><td>||||||||[||||||||||||]PAM</td><td>NA</td><td>Intergenic</td><td>NA</td><td>NA</td></tr>
<tr><td>T32094</td><td>GAGGCGCTACGACGGCTTCGGGG</td><td>1000</td><td></td><td></td><td></td><td></td><td></td><td></td><td></td><td></td><td></td></tr>
<tr><td>Oligo fwd</td><td>CACCGAGGCGCTACGACGGCTTCG</td><td></td><td></td><td></td><td></td><td></td><td></td><td></td><td></td><td></td><td></td></tr>
<tr><td>Oligo rev</td><td>AAACCGAAGCCGTCGTAGCGCCTC</td><td></td><td></td><td></td><td></td><td></td><td></td><td></td><td></td><td></td><td></td></tr>
<tr><td>Chromosome</td><td>start</td><td>end</td><td>strand</td><td>MM</td><td>target_seq</td><td>PAM</td><td>alignment</td><td>distance</td><td>position</td><td>gene name</td><td>gene id</td></tr>
<tr><td>GU305914.1</td><td>30223</td><td>30245</td><td>-</td><td>0</td><td>GAGGCGCTACGACGGCTTCG</td><td>GGG</td><td>||||||||[||||||||||||]PAM</td><td>NA</td><td>Intergenic</td><td>NA</td><td>NA</td></tr>
<tr><td>T32095</td><td>TACGACGGCTTCGGGGATGTTGG</td><td>1000</td><td></td><td></td><td></td><td></td><td></td><td></td><td></td><td></td><td></td></tr>
<tr><td>Oligo fwd</td><td>CACCTACGACGGCTTCGGGGATGT</td><td></td><td></td><td></td><td></td><td></td><td></td><td></td><td></td><td></td><td></td></tr>
<tr><td>Oligo rev</td><td>AAACACATCCCCGAAGCCGTCGTA</td><td></td><td></td><td></td><td></td><td></td><td></td><td></td><td></td><td></td><td></td></tr>
<tr><td>Chromosome</td><td>start</td><td>end</td><td>strand</td><td>MM</td><td>target_seq</td><td>PAM</td><td>alignment</td><td>distance</td><td>position</td><td>gene name</td><td>gene id</td></tr>
<tr><td>GU305914.1</td><td>30216</td><td>30238</td><td>-</td><td>0</td><td>TACGACGGCTTCGGGGATGT</td><td>TGG</td><td>||||||||[||||||||||||]PAM</td><td>NA</td><td>Intergenic</td><td>NA</td><td>NA</td></tr>
<tr><td>T32096</td><td>GTAACAGCCGCTACATCCCGCGG</td><td>1000</td><td></td><td></td><td></td><td></td><td></td><td></td><td></td><td></td><td></td></tr>
<tr><td>Oligo fwd</td><td>CACCGTAACAGCCGCTACATCCCG</td><td></td><td></td><td></td><td></td><td></td><td></td><td></td><td></td><td></td><td></td></tr>
<tr><td>Oligo rev</td><td>AAACCGGGATGTAGCGGCTGTTAC</td><td></td><td></td><td></td><td></td><td></td><td></td><td></td><td></td><td></td><td></td></tr>
<tr><td>Chromosome</td><td>start</td><td>end</td><td>strand</td><td>MM</td><td>target_seq</td><td>PAM</td><td>alignment</td><td>distance</td><td>position</td><td>gene name</td><td>gene id</td></tr>
<tr><td>GU305914.1</td><td>30194</td><td>30216</td><td>-</td><td>0</td><td>GTAACAGCCGCTACATCCCG</td><td>CGG</td><td>||||||||[||||||||||||]PAM</td><td>NA</td><td>Intergenic</td><td>NA</td><td>NA</td></tr>
<tr><td>T32097</td><td>CAGCCGCTACATCCCGCGGCTGG</td><td>1000</td><td></td><td></td><td></td><td></td><td></td><td></td><td></td><td></td><td></td></tr>
<tr><td>Oligo fwd</td><td>CACCCAGCCGCTACATCCCGCGGC</td><td></td><td></td><td></td><td></td><td></td><td></td><td></td><td></td><td></td><td></td></tr>
<tr><td>Oligo rev</td><td>AAACGCCGCGGGATGTAGCGGCTG</td><td></td><td></td><td></td><td></td><td></td><td></td><td></td><td></td><td></td><td></td></tr>
<tr><td>Chromosome</td><td>start</td><td>end</td><td>strand</td><td>MM</td><td>target_seq</td><td>PAM</td><td>alignment</td><td>distance</td><td>position</td><td>gene name</td><td>gene id</td></tr>
<tr><td>GU305914.1</td><td>30190</td><td>30212</td><td>-</td><td>0</td><td>CAGCCGCTACATCCCGCGGC</td><td>TGG</td><td>||||||||[||||||||||||]PAM</td><td>NA</td><td>Intergenic</td><td>NA</td><td>NA</td></tr>
<tr><td>T32098</td><td>AGCCGCTACATCCCGCGGCTGGG</td><td>1000</td><td></td><td></td><td></td><td></td><td></td><td></td><td></td><td></td><td></td></tr>
<tr><td>Oligo fwd</td><td>CACCAGCCGCTACATCCCGCGGCT</td><td></td><td></td><td></td><td></td><td></td><td></td><td></td><td></td><td></td><td></td></tr>
<tr><td>Oligo rev</td><td>AAACAGCCGCGGGATGTAGCGGCT</td><td></td><td></td><td></td><td></td><td></td><td></td><td></td><td></td><td></td><td></td></tr>
<tr><td>Chromosome</td><td>start</td><td>end</td><td>strand</td><td>MM</td><td>target_seq</td><td>PAM</td><td>alignment</td><td>distance</td><td>position</td><td>gene name</td><td>gene id</td></tr>
<tr><td>GU305914.1</td><td>30189</td><td>30211</td><td>-</td><td>0</td><td>AGCCGCTACATCCCGCGGCT</td><td>GGG</td><td>||||||||[||||||||||||]PAM</td><td>NA</td><td>Intergenic</td><td>NA</td><td>NA</td></tr>
<tr><td>T32099</td><td>CTACATCCCGCGGCTGGGTCCGG</td><td>1000</td><td></td><td></td><td></td><td></td><td></td><td></td><td></td><td></td><td></td></tr>
<tr><td>Oligo fwd</td><td>CACCCTACATCCCGCGGCTGGGTC</td><td></td><td></td><td></td><td></td><td></td><td></td><td></td><td></td><td></td><td></td></tr>
<tr><td>Oligo rev</td><td>AAACGACCCAGCCGCGGGATGTAG</td><td></td><td></td><td></td><td></td><td></td><td></td><td></td><td></td><td></td><td></td></tr>
<tr><td>Chromosome</td><td>start</td><td>end</td><td>strand</td><td>MM</td><td>target_seq</td><td>PAM</td><td>alignment</td><td>distance</td><td>position</td><td>gene name</td><td>gene id</td></tr>
<tr><td>GU305914.1</td><td>30184</td><td>30206</td><td>-</td><td>0</td><td>CTACATCCCGCGGCTGGGTC</td><td>CGG</td><td>||||||||[||||||||||||]PAM</td><td>NA</td><td>Intergenic</td><td>NA</td><td>NA</td></tr>
<tr><td>T32100</td><td>CTCCGCCACCCGCCCTCCCCAGG</td><td>1000</td><td></td><td></td><td></td><td></td><td></td><td></td><td></td><td></td><td></td></tr>
<tr><td>Oligo fwd</td><td>CACCCTCCGCCACCCGCCCTCCCC</td><td></td><td></td><td></td><td></td><td></td><td></td><td></td><td></td><td></td><td></td></tr>
<tr><td>Oligo rev</td><td>AAACGGGGAGGGCGGGTGGCGGAG</td><td></td><td></td><td></td><td></td><td></td><td></td><td></td><td></td><td></td><td></td></tr>
<tr><td>Chromosome</td><td>start</td><td>end</td><td>strand</td><td>MM</td><td>target_seq</td><td>PAM</td><td>alignment</td><td>distance</td><td>position</td><td>gene name</td><td>gene id</td></tr>
<tr><td>GU305914.1</td><td>30158</td><td>30180</td><td>-</td><td>0</td><td>CTCCGCCACCCGCCCTCCCC</td><td>AGG</td><td>||||||||[||||||||||||]PAM</td><td>NA</td><td>Intergenic</td><td>NA</td><td>NA</td></tr>
<tr><td>T32101</td><td>ACCCGCCCTCCCCAGGCCGCCGG</td><td>1000</td><td></td><td></td><td></td><td></td><td></td><td></td><td></td><td></td><td></td></tr>
<tr><td>Oligo fwd</td><td>CACCACCCGCCCTCCCCAGGCCGC</td><td></td><td></td><td></td><td></td><td></td><td></td><td></td><td></td><td></td><td></td></tr>
<tr><td>Oligo rev</td><td>AAACGCGGCCTGGGGAGGGCGGGT</td><td></td><td></td><td></td><td></td><td></td><td></td><td></td><td></td><td></td><td></td></tr>
</table>

| | | | | | | | | | | | |
|---|---|---|---|---|---|---|---|---|---|---|---|
| Chromosome | start | end | strand | MM | target_seq | PAM | alignment | distance | position | gene name | gene id |
| GU305914.1 | 30151 | 30173 | - | 0 | ACCCGCCCTCCCCAGGCCGC | CGG | ||||||||[||||||||||||]PAM | NA | Intergenic | NA | NA |
| T32102 | CCCGCCCTCCCCAGGCCGCCGGG | 1000 | | | | | | | | | |
| Oligo fwd | CACCCCCGCCCTCCCCAGGCCGCC | | | | | | | | | | |
| Oligo rev | AAACGGCGGCCTGGGGAGGGCGGG | | | | | | | | | | |
| Chromosome | start | end | strand | MM | target_seq | PAM | alignment | distance | position | gene name | gene id |
| GU305914.1 | 30150 | 30172 | - | 0 | CCCGCCCTCCCCAGGCCGCC | GGG | ||||||||[||||||||||||]PAM | NA | Intergenic | NA | NA |
| T32103 | CCGCCCTCCCCAGGCCGCCGGGG | 1000 | | | | | | | | | |
| Oligo fwd | CACCCCGCCCTCCCCAGGCCGCCG | | | | | | | | | | |
| Oligo rev | AAACCGGCGGCCTGGGGAGGGCGG | | | | | | | | | | |
| Chromosome | start | end | strand | MM | target_seq | PAM | alignment | distance | position | gene name | gene id |
| GU305914.1 | 30149 | 30171 | - | 0 | CCGCCCTCCCCAGGCCGCCG | GGG | ||||||||[||||||||||||]PAM | NA | Intergenic | NA | NA |
| T32104 | GCTGCAGCTCTCGCTACCGCTGG | 1000 | | | | | | | | | |
| Oligo fwd | CACCGCTGCAGCTCTCGCTACCGC | | | | | | | | | | |
| Oligo rev | AAACGCGGTAGCGAGAGCTGCAGC | | | | | | | | | | |
| Chromosome | start | end | strand | MM | target_seq | PAM | alignment | distance | position | gene name | gene id |
| GU305914.1 | 30127 | 30149 | - | 0 | GCTGCAGCTCTCGCTACCGC | TGG | ||||||||[||||||||||||]PAM | NA | Intergenic | NA | NA |
| T32105 | CGTGCCCGTCACCACCGCGCCGG | 1000 | | | | | | | | | |
| Oligo fwd | CACCCGTGCCCGTCACCACCGCGC | | | | | | | | | | |
| Oligo rev | AAACGCGCGGTGGTGACGGGCACG | | | | | | | | | | |
| Chromosome | start | end | strand | MM | target_seq | PAM | alignment | distance | position | gene name | gene id |
| GU305914.1 | 30055 | 30077 | - | 0 | CGTGCCCGTCACCACCGCGC | CGG | ||||||||[||||||||||||]PAM | NA | Intergenic | NA | NA |
| T32106 | CGTCACCACCGCGCCGGTGACGG | 1000 | | | | | | | | | |
| Oligo fwd | CACCCGTCACCACCGCGCCGGTGA | | | | | | | | | | |
| Oligo rev | AAACTCACCGGCGCGGTGGTGACG | | | | | | | | | | |
| Chromosome | start | end | strand | MM | target_seq | PAM | alignment | distance | position | gene name | gene id |
| GU305914.1 | 30049 | 30071 | - | 0 | CGTCACCACCGCGCCGGTGA | CGG | ||||||||[||||||||||||]PAM | NA | Intergenic | NA | NA |
| T32107 | GGTGACGGTCGCGCAGCCTCTGG | 1000 | | | | | | | | | |
| Oligo fwd | CACCGGTGACGGTCGCGCAGCCTC | | | | | | | | | | |
| Oligo rev | AAACGAGGCTGCGCGACCGTCACC | | | | | | | | | | |
| Chromosome | start | end | strand | MM | target_seq | PAM | alignment | distance | position | gene name | gene id |
| GU305914.1 | 30034 | 30056 | - | 0 | GGTGACGGTCGCGCAGCCTC | TGG | ||||||||[||||||||||||]PAM | NA | Intergenic | NA | NA |
| T32108 | GCCGCCGCCGCCGCCAGCACCGG | 1000 | | | | | | | | | |
| Oligo fwd | CACCGCCGCCGCCGCCGCCAGCAC | | | | | | | | | | |
| Oligo rev | AAACGTGCTGGCGGCGGCGGCGGC | | | | | | | | | | |
| Chromosome | start | end | strand | MM | target_seq | PAM | alignment | distance | position | gene name | gene id |
| GU305914.1 | 29998 | 30020 | - | 0 | GCCGCCGCCGCCGCCAGCAC | CGG | ||||||||[||||||||||||]PAM | NA | Intergenic | NA | NA |
| GU305914.1 | 48151 | 48173 | - | 3 | GCCGCCACCGCCGCCGCCAC | AGG | ||||||-|[|||||||--|||]PAM | NA | Intergenic | NA | NA |
| GU305914.1 | 170532 | 170554 | + | 4 | GTGGCCGCCGCCGCCACCAT | CGG | |--|||||[||||||||-||-]PAM | NA | Intergenic | NA | NA |
| GU305914.1 | 24876 | 24898 | - | 3 | GCCGCCGTCGCCGCCAACAT | CGG | |||||||-[||||||||-||-]PAM | NA | Intergenic | NA | NA |
| T32109 | CCGAAAAGTCTCCTCCTCAGCGG | 1000 | | | | | | | | | |
| Oligo fwd | CACCCCGAAAAGTCTCCTCCTCAG | | | | | | | | | | |
| Oligo rev | AAACCTGAGGAGGAGACTTTTCGG | | | | | | | | | | |
| Chromosome | start | end | strand | MM | target_seq | PAM | alignment | distance | position | gene name | gene id |
| GU305914.1 | 29972 | 29994 | - | 0 | CCGAAAAGTCTCCTCCTCAG | CGG | ||||||||[||||||||||||]PAM | NA | Intergenic | NA | NA |
| T32110 | TCAGCGGATCATCGAGCCCGAGG | 1000 | | | | | | | | | |
| Oligo fwd | CACCTCAGCGGATCATCGAGCCCG | | | | | | | | | | |
| Oligo rev | AAACCGGGCTCGATGATCCGCTGA | | | | | | | | | | |
| Chromosome | start | end | strand | MM | target_seq | PAM | alignment | distance | position | gene name | gene id |
| GU305914.1 | 29956 | 29978 | - | 0 | TCAGCGGATCATCGAGCCCG | AGG | ||||||||[||||||||||||]PAM | NA | Intergenic | NA | NA |
| T32111 | TCGAGCCCGAGGACTTCGCGCGG | 1000 | | | | | | | | | |
| Oligo fwd | CACCTCGAGCCCGAGGACTTCGCG | | | | | | | | | | |
| Oligo rev | AAACCGCGAAGTCCTCGGGCTCGA | | | | | | | | | | |
| Chromosome | start | end | strand | MM | target_seq | PAM | alignment | distance | position | gene name | gene id |
| GU305914.1 | 29945 | 29967 | - | 0 | TCGAGCCCGAGGACTTCGCG | CGG | ||||||||[||||||||||||]PAM | NA | Intergenic | NA | NA |
| T32112 | CGAGCCCGAGGACTTCGCGCGGG | 1000 | | | | | | | | | |
| Oligo fwd | CACCCGAGCCCGAGGACTTCGCGC | | | | | | | | | | |
| Oligo rev | AAACGCGCGAAGTCCTCGGGCTCG | | | | | | | | | | |
| Chromosome | start | end | strand | MM | target_seq | PAM | alignment | distance | position | gene name | gene id |
| GU305914.1 | 29944 | 29966 | - | 0 | CGAGCCCGAGGACTTCGCGC | GGG | ||||||||[||||||||||||]PAM | NA | Intergenic | NA | NA |
| GU305914.1 | 65750 | 65772 | + | 4 | GACGCCCGAGGACATCGCGC | GGG | ---|||||[|||||-||||||]PAM | NA | Intergenic | NA | NA |
| T32113 | TCAACAATCCCGATCTCTGCCGG | 1000 | | | | | | | | | |
| Oligo fwd | CACCTCAACAATCCCGATCTCTGC | | | | | | | | | | |
| Oligo rev | AAACGCAGAGATCGGGATTGTTGA | | | | | | | | | | |
| Chromosome | start | end | strand | MM | target_seq | PAM | alignment | distance | position | gene name | gene id |
| GU305914.1 | 29918 | 29940 | - | 0 | TCAACAATCCCGATCTCTGC | CGG | ||||||||[||||||||||||]PAM | NA | Intergenic | NA | NA |
| T32114 | CAACAATCCCGATCTCTGCCGGG | 1000 | | | | | | | | | |
| Oligo fwd | CACCCAACAATCCCGATCTCTGCC | | | | | | | | | | |
| Oligo rev | AAACGGCAGAGATCGGGATTGTTG | | | | | | | | | | |
| Chromosome | start | end | strand | MM | target_seq | PAM | alignment | distance | position | gene name | gene id |
| GU305914.1 | 29917 | 29939 | - | 0 | CAACAATCCCGATCTCTGCC | GGG | ||||||||[||||||||||||]PAM | NA | Intergenic | NA | NA |
| T32115 | GGAGTTCGTCAAGATCCGCAAGG | 1000 | | | | | | | | | |
| Oligo fwd | CACCGGAGTTCGTCAAGATCCGCA | | | | | | | | | | |
| Oligo rev | AAACTGCGGATCTTGACGAACTCC | | | | | | | | | | |
| Chromosome | start | end | strand | MM | target_seq | PAM | alignment | distance | position | gene name | gene id |

GU305914.1 29896 29918 - 0 GGAGTTCGTCAAGATCCGCA AGG ||||||||[||||||||||||]PAM NA Intergenic NA NA

T32116 GAGTTCGTCAAGATCCGCAAGGG 1000
Oligo fwd CACCGAGTTCGTCAAGATCCGCAA
Oligo rev AAACTTGCGGATCTTGACGAACTC
Chromosome start end strand MM target_seq PAM alignment distance position gene name gene id
GU305914.1 29895 29917 - 0 GAGTTCGTCAAGATCCGCAA GGG ||||||||[||||||||||||]PAM NA Intergenic NA NA

T32117 TCGTCAAGATCCGCAAGGGATGG 1000
Oligo fwd CACCTCGTCAAGATCCGCAAGGGA
Oligo rev AAACTCCCTTGCGGATCTTGACGA
Chromosome start end strand MM target_seq PAM alignment distance position gene name gene id
GU305914.1 29891 29913 - 0 TCGTCAAGATCCGCAAGGGA TGG ||||||||[||||||||||||]PAM NA Intergenic NA NA

T32118 TCAAGATCCGCAAGGGATGGCGG 1000
Oligo fwd CACCTCAAGATCCGCAAGGGATGG
Oligo rev AAACCCATCCCTTGCGGATCTTGA
Chromosome start end strand MM target_seq PAM alignment distance position gene name gene id
GU305914.1 29888 29910 - 0 TCAAGATCCGCAAGGGATGG CGG ||||||||[||||||||||||]PAM NA Intergenic NA NA

T32119 GATGGCGGATGCCGCTGACCTGG 1000
Oligo fwd CACCGATGGCGGATGCCGCTGACC
Oligo rev AAACGGTCAGCGGCATCCGCCATC
Chromosome start end strand MM target_seq PAM alignment distance position gene name gene id
GU305914.1 29873 29895 - 0 GATGGCGGATGCCGCTGACC TGG ||||||||[||||||||||||]PAM NA Intergenic NA NA

T32120 GGCCCAAGAATTACACACTCTGG 1000
Oligo fwd CACCGGCCCAAGAATTACACACTC
Oligo rev AAACGAGTGTGTAATTCTTGGGCC
Chromosome start end strand MM target_seq PAM alignment distance position gene name gene id
GU305914.1 29852 29874 - 0 GGCCCAAGAATTACACACTC TGG ||||||||[||||||||||||]PAM NA Intergenic NA NA

T32121 CAAGAATTACACACTCTGGTTGG 1000
Oligo fwd CACCCAAGAATTACACACTCTGGT
Oligo rev AAACACCAGAGTGTGTAATTCTTG
Chromosome start end strand MM target_seq PAM alignment distance position gene name gene id
GU305914.1 29848 29870 - 0 CAAGAATTACACACTCTGGT TGG ||||||||[||||||||||||]PAM NA Intergenic NA NA

T32122 CACACTCTGGTTGGAGACCGAGG 1000
Oligo fwd CACCCACACTCTGGTTGGAGACCG
Oligo rev AAACCGGTCTCCAACCAGAGTGTG
Chromosome start end strand MM target_seq PAM alignment distance position gene name gene id
GU305914.1 29839 29861 - 0 CACACTCTGGTTGGAGACCG AGG ||||||||[||||||||||||]PAM NA Intergenic NA NA

T32123 GCCGAACTCCGCATGCCCGTCGG 1000
Oligo fwd CACCGCCGAACTCCGCATGCCCGT
Oligo rev AAACACGGGCATGCGGAGTTCGGC
Chromosome start end strand MM target_seq PAM alignment distance position gene name gene id
GU305914.1 29817 29839 - 0 GCCGAACTCCGCATGCCCGT CGG ||||||||[||||||||||||]PAM NA Intergenic NA NA

T32124 CCGAACTCCGCATGCCCGTCGGG 1000
Oligo fwd CACCCCGAACTCCGCATGCCCGTC
Oligo rev AAACGACGGGCATGCGGAGTTCGG
Chromosome start end strand MM target_seq PAM alignment distance position gene name gene id
GU305914.1 29816 29838 - 0 CCGAACTCCGCATGCCCGTC GGG ||||||||[||||||||||||]PAM NA Intergenic NA NA
GU305914.1 139301 139323 + 4 CCGACCGCAGCGTGCCCGTC TGG ||||-|-|[-||-||||||||]PAM NA Intergenic NA NA

T32125 CGAACTCCGCATGCCCGTCGGGG 1000
Oligo fwd CACCCGAACTCCGCATGCCCGTCG
Oligo rev AAACCGACGGGCATGCGGAGTTCG
Chromosome start end strand MM target_seq PAM alignment distance position gene name gene id
GU305914.1 29815 29837 - 0 CGAACTCCGCATGCCCGTCG GGG ||||||||[||||||||||||]PAM NA Intergenic NA NA

T32126 TGCCCGTCGGGGAGATCAGTCGG 1000
Oligo fwd CACCTGCCCGTCGGGGAGATCAGT
Oligo rev AAACACTGATCTCCCCGACGGGCA
Chromosome start end strand MM target_seq PAM alignment distance position gene name gene id
GU305914.1 29804 29826 - 0 TGCCCGTCGGGGAGATCAGT CGG ||||||||[||||||||||||]PAM NA Intergenic NA NA

T32127 GGAGATCAGTCGGCTCTCGCAGG 1000
Oligo fwd CACCGGAGATCAGTCGGCTCTCGC
Oligo rev AAACGCGAGAGCCGACTGATCTCC
Chromosome start end strand MM target_seq PAM alignment distance position gene name gene id
GU305914.1 29794 29816 - 0 GGAGATCAGTCGGCTCTCGC AGG ||||||||[||||||||||||]PAM NA Intergenic NA NA

T32128 GCAGGAGTACCTCTGCTGTCCGG 1000
Oligo fwd CACCGCAGGAGTACCTCTGCTGTC
Oligo rev AAACGACAGCAGAGGTACTCCTGC
Chromosome start end strand MM target_seq PAM alignment distance position gene name gene id
GU305914.1 29776 29798 - 0 GCAGGAGTACCTCTGCTGTC CGG ||||||||[||||||||||||]PAM NA Intergenic NA NA

T32129 GGAGTACCTCTGCTGTCCGGAGG 1000
Oligo fwd CACCGGAGTACCTCTGCTGTCCGG
Oligo rev AAACCCGGACAGCAGAGGTACTCC
Chromosome start end strand MM target_seq PAM alignment distance position gene name gene id
GU305914.1 29773 29795 - 0 GGAGTACCTCTGCTGTCCGG AGG ||||||||[||||||||||||]PAM NA Intergenic NA NA

T32130 TCCGGAGGATCTCATCGTCCTGG 1000
Oligo fwd CACCTCCGGAGGATCTCATCGTCC

| | | | | | | | | | | | |
|---|---|---|---|---|---|---|---|---|---|---|---|
| Oligo rev | AAACGGACGATGAGATCCTCCGGA | | | | | | | | | | |
| Chromosome | start | end | strand | MM | target_seq | PAM | alignment | distance | position | gene name | gene id |
| GU305914.1 | 29758 | 29780 | - | 0 | TCCGGAGGATCTCATCGTCC | TGG | \|\|\|\|\|\|\|\|[\|\|\|\|\|\|\|\|\|\|\|\|]PAM | NA | Intergenic | NA | NA |
| T32131 | CCGGAGGATCTCATCGTCCTGGG | 1000 | | | | | | | | | |
| Oligo fwd | CACCCCGGAGGATCTCATCGTCCT | | | | | | | | | | |
| Oligo rev | AAACAGGACGATGAGATCCTCCGG | | | | | | | | | | |
| Chromosome | start | end | strand | MM | target_seq | PAM | alignment | distance | position | gene name | gene id |
| GU305914.1 | 29757 | 29779 | - | 0 | CCGGAGGATCTCATCGTCCT | GGG | \|\|\|\|\|\|\|\|[\|\|\|\|\|\|\|\|\|\|\|\|]PAM | NA | Intergenic | NA | NA |
| T32132 | GAGGATCTCATCGTCCTGGGCGG | 1000 | | | | | | | | | |
| Oligo fwd | CACCGAGGATCTCATCGTCCTGGG | | | | | | | | | | |
| Oligo rev | AAACCCCAGGACGATGAGATCCTC | | | | | | | | | | |
| Chromosome | start | end | strand | MM | target_seq | PAM | alignment | distance | position | gene name | gene id |
| GU305914.1 | 29754 | 29776 | - | 0 | GAGGATCTCATCGTCCTGGG | CGG | \|\|\|\|\|\|\|\|[\|\|\|\|\|\|\|\|\|\|\|\|]PAM | NA | Intergenic | NA | NA |
| T32133 | GGCGGCGTCCGCCGCTCGACCGG | 1000 | | | | | | | | | |
| Oligo fwd | CACCGGCGGCGTCCGCCGCTCGAC | | | | | | | | | | |
| Oligo rev | AAACGTCGAGCGGCGGACGCCGCC | | | | | | | | | | |
| Chromosome | start | end | strand | MM | target_seq | PAM | alignment | distance | position | gene name | gene id |
| GU305914.1 | 29736 | 29758 | - | 0 | GGCGGCGTCCGCCGCTCGAC | CGG | \|\|\|\|\|\|\|\|[\|\|\|\|\|\|\|\|\|\|\|\|]PAM | NA | Intergenic | NA | NA |
| T32134 | GACCGGCGATGACAAGAACGAGG | 1000 | | | | | | | | | |
| Oligo fwd | CACCGACCGGCGATGACAAGAACG | | | | | | | | | | |
| Oligo rev | AAACCGTTCTTGTCATCGCCGGTC | | | | | | | | | | |
| Chromosome | start | end | strand | MM | target_seq | PAM | alignment | distance | position | gene name | gene id |
| GU305914.1 | 29719 | 29741 | - | 0 | GACCGGCGATGACAAGAACG | AGG | \|\|\|\|\|\|\|\|[\|\|\|\|\|\|\|\|\|\|\|\|]PAM | NA | Intergenic | NA | NA |
| T32135 | CGATGACAAGAACGAGGACCTGG | 1000 | | | | | | | | | |
| Oligo fwd | CACCCGATGACAAGAACGAGGACC | | | | | | | | | | |
| Oligo rev | AAACGGTCCTCGTTCTTGTCATCG | | | | | | | | | | |
| Chromosome | start | end | strand | MM | target_seq | PAM | alignment | distance | position | gene name | gene id |
| GU305914.1 | 29713 | 29735 | - | 0 | CGATGACAAGAACGAGGACC | TGG | \|\|\|\|\|\|\|\|[\|\|\|\|\|\|\|\|\|\|\|\|]PAM | NA | Intergenic | NA | NA |
| T32136 | GATGACAAGAACGAGGACCTGGG | 1000 | | | | | | | | | |
| Oligo fwd | CACCGATGACAAGAACGAGGACCT | | | | | | | | | | |
| Oligo rev | AAACAGGTCCTCGTTCTTGTCATC | | | | | | | | | | |
| Chromosome | start | end | strand | MM | target_seq | PAM | alignment | distance | position | gene name | gene id |
| GU305914.1 | 29712 | 29734 | - | 0 | GATGACAAGAACGAGGACCT | GGG | \|\|\|\|\|\|\|\|[\|\|\|\|\|\|\|\|\|\|\|\|]PAM | NA | Intergenic | NA | NA |
| T32137 | GACCTGGGCGTCCTCTTCGCCGG | 1000 | | | | | | | | | |
| Oligo fwd | CACCGACCTGGGCGTCCTCTTCGC | | | | | | | | | | |
| Oligo rev | AAACGCGAAGAGGACGCCCAGGTC | | | | | | | | | | |
| Chromosome | start | end | strand | MM | target_seq | PAM | alignment | distance | position | gene name | gene id |
| GU305914.1 | 29697 | 29719 | - | 0 | GACCTGGGCGTCCTCTTCGC | CGG | \|\|\|\|\|\|\|\|[\|\|\|\|\|\|\|\|\|\|\|\|]PAM | NA | Intergenic | NA | NA |
| T32138 | GGGCGTCCTCTTCGCCGGCGAGG | 1000 | | | | | | | | | |
| Oligo fwd | CACCGGGCGTCCTCTTCGCCGGCG | | | | | | | | | | |
| Oligo rev | AAACCGCCGGCGAAGAGGACGCCC | | | | | | | | | | |
| Chromosome | start | end | strand | MM | target_seq | PAM | alignment | distance | position | gene name | gene id |
| GU305914.1 | 29692 | 29714 | - | 0 | GGGCGTCCTCTTCGCCGGCG | AGG | \|\|\|\|\|\|\|\|[\|\|\|\|\|\|\|\|\|\|\|\|]PAM | NA | Intergenic | NA | NA |
| T32139 | CGTCCTCTTCGCCGGCGAGGAGG | 1000 | | | | | | | | | |
| Oligo fwd | CACCCGTCCTCTTCGCCGGCGAGG | | | | | | | | | | |
| Oligo rev | AAACCCTCGCCGGCGAAGAGGACG | | | | | | | | | | |
| Chromosome | start | end | strand | MM | target_seq | PAM | alignment | distance | position | gene name | gene id |
| GU305914.1 | 29689 | 29711 | - | 0 | CGTCCTCTTCGCCGGCGAGG | AGG | \|\|\|\|\|\|\|\|[\|\|\|\|\|\|\|\|\|\|\|\|]PAM | NA | Intergenic | NA | NA |
| GU305914.1 | 111941 | 111963 | + | 4 | CGTCGCCTTCGCCTGCGACG | TGG | \|\|\|\|--\|\|[\|\|\|\|\|-\|\|\|\|-\|]PAM | NA | Intergenic | NA | NA |
| T32140 | GTCCTCTTCGCCGGCGAGGAGGG | 1000 | | | | | | | | | |
| Oligo fwd | CACCGTCCTCTTCGCCGGCGAGGA | | | | | | | | | | |
| Oligo rev | AAACTCCTCGCCGGCGAAGAGGAC | | | | | | | | | | |
| Chromosome | start | end | strand | MM | target_seq | PAM | alignment | distance | position | gene name | gene id |
| GU305914.1 | 29688 | 29710 | - | 0 | GTCCTCTTCGCCGGCGAGGA | GGG | \|\|\|\|\|\|\|\|[\|\|\|\|\|\|\|\|\|\|\|\|]PAM | NA | Intergenic | NA | NA |
| T32141 | CGTCGACGACGCCATCTATCTGG | 1000 | | | | | | | | | |
| Oligo fwd | CACCCGTCGACGACGCCATCTATC | | | | | | | | | | |
| Oligo rev | AAACGATAGATGGCGTCGTCGACG | | | | | | | | | | |
| Chromosome | start | end | strand | MM | target_seq | PAM | alignment | distance | position | gene name | gene id |
| GU305914.1 | 29641 | 29663 | - | 0 | CGTCGACGACGCCATCTATC | TGG | \|\|\|\|\|\|\|\|[\|\|\|\|\|\|\|\|\|\|\|\|]PAM | NA | Intergenic | NA | NA |
| GU305914.1 | 75083 | 75105 | - | 4 | GGTCGACAACGCCAGCTACC | AGG | -\|\|\|\|\|\|-[\|\|\|\|\|\|-\|\|\|-\|]PAM | NA | Intergenic | NA | NA |
| T32142 | GACGACGCCATCTATCTGGTCGG | 1000 | | | | | | | | | |
| Oligo fwd | CACCGACGACGCCATCTATCTGGT | | | | | | | | | | |
| Oligo rev | AAACACCAGATAGATGGCGTCGTC | | | | | | | | | | |
| Chromosome | start | end | strand | MM | target_seq | PAM | alignment | distance | position | gene name | gene id |
| GU305914.1 | 29637 | 29659 | - | 0 | GACGACGCCATCTATCTGGT | CGG | \|\|\|\|\|\|\|\|[\|\|\|\|\|\|\|\|\|\|\|\|]PAM | NA | Intergenic | NA | NA |
| T32143 | ACGACGCCATCTATCTGGTCGGG | 1000 | | | | | | | | | |
| Oligo fwd | CACCACGACGCCATCTATCTGGTC | | | | | | | | | | |
| Oligo rev | AAACGACCAGATAGATGGCGTCGT | | | | | | | | | | |
| Chromosome | start | end | strand | MM | target_seq | PAM | alignment | distance | position | gene name | gene id |
| GU305914.1 | 29636 | 29658 | - | 0 | ACGACGCCATCTATCTGGTC | GGG | \|\|\|\|\|\|\|\|[\|\|\|\|\|\|\|\|\|\|\|\|]PAM | NA | Intergenic | NA | NA |
| T32144 | CGACGCCATCTATCTGGTCGGGG | 1000 | | | | | | | | | |
| Oligo fwd | CACCCGACGCCATCTATCTGGTCG | | | | | | | | | | |
| Oligo rev | AAACCGACCAGATAGATGGCGTCG | | | | | | | | | | |
| Chromosome | start | end | strand | MM | target_seq | PAM | alignment | distance | position | gene name | gene id |
| GU305914.1 | 29635 | 29657 | - | 0 | CGACGCCATCTATCTGGTCG | GGG | \|\|\|\|\|\|\|\|[\|\|\|\|\|\|\|\|\|\|\|\|]PAM | NA | Intergenic | NA | NA |

| | | | | | | | | | | | |
|---|---|---|---|---|---|---|---|---|---|---|---|
| T32145 | CTATCTGGTCGGGGACTCCCCGG | 1000 | | | | | | | | | |
| Oligo fwd | CACCCTATCTGGTCGGGGACTCCC | | | | | | | | | | |
| Oligo rev | AAACGGGAGTCCCCGACCAGATAG | | | | | | | | | | |
| Chromosome | start | end | strand | MM | target_seq | PAM | alignment | distance | position | gene name | gene id |
| GU305914.1 | 29626 | 29648 | - | 0 | CTATCTGGTCGGGGACTCCC | CGG | \|\|\|\|\|\|\|\|[\|\|\|\|\|\|\|\|\|\|\|\|]PAM | NA | Intergenic | NA | NA |
| T32146 | TCTGGTCGGGGACTCCCCGGAGG | 1000 | | | | | | | | | |
| Oligo fwd | CACCTCTGGTCGGGGACTCCCCGG | | | | | | | | | | |
| Oligo rev | AAACCCGGGGAGTCCCCGACCAGA | | | | | | | | | | |
| Chromosome | start | end | strand | MM | target_seq | PAM | alignment | distance | position | gene name | gene id |
| GU305914.1 | 29623 | 29645 | - | 0 | TCTGGTCGGGGACTCCCCGG | AGG | \|\|\|\|\|\|\|\|[\|\|\|\|\|\|\|\|\|\|\|\|]PAM | NA | Intergenic | NA | NA |
| T32147 | CTGGTCGGGGACTCCCCGGAGGG | 1000 | | | | | | | | | |
| Oligo fwd | CACCCTGGTCGGGGACTCCCCGGA | | | | | | | | | | |
| Oligo rev | AAACTCCGGGGAGTCCCCGACCAG | | | | | | | | | | |
| Chromosome | start | end | strand | MM | target_seq | PAM | alignment | distance | position | gene name | gene id |
| GU305914.1 | 29622 | 29644 | - | 0 | CTGGTCGGGGACTCCCCGGA | GGG | \|\|\|\|\|\|\|\|[\|\|\|\|\|\|\|\|\|\|\|\|]PAM | NA | Intergenic | NA | NA |
| T32148 | CCCCGGAGGGCTTCATCAAGAGG | 1000 | | | | | | | | | |
| Oligo fwd | CACCCCCCGGAGGGCTTCATCAAG | | | | | | | | | | |
| Oligo rev | AAACCTTGATGAAGCCCTCCGGGG | | | | | | | | | | |
| Chromosome | start | end | strand | MM | target_seq | PAM | alignment | distance | position | gene name | gene id |
| GU305914.1 | 29609 | 29631 | - | 0 | CCCCGGAGGGCTTCATCAAG | AGG | \|\|\|\|\|\|\|\|[\|\|\|\|\|\|\|\|\|\|\|\|]PAM | NA | Intergenic | NA | NA |
| T32149 | CCCGGAGGGCTTCATCAAGAGGG | 1000 | | | | | | | | | |
| Oligo fwd | CACCCCCGGAGGGCTTCATCAAGA | | | | | | | | | | |
| Oligo rev | AAACTCTTGATGAAGCCCTCCGGG | | | | | | | | | | |
| Chromosome | start | end | strand | MM | target_seq | PAM | alignment | distance | position | gene name | gene id |
| GU305914.1 | 29608 | 29630 | - | 0 | CCCGGAGGGCTTCATCAAGA | GGG | \|\|\|\|\|\|\|\|[\|\|\|\|\|\|\|\|\|\|\|\|]PAM | NA | Intergenic | NA | NA |
| T32150 | CCGGAGGGCTTCATCAAGAGGGG | 1000 | | | | | | | | | |
| Oligo fwd | CACCCCGGAGGGCTTCATCAAGAG | | | | | | | | | | |
| Oligo rev | AAACCTCTTGATGAAGCCCTCCGG | | | | | | | | | | |
| Chromosome | start | end | strand | MM | target_seq | PAM | alignment | distance | position | gene name | gene id |
| GU305914.1 | 29607 | 29629 | - | 0 | CCGGAGGGCTTCATCAAGAG | GGG | \|\|\|\|\|\|\|\|[\|\|\|\|\|\|\|\|\|\|\|\|]PAM | NA | Intergenic | NA | NA |
| T32151 | GCTTCATCAAGAGGGGACTCAGG | 1000 | | | | | | | | | |
| Oligo fwd | CACCGCTTCATCAAGAGGGGACTC | | | | | | | | | | |
| Oligo rev | AAACGAGTCCCCTCTTGATGAAGC | | | | | | | | | | |
| Chromosome | start | end | strand | MM | target_seq | PAM | alignment | distance | position | gene name | gene id |
| GU305914.1 | 29600 | 29622 | - | 0 | GCTTCATCAAGAGGGGACTC | AGG | \|\|\|\|\|\|\|\|[\|\|\|\|\|\|\|\|\|\|\|\|]PAM | NA | Intergenic | NA | NA |
| T32152 | GGCGCTTCTACCCCATCTATCGG | 1000 | | | | | | | | | |
| Oligo fwd | CACCGGCGCTTCTACCCCATCTAT | | | | | | | | | | |
| Oligo rev | AAACATAGATGGGGTAGAAGCGCC | | | | | | | | | | |
| Chromosome | start | end | strand | MM | target_seq | PAM | alignment | distance | position | gene name | gene id |
| GU305914.1 | 29579 | 29601 | - | 0 | GGCGCTTCTACCCCATCTAT | CGG | \|\|\|\|\|\|\|\|[\|\|\|\|\|\|\|\|\|\|\|\|]PAM | NA | Intergenic | NA | NA |
| T32153 | GCGCTTCTACCCCATCTATCGGG | 1000 | | | | | | | | | |
| Oligo fwd | CACCGCGCTTCTACCCCATCTATC | | | | | | | | | | |
| Oligo rev | AAACGATAGATGGGGTAGAAGCGC | | | | | | | | | | |
| Chromosome | start | end | strand | MM | target_seq | PAM | alignment | distance | position | gene name | gene id |
| GU305914.1 | 29578 | 29600 | - | 0 | GCGCTTCTACCCCATCTATC | GGG | \|\|\|\|\|\|\|\|[\|\|\|\|\|\|\|\|\|\|\|\|]PAM | NA | Intergenic | NA | NA |
| T32154 | CATCTATCGGGAGATAGAGACGG | 1000 | | | | | | | | | |
| Oligo fwd | CACCCATCTATCGGGAGATAGAGA | | | | | | | | | | |
| Oligo rev | AAACTCTCTATCTCCCGATAGATG | | | | | | | | | | |
| Chromosome | start | end | strand | MM | target_seq | PAM | alignment | distance | position | gene name | gene id |
| GU305914.1 | 29566 | 29588 | - | 0 | CATCTATCGGGAGATAGAGA | CGG | \|\|\|\|\|\|\|\|[\|\|\|\|\|\|\|\|\|\|\|\|]PAM | NA | Intergenic | NA | NA |
| T32155 | TATCGGGAGATAGAGACGGACGG | 1000 | | | | | | | | | |
| Oligo fwd | CACCTATCGGGAGATAGAGACGGA | | | | | | | | | | |
| Oligo rev | AAACTCCGTCTCTATCTCCCGATA | | | | | | | | | | |
| Chromosome | start | end | strand | MM | target_seq | PAM | alignment | distance | position | gene name | gene id |
| GU305914.1 | 29562 | 29584 | - | 0 | TATCGGGAGATAGAGACGGA | CGG | \|\|\|\|\|\|\|\|[\|\|\|\|\|\|\|\|\|\|\|\|]PAM | NA | Intergenic | NA | NA |
| T32156 | GAGACGGACGGCTGCAGCGCCGG | 1000 | | | | | | | | | |
| Oligo fwd | CACCGAGACGGACGGCTGCAGCGC | | | | | | | | | | |
| Oligo rev | AAACGCGCTGCAGCCGTCCGTCTC | | | | | | | | | | |
| Chromosome | start | end | strand | MM | target_seq | PAM | alignment | distance | position | gene name | gene id |
| GU305914.1 | 29550 | 29572 | - | 0 | GAGACGGACGGCTGCAGCGC | CGG | \|\|\|\|\|\|\|\|[\|\|\|\|\|\|\|\|\|\|\|\|]PAM | NA | Intergenic | NA | NA |
| T32157 | CGCCGGCGTCACCGAGTACCTGG | 1000 | | | | | | | | | |
| Oligo fwd | CACCCGCCGGCGTCACCGAGTACC | | | | | | | | | | |
| Oligo rev | AAACGGTACTCGGTGACGCCGGCG | | | | | | | | | | |
| Chromosome | start | end | strand | MM | target_seq | PAM | alignment | distance | position | gene name | gene id |
| GU305914.1 | 29533 | 29555 | - | 0 | CGCCGGCGTCACCGAGTACC | TGG | \|\|\|\|\|\|\|\|[\|\|\|\|\|\|\|\|\|\|\|\|]PAM | NA | Intergenic | NA | NA |
| T32158 | AAGTTCTCCGCCACCTACCCCGG | 1000 | | | | | | | | | |
| Oligo fwd | CACCAAGTTCTCCGCCACCTACCC | | | | | | | | | | |
| Oligo rev | AAACGGGTAGGTGGCGGAGAACTT | | | | | | | | | | |
| Chromosome | start | end | strand | MM | target_seq | PAM | alignment | distance | position | gene name | gene id |
| GU305914.1 | 29475 | 29497 | - | 0 | AAGTTCTCCGCCACCTACCC | CGG | \|\|\|\|\|\|\|\|[\|\|\|\|\|\|\|\|\|\|\|\|]PAM | NA | Intergenic | NA | NA |
| T32159 | GCACCGTCTTCTCCCTGCCCTGG | 1000 | | | | | | | | | |
| Oligo fwd | CACCGCACCGTCTTCTCCCTGCCC | | | | | | | | | | |
| Oligo rev | AAACGGGCAGGGAGAAGACGGTGC | | | | | | | | | | |
| Chromosome | start | end | strand | MM | target_seq | PAM | alignment | distance | position | gene name | gene id |

<table>
<tr><td>GU305914.1</td><td>29453</td><td>29475</td><td>-</td><td>0</td><td>GCACCGTCTTCTCCCTGCCC</td><td>TGG</td><td>||||||||[||||||||||||]PAM</td><td>NA</td><td>Intergenic</td><td>NA</td><td>NA</td></tr>
<tr><td>GU305914.1</td><td>146708</td><td>146730</td><td>+</td><td>4</td><td>TCGCCGTCTACTCCGTGCCC</td><td>TGG</td><td>-|-|||||[|-||||-|||||]PAM</td><td>NA</td><td>Intergenic</td><td>NA</td><td>NA</td></tr>
<tr><td></td><td></td><td></td><td></td><td></td><td></td><td></td><td></td><td></td><td></td><td></td><td></td></tr>
<tr><td>T32160</td><td>TTCTCCCTGCCCTGGCCCACCGG</td><td>1000</td><td></td><td></td><td></td><td></td><td></td><td></td><td></td><td></td><td></td></tr>
<tr><td>Oligo fwd</td><td>CACCTTCTCCCTGCCCTGGCCCAC</td><td></td><td></td><td></td><td></td><td></td><td></td><td></td><td></td><td></td><td></td></tr>
<tr><td>Oligo rev</td><td>AAACGTGGGCCAGGGCAGGGAGAA</td><td></td><td></td><td></td><td></td><td></td><td></td><td></td><td></td><td></td><td></td></tr>
<tr><td>Chromosome</td><td>start</td><td>end</td><td>strand</td><td>MM</td><td>target_seq</td><td>PAM</td><td>alignment</td><td>distance</td><td>position</td><td>gene name</td><td>gene id</td></tr>
<tr><td>GU305914.1</td><td>29445</td><td>29467</td><td>-</td><td>0</td><td>TTCTCCCTGCCCTGGCCCAC</td><td>CGG</td><td>||||||||[||||||||||||]PAM</td><td>NA</td><td>Intergenic</td><td>NA</td><td>NA</td></tr>
<tr><td></td><td></td><td></td><td></td><td></td><td></td><td></td><td></td><td></td><td></td><td></td><td></td></tr>
<tr><td>T32161</td><td>CAGACGACGCAAGTTCGTCACGG</td><td>1000</td><td></td><td></td><td></td><td></td><td></td><td></td><td></td><td></td><td></td></tr>
<tr><td>Oligo fwd</td><td>CACCCAGACGACGCAAGTTCGTCA</td><td></td><td></td><td></td><td></td><td></td><td></td><td></td><td></td><td></td><td></td></tr>
<tr><td>Oligo rev</td><td>AAACTGACGAACTTGCGTCGTCTG</td><td></td><td></td><td></td><td></td><td></td><td></td><td></td><td></td><td></td><td></td></tr>
<tr><td>Chromosome</td><td>start</td><td>end</td><td>strand</td><td>MM</td><td>target_seq</td><td>PAM</td><td>alignment</td><td>distance</td><td>position</td><td>gene name</td><td>gene id</td></tr>
<tr><td>GU305914.1</td><td>29398</td><td>29420</td><td>-</td><td>0</td><td>CAGACGACGCAAGTTCGTCA</td><td>CGG</td><td>||||||||[||||||||||||]PAM</td><td>NA</td><td>Intergenic</td><td>NA</td><td>NA</td></tr>
<tr><td>GU305914.1</td><td>135600</td><td>135622</td><td>+</td><td>4</td><td>CAGACGCGGCGAGTTCTTCA</td><td>GGG</td><td>||||||--[||-|||||-|||]PAM</td><td>NA</td><td>Intergenic</td><td>NA</td><td>NA</td></tr>
<tr><td></td><td></td><td></td><td></td><td></td><td></td><td></td><td></td><td></td><td></td><td></td><td></td></tr>
<tr><td>T32162</td><td>AGACGACGCAAGTTCGTCACGGG</td><td>1000</td><td></td><td></td><td></td><td></td><td></td><td></td><td></td><td></td><td></td></tr>
<tr><td>Oligo fwd</td><td>CACCAGACGACGCAAGTTCGTCAC</td><td></td><td></td><td></td><td></td><td></td><td></td><td></td><td></td><td></td><td></td></tr>
<tr><td>Oligo rev</td><td>AAACGTGACGAACTTGCGTCGTCT</td><td></td><td></td><td></td><td></td><td></td><td></td><td></td><td></td><td></td><td></td></tr>
<tr><td>Chromosome</td><td>start</td><td>end</td><td>strand</td><td>MM</td><td>target_seq</td><td>PAM</td><td>alignment</td><td>distance</td><td>position</td><td>gene name</td><td>gene id</td></tr>
<tr><td>GU305914.1</td><td>29397</td><td>29419</td><td>-</td><td>0</td><td>AGACGACGCAAGTTCGTCAC</td><td>GGG</td><td>||||||||[||||||||||||]PAM</td><td>NA</td><td>Intergenic</td><td>NA</td><td>NA</td></tr>
<tr><td></td><td></td><td></td><td></td><td></td><td></td><td></td><td></td><td></td><td></td><td></td><td></td></tr>
<tr><td>T32163</td><td>CGCAAGTTCGTCACGGGACGCGG</td><td>1000</td><td></td><td></td><td></td><td></td><td></td><td></td><td></td><td></td><td></td></tr>
<tr><td>Oligo fwd</td><td>CACCCGCAAGTTCGTCACGGGACG</td><td></td><td></td><td></td><td></td><td></td><td></td><td></td><td></td><td></td><td></td></tr>
<tr><td>Oligo rev</td><td>AAACCGTCCCGTGACGAACTTGCG</td><td></td><td></td><td></td><td></td><td></td><td></td><td></td><td></td><td></td><td></td></tr>
<tr><td>Chromosome</td><td>start</td><td>end</td><td>strand</td><td>MM</td><td>target_seq</td><td>PAM</td><td>alignment</td><td>distance</td><td>position</td><td>gene name</td><td>gene id</td></tr>
<tr><td>GU305914.1</td><td>29391</td><td>29413</td><td>-</td><td>0</td><td>CGCAAGTTCGTCACGGGACG</td><td>CGG</td><td>||||||||[||||||||||||]PAM</td><td>NA</td><td>Intergenic</td><td>NA</td><td>NA</td></tr>
<tr><td></td><td></td><td></td><td></td><td></td><td></td><td></td><td></td><td></td><td></td><td></td><td></td></tr>
<tr><td>T32164</td><td>GGGACGCGGTTCTGACACCATGG</td><td>1000</td><td></td><td></td><td></td><td></td><td></td><td></td><td></td><td></td><td></td></tr>
<tr><td>Oligo fwd</td><td>CACCGGGACGCGGTTCTGACACCA</td><td></td><td></td><td></td><td></td><td></td><td></td><td></td><td></td><td></td><td></td></tr>
<tr><td>Oligo rev</td><td>AAACTGGTGTCAGAACCGCGTCCC</td><td></td><td></td><td></td><td></td><td></td><td></td><td></td><td></td><td></td><td></td></tr>
<tr><td>Chromosome</td><td>start</td><td>end</td><td>strand</td><td>MM</td><td>target_seq</td><td>PAM</td><td>alignment</td><td>distance</td><td>position</td><td>gene name</td><td>gene id</td></tr>
<tr><td>GU305914.1</td><td>29377</td><td>29399</td><td>-</td><td>0</td><td>GGGACGCGGTTCTGACACCA</td><td>TGG</td><td>||||||||[||||||||||||]PAM</td><td>NA</td><td>Intergenic</td><td>NA</td><td>NA</td></tr>
<tr><td></td><td></td><td></td><td></td><td></td><td></td><td></td><td></td><td></td><td></td><td></td><td></td></tr>
<tr><td>T32165</td><td>TGACACCATGGTCTACTTCGCGG</td><td>1000</td><td></td><td></td><td></td><td></td><td></td><td></td><td></td><td></td><td></td></tr>
<tr><td>Oligo fwd</td><td>CACCTGACACCATGGTCTACTTCG</td><td></td><td></td><td></td><td></td><td></td><td></td><td></td><td></td><td></td><td></td></tr>
<tr><td>Oligo rev</td><td>AAACCGAAGTAGACCATGGTGTCA</td><td></td><td></td><td></td><td></td><td></td><td></td><td></td><td></td><td></td><td></td></tr>
<tr><td>Chromosome</td><td>start</td><td>end</td><td>strand</td><td>MM</td><td>target_seq</td><td>PAM</td><td>alignment</td><td>distance</td><td>position</td><td>gene name</td><td>gene id</td></tr>
<tr><td>GU305914.1</td><td>29365</td><td>29387</td><td>-</td><td>0</td><td>TGACACCATGGTCTACTTCG</td><td>CGG</td><td>||||||||[||||||||||||]PAM</td><td>NA</td><td>Intergenic</td><td>NA</td><td>NA</td></tr>
<tr><td>GU305914.1</td><td>36604</td><td>36626</td><td>-</td><td>4</td><td>TGATCTCATGGTCAACTTCG</td><td>CGG</td><td>|||---||[|||||-||||||]PAM</td><td>NA</td><td>Intergenic</td><td>NA</td><td>NA</td></tr>
<tr><td></td><td></td><td></td><td></td><td></td><td></td><td></td><td></td><td></td><td></td><td></td><td></td></tr>
<tr><td>T32166</td><td>GTCTACTTCGCGGCCGTCATCGG</td><td>1000</td><td></td><td></td><td></td><td></td><td></td><td></td><td></td><td></td><td></td></tr>
<tr><td>Oligo fwd</td><td>CACCGTCTACTTCGCGGCCGTCAT</td><td></td><td></td><td></td><td></td><td></td><td></td><td></td><td></td><td></td><td></td></tr>
<tr><td>Oligo rev</td><td>AAACATGACGGCCGCGAAGTAGAC</td><td></td><td></td><td></td><td></td><td></td><td></td><td></td><td></td><td></td><td></td></tr>
<tr><td>Chromosome</td><td>start</td><td>end</td><td>strand</td><td>MM</td><td>target_seq</td><td>PAM</td><td>alignment</td><td>distance</td><td>position</td><td>gene name</td><td>gene id</td></tr>
<tr><td>GU305914.1</td><td>29355</td><td>29377</td><td>-</td><td>0</td><td>GTCTACTTCGCGGCCGTCAT</td><td>CGG</td><td>||||||||[||||||||||||]PAM</td><td>NA</td><td>Intergenic</td><td>NA</td><td>NA</td></tr>
<tr><td></td><td></td><td></td><td></td><td></td><td></td><td></td><td></td><td></td><td></td><td></td><td></td></tr>
<tr><td>T32167</td><td>GGCCGTCATCGGACGTTACTTGG</td><td>1000</td><td></td><td></td><td></td><td></td><td></td><td></td><td></td><td></td><td></td></tr>
<tr><td>Oligo fwd</td><td>CACCGGCCGTCATCGGACGTTACT</td><td></td><td></td><td></td><td></td><td></td><td></td><td></td><td></td><td></td><td></td></tr>
<tr><td>Oligo rev</td><td>AAACAGTAACGTCCGATGACGGCC</td><td></td><td></td><td></td><td></td><td></td><td></td><td></td><td></td><td></td><td></td></tr>
<tr><td>Chromosome</td><td>start</td><td>end</td><td>strand</td><td>MM</td><td>target_seq</td><td>PAM</td><td>alignment</td><td>distance</td><td>position</td><td>gene name</td><td>gene id</td></tr>
<tr><td>GU305914.1</td><td>29344</td><td>29366</td><td>-</td><td>0</td><td>GGCCGTCATCGGACGTTACT</td><td>TGG</td><td>||||||||[||||||||||||]PAM</td><td>NA</td><td>Intergenic</td><td>NA</td><td>NA</td></tr>
<tr><td></td><td></td><td></td><td></td><td></td><td></td><td></td><td></td><td></td><td></td><td></td><td></td></tr>
<tr><td>T32168</td><td>CTTGGACCCCGTCTTCCGCGAGG</td><td>1000</td><td></td><td></td><td></td><td></td><td></td><td></td><td></td><td></td><td></td></tr>
<tr><td>Oligo fwd</td><td>CACCCTTGGACCCCGTCTTCCGCG</td><td></td><td></td><td></td><td></td><td></td><td></td><td></td><td></td><td></td><td></td></tr>
<tr><td>Oligo rev</td><td>AAACCGCGGAAGACGGGGTCCAAG</td><td></td><td></td><td></td><td></td><td></td><td></td><td></td><td></td><td></td><td></td></tr>
<tr><td>Chromosome</td><td>start</td><td>end</td><td>strand</td><td>MM</td><td>target_seq</td><td>PAM</td><td>alignment</td><td>distance</td><td>position</td><td>gene name</td><td>gene id</td></tr>
<tr><td>GU305914.1</td><td>29326</td><td>29348</td><td>-</td><td>0</td><td>CTTGGACCCCGTCTTCCGCG</td><td>AGG</td><td>||||||||[||||||||||||]PAM</td><td>NA</td><td>Intergenic</td><td>NA</td><td>NA</td></tr>
<tr><td></td><td></td><td></td><td></td><td></td><td></td><td></td><td></td><td></td><td></td><td></td><td></td></tr>
<tr><td>T32169</td><td>CGTCTTCCGCGAGGTCCTCTTGG</td><td>1000</td><td></td><td></td><td></td><td></td><td></td><td></td><td></td><td></td><td></td></tr>
<tr><td>Oligo fwd</td><td>CACCCGTCTTCCGCGAGGTCCTCT</td><td></td><td></td><td></td><td></td><td></td><td></td><td></td><td></td><td></td><td></td></tr>
<tr><td>Oligo rev</td><td>AAACAGAGGACCTCGCGGAAGACG</td><td></td><td></td><td></td><td></td><td></td><td></td><td></td><td></td><td></td><td></td></tr>
<tr><td>Chromosome</td><td>start</td><td>end</td><td>strand</td><td>MM</td><td>target_seq</td><td>PAM</td><td>alignment</td><td>distance</td><td>position</td><td>gene name</td><td>gene id</td></tr>
<tr><td>GU305914.1</td><td>29317</td><td>29339</td><td>-</td><td>0</td><td>CGTCTTCCGCGAGGTCCTCT</td><td>TGG</td><td>||||||||[||||||||||||]PAM</td><td>NA</td><td>Intergenic</td><td>NA</td><td>NA</td></tr>
<tr><td></td><td></td><td></td><td></td><td></td><td></td><td></td><td></td><td></td><td></td><td></td><td></td></tr>
<tr><td>T32170</td><td>CTCTTGGCCGTCAACGAGAAAGG</td><td>1000</td><td></td><td></td><td></td><td></td><td></td><td></td><td></td><td></td><td></td></tr>
<tr><td>Oligo fwd</td><td>CACCCTCTTGGCCGTCAACGAGAA</td><td></td><td></td><td></td><td></td><td></td><td></td><td></td><td></td><td></td><td></td></tr>
<tr><td>Oligo rev</td><td>AAACTTCTCGTTGACGGCCAAGAG</td><td></td><td></td><td></td><td></td><td></td><td></td><td></td><td></td><td></td><td></td></tr>
<tr><td>Chromosome</td><td>start</td><td>end</td><td>strand</td><td>MM</td><td>target_seq</td><td>PAM</td><td>alignment</td><td>distance</td><td>position</td><td>gene name</td><td>gene id</td></tr>
<tr><td>GU305914.1</td><td>29301</td><td>29323</td><td>-</td><td>0</td><td>CTCTTGGCCGTCAACGAGAA</td><td>AGG</td><td>||||||||[||||||||||||]PAM</td><td>NA</td><td>Intergenic</td><td>NA</td><td>NA</td></tr>
<tr><td></td><td></td><td></td><td></td><td></td><td></td><td></td><td></td><td></td><td></td><td></td><td></td></tr>
<tr><td>T32171</td><td>CCATCTTTTCCTACAACCCCTGG</td><td>1000</td><td></td><td></td><td></td><td></td><td></td><td></td><td></td><td></td><td></td></tr>
<tr><td>Oligo fwd</td><td>CACCCCATCTTTTCCTACAACCCC</td><td></td><td></td><td></td><td></td><td></td><td></td><td></td><td></td><td></td><td></td></tr>
<tr><td>Oligo rev</td><td>AAACGGGGTTGTAGGAAAAGATGG</td><td></td><td></td><td></td><td></td><td></td><td></td><td></td><td></td><td></td><td></td></tr>
<tr><td>Chromosome</td><td>start</td><td>end</td><td>strand</td><td>MM</td><td>target_seq</td><td>PAM</td><td>alignment</td><td>distance</td><td>position</td><td>gene name</td><td>gene id</td></tr>
<tr><td>GU305914.1</td><td>29276</td><td>29298</td><td>-</td><td>0</td><td>CCATCTTTTCCTACAACCCC</td><td>TGG</td><td>||||||||[||||||||||||]PAM</td><td>NA</td><td>Intergenic</td><td>NA</td><td>NA</td></tr>
<tr><td></td><td></td><td></td><td></td><td></td><td></td><td></td><td></td><td></td><td></td><td></td><td></td></tr>
<tr><td>T32172</td><td>CATCTTTTCCTACAACCCCTGGG</td><td>1000</td><td></td><td></td><td></td><td></td><td></td><td></td><td></td><td></td><td></td></tr>
<tr><td>Oligo fwd</td><td>CACCCATCTTTTCCTACAACCCCT</td><td></td><td></td><td></td><td></td><td></td><td></td><td></td><td></td><td></td><td></td></tr>
<tr><td>Oligo rev</td><td>AAACAGGGGTTGTAGGAAAAGATG</td><td></td><td></td><td></td><td></td><td></td><td></td><td></td><td></td><td></td><td></td></tr>
<tr><td>Chromosome</td><td>start</td><td>end</td><td>strand</td><td>MM</td><td>target_seq</td><td>PAM</td><td>alignment</td><td>distance</td><td>position</td><td>gene name</td><td>gene id</td></tr>
<tr><td>GU305914.1</td><td>29275</td><td>29297</td><td>-</td><td>0</td><td>CATCTTTTCCTACAACCCCT</td><td>GGG</td><td>||||||||[||||||||||||]PAM</td><td>NA</td><td>Intergenic</td><td>NA</td><td>NA</td></tr>
<tr><td></td><td></td><td></td><td></td><td></td><td></td><td></td><td></td><td></td><td></td><td></td><td></td></tr>
<tr><td>T32173</td><td>CGAGATCACCAAGATCTGCGAGG</td><td>1000</td><td></td><td></td><td></td><td></td><td></td><td></td><td></td><td></td><td></td></tr>
<tr><td>Oligo fwd</td><td>CACCCGAGATCACCAAGATCTGCG</td><td></td><td></td><td></td><td></td><td></td><td></td><td></td><td></td><td></td><td></td></tr>
<tr><td>Oligo rev</td><td>AAACCGCAGATCTTGGTGATCTCG</td><td></td><td></td><td></td><td></td><td></td><td></td><td></td><td></td><td></td><td></td></tr>
<tr><td>Chromosome</td><td>start</td><td>end</td><td>strand</td><td>MM</td><td>target_seq</td><td>PAM</td><td>alignment</td><td>distance</td><td>position</td><td>gene name</td><td>gene id</td></tr>
<tr><td>GU305914.1</td><td>29248</td><td>29270</td><td>-</td><td>0</td><td>CGAGATCACCAAGATCTGCG</td><td>AGG</td><td>||||||||[||||||||||||]PAM</td><td>NA</td><td>Intergenic</td><td>NA</td><td>NA</td></tr>
</table>

| | | | | | | | | | | | |
|---|---|---|---|---|---|---|---|---|---|---|---|
| T32174 | GAGATCACCAAGATCTGCGAGGG | 1000 | | | | | | | | | |
| Oligo fwd | CACCGAGATCACCAAGATCTGCGA | | | | | | | | | | |
| Oligo rev | AAACTCGCAGATCTTGGTGATCTC | | | | | | | | | | |
| Chromosome | start | end | strand | MM | target_seq | PAM | alignment | distance | position | gene name | gene id |
| GU305914.1 | 29247 | 29269 | - | 0 | GAGATCACCAAGATCTGCGA | GGG | \|\|\|\|\|\|\|\|[\|\|\|\|\|\|\|\|\|\|\|\|]PAM | NA | Intergenic | NA | NA |

| | | | | | | | | | | | |
|---|---|---|---|---|---|---|---|---|---|---|---|
| T32175 | CGTCATGCAGCTATTCACTCTGG | 1000 | | | | | | | | | |
| Oligo fwd | CACCCGTCATGCAGCTATTCACTC | | | | | | | | | | |
| Oligo rev | AAACGAGTGAATAGCTGCATGACG | | | | | | | | | | |
| Chromosome | start | end | strand | MM | target_seq | PAM | alignment | distance | position | gene name | gene id |
| GU305914.1 | 29224 | 29246 | - | 0 | CGTCATGCAGCTATTCACTC | TGG | \|\|\|\|\|\|\|\|[\|\|\|\|\|\|\|\|\|\|\|\|]PAM | NA | Intergenic | NA | NA |

| | | | | | | | | | | | |
|---|---|---|---|---|---|---|---|---|---|---|---|
| T32176 | GTCATGCAGCTATTCACTCTGGG | 1000 | | | | | | | | | |
| Oligo fwd | CACCGTCATGCAGCTATTCACTCT | | | | | | | | | | |
| Oligo rev | AAACAGAGTGAATAGCTGCATGAC | | | | | | | | | | |
| Chromosome | start | end | strand | MM | target_seq | PAM | alignment | distance | position | gene name | gene id |
| GU305914.1 | 29223 | 29245 | - | 0 | GTCATGCAGCTATTCACTCT | GGG | \|\|\|\|\|\|\|\|[\|\|\|\|\|\|\|\|\|\|\|\|]PAM | NA | Intergenic | NA | NA |

| | | | | | | | | | | | |
|---|---|---|---|---|---|---|---|---|---|---|---|
| T32177 | AGCTATTCACTCTGGGACTCCGG | 1000 | | | | | | | | | |
| Oligo fwd | CACCAGCTATTCACTCTGGGACTC | | | | | | | | | | |
| Oligo rev | AAACGAGTCCCAGAGTGAATAGCT | | | | | | | | | | |
| Chromosome | start | end | strand | MM | target_seq | PAM | alignment | distance | position | gene name | gene id |
| GU305914.1 | 29216 | 29238 | - | 0 | AGCTATTCACTCTGGGACTC | CGG | \|\|\|\|\|\|\|\|[\|\|\|\|\|\|\|\|\|\|\|\|]PAM | NA | Intergenic | NA | NA |

| | | | | | | | | | | | |
|---|---|---|---|---|---|---|---|---|---|---|---|
| T32178 | GCTATTCACTCTGGGACTCCGGG | 1000 | | | | | | | | | |
| Oligo fwd | CACCGCTATTCACTCTGGGACTCC | | | | | | | | | | |
| Oligo rev | AAACGGAGTCCCAGAGTGAATAGC | | | | | | | | | | |
| Chromosome | start | end | strand | MM | target_seq | PAM | alignment | distance | position | gene name | gene id |
| GU305914.1 | 29215 | 29237 | - | 0 | GCTATTCACTCTGGGACTCC | GGG | \|\|\|\|\|\|\|\|[\|\|\|\|\|\|\|\|\|\|\|\|]PAM | NA | Intergenic | NA | NA |

| | | | | | | | | | | | |
|---|---|---|---|---|---|---|---|---|---|---|---|
| T32179 | CTATTCACTCTGGGACTCCGGGG | 1000 | | | | | | | | | |
| Oligo fwd | CACCCTATTCACTCTGGGACTCCG | | | | | | | | | | |
| Oligo rev | AAACCGGAGTCCCAGAGTGAATAG | | | | | | | | | | |
| Chromosome | start | end | strand | MM | target_seq | PAM | alignment | distance | position | gene name | gene id |
| GU305914.1 | 29214 | 29236 | - | 0 | CTATTCACTCTGGGACTCCG | GGG | \|\|\|\|\|\|\|\|[\|\|\|\|\|\|\|\|\|\|\|\|]PAM | NA | Intergenic | NA | NA |

| | | | | | | | | | | | |
|---|---|---|---|---|---|---|---|---|---|---|---|
| T32180 | TCAAAAAGAACTATCGATTCAGG | 1000 | | | | | | | | | |
| Oligo fwd | CACCTCAAAAAGAACTATCGATTC | | | | | | | | | | |
| Oligo rev | AAACGAATCGATAGTTCTTTTTGA | | | | | | | | | | |
| Chromosome | start | end | strand | MM | target_seq | PAM | alignment | distance | position | gene name | gene id |
| GU305914.1 | 29189 | 29211 | - | 0 | TCAAAAAGAACTATCGATTC | AGG | \|\|\|\|\|\|\|\|[\|\|\|\|\|\|\|\|\|\|\|\|]PAM | NA | Intergenic | NA | NA |

| | | | | | | | | | | | |
|---|---|---|---|---|---|---|---|---|---|---|---|
| T32181 | CAAAAAGAACTATCGATTCAGGG | 1000 | | | | | | | | | |
| Oligo fwd | CACCCAAAAAGAACTATCGATTCA | | | | | | | | | | |
| Oligo rev | AAACTGAATCGATAGTTCTTTTTG | | | | | | | | | | |
| Chromosome | start | end | strand | MM | target_seq | PAM | alignment | distance | position | gene name | gene id |
| GU305914.1 | 29188 | 29210 | - | 0 | CAAAAAGAACTATCGATTCA | GGG | \|\|\|\|\|\|\|\|[\|\|\|\|\|\|\|\|\|\|\|\|]PAM | NA | Intergenic | NA | NA |

| | | | | | | | | | | | |
|---|---|---|---|---|---|---|---|---|---|---|---|
| T32182 | AGAACTATCGATTCAGGGCTCGG | 1000 | | | | | | | | | |
| Oligo fwd | CACCAGAACTATCGATTCAGGGCT | | | | | | | | | | |
| Oligo rev | AAACAGCCCTGAATCGATAGTTCT | | | | | | | | | | |
| Chromosome | start | end | strand | MM | target_seq | PAM | alignment | distance | position | gene name | gene id |
| GU305914.1 | 29183 | 29205 | - | 0 | AGAACTATCGATTCAGGGCT | CGG | \|\|\|\|\|\|\|\|[\|\|\|\|\|\|\|\|\|\|\|\|]PAM | NA | Intergenic | NA | NA |

| | | | | | | | | | | | |
|---|---|---|---|---|---|---|---|---|---|---|---|
| T32183 | GGCTCGGCTGCCCGTCGCGCTGG | 1000 | | | | | | | | | |
| Oligo fwd | CACCGGCTCGGCTGCCCGTCGCGC | | | | | | | | | | |
| Oligo rev | AAACGCGCGACGGGCAGCCGAGCC | | | | | | | | | | |
| Chromosome | start | end | strand | MM | target_seq | PAM | alignment | distance | position | gene name | gene id |
| GU305914.1 | 29167 | 29189 | - | 0 | GGCTCGGCTGCCCGTCGCGC | TGG | \|\|\|\|\|\|\|\|[\|\|\|\|\|\|\|\|\|\|\|\|]PAM | NA | Intergenic | NA | NA |

| | | | | | | | | | | | |
|---|---|---|---|---|---|---|---|---|---|---|---|
| T32184 | GCTCGGCTGCCCGTCGCGCTGGG | 1000 | | | | | | | | | |
| Oligo fwd | CACCGCTCGGCTGCCCGTCGCGCT | | | | | | | | | | |
| Oligo rev | AAACAGCGCGACGGGCAGCCGAGC | | | | | | | | | | |
| Chromosome | start | end | strand | MM | target_seq | PAM | alignment | distance | position | gene name | gene id |
| GU305914.1 | 29166 | 29188 | - | 0 | GCTCGGCTGCCCGTCGCGCT | GGG | \|\|\|\|\|\|\|\|[\|\|\|\|\|\|\|\|\|\|\|\|]PAM | NA | Intergenic | NA | NA |

| | | | | | | | | | | | |
|---|---|---|---|---|---|---|---|---|---|---|---|
| T32185 | TGCCCGTCGCGCTGGGCGACCGG | 1000 | | | | | | | | | |
| Oligo fwd | CACCTGCCCGTCGCGCTGGGCGAC | | | | | | | | | | |
| Oligo rev | AAACGTCGCCCAGCGCGACGGGCA | | | | | | | | | | |
| Chromosome | start | end | strand | MM | target_seq | PAM | alignment | distance | position | gene name | gene id |
| GU305914.1 | 29159 | 29181 | - | 0 | TGCCCGTCGCGCTGGGCGAC | CGG | \|\|\|\|\|\|\|\|[\|\|\|\|\|\|\|\|\|\|\|\|]PAM | NA | Intergenic | NA | NA |

| | | | | | | | | | | | |
|---|---|---|---|---|---|---|---|---|---|---|---|
| T32186 | CAAGTGTCCCCACGTCAGTCCGG | 1000 | | | | | | | | | |
| Oligo fwd | CACCCAAGTGTCCCCACGTCAGTC | | | | | | | | | | |
| Oligo rev | AAACGACTGACGTGGGGACACTTG | | | | | | | | | | |
| Chromosome | start | end | strand | MM | target_seq | PAM | alignment | distance | position | gene name | gene id |
| GU305914.1 | 29131 | 29153 | - | 0 | CAAGTGTCCCCACGTCAGTC | CGG | \|\|\|\|\|\|\|\|[\|\|\|\|\|\|\|\|\|\|\|\|]PAM | NA | Intergenic | NA | NA |

| | | | | | | | | | | | |
|---|---|---|---|---|---|---|---|---|---|---|---|
| T32187 | GTCAGTCCGGTGTTCTTCACAGG | 1000 | | | | | | | | | |
| Oligo fwd | CACCGTCAGTCCGGTGTTCTTCAC | | | | | | | | | | |
| Oligo rev | AAACGTGAAGAACACCGGACTGAC | | | | | | | | | | |
| Chromosome | start | end | strand | MM | target_seq | PAM | alignment | distance | position | gene name | gene id |
| GU305914.1 | 29118 | 29140 | - | 0 | GTCAGTCCGGTGTTCTTCAC | AGG | \|\|\|\|\|\|\|\|[\|\|\|\|\|\|\|\|\|\|\|\|]PAM | NA | Intergenic | NA | NA |

| | | | | | | | | | | | |
|---|---|---|---|---|---|---|---|---|---|---|---|
| T32188 | TTCACAGGACAAGATCTCTGTGG | 1000 | | | | | | | | | |
| Oligo fwd | CACCTTCACAGGACAAGATCTCTG | | | | | | | | | | |
| Oligo rev | AAACCAGAGATCTTGTCCTGTGAA | | | | | | | | | | |
| Chromosome | start | end | strand | MM | target_seq | PAM | alignment | distance | position | gene name | gene id |
| GU305914.1 | 29103 | 29125 | - | 0 | TTCACAGGACAAGATCTCTG | TGG | \|\|\|\|\|\|\|\|[\|\|\|\|\|\|\|\|\|\|\|\|]PAM | NA | Intergenic | NA | NA |

<table>
<tr><td>T32189</td><td>TCACAGGACAAGATCTCTGTGGG</td><td>1000</td><td></td><td></td><td></td><td></td><td></td><td></td><td></td><td></td><td></td></tr>
<tr><td>Oligo fwd</td><td>CACCTCACAGGACAAGATCTCTGT</td><td></td><td></td><td></td><td></td><td></td><td></td><td></td><td></td><td></td><td></td></tr>
<tr><td>Oligo rev</td><td>AAACACAGAGATCTTGTCCTGTGA</td><td></td><td></td><td></td><td></td><td></td><td></td><td></td><td></td><td></td><td></td></tr>
<tr><td>Chromosome</td><td>start</td><td>end</td><td>strand</td><td>MM</td><td>target_seq</td><td>PAM</td><td>alignment</td><td>distance</td><td>position</td><td>gene name</td><td>gene id</td></tr>
<tr><td>GU305914.1</td><td>29102</td><td>29124</td><td>-</td><td>0</td><td>TCACAGGACAAGATCTCTGT</td><td>GGG</td><td>||||||||[||||||||||||]PAM</td><td>NA</td><td>Intergenic</td><td>NA</td><td>NA</td></tr>
<tr><td>T32190</td><td>CACAGGACAAGATCTCTGTGGGG</td><td>1000</td><td></td><td></td><td></td><td></td><td></td><td></td><td></td><td></td><td></td></tr>
<tr><td>Oligo fwd</td><td>CACCCACAGGACAAGATCTCTGTG</td><td></td><td></td><td></td><td></td><td></td><td></td><td></td><td></td><td></td><td></td></tr>
<tr><td>Oligo rev</td><td>AAACCACAGAGATCTTGTCCTGTG</td><td></td><td></td><td></td><td></td><td></td><td></td><td></td><td></td><td></td><td></td></tr>
<tr><td>Chromosome</td><td>start</td><td>end</td><td>strand</td><td>MM</td><td>target_seq</td><td>PAM</td><td>alignment</td><td>distance</td><td>position</td><td>gene name</td><td>gene id</td></tr>
<tr><td>GU305914.1</td><td>29101</td><td>29123</td><td>-</td><td>0</td><td>CACAGGACAAGATCTCTGTG</td><td>GGG</td><td>||||||||[||||||||||||]PAM</td><td>NA</td><td>Intergenic</td><td>NA</td><td>NA</td></tr>
<tr><td>T32191</td><td>AACAGCACCTTACGTTGCCTCGG</td><td>1000</td><td></td><td></td><td></td><td></td><td></td><td></td><td></td><td></td><td></td></tr>
<tr><td>Oligo fwd</td><td>CACCAACAGCACCTTACGTTGCCT</td><td></td><td></td><td></td><td></td><td></td><td></td><td></td><td></td><td></td><td></td></tr>
<tr><td>Oligo rev</td><td>AAACAGGCAACGTAAGGTGCTGTT</td><td></td><td></td><td></td><td></td><td></td><td></td><td></td><td></td><td></td><td></td></tr>
<tr><td>Chromosome</td><td>start</td><td>end</td><td>strand</td><td>MM</td><td>target_seq</td><td>PAM</td><td>alignment</td><td>distance</td><td>position</td><td>gene name</td><td>gene id</td></tr>
<tr><td>GU305914.1</td><td>29076</td><td>29098</td><td>-</td><td>0</td><td>AACAGCACCTTACGTTGCCT</td><td>CGG</td><td>||||||||[||||||||||||]PAM</td><td>NA</td><td>Intergenic</td><td>NA</td><td>NA</td></tr>
<tr><td>T32192</td><td>GCACCTTACGTTGCCTCGGTCGG</td><td>1000</td><td></td><td></td><td></td><td></td><td></td><td></td><td></td><td></td><td></td></tr>
<tr><td>Oligo fwd</td><td>CACCGCACCTTACGTTGCCTCGGT</td><td></td><td></td><td></td><td></td><td></td><td></td><td></td><td></td><td></td><td></td></tr>
<tr><td>Oligo rev</td><td>AAACACCGAGGCAACGTAAGGTGC</td><td></td><td></td><td></td><td></td><td></td><td></td><td></td><td></td><td></td><td></td></tr>
<tr><td>Chromosome</td><td>start</td><td>end</td><td>strand</td><td>MM</td><td>target_seq</td><td>PAM</td><td>alignment</td><td>distance</td><td>position</td><td>gene name</td><td>gene id</td></tr>
<tr><td>GU305914.1</td><td>29072</td><td>29094</td><td>-</td><td>0</td><td>GCACCTTACGTTGCCTCGGT</td><td>CGG</td><td>||||||||[||||||||||||]PAM</td><td>NA</td><td>Intergenic</td><td>NA</td><td>NA</td></tr>
<tr><td>T32193</td><td>CGGTCGGTCCTTCAATCTGCTGG</td><td>1000</td><td></td><td></td><td></td><td></td><td></td><td></td><td></td><td></td><td></td></tr>
<tr><td>Oligo fwd</td><td>CACCCGGTCGGTCCTTCAATCTGC</td><td></td><td></td><td></td><td></td><td></td><td></td><td></td><td></td><td></td><td></td></tr>
<tr><td>Oligo rev</td><td>AAACGCAGATTGAAGGACCGACCG</td><td></td><td></td><td></td><td></td><td></td><td></td><td></td><td></td><td></td><td></td></tr>
<tr><td>Chromosome</td><td>start</td><td>end</td><td>strand</td><td>MM</td><td>target_seq</td><td>PAM</td><td>alignment</td><td>distance</td><td>position</td><td>gene name</td><td>gene id</td></tr>
<tr><td>GU305914.1</td><td>29056</td><td>29078</td><td>-</td><td>0</td><td>CGGTCGGTCCTTCAATCTGC</td><td>TGG</td><td>||||||||[||||||||||||]PAM</td><td>NA</td><td>Intergenic</td><td>NA</td><td>NA</td></tr>
<tr><td>T32194</td><td>GTCCTTCAATCTGCTGGCTAAGG</td><td>1000</td><td></td><td></td><td></td><td></td><td></td><td></td><td></td><td></td><td></td></tr>
<tr><td>Oligo fwd</td><td>CACCGTCCTTCAATCTGCTGGCTA</td><td></td><td></td><td></td><td></td><td></td><td></td><td></td><td></td><td></td><td></td></tr>
<tr><td>Oligo rev</td><td>AAACTAGCCAGCAGATTGAAGGAC</td><td></td><td></td><td></td><td></td><td></td><td></td><td></td><td></td><td></td><td></td></tr>
<tr><td>Chromosome</td><td>start</td><td>end</td><td>strand</td><td>MM</td><td>target_seq</td><td>PAM</td><td>alignment</td><td>distance</td><td>position</td><td>gene name</td><td>gene id</td></tr>
<tr><td>GU305914.1</td><td>29050</td><td>29072</td><td>-</td><td>0</td><td>GTCCTTCAATCTGCTGGCTA</td><td>AGG</td><td>||||||||[||||||||||||]PAM</td><td>NA</td><td>Intergenic</td><td>NA</td><td>NA</td></tr>
<tr><td>T32195</td><td>TCCTTCAATCTGCTGGCTAAGGG</td><td>1000</td><td></td><td></td><td></td><td></td><td></td><td></td><td></td><td></td><td></td></tr>
<tr><td>Oligo fwd</td><td>CACCTCCTTCAATCTGCTGGCTAA</td><td></td><td></td><td></td><td></td><td></td><td></td><td></td><td></td><td></td><td></td></tr>
<tr><td>Oligo rev</td><td>AAACTTAGCCAGCAGATTGAAGGA</td><td></td><td></td><td></td><td></td><td></td><td></td><td></td><td></td><td></td><td></td></tr>
<tr><td>Chromosome</td><td>start</td><td>end</td><td>strand</td><td>MM</td><td>target_seq</td><td>PAM</td><td>alignment</td><td>distance</td><td>position</td><td>gene name</td><td>gene id</td></tr>
<tr><td>GU305914.1</td><td>29049</td><td>29071</td><td>-</td><td>0</td><td>TCCTTCAATCTGCTGGCTAA</td><td>GGG</td><td>||||||||[||||||||||||]PAM</td><td>NA</td><td>Intergenic</td><td>NA</td><td>NA</td></tr>
<tr><td>T32196</td><td>TTCAATCTGCTGGCTAAGGGAGG</td><td>1000</td><td></td><td></td><td></td><td></td><td></td><td></td><td></td><td></td><td></td></tr>
<tr><td>Oligo fwd</td><td>CACCTTCAATCTGCTGGCTAAGGG</td><td></td><td></td><td></td><td></td><td></td><td></td><td></td><td></td><td></td><td></td></tr>
<tr><td>Oligo rev</td><td>AAACCCCTTAGCCAGCAGATTGAA</td><td></td><td></td><td></td><td></td><td></td><td></td><td></td><td></td><td></td><td></td></tr>
<tr><td>Chromosome</td><td>start</td><td>end</td><td>strand</td><td>MM</td><td>target_seq</td><td>PAM</td><td>alignment</td><td>distance</td><td>position</td><td>gene name</td><td>gene id</td></tr>
<tr><td>GU305914.1</td><td>29046</td><td>29068</td><td>-</td><td>0</td><td>TTCAATCTGCTGGCTAAGGG</td><td>AGG</td><td>||||||||[||||||||||||]PAM</td><td>NA</td><td>Intergenic</td><td>NA</td><td>NA</td></tr>
<tr><td>T32197</td><td>TTATTTATTCAGTGAAATTATGG</td><td>1000</td><td></td><td></td><td></td><td></td><td></td><td></td><td></td><td></td><td></td></tr>
<tr><td>Oligo fwd</td><td>CACCTTATTTATTCAGTGAAATTA</td><td></td><td></td><td></td><td></td><td></td><td></td><td></td><td></td><td></td><td></td></tr>
<tr><td>Oligo rev</td><td>AAACTAATTTCACTGAATAAATAA</td><td></td><td></td><td></td><td></td><td></td><td></td><td></td><td></td><td></td><td></td></tr>
<tr><td>Chromosome</td><td>start</td><td>end</td><td>strand</td><td>MM</td><td>target_seq</td><td>PAM</td><td>alignment</td><td>distance</td><td>position</td><td>gene name</td><td>gene id</td></tr>
<tr><td>GU305914.1</td><td>28924</td><td>28946</td><td>-</td><td>0</td><td>TTATTTATTCAGTGAAATTA</td><td>TGG</td><td>||||||||[||||||||||||]PAM</td><td>NA</td><td>Intergenic</td><td>NA</td><td>NA</td></tr>
<tr><td>T32198</td><td>TTATGGAAGAAAGAAGTACATGG</td><td>1000</td><td></td><td></td><td></td><td></td><td></td><td></td><td></td><td></td><td></td></tr>
<tr><td>Oligo fwd</td><td>CACCTTATGGAAGAAAGAAGTACA</td><td></td><td></td><td></td><td></td><td></td><td></td><td></td><td></td><td></td><td></td></tr>
<tr><td>Oligo rev</td><td>AAACTGTACTTCTTTCTTCCATAA</td><td></td><td></td><td></td><td></td><td></td><td></td><td></td><td></td><td></td><td></td></tr>
<tr><td>Chromosome</td><td>start</td><td>end</td><td>strand</td><td>MM</td><td>target_seq</td><td>PAM</td><td>alignment</td><td>distance</td><td>position</td><td>gene name</td><td>gene id</td></tr>
<tr><td>GU305914.1</td><td>28907</td><td>28929</td><td>-</td><td>0</td><td>TTATGGAAGAAAGAAGTACA</td><td>TGG</td><td>||||||||[||||||||||||]PAM</td><td>NA</td><td>Intergenic</td><td>NA</td><td>NA</td></tr>
<tr><td>T32199</td><td>TATGGAAGAAAGAAGTACATGGG</td><td>1000</td><td></td><td></td><td></td><td></td><td></td><td></td><td></td><td></td><td></td></tr>
<tr><td>Oligo fwd</td><td>CACCTATGGAAGAAAGAAGTACAT</td><td></td><td></td><td></td><td></td><td></td><td></td><td></td><td></td><td></td><td></td></tr>
<tr><td>Oligo rev</td><td>AAACATGTACTTCTTTCTTCCATA</td><td></td><td></td><td></td><td></td><td></td><td></td><td></td><td></td><td></td><td></td></tr>
<tr><td>Chromosome</td><td>start</td><td>end</td><td>strand</td><td>MM</td><td>target_seq</td><td>PAM</td><td>alignment</td><td>distance</td><td>position</td><td>gene name</td><td>gene id</td></tr>
<tr><td>GU305914.1</td><td>28906</td><td>28928</td><td>-</td><td>0</td><td>TATGGAAGAAAGAAGTACAT</td><td>GGG</td><td>||||||||[||||||||||||]PAM</td><td>NA</td><td>Intergenic</td><td>NA</td><td>NA</td></tr>
<tr><td>T32200</td><td>ATGGAAGAAAGAAGTACATGGGG</td><td>1000</td><td></td><td></td><td></td><td></td><td></td><td></td><td></td><td></td><td></td></tr>
<tr><td>Oligo fwd</td><td>CACCATGGAAGAAAGAAGTACATG</td><td></td><td></td><td></td><td></td><td></td><td></td><td></td><td></td><td></td><td></td></tr>
<tr><td>Oligo rev</td><td>AAACCATGTACTTCTTTCTTCCAT</td><td></td><td></td><td></td><td></td><td></td><td></td><td></td><td></td><td></td><td></td></tr>
<tr><td>Chromosome</td><td>start</td><td>end</td><td>strand</td><td>MM</td><td>target_seq</td><td>PAM</td><td>alignment</td><td>distance</td><td>position</td><td>gene name</td><td>gene id</td></tr>
<tr><td>GU305914.1</td><td>28905</td><td>28927</td><td>-</td><td>0</td><td>ATGGAAGAAAGAAGTACATG</td><td>GGG</td><td>||||||||[||||||||||||]PAM</td><td>NA</td><td>Intergenic</td><td>NA</td><td>NA</td></tr>
<tr><td>T32201</td><td>TGGAAGAAAGAAGTACATGGGGG</td><td>1000</td><td></td><td></td><td></td><td></td><td></td><td></td><td></td><td></td><td></td></tr>
<tr><td>Oligo fwd</td><td>CACCTGGAAGAAAGAAGTACATGG</td><td></td><td></td><td></td><td></td><td></td><td></td><td></td><td></td><td></td><td></td></tr>
<tr><td>Oligo rev</td><td>AAACCCATGTACTTCTTTCTTCCA</td><td></td><td></td><td></td><td></td><td></td><td></td><td></td><td></td><td></td><td></td></tr>
<tr><td>Chromosome</td><td>start</td><td>end</td><td>strand</td><td>MM</td><td>target_seq</td><td>PAM</td><td>alignment</td><td>distance</td><td>position</td><td>gene name</td><td>gene id</td></tr>
<tr><td>GU305914.1</td><td>28904</td><td>28926</td><td>-</td><td>0</td><td>TGGAAGAAAGAAGTACATGG</td><td>GGG</td><td>||||||||[||||||||||||]PAM</td><td>NA</td><td>Intergenic</td><td>NA</td><td>NA</td></tr>
<tr><td>T32202</td><td>GGAAGAAAGAAGTACATGGGGGG</td><td>1000</td><td></td><td></td><td></td><td></td><td></td><td></td><td></td><td></td><td></td></tr>
<tr><td>Oligo fwd</td><td>CACCGGAAGAAAGAAGTACATGGG</td><td></td><td></td><td></td><td></td><td></td><td></td><td></td><td></td><td></td><td></td></tr>
<tr><td>Oligo rev</td><td>AAACCCCATGTACTTCTTTCTTCC</td><td></td><td></td><td></td><td></td><td></td><td></td><td></td><td></td><td></td><td></td></tr>
<tr><td>Chromosome</td><td>start</td><td>end</td><td>strand</td><td>MM</td><td>target_seq</td><td>PAM</td><td>alignment</td><td>distance</td><td>position</td><td>gene name</td><td>gene id</td></tr>
<tr><td>GU305914.1</td><td>28903</td><td>28925</td><td>-</td><td>0</td><td>GGAAGAAAGAAGTACATGGG</td><td>GGG</td><td>||||||||[||||||||||||]PAM</td><td>NA</td><td>Intergenic</td><td>NA</td><td>NA</td></tr>
<tr><td>T32203</td><td>GAAGAAAGAAGTACATGGGGGGG</td><td>1000</td><td></td><td></td><td></td><td></td><td></td><td></td><td></td><td></td><td></td></tr>
<tr><td>Oligo fwd</td><td>CACCGAAGAAAGAAGTACATGGGG</td><td></td><td></td><td></td><td></td><td></td><td></td><td></td><td></td><td></td><td></td></tr>
<tr><td>Oligo rev</td><td>AAACCCCCATGTACTTCTTTCTTC</td><td></td><td></td><td></td><td></td><td></td><td></td><td></td><td></td><td></td><td></td></tr>
<tr><td>Chromosome</td><td>start</td><td>end</td><td>strand</td><td>MM</td><td>target_seq</td><td>PAM</td><td>alignment</td><td>distance</td><td>position</td><td>gene name</td><td>gene id</td></tr>
</table>

GU305914.1 28902 28924 - 0 GAAGAAAGAAGTACATGGGG GGG ||||||||[||||||||||||]PAM NA Intergenic NA NA

T32204 AAGAAAGAAGTACATGGGGGGGG 1000
Oligo fwd CACCAAGAAAGAAGTACATGGGGG
Oligo rev AAACCCCCCATGTACTTCTTTCTT
Chromosome start end strand MM target_seq PAM alignment distance position gene name gene id
GU305914.1 28901 28923 - 0 AAGAAAGAAGTACATGGGGG GGG ||||||||[||||||||||||]PAM NA Intergenic NA NA

T32205 ATGGGGGGGGTGTATGAGATTGG 1000
Oligo fwd CACCATGGGGGGGGTGTATGAGAT
Oligo rev AAACATCTCATACACCCCCCCCAT
Chromosome start end strand MM target_seq PAM alignment distance position gene name gene id
GU305914.1 28888 28910 - 0 ATGGGGGGGGTGTATGAGAT TGG ||||||||[||||||||||||]PAM NA Intergenic NA NA

T32206 ACTTACTTACTCGATTCGTGTGG 1000
Oligo fwd CACCACTTACTTACTCGATTCGTG
Oligo rev AAACCACGAATCGAGTAAGTAAGT
Chromosome start end strand MM target_seq PAM alignment distance position gene name gene id
GU305914.1 28863 28885 - 0 ACTTACTTACTCGATTCGTG TGG ||||||||[||||||||||||]PAM NA Intergenic NA NA

T32207 GGCTAAGTGCTAAGTGCTTTTGG 1000
Oligo fwd CACCGGCTAAGTGCTAAGTGCTTT
Oligo rev AAACAAAGCACTTAGCACTTAGCC
Chromosome start end strand MM target_seq PAM alignment distance position gene name gene id
GU305914.1 28842 28864 - 0 GGCTAAGTGCTAAGTGCTTT TGG ||||||||[||||||||||||]PAM NA Intergenic NA NA

T32208 GCTAAGTGCTAAGTGCTTTTGGG 1000
Oligo fwd CACCGCTAAGTGCTAAGTGCTTTT
Oligo rev AAACAAAAGCACTTAGCACTTAGC
Chromosome start end strand MM target_seq PAM alignment distance position gene name gene id
GU305914.1 28841 28863 - 0 GCTAAGTGCTAAGTGCTTTT GGG ||||||||[||||||||||||]PAM NA Intergenic NA NA

T32209 CTAAGTGCTAAGTGCTTTTGGGG 1000
Oligo fwd CACCCTAAGTGCTAAGTGCTTTTG
Oligo rev AAACCAAAAGCACTTAGCACTTAG
Chromosome start end strand MM target_seq PAM alignment distance position gene name gene id
GU305914.1 28840 28862 - 0 CTAAGTGCTAAGTGCTTTTG GGG ||||||||[||||||||||||]PAM NA Intergenic NA NA

T32210 GTGCTAAGTGCTTTTGGGGTTGG 1000
Oligo fwd CACCGTGCTAAGTGCTTTTGGGGT
Oligo rev AAACACCCCAAAAGCACTTAGCAC
Chromosome start end strand MM target_seq PAM alignment distance position gene name gene id
GU305914.1 28836 28858 - 0 GTGCTAAGTGCTTTTGGGGT TGG ||||||||[||||||||||||]PAM NA Intergenic NA NA

T32211 ATTAATCTAAGATTACAGAAAGG 1000
Oligo fwd CACCATTAATCTAAGATTACAGAA
Oligo rev AAACTTCTGTAATCTTAGATTAAT
Chromosome start end strand MM target_seq PAM alignment distance position gene name gene id
GU305914.1 28802 28824 - 0 ATTAATCTAAGATTACAGAA AGG ||||||||[||||||||||||]PAM NA Intergenic NA NA

T32212 AGATTACAGAAAGGTACGCTTGG 1000
Oligo fwd CACCAGATTACAGAAAGGTACGCT
Oligo rev AAACAGCGTACCTTTCTGTAATCT
Chromosome start end strand MM target_seq PAM alignment distance position gene name gene id
GU305914.1 28793 28815 - 0 AGATTACAGAAAGGTACGCT TGG ||||||||[||||||||||||]PAM NA Intergenic NA NA

T32213 AAGGTACGCTTGGAGTACTTTGG 1000
Oligo fwd CACCAAGGTACGCTTGGAGTACTT
Oligo rev AAACAAGTACTCCAAGCGTACCTT
Chromosome start end strand MM target_seq PAM alignment distance position gene name gene id
GU305914.1 28783 28805 - 0 AAGGTACGCTTGGAGTACTT TGG ||||||||[||||||||||||]PAM NA Intergenic NA NA

T32214 CTTGGAGTACTTTGGCCCTCCGG 1000
Oligo fwd CACCCTTGGAGTACTTTGGCCCTC
Oligo rev AAACGAGGGCCAAAGTACTCCAAG
Chromosome start end strand MM target_seq PAM alignment distance position gene name gene id
GU305914.1 28775 28797 - 0 CTTGGAGTACTTTGGCCCTC CGG ||||||||[||||||||||||]PAM NA Intergenic NA NA

T32215 GGAGTACTTTGGCCCTCCGGCGG 1000
Oligo fwd CACCGGAGTACTTTGGCCCTCCGG
Oligo rev AAACCCGGAGGGCCAAAGTACTCC
Chromosome start end strand MM target_seq PAM alignment distance position gene name gene id
GU305914.1 28772 28794 - 0 GGAGTACTTTGGCCCTCCGG CGG ||||||||[||||||||||||]PAM NA Intergenic NA NA

T32216 TACTTTGGCCCTCCGGCGGCCGG 1000
Oligo fwd CACCTACTTTGGCCCTCCGGCGGC
Oligo rev AAACGCCGCCGGAGGGCCAAAGTA
Chromosome start end strand MM target_seq PAM alignment distance position gene name gene id
GU305914.1 28768 28790 - 0 TACTTTGGCCCTCCGGCGGC CGG ||||||||[||||||||||||]PAM NA Intergenic NA NA

T32217 ACTTTGGCCCTCCGGCGGCCGGG 1000
Oligo fwd CACCACTTTGGCCCTCCGGCGGCC
Oligo rev AAACGGCCGCCGGAGGGCCAAAGT
Chromosome start end strand MM target_seq PAM alignment distance position gene name gene id
GU305914.1 28767 28789 - 0 ACTTTGGCCCTCCGGCGGCC GGG ||||||||[||||||||||||]PAM NA Intergenic NA NA

T32218 CTTTGGCCCTCCGGCGGCCGGGG 1000
Oligo fwd CACCCTTTGGCCCTCCGGCGGCCG
Oligo rev AAACCGGCCGCCGGAGGGCCAAAG

<table>
<tr><td>Chromosome</td><td>start</td><td>end</td><td>strand</td><td>MM</td><td>target_seq</td><td>PAM</td><td>alignment</td><td>distance</td><td>position</td><td>gene name</td><td>gene id</td></tr>
<tr><td>GU305914.1</td><td>28766</td><td>28788</td><td>-</td><td>0</td><td>CTTTGGCCCTCCGGCGGCCG</td><td>GGG</td><td>||||||||[||||||||||||]PAM</td><td>NA</td><td>Intergenic</td><td>NA</td><td>NA</td></tr>
<tr><td>T32219</td><td>TTTGGCCCTCCGGCGGCCGGGGG</td><td>1000</td><td></td><td></td><td></td><td></td><td></td><td></td><td></td><td></td><td></td></tr>
<tr><td>Oligo fwd</td><td>CACCTTTGGCCCTCCGGCGGCCGG</td><td></td><td></td><td></td><td></td><td></td><td></td><td></td><td></td><td></td><td></td></tr>
<tr><td>Oligo rev</td><td>AAACCCGGCCGCCGGAGGGCCAAA</td><td></td><td></td><td></td><td></td><td></td><td></td><td></td><td></td><td></td><td></td></tr>
<tr><td>Chromosome</td><td>start</td><td>end</td><td>strand</td><td>MM</td><td>target_seq</td><td>PAM</td><td>alignment</td><td>distance</td><td>position</td><td>gene name</td><td>gene id</td></tr>
<tr><td>GU305914.1</td><td>28765</td><td>28787</td><td>-</td><td>0</td><td>TTTGGCCCTCCGGCGGCCGG</td><td>GGG</td><td>||||||||[||||||||||||]PAM</td><td>NA</td><td>Intergenic</td><td>NA</td><td>NA</td></tr>
<tr><td>T32220</td><td>GGCCCTCCGGCGGCCGGGGGAGG</td><td>1000</td><td></td><td></td><td></td><td></td><td></td><td></td><td></td><td></td><td></td></tr>
<tr><td>Oligo fwd</td><td>CACCGGCCCTCCGGCGGCCGGGGG</td><td></td><td></td><td></td><td></td><td></td><td></td><td></td><td></td><td></td><td></td></tr>
<tr><td>Oligo rev</td><td>AAACCCCCCGGCCGCCGGAGGGCC</td><td></td><td></td><td></td><td></td><td></td><td></td><td></td><td></td><td></td><td></td></tr>
<tr><td>Chromosome</td><td>start</td><td>end</td><td>strand</td><td>MM</td><td>target_seq</td><td>PAM</td><td>alignment</td><td>distance</td><td>position</td><td>gene name</td><td>gene id</td></tr>
<tr><td>GU305914.1</td><td>28762</td><td>28784</td><td>-</td><td>0</td><td>GGCCCTCCGGCGGCCGGGGG</td><td>AGG</td><td>||||||||[||||||||||||]PAM</td><td>NA</td><td>Intergenic</td><td>NA</td><td>NA</td></tr>
<tr><td>GU305914.1</td><td>33</td><td>55</td><td>-</td><td>4</td><td>CGCACTCAGACGGCCGGGGG</td><td>GGG</td><td>-||-|||-[|-||||||||||]PAM</td><td>NA</td><td>Intergenic</td><td>NA</td><td>NA</td></tr>
<tr><td>T32221</td><td>CCTCCGGCGGCCGGGGGAGGAGG</td><td>1000</td><td></td><td></td><td></td><td></td><td></td><td></td><td></td><td></td><td></td></tr>
<tr><td>Oligo fwd</td><td>CACCCCTCCGGCGGCCGGGGGAGG</td><td></td><td></td><td></td><td></td><td></td><td></td><td></td><td></td><td></td><td></td></tr>
<tr><td>Oligo rev</td><td>AAACCCTCCCCCGGCCGCCGGAGG</td><td></td><td></td><td></td><td></td><td></td><td></td><td></td><td></td><td></td><td></td></tr>
<tr><td>Chromosome</td><td>start</td><td>end</td><td>strand</td><td>MM</td><td>target_seq</td><td>PAM</td><td>alignment</td><td>distance</td><td>position</td><td>gene name</td><td>gene id</td></tr>
<tr><td>GU305914.1</td><td>28759</td><td>28781</td><td>-</td><td>0</td><td>CCTCCGGCGGCCGGGGGAGG</td><td>AGG</td><td>||||||||[||||||||||||]PAM</td><td>NA</td><td>Intergenic</td><td>NA</td><td>NA</td></tr>
<tr><td>T32222</td><td>GGCGGCCGGGGGAGGAGGAGAGG</td><td>1000</td><td></td><td></td><td></td><td></td><td></td><td></td><td></td><td></td><td></td></tr>
<tr><td>Oligo fwd</td><td>CACCGGCGGCCGGGGGAGGAGGAG</td><td></td><td></td><td></td><td></td><td></td><td></td><td></td><td></td><td></td><td></td></tr>
<tr><td>Oligo rev</td><td>AAACCTCCTCCTCCCCCGGCCGCC</td><td></td><td></td><td></td><td></td><td></td><td></td><td></td><td></td><td></td><td></td></tr>
<tr><td>Chromosome</td><td>start</td><td>end</td><td>strand</td><td>MM</td><td>target_seq</td><td>PAM</td><td>alignment</td><td>distance</td><td>position</td><td>gene name</td><td>gene id</td></tr>
<tr><td>GU305914.1</td><td>28754</td><td>28776</td><td>-</td><td>0</td><td>GGCGGCCGGGGGAGGAGGAG</td><td>AGG</td><td>||||||||[||||||||||||]PAM</td><td>NA</td><td>Intergenic</td><td>NA</td><td>NA</td></tr>
<tr><td>T32223</td><td>GGGGGAGGAGGAGAGGATGATGG</td><td>1000</td><td></td><td></td><td></td><td></td><td></td><td></td><td></td><td></td><td></td></tr>
<tr><td>Oligo fwd</td><td>CACCGGGGGAGGAGGAGAGGATGA</td><td></td><td></td><td></td><td></td><td></td><td></td><td></td><td></td><td></td><td></td></tr>
<tr><td>Oligo rev</td><td>AAACTCATCCTCTCCTCCTCCCCC</td><td></td><td></td><td></td><td></td><td></td><td></td><td></td><td></td><td></td><td></td></tr>
<tr><td>Chromosome</td><td>start</td><td>end</td><td>strand</td><td>MM</td><td>target_seq</td><td>PAM</td><td>alignment</td><td>distance</td><td>position</td><td>gene name</td><td>gene id</td></tr>
<tr><td>GU305914.1</td><td>28747</td><td>28769</td><td>-</td><td>0</td><td>GGGGGAGGAGGAGAGGATGA</td><td>TGG</td><td>||||||||[||||||||||||]PAM</td><td>NA</td><td>Intergenic</td><td>NA</td><td>NA</td></tr>
<tr><td>GU305914.1</td><td>69163</td><td>69185</td><td>+</td><td>4</td><td>CGAGGAGGTGGAGAGCATGA</td><td>CGG</td><td>-|-|||||[-||||||-||||]PAM</td><td>NA</td><td>Intergenic</td><td>NA</td><td>NA</td></tr>
<tr><td>GU305914.1</td><td>18851</td><td>18873</td><td>-</td><td>4</td><td>GGTGGGGGAGGAGGGGAAGA</td><td>CGG</td><td>||-||-||[|||||-|||-||]PAM</td><td>NA</td><td>Intergenic</td><td>NA</td><td>NA</td></tr>
<tr><td>T32224</td><td>GGAGGAGGAGAGGATGATGGTGG</td><td>1000</td><td></td><td></td><td></td><td></td><td></td><td></td><td></td><td></td><td></td></tr>
<tr><td>Oligo fwd</td><td>CACCGGAGGAGGAGAGGATGATGG</td><td></td><td></td><td></td><td></td><td></td><td></td><td></td><td></td><td></td><td></td></tr>
<tr><td>Oligo rev</td><td>AAACCCATCATCCTCTCCTCCTCC</td><td></td><td></td><td></td><td></td><td></td><td></td><td></td><td></td><td></td><td></td></tr>
<tr><td>Chromosome</td><td>start</td><td>end</td><td>strand</td><td>MM</td><td>target_seq</td><td>PAM</td><td>alignment</td><td>distance</td><td>position</td><td>gene name</td><td>gene id</td></tr>
<tr><td>GU305914.1</td><td>28744</td><td>28766</td><td>-</td><td>0</td><td>GGAGGAGGAGAGGATGATGG</td><td>TGG</td><td>||||||||[||||||||||||]PAM</td><td>NA</td><td>Intergenic</td><td>NA</td><td>NA</td></tr>
<tr><td>GU305914.1</td><td>69166</td><td>69188</td><td>+</td><td>3</td><td>GGAGGTGGAGAGCATGACGG</td><td>TGG</td><td>|||||-||[||||-||||-||]PAM</td><td>NA</td><td>Intergenic</td><td>NA</td><td>NA</td></tr>
<tr><td>GU305914.1</td><td>42840</td><td>42862</td><td>-</td><td>4</td><td>GAAGGAGAAGAGGATGACGA</td><td>AGG</td><td>|-|||||-[|||||||||-|-]PAM</td><td>NA</td><td>Intergenic</td><td>NA</td><td>NA</td></tr>
<tr><td>T32225</td><td>GGAGGAGAGGATGATGGTGGCGG</td><td>1000</td><td></td><td></td><td></td><td></td><td></td><td></td><td></td><td></td><td></td></tr>
<tr><td>Oligo fwd</td><td>CACCGGAGGAGAGGATGATGGTGG</td><td></td><td></td><td></td><td></td><td></td><td></td><td></td><td></td><td></td><td></td></tr>
<tr><td>Oligo rev</td><td>AAACCCACCATCATCCTCTCCTCC</td><td></td><td></td><td></td><td></td><td></td><td></td><td></td><td></td><td></td><td></td></tr>
<tr><td>Chromosome</td><td>start</td><td>end</td><td>strand</td><td>MM</td><td>target_seq</td><td>PAM</td><td>alignment</td><td>distance</td><td>position</td><td>gene name</td><td>gene id</td></tr>
<tr><td>GU305914.1</td><td>28741</td><td>28763</td><td>-</td><td>0</td><td>GGAGGAGAGGATGATGGTGG</td><td>CGG</td><td>||||||||[||||||||||||]PAM</td><td>NA</td><td>Intergenic</td><td>NA</td><td>NA</td></tr>
<tr><td>T32226</td><td>GAGGAGAGGATGATGGTGGCGGG</td><td>1000</td><td></td><td></td><td></td><td></td><td></td><td></td><td></td><td></td><td></td></tr>
<tr><td>Oligo fwd</td><td>CACCGAGGAGAGGATGATGGTGGC</td><td></td><td></td><td></td><td></td><td></td><td></td><td></td><td></td><td></td><td></td></tr>
<tr><td>Oligo rev</td><td>AAACGCCACCATCATCCTCTCCTC</td><td></td><td></td><td></td><td></td><td></td><td></td><td></td><td></td><td></td><td></td></tr>
<tr><td>Chromosome</td><td>start</td><td>end</td><td>strand</td><td>MM</td><td>target_seq</td><td>PAM</td><td>alignment</td><td>distance</td><td>position</td><td>gene name</td><td>gene id</td></tr>
<tr><td>GU305914.1</td><td>28740</td><td>28762</td><td>-</td><td>0</td><td>GAGGAGAGGATGATGGTGGC</td><td>GGG</td><td>||||||||[||||||||||||]PAM</td><td>NA</td><td>Intergenic</td><td>NA</td><td>NA</td></tr>
<tr><td>T32227</td><td>AGGAGAGGATGATGGTGGCGGGG</td><td>1000</td><td></td><td></td><td></td><td></td><td></td><td></td><td></td><td></td><td></td></tr>
<tr><td>Oligo fwd</td><td>CACCAGGAGAGGATGATGGTGGCG</td><td></td><td></td><td></td><td></td><td></td><td></td><td></td><td></td><td></td><td></td></tr>
<tr><td>Oligo rev</td><td>AAACCGCCACCATCATCCTCTCCT</td><td></td><td></td><td></td><td></td><td></td><td></td><td></td><td></td><td></td><td></td></tr>
<tr><td>Chromosome</td><td>start</td><td>end</td><td>strand</td><td>MM</td><td>target_seq</td><td>PAM</td><td>alignment</td><td>distance</td><td>position</td><td>gene name</td><td>gene id</td></tr>
<tr><td>GU305914.1</td><td>28739</td><td>28761</td><td>-</td><td>0</td><td>AGGAGAGGATGATGGTGGCG</td><td>GGG</td><td>||||||||[||||||||||||]PAM</td><td>NA</td><td>Intergenic</td><td>NA</td><td>NA</td></tr>
<tr><td>T32228</td><td>CTGCGTGACTGCTGTTGCTGCGG</td><td>1000</td><td></td><td></td><td></td><td></td><td></td><td></td><td></td><td></td><td></td></tr>
<tr><td>Oligo fwd</td><td>CACCCTGCGTGACTGCTGTTGCTG</td><td></td><td></td><td></td><td></td><td></td><td></td><td></td><td></td><td></td><td></td></tr>
<tr><td>Oligo rev</td><td>AAACCAGCAACAGCAGTCACGCAG</td><td></td><td></td><td></td><td></td><td></td><td></td><td></td><td></td><td></td><td></td></tr>
<tr><td>Chromosome</td><td>start</td><td>end</td><td>strand</td><td>MM</td><td>target_seq</td><td>PAM</td><td>alignment</td><td>distance</td><td>position</td><td>gene name</td><td>gene id</td></tr>
<tr><td>GU305914.1</td><td>28708</td><td>28730</td><td>-</td><td>0</td><td>CTGCGTGACTGCTGTTGCTG</td><td>CGG</td><td>||||||||[||||||||||||]PAM</td><td>NA</td><td>Intergenic</td><td>NA</td><td>NA</td></tr>
<tr><td>T32229</td><td>CGATCATGCGAGACGCTTGCAGG</td><td>1000</td><td></td><td></td><td></td><td></td><td></td><td></td><td></td><td></td><td></td></tr>
<tr><td>Oligo fwd</td><td>CACCCGATCATGCGAGACGCTTGC</td><td></td><td></td><td></td><td></td><td></td><td></td><td></td><td></td><td></td><td></td></tr>
<tr><td>Oligo rev</td><td>AAACGCAAGCGTCTCGCATGATCG</td><td></td><td></td><td></td><td></td><td></td><td></td><td></td><td></td><td></td><td></td></tr>
<tr><td>Chromosome</td><td>start</td><td>end</td><td>strand</td><td>MM</td><td>target_seq</td><td>PAM</td><td>alignment</td><td>distance</td><td>position</td><td>gene name</td><td>gene id</td></tr>
<tr><td>GU305914.1</td><td>28683</td><td>28705</td><td>-</td><td>0</td><td>CGATCATGCGAGACGCTTGC</td><td>AGG</td><td>||||||||[||||||||||||]PAM</td><td>NA</td><td>Intergenic</td><td>NA</td><td>NA</td></tr>
<tr><td>T32230</td><td>TTGCAGGAACAGCTCGTAGTTGG</td><td>1000</td><td></td><td></td><td></td><td></td><td></td><td></td><td></td><td></td><td></td></tr>
<tr><td>Oligo fwd</td><td>CACCTTGCAGGAACAGCTCGTAGT</td><td></td><td></td><td></td><td></td><td></td><td></td><td></td><td></td><td></td><td></td></tr>
<tr><td>Oligo rev</td><td>AAACACTACGAGCTGTTCCTGCAA</td><td></td><td></td><td></td><td></td><td></td><td></td><td></td><td></td><td></td><td></td></tr>
<tr><td>Chromosome</td><td>start</td><td>end</td><td>strand</td><td>MM</td><td>target_seq</td><td>PAM</td><td>alignment</td><td>distance</td><td>position</td><td>gene name</td><td>gene id</td></tr>
<tr><td>GU305914.1</td><td>28667</td><td>28689</td><td>-</td><td>0</td><td>TTGCAGGAACAGCTCGTAGT</td><td>TGG</td><td>||||||||[||||||||||||]PAM</td><td>NA</td><td>Intergenic</td><td>NA</td><td>NA</td></tr>
<tr><td>GU305914.1</td><td>128256</td><td>128278</td><td>-</td><td>4</td><td>GAGCAGGAAGAGGTCGTAGT</td><td>TGG</td><td>--||||||[|-||-|||||||]PAM</td><td>NA</td><td>Intergenic</td><td>NA</td><td>NA</td></tr>
<tr><td>T32231</td><td>AGCTCGTAGTTGGCGTGTGTTGG</td><td>1000</td><td></td><td></td><td></td><td></td><td></td><td></td><td></td><td></td><td></td></tr>
<tr><td>Oligo fwd</td><td>CACCAGCTCGTAGTTGGCGTGTGT</td><td></td><td></td><td></td><td></td><td></td><td></td><td></td><td></td><td></td><td></td></tr>
<tr><td>Oligo rev</td><td>AAACACACACGCCAACTACGAGCT</td><td></td><td></td><td></td><td></td><td></td><td></td><td></td><td></td><td></td><td></td></tr>
<tr><td>Chromosome</td><td>start</td><td>end</td><td>strand</td><td>MM</td><td>target_seq</td><td>PAM</td><td>alignment</td><td>distance</td><td>position</td><td>gene name</td><td>gene id</td></tr>
<tr><td>GU305914.1</td><td>28657</td><td>28679</td><td>-</td><td>0</td><td>AGCTCGTAGTTGGCGTGTGT</td><td>TGG</td><td>||||||||[||||||||||||]PAM</td><td>NA</td><td>Intergenic</td><td>NA</td><td>NA</td></tr>
<tr><td>T32232</td><td>GCTCGTAGTTGGCGTGTGTTGGG</td><td>1000</td><td></td><td></td><td></td><td></td><td></td><td></td><td></td><td></td><td></td></tr>
<tr><td>Oligo fwd</td><td>CACCGCTCGTAGTTGGCGTGTGTT</td><td></td><td></td><td></td><td></td><td></td><td></td><td></td><td></td><td></td><td></td></tr>
</table>

Oligo rev AAACAACACACGCCAACTACGAGC
Chromosome start end strand MM target_seq PAM alignment distance position gene name gene id
GU305914.1 28656 28678 - 0 GCTCGTAGTTGGCGTGTGTT GGG ||||||||[||||||||||||]PAM NA Intergenic NA NA

T32233 AGTTGGCGTGTGTTGGGATGTGG 1000
Oligo fwd CACCAGTTGGCGTGTGTTGGGATG
Oligo rev AAACCATCCCAACACACGCCAACT
Chromosome start end strand MM target_seq PAM alignment distance position gene name gene id
GU305914.1 28650 28672 - 0 AGTTGGCGTGTGTTGGGATG TGG ||||||||[||||||||||||]PAM NA Intergenic NA NA

T32234 GTTGGCGTGTGTTGGGATGTGGG 1000
Oligo fwd CACCGTTGGCGTGTGTTGGGATGT
Oligo rev AAACACATCCCAACACACGCCAAC
Chromosome start end strand MM target_seq PAM alignment distance position gene name gene id
GU305914.1 28649 28671 - 0 GTTGGCGTGTGTTGGGATGT GGG ||||||||[||||||||||||]PAM NA Intergenic NA NA

T32235 GTGTGTTGGGATGTGGGTCGTGG 1000
Oligo fwd CACCGTGTGTTGGGATGTGGGTCG
Oligo rev AAACCGACCCACATCCCAACACAC
Chromosome start end strand MM target_seq PAM alignment distance position gene name gene id
GU305914.1 28643 28665 - 0 GTGTGTTGGGATGTGGGTCG TGG ||||||||[||||||||||||]PAM NA Intergenic NA NA

T32236 TGTGTTGGGATGTGGGTCGTGGG 1000
Oligo fwd CACCTGTGTTGGGATGTGGGTCGT
Oligo rev AAACACGACCCACATCCCAACACA
Chromosome start end strand MM target_seq PAM alignment distance position gene name gene id
GU305914.1 28642 28664 - 0 TGTGTTGGGATGTGGGTCGT GGG ||||||||[||||||||||||]PAM NA Intergenic NA NA

T32237 GTGTTGGGATGTGGGTCGTGGGG 1000
Oligo fwd CACCGTGTTGGGATGTGGGTCGTG
Oligo rev AAACCACGACCCACATCCCAACAC
Chromosome start end strand MM target_seq PAM alignment distance position gene name gene id
GU305914.1 28641 28663 - 0 GTGTTGGGATGTGGGTCGTG GGG ||||||||[||||||||||||]PAM NA Intergenic NA NA

T32238 GTGGGTCGTGGGGATCTGTCTGG 1000
Oligo fwd CACCGTGGGTCGTGGGGATCTGTC
Oligo rev AAACGACAGATCCCCACGACCCAC
Chromosome start end strand MM target_seq PAM alignment distance position gene name gene id
GU305914.1 28631 28653 - 0 GTGGGTCGTGGGGATCTGTC TGG ||||||||[||||||||||||]PAM NA Intergenic NA NA

T32239 GGGGATCTGTCTGGAAGACGAGG 1000
Oligo fwd CACCGGGGATCTGTCTGGAAGACG
Oligo rev AAACCGTCTTCCAGACAGATCCCC
Chromosome start end strand MM target_seq PAM alignment distance position gene name gene id
GU305914.1 28622 28644 - 0 GGGGATCTGTCTGGAAGACG AGG ||||||||[||||||||||||]PAM NA Intergenic NA NA

T32240 GGGATCTGTCTGGAAGACGAGGG 1000
Oligo fwd CACCGGGATCTGTCTGGAAGACGA
Oligo rev AAACTCGTCTTCCAGACAGATCCC
Chromosome start end strand MM target_seq PAM alignment distance position gene name gene id
GU305914.1 28621 28643 - 0 GGGATCTGTCTGGAAGACGA GGG ||||||||[||||||||||||]PAM NA Intergenic NA NA

T32241 CGCGCACGAAGTGTATCGTGCGG 1000
Oligo fwd CACCCGCGCACGAAGTGTATCGTG
Oligo rev AAACCACGATACACTTCGTGCGCG
Chromosome start end strand MM target_seq PAM alignment distance position gene name gene id
GU305914.1 28587 28609 - 0 CGCGCACGAAGTGTATCGTG CGG ||||||||[||||||||||||]PAM NA Intergenic NA NA

T32242 GTATCGTGCGGCGCAGCATGCGG 1000
Oligo fwd CACCGTATCGTGCGGCGCAGCATG
Oligo rev AAACCATGCTGCGCCGCACGATAC
Chromosome start end strand MM target_seq PAM alignment distance position gene name gene id
GU305914.1 28575 28597 - 0 GTATCGTGCGGCGCAGCATG CGG ||||||||[||||||||||||]PAM NA Intergenic NA NA

T32243 TCGTGCGGCGCAGCATGCGGCGG 1000
Oligo fwd CACCTCGTGCGGCGCAGCATGCGG
Oligo rev AAACCCGCATGCTGCGCCGCACGA
Chromosome start end strand MM target_seq PAM alignment distance position gene name gene id
GU305914.1 28572 28594 - 0 TCGTGCGGCGCAGCATGCGG CGG ||||||||[||||||||||||]PAM NA Intergenic NA NA

T32244 CGTGCGGCGCAGCATGCGGCGGG 1000
Oligo fwd CACCCGTGCGGCGCAGCATGCGGC
Oligo rev AAACGCCGCATGCTGCGCCGCACG
Chromosome start end strand MM target_seq PAM alignment distance position gene name gene id
GU305914.1 28571 28593 - 0 CGTGCGGCGCAGCATGCGGC GGG ||||||||[||||||||||||]PAM NA Intergenic NA NA

T32245 CGGCGGGCGCTACTCTCGTCAGG 1000
Oligo fwd CACCCGGCGGGCGCTACTCTCGTC
Oligo rev AAACGACGAGAGTAGCGCCCGCCG
Chromosome start end strand MM target_seq PAM alignment distance position gene name gene id
GU305914.1 28555 28577 - 0 CGGCGGGCGCTACTCTCGTC AGG ||||||||[||||||||||||]PAM NA Intergenic NA NA

T32246 GCGCTACTCTCGTCAGGCAGCGG 1000
Oligo fwd CACCGCGCTACTCTCGTCAGGCAG
Oligo rev AAACCTGCCTGACGAGAGTAGCGC
Chromosome start end strand MM target_seq PAM alignment distance position gene name gene id
GU305914.1 28549 28571 - 0 GCGCTACTCTCGTCAGGCAG CGG ||||||||[||||||||||||]PAM NA Intergenic NA NA

T32247 CGCTACTCTCGTCAGGCAGCGGG 1000

<table>
<tr><td>Oligo fwd</td><td>CACCCGCTACTCTCGTCAGGCAGC</td><td></td><td></td><td></td><td></td><td></td><td></td><td></td><td></td><td></td><td></td></tr>
<tr><td>Oligo rev</td><td>AAACGCTGCCTGACGAGAGTAGCG</td><td></td><td></td><td></td><td></td><td></td><td></td><td></td><td></td><td></td><td></td></tr>
<tr><td>Chromosome</td><td>start</td><td>end</td><td>strand</td><td>MM</td><td>target_seq</td><td>PAM</td><td>alignment</td><td>distance</td><td>position</td><td>gene name</td><td>gene id</td></tr>
<tr><td>GU305914.1</td><td>28548</td><td>28570</td><td>-</td><td>0</td><td>CGCTACTCTCGTCAGGCAGC</td><td>GGG</td><td>||||||||[||||||||||||]PAM</td><td>NA</td><td>Intergenic</td><td>NA</td><td>NA</td></tr>
<tr><td></td><td></td><td></td><td></td><td></td><td></td><td></td><td></td><td></td><td></td><td></td><td></td></tr>
<tr><td>T32248</td><td>TACTCTCGTCAGGCAGCGGGCGG</td><td>1000</td><td></td><td></td><td></td><td></td><td></td><td></td><td></td><td></td><td></td></tr>
<tr><td>Oligo fwd</td><td>CACCTACTCTCGTCAGGCAGCGGG</td><td></td><td></td><td></td><td></td><td></td><td></td><td></td><td></td><td></td><td></td></tr>
<tr><td>Oligo rev</td><td>AAACCCCGCTGCCTGACGAGAGTA</td><td></td><td></td><td></td><td></td><td></td><td></td><td></td><td></td><td></td><td></td></tr>
<tr><td>Chromosome</td><td>start</td><td>end</td><td>strand</td><td>MM</td><td>target_seq</td><td>PAM</td><td>alignment</td><td>distance</td><td>position</td><td>gene name</td><td>gene id</td></tr>
<tr><td>GU305914.1</td><td>28545</td><td>28567</td><td>-</td><td>0</td><td>TACTCTCGTCAGGCAGCGGG</td><td>CGG</td><td>||||||||[||||||||||||]PAM</td><td>NA</td><td>Intergenic</td><td>NA</td><td>NA</td></tr>
<tr><td></td><td></td><td></td><td></td><td></td><td></td><td></td><td></td><td></td><td></td><td></td><td></td></tr>
<tr><td>T32249</td><td>GGCAGCGGGCGGAAGATGTCTGG</td><td>1000</td><td></td><td></td><td></td><td></td><td></td><td></td><td></td><td></td><td></td></tr>
<tr><td>Oligo fwd</td><td>CACCGGCAGCGGGCGGAAGATGTC</td><td></td><td></td><td></td><td></td><td></td><td></td><td></td><td></td><td></td><td></td></tr>
<tr><td>Oligo rev</td><td>AAACGACATCTTCCGCCCGCTGCC</td><td></td><td></td><td></td><td></td><td></td><td></td><td></td><td></td><td></td><td></td></tr>
<tr><td>Chromosome</td><td>start</td><td>end</td><td>strand</td><td>MM</td><td>target_seq</td><td>PAM</td><td>alignment</td><td>distance</td><td>position</td><td>gene name</td><td>gene id</td></tr>
<tr><td>GU305914.1</td><td>28534</td><td>28556</td><td>-</td><td>0</td><td>GGCAGCGGGCGGAAGATGTC</td><td>TGG</td><td>||||||||[||||||||||||]PAM</td><td>NA</td><td>Intergenic</td><td>NA</td><td>NA</td></tr>
<tr><td></td><td></td><td></td><td></td><td></td><td></td><td></td><td></td><td></td><td></td><td></td><td></td></tr>
<tr><td>T32250</td><td>GCAGCGGGCGGAAGATGTCTGGG</td><td>1000</td><td></td><td></td><td></td><td></td><td></td><td></td><td></td><td></td><td></td></tr>
<tr><td>Oligo fwd</td><td>CACCGCAGCGGGCGGAAGATGTCT</td><td></td><td></td><td></td><td></td><td></td><td></td><td></td><td></td><td></td><td></td></tr>
<tr><td>Oligo rev</td><td>AAACAGACATCTTCCGCCCGCTGC</td><td></td><td></td><td></td><td></td><td></td><td></td><td></td><td></td><td></td><td></td></tr>
<tr><td>Chromosome</td><td>start</td><td>end</td><td>strand</td><td>MM</td><td>target_seq</td><td>PAM</td><td>alignment</td><td>distance</td><td>position</td><td>gene name</td><td>gene id</td></tr>
<tr><td>GU305914.1</td><td>28533</td><td>28555</td><td>-</td><td>0</td><td>GCAGCGGGCGGAAGATGTCT</td><td>GGG</td><td>||||||||[||||||||||||]PAM</td><td>NA</td><td>Intergenic</td><td>NA</td><td>NA</td></tr>
<tr><td></td><td></td><td></td><td></td><td></td><td></td><td></td><td></td><td></td><td></td><td></td><td></td></tr>
<tr><td>T32251</td><td>TGGGTTCGCCAGAGATCTCGCGG</td><td>1000</td><td></td><td></td><td></td><td></td><td></td><td></td><td></td><td></td><td></td></tr>
<tr><td>Oligo fwd</td><td>CACCTGGGTTCGCCAGAGATCTCG</td><td></td><td></td><td></td><td></td><td></td><td></td><td></td><td></td><td></td><td></td></tr>
<tr><td>Oligo rev</td><td>AAACCGAGATCTCTGGCGAACCCA</td><td></td><td></td><td></td><td></td><td></td><td></td><td></td><td></td><td></td><td></td></tr>
<tr><td>Chromosome</td><td>start</td><td>end</td><td>strand</td><td>MM</td><td>target_seq</td><td>PAM</td><td>alignment</td><td>distance</td><td>position</td><td>gene name</td><td>gene id</td></tr>
<tr><td>GU305914.1</td><td>28514</td><td>28536</td><td>-</td><td>0</td><td>TGGGTTCGCCAGAGATCTCG</td><td>CGG</td><td>||||||||[||||||||||||]PAM</td><td>NA</td><td>Intergenic</td><td>NA</td><td>NA</td></tr>
<tr><td></td><td></td><td></td><td></td><td></td><td></td><td></td><td></td><td></td><td></td><td></td><td></td></tr>
<tr><td>T32252</td><td>TTCGAAGCCTGCCTCTTCCATGG</td><td>1000</td><td></td><td></td><td></td><td></td><td></td><td></td><td></td><td></td><td></td></tr>
<tr><td>Oligo fwd</td><td>CACCTTCGAAGCCTGCCTCTTCCA</td><td></td><td></td><td></td><td></td><td></td><td></td><td></td><td></td><td></td><td></td></tr>
<tr><td>Oligo rev</td><td>AAACTGGAAGAGGCAGGCTTCGAA</td><td></td><td></td><td></td><td></td><td></td><td></td><td></td><td></td><td></td><td></td></tr>
<tr><td>Chromosome</td><td>start</td><td>end</td><td>strand</td><td>MM</td><td>target_seq</td><td>PAM</td><td>alignment</td><td>distance</td><td>position</td><td>gene name</td><td>gene id</td></tr>
<tr><td>GU305914.1</td><td>28472</td><td>28494</td><td>-</td><td>0</td><td>TTCGAAGCCTGCCTCTTCCA</td><td>TGG</td><td>||||||||[||||||||||||]PAM</td><td>NA</td><td>Intergenic</td><td>NA</td><td>NA</td></tr>
<tr><td></td><td></td><td></td><td></td><td></td><td></td><td></td><td></td><td></td><td></td><td></td><td></td></tr>
<tr><td>T32253</td><td>ATCAGATACTTGACGTACGACGG</td><td>1000</td><td></td><td></td><td></td><td></td><td></td><td></td><td></td><td></td><td></td></tr>
<tr><td>Oligo fwd</td><td>CACCATCAGATACTTGACGTACGA</td><td></td><td></td><td></td><td></td><td></td><td></td><td></td><td></td><td></td><td></td></tr>
<tr><td>Oligo rev</td><td>AAACTCGTACGTCAAGTATCTGAT</td><td></td><td></td><td></td><td></td><td></td><td></td><td></td><td></td><td></td><td></td></tr>
<tr><td>Chromosome</td><td>start</td><td>end</td><td>strand</td><td>MM</td><td>target_seq</td><td>PAM</td><td>alignment</td><td>distance</td><td>position</td><td>gene name</td><td>gene id</td></tr>
<tr><td>GU305914.1</td><td>28438</td><td>28460</td><td>-</td><td>0</td><td>ATCAGATACTTGACGTACGA</td><td>CGG</td><td>||||||||[||||||||||||]PAM</td><td>NA</td><td>Intergenic</td><td>NA</td><td>NA</td></tr>
<tr><td></td><td></td><td></td><td></td><td></td><td></td><td></td><td></td><td></td><td></td><td></td><td></td></tr>
<tr><td>T32254</td><td>ACACGCCTTCGTTGTAGCAGCGG</td><td>1000</td><td></td><td></td><td></td><td></td><td></td><td></td><td></td><td></td><td></td></tr>
<tr><td>Oligo fwd</td><td>CACCACACGCCTTCGTTGTAGCAG</td><td></td><td></td><td></td><td></td><td></td><td></td><td></td><td></td><td></td><td></td></tr>
<tr><td>Oligo rev</td><td>AAACCTGCTACAACGAAGGCGTGT</td><td></td><td></td><td></td><td></td><td></td><td></td><td></td><td></td><td></td><td></td></tr>
<tr><td>Chromosome</td><td>start</td><td>end</td><td>strand</td><td>MM</td><td>target_seq</td><td>PAM</td><td>alignment</td><td>distance</td><td>position</td><td>gene name</td><td>gene id</td></tr>
<tr><td>GU305914.1</td><td>28407</td><td>28429</td><td>-</td><td>0</td><td>ACACGCCTTCGTTGTAGCAG</td><td>CGG</td><td>||||||||[||||||||||||]PAM</td><td>NA</td><td>Intergenic</td><td>NA</td><td>NA</td></tr>
<tr><td></td><td></td><td></td><td></td><td></td><td></td><td></td><td></td><td></td><td></td><td></td><td></td></tr>
<tr><td>T32255</td><td>AGCAGCGGCGCGCGAAAGCGAGG</td><td>1000</td><td></td><td></td><td></td><td></td><td></td><td></td><td></td><td></td><td></td></tr>
<tr><td>Oligo fwd</td><td>CACCAGCAGCGGCGCGCGAAAGCG</td><td></td><td></td><td></td><td></td><td></td><td></td><td></td><td></td><td></td><td></td></tr>
<tr><td>Oligo rev</td><td>AAACCGCTTTCGCGCGCCGCTGCT</td><td></td><td></td><td></td><td></td><td></td><td></td><td></td><td></td><td></td><td></td></tr>
<tr><td>Chromosome</td><td>start</td><td>end</td><td>strand</td><td>MM</td><td>target_seq</td><td>PAM</td><td>alignment</td><td>distance</td><td>position</td><td>gene name</td><td>gene id</td></tr>
<tr><td>GU305914.1</td><td>28392</td><td>28414</td><td>-</td><td>0</td><td>AGCAGCGGCGCGCGAAAGCG</td><td>AGG</td><td>||||||||[||||||||||||]PAM</td><td>NA</td><td>Intergenic</td><td>NA</td><td>NA</td></tr>
<tr><td></td><td></td><td></td><td></td><td></td><td></td><td></td><td></td><td></td><td></td><td></td><td></td></tr>
<tr><td>T32256</td><td>GGCGCGCGAAAGCGAGGCTGCGG</td><td>1000</td><td></td><td></td><td></td><td></td><td></td><td></td><td></td><td></td><td></td></tr>
<tr><td>Oligo fwd</td><td>CACCGGCGCGCGAAAGCGAGGCTG</td><td></td><td></td><td></td><td></td><td></td><td></td><td></td><td></td><td></td><td></td></tr>
<tr><td>Oligo rev</td><td>AAACCAGCCTCGCTTTCGCGCGCC</td><td></td><td></td><td></td><td></td><td></td><td></td><td></td><td></td><td></td><td></td></tr>
<tr><td>Chromosome</td><td>start</td><td>end</td><td>strand</td><td>MM</td><td>target_seq</td><td>PAM</td><td>alignment</td><td>distance</td><td>position</td><td>gene name</td><td>gene id</td></tr>
<tr><td>GU305914.1</td><td>28386</td><td>28408</td><td>-</td><td>0</td><td>GGCGCGCGAAAGCGAGGCTG</td><td>CGG</td><td>||||||||[||||||||||||]PAM</td><td>NA</td><td>Intergenic</td><td>NA</td><td>NA</td></tr>
<tr><td></td><td></td><td></td><td></td><td></td><td></td><td></td><td></td><td></td><td></td><td></td><td></td></tr>
<tr><td>T32257</td><td>GCGCGAAAGCGAGGCTGCGGCGG</td><td>1000</td><td></td><td></td><td></td><td></td><td></td><td></td><td></td><td></td><td></td></tr>
<tr><td>Oligo fwd</td><td>CACCGCGCGAAAGCGAGGCTGCGG</td><td></td><td></td><td></td><td></td><td></td><td></td><td></td><td></td><td></td><td></td></tr>
<tr><td>Oligo rev</td><td>AAACCCGCAGCCTCGCTTTCGCGC</td><td></td><td></td><td></td><td></td><td></td><td></td><td></td><td></td><td></td><td></td></tr>
<tr><td>Chromosome</td><td>start</td><td>end</td><td>strand</td><td>MM</td><td>target_seq</td><td>PAM</td><td>alignment</td><td>distance</td><td>position</td><td>gene name</td><td>gene id</td></tr>
<tr><td>GU305914.1</td><td>28383</td><td>28405</td><td>-</td><td>0</td><td>GCGCGAAAGCGAGGCTGCGG</td><td>CGG</td><td>||||||||[||||||||||||]PAM</td><td>NA</td><td>Intergenic</td><td>NA</td><td>NA</td></tr>
<tr><td></td><td></td><td></td><td></td><td></td><td></td><td></td><td></td><td></td><td></td><td></td><td></td></tr>
<tr><td>T32258</td><td>GAGGCTGCGGCGGCACAACGCGG</td><td>1000</td><td></td><td></td><td></td><td></td><td></td><td></td><td></td><td></td><td></td></tr>
<tr><td>Oligo fwd</td><td>CACCGAGGCTGCGGCGGCACAACG</td><td></td><td></td><td></td><td></td><td></td><td></td><td></td><td></td><td></td><td></td></tr>
<tr><td>Oligo rev</td><td>AAACCGTTGTGCCGCCGCAGCCTC</td><td></td><td></td><td></td><td></td><td></td><td></td><td></td><td></td><td></td><td></td></tr>
<tr><td>Chromosome</td><td>start</td><td>end</td><td>strand</td><td>MM</td><td>target_seq</td><td>PAM</td><td>alignment</td><td>distance</td><td>position</td><td>gene name</td><td>gene id</td></tr>
<tr><td>GU305914.1</td><td>28373</td><td>28395</td><td>-</td><td>0</td><td>GAGGCTGCGGCGGCACAACG</td><td>CGG</td><td>||||||||[||||||||||||]PAM</td><td>NA</td><td>Intergenic</td><td>NA</td><td>NA</td></tr>
<tr><td></td><td></td><td></td><td></td><td></td><td></td><td></td><td></td><td></td><td></td><td></td><td></td></tr>
<tr><td>T32259</td><td>TGCGGCGGCACAACGCGGACAGG</td><td>1000</td><td></td><td></td><td></td><td></td><td></td><td></td><td></td><td></td><td></td></tr>
<tr><td>Oligo fwd</td><td>CACCTGCGGCGGCACAACGCGGAC</td><td></td><td></td><td></td><td></td><td></td><td></td><td></td><td></td><td></td><td></td></tr>
<tr><td>Oligo rev</td><td>AAACGTCCGCGTTGTGCCGCCGCA</td><td></td><td></td><td></td><td></td><td></td><td></td><td></td><td></td><td></td><td></td></tr>
<tr><td>Chromosome</td><td>start</td><td>end</td><td>strand</td><td>MM</td><td>target_seq</td><td>PAM</td><td>alignment</td><td>distance</td><td>position</td><td>gene name</td><td>gene id</td></tr>
<tr><td>GU305914.1</td><td>28368</td><td>28390</td><td>-</td><td>0</td><td>TGCGGCGGCACAACGCGGAC</td><td>AGG</td><td>||||||||[||||||||||||]PAM</td><td>NA</td><td>Intergenic</td><td>NA</td><td>NA</td></tr>
<tr><td></td><td></td><td></td><td></td><td></td><td></td><td></td><td></td><td></td><td></td><td></td><td></td></tr>
<tr><td>T32260</td><td>ACAGGCCCTGTCCGTCTTCACGG</td><td>1000</td><td></td><td></td><td></td><td></td><td></td><td></td><td></td><td></td><td></td></tr>
<tr><td>Oligo fwd</td><td>CACCACAGGCCCTGTCCGTCTTCA</td><td></td><td></td><td></td><td></td><td></td><td></td><td></td><td></td><td></td><td></td></tr>
<tr><td>Oligo rev</td><td>AAACTGAAGACGGACAGGGCCTGT</td><td></td><td></td><td></td><td></td><td></td><td></td><td></td><td></td><td></td><td></td></tr>
<tr><td>Chromosome</td><td>start</td><td>end</td><td>strand</td><td>MM</td><td>target_seq</td><td>PAM</td><td>alignment</td><td>distance</td><td>position</td><td>gene name</td><td>gene id</td></tr>
<tr><td>GU305914.1</td><td>28350</td><td>28372</td><td>-</td><td>0</td><td>ACAGGCCCTGTCCGTCTTCA</td><td>CGG</td><td>||||||||[||||||||||||]PAM</td><td>NA</td><td>Intergenic</td><td>NA</td><td>NA</td></tr>
<tr><td></td><td></td><td></td><td></td><td></td><td></td><td></td><td></td><td></td><td></td><td></td><td></td></tr>
<tr><td>T32261</td><td>CAGGCCCTGTCCGTCTTCACGGG</td><td>1000</td><td></td><td></td><td></td><td></td><td></td><td></td><td></td><td></td><td></td></tr>
<tr><td>Oligo fwd</td><td>CACCCAGGCCCTGTCCGTCTTCAC</td><td></td><td></td><td></td><td></td><td></td><td></td><td></td><td></td><td></td><td></td></tr>
<tr><td>Oligo rev</td><td>AAACGTGAAGACGGACAGGGCCTG</td><td></td><td></td><td></td><td></td><td></td><td></td><td></td><td></td><td></td><td></td></tr>
<tr><td>Chromosome</td><td>start</td><td>end</td><td>strand</td><td>MM</td><td>target_seq</td><td>PAM</td><td>alignment</td><td>distance</td><td>position</td><td>gene name</td><td>gene id</td></tr>
<tr><td>GU305914.1</td><td>28349</td><td>28371</td><td>-</td><td>0</td><td>CAGGCCCTGTCCGTCTTCAC</td><td>GGG</td><td>||||||||[||||||||||||]PAM</td><td>NA</td><td>Intergenic</td><td>NA</td><td>NA</td></tr>
</table>

| | | | | | | | | | | | |
|---|---|---|---|---|---|---|---|---|---|---|---|
| T32262 | AGGCCCTGTCCGTCTTCACGGGG | 1000 | | | | | | | | | |
| Oligo fwd | CACCAGGCCCTGTCCGTCTTCACG | | | | | | | | | | |
| Oligo rev | AAACCGTGAAGACGGACAGGGCCT | | | | | | | | | | |
| Chromosome | start | end | strand | MM | target_seq | PAM | alignment | distance | position | gene name | gene id |
| GU305914.1 | 28348 | 28370 | - | 0 | AGGCCCTGTCCGTCTTCACG | GGG | \|\|\|\|\|\|\|\|[\|\|\|\|\|\|\|\|\|\|\|\|]PAM | NA | Intergenic | NA | NA |
| T32263 | CGTGATCTCGTTCGCCACCTCGG | 1000 | | | | | | | | | |
| Oligo fwd | CACCCGTGATCTCGTTCGCCACCT | | | | | | | | | | |
| Oligo rev | AAACAGGTGGCGAACGAGATCACG | | | | | | | | | | |
| Chromosome | start | end | strand | MM | target_seq | PAM | alignment | distance | position | gene name | gene id |
| GU305914.1 | 28322 | 28344 | - | 0 | CGTGATCTCGTTCGCCACCT | CGG | \|\|\|\|\|\|\|\|[\|\|\|\|\|\|\|\|\|\|\|\|]PAM | NA | Intergenic | NA | NA |
| GU305914.1 | 70416 | 70438 | - | 4 | CTCGATCTCGTTCGCCAGTT | CGG | \|--\|\|\|\|\|[\|\|\|\|\|\|\|\|\|--\|]PAM | NA | Intergenic | NA | NA |
| T32264 | CCACCTCGGAGCACAGCACGCGG | 1000 | | | | | | | | | |
| Oligo fwd | CACCCCACCTCGGAGCACAGCACG | | | | | | | | | | |
| Oligo rev | AAACCGTGCTGTGCTCCGAGGTGG | | | | | | | | | | |
| Chromosome | start | end | strand | MM | target_seq | PAM | alignment | distance | position | gene name | gene id |
| GU305914.1 | 28308 | 28330 | - | 0 | CCACCTCGGAGCACAGCACG | CGG | \|\|\|\|\|\|\|\|[\|\|\|\|\|\|\|\|\|\|\|\|]PAM | NA | Intergenic | NA | NA |
| T32265 | GCGGTAGTCGTCTCTCATTATGG | 1000 | | | | | | | | | |
| Oligo fwd | CACCGCGGTAGTCGTCTCTCATTA | | | | | | | | | | |
| Oligo rev | AAACTAATGAGAGACGACTACCGC | | | | | | | | | | |
| Chromosome | start | end | strand | MM | target_seq | PAM | alignment | distance | position | gene name | gene id |
| GU305914.1 | 28289 | 28311 | - | 0 | GCGGTAGTCGTCTCTCATTA | TGG | \|\|\|\|\|\|\|\|[\|\|\|\|\|\|\|\|\|\|\|\|]PAM | NA | Intergenic | NA | NA |
| T32266 | CGGTAGTCGTCTCTCATTATGGG | 1000 | | | | | | | | | |
| Oligo fwd | CACCCGGTAGTCGTCTCTCATTAT | | | | | | | | | | |
| Oligo rev | AAACATAATGAGAGACGACTACCG | | | | | | | | | | |
| Chromosome | start | end | strand | MM | target_seq | PAM | alignment | distance | position | gene name | gene id |
| GU305914.1 | 28288 | 28310 | - | 0 | CGGTAGTCGTCTCTCATTAT | GGG | \|\|\|\|\|\|\|\|[\|\|\|\|\|\|\|\|\|\|\|\|]PAM | NA | Intergenic | NA | NA |
| T32267 | CATTATGGGACACTGATAGTAGG | 1000 | | | | | | | | | |
| Oligo fwd | CACCCATTATGGGACACTGATAGT | | | | | | | | | | |
| Oligo rev | AAACACTATCAGTGTCCCATAATG | | | | | | | | | | |
| Chromosome | start | end | strand | MM | target_seq | PAM | alignment | distance | position | gene name | gene id |
| GU305914.1 | 28274 | 28296 | - | 0 | CATTATGGGACACTGATAGT | AGG | \|\|\|\|\|\|\|\|[\|\|\|\|\|\|\|\|\|\|\|\|]PAM | NA | Intergenic | NA | NA |
| T32268 | ACTGATAGTAGGCCGCGTACAGG | 1000 | | | | | | | | | |
| Oligo fwd | CACCACTGATAGTAGGCCGCGTAC | | | | | | | | | | |
| Oligo rev | AAACGTACGCGGCCTACTATCAGT | | | | | | | | | | |
| Chromosome | start | end | strand | MM | target_seq | PAM | alignment | distance | position | gene name | gene id |
| GU305914.1 | 28263 | 28285 | - | 0 | ACTGATAGTAGGCCGCGTAC | AGG | \|\|\|\|\|\|\|\|[\|\|\|\|\|\|\|\|\|\|\|\|]PAM | NA | Intergenic | NA | NA |
| T32269 | CTGATAGTAGGCCGCGTACAGGG | 1000 | | | | | | | | | |
| Oligo fwd | CACCCTGATAGTAGGCCGCGTACA | | | | | | | | | | |
| Oligo rev | AAACTGTACGCGGCCTACTATCAG | | | | | | | | | | |
| Chromosome | start | end | strand | MM | target_seq | PAM | alignment | distance | position | gene name | gene id |
| GU305914.1 | 28262 | 28284 | - | 0 | CTGATAGTAGGCCGCGTACA | GGG | \|\|\|\|\|\|\|\|[\|\|\|\|\|\|\|\|\|\|\|\|]PAM | NA | Intergenic | NA | NA |
| T32270 | GTAGGCCGCGTACAGGGTCTCGG | 1000 | | | | | | | | | |
| Oligo fwd | CACCGTAGGCCGCGTACAGGGTCT | | | | | | | | | | |
| Oligo rev | AAACAGACCCTGTACGCGGCCTAC | | | | | | | | | | |
| Chromosome | start | end | strand | MM | target_seq | PAM | alignment | distance | position | gene name | gene id |
| GU305914.1 | 28256 | 28278 | - | 0 | GTAGGCCGCGTACAGGGTCT | CGG | \|\|\|\|\|\|\|\|[\|\|\|\|\|\|\|\|\|\|\|\|]PAM | NA | Intergenic | NA | NA |
| T32271 | CCGCGTACAGGGTCTCGGTCAGG | 1000 | | | | | | | | | |
| Oligo fwd | CACCCCGCGTACAGGGTCTCGGTC | | | | | | | | | | |
| Oligo rev | AAACGACCGAGACCCTGTACGCGG | | | | | | | | | | |
| Chromosome | start | end | strand | MM | target_seq | PAM | alignment | distance | position | gene name | gene id |
| GU305914.1 | 28251 | 28273 | - | 0 | CCGCGTACAGGGTCTCGGTC | AGG | \|\|\|\|\|\|\|\|[\|\|\|\|\|\|\|\|\|\|\|\|]PAM | NA | Intergenic | NA | NA |
| T32272 | TCTCGGTCAGGTCCTCGAACTGG | 1000 | | | | | | | | | |
| Oligo fwd | CACCTCTCGGTCAGGTCCTCGAAC | | | | | | | | | | |
| Oligo rev | AAACGTTCGAGGACCTGACCGAGA | | | | | | | | | | |
| Chromosome | start | end | strand | MM | target_seq | PAM | alignment | distance | position | gene name | gene id |
| GU305914.1 | 28239 | 28261 | - | 0 | TCTCGGTCAGGTCCTCGAAC | TGG | \|\|\|\|\|\|\|\|[\|\|\|\|\|\|\|\|\|\|\|\|]PAM | NA | Intergenic | NA | NA |
| T32273 | ACACAAAGTTTCCCAGCGACAGG | 1000 | | | | | | | | | |
| Oligo fwd | CACCACACAAAGTTTCCCAGCGAC | | | | | | | | | | |
| Oligo rev | AAACGTCGCTGGGAAACTTTGTGT | | | | | | | | | | |
| Chromosome | start | end | strand | MM | target_seq | PAM | alignment | distance | position | gene name | gene id |
| GU305914.1 | 28170 | 28192 | - | 0 | ACACAAAGTTTCCCAGCGAC | AGG | \|\|\|\|\|\|\|\|[\|\|\|\|\|\|\|\|\|\|\|\|]PAM | NA | Intergenic | NA | NA |
| T32274 | GTTCAGCACATCTTCCCGTGTGG | 1000 | | | | | | | | | |
| Oligo fwd | CACCGTTCAGCACATCTTCCCGTG | | | | | | | | | | |
| Oligo rev | AAACCACGGGAAGATGTGCTGAAC | | | | | | | | | | |
| Chromosome | start | end | strand | MM | target_seq | PAM | alignment | distance | position | gene name | gene id |
| GU305914.1 | 28133 | 28155 | - | 0 | GTTCAGCACATCTTCCCGTG | TGG | \|\|\|\|\|\|\|\|[\|\|\|\|\|\|\|\|\|\|\|\|]PAM | NA | Intergenic | NA | NA |
| T32275 | TTCAGCACATCTTCCCGTGTGGG | 1000 | | | | | | | | | |
| Oligo fwd | CACCTTCAGCACATCTTCCCGTGT | | | | | | | | | | |
| Oligo rev | AAACACACGGGAAGATGTGCTGAA | | | | | | | | | | |
| Chromosome | start | end | strand | MM | target_seq | PAM | alignment | distance | position | gene name | gene id |
| GU305914.1 | 28132 | 28154 | - | 0 | TTCAGCACATCTTCCCGTGT | GGG | \|\|\|\|\|\|\|\|[\|\|\|\|\|\|\|\|\|\|\|\|]PAM | NA | Intergenic | NA | NA |
| T32276 | ACATCTTCCCGTGTGGGATGCGG | 1000 | | | | | | | | | |
| Oligo fwd | CACCACATCTTCCCGTGTGGGATG | | | | | | | | | | |
| Oligo rev | AAACCATCCCACACGGGAAGATGT | | | | | | | | | | |
| Chromosome | start | end | strand | MM | target_seq | PAM | alignment | distance | position | gene name | gene id |

GU305914.1 28126 28148 - 0 ACATCTTCCCGTGTGGGATG CGG ||||||||[||||||||||||]PAM NA Intergenic NA NA

T32277 CCCGTGTGGGATGCGGATTCAGG 1000
Oligo fwd CACCCCCGTGTGGGATGCGGATTC
Oligo rev AAACGAATCCGCATCCCACACGGG
Chromosome start end strand MM target_seq PAM alignment distance position gene name gene id
GU305914.1 28119 28141 - 0 CCCGTGTGGGATGCGGATTC AGG ||||||||[||||||||||||]PAM NA Intergenic NA NA

T32278 CCGTGTGGGATGCGGATTCAGGG 1000
Oligo fwd CACCCCGTGTGGGATGCGGATTCA
Oligo rev AAACTGAATCCGCATCCCACACGG
Chromosome start end strand MM target_seq PAM alignment distance position gene name gene id
GU305914.1 28118 28140 - 0 CCGTGTGGGATGCGGATTCA GGG ||||||||[||||||||||||]PAM NA Intergenic NA NA

T32279 GCGGATTCAGGGTCATCACTAGG 1000
Oligo fwd CACCGCGGATTCAGGGTCATCACT
Oligo rev AAACAGTGATGACCCTGAATCCGC
Chromosome start end strand MM target_seq PAM alignment distance position gene name gene id
GU305914.1 28107 28129 - 0 GCGGATTCAGGGTCATCACT AGG ||||||||[||||||||||||]PAM NA Intergenic NA NA

T32280 TATCCTCCTCGTCGCTGTTCAGG 1000
Oligo fwd CACCTATCCTCCTCGTCGCTGTTC
Oligo rev AAACGAACAGCGACGAGGAGGATA
Chromosome start end strand MM target_seq PAM alignment distance position gene name gene id
GU305914.1 28074 28096 - 0 TATCCTCCTCGTCGCTGTTC AGG ||||||||[||||||||||||]PAM NA Intergenic NA NA

T32281 TCTTGATGTTCCGCATGTACAGG 1000
Oligo fwd CACCTCTTGATGTTCCGCATGTAC
Oligo rev AAACGTACATGCGGAACATCAAGA
Chromosome start end strand MM target_seq PAM alignment distance position gene name gene id
GU305914.1 28050 28072 - 0 TCTTGATGTTCCGCATGTAC AGG ||||||||[||||||||||||]PAM NA Intergenic NA NA

T32282 CTTGATGTTCCGCATGTACAGGG 1000
Oligo fwd CACCCTTGATGTTCCGCATGTACA
Oligo rev AAACTGTACATGCGGAACATCAAG
Chromosome start end strand MM target_seq PAM alignment distance position gene name gene id
GU305914.1 28049 28071 - 0 CTTGATGTTCCGCATGTACA GGG ||||||||[||||||||||||]PAM NA Intergenic NA NA

T32283 ACAGGGTCTTCGCCCCCTCGCGG 1000
Oligo fwd CACCACAGGGTCTTCGCCCCCTCG
Oligo rev AAACCGAGGGGGCGAAGACCCTGT
Chromosome start end strand MM target_seq PAM alignment distance position gene name gene id
GU305914.1 28032 28054 - 0 ACAGGGTCTTCGCCCCCTCG CGG ||||||||[||||||||||||]PAM NA Intergenic NA NA

T32284 CAGGGTCTTCGCCCCCTCGCGGG 1000
Oligo fwd CACCCAGGGTCTTCGCCCCCTCGC
Oligo rev AAACGCGAGGGGGCGAAGACCCTG
Chromosome start end strand MM target_seq PAM alignment distance position gene name gene id
GU305914.1 28031 28053 - 0 CAGGGTCTTCGCCCCCTCGC GGG ||||||||[||||||||||||]PAM NA Intergenic NA NA

T32285 GCCCCCTCGCGGGTGAAACGCGG 1000
Oligo fwd CACCGCCCCCTCGCGGGTGAAACG
Oligo rev AAACCGTTTCACCCGCGAGGGGGC
Chromosome start end strand MM target_seq PAM alignment distance position gene name gene id
GU305914.1 28021 28043 - 0 GCCCCCTCGCGGGTGAAACG CGG ||||||||[||||||||||||]PAM NA Intergenic NA NA

T32286 CCCCCTCGCGGGTGAAACGCGGG 1000
Oligo fwd CACCCCCCCTCGCGGGTGAAACGC
Oligo rev AAACGCGTTTCACCCGCGAGGGGG
Chromosome start end strand MM target_seq PAM alignment distance position gene name gene id
GU305914.1 28020 28042 - 0 CCCCCTCGCGGGTGAAACGC GGG ||||||||[||||||||||||]PAM NA Intergenic NA NA

T32287 CCCCTCGCGGGTGAAACGCGGGG 1000
Oligo fwd CACCCCCCTCGCGGGTGAAACGCG
Oligo rev AAACCGCGTTTCACCCGCGAGGGG
Chromosome start end strand MM target_seq PAM alignment distance position gene name gene id
GU305914.1 28019 28041 - 0 CCCCTCGCGGGTGAAACGCG GGG ||||||||[||||||||||||]PAM NA Intergenic NA NA

T32288 CCCTCGCGGGTGAAACGCGGGGG 1000
Oligo fwd CACCCCCTCGCGGGTGAAACGCGG
Oligo rev AAACCCGCGTTTCACCCGCGAGGG
Chromosome start end strand MM target_seq PAM alignment distance position gene name gene id
GU305914.1 28018 28040 - 0 CCCTCGCGGGTGAAACGCGG GGG ||||||||[||||||||||||]PAM NA Intergenic NA NA

T32289 AACGCGGGGGACTGAACAGCAGG 1000
Oligo fwd CACCAACGCGGGGGACTGAACAGC
Oligo rev AAACGCTGTTCAGTCCCCCGCGTT
Chromosome start end strand MM target_seq PAM alignment distance position gene name gene id
GU305914.1 28005 28027 - 0 AACGCGGGGGACTGAACAGC AGG ||||||||[||||||||||||]PAM NA Intergenic NA NA

T32290 TGAACAGCAGGTTCTTGCTGCGG 1000
Oligo fwd CACCTGAACAGCAGGTTCTTGCTG
Oligo rev AAACCAGCAAGAACCTGCTGTTCA
Chromosome start end strand MM target_seq PAM alignment distance position gene name gene id
GU305914.1 27993 28015 - 0 TGAACAGCAGGTTCTTGCTG CGG ||||||||[||||||||||||]PAM NA Intergenic NA NA

T32291 GTGCGCACTGCGTCTTCTGTCGG 1000
Oligo fwd CACCGTGCGCACTGCGTCTTCTGT
Oligo rev AAACACAGAAGACGCAGTGCGCAC

<table>
<tr><td>Chromosome</td><td>start</td><td>end</td><td>strand</td><td>MM</td><td>target_seq</td><td>PAM</td><td>alignment</td><td>distance</td><td>position</td><td>gene name</td><td>gene id</td></tr>
<tr><td>GU305914.1</td><td>27963</td><td>27985</td><td>-</td><td>0</td><td>GTGCGCACTGCGTCTTCTGT</td><td>CGG</td><td>||||||||[||||||||||||]PAM</td><td>NA</td><td>Intergenic</td><td>NA</td><td>NA</td></tr>
<tr><td>T32292</td><td>CGCACTGCGTCTTCTGTCGGCGG</td><td>1000</td><td></td><td></td><td></td><td></td><td></td><td></td><td></td><td></td><td></td></tr>
<tr><td>Oligo fwd</td><td>CACCCGCACTGCGTCTTCTGTCGG</td><td></td><td></td><td></td><td></td><td></td><td></td><td></td><td></td><td></td><td></td></tr>
<tr><td>Oligo rev</td><td>AAACCCGACAGAAGACGCAGTGCG</td><td></td><td></td><td></td><td></td><td></td><td></td><td></td><td></td><td></td><td></td></tr>
<tr><td>Chromosome</td><td>start</td><td>end</td><td>strand</td><td>MM</td><td>target_seq</td><td>PAM</td><td>alignment</td><td>distance</td><td>position</td><td>gene name</td><td>gene id</td></tr>
<tr><td>GU305914.1</td><td>27960</td><td>27982</td><td>-</td><td>0</td><td>CGCACTGCGTCTTCTGTCGG</td><td>CGG</td><td>||||||||[||||||||||||]PAM</td><td>NA</td><td>Intergenic</td><td>NA</td><td>NA</td></tr>
<tr><td>T32293</td><td>CGGCGCTCGCGTTCCTGCATCGG</td><td>1000</td><td></td><td></td><td></td><td></td><td></td><td></td><td></td><td></td><td></td></tr>
<tr><td>Oligo fwd</td><td>CACCCGGCGCTCGCGTTCCTGCAT</td><td></td><td></td><td></td><td></td><td></td><td></td><td></td><td></td><td></td><td></td></tr>
<tr><td>Oligo rev</td><td>AAACATGCAGGAACGCGAGCGCCG</td><td></td><td></td><td></td><td></td><td></td><td></td><td></td><td></td><td></td><td></td></tr>
<tr><td>Chromosome</td><td>start</td><td>end</td><td>strand</td><td>MM</td><td>target_seq</td><td>PAM</td><td>alignment</td><td>distance</td><td>position</td><td>gene name</td><td>gene id</td></tr>
<tr><td>GU305914.1</td><td>27940</td><td>27962</td><td>-</td><td>0</td><td>CGGCGCTCGCGTTCCTGCAT</td><td>CGG</td><td>||||||||[||||||||||||]PAM</td><td>NA</td><td>Intergenic</td><td>NA</td><td>NA</td></tr>
<tr><td>T32294</td><td>CCTGCATCGGTTTCATCAGTTGG</td><td>1000</td><td></td><td></td><td></td><td></td><td></td><td></td><td></td><td></td><td></td></tr>
<tr><td>Oligo fwd</td><td>CACCCCTGCATCGGTTTCATCAGT</td><td></td><td></td><td></td><td></td><td></td><td></td><td></td><td></td><td></td><td></td></tr>
<tr><td>Oligo rev</td><td>AAACACTGATGAAACCGATGCAGG</td><td></td><td></td><td></td><td></td><td></td><td></td><td></td><td></td><td></td><td></td></tr>
<tr><td>Chromosome</td><td>start</td><td>end</td><td>strand</td><td>MM</td><td>target_seq</td><td>PAM</td><td>alignment</td><td>distance</td><td>position</td><td>gene name</td><td>gene id</td></tr>
<tr><td>GU305914.1</td><td>27927</td><td>27949</td><td>-</td><td>0</td><td>CCTGCATCGGTTTCATCAGT</td><td>TGG</td><td>||||||||[||||||||||||]PAM</td><td>NA</td><td>Intergenic</td><td>NA</td><td>NA</td></tr>
<tr><td>T32295</td><td>TTGGTCGATCGTATCGCAGCTGG</td><td>1000</td><td></td><td></td><td></td><td></td><td></td><td></td><td></td><td></td><td></td></tr>
<tr><td>Oligo fwd</td><td>CACCTTGGTCGATCGTATCGCAGC</td><td></td><td></td><td></td><td></td><td></td><td></td><td></td><td></td><td></td><td></td></tr>
<tr><td>Oligo rev</td><td>AAACGCTGCGATACGATCGACCAA</td><td></td><td></td><td></td><td></td><td></td><td></td><td></td><td></td><td></td><td></td></tr>
<tr><td>Chromosome</td><td>start</td><td>end</td><td>strand</td><td>MM</td><td>target_seq</td><td>PAM</td><td>alignment</td><td>distance</td><td>position</td><td>gene name</td><td>gene id</td></tr>
<tr><td>GU305914.1</td><td>27908</td><td>27930</td><td>-</td><td>0</td><td>TTGGTCGATCGTATCGCAGC</td><td>TGG</td><td>||||||||[||||||||||||]PAM</td><td>NA</td><td>Intergenic</td><td>NA</td><td>NA</td></tr>
<tr><td>T32296</td><td>GCTGGCCTTGTGCTGCGCCGCGG</td><td>1000</td><td></td><td></td><td></td><td></td><td></td><td></td><td></td><td></td><td></td></tr>
<tr><td>Oligo fwd</td><td>CACCGCTGGCCTTGTGCTGCGCCG</td><td></td><td></td><td></td><td></td><td></td><td></td><td></td><td></td><td></td><td></td></tr>
<tr><td>Oligo rev</td><td>AAACCGGCGCAGCACAAGGCCAGC</td><td></td><td></td><td></td><td></td><td></td><td></td><td></td><td></td><td></td><td></td></tr>
<tr><td>Chromosome</td><td>start</td><td>end</td><td>strand</td><td>MM</td><td>target_seq</td><td>PAM</td><td>alignment</td><td>distance</td><td>position</td><td>gene name</td><td>gene id</td></tr>
<tr><td>GU305914.1</td><td>27890</td><td>27912</td><td>-</td><td>0</td><td>GCTGGCCTTGTGCTGCGCCG</td><td>CGG</td><td>||||||||[||||||||||||]PAM</td><td>NA</td><td>Intergenic</td><td>NA</td><td>NA</td></tr>
<tr><td>T32297</td><td>GCTGCGCCGCGGTGATGTTCAGG</td><td>1000</td><td></td><td></td><td></td><td></td><td></td><td></td><td></td><td></td><td></td></tr>
<tr><td>Oligo fwd</td><td>CACCGCTGCGCCGCGGTGATGTTC</td><td></td><td></td><td></td><td></td><td></td><td></td><td></td><td></td><td></td><td></td></tr>
<tr><td>Oligo rev</td><td>AAACGAACATCACCGCGGCGCAGC</td><td></td><td></td><td></td><td></td><td></td><td></td><td></td><td></td><td></td><td></td></tr>
<tr><td>Chromosome</td><td>start</td><td>end</td><td>strand</td><td>MM</td><td>target_seq</td><td>PAM</td><td>alignment</td><td>distance</td><td>position</td><td>gene name</td><td>gene id</td></tr>
<tr><td>GU305914.1</td><td>27879</td><td>27901</td><td>-</td><td>0</td><td>GCTGCGCCGCGGTGATGTTC</td><td>AGG</td><td>||||||||[||||||||||||]PAM</td><td>NA</td><td>Intergenic</td><td>NA</td><td>NA</td></tr>
<tr><td>T32298</td><td>GCGCCGCGGTGATGTTCAGGCGG</td><td>1000</td><td></td><td></td><td></td><td></td><td></td><td></td><td></td><td></td><td></td></tr>
<tr><td>Oligo fwd</td><td>CACCGCGCCGCGGTGATGTTCAGG</td><td></td><td></td><td></td><td></td><td></td><td></td><td></td><td></td><td></td><td></td></tr>
<tr><td>Oligo rev</td><td>AAACCCTGAACATCACCGCGGCGC</td><td></td><td></td><td></td><td></td><td></td><td></td><td></td><td></td><td></td><td></td></tr>
<tr><td>Chromosome</td><td>start</td><td>end</td><td>strand</td><td>MM</td><td>target_seq</td><td>PAM</td><td>alignment</td><td>distance</td><td>position</td><td>gene name</td><td>gene id</td></tr>
<tr><td>GU305914.1</td><td>27876</td><td>27898</td><td>-</td><td>0</td><td>GCGCCGCGGTGATGTTCAGG</td><td>CGG</td><td>||||||||[||||||||||||]PAM</td><td>NA</td><td>Intergenic</td><td>NA</td><td>NA</td></tr>
<tr><td>T32299</td><td>TCAGGCGGCCGAGCACCCGCAGG</td><td>1000</td><td></td><td></td><td></td><td></td><td></td><td></td><td></td><td></td><td></td></tr>
<tr><td>Oligo fwd</td><td>CACCTCAGGCGGCCGAGCACCCGC</td><td></td><td></td><td></td><td></td><td></td><td></td><td></td><td></td><td></td><td></td></tr>
<tr><td>Oligo rev</td><td>AAACGCGGGTGCTCGGCCGCCTGA</td><td></td><td></td><td></td><td></td><td></td><td></td><td></td><td></td><td></td><td></td></tr>
<tr><td>Chromosome</td><td>start</td><td>end</td><td>strand</td><td>MM</td><td>target_seq</td><td>PAM</td><td>alignment</td><td>distance</td><td>position</td><td>gene name</td><td>gene id</td></tr>
<tr><td>GU305914.1</td><td>27861</td><td>27883</td><td>-</td><td>0</td><td>TCAGGCGGCCGAGCACCCGC</td><td>AGG</td><td>||||||||[||||||||||||]PAM</td><td>NA</td><td>Intergenic</td><td>NA</td><td>NA</td></tr>
<tr><td>T32300</td><td>CCGAGCACCCGCAGGCACATCGG</td><td>1000</td><td></td><td></td><td></td><td></td><td></td><td></td><td></td><td></td><td></td></tr>
<tr><td>Oligo fwd</td><td>CACCCCGAGCACCCGCAGGCACAT</td><td></td><td></td><td></td><td></td><td></td><td></td><td></td><td></td><td></td><td></td></tr>
<tr><td>Oligo rev</td><td>AAACATGTGCCTGCGGGTGCTCGG</td><td></td><td></td><td></td><td></td><td></td><td></td><td></td><td></td><td></td><td></td></tr>
<tr><td>Chromosome</td><td>start</td><td>end</td><td>strand</td><td>MM</td><td>target_seq</td><td>PAM</td><td>alignment</td><td>distance</td><td>position</td><td>gene name</td><td>gene id</td></tr>
<tr><td>GU305914.1</td><td>27853</td><td>27875</td><td>-</td><td>0</td><td>CCGAGCACCCGCAGGCACAT</td><td>CGG</td><td>||||||||[||||||||||||]PAM</td><td>NA</td><td>Intergenic</td><td>NA</td><td>NA</td></tr>
<tr><td>T32301</td><td>AGCACCCGCAGGCACATCGGAGG</td><td>1000</td><td></td><td></td><td></td><td></td><td></td><td></td><td></td><td></td><td></td></tr>
<tr><td>Oligo fwd</td><td>CACCAGCACCCGCAGGCACATCGG</td><td></td><td></td><td></td><td></td><td></td><td></td><td></td><td></td><td></td><td></td></tr>
<tr><td>Oligo rev</td><td>AAACCCGATGTGCCTGCGGGTGCT</td><td></td><td></td><td></td><td></td><td></td><td></td><td></td><td></td><td></td><td></td></tr>
<tr><td>Chromosome</td><td>start</td><td>end</td><td>strand</td><td>MM</td><td>target_seq</td><td>PAM</td><td>alignment</td><td>distance</td><td>position</td><td>gene name</td><td>gene id</td></tr>
<tr><td>GU305914.1</td><td>27850</td><td>27872</td><td>-</td><td>0</td><td>AGCACCCGCAGGCACATCGG</td><td>AGG</td><td>||||||||[||||||||||||]PAM</td><td>NA</td><td>Intergenic</td><td>NA</td><td>NA</td></tr>
<tr><td>T32302</td><td>GCACCCGCAGGCACATCGGAGGG</td><td>1000</td><td></td><td></td><td></td><td></td><td></td><td></td><td></td><td></td><td></td></tr>
<tr><td>Oligo fwd</td><td>CACCGCACCCGCAGGCACATCGGA</td><td></td><td></td><td></td><td></td><td></td><td></td><td></td><td></td><td></td><td></td></tr>
<tr><td>Oligo rev</td><td>AAACTCCGATGTGCCTGCGGGTGC</td><td></td><td></td><td></td><td></td><td></td><td></td><td></td><td></td><td></td><td></td></tr>
<tr><td>Chromosome</td><td>start</td><td>end</td><td>strand</td><td>MM</td><td>target_seq</td><td>PAM</td><td>alignment</td><td>distance</td><td>position</td><td>gene name</td><td>gene id</td></tr>
<tr><td>GU305914.1</td><td>27849</td><td>27871</td><td>-</td><td>0</td><td>GCACCCGCAGGCACATCGGA</td><td>GGG</td><td>||||||||[||||||||||||]PAM</td><td>NA</td><td>Intergenic</td><td>NA</td><td>NA</td></tr>
<tr><td>T32303</td><td>CCGCAGGCACATCGGAGGGCTGG</td><td>1000</td><td></td><td></td><td></td><td></td><td></td><td></td><td></td><td></td><td></td></tr>
<tr><td>Oligo fwd</td><td>CACCCCGCAGGCACATCGGAGGGC</td><td></td><td></td><td></td><td></td><td></td><td></td><td></td><td></td><td></td><td></td></tr>
<tr><td>Oligo rev</td><td>AAACGCCCTCCGATGTGCCTGCGG</td><td></td><td></td><td></td><td></td><td></td><td></td><td></td><td></td><td></td><td></td></tr>
<tr><td>Chromosome</td><td>start</td><td>end</td><td>strand</td><td>MM</td><td>target_seq</td><td>PAM</td><td>alignment</td><td>distance</td><td>position</td><td>gene name</td><td>gene id</td></tr>
<tr><td>GU305914.1</td><td>27845</td><td>27867</td><td>-</td><td>0</td><td>CCGCAGGCACATCGGAGGGC</td><td>TGG</td><td>||||||||[||||||||||||]PAM</td><td>NA</td><td>Intergenic</td><td>NA</td><td>NA</td></tr>
<tr><td>T32304</td><td>CGCAGGCACATCGGAGGGCTGGG</td><td>1000</td><td></td><td></td><td></td><td></td><td></td><td></td><td></td><td></td><td></td></tr>
<tr><td>Oligo fwd</td><td>CACCCGCAGGCACATCGGAGGGCT</td><td></td><td></td><td></td><td></td><td></td><td></td><td></td><td></td><td></td><td></td></tr>
<tr><td>Oligo rev</td><td>AAACAGCCCTCCGATGTGCCTGCG</td><td></td><td></td><td></td><td></td><td></td><td></td><td></td><td></td><td></td><td></td></tr>
<tr><td>Chromosome</td><td>start</td><td>end</td><td>strand</td><td>MM</td><td>target_seq</td><td>PAM</td><td>alignment</td><td>distance</td><td>position</td><td>gene name</td><td>gene id</td></tr>
<tr><td>GU305914.1</td><td>27844</td><td>27866</td><td>-</td><td>0</td><td>CGCAGGCACATCGGAGGGCT</td><td>GGG</td><td>||||||||[||||||||||||]PAM</td><td>NA</td><td>Intergenic</td><td>NA</td><td>NA</td></tr>
<tr><td>T32305</td><td>GAGGGCTGGGCGACGTCTGCTGG</td><td>1000</td><td></td><td></td><td></td><td></td><td></td><td></td><td></td><td></td><td></td></tr>
<tr><td>Oligo fwd</td><td>CACCGAGGGCTGGGCGACGTCTGC</td><td></td><td></td><td></td><td></td><td></td><td></td><td></td><td></td><td></td><td></td></tr>
<tr><td>Oligo rev</td><td>AAACGCAGACGTCGCCCAGCCCTC</td><td></td><td></td><td></td><td></td><td></td><td></td><td></td><td></td><td></td><td></td></tr>
<tr><td>Chromosome</td><td>start</td><td>end</td><td>strand</td><td>MM</td><td>target_seq</td><td>PAM</td><td>alignment</td><td>distance</td><td>position</td><td>gene name</td><td>gene id</td></tr>
<tr><td>GU305914.1</td><td>27831</td><td>27853</td><td>-</td><td>0</td><td>GAGGGCTGGGCGACGTCTGC</td><td>TGG</td><td>||||||||[||||||||||||]PAM</td><td>NA</td><td>Intergenic</td><td>NA</td><td>NA</td></tr>
<tr><td>T32306</td><td>AGGGCTGGGCGACGTCTGCTGGG</td><td>1000</td><td></td><td></td><td></td><td></td><td></td><td></td><td></td><td></td><td></td></tr>
<tr><td>Oligo fwd</td><td>CACCAGGGCTGGGCGACGTCTGCT</td><td></td><td></td><td></td><td></td><td></td><td></td><td></td><td></td><td></td><td></td></tr>
</table>

<table>
<tr><td>Oligo rev</td><td>AAACAGCAGACGTCGCCCAGCCCT</td><td></td><td></td><td></td><td></td><td></td><td></td><td></td><td></td><td></td><td></td></tr>
<tr><td>Chromosome</td><td>start</td><td>end</td><td>strand</td><td>MM</td><td>target_seq</td><td>PAM</td><td>alignment</td><td>distance</td><td>position</td><td>gene name</td><td>gene id</td></tr>
<tr><td>GU305914.1</td><td>27830</td><td>27852</td><td>-</td><td>0</td><td>AGGGCTGGGCGACGTCTGCT</td><td>GGG</td><td>||||||||[||||||||||||]PAM</td><td>NA</td><td>Intergenic</td><td>NA</td><td>NA</td></tr>
<tr><td>T32307</td><td>GGGCTGGGCGACGTCTGCTGGGG</td><td>1000</td><td></td><td></td><td></td><td></td><td></td><td></td><td></td><td></td><td></td></tr>
<tr><td>Oligo fwd</td><td>CACCGGGCTGGGCGACGTCTGCTG</td><td></td><td></td><td></td><td></td><td></td><td></td><td></td><td></td><td></td><td></td></tr>
<tr><td>Oligo rev</td><td>AAACCAGCAGACGTCGCCCAGCCC</td><td></td><td></td><td></td><td></td><td></td><td></td><td></td><td></td><td></td><td></td></tr>
<tr><td>Chromosome</td><td>start</td><td>end</td><td>strand</td><td>MM</td><td>target_seq</td><td>PAM</td><td>alignment</td><td>distance</td><td>position</td><td>gene name</td><td>gene id</td></tr>
<tr><td>GU305914.1</td><td>27829</td><td>27851</td><td>-</td><td>0</td><td>GGGCTGGGCGACGTCTGCTG</td><td>GGG</td><td>||||||||[||||||||||||]PAM</td><td>NA</td><td>Intergenic</td><td>NA</td><td>NA</td></tr>
<tr><td>T32308</td><td>TGGGCGACGTCTGCTGGGGCAGG</td><td>1000</td><td></td><td></td><td></td><td></td><td></td><td></td><td></td><td></td><td></td></tr>
<tr><td>Oligo fwd</td><td>CACCTGGGCGACGTCTGCTGGGGC</td><td></td><td></td><td></td><td></td><td></td><td></td><td></td><td></td><td></td><td></td></tr>
<tr><td>Oligo rev</td><td>AAACGCCCCAGCAGACGTCGCCCA</td><td></td><td></td><td></td><td></td><td></td><td></td><td></td><td></td><td></td><td></td></tr>
<tr><td>Chromosome</td><td>start</td><td>end</td><td>strand</td><td>MM</td><td>target_seq</td><td>PAM</td><td>alignment</td><td>distance</td><td>position</td><td>gene name</td><td>gene id</td></tr>
<tr><td>GU305914.1</td><td>27825</td><td>27847</td><td>-</td><td>0</td><td>TGGGCGACGTCTGCTGGGGC</td><td>AGG</td><td>||||||||[||||||||||||]PAM</td><td>NA</td><td>Intergenic</td><td>NA</td><td>NA</td></tr>
<tr><td>T32309</td><td>GCGACGTCTGCTGGGGCAGGAGG</td><td>1000</td><td></td><td></td><td></td><td></td><td></td><td></td><td></td><td></td><td></td></tr>
<tr><td>Oligo fwd</td><td>CACCGCGACGTCTGCTGGGGCAGG</td><td></td><td></td><td></td><td></td><td></td><td></td><td></td><td></td><td></td><td></td></tr>
<tr><td>Oligo rev</td><td>AAACCCTGCCCCAGCAGACGTCGC</td><td></td><td></td><td></td><td></td><td></td><td></td><td></td><td></td><td></td><td></td></tr>
<tr><td>Chromosome</td><td>start</td><td>end</td><td>strand</td><td>MM</td><td>target_seq</td><td>PAM</td><td>alignment</td><td>distance</td><td>position</td><td>gene name</td><td>gene id</td></tr>
<tr><td>GU305914.1</td><td>27822</td><td>27844</td><td>-</td><td>0</td><td>GCGACGTCTGCTGGGGCAGG</td><td>AGG</td><td>||||||||[||||||||||||]PAM</td><td>NA</td><td>Intergenic</td><td>NA</td><td>NA</td></tr>
<tr><td>T32310</td><td>GGGGCAGGAGGCCCTCCAGACGG</td><td>1000</td><td></td><td></td><td></td><td></td><td></td><td></td><td></td><td></td><td></td></tr>
<tr><td>Oligo fwd</td><td>CACCGGGGCAGGAGGCCCTCCAGA</td><td></td><td></td><td></td><td></td><td></td><td></td><td></td><td></td><td></td><td></td></tr>
<tr><td>Oligo rev</td><td>AAACTCTGGAGGGCCTCCTGCCCC</td><td></td><td></td><td></td><td></td><td></td><td></td><td></td><td></td><td></td><td></td></tr>
<tr><td>Chromosome</td><td>start</td><td>end</td><td>strand</td><td>MM</td><td>target_seq</td><td>PAM</td><td>alignment</td><td>distance</td><td>position</td><td>gene name</td><td>gene id</td></tr>
<tr><td>GU305914.1</td><td>27810</td><td>27832</td><td>-</td><td>0</td><td>GGGGCAGGAGGCCCTCCAGA</td><td>CGG</td><td>||||||||[||||||||||||]PAM</td><td>NA</td><td>Intergenic</td><td>NA</td><td>NA</td></tr>
<tr><td>T32311</td><td>GCAGGAGGCCCTCCAGACGGCGG</td><td>1000</td><td></td><td></td><td></td><td></td><td></td><td></td><td></td><td></td><td></td></tr>
<tr><td>Oligo fwd</td><td>CACCGCAGGAGGCCCTCCAGACGG</td><td></td><td></td><td></td><td></td><td></td><td></td><td></td><td></td><td></td><td></td></tr>
<tr><td>Oligo rev</td><td>AAACCCGTCTGGAGGGCCTCCTGC</td><td></td><td></td><td></td><td></td><td></td><td></td><td></td><td></td><td></td><td></td></tr>
<tr><td>Chromosome</td><td>start</td><td>end</td><td>strand</td><td>MM</td><td>target_seq</td><td>PAM</td><td>alignment</td><td>distance</td><td>position</td><td>gene name</td><td>gene id</td></tr>
<tr><td>GU305914.1</td><td>27807</td><td>27829</td><td>-</td><td>0</td><td>GCAGGAGGCCCTCCAGACGG</td><td>CGG</td><td>||||||||[||||||||||||]PAM</td><td>NA</td><td>Intergenic</td><td>NA</td><td>NA</td></tr>
<tr><td>T32312</td><td>CGGCGCAGCGTCTCGATGCGCGG</td><td>1000</td><td></td><td></td><td></td><td></td><td></td><td></td><td></td><td></td><td></td></tr>
<tr><td>Oligo fwd</td><td>CACCCGGCGCAGCGTCTCGATGCG</td><td></td><td></td><td></td><td></td><td></td><td></td><td></td><td></td><td></td><td></td></tr>
<tr><td>Oligo rev</td><td>AAACCGCATCGAGACGCTGCGCCG</td><td></td><td></td><td></td><td></td><td></td><td></td><td></td><td></td><td></td><td></td></tr>
<tr><td>Chromosome</td><td>start</td><td>end</td><td>strand</td><td>MM</td><td>target_seq</td><td>PAM</td><td>alignment</td><td>distance</td><td>position</td><td>gene name</td><td>gene id</td></tr>
<tr><td>GU305914.1</td><td>27787</td><td>27809</td><td>-</td><td>0</td><td>CGGCGCAGCGTCTCGATGCG</td><td>CGG</td><td>||||||||[||||||||||||]PAM</td><td>NA</td><td>Intergenic</td><td>NA</td><td>NA</td></tr>
<tr><td>T32313</td><td>CGCAGCGTCTCGATGCGCGGCGG</td><td>1000</td><td></td><td></td><td></td><td></td><td></td><td></td><td></td><td></td><td></td></tr>
<tr><td>Oligo fwd</td><td>CACCCGCAGCGTCTCGATGCGCGG</td><td></td><td></td><td></td><td></td><td></td><td></td><td></td><td></td><td></td><td></td></tr>
<tr><td>Oligo rev</td><td>AAACCCGCGCATCGAGACGCTGCG</td><td></td><td></td><td></td><td></td><td></td><td></td><td></td><td></td><td></td><td></td></tr>
<tr><td>Chromosome</td><td>start</td><td>end</td><td>strand</td><td>MM</td><td>target_seq</td><td>PAM</td><td>alignment</td><td>distance</td><td>position</td><td>gene name</td><td>gene id</td></tr>
<tr><td>GU305914.1</td><td>27784</td><td>27806</td><td>-</td><td>0</td><td>CGCAGCGTCTCGATGCGCGG</td><td>CGG</td><td>||||||||[||||||||||||]PAM</td><td>NA</td><td>Intergenic</td><td>NA</td><td>NA</td></tr>
<tr><td>T32314</td><td>GCAGCGTCTCGATGCGCGGCGGG</td><td>1000</td><td></td><td></td><td></td><td></td><td></td><td></td><td></td><td></td><td></td></tr>
<tr><td>Oligo fwd</td><td>CACCGCAGCGTCTCGATGCGCGGC</td><td></td><td></td><td></td><td></td><td></td><td></td><td></td><td></td><td></td><td></td></tr>
<tr><td>Oligo rev</td><td>AAACGCCGCGCATCGAGACGCTGC</td><td></td><td></td><td></td><td></td><td></td><td></td><td></td><td></td><td></td><td></td></tr>
<tr><td>Chromosome</td><td>start</td><td>end</td><td>strand</td><td>MM</td><td>target_seq</td><td>PAM</td><td>alignment</td><td>distance</td><td>position</td><td>gene name</td><td>gene id</td></tr>
<tr><td>GU305914.1</td><td>27783</td><td>27805</td><td>-</td><td>0</td><td>GCAGCGTCTCGATGCGCGGC</td><td>GGG</td><td>||||||||[||||||||||||]PAM</td><td>NA</td><td>Intergenic</td><td>NA</td><td>NA</td></tr>
<tr><td>GU305914.1</td><td>125840</td><td>125862</td><td>+</td><td>4</td><td>GCGGCGGGTCGATGCGCGCC</td><td>TGG</td><td>||-|||--[||||||||||-|]PAM</td><td>NA</td><td>Intergenic</td><td>NA</td><td>NA</td></tr>
<tr><td>T32315</td><td>CAGCGTCTCGATGCGCGGCGGGG</td><td>1000</td><td></td><td></td><td></td><td></td><td></td><td></td><td></td><td></td><td></td></tr>
<tr><td>Oligo fwd</td><td>CACCCAGCGTCTCGATGCGCGGCG</td><td></td><td></td><td></td><td></td><td></td><td></td><td></td><td></td><td></td><td></td></tr>
<tr><td>Oligo rev</td><td>AAACCGCCGCGCATCGAGACGCTG</td><td></td><td></td><td></td><td></td><td></td><td></td><td></td><td></td><td></td><td></td></tr>
<tr><td>Chromosome</td><td>start</td><td>end</td><td>strand</td><td>MM</td><td>target_seq</td><td>PAM</td><td>alignment</td><td>distance</td><td>position</td><td>gene name</td><td>gene id</td></tr>
<tr><td>GU305914.1</td><td>27782</td><td>27804</td><td>-</td><td>0</td><td>CAGCGTCTCGATGCGCGGCG</td><td>GGG</td><td>||||||||[||||||||||||]PAM</td><td>NA</td><td>Intergenic</td><td>NA</td><td>NA</td></tr>
<tr><td>T32316</td><td>CGGCGGGGTCATCAGATCTTTGG</td><td>1000</td><td></td><td></td><td></td><td></td><td></td><td></td><td></td><td></td><td></td></tr>
<tr><td>Oligo fwd</td><td>CACCCGGCGGGGTCATCAGATCTT</td><td></td><td></td><td></td><td></td><td></td><td></td><td></td><td></td><td></td><td></td></tr>
<tr><td>Oligo rev</td><td>AAACAAGATCTGATGACCCCGCCG</td><td></td><td></td><td></td><td></td><td></td><td></td><td></td><td></td><td></td><td></td></tr>
<tr><td>Chromosome</td><td>start</td><td>end</td><td>strand</td><td>MM</td><td>target_seq</td><td>PAM</td><td>alignment</td><td>distance</td><td>position</td><td>gene name</td><td>gene id</td></tr>
<tr><td>GU305914.1</td><td>27767</td><td>27789</td><td>-</td><td>0</td><td>CGGCGGGGTCATCAGATCTT</td><td>TGG</td><td>||||||||[||||||||||||]PAM</td><td>NA</td><td>Intergenic</td><td>NA</td><td>NA</td></tr>
<tr><td>T32317</td><td>CTTTGGACAGCGCGCGAGCACGG</td><td>1000</td><td></td><td></td><td></td><td></td><td></td><td></td><td></td><td></td><td></td></tr>
<tr><td>Oligo fwd</td><td>CACCCTTTGGACAGCGCGCGAGCA</td><td></td><td></td><td></td><td></td><td></td><td></td><td></td><td></td><td></td><td></td></tr>
<tr><td>Oligo rev</td><td>AAACTGCTCGCGCGCTGTCCAAAG</td><td></td><td></td><td></td><td></td><td></td><td></td><td></td><td></td><td></td><td></td></tr>
<tr><td>Chromosome</td><td>start</td><td>end</td><td>strand</td><td>MM</td><td>target_seq</td><td>PAM</td><td>alignment</td><td>distance</td><td>position</td><td>gene name</td><td>gene id</td></tr>
<tr><td>GU305914.1</td><td>27750</td><td>27772</td><td>-</td><td>0</td><td>CTTTGGACAGCGCGCGAGCA</td><td>CGG</td><td>||||||||[||||||||||||]PAM</td><td>NA</td><td>Intergenic</td><td>NA</td><td>NA</td></tr>
<tr><td>T32318</td><td>TTTGGACAGCGCGCGAGCACGGG</td><td>1000</td><td></td><td></td><td></td><td></td><td></td><td></td><td></td><td></td><td></td></tr>
<tr><td>Oligo fwd</td><td>CACCTTTGGACAGCGCGCGAGCAC</td><td></td><td></td><td></td><td></td><td></td><td></td><td></td><td></td><td></td><td></td></tr>
<tr><td>Oligo rev</td><td>AAACGTGCTCGCGCGCTGTCCAAA</td><td></td><td></td><td></td><td></td><td></td><td></td><td></td><td></td><td></td><td></td></tr>
<tr><td>Chromosome</td><td>start</td><td>end</td><td>strand</td><td>MM</td><td>target_seq</td><td>PAM</td><td>alignment</td><td>distance</td><td>position</td><td>gene name</td><td>gene id</td></tr>
<tr><td>GU305914.1</td><td>27749</td><td>27771</td><td>-</td><td>0</td><td>TTTGGACAGCGCGCGAGCAC</td><td>GGG</td><td>||||||||[||||||||||||]PAM</td><td>NA</td><td>Intergenic</td><td>NA</td><td>NA</td></tr>
<tr><td>T32319</td><td>CTACGCAGTAGTAATTCACCAGG</td><td>1000</td><td></td><td></td><td></td><td></td><td></td><td></td><td></td><td></td><td></td></tr>
<tr><td>Oligo fwd</td><td>CACCCTACGCAGTAGTAATTCACC</td><td></td><td></td><td></td><td></td><td></td><td></td><td></td><td></td><td></td><td></td></tr>
<tr><td>Oligo rev</td><td>AAACGGTGAATTACTACTGCGTAG</td><td></td><td></td><td></td><td></td><td></td><td></td><td></td><td></td><td></td><td></td></tr>
<tr><td>Chromosome</td><td>start</td><td>end</td><td>strand</td><td>MM</td><td>target_seq</td><td>PAM</td><td>alignment</td><td>distance</td><td>position</td><td>gene name</td><td>gene id</td></tr>
<tr><td>GU305914.1</td><td>27711</td><td>27733</td><td>-</td><td>0</td><td>CTACGCAGTAGTAATTCACC</td><td>AGG</td><td>||||||||[||||||||||||]PAM</td><td>NA</td><td>Intergenic</td><td>NA</td><td>NA</td></tr>
<tr><td>T32320</td><td>GCAGTAGTAATTCACCAGGATGG</td><td>1000</td><td></td><td></td><td></td><td></td><td></td><td></td><td></td><td></td><td></td></tr>
<tr><td>Oligo fwd</td><td>CACCGCAGTAGTAATTCACCAGGA</td><td></td><td></td><td></td><td></td><td></td><td></td><td></td><td></td><td></td><td></td></tr>
<tr><td>Oligo rev</td><td>AAACTCCTGGTGAATTACTACTGC</td><td></td><td></td><td></td><td></td><td></td><td></td><td></td><td></td><td></td><td></td></tr>
<tr><td>Chromosome</td><td>start</td><td>end</td><td>strand</td><td>MM</td><td>target_seq</td><td>PAM</td><td>alignment</td><td>distance</td><td>position</td><td>gene name</td><td>gene id</td></tr>
<tr><td>GU305914.1</td><td>27707</td><td>27729</td><td>-</td><td>0</td><td>GCAGTAGTAATTCACCAGGA</td><td>TGG</td><td>||||||||[||||||||||||]PAM</td><td>NA</td><td>Intergenic</td><td>NA</td><td>NA</td></tr>
</table>

T32321 CAGGATGGCTAGCTTGATGACGG 1000
Oligo fwd CACCCAGGATGGCTAGCTTGATGA
Oligo rev AAACTCATCAAGCTAGCCATCCTG
Chromosome start end strand MM target_seq PAM alignment distance position gene name gene id
GU305914.1 27692 27714 - 0 CAGGATGGCTAGCTTGATGA CGG ||||||||[||||||||||||]PAM NA Intergenic NA NA

T32322 AGGATGGCTAGCTTGATGACGGG 1000
Oligo fwd CACCAGGATGGCTAGCTTGATGAC
Oligo rev AAACGTCATCAAGCTAGCCATCCT
Chromosome start end strand MM target_seq PAM alignment distance position gene name gene id
GU305914.1 27691 27713 - 0 AGGATGGCTAGCTTGATGAC GGG ||||||||[||||||||||||]PAM NA Intergenic NA NA

T32323 CGCGCAGCGCAGCGCCTTCTTGG 1000
Oligo fwd CACCCGCGCAGCGCAGCGCCTTCT
Oligo rev AAACAGAAGGCGCTGCGCTGCGCG
Chromosome start end strand MM target_seq PAM alignment distance position gene name gene id
GU305914.1 27656 27678 - 0 CGCGCAGCGCAGCGCCTTCT TGG ||||||||[||||||||||||]PAM NA Intergenic NA NA

T32324 AGCGCAGCGCCTTCTTGGTCAGG 1000
Oligo fwd CACCAGCGCAGCGCCTTCTTGGTC
Oligo rev AAACGACCAAGAAGGCGCTGCGCT
Chromosome start end strand MM target_seq PAM alignment distance position gene name gene id
GU305914.1 27651 27673 - 0 AGCGCAGCGCCTTCTTGGTC AGG ||||||||[||||||||||||]PAM NA Intergenic NA NA

T32325 AGCGCCTTCTTGGTCAGGACCGG 1000
Oligo fwd CACCAGCGCCTTCTTGGTCAGGAC
Oligo rev AAACGTCCTGACCAAGAAGGCGCT
Chromosome start end strand MM target_seq PAM alignment distance position gene name gene id
GU305914.1 27646 27668 - 0 AGCGCCTTCTTGGTCAGGAC CGG ||||||||[||||||||||||]PAM NA Intergenic NA NA

T32326 CCGGCTCAGATAGCGCGTCTAGG 1000
Oligo fwd CACCCCGGCTCAGATAGCGCGTCT
Oligo rev AAACAGACGCGCTATCTGAGCCGG
Chromosome start end strand MM target_seq PAM alignment distance position gene name gene id
GU305914.1 27627 27649 - 0 CCGGCTCAGATAGCGCGTCT AGG ||||||||[||||||||||||]PAM NA Intergenic NA NA

T32327 CGGCTCAGATAGCGCGTCTAGGG 1000
Oligo fwd CACCCGGCTCAGATAGCGCGTCTA
Oligo rev AAACTAGACGCGCTATCTGAGCCG
Chromosome start end strand MM target_seq PAM alignment distance position gene name gene id
GU305914.1 27626 27648 - 0 CGGCTCAGATAGCGCGTCTA GGG ||||||||[||||||||||||]PAM NA Intergenic NA NA

T32328 GATAGCGCGTCTAGGGTGTACGG 1000
Oligo fwd CACCGATAGCGCGTCTAGGGTGTA
Oligo rev AAACTACACCCTAGACGCGCTATC
Chromosome start end strand MM target_seq PAM alignment distance position gene name gene id
GU305914.1 27619 27641 - 0 GATAGCGCGTCTAGGGTGTA CGG ||||||||[||||||||||||]PAM NA Intergenic NA NA

T32329 AGCGCGTCTAGGGTGTACGGCGG 1000
Oligo fwd CACCAGCGCGTCTAGGGTGTACGG
Oligo rev AAACCCGTACACCCTAGACGCGCT
Chromosome start end strand MM target_seq PAM alignment distance position gene name gene id
GU305914.1 27616 27638 - 0 AGCGCGTCTAGGGTGTACGG CGG ||||||||[||||||||||||]PAM NA Intergenic NA NA

T32330 TCTAGGGTGTACGGCGGCATCGG 1000
Oligo fwd CACCTCTAGGGTGTACGGCGGCAT
Oligo rev AAACATGCCGCCGTACACCCTAGA
Chromosome start end strand MM target_seq PAM alignment distance position gene name gene id
GU305914.1 27610 27632 - 0 TCTAGGGTGTACGGCGGCAT CGG ||||||||[||||||||||||]PAM NA Intergenic NA NA
GU305914.1 46077 46099 - 4 TCTAGCATGTACGGGGCCAT CGG |||||--|[||||||-|-|||]PAM NA Intergenic NA NA

T32331 GTACGGCGGCATCGGATAGCAGG 1000
Oligo fwd CACCGTACGGCGGCATCGGATAGC
Oligo rev AAACGCTATCCGATGCCGCCGTAC
Chromosome start end strand MM target_seq PAM alignment distance position gene name gene id
GU305914.1 27602 27624 - 0 GTACGGCGGCATCGGATAGC AGG ||||||||[||||||||||||]PAM NA Intergenic NA NA

T32332 GTGTTTATGATGTCGACCTGCGG 1000
Oligo fwd CACCGTGTTTATGATGTCGACCTG
Oligo rev AAACCAGGTCGACATCATAAACAC
Chromosome start end strand MM target_seq PAM alignment distance position gene name gene id
GU305914.1 27580 27602 - 0 GTGTTTATGATGTCGACCTG CGG ||||||||[||||||||||||]PAM NA Intergenic NA NA

T32333 CGACCTGCGGCGCGTGCGAGTGG 1000
Oligo fwd CACCCGACCTGCGGCGCGTGCGAG
Oligo rev AAACCTCGCACGCGCCGCAGGTCG
Chromosome start end strand MM target_seq PAM alignment distance position gene name gene id
GU305914.1 27567 27589 - 0 CGACCTGCGGCGCGTGCGAG TGG ||||||||[||||||||||||]PAM NA Intergenic NA NA

T32334 CCTGCGGCGCGTGCGAGTGGTGG 1000
Oligo fwd CACCCCTGCGGCGCGTGCGAGTGG
Oligo rev AAACCCACTCGCACGCGCCGCAGG
Chromosome start end strand MM target_seq PAM alignment distance position gene name gene id
GU305914.1 27564 27586 - 0 CCTGCGGCGCGTGCGAGTGG TGG ||||||||[||||||||||||]PAM NA Intergenic NA NA

T32335 GCGGCGCGTGCGAGTGGTGGAGG 1000
Oligo fwd CACCGCGGCGCGTGCGAGTGGTGG
Oligo rev AAACCCACCACTCGCACGCGCCGC
Chromosome start end strand MM target_seq PAM alignment distance position gene name gene id

GU305914.1 27561 27583 - 0 GCGGCGCGTGCGAGTGGTGG AGG ||||||||[||||||||||||]PAM NA Intergenic NA NA

T32336 GAGGCACGCCGCGATGAATCTGG 1000
Oligo fwd CACCGAGGCACGCCGCGATGAATC
Oligo rev AAACGATTCATCGCGGCGTGCCTC
Chromosome start end strand MM target_seq PAM alignment distance position gene name gene id
GU305914.1 27542 27564 - 0 GAGGCACGCCGCGATGAATC TGG ||||||||[||||||||||||]PAM NA Intergenic NA NA

T32337 GTCCTGACCGCTCACTACACCGG 1000
Oligo fwd CACCGTCCTGACCGCTCACTACAC
Oligo rev AAACGTGTAGTGAGCGGTCAGGAC
Chromosome start end strand MM target_seq PAM alignment distance position gene name gene id
GU305914.1 27518 27540 - 0 GTCCTGACCGCTCACTACAC CGG ||||||||[||||||||||||]PAM NA Intergenic NA NA

T32338 TCCTGACCGCTCACTACACCGGG 1000
Oligo fwd CACCTCCTGACCGCTCACTACACC
Oligo rev AAACGGTGTAGTGAGCGGTCAGGA
Chromosome start end strand MM target_seq PAM alignment distance position gene name gene id
GU305914.1 27517 27539 - 0 TCCTGACCGCTCACTACACC GGG ||||||||[||||||||||||]PAM NA Intergenic NA NA

T32339 GACCGCTCACTACACCGGGCAGG 1000
Oligo fwd CACCGACCGCTCACTACACCGGGC
Oligo rev AAACGCCCGGTGTAGTGAGCGGTC
Chromosome start end strand MM target_seq PAM alignment distance position gene name gene id
GU305914.1 27513 27535 - 0 GACCGCTCACTACACCGGGC AGG ||||||||[||||||||||||]PAM NA Intergenic NA NA

T32340 CCGAGCTCGCGCGTGATGCGCGG 1000
Oligo fwd CACCCCGAGCTCGCGCGTGATGCG
Oligo rev AAACCGCATCACGCGCGAGCTCGG
Chromosome start end strand MM target_seq PAM alignment distance position gene name gene id
GU305914.1 27487 27509 - 0 CCGAGCTCGCGCGTGATGCG CGG ||||||||[||||||||||||]PAM NA Intergenic NA NA

T32341 CGCGTGATGCGCGGAGACGACGG 1000
Oligo fwd CACCCGCGTGATGCGCGGAGACGA
Oligo rev AAACTCGTCTCCGCGCATCACGCG
Chromosome start end strand MM target_seq PAM alignment distance position gene name gene id
GU305914.1 27478 27500 - 0 CGCGTGATGCGCGGAGACGA CGG ||||||||[||||||||||||]PAM NA Intergenic NA NA
GU305914.1 66959 66981 + 4 CGAGTGGTACGTGGAGACGA AGG ||-|||-|[-||-||||||||]PAM NA Intergenic NA NA

T32342 GTGATGCGCGGAGACGACGGTGG 1000
Oligo fwd CACCGTGATGCGCGGAGACGACGG
Oligo rev AAACCCGTCGTCTCCGCGCATCAC
Chromosome start end strand MM target_seq PAM alignment distance position gene name gene id
GU305914.1 27475 27497 - 0 GTGATGCGCGGAGACGACGG TGG ||||||||[||||||||||||]PAM NA Intergenic NA NA

T32343 TGATGCGCGGAGACGACGGTGGG 1000
Oligo fwd CACCTGATGCGCGGAGACGACGGT
Oligo rev AAACACCGTCGTCTCCGCGCATCA
Chromosome start end strand MM target_seq PAM alignment distance position gene name gene id
GU305914.1 27474 27496 - 0 TGATGCGCGGAGACGACGGT GGG ||||||||[||||||||||||]PAM NA Intergenic NA NA

T32344 GCGCGGAGACGACGGTGGGACGG 1000
Oligo fwd CACCGCGCGGAGACGACGGTGGGA
Oligo rev AAACTCCCACCGTCGTCTCCGCGC
Chromosome start end strand MM target_seq PAM alignment distance position gene name gene id
GU305914.1 27470 27492 - 0 GCGCGGAGACGACGGTGGGA CGG ||||||||[||||||||||||]PAM NA Intergenic NA NA

T32345 CGCGGAGACGACGGTGGGACGGG 1000
Oligo fwd CACCCGCGGAGACGACGGTGGGAC
Oligo rev AAACGTCCCACCGTCGTCTCCGCG
Chromosome start end strand MM target_seq PAM alignment distance position gene name gene id
GU305914.1 27469 27491 - 0 CGCGGAGACGACGGTGGGAC GGG ||||||||[||||||||||||]PAM NA Intergenic NA NA

T32346 GACGACGGTGGGACGGGCAGAGG 1000
Oligo fwd CACCGACGACGGTGGGACGGGCAG
Oligo rev AAACCTGCCCGTCCCACCGTCGTC
Chromosome start end strand MM target_seq PAM alignment distance position gene name gene id
GU305914.1 27463 27485 - 0 GACGACGGTGGGACGGGCAG AGG ||||||||[||||||||||||]PAM NA Intergenic NA NA

T32347 ACGGTGGGACGGGCAGAGGCTGG 1000
Oligo fwd CACCACGGTGGGACGGGCAGAGGC
Oligo rev AAACGCCTCTGCCCGTCCCACCGT
Chromosome start end strand MM target_seq PAM alignment distance position gene name gene id
GU305914.1 27459 27481 - 0 ACGGTGGGACGGGCAGAGGC TGG ||||||||[||||||||||||]PAM NA Intergenic NA NA

T32348 CGGTGGGACGGGCAGAGGCTGGG 1000
Oligo fwd CACCCGGTGGGACGGGCAGAGGCT
Oligo rev AAACAGCCTCTGCCCGTCCCACCG
Chromosome start end strand MM target_seq PAM alignment distance position gene name gene id
GU305914.1 27458 27480 - 0 CGGTGGGACGGGCAGAGGCT GGG ||||||||[||||||||||||]PAM NA Intergenic NA NA

T32349 GGCAGAGGCTGGGTGCGATGAGG 1000
Oligo fwd CACCGGCAGAGGCTGGGTGCGATG
Oligo rev AAACCATCGCACCCAGCCTCTGCC
Chromosome start end strand MM target_seq PAM alignment distance position gene name gene id
GU305914.1 27448 27470 - 0 GGCAGAGGCTGGGTGCGATG AGG ||||||||[||||||||||||]PAM NA Intergenic NA NA

T32350 GCTGCTGCTGCTGCTGCTGCCGG 1000
Oligo fwd CACCGCTGCTGCTGCTGCTGCTGC

<table>
<tr><td>Oligo rev</td><td>AAACGCAGCAGCAGCAGCAGCAGC</td><td></td><td></td><td></td><td></td><td></td><td></td><td></td><td></td><td></td><td></td></tr>
<tr><td>Chromosome</td><td>start</td><td>end</td><td>strand</td><td>MM</td><td>target_seq</td><td>PAM</td><td>alignment</td><td>distance</td><td>position</td><td>gene name</td><td>gene id</td></tr>
<tr><td>GU305914.1</td><td>27406</td><td>27428</td><td>-</td><td>0</td><td>GCTGCTGCTGCTGCTGCTGC</td><td>CGG</td><td>||||||||[||||||||||||]PAM</td><td>NA</td><td>Intergenic</td><td>NA</td><td>NA</td></tr>
<tr><td>GU305914.1</td><td>27021</td><td>27043</td><td>-</td><td>4</td><td>GAGGCTGTTGTTGCTGCTGC</td><td>TGG</td><td>|--||||-[||-|||||||||]PAM</td><td>NA</td><td>Intergenic</td><td>NA</td><td>NA</td></tr>
<tr><td>GU305914.1</td><td>62684</td><td>62706</td><td>-</td><td>3</td><td>GCTGCCGCGGCTGCTGCCGC</td><td>CGG</td><td>|||||-||[-||||||||-||]PAM</td><td>NA</td><td>Intergenic</td><td>NA</td><td>NA</td></tr>
<tr><td>GU305914.1</td><td>168314</td><td>168336</td><td>+</td><td>4</td><td>GGGGCTGCTGGTGCTGCTCC</td><td>TGG</td><td>|--|||||[||-|||||||-|]PAM</td><td>NA</td><td>Intergenic</td><td>NA</td><td>NA</td></tr>
<tr><td>T32351</td><td>GACGACGATAGACGCGTCGTCGG</td><td>1000</td><td></td><td></td><td></td><td></td><td></td><td></td><td></td><td></td><td></td></tr>
<tr><td>Oligo fwd</td><td>CACCGACGACGATAGACGCGTCGT</td><td></td><td></td><td></td><td></td><td></td><td></td><td></td><td></td><td></td><td></td></tr>
<tr><td>Oligo rev</td><td>AAACACGACGCGTCTATCGTCGTC</td><td></td><td></td><td></td><td></td><td></td><td></td><td></td><td></td><td></td><td></td></tr>
<tr><td>Chromosome</td><td>start</td><td>end</td><td>strand</td><td>MM</td><td>target_seq</td><td>PAM</td><td>alignment</td><td>distance</td><td>position</td><td>gene name</td><td>gene id</td></tr>
<tr><td>GU305914.1</td><td>27323</td><td>27345</td><td>-</td><td>0</td><td>GACGACGATAGACGCGTCGT</td><td>CGG</td><td>||||||||[||||||||||||]PAM</td><td>NA</td><td>Intergenic</td><td>NA</td><td>NA</td></tr>
<tr><td>T32352</td><td>GACGATAGACGCGTCGTCGGCGG</td><td>1000</td><td></td><td></td><td></td><td></td><td></td><td></td><td></td><td></td><td></td></tr>
<tr><td>Oligo fwd</td><td>CACCGACGATAGACGCGTCGTCGG</td><td></td><td></td><td></td><td></td><td></td><td></td><td></td><td></td><td></td><td></td></tr>
<tr><td>Oligo rev</td><td>AAACCCGACGACGCGTCTATCGTC</td><td></td><td></td><td></td><td></td><td></td><td></td><td></td><td></td><td></td><td></td></tr>
<tr><td>Chromosome</td><td>start</td><td>end</td><td>strand</td><td>MM</td><td>target_seq</td><td>PAM</td><td>alignment</td><td>distance</td><td>position</td><td>gene name</td><td>gene id</td></tr>
<tr><td>GU305914.1</td><td>27320</td><td>27342</td><td>-</td><td>0</td><td>GACGATAGACGCGTCGTCGG</td><td>CGG</td><td>||||||||[||||||||||||]PAM</td><td>NA</td><td>Intergenic</td><td>NA</td><td>NA</td></tr>
<tr><td>GU305914.1</td><td>65618</td><td>65640</td><td>+</td><td>4</td><td>GACGCTGGGCGCGTCGCCGG</td><td>CGG</td><td>||||-|-|[-|||||||-|||]PAM</td><td>NA</td><td>Intergenic</td><td>NA</td><td>NA</td></tr>
<tr><td>T32353</td><td>CGCGTCGTCGGCGGCGCCGCCGG</td><td>1000</td><td></td><td></td><td></td><td></td><td></td><td></td><td></td><td></td><td></td></tr>
<tr><td>Oligo fwd</td><td>CACCCGCGTCGTCGGCGGCGCCGC</td><td></td><td></td><td></td><td></td><td></td><td></td><td></td><td></td><td></td><td></td></tr>
<tr><td>Oligo rev</td><td>AAACGCGGCGCCGCCGACGACGCG</td><td></td><td></td><td></td><td></td><td></td><td></td><td></td><td></td><td></td><td></td></tr>
<tr><td>Chromosome</td><td>start</td><td>end</td><td>strand</td><td>MM</td><td>target_seq</td><td>PAM</td><td>alignment</td><td>distance</td><td>position</td><td>gene name</td><td>gene id</td></tr>
<tr><td>GU305914.1</td><td>27311</td><td>27333</td><td>-</td><td>0</td><td>CGCGTCGTCGGCGGCGCCGC</td><td>CGG</td><td>||||||||[||||||||||||]PAM</td><td>NA</td><td>Intergenic</td><td>NA</td><td>NA</td></tr>
<tr><td>GU305914.1</td><td>1212</td><td>1234</td><td>+</td><td>4</td><td>TGTGTCGTCGGCGCCGCTGC</td><td>CGG</td><td>-|-|||||[|||||-|||-||]PAM</td><td>NA</td><td>Intergenic</td><td>NA</td><td>NA</td></tr>
<tr><td>GU305914.1</td><td>163743</td><td>163765</td><td>+</td><td>4</td><td>AGCTCCGTCGGCGGCGCCGG</td><td>TGG</td><td>-||--|||[|||||||||||-]PAM</td><td>NA</td><td>Intergenic</td><td>NA</td><td>NA</td></tr>
<tr><td>T32354</td><td>ATCTTCGATATCGATCACCATGG</td><td>1000</td><td></td><td></td><td></td><td></td><td></td><td></td><td></td><td></td><td></td></tr>
<tr><td>Oligo fwd</td><td>CACCATCTTCGATATCGATCACCA</td><td></td><td></td><td></td><td></td><td></td><td></td><td></td><td></td><td></td><td></td></tr>
<tr><td>Oligo rev</td><td>AAACTGGTGATCGATATCGAAGAT</td><td></td><td></td><td></td><td></td><td></td><td></td><td></td><td></td><td></td><td></td></tr>
<tr><td>Chromosome</td><td>start</td><td>end</td><td>strand</td><td>MM</td><td>target_seq</td><td>PAM</td><td>alignment</td><td>distance</td><td>position</td><td>gene name</td><td>gene id</td></tr>
<tr><td>GU305914.1</td><td>27272</td><td>27294</td><td>-</td><td>0</td><td>ATCTTCGATATCGATCACCA</td><td>TGG</td><td>||||||||[||||||||||||]PAM</td><td>NA</td><td>Intergenic</td><td>NA</td><td>NA</td></tr>
<tr><td>T32355</td><td>TTCGATATCGATCACCATGGCGG</td><td>1000</td><td></td><td></td><td></td><td></td><td></td><td></td><td></td><td></td><td></td></tr>
<tr><td>Oligo fwd</td><td>CACCTTCGATATCGATCACCATGG</td><td></td><td></td><td></td><td></td><td></td><td></td><td></td><td></td><td></td><td></td></tr>
<tr><td>Oligo rev</td><td>AAACCCATGGTGATCGATATCGAA</td><td></td><td></td><td></td><td></td><td></td><td></td><td></td><td></td><td></td><td></td></tr>
<tr><td>Chromosome</td><td>start</td><td>end</td><td>strand</td><td>MM</td><td>target_seq</td><td>PAM</td><td>alignment</td><td>distance</td><td>position</td><td>gene name</td><td>gene id</td></tr>
<tr><td>GU305914.1</td><td>27269</td><td>27291</td><td>-</td><td>0</td><td>TTCGATATCGATCACCATGG</td><td>CGG</td><td>||||||||[||||||||||||]PAM</td><td>NA</td><td>Intergenic</td><td>NA</td><td>NA</td></tr>
<tr><td>T32356</td><td>GATATCGATCACCATGGCGGTGG</td><td>1000</td><td></td><td></td><td></td><td></td><td></td><td></td><td></td><td></td><td></td></tr>
<tr><td>Oligo fwd</td><td>CACCGATATCGATCACCATGGCGG</td><td></td><td></td><td></td><td></td><td></td><td></td><td></td><td></td><td></td><td></td></tr>
<tr><td>Oligo rev</td><td>AAACCCGCCATGGTGATCGATATC</td><td></td><td></td><td></td><td></td><td></td><td></td><td></td><td></td><td></td><td></td></tr>
<tr><td>Chromosome</td><td>start</td><td>end</td><td>strand</td><td>MM</td><td>target_seq</td><td>PAM</td><td>alignment</td><td>distance</td><td>position</td><td>gene name</td><td>gene id</td></tr>
<tr><td>GU305914.1</td><td>27266</td><td>27288</td><td>-</td><td>0</td><td>GATATCGATCACCATGGCGG</td><td>TGG</td><td>||||||||[||||||||||||]PAM</td><td>NA</td><td>Intergenic</td><td>NA</td><td>NA</td></tr>
<tr><td>T32357</td><td>ATCGATCACCATGGCGGTGGCGG</td><td>1000</td><td></td><td></td><td></td><td></td><td></td><td></td><td></td><td></td><td></td></tr>
<tr><td>Oligo fwd</td><td>CACCATCGATCACCATGGCGGTGG</td><td></td><td></td><td></td><td></td><td></td><td></td><td></td><td></td><td></td><td></td></tr>
<tr><td>Oligo rev</td><td>AAACCCACCGCCATGGTGATCGAT</td><td></td><td></td><td></td><td></td><td></td><td></td><td></td><td></td><td></td><td></td></tr>
<tr><td>Chromosome</td><td>start</td><td>end</td><td>strand</td><td>MM</td><td>target_seq</td><td>PAM</td><td>alignment</td><td>distance</td><td>position</td><td>gene name</td><td>gene id</td></tr>
<tr><td>GU305914.1</td><td>27263</td><td>27285</td><td>-</td><td>0</td><td>ATCGATCACCATGGCGGTGG</td><td>CGG</td><td>||||||||[||||||||||||]PAM</td><td>NA</td><td>Intergenic</td><td>NA</td><td>NA</td></tr>
<tr><td>T32358</td><td>ACCATGGCGGTGGCGGATATCGG</td><td>1000</td><td></td><td></td><td></td><td></td><td></td><td></td><td></td><td></td><td></td></tr>
<tr><td>Oligo fwd</td><td>CACCACCATGGCGGTGGCGGATAT</td><td></td><td></td><td></td><td></td><td></td><td></td><td></td><td></td><td></td><td></td></tr>
<tr><td>Oligo rev</td><td>AAACATATCCGCCACCGCCATGGT</td><td></td><td></td><td></td><td></td><td></td><td></td><td></td><td></td><td></td><td></td></tr>
<tr><td>Chromosome</td><td>start</td><td>end</td><td>strand</td><td>MM</td><td>target_seq</td><td>PAM</td><td>alignment</td><td>distance</td><td>position</td><td>gene name</td><td>gene id</td></tr>
<tr><td>GU305914.1</td><td>27256</td><td>27278</td><td>-</td><td>0</td><td>ACCATGGCGGTGGCGGATAT</td><td>CGG</td><td>||||||||[||||||||||||]PAM</td><td>NA</td><td>Intergenic</td><td>NA</td><td>NA</td></tr>
<tr><td>T32359</td><td>CCATGGCGGTGGCGGATATCGGG</td><td>1000</td><td></td><td></td><td></td><td></td><td></td><td></td><td></td><td></td><td></td></tr>
<tr><td>Oligo fwd</td><td>CACCCCATGGCGGTGGCGGATATC</td><td></td><td></td><td></td><td></td><td></td><td></td><td></td><td></td><td></td><td></td></tr>
<tr><td>Oligo rev</td><td>AAACGATATCCGCCACCGCCATGG</td><td></td><td></td><td></td><td></td><td></td><td></td><td></td><td></td><td></td><td></td></tr>
<tr><td>Chromosome</td><td>start</td><td>end</td><td>strand</td><td>MM</td><td>target_seq</td><td>PAM</td><td>alignment</td><td>distance</td><td>position</td><td>gene name</td><td>gene id</td></tr>
<tr><td>GU305914.1</td><td>27255</td><td>27277</td><td>-</td><td>0</td><td>CCATGGCGGTGGCGGATATC</td><td>GGG</td><td>||||||||[||||||||||||]PAM</td><td>NA</td><td>Intergenic</td><td>NA</td><td>NA</td></tr>
<tr><td>T32360</td><td>CATGGCGGTGGCGGATATCGGGG</td><td>1000</td><td></td><td></td><td></td><td></td><td></td><td></td><td></td><td></td><td></td></tr>
<tr><td>Oligo fwd</td><td>CACCCATGGCGGTGGCGGATATCG</td><td></td><td></td><td></td><td></td><td></td><td></td><td></td><td></td><td></td><td></td></tr>
<tr><td>Oligo rev</td><td>AAACCGATATCCGCCACCGCCATG</td><td></td><td></td><td></td><td></td><td></td><td></td><td></td><td></td><td></td><td></td></tr>
<tr><td>Chromosome</td><td>start</td><td>end</td><td>strand</td><td>MM</td><td>target_seq</td><td>PAM</td><td>alignment</td><td>distance</td><td>position</td><td>gene name</td><td>gene id</td></tr>
<tr><td>GU305914.1</td><td>27254</td><td>27276</td><td>-</td><td>0</td><td>CATGGCGGTGGCGGATATCG</td><td>GGG</td><td>||||||||[||||||||||||]PAM</td><td>NA</td><td>Intergenic</td><td>NA</td><td>NA</td></tr>
<tr><td>T32361</td><td>TCTCCGTGTGATCGACACGACGG</td><td>1000</td><td></td><td></td><td></td><td></td><td></td><td></td><td></td><td></td><td></td></tr>
<tr><td>Oligo fwd</td><td>CACCTCTCCGTGTGATCGACACGA</td><td></td><td></td><td></td><td></td><td></td><td></td><td></td><td></td><td></td><td></td></tr>
<tr><td>Oligo rev</td><td>AAACTCGTGTCGATCACACGGAGA</td><td></td><td></td><td></td><td></td><td></td><td></td><td></td><td></td><td></td><td></td></tr>
<tr><td>Chromosome</td><td>start</td><td>end</td><td>strand</td><td>MM</td><td>target_seq</td><td>PAM</td><td>alignment</td><td>distance</td><td>position</td><td>gene name</td><td>gene id</td></tr>
<tr><td>GU305914.1</td><td>27213</td><td>27235</td><td>-</td><td>0</td><td>TCTCCGTGTGATCGACACGA</td><td>CGG</td><td>||||||||[||||||||||||]PAM</td><td>NA</td><td>Intergenic</td><td>NA</td><td>NA</td></tr>
<tr><td>T32362</td><td>CGTGTGATCGACACGACGGCAGG</td><td>1000</td><td></td><td></td><td></td><td></td><td></td><td></td><td></td><td></td><td></td></tr>
<tr><td>Oligo fwd</td><td>CACCCGTGTGATCGACACGACGGC</td><td></td><td></td><td></td><td></td><td></td><td></td><td></td><td></td><td></td><td></td></tr>
<tr><td>Oligo rev</td><td>AAACGCCGTCGTGTCGATCACACG</td><td></td><td></td><td></td><td></td><td></td><td></td><td></td><td></td><td></td><td></td></tr>
<tr><td>Chromosome</td><td>start</td><td>end</td><td>strand</td><td>MM</td><td>target_seq</td><td>PAM</td><td>alignment</td><td>distance</td><td>position</td><td>gene name</td><td>gene id</td></tr>
<tr><td>GU305914.1</td><td>27209</td><td>27231</td><td>-</td><td>0</td><td>CGTGTGATCGACACGACGGC</td><td>AGG</td><td>||||||||[||||||||||||]PAM</td><td>NA</td><td>Intergenic</td><td>NA</td><td>NA</td></tr>
<tr><td>T32363</td><td>GTGTGATCGACACGACGGCAGGG</td><td>1000</td><td></td><td></td><td></td><td></td><td></td><td></td><td></td><td></td><td></td></tr>
<tr><td>Oligo fwd</td><td>CACCGTGTGATCGACACGACGGCA</td><td></td><td></td><td></td><td></td><td></td><td></td><td></td><td></td><td></td><td></td></tr>
<tr><td>Oligo rev</td><td>AAACTGCCGTCGTGTCGATCACAC</td><td></td><td></td><td></td><td></td><td></td><td></td><td></td><td></td><td></td><td></td></tr>
<tr><td>Chromosome</td><td>start</td><td>end</td><td>strand</td><td>MM</td><td>target_seq</td><td>PAM</td><td>alignment</td><td>distance</td><td>position</td><td>gene name</td><td>gene id</td></tr>
<tr><td>GU305914.1</td><td>27208</td><td>27230</td><td>-</td><td>0</td><td>GTGTGATCGACACGACGGCA</td><td>GGG</td><td>||||||||[||||||||||||]PAM</td><td>NA</td><td>Intergenic</td><td>NA</td><td>NA</td></tr>
<tr><td>T32364</td><td>GACGGCAGGGCGTTTTGCCGCGG</td><td>1000</td><td></td><td></td><td></td><td></td><td></td><td></td><td></td><td></td><td></td></tr>
</table>

Oligo fwd CACCGACGGCAGGGCGTTTTGCCG
Oligo rev AAACCGGCAAAACGCCCTGCCGTC
Chromosome start end strand MM target_seq PAM alignment distance position gene name gene id
GU305914.1 27195 27217 - 0 GACGGCAGGGCGTTTTGCCG CGG ||||||||[||||||||||||]PAM NA Intergenic NA NA

T32365 TTGCCGCGGCGTGTGTTGCTCGG 1000
Oligo fwd CACCTTGCCGCGGCGTGTGTTGCT
Oligo rev AAACAGCAACACACGCCGCGGCAA
Chromosome start end strand MM target_seq PAM alignment distance position gene name gene id
GU305914.1 27181 27203 - 0 TTGCCGCGGCGTGTGTTGCT CGG ||||||||[||||||||||||]PAM NA Intergenic NA NA

T32366 CGGCGTGTGTTGCTCGGACCCGG 1000
Oligo fwd CACCCGGCGTGTGTTGCTCGGACC
Oligo rev AAACGGTCCGAGCAACACACGCCG
Chromosome start end strand MM target_seq PAM alignment distance position gene name gene id
GU305914.1 27175 27197 - 0 CGGCGTGTGTTGCTCGGACC CGG ||||||||[||||||||||||]PAM NA Intergenic NA NA

T32367 ATCGTCGTCATCGTCGTCGTTGG 1000
Oligo fwd CACCATCGTCGTCATCGTCGTCGT
Oligo rev AAACACGACGACGATGACGACGAT
Chromosome start end strand MM target_seq PAM alignment distance position gene name gene id
GU305914.1 27140 27162 - 0 ATCGTCGTCATCGTCGTCGT TGG ||||||||[||||||||||||]PAM NA Intergenic NA NA
GU305914.1 112237 112259 - 4 CTTGTCGTCGTCGTCCTCGT CGG -|-|||||[|-|||||-||||]PAM NA Intergenic NA NA
GU305914.1 73071 73093 - 3 ATCGTCGCCCTCGTCCTCGT GGG |||||||-[|-|||||-||||]PAM NA Intergenic NA NA

T32368 TCATCGTCGTCGTTGGCGTTCGG 1000
Oligo fwd CACCTCATCGTCGTCGTTGGCGTT
Oligo rev AAACAACGCCAACGACGACGATGA
Chromosome start end strand MM target_seq PAM alignment distance position gene name gene id
GU305914.1 27133 27155 - 0 TCATCGTCGTCGTTGGCGTT CGG ||||||||[||||||||||||]PAM NA Intergenic NA NA

T32369 CATCGTCGTCGTTGGCGTTCGGG 1000
Oligo fwd CACCCATCGTCGTCGTTGGCGTTC
Oligo rev AAACGAACGCCAACGACGACGATG
Chromosome start end strand MM target_seq PAM alignment distance position gene name gene id
GU305914.1 27132 27154 - 0 CATCGTCGTCGTTGGCGTTC GGG ||||||||[||||||||||||]PAM NA Intergenic NA NA

T32370 ATCGTCGTCGTTGGCGTTCGGGG 1000
Oligo fwd CACCATCGTCGTCGTTGGCGTTCG
Oligo rev AAACCGAACGCCAACGACGACGAT
Chromosome start end strand MM target_seq PAM alignment distance position gene name gene id
GU305914.1 27131 27153 - 0 ATCGTCGTCGTTGGCGTTCG GGG ||||||||[||||||||||||]PAM NA Intergenic NA NA

T32371 GTCGTTGGCGTTCGGGGTCCTGG 1000
Oligo fwd CACCGTCGTTGGCGTTCGGGGTCC
Oligo rev AAACGGACCCCGAACGCCAACGAC
Chromosome start end strand MM target_seq PAM alignment distance position gene name gene id
GU305914.1 27125 27147 - 0 GTCGTTGGCGTTCGGGGTCC TGG ||||||||[||||||||||||]PAM NA Intergenic NA NA

T32372 TCGTTGGCGTTCGGGGTCCTGGG 1000
Oligo fwd CACCTCGTTGGCGTTCGGGGTCCT
Oligo rev AAACAGGACCCCGAACGCCAACGA
Chromosome start end strand MM target_seq PAM alignment distance position gene name gene id
GU305914.1 27124 27146 - 0 TCGTTGGCGTTCGGGGTCCT GGG ||||||||[||||||||||||]PAM NA Intergenic NA NA

T32373 CGTTGGCGTTCGGGGTCCTGGGG 1000
Oligo fwd CACCCGTTGGCGTTCGGGGTCCTG
Oligo rev AAACCAGGACCCCGAACGCCAACG
Chromosome start end strand MM target_seq PAM alignment distance position gene name gene id
GU305914.1 27123 27145 - 0 CGTTGGCGTTCGGGGTCCTG GGG ||||||||[||||||||||||]PAM NA Intergenic NA NA

T32374 GCGTTCGGGGTCCTGGGGAGAGG 1000
Oligo fwd CACCGCGTTCGGGGTCCTGGGGAG
Oligo rev AAACCTCCCCAGGACCCCGAACGC
Chromosome start end strand MM target_seq PAM alignment distance position gene name gene id
GU305914.1 27118 27140 - 0 GCGTTCGGGGTCCTGGGGAG AGG ||||||||[||||||||||||]PAM NA Intergenic NA NA

T32375 ACGCGACACAGTCGCCTGTACGG 1000
Oligo fwd CACCACGCGACACAGTCGCCTGTA
Oligo rev AAACTACAGGCGACTGTGTCGCGT
Chromosome start end strand MM target_seq PAM alignment distance position gene name gene id
GU305914.1 27095 27117 - 0 ACGCGACACAGTCGCCTGTA CGG ||||||||[||||||||||||]PAM NA Intergenic NA NA

T32376 CGCGACACAGTCGCCTGTACGGG 1000
Oligo fwd CACCCGCGACACAGTCGCCTGTAC
Oligo rev AAACGTACAGGCGACTGTGTCGCG
Chromosome start end strand MM target_seq PAM alignment distance position gene name gene id
GU305914.1 27094 27116 - 0 CGCGACACAGTCGCCTGTAC GGG ||||||||[||||||||||||]PAM NA Intergenic NA NA

T32377 GACACAGTCGCCTGTACGGGCGG 1000
Oligo fwd CACCGACACAGTCGCCTGTACGGG
Oligo rev AAACCCCGTACAGGCGACTGTGTC
Chromosome start end strand MM target_seq PAM alignment distance position gene name gene id
GU305914.1 27091 27113 - 0 GACACAGTCGCCTGTACGGG CGG ||||||||[||||||||||||]PAM NA Intergenic NA NA

T32378 ACAGTCGCCTGTACGGGCGGAGG 1000
Oligo fwd CACCACAGTCGCCTGTACGGGCGG
Oligo rev AAACCCGCCCGTACAGGCGACTGT
Chromosome start end strand MM target_seq PAM alignment distance position gene name gene id

GU305914.1 27088 27110 - 0 ACAGTCGCCTGTACGGGCGG AGG ||||||||[||||||||||||]PAM NA Intergenic NA NA

T32379 GTCGCCTGTACGGGCGGAGGTGG 1000
Oligo fwd CACCGTCGCCTGTACGGGCGGAGG
Oligo rev AAACCCTCCGCCCGTACAGGCGAC
Chromosome start end strand MM target_seq PAM alignment distance position gene name gene id
GU305914.1 27085 27107 - 0 GTCGCCTGTACGGGCGGAGG TGG ||||||||[||||||||||||]PAM NA Intergenic NA NA

T32380 TCGCCTGTACGGGCGGAGGTGGG 1000
Oligo fwd CACCTCGCCTGTACGGGCGGAGGT
Oligo rev AAACACCTCCGCCCGTACAGGCGA
Chromosome start end strand MM target_seq PAM alignment distance position gene name gene id
GU305914.1 27084 27106 - 0 TCGCCTGTACGGGCGGAGGT GGG ||||||||[||||||||||||]PAM NA Intergenic NA NA

T32381 CGCCTGTACGGGCGGAGGTGGGG 1000
Oligo fwd CACCCGCCTGTACGGGCGGAGGTG
Oligo rev AAACCACCTCCGCCCGTACAGGCG
Chromosome start end strand MM target_seq PAM alignment distance position gene name gene id
GU305914.1 27083 27105 - 0 CGCCTGTACGGGCGGAGGTG GGG ||||||||[||||||||||||]PAM NA Intergenic NA NA

T32382 GCCTGTACGGGCGGAGGTGGGGG 1000
Oligo fwd CACCGCCTGTACGGGCGGAGGTGG
Oligo rev AAACCCACCTCCGCCCGTACAGGC
Chromosome start end strand MM target_seq PAM alignment distance position gene name gene id
GU305914.1 27082 27104 - 0 GCCTGTACGGGCGGAGGTGG GGG ||||||||[||||||||||||]PAM NA Intergenic NA NA

T32383 TGTACGGGCGGAGGTGGGGGAGG 1000
Oligo fwd CACCTGTACGGGCGGAGGTGGGGG
Oligo rev AAACCCCCCACCTCCGCCCGTACA
Chromosome start end strand MM target_seq PAM alignment distance position gene name gene id
GU305914.1 27079 27101 - 0 TGTACGGGCGGAGGTGGGGG AGG ||||||||[||||||||||||]PAM NA Intergenic NA NA

T32384 CGGAGGTGGGGGAGGATAATCGG 1000
Oligo fwd CACCCGGAGGTGGGGGAGGATAAT
Oligo rev AAACATTATCCTCCCCCACCTCCG
Chromosome start end strand MM target_seq PAM alignment distance position gene name gene id
GU305914.1 27071 27093 - 0 CGGAGGTGGGGGAGGATAAT CGG ||||||||[||||||||||||]PAM NA Intergenic NA NA

T32385 GGAGGTGGGGGAGGATAATCGGG 1000
Oligo fwd CACCGGAGGTGGGGGAGGATAATC
Oligo rev AAACGATTATCCTCCCCCACCTCC
Chromosome start end strand MM target_seq PAM alignment distance position gene name gene id
GU305914.1 27070 27092 - 0 GGAGGTGGGGGAGGATAATC GGG ||||||||[||||||||||||]PAM NA Intergenic NA NA

T32386 GGTGGGGGAGGATAATCGGGTGG 1000
Oligo fwd CACCGGTGGGGGAGGATAATCGGG
Oligo rev AAACCCCGATTATCCTCCCCCACC
Chromosome start end strand MM target_seq PAM alignment distance position gene name gene id
GU305914.1 27067 27089 - 0 GGTGGGGGAGGATAATCGGG TGG ||||||||[||||||||||||]PAM NA Intergenic NA NA

T32387 GGGGGAGGATAATCGGGTGGAGG 1000
Oligo fwd CACCGGGGGAGGATAATCGGGTGG
Oligo rev AAACCCACCCGATTATCCTCCCCC
Chromosome start end strand MM target_seq PAM alignment distance position gene name gene id
GU305914.1 27064 27086 - 0 GGGGGAGGATAATCGGGTGG AGG ||||||||[||||||||||||]PAM NA Intergenic NA NA

T32388 GGGGAGGATAATCGGGTGGAGGG 1000
Oligo fwd CACCGGGGAGGATAATCGGGTGGA
Oligo rev AAACTCCACCCGATTATCCTCCCC
Chromosome start end strand MM target_seq PAM alignment distance position gene name gene id
GU305914.1 27063 27085 - 0 GGGGAGGATAATCGGGTGGA GGG ||||||||[||||||||||||]PAM NA Intergenic NA NA

T32389 GAGGATAATCGGGTGGAGGGTGG 1000
Oligo fwd CACCGAGGATAATCGGGTGGAGGG
Oligo rev AAACCCCTCCACCCGATTATCCTC
Chromosome start end strand MM target_seq PAM alignment distance position gene name gene id
GU305914.1 27060 27082 - 0 GAGGATAATCGGGTGGAGGG TGG ||||||||[||||||||||||]PAM NA Intergenic NA NA

T32390 CGGGTGGAGGGTGGTAAGATTGG 1000
Oligo fwd CACCCGGGTGGAGGGTGGTAAGAT
Oligo rev AAACATCTTACCACCCTCCACCCG
Chromosome start end strand MM target_seq PAM alignment distance position gene name gene id
GU305914.1 27051 27073 - 0 CGGGTGGAGGGTGGTAAGAT TGG ||||||||[||||||||||||]PAM NA Intergenic NA NA

T32391 GTGGAGGGTGGTAAGATTGGCGG 1000
Oligo fwd CACCGTGGAGGGTGGTAAGATTGG
Oligo rev AAACCCAATCTTACCACCCTCCAC
Chromosome start end strand MM target_seq PAM alignment distance position gene name gene id
GU305914.1 27048 27070 - 0 GTGGAGGGTGGTAAGATTGG CGG ||||||||[||||||||||||]PAM NA Intergenic NA NA

T32392 GGGTGGTAAGATTGGCGGCGTGG 1000
Oligo fwd CACCGGGTGGTAAGATTGGCGGCG
Oligo rev AAACCGCCGCCAATCTTACCACCC
Chromosome start end strand MM target_seq PAM alignment distance position gene name gene id
GU305914.1 27043 27065 - 0 GGGTGGTAAGATTGGCGGCG TGG ||||||||[||||||||||||]PAM NA Intergenic NA NA

T32393 TGGTAAGATTGGCGGCGTGGAGG 1000
Oligo fwd CACCTGGTAAGATTGGCGGCGTGG
Oligo rev AAACCCACGCCGCCAATCTTACCA

<table>
<tr><td>Chromosome</td><td>start</td><td>end</td><td>strand</td><td>MM</td><td>target_seq</td><td>PAM</td><td>alignment</td><td>distance</td><td>position</td><td>gene name</td><td>gene id</td></tr>
<tr><td>GU305914.1</td><td>27040</td><td>27062</td><td>-</td><td>0</td><td>TGGTAAGATTGGCGGCGTGG</td><td>AGG</td><td>||||||||[||||||||||||]PAM</td><td>NA</td><td>Intergenic</td><td>NA</td><td>NA</td></tr>
<tr><td>T32394</td><td>GAGGCTGTTGTTGCTGCTGCTGG</td><td>1000</td><td></td><td></td><td></td><td></td><td></td><td></td><td></td><td></td><td></td></tr>
<tr><td>Oligo fwd</td><td>CACCGAGGCTGTTGTTGCTGCTGC</td><td></td><td></td><td></td><td></td><td></td><td></td><td></td><td></td><td></td><td></td></tr>
<tr><td>Oligo rev</td><td>AAACGCAGCAGCAACAACAGCCTC</td><td></td><td></td><td></td><td></td><td></td><td></td><td></td><td></td><td></td><td></td></tr>
<tr><td>Chromosome</td><td>start</td><td>end</td><td>strand</td><td>MM</td><td>target_seq</td><td>PAM</td><td>alignment</td><td>distance</td><td>position</td><td>gene name</td><td>gene id</td></tr>
<tr><td>GU305914.1</td><td>27021</td><td>27043</td><td>-</td><td>0</td><td>GAGGCTGTTGTTGCTGCTGC</td><td>TGG</td><td>||||||||[||||||||||||]PAM</td><td>NA</td><td>Intergenic</td><td>NA</td><td>NA</td></tr>
<tr><td>GU305914.1</td><td>27406</td><td>27428</td><td>-</td><td>4</td><td>GCTGCTGCTGCTGCTGCTGC</td><td>CGG</td><td>|--||||-[||-|||||||||]PAM</td><td>NA</td><td>Intergenic</td><td>NA</td><td>NA</td></tr>
<tr><td>GU305914.1</td><td>168314</td><td>168336</td><td>+</td><td>4</td><td>GGGGCTGCTGGTGCTGCTCC</td><td>TGG</td><td>|-|||||-[||-|||||||-|]PAM</td><td>NA</td><td>Intergenic</td><td>NA</td><td>NA</td></tr>
<tr><td>T32395</td><td>GCTGCTGCTGGCGTTGTTTTTGG</td><td>1000</td><td></td><td></td><td></td><td></td><td></td><td></td><td></td><td></td><td></td></tr>
<tr><td>Oligo fwd</td><td>CACCGCTGCTGCTGGCGTTGTTTT</td><td></td><td></td><td></td><td></td><td></td><td></td><td></td><td></td><td></td><td></td></tr>
<tr><td>Oligo rev</td><td>AAACAAAACAACGCCAGCAGCAGC</td><td></td><td></td><td></td><td></td><td></td><td></td><td></td><td></td><td></td><td></td></tr>
<tr><td>Chromosome</td><td>start</td><td>end</td><td>strand</td><td>MM</td><td>target_seq</td><td>PAM</td><td>alignment</td><td>distance</td><td>position</td><td>gene name</td><td>gene id</td></tr>
<tr><td>GU305914.1</td><td>27009</td><td>27031</td><td>-</td><td>0</td><td>GCTGCTGCTGGCGTTGTTTT</td><td>TGG</td><td>||||||||[||||||||||||]PAM</td><td>NA</td><td>Intergenic</td><td>NA</td><td>NA</td></tr>
<tr><td>T32396</td><td>GCGTTGTTTTTGGCGAGCTCCGG</td><td>1000</td><td></td><td></td><td></td><td></td><td></td><td></td><td></td><td></td><td></td></tr>
<tr><td>Oligo fwd</td><td>CACCGCGTTGTTTTTGGCGAGCTC</td><td></td><td></td><td></td><td></td><td></td><td></td><td></td><td></td><td></td><td></td></tr>
<tr><td>Oligo rev</td><td>AAACGAGCTCGCCAAAAACAACGC</td><td></td><td></td><td></td><td></td><td></td><td></td><td></td><td></td><td></td><td></td></tr>
<tr><td>Chromosome</td><td>start</td><td>end</td><td>strand</td><td>MM</td><td>target_seq</td><td>PAM</td><td>alignment</td><td>distance</td><td>position</td><td>gene name</td><td>gene id</td></tr>
<tr><td>GU305914.1</td><td>26999</td><td>27021</td><td>-</td><td>0</td><td>GCGTTGTTTTTGGCGAGCTC</td><td>CGG</td><td>||||||||[||||||||||||]PAM</td><td>NA</td><td>Intergenic</td><td>NA</td><td>NA</td></tr>
<tr><td>T32397</td><td>GCGAGCTCCGGAGCCCGCCGCGG</td><td>1000</td><td></td><td></td><td></td><td></td><td></td><td></td><td></td><td></td><td></td></tr>
<tr><td>Oligo fwd</td><td>CACCGCGAGCTCCGGAGCCCGCCG</td><td></td><td></td><td></td><td></td><td></td><td></td><td></td><td></td><td></td><td></td></tr>
<tr><td>Oligo rev</td><td>AAACCGGCGGGCTCCGGAGCTCGC</td><td></td><td></td><td></td><td></td><td></td><td></td><td></td><td></td><td></td><td></td></tr>
<tr><td>Chromosome</td><td>start</td><td>end</td><td>strand</td><td>MM</td><td>target_seq</td><td>PAM</td><td>alignment</td><td>distance</td><td>position</td><td>gene name</td><td>gene id</td></tr>
<tr><td>GU305914.1</td><td>26987</td><td>27009</td><td>-</td><td>0</td><td>GCGAGCTCCGGAGCCCGCCG</td><td>CGG</td><td>||||||||[||||||||||||]PAM</td><td>NA</td><td>Intergenic</td><td>NA</td><td>NA</td></tr>
<tr><td>T32398</td><td>CCGGAGCCCGCCGCGGCCGCCGG</td><td>1000</td><td></td><td></td><td></td><td></td><td></td><td></td><td></td><td></td><td></td></tr>
<tr><td>Oligo fwd</td><td>CACCCCGGAGCCCGCCGCGGCCGC</td><td></td><td></td><td></td><td></td><td></td><td></td><td></td><td></td><td></td><td></td></tr>
<tr><td>Oligo rev</td><td>AAACGCGGCCGCGGCGGGCTCCGG</td><td></td><td></td><td></td><td></td><td></td><td></td><td></td><td></td><td></td><td></td></tr>
<tr><td>Chromosome</td><td>start</td><td>end</td><td>strand</td><td>MM</td><td>target_seq</td><td>PAM</td><td>alignment</td><td>distance</td><td>position</td><td>gene name</td><td>gene id</td></tr>
<tr><td>GU305914.1</td><td>26980</td><td>27002</td><td>-</td><td>0</td><td>CCGGAGCCCGCCGCGGCCGC</td><td>CGG</td><td>||||||||[||||||||||||]PAM</td><td>NA</td><td>Intergenic</td><td>NA</td><td>NA</td></tr>
<tr><td>T32399</td><td>GCCCGCCGCGGCCGCCGGAGCGG</td><td>1000</td><td></td><td></td><td></td><td></td><td></td><td></td><td></td><td></td><td></td></tr>
<tr><td>Oligo fwd</td><td>CACCGCCCGCCGCGGCCGCCGGAG</td><td></td><td></td><td></td><td></td><td></td><td></td><td></td><td></td><td></td><td></td></tr>
<tr><td>Oligo rev</td><td>AAACCTCCGGCGGCCGCGGCGGGC</td><td></td><td></td><td></td><td></td><td></td><td></td><td></td><td></td><td></td><td></td></tr>
<tr><td>Chromosome</td><td>start</td><td>end</td><td>strand</td><td>MM</td><td>target_seq</td><td>PAM</td><td>alignment</td><td>distance</td><td>position</td><td>gene name</td><td>gene id</td></tr>
<tr><td>GU305914.1</td><td>26975</td><td>26997</td><td>-</td><td>0</td><td>GCCCGCCGCGGCCGCCGGAG</td><td>CGG</td><td>||||||||[||||||||||||]PAM</td><td>NA</td><td>Intergenic</td><td>NA</td><td>NA</td></tr>
<tr><td>GU305914.1</td><td>110387</td><td>110409</td><td>-</td><td>3</td><td>GCGCGTCGCGGCCGCCGCAG</td><td>AGG</td><td>||-||-||[|||||||||-||]PAM</td><td>NA</td><td>Intergenic</td><td>NA</td><td>NA</td></tr>
<tr><td>T32400</td><td>GCGGTCTCTTGAGCCTTCTTCGG</td><td>1000</td><td></td><td></td><td></td><td></td><td></td><td></td><td></td><td></td><td></td></tr>
<tr><td>Oligo fwd</td><td>CACCGCGGTCTCTTGAGCCTTCTT</td><td></td><td></td><td></td><td></td><td></td><td></td><td></td><td></td><td></td><td></td></tr>
<tr><td>Oligo rev</td><td>AAACAAGAAGGCTCAAGAGACCGC</td><td></td><td></td><td></td><td></td><td></td><td></td><td></td><td></td><td></td><td></td></tr>
<tr><td>Chromosome</td><td>start</td><td>end</td><td>strand</td><td>MM</td><td>target_seq</td><td>PAM</td><td>alignment</td><td>distance</td><td>position</td><td>gene name</td><td>gene id</td></tr>
<tr><td>GU305914.1</td><td>26956</td><td>26978</td><td>-</td><td>0</td><td>GCGGTCTCTTGAGCCTTCTT</td><td>CGG</td><td>||||||||[||||||||||||]PAM</td><td>NA</td><td>Intergenic</td><td>NA</td><td>NA</td></tr>
<tr><td>T32401</td><td>CGGTCTCTTGAGCCTTCTTCGGG</td><td>1000</td><td></td><td></td><td></td><td></td><td></td><td></td><td></td><td></td><td></td></tr>
<tr><td>Oligo fwd</td><td>CACCCGGTCTCTTGAGCCTTCTTC</td><td></td><td></td><td></td><td></td><td></td><td></td><td></td><td></td><td></td><td></td></tr>
<tr><td>Oligo rev</td><td>AAACGAAGAAGGCTCAAGAGACCG</td><td></td><td></td><td></td><td></td><td></td><td></td><td></td><td></td><td></td><td></td></tr>
<tr><td>Chromosome</td><td>start</td><td>end</td><td>strand</td><td>MM</td><td>target_seq</td><td>PAM</td><td>alignment</td><td>distance</td><td>position</td><td>gene name</td><td>gene id</td></tr>
<tr><td>GU305914.1</td><td>26955</td><td>26977</td><td>-</td><td>0</td><td>CGGTCTCTTGAGCCTTCTTC</td><td>GGG</td><td>||||||||[||||||||||||]PAM</td><td>NA</td><td>Intergenic</td><td>NA</td><td>NA</td></tr>
<tr><td>T32402</td><td>GGTCTCTTGAGCCTTCTTCGGGG</td><td>1000</td><td></td><td></td><td></td><td></td><td></td><td></td><td></td><td></td><td></td></tr>
<tr><td>Oligo fwd</td><td>CACCGGTCTCTTGAGCCTTCTTCG</td><td></td><td></td><td></td><td></td><td></td><td></td><td></td><td></td><td></td><td></td></tr>
<tr><td>Oligo rev</td><td>AAACCGAAGAAGGCTCAAGAGACC</td><td></td><td></td><td></td><td></td><td></td><td></td><td></td><td></td><td></td><td></td></tr>
<tr><td>Chromosome</td><td>start</td><td>end</td><td>strand</td><td>MM</td><td>target_seq</td><td>PAM</td><td>alignment</td><td>distance</td><td>position</td><td>gene name</td><td>gene id</td></tr>
<tr><td>GU305914.1</td><td>26954</td><td>26976</td><td>-</td><td>0</td><td>GGTCTCTTGAGCCTTCTTCG</td><td>GGG</td><td>||||||||[||||||||||||]PAM</td><td>NA</td><td>Intergenic</td><td>NA</td><td>NA</td></tr>
<tr><td>T32403</td><td>CTTGAGCCTTCTTCGGGGTCCGG</td><td>1000</td><td></td><td></td><td></td><td></td><td></td><td></td><td></td><td></td><td></td></tr>
<tr><td>Oligo fwd</td><td>CACCCTTGAGCCTTCTTCGGGGTC</td><td></td><td></td><td></td><td></td><td></td><td></td><td></td><td></td><td></td><td></td></tr>
<tr><td>Oligo rev</td><td>AAACGACCCCGAAGAAGGCTCAAG</td><td></td><td></td><td></td><td></td><td></td><td></td><td></td><td></td><td></td><td></td></tr>
<tr><td>Chromosome</td><td>start</td><td>end</td><td>strand</td><td>MM</td><td>target_seq</td><td>PAM</td><td>alignment</td><td>distance</td><td>position</td><td>gene name</td><td>gene id</td></tr>
<tr><td>GU305914.1</td><td>26949</td><td>26971</td><td>-</td><td>0</td><td>CTTGAGCCTTCTTCGGGGTC</td><td>CGG</td><td>||||||||[||||||||||||]PAM</td><td>NA</td><td>Intergenic</td><td>NA</td><td>NA</td></tr>
<tr><td>GU305914.1</td><td>94739</td><td>94761</td><td>-</td><td>4</td><td>TTTAAGCCTTTTTCGAGGTC</td><td>TGG</td><td>-||-||||[||-||||-||||]PAM</td><td>NA</td><td>Intergenic</td><td>NA</td><td>NA</td></tr>
<tr><td>T32404</td><td>CTTCTTCGGGGTCCGGCGTGCGG</td><td>1000</td><td></td><td></td><td></td><td></td><td></td><td></td><td></td><td></td><td></td></tr>
<tr><td>Oligo fwd</td><td>CACCCTTCTTCGGGGTCCGGCGTG</td><td></td><td></td><td></td><td></td><td></td><td></td><td></td><td></td><td></td><td></td></tr>
<tr><td>Oligo rev</td><td>AAACCACGCCGGACCCCGAAGAAG</td><td></td><td></td><td></td><td></td><td></td><td></td><td></td><td></td><td></td><td></td></tr>
<tr><td>Chromosome</td><td>start</td><td>end</td><td>strand</td><td>MM</td><td>target_seq</td><td>PAM</td><td>alignment</td><td>distance</td><td>position</td><td>gene name</td><td>gene id</td></tr>
<tr><td>GU305914.1</td><td>26942</td><td>26964</td><td>-</td><td>0</td><td>CTTCTTCGGGGTCCGGCGTG</td><td>CGG</td><td>||||||||[||||||||||||]PAM</td><td>NA</td><td>Intergenic</td><td>NA</td><td>NA</td></tr>
<tr><td>T32405</td><td>GGGTCCGGCGTGCGGTCGAGCGG</td><td>1000</td><td></td><td></td><td></td><td></td><td></td><td></td><td></td><td></td><td></td></tr>
<tr><td>Oligo fwd</td><td>CACCGGGTCCGGCGTGCGGTCGAG</td><td></td><td></td><td></td><td></td><td></td><td></td><td></td><td></td><td></td><td></td></tr>
<tr><td>Oligo rev</td><td>AAACCTCGACCGCACGCCGGACCC</td><td></td><td></td><td></td><td></td><td></td><td></td><td></td><td></td><td></td><td></td></tr>
<tr><td>Chromosome</td><td>start</td><td>end</td><td>strand</td><td>MM</td><td>target_seq</td><td>PAM</td><td>alignment</td><td>distance</td><td>position</td><td>gene name</td><td>gene id</td></tr>
<tr><td>GU305914.1</td><td>26934</td><td>26956</td><td>-</td><td>0</td><td>GGGTCCGGCGTGCGGTCGAG</td><td>CGG</td><td>||||||||[||||||||||||]PAM</td><td>NA</td><td>Intergenic</td><td>NA</td><td>NA</td></tr>
<tr><td>T32406</td><td>GCGTGCGGTCGAGCGGCCTCTGG</td><td>1000</td><td></td><td></td><td></td><td></td><td></td><td></td><td></td><td></td><td></td></tr>
<tr><td>Oligo fwd</td><td>CACCGCGTGCGGTCGAGCGGCCTC</td><td></td><td></td><td></td><td></td><td></td><td></td><td></td><td></td><td></td><td></td></tr>
<tr><td>Oligo rev</td><td>AAACGAGGCCGCTCGACCGCACGC</td><td></td><td></td><td></td><td></td><td></td><td></td><td></td><td></td><td></td><td></td></tr>
<tr><td>Chromosome</td><td>start</td><td>end</td><td>strand</td><td>MM</td><td>target_seq</td><td>PAM</td><td>alignment</td><td>distance</td><td>position</td><td>gene name</td><td>gene id</td></tr>
<tr><td>GU305914.1</td><td>26927</td><td>26949</td><td>-</td><td>0</td><td>GCGTGCGGTCGAGCGGCCTC</td><td>TGG</td><td>||||||||[||||||||||||]PAM</td><td>NA</td><td>Intergenic</td><td>NA</td><td>NA</td></tr>
<tr><td>T32407</td><td>CGGTCGAGCGGCCTCTGGACCGG</td><td>1000</td><td></td><td></td><td></td><td></td><td></td><td></td><td></td><td></td><td></td></tr>
<tr><td>Oligo fwd</td><td>CACCCGGTCGAGCGGCCTCTGGAC</td><td></td><td></td><td></td><td></td><td></td><td></td><td></td><td></td><td></td><td></td></tr>
<tr><td>Oligo rev</td><td>AAACGTCCAGAGGCCGCTCGACCG</td><td></td><td></td><td></td><td></td><td></td><td></td><td></td><td></td><td></td><td></td></tr>
<tr><td>Chromosome</td><td>start</td><td>end</td><td>strand</td><td>MM</td><td>target_seq</td><td>PAM</td><td>alignment</td><td>distance</td><td>position</td><td>gene name</td><td>gene id</td></tr>
</table>

<table>
<tr><td>GU305914.1</td><td>26922</td><td>26944</td><td>-</td><td>0</td><td>CGGTCGAGCGGCCTCTGGAC</td><td>CGG</td><td>||||||||[||||||||||||]PAM</td><td>NA</td><td>Intergenic</td><td>NA</td><td>NA</td></tr>
<tr><td>T32408</td><td>CGAGCGGCCTCTGGACCGGCCGG</td><td>1000</td><td></td><td></td><td></td><td></td><td></td><td></td><td></td><td></td><td></td></tr>
<tr><td>Oligo fwd</td><td>CACCCGAGCGGCCTCTGGACCGGC</td><td></td><td></td><td></td><td></td><td></td><td></td><td></td><td></td><td></td><td></td></tr>
<tr><td>Oligo rev</td><td>AAACGCCGGTCCAGAGGCCGCTCG</td><td></td><td></td><td></td><td></td><td></td><td></td><td></td><td></td><td></td><td></td></tr>
<tr><td>Chromosome</td><td>start</td><td>end</td><td>strand</td><td>MM</td><td>target_seq</td><td>PAM</td><td>alignment</td><td>distance</td><td>position</td><td>gene name</td><td>gene id</td></tr>
<tr><td>GU305914.1</td><td>26918</td><td>26940</td><td>-</td><td>0</td><td>CGAGCGGCCTCTGGACCGGC</td><td>CGG</td><td>||||||||[||||||||||||]PAM</td><td>NA</td><td>Intergenic</td><td>NA</td><td>NA</td></tr>
<tr><td>T32409</td><td>GCGGCCTCTGGACCGGCCGGCGG</td><td>1000</td><td></td><td></td><td></td><td></td><td></td><td></td><td></td><td></td><td></td></tr>
<tr><td>Oligo fwd</td><td>CACCGCGGCCTCTGGACCGGCCGG</td><td></td><td></td><td></td><td></td><td></td><td></td><td></td><td></td><td></td><td></td></tr>
<tr><td>Oligo rev</td><td>AAACCCGGCCGGTCCAGAGGCCGC</td><td></td><td></td><td></td><td></td><td></td><td></td><td></td><td></td><td></td><td></td></tr>
<tr><td>Chromosome</td><td>start</td><td>end</td><td>strand</td><td>MM</td><td>target_seq</td><td>PAM</td><td>alignment</td><td>distance</td><td>position</td><td>gene name</td><td>gene id</td></tr>
<tr><td>GU305914.1</td><td>26915</td><td>26937</td><td>-</td><td>0</td><td>GCGGCCTCTGGACCGGCCGG</td><td>CGG</td><td>||||||||[||||||||||||]PAM</td><td>NA</td><td>Intergenic</td><td>NA</td><td>NA</td></tr>
<tr><td>T32410</td><td>GCCTCTGGACCGGCCGGCGGTGG</td><td>1000</td><td></td><td></td><td></td><td></td><td></td><td></td><td></td><td></td><td></td></tr>
<tr><td>Oligo fwd</td><td>CACCGCCTCTGGACCGGCCGGCGG</td><td></td><td></td><td></td><td></td><td></td><td></td><td></td><td></td><td></td><td></td></tr>
<tr><td>Oligo rev</td><td>AAACCCGCCGGCCGGTCCAGAGGC</td><td></td><td></td><td></td><td></td><td></td><td></td><td></td><td></td><td></td><td></td></tr>
<tr><td>Chromosome</td><td>start</td><td>end</td><td>strand</td><td>MM</td><td>target_seq</td><td>PAM</td><td>alignment</td><td>distance</td><td>position</td><td>gene name</td><td>gene id</td></tr>
<tr><td>GU305914.1</td><td>26912</td><td>26934</td><td>-</td><td>0</td><td>GCCTCTGGACCGGCCGGCGG</td><td>TGG</td><td>||||||||[||||||||||||]PAM</td><td>NA</td><td>Intergenic</td><td>NA</td><td>NA</td></tr>
<tr><td>GU305914.1</td><td>19315</td><td>19337</td><td>+</td><td>4</td><td>GGATCTGGACCGTCCGGCGA</td><td>CGG</td><td>|--|||||[||||-||||||-]PAM</td><td>NA</td><td>Intergenic</td><td>NA</td><td>NA</td></tr>
<tr><td>T32411</td><td>CTTTTCCTCCGCCGATTTTTTGG</td><td>1000</td><td></td><td></td><td></td><td></td><td></td><td></td><td></td><td></td><td></td></tr>
<tr><td>Oligo fwd</td><td>CACCCTTTTCCTCCGCCGATTTTT</td><td></td><td></td><td></td><td></td><td></td><td></td><td></td><td></td><td></td><td></td></tr>
<tr><td>Oligo rev</td><td>AAACAAAAATCGGCGGAGGAAAAG</td><td></td><td></td><td></td><td></td><td></td><td></td><td></td><td></td><td></td><td></td></tr>
<tr><td>Chromosome</td><td>start</td><td>end</td><td>strand</td><td>MM</td><td>target_seq</td><td>PAM</td><td>alignment</td><td>distance</td><td>position</td><td>gene name</td><td>gene id</td></tr>
<tr><td>GU305914.1</td><td>26882</td><td>26904</td><td>-</td><td>0</td><td>CTTTTCCTCCGCCGATTTTT</td><td>TGG</td><td>||||||||[||||||||||||]PAM</td><td>NA</td><td>Intergenic</td><td>NA</td><td>NA</td></tr>
<tr><td>T32412</td><td>TCCGCCGATTTTTTGGTCGAAGG</td><td>1000</td><td></td><td></td><td></td><td></td><td></td><td></td><td></td><td></td><td></td></tr>
<tr><td>Oligo fwd</td><td>CACCTCCGCCGATTTTTTGGTCGA</td><td></td><td></td><td></td><td></td><td></td><td></td><td></td><td></td><td></td><td></td></tr>
<tr><td>Oligo rev</td><td>AAACTCGACCAAAAAATCGGCGGA</td><td></td><td></td><td></td><td></td><td></td><td></td><td></td><td></td><td></td><td></td></tr>
<tr><td>Chromosome</td><td>start</td><td>end</td><td>strand</td><td>MM</td><td>target_seq</td><td>PAM</td><td>alignment</td><td>distance</td><td>position</td><td>gene name</td><td>gene id</td></tr>
<tr><td>GU305914.1</td><td>26875</td><td>26897</td><td>-</td><td>0</td><td>TCCGCCGATTTTTTGGTCGA</td><td>AGG</td><td>||||||||[||||||||||||]PAM</td><td>NA</td><td>Intergenic</td><td>NA</td><td>NA</td></tr>
<tr><td>T32413</td><td>GTCGAAGGTCGACTCTTGCGTGG</td><td>1000</td><td></td><td></td><td></td><td></td><td></td><td></td><td></td><td></td><td></td></tr>
<tr><td>Oligo fwd</td><td>CACCGTCGAAGGTCGACTCTTGCG</td><td></td><td></td><td></td><td></td><td></td><td></td><td></td><td></td><td></td><td></td></tr>
<tr><td>Oligo rev</td><td>AAACCGCAAGAGTCGACCTTCGAC</td><td></td><td></td><td></td><td></td><td></td><td></td><td></td><td></td><td></td><td></td></tr>
<tr><td>Chromosome</td><td>start</td><td>end</td><td>strand</td><td>MM</td><td>target_seq</td><td>PAM</td><td>alignment</td><td>distance</td><td>position</td><td>gene name</td><td>gene id</td></tr>
<tr><td>GU305914.1</td><td>26860</td><td>26882</td><td>-</td><td>0</td><td>GTCGAAGGTCGACTCTTGCG</td><td>TGG</td><td>||||||||[||||||||||||]PAM</td><td>NA</td><td>Intergenic</td><td>NA</td><td>NA</td></tr>
<tr><td>T32414</td><td>TCGACTCTTGCGTGGCAGTTTGG</td><td>1000</td><td></td><td></td><td></td><td></td><td></td><td></td><td></td><td></td><td></td></tr>
<tr><td>Oligo fwd</td><td>CACCTCGACTCTTGCGTGGCAGTT</td><td></td><td></td><td></td><td></td><td></td><td></td><td></td><td></td><td></td><td></td></tr>
<tr><td>Oligo rev</td><td>AAACAACTGCCACGCAAGAGTCGA</td><td></td><td></td><td></td><td></td><td></td><td></td><td></td><td></td><td></td><td></td></tr>
<tr><td>Chromosome</td><td>start</td><td>end</td><td>strand</td><td>MM</td><td>target_seq</td><td>PAM</td><td>alignment</td><td>distance</td><td>position</td><td>gene name</td><td>gene id</td></tr>
<tr><td>GU305914.1</td><td>26852</td><td>26874</td><td>-</td><td>0</td><td>TCGACTCTTGCGTGGCAGTT</td><td>TGG</td><td>||||||||[||||||||||||]PAM</td><td>NA</td><td>Intergenic</td><td>NA</td><td>NA</td></tr>
<tr><td>T32415</td><td>CGACTCTTGCGTGGCAGTTTGGG</td><td>1000</td><td></td><td></td><td></td><td></td><td></td><td></td><td></td><td></td><td></td></tr>
<tr><td>Oligo fwd</td><td>CACCCGACTCTTGCGTGGCAGTTT</td><td></td><td></td><td></td><td></td><td></td><td></td><td></td><td></td><td></td><td></td></tr>
<tr><td>Oligo rev</td><td>AAACAAACTGCCACGCAAGAGTCG</td><td></td><td></td><td></td><td></td><td></td><td></td><td></td><td></td><td></td><td></td></tr>
<tr><td>Chromosome</td><td>start</td><td>end</td><td>strand</td><td>MM</td><td>target_seq</td><td>PAM</td><td>alignment</td><td>distance</td><td>position</td><td>gene name</td><td>gene id</td></tr>
<tr><td>GU305914.1</td><td>26851</td><td>26873</td><td>-</td><td>0</td><td>CGACTCTTGCGTGGCAGTTT</td><td>GGG</td><td>||||||||[||||||||||||]PAM</td><td>NA</td><td>Intergenic</td><td>NA</td><td>NA</td></tr>
<tr><td>T32416</td><td>CTCTTGCGTGGCAGTTTGGGTGG</td><td>1000</td><td></td><td></td><td></td><td></td><td></td><td></td><td></td><td></td><td></td></tr>
<tr><td>Oligo fwd</td><td>CACCCTCTTGCGTGGCAGTTTGGG</td><td></td><td></td><td></td><td></td><td></td><td></td><td></td><td></td><td></td><td></td></tr>
<tr><td>Oligo rev</td><td>AAACCCCAAACTGCCACGCAAGAG</td><td></td><td></td><td></td><td></td><td></td><td></td><td></td><td></td><td></td><td></td></tr>
<tr><td>Chromosome</td><td>start</td><td>end</td><td>strand</td><td>MM</td><td>target_seq</td><td>PAM</td><td>alignment</td><td>distance</td><td>position</td><td>gene name</td><td>gene id</td></tr>
<tr><td>GU305914.1</td><td>26848</td><td>26870</td><td>-</td><td>0</td><td>CTCTTGCGTGGCAGTTTGGG</td><td>TGG</td><td>||||||||[||||||||||||]PAM</td><td>NA</td><td>Intergenic</td><td>NA</td><td>NA</td></tr>
<tr><td>T32417</td><td>TCTTGCGTGGCAGTTTGGGTGGG</td><td>1000</td><td></td><td></td><td></td><td></td><td></td><td></td><td></td><td></td><td></td></tr>
<tr><td>Oligo fwd</td><td>CACCTCTTGCGTGGCAGTTTGGGT</td><td></td><td></td><td></td><td></td><td></td><td></td><td></td><td></td><td></td><td></td></tr>
<tr><td>Oligo rev</td><td>AAACACCCAAACTGCCACGCAAGA</td><td></td><td></td><td></td><td></td><td></td><td></td><td></td><td></td><td></td><td></td></tr>
<tr><td>Chromosome</td><td>start</td><td>end</td><td>strand</td><td>MM</td><td>target_seq</td><td>PAM</td><td>alignment</td><td>distance</td><td>position</td><td>gene name</td><td>gene id</td></tr>
<tr><td>GU305914.1</td><td>26847</td><td>26869</td><td>-</td><td>0</td><td>TCTTGCGTGGCAGTTTGGGT</td><td>GGG</td><td>||||||||[||||||||||||]PAM</td><td>NA</td><td>Intergenic</td><td>NA</td><td>NA</td></tr>
<tr><td>T32418</td><td>CGTGGCAGTTTGGGTGGGCTCGG</td><td>1000</td><td></td><td></td><td></td><td></td><td></td><td></td><td></td><td></td><td></td></tr>
<tr><td>Oligo fwd</td><td>CACCCGTGGCAGTTTGGGTGGGCT</td><td></td><td></td><td></td><td></td><td></td><td></td><td></td><td></td><td></td><td></td></tr>
<tr><td>Oligo rev</td><td>AAACAGCCCACCCAAACTGCCACG</td><td></td><td></td><td></td><td></td><td></td><td></td><td></td><td></td><td></td><td></td></tr>
<tr><td>Chromosome</td><td>start</td><td>end</td><td>strand</td><td>MM</td><td>target_seq</td><td>PAM</td><td>alignment</td><td>distance</td><td>position</td><td>gene name</td><td>gene id</td></tr>
<tr><td>GU305914.1</td><td>26842</td><td>26864</td><td>-</td><td>0</td><td>CGTGGCAGTTTGGGTGGGCT</td><td>CGG</td><td>||||||||[||||||||||||]PAM</td><td>NA</td><td>Intergenic</td><td>NA</td><td>NA</td></tr>
<tr><td>T32419</td><td>CGCGAATCGTCGCGCGAAGTAGG</td><td>1000</td><td></td><td></td><td></td><td></td><td></td><td></td><td></td><td></td><td></td></tr>
<tr><td>Oligo fwd</td><td>CACCCGCGAATCGTCGCGCGAAGT</td><td></td><td></td><td></td><td></td><td></td><td></td><td></td><td></td><td></td><td></td></tr>
<tr><td>Oligo rev</td><td>AAACACTTCGCGCGACGATTCGCG</td><td></td><td></td><td></td><td></td><td></td><td></td><td></td><td></td><td></td><td></td></tr>
<tr><td>Chromosome</td><td>start</td><td>end</td><td>strand</td><td>MM</td><td>target_seq</td><td>PAM</td><td>alignment</td><td>distance</td><td>position</td><td>gene name</td><td>gene id</td></tr>
<tr><td>GU305914.1</td><td>26752</td><td>26774</td><td>-</td><td>0</td><td>CGCGAATCGTCGCGCGAAGT</td><td>AGG</td><td>||||||||[||||||||||||]PAM</td><td>NA</td><td>Intergenic</td><td>NA</td><td>NA</td></tr>
<tr><td>T32420</td><td>GAATCGTCGCGCGAAGTAGGTGG</td><td>1000</td><td></td><td></td><td></td><td></td><td></td><td></td><td></td><td></td><td></td></tr>
<tr><td>Oligo fwd</td><td>CACCGAATCGTCGCGCGAAGTAGG</td><td></td><td></td><td></td><td></td><td></td><td></td><td></td><td></td><td></td><td></td></tr>
<tr><td>Oligo rev</td><td>AAACCCTACTTCGCGCGACGATTC</td><td></td><td></td><td></td><td></td><td></td><td></td><td></td><td></td><td></td><td></td></tr>
<tr><td>Chromosome</td><td>start</td><td>end</td><td>strand</td><td>MM</td><td>target_seq</td><td>PAM</td><td>alignment</td><td>distance</td><td>position</td><td>gene name</td><td>gene id</td></tr>
<tr><td>GU305914.1</td><td>26749</td><td>26771</td><td>-</td><td>0</td><td>GAATCGTCGCGCGAAGTAGG</td><td>TGG</td><td>||||||||[||||||||||||]PAM</td><td>NA</td><td>Intergenic</td><td>NA</td><td>NA</td></tr>
<tr><td>T32421</td><td>AATCGTCGCGCGAAGTAGGTGGG</td><td>1000</td><td></td><td></td><td></td><td></td><td></td><td></td><td></td><td></td><td></td></tr>
<tr><td>Oligo fwd</td><td>CACCAATCGTCGCGCGAAGTAGGT</td><td></td><td></td><td></td><td></td><td></td><td></td><td></td><td></td><td></td><td></td></tr>
<tr><td>Oligo rev</td><td>AAACACCTACTTCGCGCGACGATT</td><td></td><td></td><td></td><td></td><td></td><td></td><td></td><td></td><td></td><td></td></tr>
<tr><td>Chromosome</td><td>start</td><td>end</td><td>strand</td><td>MM</td><td>target_seq</td><td>PAM</td><td>alignment</td><td>distance</td><td>position</td><td>gene name</td><td>gene id</td></tr>
<tr><td>GU305914.1</td><td>26748</td><td>26770</td><td>-</td><td>0</td><td>AATCGTCGCGCGAAGTAGGT</td><td>GGG</td><td>||||||||[||||||||||||]PAM</td><td>NA</td><td>Intergenic</td><td>NA</td><td>NA</td></tr>
<tr><td>T32422</td><td>GTCGCGCGAAGTAGGTGGGTCGG</td><td>1000</td><td></td><td></td><td></td><td></td><td></td><td></td><td></td><td></td><td></td></tr>
<tr><td>Oligo fwd</td><td>CACCGTCGCGCGAAGTAGGTGGGT</td><td></td><td></td><td></td><td></td><td></td><td></td><td></td><td></td><td></td><td></td></tr>
</table>

| | | | | | | | | | | | | |
|---|---|---|---|---|---|---|---|---|---|---|---|---|
| Oligo rev | AAACACCCACCTACTTCGCGCGAC | | | | | | | | | | | |
| Chromosome | start | | end | strand | MM | target_seq | PAM | alignment | distance | position | gene name | gene id |
| GU305914.1 | | 26744 | 26766 | - | 0 | GTCGCGCGAAGTAGGTGGGT | CGG | \|\|\|\|\|\|\|\|[\|\|\|\|\|\|\|\|\|\|\|\|]PAM | NA | Intergenic | NA | NA |
| T32423 | TCGCGCGAAGTAGGTGGGTCGGG | | 1000 | | | | | | | | | |
| Oligo fwd | CACCTCGCGCGAAGTAGGTGGGTC | | | | | | | | | | | |
| Oligo rev | AAACGACCCACCTACTTCGCGCGA | | | | | | | | | | | |
| Chromosome | start | | end | strand | MM | target_seq | PAM | alignment | distance | position | gene name | gene id |
| GU305914.1 | | 26743 | 26765 | - | 0 | TCGCGCGAAGTAGGTGGGTC | GGG | \|\|\|\|\|\|\|\|[\|\|\|\|\|\|\|\|\|\|\|\|]PAM | NA | Intergenic | NA | NA |
| T32424 | CGCGCGAAGTAGGTGGGTCGGGG | | 1000 | | | | | | | | | |
| Oligo fwd | CACCCGCGCGAAGTAGGTGGGTCG | | | | | | | | | | | |
| Oligo rev | AAACCGACCCACCTACTTCGCGCG | | | | | | | | | | | |
| Chromosome | start | | end | strand | MM | target_seq | PAM | alignment | distance | position | gene name | gene id |
| GU305914.1 | | 26742 | 26764 | - | 0 | CGCGCGAAGTAGGTGGGTCG | GGG | \|\|\|\|\|\|\|\|[\|\|\|\|\|\|\|\|\|\|\|\|]PAM | NA | Intergenic | NA | NA |
| T32425 | CGTATCCGTCCTCATCAAAAAGG | | 1000 | | | | | | | | | |
| Oligo fwd | CACCCGTATCCGTCCTCATCAAAA | | | | | | | | | | | |
| Oligo rev | AAACTTTTGATGAGGACGGATACG | | | | | | | | | | | |
| Chromosome | start | | end | strand | MM | target_seq | PAM | alignment | distance | position | gene name | gene id |
| GU305914.1 | | 26712 | 26734 | - | 0 | CGTATCCGTCCTCATCAAAA | AGG | \|\|\|\|\|\|\|\|[\|\|\|\|\|\|\|\|\|\|\|\|]PAM | NA | Intergenic | NA | NA |
| GU305914.1 | | 115581 | 115603 | + | 4 | CGTGTCCCTCCTCAATAAAA | TGG | \|\|\|-\|\|\|-[\|\|\|\|\|\|--\|\|\|\|]PAM | NA | Intergenic | NA | NA |
| T32426 | GTATCCGTCCTCATCAAAAAGGG | | 1000 | | | | | | | | | |
| Oligo fwd | CACCGTATCCGTCCTCATCAAAAA | | | | | | | | | | | |
| Oligo rev | AAACTTTTTGATGAGGACGGATAC | | | | | | | | | | | |
| Chromosome | start | | end | strand | MM | target_seq | PAM | alignment | distance | position | gene name | gene id |
| GU305914.1 | | 26711 | 26733 | - | 0 | GTATCCGTCCTCATCAAAAA | GGG | \|\|\|\|\|\|\|\|[\|\|\|\|\|\|\|\|\|\|\|\|]PAM | NA | Intergenic | NA | NA |
| T32427 | GTCCTCATCAAAAAGGGTCACGG | | 1000 | | | | | | | | | |
| Oligo fwd | CACCGTCCTCATCAAAAAGGGTCA | | | | | | | | | | | |
| Oligo rev | AAACTGACCCTTTTTGATGAGGAC | | | | | | | | | | | |
| Chromosome | start | | end | strand | MM | target_seq | PAM | alignment | distance | position | gene name | gene id |
| GU305914.1 | | 26705 | 26727 | - | 0 | GTCCTCATCAAAAAGGGTCA | CGG | \|\|\|\|\|\|\|\|[\|\|\|\|\|\|\|\|\|\|\|\|]PAM | NA | Intergenic | NA | NA |
| T32428 | TCCTCATCAAAAAGGGTCACGGG | | 1000 | | | | | | | | | |
| Oligo fwd | CACCTCCTCATCAAAAAGGGTCAC | | | | | | | | | | | |
| Oligo rev | AAACGTGACCCTTTTTGATGAGGA | | | | | | | | | | | |
| Chromosome | start | | end | strand | MM | target_seq | PAM | alignment | distance | position | gene name | gene id |
| GU305914.1 | | 26704 | 26726 | - | 0 | TCCTCATCAAAAAGGGTCAC | GGG | \|\|\|\|\|\|\|\|[\|\|\|\|\|\|\|\|\|\|\|\|]PAM | NA | Intergenic | NA | NA |
| T32429 | ACACGCTTCCACTATCATTCCGG | | 1000 | | | | | | | | | |
| Oligo fwd | CACCACACGCTTCCACTATCATTC | | | | | | | | | | | |
| Oligo rev | AAACGAATGATAGTGGAAGCGTGT | | | | | | | | | | | |
| Chromosome | start | | end | strand | MM | target_seq | PAM | alignment | distance | position | gene name | gene id |
| GU305914.1 | | 26679 | 26701 | - | 0 | ACACGCTTCCACTATCATTC | CGG | \|\|\|\|\|\|\|\|[\|\|\|\|\|\|\|\|\|\|\|\|]PAM | NA | Intergenic | NA | NA |
| T32430 | CACGCTTCCACTATCATTCCGGG | | 1000 | | | | | | | | | |
| Oligo fwd | CACCCACGCTTCCACTATCATTCC | | | | | | | | | | | |
| Oligo rev | AAACGGAATGATAGTGGAAGCGTG | | | | | | | | | | | |
| Chromosome | start | | end | strand | MM | target_seq | PAM | alignment | distance | position | gene name | gene id |
| GU305914.1 | | 26678 | 26700 | - | 0 | CACGCTTCCACTATCATTCC | GGG | \|\|\|\|\|\|\|\|[\|\|\|\|\|\|\|\|\|\|\|\|]PAM | NA | Intergenic | NA | NA |
| T32431 | CATCGCCCGCAGTCTCACATCGG | | 1000 | | | | | | | | | |
| Oligo fwd | CACCCATCGCCCGCAGTCTCACAT | | | | | | | | | | | |
| Oligo rev | AAACATGTGAGACTGCGGGCGATG | | | | | | | | | | | |
| Chromosome | start | | end | strand | MM | target_seq | PAM | alignment | distance | position | gene name | gene id |
| GU305914.1 | | 26600 | 26622 | - | 0 | CATCGCCCGCAGTCTCACAT | CGG | \|\|\|\|\|\|\|\|[\|\|\|\|\|\|\|\|\|\|\|\|]PAM | NA | Intergenic | NA | NA |
| T32432 | GCCCGCAGTCTCACATCGGCCGG | | 1000 | | | | | | | | | |
| Oligo fwd | CACCGCCCGCAGTCTCACATCGGC | | | | | | | | | | | |
| Oligo rev | AAACGCCGATGTGAGACTGCGGGC | | | | | | | | | | | |
| Chromosome | start | | end | strand | MM | target_seq | PAM | alignment | distance | position | gene name | gene id |
| GU305914.1 | | 26596 | 26618 | - | 0 | GCCCGCAGTCTCACATCGGC | CGG | \|\|\|\|\|\|\|\|[\|\|\|\|\|\|\|\|\|\|\|\|]PAM | NA | Intergenic | NA | NA |
| T32433 | CCCGCAGTCTCACATCGGCCGGG | | 1000 | | | | | | | | | |
| Oligo fwd | CACCCCCGCAGTCTCACATCGGCC | | | | | | | | | | | |
| Oligo rev | AAACGGCCGATGTGAGACTGCGGG | | | | | | | | | | | |
| Chromosome | start | | end | strand | MM | target_seq | PAM | alignment | distance | position | gene name | gene id |
| GU305914.1 | | 26595 | 26617 | - | 0 | CCCGCAGTCTCACATCGGCC | GGG | \|\|\|\|\|\|\|\|[\|\|\|\|\|\|\|\|\|\|\|\|]PAM | NA | Intergenic | NA | NA |
| T32434 | CATAAGTAAAGTCGTCGTCCTGG | | 1000 | | | | | | | | | |
| Oligo fwd | CACCCATAAGTAAAGTCGTCGTCC | | | | | | | | | | | |
| Oligo rev | AAACGGACGACGACTTTACTTATG | | | | | | | | | | | |
| Chromosome | start | | end | strand | MM | target_seq | PAM | alignment | distance | position | gene name | gene id |
| GU305914.1 | | 26568 | 26590 | - | 0 | CATAAGTAAAGTCGTCGTCC | TGG | \|\|\|\|\|\|\|\|[\|\|\|\|\|\|\|\|\|\|\|\|]PAM | NA | Intergenic | NA | NA |
| T32435 | AGTCGTCGTCCTGGCTGTGCCGG | | 1000 | | | | | | | | | |
| Oligo fwd | CACCAGTCGTCGTCCTGGCTGTGC | | | | | | | | | | | |
| Oligo rev | AAACGCACAGCCAGGACGACGACT | | | | | | | | | | | |
| Chromosome | start | | end | strand | MM | target_seq | PAM | alignment | distance | position | gene name | gene id |
| GU305914.1 | | 26559 | 26581 | - | 0 | AGTCGTCGTCCTGGCTGTGC | CGG | \|\|\|\|\|\|\|\|[\|\|\|\|\|\|\|\|\|\|\|\|]PAM | NA | Intergenic | NA | NA |
| T32436 | CGTCGTCCTGGCTGTGCCGGCGG | | 1000 | | | | | | | | | |
| Oligo fwd | CACCCGTCGTCCTGGCTGTGCCGG | | | | | | | | | | | |
| Oligo rev | AAACCCGGCACAGCCAGGACGACG | | | | | | | | | | | |
| Chromosome | start | | end | strand | MM | target_seq | PAM | alignment | distance | position | gene name | gene id |
| GU305914.1 | | 26556 | 26578 | - | 0 | CGTCGTCCTGGCTGTGCCGG | CGG | \|\|\|\|\|\|\|\|[\|\|\|\|\|\|\|\|\|\|\|\|]PAM | NA | Intergenic | NA | NA |

<table>
<tr><td>T32437</td><td>GTCGTCCTGGCTGTGCCGGCGGG</td><td></td><td>1000</td><td></td><td></td><td></td><td></td><td></td><td></td><td></td><td></td><td></td></tr>
<tr><td>Oligo fwd</td><td>CACCGTCGTCCTGGCTGTGCCGGC</td><td></td><td></td><td></td><td></td><td></td><td></td><td></td><td></td><td></td><td></td><td></td></tr>
<tr><td>Oligo rev</td><td>AAACGCCGGCACAGCCAGGACGAC</td><td></td><td></td><td></td><td></td><td></td><td></td><td></td><td></td><td></td><td></td><td></td></tr>
<tr><td>Chromosome</td><td>start</td><td></td><td>end</td><td>strand</td><td>MM</td><td>target_seq</td><td>PAM</td><td>alignment</td><td>distance</td><td>position</td><td>gene name</td><td>gene id</td></tr>
<tr><td>GU305914.1</td><td></td><td>26555</td><td>26577</td><td>-</td><td>0</td><td>GTCGTCCTGGCTGTGCCGGC</td><td>GGG</td><td>||||||||[||||||||||||]PAM</td><td>NA</td><td>Intergenic</td><td>NA</td><td>NA</td></tr>
<tr><td>T32438</td><td>TCGTCCTGGCTGTGCCGGCGGGG</td><td></td><td>1000</td><td></td><td></td><td></td><td></td><td></td><td></td><td></td><td></td><td></td></tr>
<tr><td>Oligo fwd</td><td>CACCTCGTCCTGGCTGTGCCGGCG</td><td></td><td></td><td></td><td></td><td></td><td></td><td></td><td></td><td></td><td></td><td></td></tr>
<tr><td>Oligo rev</td><td>AAACCGCCGGCACAGCCAGGACGA</td><td></td><td></td><td></td><td></td><td></td><td></td><td></td><td></td><td></td><td></td><td></td></tr>
<tr><td>Chromosome</td><td>start</td><td></td><td>end</td><td>strand</td><td>MM</td><td>target_seq</td><td>PAM</td><td>alignment</td><td>distance</td><td>position</td><td>gene name</td><td>gene id</td></tr>
<tr><td>GU305914.1</td><td></td><td>26554</td><td>26576</td><td>-</td><td>0</td><td>TCGTCCTGGCTGTGCCGGCG</td><td>GGG</td><td>||||||||[||||||||||||]PAM</td><td>NA</td><td>Intergenic</td><td>NA</td><td>NA</td></tr>
<tr><td>T32439</td><td>TCGCCGTTTAGTTCCTCAGACGG</td><td></td><td>1000</td><td></td><td></td><td></td><td></td><td></td><td></td><td></td><td></td><td></td></tr>
<tr><td>Oligo fwd</td><td>CACCTCGCCGTTTAGTTCCTCAGA</td><td></td><td></td><td></td><td></td><td></td><td></td><td></td><td></td><td></td><td></td><td></td></tr>
<tr><td>Oligo rev</td><td>AAACTCTGAGGAACTAAACGGCGA</td><td></td><td></td><td></td><td></td><td></td><td></td><td></td><td></td><td></td><td></td><td></td></tr>
<tr><td>Chromosome</td><td>start</td><td></td><td>end</td><td>strand</td><td>MM</td><td>target_seq</td><td>PAM</td><td>alignment</td><td>distance</td><td>position</td><td>gene name</td><td>gene id</td></tr>
<tr><td>GU305914.1</td><td></td><td>26491</td><td>26513</td><td>-</td><td>0</td><td>TCGCCGTTTAGTTCCTCAGA</td><td>CGG</td><td>||||||||[||||||||||||]PAM</td><td>NA</td><td>Intergenic</td><td>NA</td><td>NA</td></tr>
<tr><td>T32440</td><td>CGCCGTTTAGTTCCTCAGACGGG</td><td></td><td>1000</td><td></td><td></td><td></td><td></td><td></td><td></td><td></td><td></td><td></td></tr>
<tr><td>Oligo fwd</td><td>CACCCGCCGTTTAGTTCCTCAGAC</td><td></td><td></td><td></td><td></td><td></td><td></td><td></td><td></td><td></td><td></td><td></td></tr>
<tr><td>Oligo rev</td><td>AAACGTCTGAGGAACTAAACGGCG</td><td></td><td></td><td></td><td></td><td></td><td></td><td></td><td></td><td></td><td></td><td></td></tr>
<tr><td>Chromosome</td><td>start</td><td></td><td>end</td><td>strand</td><td>MM</td><td>target_seq</td><td>PAM</td><td>alignment</td><td>distance</td><td>position</td><td>gene name</td><td>gene id</td></tr>
<tr><td>GU305914.1</td><td></td><td>26490</td><td>26512</td><td>-</td><td>0</td><td>CGCCGTTTAGTTCCTCAGAC</td><td>GGG</td><td>||||||||[||||||||||||]PAM</td><td>NA</td><td>Intergenic</td><td>NA</td><td>NA</td></tr>
<tr><td>T32441</td><td>TCCTCAGACGGGATGCTCAATGG</td><td></td><td>1000</td><td></td><td></td><td></td><td></td><td></td><td></td><td></td><td></td><td></td></tr>
<tr><td>Oligo fwd</td><td>CACCTCCTCAGACGGGATGCTCAA</td><td></td><td></td><td></td><td></td><td></td><td></td><td></td><td></td><td></td><td></td><td></td></tr>
<tr><td>Oligo rev</td><td>AAACTTGAGCATCCCGTCTGAGGA</td><td></td><td></td><td></td><td></td><td></td><td></td><td></td><td></td><td></td><td></td><td></td></tr>
<tr><td>Chromosome</td><td>start</td><td></td><td>end</td><td>strand</td><td>MM</td><td>target_seq</td><td>PAM</td><td>alignment</td><td>distance</td><td>position</td><td>gene name</td><td>gene id</td></tr>
<tr><td>GU305914.1</td><td></td><td>26479</td><td>26501</td><td>-</td><td>0</td><td>TCCTCAGACGGGATGCTCAA</td><td>TGG</td><td>||||||||[||||||||||||]PAM</td><td>NA</td><td>Intergenic</td><td>NA</td><td>NA</td></tr>
<tr><td>T32442</td><td>CAATGGCTTGTATGTGCTCTCGG</td><td></td><td>1000</td><td></td><td></td><td></td><td></td><td></td><td></td><td></td><td></td><td></td></tr>
<tr><td>Oligo fwd</td><td>CACCCAATGGCTTGTATGTGCTCT</td><td></td><td></td><td></td><td></td><td></td><td></td><td></td><td></td><td></td><td></td><td></td></tr>
<tr><td>Oligo rev</td><td>AAACAGAGCACATACAAGCCATTG</td><td></td><td></td><td></td><td></td><td></td><td></td><td></td><td></td><td></td><td></td><td></td></tr>
<tr><td>Chromosome</td><td>start</td><td></td><td>end</td><td>strand</td><td>MM</td><td>target_seq</td><td>PAM</td><td>alignment</td><td>distance</td><td>position</td><td>gene name</td><td>gene id</td></tr>
<tr><td>GU305914.1</td><td></td><td>26462</td><td>26484</td><td>-</td><td>0</td><td>CAATGGCTTGTATGTGCTCT</td><td>CGG</td><td>||||||||[||||||||||||]PAM</td><td>NA</td><td>Intergenic</td><td>NA</td><td>NA</td></tr>
<tr><td>T32443</td><td>TGGCTTGTATGTGCTCTCGGAGG</td><td></td><td>1000</td><td></td><td></td><td></td><td></td><td></td><td></td><td></td><td></td><td></td></tr>
<tr><td>Oligo fwd</td><td>CACCTGGCTTGTATGTGCTCTCGG</td><td></td><td></td><td></td><td></td><td></td><td></td><td></td><td></td><td></td><td></td><td></td></tr>
<tr><td>Oligo rev</td><td>AAACCCGAGAGCACATACAAGCCA</td><td></td><td></td><td></td><td></td><td></td><td></td><td></td><td></td><td></td><td></td><td></td></tr>
<tr><td>Chromosome</td><td>start</td><td></td><td>end</td><td>strand</td><td>MM</td><td>target_seq</td><td>PAM</td><td>alignment</td><td>distance</td><td>position</td><td>gene name</td><td>gene id</td></tr>
<tr><td>GU305914.1</td><td></td><td>26459</td><td>26481</td><td>-</td><td>0</td><td>TGGCTTGTATGTGCTCTCGG</td><td>AGG</td><td>||||||||[||||||||||||]PAM</td><td>NA</td><td>Intergenic</td><td>NA</td><td>NA</td></tr>
<tr><td>T32444</td><td>CTCGGAGGCCGCCGCCGCCGTGG</td><td></td><td>1000</td><td></td><td></td><td></td><td></td><td></td><td></td><td></td><td></td><td></td></tr>
<tr><td>Oligo fwd</td><td>CACCCTCGGAGGCCGCCGCCGCCG</td><td></td><td></td><td></td><td></td><td></td><td></td><td></td><td></td><td></td><td></td><td></td></tr>
<tr><td>Oligo rev</td><td>AAACCGGCGGCGGCGGCCTCCGAG</td><td></td><td></td><td></td><td></td><td></td><td></td><td></td><td></td><td></td><td></td><td></td></tr>
<tr><td>Chromosome</td><td>start</td><td></td><td>end</td><td>strand</td><td>MM</td><td>target_seq</td><td>PAM</td><td>alignment</td><td>distance</td><td>position</td><td>gene name</td><td>gene id</td></tr>
<tr><td>GU305914.1</td><td></td><td>26444</td><td>26466</td><td>-</td><td>0</td><td>CTCGGAGGCCGCCGCCGCCG</td><td>TGG</td><td>||||||||[||||||||||||]PAM</td><td>NA</td><td>Intergenic</td><td>NA</td><td>NA</td></tr>
<tr><td>GU305914.1</td><td></td><td>45592</td><td>45614</td><td>-</td><td>4</td><td>GGCGGCTGCCGCCGCCGCCG</td><td>CGG</td><td>--|||--|[||||||||||||]PAM</td><td>NA</td><td>Intergenic</td><td>NA</td><td>NA</td></tr>
<tr><td>GU305914.1</td><td></td><td>35655</td><td>35677</td><td>-</td><td>4</td><td>CCCCGCCGCCGCCGCCGCCG</td><td>TGG</td><td>|-|-|--|[||||||||||||]PAM</td><td>NA</td><td>Intergenic</td><td>NA</td><td>NA</td></tr>
<tr><td>T32445</td><td>GGAGGCCGCCGCCGCCGTGGCGG</td><td></td><td>1000</td><td></td><td></td><td></td><td></td><td></td><td></td><td></td><td></td><td></td></tr>
<tr><td>Oligo fwd</td><td>CACCGGAGGCCGCCGCCGCCGTGG</td><td></td><td></td><td></td><td></td><td></td><td></td><td></td><td></td><td></td><td></td><td></td></tr>
<tr><td>Oligo rev</td><td>AAACCCACGGCGGCGGCGGCCTCC</td><td></td><td></td><td></td><td></td><td></td><td></td><td></td><td></td><td></td><td></td><td></td></tr>
<tr><td>Chromosome</td><td>start</td><td></td><td>end</td><td>strand</td><td>MM</td><td>target_seq</td><td>PAM</td><td>alignment</td><td>distance</td><td>position</td><td>gene name</td><td>gene id</td></tr>
<tr><td>GU305914.1</td><td></td><td>26441</td><td>26463</td><td>-</td><td>0</td><td>GGAGGCCGCCGCCGCCGTGG</td><td>CGG</td><td>||||||||[||||||||||||]PAM</td><td>NA</td><td>Intergenic</td><td>NA</td><td>NA</td></tr>
<tr><td>GU305914.1</td><td></td><td>45592</td><td>45614</td><td>-</td><td>4</td><td>GGCGGCTGCCGCCGCCGCCG</td><td>CGG</td><td>||-|||-|[|||||||||--|]PAM</td><td>NA</td><td>Intergenic</td><td>NA</td><td>NA</td></tr>
<tr><td>T32446</td><td>CGCCGCCGCCGTGGCGGAGTTGG</td><td></td><td>1000</td><td></td><td></td><td></td><td></td><td></td><td></td><td></td><td></td><td></td></tr>
<tr><td>Oligo fwd</td><td>CACCCGCCGCCGCCGTGGCGGAGT</td><td></td><td></td><td></td><td></td><td></td><td></td><td></td><td></td><td></td><td></td><td></td></tr>
<tr><td>Oligo rev</td><td>AAACACTCCGCCACGGCGGCGGCG</td><td></td><td></td><td></td><td></td><td></td><td></td><td></td><td></td><td></td><td></td><td></td></tr>
<tr><td>Chromosome</td><td>start</td><td></td><td>end</td><td>strand</td><td>MM</td><td>target_seq</td><td>PAM</td><td>alignment</td><td>distance</td><td>position</td><td>gene name</td><td>gene id</td></tr>
<tr><td>GU305914.1</td><td></td><td>26435</td><td>26457</td><td>-</td><td>0</td><td>CGCCGCCGCCGTGGCGGAGT</td><td>TGG</td><td>||||||||[||||||||||||]PAM</td><td>NA</td><td>Intergenic</td><td>NA</td><td>NA</td></tr>
<tr><td>T32447</td><td>CGCCGCCGTGGCGGAGTTGGAGG</td><td></td><td>1000</td><td></td><td></td><td></td><td></td><td></td><td></td><td></td><td></td><td></td></tr>
<tr><td>Oligo fwd</td><td>CACCCGCCGCCGTGGCGGAGTTGG</td><td></td><td></td><td></td><td></td><td></td><td></td><td></td><td></td><td></td><td></td><td></td></tr>
<tr><td>Oligo rev</td><td>AAACCCAACTCCGCCACGGCGGCG</td><td></td><td></td><td></td><td></td><td></td><td></td><td></td><td></td><td></td><td></td><td></td></tr>
<tr><td>Chromosome</td><td>start</td><td></td><td>end</td><td>strand</td><td>MM</td><td>target_seq</td><td>PAM</td><td>alignment</td><td>distance</td><td>position</td><td>gene name</td><td>gene id</td></tr>
<tr><td>GU305914.1</td><td></td><td>26432</td><td>26454</td><td>-</td><td>0</td><td>CGCCGCCGTGGCGGAGTTGG</td><td>AGG</td><td>||||||||[||||||||||||]PAM</td><td>NA</td><td>Intergenic</td><td>NA</td><td>NA</td></tr>
<tr><td>T32448</td><td>CGCCGTGGCGGAGTTGGAGGCGG</td><td></td><td>1000</td><td></td><td></td><td></td><td></td><td></td><td></td><td></td><td></td><td></td></tr>
<tr><td>Oligo fwd</td><td>CACCCGCCGTGGCGGAGTTGGAGG</td><td></td><td></td><td></td><td></td><td></td><td></td><td></td><td></td><td></td><td></td><td></td></tr>
<tr><td>Oligo rev</td><td>AAACCCTCCAACTCCGCCACGGCG</td><td></td><td></td><td></td><td></td><td></td><td></td><td></td><td></td><td></td><td></td><td></td></tr>
<tr><td>Chromosome</td><td>start</td><td></td><td>end</td><td>strand</td><td>MM</td><td>target_seq</td><td>PAM</td><td>alignment</td><td>distance</td><td>position</td><td>gene name</td><td>gene id</td></tr>
<tr><td>GU305914.1</td><td></td><td>26429</td><td>26451</td><td>-</td><td>0</td><td>CGCCGTGGCGGAGTTGGAGG</td><td>CGG</td><td>||||||||[||||||||||||]PAM</td><td>NA</td><td>Intergenic</td><td>NA</td><td>NA</td></tr>
<tr><td>T32449</td><td>GCCGTGGCGGAGTTGGAGGCGGG</td><td></td><td>1000</td><td></td><td></td><td></td><td></td><td></td><td></td><td></td><td></td><td></td></tr>
<tr><td>Oligo fwd</td><td>CACCGCCGTGGCGGAGTTGGAGGC</td><td></td><td></td><td></td><td></td><td></td><td></td><td></td><td></td><td></td><td></td><td></td></tr>
<tr><td>Oligo rev</td><td>AAACGCCTCCAACTCCGCCACGGC</td><td></td><td></td><td></td><td></td><td></td><td></td><td></td><td></td><td></td><td></td><td></td></tr>
<tr><td>Chromosome</td><td>start</td><td></td><td>end</td><td>strand</td><td>MM</td><td>target_seq</td><td>PAM</td><td>alignment</td><td>distance</td><td>position</td><td>gene name</td><td>gene id</td></tr>
<tr><td>GU305914.1</td><td></td><td>26428</td><td>26450</td><td>-</td><td>0</td><td>GCCGTGGCGGAGTTGGAGGC</td><td>GGG</td><td>||||||||[||||||||||||]PAM</td><td>NA</td><td>Intergenic</td><td>NA</td><td>NA</td></tr>
<tr><td>T32450</td><td>CCGTGGCGGAGTTGGAGGCGGGG</td><td></td><td>1000</td><td></td><td></td><td></td><td></td><td></td><td></td><td></td><td></td><td></td></tr>
<tr><td>Oligo fwd</td><td>CACCCCGTGGCGGAGTTGGAGGCG</td><td></td><td></td><td></td><td></td><td></td><td></td><td></td><td></td><td></td><td></td><td></td></tr>
<tr><td>Oligo rev</td><td>AAACCGCCTCCAACTCCGCCACGG</td><td></td><td></td><td></td><td></td><td></td><td></td><td></td><td></td><td></td><td></td><td></td></tr>
<tr><td>Chromosome</td><td>start</td><td></td><td>end</td><td>strand</td><td>MM</td><td>target_seq</td><td>PAM</td><td>alignment</td><td>distance</td><td>position</td><td>gene name</td><td>gene id</td></tr>
<tr><td>GU305914.1</td><td></td><td>26427</td><td>26449</td><td>-</td><td>0</td><td>CCGTGGCGGAGTTGGAGGCG</td><td>GGG</td><td>||||||||[||||||||||||]PAM</td><td>NA</td><td>Intergenic</td><td>NA</td><td>NA</td></tr>
<tr><td>T32451</td><td>CGTGGCGGAGTTGGAGGCGGGGG</td><td></td><td>1000</td><td></td><td></td><td></td><td></td><td></td><td></td><td></td><td></td><td></td></tr>
<tr><td>Oligo fwd</td><td>CACCCGTGGCGGAGTTGGAGGCGG</td><td></td><td></td><td></td><td></td><td></td><td></td><td></td><td></td><td></td><td></td><td></td></tr>
</table>

<table>
<tr><td>Oligo rev</td><td>AAACCCGCCTCCAACTCCGCCACG</td><td></td><td></td><td></td><td></td><td></td><td></td><td></td><td></td><td></td><td></td><td></td></tr>
<tr><td>Chromosome</td><td>start</td><td></td><td>end</td><td>strand</td><td>MM</td><td>target_seq</td><td>PAM</td><td>alignment</td><td>distance</td><td>position</td><td>gene name</td><td>gene id</td></tr>
<tr><td>GU305914.1</td><td></td><td>26426</td><td>26448</td><td>-</td><td>0</td><td>CGTGGCGGAGTTGGAGGCGG</td><td>GGG</td><td>||||||||[||||||||||||]PAM</td><td>NA</td><td>Intergenic</td><td>NA</td><td>NA</td></tr>
<tr><td>T32452</td><td>CGGAGTTGGAGGCGGGGGCATGG</td><td></td><td>1000</td><td></td><td></td><td></td><td></td><td></td><td></td><td></td><td></td><td></td></tr>
<tr><td>Oligo fwd</td><td>CACCCGGAGTTGGAGGCGGGGGCA</td><td></td><td></td><td></td><td></td><td></td><td></td><td></td><td></td><td></td><td></td><td></td></tr>
<tr><td>Oligo rev</td><td>AAACTGCCCCCGCCTCCAACTCCG</td><td></td><td></td><td></td><td></td><td></td><td></td><td></td><td></td><td></td><td></td><td></td></tr>
<tr><td>Chromosome</td><td>start</td><td></td><td>end</td><td>strand</td><td>MM</td><td>target_seq</td><td>PAM</td><td>alignment</td><td>distance</td><td>position</td><td>gene name</td><td>gene id</td></tr>
<tr><td>GU305914.1</td><td></td><td>26421</td><td>26443</td><td>-</td><td>0</td><td>CGGAGTTGGAGGCGGGGGCA</td><td>TGG</td><td>||||||||[||||||||||||]PAM</td><td>NA</td><td>Intergenic</td><td>NA</td><td>NA</td></tr>
<tr><td>T32453</td><td>GGAGTTGGAGGCGGGGGCATGGG</td><td></td><td>1000</td><td></td><td></td><td></td><td></td><td></td><td></td><td></td><td></td><td></td></tr>
<tr><td>Oligo fwd</td><td>CACCGGAGTTGGAGGCGGGGGCAT</td><td></td><td></td><td></td><td></td><td></td><td></td><td></td><td></td><td></td><td></td><td></td></tr>
<tr><td>Oligo rev</td><td>AAACATGCCCCCGCCTCCAACTCC</td><td></td><td></td><td></td><td></td><td></td><td></td><td></td><td></td><td></td><td></td><td></td></tr>
<tr><td>Chromosome</td><td>start</td><td></td><td>end</td><td>strand</td><td>MM</td><td>target_seq</td><td>PAM</td><td>alignment</td><td>distance</td><td>position</td><td>gene name</td><td>gene id</td></tr>
<tr><td>GU305914.1</td><td></td><td>26420</td><td>26442</td><td>-</td><td>0</td><td>GGAGTTGGAGGCGGGGGCAT</td><td>GGG</td><td>||||||||[||||||||||||]PAM</td><td>NA</td><td>Intergenic</td><td>NA</td><td>NA</td></tr>
<tr><td>T32454</td><td>GAGTTGGAGGCGGGGGCATGGGG</td><td></td><td>1000</td><td></td><td></td><td></td><td></td><td></td><td></td><td></td><td></td><td></td></tr>
<tr><td>Oligo fwd</td><td>CACCGAGTTGGAGGCGGGGGCATG</td><td></td><td></td><td></td><td></td><td></td><td></td><td></td><td></td><td></td><td></td><td></td></tr>
<tr><td>Oligo rev</td><td>AAACCATGCCCCCGCCTCCAACTC</td><td></td><td></td><td></td><td></td><td></td><td></td><td></td><td></td><td></td><td></td><td></td></tr>
<tr><td>Chromosome</td><td>start</td><td></td><td>end</td><td>strand</td><td>MM</td><td>target_seq</td><td>PAM</td><td>alignment</td><td>distance</td><td>position</td><td>gene name</td><td>gene id</td></tr>
<tr><td>GU305914.1</td><td></td><td>26419</td><td>26441</td><td>-</td><td>0</td><td>GAGTTGGAGGCGGGGGCATG</td><td>GGG</td><td>||||||||[||||||||||||]PAM</td><td>NA</td><td>Intergenic</td><td>NA</td><td>NA</td></tr>
<tr><td>T32455</td><td>GACGACGACGACGTCGTCTCCGG</td><td></td><td>1000</td><td></td><td></td><td></td><td></td><td></td><td></td><td></td><td></td><td></td></tr>
<tr><td>Oligo fwd</td><td>CACCGACGACGACGACGTCGTCTC</td><td></td><td></td><td></td><td></td><td></td><td></td><td></td><td></td><td></td><td></td><td></td></tr>
<tr><td>Oligo rev</td><td>AAACGAGACGACGTCGTCGTCGTC</td><td></td><td></td><td></td><td></td><td></td><td></td><td></td><td></td><td></td><td></td><td></td></tr>
<tr><td>Chromosome</td><td>start</td><td></td><td>end</td><td>strand</td><td>MM</td><td>target_seq</td><td>PAM</td><td>alignment</td><td>distance</td><td>position</td><td>gene name</td><td>gene id</td></tr>
<tr><td>GU305914.1</td><td></td><td>26359</td><td>26381</td><td>-</td><td>0</td><td>GACGACGACGACGTCGTCTC</td><td>CGG</td><td>||||||||[||||||||||||]PAM</td><td>NA</td><td>Intergenic</td><td>NA</td><td>NA</td></tr>
<tr><td>T32456</td><td>ACGACGACGACGTCGTCTCCGGG</td><td></td><td>1000</td><td></td><td></td><td></td><td></td><td></td><td></td><td></td><td></td><td></td></tr>
<tr><td>Oligo fwd</td><td>CACCACGACGACGACGTCGTCTCC</td><td></td><td></td><td></td><td></td><td></td><td></td><td></td><td></td><td></td><td></td><td></td></tr>
<tr><td>Oligo rev</td><td>AAACGGAGACGACGTCGTCGTCGT</td><td></td><td></td><td></td><td></td><td></td><td></td><td></td><td></td><td></td><td></td><td></td></tr>
<tr><td>Chromosome</td><td>start</td><td></td><td>end</td><td>strand</td><td>MM</td><td>target_seq</td><td>PAM</td><td>alignment</td><td>distance</td><td>position</td><td>gene name</td><td>gene id</td></tr>
<tr><td>GU305914.1</td><td></td><td>26358</td><td>26380</td><td>-</td><td>0</td><td>ACGACGACGACGTCGTCTCC</td><td>GGG</td><td>||||||||[||||||||||||]PAM</td><td>NA</td><td>Intergenic</td><td>NA</td><td>NA</td></tr>
<tr><td>T32457</td><td>CGACGACGACGTCGTCTCCGGGG</td><td></td><td>1000</td><td></td><td></td><td></td><td></td><td></td><td></td><td></td><td></td><td></td></tr>
<tr><td>Oligo fwd</td><td>CACCCGACGACGACGTCGTCTCCG</td><td></td><td></td><td></td><td></td><td></td><td></td><td></td><td></td><td></td><td></td><td></td></tr>
<tr><td>Oligo rev</td><td>AAACCGGAGACGACGTCGTCGTCG</td><td></td><td></td><td></td><td></td><td></td><td></td><td></td><td></td><td></td><td></td><td></td></tr>
<tr><td>Chromosome</td><td>start</td><td></td><td>end</td><td>strand</td><td>MM</td><td>target_seq</td><td>PAM</td><td>alignment</td><td>distance</td><td>position</td><td>gene name</td><td>gene id</td></tr>
<tr><td>GU305914.1</td><td></td><td>26357</td><td>26379</td><td>-</td><td>0</td><td>CGACGACGACGTCGTCTCCG</td><td>GGG</td><td>||||||||[||||||||||||]PAM</td><td>NA</td><td>Intergenic</td><td>NA</td><td>NA</td></tr>
<tr><td>T32458</td><td>GACGACGACGTCGTCTCCGGGGG</td><td></td><td>1000</td><td></td><td></td><td></td><td></td><td></td><td></td><td></td><td></td><td></td></tr>
<tr><td>Oligo fwd</td><td>CACCGACGACGACGTCGTCTCCGG</td><td></td><td></td><td></td><td></td><td></td><td></td><td></td><td></td><td></td><td></td><td></td></tr>
<tr><td>Oligo rev</td><td>AAACCCGGAGACGACGTCGTCGTC</td><td></td><td></td><td></td><td></td><td></td><td></td><td></td><td></td><td></td><td></td><td></td></tr>
<tr><td>Chromosome</td><td>start</td><td></td><td>end</td><td>strand</td><td>MM</td><td>target_seq</td><td>PAM</td><td>alignment</td><td>distance</td><td>position</td><td>gene name</td><td>gene id</td></tr>
<tr><td>GU305914.1</td><td></td><td>26356</td><td>26378</td><td>-</td><td>0</td><td>GACGACGACGTCGTCTCCGG</td><td>GGG</td><td>||||||||[||||||||||||]PAM</td><td>NA</td><td>Intergenic</td><td>NA</td><td>NA</td></tr>
<tr><td>GU305914.1</td><td></td><td>60614</td><td>60636</td><td>-</td><td>4</td><td>GGCGTCTCCGTCGTCTCCGG</td><td>CGG</td><td>|-||-|--[||||||||||||]PAM</td><td>NA</td><td>Intergenic</td><td>NA</td><td>NA</td></tr>
<tr><td>T32459</td><td>ACGACGACGTCGTCTCCGGGGGG</td><td></td><td>1000</td><td></td><td></td><td></td><td></td><td></td><td></td><td></td><td></td><td></td></tr>
<tr><td>Oligo fwd</td><td>CACCACGACGACGTCGTCTCCGGG</td><td></td><td></td><td></td><td></td><td></td><td></td><td></td><td></td><td></td><td></td><td></td></tr>
<tr><td>Oligo rev</td><td>AAACCCCGGAGACGACGTCGTCGT</td><td></td><td></td><td></td><td></td><td></td><td></td><td></td><td></td><td></td><td></td><td></td></tr>
<tr><td>Chromosome</td><td>start</td><td></td><td>end</td><td>strand</td><td>MM</td><td>target_seq</td><td>PAM</td><td>alignment</td><td>distance</td><td>position</td><td>gene name</td><td>gene id</td></tr>
<tr><td>GU305914.1</td><td></td><td>26355</td><td>26377</td><td>-</td><td>0</td><td>ACGACGACGTCGTCTCCGGG</td><td>GGG</td><td>||||||||[||||||||||||]PAM</td><td>NA</td><td>Intergenic</td><td>NA</td><td>NA</td></tr>
<tr><td>T32460</td><td>GTCATCATCCGACGAGTCGTCGG</td><td></td><td>1000</td><td></td><td></td><td></td><td></td><td></td><td></td><td></td><td></td><td></td></tr>
<tr><td>Oligo fwd</td><td>CACCGTCATCATCCGACGAGTCGT</td><td></td><td></td><td></td><td></td><td></td><td></td><td></td><td></td><td></td><td></td><td></td></tr>
<tr><td>Oligo rev</td><td>AAACACGACTCGTCGGATGATGAC</td><td></td><td></td><td></td><td></td><td></td><td></td><td></td><td></td><td></td><td></td><td></td></tr>
<tr><td>Chromosome</td><td>start</td><td></td><td>end</td><td>strand</td><td>MM</td><td>target_seq</td><td>PAM</td><td>alignment</td><td>distance</td><td>position</td><td>gene name</td><td>gene id</td></tr>
<tr><td>GU305914.1</td><td></td><td>26309</td><td>26331</td><td>-</td><td>0</td><td>GTCATCATCCGACGAGTCGT</td><td>CGG</td><td>||||||||[||||||||||||]PAM</td><td>NA</td><td>Intergenic</td><td>NA</td><td>NA</td></tr>
<tr><td>T32461</td><td>CGTCGGTGACGAACATACGACGG</td><td></td><td>1000</td><td></td><td></td><td></td><td></td><td></td><td></td><td></td><td></td><td></td></tr>
<tr><td>Oligo fwd</td><td>CACCCGTCGGTGACGAACATACGA</td><td></td><td></td><td></td><td></td><td></td><td></td><td></td><td></td><td></td><td></td><td></td></tr>
<tr><td>Oligo rev</td><td>AAACTCGTATGTTCGTCACCGACG</td><td></td><td></td><td></td><td></td><td></td><td></td><td></td><td></td><td></td><td></td><td></td></tr>
<tr><td>Chromosome</td><td>start</td><td></td><td>end</td><td>strand</td><td>MM</td><td>target_seq</td><td>PAM</td><td>alignment</td><td>distance</td><td>position</td><td>gene name</td><td>gene id</td></tr>
<tr><td>GU305914.1</td><td></td><td>26292</td><td>26314</td><td>-</td><td>0</td><td>CGTCGGTGACGAACATACGA</td><td>CGG</td><td>||||||||[||||||||||||]PAM</td><td>NA</td><td>Intergenic</td><td>NA</td><td>NA</td></tr>
<tr><td>T32462</td><td>GACGAACATACGACGGTCCTTGG</td><td></td><td>1000</td><td></td><td></td><td></td><td></td><td></td><td></td><td></td><td></td><td></td></tr>
<tr><td>Oligo fwd</td><td>CACCGACGAACATACGACGGTCCT</td><td></td><td></td><td></td><td></td><td></td><td></td><td></td><td></td><td></td><td></td><td></td></tr>
<tr><td>Oligo rev</td><td>AAACAGGACCGTCGTATGTTCGTC</td><td></td><td></td><td></td><td></td><td></td><td></td><td></td><td></td><td></td><td></td><td></td></tr>
<tr><td>Chromosome</td><td>start</td><td></td><td>end</td><td>strand</td><td>MM</td><td>target_seq</td><td>PAM</td><td>alignment</td><td>distance</td><td>position</td><td>gene name</td><td>gene id</td></tr>
<tr><td>GU305914.1</td><td></td><td>26285</td><td>26307</td><td>-</td><td>0</td><td>GACGAACATACGACGGTCCT</td><td>TGG</td><td>||||||||[||||||||||||]PAM</td><td>NA</td><td>Intergenic</td><td>NA</td><td>NA</td></tr>
<tr><td>T32463</td><td>GAACATACGACGGTCCTTGGAGG</td><td></td><td>1000</td><td></td><td></td><td></td><td></td><td></td><td></td><td></td><td></td><td></td></tr>
<tr><td>Oligo fwd</td><td>CACCGAACATACGACGGTCCTTGG</td><td></td><td></td><td></td><td></td><td></td><td></td><td></td><td></td><td></td><td></td><td></td></tr>
<tr><td>Oligo rev</td><td>AAACCCAAGGACCGTCGTATGTTC</td><td></td><td></td><td></td><td></td><td></td><td></td><td></td><td></td><td></td><td></td><td></td></tr>
<tr><td>Chromosome</td><td>start</td><td></td><td>end</td><td>strand</td><td>MM</td><td>target_seq</td><td>PAM</td><td>alignment</td><td>distance</td><td>position</td><td>gene name</td><td>gene id</td></tr>
<tr><td>GU305914.1</td><td></td><td>26282</td><td>26304</td><td>-</td><td>0</td><td>GAACATACGACGGTCCTTGG</td><td>AGG</td><td>||||||||[||||||||||||]PAM</td><td>NA</td><td>Intergenic</td><td>NA</td><td>NA</td></tr>
<tr><td>T32464</td><td>GACGGTCCTTGGAGGATCGACGG</td><td></td><td>1000</td><td></td><td></td><td></td><td></td><td></td><td></td><td></td><td></td><td></td></tr>
<tr><td>Oligo fwd</td><td>CACCGACGGTCCTTGGAGGATCGA</td><td></td><td></td><td></td><td></td><td></td><td></td><td></td><td></td><td></td><td></td><td></td></tr>
<tr><td>Oligo rev</td><td>AAACTCGATCCTCCAAGGACCGTC</td><td></td><td></td><td></td><td></td><td></td><td></td><td></td><td></td><td></td><td></td><td></td></tr>
<tr><td>Chromosome</td><td>start</td><td></td><td>end</td><td>strand</td><td>MM</td><td>target_seq</td><td>PAM</td><td>alignment</td><td>distance</td><td>position</td><td>gene name</td><td>gene id</td></tr>
<tr><td>GU305914.1</td><td></td><td>26274</td><td>26296</td><td>-</td><td>0</td><td>GACGGTCCTTGGAGGATCGA</td><td>CGG</td><td>||||||||[||||||||||||]PAM</td><td>NA</td><td>Intergenic</td><td>NA</td><td>NA</td></tr>
<tr><td>T32465</td><td>AGGATCGACGGTTCATCGTTCGG</td><td></td><td>1000</td><td></td><td></td><td></td><td></td><td></td><td></td><td></td><td></td><td></td></tr>
<tr><td>Oligo fwd</td><td>CACCAGGATCGACGGTTCATCGTT</td><td></td><td></td><td></td><td></td><td></td><td></td><td></td><td></td><td></td><td></td><td></td></tr>
<tr><td>Oligo rev</td><td>AAACAACGATGAACCGTCGATCCT</td><td></td><td></td><td></td><td></td><td></td><td></td><td></td><td></td><td></td><td></td><td></td></tr>
<tr><td>Chromosome</td><td>start</td><td></td><td>end</td><td>strand</td><td>MM</td><td>target_seq</td><td>PAM</td><td>alignment</td><td>distance</td><td>position</td><td>gene name</td><td>gene id</td></tr>
<tr><td>GU305914.1</td><td></td><td>26262</td><td>26284</td><td>-</td><td>0</td><td>AGGATCGACGGTTCATCGTT</td><td>CGG</td><td>||||||||[||||||||||||]PAM</td><td>NA</td><td>Intergenic</td><td>NA</td><td>NA</td></tr>
</table>

| | | | | | | | | | | | | |
|---|---|---|---|---|---|---|---|---|---|---|---|---|
| T32466 | GGATCGACGGTTCATCGTTCGGG | | 1000 | | | | | | | | | |
| Oligo fwd | CACCGGATCGACGGTTCATCGTTC | | | | | | | | | | | |
| Oligo rev | AAACGAACGATGAACCGTCGATCC | | | | | | | | | | | |
| Chromosome | start | | end | strand | MM | target_seq | PAM | alignment | distance | position | gene name | gene id |
| GU305914.1 | | 26261 | 26283 | - | 0 | GGATCGACGGTTCATCGTTC | GGG | \|\|\|\|\|\|\|\|[\|\|\|\|\|\|\|\|\|\|\|\|]PAM | NA | Intergenic | NA | NA |
| T32467 | CGGTTCATCGTTCGGGTGAGTGG | | 1000 | | | | | | | | | |
| Oligo fwd | CACCCGGTTCATCGTTCGGGTGAG | | | | | | | | | | | |
| Oligo rev | AAACCTCACCCGAACGATGAACCG | | | | | | | | | | | |
| Chromosome | start | | end | strand | MM | target_seq | PAM | alignment | distance | position | gene name | gene id |
| GU305914.1 | | 26254 | 26276 | - | 0 | CGGTTCATCGTTCGGGTGAG | TGG | \|\|\|\|\|\|\|\|[\|\|\|\|\|\|\|\|\|\|\|\|]PAM | NA | Intergenic | NA | NA |
| T32468 | GGTTCATCGTTCGGGTGAGTGGG | | 1000 | | | | | | | | | |
| Oligo fwd | CACCGGTTCATCGTTCGGGTGAGT | | | | | | | | | | | |
| Oligo rev | AAACACTCACCCGAACGATGAACC | | | | | | | | | | | |
| Chromosome | start | | end | strand | MM | target_seq | PAM | alignment | distance | position | gene name | gene id |
| GU305914.1 | | 26253 | 26275 | - | 0 | GGTTCATCGTTCGGGTGAGT | GGG | \|\|\|\|\|\|\|\|[\|\|\|\|\|\|\|\|\|\|\|\|]PAM | NA | Intergenic | NA | NA |
| T32469 | GTTCATCGTTCGGGTGAGTGGGG | | 1000 | | | | | | | | | |
| Oligo fwd | CACCGTTCATCGTTCGGGTGAGTG | | | | | | | | | | | |
| Oligo rev | AAACCACTCACCCGAACGATGAAC | | | | | | | | | | | |
| Chromosome | start | | end | strand | MM | target_seq | PAM | alignment | distance | position | gene name | gene id |
| GU305914.1 | | 26252 | 26274 | - | 0 | GTTCATCGTTCGGGTGAGTG | GGG | \|\|\|\|\|\|\|\|[\|\|\|\|\|\|\|\|\|\|\|\|]PAM | NA | Intergenic | NA | NA |
| T32470 | TTCATCGTTCGGGTGAGTGGGGG | | 1000 | | | | | | | | | |
| Oligo fwd | CACCTTCATCGTTCGGGTGAGTGG | | | | | | | | | | | |
| Oligo rev | AAACCCACTCACCCGAACGATGAA | | | | | | | | | | | |
| Chromosome | start | | end | strand | MM | target_seq | PAM | alignment | distance | position | gene name | gene id |
| GU305914.1 | | 26251 | 26273 | - | 0 | TTCATCGTTCGGGTGAGTGG | GGG | \|\|\|\|\|\|\|\|[\|\|\|\|\|\|\|\|\|\|\|\|]PAM | NA | Intergenic | NA | NA |
| T32471 | TTCGGGTGAGTGGGGGTTCTCGG | | 1000 | | | | | | | | | |
| Oligo fwd | CACCTTCGGGTGAGTGGGGGTTCT | | | | | | | | | | | |
| Oligo rev | AAACAGAACCCCCACTCACCCGAA | | | | | | | | | | | |
| Chromosome | start | | end | strand | MM | target_seq | PAM | alignment | distance | position | gene name | gene id |
| GU305914.1 | | 26244 | 26266 | - | 0 | TTCGGGTGAGTGGGGGTTCT | CGG | \|\|\|\|\|\|\|\|[\|\|\|\|\|\|\|\|\|\|\|\|]PAM | NA | Intergenic | NA | NA |
| T32472 | GTGAGTGGGGGTTCTCGGACAGG | | 1000 | | | | | | | | | |
| Oligo fwd | CACCGTGAGTGGGGGTTCTCGGAC | | | | | | | | | | | |
| Oligo rev | AAACGTCCGAGAACCCCCACTCAC | | | | | | | | | | | |
| Chromosome | start | | end | strand | MM | target_seq | PAM | alignment | distance | position | gene name | gene id |
| GU305914.1 | | 26239 | 26261 | - | 0 | GTGAGTGGGGGTTCTCGGAC | AGG | \|\|\|\|\|\|\|\|[\|\|\|\|\|\|\|\|\|\|\|\|]PAM | NA | Intergenic | NA | NA |
| T32473 | GGTTCTCGGACAGGCGTTCGAGG | | 1000 | | | | | | | | | |
| Oligo fwd | CACCGGTTCTCGGACAGGCGTTCG | | | | | | | | | | | |
| Oligo rev | AAACCGAACGCCTGTCCGAGAACC | | | | | | | | | | | |
| Chromosome | start | | end | strand | MM | target_seq | PAM | alignment | distance | position | gene name | gene id |
| GU305914.1 | | 26230 | 26252 | - | 0 | GGTTCTCGGACAGGCGTTCG | AGG | \|\|\|\|\|\|\|\|[\|\|\|\|\|\|\|\|\|\|\|\|]PAM | NA | Intergenic | NA | NA |
| T32474 | GACAGGCGTTCGAGGTCGAAAGG | | 1000 | | | | | | | | | |
| Oligo fwd | CACCGACAGGCGTTCGAGGTCGAA | | | | | | | | | | | |
| Oligo rev | AAACTTCGACCTCGAACGCCTGTC | | | | | | | | | | | |
| Chromosome | start | | end | strand | MM | target_seq | PAM | alignment | distance | position | gene name | gene id |
| GU305914.1 | | 26222 | 26244 | - | 0 | GACAGGCGTTCGAGGTCGAA | AGG | \|\|\|\|\|\|\|\|[\|\|\|\|\|\|\|\|\|\|\|\|]PAM | NA | Intergenic | NA | NA |
| T32475 | ACAGGCGTTCGAGGTCGAAAGGG | | 1000 | | | | | | | | | |
| Oligo fwd | CACCACAGGCGTTCGAGGTCGAAA | | | | | | | | | | | |
| Oligo rev | AAACTTTCGACCTCGAACGCCTGT | | | | | | | | | | | |
| Chromosome | start | | end | strand | MM | target_seq | PAM | alignment | distance | position | gene name | gene id |
| GU305914.1 | | 26221 | 26243 | - | 0 | ACAGGCGTTCGAGGTCGAAA | GGG | \|\|\|\|\|\|\|\|[\|\|\|\|\|\|\|\|\|\|\|\|]PAM | NA | Intergenic | NA | NA |
| T32476 | CGAGGTCGAAAGGGACGCGCCGG | | 1000 | | | | | | | | | |
| Oligo fwd | CACCCGAGGTCGAAAGGGACGCGC | | | | | | | | | | | |
| Oligo rev | AAACGCGCGTCCCTTTCGACCTCG | | | | | | | | | | | |
| Chromosome | start | | end | strand | MM | target_seq | PAM | alignment | distance | position | gene name | gene id |
| GU305914.1 | | 26212 | 26234 | - | 0 | CGAGGTCGAAAGGGACGCGC | CGG | \|\|\|\|\|\|\|\|[\|\|\|\|\|\|\|\|\|\|\|\|]PAM | NA | Intergenic | NA | NA |
| T32477 | TCGAAAGGGACGCGCCGGCACGG | | 1000 | | | | | | | | | |
| Oligo fwd | CACCTCGAAAGGGACGCGCCGGCA | | | | | | | | | | | |
| Oligo rev | AAACTGCCGGCGCGTCCCTTTCGA | | | | | | | | | | | |
| Chromosome | start | | end | strand | MM | target_seq | PAM | alignment | distance | position | gene name | gene id |
| GU305914.1 | | 26207 | 26229 | - | 0 | TCGAAAGGGACGCGCCGGCA | CGG | \|\|\|\|\|\|\|\|[\|\|\|\|\|\|\|\|\|\|\|\|]PAM | NA | Intergenic | NA | NA |
| T32478 | CGAAAGGGACGCGCCGGCACGGG | | 1000 | | | | | | | | | |
| Oligo fwd | CACCCGAAAGGGACGCGCCGGCAC | | | | | | | | | | | |
| Oligo rev | AAACGTGCCGGCGCGTCCCTTTCG | | | | | | | | | | | |
| Chromosome | start | | end | strand | MM | target_seq | PAM | alignment | distance | position | gene name | gene id |
| GU305914.1 | | 26206 | 26228 | - | 0 | CGAAAGGGACGCGCCGGCAC | GGG | \|\|\|\|\|\|\|\|[\|\|\|\|\|\|\|\|\|\|\|\|]PAM | NA | Intergenic | NA | NA |
| T32479 | GAAAGGGACGCGCCGGCACGGGG | | 1000 | | | | | | | | | |
| Oligo fwd | CACCGAAAGGGACGCGCCGGCACG | | | | | | | | | | | |
| Oligo rev | AAACCGTGCCGGCGCGTCCCTTTC | | | | | | | | | | | |
| Chromosome | start | | end | strand | MM | target_seq | PAM | alignment | distance | position | gene name | gene id |
| GU305914.1 | | 26205 | 26227 | - | 0 | GAAAGGGACGCGCCGGCACG | GGG | \|\|\|\|\|\|\|\|[\|\|\|\|\|\|\|\|\|\|\|\|]PAM | NA | Intergenic | NA | NA |
| T32480 | GACGCGCCGGCACGGGGAATAGG | | 1000 | | | | | | | | | |
| Oligo fwd | CACCGACGCGCCGGCACGGGGAAT | | | | | | | | | | | |
| Oligo rev | AAACATTCCCCGTGCCGGCGCGTC | | | | | | | | | | | |
| Chromosome | start | | end | strand | MM | target_seq | PAM | alignment | distance | position | gene name | gene id |
| GU305914.1 | | 26199 | 26221 | - | 0 | GACGCGCCGGCACGGGGAAT | AGG | \|\|\|\|\|\|\|\|[\|\|\|\|\|\|\|\|\|\|\|\|]PAM | NA | Intergenic | NA | NA |

T32481 CGCCGGCACGGGGAATAGGACGG 1000
Oligo fwd CACCCGCCGGCACGGGGAATAGGA
Oligo rev AAACTCCTATTCCCCGTGCCGGCG
Chromosome start end strand MM target_seq PAM alignment distance position gene name gene id
GU305914.1 26195 26217 - 0 CGCCGGCACGGGGAATAGGA CGG ||||||||[||||||||||||]PAM NA Intergenic NA NA

T32482 GGCACGGGGAATAGGACGGCTGG 1000
Oligo fwd CACCGGCACGGGGAATAGGACGGC
Oligo rev AAACGCCGTCCTATTCCCCGTGCC
Chromosome start end strand MM target_seq PAM alignment distance position gene name gene id
GU305914.1 26191 26213 - 0 GGCACGGGGAATAGGACGGC TGG ||||||||[||||||||||||]PAM NA Intergenic NA NA

T32483 GACGGCTGGCACGCAGACCCCGG 1000
Oligo fwd CACCGACGGCTGGCACGCAGACCC
Oligo rev AAACGGGTCTGCGTGCCAGCCGTC
Chromosome start end strand MM target_seq PAM alignment distance position gene name gene id
GU305914.1 26177 26199 - 0 GACGGCTGGCACGCAGACCC CGG ||||||||[||||||||||||]PAM NA Intergenic NA NA

T32484 TAGTTGCACGTCTCCATCTCTGG 1000
Oligo fwd CACCTAGTTGCACGTCTCCATCTC
Oligo rev AAACGAGATGGAGACGTGCAACTA
Chromosome start end strand MM target_seq PAM alignment distance position gene name gene id
GU305914.1 26138 26160 - 0 TAGTTGCACGTCTCCATCTC TGG ||||||||[||||||||||||]PAM NA Intergenic NA NA

T32485 AGTTGCACGTCTCCATCTCTGGG 1000
Oligo fwd CACCAGTTGCACGTCTCCATCTCT
Oligo rev AAACAGAGATGGAGACGTGCAACT
Chromosome start end strand MM target_seq PAM alignment distance position gene name gene id
GU305914.1 26137 26159 - 0 AGTTGCACGTCTCCATCTCT GGG ||||||||[||||||||||||]PAM NA Intergenic NA NA

T32486 GCACGTCTCCATCTCTGGGCAGG 1000
Oligo fwd CACCGCACGTCTCCATCTCTGGGC
Oligo rev AAACGCCCAGAGATGGAGACGTGC
Chromosome start end strand MM target_seq PAM alignment distance position gene name gene id
GU305914.1 26133 26155 - 0 GCACGTCTCCATCTCTGGGC AGG ||||||||[||||||||||||]PAM NA Intergenic NA NA

T32487 CACGTCTCCATCTCTGGGCAGGG 1000
Oligo fwd CACCCACGTCTCCATCTCTGGGCA
Oligo rev AAACTGCCCAGAGATGGAGACGTG
Chromosome start end strand MM target_seq PAM alignment distance position gene name gene id
GU305914.1 26132 26154 - 0 CACGTCTCCATCTCTGGGCA GGG ||||||||[||||||||||||]PAM NA Intergenic NA NA

T32488 CTCCATCTCTGGGCAGGGTTCGG 1000
Oligo fwd CACCCTCCATCTCTGGGCAGGGTT
Oligo rev AAACAACCCTGCCCAGAGATGGAG
Chromosome start end strand MM target_seq PAM alignment distance position gene name gene id
GU305914.1 26127 26149 - 0 CTCCATCTCTGGGCAGGGTT CGG ||||||||[||||||||||||]PAM NA Intergenic NA NA

T32489 CTCTGGGCAGGGTTCGGATGCGG 1000
Oligo fwd CACCCTCTGGGCAGGGTTCGGATG
Oligo rev AAACCATCCGAACCCTGCCCAGAG
Chromosome start end strand MM target_seq PAM alignment distance position gene name gene id
GU305914.1 26121 26143 - 0 CTCTGGGCAGGGTTCGGATG CGG ||||||||[||||||||||||]PAM NA Intergenic NA NA

T32490 GCAGGGTTCGGATGCGGAGATGG 1000
Oligo fwd CACCGCAGGGTTCGGATGCGGAGA
Oligo rev AAACTCTCCGCATCCGAACCCTGC
Chromosome start end strand MM target_seq PAM alignment distance position gene name gene id
GU305914.1 26115 26137 - 0 GCAGGGTTCGGATGCGGAGA TGG ||||||||[||||||||||||]PAM NA Intergenic NA NA
GU305914.1 170246 170268 + 4 GACGGGTTCGGATTCGGGGA CGG |--|||||[|||||-|||-||]PAM NA Intergenic NA NA

T32491 GGATGCGGAGATGGATTCGACGG 1000
Oligo fwd CACCGGATGCGGAGATGGATTCGA
Oligo rev AAACTCGAATCCATCTCCGCATCC
Chromosome start end strand MM target_seq PAM alignment distance position gene name gene id
GU305914.1 26106 26128 - 0 GGATGCGGAGATGGATTCGA CGG ||||||||[||||||||||||]PAM NA Intergenic NA NA

T32492 GATGCGGAGATGGATTCGACGGG 1000
Oligo fwd CACCGATGCGGAGATGGATTCGAC
Oligo rev AAACGTCGAATCCATCTCCGCATC
Chromosome start end strand MM target_seq PAM alignment distance position gene name gene id
GU305914.1 26105 26127 - 0 GATGCGGAGATGGATTCGAC GGG ||||||||[||||||||||||]PAM NA Intergenic NA NA

T32493 ATGCGGAGATGGATTCGACGGGG 1000
Oligo fwd CACCATGCGGAGATGGATTCGACG
Oligo rev AAACCGTCGAATCCATCTCCGCAT
Chromosome start end strand MM target_seq PAM alignment distance position gene name gene id
GU305914.1 26104 26126 - 0 ATGCGGAGATGGATTCGACG GGG ||||||||[||||||||||||]PAM NA Intergenic NA NA

T32494 ATGGATTCGACGGGGAAAATCGG 1000
Oligo fwd CACCATGGATTCGACGGGGAAAAT
Oligo rev AAACATTTTCCCCGTCGAATCCAT
Chromosome start end strand MM target_seq PAM alignment distance position gene name gene id
GU305914.1 26096 26118 - 0 ATGGATTCGACGGGGAAAAT CGG ||||||||[||||||||||||]PAM NA Intergenic NA NA

T32495 CGACGGGGAAAATCGGCTCGCGG 1000
Oligo fwd CACCCGACGGGGAAAATCGGCTCG
Oligo rev AAACCGAGCCGATTTTCCCCGTCG

| | | | | | | | | | | | |
|---|---|---|---|---|---|---|---|---|---|---|---|
| Chromosome | start | end | strand | MM | target_seq | PAM | alignment | distance | position | gene name | gene id |
| GU305914.1 | 26089 | 26111 | - | 0 | CGACGGGGAAAATCGGCTCG | CGG | \|\|\|\|\|\|\|\|[\|\|\|\|\|\|\|\|\|\|\|\|]PAM | NA | Intergenic | NA | NA |
| T32496 | CGACGCCGCGTGCGAGCCGCTGG | 1000 | | | | | | | | | |
| Oligo fwd | CACCCGACGCCGCGTGCGAGCCGC | | | | | | | | | | |
| Oligo rev | AAACGCGGCTCGCACGCGGCGTCG | | | | | | | | | | |
| Chromosome | start | end | strand | MM | target_seq | PAM | alignment | distance | position | gene name | gene id |
| GU305914.1 | 26064 | 26086 | - | 0 | CGACGCCGCGTGCGAGCCGC | TGG | \|\|\|\|\|\|\|\|[\|\|\|\|\|\|\|\|\|\|\|\|]PAM | NA | Intergenic | NA | NA |
| GU305914.1 | 113607 | 113629 | - | 3 | CGACGTCGCGAGGGAGCCGC | AGG | \|\|\|\|\|-\|\|[\|\|-\|-\|\|\|\|\|\|\|]PAM | NA | Intergenic | NA | NA |
| T32497 | CGCCGCGTGCGAGCCGCTGGTGG | 1000 | | | | | | | | | |
| Oligo fwd | CACCCGCCGCGTGCGAGCCGCTGG | | | | | | | | | | |
| Oligo rev | AAACCCAGCGGCTCGCACGCGGCG | | | | | | | | | | |
| Chromosome | start | end | strand | MM | target_seq | PAM | alignment | distance | position | gene name | gene id |
| GU305914.1 | 26061 | 26083 | - | 0 | CGCCGCGTGCGAGCCGCTGG | TGG | \|\|\|\|\|\|\|\|[\|\|\|\|\|\|\|\|\|\|\|\|]PAM | NA | Intergenic | NA | NA |
| GU305914.1 | 64532 | 64554 | + | 4 | CGCCGTGGCCGAGCTGCTGG | AGG | \|\|\|\|\|-\|-[-\|\|\|\|\|-\|\|\|\|\|]PAM | NA | Intergenic | NA | NA |
| T32498 | GCCGCTGGTGGACGACGCCGAGG | 1000 | | | | | | | | | |
| Oligo fwd | CACCGCCGCTGGTGGACGACGCCG | | | | | | | | | | |
| Oligo rev | AAACCGGCGTCGTCCACCAGCGGC | | | | | | | | | | |
| Chromosome | start | end | strand | MM | target_seq | PAM | alignment | distance | position | gene name | gene id |
| GU305914.1 | 26049 | 26071 | - | 0 | GCCGCTGGTGGACGACGCCG | AGG | \|\|\|\|\|\|\|\|[\|\|\|\|\|\|\|\|\|\|\|\|]PAM | NA | Intergenic | NA | NA |
| GU305914.1 | 77515 | 77537 | + | 4 | GCTGCTGACGGACGACGACG | CGG | \|\|-\|\|\|\|-[-\|\|\|\|\|\|\|\|-\|\|]PAM | NA | Intergenic | NA | NA |
| T32499 | GGTGGACGACGCCGAGGCCACGG | 1000 | | | | | | | | | |
| Oligo fwd | CACCGGTGGACGACGCCGAGGCCA | | | | | | | | | | |
| Oligo rev | AAACTGGCCTCGGCGTCGTCCACC | | | | | | | | | | |
| Chromosome | start | end | strand | MM | target_seq | PAM | alignment | distance | position | gene name | gene id |
| GU305914.1 | 26043 | 26065 | - | 0 | GGTGGACGACGCCGAGGCCA | CGG | \|\|\|\|\|\|\|\|[\|\|\|\|\|\|\|\|\|\|\|\|]PAM | NA | Intergenic | NA | NA |
| T32500 | CAGCGACGTGCTGTACGAACTGG | 1000 | | | | | | | | | |
| Oligo fwd | CACCCAGCGACGTGCTGTACGAAC | | | | | | | | | | |
| Oligo rev | AAACGTTCGTACAGCACGTCGCTG | | | | | | | | | | |
| Chromosome | start | end | strand | MM | target_seq | PAM | alignment | distance | position | gene name | gene id |
| GU305914.1 | 26019 | 26041 | - | 0 | CAGCGACGTGCTGTACGAAC | TGG | \|\|\|\|\|\|\|\|[\|\|\|\|\|\|\|\|\|\|\|\|]PAM | NA | Intergenic | NA | NA |
| T32501 | TACGAACTGGCTCATGCCGCCGG | 1000 | | | | | | | | | |
| Oligo fwd | CACCTACGAACTGGCTCATGCCGC | | | | | | | | | | |
| Oligo rev | AAACGCGGCATGAGCCAGTTCGTA | | | | | | | | | | |
| Chromosome | start | end | strand | MM | target_seq | PAM | alignment | distance | position | gene name | gene id |
| GU305914.1 | 26006 | 26028 | - | 0 | TACGAACTGGCTCATGCCGC | CGG | \|\|\|\|\|\|\|\|[\|\|\|\|\|\|\|\|\|\|\|\|]PAM | NA | Intergenic | NA | NA |
| T32502 | AACTGGCTCATGCCGCCGGCCGG | 1000 | | | | | | | | | |
| Oligo fwd | CACCAACTGGCTCATGCCGCCGGC | | | | | | | | | | |
| Oligo rev | AAACGCCGGCGGCATGAGCCAGTT | | | | | | | | | | |
| Chromosome | start | end | strand | MM | target_seq | PAM | alignment | distance | position | gene name | gene id |
| GU305914.1 | 26002 | 26024 | - | 0 | AACTGGCTCATGCCGCCGGC | CGG | \|\|\|\|\|\|\|\|[\|\|\|\|\|\|\|\|\|\|\|\|]PAM | NA | Intergenic | NA | NA |
| T32503 | ACTGGCTCATGCCGCCGGCCGGG | 1000 | | | | | | | | | |
| Oligo fwd | CACCACTGGCTCATGCCGCCGGCC | | | | | | | | | | |
| Oligo rev | AAACGGCCGGCGGCATGAGCCAGT | | | | | | | | | | |
| Chromosome | start | end | strand | MM | target_seq | PAM | alignment | distance | position | gene name | gene id |
| GU305914.1 | 26001 | 26023 | - | 0 | ACTGGCTCATGCCGCCGGCC | GGG | \|\|\|\|\|\|\|\|[\|\|\|\|\|\|\|\|\|\|\|\|]PAM | NA | Intergenic | NA | NA |
| T32504 | CTGGCTCATGCCGCCGGCCGGGG | 1000 | | | | | | | | | |
| Oligo fwd | CACCCTGGCTCATGCCGCCGGCCG | | | | | | | | | | |
| Oligo rev | AAACCGGCCGGCGGCATGAGCCAG | | | | | | | | | | |
| Chromosome | start | end | strand | MM | target_seq | PAM | alignment | distance | position | gene name | gene id |
| GU305914.1 | 26000 | 26022 | - | 0 | CTGGCTCATGCCGCCGGCCG | GGG | \|\|\|\|\|\|\|\|[\|\|\|\|\|\|\|\|\|\|\|\|]PAM | NA | Intergenic | NA | NA |
| T32505 | GAGAAACTCTTCCTCTTCGCCGG | 1000 | | | | | | | | | |
| Oligo fwd | CACCGAGAAACTCTTCCTCTTCGC | | | | | | | | | | |
| Oligo rev | AAACGCGAAGAGGAAGAGTTTCTC | | | | | | | | | | |
| Chromosome | start | end | strand | MM | target_seq | PAM | alignment | distance | position | gene name | gene id |
| GU305914.1 | 25961 | 25983 | - | 0 | GAGAAACTCTTCCTCTTCGC | CGG | \|\|\|\|\|\|\|\|[\|\|\|\|\|\|\|\|\|\|\|\|]PAM | NA | Intergenic | NA | NA |
| T32506 | AAACTCTTCCTCTTCGCCGGCGG | 1000 | | | | | | | | | |
| Oligo fwd | CACCAAACTCTTCCTCTTCGCCGG | | | | | | | | | | |
| Oligo rev | AAACCCGGCGAAGAGGAAGAGTTT | | | | | | | | | | |
| Chromosome | start | end | strand | MM | target_seq | PAM | alignment | distance | position | gene name | gene id |
| GU305914.1 | 25958 | 25980 | - | 0 | AAACTCTTCCTCTTCGCCGG | CGG | \|\|\|\|\|\|\|\|[\|\|\|\|\|\|\|\|\|\|\|\|]PAM | NA | Intergenic | NA | NA |
| T32507 | AACTCTTCCTCTTCGCCGGCGGG | 1000 | | | | | | | | | |
| Oligo fwd | CACCAACTCTTCCTCTTCGCCGGC | | | | | | | | | | |
| Oligo rev | AAACGCCGGCGAAGAGGAAGAGTT | | | | | | | | | | |
| Chromosome | start | end | strand | MM | target_seq | PAM | alignment | distance | position | gene name | gene id |
| GU305914.1 | 25957 | 25979 | - | 0 | AACTCTTCCTCTTCGCCGGC | GGG | \|\|\|\|\|\|\|\|[\|\|\|\|\|\|\|\|\|\|\|\|]PAM | NA | Intergenic | NA | NA |
| T32508 | CGCCGGCGGGAACCTGCGCATGG | 1000 | | | | | | | | | |
| Oligo fwd | CACCCGCCGGCGGGAACCTGCGCA | | | | | | | | | | |
| Oligo rev | AAACTGCGCAGGTTCCCGCCGGCG | | | | | | | | | | |
| Chromosome | start | end | strand | MM | target_seq | PAM | alignment | distance | position | gene name | gene id |
| GU305914.1 | 25944 | 25966 | - | 0 | CGCCGGCGGGAACCTGCGCA | TGG | \|\|\|\|\|\|\|\|[\|\|\|\|\|\|\|\|\|\|\|\|]PAM | NA | Intergenic | NA | NA |
| T32509 | GCGGGAACCTGCGCATGGCCTGG | 1000 | | | | | | | | | |
| Oligo fwd | CACCGCGGGAACCTGCGCATGGCC | | | | | | | | | | |
| Oligo rev | AAACGGCCATGCGCAGGTTCCCGC | | | | | | | | | | |
| Chromosome | start | end | strand | MM | target_seq | PAM | alignment | distance | position | gene name | gene id |
| GU305914.1 | 25939 | 25961 | - | 0 | GCGGGAACCTGCGCATGGCC | TGG | \|\|\|\|\|\|\|\|[\|\|\|\|\|\|\|\|\|\|\|\|]PAM | NA | Intergenic | NA | NA |

T32510 CTGCGCATGGCCTGGCCCGCCGG 1000
Oligo fwd CACCCTGCGCATGGCCTGGCCCGC
Oligo rev AAACGCGGGCCAGGCCATGCGCAG
Chromosome start end strand MM target_seq PAM alignment distance position gene name gene id
GU305914.1 25931 25953 - 0 CTGCGCATGGCCTGGCCCGC CGG ||||||||[||||||||||||]PAM NA Intergenic NA NA

T32511 TGCGCATGGCCTGGCCCGCCGGG 1000
Oligo fwd CACCTGCGCATGGCCTGGCCCGCC
Oligo rev AAACGGCGGGCCAGGCCATGCGCA
Chromosome start end strand MM target_seq PAM alignment distance position gene name gene id
GU305914.1 25930 25952 - 0 TGCGCATGGCCTGGCCCGCC GGG ||||||||[||||||||||||]PAM NA Intergenic NA NA

T32512 GCGCATGGCCTGGCCCGCCGGGG 1000
Oligo fwd CACCGCGCATGGCCTGGCCCGCCG
Oligo rev AAACCGGCGGGCCAGGCCATGCGC
Chromosome start end strand MM target_seq PAM alignment distance position gene name gene id
GU305914.1 25929 25951 - 0 GCGCATGGCCTGGCCCGCCG GGG ||||||||[||||||||||||]PAM NA Intergenic NA NA

T32513 AACCTTCGTCCCCGATATCCAGG 1000
Oligo fwd CACCAACCTTCGTCCCCGATATCC
Oligo rev AAACGGATATCGGGGACGAAGGTT
Chromosome start end strand MM target_seq PAM alignment distance position gene name gene id
GU305914.1 25899 25921 - 0 AACCTTCGTCCCCGATATCC AGG ||||||||[||||||||||||]PAM NA Intergenic NA NA

T32514 ACCTTCGTCCCCGATATCCAGGG 1000
Oligo fwd CACCACCTTCGTCCCCGATATCCA
Oligo rev AAACTGGATATCGGGGACGAAGGT
Chromosome start end strand MM target_seq PAM alignment distance position gene name gene id
GU305914.1 25898 25920 - 0 ACCTTCGTCCCCGATATCCA GGG ||||||||[||||||||||||]PAM NA Intergenic NA NA

T32515 TCGTCCCCGATATCCAGGGTTGG 1000
Oligo fwd CACCTCGTCCCCGATATCCAGGGT
Oligo rev AAACACCCTGGATATCGGGGACGA
Chromosome start end strand MM target_seq PAM alignment distance position gene name gene id
GU305914.1 25894 25916 - 0 TCGTCCCCGATATCCAGGGT TGG ||||||||[||||||||||||]PAM NA Intergenic NA NA

T32516 CGTCCCCGATATCCAGGGTTGGG 1000
Oligo fwd CACCCGTCCCCGATATCCAGGGTT
Oligo rev AAACAACCCTGGATATCGGGGACG
Chromosome start end strand MM target_seq PAM alignment distance position gene name gene id
GU305914.1 25893 25915 - 0 CGTCCCCGATATCCAGGGTT GGG ||||||||[||||||||||||]PAM NA Intergenic NA NA

T32517 CAGGGTTGGGAGCCGCAGTCTGG 1000
Oligo fwd CACCCAGGGTTGGGAGCCGCAGTC
Oligo rev AAACGACTGCGGCTCCCAACCCTG
Chromosome start end strand MM target_seq PAM alignment distance position gene name gene id
GU305914.1 25880 25902 - 0 CAGGGTTGGGAGCCGCAGTC TGG ||||||||[||||||||||||]PAM NA Intergenic NA NA

T32518 GACGTCTTCGTGTATCCCATCGG 1000
Oligo fwd CACCGACGTCTTCGTGTATCCCAT
Oligo rev AAACATGGGATACACGAAGACGTC
Chromosome start end strand MM target_seq PAM alignment distance position gene name gene id
GU305914.1 25823 25845 - 0 GACGTCTTCGTGTATCCCAT CGG ||||||||[||||||||||||]PAM NA Intergenic NA NA

T32519 TATCCCATCGGCTATGCCAGCGG 1000
Oligo fwd CACCTATCCCATCGGCTATGCCAG
Oligo rev AAACCTGGCATAGCCGATGGGATA
Chromosome start end strand MM target_seq PAM alignment distance position gene name gene id
GU305914.1 25811 25833 - 0 TATCCCATCGGCTATGCCAG CGG ||||||||[||||||||||||]PAM NA Intergenic NA NA

T32520 CGGCTATGCCAGCGGACTCAAGG 1000
Oligo fwd CACCCGGCTATGCCAGCGGACTCA
Oligo rev AAACTGAGTCCGCTGGCATAGCCG
Chromosome start end strand MM target_seq PAM alignment distance position gene name gene id
GU305914.1 25803 25825 - 0 CGGCTATGCCAGCGGACTCA AGG ||||||||[||||||||||||]PAM NA Intergenic NA NA

T32521 CAAGGTCGCCGACGACACCCTGG 1000
Oligo fwd CACCCAAGGTCGCCGACGACACCC
Oligo rev AAACGGGTGTCGTCGGCGACCTTG
Chromosome start end strand MM target_seq PAM alignment distance position gene name gene id
GU305914.1 25785 25807 - 0 CAAGGTCGCCGACGACACCC TGG ||||||||[||||||||||||]PAM NA Intergenic NA NA

T32522 CTGGTCCTCGTCGACGTCTTCGG 1000
Oligo fwd CACCCTGGTCCTCGTCGACGTCTT
Oligo rev AAACAAGACGTCGACGAGGACCAG
Chromosome start end strand MM target_seq PAM alignment distance position gene name gene id
GU305914.1 25766 25788 - 0 CTGGTCCTCGTCGACGTCTT CGG ||||||||[||||||||||||]PAM NA Intergenic NA NA

T32523 GGCCGTTTCTACTGTTATCGCGG 1000
Oligo fwd CACCGGCCGTTTCTACTGTTATCG
Oligo rev AAACCGATAACAGTAGAAACGGCC
Chromosome start end strand MM target_seq PAM alignment distance position gene name gene id
GU305914.1 25745 25767 - 0 GGCCGTTTCTACTGTTATCG CGG ||||||||[||||||||||||]PAM NA Intergenic NA NA

T32524 GCCGTTTCTACTGTTATCGCGGG 1000
Oligo fwd CACCGCCGTTTCTACTGTTATCGC
Oligo rev AAACGCGATAACAGTAGAAACGGC
Chromosome start end strand MM target_seq PAM alignment distance position gene name gene id

<table>
<tr><td>GU305914.1</td><td></td><td>25744</td><td>25766</td><td>-</td><td>0</td><td>GCCGTTTCTACTGTTATCGC</td><td>GGG</td><td>||||||||[||||||||||||]PAM</td><td>NA</td><td>Intergenic</td><td>NA</td><td>NA</td></tr>
<tr><td>T32525</td><td>CTACTGTTATCGCGGGCCCCCGG</td><td></td><td>1000</td><td></td><td></td><td></td><td></td><td></td><td></td><td></td><td></td><td></td></tr>
<tr><td>Oligo fwd</td><td>CACCCTACTGTTATCGCGGGCCCC</td><td></td><td></td><td></td><td></td><td></td><td></td><td></td><td></td><td></td><td></td><td></td></tr>
<tr><td>Oligo rev</td><td>AAACGGGGCCCGCGATAACAGTAG</td><td></td><td></td><td></td><td></td><td></td><td></td><td></td><td></td><td></td><td></td><td></td></tr>
<tr><td>Chromosome</td><td>start</td><td></td><td>end</td><td>strand</td><td>MM</td><td>target_seq</td><td>PAM</td><td>alignment</td><td>distance</td><td>position</td><td>gene name</td><td>gene id</td></tr>
<tr><td>GU305914.1</td><td></td><td>25737</td><td>25759</td><td>-</td><td>0</td><td>CTACTGTTATCGCGGGCCCC</td><td>CGG</td><td>||||||||[||||||||||||]PAM</td><td>NA</td><td>Intergenic</td><td>NA</td><td>NA</td></tr>
<tr><td>T32526</td><td>CTACTACCTCGCCGCCTCGCCGG</td><td></td><td>1000</td><td></td><td></td><td></td><td></td><td></td><td></td><td></td><td></td><td></td></tr>
<tr><td>Oligo fwd</td><td>CACCCTACTACCTCGCCGCCTCGC</td><td></td><td></td><td></td><td></td><td></td><td></td><td></td><td></td><td></td><td></td><td></td></tr>
<tr><td>Oligo rev</td><td>AAACGCGAGGCGGCGAGGTAGTAG</td><td></td><td></td><td></td><td></td><td></td><td></td><td></td><td></td><td></td><td></td><td></td></tr>
<tr><td>Chromosome</td><td>start</td><td></td><td>end</td><td>strand</td><td>MM</td><td>target_seq</td><td>PAM</td><td>alignment</td><td>distance</td><td>position</td><td>gene name</td><td>gene id</td></tr>
<tr><td>GU305914.1</td><td></td><td>25704</td><td>25726</td><td>-</td><td>0</td><td>CTACTACCTCGCCGCCTCGC</td><td>CGG</td><td>||||||||[||||||||||||]PAM</td><td>NA</td><td>Intergenic</td><td>NA</td><td>NA</td></tr>
<tr><td>T32527</td><td>CCGGAGCAGTTCTCCGTCATCGG</td><td></td><td>1000</td><td></td><td></td><td></td><td></td><td></td><td></td><td></td><td></td><td></td></tr>
<tr><td>Oligo fwd</td><td>CACCCCGGAGCAGTTCTCCGTCAT</td><td></td><td></td><td></td><td></td><td></td><td></td><td></td><td></td><td></td><td></td><td></td></tr>
<tr><td>Oligo rev</td><td>AAACATGACGGAGAACTGCTCCGG</td><td></td><td></td><td></td><td></td><td></td><td></td><td></td><td></td><td></td><td></td><td></td></tr>
<tr><td>Chromosome</td><td>start</td><td></td><td>end</td><td>strand</td><td>MM</td><td>target_seq</td><td>PAM</td><td>alignment</td><td>distance</td><td>position</td><td>gene name</td><td>gene id</td></tr>
<tr><td>GU305914.1</td><td></td><td>25685</td><td>25707</td><td>-</td><td>0</td><td>CCGGAGCAGTTCTCCGTCAT</td><td>CGG</td><td>||||||||[||||||||||||]PAM</td><td>NA</td><td>Intergenic</td><td>NA</td><td>NA</td></tr>
<tr><td>T32528</td><td>CGGAGCAGTTCTCCGTCATCGGG</td><td></td><td>1000</td><td></td><td></td><td></td><td></td><td></td><td></td><td></td><td></td><td></td></tr>
<tr><td>Oligo fwd</td><td>CACCCGGAGCAGTTCTCCGTCATC</td><td></td><td></td><td></td><td></td><td></td><td></td><td></td><td></td><td></td><td></td><td></td></tr>
<tr><td>Oligo rev</td><td>AAACGATGACGGAGAACTGCTCCG</td><td></td><td></td><td></td><td></td><td></td><td></td><td></td><td></td><td></td><td></td><td></td></tr>
<tr><td>Chromosome</td><td>start</td><td></td><td>end</td><td>strand</td><td>MM</td><td>target_seq</td><td>PAM</td><td>alignment</td><td>distance</td><td>position</td><td>gene name</td><td>gene id</td></tr>
<tr><td>GU305914.1</td><td></td><td>25684</td><td>25706</td><td>-</td><td>0</td><td>CGGAGCAGTTCTCCGTCATC</td><td>GGG</td><td>||||||||[||||||||||||]PAM</td><td>NA</td><td>Intergenic</td><td>NA</td><td>NA</td></tr>
<tr><td>T32529</td><td>TTTTTATCCGATTCACTGCACGG</td><td></td><td>1000</td><td></td><td></td><td></td><td></td><td></td><td></td><td></td><td></td><td></td></tr>
<tr><td>Oligo fwd</td><td>CACCTTTTTATCCGATTCACTGCA</td><td></td><td></td><td></td><td></td><td></td><td></td><td></td><td></td><td></td><td></td><td></td></tr>
<tr><td>Oligo rev</td><td>AAACTGCAGTGAATCGGATAAAAA</td><td></td><td></td><td></td><td></td><td></td><td></td><td></td><td></td><td></td><td></td><td></td></tr>
<tr><td>Chromosome</td><td>start</td><td></td><td>end</td><td>strand</td><td>MM</td><td>target_seq</td><td>PAM</td><td>alignment</td><td>distance</td><td>position</td><td>gene name</td><td>gene id</td></tr>
<tr><td>GU305914.1</td><td></td><td>25653</td><td>25675</td><td>-</td><td>0</td><td>TTTTTATCCGATTCACTGCA</td><td>CGG</td><td>||||||||[||||||||||||]PAM</td><td>NA</td><td>Intergenic</td><td>NA</td><td>NA</td></tr>
<tr><td>T32530</td><td>TATCCGATTCACTGCACGGCAGG</td><td></td><td>1000</td><td></td><td></td><td></td><td></td><td></td><td></td><td></td><td></td><td></td></tr>
<tr><td>Oligo fwd</td><td>CACCTATCCGATTCACTGCACGGC</td><td></td><td></td><td></td><td></td><td></td><td></td><td></td><td></td><td></td><td></td><td></td></tr>
<tr><td>Oligo rev</td><td>AAACGCCGTGCAGTGAATCGGATA</td><td></td><td></td><td></td><td></td><td></td><td></td><td></td><td></td><td></td><td></td><td></td></tr>
<tr><td>Chromosome</td><td>start</td><td></td><td>end</td><td>strand</td><td>MM</td><td>target_seq</td><td>PAM</td><td>alignment</td><td>distance</td><td>position</td><td>gene name</td><td>gene id</td></tr>
<tr><td>GU305914.1</td><td></td><td>25649</td><td>25671</td><td>-</td><td>0</td><td>TATCCGATTCACTGCACGGC</td><td>AGG</td><td>||||||||[||||||||||||]PAM</td><td>NA</td><td>Intergenic</td><td>NA</td><td>NA</td></tr>
<tr><td>T32531</td><td>ACGGCAGGCCCCATCGAGATCGG</td><td></td><td>1000</td><td></td><td></td><td></td><td></td><td></td><td></td><td></td><td></td><td></td></tr>
<tr><td>Oligo fwd</td><td>CACCACGGCAGGCCCCATCGAGAT</td><td></td><td></td><td></td><td></td><td></td><td></td><td></td><td></td><td></td><td></td><td></td></tr>
<tr><td>Oligo rev</td><td>AAACATCTCGATGGGGCCTGCCGT</td><td></td><td></td><td></td><td></td><td></td><td></td><td></td><td></td><td></td><td></td><td></td></tr>
<tr><td>Chromosome</td><td>start</td><td></td><td>end</td><td>strand</td><td>MM</td><td>target_seq</td><td>PAM</td><td>alignment</td><td>distance</td><td>position</td><td>gene name</td><td>gene id</td></tr>
<tr><td>GU305914.1</td><td></td><td>25634</td><td>25656</td><td>-</td><td>0</td><td>ACGGCAGGCCCCATCGAGAT</td><td>CGG</td><td>||||||||[||||||||||||]PAM</td><td>NA</td><td>Intergenic</td><td>NA</td><td>NA</td></tr>
<tr><td>T32532</td><td>GCCTGCTGCTCAGCCGCCTCTGG</td><td></td><td>1000</td><td></td><td></td><td></td><td></td><td></td><td></td><td></td><td></td><td></td></tr>
<tr><td>Oligo fwd</td><td>CACCGCCTGCTGCTCAGCCGCCTC</td><td></td><td></td><td></td><td></td><td></td><td></td><td></td><td></td><td></td><td></td><td></td></tr>
<tr><td>Oligo rev</td><td>AAACGAGGCGGCTGAGCAGCAGGC</td><td></td><td></td><td></td><td></td><td></td><td></td><td></td><td></td><td></td><td></td><td></td></tr>
<tr><td>Chromosome</td><td>start</td><td></td><td>end</td><td>strand</td><td>MM</td><td>target_seq</td><td>PAM</td><td>alignment</td><td>distance</td><td>position</td><td>gene name</td><td>gene id</td></tr>
<tr><td>GU305914.1</td><td></td><td>25612</td><td>25634</td><td>-</td><td>0</td><td>GCCTGCTGCTCAGCCGCCTC</td><td>TGG</td><td>||||||||[||||||||||||]PAM</td><td>NA</td><td>Intergenic</td><td>NA</td><td>NA</td></tr>
<tr><td>T32533</td><td>GCCGCCTCTGGATGATGATCAGG</td><td></td><td>1000</td><td></td><td></td><td></td><td></td><td></td><td></td><td></td><td></td><td></td></tr>
<tr><td>Oligo fwd</td><td>CACCGCCGCCTCTGGATGATGATC</td><td></td><td></td><td></td><td></td><td></td><td></td><td></td><td></td><td></td><td></td><td></td></tr>
<tr><td>Oligo rev</td><td>AAACGATCATCATCCAGAGGCGGC</td><td></td><td></td><td></td><td></td><td></td><td></td><td></td><td></td><td></td><td></td><td></td></tr>
<tr><td>Chromosome</td><td>start</td><td></td><td>end</td><td>strand</td><td>MM</td><td>target_seq</td><td>PAM</td><td>alignment</td><td>distance</td><td>position</td><td>gene name</td><td>gene id</td></tr>
<tr><td>GU305914.1</td><td></td><td>25600</td><td>25622</td><td>-</td><td>0</td><td>GCCGCCTCTGGATGATGATC</td><td>AGG</td><td>||||||||[||||||||||||]PAM</td><td>NA</td><td>Intergenic</td><td>NA</td><td>NA</td></tr>
<tr><td>T32534</td><td>CTCTGGATGATGATCAGGCGCGG</td><td></td><td>1000</td><td></td><td></td><td></td><td></td><td></td><td></td><td></td><td></td><td></td></tr>
<tr><td>Oligo fwd</td><td>CACCCTCTGGATGATGATCAGGCG</td><td></td><td></td><td></td><td></td><td></td><td></td><td></td><td></td><td></td><td></td><td></td></tr>
<tr><td>Oligo rev</td><td>AAACCGCCTGATCATCATCCAGAG</td><td></td><td></td><td></td><td></td><td></td><td></td><td></td><td></td><td></td><td></td><td></td></tr>
<tr><td>Chromosome</td><td>start</td><td></td><td>end</td><td>strand</td><td>MM</td><td>target_seq</td><td>PAM</td><td>alignment</td><td>distance</td><td>position</td><td>gene name</td><td>gene id</td></tr>
<tr><td>GU305914.1</td><td></td><td>25595</td><td>25617</td><td>-</td><td>0</td><td>CTCTGGATGATGATCAGGCG</td><td>CGG</td><td>||||||||[||||||||||||]PAM</td><td>NA</td><td>Intergenic</td><td>NA</td><td>NA</td></tr>
<tr><td>T32535</td><td>TCTGGATGATGATCAGGCGCGGG</td><td></td><td>1000</td><td></td><td></td><td></td><td></td><td></td><td></td><td></td><td></td><td></td></tr>
<tr><td>Oligo fwd</td><td>CACCTCTGGATGATGATCAGGCGC</td><td></td><td></td><td></td><td></td><td></td><td></td><td></td><td></td><td></td><td></td><td></td></tr>
<tr><td>Oligo rev</td><td>AAACGCGCCTGATCATCATCCAGA</td><td></td><td></td><td></td><td></td><td></td><td></td><td></td><td></td><td></td><td></td><td></td></tr>
<tr><td>Chromosome</td><td>start</td><td></td><td>end</td><td>strand</td><td>MM</td><td>target_seq</td><td>PAM</td><td>alignment</td><td>distance</td><td>position</td><td>gene name</td><td>gene id</td></tr>
<tr><td>GU305914.1</td><td></td><td>25594</td><td>25616</td><td>-</td><td>0</td><td>TCTGGATGATGATCAGGCGC</td><td>GGG</td><td>||||||||[||||||||||||]PAM</td><td>NA</td><td>Intergenic</td><td>NA</td><td>NA</td></tr>
<tr><td>T32536</td><td>CTGGATGATGATCAGGCGCGGGG</td><td></td><td>1000</td><td></td><td></td><td></td><td></td><td></td><td></td><td></td><td></td><td></td></tr>
<tr><td>Oligo fwd</td><td>CACCCTGGATGATGATCAGGCGCG</td><td></td><td></td><td></td><td></td><td></td><td></td><td></td><td></td><td></td><td></td><td></td></tr>
<tr><td>Oligo rev</td><td>AAACCGCGCCTGATCATCATCCAG</td><td></td><td></td><td></td><td></td><td></td><td></td><td></td><td></td><td></td><td></td><td></td></tr>
<tr><td>Chromosome</td><td>start</td><td></td><td>end</td><td>strand</td><td>MM</td><td>target_seq</td><td>PAM</td><td>alignment</td><td>distance</td><td>position</td><td>gene name</td><td>gene id</td></tr>
<tr><td>GU305914.1</td><td></td><td>25593</td><td>25615</td><td>-</td><td>0</td><td>CTGGATGATGATCAGGCGCG</td><td>GGG</td><td>||||||||[||||||||||||]PAM</td><td>NA</td><td>Intergenic</td><td>NA</td><td>NA</td></tr>
<tr><td>T32537</td><td>CGCTTTGTCGTGCGCGCGCACGG</td><td></td><td>1000</td><td></td><td></td><td></td><td></td><td></td><td></td><td></td><td></td><td></td></tr>
<tr><td>Oligo fwd</td><td>CACCCGCTTTGTCGTGCGCGCGCA</td><td></td><td></td><td></td><td></td><td></td><td></td><td></td><td></td><td></td><td></td><td></td></tr>
<tr><td>Oligo rev</td><td>AAACTGCGCGCGCACGACAAAGCG</td><td></td><td></td><td></td><td></td><td></td><td></td><td></td><td></td><td></td><td></td><td></td></tr>
<tr><td>Chromosome</td><td>start</td><td></td><td>end</td><td>strand</td><td>MM</td><td>target_seq</td><td>PAM</td><td>alignment</td><td>distance</td><td>position</td><td>gene name</td><td>gene id</td></tr>
<tr><td>GU305914.1</td><td></td><td>25550</td><td>25572</td><td>-</td><td>0</td><td>CGCTTTGTCGTGCGCGCGCA</td><td>CGG</td><td>||||||||[||||||||||||]PAM</td><td>NA</td><td>Intergenic</td><td>NA</td><td>NA</td></tr>
<tr><td>T32538</td><td>CGCGCACGGCGAGACCGTCGTGG</td><td></td><td>1000</td><td></td><td></td><td></td><td></td><td></td><td></td><td></td><td></td><td></td></tr>
<tr><td>Oligo fwd</td><td>CACCCGCGCACGGCGAGACCGTCG</td><td></td><td></td><td></td><td></td><td></td><td></td><td></td><td></td><td></td><td></td><td></td></tr>
<tr><td>Oligo rev</td><td>AAACCGACGGTCTCGCCGTGCGCG</td><td></td><td></td><td></td><td></td><td></td><td></td><td></td><td></td><td></td><td></td><td></td></tr>
<tr><td>Chromosome</td><td>start</td><td></td><td>end</td><td>strand</td><td>MM</td><td>target_seq</td><td>PAM</td><td>alignment</td><td>distance</td><td>position</td><td>gene name</td><td>gene id</td></tr>
<tr><td>GU305914.1</td><td></td><td>25536</td><td>25558</td><td>-</td><td>0</td><td>CGCGCACGGCGAGACCGTCG</td><td>TGG</td><td>||||||||[||||||||||||]PAM</td><td>NA</td><td>Intergenic</td><td>NA</td><td>NA</td></tr>
<tr><td>T32539</td><td>GTGGTCGCCAGACGCCTCGTCGG</td><td></td><td>1000</td><td></td><td></td><td></td><td></td><td></td><td></td><td></td><td></td><td></td></tr>
<tr><td>Oligo fwd</td><td>CACCGTGGTCGCCAGACGCCTCGT</td><td></td><td></td><td></td><td></td><td></td><td></td><td></td><td></td><td></td><td></td><td></td></tr>
<tr><td>Oligo rev</td><td>AAACACGAGGCGTCTGGCGACCAC</td><td></td><td></td><td></td><td></td><td></td><td></td><td></td><td></td><td></td><td></td><td></td></tr>
</table>

| | | | | | | | | | | |
|---|---|---|---|---|---|---|---|---|---|---|
| Chromosome | start | end | strand | MM | target_seq | PAM | alignment | distance | position | gene name | gene id |
| GU305914.1 | 25517 | 25539 | - | 0 | GTGGTCGCCAGACGCCTCGT | CGG | \|\|\|\|\|\|\|\|[\|\|\|\|\|\|\|\|\|\|\|\|]PAM | NA | Intergenic | NA | NA |
| T32540 | TGGTCGCCAGACGCCTCGTCGGG | 1000 | | | | | | | | | |
| Oligo fwd | CACCTGGTCGCCAGACGCCTCGTC | | | | | | | | | | |
| Oligo rev | AAACGACGAGGCGTCTGGCGACCA | | | | | | | | | | |
| Chromosome | start | end | strand | MM | target_seq | PAM | alignment | distance | position | gene name | gene id |
| GU305914.1 | 25516 | 25538 | - | 0 | TGGTCGCCAGACGCCTCGTC | GGG | \|\|\|\|\|\|\|\|[\|\|\|\|\|\|\|\|\|\|\|\|]PAM | NA | Intergenic | NA | NA |
| T32541 | TCGCCAGACGCCTCGTCGGGCGG | 1000 | | | | | | | | | |
| Oligo fwd | CACCTCGCCAGACGCCTCGTCGGG | | | | | | | | | | |
| Oligo rev | AAACCCCGACGAGGCGTCTGGCGA | | | | | | | | | | |
| Chromosome | start | end | strand | MM | target_seq | PAM | alignment | distance | position | gene name | gene id |
| GU305914.1 | 25513 | 25535 | - | 0 | TCGCCAGACGCCTCGTCGGG | CGG | \|\|\|\|\|\|\|\|[\|\|\|\|\|\|\|\|\|\|\|\|]PAM | NA | Intergenic | NA | NA |
| T32542 | CCAGACGCCTCGTCGGGCGGCGG | 1000 | | | | | | | | | |
| Oligo fwd | CACCCCAGACGCCTCGTCGGGCGG | | | | | | | | | | |
| Oligo rev | AAACCCGCCCGACGAGGCGTCTGG | | | | | | | | | | |
| Chromosome | start | end | strand | MM | target_seq | PAM | alignment | distance | position | gene name | gene id |
| GU305914.1 | 25510 | 25532 | - | 0 | CCAGACGCCTCGTCGGGCGG | CGG | \|\|\|\|\|\|\|\|[\|\|\|\|\|\|\|\|\|\|\|\|]PAM | NA | Intergenic | NA | NA |
| T32543 | CAGACGCCTCGTCGGGCGGCGGG | 1000 | | | | | | | | | |
| Oligo fwd | CACCCAGACGCCTCGTCGGGCGGC | | | | | | | | | | |
| Oligo rev | AAACGCCGCCCGACGAGGCGTCTG | | | | | | | | | | |
| Chromosome | start | end | strand | MM | target_seq | PAM | alignment | distance | position | gene name | gene id |
| GU305914.1 | 25509 | 25531 | - | 0 | CAGACGCCTCGTCGGGCGGC | GGG | \|\|\|\|\|\|\|\|[\|\|\|\|\|\|\|\|\|\|\|\|]PAM | NA | Intergenic | NA | NA |
| T32544 | ACGCCTCGTCGGGCGGCGGGAGG | 1000 | | | | | | | | | |
| Oligo fwd | CACCACGCCTCGTCGGGCGGCGGG | | | | | | | | | | |
| Oligo rev | AAACCCCGCCGCCCGACGAGGCGT | | | | | | | | | | |
| Chromosome | start | end | strand | MM | target_seq | PAM | alignment | distance | position | gene name | gene id |
| GU305914.1 | 25506 | 25528 | - | 0 | ACGCCTCGTCGGGCGGCGGG | AGG | \|\|\|\|\|\|\|\|[\|\|\|\|\|\|\|\|\|\|\|\|]PAM | NA | Intergenic | NA | NA |
| T32545 | GGTGCTGCGCGTCTGCTCTCTGG | 1000 | | | | | | | | | |
| Oligo fwd | CACCGGTGCTGCGCGTCTGCTCTC | | | | | | | | | | |
| Oligo rev | AAACGAGAGCAGACGCGCAGCACC | | | | | | | | | | |
| Chromosome | start | end | strand | MM | target_seq | PAM | alignment | distance | position | gene name | gene id |
| GU305914.1 | 25485 | 25507 | - | 0 | GGTGCTGCGCGTCTGCTCTC | TGG | \|\|\|\|\|\|\|\|[\|\|\|\|\|\|\|\|\|\|\|\|]PAM | NA | Intergenic | NA | NA |
| T32546 | GTGCTGCGCGTCTGCTCTCTGGG | 1000 | | | | | | | | | |
| Oligo fwd | CACCGTGCTGCGCGTCTGCTCTCT | | | | | | | | | | |
| Oligo rev | AAACAGAGAGCAGACGCGCAGCAC | | | | | | | | | | |
| Chromosome | start | end | strand | MM | target_seq | PAM | alignment | distance | position | gene name | gene id |
| GU305914.1 | 25484 | 25506 | - | 0 | GTGCTGCGCGTCTGCTCTCT | GGG | \|\|\|\|\|\|\|\|[\|\|\|\|\|\|\|\|\|\|\|\|]PAM | NA | Intergenic | NA | NA |
| T32547 | GCTCTCTGGGTTGTAGTCGCTGG | 1000 | | | | | | | | | |
| Oligo fwd | CACCGCTCTCTGGGTTGTAGTCGC | | | | | | | | | | |
| Oligo rev | AAACGCGACTACAACCCAGAGAGC | | | | | | | | | | |
| Chromosome | start | end | strand | MM | target_seq | PAM | alignment | distance | position | gene name | gene id |
| GU305914.1 | 25471 | 25493 | - | 0 | GCTCTCTGGGTTGTAGTCGC | TGG | \|\|\|\|\|\|\|\|[\|\|\|\|\|\|\|\|\|\|\|\|]PAM | NA | Intergenic | NA | NA |
| T32548 | GTTGTAGTCGCTGGCGCAAGAGG | 1000 | | | | | | | | | |
| Oligo fwd | CACCGTTGTAGTCGCTGGCGCAAG | | | | | | | | | | |
| Oligo rev | AAACCTTGCGCCAGCGACTACAAC | | | | | | | | | | |
| Chromosome | start | end | strand | MM | target_seq | PAM | alignment | distance | position | gene name | gene id |
| GU305914.1 | 25462 | 25484 | - | 0 | GTTGTAGTCGCTGGCGCAAG | AGG | \|\|\|\|\|\|\|\|[\|\|\|\|\|\|\|\|\|\|\|\|]PAM | NA | Intergenic | NA | NA |
| T32549 | TTGTAGTCGCTGGCGCAAGAGGG | 1000 | | | | | | | | | |
| Oligo fwd | CACCTTGTAGTCGCTGGCGCAAGA | | | | | | | | | | |
| Oligo rev | AAACTCTTGCGCCAGCGACTACAA | | | | | | | | | | |
| Chromosome | start | end | strand | MM | target_seq | PAM | alignment | distance | position | gene name | gene id |
| GU305914.1 | 25461 | 25483 | - | 0 | TTGTAGTCGCTGGCGCAAGA | GGG | \|\|\|\|\|\|\|\|[\|\|\|\|\|\|\|\|\|\|\|\|]PAM | NA | Intergenic | NA | NA |
| T32550 | CTGGCGCAAGAGGGACTCTTCGG | 1000 | | | | | | | | | |
| Oligo fwd | CACCCTGGCGCAAGAGGGACTCTT | | | | | | | | | | |
| Oligo rev | AAACAAGAGTCCCTCTTGCGCCAG | | | | | | | | | | |
| Chromosome | start | end | strand | MM | target_seq | PAM | alignment | distance | position | gene name | gene id |
| GU305914.1 | 25452 | 25474 | - | 0 | CTGGCGCAAGAGGGACTCTT | CGG | \|\|\|\|\|\|\|\|[\|\|\|\|\|\|\|\|\|\|\|\|]PAM | NA | Intergenic | NA | NA |
| T32551 | TGGCGCAAGAGGGACTCTTCGGG | 1000 | | | | | | | | | |
| Oligo fwd | CACCTGGCGCAAGAGGGACTCTTC | | | | | | | | | | |
| Oligo rev | AAACGAAGAGTCCCTCTTGCGCCA | | | | | | | | | | |
| Chromosome | start | end | strand | MM | target_seq | PAM | alignment | distance | position | gene name | gene id |
| GU305914.1 | 25451 | 25473 | - | 0 | TGGCGCAAGAGGGACTCTTC | GGG | \|\|\|\|\|\|\|\|[\|\|\|\|\|\|\|\|\|\|\|\|]PAM | NA | Intergenic | NA | NA |
| T32552 | GGCGCAAGAGGGACTCTTCGGGG | 1000 | | | | | | | | | |
| Oligo fwd | CACCGGCGCAAGAGGGACTCTTCG | | | | | | | | | | |
| Oligo rev | AAACCGAAGAGTCCCTCTTGCGCC | | | | | | | | | | |
| Chromosome | start | end | strand | MM | target_seq | PAM | alignment | distance | position | gene name | gene id |
| GU305914.1 | 25450 | 25472 | - | 0 | GGCGCAAGAGGGACTCTTCG | GGG | \|\|\|\|\|\|\|\|[\|\|\|\|\|\|\|\|\|\|\|\|]PAM | NA | Intergenic | NA | NA |
| T32553 | GACTCTTCGGGGCGTCTGCATGG | 1000 | | | | | | | | | |
| Oligo fwd | CACCGACTCTTCGGGGCGTCTGCA | | | | | | | | | | |
| Oligo rev | AAACTGCAGACGCCCCGAAGAGTC | | | | | | | | | | |
| Chromosome | start | end | strand | MM | target_seq | PAM | alignment | distance | position | gene name | gene id |
| GU305914.1 | 25439 | 25461 | - | 0 | GACTCTTCGGGGCGTCTGCA | TGG | \|\|\|\|\|\|\|\|[\|\|\|\|\|\|\|\|\|\|\|\|]PAM | NA | Intergenic | NA | NA |
| T32554 | TGCGCCCTGTCCGCCCCCACTGG | 1000 | | | | | | | | | |
| Oligo fwd | CACCTGCGCCCTGTCCGCCCCCAC | | | | | | | | | | |

<table>
<tr><td>Oligo rev</td><td>AAACGTGGGGGCGGACAGGGCGCA</td><td></td><td></td><td></td><td></td><td></td><td></td><td></td><td></td><td></td><td></td><td></td></tr>
<tr><td>Chromosome</td><td>start</td><td></td><td>end</td><td>strand</td><td>MM</td><td>target_seq</td><td>PAM</td><td>alignment</td><td>distance</td><td>position</td><td>gene name</td><td>gene id</td></tr>
<tr><td>GU305914.1</td><td></td><td>25413</td><td>25435</td><td>-</td><td>0</td><td>TGCGCCCTGTCCGCCCCCAC</td><td>TGG</td><td>||||||||[||||||||||||]PAM</td><td>NA</td><td>Intergenic</td><td>NA</td><td>NA</td></tr>
<tr><td></td><td></td><td></td><td></td><td></td><td></td><td></td><td></td><td></td><td></td><td></td><td></td><td></td></tr>
<tr><td>T32555</td><td>GCGCCCTGTCCGCCCCCACTGGG</td><td></td><td>1000</td><td></td><td></td><td></td><td></td><td></td><td></td><td></td><td></td><td></td></tr>
<tr><td>Oligo fwd</td><td>CACCGCGCCCTGTCCGCCCCCACT</td><td></td><td></td><td></td><td></td><td></td><td></td><td></td><td></td><td></td><td></td><td></td></tr>
<tr><td>Oligo rev</td><td>AAACAGTGGGGGCGGACAGGGCGC</td><td></td><td></td><td></td><td></td><td></td><td></td><td></td><td></td><td></td><td></td><td></td></tr>
<tr><td>Chromosome</td><td>start</td><td></td><td>end</td><td>strand</td><td>MM</td><td>target_seq</td><td>PAM</td><td>alignment</td><td>distance</td><td>position</td><td>gene name</td><td>gene id</td></tr>
<tr><td>GU305914.1</td><td></td><td>25412</td><td>25434</td><td>-</td><td>0</td><td>GCGCCCTGTCCGCCCCCACT</td><td>GGG</td><td>||||||||[||||||||||||]PAM</td><td>NA</td><td>Intergenic</td><td>NA</td><td>NA</td></tr>
<tr><td></td><td></td><td></td><td></td><td></td><td></td><td></td><td></td><td></td><td></td><td></td><td></td><td></td></tr>
<tr><td>T32556</td><td>GAATATAGCACCGTTCCGCTTGG</td><td></td><td>1000</td><td></td><td></td><td></td><td></td><td></td><td></td><td></td><td></td><td></td></tr>
<tr><td>Oligo fwd</td><td>CACCGAATATAGCACCGTTCCGCT</td><td></td><td></td><td></td><td></td><td></td><td></td><td></td><td></td><td></td><td></td><td></td></tr>
<tr><td>Oligo rev</td><td>AAACAGCGGAACGGTGCTATATTC</td><td></td><td></td><td></td><td></td><td></td><td></td><td></td><td></td><td></td><td></td><td></td></tr>
<tr><td>Chromosome</td><td>start</td><td></td><td>end</td><td>strand</td><td>MM</td><td>target_seq</td><td>PAM</td><td>alignment</td><td>distance</td><td>position</td><td>gene name</td><td>gene id</td></tr>
<tr><td>GU305914.1</td><td></td><td>25388</td><td>25410</td><td>-</td><td>0</td><td>GAATATAGCACCGTTCCGCT</td><td>TGG</td><td>||||||||[||||||||||||]PAM</td><td>NA</td><td>Intergenic</td><td>NA</td><td>NA</td></tr>
<tr><td></td><td></td><td></td><td></td><td></td><td></td><td></td><td></td><td></td><td></td><td></td><td></td><td></td></tr>
<tr><td>T32557</td><td>TCCGCTTGGTCGAGTCAAAAAGG</td><td></td><td>1000</td><td></td><td></td><td></td><td></td><td></td><td></td><td></td><td></td><td></td></tr>
<tr><td>Oligo fwd</td><td>CACCTCCGCTTGGTCGAGTCAAAA</td><td></td><td></td><td></td><td></td><td></td><td></td><td></td><td></td><td></td><td></td><td></td></tr>
<tr><td>Oligo rev</td><td>AAACTTTTGACTCGACCAAGCGGA</td><td></td><td></td><td></td><td></td><td></td><td></td><td></td><td></td><td></td><td></td><td></td></tr>
<tr><td>Chromosome</td><td>start</td><td></td><td>end</td><td>strand</td><td>MM</td><td>target_seq</td><td>PAM</td><td>alignment</td><td>distance</td><td>position</td><td>gene name</td><td>gene id</td></tr>
<tr><td>GU305914.1</td><td></td><td>25374</td><td>25396</td><td>-</td><td>0</td><td>TCCGCTTGGTCGAGTCAAAA</td><td>AGG</td><td>||||||||[||||||||||||]PAM</td><td>NA</td><td>Intergenic</td><td>NA</td><td>NA</td></tr>
<tr><td></td><td></td><td></td><td></td><td></td><td></td><td></td><td></td><td></td><td></td><td></td><td></td><td></td></tr>
<tr><td>T32558</td><td>CCGCTTGGTCGAGTCAAAAAGGG</td><td></td><td>1000</td><td></td><td></td><td></td><td></td><td></td><td></td><td></td><td></td><td></td></tr>
<tr><td>Oligo fwd</td><td>CACCCCGCTTGGTCGAGTCAAAAA</td><td></td><td></td><td></td><td></td><td></td><td></td><td></td><td></td><td></td><td></td><td></td></tr>
<tr><td>Oligo rev</td><td>AAACTTTTTGACTCGACCAAGCGG</td><td></td><td></td><td></td><td></td><td></td><td></td><td></td><td></td><td></td><td></td><td></td></tr>
<tr><td>Chromosome</td><td>start</td><td></td><td>end</td><td>strand</td><td>MM</td><td>target_seq</td><td>PAM</td><td>alignment</td><td>distance</td><td>position</td><td>gene name</td><td>gene id</td></tr>
<tr><td>GU305914.1</td><td></td><td>25373</td><td>25395</td><td>-</td><td>0</td><td>CCGCTTGGTCGAGTCAAAAA</td><td>GGG</td><td>||||||||[||||||||||||]PAM</td><td>NA</td><td>Intergenic</td><td>NA</td><td>NA</td></tr>
<tr><td></td><td></td><td></td><td></td><td></td><td></td><td></td><td></td><td></td><td></td><td></td><td></td><td></td></tr>
<tr><td>T32559</td><td>AGTCAAAAAGGGACTGCCGCAGG</td><td></td><td>1000</td><td></td><td></td><td></td><td></td><td></td><td></td><td></td><td></td><td></td></tr>
<tr><td>Oligo fwd</td><td>CACCAGTCAAAAAGGGACTGCCGC</td><td></td><td></td><td></td><td></td><td></td><td></td><td></td><td></td><td></td><td></td><td></td></tr>
<tr><td>Oligo rev</td><td>AAACGCGGCAGTCCCTTTTTGACT</td><td></td><td></td><td></td><td></td><td></td><td></td><td></td><td></td><td></td><td></td><td></td></tr>
<tr><td>Chromosome</td><td>start</td><td></td><td>end</td><td>strand</td><td>MM</td><td>target_seq</td><td>PAM</td><td>alignment</td><td>distance</td><td>position</td><td>gene name</td><td>gene id</td></tr>
<tr><td>GU305914.1</td><td></td><td>25362</td><td>25384</td><td>-</td><td>0</td><td>AGTCAAAAAGGGACTGCCGC</td><td>AGG</td><td>||||||||[||||||||||||]PAM</td><td>NA</td><td>Intergenic</td><td>NA</td><td>NA</td></tr>
<tr><td></td><td></td><td></td><td></td><td></td><td></td><td></td><td></td><td></td><td></td><td></td><td></td><td></td></tr>
<tr><td>T32560</td><td>AAGGGACTGCCGCAGGACCCTGG</td><td></td><td>1000</td><td></td><td></td><td></td><td></td><td></td><td></td><td></td><td></td><td></td></tr>
<tr><td>Oligo fwd</td><td>CACCAAGGGACTGCCGCAGGACCC</td><td></td><td></td><td></td><td></td><td></td><td></td><td></td><td></td><td></td><td></td><td></td></tr>
<tr><td>Oligo rev</td><td>AAACGGGTCCTGCGGCAGTCCCTT</td><td></td><td></td><td></td><td></td><td></td><td></td><td></td><td></td><td></td><td></td><td></td></tr>
<tr><td>Chromosome</td><td>start</td><td></td><td>end</td><td>strand</td><td>MM</td><td>target_seq</td><td>PAM</td><td>alignment</td><td>distance</td><td>position</td><td>gene name</td><td>gene id</td></tr>
<tr><td>GU305914.1</td><td></td><td>25355</td><td>25377</td><td>-</td><td>0</td><td>AAGGGACTGCCGCAGGACCC</td><td>TGG</td><td>||||||||[||||||||||||]PAM</td><td>NA</td><td>Intergenic</td><td>NA</td><td>NA</td></tr>
<tr><td></td><td></td><td></td><td></td><td></td><td></td><td></td><td></td><td></td><td></td><td></td><td></td><td></td></tr>
<tr><td>T32561</td><td>AGGGACTGCCGCAGGACCCTGGG</td><td></td><td>1000</td><td></td><td></td><td></td><td></td><td></td><td></td><td></td><td></td><td></td></tr>
<tr><td>Oligo fwd</td><td>CACCAGGGACTGCCGCAGGACCCT</td><td></td><td></td><td></td><td></td><td></td><td></td><td></td><td></td><td></td><td></td><td></td></tr>
<tr><td>Oligo rev</td><td>AAACAGGGTCCTGCGGCAGTCCCT</td><td></td><td></td><td></td><td></td><td></td><td></td><td></td><td></td><td></td><td></td><td></td></tr>
<tr><td>Chromosome</td><td>start</td><td></td><td>end</td><td>strand</td><td>MM</td><td>target_seq</td><td>PAM</td><td>alignment</td><td>distance</td><td>position</td><td>gene name</td><td>gene id</td></tr>
<tr><td>GU305914.1</td><td></td><td>25354</td><td>25376</td><td>-</td><td>0</td><td>AGGGACTGCCGCAGGACCCT</td><td>GGG</td><td>||||||||[||||||||||||]PAM</td><td>NA</td><td>Intergenic</td><td>NA</td><td>NA</td></tr>
<tr><td></td><td></td><td></td><td></td><td></td><td></td><td></td><td></td><td></td><td></td><td></td><td></td><td></td></tr>
<tr><td>T32562</td><td>CCCTGGGATCGCTGTGTTCGTGG</td><td></td><td>1000</td><td></td><td></td><td></td><td></td><td></td><td></td><td></td><td></td><td></td></tr>
<tr><td>Oligo fwd</td><td>CACCCCCTGGGATCGCTGTGTTCG</td><td></td><td></td><td></td><td></td><td></td><td></td><td></td><td></td><td></td><td></td><td></td></tr>
<tr><td>Oligo rev</td><td>AAACCGAACACAGCGATCCCAGGG</td><td></td><td></td><td></td><td></td><td></td><td></td><td></td><td></td><td></td><td></td><td></td></tr>
<tr><td>Chromosome</td><td>start</td><td></td><td>end</td><td>strand</td><td>MM</td><td>target_seq</td><td>PAM</td><td>alignment</td><td>distance</td><td>position</td><td>gene name</td><td>gene id</td></tr>
<tr><td>GU305914.1</td><td></td><td>25338</td><td>25360</td><td>-</td><td>0</td><td>CCCTGGGATCGCTGTGTTCG</td><td>TGG</td><td>||||||||[||||||||||||]PAM</td><td>NA</td><td>Intergenic</td><td>NA</td><td>NA</td></tr>
<tr><td></td><td></td><td></td><td></td><td></td><td></td><td></td><td></td><td></td><td></td><td></td><td></td><td></td></tr>
<tr><td>T32563</td><td>CCTGGGATCGCTGTGTTCGTGGG</td><td></td><td>1000</td><td></td><td></td><td></td><td></td><td></td><td></td><td></td><td></td><td></td></tr>
<tr><td>Oligo fwd</td><td>CACCCCTGGGATCGCTGTGTTCGT</td><td></td><td></td><td></td><td></td><td></td><td></td><td></td><td></td><td></td><td></td><td></td></tr>
<tr><td>Oligo rev</td><td>AAACACGAACACAGCGATCCCAGG</td><td></td><td></td><td></td><td></td><td></td><td></td><td></td><td></td><td></td><td></td><td></td></tr>
<tr><td>Chromosome</td><td>start</td><td></td><td>end</td><td>strand</td><td>MM</td><td>target_seq</td><td>PAM</td><td>alignment</td><td>distance</td><td>position</td><td>gene name</td><td>gene id</td></tr>
<tr><td>GU305914.1</td><td></td><td>25337</td><td>25359</td><td>-</td><td>0</td><td>CCTGGGATCGCTGTGTTCGT</td><td>GGG</td><td>||||||||[||||||||||||]PAM</td><td>NA</td><td>Intergenic</td><td>NA</td><td>NA</td></tr>
<tr><td></td><td></td><td></td><td></td><td></td><td></td><td></td><td></td><td></td><td></td><td></td><td></td><td></td></tr>
<tr><td>T32564</td><td>GTGTTCGTGGGCTTGCCCTCCGG</td><td></td><td>1000</td><td></td><td></td><td></td><td></td><td></td><td></td><td></td><td></td><td></td></tr>
<tr><td>Oligo fwd</td><td>CACCGTGTTCGTGGGCTTGCCCTC</td><td></td><td></td><td></td><td></td><td></td><td></td><td></td><td></td><td></td><td></td><td></td></tr>
<tr><td>Oligo rev</td><td>AAACGAGGGCAAGCCCACGAACAC</td><td></td><td></td><td></td><td></td><td></td><td></td><td></td><td></td><td></td><td></td><td></td></tr>
<tr><td>Chromosome</td><td>start</td><td></td><td>end</td><td>strand</td><td>MM</td><td>target_seq</td><td>PAM</td><td>alignment</td><td>distance</td><td>position</td><td>gene name</td><td>gene id</td></tr>
<tr><td>GU305914.1</td><td></td><td>25325</td><td>25347</td><td>-</td><td>0</td><td>GTGTTCGTGGGCTTGCCCTC</td><td>CGG</td><td>||||||||[||||||||||||]PAM</td><td>NA</td><td>Intergenic</td><td>NA</td><td>NA</td></tr>
<tr><td></td><td></td><td></td><td></td><td></td><td></td><td></td><td></td><td></td><td></td><td></td><td></td><td></td></tr>
<tr><td>T32565</td><td>TCGTGGGCTTGCCCTCCGGTAGG</td><td></td><td>1000</td><td></td><td></td><td></td><td></td><td></td><td></td><td></td><td></td><td></td></tr>
<tr><td>Oligo fwd</td><td>CACCTCGTGGGCTTGCCCTCCGGT</td><td></td><td></td><td></td><td></td><td></td><td></td><td></td><td></td><td></td><td></td><td></td></tr>
<tr><td>Oligo rev</td><td>AAACACCGGAGGGCAAGCCCACGA</td><td></td><td></td><td></td><td></td><td></td><td></td><td></td><td></td><td></td><td></td><td></td></tr>
<tr><td>Chromosome</td><td>start</td><td></td><td>end</td><td>strand</td><td>MM</td><td>target_seq</td><td>PAM</td><td>alignment</td><td>distance</td><td>position</td><td>gene name</td><td>gene id</td></tr>
<tr><td>GU305914.1</td><td></td><td>25321</td><td>25343</td><td>-</td><td>0</td><td>TCGTGGGCTTGCCCTCCGGT</td><td>AGG</td><td>||||||||[||||||||||||]PAM</td><td>NA</td><td>Intergenic</td><td>NA</td><td>NA</td></tr>
<tr><td></td><td></td><td></td><td></td><td></td><td></td><td></td><td></td><td></td><td></td><td></td><td></td><td></td></tr>
<tr><td>T32566</td><td>CGTGGGCTTGCCCTCCGGTAGGG</td><td></td><td>1000</td><td></td><td></td><td></td><td></td><td></td><td></td><td></td><td></td><td></td></tr>
<tr><td>Oligo fwd</td><td>CACCCGTGGGCTTGCCCTCCGGTA</td><td></td><td></td><td></td><td></td><td></td><td></td><td></td><td></td><td></td><td></td><td></td></tr>
<tr><td>Oligo rev</td><td>AAACTACCGGAGGGCAAGCCCACG</td><td></td><td></td><td></td><td></td><td></td><td></td><td></td><td></td><td></td><td></td><td></td></tr>
<tr><td>Chromosome</td><td>start</td><td></td><td>end</td><td>strand</td><td>MM</td><td>target_seq</td><td>PAM</td><td>alignment</td><td>distance</td><td>position</td><td>gene name</td><td>gene id</td></tr>
<tr><td>GU305914.1</td><td></td><td>25320</td><td>25342</td><td>-</td><td>0</td><td>CGTGGGCTTGCCCTCCGGTA</td><td>GGG</td><td>||||||||[||||||||||||]PAM</td><td>NA</td><td>Intergenic</td><td>NA</td><td>NA</td></tr>
<tr><td></td><td></td><td></td><td></td><td></td><td></td><td></td><td></td><td></td><td></td><td></td><td></td><td></td></tr>
<tr><td>T32567</td><td>CGGTAGGGTGTACGCCCGACAGG</td><td></td><td>1000</td><td></td><td></td><td></td><td></td><td></td><td></td><td></td><td></td><td></td></tr>
<tr><td>Oligo fwd</td><td>CACCCGGTAGGGTGTACGCCCGAC</td><td></td><td></td><td></td><td></td><td></td><td></td><td></td><td></td><td></td><td></td><td></td></tr>
<tr><td>Oligo rev</td><td>AAACGTCGGGCGTACACCCTACCG</td><td></td><td></td><td></td><td></td><td></td><td></td><td></td><td></td><td></td><td></td><td></td></tr>
<tr><td>Chromosome</td><td>start</td><td></td><td>end</td><td>strand</td><td>MM</td><td>target_seq</td><td>PAM</td><td>alignment</td><td>distance</td><td>position</td><td>gene name</td><td>gene id</td></tr>
<tr><td>GU305914.1</td><td></td><td>25305</td><td>25327</td><td>-</td><td>0</td><td>CGGTAGGGTGTACGCCCGAC</td><td>AGG</td><td>||||||||[||||||||||||]PAM</td><td>NA</td><td>Intergenic</td><td>NA</td><td>NA</td></tr>
<tr><td></td><td></td><td></td><td></td><td></td><td></td><td></td><td></td><td></td><td></td><td></td><td></td><td></td></tr>
<tr><td>T32568</td><td>AGGGTGTACGCCCGACAGGCCGG</td><td></td><td>1000</td><td></td><td></td><td></td><td></td><td></td><td></td><td></td><td></td><td></td></tr>
<tr><td>Oligo fwd</td><td>CACCAGGGTGTACGCCCGACAGGC</td><td></td><td></td><td></td><td></td><td></td><td></td><td></td><td></td><td></td><td></td><td></td></tr>
<tr><td>Oligo rev</td><td>AAACGCCTGTCGGGCGTACACCCT</td><td></td><td></td><td></td><td></td><td></td><td></td><td></td><td></td><td></td><td></td><td></td></tr>
<tr><td>Chromosome</td><td>start</td><td></td><td>end</td><td>strand</td><td>MM</td><td>target_seq</td><td>PAM</td><td>alignment</td><td>distance</td><td>position</td><td>gene name</td><td>gene id</td></tr>
<tr><td>GU305914.1</td><td></td><td>25301</td><td>25323</td><td>-</td><td>0</td><td>AGGGTGTACGCCCGACAGGC</td><td>CGG</td><td>||||||||[||||||||||||]PAM</td><td>NA</td><td>Intergenic</td><td>NA</td><td>NA</td></tr>
<tr><td></td><td></td><td></td><td></td><td></td><td></td><td></td><td></td><td></td><td></td><td></td><td></td><td></td></tr>
<tr><td>T32569</td><td>GGGTGTACGCCCGACAGGCCGGG</td><td></td><td>1000</td><td></td><td></td><td></td><td></td><td></td><td></td><td></td><td></td><td></td></tr>
</table>

<table>
<tr><td>Oligo fwd</td><td>CACCGGGTGTACGCCCGACAGGCC</td><td></td><td></td><td></td><td></td><td></td><td></td><td></td><td></td><td></td><td></td></tr>
<tr><td>Oligo rev</td><td>AAACGGCCTGTCGGGCGTACACCC</td><td></td><td></td><td></td><td></td><td></td><td></td><td></td><td></td><td></td><td></td></tr>
<tr><td>Chromosome</td><td>start</td><td>end</td><td>strand</td><td>MM</td><td>target_seq</td><td>PAM</td><td>alignment</td><td>distance</td><td>position</td><td>gene name</td><td>gene id</td></tr>
<tr><td>GU305914.1</td><td>25300</td><td>25322</td><td>-</td><td>0</td><td>GGGTGTACGCCCGACAGGCC</td><td>GGG</td><td>||||||||[||||||||||||]PAM</td><td>NA</td><td>Intergenic</td><td>NA</td><td>NA</td></tr>
<tr><td>T32570</td><td>GGTGTACGCCCGACAGGCCGGGG</td><td>1000</td><td></td><td></td><td></td><td></td><td></td><td></td><td></td><td></td><td></td></tr>
<tr><td>Oligo fwd</td><td>CACCGGTGTACGCCCGACAGGCCG</td><td></td><td></td><td></td><td></td><td></td><td></td><td></td><td></td><td></td><td></td></tr>
<tr><td>Oligo rev</td><td>AAACCGGCCTGTCGGGCGTACACC</td><td></td><td></td><td></td><td></td><td></td><td></td><td></td><td></td><td></td><td></td></tr>
<tr><td>Chromosome</td><td>start</td><td>end</td><td>strand</td><td>MM</td><td>target_seq</td><td>PAM</td><td>alignment</td><td>distance</td><td>position</td><td>gene name</td><td>gene id</td></tr>
<tr><td>GU305914.1</td><td>25299</td><td>25321</td><td>-</td><td>0</td><td>GGTGTACGCCCGACAGGCCG</td><td>GGG</td><td>||||||||[||||||||||||]PAM</td><td>NA</td><td>Intergenic</td><td>NA</td><td>NA</td></tr>
<tr><td>T32571</td><td>TCTCATAGCCGACTCGATCCCGG</td><td>1000</td><td></td><td></td><td></td><td></td><td></td><td></td><td></td><td></td><td></td></tr>
<tr><td>Oligo fwd</td><td>CACCTCTCATAGCCGACTCGATCC</td><td></td><td></td><td></td><td></td><td></td><td></td><td></td><td></td><td></td><td></td></tr>
<tr><td>Oligo rev</td><td>AAACGGATCGAGTCGGCTATGAGA</td><td></td><td></td><td></td><td></td><td></td><td></td><td></td><td></td><td></td><td></td></tr>
<tr><td>Chromosome</td><td>start</td><td>end</td><td>strand</td><td>MM</td><td>target_seq</td><td>PAM</td><td>alignment</td><td>distance</td><td>position</td><td>gene name</td><td>gene id</td></tr>
<tr><td>GU305914.1</td><td>25263</td><td>25285</td><td>-</td><td>0</td><td>TCTCATAGCCGACTCGATCC</td><td>CGG</td><td>||||||||[||||||||||||]PAM</td><td>NA</td><td>Intergenic</td><td>NA</td><td>NA</td></tr>
<tr><td>T32572</td><td>CTCATAGCCGACTCGATCCCGGG</td><td>1000</td><td></td><td></td><td></td><td></td><td></td><td></td><td></td><td></td><td></td></tr>
<tr><td>Oligo fwd</td><td>CACCCTCATAGCCGACTCGATCCC</td><td></td><td></td><td></td><td></td><td></td><td></td><td></td><td></td><td></td><td></td></tr>
<tr><td>Oligo rev</td><td>AAACGGGATCGAGTCGGCTATGAG</td><td></td><td></td><td></td><td></td><td></td><td></td><td></td><td></td><td></td><td></td></tr>
<tr><td>Chromosome</td><td>start</td><td>end</td><td>strand</td><td>MM</td><td>target_seq</td><td>PAM</td><td>alignment</td><td>distance</td><td>position</td><td>gene name</td><td>gene id</td></tr>
<tr><td>GU305914.1</td><td>25262</td><td>25284</td><td>-</td><td>0</td><td>CTCATAGCCGACTCGATCCC</td><td>GGG</td><td>||||||||[||||||||||||]PAM</td><td>NA</td><td>Intergenic</td><td>NA</td><td>NA</td></tr>
<tr><td>T32573</td><td>GACTCGATCCCGGGCTTCGTCGG</td><td>1000</td><td></td><td></td><td></td><td></td><td></td><td></td><td></td><td></td><td></td></tr>
<tr><td>Oligo fwd</td><td>CACCGACTCGATCCCGGGCTTCGT</td><td></td><td></td><td></td><td></td><td></td><td></td><td></td><td></td><td></td><td></td></tr>
<tr><td>Oligo rev</td><td>AAACACGAAGCCCGGGATCGAGTC</td><td></td><td></td><td></td><td></td><td></td><td></td><td></td><td></td><td></td><td></td></tr>
<tr><td>Chromosome</td><td>start</td><td>end</td><td>strand</td><td>MM</td><td>target_seq</td><td>PAM</td><td>alignment</td><td>distance</td><td>position</td><td>gene name</td><td>gene id</td></tr>
<tr><td>GU305914.1</td><td>25253</td><td>25275</td><td>-</td><td>0</td><td>GACTCGATCCCGGGCTTCGT</td><td>CGG</td><td>||||||||[||||||||||||]PAM</td><td>NA</td><td>Intergenic</td><td>NA</td><td>NA</td></tr>
<tr><td>T32574</td><td>ATCCCGGGCTTCGTCGGCATCGG</td><td>1000</td><td></td><td></td><td></td><td></td><td></td><td></td><td></td><td></td><td></td></tr>
<tr><td>Oligo fwd</td><td>CACCATCCCGGGCTTCGTCGGCAT</td><td></td><td></td><td></td><td></td><td></td><td></td><td></td><td></td><td></td><td></td></tr>
<tr><td>Oligo rev</td><td>AAACATGCCGACGAAGCCCGGGAT</td><td></td><td></td><td></td><td></td><td></td><td></td><td></td><td></td><td></td><td></td></tr>
<tr><td>Chromosome</td><td>start</td><td>end</td><td>strand</td><td>MM</td><td>target_seq</td><td>PAM</td><td>alignment</td><td>distance</td><td>position</td><td>gene name</td><td>gene id</td></tr>
<tr><td>GU305914.1</td><td>25247</td><td>25269</td><td>-</td><td>0</td><td>ATCCCGGGCTTCGTCGGCAT</td><td>CGG</td><td>||||||||[||||||||||||]PAM</td><td>NA</td><td>Intergenic</td><td>NA</td><td>NA</td></tr>
<tr><td>T32575</td><td>TCGTCGGCATCGGCGCCACCAGG</td><td>1000</td><td></td><td></td><td></td><td></td><td></td><td></td><td></td><td></td><td></td></tr>
<tr><td>Oligo fwd</td><td>CACCTCGTCGGCATCGGCGCCACC</td><td></td><td></td><td></td><td></td><td></td><td></td><td></td><td></td><td></td><td></td></tr>
<tr><td>Oligo rev</td><td>AAACGGTGGCGCCGATGCCGACGA</td><td></td><td></td><td></td><td></td><td></td><td></td><td></td><td></td><td></td><td></td></tr>
<tr><td>Chromosome</td><td>start</td><td>end</td><td>strand</td><td>MM</td><td>target_seq</td><td>PAM</td><td>alignment</td><td>distance</td><td>position</td><td>gene name</td><td>gene id</td></tr>
<tr><td>GU305914.1</td><td>25237</td><td>25259</td><td>-</td><td>0</td><td>TCGTCGGCATCGGCGCCACC</td><td>AGG</td><td>||||||||[||||||||||||]PAM</td><td>NA</td><td>Intergenic</td><td>NA</td><td>NA</td></tr>
<tr><td>T32576</td><td>TACTTTGAGAATCGACGTTTCGG</td><td>1000</td><td></td><td></td><td></td><td></td><td></td><td></td><td></td><td></td><td></td></tr>
<tr><td>Oligo fwd</td><td>CACCTACTTTGAGAATCGACGTTT</td><td></td><td></td><td></td><td></td><td></td><td></td><td></td><td></td><td></td><td></td></tr>
<tr><td>Oligo rev</td><td>AAACAAACGTCGATTCTCAAAGTA</td><td></td><td></td><td></td><td></td><td></td><td></td><td></td><td></td><td></td><td></td></tr>
<tr><td>Chromosome</td><td>start</td><td>end</td><td>strand</td><td>MM</td><td>target_seq</td><td>PAM</td><td>alignment</td><td>distance</td><td>position</td><td>gene name</td><td>gene id</td></tr>
<tr><td>GU305914.1</td><td>25214</td><td>25236</td><td>-</td><td>0</td><td>TACTTTGAGAATCGACGTTT</td><td>CGG</td><td>||||||||[||||||||||||]PAM</td><td>NA</td><td>Intergenic</td><td>NA</td><td>NA</td></tr>
<tr><td>T32577</td><td>GTTTCGGCCTCATCAGCGAGCGG</td><td>1000</td><td></td><td></td><td></td><td></td><td></td><td></td><td></td><td></td><td></td></tr>
<tr><td>Oligo fwd</td><td>CACCGTTTCGGCCTCATCAGCGAG</td><td></td><td></td><td></td><td></td><td></td><td></td><td></td><td></td><td></td><td></td></tr>
<tr><td>Oligo rev</td><td>AAACCTCGCTGATGAGGCCGAAAC</td><td></td><td></td><td></td><td></td><td></td><td></td><td></td><td></td><td></td><td></td></tr>
<tr><td>Chromosome</td><td>start</td><td>end</td><td>strand</td><td>MM</td><td>target_seq</td><td>PAM</td><td>alignment</td><td>distance</td><td>position</td><td>gene name</td><td>gene id</td></tr>
<tr><td>GU305914.1</td><td>25198</td><td>25220</td><td>-</td><td>0</td><td>GTTTCGGCCTCATCAGCGAG</td><td>CGG</td><td>||||||||[||||||||||||]PAM</td><td>NA</td><td>Intergenic</td><td>NA</td><td>NA</td></tr>
<tr><td>T32578</td><td>TTTCGGCCTCATCAGCGAGCGGG</td><td>1000</td><td></td><td></td><td></td><td></td><td></td><td></td><td></td><td></td><td></td></tr>
<tr><td>Oligo fwd</td><td>CACCTTTCGGCCTCATCAGCGAGC</td><td></td><td></td><td></td><td></td><td></td><td></td><td></td><td></td><td></td><td></td></tr>
<tr><td>Oligo rev</td><td>AAACGCTCGCTGATGAGGCCGAAA</td><td></td><td></td><td></td><td></td><td></td><td></td><td></td><td></td><td></td><td></td></tr>
<tr><td>Chromosome</td><td>start</td><td>end</td><td>strand</td><td>MM</td><td>target_seq</td><td>PAM</td><td>alignment</td><td>distance</td><td>position</td><td>gene name</td><td>gene id</td></tr>
<tr><td>GU305914.1</td><td>25197</td><td>25219</td><td>-</td><td>0</td><td>TTTCGGCCTCATCAGCGAGC</td><td>GGG</td><td>||||||||[||||||||||||]PAM</td><td>NA</td><td>Intergenic</td><td>NA</td><td>NA</td></tr>
<tr><td>T32579</td><td>CTCATCAGCGAGCGGGACTCCGG</td><td>1000</td><td></td><td></td><td></td><td></td><td></td><td></td><td></td><td></td><td></td></tr>
<tr><td>Oligo fwd</td><td>CACCCTCATCAGCGAGCGGGACTC</td><td></td><td></td><td></td><td></td><td></td><td></td><td></td><td></td><td></td><td></td></tr>
<tr><td>Oligo rev</td><td>AAACGAGTCCCGCTCGCTGATGAG</td><td></td><td></td><td></td><td></td><td></td><td></td><td></td><td></td><td></td><td></td></tr>
<tr><td>Chromosome</td><td>start</td><td>end</td><td>strand</td><td>MM</td><td>target_seq</td><td>PAM</td><td>alignment</td><td>distance</td><td>position</td><td>gene name</td><td>gene id</td></tr>
<tr><td>GU305914.1</td><td>25190</td><td>25212</td><td>-</td><td>0</td><td>CTCATCAGCGAGCGGGACTC</td><td>CGG</td><td>||||||||[||||||||||||]PAM</td><td>NA</td><td>Intergenic</td><td>NA</td><td>NA</td></tr>
<tr><td>T32580</td><td>TCATCAGCGAGCGGGACTCCGGG</td><td>1000</td><td></td><td></td><td></td><td></td><td></td><td></td><td></td><td></td><td></td></tr>
<tr><td>Oligo fwd</td><td>CACCTCATCAGCGAGCGGGACTCC</td><td></td><td></td><td></td><td></td><td></td><td></td><td></td><td></td><td></td><td></td></tr>
<tr><td>Oligo rev</td><td>AAACGGAGTCCCGCTCGCTGATGA</td><td></td><td></td><td></td><td></td><td></td><td></td><td></td><td></td><td></td><td></td></tr>
<tr><td>Chromosome</td><td>start</td><td>end</td><td>strand</td><td>MM</td><td>target_seq</td><td>PAM</td><td>alignment</td><td>distance</td><td>position</td><td>gene name</td><td>gene id</td></tr>
<tr><td>GU305914.1</td><td>25189</td><td>25211</td><td>-</td><td>0</td><td>TCATCAGCGAGCGGGACTCC</td><td>GGG</td><td>||||||||[||||||||||||]PAM</td><td>NA</td><td>Intergenic</td><td>NA</td><td>NA</td></tr>
<tr><td>T32581</td><td>TCAGCGAGCGGGACTCCGGGAGG</td><td>1000</td><td></td><td></td><td></td><td></td><td></td><td></td><td></td><td></td><td></td></tr>
<tr><td>Oligo fwd</td><td>CACCTCAGCGAGCGGGACTCCGGG</td><td></td><td></td><td></td><td></td><td></td><td></td><td></td><td></td><td></td><td></td></tr>
<tr><td>Oligo rev</td><td>AAACCCCGGAGTCCCGCTCGCTGA</td><td></td><td></td><td></td><td></td><td></td><td></td><td></td><td></td><td></td><td></td></tr>
<tr><td>Chromosome</td><td>start</td><td>end</td><td>strand</td><td>MM</td><td>target_seq</td><td>PAM</td><td>alignment</td><td>distance</td><td>position</td><td>gene name</td><td>gene id</td></tr>
<tr><td>GU305914.1</td><td>25186</td><td>25208</td><td>-</td><td>0</td><td>TCAGCGAGCGGGACTCCGGG</td><td>AGG</td><td>||||||||[||||||||||||]PAM</td><td>NA</td><td>Intergenic</td><td>NA</td><td>NA</td></tr>
<tr><td>T32582</td><td>ACTCCGGGAGGAACTGTGAGAGG</td><td>1000</td><td></td><td></td><td></td><td></td><td></td><td></td><td></td><td></td><td></td></tr>
<tr><td>Oligo fwd</td><td>CACCACTCCGGGAGGAACTGTGAG</td><td></td><td></td><td></td><td></td><td></td><td></td><td></td><td></td><td></td><td></td></tr>
<tr><td>Oligo rev</td><td>AAACCTCACAGTTCCTCCCGGAGT</td><td></td><td></td><td></td><td></td><td></td><td></td><td></td><td></td><td></td><td></td></tr>
<tr><td>Chromosome</td><td>start</td><td>end</td><td>strand</td><td>MM</td><td>target_seq</td><td>PAM</td><td>alignment</td><td>distance</td><td>position</td><td>gene name</td><td>gene id</td></tr>
<tr><td>GU305914.1</td><td>25174</td><td>25196</td><td>-</td><td>0</td><td>ACTCCGGGAGGAACTGTGAG</td><td>AGG</td><td>||||||||[||||||||||||]PAM</td><td>NA</td><td>Intergenic</td><td>NA</td><td>NA</td></tr>
<tr><td>T32583</td><td>TTCCTCCTTTCTCTGTCGATCGG</td><td>1000</td><td></td><td></td><td></td><td></td><td></td><td></td><td></td><td></td><td></td></tr>
<tr><td>Oligo fwd</td><td>CACCTTCCTCCTTTCTCTGTCGAT</td><td></td><td></td><td></td><td></td><td></td><td></td><td></td><td></td><td></td><td></td></tr>
<tr><td>Oligo rev</td><td>AAACATCGACAGAGAAAGGAGGAA</td><td></td><td></td><td></td><td></td><td></td><td></td><td></td><td></td><td></td><td></td></tr>
<tr><td>Chromosome</td><td>start</td><td>end</td><td>strand</td><td>MM</td><td>target_seq</td><td>PAM</td><td>alignment</td><td>distance</td><td>position</td><td>gene name</td><td>gene id</td></tr>
<tr><td>GU305914.1</td><td>25106</td><td>25128</td><td>-</td><td>0</td><td>TTCCTCCTTTCTCTGTCGAT</td><td>CGG</td><td>||||||||[||||||||||||]PAM</td><td>NA</td><td>Intergenic</td><td>NA</td><td>NA</td></tr>
</table>

T32584 TCGATCGGTCGATCGTCAGACGG 1000
Oligo fwd CACCTCGATCGGTCGATCGTCAGA
Oligo rev AAACTCTGACGATCGACCGATCGA
Chromosome start end strand MM target_seq PAM alignment distance position gene name gene id
GU305914.1 25091 25113 - 0 TCGATCGGTCGATCGTCAGA CGG ||||||||[||||||||||||]PAM NA Intergenic NA NA

T32585 TCGATCGTCAGACGGCTACGTGG 1000
Oligo fwd CACCTCGATCGTCAGACGGCTACG
Oligo rev AAACCGTAGCCGTCTGACGATCGA
Chromosome start end strand MM target_seq PAM alignment distance position gene name gene id
GU305914.1 25083 25105 - 0 TCGATCGTCAGACGGCTACG TGG ||||||||[||||||||||||]PAM NA Intergenic NA NA

T32586 CGATCGTCAGACGGCTACGTGGG 1000
Oligo fwd CACCCGATCGTCAGACGGCTACGT
Oligo rev AAACACGTAGCCGTCTGACGATCG
Chromosome start end strand MM target_seq PAM alignment distance position gene name gene id
GU305914.1 25082 25104 - 0 CGATCGTCAGACGGCTACGT GGG ||||||||[||||||||||||]PAM NA Intergenic NA NA

T32587 CGGCTACGTGGGACACACGTCGG 1000
Oligo fwd CACCCGGCTACGTGGGACACACGT
Oligo rev AAACACGTGTGTCCCACGTAGCCG
Chromosome start end strand MM target_seq PAM alignment distance position gene name gene id
GU305914.1 25071 25093 - 0 CGGCTACGTGGGACACACGT CGG ||||||||[||||||||||||]PAM NA Intergenic NA NA

T32588 CTACGTGGGACACACGTCGGAGG 1000
Oligo fwd CACCCTACGTGGGACACACGTCGG
Oligo rev AAACCCGACGTGTGTCCCACGTAG
Chromosome start end strand MM target_seq PAM alignment distance position gene name gene id
GU305914.1 25068 25090 - 0 CTACGTGGGACACACGTCGG AGG ||||||||[||||||||||||]PAM NA Intergenic NA NA

T32589 TACGTGGGACACACGTCGGAGGG 1000
Oligo fwd CACCTACGTGGGACACACGTCGGA
Oligo rev AAACTCCGACGTGTGTCCCACGTA
Chromosome start end strand MM target_seq PAM alignment distance position gene name gene id
GU305914.1 25067 25089 - 0 TACGTGGGACACACGTCGGA GGG ||||||||[||||||||||||]PAM NA Intergenic NA NA

T32590 TGGGACACACGTCGGAGGGTCGG 1000
Oligo fwd CACCTGGGACACACGTCGGAGGGT
Oligo rev AAACACCCTCCGACGTGTGTCCCA
Chromosome start end strand MM target_seq PAM alignment distance position gene name gene id
GU305914.1 25063 25085 - 0 TGGGACACACGTCGGAGGGT CGG ||||||||[||||||||||||]PAM NA Intergenic NA NA

T32591 GGGACACACGTCGGAGGGTCGGG 1000
Oligo fwd CACCGGGACACACGTCGGAGGGTC
Oligo rev AAACGACCCTCCGACGTGTGTCCC
Chromosome start end strand MM target_seq PAM alignment distance position gene name gene id
GU305914.1 25062 25084 - 0 GGGACACACGTCGGAGGGTC GGG ||||||||[||||||||||||]PAM NA Intergenic NA NA

T32592 CACGTCGGAGGGTCGGGACACGG 1000
Oligo fwd CACCCACGTCGGAGGGTCGGGACA
Oligo rev AAACTGTCCCGACCCTCCGACGTG
Chromosome start end strand MM target_seq PAM alignment distance position gene name gene id
GU305914.1 25056 25078 - 0 CACGTCGGAGGGTCGGGACA CGG ||||||||[||||||||||||]PAM NA Intergenic NA NA

T32593 ACGGCCCCTTGCCGCGCCCACGG 1000
Oligo fwd CACCACGGCCCCTTGCCGCGCCCA
Oligo rev AAACTGGGCGCGGCAAGGGGCCGT
Chromosome start end strand MM target_seq PAM alignment distance position gene name gene id
GU305914.1 25037 25059 - 0 ACGGCCCCTTGCCGCGCCCA CGG ||||||||[||||||||||||]PAM NA Intergenic NA NA

T32594 CACGGACGCCCACACACAAAAGG 1000
Oligo fwd CACCCACGGACGCCCACACACAAA
Oligo rev AAACTTTGTGTGTGGGCGTCCGTG
Chromosome start end strand MM target_seq PAM alignment distance position gene name gene id
GU305914.1 25019 25041 - 0 CACGGACGCCCACACACAAA AGG ||||||||[||||||||||||]PAM NA Intergenic NA NA

T32595 CGCCCACACACAAAAGGCCGAGG 1000
Oligo fwd CACCCGCCCACACACAAAAGGCCG
Oligo rev AAACCGGCCTTTTGTGTGTGGGCG
Chromosome start end strand MM target_seq PAM alignment distance position gene name gene id
GU305914.1 25013 25035 - 0 CGCCCACACACAAAAGGCCG AGG ||||||||[||||||||||||]PAM NA Intergenic NA NA

T32596 GCCCACACACAAAAGGCCGAGGG 1000
Oligo fwd CACCGCCCACACACAAAAGGCCGA
Oligo rev AAACTCGGCCTTTTGTGTGTGGGC
Chromosome start end strand MM target_seq PAM alignment distance position gene name gene id
GU305914.1 25012 25034 - 0 GCCCACACACAAAAGGCCGA GGG ||||||||[||||||||||||]PAM NA Intergenic NA NA

T32597 ACGCTGTCGATCGCTCGATATGG 1000
Oligo fwd CACCACGCTGTCGATCGCTCGATA
Oligo rev AAACTATCGAGCGATCGACAGCGT
Chromosome start end strand MM target_seq PAM alignment distance position gene name gene id
GU305914.1 24949 24971 - 0 ACGCTGTCGATCGCTCGATA TGG ||||||||[||||||||||||]PAM NA Intergenic NA NA

T32598 ATCGCTCGATATGGCGCCTCCGG 1000
Oligo fwd CACCATCGCTCGATATGGCGCCTC
Oligo rev AAACGAGGCGCCATATCGAGCGAT
Chromosome start end strand MM target_seq PAM alignment distance position gene name gene id
GU305914.1 24940 24962 - 0 ATCGCTCGATATGGCGCCTC CGG ||||||||[||||||||||||]PAM NA Intergenic NA NA

| | | | | | | | | | | | |
|---|---|---|---|---|---|---|---|---|---|---|---|
| T32599 | CAGCGACTTTGCCACCGTCGAGG | 1000 | | | | | | | | | |
| Oligo fwd | CACCCAGCGACTTTGCCACCGTCG | | | | | | | | | | |
| Oligo rev | AAACCGACGGTGGCAAAGTCGCTG | | | | | | | | | | |
| Chromosome | start | end | strand | MM | target_seq | PAM | alignment | distance | position | gene name | gene id |
| GU305914.1 | 24898 | 24920 | - | 0 | CAGCGACTTTGCCACCGTCG | AGG | \|\|\|\|\|\|\|\|[\|\|\|\|\|\|\|\|\|\|\|\|]PAM | NA | Intergenic | NA | NA |
| T32600 | GCCGCCGTCGCCGCCAACATCGG | 1000 | | | | | | | | | |
| Oligo fwd | CACCGCCGCCGTCGCCGCCAACAT | | | | | | | | | | |
| Oligo rev | AAACATGTTGGCGGCGACGGCGGC | | | | | | | | | | |
| Chromosome | start | end | strand | MM | target_seq | PAM | alignment | distance | position | gene name | gene id |
| GU305914.1 | 24876 | 24898 | - | 0 | GCCGCCGTCGCCGCCAACAT | CGG | \|\|\|\|\|\|\|\|[\|\|\|\|\|\|\|\|\|\|\|\|]PAM | NA | Intergenic | NA | NA |
| GU305914.1 | 170532 | 170554 | + | 4 | GTGGCCGCCGCCGCCACCAT | CGG | \|--\|\|\|\|-[\|\|\|\|\|\|\|\|-\|\|\|]PAM | NA | Intergenic | NA | NA |
| GU305914.1 | 29998 | 30020 | - | 3 | GCCGCCGCCGCCGCCAGCAC | CGG | \|\|\|\|\|\|\|-[\|\|\|\|\|\|\|\|-\|\|-]PAM | NA | Intergenic | NA | NA |
| T32601 | CCGCCGTCGCCGCCAACATCGGG | 1000 | | | | | | | | | |
| Oligo fwd | CACCCCGCCGTCGCCGCCAACATC | | | | | | | | | | |
| Oligo rev | AAACGATGTTGGCGGCGACGGCGG | | | | | | | | | | |
| Chromosome | start | end | strand | MM | target_seq | PAM | alignment | distance | position | gene name | gene id |
| GU305914.1 | 24875 | 24897 | - | 0 | CCGCCGTCGCCGCCAACATC | GGG | \|\|\|\|\|\|\|\|[\|\|\|\|\|\|\|\|\|\|\|\|]PAM | NA | Intergenic | NA | NA |
| T32602 | GTGCGTCTCCACCACCCGAAAGG | 1000 | | | | | | | | | |
| Oligo fwd | CACCGTGCGTCTCCACCACCCGAA | | | | | | | | | | |
| Oligo rev | AAACTTCGGGTGGTGGAGACGCAC | | | | | | | | | | |
| Chromosome | start | end | strand | MM | target_seq | PAM | alignment | distance | position | gene name | gene id |
| GU305914.1 | 24846 | 24868 | - | 0 | GTGCGTCTCCACCACCCGAA | AGG | \|\|\|\|\|\|\|\|[\|\|\|\|\|\|\|\|\|\|\|\|]PAM | NA | Intergenic | NA | NA |
| T32603 | TGCGTCTCCACCACCCGAAAGGG | 1000 | | | | | | | | | |
| Oligo fwd | CACCTGCGTCTCCACCACCCGAAA | | | | | | | | | | |
| Oligo rev | AAACTTTCGGGTGGTGGAGACGCA | | | | | | | | | | |
| Chromosome | start | end | strand | MM | target_seq | PAM | alignment | distance | position | gene name | gene id |
| GU305914.1 | 24845 | 24867 | - | 0 | TGCGTCTCCACCACCCGAAA | GGG | \|\|\|\|\|\|\|\|[\|\|\|\|\|\|\|\|\|\|\|\|]PAM | NA | Intergenic | NA | NA |
| T32604 | AAAGGGCTCTCGCTCTGCATCGG | 1000 | | | | | | | | | |
| Oligo fwd | CACCAAAGGGCTCTCGCTCTGCAT | | | | | | | | | | |
| Oligo rev | AAACATGCAGAGCGAGAGCCCTTT | | | | | | | | | | |
| Chromosome | start | end | strand | MM | target_seq | PAM | alignment | distance | position | gene name | gene id |
| GU305914.1 | 24828 | 24850 | - | 0 | AAAGGGCTCTCGCTCTGCAT | CGG | \|\|\|\|\|\|\|\|[\|\|\|\|\|\|\|\|\|\|\|\|]PAM | NA | Intergenic | NA | NA |
| T32605 | CGCTCTGCATCGGCATGCGCTGG | 1000 | | | | | | | | | |
| Oligo fwd | CACCCGCTCTGCATCGGCATGCGC | | | | | | | | | | |
| Oligo rev | AAACGCGCATGCCGATGCAGAGCG | | | | | | | | | | |
| Chromosome | start | end | strand | MM | target_seq | PAM | alignment | distance | position | gene name | gene id |
| GU305914.1 | 24818 | 24840 | - | 0 | CGCTCTGCATCGGCATGCGC | TGG | \|\|\|\|\|\|\|\|[\|\|\|\|\|\|\|\|\|\|\|\|]PAM | NA | Intergenic | NA | NA |
| T32606 | CTGGCGCGAGATACCCGCGCCGG | 1000 | | | | | | | | | |
| Oligo fwd | CACCCTGGCGCGAGATACCCGCGC | | | | | | | | | | |
| Oligo rev | AAACGCGCGGGTATCTCGCGCCAG | | | | | | | | | | |
| Chromosome | start | end | strand | MM | target_seq | PAM | alignment | distance | position | gene name | gene id |
| GU305914.1 | 24799 | 24821 | - | 0 | CTGGCGCGAGATACCCGCGC | CGG | \|\|\|\|\|\|\|\|[\|\|\|\|\|\|\|\|\|\|\|\|]PAM | NA | Intergenic | NA | NA |
| T32607 | GAGATACCCGCGCCGGCCAACGG | 1000 | | | | | | | | | |
| Oligo fwd | CACCGAGATACCCGCGCCGGCCAA | | | | | | | | | | |
| Oligo rev | AAACTTGGCCGGCGCGGGTATCTC | | | | | | | | | | |
| Chromosome | start | end | strand | MM | target_seq | PAM | alignment | distance | position | gene name | gene id |
| GU305914.1 | 24792 | 24814 | - | 0 | GAGATACCCGCGCCGGCCAA | CGG | \|\|\|\|\|\|\|\|[\|\|\|\|\|\|\|\|\|\|\|\|]PAM | NA | Intergenic | NA | NA |
| T32608 | AGATACCCGCGCCGGCCAACGGG | 1000 | | | | | | | | | |
| Oligo fwd | CACCAGATACCCGCGCCGGCCAAC | | | | | | | | | | |
| Oligo rev | AAACGTTGGCCGGCGCGGGTATCT | | | | | | | | | | |
| Chromosome | start | end | strand | MM | target_seq | PAM | alignment | distance | position | gene name | gene id |
| GU305914.1 | 24791 | 24813 | - | 0 | AGATACCCGCGCCGGCCAAC | GGG | \|\|\|\|\|\|\|\|[\|\|\|\|\|\|\|\|\|\|\|\|]PAM | NA | Intergenic | NA | NA |
| T32609 | GATACCCGCGCCGGCCAACGGGG | 1000 | | | | | | | | | |
| Oligo fwd | CACCGATACCCGCGCCGGCCAACG | | | | | | | | | | |
| Oligo rev | AAACCGTTGGCCGGCGCGGGTATC | | | | | | | | | | |
| Chromosome | start | end | strand | MM | target_seq | PAM | alignment | distance | position | gene name | gene id |
| GU305914.1 | 24790 | 24812 | - | 0 | GATACCCGCGCCGGCCAACG | GGG | \|\|\|\|\|\|\|\|[\|\|\|\|\|\|\|\|\|\|\|\|]PAM | NA | Intergenic | NA | NA |
| T32610 | ATACCCGCGCCGGCCAACGGGGG | 1000 | | | | | | | | | |
| Oligo fwd | CACCATACCCGCGCCGGCCAACGG | | | | | | | | | | |
| Oligo rev | AAACCCGTTGGCCGGCGCGGGTAT | | | | | | | | | | |
| Chromosome | start | end | strand | MM | target_seq | PAM | alignment | distance | position | gene name | gene id |
| GU305914.1 | 24789 | 24811 | - | 0 | ATACCCGCGCCGGCCAACGG | GGG | \|\|\|\|\|\|\|\|[\|\|\|\|\|\|\|\|\|\|\|\|]PAM | NA | Intergenic | NA | NA |
| T32611 | TACCCGCGCCGGCCAACGGGGGG | 1000 | | | | | | | | | |
| Oligo fwd | CACCTACCCGCGCCGGCCAACGGG | | | | | | | | | | |
| Oligo rev | AAACCCCGTTGGCCGGCGCGGGTA | | | | | | | | | | |
| Chromosome | start | end | strand | MM | target_seq | PAM | alignment | distance | position | gene name | gene id |
| GU305914.1 | 24788 | 24810 | - | 0 | TACCCGCGCCGGCCAACGGG | GGG | \|\|\|\|\|\|\|\|[\|\|\|\|\|\|\|\|\|\|\|\|]PAM | NA | Intergenic | NA | NA |
| T32612 | CACGAGAACATCAAGTTCATCGG | 1000 | | | | | | | | | |
| Oligo fwd | CACCCACGAGAACATCAAGTTCAT | | | | | | | | | | |
| Oligo rev | AAACATGAACTTGATGTTCTCGTG | | | | | | | | | | |
| Chromosome | start | end | strand | MM | target_seq | PAM | alignment | distance | position | gene name | gene id |
| GU305914.1 | 24723 | 24745 | - | 0 | CACGAGAACATCAAGTTCAT | CGG | \|\|\|\|\|\|\|\|[\|\|\|\|\|\|\|\|\|\|\|\|]PAM | NA | Intergenic | NA | NA |
| T32613 | CGCCTCCGATTCTAACCTCGAGG | 1000 | | | | | | | | | |
| Oligo fwd | CACCCGCCTCCGATTCTAACCTCG | | | | | | | | | | |

<table>
<tr><td>Oligo rev</td><td>AAACCGAGGTTAGAATCGGAGGCG</td><td></td><td></td><td></td><td></td><td></td><td></td><td></td><td></td><td></td><td></td></tr>
<tr><td>Chromosome</td><td>start</td><td>end</td><td>strand</td><td>MM</td><td>target_seq</td><td>PAM</td><td>alignment</td><td>distance</td><td>position</td><td>gene name</td><td>gene id</td></tr>
<tr><td>GU305914.1</td><td>24691</td><td>24713</td><td>-</td><td>0</td><td>CGCCTCCGATTCTAACCTCG</td><td>AGG</td><td>||||||||[||||||||||||]PAM</td><td>NA</td><td>Intergenic</td><td>NA</td><td>NA</td></tr>
<tr><td></td><td></td><td></td><td></td><td></td><td></td><td></td><td></td><td></td><td></td><td></td><td></td></tr>
<tr><td>T32614</td><td>GCCTCCGATTCTAACCTCGAGGG</td><td>1000</td><td></td><td></td><td></td><td></td><td></td><td></td><td></td><td></td><td></td></tr>
<tr><td>Oligo fwd</td><td>CACCGCCTCCGATTCTAACCTCGA</td><td></td><td></td><td></td><td></td><td></td><td></td><td></td><td></td><td></td><td></td></tr>
<tr><td>Oligo rev</td><td>AAACTCGAGGTTAGAATCGGAGGC</td><td></td><td></td><td></td><td></td><td></td><td></td><td></td><td></td><td></td><td></td></tr>
<tr><td>Chromosome</td><td>start</td><td>end</td><td>strand</td><td>MM</td><td>target_seq</td><td>PAM</td><td>alignment</td><td>distance</td><td>position</td><td>gene name</td><td>gene id</td></tr>
<tr><td>GU305914.1</td><td>24690</td><td>24712</td><td>-</td><td>0</td><td>GCCTCCGATTCTAACCTCGA</td><td>GGG</td><td>||||||||[||||||||||||]PAM</td><td>NA</td><td>Intergenic</td><td>NA</td><td>NA</td></tr>
<tr><td></td><td></td><td></td><td></td><td></td><td></td><td></td><td></td><td></td><td></td><td></td><td></td></tr>
<tr><td>T32615</td><td>CCAGATCGCGCCTCCTCAACTGG</td><td>1000</td><td></td><td></td><td></td><td></td><td></td><td></td><td></td><td></td><td></td></tr>
<tr><td>Oligo fwd</td><td>CACCCCAGATCGCGCCTCCTCAAC</td><td></td><td></td><td></td><td></td><td></td><td></td><td></td><td></td><td></td><td></td></tr>
<tr><td>Oligo rev</td><td>AAACGTTGAGGAGGCGCGATCTGG</td><td></td><td></td><td></td><td></td><td></td><td></td><td></td><td></td><td></td><td></td></tr>
<tr><td>Chromosome</td><td>start</td><td>end</td><td>strand</td><td>MM</td><td>target_seq</td><td>PAM</td><td>alignment</td><td>distance</td><td>position</td><td>gene name</td><td>gene id</td></tr>
<tr><td>GU305914.1</td><td>24656</td><td>24678</td><td>-</td><td>0</td><td>CCAGATCGCGCCTCCTCAAC</td><td>TGG</td><td>||||||||[||||||||||||]PAM</td><td>NA</td><td>Intergenic</td><td>NA</td><td>NA</td></tr>
<tr><td></td><td></td><td></td><td></td><td></td><td></td><td></td><td></td><td></td><td></td><td></td><td></td></tr>
<tr><td>T32616</td><td>AACTGGCGCTTTCGCAACACCGG</td><td>1000</td><td></td><td></td><td></td><td></td><td></td><td></td><td></td><td></td><td></td></tr>
<tr><td>Oligo fwd</td><td>CACCAACTGGCGCTTTCGCAACAC</td><td></td><td></td><td></td><td></td><td></td><td></td><td></td><td></td><td></td><td></td></tr>
<tr><td>Oligo rev</td><td>AAACGTGTTGCGAAAGCGCCAGTT</td><td></td><td></td><td></td><td></td><td></td><td></td><td></td><td></td><td></td><td></td></tr>
<tr><td>Chromosome</td><td>start</td><td>end</td><td>strand</td><td>MM</td><td>target_seq</td><td>PAM</td><td>alignment</td><td>distance</td><td>position</td><td>gene name</td><td>gene id</td></tr>
<tr><td>GU305914.1</td><td>24639</td><td>24661</td><td>-</td><td>0</td><td>AACTGGCGCTTTCGCAACAC</td><td>CGG</td><td>||||||||[||||||||||||]PAM</td><td>NA</td><td>Intergenic</td><td>NA</td><td>NA</td></tr>
<tr><td></td><td></td><td></td><td></td><td></td><td></td><td></td><td></td><td></td><td></td><td></td><td></td></tr>
<tr><td>T32617</td><td>TGGCGCTTTCGCAACACCGGCGG</td><td>1000</td><td></td><td></td><td></td><td></td><td></td><td></td><td></td><td></td><td></td></tr>
<tr><td>Oligo fwd</td><td>CACCTGGCGCTTTCGCAACACCGG</td><td></td><td></td><td></td><td></td><td></td><td></td><td></td><td></td><td></td><td></td></tr>
<tr><td>Oligo rev</td><td>AAACCCGGTGTTGCGAAAGCGCCA</td><td></td><td></td><td></td><td></td><td></td><td></td><td></td><td></td><td></td><td></td></tr>
<tr><td>Chromosome</td><td>start</td><td>end</td><td>strand</td><td>MM</td><td>target_seq</td><td>PAM</td><td>alignment</td><td>distance</td><td>position</td><td>gene name</td><td>gene id</td></tr>
<tr><td>GU305914.1</td><td>24636</td><td>24658</td><td>-</td><td>0</td><td>TGGCGCTTTCGCAACACCGG</td><td>CGG</td><td>||||||||[||||||||||||]PAM</td><td>NA</td><td>Intergenic</td><td>NA</td><td>NA</td></tr>
<tr><td></td><td></td><td></td><td></td><td></td><td></td><td></td><td></td><td></td><td></td><td></td><td></td></tr>
<tr><td>T32618</td><td>CGCTTTCGCAACACCGGCGGCGG</td><td>1000</td><td></td><td></td><td></td><td></td><td></td><td></td><td></td><td></td><td></td></tr>
<tr><td>Oligo fwd</td><td>CACCCGCTTTCGCAACACCGGCGG</td><td></td><td></td><td></td><td></td><td></td><td></td><td></td><td></td><td></td><td></td></tr>
<tr><td>Oligo rev</td><td>AAACCCGCCGGTGTTGCGAAAGCG</td><td></td><td></td><td></td><td></td><td></td><td></td><td></td><td></td><td></td><td></td></tr>
<tr><td>Chromosome</td><td>start</td><td>end</td><td>strand</td><td>MM</td><td>target_seq</td><td>PAM</td><td>alignment</td><td>distance</td><td>position</td><td>gene name</td><td>gene id</td></tr>
<tr><td>GU305914.1</td><td>24633</td><td>24655</td><td>-</td><td>0</td><td>CGCTTTCGCAACACCGGCGG</td><td>CGG</td><td>||||||||[||||||||||||]PAM</td><td>NA</td><td>Intergenic</td><td>NA</td><td>NA</td></tr>
<tr><td></td><td></td><td></td><td></td><td></td><td></td><td></td><td></td><td></td><td></td><td></td><td></td></tr>
<tr><td>T32619</td><td>TTTCGCAACACCGGCGGCGGCGG</td><td>1000</td><td></td><td></td><td></td><td></td><td></td><td></td><td></td><td></td><td></td></tr>
<tr><td>Oligo fwd</td><td>CACCTTTCGCAACACCGGCGGCGG</td><td></td><td></td><td></td><td></td><td></td><td></td><td></td><td></td><td></td><td></td></tr>
<tr><td>Oligo rev</td><td>AAACCCGCCGCCGGTGTTGCGAAA</td><td></td><td></td><td></td><td></td><td></td><td></td><td></td><td></td><td></td><td></td></tr>
<tr><td>Chromosome</td><td>start</td><td>end</td><td>strand</td><td>MM</td><td>target_seq</td><td>PAM</td><td>alignment</td><td>distance</td><td>position</td><td>gene name</td><td>gene id</td></tr>
<tr><td>GU305914.1</td><td>24630</td><td>24652</td><td>-</td><td>0</td><td>TTTCGCAACACCGGCGGCGG</td><td>CGG</td><td>||||||||[||||||||||||]PAM</td><td>NA</td><td>Intergenic</td><td>NA</td><td>NA</td></tr>
<tr><td></td><td></td><td></td><td></td><td></td><td></td><td></td><td></td><td></td><td></td><td></td><td></td></tr>
<tr><td>T32620</td><td>CGCAACACCGGCGGCGGCGGTGG</td><td>1000</td><td></td><td></td><td></td><td></td><td></td><td></td><td></td><td></td><td></td></tr>
<tr><td>Oligo fwd</td><td>CACCCGCAACACCGGCGGCGGCGG</td><td></td><td></td><td></td><td></td><td></td><td></td><td></td><td></td><td></td><td></td></tr>
<tr><td>Oligo rev</td><td>AAACCCGCCGCCGCCGGTGTTGCG</td><td></td><td></td><td></td><td></td><td></td><td></td><td></td><td></td><td></td><td></td></tr>
<tr><td>Chromosome</td><td>start</td><td>end</td><td>strand</td><td>MM</td><td>target_seq</td><td>PAM</td><td>alignment</td><td>distance</td><td>position</td><td>gene name</td><td>gene id</td></tr>
<tr><td>GU305914.1</td><td>24627</td><td>24649</td><td>-</td><td>0</td><td>CGCAACACCGGCGGCGGCGG</td><td>TGG</td><td>||||||||[||||||||||||]PAM</td><td>NA</td><td>Intergenic</td><td>NA</td><td>NA</td></tr>
<tr><td>GU305914.1</td><td>56773</td><td>56795</td><td>+</td><td>3</td><td>CGCCGCAGCGGCGGCGGCGG</td><td>CGG</td><td>|||--||-[||||||||||||]PAM</td><td>NA</td><td>Intergenic</td><td>NA</td><td>NA</td></tr>
<tr><td>GU305914.1</td><td>149482</td><td>149504</td><td>-</td><td>4</td><td>GGCGACGGCGGCGGCGGCGG</td><td>CGG</td><td>-||-||--[||||||||||||]PAM</td><td>NA</td><td>Intergenic</td><td>NA</td><td>NA</td></tr>
<tr><td>GU305914.1</td><td>58135</td><td>58157</td><td>-</td><td>4</td><td>CGCAAGGCGGGCGGTGGCGG</td><td>CGG</td><td>|||||--|[-|||||-|||||]PAM</td><td>NA</td><td>Intergenic</td><td>NA</td><td>NA</td></tr>
<tr><td>GU305914.1</td><td>70181</td><td>70203</td><td>+</td><td>4</td><td>AGCAGCACCGGCAGCGCCGG</td><td>CGG</td><td>-|||-|||[||||-|||-|||]PAM</td><td>NA</td><td>Intergenic</td><td>NA</td><td>NA</td></tr>
<tr><td></td><td></td><td></td><td></td><td></td><td></td><td></td><td></td><td></td><td></td><td></td><td></td></tr>
<tr><td>T32621</td><td>AACACCGGCGGCGGCGGTGGCGG</td><td>1000</td><td></td><td></td><td></td><td></td><td></td><td></td><td></td><td></td><td></td></tr>
<tr><td>Oligo fwd</td><td>CACCAACACCGGCGGCGGCGGTGG</td><td></td><td></td><td></td><td></td><td></td><td></td><td></td><td></td><td></td><td></td></tr>
<tr><td>Oligo rev</td><td>AAACCCACCGCCGCCGCCGGTGTT</td><td></td><td></td><td></td><td></td><td></td><td></td><td></td><td></td><td></td><td></td></tr>
<tr><td>Chromosome</td><td>start</td><td>end</td><td>strand</td><td>MM</td><td>target_seq</td><td>PAM</td><td>alignment</td><td>distance</td><td>position</td><td>gene name</td><td>gene id</td></tr>
<tr><td>GU305914.1</td><td>24624</td><td>24646</td><td>-</td><td>0</td><td>AACACCGGCGGCGGCGGTGG</td><td>CGG</td><td>||||||||[||||||||||||]PAM</td><td>NA</td><td>Intergenic</td><td>NA</td><td>NA</td></tr>
<tr><td>GU305914.1</td><td>48150</td><td>48172</td><td>+</td><td>4</td><td>ACCTGTGGCGGCGGCGGTGG</td><td>CGG</td><td>|-|---||[||||||||||||]PAM</td><td>NA</td><td>Intergenic</td><td>NA</td><td>NA</td></tr>
<tr><td>GU305914.1</td><td>35653</td><td>35675</td><td>+</td><td>4</td><td>ATCCACGGCGGCGGCGGCGG</td><td>CGG</td><td>|-|--|||[|||||||||-||]PAM</td><td>NA</td><td>Intergenic</td><td>NA</td><td>NA</td></tr>
<tr><td>GU305914.1</td><td>73786</td><td>73808</td><td>-</td><td>4</td><td>AGCAGCGGCGGCGGAGGAGG</td><td>AGG</td><td>|-||-|||[||||||-||-||]PAM</td><td>NA</td><td>Intergenic</td><td>NA</td><td>NA</td></tr>
<tr><td></td><td></td><td></td><td></td><td></td><td></td><td></td><td></td><td></td><td></td><td></td><td></td></tr>
<tr><td>T32622</td><td>GGTGGCGGCGACGACACGCGCGG</td><td>1000</td><td></td><td></td><td></td><td></td><td></td><td></td><td></td><td></td><td></td></tr>
<tr><td>Oligo fwd</td><td>CACCGGTGGCGGCGACGACACGCG</td><td></td><td></td><td></td><td></td><td></td><td></td><td></td><td></td><td></td><td></td></tr>
<tr><td>Oligo rev</td><td>AAACCGCGTGTCGTCGCCGCCACC</td><td></td><td></td><td></td><td></td><td></td><td></td><td></td><td></td><td></td><td></td></tr>
<tr><td>Chromosome</td><td>start</td><td>end</td><td>strand</td><td>MM</td><td>target_seq</td><td>PAM</td><td>alignment</td><td>distance</td><td>position</td><td>gene name</td><td>gene id</td></tr>
<tr><td>GU305914.1</td><td>24609</td><td>24631</td><td>-</td><td>0</td><td>GGTGGCGGCGACGACACGCG</td><td>CGG</td><td>||||||||[||||||||||||]PAM</td><td>NA</td><td>Intergenic</td><td>NA</td><td>NA</td></tr>
<tr><td></td><td></td><td></td><td></td><td></td><td></td><td></td><td></td><td></td><td></td><td></td><td></td></tr>
<tr><td>T32623</td><td>CGGACCGCGCGATCCCGACGAGG</td><td>1000</td><td></td><td></td><td></td><td></td><td></td><td></td><td></td><td></td><td></td></tr>
<tr><td>Oligo fwd</td><td>CACCCGGACCGCGCGATCCCGACG</td><td></td><td></td><td></td><td></td><td></td><td></td><td></td><td></td><td></td><td></td></tr>
<tr><td>Oligo rev</td><td>AAACCGTCGGGATCGCGCGGTCCG</td><td></td><td></td><td></td><td></td><td></td><td></td><td></td><td></td><td></td><td></td></tr>
<tr><td>Chromosome</td><td>start</td><td>end</td><td>strand</td><td>MM</td><td>target_seq</td><td>PAM</td><td>alignment</td><td>distance</td><td>position</td><td>gene name</td><td>gene id</td></tr>
<tr><td>GU305914.1</td><td>24589</td><td>24611</td><td>-</td><td>0</td><td>CGGACCGCGCGATCCCGACG</td><td>AGG</td><td>||||||||[||||||||||||]PAM</td><td>NA</td><td>Intergenic</td><td>NA</td><td>NA</td></tr>
<tr><td></td><td></td><td></td><td></td><td></td><td></td><td></td><td></td><td></td><td></td><td></td><td></td></tr>
<tr><td>T32624</td><td>GGACCGCGCGATCCCGACGAGGG</td><td>1000</td><td></td><td></td><td></td><td></td><td></td><td></td><td></td><td></td><td></td></tr>
<tr><td>Oligo fwd</td><td>CACCGGACCGCGCGATCCCGACGA</td><td></td><td></td><td></td><td></td><td></td><td></td><td></td><td></td><td></td><td></td></tr>
<tr><td>Oligo rev</td><td>AAACTCGTCGGGATCGCGCGGTCC</td><td></td><td></td><td></td><td></td><td></td><td></td><td></td><td></td><td></td><td></td></tr>
<tr><td>Chromosome</td><td>start</td><td>end</td><td>strand</td><td>MM</td><td>target_seq</td><td>PAM</td><td>alignment</td><td>distance</td><td>position</td><td>gene name</td><td>gene id</td></tr>
<tr><td>GU305914.1</td><td>24588</td><td>24610</td><td>-</td><td>0</td><td>GGACCGCGCGATCCCGACGA</td><td>GGG</td><td>||||||||[||||||||||||]PAM</td><td>NA</td><td>Intergenic</td><td>NA</td><td>NA</td></tr>
<tr><td></td><td></td><td></td><td></td><td></td><td></td><td></td><td></td><td></td><td></td><td></td><td></td></tr>
<tr><td>T32625</td><td>GACCGCGCGATCCCGACGAGGGG</td><td>1000</td><td></td><td></td><td></td><td></td><td></td><td></td><td></td><td></td><td></td></tr>
<tr><td>Oligo fwd</td><td>CACCGACCGCGCGATCCCGACGAG</td><td></td><td></td><td></td><td></td><td></td><td></td><td></td><td></td><td></td><td></td></tr>
<tr><td>Oligo rev</td><td>AAACCTCGTCGGGATCGCGCGGTC</td><td></td><td></td><td></td><td></td><td></td><td></td><td></td><td></td><td></td><td></td></tr>
<tr><td>Chromosome</td><td>start</td><td>end</td><td>strand</td><td>MM</td><td>target_seq</td><td>PAM</td><td>alignment</td><td>distance</td><td>position</td><td>gene name</td><td>gene id</td></tr>
<tr><td>GU305914.1</td><td>24587</td><td>24609</td><td>-</td><td>0</td><td>GACCGCGCGATCCCGACGAG</td><td>GGG</td><td>||||||||[||||||||||||]PAM</td><td>NA</td><td>Intergenic</td><td>NA</td><td>NA</td></tr>
<tr><td></td><td></td><td></td><td></td><td></td><td></td><td></td><td></td><td></td><td></td><td></td><td></td></tr>
<tr><td>T32626</td><td>CGTCGCTGCCGCCGCCATCTTGG</td><td>1000</td><td></td><td></td><td></td><td></td><td></td><td></td><td></td><td></td><td></td></tr>
<tr><td>Oligo fwd</td><td>CACCCGTCGCTGCCGCCGCCATCT</td><td></td><td></td><td></td><td></td><td></td><td></td><td></td><td></td><td></td><td></td></tr>
<tr><td>Oligo rev</td><td>AAACAGATGGCGGCGGCAGCGACG</td><td></td><td></td><td></td><td></td><td></td><td></td><td></td><td></td><td></td><td></td></tr>
<tr><td>Chromosome</td><td>start</td><td>end</td><td>strand</td><td>MM</td><td>target_seq</td><td>PAM</td><td>alignment</td><td>distance</td><td>position</td><td>gene name</td><td>gene id</td></tr>
<tr><td>GU305914.1</td><td>24547</td><td>24569</td><td>-</td><td>0</td><td>CGTCGCTGCCGCCGCCATCT</td><td>TGG</td><td>||||||||[||||||||||||]PAM</td><td>NA</td><td>Intergenic</td><td>NA</td><td>NA</td></tr>
</table>

<table>
<tr><td>T32627</td><td>GTCGCTGCCGCCGCCATCTTGGG</td><td>1000</td><td></td><td></td><td></td><td></td><td></td><td></td><td></td><td></td><td></td></tr>
<tr><td>Oligo fwd</td><td>CACCGTCGCTGCCGCCGCCATCTT</td><td></td><td></td><td></td><td></td><td></td><td></td><td></td><td></td><td></td><td></td></tr>
<tr><td>Oligo rev</td><td>AAACAAGATGGCGGCGGCAGCGAC</td><td></td><td></td><td></td><td></td><td></td><td></td><td></td><td></td><td></td><td></td></tr>
<tr><td>Chromosome</td><td>start</td><td>end</td><td>strand</td><td>MM</td><td>target_seq</td><td>PAM</td><td>alignment</td><td>distance</td><td>position</td><td>gene name</td><td>gene id</td></tr>
<tr><td>GU305914.1</td><td>24546</td><td>24568</td><td>-</td><td>0</td><td>GTCGCTGCCGCCGCCATCTT</td><td>GGG</td><td>||||||||[||||||||||||]PAM</td><td>NA</td><td>Intergenic</td><td>NA</td><td>NA</td></tr>
<tr><td>GU305914.1</td><td>170532</td><td>170554</td><td>+</td><td>4</td><td>GTGGCCGCCGCCGCCACCAT</td><td>CGG</td><td>||-||-||[||||||||-|-|]PAM</td><td>NA</td><td>Intergenic</td><td>NA</td><td>NA</td></tr>
<tr><td>T32628</td><td>TCGCTGCCGCCGCCATCTTGGGG</td><td>1000</td><td></td><td></td><td></td><td></td><td></td><td></td><td></td><td></td><td></td></tr>
<tr><td>Oligo fwd</td><td>CACCTCGCTGCCGCCGCCATCTTG</td><td></td><td></td><td></td><td></td><td></td><td></td><td></td><td></td><td></td><td></td></tr>
<tr><td>Oligo rev</td><td>AAACCAAGATGGCGGCGGCAGCGA</td><td></td><td></td><td></td><td></td><td></td><td></td><td></td><td></td><td></td><td></td></tr>
<tr><td>Chromosome</td><td>start</td><td>end</td><td>strand</td><td>MM</td><td>target_seq</td><td>PAM</td><td>alignment</td><td>distance</td><td>position</td><td>gene name</td><td>gene id</td></tr>
<tr><td>GU305914.1</td><td>24545</td><td>24567</td><td>-</td><td>0</td><td>TCGCTGCCGCCGCCATCTTG</td><td>GGG</td><td>||||||||[||||||||||||]PAM</td><td>NA</td><td>Intergenic</td><td>NA</td><td>NA</td></tr>
<tr><td>T32629</td><td>TGCCGCCGCCATCTTGGGGCCGG</td><td>1000</td><td></td><td></td><td></td><td></td><td></td><td></td><td></td><td></td><td></td></tr>
<tr><td>Oligo fwd</td><td>CACCTGCCGCCGCCATCTTGGGGC</td><td></td><td></td><td></td><td></td><td></td><td></td><td></td><td></td><td></td><td></td></tr>
<tr><td>Oligo rev</td><td>AAACGCCCCAAGATGGCGGCGGCA</td><td></td><td></td><td></td><td></td><td></td><td></td><td></td><td></td><td></td><td></td></tr>
<tr><td>Chromosome</td><td>start</td><td>end</td><td>strand</td><td>MM</td><td>target_seq</td><td>PAM</td><td>alignment</td><td>distance</td><td>position</td><td>gene name</td><td>gene id</td></tr>
<tr><td>GU305914.1</td><td>24541</td><td>24563</td><td>-</td><td>0</td><td>TGCCGCCGCCATCTTGGGGC</td><td>CGG</td><td>||||||||[||||||||||||]PAM</td><td>NA</td><td>Intergenic</td><td>NA</td><td>NA</td></tr>
<tr><td>T32630</td><td>CCATCTTGGGGCCGGCGCTCAGG</td><td>1000</td><td></td><td></td><td></td><td></td><td></td><td></td><td></td><td></td><td></td></tr>
<tr><td>Oligo fwd</td><td>CACCCCATCTTGGGGCCGGCGCTC</td><td></td><td></td><td></td><td></td><td></td><td></td><td></td><td></td><td></td><td></td></tr>
<tr><td>Oligo rev</td><td>AAACGAGCGCCGGCCCCAAGATGG</td><td></td><td></td><td></td><td></td><td></td><td></td><td></td><td></td><td></td><td></td></tr>
<tr><td>Chromosome</td><td>start</td><td>end</td><td>strand</td><td>MM</td><td>target_seq</td><td>PAM</td><td>alignment</td><td>distance</td><td>position</td><td>gene name</td><td>gene id</td></tr>
<tr><td>GU305914.1</td><td>24533</td><td>24555</td><td>-</td><td>0</td><td>CCATCTTGGGGCCGGCGCTC</td><td>AGG</td><td>||||||||[||||||||||||]PAM</td><td>NA</td><td>Intergenic</td><td>NA</td><td>NA</td></tr>
<tr><td>T32631</td><td>CATCTTGGGGCCGGCGCTCAGGG</td><td>1000</td><td></td><td></td><td></td><td></td><td></td><td></td><td></td><td></td><td></td></tr>
<tr><td>Oligo fwd</td><td>CACCCATCTTGGGGCCGGCGCTCA</td><td></td><td></td><td></td><td></td><td></td><td></td><td></td><td></td><td></td><td></td></tr>
<tr><td>Oligo rev</td><td>AAACTGAGCGCCGGCCCCAAGATG</td><td></td><td></td><td></td><td></td><td></td><td></td><td></td><td></td><td></td><td></td></tr>
<tr><td>Chromosome</td><td>start</td><td>end</td><td>strand</td><td>MM</td><td>target_seq</td><td>PAM</td><td>alignment</td><td>distance</td><td>position</td><td>gene name</td><td>gene id</td></tr>
<tr><td>GU305914.1</td><td>24532</td><td>24554</td><td>-</td><td>0</td><td>CATCTTGGGGCCGGCGCTCA</td><td>GGG</td><td>||||||||[||||||||||||]PAM</td><td>NA</td><td>Intergenic</td><td>NA</td><td>NA</td></tr>
<tr><td>T32632</td><td>TTGGGGCCGGCGCTCAGGGCCGG</td><td>1000</td><td></td><td></td><td></td><td></td><td></td><td></td><td></td><td></td><td></td></tr>
<tr><td>Oligo fwd</td><td>CACCTTGGGGCCGGCGCTCAGGGC</td><td></td><td></td><td></td><td></td><td></td><td></td><td></td><td></td><td></td><td></td></tr>
<tr><td>Oligo rev</td><td>AAACGCCCTGAGCGCCGGCCCCAA</td><td></td><td></td><td></td><td></td><td></td><td></td><td></td><td></td><td></td><td></td></tr>
<tr><td>Chromosome</td><td>start</td><td>end</td><td>strand</td><td>MM</td><td>target_seq</td><td>PAM</td><td>alignment</td><td>distance</td><td>position</td><td>gene name</td><td>gene id</td></tr>
<tr><td>GU305914.1</td><td>24528</td><td>24550</td><td>-</td><td>0</td><td>TTGGGGCCGGCGCTCAGGGC</td><td>CGG</td><td>||||||||[||||||||||||]PAM</td><td>NA</td><td>Intergenic</td><td>NA</td><td>NA</td></tr>
<tr><td>T32633</td><td>GGGCCGGCGCTCAGGGCCGGCGG</td><td>1000</td><td></td><td></td><td></td><td></td><td></td><td></td><td></td><td></td><td></td></tr>
<tr><td>Oligo fwd</td><td>CACCGGGCCGGCGCTCAGGGCCGG</td><td></td><td></td><td></td><td></td><td></td><td></td><td></td><td></td><td></td><td></td></tr>
<tr><td>Oligo rev</td><td>AAACCCGGCCCTGAGCGCCGGCCC</td><td></td><td></td><td></td><td></td><td></td><td></td><td></td><td></td><td></td><td></td></tr>
<tr><td>Chromosome</td><td>start</td><td>end</td><td>strand</td><td>MM</td><td>target_seq</td><td>PAM</td><td>alignment</td><td>distance</td><td>position</td><td>gene name</td><td>gene id</td></tr>
<tr><td>GU305914.1</td><td>24525</td><td>24547</td><td>-</td><td>0</td><td>GGGCCGGCGCTCAGGGCCGG</td><td>CGG</td><td>||||||||[||||||||||||]PAM</td><td>NA</td><td>Intergenic</td><td>NA</td><td>NA</td></tr>
<tr><td>T32634</td><td>CCGGCGCTCAGGGCCGGCGGCGG</td><td>1000</td><td></td><td></td><td></td><td></td><td></td><td></td><td></td><td></td><td></td></tr>
<tr><td>Oligo fwd</td><td>CACCCCGGCGCTCAGGGCCGGCGG</td><td></td><td></td><td></td><td></td><td></td><td></td><td></td><td></td><td></td><td></td></tr>
<tr><td>Oligo rev</td><td>AAACCCGCCGGCCCTGAGCGCCGG</td><td></td><td></td><td></td><td></td><td></td><td></td><td></td><td></td><td></td><td></td></tr>
<tr><td>Chromosome</td><td>start</td><td>end</td><td>strand</td><td>MM</td><td>target_seq</td><td>PAM</td><td>alignment</td><td>distance</td><td>position</td><td>gene name</td><td>gene id</td></tr>
<tr><td>GU305914.1</td><td>24522</td><td>24544</td><td>-</td><td>0</td><td>CCGGCGCTCAGGGCCGGCGG</td><td>CGG</td><td>||||||||[||||||||||||]PAM</td><td>NA</td><td>Intergenic</td><td>NA</td><td>NA</td></tr>
<tr><td>T32635</td><td>CCACTACCGCCAGCCTCCCGAGG</td><td>1000</td><td></td><td></td><td></td><td></td><td></td><td></td><td></td><td></td><td></td></tr>
<tr><td>Oligo fwd</td><td>CACCCCACTACCGCCAGCCTCCCG</td><td></td><td></td><td></td><td></td><td></td><td></td><td></td><td></td><td></td><td></td></tr>
<tr><td>Oligo rev</td><td>AAACCGGGAGGCTGGCGGTAGTGG</td><td></td><td></td><td></td><td></td><td></td><td></td><td></td><td></td><td></td><td></td></tr>
<tr><td>Chromosome</td><td>start</td><td>end</td><td>strand</td><td>MM</td><td>target_seq</td><td>PAM</td><td>alignment</td><td>distance</td><td>position</td><td>gene name</td><td>gene id</td></tr>
<tr><td>GU305914.1</td><td>24469</td><td>24491</td><td>-</td><td>0</td><td>CCACTACCGCCAGCCTCCCG</td><td>AGG</td><td>||||||||[||||||||||||]PAM</td><td>NA</td><td>Intergenic</td><td>NA</td><td>NA</td></tr>
<tr><td>T32636</td><td>CAGCCTCCCGAGGTCTACGTCGG</td><td>1000</td><td></td><td></td><td></td><td></td><td></td><td></td><td></td><td></td><td></td></tr>
<tr><td>Oligo fwd</td><td>CACCCAGCCTCCCGAGGTCTACGT</td><td></td><td></td><td></td><td></td><td></td><td></td><td></td><td></td><td></td><td></td></tr>
<tr><td>Oligo rev</td><td>AAACACGTAGACCTCGGGAGGCTG</td><td></td><td></td><td></td><td></td><td></td><td></td><td></td><td></td><td></td><td></td></tr>
<tr><td>Chromosome</td><td>start</td><td>end</td><td>strand</td><td>MM</td><td>target_seq</td><td>PAM</td><td>alignment</td><td>distance</td><td>position</td><td>gene name</td><td>gene id</td></tr>
<tr><td>GU305914.1</td><td>24459</td><td>24481</td><td>-</td><td>0</td><td>CAGCCTCCCGAGGTCTACGT</td><td>CGG</td><td>||||||||[||||||||||||]PAM</td><td>NA</td><td>Intergenic</td><td>NA</td><td>NA</td></tr>
<tr><td>T32637</td><td>GAGGTCTACGTCGGCAGCAGCGG</td><td>1000</td><td></td><td></td><td></td><td></td><td></td><td></td><td></td><td></td><td></td></tr>
<tr><td>Oligo fwd</td><td>CACCGAGGTCTACGTCGGCAGCAG</td><td></td><td></td><td></td><td></td><td></td><td></td><td></td><td></td><td></td><td></td></tr>
<tr><td>Oligo rev</td><td>AAACCTGCTGCCGACGTAGACCTC</td><td></td><td></td><td></td><td></td><td></td><td></td><td></td><td></td><td></td><td></td></tr>
<tr><td>Chromosome</td><td>start</td><td>end</td><td>strand</td><td>MM</td><td>target_seq</td><td>PAM</td><td>alignment</td><td>distance</td><td>position</td><td>gene name</td><td>gene id</td></tr>
<tr><td>GU305914.1</td><td>24450</td><td>24472</td><td>-</td><td>0</td><td>GAGGTCTACGTCGGCAGCAG</td><td>CGG</td><td>||||||||[||||||||||||]PAM</td><td>NA</td><td>Intergenic</td><td>NA</td><td>NA</td></tr>
<tr><td>T32638</td><td>CCTGCTTCTCACGCGCAGCGAGG</td><td>1000</td><td></td><td></td><td></td><td></td><td></td><td></td><td></td><td></td><td></td></tr>
<tr><td>Oligo fwd</td><td>CACCCCTGCTTCTCACGCGCAGCG</td><td></td><td></td><td></td><td></td><td></td><td></td><td></td><td></td><td></td><td></td></tr>
<tr><td>Oligo rev</td><td>AAACCGCTGCGCGTGAGAAGCAGG</td><td></td><td></td><td></td><td></td><td></td><td></td><td></td><td></td><td></td><td></td></tr>
<tr><td>Chromosome</td><td>start</td><td>end</td><td>strand</td><td>MM</td><td>target_seq</td><td>PAM</td><td>alignment</td><td>distance</td><td>position</td><td>gene name</td><td>gene id</td></tr>
<tr><td>GU305914.1</td><td>24412</td><td>24434</td><td>-</td><td>0</td><td>CCTGCTTCTCACGCGCAGCG</td><td>AGG</td><td>||||||||[||||||||||||]PAM</td><td>NA</td><td>Intergenic</td><td>NA</td><td>NA</td></tr>
<tr><td>T32639</td><td>CAGCATGCGCGAGATCGTCAAGG</td><td>1000</td><td></td><td></td><td></td><td></td><td></td><td></td><td></td><td></td><td></td></tr>
<tr><td>Oligo fwd</td><td>CACCCAGCATGCGCGAGATCGTCA</td><td></td><td></td><td></td><td></td><td></td><td></td><td></td><td></td><td></td><td></td></tr>
<tr><td>Oligo rev</td><td>AAACTGACGATCTCGCGCATGCTG</td><td></td><td></td><td></td><td></td><td></td><td></td><td></td><td></td><td></td><td></td></tr>
<tr><td>Chromosome</td><td>start</td><td>end</td><td>strand</td><td>MM</td><td>target_seq</td><td>PAM</td><td>alignment</td><td>distance</td><td>position</td><td>gene name</td><td>gene id</td></tr>
<tr><td>GU305914.1</td><td>24370</td><td>24392</td><td>-</td><td>0</td><td>CAGCATGCGCGAGATCGTCA</td><td>AGG</td><td>||||||||[||||||||||||]PAM</td><td>NA</td><td>Intergenic</td><td>NA</td><td>NA</td></tr>
<tr><td>T32640</td><td>ATGCGCGAGATCGTCAAGGATGG</td><td>1000</td><td></td><td></td><td></td><td></td><td></td><td></td><td></td><td></td><td></td></tr>
<tr><td>Oligo fwd</td><td>CACCATGCGCGAGATCGTCAAGGA</td><td></td><td></td><td></td><td></td><td></td><td></td><td></td><td></td><td></td><td></td></tr>
<tr><td>Oligo rev</td><td>AAACTCCTTGACGATCTCGCGCAT</td><td></td><td></td><td></td><td></td><td></td><td></td><td></td><td></td><td></td><td></td></tr>
<tr><td>Chromosome</td><td>start</td><td>end</td><td>strand</td><td>MM</td><td>target_seq</td><td>PAM</td><td>alignment</td><td>distance</td><td>position</td><td>gene name</td><td>gene id</td></tr>
<tr><td>GU305914.1</td><td>24366</td><td>24388</td><td>-</td><td>0</td><td>ATGCGCGAGATCGTCAAGGA</td><td>TGG</td><td>||||||||[||||||||||||]PAM</td><td>NA</td><td>Intergenic</td><td>NA</td><td>NA</td></tr>
<tr><td>T32641</td><td>CATCAAGAATATCTACCTGCCGG</td><td>1000</td><td></td><td></td><td></td><td></td><td></td><td></td><td></td><td></td><td></td></tr>
<tr><td>Oligo fwd</td><td>CACCCATCAAGAATATCTACCTGC</td><td></td><td></td><td></td><td></td><td></td><td></td><td></td><td></td><td></td><td></td></tr>
<tr><td>Oligo rev</td><td>AAACGCAGGTAGATATTCTTGATG</td><td></td><td></td><td></td><td></td><td></td><td></td><td></td><td></td><td></td><td></td></tr>
<tr><td>Chromosome</td><td>start</td><td>end</td><td>strand</td><td>MM</td><td>target_seq</td><td>PAM</td><td>alignment</td><td>distance</td><td>position</td><td>gene name</td><td>gene id</td></tr>
</table>

GU305914.1 24343 24365 - 0 CATCAAGAATATCTACCTGC CGG ||||||||[||||||||||||]PAM NA Intergenic NA NA

T32642 CGTCGCCCCATCATCTGTCGCGG 1000
Oligo fwd CACCCGTCGCCCCATCATCTGTCG
Oligo rev AAACCGACAGATGATGGGGCGACG
Chromosome start end strand MM target_seq PAM alignment distance position gene name gene id
GU305914.1 24318 24340 - 0 CGTCGCCCCATCATCTGTCG CGG ||||||||[||||||||||||]PAM NA Intergenic NA NA

T32643 GTCGCCCCATCATCTGTCGCGGG 1000
Oligo fwd CACCGTCGCCCCATCATCTGTCGC
Oligo rev AAACGCGACAGATGATGGGGCGAC
Chromosome start end strand MM target_seq PAM alignment distance position gene name gene id
GU305914.1 24317 24339 - 0 GTCGCCCCATCATCTGTCGC GGG ||||||||[||||||||||||]PAM NA Intergenic NA NA

T32644 TCGCCCCATCATCTGTCGCGGGG 1000
Oligo fwd CACCTCGCCCCATCATCTGTCGCG
Oligo rev AAACCGCGACAGATGATGGGGCGA
Chromosome start end strand MM target_seq PAM alignment distance position gene name gene id
GU305914.1 24316 24338 - 0 TCGCCCCATCATCTGTCGCG GGG ||||||||[||||||||||||]PAM NA Intergenic NA NA

T32645 CCAACAACGACATCCTGTCCTGG 1000
Oligo fwd CACCCCAACAACGACATCCTGTCC
Oligo rev AAACGGACAGGATGTCGTTGTTGG
Chromosome start end strand MM target_seq PAM alignment distance position gene name gene id
GU305914.1 24263 24285 - 0 CCAACAACGACATCCTGTCC TGG ||||||||[||||||||||||]PAM NA Intergenic NA NA

T32646 CCTCGCCAACCGCGTCTTCTTGG 1000
Oligo fwd CACCCCTCGCCAACCGCGTCTTCT
Oligo rev AAACAGAAGACGCGGTTGGCGAGG
Chromosome start end strand MM target_seq PAM alignment distance position gene name gene id
GU305914.1 24229 24251 - 0 CCTCGCCAACCGCGTCTTCT TGG ||||||||[||||||||||||]PAM NA Intergenic NA NA

T32647 CTCGCCAACCGCGTCTTCTTGGG 1000
Oligo fwd CACCCTCGCCAACCGCGTCTTCTT
Oligo rev AAACAAGAAGACGCGGTTGGCGAG
Chromosome start end strand MM target_seq PAM alignment distance position gene name gene id
GU305914.1 24228 24250 - 0 CTCGCCAACCGCGTCTTCTT GGG ||||||||[||||||||||||]PAM NA Intergenic NA NA

T32648 TCGCCAACCGCGTCTTCTTGGGG 1000
Oligo fwd CACCTCGCCAACCGCGTCTTCTTG
Oligo rev AAACCAAGAAGACGCGGTTGGCGA
Chromosome start end strand MM target_seq PAM alignment distance position gene name gene id
GU305914.1 24227 24249 - 0 TCGCCAACCGCGTCTTCTTG GGG ||||||||[||||||||||||]PAM NA Intergenic NA NA

T32649 CGCCAACCGCGTCTTCTTGGGGG 1000
Oligo fwd CACCCGCCAACCGCGTCTTCTTGG
Oligo rev AAACCCAAGAAGACGCGGTTGGCG
Chromosome start end strand MM target_seq PAM alignment distance position gene name gene id
GU305914.1 24226 24248 - 0 CGCCAACCGCGTCTTCTTGG GGG ||||||||[||||||||||||]PAM NA Intergenic NA NA

T32650 CGCCTGTGACGAGTTCTTCGTGG 1000
Oligo fwd CACCCGCCTGTGACGAGTTCTTCG
Oligo rev AAACCGAAGAACTCGTCACAGGCG
Chromosome start end strand MM target_seq PAM alignment distance position gene name gene id
GU305914.1 24187 24209 - 0 CGCCTGTGACGAGTTCTTCG TGG ||||||||[||||||||||||]PAM NA Intergenic NA NA

T32651 GCCTGTGACGAGTTCTTCGTGGG 1000
Oligo fwd CACCGCCTGTGACGAGTTCTTCGT
Oligo rev AAACACGAAGAACTCGTCACAGGC
Chromosome start end strand MM target_seq PAM alignment distance position gene name gene id
GU305914.1 24186 24208 - 0 GCCTGTGACGAGTTCTTCGT GGG ||||||||[||||||||||||]PAM NA Intergenic NA NA

T32652 TACGACGTCGTCGCCCTTGACGG 1000
Oligo fwd CACCTACGACGTCGTCGCCCTTGA
Oligo rev AAACTCAAGGGCGACGACGTCGTA
Chromosome start end strand MM target_seq PAM alignment distance position gene name gene id
GU305914.1 24162 24184 - 0 TACGACGTCGTCGCCCTTGA CGG ||||||||[||||||||||||]PAM NA Intergenic NA NA

T32653 ACGTCGTCGCCCTTGACGGATGG 1000
Oligo fwd CACCACGTCGTCGCCCTTGACGGA
Oligo rev AAACTCCGTCAAGGGCGACGACGT
Chromosome start end strand MM target_seq PAM alignment distance position gene name gene id
GU305914.1 24158 24180 - 0 ACGTCGTCGCCCTTGACGGA TGG ||||||||[||||||||||||]PAM NA Intergenic NA NA

T32654 CGTCGTCGCCCTTGACGGATGGG 1000
Oligo fwd CACCCGTCGTCGCCCTTGACGGAT
Oligo rev AAACATCCGTCAAGGGCGACGACG
Chromosome start end strand MM target_seq PAM alignment distance position gene name gene id
GU305914.1 24157 24179 - 0 CGTCGTCGCCCTTGACGGAT GGG ||||||||[||||||||||||]PAM NA Intergenic NA NA

T32655 CCTTGACGGATGGGCCGCCGAGG 1000
Oligo fwd CACCCCTTGACGGATGGGCCGCCG
Oligo rev AAACCGGCGGCCCATCCGTCAAGG
Chromosome start end strand MM target_seq PAM alignment distance position gene name gene id
GU305914.1 24148 24170 - 0 CCTTGACGGATGGGCCGCCG AGG ||||||||[||||||||||||]PAM NA Intergenic NA NA

T32656 TGGGCCGCCGAGGCCGACGTCGG 1000
Oligo fwd CACCTGGGCCGCCGAGGCCGACGT
Oligo rev AAACACGTCGGCCTCGGCGGCCCA

| Chromosome | start | end | strand | MM | target_seq | PAM | alignment | distance | position | gene name | gene id |
|---|---|---|---|---|---|---|---|---|---|---|---|
| GU305914.1 | 24138 | 24160 | - | 0 | TGGGCCGCCGAGGCCGACGT | CGG | ||||||||[||||||||||||]PAM | NA | Intergenic | NA | NA |
| GU305914.1 | 128969 | 128991 | + | 4 | CTGGCCGCCGAGGCAGCCGT | CGG | --||||||[||||||-|-|||]PAM | NA | Intergenic | NA | NA |
| T32657 | CGAGGCCGACGTCGGCGCCATGG | 1000 | | | | | | | | | |
| Oligo fwd | CACCCGAGGCCGACGTCGGCGCCA | | | | | | | | | | |
| Oligo rev | AAACTGGCGCCGACGTCGGCCTCG | | | | | | | | | | |
| Chromosome | start | end | strand | MM | target_seq | PAM | alignment | distance | position | gene name | gene id |
| GU305914.1 | 24130 | 24152 | - | 0 | CGAGGCCGACGTCGGCGCCA | TGG | ||||||||[||||||||||||]PAM | NA | Intergenic | NA | NA |
| GU305914.1 | 118569 | 118591 | - | 4 | AGACGCCGACGCCGACGCCA | TGG | -||-||||[|||-||-|||||]PAM | NA | Intergenic | NA | NA |
| T32658 | GTCGGCGCCATGGACGTCCTCGG | 1000 | | | | | | | | | |
| Oligo fwd | CACCGTCGGCGCCATGGACGTCCT | | | | | | | | | | |
| Oligo rev | AAACAGGACGTCCATGGCGCCGAC | | | | | | | | | | |
| Chromosome | start | end | strand | MM | target_seq | PAM | alignment | distance | position | gene name | gene id |
| GU305914.1 | 24120 | 24142 | - | 0 | GTCGGCGCCATGGACGTCCT | CGG | ||||||||[||||||||||||]PAM | NA | Intergenic | NA | NA |
| GU305914.1 | 118559 | 118581 | - | 4 | GCCGACGCCATGGACGTCGG | CGG | |-||-|||[||||||||||--]PAM | NA | Intergenic | NA | NA |
| T32659 | CGACCTCTGTCGCGTCTACGTGG | 1000 | | | | | | | | | |
| Oligo fwd | CACCCGACCTCTGTCGCGTCTACG | | | | | | | | | | |
| Oligo rev | AAACCGTAGACGCGACAGAGGTCG | | | | | | | | | | |
| Chromosome | start | end | strand | MM | target_seq | PAM | alignment | distance | position | gene name | gene id |
| GU305914.1 | 24049 | 24071 | - | 0 | CGACCTCTGTCGCGTCTACG | TGG | ||||||||[||||||||||||]PAM | NA | Intergenic | NA | NA |
| T32660 | CTGTCGCGTCTACGTGGTTCAGG | 1000 | | | | | | | | | |
| Oligo fwd | CACCCTGTCGCGTCTACGTGGTTC | | | | | | | | | | |
| Oligo rev | AAACGAACCACGTAGACGCGACAG | | | | | | | | | | |
| Chromosome | start | end | strand | MM | target_seq | PAM | alignment | distance | position | gene name | gene id |
| GU305914.1 | 24043 | 24065 | - | 0 | CTGTCGCGTCTACGTGGTTC | AGG | ||||||||[||||||||||||]PAM | NA | Intergenic | NA | NA |
| T32661 | TGTCGCGTCTACGTGGTTCAGGG | 1000 | | | | | | | | | |
| Oligo fwd | CACCTGTCGCGTCTACGTGGTTCA | | | | | | | | | | |
| Oligo rev | AAACTGAACCACGTAGACGCGACA | | | | | | | | | | |
| Chromosome | start | end | strand | MM | target_seq | PAM | alignment | distance | position | gene name | gene id |
| GU305914.1 | 24042 | 24064 | - | 0 | TGTCGCGTCTACGTGGTTCA | GGG | ||||||||[||||||||||||]PAM | NA | Intergenic | NA | NA |
| T32662 | GTCGCTCTGTCTCATCGCCGAGG | 1000 | | | | | | | | | |
| Oligo fwd | CACCGTCGCTCTGTCTCATCGCCG | | | | | | | | | | |
| Oligo rev | AAACCGGCGATGAGACAGAGCGAC | | | | | | | | | | |
| Chromosome | start | end | strand | MM | target_seq | PAM | alignment | distance | position | gene name | gene id |
| GU305914.1 | 24013 | 24035 | - | 0 | GTCGCTCTGTCTCATCGCCG | AGG | ||||||||[||||||||||||]PAM | NA | Intergenic | NA | NA |
| T32663 | TACTTTGAGTTCGTCGAGCGAGG | 1000 | | | | | | | | | |
| Oligo fwd | CACCTACTTTGAGTTCGTCGAGCG | | | | | | | | | | |
| Oligo rev | AAACCGCTCGACGAACTCAAAGTA | | | | | | | | | | |
| Chromosome | start | end | strand | MM | target_seq | PAM | alignment | distance | position | gene name | gene id |
| GU305914.1 | 23988 | 24010 | - | 0 | TACTTTGAGTTCGTCGAGCG | AGG | ||||||||[||||||||||||]PAM | NA | Intergenic | NA | NA |
| T32664 | ACTTTGAGTTCGTCGAGCGAGGG | 1000 | | | | | | | | | |
| Oligo fwd | CACCACTTTGAGTTCGTCGAGCGA | | | | | | | | | | |
| Oligo rev | AAACTCGCTCGACGAACTCAAAGT | | | | | | | | | | |
| Chromosome | start | end | strand | MM | target_seq | PAM | alignment | distance | position | gene name | gene id |
| GU305914.1 | 23987 | 24009 | - | 0 | ACTTTGAGTTCGTCGAGCGA | GGG | ||||||||[||||||||||||]PAM | NA | Intergenic | NA | NA |
| T32665 | TCAGCCGCTATCACAAGAACAGG | 1000 | | | | | | | | | |
| Oligo fwd | CACCTCAGCCGCTATCACAAGAAC | | | | | | | | | | |
| Oligo rev | AAACGTTCTTGTGATAGCGGCTGA | | | | | | | | | | |
| Chromosome | start | end | strand | MM | target_seq | PAM | alignment | distance | position | gene name | gene id |
| GU305914.1 | 23963 | 23985 | - | 0 | TCAGCCGCTATCACAAGAAC | AGG | ||||||||[||||||||||||]PAM | NA | Intergenic | NA | NA |
| T32666 | CTTCATCATCGATCCGCTCGAGG | 1000 | | | | | | | | | |
| Oligo fwd | CACCCTTCATCATCGATCCGCTCG | | | | | | | | | | |
| Oligo rev | AAACCGAGCGGATCGATGATGAAG | | | | | | | | | | |
| Chromosome | start | end | strand | MM | target_seq | PAM | alignment | distance | position | gene name | gene id |
| GU305914.1 | 23938 | 23960 | - | 0 | CTTCATCATCGATCCGCTCG | AGG | ||||||||[||||||||||||]PAM | NA | Intergenic | NA | NA |
| GU305914.1 | 39777 | 39799 | - | 4 | CTTCGTCGGCGATCCGGTCG | CGG | ||||-||-[-|||||||-|||]PAM | NA | Intergenic | NA | NA |
| T32667 | CCAACAACTTCTTACACTCCAGG | 1000 | | | | | | | | | |
| Oligo fwd | CACCCCAACAACTTCTTACACTCC | | | | | | | | | | |
| Oligo rev | AAACGGAGTGTAAGAAGTTGTTGG | | | | | | | | | | |
| Chromosome | start | end | strand | MM | target_seq | PAM | alignment | distance | position | gene name | gene id |
| GU305914.1 | 23888 | 23910 | - | 0 | CCAACAACTTCTTACACTCC | AGG | ||||||||[||||||||||||]PAM | NA | Intergenic | NA | NA |
| T32668 | CAACAACTTCTTACACTCCAGGG | 1000 | | | | | | | | | |
| Oligo fwd | CACCCAACAACTTCTTACACTCCA | | | | | | | | | | |
| Oligo rev | AAACTGGAGTGTAAGAAGTTGTTG | | | | | | | | | | |
| Chromosome | start | end | strand | MM | target_seq | PAM | alignment | distance | position | gene name | gene id |
| GU305914.1 | 23887 | 23909 | - | 0 | CAACAACTTCTTACACTCCA | GGG | ||||||||[||||||||||||]PAM | NA | Intergenic | NA | NA |
| T32669 | TTACACTCCAGGGCGACCAATGG | 1000 | | | | | | | | | |
| Oligo fwd | CACCTTACACTCCAGGGCGACCAA | | | | | | | | | | |
| Oligo rev | AAACTTGGTCGCCCTGGAGTGTAA | | | | | | | | | | |
| Chromosome | start | end | strand | MM | target_seq | PAM | alignment | distance | position | gene name | gene id |
| GU305914.1 | 23877 | 23899 | - | 0 | TTACACTCCAGGGCGACCAA | TGG | ||||||||[||||||||||||]PAM | NA | Intergenic | NA | NA |
| T32670 | TACACTCCAGGGCGACCAATGGG | 1000 | | | | | | | | | |
| Oligo fwd | CACCTACACTCCAGGGCGACCAAT | | | | | | | | | | |
| Oligo rev | AAACATTGGTCGCCCTGGAGTGTA | | | | | | | | | | |
| Chromosome | start | end | strand | MM | target_seq | PAM | alignment | distance | position | gene name | gene id |

| | | | | | | | | | | | |
|---|---|---|---|---|---|---|---|---|---|---|---|
| GU305914.1 | | 23876 | 23898 | - | 0 | TACACTCCAGGGCGACCAAT | GGG | \|\|\|\|\|\|\|\|[\|\|\|\|\|\|\|\|\|\|\|\|]PAM | NA | Intergenic | NA | NA |
| T32671 | ACACTCCAGGGCGACCAATGGGG | | 1000 | | | | | | | | | |
| Oligo fwd | CACCACACTCCAGGGCGACCAATG | | | | | | | | | | | |
| Oligo rev | AAACCATTGGTCGCCCTGGAGTGT | | | | | | | | | | | |
| Chromosome | start | | end | strand | MM | target_seq | PAM | alignment | distance | position | gene name | gene id |
| GU305914.1 | | 23875 | 23897 | - | 0 | ACACTCCAGGGCGACCAATG | GGG | \|\|\|\|\|\|\|\|[\|\|\|\|\|\|\|\|\|\|\|\|]PAM | NA | Intergenic | NA | NA |
| T32672 | GCGACCAATGGGGTGCCGCATGG | | 1000 | | | | | | | | | |
| Oligo fwd | CACCGCGACCAATGGGGTGCCGCA | | | | | | | | | | | |
| Oligo rev | AAACTGCGGCACCCCATTGGTCGC | | | | | | | | | | | |
| Chromosome | start | | end | strand | MM | target_seq | PAM | alignment | distance | position | gene name | gene id |
| GU305914.1 | | 23865 | 23887 | - | 0 | GCGACCAATGGGGTGCCGCA | TGG | \|\|\|\|\|\|\|\|[\|\|\|\|\|\|\|\|\|\|\|\|]PAM | NA | Intergenic | NA | NA |
| T32673 | CGCATGGTAGCGAACCAACCTGG | | 1000 | | | | | | | | | |
| Oligo fwd | CACCCGCATGGTAGCGAACCAACC | | | | | | | | | | | |
| Oligo rev | AAACGGTTGGTTCGCTACCATGCG | | | | | | | | | | | |
| Chromosome | start | | end | strand | MM | target_seq | PAM | alignment | distance | position | gene name | gene id |
| GU305914.1 | | 23849 | 23871 | - | 0 | CGCATGGTAGCGAACCAACC | TGG | \|\|\|\|\|\|\|\|[\|\|\|\|\|\|\|\|\|\|\|\|]PAM | NA | Intergenic | NA | NA |
| T32674 | GCATGGTAGCGAACCAACCTGGG | | 1000 | | | | | | | | | |
| Oligo fwd | CACCGCATGGTAGCGAACCAACCT | | | | | | | | | | | |
| Oligo rev | AAACAGGTTGGTTCGCTACCATGC | | | | | | | | | | | |
| Chromosome | start | | end | strand | MM | target_seq | PAM | alignment | distance | position | gene name | gene id |
| GU305914.1 | | 23848 | 23870 | - | 0 | GCATGGTAGCGAACCAACCT | GGG | \|\|\|\|\|\|\|\|[\|\|\|\|\|\|\|\|\|\|\|\|]PAM | NA | Intergenic | NA | NA |
| T32675 | CCAACCTGGGAGAAGAAAACCGG | | 1000 | | | | | | | | | |
| Oligo fwd | CACCCCAACCTGGGAGAAGAAAAC | | | | | | | | | | | |
| Oligo rev | AAACGTTTTCTTCTCCCAGGTTGG | | | | | | | | | | | |
| Chromosome | start | | end | strand | MM | target_seq | PAM | alignment | distance | position | gene name | gene id |
| GU305914.1 | | 23835 | 23857 | - | 0 | CCAACCTGGGAGAAGAAAAC | CGG | \|\|\|\|\|\|\|\|[\|\|\|\|\|\|\|\|\|\|\|\|]PAM | NA | Intergenic | NA | NA |
| T32676 | CAACCTGGGAGAAGAAAACCGGG | | 1000 | | | | | | | | | |
| Oligo fwd | CACCCAACCTGGGAGAAGAAAACC | | | | | | | | | | | |
| Oligo rev | AAACGGTTTTCTTCTCCCAGGTTG | | | | | | | | | | | |
| Chromosome | start | | end | strand | MM | target_seq | PAM | alignment | distance | position | gene name | gene id |
| GU305914.1 | | 23834 | 23856 | - | 0 | CAACCTGGGAGAAGAAAACC | GGG | \|\|\|\|\|\|\|\|[\|\|\|\|\|\|\|\|\|\|\|\|]PAM | NA | Intergenic | NA | NA |
| T32677 | CCTGGGAGAAGAAAACCGGGAGG | | 1000 | | | | | | | | | |
| Oligo fwd | CACCCCTGGGAGAAGAAAACCGGG | | | | | | | | | | | |
| Oligo rev | AAACCCCGGTTTTCTTCTCCCAGG | | | | | | | | | | | |
| Chromosome | start | | end | strand | MM | target_seq | PAM | alignment | distance | position | gene name | gene id |
| GU305914.1 | | 23831 | 23853 | - | 0 | CCTGGGAGAAGAAAACCGGG | AGG | \|\|\|\|\|\|\|\|[\|\|\|\|\|\|\|\|\|\|\|\|]PAM | NA | Intergenic | NA | NA |
| T32678 | CGGGAGGAACAGCATGAGAAAGG | | 1000 | | | | | | | | | |
| Oligo fwd | CACCCGGGAGGAACAGCATGAGAA | | | | | | | | | | | |
| Oligo rev | AAACTTCTCATGCTGTTCCTCCCG | | | | | | | | | | | |
| Chromosome | start | | end | strand | MM | target_seq | PAM | alignment | distance | position | gene name | gene id |
| GU305914.1 | | 23815 | 23837 | - | 0 | CGGGAGGAACAGCATGAGAA | AGG | \|\|\|\|\|\|\|\|[\|\|\|\|\|\|\|\|\|\|\|\|]PAM | NA | Intergenic | NA | NA |
| T32679 | GACTGATGCGCTCCATCAAAAGG | | 1000 | | | | | | | | | |
| Oligo fwd | CACCGACTGATGCGCTCCATCAAA | | | | | | | | | | | |
| Oligo rev | AAACTTTGATGGAGCGCATCAGTC | | | | | | | | | | | |
| Chromosome | start | | end | strand | MM | target_seq | PAM | alignment | distance | position | gene name | gene id |
| GU305914.1 | | 23783 | 23805 | - | 0 | GACTGATGCGCTCCATCAAA | AGG | \|\|\|\|\|\|\|\|[\|\|\|\|\|\|\|\|\|\|\|\|]PAM | NA | Intergenic | NA | NA |
| T32680 | TGATGCGCTCCATCAAAAGGAGG | | 1000 | | | | | | | | | |
| Oligo fwd | CACCTGATGCGCTCCATCAAAAGG | | | | | | | | | | | |
| Oligo rev | AAACCCTTTTGATGGAGCGCATCA | | | | | | | | | | | |
| Chromosome | start | | end | strand | MM | target_seq | PAM | alignment | distance | position | gene name | gene id |
| GU305914.1 | | 23780 | 23802 | - | 0 | TGATGCGCTCCATCAAAAGG | AGG | \|\|\|\|\|\|\|\|[\|\|\|\|\|\|\|\|\|\|\|\|]PAM | NA | Intergenic | NA | NA |
| T32681 | CTCCATCAAAAGGAGGTGATCGG | | 1000 | | | | | | | | | |
| Oligo fwd | CACCCTCCATCAAAAGGAGGTGAT | | | | | | | | | | | |
| Oligo rev | AAACATCACCTCCTTTTGATGGAG | | | | | | | | | | | |
| Chromosome | start | | end | strand | MM | target_seq | PAM | alignment | distance | position | gene name | gene id |
| GU305914.1 | | 23773 | 23795 | - | 0 | CTCCATCAAAAGGAGGTGAT | CGG | \|\|\|\|\|\|\|\|[\|\|\|\|\|\|\|\|\|\|\|\|]PAM | NA | Intergenic | NA | NA |
| T32682 | TCAAAAGGAGGTGATCGGATCGG | | 1000 | | | | | | | | | |
| Oligo fwd | CACCTCAAAAGGAGGTGATCGGAT | | | | | | | | | | | |
| Oligo rev | AAACATCCGATCACCTCCTTTTGA | | | | | | | | | | | |
| Chromosome | start | | end | strand | MM | target_seq | PAM | alignment | distance | position | gene name | gene id |
| GU305914.1 | | 23768 | 23790 | - | 0 | TCAAAAGGAGGTGATCGGAT | CGG | \|\|\|\|\|\|\|\|[\|\|\|\|\|\|\|\|\|\|\|\|]PAM | NA | Intergenic | NA | NA |
| T32683 | AAGGAGGTGATCGGATCGGACGG | | 1000 | | | | | | | | | |
| Oligo fwd | CACCAAGGAGGTGATCGGATCGGA | | | | | | | | | | | |
| Oligo rev | AAACTCCGATCCGATCACCTCCTT | | | | | | | | | | | |
| Chromosome | start | | end | strand | MM | target_seq | PAM | alignment | distance | position | gene name | gene id |
| GU305914.1 | | 23764 | 23786 | - | 0 | AAGGAGGTGATCGGATCGGA | CGG | \|\|\|\|\|\|\|\|[\|\|\|\|\|\|\|\|\|\|\|\|]PAM | NA | Intergenic | NA | NA |
| T32684 | GGTGATCGGATCGGACGGACCGG | | 1000 | | | | | | | | | |
| Oligo fwd | CACCGGTGATCGGATCGGACGGAC | | | | | | | | | | | |
| Oligo rev | AAACGTCCGTCCGATCCGATCACC | | | | | | | | | | | |
| Chromosome | start | | end | strand | MM | target_seq | PAM | alignment | distance | position | gene name | gene id |
| GU305914.1 | | 23759 | 23781 | - | 0 | GGTGATCGGATCGGACGGAC | CGG | \|\|\|\|\|\|\|\|[\|\|\|\|\|\|\|\|\|\|\|\|]PAM | NA | Intergenic | NA | NA |
| T32685 | ATCGGATCGGACGGACCGGACGG | | 1000 | | | | | | | | | |
| Oligo fwd | CACCATCGGATCGGACGGACCGGA | | | | | | | | | | | |
| Oligo rev | AAACTCCGGTCCGTCCGATCCGAT | | | | | | | | | | | |

<table>
<tr><td>Chromosome</td><td>start</td><td>end</td><td>strand</td><td>MM</td><td>target_seq</td><td>PAM</td><td>alignment</td><td>distance</td><td>position</td><td>gene name</td><td>gene id</td></tr>
<tr><td>GU305914.1</td><td>23755</td><td>23777</td><td>-</td><td>0</td><td>ATCGGATCGGACGGACCGGA</td><td>CGG</td><td>||||||||[||||||||||||]PAM</td><td>NA</td><td>Intergenic</td><td>NA</td><td>NA</td></tr>
<tr><td></td><td></td><td></td><td></td><td></td><td></td><td></td><td></td><td></td><td></td><td></td><td></td></tr>
<tr><td>T32686</td><td>GACGGACCGCGACTGCTTGTCGG</td><td>1000</td><td></td><td></td><td></td><td></td><td></td><td></td><td></td><td></td><td></td></tr>
<tr><td>Oligo fwd</td><td>CACCGACGGACCGCGACTGCTTGT</td><td></td><td></td><td></td><td></td><td></td><td></td><td></td><td></td><td></td><td></td></tr>
<tr><td>Oligo rev</td><td>AAACACAAGCAGTCGCGGTCCGTC</td><td></td><td></td><td></td><td></td><td></td><td></td><td></td><td></td><td></td><td></td></tr>
<tr><td>Chromosome</td><td>start</td><td>end</td><td>strand</td><td>MM</td><td>target_seq</td><td>PAM</td><td>alignment</td><td>distance</td><td>position</td><td>gene name</td><td>gene id</td></tr>
<tr><td>GU305914.1</td><td>23737</td><td>23759</td><td>-</td><td>0</td><td>GACGGACCGCGACTGCTTGT</td><td>CGG</td><td>||||||||[||||||||||||]PAM</td><td>NA</td><td>Intergenic</td><td>NA</td><td>NA</td></tr>
<tr><td></td><td></td><td></td><td></td><td></td><td></td><td></td><td></td><td></td><td></td><td></td><td></td></tr>
<tr><td>T32687</td><td>ACGGACCGCGACTGCTTGTCGGG</td><td>1000</td><td></td><td></td><td></td><td></td><td></td><td></td><td></td><td></td><td></td></tr>
<tr><td>Oligo fwd</td><td>CACCACGGACCGCGACTGCTTGTC</td><td></td><td></td><td></td><td></td><td></td><td></td><td></td><td></td><td></td><td></td></tr>
<tr><td>Oligo rev</td><td>AAACGACAAGCAGTCGCGGTCCGT</td><td></td><td></td><td></td><td></td><td></td><td></td><td></td><td></td><td></td><td></td></tr>
<tr><td>Chromosome</td><td>start</td><td>end</td><td>strand</td><td>MM</td><td>target_seq</td><td>PAM</td><td>alignment</td><td>distance</td><td>position</td><td>gene name</td><td>gene id</td></tr>
<tr><td>GU305914.1</td><td>23736</td><td>23758</td><td>-</td><td>0</td><td>ACGGACCGCGACTGCTTGTC</td><td>GGG</td><td>||||||||[||||||||||||]PAM</td><td>NA</td><td>Intergenic</td><td>NA</td><td>NA</td></tr>
<tr><td></td><td></td><td></td><td></td><td></td><td></td><td></td><td></td><td></td><td></td><td></td><td></td></tr>
<tr><td>T32688</td><td>GACCGCGACTGCTTGTCGGGCGG</td><td>1000</td><td></td><td></td><td></td><td></td><td></td><td></td><td></td><td></td><td></td></tr>
<tr><td>Oligo fwd</td><td>CACCGACCGCGACTGCTTGTCGGG</td><td></td><td></td><td></td><td></td><td></td><td></td><td></td><td></td><td></td><td></td></tr>
<tr><td>Oligo rev</td><td>AAACCCCGACAAGCAGTCGCGGTC</td><td></td><td></td><td></td><td></td><td></td><td></td><td></td><td></td><td></td><td></td></tr>
<tr><td>Chromosome</td><td>start</td><td>end</td><td>strand</td><td>MM</td><td>target_seq</td><td>PAM</td><td>alignment</td><td>distance</td><td>position</td><td>gene name</td><td>gene id</td></tr>
<tr><td>GU305914.1</td><td>23733</td><td>23755</td><td>-</td><td>0</td><td>GACCGCGACTGCTTGTCGGG</td><td>CGG</td><td>||||||||[||||||||||||]PAM</td><td>NA</td><td>Intergenic</td><td>NA</td><td>NA</td></tr>
<tr><td></td><td></td><td></td><td></td><td></td><td></td><td></td><td></td><td></td><td></td><td></td><td></td></tr>
<tr><td>T32689</td><td>ACCGCGACTGCTTGTCGGGCGGG</td><td>1000</td><td></td><td></td><td></td><td></td><td></td><td></td><td></td><td></td><td></td></tr>
<tr><td>Oligo fwd</td><td>CACCACCGCGACTGCTTGTCGGGC</td><td></td><td></td><td></td><td></td><td></td><td></td><td></td><td></td><td></td><td></td></tr>
<tr><td>Oligo rev</td><td>AAACGCCCGACAAGCAGTCGCGGT</td><td></td><td></td><td></td><td></td><td></td><td></td><td></td><td></td><td></td><td></td></tr>
<tr><td>Chromosome</td><td>start</td><td>end</td><td>strand</td><td>MM</td><td>target_seq</td><td>PAM</td><td>alignment</td><td>distance</td><td>position</td><td>gene name</td><td>gene id</td></tr>
<tr><td>GU305914.1</td><td>23732</td><td>23754</td><td>-</td><td>0</td><td>ACCGCGACTGCTTGTCGGGC</td><td>GGG</td><td>||||||||[||||||||||||]PAM</td><td>NA</td><td>Intergenic</td><td>NA</td><td>NA</td></tr>
<tr><td></td><td></td><td></td><td></td><td></td><td></td><td></td><td></td><td></td><td></td><td></td><td></td></tr>
<tr><td>T32690</td><td>CTGCTTGTCGGGCGGGTGTCTGG</td><td>1000</td><td></td><td></td><td></td><td></td><td></td><td></td><td></td><td></td><td></td></tr>
<tr><td>Oligo fwd</td><td>CACCCTGCTTGTCGGGCGGGTGTC</td><td></td><td></td><td></td><td></td><td></td><td></td><td></td><td></td><td></td><td></td></tr>
<tr><td>Oligo rev</td><td>AAACGACACCCGCCCGACAAGCAG</td><td></td><td></td><td></td><td></td><td></td><td></td><td></td><td></td><td></td><td></td></tr>
<tr><td>Chromosome</td><td>start</td><td>end</td><td>strand</td><td>MM</td><td>target_seq</td><td>PAM</td><td>alignment</td><td>distance</td><td>position</td><td>gene name</td><td>gene id</td></tr>
<tr><td>GU305914.1</td><td>23725</td><td>23747</td><td>-</td><td>0</td><td>CTGCTTGTCGGGCGGGTGTC</td><td>TGG</td><td>||||||||[||||||||||||]PAM</td><td>NA</td><td>Intergenic</td><td>NA</td><td>NA</td></tr>
<tr><td></td><td></td><td></td><td></td><td></td><td></td><td></td><td></td><td></td><td></td><td></td><td></td></tr>
<tr><td>T32691</td><td>GTCGGGCGGGTGTCTGGCCTCGG</td><td>1000</td><td></td><td></td><td></td><td></td><td></td><td></td><td></td><td></td><td></td></tr>
<tr><td>Oligo fwd</td><td>CACCGTCGGGCGGGTGTCTGGCCT</td><td></td><td></td><td></td><td></td><td></td><td></td><td></td><td></td><td></td><td></td></tr>
<tr><td>Oligo rev</td><td>AAACAGGCCAGACACCCGCCCGAC</td><td></td><td></td><td></td><td></td><td></td><td></td><td></td><td></td><td></td><td></td></tr>
<tr><td>Chromosome</td><td>start</td><td>end</td><td>strand</td><td>MM</td><td>target_seq</td><td>PAM</td><td>alignment</td><td>distance</td><td>position</td><td>gene name</td><td>gene id</td></tr>
<tr><td>GU305914.1</td><td>23719</td><td>23741</td><td>-</td><td>0</td><td>GTCGGGCGGGTGTCTGGCCT</td><td>CGG</td><td>||||||||[||||||||||||]PAM</td><td>NA</td><td>Intergenic</td><td>NA</td><td>NA</td></tr>
<tr><td></td><td></td><td></td><td></td><td></td><td></td><td></td><td></td><td></td><td></td><td></td><td></td></tr>
<tr><td>T32692</td><td>GCGGGTGTCTGGCCTCGGTACGG</td><td>1000</td><td></td><td></td><td></td><td></td><td></td><td></td><td></td><td></td><td></td></tr>
<tr><td>Oligo fwd</td><td>CACCGCGGGTGTCTGGCCTCGGTA</td><td></td><td></td><td></td><td></td><td></td><td></td><td></td><td></td><td></td><td></td></tr>
<tr><td>Oligo rev</td><td>AAACTACCGAGGCCAGACACCCGC</td><td></td><td></td><td></td><td></td><td></td><td></td><td></td><td></td><td></td><td></td></tr>
<tr><td>Chromosome</td><td>start</td><td>end</td><td>strand</td><td>MM</td><td>target_seq</td><td>PAM</td><td>alignment</td><td>distance</td><td>position</td><td>gene name</td><td>gene id</td></tr>
<tr><td>GU305914.1</td><td>23714</td><td>23736</td><td>-</td><td>0</td><td>GCGGGTGTCTGGCCTCGGTA</td><td>CGG</td><td>||||||||[||||||||||||]PAM</td><td>NA</td><td>Intergenic</td><td>NA</td><td>NA</td></tr>
<tr><td></td><td></td><td></td><td></td><td></td><td></td><td></td><td></td><td></td><td></td><td></td><td></td></tr>
<tr><td>T32693</td><td>GTGTCTGGCCTCGGTACGGACGG</td><td>1000</td><td></td><td></td><td></td><td></td><td></td><td></td><td></td><td></td><td></td></tr>
<tr><td>Oligo fwd</td><td>CACCGTGTCTGGCCTCGGTACGGA</td><td></td><td></td><td></td><td></td><td></td><td></td><td></td><td></td><td></td><td></td></tr>
<tr><td>Oligo rev</td><td>AAACTCCGTACCGAGGCCAGACAC</td><td></td><td></td><td></td><td></td><td></td><td></td><td></td><td></td><td></td><td></td></tr>
<tr><td>Chromosome</td><td>start</td><td>end</td><td>strand</td><td>MM</td><td>target_seq</td><td>PAM</td><td>alignment</td><td>distance</td><td>position</td><td>gene name</td><td>gene id</td></tr>
<tr><td>GU305914.1</td><td>23710</td><td>23732</td><td>-</td><td>0</td><td>GTGTCTGGCCTCGGTACGGA</td><td>CGG</td><td>||||||||[||||||||||||]PAM</td><td>NA</td><td>Intergenic</td><td>NA</td><td>NA</td></tr>
<tr><td></td><td></td><td></td><td></td><td></td><td></td><td></td><td></td><td></td><td></td><td></td><td></td></tr>
<tr><td>T32694</td><td>TGTCTGGCCTCGGTACGGACGGG</td><td>1000</td><td></td><td></td><td></td><td></td><td></td><td></td><td></td><td></td><td></td></tr>
<tr><td>Oligo fwd</td><td>CACCTGTCTGGCCTCGGTACGGAC</td><td></td><td></td><td></td><td></td><td></td><td></td><td></td><td></td><td></td><td></td></tr>
<tr><td>Oligo rev</td><td>AAACGTCCGTACCGAGGCCAGACA</td><td></td><td></td><td></td><td></td><td></td><td></td><td></td><td></td><td></td><td></td></tr>
<tr><td>Chromosome</td><td>start</td><td>end</td><td>strand</td><td>MM</td><td>target_seq</td><td>PAM</td><td>alignment</td><td>distance</td><td>position</td><td>gene name</td><td>gene id</td></tr>
<tr><td>GU305914.1</td><td>23709</td><td>23731</td><td>-</td><td>0</td><td>TGTCTGGCCTCGGTACGGAC</td><td>GGG</td><td>||||||||[||||||||||||]PAM</td><td>NA</td><td>Intergenic</td><td>NA</td><td>NA</td></tr>
<tr><td></td><td></td><td></td><td></td><td></td><td></td><td></td><td></td><td></td><td></td><td></td><td></td></tr>
<tr><td>T32695</td><td>GTCTGGCCTCGGTACGGACGGGG</td><td>1000</td><td></td><td></td><td></td><td></td><td></td><td></td><td></td><td></td><td></td></tr>
<tr><td>Oligo fwd</td><td>CACCGTCTGGCCTCGGTACGGACG</td><td></td><td></td><td></td><td></td><td></td><td></td><td></td><td></td><td></td><td></td></tr>
<tr><td>Oligo rev</td><td>AAACCGTCCGTACCGAGGCCAGAC</td><td></td><td></td><td></td><td></td><td></td><td></td><td></td><td></td><td></td><td></td></tr>
<tr><td>Chromosome</td><td>start</td><td>end</td><td>strand</td><td>MM</td><td>target_seq</td><td>PAM</td><td>alignment</td><td>distance</td><td>position</td><td>gene name</td><td>gene id</td></tr>
<tr><td>GU305914.1</td><td>23708</td><td>23730</td><td>-</td><td>0</td><td>GTCTGGCCTCGGTACGGACG</td><td>GGG</td><td>||||||||[||||||||||||]PAM</td><td>NA</td><td>Intergenic</td><td>NA</td><td>NA</td></tr>
<tr><td></td><td></td><td></td><td></td><td></td><td></td><td></td><td></td><td></td><td></td><td></td><td></td></tr>
<tr><td>T32696</td><td>GGAACCGTGTGTATTGTGAACGG</td><td>1000</td><td></td><td></td><td></td><td></td><td></td><td></td><td></td><td></td><td></td></tr>
<tr><td>Oligo fwd</td><td>CACCGGAACCGTGTGTATTGTGAA</td><td></td><td></td><td></td><td></td><td></td><td></td><td></td><td></td><td></td><td></td></tr>
<tr><td>Oligo rev</td><td>AAACTTCACAATACACACGGTTCC</td><td></td><td></td><td></td><td></td><td></td><td></td><td></td><td></td><td></td><td></td></tr>
<tr><td>Chromosome</td><td>start</td><td>end</td><td>strand</td><td>MM</td><td>target_seq</td><td>PAM</td><td>alignment</td><td>distance</td><td>position</td><td>gene name</td><td>gene id</td></tr>
<tr><td>GU305914.1</td><td>23687</td><td>23709</td><td>-</td><td>0</td><td>GGAACCGTGTGTATTGTGAA</td><td>CGG</td><td>||||||||[||||||||||||]PAM</td><td>NA</td><td>Intergenic</td><td>NA</td><td>NA</td></tr>
<tr><td></td><td></td><td></td><td></td><td></td><td></td><td></td><td></td><td></td><td></td><td></td><td></td></tr>
<tr><td>T32697</td><td>GAACCGTGTGTATTGTGAACGGG</td><td>1000</td><td></td><td></td><td></td><td></td><td></td><td></td><td></td><td></td><td></td></tr>
<tr><td>Oligo fwd</td><td>CACCGAACCGTGTGTATTGTGAAC</td><td></td><td></td><td></td><td></td><td></td><td></td><td></td><td></td><td></td><td></td></tr>
<tr><td>Oligo rev</td><td>AAACGTTCACAATACACACGGTTC</td><td></td><td></td><td></td><td></td><td></td><td></td><td></td><td></td><td></td><td></td></tr>
<tr><td>Chromosome</td><td>start</td><td>end</td><td>strand</td><td>MM</td><td>target_seq</td><td>PAM</td><td>alignment</td><td>distance</td><td>position</td><td>gene name</td><td>gene id</td></tr>
<tr><td>GU305914.1</td><td>23686</td><td>23708</td><td>-</td><td>0</td><td>GAACCGTGTGTATTGTGAAC</td><td>GGG</td><td>||||||||[||||||||||||]PAM</td><td>NA</td><td>Intergenic</td><td>NA</td><td>NA</td></tr>
<tr><td></td><td></td><td></td><td></td><td></td><td></td><td></td><td></td><td></td><td></td><td></td><td></td></tr>
<tr><td>T32698</td><td>TTGTGAACGGGCTCCTGCGTCGG</td><td>1000</td><td></td><td></td><td></td><td></td><td></td><td></td><td></td><td></td><td></td></tr>
<tr><td>Oligo fwd</td><td>CACCTTGTGAACGGGCTCCTGCGT</td><td></td><td></td><td></td><td></td><td></td><td></td><td></td><td></td><td></td><td></td></tr>
<tr><td>Oligo rev</td><td>AAACACGCAGGAGCCCGTTCACAA</td><td></td><td></td><td></td><td></td><td></td><td></td><td></td><td></td><td></td><td></td></tr>
<tr><td>Chromosome</td><td>start</td><td>end</td><td>strand</td><td>MM</td><td>target_seq</td><td>PAM</td><td>alignment</td><td>distance</td><td>position</td><td>gene name</td><td>gene id</td></tr>
<tr><td>GU305914.1</td><td>23674</td><td>23696</td><td>-</td><td>0</td><td>TTGTGAACGGGCTCCTGCGT</td><td>CGG</td><td>||||||||[||||||||||||]PAM</td><td>NA</td><td>Intergenic</td><td>NA</td><td>NA</td></tr>
<tr><td></td><td></td><td></td><td></td><td></td><td></td><td></td><td></td><td></td><td></td><td></td><td></td></tr>
<tr><td>T32699</td><td>CGGGCTCCTGCGTCGGCCCGAGG</td><td>1000</td><td></td><td></td><td></td><td></td><td></td><td></td><td></td><td></td><td></td></tr>
<tr><td>Oligo fwd</td><td>CACCCGGGCTCCTGCGTCGGCCCG</td><td></td><td></td><td></td><td></td><td></td><td></td><td></td><td></td><td></td><td></td></tr>
<tr><td>Oligo rev</td><td>AAACCGGGCCGACGCAGGAGCCCG</td><td></td><td></td><td></td><td></td><td></td><td></td><td></td><td></td><td></td><td></td></tr>
<tr><td>Chromosome</td><td>start</td><td>end</td><td>strand</td><td>MM</td><td>target_seq</td><td>PAM</td><td>alignment</td><td>distance</td><td>position</td><td>gene name</td><td>gene id</td></tr>
<tr><td>GU305914.1</td><td>23667</td><td>23689</td><td>-</td><td>0</td><td>CGGGCTCCTGCGTCGGCCCG</td><td>AGG</td><td>||||||||[||||||||||||]PAM</td><td>NA</td><td>Intergenic</td><td>NA</td><td>NA</td></tr>
<tr><td>GU305914.1</td><td>116360</td><td>116382</td><td>+</td><td>4</td><td>CGGCGTCCTGCGGCGGCTCG</td><td>GGG</td><td>|||--|||[||||-||||-||]PAM</td><td>NA</td><td>Intergenic</td><td>NA</td><td>NA</td></tr>
<tr><td></td><td></td><td></td><td></td><td></td><td></td><td></td><td></td><td></td><td></td><td></td><td></td></tr>
<tr><td>T32700</td><td>CCGCCCGCATCTTCTTCTCCCGG</td><td>1000</td><td></td><td></td><td></td><td></td><td></td><td></td><td></td><td></td><td></td></tr>
</table>

<table>
<tr><td>Oligo fwd</td><td>CACCCCGCCCGCATCTTCTTCTCC</td><td></td><td></td><td></td><td></td><td></td><td></td><td></td><td></td><td></td><td></td></tr>
<tr><td>Oligo rev</td><td>AAACGGAGAAGAAGATGCGGGCGG</td><td></td><td></td><td></td><td></td><td></td><td></td><td></td><td></td><td></td><td></td></tr>
<tr><td>Chromosome</td><td>start</td><td>end</td><td>strand</td><td>MM</td><td>target_seq</td><td>PAM</td><td>alignment</td><td>distance</td><td>position</td><td>gene name</td><td>gene id</td></tr>
<tr><td>GU305914.1</td><td>23572</td><td>23594</td><td>-</td><td>0</td><td>CCGCCCGCATCTTCTTCTCC</td><td>CGG</td><td>||||||||[||||||||||||]PAM</td><td>NA</td><td>Intergenic</td><td>NA</td><td>NA</td></tr>
<tr><td>T32701</td><td>TGTCCGACGCTCGTCGATCGAGG</td><td>1000</td><td></td><td></td><td></td><td></td><td></td><td></td><td></td><td></td><td></td></tr>
<tr><td>Oligo fwd</td><td>CACCTGTCCGACGCTCGTCGATCG</td><td></td><td></td><td></td><td></td><td></td><td></td><td></td><td></td><td></td><td></td></tr>
<tr><td>Oligo rev</td><td>AAACCGATCGACGAGCGTCGGACA</td><td></td><td></td><td></td><td></td><td></td><td></td><td></td><td></td><td></td><td></td></tr>
<tr><td>Chromosome</td><td>start</td><td>end</td><td>strand</td><td>MM</td><td>target_seq</td><td>PAM</td><td>alignment</td><td>distance</td><td>position</td><td>gene name</td><td>gene id</td></tr>
<tr><td>GU305914.1</td><td>23543</td><td>23565</td><td>-</td><td>0</td><td>TGTCCGACGCTCGTCGATCG</td><td>AGG</td><td>||||||||[||||||||||||]PAM</td><td>NA</td><td>Intergenic</td><td>NA</td><td>NA</td></tr>
<tr><td>T32702</td><td>CGTCGATCGAGGCGCTACCCCGG</td><td>1000</td><td></td><td></td><td></td><td></td><td></td><td></td><td></td><td></td><td></td></tr>
<tr><td>Oligo fwd</td><td>CACCCGTCGATCGAGGCGCTACCC</td><td></td><td></td><td></td><td></td><td></td><td></td><td></td><td></td><td></td><td></td></tr>
<tr><td>Oligo rev</td><td>AAACGGGTAGCGCCTCGATCGACG</td><td></td><td></td><td></td><td></td><td></td><td></td><td></td><td></td><td></td><td></td></tr>
<tr><td>Chromosome</td><td>start</td><td>end</td><td>strand</td><td>MM</td><td>target_seq</td><td>PAM</td><td>alignment</td><td>distance</td><td>position</td><td>gene name</td><td>gene id</td></tr>
<tr><td>GU305914.1</td><td>23532</td><td>23554</td><td>-</td><td>0</td><td>CGTCGATCGAGGCGCTACCC</td><td>CGG</td><td>||||||||[||||||||||||]PAM</td><td>NA</td><td>Intergenic</td><td>NA</td><td>NA</td></tr>
<tr><td>T32703</td><td>GGCTTGAACGTGTCCCCTATCGG</td><td>1000</td><td></td><td></td><td></td><td></td><td></td><td></td><td></td><td></td><td></td></tr>
<tr><td>Oligo fwd</td><td>CACCGGCTTGAACGTGTCCCCTAT</td><td></td><td></td><td></td><td></td><td></td><td></td><td></td><td></td><td></td><td></td></tr>
<tr><td>Oligo rev</td><td>AAACATAGGGGACACGTTCAAGCC</td><td></td><td></td><td></td><td></td><td></td><td></td><td></td><td></td><td></td><td></td></tr>
<tr><td>Chromosome</td><td>start</td><td>end</td><td>strand</td><td>MM</td><td>target_seq</td><td>PAM</td><td>alignment</td><td>distance</td><td>position</td><td>gene name</td><td>gene id</td></tr>
<tr><td>GU305914.1</td><td>23511</td><td>23533</td><td>-</td><td>0</td><td>GGCTTGAACGTGTCCCCTAT</td><td>CGG</td><td>||||||||[||||||||||||]PAM</td><td>NA</td><td>Intergenic</td><td>NA</td><td>NA</td></tr>
<tr><td>T32704</td><td>TTGAACGTGTCCCCTATCGGTGG</td><td>1000</td><td></td><td></td><td></td><td></td><td></td><td></td><td></td><td></td><td></td></tr>
<tr><td>Oligo fwd</td><td>CACCTTGAACGTGTCCCCTATCGG</td><td></td><td></td><td></td><td></td><td></td><td></td><td></td><td></td><td></td><td></td></tr>
<tr><td>Oligo rev</td><td>AAACCCGATAGGGGACACGTTCAA</td><td></td><td></td><td></td><td></td><td></td><td></td><td></td><td></td><td></td><td></td></tr>
<tr><td>Chromosome</td><td>start</td><td>end</td><td>strand</td><td>MM</td><td>target_seq</td><td>PAM</td><td>alignment</td><td>distance</td><td>position</td><td>gene name</td><td>gene id</td></tr>
<tr><td>GU305914.1</td><td>23508</td><td>23530</td><td>-</td><td>0</td><td>TTGAACGTGTCCCCTATCGG</td><td>TGG</td><td>||||||||[||||||||||||]PAM</td><td>NA</td><td>Intergenic</td><td>NA</td><td>NA</td></tr>
<tr><td>T32705</td><td>AGTTTACTGCGTGTGCACGATGG</td><td>1000</td><td></td><td></td><td></td><td></td><td></td><td></td><td></td><td></td><td></td></tr>
<tr><td>Oligo fwd</td><td>CACCAGTTTACTGCGTGTGCACGA</td><td></td><td></td><td></td><td></td><td></td><td></td><td></td><td></td><td></td><td></td></tr>
<tr><td>Oligo rev</td><td>AAACTCGTGCACACGCAGTAAACT</td><td></td><td></td><td></td><td></td><td></td><td></td><td></td><td></td><td></td><td></td></tr>
<tr><td>Chromosome</td><td>start</td><td>end</td><td>strand</td><td>MM</td><td>target_seq</td><td>PAM</td><td>alignment</td><td>distance</td><td>position</td><td>gene name</td><td>gene id</td></tr>
<tr><td>GU305914.1</td><td>23484</td><td>23506</td><td>-</td><td>0</td><td>AGTTTACTGCGTGTGCACGA</td><td>TGG</td><td>||||||||[||||||||||||]PAM</td><td>NA</td><td>Intergenic</td><td>NA</td><td>NA</td></tr>
<tr><td>T32706</td><td>GTTTACTGCGTGTGCACGATGGG</td><td>1000</td><td></td><td></td><td></td><td></td><td></td><td></td><td></td><td></td><td></td></tr>
<tr><td>Oligo fwd</td><td>CACCGTTTACTGCGTGTGCACGAT</td><td></td><td></td><td></td><td></td><td></td><td></td><td></td><td></td><td></td><td></td></tr>
<tr><td>Oligo rev</td><td>AAACATCGTGCACACGCAGTAAAC</td><td></td><td></td><td></td><td></td><td></td><td></td><td></td><td></td><td></td><td></td></tr>
<tr><td>Chromosome</td><td>start</td><td>end</td><td>strand</td><td>MM</td><td>target_seq</td><td>PAM</td><td>alignment</td><td>distance</td><td>position</td><td>gene name</td><td>gene id</td></tr>
<tr><td>GU305914.1</td><td>23483</td><td>23505</td><td>-</td><td>0</td><td>GTTTACTGCGTGTGCACGAT</td><td>GGG</td><td>||||||||[||||||||||||]PAM</td><td>NA</td><td>Intergenic</td><td>NA</td><td>NA</td></tr>
<tr><td>T32707</td><td>TTTACTGCGTGTGCACGATGGGG</td><td>1000</td><td></td><td></td><td></td><td></td><td></td><td></td><td></td><td></td><td></td></tr>
<tr><td>Oligo fwd</td><td>CACCTTTACTGCGTGTGCACGATG</td><td></td><td></td><td></td><td></td><td></td><td></td><td></td><td></td><td></td><td></td></tr>
<tr><td>Oligo rev</td><td>AAACCATCGTGCACACGCAGTAAA</td><td></td><td></td><td></td><td></td><td></td><td></td><td></td><td></td><td></td><td></td></tr>
<tr><td>Chromosome</td><td>start</td><td>end</td><td>strand</td><td>MM</td><td>target_seq</td><td>PAM</td><td>alignment</td><td>distance</td><td>position</td><td>gene name</td><td>gene id</td></tr>
<tr><td>GU305914.1</td><td>23482</td><td>23504</td><td>-</td><td>0</td><td>TTTACTGCGTGTGCACGATG</td><td>GGG</td><td>||||||||[||||||||||||]PAM</td><td>NA</td><td>Intergenic</td><td>NA</td><td>NA</td></tr>
<tr><td>T32708</td><td>TTACTGCGTGTGCACGATGGGGG</td><td>1000</td><td></td><td></td><td></td><td></td><td></td><td></td><td></td><td></td><td></td></tr>
<tr><td>Oligo fwd</td><td>CACCTTACTGCGTGTGCACGATGG</td><td></td><td></td><td></td><td></td><td></td><td></td><td></td><td></td><td></td><td></td></tr>
<tr><td>Oligo rev</td><td>AAACCCATCGTGCACACGCAGTAA</td><td></td><td></td><td></td><td></td><td></td><td></td><td></td><td></td><td></td><td></td></tr>
<tr><td>Chromosome</td><td>start</td><td>end</td><td>strand</td><td>MM</td><td>target_seq</td><td>PAM</td><td>alignment</td><td>distance</td><td>position</td><td>gene name</td><td>gene id</td></tr>
<tr><td>GU305914.1</td><td>23481</td><td>23503</td><td>-</td><td>0</td><td>TTACTGCGTGTGCACGATGG</td><td>GGG</td><td>||||||||[||||||||||||]PAM</td><td>NA</td><td>Intergenic</td><td>NA</td><td>NA</td></tr>
<tr><td>T32709</td><td>CGTGTGCACGATGGGGGCCTCGG</td><td>1000</td><td></td><td></td><td></td><td></td><td></td><td></td><td></td><td></td><td></td></tr>
<tr><td>Oligo fwd</td><td>CACCCGTGTGCACGATGGGGGCCT</td><td></td><td></td><td></td><td></td><td></td><td></td><td></td><td></td><td></td><td></td></tr>
<tr><td>Oligo rev</td><td>AAACAGGCCCCCATCGTGCACACG</td><td></td><td></td><td></td><td></td><td></td><td></td><td></td><td></td><td></td><td></td></tr>
<tr><td>Chromosome</td><td>start</td><td>end</td><td>strand</td><td>MM</td><td>target_seq</td><td>PAM</td><td>alignment</td><td>distance</td><td>position</td><td>gene name</td><td>gene id</td></tr>
<tr><td>GU305914.1</td><td>23475</td><td>23497</td><td>-</td><td>0</td><td>CGTGTGCACGATGGGGGCCT</td><td>CGG</td><td>||||||||[||||||||||||]PAM</td><td>NA</td><td>Intergenic</td><td>NA</td><td>NA</td></tr>
<tr><td>T32710</td><td>CGATGGGGGCCTCGGTCAAGCGG</td><td>1000</td><td></td><td></td><td></td><td></td><td></td><td></td><td></td><td></td><td></td></tr>
<tr><td>Oligo fwd</td><td>CACCCGATGGGGGCCTCGGTCAAG</td><td></td><td></td><td></td><td></td><td></td><td></td><td></td><td></td><td></td><td></td></tr>
<tr><td>Oligo rev</td><td>AAACCTTGACCGAGGCCCCCATCG</td><td></td><td></td><td></td><td></td><td></td><td></td><td></td><td></td><td></td><td></td></tr>
<tr><td>Chromosome</td><td>start</td><td>end</td><td>strand</td><td>MM</td><td>target_seq</td><td>PAM</td><td>alignment</td><td>distance</td><td>position</td><td>gene name</td><td>gene id</td></tr>
<tr><td>GU305914.1</td><td>23467</td><td>23489</td><td>-</td><td>0</td><td>CGATGGGGGCCTCGGTCAAG</td><td>CGG</td><td>||||||||[||||||||||||]PAM</td><td>NA</td><td>Intergenic</td><td>NA</td><td>NA</td></tr>
<tr><td>T32711</td><td>GATGGGGGCCTCGGTCAAGCGGG</td><td>1000</td><td></td><td></td><td></td><td></td><td></td><td></td><td></td><td></td><td></td></tr>
<tr><td>Oligo fwd</td><td>CACCGATGGGGGCCTCGGTCAAGC</td><td></td><td></td><td></td><td></td><td></td><td></td><td></td><td></td><td></td><td></td></tr>
<tr><td>Oligo rev</td><td>AAACGCTTGACCGAGGCCCCCATC</td><td></td><td></td><td></td><td></td><td></td><td></td><td></td><td></td><td></td><td></td></tr>
<tr><td>Chromosome</td><td>start</td><td>end</td><td>strand</td><td>MM</td><td>target_seq</td><td>PAM</td><td>alignment</td><td>distance</td><td>position</td><td>gene name</td><td>gene id</td></tr>
<tr><td>GU305914.1</td><td>23466</td><td>23488</td><td>-</td><td>0</td><td>GATGGGGGCCTCGGTCAAGC</td><td>GGG</td><td>||||||||[||||||||||||]PAM</td><td>NA</td><td>Intergenic</td><td>NA</td><td>NA</td></tr>
<tr><td>T32712</td><td>ATGGGGGCCTCGGTCAAGCGGGG</td><td>1000</td><td></td><td></td><td></td><td></td><td></td><td></td><td></td><td></td><td></td></tr>
<tr><td>Oligo fwd</td><td>CACCATGGGGGCCTCGGTCAAGCG</td><td></td><td></td><td></td><td></td><td></td><td></td><td></td><td></td><td></td><td></td></tr>
<tr><td>Oligo rev</td><td>AAACCGCTTGACCGAGGCCCCCAT</td><td></td><td></td><td></td><td></td><td></td><td></td><td></td><td></td><td></td><td></td></tr>
<tr><td>Chromosome</td><td>start</td><td>end</td><td>strand</td><td>MM</td><td>target_seq</td><td>PAM</td><td>alignment</td><td>distance</td><td>position</td><td>gene name</td><td>gene id</td></tr>
<tr><td>GU305914.1</td><td>23465</td><td>23487</td><td>-</td><td>0</td><td>ATGGGGGCCTCGGTCAAGCG</td><td>GGG</td><td>||||||||[||||||||||||]PAM</td><td>NA</td><td>Intergenic</td><td>NA</td><td>NA</td></tr>
<tr><td>T32713</td><td>TGGGGGCCTCGGTCAAGCGGGGG</td><td>1000</td><td></td><td></td><td></td><td></td><td></td><td></td><td></td><td></td><td></td></tr>
<tr><td>Oligo fwd</td><td>CACCTGGGGGCCTCGGTCAAGCGG</td><td></td><td></td><td></td><td></td><td></td><td></td><td></td><td></td><td></td><td></td></tr>
<tr><td>Oligo rev</td><td>AAACCCGCTTGACCGAGGCCCCCA</td><td></td><td></td><td></td><td></td><td></td><td></td><td></td><td></td><td></td><td></td></tr>
<tr><td>Chromosome</td><td>start</td><td>end</td><td>strand</td><td>MM</td><td>target_seq</td><td>PAM</td><td>alignment</td><td>distance</td><td>position</td><td>gene name</td><td>gene id</td></tr>
<tr><td>GU305914.1</td><td>23464</td><td>23486</td><td>-</td><td>0</td><td>TGGGGGCCTCGGTCAAGCGG</td><td>GGG</td><td>||||||||[||||||||||||]PAM</td><td>NA</td><td>Intergenic</td><td>NA</td><td>NA</td></tr>
<tr><td>T32714</td><td>GGGGGCCTCGGTCAAGCGGGGGG</td><td>1000</td><td></td><td></td><td></td><td></td><td></td><td></td><td></td><td></td><td></td></tr>
<tr><td>Oligo fwd</td><td>CACCGGGGGCCTCGGTCAAGCGGG</td><td></td><td></td><td></td><td></td><td></td><td></td><td></td><td></td><td></td><td></td></tr>
<tr><td>Oligo rev</td><td>AAACCCCGCTTGACCGAGGCCCCC</td><td></td><td></td><td></td><td></td><td></td><td></td><td></td><td></td><td></td><td></td></tr>
<tr><td>Chromosome</td><td>start</td><td>end</td><td>strand</td><td>MM</td><td>target_seq</td><td>PAM</td><td>alignment</td><td>distance</td><td>position</td><td>gene name</td><td>gene id</td></tr>
<tr><td>GU305914.1</td><td>23463</td><td>23485</td><td>-</td><td>0</td><td>GGGGGCCTCGGTCAAGCGGG</td><td>GGG</td><td>||||||||[||||||||||||]PAM</td><td>NA</td><td>Intergenic</td><td>NA</td><td>NA</td></tr>
</table>

<table>
<tr><td>T32715</td><td>GCCTCGGTCAAGCGGGGGGATGG</td><td></td><td>1000</td><td></td><td></td><td></td><td></td><td></td><td></td><td></td><td></td><td></td></tr>
<tr><td>Oligo fwd</td><td>CACCGCCTCGGTCAAGCGGGGGGA</td><td></td><td></td><td></td><td></td><td></td><td></td><td></td><td></td><td></td><td></td><td></td></tr>
<tr><td>Oligo rev</td><td>AAACTCCCCCCGCTTGACCGAGGC</td><td></td><td></td><td></td><td></td><td></td><td></td><td></td><td></td><td></td><td></td><td></td></tr>
<tr><td>Chromosome</td><td>start</td><td></td><td>end</td><td>strand</td><td>MM</td><td>target_seq</td><td>PAM</td><td>alignment</td><td>distance</td><td>position</td><td>gene name</td><td>gene id</td></tr>
<tr><td>GU305914.1</td><td></td><td>23459</td><td>23481</td><td>-</td><td>0</td><td>GCCTCGGTCAAGCGGGGGGA</td><td>TGG</td><td>||||||||[||||||||||||]PAM</td><td>NA</td><td>Intergenic</td><td>NA</td><td>NA</td></tr>
<tr><td>T32716</td><td>TCAAGCGGGGGGATGGCCGACGG</td><td></td><td>1000</td><td></td><td></td><td></td><td></td><td></td><td></td><td></td><td></td><td></td></tr>
<tr><td>Oligo fwd</td><td>CACCTCAAGCGGGGGGATGGCCGA</td><td></td><td></td><td></td><td></td><td></td><td></td><td></td><td></td><td></td><td></td><td></td></tr>
<tr><td>Oligo rev</td><td>AAACTCGGCCATCCCCCCGCTTGA</td><td></td><td></td><td></td><td></td><td></td><td></td><td></td><td></td><td></td><td></td><td></td></tr>
<tr><td>Chromosome</td><td>start</td><td></td><td>end</td><td>strand</td><td>MM</td><td>target_seq</td><td>PAM</td><td>alignment</td><td>distance</td><td>position</td><td>gene name</td><td>gene id</td></tr>
<tr><td>GU305914.1</td><td></td><td>23452</td><td>23474</td><td>-</td><td>0</td><td>TCAAGCGGGGGGATGGCCGA</td><td>CGG</td><td>||||||||[||||||||||||]PAM</td><td>NA</td><td>Intergenic</td><td>NA</td><td>NA</td></tr>
<tr><td>T32717</td><td>TATCTGTCTGTTTGTCTGTCCGG</td><td></td><td>1000</td><td></td><td></td><td></td><td></td><td></td><td></td><td></td><td></td><td></td></tr>
<tr><td>Oligo fwd</td><td>CACCTATCTGTCTGTTTGTCTGTC</td><td></td><td></td><td></td><td></td><td></td><td></td><td></td><td></td><td></td><td></td><td></td></tr>
<tr><td>Oligo rev</td><td>AAACGACAGACAAACAGACAGATA</td><td></td><td></td><td></td><td></td><td></td><td></td><td></td><td></td><td></td><td></td><td></td></tr>
<tr><td>Chromosome</td><td>start</td><td></td><td>end</td><td>strand</td><td>MM</td><td>target_seq</td><td>PAM</td><td>alignment</td><td>distance</td><td>position</td><td>gene name</td><td>gene id</td></tr>
<tr><td>GU305914.1</td><td></td><td>23407</td><td>23429</td><td>-</td><td>0</td><td>TATCTGTCTGTTTGTCTGTC</td><td>CGG</td><td>||||||||[||||||||||||]PAM</td><td>NA</td><td>Intergenic</td><td>NA</td><td>NA</td></tr>
<tr><td>T32718</td><td>GTCTGTTTGTCTGTCCGGAGAGG</td><td></td><td>1000</td><td></td><td></td><td></td><td></td><td></td><td></td><td></td><td></td><td></td></tr>
<tr><td>Oligo fwd</td><td>CACCGTCTGTTTGTCTGTCCGGAG</td><td></td><td></td><td></td><td></td><td></td><td></td><td></td><td></td><td></td><td></td><td></td></tr>
<tr><td>Oligo rev</td><td>AAACCTCCGGACAGACAAACAGAC</td><td></td><td></td><td></td><td></td><td></td><td></td><td></td><td></td><td></td><td></td><td></td></tr>
<tr><td>Chromosome</td><td>start</td><td></td><td>end</td><td>strand</td><td>MM</td><td>target_seq</td><td>PAM</td><td>alignment</td><td>distance</td><td>position</td><td>gene name</td><td>gene id</td></tr>
<tr><td>GU305914.1</td><td></td><td>23402</td><td>23424</td><td>-</td><td>0</td><td>GTCTGTTTGTCTGTCCGGAG</td><td>AGG</td><td>||||||||[||||||||||||]PAM</td><td>NA</td><td>Intergenic</td><td>NA</td><td>NA</td></tr>
<tr><td>T32719</td><td>TCTGTTTGTCTGTCCGGAGAGGG</td><td></td><td>1000</td><td></td><td></td><td></td><td></td><td></td><td></td><td></td><td></td><td></td></tr>
<tr><td>Oligo fwd</td><td>CACCTCTGTTTGTCTGTCCGGAGA</td><td></td><td></td><td></td><td></td><td></td><td></td><td></td><td></td><td></td><td></td><td></td></tr>
<tr><td>Oligo rev</td><td>AAACTCTCCGGACAGACAAACAGA</td><td></td><td></td><td></td><td></td><td></td><td></td><td></td><td></td><td></td><td></td><td></td></tr>
<tr><td>Chromosome</td><td>start</td><td></td><td>end</td><td>strand</td><td>MM</td><td>target_seq</td><td>PAM</td><td>alignment</td><td>distance</td><td>position</td><td>gene name</td><td>gene id</td></tr>
<tr><td>GU305914.1</td><td></td><td>23401</td><td>23423</td><td>-</td><td>0</td><td>TCTGTTTGTCTGTCCGGAGA</td><td>GGG</td><td>||||||||[||||||||||||]PAM</td><td>NA</td><td>Intergenic</td><td>NA</td><td>NA</td></tr>
<tr><td>T32720</td><td>CTGTTTGTCTGTCCGGAGAGGGG</td><td></td><td>1000</td><td></td><td></td><td></td><td></td><td></td><td></td><td></td><td></td><td></td></tr>
<tr><td>Oligo fwd</td><td>CACCCTGTTTGTCTGTCCGGAGAG</td><td></td><td></td><td></td><td></td><td></td><td></td><td></td><td></td><td></td><td></td><td></td></tr>
<tr><td>Oligo rev</td><td>AAACCTCTCCGGACAGACAAACAG</td><td></td><td></td><td></td><td></td><td></td><td></td><td></td><td></td><td></td><td></td><td></td></tr>
<tr><td>Chromosome</td><td>start</td><td></td><td>end</td><td>strand</td><td>MM</td><td>target_seq</td><td>PAM</td><td>alignment</td><td>distance</td><td>position</td><td>gene name</td><td>gene id</td></tr>
<tr><td>GU305914.1</td><td></td><td>23400</td><td>23422</td><td>-</td><td>0</td><td>CTGTTTGTCTGTCCGGAGAG</td><td>GGG</td><td>||||||||[||||||||||||]PAM</td><td>NA</td><td>Intergenic</td><td>NA</td><td>NA</td></tr>
<tr><td>T32721</td><td>TTTGTCTGTCCGGAGAGGGGTGG</td><td></td><td>1000</td><td></td><td></td><td></td><td></td><td></td><td></td><td></td><td></td><td></td></tr>
<tr><td>Oligo fwd</td><td>CACCTTTGTCTGTCCGGAGAGGGG</td><td></td><td></td><td></td><td></td><td></td><td></td><td></td><td></td><td></td><td></td><td></td></tr>
<tr><td>Oligo rev</td><td>AAACCCCCTCTCCGGACAGACAAA</td><td></td><td></td><td></td><td></td><td></td><td></td><td></td><td></td><td></td><td></td><td></td></tr>
<tr><td>Chromosome</td><td>start</td><td></td><td>end</td><td>strand</td><td>MM</td><td>target_seq</td><td>PAM</td><td>alignment</td><td>distance</td><td>position</td><td>gene name</td><td>gene id</td></tr>
<tr><td>GU305914.1</td><td></td><td>23397</td><td>23419</td><td>-</td><td>0</td><td>TTTGTCTGTCCGGAGAGGGG</td><td>TGG</td><td>||||||||[||||||||||||]PAM</td><td>NA</td><td>Intergenic</td><td>NA</td><td>NA</td></tr>
<tr><td>T32722</td><td>TTGACCCGCTGTCATGTAAATGG</td><td></td><td>1000</td><td></td><td></td><td></td><td></td><td></td><td></td><td></td><td></td><td></td></tr>
<tr><td>Oligo fwd</td><td>CACCTTGACCCGCTGTCATGTAAA</td><td></td><td></td><td></td><td></td><td></td><td></td><td></td><td></td><td></td><td></td><td></td></tr>
<tr><td>Oligo rev</td><td>AAACTTTACATGACAGCGGGTCAA</td><td></td><td></td><td></td><td></td><td></td><td></td><td></td><td></td><td></td><td></td><td></td></tr>
<tr><td>Chromosome</td><td>start</td><td></td><td>end</td><td>strand</td><td>MM</td><td>target_seq</td><td>PAM</td><td>alignment</td><td>distance</td><td>position</td><td>gene name</td><td>gene id</td></tr>
<tr><td>GU305914.1</td><td></td><td>23368</td><td>23390</td><td>-</td><td>0</td><td>TTGACCCGCTGTCATGTAAA</td><td>TGG</td><td>||||||||[||||||||||||]PAM</td><td>NA</td><td>Intergenic</td><td>NA</td><td>NA</td></tr>
<tr><td>T32723</td><td>CCCGCTGTCATGTAAATGGACGG</td><td></td><td>1000</td><td></td><td></td><td></td><td></td><td></td><td></td><td></td><td></td><td></td></tr>
<tr><td>Oligo fwd</td><td>CACCCCCGCTGTCATGTAAATGGA</td><td></td><td></td><td></td><td></td><td></td><td></td><td></td><td></td><td></td><td></td><td></td></tr>
<tr><td>Oligo rev</td><td>AAACTCCATTTACATGACAGCGGG</td><td></td><td></td><td></td><td></td><td></td><td></td><td></td><td></td><td></td><td></td><td></td></tr>
<tr><td>Chromosome</td><td>start</td><td></td><td>end</td><td>strand</td><td>MM</td><td>target_seq</td><td>PAM</td><td>alignment</td><td>distance</td><td>position</td><td>gene name</td><td>gene id</td></tr>
<tr><td>GU305914.1</td><td></td><td>23364</td><td>23386</td><td>-</td><td>0</td><td>CCCGCTGTCATGTAAATGGA</td><td>CGG</td><td>||||||||[||||||||||||]PAM</td><td>NA</td><td>Intergenic</td><td>NA</td><td>NA</td></tr>
<tr><td>T32724</td><td>TGGACGGTTATTAAAAGATGAGG</td><td></td><td>1000</td><td></td><td></td><td></td><td></td><td></td><td></td><td></td><td></td><td></td></tr>
<tr><td>Oligo fwd</td><td>CACCTGGACGGTTATTAAAAGATG</td><td></td><td></td><td></td><td></td><td></td><td></td><td></td><td></td><td></td><td></td><td></td></tr>
<tr><td>Oligo rev</td><td>AAACCATCTTTTAATAACCGTCCA</td><td></td><td></td><td></td><td></td><td></td><td></td><td></td><td></td><td></td><td></td><td></td></tr>
<tr><td>Chromosome</td><td>start</td><td></td><td>end</td><td>strand</td><td>MM</td><td>target_seq</td><td>PAM</td><td>alignment</td><td>distance</td><td>position</td><td>gene name</td><td>gene id</td></tr>
<tr><td>GU305914.1</td><td></td><td>23348</td><td>23370</td><td>-</td><td>0</td><td>TGGACGGTTATTAAAAGATG</td><td>AGG</td><td>||||||||[||||||||||||]PAM</td><td>NA</td><td>Intergenic</td><td>NA</td><td>NA</td></tr>
<tr><td>T32725</td><td>GCTCGTTTGTCAACCGCCGTTGG</td><td></td><td>1000</td><td></td><td></td><td></td><td></td><td></td><td></td><td></td><td></td><td></td></tr>
<tr><td>Oligo fwd</td><td>CACCGCTCGTTTGTCAACCGCCGT</td><td></td><td></td><td></td><td></td><td></td><td></td><td></td><td></td><td></td><td></td><td></td></tr>
<tr><td>Oligo rev</td><td>AAACACGGCGGTTGACAAACGAGC</td><td></td><td></td><td></td><td></td><td></td><td></td><td></td><td></td><td></td><td></td><td></td></tr>
<tr><td>Chromosome</td><td>start</td><td></td><td>end</td><td>strand</td><td>MM</td><td>target_seq</td><td>PAM</td><td>alignment</td><td>distance</td><td>position</td><td>gene name</td><td>gene id</td></tr>
<tr><td>GU305914.1</td><td></td><td>23283</td><td>23305</td><td>-</td><td>0</td><td>GCTCGTTTGTCAACCGCCGT</td><td>TGG</td><td>||||||||[||||||||||||]PAM</td><td>NA</td><td>Intergenic</td><td>NA</td><td>NA</td></tr>
<tr><td>T32726</td><td>CGTTTGTCAACCGCCGTTGGCGG</td><td></td><td>1000</td><td></td><td></td><td></td><td></td><td></td><td></td><td></td><td></td><td></td></tr>
<tr><td>Oligo fwd</td><td>CACCCGTTTGTCAACCGCCGTTGG</td><td></td><td></td><td></td><td></td><td></td><td></td><td></td><td></td><td></td><td></td><td></td></tr>
<tr><td>Oligo rev</td><td>AAACCCAACGGCGGTTGACAAACG</td><td></td><td></td><td></td><td></td><td></td><td></td><td></td><td></td><td></td><td></td><td></td></tr>
<tr><td>Chromosome</td><td>start</td><td></td><td>end</td><td>strand</td><td>MM</td><td>target_seq</td><td>PAM</td><td>alignment</td><td>distance</td><td>position</td><td>gene name</td><td>gene id</td></tr>
<tr><td>GU305914.1</td><td></td><td>23280</td><td>23302</td><td>-</td><td>0</td><td>CGTTTGTCAACCGCCGTTGG</td><td>CGG</td><td>||||||||[||||||||||||]PAM</td><td>NA</td><td>Intergenic</td><td>NA</td><td>NA</td></tr>
<tr><td>T32727</td><td>TTGTCAACCGCCGTTGGCGGAGG</td><td></td><td>1000</td><td></td><td></td><td></td><td></td><td></td><td></td><td></td><td></td><td></td></tr>
<tr><td>Oligo fwd</td><td>CACCTTGTCAACCGCCGTTGGCGG</td><td></td><td></td><td></td><td></td><td></td><td></td><td></td><td></td><td></td><td></td><td></td></tr>
<tr><td>Oligo rev</td><td>AAACCCGCCAACGGCGGTTGACAA</td><td></td><td></td><td></td><td></td><td></td><td></td><td></td><td></td><td></td><td></td><td></td></tr>
<tr><td>Chromosome</td><td>start</td><td></td><td>end</td><td>strand</td><td>MM</td><td>target_seq</td><td>PAM</td><td>alignment</td><td>distance</td><td>position</td><td>gene name</td><td>gene id</td></tr>
<tr><td>GU305914.1</td><td></td><td>23277</td><td>23299</td><td>-</td><td>0</td><td>TTGTCAACCGCCGTTGGCGG</td><td>AGG</td><td>||||||||[||||||||||||]PAM</td><td>NA</td><td>Intergenic</td><td>NA</td><td>NA</td></tr>
<tr><td>T32728</td><td>TGTCAACCGCCGTTGGCGGAGGG</td><td></td><td>1000</td><td></td><td></td><td></td><td></td><td></td><td></td><td></td><td></td><td></td></tr>
<tr><td>Oligo fwd</td><td>CACCTGTCAACCGCCGTTGGCGGA</td><td></td><td></td><td></td><td></td><td></td><td></td><td></td><td></td><td></td><td></td><td></td></tr>
<tr><td>Oligo rev</td><td>AAACTCCGCCAACGGCGGTTGACA</td><td></td><td></td><td></td><td></td><td></td><td></td><td></td><td></td><td></td><td></td><td></td></tr>
<tr><td>Chromosome</td><td>start</td><td></td><td>end</td><td>strand</td><td>MM</td><td>target_seq</td><td>PAM</td><td>alignment</td><td>distance</td><td>position</td><td>gene name</td><td>gene id</td></tr>
<tr><td>GU305914.1</td><td></td><td>23276</td><td>23298</td><td>-</td><td>0</td><td>TGTCAACCGCCGTTGGCGGA</td><td>GGG</td><td>||||||||[||||||||||||]PAM</td><td>NA</td><td>Intergenic</td><td>NA</td><td>NA</td></tr>
<tr><td>T32729</td><td>CAACCGCCGTTGGCGGAGGGAGG</td><td></td><td>1000</td><td></td><td></td><td></td><td></td><td></td><td></td><td></td><td></td><td></td></tr>
<tr><td>Oligo fwd</td><td>CACCCAACCGCCGTTGGCGGAGGG</td><td></td><td></td><td></td><td></td><td></td><td></td><td></td><td></td><td></td><td></td><td></td></tr>
<tr><td>Oligo rev</td><td>AAACCCCTCCGCCAACGGCGGTTG</td><td></td><td></td><td></td><td></td><td></td><td></td><td></td><td></td><td></td><td></td><td></td></tr>
<tr><td>Chromosome</td><td>start</td><td></td><td>end</td><td>strand</td><td>MM</td><td>target_seq</td><td>PAM</td><td>alignment</td><td>distance</td><td>position</td><td>gene name</td><td>gene id</td></tr>
<tr><td>GU305914.1</td><td></td><td>23273</td><td>23295</td><td>-</td><td>0</td><td>CAACCGCCGTTGGCGGAGGG</td><td>AGG</td><td>||||||||[||||||||||||]PAM</td><td>NA</td><td>Intergenic</td><td>NA</td><td>NA</td></tr>
</table>

T32730 AACCGCCGTTGGCGGAGGGAGGG 1000
Oligo fwd CACCAACCGCCGTTGGCGGAGGGA
Oligo rev AAACTCCCTCCGCCAACGGCGGTT
Chromosome start end strand MM target_seq PAM alignment distance position gene name gene id
GU305914.1 23272 23294 - 0 AACCGCCGTTGGCGGAGGGA GGG ||||||||[||||||||||||]PAM NA Intergenic NA NA

T32731 CGTTGGCGGAGGGAGGGTGTCGG 1000
Oligo fwd CACCCGTTGGCGGAGGGAGGGTGT
Oligo rev AAACACACCCTCCCTCCGCCAACG
Chromosome start end strand MM target_seq PAM alignment distance position gene name gene id
GU305914.1 23266 23288 - 0 CGTTGGCGGAGGGAGGGTGT CGG ||||||||[||||||||||||]PAM NA Intergenic NA NA

T32732 GTTGGCGGAGGGAGGGTGTCGGG 1000
Oligo fwd CACCGTTGGCGGAGGGAGGGTGTC
Oligo rev AAACGACACCCTCCCTCCGCCAAC
Chromosome start end strand MM target_seq PAM alignment distance position gene name gene id
GU305914.1 23265 23287 - 0 GTTGGCGGAGGGAGGGTGTC GGG ||||||||[||||||||||||]PAM NA Intergenic NA NA

T32733 TCTGTCGCCTGCTGCGAATTCGG 1000
Oligo fwd CACCTCTGTCGCCTGCTGCGAATT
Oligo rev AAACAATTCGCAGCAGGCGACAGA
Chromosome start end strand MM target_seq PAM alignment distance position gene name gene id
GU305914.1 23235 23257 - 0 TCTGTCGCCTGCTGCGAATT CGG ||||||||[||||||||||||]PAM NA Intergenic NA NA

T32734 CTGTCGCCTGCTGCGAATTCGGG 1000
Oligo fwd CACCCTGTCGCCTGCTGCGAATTC
Oligo rev AAACGAATTCGCAGCAGGCGACAG
Chromosome start end strand MM target_seq PAM alignment distance position gene name gene id
GU305914.1 23234 23256 - 0 CTGTCGCCTGCTGCGAATTC GGG ||||||||[||||||||||||]PAM NA Intergenic NA NA

T32735 TCGCCTGCTGCGAATTCGGGCGG 1000
Oligo fwd CACCTCGCCTGCTGCGAATTCGGG
Oligo rev AAACCCCGAATTCGCAGCAGGCGA
Chromosome start end strand MM target_seq PAM alignment distance position gene name gene id
GU305914.1 23231 23253 - 0 TCGCCTGCTGCGAATTCGGG CGG ||||||||[||||||||||||]PAM NA Intergenic NA NA

T32736 CGCCTGCTGCGAATTCGGGCGGG 1000
Oligo fwd CACCCGCCTGCTGCGAATTCGGGC
Oligo rev AAACGCCCGAATTCGCAGCAGGCG
Chromosome start end strand MM target_seq PAM alignment distance position gene name gene id
GU305914.1 23230 23252 - 0 CGCCTGCTGCGAATTCGGGC GGG ||||||||[||||||||||||]PAM NA Intergenic NA NA

T32737 GCTGCGAATTCGGGCGGGATCGG 1000
Oligo fwd CACCGCTGCGAATTCGGGCGGGAT
Oligo rev AAACATCCCGCCCGAATTCGCAGC
Chromosome start end strand MM target_seq PAM alignment distance position gene name gene id
GU305914.1 23225 23247 - 0 GCTGCGAATTCGGGCGGGAT CGG ||||||||[||||||||||||]PAM NA Intergenic NA NA

T32738 GCGAATTCGGGCGGGATCGGCGG 1000
Oligo fwd CACCGCGAATTCGGGCGGGATCGG
Oligo rev AAACCCGATCCCGCCCGAATTCGC
Chromosome start end strand MM target_seq PAM alignment distance position gene name gene id
GU305914.1 23222 23244 - 0 GCGAATTCGGGCGGGATCGG CGG ||||||||[||||||||||||]PAM NA Intergenic NA NA

T32739 CGAATTCGGGCGGGATCGGCGGG 1000
Oligo fwd CACCCGAATTCGGGCGGGATCGGC
Oligo rev AAACGCCGATCCCGCCCGAATTCG
Chromosome start end strand MM target_seq PAM alignment distance position gene name gene id
GU305914.1 23221 23243 - 0 CGAATTCGGGCGGGATCGGC GGG ||||||||[||||||||||||]PAM NA Intergenic NA NA

T32740 GAATTCGGGCGGGATCGGCGGGG 1000
Oligo fwd CACCGAATTCGGGCGGGATCGGCG
Oligo rev AAACCGCCGATCCCGCCCGAATTC
Chromosome start end strand MM target_seq PAM alignment distance position gene name gene id
GU305914.1 23220 23242 - 0 GAATTCGGGCGGGATCGGCG GGG ||||||||[||||||||||||]PAM NA Intergenic NA NA

T32741 CCGACCTGCTCGAGAATCGCTGG 1000
Oligo fwd CACCCCGACCTGCTCGAGAATCGC
Oligo rev AAACGCGATTCTCGAGCAGGTCGG
Chromosome start end strand MM target_seq PAM alignment distance position gene name gene id
GU305914.1 23197 23219 - 0 CCGACCTGCTCGAGAATCGC TGG ||||||||[||||||||||||]PAM NA Intergenic NA NA

T32742 AATCGCTGGTTACGAGTCGCCGG 1000
Oligo fwd CACCAATCGCTGGTTACGAGTCGC
Oligo rev AAACGCGACTCGTAACCAGCGATT
Chromosome start end strand MM target_seq PAM alignment distance position gene name gene id
GU305914.1 23183 23205 - 0 AATCGCTGGTTACGAGTCGC CGG ||||||||[||||||||||||]PAM NA Intergenic NA NA

T32743 GGTTACGAGTCGCCGGTGAGTGG 1000
Oligo fwd CACCGGTTACGAGTCGCCGGTGAG
Oligo rev AAACCTCACCGGCGACTCGTAACC
Chromosome start end strand MM target_seq PAM alignment distance position gene name gene id
GU305914.1 23176 23198 - 0 GGTTACGAGTCGCCGGTGAG TGG ||||||||[||||||||||||]PAM NA Intergenic NA NA

T32744 GTTACGAGTCGCCGGTGAGTGGG 1000
Oligo fwd CACCGTTACGAGTCGCCGGTGAGT
Oligo rev AAACACTCACCGGCGACTCGTAAC
Chromosome start end strand MM target_seq PAM alignment distance position gene name gene id

| | | | | | | | | | | | |
|---|---|---|---|---|---|---|---|---|---|---|---|
| GU305914.1 | 23175 | 23197 | - | 0 | GTTACGAGTCGCCGGTGAGT | GGG | \|\|\|\|\|\|\|\|[\|\|\|\|\|\|\|\|\|\|\|\|]PAM | NA | Intergenic | NA | NA |
| T32745 | GCCGGTGAGTGGGATTCTCCCGG | 1000 | | | | | | | | | |
| Oligo fwd | CACCGCCGGTGAGTGGGATTCTCC | | | | | | | | | | |
| Oligo rev | AAACGGAGAATCCCACTCACCGGC | | | | | | | | | | |
| Chromosome | start | end | strand | MM | target_seq | PAM | alignment | distance | position | gene name | gene id |
| GU305914.1 | 23165 | 23187 | - | 0 | GCCGGTGAGTGGGATTCTCC | CGG | \|\|\|\|\|\|\|\|[\|\|\|\|\|\|\|\|\|\|\|\|]PAM | NA | Intergenic | NA | NA |
| T32746 | CCGGTGAGTGGGATTCTCCCGGG | 1000 | | | | | | | | | |
| Oligo fwd | CACCCCGGTGAGTGGGATTCTCCC | | | | | | | | | | |
| Oligo rev | AAACGGGAGAATCCCACTCACCGG | | | | | | | | | | |
| Chromosome | start | end | strand | MM | target_seq | PAM | alignment | distance | position | gene name | gene id |
| GU305914.1 | 23164 | 23186 | - | 0 | CCGGTGAGTGGGATTCTCCC | GGG | \|\|\|\|\|\|\|\|[\|\|\|\|\|\|\|\|\|\|\|\|]PAM | NA | Intergenic | NA | NA |
| T32747 | GATGACTGACAACGCTCAAACGG | 1000 | | | | | | | | | |
| Oligo fwd | CACCGATGACTGACAACGCTCAAA | | | | | | | | | | |
| Oligo rev | AAACTTTGAGCGTTGTCAGTCATC | | | | | | | | | | |
| Chromosome | start | end | strand | MM | target_seq | PAM | alignment | distance | position | gene name | gene id |
| GU305914.1 | 23129 | 23151 | - | 0 | GATGACTGACAACGCTCAAA | CGG | \|\|\|\|\|\|\|\|[\|\|\|\|\|\|\|\|\|\|\|\|]PAM | NA | Intergenic | NA | NA |
| T32748 | ACGCTCAAACGGAATAAAAGCGG | 1000 | | | | | | | | | |
| Oligo fwd | CACCACGCTCAAACGGAATAAAAG | | | | | | | | | | |
| Oligo rev | AAACCTTTTATTCCGTTTGAGCGT | | | | | | | | | | |
| Chromosome | start | end | strand | MM | target_seq | PAM | alignment | distance | position | gene name | gene id |
| GU305914.1 | 23118 | 23140 | - | 0 | ACGCTCAAACGGAATAAAAG | CGG | \|\|\|\|\|\|\|\|[\|\|\|\|\|\|\|\|\|\|\|\|]PAM | NA | Intergenic | NA | NA |
| T32749 | AACATGTCACCAGAATTTTGAGG | 1000 | | | | | | | | | |
| Oligo fwd | CACCAACATGTCACCAGAATTTTG | | | | | | | | | | |
| Oligo rev | AAACCAAAATTCTGGTGACATGTT | | | | | | | | | | |
| Chromosome | start | end | strand | MM | target_seq | PAM | alignment | distance | position | gene name | gene id |
| GU305914.1 | 23084 | 23106 | - | 0 | AACATGTCACCAGAATTTTG | AGG | \|\|\|\|\|\|\|\|[\|\|\|\|\|\|\|\|\|\|\|\|]PAM | NA | Intergenic | NA | NA |
| T32750 | CACCAGAATTTTGAGGTCTCCGG | 1000 | | | | | | | | | |
| Oligo fwd | CACCCACCAGAATTTTGAGGTCTC | | | | | | | | | | |
| Oligo rev | AAACGAGACCTCAAAATTCTGGTG | | | | | | | | | | |
| Chromosome | start | end | strand | MM | target_seq | PAM | alignment | distance | position | gene name | gene id |
| GU305914.1 | 23077 | 23099 | - | 0 | CACCAGAATTTTGAGGTCTC | CGG | \|\|\|\|\|\|\|\|[\|\|\|\|\|\|\|\|\|\|\|\|]PAM | NA | Intergenic | NA | NA |
| T32751 | TCGTCTCGAACCGTTCGCTATGG | 1000 | | | | | | | | | |
| Oligo fwd | CACCTCGTCTCGAACCGTTCGCTA | | | | | | | | | | |
| Oligo rev | AAACTAGCGAACGGTTCGAGACGA | | | | | | | | | | |
| Chromosome | start | end | strand | MM | target_seq | PAM | alignment | distance | position | gene name | gene id |
| GU305914.1 | 23041 | 23063 | - | 0 | TCGTCTCGAACCGTTCGCTA | TGG | \|\|\|\|\|\|\|\|[\|\|\|\|\|\|\|\|\|\|\|\|]PAM | NA | Intergenic | NA | NA |
| T32752 | CGTTCGCTATGGCTCGAGTGTGG | 1000 | | | | | | | | | |
| Oligo fwd | CACCCGTTCGCTATGGCTCGAGTG | | | | | | | | | | |
| Oligo rev | AAACCACTCGAGCCATAGCGAACG | | | | | | | | | | |
| Chromosome | start | end | strand | MM | target_seq | PAM | alignment | distance | position | gene name | gene id |
| GU305914.1 | 23030 | 23052 | - | 0 | CGTTCGCTATGGCTCGAGTG | TGG | \|\|\|\|\|\|\|\|[\|\|\|\|\|\|\|\|\|\|\|\|]PAM | NA | Intergenic | NA | NA |
| T32753 | GTGTGGTTTCTGTTCGCGATCGG | 1000 | | | | | | | | | |
| Oligo fwd | CACCGTGTGGTTTCTGTTCGCGAT | | | | | | | | | | |
| Oligo rev | AAACATCGCGAACAGAAACCACAC | | | | | | | | | | |
| Chromosome | start | end | strand | MM | target_seq | PAM | alignment | distance | position | gene name | gene id |
| GU305914.1 | 23013 | 23035 | - | 0 | GTGTGGTTTCTGTTCGCGAT | CGG | \|\|\|\|\|\|\|\|[\|\|\|\|\|\|\|\|\|\|\|\|]PAM | NA | Intergenic | NA | NA |
| T32754 | TGTGGTTTCTGTTCGCGATCGGG | 1000 | | | | | | | | | |
| Oligo fwd | CACCTGTGGTTTCTGTTCGCGATC | | | | | | | | | | |
| Oligo rev | AAACGATCGCGAACAGAAACCACA | | | | | | | | | | |
| Chromosome | start | end | strand | MM | target_seq | PAM | alignment | distance | position | gene name | gene id |
| GU305914.1 | 23012 | 23034 | - | 0 | TGTGGTTTCTGTTCGCGATC | GGG | \|\|\|\|\|\|\|\|[\|\|\|\|\|\|\|\|\|\|\|\|]PAM | NA | Intergenic | NA | NA |
| T32755 | GTGGTTTCTGTTCGCGATCGGGG | 1000 | | | | | | | | | |
| Oligo fwd | CACCGTGGTTTCTGTTCGCGATCG | | | | | | | | | | |
| Oligo rev | AAACCGATCGCGAACAGAAACCAC | | | | | | | | | | |
| Chromosome | start | end | strand | MM | target_seq | PAM | alignment | distance | position | gene name | gene id |
| GU305914.1 | 23011 | 23033 | - | 0 | GTGGTTTCTGTTCGCGATCG | GGG | \|\|\|\|\|\|\|\|[\|\|\|\|\|\|\|\|\|\|\|\|]PAM | NA | Intergenic | NA | NA |
| T32756 | GTTCGCGATCGGGGCGATCTCGG | 1000 | | | | | | | | | |
| Oligo fwd | CACCGTTCGCGATCGGGGCGATCT | | | | | | | | | | |
| Oligo rev | AAACAGATCGCCCCGATCGCGAAC | | | | | | | | | | |
| Chromosome | start | end | strand | MM | target_seq | PAM | alignment | distance | position | gene name | gene id |
| GU305914.1 | 23002 | 23024 | - | 0 | GTTCGCGATCGGGGCGATCT | CGG | \|\|\|\|\|\|\|\|[\|\|\|\|\|\|\|\|\|\|\|\|]PAM | NA | Intergenic | NA | NA |
| T32757 | TTCGCGATCGGGGCGATCTCGGG | 1000 | | | | | | | | | |
| Oligo fwd | CACCTTCGCGATCGGGGCGATCTC | | | | | | | | | | |
| Oligo rev | AAACGAGATCGCCCCGATCGCGAA | | | | | | | | | | |
| Chromosome | start | end | strand | MM | target_seq | PAM | alignment | distance | position | gene name | gene id |
| GU305914.1 | 23001 | 23023 | - | 0 | TTCGCGATCGGGGCGATCTC | GGG | \|\|\|\|\|\|\|\|[\|\|\|\|\|\|\|\|\|\|\|\|]PAM | NA | Intergenic | NA | NA |
| T32758 | TCGCGATCGGGGCGATCTCGGGG | 1000 | | | | | | | | | |
| Oligo fwd | CACCTCGCGATCGGGGCGATCTCG | | | | | | | | | | |
| Oligo rev | AAACCGAGATCGCCCCGATCGCGA | | | | | | | | | | |
| Chromosome | start | end | strand | MM | target_seq | PAM | alignment | distance | position | gene name | gene id |
| GU305914.1 | 23000 | 23022 | - | 0 | TCGCGATCGGGGCGATCTCG | GGG | \|\|\|\|\|\|\|\|[\|\|\|\|\|\|\|\|\|\|\|\|]PAM | NA | Intergenic | NA | NA |
| T32759 | CGCGATCGGGGCGATCTCGGGGG | 1000 | | | | | | | | | |
| Oligo fwd | CACCCGCGATCGGGGCGATCTCGG | | | | | | | | | | |
| Oligo rev | AAACCCGAGATCGCCCCGATCGCG | | | | | | | | | | |

Chromosome start end strand MM target_seq PAM alignment distance position gene name gene id
GU305914.1 22999 23021 - 0 CGCGATCGGGGCGATCTCGG GGG ||||||||[||||||||||||]PAM NA Intergenic NA NA

T32760 GGGGGTTTCCGCCTCGATGCCGG 1000
Oligo fwd CACCGGGGGTTTCCGCCTCGATGC
Oligo rev AAACGCATCGAGGCGGAAACCCCC
Chromosome start end strand MM target_seq PAM alignment distance position gene name gene id
GU305914.1 22981 23003 - 0 GGGGGTTTCCGCCTCGATGC CGG ||||||||[||||||||||||]PAM NA Intergenic NA NA

T32761 CGACAAATCTGACGACGAGACGG 1000
Oligo fwd CACCCGACAAATCTGACGACGAGA
Oligo rev AAACTCTCGTCGTCAGATTTGTCG
Chromosome start end strand MM target_seq PAM alignment distance position gene name gene id
GU305914.1 22958 22980 - 0 CGACAAATCTGACGACGAGA CGG ||||||||[||||||||||||]PAM NA Intergenic NA NA

T32762 ACCGCCGCCACTGACCGCGTTGG 1000
Oligo fwd CACCACCGCCGCCACTGACCGCGT
Oligo rev AAACACGCGGTCAGTGGCGGCGGT
Chromosome start end strand MM target_seq PAM alignment distance position gene name gene id
GU305914.1 22927 22949 - 0 ACCGCCGCCACTGACCGCGT TGG ||||||||[||||||||||||]PAM NA Intergenic NA NA

T32763 CCGCCGCCACTGACCGCGTTGGG 1000
Oligo fwd CACCCCGCCGCCACTGACCGCGTT
Oligo rev AAACAACGCGGTCAGTGGCGGCGG
Chromosome start end strand MM target_seq PAM alignment distance position gene name gene id
GU305914.1 22926 22948 - 0 CCGCCGCCACTGACCGCGTT GGG ||||||||[||||||||||||]PAM NA Intergenic NA NA

T32764 CCGCCACTGACCGCGTTGGGAGG 1000
Oligo fwd CACCCCGCCACTGACCGCGTTGGG
Oligo rev AAACCCCAACGCGGTCAGTGGCGG
Chromosome start end strand MM target_seq PAM alignment distance position gene name gene id
GU305914.1 22923 22945 - 0 CCGCCACTGACCGCGTTGGG AGG ||||||||[||||||||||||]PAM NA Intergenic NA NA

T32765 CAGCGTGCTGCGAAACGCTTCGG 1000
Oligo fwd CACCCAGCGTGCTGCGAAACGCTT
Oligo rev AAACAAGCGTTTCGCAGCACGCTG
Chromosome start end strand MM target_seq PAM alignment distance position gene name gene id
GU305914.1 22891 22913 - 0 CAGCGTGCTGCGAAACGCTT CGG ||||||||[||||||||||||]PAM NA Intergenic NA NA

T32766 CCTCGCGCACGTTTTGAGCGTGG 1000
Oligo fwd CACCCCTCGCGCACGTTTTGAGCG
Oligo rev AAACCGCTCAAAACGTGCGCGAGG
Chromosome start end strand MM target_seq PAM alignment distance position gene name gene id
GU305914.1 22861 22883 - 0 CCTCGCGCACGTTTTGAGCG TGG ||||||||[||||||||||||]PAM NA Intergenic NA NA

T32767 CTCGCGCACGTTTTGAGCGTGGG 1000
Oligo fwd CACCCTCGCGCACGTTTTGAGCGT
Oligo rev AAACACGCTCAAAACGTGCGCGAG
Chromosome start end strand MM target_seq PAM alignment distance position gene name gene id
GU305914.1 22860 22882 - 0 CTCGCGCACGTTTTGAGCGT GGG ||||||||[||||||||||||]PAM NA Intergenic NA NA

T32768 TTGAGCGTGGGTATGTTGAAAGG 1000
Oligo fwd CACCTTGAGCGTGGGTATGTTGAA
Oligo rev AAACTTCAACATACCCACGCTCAA
Chromosome start end strand MM target_seq PAM alignment distance position gene name gene id
GU305914.1 22848 22870 - 0 TTGAGCGTGGGTATGTTGAA AGG ||||||||[||||||||||||]PAM NA Intergenic NA NA

T32769 GAAAGGCACTAAGATAGCGACGG 1000
Oligo fwd CACCGAAAGGCACTAAGATAGCGA
Oligo rev AAACTCGCTATCTTAGTGCCTTTC
Chromosome start end strand MM target_seq PAM alignment distance position gene name gene id
GU305914.1 22831 22853 - 0 GAAAGGCACTAAGATAGCGA CGG ||||||||[||||||||||||]PAM NA Intergenic NA NA

T32770 AGGCACTAAGATAGCGACGGTGG 1000
Oligo fwd CACCAGGCACTAAGATAGCGACGG
Oligo rev AAACCCGTCGCTATCTTAGTGCCT
Chromosome start end strand MM target_seq PAM alignment distance position gene name gene id
GU305914.1 22828 22850 - 0 AGGCACTAAGATAGCGACGG TGG ||||||||[||||||||||||]PAM NA Intergenic NA NA

T32771 ACTAAGATAGCGACGGTGGCCGG 1000
Oligo fwd CACCACTAAGATAGCGACGGTGGC
Oligo rev AAACGCCACCGTCGCTATCTTAGT
Chromosome start end strand MM target_seq PAM alignment distance position gene name gene id
GU305914.1 22824 22846 - 0 ACTAAGATAGCGACGGTGGC CGG ||||||||[||||||||||||]PAM NA Intergenic NA NA

T32772 AGCGACGGTGGCCGGTCGTGAGG 1000
Oligo fwd CACCAGCGACGGTGGCCGGTCGTG
Oligo rev AAACCACGACCGGCCACCGTCGCT
Chromosome start end strand MM target_seq PAM alignment distance position gene name gene id
GU305914.1 22816 22838 - 0 AGCGACGGTGGCCGGTCGTG AGG ||||||||[||||||||||||]PAM NA Intergenic NA NA

T32773 GCCGGTCGTGAGGACGACGATGG 1000
Oligo fwd CACCGCCGGTCGTGAGGACGACGA
Oligo rev AAACTCGTCGTCCTCACGACCGGC
Chromosome start end strand MM target_seq PAM alignment distance position gene name gene id
GU305914.1 22806 22828 - 0 GCCGGTCGTGAGGACGACGA TGG ||||||||[||||||||||||]PAM NA Intergenic NA NA

T32774 GGACGACGATGGTATATTTTTGG 1000
Oligo fwd CACCGGACGACGATGGTATATTTT

<table>
<tr><td>Oligo rev</td><td>AAACAAAATATACCATCGTCGTCC</td><td></td><td></td><td></td><td></td><td></td><td></td><td></td><td></td><td></td><td></td><td></td></tr>
<tr><td>Chromosome</td><td>start</td><td></td><td>end</td><td>strand</td><td>MM</td><td>target_seq</td><td>PAM</td><td>alignment</td><td>distance</td><td>position</td><td>gene name</td><td>gene id</td></tr>
<tr><td>GU305914.1</td><td></td><td>22795</td><td>22817</td><td>-</td><td>0</td><td>GGACGACGATGGTATATTTT</td><td>TGG</td><td>||||||||[||||||||||||]PAM</td><td>NA</td><td>Intergenic</td><td>NA</td><td>NA</td></tr>
<tr><td></td><td></td><td></td><td></td><td></td><td></td><td></td><td></td><td></td><td></td><td></td><td></td><td></td></tr>
<tr><td>T32775</td><td>CGATGGTATATTTTTGGCTCTGG</td><td></td><td>1000</td><td></td><td></td><td></td><td></td><td></td><td></td><td></td><td></td><td></td></tr>
<tr><td>Oligo fwd</td><td>CACCCGATGGTATATTTTTGGCTC</td><td></td><td></td><td></td><td></td><td></td><td></td><td></td><td></td><td></td><td></td><td></td></tr>
<tr><td>Oligo rev</td><td>AAACGAGCCAAAAATATACCATCG</td><td></td><td></td><td></td><td></td><td></td><td></td><td></td><td></td><td></td><td></td><td></td></tr>
<tr><td>Chromosome</td><td>start</td><td></td><td>end</td><td>strand</td><td>MM</td><td>target_seq</td><td>PAM</td><td>alignment</td><td>distance</td><td>position</td><td>gene name</td><td>gene id</td></tr>
<tr><td>GU305914.1</td><td></td><td>22789</td><td>22811</td><td>-</td><td>0</td><td>CGATGGTATATTTTTGGCTC</td><td>TGG</td><td>||||||||[||||||||||||]PAM</td><td>NA</td><td>Intergenic</td><td>NA</td><td>NA</td></tr>
<tr><td></td><td></td><td></td><td></td><td></td><td></td><td></td><td></td><td></td><td></td><td></td><td></td><td></td></tr>
<tr><td>T32776</td><td>TATATTTTTGGCTCTGGTCTTGG</td><td></td><td>1000</td><td></td><td></td><td></td><td></td><td></td><td></td><td></td><td></td><td></td></tr>
<tr><td>Oligo fwd</td><td>CACCTATATTTTTGGCTCTGGTCT</td><td></td><td></td><td></td><td></td><td></td><td></td><td></td><td></td><td></td><td></td><td></td></tr>
<tr><td>Oligo rev</td><td>AAACAGACCAGAGCCAAAAATATA</td><td></td><td></td><td></td><td></td><td></td><td></td><td></td><td></td><td></td><td></td><td></td></tr>
<tr><td>Chromosome</td><td>start</td><td></td><td>end</td><td>strand</td><td>MM</td><td>target_seq</td><td>PAM</td><td>alignment</td><td>distance</td><td>position</td><td>gene name</td><td>gene id</td></tr>
<tr><td>GU305914.1</td><td></td><td>22783</td><td>22805</td><td>-</td><td>0</td><td>TATATTTTTGGCTCTGGTCT</td><td>TGG</td><td>||||||||[||||||||||||]PAM</td><td>NA</td><td>Intergenic</td><td>NA</td><td>NA</td></tr>
<tr><td></td><td></td><td></td><td></td><td></td><td></td><td></td><td></td><td></td><td></td><td></td><td></td><td></td></tr>
<tr><td>T32777</td><td>ATATTTTTGGCTCTGGTCTTGGG</td><td></td><td>1000</td><td></td><td></td><td></td><td></td><td></td><td></td><td></td><td></td><td></td></tr>
<tr><td>Oligo fwd</td><td>CACCATATTTTTGGCTCTGGTCTT</td><td></td><td></td><td></td><td></td><td></td><td></td><td></td><td></td><td></td><td></td><td></td></tr>
<tr><td>Oligo rev</td><td>AAACAAGACCAGAGCCAAAAATAT</td><td></td><td></td><td></td><td></td><td></td><td></td><td></td><td></td><td></td><td></td><td></td></tr>
<tr><td>Chromosome</td><td>start</td><td></td><td>end</td><td>strand</td><td>MM</td><td>target_seq</td><td>PAM</td><td>alignment</td><td>distance</td><td>position</td><td>gene name</td><td>gene id</td></tr>
<tr><td>GU305914.1</td><td></td><td>22782</td><td>22804</td><td>-</td><td>0</td><td>ATATTTTTGGCTCTGGTCTT</td><td>GGG</td><td>||||||||[||||||||||||]PAM</td><td>NA</td><td>Intergenic</td><td>NA</td><td>NA</td></tr>
<tr><td></td><td></td><td></td><td></td><td></td><td></td><td></td><td></td><td></td><td></td><td></td><td></td><td></td></tr>
<tr><td>T32778</td><td>TATTTTTGGCTCTGGTCTTGGGG</td><td></td><td>1000</td><td></td><td></td><td></td><td></td><td></td><td></td><td></td><td></td><td></td></tr>
<tr><td>Oligo fwd</td><td>CACCTATTTTTGGCTCTGGTCTTG</td><td></td><td></td><td></td><td></td><td></td><td></td><td></td><td></td><td></td><td></td><td></td></tr>
<tr><td>Oligo rev</td><td>AAACCAAGACCAGAGCCAAAAATA</td><td></td><td></td><td></td><td></td><td></td><td></td><td></td><td></td><td></td><td></td><td></td></tr>
<tr><td>Chromosome</td><td>start</td><td></td><td>end</td><td>strand</td><td>MM</td><td>target_seq</td><td>PAM</td><td>alignment</td><td>distance</td><td>position</td><td>gene name</td><td>gene id</td></tr>
<tr><td>GU305914.1</td><td></td><td>22781</td><td>22803</td><td>-</td><td>0</td><td>TATTTTTGGCTCTGGTCTTG</td><td>GGG</td><td>||||||||[||||||||||||]PAM</td><td>NA</td><td>Intergenic</td><td>NA</td><td>NA</td></tr>
<tr><td></td><td></td><td></td><td></td><td></td><td></td><td></td><td></td><td></td><td></td><td></td><td></td><td></td></tr>
<tr><td>T32779</td><td>ATTTTTGGCTCTGGTCTTGGGGG</td><td></td><td>1000</td><td></td><td></td><td></td><td></td><td></td><td></td><td></td><td></td><td></td></tr>
<tr><td>Oligo fwd</td><td>CACCATTTTTGGCTCTGGTCTTGG</td><td></td><td></td><td></td><td></td><td></td><td></td><td></td><td></td><td></td><td></td><td></td></tr>
<tr><td>Oligo rev</td><td>AAACCCAAGACCAGAGCCAAAAAT</td><td></td><td></td><td></td><td></td><td></td><td></td><td></td><td></td><td></td><td></td><td></td></tr>
<tr><td>Chromosome</td><td>start</td><td></td><td>end</td><td>strand</td><td>MM</td><td>target_seq</td><td>PAM</td><td>alignment</td><td>distance</td><td>position</td><td>gene name</td><td>gene id</td></tr>
<tr><td>GU305914.1</td><td></td><td>22780</td><td>22802</td><td>-</td><td>0</td><td>ATTTTTGGCTCTGGTCTTGG</td><td>GGG</td><td>||||||||[||||||||||||]PAM</td><td>NA</td><td>Intergenic</td><td>NA</td><td>NA</td></tr>
<tr><td></td><td></td><td></td><td></td><td></td><td></td><td></td><td></td><td></td><td></td><td></td><td></td><td></td></tr>
<tr><td>T32780</td><td>GGCTCTGGTCTTGGGGGTGACGG</td><td></td><td>1000</td><td></td><td></td><td></td><td></td><td></td><td></td><td></td><td></td><td></td></tr>
<tr><td>Oligo fwd</td><td>CACCGGCTCTGGTCTTGGGGGTGA</td><td></td><td></td><td></td><td></td><td></td><td></td><td></td><td></td><td></td><td></td><td></td></tr>
<tr><td>Oligo rev</td><td>AAACTCACCCCCAAGACCAGAGCC</td><td></td><td></td><td></td><td></td><td></td><td></td><td></td><td></td><td></td><td></td><td></td></tr>
<tr><td>Chromosome</td><td>start</td><td></td><td>end</td><td>strand</td><td>MM</td><td>target_seq</td><td>PAM</td><td>alignment</td><td>distance</td><td>position</td><td>gene name</td><td>gene id</td></tr>
<tr><td>GU305914.1</td><td></td><td>22774</td><td>22796</td><td>-</td><td>0</td><td>GGCTCTGGTCTTGGGGGTGA</td><td>CGG</td><td>||||||||[||||||||||||]PAM</td><td>NA</td><td>Intergenic</td><td>NA</td><td>NA</td></tr>
<tr><td></td><td></td><td></td><td></td><td></td><td></td><td></td><td></td><td></td><td></td><td></td><td></td><td></td></tr>
<tr><td>T32781</td><td>GGAGCAATTCGTGAGCCAGACGG</td><td></td><td>1000</td><td></td><td></td><td></td><td></td><td></td><td></td><td></td><td></td><td></td></tr>
<tr><td>Oligo fwd</td><td>CACCGGAGCAATTCGTGAGCCAGA</td><td></td><td></td><td></td><td></td><td></td><td></td><td></td><td></td><td></td><td></td><td></td></tr>
<tr><td>Oligo rev</td><td>AAACTCTGGCTCACGAATTGCTCC</td><td></td><td></td><td></td><td></td><td></td><td></td><td></td><td></td><td></td><td></td><td></td></tr>
<tr><td>Chromosome</td><td>start</td><td></td><td>end</td><td>strand</td><td>MM</td><td>target_seq</td><td>PAM</td><td>alignment</td><td>distance</td><td>position</td><td>gene name</td><td>gene id</td></tr>
<tr><td>GU305914.1</td><td></td><td>22753</td><td>22775</td><td>-</td><td>0</td><td>GGAGCAATTCGTGAGCCAGA</td><td>CGG</td><td>||||||||[||||||||||||]PAM</td><td>NA</td><td>Intergenic</td><td>NA</td><td>NA</td></tr>
<tr><td>GU305914.1</td><td></td><td>66023</td><td>66045</td><td>+</td><td>4</td><td>GCTGCACTACGTGAGCCAGA</td><td>CGG</td><td>|--|||-|[-|||||||||||]PAM</td><td>NA</td><td>Intergenic</td><td>NA</td><td>NA</td></tr>
<tr><td></td><td></td><td></td><td></td><td></td><td></td><td></td><td></td><td></td><td></td><td></td><td></td><td></td></tr>
<tr><td>T32782</td><td>ATTCGTGAGCCAGACGGTTTCGG</td><td></td><td>1000</td><td></td><td></td><td></td><td></td><td></td><td></td><td></td><td></td><td></td></tr>
<tr><td>Oligo fwd</td><td>CACCATTCGTGAGCCAGACGGTTT</td><td></td><td></td><td></td><td></td><td></td><td></td><td></td><td></td><td></td><td></td><td></td></tr>
<tr><td>Oligo rev</td><td>AAACAAACCGTCTGGCTCACGAAT</td><td></td><td></td><td></td><td></td><td></td><td></td><td></td><td></td><td></td><td></td><td></td></tr>
<tr><td>Chromosome</td><td>start</td><td></td><td>end</td><td>strand</td><td>MM</td><td>target_seq</td><td>PAM</td><td>alignment</td><td>distance</td><td>position</td><td>gene name</td><td>gene id</td></tr>
<tr><td>GU305914.1</td><td></td><td>22747</td><td>22769</td><td>-</td><td>0</td><td>ATTCGTGAGCCAGACGGTTT</td><td>CGG</td><td>||||||||[||||||||||||]PAM</td><td>NA</td><td>Intergenic</td><td>NA</td><td>NA</td></tr>
<tr><td></td><td></td><td></td><td></td><td></td><td></td><td></td><td></td><td></td><td></td><td></td><td></td><td></td></tr>
<tr><td>T32783</td><td>TTCGTGAGCCAGACGGTTTCGGG</td><td></td><td>1000</td><td></td><td></td><td></td><td></td><td></td><td></td><td></td><td></td><td></td></tr>
<tr><td>Oligo fwd</td><td>CACCTTCGTGAGCCAGACGGTTTC</td><td></td><td></td><td></td><td></td><td></td><td></td><td></td><td></td><td></td><td></td><td></td></tr>
<tr><td>Oligo rev</td><td>AAACGAAACCGTCTGGCTCACGAA</td><td></td><td></td><td></td><td></td><td></td><td></td><td></td><td></td><td></td><td></td><td></td></tr>
<tr><td>Chromosome</td><td>start</td><td></td><td>end</td><td>strand</td><td>MM</td><td>target_seq</td><td>PAM</td><td>alignment</td><td>distance</td><td>position</td><td>gene name</td><td>gene id</td></tr>
<tr><td>GU305914.1</td><td></td><td>22746</td><td>22768</td><td>-</td><td>0</td><td>TTCGTGAGCCAGACGGTTTC</td><td>GGG</td><td>||||||||[||||||||||||]PAM</td><td>NA</td><td>Intergenic</td><td>NA</td><td>NA</td></tr>
<tr><td></td><td></td><td></td><td></td><td></td><td></td><td></td><td></td><td></td><td></td><td></td><td></td><td></td></tr>
<tr><td>T32784</td><td>TGAGCCAGACGGTTTCGGGACGG</td><td></td><td>1000</td><td></td><td></td><td></td><td></td><td></td><td></td><td></td><td></td><td></td></tr>
<tr><td>Oligo fwd</td><td>CACCTGAGCCAGACGGTTTCGGGA</td><td></td><td></td><td></td><td></td><td></td><td></td><td></td><td></td><td></td><td></td><td></td></tr>
<tr><td>Oligo rev</td><td>AAACTCCCGAAACCGTCTGGCTCA</td><td></td><td></td><td></td><td></td><td></td><td></td><td></td><td></td><td></td><td></td><td></td></tr>
<tr><td>Chromosome</td><td>start</td><td></td><td>end</td><td>strand</td><td>MM</td><td>target_seq</td><td>PAM</td><td>alignment</td><td>distance</td><td>position</td><td>gene name</td><td>gene id</td></tr>
<tr><td>GU305914.1</td><td></td><td>22742</td><td>22764</td><td>-</td><td>0</td><td>TGAGCCAGACGGTTTCGGGA</td><td>CGG</td><td>||||||||[||||||||||||]PAM</td><td>NA</td><td>Intergenic</td><td>NA</td><td>NA</td></tr>
<tr><td></td><td></td><td></td><td></td><td></td><td></td><td></td><td></td><td></td><td></td><td></td><td></td><td></td></tr>
<tr><td>T32785</td><td>GCCAGACGGTTTCGGGACGGTGG</td><td></td><td>1000</td><td></td><td></td><td></td><td></td><td></td><td></td><td></td><td></td><td></td></tr>
<tr><td>Oligo fwd</td><td>CACCGCCAGACGGTTTCGGGACGG</td><td></td><td></td><td></td><td></td><td></td><td></td><td></td><td></td><td></td><td></td><td></td></tr>
<tr><td>Oligo rev</td><td>AAACCCGTCCCGAAACCGTCTGGC</td><td></td><td></td><td></td><td></td><td></td><td></td><td></td><td></td><td></td><td></td><td></td></tr>
<tr><td>Chromosome</td><td>start</td><td></td><td>end</td><td>strand</td><td>MM</td><td>target_seq</td><td>PAM</td><td>alignment</td><td>distance</td><td>position</td><td>gene name</td><td>gene id</td></tr>
<tr><td>GU305914.1</td><td></td><td>22739</td><td>22761</td><td>-</td><td>0</td><td>GCCAGACGGTTTCGGGACGG</td><td>TGG</td><td>||||||||[||||||||||||]PAM</td><td>NA</td><td>Intergenic</td><td>NA</td><td>NA</td></tr>
<tr><td></td><td></td><td></td><td></td><td></td><td></td><td></td><td></td><td></td><td></td><td></td><td></td><td></td></tr>
<tr><td>T32786</td><td>GGTTTCGGGACGGTGGACCCTGG</td><td></td><td>1000</td><td></td><td></td><td></td><td></td><td></td><td></td><td></td><td></td><td></td></tr>
<tr><td>Oligo fwd</td><td>CACCGGTTTCGGGACGGTGGACCC</td><td></td><td></td><td></td><td></td><td></td><td></td><td></td><td></td><td></td><td></td><td></td></tr>
<tr><td>Oligo rev</td><td>AAACGGGTCCACCGTCCCGAAACC</td><td></td><td></td><td></td><td></td><td></td><td></td><td></td><td></td><td></td><td></td><td></td></tr>
<tr><td>Chromosome</td><td>start</td><td></td><td>end</td><td>strand</td><td>MM</td><td>target_seq</td><td>PAM</td><td>alignment</td><td>distance</td><td>position</td><td>gene name</td><td>gene id</td></tr>
<tr><td>GU305914.1</td><td></td><td>22732</td><td>22754</td><td>-</td><td>0</td><td>GGTTTCGGGACGGTGGACCC</td><td>TGG</td><td>||||||||[||||||||||||]PAM</td><td>NA</td><td>Intergenic</td><td>NA</td><td>NA</td></tr>
<tr><td></td><td></td><td></td><td></td><td></td><td></td><td></td><td></td><td></td><td></td><td></td><td></td><td></td></tr>
<tr><td>T32787</td><td>GGTGGACCCTGGATTTGATCTGG</td><td></td><td>1000</td><td></td><td></td><td></td><td></td><td></td><td></td><td></td><td></td><td></td></tr>
<tr><td>Oligo fwd</td><td>CACCGGTGGACCCTGGATTTGATC</td><td></td><td></td><td></td><td></td><td></td><td></td><td></td><td></td><td></td><td></td><td></td></tr>
<tr><td>Oligo rev</td><td>AAACGATCAAATCCAGGGTCCACC</td><td></td><td></td><td></td><td></td><td></td><td></td><td></td><td></td><td></td><td></td><td></td></tr>
<tr><td>Chromosome</td><td>start</td><td></td><td>end</td><td>strand</td><td>MM</td><td>target_seq</td><td>PAM</td><td>alignment</td><td>distance</td><td>position</td><td>gene name</td><td>gene id</td></tr>
<tr><td>GU305914.1</td><td></td><td>22721</td><td>22743</td><td>-</td><td>0</td><td>GGTGGACCCTGGATTTGATC</td><td>TGG</td><td>||||||||[||||||||||||]PAM</td><td>NA</td><td>Intergenic</td><td>NA</td><td>NA</td></tr>
<tr><td></td><td></td><td></td><td></td><td></td><td></td><td></td><td></td><td></td><td></td><td></td><td></td><td></td></tr>
<tr><td>T32788</td><td>GTGGACCCTGGATTTGATCTGGG</td><td></td><td>1000</td><td></td><td></td><td></td><td></td><td></td><td></td><td></td><td></td><td></td></tr>
<tr><td>Oligo fwd</td><td>CACCGTGGACCCTGGATTTGATCT</td><td></td><td></td><td></td><td></td><td></td><td></td><td></td><td></td><td></td><td></td><td></td></tr>
<tr><td>Oligo rev</td><td>AAACAGATCAAATCCAGGGTCCAC</td><td></td><td></td><td></td><td></td><td></td><td></td><td></td><td></td><td></td><td></td><td></td></tr>
<tr><td>Chromosome</td><td>start</td><td></td><td>end</td><td>strand</td><td>MM</td><td>target_seq</td><td>PAM</td><td>alignment</td><td>distance</td><td>position</td><td>gene name</td><td>gene id</td></tr>
<tr><td>GU305914.1</td><td></td><td>22720</td><td>22742</td><td>-</td><td>0</td><td>GTGGACCCTGGATTTGATCT</td><td>GGG</td><td>||||||||[||||||||||||]PAM</td><td>NA</td><td>Intergenic</td><td>NA</td><td>NA</td></tr>
</table>

T32789 GACCCTGGATTTGATCTGGGAGG 1000
Oligo fwd CACCGACCCTGGATTTGATCTGGG
Oligo rev AAACCCCAGATCAAATCCAGGGTC
Chromosome start end strand MM target_seq PAM alignment distance position gene name gene id
GU305914.1 22717 22739 - 0 GACCCTGGATTTGATCTGGG AGG ||||||||[||||||||||||]PAM NA Intergenic NA NA

T32790 CCTGGATTTGATCTGGGAGGTGG 1000
Oligo fwd CACCCCTGGATTTGATCTGGGAGG
Oligo rev AAACCCTCCCAGATCAAATCCAGG
Chromosome start end strand MM target_seq PAM alignment distance position gene name gene id
GU305914.1 22714 22736 - 0 CCTGGATTTGATCTGGGAGG TGG ||||||||[||||||||||||]PAM NA Intergenic NA NA

T32791 GAATATAACGATACCGCGACCGG 1000
Oligo fwd CACCGAATATAACGATACCGCGAC
Oligo rev AAACGTCGCGGTATCGTTATATTC
Chromosome start end strand MM target_seq PAM alignment distance position gene name gene id
GU305914.1 22692 22714 - 0 GAATATAACGATACCGCGAC CGG ||||||||[||||||||||||]PAM NA Intergenic NA NA

T32792 ATAACGATACCGCGACCGGCAGG 1000
Oligo fwd CACCATAACGATACCGCGACCGGC
Oligo rev AAACGCCGGTCGCGGTATCGTTAT
Chromosome start end strand MM target_seq PAM alignment distance position gene name gene id
GU305914.1 22688 22710 - 0 ATAACGATACCGCGACCGGC AGG ||||||||[||||||||||||]PAM NA Intergenic NA NA

T32793 GTCGTACAGCAGAATGCTCCTGG 1000
Oligo fwd CACCGTCGTACAGCAGAATGCTCC
Oligo rev AAACGGAGCATTCTGCTGTACGAC
Chromosome start end strand MM target_seq PAM alignment distance position gene name gene id
GU305914.1 22666 22688 - 0 GTCGTACAGCAGAATGCTCC TGG ||||||||[||||||||||||]PAM NA Intergenic NA NA

T32794 TCGTACAGCAGAATGCTCCTGGG 1000
Oligo fwd CACCTCGTACAGCAGAATGCTCCT
Oligo rev AAACAGGAGCATTCTGCTGTACGA
Chromosome start end strand MM target_seq PAM alignment distance position gene name gene id
GU305914.1 22665 22687 - 0 TCGTACAGCAGAATGCTCCT GGG ||||||||[||||||||||||]PAM NA Intergenic NA NA

T32795 GAATGCTCCTGGGTCCGTATCGG 1000
Oligo fwd CACCGAATGCTCCTGGGTCCGTAT
Oligo rev AAACATACGGACCCAGGAGCATTC
Chromosome start end strand MM target_seq PAM alignment distance position gene name gene id
GU305914.1 22655 22677 - 0 GAATGCTCCTGGGTCCGTAT CGG ||||||||[||||||||||||]PAM NA Intergenic NA NA

T32796 TGTCATGAAAACCCCTTTACCGG 1000
Oligo fwd CACCTGTCATGAAAACCCCTTTAC
Oligo rev AAACGTAAAGGGGTTTTCATGACA
Chromosome start end strand MM target_seq PAM alignment distance position gene name gene id
GU305914.1 22627 22649 - 0 TGTCATGAAAACCCCTTTAC CGG ||||||||[||||||||||||]PAM NA Intergenic NA NA

T32797 GTCATGAAAACCCCTTTACCGGG 1000
Oligo fwd CACCGTCATGAAAACCCCTTTACC
Oligo rev AAACGGTAAAGGGGTTTTCATGAC
Chromosome start end strand MM target_seq PAM alignment distance position gene name gene id
GU305914.1 22626 22648 - 0 GTCATGAAAACCCCTTTACC GGG ||||||||[||||||||||||]PAM NA Intergenic NA NA

T32798 GCACGAATTACACCACGAGTTGG 1000
Oligo fwd CACCGCACGAATTACACCACGAGT
Oligo rev AAACACTCGTGGTGTAATTCGTGC
Chromosome start end strand MM target_seq PAM alignment distance position gene name gene id
GU305914.1 22604 22626 - 0 GCACGAATTACACCACGAGT TGG ||||||||[||||||||||||]PAM NA Intergenic NA NA

T32799 CACGAATTACACCACGAGTTGGG 1000
Oligo fwd CACCCACGAATTACACCACGAGTT
Oligo rev AAACAACTCGTGGTGTAATTCGTG
Chromosome start end strand MM target_seq PAM alignment distance position gene name gene id
GU305914.1 22603 22625 - 0 CACGAATTACACCACGAGTT GGG ||||||||[||||||||||||]PAM NA Intergenic NA NA

T32800 ACGAATTACACCACGAGTTGGGG 1000
Oligo fwd CACCACGAATTACACCACGAGTTG
Oligo rev AAACCAACTCGTGGTGTAATTCGT
Chromosome start end strand MM target_seq PAM alignment distance position gene name gene id
GU305914.1 22602 22624 - 0 ACGAATTACACCACGAGTTG GGG ||||||||[||||||||||||]PAM NA Intergenic NA NA

T32801 AGTTGGGGTGAGCCGTGCGCTGG 1000
Oligo fwd CACCAGTTGGGGTGAGCCGTGCGC
Oligo rev AAACGCGCACGGCTCACCCCAACT
Chromosome start end strand MM target_seq PAM alignment distance position gene name gene id
GU305914.1 22587 22609 - 0 AGTTGGGGTGAGCCGTGCGC TGG ||||||||[||||||||||||]PAM NA Intergenic NA NA

T32802 GTTGGGGTGAGCCGTGCGCTGGG 1000
Oligo fwd CACCGTTGGGGTGAGCCGTGCGCT
Oligo rev AAACAGCGCACGGCTCACCCCAAC
Chromosome start end strand MM target_seq PAM alignment distance position gene name gene id
GU305914.1 22586 22608 - 0 GTTGGGGTGAGCCGTGCGCT GGG ||||||||[||||||||||||]PAM NA Intergenic NA NA

T32803 TTGGGGTGAGCCGTGCGCTGGGG 1000
Oligo fwd CACCTTGGGGTGAGCCGTGCGCTG
Oligo rev AAACCAGCGCACGGCTCACCCCAA
Chromosome start end strand MM target_seq PAM alignment distance position gene name gene id
GU305914.1 22585 22607 - 0 TTGGGGTGAGCCGTGCGCTG GGG ||||||||[||||||||||||]PAM NA Intergenic NA NA

T32804 TGAGCCGTGCGCTGGGGTTGAGG 1000
Oligo fwd CACCTGAGCCGTGCGCTGGGGTTG
Oligo rev AAACCAACCCCAGCGCACGGCTCA
Chromosome start end strand MM target_seq PAM alignment distance position gene name gene id
GU305914.1 22579 22601 - 0 TGAGCCGTGCGCTGGGGTTG AGG ||||||||[||||||||||||]PAM NA Intergenic NA NA

T32805 GCCGTGCGCTGGGGTTGAGGTGG 1000
Oligo fwd CACCGCCGTGCGCTGGGGTTGAGG
Oligo rev AAACCCTCAACCCCAGCGCACGGC
Chromosome start end strand MM target_seq PAM alignment distance position gene name gene id
GU305914.1 22576 22598 - 0 GCCGTGCGCTGGGGTTGAGG TGG ||||||||[||||||||||||]PAM NA Intergenic NA NA

T32806 CCGTGCGCTGGGGTTGAGGTGGG 1000
Oligo fwd CACCCCGTGCGCTGGGGTTGAGGT
Oligo rev AAACACCTCAACCCCAGCGCACGG
Chromosome start end strand MM target_seq PAM alignment distance position gene name gene id
GU305914.1 22575 22597 - 0 CCGTGCGCTGGGGTTGAGGT GGG ||||||||[||||||||||||]PAM NA Intergenic NA NA

T32807 GCTGGGGTTGAGGTGGGATCCGG 1000
Oligo fwd CACCGCTGGGGTTGAGGTGGGATC
Oligo rev AAACGATCCCACCTCAACCCCAGC
Chromosome start end strand MM target_seq PAM alignment distance position gene name gene id
GU305914.1 22569 22591 - 0 GCTGGGGTTGAGGTGGGATC CGG ||||||||[||||||||||||]PAM NA Intergenic NA NA

T32808 GGTTGAGGTGGGATCCGGAACGG 1000
Oligo fwd CACCGGTTGAGGTGGGATCCGGAA
Oligo rev AAACTTCCGGATCCCACCTCAACC
Chromosome start end strand MM target_seq PAM alignment distance position gene name gene id
GU305914.1 22564 22586 - 0 GGTTGAGGTGGGATCCGGAA CGG ||||||||[||||||||||||]PAM NA Intergenic NA NA

T32809 GTTGAGGTGGGATCCGGAACGGG 1000
Oligo fwd CACCGTTGAGGTGGGATCCGGAAC
Oligo rev AAACGTTCCGGATCCCACCTCAAC
Chromosome start end strand MM target_seq PAM alignment distance position gene name gene id
GU305914.1 22563 22585 - 0 GTTGAGGTGGGATCCGGAAC GGG ||||||||[||||||||||||]PAM NA Intergenic NA NA

T32810 TTGAGGTGGGATCCGGAACGGGG 1000
Oligo fwd CACCTTGAGGTGGGATCCGGAACG
Oligo rev AAACCGTTCCGGATCCCACCTCAA
Chromosome start end strand MM target_seq PAM alignment distance position gene name gene id
GU305914.1 22562 22584 - 0 TTGAGGTGGGATCCGGAACG GGG ||||||||[||||||||||||]PAM NA Intergenic NA NA

T32811 GAACGGGGAATACCGTTGTCCGG 1000
Oligo fwd CACCGAACGGGGAATACCGTTGTC
Oligo rev AAACGACAACGGTATTCCCCGTTC
Chromosome start end strand MM target_seq PAM alignment distance position gene name gene id
GU305914.1 22547 22569 - 0 GAACGGGGAATACCGTTGTC CGG ||||||||[||||||||||||]PAM NA Intergenic NA NA

T32812 AACGGGGAATACCGTTGTCCGGG 1000
Oligo fwd CACCAACGGGGAATACCGTTGTCC
Oligo rev AAACGGACAACGGTATTCCCCGTT
Chromosome start end strand MM target_seq PAM alignment distance position gene name gene id
GU305914.1 22546 22568 - 0 AACGGGGAATACCGTTGTCC GGG ||||||||[||||||||||||]PAM NA Intergenic NA NA

T32813 GGGGAATACCGTTGTCCGGGTGG 1000
Oligo fwd CACCGGGGAATACCGTTGTCCGGG
Oligo rev AAACCCCGGACAACGGTATTCCCC
Chromosome start end strand MM target_seq PAM alignment distance position gene name gene id
GU305914.1 22543 22565 - 0 GGGGAATACCGTTGTCCGGG TGG ||||||||[||||||||||||]PAM NA Intergenic NA NA

T32814 GGGAATACCGTTGTCCGGGTGGG 1000
Oligo fwd CACCGGGAATACCGTTGTCCGGGT
Oligo rev AAACACCCGGACAACGGTATTCCC
Chromosome start end strand MM target_seq PAM alignment distance position gene name gene id
GU305914.1 22542 22564 - 0 GGGAATACCGTTGTCCGGGT GGG ||||||||[||||||||||||]PAM NA Intergenic NA NA

T32815 GGAATACCGTTGTCCGGGTGGGG 1000
Oligo fwd CACCGGAATACCGTTGTCCGGGTG
Oligo rev AAACCACCCGGACAACGGTATTCC
Chromosome start end strand MM target_seq PAM alignment distance position gene name gene id
GU305914.1 22541 22563 - 0 GGAATACCGTTGTCCGGGTG GGG ||||||||[||||||||||||]PAM NA Intergenic NA NA

T32816 TAAAGTCGCACGCCCTCCAAAGG 1000
Oligo fwd CACCTAAAGTCGCACGCCCTCCAA
Oligo rev AAACTTGGAGGGCGTGCGACTTTA
Chromosome start end strand MM target_seq PAM alignment distance position gene name gene id
GU305914.1 22516 22538 - 0 TAAAGTCGCACGCCCTCCAA AGG ||||||||[||||||||||||]PAM NA Intergenic NA NA

T32817 CGTCGACTACAAGAAATTCCAGG 1000
Oligo fwd CACCCGTCGACTACAAGAAATTCC
Oligo rev AAACGGAATTTCTTGTAGTCGACG
Chromosome start end strand MM target_seq PAM alignment distance position gene name gene id
GU305914.1 22492 22514 - 0 CGTCGACTACAAGAAATTCC AGG ||||||||[||||||||||||]PAM NA Intergenic NA NA

T32818 ACAGTGAGAGAGCACGCGACTGG 1000
Oligo fwd CACCACAGTGAGAGAGCACGCGAC
Oligo rev AAACGTCGCGTGCTCTCTCACTGT
Chromosome start end strand MM target_seq PAM alignment distance position gene name gene id

GU305914.1 22452 22474 - 0 ACAGTGAGAGAGCACGCGAC TGG ||||||||[||||||||||||]PAM NA Intergenic NA NA

T32819 CGCGACTGGAGACAAAGCCGTGG 1000
Oligo fwd CACCCGCGACTGGAGACAAAGCCG
Oligo rev AAACCGGCTTTGTCTCCAGTCGCG
Chromosome start end strand MM target_seq PAM alignment distance position gene name gene id
GU305914.1 22438 22460 - 0 CGCGACTGGAGACAAAGCCG TGG ||||||||[||||||||||||]PAM NA Intergenic NA NA

T32820 GGAGACAAAGCCGTGGCGCTCGG 1000
Oligo fwd CACCGGAGACAAAGCCGTGGCGCT
Oligo rev AAACAGCGCCACGGCTTTGTCTCC
Chromosome start end strand MM target_seq PAM alignment distance position gene name gene id
GU305914.1 22431 22453 - 0 GGAGACAAAGCCGTGGCGCT CGG ||||||||[||||||||||||]PAM NA Intergenic NA NA

T32821 GGCGCTCGGCATCTTTCCCATGG 1000
Oligo fwd CACCGGCGCTCGGCATCTTTCCCA
Oligo rev AAACTGGGAAAGATGCCGAGCGCC
Chromosome start end strand MM target_seq PAM alignment distance position gene name gene id
GU305914.1 22417 22439 - 0 GGCGCTCGGCATCTTTCCCA TGG ||||||||[||||||||||||]PAM NA Intergenic NA NA

T32822 TCTTGTCGAACCGCATCTCTCGG 1000
Oligo fwd CACCTCTTGTCGAACCGCATCTCT
Oligo rev AAACAGAGATGCGGTTCGACAAGA
Chromosome start end strand MM target_seq PAM alignment distance position gene name gene id
GU305914.1 22391 22413 - 0 TCTTGTCGAACCGCATCTCT CGG ||||||||[||||||||||||]PAM NA Intergenic NA NA

T32823 CTTGTCGAACCGCATCTCTCGGG 1000
Oligo fwd CACCCTTGTCGAACCGCATCTCTC
Oligo rev AAACGAGAGATGCGGTTCGACAAG
Chromosome start end strand MM target_seq PAM alignment distance position gene name gene id
GU305914.1 22390 22412 - 0 CTTGTCGAACCGCATCTCTC GGG ||||||||[||||||||||||]PAM NA Intergenic NA NA

T32824 GTCGAACCGCATCTCTCGGGTGG 1000
Oligo fwd CACCGTCGAACCGCATCTCTCGGG
Oligo rev AAACCCCGAGAGATGCGGTTCGAC
Chromosome start end strand MM target_seq PAM alignment distance position gene name gene id
GU305914.1 22387 22409 - 0 GTCGAACCGCATCTCTCGGG TGG ||||||||[||||||||||||]PAM NA Intergenic NA NA

T32825 TCGAACCGCATCTCTCGGGTGGG 1000
Oligo fwd CACCTCGAACCGCATCTCTCGGGT
Oligo rev AAACACCCGAGAGATGCGGTTCGA
Chromosome start end strand MM target_seq PAM alignment distance position gene name gene id
GU305914.1 22386 22408 - 0 TCGAACCGCATCTCTCGGGT GGG ||||||||[||||||||||||]PAM NA Intergenic NA NA

T32826 CTCGGGTGGGACTACATCACTGG 1000
Oligo fwd CACCCTCGGGTGGGACTACATCAC
Oligo rev AAACGTGATGTAGTCCCACCCGAG
Chromosome start end strand MM target_seq PAM alignment distance position gene name gene id
GU305914.1 22373 22395 - 0 CTCGGGTGGGACTACATCAC TGG ||||||||[||||||||||||]PAM NA Intergenic NA NA

T32827 GGTGGGACTACATCACTGGACGG 1000
Oligo fwd CACCGGTGGGACTACATCACTGGA
Oligo rev AAACTCCAGTGATGTAGTCCCACC
Chromosome start end strand MM target_seq PAM alignment distance position gene name gene id
GU305914.1 22369 22391 - 0 GGTGGGACTACATCACTGGA CGG ||||||||[||||||||||||]PAM NA Intergenic NA NA

T32828 TTTCACCCCGCTGATGTCGCTGG 1000
Oligo fwd CACCTTTCACCCCGCTGATGTCGC
Oligo rev AAACGCGACATCAGCGGGGTGAAA
Chromosome start end strand MM target_seq PAM alignment distance position gene name gene id
GU305914.1 22341 22363 - 0 TTTCACCCCGCTGATGTCGC TGG ||||||||[||||||||||||]PAM NA Intergenic NA NA

T32829 TTCACCCCGCTGATGTCGCTGGG 1000
Oligo fwd CACCTTCACCCCGCTGATGTCGCT
Oligo rev AAACAGCGACATCAGCGGGGTGAA
Chromosome start end strand MM target_seq PAM alignment distance position gene name gene id
GU305914.1 22340 22362 - 0 TTCACCCCGCTGATGTCGCT GGG ||||||||[||||||||||||]PAM NA Intergenic NA NA

T32830 GCTGATGTCGCTGGGAAAAACGG 1000
Oligo fwd CACCGCTGATGTCGCTGGGAAAAA
Oligo rev AAACTTTTTCCCAGCGACATCAGC
Chromosome start end strand MM target_seq PAM alignment distance position gene name gene id
GU305914.1 22332 22354 - 0 GCTGATGTCGCTGGGAAAAA CGG ||||||||[||||||||||||]PAM NA Intergenic NA NA

T32831 TGTCGCTGGGAAAAACGGAAAGG 1000
Oligo fwd CACCTGTCGCTGGGAAAAACGGAA
Oligo rev AAACTTCCGTTTTTCCCAGCGACA
Chromosome start end strand MM target_seq PAM alignment distance position gene name gene id
GU305914.1 22327 22349 - 0 TGTCGCTGGGAAAAACGGAA AGG ||||||||[||||||||||||]PAM NA Intergenic NA NA

T32832 CCGAGATTCGAGAGCTCTCCCGG 1000
Oligo fwd CACCCCGAGATTCGAGAGCTCTCC
Oligo rev AAACGGAGAGCTCTCGAATCTCGG
Chromosome start end strand MM target_seq PAM alignment distance position gene name gene id
GU305914.1 22302 22324 - 0 CCGAGATTCGAGAGCTCTCC CGG ||||||||[||||||||||||]PAM NA Intergenic NA NA

T32833 CGAGATTCGAGAGCTCTCCCGGG 1000
Oligo fwd CACCCGAGATTCGAGAGCTCTCCC
Oligo rev AAACGGGAGAGCTCTCGAATCTCG

Chromosome start end strand MM target_seq PAM alignment distance position gene name gene id
GU305914.1 22301 22323 - 0 CGAGATTCGAGAGCTCTCCC GGG ||||||||[||||||||||||]PAM NA Intergenic NA NA

T32834 TCGAGAGCTCTCCCGGGTTCCGG 1000
Oligo fwd CACCTCGAGAGCTCTCCCGGGTTC
Oligo rev AAACGAACCCGGGAGAGCTCTCGA
Chromosome start end strand MM target_seq PAM alignment distance position gene name gene id
GU305914.1 22295 22317 - 0 TCGAGAGCTCTCCCGGGTTC CGG ||||||||[||||||||||||]PAM NA Intergenic NA NA

T32835 CCCGGGTTCCGGTTTCTCGTCGG 1000
Oligo fwd CACCCCCGGGTTCCGGTTTCTCGT
Oligo rev AAACACGAGAAACCGGAACCCGGG
Chromosome start end strand MM target_seq PAM alignment distance position gene name gene id
GU305914.1 22284 22306 - 0 CCCGGGTTCCGGTTTCTCGT CGG ||||||||[||||||||||||]PAM NA Intergenic NA NA

T32836 CCGGGTTCCGGTTTCTCGTCGGG 1000
Oligo fwd CACCCCGGGTTCCGGTTTCTCGTC
Oligo rev AAACGACGAGAAACCGGAACCCGG
Chromosome start end strand MM target_seq PAM alignment distance position gene name gene id
GU305914.1 22283 22305 - 0 CCGGGTTCCGGTTTCTCGTC GGG ||||||||[||||||||||||]PAM NA Intergenic NA NA

T32837 GTTTCTCGTCGGGACCATGTTGG 1000
Oligo fwd CACCGTTTCTCGTCGGGACCATGT
Oligo rev AAACACATGGTCCCGACGAGAAAC
Chromosome start end strand MM target_seq PAM alignment distance position gene name gene id
GU305914.1 22273 22295 - 0 GTTTCTCGTCGGGACCATGT TGG ||||||||[||||||||||||]PAM NA Intergenic NA NA

T32838 TTTCTCGTCGGGACCATGTTGGG 1000
Oligo fwd CACCTTTCTCGTCGGGACCATGTT
Oligo rev AAACAACATGGTCCCGACGAGAAA
Chromosome start end strand MM target_seq PAM alignment distance position gene name gene id
GU305914.1 22272 22294 - 0 TTTCTCGTCGGGACCATGTT GGG ||||||||[||||||||||||]PAM NA Intergenic NA NA

T32839 GTCGGGACCATGTTGGGCGCCGG 1000
Oligo fwd CACCGTCGGGACCATGTTGGGCGC
Oligo rev AAACGCGCCCAACATGGTCCCGAC
Chromosome start end strand MM target_seq PAM alignment distance position gene name gene id
GU305914.1 22266 22288 - 0 GTCGGGACCATGTTGGGCGC CGG ||||||||[||||||||||||]PAM NA Intergenic NA NA

T32840 GTTGGGCGCCGGCACGATGACGG 1000
Oligo fwd CACCGTTGGGCGCCGGCACGATGA
Oligo rev AAACTCATCGTGCCGGCGCCCAAC
Chromosome start end strand MM target_seq PAM alignment distance position gene name gene id
GU305914.1 22255 22277 - 0 GTTGGGCGCCGGCACGATGA CGG ||||||||[||||||||||||]PAM NA Intergenic NA NA

T32841 ATGACGGTCATCGCGACCGTCGG 1000
Oligo fwd CACCATGACGGTCATCGCGACCGT
Oligo rev AAACACGGTCGCGATGACCGTCAT
Chromosome start end strand MM target_seq PAM alignment distance position gene name gene id
GU305914.1 22239 22261 - 0 ATGACGGTCATCGCGACCGT CGG ||||||||[||||||||||||]PAM NA Intergenic NA NA

T32842 TCAAGATCGTGACGACCCTGTGG 1000
Oligo fwd CACCTCAAGATCGTGACGACCCTG
Oligo rev AAACCAGGGTCGTCACGATCTTGA
Chromosome start end strand MM target_seq PAM alignment distance position gene name gene id
GU305914.1 22196 22218 - 0 TCAAGATCGTGACGACCCTG TGG ||||||||[||||||||||||]PAM NA Intergenic NA NA

T32843 CAAGATCGTGACGACCCTGTGGG 1000
Oligo fwd CACCCAAGATCGTGACGACCCTGT
Oligo rev AAACACAGGGTCGTCACGATCTTG
Chromosome start end strand MM target_seq PAM alignment distance position gene name gene id
GU305914.1 22195 22217 - 0 CAAGATCGTGACGACCCTGT GGG ||||||||[||||||||||||]PAM NA Intergenic NA NA

T32844 TGACGACCCTGTGGGAATCGCGG 1000
Oligo fwd CACCTGACGACCCTGTGGGAATCG
Oligo rev AAACCGATTCCCACAGGGTCGTCA
Chromosome start end strand MM target_seq PAM alignment distance position gene name gene id
GU305914.1 22187 22209 - 0 TGACGACCCTGTGGGAATCG CGG ||||||||[||||||||||||]PAM NA Intergenic NA NA

T32845 ACCCTGTGGGAATCGCGGTACGG 1000
Oligo fwd CACCACCCTGTGGGAATCGCGGTA
Oligo rev AAACTACCGCGATTCCCACAGGGT
Chromosome start end strand MM target_seq PAM alignment distance position gene name gene id
GU305914.1 22182 22204 - 0 ACCCTGTGGGAATCGCGGTA CGG ||||||||[||||||||||||]PAM NA Intergenic NA NA

T32846 GGTACGGTTCTGATTCGAACCGG 1000
Oligo fwd CACCGGTACGGTTCTGATTCGAAC
Oligo rev AAACGTTCGAATCAGAACCGTACC
Chromosome start end strand MM target_seq PAM alignment distance position gene name gene id
GU305914.1 22166 22188 - 0 GGTACGGTTCTGATTCGAAC CGG ||||||||[||||||||||||]PAM NA Intergenic NA NA

T32847 ATTCGAACCGGCATTTTCAACGG 1000
Oligo fwd CACCATTCGAACCGGCATTTTCAA
Oligo rev AAACTTGAAAATGCCGGTTCGAAT
Chromosome start end strand MM target_seq PAM alignment distance position gene name gene id
GU305914.1 22154 22176 - 0 ATTCGAACCGGCATTTTCAA CGG ||||||||[||||||||||||]PAM NA Intergenic NA NA

T32848 GTCGACGCGCATATGATCTTCGG 1000
Oligo fwd CACCGTCGACGCGCATATGATCTT

| | | | | | | | | | | | | |
|---|---|---|---|---|---|---|---|---|---|---|---|---|
| Oligo rev | AAACAAGATCATATGCGCGTCGAC | | | | | | | | | | | |
| Chromosome | start | | end | strand | MM | target_seq | PAM | alignment | distance | position | gene name | gene id |
| GU305914.1 | | 22119 | 22141 | - | 0 | GTCGACGCGCATATGATCTT | CGG | \|\|\|\|\|\|\|\|[\|\|\|\|\|\|\|\|\|\|\|\|]PAM | NA | Intergenic | NA | NA |
| T32849 | CATATGATCTTCGGAGCGCGCGG | | 1000 | | | | | | | | | |
| Oligo fwd | CACCCATATGATCTTCGGAGCGCG | | | | | | | | | | | |
| Oligo rev | AAACCGCGCTCCGAAGATCATATG | | | | | | | | | | | |
| Chromosome | start | | end | strand | MM | target_seq | PAM | alignment | distance | position | gene name | gene id |
| GU305914.1 | | 22110 | 22132 | - | 0 | CATATGATCTTCGGAGCGCG | CGG | \|\|\|\|\|\|\|\|[\|\|\|\|\|\|\|\|\|\|\|\|]PAM | NA | Intergenic | NA | NA |
| T32850 | ATATGATCTTCGGAGCGCGCGGG | | 1000 | | | | | | | | | |
| Oligo fwd | CACCATATGATCTTCGGAGCGCGC | | | | | | | | | | | |
| Oligo rev | AAACGCGCGCTCCGAAGATCATAT | | | | | | | | | | | |
| Chromosome | start | | end | strand | MM | target_seq | PAM | alignment | distance | position | gene name | gene id |
| GU305914.1 | | 22109 | 22131 | - | 0 | ATATGATCTTCGGAGCGCGC | GGG | \|\|\|\|\|\|\|\|[\|\|\|\|\|\|\|\|\|\|\|\|]PAM | NA | Intergenic | NA | NA |
| T32851 | GATCTTCGGAGCGCGCGGGTTGG | | 1000 | | | | | | | | | |
| Oligo fwd | CACCGATCTTCGGAGCGCGCGGGT | | | | | | | | | | | |
| Oligo rev | AAACACCCGCGCGCTCCGAAGATC | | | | | | | | | | | |
| Chromosome | start | | end | strand | MM | target_seq | PAM | alignment | distance | position | gene name | gene id |
| GU305914.1 | | 22105 | 22127 | - | 0 | GATCTTCGGAGCGCGCGGGT | TGG | \|\|\|\|\|\|\|\|[\|\|\|\|\|\|\|\|\|\|\|\|]PAM | NA | Intergenic | NA | NA |
| T32852 | ATCTTCGGAGCGCGCGGGTTGGG | | 1000 | | | | | | | | | |
| Oligo fwd | CACCATCTTCGGAGCGCGCGGGTT | | | | | | | | | | | |
| Oligo rev | AAACAACCCGCGCGCTCCGAAGAT | | | | | | | | | | | |
| Chromosome | start | | end | strand | MM | target_seq | PAM | alignment | distance | position | gene name | gene id |
| GU305914.1 | | 22104 | 22126 | - | 0 | ATCTTCGGAGCGCGCGGGTT | GGG | \|\|\|\|\|\|\|\|[\|\|\|\|\|\|\|\|\|\|\|\|]PAM | NA | Intergenic | NA | NA |
| T32853 | GTTGGGCCGCCGCGACCGACCGG | | 1000 | | | | | | | | | |
| Oligo fwd | CACCGTTGGGCCGCCGCGACCGAC | | | | | | | | | | | |
| Oligo rev | AAACGTCGGTCGCGGCGGCCCAAC | | | | | | | | | | | |
| Chromosome | start | | end | strand | MM | target_seq | PAM | alignment | distance | position | gene name | gene id |
| GU305914.1 | | 22087 | 22109 | - | 0 | GTTGGGCCGCCGCGACCGAC | CGG | \|\|\|\|\|\|\|\|[\|\|\|\|\|\|\|\|\|\|\|\|]PAM | NA | Intergenic | NA | NA |
| T32854 | GGGCCGCCGCGACCGACCGGAGG | | 1000 | | | | | | | | | |
| Oligo fwd | CACCGGGCCGCCGCGACCGACCGG | | | | | | | | | | | |
| Oligo rev | AAACCCGGTCGGTCGCGGCGGCCC | | | | | | | | | | | |
| Chromosome | start | | end | strand | MM | target_seq | PAM | alignment | distance | position | gene name | gene id |
| GU305914.1 | | 22084 | 22106 | - | 0 | GGGCCGCCGCGACCGACCGG | AGG | \|\|\|\|\|\|\|\|[\|\|\|\|\|\|\|\|\|\|\|\|]PAM | NA | Intergenic | NA | NA |
| T32855 | GGCCGCCGCGACCGACCGGAGGG | | 1000 | | | | | | | | | |
| Oligo fwd | CACCGGCCGCCGCGACCGACCGGA | | | | | | | | | | | |
| Oligo rev | AAACTCCGGTCGGTCGCGGCGGCC | | | | | | | | | | | |
| Chromosome | start | | end | strand | MM | target_seq | PAM | alignment | distance | position | gene name | gene id |
| GU305914.1 | | 22083 | 22105 | - | 0 | GGCCGCCGCGACCGACCGGA | GGG | \|\|\|\|\|\|\|\|[\|\|\|\|\|\|\|\|\|\|\|\|]PAM | NA | Intergenic | NA | NA |
| T32856 | GACCGACCGGAGGGATACCGCGG | | 1000 | | | | | | | | | |
| Oligo fwd | CACCGACCGACCGGAGGGATACCG | | | | | | | | | | | |
| Oligo rev | AAACCGGTATCCCTCCGGTCGGTC | | | | | | | | | | | |
| Chromosome | start | | end | strand | MM | target_seq | PAM | alignment | distance | position | gene name | gene id |
| GU305914.1 | | 22074 | 22096 | - | 0 | GACCGACCGGAGGGATACCG | CGG | \|\|\|\|\|\|\|\|[\|\|\|\|\|\|\|\|\|\|\|\|]PAM | NA | Intergenic | NA | NA |
| T32857 | GAGGGATACCGCGGCCTGCCCGG | | 1000 | | | | | | | | | |
| Oligo fwd | CACCGAGGGATACCGCGGCCTGCC | | | | | | | | | | | |
| Oligo rev | AAACGGCAGGCCGCGGTATCCCTC | | | | | | | | | | | |
| Chromosome | start | | end | strand | MM | target_seq | PAM | alignment | distance | position | gene name | gene id |
| GU305914.1 | | 22065 | 22087 | - | 0 | GAGGGATACCGCGGCCTGCC | CGG | \|\|\|\|\|\|\|\|[\|\|\|\|\|\|\|\|\|\|\|\|]PAM | NA | Intergenic | NA | NA |
| T32858 | AGGGATACCGCGGCCTGCCCGGG | | 1000 | | | | | | | | | |
| Oligo fwd | CACCAGGGATACCGCGGCCTGCCC | | | | | | | | | | | |
| Oligo rev | AAACGGGCAGGCCGCGGTATCCCT | | | | | | | | | | | |
| Chromosome | start | | end | strand | MM | target_seq | PAM | alignment | distance | position | gene name | gene id |
| GU305914.1 | | 22064 | 22086 | - | 0 | AGGGATACCGCGGCCTGCCC | GGG | \|\|\|\|\|\|\|\|[\|\|\|\|\|\|\|\|\|\|\|\|]PAM | NA | Intergenic | NA | NA |
| T32859 | GGGATACCGCGGCCTGCCCGGGG | | 1000 | | | | | | | | | |
| Oligo fwd | CACCGGGATACCGCGGCCTGCCCG | | | | | | | | | | | |
| Oligo rev | AAACCGGGCAGGCCGCGGTATCCC | | | | | | | | | | | |
| Chromosome | start | | end | strand | MM | target_seq | PAM | alignment | distance | position | gene name | gene id |
| GU305914.1 | | 22063 | 22085 | - | 0 | GGGATACCGCGGCCTGCCCG | GGG | \|\|\|\|\|\|\|\|[\|\|\|\|\|\|\|\|\|\|\|\|]PAM | NA | Intergenic | NA | NA |
| T32860 | CCGCGGCCTGCCCGGGGCCGTGG | | 1000 | | | | | | | | | |
| Oligo fwd | CACCCCGCGGCCTGCCCGGGGCCG | | | | | | | | | | | |
| Oligo rev | AAACCGGCCCCGGGCAGGCCGCGG | | | | | | | | | | | |
| Chromosome | start | | end | strand | MM | target_seq | PAM | alignment | distance | position | gene name | gene id |
| GU305914.1 | | 22057 | 22079 | - | 0 | CCGCGGCCTGCCCGGGGCCG | TGG | \|\|\|\|\|\|\|\|[\|\|\|\|\|\|\|\|\|\|\|\|]PAM | NA | Intergenic | NA | NA |
| T32861 | GGCCTGCCCGGGGCCGTGGCCGG | | 1000 | | | | | | | | | |
| Oligo fwd | CACCGGCCTGCCCGGGGCCGTGGC | | | | | | | | | | | |
| Oligo rev | AAACGCCACGGCCCCGGGCAGGCC | | | | | | | | | | | |
| Chromosome | start | | end | strand | MM | target_seq | PAM | alignment | distance | position | gene name | gene id |
| GU305914.1 | | 22053 | 22075 | - | 0 | GGCCTGCCCGGGGCCGTGGC | CGG | \|\|\|\|\|\|\|\|[\|\|\|\|\|\|\|\|\|\|\|\|]PAM | NA | Intergenic | NA | NA |
| T32862 | GCCTGCCCGGGGCCGTGGCCGGG | | 1000 | | | | | | | | | |
| Oligo fwd | CACCGCCTGCCCGGGGCCGTGGCC | | | | | | | | | | | |
| Oligo rev | AAACGGCCACGGCCCCGGGCAGGC | | | | | | | | | | | |
| Chromosome | start | | end | strand | MM | target_seq | PAM | alignment | distance | position | gene name | gene id |
| GU305914.1 | | 22052 | 22074 | - | 0 | GCCTGCCCGGGGCCGTGGCC | GGG | \|\|\|\|\|\|\|\|[\|\|\|\|\|\|\|\|\|\|\|\|]PAM | NA | Intergenic | NA | NA |
| T32863 | GGCCGTGGCCGGGTTAGTGCCGG | | 1000 | | | | | | | | | |

Oligo fwd CACCGGCCGTGGCCGGGTTAGTGC
Oligo rev AAACGCACTAACCCGGCCACGGCC
Chromosome start end strand MM target_seq PAM alignment distance position gene name gene id
GU305914.1 22042 22064 - 0 GGCCGTGGCCGGGTTAGTGC CGG ||||||||[||||||||||||]PAM NA Intergenic NA NA

T32864 CGTGGCCGGGTTAGTGCCGGCGG 1000
Oligo fwd CACCCGTGGCCGGGTTAGTGCCGG
Oligo rev AAACCCGGCACTAACCCGGCCACG
Chromosome start end strand MM target_seq PAM alignment distance position gene name gene id
GU305914.1 22039 22061 - 0 CGTGGCCGGGTTAGTGCCGG CGG ||||||||[||||||||||||]PAM NA Intergenic NA NA

T32865 GTGCCGGCGGCGATGACCGTCGG 1000
Oligo fwd CACCGTGCCGGCGGCGATGACCGT
Oligo rev AAACACGGTCATCGCCGCCGGCAC
Chromosome start end strand MM target_seq PAM alignment distance position gene name gene id
GU305914.1 22026 22048 - 0 GTGCCGGCGGCGATGACCGT CGG ||||||||[||||||||||||]PAM NA Intergenic NA NA
GU305914.1 139830 139852 + 4 GTGGCCGCGGCGATCGCCGT TGG |||-|-||[||||||--||||]PAM NA Intergenic NA NA

T32866 GGCGGCGATGACCGTCGGCGTGG 1000
Oligo fwd CACCGGCGGCGATGACCGTCGGCG
Oligo rev AAACCGCCGACGGTCATCGCCGCC
Chromosome start end strand MM target_seq PAM alignment distance position gene name gene id
GU305914.1 22021 22043 - 0 GGCGGCGATGACCGTCGGCG TGG ||||||||[||||||||||||]PAM NA Intergenic NA NA

T32867 GATGACCGTCGGCGTGGCCGTGG 1000
Oligo fwd CACCGATGACCGTCGGCGTGGCCG
Oligo rev AAACCGGCCACGCCGACGGTCATC
Chromosome start end strand MM target_seq PAM alignment distance position gene name gene id
GU305914.1 22015 22037 - 0 GATGACCGTCGGCGTGGCCG TGG ||||||||[||||||||||||]PAM NA Intergenic NA NA

T32868 GACCGTCGGCGTGGCCGTGGCGG 1000
Oligo fwd CACCGACCGTCGGCGTGGCCGTGG
Oligo rev AAACCCACGGCCACGCCGACGGTC
Chromosome start end strand MM target_seq PAM alignment distance position gene name gene id
GU305914.1 22012 22034 - 0 GACCGTCGGCGTGGCCGTGG CGG ||||||||[||||||||||||]PAM NA Intergenic NA NA

T32869 CCTCACCGTCCTGCCGACGACGG 1000
Oligo fwd CACCCCTCACCGTCCTGCCGACGA
Oligo rev AAACTCGTCGGCAGGACGGTGAGG
Chromosome start end strand MM target_seq PAM alignment distance position gene name gene id
GU305914.1 21988 22010 - 0 CCTCACCGTCCTGCCGACGA CGG ||||||||[||||||||||||]PAM NA Intergenic NA NA
GU305914.1 145031 145053 - 4 CTTCACCTTCGTGCTGACGA TGG |-|||||-[||-|||-|||||]PAM NA Intergenic NA NA

T32870 CGTCCTGCCGACGACGGCCCTGG 1000
Oligo fwd CACCCGTCCTGCCGACGACGGCCC
Oligo rev AAACGGGCCGTCGTCGGCAGGACG
Chromosome start end strand MM target_seq PAM alignment distance position gene name gene id
GU305914.1 21982 22004 - 0 CGTCCTGCCGACGACGGCCC TGG ||||||||[||||||||||||]PAM NA Intergenic NA NA

T32871 CAGCATCAGCATCAGCACAGCGG 1000
Oligo fwd CACCCAGCATCAGCATCAGCACAG
Oligo rev AAACCTGTGCTGATGCTGATGCTG
Chromosome start end strand MM target_seq PAM alignment distance position gene name gene id
GU305914.1 21945 21967 - 0 CAGCATCAGCATCAGCACAG CGG ||||||||[||||||||||||]PAM NA Intergenic NA NA

T32872 ATTCGCACGAAACCTCCAAGTGG 1000
Oligo fwd CACCATTCGCACGAAACCTCCAAG
Oligo rev AAACCTTGGAGGTTTCGTGCGAAT
Chromosome start end strand MM target_seq PAM alignment distance position gene name gene id
GU305914.1 21920 21942 - 0 ATTCGCACGAAACCTCCAAG TGG ||||||||[||||||||||||]PAM NA Intergenic NA NA

T32873 CGCACGAAACCTCCAAGTGGTGG 1000
Oligo fwd CACCCGCACGAAACCTCCAAGTGG
Oligo rev AAACCCACTTGGAGGTTTCGTGCG
Chromosome start end strand MM target_seq PAM alignment distance position gene name gene id
GU305914.1 21917 21939 - 0 CGCACGAAACCTCCAAGTGG TGG ||||||||[||||||||||||]PAM NA Intergenic NA NA

T32874 GAAACCTCCAAGTGGTGGAGTGG 1000
Oligo fwd CACCGAAACCTCCAAGTGGTGGAG
Oligo rev AAACCTCCACCACTTGGAGGTTTC
Chromosome start end strand MM target_seq PAM alignment distance position gene name gene id
GU305914.1 21912 21934 - 0 GAAACCTCCAAGTGGTGGAG TGG ||||||||[||||||||||||]PAM NA Intergenic NA NA

T32875 AAACCTCCAAGTGGTGGAGTGGG 1000
Oligo fwd CACCAAACCTCCAAGTGGTGGAGT
Oligo rev AAACACTCCACCACTTGGAGGTTT
Chromosome start end strand MM target_seq PAM alignment distance position gene name gene id
GU305914.1 21911 21933 - 0 AAACCTCCAAGTGGTGGAGT GGG ||||||||[||||||||||||]PAM NA Intergenic NA NA

T32876 AACCTCCAAGTGGTGGAGTGGGG 1000
Oligo fwd CACCAACCTCCAAGTGGTGGAGTG
Oligo rev AAACCACTCCACCACTTGGAGGTT
Chromosome start end strand MM target_seq PAM alignment distance position gene name gene id
GU305914.1 21910 21932 - 0 AACCTCCAAGTGGTGGAGTG GGG ||||||||[||||||||||||]PAM NA Intergenic NA NA

T32877 ACCTCCAAGTGGTGGAGTGGGGG 1000
Oligo fwd CACCACCTCCAAGTGGTGGAGTGG
Oligo rev AAACCCACTCCACCACTTGGAGGT
Chromosome start end strand MM target_seq PAM alignment distance position gene name gene id

| | | | | | | | | | | | |
|---|---|---|---|---|---|---|---|---|---|---|---|
| GU305914.1 | 21909 | 21931 | - | 0 | ACCTCCAAGTGGTGGAGTGG | GGG | \|\|\|\|\|\|\|\|[\|\|\|\|\|\|\|\|\|\|\|\|]PAM | NA | Intergenic | NA | NA |
| T32878 | CAAGTGGTGGAGTGGGGGAATGG | 1000 | | | | | | | | | |
| Oligo fwd | CACCCAAGTGGTGGAGTGGGGGAA | | | | | | | | | | |
| Oligo rev | AAACTTCCCCCACTCCACCACTTG | | | | | | | | | | |
| Chromosome | start | end | strand | MM | target_seq | PAM | alignment | distance | position | gene name | gene id |
| GU305914.1 | 21904 | 21926 | - | 0 | CAAGTGGTGGAGTGGGGGAA | TGG | \|\|\|\|\|\|\|\|[\|\|\|\|\|\|\|\|\|\|\|\|]PAM | NA | Intergenic | NA | NA |
| T32879 | AAGTGGTGGAGTGGGGGAATGGG | 1000 | | | | | | | | | |
| Oligo fwd | CACCAAGTGGTGGAGTGGGGGAAT | | | | | | | | | | |
| Oligo rev | AAACATTCCCCCACTCCACCACTT | | | | | | | | | | |
| Chromosome | start | end | strand | MM | target_seq | PAM | alignment | distance | position | gene name | gene id |
| GU305914.1 | 21903 | 21925 | - | 0 | AAGTGGTGGAGTGGGGGAAT | GGG | \|\|\|\|\|\|\|\|[\|\|\|\|\|\|\|\|\|\|\|\|]PAM | NA | Intergenic | NA | NA |
| T32880 | CTTCCGAGTTCGATAAGCCGCGG | 1000 | | | | | | | | | |
| Oligo fwd | CACCCTTCCGAGTTCGATAAGCCG | | | | | | | | | | |
| Oligo rev | AAACCGGCTTATCGAACTCGGAAG | | | | | | | | | | |
| Chromosome | start | end | strand | MM | target_seq | PAM | alignment | distance | position | gene name | gene id |
| GU305914.1 | 21872 | 21894 | - | 0 | CTTCCGAGTTCGATAAGCCG | CGG | \|\|\|\|\|\|\|\|[\|\|\|\|\|\|\|\|\|\|\|\|]PAM | NA | Intergenic | NA | NA |
| T32881 | CGATAAGCCGCGGTATCCCCCGG | 1000 | | | | | | | | | |
| Oligo fwd | CACCCGATAAGCCGCGGTATCCCC | | | | | | | | | | |
| Oligo rev | AAACGGGGATACCGCGGCTTATCG | | | | | | | | | | |
| Chromosome | start | end | strand | MM | target_seq | PAM | alignment | distance | position | gene name | gene id |
| GU305914.1 | 21862 | 21884 | - | 0 | CGATAAGCCGCGGTATCCCC | CGG | \|\|\|\|\|\|\|\|[\|\|\|\|\|\|\|\|\|\|\|\|]PAM | NA | Intergenic | NA | NA |
| T32882 | GATAAGCCGCGGTATCCCCCGGG | 1000 | | | | | | | | | |
| Oligo fwd | CACCGATAAGCCGCGGTATCCCCC | | | | | | | | | | |
| Oligo rev | AAACGGGGGATACCGCGGCTTATC | | | | | | | | | | |
| Chromosome | start | end | strand | MM | target_seq | PAM | alignment | distance | position | gene name | gene id |
| GU305914.1 | 21861 | 21883 | - | 0 | GATAAGCCGCGGTATCCCCC | GGG | \|\|\|\|\|\|\|\|[\|\|\|\|\|\|\|\|\|\|\|\|]PAM | NA | Intergenic | NA | NA |
| GU305914.1 | 22068 | 22090 | + | 4 | GGCAGGCCGCGGTATCCCTC | CGG | \|--\|-\|\|\|[\|\|\|\|\|\|\|\|\|\|-\|]PAM | NA | Intergenic | NA | NA |
| T32883 | AAGCCGCGGTATCCCCCGGGCGG | 1000 | | | | | | | | | |
| Oligo fwd | CACCAAGCCGCGGTATCCCCCGGG | | | | | | | | | | |
| Oligo rev | AAACCCCGGGGGATACCGCGGCTT | | | | | | | | | | |
| Chromosome | start | end | strand | MM | target_seq | PAM | alignment | distance | position | gene name | gene id |
| GU305914.1 | 21858 | 21880 | - | 0 | AAGCCGCGGTATCCCCCGGG | CGG | \|\|\|\|\|\|\|\|[\|\|\|\|\|\|\|\|\|\|\|\|]PAM | NA | Intergenic | NA | NA |
| T32884 | AGCCGCGGTATCCCCCGGGCGGG | 1000 | | | | | | | | | |
| Oligo fwd | CACCAGCCGCGGTATCCCCCGGGC | | | | | | | | | | |
| Oligo rev | AAACGCCCGGGGGATACCGCGGCT | | | | | | | | | | |
| Chromosome | start | end | strand | MM | target_seq | PAM | alignment | distance | position | gene name | gene id |
| GU305914.1 | 21857 | 21879 | - | 0 | AGCCGCGGTATCCCCCGGGC | GGG | \|\|\|\|\|\|\|\|[\|\|\|\|\|\|\|\|\|\|\|\|]PAM | NA | Intergenic | NA | NA |
| T32885 | CGGTATCCCCCGGGCGGGCGAGG | 1000 | | | | | | | | | |
| Oligo fwd | CACCCGGTATCCCCCGGGCGGGCG | | | | | | | | | | |
| Oligo rev | AAACCGCCCGCCCGGGGGATACCG | | | | | | | | | | |
| Chromosome | start | end | strand | MM | target_seq | PAM | alignment | distance | position | gene name | gene id |
| GU305914.1 | 21852 | 21874 | - | 0 | CGGTATCCCCCGGGCGGGCG | AGG | \|\|\|\|\|\|\|\|[\|\|\|\|\|\|\|\|\|\|\|\|]PAM | NA | Intergenic | NA | NA |
| T32886 | CGGGCGGGCGAGGATTCAAGAGG | 1000 | | | | | | | | | |
| Oligo fwd | CACCCGGGCGGGCGAGGATTCAAG | | | | | | | | | | |
| Oligo rev | AAACCTTGAATCCTCGCCCGCCCG | | | | | | | | | | |
| Chromosome | start | end | strand | MM | target_seq | PAM | alignment | distance | position | gene name | gene id |
| GU305914.1 | 21842 | 21864 | - | 0 | CGGGCGGGCGAGGATTCAAG | AGG | \|\|\|\|\|\|\|\|[\|\|\|\|\|\|\|\|\|\|\|\|]PAM | NA | Intergenic | NA | NA |
| T32887 | GAGGTGCACGTCGACCACGCCGG | 1000 | | | | | | | | | |
| Oligo fwd | CACCGAGGTGCACGTCGACCACGC | | | | | | | | | | |
| Oligo rev | AAACGCGTGGTCGACGTGCACCTC | | | | | | | | | | |
| Chromosome | start | end | strand | MM | target_seq | PAM | alignment | distance | position | gene name | gene id |
| GU305914.1 | 21823 | 21845 | - | 0 | GAGGTGCACGTCGACCACGC | CGG | \|\|\|\|\|\|\|\|[\|\|\|\|\|\|\|\|\|\|\|\|]PAM | NA | Intergenic | NA | NA |
| T32888 | CGCTTCTTTCTACGACGCCGAGG | 1000 | | | | | | | | | |
| Oligo fwd | CACCCGCTTCTTTCTACGACGCCG | | | | | | | | | | |
| Oligo rev | AAACCGGCGTCGTAGAAAGAAGCG | | | | | | | | | | |
| Chromosome | start | end | strand | MM | target_seq | PAM | alignment | distance | position | gene name | gene id |
| GU305914.1 | 21790 | 21812 | - | 0 | CGCTTCTTTCTACGACGCCG | AGG | \|\|\|\|\|\|\|\|[\|\|\|\|\|\|\|\|\|\|\|\|]PAM | NA | Intergenic | NA | NA |
| T32889 | TTCTTTCTACGACGCCGAGGAGG | 1000 | | | | | | | | | |
| Oligo fwd | CACCTTCTTTCTACGACGCCGAGG | | | | | | | | | | |
| Oligo rev | AAACCCTCGGCGTCGTAGAAAGAA | | | | | | | | | | |
| Chromosome | start | end | strand | MM | target_seq | PAM | alignment | distance | position | gene name | gene id |
| GU305914.1 | 21787 | 21809 | - | 0 | TTCTTTCTACGACGCCGAGG | AGG | \|\|\|\|\|\|\|\|[\|\|\|\|\|\|\|\|\|\|\|\|]PAM | NA | Intergenic | NA | NA |
| T32890 | CTGACGACGTACCAAGATAGCGG | 1000 | | | | | | | | | |
| Oligo fwd | CACCCTGACGACGTACCAAGATAG | | | | | | | | | | |
| Oligo rev | AAACCTATCTTGGTACGTCGTCAG | | | | | | | | | | |
| Chromosome | start | end | strand | MM | target_seq | PAM | alignment | distance | position | gene name | gene id |
| GU305914.1 | 21762 | 21784 | - | 0 | CTGACGACGTACCAAGATAG | CGG | \|\|\|\|\|\|\|\|[\|\|\|\|\|\|\|\|\|\|\|\|]PAM | NA | Intergenic | NA | NA |
| T32891 | CTTCCTGACCTTCGATTGTCCGG | 1000 | | | | | | | | | |
| Oligo fwd | CACCCTTCCTGACCTTCGATTGTC | | | | | | | | | | |
| Oligo rev | AAACGACAATCGAAGGTCAGGAAG | | | | | | | | | | |
| Chromosome | start | end | strand | MM | target_seq | PAM | alignment | distance | position | gene name | gene id |
| GU305914.1 | 21730 | 21752 | - | 0 | CTTCCTGACCTTCGATTGTC | CGG | \|\|\|\|\|\|\|\|[\|\|\|\|\|\|\|\|\|\|\|\|]PAM | NA | Intergenic | NA | NA |
| T32892 | CTGACCTTCGATTGTCCGGACGG | 1000 | | | | | | | | | |
| Oligo fwd | CACCCTGACCTTCGATTGTCCGGA | | | | | | | | | | |

| | | | | | | | | | | | | |
|---|---|---|---|---|---|---|---|---|---|---|---|---|
| Oligo rev | AAACTCCGGACAATCGAAGGTCAG | | | | | | | | | | | |
| Chromosome | start | | end | strand | MM | target_seq | PAM | alignment | distance | position | gene name | gene id |
| GU305914.1 | | 21726 | 21748 | - | 0 | CTGACCTTCGATTGTCCGGA | CGG | \|\|\|\|\|\|\|\|[\|\|\|\|\|\|\|\|\|\|\|\|]PAM | NA | Intergenic | NA | NA |
| T32893 | ACCTGCCGTTTCAAGACCCGCGG | | 1000 | | | | | | | | | |
| Oligo fwd | CACCACCTGCCGTTTCAAGACCCG | | | | | | | | | | | |
| Oligo rev | AAACCGGGTCTTGAAACGGCAGGT | | | | | | | | | | | |
| Chromosome | start | | end | strand | MM | target_seq | PAM | alignment | distance | position | gene name | gene id |
| GU305914.1 | | 21690 | 21712 | - | 0 | ACCTGCCGTTTCAAGACCCG | CGG | \|\|\|\|\|\|\|\|[\|\|\|\|\|\|\|\|\|\|\|\|]PAM | NA | Intergenic | NA | NA |
| T32894 | CCGTTTCAAGACCCGCGGACAGG | | 1000 | | | | | | | | | |
| Oligo fwd | CACCCCGTTTCAAGACCCGCGGAC | | | | | | | | | | | |
| Oligo rev | AAACGTCCGCGGGTCTTGAAACGG | | | | | | | | | | | |
| Chromosome | start | | end | strand | MM | target_seq | PAM | alignment | distance | position | gene name | gene id |
| GU305914.1 | | 21685 | 21707 | - | 0 | CCGTTTCAAGACCCGCGGAC | AGG | \|\|\|\|\|\|\|\|[\|\|\|\|\|\|\|\|\|\|\|\|]PAM | NA | Intergenic | NA | NA |
| T32895 | CAAGACCCGCGGACAGGTCTTGG | | 1000 | | | | | | | | | |
| Oligo fwd | CACCCAAGACCCGCGGACAGGTCT | | | | | | | | | | | |
| Oligo rev | AAACAGACCTGTCCGCGGGTCTTG | | | | | | | | | | | |
| Chromosome | start | | end | strand | MM | target_seq | PAM | alignment | distance | position | gene name | gene id |
| GU305914.1 | | 21679 | 21701 | - | 0 | CAAGACCCGCGGACAGGTCT | TGG | \|\|\|\|\|\|\|\|[\|\|\|\|\|\|\|\|\|\|\|\|]PAM | NA | Intergenic | NA | NA |
| T32896 | CAGGTCTTGGACGTGATTATCGG | | 1000 | | | | | | | | | |
| Oligo fwd | CACCCAGGTCTTGGACGTGATTAT | | | | | | | | | | | |
| Oligo rev | AAACATAATCACGTCCAAGACCTG | | | | | | | | | | | |
| Chromosome | start | | end | strand | MM | target_seq | PAM | alignment | distance | position | gene name | gene id |
| GU305914.1 | | 21666 | 21688 | - | 0 | CAGGTCTTGGACGTGATTAT | CGG | \|\|\|\|\|\|\|\|[\|\|\|\|\|\|\|\|\|\|\|\|]PAM | NA | Intergenic | NA | NA |
| T32897 | TCTTGGACGTGATTATCGGCTGG | | 1000 | | | | | | | | | |
| Oligo fwd | CACCTCTTGGACGTGATTATCGGC | | | | | | | | | | | |
| Oligo rev | AAACGCCGATAATCACGTCCAAGA | | | | | | | | | | | |
| Chromosome | start | | end | strand | MM | target_seq | PAM | alignment | distance | position | gene name | gene id |
| GU305914.1 | | 21662 | 21684 | - | 0 | TCTTGGACGTGATTATCGGC | TGG | \|\|\|\|\|\|\|\|[\|\|\|\|\|\|\|\|\|\|\|\|]PAM | NA | Intergenic | NA | NA |
| T32898 | CTTGGACGTGATTATCGGCTGGG | | 1000 | | | | | | | | | |
| Oligo fwd | CACCCTTGGACGTGATTATCGGCT | | | | | | | | | | | |
| Oligo rev | AAACAGCCGATAATCACGTCCAAG | | | | | | | | | | | |
| Chromosome | start | | end | strand | MM | target_seq | PAM | alignment | distance | position | gene name | gene id |
| GU305914.1 | | 21661 | 21683 | - | 0 | CTTGGACGTGATTATCGGCT | GGG | \|\|\|\|\|\|\|\|[\|\|\|\|\|\|\|\|\|\|\|\|]PAM | NA | Intergenic | NA | NA |
| T32899 | ATCGGCTGGGCCTCCCAGAGCGG | | 1000 | | | | | | | | | |
| Oligo fwd | CACCATCGGCTGGGCCTCCCAGAG | | | | | | | | | | | |
| Oligo rev | AAACCTCTGGGAGGCCCAGCCGAT | | | | | | | | | | | |
| Chromosome | start | | end | strand | MM | target_seq | PAM | alignment | distance | position | gene name | gene id |
| GU305914.1 | | 21648 | 21670 | - | 0 | ATCGGCTGGGCCTCCCAGAG | CGG | \|\|\|\|\|\|\|\|[\|\|\|\|\|\|\|\|\|\|\|\|]PAM | NA | Intergenic | NA | NA |
| T32900 | CCAGAGCGGCTACGAGTTGCAGG | | 1000 | | | | | | | | | |
| Oligo fwd | CACCCCAGAGCGGCTACGAGTTGC | | | | | | | | | | | |
| Oligo rev | AAACGCAACTCGTAGCCGCTCTGG | | | | | | | | | | | |
| Chromosome | start | | end | strand | MM | target_seq | PAM | alignment | distance | position | gene name | gene id |
| GU305914.1 | | 21634 | 21656 | - | 0 | CCAGAGCGGCTACGAGTTGC | AGG | \|\|\|\|\|\|\|\|[\|\|\|\|\|\|\|\|\|\|\|\|]PAM | NA | Intergenic | NA | NA |
| T32901 | ACGAGTTGCAGGACCTGCACAGG | | 1000 | | | | | | | | | |
| Oligo fwd | CACCACGAGTTGCAGGACCTGCAC | | | | | | | | | | | |
| Oligo rev | AAACGTGCAGGTCCTGCAACTCGT | | | | | | | | | | | |
| Chromosome | start | | end | strand | MM | target_seq | PAM | alignment | distance | position | gene name | gene id |
| GU305914.1 | | 21623 | 21645 | - | 0 | ACGAGTTGCAGGACCTGCAC | AGG | \|\|\|\|\|\|\|\|[\|\|\|\|\|\|\|\|\|\|\|\|]PAM | NA | Intergenic | NA | NA |
| T32902 | CGAGTTGCAGGACCTGCACAGGG | | 1000 | | | | | | | | | |
| Oligo fwd | CACCCGAGTTGCAGGACCTGCACA | | | | | | | | | | | |
| Oligo rev | AAACTGTGCAGGTCCTGCAACTCG | | | | | | | | | | | |
| Chromosome | start | | end | strand | MM | target_seq | PAM | alignment | distance | position | gene name | gene id |
| GU305914.1 | | 21622 | 21644 | - | 0 | CGAGTTGCAGGACCTGCACA | GGG | \|\|\|\|\|\|\|\|[\|\|\|\|\|\|\|\|\|\|\|\|]PAM | NA | Intergenic | NA | NA |
| T32903 | TGCAGGACCTGCACAGGGAATGG | | 1000 | | | | | | | | | |
| Oligo fwd | CACCTGCAGGACCTGCACAGGGAA | | | | | | | | | | | |
| Oligo rev | AAACTTCCCTGTGCAGGTCCTGCA | | | | | | | | | | | |
| Chromosome | start | | end | strand | MM | target_seq | PAM | alignment | distance | position | gene name | gene id |
| GU305914.1 | | 21617 | 21639 | - | 0 | TGCAGGACCTGCACAGGGAA | TGG | \|\|\|\|\|\|\|\|[\|\|\|\|\|\|\|\|\|\|\|\|]PAM | NA | Intergenic | NA | NA |
| T32904 | ACCAGCGTTGCAACATACGCAGG | | 1000 | | | | | | | | | |
| Oligo fwd | CACCACCAGCGTTGCAACATACGC | | | | | | | | | | | |
| Oligo rev | AAACGCGTATGTTGCAACGCTGGT | | | | | | | | | | | |
| Chromosome | start | | end | strand | MM | target_seq | PAM | alignment | distance | position | gene name | gene id |
| GU305914.1 | | 21590 | 21612 | - | 0 | ACCAGCGTTGCAACATACGC | AGG | \|\|\|\|\|\|\|\|[\|\|\|\|\|\|\|\|\|\|\|\|]PAM | NA | Intergenic | NA | NA |
| T32905 | CCAGCGTTGCAACATACGCAGGG | | 1000 | | | | | | | | | |
| Oligo fwd | CACCCCAGCGTTGCAACATACGCA | | | | | | | | | | | |
| Oligo rev | AAACTGCGTATGTTGCAACGCTGG | | | | | | | | | | | |
| Chromosome | start | | end | strand | MM | target_seq | PAM | alignment | distance | position | gene name | gene id |
| GU305914.1 | | 21589 | 21611 | - | 0 | CCAGCGTTGCAACATACGCA | GGG | \|\|\|\|\|\|\|\|[\|\|\|\|\|\|\|\|\|\|\|\|]PAM | NA | Intergenic | NA | NA |
| T32906 | GAACACTACATCAACCCCTACGG | | 1000 | | | | | | | | | |
| Oligo fwd | CACCGAACACTACATCAACCCCTA | | | | | | | | | | | |
| Oligo rev | AAACTAGGGGTTGATGTAGTGTTC | | | | | | | | | | | |
| Chromosome | start | | end | strand | MM | target_seq | PAM | alignment | distance | position | gene name | gene id |
| GU305914.1 | | 21555 | 21577 | - | 0 | GAACACTACATCAACCCCTA | CGG | \|\|\|\|\|\|\|\|[\|\|\|\|\|\|\|\|\|\|\|\|]PAM | NA | Intergenic | NA | NA |
| T32907 | CCTACGGACAGCGCGACAAGAGG | | 1000 | | | | | | | | | |

<table>
<tr><td>Oligo fwd</td><td>CACCCCTACGGACAGCGCGACAAG</td><td></td><td></td><td></td><td></td><td></td><td></td><td></td><td></td><td></td><td></td></tr>
<tr><td>Oligo rev</td><td>AAACCTTGTCGCGCTGTCCGTAGG</td><td></td><td></td><td></td><td></td><td></td><td></td><td></td><td></td><td></td><td></td></tr>
<tr><td>Chromosome</td><td>start</td><td>end</td><td>strand</td><td>MM</td><td>target_seq</td><td>PAM</td><td>alignment</td><td>distance</td><td>position</td><td>gene name</td><td>gene id</td></tr>
<tr><td>GU305914.1</td><td>21539</td><td>21561</td><td>-</td><td>0</td><td>CCTACGGACAGCGCGACAAG</td><td>AGG</td><td>||||||||[||||||||||||]PAM</td><td>NA</td><td>Intergenic</td><td>NA</td><td>NA</td></tr>
<tr><td></td><td></td><td></td><td></td><td></td><td></td><td></td><td></td><td></td><td></td><td></td><td></td></tr>
<tr><td>T32908</td><td>CGCGACAAGAGGATCATGTGCGG</td><td>1000</td><td></td><td></td><td></td><td></td><td></td><td></td><td></td><td></td><td></td></tr>
<tr><td>Oligo fwd</td><td>CACCCGCGACAAGAGGATCATGTG</td><td></td><td></td><td></td><td></td><td></td><td></td><td></td><td></td><td></td><td></td></tr>
<tr><td>Oligo rev</td><td>AAACCACATGATCCTCTTGTCGCG</td><td></td><td></td><td></td><td></td><td></td><td></td><td></td><td></td><td></td><td></td></tr>
<tr><td>Chromosome</td><td>start</td><td>end</td><td>strand</td><td>MM</td><td>target_seq</td><td>PAM</td><td>alignment</td><td>distance</td><td>position</td><td>gene name</td><td>gene id</td></tr>
<tr><td>GU305914.1</td><td>21528</td><td>21550</td><td>-</td><td>0</td><td>CGCGACAAGAGGATCATGTG</td><td>CGG</td><td>||||||||[||||||||||||]PAM</td><td>NA</td><td>Intergenic</td><td>NA</td><td>NA</td></tr>
<tr><td></td><td></td><td></td><td></td><td></td><td></td><td></td><td></td><td></td><td></td><td></td><td></td></tr>
<tr><td>T32909</td><td>AGGATCATGTGCGGCAACAGCGG</td><td>1000</td><td></td><td></td><td></td><td></td><td></td><td></td><td></td><td></td><td></td></tr>
<tr><td>Oligo fwd</td><td>CACCAGGATCATGTGCGGCAACAG</td><td></td><td></td><td></td><td></td><td></td><td></td><td></td><td></td><td></td><td></td></tr>
<tr><td>Oligo rev</td><td>AAACCTGTTGCCGCACATGATCCT</td><td></td><td></td><td></td><td></td><td></td><td></td><td></td><td></td><td></td><td></td></tr>
<tr><td>Chromosome</td><td>start</td><td>end</td><td>strand</td><td>MM</td><td>target_seq</td><td>PAM</td><td>alignment</td><td>distance</td><td>position</td><td>gene name</td><td>gene id</td></tr>
<tr><td>GU305914.1</td><td>21519</td><td>21541</td><td>-</td><td>0</td><td>AGGATCATGTGCGGCAACAG</td><td>CGG</td><td>||||||||[||||||||||||]PAM</td><td>NA</td><td>Intergenic</td><td>NA</td><td>NA</td></tr>
<tr><td></td><td></td><td></td><td></td><td></td><td></td><td></td><td></td><td></td><td></td><td></td><td></td></tr>
<tr><td>T32910</td><td>GCAACAGCGGCACCTGCGTGTGG</td><td>1000</td><td></td><td></td><td></td><td></td><td></td><td></td><td></td><td></td><td></td></tr>
<tr><td>Oligo fwd</td><td>CACCGCAACAGCGGCACCTGCGTG</td><td></td><td></td><td></td><td></td><td></td><td></td><td></td><td></td><td></td><td></td></tr>
<tr><td>Oligo rev</td><td>AAACCACGCAGGTGCCGCTGTTGC</td><td></td><td></td><td></td><td></td><td></td><td></td><td></td><td></td><td></td><td></td></tr>
<tr><td>Chromosome</td><td>start</td><td>end</td><td>strand</td><td>MM</td><td>target_seq</td><td>PAM</td><td>alignment</td><td>distance</td><td>position</td><td>gene name</td><td>gene id</td></tr>
<tr><td>GU305914.1</td><td>21506</td><td>21528</td><td>-</td><td>0</td><td>GCAACAGCGGCACCTGCGTG</td><td>TGG</td><td>||||||||[||||||||||||]PAM</td><td>NA</td><td>Intergenic</td><td>NA</td><td>NA</td></tr>
<tr><td>GU305914.1</td><td>31558</td><td>31580</td><td>-</td><td>4</td><td>GGTACAGCGCGACCTGCGTG</td><td>AGG</td><td>|--|||||[|--|||||||||]PAM</td><td>NA</td><td>Intergenic</td><td>NA</td><td>NA</td></tr>
<tr><td></td><td></td><td></td><td></td><td></td><td></td><td></td><td></td><td></td><td></td><td></td><td></td></tr>
<tr><td>T32911</td><td>GGCACCTGCGTGTGGAACTTCGG</td><td>1000</td><td></td><td></td><td></td><td></td><td></td><td></td><td></td><td></td><td></td></tr>
<tr><td>Oligo fwd</td><td>CACCGGCACCTGCGTGTGGAACTT</td><td></td><td></td><td></td><td></td><td></td><td></td><td></td><td></td><td></td><td></td></tr>
<tr><td>Oligo rev</td><td>AAACAAGTTCCACACGCAGGTGCC</td><td></td><td></td><td></td><td></td><td></td><td></td><td></td><td></td><td></td><td></td></tr>
<tr><td>Chromosome</td><td>start</td><td>end</td><td>strand</td><td>MM</td><td>target_seq</td><td>PAM</td><td>alignment</td><td>distance</td><td>position</td><td>gene name</td><td>gene id</td></tr>
<tr><td>GU305914.1</td><td>21498</td><td>21520</td><td>-</td><td>0</td><td>GGCACCTGCGTGTGGAACTT</td><td>CGG</td><td>||||||||[||||||||||||]PAM</td><td>NA</td><td>Intergenic</td><td>NA</td><td>NA</td></tr>
<tr><td></td><td></td><td></td><td></td><td></td><td></td><td></td><td></td><td></td><td></td><td></td><td></td></tr>
<tr><td>T32912</td><td>GCGTGTGGAACTTCGGAAAATGG</td><td>1000</td><td></td><td></td><td></td><td></td><td></td><td></td><td></td><td></td><td></td></tr>
<tr><td>Oligo fwd</td><td>CACCGCGTGTGGAACTTCGGAAAA</td><td></td><td></td><td></td><td></td><td></td><td></td><td></td><td></td><td></td><td></td></tr>
<tr><td>Oligo rev</td><td>AAACTTTTCCGAAGTTCCACACGC</td><td></td><td></td><td></td><td></td><td></td><td></td><td></td><td></td><td></td><td></td></tr>
<tr><td>Chromosome</td><td>start</td><td>end</td><td>strand</td><td>MM</td><td>target_seq</td><td>PAM</td><td>alignment</td><td>distance</td><td>position</td><td>gene name</td><td>gene id</td></tr>
<tr><td>GU305914.1</td><td>21491</td><td>21513</td><td>-</td><td>0</td><td>GCGTGTGGAACTTCGGAAAA</td><td>TGG</td><td>||||||||[||||||||||||]PAM</td><td>NA</td><td>Intergenic</td><td>NA</td><td>NA</td></tr>
<tr><td></td><td></td><td></td><td></td><td></td><td></td><td></td><td></td><td></td><td></td><td></td><td></td></tr>
<tr><td>T32913</td><td>CGTGTGGAACTTCGGAAAATGGG</td><td>1000</td><td></td><td></td><td></td><td></td><td></td><td></td><td></td><td></td><td></td></tr>
<tr><td>Oligo fwd</td><td>CACCCGTGTGGAACTTCGGAAAAT</td><td></td><td></td><td></td><td></td><td></td><td></td><td></td><td></td><td></td><td></td></tr>
<tr><td>Oligo rev</td><td>AAACATTTTCCGAAGTTCCACACG</td><td></td><td></td><td></td><td></td><td></td><td></td><td></td><td></td><td></td><td></td></tr>
<tr><td>Chromosome</td><td>start</td><td>end</td><td>strand</td><td>MM</td><td>target_seq</td><td>PAM</td><td>alignment</td><td>distance</td><td>position</td><td>gene name</td><td>gene id</td></tr>
<tr><td>GU305914.1</td><td>21490</td><td>21512</td><td>-</td><td>0</td><td>CGTGTGGAACTTCGGAAAAT</td><td>GGG</td><td>||||||||[||||||||||||]PAM</td><td>NA</td><td>Intergenic</td><td>NA</td><td>NA</td></tr>
<tr><td></td><td></td><td></td><td></td><td></td><td></td><td></td><td></td><td></td><td></td><td></td><td></td></tr>
<tr><td>T32914</td><td>CGGAAAATGGGCCAACGATCTGG</td><td>1000</td><td></td><td></td><td></td><td></td><td></td><td></td><td></td><td></td><td></td></tr>
<tr><td>Oligo fwd</td><td>CACCCGGAAAATGGGCCAACGATC</td><td></td><td></td><td></td><td></td><td></td><td></td><td></td><td></td><td></td><td></td></tr>
<tr><td>Oligo rev</td><td>AAACGATCGTTGGCCCATTTTCCG</td><td></td><td></td><td></td><td></td><td></td><td></td><td></td><td></td><td></td><td></td></tr>
<tr><td>Chromosome</td><td>start</td><td>end</td><td>strand</td><td>MM</td><td>target_seq</td><td>PAM</td><td>alignment</td><td>distance</td><td>position</td><td>gene name</td><td>gene id</td></tr>
<tr><td>GU305914.1</td><td>21478</td><td>21500</td><td>-</td><td>0</td><td>CGGAAAATGGGCCAACGATC</td><td>TGG</td><td>||||||||[||||||||||||]PAM</td><td>NA</td><td>Intergenic</td><td>NA</td><td>NA</td></tr>
<tr><td></td><td></td><td></td><td></td><td></td><td></td><td></td><td></td><td></td><td></td><td></td><td></td></tr>
<tr><td>T32915</td><td>AAATGGGCCAACGATCTGGCCGG</td><td>1000</td><td></td><td></td><td></td><td></td><td></td><td></td><td></td><td></td><td></td></tr>
<tr><td>Oligo fwd</td><td>CACCAAATGGGCCAACGATCTGGC</td><td></td><td></td><td></td><td></td><td></td><td></td><td></td><td></td><td></td><td></td></tr>
<tr><td>Oligo rev</td><td>AAACGCCAGATCGTTGGCCCATTT</td><td></td><td></td><td></td><td></td><td></td><td></td><td></td><td></td><td></td><td></td></tr>
<tr><td>Chromosome</td><td>start</td><td>end</td><td>strand</td><td>MM</td><td>target_seq</td><td>PAM</td><td>alignment</td><td>distance</td><td>position</td><td>gene name</td><td>gene id</td></tr>
<tr><td>GU305914.1</td><td>21474</td><td>21496</td><td>-</td><td>0</td><td>AAATGGGCCAACGATCTGGC</td><td>CGG</td><td>||||||||[||||||||||||]PAM</td><td>NA</td><td>Intergenic</td><td>NA</td><td>NA</td></tr>
<tr><td></td><td></td><td></td><td></td><td></td><td></td><td></td><td></td><td></td><td></td><td></td><td></td></tr>
<tr><td>T32916</td><td>CTACTTCACCGCCATGTTCTCGG</td><td>1000</td><td></td><td></td><td></td><td></td><td></td><td></td><td></td><td></td><td></td></tr>
<tr><td>Oligo fwd</td><td>CACCCTACTTCACCGCCATGTTCT</td><td></td><td></td><td></td><td></td><td></td><td></td><td></td><td></td><td></td><td></td></tr>
<tr><td>Oligo rev</td><td>AAACAGAACATGGCGGTGAAGTAG</td><td></td><td></td><td></td><td></td><td></td><td></td><td></td><td></td><td></td><td></td></tr>
<tr><td>Chromosome</td><td>start</td><td>end</td><td>strand</td><td>MM</td><td>target_seq</td><td>PAM</td><td>alignment</td><td>distance</td><td>position</td><td>gene name</td><td>gene id</td></tr>
<tr><td>GU305914.1</td><td>21433</td><td>21455</td><td>-</td><td>0</td><td>CTACTTCACCGCCATGTTCT</td><td>CGG</td><td>||||||||[||||||||||||]PAM</td><td>NA</td><td>Intergenic</td><td>NA</td><td>NA</td></tr>
<tr><td></td><td></td><td></td><td></td><td></td><td></td><td></td><td></td><td></td><td></td><td></td><td></td></tr>
<tr><td>T32917</td><td>ATGTTCTCGGACCTGCACAACGG</td><td>1000</td><td></td><td></td><td></td><td></td><td></td><td></td><td></td><td></td><td></td></tr>
<tr><td>Oligo fwd</td><td>CACCATGTTCTCGGACCTGCACAA</td><td></td><td></td><td></td><td></td><td></td><td></td><td></td><td></td><td></td><td></td></tr>
<tr><td>Oligo rev</td><td>AAACTTGTGCAGGTCCGAGAACAT</td><td></td><td></td><td></td><td></td><td></td><td></td><td></td><td></td><td></td><td></td></tr>
<tr><td>Chromosome</td><td>start</td><td>end</td><td>strand</td><td>MM</td><td>target_seq</td><td>PAM</td><td>alignment</td><td>distance</td><td>position</td><td>gene name</td><td>gene id</td></tr>
<tr><td>GU305914.1</td><td>21420</td><td>21442</td><td>-</td><td>0</td><td>ATGTTCTCGGACCTGCACAA</td><td>CGG</td><td>||||||||[||||||||||||]PAM</td><td>NA</td><td>Intergenic</td><td>NA</td><td>NA</td></tr>
<tr><td></td><td></td><td></td><td></td><td></td><td></td><td></td><td></td><td></td><td></td><td></td><td></td></tr>
<tr><td>T32918</td><td>CACAACGGCAACGTGAGTATAGG</td><td>1000</td><td></td><td></td><td></td><td></td><td></td><td></td><td></td><td></td><td></td></tr>
<tr><td>Oligo fwd</td><td>CACCCACAACGGCAACGTGAGTAT</td><td></td><td></td><td></td><td></td><td></td><td></td><td></td><td></td><td></td><td></td></tr>
<tr><td>Oligo rev</td><td>AAACATACTCACGTTGCCGTTGTG</td><td></td><td></td><td></td><td></td><td></td><td></td><td></td><td></td><td></td><td></td></tr>
<tr><td>Chromosome</td><td>start</td><td>end</td><td>strand</td><td>MM</td><td>target_seq</td><td>PAM</td><td>alignment</td><td>distance</td><td>position</td><td>gene name</td><td>gene id</td></tr>
<tr><td>GU305914.1</td><td>21405</td><td>21427</td><td>-</td><td>0</td><td>CACAACGGCAACGTGAGTAT</td><td>AGG</td><td>||||||||[||||||||||||]PAM</td><td>NA</td><td>Intergenic</td><td>NA</td><td>NA</td></tr>
<tr><td></td><td></td><td></td><td></td><td></td><td></td><td></td><td></td><td></td><td></td><td></td><td></td></tr>
<tr><td>T32919</td><td>CGGCAACGTGAGTATAGGCCAGG</td><td>1000</td><td></td><td></td><td></td><td></td><td></td><td></td><td></td><td></td><td></td></tr>
<tr><td>Oligo fwd</td><td>CACCCGGCAACGTGAGTATAGGCC</td><td></td><td></td><td></td><td></td><td></td><td></td><td></td><td></td><td></td><td></td></tr>
<tr><td>Oligo rev</td><td>AAACGGCCTATACTCACGTTGCCG</td><td></td><td></td><td></td><td></td><td></td><td></td><td></td><td></td><td></td><td></td></tr>
<tr><td>Chromosome</td><td>start</td><td>end</td><td>strand</td><td>MM</td><td>target_seq</td><td>PAM</td><td>alignment</td><td>distance</td><td>position</td><td>gene name</td><td>gene id</td></tr>
<tr><td>GU305914.1</td><td>21400</td><td>21422</td><td>-</td><td>0</td><td>CGGCAACGTGAGTATAGGCC</td><td>AGG</td><td>||||||||[||||||||||||]PAM</td><td>NA</td><td>Intergenic</td><td>NA</td><td>NA</td></tr>
<tr><td></td><td></td><td></td><td></td><td></td><td></td><td></td><td></td><td></td><td></td><td></td><td></td></tr>
<tr><td>T32920</td><td>CGTGAGTATAGGCCAGGTGTTGG</td><td>1000</td><td></td><td></td><td></td><td></td><td></td><td></td><td></td><td></td><td></td></tr>
<tr><td>Oligo fwd</td><td>CACCCGTGAGTATAGGCCAGGTGT</td><td></td><td></td><td></td><td></td><td></td><td></td><td></td><td></td><td></td><td></td></tr>
<tr><td>Oligo rev</td><td>AAACACACCTGGCCTATACTCACG</td><td></td><td></td><td></td><td></td><td></td><td></td><td></td><td></td><td></td><td></td></tr>
<tr><td>Chromosome</td><td>start</td><td>end</td><td>strand</td><td>MM</td><td>target_seq</td><td>PAM</td><td>alignment</td><td>distance</td><td>position</td><td>gene name</td><td>gene id</td></tr>
<tr><td>GU305914.1</td><td>21394</td><td>21416</td><td>-</td><td>0</td><td>CGTGAGTATAGGCCAGGTGT</td><td>TGG</td><td>||||||||[||||||||||||]PAM</td><td>NA</td><td>Intergenic</td><td>NA</td><td>NA</td></tr>
<tr><td></td><td></td><td></td><td></td><td></td><td></td><td></td><td></td><td></td><td></td><td></td><td></td></tr>
<tr><td>T32921</td><td>GTTGACCGCCGCGCACGCCGAGG</td><td>1000</td><td></td><td></td><td></td><td></td><td></td><td></td><td></td><td></td><td></td></tr>
<tr><td>Oligo fwd</td><td>CACCGTTGACCGCCGCGCACGCCG</td><td></td><td></td><td></td><td></td><td></td><td></td><td></td><td></td><td></td><td></td></tr>
<tr><td>Oligo rev</td><td>AAACCGGCGTGCGCGGCGGTCAAC</td><td></td><td></td><td></td><td></td><td></td><td></td><td></td><td></td><td></td><td></td></tr>
<tr><td>Chromosome</td><td>start</td><td>end</td><td>strand</td><td>MM</td><td>target_seq</td><td>PAM</td><td>alignment</td><td>distance</td><td>position</td><td>gene name</td><td>gene id</td></tr>
<tr><td>GU305914.1</td><td>21343</td><td>21365</td><td>-</td><td>0</td><td>GTTGACCGCCGCGCACGCCG</td><td>AGG</td><td>||||||||[||||||||||||]PAM</td><td>NA</td><td>Intergenic</td><td>NA</td><td>NA</td></tr>
</table>

<table>
<tr><td>T32922</td><td>GACCGCCGCGCACGCCGAGGCGG</td><td>1000</td><td></td><td></td><td></td><td></td><td></td><td></td><td></td><td></td><td></td></tr>
<tr><td>Oligo fwd</td><td>CACCGACCGCCGCGCACGCCGAGG</td><td></td><td></td><td></td><td></td><td></td><td></td><td></td><td></td><td></td><td></td></tr>
<tr><td>Oligo rev</td><td>AAACCCTCGGCGTGCGCGGCGGTC</td><td></td><td></td><td></td><td></td><td></td><td></td><td></td><td></td><td></td><td></td></tr>
<tr><td>Chromosome</td><td>start</td><td>end</td><td>strand</td><td>MM</td><td>target_seq</td><td>PAM</td><td>alignment</td><td>distance</td><td>position</td><td>gene name</td><td>gene id</td></tr>
<tr><td>GU305914.1</td><td>21340</td><td>21362</td><td>-</td><td>0</td><td>GACCGCCGCGCACGCCGAGG</td><td>CGG</td><td>||||||||[||||||||||||]PAM</td><td>NA</td><td>Intergenic</td><td>NA</td><td>NA</td></tr>
<tr><td>T32923</td><td>CACGCCGAGGCGGTCGTGCACGG</td><td>1000</td><td></td><td></td><td></td><td></td><td></td><td></td><td></td><td></td><td></td></tr>
<tr><td>Oligo fwd</td><td>CACCCACGCCGAGGCGGTCGTGCA</td><td></td><td></td><td></td><td></td><td></td><td></td><td></td><td></td><td></td><td></td></tr>
<tr><td>Oligo rev</td><td>AAACTGCACGACCGCCTCGGCGTG</td><td></td><td></td><td></td><td></td><td></td><td></td><td></td><td></td><td></td><td></td></tr>
<tr><td>Chromosome</td><td>start</td><td>end</td><td>strand</td><td>MM</td><td>target_seq</td><td>PAM</td><td>alignment</td><td>distance</td><td>position</td><td>gene name</td><td>gene id</td></tr>
<tr><td>GU305914.1</td><td>21330</td><td>21352</td><td>-</td><td>0</td><td>CACGCCGAGGCGGTCGTGCA</td><td>CGG</td><td>||||||||[||||||||||||]PAM</td><td>NA</td><td>Intergenic</td><td>NA</td><td>NA</td></tr>
<tr><td>T32924</td><td>GGCGGTCGTGCACGGATGTTCGG</td><td>1000</td><td></td><td></td><td></td><td></td><td></td><td></td><td></td><td></td><td></td></tr>
<tr><td>Oligo fwd</td><td>CACCGGCGGTCGTGCACGGATGTT</td><td></td><td></td><td></td><td></td><td></td><td></td><td></td><td></td><td></td><td></td></tr>
<tr><td>Oligo rev</td><td>AAACAACATCCGTGCACGACCGCC</td><td></td><td></td><td></td><td></td><td></td><td></td><td></td><td></td><td></td><td></td></tr>
<tr><td>Chromosome</td><td>start</td><td>end</td><td>strand</td><td>MM</td><td>target_seq</td><td>PAM</td><td>alignment</td><td>distance</td><td>position</td><td>gene name</td><td>gene id</td></tr>
<tr><td>GU305914.1</td><td>21322</td><td>21344</td><td>-</td><td>0</td><td>GGCGGTCGTGCACGGATGTT</td><td>CGG</td><td>||||||||[||||||||||||]PAM</td><td>NA</td><td>Intergenic</td><td>NA</td><td>NA</td></tr>
<tr><td>T32925</td><td>GTCCATCATCGACATTCCTCTGG</td><td>1000</td><td></td><td></td><td></td><td></td><td></td><td></td><td></td><td></td><td></td></tr>
<tr><td>Oligo fwd</td><td>CACCGTCCATCATCGACATTCCTC</td><td></td><td></td><td></td><td></td><td></td><td></td><td></td><td></td><td></td><td></td></tr>
<tr><td>Oligo rev</td><td>AAACGAGGAATGTCGATGATGGAC</td><td></td><td></td><td></td><td></td><td></td><td></td><td></td><td></td><td></td><td></td></tr>
<tr><td>Chromosome</td><td>start</td><td>end</td><td>strand</td><td>MM</td><td>target_seq</td><td>PAM</td><td>alignment</td><td>distance</td><td>position</td><td>gene name</td><td>gene id</td></tr>
<tr><td>GU305914.1</td><td>21274</td><td>21296</td><td>-</td><td>0</td><td>GTCCATCATCGACATTCCTC</td><td>TGG</td><td>||||||||[||||||||||||]PAM</td><td>NA</td><td>Intergenic</td><td>NA</td><td>NA</td></tr>
<tr><td>T32926</td><td>CCTCTGGTCTATTACCGCATCGG</td><td>1000</td><td></td><td></td><td></td><td></td><td></td><td></td><td></td><td></td><td></td></tr>
<tr><td>Oligo fwd</td><td>CACCCCTCTGGTCTATTACCGCAT</td><td></td><td></td><td></td><td></td><td></td><td></td><td></td><td></td><td></td><td></td></tr>
<tr><td>Oligo rev</td><td>AAACATGCGGTAATAGACCAGAGG</td><td></td><td></td><td></td><td></td><td></td><td></td><td></td><td></td><td></td><td></td></tr>
<tr><td>Chromosome</td><td>start</td><td>end</td><td>strand</td><td>MM</td><td>target_seq</td><td>PAM</td><td>alignment</td><td>distance</td><td>position</td><td>gene name</td><td>gene id</td></tr>
<tr><td>GU305914.1</td><td>21258</td><td>21280</td><td>-</td><td>0</td><td>CCTCTGGTCTATTACCGCAT</td><td>CGG</td><td>||||||||[||||||||||||]PAM</td><td>NA</td><td>Intergenic</td><td>NA</td><td>NA</td></tr>
<tr><td>T32927</td><td>CTCTGGTCTATTACCGCATCGGG</td><td>1000</td><td></td><td></td><td></td><td></td><td></td><td></td><td></td><td></td><td></td></tr>
<tr><td>Oligo fwd</td><td>CACCCTCTGGTCTATTACCGCATC</td><td></td><td></td><td></td><td></td><td></td><td></td><td></td><td></td><td></td><td></td></tr>
<tr><td>Oligo rev</td><td>AAACGATGCGGTAATAGACCAGAG</td><td></td><td></td><td></td><td></td><td></td><td></td><td></td><td></td><td></td><td></td></tr>
<tr><td>Chromosome</td><td>start</td><td>end</td><td>strand</td><td>MM</td><td>target_seq</td><td>PAM</td><td>alignment</td><td>distance</td><td>position</td><td>gene name</td><td>gene id</td></tr>
<tr><td>GU305914.1</td><td>21257</td><td>21279</td><td>-</td><td>0</td><td>CTCTGGTCTATTACCGCATC</td><td>GGG</td><td>||||||||[||||||||||||]PAM</td><td>NA</td><td>Intergenic</td><td>NA</td><td>NA</td></tr>
<tr><td>T32928</td><td>ACCGCATCGGGATTTTCAACTGG</td><td>1000</td><td></td><td></td><td></td><td></td><td></td><td></td><td></td><td></td><td></td></tr>
<tr><td>Oligo fwd</td><td>CACCACCGCATCGGGATTTTCAAC</td><td></td><td></td><td></td><td></td><td></td><td></td><td></td><td></td><td></td><td></td></tr>
<tr><td>Oligo rev</td><td>AAACGTTGAAAATCCCGATGCGGT</td><td></td><td></td><td></td><td></td><td></td><td></td><td></td><td></td><td></td><td></td></tr>
<tr><td>Chromosome</td><td>start</td><td>end</td><td>strand</td><td>MM</td><td>target_seq</td><td>PAM</td><td>alignment</td><td>distance</td><td>position</td><td>gene name</td><td>gene id</td></tr>
<tr><td>GU305914.1</td><td>21245</td><td>21267</td><td>-</td><td>0</td><td>ACCGCATCGGGATTTTCAAC</td><td>TGG</td><td>||||||||[||||||||||||]PAM</td><td>NA</td><td>Intergenic</td><td>NA</td><td>NA</td></tr>
<tr><td>T32929</td><td>CTTCGATTCCCCCGATCGCACGG</td><td>1000</td><td></td><td></td><td></td><td></td><td></td><td></td><td></td><td></td><td></td></tr>
<tr><td>Oligo fwd</td><td>CACCCTTCGATTCCCCCGATCGCA</td><td></td><td></td><td></td><td></td><td></td><td></td><td></td><td></td><td></td><td></td></tr>
<tr><td>Oligo rev</td><td>AAACTGCGATCGGGGGAATCGAAG</td><td></td><td></td><td></td><td></td><td></td><td></td><td></td><td></td><td></td><td></td></tr>
<tr><td>Chromosome</td><td>start</td><td>end</td><td>strand</td><td>MM</td><td>target_seq</td><td>PAM</td><td>alignment</td><td>distance</td><td>position</td><td>gene name</td><td>gene id</td></tr>
<tr><td>GU305914.1</td><td>21220</td><td>21242</td><td>-</td><td>0</td><td>CTTCGATTCCCCCGATCGCA</td><td>CGG</td><td>||||||||[||||||||||||]PAM</td><td>NA</td><td>Intergenic</td><td>NA</td><td>NA</td></tr>
<tr><td>T32930</td><td>TTCGATTCCCCCGATCGCACGGG</td><td>1000</td><td></td><td></td><td></td><td></td><td></td><td></td><td></td><td></td><td></td></tr>
<tr><td>Oligo fwd</td><td>CACCTTCGATTCCCCCGATCGCAC</td><td></td><td></td><td></td><td></td><td></td><td></td><td></td><td></td><td></td><td></td></tr>
<tr><td>Oligo rev</td><td>AAACGTGCGATCGGGGGAATCGAA</td><td></td><td></td><td></td><td></td><td></td><td></td><td></td><td></td><td></td><td></td></tr>
<tr><td>Chromosome</td><td>start</td><td>end</td><td>strand</td><td>MM</td><td>target_seq</td><td>PAM</td><td>alignment</td><td>distance</td><td>position</td><td>gene name</td><td>gene id</td></tr>
<tr><td>GU305914.1</td><td>21219</td><td>21241</td><td>-</td><td>0</td><td>TTCGATTCCCCCGATCGCAC</td><td>GGG</td><td>||||||||[||||||||||||]PAM</td><td>NA</td><td>Intergenic</td><td>NA</td><td>NA</td></tr>
<tr><td>T32931</td><td>CCCGATCGCACGGGAGATAATGG</td><td>1000</td><td></td><td></td><td></td><td></td><td></td><td></td><td></td><td></td><td></td></tr>
<tr><td>Oligo fwd</td><td>CACCCCCGATCGCACGGGAGATAA</td><td></td><td></td><td></td><td></td><td></td><td></td><td></td><td></td><td></td><td></td></tr>
<tr><td>Oligo rev</td><td>AAACTTATCTCCCGTGCGATCGGG</td><td></td><td></td><td></td><td></td><td></td><td></td><td></td><td></td><td></td><td></td></tr>
<tr><td>Chromosome</td><td>start</td><td>end</td><td>strand</td><td>MM</td><td>target_seq</td><td>PAM</td><td>alignment</td><td>distance</td><td>position</td><td>gene name</td><td>gene id</td></tr>
<tr><td>GU305914.1</td><td>21210</td><td>21232</td><td>-</td><td>0</td><td>CCCGATCGCACGGGAGATAA</td><td>TGG</td><td>||||||||[||||||||||||]PAM</td><td>NA</td><td>Intergenic</td><td>NA</td><td>NA</td></tr>
<tr><td>T32932</td><td>GAGATAATGGAGAGCCGTTGCGG</td><td>1000</td><td></td><td></td><td></td><td></td><td></td><td></td><td></td><td></td><td></td></tr>
<tr><td>Oligo fwd</td><td>CACCGAGATAATGGAGAGCCGTTG</td><td></td><td></td><td></td><td></td><td></td><td></td><td></td><td></td><td></td><td></td></tr>
<tr><td>Oligo rev</td><td>AAACCAACGGCTCTCCATTATCTC</td><td></td><td></td><td></td><td></td><td></td><td></td><td></td><td></td><td></td><td></td></tr>
<tr><td>Chromosome</td><td>start</td><td>end</td><td>strand</td><td>MM</td><td>target_seq</td><td>PAM</td><td>alignment</td><td>distance</td><td>position</td><td>gene name</td><td>gene id</td></tr>
<tr><td>GU305914.1</td><td>21197</td><td>21219</td><td>-</td><td>0</td><td>GAGATAATGGAGAGCCGTTG</td><td>CGG</td><td>||||||||[||||||||||||]PAM</td><td>NA</td><td>Intergenic</td><td>NA</td><td>NA</td></tr>
<tr><td>T32933</td><td>GAGCCGTTGCGGATCCTCGCCGG</td><td>1000</td><td></td><td></td><td></td><td></td><td></td><td></td><td></td><td></td><td></td></tr>
<tr><td>Oligo fwd</td><td>CACCGAGCCGTTGCGGATCCTCGC</td><td></td><td></td><td></td><td></td><td></td><td></td><td></td><td></td><td></td><td></td></tr>
<tr><td>Oligo rev</td><td>AAACGCGAGGATCCGCAACGGCTC</td><td></td><td></td><td></td><td></td><td></td><td></td><td></td><td></td><td></td><td></td></tr>
<tr><td>Chromosome</td><td>start</td><td>end</td><td>strand</td><td>MM</td><td>target_seq</td><td>PAM</td><td>alignment</td><td>distance</td><td>position</td><td>gene name</td><td>gene id</td></tr>
<tr><td>GU305914.1</td><td>21186</td><td>21208</td><td>-</td><td>0</td><td>GAGCCGTTGCGGATCCTCGC</td><td>CGG</td><td>||||||||[||||||||||||]PAM</td><td>NA</td><td>Intergenic</td><td>NA</td><td>NA</td></tr>
<tr><td>T32934</td><td>AGCCGTTGCGGATCCTCGCCGGG</td><td>1000</td><td></td><td></td><td></td><td></td><td></td><td></td><td></td><td></td><td></td></tr>
<tr><td>Oligo fwd</td><td>CACCAGCCGTTGCGGATCCTCGCC</td><td></td><td></td><td></td><td></td><td></td><td></td><td></td><td></td><td></td><td></td></tr>
<tr><td>Oligo rev</td><td>AAACGGCGAGGATCCGCAACGGCT</td><td></td><td></td><td></td><td></td><td></td><td></td><td></td><td></td><td></td><td></td></tr>
<tr><td>Chromosome</td><td>start</td><td>end</td><td>strand</td><td>MM</td><td>target_seq</td><td>PAM</td><td>alignment</td><td>distance</td><td>position</td><td>gene name</td><td>gene id</td></tr>
<tr><td>GU305914.1</td><td>21185</td><td>21207</td><td>-</td><td>0</td><td>AGCCGTTGCGGATCCTCGCC</td><td>GGG</td><td>||||||||[||||||||||||]PAM</td><td>NA</td><td>Intergenic</td><td>NA</td><td>NA</td></tr>
<tr><td>T32935</td><td>GCCGTTGCGGATCCTCGCCGGGG</td><td>1000</td><td></td><td></td><td></td><td></td><td></td><td></td><td></td><td></td><td></td></tr>
<tr><td>Oligo fwd</td><td>CACCGCCGTTGCGGATCCTCGCCG</td><td></td><td></td><td></td><td></td><td></td><td></td><td></td><td></td><td></td><td></td></tr>
<tr><td>Oligo rev</td><td>AAACCGGCGAGGATCCGCAACGGC</td><td></td><td></td><td></td><td></td><td></td><td></td><td></td><td></td><td></td><td></td></tr>
<tr><td>Chromosome</td><td>start</td><td>end</td><td>strand</td><td>MM</td><td>target_seq</td><td>PAM</td><td>alignment</td><td>distance</td><td>position</td><td>gene name</td><td>gene id</td></tr>
<tr><td>GU305914.1</td><td>21184</td><td>21206</td><td>-</td><td>0</td><td>GCCGTTGCGGATCCTCGCCG</td><td>GGG</td><td>||||||||[||||||||||||]PAM</td><td>NA</td><td>Intergenic</td><td>NA</td><td>NA</td></tr>
<tr><td>T32936</td><td>CCGGGGCTGTGTTGTTTAACAGG</td><td>1000</td><td></td><td></td><td></td><td></td><td></td><td></td><td></td><td></td><td></td></tr>
<tr><td>Oligo fwd</td><td>CACCCCGGGGCTGTGTTGTTTAAC</td><td></td><td></td><td></td><td></td><td></td><td></td><td></td><td></td><td></td><td></td></tr>
<tr><td>Oligo rev</td><td>AAACGTTAAACAACACAGCCCCGG</td><td></td><td></td><td></td><td></td><td></td><td></td><td></td><td></td><td></td><td></td></tr>
<tr><td>Chromosome</td><td>start</td><td>end</td><td>strand</td><td>MM</td><td>target_seq</td><td>PAM</td><td>alignment</td><td>distance</td><td>position</td><td>gene name</td><td>gene id</td></tr>
</table>

GU305914.1 21167 21189 - 0 CCGGGGCTGTGTTGTTTAAC AGG ||||||||[||||||||||||]PAM NA Intergenic NA NA

T32937 CGGGGCTGTGTTGTTTAACAGGG 1000
Oligo fwd CACCCGGGGCTGTGTTGTTTAACA
Oligo rev AAACTGTTAAACAACACAGCCCCG
Chromosome start end strand MM target_seq PAM alignment distance position gene name gene id
GU305914.1 21166 21188 - 0 CGGGGCTGTGTTGTTTAACA GGG ||||||||[||||||||||||]PAM NA Intergenic NA NA

T32938 GACATGTACGACCTCATACACGG 1000
Oligo fwd CACCGACATGTACGACCTCATACA
Oligo rev AAACTGTATGAGGTCGTACATGTC
Chromosome start end strand MM target_seq PAM alignment distance position gene name gene id
GU305914.1 21144 21166 - 0 GACATGTACGACCTCATACA CGG ||||||||[||||||||||||]PAM NA Intergenic NA NA

T32939 GTACGACCTCATACACGGAACGG 1000
Oligo fwd CACCGTACGACCTCATACACGGAA
Oligo rev AAACTTCCGTGTATGAGGTCGTAC
Chromosome start end strand MM target_seq PAM alignment distance position gene name gene id
GU305914.1 21139 21161 - 0 GTACGACCTCATACACGGAA CGG ||||||||[||||||||||||]PAM NA Intergenic NA NA

T32940 CGACCTCATACACGGAACGGAGG 1000
Oligo fwd CACCCGACCTCATACACGGAACGG
Oligo rev AAACCCGTTCCGTGTATGAGGTCG
Chromosome start end strand MM target_seq PAM alignment distance position gene name gene id
GU305914.1 21136 21158 - 0 CGACCTCATACACGGAACGG AGG ||||||||[||||||||||||]PAM NA Intergenic NA NA

T32941 GACCTCATACACGGAACGGAGGG 1000
Oligo fwd CACCGACCTCATACACGGAACGGA
Oligo rev AAACTCCGTTCCGTGTATGAGGTC
Chromosome start end strand MM target_seq PAM alignment distance position gene name gene id
GU305914.1 21135 21157 - 0 GACCTCATACACGGAACGGA GGG ||||||||[||||||||||||]PAM NA Intergenic NA NA

T32942 AGCGTACGTCATGTCCCCTACGG 1000
Oligo fwd CACCAGCGTACGTCATGTCCCCTA
Oligo rev AAACTAGGGGACATGACGTACGCT
Chromosome start end strand MM target_seq PAM alignment distance position gene name gene id
GU305914.1 21103 21125 - 0 AGCGTACGTCATGTCCCCTA CGG ||||||||[||||||||||||]PAM NA Intergenic NA NA

T32943 CGTCATGTCCCCTACGGACACGG 1000
Oligo fwd CACCCGTCATGTCCCCTACGGACA
Oligo rev AAACTGTCCGTAGGGGACATGACG
Chromosome start end strand MM target_seq PAM alignment distance position gene name gene id
GU305914.1 21097 21119 - 0 CGTCATGTCCCCTACGGACA CGG ||||||||[||||||||||||]PAM NA Intergenic NA NA

T32944 TGTCCCCTACGGACACGGTGTGG 1000
Oligo fwd CACCTGTCCCCTACGGACACGGTG
Oligo rev AAACCACCGTGTCCGTAGGGGACA
Chromosome start end strand MM target_seq PAM alignment distance position gene name gene id
GU305914.1 21092 21114 - 0 TGTCCCCTACGGACACGGTG TGG ||||||||[||||||||||||]PAM NA Intergenic NA NA

T32945 ACGGACACGGTGTGGCGAGACGG 1000
Oligo fwd CACCACGGACACGGTGTGGCGAGA
Oligo rev AAACTCTCGCCACACCGTGTCCGT
Chromosome start end strand MM target_seq PAM alignment distance position gene name gene id
GU305914.1 21084 21106 - 0 ACGGACACGGTGTGGCGAGA CGG ||||||||[||||||||||||]PAM NA Intergenic NA NA

T32946 CGGACACGGTGTGGCGAGACGGG 1000
Oligo fwd CACCCGGACACGGTGTGGCGAGAC
Oligo rev AAACGTCTCGCCACACCGTGTCCG
Chromosome start end strand MM target_seq PAM alignment distance position gene name gene id
GU305914.1 21083 21105 - 0 CGGACACGGTGTGGCGAGAC GGG ||||||||[||||||||||||]PAM NA Intergenic NA NA

T32947 GGACACGGTGTGGCGAGACGGGG 1000
Oligo fwd CACCGGACACGGTGTGGCGAGACG
Oligo rev AAACCGTCTCGCCACACCGTGTCC
Chromosome start end strand MM target_seq PAM alignment distance position gene name gene id
GU305914.1 21082 21104 - 0 GGACACGGTGTGGCGAGACG GGG ||||||||[||||||||||||]PAM NA Intergenic NA NA

T32948 GACACGGTGTGGCGAGACGGGGG 1000
Oligo fwd CACCGACACGGTGTGGCGAGACGG
Oligo rev AAACCCGTCTCGCCACACCGTGTC
Chromosome start end strand MM target_seq PAM alignment distance position gene name gene id
GU305914.1 21081 21103 - 0 GACACGGTGTGGCGAGACGG GGG ||||||||[||||||||||||]PAM NA Intergenic NA NA
GU305914.1 195212 195234 - 4 GAGCCGGTGTATCGAGACGG CGG ||--||||[||--||||||||]PAM NA Intergenic NA NA

T32949 ACGGTGTGGCGAGACGGGGGAGG 1000
Oligo fwd CACCACGGTGTGGCGAGACGGGGG
Oligo rev AAACCCCCCGTCTCGCCACACCGT
Chromosome start end strand MM target_seq PAM alignment distance position gene name gene id
GU305914.1 21078 21100 - 0 ACGGTGTGGCGAGACGGGGG AGG ||||||||[||||||||||||]PAM NA Intergenic NA NA

T32950 CAACCTCACCTTCAGATTCGTGG 1000
Oligo fwd CACCCAACCTCACCTTCAGATTCG
Oligo rev AAACCGAATCTGAAGGTGAGGTTG
Chromosome start end strand MM target_seq PAM alignment distance position gene name gene id
GU305914.1 21055 21077 - 0 CAACCTCACCTTCAGATTCG TGG ||||||||[||||||||||||]PAM NA Intergenic NA NA

T32951 CTTCAGATTCGTGGTTAGAAAGG 1000
Oligo fwd CACCCTTCAGATTCGTGGTTAGAA

Oligo rev AAACTTCTAACCACGAATCTGAAG
Chromosome start end strand MM target_seq PAM alignment distance position gene name gene id
GU305914.1 21046 21068 - 0 CTTCAGATTCGTGGTTAGAA AGG ||||||||[||||||||||||]PAM NA Intergenic NA NA

T32952 AGATTCGTGGTTAGAAAGGCCGG 1000
Oligo fwd CACCAGATTCGTGGTTAGAAAGGC
Oligo rev AAACGCCTTTCTAACCACGAATCT
Chromosome start end strand MM target_seq PAM alignment distance position gene name gene id
GU305914.1 21042 21064 - 0 AGATTCGTGGTTAGAAAGGC CGG ||||||||[||||||||||||]PAM NA Intergenic NA NA

T32953 AGAAAGGCCGGCGTATACCGAGG 1000
Oligo fwd CACCAGAAAGGCCGGCGTATACCG
Oligo rev AAACCGGTATACGCCGGCCTTTCT
Chromosome start end strand MM target_seq PAM alignment distance position gene name gene id
GU305914.1 21030 21052 - 0 AGAAAGGCCGGCGTATACCG AGG ||||||||[||||||||||||]PAM NA Intergenic NA NA

T32954 CGAGGTCTCCTCACCATCGACGG 1000
Oligo fwd CACCCGAGGTCTCCTCACCATCGA
Oligo rev AAACTCGATGGTGAGGAGACCTCG
Chromosome start end strand MM target_seq PAM alignment distance position gene name gene id
GU305914.1 21012 21034 - 0 CGAGGTCTCCTCACCATCGA CGG ||||||||[||||||||||||]PAM NA Intergenic NA NA

T32955 GAGGTCTCCTCACCATCGACGGG 1000
Oligo fwd CACCGAGGTCTCCTCACCATCGAC
Oligo rev AAACGTCGATGGTGAGGAGACCTC
Chromosome start end strand MM target_seq PAM alignment distance position gene name gene id
GU305914.1 21011 21033 - 0 GAGGTCTCCTCACCATCGAC GGG ||||||||[||||||||||||]PAM NA Intergenic NA NA

T32956 AGGTCTCCTCACCATCGACGGGG 1000
Oligo fwd CACCAGGTCTCCTCACCATCGACG
Oligo rev AAACCGTCGATGGTGAGGAGACCT
Chromosome start end strand MM target_seq PAM alignment distance position gene name gene id
GU305914.1 21010 21032 - 0 AGGTCTCCTCACCATCGACG GGG ||||||||[||||||||||||]PAM NA Intergenic NA NA

T32957 CTACTCCTGCGTCTTCACAGTGG 1000
Oligo fwd CACCCTACTCCTGCGTCTTCACAG
Oligo rev AAACCTGTGAAGACGCAGGAGTAG
Chromosome start end strand MM target_seq PAM alignment distance position gene name gene id
GU305914.1 20980 21002 - 0 CTACTCCTGCGTCTTCACAG TGG ||||||||[||||||||||||]PAM NA Intergenic NA NA

T32958 CTGCGTCTTCACAGTGGTCGAGG 1000
Oligo fwd CACCCTGCGTCTTCACAGTGGTCG
Oligo rev AAACCGACCACTGTGAAGACGCAG
Chromosome start end strand MM target_seq PAM alignment distance position gene name gene id
GU305914.1 20974 20996 - 0 CTGCGTCTTCACAGTGGTCG AGG ||||||||[||||||||||||]PAM NA Intergenic NA NA

T32959 CCAGTGCGCCCGCCTCAAAGCGG 1000
Oligo fwd CACCCCAGTGCGCCCGCCTCAAAG
Oligo rev AAACCTTTGAGGCGGGCGCACTGG
Chromosome start end strand MM target_seq PAM alignment distance position gene name gene id
GU305914.1 20950 20972 - 0 CCAGTGCGCCCGCCTCAAAG CGG ||||||||[||||||||||||]PAM NA Intergenic NA NA

T32960 CCGCCTCAAAGCGGACAACGAGG 1000
Oligo fwd CACCCCGCCTCAAAGCGGACAACG
Oligo rev AAACCGTTGTCCGCTTTGAGGCGG
Chromosome start end strand MM target_seq PAM alignment distance position gene name gene id
GU305914.1 20941 20963 - 0 CCGCCTCAAAGCGGACAACG AGG ||||||||[||||||||||||]PAM NA Intergenic NA NA

T32961 GACAACGAGGCGAAAGCGAGCGG 1000
Oligo fwd CACCGACAACGAGGCGAAAGCGAG
Oligo rev AAACCTCGCTTTCGCCTCGTTGTC
Chromosome start end strand MM target_seq PAM alignment distance position gene name gene id
GU305914.1 20928 20950 - 0 GACAACGAGGCGAAAGCGAG CGG ||||||||[||||||||||||]PAM NA Intergenic NA NA

T32962 GAGCGGATTCCCCAGCGCCGCGG 1000
Oligo fwd CACCGAGCGGATTCCCCAGCGCCG
Oligo rev AAACCGGCGCTGGGGAATCCGCTC
Chromosome start end strand MM target_seq PAM alignment distance position gene name gene id
GU305914.1 20911 20933 - 0 GAGCGGATTCCCCAGCGCCG CGG ||||||||[||||||||||||]PAM NA Intergenic NA NA

T32963 GCCTCTCTACCTACCGCTCGCGG 1000
Oligo fwd CACCGCCTCTCTACCTACCGCTCG
Oligo rev AAACCGAGCGGTAGGTAGAGAGGC
Chromosome start end strand MM target_seq PAM alignment distance position gene name gene id
GU305914.1 20877 20899 - 0 GCCTCTCTACCTACCGCTCG CGG ||||||||[||||||||||||]PAM NA Intergenic NA NA

T32964 TCTACCTACCGCTCGCGGACCGG 1000
Oligo fwd CACCTCTACCTACCGCTCGCGGAC
Oligo rev AAACGTCCGCGAGCGGTAGGTAGA
Chromosome start end strand MM target_seq PAM alignment distance position gene name gene id
GU305914.1 20872 20894 - 0 TCTACCTACCGCTCGCGGAC CGG ||||||||[||||||||||||]PAM NA Intergenic NA NA

T32965 TTCCTCGACGACCCTCGAGAAGG 1000
Oligo fwd CACCTTCCTCGACGACCCTCGAGA
Oligo rev AAACTCTCGAGGGTCGTCGAGGAA
Chromosome start end strand MM target_seq PAM alignment distance position gene name gene id
GU305914.1 20845 20867 - 0 TTCCTCGACGACCCTCGAGA AGG ||||||||[||||||||||||]PAM NA Intergenic NA NA

T32966 ACCCTCGAGAAGGACGATGTTGG 1000

<table>
<tr><td>Oligo fwd</td><td>CACCACCCTCGAGAAGGACGATGT</td><td></td><td></td><td></td><td></td><td></td><td></td><td></td><td></td><td></td><td></td></tr>
<tr><td>Oligo rev</td><td>AAACACATCGTCCTTCTCGAGGGT</td><td></td><td></td><td></td><td></td><td></td><td></td><td></td><td></td><td></td><td></td></tr>
<tr><td>Chromosome</td><td>start</td><td>end</td><td>strand</td><td>MM</td><td>target_seq</td><td>PAM</td><td>alignment</td><td>distance</td><td>position</td><td>gene name</td><td>gene id</td></tr>
<tr><td>GU305914.1</td><td>20835</td><td>20857</td><td>-</td><td>0</td><td>ACCCTCGAGAAGGACGATGT</td><td>TGG</td><td>||||||||[||||||||||||]PAM</td><td>NA</td><td>Intergenic</td><td>NA</td><td>NA</td></tr>
<tr><td></td><td></td><td></td><td></td><td></td><td></td><td></td><td></td><td></td><td></td><td></td><td></td></tr>
<tr><td>T32967</td><td>AAGGACGATGTTGGAACTCGAGG</td><td>1000</td><td></td><td></td><td></td><td></td><td></td><td></td><td></td><td></td><td></td></tr>
<tr><td>Oligo fwd</td><td>CACCAAGGACGATGTTGGAACTCG</td><td></td><td></td><td></td><td></td><td></td><td></td><td></td><td></td><td></td><td></td></tr>
<tr><td>Oligo rev</td><td>AAACCGAGTTCCAACATCGTCCTT</td><td></td><td></td><td></td><td></td><td></td><td></td><td></td><td></td><td></td><td></td></tr>
<tr><td>Chromosome</td><td>start</td><td>end</td><td>strand</td><td>MM</td><td>target_seq</td><td>PAM</td><td>alignment</td><td>distance</td><td>position</td><td>gene name</td><td>gene id</td></tr>
<tr><td>GU305914.1</td><td>20826</td><td>20848</td><td>-</td><td>0</td><td>AAGGACGATGTTGGAACTCG</td><td>AGG</td><td>||||||||[||||||||||||]PAM</td><td>NA</td><td>Intergenic</td><td>NA</td><td>NA</td></tr>
<tr><td></td><td></td><td></td><td></td><td></td><td></td><td></td><td></td><td></td><td></td><td></td><td></td></tr>
<tr><td>T32968</td><td>TGGAACTCGAGGCGAGTCTTCGG</td><td>1000</td><td></td><td></td><td></td><td></td><td></td><td></td><td></td><td></td><td></td></tr>
<tr><td>Oligo fwd</td><td>CACCTGGAACTCGAGGCGAGTCTT</td><td></td><td></td><td></td><td></td><td></td><td></td><td></td><td></td><td></td><td></td></tr>
<tr><td>Oligo rev</td><td>AAACAAGACTCGCCTCGAGTTCCA</td><td></td><td></td><td></td><td></td><td></td><td></td><td></td><td></td><td></td><td></td></tr>
<tr><td>Chromosome</td><td>start</td><td>end</td><td>strand</td><td>MM</td><td>target_seq</td><td>PAM</td><td>alignment</td><td>distance</td><td>position</td><td>gene name</td><td>gene id</td></tr>
<tr><td>GU305914.1</td><td>20815</td><td>20837</td><td>-</td><td>0</td><td>TGGAACTCGAGGCGAGTCTT</td><td>CGG</td><td>||||||||[||||||||||||]PAM</td><td>NA</td><td>Intergenic</td><td>NA</td><td>NA</td></tr>
<tr><td></td><td></td><td></td><td></td><td></td><td></td><td></td><td></td><td></td><td></td><td></td><td></td></tr>
<tr><td>T32969</td><td>AGTCTTCGGAGCTGTACGCTAGG</td><td>1000</td><td></td><td></td><td></td><td></td><td></td><td></td><td></td><td></td><td></td></tr>
<tr><td>Oligo fwd</td><td>CACCAGTCTTCGGAGCTGTACGCT</td><td></td><td></td><td></td><td></td><td></td><td></td><td></td><td></td><td></td><td></td></tr>
<tr><td>Oligo rev</td><td>AAACAGCGTACAGCTCCGAAGACT</td><td></td><td></td><td></td><td></td><td></td><td></td><td></td><td></td><td></td><td></td></tr>
<tr><td>Chromosome</td><td>start</td><td>end</td><td>strand</td><td>MM</td><td>target_seq</td><td>PAM</td><td>alignment</td><td>distance</td><td>position</td><td>gene name</td><td>gene id</td></tr>
<tr><td>GU305914.1</td><td>20801</td><td>20823</td><td>-</td><td>0</td><td>AGTCTTCGGAGCTGTACGCT</td><td>AGG</td><td>||||||||[||||||||||||]PAM</td><td>NA</td><td>Intergenic</td><td>NA</td><td>NA</td></tr>
<tr><td></td><td></td><td></td><td></td><td></td><td></td><td></td><td></td><td></td><td></td><td></td><td></td></tr>
<tr><td>T32970</td><td>GTCTTCGGAGCTGTACGCTAGGG</td><td>1000</td><td></td><td></td><td></td><td></td><td></td><td></td><td></td><td></td><td></td></tr>
<tr><td>Oligo fwd</td><td>CACCGTCTTCGGAGCTGTACGCTA</td><td></td><td></td><td></td><td></td><td></td><td></td><td></td><td></td><td></td><td></td></tr>
<tr><td>Oligo rev</td><td>AAACTAGCGTACAGCTCCGAAGAC</td><td></td><td></td><td></td><td></td><td></td><td></td><td></td><td></td><td></td><td></td></tr>
<tr><td>Chromosome</td><td>start</td><td>end</td><td>strand</td><td>MM</td><td>target_seq</td><td>PAM</td><td>alignment</td><td>distance</td><td>position</td><td>gene name</td><td>gene id</td></tr>
<tr><td>GU305914.1</td><td>20800</td><td>20822</td><td>-</td><td>0</td><td>GTCTTCGGAGCTGTACGCTA</td><td>GGG</td><td>||||||||[||||||||||||]PAM</td><td>NA</td><td>Intergenic</td><td>NA</td><td>NA</td></tr>
<tr><td></td><td></td><td></td><td></td><td></td><td></td><td></td><td></td><td></td><td></td><td></td><td></td></tr>
<tr><td>T32971</td><td>GTGATCGCCGTCGTCGCCGTCGG</td><td>1000</td><td></td><td></td><td></td><td></td><td></td><td></td><td></td><td></td><td></td></tr>
<tr><td>Oligo fwd</td><td>CACCGTGATCGCCGTCGTCGCCGT</td><td></td><td></td><td></td><td></td><td></td><td></td><td></td><td></td><td></td><td></td></tr>
<tr><td>Oligo rev</td><td>AAACACGGCGACGACGGCGATCAC</td><td></td><td></td><td></td><td></td><td></td><td></td><td></td><td></td><td></td><td></td></tr>
<tr><td>Chromosome</td><td>start</td><td>end</td><td>strand</td><td>MM</td><td>target_seq</td><td>PAM</td><td>alignment</td><td>distance</td><td>position</td><td>gene name</td><td>gene id</td></tr>
<tr><td>GU305914.1</td><td>20748</td><td>20770</td><td>-</td><td>0</td><td>GTGATCGCCGTCGTCGCCGT</td><td>CGG</td><td>||||||||[||||||||||||]PAM</td><td>NA</td><td>Intergenic</td><td>NA</td><td>NA</td></tr>
<tr><td>GU305914.1</td><td>31683</td><td>31705</td><td>-</td><td>4</td><td>GTGGTCGGCGTCAACGCCGT</td><td>GGG</td><td>|||-|||-[||||--||||||]PAM</td><td>NA</td><td>Intergenic</td><td>NA</td><td>NA</td></tr>
<tr><td></td><td></td><td></td><td></td><td></td><td></td><td></td><td></td><td></td><td></td><td></td><td></td></tr>
<tr><td>T32972</td><td>TCGGTGCCGTCGTGTGCTACTGG</td><td>1000</td><td></td><td></td><td></td><td></td><td></td><td></td><td></td><td></td><td></td></tr>
<tr><td>Oligo fwd</td><td>CACCTCGGTGCCGTCGTGTGCTAC</td><td></td><td></td><td></td><td></td><td></td><td></td><td></td><td></td><td></td><td></td></tr>
<tr><td>Oligo rev</td><td>AAACGTAGCACACGACGGCACCGA</td><td></td><td></td><td></td><td></td><td></td><td></td><td></td><td></td><td></td><td></td></tr>
<tr><td>Chromosome</td><td>start</td><td>end</td><td>strand</td><td>MM</td><td>target_seq</td><td>PAM</td><td>alignment</td><td>distance</td><td>position</td><td>gene name</td><td>gene id</td></tr>
<tr><td>GU305914.1</td><td>20729</td><td>20751</td><td>-</td><td>0</td><td>TCGGTGCCGTCGTGTGCTAC</td><td>TGG</td><td>||||||||[||||||||||||]PAM</td><td>NA</td><td>Intergenic</td><td>NA</td><td>NA</td></tr>
<tr><td></td><td></td><td></td><td></td><td></td><td></td><td></td><td></td><td></td><td></td><td></td><td></td></tr>
<tr><td>T32973</td><td>CGGTGCCGTCGTGTGCTACTGGG</td><td>1000</td><td></td><td></td><td></td><td></td><td></td><td></td><td></td><td></td><td></td></tr>
<tr><td>Oligo fwd</td><td>CACCCGGTGCCGTCGTGTGCTACT</td><td></td><td></td><td></td><td></td><td></td><td></td><td></td><td></td><td></td><td></td></tr>
<tr><td>Oligo rev</td><td>AAACAGTAGCACACGACGGCACCG</td><td></td><td></td><td></td><td></td><td></td><td></td><td></td><td></td><td></td><td></td></tr>
<tr><td>Chromosome</td><td>start</td><td>end</td><td>strand</td><td>MM</td><td>target_seq</td><td>PAM</td><td>alignment</td><td>distance</td><td>position</td><td>gene name</td><td>gene id</td></tr>
<tr><td>GU305914.1</td><td>20728</td><td>20750</td><td>-</td><td>0</td><td>CGGTGCCGTCGTGTGCTACT</td><td>GGG</td><td>||||||||[||||||||||||]PAM</td><td>NA</td><td>Intergenic</td><td>NA</td><td>NA</td></tr>
<tr><td></td><td></td><td></td><td></td><td></td><td></td><td></td><td></td><td></td><td></td><td></td><td></td></tr>
<tr><td>T32974</td><td>GGTGCCGTCGTGTGCTACTGGGG</td><td>1000</td><td></td><td></td><td></td><td></td><td></td><td></td><td></td><td></td><td></td></tr>
<tr><td>Oligo fwd</td><td>CACCGGTGCCGTCGTGTGCTACTG</td><td></td><td></td><td></td><td></td><td></td><td></td><td></td><td></td><td></td><td></td></tr>
<tr><td>Oligo rev</td><td>AAACCAGTAGCACACGACGGCACC</td><td></td><td></td><td></td><td></td><td></td><td></td><td></td><td></td><td></td><td></td></tr>
<tr><td>Chromosome</td><td>start</td><td>end</td><td>strand</td><td>MM</td><td>target_seq</td><td>PAM</td><td>alignment</td><td>distance</td><td>position</td><td>gene name</td><td>gene id</td></tr>
<tr><td>GU305914.1</td><td>20727</td><td>20749</td><td>-</td><td>0</td><td>GGTGCCGTCGTGTGCTACTG</td><td>GGG</td><td>||||||||[||||||||||||]PAM</td><td>NA</td><td>Intergenic</td><td>NA</td><td>NA</td></tr>
<tr><td></td><td></td><td></td><td></td><td></td><td></td><td></td><td></td><td></td><td></td><td></td><td></td></tr>
<tr><td>T32975</td><td>GTGTGCTACTGGGGACTGCCCGG</td><td>1000</td><td></td><td></td><td></td><td></td><td></td><td></td><td></td><td></td><td></td></tr>
<tr><td>Oligo fwd</td><td>CACCGTGTGCTACTGGGGACTGCC</td><td></td><td></td><td></td><td></td><td></td><td></td><td></td><td></td><td></td><td></td></tr>
<tr><td>Oligo rev</td><td>AAACGGCAGTCCCCAGTAGCACAC</td><td></td><td></td><td></td><td></td><td></td><td></td><td></td><td></td><td></td><td></td></tr>
<tr><td>Chromosome</td><td>start</td><td>end</td><td>strand</td><td>MM</td><td>target_seq</td><td>PAM</td><td>alignment</td><td>distance</td><td>position</td><td>gene name</td><td>gene id</td></tr>
<tr><td>GU305914.1</td><td>20718</td><td>20740</td><td>-</td><td>0</td><td>GTGTGCTACTGGGGACTGCC</td><td>CGG</td><td>||||||||[||||||||||||]PAM</td><td>NA</td><td>Intergenic</td><td>NA</td><td>NA</td></tr>
<tr><td></td><td></td><td></td><td></td><td></td><td></td><td></td><td></td><td></td><td></td><td></td><td></td></tr>
<tr><td>T32976</td><td>TGGGGACTGCCCGGTGCCAGCGG</td><td>1000</td><td></td><td></td><td></td><td></td><td></td><td></td><td></td><td></td><td></td></tr>
<tr><td>Oligo fwd</td><td>CACCTGGGGACTGCCCGGTGCCAG</td><td></td><td></td><td></td><td></td><td></td><td></td><td></td><td></td><td></td><td></td></tr>
<tr><td>Oligo rev</td><td>AAACCTGGCACCGGGCAGTCCCCA</td><td></td><td></td><td></td><td></td><td></td><td></td><td></td><td></td><td></td><td></td></tr>
<tr><td>Chromosome</td><td>start</td><td>end</td><td>strand</td><td>MM</td><td>target_seq</td><td>PAM</td><td>alignment</td><td>distance</td><td>position</td><td>gene name</td><td>gene id</td></tr>
<tr><td>GU305914.1</td><td>20709</td><td>20731</td><td>-</td><td>0</td><td>TGGGGACTGCCCGGTGCCAG</td><td>CGG</td><td>||||||||[||||||||||||]PAM</td><td>NA</td><td>Intergenic</td><td>NA</td><td>NA</td></tr>
<tr><td></td><td></td><td></td><td></td><td></td><td></td><td></td><td></td><td></td><td></td><td></td><td></td></tr>
<tr><td>T32977</td><td>ACTGCCCGGTGCCAGCGGCGTGG</td><td>1000</td><td></td><td></td><td></td><td></td><td></td><td></td><td></td><td></td><td></td></tr>
<tr><td>Oligo fwd</td><td>CACCACTGCCCGGTGCCAGCGGCG</td><td></td><td></td><td></td><td></td><td></td><td></td><td></td><td></td><td></td><td></td></tr>
<tr><td>Oligo rev</td><td>AAACCGCCGCTGGCACCGGGCAGT</td><td></td><td></td><td></td><td></td><td></td><td></td><td></td><td></td><td></td><td></td></tr>
<tr><td>Chromosome</td><td>start</td><td>end</td><td>strand</td><td>MM</td><td>target_seq</td><td>PAM</td><td>alignment</td><td>distance</td><td>position</td><td>gene name</td><td>gene id</td></tr>
<tr><td>GU305914.1</td><td>20704</td><td>20726</td><td>-</td><td>0</td><td>ACTGCCCGGTGCCAGCGGCG</td><td>TGG</td><td>||||||||[||||||||||||]PAM</td><td>NA</td><td>Intergenic</td><td>NA</td><td>NA</td></tr>
<tr><td></td><td></td><td></td><td></td><td></td><td></td><td></td><td></td><td></td><td></td><td></td><td></td></tr>
<tr><td>T32978</td><td>GCCCGGTGCCAGCGGCGTGGTGG</td><td>1000</td><td></td><td></td><td></td><td></td><td></td><td></td><td></td><td></td><td></td></tr>
<tr><td>Oligo fwd</td><td>CACCGCCCGGTGCCAGCGGCGTGG</td><td></td><td></td><td></td><td></td><td></td><td></td><td></td><td></td><td></td><td></td></tr>
<tr><td>Oligo rev</td><td>AAACCCACGCCGCTGGCACCGGGC</td><td></td><td></td><td></td><td></td><td></td><td></td><td></td><td></td><td></td><td></td></tr>
<tr><td>Chromosome</td><td>start</td><td>end</td><td>strand</td><td>MM</td><td>target_seq</td><td>PAM</td><td>alignment</td><td>distance</td><td>position</td><td>gene name</td><td>gene id</td></tr>
<tr><td>GU305914.1</td><td>20701</td><td>20723</td><td>-</td><td>0</td><td>GCCCGGTGCCAGCGGCGTGG</td><td>TGG</td><td>||||||||[||||||||||||]PAM</td><td>NA</td><td>Intergenic</td><td>NA</td><td>NA</td></tr>
<tr><td></td><td></td><td></td><td></td><td></td><td></td><td></td><td></td><td></td><td></td><td></td><td></td></tr>
<tr><td>T32979</td><td>GGTGGAAGACGCGTACTCCGAGG</td><td>1000</td><td></td><td></td><td></td><td></td><td></td><td></td><td></td><td></td><td></td></tr>
<tr><td>Oligo fwd</td><td>CACCGGTGGAAGACGCGTACTCCG</td><td></td><td></td><td></td><td></td><td></td><td></td><td></td><td></td><td></td><td></td></tr>
<tr><td>Oligo rev</td><td>AAACCGGAGTACGCGTCTTCCACC</td><td></td><td></td><td></td><td></td><td></td><td></td><td></td><td></td><td></td><td></td></tr>
<tr><td>Chromosome</td><td>start</td><td>end</td><td>strand</td><td>MM</td><td>target_seq</td><td>PAM</td><td>alignment</td><td>distance</td><td>position</td><td>gene name</td><td>gene id</td></tr>
<tr><td>GU305914.1</td><td>20683</td><td>20705</td><td>-</td><td>0</td><td>GGTGGAAGACGCGTACTCCG</td><td>AGG</td><td>||||||||[||||||||||||]PAM</td><td>NA</td><td>Intergenic</td><td>NA</td><td>NA</td></tr>
<tr><td></td><td></td><td></td><td></td><td></td><td></td><td></td><td></td><td></td><td></td><td></td><td></td></tr>
<tr><td>T32980</td><td>GCGTACTCCGAGGCCTGCGACGG</td><td>1000</td><td></td><td></td><td></td><td></td><td></td><td></td><td></td><td></td><td></td></tr>
<tr><td>Oligo fwd</td><td>CACCGCGTACTCCGAGGCCTGCGA</td><td></td><td></td><td></td><td></td><td></td><td></td><td></td><td></td><td></td><td></td></tr>
<tr><td>Oligo rev</td><td>AAACTCGCAGGCCTCGGAGTACGC</td><td></td><td></td><td></td><td></td><td></td><td></td><td></td><td></td><td></td><td></td></tr>
<tr><td>Chromosome</td><td>start</td><td>end</td><td>strand</td><td>MM</td><td>target_seq</td><td>PAM</td><td>alignment</td><td>distance</td><td>position</td><td>gene name</td><td>gene id</td></tr>
<tr><td>GU305914.1</td><td>20673</td><td>20695</td><td>-</td><td>0</td><td>GCGTACTCCGAGGCCTGCGA</td><td>CGG</td><td>||||||||[||||||||||||]PAM</td><td>NA</td><td>Intergenic</td><td>NA</td><td>NA</td></tr>
</table>

| | | | | | | | | | | | |
|---|---|---|---|---|---|---|---|---|---|---|---|
| T32981 | TCCGAGGCCTGCGACGGAGACGG | 1000 | | | | | | | | | |
| Oligo fwd | CACCTCCGAGGCCTGCGACGGAGA | | | | | | | | | | |
| Oligo rev | AAACTCTCCGTCGCAGGCCTCGGA | | | | | | | | | | |
| Chromosome | start | end | strand | MM | target_seq | PAM | alignment | distance | position | gene name | gene id |
| GU305914.1 | 20667 | 20689 | - | 0 | TCCGAGGCCTGCGACGGAGA | CGG | \|\|\|\|\|\|\|\|[\|\|\|\|\|\|\|\|\|\|\|\|]PAM | NA | Intergenic | NA | NA |
| T32982 | CTGCGACGGAGACGGCGATACGG | 1000 | | | | | | | | | |
| Oligo fwd | CACCCTGCGACGGAGACGGCGATA | | | | | | | | | | |
| Oligo rev | AAACTATCGCCGTCTCCGTCGCAG | | | | | | | | | | |
| Chromosome | start | end | strand | MM | target_seq | PAM | alignment | distance | position | gene name | gene id |
| GU305914.1 | 20659 | 20681 | - | 0 | CTGCGACGGAGACGGCGATA | CGG | \|\|\|\|\|\|\|\|[\|\|\|\|\|\|\|\|\|\|\|\|]PAM | NA | Intergenic | NA | NA |
| T32983 | CGGAGACGGCGATACGGTCGCGG | 1000 | | | | | | | | | |
| Oligo fwd | CACCCGGAGACGGCGATACGGTCG | | | | | | | | | | |
| Oligo rev | AAACCGACCGTATCGCCGTCTCCG | | | | | | | | | | |
| Chromosome | start | end | strand | MM | target_seq | PAM | alignment | distance | position | gene name | gene id |
| GU305914.1 | 20653 | 20675 | - | 0 | CGGAGACGGCGATACGGTCG | CGG | \|\|\|\|\|\|\|\|[\|\|\|\|\|\|\|\|\|\|\|\|]PAM | NA | Intergenic | NA | NA |
| T32984 | TACGGTCGCGGAACTCGTCCTGG | 1000 | | | | | | | | | |
| Oligo fwd | CACCTACGGTCGCGGAACTCGTCC | | | | | | | | | | |
| Oligo rev | AAACGGACGAGTTCCGCGACCGTA | | | | | | | | | | |
| Chromosome | start | end | strand | MM | target_seq | PAM | alignment | distance | position | gene name | gene id |
| GU305914.1 | 20641 | 20663 | - | 0 | TACGGTCGCGGAACTCGTCC | TGG | \|\|\|\|\|\|\|\|[\|\|\|\|\|\|\|\|\|\|\|\|]PAM | NA | Intergenic | NA | NA |
| GU305914.1 | 164662 | 164684 | - | 4 | TACGGCGGCGGAACTCGAGC | TGG | \|\|\|\|\|--\|[\|\|\|\|\|\|\|\|\|--\|]PAM | NA | Intergenic | NA | NA |
| T32985 | CGCGGAACTCGTCCTGGAAGAGG | 1000 | | | | | | | | | |
| Oligo fwd | CACCCGCGGAACTCGTCCTGGAAG | | | | | | | | | | |
| Oligo rev | AAACCTTCCAGGACGAGTTCCGCG | | | | | | | | | | |
| Chromosome | start | end | strand | MM | target_seq | PAM | alignment | distance | position | gene name | gene id |
| GU305914.1 | 20635 | 20657 | - | 0 | CGCGGAACTCGTCCTGGAAG | AGG | \|\|\|\|\|\|\|\|[\|\|\|\|\|\|\|\|\|\|\|\|]PAM | NA | Intergenic | NA | NA |
| T32986 | CGTCCTGGAAGAGGAACAGATGG | 1000 | | | | | | | | | |
| Oligo fwd | CACCCGTCCTGGAAGAGGAACAGA | | | | | | | | | | |
| Oligo rev | AAACTCTGTTCCTCTTCCAGGACG | | | | | | | | | | |
| Chromosome | start | end | strand | MM | target_seq | PAM | alignment | distance | position | gene name | gene id |
| GU305914.1 | 20626 | 20648 | - | 0 | CGTCCTGGAAGAGGAACAGA | TGG | \|\|\|\|\|\|\|\|[\|\|\|\|\|\|\|\|\|\|\|\|]PAM | NA | Intergenic | NA | NA |
| T32987 | AGAGGAACAGATGGCCGTCGAGG | 1000 | | | | | | | | | |
| Oligo fwd | CACCAGAGGAACAGATGGCCGTCG | | | | | | | | | | |
| Oligo rev | AAACCGACGGCCATCTGTTCCTCT | | | | | | | | | | |
| Chromosome | start | end | strand | MM | target_seq | PAM | alignment | distance | position | gene name | gene id |
| GU305914.1 | 20617 | 20639 | - | 0 | AGAGGAACAGATGGCCGTCG | AGG | \|\|\|\|\|\|\|\|[\|\|\|\|\|\|\|\|\|\|\|\|]PAM | NA | Intergenic | NA | NA |
| T32988 | CGTCGAGGAGCAGTCGCCCAAGG | 1000 | | | | | | | | | |
| Oligo fwd | CACCCGTCGAGGAGCAGTCGCCCA | | | | | | | | | | |
| Oligo rev | AAACTGGGCGACTGCTCCTCGACG | | | | | | | | | | |
| Chromosome | start | end | strand | MM | target_seq | PAM | alignment | distance | position | gene name | gene id |
| GU305914.1 | 20602 | 20624 | - | 0 | CGTCGAGGAGCAGTCGCCCA | AGG | \|\|\|\|\|\|\|\|[\|\|\|\|\|\|\|\|\|\|\|\|]PAM | NA | Intergenic | NA | NA |
| T32989 | TAACTAGAACATCCCGACCGCGG | 1000 | | | | | | | | | |
| Oligo fwd | CACCTAACTAGAACATCCCGACCG | | | | | | | | | | |
| Oligo rev | AAACCGGTCGGGATGTTCTAGTTA | | | | | | | | | | |
| Chromosome | start | end | strand | MM | target_seq | PAM | alignment | distance | position | gene name | gene id |
| GU305914.1 | 20514 | 20536 | - | 0 | TAACTAGAACATCCCGACCG | CGG | \|\|\|\|\|\|\|\|[\|\|\|\|\|\|\|\|\|\|\|\|]PAM | NA | Intergenic | NA | NA |
| T32990 | AACTAGAACATCCCGACCGCGGG | 1000 | | | | | | | | | |
| Oligo fwd | CACCAACTAGAACATCCCGACCGC | | | | | | | | | | |
| Oligo rev | AAACGCGGTCGGGATGTTCTAGTT | | | | | | | | | | |
| Chromosome | start | end | strand | MM | target_seq | PAM | alignment | distance | position | gene name | gene id |
| GU305914.1 | 20513 | 20535 | - | 0 | AACTAGAACATCCCGACCGC | GGG | \|\|\|\|\|\|\|\|[\|\|\|\|\|\|\|\|\|\|\|\|]PAM | NA | Intergenic | NA | NA |
| T32991 | CATCCCGACCGCGGGACTGAAGG | 1000 | | | | | | | | | |
| Oligo fwd | CACCCATCCCGACCGCGGGACTGA | | | | | | | | | | |
| Oligo rev | AAACTCAGTCCCGCGGTCGGGATG | | | | | | | | | | |
| Chromosome | start | end | strand | MM | target_seq | PAM | alignment | distance | position | gene name | gene id |
| GU305914.1 | 20505 | 20527 | - | 0 | CATCCCGACCGCGGGACTGA | AGG | \|\|\|\|\|\|\|\|[\|\|\|\|\|\|\|\|\|\|\|\|]PAM | NA | Intergenic | NA | NA |
| T32992 | CCCGACCGCGGGACTGAAGGAGG | 1000 | | | | | | | | | |
| Oligo fwd | CACCCCCGACCGCGGGACTGAAGG | | | | | | | | | | |
| Oligo rev | AAACCCTTCAGTCCCGCGGTCGGG | | | | | | | | | | |
| Chromosome | start | end | strand | MM | target_seq | PAM | alignment | distance | position | gene name | gene id |
| GU305914.1 | 20502 | 20524 | - | 0 | CCCGACCGCGGGACTGAAGG | AGG | \|\|\|\|\|\|\|\|[\|\|\|\|\|\|\|\|\|\|\|\|]PAM | NA | Intergenic | NA | NA |
| GU305914.1 | 121707 | 121729 | + | 4 | CGCGTCCGCGGGCGTGAAGG | TGG | \|-\|\|-\|\|\|[\|\|\|\|--\|\|\|\|\|\|]PAM | NA | Intergenic | NA | NA |
| T32993 | CCGCGGGACTGAAGGAGGCCCGG | 1000 | | | | | | | | | |
| Oligo fwd | CACCCCGCGGGACTGAAGGAGGCC | | | | | | | | | | |
| Oligo rev | AAACGGCCTCCTTCAGTCCCGCGG | | | | | | | | | | |
| Chromosome | start | end | strand | MM | target_seq | PAM | alignment | distance | position | gene name | gene id |
| GU305914.1 | 20497 | 20519 | - | 0 | CCGCGGGACTGAAGGAGGCC | CGG | \|\|\|\|\|\|\|\|[\|\|\|\|\|\|\|\|\|\|\|\|]PAM | NA | Intergenic | NA | NA |
| T32994 | GGGACTGAAGGAGGCCCGGCCGG | 1000 | | | | | | | | | |
| Oligo fwd | CACCGGGACTGAAGGAGGCCCGGC | | | | | | | | | | |
| Oligo rev | AAACGCCGGGCCTCCTTCAGTCCC | | | | | | | | | | |
| Chromosome | start | end | strand | MM | target_seq | PAM | alignment | distance | position | gene name | gene id |
| GU305914.1 | 20493 | 20515 | - | 0 | GGGACTGAAGGAGGCCCGGC | CGG | \|\|\|\|\|\|\|\|[\|\|\|\|\|\|\|\|\|\|\|\|]PAM | NA | Intergenic | NA | NA |
| T32995 | CTGAAGGAGGCCCGGCCGGCTGG | 1000 | | | | | | | | | |
| Oligo fwd | CACCCTGAAGGAGGCCCGGCCGGC | | | | | | | | | | |

<table>
<tr><td>Oligo rev</td><td>AAACGCCGGCCGGGCCTCCTTCAG</td></tr>
<tr><td>Chromosome</td><td>start</td><td>end</td><td>strand</td><td>MM</td><td>target_seq</td><td>PAM</td><td>alignment</td><td>distance</td><td>position</td><td>gene name</td><td>gene id</td></tr>
<tr><td>GU305914.1</td><td>20489</td><td>20511</td><td>-</td><td>0</td><td>CTGAAGGAGGCCCGGCCGGC</td><td>TGG</td><td>||||||||[||||||||||||]PAM</td><td>NA</td><td>Intergenic</td><td>NA</td><td>NA</td></tr>
<tr><td>T32996</td><td>AGGAGGCCCGGCCGGCTGGCTGG</td><td>1000</td></tr>
<tr><td>Oligo fwd</td><td>CACCAGGAGGCCCGGCCGGCTGGC</td></tr>
<tr><td>Oligo rev</td><td>AAACGCCAGCCGGCCGGGCCTCCT</td></tr>
<tr><td>Chromosome</td><td>start</td><td>end</td><td>strand</td><td>MM</td><td>target_seq</td><td>PAM</td><td>alignment</td><td>distance</td><td>position</td><td>gene name</td><td>gene id</td></tr>
<tr><td>GU305914.1</td><td>20485</td><td>20507</td><td>-</td><td>0</td><td>AGGAGGCCCGGCCGGCTGGC</td><td>TGG</td><td>||||||||[||||||||||||]PAM</td><td>NA</td><td>Intergenic</td><td>NA</td><td>NA</td></tr>
<tr><td>T32997</td><td>AGGCCCGGCCGGCTGGCTGGCGG</td><td>1000</td></tr>
<tr><td>Oligo fwd</td><td>CACCAGGCCCGGCCGGCTGGCTGG</td></tr>
<tr><td>Oligo rev</td><td>AAACCCAGCCAGCCGGCCGGGCCT</td></tr>
<tr><td>Chromosome</td><td>start</td><td>end</td><td>strand</td><td>MM</td><td>target_seq</td><td>PAM</td><td>alignment</td><td>distance</td><td>position</td><td>gene name</td><td>gene id</td></tr>
<tr><td>GU305914.1</td><td>20482</td><td>20504</td><td>-</td><td>0</td><td>AGGCCCGGCCGGCTGGCTGG</td><td>CGG</td><td>||||||||[||||||||||||]PAM</td><td>NA</td><td>Intergenic</td><td>NA</td><td>NA</td></tr>
<tr><td>T32998</td><td>GGCCCGGCCGGCTGGCTGGCGGG</td><td>1000</td></tr>
<tr><td>Oligo fwd</td><td>CACCGGCCCGGCCGGCTGGCTGGC</td></tr>
<tr><td>Oligo rev</td><td>AAACGCCAGCCAGCCGGCCGGGCC</td></tr>
<tr><td>Chromosome</td><td>start</td><td>end</td><td>strand</td><td>MM</td><td>target_seq</td><td>PAM</td><td>alignment</td><td>distance</td><td>position</td><td>gene name</td><td>gene id</td></tr>
<tr><td>GU305914.1</td><td>20481</td><td>20503</td><td>-</td><td>0</td><td>GGCCCGGCCGGCTGGCTGGC</td><td>GGG</td><td>||||||||[||||||||||||]PAM</td><td>NA</td><td>Intergenic</td><td>NA</td><td>NA</td></tr>
<tr><td>T32999</td><td>CCGGCTGGCTGGCGGGCGTGCGG</td><td>1000</td></tr>
<tr><td>Oligo fwd</td><td>CACCCCGGCTGGCTGGCGGGCGTG</td></tr>
<tr><td>Oligo rev</td><td>AAACCACGCCCGCCAGCCAGCCGG</td></tr>
<tr><td>Chromosome</td><td>start</td><td>end</td><td>strand</td><td>MM</td><td>target_seq</td><td>PAM</td><td>alignment</td><td>distance</td><td>position</td><td>gene name</td><td>gene id</td></tr>
<tr><td>GU305914.1</td><td>20474</td><td>20496</td><td>-</td><td>0</td><td>CCGGCTGGCTGGCGGGCGTG</td><td>CGG</td><td>||||||||[||||||||||||]PAM</td><td>NA</td><td>Intergenic</td><td>NA</td><td>NA</td></tr>
<tr><td>T33000</td><td>TGGCTGGCGGGCGTGCGGTCAGG</td><td>1000</td></tr>
<tr><td>Oligo fwd</td><td>CACCTGGCTGGCGGGCGTGCGGTC</td></tr>
<tr><td>Oligo rev</td><td>AAACGACCGCACGCCCGCCAGCCA</td></tr>
<tr><td>Chromosome</td><td>start</td><td>end</td><td>strand</td><td>MM</td><td>target_seq</td><td>PAM</td><td>alignment</td><td>distance</td><td>position</td><td>gene name</td><td>gene id</td></tr>
<tr><td>GU305914.1</td><td>20469</td><td>20491</td><td>-</td><td>0</td><td>TGGCTGGCGGGCGTGCGGTC</td><td>AGG</td><td>||||||||[||||||||||||]PAM</td><td>NA</td><td>Intergenic</td><td>NA</td><td>NA</td></tr>
<tr><td>T33001</td><td>CGATGTCCGTGTCTCACTTCTGG</td><td>1000</td></tr>
<tr><td>Oligo fwd</td><td>CACCCGATGTCCGTGTCTCACTTC</td></tr>
<tr><td>Oligo rev</td><td>AAACGAAGTGAGACACGGACATCG</td></tr>
<tr><td>Chromosome</td><td>start</td><td>end</td><td>strand</td><td>MM</td><td>target_seq</td><td>PAM</td><td>alignment</td><td>distance</td><td>position</td><td>gene name</td><td>gene id</td></tr>
<tr><td>GU305914.1</td><td>20408</td><td>20430</td><td>-</td><td>0</td><td>CGATGTCCGTGTCTCACTTC</td><td>TGG</td><td>||||||||[||||||||||||]PAM</td><td>NA</td><td>Intergenic</td><td>NA</td><td>NA</td></tr>
<tr><td>T33002</td><td>TCTCACTTCTGGTTTCTTTGAGG</td><td>1000</td></tr>
<tr><td>Oligo fwd</td><td>CACCTCTCACTTCTGGTTTCTTTG</td></tr>
<tr><td>Oligo rev</td><td>AAACCAAAGAAACCAGAAGTGAGA</td></tr>
<tr><td>Chromosome</td><td>start</td><td>end</td><td>strand</td><td>MM</td><td>target_seq</td><td>PAM</td><td>alignment</td><td>distance</td><td>position</td><td>gene name</td><td>gene id</td></tr>
<tr><td>GU305914.1</td><td>20397</td><td>20419</td><td>-</td><td>0</td><td>TCTCACTTCTGGTTTCTTTG</td><td>AGG</td><td>||||||||[||||||||||||]PAM</td><td>NA</td><td>Intergenic</td><td>NA</td><td>NA</td></tr>
<tr><td>T33003</td><td>TTGAGGATCTCGAGATGAACCGG</td><td>1000</td></tr>
<tr><td>Oligo fwd</td><td>CACCTTGAGGATCTCGAGATGAAC</td></tr>
<tr><td>Oligo rev</td><td>AAACGTTCATCTCGAGATCCTCAA</td></tr>
<tr><td>Chromosome</td><td>start</td><td>end</td><td>strand</td><td>MM</td><td>target_seq</td><td>PAM</td><td>alignment</td><td>distance</td><td>position</td><td>gene name</td><td>gene id</td></tr>
<tr><td>GU305914.1</td><td>20380</td><td>20402</td><td>-</td><td>0</td><td>TTGAGGATCTCGAGATGAAC</td><td>CGG</td><td>||||||||[||||||||||||]PAM</td><td>NA</td><td>Intergenic</td><td>NA</td><td>NA</td></tr>
<tr><td>T33004</td><td>TGAGGATCTCGAGATGAACCGGG</td><td>1000</td></tr>
<tr><td>Oligo fwd</td><td>CACCTGAGGATCTCGAGATGAACC</td></tr>
<tr><td>Oligo rev</td><td>AAACGGTTCATCTCGAGATCCTCA</td></tr>
<tr><td>Chromosome</td><td>start</td><td>end</td><td>strand</td><td>MM</td><td>target_seq</td><td>PAM</td><td>alignment</td><td>distance</td><td>position</td><td>gene name</td><td>gene id</td></tr>
<tr><td>GU305914.1</td><td>20379</td><td>20401</td><td>-</td><td>0</td><td>TGAGGATCTCGAGATGAACC</td><td>GGG</td><td>||||||||[||||||||||||]PAM</td><td>NA</td><td>Intergenic</td><td>NA</td><td>NA</td></tr>
<tr><td>T33005</td><td>GAGGATCTCGAGATGAACCGGGG</td><td>1000</td></tr>
<tr><td>Oligo fwd</td><td>CACCGAGGATCTCGAGATGAACCG</td></tr>
<tr><td>Oligo rev</td><td>AAACCGGTTCATCTCGAGATCCTC</td></tr>
<tr><td>Chromosome</td><td>start</td><td>end</td><td>strand</td><td>MM</td><td>target_seq</td><td>PAM</td><td>alignment</td><td>distance</td><td>position</td><td>gene name</td><td>gene id</td></tr>
<tr><td>GU305914.1</td><td>20378</td><td>20400</td><td>-</td><td>0</td><td>GAGGATCTCGAGATGAACCG</td><td>GGG</td><td>||||||||[||||||||||||]PAM</td><td>NA</td><td>Intergenic</td><td>NA</td><td>NA</td></tr>
<tr><td>T33006</td><td>GATGAACCGGGGAAACATAATGG</td><td>1000</td></tr>
<tr><td>Oligo fwd</td><td>CACCGATGAACCGGGGAAACATAA</td></tr>
<tr><td>Oligo rev</td><td>AAACTTATGTTTCCCCGGTTCATC</td></tr>
<tr><td>Chromosome</td><td>start</td><td>end</td><td>strand</td><td>MM</td><td>target_seq</td><td>PAM</td><td>alignment</td><td>distance</td><td>position</td><td>gene name</td><td>gene id</td></tr>
<tr><td>GU305914.1</td><td>20367</td><td>20389</td><td>-</td><td>0</td><td>GATGAACCGGGGAAACATAA</td><td>TGG</td><td>||||||||[||||||||||||]PAM</td><td>NA</td><td>Intergenic</td><td>NA</td><td>NA</td></tr>
<tr><td>T33007</td><td>AACCGGGGAAACATAATGGTAGG</td><td>1000</td></tr>
<tr><td>Oligo fwd</td><td>CACCAACCGGGGAAACATAATGGT</td></tr>
<tr><td>Oligo rev</td><td>AAACACCATTATGTTTCCCCGGTT</td></tr>
<tr><td>Chromosome</td><td>start</td><td>end</td><td>strand</td><td>MM</td><td>target_seq</td><td>PAM</td><td>alignment</td><td>distance</td><td>position</td><td>gene name</td><td>gene id</td></tr>
<tr><td>GU305914.1</td><td>20363</td><td>20385</td><td>-</td><td>0</td><td>AACCGGGGAAACATAATGGT</td><td>AGG</td><td>||||||||[||||||||||||]PAM</td><td>NA</td><td>Intergenic</td><td>NA</td><td>NA</td></tr>
<tr><td>T33008</td><td>ACCGGGGAAACATAATGGTAGGG</td><td>1000</td></tr>
<tr><td>Oligo fwd</td><td>CACCACCGGGGAAACATAATGGTA</td></tr>
<tr><td>Oligo rev</td><td>AAACTACCATTATGTTTCCCCGGT</td></tr>
<tr><td>Chromosome</td><td>start</td><td>end</td><td>strand</td><td>MM</td><td>target_seq</td><td>PAM</td><td>alignment</td><td>distance</td><td>position</td><td>gene name</td><td>gene id</td></tr>
<tr><td>GU305914.1</td><td>20362</td><td>20384</td><td>-</td><td>0</td><td>ACCGGGGAAACATAATGGTA</td><td>GGG</td><td>||||||||[||||||||||||]PAM</td><td>NA</td><td>Intergenic</td><td>NA</td><td>NA</td></tr>
<tr><td>T33009</td><td>CCGGGGAAACATAATGGTAGGGG</td><td>1000</td></tr>
<tr><td>Oligo fwd</td><td>CACCCCGGGGAAACATAATGGTAG</td></tr>
<tr><td>Oligo rev</td><td>AAACCTACCATTATGTTTCCCCGG</td></tr>
<tr><td>Chromosome</td><td>start</td><td>end</td><td>strand</td><td>MM</td><td>target_seq</td><td>PAM</td><td>alignment</td><td>distance</td><td>position</td><td>gene name</td><td>gene id</td></tr>
<tr><td>GU305914.1</td><td>20361</td><td>20383</td><td>-</td><td>0</td><td>CCGGGGAAACATAATGGTAG</td><td>GGG</td><td>||||||||[||||||||||||]PAM</td><td>NA</td><td>Intergenic</td><td>NA</td><td>NA</td></tr>
<tr><td>T33010</td><td>GGAAACATAATGGTAGGGGATGG</td><td>1000</td></tr>
</table>

<table>
<tr><td>Oligo fwd</td><td>CACCGGAAACATAATGGTAGGGGA</td><td></td><td></td><td></td><td></td><td></td><td></td><td></td><td></td><td></td><td></td><td></td></tr>
<tr><td>Oligo rev</td><td>AAACTCCCCTACCATTATGTTTCC</td><td></td><td></td><td></td><td></td><td></td><td></td><td></td><td></td><td></td><td></td><td></td></tr>
<tr><td>Chromosome</td><td>start</td><td></td><td>end</td><td>strand</td><td>MM</td><td>target_seq</td><td>PAM</td><td>alignment</td><td>distance</td><td>position</td><td>gene name</td><td>gene id</td></tr>
<tr><td>GU305914.1</td><td></td><td>20357</td><td>20379</td><td>-</td><td>0</td><td>GGAAACATAATGGTAGGGGA</td><td>TGG</td><td>||||||||[||||||||||||]PAM</td><td>NA</td><td>Intergenic</td><td>NA</td><td>NA</td></tr>
<tr><td>T33011</td><td>GAAACATAATGGTAGGGGATGGG</td><td></td><td>1000</td><td></td><td></td><td></td><td></td><td></td><td></td><td></td><td></td><td></td></tr>
<tr><td>Oligo fwd</td><td>CACCGAAACATAATGGTAGGGGAT</td><td></td><td></td><td></td><td></td><td></td><td></td><td></td><td></td><td></td><td></td><td></td></tr>
<tr><td>Oligo rev</td><td>AAACATCCCCTACCATTATGTTTC</td><td></td><td></td><td></td><td></td><td></td><td></td><td></td><td></td><td></td><td></td><td></td></tr>
<tr><td>Chromosome</td><td>start</td><td></td><td>end</td><td>strand</td><td>MM</td><td>target_seq</td><td>PAM</td><td>alignment</td><td>distance</td><td>position</td><td>gene name</td><td>gene id</td></tr>
<tr><td>GU305914.1</td><td></td><td>20356</td><td>20378</td><td>-</td><td>0</td><td>GAAACATAATGGTAGGGGAT</td><td>GGG</td><td>||||||||[||||||||||||]PAM</td><td>NA</td><td>Intergenic</td><td>NA</td><td>NA</td></tr>
<tr><td>T33012</td><td>ATAATGGTAGGGGATGGGTGTGG</td><td></td><td>1000</td><td></td><td></td><td></td><td></td><td></td><td></td><td></td><td></td><td></td></tr>
<tr><td>Oligo fwd</td><td>CACCATAATGGTAGGGGATGGGTG</td><td></td><td></td><td></td><td></td><td></td><td></td><td></td><td></td><td></td><td></td><td></td></tr>
<tr><td>Oligo rev</td><td>AAACCACCCATCCCCTACCATTAT</td><td></td><td></td><td></td><td></td><td></td><td></td><td></td><td></td><td></td><td></td><td></td></tr>
<tr><td>Chromosome</td><td>start</td><td></td><td>end</td><td>strand</td><td>MM</td><td>target_seq</td><td>PAM</td><td>alignment</td><td>distance</td><td>position</td><td>gene name</td><td>gene id</td></tr>
<tr><td>GU305914.1</td><td></td><td>20351</td><td>20373</td><td>-</td><td>0</td><td>ATAATGGTAGGGGATGGGTG</td><td>TGG</td><td>||||||||[||||||||||||]PAM</td><td>NA</td><td>Intergenic</td><td>NA</td><td>NA</td></tr>
<tr><td>T33013</td><td>ATGATACTCATATCCTGAGTCGG</td><td></td><td>1000</td><td></td><td></td><td></td><td></td><td></td><td></td><td></td><td></td><td></td></tr>
<tr><td>Oligo fwd</td><td>CACCATGATACTCATATCCTGAGT</td><td></td><td></td><td></td><td></td><td></td><td></td><td></td><td></td><td></td><td></td><td></td></tr>
<tr><td>Oligo rev</td><td>AAACACTCAGGATATGAGTATCAT</td><td></td><td></td><td></td><td></td><td></td><td></td><td></td><td></td><td></td><td></td><td></td></tr>
<tr><td>Chromosome</td><td>start</td><td></td><td>end</td><td>strand</td><td>MM</td><td>target_seq</td><td>PAM</td><td>alignment</td><td>distance</td><td>position</td><td>gene name</td><td>gene id</td></tr>
<tr><td>GU305914.1</td><td></td><td>20326</td><td>20348</td><td>-</td><td>0</td><td>ATGATACTCATATCCTGAGT</td><td>CGG</td><td>||||||||[||||||||||||]PAM</td><td>NA</td><td>Intergenic</td><td>NA</td><td>NA</td></tr>
<tr><td>T33014</td><td>GAGTCGGAGTAAACGTCAGTCGG</td><td></td><td>1000</td><td></td><td></td><td></td><td></td><td></td><td></td><td></td><td></td><td></td></tr>
<tr><td>Oligo fwd</td><td>CACCGAGTCGGAGTAAACGTCAGT</td><td></td><td></td><td></td><td></td><td></td><td></td><td></td><td></td><td></td><td></td><td></td></tr>
<tr><td>Oligo rev</td><td>AAACACTGACGTTTACTCCGACTC</td><td></td><td></td><td></td><td></td><td></td><td></td><td></td><td></td><td></td><td></td><td></td></tr>
<tr><td>Chromosome</td><td>start</td><td></td><td>end</td><td>strand</td><td>MM</td><td>target_seq</td><td>PAM</td><td>alignment</td><td>distance</td><td>position</td><td>gene name</td><td>gene id</td></tr>
<tr><td>GU305914.1</td><td></td><td>20310</td><td>20332</td><td>-</td><td>0</td><td>GAGTCGGAGTAAACGTCAGT</td><td>CGG</td><td>||||||||[||||||||||||]PAM</td><td>NA</td><td>Intergenic</td><td>NA</td><td>NA</td></tr>
<tr><td>T33015</td><td>GTGTTATAAGACGAGCGTCTTGG</td><td></td><td>1000</td><td></td><td></td><td></td><td></td><td></td><td></td><td></td><td></td><td></td></tr>
<tr><td>Oligo fwd</td><td>CACCGTGTTATAAGACGAGCGTCT</td><td></td><td></td><td></td><td></td><td></td><td></td><td></td><td></td><td></td><td></td><td></td></tr>
<tr><td>Oligo rev</td><td>AAACAGACGCTCGTCTTATAACAC</td><td></td><td></td><td></td><td></td><td></td><td></td><td></td><td></td><td></td><td></td><td></td></tr>
<tr><td>Chromosome</td><td>start</td><td></td><td>end</td><td>strand</td><td>MM</td><td>target_seq</td><td>PAM</td><td>alignment</td><td>distance</td><td>position</td><td>gene name</td><td>gene id</td></tr>
<tr><td>GU305914.1</td><td></td><td>20288</td><td>20310</td><td>-</td><td>0</td><td>GTGTTATAAGACGAGCGTCT</td><td>TGG</td><td>||||||||[||||||||||||]PAM</td><td>NA</td><td>Intergenic</td><td>NA</td><td>NA</td></tr>
<tr><td>T33016</td><td>TTGGCAAGCACCACTGCTGTCGG</td><td></td><td>1000</td><td></td><td></td><td></td><td></td><td></td><td></td><td></td><td></td><td></td></tr>
<tr><td>Oligo fwd</td><td>CACCTTGGCAAGCACCACTGCTGT</td><td></td><td></td><td></td><td></td><td></td><td></td><td></td><td></td><td></td><td></td><td></td></tr>
<tr><td>Oligo rev</td><td>AAACACAGCAGTGGTGCTTGCCAA</td><td></td><td></td><td></td><td></td><td></td><td></td><td></td><td></td><td></td><td></td><td></td></tr>
<tr><td>Chromosome</td><td>start</td><td></td><td>end</td><td>strand</td><td>MM</td><td>target_seq</td><td>PAM</td><td>alignment</td><td>distance</td><td>position</td><td>gene name</td><td>gene id</td></tr>
<tr><td>GU305914.1</td><td></td><td>20269</td><td>20291</td><td>-</td><td>0</td><td>TTGGCAAGCACCACTGCTGT</td><td>CGG</td><td>||||||||[||||||||||||]PAM</td><td>NA</td><td>Intergenic</td><td>NA</td><td>NA</td></tr>
<tr><td>T33017</td><td>GTTTCAGAGCGACATACACGAGG</td><td></td><td>1000</td><td></td><td></td><td></td><td></td><td></td><td></td><td></td><td></td><td></td></tr>
<tr><td>Oligo fwd</td><td>CACCGTTTCAGAGCGACATACACG</td><td></td><td></td><td></td><td></td><td></td><td></td><td></td><td></td><td></td><td></td><td></td></tr>
<tr><td>Oligo rev</td><td>AAACCGTGTATGTCGCTCTGAAAC</td><td></td><td></td><td></td><td></td><td></td><td></td><td></td><td></td><td></td><td></td><td></td></tr>
<tr><td>Chromosome</td><td>start</td><td></td><td>end</td><td>strand</td><td>MM</td><td>target_seq</td><td>PAM</td><td>alignment</td><td>distance</td><td>position</td><td>gene name</td><td>gene id</td></tr>
<tr><td>GU305914.1</td><td></td><td>20240</td><td>20262</td><td>-</td><td>0</td><td>GTTTCAGAGCGACATACACG</td><td>AGG</td><td>||||||||[||||||||||||]PAM</td><td>NA</td><td>Intergenic</td><td>NA</td><td>NA</td></tr>
<tr><td>T33018</td><td>TTTCAGAGCGACATACACGAGGG</td><td></td><td>1000</td><td></td><td></td><td></td><td></td><td></td><td></td><td></td><td></td><td></td></tr>
<tr><td>Oligo fwd</td><td>CACCTTTCAGAGCGACATACACGA</td><td></td><td></td><td></td><td></td><td></td><td></td><td></td><td></td><td></td><td></td><td></td></tr>
<tr><td>Oligo rev</td><td>AAACTCGTGTATGTCGCTCTGAAA</td><td></td><td></td><td></td><td></td><td></td><td></td><td></td><td></td><td></td><td></td><td></td></tr>
<tr><td>Chromosome</td><td>start</td><td></td><td>end</td><td>strand</td><td>MM</td><td>target_seq</td><td>PAM</td><td>alignment</td><td>distance</td><td>position</td><td>gene name</td><td>gene id</td></tr>
<tr><td>GU305914.1</td><td></td><td>20239</td><td>20261</td><td>-</td><td>0</td><td>TTTCAGAGCGACATACACGA</td><td>GGG</td><td>||||||||[||||||||||||]PAM</td><td>NA</td><td>Intergenic</td><td>NA</td><td>NA</td></tr>
<tr><td>T33019</td><td>TTCAGAGCGACATACACGAGGGG</td><td></td><td>1000</td><td></td><td></td><td></td><td></td><td></td><td></td><td></td><td></td><td></td></tr>
<tr><td>Oligo fwd</td><td>CACCTTCAGAGCGACATACACGAG</td><td></td><td></td><td></td><td></td><td></td><td></td><td></td><td></td><td></td><td></td><td></td></tr>
<tr><td>Oligo rev</td><td>AAACCTCGTGTATGTCGCTCTGAA</td><td></td><td></td><td></td><td></td><td></td><td></td><td></td><td></td><td></td><td></td><td></td></tr>
<tr><td>Chromosome</td><td>start</td><td></td><td>end</td><td>strand</td><td>MM</td><td>target_seq</td><td>PAM</td><td>alignment</td><td>distance</td><td>position</td><td>gene name</td><td>gene id</td></tr>
<tr><td>GU305914.1</td><td></td><td>20238</td><td>20260</td><td>-</td><td>0</td><td>TTCAGAGCGACATACACGAG</td><td>GGG</td><td>||||||||[||||||||||||]PAM</td><td>NA</td><td>Intergenic</td><td>NA</td><td>NA</td></tr>
<tr><td>T33020</td><td>GAGCGACATACACGAGGGGTTGG</td><td></td><td>1000</td><td></td><td></td><td></td><td></td><td></td><td></td><td></td><td></td><td></td></tr>
<tr><td>Oligo fwd</td><td>CACCGAGCGACATACACGAGGGGT</td><td></td><td></td><td></td><td></td><td></td><td></td><td></td><td></td><td></td><td></td><td></td></tr>
<tr><td>Oligo rev</td><td>AAACACCCCTCGTGTATGTCGCTC</td><td></td><td></td><td></td><td></td><td></td><td></td><td></td><td></td><td></td><td></td><td></td></tr>
<tr><td>Chromosome</td><td>start</td><td></td><td>end</td><td>strand</td><td>MM</td><td>target_seq</td><td>PAM</td><td>alignment</td><td>distance</td><td>position</td><td>gene name</td><td>gene id</td></tr>
<tr><td>GU305914.1</td><td></td><td>20234</td><td>20256</td><td>-</td><td>0</td><td>GAGCGACATACACGAGGGGT</td><td>TGG</td><td>||||||||[||||||||||||]PAM</td><td>NA</td><td>Intergenic</td><td>NA</td><td>NA</td></tr>
<tr><td>T33021</td><td>ACATACACGAGGGGTTGGTACGG</td><td></td><td>1000</td><td></td><td></td><td></td><td></td><td></td><td></td><td></td><td></td><td></td></tr>
<tr><td>Oligo fwd</td><td>CACCACATACACGAGGGGTTGGTA</td><td></td><td></td><td></td><td></td><td></td><td></td><td></td><td></td><td></td><td></td><td></td></tr>
<tr><td>Oligo rev</td><td>AAACTACCAACCCCTCGTGTATGT</td><td></td><td></td><td></td><td></td><td></td><td></td><td></td><td></td><td></td><td></td><td></td></tr>
<tr><td>Chromosome</td><td>start</td><td></td><td>end</td><td>strand</td><td>MM</td><td>target_seq</td><td>PAM</td><td>alignment</td><td>distance</td><td>position</td><td>gene name</td><td>gene id</td></tr>
<tr><td>GU305914.1</td><td></td><td>20229</td><td>20251</td><td>-</td><td>0</td><td>ACATACACGAGGGGTTGGTA</td><td>CGG</td><td>||||||||[||||||||||||]PAM</td><td>NA</td><td>Intergenic</td><td>NA</td><td>NA</td></tr>
<tr><td>T33022</td><td>TGGTACGGTTCGAGCGATTTTGG</td><td></td><td>1000</td><td></td><td></td><td></td><td></td><td></td><td></td><td></td><td></td><td></td></tr>
<tr><td>Oligo fwd</td><td>CACCTGGTACGGTTCGAGCGATTT</td><td></td><td></td><td></td><td></td><td></td><td></td><td></td><td></td><td></td><td></td><td></td></tr>
<tr><td>Oligo rev</td><td>AAACAAATCGCTCGAACCGTACCA</td><td></td><td></td><td></td><td></td><td></td><td></td><td></td><td></td><td></td><td></td><td></td></tr>
<tr><td>Chromosome</td><td>start</td><td></td><td>end</td><td>strand</td><td>MM</td><td>target_seq</td><td>PAM</td><td>alignment</td><td>distance</td><td>position</td><td>gene name</td><td>gene id</td></tr>
<tr><td>GU305914.1</td><td></td><td>20214</td><td>20236</td><td>-</td><td>0</td><td>TGGTACGGTTCGAGCGATTT</td><td>TGG</td><td>||||||||[||||||||||||]PAM</td><td>NA</td><td>Intergenic</td><td>NA</td><td>NA</td></tr>
<tr><td>T33023</td><td>AGTCCGAGACGTCCGCCGCATGG</td><td></td><td>1000</td><td></td><td></td><td></td><td></td><td></td><td></td><td></td><td></td><td></td></tr>
<tr><td>Oligo fwd</td><td>CACCAGTCCGAGACGTCCGCCGCA</td><td></td><td></td><td></td><td></td><td></td><td></td><td></td><td></td><td></td><td></td><td></td></tr>
<tr><td>Oligo rev</td><td>AAACTGCGGCGGACGTCTCGGACT</td><td></td><td></td><td></td><td></td><td></td><td></td><td></td><td></td><td></td><td></td><td></td></tr>
<tr><td>Chromosome</td><td>start</td><td></td><td>end</td><td>strand</td><td>MM</td><td>target_seq</td><td>PAM</td><td>alignment</td><td>distance</td><td>position</td><td>gene name</td><td>gene id</td></tr>
<tr><td>GU305914.1</td><td></td><td>20190</td><td>20212</td><td>-</td><td>0</td><td>AGTCCGAGACGTCCGCCGCA</td><td>TGG</td><td>||||||||[||||||||||||]PAM</td><td>NA</td><td>Intergenic</td><td>NA</td><td>NA</td></tr>
<tr><td>T33024</td><td>GTCCGCCGCATGGCTGACACTGG</td><td></td><td>1000</td><td></td><td></td><td></td><td></td><td></td><td></td><td></td><td></td><td></td></tr>
<tr><td>Oligo fwd</td><td>CACCGTCCGCCGCATGGCTGACAC</td><td></td><td></td><td></td><td></td><td></td><td></td><td></td><td></td><td></td><td></td><td></td></tr>
<tr><td>Oligo rev</td><td>AAACGTGTCAGCCATGCGGCGGAC</td><td></td><td></td><td></td><td></td><td></td><td></td><td></td><td></td><td></td><td></td><td></td></tr>
<tr><td>Chromosome</td><td>start</td><td></td><td>end</td><td>strand</td><td>MM</td><td>target_seq</td><td>PAM</td><td>alignment</td><td>distance</td><td>position</td><td>gene name</td><td>gene id</td></tr>
<tr><td>GU305914.1</td><td></td><td>20180</td><td>20202</td><td>-</td><td>0</td><td>GTCCGCCGCATGGCTGACAC</td><td>TGG</td><td>||||||||[||||||||||||]PAM</td><td>NA</td><td>Intergenic</td><td>NA</td><td>NA</td></tr>
</table>

<table>
<tr><td>T33025</td><td>TCCGCCGCATGGCTGACACTGGG</td><td>1000</td><td></td><td></td><td></td><td></td><td></td><td></td><td></td><td></td><td></td></tr>
<tr><td>Oligo fwd</td><td>CACCTCCGCCGCATGGCTGACACT</td><td></td><td></td><td></td><td></td><td></td><td></td><td></td><td></td><td></td><td></td></tr>
<tr><td>Oligo rev</td><td>AAACAGTGTCAGCCATGCGGCGGA</td><td></td><td></td><td></td><td></td><td></td><td></td><td></td><td></td><td></td><td></td></tr>
<tr><td>Chromosome</td><td>start</td><td>end</td><td>strand</td><td>MM</td><td>target_seq</td><td>PAM</td><td>alignment</td><td>distance</td><td>position</td><td>gene name</td><td>gene id</td></tr>
<tr><td>GU305914.1</td><td>20179</td><td>20201</td><td>-</td><td>0</td><td>TCCGCCGCATGGCTGACACT</td><td>GGG</td><td>||||||||[||||||||||||]PAM</td><td>NA</td><td>Intergenic</td><td>NA</td><td>NA</td></tr>
<tr><td>T33026</td><td>CGCATGGCTGACACTGGGCACGG</td><td>1000</td><td></td><td></td><td></td><td></td><td></td><td></td><td></td><td></td><td></td></tr>
<tr><td>Oligo fwd</td><td>CACCCGCATGGCTGACACTGGGCA</td><td></td><td></td><td></td><td></td><td></td><td></td><td></td><td></td><td></td><td></td></tr>
<tr><td>Oligo rev</td><td>AAACTGCCCAGTGTCAGCCATGCG</td><td></td><td></td><td></td><td></td><td></td><td></td><td></td><td></td><td></td><td></td></tr>
<tr><td>Chromosome</td><td>start</td><td>end</td><td>strand</td><td>MM</td><td>target_seq</td><td>PAM</td><td>alignment</td><td>distance</td><td>position</td><td>gene name</td><td>gene id</td></tr>
<tr><td>GU305914.1</td><td>20174</td><td>20196</td><td>-</td><td>0</td><td>CGCATGGCTGACACTGGGCA</td><td>CGG</td><td>||||||||[||||||||||||]PAM</td><td>NA</td><td>Intergenic</td><td>NA</td><td>NA</td></tr>
<tr><td>T33027</td><td>GGGCACGGTACCCGAACGAGAGG</td><td>1000</td><td></td><td></td><td></td><td></td><td></td><td></td><td></td><td></td><td></td></tr>
<tr><td>Oligo fwd</td><td>CACCGGGCACGGTACCCGAACGAG</td><td></td><td></td><td></td><td></td><td></td><td></td><td></td><td></td><td></td><td></td></tr>
<tr><td>Oligo rev</td><td>AAACCTCGTTCGGGTACCGTGCCC</td><td></td><td></td><td></td><td></td><td></td><td></td><td></td><td></td><td></td><td></td></tr>
<tr><td>Chromosome</td><td>start</td><td>end</td><td>strand</td><td>MM</td><td>target_seq</td><td>PAM</td><td>alignment</td><td>distance</td><td>position</td><td>gene name</td><td>gene id</td></tr>
<tr><td>GU305914.1</td><td>20159</td><td>20181</td><td>-</td><td>0</td><td>GGGCACGGTACCCGAACGAG</td><td>AGG</td><td>||||||||[||||||||||||]PAM</td><td>NA</td><td>Intergenic</td><td>NA</td><td>NA</td></tr>
<tr><td>T33028</td><td>GTACCCGAACGAGAGGTTCGAGG</td><td>1000</td><td></td><td></td><td></td><td></td><td></td><td></td><td></td><td></td><td></td></tr>
<tr><td>Oligo fwd</td><td>CACCGTACCCGAACGAGAGGTTCG</td><td></td><td></td><td></td><td></td><td></td><td></td><td></td><td></td><td></td><td></td></tr>
<tr><td>Oligo rev</td><td>AAACCGAACCTCTCGTTCGGGTAC</td><td></td><td></td><td></td><td></td><td></td><td></td><td></td><td></td><td></td><td></td></tr>
<tr><td>Chromosome</td><td>start</td><td>end</td><td>strand</td><td>MM</td><td>target_seq</td><td>PAM</td><td>alignment</td><td>distance</td><td>position</td><td>gene name</td><td>gene id</td></tr>
<tr><td>GU305914.1</td><td>20152</td><td>20174</td><td>-</td><td>0</td><td>GTACCCGAACGAGAGGTTCG</td><td>AGG</td><td>||||||||[||||||||||||]PAM</td><td>NA</td><td>Intergenic</td><td>NA</td><td>NA</td></tr>
<tr><td>T33029</td><td>TACCCGAACGAGAGGTTCGAGGG</td><td>1000</td><td></td><td></td><td></td><td></td><td></td><td></td><td></td><td></td><td></td></tr>
<tr><td>Oligo fwd</td><td>CACCTACCCGAACGAGAGGTTCGA</td><td></td><td></td><td></td><td></td><td></td><td></td><td></td><td></td><td></td><td></td></tr>
<tr><td>Oligo rev</td><td>AAACTCGAACCTCTCGTTCGGGTA</td><td></td><td></td><td></td><td></td><td></td><td></td><td></td><td></td><td></td><td></td></tr>
<tr><td>Chromosome</td><td>start</td><td>end</td><td>strand</td><td>MM</td><td>target_seq</td><td>PAM</td><td>alignment</td><td>distance</td><td>position</td><td>gene name</td><td>gene id</td></tr>
<tr><td>GU305914.1</td><td>20151</td><td>20173</td><td>-</td><td>0</td><td>TACCCGAACGAGAGGTTCGA</td><td>GGG</td><td>||||||||[||||||||||||]PAM</td><td>NA</td><td>Intergenic</td><td>NA</td><td>NA</td></tr>
<tr><td>T33030</td><td>CCCGTCGATATGCCGTCAGACGG</td><td>1000</td><td></td><td></td><td></td><td></td><td></td><td></td><td></td><td></td><td></td></tr>
<tr><td>Oligo fwd</td><td>CACCCCCGTCGATATGCCGTCAGA</td><td></td><td></td><td></td><td></td><td></td><td></td><td></td><td></td><td></td><td></td></tr>
<tr><td>Oligo rev</td><td>AAACTCTGACGGCATATCGACGGG</td><td></td><td></td><td></td><td></td><td></td><td></td><td></td><td></td><td></td><td></td></tr>
<tr><td>Chromosome</td><td>start</td><td>end</td><td>strand</td><td>MM</td><td>target_seq</td><td>PAM</td><td>alignment</td><td>distance</td><td>position</td><td>gene name</td><td>gene id</td></tr>
<tr><td>GU305914.1</td><td>20108</td><td>20130</td><td>-</td><td>0</td><td>CCCGTCGATATGCCGTCAGA</td><td>CGG</td><td>||||||||[||||||||||||]PAM</td><td>NA</td><td>Intergenic</td><td>NA</td><td>NA</td></tr>
<tr><td>T33031</td><td>GCCGTCAGACGGCAGTGCCGCGG</td><td>1000</td><td></td><td></td><td></td><td></td><td></td><td></td><td></td><td></td><td></td></tr>
<tr><td>Oligo fwd</td><td>CACCGCCGTCAGACGGCAGTGCCG</td><td></td><td></td><td></td><td></td><td></td><td></td><td></td><td></td><td></td><td></td></tr>
<tr><td>Oligo rev</td><td>AAACCGGCACTGCCGTCTGACGGC</td><td></td><td></td><td></td><td></td><td></td><td></td><td></td><td></td><td></td><td></td></tr>
<tr><td>Chromosome</td><td>start</td><td>end</td><td>strand</td><td>MM</td><td>target_seq</td><td>PAM</td><td>alignment</td><td>distance</td><td>position</td><td>gene name</td><td>gene id</td></tr>
<tr><td>GU305914.1</td><td>20097</td><td>20119</td><td>-</td><td>0</td><td>GCCGTCAGACGGCAGTGCCG</td><td>CGG</td><td>||||||||[||||||||||||]PAM</td><td>NA</td><td>Intergenic</td><td>NA</td><td>NA</td></tr>
<tr><td>T33032</td><td>CGGCAGTGCCGCGGCAGAGCCGG</td><td>1000</td><td></td><td></td><td></td><td></td><td></td><td></td><td></td><td></td><td></td></tr>
<tr><td>Oligo fwd</td><td>CACCCGGCAGTGCCGCGGCAGAGC</td><td></td><td></td><td></td><td></td><td></td><td></td><td></td><td></td><td></td><td></td></tr>
<tr><td>Oligo rev</td><td>AAACGCTCTGCCGCGGCACTGCCG</td><td></td><td></td><td></td><td></td><td></td><td></td><td></td><td></td><td></td><td></td></tr>
<tr><td>Chromosome</td><td>start</td><td>end</td><td>strand</td><td>MM</td><td>target_seq</td><td>PAM</td><td>alignment</td><td>distance</td><td>position</td><td>gene name</td><td>gene id</td></tr>
<tr><td>GU305914.1</td><td>20088</td><td>20110</td><td>-</td><td>0</td><td>CGGCAGTGCCGCGGCAGAGC</td><td>CGG</td><td>||||||||[||||||||||||]PAM</td><td>NA</td><td>Intergenic</td><td>NA</td><td>NA</td></tr>
<tr><td>T33033</td><td>AGTGCCGCGGCAGAGCCGGTAGG</td><td>1000</td><td></td><td></td><td></td><td></td><td></td><td></td><td></td><td></td><td></td></tr>
<tr><td>Oligo fwd</td><td>CACCAGTGCCGCGGCAGAGCCGGT</td><td></td><td></td><td></td><td></td><td></td><td></td><td></td><td></td><td></td><td></td></tr>
<tr><td>Oligo rev</td><td>AAACACCGGCTCTGCCGCGGCACT</td><td></td><td></td><td></td><td></td><td></td><td></td><td></td><td></td><td></td><td></td></tr>
<tr><td>Chromosome</td><td>start</td><td>end</td><td>strand</td><td>MM</td><td>target_seq</td><td>PAM</td><td>alignment</td><td>distance</td><td>position</td><td>gene name</td><td>gene id</td></tr>
<tr><td>GU305914.1</td><td>20084</td><td>20106</td><td>-</td><td>0</td><td>AGTGCCGCGGCAGAGCCGGT</td><td>AGG</td><td>||||||||[||||||||||||]PAM</td><td>NA</td><td>Intergenic</td><td>NA</td><td>NA</td></tr>
<tr><td>T33034</td><td>GCCGCGGCAGAGCCGGTAGGTGG</td><td>1000</td><td></td><td></td><td></td><td></td><td></td><td></td><td></td><td></td><td></td></tr>
<tr><td>Oligo fwd</td><td>CACCGCCGCGGCAGAGCCGGTAGG</td><td></td><td></td><td></td><td></td><td></td><td></td><td></td><td></td><td></td><td></td></tr>
<tr><td>Oligo rev</td><td>AAACCCTACCGGCTCTGCCGCGGC</td><td></td><td></td><td></td><td></td><td></td><td></td><td></td><td></td><td></td><td></td></tr>
<tr><td>Chromosome</td><td>start</td><td>end</td><td>strand</td><td>MM</td><td>target_seq</td><td>PAM</td><td>alignment</td><td>distance</td><td>position</td><td>gene name</td><td>gene id</td></tr>
<tr><td>GU305914.1</td><td>20081</td><td>20103</td><td>-</td><td>0</td><td>GCCGCGGCAGAGCCGGTAGG</td><td>TGG</td><td>||||||||[||||||||||||]PAM</td><td>NA</td><td>Intergenic</td><td>NA</td><td>NA</td></tr>
<tr><td>T33035</td><td>CCGCGGCAGAGCCGGTAGGTGGG</td><td>1000</td><td></td><td></td><td></td><td></td><td></td><td></td><td></td><td></td><td></td></tr>
<tr><td>Oligo fwd</td><td>CACCCCGCGGCAGAGCCGGTAGGT</td><td></td><td></td><td></td><td></td><td></td><td></td><td></td><td></td><td></td><td></td></tr>
<tr><td>Oligo rev</td><td>AAACACCTACCGGCTCTGCCGCGG</td><td></td><td></td><td></td><td></td><td></td><td></td><td></td><td></td><td></td><td></td></tr>
<tr><td>Chromosome</td><td>start</td><td>end</td><td>strand</td><td>MM</td><td>target_seq</td><td>PAM</td><td>alignment</td><td>distance</td><td>position</td><td>gene name</td><td>gene id</td></tr>
<tr><td>GU305914.1</td><td>20080</td><td>20102</td><td>-</td><td>0</td><td>CCGCGGCAGAGCCGGTAGGT</td><td>GGG</td><td>||||||||[||||||||||||]PAM</td><td>NA</td><td>Intergenic</td><td>NA</td><td>NA</td></tr>
<tr><td>T33036</td><td>ATTCAACCGCCCGATCTCAAAGG</td><td>1000</td><td></td><td></td><td></td><td></td><td></td><td></td><td></td><td></td><td></td></tr>
<tr><td>Oligo fwd</td><td>CACCATTCAACCGCCCGATCTCAA</td><td></td><td></td><td></td><td></td><td></td><td></td><td></td><td></td><td></td><td></td></tr>
<tr><td>Oligo rev</td><td>AAACTTGAGATCGGGCGGTTGAAT</td><td></td><td></td><td></td><td></td><td></td><td></td><td></td><td></td><td></td><td></td></tr>
<tr><td>Chromosome</td><td>start</td><td>end</td><td>strand</td><td>MM</td><td>target_seq</td><td>PAM</td><td>alignment</td><td>distance</td><td>position</td><td>gene name</td><td>gene id</td></tr>
<tr><td>GU305914.1</td><td>20053</td><td>20075</td><td>-</td><td>0</td><td>ATTCAACCGCCCGATCTCAA</td><td>AGG</td><td>||||||||[||||||||||||]PAM</td><td>NA</td><td>Intergenic</td><td>NA</td><td>NA</td></tr>
<tr><td>T33037</td><td>AGCGTCAGCAACAGCAGTCGCGG</td><td>1000</td><td></td><td></td><td></td><td></td><td></td><td></td><td></td><td></td><td></td></tr>
<tr><td>Oligo fwd</td><td>CACCAGCGTCAGCAACAGCAGTCG</td><td></td><td></td><td></td><td></td><td></td><td></td><td></td><td></td><td></td><td></td></tr>
<tr><td>Oligo rev</td><td>AAACCGACTGCTGTTGCTGACGCT</td><td></td><td></td><td></td><td></td><td></td><td></td><td></td><td></td><td></td><td></td></tr>
<tr><td>Chromosome</td><td>start</td><td>end</td><td>strand</td><td>MM</td><td>target_seq</td><td>PAM</td><td>alignment</td><td>distance</td><td>position</td><td>gene name</td><td>gene id</td></tr>
<tr><td>GU305914.1</td><td>20026</td><td>20048</td><td>-</td><td>0</td><td>AGCGTCAGCAACAGCAGTCG</td><td>CGG</td><td>||||||||[||||||||||||]PAM</td><td>NA</td><td>Intergenic</td><td>NA</td><td>NA</td></tr>
<tr><td>T33038</td><td>AGAGCAGCAACAAGTGCCGACGG</td><td>1000</td><td></td><td></td><td></td><td></td><td></td><td></td><td></td><td></td><td></td></tr>
<tr><td>Oligo fwd</td><td>CACCAGAGCAGCAACAAGTGCCGA</td><td></td><td></td><td></td><td></td><td></td><td></td><td></td><td></td><td></td><td></td></tr>
<tr><td>Oligo rev</td><td>AAACTCGGCACTTGTTGCTGCTCT</td><td></td><td></td><td></td><td></td><td></td><td></td><td></td><td></td><td></td><td></td></tr>
<tr><td>Chromosome</td><td>start</td><td>end</td><td>strand</td><td>MM</td><td>target_seq</td><td>PAM</td><td>alignment</td><td>distance</td><td>position</td><td>gene name</td><td>gene id</td></tr>
<tr><td>GU305914.1</td><td>19971</td><td>19993</td><td>-</td><td>0</td><td>AGAGCAGCAACAAGTGCCGA</td><td>CGG</td><td>||||||||[||||||||||||]PAM</td><td>NA</td><td>Intergenic</td><td>NA</td><td>NA</td></tr>
<tr><td>T33039</td><td>CGCTACCGTCTTCGATTCCTCGG</td><td>1000</td><td></td><td></td><td></td><td></td><td></td><td></td><td></td><td></td><td></td></tr>
<tr><td>Oligo fwd</td><td>CACCCGCTACCGTCTTCGATTCCT</td><td></td><td></td><td></td><td></td><td></td><td></td><td></td><td></td><td></td><td></td></tr>
<tr><td>Oligo rev</td><td>AAACAGGAATCGAAGACGGTAGCG</td><td></td><td></td><td></td><td></td><td></td><td></td><td></td><td></td><td></td><td></td></tr>
<tr><td>Chromosome</td><td>start</td><td>end</td><td>strand</td><td>MM</td><td>target_seq</td><td>PAM</td><td>alignment</td><td>distance</td><td>position</td><td>gene name</td><td>gene id</td></tr>
<tr><td>GU305914.1</td><td>19923</td><td>19945</td><td>-</td><td>0</td><td>CGCTACCGTCTTCGATTCCT</td><td>CGG</td><td>||||||||[||||||||||||]PAM</td><td>NA</td><td>Intergenic</td><td>NA</td><td>NA</td></tr>
</table>

<table>
<tr><td>T33040</td><td>TACCGTCTTCGATTCCTCGGCGG</td><td>1000</td><td></td><td></td><td></td><td></td><td></td><td></td><td></td><td></td><td></td></tr>
<tr><td>Oligo fwd</td><td>CACCTACCGTCTTCGATTCCTCGG</td><td></td><td></td><td></td><td></td><td></td><td></td><td></td><td></td><td></td><td></td></tr>
<tr><td>Oligo rev</td><td>AAACCCGAGGAATCGAAGACGGTA</td><td></td><td></td><td></td><td></td><td></td><td></td><td></td><td></td><td></td><td></td></tr>
<tr><td>Chromosome</td><td>start</td><td>end</td><td>strand</td><td>MM</td><td>target_seq</td><td>PAM</td><td>alignment</td><td>distance</td><td>position</td><td>gene name</td><td>gene id</td></tr>
<tr><td>GU305914.1</td><td>19920</td><td>19942</td><td>-</td><td>0</td><td>TACCGTCTTCGATTCCTCGG</td><td>CGG</td><td>||||||||[||||||||||||]PAM</td><td>NA</td><td>Intergenic</td><td>NA</td><td>NA</td></tr>
<tr><td>T33041</td><td>CTTCGATTCCTCGGCGGCGACGG</td><td>1000</td><td></td><td></td><td></td><td></td><td></td><td></td><td></td><td></td><td></td></tr>
<tr><td>Oligo fwd</td><td>CACCCTTCGATTCCTCGGCGGCGA</td><td></td><td></td><td></td><td></td><td></td><td></td><td></td><td></td><td></td><td></td></tr>
<tr><td>Oligo rev</td><td>AAACTCGCCGCCGAGGAATCGAAG</td><td></td><td></td><td></td><td></td><td></td><td></td><td></td><td></td><td></td><td></td></tr>
<tr><td>Chromosome</td><td>start</td><td>end</td><td>strand</td><td>MM</td><td>target_seq</td><td>PAM</td><td>alignment</td><td>distance</td><td>position</td><td>gene name</td><td>gene id</td></tr>
<tr><td>GU305914.1</td><td>19914</td><td>19936</td><td>-</td><td>0</td><td>CTTCGATTCCTCGGCGGCGA</td><td>CGG</td><td>||||||||[||||||||||||]PAM</td><td>NA</td><td>Intergenic</td><td>NA</td><td>NA</td></tr>
<tr><td>T33042</td><td>TTCGATTCCTCGGCGGCGACGGG</td><td>1000</td><td></td><td></td><td></td><td></td><td></td><td></td><td></td><td></td><td></td></tr>
<tr><td>Oligo fwd</td><td>CACCTTCGATTCCTCGGCGGCGAC</td><td></td><td></td><td></td><td></td><td></td><td></td><td></td><td></td><td></td><td></td></tr>
<tr><td>Oligo rev</td><td>AAACGTCGCCGCCGAGGAATCGAA</td><td></td><td></td><td></td><td></td><td></td><td></td><td></td><td></td><td></td><td></td></tr>
<tr><td>Chromosome</td><td>start</td><td>end</td><td>strand</td><td>MM</td><td>target_seq</td><td>PAM</td><td>alignment</td><td>distance</td><td>position</td><td>gene name</td><td>gene id</td></tr>
<tr><td>GU305914.1</td><td>19913</td><td>19935</td><td>-</td><td>0</td><td>TTCGATTCCTCGGCGGCGAC</td><td>GGG</td><td>||||||||[||||||||||||]PAM</td><td>NA</td><td>Intergenic</td><td>NA</td><td>NA</td></tr>
<tr><td>GU305914.1</td><td>76808</td><td>76830</td><td>+</td><td>4</td><td>TTGCACTCCTCGGCGGGGAC</td><td>GGG</td><td>||--|-||[||||||||-|||]PAM</td><td>NA</td><td>Intergenic</td><td>NA</td><td>NA</td></tr>
<tr><td>T33043</td><td>TCGATTCCTCGGCGGCGACGGGG</td><td>1000</td><td></td><td></td><td></td><td></td><td></td><td></td><td></td><td></td><td></td></tr>
<tr><td>Oligo fwd</td><td>CACCTCGATTCCTCGGCGGCGACG</td><td></td><td></td><td></td><td></td><td></td><td></td><td></td><td></td><td></td><td></td></tr>
<tr><td>Oligo rev</td><td>AAACCGTCGCCGCCGAGGAATCGA</td><td></td><td></td><td></td><td></td><td></td><td></td><td></td><td></td><td></td><td></td></tr>
<tr><td>Chromosome</td><td>start</td><td>end</td><td>strand</td><td>MM</td><td>target_seq</td><td>PAM</td><td>alignment</td><td>distance</td><td>position</td><td>gene name</td><td>gene id</td></tr>
<tr><td>GU305914.1</td><td>19912</td><td>19934</td><td>-</td><td>0</td><td>TCGATTCCTCGGCGGCGACG</td><td>GGG</td><td>||||||||[||||||||||||]PAM</td><td>NA</td><td>Intergenic</td><td>NA</td><td>NA</td></tr>
<tr><td>T33044</td><td>ATTCCTCGGCGGCGACGGGGTGG</td><td>1000</td><td></td><td></td><td></td><td></td><td></td><td></td><td></td><td></td><td></td></tr>
<tr><td>Oligo fwd</td><td>CACCATTCCTCGGCGGCGACGGGG</td><td></td><td></td><td></td><td></td><td></td><td></td><td></td><td></td><td></td><td></td></tr>
<tr><td>Oligo rev</td><td>AAACCCCCGTCGCCGCCGAGGAAT</td><td></td><td></td><td></td><td></td><td></td><td></td><td></td><td></td><td></td><td></td></tr>
<tr><td>Chromosome</td><td>start</td><td>end</td><td>strand</td><td>MM</td><td>target_seq</td><td>PAM</td><td>alignment</td><td>distance</td><td>position</td><td>gene name</td><td>gene id</td></tr>
<tr><td>GU305914.1</td><td>19909</td><td>19931</td><td>-</td><td>0</td><td>ATTCCTCGGCGGCGACGGGG</td><td>TGG</td><td>||||||||[||||||||||||]PAM</td><td>NA</td><td>Intergenic</td><td>NA</td><td>NA</td></tr>
<tr><td>T33045</td><td>CGGCGGCGACGGGGTGGATCAGG</td><td>1000</td><td></td><td></td><td></td><td></td><td></td><td></td><td></td><td></td><td></td></tr>
<tr><td>Oligo fwd</td><td>CACCCGGCGGCGACGGGGTGGATC</td><td></td><td></td><td></td><td></td><td></td><td></td><td></td><td></td><td></td><td></td></tr>
<tr><td>Oligo rev</td><td>AAACGATCCACCCCGTCGCCGCCG</td><td></td><td></td><td></td><td></td><td></td><td></td><td></td><td></td><td></td><td></td></tr>
<tr><td>Chromosome</td><td>start</td><td>end</td><td>strand</td><td>MM</td><td>target_seq</td><td>PAM</td><td>alignment</td><td>distance</td><td>position</td><td>gene name</td><td>gene id</td></tr>
<tr><td>GU305914.1</td><td>19903</td><td>19925</td><td>-</td><td>0</td><td>CGGCGGCGACGGGGTGGATC</td><td>AGG</td><td>||||||||[||||||||||||]PAM</td><td>NA</td><td>Intergenic</td><td>NA</td><td>NA</td></tr>
<tr><td>T33046</td><td>GGCGGCGACGGGGTGGATCAGGG</td><td>1000</td><td></td><td></td><td></td><td></td><td></td><td></td><td></td><td></td><td></td></tr>
<tr><td>Oligo fwd</td><td>CACCGGCGGCGACGGGGTGGATCA</td><td></td><td></td><td></td><td></td><td></td><td></td><td></td><td></td><td></td><td></td></tr>
<tr><td>Oligo rev</td><td>AAACTGATCCACCCCGTCGCCGCC</td><td></td><td></td><td></td><td></td><td></td><td></td><td></td><td></td><td></td><td></td></tr>
<tr><td>Chromosome</td><td>start</td><td>end</td><td>strand</td><td>MM</td><td>target_seq</td><td>PAM</td><td>alignment</td><td>distance</td><td>position</td><td>gene name</td><td>gene id</td></tr>
<tr><td>GU305914.1</td><td>19902</td><td>19924</td><td>-</td><td>0</td><td>GGCGGCGACGGGGTGGATCA</td><td>GGG</td><td>||||||||[||||||||||||]PAM</td><td>NA</td><td>Intergenic</td><td>NA</td><td>NA</td></tr>
<tr><td>T33047</td><td>AGGGTCTCTAGCACGCCATTCGG</td><td>1000</td><td></td><td></td><td></td><td></td><td></td><td></td><td></td><td></td><td></td></tr>
<tr><td>Oligo fwd</td><td>CACCAGGGTCTCTAGCACGCCATT</td><td></td><td></td><td></td><td></td><td></td><td></td><td></td><td></td><td></td><td></td></tr>
<tr><td>Oligo rev</td><td>AAACAATGGCGTGCTAGAGACCCT</td><td></td><td></td><td></td><td></td><td></td><td></td><td></td><td></td><td></td><td></td></tr>
<tr><td>Chromosome</td><td>start</td><td>end</td><td>strand</td><td>MM</td><td>target_seq</td><td>PAM</td><td>alignment</td><td>distance</td><td>position</td><td>gene name</td><td>gene id</td></tr>
<tr><td>GU305914.1</td><td>19883</td><td>19905</td><td>-</td><td>0</td><td>AGGGTCTCTAGCACGCCATT</td><td>CGG</td><td>||||||||[||||||||||||]PAM</td><td>NA</td><td>Intergenic</td><td>NA</td><td>NA</td></tr>
<tr><td>T33048</td><td>AATCACCTGCTCCTACCTCGTGG</td><td>1000</td><td></td><td></td><td></td><td></td><td></td><td></td><td></td><td></td><td></td></tr>
<tr><td>Oligo fwd</td><td>CACCAATCACCTGCTCCTACCTCG</td><td></td><td></td><td></td><td></td><td></td><td></td><td></td><td></td><td></td><td></td></tr>
<tr><td>Oligo rev</td><td>AAACCGAGGTAGGAGCAGGTGATT</td><td></td><td></td><td></td><td></td><td></td><td></td><td></td><td></td><td></td><td></td></tr>
<tr><td>Chromosome</td><td>start</td><td>end</td><td>strand</td><td>MM</td><td>target_seq</td><td>PAM</td><td>alignment</td><td>distance</td><td>position</td><td>gene name</td><td>gene id</td></tr>
<tr><td>GU305914.1</td><td>19825</td><td>19847</td><td>-</td><td>0</td><td>AATCACCTGCTCCTACCTCG</td><td>TGG</td><td>||||||||[||||||||||||]PAM</td><td>NA</td><td>Intergenic</td><td>NA</td><td>NA</td></tr>
<tr><td>T33049</td><td>GTGGACGACTCTTTTGACTCCGG</td><td>1000</td><td></td><td></td><td></td><td></td><td></td><td></td><td></td><td></td><td></td></tr>
<tr><td>Oligo fwd</td><td>CACCGTGGACGACTCTTTTGACTC</td><td></td><td></td><td></td><td></td><td></td><td></td><td></td><td></td><td></td><td></td></tr>
<tr><td>Oligo rev</td><td>AAACGAGTCAAAAGAGTCGTCCAC</td><td></td><td></td><td></td><td></td><td></td><td></td><td></td><td></td><td></td><td></td></tr>
<tr><td>Chromosome</td><td>start</td><td>end</td><td>strand</td><td>MM</td><td>target_seq</td><td>PAM</td><td>alignment</td><td>distance</td><td>position</td><td>gene name</td><td>gene id</td></tr>
<tr><td>GU305914.1</td><td>19806</td><td>19828</td><td>-</td><td>0</td><td>GTGGACGACTCTTTTGACTC</td><td>CGG</td><td>||||||||[||||||||||||]PAM</td><td>NA</td><td>Intergenic</td><td>NA</td><td>NA</td></tr>
<tr><td>T33050</td><td>CCCGCACGCCCCCGTCTTCCTGG</td><td>1000</td><td></td><td></td><td></td><td></td><td></td><td></td><td></td><td></td><td></td></tr>
<tr><td>Oligo fwd</td><td>CACCCCCGCACGCCCCCGTCTTCC</td><td></td><td></td><td></td><td></td><td></td><td></td><td></td><td></td><td></td><td></td></tr>
<tr><td>Oligo rev</td><td>AAACGGAAGACGGGGGCGTGCGGG</td><td></td><td></td><td></td><td></td><td></td><td></td><td></td><td></td><td></td><td></td></tr>
<tr><td>Chromosome</td><td>start</td><td>end</td><td>strand</td><td>MM</td><td>target_seq</td><td>PAM</td><td>alignment</td><td>distance</td><td>position</td><td>gene name</td><td>gene id</td></tr>
<tr><td>GU305914.1</td><td>19767</td><td>19789</td><td>-</td><td>0</td><td>CCCGCACGCCCCCGTCTTCC</td><td>TGG</td><td>||||||||[||||||||||||]PAM</td><td>NA</td><td>Intergenic</td><td>NA</td><td>NA</td></tr>
<tr><td>T33051</td><td>CCGCACGCCCCCGTCTTCCTGGG</td><td>1000</td><td></td><td></td><td></td><td></td><td></td><td></td><td></td><td></td><td></td></tr>
<tr><td>Oligo fwd</td><td>CACCCCGCACGCCCCCGTCTTCCT</td><td></td><td></td><td></td><td></td><td></td><td></td><td></td><td></td><td></td><td></td></tr>
<tr><td>Oligo rev</td><td>AAACAGGAAGACGGGGGCGTGCGG</td><td></td><td></td><td></td><td></td><td></td><td></td><td></td><td></td><td></td><td></td></tr>
<tr><td>Chromosome</td><td>start</td><td>end</td><td>strand</td><td>MM</td><td>target_seq</td><td>PAM</td><td>alignment</td><td>distance</td><td>position</td><td>gene name</td><td>gene id</td></tr>
<tr><td>GU305914.1</td><td>19766</td><td>19788</td><td>-</td><td>0</td><td>CCGCACGCCCCCGTCTTCCT</td><td>GGG</td><td>||||||||[||||||||||||]PAM</td><td>NA</td><td>Intergenic</td><td>NA</td><td>NA</td></tr>
<tr><td>T33052</td><td>CTGGGATACGCGCCCCATCTCGG</td><td>1000</td><td></td><td></td><td></td><td></td><td></td><td></td><td></td><td></td><td></td></tr>
<tr><td>Oligo fwd</td><td>CACCCTGGGATACGCGCCCCATCT</td><td></td><td></td><td></td><td></td><td></td><td></td><td></td><td></td><td></td><td></td></tr>
<tr><td>Oligo rev</td><td>AAACAGATGGGGCGCGTATCCCAG</td><td></td><td></td><td></td><td></td><td></td><td></td><td></td><td></td><td></td><td></td></tr>
<tr><td>Chromosome</td><td>start</td><td>end</td><td>strand</td><td>MM</td><td>target_seq</td><td>PAM</td><td>alignment</td><td>distance</td><td>position</td><td>gene name</td><td>gene id</td></tr>
<tr><td>GU305914.1</td><td>19748</td><td>19770</td><td>-</td><td>0</td><td>CTGGGATACGCGCCCCATCT</td><td>CGG</td><td>||||||||[||||||||||||]PAM</td><td>NA</td><td>Intergenic</td><td>NA</td><td>NA</td></tr>
<tr><td>T33053</td><td>GATACGCGCCCCATCTCGGATGG</td><td>1000</td><td></td><td></td><td></td><td></td><td></td><td></td><td></td><td></td><td></td></tr>
<tr><td>Oligo fwd</td><td>CACCGATACGCGCCCCATCTCGGA</td><td></td><td></td><td></td><td></td><td></td><td></td><td></td><td></td><td></td><td></td></tr>
<tr><td>Oligo rev</td><td>AAACTCCGAGATGGGGCGCGTATC</td><td></td><td></td><td></td><td></td><td></td><td></td><td></td><td></td><td></td><td></td></tr>
<tr><td>Chromosome</td><td>start</td><td>end</td><td>strand</td><td>MM</td><td>target_seq</td><td>PAM</td><td>alignment</td><td>distance</td><td>position</td><td>gene name</td><td>gene id</td></tr>
<tr><td>GU305914.1</td><td>19744</td><td>19766</td><td>-</td><td>0</td><td>GATACGCGCCCCATCTCGGA</td><td>TGG</td><td>||||||||[||||||||||||]PAM</td><td>NA</td><td>Intergenic</td><td>NA</td><td>NA</td></tr>
<tr><td>T33054</td><td>TCCGCGTTTCAGCCGTTACCCGG</td><td>1000</td><td></td><td></td><td></td><td></td><td></td><td></td><td></td><td></td><td></td></tr>
<tr><td>Oligo fwd</td><td>CACCTCCGCGTTTCAGCCGTTACC</td><td></td><td></td><td></td><td></td><td></td><td></td><td></td><td></td><td></td><td></td></tr>
<tr><td>Oligo rev</td><td>AAACGGTAACGGCTGAAACGCGGA</td><td></td><td></td><td></td><td></td><td></td><td></td><td></td><td></td><td></td><td></td></tr>
</table>

<table>
<tr><td>Chromosome</td><td>start</td><td>end</td><td>strand</td><td>MM</td><td>target_seq</td><td>PAM</td><td>alignment</td><td>distance</td><td>position</td><td>gene name</td><td>gene id</td></tr>
<tr><td>GU305914.1</td><td>19712</td><td>19734</td><td>-</td><td>0</td><td>TCCGCGTTTCAGCCGTTACC</td><td>CGG</td><td>||||||||[||||||||||||]PAM</td><td>NA</td><td>Intergenic</td><td>NA</td><td>NA</td></tr>
<tr><td>T33055</td><td>GTTTCAGCCGTTACCCGGCCCGG</td><td>1000</td><td></td><td></td><td></td><td></td><td></td><td></td><td></td><td></td><td></td></tr>
<tr><td>Oligo fwd</td><td>CACCGTTTCAGCCGTTACCCGGCC</td><td></td><td></td><td></td><td></td><td></td><td></td><td></td><td></td><td></td><td></td></tr>
<tr><td>Oligo rev</td><td>AAACGGCCGGGTAACGGCTGAAAC</td><td></td><td></td><td></td><td></td><td></td><td></td><td></td><td></td><td></td><td></td></tr>
<tr><td>Chromosome</td><td>start</td><td>end</td><td>strand</td><td>MM</td><td>target_seq</td><td>PAM</td><td>alignment</td><td>distance</td><td>position</td><td>gene name</td><td>gene id</td></tr>
<tr><td>GU305914.1</td><td>19707</td><td>19729</td><td>-</td><td>0</td><td>GTTTCAGCCGTTACCCGGCC</td><td>CGG</td><td>||||||||[||||||||||||]PAM</td><td>NA</td><td>Intergenic</td><td>NA</td><td>NA</td></tr>
<tr><td>T33056</td><td>CCACCTTCCAACCCTTCCCTTGG</td><td>1000</td><td></td><td></td><td></td><td></td><td></td><td></td><td></td><td></td><td></td></tr>
<tr><td>Oligo fwd</td><td>CACCCCACCTTCCAACCCTTCCCT</td><td></td><td></td><td></td><td></td><td></td><td></td><td></td><td></td><td></td><td></td></tr>
<tr><td>Oligo rev</td><td>AAACAGGGAAGGGTTGGAAGGTGG</td><td></td><td></td><td></td><td></td><td></td><td></td><td></td><td></td><td></td><td></td></tr>
<tr><td>Chromosome</td><td>start</td><td>end</td><td>strand</td><td>MM</td><td>target_seq</td><td>PAM</td><td>alignment</td><td>distance</td><td>position</td><td>gene name</td><td>gene id</td></tr>
<tr><td>GU305914.1</td><td>19678</td><td>19700</td><td>-</td><td>0</td><td>CCACCTTCCAACCCTTCCCT</td><td>TGG</td><td>||||||||[||||||||||||]PAM</td><td>NA</td><td>Intergenic</td><td>NA</td><td>NA</td></tr>
<tr><td>T33057</td><td>ACGCAGTTCATCCTGCCGTTCGG</td><td>1000</td><td></td><td></td><td></td><td></td><td></td><td></td><td></td><td></td><td></td></tr>
<tr><td>Oligo fwd</td><td>CACCACGCAGTTCATCCTGCCGTT</td><td></td><td></td><td></td><td></td><td></td><td></td><td></td><td></td><td></td><td></td></tr>
<tr><td>Oligo rev</td><td>AAACAACGGCAGGATGAACTGCGT</td><td></td><td></td><td></td><td></td><td></td><td></td><td></td><td></td><td></td><td></td></tr>
<tr><td>Chromosome</td><td>start</td><td>end</td><td>strand</td><td>MM</td><td>target_seq</td><td>PAM</td><td>alignment</td><td>distance</td><td>position</td><td>gene name</td><td>gene id</td></tr>
<tr><td>GU305914.1</td><td>19562</td><td>19584</td><td>-</td><td>0</td><td>ACGCAGTTCATCCTGCCGTT</td><td>CGG</td><td>||||||||[||||||||||||]PAM</td><td>NA</td><td>Intergenic</td><td>NA</td><td>NA</td></tr>
<tr><td>T33058</td><td>TTCGGAATCGTATCGCCGTTCGG</td><td>1000</td><td></td><td></td><td></td><td></td><td></td><td></td><td></td><td></td><td></td></tr>
<tr><td>Oligo fwd</td><td>CACCTTCGGAATCGTATCGCCGTT</td><td></td><td></td><td></td><td></td><td></td><td></td><td></td><td></td><td></td><td></td></tr>
<tr><td>Oligo rev</td><td>AAACAACGGCGATACGATTCCGAA</td><td></td><td></td><td></td><td></td><td></td><td></td><td></td><td></td><td></td><td></td></tr>
<tr><td>Chromosome</td><td>start</td><td>end</td><td>strand</td><td>MM</td><td>target_seq</td><td>PAM</td><td>alignment</td><td>distance</td><td>position</td><td>gene name</td><td>gene id</td></tr>
<tr><td>GU305914.1</td><td>19544</td><td>19566</td><td>-</td><td>0</td><td>TTCGGAATCGTATCGCCGTT</td><td>CGG</td><td>||||||||[||||||||||||]PAM</td><td>NA</td><td>Intergenic</td><td>NA</td><td>NA</td></tr>
<tr><td>T33059</td><td>ATCGTATCGCCGTTCGGCAGCGG</td><td>1000</td><td></td><td></td><td></td><td></td><td></td><td></td><td></td><td></td><td></td></tr>
<tr><td>Oligo fwd</td><td>CACCATCGTATCGCCGTTCGGCAG</td><td></td><td></td><td></td><td></td><td></td><td></td><td></td><td></td><td></td><td></td></tr>
<tr><td>Oligo rev</td><td>AAACCTGCCGAACGGCGATACGAT</td><td></td><td></td><td></td><td></td><td></td><td></td><td></td><td></td><td></td><td></td></tr>
<tr><td>Chromosome</td><td>start</td><td>end</td><td>strand</td><td>MM</td><td>target_seq</td><td>PAM</td><td>alignment</td><td>distance</td><td>position</td><td>gene name</td><td>gene id</td></tr>
<tr><td>GU305914.1</td><td>19538</td><td>19560</td><td>-</td><td>0</td><td>ATCGTATCGCCGTTCGGCAG</td><td>CGG</td><td>||||||||[||||||||||||]PAM</td><td>NA</td><td>Intergenic</td><td>NA</td><td>NA</td></tr>
<tr><td>T33060</td><td>CGCCGTCCCCATCGCGACGATGG</td><td>1000</td><td></td><td></td><td></td><td></td><td></td><td></td><td></td><td></td><td></td></tr>
<tr><td>Oligo fwd</td><td>CACCCGCCGTCCCCATCGCGACGA</td><td></td><td></td><td></td><td></td><td></td><td></td><td></td><td></td><td></td><td></td></tr>
<tr><td>Oligo rev</td><td>AAACTCGTCGCGATGGGGACGGCG</td><td></td><td></td><td></td><td></td><td></td><td></td><td></td><td></td><td></td><td></td></tr>
<tr><td>Chromosome</td><td>start</td><td>end</td><td>strand</td><td>MM</td><td>target_seq</td><td>PAM</td><td>alignment</td><td>distance</td><td>position</td><td>gene name</td><td>gene id</td></tr>
<tr><td>GU305914.1</td><td>19512</td><td>19534</td><td>-</td><td>0</td><td>CGCCGTCCCCATCGCGACGA</td><td>TGG</td><td>||||||||[||||||||||||]PAM</td><td>NA</td><td>Intergenic</td><td>NA</td><td>NA</td></tr>
<tr><td>GU305914.1</td><td>194496</td><td>194518</td><td>-</td><td>4</td><td>CCCCGTCTCCATCGTGACGG</td><td>CGG</td><td>|-|||||-[||||||-||||-]PAM</td><td>NA</td><td>Intergenic</td><td>NA</td><td>NA</td></tr>
<tr><td>T33061</td><td>GCCGTCCCCATCGCGACGATGGG</td><td>1000</td><td></td><td></td><td></td><td></td><td></td><td></td><td></td><td></td><td></td></tr>
<tr><td>Oligo fwd</td><td>CACCGCCGTCCCCATCGCGACGAT</td><td></td><td></td><td></td><td></td><td></td><td></td><td></td><td></td><td></td><td></td></tr>
<tr><td>Oligo rev</td><td>AAACATCGTCGCGATGGGGACGGC</td><td></td><td></td><td></td><td></td><td></td><td></td><td></td><td></td><td></td><td></td></tr>
<tr><td>Chromosome</td><td>start</td><td>end</td><td>strand</td><td>MM</td><td>target_seq</td><td>PAM</td><td>alignment</td><td>distance</td><td>position</td><td>gene name</td><td>gene id</td></tr>
<tr><td>GU305914.1</td><td>19511</td><td>19533</td><td>-</td><td>0</td><td>GCCGTCCCCATCGCGACGAT</td><td>GGG</td><td>||||||||[||||||||||||]PAM</td><td>NA</td><td>Intergenic</td><td>NA</td><td>NA</td></tr>
<tr><td>T33062</td><td>CACGCTCATCGCCAGCAGCGCGG</td><td>1000</td><td></td><td></td><td></td><td></td><td></td><td></td><td></td><td></td><td></td></tr>
<tr><td>Oligo fwd</td><td>CACCCACGCTCATCGCCAGCAGCG</td><td></td><td></td><td></td><td></td><td></td><td></td><td></td><td></td><td></td><td></td></tr>
<tr><td>Oligo rev</td><td>AAACCGCTGCTGGCGATGAGCGTG</td><td></td><td></td><td></td><td></td><td></td><td></td><td></td><td></td><td></td><td></td></tr>
<tr><td>Chromosome</td><td>start</td><td>end</td><td>strand</td><td>MM</td><td>target_seq</td><td>PAM</td><td>alignment</td><td>distance</td><td>position</td><td>gene name</td><td>gene id</td></tr>
<tr><td>GU305914.1</td><td>19464</td><td>19486</td><td>-</td><td>0</td><td>CACGCTCATCGCCAGCAGCG</td><td>CGG</td><td>||||||||[||||||||||||]PAM</td><td>NA</td><td>Intergenic</td><td>NA</td><td>NA</td></tr>
<tr><td>T33063</td><td>CGCCAGCAGCGCGGTGCCTCCGG</td><td>1000</td><td></td><td></td><td></td><td></td><td></td><td></td><td></td><td></td><td></td></tr>
<tr><td>Oligo fwd</td><td>CACCCGCCAGCAGCGCGGTGCCTC</td><td></td><td></td><td></td><td></td><td></td><td></td><td></td><td></td><td></td><td></td></tr>
<tr><td>Oligo rev</td><td>AAACGAGGCACCGCGCTGCTGGCG</td><td></td><td></td><td></td><td></td><td></td><td></td><td></td><td></td><td></td><td></td></tr>
<tr><td>Chromosome</td><td>start</td><td>end</td><td>strand</td><td>MM</td><td>target_seq</td><td>PAM</td><td>alignment</td><td>distance</td><td>position</td><td>gene name</td><td>gene id</td></tr>
<tr><td>GU305914.1</td><td>19455</td><td>19477</td><td>-</td><td>0</td><td>CGCCAGCAGCGCGGTGCCTC</td><td>CGG</td><td>||||||||[||||||||||||]PAM</td><td>NA</td><td>Intergenic</td><td>NA</td><td>NA</td></tr>
<tr><td>T33064</td><td>TCCGGCACCTCCGCCGACGCCGG</td><td>1000</td><td></td><td></td><td></td><td></td><td></td><td></td><td></td><td></td><td></td></tr>
<tr><td>Oligo fwd</td><td>CACCTCCGGCACCTCCGCCGACGC</td><td></td><td></td><td></td><td></td><td></td><td></td><td></td><td></td><td></td><td></td></tr>
<tr><td>Oligo rev</td><td>AAACGCGTCGGCGGAGGTGCCGGA</td><td></td><td></td><td></td><td></td><td></td><td></td><td></td><td></td><td></td><td></td></tr>
<tr><td>Chromosome</td><td>start</td><td>end</td><td>strand</td><td>MM</td><td>target_seq</td><td>PAM</td><td>alignment</td><td>distance</td><td>position</td><td>gene name</td><td>gene id</td></tr>
<tr><td>GU305914.1</td><td>19437</td><td>19459</td><td>-</td><td>0</td><td>TCCGGCACCTCCGCCGACGC</td><td>CGG</td><td>||||||||[||||||||||||]PAM</td><td>NA</td><td>Intergenic</td><td>NA</td><td>NA</td></tr>
<tr><td>GU305914.1</td><td>152551</td><td>152573</td><td>-</td><td>3</td><td>GCCGGCACCGCCGCCGCCGC</td><td>AGG</td><td>-|||||||[|-||||||-|||]PAM</td><td>NA</td><td>Intergenic</td><td>NA</td><td>NA</td></tr>
<tr><td>T33065</td><td>GGCACCTCCGCCGACGCCGGCGG</td><td>1000</td><td></td><td></td><td></td><td></td><td></td><td></td><td></td><td></td><td></td></tr>
<tr><td>Oligo fwd</td><td>CACCGGCACCTCCGCCGACGCCGG</td><td></td><td></td><td></td><td></td><td></td><td></td><td></td><td></td><td></td><td></td></tr>
<tr><td>Oligo rev</td><td>AAACCCGGCGTCGGCGGAGGTGCC</td><td></td><td></td><td></td><td></td><td></td><td></td><td></td><td></td><td></td><td></td></tr>
<tr><td>Chromosome</td><td>start</td><td>end</td><td>strand</td><td>MM</td><td>target_seq</td><td>PAM</td><td>alignment</td><td>distance</td><td>position</td><td>gene name</td><td>gene id</td></tr>
<tr><td>GU305914.1</td><td>19434</td><td>19456</td><td>-</td><td>0</td><td>GGCACCTCCGCCGACGCCGG</td><td>CGG</td><td>||||||||[||||||||||||]PAM</td><td>NA</td><td>Intergenic</td><td>NA</td><td>NA</td></tr>
<tr><td>T33066</td><td>GCACCTCCGCCGACGCCGGCGGG</td><td>1000</td><td></td><td></td><td></td><td></td><td></td><td></td><td></td><td></td><td></td></tr>
<tr><td>Oligo fwd</td><td>CACCGCACCTCCGCCGACGCCGGC</td><td></td><td></td><td></td><td></td><td></td><td></td><td></td><td></td><td></td><td></td></tr>
<tr><td>Oligo rev</td><td>AAACGCCGGCGTCGGCGGAGGTGC</td><td></td><td></td><td></td><td></td><td></td><td></td><td></td><td></td><td></td><td></td></tr>
<tr><td>Chromosome</td><td>start</td><td>end</td><td>strand</td><td>MM</td><td>target_seq</td><td>PAM</td><td>alignment</td><td>distance</td><td>position</td><td>gene name</td><td>gene id</td></tr>
<tr><td>GU305914.1</td><td>19433</td><td>19455</td><td>-</td><td>0</td><td>GCACCTCCGCCGACGCCGGC</td><td>GGG</td><td>||||||||[||||||||||||]PAM</td><td>NA</td><td>Intergenic</td><td>NA</td><td>NA</td></tr>
<tr><td>T33067</td><td>GTAACGTCCCCATACTACCCAGG</td><td>1000</td><td></td><td></td><td></td><td></td><td></td><td></td><td></td><td></td><td></td></tr>
<tr><td>Oligo fwd</td><td>CACCGTAACGTCCCCATACTACCC</td><td></td><td></td><td></td><td></td><td></td><td></td><td></td><td></td><td></td><td></td></tr>
<tr><td>Oligo rev</td><td>AAACGGGTAGTATGGGGACGTTAC</td><td></td><td></td><td></td><td></td><td></td><td></td><td></td><td></td><td></td><td></td></tr>
<tr><td>Chromosome</td><td>start</td><td>end</td><td>strand</td><td>MM</td><td>target_seq</td><td>PAM</td><td>alignment</td><td>distance</td><td>position</td><td>gene name</td><td>gene id</td></tr>
<tr><td>GU305914.1</td><td>19411</td><td>19433</td><td>-</td><td>0</td><td>GTAACGTCCCCATACTACCC</td><td>AGG</td><td>||||||||[||||||||||||]PAM</td><td>NA</td><td>Intergenic</td><td>NA</td><td>NA</td></tr>
<tr><td>T33068</td><td>GCGATGTAGCGTGCGTCCTCCGG</td><td>1000</td><td></td><td></td><td></td><td></td><td></td><td></td><td></td><td></td><td></td></tr>
<tr><td>Oligo fwd</td><td>CACCGCGATGTAGCGTGCGTCCTC</td><td></td><td></td><td></td><td></td><td></td><td></td><td></td><td></td><td></td><td></td></tr>
<tr><td>Oligo rev</td><td>AAACGAGGACGCACGCTACATCGC</td><td></td><td></td><td></td><td></td><td></td><td></td><td></td><td></td><td></td><td></td></tr>
<tr><td>Chromosome</td><td>start</td><td>end</td><td>strand</td><td>MM</td><td>target_seq</td><td>PAM</td><td>alignment</td><td>distance</td><td>position</td><td>gene name</td><td>gene id</td></tr>
<tr><td>GU305914.1</td><td>19350</td><td>19372</td><td>-</td><td>0</td><td>GCGATGTAGCGTGCGTCCTC</td><td>CGG</td><td>||||||||[||||||||||||]PAM</td><td>NA</td><td>Intergenic</td><td>NA</td><td>NA</td></tr>
</table>

<table>
<tr><td>T33069</td><td>CGATGTAGCGTGCGTCCTCCGGG</td><td></td><td>1000</td><td></td><td></td><td></td><td></td><td></td><td></td><td></td><td></td><td></td></tr>
<tr><td>Oligo fwd</td><td>CACCCGATGTAGCGTGCGTCCTCC</td><td></td><td></td><td></td><td></td><td></td><td></td><td></td><td></td><td></td><td></td><td></td></tr>
<tr><td>Oligo rev</td><td>AAACGGAGGACGCACGCTACATCG</td><td></td><td></td><td></td><td></td><td></td><td></td><td></td><td></td><td></td><td></td><td></td></tr>
<tr><td>Chromosome</td><td>start</td><td></td><td>end</td><td>strand</td><td>MM</td><td>target_seq</td><td>PAM</td><td>alignment</td><td>distance</td><td>position</td><td>gene name</td><td>gene id</td></tr>
<tr><td>GU305914.1</td><td></td><td>19349</td><td>19371</td><td>-</td><td>0</td><td>CGATGTAGCGTGCGTCCTCC</td><td>GGG</td><td>||||||||[||||||||||||]PAM</td><td>NA</td><td>Intergenic</td><td>NA</td><td>NA</td></tr>
<tr><td>T33070</td><td>GTAGCGTGCGTCCTCCGGGTAGG</td><td></td><td>1000</td><td></td><td></td><td></td><td></td><td></td><td></td><td></td><td></td><td></td></tr>
<tr><td>Oligo fwd</td><td>CACCGTAGCGTGCGTCCTCCGGGT</td><td></td><td></td><td></td><td></td><td></td><td></td><td></td><td></td><td></td><td></td><td></td></tr>
<tr><td>Oligo rev</td><td>AAACACCCGGAGGACGCACGCTAC</td><td></td><td></td><td></td><td></td><td></td><td></td><td></td><td></td><td></td><td></td><td></td></tr>
<tr><td>Chromosome</td><td>start</td><td></td><td>end</td><td>strand</td><td>MM</td><td>target_seq</td><td>PAM</td><td>alignment</td><td>distance</td><td>position</td><td>gene name</td><td>gene id</td></tr>
<tr><td>GU305914.1</td><td></td><td>19345</td><td>19367</td><td>-</td><td>0</td><td>GTAGCGTGCGTCCTCCGGGT</td><td>AGG</td><td>||||||||[||||||||||||]PAM</td><td>NA</td><td>Intergenic</td><td>NA</td><td>NA</td></tr>
<tr><td>T33071</td><td>GGTAGGATGACGCCCGTCGCCGG</td><td></td><td>1000</td><td></td><td></td><td></td><td></td><td></td><td></td><td></td><td></td><td></td></tr>
<tr><td>Oligo fwd</td><td>CACCGGTAGGATGACGCCCGTCGC</td><td></td><td></td><td></td><td></td><td></td><td></td><td></td><td></td><td></td><td></td><td></td></tr>
<tr><td>Oligo rev</td><td>AAACGCGACGGGCGTCATCCTACC</td><td></td><td></td><td></td><td></td><td></td><td></td><td></td><td></td><td></td><td></td><td></td></tr>
<tr><td>Chromosome</td><td>start</td><td></td><td>end</td><td>strand</td><td>MM</td><td>target_seq</td><td>PAM</td><td>alignment</td><td>distance</td><td>position</td><td>gene name</td><td>gene id</td></tr>
<tr><td>GU305914.1</td><td></td><td>19328</td><td>19350</td><td>-</td><td>0</td><td>GGTAGGATGACGCCCGTCGC</td><td>CGG</td><td>||||||||[||||||||||||]PAM</td><td>NA</td><td>Intergenic</td><td>NA</td><td>NA</td></tr>
<tr><td>T33072</td><td>GGATGACGCCCGTCGCCGGACGG</td><td></td><td>1000</td><td></td><td></td><td></td><td></td><td></td><td></td><td></td><td></td><td></td></tr>
<tr><td>Oligo fwd</td><td>CACCGGATGACGCCCGTCGCCGGA</td><td></td><td></td><td></td><td></td><td></td><td></td><td></td><td></td><td></td><td></td><td></td></tr>
<tr><td>Oligo rev</td><td>AAACTCCGGCGACGGGCGTCATCC</td><td></td><td></td><td></td><td></td><td></td><td></td><td></td><td></td><td></td><td></td><td></td></tr>
<tr><td>Chromosome</td><td>start</td><td></td><td>end</td><td>strand</td><td>MM</td><td>target_seq</td><td>PAM</td><td>alignment</td><td>distance</td><td>position</td><td>gene name</td><td>gene id</td></tr>
<tr><td>GU305914.1</td><td></td><td>19324</td><td>19346</td><td>-</td><td>0</td><td>GGATGACGCCCGTCGCCGGA</td><td>CGG</td><td>||||||||[||||||||||||]PAM</td><td>NA</td><td>Intergenic</td><td>NA</td><td>NA</td></tr>
<tr><td>T33073</td><td>TCCAGATCCCCCGTCGCGCGAGG</td><td></td><td>1000</td><td></td><td></td><td></td><td></td><td></td><td></td><td></td><td></td><td></td></tr>
<tr><td>Oligo fwd</td><td>CACCTCCAGATCCCCCGTCGCGCG</td><td></td><td></td><td></td><td></td><td></td><td></td><td></td><td></td><td></td><td></td><td></td></tr>
<tr><td>Oligo rev</td><td>AAACCGCGCGACGGGGGATCTGGA</td><td></td><td></td><td></td><td></td><td></td><td></td><td></td><td></td><td></td><td></td><td></td></tr>
<tr><td>Chromosome</td><td>start</td><td></td><td>end</td><td>strand</td><td>MM</td><td>target_seq</td><td>PAM</td><td>alignment</td><td>distance</td><td>position</td><td>gene name</td><td>gene id</td></tr>
<tr><td>GU305914.1</td><td></td><td>19301</td><td>19323</td><td>-</td><td>0</td><td>TCCAGATCCCCCGTCGCGCG</td><td>AGG</td><td>||||||||[||||||||||||]PAM</td><td>NA</td><td>Intergenic</td><td>NA</td><td>NA</td></tr>
<tr><td>T33074</td><td>CGTCGCGCGAGGAAAGAAAAAGG</td><td></td><td>1000</td><td></td><td></td><td></td><td></td><td></td><td></td><td></td><td></td><td></td></tr>
<tr><td>Oligo fwd</td><td>CACCCGTCGCGCGAGGAAAGAAAA</td><td></td><td></td><td></td><td></td><td></td><td></td><td></td><td></td><td></td><td></td><td></td></tr>
<tr><td>Oligo rev</td><td>AAACTTTTCTTTCCTCGCGCGACG</td><td></td><td></td><td></td><td></td><td></td><td></td><td></td><td></td><td></td><td></td><td></td></tr>
<tr><td>Chromosome</td><td>start</td><td></td><td>end</td><td>strand</td><td>MM</td><td>target_seq</td><td>PAM</td><td>alignment</td><td>distance</td><td>position</td><td>gene name</td><td>gene id</td></tr>
<tr><td>GU305914.1</td><td></td><td>19290</td><td>19312</td><td>-</td><td>0</td><td>CGTCGCGCGAGGAAAGAAAA</td><td>AGG</td><td>||||||||[||||||||||||]PAM</td><td>NA</td><td>Intergenic</td><td>NA</td><td>NA</td></tr>
<tr><td>T33075</td><td>CGCGCGAGGAAAGAAAAAGGCGG</td><td></td><td>1000</td><td></td><td></td><td></td><td></td><td></td><td></td><td></td><td></td><td></td></tr>
<tr><td>Oligo fwd</td><td>CACCCGCGCGAGGAAAGAAAAAGG</td><td></td><td></td><td></td><td></td><td></td><td></td><td></td><td></td><td></td><td></td><td></td></tr>
<tr><td>Oligo rev</td><td>AAACCCTTTTTCTTTCCTCGCGCG</td><td></td><td></td><td></td><td></td><td></td><td></td><td></td><td></td><td></td><td></td><td></td></tr>
<tr><td>Chromosome</td><td>start</td><td></td><td>end</td><td>strand</td><td>MM</td><td>target_seq</td><td>PAM</td><td>alignment</td><td>distance</td><td>position</td><td>gene name</td><td>gene id</td></tr>
<tr><td>GU305914.1</td><td></td><td>19287</td><td>19309</td><td>-</td><td>0</td><td>CGCGCGAGGAAAGAAAAAGG</td><td>CGG</td><td>||||||||[||||||||||||]PAM</td><td>NA</td><td>Intergenic</td><td>NA</td><td>NA</td></tr>
<tr><td>T33076</td><td>AGTAACAACAATTTATTGAACGG</td><td></td><td>1000</td><td></td><td></td><td></td><td></td><td></td><td></td><td></td><td></td><td></td></tr>
<tr><td>Oligo fwd</td><td>CACCAGTAACAACAATTTATTGAA</td><td></td><td></td><td></td><td></td><td></td><td></td><td></td><td></td><td></td><td></td><td></td></tr>
<tr><td>Oligo rev</td><td>AAACTTCAATAAATTGTTGTTACT</td><td></td><td></td><td></td><td></td><td></td><td></td><td></td><td></td><td></td><td></td><td></td></tr>
<tr><td>Chromosome</td><td>start</td><td></td><td>end</td><td>strand</td><td>MM</td><td>target_seq</td><td>PAM</td><td>alignment</td><td>distance</td><td>position</td><td>gene name</td><td>gene id</td></tr>
<tr><td>GU305914.1</td><td></td><td>19262</td><td>19284</td><td>-</td><td>0</td><td>AGTAACAACAATTTATTGAA</td><td>CGG</td><td>||||||||[||||||||||||]PAM</td><td>NA</td><td>Intergenic</td><td>NA</td><td>NA</td></tr>
<tr><td>T33077</td><td>AACAACAATTTATTGAACGGAGG</td><td></td><td>1000</td><td></td><td></td><td></td><td></td><td></td><td></td><td></td><td></td><td></td></tr>
<tr><td>Oligo fwd</td><td>CACCAACAACAATTTATTGAACGG</td><td></td><td></td><td></td><td></td><td></td><td></td><td></td><td></td><td></td><td></td><td></td></tr>
<tr><td>Oligo rev</td><td>AAACCCGTTCAATAAATTGTTGTT</td><td></td><td></td><td></td><td></td><td></td><td></td><td></td><td></td><td></td><td></td><td></td></tr>
<tr><td>Chromosome</td><td>start</td><td></td><td>end</td><td>strand</td><td>MM</td><td>target_seq</td><td>PAM</td><td>alignment</td><td>distance</td><td>position</td><td>gene name</td><td>gene id</td></tr>
<tr><td>GU305914.1</td><td></td><td>19259</td><td>19281</td><td>-</td><td>0</td><td>AACAACAATTTATTGAACGG</td><td>AGG</td><td>||||||||[||||||||||||]PAM</td><td>NA</td><td>Intergenic</td><td>NA</td><td>NA</td></tr>
<tr><td>T33078</td><td>ACAACAATTTATTGAACGGAGGG</td><td></td><td>1000</td><td></td><td></td><td></td><td></td><td></td><td></td><td></td><td></td><td></td></tr>
<tr><td>Oligo fwd</td><td>CACCACAACAATTTATTGAACGGA</td><td></td><td></td><td></td><td></td><td></td><td></td><td></td><td></td><td></td><td></td><td></td></tr>
<tr><td>Oligo rev</td><td>AAACTCCGTTCAATAAATTGTTGT</td><td></td><td></td><td></td><td></td><td></td><td></td><td></td><td></td><td></td><td></td><td></td></tr>
<tr><td>Chromosome</td><td>start</td><td></td><td>end</td><td>strand</td><td>MM</td><td>target_seq</td><td>PAM</td><td>alignment</td><td>distance</td><td>position</td><td>gene name</td><td>gene id</td></tr>
<tr><td>GU305914.1</td><td></td><td>19258</td><td>19280</td><td>-</td><td>0</td><td>ACAACAATTTATTGAACGGA</td><td>GGG</td><td>||||||||[||||||||||||]PAM</td><td>NA</td><td>Intergenic</td><td>NA</td><td>NA</td></tr>
<tr><td>T33079</td><td>TTTATTGAACGGAGGGCCTGTGG</td><td></td><td>1000</td><td></td><td></td><td></td><td></td><td></td><td></td><td></td><td></td><td></td></tr>
<tr><td>Oligo fwd</td><td>CACCTTTATTGAACGGAGGGCCTG</td><td></td><td></td><td></td><td></td><td></td><td></td><td></td><td></td><td></td><td></td><td></td></tr>
<tr><td>Oligo rev</td><td>AAACCAGGCCCTCCGTTCAATAAA</td><td></td><td></td><td></td><td></td><td></td><td></td><td></td><td></td><td></td><td></td><td></td></tr>
<tr><td>Chromosome</td><td>start</td><td></td><td>end</td><td>strand</td><td>MM</td><td>target_seq</td><td>PAM</td><td>alignment</td><td>distance</td><td>position</td><td>gene name</td><td>gene id</td></tr>
<tr><td>GU305914.1</td><td></td><td>19251</td><td>19273</td><td>-</td><td>0</td><td>TTTATTGAACGGAGGGCCTG</td><td>TGG</td><td>||||||||[||||||||||||]PAM</td><td>NA</td><td>Intergenic</td><td>NA</td><td>NA</td></tr>
<tr><td>T33080</td><td>GGGCCTGTGGAGACGACCACCGG</td><td></td><td>1000</td><td></td><td></td><td></td><td></td><td></td><td></td><td></td><td></td><td></td></tr>
<tr><td>Oligo fwd</td><td>CACCGGGCCTGTGGAGACGACCAC</td><td></td><td></td><td></td><td></td><td></td><td></td><td></td><td></td><td></td><td></td><td></td></tr>
<tr><td>Oligo rev</td><td>AAACGTGGTCGTCTCCACAGGCCC</td><td></td><td></td><td></td><td></td><td></td><td></td><td></td><td></td><td></td><td></td><td></td></tr>
<tr><td>Chromosome</td><td>start</td><td></td><td>end</td><td>strand</td><td>MM</td><td>target_seq</td><td>PAM</td><td>alignment</td><td>distance</td><td>position</td><td>gene name</td><td>gene id</td></tr>
<tr><td>GU305914.1</td><td></td><td>19238</td><td>19260</td><td>-</td><td>0</td><td>GGGCCTGTGGAGACGACCAC</td><td>CGG</td><td>||||||||[||||||||||||]PAM</td><td>NA</td><td>Intergenic</td><td>NA</td><td>NA</td></tr>
<tr><td>T33081</td><td>GACCACCGGCTCGCGCAGAGCGG</td><td></td><td>1000</td><td></td><td></td><td></td><td></td><td></td><td></td><td></td><td></td><td></td></tr>
<tr><td>Oligo fwd</td><td>CACCGACCACCGGCTCGCGCAGAG</td><td></td><td></td><td></td><td></td><td></td><td></td><td></td><td></td><td></td><td></td><td></td></tr>
<tr><td>Oligo rev</td><td>AAACCTCTGCGCGAGCCGGTGGTC</td><td></td><td></td><td></td><td></td><td></td><td></td><td></td><td></td><td></td><td></td><td></td></tr>
<tr><td>Chromosome</td><td>start</td><td></td><td>end</td><td>strand</td><td>MM</td><td>target_seq</td><td>PAM</td><td>alignment</td><td>distance</td><td>position</td><td>gene name</td><td>gene id</td></tr>
<tr><td>GU305914.1</td><td></td><td>19224</td><td>19246</td><td>-</td><td>0</td><td>GACCACCGGCTCGCGCAGAG</td><td>CGG</td><td>||||||||[||||||||||||]PAM</td><td>NA</td><td>Intergenic</td><td>NA</td><td>NA</td></tr>
<tr><td>T33082</td><td>ACCGGCTCGCGCAGAGCGGCCGG</td><td></td><td>1000</td><td></td><td></td><td></td><td></td><td></td><td></td><td></td><td></td><td></td></tr>
<tr><td>Oligo fwd</td><td>CACCACCGGCTCGCGCAGAGCGGC</td><td></td><td></td><td></td><td></td><td></td><td></td><td></td><td></td><td></td><td></td><td></td></tr>
<tr><td>Oligo rev</td><td>AAACGCCGCTCTGCGCGAGCCGGT</td><td></td><td></td><td></td><td></td><td></td><td></td><td></td><td></td><td></td><td></td><td></td></tr>
<tr><td>Chromosome</td><td>start</td><td></td><td>end</td><td>strand</td><td>MM</td><td>target_seq</td><td>PAM</td><td>alignment</td><td>distance</td><td>position</td><td>gene name</td><td>gene id</td></tr>
<tr><td>GU305914.1</td><td></td><td>19220</td><td>19242</td><td>-</td><td>0</td><td>ACCGGCTCGCGCAGAGCGGC</td><td>CGG</td><td>||||||||[||||||||||||]PAM</td><td>NA</td><td>Intergenic</td><td>NA</td><td>NA</td></tr>
<tr><td>T33083</td><td>GGCTCGCGCAGAGCGGCCGGAGG</td><td></td><td>1000</td><td></td><td></td><td></td><td></td><td></td><td></td><td></td><td></td><td></td></tr>
<tr><td>Oligo fwd</td><td>CACCGGCTCGCGCAGAGCGGCCGG</td><td></td><td></td><td></td><td></td><td></td><td></td><td></td><td></td><td></td><td></td><td></td></tr>
<tr><td>Oligo rev</td><td>AAACCCGGCCGCTCTGCGCGAGCC</td><td></td><td></td><td></td><td></td><td></td><td></td><td></td><td></td><td></td><td></td><td></td></tr>
<tr><td>Chromosome</td><td>start</td><td></td><td>end</td><td>strand</td><td>MM</td><td>target_seq</td><td>PAM</td><td>alignment</td><td>distance</td><td>position</td><td>gene name</td><td>gene id</td></tr>
<tr><td>GU305914.1</td><td></td><td>19217</td><td>19239</td><td>-</td><td>0</td><td>GGCTCGCGCAGAGCGGCCGG</td><td>AGG</td><td>||||||||[||||||||||||]PAM</td><td>NA</td><td>Intergenic</td><td>NA</td><td>NA</td></tr>
</table>

<table>
<tr><td>T33084</td><td>CGGCCGGAGGTGTCCCGCCTCGG</td><td>1000</td><td></td><td></td><td></td><td></td><td></td><td></td><td></td><td></td><td></td></tr>
<tr><td>Oligo fwd</td><td>CACCCGGCCGGAGGTGTCCCGCCT</td><td></td><td></td><td></td><td></td><td></td><td></td><td></td><td></td><td></td><td></td></tr>
<tr><td>Oligo rev</td><td>AAACAGGCGGGACACCTCCGGCCG</td><td></td><td></td><td></td><td></td><td></td><td></td><td></td><td></td><td></td><td></td></tr>
<tr><td>Chromosome</td><td>start</td><td>end</td><td>strand</td><td>MM</td><td>target_seq</td><td>PAM</td><td>alignment</td><td>distance</td><td>position</td><td>gene name</td><td>gene id</td></tr>
<tr><td>GU305914.1</td><td>19204</td><td>19226</td><td>-</td><td>0</td><td>CGGCCGGAGGTGTCCCGCCT</td><td>CGG</td><td>||||||||[||||||||||||]PAM</td><td>NA</td><td>Intergenic</td><td>NA</td><td>NA</td></tr>
<tr><td>T33085</td><td>TGTCCCGCCTCGGCAGAAAGCGG</td><td>1000</td><td></td><td></td><td></td><td></td><td></td><td></td><td></td><td></td><td></td></tr>
<tr><td>Oligo fwd</td><td>CACCTGTCCCGCCTCGGCAGAAAG</td><td></td><td></td><td></td><td></td><td></td><td></td><td></td><td></td><td></td><td></td></tr>
<tr><td>Oligo rev</td><td>AAACCTTTCTGCCGAGGCGGGACA</td><td></td><td></td><td></td><td></td><td></td><td></td><td></td><td></td><td></td><td></td></tr>
<tr><td>Chromosome</td><td>start</td><td>end</td><td>strand</td><td>MM</td><td>target_seq</td><td>PAM</td><td>alignment</td><td>distance</td><td>position</td><td>gene name</td><td>gene id</td></tr>
<tr><td>GU305914.1</td><td>19194</td><td>19216</td><td>-</td><td>0</td><td>TGTCCCGCCTCGGCAGAAAG</td><td>CGG</td><td>||||||||[||||||||||||]PAM</td><td>NA</td><td>Intergenic</td><td>NA</td><td>NA</td></tr>
<tr><td>T33086</td><td>GCCTCGGCAGAAAGCGGCATCGG</td><td>1000</td><td></td><td></td><td></td><td></td><td></td><td></td><td></td><td></td><td></td></tr>
<tr><td>Oligo fwd</td><td>CACCGCCTCGGCAGAAAGCGGCAT</td><td></td><td></td><td></td><td></td><td></td><td></td><td></td><td></td><td></td><td></td></tr>
<tr><td>Oligo rev</td><td>AAACATGCCGCTTTCTGCCGAGGC</td><td></td><td></td><td></td><td></td><td></td><td></td><td></td><td></td><td></td><td></td></tr>
<tr><td>Chromosome</td><td>start</td><td>end</td><td>strand</td><td>MM</td><td>target_seq</td><td>PAM</td><td>alignment</td><td>distance</td><td>position</td><td>gene name</td><td>gene id</td></tr>
<tr><td>GU305914.1</td><td>19188</td><td>19210</td><td>-</td><td>0</td><td>GCCTCGGCAGAAAGCGGCAT</td><td>CGG</td><td>||||||||[||||||||||||]PAM</td><td>NA</td><td>Intergenic</td><td>NA</td><td>NA</td></tr>
<tr><td>T33087</td><td>GGCAGCATCGTCGTCGTCCTCGG</td><td>1000</td><td></td><td></td><td></td><td></td><td></td><td></td><td></td><td></td><td></td></tr>
<tr><td>Oligo fwd</td><td>CACCGGCAGCATCGTCGTCGTCCT</td><td></td><td></td><td></td><td></td><td></td><td></td><td></td><td></td><td></td><td></td></tr>
<tr><td>Oligo rev</td><td>AAACAGGACGACGACGATGCTGCC</td><td></td><td></td><td></td><td></td><td></td><td></td><td></td><td></td><td></td><td></td></tr>
<tr><td>Chromosome</td><td>start</td><td>end</td><td>strand</td><td>MM</td><td>target_seq</td><td>PAM</td><td>alignment</td><td>distance</td><td>position</td><td>gene name</td><td>gene id</td></tr>
<tr><td>GU305914.1</td><td>19167</td><td>19189</td><td>-</td><td>0</td><td>GGCAGCATCGTCGTCGTCCT</td><td>CGG</td><td>||||||||[||||||||||||]PAM</td><td>NA</td><td>Intergenic</td><td>NA</td><td>NA</td></tr>
<tr><td>T33088</td><td>TCGTCCTCGGCATCTTCTTCCGG</td><td>1000</td><td></td><td></td><td></td><td></td><td></td><td></td><td></td><td></td><td></td></tr>
<tr><td>Oligo fwd</td><td>CACCTCGTCCTCGGCATCTTCTTC</td><td></td><td></td><td></td><td></td><td></td><td></td><td></td><td></td><td></td><td></td></tr>
<tr><td>Oligo rev</td><td>AAACGAAGAAGATGCCGAGGACGA</td><td></td><td></td><td></td><td></td><td></td><td></td><td></td><td></td><td></td><td></td></tr>
<tr><td>Chromosome</td><td>start</td><td>end</td><td>strand</td><td>MM</td><td>target_seq</td><td>PAM</td><td>alignment</td><td>distance</td><td>position</td><td>gene name</td><td>gene id</td></tr>
<tr><td>GU305914.1</td><td>19154</td><td>19176</td><td>-</td><td>0</td><td>TCGTCCTCGGCATCTTCTTC</td><td>CGG</td><td>||||||||[||||||||||||]PAM</td><td>NA</td><td>Intergenic</td><td>NA</td><td>NA</td></tr>
<tr><td>T33089</td><td>TCTTCTTCCGGACCGTCTCGTGG</td><td>1000</td><td></td><td></td><td></td><td></td><td></td><td></td><td></td><td></td><td></td></tr>
<tr><td>Oligo fwd</td><td>CACCTCTTCTTCCGGACCGTCTCG</td><td></td><td></td><td></td><td></td><td></td><td></td><td></td><td></td><td></td><td></td></tr>
<tr><td>Oligo rev</td><td>AAACCGAGACGGTCCGGAAGAAGA</td><td></td><td></td><td></td><td></td><td></td><td></td><td></td><td></td><td></td><td></td></tr>
<tr><td>Chromosome</td><td>start</td><td>end</td><td>strand</td><td>MM</td><td>target_seq</td><td>PAM</td><td>alignment</td><td>distance</td><td>position</td><td>gene name</td><td>gene id</td></tr>
<tr><td>GU305914.1</td><td>19142</td><td>19164</td><td>-</td><td>0</td><td>TCTTCTTCCGGACCGTCTCG</td><td>TGG</td><td>||||||||[||||||||||||]PAM</td><td>NA</td><td>Intergenic</td><td>NA</td><td>NA</td></tr>
<tr><td>T33090</td><td>CTTCCGGACCGTCTCGTGGCCGG</td><td>1000</td><td></td><td></td><td></td><td></td><td></td><td></td><td></td><td></td><td></td></tr>
<tr><td>Oligo fwd</td><td>CACCCTTCCGGACCGTCTCGTGGC</td><td></td><td></td><td></td><td></td><td></td><td></td><td></td><td></td><td></td><td></td></tr>
<tr><td>Oligo rev</td><td>AAACGCCACGAGACGGTCCGGAAG</td><td></td><td></td><td></td><td></td><td></td><td></td><td></td><td></td><td></td><td></td></tr>
<tr><td>Chromosome</td><td>start</td><td>end</td><td>strand</td><td>MM</td><td>target_seq</td><td>PAM</td><td>alignment</td><td>distance</td><td>position</td><td>gene name</td><td>gene id</td></tr>
<tr><td>GU305914.1</td><td>19138</td><td>19160</td><td>-</td><td>0</td><td>CTTCCGGACCGTCTCGTGGC</td><td>CGG</td><td>||||||||[||||||||||||]PAM</td><td>NA</td><td>Intergenic</td><td>NA</td><td>NA</td></tr>
<tr><td>T33091</td><td>GTCTCGTGGCCGGATCATCTCGG</td><td>1000</td><td></td><td></td><td></td><td></td><td></td><td></td><td></td><td></td><td></td></tr>
<tr><td>Oligo fwd</td><td>CACCGTCTCGTGGCCGGATCATCT</td><td></td><td></td><td></td><td></td><td></td><td></td><td></td><td></td><td></td><td></td></tr>
<tr><td>Oligo rev</td><td>AAACAGATGATCCGGCCACGAGAC</td><td></td><td></td><td></td><td></td><td></td><td></td><td></td><td></td><td></td><td></td></tr>
<tr><td>Chromosome</td><td>start</td><td>end</td><td>strand</td><td>MM</td><td>target_seq</td><td>PAM</td><td>alignment</td><td>distance</td><td>position</td><td>gene name</td><td>gene id</td></tr>
<tr><td>GU305914.1</td><td>19128</td><td>19150</td><td>-</td><td>0</td><td>GTCTCGTGGCCGGATCATCT</td><td>CGG</td><td>||||||||[||||||||||||]PAM</td><td>NA</td><td>Intergenic</td><td>NA</td><td>NA</td></tr>
<tr><td>T33092</td><td>CTCGGTCAGTGACCCGAACCTGG</td><td>1000</td><td></td><td></td><td></td><td></td><td></td><td></td><td></td><td></td><td></td></tr>
<tr><td>Oligo fwd</td><td>CACCCTCGGTCAGTGACCCGAACC</td><td></td><td></td><td></td><td></td><td></td><td></td><td></td><td></td><td></td><td></td></tr>
<tr><td>Oligo rev</td><td>AAACGGTTCGGGTCACTGACCGAG</td><td></td><td></td><td></td><td></td><td></td><td></td><td></td><td></td><td></td><td></td></tr>
<tr><td>Chromosome</td><td>start</td><td>end</td><td>strand</td><td>MM</td><td>target_seq</td><td>PAM</td><td>alignment</td><td>distance</td><td>position</td><td>gene name</td><td>gene id</td></tr>
<tr><td>GU305914.1</td><td>19110</td><td>19132</td><td>-</td><td>0</td><td>CTCGGTCAGTGACCCGAACC</td><td>TGG</td><td>||||||||[||||||||||||]PAM</td><td>NA</td><td>Intergenic</td><td>NA</td><td>NA</td></tr>
<tr><td>T33093</td><td>TCGGTCAGTGACCCGAACCTGGG</td><td>1000</td><td></td><td></td><td></td><td></td><td></td><td></td><td></td><td></td><td></td></tr>
<tr><td>Oligo fwd</td><td>CACCTCGGTCAGTGACCCGAACCT</td><td></td><td></td><td></td><td></td><td></td><td></td><td></td><td></td><td></td><td></td></tr>
<tr><td>Oligo rev</td><td>AAACAGGTTCGGGTCACTGACCGA</td><td></td><td></td><td></td><td></td><td></td><td></td><td></td><td></td><td></td><td></td></tr>
<tr><td>Chromosome</td><td>start</td><td>end</td><td>strand</td><td>MM</td><td>target_seq</td><td>PAM</td><td>alignment</td><td>distance</td><td>position</td><td>gene name</td><td>gene id</td></tr>
<tr><td>GU305914.1</td><td>19109</td><td>19131</td><td>-</td><td>0</td><td>TCGGTCAGTGACCCGAACCT</td><td>GGG</td><td>||||||||[||||||||||||]PAM</td><td>NA</td><td>Intergenic</td><td>NA</td><td>NA</td></tr>
<tr><td>T33094</td><td>CAGTGACCCGAACCTGGGCGCGG</td><td>1000</td><td></td><td></td><td></td><td></td><td></td><td></td><td></td><td></td><td></td></tr>
<tr><td>Oligo fwd</td><td>CACCCAGTGACCCGAACCTGGGCG</td><td></td><td></td><td></td><td></td><td></td><td></td><td></td><td></td><td></td><td></td></tr>
<tr><td>Oligo rev</td><td>AAACCGCCCAGGTTCGGGTCACTG</td><td></td><td></td><td></td><td></td><td></td><td></td><td></td><td></td><td></td><td></td></tr>
<tr><td>Chromosome</td><td>start</td><td>end</td><td>strand</td><td>MM</td><td>target_seq</td><td>PAM</td><td>alignment</td><td>distance</td><td>position</td><td>gene name</td><td>gene id</td></tr>
<tr><td>GU305914.1</td><td>19104</td><td>19126</td><td>-</td><td>0</td><td>CAGTGACCCGAACCTGGGCG</td><td>CGG</td><td>||||||||[||||||||||||]PAM</td><td>NA</td><td>Intergenic</td><td>NA</td><td>NA</td></tr>
<tr><td>T33095</td><td>CCTCCTCGTCGTCACAGATGTGG</td><td>1000</td><td></td><td></td><td></td><td></td><td></td><td></td><td></td><td></td><td></td></tr>
<tr><td>Oligo fwd</td><td>CACCCCTCCTCGTCGTCACAGATG</td><td></td><td></td><td></td><td></td><td></td><td></td><td></td><td></td><td></td><td></td></tr>
<tr><td>Oligo rev</td><td>AAACCATCTGTGACGACGAGGAGG</td><td></td><td></td><td></td><td></td><td></td><td></td><td></td><td></td><td></td><td></td></tr>
<tr><td>Chromosome</td><td>start</td><td>end</td><td>strand</td><td>MM</td><td>target_seq</td><td>PAM</td><td>alignment</td><td>distance</td><td>position</td><td>gene name</td><td>gene id</td></tr>
<tr><td>GU305914.1</td><td>19078</td><td>19100</td><td>-</td><td>0</td><td>CCTCCTCGTCGTCACAGATG</td><td>TGG</td><td>||||||||[||||||||||||]PAM</td><td>NA</td><td>Intergenic</td><td>NA</td><td>NA</td></tr>
<tr><td>T33096</td><td>CTCCTCGTCGTCACAGATGTGGG</td><td>1000</td><td></td><td></td><td></td><td></td><td></td><td></td><td></td><td></td><td></td></tr>
<tr><td>Oligo fwd</td><td>CACCCTCCTCGTCGTCACAGATGT</td><td></td><td></td><td></td><td></td><td></td><td></td><td></td><td></td><td></td><td></td></tr>
<tr><td>Oligo rev</td><td>AAACACATCTGTGACGACGAGGAG</td><td></td><td></td><td></td><td></td><td></td><td></td><td></td><td></td><td></td><td></td></tr>
<tr><td>Chromosome</td><td>start</td><td>end</td><td>strand</td><td>MM</td><td>target_seq</td><td>PAM</td><td>alignment</td><td>distance</td><td>position</td><td>gene name</td><td>gene id</td></tr>
<tr><td>GU305914.1</td><td>19077</td><td>19099</td><td>-</td><td>0</td><td>CTCCTCGTCGTCACAGATGT</td><td>GGG</td><td>||||||||[||||||||||||]PAM</td><td>NA</td><td>Intergenic</td><td>NA</td><td>NA</td></tr>
<tr><td>T33097</td><td>GCCGCCGTCGTCCAACCCTTCGG</td><td>1000</td><td></td><td></td><td></td><td></td><td></td><td></td><td></td><td></td><td></td></tr>
<tr><td>Oligo fwd</td><td>CACCGCCGCCGTCGTCCAACCCTT</td><td></td><td></td><td></td><td></td><td></td><td></td><td></td><td></td><td></td><td></td></tr>
<tr><td>Oligo rev</td><td>AAACAAGGGTTGGACGACGGCGGC</td><td></td><td></td><td></td><td></td><td></td><td></td><td></td><td></td><td></td><td></td></tr>
<tr><td>Chromosome</td><td>start</td><td>end</td><td>strand</td><td>MM</td><td>target_seq</td><td>PAM</td><td>alignment</td><td>distance</td><td>position</td><td>gene name</td><td>gene id</td></tr>
<tr><td>GU305914.1</td><td>19049</td><td>19071</td><td>-</td><td>0</td><td>GCCGCCGTCGTCCAACCCTT</td><td>CGG</td><td>||||||||[||||||||||||]PAM</td><td>NA</td><td>Intergenic</td><td>NA</td><td>NA</td></tr>
<tr><td>T33098</td><td>AGTCGTCGATCAGTTTGCGCCGG</td><td>1000</td><td></td><td></td><td></td><td></td><td></td><td></td><td></td><td></td><td></td></tr>
<tr><td>Oligo fwd</td><td>CACCAGTCGTCGATCAGTTTGCGC</td><td></td><td></td><td></td><td></td><td></td><td></td><td></td><td></td><td></td><td></td></tr>
<tr><td>Oligo rev</td><td>AAACGCGCAAACTGATCGACGACT</td><td></td><td></td><td></td><td></td><td></td><td></td><td></td><td></td><td></td><td></td></tr>
<tr><td>Chromosome</td><td>start</td><td>end</td><td>strand</td><td>MM</td><td>target_seq</td><td>PAM</td><td>alignment</td><td>distance</td><td>position</td><td>gene name</td><td>gene id</td></tr>
</table>

| | | | | | | | | | | | | |
|---|---|---|---|---|---|---|---|---|---|---|---|---|
| GU305914.1 | | 19026 | 19048 | - | 0 | AGTCGTCGATCAGTTTGCGC | CGG | \|\|\|\|\|\|\|\|[\|\|\|\|\|\|\|\|\|\|\|\|]PAM | NA | Intergenic | NA | NA |
| T33099 | GTCGTCGATCAGTTTGCGCCGGG | | 1000 | | | | | | | | | |
| Oligo fwd | CACCGTCGTCGATCAGTTTGCGCC | | | | | | | | | | | |
| Oligo rev | AAACGGCGCAAACTGATCGACGAC | | | | | | | | | | | |
| Chromosome | start | | end | strand | MM | target_seq | PAM | alignment | distance | position | gene name | gene id |
| GU305914.1 | | 19025 | 19047 | - | 0 | GTCGTCGATCAGTTTGCGCC | GGG | \|\|\|\|\|\|\|\|[\|\|\|\|\|\|\|\|\|\|\|\|]PAM | NA | Intergenic | NA | NA |
| T33100 | TCAGTTTGCGCCGGGACAGCAGG | | 1000 | | | | | | | | | |
| Oligo fwd | CACCTCAGTTTGCGCCGGGACAGC | | | | | | | | | | | |
| Oligo rev | AAACGCTGTCCCGGCGCAAACTGA | | | | | | | | | | | |
| Chromosome | start | | end | strand | MM | target_seq | PAM | alignment | distance | position | gene name | gene id |
| GU305914.1 | | 19017 | 19039 | - | 0 | TCAGTTTGCGCCGGGACAGC | AGG | \|\|\|\|\|\|\|\|[\|\|\|\|\|\|\|\|\|\|\|\|]PAM | NA | Intergenic | NA | NA |
| T33101 | CAGTTTGCGCCGGGACAGCAGGG | | 1000 | | | | | | | | | |
| Oligo fwd | CACCCAGTTTGCGCCGGGACAGCA | | | | | | | | | | | |
| Oligo rev | AAACTGCTGTCCCGGCGCAAACTG | | | | | | | | | | | |
| Chromosome | start | | end | strand | MM | target_seq | PAM | alignment | distance | position | gene name | gene id |
| GU305914.1 | | 19016 | 19038 | - | 0 | CAGTTTGCGCCGGGACAGCA | GGG | \|\|\|\|\|\|\|\|[\|\|\|\|\|\|\|\|\|\|\|\|]PAM | NA | Intergenic | NA | NA |
| T33102 | GCAGCACAGCCACCACACCCAGG | | 1000 | | | | | | | | | |
| Oligo fwd | CACCGCAGCACAGCCACCACACCC | | | | | | | | | | | |
| Oligo rev | AAACGGGTGTGGTGGCTGTGCTGC | | | | | | | | | | | |
| Chromosome | start | | end | strand | MM | target_seq | PAM | alignment | distance | position | gene name | gene id |
| GU305914.1 | | 18987 | 19009 | - | 0 | GCAGCACAGCCACCACACCC | AGG | \|\|\|\|\|\|\|\|[\|\|\|\|\|\|\|\|\|\|\|\|]PAM | NA | Intergenic | NA | NA |
| T33103 | CACACGCGACATCGAGTAACAGG | | 1000 | | | | | | | | | |
| Oligo fwd | CACCCACACGCGACATCGAGTAAC | | | | | | | | | | | |
| Oligo rev | AAACGTTACTCGATGTCGCGTGTG | | | | | | | | | | | |
| Chromosome | start | | end | strand | MM | target_seq | PAM | alignment | distance | position | gene name | gene id |
| GU305914.1 | | 18926 | 18948 | - | 0 | CACACGCGACATCGAGTAAC | AGG | \|\|\|\|\|\|\|\|[\|\|\|\|\|\|\|\|\|\|\|\|]PAM | NA | Intergenic | NA | NA |
| T33104 | ACACGCGACATCGAGTAACAGGG | | 1000 | | | | | | | | | |
| Oligo fwd | CACCACACGCGACATCGAGTAACA | | | | | | | | | | | |
| Oligo rev | AAACTGTTACTCGATGTCGCGTGT | | | | | | | | | | | |
| Chromosome | start | | end | strand | MM | target_seq | PAM | alignment | distance | position | gene name | gene id |
| GU305914.1 | | 18925 | 18947 | - | 0 | ACACGCGACATCGAGTAACA | GGG | \|\|\|\|\|\|\|\|[\|\|\|\|\|\|\|\|\|\|\|\|]PAM | NA | Intergenic | NA | NA |
| T33105 | CGACATCGAGTAACAGGGTCTGG | | 1000 | | | | | | | | | |
| Oligo fwd | CACCCGACATCGAGTAACAGGGTC | | | | | | | | | | | |
| Oligo rev | AAACGACCCTGTTACTCGATGTCG | | | | | | | | | | | |
| Chromosome | start | | end | strand | MM | target_seq | PAM | alignment | distance | position | gene name | gene id |
| GU305914.1 | | 18920 | 18942 | - | 0 | CGACATCGAGTAACAGGGTC | TGG | \|\|\|\|\|\|\|\|[\|\|\|\|\|\|\|\|\|\|\|\|]PAM | NA | Intergenic | NA | NA |
| T33106 | AGTAACAGGGTCTGGCCGTTCGG | | 1000 | | | | | | | | | |
| Oligo fwd | CACCAGTAACAGGGTCTGGCCGTT | | | | | | | | | | | |
| Oligo rev | AAACAACGGCCAGACCCTGTTACT | | | | | | | | | | | |
| Chromosome | start | | end | strand | MM | target_seq | PAM | alignment | distance | position | gene name | gene id |
| GU305914.1 | | 18912 | 18934 | - | 0 | AGTAACAGGGTCTGGCCGTT | CGG | \|\|\|\|\|\|\|\|[\|\|\|\|\|\|\|\|\|\|\|\|]PAM | NA | Intergenic | NA | NA |
| T33107 | GTAACAGGGTCTGGCCGTTCGGG | | 1000 | | | | | | | | | |
| Oligo fwd | CACCGTAACAGGGTCTGGCCGTTC | | | | | | | | | | | |
| Oligo rev | AAACGAACGGCCAGACCCTGTTAC | | | | | | | | | | | |
| Chromosome | start | | end | strand | MM | target_seq | PAM | alignment | distance | position | gene name | gene id |
| GU305914.1 | | 18911 | 18933 | - | 0 | GTAACAGGGTCTGGCCGTTC | GGG | \|\|\|\|\|\|\|\|[\|\|\|\|\|\|\|\|\|\|\|\|]PAM | NA | Intergenic | NA | NA |
| T33108 | TAACAGGGTCTGGCCGTTCGGGG | | 1000 | | | | | | | | | |
| Oligo fwd | CACCTAACAGGGTCTGGCCGTTCG | | | | | | | | | | | |
| Oligo rev | AAACCGAACGGCCAGACCCTGTTA | | | | | | | | | | | |
| Chromosome | start | | end | strand | MM | target_seq | PAM | alignment | distance | position | gene name | gene id |
| GU305914.1 | | 18910 | 18932 | - | 0 | TAACAGGGTCTGGCCGTTCG | GGG | \|\|\|\|\|\|\|\|[\|\|\|\|\|\|\|\|\|\|\|\|]PAM | NA | Intergenic | NA | NA |
| T33109 | AACAGGGTCTGGCCGTTCGGGGG | | 1000 | | | | | | | | | |
| Oligo fwd | CACCAACAGGGTCTGGCCGTTCGG | | | | | | | | | | | |
| Oligo rev | AAACCCGAACGGCCAGACCCTGTT | | | | | | | | | | | |
| Chromosome | start | | end | strand | MM | target_seq | PAM | alignment | distance | position | gene name | gene id |
| GU305914.1 | | 18909 | 18931 | - | 0 | AACAGGGTCTGGCCGTTCGG | GGG | \|\|\|\|\|\|\|\|[\|\|\|\|\|\|\|\|\|\|\|\|]PAM | NA | Intergenic | NA | NA |
| T33110 | ACAGGGTCTGGCCGTTCGGGGGG | | 1000 | | | | | | | | | |
| Oligo fwd | CACCACAGGGTCTGGCCGTTCGGG | | | | | | | | | | | |
| Oligo rev | AAACCCCGAACGGCCAGACCCTGT | | | | | | | | | | | |
| Chromosome | start | | end | strand | MM | target_seq | PAM | alignment | distance | position | gene name | gene id |
| GU305914.1 | | 18908 | 18930 | - | 0 | ACAGGGTCTGGCCGTTCGGG | GGG | \|\|\|\|\|\|\|\|[\|\|\|\|\|\|\|\|\|\|\|\|]PAM | NA | Intergenic | NA | NA |
| T33111 | CAGGGTCTGGCCGTTCGGGGGGG | | 1000 | | | | | | | | | |
| Oligo fwd | CACCCAGGGTCTGGCCGTTCGGGG | | | | | | | | | | | |
| Oligo rev | AAACCCCCGAACGGCCAGACCCTG | | | | | | | | | | | |
| Chromosome | start | | end | strand | MM | target_seq | PAM | alignment | distance | position | gene name | gene id |
| GU305914.1 | | 18907 | 18929 | - | 0 | CAGGGTCTGGCCGTTCGGGG | GGG | \|\|\|\|\|\|\|\|[\|\|\|\|\|\|\|\|\|\|\|\|]PAM | NA | Intergenic | NA | NA |
| T33112 | TCTGGCCGTTCGGGGGGGAGCGG | | 1000 | | | | | | | | | |
| Oligo fwd | CACCTCTGGCCGTTCGGGGGGGAG | | | | | | | | | | | |
| Oligo rev | AAACCTCCCCCCCGAACGGCCAGA | | | | | | | | | | | |
| Chromosome | start | | end | strand | MM | target_seq | PAM | alignment | distance | position | gene name | gene id |
| GU305914.1 | | 18902 | 18924 | - | 0 | TCTGGCCGTTCGGGGGGGAG | CGG | \|\|\|\|\|\|\|\|[\|\|\|\|\|\|\|\|\|\|\|\|]PAM | NA | Intergenic | NA | NA |
| T33113 | CTGGCCGTTCGGGGGGGAGCGGG | | 1000 | | | | | | | | | |
| Oligo fwd | CACCCTGGCCGTTCGGGGGGGAGC | | | | | | | | | | | |
| Oligo rev | AAACGCTCCCCCCCGAACGGCCAG | | | | | | | | | | | |

<table>
<tr><td>Chromosome</td><td>start</td><td>end</td><td>strand</td><td>MM</td><td>target_seq</td><td>PAM</td><td>alignment</td><td>distance</td><td>position</td><td>gene name</td><td>gene id</td></tr>
<tr><td>GU305914.1</td><td>18901</td><td>18923</td><td>-</td><td>0</td><td>CTGGCCGTTCGGGGGGGAGC</td><td>GGG</td><td>||||||||[||||||||||||]PAM</td><td>NA</td><td>Intergenic</td><td>NA</td><td>NA</td></tr>
<tr><td>T33114</td><td>TCGGGGGGGAGCGGGCGTGCCGG</td><td>1000</td><td></td><td></td><td></td><td></td><td></td><td></td><td></td><td></td><td></td></tr>
<tr><td>Oligo fwd</td><td>CACCTCGGGGGGGAGCGGGCGTGC</td><td></td><td></td><td></td><td></td><td></td><td></td><td></td><td></td><td></td><td></td></tr>
<tr><td>Oligo rev</td><td>AAACGCACGCCCGCTCCCCCCCGA</td><td></td><td></td><td></td><td></td><td></td><td></td><td></td><td></td><td></td><td></td></tr>
<tr><td>Chromosome</td><td>start</td><td>end</td><td>strand</td><td>MM</td><td>target_seq</td><td>PAM</td><td>alignment</td><td>distance</td><td>position</td><td>gene name</td><td>gene id</td></tr>
<tr><td>GU305914.1</td><td>18893</td><td>18915</td><td>-</td><td>0</td><td>TCGGGGGGGAGCGGGCGTGC</td><td>CGG</td><td>||||||||[||||||||||||]PAM</td><td>NA</td><td>Intergenic</td><td>NA</td><td>NA</td></tr>
<tr><td>T33115</td><td>CGGGGGGGAGCGGGCGTGCCGGG</td><td>1000</td><td></td><td></td><td></td><td></td><td></td><td></td><td></td><td></td><td></td></tr>
<tr><td>Oligo fwd</td><td>CACCCGGGGGGGAGCGGGCGTGCC</td><td></td><td></td><td></td><td></td><td></td><td></td><td></td><td></td><td></td><td></td></tr>
<tr><td>Oligo rev</td><td>AAACGGCACGCCCGCTCCCCCCCG</td><td></td><td></td><td></td><td></td><td></td><td></td><td></td><td></td><td></td><td></td></tr>
<tr><td>Chromosome</td><td>start</td><td>end</td><td>strand</td><td>MM</td><td>target_seq</td><td>PAM</td><td>alignment</td><td>distance</td><td>position</td><td>gene name</td><td>gene id</td></tr>
<tr><td>GU305914.1</td><td>18892</td><td>18914</td><td>-</td><td>0</td><td>CGGGGGGGAGCGGGCGTGCC</td><td>GGG</td><td>||||||||[||||||||||||]PAM</td><td>NA</td><td>Intergenic</td><td>NA</td><td>NA</td></tr>
<tr><td>GU305914.1</td><td>197469</td><td>197491</td><td>-</td><td>4</td><td>CCGCGGGGAGCTGGCGTCCC</td><td>TGG</td><td>|-|-||||[|||-|||||-||]PAM</td><td>NA</td><td>Intergenic</td><td>NA</td><td>NA</td></tr>
<tr><td>T33116</td><td>GGGATGTCGCACTCGTCAGTCGG</td><td>1000</td><td></td><td></td><td></td><td></td><td></td><td></td><td></td><td></td><td></td></tr>
<tr><td>Oligo fwd</td><td>CACCGGGATGTCGCACTCGTCAGT</td><td></td><td></td><td></td><td></td><td></td><td></td><td></td><td></td><td></td><td></td></tr>
<tr><td>Oligo rev</td><td>AAACACTGACGAGTGCGACATCCC</td><td></td><td></td><td></td><td></td><td></td><td></td><td></td><td></td><td></td><td></td></tr>
<tr><td>Chromosome</td><td>start</td><td>end</td><td>strand</td><td>MM</td><td>target_seq</td><td>PAM</td><td>alignment</td><td>distance</td><td>position</td><td>gene name</td><td>gene id</td></tr>
<tr><td>GU305914.1</td><td>18872</td><td>18894</td><td>-</td><td>0</td><td>GGGATGTCGCACTCGTCAGT</td><td>CGG</td><td>||||||||[||||||||||||]PAM</td><td>NA</td><td>Intergenic</td><td>NA</td><td>NA</td></tr>
<tr><td>T33117</td><td>ATGTCGCACTCGTCAGTCGGTGG</td><td>1000</td><td></td><td></td><td></td><td></td><td></td><td></td><td></td><td></td><td></td></tr>
<tr><td>Oligo fwd</td><td>CACCATGTCGCACTCGTCAGTCGG</td><td></td><td></td><td></td><td></td><td></td><td></td><td></td><td></td><td></td><td></td></tr>
<tr><td>Oligo rev</td><td>AAACCCGACTGACGAGTGCGACAT</td><td></td><td></td><td></td><td></td><td></td><td></td><td></td><td></td><td></td><td></td></tr>
<tr><td>Chromosome</td><td>start</td><td>end</td><td>strand</td><td>MM</td><td>target_seq</td><td>PAM</td><td>alignment</td><td>distance</td><td>position</td><td>gene name</td><td>gene id</td></tr>
<tr><td>GU305914.1</td><td>18869</td><td>18891</td><td>-</td><td>0</td><td>ATGTCGCACTCGTCAGTCGG</td><td>TGG</td><td>||||||||[||||||||||||]PAM</td><td>NA</td><td>Intergenic</td><td>NA</td><td>NA</td></tr>
<tr><td>T33118</td><td>TGTCGCACTCGTCAGTCGGTGGG</td><td>1000</td><td></td><td></td><td></td><td></td><td></td><td></td><td></td><td></td><td></td></tr>
<tr><td>Oligo fwd</td><td>CACCTGTCGCACTCGTCAGTCGGT</td><td></td><td></td><td></td><td></td><td></td><td></td><td></td><td></td><td></td><td></td></tr>
<tr><td>Oligo rev</td><td>AAACACCGACTGACGAGTGCGACA</td><td></td><td></td><td></td><td></td><td></td><td></td><td></td><td></td><td></td><td></td></tr>
<tr><td>Chromosome</td><td>start</td><td>end</td><td>strand</td><td>MM</td><td>target_seq</td><td>PAM</td><td>alignment</td><td>distance</td><td>position</td><td>gene name</td><td>gene id</td></tr>
<tr><td>GU305914.1</td><td>18868</td><td>18890</td><td>-</td><td>0</td><td>TGTCGCACTCGTCAGTCGGT</td><td>GGG</td><td>||||||||[||||||||||||]PAM</td><td>NA</td><td>Intergenic</td><td>NA</td><td>NA</td></tr>
<tr><td>T33119</td><td>GTCGCACTCGTCAGTCGGTGGGG</td><td>1000</td><td></td><td></td><td></td><td></td><td></td><td></td><td></td><td></td><td></td></tr>
<tr><td>Oligo fwd</td><td>CACCGTCGCACTCGTCAGTCGGTG</td><td></td><td></td><td></td><td></td><td></td><td></td><td></td><td></td><td></td><td></td></tr>
<tr><td>Oligo rev</td><td>AAACCACCGACTGACGAGTGCGAC</td><td></td><td></td><td></td><td></td><td></td><td></td><td></td><td></td><td></td><td></td></tr>
<tr><td>Chromosome</td><td>start</td><td>end</td><td>strand</td><td>MM</td><td>target_seq</td><td>PAM</td><td>alignment</td><td>distance</td><td>position</td><td>gene name</td><td>gene id</td></tr>
<tr><td>GU305914.1</td><td>18867</td><td>18889</td><td>-</td><td>0</td><td>GTCGCACTCGTCAGTCGGTG</td><td>GGG</td><td>||||||||[||||||||||||]PAM</td><td>NA</td><td>Intergenic</td><td>NA</td><td>NA</td></tr>
<tr><td>GU305914.1</td><td>73157</td><td>73179</td><td>-</td><td>4</td><td>GACGCTCTCTTCAGTCGGGG</td><td>AGG</td><td>|-|||-||[|-||||||||-|]PAM</td><td>NA</td><td>Intergenic</td><td>NA</td><td>NA</td></tr>
<tr><td>T33120</td><td>TCGCACTCGTCAGTCGGTGGGGG</td><td>1000</td><td></td><td></td><td></td><td></td><td></td><td></td><td></td><td></td><td></td></tr>
<tr><td>Oligo fwd</td><td>CACCTCGCACTCGTCAGTCGGTGG</td><td></td><td></td><td></td><td></td><td></td><td></td><td></td><td></td><td></td><td></td></tr>
<tr><td>Oligo rev</td><td>AAACCCACCGACTGACGAGTGCGA</td><td></td><td></td><td></td><td></td><td></td><td></td><td></td><td></td><td></td><td></td></tr>
<tr><td>Chromosome</td><td>start</td><td>end</td><td>strand</td><td>MM</td><td>target_seq</td><td>PAM</td><td>alignment</td><td>distance</td><td>position</td><td>gene name</td><td>gene id</td></tr>
<tr><td>GU305914.1</td><td>18866</td><td>18888</td><td>-</td><td>0</td><td>TCGCACTCGTCAGTCGGTGG</td><td>GGG</td><td>||||||||[||||||||||||]PAM</td><td>NA</td><td>Intergenic</td><td>NA</td><td>NA</td></tr>
<tr><td>T33121</td><td>CACTCGTCAGTCGGTGGGGGAGG</td><td>1000</td><td></td><td></td><td></td><td></td><td></td><td></td><td></td><td></td><td></td></tr>
<tr><td>Oligo fwd</td><td>CACCCACTCGTCAGTCGGTGGGGG</td><td></td><td></td><td></td><td></td><td></td><td></td><td></td><td></td><td></td><td></td></tr>
<tr><td>Oligo rev</td><td>AAACCCCCCACCGACTGACGAGTG</td><td></td><td></td><td></td><td></td><td></td><td></td><td></td><td></td><td></td><td></td></tr>
<tr><td>Chromosome</td><td>start</td><td>end</td><td>strand</td><td>MM</td><td>target_seq</td><td>PAM</td><td>alignment</td><td>distance</td><td>position</td><td>gene name</td><td>gene id</td></tr>
<tr><td>GU305914.1</td><td>18863</td><td>18885</td><td>-</td><td>0</td><td>CACTCGTCAGTCGGTGGGGG</td><td>AGG</td><td>||||||||[||||||||||||]PAM</td><td>NA</td><td>Intergenic</td><td>NA</td><td>NA</td></tr>
<tr><td>T33122</td><td>TCGTCAGTCGGTGGGGGAGGAGG</td><td>1000</td><td></td><td></td><td></td><td></td><td></td><td></td><td></td><td></td><td></td></tr>
<tr><td>Oligo fwd</td><td>CACCTCGTCAGTCGGTGGGGGAGG</td><td></td><td></td><td></td><td></td><td></td><td></td><td></td><td></td><td></td><td></td></tr>
<tr><td>Oligo rev</td><td>AAACCCTCCCCCACCGACTGACGA</td><td></td><td></td><td></td><td></td><td></td><td></td><td></td><td></td><td></td><td></td></tr>
<tr><td>Chromosome</td><td>start</td><td>end</td><td>strand</td><td>MM</td><td>target_seq</td><td>PAM</td><td>alignment</td><td>distance</td><td>position</td><td>gene name</td><td>gene id</td></tr>
<tr><td>GU305914.1</td><td>18860</td><td>18882</td><td>-</td><td>0</td><td>TCGTCAGTCGGTGGGGGAGG</td><td>AGG</td><td>||||||||[||||||||||||]PAM</td><td>NA</td><td>Intergenic</td><td>NA</td><td>NA</td></tr>
<tr><td>T33123</td><td>CGTCAGTCGGTGGGGGAGGAGGG</td><td>1000</td><td></td><td></td><td></td><td></td><td></td><td></td><td></td><td></td><td></td></tr>
<tr><td>Oligo fwd</td><td>CACCCGTCAGTCGGTGGGGGAGGA</td><td></td><td></td><td></td><td></td><td></td><td></td><td></td><td></td><td></td><td></td></tr>
<tr><td>Oligo rev</td><td>AAACTCCTCCCCCACCGACTGACG</td><td></td><td></td><td></td><td></td><td></td><td></td><td></td><td></td><td></td><td></td></tr>
<tr><td>Chromosome</td><td>start</td><td>end</td><td>strand</td><td>MM</td><td>target_seq</td><td>PAM</td><td>alignment</td><td>distance</td><td>position</td><td>gene name</td><td>gene id</td></tr>
<tr><td>GU305914.1</td><td>18859</td><td>18881</td><td>-</td><td>0</td><td>CGTCAGTCGGTGGGGGAGGA</td><td>GGG</td><td>||||||||[||||||||||||]PAM</td><td>NA</td><td>Intergenic</td><td>NA</td><td>NA</td></tr>
<tr><td>T33124</td><td>GTCAGTCGGTGGGGGAGGAGGGG</td><td>1000</td><td></td><td></td><td></td><td></td><td></td><td></td><td></td><td></td><td></td></tr>
<tr><td>Oligo fwd</td><td>CACCGTCAGTCGGTGGGGGAGGAG</td><td></td><td></td><td></td><td></td><td></td><td></td><td></td><td></td><td></td><td></td></tr>
<tr><td>Oligo rev</td><td>AAACCTCCTCCCCCACCGACTGAC</td><td></td><td></td><td></td><td></td><td></td><td></td><td></td><td></td><td></td><td></td></tr>
<tr><td>Chromosome</td><td>start</td><td>end</td><td>strand</td><td>MM</td><td>target_seq</td><td>PAM</td><td>alignment</td><td>distance</td><td>position</td><td>gene name</td><td>gene id</td></tr>
<tr><td>GU305914.1</td><td>18858</td><td>18880</td><td>-</td><td>0</td><td>GTCAGTCGGTGGGGGAGGAG</td><td>GGG</td><td>||||||||[||||||||||||]PAM</td><td>NA</td><td>Intergenic</td><td>NA</td><td>NA</td></tr>
<tr><td>T33125</td><td>GGTGGGGGAGGAGGGGAAGACGG</td><td>1000</td><td></td><td></td><td></td><td></td><td></td><td></td><td></td><td></td><td></td></tr>
<tr><td>Oligo fwd</td><td>CACCGGTGGGGGAGGAGGGGAAGA</td><td></td><td></td><td></td><td></td><td></td><td></td><td></td><td></td><td></td><td></td></tr>
<tr><td>Oligo rev</td><td>AAACTCTTCCCCTCCTCCCCCACC</td><td></td><td></td><td></td><td></td><td></td><td></td><td></td><td></td><td></td><td></td></tr>
<tr><td>Chromosome</td><td>start</td><td>end</td><td>strand</td><td>MM</td><td>target_seq</td><td>PAM</td><td>alignment</td><td>distance</td><td>position</td><td>gene name</td><td>gene id</td></tr>
<tr><td>GU305914.1</td><td>18851</td><td>18873</td><td>-</td><td>0</td><td>GGTGGGGGAGGAGGGGAAGA</td><td>CGG</td><td>||||||||[||||||||||||]PAM</td><td>NA</td><td>Intergenic</td><td>NA</td><td>NA</td></tr>
<tr><td>GU305914.1</td><td>86798</td><td>86820</td><td>-</td><td>3</td><td>GCCGGGGCAGGAGGGGAAGA</td><td>GGG</td><td>|--||||-[||||||||||||]PAM</td><td>NA</td><td>Intergenic</td><td>NA</td><td>NA</td></tr>
<tr><td>GU305914.1</td><td>82326</td><td>82348</td><td>-</td><td>4</td><td>GATGCTGGAGGAGGGGAGGA</td><td>AGG</td><td>|-||--||[|||||||||-||]PAM</td><td>NA</td><td>Intergenic</td><td>NA</td><td>NA</td></tr>
<tr><td>GU305914.1</td><td>45315</td><td>45337</td><td>-</td><td>4</td><td>AGTGGAGGAGGAGGAGGAGA</td><td>AGG</td><td>-||||-||[||||||-|-|||]PAM</td><td>NA</td><td>Intergenic</td><td>NA</td><td>NA</td></tr>
<tr><td>GU305914.1</td><td>28747</td><td>28769</td><td>-</td><td>4</td><td>GGGGGAGGAGGAGAGGATGA</td><td>TGG</td><td>||-||-||[|||||-|||-||]PAM</td><td>NA</td><td>Intergenic</td><td>NA</td><td>NA</td></tr>
<tr><td>T33126</td><td>TCCTGCGATAATCTTTTCTTAGG</td><td>1000</td><td></td><td></td><td></td><td></td><td></td><td></td><td></td><td></td><td></td></tr>
<tr><td>Oligo fwd</td><td>CACCTCCTGCGATAATCTTTTCTT</td><td></td><td></td><td></td><td></td><td></td><td></td><td></td><td></td><td></td><td></td></tr>
<tr><td>Oligo rev</td><td>AAACAAGAAAAGATTATCGCAGGA</td><td></td><td></td><td></td><td></td><td></td><td></td><td></td><td></td><td></td><td></td></tr>
<tr><td>Chromosome</td><td>start</td><td>end</td><td>strand</td><td>MM</td><td>target_seq</td><td>PAM</td><td>alignment</td><td>distance</td><td>position</td><td>gene name</td><td>gene id</td></tr>
<tr><td>GU305914.1</td><td>18827</td><td>18849</td><td>-</td><td>0</td><td>TCCTGCGATAATCTTTTCTT</td><td>AGG</td><td>||||||||[||||||||||||]PAM</td><td>NA</td><td>Intergenic</td><td>NA</td><td>NA</td></tr>
<tr><td>T33127</td><td>CTTAGGTTCGCCAATCTCCTCGG</td><td>1000</td><td></td><td></td><td></td><td></td><td></td><td></td><td></td><td></td><td></td></tr>
<tr><td>Oligo fwd</td><td>CACCCTTAGGTTCGCCAATCTCCT</td><td></td><td></td><td></td><td></td><td></td><td></td><td></td><td></td><td></td><td></td></tr>
</table>

| | | | | | | | | | | | |
|---|---|---|---|---|---|---|---|---|---|---|---|
| Oligo rev | AAACAGGAGATTGGCGAACCTAAG | | | | | | | | | | |
| Chromosome | start | end | strand | MM | target_seq | PAM | alignment | distance | position | gene name | gene id |
| GU305914.1 | 18810 | 18832 | - | 0 | CTTAGGTTCGCCAATCTCCT | CGG | \|\|\|\|\|\|\|\|[\|\|\|\|\|\|\|\|\|\|\|\|]PAM | NA | Intergenic | NA | NA |
| T33128 | CCCGCTACTGTCACTGCTGCTGG | 1000 | | | | | | | | | |
| Oligo fwd | CACCCCCGCTACTGTCACTGCTGC | | | | | | | | | | |
| Oligo rev | AAACGCAGCAGTGACAGTAGCGGG | | | | | | | | | | |
| Chromosome | start | end | strand | MM | target_seq | PAM | alignment | distance | position | gene name | gene id |
| GU305914.1 | 18756 | 18778 | - | 0 | CCCGCTACTGTCACTGCTGC | TGG | \|\|\|\|\|\|\|\|[\|\|\|\|\|\|\|\|\|\|\|\|]PAM | NA | Intergenic | NA | NA |
| GU305914.1 | 132168 | 132190 | - | 4 | CCCGTTCCTGTCACAGGTGC | GGG | \|\|\|\|-\|-\|[\|\|\|\|\|\|-\|-\|\|\|]PAM | NA | Intergenic | NA | NA |
| T33129 | CTGGACTCTGTGATCGACAGCGG | 1000 | | | | | | | | | |
| Oligo fwd | CACCCTGGACTCTGTGATCGACAG | | | | | | | | | | |
| Oligo rev | AAACCTGTCGATCACAGAGTCCAG | | | | | | | | | | |
| Chromosome | start | end | strand | MM | target_seq | PAM | alignment | distance | position | gene name | gene id |
| GU305914.1 | 18737 | 18759 | - | 0 | CTGGACTCTGTGATCGACAG | CGG | \|\|\|\|\|\|\|\|[\|\|\|\|\|\|\|\|\|\|\|\|]PAM | NA | Intergenic | NA | NA |
| T33130 | CAGCGGCACCTTCGACGACTCGG | 1000 | | | | | | | | | |
| Oligo fwd | CACCCAGCGGCACCTTCGACGACT | | | | | | | | | | |
| Oligo rev | AAACAGTCGTCGAAGGTGCCGCTG | | | | | | | | | | |
| Chromosome | start | end | strand | MM | target_seq | PAM | alignment | distance | position | gene name | gene id |
| GU305914.1 | 18720 | 18742 | - | 0 | CAGCGGCACCTTCGACGACT | CGG | \|\|\|\|\|\|\|\|[\|\|\|\|\|\|\|\|\|\|\|\|]PAM | NA | Intergenic | NA | NA |
| T33131 | CGGCACCTTCGACGACTCGGTGG | 1000 | | | | | | | | | |
| Oligo fwd | CACCCGGCACCTTCGACGACTCGG | | | | | | | | | | |
| Oligo rev | AAACCCGAGTCGTCGAAGGTGCCG | | | | | | | | | | |
| Chromosome | start | end | strand | MM | target_seq | PAM | alignment | distance | position | gene name | gene id |
| GU305914.1 | 18717 | 18739 | - | 0 | CGGCACCTTCGACGACTCGG | TGG | \|\|\|\|\|\|\|\|[\|\|\|\|\|\|\|\|\|\|\|\|]PAM | NA | Intergenic | NA | NA |
| T33132 | CACCTTCGACGACTCGGTGGTGG | 1000 | | | | | | | | | |
| Oligo fwd | CACCCACCTTCGACGACTCGGTGG | | | | | | | | | | |
| Oligo rev | AAACCCACCGAGTCGTCGAAGGTG | | | | | | | | | | |
| Chromosome | start | end | strand | MM | target_seq | PAM | alignment | distance | position | gene name | gene id |
| GU305914.1 | 18714 | 18736 | - | 0 | CACCTTCGACGACTCGGTGG | TGG | \|\|\|\|\|\|\|\|[\|\|\|\|\|\|\|\|\|\|\|\|]PAM | NA | Intergenic | NA | NA |
| T33133 | ACCTTCGACGACTCGGTGGTGGG | 1000 | | | | | | | | | |
| Oligo fwd | CACCACCTTCGACGACTCGGTGGT | | | | | | | | | | |
| Oligo rev | AAACACCACCGAGTCGTCGAAGGT | | | | | | | | | | |
| Chromosome | start | end | strand | MM | target_seq | PAM | alignment | distance | position | gene name | gene id |
| GU305914.1 | 18713 | 18735 | - | 0 | ACCTTCGACGACTCGGTGGT | GGG | \|\|\|\|\|\|\|\|[\|\|\|\|\|\|\|\|\|\|\|\|]PAM | NA | Intergenic | NA | NA |
| T33134 | GACTCGGTGGTGGGCGAATCCGG | 1000 | | | | | | | | | |
| Oligo fwd | CACCGACTCGGTGGTGGGCGAATC | | | | | | | | | | |
| Oligo rev | AAACGATTCGCCCACCACCGAGTC | | | | | | | | | | |
| Chromosome | start | end | strand | MM | target_seq | PAM | alignment | distance | position | gene name | gene id |
| GU305914.1 | 18704 | 18726 | - | 0 | GACTCGGTGGTGGGCGAATC | CGG | \|\|\|\|\|\|\|\|[\|\|\|\|\|\|\|\|\|\|\|\|]PAM | NA | Intergenic | NA | NA |
| T33135 | ACTCGGTGGTGGGCGAATCCGGG | 1000 | | | | | | | | | |
| Oligo fwd | CACCACTCGGTGGTGGGCGAATCC | | | | | | | | | | |
| Oligo rev | AAACGGATTCGCCCACCACCGAGT | | | | | | | | | | |
| Chromosome | start | end | strand | MM | target_seq | PAM | alignment | distance | position | gene name | gene id |
| GU305914.1 | 18703 | 18725 | - | 0 | ACTCGGTGGTGGGCGAATCC | GGG | \|\|\|\|\|\|\|\|[\|\|\|\|\|\|\|\|\|\|\|\|]PAM | NA | Intergenic | NA | NA |
| T33136 | GTGGGCGAATCCGGGACCGCTGG | 1000 | | | | | | | | | |
| Oligo fwd | CACCGTGGGCGAATCCGGGACCGC | | | | | | | | | | |
| Oligo rev | AAACGCGGTCCCGGATTCGCCCAC | | | | | | | | | | |
| Chromosome | start | end | strand | MM | target_seq | PAM | alignment | distance | position | gene name | gene id |
| GU305914.1 | 18695 | 18717 | - | 0 | GTGGGCGAATCCGGGACCGC | TGG | \|\|\|\|\|\|\|\|[\|\|\|\|\|\|\|\|\|\|\|\|]PAM | NA | Intergenic | NA | NA |
| T33137 | GGCGAAGATGCTACAAGAGACGG | 1000 | | | | | | | | | |
| Oligo fwd | CACCGGCGAAGATGCTACAAGAGA | | | | | | | | | | |
| Oligo rev | AAACTCTCTTGTAGCATCTTCGCC | | | | | | | | | | |
| Chromosome | start | end | strand | MM | target_seq | PAM | alignment | distance | position | gene name | gene id |
| GU305914.1 | 18674 | 18696 | - | 0 | GGCGAAGATGCTACAAGAGA | CGG | \|\|\|\|\|\|\|\|[\|\|\|\|\|\|\|\|\|\|\|\|]PAM | NA | Intergenic | NA | NA |
| T33138 | GATGCTACAAGAGACGGCAGAGG | 1000 | | | | | | | | | |
| Oligo fwd | CACCGATGCTACAAGAGACGGCAG | | | | | | | | | | |
| Oligo rev | AAACCTGCCGTCTCTTGTAGCATC | | | | | | | | | | |
| Chromosome | start | end | strand | MM | target_seq | PAM | alignment | distance | position | gene name | gene id |
| GU305914.1 | 18668 | 18690 | - | 0 | GATGCTACAAGAGACGGCAG | AGG | \|\|\|\|\|\|\|\|[\|\|\|\|\|\|\|\|\|\|\|\|]PAM | NA | Intergenic | NA | NA |
| T33139 | GCTACAAGAGACGGCAGAGGTGG | 1000 | | | | | | | | | |
| Oligo fwd | CACCGCTACAAGAGACGGCAGAGG | | | | | | | | | | |
| Oligo rev | AAACCCTCTGCCGTCTCTTGTAGC | | | | | | | | | | |
| Chromosome | start | end | strand | MM | target_seq | PAM | alignment | distance | position | gene name | gene id |
| GU305914.1 | 18665 | 18687 | - | 0 | GCTACAAGAGACGGCAGAGG | TGG | \|\|\|\|\|\|\|\|[\|\|\|\|\|\|\|\|\|\|\|\|]PAM | NA | Intergenic | NA | NA |
| T33140 | ACAAGAGACGGCAGAGGTGGCGG | 1000 | | | | | | | | | |
| Oligo fwd | CACCACAAGAGACGGCAGAGGTGG | | | | | | | | | | |
| Oligo rev | AAACCCACCTCTGCCGTCTCTTGT | | | | | | | | | | |
| Chromosome | start | end | strand | MM | target_seq | PAM | alignment | distance | position | gene name | gene id |
| GU305914.1 | 18662 | 18684 | - | 0 | ACAAGAGACGGCAGAGGTGG | CGG | \|\|\|\|\|\|\|\|[\|\|\|\|\|\|\|\|\|\|\|\|]PAM | NA | Intergenic | NA | NA |
| T33141 | AGAGACGGCAGAGGTGGCGGAGG | 1000 | | | | | | | | | |
| Oligo fwd | CACCAGAGACGGCAGAGGTGGCGG | | | | | | | | | | |
| Oligo rev | AAACCCGCCACCTCTGCCGTCTCT | | | | | | | | | | |
| Chromosome | start | end | strand | MM | target_seq | PAM | alignment | distance | position | gene name | gene id |
| GU305914.1 | 18659 | 18681 | - | 0 | AGAGACGGCAGAGGTGGCGG | AGG | \|\|\|\|\|\|\|\|[\|\|\|\|\|\|\|\|\|\|\|\|]PAM | NA | Intergenic | NA | NA |

T33142 GACGGCAGAGGTGGCGGAGGCGG 1000
Oligo fwd CACCGACGGCAGAGGTGGCGGAGG
Oligo rev AAACCCTCCGCCACCTCTGCCGTC
Chromosome start end strand MM target_seq PAM alignment distance position gene name gene id
GU305914.1 18656 18678 - 0 GACGGCAGAGGTGGCGGAGG CGG ||||||||[||||||||||||]PAM NA Intergenic NA NA
GU305914.1 170649 170671 + 3 AAAGGCAGAGGTAGCGGAGG CGG -|-|||||[||||-|||||||]PAM NA Intergenic NA NA
GU305914.1 73789 73811 - 4 GCCAGCAGCGGCGGCGGAGG AGG |-|-||||[-||-||||||||]PAM NA Intergenic NA NA
GU305914.1 74622 74644 + 4 CACGACCGAGGTGGCGAAGG TGG -|||-|-|[||||||||-|||]PAM NA Intergenic NA NA
GU305914.1 48156 48178 + 4 GGCGGCGGCGGTGGCGGCGG CGG |-||||-|[-||||||||-||]PAM NA Intergenic NA NA
GU305914.1 90253 90275 - 3 GACGGCGGAGGCGGCGGAGC CGG ||||||-|[|||-|||||||-]PAM NA Intergenic NA NA

T33143 ACGGCAGAGGTGGCGGAGGCGGG 1000
Oligo fwd CACCACGGCAGAGGTGGCGGAGGC
Oligo rev AAACGCCTCCGCCACCTCTGCCGT
Chromosome start end strand MM target_seq PAM alignment distance position gene name gene id
GU305914.1 18655 18677 - 0 ACGGCAGAGGTGGCGGAGGC GGG ||||||||[||||||||||||]PAM NA Intergenic NA NA

T33144 CGGCAGAGGTGGCGGAGGCGGGG 1000
Oligo fwd CACCCGGCAGAGGTGGCGGAGGCG
Oligo rev AAACCGCCTCCGCCACCTCTGCCG
Chromosome start end strand MM target_seq PAM alignment distance position gene name gene id
GU305914.1 18654 18676 - 0 CGGCAGAGGTGGCGGAGGCG GGG ||||||||[||||||||||||]PAM NA Intergenic NA NA
GU305914.1 62445 62467 + 4 CGGCGGGGGTGGTGGCGGCG GGG ||||-|-|[||||-||-||||]PAM NA Intergenic NA NA

T33145 GGCAGAGGTGGCGGAGGCGGGGG 1000
Oligo fwd CACCGGCAGAGGTGGCGGAGGCGG
Oligo rev AAACCCGCCTCCGCCACCTCTGCC
Chromosome start end strand MM target_seq PAM alignment distance position gene name gene id
GU305914.1 18653 18675 - 0 GGCAGAGGTGGCGGAGGCGG GGG ||||||||[||||||||||||]PAM NA Intergenic NA NA
GU305914.1 48159 48181 + 3 GGCGGCGGTGGCGGCGGCGG TGG |||-|-||[||||||-|||||]PAM NA Intergenic NA NA
GU305914.1 86784 86806 - 3 GGAAGAGGGGGCGGGGGCGG CGG ||-|||||[-|||||-|||||]PAM NA Intergenic NA NA
GU305914.1 161754 161776 - 3 GGCCGAGGAGGCGGCGGCGG AGG |||-||||[-|||||-|||||]PAM NA Intergenic NA NA
GU305914.1 67742 67764 + 4 GGGAGCGGCGGCGGCGGCGG CGG ||-||-||[-|||||-|||||]PAM NA Intergenic NA NA
GU305914.1 154404 154426 + 4 GGCGGTGGCGGCGGCGGCGG CGG |||-|-||[-|||||-|||||]PAM NA Intergenic NA NA
GU305914.1 58126 58148 - 4 GGCGGTGGCGGCGGCGGCGG CGG |||-|-||[-|||||-|||||]PAM NA Intergenic NA NA
GU305914.1 62446 62468 + 4 GGCGGGGGTGGTGGCGGCGG GGG |||-|-||[|||-||-|||||]PAM NA Intergenic NA NA
GU305914.1 114658 114680 - 4 GGCGGTGGTGGTGGCGGCGG CGG |||-|-||[|||-||-|||||]PAM NA Intergenic NA NA
GU305914.1 90250 90272 - 3 GGCGGAGGCGGCGGAGCCGG CGG |||-||||[-|||||||-|||]PAM NA Intergenic NA NA
GU305914.1 168817 168839 + 4 GGCGGTGGTGGCAGCGGCGG CGG |||-|-||[||||-|-|||||]PAM NA Intergenic NA NA
GU305914.1 73786 73808 - 4 AGCAGCGGCGGCGGAGGAGG AGG -||||-||[-||||||||-||]PAM NA Intergenic NA NA
GU305914.1 109328 109350 + 4 GACGGAGGTGGCGGGGACGG CGG |-|-||||[||||||-|-|||]PAM NA Intergenic NA NA

T33146 TGGCGGAGGCGGGGGAAACGTGG 1000
Oligo fwd CACCTGGCGGAGGCGGGGGAAACG
Oligo rev AAACCGTTTCCCCCGCCTCCGCCA
Chromosome start end strand MM target_seq PAM alignment distance position gene name gene id
GU305914.1 18645 18667 - 0 TGGCGGAGGCGGGGGAAACG TGG ||||||||[||||||||||||]PAM NA Intergenic NA NA

T33147 GGCGGAGGCGGGGGAAACGTGGG 1000
Oligo fwd CACCGGCGGAGGCGGGGGAAACGT
Oligo rev AAACACGTTTCCCCCGCCTCCGCC
Chromosome start end strand MM target_seq PAM alignment distance position gene name gene id
GU305914.1 18644 18666 - 0 GGCGGAGGCGGGGGAAACGT GGG ||||||||[||||||||||||]PAM NA Intergenic NA NA

T33148 AGGCGGGGGAAACGTGGGCCTGG 1000
Oligo fwd CACCAGGCGGGGGAAACGTGGGCC
Oligo rev AAACGGCCCACGTTTCCCCCGCCT
Chromosome start end strand MM target_seq PAM alignment distance position gene name gene id
GU305914.1 18639 18661 - 0 AGGCGGGGGAAACGTGGGCC TGG ||||||||[||||||||||||]PAM NA Intergenic NA NA

T33149 GGCGGGGGAAACGTGGGCCTGGG 1000
Oligo fwd CACCGGCGGGGGAAACGTGGGCCT
Oligo rev AAACAGGCCCACGTTTCCCCCGCC
Chromosome start end strand MM target_seq PAM alignment distance position gene name gene id
GU305914.1 18638 18660 - 0 GGCGGGGGAAACGTGGGCCT GGG ||||||||[||||||||||||]PAM NA Intergenic NA NA

T33150 GGGGGAAACGTGGGCCTGGGCGG 1000
Oligo fwd CACCGGGGGAAACGTGGGCCTGGG
Oligo rev AAACCCCAGGCCCACGTTTCCCCC
Chromosome start end strand MM target_seq PAM alignment distance position gene name gene id
GU305914.1 18635 18657 - 0 GGGGGAAACGTGGGCCTGGG CGG ||||||||[||||||||||||]PAM NA Intergenic NA NA

T33151 CGGTGACGACAACACCAACGAGG 1000
Oligo fwd CACCCGGTGACGACAACACCAACG
Oligo rev AAACCGTTGGTGTTGTCGTCACCG
Chromosome start end strand MM target_seq PAM alignment distance position gene name gene id
GU305914.1 18615 18637 - 0 CGGTGACGACAACACCAACG AGG ||||||||[||||||||||||]PAM NA Intergenic NA NA

T33152 AACGAGGCCCCTCCGCTCCTCGG 1000
Oligo fwd CACCAACGAGGCCCCTCCGCTCCT
Oligo rev AAACAGGAGCGGAGGGGCCTCGTT
Chromosome start end strand MM target_seq PAM alignment distance position gene name gene id
GU305914.1 18599 18621 - 0 AACGAGGCCCCTCCGCTCCT CGG ||||||||[||||||||||||]PAM NA Intergenic NA NA

T33153 GAGGCCCCTCCGCTCCTCGGAGG 1000
Oligo fwd CACCGAGGCCCCTCCGCTCCTCGG
Oligo rev AAACCCGAGGAGCGGAGGGGCCTC
Chromosome start end strand MM target_seq PAM alignment distance position gene name gene id
GU305914.1 18596 18618 - 0 GAGGCCCCTCCGCTCCTCGG AGG ||||||||[||||||||||||]PAM NA Intergenic NA NA

| | | | | | | | | | | | |
|---|---|---|---|---|---|---|---|---|---|---|---|
| T33154 | CCGCTCCTCGGAGGAGTCGCCGG | 1000 | | | | | | | | | |
| Oligo fwd | CACCCCGCTCCTCGGAGGAGTCGC | | | | | | | | | | |
| Oligo rev | AAACGCGACTCCTCCGAGGAGCGG | | | | | | | | | | |
| Chromosome | start | end | strand | MM | target_seq | PAM | alignment | distance | position | gene name | gene id |
| GU305914.1 | 18587 | 18609 | - | 0 | CCGCTCCTCGGAGGAGTCGC | CGG | \|\|\|\|\|\|\|\|[\|\|\|\|\|\|\|\|\|\|\|\|]PAM | NA | Intergenic | NA | NA |
| T33155 | CCTCGGAGGAGTCGCCGGAGAGG | 1000 | | | | | | | | | |
| Oligo fwd | CACCCCTCGGAGGAGTCGCCGGAG | | | | | | | | | | |
| Oligo rev | AAACCTCCGGCGACTCCTCCGAGG | | | | | | | | | | |
| Chromosome | start | end | strand | MM | target_seq | PAM | alignment | distance | position | gene name | gene id |
| GU305914.1 | 18582 | 18604 | - | 0 | CCTCGGAGGAGTCGCCGGAG | AGG | \|\|\|\|\|\|\|\|[\|\|\|\|\|\|\|\|\|\|\|\|]PAM | NA | Intergenic | NA | NA |
| T33156 | GGAGTCGCCGGAGAGGTCGTCGG | 1000 | | | | | | | | | |
| Oligo fwd | CACCGGAGTCGCCGGAGAGGTCGT | | | | | | | | | | |
| Oligo rev | AAACACGACCTCTCCGGCGACTCC | | | | | | | | | | |
| Chromosome | start | end | strand | MM | target_seq | PAM | alignment | distance | position | gene name | gene id |
| GU305914.1 | 18575 | 18597 | - | 0 | GGAGTCGCCGGAGAGGTCGT | CGG | \|\|\|\|\|\|\|\|[\|\|\|\|\|\|\|\|\|\|\|\|]PAM | NA | Intergenic | NA | NA |
| T33157 | CGGCGTCGAGTCCGAACCGTCGG | 1000 | | | | | | | | | |
| Oligo fwd | CACCCGGCGTCGAGTCCGAACCGT | | | | | | | | | | |
| Oligo rev | AAACACGGTTCGGACTCGACGCCG | | | | | | | | | | |
| Chromosome | start | end | strand | MM | target_seq | PAM | alignment | distance | position | gene name | gene id |
| GU305914.1 | 18555 | 18577 | - | 0 | CGGCGTCGAGTCCGAACCGT | CGG | \|\|\|\|\|\|\|\|[\|\|\|\|\|\|\|\|\|\|\|\|]PAM | NA | Intergenic | NA | NA |
| T33158 | GGCGTCGAGTCCGAACCGTCGGG | 1000 | | | | | | | | | |
| Oligo fwd | CACCGGCGTCGAGTCCGAACCGTC | | | | | | | | | | |
| Oligo rev | AAACGACGGTTCGGACTCGACGCC | | | | | | | | | | |
| Chromosome | start | end | strand | MM | target_seq | PAM | alignment | distance | position | gene name | gene id |
| GU305914.1 | 18554 | 18576 | - | 0 | GGCGTCGAGTCCGAACCGTC | GGG | \|\|\|\|\|\|\|\|[\|\|\|\|\|\|\|\|\|\|\|\|]PAM | NA | Intergenic | NA | NA |
| T33159 | CGAGTCCGAACCGTCGGGTTCGG | 1000 | | | | | | | | | |
| Oligo fwd | CACCCGAGTCCGAACCGTCGGGTT | | | | | | | | | | |
| Oligo rev | AAACAACCCGACGGTTCGGACTCG | | | | | | | | | | |
| Chromosome | start | end | strand | MM | target_seq | PAM | alignment | distance | position | gene name | gene id |
| GU305914.1 | 18549 | 18571 | - | 0 | CGAGTCCGAACCGTCGGGTT | CGG | \|\|\|\|\|\|\|\|[\|\|\|\|\|\|\|\|\|\|\|\|]PAM | NA | Intergenic | NA | NA |
| T33160 | AACCGTCGGGTTCGGCTCTGAGG | 1000 | | | | | | | | | |
| Oligo fwd | CACCAACCGTCGGGTTCGGCTCTG | | | | | | | | | | |
| Oligo rev | AAACCAGAGCCGAACCCGACGGTT | | | | | | | | | | |
| Chromosome | start | end | strand | MM | target_seq | PAM | alignment | distance | position | gene name | gene id |
| GU305914.1 | 18541 | 18563 | - | 0 | AACCGTCGGGTTCGGCTCTG | AGG | \|\|\|\|\|\|\|\|[\|\|\|\|\|\|\|\|\|\|\|\|]PAM | NA | Intergenic | NA | NA |
| T33161 | ACCGTCGGGTTCGGCTCTGAGGG | 1000 | | | | | | | | | |
| Oligo fwd | CACCACCGTCGGGTTCGGCTCTGA | | | | | | | | | | |
| Oligo rev | AAACTCAGAGCCGAACCCGACGGT | | | | | | | | | | |
| Chromosome | start | end | strand | MM | target_seq | PAM | alignment | distance | position | gene name | gene id |
| GU305914.1 | 18540 | 18562 | - | 0 | ACCGTCGGGTTCGGCTCTGA | GGG | \|\|\|\|\|\|\|\|[\|\|\|\|\|\|\|\|\|\|\|\|]PAM | NA | Intergenic | NA | NA |
| T33162 | GGTTCGGCTCTGAGGGTCGTCGG | 1000 | | | | | | | | | |
| Oligo fwd | CACCGGTTCGGCTCTGAGGGTCGT | | | | | | | | | | |
| Oligo rev | AAACACGACCCTCAGAGCCGAACC | | | | | | | | | | |
| Chromosome | start | end | strand | MM | target_seq | PAM | alignment | distance | position | gene name | gene id |
| GU305914.1 | 18533 | 18555 | - | 0 | GGTTCGGCTCTGAGGGTCGT | CGG | \|\|\|\|\|\|\|\|[\|\|\|\|\|\|\|\|\|\|\|\|]PAM | NA | Intergenic | NA | NA |
| T33163 | TCGGCTCTGAGGGTCGTCGGTGG | 1000 | | | | | | | | | |
| Oligo fwd | CACCTCGGCTCTGAGGGTCGTCGG | | | | | | | | | | |
| Oligo rev | AAACCCGACGACCCTCAGAGCCGA | | | | | | | | | | |
| Chromosome | start | end | strand | MM | target_seq | PAM | alignment | distance | position | gene name | gene id |
| GU305914.1 | 18530 | 18552 | - | 0 | TCGGCTCTGAGGGTCGTCGG | TGG | \|\|\|\|\|\|\|\|[\|\|\|\|\|\|\|\|\|\|\|\|]PAM | NA | Intergenic | NA | NA |
| T33164 | TGGCGACGACGCCCCGTCGTTGG | 1000 | | | | | | | | | |
| Oligo fwd | CACCTGGCGACGACGCCCCGTCGT | | | | | | | | | | |
| Oligo rev | AAACACGACGGGGCGTCGTCGCCA | | | | | | | | | | |
| Chromosome | start | end | strand | MM | target_seq | PAM | alignment | distance | position | gene name | gene id |
| GU305914.1 | 18510 | 18532 | - | 0 | TGGCGACGACGCCCCGTCGT | TGG | \|\|\|\|\|\|\|\|[\|\|\|\|\|\|\|\|\|\|\|\|]PAM | NA | Intergenic | NA | NA |
| GU305914.1 | 118882 | 118904 | + | 4 | GGACGACGACGCCTCGTCCT | GGG | -\|-\|\|\|\|\|[\|\|\|\|\|-\|\|\|\|-\|]PAM | NA | Intergenic | NA | NA |
| T33165 | CCGTCGTTGGTACCTTCTTCTGG | 1000 | | | | | | | | | |
| Oligo fwd | CACCCCGTCGTTGGTACCTTCTTC | | | | | | | | | | |
| Oligo rev | AAACGAAGAAGGTACCAACGACGG | | | | | | | | | | |
| Chromosome | start | end | strand | MM | target_seq | PAM | alignment | distance | position | gene name | gene id |
| GU305914.1 | 18497 | 18519 | - | 0 | CCGTCGTTGGTACCTTCTTC | TGG | \|\|\|\|\|\|\|\|[\|\|\|\|\|\|\|\|\|\|\|\|]PAM | NA | Intergenic | NA | NA |
| T33166 | ACCTTCTTCTGGCCCGTTCGAGG | 1000 | | | | | | | | | |
| Oligo fwd | CACCACCTTCTTCTGGCCCGTTCG | | | | | | | | | | |
| Oligo rev | AAACCGAACGGGCCAGAAGAAGGT | | | | | | | | | | |
| Chromosome | start | end | strand | MM | target_seq | PAM | alignment | distance | position | gene name | gene id |
| GU305914.1 | 18486 | 18508 | - | 0 | ACCTTCTTCTGGCCCGTTCG | AGG | \|\|\|\|\|\|\|\|[\|\|\|\|\|\|\|\|\|\|\|\|]PAM | NA | Intergenic | NA | NA |
| T33167 | TCTGGCCCGTTCGAGGAATCCGG | 1000 | | | | | | | | | |
| Oligo fwd | CACCTCTGGCCCGTTCGAGGAATC | | | | | | | | | | |
| Oligo rev | AAACGATTCCTCGAACGGGCCAGA | | | | | | | | | | |
| Chromosome | start | end | strand | MM | target_seq | PAM | alignment | distance | position | gene name | gene id |
| GU305914.1 | 18479 | 18501 | - | 0 | TCTGGCCCGTTCGAGGAATC | CGG | \|\|\|\|\|\|\|\|[\|\|\|\|\|\|\|\|\|\|\|\|]PAM | NA | Intergenic | NA | NA |
| T33168 | GGCCCGTTCGAGGAATCCGGCGG | 1000 | | | | | | | | | |
| Oligo fwd | CACCGGCCCGTTCGAGGAATCCGG | | | | | | | | | | |
| Oligo rev | AAACCCGGATTCCTCGAACGGGCC | | | | | | | | | | |

Chromosome start end strand MM target_seq PAM alignment distance position gene name gene id
GU305914.1 18476 18498 - 0 GGCCCGTTCGAGGAATCCGG CGG ||||||||[||||||||||||]PAM NA Intergenic NA NA

T33169 AGGAATCCGGCGGCTGTCCGAGG 1000
Oligo fwd CACCAGGAATCCGGCGGCTGTCCG
Oligo rev AAACCGGACAGCCGCCGGATTCCT
Chromosome start end strand MM target_seq PAM alignment distance position gene name gene id
GU305914.1 18466 18488 - 0 AGGAATCCGGCGGCTGTCCG AGG ||||||||[||||||||||||]PAM NA Intergenic NA NA

T33170 CGCTTCAGATTCCACCACCACGG 1000
Oligo fwd CACCCGCTTCAGATTCCACCACCA
Oligo rev AAACTGGTGGTGGAATCTGAAGCG
Chromosome start end strand MM target_seq PAM alignment distance position gene name gene id
GU305914.1 18426 18448 - 0 CGCTTCAGATTCCACCACCA CGG ||||||||[||||||||||||]PAM NA Intergenic NA NA

T33171 AGATTCCACCACCACGGACTCGG 1000
Oligo fwd CACCAGATTCCACCACCACGGACT
Oligo rev AAACAGTCCGTGGTGGTGGAATCT
Chromosome start end strand MM target_seq PAM alignment distance position gene name gene id
GU305914.1 18420 18442 - 0 AGATTCCACCACCACGGACT CGG ||||||||[||||||||||||]PAM NA Intergenic NA NA

T33172 CCACGGACTCGGAGTCGAAATGG 1000
Oligo fwd CACCCCACGGACTCGGAGTCGAAA
Oligo rev AAACTTTCGACTCCGAGTCCGTGG
Chromosome start end strand MM target_seq PAM alignment distance position gene name gene id
GU305914.1 18409 18431 - 0 CCACGGACTCGGAGTCGAAA TGG ||||||||[||||||||||||]PAM NA Intergenic NA NA

T33173 CACGGACTCGGAGTCGAAATGGG 1000
Oligo fwd CACCCACGGACTCGGAGTCGAAAT
Oligo rev AAACATTTCGACTCCGAGTCCGTG
Chromosome start end strand MM target_seq PAM alignment distance position gene name gene id
GU305914.1 18408 18430 - 0 CACGGACTCGGAGTCGAAAT GGG ||||||||[||||||||||||]PAM NA Intergenic NA NA

T33174 AAATGGGCGCTTCGTTCGTGCGG 1000
Oligo fwd CACCAAATGGGCGCTTCGTTCGTG
Oligo rev AAACCACGAACGAAGCGCCCATTT
Chromosome start end strand MM target_seq PAM alignment distance position gene name gene id
GU305914.1 18392 18414 - 0 AAATGGGCGCTTCGTTCGTG CGG ||||||||[||||||||||||]PAM NA Intergenic NA NA

T33175 TCGTTCGTGCGGATCGTTCGAGG 1000
Oligo fwd CACCTCGTTCGTGCGGATCGTTCG
Oligo rev AAACCGAACGATCCGCACGAACGA
Chromosome start end strand MM target_seq PAM alignment distance position gene name gene id
GU305914.1 18381 18403 - 0 TCGTTCGTGCGGATCGTTCG AGG ||||||||[||||||||||||]PAM NA Intergenic NA NA

T33176 CGTTCGTGCGGATCGTTCGAGGG 1000
Oligo fwd CACCCGTTCGTGCGGATCGTTCGA
Oligo rev AAACTCGAACGATCCGCACGAACG
Chromosome start end strand MM target_seq PAM alignment distance position gene name gene id
GU305914.1 18380 18402 - 0 CGTTCGTGCGGATCGTTCGA GGG ||||||||[||||||||||||]PAM NA Intergenic NA NA

T33177 GTTCGTGCGGATCGTTCGAGGGG 1000
Oligo fwd CACCGTTCGTGCGGATCGTTCGAG
Oligo rev AAACCTCGAACGATCCGCACGAAC
Chromosome start end strand MM target_seq PAM alignment distance position gene name gene id
GU305914.1 18379 18401 - 0 GTTCGTGCGGATCGTTCGAG GGG ||||||||[||||||||||||]PAM NA Intergenic NA NA

T33178 TTCGTGCGGATCGTTCGAGGGGG 1000
Oligo fwd CACCTTCGTGCGGATCGTTCGAGG
Oligo rev AAACCCTCGAACGATCCGCACGAA
Chromosome start end strand MM target_seq PAM alignment distance position gene name gene id
GU305914.1 18378 18400 - 0 TTCGTGCGGATCGTTCGAGG GGG ||||||||[||||||||||||]PAM NA Intergenic NA NA

T33179 TCGAACGATCAGAAGAGATCTGG 1000
Oligo fwd CACCTCGAACGATCAGAAGAGATC
Oligo rev AAACGATCTCTTCTGATCGTTCGA
Chromosome start end strand MM target_seq PAM alignment distance position gene name gene id
GU305914.1 18350 18372 - 0 TCGAACGATCAGAAGAGATC TGG ||||||||[||||||||||||]PAM NA Intergenic NA NA

T33180 AGAAGAGATCTGGAATCGAGAGG 1000
Oligo fwd CACCAGAAGAGATCTGGAATCGAG
Oligo rev AAACCTCGATTCCAGATCTCTTCT
Chromosome start end strand MM target_seq PAM alignment distance position gene name gene id
GU305914.1 18340 18362 - 0 AGAAGAGATCTGGAATCGAG AGG ||||||||[||||||||||||]PAM NA Intergenic NA NA

T33181 GAAGAGATCTGGAATCGAGAGGG 1000
Oligo fwd CACCGAAGAGATCTGGAATCGAGA
Oligo rev AAACTCTCGATTCCAGATCTCTTC
Chromosome start end strand MM target_seq PAM alignment distance position gene name gene id
GU305914.1 18339 18361 - 0 GAAGAGATCTGGAATCGAGA GGG ||||||||[||||||||||||]PAM NA Intergenic NA NA

T33182 TGGAATCGAGAGGGCGAGTCCGG 1000
Oligo fwd CACCTGGAATCGAGAGGGCGAGTC
Oligo rev AAACGACTCGCCCTCTCGATTCCA
Chromosome start end strand MM target_seq PAM alignment distance position gene name gene id
GU305914.1 18330 18352 - 0 TGGAATCGAGAGGGCGAGTC CGG ||||||||[||||||||||||]PAM NA Intergenic NA NA

T33183 AGGGCGAGTCCGGCTCGTGTCGG 1000
Oligo fwd CACCAGGGCGAGTCCGGCTCGTGT

<table>
<tr><td>Oligo rev</td><td>AAACACACGAGCCGGACTCGCCCT</td><td></td><td></td><td></td><td></td><td></td><td></td><td></td><td></td><td></td><td></td><td></td></tr>
<tr><td>Chromosome</td><td>start</td><td></td><td>end</td><td>strand</td><td>MM</td><td>target_seq</td><td>PAM</td><td>alignment</td><td>distance</td><td>position</td><td>gene name</td><td>gene id</td></tr>
<tr><td>GU305914.1</td><td></td><td>18320</td><td>18342</td><td>-</td><td>0</td><td>AGGGCGAGTCCGGCTCGTGT</td><td>CGG</td><td>||||||||[||||||||||||]PAM</td><td>NA</td><td>Intergenic</td><td>NA</td><td>NA</td></tr>
<tr><td>T33184</td><td>CGTGTCGGCTTTCGTCCTATCGG</td><td></td><td>1000</td><td></td><td></td><td></td><td></td><td></td><td></td><td></td><td></td><td></td></tr>
<tr><td>Oligo fwd</td><td>CACCCGTGTCGGCTTTCGTCCTAT</td><td></td><td></td><td></td><td></td><td></td><td></td><td></td><td></td><td></td><td></td><td></td></tr>
<tr><td>Oligo rev</td><td>AAACATAGGACGAAAGCCGACACG</td><td></td><td></td><td></td><td></td><td></td><td></td><td></td><td></td><td></td><td></td><td></td></tr>
<tr><td>Chromosome</td><td>start</td><td></td><td>end</td><td>strand</td><td>MM</td><td>target_seq</td><td>PAM</td><td>alignment</td><td>distance</td><td>position</td><td>gene name</td><td>gene id</td></tr>
<tr><td>GU305914.1</td><td></td><td>18305</td><td>18327</td><td>-</td><td>0</td><td>CGTGTCGGCTTTCGTCCTAT</td><td>CGG</td><td>||||||||[||||||||||||]PAM</td><td>NA</td><td>Intergenic</td><td>NA</td><td>NA</td></tr>
<tr><td>T33185</td><td>CTCCGAGCGACGCCTCCAGTCGG</td><td></td><td>1000</td><td></td><td></td><td></td><td></td><td></td><td></td><td></td><td></td><td></td></tr>
<tr><td>Oligo fwd</td><td>CACCCTCCGAGCGACGCCTCCAGT</td><td></td><td></td><td></td><td></td><td></td><td></td><td></td><td></td><td></td><td></td><td></td></tr>
<tr><td>Oligo rev</td><td>AAACACTGGAGGCGTCGCTCGGAG</td><td></td><td></td><td></td><td></td><td></td><td></td><td></td><td></td><td></td><td></td><td></td></tr>
<tr><td>Chromosome</td><td>start</td><td></td><td>end</td><td>strand</td><td>MM</td><td>target_seq</td><td>PAM</td><td>alignment</td><td>distance</td><td>position</td><td>gene name</td><td>gene id</td></tr>
<tr><td>GU305914.1</td><td></td><td>18277</td><td>18299</td><td>-</td><td>0</td><td>CTCCGAGCGACGCCTCCAGT</td><td>CGG</td><td>||||||||[||||||||||||]PAM</td><td>NA</td><td>Intergenic</td><td>NA</td><td>NA</td></tr>
<tr><td>T33186</td><td>TCCGAGCGACGCCTCCAGTCGGG</td><td></td><td>1000</td><td></td><td></td><td></td><td></td><td></td><td></td><td></td><td></td><td></td></tr>
<tr><td>Oligo fwd</td><td>CACCTCCGAGCGACGCCTCCAGTC</td><td></td><td></td><td></td><td></td><td></td><td></td><td></td><td></td><td></td><td></td><td></td></tr>
<tr><td>Oligo rev</td><td>AAACGACTGGAGGCGTCGCTCGGA</td><td></td><td></td><td></td><td></td><td></td><td></td><td></td><td></td><td></td><td></td><td></td></tr>
<tr><td>Chromosome</td><td>start</td><td></td><td>end</td><td>strand</td><td>MM</td><td>target_seq</td><td>PAM</td><td>alignment</td><td>distance</td><td>position</td><td>gene name</td><td>gene id</td></tr>
<tr><td>GU305914.1</td><td></td><td>18276</td><td>18298</td><td>-</td><td>0</td><td>TCCGAGCGACGCCTCCAGTC</td><td>GGG</td><td>||||||||[||||||||||||]PAM</td><td>NA</td><td>Intergenic</td><td>NA</td><td>NA</td></tr>
<tr><td>T33187</td><td>CTCCAGTCGGGAGACGACTCCGG</td><td></td><td>1000</td><td></td><td></td><td></td><td></td><td></td><td></td><td></td><td></td><td></td></tr>
<tr><td>Oligo fwd</td><td>CACCCTCCAGTCGGGAGACGACTC</td><td></td><td></td><td></td><td></td><td></td><td></td><td></td><td></td><td></td><td></td><td></td></tr>
<tr><td>Oligo rev</td><td>AAACGAGTCGTCTCCCGACTGGAG</td><td></td><td></td><td></td><td></td><td></td><td></td><td></td><td></td><td></td><td></td><td></td></tr>
<tr><td>Chromosome</td><td>start</td><td></td><td>end</td><td>strand</td><td>MM</td><td>target_seq</td><td>PAM</td><td>alignment</td><td>distance</td><td>position</td><td>gene name</td><td>gene id</td></tr>
<tr><td>GU305914.1</td><td></td><td>18264</td><td>18286</td><td>-</td><td>0</td><td>CTCCAGTCGGGAGACGACTC</td><td>CGG</td><td>||||||||[||||||||||||]PAM</td><td>NA</td><td>Intergenic</td><td>NA</td><td>NA</td></tr>
<tr><td>GU305914.1</td><td></td><td>77062</td><td>77084</td><td>+</td><td>4</td><td>CTCCGGGCGGGAGACGATGC</td><td>TGG</td><td>||||-|-|[|||||||||--|]PAM</td><td>NA</td><td>Intergenic</td><td>NA</td><td>NA</td></tr>
<tr><td>T33188</td><td>GTCGGGAGACGACTCCGGAGAGG</td><td></td><td>1000</td><td></td><td></td><td></td><td></td><td></td><td></td><td></td><td></td><td></td></tr>
<tr><td>Oligo fwd</td><td>CACCGTCGGGAGACGACTCCGGAG</td><td></td><td></td><td></td><td></td><td></td><td></td><td></td><td></td><td></td><td></td><td></td></tr>
<tr><td>Oligo rev</td><td>AAACCTCCGGAGTCGTCTCCCGAC</td><td></td><td></td><td></td><td></td><td></td><td></td><td></td><td></td><td></td><td></td><td></td></tr>
<tr><td>Chromosome</td><td>start</td><td></td><td>end</td><td>strand</td><td>MM</td><td>target_seq</td><td>PAM</td><td>alignment</td><td>distance</td><td>position</td><td>gene name</td><td>gene id</td></tr>
<tr><td>GU305914.1</td><td></td><td>18259</td><td>18281</td><td>-</td><td>0</td><td>GTCGGGAGACGACTCCGGAG</td><td>AGG</td><td>||||||||[||||||||||||]PAM</td><td>NA</td><td>Intergenic</td><td>NA</td><td>NA</td></tr>
<tr><td>T33189</td><td>GAGACGACTCCGGAGAGGCGCGG</td><td></td><td>1000</td><td></td><td></td><td></td><td></td><td></td><td></td><td></td><td></td><td></td></tr>
<tr><td>Oligo fwd</td><td>CACCGAGACGACTCCGGAGAGGCG</td><td></td><td></td><td></td><td></td><td></td><td></td><td></td><td></td><td></td><td></td><td></td></tr>
<tr><td>Oligo rev</td><td>AAACCGCCTCTCCGGAGTCGTCTC</td><td></td><td></td><td></td><td></td><td></td><td></td><td></td><td></td><td></td><td></td><td></td></tr>
<tr><td>Chromosome</td><td>start</td><td></td><td>end</td><td>strand</td><td>MM</td><td>target_seq</td><td>PAM</td><td>alignment</td><td>distance</td><td>position</td><td>gene name</td><td>gene id</td></tr>
<tr><td>GU305914.1</td><td></td><td>18254</td><td>18276</td><td>-</td><td>0</td><td>GAGACGACTCCGGAGAGGCG</td><td>CGG</td><td>||||||||[||||||||||||]PAM</td><td>NA</td><td>Intergenic</td><td>NA</td><td>NA</td></tr>
<tr><td>T33190</td><td>AGACGACTCCGGAGAGGCGCGGG</td><td></td><td>1000</td><td></td><td></td><td></td><td></td><td></td><td></td><td></td><td></td><td></td></tr>
<tr><td>Oligo fwd</td><td>CACCAGACGACTCCGGAGAGGCGC</td><td></td><td></td><td></td><td></td><td></td><td></td><td></td><td></td><td></td><td></td><td></td></tr>
<tr><td>Oligo rev</td><td>AAACGCGCCTCTCCGGAGTCGTCT</td><td></td><td></td><td></td><td></td><td></td><td></td><td></td><td></td><td></td><td></td><td></td></tr>
<tr><td>Chromosome</td><td>start</td><td></td><td>end</td><td>strand</td><td>MM</td><td>target_seq</td><td>PAM</td><td>alignment</td><td>distance</td><td>position</td><td>gene name</td><td>gene id</td></tr>
<tr><td>GU305914.1</td><td></td><td>18253</td><td>18275</td><td>-</td><td>0</td><td>AGACGACTCCGGAGAGGCGC</td><td>GGG</td><td>||||||||[||||||||||||]PAM</td><td>NA</td><td>Intergenic</td><td>NA</td><td>NA</td></tr>
<tr><td>T33191</td><td>CGACTCCGGAGAGGCGCGGGAGG</td><td></td><td>1000</td><td></td><td></td><td></td><td></td><td></td><td></td><td></td><td></td><td></td></tr>
<tr><td>Oligo fwd</td><td>CACCCGACTCCGGAGAGGCGCGGG</td><td></td><td></td><td></td><td></td><td></td><td></td><td></td><td></td><td></td><td></td><td></td></tr>
<tr><td>Oligo rev</td><td>AAACCCCGCGCCTCTCCGGAGTCG</td><td></td><td></td><td></td><td></td><td></td><td></td><td></td><td></td><td></td><td></td><td></td></tr>
<tr><td>Chromosome</td><td>start</td><td></td><td>end</td><td>strand</td><td>MM</td><td>target_seq</td><td>PAM</td><td>alignment</td><td>distance</td><td>position</td><td>gene name</td><td>gene id</td></tr>
<tr><td>GU305914.1</td><td></td><td>18250</td><td>18272</td><td>-</td><td>0</td><td>CGACTCCGGAGAGGCGCGGG</td><td>AGG</td><td>||||||||[||||||||||||]PAM</td><td>NA</td><td>Intergenic</td><td>NA</td><td>NA</td></tr>
<tr><td>T33192</td><td>TTCGTCCCGAGAACGCCCCGCGG</td><td></td><td>1000</td><td></td><td></td><td></td><td></td><td></td><td></td><td></td><td></td><td></td></tr>
<tr><td>Oligo fwd</td><td>CACCTTCGTCCCGAGAACGCCCCG</td><td></td><td></td><td></td><td></td><td></td><td></td><td></td><td></td><td></td><td></td><td></td></tr>
<tr><td>Oligo rev</td><td>AAACCGGGGCGTTCTCGGGACGAA</td><td></td><td></td><td></td><td></td><td></td><td></td><td></td><td></td><td></td><td></td><td></td></tr>
<tr><td>Chromosome</td><td>start</td><td></td><td>end</td><td>strand</td><td>MM</td><td>target_seq</td><td>PAM</td><td>alignment</td><td>distance</td><td>position</td><td>gene name</td><td>gene id</td></tr>
<tr><td>GU305914.1</td><td></td><td>18186</td><td>18208</td><td>-</td><td>0</td><td>TTCGTCCCGAGAACGCCCCG</td><td>CGG</td><td>||||||||[||||||||||||]PAM</td><td>NA</td><td>Intergenic</td><td>NA</td><td>NA</td></tr>
<tr><td>T33193</td><td>ACCGCCAACATCCCGTCCCGCGG</td><td></td><td>1000</td><td></td><td></td><td></td><td></td><td></td><td></td><td></td><td></td><td></td></tr>
<tr><td>Oligo fwd</td><td>CACCACCGCCAACATCCCGTCCCG</td><td></td><td></td><td></td><td></td><td></td><td></td><td></td><td></td><td></td><td></td><td></td></tr>
<tr><td>Oligo rev</td><td>AAACCGGGACGGGATGTTGGCGGT</td><td></td><td></td><td></td><td></td><td></td><td></td><td></td><td></td><td></td><td></td><td></td></tr>
<tr><td>Chromosome</td><td>start</td><td></td><td>end</td><td>strand</td><td>MM</td><td>target_seq</td><td>PAM</td><td>alignment</td><td>distance</td><td>position</td><td>gene name</td><td>gene id</td></tr>
<tr><td>GU305914.1</td><td></td><td>18158</td><td>18180</td><td>-</td><td>0</td><td>ACCGCCAACATCCCGTCCCG</td><td>CGG</td><td>||||||||[||||||||||||]PAM</td><td>NA</td><td>Intergenic</td><td>NA</td><td>NA</td></tr>
<tr><td>T33194</td><td>GCCAACATCCCGTCCCGCGGCGG</td><td></td><td>1000</td><td></td><td></td><td></td><td></td><td></td><td></td><td></td><td></td><td></td></tr>
<tr><td>Oligo fwd</td><td>CACCGCCAACATCCCGTCCCGCGG</td><td></td><td></td><td></td><td></td><td></td><td></td><td></td><td></td><td></td><td></td><td></td></tr>
<tr><td>Oligo rev</td><td>AAACCCGCGGGACGGGATGTTGGC</td><td></td><td></td><td></td><td></td><td></td><td></td><td></td><td></td><td></td><td></td><td></td></tr>
<tr><td>Chromosome</td><td>start</td><td></td><td>end</td><td>strand</td><td>MM</td><td>target_seq</td><td>PAM</td><td>alignment</td><td>distance</td><td>position</td><td>gene name</td><td>gene id</td></tr>
<tr><td>GU305914.1</td><td></td><td>18155</td><td>18177</td><td>-</td><td>0</td><td>GCCAACATCCCGTCCCGCGG</td><td>CGG</td><td>||||||||[||||||||||||]PAM</td><td>NA</td><td>Intergenic</td><td>NA</td><td>NA</td></tr>
<tr><td>T33195</td><td>CCAACATCCCGTCCCGCGGCGGG</td><td></td><td>1000</td><td></td><td></td><td></td><td></td><td></td><td></td><td></td><td></td><td></td></tr>
<tr><td>Oligo fwd</td><td>CACCCCAACATCCCGTCCCGCGGC</td><td></td><td></td><td></td><td></td><td></td><td></td><td></td><td></td><td></td><td></td><td></td></tr>
<tr><td>Oligo rev</td><td>AAACGCCGCGGGACGGGATGTTGG</td><td></td><td></td><td></td><td></td><td></td><td></td><td></td><td></td><td></td><td></td><td></td></tr>
<tr><td>Chromosome</td><td>start</td><td></td><td>end</td><td>strand</td><td>MM</td><td>target_seq</td><td>PAM</td><td>alignment</td><td>distance</td><td>position</td><td>gene name</td><td>gene id</td></tr>
<tr><td>GU305914.1</td><td></td><td>18154</td><td>18176</td><td>-</td><td>0</td><td>CCAACATCCCGTCCCGCGGC</td><td>GGG</td><td>||||||||[||||||||||||]PAM</td><td>NA</td><td>Intergenic</td><td>NA</td><td>NA</td></tr>
<tr><td>T33196</td><td>ATCCCGTCCCGCGGCGGGCCCGG</td><td></td><td>1000</td><td></td><td></td><td></td><td></td><td></td><td></td><td></td><td></td><td></td></tr>
<tr><td>Oligo fwd</td><td>CACCATCCCGTCCCGCGGCGGGCC</td><td></td><td></td><td></td><td></td><td></td><td></td><td></td><td></td><td></td><td></td><td></td></tr>
<tr><td>Oligo rev</td><td>AAACGGCCCGCCGCGGGACGGGAT</td><td></td><td></td><td></td><td></td><td></td><td></td><td></td><td></td><td></td><td></td><td></td></tr>
<tr><td>Chromosome</td><td>start</td><td></td><td>end</td><td>strand</td><td>MM</td><td>target_seq</td><td>PAM</td><td>alignment</td><td>distance</td><td>position</td><td>gene name</td><td>gene id</td></tr>
<tr><td>GU305914.1</td><td></td><td>18149</td><td>18171</td><td>-</td><td>0</td><td>ATCCCGTCCCGCGGCGGGCC</td><td>CGG</td><td>||||||||[||||||||||||]PAM</td><td>NA</td><td>Intergenic</td><td>NA</td><td>NA</td></tr>
<tr><td>T33197</td><td>CCGTCCCGCGGCGGGCCCGGAGG</td><td></td><td>1000</td><td></td><td></td><td></td><td></td><td></td><td></td><td></td><td></td><td></td></tr>
<tr><td>Oligo fwd</td><td>CACCCCGTCCCGCGGCGGGCCCGG</td><td></td><td></td><td></td><td></td><td></td><td></td><td></td><td></td><td></td><td></td><td></td></tr>
<tr><td>Oligo rev</td><td>AAACCCGGGCCCGCCGCGGGACGG</td><td></td><td></td><td></td><td></td><td></td><td></td><td></td><td></td><td></td><td></td><td></td></tr>
<tr><td>Chromosome</td><td>start</td><td></td><td>end</td><td>strand</td><td>MM</td><td>target_seq</td><td>PAM</td><td>alignment</td><td>distance</td><td>position</td><td>gene name</td><td>gene id</td></tr>
<tr><td>GU305914.1</td><td></td><td>18146</td><td>18168</td><td>-</td><td>0</td><td>CCGTCCCGCGGCGGGCCCGG</td><td>AGG</td><td>||||||||[||||||||||||]PAM</td><td>NA</td><td>Intergenic</td><td>NA</td><td>NA</td></tr>
</table>

<table>
<tr><td>T33198</td><td>TTCTTCCGCGACAGAGATGCCGG</td><td>1000</td><td></td><td></td><td></td><td></td><td></td><td></td><td></td><td></td><td></td></tr>
<tr><td>Oligo fwd</td><td>CACCTTCTTCCGCGACAGAGATGC</td><td></td><td></td><td></td><td></td><td></td><td></td><td></td><td></td><td></td><td></td></tr>
<tr><td>Oligo rev</td><td>AAACGCATCTCTGTCGCGGAAGAA</td><td></td><td></td><td></td><td></td><td></td><td></td><td></td><td></td><td></td><td></td></tr>
<tr><td>Chromosome</td><td>start</td><td>end</td><td>strand</td><td>MM</td><td>target_seq</td><td>PAM</td><td>alignment</td><td>distance</td><td>position</td><td>gene name</td><td>gene id</td></tr>
<tr><td>GU305914.1</td><td>18123</td><td>18145</td><td>-</td><td>0</td><td>TTCTTCCGCGACAGAGATGC</td><td>CGG</td><td>||||||||[||||||||||||]PAM</td><td>NA</td><td>Intergenic</td><td>NA</td><td>NA</td></tr>
<tr><td>T33199</td><td>CGACAGAGATGCCGGCCACGAGG</td><td>1000</td><td></td><td></td><td></td><td></td><td></td><td></td><td></td><td></td><td></td></tr>
<tr><td>Oligo fwd</td><td>CACCCGACAGAGATGCCGGCCACG</td><td></td><td></td><td></td><td></td><td></td><td></td><td></td><td></td><td></td><td></td></tr>
<tr><td>Oligo rev</td><td>AAACCGTGGCCGGCATCTCTGTCG</td><td></td><td></td><td></td><td></td><td></td><td></td><td></td><td></td><td></td><td></td></tr>
<tr><td>Chromosome</td><td>start</td><td>end</td><td>strand</td><td>MM</td><td>target_seq</td><td>PAM</td><td>alignment</td><td>distance</td><td>position</td><td>gene name</td><td>gene id</td></tr>
<tr><td>GU305914.1</td><td>18115</td><td>18137</td><td>-</td><td>0</td><td>CGACAGAGATGCCGGCCACG</td><td>AGG</td><td>||||||||[||||||||||||]PAM</td><td>NA</td><td>Intergenic</td><td>NA</td><td>NA</td></tr>
<tr><td>T33200</td><td>AGATGCCGGCCACGAGGCTGCGG</td><td>1000</td><td></td><td></td><td></td><td></td><td></td><td></td><td></td><td></td><td></td></tr>
<tr><td>Oligo fwd</td><td>CACCAGATGCCGGCCACGAGGCTG</td><td></td><td></td><td></td><td></td><td></td><td></td><td></td><td></td><td></td><td></td></tr>
<tr><td>Oligo rev</td><td>AAACCAGCCTCGTGGCCGGCATCT</td><td></td><td></td><td></td><td></td><td></td><td></td><td></td><td></td><td></td><td></td></tr>
<tr><td>Chromosome</td><td>start</td><td>end</td><td>strand</td><td>MM</td><td>target_seq</td><td>PAM</td><td>alignment</td><td>distance</td><td>position</td><td>gene name</td><td>gene id</td></tr>
<tr><td>GU305914.1</td><td>18109</td><td>18131</td><td>-</td><td>0</td><td>AGATGCCGGCCACGAGGCTG</td><td>CGG</td><td>||||||||[||||||||||||]PAM</td><td>NA</td><td>Intergenic</td><td>NA</td><td>NA</td></tr>
<tr><td>T33201</td><td>TGCCGGCCACGAGGCTGCGGCGG</td><td>1000</td><td></td><td></td><td></td><td></td><td></td><td></td><td></td><td></td><td></td></tr>
<tr><td>Oligo fwd</td><td>CACCTGCCGGCCACGAGGCTGCGG</td><td></td><td></td><td></td><td></td><td></td><td></td><td></td><td></td><td></td><td></td></tr>
<tr><td>Oligo rev</td><td>AAACCCGCAGCCTCGTGGCCGGCA</td><td></td><td></td><td></td><td></td><td></td><td></td><td></td><td></td><td></td><td></td></tr>
<tr><td>Chromosome</td><td>start</td><td>end</td><td>strand</td><td>MM</td><td>target_seq</td><td>PAM</td><td>alignment</td><td>distance</td><td>position</td><td>gene name</td><td>gene id</td></tr>
<tr><td>GU305914.1</td><td>18106</td><td>18128</td><td>-</td><td>0</td><td>TGCCGGCCACGAGGCTGCGG</td><td>CGG</td><td>||||||||[||||||||||||]PAM</td><td>NA</td><td>Intergenic</td><td>NA</td><td>NA</td></tr>
<tr><td>T33202</td><td>CCAGCTCCCCGACACCCAAGAGG</td><td>1000</td><td></td><td></td><td></td><td></td><td></td><td></td><td></td><td></td><td></td></tr>
<tr><td>Oligo fwd</td><td>CACCCCAGCTCCCCGACACCCAAG</td><td></td><td></td><td></td><td></td><td></td><td></td><td></td><td></td><td></td><td></td></tr>
<tr><td>Oligo rev</td><td>AAACCTTGGGTGTCGGGGAGCTGG</td><td></td><td></td><td></td><td></td><td></td><td></td><td></td><td></td><td></td><td></td></tr>
<tr><td>Chromosome</td><td>start</td><td>end</td><td>strand</td><td>MM</td><td>target_seq</td><td>PAM</td><td>alignment</td><td>distance</td><td>position</td><td>gene name</td><td>gene id</td></tr>
<tr><td>GU305914.1</td><td>18007</td><td>18029</td><td>-</td><td>0</td><td>CCAGCTCCCCGACACCCAAG</td><td>AGG</td><td>||||||||[||||||||||||]PAM</td><td>NA</td><td>Intergenic</td><td>NA</td><td>NA</td></tr>
<tr><td>T33203</td><td>CCGACACCCAAGAGGTCGCGCGG</td><td>1000</td><td></td><td></td><td></td><td></td><td></td><td></td><td></td><td></td><td></td></tr>
<tr><td>Oligo fwd</td><td>CACCCCGACACCCAAGAGGTCGCG</td><td></td><td></td><td></td><td></td><td></td><td></td><td></td><td></td><td></td><td></td></tr>
<tr><td>Oligo rev</td><td>AAACCGCGACCTCTTGGGTGTCGG</td><td></td><td></td><td></td><td></td><td></td><td></td><td></td><td></td><td></td><td></td></tr>
<tr><td>Chromosome</td><td>start</td><td>end</td><td>strand</td><td>MM</td><td>target_seq</td><td>PAM</td><td>alignment</td><td>distance</td><td>position</td><td>gene name</td><td>gene id</td></tr>
<tr><td>GU305914.1</td><td>17999</td><td>18021</td><td>-</td><td>0</td><td>CCGACACCCAAGAGGTCGCG</td><td>CGG</td><td>||||||||[||||||||||||]PAM</td><td>NA</td><td>Intergenic</td><td>NA</td><td>NA</td></tr>
<tr><td>T33204</td><td>CCAAGAGGTCGCGCGGCAGACGG</td><td>1000</td><td></td><td></td><td></td><td></td><td></td><td></td><td></td><td></td><td></td></tr>
<tr><td>Oligo fwd</td><td>CACCCCAAGAGGTCGCGCGGCAGA</td><td></td><td></td><td></td><td></td><td></td><td></td><td></td><td></td><td></td><td></td></tr>
<tr><td>Oligo rev</td><td>AAACTCTGCCGCGCGACCTCTTGG</td><td></td><td></td><td></td><td></td><td></td><td></td><td></td><td></td><td></td><td></td></tr>
<tr><td>Chromosome</td><td>start</td><td>end</td><td>strand</td><td>MM</td><td>target_seq</td><td>PAM</td><td>alignment</td><td>distance</td><td>position</td><td>gene name</td><td>gene id</td></tr>
<tr><td>GU305914.1</td><td>17992</td><td>18014</td><td>-</td><td>0</td><td>CCAAGAGGTCGCGCGGCAGA</td><td>CGG</td><td>||||||||[||||||||||||]PAM</td><td>NA</td><td>Intergenic</td><td>NA</td><td>NA</td></tr>
<tr><td>T33205</td><td>CGGCAGACGGACGACCGTCACGG</td><td>1000</td><td></td><td></td><td></td><td></td><td></td><td></td><td></td><td></td><td></td></tr>
<tr><td>Oligo fwd</td><td>CACCCGGCAGACGGACGACCGTCA</td><td></td><td></td><td></td><td></td><td></td><td></td><td></td><td></td><td></td><td></td></tr>
<tr><td>Oligo rev</td><td>AAACTGACGGTCGTCCGTCTGCCG</td><td></td><td></td><td></td><td></td><td></td><td></td><td></td><td></td><td></td><td></td></tr>
<tr><td>Chromosome</td><td>start</td><td>end</td><td>strand</td><td>MM</td><td>target_seq</td><td>PAM</td><td>alignment</td><td>distance</td><td>position</td><td>gene name</td><td>gene id</td></tr>
<tr><td>GU305914.1</td><td>17979</td><td>18001</td><td>-</td><td>0</td><td>CGGCAGACGGACGACCGTCA</td><td>CGG</td><td>||||||||[||||||||||||]PAM</td><td>NA</td><td>Intergenic</td><td>NA</td><td>NA</td></tr>
<tr><td>T33206</td><td>TCGTACCTGCCCCGATATCCAGG</td><td>1000</td><td></td><td></td><td></td><td></td><td></td><td></td><td></td><td></td><td></td></tr>
<tr><td>Oligo fwd</td><td>CACCTCGTACCTGCCCCGATATCC</td><td></td><td></td><td></td><td></td><td></td><td></td><td></td><td></td><td></td><td></td></tr>
<tr><td>Oligo rev</td><td>AAACGGATATCGGGGCAGGTACGA</td><td></td><td></td><td></td><td></td><td></td><td></td><td></td><td></td><td></td><td></td></tr>
<tr><td>Chromosome</td><td>start</td><td>end</td><td>strand</td><td>MM</td><td>target_seq</td><td>PAM</td><td>alignment</td><td>distance</td><td>position</td><td>gene name</td><td>gene id</td></tr>
<tr><td>GU305914.1</td><td>17941</td><td>17963</td><td>-</td><td>0</td><td>TCGTACCTGCCCCGATATCC</td><td>AGG</td><td>||||||||[||||||||||||]PAM</td><td>NA</td><td>Intergenic</td><td>NA</td><td>NA</td></tr>
<tr><td>T33207</td><td>TACCTGCCCCGATATCCAGGAGG</td><td>1000</td><td></td><td></td><td></td><td></td><td></td><td></td><td></td><td></td><td></td></tr>
<tr><td>Oligo fwd</td><td>CACCTACCTGCCCCGATATCCAGG</td><td></td><td></td><td></td><td></td><td></td><td></td><td></td><td></td><td></td><td></td></tr>
<tr><td>Oligo rev</td><td>AAACCCTGGATATCGGGGCAGGTA</td><td></td><td></td><td></td><td></td><td></td><td></td><td></td><td></td><td></td><td></td></tr>
<tr><td>Chromosome</td><td>start</td><td>end</td><td>strand</td><td>MM</td><td>target_seq</td><td>PAM</td><td>alignment</td><td>distance</td><td>position</td><td>gene name</td><td>gene id</td></tr>
<tr><td>GU305914.1</td><td>17938</td><td>17960</td><td>-</td><td>0</td><td>TACCTGCCCCGATATCCAGG</td><td>AGG</td><td>||||||||[||||||||||||]PAM</td><td>NA</td><td>Intergenic</td><td>NA</td><td>NA</td></tr>
<tr><td>T33208</td><td>AGGAGGCGACAGTCTCTCGACGG</td><td>1000</td><td></td><td></td><td></td><td></td><td></td><td></td><td></td><td></td><td></td></tr>
<tr><td>Oligo fwd</td><td>CACCAGGAGGCGACAGTCTCTCGA</td><td></td><td></td><td></td><td></td><td></td><td></td><td></td><td></td><td></td><td></td></tr>
<tr><td>Oligo rev</td><td>AAACTCGAGAGACTGTCGCCTCCT</td><td></td><td></td><td></td><td></td><td></td><td></td><td></td><td></td><td></td><td></td></tr>
<tr><td>Chromosome</td><td>start</td><td>end</td><td>strand</td><td>MM</td><td>target_seq</td><td>PAM</td><td>alignment</td><td>distance</td><td>position</td><td>gene name</td><td>gene id</td></tr>
<tr><td>GU305914.1</td><td>17921</td><td>17943</td><td>-</td><td>0</td><td>AGGAGGCGACAGTCTCTCGA</td><td>CGG</td><td>||||||||[||||||||||||]PAM</td><td>NA</td><td>Intergenic</td><td>NA</td><td>NA</td></tr>
<tr><td>T33209</td><td>GGAGGCGACAGTCTCTCGACGGG</td><td>1000</td><td></td><td></td><td></td><td></td><td></td><td></td><td></td><td></td><td></td></tr>
<tr><td>Oligo fwd</td><td>CACCGGAGGCGACAGTCTCTCGAC</td><td></td><td></td><td></td><td></td><td></td><td></td><td></td><td></td><td></td><td></td></tr>
<tr><td>Oligo rev</td><td>AAACGTCGAGAGACTGTCGCCTCC</td><td></td><td></td><td></td><td></td><td></td><td></td><td></td><td></td><td></td><td></td></tr>
<tr><td>Chromosome</td><td>start</td><td>end</td><td>strand</td><td>MM</td><td>target_seq</td><td>PAM</td><td>alignment</td><td>distance</td><td>position</td><td>gene name</td><td>gene id</td></tr>
<tr><td>GU305914.1</td><td>17920</td><td>17942</td><td>-</td><td>0</td><td>GGAGGCGACAGTCTCTCGAC</td><td>GGG</td><td>||||||||[||||||||||||]PAM</td><td>NA</td><td>Intergenic</td><td>NA</td><td>NA</td></tr>
<tr><td>T33210</td><td>CGCTACTTTTCCTGAAAGCTCGG</td><td>1000</td><td></td><td></td><td></td><td></td><td></td><td></td><td></td><td></td><td></td></tr>
<tr><td>Oligo fwd</td><td>CACCCGCTACTTTTCCTGAAAGCT</td><td></td><td></td><td></td><td></td><td></td><td></td><td></td><td></td><td></td><td></td></tr>
<tr><td>Oligo rev</td><td>AAACAGCTTTCAGGAAAAGTAGCG</td><td></td><td></td><td></td><td></td><td></td><td></td><td></td><td></td><td></td><td></td></tr>
<tr><td>Chromosome</td><td>start</td><td>end</td><td>strand</td><td>MM</td><td>target_seq</td><td>PAM</td><td>alignment</td><td>distance</td><td>position</td><td>gene name</td><td>gene id</td></tr>
<tr><td>GU305914.1</td><td>17886</td><td>17908</td><td>-</td><td>0</td><td>CGCTACTTTTCCTGAAAGCT</td><td>CGG</td><td>||||||||[||||||||||||]PAM</td><td>NA</td><td>Intergenic</td><td>NA</td><td>NA</td></tr>
<tr><td>T33211</td><td>GAAAGCTCGGCTGCTCGACTCGG</td><td>1000</td><td></td><td></td><td></td><td></td><td></td><td></td><td></td><td></td><td></td></tr>
<tr><td>Oligo fwd</td><td>CACCGAAAGCTCGGCTGCTCGACT</td><td></td><td></td><td></td><td></td><td></td><td></td><td></td><td></td><td></td><td></td></tr>
<tr><td>Oligo rev</td><td>AAACAGTCGAGCAGCCGAGCTTTC</td><td></td><td></td><td></td><td></td><td></td><td></td><td></td><td></td><td></td><td></td></tr>
<tr><td>Chromosome</td><td>start</td><td>end</td><td>strand</td><td>MM</td><td>target_seq</td><td>PAM</td><td>alignment</td><td>distance</td><td>position</td><td>gene name</td><td>gene id</td></tr>
<tr><td>GU305914.1</td><td>17873</td><td>17895</td><td>-</td><td>0</td><td>GAAAGCTCGGCTGCTCGACT</td><td>CGG</td><td>||||||||[||||||||||||]PAM</td><td>NA</td><td>Intergenic</td><td>NA</td><td>NA</td></tr>
<tr><td>T33212</td><td>AGCTCGGCTGCTCGACTCGGCGG</td><td>1000</td><td></td><td></td><td></td><td></td><td></td><td></td><td></td><td></td><td></td></tr>
<tr><td>Oligo fwd</td><td>CACCAGCTCGGCTGCTCGACTCGG</td><td></td><td></td><td></td><td></td><td></td><td></td><td></td><td></td><td></td><td></td></tr>
<tr><td>Oligo rev</td><td>AAACCCGAGTCGAGCAGCCGAGCT</td><td></td><td></td><td></td><td></td><td></td><td></td><td></td><td></td><td></td><td></td></tr>
<tr><td>Chromosome</td><td>start</td><td>end</td><td>strand</td><td>MM</td><td>target_seq</td><td>PAM</td><td>alignment</td><td>distance</td><td>position</td><td>gene name</td><td>gene id</td></tr>
<tr><td>GU305914.1</td><td>17870</td><td>17892</td><td>-</td><td>0</td><td>AGCTCGGCTGCTCGACTCGG</td><td>CGG</td><td>||||||||[||||||||||||]PAM</td><td>NA</td><td>Intergenic</td><td>NA</td><td>NA</td></tr>
</table>

<table>
<tr><td>T33213</td><td>CGGCGGCAATCCCAGACCTGTGG</td><td>1000</td><td></td><td></td><td></td><td></td><td></td><td></td><td></td><td></td><td></td></tr>
<tr><td>Oligo fwd</td><td>CACCCGGCGGCAATCCCAGACCTG</td><td></td><td></td><td></td><td></td><td></td><td></td><td></td><td></td><td></td><td></td></tr>
<tr><td>Oligo rev</td><td>AAACCAGGTCTGGGATTGCCGCCG</td><td></td><td></td><td></td><td></td><td></td><td></td><td></td><td></td><td></td><td></td></tr>
<tr><td>Chromosome</td><td>start</td><td>end</td><td>strand</td><td>MM</td><td>target_seq</td><td>PAM</td><td>alignment</td><td>distance</td><td>position</td><td>gene name</td><td>gene id</td></tr>
<tr><td>GU305914.1</td><td>17853</td><td>17875</td><td>-</td><td>0</td><td>CGGCGGCAATCCCAGACCTG</td><td>TGG</td><td>||||||||[||||||||||||]PAM</td><td>NA</td><td>Intergenic</td><td>NA</td><td>NA</td></tr>
<tr><td>T33214</td><td>AATCCCAGACCTGTGGAAAACGG</td><td>1000</td><td></td><td></td><td></td><td></td><td></td><td></td><td></td><td></td><td></td></tr>
<tr><td>Oligo fwd</td><td>CACCAATCCCAGACCTGTGGAAAA</td><td></td><td></td><td></td><td></td><td></td><td></td><td></td><td></td><td></td><td></td></tr>
<tr><td>Oligo rev</td><td>AAACTTTTCCACAGGTCTGGGATT</td><td></td><td></td><td></td><td></td><td></td><td></td><td></td><td></td><td></td><td></td></tr>
<tr><td>Chromosome</td><td>start</td><td>end</td><td>strand</td><td>MM</td><td>target_seq</td><td>PAM</td><td>alignment</td><td>distance</td><td>position</td><td>gene name</td><td>gene id</td></tr>
<tr><td>GU305914.1</td><td>17846</td><td>17868</td><td>-</td><td>0</td><td>AATCCCAGACCTGTGGAAAA</td><td>CGG</td><td>||||||||[||||||||||||]PAM</td><td>NA</td><td>Intergenic</td><td>NA</td><td>NA</td></tr>
<tr><td>T33215</td><td>ATCCCAGACCTGTGGAAAACGGG</td><td>1000</td><td></td><td></td><td></td><td></td><td></td><td></td><td></td><td></td><td></td></tr>
<tr><td>Oligo fwd</td><td>CACCATCCCAGACCTGTGGAAAAC</td><td></td><td></td><td></td><td></td><td></td><td></td><td></td><td></td><td></td><td></td></tr>
<tr><td>Oligo rev</td><td>AAACGTTTTCCACAGGTCTGGGAT</td><td></td><td></td><td></td><td></td><td></td><td></td><td></td><td></td><td></td><td></td></tr>
<tr><td>Chromosome</td><td>start</td><td>end</td><td>strand</td><td>MM</td><td>target_seq</td><td>PAM</td><td>alignment</td><td>distance</td><td>position</td><td>gene name</td><td>gene id</td></tr>
<tr><td>GU305914.1</td><td>17845</td><td>17867</td><td>-</td><td>0</td><td>ATCCCAGACCTGTGGAAAAC</td><td>GGG</td><td>||||||||[||||||||||||]PAM</td><td>NA</td><td>Intergenic</td><td>NA</td><td>NA</td></tr>
<tr><td>T33216</td><td>TCCCAGACCTGTGGAAAACGGGG</td><td>1000</td><td></td><td></td><td></td><td></td><td></td><td></td><td></td><td></td><td></td></tr>
<tr><td>Oligo fwd</td><td>CACCTCCCAGACCTGTGGAAAACG</td><td></td><td></td><td></td><td></td><td></td><td></td><td></td><td></td><td></td><td></td></tr>
<tr><td>Oligo rev</td><td>AAACCGTTTTCCACAGGTCTGGGA</td><td></td><td></td><td></td><td></td><td></td><td></td><td></td><td></td><td></td><td></td></tr>
<tr><td>Chromosome</td><td>start</td><td>end</td><td>strand</td><td>MM</td><td>target_seq</td><td>PAM</td><td>alignment</td><td>distance</td><td>position</td><td>gene name</td><td>gene id</td></tr>
<tr><td>GU305914.1</td><td>17844</td><td>17866</td><td>-</td><td>0</td><td>TCCCAGACCTGTGGAAAACG</td><td>GGG</td><td>||||||||[||||||||||||]PAM</td><td>NA</td><td>Intergenic</td><td>NA</td><td>NA</td></tr>
<tr><td>T33217</td><td>TGGAAAACGGGGTGATCACGCGG</td><td>1000</td><td></td><td></td><td></td><td></td><td></td><td></td><td></td><td></td><td></td></tr>
<tr><td>Oligo fwd</td><td>CACCTGGAAAACGGGGTGATCACG</td><td></td><td></td><td></td><td></td><td></td><td></td><td></td><td></td><td></td><td></td></tr>
<tr><td>Oligo rev</td><td>AAACCGTGATCACCCCGTTTTCCA</td><td></td><td></td><td></td><td></td><td></td><td></td><td></td><td></td><td></td><td></td></tr>
<tr><td>Chromosome</td><td>start</td><td>end</td><td>strand</td><td>MM</td><td>target_seq</td><td>PAM</td><td>alignment</td><td>distance</td><td>position</td><td>gene name</td><td>gene id</td></tr>
<tr><td>GU305914.1</td><td>17833</td><td>17855</td><td>-</td><td>0</td><td>TGGAAAACGGGGTGATCACG</td><td>CGG</td><td>||||||||[||||||||||||]PAM</td><td>NA</td><td>Intergenic</td><td>NA</td><td>NA</td></tr>
<tr><td>T33218</td><td>GGAAAACGGGGTGATCACGCGGG</td><td>1000</td><td></td><td></td><td></td><td></td><td></td><td></td><td></td><td></td><td></td></tr>
<tr><td>Oligo fwd</td><td>CACCGGAAAACGGGGTGATCACGC</td><td></td><td></td><td></td><td></td><td></td><td></td><td></td><td></td><td></td><td></td></tr>
<tr><td>Oligo rev</td><td>AAACGCGTGATCACCCCGTTTTCC</td><td></td><td></td><td></td><td></td><td></td><td></td><td></td><td></td><td></td><td></td></tr>
<tr><td>Chromosome</td><td>start</td><td>end</td><td>strand</td><td>MM</td><td>target_seq</td><td>PAM</td><td>alignment</td><td>distance</td><td>position</td><td>gene name</td><td>gene id</td></tr>
<tr><td>GU305914.1</td><td>17832</td><td>17854</td><td>-</td><td>0</td><td>GGAAAACGGGGTGATCACGC</td><td>GGG</td><td>||||||||[||||||||||||]PAM</td><td>NA</td><td>Intergenic</td><td>NA</td><td>NA</td></tr>
<tr><td>T33219</td><td>CGGGATCGCGCCAAACGCGACGG</td><td>1000</td><td></td><td></td><td></td><td></td><td></td><td></td><td></td><td></td><td></td></tr>
<tr><td>Oligo fwd</td><td>CACCCGGGATCGCGCCAAACGCGA</td><td></td><td></td><td></td><td></td><td></td><td></td><td></td><td></td><td></td><td></td></tr>
<tr><td>Oligo rev</td><td>AAACTCGCGTTTGGCGCGATCCCG</td><td></td><td></td><td></td><td></td><td></td><td></td><td></td><td></td><td></td><td></td></tr>
<tr><td>Chromosome</td><td>start</td><td>end</td><td>strand</td><td>MM</td><td>target_seq</td><td>PAM</td><td>alignment</td><td>distance</td><td>position</td><td>gene name</td><td>gene id</td></tr>
<tr><td>GU305914.1</td><td>17813</td><td>17835</td><td>-</td><td>0</td><td>CGGGATCGCGCCAAACGCGA</td><td>CGG</td><td>||||||||[||||||||||||]PAM</td><td>NA</td><td>Intergenic</td><td>NA</td><td>NA</td></tr>
<tr><td>T33220</td><td>AGCCGCTGCACACTCCTCCGAGG</td><td>1000</td><td></td><td></td><td></td><td></td><td></td><td></td><td></td><td></td><td></td></tr>
<tr><td>Oligo fwd</td><td>CACCAGCCGCTGCACACTCCTCCG</td><td></td><td></td><td></td><td></td><td></td><td></td><td></td><td></td><td></td><td></td></tr>
<tr><td>Oligo rev</td><td>AAACCGGAGGAGTGTGCAGCGGCT</td><td></td><td></td><td></td><td></td><td></td><td></td><td></td><td></td><td></td><td></td></tr>
<tr><td>Chromosome</td><td>start</td><td>end</td><td>strand</td><td>MM</td><td>target_seq</td><td>PAM</td><td>alignment</td><td>distance</td><td>position</td><td>gene name</td><td>gene id</td></tr>
<tr><td>GU305914.1</td><td>17776</td><td>17798</td><td>-</td><td>0</td><td>AGCCGCTGCACACTCCTCCG</td><td>AGG</td><td>||||||||[||||||||||||]PAM</td><td>NA</td><td>Intergenic</td><td>NA</td><td>NA</td></tr>
<tr><td>T33221</td><td>AAAGCCTCCCAGATCGCCTGAGG</td><td>1000</td><td></td><td></td><td></td><td></td><td></td><td></td><td></td><td></td><td></td></tr>
<tr><td>Oligo fwd</td><td>CACCAAAGCCTCCCAGATCGCCTG</td><td></td><td></td><td></td><td></td><td></td><td></td><td></td><td></td><td></td><td></td></tr>
<tr><td>Oligo rev</td><td>AAACCAGGCGATCTGGGAGGCTTT</td><td></td><td></td><td></td><td></td><td></td><td></td><td></td><td></td><td></td><td></td></tr>
<tr><td>Chromosome</td><td>start</td><td>end</td><td>strand</td><td>MM</td><td>target_seq</td><td>PAM</td><td>alignment</td><td>distance</td><td>position</td><td>gene name</td><td>gene id</td></tr>
<tr><td>GU305914.1</td><td>17748</td><td>17770</td><td>-</td><td>0</td><td>AAAGCCTCCCAGATCGCCTG</td><td>AGG</td><td>||||||||[||||||||||||]PAM</td><td>NA</td><td>Intergenic</td><td>NA</td><td>NA</td></tr>
<tr><td>T33222</td><td>CAACAACAAAGCGACGACGACGG</td><td>1000</td><td></td><td></td><td></td><td></td><td></td><td></td><td></td><td></td><td></td></tr>
<tr><td>Oligo fwd</td><td>CACCCAACAACAAAGCGACGACGA</td><td></td><td></td><td></td><td></td><td></td><td></td><td></td><td></td><td></td><td></td></tr>
<tr><td>Oligo rev</td><td>AAACTCGTCGTCGCTTTGTTGTTG</td><td></td><td></td><td></td><td></td><td></td><td></td><td></td><td></td><td></td><td></td></tr>
<tr><td>Chromosome</td><td>start</td><td>end</td><td>strand</td><td>MM</td><td>target_seq</td><td>PAM</td><td>alignment</td><td>distance</td><td>position</td><td>gene name</td><td>gene id</td></tr>
<tr><td>GU305914.1</td><td>17723</td><td>17745</td><td>-</td><td>0</td><td>CAACAACAAAGCGACGACGA</td><td>CGG</td><td>||||||||[||||||||||||]PAM</td><td>NA</td><td>Intergenic</td><td>NA</td><td>NA</td></tr>
<tr><td>T33223</td><td>CAAAGCGACGACGACGGCCACGG</td><td>1000</td><td></td><td></td><td></td><td></td><td></td><td></td><td></td><td></td><td></td></tr>
<tr><td>Oligo fwd</td><td>CACCCAAAGCGACGACGACGGCCA</td><td></td><td></td><td></td><td></td><td></td><td></td><td></td><td></td><td></td><td></td></tr>
<tr><td>Oligo rev</td><td>AAACTGGCCGTCGTCGTCGCTTTG</td><td></td><td></td><td></td><td></td><td></td><td></td><td></td><td></td><td></td><td></td></tr>
<tr><td>Chromosome</td><td>start</td><td>end</td><td>strand</td><td>MM</td><td>target_seq</td><td>PAM</td><td>alignment</td><td>distance</td><td>position</td><td>gene name</td><td>gene id</td></tr>
<tr><td>GU305914.1</td><td>17717</td><td>17739</td><td>-</td><td>0</td><td>CAAAGCGACGACGACGGCCA</td><td>CGG</td><td>||||||||[||||||||||||]PAM</td><td>NA</td><td>Intergenic</td><td>NA</td><td>NA</td></tr>
<tr><td>T33224</td><td>GACGACGACGGCCACGGCGATGG</td><td>1000</td><td></td><td></td><td></td><td></td><td></td><td></td><td></td><td></td><td></td></tr>
<tr><td>Oligo fwd</td><td>CACCGACGACGACGGCCACGGCGA</td><td></td><td></td><td></td><td></td><td></td><td></td><td></td><td></td><td></td><td></td></tr>
<tr><td>Oligo rev</td><td>AAACTCGCCGTGGCCGTCGTCGTC</td><td></td><td></td><td></td><td></td><td></td><td></td><td></td><td></td><td></td><td></td></tr>
<tr><td>Chromosome</td><td>start</td><td>end</td><td>strand</td><td>MM</td><td>target_seq</td><td>PAM</td><td>alignment</td><td>distance</td><td>position</td><td>gene name</td><td>gene id</td></tr>
<tr><td>GU305914.1</td><td>17711</td><td>17733</td><td>-</td><td>0</td><td>GACGACGACGGCCACGGCGA</td><td>TGG</td><td>||||||||[||||||||||||]PAM</td><td>NA</td><td>Intergenic</td><td>NA</td><td>NA</td></tr>
<tr><td>GU305914.1</td><td>20756</td><td>20778</td><td>+</td><td>3</td><td>GACGACGGCGATCACGGCGA</td><td>GGG</td><td>|||||||-[||--||||||||]PAM</td><td>NA</td><td>Intergenic</td><td>NA</td><td>NA</td></tr>
<tr><td>T33225</td><td>CGGCCACGGCGATGGCGCGACGG</td><td>1000</td><td></td><td></td><td></td><td></td><td></td><td></td><td></td><td></td><td></td></tr>
<tr><td>Oligo fwd</td><td>CACCCGGCCACGGCGATGGCGCGA</td><td></td><td></td><td></td><td></td><td></td><td></td><td></td><td></td><td></td><td></td></tr>
<tr><td>Oligo rev</td><td>AAACTCGCGCCATCGCCGTGGCCG</td><td></td><td></td><td></td><td></td><td></td><td></td><td></td><td></td><td></td><td></td></tr>
<tr><td>Chromosome</td><td>start</td><td>end</td><td>strand</td><td>MM</td><td>target_seq</td><td>PAM</td><td>alignment</td><td>distance</td><td>position</td><td>gene name</td><td>gene id</td></tr>
<tr><td>GU305914.1</td><td>17703</td><td>17725</td><td>-</td><td>0</td><td>CGGCCACGGCGATGGCGCGA</td><td>CGG</td><td>||||||||[||||||||||||]PAM</td><td>NA</td><td>Intergenic</td><td>NA</td><td>NA</td></tr>
<tr><td>T33226</td><td>ATGGCGCGACGGACGACCGTCGG</td><td>1000</td><td></td><td></td><td></td><td></td><td></td><td></td><td></td><td></td><td></td></tr>
<tr><td>Oligo fwd</td><td>CACCATGGCGCGACGGACGACCGT</td><td></td><td></td><td></td><td></td><td></td><td></td><td></td><td></td><td></td><td></td></tr>
<tr><td>Oligo rev</td><td>AAACACGGTCGTCCGTCGCGCCAT</td><td></td><td></td><td></td><td></td><td></td><td></td><td></td><td></td><td></td><td></td></tr>
<tr><td>Chromosome</td><td>start</td><td>end</td><td>strand</td><td>MM</td><td>target_seq</td><td>PAM</td><td>alignment</td><td>distance</td><td>position</td><td>gene name</td><td>gene id</td></tr>
<tr><td>GU305914.1</td><td>17692</td><td>17714</td><td>-</td><td>0</td><td>ATGGCGCGACGGACGACCGT</td><td>CGG</td><td>||||||||[||||||||||||]PAM</td><td>NA</td><td>Intergenic</td><td>NA</td><td>NA</td></tr>
<tr><td>T33227</td><td>CGTCGGTCGTCATCCGACTCCGG</td><td>1000</td><td></td><td></td><td></td><td></td><td></td><td></td><td></td><td></td><td></td></tr>
<tr><td>Oligo fwd</td><td>CACCCGTCGGTCGTCATCCGACTC</td><td></td><td></td><td></td><td></td><td></td><td></td><td></td><td></td><td></td><td></td></tr>
<tr><td>Oligo rev</td><td>AAACGAGTCGGATGACGACCGACG</td><td></td><td></td><td></td><td></td><td></td><td></td><td></td><td></td><td></td><td></td></tr>
</table>

<table>
<tr><td>Chromosome</td><td>start</td><td>end</td><td>strand</td><td>MM</td><td>target_seq</td><td>PAM</td><td>alignment</td><td>distance</td><td>position</td><td>gene name</td><td>gene id</td></tr>
<tr><td>GU305914.1</td><td>17675</td><td>17697</td><td>-</td><td>0</td><td>CGTCGGTCGTCATCCGACTC</td><td>CGG</td><td>||||||||[||||||||||||]PAM</td><td>NA</td><td>Intergenic</td><td>NA</td><td>NA</td></tr>
<tr><td>T33228</td><td>GCGCCGCGACCACAAAATATCGG</td><td>1000</td><td></td><td></td><td></td><td></td><td></td><td></td><td></td><td></td><td></td></tr>
<tr><td>Oligo fwd</td><td>CACCGCGCCGCGACCACAAAATAT</td><td></td><td></td><td></td><td></td><td></td><td></td><td></td><td></td><td></td><td></td></tr>
<tr><td>Oligo rev</td><td>AAACATATTTTGTGGTCGCGGCGC</td><td></td><td></td><td></td><td></td><td></td><td></td><td></td><td></td><td></td><td></td></tr>
<tr><td>Chromosome</td><td>start</td><td>end</td><td>strand</td><td>MM</td><td>target_seq</td><td>PAM</td><td>alignment</td><td>distance</td><td>position</td><td>gene name</td><td>gene id</td></tr>
<tr><td>GU305914.1</td><td>17631</td><td>17653</td><td>-</td><td>0</td><td>GCGCCGCGACCACAAAATAT</td><td>CGG</td><td>||||||||[||||||||||||]PAM</td><td>NA</td><td>Intergenic</td><td>NA</td><td>NA</td></tr>
<tr><td>T33229</td><td>CCACAAAATATCGGTCGATGAGG</td><td>1000</td><td></td><td></td><td></td><td></td><td></td><td></td><td></td><td></td><td></td></tr>
<tr><td>Oligo fwd</td><td>CACCCCACAAAATATCGGTCGATG</td><td></td><td></td><td></td><td></td><td></td><td></td><td></td><td></td><td></td><td></td></tr>
<tr><td>Oligo rev</td><td>AAACCATCGACCGATATTTTGTGG</td><td></td><td></td><td></td><td></td><td></td><td></td><td></td><td></td><td></td><td></td></tr>
<tr><td>Chromosome</td><td>start</td><td>end</td><td>strand</td><td>MM</td><td>target_seq</td><td>PAM</td><td>alignment</td><td>distance</td><td>position</td><td>gene name</td><td>gene id</td></tr>
<tr><td>GU305914.1</td><td>17622</td><td>17644</td><td>-</td><td>0</td><td>CCACAAAATATCGGTCGATG</td><td>AGG</td><td>||||||||[||||||||||||]PAM</td><td>NA</td><td>Intergenic</td><td>NA</td><td>NA</td></tr>
<tr><td>T33230</td><td>CACAAAATATCGGTCGATGAGGG</td><td>1000</td><td></td><td></td><td></td><td></td><td></td><td></td><td></td><td></td><td></td></tr>
<tr><td>Oligo fwd</td><td>CACCCACAAAATATCGGTCGATGA</td><td></td><td></td><td></td><td></td><td></td><td></td><td></td><td></td><td></td><td></td></tr>
<tr><td>Oligo rev</td><td>AAACTCATCGACCGATATTTTGTG</td><td></td><td></td><td></td><td></td><td></td><td></td><td></td><td></td><td></td><td></td></tr>
<tr><td>Chromosome</td><td>start</td><td>end</td><td>strand</td><td>MM</td><td>target_seq</td><td>PAM</td><td>alignment</td><td>distance</td><td>position</td><td>gene name</td><td>gene id</td></tr>
<tr><td>GU305914.1</td><td>17621</td><td>17643</td><td>-</td><td>0</td><td>CACAAAATATCGGTCGATGA</td><td>GGG</td><td>||||||||[||||||||||||]PAM</td><td>NA</td><td>Intergenic</td><td>NA</td><td>NA</td></tr>
<tr><td>T33231</td><td>ACAAAATATCGGTCGATGAGGGG</td><td>1000</td><td></td><td></td><td></td><td></td><td></td><td></td><td></td><td></td><td></td></tr>
<tr><td>Oligo fwd</td><td>CACCACAAAATATCGGTCGATGAG</td><td></td><td></td><td></td><td></td><td></td><td></td><td></td><td></td><td></td><td></td></tr>
<tr><td>Oligo rev</td><td>AAACCTCATCGACCGATATTTTGT</td><td></td><td></td><td></td><td></td><td></td><td></td><td></td><td></td><td></td><td></td></tr>
<tr><td>Chromosome</td><td>start</td><td>end</td><td>strand</td><td>MM</td><td>target_seq</td><td>PAM</td><td>alignment</td><td>distance</td><td>position</td><td>gene name</td><td>gene id</td></tr>
<tr><td>GU305914.1</td><td>17620</td><td>17642</td><td>-</td><td>0</td><td>ACAAAATATCGGTCGATGAG</td><td>GGG</td><td>||||||||[||||||||||||]PAM</td><td>NA</td><td>Intergenic</td><td>NA</td><td>NA</td></tr>
<tr><td>T33232</td><td>CTCAGAAGAAACATAAGACTCGG</td><td>1000</td><td></td><td></td><td></td><td></td><td></td><td></td><td></td><td></td><td></td></tr>
<tr><td>Oligo fwd</td><td>CACCCTCAGAAGAAACATAAGACT</td><td></td><td></td><td></td><td></td><td></td><td></td><td></td><td></td><td></td><td></td></tr>
<tr><td>Oligo rev</td><td>AAACAGTCTTATGTTTCTTCTGAG</td><td></td><td></td><td></td><td></td><td></td><td></td><td></td><td></td><td></td><td></td></tr>
<tr><td>Chromosome</td><td>start</td><td>end</td><td>strand</td><td>MM</td><td>target_seq</td><td>PAM</td><td>alignment</td><td>distance</td><td>position</td><td>gene name</td><td>gene id</td></tr>
<tr><td>GU305914.1</td><td>17596</td><td>17618</td><td>-</td><td>0</td><td>CTCAGAAGAAACATAAGACT</td><td>CGG</td><td>||||||||[||||||||||||]PAM</td><td>NA</td><td>Intergenic</td><td>NA</td><td>NA</td></tr>
<tr><td>T33233</td><td>TAAGACTCGGAGCCCGAGCAAGG</td><td>1000</td><td></td><td></td><td></td><td></td><td></td><td></td><td></td><td></td><td></td></tr>
<tr><td>Oligo fwd</td><td>CACCTAAGACTCGGAGCCCGAGCA</td><td></td><td></td><td></td><td></td><td></td><td></td><td></td><td></td><td></td><td></td></tr>
<tr><td>Oligo rev</td><td>AAACTGCTCGGGCTCCGAGTCTTA</td><td></td><td></td><td></td><td></td><td></td><td></td><td></td><td></td><td></td><td></td></tr>
<tr><td>Chromosome</td><td>start</td><td>end</td><td>strand</td><td>MM</td><td>target_seq</td><td>PAM</td><td>alignment</td><td>distance</td><td>position</td><td>gene name</td><td>gene id</td></tr>
<tr><td>GU305914.1</td><td>17583</td><td>17605</td><td>-</td><td>0</td><td>TAAGACTCGGAGCCCGAGCA</td><td>AGG</td><td>||||||||[||||||||||||]PAM</td><td>NA</td><td>Intergenic</td><td>NA</td><td>NA</td></tr>
<tr><td>T33234</td><td>AAGACTCGGAGCCCGAGCAAGGG</td><td>1000</td><td></td><td></td><td></td><td></td><td></td><td></td><td></td><td></td><td></td></tr>
<tr><td>Oligo fwd</td><td>CACCAAGACTCGGAGCCCGAGCAA</td><td></td><td></td><td></td><td></td><td></td><td></td><td></td><td></td><td></td><td></td></tr>
<tr><td>Oligo rev</td><td>AAACTTGCTCGGGCTCCGAGTCTT</td><td></td><td></td><td></td><td></td><td></td><td></td><td></td><td></td><td></td><td></td></tr>
<tr><td>Chromosome</td><td>start</td><td>end</td><td>strand</td><td>MM</td><td>target_seq</td><td>PAM</td><td>alignment</td><td>distance</td><td>position</td><td>gene name</td><td>gene id</td></tr>
<tr><td>GU305914.1</td><td>17582</td><td>17604</td><td>-</td><td>0</td><td>AAGACTCGGAGCCCGAGCAA</td><td>GGG</td><td>||||||||[||||||||||||]PAM</td><td>NA</td><td>Intergenic</td><td>NA</td><td>NA</td></tr>
<tr><td>T33235</td><td>AGACTCGGAGCCCGAGCAAGGGG</td><td>1000</td><td></td><td></td><td></td><td></td><td></td><td></td><td></td><td></td><td></td></tr>
<tr><td>Oligo fwd</td><td>CACCAGACTCGGAGCCCGAGCAAG</td><td></td><td></td><td></td><td></td><td></td><td></td><td></td><td></td><td></td><td></td></tr>
<tr><td>Oligo rev</td><td>AAACCTTGCTCGGGCTCCGAGTCT</td><td></td><td></td><td></td><td></td><td></td><td></td><td></td><td></td><td></td><td></td></tr>
<tr><td>Chromosome</td><td>start</td><td>end</td><td>strand</td><td>MM</td><td>target_seq</td><td>PAM</td><td>alignment</td><td>distance</td><td>position</td><td>gene name</td><td>gene id</td></tr>
<tr><td>GU305914.1</td><td>17581</td><td>17603</td><td>-</td><td>0</td><td>AGACTCGGAGCCCGAGCAAG</td><td>GGG</td><td>||||||||[||||||||||||]PAM</td><td>NA</td><td>Intergenic</td><td>NA</td><td>NA</td></tr>
<tr><td>T33236</td><td>CAAATCGCCCGCTGCGATGACGG</td><td>1000</td><td></td><td></td><td></td><td></td><td></td><td></td><td></td><td></td><td></td></tr>
<tr><td>Oligo fwd</td><td>CACCCAAATCGCCCGCTGCGATGA</td><td></td><td></td><td></td><td></td><td></td><td></td><td></td><td></td><td></td><td></td></tr>
<tr><td>Oligo rev</td><td>AAACTCATCGCAGCGGGCGATTTG</td><td></td><td></td><td></td><td></td><td></td><td></td><td></td><td></td><td></td><td></td></tr>
<tr><td>Chromosome</td><td>start</td><td>end</td><td>strand</td><td>MM</td><td>target_seq</td><td>PAM</td><td>alignment</td><td>distance</td><td>position</td><td>gene name</td><td>gene id</td></tr>
<tr><td>GU305914.1</td><td>17547</td><td>17569</td><td>-</td><td>0</td><td>CAAATCGCCCGCTGCGATGA</td><td>CGG</td><td>||||||||[||||||||||||]PAM</td><td>NA</td><td>Intergenic</td><td>NA</td><td>NA</td></tr>
<tr><td>GU305914.1</td><td>112175</td><td>112197</td><td>+</td><td>4</td><td>CAAGTCACCGGCCGCGATGA</td><td>AGG</td><td>|||-||-|[|-||-|||||||]PAM</td><td>NA</td><td>Intergenic</td><td>NA</td><td>NA</td></tr>
<tr><td>T33237</td><td>GACGAACCCAAAGATACTCGCGG</td><td>1000</td><td></td><td></td><td></td><td></td><td></td><td></td><td></td><td></td><td></td></tr>
<tr><td>Oligo fwd</td><td>CACCGACGAACCCAAAGATACTCG</td><td></td><td></td><td></td><td></td><td></td><td></td><td></td><td></td><td></td><td></td></tr>
<tr><td>Oligo rev</td><td>AAACCGAGTATCTTTGGGTTCGTC</td><td></td><td></td><td></td><td></td><td></td><td></td><td></td><td></td><td></td><td></td></tr>
<tr><td>Chromosome</td><td>start</td><td>end</td><td>strand</td><td>MM</td><td>target_seq</td><td>PAM</td><td>alignment</td><td>distance</td><td>position</td><td>gene name</td><td>gene id</td></tr>
<tr><td>GU305914.1</td><td>17519</td><td>17541</td><td>-</td><td>0</td><td>GACGAACCCAAAGATACTCG</td><td>CGG</td><td>||||||||[||||||||||||]PAM</td><td>NA</td><td>Intergenic</td><td>NA</td><td>NA</td></tr>
<tr><td>T33238</td><td>CCCAAAGATACTCGCGGACCCGG</td><td>1000</td><td></td><td></td><td></td><td></td><td></td><td></td><td></td><td></td><td></td></tr>
<tr><td>Oligo fwd</td><td>CACCCCCAAAGATACTCGCGGACC</td><td></td><td></td><td></td><td></td><td></td><td></td><td></td><td></td><td></td><td></td></tr>
<tr><td>Oligo rev</td><td>AAACGGTCCGCGAGTATCTTTGGG</td><td></td><td></td><td></td><td></td><td></td><td></td><td></td><td></td><td></td><td></td></tr>
<tr><td>Chromosome</td><td>start</td><td>end</td><td>strand</td><td>MM</td><td>target_seq</td><td>PAM</td><td>alignment</td><td>distance</td><td>position</td><td>gene name</td><td>gene id</td></tr>
<tr><td>GU305914.1</td><td>17513</td><td>17535</td><td>-</td><td>0</td><td>CCCAAAGATACTCGCGGACC</td><td>CGG</td><td>||||||||[||||||||||||]PAM</td><td>NA</td><td>Intergenic</td><td>NA</td><td>NA</td></tr>
<tr><td>T33239</td><td>GTTTTGTGCAGAGAAACTTGAGG</td><td>1000</td><td></td><td></td><td></td><td></td><td></td><td></td><td></td><td></td><td></td></tr>
<tr><td>Oligo fwd</td><td>CACCGTTTTGTGCAGAGAAACTTG</td><td></td><td></td><td></td><td></td><td></td><td></td><td></td><td></td><td></td><td></td></tr>
<tr><td>Oligo rev</td><td>AAACCAAGTTTCTCTGCACAAAAC</td><td></td><td></td><td></td><td></td><td></td><td></td><td></td><td></td><td></td><td></td></tr>
<tr><td>Chromosome</td><td>start</td><td>end</td><td>strand</td><td>MM</td><td>target_seq</td><td>PAM</td><td>alignment</td><td>distance</td><td>position</td><td>gene name</td><td>gene id</td></tr>
<tr><td>GU305914.1</td><td>17491</td><td>17513</td><td>-</td><td>0</td><td>GTTTTGTGCAGAGAAACTTG</td><td>AGG</td><td>||||||||[||||||||||||]PAM</td><td>NA</td><td>Intergenic</td><td>NA</td><td>NA</td></tr>
<tr><td>T33240</td><td>TTGTGCAGAGAAACTTGAGGAGG</td><td>1000</td><td></td><td></td><td></td><td></td><td></td><td></td><td></td><td></td><td></td></tr>
<tr><td>Oligo fwd</td><td>CACCTTGTGCAGAGAAACTTGAGG</td><td></td><td></td><td></td><td></td><td></td><td></td><td></td><td></td><td></td><td></td></tr>
<tr><td>Oligo rev</td><td>AAACCCTCAAGTTTCTCTGCACAA</td><td></td><td></td><td></td><td></td><td></td><td></td><td></td><td></td><td></td><td></td></tr>
<tr><td>Chromosome</td><td>start</td><td>end</td><td>strand</td><td>MM</td><td>target_seq</td><td>PAM</td><td>alignment</td><td>distance</td><td>position</td><td>gene name</td><td>gene id</td></tr>
<tr><td>GU305914.1</td><td>17488</td><td>17510</td><td>-</td><td>0</td><td>TTGTGCAGAGAAACTTGAGG</td><td>AGG</td><td>||||||||[||||||||||||]PAM</td><td>NA</td><td>Intergenic</td><td>NA</td><td>NA</td></tr>
<tr><td>T33241</td><td>TGTGCAGAGAAACTTGAGGAGGG</td><td>1000</td><td></td><td></td><td></td><td></td><td></td><td></td><td></td><td></td><td></td></tr>
<tr><td>Oligo fwd</td><td>CACCTGTGCAGAGAAACTTGAGGA</td><td></td><td></td><td></td><td></td><td></td><td></td><td></td><td></td><td></td><td></td></tr>
<tr><td>Oligo rev</td><td>AAACTCCTCAAGTTTCTCTGCACA</td><td></td><td></td><td></td><td></td><td></td><td></td><td></td><td></td><td></td><td></td></tr>
<tr><td>Chromosome</td><td>start</td><td>end</td><td>strand</td><td>MM</td><td>target_seq</td><td>PAM</td><td>alignment</td><td>distance</td><td>position</td><td>gene name</td><td>gene id</td></tr>
<tr><td>GU305914.1</td><td>17487</td><td>17509</td><td>-</td><td>0</td><td>TGTGCAGAGAAACTTGAGGA</td><td>GGG</td><td>||||||||[||||||||||||]PAM</td><td>NA</td><td>Intergenic</td><td>NA</td><td>NA</td></tr>
<tr><td>T33242</td><td>GCAGAGAAACTTGAGGAGGGTGG</td><td>1000</td><td></td><td></td><td></td><td></td><td></td><td></td><td></td><td></td><td></td></tr>
</table>

| | | | | | | | | | | | |
|---|---|---|---|---|---|---|---|---|---|---|---|
| Oligo fwd | CACCGCAGAGAAACTTGAGGAGGG | | | | | | | | | | |
| Oligo rev | AAACCCCTCCTCAAGTTTCTCTGC | | | | | | | | | | |
| Chromosome | start | end | strand | MM | target_seq | PAM | alignment | distance | position | gene name | gene id |
| GU305914.1 | 17484 | 17506 | - | 0 | GCAGAGAAACTTGAGGAGGG | TGG | \|\|\|\|\|\|\|\|[\|\|\|\|\|\|\|\|\|\|\|\|]PAM | NA | Intergenic | NA | NA |

| | | | | | | | | | | | |
|---|---|---|---|---|---|---|---|---|---|---|---|
| T33243 | GGCTTCACGTGTCGCCAGAGAGG | 1000 | | | | | | | | | |
| Oligo fwd | CACCGGCTTCACGTGTCGCCAGAG | | | | | | | | | | |
| Oligo rev | AAACCTCTGGCGACACGTGAAGCC | | | | | | | | | | |
| Chromosome | start | end | strand | MM | target_seq | PAM | alignment | distance | position | gene name | gene id |
| GU305914.1 | 17463 | 17485 | - | 0 | GGCTTCACGTGTCGCCAGAG | AGG | \|\|\|\|\|\|\|\|[\|\|\|\|\|\|\|\|\|\|\|\|]PAM | NA | Intergenic | NA | NA |

| | | | | | | | | | | | |
|---|---|---|---|---|---|---|---|---|---|---|---|
| T33244 | ACCCCAACAACAGTCTCTCGCGG | 1000 | | | | | | | | | |
| Oligo fwd | CACCACCCCAACAACAGTCTCTCG | | | | | | | | | | |
| Oligo rev | AAACCGAGAGACTGTTGTTGGGGT | | | | | | | | | | |
| Chromosome | start | end | strand | MM | target_seq | PAM | alignment | distance | position | gene name | gene id |
| GU305914.1 | 17374 | 17396 | - | 0 | ACCCCAACAACAGTCTCTCG | CGG | \|\|\|\|\|\|\|\|[\|\|\|\|\|\|\|\|\|\|\|\|]PAM | NA | Intergenic | NA | NA |

| | | | | | | | | | | | |
|---|---|---|---|---|---|---|---|---|---|---|---|
| T33245 | TCTCTCGCGGCATCAGACCGAGG | 1000 | | | | | | | | | |
| Oligo fwd | CACCTCTCTCGCGGCATCAGACCG | | | | | | | | | | |
| Oligo rev | AAACCGGTCTGATGCCGCGAGAGA | | | | | | | | | | |
| Chromosome | start | end | strand | MM | target_seq | PAM | alignment | distance | position | gene name | gene id |
| GU305914.1 | 17361 | 17383 | - | 0 | TCTCTCGCGGCATCAGACCG | AGG | \|\|\|\|\|\|\|\|[\|\|\|\|\|\|\|\|\|\|\|\|]PAM | NA | Intergenic | NA | NA |

| | | | | | | | | | | | |
|---|---|---|---|---|---|---|---|---|---|---|---|
| T33246 | CGGCATCAGACCGAGGACACCGG | 1000 | | | | | | | | | |
| Oligo fwd | CACCCGGCATCAGACCGAGGACAC | | | | | | | | | | |
| Oligo rev | AAACGTGTCCTCGGTCTGATGCCG | | | | | | | | | | |
| Chromosome | start | end | strand | MM | target_seq | PAM | alignment | distance | position | gene name | gene id |
| GU305914.1 | 17354 | 17376 | - | 0 | CGGCATCAGACCGAGGACAC | CGG | \|\|\|\|\|\|\|\|[\|\|\|\|\|\|\|\|\|\|\|\|]PAM | NA | Intergenic | NA | NA |

| | | | | | | | | | | | |
|---|---|---|---|---|---|---|---|---|---|---|---|
| T33247 | ACCGAGGACACCGGACTCGAAGG | 1000 | | | | | | | | | |
| Oligo fwd | CACCACCGAGGACACCGGACTCGA | | | | | | | | | | |
| Oligo rev | AAACTCGAGTCCGGTGTCCTCGGT | | | | | | | | | | |
| Chromosome | start | end | strand | MM | target_seq | PAM | alignment | distance | position | gene name | gene id |
| GU305914.1 | 17345 | 17367 | - | 0 | ACCGAGGACACCGGACTCGA | AGG | \|\|\|\|\|\|\|\|[\|\|\|\|\|\|\|\|\|\|\|\|]PAM | NA | Intergenic | NA | NA |

| | | | | | | | | | | | |
|---|---|---|---|---|---|---|---|---|---|---|---|
| T33248 | GAGGACACCGGACTCGAAGGCGG | 1000 | | | | | | | | | |
| Oligo fwd | CACCGAGGACACCGGACTCGAAGG | | | | | | | | | | |
| Oligo rev | AAACCCTTCGAGTCCGGTGTCCTC | | | | | | | | | | |
| Chromosome | start | end | strand | MM | target_seq | PAM | alignment | distance | position | gene name | gene id |
| GU305914.1 | 17342 | 17364 | - | 0 | GAGGACACCGGACTCGAAGG | CGG | \|\|\|\|\|\|\|\|[\|\|\|\|\|\|\|\|\|\|\|\|]PAM | NA | Intergenic | NA | NA |

| | | | | | | | | | | | |
|---|---|---|---|---|---|---|---|---|---|---|---|
| T33249 | CGGACTCGAAGGCGGCTCCGAGG | 1000 | | | | | | | | | |
| Oligo fwd | CACCCGGACTCGAAGGCGGCTCCG | | | | | | | | | | |
| Oligo rev | AAACCGGAGCCGCCTTCGAGTCCG | | | | | | | | | | |
| Chromosome | start | end | strand | MM | target_seq | PAM | alignment | distance | position | gene name | gene id |
| GU305914.1 | 17334 | 17356 | - | 0 | CGGACTCGAAGGCGGCTCCG | AGG | \|\|\|\|\|\|\|\|[\|\|\|\|\|\|\|\|\|\|\|\|]PAM | NA | Intergenic | NA | NA |

| | | | | | | | | | | | |
|---|---|---|---|---|---|---|---|---|---|---|---|
| T33250 | GCCGCACCCGACACCACCTCCGG | 1000 | | | | | | | | | |
| Oligo fwd | CACCGCCGCACCCGACACCACCTC | | | | | | | | | | |
| Oligo rev | AAACGAGGTGGTGTCGGGTGCGGC | | | | | | | | | | |
| Chromosome | start | end | strand | MM | target_seq | PAM | alignment | distance | position | gene name | gene id |
| GU305914.1 | 17300 | 17322 | - | 0 | GCCGCACCCGACACCACCTC | CGG | \|\|\|\|\|\|\|\|[\|\|\|\|\|\|\|\|\|\|\|\|]PAM | NA | Intergenic | NA | NA |

| | | | | | | | | | | | |
|---|---|---|---|---|---|---|---|---|---|---|---|
| T33251 | GCACCCGACACCACCTCCGGCGG | 1000 | | | | | | | | | |
| Oligo fwd | CACCGCACCCGACACCACCTCCGG | | | | | | | | | | |
| Oligo rev | AAACCCGGAGGTGGTGTCGGGTGC | | | | | | | | | | |
| Chromosome | start | end | strand | MM | target_seq | PAM | alignment | distance | position | gene name | gene id |
| GU305914.1 | 17297 | 17319 | - | 0 | GCACCCGACACCACCTCCGG | CGG | \|\|\|\|\|\|\|\|[\|\|\|\|\|\|\|\|\|\|\|\|]PAM | NA | Intergenic | NA | NA |

| | | | | | | | | | | | |
|---|---|---|---|---|---|---|---|---|---|---|---|
| T33252 | CACCTCCGGCGGTGAAGAAGAGG | 1000 | | | | | | | | | |
| Oligo fwd | CACCCACCTCCGGCGGTGAAGAAG | | | | | | | | | | |
| Oligo rev | AAACCTTCTTCACCGCCGGAGGTG | | | | | | | | | | |
| Chromosome | start | end | strand | MM | target_seq | PAM | alignment | distance | position | gene name | gene id |
| GU305914.1 | 17286 | 17308 | - | 0 | CACCTCCGGCGGTGAAGAAG | AGG | \|\|\|\|\|\|\|\|[\|\|\|\|\|\|\|\|\|\|\|\|]PAM | NA | Intergenic | NA | NA |

| | | | | | | | | | | | |
|---|---|---|---|---|---|---|---|---|---|---|---|
| T33253 | CTCCGGCGGTGAAGAAGAGGAGG | 1000 | | | | | | | | | |
| Oligo fwd | CACCCTCCGGCGGTGAAGAAGAGG | | | | | | | | | | |
| Oligo rev | AAACCCTCTTCTTCACCGCCGGAG | | | | | | | | | | |
| Chromosome | start | end | strand | MM | target_seq | PAM | alignment | distance | position | gene name | gene id |
| GU305914.1 | 17283 | 17305 | - | 0 | CTCCGGCGGTGAAGAAGAGG | AGG | \|\|\|\|\|\|\|\|[\|\|\|\|\|\|\|\|\|\|\|\|]PAM | NA | Intergenic | NA | NA |
| GU305914.1 | 27387 | 27409 | + | 4 | CTGCGACGGGGAAGAAGAGC | CGG | \|\|-\|\|-\|\|[\|-\|\|\|\|\|\|\|\|\|-]PAM | NA | Intergenic | NA | NA |

| | | | | | | | | | | | |
|---|---|---|---|---|---|---|---|---|---|---|---|
| T33254 | AGAGGAGGTCGACGAGATCGAGG | 1000 | | | | | | | | | |
| Oligo fwd | CACCAGAGGAGGTCGACGAGATCG | | | | | | | | | | |
| Oligo rev | AAACCGATCTCGTCGACCTCCTCT | | | | | | | | | | |
| Chromosome | start | end | strand | MM | target_seq | PAM | alignment | distance | position | gene name | gene id |
| GU305914.1 | 17268 | 17290 | - | 0 | AGAGGAGGTCGACGAGATCG | AGG | \|\|\|\|\|\|\|\|[\|\|\|\|\|\|\|\|\|\|\|\|]PAM | NA | Intergenic | NA | NA |

| | | | | | | | | | | | |
|---|---|---|---|---|---|---|---|---|---|---|---|
| T33255 | CGACGAGATCGAGGCAGCTGAGG | 1000 | | | | | | | | | |
| Oligo fwd | CACCCGACGAGATCGAGGCAGCTG | | | | | | | | | | |
| Oligo rev | AAACCAGCTGCCTCGATCTCGTCG | | | | | | | | | | |
| Chromosome | start | end | strand | MM | target_seq | PAM | alignment | distance | position | gene name | gene id |
| GU305914.1 | 17259 | 17281 | - | 0 | CGACGAGATCGAGGCAGCTG | AGG | \|\|\|\|\|\|\|\|[\|\|\|\|\|\|\|\|\|\|\|\|]PAM | NA | Intergenic | NA | NA |

| | | | | | | | | | | | |
|---|---|---|---|---|---|---|---|---|---|---|---|
| T33256 | CGAGGCAGCTGAGGCCCTTTCGG | 1000 | | | | | | | | | |
| Oligo fwd | CACCCGAGGCAGCTGAGGCCCTTT | | | | | | | | | | |
| Oligo rev | AAACAAAGGGCCTCAGCTGCCTCG | | | | | | | | | | |
| Chromosome | start | end | strand | MM | target_seq | PAM | alignment | distance | position | gene name | gene id |
| GU305914.1 | 17250 | 17272 | - | 0 | CGAGGCAGCTGAGGCCCTTT | CGG | \|\|\|\|\|\|\|\|[\|\|\|\|\|\|\|\|\|\|\|\|]PAM | NA | Intergenic | NA | NA |

| | | | | | | | | | | | |
|---|---|---|---|---|---|---|---|---|---|---|---|
| T33257 | TGAGGCCCTTTCGGACGCTTTGG | 1000 | | | | | | | | | |
| Oligo fwd | CACCTGAGGCCCTTTCGGACGCTT | | | | | | | | | | |
| Oligo rev | AAACAAGCGTCCGAAAGGGCCTCA | | | | | | | | | | |
| Chromosome | start | end | strand | MM | target_seq | PAM | alignment | distance | position | gene name | gene id |
| GU305914.1 | 17241 | 17263 | - | 0 | TGAGGCCCTTTCGGACGCTT | TGG | \|\|\|\|\|\|\|\|[\|\|\|\|\|\|\|\|\|\|\|\|]PAM | NA | Intergenic | NA | NA |

| | | | | | | | | | | | |
|---|---|---|---|---|---|---|---|---|---|---|---|
| T33258 | GGACGCTTTGGCGCTGCTCAAGG | 1000 | | | | | | | | | |
| Oligo fwd | CACCGGACGCTTTGGCGCTGCTCA | | | | | | | | | | |
| Oligo rev | AAACTGAGCAGCGCCAAAGCGTCC | | | | | | | | | | |
| Chromosome | start | end | strand | MM | target_seq | PAM | alignment | distance | position | gene name | gene id |
| GU305914.1 | 17229 | 17251 | - | 0 | GGACGCTTTGGCGCTGCTCA | AGG | \|\|\|\|\|\|\|\|[\|\|\|\|\|\|\|\|\|\|\|\|]PAM | NA | Intergenic | NA | NA |

| | | | | | | | | | | | |
|---|---|---|---|---|---|---|---|---|---|---|---|
| T33259 | GACGCTTTGGCGCTGCTCAAGGG | 1000 | | | | | | | | | |
| Oligo fwd | CACCGACGCTTTGGCGCTGCTCAA | | | | | | | | | | |
| Oligo rev | AAACTTGAGCAGCGCCAAAGCGTC | | | | | | | | | | |
| Chromosome | start | end | strand | MM | target_seq | PAM | alignment | distance | position | gene name | gene id |
| GU305914.1 | 17228 | 17250 | - | 0 | GACGCTTTGGCGCTGCTCAA | GGG | \|\|\|\|\|\|\|\|[\|\|\|\|\|\|\|\|\|\|\|\|]PAM | NA | Intergenic | NA | NA |

| | | | | | | | | | | | |
|---|---|---|---|---|---|---|---|---|---|---|---|
| T33260 | TTGGCGCTGCTCAAGGGACCTGG | 1000 | | | | | | | | | |
| Oligo fwd | CACCTTGGCGCTGCTCAAGGGACC | | | | | | | | | | |
| Oligo rev | AAACGGTCCCTTGAGCAGCGCCAA | | | | | | | | | | |
| Chromosome | start | end | strand | MM | target_seq | PAM | alignment | distance | position | gene name | gene id |
| GU305914.1 | 17222 | 17244 | - | 0 | TTGGCGCTGCTCAAGGGACC | TGG | \|\|\|\|\|\|\|\|[\|\|\|\|\|\|\|\|\|\|\|\|]PAM | NA | Intergenic | NA | NA |

| | | | | | | | | | | | |
|---|---|---|---|---|---|---|---|---|---|---|---|
| T33261 | TGGCGCTGCTCAAGGGACCTGGG | 1000 | | | | | | | | | |
| Oligo fwd | CACCTGGCGCTGCTCAAGGGACCT | | | | | | | | | | |
| Oligo rev | AAACAGGTCCCTTGAGCAGCGCCA | | | | | | | | | | |
| Chromosome | start | end | strand | MM | target_seq | PAM | alignment | distance | position | gene name | gene id |
| GU305914.1 | 17221 | 17243 | - | 0 | TGGCGCTGCTCAAGGGACCT | GGG | \|\|\|\|\|\|\|\|[\|\|\|\|\|\|\|\|\|\|\|\|]PAM | NA | Intergenic | NA | NA |

| | | | | | | | | | | | |
|---|---|---|---|---|---|---|---|---|---|---|---|
| T33262 | GGCGCTGCTCAAGGGACCTGGGG | 1000 | | | | | | | | | |
| Oligo fwd | CACCGGCGCTGCTCAAGGGACCTG | | | | | | | | | | |
| Oligo rev | AAACCAGGTCCCTTGAGCAGCGCC | | | | | | | | | | |
| Chromosome | start | end | strand | MM | target_seq | PAM | alignment | distance | position | gene name | gene id |
| GU305914.1 | 17220 | 17242 | - | 0 | GGCGCTGCTCAAGGGACCTG | GGG | \|\|\|\|\|\|\|\|[\|\|\|\|\|\|\|\|\|\|\|\|]PAM | NA | Intergenic | NA | NA |

| | | | | | | | | | | | |
|---|---|---|---|---|---|---|---|---|---|---|---|
| T33263 | GCTCAAGGGACCTGGGGACGAGG | 1000 | | | | | | | | | |
| Oligo fwd | CACCGCTCAAGGGACCTGGGGACG | | | | | | | | | | |
| Oligo rev | AAACCGTCCCCAGGTCCCTTGAGC | | | | | | | | | | |
| Chromosome | start | end | strand | MM | target_seq | PAM | alignment | distance | position | gene name | gene id |
| GU305914.1 | 17214 | 17236 | - | 0 | GCTCAAGGGACCTGGGGACG | AGG | \|\|\|\|\|\|\|\|[\|\|\|\|\|\|\|\|\|\|\|\|]PAM | NA | Intergenic | NA | NA |

| | | | | | | | | | | | |
|---|---|---|---|---|---|---|---|---|---|---|---|
| T33264 | TGGGGACGAGGACGATCCTTTGG | 1000 | | | | | | | | | |
| Oligo fwd | CACCTGGGGACGAGGACGATCCTT | | | | | | | | | | |
| Oligo rev | AAACAAGGATCGTCCTCGTCCCCA | | | | | | | | | | |
| Chromosome | start | end | strand | MM | target_seq | PAM | alignment | distance | position | gene name | gene id |
| GU305914.1 | 17202 | 17224 | - | 0 | TGGGGACGAGGACGATCCTT | TGG | \|\|\|\|\|\|\|\|[\|\|\|\|\|\|\|\|\|\|\|\|]PAM | NA | Intergenic | NA | NA |

| | | | | | | | | | | | |
|---|---|---|---|---|---|---|---|---|---|---|---|
| T33265 | GGGGACGAGGACGATCCTTTGGG | 1000 | | | | | | | | | |
| Oligo fwd | CACCGGGGACGAGGACGATCCTTT | | | | | | | | | | |
| Oligo rev | AAACAAAGGATCGTCCTCGTCCCC | | | | | | | | | | |
| Chromosome | start | end | strand | MM | target_seq | PAM | alignment | distance | position | gene name | gene id |
| GU305914.1 | 17201 | 17223 | - | 0 | GGGGACGAGGACGATCCTTT | GGG | \|\|\|\|\|\|\|\|[\|\|\|\|\|\|\|\|\|\|\|\|]PAM | NA | Intergenic | NA | NA |

| | | | | | | | | | | | |
|---|---|---|---|---|---|---|---|---|---|---|---|
| T33266 | GGGACGAGGACGATCCTTTGGGG | 1000 | | | | | | | | | |
| Oligo fwd | CACCGGGACGAGGACGATCCTTTG | | | | | | | | | | |
| Oligo rev | AAACCAAAGGATCGTCCTCGTCCC | | | | | | | | | | |
| Chromosome | start | end | strand | MM | target_seq | PAM | alignment | distance | position | gene name | gene id |
| GU305914.1 | 17200 | 17222 | - | 0 | GGGACGAGGACGATCCTTTG | GGG | \|\|\|\|\|\|\|\|[\|\|\|\|\|\|\|\|\|\|\|\|]PAM | NA | Intergenic | NA | NA |

| | | | | | | | | | | | |
|---|---|---|---|---|---|---|---|---|---|---|---|
| T33267 | GGACGAGGACGATCCTTTGGGGG | 1000 | | | | | | | | | |
| Oligo fwd | CACCGGACGAGGACGATCCTTTGG | | | | | | | | | | |
| Oligo rev | AAACCCAAAGGATCGTCCTCGTCC | | | | | | | | | | |
| Chromosome | start | end | strand | MM | target_seq | PAM | alignment | distance | position | gene name | gene id |
| GU305914.1 | 17199 | 17221 | - | 0 | GGACGAGGACGATCCTTTGG | GGG | \|\|\|\|\|\|\|\|[\|\|\|\|\|\|\|\|\|\|\|\|]PAM | NA | Intergenic | NA | NA |

| | | | | | | | | | | | |
|---|---|---|---|---|---|---|---|---|---|---|---|
| T33268 | GGGGATGCCGTCGTACCTGATGG | 1000 | | | | | | | | | |
| Oligo fwd | CACCGGGGATGCCGTCGTACCTGA | | | | | | | | | | |
| Oligo rev | AAACTCAGGTACGACGGCATCCCC | | | | | | | | | | |
| Chromosome | start | end | strand | MM | target_seq | PAM | alignment | distance | position | gene name | gene id |
| GU305914.1 | 17180 | 17202 | - | 0 | GGGGATGCCGTCGTACCTGA | TGG | \|\|\|\|\|\|\|\|[\|\|\|\|\|\|\|\|\|\|\|\|]PAM | NA | Intergenic | NA | NA |
| GU305914.1 | 222401 | 222423 | - | 4 | GGGAACGCCTTCGTACCCGA | CGG | \|\|\|-\|-\|\|[\|-\|\|\|\|\|\|\|-\|\|]PAM | NA | Intergenic | NA | NA |

| | | | | | | | | | | | |
|---|---|---|---|---|---|---|---|---|---|---|---|
| T33269 | CGTCGTACCTGATGGCATGACGG | 1000 | | | | | | | | | |
| Oligo fwd | CACCCGTCGTACCTGATGGCATGA | | | | | | | | | | |
| Oligo rev | AAACTCATGCCATCAGGTACGACG | | | | | | | | | | |
| Chromosome | start | end | strand | MM | target_seq | PAM | alignment | distance | position | gene name | gene id |
| GU305914.1 | 17172 | 17194 | - | 0 | CGTCGTACCTGATGGCATGA | CGG | \|\|\|\|\|\|\|\|[\|\|\|\|\|\|\|\|\|\|\|\|]PAM | NA | Intergenic | NA | NA |

| | | | | | | | | | | | |
|---|---|---|---|---|---|---|---|---|---|---|---|
| T33270 | GTCGTACCTGATGGCATGACGGG | 1000 | | | | | | | | | |
| Oligo fwd | CACCGTCGTACCTGATGGCATGAC | | | | | | | | | | |
| Oligo rev | AAACGTCATGCCATCAGGTACGAC | | | | | | | | | | |
| Chromosome | start | end | strand | MM | target_seq | PAM | alignment | distance | position | gene name | gene id |
| GU305914.1 | 17171 | 17193 | - | 0 | GTCGTACCTGATGGCATGAC | GGG | \|\|\|\|\|\|\|\|[\|\|\|\|\|\|\|\|\|\|\|\|]PAM | NA | Intergenic | NA | NA |

| | | |
|---|---|---|
| T33271 | ATGGCATGACGGGAGTTGCCAGG | 1000 |
| Oligo fwd | CACCATGGCATGACGGGAGTTGCC | |
| Oligo rev | AAACGGCAACTCCCGTCATGCCAT | |

Chromosome start end strand MM target_seq PAM alignment distance position gene name gene id
GU305914.1 17161 17183 - 0 ATGGCATGACGGGAGTTGCC AGG ||||||||[||||||||||||]PAM NA Intergenic NA NA

T33272 TGGCATGACGGGAGTTGCCAGGG 1000
Oligo fwd CACCTGGCATGACGGGAGTTGCCA
Oligo rev AAACTGGCAACTCCCGTCATGCCA
Chromosome start end strand MM target_seq PAM alignment distance position gene name gene id
GU305914.1 17160 17182 - 0 TGGCATGACGGGAGTTGCCA GGG ||||||||[||||||||||||]PAM NA Intergenic NA NA

T33273 GGGAGTTGCCAGGGTCGTCGAGG 1000
Oligo fwd CACCGGGAGTTGCCAGGGTCGTCG
Oligo rev AAACCGACGACCCTGGCAACTCCC
Chromosome start end strand MM target_seq PAM alignment distance position gene name gene id
GU305914.1 17151 17173 - 0 GGGAGTTGCCAGGGTCGTCG AGG ||||||||[||||||||||||]PAM NA Intergenic NA NA

T33274 GAAATACTTAACGACGAGATCGG 1000
Oligo fwd CACCGAAATACTTAACGACGAGAT
Oligo rev AAACATCTCGTCGTTAAGTATTTC
Chromosome start end strand MM target_seq PAM alignment distance position gene name gene id
GU305914.1 17096 17118 - 0 GAAATACTTAACGACGAGAT CGG ||||||||[||||||||||||]PAM NA Intergenic NA NA

T33275 ATACTTAACGACGAGATCGGTGG 1000
Oligo fwd CACCATACTTAACGACGAGATCGG
Oligo rev AAACCCGATCTCGTCGTTAAGTAT
Chromosome start end strand MM target_seq PAM alignment distance position gene name gene id
GU305914.1 17093 17115 - 0 ATACTTAACGACGAGATCGG TGG ||||||||[||||||||||||]PAM NA Intergenic NA NA

T33276 ACGAGATCGGTGGACCTCTCTGG 1000
Oligo fwd CACCACGAGATCGGTGGACCTCTC
Oligo rev AAACGAGAGGTCCACCGATCTCGT
Chromosome start end strand MM target_seq PAM alignment distance position gene name gene id
GU305914.1 17083 17105 - 0 ACGAGATCGGTGGACCTCTC TGG ||||||||[||||||||||||]PAM NA Intergenic NA NA

T33277 AGATCGGTGGACCTCTCTGGTGG 1000
Oligo fwd CACCAGATCGGTGGACCTCTCTGG
Oligo rev AAACCCAGAGAGGTCCACCGATCT
Chromosome start end strand MM target_seq PAM alignment distance position gene name gene id
GU305914.1 17080 17102 - 0 AGATCGGTGGACCTCTCTGG TGG ||||||||[||||||||||||]PAM NA Intergenic NA NA

T33278 GATCGGTGGACCTCTCTGGTGGG 1000
Oligo fwd CACCGATCGGTGGACCTCTCTGGT
Oligo rev AAACACCAGAGAGGTCCACCGATC
Chromosome start end strand MM target_seq PAM alignment distance position gene name gene id
GU305914.1 17079 17101 - 0 GATCGGTGGACCTCTCTGGT GGG ||||||||[||||||||||||]PAM NA Intergenic NA NA

T33279 CCTCTCTGGTGGGATGATTGAGG 1000
Oligo fwd CACCCCTCTCTGGTGGGATGATTG
Oligo rev AAACCAATCATCCCACCAGAGAGG
Chromosome start end strand MM target_seq PAM alignment distance position gene name gene id
GU305914.1 17069 17091 - 0 CCTCTCTGGTGGGATGATTG AGG ||||||||[||||||||||||]PAM NA Intergenic NA NA

T33280 CTCTGGTGGGATGATTGAGGTGG 1000
Oligo fwd CACCCTCTGGTGGGATGATTGAGG
Oligo rev AAACCCTCAATCATCCCACCAGAG
Chromosome start end strand MM target_seq PAM alignment distance position gene name gene id
GU305914.1 17066 17088 - 0 CTCTGGTGGGATGATTGAGG TGG ||||||||[||||||||||||]PAM NA Intergenic NA NA

T33281 TCTGGTGGGATGATTGAGGTGGG 1000
Oligo fwd CACCTCTGGTGGGATGATTGAGGT
Oligo rev AAACACCTCAATCATCCCACCAGA
Chromosome start end strand MM target_seq PAM alignment distance position gene name gene id
GU305914.1 17065 17087 - 0 TCTGGTGGGATGATTGAGGT GGG ||||||||[||||||||||||]PAM NA Intergenic NA NA

T33282 CTGGTGGGATGATTGAGGTGGGG 1000
Oligo fwd CACCCTGGTGGGATGATTGAGGTG
Oligo rev AAACCACCTCAATCATCCCACCAG
Chromosome start end strand MM target_seq PAM alignment distance position gene name gene id
GU305914.1 17064 17086 - 0 CTGGTGGGATGATTGAGGTG GGG ||||||||[||||||||||||]PAM NA Intergenic NA NA

T33283 TGGTGGGATGATTGAGGTGGGGG 1000
Oligo fwd CACCTGGTGGGATGATTGAGGTGG
Oligo rev AAACCCACCTCAATCATCCCACCA
Chromosome start end strand MM target_seq PAM alignment distance position gene name gene id
GU305914.1 17063 17085 - 0 TGGTGGGATGATTGAGGTGG GGG ||||||||[||||||||||||]PAM NA Intergenic NA NA

T33284 GGTGGGATGATTGAGGTGGGGGG 1000
Oligo fwd CACCGGTGGGATGATTGAGGTGGG
Oligo rev AAACCCCACCTCAATCATCCCACC
Chromosome start end strand MM target_seq PAM alignment distance position gene name gene id
GU305914.1 17062 17084 - 0 GGTGGGATGATTGAGGTGGG GGG ||||||||[||||||||||||]PAM NA Intergenic NA NA

T33285 GTGGGATGATTGAGGTGGGGGGG 1000
Oligo fwd CACCGTGGGATGATTGAGGTGGGG
Oligo rev AAACCCCCACCTCAATCATCCCAC
Chromosome start end strand MM target_seq PAM alignment distance position gene name gene id
GU305914.1 17061 17083 - 0 GTGGGATGATTGAGGTGGGG GGG ||||||||[||||||||||||]PAM NA Intergenic NA NA

T33286 CTGTCTGCTTTCTTTGAAATCGG 1000
Oligo fwd CACCCTGTCTGCTTTCTTTGAAAT

Oligo rev AAACATTTCAAAGAAAGCAGACAG
Chromosome start end strand MM target_seq PAM alignment distance position gene name gene id
GU305914.1 16981 17003 - 0 CTGTCTGCTTTCTTTGAAAT CGG ||||||||[||||||||||||]PAM NA Intergenic NA NA

T33287 AGAACGTCCGCCTTCGAGAATGG 1000
Oligo fwd CACCAGAACGTCCGCCTTCGAGAA
Oligo rev AAACTTCTCGAAGGCGGACGTTCT
Chromosome start end strand MM target_seq PAM alignment distance position gene name gene id
GU305914.1 16948 16970 - 0 AGAACGTCCGCCTTCGAGAA TGG ||||||||[||||||||||||]PAM NA Intergenic NA NA

T33288 GAACGTCCGCCTTCGAGAATGGG 1000
Oligo fwd CACCGAACGTCCGCCTTCGAGAAT
Oligo rev AAACATTCTCGAAGGCGGACGTTC
Chromosome start end strand MM target_seq PAM alignment distance position gene name gene id
GU305914.1 16947 16969 - 0 GAACGTCCGCCTTCGAGAAT GGG ||||||||[||||||||||||]PAM NA Intergenic NA NA

T33289 AACGTCCGCCTTCGAGAATGGGG 1000
Oligo fwd CACCAACGTCCGCCTTCGAGAATG
Oligo rev AAACCATTCTCGAAGGCGGACGTT
Chromosome start end strand MM target_seq PAM alignment distance position gene name gene id
GU305914.1 16946 16968 - 0 AACGTCCGCCTTCGAGAATG GGG ||||||||[||||||||||||]PAM NA Intergenic NA NA

T33290 TCCGCCTTCGAGAATGGGGACGG 1000
Oligo fwd CACCTCCGCCTTCGAGAATGGGGA
Oligo rev AAACTCCCCATTCTCGAAGGCGGA
Chromosome start end strand MM target_seq PAM alignment distance position gene name gene id
GU305914.1 16942 16964 - 0 TCCGCCTTCGAGAATGGGGA CGG ||||||||[||||||||||||]PAM NA Intergenic NA NA

T33291 CCGCCTTCGAGAATGGGGACGGG 1000
Oligo fwd CACCCCGCCTTCGAGAATGGGGAC
Oligo rev AAACGTCCCCATTCTCGAAGGCGG
Chromosome start end strand MM target_seq PAM alignment distance position gene name gene id
GU305914.1 16941 16963 - 0 CCGCCTTCGAGAATGGGGAC GGG ||||||||[||||||||||||]PAM NA Intergenic NA NA

T33292 CCTTCGAGAATGGGGACGGGAGG 1000
Oligo fwd CACCCCTTCGAGAATGGGGACGGG
Oligo rev AAACCCCGTCCCCATTCTCGAAGG
Chromosome start end strand MM target_seq PAM alignment distance position gene name gene id
GU305914.1 16938 16960 - 0 CCTTCGAGAATGGGGACGGG AGG ||||||||[||||||||||||]PAM NA Intergenic NA NA

T33293 GAGAATGGGGACGGGAGGAAAGG 1000
Oligo fwd CACCGAGAATGGGGACGGGAGGAA
Oligo rev AAACTTCCTCCCGTCCCCATTCTC
Chromosome start end strand MM target_seq PAM alignment distance position gene name gene id
GU305914.1 16933 16955 - 0 GAGAATGGGGACGGGAGGAA AGG ||||||||[||||||||||||]PAM NA Intergenic NA NA

T33294 AATGGGGACGGGAGGAAAGGCGG 1000
Oligo fwd CACCAATGGGGACGGGAGGAAAGG
Oligo rev AAACCCTTTCCTCCCGTCCCCATT
Chromosome start end strand MM target_seq PAM alignment distance position gene name gene id
GU305914.1 16930 16952 - 0 AATGGGGACGGGAGGAAAGG CGG ||||||||[||||||||||||]PAM NA Intergenic NA NA

T33295 GGGACGGGAGGAAAGGCGGACGG 1000
Oligo fwd CACCGGGACGGGAGGAAAGGCGGA
Oligo rev AAACTCCGCCTTTCCTCCCGTCCC
Chromosome start end strand MM target_seq PAM alignment distance position gene name gene id
GU305914.1 16926 16948 - 0 GGGACGGGAGGAAAGGCGGA CGG ||||||||[||||||||||||]PAM NA Intergenic NA NA

T33296 AAAGGCGGACGGTACGTTCTCGG 1000
Oligo fwd CACCAAAGGCGGACGGTACGTTCT
Oligo rev AAACAGAACGTACCGTCCGCCTTT
Chromosome start end strand MM target_seq PAM alignment distance position gene name gene id
GU305914.1 16915 16937 - 0 AAAGGCGGACGGTACGTTCT CGG ||||||||[||||||||||||]PAM NA Intergenic NA NA

T33297 AAGGCGGACGGTACGTTCTCGGG 1000
Oligo fwd CACCAAGGCGGACGGTACGTTCTC
Oligo rev AAACGAGAACGTACCGTCCGCCTT
Chromosome start end strand MM target_seq PAM alignment distance position gene name gene id
GU305914.1 16914 16936 - 0 AAGGCGGACGGTACGTTCTC GGG ||||||||[||||||||||||]PAM NA Intergenic NA NA

T33298 CGGGTTTCTATCTGCTGTTCTGG 1000
Oligo fwd CACCCGGGTTTCTATCTGCTGTTC
Oligo rev AAACGAACAGCAGATAGAAACCCG
Chromosome start end strand MM target_seq PAM alignment distance position gene name gene id
GU305914.1 16895 16917 - 0 CGGGTTTCTATCTGCTGTTC TGG ||||||||[||||||||||||]PAM NA Intergenic NA NA

T33299 GCCGTGCGTGCGCAGAACTGCGG 1000
Oligo fwd CACCGCCGTGCGTGCGCAGAACTG
Oligo rev AAACCAGTTCTGCGCACGCACGGC
Chromosome start end strand MM target_seq PAM alignment distance position gene name gene id
GU305914.1 16863 16885 - 0 GCCGTGCGTGCGCAGAACTG CGG ||||||||[||||||||||||]PAM NA Intergenic NA NA

T33300 CAGAACTGCGGCGTCAGTTTCGG 1000
Oligo fwd CACCCAGAACTGCGGCGTCAGTTT
Oligo rev AAACAAACTGACGCCGCAGTTCTG
Chromosome start end strand MM target_seq PAM alignment distance position gene name gene id
GU305914.1 16851 16873 - 0 CAGAACTGCGGCGTCAGTTT CGG ||||||||[||||||||||||]PAM NA Intergenic NA NA
GU305914.1 82034 82056 - 4 CAGATCTCCTGCGTCTGTTT CGG ||||-||-[|-|||||-||||]PAM NA Intergenic NA NA

<table>
<tr><td>T33301</td><td>TGATGAATATCTCACCGCCACGG</td><td>1000</td></tr>
<tr><td>Oligo fwd</td><td>CACCTGATGAATATCTCACCGCCA</td></tr>
<tr><td>Oligo rev</td><td>AAACTGGCGGTGAGATATTCATCA</td></tr>
<tr><td>Chromosome</td><td>start</td><td>end</td><td>strand</td><td>MM</td><td>target_seq</td><td>PAM</td><td>alignment</td><td>distance</td><td>position</td><td>gene name</td><td>gene id</td></tr>
<tr><td>GU305914.1</td><td>16810</td><td>16832</td><td>-</td><td>0</td><td>TGATGAATATCTCACCGCCA</td><td>CGG</td><td>||||||||[||||||||||||]PAM</td><td>NA</td><td>Intergenic</td><td>NA</td><td>NA</td></tr>
<tr><td>T33302</td><td>AATATCTCACCGCCACGGTGCGG</td><td>1000</td></tr>
<tr><td>Oligo fwd</td><td>CACCAATATCTCACCGCCACGGTG</td></tr>
<tr><td>Oligo rev</td><td>AAACCACCGTGGCGGTGAGATATT</td></tr>
<tr><td>Chromosome</td><td>start</td><td>end</td><td>strand</td><td>MM</td><td>target_seq</td><td>PAM</td><td>alignment</td><td>distance</td><td>position</td><td>gene name</td><td>gene id</td></tr>
<tr><td>GU305914.1</td><td>16805</td><td>16827</td><td>-</td><td>0</td><td>AATATCTCACCGCCACGGTG</td><td>CGG</td><td>||||||||[||||||||||||]PAM</td><td>NA</td><td>Intergenic</td><td>NA</td><td>NA</td></tr>
<tr><td>T33303</td><td>GCCACGGTGCGGAGCGAGTGCGG</td><td>1000</td></tr>
<tr><td>Oligo fwd</td><td>CACCGCCACGGTGCGGAGCGAGTG</td></tr>
<tr><td>Oligo rev</td><td>AAACCACTCGCTCCGCACCGTGGC</td></tr>
<tr><td>Chromosome</td><td>start</td><td>end</td><td>strand</td><td>MM</td><td>target_seq</td><td>PAM</td><td>alignment</td><td>distance</td><td>position</td><td>gene name</td><td>gene id</td></tr>
<tr><td>GU305914.1</td><td>16794</td><td>16816</td><td>-</td><td>0</td><td>GCCACGGTGCGGAGCGAGTG</td><td>CGG</td><td>||||||||[||||||||||||]PAM</td><td>NA</td><td>Intergenic</td><td>NA</td><td>NA</td></tr>
<tr><td>T33304</td><td>ATCAACATAGAGTTCTACGCCGG</td><td>1000</td></tr>
<tr><td>Oligo fwd</td><td>CACCATCAACATAGAGTTCTACGC</td></tr>
<tr><td>Oligo rev</td><td>AAACGCGTAGAACTCTATGTTGAT</td></tr>
<tr><td>Chromosome</td><td>start</td><td>end</td><td>strand</td><td>MM</td><td>target_seq</td><td>PAM</td><td>alignment</td><td>distance</td><td>position</td><td>gene name</td><td>gene id</td></tr>
<tr><td>GU305914.1</td><td>16764</td><td>16786</td><td>-</td><td>0</td><td>ATCAACATAGAGTTCTACGC</td><td>CGG</td><td>||||||||[||||||||||||]PAM</td><td>NA</td><td>Intergenic</td><td>NA</td><td>NA</td></tr>
<tr><td>T33305</td><td>CATAGAGTTCTACGCCGGCATGG</td><td>1000</td></tr>
<tr><td>Oligo fwd</td><td>CACCCATAGAGTTCTACGCCGGCA</td></tr>
<tr><td>Oligo rev</td><td>AAACTGCCGGCGTAGAACTCTATG</td></tr>
<tr><td>Chromosome</td><td>start</td><td>end</td><td>strand</td><td>MM</td><td>target_seq</td><td>PAM</td><td>alignment</td><td>distance</td><td>position</td><td>gene name</td><td>gene id</td></tr>
<tr><td>GU305914.1</td><td>16759</td><td>16781</td><td>-</td><td>0</td><td>CATAGAGTTCTACGCCGGCA</td><td>TGG</td><td>||||||||[||||||||||||]PAM</td><td>NA</td><td>Intergenic</td><td>NA</td><td>NA</td></tr>
<tr><td>T33306</td><td>CATGGTTCTCAACACGAGCGAGG</td><td>1000</td></tr>
<tr><td>Oligo fwd</td><td>CACCCATGGTTCTCAACACGAGCG</td></tr>
<tr><td>Oligo rev</td><td>AAACCGCTCGTGTTGAGAACCATG</td></tr>
<tr><td>Chromosome</td><td>start</td><td>end</td><td>strand</td><td>MM</td><td>target_seq</td><td>PAM</td><td>alignment</td><td>distance</td><td>position</td><td>gene name</td><td>gene id</td></tr>
<tr><td>GU305914.1</td><td>16741</td><td>16763</td><td>-</td><td>0</td><td>CATGGTTCTCAACACGAGCG</td><td>AGG</td><td>||||||||[||||||||||||]PAM</td><td>NA</td><td>Intergenic</td><td>NA</td><td>NA</td></tr>
<tr><td>T33307</td><td>TTCTCAACACGAGCGAGGTTTGG</td><td>1000</td></tr>
<tr><td>Oligo fwd</td><td>CACCTTCTCAACACGAGCGAGGTT</td></tr>
<tr><td>Oligo rev</td><td>AAACAACCTCGCTCGTGTTGAGAA</td></tr>
<tr><td>Chromosome</td><td>start</td><td>end</td><td>strand</td><td>MM</td><td>target_seq</td><td>PAM</td><td>alignment</td><td>distance</td><td>position</td><td>gene name</td><td>gene id</td></tr>
<tr><td>GU305914.1</td><td>16736</td><td>16758</td><td>-</td><td>0</td><td>TTCTCAACACGAGCGAGGTT</td><td>TGG</td><td>||||||||[||||||||||||]PAM</td><td>NA</td><td>Intergenic</td><td>NA</td><td>NA</td></tr>
<tr><td>T33308</td><td>CACGAGCGAGGTTTGGATGATGG</td><td>1000</td></tr>
<tr><td>Oligo fwd</td><td>CACCCACGAGCGAGGTTTGGATGA</td></tr>
<tr><td>Oligo rev</td><td>AAACTCATCCAAACCTCGCTCGTG</td></tr>
<tr><td>Chromosome</td><td>start</td><td>end</td><td>strand</td><td>MM</td><td>target_seq</td><td>PAM</td><td>alignment</td><td>distance</td><td>position</td><td>gene name</td><td>gene id</td></tr>
<tr><td>GU305914.1</td><td>16729</td><td>16751</td><td>-</td><td>0</td><td>CACGAGCGAGGTTTGGATGA</td><td>TGG</td><td>||||||||[||||||||||||]PAM</td><td>NA</td><td>Intergenic</td><td>NA</td><td>NA</td></tr>
<tr><td>T33309</td><td>ACGAGCGAGGTTTGGATGATGGG</td><td>1000</td></tr>
<tr><td>Oligo fwd</td><td>CACCACGAGCGAGGTTTGGATGAT</td></tr>
<tr><td>Oligo rev</td><td>AAACATCATCCAAACCTCGCTCGT</td></tr>
<tr><td>Chromosome</td><td>start</td><td>end</td><td>strand</td><td>MM</td><td>target_seq</td><td>PAM</td><td>alignment</td><td>distance</td><td>position</td><td>gene name</td><td>gene id</td></tr>
<tr><td>GU305914.1</td><td>16728</td><td>16750</td><td>-</td><td>0</td><td>ACGAGCGAGGTTTGGATGAT</td><td>GGG</td><td>||||||||[||||||||||||]PAM</td><td>NA</td><td>Intergenic</td><td>NA</td><td>NA</td></tr>
<tr><td>T33310</td><td>AGCGAGGTTTGGATGATGGGAGG</td><td>1000</td></tr>
<tr><td>Oligo fwd</td><td>CACCAGCGAGGTTTGGATGATGGG</td></tr>
<tr><td>Oligo rev</td><td>AAACCCCATCATCCAAACCTCGCT</td></tr>
<tr><td>Chromosome</td><td>start</td><td>end</td><td>strand</td><td>MM</td><td>target_seq</td><td>PAM</td><td>alignment</td><td>distance</td><td>position</td><td>gene name</td><td>gene id</td></tr>
<tr><td>GU305914.1</td><td>16725</td><td>16747</td><td>-</td><td>0</td><td>AGCGAGGTTTGGATGATGGG</td><td>AGG</td><td>||||||||[||||||||||||]PAM</td><td>NA</td><td>Intergenic</td><td>NA</td><td>NA</td></tr>
<tr><td>T33311</td><td>CTGTTCGAAGTCTTGCCGAACGG</td><td>1000</td></tr>
<tr><td>Oligo fwd</td><td>CACCCTGTTCGAAGTCTTGCCGAA</td></tr>
<tr><td>Oligo rev</td><td>AAACTTCGGCAAGACTTCGAACAG</td></tr>
<tr><td>Chromosome</td><td>start</td><td>end</td><td>strand</td><td>MM</td><td>target_seq</td><td>PAM</td><td>alignment</td><td>distance</td><td>position</td><td>gene name</td><td>gene id</td></tr>
<tr><td>GU305914.1</td><td>16683</td><td>16705</td><td>-</td><td>0</td><td>CTGTTCGAAGTCTTGCCGAA</td><td>CGG</td><td>||||||||[||||||||||||]PAM</td><td>NA</td><td>Intergenic</td><td>NA</td><td>NA</td></tr>
<tr><td>T33312</td><td>CCATCAACAACGACGTTAGATGG</td><td>1000</td></tr>
<tr><td>Oligo fwd</td><td>CACCCCATCAACAACGACGTTAGA</td></tr>
<tr><td>Oligo rev</td><td>AAACTCTAACGTCGTTGTTGATGG</td></tr>
<tr><td>Chromosome</td><td>start</td><td>end</td><td>strand</td><td>MM</td><td>target_seq</td><td>PAM</td><td>alignment</td><td>distance</td><td>position</td><td>gene name</td><td>gene id</td></tr>
<tr><td>GU305914.1</td><td>16658</td><td>16680</td><td>-</td><td>0</td><td>CCATCAACAACGACGTTAGA</td><td>TGG</td><td>||||||||[||||||||||||]PAM</td><td>NA</td><td>Intergenic</td><td>NA</td><td>NA</td></tr>
<tr><td>T33313</td><td>CATCAACAACGACGTTAGATGGG</td><td>1000</td></tr>
<tr><td>Oligo fwd</td><td>CACCCATCAACAACGACGTTAGAT</td></tr>
<tr><td>Oligo rev</td><td>AAACATCTAACGTCGTTGTTGATG</td></tr>
<tr><td>Chromosome</td><td>start</td><td>end</td><td>strand</td><td>MM</td><td>target_seq</td><td>PAM</td><td>alignment</td><td>distance</td><td>position</td><td>gene name</td><td>gene id</td></tr>
<tr><td>GU305914.1</td><td>16657</td><td>16679</td><td>-</td><td>0</td><td>CATCAACAACGACGTTAGAT</td><td>GGG</td><td>||||||||[||||||||||||]PAM</td><td>NA</td><td>Intergenic</td><td>NA</td><td>NA</td></tr>
<tr><td>T33314</td><td>CGTTAGATGGGTGAATATCACGG</td><td>1000</td></tr>
<tr><td>Oligo fwd</td><td>CACCCGTTAGATGGGTGAATATCA</td></tr>
<tr><td>Oligo rev</td><td>AAACTGATATTCACCCATCTAACG</td></tr>
<tr><td>Chromosome</td><td>start</td><td>end</td><td>strand</td><td>MM</td><td>target_seq</td><td>PAM</td><td>alignment</td><td>distance</td><td>position</td><td>gene name</td><td>gene id</td></tr>
<tr><td>GU305914.1</td><td>16645</td><td>16667</td><td>-</td><td>0</td><td>CGTTAGATGGGTGAATATCA</td><td>CGG</td><td>||||||||[||||||||||||]PAM</td><td>NA</td><td>Intergenic</td><td>NA</td><td>NA</td></tr>
<tr><td>T33315</td><td>GTGAATATCACGGAGCGTGACGG</td><td>1000</td></tr>
<tr><td>Oligo fwd</td><td>CACCGTGAATATCACGGAGCGTGA</td></tr>
<tr><td>Oligo rev</td><td>AAACTCACGCTCCGTGATATTCAC</td></tr>
<tr><td>Chromosome</td><td>start</td><td>end</td><td>strand</td><td>MM</td><td>target_seq</td><td>PAM</td><td>alignment</td><td>distance</td><td>position</td><td>gene name</td><td>gene id</td></tr>
<tr><td>GU305914.1</td><td>16635</td><td>16657</td><td>-</td><td>0</td><td>GTGAATATCACGGAGCGTGA</td><td>CGG</td><td>||||||||[||||||||||||]PAM</td><td>NA</td><td>Intergenic</td><td>NA</td><td>NA</td></tr>
</table>

<table>
<tr><td>T33316</td><td>TGAATATCACGGAGCGTGACGGG</td><td></td><td>1000</td><td></td><td></td><td></td><td></td><td></td><td></td><td></td><td></td><td></td></tr>
<tr><td>Oligo fwd</td><td>CACCTGAATATCACGGAGCGTGAC</td><td></td><td></td><td></td><td></td><td></td><td></td><td></td><td></td><td></td><td></td><td></td></tr>
<tr><td>Oligo rev</td><td>AAACGTCACGCTCCGTGATATTCA</td><td></td><td></td><td></td><td></td><td></td><td></td><td></td><td></td><td></td><td></td><td></td></tr>
<tr><td>Chromosome</td><td>start</td><td></td><td>end</td><td>strand</td><td>MM</td><td>target_seq</td><td>PAM</td><td>alignment</td><td>distance</td><td>position</td><td>gene name</td><td>gene id</td></tr>
<tr><td>GU305914.1</td><td></td><td>16634</td><td>16656</td><td>-</td><td>0</td><td>TGAATATCACGGAGCGTGAC</td><td>GGG</td><td>||||||||[||||||||||||]PAM</td><td>NA</td><td>Intergenic</td><td>NA</td><td>NA</td></tr>
<tr><td>T33317</td><td>GAATATCACGGAGCGTGACGGGG</td><td></td><td>1000</td><td></td><td></td><td></td><td></td><td></td><td></td><td></td><td></td><td></td></tr>
<tr><td>Oligo fwd</td><td>CACCGAATATCACGGAGCGTGACG</td><td></td><td></td><td></td><td></td><td></td><td></td><td></td><td></td><td></td><td></td><td></td></tr>
<tr><td>Oligo rev</td><td>AAACCGTCACGCTCCGTGATATTC</td><td></td><td></td><td></td><td></td><td></td><td></td><td></td><td></td><td></td><td></td><td></td></tr>
<tr><td>Chromosome</td><td>start</td><td></td><td>end</td><td>strand</td><td>MM</td><td>target_seq</td><td>PAM</td><td>alignment</td><td>distance</td><td>position</td><td>gene name</td><td>gene id</td></tr>
<tr><td>GU305914.1</td><td></td><td>16633</td><td>16655</td><td>-</td><td>0</td><td>GAATATCACGGAGCGTGACG</td><td>GGG</td><td>||||||||[||||||||||||]PAM</td><td>NA</td><td>Intergenic</td><td>NA</td><td>NA</td></tr>
<tr><td>T33318</td><td>TAAATTTTTGATGTCCCAGACGG</td><td></td><td>1000</td><td></td><td></td><td></td><td></td><td></td><td></td><td></td><td></td><td></td></tr>
<tr><td>Oligo fwd</td><td>CACCTAAATTTTTGATGTCCCAGA</td><td></td><td></td><td></td><td></td><td></td><td></td><td></td><td></td><td></td><td></td><td></td></tr>
<tr><td>Oligo rev</td><td>AAACTCTGGGACATCAAAAATTTA</td><td></td><td></td><td></td><td></td><td></td><td></td><td></td><td></td><td></td><td></td><td></td></tr>
<tr><td>Chromosome</td><td>start</td><td></td><td>end</td><td>strand</td><td>MM</td><td>target_seq</td><td>PAM</td><td>alignment</td><td>distance</td><td>position</td><td>gene name</td><td>gene id</td></tr>
<tr><td>GU305914.1</td><td></td><td>16606</td><td>16628</td><td>-</td><td>0</td><td>TAAATTTTTGATGTCCCAGA</td><td>CGG</td><td>||||||||[||||||||||||]PAM</td><td>NA</td><td>Intergenic</td><td>NA</td><td>NA</td></tr>
<tr><td>T33319</td><td>ACATGTAAAGAAGTTTCATGAGG</td><td></td><td>1000</td><td></td><td></td><td></td><td></td><td></td><td></td><td></td><td></td><td></td></tr>
<tr><td>Oligo fwd</td><td>CACCACATGTAAAGAAGTTTCATG</td><td></td><td></td><td></td><td></td><td></td><td></td><td></td><td></td><td></td><td></td><td></td></tr>
<tr><td>Oligo rev</td><td>AAACCATGAAACTTCTTTACATGT</td><td></td><td></td><td></td><td></td><td></td><td></td><td></td><td></td><td></td><td></td><td></td></tr>
<tr><td>Chromosome</td><td>start</td><td></td><td>end</td><td>strand</td><td>MM</td><td>target_seq</td><td>PAM</td><td>alignment</td><td>distance</td><td>position</td><td>gene name</td><td>gene id</td></tr>
<tr><td>GU305914.1</td><td></td><td>16582</td><td>16604</td><td>-</td><td>0</td><td>ACATGTAAAGAAGTTTCATG</td><td>AGG</td><td>||||||||[||||||||||||]PAM</td><td>NA</td><td>Intergenic</td><td>NA</td><td>NA</td></tr>
<tr><td>T33320</td><td>TAAAGAAGTTTCATGAGGTGTGG</td><td></td><td>1000</td><td></td><td></td><td></td><td></td><td></td><td></td><td></td><td></td><td></td></tr>
<tr><td>Oligo fwd</td><td>CACCTAAAGAAGTTTCATGAGGTG</td><td></td><td></td><td></td><td></td><td></td><td></td><td></td><td></td><td></td><td></td><td></td></tr>
<tr><td>Oligo rev</td><td>AAACCACCTCATGAAACTTCTTTA</td><td></td><td></td><td></td><td></td><td></td><td></td><td></td><td></td><td></td><td></td><td></td></tr>
<tr><td>Chromosome</td><td>start</td><td></td><td>end</td><td>strand</td><td>MM</td><td>target_seq</td><td>PAM</td><td>alignment</td><td>distance</td><td>position</td><td>gene name</td><td>gene id</td></tr>
<tr><td>GU305914.1</td><td></td><td>16577</td><td>16599</td><td>-</td><td>0</td><td>TAAAGAAGTTTCATGAGGTG</td><td>TGG</td><td>||||||||[||||||||||||]PAM</td><td>NA</td><td>Intergenic</td><td>NA</td><td>NA</td></tr>
<tr><td>T33321</td><td>AGTTTCATGAGGTGTGGAAACGG</td><td></td><td>1000</td><td></td><td></td><td></td><td></td><td></td><td></td><td></td><td></td><td></td></tr>
<tr><td>Oligo fwd</td><td>CACCAGTTTCATGAGGTGTGGAAA</td><td></td><td></td><td></td><td></td><td></td><td></td><td></td><td></td><td></td><td></td><td></td></tr>
<tr><td>Oligo rev</td><td>AAACTTTCCACACCTCATGAAACT</td><td></td><td></td><td></td><td></td><td></td><td></td><td></td><td></td><td></td><td></td><td></td></tr>
<tr><td>Chromosome</td><td>start</td><td></td><td>end</td><td>strand</td><td>MM</td><td>target_seq</td><td>PAM</td><td>alignment</td><td>distance</td><td>position</td><td>gene name</td><td>gene id</td></tr>
<tr><td>GU305914.1</td><td></td><td>16571</td><td>16593</td><td>-</td><td>0</td><td>AGTTTCATGAGGTGTGGAAA</td><td>CGG</td><td>||||||||[||||||||||||]PAM</td><td>NA</td><td>Intergenic</td><td>NA</td><td>NA</td></tr>
<tr><td>T33322</td><td>GTTTCATGAGGTGTGGAAACGGG</td><td></td><td>1000</td><td></td><td></td><td></td><td></td><td></td><td></td><td></td><td></td><td></td></tr>
<tr><td>Oligo fwd</td><td>CACCGTTTCATGAGGTGTGGAAAC</td><td></td><td></td><td></td><td></td><td></td><td></td><td></td><td></td><td></td><td></td><td></td></tr>
<tr><td>Oligo rev</td><td>AAACGTTTCCACACCTCATGAAAC</td><td></td><td></td><td></td><td></td><td></td><td></td><td></td><td></td><td></td><td></td><td></td></tr>
<tr><td>Chromosome</td><td>start</td><td></td><td>end</td><td>strand</td><td>MM</td><td>target_seq</td><td>PAM</td><td>alignment</td><td>distance</td><td>position</td><td>gene name</td><td>gene id</td></tr>
<tr><td>GU305914.1</td><td></td><td>16570</td><td>16592</td><td>-</td><td>0</td><td>GTTTCATGAGGTGTGGAAAC</td><td>GGG</td><td>||||||||[||||||||||||]PAM</td><td>NA</td><td>Intergenic</td><td>NA</td><td>NA</td></tr>
<tr><td>T33323</td><td>TCATGAGGTGTGGAAACGGGCGG</td><td></td><td>1000</td><td></td><td></td><td></td><td></td><td></td><td></td><td></td><td></td><td></td></tr>
<tr><td>Oligo fwd</td><td>CACCTCATGAGGTGTGGAAACGGG</td><td></td><td></td><td></td><td></td><td></td><td></td><td></td><td></td><td></td><td></td><td></td></tr>
<tr><td>Oligo rev</td><td>AAACCCCGTTTCCACACCTCATGA</td><td></td><td></td><td></td><td></td><td></td><td></td><td></td><td></td><td></td><td></td><td></td></tr>
<tr><td>Chromosome</td><td>start</td><td></td><td>end</td><td>strand</td><td>MM</td><td>target_seq</td><td>PAM</td><td>alignment</td><td>distance</td><td>position</td><td>gene name</td><td>gene id</td></tr>
<tr><td>GU305914.1</td><td></td><td>16567</td><td>16589</td><td>-</td><td>0</td><td>TCATGAGGTGTGGAAACGGG</td><td>CGG</td><td>||||||||[||||||||||||]PAM</td><td>NA</td><td>Intergenic</td><td>NA</td><td>NA</td></tr>
<tr><td>T33324</td><td>CATGAGGTGTGGAAACGGGCGGG</td><td></td><td>1000</td><td></td><td></td><td></td><td></td><td></td><td></td><td></td><td></td><td></td></tr>
<tr><td>Oligo fwd</td><td>CACCCATGAGGTGTGGAAACGGGC</td><td></td><td></td><td></td><td></td><td></td><td></td><td></td><td></td><td></td><td></td><td></td></tr>
<tr><td>Oligo rev</td><td>AAACGCCCGTTTCCACACCTCATG</td><td></td><td></td><td></td><td></td><td></td><td></td><td></td><td></td><td></td><td></td><td></td></tr>
<tr><td>Chromosome</td><td>start</td><td></td><td>end</td><td>strand</td><td>MM</td><td>target_seq</td><td>PAM</td><td>alignment</td><td>distance</td><td>position</td><td>gene name</td><td>gene id</td></tr>
<tr><td>GU305914.1</td><td></td><td>16566</td><td>16588</td><td>-</td><td>0</td><td>CATGAGGTGTGGAAACGGGC</td><td>GGG</td><td>||||||||[||||||||||||]PAM</td><td>NA</td><td>Intergenic</td><td>NA</td><td>NA</td></tr>
<tr><td>T33325</td><td>CCGCAAGATTTTACTCACATCGG</td><td></td><td>1000</td><td></td><td></td><td></td><td></td><td></td><td></td><td></td><td></td><td></td></tr>
<tr><td>Oligo fwd</td><td>CACCCCGCAAGATTTTACTCACAT</td><td></td><td></td><td></td><td></td><td></td><td></td><td></td><td></td><td></td><td></td><td></td></tr>
<tr><td>Oligo rev</td><td>AAACATGTGAGTAAAATCTTGCGG</td><td></td><td></td><td></td><td></td><td></td><td></td><td></td><td></td><td></td><td></td><td></td></tr>
<tr><td>Chromosome</td><td>start</td><td></td><td>end</td><td>strand</td><td>MM</td><td>target_seq</td><td>PAM</td><td>alignment</td><td>distance</td><td>position</td><td>gene name</td><td>gene id</td></tr>
<tr><td>GU305914.1</td><td></td><td>16530</td><td>16552</td><td>-</td><td>0</td><td>CCGCAAGATTTTACTCACAT</td><td>CGG</td><td>||||||||[||||||||||||]PAM</td><td>NA</td><td>Intergenic</td><td>NA</td><td>NA</td></tr>
<tr><td>T33326</td><td>CTGCCGCGTGAGCCACGTGACGG</td><td></td><td>1000</td><td></td><td></td><td></td><td></td><td></td><td></td><td></td><td></td><td></td></tr>
<tr><td>Oligo fwd</td><td>CACCCTGCCGCGTGAGCCACGTGA</td><td></td><td></td><td></td><td></td><td></td><td></td><td></td><td></td><td></td><td></td><td></td></tr>
<tr><td>Oligo rev</td><td>AAACTCACGTGGCTCACGCGGCAG</td><td></td><td></td><td></td><td></td><td></td><td></td><td></td><td></td><td></td><td></td><td></td></tr>
<tr><td>Chromosome</td><td>start</td><td></td><td>end</td><td>strand</td><td>MM</td><td>target_seq</td><td>PAM</td><td>alignment</td><td>distance</td><td>position</td><td>gene name</td><td>gene id</td></tr>
<tr><td>GU305914.1</td><td></td><td>16495</td><td>16517</td><td>-</td><td>0</td><td>CTGCCGCGTGAGCCACGTGA</td><td>CGG</td><td>||||||||[||||||||||||]PAM</td><td>NA</td><td>Intergenic</td><td>NA</td><td>NA</td></tr>
<tr><td>T33327</td><td>CGAGCGAAACTTTCTTCAGTTGG</td><td></td><td>1000</td><td></td><td></td><td></td><td></td><td></td><td></td><td></td><td></td><td></td></tr>
<tr><td>Oligo fwd</td><td>CACCCGAGCGAAACTTTCTTCAGT</td><td></td><td></td><td></td><td></td><td></td><td></td><td></td><td></td><td></td><td></td><td></td></tr>
<tr><td>Oligo rev</td><td>AAACACTGAAGAAAGTTTCGCTCG</td><td></td><td></td><td></td><td></td><td></td><td></td><td></td><td></td><td></td><td></td><td></td></tr>
<tr><td>Chromosome</td><td>start</td><td></td><td>end</td><td>strand</td><td>MM</td><td>target_seq</td><td>PAM</td><td>alignment</td><td>distance</td><td>position</td><td>gene name</td><td>gene id</td></tr>
<tr><td>GU305914.1</td><td></td><td>16453</td><td>16475</td><td>-</td><td>0</td><td>CGAGCGAAACTTTCTTCAGT</td><td>TGG</td><td>||||||||[||||||||||||]PAM</td><td>NA</td><td>Intergenic</td><td>NA</td><td>NA</td></tr>
<tr><td>T33328</td><td>TTACAGAAACCATACGTACCCGG</td><td></td><td>1000</td><td></td><td></td><td></td><td></td><td></td><td></td><td></td><td></td><td></td></tr>
<tr><td>Oligo fwd</td><td>CACCTTACAGAAACCATACGTACC</td><td></td><td></td><td></td><td></td><td></td><td></td><td></td><td></td><td></td><td></td><td></td></tr>
<tr><td>Oligo rev</td><td>AAACGGTACGTATGGTTTCTGTAA</td><td></td><td></td><td></td><td></td><td></td><td></td><td></td><td></td><td></td><td></td><td></td></tr>
<tr><td>Chromosome</td><td>start</td><td></td><td>end</td><td>strand</td><td>MM</td><td>target_seq</td><td>PAM</td><td>alignment</td><td>distance</td><td>position</td><td>gene name</td><td>gene id</td></tr>
<tr><td>GU305914.1</td><td></td><td>16414</td><td>16436</td><td>-</td><td>0</td><td>TTACAGAAACCATACGTACC</td><td>CGG</td><td>||||||||[||||||||||||]PAM</td><td>NA</td><td>Intergenic</td><td>NA</td><td>NA</td></tr>
<tr><td>T33329</td><td>CAGAAACCATACGTACCCGGTGG</td><td></td><td>1000</td><td></td><td></td><td></td><td></td><td></td><td></td><td></td><td></td><td></td></tr>
<tr><td>Oligo fwd</td><td>CACCCAGAAACCATACGTACCCGG</td><td></td><td></td><td></td><td></td><td></td><td></td><td></td><td></td><td></td><td></td><td></td></tr>
<tr><td>Oligo rev</td><td>AAACCCGGGTACGTATGGTTTCTG</td><td></td><td></td><td></td><td></td><td></td><td></td><td></td><td></td><td></td><td></td><td></td></tr>
<tr><td>Chromosome</td><td>start</td><td></td><td>end</td><td>strand</td><td>MM</td><td>target_seq</td><td>PAM</td><td>alignment</td><td>distance</td><td>position</td><td>gene name</td><td>gene id</td></tr>
<tr><td>GU305914.1</td><td></td><td>16411</td><td>16433</td><td>-</td><td>0</td><td>CAGAAACCATACGTACCCGG</td><td>TGG</td><td>||||||||[||||||||||||]PAM</td><td>NA</td><td>Intergenic</td><td>NA</td><td>NA</td></tr>
<tr><td>T33330</td><td>AGAAACCATACGTACCCGGTGGG</td><td></td><td>1000</td><td></td><td></td><td></td><td></td><td></td><td></td><td></td><td></td><td></td></tr>
<tr><td>Oligo fwd</td><td>CACCAGAAACCATACGTACCCGGT</td><td></td><td></td><td></td><td></td><td></td><td></td><td></td><td></td><td></td><td></td><td></td></tr>
<tr><td>Oligo rev</td><td>AAACACCGGGTACGTATGGTTTCT</td><td></td><td></td><td></td><td></td><td></td><td></td><td></td><td></td><td></td><td></td><td></td></tr>
<tr><td>Chromosome</td><td>start</td><td></td><td>end</td><td>strand</td><td>MM</td><td>target_seq</td><td>PAM</td><td>alignment</td><td>distance</td><td>position</td><td>gene name</td><td>gene id</td></tr>
</table>

<table>
<tr><td>GU305914.1</td><td>16410</td><td>16432</td><td>-</td><td>0</td><td>AGAAACCATACGTACCCGGT</td><td>GGG</td><td>||||||||[||||||||||||]PAM</td><td>NA</td><td>Intergenic</td><td>NA</td><td>NA</td></tr>
<tr><td>T33331</td><td>ACGTACCCGGTGGGAGACGTTGG</td><td>1000</td><td></td><td></td><td></td><td></td><td></td><td></td><td></td><td></td><td></td></tr>
<tr><td>Oligo fwd</td><td>CACCACGTACCCGGTGGGAGACGT</td><td></td><td></td><td></td><td></td><td></td><td></td><td></td><td></td><td></td><td></td></tr>
<tr><td>Oligo rev</td><td>AAACACGTCTCCCACCGGGTACGT</td><td></td><td></td><td></td><td></td><td></td><td></td><td></td><td></td><td></td><td></td></tr>
<tr><td>Chromosome</td><td>start</td><td>end</td><td>strand</td><td>MM</td><td>target_seq</td><td>PAM</td><td>alignment</td><td>distance</td><td>position</td><td>gene name</td><td>gene id</td></tr>
<tr><td>GU305914.1</td><td>16401</td><td>16423</td><td>-</td><td>0</td><td>ACGTACCCGGTGGGAGACGT</td><td>TGG</td><td>||||||||[||||||||||||]PAM</td><td>NA</td><td>Intergenic</td><td>NA</td><td>NA</td></tr>
<tr><td>T33332</td><td>TGGAAAACTGTTCGAGTCCGTGG</td><td>1000</td><td></td><td></td><td></td><td></td><td></td><td></td><td></td><td></td><td></td></tr>
<tr><td>Oligo fwd</td><td>CACCTGGAAAACTGTTCGAGTCCG</td><td></td><td></td><td></td><td></td><td></td><td></td><td></td><td></td><td></td><td></td></tr>
<tr><td>Oligo rev</td><td>AAACCGGACTCGAACAGTTTTCCA</td><td></td><td></td><td></td><td></td><td></td><td></td><td></td><td></td><td></td><td></td></tr>
<tr><td>Chromosome</td><td>start</td><td>end</td><td>strand</td><td>MM</td><td>target_seq</td><td>PAM</td><td>alignment</td><td>distance</td><td>position</td><td>gene name</td><td>gene id</td></tr>
<tr><td>GU305914.1</td><td>16381</td><td>16403</td><td>-</td><td>0</td><td>TGGAAAACTGTTCGAGTCCG</td><td>TGG</td><td>||||||||[||||||||||||]PAM</td><td>NA</td><td>Intergenic</td><td>NA</td><td>NA</td></tr>
<tr><td>T33333</td><td>CTGTTCGAGTCCGTGGCCCGTGG</td><td>1000</td><td></td><td></td><td></td><td></td><td></td><td></td><td></td><td></td><td></td></tr>
<tr><td>Oligo fwd</td><td>CACCCTGTTCGAGTCCGTGGCCCG</td><td></td><td></td><td></td><td></td><td></td><td></td><td></td><td></td><td></td><td></td></tr>
<tr><td>Oligo rev</td><td>AAACCGGGCCACGGACTCGAACAG</td><td></td><td></td><td></td><td></td><td></td><td></td><td></td><td></td><td></td><td></td></tr>
<tr><td>Chromosome</td><td>start</td><td>end</td><td>strand</td><td>MM</td><td>target_seq</td><td>PAM</td><td>alignment</td><td>distance</td><td>position</td><td>gene name</td><td>gene id</td></tr>
<tr><td>GU305914.1</td><td>16374</td><td>16396</td><td>-</td><td>0</td><td>CTGTTCGAGTCCGTGGCCCG</td><td>TGG</td><td>||||||||[||||||||||||]PAM</td><td>NA</td><td>Intergenic</td><td>NA</td><td>NA</td></tr>
<tr><td>T33334</td><td>TGTTCGAGTCCGTGGCCCGTGGG</td><td>1000</td><td></td><td></td><td></td><td></td><td></td><td></td><td></td><td></td><td></td></tr>
<tr><td>Oligo fwd</td><td>CACCTGTTCGAGTCCGTGGCCCGT</td><td></td><td></td><td></td><td></td><td></td><td></td><td></td><td></td><td></td><td></td></tr>
<tr><td>Oligo rev</td><td>AAACACGGGCCACGGACTCGAACA</td><td></td><td></td><td></td><td></td><td></td><td></td><td></td><td></td><td></td><td></td></tr>
<tr><td>Chromosome</td><td>start</td><td>end</td><td>strand</td><td>MM</td><td>target_seq</td><td>PAM</td><td>alignment</td><td>distance</td><td>position</td><td>gene name</td><td>gene id</td></tr>
<tr><td>GU305914.1</td><td>16373</td><td>16395</td><td>-</td><td>0</td><td>TGTTCGAGTCCGTGGCCCGT</td><td>GGG</td><td>||||||||[||||||||||||]PAM</td><td>NA</td><td>Intergenic</td><td>NA</td><td>NA</td></tr>
<tr><td>T33335</td><td>GTTCGAGTCCGTGGCCCGTGGGG</td><td>1000</td><td></td><td></td><td></td><td></td><td></td><td></td><td></td><td></td><td></td></tr>
<tr><td>Oligo fwd</td><td>CACCGTTCGAGTCCGTGGCCCGTG</td><td></td><td></td><td></td><td></td><td></td><td></td><td></td><td></td><td></td><td></td></tr>
<tr><td>Oligo rev</td><td>AAACCACGGGCCACGGACTCGAAC</td><td></td><td></td><td></td><td></td><td></td><td></td><td></td><td></td><td></td><td></td></tr>
<tr><td>Chromosome</td><td>start</td><td>end</td><td>strand</td><td>MM</td><td>target_seq</td><td>PAM</td><td>alignment</td><td>distance</td><td>position</td><td>gene name</td><td>gene id</td></tr>
<tr><td>GU305914.1</td><td>16372</td><td>16394</td><td>-</td><td>0</td><td>GTTCGAGTCCGTGGCCCGTG</td><td>GGG</td><td>||||||||[||||||||||||]PAM</td><td>NA</td><td>Intergenic</td><td>NA</td><td>NA</td></tr>
<tr><td>T33336</td><td>GAGTCCGTGGCCCGTGGGGTAGG</td><td>1000</td><td></td><td></td><td></td><td></td><td></td><td></td><td></td><td></td><td></td></tr>
<tr><td>Oligo fwd</td><td>CACCGAGTCCGTGGCCCGTGGGGT</td><td></td><td></td><td></td><td></td><td></td><td></td><td></td><td></td><td></td><td></td></tr>
<tr><td>Oligo rev</td><td>AAACACCCCACGGGCCACGGACTC</td><td></td><td></td><td></td><td></td><td></td><td></td><td></td><td></td><td></td><td></td></tr>
<tr><td>Chromosome</td><td>start</td><td>end</td><td>strand</td><td>MM</td><td>target_seq</td><td>PAM</td><td>alignment</td><td>distance</td><td>position</td><td>gene name</td><td>gene id</td></tr>
<tr><td>GU305914.1</td><td>16368</td><td>16390</td><td>-</td><td>0</td><td>GAGTCCGTGGCCCGTGGGGT</td><td>AGG</td><td>||||||||[||||||||||||]PAM</td><td>NA</td><td>Intergenic</td><td>NA</td><td>NA</td></tr>
<tr><td>T33337</td><td>ATAACAGCCACATATACCAACGG</td><td>1000</td><td></td><td></td><td></td><td></td><td></td><td></td><td></td><td></td><td></td></tr>
<tr><td>Oligo fwd</td><td>CACCATAACAGCCACATATACCAA</td><td></td><td></td><td></td><td></td><td></td><td></td><td></td><td></td><td></td><td></td></tr>
<tr><td>Oligo rev</td><td>AAACTTGGTATATGTGGCTGTTAT</td><td></td><td></td><td></td><td></td><td></td><td></td><td></td><td></td><td></td><td></td></tr>
<tr><td>Chromosome</td><td>start</td><td>end</td><td>strand</td><td>MM</td><td>target_seq</td><td>PAM</td><td>alignment</td><td>distance</td><td>position</td><td>gene name</td><td>gene id</td></tr>
<tr><td>GU305914.1</td><td>16325</td><td>16347</td><td>-</td><td>0</td><td>ATAACAGCCACATATACCAA</td><td>CGG</td><td>||||||||[||||||||||||]PAM</td><td>NA</td><td>Intergenic</td><td>NA</td><td>NA</td></tr>
<tr><td>T33338</td><td>ACAGCCACATATACCAACGGTGG</td><td>1000</td><td></td><td></td><td></td><td></td><td></td><td></td><td></td><td></td><td></td></tr>
<tr><td>Oligo fwd</td><td>CACCACAGCCACATATACCAACGG</td><td></td><td></td><td></td><td></td><td></td><td></td><td></td><td></td><td></td><td></td></tr>
<tr><td>Oligo rev</td><td>AAACCCGTTGGTATATGTGGCTGT</td><td></td><td></td><td></td><td></td><td></td><td></td><td></td><td></td><td></td><td></td></tr>
<tr><td>Chromosome</td><td>start</td><td>end</td><td>strand</td><td>MM</td><td>target_seq</td><td>PAM</td><td>alignment</td><td>distance</td><td>position</td><td>gene name</td><td>gene id</td></tr>
<tr><td>GU305914.1</td><td>16322</td><td>16344</td><td>-</td><td>0</td><td>ACAGCCACATATACCAACGG</td><td>TGG</td><td>||||||||[||||||||||||]PAM</td><td>NA</td><td>Intergenic</td><td>NA</td><td>NA</td></tr>
<tr><td>T33339</td><td>CCACATATACCAACGGTGGAAGG</td><td>1000</td><td></td><td></td><td></td><td></td><td></td><td></td><td></td><td></td><td></td></tr>
<tr><td>Oligo fwd</td><td>CACCCCACATATACCAACGGTGGA</td><td></td><td></td><td></td><td></td><td></td><td></td><td></td><td></td><td></td><td></td></tr>
<tr><td>Oligo rev</td><td>AAACTCCACCGTTGGTATATGTGG</td><td></td><td></td><td></td><td></td><td></td><td></td><td></td><td></td><td></td><td></td></tr>
<tr><td>Chromosome</td><td>start</td><td>end</td><td>strand</td><td>MM</td><td>target_seq</td><td>PAM</td><td>alignment</td><td>distance</td><td>position</td><td>gene name</td><td>gene id</td></tr>
<tr><td>GU305914.1</td><td>16318</td><td>16340</td><td>-</td><td>0</td><td>CCACATATACCAACGGTGGA</td><td>AGG</td><td>||||||||[||||||||||||]PAM</td><td>NA</td><td>Intergenic</td><td>NA</td><td>NA</td></tr>
<tr><td>T33340</td><td>CATATACCAACGGTGGAAGGAGG</td><td>1000</td><td></td><td></td><td></td><td></td><td></td><td></td><td></td><td></td><td></td></tr>
<tr><td>Oligo fwd</td><td>CACCCATATACCAACGGTGGAAGG</td><td></td><td></td><td></td><td></td><td></td><td></td><td></td><td></td><td></td><td></td></tr>
<tr><td>Oligo rev</td><td>AAACCCTTCCACCGTTGGTATATG</td><td></td><td></td><td></td><td></td><td></td><td></td><td></td><td></td><td></td><td></td></tr>
<tr><td>Chromosome</td><td>start</td><td>end</td><td>strand</td><td>MM</td><td>target_seq</td><td>PAM</td><td>alignment</td><td>distance</td><td>position</td><td>gene name</td><td>gene id</td></tr>
<tr><td>GU305914.1</td><td>16315</td><td>16337</td><td>-</td><td>0</td><td>CATATACCAACGGTGGAAGG</td><td>AGG</td><td>||||||||[||||||||||||]PAM</td><td>NA</td><td>Intergenic</td><td>NA</td><td>NA</td></tr>
<tr><td>T33341</td><td>GGTGGAAGGAGGTCTGTGTGAGG</td><td>1000</td><td></td><td></td><td></td><td></td><td></td><td></td><td></td><td></td><td></td></tr>
<tr><td>Oligo fwd</td><td>CACCGGTGGAAGGAGGTCTGTGTG</td><td></td><td></td><td></td><td></td><td></td><td></td><td></td><td></td><td></td><td></td></tr>
<tr><td>Oligo rev</td><td>AAACCACACAGACCTCCTTCCACC</td><td></td><td></td><td></td><td></td><td></td><td></td><td></td><td></td><td></td><td></td></tr>
<tr><td>Chromosome</td><td>start</td><td>end</td><td>strand</td><td>MM</td><td>target_seq</td><td>PAM</td><td>alignment</td><td>distance</td><td>position</td><td>gene name</td><td>gene id</td></tr>
<tr><td>GU305914.1</td><td>16304</td><td>16326</td><td>-</td><td>0</td><td>GGTGGAAGGAGGTCTGTGTG</td><td>AGG</td><td>||||||||[||||||||||||]PAM</td><td>NA</td><td>Intergenic</td><td>NA</td><td>NA</td></tr>
<tr><td>T33342</td><td>GTGGAAGGAGGTCTGTGTGAGGG</td><td>1000</td><td></td><td></td><td></td><td></td><td></td><td></td><td></td><td></td><td></td></tr>
<tr><td>Oligo fwd</td><td>CACCGTGGAAGGAGGTCTGTGTGA</td><td></td><td></td><td></td><td></td><td></td><td></td><td></td><td></td><td></td><td></td></tr>
<tr><td>Oligo rev</td><td>AAACTCACACAGACCTCCTTCCAC</td><td></td><td></td><td></td><td></td><td></td><td></td><td></td><td></td><td></td><td></td></tr>
<tr><td>Chromosome</td><td>start</td><td>end</td><td>strand</td><td>MM</td><td>target_seq</td><td>PAM</td><td>alignment</td><td>distance</td><td>position</td><td>gene name</td><td>gene id</td></tr>
<tr><td>GU305914.1</td><td>16303</td><td>16325</td><td>-</td><td>0</td><td>GTGGAAGGAGGTCTGTGTGA</td><td>GGG</td><td>||||||||[||||||||||||]PAM</td><td>NA</td><td>Intergenic</td><td>NA</td><td>NA</td></tr>
<tr><td>T33343</td><td>GAAGGAGGTCTGTGTGAGGGTGG</td><td>1000</td><td></td><td></td><td></td><td></td><td></td><td></td><td></td><td></td><td></td></tr>
<tr><td>Oligo fwd</td><td>CACCGAAGGAGGTCTGTGTGAGGG</td><td></td><td></td><td></td><td></td><td></td><td></td><td></td><td></td><td></td><td></td></tr>
<tr><td>Oligo rev</td><td>AAACCCCTCACACAGACCTCCTTC</td><td></td><td></td><td></td><td></td><td></td><td></td><td></td><td></td><td></td><td></td></tr>
<tr><td>Chromosome</td><td>start</td><td>end</td><td>strand</td><td>MM</td><td>target_seq</td><td>PAM</td><td>alignment</td><td>distance</td><td>position</td><td>gene name</td><td>gene id</td></tr>
<tr><td>GU305914.1</td><td>16300</td><td>16322</td><td>-</td><td>0</td><td>GAAGGAGGTCTGTGTGAGGG</td><td>TGG</td><td>||||||||[||||||||||||]PAM</td><td>NA</td><td>Intergenic</td><td>NA</td><td>NA</td></tr>
<tr><td>T33344</td><td>GGGTGGCGCACGCCTCGATGCGG</td><td>1000</td><td></td><td></td><td></td><td></td><td></td><td></td><td></td><td></td><td></td></tr>
<tr><td>Oligo fwd</td><td>CACCGGGTGGCGCACGCCTCGATG</td><td></td><td></td><td></td><td></td><td></td><td></td><td></td><td></td><td></td><td></td></tr>
<tr><td>Oligo rev</td><td>AAACCATCGAGGCGTGCGCCACCC</td><td></td><td></td><td></td><td></td><td></td><td></td><td></td><td></td><td></td><td></td></tr>
<tr><td>Chromosome</td><td>start</td><td>end</td><td>strand</td><td>MM</td><td>target_seq</td><td>PAM</td><td>alignment</td><td>distance</td><td>position</td><td>gene name</td><td>gene id</td></tr>
<tr><td>GU305914.1</td><td>16283</td><td>16305</td><td>-</td><td>0</td><td>GGGTGGCGCACGCCTCGATG</td><td>CGG</td><td>||||||||[||||||||||||]PAM</td><td>NA</td><td>Intergenic</td><td>NA</td><td>NA</td></tr>
<tr><td>T33345</td><td>TTCTCATACGAGAGATCGAACGG</td><td>1000</td><td></td><td></td><td></td><td></td><td></td><td></td><td></td><td></td><td></td></tr>
<tr><td>Oligo fwd</td><td>CACCTTCTCATACGAGAGATCGAA</td><td></td><td></td><td></td><td></td><td></td><td></td><td></td><td></td><td></td><td></td></tr>
<tr><td>Oligo rev</td><td>AAACTTCGATCTCTCGTATGAGAA</td><td></td><td></td><td></td><td></td><td></td><td></td><td></td><td></td><td></td><td></td></tr>
</table>

Chromosome start end strand MM target_seq PAM alignment distance position gene name gene id
GU305914.1 16248 16270 - 0 TTCTCATACGAGAGATCGAA CGG ||||||||[||||||||||||]PAM NA Intergenic NA NA

T33346 CGAACGGTACCGTCAGATGTCGG 1000
Oligo fwd CACCCGAACGGTACCGTCAGATGT
Oligo rev AAACACATCTGACGGTACCGTTCG
Chromosome start end strand MM target_seq PAM alignment distance position gene name gene id
GU305914.1 16232 16254 - 0 CGAACGGTACCGTCAGATGT CGG ||||||||[||||||||||||]PAM NA Intergenic NA NA

T33347 ACCATCACCCGCTAGATTCATGG 1000
Oligo fwd CACCACCATCACCCGCTAGATTCA
Oligo rev AAACTGAATCTAGCGGGTGATGGT
Chromosome start end strand MM target_seq PAM alignment distance position gene name gene id
GU305914.1 16201 16223 - 0 ACCATCACCCGCTAGATTCA TGG ||||||||[||||||||||||]PAM NA Intergenic NA NA

T33348 ATCACCCGCTAGATTCATGGCGG 1000
Oligo fwd CACCATCACCCGCTAGATTCATGG
Oligo rev AAACCCATGAATCTAGCGGGTGAT
Chromosome start end strand MM target_seq PAM alignment distance position gene name gene id
GU305914.1 16198 16220 - 0 ATCACCCGCTAGATTCATGG CGG ||||||||[||||||||||||]PAM NA Intergenic NA NA

T33349 TCACCCGCTAGATTCATGGCGGG 1000
Oligo fwd CACCTCACCCGCTAGATTCATGGC
Oligo rev AAACGCCATGAATCTAGCGGGTGA
Chromosome start end strand MM target_seq PAM alignment distance position gene name gene id
GU305914.1 16197 16219 - 0 TCACCCGCTAGATTCATGGC GGG ||||||||[||||||||||||]PAM NA Intergenic NA NA

T33350 GCGGGTATGAATACCCCTTACGG 1000
Oligo fwd CACCGCGGGTATGAATACCCCTTA
Oligo rev AAACTAAGGGGTATTCATACCCGC
Chromosome start end strand MM target_seq PAM alignment distance position gene name gene id
GU305914.1 16179 16201 - 0 GCGGGTATGAATACCCCTTA CGG ||||||||[||||||||||||]PAM NA Intergenic NA NA

T33351 ACCAGATACAACTACACCACCGG 1000
Oligo fwd CACCACCAGATACAACTACACCAC
Oligo rev AAACGTGGTGTAGTTGTATCTGGT
Chromosome start end strand MM target_seq PAM alignment distance position gene name gene id
GU305914.1 16149 16171 - 0 ACCAGATACAACTACACCAC CGG ||||||||[||||||||||||]PAM NA Intergenic NA NA

T33352 CACCACCGGCGTGTCGAGCACGG 1000
Oligo fwd CACCCACCACCGGCGTGTCGAGCA
Oligo rev AAACTGCTCGACACGCCGGTGGTG
Chromosome start end strand MM target_seq PAM alignment distance position gene name gene id
GU305914.1 16135 16157 - 0 CACCACCGGCGTGTCGAGCA CGG ||||||||[||||||||||||]PAM NA Intergenic NA NA

T33353 CGTGTCGAGCACGGTCAACGTGG 1000
Oligo fwd CACCCGTGTCGAGCACGGTCAACG
Oligo rev AAACCGTTGACCGTGCTCGACACG
Chromosome start end strand MM target_seq PAM alignment distance position gene name gene id
GU305914.1 16126 16148 - 0 CGTGTCGAGCACGGTCAACG TGG ||||||||[||||||||||||]PAM NA Intergenic NA NA

T33354 TGCGATTTTCAGACCCTGTTCGG 1000
Oligo fwd CACCTGCGATTTTCAGACCCTGTT
Oligo rev AAACAACAGGGTCTGAAAATCGCA
Chromosome start end strand MM target_seq PAM alignment distance position gene name gene id
GU305914.1 16077 16099 - 0 TGCGATTTTCAGACCCTGTT CGG ||||||||[||||||||||||]PAM NA Intergenic NA NA

T33355 TTCAGACCCTGTTCGGCCGCTGG 1000
Oligo fwd CACCTTCAGACCCTGTTCGGCCGC
Oligo rev AAACGCGGCCGAACAGGGTCTGAA
Chromosome start end strand MM target_seq PAM alignment distance position gene name gene id
GU305914.1 16070 16092 - 0 TTCAGACCCTGTTCGGCCGC TGG ||||||||[||||||||||||]PAM NA Intergenic NA NA

T33356 AGCTATGAACTGCCAACACCCGG 1000
Oligo fwd CACCAGCTATGAACTGCCAACACC
Oligo rev AAACGGTGTTGGCAGTTCATAGCT
Chromosome start end strand MM target_seq PAM alignment distance position gene name gene id
GU305914.1 16047 16069 - 0 AGCTATGAACTGCCAACACC CGG ||||||||[||||||||||||]PAM NA Intergenic NA NA

T33357 CCAACACCCGGCCATGTGCGTGG 1000
Oligo fwd CACCCCAACACCCGGCCATGTGCG
Oligo rev AAACCGCACATGGCCGGGTGTTGG
Chromosome start end strand MM target_seq PAM alignment distance position gene name gene id
GU305914.1 16035 16057 - 0 CCAACACCCGGCCATGTGCG TGG ||||||||[||||||||||||]PAM NA Intergenic NA NA

T33358 ACGTCGAGATCGACGTCCGAAGG 1000
Oligo fwd CACCACGTCGAGATCGACGTCCGA
Oligo rev AAACTCGGACGTCGATCTCGACGT
Chromosome start end strand MM target_seq PAM alignment distance position gene name gene id
GU305914.1 15998 16020 - 0 ACGTCGAGATCGACGTCCGA AGG ||||||||[||||||||||||]PAM NA Intergenic NA NA
GU305914.1 135773 135795 + 4 CCGTCGACGTCGACGTCGGA CGG -||||||-[-||||||||-||]PAM NA Intergenic NA NA

T33359 AAAACAAAACGCTGCGCGAGCGG 1000
Oligo fwd CACCAAAACAAAACGCTGCGCGAG
Oligo rev AAACCTCGCGCAGCGTTTTGTTTT
Chromosome start end strand MM target_seq PAM alignment distance position gene name gene id
GU305914.1 15957 15979 - 0 AAAACAAAACGCTGCGCGAG CGG ||||||||[||||||||||||]PAM NA Intergenic NA NA

T33360 GGACAGTTGTGTGTGCACCTCGG 1000

<table>
<tr><td>Oligo fwd</td><td>CACCGGACAGTTGTGTGTGCACCT</td><td></td><td></td><td></td><td></td><td></td><td></td><td></td><td></td><td></td><td></td><td></td></tr>
<tr><td>Oligo rev</td><td>AAACAGGTGCACACACAACTGTCC</td><td></td><td></td><td></td><td></td><td></td><td></td><td></td><td></td><td></td><td></td><td></td></tr>
<tr><td>Chromosome</td><td>start</td><td></td><td>end</td><td>strand</td><td>MM</td><td>target_seq</td><td>PAM</td><td>alignment</td><td>distance</td><td>position</td><td>gene name</td><td>gene id</td></tr>
<tr><td>GU305914.1</td><td></td><td>15936</td><td>15958</td><td>-</td><td>0</td><td>GGACAGTTGTGTGTGCACCT</td><td>CGG</td><td>||||||||[||||||||||||]PAM</td><td>NA</td><td>Intergenic</td><td>NA</td><td>NA</td></tr>
<tr><td></td><td></td><td></td><td></td><td></td><td></td><td></td><td></td><td></td><td></td><td></td><td></td><td></td></tr>
<tr><td>T33361</td><td>TTGATCCTGAACTTATGCATAGG</td><td></td><td>1000</td><td></td><td></td><td></td><td></td><td></td><td></td><td></td><td></td><td></td></tr>
<tr><td>Oligo fwd</td><td>CACCTTGATCCTGAACTTATGCAT</td><td></td><td></td><td></td><td></td><td></td><td></td><td></td><td></td><td></td><td></td><td></td></tr>
<tr><td>Oligo rev</td><td>AAACATGCATAAGTTCAGGATCAA</td><td></td><td></td><td></td><td></td><td></td><td></td><td></td><td></td><td></td><td></td><td></td></tr>
<tr><td>Chromosome</td><td>start</td><td></td><td>end</td><td>strand</td><td>MM</td><td>target_seq</td><td>PAM</td><td>alignment</td><td>distance</td><td>position</td><td>gene name</td><td>gene id</td></tr>
<tr><td>GU305914.1</td><td></td><td>15864</td><td>15886</td><td>-</td><td>0</td><td>TTGATCCTGAACTTATGCAT</td><td>AGG</td><td>||||||||[||||||||||||]PAM</td><td>NA</td><td>Intergenic</td><td>NA</td><td>NA</td></tr>
<tr><td></td><td></td><td></td><td></td><td></td><td></td><td></td><td></td><td></td><td></td><td></td><td></td><td></td></tr>
<tr><td>T33362</td><td>TGACAGATTGATGTATCACAAGG</td><td></td><td>1000</td><td></td><td></td><td></td><td></td><td></td><td></td><td></td><td></td><td></td></tr>
<tr><td>Oligo fwd</td><td>CACCTGACAGATTGATGTATCACA</td><td></td><td></td><td></td><td></td><td></td><td></td><td></td><td></td><td></td><td></td><td></td></tr>
<tr><td>Oligo rev</td><td>AAACTGTGATACATCAATCTGTCA</td><td></td><td></td><td></td><td></td><td></td><td></td><td></td><td></td><td></td><td></td><td></td></tr>
<tr><td>Chromosome</td><td>start</td><td></td><td>end</td><td>strand</td><td>MM</td><td>target_seq</td><td>PAM</td><td>alignment</td><td>distance</td><td>position</td><td>gene name</td><td>gene id</td></tr>
<tr><td>GU305914.1</td><td></td><td>15823</td><td>15845</td><td>-</td><td>0</td><td>TGACAGATTGATGTATCACA</td><td>AGG</td><td>||||||||[||||||||||||]PAM</td><td>NA</td><td>Intergenic</td><td>NA</td><td>NA</td></tr>
<tr><td></td><td></td><td></td><td></td><td></td><td></td><td></td><td></td><td></td><td></td><td></td><td></td><td></td></tr>
<tr><td>T33363</td><td>AGATTGATGTATCACAAGGAAGG</td><td></td><td>1000</td><td></td><td></td><td></td><td></td><td></td><td></td><td></td><td></td><td></td></tr>
<tr><td>Oligo fwd</td><td>CACCAGATTGATGTATCACAAGGA</td><td></td><td></td><td></td><td></td><td></td><td></td><td></td><td></td><td></td><td></td><td></td></tr>
<tr><td>Oligo rev</td><td>AAACTCCTTGTGATACATCAATCT</td><td></td><td></td><td></td><td></td><td></td><td></td><td></td><td></td><td></td><td></td><td></td></tr>
<tr><td>Chromosome</td><td>start</td><td></td><td>end</td><td>strand</td><td>MM</td><td>target_seq</td><td>PAM</td><td>alignment</td><td>distance</td><td>position</td><td>gene name</td><td>gene id</td></tr>
<tr><td>GU305914.1</td><td></td><td>15819</td><td>15841</td><td>-</td><td>0</td><td>AGATTGATGTATCACAAGGA</td><td>AGG</td><td>||||||||[||||||||||||]PAM</td><td>NA</td><td>Intergenic</td><td>NA</td><td>NA</td></tr>
<tr><td></td><td></td><td></td><td></td><td></td><td></td><td></td><td></td><td></td><td></td><td></td><td></td><td></td></tr>
<tr><td>T33364</td><td>GATTGATGTATCACAAGGAAGGG</td><td></td><td>1000</td><td></td><td></td><td></td><td></td><td></td><td></td><td></td><td></td><td></td></tr>
<tr><td>Oligo fwd</td><td>CACCGATTGATGTATCACAAGGAA</td><td></td><td></td><td></td><td></td><td></td><td></td><td></td><td></td><td></td><td></td><td></td></tr>
<tr><td>Oligo rev</td><td>AAACTTCCTTGTGATACATCAATC</td><td></td><td></td><td></td><td></td><td></td><td></td><td></td><td></td><td></td><td></td><td></td></tr>
<tr><td>Chromosome</td><td>start</td><td></td><td>end</td><td>strand</td><td>MM</td><td>target_seq</td><td>PAM</td><td>alignment</td><td>distance</td><td>position</td><td>gene name</td><td>gene id</td></tr>
<tr><td>GU305914.1</td><td></td><td>15818</td><td>15840</td><td>-</td><td>0</td><td>GATTGATGTATCACAAGGAA</td><td>GGG</td><td>||||||||[||||||||||||]PAM</td><td>NA</td><td>Intergenic</td><td>NA</td><td>NA</td></tr>
<tr><td></td><td></td><td></td><td></td><td></td><td></td><td></td><td></td><td></td><td></td><td></td><td></td><td></td></tr>
<tr><td>T33365</td><td>ATATCCCGCTCACGTTACAGAGG</td><td></td><td>1000</td><td></td><td></td><td></td><td></td><td></td><td></td><td></td><td></td><td></td></tr>
<tr><td>Oligo fwd</td><td>CACCATATCCCGCTCACGTTACAG</td><td></td><td></td><td></td><td></td><td></td><td></td><td></td><td></td><td></td><td></td><td></td></tr>
<tr><td>Oligo rev</td><td>AAACCTGTAACGTGAGCGGGATAT</td><td></td><td></td><td></td><td></td><td></td><td></td><td></td><td></td><td></td><td></td><td></td></tr>
<tr><td>Chromosome</td><td>start</td><td></td><td>end</td><td>strand</td><td>MM</td><td>target_seq</td><td>PAM</td><td>alignment</td><td>distance</td><td>position</td><td>gene name</td><td>gene id</td></tr>
<tr><td>GU305914.1</td><td></td><td>15795</td><td>15817</td><td>-</td><td>0</td><td>ATATCCCGCTCACGTTACAG</td><td>AGG</td><td>||||||||[||||||||||||]PAM</td><td>NA</td><td>Intergenic</td><td>NA</td><td>NA</td></tr>
<tr><td></td><td></td><td></td><td></td><td></td><td></td><td></td><td></td><td></td><td></td><td></td><td></td><td></td></tr>
<tr><td>T33366</td><td>CGCTCACGTTACAGAGGAATAGG</td><td></td><td>1000</td><td></td><td></td><td></td><td></td><td></td><td></td><td></td><td></td><td></td></tr>
<tr><td>Oligo fwd</td><td>CACCCGCTCACGTTACAGAGGAAT</td><td></td><td></td><td></td><td></td><td></td><td></td><td></td><td></td><td></td><td></td><td></td></tr>
<tr><td>Oligo rev</td><td>AAACATTCCTCTGTAACGTGAGCG</td><td></td><td></td><td></td><td></td><td></td><td></td><td></td><td></td><td></td><td></td><td></td></tr>
<tr><td>Chromosome</td><td>start</td><td></td><td>end</td><td>strand</td><td>MM</td><td>target_seq</td><td>PAM</td><td>alignment</td><td>distance</td><td>position</td><td>gene name</td><td>gene id</td></tr>
<tr><td>GU305914.1</td><td></td><td>15789</td><td>15811</td><td>-</td><td>0</td><td>CGCTCACGTTACAGAGGAAT</td><td>AGG</td><td>||||||||[||||||||||||]PAM</td><td>NA</td><td>Intergenic</td><td>NA</td><td>NA</td></tr>
<tr><td></td><td></td><td></td><td></td><td></td><td></td><td></td><td></td><td></td><td></td><td></td><td></td><td></td></tr>
<tr><td>T33367</td><td>CTGAGCTCCCCTTATTTCGATGG</td><td></td><td>1000</td><td></td><td></td><td></td><td></td><td></td><td></td><td></td><td></td><td></td></tr>
<tr><td>Oligo fwd</td><td>CACCCTGAGCTCCCCTTATTTCGA</td><td></td><td></td><td></td><td></td><td></td><td></td><td></td><td></td><td></td><td></td><td></td></tr>
<tr><td>Oligo rev</td><td>AAACTCGAAATAAGGGGAGCTCAG</td><td></td><td></td><td></td><td></td><td></td><td></td><td></td><td></td><td></td><td></td><td></td></tr>
<tr><td>Chromosome</td><td>start</td><td></td><td>end</td><td>strand</td><td>MM</td><td>target_seq</td><td>PAM</td><td>alignment</td><td>distance</td><td>position</td><td>gene name</td><td>gene id</td></tr>
<tr><td>GU305914.1</td><td></td><td>15730</td><td>15752</td><td>-</td><td>0</td><td>CTGAGCTCCCCTTATTTCGA</td><td>TGG</td><td>||||||||[||||||||||||]PAM</td><td>NA</td><td>Intergenic</td><td>NA</td><td>NA</td></tr>
<tr><td></td><td></td><td></td><td></td><td></td><td></td><td></td><td></td><td></td><td></td><td></td><td></td><td></td></tr>
<tr><td>T33368</td><td>AAGATTTCAGACAAAAAGTATGG</td><td></td><td>1000</td><td></td><td></td><td></td><td></td><td></td><td></td><td></td><td></td><td></td></tr>
<tr><td>Oligo fwd</td><td>CACCAAGATTTCAGACAAAAAGTA</td><td></td><td></td><td></td><td></td><td></td><td></td><td></td><td></td><td></td><td></td><td></td></tr>
<tr><td>Oligo rev</td><td>AAACTACTTTTTGTCTGAAATCTT</td><td></td><td></td><td></td><td></td><td></td><td></td><td></td><td></td><td></td><td></td><td></td></tr>
<tr><td>Chromosome</td><td>start</td><td></td><td>end</td><td>strand</td><td>MM</td><td>target_seq</td><td>PAM</td><td>alignment</td><td>distance</td><td>position</td><td>gene name</td><td>gene id</td></tr>
<tr><td>GU305914.1</td><td></td><td>15698</td><td>15720</td><td>-</td><td>0</td><td>AAGATTTCAGACAAAAAGTA</td><td>TGG</td><td>||||||||[||||||||||||]PAM</td><td>NA</td><td>Intergenic</td><td>NA</td><td>NA</td></tr>
<tr><td></td><td></td><td></td><td></td><td></td><td></td><td></td><td></td><td></td><td></td><td></td><td></td><td></td></tr>
<tr><td>T33369</td><td>TTATAATTGTCATGATTAAGAGG</td><td></td><td>1000</td><td></td><td></td><td></td><td></td><td></td><td></td><td></td><td></td><td></td></tr>
<tr><td>Oligo fwd</td><td>CACCTTATAATTGTCATGATTAAG</td><td></td><td></td><td></td><td></td><td></td><td></td><td></td><td></td><td></td><td></td><td></td></tr>
<tr><td>Oligo rev</td><td>AAACCTTAATCATGACAATTATAA</td><td></td><td></td><td></td><td></td><td></td><td></td><td></td><td></td><td></td><td></td><td></td></tr>
<tr><td>Chromosome</td><td>start</td><td></td><td>end</td><td>strand</td><td>MM</td><td>target_seq</td><td>PAM</td><td>alignment</td><td>distance</td><td>position</td><td>gene name</td><td>gene id</td></tr>
<tr><td>GU305914.1</td><td></td><td>15625</td><td>15647</td><td>-</td><td>0</td><td>TTATAATTGTCATGATTAAG</td><td>AGG</td><td>||||||||[||||||||||||]PAM</td><td>NA</td><td>Intergenic</td><td>NA</td><td>NA</td></tr>
<tr><td></td><td></td><td></td><td></td><td></td><td></td><td></td><td></td><td></td><td></td><td></td><td></td><td></td></tr>
<tr><td>T33370</td><td>TATAATTGTCATGATTAAGAGGG</td><td></td><td>1000</td><td></td><td></td><td></td><td></td><td></td><td></td><td></td><td></td><td></td></tr>
<tr><td>Oligo fwd</td><td>CACCTATAATTGTCATGATTAAGA</td><td></td><td></td><td></td><td></td><td></td><td></td><td></td><td></td><td></td><td></td><td></td></tr>
<tr><td>Oligo rev</td><td>AAACTCTTAATCATGACAATTATA</td><td></td><td></td><td></td><td></td><td></td><td></td><td></td><td></td><td></td><td></td><td></td></tr>
<tr><td>Chromosome</td><td>start</td><td></td><td>end</td><td>strand</td><td>MM</td><td>target_seq</td><td>PAM</td><td>alignment</td><td>distance</td><td>position</td><td>gene name</td><td>gene id</td></tr>
<tr><td>GU305914.1</td><td></td><td>15624</td><td>15646</td><td>-</td><td>0</td><td>TATAATTGTCATGATTAAGA</td><td>GGG</td><td>||||||||[||||||||||||]PAM</td><td>NA</td><td>Intergenic</td><td>NA</td><td>NA</td></tr>
<tr><td></td><td></td><td></td><td></td><td></td><td></td><td></td><td></td><td></td><td></td><td></td><td></td><td></td></tr>
<tr><td>T33371</td><td>ATGATTAAGAGGGATATTATCGG</td><td></td><td>1000</td><td></td><td></td><td></td><td></td><td></td><td></td><td></td><td></td><td></td></tr>
<tr><td>Oligo fwd</td><td>CACCATGATTAAGAGGGATATTAT</td><td></td><td></td><td></td><td></td><td></td><td></td><td></td><td></td><td></td><td></td><td></td></tr>
<tr><td>Oligo rev</td><td>AAACATAATATCCCTCTTAATCAT</td><td></td><td></td><td></td><td></td><td></td><td></td><td></td><td></td><td></td><td></td><td></td></tr>
<tr><td>Chromosome</td><td>start</td><td></td><td>end</td><td>strand</td><td>MM</td><td>target_seq</td><td>PAM</td><td>alignment</td><td>distance</td><td>position</td><td>gene name</td><td>gene id</td></tr>
<tr><td>GU305914.1</td><td></td><td>15614</td><td>15636</td><td>-</td><td>0</td><td>ATGATTAAGAGGGATATTAT</td><td>CGG</td><td>||||||||[||||||||||||]PAM</td><td>NA</td><td>Intergenic</td><td>NA</td><td>NA</td></tr>
<tr><td></td><td></td><td></td><td></td><td></td><td></td><td></td><td></td><td></td><td></td><td></td><td></td><td></td></tr>
<tr><td>T33372</td><td>ATTATCGGATTAGATAACTTCGG</td><td></td><td>1000</td><td></td><td></td><td></td><td></td><td></td><td></td><td></td><td></td><td></td></tr>
<tr><td>Oligo fwd</td><td>CACCATTATCGGATTAGATAACTT</td><td></td><td></td><td></td><td></td><td></td><td></td><td></td><td></td><td></td><td></td><td></td></tr>
<tr><td>Oligo rev</td><td>AAACAAGTTATCTAATCCGATAAT</td><td></td><td></td><td></td><td></td><td></td><td></td><td></td><td></td><td></td><td></td><td></td></tr>
<tr><td>Chromosome</td><td>start</td><td></td><td>end</td><td>strand</td><td>MM</td><td>target_seq</td><td>PAM</td><td>alignment</td><td>distance</td><td>position</td><td>gene name</td><td>gene id</td></tr>
<tr><td>GU305914.1</td><td></td><td>15599</td><td>15621</td><td>-</td><td>0</td><td>ATTATCGGATTAGATAACTT</td><td>CGG</td><td>||||||||[||||||||||||]PAM</td><td>NA</td><td>Intergenic</td><td>NA</td><td>NA</td></tr>
<tr><td></td><td></td><td></td><td></td><td></td><td></td><td></td><td></td><td></td><td></td><td></td><td></td><td></td></tr>
<tr><td>T33373</td><td>ATTAGATAACTTCGGAATCGTGG</td><td></td><td>1000</td><td></td><td></td><td></td><td></td><td></td><td></td><td></td><td></td><td></td></tr>
<tr><td>Oligo fwd</td><td>CACCATTAGATAACTTCGGAATCG</td><td></td><td></td><td></td><td></td><td></td><td></td><td></td><td></td><td></td><td></td><td></td></tr>
<tr><td>Oligo rev</td><td>AAACCGATTCCGAAGTTATCTAAT</td><td></td><td></td><td></td><td></td><td></td><td></td><td></td><td></td><td></td><td></td><td></td></tr>
<tr><td>Chromosome</td><td>start</td><td></td><td>end</td><td>strand</td><td>MM</td><td>target_seq</td><td>PAM</td><td>alignment</td><td>distance</td><td>position</td><td>gene name</td><td>gene id</td></tr>
<tr><td>GU305914.1</td><td></td><td>15591</td><td>15613</td><td>-</td><td>0</td><td>ATTAGATAACTTCGGAATCG</td><td>TGG</td><td>||||||||[||||||||||||]PAM</td><td>NA</td><td>Intergenic</td><td>NA</td><td>NA</td></tr>
<tr><td></td><td></td><td></td><td></td><td></td><td></td><td></td><td></td><td></td><td></td><td></td><td></td><td></td></tr>
<tr><td>T33374</td><td>GCACTCGATCTTAATAGCGTAGG</td><td></td><td>1000</td><td></td><td></td><td></td><td></td><td></td><td></td><td></td><td></td><td></td></tr>
<tr><td>Oligo fwd</td><td>CACCGCACTCGATCTTAATAGCGT</td><td></td><td></td><td></td><td></td><td></td><td></td><td></td><td></td><td></td><td></td><td></td></tr>
<tr><td>Oligo rev</td><td>AAACACGCTATTAAGATCGAGTGC</td><td></td><td></td><td></td><td></td><td></td><td></td><td></td><td></td><td></td><td></td><td></td></tr>
<tr><td>Chromosome</td><td>start</td><td></td><td>end</td><td>strand</td><td>MM</td><td>target_seq</td><td>PAM</td><td>alignment</td><td>distance</td><td>position</td><td>gene name</td><td>gene id</td></tr>
<tr><td>GU305914.1</td><td></td><td>15569</td><td>15591</td><td>-</td><td>0</td><td>GCACTCGATCTTAATAGCGT</td><td>AGG</td><td>||||||||[||||||||||||]PAM</td><td>NA</td><td>Intergenic</td><td>NA</td><td>NA</td></tr>
</table>

<table>
<tr><td>T33375</td><td>CTCGATCTTAATAGCGTAGGCGG</td><td>1000</td></tr>
<tr><td>Oligo fwd</td><td>CACCCTCGATCTTAATAGCGTAGG</td></tr>
<tr><td>Oligo rev</td><td>AAACCCTACGCTATTAAGATCGAG</td></tr>
<tr><td>Chromosome</td><td>start</td><td>end</td><td>strand</td><td>MM</td><td>target_seq</td><td>PAM</td><td>alignment</td><td>distance</td><td>position</td><td>gene name</td><td>gene id</td></tr>
<tr><td>GU305914.1</td><td>15566</td><td>15588</td><td>-</td><td>0</td><td>CTCGATCTTAATAGCGTAGG</td><td>CGG</td><td>||||||||[||||||||||||]PAM</td><td>NA</td><td>Intergenic</td><td>NA</td><td>NA</td></tr>
<tr><td>T33376</td><td>TCGATCTTAATAGCGTAGGCGGG</td><td>1000</td></tr>
<tr><td>Oligo fwd</td><td>CACCTCGATCTTAATAGCGTAGGC</td></tr>
<tr><td>Oligo rev</td><td>AAACGCCTACGCTATTAAGATCGA</td></tr>
<tr><td>Chromosome</td><td>start</td><td>end</td><td>strand</td><td>MM</td><td>target_seq</td><td>PAM</td><td>alignment</td><td>distance</td><td>position</td><td>gene name</td><td>gene id</td></tr>
<tr><td>GU305914.1</td><td>15565</td><td>15587</td><td>-</td><td>0</td><td>TCGATCTTAATAGCGTAGGC</td><td>GGG</td><td>||||||||[||||||||||||]PAM</td><td>NA</td><td>Intergenic</td><td>NA</td><td>NA</td></tr>
<tr><td>T33377</td><td>CTTAATAGCGTAGGCGGGTGCGG</td><td>1000</td></tr>
<tr><td>Oligo fwd</td><td>CACCCTTAATAGCGTAGGCGGGTG</td></tr>
<tr><td>Oligo rev</td><td>AAACCACCCGCCTACGCTATTAAG</td></tr>
<tr><td>Chromosome</td><td>start</td><td>end</td><td>strand</td><td>MM</td><td>target_seq</td><td>PAM</td><td>alignment</td><td>distance</td><td>position</td><td>gene name</td><td>gene id</td></tr>
<tr><td>GU305914.1</td><td>15560</td><td>15582</td><td>-</td><td>0</td><td>CTTAATAGCGTAGGCGGGTG</td><td>CGG</td><td>||||||||[||||||||||||]PAM</td><td>NA</td><td>Intergenic</td><td>NA</td><td>NA</td></tr>
<tr><td>T33378</td><td>TTAATAGCGTAGGCGGGTGCGGG</td><td>1000</td></tr>
<tr><td>Oligo fwd</td><td>CACCTTAATAGCGTAGGCGGGTGC</td></tr>
<tr><td>Oligo rev</td><td>AAACGCACCCGCCTACGCTATTAA</td></tr>
<tr><td>Chromosome</td><td>start</td><td>end</td><td>strand</td><td>MM</td><td>target_seq</td><td>PAM</td><td>alignment</td><td>distance</td><td>position</td><td>gene name</td><td>gene id</td></tr>
<tr><td>GU305914.1</td><td>15559</td><td>15581</td><td>-</td><td>0</td><td>TTAATAGCGTAGGCGGGTGC</td><td>GGG</td><td>||||||||[||||||||||||]PAM</td><td>NA</td><td>Intergenic</td><td>NA</td><td>NA</td></tr>
<tr><td>T33379</td><td>TAGCGTAGGCGGGTGCGGGACGG</td><td>1000</td></tr>
<tr><td>Oligo fwd</td><td>CACCTAGCGTAGGCGGGTGCGGGA</td></tr>
<tr><td>Oligo rev</td><td>AAACTCCCGCACCCGCCTACGCTA</td></tr>
<tr><td>Chromosome</td><td>start</td><td>end</td><td>strand</td><td>MM</td><td>target_seq</td><td>PAM</td><td>alignment</td><td>distance</td><td>position</td><td>gene name</td><td>gene id</td></tr>
<tr><td>GU305914.1</td><td>15555</td><td>15577</td><td>-</td><td>0</td><td>TAGCGTAGGCGGGTGCGGGA</td><td>CGG</td><td>||||||||[||||||||||||]PAM</td><td>NA</td><td>Intergenic</td><td>NA</td><td>NA</td></tr>
<tr><td>T33380</td><td>GTAGGCGGGTGCGGGACGGTCGG</td><td>1000</td></tr>
<tr><td>Oligo fwd</td><td>CACCGTAGGCGGGTGCGGGACGGT</td></tr>
<tr><td>Oligo rev</td><td>AAACACCGTCCCGCACCCGCCTAC</td></tr>
<tr><td>Chromosome</td><td>start</td><td>end</td><td>strand</td><td>MM</td><td>target_seq</td><td>PAM</td><td>alignment</td><td>distance</td><td>position</td><td>gene name</td><td>gene id</td></tr>
<tr><td>GU305914.1</td><td>15551</td><td>15573</td><td>-</td><td>0</td><td>GTAGGCGGGTGCGGGACGGT</td><td>CGG</td><td>||||||||[||||||||||||]PAM</td><td>NA</td><td>Intergenic</td><td>NA</td><td>NA</td></tr>
<tr><td>T33381</td><td>GGCGGGTGCGGGACGGTCGGTGG</td><td>1000</td></tr>
<tr><td>Oligo fwd</td><td>CACCGGCGGGTGCGGGACGGTCGG</td></tr>
<tr><td>Oligo rev</td><td>AAACCCGACCGTCCCGCACCCGCC</td></tr>
<tr><td>Chromosome</td><td>start</td><td>end</td><td>strand</td><td>MM</td><td>target_seq</td><td>PAM</td><td>alignment</td><td>distance</td><td>position</td><td>gene name</td><td>gene id</td></tr>
<tr><td>GU305914.1</td><td>15548</td><td>15570</td><td>-</td><td>0</td><td>GGCGGGTGCGGGACGGTCGG</td><td>TGG</td><td>||||||||[||||||||||||]PAM</td><td>NA</td><td>Intergenic</td><td>NA</td><td>NA</td></tr>
<tr><td>T33382</td><td>TGCGGGACGGTCGGTGGAAACGG</td><td>1000</td></tr>
<tr><td>Oligo fwd</td><td>CACCTGCGGGACGGTCGGTGGAAA</td></tr>
<tr><td>Oligo rev</td><td>AAACTTTCCACCGACCGTCCCGCA</td></tr>
<tr><td>Chromosome</td><td>start</td><td>end</td><td>strand</td><td>MM</td><td>target_seq</td><td>PAM</td><td>alignment</td><td>distance</td><td>position</td><td>gene name</td><td>gene id</td></tr>
<tr><td>GU305914.1</td><td>15542</td><td>15564</td><td>-</td><td>0</td><td>TGCGGGACGGTCGGTGGAAA</td><td>CGG</td><td>||||||||[||||||||||||]PAM</td><td>NA</td><td>Intergenic</td><td>NA</td><td>NA</td></tr>
<tr><td>T33383</td><td>CGGTGGAAACGGTGTTGTTGAGG</td><td>1000</td></tr>
<tr><td>Oligo fwd</td><td>CACCCGGTGGAAACGGTGTTGTTG</td></tr>
<tr><td>Oligo rev</td><td>AAACCAACAACACCGTTTCCACCG</td></tr>
<tr><td>Chromosome</td><td>start</td><td>end</td><td>strand</td><td>MM</td><td>target_seq</td><td>PAM</td><td>alignment</td><td>distance</td><td>position</td><td>gene name</td><td>gene id</td></tr>
<tr><td>GU305914.1</td><td>15531</td><td>15553</td><td>-</td><td>0</td><td>CGGTGGAAACGGTGTTGTTG</td><td>AGG</td><td>||||||||[||||||||||||]PAM</td><td>NA</td><td>Intergenic</td><td>NA</td><td>NA</td></tr>
<tr><td>T33384</td><td>CGGTGTTGTTGAGGATCGACAGG</td><td>1000</td></tr>
<tr><td>Oligo fwd</td><td>CACCCGGTGTTGTTGAGGATCGAC</td></tr>
<tr><td>Oligo rev</td><td>AAACGTCGATCCTCAACAACACCG</td></tr>
<tr><td>Chromosome</td><td>start</td><td>end</td><td>strand</td><td>MM</td><td>target_seq</td><td>PAM</td><td>alignment</td><td>distance</td><td>position</td><td>gene name</td><td>gene id</td></tr>
<tr><td>GU305914.1</td><td>15522</td><td>15544</td><td>-</td><td>0</td><td>CGGTGTTGTTGAGGATCGAC</td><td>AGG</td><td>||||||||[||||||||||||]PAM</td><td>NA</td><td>Intergenic</td><td>NA</td><td>NA</td></tr>
<tr><td>T33385</td><td>TATCCACTGCTCGAACATGAAGG</td><td>1000</td></tr>
<tr><td>Oligo fwd</td><td>CACCTATCCACTGCTCGAACATGA</td></tr>
<tr><td>Oligo rev</td><td>AAACTCATGTTCGAGCAGTGGATA</td></tr>
<tr><td>Chromosome</td><td>start</td><td>end</td><td>strand</td><td>MM</td><td>target_seq</td><td>PAM</td><td>alignment</td><td>distance</td><td>position</td><td>gene name</td><td>gene id</td></tr>
<tr><td>GU305914.1</td><td>15476</td><td>15498</td><td>-</td><td>0</td><td>TATCCACTGCTCGAACATGA</td><td>AGG</td><td>||||||||[||||||||||||]PAM</td><td>NA</td><td>Intergenic</td><td>NA</td><td>NA</td></tr>
<tr><td>T33386</td><td>TCGAACATGAAGGAGATACATGG</td><td>1000</td></tr>
<tr><td>Oligo fwd</td><td>CACCTCGAACATGAAGGAGATACA</td></tr>
<tr><td>Oligo rev</td><td>AAACTGTATCTCCTTCATGTTCGA</td></tr>
<tr><td>Chromosome</td><td>start</td><td>end</td><td>strand</td><td>MM</td><td>target_seq</td><td>PAM</td><td>alignment</td><td>distance</td><td>position</td><td>gene name</td><td>gene id</td></tr>
<tr><td>GU305914.1</td><td>15466</td><td>15488</td><td>-</td><td>0</td><td>TCGAACATGAAGGAGATACA</td><td>TGG</td><td>||||||||[||||||||||||]PAM</td><td>NA</td><td>Intergenic</td><td>NA</td><td>NA</td></tr>
<tr><td>T33387</td><td>AGGAGATACATGGACTTCTCCGG</td><td>1000</td></tr>
<tr><td>Oligo fwd</td><td>CACCAGGAGATACATGGACTTCTC</td></tr>
<tr><td>Oligo rev</td><td>AAACGAGAAGTCCATGTATCTCCT</td></tr>
<tr><td>Chromosome</td><td>start</td><td>end</td><td>strand</td><td>MM</td><td>target_seq</td><td>PAM</td><td>alignment</td><td>distance</td><td>position</td><td>gene name</td><td>gene id</td></tr>
<tr><td>GU305914.1</td><td>15456</td><td>15478</td><td>-</td><td>0</td><td>AGGAGATACATGGACTTCTC</td><td>CGG</td><td>||||||||[||||||||||||]PAM</td><td>NA</td><td>Intergenic</td><td>NA</td><td>NA</td></tr>
<tr><td>T33388</td><td>GTCGAGTTTGAGTTTATCGATGG</td><td>1000</td></tr>
<tr><td>Oligo fwd</td><td>CACCGTCGAGTTTGAGTTTATCGA</td></tr>
<tr><td>Oligo rev</td><td>AAACTCGATAAACTCAAACTCGAC</td></tr>
<tr><td>Chromosome</td><td>start</td><td>end</td><td>strand</td><td>MM</td><td>target_seq</td><td>PAM</td><td>alignment</td><td>distance</td><td>position</td><td>gene name</td><td>gene id</td></tr>
<tr><td>GU305914.1</td><td>15426</td><td>15448</td><td>-</td><td>0</td><td>GTCGAGTTTGAGTTTATCGA</td><td>TGG</td><td>||||||||[||||||||||||]PAM</td><td>NA</td><td>Intergenic</td><td>NA</td><td>NA</td></tr>
<tr><td>T33389</td><td>AGTTTATCGATGGTTAGACGCGG</td><td>1000</td></tr>
<tr><td>Oligo fwd</td><td>CACCAGTTTATCGATGGTTAGACG</td></tr>
<tr><td>Oligo rev</td><td>AAACCGTCTAACCATCGATAAACT</td></tr>
<tr><td>Chromosome</td><td>start</td><td>end</td><td>strand</td><td>MM</td><td>target_seq</td><td>PAM</td><td>alignment</td><td>distance</td><td>position</td><td>gene name</td><td>gene id</td></tr>
<tr><td>GU305914.1</td><td>15416</td><td>15438</td><td>-</td><td>0</td><td>AGTTTATCGATGGTTAGACG</td><td>CGG</td><td>||||||||[||||||||||||]PAM</td><td>NA</td><td>Intergenic</td><td>NA</td><td>NA</td></tr>
</table>

```
T33390	TGTCCGAAGAATTTGTCCACAGG	1000
Oligo fwd	CACCTGTCCGAAGAATTTGTCCAC
Oligo rev	AAACGTGGACAAATTCTTCGGACA
Chromosome	start	end	strand	MM	target_seq	PAM	alignment	distance	position	gene name	gene id
GU305914.1	15388	15410	-	0	TGTCCGAAGAATTTGTCCAC	AGG	||||||||[||||||||||||]PAM	NA	Intergenic	NA	NA

T33391	CGAAGAATTTGTCCACAGGTCGG	1000
Oligo fwd	CACCCGAAGAATTTGTCCACAGGT
Oligo rev	AAACACCTGTGGACAAATTCTTCG
Chromosome	start	end	strand	MM	target_seq	PAM	alignment	distance	position	gene name	gene id
GU305914.1	15384	15406	-	0	CGAAGAATTTGTCCACAGGT	CGG	||||||||[||||||||||||]PAM	NA	Intergenic	NA	NA

T33392	CAGGTCGGTGAATGTCGCGCCGG	1000
Oligo fwd	CACCCAGGTCGGTGAATGTCGCGC
Oligo rev	AAACGCGCGACATTCACCGACCTG
Chromosome	start	end	strand	MM	target_seq	PAM	alignment	distance	position	gene name	gene id
GU305914.1	15369	15391	-	0	CAGGTCGGTGAATGTCGCGC	CGG	||||||||[||||||||||||]PAM	NA	Intergenic	NA	NA

T33393	ACAGCTGAATATACACTTATCGG	1000
Oligo fwd	CACCACAGCTGAATATACACTTAT
Oligo rev	AAACATAAGTGTATATTCAGCTGT
Chromosome	start	end	strand	MM	target_seq	PAM	alignment	distance	position	gene name	gene id
GU305914.1	15234	15256	-	0	ACAGCTGAATATACACTTAT	CGG	||||||||[||||||||||||]PAM	NA	Intergenic	NA	NA

T33394	AAGCGTCCAGTGCGATATCAAGG	1000
Oligo fwd	CACCAAGCGTCCAGTGCGATATCA
Oligo rev	AAACTGATATCGCACTGGACGCTT
Chromosome	start	end	strand	MM	target_seq	PAM	alignment	distance	position	gene name	gene id
GU305914.1	15172	15194	-	0	AAGCGTCCAGTGCGATATCA	AGG	||||||||[||||||||||||]PAM	NA	Intergenic	NA	NA

T33395	ATATTCACTGTCATCTTGTAAGG	1000
Oligo fwd	CACCATATTCACTGTCATCTTGTA
Oligo rev	AAACTACAAGATGACAGTGAATAT
Chromosome	start	end	strand	MM	target_seq	PAM	alignment	distance	position	gene name	gene id
GU305914.1	15086	15108	-	0	ATATTCACTGTCATCTTGTA	AGG	||||||||[||||||||||||]PAM	NA	Intergenic	NA	NA

T33396	ACTGTCATCTTGTAAGGTCGAGG	1000
Oligo fwd	CACCACTGTCATCTTGTAAGGTCG
Oligo rev	AAACCGACCTTACAAGATGACAGT
Chromosome	start	end	strand	MM	target_seq	PAM	alignment	distance	position	gene name	gene id
GU305914.1	15080	15102	-	0	ACTGTCATCTTGTAAGGTCG	AGG	||||||||[||||||||||||]PAM	NA	Intergenic	NA	NA

T33397	CTGTCATCTTGTAAGGTCGAGGG	1000
Oligo fwd	CACCCTGTCATCTTGTAAGGTCGA
Oligo rev	AAACTCGACCTTACAAGATGACAG
Chromosome	start	end	strand	MM	target_seq	PAM	alignment	distance	position	gene name	gene id
GU305914.1	15079	15101	-	0	CTGTCATCTTGTAAGGTCGA	GGG	||||||||[||||||||||||]PAM	NA	Intergenic	NA	NA

T33398	AGATGAATATATAGATCCATAGG	1000
Oligo fwd	CACCAGATGAATATATAGATCCAT
Oligo rev	AAACATGGATCTATATATTCATCT
Chromosome	start	end	strand	MM	target_seq	PAM	alignment	distance	position	gene name	gene id
GU305914.1	15041	15063	-	0	AGATGAATATATAGATCCAT	AGG	||||||||[||||||||||||]PAM	NA	Intergenic	NA	NA

T33399	TGAATATATAGATCCATAGGAGG	1000
Oligo fwd	CACCTGAATATATAGATCCATAGG
Oligo rev	AAACCCTATGGATCTATATATTCA
Chromosome	start	end	strand	MM	target_seq	PAM	alignment	distance	position	gene name	gene id
GU305914.1	15038	15060	-	0	TGAATATATAGATCCATAGG	AGG	||||||||[||||||||||||]PAM	NA	Intergenic	NA	NA

T33400	GTATAATTTCGTCTTCTGTGTGG	1000
Oligo fwd	CACCGTATAATTTCGTCTTCTGTG
Oligo rev	AAACCACAGAAGACGAAATTATAC
Chromosome	start	end	strand	MM	target_seq	PAM	alignment	distance	position	gene name	gene id
GU305914.1	14921	14943	-	0	GTATAATTTCGTCTTCTGTG	TGG	||||||||[||||||||||||]PAM	NA	Intergenic	NA	NA
GU305914.1	136537	136559	+	4	CGGTAATTTCGTCTTCTGTC	TGG	---|||||[|||||||||||-]PAM	NA	Intergenic	NA	NA

T33401	TCTGTGTGGCTTGTAGTAGATGG	1000
Oligo fwd	CACCTCTGTGTGGCTTGTAGTAGA
Oligo rev	AAACTCTACTACAAGCCACACAGA
Chromosome	start	end	strand	MM	target_seq	PAM	alignment	distance	position	gene name	gene id
GU305914.1	14907	14929	-	0	TCTGTGTGGCTTGTAGTAGA	TGG	||||||||[||||||||||||]PAM	NA	Intergenic	NA	NA

T33402	GCTTGTAGTAGATGGAGCAATGG	1000
Oligo fwd	CACCGCTTGTAGTAGATGGAGCAA
Oligo rev	AAACTTGCTCCATCTACTACAAGC
Chromosome	start	end	strand	MM	target_seq	PAM	alignment	distance	position	gene name	gene id
GU305914.1	14899	14921	-	0	GCTTGTAGTAGATGGAGCAA	TGG	||||||||[||||||||||||]PAM	NA	Intergenic	NA	NA

T33403	TAGTGCGCGATGTGACAACATGG	1000
Oligo fwd	CACCTAGTGCGCGATGTGACAACA
Oligo rev	AAACTGTTGTCACATCGCGCACTA
Chromosome	start	end	strand	MM	target_seq	PAM	alignment	distance	position	gene name	gene id
GU305914.1	14867	14889	-	0	TAGTGCGCGATGTGACAACA	TGG	||||||||[||||||||||||]PAM	NA	Intergenic	NA	NA

T33404	AGTGCGCGATGTGACAACATGGG	1000
Oligo fwd	CACCAGTGCGCGATGTGACAACAT
Oligo rev	AAACATGTTGTCACATCGCGCACT
```

Chromosome start end strand MM target_seq PAM alignment distance position gene name gene id
GU305914.1 14866 14888 - 0 AGTGCGCGATGTGACAACAT GGG ||||||||[||||||||||||]PAM NA Intergenic NA NA

T33405 CGTTGTATTCGAATATATCTTGG 1000
Oligo fwd CACCCGTTGTATTCGAATATATCT
Oligo rev AAACAGATATATTCGAATACAACG
Chromosome start end strand MM target_seq PAM alignment distance position gene name gene id
GU305914.1 14812 14834 - 0 CGTTGTATTCGAATATATCT TGG ||||||||[||||||||||||]PAM NA Intergenic NA NA

T33406 TGATGTCGTTTGCGATTCTGTGG 1000
Oligo fwd CACCTGATGTCGTTTGCGATTCTG
Oligo rev AAACCAGAATCGCAAACGACATCA
Chromosome start end strand MM target_seq PAM alignment distance position gene name gene id
GU305914.1 14758 14780 - 0 TGATGTCGTTTGCGATTCTG TGG ||||||||[||||||||||||]PAM NA Intergenic NA NA

T33407 TTGCGATTCTGTGGATTTGCTGG 1000
Oligo fwd CACCTTGCGATTCTGTGGATTTGC
Oligo rev AAACGCAAATCCACAGAATCGCAA
Chromosome start end strand MM target_seq PAM alignment distance position gene name gene id
GU305914.1 14749 14771 - 0 TTGCGATTCTGTGGATTTGC TGG ||||||||[||||||||||||]PAM NA Intergenic NA NA

T33408 TGCGATTCTGTGGATTTGCTGGG 1000
Oligo fwd CACCTGCGATTCTGTGGATTTGCT
Oligo rev AAACAGCAAATCCACAGAATCGCA
Chromosome start end strand MM target_seq PAM alignment distance position gene name gene id
GU305914.1 14748 14770 - 0 TGCGATTCTGTGGATTTGCT GGG ||||||||[||||||||||||]PAM NA Intergenic NA NA

T33409 TCTGTGGATTTGCTGGGTGACGG 1000
Oligo fwd CACCTCTGTGGATTTGCTGGGTGA
Oligo rev AAACTCACCCAGCAAATCCACAGA
Chromosome start end strand MM target_seq PAM alignment distance position gene name gene id
GU305914.1 14742 14764 - 0 TCTGTGGATTTGCTGGGTGA CGG ||||||||[||||||||||||]PAM NA Intergenic NA NA

T33410 GGATTTGCTGGGTGACGGCGTGG 1000
Oligo fwd CACCGGATTTGCTGGGTGACGGCG
Oligo rev AAACCGCCGTCACCCAGCAAATCC
Chromosome start end strand MM target_seq PAM alignment distance position gene name gene id
GU305914.1 14737 14759 - 0 GGATTTGCTGGGTGACGGCG TGG ||||||||[||||||||||||]PAM NA Intergenic NA NA
GU305914.1 14716 14738 - 4 GGATCTGATGGGTGGCGGTG TGG ||||-||-[||||||-|||-|]PAM NA Intergenic NA NA

T33411 GGGTGACGGCGTGGATCTGATGG 1000
Oligo fwd CACCGGGTGACGGCGTGGATCTGA
Oligo rev AAACTCAGATCCACGCCGTCACCC
Chromosome start end strand MM target_seq PAM alignment distance position gene name gene id
GU305914.1 14728 14750 - 0 GGGTGACGGCGTGGATCTGA TGG ||||||||[||||||||||||]PAM NA Intergenic NA NA

T33412 GGTGACGGCGTGGATCTGATGGG 1000
Oligo fwd CACCGGTGACGGCGTGGATCTGAT
Oligo rev AAACATCAGATCCACGCCGTCACC
Chromosome start end strand MM target_seq PAM alignment distance position gene name gene id
GU305914.1 14727 14749 - 0 GGTGACGGCGTGGATCTGAT GGG ||||||||[||||||||||||]PAM NA Intergenic NA NA

T33413 GACGGCGTGGATCTGATGGGTGG 1000
Oligo fwd CACCGACGGCGTGGATCTGATGGG
Oligo rev AAACCCCATCAGATCCACGCCGTC
Chromosome start end strand MM target_seq PAM alignment distance position gene name gene id
GU305914.1 14724 14746 - 0 GACGGCGTGGATCTGATGGG TGG ||||||||[||||||||||||]PAM NA Intergenic NA NA
GU305914.1 14703 14725 - 4 GGCGGTGTGGATCCGTTGGG TGG |-|||-||[|||||-|-||||]PAM NA Intergenic NA NA

T33414 GGCGTGGATCTGATGGGTGGCGG 1000
Oligo fwd CACCGGCGTGGATCTGATGGGTGG
Oligo rev AAACCCACCCATCAGATCCACGCC
Chromosome start end strand MM target_seq PAM alignment distance position gene name gene id
GU305914.1 14721 14743 - 0 GGCGTGGATCTGATGGGTGG CGG ||||||||[||||||||||||]PAM NA Intergenic NA NA
GU305914.1 14700 14722 - 3 GGTGTGGATCCGTTGGGTGG TGG ||-|||||[||-|-|||||||]PAM NA Intergenic NA NA

T33415 GGATCTGATGGGTGGCGGTGTGG 1000
Oligo fwd CACCGGATCTGATGGGTGGCGGTG
Oligo rev AAACCACCGCCACCCATCAGATCC
Chromosome start end strand MM target_seq PAM alignment distance position gene name gene id
GU305914.1 14716 14738 - 0 GGATCTGATGGGTGGCGGTG TGG ||||||||[||||||||||||]PAM NA Intergenic NA NA
GU305914.1 14737 14759 - 4 GGATTTGCTGGGTGACGGCG TGG ||||-||-[||||||-|||-|]PAM NA Intergenic NA NA

T33416 GGGTGGCGGTGTGGATCCGTTGG 1000
Oligo fwd CACCGGGTGGCGGTGTGGATCCGT
Oligo rev AAACACGGATCCACACCGCCACCC
Chromosome start end strand MM target_seq PAM alignment distance position gene name gene id
GU305914.1 14707 14729 - 0 GGGTGGCGGTGTGGATCCGT TGG ||||||||[||||||||||||]PAM NA Intergenic NA NA

T33417 GGTGGCGGTGTGGATCCGTTGGG 1000
Oligo fwd CACCGGTGGCGGTGTGGATCCGTT
Oligo rev AAACAACGGATCCACACCGCCACC
Chromosome start end strand MM target_seq PAM alignment distance position gene name gene id
GU305914.1 14706 14728 - 0 GGTGGCGGTGTGGATCCGTT GGG ||||||||[||||||||||||]PAM NA Intergenic NA NA

T33418 GGCGGTGTGGATCCGTTGGGTGG 1000
Oligo fwd CACCGGCGGTGTGGATCCGTTGGG
Oligo rev AAACCCCAACGGATCCACACCGCC
Chromosome start end strand MM target_seq PAM alignment distance position gene name gene id

<table>
<tr><td>GU305914.1</td><td></td><td>14703</td><td>14725</td><td>-</td><td>0</td><td>GGCGGTGTGGATCCGTTGGG</td><td>TGG</td><td>||||||||[||||||||||||]PAM</td><td>NA</td><td>Intergenic</td><td>NA</td><td>NA</td></tr>
<tr><td>GU305914.1</td><td></td><td>14724</td><td>14746</td><td>-</td><td>4</td><td>GACGGCGTGGATCTGATGGG</td><td>TGG</td><td>|-|||-||[|||||-|-||||]PAM</td><td>NA</td><td>Intergenic</td><td>NA</td><td>NA</td></tr>
<tr><td>T33419</td><td>GGTGTGGATCCGTTGGGTGGTGG</td><td></td><td>1000</td><td></td><td></td><td></td><td></td><td></td><td></td><td></td><td></td><td></td></tr>
<tr><td>Oligo fwd</td><td>CACCGGTGTGGATCCGTTGGGTGG</td><td></td><td></td><td></td><td></td><td></td><td></td><td></td><td></td><td></td><td></td><td></td></tr>
<tr><td>Oligo rev</td><td>AAACCCACCCAACGGATCCACACC</td><td></td><td></td><td></td><td></td><td></td><td></td><td></td><td></td><td></td><td></td><td></td></tr>
<tr><td>Chromosome</td><td>start</td><td></td><td>end</td><td>strand</td><td>MM</td><td>target_seq</td><td>PAM</td><td>alignment</td><td>distance</td><td>position</td><td>gene name</td><td>gene id</td></tr>
<tr><td>GU305914.1</td><td></td><td>14700</td><td>14722</td><td>-</td><td>0</td><td>GGTGTGGATCCGTTGGGTGG</td><td>TGG</td><td>||||||||[||||||||||||]PAM</td><td>NA</td><td>Intergenic</td><td>NA</td><td>NA</td></tr>
<tr><td>GU305914.1</td><td></td><td>14721</td><td>14743</td><td>-</td><td>3</td><td>GGCGTGGATCTGATGGGTGG</td><td>CGG</td><td>||-|||||[||-|-|||||||]PAM</td><td>NA</td><td>Intergenic</td><td>NA</td><td>NA</td></tr>
<tr><td>T33420</td><td>CCGTTGGGTGGTGGAGTCGACGG</td><td></td><td>1000</td><td></td><td></td><td></td><td></td><td></td><td></td><td></td><td></td><td></td></tr>
<tr><td>Oligo fwd</td><td>CACCCCGTTGGGTGGTGGAGTCGA</td><td></td><td></td><td></td><td></td><td></td><td></td><td></td><td></td><td></td><td></td><td></td></tr>
<tr><td>Oligo rev</td><td>AAACTCGACTCCACCACCCAACGG</td><td></td><td></td><td></td><td></td><td></td><td></td><td></td><td></td><td></td><td></td><td></td></tr>
<tr><td>Chromosome</td><td>start</td><td></td><td>end</td><td>strand</td><td>MM</td><td>target_seq</td><td>PAM</td><td>alignment</td><td>distance</td><td>position</td><td>gene name</td><td>gene id</td></tr>
<tr><td>GU305914.1</td><td></td><td>14691</td><td>14713</td><td>-</td><td>0</td><td>CCGTTGGGTGGTGGAGTCGA</td><td>CGG</td><td>||||||||[||||||||||||]PAM</td><td>NA</td><td>Intergenic</td><td>NA</td><td>NA</td></tr>
<tr><td>T33421</td><td>CTGTGACGATGTCGTTGTTGTGG</td><td></td><td>1000</td><td></td><td></td><td></td><td></td><td></td><td></td><td></td><td></td><td></td></tr>
<tr><td>Oligo fwd</td><td>CACCCTGTGACGATGTCGTTGTTG</td><td></td><td></td><td></td><td></td><td></td><td></td><td></td><td></td><td></td><td></td><td></td></tr>
<tr><td>Oligo rev</td><td>AAACCAACAACGACATCGTCACAG</td><td></td><td></td><td></td><td></td><td></td><td></td><td></td><td></td><td></td><td></td><td></td></tr>
<tr><td>Chromosome</td><td>start</td><td></td><td>end</td><td>strand</td><td>MM</td><td>target_seq</td><td>PAM</td><td>alignment</td><td>distance</td><td>position</td><td>gene name</td><td>gene id</td></tr>
<tr><td>GU305914.1</td><td></td><td>14659</td><td>14681</td><td>-</td><td>0</td><td>CTGTGACGATGTCGTTGTTG</td><td>TGG</td><td>||||||||[||||||||||||]PAM</td><td>NA</td><td>Intergenic</td><td>NA</td><td>NA</td></tr>
<tr><td>T33422</td><td>TGTGACGATGTCGTTGTTGTGGG</td><td></td><td>1000</td><td></td><td></td><td></td><td></td><td></td><td></td><td></td><td></td><td></td></tr>
<tr><td>Oligo fwd</td><td>CACCTGTGACGATGTCGTTGTTGT</td><td></td><td></td><td></td><td></td><td></td><td></td><td></td><td></td><td></td><td></td><td></td></tr>
<tr><td>Oligo rev</td><td>AAACACAACAACGACATCGTCACA</td><td></td><td></td><td></td><td></td><td></td><td></td><td></td><td></td><td></td><td></td><td></td></tr>
<tr><td>Chromosome</td><td>start</td><td></td><td>end</td><td>strand</td><td>MM</td><td>target_seq</td><td>PAM</td><td>alignment</td><td>distance</td><td>position</td><td>gene name</td><td>gene id</td></tr>
<tr><td>GU305914.1</td><td></td><td>14658</td><td>14680</td><td>-</td><td>0</td><td>TGTGACGATGTCGTTGTTGT</td><td>GGG</td><td>||||||||[||||||||||||]PAM</td><td>NA</td><td>Intergenic</td><td>NA</td><td>NA</td></tr>
<tr><td>T33423</td><td>GATATTTGAGTAACGAATTGTGG</td><td></td><td>1000</td><td></td><td></td><td></td><td></td><td></td><td></td><td></td><td></td><td></td></tr>
<tr><td>Oligo fwd</td><td>CACCGATATTTGAGTAACGAATTG</td><td></td><td></td><td></td><td></td><td></td><td></td><td></td><td></td><td></td><td></td><td></td></tr>
<tr><td>Oligo rev</td><td>AAACCAATTCGTTACTCAAATATC</td><td></td><td></td><td></td><td></td><td></td><td></td><td></td><td></td><td></td><td></td><td></td></tr>
<tr><td>Chromosome</td><td>start</td><td></td><td>end</td><td>strand</td><td>MM</td><td>target_seq</td><td>PAM</td><td>alignment</td><td>distance</td><td>position</td><td>gene name</td><td>gene id</td></tr>
<tr><td>GU305914.1</td><td></td><td>14600</td><td>14622</td><td>-</td><td>0</td><td>GATATTTGAGTAACGAATTG</td><td>TGG</td><td>||||||||[||||||||||||]PAM</td><td>NA</td><td>Intergenic</td><td>NA</td><td>NA</td></tr>
<tr><td>T33424</td><td>ATATTTGAGTAACGAATTGTGGG</td><td></td><td>1000</td><td></td><td></td><td></td><td></td><td></td><td></td><td></td><td></td><td></td></tr>
<tr><td>Oligo fwd</td><td>CACCATATTTGAGTAACGAATTGT</td><td></td><td></td><td></td><td></td><td></td><td></td><td></td><td></td><td></td><td></td><td></td></tr>
<tr><td>Oligo rev</td><td>AAACACAATTCGTTACTCAAATAT</td><td></td><td></td><td></td><td></td><td></td><td></td><td></td><td></td><td></td><td></td><td></td></tr>
<tr><td>Chromosome</td><td>start</td><td></td><td>end</td><td>strand</td><td>MM</td><td>target_seq</td><td>PAM</td><td>alignment</td><td>distance</td><td>position</td><td>gene name</td><td>gene id</td></tr>
<tr><td>GU305914.1</td><td></td><td>14599</td><td>14621</td><td>-</td><td>0</td><td>ATATTTGAGTAACGAATTGT</td><td>GGG</td><td>||||||||[||||||||||||]PAM</td><td>NA</td><td>Intergenic</td><td>NA</td><td>NA</td></tr>
<tr><td>T33425</td><td>TATTTGAGTAACGAATTGTGGGG</td><td></td><td>1000</td><td></td><td></td><td></td><td></td><td></td><td></td><td></td><td></td><td></td></tr>
<tr><td>Oligo fwd</td><td>CACCTATTTGAGTAACGAATTGTG</td><td></td><td></td><td></td><td></td><td></td><td></td><td></td><td></td><td></td><td></td><td></td></tr>
<tr><td>Oligo rev</td><td>AAACCACAATTCGTTACTCAAATA</td><td></td><td></td><td></td><td></td><td></td><td></td><td></td><td></td><td></td><td></td><td></td></tr>
<tr><td>Chromosome</td><td>start</td><td></td><td>end</td><td>strand</td><td>MM</td><td>target_seq</td><td>PAM</td><td>alignment</td><td>distance</td><td>position</td><td>gene name</td><td>gene id</td></tr>
<tr><td>GU305914.1</td><td></td><td>14598</td><td>14620</td><td>-</td><td>0</td><td>TATTTGAGTAACGAATTGTG</td><td>GGG</td><td>||||||||[||||||||||||]PAM</td><td>NA</td><td>Intergenic</td><td>NA</td><td>NA</td></tr>
<tr><td>T33426</td><td>GTTGATTGAAGTGTATCCCTAGG</td><td></td><td>1000</td><td></td><td></td><td></td><td></td><td></td><td></td><td></td><td></td><td></td></tr>
<tr><td>Oligo fwd</td><td>CACCGTTGATTGAAGTGTATCCCT</td><td></td><td></td><td></td><td></td><td></td><td></td><td></td><td></td><td></td><td></td><td></td></tr>
<tr><td>Oligo rev</td><td>AAACAGGGATACACTTCAATCAAC</td><td></td><td></td><td></td><td></td><td></td><td></td><td></td><td></td><td></td><td></td><td></td></tr>
<tr><td>Chromosome</td><td>start</td><td></td><td>end</td><td>strand</td><td>MM</td><td>target_seq</td><td>PAM</td><td>alignment</td><td>distance</td><td>position</td><td>gene name</td><td>gene id</td></tr>
<tr><td>GU305914.1</td><td></td><td>14555</td><td>14577</td><td>-</td><td>0</td><td>GTTGATTGAAGTGTATCCCT</td><td>AGG</td><td>||||||||[||||||||||||]PAM</td><td>NA</td><td>Intergenic</td><td>NA</td><td>NA</td></tr>
<tr><td>T33427</td><td>GATTGAAGTGTATCCCTAGGAGG</td><td></td><td>1000</td><td></td><td></td><td></td><td></td><td></td><td></td><td></td><td></td><td></td></tr>
<tr><td>Oligo fwd</td><td>CACCGATTGAAGTGTATCCCTAGG</td><td></td><td></td><td></td><td></td><td></td><td></td><td></td><td></td><td></td><td></td><td></td></tr>
<tr><td>Oligo rev</td><td>AAACCCTAGGGATACACTTCAATC</td><td></td><td></td><td></td><td></td><td></td><td></td><td></td><td></td><td></td><td></td><td></td></tr>
<tr><td>Chromosome</td><td>start</td><td></td><td>end</td><td>strand</td><td>MM</td><td>target_seq</td><td>PAM</td><td>alignment</td><td>distance</td><td>position</td><td>gene name</td><td>gene id</td></tr>
<tr><td>GU305914.1</td><td></td><td>14552</td><td>14574</td><td>-</td><td>0</td><td>GATTGAAGTGTATCCCTAGG</td><td>AGG</td><td>||||||||[||||||||||||]PAM</td><td>NA</td><td>Intergenic</td><td>NA</td><td>NA</td></tr>
<tr><td>T33428</td><td>CTAGGAGGCTCTGATTCACATGG</td><td></td><td>1000</td><td></td><td></td><td></td><td></td><td></td><td></td><td></td><td></td><td></td></tr>
<tr><td>Oligo fwd</td><td>CACCCTAGGAGGCTCTGATTCACA</td><td></td><td></td><td></td><td></td><td></td><td></td><td></td><td></td><td></td><td></td><td></td></tr>
<tr><td>Oligo rev</td><td>AAACTGTGAATCAGAGCCTCCTAG</td><td></td><td></td><td></td><td></td><td></td><td></td><td></td><td></td><td></td><td></td><td></td></tr>
<tr><td>Chromosome</td><td>start</td><td></td><td>end</td><td>strand</td><td>MM</td><td>target_seq</td><td>PAM</td><td>alignment</td><td>distance</td><td>position</td><td>gene name</td><td>gene id</td></tr>
<tr><td>GU305914.1</td><td></td><td>14537</td><td>14559</td><td>-</td><td>0</td><td>CTAGGAGGCTCTGATTCACA</td><td>TGG</td><td>||||||||[||||||||||||]PAM</td><td>NA</td><td>Intergenic</td><td>NA</td><td>NA</td></tr>
<tr><td>T33429</td><td>TTCCAAGTGCCAGTGTAATACGG</td><td></td><td>1000</td><td></td><td></td><td></td><td></td><td></td><td></td><td></td><td></td><td></td></tr>
<tr><td>Oligo fwd</td><td>CACCTTCCAAGTGCCAGTGTAATA</td><td></td><td></td><td></td><td></td><td></td><td></td><td></td><td></td><td></td><td></td><td></td></tr>
<tr><td>Oligo rev</td><td>AAACTATTACACTGGCACTTGGAA</td><td></td><td></td><td></td><td></td><td></td><td></td><td></td><td></td><td></td><td></td><td></td></tr>
<tr><td>Chromosome</td><td>start</td><td></td><td>end</td><td>strand</td><td>MM</td><td>target_seq</td><td>PAM</td><td>alignment</td><td>distance</td><td>position</td><td>gene name</td><td>gene id</td></tr>
<tr><td>GU305914.1</td><td></td><td>14493</td><td>14515</td><td>-</td><td>0</td><td>TTCCAAGTGCCAGTGTAATA</td><td>CGG</td><td>||||||||[||||||||||||]PAM</td><td>NA</td><td>Intergenic</td><td>NA</td><td>NA</td></tr>
<tr><td>T33430</td><td>GCCAGTGTAATACGGTATTGAGG</td><td></td><td>1000</td><td></td><td></td><td></td><td></td><td></td><td></td><td></td><td></td><td></td></tr>
<tr><td>Oligo fwd</td><td>CACCGCCAGTGTAATACGGTATTG</td><td></td><td></td><td></td><td></td><td></td><td></td><td></td><td></td><td></td><td></td><td></td></tr>
<tr><td>Oligo rev</td><td>AAACCAATACCGTATTACACTGGC</td><td></td><td></td><td></td><td></td><td></td><td></td><td></td><td></td><td></td><td></td><td></td></tr>
<tr><td>Chromosome</td><td>start</td><td></td><td>end</td><td>strand</td><td>MM</td><td>target_seq</td><td>PAM</td><td>alignment</td><td>distance</td><td>position</td><td>gene name</td><td>gene id</td></tr>
<tr><td>GU305914.1</td><td></td><td>14485</td><td>14507</td><td>-</td><td>0</td><td>GCCAGTGTAATACGGTATTG</td><td>AGG</td><td>||||||||[||||||||||||]PAM</td><td>NA</td><td>Intergenic</td><td>NA</td><td>NA</td></tr>
<tr><td>T33431</td><td>AAAATCTATAATCATGATATTGG</td><td></td><td>1000</td><td></td><td></td><td></td><td></td><td></td><td></td><td></td><td></td><td></td></tr>
<tr><td>Oligo fwd</td><td>CACCAAAATCTATAATCATGATAT</td><td></td><td></td><td></td><td></td><td></td><td></td><td></td><td></td><td></td><td></td><td></td></tr>
<tr><td>Oligo rev</td><td>AAACATATCATGATTATAGATTTT</td><td></td><td></td><td></td><td></td><td></td><td></td><td></td><td></td><td></td><td></td><td></td></tr>
<tr><td>Chromosome</td><td>start</td><td></td><td>end</td><td>strand</td><td>MM</td><td>target_seq</td><td>PAM</td><td>alignment</td><td>distance</td><td>position</td><td>gene name</td><td>gene id</td></tr>
<tr><td>GU305914.1</td><td></td><td>14440</td><td>14462</td><td>-</td><td>0</td><td>AAAATCTATAATCATGATAT</td><td>TGG</td><td>||||||||[||||||||||||]PAM</td><td>NA</td><td>Intergenic</td><td>NA</td><td>NA</td></tr>
<tr><td>T33432</td><td>AAATATGCGCCATACGATCTTGG</td><td></td><td>1000</td><td></td><td></td><td></td><td></td><td></td><td></td><td></td><td></td><td></td></tr>
<tr><td>Oligo fwd</td><td>CACCAAATATGCGCCATACGATCT</td><td></td><td></td><td></td><td></td><td></td><td></td><td></td><td></td><td></td><td></td><td></td></tr>
<tr><td>Oligo rev</td><td>AAACAGATCGTATGGCGCATATTT</td><td></td><td></td><td></td><td></td><td></td><td></td><td></td><td></td><td></td><td></td><td></td></tr>
<tr><td>Chromosome</td><td>start</td><td></td><td>end</td><td>strand</td><td>MM</td><td>target_seq</td><td>PAM</td><td>alignment</td><td>distance</td><td>position</td><td>gene name</td><td>gene id</td></tr>
<tr><td>GU305914.1</td><td></td><td>14409</td><td>14431</td><td>-</td><td>0</td><td>AAATATGCGCCATACGATCT</td><td>TGG</td><td>||||||||[||||||||||||]PAM</td><td>NA</td><td>Intergenic</td><td>NA</td><td>NA</td></tr>
<tr><td>T33433</td><td>GATCTTGGTTGCCGATGTTTTGG</td><td></td><td>1000</td><td></td><td></td><td></td><td></td><td></td><td></td><td></td><td></td><td></td></tr>
</table>

<table>
<tr><td>Oligo fwd</td><td>CACCGATCTTGGTTGCCGATGTTT</td><td></td><td></td><td></td><td></td><td></td><td></td><td></td><td></td><td></td><td></td><td></td></tr>
<tr><td>Oligo rev</td><td>AAACAAACATCGGCAACCAAGATC</td><td></td><td></td><td></td><td></td><td></td><td></td><td></td><td></td><td></td><td></td><td></td></tr>
<tr><td>Chromosome</td><td>start</td><td></td><td>end</td><td>strand</td><td>MM</td><td>target_seq</td><td>PAM</td><td>alignment</td><td>distance</td><td>position</td><td>gene name</td><td>gene id</td></tr>
<tr><td>GU305914.1</td><td></td><td>14394</td><td>14416</td><td>-</td><td>0</td><td>GATCTTGGTTGCCGATGTTT</td><td>TGG</td><td>||||||||[||||||||||||]PAM</td><td>NA</td><td>Intergenic</td><td>NA</td><td>NA</td></tr>
<tr><td>T33434</td><td>CTTGGTTGCCGATGTTTTGGTGG</td><td></td><td>1000</td><td></td><td></td><td></td><td></td><td></td><td></td><td></td><td></td><td></td></tr>
<tr><td>Oligo fwd</td><td>CACCCTTGGTTGCCGATGTTTTGG</td><td></td><td></td><td></td><td></td><td></td><td></td><td></td><td></td><td></td><td></td><td></td></tr>
<tr><td>Oligo rev</td><td>AAACCCAAAACATCGGCAACCAAG</td><td></td><td></td><td></td><td></td><td></td><td></td><td></td><td></td><td></td><td></td><td></td></tr>
<tr><td>Chromosome</td><td>start</td><td></td><td>end</td><td>strand</td><td>MM</td><td>target_seq</td><td>PAM</td><td>alignment</td><td>distance</td><td>position</td><td>gene name</td><td>gene id</td></tr>
<tr><td>GU305914.1</td><td></td><td>14391</td><td>14413</td><td>-</td><td>0</td><td>CTTGGTTGCCGATGTTTTGG</td><td>TGG</td><td>||||||||[||||||||||||]PAM</td><td>NA</td><td>Intergenic</td><td>NA</td><td>NA</td></tr>
<tr><td>T33435</td><td>TCAGAAATCGAAGTGATGTCCGG</td><td></td><td>1000</td><td></td><td></td><td></td><td></td><td></td><td></td><td></td><td></td><td></td></tr>
<tr><td>Oligo fwd</td><td>CACCTCAGAAATCGAAGTGATGTC</td><td></td><td></td><td></td><td></td><td></td><td></td><td></td><td></td><td></td><td></td><td></td></tr>
<tr><td>Oligo rev</td><td>AAACGACATCACTTCGATTTCTGA</td><td></td><td></td><td></td><td></td><td></td><td></td><td></td><td></td><td></td><td></td><td></td></tr>
<tr><td>Chromosome</td><td>start</td><td></td><td>end</td><td>strand</td><td>MM</td><td>target_seq</td><td>PAM</td><td>alignment</td><td>distance</td><td>position</td><td>gene name</td><td>gene id</td></tr>
<tr><td>GU305914.1</td><td></td><td>14337</td><td>14359</td><td>-</td><td>0</td><td>TCAGAAATCGAAGTGATGTC</td><td>CGG</td><td>||||||||[||||||||||||]PAM</td><td>NA</td><td>Intergenic</td><td>NA</td><td>NA</td></tr>
<tr><td>T33436</td><td>AACAGATGTAGAAGTTCTTACGG</td><td></td><td>1000</td><td></td><td></td><td></td><td></td><td></td><td></td><td></td><td></td><td></td></tr>
<tr><td>Oligo fwd</td><td>CACCAACAGATGTAGAAGTTCTTA</td><td></td><td></td><td></td><td></td><td></td><td></td><td></td><td></td><td></td><td></td><td></td></tr>
<tr><td>Oligo rev</td><td>AAACTAAGAACTTCTACATCTGTT</td><td></td><td></td><td></td><td></td><td></td><td></td><td></td><td></td><td></td><td></td><td></td></tr>
<tr><td>Chromosome</td><td>start</td><td></td><td>end</td><td>strand</td><td>MM</td><td>target_seq</td><td>PAM</td><td>alignment</td><td>distance</td><td>position</td><td>gene name</td><td>gene id</td></tr>
<tr><td>GU305914.1</td><td></td><td>14314</td><td>14336</td><td>-</td><td>0</td><td>AACAGATGTAGAAGTTCTTA</td><td>CGG</td><td>||||||||[||||||||||||]PAM</td><td>NA</td><td>Intergenic</td><td>NA</td><td>NA</td></tr>
<tr><td>T33437</td><td>GGTCCATGTTGCGTACCATATGG</td><td></td><td>1000</td><td></td><td></td><td></td><td></td><td></td><td></td><td></td><td></td><td></td></tr>
<tr><td>Oligo fwd</td><td>CACCGGTCCATGTTGCGTACCATA</td><td></td><td></td><td></td><td></td><td></td><td></td><td></td><td></td><td></td><td></td><td></td></tr>
<tr><td>Oligo rev</td><td>AAACTATGGTACGCAACATGGACC</td><td></td><td></td><td></td><td></td><td></td><td></td><td></td><td></td><td></td><td></td><td></td></tr>
<tr><td>Chromosome</td><td>start</td><td></td><td>end</td><td>strand</td><td>MM</td><td>target_seq</td><td>PAM</td><td>alignment</td><td>distance</td><td>position</td><td>gene name</td><td>gene id</td></tr>
<tr><td>GU305914.1</td><td></td><td>14293</td><td>14315</td><td>-</td><td>0</td><td>GGTCCATGTTGCGTACCATA</td><td>TGG</td><td>||||||||[||||||||||||]PAM</td><td>NA</td><td>Intergenic</td><td>NA</td><td>NA</td></tr>
<tr><td>T33438</td><td>CATGTTGCGTACCATATGGCAGG</td><td></td><td>1000</td><td></td><td></td><td></td><td></td><td></td><td></td><td></td><td></td><td></td></tr>
<tr><td>Oligo fwd</td><td>CACCCATGTTGCGTACCATATGGC</td><td></td><td></td><td></td><td></td><td></td><td></td><td></td><td></td><td></td><td></td><td></td></tr>
<tr><td>Oligo rev</td><td>AAACGCCATATGGTACGCAACATG</td><td></td><td></td><td></td><td></td><td></td><td></td><td></td><td></td><td></td><td></td><td></td></tr>
<tr><td>Chromosome</td><td>start</td><td></td><td>end</td><td>strand</td><td>MM</td><td>target_seq</td><td>PAM</td><td>alignment</td><td>distance</td><td>position</td><td>gene name</td><td>gene id</td></tr>
<tr><td>GU305914.1</td><td></td><td>14289</td><td>14311</td><td>-</td><td>0</td><td>CATGTTGCGTACCATATGGC</td><td>AGG</td><td>||||||||[||||||||||||]PAM</td><td>NA</td><td>Intergenic</td><td>NA</td><td>NA</td></tr>
<tr><td>T33439</td><td>CATATGGCAGGAATACATACAGG</td><td></td><td>1000</td><td></td><td></td><td></td><td></td><td></td><td></td><td></td><td></td><td></td></tr>
<tr><td>Oligo fwd</td><td>CACCCATATGGCAGGAATACATAC</td><td></td><td></td><td></td><td></td><td></td><td></td><td></td><td></td><td></td><td></td><td></td></tr>
<tr><td>Oligo rev</td><td>AAACGTATGTATTCCTGCCATATG</td><td></td><td></td><td></td><td></td><td></td><td></td><td></td><td></td><td></td><td></td><td></td></tr>
<tr><td>Chromosome</td><td>start</td><td></td><td>end</td><td>strand</td><td>MM</td><td>target_seq</td><td>PAM</td><td>alignment</td><td>distance</td><td>position</td><td>gene name</td><td>gene id</td></tr>
<tr><td>GU305914.1</td><td></td><td>14277</td><td>14299</td><td>-</td><td>0</td><td>CATATGGCAGGAATACATAC</td><td>AGG</td><td>||||||||[||||||||||||]PAM</td><td>NA</td><td>Intergenic</td><td>NA</td><td>NA</td></tr>
<tr><td>T33440</td><td>GCGTGATGCGCTTGCAACGCTGG</td><td></td><td>1000</td><td></td><td></td><td></td><td></td><td></td><td></td><td></td><td></td><td></td></tr>
<tr><td>Oligo fwd</td><td>CACCGCGTGATGCGCTTGCAACGC</td><td></td><td></td><td></td><td></td><td></td><td></td><td></td><td></td><td></td><td></td><td></td></tr>
<tr><td>Oligo rev</td><td>AAACGCGTTGCAAGCGCATCACGC</td><td></td><td></td><td></td><td></td><td></td><td></td><td></td><td></td><td></td><td></td><td></td></tr>
<tr><td>Chromosome</td><td>start</td><td></td><td>end</td><td>strand</td><td>MM</td><td>target_seq</td><td>PAM</td><td>alignment</td><td>distance</td><td>position</td><td>gene name</td><td>gene id</td></tr>
<tr><td>GU305914.1</td><td></td><td>14221</td><td>14243</td><td>-</td><td>0</td><td>GCGTGATGCGCTTGCAACGC</td><td>TGG</td><td>||||||||[||||||||||||]PAM</td><td>NA</td><td>Intergenic</td><td>NA</td><td>NA</td></tr>
<tr><td>T33441</td><td>GCGCTTGCAACGCTGGCGAATGG</td><td></td><td>1000</td><td></td><td></td><td></td><td></td><td></td><td></td><td></td><td></td><td></td></tr>
<tr><td>Oligo fwd</td><td>CACCGCGCTTGCAACGCTGGCGAA</td><td></td><td></td><td></td><td></td><td></td><td></td><td></td><td></td><td></td><td></td><td></td></tr>
<tr><td>Oligo rev</td><td>AAACTTCGCCAGCGTTGCAAGCGC</td><td></td><td></td><td></td><td></td><td></td><td></td><td></td><td></td><td></td><td></td><td></td></tr>
<tr><td>Chromosome</td><td>start</td><td></td><td>end</td><td>strand</td><td>MM</td><td>target_seq</td><td>PAM</td><td>alignment</td><td>distance</td><td>position</td><td>gene name</td><td>gene id</td></tr>
<tr><td>GU305914.1</td><td></td><td>14214</td><td>14236</td><td>-</td><td>0</td><td>GCGCTTGCAACGCTGGCGAA</td><td>TGG</td><td>||||||||[||||||||||||]PAM</td><td>NA</td><td>Intergenic</td><td>NA</td><td>NA</td></tr>
<tr><td>T33442</td><td>GGTACCCGTACACACATACCTGG</td><td></td><td>1000</td><td></td><td></td><td></td><td></td><td></td><td></td><td></td><td></td><td></td></tr>
<tr><td>Oligo fwd</td><td>CACCGGTACCCGTACACACATACC</td><td></td><td></td><td></td><td></td><td></td><td></td><td></td><td></td><td></td><td></td><td></td></tr>
<tr><td>Oligo rev</td><td>AAACGGTATGTGTGTACGGGTACC</td><td></td><td></td><td></td><td></td><td></td><td></td><td></td><td></td><td></td><td></td><td></td></tr>
<tr><td>Chromosome</td><td>start</td><td></td><td>end</td><td>strand</td><td>MM</td><td>target_seq</td><td>PAM</td><td>alignment</td><td>distance</td><td>position</td><td>gene name</td><td>gene id</td></tr>
<tr><td>GU305914.1</td><td></td><td>14193</td><td>14215</td><td>-</td><td>0</td><td>GGTACCCGTACACACATACC</td><td>TGG</td><td>||||||||[||||||||||||]PAM</td><td>NA</td><td>Intergenic</td><td>NA</td><td>NA</td></tr>
<tr><td>T33443</td><td>ACCTGGAGTAAAATCCATACCGG</td><td></td><td>1000</td><td></td><td></td><td></td><td></td><td></td><td></td><td></td><td></td><td></td></tr>
<tr><td>Oligo fwd</td><td>CACCACCTGGAGTAAAATCCATAC</td><td></td><td></td><td></td><td></td><td></td><td></td><td></td><td></td><td></td><td></td><td></td></tr>
<tr><td>Oligo rev</td><td>AAACGTATGGATTTTACTCCAGGT</td><td></td><td></td><td></td><td></td><td></td><td></td><td></td><td></td><td></td><td></td><td></td></tr>
<tr><td>Chromosome</td><td>start</td><td></td><td>end</td><td>strand</td><td>MM</td><td>target_seq</td><td>PAM</td><td>alignment</td><td>distance</td><td>position</td><td>gene name</td><td>gene id</td></tr>
<tr><td>GU305914.1</td><td></td><td>14176</td><td>14198</td><td>-</td><td>0</td><td>ACCTGGAGTAAAATCCATAC</td><td>CGG</td><td>||||||||[||||||||||||]PAM</td><td>NA</td><td>Intergenic</td><td>NA</td><td>NA</td></tr>
<tr><td>T33444</td><td>GTAAAATCCATACCGGAAGACGG</td><td></td><td>1000</td><td></td><td></td><td></td><td></td><td></td><td></td><td></td><td></td><td></td></tr>
<tr><td>Oligo fwd</td><td>CACCGTAAAATCCATACCGGAAGA</td><td></td><td></td><td></td><td></td><td></td><td></td><td></td><td></td><td></td><td></td><td></td></tr>
<tr><td>Oligo rev</td><td>AAACTCTTCCGGTATGGATTTTAC</td><td></td><td></td><td></td><td></td><td></td><td></td><td></td><td></td><td></td><td></td><td></td></tr>
<tr><td>Chromosome</td><td>start</td><td></td><td>end</td><td>strand</td><td>MM</td><td>target_seq</td><td>PAM</td><td>alignment</td><td>distance</td><td>position</td><td>gene name</td><td>gene id</td></tr>
<tr><td>GU305914.1</td><td></td><td>14169</td><td>14191</td><td>-</td><td>0</td><td>GTAAAATCCATACCGGAAGA</td><td>CGG</td><td>||||||||[||||||||||||]PAM</td><td>NA</td><td>Intergenic</td><td>NA</td><td>NA</td></tr>
<tr><td>T33445</td><td>TAAAATCCATACCGGAAGACGGG</td><td></td><td>1000</td><td></td><td></td><td></td><td></td><td></td><td></td><td></td><td></td><td></td></tr>
<tr><td>Oligo fwd</td><td>CACCTAAAATCCATACCGGAAGAC</td><td></td><td></td><td></td><td></td><td></td><td></td><td></td><td></td><td></td><td></td><td></td></tr>
<tr><td>Oligo rev</td><td>AAACGTCTTCCGGTATGGATTTTA</td><td></td><td></td><td></td><td></td><td></td><td></td><td></td><td></td><td></td><td></td><td></td></tr>
<tr><td>Chromosome</td><td>start</td><td></td><td>end</td><td>strand</td><td>MM</td><td>target_seq</td><td>PAM</td><td>alignment</td><td>distance</td><td>position</td><td>gene name</td><td>gene id</td></tr>
<tr><td>GU305914.1</td><td></td><td>14168</td><td>14190</td><td>-</td><td>0</td><td>TAAAATCCATACCGGAAGAC</td><td>GGG</td><td>||||||||[||||||||||||]PAM</td><td>NA</td><td>Intergenic</td><td>NA</td><td>NA</td></tr>
<tr><td>T33446</td><td>CCATACCGGAAGACGGGCTTTGG</td><td></td><td>1000</td><td></td><td></td><td></td><td></td><td></td><td></td><td></td><td></td><td></td></tr>
<tr><td>Oligo fwd</td><td>CACCCCATACCGGAAGACGGGCTT</td><td></td><td></td><td></td><td></td><td></td><td></td><td></td><td></td><td></td><td></td><td></td></tr>
<tr><td>Oligo rev</td><td>AAACAAGCCCGTCTTCCGGTATGG</td><td></td><td></td><td></td><td></td><td></td><td></td><td></td><td></td><td></td><td></td><td></td></tr>
<tr><td>Chromosome</td><td>start</td><td></td><td>end</td><td>strand</td><td>MM</td><td>target_seq</td><td>PAM</td><td>alignment</td><td>distance</td><td>position</td><td>gene name</td><td>gene id</td></tr>
<tr><td>GU305914.1</td><td></td><td>14162</td><td>14184</td><td>-</td><td>0</td><td>CCATACCGGAAGACGGGCTT</td><td>TGG</td><td>||||||||[||||||||||||]PAM</td><td>NA</td><td>Intergenic</td><td>NA</td><td>NA</td></tr>
<tr><td>T33447</td><td>GTAAGATATATCCGACGTGACGG</td><td></td><td>1000</td><td></td><td></td><td></td><td></td><td></td><td></td><td></td><td></td><td></td></tr>
<tr><td>Oligo fwd</td><td>CACCGTAAGATATATCCGACGTGA</td><td></td><td></td><td></td><td></td><td></td><td></td><td></td><td></td><td></td><td></td><td></td></tr>
<tr><td>Oligo rev</td><td>AAACTCACGTCGGATATATCTTAC</td><td></td><td></td><td></td><td></td><td></td><td></td><td></td><td></td><td></td><td></td><td></td></tr>
<tr><td>Chromosome</td><td>start</td><td></td><td>end</td><td>strand</td><td>MM</td><td>target_seq</td><td>PAM</td><td>alignment</td><td>distance</td><td>position</td><td>gene name</td><td>gene id</td></tr>
<tr><td>GU305914.1</td><td></td><td>14090</td><td>14112</td><td>-</td><td>0</td><td>GTAAGATATATCCGACGTGA</td><td>CGG</td><td>||||||||[||||||||||||]PAM</td><td>NA</td><td>Intergenic</td><td>NA</td><td>NA</td></tr>
</table>

T33448 GACGGTACATCTTCTTGATACGG 1000
Oligo fwd CACCGACGGTACATCTTCTTGATA
Oligo rev AAACTATCAAGAAGATGTACCGTC
Chromosome start end strand MM target_seq PAM alignment distance position gene name gene id
GU305914.1 14072 14094 - 0 GACGGTACATCTTCTTGATA CGG ||||||||[||||||||||||]PAM NA Intergenic NA NA

T33449 CGGTAAAAGACTCGAAAGTGTGG 1000
Oligo fwd CACCCGGTAAAAGACTCGAAAGTG
Oligo rev AAACCACTTTCGAGTCTTTTACCG
Chromosome start end strand MM target_seq PAM alignment distance position gene name gene id
GU305914.1 14052 14074 - 0 CGGTAAAAGACTCGAAAGTG TGG ||||||||[||||||||||||]PAM NA Intergenic NA NA

T33450 AGACTCGAAAGTGTGGTCGAAGG 1000
Oligo fwd CACCAGACTCGAAAGTGTGGTCGA
Oligo rev AAACTCGACCACACTTTCGAGTCT
Chromosome start end strand MM target_seq PAM alignment distance position gene name gene id
GU305914.1 14045 14067 - 0 AGACTCGAAAGTGTGGTCGA AGG ||||||||[||||||||||||]PAM NA Intergenic NA NA

T33451 GAAGGATAGCGAGTGTGAAATGG 1000
Oligo fwd CACCGAAGGATAGCGAGTGTGAAA
Oligo rev AAACTTTCACACTCGCTATCCTTC
Chromosome start end strand MM target_seq PAM alignment distance position gene name gene id
GU305914.1 14027 14049 - 0 GAAGGATAGCGAGTGTGAAA TGG ||||||||[||||||||||||]PAM NA Intergenic NA NA

T33452 AAGGATAGCGAGTGTGAAATGGG 1000
Oligo fwd CACCAAGGATAGCGAGTGTGAAAT
Oligo rev AAACATTTCACACTCGCTATCCTT
Chromosome start end strand MM target_seq PAM alignment distance position gene name gene id
GU305914.1 14026 14048 - 0 AAGGATAGCGAGTGTGAAAT GGG ||||||||[||||||||||||]PAM NA Intergenic NA NA

T33453 AGGATAGCGAGTGTGAAATGGGG 1000
Oligo fwd CACCAGGATAGCGAGTGTGAAATG
Oligo rev AAACCATTTCACACTCGCTATCCT
Chromosome start end strand MM target_seq PAM alignment distance position gene name gene id
GU305914.1 14025 14047 - 0 AGGATAGCGAGTGTGAAATG GGG ||||||||[||||||||||||]PAM NA Intergenic NA NA

T33454 CGAGTGTGAAATGGGGAAACTGG 1000
Oligo fwd CACCCGAGTGTGAAATGGGGAAAC
Oligo rev AAACGTTTCCCCATTTCACACTCG
Chromosome start end strand MM target_seq PAM alignment distance position gene name gene id
GU305914.1 14018 14040 - 0 CGAGTGTGAAATGGGGAAAC TGG ||||||||[||||||||||||]PAM NA Intergenic NA NA

T33455 ACGAATGCTGACACATTTTCCGG 1000
Oligo fwd CACCACGAATGCTGACACATTTTC
Oligo rev AAACGAAAATGTGTCAGCATTCGT
Chromosome start end strand MM target_seq PAM alignment distance position gene name gene id
GU305914.1 13963 13985 - 0 ACGAATGCTGACACATTTTC CGG ||||||||[||||||||||||]PAM NA Intergenic NA NA

T33456 CGATCGTGCGCTGATCCTGACGG 1000
Oligo fwd CACCCGATCGTGCGCTGATCCTGA
Oligo rev AAACTCAGGATCAGCGCACGATCG
Chromosome start end strand MM target_seq PAM alignment distance position gene name gene id
GU305914.1 13939 13961 - 0 CGATCGTGCGCTGATCCTGA CGG ||||||||[||||||||||||]PAM NA Intergenic NA NA

T33457 GATCGTGCGCTGATCCTGACGGG 1000
Oligo fwd CACCGATCGTGCGCTGATCCTGAC
Oligo rev AAACGTCAGGATCAGCGCACGATC
Chromosome start end strand MM target_seq PAM alignment distance position gene name gene id
GU305914.1 13938 13960 - 0 GATCGTGCGCTGATCCTGAC GGG ||||||||[||||||||||||]PAM NA Intergenic NA NA

T33458 ATCGTGCGCTGATCCTGACGGGG 1000
Oligo fwd CACCATCGTGCGCTGATCCTGACG
Oligo rev AAACCGTCAGGATCAGCGCACGAT
Chromosome start end strand MM target_seq PAM alignment distance position gene name gene id
GU305914.1 13937 13959 - 0 ATCGTGCGCTGATCCTGACG GGG ||||||||[||||||||||||]PAM NA Intergenic NA NA

T33459 GCGCTGATCCTGACGGGGATCGG 1000
Oligo fwd CACCGCGCTGATCCTGACGGGGAT
Oligo rev AAACATCCCCGTCAGGATCAGCGC
Chromosome start end strand MM target_seq PAM alignment distance position gene name gene id
GU305914.1 13932 13954 - 0 GCGCTGATCCTGACGGGGAT CGG ||||||||[||||||||||||]PAM NA Intergenic NA NA

T33460 CACATAGCGAAAATCTGTCAAGG 1000
Oligo fwd CACCCACATAGCGAAAATCTGTCA
Oligo rev AAACTGACAGATTTTCGCTATGTG
Chromosome start end strand MM target_seq PAM alignment distance position gene name gene id
GU305914.1 13905 13927 - 0 CACATAGCGAAAATCTGTCA AGG ||||||||[||||||||||||]PAM NA Intergenic NA NA

T33461 GCTTGTGCCGCGTGACGTAAAGG 1000
Oligo fwd CACCGCTTGTGCCGCGTGACGTAA
Oligo rev AAACTTACGTCACGCGGCACAAGC
Chromosome start end strand MM target_seq PAM alignment distance position gene name gene id
GU305914.1 13835 13857 - 0 GCTTGTGCCGCGTGACGTAA AGG ||||||||[||||||||||||]PAM NA Intergenic NA NA

T33462 TGCCGCGTGACGTAAAGGCGCGG 1000
Oligo fwd CACCTGCCGCGTGACGTAAAGGCG
Oligo rev AAACCGCCTTTACGTCACGCGGCA
Chromosome start end strand MM target_seq PAM alignment distance position gene name gene id
GU305914.1 13830 13852 - 0 TGCCGCGTGACGTAAAGGCG CGG ||||||||[||||||||||||]PAM NA Intergenic NA NA

<table>
<tr><td>T33463</td><td>AAAGGCGCGGCGCTACTAGTAGG</td><td>1000</td><td></td><td></td><td></td><td></td><td></td><td></td><td></td><td></td><td></td></tr>
<tr><td>Oligo fwd</td><td>CACCAAAGGCGCGGCGCTACTAGT</td><td></td><td></td><td></td><td></td><td></td><td></td><td></td><td></td><td></td><td></td></tr>
<tr><td>Oligo rev</td><td>AAACACTAGTAGCGCCGCGCCTTT</td><td></td><td></td><td></td><td></td><td></td><td></td><td></td><td></td><td></td><td></td></tr>
<tr><td>Chromosome</td><td>start</td><td>end</td><td>strand</td><td>MM</td><td>target_seq</td><td>PAM</td><td>alignment</td><td>distance</td><td>position</td><td>gene name</td><td>gene id</td></tr>
<tr><td>GU305914.1</td><td>13817</td><td>13839</td><td>-</td><td>0</td><td>AAAGGCGCGGCGCTACTAGT</td><td>AGG</td><td>||||||||[||||||||||||]PAM</td><td>NA</td><td>Intergenic</td><td>NA</td><td>NA</td></tr>
<tr><td>T33464</td><td>AGGATTCTGAGCCGACGACACGG</td><td>1000</td><td></td><td></td><td></td><td></td><td></td><td></td><td></td><td></td><td></td></tr>
<tr><td>Oligo fwd</td><td>CACCAGGATTCTGAGCCGACGACA</td><td></td><td></td><td></td><td></td><td></td><td></td><td></td><td></td><td></td><td></td></tr>
<tr><td>Oligo rev</td><td>AAACTGTCGTCGGCTCAGAATCCT</td><td></td><td></td><td></td><td></td><td></td><td></td><td></td><td></td><td></td><td></td></tr>
<tr><td>Chromosome</td><td>start</td><td>end</td><td>strand</td><td>MM</td><td>target_seq</td><td>PAM</td><td>alignment</td><td>distance</td><td>position</td><td>gene name</td><td>gene id</td></tr>
<tr><td>GU305914.1</td><td>13797</td><td>13819</td><td>-</td><td>0</td><td>AGGATTCTGAGCCGACGACA</td><td>CGG</td><td>||||||||[||||||||||||]PAM</td><td>NA</td><td>Intergenic</td><td>NA</td><td>NA</td></tr>
<tr><td>T33465</td><td>AGCCGACGACACGGTGATCTCGG</td><td>1000</td><td></td><td></td><td></td><td></td><td></td><td></td><td></td><td></td><td></td></tr>
<tr><td>Oligo fwd</td><td>CACCAGCCGACGACACGGTGATCT</td><td></td><td></td><td></td><td></td><td></td><td></td><td></td><td></td><td></td><td></td></tr>
<tr><td>Oligo rev</td><td>AAACAGATCACCGTGTCGTCGGCT</td><td></td><td></td><td></td><td></td><td></td><td></td><td></td><td></td><td></td><td></td></tr>
<tr><td>Chromosome</td><td>start</td><td>end</td><td>strand</td><td>MM</td><td>target_seq</td><td>PAM</td><td>alignment</td><td>distance</td><td>position</td><td>gene name</td><td>gene id</td></tr>
<tr><td>GU305914.1</td><td>13788</td><td>13810</td><td>-</td><td>0</td><td>AGCCGACGACACGGTGATCT</td><td>CGG</td><td>||||||||[||||||||||||]PAM</td><td>NA</td><td>Intergenic</td><td>NA</td><td>NA</td></tr>
<tr><td>T33466</td><td>GCCGACGACACGGTGATCTCGGG</td><td>1000</td><td></td><td></td><td></td><td></td><td></td><td></td><td></td><td></td><td></td></tr>
<tr><td>Oligo fwd</td><td>CACCGCCGACGACACGGTGATCTC</td><td></td><td></td><td></td><td></td><td></td><td></td><td></td><td></td><td></td><td></td></tr>
<tr><td>Oligo rev</td><td>AAACGAGATCACCGTGTCGTCGGC</td><td></td><td></td><td></td><td></td><td></td><td></td><td></td><td></td><td></td><td></td></tr>
<tr><td>Chromosome</td><td>start</td><td>end</td><td>strand</td><td>MM</td><td>target_seq</td><td>PAM</td><td>alignment</td><td>distance</td><td>position</td><td>gene name</td><td>gene id</td></tr>
<tr><td>GU305914.1</td><td>13787</td><td>13809</td><td>-</td><td>0</td><td>GCCGACGACACGGTGATCTC</td><td>GGG</td><td>||||||||[||||||||||||]PAM</td><td>NA</td><td>Intergenic</td><td>NA</td><td>NA</td></tr>
<tr><td>T33467</td><td>ACCGATGTTCCTCTTGTTGTAGG</td><td>1000</td><td></td><td></td><td></td><td></td><td></td><td></td><td></td><td></td><td></td></tr>
<tr><td>Oligo fwd</td><td>CACCACCGATGTTCCTCTTGTTGT</td><td></td><td></td><td></td><td></td><td></td><td></td><td></td><td></td><td></td><td></td></tr>
<tr><td>Oligo rev</td><td>AAACACAACAAGAGGAACATCGGT</td><td></td><td></td><td></td><td></td><td></td><td></td><td></td><td></td><td></td><td></td></tr>
<tr><td>Chromosome</td><td>start</td><td>end</td><td>strand</td><td>MM</td><td>target_seq</td><td>PAM</td><td>alignment</td><td>distance</td><td>position</td><td>gene name</td><td>gene id</td></tr>
<tr><td>GU305914.1</td><td>13754</td><td>13776</td><td>-</td><td>0</td><td>ACCGATGTTCCTCTTGTTGT</td><td>AGG</td><td>||||||||[||||||||||||]PAM</td><td>NA</td><td>Intergenic</td><td>NA</td><td>NA</td></tr>
<tr><td>T33468</td><td>CTGATCATCTGTAGACTTTTCGG</td><td>1000</td><td></td><td></td><td></td><td></td><td></td><td></td><td></td><td></td><td></td></tr>
<tr><td>Oligo fwd</td><td>CACCCTGATCATCTGTAGACTTTT</td><td></td><td></td><td></td><td></td><td></td><td></td><td></td><td></td><td></td><td></td></tr>
<tr><td>Oligo rev</td><td>AAACAAAAGTCTACAGATGATCAG</td><td></td><td></td><td></td><td></td><td></td><td></td><td></td><td></td><td></td><td></td></tr>
<tr><td>Chromosome</td><td>start</td><td>end</td><td>strand</td><td>MM</td><td>target_seq</td><td>PAM</td><td>alignment</td><td>distance</td><td>position</td><td>gene name</td><td>gene id</td></tr>
<tr><td>GU305914.1</td><td>13730</td><td>13752</td><td>-</td><td>0</td><td>CTGATCATCTGTAGACTTTT</td><td>CGG</td><td>||||||||[||||||||||||]PAM</td><td>NA</td><td>Intergenic</td><td>NA</td><td>NA</td></tr>
<tr><td>T33469</td><td>ATCTGTAGACTTTTCGGCGTCGG</td><td>1000</td><td></td><td></td><td></td><td></td><td></td><td></td><td></td><td></td><td></td></tr>
<tr><td>Oligo fwd</td><td>CACCATCTGTAGACTTTTCGGCGT</td><td></td><td></td><td></td><td></td><td></td><td></td><td></td><td></td><td></td><td></td></tr>
<tr><td>Oligo rev</td><td>AAACACGCCGAAAAGTCTACAGAT</td><td></td><td></td><td></td><td></td><td></td><td></td><td></td><td></td><td></td><td></td></tr>
<tr><td>Chromosome</td><td>start</td><td>end</td><td>strand</td><td>MM</td><td>target_seq</td><td>PAM</td><td>alignment</td><td>distance</td><td>position</td><td>gene name</td><td>gene id</td></tr>
<tr><td>GU305914.1</td><td>13724</td><td>13746</td><td>-</td><td>0</td><td>ATCTGTAGACTTTTCGGCGT</td><td>CGG</td><td>||||||||[||||||||||||]PAM</td><td>NA</td><td>Intergenic</td><td>NA</td><td>NA</td></tr>
<tr><td>T33470</td><td>AGACTTTTCGGCGTCGGTGACGG</td><td>1000</td><td></td><td></td><td></td><td></td><td></td><td></td><td></td><td></td><td></td></tr>
<tr><td>Oligo fwd</td><td>CACCAGACTTTTCGGCGTCGGTGA</td><td></td><td></td><td></td><td></td><td></td><td></td><td></td><td></td><td></td><td></td></tr>
<tr><td>Oligo rev</td><td>AAACTCACCGACGCCGAAAAGTCT</td><td></td><td></td><td></td><td></td><td></td><td></td><td></td><td></td><td></td><td></td></tr>
<tr><td>Chromosome</td><td>start</td><td>end</td><td>strand</td><td>MM</td><td>target_seq</td><td>PAM</td><td>alignment</td><td>distance</td><td>position</td><td>gene name</td><td>gene id</td></tr>
<tr><td>GU305914.1</td><td>13718</td><td>13740</td><td>-</td><td>0</td><td>AGACTTTTCGGCGTCGGTGA</td><td>CGG</td><td>||||||||[||||||||||||]PAM</td><td>NA</td><td>Intergenic</td><td>NA</td><td>NA</td></tr>
<tr><td>T33471</td><td>TCGATACGACGACTCGACGTCGG</td><td>1000</td><td></td><td></td><td></td><td></td><td></td><td></td><td></td><td></td><td></td></tr>
<tr><td>Oligo fwd</td><td>CACCTCGATACGACGACTCGACGT</td><td></td><td></td><td></td><td></td><td></td><td></td><td></td><td></td><td></td><td></td></tr>
<tr><td>Oligo rev</td><td>AAACACGTCGAGTCGTCGTATCGA</td><td></td><td></td><td></td><td></td><td></td><td></td><td></td><td></td><td></td><td></td></tr>
<tr><td>Chromosome</td><td>start</td><td>end</td><td>strand</td><td>MM</td><td>target_seq</td><td>PAM</td><td>alignment</td><td>distance</td><td>position</td><td>gene name</td><td>gene id</td></tr>
<tr><td>GU305914.1</td><td>13691</td><td>13713</td><td>-</td><td>0</td><td>TCGATACGACGACTCGACGT</td><td>CGG</td><td>||||||||[||||||||||||]PAM</td><td>NA</td><td>Intergenic</td><td>NA</td><td>NA</td></tr>
<tr><td>T33472</td><td>ACGACGACTCGACGTCGGTTTGG</td><td>1000</td><td></td><td></td><td></td><td></td><td></td><td></td><td></td><td></td><td></td></tr>
<tr><td>Oligo fwd</td><td>CACCACGACGACTCGACGTCGGTT</td><td></td><td></td><td></td><td></td><td></td><td></td><td></td><td></td><td></td><td></td></tr>
<tr><td>Oligo rev</td><td>AAACAACCGACGTCGAGTCGTCGT</td><td></td><td></td><td></td><td></td><td></td><td></td><td></td><td></td><td></td><td></td></tr>
<tr><td>Chromosome</td><td>start</td><td>end</td><td>strand</td><td>MM</td><td>target_seq</td><td>PAM</td><td>alignment</td><td>distance</td><td>position</td><td>gene name</td><td>gene id</td></tr>
<tr><td>GU305914.1</td><td>13686</td><td>13708</td><td>-</td><td>0</td><td>ACGACGACTCGACGTCGGTT</td><td>TGG</td><td>||||||||[||||||||||||]PAM</td><td>NA</td><td>Intergenic</td><td>NA</td><td>NA</td></tr>
<tr><td>T33473</td><td>CAACTCACTTTGTTTATCCGCGG</td><td>1000</td><td></td><td></td><td></td><td></td><td></td><td></td><td></td><td></td><td></td></tr>
<tr><td>Oligo fwd</td><td>CACCCAACTCACTTTGTTTATCCG</td><td></td><td></td><td></td><td></td><td></td><td></td><td></td><td></td><td></td><td></td></tr>
<tr><td>Oligo rev</td><td>AAACCGGATAAACAAAGTGAGTTG</td><td></td><td></td><td></td><td></td><td></td><td></td><td></td><td></td><td></td><td></td></tr>
<tr><td>Chromosome</td><td>start</td><td>end</td><td>strand</td><td>MM</td><td>target_seq</td><td>PAM</td><td>alignment</td><td>distance</td><td>position</td><td>gene name</td><td>gene id</td></tr>
<tr><td>GU305914.1</td><td>13647</td><td>13669</td><td>-</td><td>0</td><td>CAACTCACTTTGTTTATCCG</td><td>CGG</td><td>||||||||[||||||||||||]PAM</td><td>NA</td><td>Intergenic</td><td>NA</td><td>NA</td></tr>
<tr><td>T33474</td><td>AACTCACTTTGTTTATCCGCGGG</td><td>1000</td><td></td><td></td><td></td><td></td><td></td><td></td><td></td><td></td><td></td></tr>
<tr><td>Oligo fwd</td><td>CACCAACTCACTTTGTTTATCCGC</td><td></td><td></td><td></td><td></td><td></td><td></td><td></td><td></td><td></td><td></td></tr>
<tr><td>Oligo rev</td><td>AAACGCGGATAAACAAAGTGAGTT</td><td></td><td></td><td></td><td></td><td></td><td></td><td></td><td></td><td></td><td></td></tr>
<tr><td>Chromosome</td><td>start</td><td>end</td><td>strand</td><td>MM</td><td>target_seq</td><td>PAM</td><td>alignment</td><td>distance</td><td>position</td><td>gene name</td><td>gene id</td></tr>
<tr><td>GU305914.1</td><td>13646</td><td>13668</td><td>-</td><td>0</td><td>AACTCACTTTGTTTATCCGC</td><td>GGG</td><td>||||||||[||||||||||||]PAM</td><td>NA</td><td>Intergenic</td><td>NA</td><td>NA</td></tr>
<tr><td>T33475</td><td>TACCGATCGCAATTGCCGTCTGG</td><td>1000</td><td></td><td></td><td></td><td></td><td></td><td></td><td></td><td></td><td></td></tr>
<tr><td>Oligo fwd</td><td>CACCTACCGATCGCAATTGCCGTC</td><td></td><td></td><td></td><td></td><td></td><td></td><td></td><td></td><td></td><td></td></tr>
<tr><td>Oligo rev</td><td>AAACGACGGCAATTGCGATCGGTA</td><td></td><td></td><td></td><td></td><td></td><td></td><td></td><td></td><td></td><td></td></tr>
<tr><td>Chromosome</td><td>start</td><td>end</td><td>strand</td><td>MM</td><td>target_seq</td><td>PAM</td><td>alignment</td><td>distance</td><td>position</td><td>gene name</td><td>gene id</td></tr>
<tr><td>GU305914.1</td><td>13621</td><td>13643</td><td>-</td><td>0</td><td>TACCGATCGCAATTGCCGTC</td><td>TGG</td><td>||||||||[||||||||||||]PAM</td><td>NA</td><td>Intergenic</td><td>NA</td><td>NA</td></tr>
<tr><td>T33476</td><td>ACCGATCGCAATTGCCGTCTGGG</td><td>1000</td><td></td><td></td><td></td><td></td><td></td><td></td><td></td><td></td><td></td></tr>
<tr><td>Oligo fwd</td><td>CACCACCGATCGCAATTGCCGTCT</td><td></td><td></td><td></td><td></td><td></td><td></td><td></td><td></td><td></td><td></td></tr>
<tr><td>Oligo rev</td><td>AAACAGACGGCAATTGCGATCGGT</td><td></td><td></td><td></td><td></td><td></td><td></td><td></td><td></td><td></td><td></td></tr>
<tr><td>Chromosome</td><td>start</td><td>end</td><td>strand</td><td>MM</td><td>target_seq</td><td>PAM</td><td>alignment</td><td>distance</td><td>position</td><td>gene name</td><td>gene id</td></tr>
<tr><td>GU305914.1</td><td>13620</td><td>13642</td><td>-</td><td>0</td><td>ACCGATCGCAATTGCCGTCT</td><td>GGG</td><td>||||||||[||||||||||||]PAM</td><td>NA</td><td>Intergenic</td><td>NA</td><td>NA</td></tr>
<tr><td>T33477</td><td>TCTGGGTCTCGATTTTCTCGCGG</td><td>1000</td><td></td><td></td><td></td><td></td><td></td><td></td><td></td><td></td><td></td></tr>
<tr><td>Oligo fwd</td><td>CACCTCTGGGTCTCGATTTTCTCG</td><td></td><td></td><td></td><td></td><td></td><td></td><td></td><td></td><td></td><td></td></tr>
<tr><td>Oligo rev</td><td>AAACCGAGAAAATCGAGACCCAGA</td><td></td><td></td><td></td><td></td><td></td><td></td><td></td><td></td><td></td><td></td></tr>
<tr><td>Chromosome</td><td>start</td><td>end</td><td>strand</td><td>MM</td><td>target_seq</td><td>PAM</td><td>alignment</td><td>distance</td><td>position</td><td>gene name</td><td>gene id</td></tr>
</table>

| | | | | | | | | | | | | |
|---|---|---|---|---|---|---|---|---|---|---|---|---|
| GU305914.1 | | 13603 | 13625 | - | 0 | TCTGGGTCTCGATTTTCTCG | CGG | \|\|\|\|\|\|\|\|[\|\|\|\|\|\|\|\|\|\|\|\|]PAM | NA | Intergenic | NA | NA |
| T33478 | CTGGGTCTCGATTTTCTCGCGGG | | 1000 | | | | | | | | | |
| Oligo fwd | CACCCTGGGTCTCGATTTTCTCGC | | | | | | | | | | | |
| Oligo rev | AAACGCGAGAAAATCGAGACCCAG | | | | | | | | | | | |
| Chromosome | start | | end | strand | MM | target_seq | PAM | alignment | distance | position | gene name | gene id |
| GU305914.1 | | 13602 | 13624 | - | 0 | CTGGGTCTCGATTTTCTCGC | GGG | \|\|\|\|\|\|\|\|[\|\|\|\|\|\|\|\|\|\|\|\|]PAM | NA | Intergenic | NA | NA |
| T33479 | ATTTTCTCGCGGGCGAACGCTGG | | 1000 | | | | | | | | | |
| Oligo fwd | CACCATTTTCTCGCGGGCGAACGC | | | | | | | | | | | |
| Oligo rev | AAACGCGTTCGCCCGCGAGAAAAT | | | | | | | | | | | |
| Chromosome | start | | end | strand | MM | target_seq | PAM | alignment | distance | position | gene name | gene id |
| GU305914.1 | | 13592 | 13614 | - | 0 | ATTTTCTCGCGGGCGAACGC | TGG | \|\|\|\|\|\|\|\|[\|\|\|\|\|\|\|\|\|\|\|\|]PAM | NA | Intergenic | NA | NA |
| T33480 | CTCGCGGGCGAACGCTGGATAGG | | 1000 | | | | | | | | | |
| Oligo fwd | CACCCTCGCGGGCGAACGCTGGAT | | | | | | | | | | | |
| Oligo rev | AAACATCCAGCGTTCGCCCGCGAG | | | | | | | | | | | |
| Chromosome | start | | end | strand | MM | target_seq | PAM | alignment | distance | position | gene name | gene id |
| GU305914.1 | | 13587 | 13609 | - | 0 | CTCGCGGGCGAACGCTGGAT | AGG | \|\|\|\|\|\|\|\|[\|\|\|\|\|\|\|\|\|\|\|\|]PAM | NA | Intergenic | NA | NA |
| T33481 | CATCTTCGAGTTTAGACAGCCGG | | 1000 | | | | | | | | | |
| Oligo fwd | CACCCATCTTCGAGTTTAGACAGC | | | | | | | | | | | |
| Oligo rev | AAACGCTGTCTAAACTCGAAGATG | | | | | | | | | | | |
| Chromosome | start | | end | strand | MM | target_seq | PAM | alignment | distance | position | gene name | gene id |
| GU305914.1 | | 13557 | 13579 | - | 0 | CATCTTCGAGTTTAGACAGC | CGG | \|\|\|\|\|\|\|\|[\|\|\|\|\|\|\|\|\|\|\|\|]PAM | NA | Intergenic | NA | NA |
| T33482 | GTTGACAGATTAGTCCCAATTGG | | 1000 | | | | | | | | | |
| Oligo fwd | CACCGTTGACAGATTAGTCCCAAT | | | | | | | | | | | |
| Oligo rev | AAACATTGGGACTAATCTGTCAAC | | | | | | | | | | | |
| Chromosome | start | | end | strand | MM | target_seq | PAM | alignment | distance | position | gene name | gene id |
| GU305914.1 | | 13531 | 13553 | - | 0 | GTTGACAGATTAGTCCCAAT | TGG | \|\|\|\|\|\|\|\|[\|\|\|\|\|\|\|\|\|\|\|\|]PAM | NA | Intergenic | NA | NA |
| T33483 | TGATGAGATATAGATGTTGTCGG | | 1000 | | | | | | | | | |
| Oligo fwd | CACCTGATGAGATATAGATGTTGT | | | | | | | | | | | |
| Oligo rev | AAACACAACATCTATATCTCATCA | | | | | | | | | | | |
| Chromosome | start | | end | strand | MM | target_seq | PAM | alignment | distance | position | gene name | gene id |
| GU305914.1 | | 13494 | 13516 | - | 0 | TGATGAGATATAGATGTTGT | CGG | \|\|\|\|\|\|\|\|[\|\|\|\|\|\|\|\|\|\|\|\|]PAM | NA | Intergenic | NA | NA |
| T33484 | GATATAGATGTTGTCGGTAACGG | | 1000 | | | | | | | | | |
| Oligo fwd | CACCGATATAGATGTTGTCGGTAA | | | | | | | | | | | |
| Oligo rev | AAACTTACCGACAACATCTATATC | | | | | | | | | | | |
| Chromosome | start | | end | strand | MM | target_seq | PAM | alignment | distance | position | gene name | gene id |
| GU305914.1 | | 13488 | 13510 | - | 0 | GATATAGATGTTGTCGGTAA | CGG | \|\|\|\|\|\|\|\|[\|\|\|\|\|\|\|\|\|\|\|\|]PAM | NA | Intergenic | NA | NA |
| T33485 | ATATAGATGTTGTCGGTAACGGG | | 1000 | | | | | | | | | |
| Oligo fwd | CACCATATAGATGTTGTCGGTAAC | | | | | | | | | | | |
| Oligo rev | AAACGTTACCGACAACATCTATAT | | | | | | | | | | | |
| Chromosome | start | | end | strand | MM | target_seq | PAM | alignment | distance | position | gene name | gene id |
| GU305914.1 | | 13487 | 13509 | - | 0 | ATATAGATGTTGTCGGTAAC | GGG | \|\|\|\|\|\|\|\|[\|\|\|\|\|\|\|\|\|\|\|\|]PAM | NA | Intergenic | NA | NA |
| T33486 | CCGCAGAACTGTTATGTGAAAGG | | 1000 | | | | | | | | | |
| Oligo fwd | CACCCCGCAGAACTGTTATGTGAA | | | | | | | | | | | |
| Oligo rev | AAACTTCACATAACAGTTCTGCGG | | | | | | | | | | | |
| Chromosome | start | | end | strand | MM | target_seq | PAM | alignment | distance | position | gene name | gene id |
| GU305914.1 | | 13450 | 13472 | - | 0 | CCGCAGAACTGTTATGTGAA | AGG | \|\|\|\|\|\|\|\|[\|\|\|\|\|\|\|\|\|\|\|\|]PAM | NA | Intergenic | NA | NA |
| T33487 | TTCCAGAAGCCGACTAATGTTGG | | 1000 | | | | | | | | | |
| Oligo fwd | CACCTTCCAGAAGCCGACTAATGT | | | | | | | | | | | |
| Oligo rev | AAACACATTAGTCGGCTTCTGGAA | | | | | | | | | | | |
| Chromosome | start | | end | strand | MM | target_seq | PAM | alignment | distance | position | gene name | gene id |
| GU305914.1 | | 13400 | 13422 | - | 0 | TTCCAGAAGCCGACTAATGT | TGG | \|\|\|\|\|\|\|\|[\|\|\|\|\|\|\|\|\|\|\|\|]PAM | NA | Intergenic | NA | NA |
| T33488 | CAGAAGCCGACTAATGTTGGTGG | | 1000 | | | | | | | | | |
| Oligo fwd | CACCCAGAAGCCGACTAATGTTGG | | | | | | | | | | | |
| Oligo rev | AAACCCAACATTAGTCGGCTTCTG | | | | | | | | | | | |
| Chromosome | start | | end | strand | MM | target_seq | PAM | alignment | distance | position | gene name | gene id |
| GU305914.1 | | 13397 | 13419 | - | 0 | CAGAAGCCGACTAATGTTGG | TGG | \|\|\|\|\|\|\|\|[\|\|\|\|\|\|\|\|\|\|\|\|]PAM | NA | Intergenic | NA | NA |
| T33489 | ACTAATGTTGGTGGAGATGTAGG | | 1000 | | | | | | | | | |
| Oligo fwd | CACCACTAATGTTGGTGGAGATGT | | | | | | | | | | | |
| Oligo rev | AAACACATCTCCACCAACATTAGT | | | | | | | | | | | |
| Chromosome | start | | end | strand | MM | target_seq | PAM | alignment | distance | position | gene name | gene id |
| GU305914.1 | | 13388 | 13410 | - | 0 | ACTAATGTTGGTGGAGATGT | AGG | \|\|\|\|\|\|\|\|[\|\|\|\|\|\|\|\|\|\|\|\|]PAM | NA | Intergenic | NA | NA |
| T33490 | GTGTAGACATCAACTGAACGTGG | | 1000 | | | | | | | | | |
| Oligo fwd | CACCGTGTAGACATCAACTGAACG | | | | | | | | | | | |
| Oligo rev | AAACCGTTCAGTTGATGTCTACAC | | | | | | | | | | | |
| Chromosome | start | | end | strand | MM | target_seq | PAM | alignment | distance | position | gene name | gene id |
| GU305914.1 | | 13366 | 13388 | - | 0 | GTGTAGACATCAACTGAACG | TGG | \|\|\|\|\|\|\|\|[\|\|\|\|\|\|\|\|\|\|\|\|]PAM | NA | Intergenic | NA | NA |
| T33491 | TGTAGACATCAACTGAACGTGGG | | 1000 | | | | | | | | | |
| Oligo fwd | CACCTGTAGACATCAACTGAACGT | | | | | | | | | | | |
| Oligo rev | AAACACGTTCAGTTGATGTCTACA | | | | | | | | | | | |
| Chromosome | start | | end | strand | MM | target_seq | PAM | alignment | distance | position | gene name | gene id |
| GU305914.1 | | 13365 | 13387 | - | 0 | TGTAGACATCAACTGAACGT | GGG | \|\|\|\|\|\|\|\|[\|\|\|\|\|\|\|\|\|\|\|\|]PAM | NA | Intergenic | NA | NA |
| T33492 | TGAACGTGGGCGAGTACTCGAGG | | 1000 | | | | | | | | | |
| Oligo fwd | CACCTGAACGTGGGCGAGTACTCG | | | | | | | | | | | |
| Oligo rev | AAACCGAGTACTCGCCCACGTTCA | | | | | | | | | | | |

Chromosome start end strand MM target_seq PAM alignment distance position gene name gene id
GU305914.1 13352 13374 - 0 TGAACGTGGGCGAGTACTCG AGG ||||||||[||||||||||||]PAM NA Intergenic NA NA

T33493 TGGGCGAGTACTCGAGGTGACGG 1000
Oligo fwd CACCTGGGCGAGTACTCGAGGTGA
Oligo rev AAACTCACCTCGAGTACTCGCCCA
Chromosome start end strand MM target_seq PAM alignment distance position gene name gene id
GU305914.1 13346 13368 - 0 TGGGCGAGTACTCGAGGTGA CGG ||||||||[||||||||||||]PAM NA Intergenic NA NA

T33494 GGGCGAGTACTCGAGGTGACGGG 1000
Oligo fwd CACCGGGCGAGTACTCGAGGTGAC
Oligo rev AAACGTCACCTCGAGTACTCGCCC
Chromosome start end strand MM target_seq PAM alignment distance position gene name gene id
GU305914.1 13345 13367 - 0 GGGCGAGTACTCGAGGTGAC GGG ||||||||[||||||||||||]PAM NA Intergenic NA NA

T33495 ACGTCCCGTACCACGTTGCCTGG 1000
Oligo fwd CACCACGTCCCGTACCACGTTGCC
Oligo rev AAACGGCAACGTGGTACGGGACGT
Chromosome start end strand MM target_seq PAM alignment distance position gene name gene id
GU305914.1 13291 13313 - 0 ACGTCCCGTACCACGTTGCC TGG ||||||||[||||||||||||]PAM NA Intergenic NA NA

T33496 CGTCCCGTACCACGTTGCCTGGG 1000
Oligo fwd CACCCGTCCCGTACCACGTTGCCT
Oligo rev AAACAGGCAACGTGGTACGGGACG
Chromosome start end strand MM target_seq PAM alignment distance position gene name gene id
GU305914.1 13290 13312 - 0 CGTCCCGTACCACGTTGCCT GGG ||||||||[||||||||||||]PAM NA Intergenic NA NA

T33497 ATTTGAATTGACATGAGTAGTGG 1000
Oligo fwd CACCATTTGAATTGACATGAGTAG
Oligo rev AAACCTACTCATGTCAATTCAAAT
Chromosome start end strand MM target_seq PAM alignment distance position gene name gene id
GU305914.1 13255 13277 - 0 ATTTGAATTGACATGAGTAG TGG ||||||||[||||||||||||]PAM NA Intergenic NA NA

T33498 ACATGAGTAGTGGACCCTGTTGG 1000
Oligo fwd CACCACATGAGTAGTGGACCCTGT
Oligo rev AAACACAGGGTCCACTACTCATGT
Chromosome start end strand MM target_seq PAM alignment distance position gene name gene id
GU305914.1 13245 13267 - 0 ACATGAGTAGTGGACCCTGT TGG ||||||||[||||||||||||]PAM NA Intergenic NA NA

T33499 CTTTCTCGTCGACACGTACGCGG 1000
Oligo fwd CACCCTTTCTCGTCGACACGTACG
Oligo rev AAACCGTACGTGTCGACGAGAAAG
Chromosome start end strand MM target_seq PAM alignment distance position gene name gene id
GU305914.1 13219 13241 - 0 CTTTCTCGTCGACACGTACG CGG ||||||||[||||||||||||]PAM NA Intergenic NA NA
GU305914.1 108133 108155 - 4 CTGTCTCTTCAACACGCACG CGG ||-||||-[||-|||||-|||]PAM NA Intergenic NA NA

T33500 TCGAAAAAAATCACACGTTTCTGG 1000
Oligo fwd CACCTCGAAAAAATCACACGTTTC
Oligo rev AAACGAAACGTGTGATTTTTTCGA
Chromosome start end strand MM target_seq PAM alignment distance position gene name gene id
GU305914.1 13181 13203 - 0 TCGAAAAAATCACACGTTTC TGG ||||||||[||||||||||||]PAM NA Intergenic NA NA

T33501 CTGTAGATTCAACAGCGCTCAGG 1000
Oligo fwd CACCCTGTAGATTCAACAGCGCTC
Oligo rev AAACGAGCGCTGTTGAATCTACAG
Chromosome start end strand MM target_seq PAM alignment distance position gene name gene id
GU305914.1 13123 13145 - 0 CTGTAGATTCAACAGCGCTC AGG ||||||||[||||||||||||]PAM NA Intergenic NA NA

T33502 ATTAAAAAAAATACCAAACTCTGG 1000
Oligo fwd CACCATTAAAAAAAATACCAAACTC
Oligo rev AAACGAGTTTGGTATTTTTTTAAT
Chromosome start end strand MM target_seq PAM alignment distance position gene name gene id
GU305914.1 13057 13079 - 0 ATTAAAAAAAATACCAAACTC TGG ||||||||[||||||||||||]PAM NA Intergenic NA NA

T33503 TGATAAGAATTATACTTTAATGG 1000
Oligo fwd CACCTGATAAGAATTATACTTTAA
Oligo rev AAACTTAAAGTATAATTCTTATCA
Chromosome start end strand MM target_seq PAM alignment distance position gene name gene id
GU305914.1 13028 13050 - 0 TGATAAGAATTATACTTTAA TGG ||||||||[||||||||||||]PAM NA Intergenic NA NA

T33504 GATAAGAATTATACTTTAATGGG 1000
Oligo fwd CACCGATAAGAATTATACTTTAAT
Oligo rev AAACATTAAAGTATAATTCTTATC
Chromosome start end strand MM target_seq PAM alignment distance position gene name gene id
GU305914.1 13027 13049 - 0 GATAAGAATTATACTTTAAT GGG ||||||||[||||||||||||]PAM NA Intergenic NA NA

T33505 ATAAGAATTATACTTTAATGGGG 1000
Oligo fwd CACCATAAGAATTATACTTTAATG
Oligo rev AAACCATTAAAGTATAATTCTTAT
Chromosome start end strand MM target_seq PAM alignment distance position gene name gene id
GU305914.1 13026 13048 - 0 ATAAGAATTATACTTTAATG GGG ||||||||[||||||||||||]PAM NA Intergenic NA NA

T33506 TAAGAATTATACTTTAATGGGGG 1000
Oligo fwd CACCTAAGAATTATACTTTAATGG
Oligo rev AAACCCATTAAAGTATAATTCTTA
Chromosome start end strand MM target_seq PAM alignment distance position gene name gene id
GU305914.1 13025 13047 - 0 TAAGAATTATACTTTAATGG GGG ||||||||[||||||||||||]PAM NA Intergenic NA NA

T33507 TAGAACACGATAAACTCTAGCGG 1000

| | | | | | | | | | | | |
|---|---|---|---|---|---|---|---|---|---|---|---|
| Oligo fwd | CACCTAGAACACGATAAACTCTAG | | | | | | | | | | |
| Oligo rev | AAACCTAGAGTTTATCGTGTTCTA | | | | | | | | | | |
| Chromosome | start | end | strand | MM | target_seq | PAM | alignment | distance | position | gene name | gene id |
| GU305914.1 | 12994 | 13016 | - | 0 | TAGAACACGATAAACTCTAG | CGG | \|\|\|\|\|\|\|\|[\|\|\|\|\|\|\|\|\|\|\|\|]PAM | NA | Intergenic | NA | NA |
| T33508 | ACGATAAACTCTAGCGGTTGCGG | 1000 | | | | | | | | | |
| Oligo fwd | CACCACGATAAACTCTAGCGGTTG | | | | | | | | | | |
| Oligo rev | AAACCAACCGCTAGAGTTTATCGT | | | | | | | | | | |
| Chromosome | start | end | strand | MM | target_seq | PAM | alignment | distance | position | gene name | gene id |
| GU305914.1 | 12988 | 13010 | - | 0 | ACGATAAACTCTAGCGGTTG | CGG | \|\|\|\|\|\|\|\|[\|\|\|\|\|\|\|\|\|\|\|\|]PAM | NA | Intergenic | NA | NA |
| T33509 | ACTCTAGCGGTTGCGGATTGCGG | 1000 | | | | | | | | | |
| Oligo fwd | CACCACTCTAGCGGTTGCGGATTG | | | | | | | | | | |
| Oligo rev | AAACCAATCCGCAACCGCTAGAGT | | | | | | | | | | |
| Chromosome | start | end | strand | MM | target_seq | PAM | alignment | distance | position | gene name | gene id |
| GU305914.1 | 12981 | 13003 | - | 0 | ACTCTAGCGGTTGCGGATTG | CGG | \|\|\|\|\|\|\|\|[\|\|\|\|\|\|\|\|\|\|\|\|]PAM | NA | Intergenic | NA | NA |
| T33510 | ACCGAAAGAATAGATCGTCGCGG | 1000 | | | | | | | | | |
| Oligo fwd | CACCACCGAAAGAATAGATCGTCG | | | | | | | | | | |
| Oligo rev | AAACCGACGATCTATTCTTTCGGT | | | | | | | | | | |
| Chromosome | start | end | strand | MM | target_seq | PAM | alignment | distance | position | gene name | gene id |
| GU305914.1 | 12957 | 12979 | - | 0 | ACCGAAAGAATAGATCGTCG | CGG | \|\|\|\|\|\|\|\|[\|\|\|\|\|\|\|\|\|\|\|\|]PAM | NA | Intergenic | NA | NA |
| T33511 | CGGCGAACAGCAGACCCAGCAGG | 1000 | | | | | | | | | |
| Oligo fwd | CACCCGGCGAACAGCAGACCCAGC | | | | | | | | | | |
| Oligo rev | AAACGCTGGGTCTGCTGTTCGCCG | | | | | | | | | | |
| Chromosome | start | end | strand | MM | target_seq | PAM | alignment | distance | position | gene name | gene id |
| GU305914.1 | 12937 | 12959 | - | 0 | CGGCGAACAGCAGACCCAGC | AGG | \|\|\|\|\|\|\|\|[\|\|\|\|\|\|\|\|\|\|\|\|]PAM | NA | Intergenic | NA | NA |
| T33512 | CCAGCAGGAGCCCTATCGTCAGG | 1000 | | | | | | | | | |
| Oligo fwd | CACCCCAGCAGGAGCCCTATCGTC | | | | | | | | | | |
| Oligo rev | AAACGACGATAGGGCTCCTGCTGG | | | | | | | | | | |
| Chromosome | start | end | strand | MM | target_seq | PAM | alignment | distance | position | gene name | gene id |
| GU305914.1 | 12922 | 12944 | - | 0 | CCAGCAGGAGCCCTATCGTC | AGG | \|\|\|\|\|\|\|\|[\|\|\|\|\|\|\|\|\|\|\|\|]PAM | NA | Intergenic | NA | NA |
| T33513 | TATCGTCAGGACGATGATAGAGG | 1000 | | | | | | | | | |
| Oligo fwd | CACCTATCGTCAGGACGATGATAG | | | | | | | | | | |
| Oligo rev | AAACCTATCATCGTCCTGACGATA | | | | | | | | | | |
| Chromosome | start | end | strand | MM | target_seq | PAM | alignment | distance | position | gene name | gene id |
| GU305914.1 | 12909 | 12931 | - | 0 | TATCGTCAGGACGATGATAG | AGG | \|\|\|\|\|\|\|\|[\|\|\|\|\|\|\|\|\|\|\|\|]PAM | NA | Intergenic | NA | NA |
| T33514 | TCTCGTGTGTGCTCTCGTCGAGG | 1000 | | | | | | | | | |
| Oligo fwd | CACCTCTCGTGTGTGCTCTCGTCG | | | | | | | | | | |
| Oligo rev | AAACCGACGAGAGCACACACGAGA | | | | | | | | | | |
| Chromosome | start | end | strand | MM | target_seq | PAM | alignment | distance | position | gene name | gene id |
| GU305914.1 | 12870 | 12892 | - | 0 | TCTCGTGTGTGCTCTCGTCG | AGG | \|\|\|\|\|\|\|\|[\|\|\|\|\|\|\|\|\|\|\|\|]PAM | NA | Intergenic | NA | NA |
| T33515 | CGTGTGTGCTCTCGTCGAGGTGG | 1000 | | | | | | | | | |
| Oligo fwd | CACCCGTGTGTGCTCTCGTCGAGG | | | | | | | | | | |
| Oligo rev | AAACCCTCGACGAGAGCACACACG | | | | | | | | | | |
| Chromosome | start | end | strand | MM | target_seq | PAM | alignment | distance | position | gene name | gene id |
| GU305914.1 | 12867 | 12889 | - | 0 | CGTGTGTGCTCTCGTCGAGG | TGG | \|\|\|\|\|\|\|\|[\|\|\|\|\|\|\|\|\|\|\|\|]PAM | NA | Intergenic | NA | NA |
| T33516 | TGTGCTCTCGTCGAGGTGGTTGG | 1000 | | | | | | | | | |
| Oligo fwd | CACCTGTGCTCTCGTCGAGGTGGT | | | | | | | | | | |
| Oligo rev | AAACACCACCTCGACGAGAGCACA | | | | | | | | | | |
| Chromosome | start | end | strand | MM | target_seq | PAM | alignment | distance | position | gene name | gene id |
| GU305914.1 | 12863 | 12885 | - | 0 | TGTGCTCTCGTCGAGGTGGT | TGG | \|\|\|\|\|\|\|\|[\|\|\|\|\|\|\|\|\|\|\|\|]PAM | NA | Intergenic | NA | NA |
| T33517 | GTGCTCTCGTCGAGGTGGTTGGG | 1000 | | | | | | | | | |
| Oligo fwd | CACCGTGCTCTCGTCGAGGTGGTT | | | | | | | | | | |
| Oligo rev | AAACAACCACCTCGACGAGAGCAC | | | | | | | | | | |
| Chromosome | start | end | strand | MM | target_seq | PAM | alignment | distance | position | gene name | gene id |
| GU305914.1 | 12862 | 12884 | - | 0 | GTGCTCTCGTCGAGGTGGTT | GGG | \|\|\|\|\|\|\|\|[\|\|\|\|\|\|\|\|\|\|\|\|]PAM | NA | Intergenic | NA | NA |
| T33518 | GAGGTGGTTGGGCGATCGTTAGG | 1000 | | | | | | | | | |
| Oligo fwd | CACCGAGGTGGTTGGGCGATCGTT | | | | | | | | | | |
| Oligo rev | AAACAACGATCGCCCAACCACCTC | | | | | | | | | | |
| Chromosome | start | end | strand | MM | target_seq | PAM | alignment | distance | position | gene name | gene id |
| GU305914.1 | 12851 | 12873 | - | 0 | GAGGTGGTTGGGCGATCGTT | AGG | \|\|\|\|\|\|\|\|[\|\|\|\|\|\|\|\|\|\|\|\|]PAM | NA | Intergenic | NA | NA |
| T33519 | CGATCGTTAGGCGCGACAATTGG | 1000 | | | | | | | | | |
| Oligo fwd | CACCCGATCGTTAGGCGCGACAAT | | | | | | | | | | |
| Oligo rev | AAACATTGTCGCGCCTAACGATCG | | | | | | | | | | |
| Chromosome | start | end | strand | MM | target_seq | PAM | alignment | distance | position | gene name | gene id |
| GU305914.1 | 12839 | 12861 | - | 0 | CGATCGTTAGGCGCGACAAT | TGG | \|\|\|\|\|\|\|\|[\|\|\|\|\|\|\|\|\|\|\|\|]PAM | NA | Intergenic | NA | NA |
| T33520 | ACCGTCACCATTCAAATAAATGG | 1000 | | | | | | | | | |
| Oligo fwd | CACCACCGTCACCATTCAAATAAA | | | | | | | | | | |
| Oligo rev | AAACTTTATTTGAATGGTGACGGT | | | | | | | | | | |
| Chromosome | start | end | strand | MM | target_seq | PAM | alignment | distance | position | gene name | gene id |
| GU305914.1 | 12807 | 12829 | - | 0 | ACCGTCACCATTCAAATAAA | TGG | \|\|\|\|\|\|\|\|[\|\|\|\|\|\|\|\|\|\|\|\|]PAM | NA | Intergenic | NA | NA |
| T33521 | CCATTCAAATAAATGGTCGTTGG | 1000 | | | | | | | | | |
| Oligo fwd | CACCCCATTCAAATAAATGGTCGT | | | | | | | | | | |
| Oligo rev | AAACACGACCATTTATTTGAATGG | | | | | | | | | | |
| Chromosome | start | end | strand | MM | target_seq | PAM | alignment | distance | position | gene name | gene id |
| GU305914.1 | 12800 | 12822 | - | 0 | CCATTCAAATAAATGGTCGT | TGG | \|\|\|\|\|\|\|\|[\|\|\|\|\|\|\|\|\|\|\|\|]PAM | NA | Intergenic | NA | NA |

<table>
<tr><td>T33522</td><td>GTCGTTGGCGTATCGCTGTTTGG</td><td>1000</td><td></td><td></td><td></td><td></td><td></td><td></td><td></td><td></td><td></td></tr>
<tr><td>Oligo fwd</td><td>CACCGTCGTTGGCGTATCGCTGTT</td><td></td><td></td><td></td><td></td><td></td><td></td><td></td><td></td><td></td><td></td></tr>
<tr><td>Oligo rev</td><td>AAACAACAGCGATACGCCAACGAC</td><td></td><td></td><td></td><td></td><td></td><td></td><td></td><td></td><td></td><td></td></tr>
<tr><td>Chromosome</td><td>start</td><td>end</td><td>strand</td><td>MM</td><td>target_seq</td><td>PAM</td><td>alignment</td><td>distance</td><td>position</td><td>gene name</td><td>gene id</td></tr>
<tr><td>GU305914.1</td><td>12785</td><td>12807</td><td>-</td><td>0</td><td>GTCGTTGGCGTATCGCTGTT</td><td>TGG</td><td>||||||||[||||||||||||]PAM</td><td>NA</td><td>Intergenic</td><td>NA</td><td>NA</td></tr>
<tr><td>T33523</td><td>TTCGTGTCCGTGCTCTTACGTGG</td><td>1000</td><td></td><td></td><td></td><td></td><td></td><td></td><td></td><td></td><td></td></tr>
<tr><td>Oligo fwd</td><td>CACCTTCGTGTCCGTGCTCTTACG</td><td></td><td></td><td></td><td></td><td></td><td></td><td></td><td></td><td></td><td></td></tr>
<tr><td>Oligo rev</td><td>AAACCGTAAGAGCACGGACACGAA</td><td></td><td></td><td></td><td></td><td></td><td></td><td></td><td></td><td></td><td></td></tr>
<tr><td>Chromosome</td><td>start</td><td>end</td><td>strand</td><td>MM</td><td>target_seq</td><td>PAM</td><td>alignment</td><td>distance</td><td>position</td><td>gene name</td><td>gene id</td></tr>
<tr><td>GU305914.1</td><td>12746</td><td>12768</td><td>-</td><td>0</td><td>TTCGTGTCCGTGCTCTTACG</td><td>TGG</td><td>||||||||[||||||||||||]PAM</td><td>NA</td><td>Intergenic</td><td>NA</td><td>NA</td></tr>
<tr><td>T33524</td><td>GAGTGATTTGTTCATATCTCCGG</td><td>1000</td><td></td><td></td><td></td><td></td><td></td><td></td><td></td><td></td><td></td></tr>
<tr><td>Oligo fwd</td><td>CACCGAGTGATTTGTTCATATCTC</td><td></td><td></td><td></td><td></td><td></td><td></td><td></td><td></td><td></td><td></td></tr>
<tr><td>Oligo rev</td><td>AAACGAGATATGAACAAATCACTC</td><td></td><td></td><td></td><td></td><td></td><td></td><td></td><td></td><td></td><td></td></tr>
<tr><td>Chromosome</td><td>start</td><td>end</td><td>strand</td><td>MM</td><td>target_seq</td><td>PAM</td><td>alignment</td><td>distance</td><td>position</td><td>gene name</td><td>gene id</td></tr>
<tr><td>GU305914.1</td><td>12678</td><td>12700</td><td>-</td><td>0</td><td>GAGTGATTTGTTCATATCTC</td><td>CGG</td><td>||||||||[||||||||||||]PAM</td><td>NA</td><td>Intergenic</td><td>NA</td><td>NA</td></tr>
<tr><td>T33525</td><td>GATTTGTTCATATCTCCGGCTGG</td><td>1000</td><td></td><td></td><td></td><td></td><td></td><td></td><td></td><td></td><td></td></tr>
<tr><td>Oligo fwd</td><td>CACCGATTTGTTCATATCTCCGGC</td><td></td><td></td><td></td><td></td><td></td><td></td><td></td><td></td><td></td><td></td></tr>
<tr><td>Oligo rev</td><td>AAACGCCGGAGATATGAACAAATC</td><td></td><td></td><td></td><td></td><td></td><td></td><td></td><td></td><td></td><td></td></tr>
<tr><td>Chromosome</td><td>start</td><td>end</td><td>strand</td><td>MM</td><td>target_seq</td><td>PAM</td><td>alignment</td><td>distance</td><td>position</td><td>gene name</td><td>gene id</td></tr>
<tr><td>GU305914.1</td><td>12674</td><td>12696</td><td>-</td><td>0</td><td>GATTTGTTCATATCTCCGGC</td><td>TGG</td><td>||||||||[||||||||||||]PAM</td><td>NA</td><td>Intergenic</td><td>NA</td><td>NA</td></tr>
<tr><td>T33526</td><td>GTTCATATCTCCGGCTGGTGCGG</td><td>1000</td><td></td><td></td><td></td><td></td><td></td><td></td><td></td><td></td><td></td></tr>
<tr><td>Oligo fwd</td><td>CACCGTTCATATCTCCGGCTGGTG</td><td></td><td></td><td></td><td></td><td></td><td></td><td></td><td></td><td></td><td></td></tr>
<tr><td>Oligo rev</td><td>AAACCACCAGCCGGAGATATGAAC</td><td></td><td></td><td></td><td></td><td></td><td></td><td></td><td></td><td></td><td></td></tr>
<tr><td>Chromosome</td><td>start</td><td>end</td><td>strand</td><td>MM</td><td>target_seq</td><td>PAM</td><td>alignment</td><td>distance</td><td>position</td><td>gene name</td><td>gene id</td></tr>
<tr><td>GU305914.1</td><td>12669</td><td>12691</td><td>-</td><td>0</td><td>GTTCATATCTCCGGCTGGTG</td><td>CGG</td><td>||||||||[||||||||||||]PAM</td><td>NA</td><td>Intergenic</td><td>NA</td><td>NA</td></tr>
<tr><td>T33527</td><td>GTATGTAGCTGAATGAGAGATGG</td><td>1000</td><td></td><td></td><td></td><td></td><td></td><td></td><td></td><td></td><td></td></tr>
<tr><td>Oligo fwd</td><td>CACCGTATGTAGCTGAATGAGAGA</td><td></td><td></td><td></td><td></td><td></td><td></td><td></td><td></td><td></td><td></td></tr>
<tr><td>Oligo rev</td><td>AAACTCTCTCATTCAGCTACATAC</td><td></td><td></td><td></td><td></td><td></td><td></td><td></td><td></td><td></td><td></td></tr>
<tr><td>Chromosome</td><td>start</td><td>end</td><td>strand</td><td>MM</td><td>target_seq</td><td>PAM</td><td>alignment</td><td>distance</td><td>position</td><td>gene name</td><td>gene id</td></tr>
<tr><td>GU305914.1</td><td>12565</td><td>12587</td><td>-</td><td>0</td><td>GTATGTAGCTGAATGAGAGA</td><td>TGG</td><td>||||||||[||||||||||||]PAM</td><td>NA</td><td>Intergenic</td><td>NA</td><td>NA</td></tr>
<tr><td>T33528</td><td>TACATATTGTTCTAAGTAAAAGG</td><td>1000</td><td></td><td></td><td></td><td></td><td></td><td></td><td></td><td></td><td></td></tr>
<tr><td>Oligo fwd</td><td>CACCTACATATTGTTCTAAGTAAA</td><td></td><td></td><td></td><td></td><td></td><td></td><td></td><td></td><td></td><td></td></tr>
<tr><td>Oligo rev</td><td>AAACTTTACTTAGAACAATATGTA</td><td></td><td></td><td></td><td></td><td></td><td></td><td></td><td></td><td></td><td></td></tr>
<tr><td>Chromosome</td><td>start</td><td>end</td><td>strand</td><td>MM</td><td>target_seq</td><td>PAM</td><td>alignment</td><td>distance</td><td>position</td><td>gene name</td><td>gene id</td></tr>
<tr><td>GU305914.1</td><td>12540</td><td>12562</td><td>-</td><td>0</td><td>TACATATTGTTCTAAGTAAA</td><td>AGG</td><td>||||||||[||||||||||||]PAM</td><td>NA</td><td>Intergenic</td><td>NA</td><td>NA</td></tr>
<tr><td>T33529</td><td>ACATATTGTTCTAAGTAAAAGGG</td><td>1000</td><td></td><td></td><td></td><td></td><td></td><td></td><td></td><td></td><td></td></tr>
<tr><td>Oligo fwd</td><td>CACCACATATTGTTCTAAGTAAAA</td><td></td><td></td><td></td><td></td><td></td><td></td><td></td><td></td><td></td><td></td></tr>
<tr><td>Oligo rev</td><td>AAACTTTTACTTAGAACAATATGT</td><td></td><td></td><td></td><td></td><td></td><td></td><td></td><td></td><td></td><td></td></tr>
<tr><td>Chromosome</td><td>start</td><td>end</td><td>strand</td><td>MM</td><td>target_seq</td><td>PAM</td><td>alignment</td><td>distance</td><td>position</td><td>gene name</td><td>gene id</td></tr>
<tr><td>GU305914.1</td><td>12539</td><td>12561</td><td>-</td><td>0</td><td>ACATATTGTTCTAAGTAAAA</td><td>GGG</td><td>||||||||[||||||||||||]PAM</td><td>NA</td><td>Intergenic</td><td>NA</td><td>NA</td></tr>
<tr><td>T33530</td><td>CATATTGTTCTAAGTAAAAGGGG</td><td>1000</td><td></td><td></td><td></td><td></td><td></td><td></td><td></td><td></td><td></td></tr>
<tr><td>Oligo fwd</td><td>CACCCATATTGTTCTAAGTAAAAG</td><td></td><td></td><td></td><td></td><td></td><td></td><td></td><td></td><td></td><td></td></tr>
<tr><td>Oligo rev</td><td>AAACCTTTTACTTAGAACAATATG</td><td></td><td></td><td></td><td></td><td></td><td></td><td></td><td></td><td></td><td></td></tr>
<tr><td>Chromosome</td><td>start</td><td>end</td><td>strand</td><td>MM</td><td>target_seq</td><td>PAM</td><td>alignment</td><td>distance</td><td>position</td><td>gene name</td><td>gene id</td></tr>
<tr><td>GU305914.1</td><td>12538</td><td>12560</td><td>-</td><td>0</td><td>CATATTGTTCTAAGTAAAAG</td><td>GGG</td><td>||||||||[||||||||||||]PAM</td><td>NA</td><td>Intergenic</td><td>NA</td><td>NA</td></tr>
<tr><td>T33531</td><td>GTTCTAAGTAAAAGGGGATACGG</td><td>1000</td><td></td><td></td><td></td><td></td><td></td><td></td><td></td><td></td><td></td></tr>
<tr><td>Oligo fwd</td><td>CACCGTTCTAAGTAAAAGGGGATA</td><td></td><td></td><td></td><td></td><td></td><td></td><td></td><td></td><td></td><td></td></tr>
<tr><td>Oligo rev</td><td>AAACTATCCCCTTTTACTTAGAAC</td><td></td><td></td><td></td><td></td><td></td><td></td><td></td><td></td><td></td><td></td></tr>
<tr><td>Chromosome</td><td>start</td><td>end</td><td>strand</td><td>MM</td><td>target_seq</td><td>PAM</td><td>alignment</td><td>distance</td><td>position</td><td>gene name</td><td>gene id</td></tr>
<tr><td>GU305914.1</td><td>12532</td><td>12554</td><td>-</td><td>0</td><td>GTTCTAAGTAAAAGGGGATA</td><td>CGG</td><td>||||||||[||||||||||||]PAM</td><td>NA</td><td>Intergenic</td><td>NA</td><td>NA</td></tr>
<tr><td>T33532</td><td>TTCTAAGTAAAAGGGGATACGGG</td><td>1000</td><td></td><td></td><td></td><td></td><td></td><td></td><td></td><td></td><td></td></tr>
<tr><td>Oligo fwd</td><td>CACCTTCTAAGTAAAAGGGGATAC</td><td></td><td></td><td></td><td></td><td></td><td></td><td></td><td></td><td></td><td></td></tr>
<tr><td>Oligo rev</td><td>AAACGTATCCCCTTTTACTTAGAA</td><td></td><td></td><td></td><td></td><td></td><td></td><td></td><td></td><td></td><td></td></tr>
<tr><td>Chromosome</td><td>start</td><td>end</td><td>strand</td><td>MM</td><td>target_seq</td><td>PAM</td><td>alignment</td><td>distance</td><td>position</td><td>gene name</td><td>gene id</td></tr>
<tr><td>GU305914.1</td><td>12531</td><td>12553</td><td>-</td><td>0</td><td>TTCTAAGTAAAAGGGGATAC</td><td>GGG</td><td>||||||||[||||||||||||]PAM</td><td>NA</td><td>Intergenic</td><td>NA</td><td>NA</td></tr>
<tr><td>T33533</td><td>TAAGTAAAAGGGGATACGGGCGG</td><td>1000</td><td></td><td></td><td></td><td></td><td></td><td></td><td></td><td></td><td></td></tr>
<tr><td>Oligo fwd</td><td>CACCTAAGTAAAAGGGGATACGGG</td><td></td><td></td><td></td><td></td><td></td><td></td><td></td><td></td><td></td><td></td></tr>
<tr><td>Oligo rev</td><td>AAACCCCGTATCCCCTTTTACTTA</td><td></td><td></td><td></td><td></td><td></td><td></td><td></td><td></td><td></td><td></td></tr>
<tr><td>Chromosome</td><td>start</td><td>end</td><td>strand</td><td>MM</td><td>target_seq</td><td>PAM</td><td>alignment</td><td>distance</td><td>position</td><td>gene name</td><td>gene id</td></tr>
<tr><td>GU305914.1</td><td>12528</td><td>12550</td><td>-</td><td>0</td><td>TAAGTAAAAGGGGATACGGG</td><td>CGG</td><td>||||||||[||||||||||||]PAM</td><td>NA</td><td>Intergenic</td><td>NA</td><td>NA</td></tr>
<tr><td>T33534</td><td>AAGTAAAAGGGGATACGGGCGGG</td><td>1000</td><td></td><td></td><td></td><td></td><td></td><td></td><td></td><td></td><td></td></tr>
<tr><td>Oligo fwd</td><td>CACCAAGTAAAAGGGGATACGGGC</td><td></td><td></td><td></td><td></td><td></td><td></td><td></td><td></td><td></td><td></td></tr>
<tr><td>Oligo rev</td><td>AAACGCCCGTATCCCCTTTTACTT</td><td></td><td></td><td></td><td></td><td></td><td></td><td></td><td></td><td></td><td></td></tr>
<tr><td>Chromosome</td><td>start</td><td>end</td><td>strand</td><td>MM</td><td>target_seq</td><td>PAM</td><td>alignment</td><td>distance</td><td>position</td><td>gene name</td><td>gene id</td></tr>
<tr><td>GU305914.1</td><td>12527</td><td>12549</td><td>-</td><td>0</td><td>AAGTAAAAGGGGATACGGGC</td><td>GGG</td><td>||||||||[||||||||||||]PAM</td><td>NA</td><td>Intergenic</td><td>NA</td><td>NA</td></tr>
<tr><td>T33535</td><td>ACGAACCCAATCATTTGAAGCGG</td><td>1000</td><td></td><td></td><td></td><td></td><td></td><td></td><td></td><td></td><td></td></tr>
<tr><td>Oligo fwd</td><td>CACCACGAACCCAATCATTTGAAG</td><td></td><td></td><td></td><td></td><td></td><td></td><td></td><td></td><td></td><td></td></tr>
<tr><td>Oligo rev</td><td>AAACCTTCAAATGATTGGGTTCGT</td><td></td><td></td><td></td><td></td><td></td><td></td><td></td><td></td><td></td><td></td></tr>
<tr><td>Chromosome</td><td>start</td><td>end</td><td>strand</td><td>MM</td><td>target_seq</td><td>PAM</td><td>alignment</td><td>distance</td><td>position</td><td>gene name</td><td>gene id</td></tr>
<tr><td>GU305914.1</td><td>12492</td><td>12514</td><td>-</td><td>0</td><td>ACGAACCCAATCATTTGAAG</td><td>CGG</td><td>||||||||[||||||||||||]PAM</td><td>NA</td><td>Intergenic</td><td>NA</td><td>NA</td></tr>
<tr><td>T33536</td><td>TGAAGCGGTCGAAACCTCTGCGG</td><td>1000</td><td></td><td></td><td></td><td></td><td></td><td></td><td></td><td></td><td></td></tr>
<tr><td>Oligo fwd</td><td>CACCTGAAGCGGTCGAAACCTCTG</td><td></td><td></td><td></td><td></td><td></td><td></td><td></td><td></td><td></td><td></td></tr>
<tr><td>Oligo rev</td><td>AAACCAGAGGTTTCGACCGCTTCA</td><td></td><td></td><td></td><td></td><td></td><td></td><td></td><td></td><td></td><td></td></tr>
<tr><td>Chromosome</td><td>start</td><td>end</td><td>strand</td><td>MM</td><td>target_seq</td><td>PAM</td><td>alignment</td><td>distance</td><td>position</td><td>gene name</td><td>gene id</td></tr>
<tr><td>GU305914.1</td><td>12477</td><td>12499</td><td>-</td><td>0</td><td>TGAAGCGGTCGAAACCTCTG</td><td>CGG</td><td>||||||||[||||||||||||]PAM</td><td>NA</td><td>Intergenic</td><td>NA</td><td>NA</td></tr>
</table>

<table>
<tr><td>GU305914.1</td><td>81222</td><td>81244</td><td>-</td><td>4</td><td>TGAGGGGGACGAAACCTCGG</td><td>AGG</td><td>|||-|-||[-|||||||||-|]PAM</td><td>NA</td><td>Intergenic</td><td>NA</td><td>NA</td></tr>
<tr><td></td><td></td><td></td><td></td><td></td><td></td><td></td><td></td><td></td><td></td><td></td><td></td></tr>
<tr><td>T33537</td><td>AGCAGATACGAGCACGATCGCGG</td><td>1000</td><td></td><td></td><td></td><td></td><td></td><td></td><td></td><td></td><td></td></tr>
<tr><td>Oligo fwd</td><td>CACCAGCAGATACGAGCACGATCG</td><td></td><td></td><td></td><td></td><td></td><td></td><td></td><td></td><td></td><td></td></tr>
<tr><td>Oligo rev</td><td>AAACCGATCGTGCTCGTATCTGCT</td><td></td><td></td><td></td><td></td><td></td><td></td><td></td><td></td><td></td><td></td></tr>
<tr><td>Chromosome</td><td>start</td><td>end</td><td>strand</td><td>MM</td><td>target_seq</td><td>PAM</td><td>alignment</td><td>distance</td><td>position</td><td>gene name</td><td>gene id</td></tr>
<tr><td>GU305914.1</td><td>12416</td><td>12438</td><td>-</td><td>0</td><td>AGCAGATACGAGCACGATCG</td><td>CGG</td><td>||||||||[||||||||||||]PAM</td><td>NA</td><td>Intergenic</td><td>NA</td><td>NA</td></tr>
<tr><td>GU305914.1</td><td>12880</td><td>12902</td><td>+</td><td>4</td><td>AGCACACACGAGAACGAGCG</td><td>CGG</td><td>||||-|-|[||||-||||-||]PAM</td><td>NA</td><td>Intergenic</td><td>NA</td><td>NA</td></tr>
<tr><td></td><td></td><td></td><td></td><td></td><td></td><td></td><td></td><td></td><td></td><td></td><td></td></tr>
<tr><td>T33538</td><td>TCTACTGTTCGTGTCGATGTTGG</td><td>1000</td><td></td><td></td><td></td><td></td><td></td><td></td><td></td><td></td><td></td></tr>
<tr><td>Oligo fwd</td><td>CACCTCTACTGTTCGTGTCGATGT</td><td></td><td></td><td></td><td></td><td></td><td></td><td></td><td></td><td></td><td></td></tr>
<tr><td>Oligo rev</td><td>AAACACATCGACACGAACAGTAGA</td><td></td><td></td><td></td><td></td><td></td><td></td><td></td><td></td><td></td><td></td></tr>
<tr><td>Chromosome</td><td>start</td><td>end</td><td>strand</td><td>MM</td><td>target_seq</td><td>PAM</td><td>alignment</td><td>distance</td><td>position</td><td>gene name</td><td>gene id</td></tr>
<tr><td>GU305914.1</td><td>12379</td><td>12401</td><td>-</td><td>0</td><td>TCTACTGTTCGTGTCGATGT</td><td>TGG</td><td>||||||||[||||||||||||]PAM</td><td>NA</td><td>Intergenic</td><td>NA</td><td>NA</td></tr>
<tr><td></td><td></td><td></td><td></td><td></td><td></td><td></td><td></td><td></td><td></td><td></td><td></td></tr>
<tr><td>T33539</td><td>GCAAAGTTAGCACGTTGAACCGG</td><td>1000</td><td></td><td></td><td></td><td></td><td></td><td></td><td></td><td></td><td></td></tr>
<tr><td>Oligo fwd</td><td>CACCGCAAAGTTAGCACGTTGAAC</td><td></td><td></td><td></td><td></td><td></td><td></td><td></td><td></td><td></td><td></td></tr>
<tr><td>Oligo rev</td><td>AAACGTTCAACGTGCTAACTTTGC</td><td></td><td></td><td></td><td></td><td></td><td></td><td></td><td></td><td></td><td></td></tr>
<tr><td>Chromosome</td><td>start</td><td>end</td><td>strand</td><td>MM</td><td>target_seq</td><td>PAM</td><td>alignment</td><td>distance</td><td>position</td><td>gene name</td><td>gene id</td></tr>
<tr><td>GU305914.1</td><td>12325</td><td>12347</td><td>-</td><td>0</td><td>GCAAAGTTAGCACGTTGAAC</td><td>CGG</td><td>||||||||[||||||||||||]PAM</td><td>NA</td><td>Intergenic</td><td>NA</td><td>NA</td></tr>
<tr><td></td><td></td><td></td><td></td><td></td><td></td><td></td><td></td><td></td><td></td><td></td><td></td></tr>
<tr><td>T33540</td><td>CAAAGTTAGCACGTTGAACCGGG</td><td>1000</td><td></td><td></td><td></td><td></td><td></td><td></td><td></td><td></td><td></td></tr>
<tr><td>Oligo fwd</td><td>CACCCAAAGTTAGCACGTTGAACC</td><td></td><td></td><td></td><td></td><td></td><td></td><td></td><td></td><td></td><td></td></tr>
<tr><td>Oligo rev</td><td>AAACGGTTCAACGTGCTAACTTTG</td><td></td><td></td><td></td><td></td><td></td><td></td><td></td><td></td><td></td><td></td></tr>
<tr><td>Chromosome</td><td>start</td><td>end</td><td>strand</td><td>MM</td><td>target_seq</td><td>PAM</td><td>alignment</td><td>distance</td><td>position</td><td>gene name</td><td>gene id</td></tr>
<tr><td>GU305914.1</td><td>12324</td><td>12346</td><td>-</td><td>0</td><td>CAAAGTTAGCACGTTGAACC</td><td>GGG</td><td>||||||||[||||||||||||]PAM</td><td>NA</td><td>Intergenic</td><td>NA</td><td>NA</td></tr>
<tr><td></td><td></td><td></td><td></td><td></td><td></td><td></td><td></td><td></td><td></td><td></td><td></td></tr>
<tr><td>T33541</td><td>GGATATGCGCATTTTTTCTGAGG</td><td>1000</td><td></td><td></td><td></td><td></td><td></td><td></td><td></td><td></td><td></td></tr>
<tr><td>Oligo fwd</td><td>CACCGGATATGCGCATTTTTTCTG</td><td></td><td></td><td></td><td></td><td></td><td></td><td></td><td></td><td></td><td></td></tr>
<tr><td>Oligo rev</td><td>AAACCAGAAAAAATGCGCATATCC</td><td></td><td></td><td></td><td></td><td></td><td></td><td></td><td></td><td></td><td></td></tr>
<tr><td>Chromosome</td><td>start</td><td>end</td><td>strand</td><td>MM</td><td>target_seq</td><td>PAM</td><td>alignment</td><td>distance</td><td>position</td><td>gene name</td><td>gene id</td></tr>
<tr><td>GU305914.1</td><td>12303</td><td>12325</td><td>-</td><td>0</td><td>GGATATGCGCATTTTTTCTG</td><td>AGG</td><td>||||||||[||||||||||||]PAM</td><td>NA</td><td>Intergenic</td><td>NA</td><td>NA</td></tr>
<tr><td></td><td></td><td></td><td></td><td></td><td></td><td></td><td></td><td></td><td></td><td></td><td></td></tr>
<tr><td>T33542</td><td>CAAAGCGTAATATAACATTTTGG</td><td>1000</td><td></td><td></td><td></td><td></td><td></td><td></td><td></td><td></td><td></td></tr>
<tr><td>Oligo fwd</td><td>CACCCAAAGCGTAATATAACATTT</td><td></td><td></td><td></td><td></td><td></td><td></td><td></td><td></td><td></td><td></td></tr>
<tr><td>Oligo rev</td><td>AAACAAATGTTATATTACGCTTTG</td><td></td><td></td><td></td><td></td><td></td><td></td><td></td><td></td><td></td><td></td></tr>
<tr><td>Chromosome</td><td>start</td><td>end</td><td>strand</td><td>MM</td><td>target_seq</td><td>PAM</td><td>alignment</td><td>distance</td><td>position</td><td>gene name</td><td>gene id</td></tr>
<tr><td>GU305914.1</td><td>12277</td><td>12299</td><td>-</td><td>0</td><td>CAAAGCGTAATATAACATTT</td><td>TGG</td><td>||||||||[||||||||||||]PAM</td><td>NA</td><td>Intergenic</td><td>NA</td><td>NA</td></tr>
<tr><td></td><td></td><td></td><td></td><td></td><td></td><td></td><td></td><td></td><td></td><td></td><td></td></tr>
<tr><td>T33543</td><td>AAAGCGTAATATAACATTTTGGG</td><td>1000</td><td></td><td></td><td></td><td></td><td></td><td></td><td></td><td></td><td></td></tr>
<tr><td>Oligo fwd</td><td>CACCAAAGCGTAATATAACATTTT</td><td></td><td></td><td></td><td></td><td></td><td></td><td></td><td></td><td></td><td></td></tr>
<tr><td>Oligo rev</td><td>AAACAAAATGTTATATTACGCTTT</td><td></td><td></td><td></td><td></td><td></td><td></td><td></td><td></td><td></td><td></td></tr>
<tr><td>Chromosome</td><td>start</td><td>end</td><td>strand</td><td>MM</td><td>target_seq</td><td>PAM</td><td>alignment</td><td>distance</td><td>position</td><td>gene name</td><td>gene id</td></tr>
<tr><td>GU305914.1</td><td>12276</td><td>12298</td><td>-</td><td>0</td><td>AAAGCGTAATATAACATTTT</td><td>GGG</td><td>||||||||[||||||||||||]PAM</td><td>NA</td><td>Intergenic</td><td>NA</td><td>NA</td></tr>
<tr><td></td><td></td><td></td><td></td><td></td><td></td><td></td><td></td><td></td><td></td><td></td><td></td></tr>
<tr><td>T33544</td><td>CGTAATATAACATTTTGGGTCGG</td><td>1000</td><td></td><td></td><td></td><td></td><td></td><td></td><td></td><td></td><td></td></tr>
<tr><td>Oligo fwd</td><td>CACCCGTAATATAACATTTTGGGT</td><td></td><td></td><td></td><td></td><td></td><td></td><td></td><td></td><td></td><td></td></tr>
<tr><td>Oligo rev</td><td>AAACACCCAAAATGTTATATTACG</td><td></td><td></td><td></td><td></td><td></td><td></td><td></td><td></td><td></td><td></td></tr>
<tr><td>Chromosome</td><td>start</td><td>end</td><td>strand</td><td>MM</td><td>target_seq</td><td>PAM</td><td>alignment</td><td>distance</td><td>position</td><td>gene name</td><td>gene id</td></tr>
<tr><td>GU305914.1</td><td>12272</td><td>12294</td><td>-</td><td>0</td><td>CGTAATATAACATTTTGGGT</td><td>CGG</td><td>||||||||[||||||||||||]PAM</td><td>NA</td><td>Intergenic</td><td>NA</td><td>NA</td></tr>
<tr><td></td><td></td><td></td><td></td><td></td><td></td><td></td><td></td><td></td><td></td><td></td><td></td></tr>
<tr><td>T33545</td><td>ATATATCACCAGTACCTGAATGG</td><td>1000</td><td></td><td></td><td></td><td></td><td></td><td></td><td></td><td></td><td></td></tr>
<tr><td>Oligo fwd</td><td>CACCATATATCACCAGTACCTGAA</td><td></td><td></td><td></td><td></td><td></td><td></td><td></td><td></td><td></td><td></td></tr>
<tr><td>Oligo rev</td><td>AAACTTCAGGTACTGGTGATATAT</td><td></td><td></td><td></td><td></td><td></td><td></td><td></td><td></td><td></td><td></td></tr>
<tr><td>Chromosome</td><td>start</td><td>end</td><td>strand</td><td>MM</td><td>target_seq</td><td>PAM</td><td>alignment</td><td>distance</td><td>position</td><td>gene name</td><td>gene id</td></tr>
<tr><td>GU305914.1</td><td>12225</td><td>12247</td><td>-</td><td>0</td><td>ATATATCACCAGTACCTGAA</td><td>TGG</td><td>||||||||[||||||||||||]PAM</td><td>NA</td><td>Intergenic</td><td>NA</td><td>NA</td></tr>
<tr><td></td><td></td><td></td><td></td><td></td><td></td><td></td><td></td><td></td><td></td><td></td><td></td></tr>
<tr><td>T33546</td><td>ATCCACAAGTCTGATGATAAAGG</td><td>1000</td><td></td><td></td><td></td><td></td><td></td><td></td><td></td><td></td><td></td></tr>
<tr><td>Oligo fwd</td><td>CACCATCCACAAGTCTGATGATAA</td><td></td><td></td><td></td><td></td><td></td><td></td><td></td><td></td><td></td><td></td></tr>
<tr><td>Oligo rev</td><td>AAACTTATCATCAGACTTGTGGAT</td><td></td><td></td><td></td><td></td><td></td><td></td><td></td><td></td><td></td><td></td></tr>
<tr><td>Chromosome</td><td>start</td><td>end</td><td>strand</td><td>MM</td><td>target_seq</td><td>PAM</td><td>alignment</td><td>distance</td><td>position</td><td>gene name</td><td>gene id</td></tr>
<tr><td>GU305914.1</td><td>12202</td><td>12224</td><td>-</td><td>0</td><td>ATCCACAAGTCTGATGATAA</td><td>AGG</td><td>||||||||[||||||||||||]PAM</td><td>NA</td><td>Intergenic</td><td>NA</td><td>NA</td></tr>
<tr><td></td><td></td><td></td><td></td><td></td><td></td><td></td><td></td><td></td><td></td><td></td><td></td></tr>
<tr><td>T33547</td><td>AAGTCTGATGATAAAGGCCATGG</td><td>1000</td><td></td><td></td><td></td><td></td><td></td><td></td><td></td><td></td><td></td></tr>
<tr><td>Oligo fwd</td><td>CACCAAGTCTGATGATAAAGGCCA</td><td></td><td></td><td></td><td></td><td></td><td></td><td></td><td></td><td></td><td></td></tr>
<tr><td>Oligo rev</td><td>AAACTGGCCTTTATCATCAGACTT</td><td></td><td></td><td></td><td></td><td></td><td></td><td></td><td></td><td></td><td></td></tr>
<tr><td>Chromosome</td><td>start</td><td>end</td><td>strand</td><td>MM</td><td>target_seq</td><td>PAM</td><td>alignment</td><td>distance</td><td>position</td><td>gene name</td><td>gene id</td></tr>
<tr><td>GU305914.1</td><td>12196</td><td>12218</td><td>-</td><td>0</td><td>AAGTCTGATGATAAAGGCCA</td><td>TGG</td><td>||||||||[||||||||||||]PAM</td><td>NA</td><td>Intergenic</td><td>NA</td><td>NA</td></tr>
<tr><td></td><td></td><td></td><td></td><td></td><td></td><td></td><td></td><td></td><td></td><td></td><td></td></tr>
<tr><td>T33548</td><td>TGATAAAGGCCATGGTTTTAAGG</td><td>1000</td><td></td><td></td><td></td><td></td><td></td><td></td><td></td><td></td><td></td></tr>
<tr><td>Oligo fwd</td><td>CACCTGATAAAGGCCATGGTTTTA</td><td></td><td></td><td></td><td></td><td></td><td></td><td></td><td></td><td></td><td></td></tr>
<tr><td>Oligo rev</td><td>AAACTAAAACCATGGCCTTTATCA</td><td></td><td></td><td></td><td></td><td></td><td></td><td></td><td></td><td></td><td></td></tr>
<tr><td>Chromosome</td><td>start</td><td>end</td><td>strand</td><td>MM</td><td>target_seq</td><td>PAM</td><td>alignment</td><td>distance</td><td>position</td><td>gene name</td><td>gene id</td></tr>
<tr><td>GU305914.1</td><td>12188</td><td>12210</td><td>-</td><td>0</td><td>TGATAAAGGCCATGGTTTTA</td><td>AGG</td><td>||||||||[||||||||||||]PAM</td><td>NA</td><td>Intergenic</td><td>NA</td><td>NA</td></tr>
<tr><td></td><td></td><td></td><td></td><td></td><td></td><td></td><td></td><td></td><td></td><td></td><td></td></tr>
<tr><td>T33549</td><td>TGGTTTTAAGGTAGATATTCCGG</td><td>1000</td><td></td><td></td><td></td><td></td><td></td><td></td><td></td><td></td><td></td></tr>
<tr><td>Oligo fwd</td><td>CACCTGGTTTTAAGGTAGATATTC</td><td></td><td></td><td></td><td></td><td></td><td></td><td></td><td></td><td></td><td></td></tr>
<tr><td>Oligo rev</td><td>AAACGAATATCTACCTTAAAACCA</td><td></td><td></td><td></td><td></td><td></td><td></td><td></td><td></td><td></td><td></td></tr>
<tr><td>Chromosome</td><td>start</td><td>end</td><td>strand</td><td>MM</td><td>target_seq</td><td>PAM</td><td>alignment</td><td>distance</td><td>position</td><td>gene name</td><td>gene id</td></tr>
<tr><td>GU305914.1</td><td>12176</td><td>12198</td><td>-</td><td>0</td><td>TGGTTTTAAGGTAGATATTC</td><td>CGG</td><td>||||||||[||||||||||||]PAM</td><td>NA</td><td>Intergenic</td><td>NA</td><td>NA</td></tr>
<tr><td></td><td></td><td></td><td></td><td></td><td></td><td></td><td></td><td></td><td></td><td></td><td></td></tr>
<tr><td>T33550</td><td>GGTAGATATTCCGGTTAACGTGG</td><td>1000</td><td></td><td></td><td></td><td></td><td></td><td></td><td></td><td></td><td></td></tr>
<tr><td>Oligo fwd</td><td>CACCGGTAGATATTCCGGTTAACG</td><td></td><td></td><td></td><td></td><td></td><td></td><td></td><td></td><td></td><td></td></tr>
<tr><td>Oligo rev</td><td>AAACCGTTAACCGGAATATCTACC</td><td></td><td></td><td></td><td></td><td></td><td></td><td></td><td></td><td></td><td></td></tr>
<tr><td>Chromosome</td><td>start</td><td>end</td><td>strand</td><td>MM</td><td>target_seq</td><td>PAM</td><td>alignment</td><td>distance</td><td>position</td><td>gene name</td><td>gene id</td></tr>
<tr><td>GU305914.1</td><td>12167</td><td>12189</td><td>-</td><td>0</td><td>GGTAGATATTCCGGTTAACG</td><td>TGG</td><td>||||||||[||||||||||||]PAM</td><td>NA</td><td>Intergenic</td><td>NA</td><td>NA</td></tr>
<tr><td></td><td></td><td></td><td></td><td></td><td></td><td></td><td></td><td></td><td></td><td></td><td></td></tr>
<tr><td>T33551</td><td>GATATTCCGGTTAACGTGGTAGG</td><td>1000</td><td></td><td></td><td></td><td></td><td></td><td></td><td></td><td></td><td></td></tr>
<tr><td>Oligo fwd</td><td>CACCGATATTCCGGTTAACGTGGT</td><td></td><td></td><td></td><td></td><td></td><td></td><td></td><td></td><td></td><td></td></tr>
</table>

<table>
<tr><td>Oligo rev</td><td>AAACACCACGTTAACCGGAATATC</td><td></td><td></td><td></td><td></td><td></td><td></td><td></td><td></td><td></td><td></td><td></td></tr>
<tr><td>Chromosome</td><td>start</td><td></td><td>end</td><td>strand</td><td>MM</td><td>target_seq</td><td>PAM</td><td>alignment</td><td>distance</td><td>position</td><td>gene name</td><td>gene id</td></tr>
<tr><td>GU305914.1</td><td></td><td>12163</td><td>12185</td><td>-</td><td>0</td><td>GATATTCCGGTTAACGTGGT</td><td>AGG</td><td>||||||||[||||||||||||]PAM</td><td>NA</td><td>Intergenic</td><td>NA</td><td>NA</td></tr>
<tr><td>T33552</td><td>TGTGCGTACAAACAATCGACCGG</td><td></td><td>1000</td><td></td><td></td><td></td><td></td><td></td><td></td><td></td><td></td><td></td></tr>
<tr><td>Oligo fwd</td><td>CACCTGTGCGTACAAACAATCGAC</td><td></td><td></td><td></td><td></td><td></td><td></td><td></td><td></td><td></td><td></td><td></td></tr>
<tr><td>Oligo rev</td><td>AAACGTCGATTGTTTGTACGCACA</td><td></td><td></td><td></td><td></td><td></td><td></td><td></td><td></td><td></td><td></td><td></td></tr>
<tr><td>Chromosome</td><td>start</td><td></td><td>end</td><td>strand</td><td>MM</td><td>target_seq</td><td>PAM</td><td>alignment</td><td>distance</td><td>position</td><td>gene name</td><td>gene id</td></tr>
<tr><td>GU305914.1</td><td></td><td>12083</td><td>12105</td><td>-</td><td>0</td><td>TGTGCGTACAAACAATCGAC</td><td>CGG</td><td>||||||||[||||||||||||]PAM</td><td>NA</td><td>Intergenic</td><td>NA</td><td>NA</td></tr>
<tr><td>T33553</td><td>ACCAAGTTGCGTTCATGCAAAGG</td><td></td><td>1000</td><td></td><td></td><td></td><td></td><td></td><td></td><td></td><td></td><td></td></tr>
<tr><td>Oligo fwd</td><td>CACCACCAAGTTGCGTTCATGCAA</td><td></td><td></td><td></td><td></td><td></td><td></td><td></td><td></td><td></td><td></td><td></td></tr>
<tr><td>Oligo rev</td><td>AAACTTGCATGAACGCAACTTGGT</td><td></td><td></td><td></td><td></td><td></td><td></td><td></td><td></td><td></td><td></td><td></td></tr>
<tr><td>Chromosome</td><td>start</td><td></td><td>end</td><td>strand</td><td>MM</td><td>target_seq</td><td>PAM</td><td>alignment</td><td>distance</td><td>position</td><td>gene name</td><td>gene id</td></tr>
<tr><td>GU305914.1</td><td></td><td>12060</td><td>12082</td><td>-</td><td>0</td><td>ACCAAGTTGCGTTCATGCAA</td><td>AGG</td><td>||||||||[||||||||||||]PAM</td><td>NA</td><td>Intergenic</td><td>NA</td><td>NA</td></tr>
<tr><td>T33554</td><td>CATGCAAAGGTATTCTTCGACGG</td><td></td><td>1000</td><td></td><td></td><td></td><td></td><td></td><td></td><td></td><td></td><td></td></tr>
<tr><td>Oligo fwd</td><td>CACCCATGCAAAGGTATTCTTCGA</td><td></td><td></td><td></td><td></td><td></td><td></td><td></td><td></td><td></td><td></td><td></td></tr>
<tr><td>Oligo rev</td><td>AAACTCGAAGAATACCTTTGCATG</td><td></td><td></td><td></td><td></td><td></td><td></td><td></td><td></td><td></td><td></td><td></td></tr>
<tr><td>Chromosome</td><td>start</td><td></td><td>end</td><td>strand</td><td>MM</td><td>target_seq</td><td>PAM</td><td>alignment</td><td>distance</td><td>position</td><td>gene name</td><td>gene id</td></tr>
<tr><td>GU305914.1</td><td></td><td>12047</td><td>12069</td><td>-</td><td>0</td><td>CATGCAAAGGTATTCTTCGA</td><td>CGG</td><td>||||||||[||||||||||||]PAM</td><td>NA</td><td>Intergenic</td><td>NA</td><td>NA</td></tr>
<tr><td>T33555</td><td>ATGCAAAGGTATTCTTCGACGGG</td><td></td><td>1000</td><td></td><td></td><td></td><td></td><td></td><td></td><td></td><td></td><td></td></tr>
<tr><td>Oligo fwd</td><td>CACCATGCAAAGGTATTCTTCGAC</td><td></td><td></td><td></td><td></td><td></td><td></td><td></td><td></td><td></td><td></td><td></td></tr>
<tr><td>Oligo rev</td><td>AAACGTCGAAGAATACCTTTGCAT</td><td></td><td></td><td></td><td></td><td></td><td></td><td></td><td></td><td></td><td></td><td></td></tr>
<tr><td>Chromosome</td><td>start</td><td></td><td>end</td><td>strand</td><td>MM</td><td>target_seq</td><td>PAM</td><td>alignment</td><td>distance</td><td>position</td><td>gene name</td><td>gene id</td></tr>
<tr><td>GU305914.1</td><td></td><td>12046</td><td>12068</td><td>-</td><td>0</td><td>ATGCAAAGGTATTCTTCGAC</td><td>GGG</td><td>||||||||[||||||||||||]PAM</td><td>NA</td><td>Intergenic</td><td>NA</td><td>NA</td></tr>
<tr><td>T33556</td><td>CGGGACATCTAAATATTTTGAGG</td><td></td><td>1000</td><td></td><td></td><td></td><td></td><td></td><td></td><td></td><td></td><td></td></tr>
<tr><td>Oligo fwd</td><td>CACCCGGGACATCTAAATATTTTG</td><td></td><td></td><td></td><td></td><td></td><td></td><td></td><td></td><td></td><td></td><td></td></tr>
<tr><td>Oligo rev</td><td>AAACCAAAATATTTAGATGTCCCG</td><td></td><td></td><td></td><td></td><td></td><td></td><td></td><td></td><td></td><td></td><td></td></tr>
<tr><td>Chromosome</td><td>start</td><td></td><td>end</td><td>strand</td><td>MM</td><td>target_seq</td><td>PAM</td><td>alignment</td><td>distance</td><td>position</td><td>gene name</td><td>gene id</td></tr>
<tr><td>GU305914.1</td><td></td><td>12027</td><td>12049</td><td>-</td><td>0</td><td>CGGGACATCTAAATATTTTG</td><td>AGG</td><td>||||||||[||||||||||||]PAM</td><td>NA</td><td>Intergenic</td><td>NA</td><td>NA</td></tr>
<tr><td>T33557</td><td>AATATTTTGAGGATAGTAACTGG</td><td></td><td>1000</td><td></td><td></td><td></td><td></td><td></td><td></td><td></td><td></td><td></td></tr>
<tr><td>Oligo fwd</td><td>CACCAATATTTTGAGGATAGTAAC</td><td></td><td></td><td></td><td></td><td></td><td></td><td></td><td></td><td></td><td></td><td></td></tr>
<tr><td>Oligo rev</td><td>AAACGTTACTATCCTCAAAATATT</td><td></td><td></td><td></td><td></td><td></td><td></td><td></td><td></td><td></td><td></td><td></td></tr>
<tr><td>Chromosome</td><td>start</td><td></td><td>end</td><td>strand</td><td>MM</td><td>target_seq</td><td>PAM</td><td>alignment</td><td>distance</td><td>position</td><td>gene name</td><td>gene id</td></tr>
<tr><td>GU305914.1</td><td></td><td>12016</td><td>12038</td><td>-</td><td>0</td><td>AATATTTTGAGGATAGTAAC</td><td>TGG</td><td>||||||||[||||||||||||]PAM</td><td>NA</td><td>Intergenic</td><td>NA</td><td>NA</td></tr>
<tr><td>T33558</td><td>TTTATCCACTTGTTTATTCGCGG</td><td></td><td>1000</td><td></td><td></td><td></td><td></td><td></td><td></td><td></td><td></td><td></td></tr>
<tr><td>Oligo fwd</td><td>CACCTTTATCCACTTGTTTATTCG</td><td></td><td></td><td></td><td></td><td></td><td></td><td></td><td></td><td></td><td></td><td></td></tr>
<tr><td>Oligo rev</td><td>AAACCGAATAAACAAGTGGATAAA</td><td></td><td></td><td></td><td></td><td></td><td></td><td></td><td></td><td></td><td></td><td></td></tr>
<tr><td>Chromosome</td><td>start</td><td></td><td>end</td><td>strand</td><td>MM</td><td>target_seq</td><td>PAM</td><td>alignment</td><td>distance</td><td>position</td><td>gene name</td><td>gene id</td></tr>
<tr><td>GU305914.1</td><td></td><td>11993</td><td>12015</td><td>-</td><td>0</td><td>TTTATCCACTTGTTTATTCG</td><td>CGG</td><td>||||||||[||||||||||||]PAM</td><td>NA</td><td>Intergenic</td><td>NA</td><td>NA</td></tr>
<tr><td>T33559</td><td>AACAGCGACATGCATTTTCCTGG</td><td></td><td>1000</td><td></td><td></td><td></td><td></td><td></td><td></td><td></td><td></td><td></td></tr>
<tr><td>Oligo fwd</td><td>CACCAACAGCGACATGCATTTTCC</td><td></td><td></td><td></td><td></td><td></td><td></td><td></td><td></td><td></td><td></td><td></td></tr>
<tr><td>Oligo rev</td><td>AAACGGAAAATGCATGTCGCTGTT</td><td></td><td></td><td></td><td></td><td></td><td></td><td></td><td></td><td></td><td></td><td></td></tr>
<tr><td>Chromosome</td><td>start</td><td></td><td>end</td><td>strand</td><td>MM</td><td>target_seq</td><td>PAM</td><td>alignment</td><td>distance</td><td>position</td><td>gene name</td><td>gene id</td></tr>
<tr><td>GU305914.1</td><td></td><td>11944</td><td>11966</td><td>-</td><td>0</td><td>AACAGCGACATGCATTTTCC</td><td>TGG</td><td>||||||||[||||||||||||]PAM</td><td>NA</td><td>Intergenic</td><td>NA</td><td>NA</td></tr>
<tr><td>T33560</td><td>CGACATGCATTTTCCTGGTTCGG</td><td></td><td>1000</td><td></td><td></td><td></td><td></td><td></td><td></td><td></td><td></td><td></td></tr>
<tr><td>Oligo fwd</td><td>CACCCGACATGCATTTTCCTGGTT</td><td></td><td></td><td></td><td></td><td></td><td></td><td></td><td></td><td></td><td></td><td></td></tr>
<tr><td>Oligo rev</td><td>AAACAACCAGGAAAATGCATGTCG</td><td></td><td></td><td></td><td></td><td></td><td></td><td></td><td></td><td></td><td></td><td></td></tr>
<tr><td>Chromosome</td><td>start</td><td></td><td>end</td><td>strand</td><td>MM</td><td>target_seq</td><td>PAM</td><td>alignment</td><td>distance</td><td>position</td><td>gene name</td><td>gene id</td></tr>
<tr><td>GU305914.1</td><td></td><td>11939</td><td>11961</td><td>-</td><td>0</td><td>CGACATGCATTTTCCTGGTT</td><td>CGG</td><td>||||||||[||||||||||||]PAM</td><td>NA</td><td>Intergenic</td><td>NA</td><td>NA</td></tr>
<tr><td>T33561</td><td>GACATGCATTTTCCTGGTTCGGG</td><td></td><td>1000</td><td></td><td></td><td></td><td></td><td></td><td></td><td></td><td></td><td></td></tr>
<tr><td>Oligo fwd</td><td>CACCGACATGCATTTTCCTGGTTC</td><td></td><td></td><td></td><td></td><td></td><td></td><td></td><td></td><td></td><td></td><td></td></tr>
<tr><td>Oligo rev</td><td>AAACGAACCAGGAAAATGCATGTC</td><td></td><td></td><td></td><td></td><td></td><td></td><td></td><td></td><td></td><td></td><td></td></tr>
<tr><td>Chromosome</td><td>start</td><td></td><td>end</td><td>strand</td><td>MM</td><td>target_seq</td><td>PAM</td><td>alignment</td><td>distance</td><td>position</td><td>gene name</td><td>gene id</td></tr>
<tr><td>GU305914.1</td><td></td><td>11938</td><td>11960</td><td>-</td><td>0</td><td>GACATGCATTTTCCTGGTTC</td><td>GGG</td><td>||||||||[||||||||||||]PAM</td><td>NA</td><td>Intergenic</td><td>NA</td><td>NA</td></tr>
<tr><td>T33562</td><td>ATGCATTTTCCTGGTTCGGGAGG</td><td></td><td>1000</td><td></td><td></td><td></td><td></td><td></td><td></td><td></td><td></td><td></td></tr>
<tr><td>Oligo fwd</td><td>CACCATGCATTTTCCTGGTTCGGG</td><td></td><td></td><td></td><td></td><td></td><td></td><td></td><td></td><td></td><td></td><td></td></tr>
<tr><td>Oligo rev</td><td>AAACCCCGAACCAGGAAAATGCAT</td><td></td><td></td><td></td><td></td><td></td><td></td><td></td><td></td><td></td><td></td><td></td></tr>
<tr><td>Chromosome</td><td>start</td><td></td><td>end</td><td>strand</td><td>MM</td><td>target_seq</td><td>PAM</td><td>alignment</td><td>distance</td><td>position</td><td>gene name</td><td>gene id</td></tr>
<tr><td>GU305914.1</td><td></td><td>11935</td><td>11957</td><td>-</td><td>0</td><td>ATGCATTTTCCTGGTTCGGG</td><td>AGG</td><td>||||||||[||||||||||||]PAM</td><td>NA</td><td>Intergenic</td><td>NA</td><td>NA</td></tr>
<tr><td>T33563</td><td>CATTTTCCTGGTTCGGGAGGAGG</td><td></td><td>1000</td><td></td><td></td><td></td><td></td><td></td><td></td><td></td><td></td><td></td></tr>
<tr><td>Oligo fwd</td><td>CACCCATTTTCCTGGTTCGGGAGG</td><td></td><td></td><td></td><td></td><td></td><td></td><td></td><td></td><td></td><td></td><td></td></tr>
<tr><td>Oligo rev</td><td>AAACCCTCCCGAACCAGGAAAATG</td><td></td><td></td><td></td><td></td><td></td><td></td><td></td><td></td><td></td><td></td><td></td></tr>
<tr><td>Chromosome</td><td>start</td><td></td><td>end</td><td>strand</td><td>MM</td><td>target_seq</td><td>PAM</td><td>alignment</td><td>distance</td><td>position</td><td>gene name</td><td>gene id</td></tr>
<tr><td>GU305914.1</td><td></td><td>11932</td><td>11954</td><td>-</td><td>0</td><td>CATTTTCCTGGTTCGGGAGG</td><td>AGG</td><td>||||||||[||||||||||||]PAM</td><td>NA</td><td>Intergenic</td><td>NA</td><td>NA</td></tr>
<tr><td>T33564</td><td>TTCGGGAGGAGGACGTCTTTCGG</td><td></td><td>1000</td><td></td><td></td><td></td><td></td><td></td><td></td><td></td><td></td><td></td></tr>
<tr><td>Oligo fwd</td><td>CACCTTCGGGAGGAGGACGTCTTT</td><td></td><td></td><td></td><td></td><td></td><td></td><td></td><td></td><td></td><td></td><td></td></tr>
<tr><td>Oligo rev</td><td>AAACAAAGACGTCCTCCTCCCGAA</td><td></td><td></td><td></td><td></td><td></td><td></td><td></td><td></td><td></td><td></td><td></td></tr>
<tr><td>Chromosome</td><td>start</td><td></td><td>end</td><td>strand</td><td>MM</td><td>target_seq</td><td>PAM</td><td>alignment</td><td>distance</td><td>position</td><td>gene name</td><td>gene id</td></tr>
<tr><td>GU305914.1</td><td></td><td>11921</td><td>11943</td><td>-</td><td>0</td><td>TTCGGGAGGAGGACGTCTTT</td><td>CGG</td><td>||||||||[||||||||||||]PAM</td><td>NA</td><td>Intergenic</td><td>NA</td><td>NA</td></tr>
<tr><td>T33565</td><td>AGGAGGACGTCTTTCGGAGACGG</td><td></td><td>1000</td><td></td><td></td><td></td><td></td><td></td><td></td><td></td><td></td><td></td></tr>
<tr><td>Oligo fwd</td><td>CACCAGGAGGACGTCTTTCGGAGA</td><td></td><td></td><td></td><td></td><td></td><td></td><td></td><td></td><td></td><td></td><td></td></tr>
<tr><td>Oligo rev</td><td>AAACTCTCCGAAAGACGTCCTCCT</td><td></td><td></td><td></td><td></td><td></td><td></td><td></td><td></td><td></td><td></td><td></td></tr>
<tr><td>Chromosome</td><td>start</td><td></td><td>end</td><td>strand</td><td>MM</td><td>target_seq</td><td>PAM</td><td>alignment</td><td>distance</td><td>position</td><td>gene name</td><td>gene id</td></tr>
<tr><td>GU305914.1</td><td></td><td>11915</td><td>11937</td><td>-</td><td>0</td><td>AGGAGGACGTCTTTCGGAGA</td><td>CGG</td><td>||||||||[||||||||||||]PAM</td><td>NA</td><td>Intergenic</td><td>NA</td><td>NA</td></tr>
<tr><td>T33566</td><td>TTAACACACGACGTATTTTTTGG</td><td></td><td>1000</td><td></td><td></td><td></td><td></td><td></td><td></td><td></td><td></td><td></td></tr>
</table>

<table>
<tr><td>Oligo fwd</td><td>CACCTTAACACACGACGTATTTTT</td><td></td><td></td><td></td><td></td><td></td><td></td><td></td><td></td><td></td><td></td><td></td></tr>
<tr><td>Oligo rev</td><td>AAACAAAAATACGTCGTGTGTTAA</td><td></td><td></td><td></td><td></td><td></td><td></td><td></td><td></td><td></td><td></td><td></td></tr>
<tr><td>Chromosome</td><td>start</td><td></td><td>end</td><td>strand</td><td>MM</td><td>target_seq</td><td>PAM</td><td>alignment</td><td>distance</td><td>position</td><td>gene name</td><td>gene id</td></tr>
<tr><td>GU305914.1</td><td></td><td>11884</td><td>11906</td><td>-</td><td>0</td><td>TTAACACACGACGTATTTTT</td><td>TGG</td><td>||||||||[||||||||||||]PAM</td><td>NA</td><td>Intergenic</td><td>NA</td><td>NA</td></tr>
<tr><td>T33567</td><td>ACATTCGTTGCCTGTGCTAGCGG</td><td></td><td>1000</td><td></td><td></td><td></td><td></td><td></td><td></td><td></td><td></td><td></td></tr>
<tr><td>Oligo fwd</td><td>CACCACATTCGTTGCCTGTGCTAG</td><td></td><td></td><td></td><td></td><td></td><td></td><td></td><td></td><td></td><td></td><td></td></tr>
<tr><td>Oligo rev</td><td>AAACCTAGCACAGGCAACGAATGT</td><td></td><td></td><td></td><td></td><td></td><td></td><td></td><td></td><td></td><td></td><td></td></tr>
<tr><td>Chromosome</td><td>start</td><td></td><td>end</td><td>strand</td><td>MM</td><td>target_seq</td><td>PAM</td><td>alignment</td><td>distance</td><td>position</td><td>gene name</td><td>gene id</td></tr>
<tr><td>GU305914.1</td><td></td><td>11851</td><td>11873</td><td>-</td><td>0</td><td>ACATTCGTTGCCTGTGCTAG</td><td>CGG</td><td>||||||||[||||||||||||]PAM</td><td>NA</td><td>Intergenic</td><td>NA</td><td>NA</td></tr>
<tr><td>T33568</td><td>CGATGCGAGCGAGAATGTGACGG</td><td></td><td>1000</td><td></td><td></td><td></td><td></td><td></td><td></td><td></td><td></td><td></td></tr>
<tr><td>Oligo fwd</td><td>CACCCGATGCGAGCGAGAATGTGA</td><td></td><td></td><td></td><td></td><td></td><td></td><td></td><td></td><td></td><td></td><td></td></tr>
<tr><td>Oligo rev</td><td>AAACTCACATTCTCGCTCGCATCG</td><td></td><td></td><td></td><td></td><td></td><td></td><td></td><td></td><td></td><td></td><td></td></tr>
<tr><td>Chromosome</td><td>start</td><td></td><td>end</td><td>strand</td><td>MM</td><td>target_seq</td><td>PAM</td><td>alignment</td><td>distance</td><td>position</td><td>gene name</td><td>gene id</td></tr>
<tr><td>GU305914.1</td><td></td><td>11813</td><td>11835</td><td>-</td><td>0</td><td>CGATGCGAGCGAGAATGTGA</td><td>CGG</td><td>||||||||[||||||||||||]PAM</td><td>NA</td><td>Intergenic</td><td>NA</td><td>NA</td></tr>
<tr><td>T33569</td><td>TGACGGTGATAAACGACGTCAGG</td><td></td><td>1000</td><td></td><td></td><td></td><td></td><td></td><td></td><td></td><td></td><td></td></tr>
<tr><td>Oligo fwd</td><td>CACCTGACGGTGATAAACGACGTC</td><td></td><td></td><td></td><td></td><td></td><td></td><td></td><td></td><td></td><td></td><td></td></tr>
<tr><td>Oligo rev</td><td>AAACGACGTCGTTTATCACCGTCA</td><td></td><td></td><td></td><td></td><td></td><td></td><td></td><td></td><td></td><td></td><td></td></tr>
<tr><td>Chromosome</td><td>start</td><td></td><td>end</td><td>strand</td><td>MM</td><td>target_seq</td><td>PAM</td><td>alignment</td><td>distance</td><td>position</td><td>gene name</td><td>gene id</td></tr>
<tr><td>GU305914.1</td><td></td><td>11796</td><td>11818</td><td>-</td><td>0</td><td>TGACGGTGATAAACGACGTC</td><td>AGG</td><td>||||||||[||||||||||||]PAM</td><td>NA</td><td>Intergenic</td><td>NA</td><td>NA</td></tr>
<tr><td>T33570</td><td>CGGTGATAAACGACGTCAGGTGG</td><td></td><td>1000</td><td></td><td></td><td></td><td></td><td></td><td></td><td></td><td></td><td></td></tr>
<tr><td>Oligo fwd</td><td>CACCCGGTGATAAACGACGTCAGG</td><td></td><td></td><td></td><td></td><td></td><td></td><td></td><td></td><td></td><td></td><td></td></tr>
<tr><td>Oligo rev</td><td>AAACCCTGACGTCGTTTATCACCG</td><td></td><td></td><td></td><td></td><td></td><td></td><td></td><td></td><td></td><td></td><td></td></tr>
<tr><td>Chromosome</td><td>start</td><td></td><td>end</td><td>strand</td><td>MM</td><td>target_seq</td><td>PAM</td><td>alignment</td><td>distance</td><td>position</td><td>gene name</td><td>gene id</td></tr>
<tr><td>GU305914.1</td><td></td><td>11793</td><td>11815</td><td>-</td><td>0</td><td>CGGTGATAAACGACGTCAGG</td><td>TGG</td><td>||||||||[||||||||||||]PAM</td><td>NA</td><td>Intergenic</td><td>NA</td><td>NA</td></tr>
<tr><td>T33571</td><td>TAAACGACGTCAGGTGGTGACGG</td><td></td><td>1000</td><td></td><td></td><td></td><td></td><td></td><td></td><td></td><td></td><td></td></tr>
<tr><td>Oligo fwd</td><td>CACCTAAACGACGTCAGGTGGTGA</td><td></td><td></td><td></td><td></td><td></td><td></td><td></td><td></td><td></td><td></td><td></td></tr>
<tr><td>Oligo rev</td><td>AAACTCACCACCTGACGTCGTTTA</td><td></td><td></td><td></td><td></td><td></td><td></td><td></td><td></td><td></td><td></td><td></td></tr>
<tr><td>Chromosome</td><td>start</td><td></td><td>end</td><td>strand</td><td>MM</td><td>target_seq</td><td>PAM</td><td>alignment</td><td>distance</td><td>position</td><td>gene name</td><td>gene id</td></tr>
<tr><td>GU305914.1</td><td></td><td>11787</td><td>11809</td><td>-</td><td>0</td><td>TAAACGACGTCAGGTGGTGA</td><td>CGG</td><td>||||||||[||||||||||||]PAM</td><td>NA</td><td>Intergenic</td><td>NA</td><td>NA</td></tr>
<tr><td>T33572</td><td>CAGGTGGTGACGGAACATTTCGG</td><td></td><td>1000</td><td></td><td></td><td></td><td></td><td></td><td></td><td></td><td></td><td></td></tr>
<tr><td>Oligo fwd</td><td>CACCCAGGTGGTGACGGAACATTT</td><td></td><td></td><td></td><td></td><td></td><td></td><td></td><td></td><td></td><td></td><td></td></tr>
<tr><td>Oligo rev</td><td>AAACAAATGTTCCGTCACCACCTG</td><td></td><td></td><td></td><td></td><td></td><td></td><td></td><td></td><td></td><td></td><td></td></tr>
<tr><td>Chromosome</td><td>start</td><td></td><td>end</td><td>strand</td><td>MM</td><td>target_seq</td><td>PAM</td><td>alignment</td><td>distance</td><td>position</td><td>gene name</td><td>gene id</td></tr>
<tr><td>GU305914.1</td><td></td><td>11777</td><td>11799</td><td>-</td><td>0</td><td>CAGGTGGTGACGGAACATTT</td><td>CGG</td><td>||||||||[||||||||||||]PAM</td><td>NA</td><td>Intergenic</td><td>NA</td><td>NA</td></tr>
<tr><td>T33573</td><td>TCCGAGTTGAGTGTAATGTGAGG</td><td></td><td>1000</td><td></td><td></td><td></td><td></td><td></td><td></td><td></td><td></td><td></td></tr>
<tr><td>Oligo fwd</td><td>CACCTCCGAGTTGAGTGTAATGTG</td><td></td><td></td><td></td><td></td><td></td><td></td><td></td><td></td><td></td><td></td><td></td></tr>
<tr><td>Oligo rev</td><td>AAACCACATTACACTCAACTCGGA</td><td></td><td></td><td></td><td></td><td></td><td></td><td></td><td></td><td></td><td></td><td></td></tr>
<tr><td>Chromosome</td><td>start</td><td></td><td>end</td><td>strand</td><td>MM</td><td>target_seq</td><td>PAM</td><td>alignment</td><td>distance</td><td>position</td><td>gene name</td><td>gene id</td></tr>
<tr><td>GU305914.1</td><td></td><td>11736</td><td>11758</td><td>-</td><td>0</td><td>TCCGAGTTGAGTGTAATGTG</td><td>AGG</td><td>||||||||[||||||||||||]PAM</td><td>NA</td><td>Intergenic</td><td>NA</td><td>NA</td></tr>
<tr><td>T33574</td><td>GTACGACGACCATCTAAAAGTGG</td><td></td><td>1000</td><td></td><td></td><td></td><td></td><td></td><td></td><td></td><td></td><td></td></tr>
<tr><td>Oligo fwd</td><td>CACCGTACGACGACCATCTAAAAG</td><td></td><td></td><td></td><td></td><td></td><td></td><td></td><td></td><td></td><td></td><td></td></tr>
<tr><td>Oligo rev</td><td>AAACCTTTTAGATGGTCGTCGTAC</td><td></td><td></td><td></td><td></td><td></td><td></td><td></td><td></td><td></td><td></td><td></td></tr>
<tr><td>Chromosome</td><td>start</td><td></td><td>end</td><td>strand</td><td>MM</td><td>target_seq</td><td>PAM</td><td>alignment</td><td>distance</td><td>position</td><td>gene name</td><td>gene id</td></tr>
<tr><td>GU305914.1</td><td></td><td>11690</td><td>11712</td><td>-</td><td>0</td><td>GTACGACGACCATCTAAAAG</td><td>TGG</td><td>||||||||[||||||||||||]PAM</td><td>NA</td><td>Intergenic</td><td>NA</td><td>NA</td></tr>
<tr><td>T33575</td><td>ACAATTTTTATTTGTTGAACCGG</td><td></td><td>1000</td><td></td><td></td><td></td><td></td><td></td><td></td><td></td><td></td><td></td></tr>
<tr><td>Oligo fwd</td><td>CACCACAATTTTTATTTGTTGAAC</td><td></td><td></td><td></td><td></td><td></td><td></td><td></td><td></td><td></td><td></td><td></td></tr>
<tr><td>Oligo rev</td><td>AAACGTTCAACAAATAAAAATTGT</td><td></td><td></td><td></td><td></td><td></td><td></td><td></td><td></td><td></td><td></td><td></td></tr>
<tr><td>Chromosome</td><td>start</td><td></td><td>end</td><td>strand</td><td>MM</td><td>target_seq</td><td>PAM</td><td>alignment</td><td>distance</td><td>position</td><td>gene name</td><td>gene id</td></tr>
<tr><td>GU305914.1</td><td></td><td>11630</td><td>11652</td><td>-</td><td>0</td><td>ACAATTTTTATTTGTTGAAC</td><td>CGG</td><td>||||||||[||||||||||||]PAM</td><td>NA</td><td>Intergenic</td><td>NA</td><td>NA</td></tr>
<tr><td>T33576</td><td>CAATTTTTATTTGTTGAACCGGG</td><td></td><td>1000</td><td></td><td></td><td></td><td></td><td></td><td></td><td></td><td></td><td></td></tr>
<tr><td>Oligo fwd</td><td>CACCCAATTTTTATTTGTTGAACC</td><td></td><td></td><td></td><td></td><td></td><td></td><td></td><td></td><td></td><td></td><td></td></tr>
<tr><td>Oligo rev</td><td>AAACGGTTCAACAAATAAAAATTG</td><td></td><td></td><td></td><td></td><td></td><td></td><td></td><td></td><td></td><td></td><td></td></tr>
<tr><td>Chromosome</td><td>start</td><td></td><td>end</td><td>strand</td><td>MM</td><td>target_seq</td><td>PAM</td><td>alignment</td><td>distance</td><td>position</td><td>gene name</td><td>gene id</td></tr>
<tr><td>GU305914.1</td><td></td><td>11629</td><td>11651</td><td>-</td><td>0</td><td>CAATTTTTATTTGTTGAACC</td><td>GGG</td><td>||||||||[||||||||||||]PAM</td><td>NA</td><td>Intergenic</td><td>NA</td><td>NA</td></tr>
<tr><td>T33577</td><td>AATTTTTATTTGTTGAACCGGGG</td><td></td><td>1000</td><td></td><td></td><td></td><td></td><td></td><td></td><td></td><td></td><td></td></tr>
<tr><td>Oligo fwd</td><td>CACCAATTTTTATTTGTTGAACCG</td><td></td><td></td><td></td><td></td><td></td><td></td><td></td><td></td><td></td><td></td><td></td></tr>
<tr><td>Oligo rev</td><td>AAACCGGTTCAACAAATAAAAATT</td><td></td><td></td><td></td><td></td><td></td><td></td><td></td><td></td><td></td><td></td><td></td></tr>
<tr><td>Chromosome</td><td>start</td><td></td><td>end</td><td>strand</td><td>MM</td><td>target_seq</td><td>PAM</td><td>alignment</td><td>distance</td><td>position</td><td>gene name</td><td>gene id</td></tr>
<tr><td>GU305914.1</td><td></td><td>11628</td><td>11650</td><td>-</td><td>0</td><td>AATTTTTATTTGTTGAACCG</td><td>GGG</td><td>||||||||[||||||||||||]PAM</td><td>NA</td><td>Intergenic</td><td>NA</td><td>NA</td></tr>
<tr><td>T33578</td><td>AAATACTGCTTATGCAGCACTGG</td><td></td><td>1000</td><td></td><td></td><td></td><td></td><td></td><td></td><td></td><td></td><td></td></tr>
<tr><td>Oligo fwd</td><td>CACCAAATACTGCTTATGCAGCAC</td><td></td><td></td><td></td><td></td><td></td><td></td><td></td><td></td><td></td><td></td><td></td></tr>
<tr><td>Oligo rev</td><td>AAACGTGCTGCATAAGCAGTATTT</td><td></td><td></td><td></td><td></td><td></td><td></td><td></td><td></td><td></td><td></td><td></td></tr>
<tr><td>Chromosome</td><td>start</td><td></td><td>end</td><td>strand</td><td>MM</td><td>target_seq</td><td>PAM</td><td>alignment</td><td>distance</td><td>position</td><td>gene name</td><td>gene id</td></tr>
<tr><td>GU305914.1</td><td></td><td>11601</td><td>11623</td><td>-</td><td>0</td><td>AAATACTGCTTATGCAGCAC</td><td>TGG</td><td>||||||||[||||||||||||]PAM</td><td>NA</td><td>Intergenic</td><td>NA</td><td>NA</td></tr>
<tr><td>T33579</td><td>GCACTGGTTGATATGTGTAACGG</td><td></td><td>1000</td><td></td><td></td><td></td><td></td><td></td><td></td><td></td><td></td><td></td></tr>
<tr><td>Oligo fwd</td><td>CACCGCACTGGTTGATATGTGTAA</td><td></td><td></td><td></td><td></td><td></td><td></td><td></td><td></td><td></td><td></td><td></td></tr>
<tr><td>Oligo rev</td><td>AAACTTACACATATCAACCAGTGC</td><td></td><td></td><td></td><td></td><td></td><td></td><td></td><td></td><td></td><td></td><td></td></tr>
<tr><td>Chromosome</td><td>start</td><td></td><td>end</td><td>strand</td><td>MM</td><td>target_seq</td><td>PAM</td><td>alignment</td><td>distance</td><td>position</td><td>gene name</td><td>gene id</td></tr>
<tr><td>GU305914.1</td><td></td><td>11585</td><td>11607</td><td>-</td><td>0</td><td>GCACTGGTTGATATGTGTAA</td><td>CGG</td><td>||||||||[||||||||||||]PAM</td><td>NA</td><td>Intergenic</td><td>NA</td><td>NA</td></tr>
<tr><td>T33580</td><td>CTGGTTGATATGTGTAACGGTGG</td><td></td><td>1000</td><td></td><td></td><td></td><td></td><td></td><td></td><td></td><td></td><td></td></tr>
<tr><td>Oligo fwd</td><td>CACCCTGGTTGATATGTGTAACGG</td><td></td><td></td><td></td><td></td><td></td><td></td><td></td><td></td><td></td><td></td><td></td></tr>
<tr><td>Oligo rev</td><td>AAACCCGTTACACATATCAACCAG</td><td></td><td></td><td></td><td></td><td></td><td></td><td></td><td></td><td></td><td></td><td></td></tr>
<tr><td>Chromosome</td><td>start</td><td></td><td>end</td><td>strand</td><td>MM</td><td>target_seq</td><td>PAM</td><td>alignment</td><td>distance</td><td>position</td><td>gene name</td><td>gene id</td></tr>
<tr><td>GU305914.1</td><td></td><td>11582</td><td>11604</td><td>-</td><td>0</td><td>CTGGTTGATATGTGTAACGG</td><td>TGG</td><td>||||||||[||||||||||||]PAM</td><td>NA</td><td>Intergenic</td><td>NA</td><td>NA</td></tr>
</table>

| | | | | | | | | | | | |
|---|---|---|---|---|---|---|---|---|---|---|---|
| T33581 | AACGGTGGAAGAAAATACGTAGG | 1000 | | | | | | | | | |
| Oligo fwd | CACCAACGGTGGAAGAAAATACGT | | | | | | | | | | |
| Oligo rev | AAACACGTATTTTCTTCCACCGTT | | | | | | | | | | |
| Chromosome | start | end | strand | MM | target_seq | PAM | alignment | distance | position | gene name | gene id |
| GU305914.1 | 11567 | 11589 | - | 0 | AACGGTGGAAGAAAATACGT | AGG | \|\|\|\|\|\|\|\|[\|\|\|\|\|\|\|\|\|\|\|\|]PAM | NA | Intergenic | NA | NA |
| T33582 | TGGAAGAAAATACGTAGGATCGG | 1000 | | | | | | | | | |
| Oligo fwd | CACCTGGAAGAAAATACGTAGGAT | | | | | | | | | | |
| Oligo rev | AAACATCCTACGTATTTTCTTCCA | | | | | | | | | | |
| Chromosome | start | end | strand | MM | target_seq | PAM | alignment | distance | position | gene name | gene id |
| GU305914.1 | 11562 | 11584 | - | 0 | TGGAAGAAAATACGTAGGAT | CGG | \|\|\|\|\|\|\|\|[\|\|\|\|\|\|\|\|\|\|\|\|]PAM | NA | Intergenic | NA | NA |
| T33583 | AGAAAATACGTAGGATCGGAAGG | 1000 | | | | | | | | | |
| Oligo fwd | CACCAGAAAATACGTAGGATCGGA | | | | | | | | | | |
| Oligo rev | AAACTCCGATCCTACGTATTTTCT | | | | | | | | | | |
| Chromosome | start | end | strand | MM | target_seq | PAM | alignment | distance | position | gene name | gene id |
| GU305914.1 | 11558 | 11580 | - | 0 | AGAAAATACGTAGGATCGGA | AGG | \|\|\|\|\|\|\|\|[\|\|\|\|\|\|\|\|\|\|\|\|]PAM | NA | Intergenic | NA | NA |
| T33584 | GAAAATACGTAGGATCGGAAGGG | 1000 | | | | | | | | | |
| Oligo fwd | CACCGAAAATACGTAGGATCGGAA | | | | | | | | | | |
| Oligo rev | AAACTTCCGATCCTACGTATTTTC | | | | | | | | | | |
| Chromosome | start | end | strand | MM | target_seq | PAM | alignment | distance | position | gene name | gene id |
| GU305914.1 | 11557 | 11579 | - | 0 | GAAAATACGTAGGATCGGAA | GGG | \|\|\|\|\|\|\|\|[\|\|\|\|\|\|\|\|\|\|\|\|]PAM | NA | Intergenic | NA | NA |
| T33585 | ACGTAGGATCGGAAGGGTCGTGG | 1000 | | | | | | | | | |
| Oligo fwd | CACCACGTAGGATCGGAAGGGTCG | | | | | | | | | | |
| Oligo rev | AAACCGACCCTTCCGATCCTACGT | | | | | | | | | | |
| Chromosome | start | end | strand | MM | target_seq | PAM | alignment | distance | position | gene name | gene id |
| GU305914.1 | 11551 | 11573 | - | 0 | ACGTAGGATCGGAAGGGTCG | TGG | \|\|\|\|\|\|\|\|[\|\|\|\|\|\|\|\|\|\|\|\|]PAM | NA | Intergenic | NA | NA |
| T33586 | TGTCTATCGATATATGTGCTCGG | 1000 | | | | | | | | | |
| Oligo fwd | CACCTGTCTATCGATATATGTGCT | | | | | | | | | | |
| Oligo rev | AAACAGCACATATATCGATAGACA | | | | | | | | | | |
| Chromosome | start | end | strand | MM | target_seq | PAM | alignment | distance | position | gene name | gene id |
| GU305914.1 | 11528 | 11550 | - | 0 | TGTCTATCGATATATGTGCT | CGG | \|\|\|\|\|\|\|\|[\|\|\|\|\|\|\|\|\|\|\|\|]PAM | NA | Intergenic | NA | NA |
| T33587 | GATATATGTGCTCGGATATCTGG | 1000 | | | | | | | | | |
| Oligo fwd | CACCGATATATGTGCTCGGATATC | | | | | | | | | | |
| Oligo rev | AAACGATATCCGAGCACATATATC | | | | | | | | | | |
| Chromosome | start | end | strand | MM | target_seq | PAM | alignment | distance | position | gene name | gene id |
| GU305914.1 | 11520 | 11542 | - | 0 | GATATATGTGCTCGGATATC | TGG | \|\|\|\|\|\|\|\|[\|\|\|\|\|\|\|\|\|\|\|\|]PAM | NA | Intergenic | NA | NA |
| T33588 | CGGATATCTGGTCGAATAGCTGG | 1000 | | | | | | | | | |
| Oligo fwd | CACCCGGATATCTGGTCGAATAGC | | | | | | | | | | |
| Oligo rev | AAACGCTATTCGACCAGATATCCG | | | | | | | | | | |
| Chromosome | start | end | strand | MM | target_seq | PAM | alignment | distance | position | gene name | gene id |
| GU305914.1 | 11508 | 11530 | - | 0 | CGGATATCTGGTCGAATAGC | TGG | \|\|\|\|\|\|\|\|[\|\|\|\|\|\|\|\|\|\|\|\|]PAM | NA | Intergenic | NA | NA |
| T33589 | TATCTGGTCGAATAGCTGGTTGG | 1000 | | | | | | | | | |
| Oligo fwd | CACCTATCTGGTCGAATAGCTGGT | | | | | | | | | | |
| Oligo rev | AAACACCAGCTATTCGACCAGATA | | | | | | | | | | |
| Chromosome | start | end | strand | MM | target_seq | PAM | alignment | distance | position | gene name | gene id |
| GU305914.1 | 11504 | 11526 | - | 0 | TATCTGGTCGAATAGCTGGT | TGG | \|\|\|\|\|\|\|\|[\|\|\|\|\|\|\|\|\|\|\|\|]PAM | NA | Intergenic | NA | NA |
| T33590 | AGCTGGTTGGATAACTGCTATGG | 1000 | | | | | | | | | |
| Oligo fwd | CACCAGCTGGTTGGATAACTGCTA | | | | | | | | | | |
| Oligo rev | AAACTAGCAGTTATCCAACCAGCT | | | | | | | | | | |
| Chromosome | start | end | strand | MM | target_seq | PAM | alignment | distance | position | gene name | gene id |
| GU305914.1 | 11491 | 11513 | - | 0 | AGCTGGTTGGATAACTGCTA | TGG | \|\|\|\|\|\|\|\|[\|\|\|\|\|\|\|\|\|\|\|\|]PAM | NA | Intergenic | NA | NA |
| T33591 | GCTGGTTGGATAACTGCTATGGG | 1000 | | | | | | | | | |
| Oligo fwd | CACCGCTGGTTGGATAACTGCTAT | | | | | | | | | | |
| Oligo rev | AAACATAGCAGTTATCCAACCAGC | | | | | | | | | | |
| Chromosome | start | end | strand | MM | target_seq | PAM | alignment | distance | position | gene name | gene id |
| GU305914.1 | 11490 | 11512 | - | 0 | GCTGGTTGGATAACTGCTAT | GGG | \|\|\|\|\|\|\|\|[\|\|\|\|\|\|\|\|\|\|\|\|]PAM | NA | Intergenic | NA | NA |
| T33592 | TGCTATGGGACACTAGAAAACGG | 1000 | | | | | | | | | |
| Oligo fwd | CACCTGCTATGGGACACTAGAAAA | | | | | | | | | | |
| Oligo rev | AAACTTTTCTAGTGTCCCATAGCA | | | | | | | | | | |
| Chromosome | start | end | strand | MM | target_seq | PAM | alignment | distance | position | gene name | gene id |
| GU305914.1 | 11476 | 11498 | - | 0 | TGCTATGGGACACTAGAAAA | CGG | \|\|\|\|\|\|\|\|[\|\|\|\|\|\|\|\|\|\|\|\|]PAM | NA | Intergenic | NA | NA |
| T33593 | GCTATGGGACACTAGAAAACGGG | 1000 | | | | | | | | | |
| Oligo fwd | CACCGCTATGGGACACTAGAAAAC | | | | | | | | | | |
| Oligo rev | AAACGTTTTCTAGTGTCCCATAGC | | | | | | | | | | |
| Chromosome | start | end | strand | MM | target_seq | PAM | alignment | distance | position | gene name | gene id |
| GU305914.1 | 11475 | 11497 | - | 0 | GCTATGGGACACTAGAAAAC | GGG | \|\|\|\|\|\|\|\|[\|\|\|\|\|\|\|\|\|\|\|\|]PAM | NA | Intergenic | NA | NA |
| T33594 | GGGACACTAGAAAACGGGCGCGG | 1000 | | | | | | | | | |
| Oligo fwd | CACCGGGACACTAGAAAACGGGCG | | | | | | | | | | |
| Oligo rev | AAACCGCCCGTTTTCTAGTGTCCC | | | | | | | | | | |
| Chromosome | start | end | strand | MM | target_seq | PAM | alignment | distance | position | gene name | gene id |
| GU305914.1 | 11470 | 11492 | - | 0 | GGGACACTAGAAAACGGGCG | CGG | \|\|\|\|\|\|\|\|[\|\|\|\|\|\|\|\|\|\|\|\|]PAM | NA | Intergenic | NA | NA |
| T33595 | ACACTAGAAAACGGGCGCGGAGG | 1000 | | | | | | | | | |
| Oligo fwd | CACCACACTAGAAAACGGGCGCGG | | | | | | | | | | |
| Oligo rev | AAACCCGCGCCCGTTTTCTAGTGT | | | | | | | | | | |
| Chromosome | start | end | strand | MM | target_seq | PAM | alignment | distance | position | gene name | gene id |
| GU305914.1 | 11467 | 11489 | - | 0 | ACACTAGAAAACGGGCGCGG | AGG | \|\|\|\|\|\|\|\|[\|\|\|\|\|\|\|\|\|\|\|\|]PAM | NA | Intergenic | NA | NA |

<table>
<tr><td>T33596</td><td>TCTATACAGATTCAGTCGTCTGG</td><td>1000</td><td></td><td></td><td></td><td></td><td></td><td></td><td></td><td></td><td></td></tr>
<tr><td>Oligo fwd</td><td>CACCTCTATACAGATTCAGTCGTC</td><td></td><td></td><td></td><td></td><td></td><td></td><td></td><td></td><td></td><td></td></tr>
<tr><td>Oligo rev</td><td>AAACGACGACTGAATCTGTATAGA</td><td></td><td></td><td></td><td></td><td></td><td></td><td></td><td></td><td></td><td></td></tr>
<tr><td>Chromosome</td><td>start</td><td>end</td><td>strand</td><td>MM</td><td>target_seq</td><td>PAM</td><td>alignment</td><td>distance</td><td>position</td><td>gene name</td><td>gene id</td></tr>
<tr><td>GU305914.1</td><td>11439</td><td>11461</td><td>-</td><td>0</td><td>TCTATACAGATTCAGTCGTC</td><td>TGG</td><td>||||||||[||||||||||||]PAM</td><td>NA</td><td>Intergenic</td><td>NA</td><td>NA</td></tr>
<tr><td></td><td></td><td></td><td></td><td></td><td></td><td></td><td></td><td></td><td></td><td></td><td></td></tr>
<tr><td>T33597</td><td>GACTATTACGATCACGATCGCGG</td><td>1000</td><td></td><td></td><td></td><td></td><td></td><td></td><td></td><td></td><td></td></tr>
<tr><td>Oligo fwd</td><td>CACCGACTATTACGATCACGATCG</td><td></td><td></td><td></td><td></td><td></td><td></td><td></td><td></td><td></td><td></td></tr>
<tr><td>Oligo rev</td><td>AAACCGATCGTGATCGTAATAGTC</td><td></td><td></td><td></td><td></td><td></td><td></td><td></td><td></td><td></td><td></td></tr>
<tr><td>Chromosome</td><td>start</td><td>end</td><td>strand</td><td>MM</td><td>target_seq</td><td>PAM</td><td>alignment</td><td>distance</td><td>position</td><td>gene name</td><td>gene id</td></tr>
<tr><td>GU305914.1</td><td>11358</td><td>11380</td><td>-</td><td>0</td><td>GACTATTACGATCACGATCG</td><td>CGG</td><td>||||||||[||||||||||||]PAM</td><td>NA</td><td>Intergenic</td><td>NA</td><td>NA</td></tr>
<tr><td></td><td></td><td></td><td></td><td></td><td></td><td></td><td></td><td></td><td></td><td></td><td></td></tr>
<tr><td>T33598</td><td>GTGCCGCTACTTGCTGTCTGTGG</td><td>1000</td><td></td><td></td><td></td><td></td><td></td><td></td><td></td><td></td><td></td></tr>
<tr><td>Oligo fwd</td><td>CACCGTGCCGCTACTTGCTGTCTG</td><td></td><td></td><td></td><td></td><td></td><td></td><td></td><td></td><td></td><td></td></tr>
<tr><td>Oligo rev</td><td>AAACCAGACAGCAAGTAGCGGCAC</td><td></td><td></td><td></td><td></td><td></td><td></td><td></td><td></td><td></td><td></td></tr>
<tr><td>Chromosome</td><td>start</td><td>end</td><td>strand</td><td>MM</td><td>target_seq</td><td>PAM</td><td>alignment</td><td>distance</td><td>position</td><td>gene name</td><td>gene id</td></tr>
<tr><td>GU305914.1</td><td>11330</td><td>11352</td><td>-</td><td>0</td><td>GTGCCGCTACTTGCTGTCTG</td><td>TGG</td><td>||||||||[||||||||||||]PAM</td><td>NA</td><td>Intergenic</td><td>NA</td><td>NA</td></tr>
<tr><td></td><td></td><td></td><td></td><td></td><td></td><td></td><td></td><td></td><td></td><td></td><td></td></tr>
<tr><td>T33599</td><td>TACCCCGAAGTTTTCACCCGCGG</td><td>1000</td><td></td><td></td><td></td><td></td><td></td><td></td><td></td><td></td><td></td></tr>
<tr><td>Oligo fwd</td><td>CACCTACCCCGAAGTTTTCACCCG</td><td></td><td></td><td></td><td></td><td></td><td></td><td></td><td></td><td></td><td></td></tr>
<tr><td>Oligo rev</td><td>AAACCGGGTGAAAACTTCGGGGTA</td><td></td><td></td><td></td><td></td><td></td><td></td><td></td><td></td><td></td><td></td></tr>
<tr><td>Chromosome</td><td>start</td><td>end</td><td>strand</td><td>MM</td><td>target_seq</td><td>PAM</td><td>alignment</td><td>distance</td><td>position</td><td>gene name</td><td>gene id</td></tr>
<tr><td>GU305914.1</td><td>11298</td><td>11320</td><td>-</td><td>0</td><td>TACCCCGAAGTTTTCACCCG</td><td>CGG</td><td>||||||||[||||||||||||]PAM</td><td>NA</td><td>Intergenic</td><td>NA</td><td>NA</td></tr>
<tr><td></td><td></td><td></td><td></td><td></td><td></td><td></td><td></td><td></td><td></td><td></td><td></td></tr>
<tr><td>T33600</td><td>TTTTCACCCGCGGATACAAAAGG</td><td>1000</td><td></td><td></td><td></td><td></td><td></td><td></td><td></td><td></td><td></td></tr>
<tr><td>Oligo fwd</td><td>CACCTTTTCACCCGCGGATACAAA</td><td></td><td></td><td></td><td></td><td></td><td></td><td></td><td></td><td></td><td></td></tr>
<tr><td>Oligo rev</td><td>AAACTTTGTATCCGCGGGTGAAAA</td><td></td><td></td><td></td><td></td><td></td><td></td><td></td><td></td><td></td><td></td></tr>
<tr><td>Chromosome</td><td>start</td><td>end</td><td>strand</td><td>MM</td><td>target_seq</td><td>PAM</td><td>alignment</td><td>distance</td><td>position</td><td>gene name</td><td>gene id</td></tr>
<tr><td>GU305914.1</td><td>11288</td><td>11310</td><td>-</td><td>0</td><td>TTTTCACCCGCGGATACAAA</td><td>AGG</td><td>||||||||[||||||||||||]PAM</td><td>NA</td><td>Intergenic</td><td>NA</td><td>NA</td></tr>
<tr><td></td><td></td><td></td><td></td><td></td><td></td><td></td><td></td><td></td><td></td><td></td><td></td></tr>
<tr><td>T33601</td><td>CTCGACGATCGTGTAGTAGTCGG</td><td>1000</td><td></td><td></td><td></td><td></td><td></td><td></td><td></td><td></td><td></td></tr>
<tr><td>Oligo fwd</td><td>CACCCTCGACGATCGTGTAGTAGT</td><td></td><td></td><td></td><td></td><td></td><td></td><td></td><td></td><td></td><td></td></tr>
<tr><td>Oligo rev</td><td>AAACACTACTACACGATCGTCGAG</td><td></td><td></td><td></td><td></td><td></td><td></td><td></td><td></td><td></td><td></td></tr>
<tr><td>Chromosome</td><td>start</td><td>end</td><td>strand</td><td>MM</td><td>target_seq</td><td>PAM</td><td>alignment</td><td>distance</td><td>position</td><td>gene name</td><td>gene id</td></tr>
<tr><td>GU305914.1</td><td>11204</td><td>11226</td><td>-</td><td>0</td><td>CTCGACGATCGTGTAGTAGT</td><td>CGG</td><td>||||||||[||||||||||||]PAM</td><td>NA</td><td>Intergenic</td><td>NA</td><td>NA</td></tr>
<tr><td></td><td></td><td></td><td></td><td></td><td></td><td></td><td></td><td></td><td></td><td></td><td></td></tr>
<tr><td>T33602</td><td>CGGTCGTCGTGTCGTAATCGCGG</td><td>1000</td><td></td><td></td><td></td><td></td><td></td><td></td><td></td><td></td><td></td></tr>
<tr><td>Oligo fwd</td><td>CACCCGGTCGTCGTGTCGTAATCG</td><td></td><td></td><td></td><td></td><td></td><td></td><td></td><td></td><td></td><td></td></tr>
<tr><td>Oligo rev</td><td>AAACCGATTACGACACGACGACCG</td><td></td><td></td><td></td><td></td><td></td><td></td><td></td><td></td><td></td><td></td></tr>
<tr><td>Chromosome</td><td>start</td><td>end</td><td>strand</td><td>MM</td><td>target_seq</td><td>PAM</td><td>alignment</td><td>distance</td><td>position</td><td>gene name</td><td>gene id</td></tr>
<tr><td>GU305914.1</td><td>11184</td><td>11206</td><td>-</td><td>0</td><td>CGGTCGTCGTGTCGTAATCG</td><td>CGG</td><td>||||||||[||||||||||||]PAM</td><td>NA</td><td>Intergenic</td><td>NA</td><td>NA</td></tr>
<tr><td></td><td></td><td></td><td></td><td></td><td></td><td></td><td></td><td></td><td></td><td></td><td></td></tr>
<tr><td>T33603</td><td>GTACAATAGATTTCTTTCTCCGG</td><td>1000</td><td></td><td></td><td></td><td></td><td></td><td></td><td></td><td></td><td></td></tr>
<tr><td>Oligo fwd</td><td>CACCGTACAATAGATTTCTTTCTC</td><td></td><td></td><td></td><td></td><td></td><td></td><td></td><td></td><td></td><td></td></tr>
<tr><td>Oligo rev</td><td>AAACGAGAAAGAAATCTATTGTAC</td><td></td><td></td><td></td><td></td><td></td><td></td><td></td><td></td><td></td><td></td></tr>
<tr><td>Chromosome</td><td>start</td><td>end</td><td>strand</td><td>MM</td><td>target_seq</td><td>PAM</td><td>alignment</td><td>distance</td><td>position</td><td>gene name</td><td>gene id</td></tr>
<tr><td>GU305914.1</td><td>11144</td><td>11166</td><td>-</td><td>0</td><td>GTACAATAGATTTCTTTCTC</td><td>CGG</td><td>||||||||[||||||||||||]PAM</td><td>NA</td><td>Intergenic</td><td>NA</td><td>NA</td></tr>
<tr><td></td><td></td><td></td><td></td><td></td><td></td><td></td><td></td><td></td><td></td><td></td><td></td></tr>
<tr><td>T33604</td><td>TTCTTTCTCCGGTTTCTTCGTGG</td><td>1000</td><td></td><td></td><td></td><td></td><td></td><td></td><td></td><td></td><td></td></tr>
<tr><td>Oligo fwd</td><td>CACCTTCTTTCTCCGGTTTCTTCG</td><td></td><td></td><td></td><td></td><td></td><td></td><td></td><td></td><td></td><td></td></tr>
<tr><td>Oligo rev</td><td>AAACCGAAGAAACCGGAGAAAGAA</td><td></td><td></td><td></td><td></td><td></td><td></td><td></td><td></td><td></td><td></td></tr>
<tr><td>Chromosome</td><td>start</td><td>end</td><td>strand</td><td>MM</td><td>target_seq</td><td>PAM</td><td>alignment</td><td>distance</td><td>position</td><td>gene name</td><td>gene id</td></tr>
<tr><td>GU305914.1</td><td>11133</td><td>11155</td><td>-</td><td>0</td><td>TTCTTTCTCCGGTTTCTTCG</td><td>TGG</td><td>||||||||[||||||||||||]PAM</td><td>NA</td><td>Intergenic</td><td>NA</td><td>NA</td></tr>
<tr><td></td><td></td><td></td><td></td><td></td><td></td><td></td><td></td><td></td><td></td><td></td><td></td></tr>
<tr><td>T33605</td><td>TCCGGTTTCTTCGTGGCGTTAGG</td><td>1000</td><td></td><td></td><td></td><td></td><td></td><td></td><td></td><td></td><td></td></tr>
<tr><td>Oligo fwd</td><td>CACCTCCGGTTTCTTCGTGGCGTT</td><td></td><td></td><td></td><td></td><td></td><td></td><td></td><td></td><td></td><td></td></tr>
<tr><td>Oligo rev</td><td>AAACAACGCCACGAAGAAACCGGA</td><td></td><td></td><td></td><td></td><td></td><td></td><td></td><td></td><td></td><td></td></tr>
<tr><td>Chromosome</td><td>start</td><td>end</td><td>strand</td><td>MM</td><td>target_seq</td><td>PAM</td><td>alignment</td><td>distance</td><td>position</td><td>gene name</td><td>gene id</td></tr>
<tr><td>GU305914.1</td><td>11126</td><td>11148</td><td>-</td><td>0</td><td>TCCGGTTTCTTCGTGGCGTT</td><td>AGG</td><td>||||||||[||||||||||||]PAM</td><td>NA</td><td>Intergenic</td><td>NA</td><td>NA</td></tr>
<tr><td></td><td></td><td></td><td></td><td></td><td></td><td></td><td></td><td></td><td></td><td></td><td></td></tr>
<tr><td>T33606</td><td>CCGGTTTCTTCGTGGCGTTAGGG</td><td>1000</td><td></td><td></td><td></td><td></td><td></td><td></td><td></td><td></td><td></td></tr>
<tr><td>Oligo fwd</td><td>CACCCCGGTTTCTTCGTGGCGTTA</td><td></td><td></td><td></td><td></td><td></td><td></td><td></td><td></td><td></td><td></td></tr>
<tr><td>Oligo rev</td><td>AAACTAACGCCACGAAGAAACCGG</td><td></td><td></td><td></td><td></td><td></td><td></td><td></td><td></td><td></td><td></td></tr>
<tr><td>Chromosome</td><td>start</td><td>end</td><td>strand</td><td>MM</td><td>target_seq</td><td>PAM</td><td>alignment</td><td>distance</td><td>position</td><td>gene name</td><td>gene id</td></tr>
<tr><td>GU305914.1</td><td>11125</td><td>11147</td><td>-</td><td>0</td><td>CCGGTTTCTTCGTGGCGTTA</td><td>GGG</td><td>||||||||[||||||||||||]PAM</td><td>NA</td><td>Intergenic</td><td>NA</td><td>NA</td></tr>
<tr><td></td><td></td><td></td><td></td><td></td><td></td><td></td><td></td><td></td><td></td><td></td><td></td></tr>
<tr><td>T33607</td><td>AGGGTAGACGTAAAGATAGCCGG</td><td>1000</td><td></td><td></td><td></td><td></td><td></td><td></td><td></td><td></td><td></td></tr>
<tr><td>Oligo fwd</td><td>CACCAGGGTAGACGTAAAGATAGC</td><td></td><td></td><td></td><td></td><td></td><td></td><td></td><td></td><td></td><td></td></tr>
<tr><td>Oligo rev</td><td>AAACGCTATCTTTACGTCTACCCT</td><td></td><td></td><td></td><td></td><td></td><td></td><td></td><td></td><td></td><td></td></tr>
<tr><td>Chromosome</td><td>start</td><td>end</td><td>strand</td><td>MM</td><td>target_seq</td><td>PAM</td><td>alignment</td><td>distance</td><td>position</td><td>gene name</td><td>gene id</td></tr>
<tr><td>GU305914.1</td><td>11106</td><td>11128</td><td>-</td><td>0</td><td>AGGGTAGACGTAAAGATAGC</td><td>CGG</td><td>||||||||[||||||||||||]PAM</td><td>NA</td><td>Intergenic</td><td>NA</td><td>NA</td></tr>
<tr><td></td><td></td><td></td><td></td><td></td><td></td><td></td><td></td><td></td><td></td><td></td><td></td></tr>
<tr><td>T33608</td><td>CGACCCATAAACCAGATAATCGG</td><td>1000</td><td></td><td></td><td></td><td></td><td></td><td></td><td></td><td></td><td></td></tr>
<tr><td>Oligo fwd</td><td>CACCCGACCCATAAACCAGATAAT</td><td></td><td></td><td></td><td></td><td></td><td></td><td></td><td></td><td></td><td></td></tr>
<tr><td>Oligo rev</td><td>AAACATTATCTGGTTTATGGGTCG</td><td></td><td></td><td></td><td></td><td></td><td></td><td></td><td></td><td></td><td></td></tr>
<tr><td>Chromosome</td><td>start</td><td>end</td><td>strand</td><td>MM</td><td>target_seq</td><td>PAM</td><td>alignment</td><td>distance</td><td>position</td><td>gene name</td><td>gene id</td></tr>
<tr><td>GU305914.1</td><td>11074</td><td>11096</td><td>-</td><td>0</td><td>CGACCCATAAACCAGATAAT</td><td>CGG</td><td>||||||||[||||||||||||]PAM</td><td>NA</td><td>Intergenic</td><td>NA</td><td>NA</td></tr>
<tr><td></td><td></td><td></td><td></td><td></td><td></td><td></td><td></td><td></td><td></td><td></td><td></td></tr>
<tr><td>T33609</td><td>CCCATAAACCAGATAATCGGCGG</td><td>1000</td><td></td><td></td><td></td><td></td><td></td><td></td><td></td><td></td><td></td></tr>
<tr><td>Oligo fwd</td><td>CACCCCCATAAACCAGATAATCGG</td><td></td><td></td><td></td><td></td><td></td><td></td><td></td><td></td><td></td><td></td></tr>
<tr><td>Oligo rev</td><td>AAACCCGATTATCTGGTTTATGGG</td><td></td><td></td><td></td><td></td><td></td><td></td><td></td><td></td><td></td><td></td></tr>
<tr><td>Chromosome</td><td>start</td><td>end</td><td>strand</td><td>MM</td><td>target_seq</td><td>PAM</td><td>alignment</td><td>distance</td><td>position</td><td>gene name</td><td>gene id</td></tr>
<tr><td>GU305914.1</td><td>11071</td><td>11093</td><td>-</td><td>0</td><td>CCCATAAACCAGATAATCGG</td><td>CGG</td><td>||||||||[||||||||||||]PAM</td><td>NA</td><td>Intergenic</td><td>NA</td><td>NA</td></tr>
<tr><td></td><td></td><td></td><td></td><td></td><td></td><td></td><td></td><td></td><td></td><td></td><td></td></tr>
<tr><td>T33610</td><td>GATAATCGGCGGTGATGTGTAGG</td><td>1000</td><td></td><td></td><td></td><td></td><td></td><td></td><td></td><td></td><td></td></tr>
<tr><td>Oligo fwd</td><td>CACCGATAATCGGCGGTGATGTGT</td><td></td><td></td><td></td><td></td><td></td><td></td><td></td><td></td><td></td><td></td></tr>
<tr><td>Oligo rev</td><td>AAACACACATCACCGCCGATTATC</td><td></td><td></td><td></td><td></td><td></td><td></td><td></td><td></td><td></td><td></td></tr>
<tr><td>Chromosome</td><td>start</td><td>end</td><td>strand</td><td>MM</td><td>target_seq</td><td>PAM</td><td>alignment</td><td>distance</td><td>position</td><td>gene name</td><td>gene id</td></tr>
</table>

<table>
<tr><td>GU305914.1</td><td>11060</td><td>11082</td><td>-</td><td>0</td><td>GATAATCGGCGGTGATGTGT</td><td>AGG</td><td>||||||||[||||||||||||]PAM</td><td>NA</td><td>Intergenic</td><td>NA</td><td>NA</td></tr>
<tr><td>T33611</td><td>ATAATCGGCGGTGATGTGTAGGG</td><td>1000</td><td></td><td></td><td></td><td></td><td></td><td></td><td></td><td></td><td></td></tr>
<tr><td>Oligo fwd</td><td>CACCATAATCGGCGGTGATGTGTA</td><td></td><td></td><td></td><td></td><td></td><td></td><td></td><td></td><td></td><td></td></tr>
<tr><td>Oligo rev</td><td>AAACTACACATCACCGCCGATTAT</td><td></td><td></td><td></td><td></td><td></td><td></td><td></td><td></td><td></td><td></td></tr>
<tr><td>Chromosome</td><td>start</td><td>end</td><td>strand</td><td>MM</td><td>target_seq</td><td>PAM</td><td>alignment</td><td>distance</td><td>position</td><td>gene name</td><td>gene id</td></tr>
<tr><td>GU305914.1</td><td>11059</td><td>11081</td><td>-</td><td>0</td><td>ATAATCGGCGGTGATGTGTA</td><td>GGG</td><td>||||||||[||||||||||||]PAM</td><td>NA</td><td>Intergenic</td><td>NA</td><td>NA</td></tr>
<tr><td>T33612</td><td>TAATCGGCGGTGATGTGTAGGGG</td><td>1000</td><td></td><td></td><td></td><td></td><td></td><td></td><td></td><td></td><td></td></tr>
<tr><td>Oligo fwd</td><td>CACCTAATCGGCGGTGATGTGTAG</td><td></td><td></td><td></td><td></td><td></td><td></td><td></td><td></td><td></td><td></td></tr>
<tr><td>Oligo rev</td><td>AAACCTACACATCACCGCCGATTA</td><td></td><td></td><td></td><td></td><td></td><td></td><td></td><td></td><td></td><td></td></tr>
<tr><td>Chromosome</td><td>start</td><td>end</td><td>strand</td><td>MM</td><td>target_seq</td><td>PAM</td><td>alignment</td><td>distance</td><td>position</td><td>gene name</td><td>gene id</td></tr>
<tr><td>GU305914.1</td><td>11058</td><td>11080</td><td>-</td><td>0</td><td>TAATCGGCGGTGATGTGTAG</td><td>GGG</td><td>||||||||[||||||||||||]PAM</td><td>NA</td><td>Intergenic</td><td>NA</td><td>NA</td></tr>
<tr><td>T33613</td><td>GGGGTGAAACCATGAAACATAGG</td><td>1000</td><td></td><td></td><td></td><td></td><td></td><td></td><td></td><td></td><td></td></tr>
<tr><td>Oligo fwd</td><td>CACCGGGGTGAAACCATGAAACAT</td><td></td><td></td><td></td><td></td><td></td><td></td><td></td><td></td><td></td><td></td></tr>
<tr><td>Oligo rev</td><td>AAACATGTTTCATGGTTTCACCCC</td><td></td><td></td><td></td><td></td><td></td><td></td><td></td><td></td><td></td><td></td></tr>
<tr><td>Chromosome</td><td>start</td><td>end</td><td>strand</td><td>MM</td><td>target_seq</td><td>PAM</td><td>alignment</td><td>distance</td><td>position</td><td>gene name</td><td>gene id</td></tr>
<tr><td>GU305914.1</td><td>11039</td><td>11061</td><td>-</td><td>0</td><td>GGGGTGAAACCATGAAACAT</td><td>AGG</td><td>||||||||[||||||||||||]PAM</td><td>NA</td><td>Intergenic</td><td>NA</td><td>NA</td></tr>
<tr><td>T33614</td><td>GTGAAACCATGAAACATAGGAGG</td><td>1000</td><td></td><td></td><td></td><td></td><td></td><td></td><td></td><td></td><td></td></tr>
<tr><td>Oligo fwd</td><td>CACCGTGAAACCATGAAACATAGG</td><td></td><td></td><td></td><td></td><td></td><td></td><td></td><td></td><td></td><td></td></tr>
<tr><td>Oligo rev</td><td>AAACCCTATGTTTCATGGTTTCAC</td><td></td><td></td><td></td><td></td><td></td><td></td><td></td><td></td><td></td><td></td></tr>
<tr><td>Chromosome</td><td>start</td><td>end</td><td>strand</td><td>MM</td><td>target_seq</td><td>PAM</td><td>alignment</td><td>distance</td><td>position</td><td>gene name</td><td>gene id</td></tr>
<tr><td>GU305914.1</td><td>11036</td><td>11058</td><td>-</td><td>0</td><td>GTGAAACCATGAAACATAGG</td><td>AGG</td><td>||||||||[||||||||||||]PAM</td><td>NA</td><td>Intergenic</td><td>NA</td><td>NA</td></tr>
<tr><td>T33615</td><td>AGTAAAGAACAGACTACCGATGG</td><td>1000</td><td></td><td></td><td></td><td></td><td></td><td></td><td></td><td></td><td></td></tr>
<tr><td>Oligo fwd</td><td>CACCAGTAAAGAACAGACTACCGA</td><td></td><td></td><td></td><td></td><td></td><td></td><td></td><td></td><td></td><td></td></tr>
<tr><td>Oligo rev</td><td>AAACTCGGTAGTCTGTTCTTTACT</td><td></td><td></td><td></td><td></td><td></td><td></td><td></td><td></td><td></td><td></td></tr>
<tr><td>Chromosome</td><td>start</td><td>end</td><td>strand</td><td>MM</td><td>target_seq</td><td>PAM</td><td>alignment</td><td>distance</td><td>position</td><td>gene name</td><td>gene id</td></tr>
<tr><td>GU305914.1</td><td>10998</td><td>11020</td><td>-</td><td>0</td><td>AGTAAAGAACAGACTACCGA</td><td>TGG</td><td>||||||||[||||||||||||]PAM</td><td>NA</td><td>Intergenic</td><td>NA</td><td>NA</td></tr>
<tr><td>T33616</td><td>ATTATTTACGCACCAGACACCGG</td><td>1000</td><td></td><td></td><td></td><td></td><td></td><td></td><td></td><td></td><td></td></tr>
<tr><td>Oligo fwd</td><td>CACCATTATTTACGCACCAGACAC</td><td></td><td></td><td></td><td></td><td></td><td></td><td></td><td></td><td></td><td></td></tr>
<tr><td>Oligo rev</td><td>AAACGTGTCTGGTGCGTAAATAAT</td><td></td><td></td><td></td><td></td><td></td><td></td><td></td><td></td><td></td><td></td></tr>
<tr><td>Chromosome</td><td>start</td><td>end</td><td>strand</td><td>MM</td><td>target_seq</td><td>PAM</td><td>alignment</td><td>distance</td><td>position</td><td>gene name</td><td>gene id</td></tr>
<tr><td>GU305914.1</td><td>10971</td><td>10993</td><td>-</td><td>0</td><td>ATTATTTACGCACCAGACAC</td><td>CGG</td><td>||||||||[||||||||||||]PAM</td><td>NA</td><td>Intergenic</td><td>NA</td><td>NA</td></tr>
<tr><td>T33617</td><td>ACACATATGTCTGTAAGTAAAGG</td><td>1000</td><td></td><td></td><td></td><td></td><td></td><td></td><td></td><td></td><td></td></tr>
<tr><td>Oligo fwd</td><td>CACCACACATATGTCTGTAAGTAA</td><td></td><td></td><td></td><td></td><td></td><td></td><td></td><td></td><td></td><td></td></tr>
<tr><td>Oligo rev</td><td>AAACTTACTTACAGACATATGTGT</td><td></td><td></td><td></td><td></td><td></td><td></td><td></td><td></td><td></td><td></td></tr>
<tr><td>Chromosome</td><td>start</td><td>end</td><td>strand</td><td>MM</td><td>target_seq</td><td>PAM</td><td>alignment</td><td>distance</td><td>position</td><td>gene name</td><td>gene id</td></tr>
<tr><td>GU305914.1</td><td>10934</td><td>10956</td><td>-</td><td>0</td><td>ACACATATGTCTGTAAGTAA</td><td>AGG</td><td>||||||||[||||||||||||]PAM</td><td>NA</td><td>Intergenic</td><td>NA</td><td>NA</td></tr>
<tr><td>T33618</td><td>CACATATGTCTGTAAGTAAAGGG</td><td>1000</td><td></td><td></td><td></td><td></td><td></td><td></td><td></td><td></td><td></td></tr>
<tr><td>Oligo fwd</td><td>CACCCACATATGTCTGTAAGTAAA</td><td></td><td></td><td></td><td></td><td></td><td></td><td></td><td></td><td></td><td></td></tr>
<tr><td>Oligo rev</td><td>AAACTTTACTTACAGACATATGTG</td><td></td><td></td><td></td><td></td><td></td><td></td><td></td><td></td><td></td><td></td></tr>
<tr><td>Chromosome</td><td>start</td><td>end</td><td>strand</td><td>MM</td><td>target_seq</td><td>PAM</td><td>alignment</td><td>distance</td><td>position</td><td>gene name</td><td>gene id</td></tr>
<tr><td>GU305914.1</td><td>10933</td><td>10955</td><td>-</td><td>0</td><td>CACATATGTCTGTAAGTAAA</td><td>GGG</td><td>||||||||[||||||||||||]PAM</td><td>NA</td><td>Intergenic</td><td>NA</td><td>NA</td></tr>
<tr><td>T33619</td><td>CGTCTTTCTGTACAGTATCCCGG</td><td>1000</td><td></td><td></td><td></td><td></td><td></td><td></td><td></td><td></td><td></td></tr>
<tr><td>Oligo fwd</td><td>CACCCGTCTTTCTGTACAGTATCC</td><td></td><td></td><td></td><td></td><td></td><td></td><td></td><td></td><td></td><td></td></tr>
<tr><td>Oligo rev</td><td>AAACGGATACTGTACAGAAAGACG</td><td></td><td></td><td></td><td></td><td></td><td></td><td></td><td></td><td></td><td></td></tr>
<tr><td>Chromosome</td><td>start</td><td>end</td><td>strand</td><td>MM</td><td>target_seq</td><td>PAM</td><td>alignment</td><td>distance</td><td>position</td><td>gene name</td><td>gene id</td></tr>
<tr><td>GU305914.1</td><td>10859</td><td>10881</td><td>-</td><td>0</td><td>CGTCTTTCTGTACAGTATCC</td><td>CGG</td><td>||||||||[||||||||||||]PAM</td><td>NA</td><td>Intergenic</td><td>NA</td><td>NA</td></tr>
<tr><td>T33620</td><td>CAGTATCCCGGACTGACGCCCGG</td><td>1000</td><td></td><td></td><td></td><td></td><td></td><td></td><td></td><td></td><td></td></tr>
<tr><td>Oligo fwd</td><td>CACCCAGTATCCCGGACTGACGCC</td><td></td><td></td><td></td><td></td><td></td><td></td><td></td><td></td><td></td><td></td></tr>
<tr><td>Oligo rev</td><td>AAACGGCGTCAGTCCGGGATACTG</td><td></td><td></td><td></td><td></td><td></td><td></td><td></td><td></td><td></td><td></td></tr>
<tr><td>Chromosome</td><td>start</td><td>end</td><td>strand</td><td>MM</td><td>target_seq</td><td>PAM</td><td>alignment</td><td>distance</td><td>position</td><td>gene name</td><td>gene id</td></tr>
<tr><td>GU305914.1</td><td>10847</td><td>10869</td><td>-</td><td>0</td><td>CAGTATCCCGGACTGACGCC</td><td>CGG</td><td>||||||||[||||||||||||]PAM</td><td>NA</td><td>Intergenic</td><td>NA</td><td>NA</td></tr>
<tr><td>T33621</td><td>TGAAAACTTCGTACAGTTATTGG</td><td>1000</td><td></td><td></td><td></td><td></td><td></td><td></td><td></td><td></td><td></td></tr>
<tr><td>Oligo fwd</td><td>CACCTGAAAACTTCGTACAGTTAT</td><td></td><td></td><td></td><td></td><td></td><td></td><td></td><td></td><td></td><td></td></tr>
<tr><td>Oligo rev</td><td>AAACATAACTGTACGAAGTTTTCA</td><td></td><td></td><td></td><td></td><td></td><td></td><td></td><td></td><td></td><td></td></tr>
<tr><td>Chromosome</td><td>start</td><td>end</td><td>strand</td><td>MM</td><td>target_seq</td><td>PAM</td><td>alignment</td><td>distance</td><td>position</td><td>gene name</td><td>gene id</td></tr>
<tr><td>GU305914.1</td><td>10821</td><td>10843</td><td>-</td><td>0</td><td>TGAAAACTTCGTACAGTTAT</td><td>TGG</td><td>||||||||[||||||||||||]PAM</td><td>NA</td><td>Intergenic</td><td>NA</td><td>NA</td></tr>
<tr><td>T33622</td><td>TTCGTACAGTTATTGGTTTTCGG</td><td>1000</td><td></td><td></td><td></td><td></td><td></td><td></td><td></td><td></td><td></td></tr>
<tr><td>Oligo fwd</td><td>CACCTTCGTACAGTTATTGGTTTT</td><td></td><td></td><td></td><td></td><td></td><td></td><td></td><td></td><td></td><td></td></tr>
<tr><td>Oligo rev</td><td>AAACAAAACCAATAACTGTACGAA</td><td></td><td></td><td></td><td></td><td></td><td></td><td></td><td></td><td></td><td></td></tr>
<tr><td>Chromosome</td><td>start</td><td>end</td><td>strand</td><td>MM</td><td>target_seq</td><td>PAM</td><td>alignment</td><td>distance</td><td>position</td><td>gene name</td><td>gene id</td></tr>
<tr><td>GU305914.1</td><td>10814</td><td>10836</td><td>-</td><td>0</td><td>TTCGTACAGTTATTGGTTTT</td><td>CGG</td><td>||||||||[||||||||||||]PAM</td><td>NA</td><td>Intergenic</td><td>NA</td><td>NA</td></tr>
<tr><td>T33623</td><td>CAGTTATTGGTTTTCGGAGCTGG</td><td>1000</td><td></td><td></td><td></td><td></td><td></td><td></td><td></td><td></td><td></td></tr>
<tr><td>Oligo fwd</td><td>CACCCAGTTATTGGTTTTCGGAGC</td><td></td><td></td><td></td><td></td><td></td><td></td><td></td><td></td><td></td><td></td></tr>
<tr><td>Oligo rev</td><td>AAACGCTCCGAAAACCAATAACTG</td><td></td><td></td><td></td><td></td><td></td><td></td><td></td><td></td><td></td><td></td></tr>
<tr><td>Chromosome</td><td>start</td><td>end</td><td>strand</td><td>MM</td><td>target_seq</td><td>PAM</td><td>alignment</td><td>distance</td><td>position</td><td>gene name</td><td>gene id</td></tr>
<tr><td>GU305914.1</td><td>10808</td><td>10830</td><td>-</td><td>0</td><td>CAGTTATTGGTTTTCGGAGC</td><td>TGG</td><td>||||||||[||||||||||||]PAM</td><td>NA</td><td>Intergenic</td><td>NA</td><td>NA</td></tr>
<tr><td>T33624</td><td>GGAGCTGGAGCACGCGCCGCCGG</td><td>1000</td><td></td><td></td><td></td><td></td><td></td><td></td><td></td><td></td><td></td></tr>
<tr><td>Oligo fwd</td><td>CACCGGAGCTGGAGCACGCGCCGC</td><td></td><td></td><td></td><td></td><td></td><td></td><td></td><td></td><td></td><td></td></tr>
<tr><td>Oligo rev</td><td>AAACGCGGCGCGTGCTCCAGCTCC</td><td></td><td></td><td></td><td></td><td></td><td></td><td></td><td></td><td></td><td></td></tr>
<tr><td>Chromosome</td><td>start</td><td>end</td><td>strand</td><td>MM</td><td>target_seq</td><td>PAM</td><td>alignment</td><td>distance</td><td>position</td><td>gene name</td><td>gene id</td></tr>
<tr><td>GU305914.1</td><td>10793</td><td>10815</td><td>-</td><td>0</td><td>GGAGCTGGAGCACGCGCCGC</td><td>CGG</td><td>||||||||[||||||||||||]PAM</td><td>NA</td><td>Intergenic</td><td>NA</td><td>NA</td></tr>
<tr><td>T33625</td><td>GAGCTACGAAGTAGATTACAAGG</td><td>1000</td><td></td><td></td><td></td><td></td><td></td><td></td><td></td><td></td><td></td></tr>
<tr><td>Oligo fwd</td><td>CACCGAGCTACGAAGTAGATTACA</td><td></td><td></td><td></td><td></td><td></td><td></td><td></td><td></td><td></td><td></td></tr>
<tr><td>Oligo rev</td><td>AAACTGTAATCTACTTCGTAGCTC</td><td></td><td></td><td></td><td></td><td></td><td></td><td></td><td></td><td></td><td></td></tr>
</table>

<table>
<tr><td>Chromosome</td><td>start</td><td></td><td>end</td><td>strand</td><td>MM</td><td>target_seq</td><td>PAM</td><td>alignment</td><td>distance</td><td>position</td><td>gene name</td><td>gene id</td></tr>
<tr><td>GU305914.1</td><td></td><td>10750</td><td>10772</td><td>-</td><td>0</td><td>GAGCTACGAAGTAGATTACA</td><td>AGG</td><td>||||||||[||||||||||||]PAM</td><td>NA</td><td>Intergenic</td><td>NA</td><td>NA</td></tr>
<tr><td>T33626</td><td>GGATGAAAAGCGATCTCCCCAGG</td><td></td><td>1000</td><td></td><td></td><td></td><td></td><td></td><td></td><td></td><td></td><td></td></tr>
<tr><td>Oligo fwd</td><td>CACCGGATGAAAAGCGATCTCCCC</td><td></td><td></td><td></td><td></td><td></td><td></td><td></td><td></td><td></td><td></td><td></td></tr>
<tr><td>Oligo rev</td><td>AAACGGGGAGATCGCTTTTCATCC</td><td></td><td></td><td></td><td></td><td></td><td></td><td></td><td></td><td></td><td></td><td></td></tr>
<tr><td>Chromosome</td><td>start</td><td></td><td>end</td><td>strand</td><td>MM</td><td>target_seq</td><td>PAM</td><td>alignment</td><td>distance</td><td>position</td><td>gene name</td><td>gene id</td></tr>
<tr><td>GU305914.1</td><td></td><td>10729</td><td>10751</td><td>-</td><td>0</td><td>GGATGAAAAGCGATCTCCCC</td><td>AGG</td><td>||||||||[||||||||||||]PAM</td><td>NA</td><td>Intergenic</td><td>NA</td><td>NA</td></tr>
<tr><td>T33627</td><td>AAAGCGATCTCCCCAGGTAGCGG</td><td></td><td>1000</td><td></td><td></td><td></td><td></td><td></td><td></td><td></td><td></td><td></td></tr>
<tr><td>Oligo fwd</td><td>CACCAAAGCGATCTCCCCAGGTAG</td><td></td><td></td><td></td><td></td><td></td><td></td><td></td><td></td><td></td><td></td><td></td></tr>
<tr><td>Oligo rev</td><td>AAACCTACCTGGGGAGATCGCTTT</td><td></td><td></td><td></td><td></td><td></td><td></td><td></td><td></td><td></td><td></td><td></td></tr>
<tr><td>Chromosome</td><td>start</td><td></td><td>end</td><td>strand</td><td>MM</td><td>target_seq</td><td>PAM</td><td>alignment</td><td>distance</td><td>position</td><td>gene name</td><td>gene id</td></tr>
<tr><td>GU305914.1</td><td></td><td>10723</td><td>10745</td><td>-</td><td>0</td><td>AAAGCGATCTCCCCAGGTAG</td><td>CGG</td><td>||||||||[||||||||||||]PAM</td><td>NA</td><td>Intergenic</td><td>NA</td><td>NA</td></tr>
<tr><td>T33628</td><td>CAGGTAGCGGTGAGCCATCTCGG</td><td></td><td>1000</td><td></td><td></td><td></td><td></td><td></td><td></td><td></td><td></td><td></td></tr>
<tr><td>Oligo fwd</td><td>CACCCAGGTAGCGGTGAGCCATCT</td><td></td><td></td><td></td><td></td><td></td><td></td><td></td><td></td><td></td><td></td><td></td></tr>
<tr><td>Oligo rev</td><td>AAACAGATGGCTCACCGCTACCTG</td><td></td><td></td><td></td><td></td><td></td><td></td><td></td><td></td><td></td><td></td><td></td></tr>
<tr><td>Chromosome</td><td>start</td><td></td><td>end</td><td>strand</td><td>MM</td><td>target_seq</td><td>PAM</td><td>alignment</td><td>distance</td><td>position</td><td>gene name</td><td>gene id</td></tr>
<tr><td>GU305914.1</td><td></td><td>10710</td><td>10732</td><td>-</td><td>0</td><td>CAGGTAGCGGTGAGCCATCT</td><td>CGG</td><td>||||||||[||||||||||||]PAM</td><td>NA</td><td>Intergenic</td><td>NA</td><td>NA</td></tr>
<tr><td>T33629</td><td>TGAGCCATCTCGGTGCTGTACGG</td><td></td><td>1000</td><td></td><td></td><td></td><td></td><td></td><td></td><td></td><td></td><td></td></tr>
<tr><td>Oligo fwd</td><td>CACCTGAGCCATCTCGGTGCTGTA</td><td></td><td></td><td></td><td></td><td></td><td></td><td></td><td></td><td></td><td></td><td></td></tr>
<tr><td>Oligo rev</td><td>AAACTACAGCACCGAGATGGCTCA</td><td></td><td></td><td></td><td></td><td></td><td></td><td></td><td></td><td></td><td></td><td></td></tr>
<tr><td>Chromosome</td><td>start</td><td></td><td>end</td><td>strand</td><td>MM</td><td>target_seq</td><td>PAM</td><td>alignment</td><td>distance</td><td>position</td><td>gene name</td><td>gene id</td></tr>
<tr><td>GU305914.1</td><td></td><td>10700</td><td>10722</td><td>-</td><td>0</td><td>TGAGCCATCTCGGTGCTGTA</td><td>CGG</td><td>||||||||[||||||||||||]PAM</td><td>NA</td><td>Intergenic</td><td>NA</td><td>NA</td></tr>
<tr><td>GU305914.1</td><td></td><td>33733</td><td>33755</td><td>-</td><td>3</td><td>TACGCCATCTCGCTGCTGTA</td><td>CGG</td><td>|--|||||[||||-|||||||]PAM</td><td>NA</td><td>Intergenic</td><td>NA</td><td>NA</td></tr>
<tr><td>T33630</td><td>GAGCCATCTCGGTGCTGTACGGG</td><td></td><td>1000</td><td></td><td></td><td></td><td></td><td></td><td></td><td></td><td></td><td></td></tr>
<tr><td>Oligo fwd</td><td>CACCGAGCCATCTCGGTGCTGTAC</td><td></td><td></td><td></td><td></td><td></td><td></td><td></td><td></td><td></td><td></td><td></td></tr>
<tr><td>Oligo rev</td><td>AAACGTACAGCACCGAGATGGCTC</td><td></td><td></td><td></td><td></td><td></td><td></td><td></td><td></td><td></td><td></td><td></td></tr>
<tr><td>Chromosome</td><td>start</td><td></td><td>end</td><td>strand</td><td>MM</td><td>target_seq</td><td>PAM</td><td>alignment</td><td>distance</td><td>position</td><td>gene name</td><td>gene id</td></tr>
<tr><td>GU305914.1</td><td></td><td>10699</td><td>10721</td><td>-</td><td>0</td><td>GAGCCATCTCGGTGCTGTAC</td><td>GGG</td><td>||||||||[||||||||||||]PAM</td><td>NA</td><td>Intergenic</td><td>NA</td><td>NA</td></tr>
<tr><td>T33631</td><td>CAGAGTGTGATCTACCGTACAGG</td><td></td><td>1000</td><td></td><td></td><td></td><td></td><td></td><td></td><td></td><td></td><td></td></tr>
<tr><td>Oligo fwd</td><td>CACCCAGAGTGTGATCTACCGTAC</td><td></td><td></td><td></td><td></td><td></td><td></td><td></td><td></td><td></td><td></td><td></td></tr>
<tr><td>Oligo rev</td><td>AAACGTACGGTAGATCACACTCTG</td><td></td><td></td><td></td><td></td><td></td><td></td><td></td><td></td><td></td><td></td><td></td></tr>
<tr><td>Chromosome</td><td>start</td><td></td><td>end</td><td>strand</td><td>MM</td><td>target_seq</td><td>PAM</td><td>alignment</td><td>distance</td><td>position</td><td>gene name</td><td>gene id</td></tr>
<tr><td>GU305914.1</td><td></td><td>10659</td><td>10681</td><td>-</td><td>0</td><td>CAGAGTGTGATCTACCGTAC</td><td>AGG</td><td>||||||||[||||||||||||]PAM</td><td>NA</td><td>Intergenic</td><td>NA</td><td>NA</td></tr>
<tr><td>T33632</td><td>TCCGCAGCTACATATACGCCAGG</td><td></td><td>1000</td><td></td><td></td><td></td><td></td><td></td><td></td><td></td><td></td><td></td></tr>
<tr><td>Oligo fwd</td><td>CACCTCCGCAGCTACATATACGCC</td><td></td><td></td><td></td><td></td><td></td><td></td><td></td><td></td><td></td><td></td><td></td></tr>
<tr><td>Oligo rev</td><td>AAACGGCGTATATGTAGCTGCGGA</td><td></td><td></td><td></td><td></td><td></td><td></td><td></td><td></td><td></td><td></td><td></td></tr>
<tr><td>Chromosome</td><td>start</td><td></td><td>end</td><td>strand</td><td>MM</td><td>target_seq</td><td>PAM</td><td>alignment</td><td>distance</td><td>position</td><td>gene name</td><td>gene id</td></tr>
<tr><td>GU305914.1</td><td></td><td>10629</td><td>10651</td><td>-</td><td>0</td><td>TCCGCAGCTACATATACGCC</td><td>AGG</td><td>||||||||[||||||||||||]PAM</td><td>NA</td><td>Intergenic</td><td>NA</td><td>NA</td></tr>
<tr><td>T33633</td><td>CTACATATACGCCAGGTACGTGG</td><td></td><td>1000</td><td></td><td></td><td></td><td></td><td></td><td></td><td></td><td></td><td></td></tr>
<tr><td>Oligo fwd</td><td>CACCCTACATATACGCCAGGTACG</td><td></td><td></td><td></td><td></td><td></td><td></td><td></td><td></td><td></td><td></td><td></td></tr>
<tr><td>Oligo rev</td><td>AAACCGTACCTGGCGTATATGTAG</td><td></td><td></td><td></td><td></td><td></td><td></td><td></td><td></td><td></td><td></td><td></td></tr>
<tr><td>Chromosome</td><td>start</td><td></td><td>end</td><td>strand</td><td>MM</td><td>target_seq</td><td>PAM</td><td>alignment</td><td>distance</td><td>position</td><td>gene name</td><td>gene id</td></tr>
<tr><td>GU305914.1</td><td></td><td>10622</td><td>10644</td><td>-</td><td>0</td><td>CTACATATACGCCAGGTACG</td><td>TGG</td><td>||||||||[||||||||||||]PAM</td><td>NA</td><td>Intergenic</td><td>NA</td><td>NA</td></tr>
<tr><td>T33634</td><td>TACATATACGCCAGGTACGTGGG</td><td></td><td>1000</td><td></td><td></td><td></td><td></td><td></td><td></td><td></td><td></td><td></td></tr>
<tr><td>Oligo fwd</td><td>CACCTACATATACGCCAGGTACGT</td><td></td><td></td><td></td><td></td><td></td><td></td><td></td><td></td><td></td><td></td><td></td></tr>
<tr><td>Oligo rev</td><td>AAACACGTACCTGGCGTATATGTA</td><td></td><td></td><td></td><td></td><td></td><td></td><td></td><td></td><td></td><td></td><td></td></tr>
<tr><td>Chromosome</td><td>start</td><td></td><td>end</td><td>strand</td><td>MM</td><td>target_seq</td><td>PAM</td><td>alignment</td><td>distance</td><td>position</td><td>gene name</td><td>gene id</td></tr>
<tr><td>GU305914.1</td><td></td><td>10621</td><td>10643</td><td>-</td><td>0</td><td>TACATATACGCCAGGTACGT</td><td>GGG</td><td>||||||||[||||||||||||]PAM</td><td>NA</td><td>Intergenic</td><td>NA</td><td>NA</td></tr>
<tr><td>T33635</td><td>ATATACGCCAGGTACGTGGGCGG</td><td></td><td>1000</td><td></td><td></td><td></td><td></td><td></td><td></td><td></td><td></td><td></td></tr>
<tr><td>Oligo fwd</td><td>CACCATATACGCCAGGTACGTGGG</td><td></td><td></td><td></td><td></td><td></td><td></td><td></td><td></td><td></td><td></td><td></td></tr>
<tr><td>Oligo rev</td><td>AAACCCCACGTACCTGGCGTATAT</td><td></td><td></td><td></td><td></td><td></td><td></td><td></td><td></td><td></td><td></td><td></td></tr>
<tr><td>Chromosome</td><td>start</td><td></td><td>end</td><td>strand</td><td>MM</td><td>target_seq</td><td>PAM</td><td>alignment</td><td>distance</td><td>position</td><td>gene name</td><td>gene id</td></tr>
<tr><td>GU305914.1</td><td></td><td>10618</td><td>10640</td><td>-</td><td>0</td><td>ATATACGCCAGGTACGTGGG</td><td>CGG</td><td>||||||||[||||||||||||]PAM</td><td>NA</td><td>Intergenic</td><td>NA</td><td>NA</td></tr>
<tr><td>T33636</td><td>TATACGCCAGGTACGTGGGCGGG</td><td></td><td>1000</td><td></td><td></td><td></td><td></td><td></td><td></td><td></td><td></td><td></td></tr>
<tr><td>Oligo fwd</td><td>CACCTATACGCCAGGTACGTGGGC</td><td></td><td></td><td></td><td></td><td></td><td></td><td></td><td></td><td></td><td></td><td></td></tr>
<tr><td>Oligo rev</td><td>AAACGCCCACGTACCTGGCGTATA</td><td></td><td></td><td></td><td></td><td></td><td></td><td></td><td></td><td></td><td></td><td></td></tr>
<tr><td>Chromosome</td><td>start</td><td></td><td>end</td><td>strand</td><td>MM</td><td>target_seq</td><td>PAM</td><td>alignment</td><td>distance</td><td>position</td><td>gene name</td><td>gene id</td></tr>
<tr><td>GU305914.1</td><td></td><td>10617</td><td>10639</td><td>-</td><td>0</td><td>TATACGCCAGGTACGTGGGC</td><td>GGG</td><td>||||||||[||||||||||||]PAM</td><td>NA</td><td>Intergenic</td><td>NA</td><td>NA</td></tr>
<tr><td>T33637</td><td>AGGTACGTGGGCGGGATCTACGG</td><td></td><td>1000</td><td></td><td></td><td></td><td></td><td></td><td></td><td></td><td></td><td></td></tr>
<tr><td>Oligo fwd</td><td>CACCAGGTACGTGGGCGGGATCTA</td><td></td><td></td><td></td><td></td><td></td><td></td><td></td><td></td><td></td><td></td><td></td></tr>
<tr><td>Oligo rev</td><td>AAACTAGATCCCGCCCACGTACCT</td><td></td><td></td><td></td><td></td><td></td><td></td><td></td><td></td><td></td><td></td><td></td></tr>
<tr><td>Chromosome</td><td>start</td><td></td><td>end</td><td>strand</td><td>MM</td><td>target_seq</td><td>PAM</td><td>alignment</td><td>distance</td><td>position</td><td>gene name</td><td>gene id</td></tr>
<tr><td>GU305914.1</td><td></td><td>10609</td><td>10631</td><td>-</td><td>0</td><td>AGGTACGTGGGCGGGATCTA</td><td>CGG</td><td>||||||||[||||||||||||]PAM</td><td>NA</td><td>Intergenic</td><td>NA</td><td>NA</td></tr>
<tr><td>T33638</td><td>GGTACGTGGGCGGGATCTACGGG</td><td></td><td>1000</td><td></td><td></td><td></td><td></td><td></td><td></td><td></td><td></td><td></td></tr>
<tr><td>Oligo fwd</td><td>CACCGGTACGTGGGCGGGATCTAC</td><td></td><td></td><td></td><td></td><td></td><td></td><td></td><td></td><td></td><td></td><td></td></tr>
<tr><td>Oligo rev</td><td>AAACGTAGATCCCGCCCACGTACC</td><td></td><td></td><td></td><td></td><td></td><td></td><td></td><td></td><td></td><td></td><td></td></tr>
<tr><td>Chromosome</td><td>start</td><td></td><td>end</td><td>strand</td><td>MM</td><td>target_seq</td><td>PAM</td><td>alignment</td><td>distance</td><td>position</td><td>gene name</td><td>gene id</td></tr>
<tr><td>GU305914.1</td><td></td><td>10608</td><td>10630</td><td>-</td><td>0</td><td>GGTACGTGGGCGGGATCTAC</td><td>GGG</td><td>||||||||[||||||||||||]PAM</td><td>NA</td><td>Intergenic</td><td>NA</td><td>NA</td></tr>
<tr><td>T33639</td><td>GGCGGGATCTACGGGAACTCAGG</td><td></td><td>1000</td><td></td><td></td><td></td><td></td><td></td><td></td><td></td><td></td><td></td></tr>
<tr><td>Oligo fwd</td><td>CACCGGCGGGATCTACGGGAACTC</td><td></td><td></td><td></td><td></td><td></td><td></td><td></td><td></td><td></td><td></td><td></td></tr>
<tr><td>Oligo rev</td><td>AAACGAGTTCCCGTAGATCCCGCC</td><td></td><td></td><td></td><td></td><td></td><td></td><td></td><td></td><td></td><td></td><td></td></tr>
<tr><td>Chromosome</td><td>start</td><td></td><td>end</td><td>strand</td><td>MM</td><td>target_seq</td><td>PAM</td><td>alignment</td><td>distance</td><td>position</td><td>gene name</td><td>gene id</td></tr>
<tr><td>GU305914.1</td><td></td><td>10600</td><td>10622</td><td>-</td><td>0</td><td>GGCGGGATCTACGGGAACTC</td><td>AGG</td><td>||||||||[||||||||||||]PAM</td><td>NA</td><td>Intergenic</td><td>NA</td><td>NA</td></tr>
<tr><td>T33640</td><td>ATCTACGGGAACTCAGGTGACGG</td><td></td><td>1000</td><td></td><td></td><td></td><td></td><td></td><td></td><td></td><td></td><td></td></tr>
</table>

<table>
<tr><td>Oligo fwd</td><td>CACCATCTACGGGAACTCAGGTGA</td><td></td><td></td><td></td><td></td><td></td><td></td><td></td><td></td><td></td><td></td></tr>
<tr><td>Oligo rev</td><td>AAACTCACCTGAGTTCCCGTAGAT</td><td></td><td></td><td></td><td></td><td></td><td></td><td></td><td></td><td></td><td></td></tr>
<tr><td>Chromosome</td><td>start</td><td>end</td><td>strand</td><td>MM</td><td>target_seq</td><td>PAM</td><td>alignment</td><td>distance</td><td>position</td><td>gene name</td><td>gene id</td></tr>
<tr><td>GU305914.1</td><td>10594</td><td>10616</td><td>-</td><td>0</td><td>ATCTACGGGAACTCAGGTGA</td><td>CGG</td><td>||||||||[||||||||||||]PAM</td><td>NA</td><td>Intergenic</td><td>NA</td><td>NA</td></tr>
<tr><td></td><td></td><td></td><td></td><td></td><td></td><td></td><td></td><td></td><td></td><td></td><td></td></tr>
<tr><td>T33641</td><td>TCTACGGGAACTCAGGTGACGGG</td><td>1000</td><td></td><td></td><td></td><td></td><td></td><td></td><td></td><td></td><td></td></tr>
<tr><td>Oligo fwd</td><td>CACCTCTACGGGAACTCAGGTGAC</td><td></td><td></td><td></td><td></td><td></td><td></td><td></td><td></td><td></td><td></td></tr>
<tr><td>Oligo rev</td><td>AAACGTCACCTGAGTTCCCGTAGA</td><td></td><td></td><td></td><td></td><td></td><td></td><td></td><td></td><td></td><td></td></tr>
<tr><td>Chromosome</td><td>start</td><td>end</td><td>strand</td><td>MM</td><td>target_seq</td><td>PAM</td><td>alignment</td><td>distance</td><td>position</td><td>gene name</td><td>gene id</td></tr>
<tr><td>GU305914.1</td><td>10593</td><td>10615</td><td>-</td><td>0</td><td>TCTACGGGAACTCAGGTGAC</td><td>GGG</td><td>||||||||[||||||||||||]PAM</td><td>NA</td><td>Intergenic</td><td>NA</td><td>NA</td></tr>
<tr><td></td><td></td><td></td><td></td><td></td><td></td><td></td><td></td><td></td><td></td><td></td><td></td></tr>
<tr><td>T33642</td><td>CTACGGGAACTCAGGTGACGGGG</td><td>1000</td><td></td><td></td><td></td><td></td><td></td><td></td><td></td><td></td><td></td></tr>
<tr><td>Oligo fwd</td><td>CACCCTACGGGAACTCAGGTGACG</td><td></td><td></td><td></td><td></td><td></td><td></td><td></td><td></td><td></td><td></td></tr>
<tr><td>Oligo rev</td><td>AAACCGTCACCTGAGTTCCCGTAG</td><td></td><td></td><td></td><td></td><td></td><td></td><td></td><td></td><td></td><td></td></tr>
<tr><td>Chromosome</td><td>start</td><td>end</td><td>strand</td><td>MM</td><td>target_seq</td><td>PAM</td><td>alignment</td><td>distance</td><td>position</td><td>gene name</td><td>gene id</td></tr>
<tr><td>GU305914.1</td><td>10592</td><td>10614</td><td>-</td><td>0</td><td>CTACGGGAACTCAGGTGACG</td><td>GGG</td><td>||||||||[||||||||||||]PAM</td><td>NA</td><td>Intergenic</td><td>NA</td><td>NA</td></tr>
<tr><td></td><td></td><td></td><td></td><td></td><td></td><td></td><td></td><td></td><td></td><td></td><td></td></tr>
<tr><td>T33643</td><td>AGGTGACGGGGAAATCACCTAGG</td><td>1000</td><td></td><td></td><td></td><td></td><td></td><td></td><td></td><td></td><td></td></tr>
<tr><td>Oligo fwd</td><td>CACCAGGTGACGGGGAAATCACCT</td><td></td><td></td><td></td><td></td><td></td><td></td><td></td><td></td><td></td><td></td></tr>
<tr><td>Oligo rev</td><td>AAACAGGTGATTTCCCCGTCACCT</td><td></td><td></td><td></td><td></td><td></td><td></td><td></td><td></td><td></td><td></td></tr>
<tr><td>Chromosome</td><td>start</td><td>end</td><td>strand</td><td>MM</td><td>target_seq</td><td>PAM</td><td>alignment</td><td>distance</td><td>position</td><td>gene name</td><td>gene id</td></tr>
<tr><td>GU305914.1</td><td>10580</td><td>10602</td><td>-</td><td>0</td><td>AGGTGACGGGGAAATCACCT</td><td>AGG</td><td>||||||||[||||||||||||]PAM</td><td>NA</td><td>Intergenic</td><td>NA</td><td>NA</td></tr>
<tr><td>GU305914.1</td><td>179328</td><td>179350</td><td>+</td><td>4</td><td>AGGAGAGGAGGAGATCACCT</td><td>GGG</td><td>|||-||-|[-|||-|||||||]PAM</td><td>NA</td><td>Intergenic</td><td>NA</td><td>NA</td></tr>
<tr><td></td><td></td><td></td><td></td><td></td><td></td><td></td><td></td><td></td><td></td><td></td><td></td></tr>
<tr><td>T33644</td><td>GGGGAAATCACCTAGGTAAACGG</td><td>1000</td><td></td><td></td><td></td><td></td><td></td><td></td><td></td><td></td><td></td></tr>
<tr><td>Oligo fwd</td><td>CACCGGGGAAATCACCTAGGTAAA</td><td></td><td></td><td></td><td></td><td></td><td></td><td></td><td></td><td></td><td></td></tr>
<tr><td>Oligo rev</td><td>AAACTTTACCTAGGTGATTTCCCC</td><td></td><td></td><td></td><td></td><td></td><td></td><td></td><td></td><td></td><td></td></tr>
<tr><td>Chromosome</td><td>start</td><td>end</td><td>strand</td><td>MM</td><td>target_seq</td><td>PAM</td><td>alignment</td><td>distance</td><td>position</td><td>gene name</td><td>gene id</td></tr>
<tr><td>GU305914.1</td><td>10573</td><td>10595</td><td>-</td><td>0</td><td>GGGGAAATCACCTAGGTAAA</td><td>CGG</td><td>||||||||[||||||||||||]PAM</td><td>NA</td><td>Intergenic</td><td>NA</td><td>NA</td></tr>
<tr><td></td><td></td><td></td><td></td><td></td><td></td><td></td><td></td><td></td><td></td><td></td><td></td></tr>
<tr><td>T33645</td><td>GAAATCACCTAGGTAAACGGAGG</td><td>1000</td><td></td><td></td><td></td><td></td><td></td><td></td><td></td><td></td><td></td></tr>
<tr><td>Oligo fwd</td><td>CACCGAAATCACCTAGGTAAACGG</td><td></td><td></td><td></td><td></td><td></td><td></td><td></td><td></td><td></td><td></td></tr>
<tr><td>Oligo rev</td><td>AAACCCGTTTACCTAGGTGATTTC</td><td></td><td></td><td></td><td></td><td></td><td></td><td></td><td></td><td></td><td></td></tr>
<tr><td>Chromosome</td><td>start</td><td>end</td><td>strand</td><td>MM</td><td>target_seq</td><td>PAM</td><td>alignment</td><td>distance</td><td>position</td><td>gene name</td><td>gene id</td></tr>
<tr><td>GU305914.1</td><td>10570</td><td>10592</td><td>-</td><td>0</td><td>GAAATCACCTAGGTAAACGG</td><td>AGG</td><td>||||||||[||||||||||||]PAM</td><td>NA</td><td>Intergenic</td><td>NA</td><td>NA</td></tr>
<tr><td></td><td></td><td></td><td></td><td></td><td></td><td></td><td></td><td></td><td></td><td></td><td></td></tr>
<tr><td>T33646</td><td>ATCACCTAGGTAAACGGAGGCGG</td><td>1000</td><td></td><td></td><td></td><td></td><td></td><td></td><td></td><td></td><td></td></tr>
<tr><td>Oligo fwd</td><td>CACCATCACCTAGGTAAACGGAGG</td><td></td><td></td><td></td><td></td><td></td><td></td><td></td><td></td><td></td><td></td></tr>
<tr><td>Oligo rev</td><td>AAACCCTCCGTTTACCTAGGTGAT</td><td></td><td></td><td></td><td></td><td></td><td></td><td></td><td></td><td></td><td></td></tr>
<tr><td>Chromosome</td><td>start</td><td>end</td><td>strand</td><td>MM</td><td>target_seq</td><td>PAM</td><td>alignment</td><td>distance</td><td>position</td><td>gene name</td><td>gene id</td></tr>
<tr><td>GU305914.1</td><td>10567</td><td>10589</td><td>-</td><td>0</td><td>ATCACCTAGGTAAACGGAGG</td><td>CGG</td><td>||||||||[||||||||||||]PAM</td><td>NA</td><td>Intergenic</td><td>NA</td><td>NA</td></tr>
<tr><td></td><td></td><td></td><td></td><td></td><td></td><td></td><td></td><td></td><td></td><td></td><td></td></tr>
<tr><td>T33647</td><td>TGAGTCATATCACAACACACTGG</td><td>1000</td><td></td><td></td><td></td><td></td><td></td><td></td><td></td><td></td><td></td></tr>
<tr><td>Oligo fwd</td><td>CACCTGAGTCATATCACAACACAC</td><td></td><td></td><td></td><td></td><td></td><td></td><td></td><td></td><td></td><td></td></tr>
<tr><td>Oligo rev</td><td>AAACGTGTGTTGTGATATGACTCA</td><td></td><td></td><td></td><td></td><td></td><td></td><td></td><td></td><td></td><td></td></tr>
<tr><td>Chromosome</td><td>start</td><td>end</td><td>strand</td><td>MM</td><td>target_seq</td><td>PAM</td><td>alignment</td><td>distance</td><td>position</td><td>gene name</td><td>gene id</td></tr>
<tr><td>GU305914.1</td><td>10486</td><td>10508</td><td>-</td><td>0</td><td>TGAGTCATATCACAACACAC</td><td>TGG</td><td>||||||||[||||||||||||]PAM</td><td>NA</td><td>Intergenic</td><td>NA</td><td>NA</td></tr>
<tr><td></td><td></td><td></td><td></td><td></td><td></td><td></td><td></td><td></td><td></td><td></td><td></td></tr>
<tr><td>T33648</td><td>TCATATCACAACACACTGGTTGG</td><td>1000</td><td></td><td></td><td></td><td></td><td></td><td></td><td></td><td></td><td></td></tr>
<tr><td>Oligo fwd</td><td>CACCTCATATCACAACACACTGGT</td><td></td><td></td><td></td><td></td><td></td><td></td><td></td><td></td><td></td><td></td></tr>
<tr><td>Oligo rev</td><td>AAACACCAGTGTGTTGTGATATGA</td><td></td><td></td><td></td><td></td><td></td><td></td><td></td><td></td><td></td><td></td></tr>
<tr><td>Chromosome</td><td>start</td><td>end</td><td>strand</td><td>MM</td><td>target_seq</td><td>PAM</td><td>alignment</td><td>distance</td><td>position</td><td>gene name</td><td>gene id</td></tr>
<tr><td>GU305914.1</td><td>10482</td><td>10504</td><td>-</td><td>0</td><td>TCATATCACAACACACTGGT</td><td>TGG</td><td>||||||||[||||||||||||]PAM</td><td>NA</td><td>Intergenic</td><td>NA</td><td>NA</td></tr>
<tr><td></td><td></td><td></td><td></td><td></td><td></td><td></td><td></td><td></td><td></td><td></td><td></td></tr>
<tr><td>T33649</td><td>TATCACAACACACTGGTTGGTGG</td><td>1000</td><td></td><td></td><td></td><td></td><td></td><td></td><td></td><td></td><td></td></tr>
<tr><td>Oligo fwd</td><td>CACCTATCACAACACACTGGTTGG</td><td></td><td></td><td></td><td></td><td></td><td></td><td></td><td></td><td></td><td></td></tr>
<tr><td>Oligo rev</td><td>AAACCCAACCAGTGTGTTGTGATA</td><td></td><td></td><td></td><td></td><td></td><td></td><td></td><td></td><td></td><td></td></tr>
<tr><td>Chromosome</td><td>start</td><td>end</td><td>strand</td><td>MM</td><td>target_seq</td><td>PAM</td><td>alignment</td><td>distance</td><td>position</td><td>gene name</td><td>gene id</td></tr>
<tr><td>GU305914.1</td><td>10479</td><td>10501</td><td>-</td><td>0</td><td>TATCACAACACACTGGTTGG</td><td>TGG</td><td>||||||||[||||||||||||]PAM</td><td>NA</td><td>Intergenic</td><td>NA</td><td>NA</td></tr>
<tr><td></td><td></td><td></td><td></td><td></td><td></td><td></td><td></td><td></td><td></td><td></td><td></td></tr>
<tr><td>T33650</td><td>ATCACAACACACTGGTTGGTGGG</td><td>1000</td><td></td><td></td><td></td><td></td><td></td><td></td><td></td><td></td><td></td></tr>
<tr><td>Oligo fwd</td><td>CACCATCACAACACACTGGTTGGT</td><td></td><td></td><td></td><td></td><td></td><td></td><td></td><td></td><td></td><td></td></tr>
<tr><td>Oligo rev</td><td>AAACACCAACCAGTGTGTTGTGAT</td><td></td><td></td><td></td><td></td><td></td><td></td><td></td><td></td><td></td><td></td></tr>
<tr><td>Chromosome</td><td>start</td><td>end</td><td>strand</td><td>MM</td><td>target_seq</td><td>PAM</td><td>alignment</td><td>distance</td><td>position</td><td>gene name</td><td>gene id</td></tr>
<tr><td>GU305914.1</td><td>10478</td><td>10500</td><td>-</td><td>0</td><td>ATCACAACACACTGGTTGGT</td><td>GGG</td><td>||||||||[||||||||||||]PAM</td><td>NA</td><td>Intergenic</td><td>NA</td><td>NA</td></tr>
<tr><td></td><td></td><td></td><td></td><td></td><td></td><td></td><td></td><td></td><td></td><td></td><td></td></tr>
<tr><td>T33651</td><td>TCACAACACACTGGTTGGTGGGG</td><td>1000</td><td></td><td></td><td></td><td></td><td></td><td></td><td></td><td></td><td></td></tr>
<tr><td>Oligo fwd</td><td>CACCTCACAACACACTGGTTGGTG</td><td></td><td></td><td></td><td></td><td></td><td></td><td></td><td></td><td></td><td></td></tr>
<tr><td>Oligo rev</td><td>AAACCACCAACCAGTGTGTTGTGA</td><td></td><td></td><td></td><td></td><td></td><td></td><td></td><td></td><td></td><td></td></tr>
<tr><td>Chromosome</td><td>start</td><td>end</td><td>strand</td><td>MM</td><td>target_seq</td><td>PAM</td><td>alignment</td><td>distance</td><td>position</td><td>gene name</td><td>gene id</td></tr>
<tr><td>GU305914.1</td><td>10477</td><td>10499</td><td>-</td><td>0</td><td>TCACAACACACTGGTTGGTG</td><td>GGG</td><td>||||||||[||||||||||||]PAM</td><td>NA</td><td>Intergenic</td><td>NA</td><td>NA</td></tr>
<tr><td></td><td></td><td></td><td></td><td></td><td></td><td></td><td></td><td></td><td></td><td></td><td></td></tr>
<tr><td>T33652</td><td>CTGGTTGGTGGGGAATTCAGCGG</td><td>1000</td><td></td><td></td><td></td><td></td><td></td><td></td><td></td><td></td><td></td></tr>
<tr><td>Oligo fwd</td><td>CACCCTGGTTGGTGGGGAATTCAG</td><td></td><td></td><td></td><td></td><td></td><td></td><td></td><td></td><td></td><td></td></tr>
<tr><td>Oligo rev</td><td>AAACCTGAATTCCCCACCAACCAG</td><td></td><td></td><td></td><td></td><td></td><td></td><td></td><td></td><td></td><td></td></tr>
<tr><td>Chromosome</td><td>start</td><td>end</td><td>strand</td><td>MM</td><td>target_seq</td><td>PAM</td><td>alignment</td><td>distance</td><td>position</td><td>gene name</td><td>gene id</td></tr>
<tr><td>GU305914.1</td><td>10467</td><td>10489</td><td>-</td><td>0</td><td>CTGGTTGGTGGGGAATTCAG</td><td>CGG</td><td>||||||||[||||||||||||]PAM</td><td>NA</td><td>Intergenic</td><td>NA</td><td>NA</td></tr>
<tr><td></td><td></td><td></td><td></td><td></td><td></td><td></td><td></td><td></td><td></td><td></td><td></td></tr>
<tr><td>T33653</td><td>TGGTGGGGAATTCAGCGGATCGG</td><td>1000</td><td></td><td></td><td></td><td></td><td></td><td></td><td></td><td></td><td></td></tr>
<tr><td>Oligo fwd</td><td>CACCTGGTGGGGAATTCAGCGGAT</td><td></td><td></td><td></td><td></td><td></td><td></td><td></td><td></td><td></td><td></td></tr>
<tr><td>Oligo rev</td><td>AAACATCCGCTGAATTCCCCACCA</td><td></td><td></td><td></td><td></td><td></td><td></td><td></td><td></td><td></td><td></td></tr>
<tr><td>Chromosome</td><td>start</td><td>end</td><td>strand</td><td>MM</td><td>target_seq</td><td>PAM</td><td>alignment</td><td>distance</td><td>position</td><td>gene name</td><td>gene id</td></tr>
<tr><td>GU305914.1</td><td>10462</td><td>10484</td><td>-</td><td>0</td><td>TGGTGGGGAATTCAGCGGAT</td><td>CGG</td><td>||||||||[||||||||||||]PAM</td><td>NA</td><td>Intergenic</td><td>NA</td><td>NA</td></tr>
<tr><td></td><td></td><td></td><td></td><td></td><td></td><td></td><td></td><td></td><td></td><td></td><td></td></tr>
<tr><td>T33654</td><td>GAATTCAGCGGATCGGATTCAGG</td><td>1000</td><td></td><td></td><td></td><td></td><td></td><td></td><td></td><td></td><td></td></tr>
<tr><td>Oligo fwd</td><td>CACCGAATTCAGCGGATCGGATTC</td><td></td><td></td><td></td><td></td><td></td><td></td><td></td><td></td><td></td><td></td></tr>
<tr><td>Oligo rev</td><td>AAACGAATCCGATCCGCTGAATTC</td><td></td><td></td><td></td><td></td><td></td><td></td><td></td><td></td><td></td><td></td></tr>
<tr><td>Chromosome</td><td>start</td><td>end</td><td>strand</td><td>MM</td><td>target_seq</td><td>PAM</td><td>alignment</td><td>distance</td><td>position</td><td>gene name</td><td>gene id</td></tr>
<tr><td>GU305914.1</td><td>10455</td><td>10477</td><td>-</td><td>0</td><td>GAATTCAGCGGATCGGATTC</td><td>AGG</td><td>||||||||[||||||||||||]PAM</td><td>NA</td><td>Intergenic</td><td>NA</td><td>NA</td></tr>
</table>

<table>
<tr><td>T33655</td><td>CAGCGGATCGGATTCAGGTCTGG</td><td></td><td>1000</td><td></td><td></td><td></td><td></td><td></td><td></td><td></td><td></td><td></td></tr>
<tr><td>Oligo fwd</td><td>CACCCAGCGGATCGGATTCAGGTC</td><td></td><td></td><td></td><td></td><td></td><td></td><td></td><td></td><td></td><td></td><td></td></tr>
<tr><td>Oligo rev</td><td>AAACGACCTGAATCCGATCCGCTG</td><td></td><td></td><td></td><td></td><td></td><td></td><td></td><td></td><td></td><td></td><td></td></tr>
<tr><td>Chromosome</td><td>start</td><td></td><td>end</td><td>strand</td><td>MM</td><td>target_seq</td><td>PAM</td><td>alignment</td><td>distance</td><td>position</td><td>gene name</td><td>gene id</td></tr>
<tr><td>GU305914.1</td><td></td><td>10450</td><td>10472</td><td>-</td><td>0</td><td>CAGCGGATCGGATTCAGGTC</td><td>TGG</td><td>||||||||[||||||||||||]PAM</td><td>NA</td><td>Intergenic</td><td>NA</td><td>NA</td></tr>
<tr><td>T33656</td><td>GGATTCAGGTCTGGAACTCTCGG</td><td></td><td>1000</td><td></td><td></td><td></td><td></td><td></td><td></td><td></td><td></td><td></td></tr>
<tr><td>Oligo fwd</td><td>CACCGGATTCAGGTCTGGAACTCT</td><td></td><td></td><td></td><td></td><td></td><td></td><td></td><td></td><td></td><td></td><td></td></tr>
<tr><td>Oligo rev</td><td>AAACAGAGTTCCAGACCTGAATCC</td><td></td><td></td><td></td><td></td><td></td><td></td><td></td><td></td><td></td><td></td><td></td></tr>
<tr><td>Chromosome</td><td>start</td><td></td><td>end</td><td>strand</td><td>MM</td><td>target_seq</td><td>PAM</td><td>alignment</td><td>distance</td><td>position</td><td>gene name</td><td>gene id</td></tr>
<tr><td>GU305914.1</td><td></td><td>10441</td><td>10463</td><td>-</td><td>0</td><td>GGATTCAGGTCTGGAACTCT</td><td>CGG</td><td>||||||||[||||||||||||]PAM</td><td>NA</td><td>Intergenic</td><td>NA</td><td>NA</td></tr>
<tr><td>T33657</td><td>GATTCAGGTCTGGAACTCTCGGG</td><td></td><td>1000</td><td></td><td></td><td></td><td></td><td></td><td></td><td></td><td></td><td></td></tr>
<tr><td>Oligo fwd</td><td>CACCGATTCAGGTCTGGAACTCTC</td><td></td><td></td><td></td><td></td><td></td><td></td><td></td><td></td><td></td><td></td><td></td></tr>
<tr><td>Oligo rev</td><td>AAACGAGAGTTCCAGACCTGAATC</td><td></td><td></td><td></td><td></td><td></td><td></td><td></td><td></td><td></td><td></td><td></td></tr>
<tr><td>Chromosome</td><td>start</td><td></td><td>end</td><td>strand</td><td>MM</td><td>target_seq</td><td>PAM</td><td>alignment</td><td>distance</td><td>position</td><td>gene name</td><td>gene id</td></tr>
<tr><td>GU305914.1</td><td></td><td>10440</td><td>10462</td><td>-</td><td>0</td><td>GATTCAGGTCTGGAACTCTC</td><td>GGG</td><td>||||||||[||||||||||||]PAM</td><td>NA</td><td>Intergenic</td><td>NA</td><td>NA</td></tr>
<tr><td>T33658</td><td>TCTGGAACTCTCGGGATCGCTGG</td><td></td><td>1000</td><td></td><td></td><td></td><td></td><td></td><td></td><td></td><td></td><td></td></tr>
<tr><td>Oligo fwd</td><td>CACCTCTGGAACTCTCGGGATCGC</td><td></td><td></td><td></td><td></td><td></td><td></td><td></td><td></td><td></td><td></td><td></td></tr>
<tr><td>Oligo rev</td><td>AAACGCGATCCCGAGAGTTCCAGA</td><td></td><td></td><td></td><td></td><td></td><td></td><td></td><td></td><td></td><td></td><td></td></tr>
<tr><td>Chromosome</td><td>start</td><td></td><td>end</td><td>strand</td><td>MM</td><td>target_seq</td><td>PAM</td><td>alignment</td><td>distance</td><td>position</td><td>gene name</td><td>gene id</td></tr>
<tr><td>GU305914.1</td><td></td><td>10432</td><td>10454</td><td>-</td><td>0</td><td>TCTGGAACTCTCGGGATCGC</td><td>TGG</td><td>||||||||[||||||||||||]PAM</td><td>NA</td><td>Intergenic</td><td>NA</td><td>NA</td></tr>
<tr><td>T33659</td><td>GGAACTCTCGGGATCGCTGGTGG</td><td></td><td>1000</td><td></td><td></td><td></td><td></td><td></td><td></td><td></td><td></td><td></td></tr>
<tr><td>Oligo fwd</td><td>CACCGGAACTCTCGGGATCGCTGG</td><td></td><td></td><td></td><td></td><td></td><td></td><td></td><td></td><td></td><td></td><td></td></tr>
<tr><td>Oligo rev</td><td>AAACCCAGCGATCCCGAGAGTTCC</td><td></td><td></td><td></td><td></td><td></td><td></td><td></td><td></td><td></td><td></td><td></td></tr>
<tr><td>Chromosome</td><td>start</td><td></td><td>end</td><td>strand</td><td>MM</td><td>target_seq</td><td>PAM</td><td>alignment</td><td>distance</td><td>position</td><td>gene name</td><td>gene id</td></tr>
<tr><td>GU305914.1</td><td></td><td>10429</td><td>10451</td><td>-</td><td>0</td><td>GGAACTCTCGGGATCGCTGG</td><td>TGG</td><td>||||||||[||||||||||||]PAM</td><td>NA</td><td>Intergenic</td><td>NA</td><td>NA</td></tr>
<tr><td>T33660</td><td>AATGACCGCGCACACGCGCGAGG</td><td></td><td>1000</td><td></td><td></td><td></td><td></td><td></td><td></td><td></td><td></td><td></td></tr>
<tr><td>Oligo fwd</td><td>CACCAATGACCGCGCACACGCGCG</td><td></td><td></td><td></td><td></td><td></td><td></td><td></td><td></td><td></td><td></td><td></td></tr>
<tr><td>Oligo rev</td><td>AAACCGCGCGTGTGCGCGGTCATT</td><td></td><td></td><td></td><td></td><td></td><td></td><td></td><td></td><td></td><td></td><td></td></tr>
<tr><td>Chromosome</td><td>start</td><td></td><td>end</td><td>strand</td><td>MM</td><td>target_seq</td><td>PAM</td><td>alignment</td><td>distance</td><td>position</td><td>gene name</td><td>gene id</td></tr>
<tr><td>GU305914.1</td><td></td><td>10403</td><td>10425</td><td>-</td><td>0</td><td>AATGACCGCGCACACGCGCG</td><td>AGG</td><td>||||||||[||||||||||||]PAM</td><td>NA</td><td>Intergenic</td><td>NA</td><td>NA</td></tr>
<tr><td>T33661</td><td>CACGCGCGAGGTACGAGTCGAGG</td><td></td><td>1000</td><td></td><td></td><td></td><td></td><td></td><td></td><td></td><td></td><td></td></tr>
<tr><td>Oligo fwd</td><td>CACCCACGCGCGAGGTACGAGTCG</td><td></td><td></td><td></td><td></td><td></td><td></td><td></td><td></td><td></td><td></td><td></td></tr>
<tr><td>Oligo rev</td><td>AAACCGACTCGTACCTCGCGCGTG</td><td></td><td></td><td></td><td></td><td></td><td></td><td></td><td></td><td></td><td></td><td></td></tr>
<tr><td>Chromosome</td><td>start</td><td></td><td>end</td><td>strand</td><td>MM</td><td>target_seq</td><td>PAM</td><td>alignment</td><td>distance</td><td>position</td><td>gene name</td><td>gene id</td></tr>
<tr><td>GU305914.1</td><td></td><td>10391</td><td>10413</td><td>-</td><td>0</td><td>CACGCGCGAGGTACGAGTCG</td><td>AGG</td><td>||||||||[||||||||||||]PAM</td><td>NA</td><td>Intergenic</td><td>NA</td><td>NA</td></tr>
<tr><td>T33662</td><td>CGAGGAGAGCCCATTTTCTCAGG</td><td></td><td>1000</td><td></td><td></td><td></td><td></td><td></td><td></td><td></td><td></td><td></td></tr>
<tr><td>Oligo fwd</td><td>CACCCGAGGAGAGCCCATTTTCTC</td><td></td><td></td><td></td><td></td><td></td><td></td><td></td><td></td><td></td><td></td><td></td></tr>
<tr><td>Oligo rev</td><td>AAACGAGAAAATGGGCTCTCCTCG</td><td></td><td></td><td></td><td></td><td></td><td></td><td></td><td></td><td></td><td></td><td></td></tr>
<tr><td>Chromosome</td><td>start</td><td></td><td>end</td><td>strand</td><td>MM</td><td>target_seq</td><td>PAM</td><td>alignment</td><td>distance</td><td>position</td><td>gene name</td><td>gene id</td></tr>
<tr><td>GU305914.1</td><td></td><td>10373</td><td>10395</td><td>-</td><td>0</td><td>CGAGGAGAGCCCATTTTCTC</td><td>AGG</td><td>||||||||[||||||||||||]PAM</td><td>NA</td><td>Intergenic</td><td>NA</td><td>NA</td></tr>
<tr><td>T33663</td><td>GCTACTAAGATGACCATTAACGG</td><td></td><td>1000</td><td></td><td></td><td></td><td></td><td></td><td></td><td></td><td></td><td></td></tr>
<tr><td>Oligo fwd</td><td>CACCGCTACTAAGATGACCATTAA</td><td></td><td></td><td></td><td></td><td></td><td></td><td></td><td></td><td></td><td></td><td></td></tr>
<tr><td>Oligo rev</td><td>AAACTTAATGGTCATCTTAGTAGC</td><td></td><td></td><td></td><td></td><td></td><td></td><td></td><td></td><td></td><td></td><td></td></tr>
<tr><td>Chromosome</td><td>start</td><td></td><td>end</td><td>strand</td><td>MM</td><td>target_seq</td><td>PAM</td><td>alignment</td><td>distance</td><td>position</td><td>gene name</td><td>gene id</td></tr>
<tr><td>GU305914.1</td><td></td><td>10311</td><td>10333</td><td>-</td><td>0</td><td>GCTACTAAGATGACCATTAA</td><td>CGG</td><td>||||||||[||||||||||||]PAM</td><td>NA</td><td>Intergenic</td><td>NA</td><td>NA</td></tr>
<tr><td>T33664</td><td>GATCGCGACCGTCGTATTCGAGG</td><td></td><td>1000</td><td></td><td></td><td></td><td></td><td></td><td></td><td></td><td></td><td></td></tr>
<tr><td>Oligo fwd</td><td>CACCGATCGCGACCGTCGTATTCG</td><td></td><td></td><td></td><td></td><td></td><td></td><td></td><td></td><td></td><td></td><td></td></tr>
<tr><td>Oligo rev</td><td>AAACCGAATACGACGGTCGCGATC</td><td></td><td></td><td></td><td></td><td></td><td></td><td></td><td></td><td></td><td></td><td></td></tr>
<tr><td>Chromosome</td><td>start</td><td></td><td>end</td><td>strand</td><td>MM</td><td>target_seq</td><td>PAM</td><td>alignment</td><td>distance</td><td>position</td><td>gene name</td><td>gene id</td></tr>
<tr><td>GU305914.1</td><td></td><td>10282</td><td>10304</td><td>-</td><td>0</td><td>GATCGCGACCGTCGTATTCG</td><td>AGG</td><td>||||||||[||||||||||||]PAM</td><td>NA</td><td>Intergenic</td><td>NA</td><td>NA</td></tr>
<tr><td>T33665</td><td>CGCGACGATAACAGCGTTATCGG</td><td></td><td>1000</td><td></td><td></td><td></td><td></td><td></td><td></td><td></td><td></td><td></td></tr>
<tr><td>Oligo fwd</td><td>CACCCGCGACGATAACAGCGTTAT</td><td></td><td></td><td></td><td></td><td></td><td></td><td></td><td></td><td></td><td></td><td></td></tr>
<tr><td>Oligo rev</td><td>AAACATAACGCTGTTATCGTCGCG</td><td></td><td></td><td></td><td></td><td></td><td></td><td></td><td></td><td></td><td></td><td></td></tr>
<tr><td>Chromosome</td><td>start</td><td></td><td>end</td><td>strand</td><td>MM</td><td>target_seq</td><td>PAM</td><td>alignment</td><td>distance</td><td>position</td><td>gene name</td><td>gene id</td></tr>
<tr><td>GU305914.1</td><td></td><td>10243</td><td>10265</td><td>-</td><td>0</td><td>CGCGACGATAACAGCGTTAT</td><td>CGG</td><td>||||||||[||||||||||||]PAM</td><td>NA</td><td>Intergenic</td><td>NA</td><td>NA</td></tr>
<tr><td>T33666</td><td>AGCGTTATCGGTGCTTGACGCGG</td><td></td><td>1000</td><td></td><td></td><td></td><td></td><td></td><td></td><td></td><td></td><td></td></tr>
<tr><td>Oligo fwd</td><td>CACCAGCGTTATCGGTGCTTGACG</td><td></td><td></td><td></td><td></td><td></td><td></td><td></td><td></td><td></td><td></td><td></td></tr>
<tr><td>Oligo rev</td><td>AAACCGTCAAGCACCGATAACGCT</td><td></td><td></td><td></td><td></td><td></td><td></td><td></td><td></td><td></td><td></td><td></td></tr>
<tr><td>Chromosome</td><td>start</td><td></td><td>end</td><td>strand</td><td>MM</td><td>target_seq</td><td>PAM</td><td>alignment</td><td>distance</td><td>position</td><td>gene name</td><td>gene id</td></tr>
<tr><td>GU305914.1</td><td></td><td>10231</td><td>10253</td><td>-</td><td>0</td><td>AGCGTTATCGGTGCTTGACG</td><td>CGG</td><td>||||||||[||||||||||||]PAM</td><td>NA</td><td>Intergenic</td><td>NA</td><td>NA</td></tr>
<tr><td>T33667</td><td>GCTTGACGCGGTGTTCGAAACGG</td><td></td><td>1000</td><td></td><td></td><td></td><td></td><td></td><td></td><td></td><td></td><td></td></tr>
<tr><td>Oligo fwd</td><td>CACCGCTTGACGCGGTGTTCGAAA</td><td></td><td></td><td></td><td></td><td></td><td></td><td></td><td></td><td></td><td></td><td></td></tr>
<tr><td>Oligo rev</td><td>AAACTTTCGAACACCGCGTCAAGC</td><td></td><td></td><td></td><td></td><td></td><td></td><td></td><td></td><td></td><td></td><td></td></tr>
<tr><td>Chromosome</td><td>start</td><td></td><td>end</td><td>strand</td><td>MM</td><td>target_seq</td><td>PAM</td><td>alignment</td><td>distance</td><td>position</td><td>gene name</td><td>gene id</td></tr>
<tr><td>GU305914.1</td><td></td><td>10219</td><td>10241</td><td>-</td><td>0</td><td>GCTTGACGCGGTGTTCGAAA</td><td>CGG</td><td>||||||||[||||||||||||]PAM</td><td>NA</td><td>Intergenic</td><td>NA</td><td>NA</td></tr>
<tr><td>T33668</td><td>TGACGCGGTGTTCGAAACGGTGG</td><td></td><td>1000</td><td></td><td></td><td></td><td></td><td></td><td></td><td></td><td></td><td></td></tr>
<tr><td>Oligo fwd</td><td>CACCTGACGCGGTGTTCGAAACGG</td><td></td><td></td><td></td><td></td><td></td><td></td><td></td><td></td><td></td><td></td><td></td></tr>
<tr><td>Oligo rev</td><td>AAACCCGTTTCGAACACCGCGTCA</td><td></td><td></td><td></td><td></td><td></td><td></td><td></td><td></td><td></td><td></td><td></td></tr>
<tr><td>Chromosome</td><td>start</td><td></td><td>end</td><td>strand</td><td>MM</td><td>target_seq</td><td>PAM</td><td>alignment</td><td>distance</td><td>position</td><td>gene name</td><td>gene id</td></tr>
<tr><td>GU305914.1</td><td></td><td>10216</td><td>10238</td><td>-</td><td>0</td><td>TGACGCGGTGTTCGAAACGG</td><td>TGG</td><td>||||||||[||||||||||||]PAM</td><td>NA</td><td>Intergenic</td><td>NA</td><td>NA</td></tr>
<tr><td>T33669</td><td>GGTGTTCGAAACGGTGGTATCGG</td><td></td><td>1000</td><td></td><td></td><td></td><td></td><td></td><td></td><td></td><td></td><td></td></tr>
<tr><td>Oligo fwd</td><td>CACCGGTGTTCGAAACGGTGGTAT</td><td></td><td></td><td></td><td></td><td></td><td></td><td></td><td></td><td></td><td></td><td></td></tr>
<tr><td>Oligo rev</td><td>AAACATACCACCGTTTCGAACACC</td><td></td><td></td><td></td><td></td><td></td><td></td><td></td><td></td><td></td><td></td><td></td></tr>
<tr><td>Chromosome</td><td>start</td><td></td><td>end</td><td>strand</td><td>MM</td><td>target_seq</td><td>PAM</td><td>alignment</td><td>distance</td><td>position</td><td>gene name</td><td>gene id</td></tr>
</table>

GU305914.1 10210 10232 - 0 GGTGTTCGAAACGGTGGTAT CGG ||||||||[||||||||||||]PAM NA Intergenic NA NA

T33670 GTTCGAAACGGTGGTATCGGTGG 1000
Oligo fwd CACCGTTCGAAACGGTGGTATCGG
Oligo rev AAACCCGATACCACCGTTTCGAAC
Chromosome start end strand MM target_seq PAM alignment distance position gene name gene id
GU305914.1 10207 10229 - 0 GTTCGAAACGGTGGTATCGG TGG ||||||||[||||||||||||]PAM NA Intergenic NA NA

T33671 CGGTGGTATCGGTGGCCTGTCGG 1000
Oligo fwd CACCCGGTGGTATCGGTGGCCTGT
Oligo rev AAACACAGGCCACCGATACCACCG
Chromosome start end strand MM target_seq PAM alignment distance position gene name gene id
GU305914.1 10199 10221 - 0 CGGTGGTATCGGTGGCCTGT CGG ||||||||[||||||||||||]PAM NA Intergenic NA NA

T33672 GGTGGTATCGGTGGCCTGTCGGG 1000
Oligo fwd CACCGGTGGTATCGGTGGCCTGTC
Oligo rev AAACGACAGGCCACCGATACCACC
Chromosome start end strand MM target_seq PAM alignment distance position gene name gene id
GU305914.1 10198 10220 - 0 GGTGGTATCGGTGGCCTGTC GGG ||||||||[||||||||||||]PAM NA Intergenic NA NA

T33673 ATCGGTGGCCTGTCGGGTGACGG 1000
Oligo fwd CACCATCGGTGGCCTGTCGGGTGA
Oligo rev AAACTCACCCGACAGGCCACCGAT
Chromosome start end strand MM target_seq PAM alignment distance position gene name gene id
GU305914.1 10192 10214 - 0 ATCGGTGGCCTGTCGGGTGA CGG ||||||||[||||||||||||]PAM NA Intergenic NA NA

T33674 TCGGTGGCCTGTCGGGTGACGGG 1000
Oligo fwd CACCTCGGTGGCCTGTCGGGTGAC
Oligo rev AAACGTCACCCGACAGGCCACCGA
Chromosome start end strand MM target_seq PAM alignment distance position gene name gene id
GU305914.1 10191 10213 - 0 TCGGTGGCCTGTCGGGTGAC GGG ||||||||[||||||||||||]PAM NA Intergenic NA NA

T33675 TGGCCTGTCGGGTGACGGGCTGG 1000
Oligo fwd CACCTGGCCTGTCGGGTGACGGGC
Oligo rev AAACGCCCGTCACCCGACAGGCCA
Chromosome start end strand MM target_seq PAM alignment distance position gene name gene id
GU305914.1 10187 10209 - 0 TGGCCTGTCGGGTGACGGGC TGG ||||||||[||||||||||||]PAM NA Intergenic NA NA

T33676 GGCCTGTCGGGTGACGGGCTGGG 1000
Oligo fwd CACCGGCCTGTCGGGTGACGGGCT
Oligo rev AAACAGCCCGTCACCCGACAGGCC
Chromosome start end strand MM target_seq PAM alignment distance position gene name gene id
GU305914.1 10186 10208 - 0 GGCCTGTCGGGTGACGGGCT GGG ||||||||[||||||||||||]PAM NA Intergenic NA NA
GU305914.1 9212 9234 - 4 GACCTGCGGGGTGACGGACT GGG |-||||--[|||||||||-||]PAM NA Intergenic NA NA

T33677 CGGGCTGGGATCGTGATGATAGG 1000
Oligo fwd CACCCGGGCTGGGATCGTGATGAT
Oligo rev AAACATCATCACGATCCCAGCCCG
Chromosome start end strand MM target_seq PAM alignment distance position gene name gene id
GU305914.1 10172 10194 - 0 CGGGCTGGGATCGTGATGAT AGG ||||||||[||||||||||||]PAM NA Intergenic NA NA

T33678 GGGCTGGGATCGTGATGATAGGG 1000
Oligo fwd CACCGGGCTGGGATCGTGATGATA
Oligo rev AAACTATCATCACGATCCCAGCCC
Chromosome start end strand MM target_seq PAM alignment distance position gene name gene id
GU305914.1 10171 10193 - 0 GGGCTGGGATCGTGATGATA GGG ||||||||[||||||||||||]PAM NA Intergenic NA NA

T33679 TCGTTTAGTTCGTGAGATCGTGG 1000
Oligo fwd CACCTCGTTTAGTTCGTGAGATCG
Oligo rev AAACCGATCTCACGAACTAAACGA
Chromosome start end strand MM target_seq PAM alignment distance position gene name gene id
GU305914.1 10129 10151 - 0 TCGTTTAGTTCGTGAGATCG TGG ||||||||[||||||||||||]PAM NA Intergenic NA NA

T33680 TTAGTTCGTGAGATCGTGGTCGG 1000
Oligo fwd CACCTTAGTTCGTGAGATCGTGGT
Oligo rev AAACACCACGATCTCACGAACTAA
Chromosome start end strand MM target_seq PAM alignment distance position gene name gene id
GU305914.1 10125 10147 - 0 TTAGTTCGTGAGATCGTGGT CGG ||||||||[||||||||||||]PAM NA Intergenic NA NA

T33681 GTTCGTGAGATCGTGGTCGGAGG 1000
Oligo fwd CACCGTTCGTGAGATCGTGGTCGG
Oligo rev AAACCCGACCACGATCTCACGAAC
Chromosome start end strand MM target_seq PAM alignment distance position gene name gene id
GU305914.1 10122 10144 - 0 GTTCGTGAGATCGTGGTCGG AGG ||||||||[||||||||||||]PAM NA Intergenic NA NA

T33682 AGGATTTGTAGTTACATGTATGG 1000
Oligo fwd CACCAGGATTTGTAGTTACATGTA
Oligo rev AAACTACATGTAACTACAAATCCT
Chromosome start end strand MM target_seq PAM alignment distance position gene name gene id
GU305914.1 10102 10124 - 0 AGGATTTGTAGTTACATGTA TGG ||||||||[||||||||||||]PAM NA Intergenic NA NA

T33683 GTTACATGTATGGTTTTATTCGG 1000
Oligo fwd CACCGTTACATGTATGGTTTTATT
Oligo rev AAACAATAAAACCATACATGTAAC
Chromosome start end strand MM target_seq PAM alignment distance position gene name gene id
GU305914.1 10092 10114 - 0 GTTACATGTATGGTTTTATT CGG ||||||||[||||||||||||]PAM NA Intergenic NA NA

T33684 TGTACAGAAACAGTTTCCTCTGG 1000
Oligo fwd CACCTGTACAGAAACAGTTTCCTC

<table>
<tr><td>Oligo rev</td><td>AAACGAGGAAACTGTTTCTGTACA</td><td></td><td></td><td></td><td></td><td></td><td></td><td></td><td></td><td></td><td></td></tr>
<tr><td>Chromosome</td><td>start</td><td>end</td><td>strand</td><td>MM</td><td>target_seq</td><td>PAM</td><td>alignment</td><td>distance</td><td>position</td><td>gene name</td><td>gene id</td></tr>
<tr><td>GU305914.1</td><td>10041</td><td>10063</td><td>-</td><td>0</td><td>TGTACAGAAACAGTTTCCTC</td><td>TGG</td><td>||||||||[||||||||||||]PAM</td><td>NA</td><td>Intergenic</td><td>NA</td><td>NA</td></tr>
<tr><td>T33685</td><td>GTACAGAAACAGTTTCCTCTGGG</td><td>1000</td><td></td><td></td><td></td><td></td><td></td><td></td><td></td><td></td><td></td></tr>
<tr><td>Oligo fwd</td><td>CACCGTACAGAAACAGTTTCCTCT</td><td></td><td></td><td></td><td></td><td></td><td></td><td></td><td></td><td></td><td></td></tr>
<tr><td>Oligo rev</td><td>AAACAGAGGAAACTGTTTCTGTAC</td><td></td><td></td><td></td><td></td><td></td><td></td><td></td><td></td><td></td><td></td></tr>
<tr><td>Chromosome</td><td>start</td><td>end</td><td>strand</td><td>MM</td><td>target_seq</td><td>PAM</td><td>alignment</td><td>distance</td><td>position</td><td>gene name</td><td>gene id</td></tr>
<tr><td>GU305914.1</td><td>10040</td><td>10062</td><td>-</td><td>0</td><td>GTACAGAAACAGTTTCCTCT</td><td>GGG</td><td>||||||||[||||||||||||]PAM</td><td>NA</td><td>Intergenic</td><td>NA</td><td>NA</td></tr>
<tr><td>T33686</td><td>TACAGAAACAGTTTCCTCTGGGG</td><td>1000</td><td></td><td></td><td></td><td></td><td></td><td></td><td></td><td></td><td></td></tr>
<tr><td>Oligo fwd</td><td>CACCTACAGAAACAGTTTCCTCTG</td><td></td><td></td><td></td><td></td><td></td><td></td><td></td><td></td><td></td><td></td></tr>
<tr><td>Oligo rev</td><td>AAACCAGAGGAAACTGTTTCTGTA</td><td></td><td></td><td></td><td></td><td></td><td></td><td></td><td></td><td></td><td></td></tr>
<tr><td>Chromosome</td><td>start</td><td>end</td><td>strand</td><td>MM</td><td>target_seq</td><td>PAM</td><td>alignment</td><td>distance</td><td>position</td><td>gene name</td><td>gene id</td></tr>
<tr><td>GU305914.1</td><td>10039</td><td>10061</td><td>-</td><td>0</td><td>TACAGAAACAGTTTCCTCTG</td><td>GGG</td><td>||||||||[||||||||||||]PAM</td><td>NA</td><td>Intergenic</td><td>NA</td><td>NA</td></tr>
<tr><td>T33687</td><td>CTGGGGACGAAACGAGATACAGG</td><td>1000</td><td></td><td></td><td></td><td></td><td></td><td></td><td></td><td></td><td></td></tr>
<tr><td>Oligo fwd</td><td>CACCCTGGGGACGAAACGAGATAC</td><td></td><td></td><td></td><td></td><td></td><td></td><td></td><td></td><td></td><td></td></tr>
<tr><td>Oligo rev</td><td>AAACGTATCTCGTTTCGTCCCCAG</td><td></td><td></td><td></td><td></td><td></td><td></td><td></td><td></td><td></td><td></td></tr>
<tr><td>Chromosome</td><td>start</td><td>end</td><td>strand</td><td>MM</td><td>target_seq</td><td>PAM</td><td>alignment</td><td>distance</td><td>position</td><td>gene name</td><td>gene id</td></tr>
<tr><td>GU305914.1</td><td>10022</td><td>10044</td><td>-</td><td>0</td><td>CTGGGGACGAAACGAGATAC</td><td>AGG</td><td>||||||||[||||||||||||]PAM</td><td>NA</td><td>Intergenic</td><td>NA</td><td>NA</td></tr>
<tr><td>T33688</td><td>AGATACAGGTAACCGTCATGCGG</td><td>1000</td><td></td><td></td><td></td><td></td><td></td><td></td><td></td><td></td><td></td></tr>
<tr><td>Oligo fwd</td><td>CACCAGATACAGGTAACCGTCATG</td><td></td><td></td><td></td><td></td><td></td><td></td><td></td><td></td><td></td><td></td></tr>
<tr><td>Oligo rev</td><td>AAACCATGACGGTTACCTGTATCT</td><td></td><td></td><td></td><td></td><td></td><td></td><td></td><td></td><td></td><td></td></tr>
<tr><td>Chromosome</td><td>start</td><td>end</td><td>strand</td><td>MM</td><td>target_seq</td><td>PAM</td><td>alignment</td><td>distance</td><td>position</td><td>gene name</td><td>gene id</td></tr>
<tr><td>GU305914.1</td><td>10008</td><td>10030</td><td>-</td><td>0</td><td>AGATACAGGTAACCGTCATG</td><td>CGG</td><td>||||||||[||||||||||||]PAM</td><td>NA</td><td>Intergenic</td><td>NA</td><td>NA</td></tr>
<tr><td>T33689</td><td>GGATTCACCGTGAACCCCTTTGG</td><td>1000</td><td></td><td></td><td></td><td></td><td></td><td></td><td></td><td></td><td></td></tr>
<tr><td>Oligo fwd</td><td>CACCGGATTCACCGTGAACCCCTT</td><td></td><td></td><td></td><td></td><td></td><td></td><td></td><td></td><td></td><td></td></tr>
<tr><td>Oligo rev</td><td>AAACAAGGGGTTCACGGTGAATCC</td><td></td><td></td><td></td><td></td><td></td><td></td><td></td><td></td><td></td><td></td></tr>
<tr><td>Chromosome</td><td>start</td><td>end</td><td>strand</td><td>MM</td><td>target_seq</td><td>PAM</td><td>alignment</td><td>distance</td><td>position</td><td>gene name</td><td>gene id</td></tr>
<tr><td>GU305914.1</td><td>9987</td><td>10009</td><td>-</td><td>0</td><td>GGATTCACCGTGAACCCCTT</td><td>TGG</td><td>||||||||[||||||||||||]PAM</td><td>NA</td><td>Intergenic</td><td>NA</td><td>NA</td></tr>
<tr><td>T33690</td><td>TTCACCGTGAACCCCTTTGGAGG</td><td>1000</td><td></td><td></td><td></td><td></td><td></td><td></td><td></td><td></td><td></td></tr>
<tr><td>Oligo fwd</td><td>CACCTTCACCGTGAACCCCTTTGG</td><td></td><td></td><td></td><td></td><td></td><td></td><td></td><td></td><td></td><td></td></tr>
<tr><td>Oligo rev</td><td>AAACCCAAAGGGGTTCACGGTGAA</td><td></td><td></td><td></td><td></td><td></td><td></td><td></td><td></td><td></td><td></td></tr>
<tr><td>Chromosome</td><td>start</td><td>end</td><td>strand</td><td>MM</td><td>target_seq</td><td>PAM</td><td>alignment</td><td>distance</td><td>position</td><td>gene name</td><td>gene id</td></tr>
<tr><td>GU305914.1</td><td>9984</td><td>10006</td><td>-</td><td>0</td><td>TTCACCGTGAACCCCTTTGG</td><td>AGG</td><td>||||||||[||||||||||||]PAM</td><td>NA</td><td>Intergenic</td><td>NA</td><td>NA</td></tr>
<tr><td>T33691</td><td>ACCGTGAACCCCTTTGGAGGTGG</td><td>1000</td><td></td><td></td><td></td><td></td><td></td><td></td><td></td><td></td><td></td></tr>
<tr><td>Oligo fwd</td><td>CACCACCGTGAACCCCTTTGGAGG</td><td></td><td></td><td></td><td></td><td></td><td></td><td></td><td></td><td></td><td></td></tr>
<tr><td>Oligo rev</td><td>AAACCCTCCAAAGGGGTTCACGGT</td><td></td><td></td><td></td><td></td><td></td><td></td><td></td><td></td><td></td><td></td></tr>
<tr><td>Chromosome</td><td>start</td><td>end</td><td>strand</td><td>MM</td><td>target_seq</td><td>PAM</td><td>alignment</td><td>distance</td><td>position</td><td>gene name</td><td>gene id</td></tr>
<tr><td>GU305914.1</td><td>9981</td><td>10003</td><td>-</td><td>0</td><td>ACCGTGAACCCCTTTGGAGG</td><td>TGG</td><td>||||||||[||||||||||||]PAM</td><td>NA</td><td>Intergenic</td><td>NA</td><td>NA</td></tr>
<tr><td>T33692</td><td>ACTAGAATTTTATCAAACTTTGG</td><td>1000</td><td></td><td></td><td></td><td></td><td></td><td></td><td></td><td></td><td></td></tr>
<tr><td>Oligo fwd</td><td>CACCACTAGAATTTTATCAAACTT</td><td></td><td></td><td></td><td></td><td></td><td></td><td></td><td></td><td></td><td></td></tr>
<tr><td>Oligo rev</td><td>AAACAAGTTTGATAAAATTCTAGT</td><td></td><td></td><td></td><td></td><td></td><td></td><td></td><td></td><td></td><td></td></tr>
<tr><td>Chromosome</td><td>start</td><td>end</td><td>strand</td><td>MM</td><td>target_seq</td><td>PAM</td><td>alignment</td><td>distance</td><td>position</td><td>gene name</td><td>gene id</td></tr>
<tr><td>GU305914.1</td><td>9939</td><td>9961</td><td>-</td><td>0</td><td>ACTAGAATTTTATCAAACTT</td><td>TGG</td><td>||||||||[||||||||||||]PAM</td><td>NA</td><td>Intergenic</td><td>NA</td><td>NA</td></tr>
<tr><td>T33693</td><td>GTCTGTCGAATTGTATAAAGCGG</td><td>1000</td><td></td><td></td><td></td><td></td><td></td><td></td><td></td><td></td><td></td></tr>
<tr><td>Oligo fwd</td><td>CACCGTCTGTCGAATTGTATAAAG</td><td></td><td></td><td></td><td></td><td></td><td></td><td></td><td></td><td></td><td></td></tr>
<tr><td>Oligo rev</td><td>AAACCTTTATACAATTCGACAGAC</td><td></td><td></td><td></td><td></td><td></td><td></td><td></td><td></td><td></td><td></td></tr>
<tr><td>Chromosome</td><td>start</td><td>end</td><td>strand</td><td>MM</td><td>target_seq</td><td>PAM</td><td>alignment</td><td>distance</td><td>position</td><td>gene name</td><td>gene id</td></tr>
<tr><td>GU305914.1</td><td>9907</td><td>9929</td><td>-</td><td>0</td><td>GTCTGTCGAATTGTATAAAG</td><td>CGG</td><td>||||||||[||||||||||||]PAM</td><td>NA</td><td>Intergenic</td><td>NA</td><td>NA</td></tr>
<tr><td>T33694</td><td>TATAAAGCGGTCGCATTCATTGG</td><td>1000</td><td></td><td></td><td></td><td></td><td></td><td></td><td></td><td></td><td></td></tr>
<tr><td>Oligo fwd</td><td>CACCTATAAAGCGGTCGCATTCAT</td><td></td><td></td><td></td><td></td><td></td><td></td><td></td><td></td><td></td><td></td></tr>
<tr><td>Oligo rev</td><td>AAACATGAATGCGACCGCTTTATA</td><td></td><td></td><td></td><td></td><td></td><td></td><td></td><td></td><td></td><td></td></tr>
<tr><td>Chromosome</td><td>start</td><td>end</td><td>strand</td><td>MM</td><td>target_seq</td><td>PAM</td><td>alignment</td><td>distance</td><td>position</td><td>gene name</td><td>gene id</td></tr>
<tr><td>GU305914.1</td><td>9894</td><td>9916</td><td>-</td><td>0</td><td>TATAAAGCGGTCGCATTCAT</td><td>TGG</td><td>||||||||[||||||||||||]PAM</td><td>NA</td><td>Intergenic</td><td>NA</td><td>NA</td></tr>
<tr><td>GU305914.1</td><td>49326</td><td>49348</td><td>-</td><td>4</td><td>TATAAGGAGGTCACATTCTT</td><td>TGG</td><td>|||||-|-[||||-|||||-|]PAM</td><td>NA</td><td>Intergenic</td><td>NA</td><td>NA</td></tr>
<tr><td>T33695</td><td>ATAAAGCGGTCGCATTCATTGGG</td><td>1000</td><td></td><td></td><td></td><td></td><td></td><td></td><td></td><td></td><td></td></tr>
<tr><td>Oligo fwd</td><td>CACCATAAAGCGGTCGCATTCATT</td><td></td><td></td><td></td><td></td><td></td><td></td><td></td><td></td><td></td><td></td></tr>
<tr><td>Oligo rev</td><td>AAACAATGAATGCGACCGCTTTAT</td><td></td><td></td><td></td><td></td><td></td><td></td><td></td><td></td><td></td><td></td></tr>
<tr><td>Chromosome</td><td>start</td><td>end</td><td>strand</td><td>MM</td><td>target_seq</td><td>PAM</td><td>alignment</td><td>distance</td><td>position</td><td>gene name</td><td>gene id</td></tr>
<tr><td>GU305914.1</td><td>9893</td><td>9915</td><td>-</td><td>0</td><td>ATAAAGCGGTCGCATTCATT</td><td>GGG</td><td>||||||||[||||||||||||]PAM</td><td>NA</td><td>Intergenic</td><td>NA</td><td>NA</td></tr>
<tr><td>T33696</td><td>ATTCATTGGGATCCGATTAACGG</td><td>1000</td><td></td><td></td><td></td><td></td><td></td><td></td><td></td><td></td><td></td></tr>
<tr><td>Oligo fwd</td><td>CACCATTCATTGGGATCCGATTAA</td><td></td><td></td><td></td><td></td><td></td><td></td><td></td><td></td><td></td><td></td></tr>
<tr><td>Oligo rev</td><td>AAACTTAATCGGATCCCAATGAAT</td><td></td><td></td><td></td><td></td><td></td><td></td><td></td><td></td><td></td><td></td></tr>
<tr><td>Chromosome</td><td>start</td><td>end</td><td>strand</td><td>MM</td><td>target_seq</td><td>PAM</td><td>alignment</td><td>distance</td><td>position</td><td>gene name</td><td>gene id</td></tr>
<tr><td>GU305914.1</td><td>9880</td><td>9902</td><td>-</td><td>0</td><td>ATTCATTGGGATCCGATTAA</td><td>CGG</td><td>||||||||[||||||||||||]PAM</td><td>NA</td><td>Intergenic</td><td>NA</td><td>NA</td></tr>
<tr><td>T33697</td><td>CTGCCATGTTGCTTCAATACAGG</td><td>1000</td><td></td><td></td><td></td><td></td><td></td><td></td><td></td><td></td><td></td></tr>
<tr><td>Oligo fwd</td><td>CACCCTGCCATGTTGCTTCAATAC</td><td></td><td></td><td></td><td></td><td></td><td></td><td></td><td></td><td></td><td></td></tr>
<tr><td>Oligo rev</td><td>AAACGTATTGAAGCAACATGGCAG</td><td></td><td></td><td></td><td></td><td></td><td></td><td></td><td></td><td></td><td></td></tr>
<tr><td>Chromosome</td><td>start</td><td>end</td><td>strand</td><td>MM</td><td>target_seq</td><td>PAM</td><td>alignment</td><td>distance</td><td>position</td><td>gene name</td><td>gene id</td></tr>
<tr><td>GU305914.1</td><td>9847</td><td>9869</td><td>-</td><td>0</td><td>CTGCCATGTTGCTTCAATAC</td><td>AGG</td><td>||||||||[||||||||||||]PAM</td><td>NA</td><td>Intergenic</td><td>NA</td><td>NA</td></tr>
<tr><td>T33698</td><td>GTTGCTTCAATACAGGCCTCAGG</td><td>1000</td><td></td><td></td><td></td><td></td><td></td><td></td><td></td><td></td><td></td></tr>
<tr><td>Oligo fwd</td><td>CACCGTTGCTTCAATACAGGCCTC</td><td></td><td></td><td></td><td></td><td></td><td></td><td></td><td></td><td></td><td></td></tr>
<tr><td>Oligo rev</td><td>AAACGAGGCCTGTATTGAAGCAAC</td><td></td><td></td><td></td><td></td><td></td><td></td><td></td><td></td><td></td><td></td></tr>
<tr><td>Chromosome</td><td>start</td><td>end</td><td>strand</td><td>MM</td><td>target_seq</td><td>PAM</td><td>alignment</td><td>distance</td><td>position</td><td>gene name</td><td>gene id</td></tr>
<tr><td>GU305914.1</td><td>9840</td><td>9862</td><td>-</td><td>0</td><td>GTTGCTTCAATACAGGCCTC</td><td>AGG</td><td>||||||||[||||||||||||]PAM</td><td>NA</td><td>Intergenic</td><td>NA</td><td>NA</td></tr>
</table>

<table>
<tr><td>T33699</td><td>ATATACAGATGCGCCTGTACAGG</td><td></td><td>1000</td><td></td><td></td><td></td><td></td><td></td><td></td><td></td><td></td><td></td></tr>
<tr><td>Oligo fwd</td><td>CACCATATACAGATGCGCCTGTAC</td><td></td><td></td><td></td><td></td><td></td><td></td><td></td><td></td><td></td><td></td><td></td></tr>
<tr><td>Oligo rev</td><td>AAACGTACAGGCGCATCTGTATAT</td><td></td><td></td><td></td><td></td><td></td><td></td><td></td><td></td><td></td><td></td><td></td></tr>
<tr><td>Chromosome</td><td>start</td><td></td><td>end</td><td>strand</td><td>MM</td><td>target_seq</td><td>PAM</td><td>alignment</td><td>distance</td><td>position</td><td>gene name</td><td>gene id</td></tr>
<tr><td>GU305914.1</td><td></td><td>9760</td><td>9782</td><td>-</td><td>0</td><td>ATATACAGATGCGCCTGTAC</td><td>AGG</td><td>||||||||[||||||||||||]PAM</td><td>NA</td><td>Intergenic</td><td>NA</td><td>NA</td></tr>
<tr><td>T33700</td><td>GATGCGCCTGTACAGGTACCTGG</td><td></td><td>1000</td><td></td><td></td><td></td><td></td><td></td><td></td><td></td><td></td><td></td></tr>
<tr><td>Oligo fwd</td><td>CACCGATGCGCCTGTACAGGTACC</td><td></td><td></td><td></td><td></td><td></td><td></td><td></td><td></td><td></td><td></td><td></td></tr>
<tr><td>Oligo rev</td><td>AAACGGTACCTGTACAGGCGCATC</td><td></td><td></td><td></td><td></td><td></td><td></td><td></td><td></td><td></td><td></td><td></td></tr>
<tr><td>Chromosome</td><td>start</td><td></td><td>end</td><td>strand</td><td>MM</td><td>target_seq</td><td>PAM</td><td>alignment</td><td>distance</td><td>position</td><td>gene name</td><td>gene id</td></tr>
<tr><td>GU305914.1</td><td></td><td>9753</td><td>9775</td><td>-</td><td>0</td><td>GATGCGCCTGTACAGGTACC</td><td>TGG</td><td>||||||||[||||||||||||]PAM</td><td>NA</td><td>Intergenic</td><td>NA</td><td>NA</td></tr>
<tr><td>T33701</td><td>GCGCCTGTACAGGTACCTGGCGG</td><td></td><td>1000</td><td></td><td></td><td></td><td></td><td></td><td></td><td></td><td></td><td></td></tr>
<tr><td>Oligo fwd</td><td>CACCGCGCCTGTACAGGTACCTGG</td><td></td><td></td><td></td><td></td><td></td><td></td><td></td><td></td><td></td><td></td><td></td></tr>
<tr><td>Oligo rev</td><td>AAACCCAGGTACCTGTACAGGCGC</td><td></td><td></td><td></td><td></td><td></td><td></td><td></td><td></td><td></td><td></td><td></td></tr>
<tr><td>Chromosome</td><td>start</td><td></td><td>end</td><td>strand</td><td>MM</td><td>target_seq</td><td>PAM</td><td>alignment</td><td>distance</td><td>position</td><td>gene name</td><td>gene id</td></tr>
<tr><td>GU305914.1</td><td></td><td>9750</td><td>9772</td><td>-</td><td>0</td><td>GCGCCTGTACAGGTACCTGG</td><td>CGG</td><td>||||||||[||||||||||||]PAM</td><td>NA</td><td>Intergenic</td><td>NA</td><td>NA</td></tr>
<tr><td>T33702</td><td>ACAGGTACCTGGCGGTGTCCTGG</td><td></td><td>1000</td><td></td><td></td><td></td><td></td><td></td><td></td><td></td><td></td><td></td></tr>
<tr><td>Oligo fwd</td><td>CACCACAGGTACCTGGCGGTGTCC</td><td></td><td></td><td></td><td></td><td></td><td></td><td></td><td></td><td></td><td></td><td></td></tr>
<tr><td>Oligo rev</td><td>AAACGGACACCGCCAGGTACCTGT</td><td></td><td></td><td></td><td></td><td></td><td></td><td></td><td></td><td></td><td></td><td></td></tr>
<tr><td>Chromosome</td><td>start</td><td></td><td>end</td><td>strand</td><td>MM</td><td>target_seq</td><td>PAM</td><td>alignment</td><td>distance</td><td>position</td><td>gene name</td><td>gene id</td></tr>
<tr><td>GU305914.1</td><td></td><td>9742</td><td>9764</td><td>-</td><td>0</td><td>ACAGGTACCTGGCGGTGTCC</td><td>TGG</td><td>||||||||[||||||||||||]PAM</td><td>NA</td><td>Intergenic</td><td>NA</td><td>NA</td></tr>
<tr><td>T33703</td><td>GGTACCTGGCGGTGTCCTGGAGG</td><td></td><td>1000</td><td></td><td></td><td></td><td></td><td></td><td></td><td></td><td></td><td></td></tr>
<tr><td>Oligo fwd</td><td>CACCGGTACCTGGCGGTGTCCTGG</td><td></td><td></td><td></td><td></td><td></td><td></td><td></td><td></td><td></td><td></td><td></td></tr>
<tr><td>Oligo rev</td><td>AAACCCAGGACACCGCCAGGTACC</td><td></td><td></td><td></td><td></td><td></td><td></td><td></td><td></td><td></td><td></td><td></td></tr>
<tr><td>Chromosome</td><td>start</td><td></td><td>end</td><td>strand</td><td>MM</td><td>target_seq</td><td>PAM</td><td>alignment</td><td>distance</td><td>position</td><td>gene name</td><td>gene id</td></tr>
<tr><td>GU305914.1</td><td></td><td>9739</td><td>9761</td><td>-</td><td>0</td><td>GGTACCTGGCGGTGTCCTGG</td><td>AGG</td><td>||||||||[||||||||||||]PAM</td><td>NA</td><td>Intergenic</td><td>NA</td><td>NA</td></tr>
<tr><td>T33704</td><td>GAGTAAGATTTCGAGATAACCGG</td><td></td><td>1000</td><td></td><td></td><td></td><td></td><td></td><td></td><td></td><td></td><td></td></tr>
<tr><td>Oligo fwd</td><td>CACCGAGTAAGATTTCGAGATAAC</td><td></td><td></td><td></td><td></td><td></td><td></td><td></td><td></td><td></td><td></td><td></td></tr>
<tr><td>Oligo rev</td><td>AAACGTTATCTCGAAATCTTACTC</td><td></td><td></td><td></td><td></td><td></td><td></td><td></td><td></td><td></td><td></td><td></td></tr>
<tr><td>Chromosome</td><td>start</td><td></td><td>end</td><td>strand</td><td>MM</td><td>target_seq</td><td>PAM</td><td>alignment</td><td>distance</td><td>position</td><td>gene name</td><td>gene id</td></tr>
<tr><td>GU305914.1</td><td></td><td>9717</td><td>9739</td><td>-</td><td>0</td><td>GAGTAAGATTTCGAGATAAC</td><td>CGG</td><td>||||||||[||||||||||||]PAM</td><td>NA</td><td>Intergenic</td><td>NA</td><td>NA</td></tr>
<tr><td>T33705</td><td>GCGACGCGAAGATCACTAAGTGG</td><td></td><td>1000</td><td></td><td></td><td></td><td></td><td></td><td></td><td></td><td></td><td></td></tr>
<tr><td>Oligo fwd</td><td>CACCGCGACGCGAAGATCACTAAG</td><td></td><td></td><td></td><td></td><td></td><td></td><td></td><td></td><td></td><td></td><td></td></tr>
<tr><td>Oligo rev</td><td>AAACCTTAGTGATCTTCGCGTCGC</td><td></td><td></td><td></td><td></td><td></td><td></td><td></td><td></td><td></td><td></td><td></td></tr>
<tr><td>Chromosome</td><td>start</td><td></td><td>end</td><td>strand</td><td>MM</td><td>target_seq</td><td>PAM</td><td>alignment</td><td>distance</td><td>position</td><td>gene name</td><td>gene id</td></tr>
<tr><td>GU305914.1</td><td></td><td>9683</td><td>9705</td><td>-</td><td>0</td><td>GCGACGCGAAGATCACTAAG</td><td>TGG</td><td>||||||||[||||||||||||]PAM</td><td>NA</td><td>Intergenic</td><td>NA</td><td>NA</td></tr>
<tr><td>T33706</td><td>CGACGCGAAGATCACTAAGTGGG</td><td></td><td>1000</td><td></td><td></td><td></td><td></td><td></td><td></td><td></td><td></td><td></td></tr>
<tr><td>Oligo fwd</td><td>CACCCGACGCGAAGATCACTAAGT</td><td></td><td></td><td></td><td></td><td></td><td></td><td></td><td></td><td></td><td></td><td></td></tr>
<tr><td>Oligo rev</td><td>AAACACTTAGTGATCTTCGCGTCG</td><td></td><td></td><td></td><td></td><td></td><td></td><td></td><td></td><td></td><td></td><td></td></tr>
<tr><td>Chromosome</td><td>start</td><td></td><td>end</td><td>strand</td><td>MM</td><td>target_seq</td><td>PAM</td><td>alignment</td><td>distance</td><td>position</td><td>gene name</td><td>gene id</td></tr>
<tr><td>GU305914.1</td><td></td><td>9682</td><td>9704</td><td>-</td><td>0</td><td>CGACGCGAAGATCACTAAGT</td><td>GGG</td><td>||||||||[||||||||||||]PAM</td><td>NA</td><td>Intergenic</td><td>NA</td><td>NA</td></tr>
<tr><td>T33707</td><td>GACGCGAAGATCACTAAGTGGGG</td><td></td><td>1000</td><td></td><td></td><td></td><td></td><td></td><td></td><td></td><td></td><td></td></tr>
<tr><td>Oligo fwd</td><td>CACCGACGCGAAGATCACTAAGTG</td><td></td><td></td><td></td><td></td><td></td><td></td><td></td><td></td><td></td><td></td><td></td></tr>
<tr><td>Oligo rev</td><td>AAACCACTTAGTGATCTTCGCGTC</td><td></td><td></td><td></td><td></td><td></td><td></td><td></td><td></td><td></td><td></td><td></td></tr>
<tr><td>Chromosome</td><td>start</td><td></td><td>end</td><td>strand</td><td>MM</td><td>target_seq</td><td>PAM</td><td>alignment</td><td>distance</td><td>position</td><td>gene name</td><td>gene id</td></tr>
<tr><td>GU305914.1</td><td></td><td>9681</td><td>9703</td><td>-</td><td>0</td><td>GACGCGAAGATCACTAAGTG</td><td>GGG</td><td>||||||||[||||||||||||]PAM</td><td>NA</td><td>Intergenic</td><td>NA</td><td>NA</td></tr>
<tr><td>T33708</td><td>CACTAAGTGGGGTGCGAAAACGG</td><td></td><td>1000</td><td></td><td></td><td></td><td></td><td></td><td></td><td></td><td></td><td></td></tr>
<tr><td>Oligo fwd</td><td>CACCCACTAAGTGGGGTGCGAAAA</td><td></td><td></td><td></td><td></td><td></td><td></td><td></td><td></td><td></td><td></td><td></td></tr>
<tr><td>Oligo rev</td><td>AAACTTTTCGCACCCCACTTAGTG</td><td></td><td></td><td></td><td></td><td></td><td></td><td></td><td></td><td></td><td></td><td></td></tr>
<tr><td>Chromosome</td><td>start</td><td></td><td>end</td><td>strand</td><td>MM</td><td>target_seq</td><td>PAM</td><td>alignment</td><td>distance</td><td>position</td><td>gene name</td><td>gene id</td></tr>
<tr><td>GU305914.1</td><td></td><td>9670</td><td>9692</td><td>-</td><td>0</td><td>CACTAAGTGGGGTGCGAAAA</td><td>CGG</td><td>||||||||[||||||||||||]PAM</td><td>NA</td><td>Intergenic</td><td>NA</td><td>NA</td></tr>
<tr><td>T33709</td><td>GAGAGTTGCAACAGACGATCTGG</td><td></td><td>1000</td><td></td><td></td><td></td><td></td><td></td><td></td><td></td><td></td><td></td></tr>
<tr><td>Oligo fwd</td><td>CACCGAGAGTTGCAACAGACGATC</td><td></td><td></td><td></td><td></td><td></td><td></td><td></td><td></td><td></td><td></td><td></td></tr>
<tr><td>Oligo rev</td><td>AAACGATCGTCTGTTGCAACTCTC</td><td></td><td></td><td></td><td></td><td></td><td></td><td></td><td></td><td></td><td></td><td></td></tr>
<tr><td>Chromosome</td><td>start</td><td></td><td>end</td><td>strand</td><td>MM</td><td>target_seq</td><td>PAM</td><td>alignment</td><td>distance</td><td>position</td><td>gene name</td><td>gene id</td></tr>
<tr><td>GU305914.1</td><td></td><td>9636</td><td>9658</td><td>-</td><td>0</td><td>GAGAGTTGCAACAGACGATC</td><td>TGG</td><td>||||||||[||||||||||||]PAM</td><td>NA</td><td>Intergenic</td><td>NA</td><td>NA</td></tr>
<tr><td>T33710</td><td>GACTTTCATGATGTATCACCCGG</td><td></td><td>1000</td><td></td><td></td><td></td><td></td><td></td><td></td><td></td><td></td><td></td></tr>
<tr><td>Oligo fwd</td><td>CACCGACTTTCATGATGTATCACC</td><td></td><td></td><td></td><td></td><td></td><td></td><td></td><td></td><td></td><td></td><td></td></tr>
<tr><td>Oligo rev</td><td>AAACGGTGATACATCATGAAAGTC</td><td></td><td></td><td></td><td></td><td></td><td></td><td></td><td></td><td></td><td></td><td></td></tr>
<tr><td>Chromosome</td><td>start</td><td></td><td>end</td><td>strand</td><td>MM</td><td>target_seq</td><td>PAM</td><td>alignment</td><td>distance</td><td>position</td><td>gene name</td><td>gene id</td></tr>
<tr><td>GU305914.1</td><td></td><td>9610</td><td>9632</td><td>-</td><td>0</td><td>GACTTTCATGATGTATCACC</td><td>CGG</td><td>||||||||[||||||||||||]PAM</td><td>NA</td><td>Intergenic</td><td>NA</td><td>NA</td></tr>
<tr><td>T33711</td><td>CACGATGATATTTACAGACTCGG</td><td></td><td>1000</td><td></td><td></td><td></td><td></td><td></td><td></td><td></td><td></td><td></td></tr>
<tr><td>Oligo fwd</td><td>CACCCACGATGATATTTACAGACT</td><td></td><td></td><td></td><td></td><td></td><td></td><td></td><td></td><td></td><td></td><td></td></tr>
<tr><td>Oligo rev</td><td>AAACAGTCTGTAAATATCATCGTG</td><td></td><td></td><td></td><td></td><td></td><td></td><td></td><td></td><td></td><td></td><td></td></tr>
<tr><td>Chromosome</td><td>start</td><td></td><td>end</td><td>strand</td><td>MM</td><td>target_seq</td><td>PAM</td><td>alignment</td><td>distance</td><td>position</td><td>gene name</td><td>gene id</td></tr>
<tr><td>GU305914.1</td><td></td><td>9558</td><td>9580</td><td>-</td><td>0</td><td>CACGATGATATTTACAGACT</td><td>CGG</td><td>||||||||[||||||||||||]PAM</td><td>NA</td><td>Intergenic</td><td>NA</td><td>NA</td></tr>
<tr><td>T33712</td><td>TGATATTTACAGACTCGGACAGG</td><td></td><td>1000</td><td></td><td></td><td></td><td></td><td></td><td></td><td></td><td></td><td></td></tr>
<tr><td>Oligo fwd</td><td>CACCTGATATTTACAGACTCGGAC</td><td></td><td></td><td></td><td></td><td></td><td></td><td></td><td></td><td></td><td></td><td></td></tr>
<tr><td>Oligo rev</td><td>AAACGTCCGAGTCTGTAAATATCA</td><td></td><td></td><td></td><td></td><td></td><td></td><td></td><td></td><td></td><td></td><td></td></tr>
<tr><td>Chromosome</td><td>start</td><td></td><td>end</td><td>strand</td><td>MM</td><td>target_seq</td><td>PAM</td><td>alignment</td><td>distance</td><td>position</td><td>gene name</td><td>gene id</td></tr>
<tr><td>GU305914.1</td><td></td><td>9553</td><td>9575</td><td>-</td><td>0</td><td>TGATATTTACAGACTCGGAC</td><td>AGG</td><td>||||||||[||||||||||||]PAM</td><td>NA</td><td>Intergenic</td><td>NA</td><td>NA</td></tr>
<tr><td>T33713</td><td>TTTACAGACTCGGACAGGTGTGG</td><td></td><td>1000</td><td></td><td></td><td></td><td></td><td></td><td></td><td></td><td></td><td></td></tr>
<tr><td>Oligo fwd</td><td>CACCTTTACAGACTCGGACAGGTG</td><td></td><td></td><td></td><td></td><td></td><td></td><td></td><td></td><td></td><td></td><td></td></tr>
<tr><td>Oligo rev</td><td>AAACCACCTGTCCGAGTCTGTAAA</td><td></td><td></td><td></td><td></td><td></td><td></td><td></td><td></td><td></td><td></td><td></td></tr>
<tr><td>Chromosome</td><td>start</td><td></td><td>end</td><td>strand</td><td>MM</td><td>target_seq</td><td>PAM</td><td>alignment</td><td>distance</td><td>position</td><td>gene name</td><td>gene id</td></tr>
<tr><td>GU305914.1</td><td></td><td>9548</td><td>9570</td><td>-</td><td>0</td><td>TTTACAGACTCGGACAGGTG</td><td>TGG</td><td>||||||||[||||||||||||]PAM</td><td>NA</td><td>Intergenic</td><td>NA</td><td>NA</td></tr>
</table>

| | | | | | | | | | | | |
|---|---|---|---|---|---|---|---|---|---|---|---|
| T33714 | TTACAGACTCGGACAGGTGTGGG | | 1000 | | | | | | | | |
| Oligo fwd | CACCTTACAGACTCGGACAGGTGT | | | | | | | | | | |
| Oligo rev | AAACACACCTGTCCGAGTCTGTAA | | | | | | | | | | |
| Chromosome | start | | end | strand | MM | target_seq | PAM | alignment | distance | position | gene name | gene id |
| GU305914.1 | | 9547 | 9569 | - | 0 | TTACAGACTCGGACAGGTGT | GGG | \|\|\|\|\|\|\|\|[\|\|\|\|\|\|\|\|\|\|\|\|]PAM | NA | Intergenic | NA | NA |

| | | | | | | | | | | | | |
|---|---|---|---|---|---|---|---|---|---|---|---|---|
| T33715 | CAGACTCGGACAGGTGTGGGCGG | | 1000 | | | | | | | | | |
| Oligo fwd | CACCCAGACTCGGACAGGTGTGGG | | | | | | | | | | | |
| Oligo rev | AAACCCCACACCTGTCCGAGTCTG | | | | | | | | | | | |
| Chromosome | start | | end | strand | MM | target_seq | PAM | alignment | distance | position | gene name | gene id |
| GU305914.1 | | 9544 | 9566 | - | 0 | CAGACTCGGACAGGTGTGGG | CGG | \|\|\|\|\|\|\|\|[\|\|\|\|\|\|\|\|\|\|\|\|]PAM | NA | Intergenic | NA | NA |

| | | | | | | | | | | | | |
|---|---|---|---|---|---|---|---|---|---|---|---|---|
| T33716 | GGACAGGTGTGGGCGGTTCCAGG | | 1000 | | | | | | | | | |
| Oligo fwd | CACCGGACAGGTGTGGGCGGTTCC | | | | | | | | | | | |
| Oligo rev | AAACGGAACCGCCCACACCTGTCC | | | | | | | | | | | |
| Chromosome | start | | end | strand | MM | target_seq | PAM | alignment | distance | position | gene name | gene id |
| GU305914.1 | | 8544 | 8566 | - | 0 | GGACAGGTGTGGGCGGTTCC | AGG | \|\|\|\|\|\|\|\|[\|\|\|\|\|\|\|\|\|\|\|\|]PAM | NA | Intergenic | NA | NA |
| GU305914.1 | | 9537 | 9559 | - | 0 | GGACAGGTGTGGGCGGTTCC | AGG | \|\|\|\|\|\|\|\|[\|\|\|\|\|\|\|\|\|\|\|\|]PAM | NA | Intergenic | NA | NA |

| | | | | | | | | | | | | |
|---|---|---|---|---|---|---|---|---|---|---|---|---|
| T33717 | CCAGGTGTACGTAGAAAATGTGG | | 1000 | | | | | | | | | |
| Oligo fwd | CACCCCAGGTGTACGTAGAAAATG | | | | | | | | | | | |
| Oligo rev | AAACCATTTTCTACGTACACCTGG | | | | | | | | | | | |
| Chromosome | start | | end | strand | MM | target_seq | PAM | alignment | distance | position | gene name | gene id |
| GU305914.1 | | 9519 | 9541 | - | 0 | CCAGGTGTACGTAGAAAATG | TGG | \|\|\|\|\|\|\|\|[\|\|\|\|\|\|\|\|\|\|\|\|]PAM | NA | Intergenic | NA | NA |

| | | | | | | | | | | | | |
|---|---|---|---|---|---|---|---|---|---|---|---|---|
| T33718 | CAGGTGTACGTAGAAAATGTGGG | | 1000 | | | | | | | | | |
| Oligo fwd | CACCCAGGTGTACGTAGAAAATGT | | | | | | | | | | | |
| Oligo rev | AAACACATTTTCTACGTACACCTG | | | | | | | | | | | |
| Chromosome | start | | end | strand | MM | target_seq | PAM | alignment | distance | position | gene name | gene id |
| GU305914.1 | | 9518 | 9540 | - | 0 | CAGGTGTACGTAGAAAATGT | GGG | \|\|\|\|\|\|\|\|[\|\|\|\|\|\|\|\|\|\|\|\|]PAM | NA | Intergenic | NA | NA |

| | | | | | | | | | | | | |
|---|---|---|---|---|---|---|---|---|---|---|---|---|
| T33719 | ACGCACGAGATGCCGTTGAACGG | | 1000 | | | | | | | | | |
| Oligo fwd | CACCACGCACGAGATGCCGTTGAA | | | | | | | | | | | |
| Oligo rev | AAACTTCAACGGCATCTCGTGCGT | | | | | | | | | | | |
| Chromosome | start | | end | strand | MM | target_seq | PAM | alignment | distance | position | gene name | gene id |
| GU305914.1 | | 9486 | 9508 | - | 0 | ACGCACGAGATGCCGTTGAA | CGG | \|\|\|\|\|\|\|\|[\|\|\|\|\|\|\|\|\|\|\|\|]PAM | NA | Intergenic | NA | NA |

| | | | | | | | | | | | | |
|---|---|---|---|---|---|---|---|---|---|---|---|---|
| T33720 | CGAGATGCCGTTGAACGGAGAGG | | 1000 | | | | | | | | | |
| Oligo fwd | CACCCGAGATGCCGTTGAACGGAG | | | | | | | | | | | |
| Oligo rev | AAACCTCCGTTCAACGGCATCTCG | | | | | | | | | | | |
| Chromosome | start | | end | strand | MM | target_seq | PAM | alignment | distance | position | gene name | gene id |
| GU305914.1 | | 9481 | 9503 | - | 0 | CGAGATGCCGTTGAACGGAG | AGG | \|\|\|\|\|\|\|\|[\|\|\|\|\|\|\|\|\|\|\|\|]PAM | NA | Intergenic | NA | NA |

| | | | | | | | | | | | | |
|---|---|---|---|---|---|---|---|---|---|---|---|---|
| T33721 | TGCCGTTGAACGGAGAGGCTCGG | | 1000 | | | | | | | | | |
| Oligo fwd | CACCTGCCGTTGAACGGAGAGGCT | | | | | | | | | | | |
| Oligo rev | AAACAGCCTCTCCGTTCAACGGCA | | | | | | | | | | | |
| Chromosome | start | | end | strand | MM | target_seq | PAM | alignment | distance | position | gene name | gene id |
| GU305914.1 | | 9476 | 9498 | - | 0 | TGCCGTTGAACGGAGAGGCT | CGG | \|\|\|\|\|\|\|\|[\|\|\|\|\|\|\|\|\|\|\|\|]PAM | NA | Intergenic | NA | NA |
| GU305914.1 | | 8492 | 8514 | - | 4 | GGATGTTGTACGGAGAGGCT | CGG | -\|--\|\|\|\|[-\|\|\|\|\|\|\|\|\|\|\|]PAM | NA | Intergenic | NA | NA |

| | | | | | | | | | | | | |
|---|---|---|---|---|---|---|---|---|---|---|---|---|
| T33722 | ACGGAGAGGCTCGGTTTCAACGG | | 1000 | | | | | | | | | |
| Oligo fwd | CACCACGGAGAGGCTCGGTTTCAA | | | | | | | | | | | |
| Oligo rev | AAACTTGAAACCGAGCCTCTCCGT | | | | | | | | | | | |
| Chromosome | start | | end | strand | MM | target_seq | PAM | alignment | distance | position | gene name | gene id |
| GU305914.1 | | 9467 | 9489 | - | 0 | ACGGAGAGGCTCGGTTTCAA | CGG | \|\|\|\|\|\|\|\|[\|\|\|\|\|\|\|\|\|\|\|\|]PAM | NA | Intergenic | NA | NA |
| GU305914.1 | | 8483 | 8505 | - | 2 | ACGGAGAGGCTCGGATTCAT | CGG | \|\|\|\|\|\|\|\|[\|\|\|\|\|\|-\|\|\|\|-]PAM | NA | Intergenic | NA | NA |

| | | | | | | | | | | | | |
|---|---|---|---|---|---|---|---|---|---|---|---|---|
| T33723 | AGAGGCTCGGTTTCAACGGTTGG | | 1000 | | | | | | | | | |
| Oligo fwd | CACCAGAGGCTCGGTTTCAACGGT | | | | | | | | | | | |
| Oligo rev | AAACACCGTTGAAACCGAGCCTCT | | | | | | | | | | | |
| Chromosome | start | | end | strand | MM | target_seq | PAM | alignment | distance | position | gene name | gene id |
| GU305914.1 | | 9463 | 9485 | - | 0 | AGAGGCTCGGTTTCAACGGT | TGG | \|\|\|\|\|\|\|\|[\|\|\|\|\|\|\|\|\|\|\|\|]PAM | NA | Intergenic | NA | NA |
| GU305914.1 | | 8479 | 8501 | - | 2 | AGAGGCTCGGATTCATCGGT | TGG | \|\|\|\|\|\|\|\|[\|\|-\|\|\|\|-\|\|\|\|]PAM | NA | Intergenic | NA | NA |

| | | | | | | | | | | | | |
|---|---|---|---|---|---|---|---|---|---|---|---|---|
| T33724 | GTTTCAACGGTTGGTCTGTCTGG | | 1000 | | | | | | | | | |
| Oligo fwd | CACCGTTTCAACGGTTGGTCTGTC | | | | | | | | | | | |
| Oligo rev | AAACGACAGACCAACCGTTGAAAC | | | | | | | | | | | |
| Chromosome | start | | end | strand | MM | target_seq | PAM | alignment | distance | position | gene name | gene id |
| GU305914.1 | | 9454 | 9476 | - | 0 | GTTTCAACGGTTGGTCTGTC | TGG | \|\|\|\|\|\|\|\|[\|\|\|\|\|\|\|\|\|\|\|\|]PAM | NA | Intergenic | NA | NA |

| | | | | | | | | | | | | |
|---|---|---|---|---|---|---|---|---|---|---|---|---|
| T33725 | CTGGACTGTTGCCCATGAAACGG | | 1000 | | | | | | | | | |
| Oligo fwd | CACCCTGGACTGTTGCCCATGAAA | | | | | | | | | | | |
| Oligo rev | AAACTTTCATGGGCAACAGTCCAG | | | | | | | | | | | |
| Chromosome | start | | end | strand | MM | target_seq | PAM | alignment | distance | position | gene name | gene id |
| GU305914.1 | | 9435 | 9457 | - | 0 | CTGGACTGTTGCCCATGAAA | CGG | \|\|\|\|\|\|\|\|[\|\|\|\|\|\|\|\|\|\|\|\|]PAM | NA | Intergenic | NA | NA |

| | | | | | | | | | | | | |
|---|---|---|---|---|---|---|---|---|---|---|---|---|
| T33726 | TGTTGCCCATGAAACGGACGCGG | | 1000 | | | | | | | | | |
| Oligo fwd | CACCTGTTGCCCATGAAACGGACG | | | | | | | | | | | |
| Oligo rev | AAACCGTCCGTTTCATGGGCAACA | | | | | | | | | | | |
| Chromosome | start | | end | strand | MM | target_seq | PAM | alignment | distance | position | gene name | gene id |
| GU305914.1 | | 9429 | 9451 | - | 0 | TGTTGCCCATGAAACGGACG | CGG | \|\|\|\|\|\|\|\|[\|\|\|\|\|\|\|\|\|\|\|\|]PAM | NA | Intergenic | NA | NA |

| | | | | | | | | | | | | |
|---|---|---|---|---|---|---|---|---|---|---|---|---|
| T33727 | CCATGAAACGGACGCGGAAACGG | | 1000 | | | | | | | | | |
| Oligo fwd | CACCCCATGAAACGGACGCGGAAA | | | | | | | | | | | |
| Oligo rev | AAACTTTCCGCGTCCGTTTCATGG | | | | | | | | | | | |
| Chromosome | start | | end | strand | MM | target_seq | PAM | alignment | distance | position | gene name | gene id |
| GU305914.1 | | 9423 | 9445 | - | 0 | CCATGAAACGGACGCGGAAA | CGG | \|\|\|\|\|\|\|\|[\|\|\|\|\|\|\|\|\|\|\|\|]PAM | NA | Intergenic | NA | NA |

<table>
<tr><td>T33728</td><td>AACGGTCTACGTATGAATCGAGG</td><td>1000</td></tr>
<tr><td>Oligo fwd</td><td>CACCAACGGTCTACGTATGAATCG</td></tr>
<tr><td>Oligo rev</td><td>AAACCGATTCATACGTAGACCGTT</td></tr>
<tr><td>Chromosome</td><td>start</td><td>end</td><td>strand</td><td>MM</td><td>target_seq</td><td>PAM</td><td>alignment</td><td>distance</td><td>position</td><td>gene name</td><td>gene id</td></tr>
<tr><td>GU305914.1</td><td>9405</td><td>9427</td><td>-</td><td>0</td><td>AACGGTCTACGTATGAATCG</td><td>AGG</td><td>||||||||[||||||||||||]PAM</td><td>NA</td><td>Intergenic</td><td>NA</td><td>NA</td></tr>
<tr><td></td></tr>
<tr><td>T33729</td><td>CAATAAACCCAGCACTATCACGG</td><td>1000</td></tr>
<tr><td>Oligo fwd</td><td>CACCCAATAAACCCAGCACTATCA</td></tr>
<tr><td>Oligo rev</td><td>AAACTGATAGTGCTGGGTTTATTG</td></tr>
<tr><td>Chromosome</td><td>start</td><td>end</td><td>strand</td><td>MM</td><td>target_seq</td><td>PAM</td><td>alignment</td><td>distance</td><td>position</td><td>gene name</td><td>gene id</td></tr>
<tr><td>GU305914.1</td><td>9350</td><td>9372</td><td>-</td><td>0</td><td>CAATAAACCCAGCACTATCA</td><td>CGG</td><td>||||||||[||||||||||||]PAM</td><td>NA</td><td>Intergenic</td><td>NA</td><td>NA</td></tr>
<tr><td></td></tr>
<tr><td>T33730</td><td>TACGATGACCACTAGCGACACGG</td><td>1000</td></tr>
<tr><td>Oligo fwd</td><td>CACCTACGATGACCACTAGCGACA</td></tr>
<tr><td>Oligo rev</td><td>AAACTGTCGCTAGTGGTCATCGTA</td></tr>
<tr><td>Chromosome</td><td>start</td><td>end</td><td>strand</td><td>MM</td><td>target_seq</td><td>PAM</td><td>alignment</td><td>distance</td><td>position</td><td>gene name</td><td>gene id</td></tr>
<tr><td>GU305914.1</td><td>9320</td><td>9342</td><td>-</td><td>0</td><td>TACGATGACCACTAGCGACA</td><td>CGG</td><td>||||||||[||||||||||||]PAM</td><td>NA</td><td>Intergenic</td><td>NA</td><td>NA</td></tr>
<tr><td></td></tr>
<tr><td>T33731</td><td>CACTAGCGACACGGCGATCGCGG</td><td>1000</td></tr>
<tr><td>Oligo fwd</td><td>CACCCACTAGCGACACGGCGATCG</td></tr>
<tr><td>Oligo rev</td><td>AAACCGATCGCCGTGTCGCTAGTG</td></tr>
<tr><td>Chromosome</td><td>start</td><td>end</td><td>strand</td><td>MM</td><td>target_seq</td><td>PAM</td><td>alignment</td><td>distance</td><td>position</td><td>gene name</td><td>gene id</td></tr>
<tr><td>GU305914.1</td><td>9311</td><td>9333</td><td>-</td><td>0</td><td>CACTAGCGACACGGCGATCG</td><td>CGG</td><td>||||||||[||||||||||||]PAM</td><td>NA</td><td>Intergenic</td><td>NA</td><td>NA</td></tr>
<tr><td></td></tr>
<tr><td>T33732</td><td>AGCGACACGGCGATCGCGGTCGG</td><td>1000</td></tr>
<tr><td>Oligo fwd</td><td>CACCAGCGACACGGCGATCGCGGT</td></tr>
<tr><td>Oligo rev</td><td>AAACACCGCGATCGCCGTGTCGCT</td></tr>
<tr><td>Chromosome</td><td>start</td><td>end</td><td>strand</td><td>MM</td><td>target_seq</td><td>PAM</td><td>alignment</td><td>distance</td><td>position</td><td>gene name</td><td>gene id</td></tr>
<tr><td>GU305914.1</td><td>9307</td><td>9329</td><td>-</td><td>0</td><td>AGCGACACGGCGATCGCGGT</td><td>CGG</td><td>||||||||[||||||||||||]PAM</td><td>NA</td><td>Intergenic</td><td>NA</td><td>NA</td></tr>
<tr><td></td></tr>
<tr><td>T33733</td><td>TAGCGTTATCGATTTCGAGTTGG</td><td>1000</td></tr>
<tr><td>Oligo fwd</td><td>CACCTAGCGTTATCGATTTCGAGT</td></tr>
<tr><td>Oligo rev</td><td>AAACACTCGAAATCGATAACGCTA</td></tr>
<tr><td>Chromosome</td><td>start</td><td>end</td><td>strand</td><td>MM</td><td>target_seq</td><td>PAM</td><td>alignment</td><td>distance</td><td>position</td><td>gene name</td><td>gene id</td></tr>
<tr><td>GU305914.1</td><td>9246</td><td>9268</td><td>-</td><td>0</td><td>TAGCGTTATCGATTTCGAGT</td><td>TGG</td><td>||||||||[||||||||||||]PAM</td><td>NA</td><td>Intergenic</td><td>NA</td><td>NA</td></tr>
<tr><td></td></tr>
<tr><td>T33734</td><td>AGCGTTATCGATTTCGAGTTGGG</td><td>1000</td></tr>
<tr><td>Oligo fwd</td><td>CACCAGCGTTATCGATTTCGAGTT</td></tr>
<tr><td>Oligo rev</td><td>AAACAACTCGAAATCGATAACGCT</td></tr>
<tr><td>Chromosome</td><td>start</td><td>end</td><td>strand</td><td>MM</td><td>target_seq</td><td>PAM</td><td>alignment</td><td>distance</td><td>position</td><td>gene name</td><td>gene id</td></tr>
<tr><td>GU305914.1</td><td>9245</td><td>9267</td><td>-</td><td>0</td><td>AGCGTTATCGATTTCGAGTT</td><td>GGG</td><td>||||||||[||||||||||||]PAM</td><td>NA</td><td>Intergenic</td><td>NA</td><td>NA</td></tr>
<tr><td></td></tr>
<tr><td>T33735</td><td>TGGGTAGTATCATCGACCTGCGG</td><td>1000</td></tr>
<tr><td>Oligo fwd</td><td>CACCTGGGTAGTATCATCGACCTG</td></tr>
<tr><td>Oligo rev</td><td>AAACCAGGTCGATGATACTACCCA</td></tr>
<tr><td>Chromosome</td><td>start</td><td>end</td><td>strand</td><td>MM</td><td>target_seq</td><td>PAM</td><td>alignment</td><td>distance</td><td>position</td><td>gene name</td><td>gene id</td></tr>
<tr><td>GU305914.1</td><td>9226</td><td>9248</td><td>-</td><td>0</td><td>TGGGTAGTATCATCGACCTG</td><td>CGG</td><td>||||||||[||||||||||||]PAM</td><td>NA</td><td>Intergenic</td><td>NA</td><td>NA</td></tr>
<tr><td></td></tr>
<tr><td>T33736</td><td>GGGTAGTATCATCGACCTGCGGG</td><td>1000</td></tr>
<tr><td>Oligo fwd</td><td>CACCGGGTAGTATCATCGACCTGC</td></tr>
<tr><td>Oligo rev</td><td>AAACGCAGGTCGATGATACTACCC</td></tr>
<tr><td>Chromosome</td><td>start</td><td>end</td><td>strand</td><td>MM</td><td>target_seq</td><td>PAM</td><td>alignment</td><td>distance</td><td>position</td><td>gene name</td><td>gene id</td></tr>
<tr><td>GU305914.1</td><td>9225</td><td>9247</td><td>-</td><td>0</td><td>GGGTAGTATCATCGACCTGC</td><td>GGG</td><td>||||||||[||||||||||||]PAM</td><td>NA</td><td>Intergenic</td><td>NA</td><td>NA</td></tr>
<tr><td></td></tr>
<tr><td>T33737</td><td>GGTAGTATCATCGACCTGCGGGG</td><td>1000</td></tr>
<tr><td>Oligo fwd</td><td>CACCGGTAGTATCATCGACCTGCG</td></tr>
<tr><td>Oligo rev</td><td>AAACCGCAGGTCGATGATACTACC</td></tr>
<tr><td>Chromosome</td><td>start</td><td>end</td><td>strand</td><td>MM</td><td>target_seq</td><td>PAM</td><td>alignment</td><td>distance</td><td>position</td><td>gene name</td><td>gene id</td></tr>
<tr><td>GU305914.1</td><td>9224</td><td>9246</td><td>-</td><td>0</td><td>GGTAGTATCATCGACCTGCG</td><td>GGG</td><td>||||||||[||||||||||||]PAM</td><td>NA</td><td>Intergenic</td><td>NA</td><td>NA</td></tr>
<tr><td></td></tr>
<tr><td>T33738</td><td>ATCATCGACCTGCGGGGTGACGG</td><td>1000</td></tr>
<tr><td>Oligo fwd</td><td>CACCATCATCGACCTGCGGGGTGA</td></tr>
<tr><td>Oligo rev</td><td>AAACTCACCCCGCAGGTCGATGAT</td></tr>
<tr><td>Chromosome</td><td>start</td><td>end</td><td>strand</td><td>MM</td><td>target_seq</td><td>PAM</td><td>alignment</td><td>distance</td><td>position</td><td>gene name</td><td>gene id</td></tr>
<tr><td>GU305914.1</td><td>9218</td><td>9240</td><td>-</td><td>0</td><td>ATCATCGACCTGCGGGGTGA</td><td>CGG</td><td>||||||||[||||||||||||]PAM</td><td>NA</td><td>Intergenic</td><td>NA</td><td>NA</td></tr>
<tr><td></td></tr>
<tr><td>T33739</td><td>CGACCTGCGGGGTGACGGACTGG</td><td>1000</td></tr>
<tr><td>Oligo fwd</td><td>CACCCGACCTGCGGGGTGACGGAC</td></tr>
<tr><td>Oligo rev</td><td>AAACGTCCGTCACCCCGCAGGTCG</td></tr>
<tr><td>Chromosome</td><td>start</td><td>end</td><td>strand</td><td>MM</td><td>target_seq</td><td>PAM</td><td>alignment</td><td>distance</td><td>position</td><td>gene name</td><td>gene id</td></tr>
<tr><td>GU305914.1</td><td>9213</td><td>9235</td><td>-</td><td>0</td><td>CGACCTGCGGGGTGACGGAC</td><td>TGG</td><td>||||||||[||||||||||||]PAM</td><td>NA</td><td>Intergenic</td><td>NA</td><td>NA</td></tr>
<tr><td></td></tr>
<tr><td>T33740</td><td>GACCTGCGGGGTGACGGACTGGG</td><td>1000</td></tr>
<tr><td>Oligo fwd</td><td>CACCGACCTGCGGGGTGACGGACT</td></tr>
<tr><td>Oligo rev</td><td>AAACAGTCCGTCACCCCGCAGGTC</td></tr>
<tr><td>Chromosome</td><td>start</td><td>end</td><td>strand</td><td>MM</td><td>target_seq</td><td>PAM</td><td>alignment</td><td>distance</td><td>position</td><td>gene name</td><td>gene id</td></tr>
<tr><td>GU305914.1</td><td>9212</td><td>9234</td><td>-</td><td>0</td><td>GACCTGCGGGGTGACGGACT</td><td>GGG</td><td>||||||||[||||||||||||]PAM</td><td>NA</td><td>Intergenic</td><td>NA</td><td>NA</td></tr>
<tr><td>GU305914.1</td><td>10186</td><td>10208</td><td>-</td><td>4</td><td>GGCCTGTCGGGTGACGGGCT</td><td>GGG</td><td>|-||||--[|||||||||-||]PAM</td><td>NA</td><td>Intergenic</td><td>NA</td><td>NA</td></tr>
<tr><td></td></tr>
<tr><td>T33741</td><td>GTGACGGACTGGGATCGTGATGG</td><td>1000</td></tr>
<tr><td>Oligo fwd</td><td>CACCGTGACGGACTGGGATCGTGA</td></tr>
<tr><td>Oligo rev</td><td>AAACTCACGATCCCAGTCCGTCAC</td></tr>
<tr><td>Chromosome</td><td>start</td><td>end</td><td>strand</td><td>MM</td><td>target_seq</td><td>PAM</td><td>alignment</td><td>distance</td><td>position</td><td>gene name</td><td>gene id</td></tr>
<tr><td>GU305914.1</td><td>9202</td><td>9224</td><td>-</td><td>0</td><td>GTGACGGACTGGGATCGTGA</td><td>TGG</td><td>||||||||[||||||||||||]PAM</td><td>NA</td><td>Intergenic</td><td>NA</td><td>NA</td></tr>
<tr><td></td></tr>
<tr><td>T33742</td><td>AAATTTTGTACCGCTTCGTGTGG</td><td>1000</td></tr>
<tr><td>Oligo fwd</td><td>CACCAAATTTTGTACCGCTTCGTG</td></tr>
<tr><td>Oligo rev</td><td>AAACCACGAAGCGGTACAAAATTT</td></tr>
<tr><td>Chromosome</td><td>start</td><td>end</td><td>strand</td><td>MM</td><td>target_seq</td><td>PAM</td><td>alignment</td><td>distance</td><td>position</td><td>gene name</td><td>gene id</td></tr>
</table>

<table>
<tr><td>GU305914.1</td><td>9161</td><td>9183</td><td>-</td><td>0</td><td>AAATTTTGTACCGCTTCGTG</td><td>TGG</td><td>||||||||[||||||||||||]PAM</td><td>NA</td><td>Intergenic</td><td>NA</td><td>NA</td></tr>
<tr><td>T33743</td><td>GCTTCGTGTGGTCCGTGTCTCGG</td><td>1000</td><td></td><td></td><td></td><td></td><td></td><td></td><td></td><td></td><td></td></tr>
<tr><td>Oligo fwd</td><td>CACCGCTTCGTGTGGTCCGTGTCT</td><td></td><td></td><td></td><td></td><td></td><td></td><td></td><td></td><td></td><td></td></tr>
<tr><td>Oligo rev</td><td>AAACAGACACGGACCACACGAAGC</td><td></td><td></td><td></td><td></td><td></td><td></td><td></td><td></td><td></td><td></td></tr>
<tr><td>Chromosome</td><td>start</td><td>end</td><td>strand</td><td>MM</td><td>target_seq</td><td>PAM</td><td>alignment</td><td>distance</td><td>position</td><td>gene name</td><td>gene id</td></tr>
<tr><td>GU305914.1</td><td>9149</td><td>9171</td><td>-</td><td>0</td><td>GCTTCGTGTGGTCCGTGTCT</td><td>CGG</td><td>||||||||[||||||||||||]PAM</td><td>NA</td><td>Intergenic</td><td>NA</td><td>NA</td></tr>
<tr><td>T33744</td><td>GGTGCTCGATCGCGATCTTGTGG</td><td>1000</td><td></td><td></td><td></td><td></td><td></td><td></td><td></td><td></td><td></td></tr>
<tr><td>Oligo fwd</td><td>CACCGGTGCTCGATCGCGATCTTG</td><td></td><td></td><td></td><td></td><td></td><td></td><td></td><td></td><td></td><td></td></tr>
<tr><td>Oligo rev</td><td>AAACCAAGATCGCGATCGAGCACC</td><td></td><td></td><td></td><td></td><td></td><td></td><td></td><td></td><td></td><td></td></tr>
<tr><td>Chromosome</td><td>start</td><td>end</td><td>strand</td><td>MM</td><td>target_seq</td><td>PAM</td><td>alignment</td><td>distance</td><td>position</td><td>gene name</td><td>gene id</td></tr>
<tr><td>GU305914.1</td><td>9128</td><td>9150</td><td>-</td><td>0</td><td>GGTGCTCGATCGCGATCTTG</td><td>TGG</td><td>||||||||[||||||||||||]PAM</td><td>NA</td><td>Intergenic</td><td>NA</td><td>NA</td></tr>
<tr><td>T33745</td><td>GATCGCGATCTTGTGGTCGAAGG</td><td>1000</td><td></td><td></td><td></td><td></td><td></td><td></td><td></td><td></td><td></td></tr>
<tr><td>Oligo fwd</td><td>CACCGATCGCGATCTTGTGGTCGA</td><td></td><td></td><td></td><td></td><td></td><td></td><td></td><td></td><td></td><td></td></tr>
<tr><td>Oligo rev</td><td>AAACTCGACCACAAGATCGCGATC</td><td></td><td></td><td></td><td></td><td></td><td></td><td></td><td></td><td></td><td></td></tr>
<tr><td>Chromosome</td><td>start</td><td>end</td><td>strand</td><td>MM</td><td>target_seq</td><td>PAM</td><td>alignment</td><td>distance</td><td>position</td><td>gene name</td><td>gene id</td></tr>
<tr><td>GU305914.1</td><td>9121</td><td>9143</td><td>-</td><td>0</td><td>GATCGCGATCTTGTGGTCGA</td><td>AGG</td><td>||||||||[||||||||||||]PAM</td><td>NA</td><td>Intergenic</td><td>NA</td><td>NA</td></tr>
<tr><td>T33746</td><td>TCTTGTGGTCGAAGGTGTCGTGG</td><td>1000</td><td></td><td></td><td></td><td></td><td></td><td></td><td></td><td></td><td></td></tr>
<tr><td>Oligo fwd</td><td>CACCTCTTGTGGTCGAAGGTGTCG</td><td></td><td></td><td></td><td></td><td></td><td></td><td></td><td></td><td></td><td></td></tr>
<tr><td>Oligo rev</td><td>AAACCGACACCTTCGACCACAAGA</td><td></td><td></td><td></td><td></td><td></td><td></td><td></td><td></td><td></td><td></td></tr>
<tr><td>Chromosome</td><td>start</td><td>end</td><td>strand</td><td>MM</td><td>target_seq</td><td>PAM</td><td>alignment</td><td>distance</td><td>position</td><td>gene name</td><td>gene id</td></tr>
<tr><td>GU305914.1</td><td>9113</td><td>9135</td><td>-</td><td>0</td><td>TCTTGTGGTCGAAGGTGTCG</td><td>TGG</td><td>||||||||[||||||||||||]PAM</td><td>NA</td><td>Intergenic</td><td>NA</td><td>NA</td></tr>
<tr><td>T33747</td><td>GTCGAAGGTGTCGTGGTTATAGG</td><td>1000</td><td></td><td></td><td></td><td></td><td></td><td></td><td></td><td></td><td></td></tr>
<tr><td>Oligo fwd</td><td>CACCGTCGAAGGTGTCGTGGTTAT</td><td></td><td></td><td></td><td></td><td></td><td></td><td></td><td></td><td></td><td></td></tr>
<tr><td>Oligo rev</td><td>AAACATAACCACGACACCTTCGAC</td><td></td><td></td><td></td><td></td><td></td><td></td><td></td><td></td><td></td><td></td></tr>
<tr><td>Chromosome</td><td>start</td><td>end</td><td>strand</td><td>MM</td><td>target_seq</td><td>PAM</td><td>alignment</td><td>distance</td><td>position</td><td>gene name</td><td>gene id</td></tr>
<tr><td>GU305914.1</td><td>9106</td><td>9128</td><td>-</td><td>0</td><td>GTCGAAGGTGTCGTGGTTAT</td><td>AGG</td><td>||||||||[||||||||||||]PAM</td><td>NA</td><td>Intergenic</td><td>NA</td><td>NA</td></tr>
<tr><td>T33748</td><td>GGTGTCGTGGTTATAGGTTTCGG</td><td>1000</td><td></td><td></td><td></td><td></td><td></td><td></td><td></td><td></td><td></td></tr>
<tr><td>Oligo fwd</td><td>CACCGGTGTCGTGGTTATAGGTTT</td><td></td><td></td><td></td><td></td><td></td><td></td><td></td><td></td><td></td><td></td></tr>
<tr><td>Oligo rev</td><td>AAACAAACCTATAACCACGACACC</td><td></td><td></td><td></td><td></td><td></td><td></td><td></td><td></td><td></td><td></td></tr>
<tr><td>Chromosome</td><td>start</td><td>end</td><td>strand</td><td>MM</td><td>target_seq</td><td>PAM</td><td>alignment</td><td>distance</td><td>position</td><td>gene name</td><td>gene id</td></tr>
<tr><td>GU305914.1</td><td>9100</td><td>9122</td><td>-</td><td>0</td><td>GGTGTCGTGGTTATAGGTTT</td><td>CGG</td><td>||||||||[||||||||||||]PAM</td><td>NA</td><td>Intergenic</td><td>NA</td><td>NA</td></tr>
<tr><td>T33749</td><td>GTCGTGGTTATAGGTTTCGGCGG</td><td>1000</td><td></td><td></td><td></td><td></td><td></td><td></td><td></td><td></td><td></td></tr>
<tr><td>Oligo fwd</td><td>CACCGTCGTGGTTATAGGTTTCGG</td><td></td><td></td><td></td><td></td><td></td><td></td><td></td><td></td><td></td><td></td></tr>
<tr><td>Oligo rev</td><td>AAACCCGAAACCTATAACCACGAC</td><td></td><td></td><td></td><td></td><td></td><td></td><td></td><td></td><td></td><td></td></tr>
<tr><td>Chromosome</td><td>start</td><td>end</td><td>strand</td><td>MM</td><td>target_seq</td><td>PAM</td><td>alignment</td><td>distance</td><td>position</td><td>gene name</td><td>gene id</td></tr>
<tr><td>GU305914.1</td><td>9097</td><td>9119</td><td>-</td><td>0</td><td>GTCGTGGTTATAGGTTTCGG</td><td>CGG</td><td>||||||||[||||||||||||]PAM</td><td>NA</td><td>Intergenic</td><td>NA</td><td>NA</td></tr>
<tr><td>T33750</td><td>TGATAGATCGAAACTGTTACCGG</td><td>1000</td><td></td><td></td><td></td><td></td><td></td><td></td><td></td><td></td><td></td></tr>
<tr><td>Oligo fwd</td><td>CACCTGATAGATCGAAACTGTTAC</td><td></td><td></td><td></td><td></td><td></td><td></td><td></td><td></td><td></td><td></td></tr>
<tr><td>Oligo rev</td><td>AAACGTAACAGTTTCGATCTATCA</td><td></td><td></td><td></td><td></td><td></td><td></td><td></td><td></td><td></td><td></td></tr>
<tr><td>Chromosome</td><td>start</td><td>end</td><td>strand</td><td>MM</td><td>target_seq</td><td>PAM</td><td>alignment</td><td>distance</td><td>position</td><td>gene name</td><td>gene id</td></tr>
<tr><td>GU305914.1</td><td>9053</td><td>9075</td><td>-</td><td>0</td><td>TGATAGATCGAAACTGTTAC</td><td>CGG</td><td>||||||||[||||||||||||]PAM</td><td>NA</td><td>Intergenic</td><td>NA</td><td>NA</td></tr>
<tr><td>T33751</td><td>TCGAAACTGTTACCGGCGCTAGG</td><td>1000</td><td></td><td></td><td></td><td></td><td></td><td></td><td></td><td></td><td></td></tr>
<tr><td>Oligo fwd</td><td>CACCTCGAAACTGTTACCGGCGCT</td><td></td><td></td><td></td><td></td><td></td><td></td><td></td><td></td><td></td><td></td></tr>
<tr><td>Oligo rev</td><td>AAACAGCGCCGGTAACAGTTTCGA</td><td></td><td></td><td></td><td></td><td></td><td></td><td></td><td></td><td></td><td></td></tr>
<tr><td>Chromosome</td><td>start</td><td>end</td><td>strand</td><td>MM</td><td>target_seq</td><td>PAM</td><td>alignment</td><td>distance</td><td>position</td><td>gene name</td><td>gene id</td></tr>
<tr><td>GU305914.1</td><td>9046</td><td>9068</td><td>-</td><td>0</td><td>TCGAAACTGTTACCGGCGCT</td><td>AGG</td><td>||||||||[||||||||||||]PAM</td><td>NA</td><td>Intergenic</td><td>NA</td><td>NA</td></tr>
<tr><td>T33752</td><td>CGGCGCTAGGATAGATGTAGAGG</td><td>1000</td><td></td><td></td><td></td><td></td><td></td><td></td><td></td><td></td><td></td></tr>
<tr><td>Oligo fwd</td><td>CACCCGGCGCTAGGATAGATGTAG</td><td></td><td></td><td></td><td></td><td></td><td></td><td></td><td></td><td></td><td></td></tr>
<tr><td>Oligo rev</td><td>AAACCTACATCTATCCTAGCGCCG</td><td></td><td></td><td></td><td></td><td></td><td></td><td></td><td></td><td></td><td></td></tr>
<tr><td>Chromosome</td><td>start</td><td>end</td><td>strand</td><td>MM</td><td>target_seq</td><td>PAM</td><td>alignment</td><td>distance</td><td>position</td><td>gene name</td><td>gene id</td></tr>
<tr><td>GU305914.1</td><td>9033</td><td>9055</td><td>-</td><td>0</td><td>CGGCGCTAGGATAGATGTAG</td><td>AGG</td><td>||||||||[||||||||||||]PAM</td><td>NA</td><td>Intergenic</td><td>NA</td><td>NA</td></tr>
<tr><td>T33753</td><td>GATGTAGAGGTAACCGTTACCGG</td><td>1000</td><td></td><td></td><td></td><td></td><td></td><td></td><td></td><td></td><td></td></tr>
<tr><td>Oligo fwd</td><td>CACCGATGTAGAGGTAACCGTTAC</td><td></td><td></td><td></td><td></td><td></td><td></td><td></td><td></td><td></td><td></td></tr>
<tr><td>Oligo rev</td><td>AAACGTAACGGTTACCTCTACATC</td><td></td><td></td><td></td><td></td><td></td><td></td><td></td><td></td><td></td><td></td></tr>
<tr><td>Chromosome</td><td>start</td><td>end</td><td>strand</td><td>MM</td><td>target_seq</td><td>PAM</td><td>alignment</td><td>distance</td><td>position</td><td>gene name</td><td>gene id</td></tr>
<tr><td>GU305914.1</td><td>9020</td><td>9042</td><td>-</td><td>0</td><td>GATGTAGAGGTAACCGTTAC</td><td>CGG</td><td>||||||||[||||||||||||]PAM</td><td>NA</td><td>Intergenic</td><td>NA</td><td>NA</td></tr>
<tr><td>T33754</td><td>GAATTTAATTCGAAACCCGTCGG</td><td>1000</td><td></td><td></td><td></td><td></td><td></td><td></td><td></td><td></td><td></td></tr>
<tr><td>Oligo fwd</td><td>CACCGAATTTAATTCGAAACCCGT</td><td></td><td></td><td></td><td></td><td></td><td></td><td></td><td></td><td></td><td></td></tr>
<tr><td>Oligo rev</td><td>AAACACGGGTTTCGAATTAAATTC</td><td></td><td></td><td></td><td></td><td></td><td></td><td></td><td></td><td></td><td></td></tr>
<tr><td>Chromosome</td><td>start</td><td>end</td><td>strand</td><td>MM</td><td>target_seq</td><td>PAM</td><td>alignment</td><td>distance</td><td>position</td><td>gene name</td><td>gene id</td></tr>
<tr><td>GU305914.1</td><td>8998</td><td>9020</td><td>-</td><td>0</td><td>GAATTTAATTCGAAACCCGT</td><td>CGG</td><td>||||||||[||||||||||||]PAM</td><td>NA</td><td>Intergenic</td><td>NA</td><td>NA</td></tr>
<tr><td>T33755</td><td>TTTAATTCGAAACCCGTCGGAGG</td><td>1000</td><td></td><td></td><td></td><td></td><td></td><td></td><td></td><td></td><td></td></tr>
<tr><td>Oligo fwd</td><td>CACCTTTAATTCGAAACCCGTCGG</td><td></td><td></td><td></td><td></td><td></td><td></td><td></td><td></td><td></td><td></td></tr>
<tr><td>Oligo rev</td><td>AAACCCGACGGGTTTCGAATTAAA</td><td></td><td></td><td></td><td></td><td></td><td></td><td></td><td></td><td></td><td></td></tr>
<tr><td>Chromosome</td><td>start</td><td>end</td><td>strand</td><td>MM</td><td>target_seq</td><td>PAM</td><td>alignment</td><td>distance</td><td>position</td><td>gene name</td><td>gene id</td></tr>
<tr><td>GU305914.1</td><td>8995</td><td>9017</td><td>-</td><td>0</td><td>TTTAATTCGAAACCCGTCGG</td><td>AGG</td><td>||||||||[||||||||||||]PAM</td><td>NA</td><td>Intergenic</td><td>NA</td><td>NA</td></tr>
<tr><td>T33756</td><td>AATTCGAAACCCGTCGGAGGTGG</td><td>1000</td><td></td><td></td><td></td><td></td><td></td><td></td><td></td><td></td><td></td></tr>
<tr><td>Oligo fwd</td><td>CACCAATTCGAAACCCGTCGGAGG</td><td></td><td></td><td></td><td></td><td></td><td></td><td></td><td></td><td></td><td></td></tr>
<tr><td>Oligo rev</td><td>AAACCCTCCGACGGGTTTCGAATT</td><td></td><td></td><td></td><td></td><td></td><td></td><td></td><td></td><td></td><td></td></tr>
<tr><td>Chromosome</td><td>start</td><td>end</td><td>strand</td><td>MM</td><td>target_seq</td><td>PAM</td><td>alignment</td><td>distance</td><td>position</td><td>gene name</td><td>gene id</td></tr>
<tr><td>GU305914.1</td><td>8992</td><td>9014</td><td>-</td><td>0</td><td>AATTCGAAACCCGTCGGAGG</td><td>TGG</td><td>||||||||[||||||||||||]PAM</td><td>NA</td><td>Intergenic</td><td>NA</td><td>NA</td></tr>
<tr><td>T33757</td><td>GAAACCCGTCGGAGGTGGTCTGG</td><td>1000</td><td></td><td></td><td></td><td></td><td></td><td></td><td></td><td></td><td></td></tr>
<tr><td>Oligo fwd</td><td>CACCGAAACCCGTCGGAGGTGGTC</td><td></td><td></td><td></td><td></td><td></td><td></td><td></td><td></td><td></td><td></td></tr>
<tr><td>Oligo rev</td><td>AAACGACCACCTCCGACGGGTTTC</td><td></td><td></td><td></td><td></td><td></td><td></td><td></td><td></td><td></td><td></td></tr>
</table>

| | | | | | | | | | | | |
|---|---|---|---|---|---|---|---|---|---|---|---|
| Chromosome | start | end | strand | MM | target_seq | PAM | alignment | distance | position | gene name | gene id |
| GU305914.1 | 8987 | 9009 | - | 0 | GAAACCCGTCGGAGGTGGTC | TGG | \|\|\|\|\|\|\|\|[\|\|\|\|\|\|\|\|\|\|\|\|]PAM | NA | Intergenic | NA | NA |
| T33758 | GTGATGAGTTTTTAAACGTAGG | 1000 | | | | | | | | | |
| Oligo fwd | CACCGTGATGAGTTTTTAAACGT | | | | | | | | | | |
| Oligo rev | AAACACGTTTAAAAACTCATCAC | | | | | | | | | | |
| Chromosome | start | end | strand | MM | target_seq | PAM | alignment | distance | position | gene name | gene id |
| GU305914.1 | 8950 | 8972 | - | 0 | GTGATGAGTTTTTAAACGT | AGG | \|\|\|\|\|\|\|\|[\|\|\|\|\|\|\|\|\|\|\|\|]PAM | NA | Intergenic | NA | NA |
| T33759 | ATGAGTTTTTAAACGTAGGCGG | 1000 | | | | | | | | | |
| Oligo fwd | CACCATGAGTTTTTAAACGTAGG | | | | | | | | | | |
| Oligo rev | AAACCCTACGTTTAAAAACTCAT | | | | | | | | | | |
| Chromosome | start | end | strand | MM | target_seq | PAM | alignment | distance | position | gene name | gene id |
| GU305914.1 | 8947 | 8969 | - | 0 | ATGAGTTTTTAAACGTAGG | CGG | \|\|\|\|\|\|\|\|[\|\|\|\|\|\|\|\|\|\|\|\|]PAM | NA | Intergenic | NA | NA |
| GU305914.1 | 7750 | 7772 | - | 4 | CTGAGTCCTTGAAACGTAGG | TGG | -\|\|\|\|\|--[\|\|-\|\|\|\|\|\|\|\|\|]PAM | NA | Intergenic | NA | NA |
| T33760 | TACAACACGCTCACGTTAATCGG | 1000 | | | | | | | | | |
| Oligo fwd | CACCTACAACACGCTCACGTTAAT | | | | | | | | | | |
| Oligo rev | AAACATTAACGTGAGCGTGTTGTA | | | | | | | | | | |
| Chromosome | start | end | strand | MM | target_seq | PAM | alignment | distance | position | gene name | gene id |
| GU305914.1 | 8905 | 8927 | - | 0 | TACAACACGCTCACGTTAAT | CGG | \|\|\|\|\|\|\|\|[\|\|\|\|\|\|\|\|\|\|\|\|]PAM | NA | Intergenic | NA | NA |
| T33761 | ACAACACGCTCACGTTAATCGGG | 1000 | | | | | | | | | |
| Oligo fwd | CACCACAACACGCTCACGTTAATC | | | | | | | | | | |
| Oligo rev | AAACGATTAACGTGAGCGTGTTGT | | | | | | | | | | |
| Chromosome | start | end | strand | MM | target_seq | PAM | alignment | distance | position | gene name | gene id |
| GU305914.1 | 8904 | 8926 | - | 0 | ACAACACGCTCACGTTAATC | GGG | \|\|\|\|\|\|\|\|[\|\|\|\|\|\|\|\|\|\|\|\|]PAM | NA | Intergenic | NA | NA |
| T33762 | GTTGCATTAATACAAAATGCAGG | 1000 | | | | | | | | | |
| Oligo fwd | CACCGTTGCATTAATACAAAATGC | | | | | | | | | | |
| Oligo rev | AAACGCATTTTGTATTAATGCAAC | | | | | | | | | | |
| Chromosome | start | end | strand | MM | target_seq | PAM | alignment | distance | position | gene name | gene id |
| GU305914.1 | 8851 | 8873 | - | 0 | GTTGCATTAATACAAAATGC | AGG | \|\|\|\|\|\|\|\|[\|\|\|\|\|\|\|\|\|\|\|\|]PAM | NA | Intergenic | NA | NA |
| T33763 | CTCACATGTATAGACCCAAACGG | 1000 | | | | | | | | | |
| Oligo fwd | CACCCTCACATGTATAGACCCAAA | | | | | | | | | | |
| Oligo rev | AAACTTTGGGTCTATACATGTGAG | | | | | | | | | | |
| Chromosome | start | end | strand | MM | target_seq | PAM | alignment | distance | position | gene name | gene id |
| GU305914.1 | 8825 | 8847 | - | 0 | CTCACATGTATAGACCCAAA | CGG | \|\|\|\|\|\|\|\|[\|\|\|\|\|\|\|\|\|\|\|\|]PAM | NA | Intergenic | NA | NA |
| T33764 | CTTATAGTTCTACATTGCCCTGG | 1000 | | | | | | | | | |
| Oligo fwd | CACCCTTATAGTTCTACATTGCCC | | | | | | | | | | |
| Oligo rev | AAACGGGCAATGTAGAACTATAAG | | | | | | | | | | |
| Chromosome | start | end | strand | MM | target_seq | PAM | alignment | distance | position | gene name | gene id |
| GU305914.1 | 8764 | 8786 | - | 0 | CTTATAGTTCTACATTGCCC | TGG | \|\|\|\|\|\|\|\|[\|\|\|\|\|\|\|\|\|\|\|\|]PAM | NA | Intergenic | NA | NA |
| T33765 | TCTACATTGCCCTGGTGTGACGG | 1000 | | | | | | | | | |
| Oligo fwd | CACCTCTACATTGCCCTGGTGTGA | | | | | | | | | | |
| Oligo rev | AAACTCACACCAGGGCAATGTAGA | | | | | | | | | | |
| Chromosome | start | end | strand | MM | target_seq | PAM | alignment | distance | position | gene name | gene id |
| GU305914.1 | 8756 | 8778 | - | 0 | TCTACATTGCCCTGGTGTGA | CGG | \|\|\|\|\|\|\|\|[\|\|\|\|\|\|\|\|\|\|\|\|]PAM | NA | Intergenic | NA | NA |
| T33766 | TATCATTTAATGTGAACAAATGG | 1000 | | | | | | | | | |
| Oligo fwd | CACCTATCATTTAATGTGAACAAA | | | | | | | | | | |
| Oligo rev | AAACTTTGTTCACATTAAATGATA | | | | | | | | | | |
| Chromosome | start | end | strand | MM | target_seq | PAM | alignment | distance | position | gene name | gene id |
| GU305914.1 | 8605 | 8627 | - | 0 | TATCATTTAATGTGAACAAA | TGG | \|\|\|\|\|\|\|\|[\|\|\|\|\|\|\|\|\|\|\|\|]PAM | NA | Intergenic | NA | NA |
| T33767 | TAATGTGAACAAATGGATACCGG | 1000 | | | | | | | | | |
| Oligo fwd | CACCTAATGTGAACAAATGGATAC | | | | | | | | | | |
| Oligo rev | AAACGTATCCATTTGTTCACATTA | | | | | | | | | | |
| Chromosome | start | end | strand | MM | target_seq | PAM | alignment | distance | position | gene name | gene id |
| GU305914.1 | 8598 | 8620 | - | 0 | TAATGTGAACAAATGGATAC | CGG | \|\|\|\|\|\|\|\|[\|\|\|\|\|\|\|\|\|\|\|\|]PAM | NA | Intergenic | NA | NA |
| T33768 | CAAATGGATACCGGCGCAACAGG | 1000 | | | | | | | | | |
| Oligo fwd | CACCCAAATGGATACCGGCGCAAC | | | | | | | | | | |
| Oligo rev | AAACGTTGCGCCGGTATCCATTTG | | | | | | | | | | |
| Chromosome | start | end | strand | MM | target_seq | PAM | alignment | distance | position | gene name | gene id |
| GU305914.1 | 8589 | 8611 | - | 0 | CAAATGGATACCGGCGCAAC | AGG | \|\|\|\|\|\|\|\|[\|\|\|\|\|\|\|\|\|\|\|\|]PAM | NA | Intergenic | NA | NA |
| T33769 | TACCGGCGCAACAGGTGCGCCGG | 1000 | | | | | | | | | |
| Oligo fwd | CACCTACCGGCGCAACAGGTGCGC | | | | | | | | | | |
| Oligo rev | AAACGCGCACCTGTTGCGCCGGTA | | | | | | | | | | |
| Chromosome | start | end | strand | MM | target_seq | PAM | alignment | distance | position | gene name | gene id |
| GU305914.1 | 8581 | 8603 | - | 0 | TACCGGCGCAACAGGTGCGC | CGG | \|\|\|\|\|\|\|\|[\|\|\|\|\|\|\|\|\|\|\|\|]PAM | NA | Intergenic | NA | NA |
| T33770 | GCGCCGGTATTTACATACTCAGG | 1000 | | | | | | | | | |
| Oligo fwd | CACCGCGCCGGTATTTACATACTC | | | | | | | | | | |
| Oligo rev | AAACGAGTATGTAAATACCGGCGC | | | | | | | | | | |
| Chromosome | start | end | strand | MM | target_seq | PAM | alignment | distance | position | gene name | gene id |
| GU305914.1 | 8565 | 8587 | - | 0 | GCGCCGGTATTTACATACTC | AGG | \|\|\|\|\|\|\|\|[\|\|\|\|\|\|\|\|\|\|\|\|]PAM | NA | Intergenic | NA | NA |
| T33771 | GGTATTTACATACTCAGGACAGG | 1000 | | | | | | | | | |
| Oligo fwd | CACCGGTATTTACATACTCAGGAC | | | | | | | | | | |
| Oligo rev | AAACGTCCTGAGTATGTAAATACC | | | | | | | | | | |
| Chromosome | start | end | strand | MM | target_seq | PAM | alignment | distance | position | gene name | gene id |
| GU305914.1 | 8560 | 8582 | - | 0 | GGTATTTACATACTCAGGAC | AGG | \|\|\|\|\|\|\|\|[\|\|\|\|\|\|\|\|\|\|\|\|]PAM | NA | Intergenic | NA | NA |
| T33772 | TTACATACTCAGGACAGGTGTGG | 1000 | | | | | | | | | |

<table>
<tr><td>Oligo fwd</td><td>CACCTTACATACTCAGGACAGGTG</td><td></td><td></td><td></td><td></td><td></td><td></td><td></td><td></td><td></td><td></td></tr>
<tr><td>Oligo rev</td><td>AAACCACCTGTCCTGAGTATGTAA</td><td></td><td></td><td></td><td></td><td></td><td></td><td></td><td></td><td></td><td></td></tr>
<tr><td>Chromosome</td><td>start</td><td>end</td><td>strand</td><td>MM</td><td>target_seq</td><td>PAM</td><td>alignment</td><td>distance</td><td>position</td><td>gene name</td><td>gene id</td></tr>
<tr><td>GU305914.1</td><td>8555</td><td>8577</td><td>-</td><td>0</td><td>TTACATACTCAGGACAGGTG</td><td>TGG</td><td>||||||||[||||||||||||]PAM</td><td>NA</td><td>Intergenic</td><td>NA</td><td>NA</td></tr>
<tr><td>T33773</td><td>TACATACTCAGGACAGGTGTGGG</td><td>1000</td><td></td><td></td><td></td><td></td><td></td><td></td><td></td><td></td><td></td></tr>
<tr><td>Oligo fwd</td><td>CACCTACATACTCAGGACAGGTGT</td><td></td><td></td><td></td><td></td><td></td><td></td><td></td><td></td><td></td><td></td></tr>
<tr><td>Oligo rev</td><td>AAACACACCTGTCCTGAGTATGTA</td><td></td><td></td><td></td><td></td><td></td><td></td><td></td><td></td><td></td><td></td></tr>
<tr><td>Chromosome</td><td>start</td><td>end</td><td>strand</td><td>MM</td><td>target_seq</td><td>PAM</td><td>alignment</td><td>distance</td><td>position</td><td>gene name</td><td>gene id</td></tr>
<tr><td>GU305914.1</td><td>8554</td><td>8576</td><td>-</td><td>0</td><td>TACATACTCAGGACAGGTGT</td><td>GGG</td><td>||||||||[||||||||||||]PAM</td><td>NA</td><td>Intergenic</td><td>NA</td><td>NA</td></tr>
<tr><td>T33774</td><td>ATACTCAGGACAGGTGTGGGCGG</td><td>1000</td><td></td><td></td><td></td><td></td><td></td><td></td><td></td><td></td><td></td></tr>
<tr><td>Oligo fwd</td><td>CACCATACTCAGGACAGGTGTGGG</td><td></td><td></td><td></td><td></td><td></td><td></td><td></td><td></td><td></td><td></td></tr>
<tr><td>Oligo rev</td><td>AAACCCCACACCTGTCCTGAGTAT</td><td></td><td></td><td></td><td></td><td></td><td></td><td></td><td></td><td></td><td></td></tr>
<tr><td>Chromosome</td><td>start</td><td>end</td><td>strand</td><td>MM</td><td>target_seq</td><td>PAM</td><td>alignment</td><td>distance</td><td>position</td><td>gene name</td><td>gene id</td></tr>
<tr><td>GU305914.1</td><td>8551</td><td>8573</td><td>-</td><td>0</td><td>ATACTCAGGACAGGTGTGGG</td><td>CGG</td><td>||||||||[||||||||||||]PAM</td><td>NA</td><td>Intergenic</td><td>NA</td><td>NA</td></tr>
<tr><td>GU305914.1</td><td>7381</td><td>7403</td><td>-</td><td>3</td><td>ATACTCACCGCAGGTGTGGG</td><td>CGG</td><td>|||||||-[--||||||||||]PAM</td><td>NA</td><td>Intergenic</td><td>NA</td><td>NA</td></tr>
<tr><td>T33775</td><td>GGACAGGTGTGGGCGGTTCCAGG</td><td>1000</td><td></td><td></td><td></td><td></td><td></td><td></td><td></td><td></td><td></td></tr>
<tr><td>Oligo fwd</td><td>CACCGGACAGGTGTGGGCGGTTCC</td><td></td><td></td><td></td><td></td><td></td><td></td><td></td><td></td><td></td><td></td></tr>
<tr><td>Oligo rev</td><td>AAACGGAACCGCCCACACCTGTCC</td><td></td><td></td><td></td><td></td><td></td><td></td><td></td><td></td><td></td><td></td></tr>
<tr><td>Chromosome</td><td>start</td><td>end</td><td>strand</td><td>MM</td><td>target_seq</td><td>PAM</td><td>alignment</td><td>distance</td><td>position</td><td>gene name</td><td>gene id</td></tr>
<tr><td>GU305914.1</td><td>8544</td><td>8566</td><td>-</td><td>0</td><td>GGACAGGTGTGGGCGGTTCC</td><td>AGG</td><td>||||||||[||||||||||||]PAM</td><td>NA</td><td>Intergenic</td><td>NA</td><td>NA</td></tr>
<tr><td>GU305914.1</td><td>9537</td><td>9559</td><td>-</td><td>0</td><td>GGACAGGTGTGGGCGGTTCC</td><td>AGG</td><td>||||||||[||||||||||||]PAM</td><td>NA</td><td>Intergenic</td><td>NA</td><td>NA</td></tr>
<tr><td>T33776</td><td>ACGTAAGCAGATTAAGACGCAGG</td><td>1000</td><td></td><td></td><td></td><td></td><td></td><td></td><td></td><td></td><td></td></tr>
<tr><td>Oligo fwd</td><td>CACCACGTAAGCAGATTAAGACGC</td><td></td><td></td><td></td><td></td><td></td><td></td><td></td><td></td><td></td><td></td></tr>
<tr><td>Oligo rev</td><td>AAACGCGTCTTAATCTGCTTACGT</td><td></td><td></td><td></td><td></td><td></td><td></td><td></td><td></td><td></td><td></td></tr>
<tr><td>Chromosome</td><td>start</td><td>end</td><td>strand</td><td>MM</td><td>target_seq</td><td>PAM</td><td>alignment</td><td>distance</td><td>position</td><td>gene name</td><td>gene id</td></tr>
<tr><td>GU305914.1</td><td>8518</td><td>8540</td><td>-</td><td>0</td><td>ACGTAAGCAGATTAAGACGC</td><td>AGG</td><td>||||||||[||||||||||||]PAM</td><td>NA</td><td>Intergenic</td><td>NA</td><td>NA</td></tr>
<tr><td>T33777</td><td>AGCAGATTAAGACGCAGGAGAGG</td><td>1000</td><td></td><td></td><td></td><td></td><td></td><td></td><td></td><td></td><td></td></tr>
<tr><td>Oligo fwd</td><td>CACCAGCAGATTAAGACGCAGGAG</td><td></td><td></td><td></td><td></td><td></td><td></td><td></td><td></td><td></td><td></td></tr>
<tr><td>Oligo rev</td><td>AAACCTCCTGCGTCTTAATCTGCT</td><td></td><td></td><td></td><td></td><td></td><td></td><td></td><td></td><td></td><td></td></tr>
<tr><td>Chromosome</td><td>start</td><td>end</td><td>strand</td><td>MM</td><td>target_seq</td><td>PAM</td><td>alignment</td><td>distance</td><td>position</td><td>gene name</td><td>gene id</td></tr>
<tr><td>GU305914.1</td><td>8513</td><td>8535</td><td>-</td><td>0</td><td>AGCAGATTAAGACGCAGGAG</td><td>AGG</td><td>||||||||[||||||||||||]PAM</td><td>NA</td><td>Intergenic</td><td>NA</td><td>NA</td></tr>
<tr><td>T33778</td><td>ACGCAGGAGAGGATGTTGTACGG</td><td>1000</td><td></td><td></td><td></td><td></td><td></td><td></td><td></td><td></td><td></td></tr>
<tr><td>Oligo fwd</td><td>CACCACGCAGGAGAGGATGTTGTA</td><td></td><td></td><td></td><td></td><td></td><td></td><td></td><td></td><td></td><td></td></tr>
<tr><td>Oligo rev</td><td>AAACTACAACATCCTCTCCTGCGT</td><td></td><td></td><td></td><td></td><td></td><td></td><td></td><td></td><td></td><td></td></tr>
<tr><td>Chromosome</td><td>start</td><td>end</td><td>strand</td><td>MM</td><td>target_seq</td><td>PAM</td><td>alignment</td><td>distance</td><td>position</td><td>gene name</td><td>gene id</td></tr>
<tr><td>GU305914.1</td><td>8502</td><td>8524</td><td>-</td><td>0</td><td>ACGCAGGAGAGGATGTTGTA</td><td>CGG</td><td>||||||||[||||||||||||]PAM</td><td>NA</td><td>Intergenic</td><td>NA</td><td>NA</td></tr>
<tr><td>T33779</td><td>GGAGAGGATGTTGTACGGAGAGG</td><td>1000</td><td></td><td></td><td></td><td></td><td></td><td></td><td></td><td></td><td></td></tr>
<tr><td>Oligo fwd</td><td>CACCGGAGAGGATGTTGTACGGAG</td><td></td><td></td><td></td><td></td><td></td><td></td><td></td><td></td><td></td><td></td></tr>
<tr><td>Oligo rev</td><td>AAACCTCCGTACAACATCCTCTCC</td><td></td><td></td><td></td><td></td><td></td><td></td><td></td><td></td><td></td><td></td></tr>
<tr><td>Chromosome</td><td>start</td><td>end</td><td>strand</td><td>MM</td><td>target_seq</td><td>PAM</td><td>alignment</td><td>distance</td><td>position</td><td>gene name</td><td>gene id</td></tr>
<tr><td>GU305914.1</td><td>8497</td><td>8519</td><td>-</td><td>0</td><td>GGAGAGGATGTTGTACGGAG</td><td>AGG</td><td>||||||||[||||||||||||]PAM</td><td>NA</td><td>Intergenic</td><td>NA</td><td>NA</td></tr>
<tr><td>T33780</td><td>GGATGTTGTACGGAGAGGCTCGG</td><td>1000</td><td></td><td></td><td></td><td></td><td></td><td></td><td></td><td></td><td></td></tr>
<tr><td>Oligo fwd</td><td>CACCGGATGTTGTACGGAGAGGCT</td><td></td><td></td><td></td><td></td><td></td><td></td><td></td><td></td><td></td><td></td></tr>
<tr><td>Oligo rev</td><td>AAACAGCCTCTCCGTACAACATCC</td><td></td><td></td><td></td><td></td><td></td><td></td><td></td><td></td><td></td><td></td></tr>
<tr><td>Chromosome</td><td>start</td><td>end</td><td>strand</td><td>MM</td><td>target_seq</td><td>PAM</td><td>alignment</td><td>distance</td><td>position</td><td>gene name</td><td>gene id</td></tr>
<tr><td>GU305914.1</td><td>8492</td><td>8514</td><td>-</td><td>0</td><td>GGATGTTGTACGGAGAGGCT</td><td>CGG</td><td>||||||||[||||||||||||]PAM</td><td>NA</td><td>Intergenic</td><td>NA</td><td>NA</td></tr>
<tr><td>GU305914.1</td><td>9476</td><td>9498</td><td>-</td><td>4</td><td>TGCCGTTGAACGGAGAGGCT</td><td>CGG</td><td>-|--||||[-|||||||||||]PAM</td><td>NA</td><td>Intergenic</td><td>NA</td><td>NA</td></tr>
<tr><td>T33781</td><td>ACGGAGAGGCTCGGATTCATCGG</td><td>1000</td><td></td><td></td><td></td><td></td><td></td><td></td><td></td><td></td><td></td></tr>
<tr><td>Oligo fwd</td><td>CACCACGGAGAGGCTCGGATTCAT</td><td></td><td></td><td></td><td></td><td></td><td></td><td></td><td></td><td></td><td></td></tr>
<tr><td>Oligo rev</td><td>AAACATGAATCCGAGCCTCTCCGT</td><td></td><td></td><td></td><td></td><td></td><td></td><td></td><td></td><td></td><td></td></tr>
<tr><td>Chromosome</td><td>start</td><td>end</td><td>strand</td><td>MM</td><td>target_seq</td><td>PAM</td><td>alignment</td><td>distance</td><td>position</td><td>gene name</td><td>gene id</td></tr>
<tr><td>GU305914.1</td><td>8483</td><td>8505</td><td>-</td><td>0</td><td>ACGGAGAGGCTCGGATTCAT</td><td>CGG</td><td>||||||||[||||||||||||]PAM</td><td>NA</td><td>Intergenic</td><td>NA</td><td>NA</td></tr>
<tr><td>GU305914.1</td><td>9467</td><td>9489</td><td>-</td><td>2</td><td>ACGGAGAGGCTCGGTTTCAA</td><td>CGG</td><td>||||||||[||||||-||||-]PAM</td><td>NA</td><td>Intergenic</td><td>NA</td><td>NA</td></tr>
<tr><td>T33782</td><td>AGAGGCTCGGATTCATCGGTTGG</td><td>1000</td><td></td><td></td><td></td><td></td><td></td><td></td><td></td><td></td><td></td></tr>
<tr><td>Oligo fwd</td><td>CACCAGAGGCTCGGATTCATCGGT</td><td></td><td></td><td></td><td></td><td></td><td></td><td></td><td></td><td></td><td></td></tr>
<tr><td>Oligo rev</td><td>AAACACCGATGAATCCGAGCCTCT</td><td></td><td></td><td></td><td></td><td></td><td></td><td></td><td></td><td></td><td></td></tr>
<tr><td>Chromosome</td><td>start</td><td>end</td><td>strand</td><td>MM</td><td>target_seq</td><td>PAM</td><td>alignment</td><td>distance</td><td>position</td><td>gene name</td><td>gene id</td></tr>
<tr><td>GU305914.1</td><td>8479</td><td>8501</td><td>-</td><td>0</td><td>AGAGGCTCGGATTCATCGGT</td><td>TGG</td><td>||||||||[||||||||||||]PAM</td><td>NA</td><td>Intergenic</td><td>NA</td><td>NA</td></tr>
<tr><td>GU305914.1</td><td>9463</td><td>9485</td><td>-</td><td>2</td><td>AGAGGCTCGGTTTCAACGGT</td><td>TGG</td><td>||||||||[||-||||-||||]PAM</td><td>NA</td><td>Intergenic</td><td>NA</td><td>NA</td></tr>
<tr><td>T33783</td><td>TCGGTTGGTCTAAAAGTCTCTGG</td><td>1000</td><td></td><td></td><td></td><td></td><td></td><td></td><td></td><td></td><td></td></tr>
<tr><td>Oligo fwd</td><td>CACCTCGGTTGGTCTAAAAGTCTC</td><td></td><td></td><td></td><td></td><td></td><td></td><td></td><td></td><td></td><td></td></tr>
<tr><td>Oligo rev</td><td>AAACGAGACTTTTAGACCAACCGA</td><td></td><td></td><td></td><td></td><td></td><td></td><td></td><td></td><td></td><td></td></tr>
<tr><td>Chromosome</td><td>start</td><td>end</td><td>strand</td><td>MM</td><td>target_seq</td><td>PAM</td><td>alignment</td><td>distance</td><td>position</td><td>gene name</td><td>gene id</td></tr>
<tr><td>GU305914.1</td><td>8464</td><td>8486</td><td>-</td><td>0</td><td>TCGGTTGGTCTAAAAGTCTC</td><td>TGG</td><td>||||||||[||||||||||||]PAM</td><td>NA</td><td>Intergenic</td><td>NA</td><td>NA</td></tr>
<tr><td>T33784</td><td>CGCGAGAACTGAAACGAGCACGG</td><td>1000</td><td></td><td></td><td></td><td></td><td></td><td></td><td></td><td></td><td></td></tr>
<tr><td>Oligo fwd</td><td>CACCCGCGAGAACTGAAACGAGCA</td><td></td><td></td><td></td><td></td><td></td><td></td><td></td><td></td><td></td><td></td></tr>
<tr><td>Oligo rev</td><td>AAACTGCTCGTTTCAGTTCTCGCG</td><td></td><td></td><td></td><td></td><td></td><td></td><td></td><td></td><td></td><td></td></tr>
<tr><td>Chromosome</td><td>start</td><td>end</td><td>strand</td><td>MM</td><td>target_seq</td><td>PAM</td><td>alignment</td><td>distance</td><td>position</td><td>gene name</td><td>gene id</td></tr>
<tr><td>GU305914.1</td><td>8439</td><td>8461</td><td>-</td><td>0</td><td>CGCGAGAACTGAAACGAGCA</td><td>CGG</td><td>||||||||[||||||||||||]PAM</td><td>NA</td><td>Intergenic</td><td>NA</td><td>NA</td></tr>
<tr><td>T33785</td><td>GAAACGAGCACGGAGACGTTCGG</td><td>1000</td><td></td><td></td><td></td><td></td><td></td><td></td><td></td><td></td><td></td></tr>
<tr><td>Oligo fwd</td><td>CACCGAAACGAGCACGGAGACGTT</td><td></td><td></td><td></td><td></td><td></td><td></td><td></td><td></td><td></td><td></td></tr>
<tr><td>Oligo rev</td><td>AAACAACGTCTCCGTGCTCGTTTC</td><td></td><td></td><td></td><td></td><td></td><td></td><td></td><td></td><td></td><td></td></tr>
<tr><td>Chromosome</td><td>start</td><td>end</td><td>strand</td><td>MM</td><td>target_seq</td><td>PAM</td><td>alignment</td><td>distance</td><td>position</td><td>gene name</td><td>gene id</td></tr>
<tr><td>GU305914.1</td><td>8429</td><td>8451</td><td>-</td><td>0</td><td>GAAACGAGCACGGAGACGTT</td><td>CGG</td><td>||||||||[||||||||||||]PAM</td><td>NA</td><td>Intergenic</td><td>NA</td><td>NA</td></tr>
<tr><td>T33786</td><td>GACGTTCGGCGTACAAATCGAGG</td><td>1000</td><td></td><td></td><td></td><td></td><td></td><td></td><td></td><td></td><td></td></tr>
</table>

<table>
<tr><td>Oligo fwd</td><td>CACCGACGTTCGGCGTACAAATCG</td><td></td><td></td><td></td><td></td><td></td><td></td><td></td><td></td><td></td><td></td></tr>
<tr><td>Oligo rev</td><td>AAACCGATTTGTACGCCGAACGTC</td><td></td><td></td><td></td><td></td><td></td><td></td><td></td><td></td><td></td><td></td></tr>
<tr><td>Chromosome</td><td>start</td><td>end</td><td>strand</td><td>MM</td><td>target_seq</td><td>PAM</td><td>alignment</td><td>distance</td><td>position</td><td>gene name</td><td>gene id</td></tr>
<tr><td>GU305914.1</td><td>8415</td><td>8437</td><td>-</td><td>0</td><td>GACGTTCGGCGTACAAATCG</td><td>AGG</td><td>||||||||[||||||||||||]PAM</td><td>NA</td><td>Intergenic</td><td>NA</td><td>NA</td></tr>
<tr><td>T33787</td><td>GATCGTCACCGCTACCGACACGG</td><td>1000</td><td></td><td></td><td></td><td></td><td></td><td></td><td></td><td></td><td></td></tr>
<tr><td>Oligo fwd</td><td>CACCGATCGTCACCGCTACCGACA</td><td></td><td></td><td></td><td></td><td></td><td></td><td></td><td></td><td></td><td></td></tr>
<tr><td>Oligo rev</td><td>AAACTGTCGGTAGCGGTGACGATC</td><td></td><td></td><td></td><td></td><td></td><td></td><td></td><td></td><td></td><td></td></tr>
<tr><td>Chromosome</td><td>start</td><td>end</td><td>strand</td><td>MM</td><td>target_seq</td><td>PAM</td><td>alignment</td><td>distance</td><td>position</td><td>gene name</td><td>gene id</td></tr>
<tr><td>GU305914.1</td><td>8327</td><td>8349</td><td>-</td><td>0</td><td>GATCGTCACCGCTACCGACA</td><td>CGG</td><td>||||||||[||||||||||||]PAM</td><td>NA</td><td>Intergenic</td><td>NA</td><td>NA</td></tr>
<tr><td>T33788</td><td>ATGCGAAGTCTGCATCTCTGCGG</td><td>1000</td><td></td><td></td><td></td><td></td><td></td><td></td><td></td><td></td><td></td></tr>
<tr><td>Oligo fwd</td><td>CACCATGCGAAGTCTGCATCTCTG</td><td></td><td></td><td></td><td></td><td></td><td></td><td></td><td></td><td></td><td></td></tr>
<tr><td>Oligo rev</td><td>AAACCAGAGATGCAGACTTCGCAT</td><td></td><td></td><td></td><td></td><td></td><td></td><td></td><td></td><td></td><td></td></tr>
<tr><td>Chromosome</td><td>start</td><td>end</td><td>strand</td><td>MM</td><td>target_seq</td><td>PAM</td><td>alignment</td><td>distance</td><td>position</td><td>gene name</td><td>gene id</td></tr>
<tr><td>GU305914.1</td><td>8291</td><td>8313</td><td>-</td><td>0</td><td>ATGCGAAGTCTGCATCTCTG</td><td>CGG</td><td>||||||||[||||||||||||]PAM</td><td>NA</td><td>Intergenic</td><td>NA</td><td>NA</td></tr>
<tr><td>T33789</td><td>AGTCTGCATCTCTGCGGTTGCGG</td><td>1000</td><td></td><td></td><td></td><td></td><td></td><td></td><td></td><td></td><td></td></tr>
<tr><td>Oligo fwd</td><td>CACCAGTCTGCATCTCTGCGGTTG</td><td></td><td></td><td></td><td></td><td></td><td></td><td></td><td></td><td></td><td></td></tr>
<tr><td>Oligo rev</td><td>AAACCAACCGCAGAGATGCAGACT</td><td></td><td></td><td></td><td></td><td></td><td></td><td></td><td></td><td></td><td></td></tr>
<tr><td>Chromosome</td><td>start</td><td>end</td><td>strand</td><td>MM</td><td>target_seq</td><td>PAM</td><td>alignment</td><td>distance</td><td>position</td><td>gene name</td><td>gene id</td></tr>
<tr><td>GU305914.1</td><td>8285</td><td>8307</td><td>-</td><td>0</td><td>AGTCTGCATCTCTGCGGTTG</td><td>CGG</td><td>||||||||[||||||||||||]PAM</td><td>NA</td><td>Intergenic</td><td>NA</td><td>NA</td></tr>
<tr><td>T33790</td><td>CATCTCTGCGGTTGCGGCGACGG</td><td>1000</td><td></td><td></td><td></td><td></td><td></td><td></td><td></td><td></td><td></td></tr>
<tr><td>Oligo fwd</td><td>CACCCATCTCTGCGGTTGCGGCGA</td><td></td><td></td><td></td><td></td><td></td><td></td><td></td><td></td><td></td><td></td></tr>
<tr><td>Oligo rev</td><td>AAACTCGCCGCAACCGCAGAGATG</td><td></td><td></td><td></td><td></td><td></td><td></td><td></td><td></td><td></td><td></td></tr>
<tr><td>Chromosome</td><td>start</td><td>end</td><td>strand</td><td>MM</td><td>target_seq</td><td>PAM</td><td>alignment</td><td>distance</td><td>position</td><td>gene name</td><td>gene id</td></tr>
<tr><td>GU305914.1</td><td>8279</td><td>8301</td><td>-</td><td>0</td><td>CATCTCTGCGGTTGCGGCGA</td><td>CGG</td><td>||||||||[||||||||||||]PAM</td><td>NA</td><td>Intergenic</td><td>NA</td><td>NA</td></tr>
<tr><td>T33791</td><td>GACGGTATCGTCTGCGTTGTCGG</td><td>1000</td><td></td><td></td><td></td><td></td><td></td><td></td><td></td><td></td><td></td></tr>
<tr><td>Oligo fwd</td><td>CACCGACGGTATCGTCTGCGTTGT</td><td></td><td></td><td></td><td></td><td></td><td></td><td></td><td></td><td></td><td></td></tr>
<tr><td>Oligo rev</td><td>AAACACAACGCAGACGATACCGTC</td><td></td><td></td><td></td><td></td><td></td><td></td><td></td><td></td><td></td><td></td></tr>
<tr><td>Chromosome</td><td>start</td><td>end</td><td>strand</td><td>MM</td><td>target_seq</td><td>PAM</td><td>alignment</td><td>distance</td><td>position</td><td>gene name</td><td>gene id</td></tr>
<tr><td>GU305914.1</td><td>8261</td><td>8283</td><td>-</td><td>0</td><td>GACGGTATCGTCTGCGTTGT</td><td>CGG</td><td>||||||||[||||||||||||]PAM</td><td>NA</td><td>Intergenic</td><td>NA</td><td>NA</td></tr>
<tr><td>T33792</td><td>ACGGTATCGTCTGCGTTGTCGGG</td><td>1000</td><td></td><td></td><td></td><td></td><td></td><td></td><td></td><td></td><td></td></tr>
<tr><td>Oligo fwd</td><td>CACCACGGTATCGTCTGCGTTGTC</td><td></td><td></td><td></td><td></td><td></td><td></td><td></td><td></td><td></td><td></td></tr>
<tr><td>Oligo rev</td><td>AAACGACAACGCAGACGATACCGT</td><td></td><td></td><td></td><td></td><td></td><td></td><td></td><td></td><td></td><td></td></tr>
<tr><td>Chromosome</td><td>start</td><td>end</td><td>strand</td><td>MM</td><td>target_seq</td><td>PAM</td><td>alignment</td><td>distance</td><td>position</td><td>gene name</td><td>gene id</td></tr>
<tr><td>GU305914.1</td><td>8260</td><td>8282</td><td>-</td><td>0</td><td>ACGGTATCGTCTGCGTTGTC</td><td>GGG</td><td>||||||||[||||||||||||]PAM</td><td>NA</td><td>Intergenic</td><td>NA</td><td>NA</td></tr>
<tr><td>T33793</td><td>CGGTATCGTCTGCGTTGTCGGGG</td><td>1000</td><td></td><td></td><td></td><td></td><td></td><td></td><td></td><td></td><td></td></tr>
<tr><td>Oligo fwd</td><td>CACCCGGTATCGTCTGCGTTGTCG</td><td></td><td></td><td></td><td></td><td></td><td></td><td></td><td></td><td></td><td></td></tr>
<tr><td>Oligo rev</td><td>AAACCGACAACGCAGACGATACCG</td><td></td><td></td><td></td><td></td><td></td><td></td><td></td><td></td><td></td><td></td></tr>
<tr><td>Chromosome</td><td>start</td><td>end</td><td>strand</td><td>MM</td><td>target_seq</td><td>PAM</td><td>alignment</td><td>distance</td><td>position</td><td>gene name</td><td>gene id</td></tr>
<tr><td>GU305914.1</td><td>8259</td><td>8281</td><td>-</td><td>0</td><td>CGGTATCGTCTGCGTTGTCG</td><td>GGG</td><td>||||||||[||||||||||||]PAM</td><td>NA</td><td>Intergenic</td><td>NA</td><td>NA</td></tr>
<tr><td>T33794</td><td>ATCGTCTGCGTTGTCGGGGCCGG</td><td>1000</td><td></td><td></td><td></td><td></td><td></td><td></td><td></td><td></td><td></td></tr>
<tr><td>Oligo fwd</td><td>CACCATCGTCTGCGTTGTCGGGGC</td><td></td><td></td><td></td><td></td><td></td><td></td><td></td><td></td><td></td><td></td></tr>
<tr><td>Oligo rev</td><td>AAACGCCCCGACAACGCAGACGAT</td><td></td><td></td><td></td><td></td><td></td><td></td><td></td><td></td><td></td><td></td></tr>
<tr><td>Chromosome</td><td>start</td><td>end</td><td>strand</td><td>MM</td><td>target_seq</td><td>PAM</td><td>alignment</td><td>distance</td><td>position</td><td>gene name</td><td>gene id</td></tr>
<tr><td>GU305914.1</td><td>8255</td><td>8277</td><td>-</td><td>0</td><td>ATCGTCTGCGTTGTCGGGGC</td><td>CGG</td><td>||||||||[||||||||||||]PAM</td><td>NA</td><td>Intergenic</td><td>NA</td><td>NA</td></tr>
<tr><td>T33795</td><td>TCGTCTGCGTTGTCGGGGCCGGG</td><td>1000</td><td></td><td></td><td></td><td></td><td></td><td></td><td></td><td></td><td></td></tr>
<tr><td>Oligo fwd</td><td>CACCTCGTCTGCGTTGTCGGGGCC</td><td></td><td></td><td></td><td></td><td></td><td></td><td></td><td></td><td></td><td></td></tr>
<tr><td>Oligo rev</td><td>AAACGGCCCCGACAACGCAGACGA</td><td></td><td></td><td></td><td></td><td></td><td></td><td></td><td></td><td></td><td></td></tr>
<tr><td>Chromosome</td><td>start</td><td>end</td><td>strand</td><td>MM</td><td>target_seq</td><td>PAM</td><td>alignment</td><td>distance</td><td>position</td><td>gene name</td><td>gene id</td></tr>
<tr><td>GU305914.1</td><td>8254</td><td>8276</td><td>-</td><td>0</td><td>TCGTCTGCGTTGTCGGGGCC</td><td>GGG</td><td>||||||||[||||||||||||]PAM</td><td>NA</td><td>Intergenic</td><td>NA</td><td>NA</td></tr>
<tr><td>T33796</td><td>GGGGCCGGGAAAAAGAGTTGTGG</td><td>1000</td><td></td><td></td><td></td><td></td><td></td><td></td><td></td><td></td><td></td></tr>
<tr><td>Oligo fwd</td><td>CACCGGGGCCGGGAAAAAGAGTTG</td><td></td><td></td><td></td><td></td><td></td><td></td><td></td><td></td><td></td><td></td></tr>
<tr><td>Oligo rev</td><td>AAACCAACTCTTTTTCCCGGCCCC</td><td></td><td></td><td></td><td></td><td></td><td></td><td></td><td></td><td></td><td></td></tr>
<tr><td>Chromosome</td><td>start</td><td>end</td><td>strand</td><td>MM</td><td>target_seq</td><td>PAM</td><td>alignment</td><td>distance</td><td>position</td><td>gene name</td><td>gene id</td></tr>
<tr><td>GU305914.1</td><td>8240</td><td>8262</td><td>-</td><td>0</td><td>GGGGCCGGGAAAAAGAGTTG</td><td>TGG</td><td>||||||||[||||||||||||]PAM</td><td>NA</td><td>Intergenic</td><td>NA</td><td>NA</td></tr>
<tr><td>T33797</td><td>GGGCCGGGAAAAAGAGTTGTGGG</td><td>1000</td><td></td><td></td><td></td><td></td><td></td><td></td><td></td><td></td><td></td></tr>
<tr><td>Oligo fwd</td><td>CACCGGGCCGGGAAAAAGAGTTGT</td><td></td><td></td><td></td><td></td><td></td><td></td><td></td><td></td><td></td><td></td></tr>
<tr><td>Oligo rev</td><td>AAACACAACTCTTTTTCCCGGCCC</td><td></td><td></td><td></td><td></td><td></td><td></td><td></td><td></td><td></td><td></td></tr>
<tr><td>Chromosome</td><td>start</td><td>end</td><td>strand</td><td>MM</td><td>target_seq</td><td>PAM</td><td>alignment</td><td>distance</td><td>position</td><td>gene name</td><td>gene id</td></tr>
<tr><td>GU305914.1</td><td>8239</td><td>8261</td><td>-</td><td>0</td><td>GGGCCGGGAAAAAGAGTTGT</td><td>GGG</td><td>||||||||[||||||||||||]PAM</td><td>NA</td><td>Intergenic</td><td>NA</td><td>NA</td></tr>
<tr><td>T33798</td><td>GGCCGGGAAAAAGAGTTGTGGGG</td><td>1000</td><td></td><td></td><td></td><td></td><td></td><td></td><td></td><td></td><td></td></tr>
<tr><td>Oligo fwd</td><td>CACCGGCCGGGAAAAAGAGTTGTG</td><td></td><td></td><td></td><td></td><td></td><td></td><td></td><td></td><td></td><td></td></tr>
<tr><td>Oligo rev</td><td>AAACCACAACTCTTTTTCCCGGCC</td><td></td><td></td><td></td><td></td><td></td><td></td><td></td><td></td><td></td><td></td></tr>
<tr><td>Chromosome</td><td>start</td><td>end</td><td>strand</td><td>MM</td><td>target_seq</td><td>PAM</td><td>alignment</td><td>distance</td><td>position</td><td>gene name</td><td>gene id</td></tr>
<tr><td>GU305914.1</td><td>8238</td><td>8260</td><td>-</td><td>0</td><td>GGCCGGGAAAAAGAGTTGTG</td><td>GGG</td><td>||||||||[||||||||||||]PAM</td><td>NA</td><td>Intergenic</td><td>NA</td><td>NA</td></tr>
<tr><td>T33799</td><td>TAGAAAACTTAACGAGCGCCTGG</td><td>1000</td><td></td><td></td><td></td><td></td><td></td><td></td><td></td><td></td><td></td></tr>
<tr><td>Oligo fwd</td><td>CACCTAGAAAACTTAACGAGCGCC</td><td></td><td></td><td></td><td></td><td></td><td></td><td></td><td></td><td></td><td></td></tr>
<tr><td>Oligo rev</td><td>AAACGGCGCTCGTTAAGTTTTCTA</td><td></td><td></td><td></td><td></td><td></td><td></td><td></td><td></td><td></td><td></td></tr>
<tr><td>Chromosome</td><td>start</td><td>end</td><td>strand</td><td>MM</td><td>target_seq</td><td>PAM</td><td>alignment</td><td>distance</td><td>position</td><td>gene name</td><td>gene id</td></tr>
<tr><td>GU305914.1</td><td>8208</td><td>8230</td><td>-</td><td>0</td><td>TAGAAAACTTAACGAGCGCC</td><td>TGG</td><td>||||||||[||||||||||||]PAM</td><td>NA</td><td>Intergenic</td><td>NA</td><td>NA</td></tr>
<tr><td>T33800</td><td>AGAAAACTTAACGAGCGCCTGGG</td><td>1000</td><td></td><td></td><td></td><td></td><td></td><td></td><td></td><td></td><td></td></tr>
<tr><td>Oligo fwd</td><td>CACCAGAAAACTTAACGAGCGCCT</td><td></td><td></td><td></td><td></td><td></td><td></td><td></td><td></td><td></td><td></td></tr>
<tr><td>Oligo rev</td><td>AAACAGGCGCTCGTTAAGTTTTCT</td><td></td><td></td><td></td><td></td><td></td><td></td><td></td><td></td><td></td><td></td></tr>
<tr><td>Chromosome</td><td>start</td><td>end</td><td>strand</td><td>MM</td><td>target_seq</td><td>PAM</td><td>alignment</td><td>distance</td><td>position</td><td>gene name</td><td>gene id</td></tr>
<tr><td>GU305914.1</td><td>8207</td><td>8229</td><td>-</td><td>0</td><td>AGAAAACTTAACGAGCGCCT</td><td>GGG</td><td>||||||||[||||||||||||]PAM</td><td>NA</td><td>Intergenic</td><td>NA</td><td>NA</td></tr>
</table>

<table>
<tr><td>T33801</td><td>AACGAGCGCCTGGGTGATCGAGG</td><td></td><td>1000</td><td></td><td></td><td></td><td></td><td></td><td></td><td></td><td></td><td></td></tr>
<tr><td>Oligo fwd</td><td>CACCAACGAGCGCCTGGGTGATCG</td><td></td><td></td><td></td><td></td><td></td><td></td><td></td><td></td><td></td><td></td><td></td></tr>
<tr><td>Oligo rev</td><td>AAACCGATCACCCAGGCGCTCGTT</td><td></td><td></td><td></td><td></td><td></td><td></td><td></td><td></td><td></td><td></td><td></td></tr>
<tr><td>Chromosome</td><td>start</td><td></td><td>end</td><td>strand</td><td>MM</td><td>target_seq</td><td>PAM</td><td>alignment</td><td>distance</td><td>position</td><td>gene name</td><td>gene id</td></tr>
<tr><td>GU305914.1</td><td></td><td>8198</td><td>8220</td><td>-</td><td>0</td><td>AACGAGCGCCTGGGTGATCG</td><td>AGG</td><td>||||||||[||||||||||||]PAM</td><td>NA</td><td>Intergenic</td><td>NA</td><td>NA</td></tr>
<tr><td>T33802</td><td>GGGTGATCGAGGTATGTTTTCGG</td><td></td><td>1000</td><td></td><td></td><td></td><td></td><td></td><td></td><td></td><td></td><td></td></tr>
<tr><td>Oligo fwd</td><td>CACCGGGTGATCGAGGTATGTTTT</td><td></td><td></td><td></td><td></td><td></td><td></td><td></td><td></td><td></td><td></td><td></td></tr>
<tr><td>Oligo rev</td><td>AAACAAAACATACCTCGATCACCC</td><td></td><td></td><td></td><td></td><td></td><td></td><td></td><td></td><td></td><td></td><td></td></tr>
<tr><td>Chromosome</td><td>start</td><td></td><td>end</td><td>strand</td><td>MM</td><td>target_seq</td><td>PAM</td><td>alignment</td><td>distance</td><td>position</td><td>gene name</td><td>gene id</td></tr>
<tr><td>GU305914.1</td><td></td><td>8187</td><td>8209</td><td>-</td><td>0</td><td>GGGTGATCGAGGTATGTTTT</td><td>CGG</td><td>||||||||[||||||||||||]PAM</td><td>NA</td><td>Intergenic</td><td>NA</td><td>NA</td></tr>
<tr><td>T33803</td><td>GGTGATCGAGGTATGTTTTCGGG</td><td></td><td>1000</td><td></td><td></td><td></td><td></td><td></td><td></td><td></td><td></td><td></td></tr>
<tr><td>Oligo fwd</td><td>CACCGGTGATCGAGGTATGTTTTC</td><td></td><td></td><td></td><td></td><td></td><td></td><td></td><td></td><td></td><td></td><td></td></tr>
<tr><td>Oligo rev</td><td>AAACGAAAACATACCTCGATCACC</td><td></td><td></td><td></td><td></td><td></td><td></td><td></td><td></td><td></td><td></td><td></td></tr>
<tr><td>Chromosome</td><td>start</td><td></td><td>end</td><td>strand</td><td>MM</td><td>target_seq</td><td>PAM</td><td>alignment</td><td>distance</td><td>position</td><td>gene name</td><td>gene id</td></tr>
<tr><td>GU305914.1</td><td></td><td>8186</td><td>8208</td><td>-</td><td>0</td><td>GGTGATCGAGGTATGTTTTC</td><td>GGG</td><td>||||||||[||||||||||||]PAM</td><td>NA</td><td>Intergenic</td><td>NA</td><td>NA</td></tr>
<tr><td>T33804</td><td>CGAGGTATGTTTTCGGGTGATGG</td><td></td><td>1000</td><td></td><td></td><td></td><td></td><td></td><td></td><td></td><td></td><td></td></tr>
<tr><td>Oligo fwd</td><td>CACCCGAGGTATGTTTTCGGGTGA</td><td></td><td></td><td></td><td></td><td></td><td></td><td></td><td></td><td></td><td></td><td></td></tr>
<tr><td>Oligo rev</td><td>AAACTCACCCGAAAACATACCTCG</td><td></td><td></td><td></td><td></td><td></td><td></td><td></td><td></td><td></td><td></td><td></td></tr>
<tr><td>Chromosome</td><td>start</td><td></td><td>end</td><td>strand</td><td>MM</td><td>target_seq</td><td>PAM</td><td>alignment</td><td>distance</td><td>position</td><td>gene name</td><td>gene id</td></tr>
<tr><td>GU305914.1</td><td></td><td>8180</td><td>8202</td><td>-</td><td>0</td><td>CGAGGTATGTTTTCGGGTGA</td><td>TGG</td><td>||||||||[||||||||||||]PAM</td><td>NA</td><td>Intergenic</td><td>NA</td><td>NA</td></tr>
<tr><td>T33805</td><td>GAGGTATGTTTTCGGGTGATGGG</td><td></td><td>1000</td><td></td><td></td><td></td><td></td><td></td><td></td><td></td><td></td><td></td></tr>
<tr><td>Oligo fwd</td><td>CACCGAGGTATGTTTTCGGGTGAT</td><td></td><td></td><td></td><td></td><td></td><td></td><td></td><td></td><td></td><td></td><td></td></tr>
<tr><td>Oligo rev</td><td>AAACATCACCCGAAAACATACCTC</td><td></td><td></td><td></td><td></td><td></td><td></td><td></td><td></td><td></td><td></td><td></td></tr>
<tr><td>Chromosome</td><td>start</td><td></td><td>end</td><td>strand</td><td>MM</td><td>target_seq</td><td>PAM</td><td>alignment</td><td>distance</td><td>position</td><td>gene name</td><td>gene id</td></tr>
<tr><td>GU305914.1</td><td></td><td>8179</td><td>8201</td><td>-</td><td>0</td><td>GAGGTATGTTTTCGGGTGAT</td><td>GGG</td><td>||||||||[||||||||||||]PAM</td><td>NA</td><td>Intergenic</td><td>NA</td><td>NA</td></tr>
<tr><td>T33806</td><td>ATGTTTTCGGGTGATGGGCTTGG</td><td></td><td>1000</td><td></td><td></td><td></td><td></td><td></td><td></td><td></td><td></td><td></td></tr>
<tr><td>Oligo fwd</td><td>CACCATGTTTTCGGGTGATGGGCT</td><td></td><td></td><td></td><td></td><td></td><td></td><td></td><td></td><td></td><td></td><td></td></tr>
<tr><td>Oligo rev</td><td>AAACAGCCCATCACCCGAAAACAT</td><td></td><td></td><td></td><td></td><td></td><td></td><td></td><td></td><td></td><td></td><td></td></tr>
<tr><td>Chromosome</td><td>start</td><td></td><td>end</td><td>strand</td><td>MM</td><td>target_seq</td><td>PAM</td><td>alignment</td><td>distance</td><td>position</td><td>gene name</td><td>gene id</td></tr>
<tr><td>GU305914.1</td><td></td><td>8174</td><td>8196</td><td>-</td><td>0</td><td>ATGTTTTCGGGTGATGGGCT</td><td>TGG</td><td>||||||||[||||||||||||]PAM</td><td>NA</td><td>Intergenic</td><td>NA</td><td>NA</td></tr>
<tr><td>T33807</td><td>TGTTTTCGGGTGATGGGCTTGGG</td><td></td><td>1000</td><td></td><td></td><td></td><td></td><td></td><td></td><td></td><td></td><td></td></tr>
<tr><td>Oligo fwd</td><td>CACCTGTTTTCGGGTGATGGGCTT</td><td></td><td></td><td></td><td></td><td></td><td></td><td></td><td></td><td></td><td></td><td></td></tr>
<tr><td>Oligo rev</td><td>AAACAAGCCCATCACCCGAAAACA</td><td></td><td></td><td></td><td></td><td></td><td></td><td></td><td></td><td></td><td></td><td></td></tr>
<tr><td>Chromosome</td><td>start</td><td></td><td>end</td><td>strand</td><td>MM</td><td>target_seq</td><td>PAM</td><td>alignment</td><td>distance</td><td>position</td><td>gene name</td><td>gene id</td></tr>
<tr><td>GU305914.1</td><td></td><td>8173</td><td>8195</td><td>-</td><td>0</td><td>TGTTTTCGGGTGATGGGCTT</td><td>GGG</td><td>||||||||[||||||||||||]PAM</td><td>NA</td><td>Intergenic</td><td>NA</td><td>NA</td></tr>
<tr><td>T33808</td><td>GTGATGGGCTTGGGTCGCGCCGG</td><td></td><td>1000</td><td></td><td></td><td></td><td></td><td></td><td></td><td></td><td></td><td></td></tr>
<tr><td>Oligo fwd</td><td>CACCGTGATGGGCTTGGGTCGCGC</td><td></td><td></td><td></td><td></td><td></td><td></td><td></td><td></td><td></td><td></td><td></td></tr>
<tr><td>Oligo rev</td><td>AAACGCGCGACCCAAGCCCATCAC</td><td></td><td></td><td></td><td></td><td></td><td></td><td></td><td></td><td></td><td></td><td></td></tr>
<tr><td>Chromosome</td><td>start</td><td></td><td>end</td><td>strand</td><td>MM</td><td>target_seq</td><td>PAM</td><td>alignment</td><td>distance</td><td>position</td><td>gene name</td><td>gene id</td></tr>
<tr><td>GU305914.1</td><td></td><td>8164</td><td>8186</td><td>-</td><td>0</td><td>GTGATGGGCTTGGGTCGCGC</td><td>CGG</td><td>||||||||[||||||||||||]PAM</td><td>NA</td><td>Intergenic</td><td>NA</td><td>NA</td></tr>
<tr><td>T33809</td><td>ATGGGCTTGGGTCGCGCCGGTGG</td><td></td><td>1000</td><td></td><td></td><td></td><td></td><td></td><td></td><td></td><td></td><td></td></tr>
<tr><td>Oligo fwd</td><td>CACCATGGGCTTGGGTCGCGCCGG</td><td></td><td></td><td></td><td></td><td></td><td></td><td></td><td></td><td></td><td></td><td></td></tr>
<tr><td>Oligo rev</td><td>AAACCCGGCGCGACCCAAGCCCAT</td><td></td><td></td><td></td><td></td><td></td><td></td><td></td><td></td><td></td><td></td><td></td></tr>
<tr><td>Chromosome</td><td>start</td><td></td><td>end</td><td>strand</td><td>MM</td><td>target_seq</td><td>PAM</td><td>alignment</td><td>distance</td><td>position</td><td>gene name</td><td>gene id</td></tr>
<tr><td>GU305914.1</td><td></td><td>8161</td><td>8183</td><td>-</td><td>0</td><td>ATGGGCTTGGGTCGCGCCGG</td><td>TGG</td><td>||||||||[||||||||||||]PAM</td><td>NA</td><td>Intergenic</td><td>NA</td><td>NA</td></tr>
<tr><td>T33810</td><td>TGGGCTTGGGTCGCGCCGGTGGG</td><td></td><td>1000</td><td></td><td></td><td></td><td></td><td></td><td></td><td></td><td></td><td></td></tr>
<tr><td>Oligo fwd</td><td>CACCTGGGCTTGGGTCGCGCCGGT</td><td></td><td></td><td></td><td></td><td></td><td></td><td></td><td></td><td></td><td></td><td></td></tr>
<tr><td>Oligo rev</td><td>AAACACCGGCGCGACCCAAGCCCA</td><td></td><td></td><td></td><td></td><td></td><td></td><td></td><td></td><td></td><td></td><td></td></tr>
<tr><td>Chromosome</td><td>start</td><td></td><td>end</td><td>strand</td><td>MM</td><td>target_seq</td><td>PAM</td><td>alignment</td><td>distance</td><td>position</td><td>gene name</td><td>gene id</td></tr>
<tr><td>GU305914.1</td><td></td><td>8160</td><td>8182</td><td>-</td><td>0</td><td>TGGGCTTGGGTCGCGCCGGT</td><td>GGG</td><td>||||||||[||||||||||||]PAM</td><td>NA</td><td>Intergenic</td><td>NA</td><td>NA</td></tr>
<tr><td>T33811</td><td>TTGGGTCGCGCCGGTGGGACTGG</td><td></td><td>1000</td><td></td><td></td><td></td><td></td><td></td><td></td><td></td><td></td><td></td></tr>
<tr><td>Oligo fwd</td><td>CACCTTGGGTCGCGCCGGTGGGAC</td><td></td><td></td><td></td><td></td><td></td><td></td><td></td><td></td><td></td><td></td><td></td></tr>
<tr><td>Oligo rev</td><td>AAACGTCCCACCGGCGCGACCCAA</td><td></td><td></td><td></td><td></td><td></td><td></td><td></td><td></td><td></td><td></td><td></td></tr>
<tr><td>Chromosome</td><td>start</td><td></td><td>end</td><td>strand</td><td>MM</td><td>target_seq</td><td>PAM</td><td>alignment</td><td>distance</td><td>position</td><td>gene name</td><td>gene id</td></tr>
<tr><td>GU305914.1</td><td></td><td>8155</td><td>8177</td><td>-</td><td>0</td><td>TTGGGTCGCGCCGGTGGGAC</td><td>TGG</td><td>||||||||[||||||||||||]PAM</td><td>NA</td><td>Intergenic</td><td>NA</td><td>NA</td></tr>
<tr><td>T33812</td><td>GCGCCGGTGGGACTGGTTTCTGG</td><td></td><td>1000</td><td></td><td></td><td></td><td></td><td></td><td></td><td></td><td></td><td></td></tr>
<tr><td>Oligo fwd</td><td>CACCGCGCCGGTGGGACTGGTTTC</td><td></td><td></td><td></td><td></td><td></td><td></td><td></td><td></td><td></td><td></td><td></td></tr>
<tr><td>Oligo rev</td><td>AAACGAAACCAGTCCCACCGGCGC</td><td></td><td></td><td></td><td></td><td></td><td></td><td></td><td></td><td></td><td></td><td></td></tr>
<tr><td>Chromosome</td><td>start</td><td></td><td>end</td><td>strand</td><td>MM</td><td>target_seq</td><td>PAM</td><td>alignment</td><td>distance</td><td>position</td><td>gene name</td><td>gene id</td></tr>
<tr><td>GU305914.1</td><td></td><td>8148</td><td>8170</td><td>-</td><td>0</td><td>GCGCCGGTGGGACTGGTTTC</td><td>TGG</td><td>||||||||[||||||||||||]PAM</td><td>NA</td><td>Intergenic</td><td>NA</td><td>NA</td></tr>
<tr><td>T33813</td><td>CGCCGGTGGGACTGGTTTCTGGG</td><td></td><td>1000</td><td></td><td></td><td></td><td></td><td></td><td></td><td></td><td></td><td></td></tr>
<tr><td>Oligo fwd</td><td>CACCCGCCGGTGGGACTGGTTTCT</td><td></td><td></td><td></td><td></td><td></td><td></td><td></td><td></td><td></td><td></td><td></td></tr>
<tr><td>Oligo rev</td><td>AAACAGAAACCAGTCCCACCGGCG</td><td></td><td></td><td></td><td></td><td></td><td></td><td></td><td></td><td></td><td></td><td></td></tr>
<tr><td>Chromosome</td><td>start</td><td></td><td>end</td><td>strand</td><td>MM</td><td>target_seq</td><td>PAM</td><td>alignment</td><td>distance</td><td>position</td><td>gene name</td><td>gene id</td></tr>
<tr><td>GU305914.1</td><td></td><td>8147</td><td>8169</td><td>-</td><td>0</td><td>CGCCGGTGGGACTGGTTTCT</td><td>GGG</td><td>||||||||[||||||||||||]PAM</td><td>NA</td><td>Intergenic</td><td>NA</td><td>NA</td></tr>
<tr><td>T33814</td><td>GGTGGGACTGGTTTCTGGGTAGG</td><td></td><td>1000</td><td></td><td></td><td></td><td></td><td></td><td></td><td></td><td></td><td></td></tr>
<tr><td>Oligo fwd</td><td>CACCGGTGGGACTGGTTTCTGGGT</td><td></td><td></td><td></td><td></td><td></td><td></td><td></td><td></td><td></td><td></td><td></td></tr>
<tr><td>Oligo rev</td><td>AAACACCCAGAAACCAGTCCCACC</td><td></td><td></td><td></td><td></td><td></td><td></td><td></td><td></td><td></td><td></td><td></td></tr>
<tr><td>Chromosome</td><td>start</td><td></td><td>end</td><td>strand</td><td>MM</td><td>target_seq</td><td>PAM</td><td>alignment</td><td>distance</td><td>position</td><td>gene name</td><td>gene id</td></tr>
<tr><td>GU305914.1</td><td></td><td>8143</td><td>8165</td><td>-</td><td>0</td><td>GGTGGGACTGGTTTCTGGGT</td><td>AGG</td><td>||||||||[||||||||||||]PAM</td><td>NA</td><td>Intergenic</td><td>NA</td><td>NA</td></tr>
<tr><td>T33815</td><td>GGACTGGTTTCTGGGTAGGATGG</td><td></td><td>1000</td><td></td><td></td><td></td><td></td><td></td><td></td><td></td><td></td><td></td></tr>
<tr><td>Oligo fwd</td><td>CACCGGACTGGTTTCTGGGTAGGA</td><td></td><td></td><td></td><td></td><td></td><td></td><td></td><td></td><td></td><td></td><td></td></tr>
<tr><td>Oligo rev</td><td>AAACTCCTACCCAGAAACCAGTCC</td><td></td><td></td><td></td><td></td><td></td><td></td><td></td><td></td><td></td><td></td><td></td></tr>
<tr><td>Chromosome</td><td>start</td><td></td><td>end</td><td>strand</td><td>MM</td><td>target_seq</td><td>PAM</td><td>alignment</td><td>distance</td><td>position</td><td>gene name</td><td>gene id</td></tr>
<tr><td>GU305914.1</td><td></td><td>8139</td><td>8161</td><td>-</td><td>0</td><td>GGACTGGTTTCTGGGTAGGA</td><td>TGG</td><td>||||||||[||||||||||||]PAM</td><td>NA</td><td>Intergenic</td><td>NA</td><td>NA</td></tr>
</table>

T33816 GACTGGTTTCTGGGTAGGATGGG 1000
Oligo fwd CACCGACTGGTTTCTGGGTAGGAT
Oligo rev AAACATCCTACCCAGAAACCAGTC
Chromosome start end strand MM target_seq PAM alignment distance position gene name gene id
GU305914.1 8138 8160 - 0 GACTGGTTTCTGGGTAGGAT GGG ||||||||[||||||||||||]PAM NA Intergenic NA NA

T33817 ACTGGTTTCTGGGTAGGATGGGG 1000
Oligo fwd CACCACTGGTTTCTGGGTAGGATG
Oligo rev AAACCATCCTACCCAGAAACCAGT
Chromosome start end strand MM target_seq PAM alignment distance position gene name gene id
GU305914.1 8137 8159 - 0 ACTGGTTTCTGGGTAGGATG GGG ||||||||[||||||||||||]PAM NA Intergenic NA NA

T33818 TTTCTGGGTAGGATGGGGTATGG 1000
Oligo fwd CACCTTTCTGGGTAGGATGGGGTA
Oligo rev AAACTACCCCATCCTACCCAGAAA
Chromosome start end strand MM target_seq PAM alignment distance position gene name gene id
GU305914.1 8132 8154 - 0 TTTCTGGGTAGGATGGGGTA TGG ||||||||[||||||||||||]PAM NA Intergenic NA NA

T33819 TTCTGGGTAGGATGGGGTATGGG 1000
Oligo fwd CACCTTCTGGGTAGGATGGGGTAT
Oligo rev AAACATACCCCATCCTACCCAGAA
Chromosome start end strand MM target_seq PAM alignment distance position gene name gene id
GU305914.1 8131 8153 - 0 TTCTGGGTAGGATGGGGTAT GGG ||||||||[||||||||||||]PAM NA Intergenic NA NA

T33820 TGGGGTATGGGTTTTGTAGTCGG 1000
Oligo fwd CACCTGGGGTATGGGTTTTGTAGT
Oligo rev AAACACTACAAAACCCATACCCCA
Chromosome start end strand MM target_seq PAM alignment distance position gene name gene id
GU305914.1 8119 8141 - 0 TGGGGTATGGGTTTTGTAGT CGG ||||||||[||||||||||||]PAM NA Intergenic NA NA

T33821 GGGGTATGGGTTTTGTAGTCGGG 1000
Oligo fwd CACCGGGGTATGGGTTTTGTAGTC
Oligo rev AAACGACTACAAAACCCATACCCC
Chromosome start end strand MM target_seq PAM alignment distance position gene name gene id
GU305914.1 8118 8140 - 0 GGGGTATGGGTTTTGTAGTC GGG ||||||||[||||||||||||]PAM NA Intergenic NA NA

T33822 GTATGGGTTTTGTAGTCGGGCGG 1000
Oligo fwd CACCGTATGGGTTTTGTAGTCGGG
Oligo rev AAACCCCGACTACAAAACCCATAC
Chromosome start end strand MM target_seq PAM alignment distance position gene name gene id
GU305914.1 8115 8137 - 0 GTATGGGTTTTGTAGTCGGG CGG ||||||||[||||||||||||]PAM NA Intergenic NA NA

T33823 TTTTGTAGTCGGGCGGACCGTGG 1000
Oligo fwd CACCTTTTGTAGTCGGGCGGACCG
Oligo rev AAACCGGTCCGCCCGACTACAAAA
Chromosome start end strand MM target_seq PAM alignment distance position gene name gene id
GU305914.1 8108 8130 - 0 TTTTGTAGTCGGGCGGACCG TGG ||||||||[||||||||||||]PAM NA Intergenic NA NA
GU305914.1 8048 8070 - 2 TCTGGTAGTCGGGCGGACCG TGG |-|-||||[||||||||||||]PAM NA Intergenic NA NA

T33824 GTCGGGCGGACCGTGGTCGTTGG 1000
Oligo fwd CACCGTCGGGCGGACCGTGGTCGT
Oligo rev AAACACGACCACGGTCCGCCCGAC
Chromosome start end strand MM target_seq PAM alignment distance position gene name gene id
GU305914.1 8101 8123 - 0 GTCGGGCGGACCGTGGTCGT TGG ||||||||[||||||||||||]PAM NA Intergenic NA NA
GU305914.1 8041 8063 - 0 GTCGGGCGGACCGTGGTCGT GGG ||||||||[||||||||||||]PAM NA Intergenic NA NA
GU305914.1 8071 8093 - 3 GTAGGACTGACCGTGGTCGT TGG ||-||-|-[||||||||||||]PAM NA Intergenic NA NA

T33825 GCGGACCGTGGTCGTTGGTCTGG 1000
Oligo fwd CACCGCGGACCGTGGTCGTTGGTC
Oligo rev AAACGACCAACGACCACGGTCCGC
Chromosome start end strand MM target_seq PAM alignment distance position gene name gene id
GU305914.1 8096 8118 - 0 GCGGACCGTGGTCGTTGGTC TGG ||||||||[||||||||||||]PAM NA Intergenic NA NA
GU305914.1 8066 8088 - 2 ACTGACCGTGGTCGTTGGTC TGG -|-|||||[||||||||||||]PAM NA Intergenic NA NA
GU305914.1 8036 8058 - 1 GCGGACCGTGGTCGTGGGTC TGG ||||||||[|||||||-||||]PAM NA Intergenic NA NA

T33826 GTGGTCGTTGGTCTGGTAGTAGG 1000
Oligo fwd CACCGTGGTCGTTGGTCTGGTAGT
Oligo rev AAACACTACCAGACCAACGACCAC
Chromosome start end strand MM target_seq PAM alignment distance position gene name gene id
GU305914.1 8059 8081 - 0 GTGGTCGTTGGTCTGGTAGT CGG ||||||||[||||||||||||]PAM NA Intergenic NA NA
GU305914.1 8089 8111 - 0 GTGGTCGTTGGTCTGGTAGT AGG ||||||||[||||||||||||]PAM NA Intergenic NA NA
GU305914.1 8029 8051 - 1 GTGGTCGTGGGTCTGGTAGT AGG ||||||||[-|||||||||||]PAM NA Intergenic NA NA

T33827 TCTGGTAGTAGGACTGACCGTGG 1000
Oligo fwd CACCTCTGGTAGTAGGACTGACCG
Oligo rev AAACCGGTCAGTCCTACTACCAGA
Chromosome start end strand MM target_seq PAM alignment distance position gene name gene id
GU305914.1 8078 8100 - 0 TCTGGTAGTAGGACTGACCG TGG ||||||||[||||||||||||]PAM NA Intergenic NA NA
GU305914.1 8018 8040 - 2 TCTGGTAGTAGGACGGACTG TGG ||||||||[||||||-|||-|]PAM NA Intergenic NA NA

T33828 GTAGGACTGACCGTGGTCGTTGG 1000
Oligo fwd CACCGTAGGACTGACCGTGGTCGT
Oligo rev AAACACGACCACGGTCAGTCCTAC
Chromosome start end strand MM target_seq PAM alignment distance position gene name gene id
GU305914.1 8071 8093 - 0 GTAGGACTGACCGTGGTCGT TGG ||||||||[||||||||||||]PAM NA Intergenic NA NA
GU305914.1 8101 8123 - 3 GTCGGGCGGACCGTGGTCGT TGG ||-||-|-[||||||||||||]PAM NA Intergenic NA NA
GU305914.1 8041 8063 - 3 GTCGGGCGGACCGTGGTCGT GGG ||-||-|-[||||||||||||]PAM NA Intergenic NA NA

<table>
<tr><td>T33829</td><td>ACTGACCGTGGTCGTTGGTCTGG</td><td>1000</td><td></td><td></td><td></td><td></td><td></td><td></td><td></td><td></td><td></td></tr>
<tr><td>Oligo fwd</td><td>CACCACTGACCGTGGTCGTTGGTC</td><td></td><td></td><td></td><td></td><td></td><td></td><td></td><td></td><td></td><td></td></tr>
<tr><td>Oligo rev</td><td>AAACGACCAACGACCACGGTCAGT</td><td></td><td></td><td></td><td></td><td></td><td></td><td></td><td></td><td></td><td></td></tr>
<tr><td>Chromosome</td><td>start</td><td>end</td><td>strand</td><td>MM</td><td>target_seq</td><td>PAM</td><td>alignment</td><td>distance</td><td>position</td><td>gene name</td><td>gene id</td></tr>
<tr><td>GU305914.1</td><td>8066</td><td>8088</td><td>-</td><td>0</td><td>ACTGACCGTGGTCGTTGGTC</td><td>TGG</td><td>||||||||[||||||||||||]PAM</td><td>NA</td><td>Intergenic</td><td>NA</td><td>NA</td></tr>
<tr><td>GU305914.1</td><td>8096</td><td>8118</td><td>-</td><td>2</td><td>GCGGACCGTGGTCGTTGGTC</td><td>TGG</td><td>-|-|||||[||||||||||||]PAM</td><td>NA</td><td>Intergenic</td><td>NA</td><td>NA</td></tr>
<tr><td>GU305914.1</td><td>8036</td><td>8058</td><td>-</td><td>3</td><td>GCGGACCGTGGTCGTGGGTC</td><td>TGG</td><td>-|-|||||[|||||||-||||]PAM</td><td>NA</td><td>Intergenic</td><td>NA</td><td>NA</td></tr>
<tr><td>GU305914.1</td><td>198552</td><td>198574</td><td>-</td><td>3</td><td>ACTGCTCGTGGTCGTTGATC</td><td>GGG</td><td>||||--||[|||||||||-||]PAM</td><td>NA</td><td>Intergenic</td><td>NA</td><td>NA</td></tr>
<tr><td>T33830</td><td>GTGGTCGTTGGTCTGGTAGTCGG</td><td>1000</td><td></td><td></td><td></td><td></td><td></td><td></td><td></td><td></td><td></td></tr>
<tr><td>Oligo fwd</td><td>CACCGTGGTCGTTGGTCTGGTAGT</td><td></td><td></td><td></td><td></td><td></td><td></td><td></td><td></td><td></td><td></td></tr>
<tr><td>Oligo rev</td><td>AAACACTACCAGACCAACGACCAC</td><td></td><td></td><td></td><td></td><td></td><td></td><td></td><td></td><td></td><td></td></tr>
<tr><td>Chromosome</td><td>start</td><td>end</td><td>strand</td><td>MM</td><td>target_seq</td><td>PAM</td><td>alignment</td><td>distance</td><td>position</td><td>gene name</td><td>gene id</td></tr>
<tr><td>GU305914.1</td><td>8059</td><td>8081</td><td>-</td><td>0</td><td>GTGGTCGTTGGTCTGGTAGT</td><td>CGG</td><td>||||||||[||||||||||||]PAM</td><td>NA</td><td>Intergenic</td><td>NA</td><td>NA</td></tr>
<tr><td>GU305914.1</td><td>8089</td><td>8111</td><td>-</td><td>0</td><td>GTGGTCGTTGGTCTGGTAGT</td><td>AGG</td><td>||||||||[||||||||||||]PAM</td><td>NA</td><td>Intergenic</td><td>NA</td><td>NA</td></tr>
<tr><td>GU305914.1</td><td>8029</td><td>8051</td><td>-</td><td>1</td><td>GTGGTCGTGGGTCTGGTAGT</td><td>AGG</td><td>||||||||[-|||||||||||]PAM</td><td>NA</td><td>Intergenic</td><td>NA</td><td>NA</td></tr>
<tr><td>T33831</td><td>TGGTCGTTGGTCTGGTAGTCGGG</td><td>1000</td><td></td><td></td><td></td><td></td><td></td><td></td><td></td><td></td><td></td></tr>
<tr><td>Oligo fwd</td><td>CACCTGGTCGTTGGTCTGGTAGTC</td><td></td><td></td><td></td><td></td><td></td><td></td><td></td><td></td><td></td><td></td></tr>
<tr><td>Oligo rev</td><td>AAACGACTACCAGACCAACGACCA</td><td></td><td></td><td></td><td></td><td></td><td></td><td></td><td></td><td></td><td></td></tr>
<tr><td>Chromosome</td><td>start</td><td>end</td><td>strand</td><td>MM</td><td>target_seq</td><td>PAM</td><td>alignment</td><td>distance</td><td>position</td><td>gene name</td><td>gene id</td></tr>
<tr><td>GU305914.1</td><td>8058</td><td>8080</td><td>-</td><td>0</td><td>TGGTCGTTGGTCTGGTAGTC</td><td>GGG</td><td>||||||||[||||||||||||]PAM</td><td>NA</td><td>Intergenic</td><td>NA</td><td>NA</td></tr>
<tr><td>T33832</td><td>TCGTTGGTCTGGTAGTCGGGCGG</td><td>1000</td><td></td><td></td><td></td><td></td><td></td><td></td><td></td><td></td><td></td></tr>
<tr><td>Oligo fwd</td><td>CACCTCGTTGGTCTGGTAGTCGGG</td><td></td><td></td><td></td><td></td><td></td><td></td><td></td><td></td><td></td><td></td></tr>
<tr><td>Oligo rev</td><td>AAACCCCGACTACCAGACCAACGA</td><td></td><td></td><td></td><td></td><td></td><td></td><td></td><td></td><td></td><td></td></tr>
<tr><td>Chromosome</td><td>start</td><td>end</td><td>strand</td><td>MM</td><td>target_seq</td><td>PAM</td><td>alignment</td><td>distance</td><td>position</td><td>gene name</td><td>gene id</td></tr>
<tr><td>GU305914.1</td><td>8055</td><td>8077</td><td>-</td><td>0</td><td>TCGTTGGTCTGGTAGTCGGG</td><td>CGG</td><td>||||||||[||||||||||||]PAM</td><td>NA</td><td>Intergenic</td><td>NA</td><td>NA</td></tr>
<tr><td>GU305914.1</td><td>8025</td><td>8047</td><td>-</td><td>3</td><td>TCGTGGGTCTGGTAGTAGGA</td><td>CGG</td><td>||||-|||[||||||||-||-]PAM</td><td>NA</td><td>Intergenic</td><td>NA</td><td>NA</td></tr>
<tr><td>T33833</td><td>TCTGGTAGTCGGGCGGACCGTGG</td><td>1000</td><td></td><td></td><td></td><td></td><td></td><td></td><td></td><td></td><td></td></tr>
<tr><td>Oligo fwd</td><td>CACCTCTGGTAGTCGGGCGGACCG</td><td></td><td></td><td></td><td></td><td></td><td></td><td></td><td></td><td></td><td></td></tr>
<tr><td>Oligo rev</td><td>AAACCGGTCCGCCCGACTACCAGA</td><td></td><td></td><td></td><td></td><td></td><td></td><td></td><td></td><td></td><td></td></tr>
<tr><td>Chromosome</td><td>start</td><td>end</td><td>strand</td><td>MM</td><td>target_seq</td><td>PAM</td><td>alignment</td><td>distance</td><td>position</td><td>gene name</td><td>gene id</td></tr>
<tr><td>GU305914.1</td><td>8048</td><td>8070</td><td>-</td><td>0</td><td>TCTGGTAGTCGGGCGGACCG</td><td>TGG</td><td>||||||||[||||||||||||]PAM</td><td>NA</td><td>Intergenic</td><td>NA</td><td>NA</td></tr>
<tr><td>GU305914.1</td><td>8108</td><td>8130</td><td>-</td><td>2</td><td>TTTTGTAGTCGGGCGGACCG</td><td>TGG</td><td>|-|-||||[||||||||||||]PAM</td><td>NA</td><td>Intergenic</td><td>NA</td><td>NA</td></tr>
<tr><td>T33834</td><td>AGTCGGGCGGACCGTGGTCGTGG</td><td>1000</td><td></td><td></td><td></td><td></td><td></td><td></td><td></td><td></td><td></td></tr>
<tr><td>Oligo fwd</td><td>CACCAGTCGGGCGGACCGTGGTCG</td><td></td><td></td><td></td><td></td><td></td><td></td><td></td><td></td><td></td><td></td></tr>
<tr><td>Oligo rev</td><td>AAACCGACCACGGTCCGCCCGACT</td><td></td><td></td><td></td><td></td><td></td><td></td><td></td><td></td><td></td><td></td></tr>
<tr><td>Chromosome</td><td>start</td><td>end</td><td>strand</td><td>MM</td><td>target_seq</td><td>PAM</td><td>alignment</td><td>distance</td><td>position</td><td>gene name</td><td>gene id</td></tr>
<tr><td>GU305914.1</td><td>8042</td><td>8064</td><td>-</td><td>0</td><td>AGTCGGGCGGACCGTGGTCG</td><td>TGG</td><td>||||||||[||||||||||||]PAM</td><td>NA</td><td>Intergenic</td><td>NA</td><td>NA</td></tr>
<tr><td>GU305914.1</td><td>8012</td><td>8034</td><td>-</td><td>4</td><td>AGTAGGACGGACTGTGGTCT</td><td>GGG</td><td>|||-||-|[||||-||||||-]PAM</td><td>NA</td><td>Intergenic</td><td>NA</td><td>NA</td></tr>
<tr><td>T33835</td><td>GTCGGGCGGACCGTGGTCGTGGG</td><td>1000</td><td></td><td></td><td></td><td></td><td></td><td></td><td></td><td></td><td></td></tr>
<tr><td>Oligo fwd</td><td>CACCGTCGGGCGGACCGTGGTCGT</td><td></td><td></td><td></td><td></td><td></td><td></td><td></td><td></td><td></td><td></td></tr>
<tr><td>Oligo rev</td><td>AAACACGACCACGGTCCGCCCGAC</td><td></td><td></td><td></td><td></td><td></td><td></td><td></td><td></td><td></td><td></td></tr>
<tr><td>Chromosome</td><td>start</td><td>end</td><td>strand</td><td>MM</td><td>target_seq</td><td>PAM</td><td>alignment</td><td>distance</td><td>position</td><td>gene name</td><td>gene id</td></tr>
<tr><td>GU305914.1</td><td>8101</td><td>8123</td><td>-</td><td>0</td><td>GTCGGGCGGACCGTGGTCGT</td><td>TGG</td><td>||||||||[||||||||||||]PAM</td><td>NA</td><td>Intergenic</td><td>NA</td><td>NA</td></tr>
<tr><td>GU305914.1</td><td>8041</td><td>8063</td><td>-</td><td>0</td><td>GTCGGGCGGACCGTGGTCGT</td><td>GGG</td><td>||||||||[||||||||||||]PAM</td><td>NA</td><td>Intergenic</td><td>NA</td><td>NA</td></tr>
<tr><td>GU305914.1</td><td>8071</td><td>8093</td><td>-</td><td>3</td><td>GTAGGACTGACCGTGGTCGT</td><td>TGG</td><td>||-||-|-[||||||||||||]PAM</td><td>NA</td><td>Intergenic</td><td>NA</td><td>NA</td></tr>
<tr><td>T33836</td><td>GCGGACCGTGGTCGTGGGTCTGG</td><td>1000</td><td></td><td></td><td></td><td></td><td></td><td></td><td></td><td></td><td></td></tr>
<tr><td>Oligo fwd</td><td>CACCGCGGACCGTGGTCGTGGGTC</td><td></td><td></td><td></td><td></td><td></td><td></td><td></td><td></td><td></td><td></td></tr>
<tr><td>Oligo rev</td><td>AAACGACCCACGACCACGGTCCGC</td><td></td><td></td><td></td><td></td><td></td><td></td><td></td><td></td><td></td><td></td></tr>
<tr><td>Chromosome</td><td>start</td><td>end</td><td>strand</td><td>MM</td><td>target_seq</td><td>PAM</td><td>alignment</td><td>distance</td><td>position</td><td>gene name</td><td>gene id</td></tr>
<tr><td>GU305914.1</td><td>8036</td><td>8058</td><td>-</td><td>0</td><td>GCGGACCGTGGTCGTGGGTC</td><td>TGG</td><td>||||||||[||||||||||||]PAM</td><td>NA</td><td>Intergenic</td><td>NA</td><td>NA</td></tr>
<tr><td>GU305914.1</td><td>8096</td><td>8118</td><td>-</td><td>1</td><td>GCGGACCGTGGTCGTTGGTC</td><td>TGG</td><td>||||||||[|||||||-||||]PAM</td><td>NA</td><td>Intergenic</td><td>NA</td><td>NA</td></tr>
<tr><td>GU305914.1</td><td>8066</td><td>8088</td><td>-</td><td>3</td><td>ACTGACCGTGGTCGTTGGTC</td><td>TGG</td><td>-|-|||||[|||||||-||||]PAM</td><td>NA</td><td>Intergenic</td><td>NA</td><td>NA</td></tr>
<tr><td>T33837</td><td>GTGGTCGTGGGTCTGGTAGTAGG</td><td>1000</td><td></td><td></td><td></td><td></td><td></td><td></td><td></td><td></td><td></td></tr>
<tr><td>Oligo fwd</td><td>CACCGTGGTCGTGGGTCTGGTAGT</td><td></td><td></td><td></td><td></td><td></td><td></td><td></td><td></td><td></td><td></td></tr>
<tr><td>Oligo rev</td><td>AAACACTACCAGACCCACGACCAC</td><td></td><td></td><td></td><td></td><td></td><td></td><td></td><td></td><td></td><td></td></tr>
<tr><td>Chromosome</td><td>start</td><td>end</td><td>strand</td><td>MM</td><td>target_seq</td><td>PAM</td><td>alignment</td><td>distance</td><td>position</td><td>gene name</td><td>gene id</td></tr>
<tr><td>GU305914.1</td><td>8029</td><td>8051</td><td>-</td><td>0</td><td>GTGGTCGTGGGTCTGGTAGT</td><td>AGG</td><td>||||||||[||||||||||||]PAM</td><td>NA</td><td>Intergenic</td><td>NA</td><td>NA</td></tr>
<tr><td>GU305914.1</td><td>8059</td><td>8081</td><td>-</td><td>1</td><td>GTGGTCGTTGGTCTGGTAGT</td><td>CGG</td><td>||||||||[-|||||||||||]PAM</td><td>NA</td><td>Intergenic</td><td>NA</td><td>NA</td></tr>
<tr><td>GU305914.1</td><td>8089</td><td>8111</td><td>-</td><td>1</td><td>GTGGTCGTTGGTCTGGTAGT</td><td>AGG</td><td>||||||||[-|||||||||||]PAM</td><td>NA</td><td>Intergenic</td><td>NA</td><td>NA</td></tr>
<tr><td>T33838</td><td>TCGTGGGTCTGGTAGTAGGACGG</td><td>1000</td><td></td><td></td><td></td><td></td><td></td><td></td><td></td><td></td><td></td></tr>
<tr><td>Oligo fwd</td><td>CACCTCGTGGGTCTGGTAGTAGGA</td><td></td><td></td><td></td><td></td><td></td><td></td><td></td><td></td><td></td><td></td></tr>
<tr><td>Oligo rev</td><td>AAACTCCTACTACCAGACCCACGA</td><td></td><td></td><td></td><td></td><td></td><td></td><td></td><td></td><td></td><td></td></tr>
<tr><td>Chromosome</td><td>start</td><td>end</td><td>strand</td><td>MM</td><td>target_seq</td><td>PAM</td><td>alignment</td><td>distance</td><td>position</td><td>gene name</td><td>gene id</td></tr>
<tr><td>GU305914.1</td><td>8025</td><td>8047</td><td>-</td><td>0</td><td>TCGTGGGTCTGGTAGTAGGA</td><td>CGG</td><td>||||||||[||||||||||||]PAM</td><td>NA</td><td>Intergenic</td><td>NA</td><td>NA</td></tr>
<tr><td>GU305914.1</td><td>8055</td><td>8077</td><td>-</td><td>3</td><td>TCGTTGGTCTGGTAGTCGGG</td><td>CGG</td><td>||||-|||[||||||||-||-]PAM</td><td>NA</td><td>Intergenic</td><td>NA</td><td>NA</td></tr>
<tr><td>T33839</td><td>TCTGGTAGTAGGACGGACTGTGG</td><td>1000</td><td></td><td></td><td></td><td></td><td></td><td></td><td></td><td></td><td></td></tr>
<tr><td>Oligo fwd</td><td>CACCTCTGGTAGTAGGACGGACTG</td><td></td><td></td><td></td><td></td><td></td><td></td><td></td><td></td><td></td><td></td></tr>
<tr><td>Oligo rev</td><td>AAACCAGTCCGTCCTACTACCAGA</td><td></td><td></td><td></td><td></td><td></td><td></td><td></td><td></td><td></td><td></td></tr>
<tr><td>Chromosome</td><td>start</td><td>end</td><td>strand</td><td>MM</td><td>target_seq</td><td>PAM</td><td>alignment</td><td>distance</td><td>position</td><td>gene name</td><td>gene id</td></tr>
<tr><td>GU305914.1</td><td>8018</td><td>8040</td><td>-</td><td>0</td><td>TCTGGTAGTAGGACGGACTG</td><td>TGG</td><td>||||||||[||||||||||||]PAM</td><td>NA</td><td>Intergenic</td><td>NA</td><td>NA</td></tr>
<tr><td>GU305914.1</td><td>8078</td><td>8100</td><td>-</td><td>2</td><td>TCTGGTAGTAGGACTGACCG</td><td>TGG</td><td>||||||||[||||||-|||-|]PAM</td><td>NA</td><td>Intergenic</td><td>NA</td><td>NA</td></tr>
<tr><td>T33840</td><td>TAGTAGGACGGACTGTGGTCTGG</td><td>1000</td><td></td><td></td><td></td><td></td><td></td><td></td><td></td><td></td><td></td></tr>
<tr><td>Oligo fwd</td><td>CACCTAGTAGGACGGACTGTGGTC</td><td></td><td></td><td></td><td></td><td></td><td></td><td></td><td></td><td></td><td></td></tr>
<tr><td>Oligo rev</td><td>AAACGACCACAGTCCGTCCTACTA</td><td></td><td></td><td></td><td></td><td></td><td></td><td></td><td></td><td></td><td></td></tr>
<tr><td>Chromosome</td><td>start</td><td>end</td><td>strand</td><td>MM</td><td>target_seq</td><td>PAM</td><td>alignment</td><td>distance</td><td>position</td><td>gene name</td><td>gene id</td></tr>
<tr><td>GU305914.1</td><td>8013</td><td>8035</td><td>-</td><td>0</td><td>TAGTAGGACGGACTGTGGTC</td><td>TGG</td><td>||||||||[||||||||||||]PAM</td><td>NA</td><td>Intergenic</td><td>NA</td><td>NA</td></tr>
<tr><td>T33841</td><td>AGTAGGACGGACTGTGGTCTGGG</td><td>1000</td><td></td><td></td><td></td><td></td><td></td><td></td><td></td><td></td><td></td></tr>
</table>

Oligo fwd CACCAGTAGGACGGACTGTGGTCT
Oligo rev AAACAGACCACAGTCCGTCCTACT
Chromosome start end strand MM target_seq PAM alignment distance position gene name gene id
GU305914.1 8012 8034 - 0 AGTAGGACGGACTGTGGTCT GGG ||||||||[||||||||||||]PAM NA Intergenic NA NA
GU305914.1 8042 8064 - 4 AGTCGGGCGGACCGTGGTCG TGG |||-||-|[||||-||||||-]PAM NA Intergenic NA NA

T33842 GTAGGACGGACTGTGGTCTGGGG 1000
Oligo fwd CACCGTAGGACGGACTGTGGTCTG
Oligo rev AAACCAGACCACAGTCCGTCCTAC
Chromosome start end strand MM target_seq PAM alignment distance position gene name gene id
GU305914.1 8011 8033 - 0 GTAGGACGGACTGTGGTCTG GGG ||||||||[||||||||||||]PAM NA Intergenic NA NA

T33843 ACGGACTGTGGTCTGGGGTTCGG 1000
Oligo fwd CACCACGGACTGTGGTCTGGGGTT
Oligo rev AAACAACCCCAGACCACAGTCCGT
Chromosome start end strand MM target_seq PAM alignment distance position gene name gene id
GU305914.1 8006 8028 - 0 ACGGACTGTGGTCTGGGGTT CGG ||||||||[||||||||||||]PAM NA Intergenic NA NA

T33844 GGTCTGGGGTTCGGATCGTGTGG 1000
Oligo fwd CACCGGTCTGGGGTTCGGATCGTG
Oligo rev AAACCACGATCCGAACCCCAGACC
Chromosome start end strand MM target_seq PAM alignment distance position gene name gene id
GU305914.1 7997 8019 - 0 GGTCTGGGGTTCGGATCGTG TGG ||||||||[||||||||||||]PAM NA Intergenic NA NA

T33845 TCGTGTGGTTCTCCGTTGTGTGG 1000
Oligo fwd CACCTCGTGTGGTTCTCCGTTGTG
Oligo rev AAACCACAACGGAGAACCACACGA
Chromosome start end strand MM target_seq PAM alignment distance position gene name gene id
GU305914.1 7982 8004 - 0 TCGTGTGGTTCTCCGTTGTG TGG ||||||||[||||||||||||]PAM NA Intergenic NA NA

T33846 TGTGGTTCTCCGTTGTGTGGTGG 1000
Oligo fwd CACCTGTGGTTCTCCGTTGTGTGG
Oligo rev AAACCCACACAACGGAGAACCACA
Chromosome start end strand MM target_seq PAM alignment distance position gene name gene id
GU305914.1 7979 8001 - 0 TGTGGTTCTCCGTTGTGTGG TGG ||||||||[||||||||||||]PAM NA Intergenic NA NA

T33847 GTGGTTCTCCGTTGTGTGGTGGG 1000
Oligo fwd CACCGTGGTTCTCCGTTGTGTGGT
Oligo rev AAACACCACACAACGGAGAACCAC
Chromosome start end strand MM target_seq PAM alignment distance position gene name gene id
GU305914.1 7978 8000 - 0 GTGGTTCTCCGTTGTGTGGT GGG ||||||||[||||||||||||]PAM NA Intergenic NA NA

T33848 CTCCGTTGTGTGGTGGGAATCGG 1000
Oligo fwd CACCCTCCGTTGTGTGGTGGGAAT
Oligo rev AAACATTCCCACCACACAACGGAG
Chromosome start end strand MM target_seq PAM alignment distance position gene name gene id
GU305914.1 7972 7994 - 0 CTCCGTTGTGTGGTGGGAAT CGG ||||||||[||||||||||||]PAM NA Intergenic NA NA

T33849 TTGTGTGGTGGGAATCGGCACGG 1000
Oligo fwd CACCTTGTGTGGTGGGAATCGGCA
Oligo rev AAACTGCCGATTCCCACCACACAA
Chromosome start end strand MM target_seq PAM alignment distance position gene name gene id
GU305914.1 7967 7989 - 0 TTGTGTGGTGGGAATCGGCA CGG ||||||||[||||||||||||]PAM NA Intergenic NA NA

T33850 TGTGTGGTGGGAATCGGCACGGG 1000
Oligo fwd CACCTGTGTGGTGGGAATCGGCAC
Oligo rev AAACGTGCCGATTCCCACCACACA
Chromosome start end strand MM target_seq PAM alignment distance position gene name gene id
GU305914.1 7966 7988 - 0 TGTGTGGTGGGAATCGGCAC GGG ||||||||[||||||||||||]PAM NA Intergenic NA NA

T33851 GGGAATCGGCACGGGTTTCGTGG 1000
Oligo fwd CACCGGGAATCGGCACGGGTTTCG
Oligo rev AAACCGAAACCCGTGCCGATTCCC
Chromosome start end strand MM target_seq PAM alignment distance position gene name gene id
GU305914.1 7958 7980 - 0 GGGAATCGGCACGGGTTTCG TGG ||||||||[||||||||||||]PAM NA Intergenic NA NA

T33852 GTACTACGTTCTTTCGTCGTAGG 1000
Oligo fwd CACCGTACTACGTTCTTTCGTCGT
Oligo rev AAACACGACGAAAGAACGTAGTAC
Chromosome start end strand MM target_seq PAM alignment distance position gene name gene id
GU305914.1 7936 7958 - 0 GTACTACGTTCTTTCGTCGT AGG ||||||||[||||||||||||]PAM NA Intergenic NA NA

T33853 GTCGTAGGTTCCTCAGTCGCTGG 1000
Oligo fwd CACCGTCGTAGGTTCCTCAGTCGC
Oligo rev AAACGCGACTGAGGAACCTACGAC
Chromosome start end strand MM target_seq PAM alignment distance position gene name gene id
GU305914.1 7921 7943 - 0 GTCGTAGGTTCCTCAGTCGC TGG ||||||||[||||||||||||]PAM NA Intergenic NA NA

T33854 CTCAGTCGCTGGACCATAATCGG 1000
Oligo fwd CACCCTCAGTCGCTGGACCATAAT
Oligo rev AAACATTATGGTCCAGCGACTGAG
Chromosome start end strand MM target_seq PAM alignment distance position gene name gene id
GU305914.1 7910 7932 - 0 CTCAGTCGCTGGACCATAAT CGG ||||||||[||||||||||||]PAM NA Intergenic NA NA

T33855 TCAGTCGCTGGACCATAATCGGG 1000
Oligo fwd CACCTCAGTCGCTGGACCATAATC
Oligo rev AAACGATTATGGTCCAGCGACTGA
Chromosome start end strand MM target_seq PAM alignment distance position gene name gene id
GU305914.1 7909 7931 - 0 TCAGTCGCTGGACCATAATC GGG ||||||||[||||||||||||]PAM NA Intergenic NA NA

| | | | | | | | | | | | |
|---|---|---|---|---|---|---|---|---|---|---|---|
| T33856 | ACCTTACAGCCCACAGTGAATGG | 1000 | | | | | | | | | |
| Oligo fwd | CACCACCTTACAGCCCACAGTGAA | | | | | | | | | | |
| Oligo rev | AAACTTCACTGTGGGCTGTAAGGT | | | | | | | | | | |
| Chromosome | start | end | strand | MM | target_seq | PAM | alignment | distance | position | gene name | gene id |
| GU305914.1 | 7867 | 7889 | - | 0 | ACCTTACAGCCCACAGTGAA | TGG | \|\|\|\|\|\|\|\|[\|\|\|\|\|\|\|\|\|\|\|\|]PAM | NA | Intergenic | NA | NA |

| | | | | | | | | | | | |
|---|---|---|---|---|---|---|---|---|---|---|---|
| T33857 | TGTTTTATATCGAACCCATCTGG | 1000 | | | | | | | | | |
| Oligo fwd | CACCTGTTTTATATCGAACCCATC | | | | | | | | | | |
| Oligo rev | AAACGATGGGTTCGATATAAAACA | | | | | | | | | | |
| Chromosome | start | end | strand | MM | target_seq | PAM | alignment | distance | position | gene name | gene id |
| GU305914.1 | 7804 | 7826 | - | 0 | TGTTTTATATCGAACCCATC | TGG | \|\|\|\|\|\|\|\|[\|\|\|\|\|\|\|\|\|\|\|\|]PAM | NA | Intergenic | NA | NA |

| | | | | | | | | | | | |
|---|---|---|---|---|---|---|---|---|---|---|---|
| T33858 | ATACTGAGTCCTTGAAACGTAGG | 1000 | | | | | | | | | |
| Oligo fwd | CACCATACTGAGTCCTTGAAACGT | | | | | | | | | | |
| Oligo rev | AAACACGTTTCAAGGACTCAGTAT | | | | | | | | | | |
| Chromosome | start | end | strand | MM | target_seq | PAM | alignment | distance | position | gene name | gene id |
| GU305914.1 | 7753 | 7775 | - | 0 | ATACTGAGTCCTTGAAACGT | AGG | \|\|\|\|\|\|\|\|[\|\|\|\|\|\|\|\|\|\|\|\|]PAM | NA | Intergenic | NA | NA |

| | | | | | | | | | | | |
|---|---|---|---|---|---|---|---|---|---|---|---|
| T33859 | CTGAGTCCTTGAAACGTAGGTGG | 1000 | | | | | | | | | |
| Oligo fwd | CACCCTGAGTCCTTGAAACGTAGG | | | | | | | | | | |
| Oligo rev | AAACCCTACGTTTCAAGGACTCAG | | | | | | | | | | |
| Chromosome | start | end | strand | MM | target_seq | PAM | alignment | distance | position | gene name | gene id |
| GU305914.1 | 7750 | 7772 | - | 0 | CTGAGTCCTTGAAACGTAGG | TGG | \|\|\|\|\|\|\|\|[\|\|\|\|\|\|\|\|\|\|\|\|]PAM | NA | Intergenic | NA | NA |
| GU305914.1 | 8947 | 8969 | - | 4 | ATGAGTTTTTTAAACGTAGG | CGG | -\|\|\|\|\|--[\|\|-\|\|\|\|\|\|\|\|\|]PAM | NA | Intergenic | NA | NA |

| | | | | | | | | | | | |
|---|---|---|---|---|---|---|---|---|---|---|---|
| T33860 | GAAACGTAGGTGGTTTTGTCAGG | 1000 | | | | | | | | | |
| Oligo fwd | CACCGAAACGTAGGTGGTTTTGTC | | | | | | | | | | |
| Oligo rev | AAACGACAAAACCACCTACGTTTC | | | | | | | | | | |
| Chromosome | start | end | strand | MM | target_seq | PAM | alignment | distance | position | gene name | gene id |
| GU305914.1 | 7740 | 7762 | - | 0 | GAAACGTAGGTGGTTTTGTC | AGG | \|\|\|\|\|\|\|\|[\|\|\|\|\|\|\|\|\|\|\|\|]PAM | NA | Intergenic | NA | NA |

| | | | | | | | | | | | |
|---|---|---|---|---|---|---|---|---|---|---|---|
| T33861 | AAACGTAGGTGGTTTTGTCAGGG | 1000 | | | | | | | | | |
| Oligo fwd | CACCAAACGTAGGTGGTTTTGTCA | | | | | | | | | | |
| Oligo rev | AAACTGACAAAACCACCTACGTTT | | | | | | | | | | |
| Chromosome | start | end | strand | MM | target_seq | PAM | alignment | distance | position | gene name | gene id |
| GU305914.1 | 7739 | 7761 | - | 0 | AAACGTAGGTGGTTTTGTCA | GGG | \|\|\|\|\|\|\|\|[\|\|\|\|\|\|\|\|\|\|\|\|]PAM | NA | Intergenic | NA | NA |

| | | | | | | | | | | | |
|---|---|---|---|---|---|---|---|---|---|---|---|
| T33862 | AAAAACCGCCCGTATCCATTTGG | 1000 | | | | | | | | | |
| Oligo fwd | CACCAAAAACCGCCCGTATCCATT | | | | | | | | | | |
| Oligo rev | AAACAATGGATACGGGCGGTTTTT | | | | | | | | | | |
| Chromosome | start | end | strand | MM | target_seq | PAM | alignment | distance | position | gene name | gene id |
| GU305914.1 | 7707 | 7729 | - | 0 | AAAAACCGCCCGTATCCATT | TGG | \|\|\|\|\|\|\|\|[\|\|\|\|\|\|\|\|\|\|\|\|]PAM | NA | Intergenic | NA | NA |

| | | | | | | | | | | | |
|---|---|---|---|---|---|---|---|---|---|---|---|
| T33863 | CATTTAACCTGCCACGATGCCGG | 1000 | | | | | | | | | |
| Oligo fwd | CACCCATTTAACCTGCCACGATGC | | | | | | | | | | |
| Oligo rev | AAACGCATCGTGGCAGGTTAAATG | | | | | | | | | | |
| Chromosome | start | end | strand | MM | target_seq | PAM | alignment | distance | position | gene name | gene id |
| GU305914.1 | 7669 | 7691 | - | 0 | CATTTAACCTGCCACGATGC | CGG | \|\|\|\|\|\|\|\|[\|\|\|\|\|\|\|\|\|\|\|\|]PAM | NA | Intergenic | NA | NA |

| | | | | | | | | | | | |
|---|---|---|---|---|---|---|---|---|---|---|---|
| T33864 | GGAATGCAGATATCACCGAGTGG | 1000 | | | | | | | | | |
| Oligo fwd | CACCGGAATGCAGATATCACCGAG | | | | | | | | | | |
| Oligo rev | AAACCTCGGTGATATCTGCATTCC | | | | | | | | | | |
| Chromosome | start | end | strand | MM | target_seq | PAM | alignment | distance | position | gene name | gene id |
| GU305914.1 | 7648 | 7670 | - | 0 | GGAATGCAGATATCACCGAG | TGG | \|\|\|\|\|\|\|\|[\|\|\|\|\|\|\|\|\|\|\|\|]PAM | NA | Intergenic | NA | NA |

| | | | | | | | | | | | |
|---|---|---|---|---|---|---|---|---|---|---|---|
| T33865 | ATCCCCAACCGTCCACACCTAGG | 1000 | | | | | | | | | |
| Oligo fwd | CACCATCCCCAACCGTCCACACCT | | | | | | | | | | |
| Oligo rev | AAACAGGTGTGGACGGTTGGGGAT | | | | | | | | | | |
| Chromosome | start | end | strand | MM | target_seq | PAM | alignment | distance | position | gene name | gene id |
| GU305914.1 | 7617 | 7639 | - | 0 | ATCCCCAACCGTCCACACCT | AGG | \|\|\|\|\|\|\|\|[\|\|\|\|\|\|\|\|\|\|\|\|]PAM | NA | Intergenic | NA | NA |

| | | | | | | | | | | | |
|---|---|---|---|---|---|---|---|---|---|---|---|
| T33866 | TAGGTTACTGTCGTTACGTACGG | 1000 | | | | | | | | | |
| Oligo fwd | CACCTAGGTTACTGTCGTTACGTA | | | | | | | | | | |
| Oligo rev | AAACTACGTAACGACAGTAACCTA | | | | | | | | | | |
| Chromosome | start | end | strand | MM | target_seq | PAM | alignment | distance | position | gene name | gene id |
| GU305914.1 | 7598 | 7620 | - | 0 | TAGGTTACTGTCGTTACGTA | CGG | \|\|\|\|\|\|\|\|[\|\|\|\|\|\|\|\|\|\|\|\|]PAM | NA | Intergenic | NA | NA |

| | | | | | | | | | | | |
|---|---|---|---|---|---|---|---|---|---|---|---|
| T33867 | AGGTTACTGTCGTTACGTACGGG | 1000 | | | | | | | | | |
| Oligo fwd | CACCAGGTTACTGTCGTTACGTAC | | | | | | | | | | |
| Oligo rev | AAACGTACGTAACGACAGTAACCT | | | | | | | | | | |
| Chromosome | start | end | strand | MM | target_seq | PAM | alignment | distance | position | gene name | gene id |
| GU305914.1 | 7597 | 7619 | - | 0 | AGGTTACTGTCGTTACGTAC | GGG | \|\|\|\|\|\|\|\|[\|\|\|\|\|\|\|\|\|\|\|\|]PAM | NA | Intergenic | NA | NA |

| | | | | | | | | | | | |
|---|---|---|---|---|---|---|---|---|---|---|---|
| T33868 | AGAATCGTACAACTTCCTTGTGG | 1000 | | | | | | | | | |
| Oligo fwd | CACCAGAATCGTACAACTTCCTTG | | | | | | | | | | |
| Oligo rev | AAACCAAGGAAGTTGTACGATTCT | | | | | | | | | | |
| Chromosome | start | end | strand | MM | target_seq | PAM | alignment | distance | position | gene name | gene id |
| GU305914.1 | 7564 | 7586 | - | 0 | AGAATCGTACAACTTCCTTG | TGG | \|\|\|\|\|\|\|\|[\|\|\|\|\|\|\|\|\|\|\|\|]PAM | NA | Intergenic | NA | NA |

| | | | | | | | | | | | |
|---|---|---|---|---|---|---|---|---|---|---|---|
| T33869 | ACAACTTCCTTGTGGTAGATTGG | 1000 | | | | | | | | | |
| Oligo fwd | CACCACAACTTCCTTGTGGTAGAT | | | | | | | | | | |
| Oligo rev | AAACATCTACCACAAGGAAGTTGT | | | | | | | | | | |
| Chromosome | start | end | strand | MM | target_seq | PAM | alignment | distance | position | gene name | gene id |
| GU305914.1 | 7556 | 7578 | - | 0 | ACAACTTCCTTGTGGTAGAT | TGG | \|\|\|\|\|\|\|\|[\|\|\|\|\|\|\|\|\|\|\|\|]PAM | NA | Intergenic | NA | NA |

| | | |
|---|---|---|
| T33870 | GCAGATCATCGATAAAACCACGG | 1000 |
| Oligo fwd | CACCGCAGATCATCGATAAAACCA | |
| Oligo rev | AAACTGGTTTTATCGATGATCTGC | |

<table>
<tr><td>Chromosome</td><td>start</td><td>end</td><td>strand</td><td>MM</td><td>target_seq</td><td>PAM</td><td>alignment</td><td>distance</td><td>position</td><td>gene name</td><td>gene id</td></tr>
<tr><td>GU305914.1</td><td>7517</td><td>7539</td><td>-</td><td>0</td><td>GCAGATCATCGATAAAACCA</td><td>CGG</td><td>||||||||[||||||||||||]PAM</td><td>NA</td><td>Intergenic</td><td>NA</td><td>NA</td></tr>
<tr><td>T33871</td><td>GATAGAGACGCATCGTGTCCAGG</td><td>1000</td><td></td><td></td><td></td><td></td><td></td><td></td><td></td><td></td><td></td></tr>
<tr><td>Oligo fwd</td><td>CACCGATAGAGACGCATCGTGTCC</td><td></td><td></td><td></td><td></td><td></td><td></td><td></td><td></td><td></td><td></td></tr>
<tr><td>Oligo rev</td><td>AAACGGACACGATGCGTCTCTATC</td><td></td><td></td><td></td><td></td><td></td><td></td><td></td><td></td><td></td><td></td></tr>
<tr><td>Chromosome</td><td>start</td><td>end</td><td>strand</td><td>MM</td><td>target_seq</td><td>PAM</td><td>alignment</td><td>distance</td><td>position</td><td>gene name</td><td>gene id</td></tr>
<tr><td>GU305914.1</td><td>7449</td><td>7471</td><td>-</td><td>0</td><td>GATAGAGACGCATCGTGTCC</td><td>AGG</td><td>||||||||[||||||||||||]PAM</td><td>NA</td><td>Intergenic</td><td>NA</td><td>NA</td></tr>
<tr><td>T33872</td><td>GATACGTACCGTCGCACCTGCGG</td><td>1000</td><td></td><td></td><td></td><td></td><td></td><td></td><td></td><td></td><td></td></tr>
<tr><td>Oligo fwd</td><td>CACCGATACGTACCGTCGCACCTG</td><td></td><td></td><td></td><td></td><td></td><td></td><td></td><td></td><td></td><td></td></tr>
<tr><td>Oligo rev</td><td>AAACCAGGTGCGACGGTACGTATC</td><td></td><td></td><td></td><td></td><td></td><td></td><td></td><td></td><td></td><td></td></tr>
<tr><td>Chromosome</td><td>start</td><td>end</td><td>strand</td><td>MM</td><td>target_seq</td><td>PAM</td><td>alignment</td><td>distance</td><td>position</td><td>gene name</td><td>gene id</td></tr>
<tr><td>GU305914.1</td><td>7419</td><td>7441</td><td>-</td><td>0</td><td>GATACGTACCGTCGCACCTG</td><td>CGG</td><td>||||||||[||||||||||||]PAM</td><td>NA</td><td>Intergenic</td><td>NA</td><td>NA</td></tr>
<tr><td>T33873</td><td>ACGTACCGTCGCACCTGCGGCGG</td><td>1000</td><td></td><td></td><td></td><td></td><td></td><td></td><td></td><td></td><td></td></tr>
<tr><td>Oligo fwd</td><td>CACCACGTACCGTCGCACCTGCGG</td><td></td><td></td><td></td><td></td><td></td><td></td><td></td><td></td><td></td><td></td></tr>
<tr><td>Oligo rev</td><td>AAACCCGCAGGTGCGACGGTACGT</td><td></td><td></td><td></td><td></td><td></td><td></td><td></td><td></td><td></td><td></td></tr>
<tr><td>Chromosome</td><td>start</td><td>end</td><td>strand</td><td>MM</td><td>target_seq</td><td>PAM</td><td>alignment</td><td>distance</td><td>position</td><td>gene name</td><td>gene id</td></tr>
<tr><td>GU305914.1</td><td>7416</td><td>7438</td><td>-</td><td>0</td><td>ACGTACCGTCGCACCTGCGG</td><td>CGG</td><td>||||||||[||||||||||||]PAM</td><td>NA</td><td>Intergenic</td><td>NA</td><td>NA</td></tr>
<tr><td>T33874</td><td>ACCGTCGCACCTGCGGCGGTCGG</td><td>1000</td><td></td><td></td><td></td><td></td><td></td><td></td><td></td><td></td><td></td></tr>
<tr><td>Oligo fwd</td><td>CACCACCGTCGCACCTGCGGCGGT</td><td></td><td></td><td></td><td></td><td></td><td></td><td></td><td></td><td></td><td></td></tr>
<tr><td>Oligo rev</td><td>AAACACCGCCGCAGGTGCGACGGT</td><td></td><td></td><td></td><td></td><td></td><td></td><td></td><td></td><td></td><td></td></tr>
<tr><td>Chromosome</td><td>start</td><td>end</td><td>strand</td><td>MM</td><td>target_seq</td><td>PAM</td><td>alignment</td><td>distance</td><td>position</td><td>gene name</td><td>gene id</td></tr>
<tr><td>GU305914.1</td><td>7412</td><td>7434</td><td>-</td><td>0</td><td>ACCGTCGCACCTGCGGCGGT</td><td>CGG</td><td>||||||||[||||||||||||]PAM</td><td>NA</td><td>Intergenic</td><td>NA</td><td>NA</td></tr>
<tr><td>T33875</td><td>GCCCTTTACATACTCACCGCAGG</td><td>1000</td><td></td><td></td><td></td><td></td><td></td><td></td><td></td><td></td><td></td></tr>
<tr><td>Oligo fwd</td><td>CACCGCCCTTTACATACTCACCGC</td><td></td><td></td><td></td><td></td><td></td><td></td><td></td><td></td><td></td><td></td></tr>
<tr><td>Oligo rev</td><td>AAACGCGGTGAGTATGTAAAGGGC</td><td></td><td></td><td></td><td></td><td></td><td></td><td></td><td></td><td></td><td></td></tr>
<tr><td>Chromosome</td><td>start</td><td>end</td><td>strand</td><td>MM</td><td>target_seq</td><td>PAM</td><td>alignment</td><td>distance</td><td>position</td><td>gene name</td><td>gene id</td></tr>
<tr><td>GU305914.1</td><td>7390</td><td>7412</td><td>-</td><td>0</td><td>GCCCTTTACATACTCACCGC</td><td>AGG</td><td>||||||||[||||||||||||]PAM</td><td>NA</td><td>Intergenic</td><td>NA</td><td>NA</td></tr>
<tr><td>T33876</td><td>TTACATACTCACCGCAGGTGTGG</td><td>1000</td><td></td><td></td><td></td><td></td><td></td><td></td><td></td><td></td><td></td></tr>
<tr><td>Oligo fwd</td><td>CACCTTACATACTCACCGCAGGTG</td><td></td><td></td><td></td><td></td><td></td><td></td><td></td><td></td><td></td><td></td></tr>
<tr><td>Oligo rev</td><td>AAACCACCTGCGGTGAGTATGTAA</td><td></td><td></td><td></td><td></td><td></td><td></td><td></td><td></td><td></td><td></td></tr>
<tr><td>Chromosome</td><td>start</td><td>end</td><td>strand</td><td>MM</td><td>target_seq</td><td>PAM</td><td>alignment</td><td>distance</td><td>position</td><td>gene name</td><td>gene id</td></tr>
<tr><td>GU305914.1</td><td>7385</td><td>7407</td><td>-</td><td>0</td><td>TTACATACTCACCGCAGGTG</td><td>TGG</td><td>||||||||[||||||||||||]PAM</td><td>NA</td><td>Intergenic</td><td>NA</td><td>NA</td></tr>
<tr><td>T33877</td><td>TACATACTCACCGCAGGTGTGGG</td><td>1000</td><td></td><td></td><td></td><td></td><td></td><td></td><td></td><td></td><td></td></tr>
<tr><td>Oligo fwd</td><td>CACCTACATACTCACCGCAGGTGT</td><td></td><td></td><td></td><td></td><td></td><td></td><td></td><td></td><td></td><td></td></tr>
<tr><td>Oligo rev</td><td>AAACACACCTGCGGTGAGTATGTA</td><td></td><td></td><td></td><td></td><td></td><td></td><td></td><td></td><td></td><td></td></tr>
<tr><td>Chromosome</td><td>start</td><td>end</td><td>strand</td><td>MM</td><td>target_seq</td><td>PAM</td><td>alignment</td><td>distance</td><td>position</td><td>gene name</td><td>gene id</td></tr>
<tr><td>GU305914.1</td><td>7384</td><td>7406</td><td>-</td><td>0</td><td>TACATACTCACCGCAGGTGT</td><td>GGG</td><td>||||||||[||||||||||||]PAM</td><td>NA</td><td>Intergenic</td><td>NA</td><td>NA</td></tr>
<tr><td>T33878</td><td>ATACTCACCGCAGGTGTGGGCGG</td><td>1000</td><td></td><td></td><td></td><td></td><td></td><td></td><td></td><td></td><td></td></tr>
<tr><td>Oligo fwd</td><td>CACCATACTCACCGCAGGTGTGGG</td><td></td><td></td><td></td><td></td><td></td><td></td><td></td><td></td><td></td><td></td></tr>
<tr><td>Oligo rev</td><td>AAACCCCACACCTGCGGTGAGTAT</td><td></td><td></td><td></td><td></td><td></td><td></td><td></td><td></td><td></td><td></td></tr>
<tr><td>Chromosome</td><td>start</td><td>end</td><td>strand</td><td>MM</td><td>target_seq</td><td>PAM</td><td>alignment</td><td>distance</td><td>position</td><td>gene name</td><td>gene id</td></tr>
<tr><td>GU305914.1</td><td>7381</td><td>7403</td><td>-</td><td>0</td><td>ATACTCACCGCAGGTGTGGG</td><td>CGG</td><td>||||||||[||||||||||||]PAM</td><td>NA</td><td>Intergenic</td><td>NA</td><td>NA</td></tr>
<tr><td>GU305914.1</td><td>8551</td><td>8573</td><td>-</td><td>3</td><td>ATACTCAGGACAGGTGTGGG</td><td>CGG</td><td>|||||||-[--||||||||||]PAM</td><td>NA</td><td>Intergenic</td><td>NA</td><td>NA</td></tr>
<tr><td>T33879</td><td>CCGCAGGTGTGGGCGGTGACAGG</td><td>1000</td><td></td><td></td><td></td><td></td><td></td><td></td><td></td><td></td><td></td></tr>
<tr><td>Oligo fwd</td><td>CACCCCGCAGGTGTGGGCGGTGAC</td><td></td><td></td><td></td><td></td><td></td><td></td><td></td><td></td><td></td><td></td></tr>
<tr><td>Oligo rev</td><td>AAACGTCACCGCCCACACCTGCGG</td><td></td><td></td><td></td><td></td><td></td><td></td><td></td><td></td><td></td><td></td></tr>
<tr><td>Chromosome</td><td>start</td><td>end</td><td>strand</td><td>MM</td><td>target_seq</td><td>PAM</td><td>alignment</td><td>distance</td><td>position</td><td>gene name</td><td>gene id</td></tr>
<tr><td>GU305914.1</td><td>7374</td><td>7396</td><td>-</td><td>0</td><td>CCGCAGGTGTGGGCGGTGAC</td><td>AGG</td><td>||||||||[||||||||||||]PAM</td><td>NA</td><td>Intergenic</td><td>NA</td><td>NA</td></tr>
<tr><td>T33880</td><td>TGACAGGTACGTAGAATTGTTGG</td><td>1000</td><td></td><td></td><td></td><td></td><td></td><td></td><td></td><td></td><td></td></tr>
<tr><td>Oligo fwd</td><td>CACCTGACAGGTACGTAGAATTGT</td><td></td><td></td><td></td><td></td><td></td><td></td><td></td><td></td><td></td><td></td></tr>
<tr><td>Oligo rev</td><td>AAACACAATTCTACGTACCTGTCA</td><td></td><td></td><td></td><td></td><td></td><td></td><td></td><td></td><td></td><td></td></tr>
<tr><td>Chromosome</td><td>start</td><td>end</td><td>strand</td><td>MM</td><td>target_seq</td><td>PAM</td><td>alignment</td><td>distance</td><td>position</td><td>gene name</td><td>gene id</td></tr>
<tr><td>GU305914.1</td><td>7358</td><td>7380</td><td>-</td><td>0</td><td>TGACAGGTACGTAGAATTGT</td><td>TGG</td><td>||||||||[||||||||||||]PAM</td><td>NA</td><td>Intergenic</td><td>NA</td><td>NA</td></tr>
<tr><td>T33881</td><td>TTGTTGGCAGATACATCAGCCGG</td><td>1000</td><td></td><td></td><td></td><td></td><td></td><td></td><td></td><td></td><td></td></tr>
<tr><td>Oligo fwd</td><td>CACCTTGTTGGCAGATACATCAGC</td><td></td><td></td><td></td><td></td><td></td><td></td><td></td><td></td><td></td><td></td></tr>
<tr><td>Oligo rev</td><td>AAACGCTGATGTATCTGCCAACAA</td><td></td><td></td><td></td><td></td><td></td><td></td><td></td><td></td><td></td><td></td></tr>
<tr><td>Chromosome</td><td>start</td><td>end</td><td>strand</td><td>MM</td><td>target_seq</td><td>PAM</td><td>alignment</td><td>distance</td><td>position</td><td>gene name</td><td>gene id</td></tr>
<tr><td>GU305914.1</td><td>7342</td><td>7364</td><td>-</td><td>0</td><td>TTGTTGGCAGATACATCAGC</td><td>CGG</td><td>||||||||[||||||||||||]PAM</td><td>NA</td><td>Intergenic</td><td>NA</td><td>NA</td></tr>
<tr><td>T33882</td><td>TTGGCAGATACATCAGCCGGCGG</td><td>1000</td><td></td><td></td><td></td><td></td><td></td><td></td><td></td><td></td><td></td></tr>
<tr><td>Oligo fwd</td><td>CACCTTGGCAGATACATCAGCCGG</td><td></td><td></td><td></td><td></td><td></td><td></td><td></td><td></td><td></td><td></td></tr>
<tr><td>Oligo rev</td><td>AAACCCGGCTGATGTATCTGCCAA</td><td></td><td></td><td></td><td></td><td></td><td></td><td></td><td></td><td></td><td></td></tr>
<tr><td>Chromosome</td><td>start</td><td>end</td><td>strand</td><td>MM</td><td>target_seq</td><td>PAM</td><td>alignment</td><td>distance</td><td>position</td><td>gene name</td><td>gene id</td></tr>
<tr><td>GU305914.1</td><td>7339</td><td>7361</td><td>-</td><td>0</td><td>TTGGCAGATACATCAGCCGG</td><td>CGG</td><td>||||||||[||||||||||||]PAM</td><td>NA</td><td>Intergenic</td><td>NA</td><td>NA</td></tr>
<tr><td>T33883</td><td>ACATCAGCCGGCGGACGTAGAGG</td><td>1000</td><td></td><td></td><td></td><td></td><td></td><td></td><td></td><td></td><td></td></tr>
<tr><td>Oligo fwd</td><td>CACCACATCAGCCGGCGGACGTAG</td><td></td><td></td><td></td><td></td><td></td><td></td><td></td><td></td><td></td><td></td></tr>
<tr><td>Oligo rev</td><td>AAACCTACGTCCGCCGGCTGATGT</td><td></td><td></td><td></td><td></td><td></td><td></td><td></td><td></td><td></td><td></td></tr>
<tr><td>Chromosome</td><td>start</td><td>end</td><td>strand</td><td>MM</td><td>target_seq</td><td>PAM</td><td>alignment</td><td>distance</td><td>position</td><td>gene name</td><td>gene id</td></tr>
<tr><td>GU305914.1</td><td>7330</td><td>7352</td><td>-</td><td>0</td><td>ACATCAGCCGGCGGACGTAG</td><td>AGG</td><td>||||||||[||||||||||||]PAM</td><td>NA</td><td>Intergenic</td><td>NA</td><td>NA</td></tr>
<tr><td>T33884</td><td>ACGTAGAGGTTCAGATTCACCGG</td><td>1000</td><td></td><td></td><td></td><td></td><td></td><td></td><td></td><td></td><td></td></tr>
<tr><td>Oligo fwd</td><td>CACCACGTAGAGGTTCAGATTCAC</td><td></td><td></td><td></td><td></td><td></td><td></td><td></td><td></td><td></td><td></td></tr>
<tr><td>Oligo rev</td><td>AAACGTGAATCTGAACCTCTACGT</td><td></td><td></td><td></td><td></td><td></td><td></td><td></td><td></td><td></td><td></td></tr>
<tr><td>Chromosome</td><td>start</td><td>end</td><td>strand</td><td>MM</td><td>target_seq</td><td>PAM</td><td>alignment</td><td>distance</td><td>position</td><td>gene name</td><td>gene id</td></tr>
<tr><td>GU305914.1</td><td>7316</td><td>7338</td><td>-</td><td>0</td><td>ACGTAGAGGTTCAGATTCAC</td><td>CGG</td><td>||||||||[||||||||||||]PAM</td><td>NA</td><td>Intergenic</td><td>NA</td><td>NA</td></tr>
<tr><td>T33885</td><td>GGTTCAGATTCACCGGAACGTGG</td><td>1000</td><td></td><td></td><td></td><td></td><td></td><td></td><td></td><td></td><td></td></tr>
</table>

<table>
<tr><td>Oligo fwd</td><td>CACCGGTTCAGATTCACCGGAACG</td><td></td><td></td><td></td><td></td><td></td><td></td><td></td><td></td><td></td><td></td></tr>
<tr><td>Oligo rev</td><td>AAACCGTTCCGGTGAATCTGAACC</td><td></td><td></td><td></td><td></td><td></td><td></td><td></td><td></td><td></td><td></td></tr>
<tr><td>Chromosome</td><td>start</td><td>end</td><td>strand</td><td>MM</td><td>target_seq</td><td>PAM</td><td>alignment</td><td>distance</td><td>position</td><td>gene name</td><td>gene id</td></tr>
<tr><td>GU305914.1</td><td>7309</td><td>7331</td><td>-</td><td>0</td><td>GGTTCAGATTCACCGGAACG</td><td>TGG</td><td>||||||||[||||||||||||]PAM</td><td>NA</td><td>Intergenic</td><td>NA</td><td>NA</td></tr>
<tr><td></td><td></td><td></td><td></td><td></td><td></td><td></td><td></td><td></td><td></td><td></td><td></td></tr>
<tr><td>T33886</td><td>AGATTCACCGGAACGTGGTTCGG</td><td>1000</td><td></td><td></td><td></td><td></td><td></td><td></td><td></td><td></td><td></td></tr>
<tr><td>Oligo fwd</td><td>CACCAGATTCACCGGAACGTGGTT</td><td></td><td></td><td></td><td></td><td></td><td></td><td></td><td></td><td></td><td></td></tr>
<tr><td>Oligo rev</td><td>AAACAACCACGTTCCGGTGAATCT</td><td></td><td></td><td></td><td></td><td></td><td></td><td></td><td></td><td></td><td></td></tr>
<tr><td>Chromosome</td><td>start</td><td>end</td><td>strand</td><td>MM</td><td>target_seq</td><td>PAM</td><td>alignment</td><td>distance</td><td>position</td><td>gene name</td><td>gene id</td></tr>
<tr><td>GU305914.1</td><td>7304</td><td>7326</td><td>-</td><td>0</td><td>AGATTCACCGGAACGTGGTT</td><td>CGG</td><td>||||||||[||||||||||||]PAM</td><td>NA</td><td>Intergenic</td><td>NA</td><td>NA</td></tr>
<tr><td></td><td></td><td></td><td></td><td></td><td></td><td></td><td></td><td></td><td></td><td></td><td></td></tr>
<tr><td>T33887</td><td>ACGTGGTTCGGATCCGTGACCGG</td><td>1000</td><td></td><td></td><td></td><td></td><td></td><td></td><td></td><td></td><td></td></tr>
<tr><td>Oligo fwd</td><td>CACCACGTGGTTCGGATCCGTGAC</td><td></td><td></td><td></td><td></td><td></td><td></td><td></td><td></td><td></td><td></td></tr>
<tr><td>Oligo rev</td><td>AAACGTCACGGATCCGAACCACGT</td><td></td><td></td><td></td><td></td><td></td><td></td><td></td><td></td><td></td><td></td></tr>
<tr><td>Chromosome</td><td>start</td><td>end</td><td>strand</td><td>MM</td><td>target_seq</td><td>PAM</td><td>alignment</td><td>distance</td><td>position</td><td>gene name</td><td>gene id</td></tr>
<tr><td>GU305914.1</td><td>7292</td><td>7314</td><td>-</td><td>0</td><td>ACGTGGTTCGGATCCGTGAC</td><td>CGG</td><td>||||||||[||||||||||||]PAM</td><td>NA</td><td>Intergenic</td><td>NA</td><td>NA</td></tr>
<tr><td></td><td></td><td></td><td></td><td></td><td></td><td></td><td></td><td></td><td></td><td></td><td></td></tr>
<tr><td>T33888</td><td>CGTGGTTCGGATCCGTGACCGGG</td><td>1000</td><td></td><td></td><td></td><td></td><td></td><td></td><td></td><td></td><td></td></tr>
<tr><td>Oligo fwd</td><td>CACCCGTGGTTCGGATCCGTGACC</td><td></td><td></td><td></td><td></td><td></td><td></td><td></td><td></td><td></td><td></td></tr>
<tr><td>Oligo rev</td><td>AAACGGTCACGGATCCGAACCACG</td><td></td><td></td><td></td><td></td><td></td><td></td><td></td><td></td><td></td><td></td></tr>
<tr><td>Chromosome</td><td>start</td><td>end</td><td>strand</td><td>MM</td><td>target_seq</td><td>PAM</td><td>alignment</td><td>distance</td><td>position</td><td>gene name</td><td>gene id</td></tr>
<tr><td>GU305914.1</td><td>7291</td><td>7313</td><td>-</td><td>0</td><td>CGTGGTTCGGATCCGTGACC</td><td>GGG</td><td>||||||||[||||||||||||]PAM</td><td>NA</td><td>Intergenic</td><td>NA</td><td>NA</td></tr>
<tr><td></td><td></td><td></td><td></td><td></td><td></td><td></td><td></td><td></td><td></td><td></td><td></td></tr>
<tr><td>T33889</td><td>CTTCTGCCGTTCCATAGACCCGG</td><td>1000</td><td></td><td></td><td></td><td></td><td></td><td></td><td></td><td></td><td></td></tr>
<tr><td>Oligo fwd</td><td>CACCCTTCTGCCGTTCCATAGACC</td><td></td><td></td><td></td><td></td><td></td><td></td><td></td><td></td><td></td><td></td></tr>
<tr><td>Oligo rev</td><td>AAACGGTCTATGGAACGGCAGAAG</td><td></td><td></td><td></td><td></td><td></td><td></td><td></td><td></td><td></td><td></td></tr>
<tr><td>Chromosome</td><td>start</td><td>end</td><td>strand</td><td>MM</td><td>target_seq</td><td>PAM</td><td>alignment</td><td>distance</td><td>position</td><td>gene name</td><td>gene id</td></tr>
<tr><td>GU305914.1</td><td>7154</td><td>7176</td><td>-</td><td>0</td><td>CTTCTGCCGTTCCATAGACC</td><td>CGG</td><td>||||||||[||||||||||||]PAM</td><td>NA</td><td>Intergenic</td><td>NA</td><td>NA</td></tr>
<tr><td></td><td></td><td></td><td></td><td></td><td></td><td></td><td></td><td></td><td></td><td></td><td></td></tr>
<tr><td>T33890</td><td>AGACCCGGTCGTCTCGTCGTCGG</td><td>1000</td><td></td><td></td><td></td><td></td><td></td><td></td><td></td><td></td><td></td></tr>
<tr><td>Oligo fwd</td><td>CACCAGACCCGGTCGTCTCGTCGT</td><td></td><td></td><td></td><td></td><td></td><td></td><td></td><td></td><td></td><td></td></tr>
<tr><td>Oligo rev</td><td>AAACACGACGAGACGACCGGGTCT</td><td></td><td></td><td></td><td></td><td></td><td></td><td></td><td></td><td></td><td></td></tr>
<tr><td>Chromosome</td><td>start</td><td>end</td><td>strand</td><td>MM</td><td>target_seq</td><td>PAM</td><td>alignment</td><td>distance</td><td>position</td><td>gene name</td><td>gene id</td></tr>
<tr><td>GU305914.1</td><td>7139</td><td>7161</td><td>-</td><td>0</td><td>AGACCCGGTCGTCTCGTCGT</td><td>CGG</td><td>||||||||[||||||||||||]PAM</td><td>NA</td><td>Intergenic</td><td>NA</td><td>NA</td></tr>
<tr><td></td><td></td><td></td><td></td><td></td><td></td><td></td><td></td><td></td><td></td><td></td><td></td></tr>
<tr><td>T33891</td><td>GGTCGTCTCGTCGTCGGTGTTGG</td><td>1000</td><td></td><td></td><td></td><td></td><td></td><td></td><td></td><td></td><td></td></tr>
<tr><td>Oligo fwd</td><td>CACCGGTCGTCTCGTCGTCGGTGT</td><td></td><td></td><td></td><td></td><td></td><td></td><td></td><td></td><td></td><td></td></tr>
<tr><td>Oligo rev</td><td>AAACACACCGACGACGAGACGACC</td><td></td><td></td><td></td><td></td><td></td><td></td><td></td><td></td><td></td><td></td></tr>
<tr><td>Chromosome</td><td>start</td><td>end</td><td>strand</td><td>MM</td><td>target_seq</td><td>PAM</td><td>alignment</td><td>distance</td><td>position</td><td>gene name</td><td>gene id</td></tr>
<tr><td>GU305914.1</td><td>7133</td><td>7155</td><td>-</td><td>0</td><td>GGTCGTCTCGTCGTCGGTGT</td><td>TGG</td><td>||||||||[||||||||||||]PAM</td><td>NA</td><td>Intergenic</td><td>NA</td><td>NA</td></tr>
<tr><td></td><td></td><td></td><td></td><td></td><td></td><td></td><td></td><td></td><td></td><td></td><td></td></tr>
<tr><td>T33892</td><td>AATCGACAGAAACGTTAACGCGG</td><td>1000</td><td></td><td></td><td></td><td></td><td></td><td></td><td></td><td></td><td></td></tr>
<tr><td>Oligo fwd</td><td>CACCAATCGACAGAAACGTTAACG</td><td></td><td></td><td></td><td></td><td></td><td></td><td></td><td></td><td></td><td></td></tr>
<tr><td>Oligo rev</td><td>AAACCGTTAACGTTTCTGTCGATT</td><td></td><td></td><td></td><td></td><td></td><td></td><td></td><td></td><td></td><td></td></tr>
<tr><td>Chromosome</td><td>start</td><td>end</td><td>strand</td><td>MM</td><td>target_seq</td><td>PAM</td><td>alignment</td><td>distance</td><td>position</td><td>gene name</td><td>gene id</td></tr>
<tr><td>GU305914.1</td><td>7110</td><td>7132</td><td>-</td><td>0</td><td>AATCGACAGAAACGTTAACG</td><td>CGG</td><td>||||||||[||||||||||||]PAM</td><td>NA</td><td>Intergenic</td><td>NA</td><td>NA</td></tr>
<tr><td></td><td></td><td></td><td></td><td></td><td></td><td></td><td></td><td></td><td></td><td></td><td></td></tr>
<tr><td>T33893</td><td>GACAGAAACGTTAACGCGGCTGG</td><td>1000</td><td></td><td></td><td></td><td></td><td></td><td></td><td></td><td></td><td></td></tr>
<tr><td>Oligo fwd</td><td>CACCGACAGAAACGTTAACGCGGC</td><td></td><td></td><td></td><td></td><td></td><td></td><td></td><td></td><td></td><td></td></tr>
<tr><td>Oligo rev</td><td>AAACGCCGCGTTAACGTTTCTGTC</td><td></td><td></td><td></td><td></td><td></td><td></td><td></td><td></td><td></td><td></td></tr>
<tr><td>Chromosome</td><td>start</td><td>end</td><td>strand</td><td>MM</td><td>target_seq</td><td>PAM</td><td>alignment</td><td>distance</td><td>position</td><td>gene name</td><td>gene id</td></tr>
<tr><td>GU305914.1</td><td>7106</td><td>7128</td><td>-</td><td>0</td><td>GACAGAAACGTTAACGCGGC</td><td>TGG</td><td>||||||||[||||||||||||]PAM</td><td>NA</td><td>Intergenic</td><td>NA</td><td>NA</td></tr>
<tr><td></td><td></td><td></td><td></td><td></td><td></td><td></td><td></td><td></td><td></td><td></td><td></td></tr>
<tr><td>T33894</td><td>ACGTTAACGCGGCTGGACGTCGG</td><td>1000</td><td></td><td></td><td></td><td></td><td></td><td></td><td></td><td></td><td></td></tr>
<tr><td>Oligo fwd</td><td>CACCACGTTAACGCGGCTGGACGT</td><td></td><td></td><td></td><td></td><td></td><td></td><td></td><td></td><td></td><td></td></tr>
<tr><td>Oligo rev</td><td>AAACACGTCCAGCCGCGTTAACGT</td><td></td><td></td><td></td><td></td><td></td><td></td><td></td><td></td><td></td><td></td></tr>
<tr><td>Chromosome</td><td>start</td><td>end</td><td>strand</td><td>MM</td><td>target_seq</td><td>PAM</td><td>alignment</td><td>distance</td><td>position</td><td>gene name</td><td>gene id</td></tr>
<tr><td>GU305914.1</td><td>7099</td><td>7121</td><td>-</td><td>0</td><td>ACGTTAACGCGGCTGGACGT</td><td>CGG</td><td>||||||||[||||||||||||]PAM</td><td>NA</td><td>Intergenic</td><td>NA</td><td>NA</td></tr>
<tr><td></td><td></td><td></td><td></td><td></td><td></td><td></td><td></td><td></td><td></td><td></td><td></td></tr>
<tr><td>T33895</td><td>ACGCGGCTGGACGTCGGTATCGG</td><td>1000</td><td></td><td></td><td></td><td></td><td></td><td></td><td></td><td></td><td></td></tr>
<tr><td>Oligo fwd</td><td>CACCACGCGGCTGGACGTCGGTAT</td><td></td><td></td><td></td><td></td><td></td><td></td><td></td><td></td><td></td><td></td></tr>
<tr><td>Oligo rev</td><td>AAACATACCGACGTCCAGCCGCGT</td><td></td><td></td><td></td><td></td><td></td><td></td><td></td><td></td><td></td><td></td></tr>
<tr><td>Chromosome</td><td>start</td><td>end</td><td>strand</td><td>MM</td><td>target_seq</td><td>PAM</td><td>alignment</td><td>distance</td><td>position</td><td>gene name</td><td>gene id</td></tr>
<tr><td>GU305914.1</td><td>7093</td><td>7115</td><td>-</td><td>0</td><td>ACGCGGCTGGACGTCGGTAT</td><td>CGG</td><td>||||||||[||||||||||||]PAM</td><td>NA</td><td>Intergenic</td><td>NA</td><td>NA</td></tr>
<tr><td></td><td></td><td></td><td></td><td></td><td></td><td></td><td></td><td></td><td></td><td></td><td></td></tr>
<tr><td>T33896</td><td>ACGTCGGTATCGGCGATGTGTGG</td><td>1000</td><td></td><td></td><td></td><td></td><td></td><td></td><td></td><td></td><td></td></tr>
<tr><td>Oligo fwd</td><td>CACCACGTCGGTATCGGCGATGTG</td><td></td><td></td><td></td><td></td><td></td><td></td><td></td><td></td><td></td><td></td></tr>
<tr><td>Oligo rev</td><td>AAACCACATCGCCGATACCGACGT</td><td></td><td></td><td></td><td></td><td></td><td></td><td></td><td></td><td></td><td></td></tr>
<tr><td>Chromosome</td><td>start</td><td>end</td><td>strand</td><td>MM</td><td>target_seq</td><td>PAM</td><td>alignment</td><td>distance</td><td>position</td><td>gene name</td><td>gene id</td></tr>
<tr><td>GU305914.1</td><td>7083</td><td>7105</td><td>-</td><td>0</td><td>ACGTCGGTATCGGCGATGTG</td><td>TGG</td><td>||||||||[||||||||||||]PAM</td><td>NA</td><td>Intergenic</td><td>NA</td><td>NA</td></tr>
<tr><td></td><td></td><td></td><td></td><td></td><td></td><td></td><td></td><td></td><td></td><td></td><td></td></tr>
<tr><td>T33897</td><td>TATCGGCGATGTGTGGTTGACGG</td><td>1000</td><td></td><td></td><td></td><td></td><td></td><td></td><td></td><td></td><td></td></tr>
<tr><td>Oligo fwd</td><td>CACCTATCGGCGATGTGTGGTTGA</td><td></td><td></td><td></td><td></td><td></td><td></td><td></td><td></td><td></td><td></td></tr>
<tr><td>Oligo rev</td><td>AAACTCAACCACACATCGCCGATA</td><td></td><td></td><td></td><td></td><td></td><td></td><td></td><td></td><td></td><td></td></tr>
<tr><td>Chromosome</td><td>start</td><td>end</td><td>strand</td><td>MM</td><td>target_seq</td><td>PAM</td><td>alignment</td><td>distance</td><td>position</td><td>gene name</td><td>gene id</td></tr>
<tr><td>GU305914.1</td><td>7076</td><td>7098</td><td>-</td><td>0</td><td>TATCGGCGATGTGTGGTTGA</td><td>CGG</td><td>||||||||[||||||||||||]PAM</td><td>NA</td><td>Intergenic</td><td>NA</td><td>NA</td></tr>
<tr><td></td><td></td><td></td><td></td><td></td><td></td><td></td><td></td><td></td><td></td><td></td><td></td></tr>
<tr><td>T33898</td><td>GGCGATGTGTGGTTGACGGTTGG</td><td>1000</td><td></td><td></td><td></td><td></td><td></td><td></td><td></td><td></td><td></td></tr>
<tr><td>Oligo fwd</td><td>CACCGGCGATGTGTGGTTGACGGT</td><td></td><td></td><td></td><td></td><td></td><td></td><td></td><td></td><td></td><td></td></tr>
<tr><td>Oligo rev</td><td>AAACACCGTCAACCACACATCGCC</td><td></td><td></td><td></td><td></td><td></td><td></td><td></td><td></td><td></td><td></td></tr>
<tr><td>Chromosome</td><td>start</td><td>end</td><td>strand</td><td>MM</td><td>target_seq</td><td>PAM</td><td>alignment</td><td>distance</td><td>position</td><td>gene name</td><td>gene id</td></tr>
<tr><td>GU305914.1</td><td>7072</td><td>7094</td><td>-</td><td>0</td><td>GGCGATGTGTGGTTGACGGT</td><td>TGG</td><td>||||||||[||||||||||||]PAM</td><td>NA</td><td>Intergenic</td><td>NA</td><td>NA</td></tr>
<tr><td></td><td></td><td></td><td></td><td></td><td></td><td></td><td></td><td></td><td></td><td></td><td></td></tr>
<tr><td>T33899</td><td>GCGATGTGTGGTTGACGGTTGGG</td><td>1000</td><td></td><td></td><td></td><td></td><td></td><td></td><td></td><td></td><td></td></tr>
<tr><td>Oligo fwd</td><td>CACCGCGATGTGTGGTTGACGGTT</td><td></td><td></td><td></td><td></td><td></td><td></td><td></td><td></td><td></td><td></td></tr>
<tr><td>Oligo rev</td><td>AAACAACCGTCAACCACACATCGC</td><td></td><td></td><td></td><td></td><td></td><td></td><td></td><td></td><td></td><td></td></tr>
<tr><td>Chromosome</td><td>start</td><td>end</td><td>strand</td><td>MM</td><td>target_seq</td><td>PAM</td><td>alignment</td><td>distance</td><td>position</td><td>gene name</td><td>gene id</td></tr>
<tr><td>GU305914.1</td><td>7071</td><td>7093</td><td>-</td><td>0</td><td>GCGATGTGTGGTTGACGGTT</td><td>GGG</td><td>||||||||[||||||||||||]PAM</td><td>NA</td><td>Intergenic</td><td>NA</td><td>NA</td></tr>
</table>

| | | | | | | | | | | | | |
|---|---|---|---|---|---|---|---|---|---|---|---|---|
| T33900 | CGATGTGTGGTTGACGGTTGGGG | | 1000 | | | | | | | | | |
| Oligo fwd | CACCCGATGTGTGGTTGACGGTTG | | | | | | | | | | | |
| Oligo rev | AAACCAACCGTCAACCACACATCG | | | | | | | | | | | |
| Chromosome | start | | end | strand | MM | target_seq | PAM | alignment | distance | position | gene name | gene id |
| GU305914.1 | | 7070 | 7092 | - | 0 | CGATGTGTGGTTGACGGTTG | GGG | \|\|\|\|\|\|\|\|[\|\|\|\|\|\|\|\|\|\|\|\|]PAM | NA | Intergenic | NA | NA |
| T33901 | GTGGTTGACGGTTGGGGATTCGG | | 1000 | | | | | | | | | |
| Oligo fwd | CACCGTGGTTGACGGTTGGGGATT | | | | | | | | | | | |
| Oligo rev | AAACAATCCCCAACCGTCAACCAC | | | | | | | | | | | |
| Chromosome | start | | end | strand | MM | target_seq | PAM | alignment | distance | position | gene name | gene id |
| GU305914.1 | | 7064 | 7086 | - | 0 | GTGGTTGACGGTTGGGGATT | CGG | \|\|\|\|\|\|\|\|[\|\|\|\|\|\|\|\|\|\|\|\|]PAM | NA | Intergenic | NA | NA |
| T33902 | CGGTTGGGGATTCGGTCGCCCGG | | 1000 | | | | | | | | | |
| Oligo fwd | CACCCGGTTGGGGATTCGGTCGCC | | | | | | | | | | | |
| Oligo rev | AAACGGCGACCGAATCCCCAACCG | | | | | | | | | | | |
| Chromosome | start | | end | strand | MM | target_seq | PAM | alignment | distance | position | gene name | gene id |
| GU305914.1 | | 7056 | 7078 | - | 0 | CGGTTGGGGATTCGGTCGCC | CGG | \|\|\|\|\|\|\|\|[\|\|\|\|\|\|\|\|\|\|\|\|]PAM | NA | Intergenic | NA | NA |
| T33903 | GGATTCGGTCGCCCGGCGTGTGG | | 1000 | | | | | | | | | |
| Oligo fwd | CACCGGATTCGGTCGCCCGGCGTG | | | | | | | | | | | |
| Oligo rev | AAACCACGCCGGGCGACCGAATCC | | | | | | | | | | | |
| Chromosome | start | | end | strand | MM | target_seq | PAM | alignment | distance | position | gene name | gene id |
| GU305914.1 | | 7049 | 7071 | - | 0 | GGATTCGGTCGCCCGGCGTG | TGG | \|\|\|\|\|\|\|\|[\|\|\|\|\|\|\|\|\|\|\|\|]PAM | NA | Intergenic | NA | NA |
| T33904 | TGTGGTGATGAGTTTTCTGATGG | | 1000 | | | | | | | | | |
| Oligo fwd | CACCTGTGGTGATGAGTTTTCTGA | | | | | | | | | | | |
| Oligo rev | AAACTCAGAAAACTCATCACCACA | | | | | | | | | | | |
| Chromosome | start | | end | strand | MM | target_seq | PAM | alignment | distance | position | gene name | gene id |
| GU305914.1 | | 7031 | 7053 | - | 0 | TGTGGTGATGAGTTTTCTGA | TGG | \|\|\|\|\|\|\|\|[\|\|\|\|\|\|\|\|\|\|\|\|]PAM | NA | Intergenic | NA | NA |
| T33905 | AGAAGTATGAGTTACAGATAAGG | | 1000 | | | | | | | | | |
| Oligo fwd | CACCAGAAGTATGAGTTACAGATA | | | | | | | | | | | |
| Oligo rev | AAACTATCTGTAACTCATACTTCT | | | | | | | | | | | |
| Chromosome | start | | end | strand | MM | target_seq | PAM | alignment | distance | position | gene name | gene id |
| GU305914.1 | | 7007 | 7029 | - | 0 | AGAAGTATGAGTTACAGATA | AGG | \|\|\|\|\|\|\|\|[\|\|\|\|\|\|\|\|\|\|\|\|]PAM | NA | Intergenic | NA | NA |
| T33906 | AGATAAGGTTGTATAACTCGTGG | | 1000 | | | | | | | | | |
| Oligo fwd | CACCAGATAAGGTTGTATAACTCG | | | | | | | | | | | |
| Oligo rev | AAACCGAGTTATACAACCTTATCT | | | | | | | | | | | |
| Chromosome | start | | end | strand | MM | target_seq | PAM | alignment | distance | position | gene name | gene id |
| GU305914.1 | | 6992 | 7014 | - | 0 | AGATAAGGTTGTATAACTCG | TGG | \|\|\|\|\|\|\|\|[\|\|\|\|\|\|\|\|\|\|\|\|]PAM | NA | Intergenic | NA | NA |
| T33907 | GTTGTATAACTCGTGGTTGTAGG | | 1000 | | | | | | | | | |
| Oligo fwd | CACCGTTGTATAACTCGTGGTTGT | | | | | | | | | | | |
| Oligo rev | AAACACAACCACGAGTTATACAAC | | | | | | | | | | | |
| Chromosome | start | | end | strand | MM | target_seq | PAM | alignment | distance | position | gene name | gene id |
| GU305914.1 | | 6985 | 7007 | - | 0 | GTTGTATAACTCGTGGTTGT | AGG | \|\|\|\|\|\|\|\|[\|\|\|\|\|\|\|\|\|\|\|\|]PAM | NA | Intergenic | NA | NA |
| T33908 | TTGTATAACTCGTGGTTGTAGGG | | 1000 | | | | | | | | | |
| Oligo fwd | CACCTTGTATAACTCGTGGTTGTA | | | | | | | | | | | |
| Oligo rev | AAACTACAACCACGAGTTATACAA | | | | | | | | | | | |
| Chromosome | start | | end | strand | MM | target_seq | PAM | alignment | distance | position | gene name | gene id |
| GU305914.1 | | 6984 | 7006 | - | 0 | TTGTATAACTCGTGGTTGTA | GGG | \|\|\|\|\|\|\|\|[\|\|\|\|\|\|\|\|\|\|\|\|]PAM | NA | Intergenic | NA | NA |
| T33909 | AACCGTCGTCGTGCGTTTCTTGG | | 1000 | | | | | | | | | |
| Oligo fwd | CACCAACCGTCGTCGTGCGTTTCT | | | | | | | | | | | |
| Oligo rev | AAACAGAAACGCACGACGACGGTT | | | | | | | | | | | |
| Chromosome | start | | end | strand | MM | target_seq | PAM | alignment | distance | position | gene name | gene id |
| GU305914.1 | | 6959 | 6981 | - | 0 | AACCGTCGTCGTGCGTTTCT | TGG | \|\|\|\|\|\|\|\|[\|\|\|\|\|\|\|\|\|\|\|\|]PAM | NA | Intergenic | NA | NA |
| T33910 | ACCGTCGTCGTGCGTTTCTTGGG | | 1000 | | | | | | | | | |
| Oligo fwd | CACCACCGTCGTCGTGCGTTTCTT | | | | | | | | | | | |
| Oligo rev | AAACAAGAAACGCACGACGACGGT | | | | | | | | | | | |
| Chromosome | start | | end | strand | MM | target_seq | PAM | alignment | distance | position | gene name | gene id |
| GU305914.1 | | 6958 | 6980 | - | 0 | ACCGTCGTCGTGCGTTTCTT | GGG | \|\|\|\|\|\|\|\|[\|\|\|\|\|\|\|\|\|\|\|\|]PAM | NA | Intergenic | NA | NA |
| T33911 | GTGCGTTTCTTGGGCCGTCTTGG | | 1000 | | | | | | | | | |
| Oligo fwd | CACCGTGCGTTTCTTGGGCCGTCT | | | | | | | | | | | |
| Oligo rev | AAACAGACGGCCCAAGAAACGCAC | | | | | | | | | | | |
| Chromosome | start | | end | strand | MM | target_seq | PAM | alignment | distance | position | gene name | gene id |
| GU305914.1 | | 6949 | 6971 | - | 0 | GTGCGTTTCTTGGGCCGTCT | TGG | \|\|\|\|\|\|\|\|[\|\|\|\|\|\|\|\|\|\|\|\|]PAM | NA | Intergenic | NA | NA |
| T33912 | CGTTTCTTGGGCCGTCTTGGTGG | | 1000 | | | | | | | | | |
| Oligo fwd | CACCCGTTTCTTGGGCCGTCTTGG | | | | | | | | | | | |
| Oligo rev | AAACCCAAGACGGCCCAAGAAACG | | | | | | | | | | | |
| Chromosome | start | | end | strand | MM | target_seq | PAM | alignment | distance | position | gene name | gene id |
| GU305914.1 | | 6946 | 6968 | - | 0 | CGTTTCTTGGGCCGTCTTGG | TGG | \|\|\|\|\|\|\|\|[\|\|\|\|\|\|\|\|\|\|\|\|]PAM | NA | Intergenic | NA | NA |
| T33913 | TTCTTGGGCCGTCTTGGTGGAGG | | 1000 | | | | | | | | | |
| Oligo fwd | CACCTTCTTGGGCCGTCTTGGTGG | | | | | | | | | | | |
| Oligo rev | AAACCCACCAAGACGGCCCAAGAA | | | | | | | | | | | |
| Chromosome | start | | end | strand | MM | target_seq | PAM | alignment | distance | position | gene name | gene id |
| GU305914.1 | | 6943 | 6965 | - | 0 | TTCTTGGGCCGTCTTGGTGG | AGG | \|\|\|\|\|\|\|\|[\|\|\|\|\|\|\|\|\|\|\|\|]PAM | NA | Intergenic | NA | NA |
| T33914 | GGCCGTCTTGGTGGAGGTTTAGG | | 1000 | | | | | | | | | |
| Oligo fwd | CACCGGCCGTCTTGGTGGAGGTTT | | | | | | | | | | | |
| Oligo rev | AAACAAACCTCCACCAAGACGGCC | | | | | | | | | | | |
| Chromosome | start | | end | strand | MM | target_seq | PAM | alignment | distance | position | gene name | gene id |
| GU305914.1 | | 6937 | 6959 | - | 0 | GGCCGTCTTGGTGGAGGTTT | AGG | \|\|\|\|\|\|\|\|[\|\|\|\|\|\|\|\|\|\|\|\|]PAM | NA | Intergenic | NA | NA |

<table>
<tr><td>T33915</td><td>AGGTTTAGGTCCGTAGCCGTCGG</td><td>1000</td><td></td><td></td><td></td><td></td><td></td><td></td><td></td><td></td><td></td></tr>
<tr><td>Oligo fwd</td><td>CACCAGGTTTAGGTCCGTAGCCGT</td><td></td><td></td><td></td><td></td><td></td><td></td><td></td><td></td><td></td><td></td></tr>
<tr><td>Oligo rev</td><td>AAACACGGCTACGGACCTAAACCT</td><td></td><td></td><td></td><td></td><td></td><td></td><td></td><td></td><td></td><td></td></tr>
<tr><td>Chromosome</td><td>start</td><td>end</td><td>strand</td><td>MM</td><td>target_seq</td><td>PAM</td><td>alignment</td><td>distance</td><td>position</td><td>gene name</td><td>gene id</td></tr>
<tr><td>GU305914.1</td><td>6923</td><td>6945</td><td>-</td><td>0</td><td>AGGTTTAGGTCCGTAGCCGT</td><td>CGG</td><td>||||||||[||||||||||||]PAM</td><td>NA</td><td>Intergenic</td><td>NA</td><td>NA</td></tr>
<tr><td>T33916</td><td>AATCGTCAGTTTACACGTGACGG</td><td>1000</td><td></td><td></td><td></td><td></td><td></td><td></td><td></td><td></td><td></td></tr>
<tr><td>Oligo fwd</td><td>CACCAATCGTCAGTTTACACGTGA</td><td></td><td></td><td></td><td></td><td></td><td></td><td></td><td></td><td></td><td></td></tr>
<tr><td>Oligo rev</td><td>AAACTCACGTGTAAACTGACGATT</td><td></td><td></td><td></td><td></td><td></td><td></td><td></td><td></td><td></td><td></td></tr>
<tr><td>Chromosome</td><td>start</td><td>end</td><td>strand</td><td>MM</td><td>target_seq</td><td>PAM</td><td>alignment</td><td>distance</td><td>position</td><td>gene name</td><td>gene id</td></tr>
<tr><td>GU305914.1</td><td>6890</td><td>6912</td><td>-</td><td>0</td><td>AATCGTCAGTTTACACGTGA</td><td>CGG</td><td>||||||||[||||||||||||]PAM</td><td>NA</td><td>Intergenic</td><td>NA</td><td>NA</td></tr>
<tr><td>T33917</td><td>TGACGGACACTGATCTAGTTAGG</td><td>1000</td><td></td><td></td><td></td><td></td><td></td><td></td><td></td><td></td><td></td></tr>
<tr><td>Oligo fwd</td><td>CACCTGACGGACACTGATCTAGTT</td><td></td><td></td><td></td><td></td><td></td><td></td><td></td><td></td><td></td><td></td></tr>
<tr><td>Oligo rev</td><td>AAACAACTAGATCAGTGTCCGTCA</td><td></td><td></td><td></td><td></td><td></td><td></td><td></td><td></td><td></td><td></td></tr>
<tr><td>Chromosome</td><td>start</td><td>end</td><td>strand</td><td>MM</td><td>target_seq</td><td>PAM</td><td>alignment</td><td>distance</td><td>position</td><td>gene name</td><td>gene id</td></tr>
<tr><td>GU305914.1</td><td>6873</td><td>6895</td><td>-</td><td>0</td><td>TGACGGACACTGATCTAGTT</td><td>AGG</td><td>||||||||[||||||||||||]PAM</td><td>NA</td><td>Intergenic</td><td>NA</td><td>NA</td></tr>
<tr><td>T33918</td><td>GGACACTGATCTAGTTAGGTTGG</td><td>1000</td><td></td><td></td><td></td><td></td><td></td><td></td><td></td><td></td><td></td></tr>
<tr><td>Oligo fwd</td><td>CACCGGACACTGATCTAGTTAGGT</td><td></td><td></td><td></td><td></td><td></td><td></td><td></td><td></td><td></td><td></td></tr>
<tr><td>Oligo rev</td><td>AAACACCTAACTAGATCAGTGTCC</td><td></td><td></td><td></td><td></td><td></td><td></td><td></td><td></td><td></td><td></td></tr>
<tr><td>Chromosome</td><td>start</td><td>end</td><td>strand</td><td>MM</td><td>target_seq</td><td>PAM</td><td>alignment</td><td>distance</td><td>position</td><td>gene name</td><td>gene id</td></tr>
<tr><td>GU305914.1</td><td>6869</td><td>6891</td><td>-</td><td>0</td><td>GGACACTGATCTAGTTAGGT</td><td>TGG</td><td>||||||||[||||||||||||]PAM</td><td>NA</td><td>Intergenic</td><td>NA</td><td>NA</td></tr>
<tr><td>T33919</td><td>TTAGGTTGGAAGAGACGATAAGG</td><td>1000</td><td></td><td></td><td></td><td></td><td></td><td></td><td></td><td></td><td></td></tr>
<tr><td>Oligo fwd</td><td>CACCTTAGGTTGGAAGAGACGATA</td><td></td><td></td><td></td><td></td><td></td><td></td><td></td><td></td><td></td><td></td></tr>
<tr><td>Oligo rev</td><td>AAACTATCGTCTCTTCCAACCTAA</td><td></td><td></td><td></td><td></td><td></td><td></td><td></td><td></td><td></td><td></td></tr>
<tr><td>Chromosome</td><td>start</td><td>end</td><td>strand</td><td>MM</td><td>target_seq</td><td>PAM</td><td>alignment</td><td>distance</td><td>position</td><td>gene name</td><td>gene id</td></tr>
<tr><td>GU305914.1</td><td>6855</td><td>6877</td><td>-</td><td>0</td><td>TTAGGTTGGAAGAGACGATA</td><td>AGG</td><td>||||||||[||||||||||||]PAM</td><td>NA</td><td>Intergenic</td><td>NA</td><td>NA</td></tr>
<tr><td>T33920</td><td>GACGATAAGGCGCCCTGACGTGG</td><td>1000</td><td></td><td></td><td></td><td></td><td></td><td></td><td></td><td></td><td></td></tr>
<tr><td>Oligo fwd</td><td>CACCGACGATAAGGCGCCCTGACG</td><td></td><td></td><td></td><td></td><td></td><td></td><td></td><td></td><td></td><td></td></tr>
<tr><td>Oligo rev</td><td>AAACCGTCAGGGCGCCTTATCGTC</td><td></td><td></td><td></td><td></td><td></td><td></td><td></td><td></td><td></td><td></td></tr>
<tr><td>Chromosome</td><td>start</td><td>end</td><td>strand</td><td>MM</td><td>target_seq</td><td>PAM</td><td>alignment</td><td>distance</td><td>position</td><td>gene name</td><td>gene id</td></tr>
<tr><td>GU305914.1</td><td>6842</td><td>6864</td><td>-</td><td>0</td><td>GACGATAAGGCGCCCTGACG</td><td>TGG</td><td>||||||||[||||||||||||]PAM</td><td>NA</td><td>Intergenic</td><td>NA</td><td>NA</td></tr>
<tr><td>T33921</td><td>ACGATAAGGCGCCCTGACGTGGG</td><td>1000</td><td></td><td></td><td></td><td></td><td></td><td></td><td></td><td></td><td></td></tr>
<tr><td>Oligo fwd</td><td>CACCACGATAAGGCGCCCTGACGT</td><td></td><td></td><td></td><td></td><td></td><td></td><td></td><td></td><td></td><td></td></tr>
<tr><td>Oligo rev</td><td>AAACACGTCAGGGCGCCTTATCGT</td><td></td><td></td><td></td><td></td><td></td><td></td><td></td><td></td><td></td><td></td></tr>
<tr><td>Chromosome</td><td>start</td><td>end</td><td>strand</td><td>MM</td><td>target_seq</td><td>PAM</td><td>alignment</td><td>distance</td><td>position</td><td>gene name</td><td>gene id</td></tr>
<tr><td>GU305914.1</td><td>6841</td><td>6863</td><td>-</td><td>0</td><td>ACGATAAGGCGCCCTGACGT</td><td>GGG</td><td>||||||||[||||||||||||]PAM</td><td>NA</td><td>Intergenic</td><td>NA</td><td>NA</td></tr>
<tr><td>T33922</td><td>TGTATAATCTGTAGCGTTGATGG</td><td>1000</td><td></td><td></td><td></td><td></td><td></td><td></td><td></td><td></td><td></td></tr>
<tr><td>Oligo fwd</td><td>CACCTGTATAATCTGTAGCGTTGA</td><td></td><td></td><td></td><td></td><td></td><td></td><td></td><td></td><td></td><td></td></tr>
<tr><td>Oligo rev</td><td>AAACTCAACGCTACAGATTATACA</td><td></td><td></td><td></td><td></td><td></td><td></td><td></td><td></td><td></td><td></td></tr>
<tr><td>Chromosome</td><td>start</td><td>end</td><td>strand</td><td>MM</td><td>target_seq</td><td>PAM</td><td>alignment</td><td>distance</td><td>position</td><td>gene name</td><td>gene id</td></tr>
<tr><td>GU305914.1</td><td>6812</td><td>6834</td><td>-</td><td>0</td><td>TGTATAATCTGTAGCGTTGA</td><td>TGG</td><td>||||||||[||||||||||||]PAM</td><td>NA</td><td>Intergenic</td><td>NA</td><td>NA</td></tr>
<tr><td>T33923</td><td>TAATCTGTAGCGTTGATGGTAGG</td><td>1000</td><td></td><td></td><td></td><td></td><td></td><td></td><td></td><td></td><td></td></tr>
<tr><td>Oligo fwd</td><td>CACCTAATCTGTAGCGTTGATGGT</td><td></td><td></td><td></td><td></td><td></td><td></td><td></td><td></td><td></td><td></td></tr>
<tr><td>Oligo rev</td><td>AAACACCATCAACGCTACAGATTA</td><td></td><td></td><td></td><td></td><td></td><td></td><td></td><td></td><td></td><td></td></tr>
<tr><td>Chromosome</td><td>start</td><td>end</td><td>strand</td><td>MM</td><td>target_seq</td><td>PAM</td><td>alignment</td><td>distance</td><td>position</td><td>gene name</td><td>gene id</td></tr>
<tr><td>GU305914.1</td><td>6808</td><td>6830</td><td>-</td><td>0</td><td>TAATCTGTAGCGTTGATGGT</td><td>AGG</td><td>||||||||[||||||||||||]PAM</td><td>NA</td><td>Intergenic</td><td>NA</td><td>NA</td></tr>
<tr><td>T33924</td><td>TCTGTAGCGTTGATGGTAGGCGG</td><td>1000</td><td></td><td></td><td></td><td></td><td></td><td></td><td></td><td></td><td></td></tr>
<tr><td>Oligo fwd</td><td>CACCTCTGTAGCGTTGATGGTAGG</td><td></td><td></td><td></td><td></td><td></td><td></td><td></td><td></td><td></td><td></td></tr>
<tr><td>Oligo rev</td><td>AAACCCTACCATCAACGCTACAGA</td><td></td><td></td><td></td><td></td><td></td><td></td><td></td><td></td><td></td><td></td></tr>
<tr><td>Chromosome</td><td>start</td><td>end</td><td>strand</td><td>MM</td><td>target_seq</td><td>PAM</td><td>alignment</td><td>distance</td><td>position</td><td>gene name</td><td>gene id</td></tr>
<tr><td>GU305914.1</td><td>6805</td><td>6827</td><td>-</td><td>0</td><td>TCTGTAGCGTTGATGGTAGG</td><td>CGG</td><td>||||||||[||||||||||||]PAM</td><td>NA</td><td>Intergenic</td><td>NA</td><td>NA</td></tr>
<tr><td>T33925</td><td>AGCGTTGATGGTAGGCGGTGTGG</td><td>1000</td><td></td><td></td><td></td><td></td><td></td><td></td><td></td><td></td><td></td></tr>
<tr><td>Oligo fwd</td><td>CACCAGCGTTGATGGTAGGCGGTG</td><td></td><td></td><td></td><td></td><td></td><td></td><td></td><td></td><td></td><td></td></tr>
<tr><td>Oligo rev</td><td>AAACCACCGCCTACCATCAACGCT</td><td></td><td></td><td></td><td></td><td></td><td></td><td></td><td></td><td></td><td></td></tr>
<tr><td>Chromosome</td><td>start</td><td>end</td><td>strand</td><td>MM</td><td>target_seq</td><td>PAM</td><td>alignment</td><td>distance</td><td>position</td><td>gene name</td><td>gene id</td></tr>
<tr><td>GU305914.1</td><td>6800</td><td>6822</td><td>-</td><td>0</td><td>AGCGTTGATGGTAGGCGGTG</td><td>TGG</td><td>||||||||[||||||||||||]PAM</td><td>NA</td><td>Intergenic</td><td>NA</td><td>NA</td></tr>
<tr><td>T33926</td><td>ATGGTAGGCGGTGTGGTAGTCGG</td><td>1000</td><td></td><td></td><td></td><td></td><td></td><td></td><td></td><td></td><td></td></tr>
<tr><td>Oligo fwd</td><td>CACCATGGTAGGCGGTGTGGTAGT</td><td></td><td></td><td></td><td></td><td></td><td></td><td></td><td></td><td></td><td></td></tr>
<tr><td>Oligo rev</td><td>AAACACTACCACACCGCCTACCAT</td><td></td><td></td><td></td><td></td><td></td><td></td><td></td><td></td><td></td><td></td></tr>
<tr><td>Chromosome</td><td>start</td><td>end</td><td>strand</td><td>MM</td><td>target_seq</td><td>PAM</td><td>alignment</td><td>distance</td><td>position</td><td>gene name</td><td>gene id</td></tr>
<tr><td>GU305914.1</td><td>6793</td><td>6815</td><td>-</td><td>0</td><td>ATGGTAGGCGGTGTGGTAGT</td><td>CGG</td><td>||||||||[||||||||||||]PAM</td><td>NA</td><td>Intergenic</td><td>NA</td><td>NA</td></tr>
<tr><td>T33927</td><td>GTGTGGTAGTCGGCTGCGTTTGG</td><td>1000</td><td></td><td></td><td></td><td></td><td></td><td></td><td></td><td></td><td></td></tr>
<tr><td>Oligo fwd</td><td>CACCGTGTGGTAGTCGGCTGCGTT</td><td></td><td></td><td></td><td></td><td></td><td></td><td></td><td></td><td></td><td></td></tr>
<tr><td>Oligo rev</td><td>AAACAACGCAGCCGACTACCACAC</td><td></td><td></td><td></td><td></td><td></td><td></td><td></td><td></td><td></td><td></td></tr>
<tr><td>Chromosome</td><td>start</td><td>end</td><td>strand</td><td>MM</td><td>target_seq</td><td>PAM</td><td>alignment</td><td>distance</td><td>position</td><td>gene name</td><td>gene id</td></tr>
<tr><td>GU305914.1</td><td>6783</td><td>6805</td><td>-</td><td>0</td><td>GTGTGGTAGTCGGCTGCGTT</td><td>TGG</td><td>||||||||[||||||||||||]PAM</td><td>NA</td><td>Intergenic</td><td>NA</td><td>NA</td></tr>
<tr><td>T33928</td><td>TTAAAAGTCATGCTAAACAAAGG</td><td>1000</td><td></td><td></td><td></td><td></td><td></td><td></td><td></td><td></td><td></td></tr>
<tr><td>Oligo fwd</td><td>CACCTTAAAAGTCATGCTAAACAA</td><td></td><td></td><td></td><td></td><td></td><td></td><td></td><td></td><td></td><td></td></tr>
<tr><td>Oligo rev</td><td>AAACTTGTTTAGCATGACTTTTAA</td><td></td><td></td><td></td><td></td><td></td><td></td><td></td><td></td><td></td><td></td></tr>
<tr><td>Chromosome</td><td>start</td><td>end</td><td>strand</td><td>MM</td><td>target_seq</td><td>PAM</td><td>alignment</td><td>distance</td><td>position</td><td>gene name</td><td>gene id</td></tr>
<tr><td>GU305914.1</td><td>6760</td><td>6782</td><td>-</td><td>0</td><td>TTAAAAGTCATGCTAAACAA</td><td>AGG</td><td>||||||||[||||||||||||]PAM</td><td>NA</td><td>Intergenic</td><td>NA</td><td>NA</td></tr>
<tr><td>T33929</td><td>AAAGTCATGCTAAACAAAGGCGG</td><td>1000</td><td></td><td></td><td></td><td></td><td></td><td></td><td></td><td></td><td></td></tr>
<tr><td>Oligo fwd</td><td>CACCAAAGTCATGCTAAACAAAGG</td><td></td><td></td><td></td><td></td><td></td><td></td><td></td><td></td><td></td><td></td></tr>
<tr><td>Oligo rev</td><td>AAACCCTTTGTTTAGCATGACTTT</td><td></td><td></td><td></td><td></td><td></td><td></td><td></td><td></td><td></td><td></td></tr>
<tr><td>Chromosome</td><td>start</td><td>end</td><td>strand</td><td>MM</td><td>target_seq</td><td>PAM</td><td>alignment</td><td>distance</td><td>position</td><td>gene name</td><td>gene id</td></tr>
</table>

<table>
<tr><td>GU305914.1</td><td></td><td>6757</td><td>6779</td><td>-</td><td>0</td><td>AAAGTCATGCTAAACAAAGG</td><td>CGG</td><td>||||||||[||||||||||||]PAM</td><td>NA</td><td>Intergenic</td><td>NA</td><td>NA</td></tr>
<tr><td>T33930</td><td>CAAAGGCGGACTGCTGTTCGAGG</td><td></td><td>1000</td><td></td><td></td><td></td><td></td><td></td><td></td><td></td><td></td><td></td></tr>
<tr><td>Oligo fwd</td><td>CACCCAAAGGCGGACTGCTGTTCG</td><td></td><td></td><td></td><td></td><td></td><td></td><td></td><td></td><td></td><td></td><td></td></tr>
<tr><td>Oligo rev</td><td>AAACCGAACAGCAGTCCGCCTTTG</td><td></td><td></td><td></td><td></td><td></td><td></td><td></td><td></td><td></td><td></td><td></td></tr>
<tr><td>Chromosome</td><td>start</td><td></td><td>end</td><td>strand</td><td>MM</td><td>target_seq</td><td>PAM</td><td>alignment</td><td>distance</td><td>position</td><td>gene name</td><td>gene id</td></tr>
<tr><td>GU305914.1</td><td></td><td>6743</td><td>6765</td><td>-</td><td>0</td><td>CAAAGGCGGACTGCTGTTCG</td><td>AGG</td><td>||||||||[||||||||||||]PAM</td><td>NA</td><td>Intergenic</td><td>NA</td><td>NA</td></tr>
<tr><td>T33931</td><td>GACTTGAAATAGACTCCGAACGG</td><td></td><td>1000</td><td></td><td></td><td></td><td></td><td></td><td></td><td></td><td></td><td></td></tr>
<tr><td>Oligo fwd</td><td>CACCGACTTGAAATAGACTCCGAA</td><td></td><td></td><td></td><td></td><td></td><td></td><td></td><td></td><td></td><td></td><td></td></tr>
<tr><td>Oligo rev</td><td>AAACTTCGGAGTCTATTTCAAGTC</td><td></td><td></td><td></td><td></td><td></td><td></td><td></td><td></td><td></td><td></td><td></td></tr>
<tr><td>Chromosome</td><td>start</td><td></td><td>end</td><td>strand</td><td>MM</td><td>target_seq</td><td>PAM</td><td>alignment</td><td>distance</td><td>position</td><td>gene name</td><td>gene id</td></tr>
<tr><td>GU305914.1</td><td></td><td>6721</td><td>6743</td><td>-</td><td>0</td><td>GACTTGAAATAGACTCCGAA</td><td>CGG</td><td>||||||||[||||||||||||]PAM</td><td>NA</td><td>Intergenic</td><td>NA</td><td>NA</td></tr>
<tr><td>T33932</td><td>AAATAGACTCCGAACGGTAACGG</td><td></td><td>1000</td><td></td><td></td><td></td><td></td><td></td><td></td><td></td><td></td><td></td></tr>
<tr><td>Oligo fwd</td><td>CACCAAATAGACTCCGAACGGTAA</td><td></td><td></td><td></td><td></td><td></td><td></td><td></td><td></td><td></td><td></td><td></td></tr>
<tr><td>Oligo rev</td><td>AAACTTACCGTTCGGAGTCTATTT</td><td></td><td></td><td></td><td></td><td></td><td></td><td></td><td></td><td></td><td></td><td></td></tr>
<tr><td>Chromosome</td><td>start</td><td></td><td>end</td><td>strand</td><td>MM</td><td>target_seq</td><td>PAM</td><td>alignment</td><td>distance</td><td>position</td><td>gene name</td><td>gene id</td></tr>
<tr><td>GU305914.1</td><td></td><td>6715</td><td>6737</td><td>-</td><td>0</td><td>AAATAGACTCCGAACGGTAA</td><td>CGG</td><td>||||||||[||||||||||||]PAM</td><td>NA</td><td>Intergenic</td><td>NA</td><td>NA</td></tr>
<tr><td>T33933</td><td>CCACGTACCTTCCCACGATGCGG</td><td></td><td>1000</td><td></td><td></td><td></td><td></td><td></td><td></td><td></td><td></td><td></td></tr>
<tr><td>Oligo fwd</td><td>CACCCCACGTACCTTCCCACGATG</td><td></td><td></td><td></td><td></td><td></td><td></td><td></td><td></td><td></td><td></td><td></td></tr>
<tr><td>Oligo rev</td><td>AAACCATCGTGGGAAGGTACGTGG</td><td></td><td></td><td></td><td></td><td></td><td></td><td></td><td></td><td></td><td></td><td></td></tr>
<tr><td>Chromosome</td><td>start</td><td></td><td>end</td><td>strand</td><td>MM</td><td>target_seq</td><td>PAM</td><td>alignment</td><td>distance</td><td>position</td><td>gene name</td><td>gene id</td></tr>
<tr><td>GU305914.1</td><td></td><td>6677</td><td>6699</td><td>-</td><td>0</td><td>CCACGTACCTTCCCACGATG</td><td>CGG</td><td>||||||||[||||||||||||]PAM</td><td>NA</td><td>Intergenic</td><td>NA</td><td>NA</td></tr>
<tr><td>T33934</td><td>CACGTACCTTCCCACGATGCGGG</td><td></td><td>1000</td><td></td><td></td><td></td><td></td><td></td><td></td><td></td><td></td><td></td></tr>
<tr><td>Oligo fwd</td><td>CACCCACGTACCTTCCCACGATGC</td><td></td><td></td><td></td><td></td><td></td><td></td><td></td><td></td><td></td><td></td><td></td></tr>
<tr><td>Oligo rev</td><td>AAACGCATCGTGGGAAGGTACGTG</td><td></td><td></td><td></td><td></td><td></td><td></td><td></td><td></td><td></td><td></td><td></td></tr>
<tr><td>Chromosome</td><td>start</td><td></td><td>end</td><td>strand</td><td>MM</td><td>target_seq</td><td>PAM</td><td>alignment</td><td>distance</td><td>position</td><td>gene name</td><td>gene id</td></tr>
<tr><td>GU305914.1</td><td></td><td>6676</td><td>6698</td><td>-</td><td>0</td><td>CACGTACCTTCCCACGATGC</td><td>GGG</td><td>||||||||[||||||||||||]PAM</td><td>NA</td><td>Intergenic</td><td>NA</td><td>NA</td></tr>
<tr><td>T33935</td><td>CCACGATGCGGGAATACAGATGG</td><td></td><td>1000</td><td></td><td></td><td></td><td></td><td></td><td></td><td></td><td></td><td></td></tr>
<tr><td>Oligo fwd</td><td>CACCCCACGATGCGGGAATACAGA</td><td></td><td></td><td></td><td></td><td></td><td></td><td></td><td></td><td></td><td></td><td></td></tr>
<tr><td>Oligo rev</td><td>AAACTCTGTATTCCCGCATCGTGG</td><td></td><td></td><td></td><td></td><td></td><td></td><td></td><td></td><td></td><td></td><td></td></tr>
<tr><td>Chromosome</td><td>start</td><td></td><td>end</td><td>strand</td><td>MM</td><td>target_seq</td><td>PAM</td><td>alignment</td><td>distance</td><td>position</td><td>gene name</td><td>gene id</td></tr>
<tr><td>GU305914.1</td><td></td><td>6665</td><td>6687</td><td>-</td><td>0</td><td>CCACGATGCGGGAATACAGA</td><td>TGG</td><td>||||||||[||||||||||||]PAM</td><td>NA</td><td>Intergenic</td><td>NA</td><td>NA</td></tr>
<tr><td>T33936</td><td>CTGTAGTTAGTAGCGAGTGACGG</td><td></td><td>1000</td><td></td><td></td><td></td><td></td><td></td><td></td><td></td><td></td><td></td></tr>
<tr><td>Oligo fwd</td><td>CACCCTGTAGTTAGTAGCGAGTGA</td><td></td><td></td><td></td><td></td><td></td><td></td><td></td><td></td><td></td><td></td><td></td></tr>
<tr><td>Oligo rev</td><td>AAACTCACTCGCTACTAACTACAG</td><td></td><td></td><td></td><td></td><td></td><td></td><td></td><td></td><td></td><td></td><td></td></tr>
<tr><td>Chromosome</td><td>start</td><td></td><td>end</td><td>strand</td><td>MM</td><td>target_seq</td><td>PAM</td><td>alignment</td><td>distance</td><td>position</td><td>gene name</td><td>gene id</td></tr>
<tr><td>GU305914.1</td><td></td><td>6559</td><td>6581</td><td>-</td><td>0</td><td>CTGTAGTTAGTAGCGAGTGA</td><td>CGG</td><td>||||||||[||||||||||||]PAM</td><td>NA</td><td>Intergenic</td><td>NA</td><td>NA</td></tr>
<tr><td>T33937</td><td>TGTAGTTAGTAGCGAGTGACGGG</td><td></td><td>1000</td><td></td><td></td><td></td><td></td><td></td><td></td><td></td><td></td><td></td></tr>
<tr><td>Oligo fwd</td><td>CACCTGTAGTTAGTAGCGAGTGAC</td><td></td><td></td><td></td><td></td><td></td><td></td><td></td><td></td><td></td><td></td><td></td></tr>
<tr><td>Oligo rev</td><td>AAACGTCACTCGCTACTAACTACA</td><td></td><td></td><td></td><td></td><td></td><td></td><td></td><td></td><td></td><td></td><td></td></tr>
<tr><td>Chromosome</td><td>start</td><td></td><td>end</td><td>strand</td><td>MM</td><td>target_seq</td><td>PAM</td><td>alignment</td><td>distance</td><td>position</td><td>gene name</td><td>gene id</td></tr>
<tr><td>GU305914.1</td><td></td><td>6558</td><td>6580</td><td>-</td><td>0</td><td>TGTAGTTAGTAGCGAGTGAC</td><td>GGG</td><td>||||||||[||||||||||||]PAM</td><td>NA</td><td>Intergenic</td><td>NA</td><td>NA</td></tr>
<tr><td>T33938</td><td>ACTCAGACAAGAGTGTCAAGAGG</td><td></td><td>1000</td><td></td><td></td><td></td><td></td><td></td><td></td><td></td><td></td><td></td></tr>
<tr><td>Oligo fwd</td><td>CACCACTCAGACAAGAGTGTCAAG</td><td></td><td></td><td></td><td></td><td></td><td></td><td></td><td></td><td></td><td></td><td></td></tr>
<tr><td>Oligo rev</td><td>AAACCTTGACACTCTTGTCTGAGT</td><td></td><td></td><td></td><td></td><td></td><td></td><td></td><td></td><td></td><td></td><td></td></tr>
<tr><td>Chromosome</td><td>start</td><td></td><td>end</td><td>strand</td><td>MM</td><td>target_seq</td><td>PAM</td><td>alignment</td><td>distance</td><td>position</td><td>gene name</td><td>gene id</td></tr>
<tr><td>GU305914.1</td><td></td><td>6407</td><td>6429</td><td>-</td><td>0</td><td>ACTCAGACAAGAGTGTCAAG</td><td>AGG</td><td>||||||||[||||||||||||]PAM</td><td>NA</td><td>Intergenic</td><td>NA</td><td>NA</td></tr>
<tr><td>T33939</td><td>CAAGAGTGTCAAGAGGACCACGG</td><td></td><td>1000</td><td></td><td></td><td></td><td></td><td></td><td></td><td></td><td></td><td></td></tr>
<tr><td>Oligo fwd</td><td>CACCCAAGAGTGTCAAGAGGACCA</td><td></td><td></td><td></td><td></td><td></td><td></td><td></td><td></td><td></td><td></td><td></td></tr>
<tr><td>Oligo rev</td><td>AAACTGGTCCTCTTGACACTCTTG</td><td></td><td></td><td></td><td></td><td></td><td></td><td></td><td></td><td></td><td></td><td></td></tr>
<tr><td>Chromosome</td><td>start</td><td></td><td>end</td><td>strand</td><td>MM</td><td>target_seq</td><td>PAM</td><td>alignment</td><td>distance</td><td>position</td><td>gene name</td><td>gene id</td></tr>
<tr><td>GU305914.1</td><td></td><td>6400</td><td>6422</td><td>-</td><td>0</td><td>CAAGAGTGTCAAGAGGACCA</td><td>CGG</td><td>||||||||[||||||||||||]PAM</td><td>NA</td><td>Intergenic</td><td>NA</td><td>NA</td></tr>
<tr><td>T33940</td><td>AACAGACACGACAGACCGTTTGG</td><td></td><td>1000</td><td></td><td></td><td></td><td></td><td></td><td></td><td></td><td></td><td></td></tr>
<tr><td>Oligo fwd</td><td>CACCAACAGACACGACAGACCGTT</td><td></td><td></td><td></td><td></td><td></td><td></td><td></td><td></td><td></td><td></td><td></td></tr>
<tr><td>Oligo rev</td><td>AAACAACGGTCTGTCGTGTCTGTT</td><td></td><td></td><td></td><td></td><td></td><td></td><td></td><td></td><td></td><td></td><td></td></tr>
<tr><td>Chromosome</td><td>start</td><td></td><td>end</td><td>strand</td><td>MM</td><td>target_seq</td><td>PAM</td><td>alignment</td><td>distance</td><td>position</td><td>gene name</td><td>gene id</td></tr>
<tr><td>GU305914.1</td><td></td><td>6364</td><td>6386</td><td>-</td><td>0</td><td>AACAGACACGACAGACCGTT</td><td>TGG</td><td>||||||||[||||||||||||]PAM</td><td>NA</td><td>Intergenic</td><td>NA</td><td>NA</td></tr>
<tr><td>T33941</td><td>CACGACAGACCGTTTGGTGTAGG</td><td></td><td>1000</td><td></td><td></td><td></td><td></td><td></td><td></td><td></td><td></td><td></td></tr>
<tr><td>Oligo fwd</td><td>CACCCACGACAGACCGTTTGGTGT</td><td></td><td></td><td></td><td></td><td></td><td></td><td></td><td></td><td></td><td></td><td></td></tr>
<tr><td>Oligo rev</td><td>AAACACACCAAACGGTCTGTCGTG</td><td></td><td></td><td></td><td></td><td></td><td></td><td></td><td></td><td></td><td></td><td></td></tr>
<tr><td>Chromosome</td><td>start</td><td></td><td>end</td><td>strand</td><td>MM</td><td>target_seq</td><td>PAM</td><td>alignment</td><td>distance</td><td>position</td><td>gene name</td><td>gene id</td></tr>
<tr><td>GU305914.1</td><td></td><td>6358</td><td>6380</td><td>-</td><td>0</td><td>CACGACAGACCGTTTGGTGT</td><td>AGG</td><td>||||||||[||||||||||||]PAM</td><td>NA</td><td>Intergenic</td><td>NA</td><td>NA</td></tr>
<tr><td>T33942</td><td>AAACAGTTTCCATTTTTATTTGG</td><td></td><td>1000</td><td></td><td></td><td></td><td></td><td></td><td></td><td></td><td></td><td></td></tr>
<tr><td>Oligo fwd</td><td>CACCAAACAGTTTCCATTTTTATT</td><td></td><td></td><td></td><td></td><td></td><td></td><td></td><td></td><td></td><td></td><td></td></tr>
<tr><td>Oligo rev</td><td>AAACAATAAAAATGGAAACTGTTT</td><td></td><td></td><td></td><td></td><td></td><td></td><td></td><td></td><td></td><td></td><td></td></tr>
<tr><td>Chromosome</td><td>start</td><td></td><td>end</td><td>strand</td><td>MM</td><td>target_seq</td><td>PAM</td><td>alignment</td><td>distance</td><td>position</td><td>gene name</td><td>gene id</td></tr>
<tr><td>GU305914.1</td><td></td><td>6330</td><td>6352</td><td>-</td><td>0</td><td>AAACAGTTTCCATTTTTATT</td><td>TGG</td><td>||||||||[||||||||||||]PAM</td><td>NA</td><td>Intergenic</td><td>NA</td><td>NA</td></tr>
<tr><td>T33943</td><td>CCATTTTTATTTGGTAAGCAAGG</td><td></td><td>1000</td><td></td><td></td><td></td><td></td><td></td><td></td><td></td><td></td><td></td></tr>
<tr><td>Oligo fwd</td><td>CACCCCATTTTTATTTGGTAAGCA</td><td></td><td></td><td></td><td></td><td></td><td></td><td></td><td></td><td></td><td></td><td></td></tr>
<tr><td>Oligo rev</td><td>AAACTGCTTACCAAATAAAAATGG</td><td></td><td></td><td></td><td></td><td></td><td></td><td></td><td></td><td></td><td></td><td></td></tr>
<tr><td>Chromosome</td><td>start</td><td></td><td>end</td><td>strand</td><td>MM</td><td>target_seq</td><td>PAM</td><td>alignment</td><td>distance</td><td>position</td><td>gene name</td><td>gene id</td></tr>
<tr><td>GU305914.1</td><td></td><td>6321</td><td>6343</td><td>-</td><td>0</td><td>CCATTTTTATTTGGTAAGCA</td><td>AGG</td><td>||||||||[||||||||||||]PAM</td><td>NA</td><td>Intergenic</td><td>NA</td><td>NA</td></tr>
<tr><td>T33944</td><td>CATTTTTATTTGGTAAGCAAGGG</td><td></td><td>1000</td><td></td><td></td><td></td><td></td><td></td><td></td><td></td><td></td><td></td></tr>
<tr><td>Oligo fwd</td><td>CACCCATTTTTATTTGGTAAGCAA</td><td></td><td></td><td></td><td></td><td></td><td></td><td></td><td></td><td></td><td></td><td></td></tr>
<tr><td>Oligo rev</td><td>AAACTTGCTTACCAAATAAAAATG</td><td></td><td></td><td></td><td></td><td></td><td></td><td></td><td></td><td></td><td></td><td></td></tr>
</table>

Chromosome start end strand MM target_seq PAM alignment distance position gene name gene id
GU305914.1 6320 6342 - 0 CATTTTTATTTGGTAAGCAA GGG ||||||||[||||||||||||]PAM NA Intergenic NA NA

T33945 ATTTTTATTTGGTAAGCAAGGGG 1000
Oligo fwd CACCATTTTTATTTGGTAAGCAAG
Oligo rev AAACCTTGCTTACCAAATAAAAAT
Chromosome start end strand MM target_seq PAM alignment distance position gene name gene id
GU305914.1 6319 6341 - 0 ATTTTTATTTGGTAAGCAAG GGG ||||||||[||||||||||||]PAM NA Intergenic NA NA

T33946 TTTTTATTTGGTAAGCAAGGGGG 1000
Oligo fwd CACCTTTTTATTTGGTAAGCAAGG
Oligo rev AAACCCTTGCTTACCAAATAAAAA
Chromosome start end strand MM target_seq PAM alignment distance position gene name gene id
GU305914.1 6318 6340 - 0 TTTTTATTTGGTAAGCAAGG GGG ||||||||[||||||||||||]PAM NA Intergenic NA NA

T33947 TGATTGATCTTTACCGTCCGCGG 1000
Oligo fwd CACCTGATTGATCTTTACCGTCCG
Oligo rev AAACCGGACGGTAAAGATCAATCA
Chromosome start end strand MM target_seq PAM alignment distance position gene name gene id
GU305914.1 6288 6310 - 0 TGATTGATCTTTACCGTCCG CGG ||||||||[||||||||||||]PAM NA Intergenic NA NA

T33948 GATTGATCTTTACCGTCCGCGGG 1000
Oligo fwd CACCGATTGATCTTTACCGTCCGC
Oligo rev AAACGCGGACGGTAAAGATCAATC
Chromosome start end strand MM target_seq PAM alignment distance position gene name gene id
GU305914.1 6287 6309 - 0 GATTGATCTTTACCGTCCGC GGG ||||||||[||||||||||||]PAM NA Intergenic NA NA

T33949 ATTGATCTTTACCGTCCGCGGGG 1000
Oligo fwd CACCATTGATCTTTACCGTCCGCG
Oligo rev AAACCGCGGACGGTAAAGATCAAT
Chromosome start end strand MM target_seq PAM alignment distance position gene name gene id
GU305914.1 6286 6308 - 0 ATTGATCTTTACCGTCCGCG GGG ||||||||[||||||||||||]PAM NA Intergenic NA NA

T33950 CTTTACCGTCCGCGGGGATGCGG 1000
Oligo fwd CACCCTTTACCGTCCGCGGGGATG
Oligo rev AAACCATCCCCGCGGACGGTAAAG
Chromosome start end strand MM target_seq PAM alignment distance position gene name gene id
GU305914.1 6280 6302 - 0 CTTTACCGTCCGCGGGGATG CGG ||||||||[||||||||||||]PAM NA Intergenic NA NA

T33951 ACCGTCCGCGGGGATGCGGACGG 1000
Oligo fwd CACCACCGTCCGCGGGGATGCGGA
Oligo rev AAACTCCGCATCCCCGCGGACGGT
Chromosome start end strand MM target_seq PAM alignment distance position gene name gene id
GU305914.1 6276 6298 - 0 ACCGTCCGCGGGGATGCGGA CGG ||||||||[||||||||||||]PAM NA Intergenic NA NA

T33952 CGCTTGAGCAGTCTAGCTAGCGG 1000
Oligo fwd CACCCGCTTGAGCAGTCTAGCTAG
Oligo rev AAACCTAGCTAGACTGCTCAAGCG
Chromosome start end strand MM target_seq PAM alignment distance position gene name gene id
GU305914.1 6233 6255 - 0 CGCTTGAGCAGTCTAGCTAG CGG ||||||||[||||||||||||]PAM NA Intergenic NA NA

T33953 GCAGTCTAGCTAGCGGCTCGCGG 1000
Oligo fwd CACCGCAGTCTAGCTAGCGGCTCG
Oligo rev AAACCGAGCCGCTAGCTAGACTGC
Chromosome start end strand MM target_seq PAM alignment distance position gene name gene id
GU305914.1 6226 6248 - 0 GCAGTCTAGCTAGCGGCTCG CGG ||||||||[||||||||||||]PAM NA Intergenic NA NA

T33954 AGCTAGCGGCTCGCGGCCGTAGG 1000
Oligo fwd CACCAGCTAGCGGCTCGCGGCCGT
Oligo rev AAACACGGCCGCGAGCCGCTAGCT
Chromosome start end strand MM target_seq PAM alignment distance position gene name gene id
GU305914.1 6219 6241 - 0 AGCTAGCGGCTCGCGGCCGT AGG ||||||||[||||||||||||]PAM NA Intergenic NA NA

T33955 GACAGCGAGCCAGAGTAGCGAGG 1000
Oligo fwd CACCGACAGCGAGCCAGAGTAGCG
Oligo rev AAACCGCTACTCTGGCTCGCTGTC
Chromosome start end strand MM target_seq PAM alignment distance position gene name gene id
GU305914.1 6177 6199 - 0 GACAGCGAGCCAGAGTAGCG AGG ||||||||[||||||||||||]PAM NA Intergenic NA NA

T33956 CCAGAGTAGCGAGGCGATCACGG 1000
Oligo fwd CACCCCAGAGTAGCGAGGCGATCA
Oligo rev AAACTGATCGCCTCGCTACTCTGG
Chromosome start end strand MM target_seq PAM alignment distance position gene name gene id
GU305914.1 6168 6190 - 0 CCAGAGTAGCGAGGCGATCA CGG ||||||||[||||||||||||]PAM NA Intergenic NA NA

T33957 CGATCACGGAGATGACGAGTAGG 1000
Oligo fwd CACCCGATCACGGAGATGACGAGT
Oligo rev AAACACTCGTCATCTCCGTGATCG
Chromosome start end strand MM target_seq PAM alignment distance position gene name gene id
GU305914.1 6154 6176 - 0 CGATCACGGAGATGACGAGT AGG ||||||||[||||||||||||]PAM NA Intergenic NA NA

T33958 CGGAGATGACGAGTAGGAGCTGG 1000
Oligo fwd CACCCGGAGATGACGAGTAGGAGC
Oligo rev AAACGCTCCTACTCGTCATCTCCG
Chromosome start end strand MM target_seq PAM alignment distance position gene name gene id
GU305914.1 6148 6170 - 0 CGGAGATGACGAGTAGGAGC TGG ||||||||[||||||||||||]PAM NA Intergenic NA NA

T33959 AGATGACGAGTAGGAGCTGGCGG 1000
Oligo fwd CACCAGATGACGAGTAGGAGCTGG

| | | | | | | | | | | | | |
|---|---|---|---|---|---|---|---|---|---|---|---|---|
| Oligo rev | AAACCCAGCTCCTACTCGTCATCT | | | | | | | | | | | |
| Chromosome | start | | end | strand | MM | target_seq | PAM | alignment | distance | position | gene name | gene id |
| GU305914.1 | | 6145 | 6167 | - | 0 | AGATGACGAGTAGGAGCTGG | CGG | \|\|\|\|\|\|\|\|[\|\|\|\|\|\|\|\|\|\|\|\|]PAM | NA | Intergenic | NA | NA |
| T33960 | TGACGAGTAGGAGCTGGCGGCGG | | 1000 | | | | | | | | | |
| Oligo fwd | CACCTGACGAGTAGGAGCTGGCGG | | | | | | | | | | | |
| Oligo rev | AAACCCGCCAGCTCCTACTCGTCA | | | | | | | | | | | |
| Chromosome | start | | end | strand | MM | target_seq | PAM | alignment | distance | position | gene name | gene id |
| GU305914.1 | | 6142 | 6164 | - | 0 | TGACGAGTAGGAGCTGGCGG | CGG | \|\|\|\|\|\|\|\|[\|\|\|\|\|\|\|\|\|\|\|\|]PAM | NA | Intergenic | NA | NA |
| GU305914.1 | | 99490 | 99512 | - | 4 | TTACGAGGAGGAGAGGGCGG | TGG | \|-\|\|\|\|\|-[\|\|\|\|\|--\|\|\|\|\|]PAM | NA | Intergenic | NA | NA |
| T33961 | CGAGTAGGAGCTGGCGGCGGCGG | | 1000 | | | | | | | | | |
| Oligo fwd | CACCCGAGTAGGAGCTGGCGGCGG | | | | | | | | | | | |
| Oligo rev | AAACCCGCCGCCAGCTCCTACTCG | | | | | | | | | | | |
| Chromosome | start | | end | strand | MM | target_seq | PAM | alignment | distance | position | gene name | gene id |
| GU305914.1 | | 6139 | 6161 | - | 0 | CGAGTAGGAGCTGGCGGCGG | CGG | \|\|\|\|\|\|\|\|[\|\|\|\|\|\|\|\|\|\|\|\|]PAM | NA | Intergenic | NA | NA |
| T33962 | AGCTGGCGGCGGCGGCGTGATGG | | 1000 | | | | | | | | | |
| Oligo fwd | CACCAGCTGGCGGCGGCGGCGTGA | | | | | | | | | | | |
| Oligo rev | AAACTCACGCCGCCGCCGCCAGCT | | | | | | | | | | | |
| Chromosome | start | | end | strand | MM | target_seq | PAM | alignment | distance | position | gene name | gene id |
| GU305914.1 | | 6131 | 6153 | - | 0 | AGCTGGCGGCGGCGGCGTGA | TGG | \|\|\|\|\|\|\|\|[\|\|\|\|\|\|\|\|\|\|\|\|]PAM | NA | Intergenic | NA | NA |
| T33963 | CCGCATTCTCTGACGCCACGCGG | | 1000 | | | | | | | | | |
| Oligo fwd | CACCCCGCATTCTCTGACGCCACG | | | | | | | | | | | |
| Oligo rev | AAACCGTGGCGTCAGAGAATGCGG | | | | | | | | | | | |
| Chromosome | start | | end | strand | MM | target_seq | PAM | alignment | distance | position | gene name | gene id |
| GU305914.1 | | 6093 | 6115 | - | 0 | CCGCATTCTCTGACGCCACG | CGG | \|\|\|\|\|\|\|\|[\|\|\|\|\|\|\|\|\|\|\|\|]PAM | NA | Intergenic | NA | NA |
| T33964 | CGCCACGCGGCGTAGTCGAATGG | | 1000 | | | | | | | | | |
| Oligo fwd | CACCCGCCACGCGGCGTAGTCGAA | | | | | | | | | | | |
| Oligo rev | AAACTTCGACTACGCCGCGTGGCG | | | | | | | | | | | |
| Chromosome | start | | end | strand | MM | target_seq | PAM | alignment | distance | position | gene name | gene id |
| GU305914.1 | | 6080 | 6102 | - | 0 | CGCCACGCGGCGTAGTCGAA | TGG | \|\|\|\|\|\|\|\|[\|\|\|\|\|\|\|\|\|\|\|\|]PAM | NA | Intergenic | NA | NA |
| T33965 | CCGTCTGTGAGCAACGTGTCCGG | | 1000 | | | | | | | | | |
| Oligo fwd | CACCCCGTCTGTGAGCAACGTGTC | | | | | | | | | | | |
| Oligo rev | AAACGACACGTTGCTCACAGACGG | | | | | | | | | | | |
| Chromosome | start | | end | strand | MM | target_seq | PAM | alignment | distance | position | gene name | gene id |
| GU305914.1 | | 6053 | 6075 | - | 0 | CCGTCTGTGAGCAACGTGTC | CGG | \|\|\|\|\|\|\|\|[\|\|\|\|\|\|\|\|\|\|\|\|]PAM | NA | Intergenic | NA | NA |
| T33966 | GTGTCCGGACCGCTGTAATCTGG | | 1000 | | | | | | | | | |
| Oligo fwd | CACCGTGTCCGGACCGCTGTAATC | | | | | | | | | | | |
| Oligo rev | AAACGATTACAGCGGTCCGGACAC | | | | | | | | | | | |
| Chromosome | start | | end | strand | MM | target_seq | PAM | alignment | distance | position | gene name | gene id |
| GU305914.1 | | 6038 | 6060 | - | 0 | GTGTCCGGACCGCTGTAATC | TGG | \|\|\|\|\|\|\|\|[\|\|\|\|\|\|\|\|\|\|\|\|]PAM | NA | Intergenic | NA | NA |
| T33967 | TGTCCGGACCGCTGTAATCTGGG | | 1000 | | | | | | | | | |
| Oligo fwd | CACCTGTCCGGACCGCTGTAATCT | | | | | | | | | | | |
| Oligo rev | AAACAGATTACAGCGGTCCGGACA | | | | | | | | | | | |
| Chromosome | start | | end | strand | MM | target_seq | PAM | alignment | distance | position | gene name | gene id |
| GU305914.1 | | 6037 | 6059 | - | 0 | TGTCCGGACCGCTGTAATCT | GGG | \|\|\|\|\|\|\|\|[\|\|\|\|\|\|\|\|\|\|\|\|]PAM | NA | Intergenic | NA | NA |
| T33968 | CCGCTCATGTTCTTCATCTGCGG | | 1000 | | | | | | | | | |
| Oligo fwd | CACCCCGCTCATGTTCTTCATCTG | | | | | | | | | | | |
| Oligo rev | AAACCAGATGAAGAACATGAGCGG | | | | | | | | | | | |
| Chromosome | start | | end | strand | MM | target_seq | PAM | alignment | distance | position | gene name | gene id |
| GU305914.1 | | 6002 | 6024 | - | 0 | CCGCTCATGTTCTTCATCTG | CGG | \|\|\|\|\|\|\|\|[\|\|\|\|\|\|\|\|\|\|\|\|]PAM | NA | Intergenic | NA | NA |
| T33969 | CGCTCATGTTCTTCATCTGCGGG | | 1000 | | | | | | | | | |
| Oligo fwd | CACCCGCTCATGTTCTTCATCTGC | | | | | | | | | | | |
| Oligo rev | AAACGCAGATGAAGAACATGAGCG | | | | | | | | | | | |
| Chromosome | start | | end | strand | MM | target_seq | PAM | alignment | distance | position | gene name | gene id |
| GU305914.1 | | 6001 | 6023 | - | 0 | CGCTCATGTTCTTCATCTGC | GGG | \|\|\|\|\|\|\|\|[\|\|\|\|\|\|\|\|\|\|\|\|]PAM | NA | Intergenic | NA | NA |
| T33970 | TTCATCTGCGGGTCGTCGTCCGG | | 1000 | | | | | | | | | |
| Oligo fwd | CACCTTCATCTGCGGGTCGTCGTC | | | | | | | | | | | |
| Oligo rev | AAACGACGACGACCCGCAGATGAA | | | | | | | | | | | |
| Chromosome | start | | end | strand | MM | target_seq | PAM | alignment | distance | position | gene name | gene id |
| GU305914.1 | | 5990 | 6012 | - | 0 | TTCATCTGCGGGTCGTCGTC | CGG | \|\|\|\|\|\|\|\|[\|\|\|\|\|\|\|\|\|\|\|\|]PAM | NA | Intergenic | NA | NA |
| T33971 | TCTGCGGGTCGTCGTCCGGCAGG | | 1000 | | | | | | | | | |
| Oligo fwd | CACCTCTGCGGGTCGTCGTCCGGC | | | | | | | | | | | |
| Oligo rev | AAACGCCGGACGACGACCCGCAGA | | | | | | | | | | | |
| Chromosome | start | | end | strand | MM | target_seq | PAM | alignment | distance | position | gene name | gene id |
| GU305914.1 | | 5986 | 6008 | - | 0 | TCTGCGGGTCGTCGTCCGGC | AGG | \|\|\|\|\|\|\|\|[\|\|\|\|\|\|\|\|\|\|\|\|]PAM | NA | Intergenic | NA | NA |
| T33972 | TCGTCCATAGACACACTATCAGG | | 1000 | | | | | | | | | |
| Oligo fwd | CACCTCGTCCATAGACACACTATC | | | | | | | | | | | |
| Oligo rev | AAACGATAGTGTGTCTATGGACGA | | | | | | | | | | | |
| Chromosome | start | | end | strand | MM | target_seq | PAM | alignment | distance | position | gene name | gene id |
| GU305914.1 | | 5944 | 5966 | - | 0 | TCGTCCATAGACACACTATC | AGG | \|\|\|\|\|\|\|\|[\|\|\|\|\|\|\|\|\|\|\|\|]PAM | NA | Intergenic | NA | NA |
| T33973 | GACACACTATCAGGCGACACCGG | | 1000 | | | | | | | | | |
| Oligo fwd | CACCGACACACTATCAGGCGACAC | | | | | | | | | | | |
| Oligo rev | AAACGTGTCGCCTGATAGTGTGTC | | | | | | | | | | | |
| Chromosome | start | | end | strand | MM | target_seq | PAM | alignment | distance | position | gene name | gene id |
| GU305914.1 | | 5935 | 5957 | - | 0 | GACACACTATCAGGCGACAC | CGG | \|\|\|\|\|\|\|\|[\|\|\|\|\|\|\|\|\|\|\|\|]PAM | NA | Intergenic | NA | NA |

<table>
<tr><td>T33974</td><td>GACACCGGACGCTGTCTGCGAGG</td><td></td><td>1000</td><td></td><td></td><td></td><td></td><td></td><td></td><td></td><td></td><td></td></tr>
<tr><td>Oligo fwd</td><td>CACCGACACCGGACGCTGTCTGCG</td><td></td><td></td><td></td><td></td><td></td><td></td><td></td><td></td><td></td><td></td><td></td></tr>
<tr><td>Oligo rev</td><td>AAACCGCAGACAGCGTCCGGTGTC</td><td></td><td></td><td></td><td></td><td></td><td></td><td></td><td></td><td></td><td></td><td></td></tr>
<tr><td>Chromosome</td><td>start</td><td></td><td>end</td><td>strand</td><td>MM</td><td>target_seq</td><td>PAM</td><td>alignment</td><td>distance</td><td>position</td><td>gene name</td><td>gene id</td></tr>
<tr><td>GU305914.1</td><td></td><td>5920</td><td>5942</td><td>-</td><td>0</td><td>GACACCGGACGCTGTCTGCG</td><td>AGG</td><td>||||||||[||||||||||||]PAM</td><td>NA</td><td>Intergenic</td><td>NA</td><td>NA</td></tr>
<tr><td>T33975</td><td>CTGCGAGGCTCGCGTTAGATCGG</td><td></td><td>1000</td><td></td><td></td><td></td><td></td><td></td><td></td><td></td><td></td><td></td></tr>
<tr><td>Oligo fwd</td><td>CACCCTGCGAGGCTCGCGTTAGAT</td><td></td><td></td><td></td><td></td><td></td><td></td><td></td><td></td><td></td><td></td><td></td></tr>
<tr><td>Oligo rev</td><td>AAACATCTAACGCGAGCCTCGCAG</td><td></td><td></td><td></td><td></td><td></td><td></td><td></td><td></td><td></td><td></td><td></td></tr>
<tr><td>Chromosome</td><td>start</td><td></td><td>end</td><td>strand</td><td>MM</td><td>target_seq</td><td>PAM</td><td>alignment</td><td>distance</td><td>position</td><td>gene name</td><td>gene id</td></tr>
<tr><td>GU305914.1</td><td></td><td>5905</td><td>5927</td><td>-</td><td>0</td><td>CTGCGAGGCTCGCGTTAGAT</td><td>CGG</td><td>||||||||[||||||||||||]PAM</td><td>NA</td><td>Intergenic</td><td>NA</td><td>NA</td></tr>
<tr><td>T33976</td><td>TGCGAGGCTCGCGTTAGATCGGG</td><td></td><td>1000</td><td></td><td></td><td></td><td></td><td></td><td></td><td></td><td></td><td></td></tr>
<tr><td>Oligo fwd</td><td>CACCTGCGAGGCTCGCGTTAGATC</td><td></td><td></td><td></td><td></td><td></td><td></td><td></td><td></td><td></td><td></td><td></td></tr>
<tr><td>Oligo rev</td><td>AAACGATCTAACGCGAGCCTCGCA</td><td></td><td></td><td></td><td></td><td></td><td></td><td></td><td></td><td></td><td></td><td></td></tr>
<tr><td>Chromosome</td><td>start</td><td></td><td>end</td><td>strand</td><td>MM</td><td>target_seq</td><td>PAM</td><td>alignment</td><td>distance</td><td>position</td><td>gene name</td><td>gene id</td></tr>
<tr><td>GU305914.1</td><td></td><td>5904</td><td>5926</td><td>-</td><td>0</td><td>TGCGAGGCTCGCGTTAGATC</td><td>GGG</td><td>||||||||[||||||||||||]PAM</td><td>NA</td><td>Intergenic</td><td>NA</td><td>NA</td></tr>
<tr><td>T33977</td><td>GTCGACTTTGAAACCGTTCGCGG</td><td></td><td>1000</td><td></td><td></td><td></td><td></td><td></td><td></td><td></td><td></td><td></td></tr>
<tr><td>Oligo fwd</td><td>CACCGTCGACTTTGAAACCGTTCG</td><td></td><td></td><td></td><td></td><td></td><td></td><td></td><td></td><td></td><td></td><td></td></tr>
<tr><td>Oligo rev</td><td>AAACCGAACGGTTTCAAAGTCGAC</td><td></td><td></td><td></td><td></td><td></td><td></td><td></td><td></td><td></td><td></td><td></td></tr>
<tr><td>Chromosome</td><td>start</td><td></td><td>end</td><td>strand</td><td>MM</td><td>target_seq</td><td>PAM</td><td>alignment</td><td>distance</td><td>position</td><td>gene name</td><td>gene id</td></tr>
<tr><td>GU305914.1</td><td></td><td>5856</td><td>5878</td><td>-</td><td>0</td><td>GTCGACTTTGAAACCGTTCG</td><td>CGG</td><td>||||||||[||||||||||||]PAM</td><td>NA</td><td>Intergenic</td><td>NA</td><td>NA</td></tr>
<tr><td>T33978</td><td>GACTTTGAAACCGTTCGCGGCGG</td><td></td><td>1000</td><td></td><td></td><td></td><td></td><td></td><td></td><td></td><td></td><td></td></tr>
<tr><td>Oligo fwd</td><td>CACCGACTTTGAAACCGTTCGCGG</td><td></td><td></td><td></td><td></td><td></td><td></td><td></td><td></td><td></td><td></td><td></td></tr>
<tr><td>Oligo rev</td><td>AAACCCGCGAACGGTTTCAAAGTC</td><td></td><td></td><td></td><td></td><td></td><td></td><td></td><td></td><td></td><td></td><td></td></tr>
<tr><td>Chromosome</td><td>start</td><td></td><td>end</td><td>strand</td><td>MM</td><td>target_seq</td><td>PAM</td><td>alignment</td><td>distance</td><td>position</td><td>gene name</td><td>gene id</td></tr>
<tr><td>GU305914.1</td><td></td><td>5853</td><td>5875</td><td>-</td><td>0</td><td>GACTTTGAAACCGTTCGCGG</td><td>CGG</td><td>||||||||[||||||||||||]PAM</td><td>NA</td><td>Intergenic</td><td>NA</td><td>NA</td></tr>
<tr><td>T33979</td><td>ACGATTTTCCGTCGATAAAAAGG</td><td></td><td>1000</td><td></td><td></td><td></td><td></td><td></td><td></td><td></td><td></td><td></td></tr>
<tr><td>Oligo fwd</td><td>CACCACGATTTTCCGTCGATAAAA</td><td></td><td></td><td></td><td></td><td></td><td></td><td></td><td></td><td></td><td></td><td></td></tr>
<tr><td>Oligo rev</td><td>AAACTTTTATCGACGGAAAATCGT</td><td></td><td></td><td></td><td></td><td></td><td></td><td></td><td></td><td></td><td></td><td></td></tr>
<tr><td>Chromosome</td><td>start</td><td></td><td>end</td><td>strand</td><td>MM</td><td>target_seq</td><td>PAM</td><td>alignment</td><td>distance</td><td>position</td><td>gene name</td><td>gene id</td></tr>
<tr><td>GU305914.1</td><td></td><td>5828</td><td>5850</td><td>-</td><td>0</td><td>ACGATTTTCCGTCGATAAAA</td><td>AGG</td><td>||||||||[||||||||||||]PAM</td><td>NA</td><td>Intergenic</td><td>NA</td><td>NA</td></tr>
<tr><td>T33980</td><td>CGATTTTCCGTCGATAAAAAGGG</td><td></td><td>1000</td><td></td><td></td><td></td><td></td><td></td><td></td><td></td><td></td><td></td></tr>
<tr><td>Oligo fwd</td><td>CACCCGATTTTCCGTCGATAAAAA</td><td></td><td></td><td></td><td></td><td></td><td></td><td></td><td></td><td></td><td></td><td></td></tr>
<tr><td>Oligo rev</td><td>AAACTTTTTATCGACGGAAAATCG</td><td></td><td></td><td></td><td></td><td></td><td></td><td></td><td></td><td></td><td></td><td></td></tr>
<tr><td>Chromosome</td><td>start</td><td></td><td>end</td><td>strand</td><td>MM</td><td>target_seq</td><td>PAM</td><td>alignment</td><td>distance</td><td>position</td><td>gene name</td><td>gene id</td></tr>
<tr><td>GU305914.1</td><td></td><td>5827</td><td>5849</td><td>-</td><td>0</td><td>CGATTTTCCGTCGATAAAAA</td><td>GGG</td><td>||||||||[||||||||||||]PAM</td><td>NA</td><td>Intergenic</td><td>NA</td><td>NA</td></tr>
<tr><td>T33981</td><td>TGCATGTTACCATCCAACGTAGG</td><td></td><td>1000</td><td></td><td></td><td></td><td></td><td></td><td></td><td></td><td></td><td></td></tr>
<tr><td>Oligo fwd</td><td>CACCTGCATGTTACCATCCAACGT</td><td></td><td></td><td></td><td></td><td></td><td></td><td></td><td></td><td></td><td></td><td></td></tr>
<tr><td>Oligo rev</td><td>AAACACGTTGGATGGTAACATGCA</td><td></td><td></td><td></td><td></td><td></td><td></td><td></td><td></td><td></td><td></td><td></td></tr>
<tr><td>Chromosome</td><td>start</td><td></td><td>end</td><td>strand</td><td>MM</td><td>target_seq</td><td>PAM</td><td>alignment</td><td>distance</td><td>position</td><td>gene name</td><td>gene id</td></tr>
<tr><td>GU305914.1</td><td></td><td>5804</td><td>5826</td><td>-</td><td>0</td><td>TGCATGTTACCATCCAACGT</td><td>AGG</td><td>||||||||[||||||||||||]PAM</td><td>NA</td><td>Intergenic</td><td>NA</td><td>NA</td></tr>
<tr><td>T33982</td><td>ATGTTACCATCCAACGTAGGCGG</td><td></td><td>1000</td><td></td><td></td><td></td><td></td><td></td><td></td><td></td><td></td><td></td></tr>
<tr><td>Oligo fwd</td><td>CACCATGTTACCATCCAACGTAGG</td><td></td><td></td><td></td><td></td><td></td><td></td><td></td><td></td><td></td><td></td><td></td></tr>
<tr><td>Oligo rev</td><td>AAACCCTACGTTGGATGGTAACAT</td><td></td><td></td><td></td><td></td><td></td><td></td><td></td><td></td><td></td><td></td><td></td></tr>
<tr><td>Chromosome</td><td>start</td><td></td><td>end</td><td>strand</td><td>MM</td><td>target_seq</td><td>PAM</td><td>alignment</td><td>distance</td><td>position</td><td>gene name</td><td>gene id</td></tr>
<tr><td>GU305914.1</td><td></td><td>5801</td><td>5823</td><td>-</td><td>0</td><td>ATGTTACCATCCAACGTAGG</td><td>CGG</td><td>||||||||[||||||||||||]PAM</td><td>NA</td><td>Intergenic</td><td>NA</td><td>NA</td></tr>
<tr><td>T33983</td><td>AGGCGGTGAGCTCGTTGTCGAGG</td><td></td><td>1000</td><td></td><td></td><td></td><td></td><td></td><td></td><td></td><td></td><td></td></tr>
<tr><td>Oligo fwd</td><td>CACCAGGCGGTGAGCTCGTTGTCG</td><td></td><td></td><td></td><td></td><td></td><td></td><td></td><td></td><td></td><td></td><td></td></tr>
<tr><td>Oligo rev</td><td>AAACCGACAACGAGCTCACCGCCT</td><td></td><td></td><td></td><td></td><td></td><td></td><td></td><td></td><td></td><td></td><td></td></tr>
<tr><td>Chromosome</td><td>start</td><td></td><td>end</td><td>strand</td><td>MM</td><td>target_seq</td><td>PAM</td><td>alignment</td><td>distance</td><td>position</td><td>gene name</td><td>gene id</td></tr>
<tr><td>GU305914.1</td><td></td><td>5784</td><td>5806</td><td>-</td><td>0</td><td>AGGCGGTGAGCTCGTTGTCG</td><td>AGG</td><td>||||||||[||||||||||||]PAM</td><td>NA</td><td>Intergenic</td><td>NA</td><td>NA</td></tr>
<tr><td>T33984</td><td>GTCGAGGTAAAGTAGACGATCGG</td><td></td><td>1000</td><td></td><td></td><td></td><td></td><td></td><td></td><td></td><td></td><td></td></tr>
<tr><td>Oligo fwd</td><td>CACCGTCGAGGTAAAGTAGACGAT</td><td></td><td></td><td></td><td></td><td></td><td></td><td></td><td></td><td></td><td></td><td></td></tr>
<tr><td>Oligo rev</td><td>AAACATCGTCTACTTTACCTCGAC</td><td></td><td></td><td></td><td></td><td></td><td></td><td></td><td></td><td></td><td></td><td></td></tr>
<tr><td>Chromosome</td><td>start</td><td></td><td>end</td><td>strand</td><td>MM</td><td>target_seq</td><td>PAM</td><td>alignment</td><td>distance</td><td>position</td><td>gene name</td><td>gene id</td></tr>
<tr><td>GU305914.1</td><td></td><td>5768</td><td>5790</td><td>-</td><td>0</td><td>GTCGAGGTAAAGTAGACGAT</td><td>CGG</td><td>||||||||[||||||||||||]PAM</td><td>NA</td><td>Intergenic</td><td>NA</td><td>NA</td></tr>
<tr><td>T33985</td><td>GTAGACGATCGGCGATACCGAGG</td><td></td><td>1000</td><td></td><td></td><td></td><td></td><td></td><td></td><td></td><td></td><td></td></tr>
<tr><td>Oligo fwd</td><td>CACCGTAGACGATCGGCGATACCG</td><td></td><td></td><td></td><td></td><td></td><td></td><td></td><td></td><td></td><td></td><td></td></tr>
<tr><td>Oligo rev</td><td>AAACCGGTATCGCCGATCGTCTAC</td><td></td><td></td><td></td><td></td><td></td><td></td><td></td><td></td><td></td><td></td><td></td></tr>
<tr><td>Chromosome</td><td>start</td><td></td><td>end</td><td>strand</td><td>MM</td><td>target_seq</td><td>PAM</td><td>alignment</td><td>distance</td><td>position</td><td>gene name</td><td>gene id</td></tr>
<tr><td>GU305914.1</td><td></td><td>5757</td><td>5779</td><td>-</td><td>0</td><td>GTAGACGATCGGCGATACCG</td><td>AGG</td><td>||||||||[||||||||||||]PAM</td><td>NA</td><td>Intergenic</td><td>NA</td><td>NA</td></tr>
<tr><td>T33986</td><td>GACCCAGTGTACGTACCACGTGG</td><td></td><td>1000</td><td></td><td></td><td></td><td></td><td></td><td></td><td></td><td></td><td></td></tr>
<tr><td>Oligo fwd</td><td>CACCGACCCAGTGTACGTACCACG</td><td></td><td></td><td></td><td></td><td></td><td></td><td></td><td></td><td></td><td></td><td></td></tr>
<tr><td>Oligo rev</td><td>AAACCGTGGTACGTACACTGGGTC</td><td></td><td></td><td></td><td></td><td></td><td></td><td></td><td></td><td></td><td></td><td></td></tr>
<tr><td>Chromosome</td><td>start</td><td></td><td>end</td><td>strand</td><td>MM</td><td>target_seq</td><td>PAM</td><td>alignment</td><td>distance</td><td>position</td><td>gene name</td><td>gene id</td></tr>
<tr><td>GU305914.1</td><td></td><td>5718</td><td>5740</td><td>-</td><td>0</td><td>GACCCAGTGTACGTACCACG</td><td>TGG</td><td>||||||||[||||||||||||]PAM</td><td>NA</td><td>Intergenic</td><td>NA</td><td>NA</td></tr>
<tr><td>T33987</td><td>GGCATTCACGTAAACGTTGTTGG</td><td></td><td>1000</td><td></td><td></td><td></td><td></td><td></td><td></td><td></td><td></td><td></td></tr>
<tr><td>Oligo fwd</td><td>CACCGGCATTCACGTAAACGTTGT</td><td></td><td></td><td></td><td></td><td></td><td></td><td></td><td></td><td></td><td></td><td></td></tr>
<tr><td>Oligo rev</td><td>AAACACAACGTTTACGTGAATGCC</td><td></td><td></td><td></td><td></td><td></td><td></td><td></td><td></td><td></td><td></td><td></td></tr>
<tr><td>Chromosome</td><td>start</td><td></td><td>end</td><td>strand</td><td>MM</td><td>target_seq</td><td>PAM</td><td>alignment</td><td>distance</td><td>position</td><td>gene name</td><td>gene id</td></tr>
<tr><td>GU305914.1</td><td></td><td>5697</td><td>5719</td><td>-</td><td>0</td><td>GGCATTCACGTAAACGTTGT</td><td>TGG</td><td>||||||||[||||||||||||]PAM</td><td>NA</td><td>Intergenic</td><td>NA</td><td>NA</td></tr>
<tr><td>T33988</td><td>GCATTCACGTAAACGTTGTTGGG</td><td></td><td>1000</td><td></td><td></td><td></td><td></td><td></td><td></td><td></td><td></td><td></td></tr>
<tr><td>Oligo fwd</td><td>CACCGCATTCACGTAAACGTTGTT</td><td></td><td></td><td></td><td></td><td></td><td></td><td></td><td></td><td></td><td></td><td></td></tr>
<tr><td>Oligo rev</td><td>AAACAACAACGTTTACGTGAATGC</td><td></td><td></td><td></td><td></td><td></td><td></td><td></td><td></td><td></td><td></td><td></td></tr>
<tr><td>Chromosome</td><td>start</td><td></td><td>end</td><td>strand</td><td>MM</td><td>target_seq</td><td>PAM</td><td>alignment</td><td>distance</td><td>position</td><td>gene name</td><td>gene id</td></tr>
<tr><td>GU305914.1</td><td></td><td>5696</td><td>5718</td><td>-</td><td>0</td><td>GCATTCACGTAAACGTTGTT</td><td>GGG</td><td>||||||||[||||||||||||]PAM</td><td>NA</td><td>Intergenic</td><td>NA</td><td>NA</td></tr>
</table>

<table>
<tr><td>T33989</td><td>AACGTTGTTGGGTATCTCACAGG</td><td>1000</td><td></td><td></td><td></td><td></td><td></td><td></td><td></td><td></td><td></td></tr>
<tr><td>Oligo fwd</td><td>CACCAACGTTGTTGGGTATCTCAC</td><td></td><td></td><td></td><td></td><td></td><td></td><td></td><td></td><td></td><td></td></tr>
<tr><td>Oligo rev</td><td>AAACGTGAGATACCCAACAACGTT</td><td></td><td></td><td></td><td></td><td></td><td></td><td></td><td></td><td></td><td></td></tr>
<tr><td>Chromosome</td><td>start</td><td>end</td><td>strand</td><td>MM</td><td>target_seq</td><td>PAM</td><td>alignment</td><td>distance</td><td>position</td><td>gene name</td><td>gene id</td></tr>
<tr><td>GU305914.1</td><td>5685</td><td>5707</td><td>-</td><td>0</td><td>AACGTTGTTGGGTATCTCAC</td><td>AGG</td><td>||||||||[||||||||||||]PAM</td><td>NA</td><td>Intergenic</td><td>NA</td><td>NA</td></tr>
<tr><td>T33990</td><td>ACAGGCGAATCTCTTCTCACCGG</td><td>1000</td><td></td><td></td><td></td><td></td><td></td><td></td><td></td><td></td><td></td></tr>
<tr><td>Oligo fwd</td><td>CACCACAGGCGAATCTCTTCTCAC</td><td></td><td></td><td></td><td></td><td></td><td></td><td></td><td></td><td></td><td></td></tr>
<tr><td>Oligo rev</td><td>AAACGTGAGAAGAGATTCGCCTGT</td><td></td><td></td><td></td><td></td><td></td><td></td><td></td><td></td><td></td><td></td></tr>
<tr><td>Chromosome</td><td>start</td><td>end</td><td>strand</td><td>MM</td><td>target_seq</td><td>PAM</td><td>alignment</td><td>distance</td><td>position</td><td>gene name</td><td>gene id</td></tr>
<tr><td>GU305914.1</td><td>5667</td><td>5689</td><td>-</td><td>0</td><td>ACAGGCGAATCTCTTCTCAC</td><td>CGG</td><td>||||||||[||||||||||||]PAM</td><td>NA</td><td>Intergenic</td><td>NA</td><td>NA</td></tr>
<tr><td>T33991</td><td>TTGAAACATTTGAATGTGTTCGG</td><td>1000</td><td></td><td></td><td></td><td></td><td></td><td></td><td></td><td></td><td></td></tr>
<tr><td>Oligo fwd</td><td>CACCTTGAAACATTTGAATGTGTT</td><td></td><td></td><td></td><td></td><td></td><td></td><td></td><td></td><td></td><td></td></tr>
<tr><td>Oligo rev</td><td>AAACAACACATTCAAATGTTTCAA</td><td></td><td></td><td></td><td></td><td></td><td></td><td></td><td></td><td></td><td></td></tr>
<tr><td>Chromosome</td><td>start</td><td>end</td><td>strand</td><td>MM</td><td>target_seq</td><td>PAM</td><td>alignment</td><td>distance</td><td>position</td><td>gene name</td><td>gene id</td></tr>
<tr><td>GU305914.1</td><td>5606</td><td>5628</td><td>-</td><td>0</td><td>TTGAAACATTTGAATGTGTT</td><td>CGG</td><td>||||||||[||||||||||||]PAM</td><td>NA</td><td>Intergenic</td><td>NA</td><td>NA</td></tr>
<tr><td>T33992</td><td>TGTTCGGTATCTTCTTCATCTGG</td><td>1000</td><td></td><td></td><td></td><td></td><td></td><td></td><td></td><td></td><td></td></tr>
<tr><td>Oligo fwd</td><td>CACCTGTTCGGTATCTTCTTCATC</td><td></td><td></td><td></td><td></td><td></td><td></td><td></td><td></td><td></td><td></td></tr>
<tr><td>Oligo rev</td><td>AAACGATGAAGAAGATACCGAACA</td><td></td><td></td><td></td><td></td><td></td><td></td><td></td><td></td><td></td><td></td></tr>
<tr><td>Chromosome</td><td>start</td><td>end</td><td>strand</td><td>MM</td><td>target_seq</td><td>PAM</td><td>alignment</td><td>distance</td><td>position</td><td>gene name</td><td>gene id</td></tr>
<tr><td>GU305914.1</td><td>5590</td><td>5612</td><td>-</td><td>0</td><td>TGTTCGGTATCTTCTTCATC</td><td>TGG</td><td>||||||||[||||||||||||]PAM</td><td>NA</td><td>Intergenic</td><td>NA</td><td>NA</td></tr>
<tr><td>T33993</td><td>TCTTCATCTGGTTTTTCAGTCGG</td><td>1000</td><td></td><td></td><td></td><td></td><td></td><td></td><td></td><td></td><td></td></tr>
<tr><td>Oligo fwd</td><td>CACCTCTTCATCTGGTTTTTCAGT</td><td></td><td></td><td></td><td></td><td></td><td></td><td></td><td></td><td></td><td></td></tr>
<tr><td>Oligo rev</td><td>AAACACTGAAAAACCAGATGAAGA</td><td></td><td></td><td></td><td></td><td></td><td></td><td></td><td></td><td></td><td></td></tr>
<tr><td>Chromosome</td><td>start</td><td>end</td><td>strand</td><td>MM</td><td>target_seq</td><td>PAM</td><td>alignment</td><td>distance</td><td>position</td><td>gene name</td><td>gene id</td></tr>
<tr><td>GU305914.1</td><td>5578</td><td>5600</td><td>-</td><td>0</td><td>TCTTCATCTGGTTTTTCAGT</td><td>CGG</td><td>||||||||[||||||||||||]PAM</td><td>NA</td><td>Intergenic</td><td>NA</td><td>NA</td></tr>
<tr><td>T33994</td><td>GTTTTTCAGTCGGTTTATCTCGG</td><td>1000</td><td></td><td></td><td></td><td></td><td></td><td></td><td></td><td></td><td></td></tr>
<tr><td>Oligo fwd</td><td>CACCGTTTTTCAGTCGGTTTATCT</td><td></td><td></td><td></td><td></td><td></td><td></td><td></td><td></td><td></td><td></td></tr>
<tr><td>Oligo rev</td><td>AAACAGATAAACCGACTGAAAAAC</td><td></td><td></td><td></td><td></td><td></td><td></td><td></td><td></td><td></td><td></td></tr>
<tr><td>Chromosome</td><td>start</td><td>end</td><td>strand</td><td>MM</td><td>target_seq</td><td>PAM</td><td>alignment</td><td>distance</td><td>position</td><td>gene name</td><td>gene id</td></tr>
<tr><td>GU305914.1</td><td>5568</td><td>5590</td><td>-</td><td>0</td><td>GTTTTTCAGTCGGTTTATCT</td><td>CGG</td><td>||||||||[||||||||||||]PAM</td><td>NA</td><td>Intergenic</td><td>NA</td><td>NA</td></tr>
<tr><td>T33995</td><td>GAAACGTTGCACCCCACGATAGG</td><td>1000</td><td></td><td></td><td></td><td></td><td></td><td></td><td></td><td></td><td></td></tr>
<tr><td>Oligo fwd</td><td>CACCGAAACGTTGCACCCCACGAT</td><td></td><td></td><td></td><td></td><td></td><td></td><td></td><td></td><td></td><td></td></tr>
<tr><td>Oligo rev</td><td>AAACATCGTGGGGTGCAACGTTTC</td><td></td><td></td><td></td><td></td><td></td><td></td><td></td><td></td><td></td><td></td></tr>
<tr><td>Chromosome</td><td>start</td><td>end</td><td>strand</td><td>MM</td><td>target_seq</td><td>PAM</td><td>alignment</td><td>distance</td><td>position</td><td>gene name</td><td>gene id</td></tr>
<tr><td>GU305914.1</td><td>5540</td><td>5562</td><td>-</td><td>0</td><td>GAAACGTTGCACCCCACGAT</td><td>AGG</td><td>||||||||[||||||||||||]PAM</td><td>NA</td><td>Intergenic</td><td>NA</td><td>NA</td></tr>
<tr><td>T33996</td><td>TCTTTCGTTTCGTTAGTCGACGG</td><td>1000</td><td></td><td></td><td></td><td></td><td></td><td></td><td></td><td></td><td></td></tr>
<tr><td>Oligo fwd</td><td>CACCTCTTTCGTTTCGTTAGTCGA</td><td></td><td></td><td></td><td></td><td></td><td></td><td></td><td></td><td></td><td></td></tr>
<tr><td>Oligo rev</td><td>AAACTCGACTAACGAAACGAAAGA</td><td></td><td></td><td></td><td></td><td></td><td></td><td></td><td></td><td></td><td></td></tr>
<tr><td>Chromosome</td><td>start</td><td>end</td><td>strand</td><td>MM</td><td>target_seq</td><td>PAM</td><td>alignment</td><td>distance</td><td>position</td><td>gene name</td><td>gene id</td></tr>
<tr><td>GU305914.1</td><td>5486</td><td>5508</td><td>-</td><td>0</td><td>TCTTTCGTTTCGTTAGTCGA</td><td>CGG</td><td>||||||||[||||||||||||]PAM</td><td>NA</td><td>Intergenic</td><td>NA</td><td>NA</td></tr>
<tr><td>T33997</td><td>TTCGTTTCGTTAGTCGACGGAGG</td><td>1000</td><td></td><td></td><td></td><td></td><td></td><td></td><td></td><td></td><td></td></tr>
<tr><td>Oligo fwd</td><td>CACCTTCGTTTCGTTAGTCGACGG</td><td></td><td></td><td></td><td></td><td></td><td></td><td></td><td></td><td></td><td></td></tr>
<tr><td>Oligo rev</td><td>AAACCCGTCGACTAACGAAACGAA</td><td></td><td></td><td></td><td></td><td></td><td></td><td></td><td></td><td></td><td></td></tr>
<tr><td>Chromosome</td><td>start</td><td>end</td><td>strand</td><td>MM</td><td>target_seq</td><td>PAM</td><td>alignment</td><td>distance</td><td>position</td><td>gene name</td><td>gene id</td></tr>
<tr><td>GU305914.1</td><td>5483</td><td>5505</td><td>-</td><td>0</td><td>TTCGTTTCGTTAGTCGACGG</td><td>AGG</td><td>||||||||[||||||||||||]PAM</td><td>NA</td><td>Intergenic</td><td>NA</td><td>NA</td></tr>
<tr><td>T33998</td><td>GTCGACGGAGGCATAATCTGAGG</td><td>1000</td><td></td><td></td><td></td><td></td><td></td><td></td><td></td><td></td><td></td></tr>
<tr><td>Oligo fwd</td><td>CACCGTCGACGGAGGCATAATCTG</td><td></td><td></td><td></td><td></td><td></td><td></td><td></td><td></td><td></td><td></td></tr>
<tr><td>Oligo rev</td><td>AAACCAGATTATGCCTCCGTCGAC</td><td></td><td></td><td></td><td></td><td></td><td></td><td></td><td></td><td></td><td></td></tr>
<tr><td>Chromosome</td><td>start</td><td>end</td><td>strand</td><td>MM</td><td>target_seq</td><td>PAM</td><td>alignment</td><td>distance</td><td>position</td><td>gene name</td><td>gene id</td></tr>
<tr><td>GU305914.1</td><td>5471</td><td>5493</td><td>-</td><td>0</td><td>GTCGACGGAGGCATAATCTG</td><td>AGG</td><td>||||||||[||||||||||||]PAM</td><td>NA</td><td>Intergenic</td><td>NA</td><td>NA</td></tr>
<tr><td>T33999</td><td>GGAGGCATAATCTGAGGCATAGG</td><td>1000</td><td></td><td></td><td></td><td></td><td></td><td></td><td></td><td></td><td></td></tr>
<tr><td>Oligo fwd</td><td>CACCGGAGGCATAATCTGAGGCAT</td><td></td><td></td><td></td><td></td><td></td><td></td><td></td><td></td><td></td><td></td></tr>
<tr><td>Oligo rev</td><td>AAACATGCCTCAGATTATGCCTCC</td><td></td><td></td><td></td><td></td><td></td><td></td><td></td><td></td><td></td><td></td></tr>
<tr><td>Chromosome</td><td>start</td><td>end</td><td>strand</td><td>MM</td><td>target_seq</td><td>PAM</td><td>alignment</td><td>distance</td><td>position</td><td>gene name</td><td>gene id</td></tr>
<tr><td>GU305914.1</td><td>5465</td><td>5487</td><td>-</td><td>0</td><td>GGAGGCATAATCTGAGGCAT</td><td>AGG</td><td>||||||||[||||||||||||]PAM</td><td>NA</td><td>Intergenic</td><td>NA</td><td>NA</td></tr>
<tr><td>T34000</td><td>GAGGCATAATCTGAGGCATAGGG</td><td>1000</td><td></td><td></td><td></td><td></td><td></td><td></td><td></td><td></td><td></td></tr>
<tr><td>Oligo fwd</td><td>CACCGAGGCATAATCTGAGGCATA</td><td></td><td></td><td></td><td></td><td></td><td></td><td></td><td></td><td></td><td></td></tr>
<tr><td>Oligo rev</td><td>AAACTATGCCTCAGATTATGCCTC</td><td></td><td></td><td></td><td></td><td></td><td></td><td></td><td></td><td></td><td></td></tr>
<tr><td>Chromosome</td><td>start</td><td>end</td><td>strand</td><td>MM</td><td>target_seq</td><td>PAM</td><td>alignment</td><td>distance</td><td>position</td><td>gene name</td><td>gene id</td></tr>
<tr><td>GU305914.1</td><td>5464</td><td>5486</td><td>-</td><td>0</td><td>GAGGCATAATCTGAGGCATA</td><td>GGG</td><td>||||||||[||||||||||||]PAM</td><td>NA</td><td>Intergenic</td><td>NA</td><td>NA</td></tr>
<tr><td>T34001</td><td>ATAATCTGAGGCATAGGGAGCGG</td><td>1000</td><td></td><td></td><td></td><td></td><td></td><td></td><td></td><td></td><td></td></tr>
<tr><td>Oligo fwd</td><td>CACCATAATCTGAGGCATAGGGAG</td><td></td><td></td><td></td><td></td><td></td><td></td><td></td><td></td><td></td><td></td></tr>
<tr><td>Oligo rev</td><td>AAACCTCCCTATGCCTCAGATTAT</td><td></td><td></td><td></td><td></td><td></td><td></td><td></td><td></td><td></td><td></td></tr>
<tr><td>Chromosome</td><td>start</td><td>end</td><td>strand</td><td>MM</td><td>target_seq</td><td>PAM</td><td>alignment</td><td>distance</td><td>position</td><td>gene name</td><td>gene id</td></tr>
<tr><td>GU305914.1</td><td>5459</td><td>5481</td><td>-</td><td>0</td><td>ATAATCTGAGGCATAGGGAG</td><td>CGG</td><td>||||||||[||||||||||||]PAM</td><td>NA</td><td>Intergenic</td><td>NA</td><td>NA</td></tr>
<tr><td>T34002</td><td>ATCATGAAACTGCTAAAATCTGG</td><td>1000</td><td></td><td></td><td></td><td></td><td></td><td></td><td></td><td></td><td></td></tr>
<tr><td>Oligo fwd</td><td>CACCATCATGAAACTGCTAAAATC</td><td></td><td></td><td></td><td></td><td></td><td></td><td></td><td></td><td></td><td></td></tr>
<tr><td>Oligo rev</td><td>AAACGATTTTAGCAGTTTCATGAT</td><td></td><td></td><td></td><td></td><td></td><td></td><td></td><td></td><td></td><td></td></tr>
<tr><td>Chromosome</td><td>start</td><td>end</td><td>strand</td><td>MM</td><td>target_seq</td><td>PAM</td><td>alignment</td><td>distance</td><td>position</td><td>gene name</td><td>gene id</td></tr>
<tr><td>GU305914.1</td><td>5432</td><td>5454</td><td>-</td><td>0</td><td>ATCATGAAACTGCTAAAATC</td><td>TGG</td><td>||||||||[||||||||||||]PAM</td><td>NA</td><td>Intergenic</td><td>NA</td><td>NA</td></tr>
<tr><td>T34003</td><td>CTGCTAAAATCTGGTATTATTGG</td><td>1000</td><td></td><td></td><td></td><td></td><td></td><td></td><td></td><td></td><td></td></tr>
<tr><td>Oligo fwd</td><td>CACCCTGCTAAAATCTGGTATTAT</td><td></td><td></td><td></td><td></td><td></td><td></td><td></td><td></td><td></td><td></td></tr>
<tr><td>Oligo rev</td><td>AAACATAATACCAGATTTTAGCAG</td><td></td><td></td><td></td><td></td><td></td><td></td><td></td><td></td><td></td><td></td></tr>
<tr><td>Chromosome</td><td>start</td><td>end</td><td>strand</td><td>MM</td><td>target_seq</td><td>PAM</td><td>alignment</td><td>distance</td><td>position</td><td>gene name</td><td>gene id</td></tr>
</table>

<table>
<tr><td>GU305914.1</td><td>5423</td><td>5445</td><td>-</td><td>0</td><td>CTGCTAAAATCTGGTATTAT</td><td>TGG</td><td>||||||||[||||||||||||]PAM</td><td>NA</td><td>Intergenic</td><td>NA</td><td>NA</td></tr>
<tr><td>T34004</td><td>GTATTAGCGATTCTCCCATCTGG</td><td>1000</td><td></td><td></td><td></td><td></td><td></td><td></td><td></td><td></td><td></td></tr>
<tr><td>Oligo fwd</td><td>CACCGTATTAGCGATTCTCCCATC</td><td></td><td></td><td></td><td></td><td></td><td></td><td></td><td></td><td></td><td></td></tr>
<tr><td>Oligo rev</td><td>AAACGATGGGAGAATCGCTAATAC</td><td></td><td></td><td></td><td></td><td></td><td></td><td></td><td></td><td></td><td></td></tr>
<tr><td>Chromosome</td><td>start</td><td>end</td><td>strand</td><td>MM</td><td>target_seq</td><td>PAM</td><td>alignment</td><td>distance</td><td>position</td><td>gene name</td><td>gene id</td></tr>
<tr><td>GU305914.1</td><td>5401</td><td>5423</td><td>-</td><td>0</td><td>GTATTAGCGATTCTCCCATC</td><td>TGG</td><td>||||||||[||||||||||||]PAM</td><td>NA</td><td>Intergenic</td><td>NA</td><td>NA</td></tr>
<tr><td>T34005</td><td>TCCCATCTGGAAAGTCATCGAGG</td><td>1000</td><td></td><td></td><td></td><td></td><td></td><td></td><td></td><td></td><td></td></tr>
<tr><td>Oligo fwd</td><td>CACCTCCCATCTGGAAAGTCATCG</td><td></td><td></td><td></td><td></td><td></td><td></td><td></td><td></td><td></td><td></td></tr>
<tr><td>Oligo rev</td><td>AAACCGATGACTTTCCAGATGGGA</td><td></td><td></td><td></td><td></td><td></td><td></td><td></td><td></td><td></td><td></td></tr>
<tr><td>Chromosome</td><td>start</td><td>end</td><td>strand</td><td>MM</td><td>target_seq</td><td>PAM</td><td>alignment</td><td>distance</td><td>position</td><td>gene name</td><td>gene id</td></tr>
<tr><td>GU305914.1</td><td>5388</td><td>5410</td><td>-</td><td>0</td><td>TCCCATCTGGAAAGTCATCG</td><td>AGG</td><td>||||||||[||||||||||||]PAM</td><td>NA</td><td>Intergenic</td><td>NA</td><td>NA</td></tr>
<tr><td>T34006</td><td>ACGAGCGCCAAGAGCTGTACCGG</td><td>1000</td><td></td><td></td><td></td><td></td><td></td><td></td><td></td><td></td><td></td></tr>
<tr><td>Oligo fwd</td><td>CACCACGAGCGCCAAGAGCTGTAC</td><td></td><td></td><td></td><td></td><td></td><td></td><td></td><td></td><td></td><td></td></tr>
<tr><td>Oligo rev</td><td>AAACGTACAGCTCTTGGCGCTCGT</td><td></td><td></td><td></td><td></td><td></td><td></td><td></td><td></td><td></td><td></td></tr>
<tr><td>Chromosome</td><td>start</td><td>end</td><td>strand</td><td>MM</td><td>target_seq</td><td>PAM</td><td>alignment</td><td>distance</td><td>position</td><td>gene name</td><td>gene id</td></tr>
<tr><td>GU305914.1</td><td>5354</td><td>5376</td><td>-</td><td>0</td><td>ACGAGCGCCAAGAGCTGTAC</td><td>CGG</td><td>||||||||[||||||||||||]PAM</td><td>NA</td><td>Intergenic</td><td>NA</td><td>NA</td></tr>
<tr><td>T34007</td><td>AGAGCTGTACCGGACGCGACCGG</td><td>1000</td><td></td><td></td><td></td><td></td><td></td><td></td><td></td><td></td><td></td></tr>
<tr><td>Oligo fwd</td><td>CACCAGAGCTGTACCGGACGCGAC</td><td></td><td></td><td></td><td></td><td></td><td></td><td></td><td></td><td></td><td></td></tr>
<tr><td>Oligo rev</td><td>AAACGTCGCGTCCGGTACAGCTCT</td><td></td><td></td><td></td><td></td><td></td><td></td><td></td><td></td><td></td><td></td></tr>
<tr><td>Chromosome</td><td>start</td><td>end</td><td>strand</td><td>MM</td><td>target_seq</td><td>PAM</td><td>alignment</td><td>distance</td><td>position</td><td>gene name</td><td>gene id</td></tr>
<tr><td>GU305914.1</td><td>5344</td><td>5366</td><td>-</td><td>0</td><td>AGAGCTGTACCGGACGCGAC</td><td>CGG</td><td>||||||||[||||||||||||]PAM</td><td>NA</td><td>Intergenic</td><td>NA</td><td>NA</td></tr>
<tr><td>T34008</td><td>GAGCTGTACCGGACGCGACCGGG</td><td>1000</td><td></td><td></td><td></td><td></td><td></td><td></td><td></td><td></td><td></td></tr>
<tr><td>Oligo fwd</td><td>CACCGAGCTGTACCGGACGCGACC</td><td></td><td></td><td></td><td></td><td></td><td></td><td></td><td></td><td></td><td></td></tr>
<tr><td>Oligo rev</td><td>AAACGGTCGCGTCCGGTACAGCTC</td><td></td><td></td><td></td><td></td><td></td><td></td><td></td><td></td><td></td><td></td></tr>
<tr><td>Chromosome</td><td>start</td><td>end</td><td>strand</td><td>MM</td><td>target_seq</td><td>PAM</td><td>alignment</td><td>distance</td><td>position</td><td>gene name</td><td>gene id</td></tr>
<tr><td>GU305914.1</td><td>5343</td><td>5365</td><td>-</td><td>0</td><td>GAGCTGTACCGGACGCGACC</td><td>GGG</td><td>||||||||[||||||||||||]PAM</td><td>NA</td><td>Intergenic</td><td>NA</td><td>NA</td></tr>
<tr><td>T34009</td><td>TGTCAACGTATCGCTCCAACTGG</td><td>1000</td><td></td><td></td><td></td><td></td><td></td><td></td><td></td><td></td><td></td></tr>
<tr><td>Oligo fwd</td><td>CACCTGTCAACGTATCGCTCCAAC</td><td></td><td></td><td></td><td></td><td></td><td></td><td></td><td></td><td></td><td></td></tr>
<tr><td>Oligo rev</td><td>AAACGTTGGAGCGATACGTTGACA</td><td></td><td></td><td></td><td></td><td></td><td></td><td></td><td></td><td></td><td></td></tr>
<tr><td>Chromosome</td><td>start</td><td>end</td><td>strand</td><td>MM</td><td>target_seq</td><td>PAM</td><td>alignment</td><td>distance</td><td>position</td><td>gene name</td><td>gene id</td></tr>
<tr><td>GU305914.1</td><td>5295</td><td>5317</td><td>-</td><td>0</td><td>TGTCAACGTATCGCTCCAAC</td><td>TGG</td><td>||||||||[||||||||||||]PAM</td><td>NA</td><td>Intergenic</td><td>NA</td><td>NA</td></tr>
<tr><td>T34010</td><td>GTCAACGTATCGCTCCAACTGGG</td><td>1000</td><td></td><td></td><td></td><td></td><td></td><td></td><td></td><td></td><td></td></tr>
<tr><td>Oligo fwd</td><td>CACCGTCAACGTATCGCTCCAACT</td><td></td><td></td><td></td><td></td><td></td><td></td><td></td><td></td><td></td><td></td></tr>
<tr><td>Oligo rev</td><td>AAACAGTTGGAGCGATACGTTGAC</td><td></td><td></td><td></td><td></td><td></td><td></td><td></td><td></td><td></td><td></td></tr>
<tr><td>Chromosome</td><td>start</td><td>end</td><td>strand</td><td>MM</td><td>target_seq</td><td>PAM</td><td>alignment</td><td>distance</td><td>position</td><td>gene name</td><td>gene id</td></tr>
<tr><td>GU305914.1</td><td>5294</td><td>5316</td><td>-</td><td>0</td><td>GTCAACGTATCGCTCCAACT</td><td>GGG</td><td>||||||||[||||||||||||]PAM</td><td>NA</td><td>Intergenic</td><td>NA</td><td>NA</td></tr>
<tr><td>T34011</td><td>CGCTCCAACTGGGCATAGATTGG</td><td>1000</td><td></td><td></td><td></td><td></td><td></td><td></td><td></td><td></td><td></td></tr>
<tr><td>Oligo fwd</td><td>CACCCGCTCCAACTGGGCATAGAT</td><td></td><td></td><td></td><td></td><td></td><td></td><td></td><td></td><td></td><td></td></tr>
<tr><td>Oligo rev</td><td>AAACATCTATGCCCAGTTGGAGCG</td><td></td><td></td><td></td><td></td><td></td><td></td><td></td><td></td><td></td><td></td></tr>
<tr><td>Chromosome</td><td>start</td><td>end</td><td>strand</td><td>MM</td><td>target_seq</td><td>PAM</td><td>alignment</td><td>distance</td><td>position</td><td>gene name</td><td>gene id</td></tr>
<tr><td>GU305914.1</td><td>5284</td><td>5306</td><td>-</td><td>0</td><td>CGCTCCAACTGGGCATAGAT</td><td>TGG</td><td>||||||||[||||||||||||]PAM</td><td>NA</td><td>Intergenic</td><td>NA</td><td>NA</td></tr>
<tr><td>T34012</td><td>CTGGGCATAGATTGGCTTCATGG</td><td>1000</td><td></td><td></td><td></td><td></td><td></td><td></td><td></td><td></td><td></td></tr>
<tr><td>Oligo fwd</td><td>CACCCTGGGCATAGATTGGCTTCA</td><td></td><td></td><td></td><td></td><td></td><td></td><td></td><td></td><td></td><td></td></tr>
<tr><td>Oligo rev</td><td>AAACTGAAGCCAATCTATGCCCAG</td><td></td><td></td><td></td><td></td><td></td><td></td><td></td><td></td><td></td><td></td></tr>
<tr><td>Chromosome</td><td>start</td><td>end</td><td>strand</td><td>MM</td><td>target_seq</td><td>PAM</td><td>alignment</td><td>distance</td><td>position</td><td>gene name</td><td>gene id</td></tr>
<tr><td>GU305914.1</td><td>5276</td><td>5298</td><td>-</td><td>0</td><td>CTGGGCATAGATTGGCTTCA</td><td>TGG</td><td>||||||||[||||||||||||]PAM</td><td>NA</td><td>Intergenic</td><td>NA</td><td>NA</td></tr>
<tr><td>T34013</td><td>TATATATGTTTGCGTGATTTTGG</td><td>1000</td><td></td><td></td><td></td><td></td><td></td><td></td><td></td><td></td><td></td></tr>
<tr><td>Oligo fwd</td><td>CACCTATATATGTTTGCGTGATTT</td><td></td><td></td><td></td><td></td><td></td><td></td><td></td><td></td><td></td><td></td></tr>
<tr><td>Oligo rev</td><td>AAACAAATCACGCAAACATATATA</td><td></td><td></td><td></td><td></td><td></td><td></td><td></td><td></td><td></td><td></td></tr>
<tr><td>Chromosome</td><td>start</td><td>end</td><td>strand</td><td>MM</td><td>target_seq</td><td>PAM</td><td>alignment</td><td>distance</td><td>position</td><td>gene name</td><td>gene id</td></tr>
<tr><td>GU305914.1</td><td>5247</td><td>5269</td><td>-</td><td>0</td><td>TATATATGTTTGCGTGATTT</td><td>TGG</td><td>||||||||[||||||||||||]PAM</td><td>NA</td><td>Intergenic</td><td>NA</td><td>NA</td></tr>
<tr><td>T34014</td><td>ATATATGTTTGCGTGATTTTGGG</td><td>1000</td><td></td><td></td><td></td><td></td><td></td><td></td><td></td><td></td><td></td></tr>
<tr><td>Oligo fwd</td><td>CACCATATATGTTTGCGTGATTTT</td><td></td><td></td><td></td><td></td><td></td><td></td><td></td><td></td><td></td><td></td></tr>
<tr><td>Oligo rev</td><td>AAACAAAATCACGCAAACATATAT</td><td></td><td></td><td></td><td></td><td></td><td></td><td></td><td></td><td></td><td></td></tr>
<tr><td>Chromosome</td><td>start</td><td>end</td><td>strand</td><td>MM</td><td>target_seq</td><td>PAM</td><td>alignment</td><td>distance</td><td>position</td><td>gene name</td><td>gene id</td></tr>
<tr><td>GU305914.1</td><td>5246</td><td>5268</td><td>-</td><td>0</td><td>ATATATGTTTGCGTGATTTT</td><td>GGG</td><td>||||||||[||||||||||||]PAM</td><td>NA</td><td>Intergenic</td><td>NA</td><td>NA</td></tr>
<tr><td>T34015</td><td>CGTGTTATCTTCATCATCACCGG</td><td>1000</td><td></td><td></td><td></td><td></td><td></td><td></td><td></td><td></td><td></td></tr>
<tr><td>Oligo fwd</td><td>CACCCGTGTTATCTTCATCATCAC</td><td></td><td></td><td></td><td></td><td></td><td></td><td></td><td></td><td></td><td></td></tr>
<tr><td>Oligo rev</td><td>AAACGTGATGATGAAGATAACACG</td><td></td><td></td><td></td><td></td><td></td><td></td><td></td><td></td><td></td><td></td></tr>
<tr><td>Chromosome</td><td>start</td><td>end</td><td>strand</td><td>MM</td><td>target_seq</td><td>PAM</td><td>alignment</td><td>distance</td><td>position</td><td>gene name</td><td>gene id</td></tr>
<tr><td>GU305914.1</td><td>5223</td><td>5245</td><td>-</td><td>0</td><td>CGTGTTATCTTCATCATCAC</td><td>CGG</td><td>||||||||[||||||||||||]PAM</td><td>NA</td><td>Intergenic</td><td>NA</td><td>NA</td></tr>
<tr><td>T34016</td><td>GATCTCGATGCTCTCCAGATCGG</td><td>1000</td><td></td><td></td><td></td><td></td><td></td><td></td><td></td><td></td><td></td></tr>
<tr><td>Oligo fwd</td><td>CACCGATCTCGATGCTCTCCAGAT</td><td></td><td></td><td></td><td></td><td></td><td></td><td></td><td></td><td></td><td></td></tr>
<tr><td>Oligo rev</td><td>AAACATCTGGAGAGCATCGAGATC</td><td></td><td></td><td></td><td></td><td></td><td></td><td></td><td></td><td></td><td></td></tr>
<tr><td>Chromosome</td><td>start</td><td>end</td><td>strand</td><td>MM</td><td>target_seq</td><td>PAM</td><td>alignment</td><td>distance</td><td>position</td><td>gene name</td><td>gene id</td></tr>
<tr><td>GU305914.1</td><td>5170</td><td>5192</td><td>-</td><td>0</td><td>GATCTCGATGCTCTCCAGAT</td><td>CGG</td><td>||||||||[||||||||||||]PAM</td><td>NA</td><td>Intergenic</td><td>NA</td><td>NA</td></tr>
<tr><td>T34017</td><td>TCGGAAGAAACAACTACAGATGG</td><td>1000</td><td></td><td></td><td></td><td></td><td></td><td></td><td></td><td></td><td></td></tr>
<tr><td>Oligo fwd</td><td>CACCTCGGAAGAAACAACTACAGA</td><td></td><td></td><td></td><td></td><td></td><td></td><td></td><td></td><td></td><td></td></tr>
<tr><td>Oligo rev</td><td>AAACTCTGTAGTTGTTTCTTCCGA</td><td></td><td></td><td></td><td></td><td></td><td></td><td></td><td></td><td></td><td></td></tr>
<tr><td>Chromosome</td><td>start</td><td>end</td><td>strand</td><td>MM</td><td>target_seq</td><td>PAM</td><td>alignment</td><td>distance</td><td>position</td><td>gene name</td><td>gene id</td></tr>
<tr><td>GU305914.1</td><td>5151</td><td>5173</td><td>-</td><td>0</td><td>TCGGAAGAAACAACTACAGA</td><td>TGG</td><td>||||||||[||||||||||||]PAM</td><td>NA</td><td>Intergenic</td><td>NA</td><td>NA</td></tr>
<tr><td>T34018</td><td>AGAAACAACTACAGATGGATCGG</td><td>1000</td><td></td><td></td><td></td><td></td><td></td><td></td><td></td><td></td><td></td></tr>
<tr><td>Oligo fwd</td><td>CACCAGAAACAACTACAGATGGAT</td><td></td><td></td><td></td><td></td><td></td><td></td><td></td><td></td><td></td><td></td></tr>
<tr><td>Oligo rev</td><td>AAACATCCATCTGTAGTTGTTTCT</td><td></td><td></td><td></td><td></td><td></td><td></td><td></td><td></td><td></td><td></td></tr>
</table>

<table>
<tr><td>Chromosome</td><td>start</td><td></td><td>end</td><td>strand</td><td>MM</td><td>target_seq</td><td>PAM</td><td>alignment</td><td>distance</td><td>position</td><td>gene name</td><td>gene id</td></tr>
<tr><td>GU305914.1</td><td></td><td>5146</td><td>5168</td><td>-</td><td>0</td><td>AGAAACAACTACAGATGGAT</td><td>CGG</td><td>||||||||[||||||||||||]PAM</td><td>NA</td><td>Intergenic</td><td>NA</td><td>NA</td></tr>
<tr><td>T34019</td><td>ATGGATCGGATTCAGTCCCTCGG</td><td></td><td>1000</td><td></td><td></td><td></td><td></td><td></td><td></td><td></td><td></td><td></td></tr>
<tr><td>Oligo fwd</td><td>CACCATGGATCGGATTCAGTCCCT</td><td></td><td></td><td></td><td></td><td></td><td></td><td></td><td></td><td></td><td></td><td></td></tr>
<tr><td>Oligo rev</td><td>AAACAGGGACTGAATCCGATCCAT</td><td></td><td></td><td></td><td></td><td></td><td></td><td></td><td></td><td></td><td></td><td></td></tr>
<tr><td>Chromosome</td><td>start</td><td></td><td>end</td><td>strand</td><td>MM</td><td>target_seq</td><td>PAM</td><td>alignment</td><td>distance</td><td>position</td><td>gene name</td><td>gene id</td></tr>
<tr><td>GU305914.1</td><td></td><td>5132</td><td>5154</td><td>-</td><td>0</td><td>ATGGATCGGATTCAGTCCCT</td><td>CGG</td><td>||||||||[||||||||||||]PAM</td><td>NA</td><td>Intergenic</td><td>NA</td><td>NA</td></tr>
<tr><td>T34020</td><td>GTATTCGACAGCAACGTCAGCGG</td><td></td><td>1000</td><td></td><td></td><td></td><td></td><td></td><td></td><td></td><td></td><td></td></tr>
<tr><td>Oligo fwd</td><td>CACCGTATTCGACAGCAACGTCAG</td><td></td><td></td><td></td><td></td><td></td><td></td><td></td><td></td><td></td><td></td><td></td></tr>
<tr><td>Oligo rev</td><td>AAACCTGACGTTGCTGTCGAATAC</td><td></td><td></td><td></td><td></td><td></td><td></td><td></td><td></td><td></td><td></td><td></td></tr>
<tr><td>Chromosome</td><td>start</td><td></td><td>end</td><td>strand</td><td>MM</td><td>target_seq</td><td>PAM</td><td>alignment</td><td>distance</td><td>position</td><td>gene name</td><td>gene id</td></tr>
<tr><td>GU305914.1</td><td></td><td>5110</td><td>5132</td><td>-</td><td>0</td><td>GTATTCGACAGCAACGTCAG</td><td>CGG</td><td>||||||||[||||||||||||]PAM</td><td>NA</td><td>Intergenic</td><td>NA</td><td>NA</td></tr>
<tr><td>T34021</td><td>TTCGACAGCAACGTCAGCGGCGG</td><td></td><td>1000</td><td></td><td></td><td></td><td></td><td></td><td></td><td></td><td></td><td></td></tr>
<tr><td>Oligo fwd</td><td>CACCTTCGACAGCAACGTCAGCGG</td><td></td><td></td><td></td><td></td><td></td><td></td><td></td><td></td><td></td><td></td><td></td></tr>
<tr><td>Oligo rev</td><td>AAACCCGCTGACGTTGCTGTCGAA</td><td></td><td></td><td></td><td></td><td></td><td></td><td></td><td></td><td></td><td></td><td></td></tr>
<tr><td>Chromosome</td><td>start</td><td></td><td>end</td><td>strand</td><td>MM</td><td>target_seq</td><td>PAM</td><td>alignment</td><td>distance</td><td>position</td><td>gene name</td><td>gene id</td></tr>
<tr><td>GU305914.1</td><td></td><td>5107</td><td>5129</td><td>-</td><td>0</td><td>TTCGACAGCAACGTCAGCGG</td><td>CGG</td><td>||||||||[||||||||||||]PAM</td><td>NA</td><td>Intergenic</td><td>NA</td><td>NA</td></tr>
<tr><td>T34022</td><td>ACTTGTGTTTGTGATTGTAATGG</td><td></td><td>1000</td><td></td><td></td><td></td><td></td><td></td><td></td><td></td><td></td><td></td></tr>
<tr><td>Oligo fwd</td><td>CACCACTTGTGTTTGTGATTGTAA</td><td></td><td></td><td></td><td></td><td></td><td></td><td></td><td></td><td></td><td></td><td></td></tr>
<tr><td>Oligo rev</td><td>AAACTTACAATCACAAACACAAGT</td><td></td><td></td><td></td><td></td><td></td><td></td><td></td><td></td><td></td><td></td><td></td></tr>
<tr><td>Chromosome</td><td>start</td><td></td><td>end</td><td>strand</td><td>MM</td><td>target_seq</td><td>PAM</td><td>alignment</td><td>distance</td><td>position</td><td>gene name</td><td>gene id</td></tr>
<tr><td>GU305914.1</td><td></td><td>5084</td><td>5106</td><td>-</td><td>0</td><td>ACTTGTGTTTGTGATTGTAA</td><td>TGG</td><td>||||||||[||||||||||||]PAM</td><td>NA</td><td>Intergenic</td><td>NA</td><td>NA</td></tr>
<tr><td>T34023</td><td>AATACACTCATCGACATTTGTGG</td><td></td><td>1000</td><td></td><td></td><td></td><td></td><td></td><td></td><td></td><td></td><td></td></tr>
<tr><td>Oligo fwd</td><td>CACCAATACACTCATCGACATTTG</td><td></td><td></td><td></td><td></td><td></td><td></td><td></td><td></td><td></td><td></td><td></td></tr>
<tr><td>Oligo rev</td><td>AAACCAAATGTCGATGAGTGTATT</td><td></td><td></td><td></td><td></td><td></td><td></td><td></td><td></td><td></td><td></td><td></td></tr>
<tr><td>Chromosome</td><td>start</td><td></td><td>end</td><td>strand</td><td>MM</td><td>target_seq</td><td>PAM</td><td>alignment</td><td>distance</td><td>position</td><td>gene name</td><td>gene id</td></tr>
<tr><td>GU305914.1</td><td></td><td>4998</td><td>5020</td><td>-</td><td>0</td><td>AATACACTCATCGACATTTG</td><td>TGG</td><td>||||||||[||||||||||||]PAM</td><td>NA</td><td>Intergenic</td><td>NA</td><td>NA</td></tr>
<tr><td>T34024</td><td>TGTGGCTGTGATGTTTCAGACGG</td><td></td><td>1000</td><td></td><td></td><td></td><td></td><td></td><td></td><td></td><td></td><td></td></tr>
<tr><td>Oligo fwd</td><td>CACCTGTGGCTGTGATGTTTCAGA</td><td></td><td></td><td></td><td></td><td></td><td></td><td></td><td></td><td></td><td></td><td></td></tr>
<tr><td>Oligo rev</td><td>AAACTCTGAAACATCACAGCCACA</td><td></td><td></td><td></td><td></td><td></td><td></td><td></td><td></td><td></td><td></td><td></td></tr>
<tr><td>Chromosome</td><td>start</td><td></td><td>end</td><td>strand</td><td>MM</td><td>target_seq</td><td>PAM</td><td>alignment</td><td>distance</td><td>position</td><td>gene name</td><td>gene id</td></tr>
<tr><td>GU305914.1</td><td></td><td>4980</td><td>5002</td><td>-</td><td>0</td><td>TGTGGCTGTGATGTTTCAGA</td><td>CGG</td><td>||||||||[||||||||||||]PAM</td><td>NA</td><td>Intergenic</td><td>NA</td><td>NA</td></tr>
<tr><td>T34025</td><td>CTCTTCTTTGTGTATTCCATTGG</td><td></td><td>1000</td><td></td><td></td><td></td><td></td><td></td><td></td><td></td><td></td><td></td></tr>
<tr><td>Oligo fwd</td><td>CACCCTCTTCTTTGTGTATTCCAT</td><td></td><td></td><td></td><td></td><td></td><td></td><td></td><td></td><td></td><td></td><td></td></tr>
<tr><td>Oligo rev</td><td>AAACATGGAATACACAAAGAAGAG</td><td></td><td></td><td></td><td></td><td></td><td></td><td></td><td></td><td></td><td></td><td></td></tr>
<tr><td>Chromosome</td><td>start</td><td></td><td>end</td><td>strand</td><td>MM</td><td>target_seq</td><td>PAM</td><td>alignment</td><td>distance</td><td>position</td><td>gene name</td><td>gene id</td></tr>
<tr><td>GU305914.1</td><td></td><td>4944</td><td>4966</td><td>-</td><td>0</td><td>CTCTTCTTTGTGTATTCCAT</td><td>TGG</td><td>||||||||[||||||||||||]PAM</td><td>NA</td><td>Intergenic</td><td>NA</td><td>NA</td></tr>
<tr><td>T34026</td><td>TATTCCATTGGATACCCGAGTGG</td><td></td><td>1000</td><td></td><td></td><td></td><td></td><td></td><td></td><td></td><td></td><td></td></tr>
<tr><td>Oligo fwd</td><td>CACCTATTCCATTGGATACCCGAG</td><td></td><td></td><td></td><td></td><td></td><td></td><td></td><td></td><td></td><td></td><td></td></tr>
<tr><td>Oligo rev</td><td>AAACCTCGGGTATCCAATGGAATA</td><td></td><td></td><td></td><td></td><td></td><td></td><td></td><td></td><td></td><td></td><td></td></tr>
<tr><td>Chromosome</td><td>start</td><td></td><td>end</td><td>strand</td><td>MM</td><td>target_seq</td><td>PAM</td><td>alignment</td><td>distance</td><td>position</td><td>gene name</td><td>gene id</td></tr>
<tr><td>GU305914.1</td><td></td><td>4932</td><td>4954</td><td>-</td><td>0</td><td>TATTCCATTGGATACCCGAG</td><td>TGG</td><td>||||||||[||||||||||||]PAM</td><td>NA</td><td>Intergenic</td><td>NA</td><td>NA</td></tr>
<tr><td>T34027</td><td>CGTCGCTATGATGCTTAGATTGG</td><td></td><td>1000</td><td></td><td></td><td></td><td></td><td></td><td></td><td></td><td></td><td></td></tr>
<tr><td>Oligo fwd</td><td>CACCCGTCGCTATGATGCTTAGAT</td><td></td><td></td><td></td><td></td><td></td><td></td><td></td><td></td><td></td><td></td><td></td></tr>
<tr><td>Oligo rev</td><td>AAACATCTAAGCATCATAGCGACG</td><td></td><td></td><td></td><td></td><td></td><td></td><td></td><td></td><td></td><td></td><td></td></tr>
<tr><td>Chromosome</td><td>start</td><td></td><td>end</td><td>strand</td><td>MM</td><td>target_seq</td><td>PAM</td><td>alignment</td><td>distance</td><td>position</td><td>gene name</td><td>gene id</td></tr>
<tr><td>GU305914.1</td><td></td><td>4891</td><td>4913</td><td>-</td><td>0</td><td>CGTCGCTATGATGCTTAGAT</td><td>TGG</td><td>||||||||[||||||||||||]PAM</td><td>NA</td><td>Intergenic</td><td>NA</td><td>NA</td></tr>
<tr><td>T34028</td><td>TATGATGCTTAGATTGGCAACGG</td><td></td><td>1000</td><td></td><td></td><td></td><td></td><td></td><td></td><td></td><td></td><td></td></tr>
<tr><td>Oligo fwd</td><td>CACCTATGATGCTTAGATTGGCAA</td><td></td><td></td><td></td><td></td><td></td><td></td><td></td><td></td><td></td><td></td><td></td></tr>
<tr><td>Oligo rev</td><td>AAACTTGCCAATCTAAGCATCATA</td><td></td><td></td><td></td><td></td><td></td><td></td><td></td><td></td><td></td><td></td><td></td></tr>
<tr><td>Chromosome</td><td>start</td><td></td><td>end</td><td>strand</td><td>MM</td><td>target_seq</td><td>PAM</td><td>alignment</td><td>distance</td><td>position</td><td>gene name</td><td>gene id</td></tr>
<tr><td>GU305914.1</td><td></td><td>4885</td><td>4907</td><td>-</td><td>0</td><td>TATGATGCTTAGATTGGCAA</td><td>CGG</td><td>||||||||[||||||||||||]PAM</td><td>NA</td><td>Intergenic</td><td>NA</td><td>NA</td></tr>
<tr><td>T34029</td><td>GCTTAGATTGGCAACGGCTGTGG</td><td></td><td>1000</td><td></td><td></td><td></td><td></td><td></td><td></td><td></td><td></td><td></td></tr>
<tr><td>Oligo fwd</td><td>CACCGCTTAGATTGGCAACGGCTG</td><td></td><td></td><td></td><td></td><td></td><td></td><td></td><td></td><td></td><td></td><td></td></tr>
<tr><td>Oligo rev</td><td>AAACCAGCCGTTGCCAATCTAAGC</td><td></td><td></td><td></td><td></td><td></td><td></td><td></td><td></td><td></td><td></td><td></td></tr>
<tr><td>Chromosome</td><td>start</td><td></td><td>end</td><td>strand</td><td>MM</td><td>target_seq</td><td>PAM</td><td>alignment</td><td>distance</td><td>position</td><td>gene name</td><td>gene id</td></tr>
<tr><td>GU305914.1</td><td></td><td>4879</td><td>4901</td><td>-</td><td>0</td><td>GCTTAGATTGGCAACGGCTG</td><td>TGG</td><td>||||||||[||||||||||||]PAM</td><td>NA</td><td>Intergenic</td><td>NA</td><td>NA</td></tr>
<tr><td>T34030</td><td>ACGAGTTTCGATGTACCCATCGG</td><td></td><td>1000</td><td></td><td></td><td></td><td></td><td></td><td></td><td></td><td></td><td></td></tr>
<tr><td>Oligo fwd</td><td>CACCACGAGTTTCGATGTACCCAT</td><td></td><td></td><td></td><td></td><td></td><td></td><td></td><td></td><td></td><td></td><td></td></tr>
<tr><td>Oligo rev</td><td>AAACATGGGTACATCGAAACTCGT</td><td></td><td></td><td></td><td></td><td></td><td></td><td></td><td></td><td></td><td></td><td></td></tr>
<tr><td>Chromosome</td><td>start</td><td></td><td>end</td><td>strand</td><td>MM</td><td>target_seq</td><td>PAM</td><td>alignment</td><td>distance</td><td>position</td><td>gene name</td><td>gene id</td></tr>
<tr><td>GU305914.1</td><td></td><td>4843</td><td>4865</td><td>-</td><td>0</td><td>ACGAGTTTCGATGTACCCAT</td><td>CGG</td><td>||||||||[||||||||||||]PAM</td><td>NA</td><td>Intergenic</td><td>NA</td><td>NA</td></tr>
<tr><td>T34031</td><td>AGTTTCGATGTACCCATCGGAGG</td><td></td><td>1000</td><td></td><td></td><td></td><td></td><td></td><td></td><td></td><td></td><td></td></tr>
<tr><td>Oligo fwd</td><td>CACCAGTTTCGATGTACCCATCGG</td><td></td><td></td><td></td><td></td><td></td><td></td><td></td><td></td><td></td><td></td><td></td></tr>
<tr><td>Oligo rev</td><td>AAACCCGATGGGTACATCGAAACT</td><td></td><td></td><td></td><td></td><td></td><td></td><td></td><td></td><td></td><td></td><td></td></tr>
<tr><td>Chromosome</td><td>start</td><td></td><td>end</td><td>strand</td><td>MM</td><td>target_seq</td><td>PAM</td><td>alignment</td><td>distance</td><td>position</td><td>gene name</td><td>gene id</td></tr>
<tr><td>GU305914.1</td><td></td><td>4840</td><td>4862</td><td>-</td><td>0</td><td>AGTTTCGATGTACCCATCGG</td><td>AGG</td><td>||||||||[||||||||||||]PAM</td><td>NA</td><td>Intergenic</td><td>NA</td><td>NA</td></tr>
<tr><td>T34032</td><td>GTGCACGTCCTTCACGAGATCGG</td><td></td><td>1000</td><td></td><td></td><td></td><td></td><td></td><td></td><td></td><td></td><td></td></tr>
<tr><td>Oligo fwd</td><td>CACCGTGCACGTCCTTCACGAGAT</td><td></td><td></td><td></td><td></td><td></td><td></td><td></td><td></td><td></td><td></td><td></td></tr>
<tr><td>Oligo rev</td><td>AAACATCTCGTGAAGGACGTGCAC</td><td></td><td></td><td></td><td></td><td></td><td></td><td></td><td></td><td></td><td></td><td></td></tr>
<tr><td>Chromosome</td><td>start</td><td></td><td>end</td><td>strand</td><td>MM</td><td>target_seq</td><td>PAM</td><td>alignment</td><td>distance</td><td>position</td><td>gene name</td><td>gene id</td></tr>
<tr><td>GU305914.1</td><td></td><td>4759</td><td>4781</td><td>-</td><td>0</td><td>GTGCACGTCCTTCACGAGAT</td><td>CGG</td><td>||||||||[||||||||||||]PAM</td><td>NA</td><td>Intergenic</td><td>NA</td><td>NA</td></tr>
<tr><td>T34033</td><td>CTTCACGAGATCGGCTCGTACGG</td><td></td><td>1000</td><td></td><td></td><td></td><td></td><td></td><td></td><td></td><td></td><td></td></tr>
<tr><td>Oligo fwd</td><td>CACCCTTCACGAGATCGGCTCGTA</td><td></td><td></td><td></td><td></td><td></td><td></td><td></td><td></td><td></td><td></td><td></td></tr>
</table>

<table>
<tr><td>Oligo rev</td><td>AAACTACGAGCCGATCTCGTGAAG</td><td></td><td></td><td></td><td></td><td></td><td></td><td></td><td></td><td></td><td></td></tr>
<tr><td>Chromosome</td><td>start</td><td>end</td><td>strand</td><td>MM</td><td>target_seq</td><td>PAM</td><td>alignment</td><td>distance</td><td>position</td><td>gene name</td><td>gene id</td></tr>
<tr><td>GU305914.1</td><td>4750</td><td>4772</td><td>-</td><td>0</td><td>CTTCACGAGATCGGCTCGTA</td><td>CGG</td><td>||||||||[||||||||||||]PAM</td><td>NA</td><td>Intergenic</td><td>NA</td><td>NA</td></tr>
<tr><td></td><td></td><td></td><td></td><td></td><td></td><td></td><td></td><td></td><td></td><td></td><td></td></tr>
<tr><td>T34034</td><td>GGCTGAAGAAATATGAAGACCGG</td><td>1000</td><td></td><td></td><td></td><td></td><td></td><td></td><td></td><td></td><td></td></tr>
<tr><td>Oligo fwd</td><td>CACCGGCTGAAGAAATATGAAGAC</td><td></td><td></td><td></td><td></td><td></td><td></td><td></td><td></td><td></td><td></td></tr>
<tr><td>Oligo rev</td><td>AAACGTCTTCATATTTCTTCAGCC</td><td></td><td></td><td></td><td></td><td></td><td></td><td></td><td></td><td></td><td></td></tr>
<tr><td>Chromosome</td><td>start</td><td>end</td><td>strand</td><td>MM</td><td>target_seq</td><td>PAM</td><td>alignment</td><td>distance</td><td>position</td><td>gene name</td><td>gene id</td></tr>
<tr><td>GU305914.1</td><td>4729</td><td>4751</td><td>-</td><td>0</td><td>GGCTGAAGAAATATGAAGAC</td><td>CGG</td><td>||||||||[||||||||||||]PAM</td><td>NA</td><td>Intergenic</td><td>NA</td><td>NA</td></tr>
<tr><td></td><td></td><td></td><td></td><td></td><td></td><td></td><td></td><td></td><td></td><td></td><td></td></tr>
<tr><td>T34035</td><td>ACGCGCAATCCTGTTTTATTTGG</td><td>1000</td><td></td><td></td><td></td><td></td><td></td><td></td><td></td><td></td><td></td></tr>
<tr><td>Oligo fwd</td><td>CACCACGCGCAATCCTGTTTTATT</td><td></td><td></td><td></td><td></td><td></td><td></td><td></td><td></td><td></td><td></td></tr>
<tr><td>Oligo rev</td><td>AAACAATAAAACAGGATTGCGCGT</td><td></td><td></td><td></td><td></td><td></td><td></td><td></td><td></td><td></td><td></td></tr>
<tr><td>Chromosome</td><td>start</td><td>end</td><td>strand</td><td>MM</td><td>target_seq</td><td>PAM</td><td>alignment</td><td>distance</td><td>position</td><td>gene name</td><td>gene id</td></tr>
<tr><td>GU305914.1</td><td>4698</td><td>4720</td><td>-</td><td>0</td><td>ACGCGCAATCCTGTTTTATT</td><td>TGG</td><td>||||||||[||||||||||||]PAM</td><td>NA</td><td>Intergenic</td><td>NA</td><td>NA</td></tr>
<tr><td></td><td></td><td></td><td></td><td></td><td></td><td></td><td></td><td></td><td></td><td></td><td></td></tr>
<tr><td>T34036</td><td>CGCGCAATCCTGTTTTATTTGGG</td><td>1000</td><td></td><td></td><td></td><td></td><td></td><td></td><td></td><td></td><td></td></tr>
<tr><td>Oligo fwd</td><td>CACCCGCGCAATCCTGTTTTATTT</td><td></td><td></td><td></td><td></td><td></td><td></td><td></td><td></td><td></td><td></td></tr>
<tr><td>Oligo rev</td><td>AAACAAATAAAACAGGATTGCGCG</td><td></td><td></td><td></td><td></td><td></td><td></td><td></td><td></td><td></td><td></td></tr>
<tr><td>Chromosome</td><td>start</td><td>end</td><td>strand</td><td>MM</td><td>target_seq</td><td>PAM</td><td>alignment</td><td>distance</td><td>position</td><td>gene name</td><td>gene id</td></tr>
<tr><td>GU305914.1</td><td>4697</td><td>4719</td><td>-</td><td>0</td><td>CGCGCAATCCTGTTTTATTT</td><td>GGG</td><td>||||||||[||||||||||||]PAM</td><td>NA</td><td>Intergenic</td><td>NA</td><td>NA</td></tr>
<tr><td></td><td></td><td></td><td></td><td></td><td></td><td></td><td></td><td></td><td></td><td></td><td></td></tr>
<tr><td>T34037</td><td>AATCCTGTTTTATTTGGGTTCGG</td><td>1000</td><td></td><td></td><td></td><td></td><td></td><td></td><td></td><td></td><td></td></tr>
<tr><td>Oligo fwd</td><td>CACCAATCCTGTTTTATTTGGGTT</td><td></td><td></td><td></td><td></td><td></td><td></td><td></td><td></td><td></td><td></td></tr>
<tr><td>Oligo rev</td><td>AAACAACCCAAATAAAACAGGATT</td><td></td><td></td><td></td><td></td><td></td><td></td><td></td><td></td><td></td><td></td></tr>
<tr><td>Chromosome</td><td>start</td><td>end</td><td>strand</td><td>MM</td><td>target_seq</td><td>PAM</td><td>alignment</td><td>distance</td><td>position</td><td>gene name</td><td>gene id</td></tr>
<tr><td>GU305914.1</td><td>4692</td><td>4714</td><td>-</td><td>0</td><td>AATCCTGTTTTATTTGGGTT</td><td>CGG</td><td>||||||||[||||||||||||]PAM</td><td>NA</td><td>Intergenic</td><td>NA</td><td>NA</td></tr>
<tr><td></td><td></td><td></td><td></td><td></td><td></td><td></td><td></td><td></td><td></td><td></td><td></td></tr>
<tr><td>T34038</td><td>CAATCTAGTCGAGTAAGCGACGG</td><td>1000</td><td></td><td></td><td></td><td></td><td></td><td></td><td></td><td></td><td></td></tr>
<tr><td>Oligo fwd</td><td>CACCCAATCTAGTCGAGTAAGCGA</td><td></td><td></td><td></td><td></td><td></td><td></td><td></td><td></td><td></td><td></td></tr>
<tr><td>Oligo rev</td><td>AAACTCGCTTACTCGACTAGATTG</td><td></td><td></td><td></td><td></td><td></td><td></td><td></td><td></td><td></td><td></td></tr>
<tr><td>Chromosome</td><td>start</td><td>end</td><td>strand</td><td>MM</td><td>target_seq</td><td>PAM</td><td>alignment</td><td>distance</td><td>position</td><td>gene name</td><td>gene id</td></tr>
<tr><td>GU305914.1</td><td>4645</td><td>4667</td><td>-</td><td>0</td><td>CAATCTAGTCGAGTAAGCGA</td><td>CGG</td><td>||||||||[||||||||||||]PAM</td><td>NA</td><td>Intergenic</td><td>NA</td><td>NA</td></tr>
<tr><td></td><td></td><td></td><td></td><td></td><td></td><td></td><td></td><td></td><td></td><td></td><td></td></tr>
<tr><td>T34039</td><td>GATAACGTGCCGTTGAACGTTGG</td><td>1000</td><td></td><td></td><td></td><td></td><td></td><td></td><td></td><td></td><td></td></tr>
<tr><td>Oligo fwd</td><td>CACCGATAACGTGCCGTTGAACGT</td><td></td><td></td><td></td><td></td><td></td><td></td><td></td><td></td><td></td><td></td></tr>
<tr><td>Oligo rev</td><td>AAACACGTTCAACGGCACGTTATC</td><td></td><td></td><td></td><td></td><td></td><td></td><td></td><td></td><td></td><td></td></tr>
<tr><td>Chromosome</td><td>start</td><td>end</td><td>strand</td><td>MM</td><td>target_seq</td><td>PAM</td><td>alignment</td><td>distance</td><td>position</td><td>gene name</td><td>gene id</td></tr>
<tr><td>GU305914.1</td><td>4623</td><td>4645</td><td>-</td><td>0</td><td>GATAACGTGCCGTTGAACGT</td><td>TGG</td><td>||||||||[||||||||||||]PAM</td><td>NA</td><td>Intergenic</td><td>NA</td><td>NA</td></tr>
<tr><td></td><td></td><td></td><td></td><td></td><td></td><td></td><td></td><td></td><td></td><td></td><td></td></tr>
<tr><td>T34040</td><td>AACGTGCCGTTGAACGTTGGTGG</td><td>1000</td><td></td><td></td><td></td><td></td><td></td><td></td><td></td><td></td><td></td></tr>
<tr><td>Oligo fwd</td><td>CACCAACGTGCCGTTGAACGTTGG</td><td></td><td></td><td></td><td></td><td></td><td></td><td></td><td></td><td></td><td></td></tr>
<tr><td>Oligo rev</td><td>AAACCCAACGTTCAACGGCACGTT</td><td></td><td></td><td></td><td></td><td></td><td></td><td></td><td></td><td></td><td></td></tr>
<tr><td>Chromosome</td><td>start</td><td>end</td><td>strand</td><td>MM</td><td>target_seq</td><td>PAM</td><td>alignment</td><td>distance</td><td>position</td><td>gene name</td><td>gene id</td></tr>
<tr><td>GU305914.1</td><td>4620</td><td>4642</td><td>-</td><td>0</td><td>AACGTGCCGTTGAACGTTGG</td><td>TGG</td><td>||||||||[||||||||||||]PAM</td><td>NA</td><td>Intergenic</td><td>NA</td><td>NA</td></tr>
<tr><td></td><td></td><td></td><td></td><td></td><td></td><td></td><td></td><td></td><td></td><td></td><td></td></tr>
<tr><td>T34041</td><td>CGTTGAACGTTGGTGGCGAACGG</td><td>1000</td><td></td><td></td><td></td><td></td><td></td><td></td><td></td><td></td><td></td></tr>
<tr><td>Oligo fwd</td><td>CACCCGTTGAACGTTGGTGGCGAA</td><td></td><td></td><td></td><td></td><td></td><td></td><td></td><td></td><td></td><td></td></tr>
<tr><td>Oligo rev</td><td>AAACTTCGCCACCAACGTTCAACG</td><td></td><td></td><td></td><td></td><td></td><td></td><td></td><td></td><td></td><td></td></tr>
<tr><td>Chromosome</td><td>start</td><td>end</td><td>strand</td><td>MM</td><td>target_seq</td><td>PAM</td><td>alignment</td><td>distance</td><td>position</td><td>gene name</td><td>gene id</td></tr>
<tr><td>GU305914.1</td><td>4613</td><td>4635</td><td>-</td><td>0</td><td>CGTTGAACGTTGGTGGCGAA</td><td>CGG</td><td>||||||||[||||||||||||]PAM</td><td>NA</td><td>Intergenic</td><td>NA</td><td>NA</td></tr>
<tr><td></td><td></td><td></td><td></td><td></td><td></td><td></td><td></td><td></td><td></td><td></td><td></td></tr>
<tr><td>T34042</td><td>GAACGTTGGTGGCGAACGGTCGG</td><td>1000</td><td></td><td></td><td></td><td></td><td></td><td></td><td></td><td></td><td></td></tr>
<tr><td>Oligo fwd</td><td>CACCGAACGTTGGTGGCGAACGGT</td><td></td><td></td><td></td><td></td><td></td><td></td><td></td><td></td><td></td><td></td></tr>
<tr><td>Oligo rev</td><td>AAACACCGTTCGCCACCAACGTTC</td><td></td><td></td><td></td><td></td><td></td><td></td><td></td><td></td><td></td><td></td></tr>
<tr><td>Chromosome</td><td>start</td><td>end</td><td>strand</td><td>MM</td><td>target_seq</td><td>PAM</td><td>alignment</td><td>distance</td><td>position</td><td>gene name</td><td>gene id</td></tr>
<tr><td>GU305914.1</td><td>4609</td><td>4631</td><td>-</td><td>0</td><td>GAACGTTGGTGGCGAACGGT</td><td>CGG</td><td>||||||||[||||||||||||]PAM</td><td>NA</td><td>Intergenic</td><td>NA</td><td>NA</td></tr>
<tr><td></td><td></td><td></td><td></td><td></td><td></td><td></td><td></td><td></td><td></td><td></td><td></td></tr>
<tr><td>T34043</td><td>GTCGGTCGACACGAATCGAATGG</td><td>1000</td><td></td><td></td><td></td><td></td><td></td><td></td><td></td><td></td><td></td></tr>
<tr><td>Oligo fwd</td><td>CACCGTCGGTCGACACGAATCGAA</td><td></td><td></td><td></td><td></td><td></td><td></td><td></td><td></td><td></td><td></td></tr>
<tr><td>Oligo rev</td><td>AAACTTCGATTCGTGTCGACCGAC</td><td></td><td></td><td></td><td></td><td></td><td></td><td></td><td></td><td></td><td></td></tr>
<tr><td>Chromosome</td><td>start</td><td>end</td><td>strand</td><td>MM</td><td>target_seq</td><td>PAM</td><td>alignment</td><td>distance</td><td>position</td><td>gene name</td><td>gene id</td></tr>
<tr><td>GU305914.1</td><td>4591</td><td>4613</td><td>-</td><td>0</td><td>GTCGGTCGACACGAATCGAA</td><td>TGG</td><td>||||||||[||||||||||||]PAM</td><td>NA</td><td>Intergenic</td><td>NA</td><td>NA</td></tr>
<tr><td></td><td></td><td></td><td></td><td></td><td></td><td></td><td></td><td></td><td></td><td></td><td></td></tr>
<tr><td>T34044</td><td>GTCGACACGAATCGAATGGCCGG</td><td>1000</td><td></td><td></td><td></td><td></td><td></td><td></td><td></td><td></td><td></td></tr>
<tr><td>Oligo fwd</td><td>CACCGTCGACACGAATCGAATGGC</td><td></td><td></td><td></td><td></td><td></td><td></td><td></td><td></td><td></td><td></td></tr>
<tr><td>Oligo rev</td><td>AAACGCCATTCGATTCGTGTCGAC</td><td></td><td></td><td></td><td></td><td></td><td></td><td></td><td></td><td></td><td></td></tr>
<tr><td>Chromosome</td><td>start</td><td>end</td><td>strand</td><td>MM</td><td>target_seq</td><td>PAM</td><td>alignment</td><td>distance</td><td>position</td><td>gene name</td><td>gene id</td></tr>
<tr><td>GU305914.1</td><td>4587</td><td>4609</td><td>-</td><td>0</td><td>GTCGACACGAATCGAATGGC</td><td>CGG</td><td>||||||||[||||||||||||]PAM</td><td>NA</td><td>Intergenic</td><td>NA</td><td>NA</td></tr>
<tr><td></td><td></td><td></td><td></td><td></td><td></td><td></td><td></td><td></td><td></td><td></td><td></td></tr>
<tr><td>T34045</td><td>ACCAAGTTGTCCCATTTCCCAGG</td><td>1000</td><td></td><td></td><td></td><td></td><td></td><td></td><td></td><td></td><td></td></tr>
<tr><td>Oligo fwd</td><td>CACCACCAAGTTGTCCCATTTCCC</td><td></td><td></td><td></td><td></td><td></td><td></td><td></td><td></td><td></td><td></td></tr>
<tr><td>Oligo rev</td><td>AAACGGGAAATGGGACAACTTGGT</td><td></td><td></td><td></td><td></td><td></td><td></td><td></td><td></td><td></td><td></td></tr>
<tr><td>Chromosome</td><td>start</td><td>end</td><td>strand</td><td>MM</td><td>target_seq</td><td>PAM</td><td>alignment</td><td>distance</td><td>position</td><td>gene name</td><td>gene id</td></tr>
<tr><td>GU305914.1</td><td>4559</td><td>4581</td><td>-</td><td>0</td><td>ACCAAGTTGTCCCATTTCCC</td><td>AGG</td><td>||||||||[||||||||||||]PAM</td><td>NA</td><td>Intergenic</td><td>NA</td><td>NA</td></tr>
<tr><td></td><td></td><td></td><td></td><td></td><td></td><td></td><td></td><td></td><td></td><td></td><td></td></tr>
<tr><td>T34046</td><td>TTTGCGTAGACGTTTTCCGTAGG</td><td>1000</td><td></td><td></td><td></td><td></td><td></td><td></td><td></td><td></td><td></td></tr>
<tr><td>Oligo fwd</td><td>CACCTTTGCGTAGACGTTTTCCGT</td><td></td><td></td><td></td><td></td><td></td><td></td><td></td><td></td><td></td><td></td></tr>
<tr><td>Oligo rev</td><td>AAACACGGAAAACGTCTACGCAAA</td><td></td><td></td><td></td><td></td><td></td><td></td><td></td><td></td><td></td><td></td></tr>
<tr><td>Chromosome</td><td>start</td><td>end</td><td>strand</td><td>MM</td><td>target_seq</td><td>PAM</td><td>alignment</td><td>distance</td><td>position</td><td>gene name</td><td>gene id</td></tr>
<tr><td>GU305914.1</td><td>4512</td><td>4534</td><td>-</td><td>0</td><td>TTTGCGTAGACGTTTTCCGT</td><td>AGG</td><td>||||||||[||||||||||||]PAM</td><td>NA</td><td>Intergenic</td><td>NA</td><td>NA</td></tr>
<tr><td></td><td></td><td></td><td></td><td></td><td></td><td></td><td></td><td></td><td></td><td></td><td></td></tr>
<tr><td>T34047</td><td>AGACTTCTTAAGAAACCGCGTGG</td><td>1000</td><td></td><td></td><td></td><td></td><td></td><td></td><td></td><td></td><td></td></tr>
<tr><td>Oligo fwd</td><td>CACCAGACTTCTTAAGAAACCGCG</td><td></td><td></td><td></td><td></td><td></td><td></td><td></td><td></td><td></td><td></td></tr>
<tr><td>Oligo rev</td><td>AAACCGCGGTTTCTTAAGAAGTCT</td><td></td><td></td><td></td><td></td><td></td><td></td><td></td><td></td><td></td><td></td></tr>
<tr><td>Chromosome</td><td>start</td><td>end</td><td>strand</td><td>MM</td><td>target_seq</td><td>PAM</td><td>alignment</td><td>distance</td><td>position</td><td>gene name</td><td>gene id</td></tr>
<tr><td>GU305914.1</td><td>4450</td><td>4472</td><td>-</td><td>0</td><td>AGACTTCTTAAGAAACCGCG</td><td>TGG</td><td>||||||||[||||||||||||]PAM</td><td>NA</td><td>Intergenic</td><td>NA</td><td>NA</td></tr>
<tr><td></td><td></td><td></td><td></td><td></td><td></td><td></td><td></td><td></td><td></td><td></td><td></td></tr>
<tr><td>T34048</td><td>GCACTTGCTTGTATACTTCCAGG</td><td>1000</td><td></td><td></td><td></td><td></td><td></td><td></td><td></td><td></td><td></td></tr>
</table>

<table>
<tr><td>Oligo fwd</td><td>CACCGCACTTGCTTGTATACTTCC</td><td></td><td></td><td></td><td></td><td></td><td></td><td></td><td></td><td></td><td></td><td></td></tr>
<tr><td>Oligo rev</td><td>AAACGGAAGTATACAAGCAAGTGC</td><td></td><td></td><td></td><td></td><td></td><td></td><td></td><td></td><td></td><td></td><td></td></tr>
<tr><td>Chromosome</td><td>start</td><td></td><td>end</td><td>strand</td><td>MM</td><td>target_seq</td><td>PAM</td><td>alignment</td><td>distance</td><td>position</td><td>gene name</td><td>gene id</td></tr>
<tr><td>GU305914.1</td><td></td><td>4411</td><td>4433</td><td>-</td><td>0</td><td>GCACTTGCTTGTATACTTCC</td><td>AGG</td><td>||||||||[||||||||||||]PAM</td><td>NA</td><td>Intergenic</td><td>NA</td><td>NA</td></tr>
<tr><td></td><td></td><td></td><td></td><td></td><td></td><td></td><td></td><td></td><td></td><td></td><td></td><td></td></tr>
<tr><td>T34049</td><td>CACTTGCTTGTATACTTCCAGGG</td><td></td><td>1000</td><td></td><td></td><td></td><td></td><td></td><td></td><td></td><td></td><td></td></tr>
<tr><td>Oligo fwd</td><td>CACCCACTTGCTTGTATACTTCCA</td><td></td><td></td><td></td><td></td><td></td><td></td><td></td><td></td><td></td><td></td><td></td></tr>
<tr><td>Oligo rev</td><td>AAACTGGAAGTATACAAGCAAGTG</td><td></td><td></td><td></td><td></td><td></td><td></td><td></td><td></td><td></td><td></td><td></td></tr>
<tr><td>Chromosome</td><td>start</td><td></td><td>end</td><td>strand</td><td>MM</td><td>target_seq</td><td>PAM</td><td>alignment</td><td>distance</td><td>position</td><td>gene name</td><td>gene id</td></tr>
<tr><td>GU305914.1</td><td></td><td>4410</td><td>4432</td><td>-</td><td>0</td><td>CACTTGCTTGTATACTTCCA</td><td>GGG</td><td>||||||||[||||||||||||]PAM</td><td>NA</td><td>Intergenic</td><td>NA</td><td>NA</td></tr>
<tr><td></td><td></td><td></td><td></td><td></td><td></td><td></td><td></td><td></td><td></td><td></td><td></td><td></td></tr>
<tr><td>T34050</td><td>TTTGTCGCCACCTAAAGCTATGG</td><td></td><td>1000</td><td></td><td></td><td></td><td></td><td></td><td></td><td></td><td></td><td></td></tr>
<tr><td>Oligo fwd</td><td>CACCTTTGTCGCCACCTAAAGCTA</td><td></td><td></td><td></td><td></td><td></td><td></td><td></td><td></td><td></td><td></td><td></td></tr>
<tr><td>Oligo rev</td><td>AAACTAGCTTTAGGTGGCGACAAA</td><td></td><td></td><td></td><td></td><td></td><td></td><td></td><td></td><td></td><td></td><td></td></tr>
<tr><td>Chromosome</td><td>start</td><td></td><td>end</td><td>strand</td><td>MM</td><td>target_seq</td><td>PAM</td><td>alignment</td><td>distance</td><td>position</td><td>gene name</td><td>gene id</td></tr>
<tr><td>GU305914.1</td><td></td><td>4381</td><td>4403</td><td>-</td><td>0</td><td>TTTGTCGCCACCTAAAGCTA</td><td>TGG</td><td>||||||||[||||||||||||]PAM</td><td>NA</td><td>Intergenic</td><td>NA</td><td>NA</td></tr>
<tr><td></td><td></td><td></td><td></td><td></td><td></td><td></td><td></td><td></td><td></td><td></td><td></td><td></td></tr>
<tr><td>T34051</td><td>TATTTATCGAATGATGCAGATGG</td><td></td><td>1000</td><td></td><td></td><td></td><td></td><td></td><td></td><td></td><td></td><td></td></tr>
<tr><td>Oligo fwd</td><td>CACCTATTTATCGAATGATGCAGA</td><td></td><td></td><td></td><td></td><td></td><td></td><td></td><td></td><td></td><td></td><td></td></tr>
<tr><td>Oligo rev</td><td>AAACTCTGCATCATTCGATAAATA</td><td></td><td></td><td></td><td></td><td></td><td></td><td></td><td></td><td></td><td></td><td></td></tr>
<tr><td>Chromosome</td><td>start</td><td></td><td>end</td><td>strand</td><td>MM</td><td>target_seq</td><td>PAM</td><td>alignment</td><td>distance</td><td>position</td><td>gene name</td><td>gene id</td></tr>
<tr><td>GU305914.1</td><td></td><td>4353</td><td>4375</td><td>-</td><td>0</td><td>TATTTATCGAATGATGCAGA</td><td>TGG</td><td>||||||||[||||||||||||]PAM</td><td>NA</td><td>Intergenic</td><td>NA</td><td>NA</td></tr>
<tr><td></td><td></td><td></td><td></td><td></td><td></td><td></td><td></td><td></td><td></td><td></td><td></td><td></td></tr>
<tr><td>T34052</td><td>TGCAGATGGAAACCGTGTTCTGG</td><td></td><td>1000</td><td></td><td></td><td></td><td></td><td></td><td></td><td></td><td></td><td></td></tr>
<tr><td>Oligo fwd</td><td>CACCTGCAGATGGAAACCGTGTTC</td><td></td><td></td><td></td><td></td><td></td><td></td><td></td><td></td><td></td><td></td><td></td></tr>
<tr><td>Oligo rev</td><td>AAACGAACACGGTTTCCATCTGCA</td><td></td><td></td><td></td><td></td><td></td><td></td><td></td><td></td><td></td><td></td><td></td></tr>
<tr><td>Chromosome</td><td>start</td><td></td><td>end</td><td>strand</td><td>MM</td><td>target_seq</td><td>PAM</td><td>alignment</td><td>distance</td><td>position</td><td>gene name</td><td>gene id</td></tr>
<tr><td>GU305914.1</td><td></td><td>4339</td><td>4361</td><td>-</td><td>0</td><td>TGCAGATGGAAACCGTGTTC</td><td>TGG</td><td>||||||||[||||||||||||]PAM</td><td>NA</td><td>Intergenic</td><td>NA</td><td>NA</td></tr>
<tr><td></td><td></td><td></td><td></td><td></td><td></td><td></td><td></td><td></td><td></td><td></td><td></td><td></td></tr>
<tr><td>T34053</td><td>TCCTATCCATGTTGTTATTCCGG</td><td></td><td>1000</td><td></td><td></td><td></td><td></td><td></td><td></td><td></td><td></td><td></td></tr>
<tr><td>Oligo fwd</td><td>CACCTCCTATCCATGTTGTTATTC</td><td></td><td></td><td></td><td></td><td></td><td></td><td></td><td></td><td></td><td></td><td></td></tr>
<tr><td>Oligo rev</td><td>AAACGAATAACAACATGGATAGGA</td><td></td><td></td><td></td><td></td><td></td><td></td><td></td><td></td><td></td><td></td><td></td></tr>
<tr><td>Chromosome</td><td>start</td><td></td><td>end</td><td>strand</td><td>MM</td><td>target_seq</td><td>PAM</td><td>alignment</td><td>distance</td><td>position</td><td>gene name</td><td>gene id</td></tr>
<tr><td>GU305914.1</td><td></td><td>4309</td><td>4331</td><td>-</td><td>0</td><td>TCCTATCCATGTTGTTATTC</td><td>CGG</td><td>||||||||[||||||||||||]PAM</td><td>NA</td><td>Intergenic</td><td>NA</td><td>NA</td></tr>
<tr><td></td><td></td><td></td><td></td><td></td><td></td><td></td><td></td><td></td><td></td><td></td><td></td><td></td></tr>
<tr><td>T34054</td><td>CCATGTTGTTATTCCGGACTTGG</td><td></td><td>1000</td><td></td><td></td><td></td><td></td><td></td><td></td><td></td><td></td><td></td></tr>
<tr><td>Oligo fwd</td><td>CACCCCATGTTGTTATTCCGGACT</td><td></td><td></td><td></td><td></td><td></td><td></td><td></td><td></td><td></td><td></td><td></td></tr>
<tr><td>Oligo rev</td><td>AAACAGTCCGGAATAACAACATGG</td><td></td><td></td><td></td><td></td><td></td><td></td><td></td><td></td><td></td><td></td><td></td></tr>
<tr><td>Chromosome</td><td>start</td><td></td><td>end</td><td>strand</td><td>MM</td><td>target_seq</td><td>PAM</td><td>alignment</td><td>distance</td><td>position</td><td>gene name</td><td>gene id</td></tr>
<tr><td>GU305914.1</td><td></td><td>4303</td><td>4325</td><td>-</td><td>0</td><td>CCATGTTGTTATTCCGGACT</td><td>TGG</td><td>||||||||[||||||||||||]PAM</td><td>NA</td><td>Intergenic</td><td>NA</td><td>NA</td></tr>
<tr><td></td><td></td><td></td><td></td><td></td><td></td><td></td><td></td><td></td><td></td><td></td><td></td><td></td></tr>
<tr><td>T34055</td><td>CATGTTGTTATTCCGGACTTGGG</td><td></td><td>1000</td><td></td><td></td><td></td><td></td><td></td><td></td><td></td><td></td><td></td></tr>
<tr><td>Oligo fwd</td><td>CACCCATGTTGTTATTCCGGACTT</td><td></td><td></td><td></td><td></td><td></td><td></td><td></td><td></td><td></td><td></td><td></td></tr>
<tr><td>Oligo rev</td><td>AAACAAGTCCGGAATAACAACATG</td><td></td><td></td><td></td><td></td><td></td><td></td><td></td><td></td><td></td><td></td><td></td></tr>
<tr><td>Chromosome</td><td>start</td><td></td><td>end</td><td>strand</td><td>MM</td><td>target_seq</td><td>PAM</td><td>alignment</td><td>distance</td><td>position</td><td>gene name</td><td>gene id</td></tr>
<tr><td>GU305914.1</td><td></td><td>4302</td><td>4324</td><td>-</td><td>0</td><td>CATGTTGTTATTCCGGACTT</td><td>GGG</td><td>||||||||[||||||||||||]PAM</td><td>NA</td><td>Intergenic</td><td>NA</td><td>NA</td></tr>
<tr><td></td><td></td><td></td><td></td><td></td><td></td><td></td><td></td><td></td><td></td><td></td><td></td><td></td></tr>
<tr><td>T34056</td><td>ATTCCGGACTTGGGTGATTGTGG</td><td></td><td>1000</td><td></td><td></td><td></td><td></td><td></td><td></td><td></td><td></td><td></td></tr>
<tr><td>Oligo fwd</td><td>CACCATTCCGGACTTGGGTGATTG</td><td></td><td></td><td></td><td></td><td></td><td></td><td></td><td></td><td></td><td></td><td></td></tr>
<tr><td>Oligo rev</td><td>AAACCAATCACCCAAGTCCGGAAT</td><td></td><td></td><td></td><td></td><td></td><td></td><td></td><td></td><td></td><td></td><td></td></tr>
<tr><td>Chromosome</td><td>start</td><td></td><td>end</td><td>strand</td><td>MM</td><td>target_seq</td><td>PAM</td><td>alignment</td><td>distance</td><td>position</td><td>gene name</td><td>gene id</td></tr>
<tr><td>GU305914.1</td><td></td><td>4293</td><td>4315</td><td>-</td><td>0</td><td>ATTCCGGACTTGGGTGATTG</td><td>TGG</td><td>||||||||[||||||||||||]PAM</td><td>NA</td><td>Intergenic</td><td>NA</td><td>NA</td></tr>
<tr><td></td><td></td><td></td><td></td><td></td><td></td><td></td><td></td><td></td><td></td><td></td><td></td><td></td></tr>
<tr><td>T34057</td><td>CCGGACTTGGGTGATTGTGGTGG</td><td></td><td>1000</td><td></td><td></td><td></td><td></td><td></td><td></td><td></td><td></td><td></td></tr>
<tr><td>Oligo fwd</td><td>CACCCCGGACTTGGGTGATTGTGG</td><td></td><td></td><td></td><td></td><td></td><td></td><td></td><td></td><td></td><td></td><td></td></tr>
<tr><td>Oligo rev</td><td>AAACCCACAATCACCCAAGTCCGG</td><td></td><td></td><td></td><td></td><td></td><td></td><td></td><td></td><td></td><td></td><td></td></tr>
<tr><td>Chromosome</td><td>start</td><td></td><td>end</td><td>strand</td><td>MM</td><td>target_seq</td><td>PAM</td><td>alignment</td><td>distance</td><td>position</td><td>gene name</td><td>gene id</td></tr>
<tr><td>GU305914.1</td><td></td><td>4290</td><td>4312</td><td>-</td><td>0</td><td>CCGGACTTGGGTGATTGTGG</td><td>TGG</td><td>||||||||[||||||||||||]PAM</td><td>NA</td><td>Intergenic</td><td>NA</td><td>NA</td></tr>
<tr><td></td><td></td><td></td><td></td><td></td><td></td><td></td><td></td><td></td><td></td><td></td><td></td><td></td></tr>
<tr><td>T34058</td><td>GGGTGATTGTGGTGGTCGTCTGG</td><td></td><td>1000</td><td></td><td></td><td></td><td></td><td></td><td></td><td></td><td></td><td></td></tr>
<tr><td>Oligo fwd</td><td>CACCGGGTGATTGTGGTGGTCGTC</td><td></td><td></td><td></td><td></td><td></td><td></td><td></td><td></td><td></td><td></td><td></td></tr>
<tr><td>Oligo rev</td><td>AAACGACGACCACCACAATCACCC</td><td></td><td></td><td></td><td></td><td></td><td></td><td></td><td></td><td></td><td></td><td></td></tr>
<tr><td>Chromosome</td><td>start</td><td></td><td>end</td><td>strand</td><td>MM</td><td>target_seq</td><td>PAM</td><td>alignment</td><td>distance</td><td>position</td><td>gene name</td><td>gene id</td></tr>
<tr><td>GU305914.1</td><td></td><td>4282</td><td>4304</td><td>-</td><td>0</td><td>GGGTGATTGTGGTGGTCGTC</td><td>TGG</td><td>||||||||[||||||||||||]PAM</td><td>NA</td><td>Intergenic</td><td>NA</td><td>NA</td></tr>
<tr><td></td><td></td><td></td><td></td><td></td><td></td><td></td><td></td><td></td><td></td><td></td><td></td><td></td></tr>
<tr><td>T34059</td><td>GGTGGTCGTCTGGATCTGTGCGG</td><td></td><td>1000</td><td></td><td></td><td></td><td></td><td></td><td></td><td></td><td></td><td></td></tr>
<tr><td>Oligo fwd</td><td>CACCGGTGGTCGTCTGGATCTGTG</td><td></td><td></td><td></td><td></td><td></td><td></td><td></td><td></td><td></td><td></td><td></td></tr>
<tr><td>Oligo rev</td><td>AAACCACAGATCCAGACGACCACC</td><td></td><td></td><td></td><td></td><td></td><td></td><td></td><td></td><td></td><td></td><td></td></tr>
<tr><td>Chromosome</td><td>start</td><td></td><td>end</td><td>strand</td><td>MM</td><td>target_seq</td><td>PAM</td><td>alignment</td><td>distance</td><td>position</td><td>gene name</td><td>gene id</td></tr>
<tr><td>GU305914.1</td><td></td><td>4272</td><td>4294</td><td>-</td><td>0</td><td>GGTGGTCGTCTGGATCTGTG</td><td>CGG</td><td>||||||||[||||||||||||]PAM</td><td>NA</td><td>Intergenic</td><td>NA</td><td>NA</td></tr>
<tr><td></td><td></td><td></td><td></td><td></td><td></td><td></td><td></td><td></td><td></td><td></td><td></td><td></td></tr>
<tr><td>T34060</td><td>GGATCTGTGCGGTGAATCTCCGG</td><td></td><td>1000</td><td></td><td></td><td></td><td></td><td></td><td></td><td></td><td></td><td></td></tr>
<tr><td>Oligo fwd</td><td>CACCGGATCTGTGCGGTGAATCTC</td><td></td><td></td><td></td><td></td><td></td><td></td><td></td><td></td><td></td><td></td><td></td></tr>
<tr><td>Oligo rev</td><td>AAACGAGATTCACCGCACAGATCC</td><td></td><td></td><td></td><td></td><td></td><td></td><td></td><td></td><td></td><td></td><td></td></tr>
<tr><td>Chromosome</td><td>start</td><td></td><td>end</td><td>strand</td><td>MM</td><td>target_seq</td><td>PAM</td><td>alignment</td><td>distance</td><td>position</td><td>gene name</td><td>gene id</td></tr>
<tr><td>GU305914.1</td><td></td><td>4261</td><td>4283</td><td>-</td><td>0</td><td>GGATCTGTGCGGTGAATCTC</td><td>CGG</td><td>||||||||[||||||||||||]PAM</td><td>NA</td><td>Intergenic</td><td>NA</td><td>NA</td></tr>
<tr><td></td><td></td><td></td><td></td><td></td><td></td><td></td><td></td><td></td><td></td><td></td><td></td><td></td></tr>
<tr><td>T34061</td><td>TAGAACCGAGCGCCACATGACGG</td><td></td><td>1000</td><td></td><td></td><td></td><td></td><td></td><td></td><td></td><td></td><td></td></tr>
<tr><td>Oligo fwd</td><td>CACCTAGAACCGAGCGCCACATGA</td><td></td><td></td><td></td><td></td><td></td><td></td><td></td><td></td><td></td><td></td><td></td></tr>
<tr><td>Oligo rev</td><td>AAACTCATGTGGCGCTCGGTTCTA</td><td></td><td></td><td></td><td></td><td></td><td></td><td></td><td></td><td></td><td></td><td></td></tr>
<tr><td>Chromosome</td><td>start</td><td></td><td>end</td><td>strand</td><td>MM</td><td>target_seq</td><td>PAM</td><td>alignment</td><td>distance</td><td>position</td><td>gene name</td><td>gene id</td></tr>
<tr><td>GU305914.1</td><td></td><td>4204</td><td>4226</td><td>-</td><td>0</td><td>TAGAACCGAGCGCCACATGA</td><td>CGG</td><td>||||||||[||||||||||||]PAM</td><td>NA</td><td>Intergenic</td><td>NA</td><td>NA</td></tr>
<tr><td></td><td></td><td></td><td></td><td></td><td></td><td></td><td></td><td></td><td></td><td></td><td></td><td></td></tr>
<tr><td>T34062</td><td>GAGCGCCACATGACGGTACATGG</td><td></td><td>1000</td><td></td><td></td><td></td><td></td><td></td><td></td><td></td><td></td><td></td></tr>
<tr><td>Oligo fwd</td><td>CACCGAGCGCCACATGACGGTACA</td><td></td><td></td><td></td><td></td><td></td><td></td><td></td><td></td><td></td><td></td><td></td></tr>
<tr><td>Oligo rev</td><td>AAACTGTACCGTCATGTGGCGCTC</td><td></td><td></td><td></td><td></td><td></td><td></td><td></td><td></td><td></td><td></td><td></td></tr>
<tr><td>Chromosome</td><td>start</td><td></td><td>end</td><td>strand</td><td>MM</td><td>target_seq</td><td>PAM</td><td>alignment</td><td>distance</td><td>position</td><td>gene name</td><td>gene id</td></tr>
<tr><td>GU305914.1</td><td></td><td>4197</td><td>4219</td><td>-</td><td>0</td><td>GAGCGCCACATGACGGTACA</td><td>TGG</td><td>||||||||[||||||||||||]PAM</td><td>NA</td><td>Intergenic</td><td>NA</td><td>NA</td></tr>
</table>

<table>
<tr><td>T34063</td><td>CTGATTCTAGACCATGTATGAGG</td><td>1000</td><td></td><td></td><td></td><td></td><td></td><td></td><td></td><td></td><td></td></tr>
<tr><td>Oligo fwd</td><td>CACCCTGATTCTAGACCATGTATG</td><td></td><td></td><td></td><td></td><td></td><td></td><td></td><td></td><td></td><td></td></tr>
<tr><td>Oligo rev</td><td>AAACCATACATGGTCTAGAATCAG</td><td></td><td></td><td></td><td></td><td></td><td></td><td></td><td></td><td></td><td></td></tr>
<tr><td>Chromosome</td><td>start</td><td>end</td><td>strand</td><td>MM</td><td>target_seq</td><td>PAM</td><td>alignment</td><td>distance</td><td>position</td><td>gene name</td><td>gene id</td></tr>
<tr><td>GU305914.1</td><td>4144</td><td>4166</td><td>-</td><td>0</td><td>CTGATTCTAGACCATGTATG</td><td>AGG</td><td>||||||||[||||||||||||]PAM</td><td>NA</td><td>Intergenic</td><td>NA</td><td>NA</td></tr>
<tr><td>T34064</td><td>ATGAGGATATCTACGTTTGACGG</td><td>1000</td><td></td><td></td><td></td><td></td><td></td><td></td><td></td><td></td><td></td></tr>
<tr><td>Oligo fwd</td><td>CACCATGAGGATATCTACGTTTGA</td><td></td><td></td><td></td><td></td><td></td><td></td><td></td><td></td><td></td><td></td></tr>
<tr><td>Oligo rev</td><td>AAACTCAAACGTAGATATCCTCAT</td><td></td><td></td><td></td><td></td><td></td><td></td><td></td><td></td><td></td><td></td></tr>
<tr><td>Chromosome</td><td>start</td><td>end</td><td>strand</td><td>MM</td><td>target_seq</td><td>PAM</td><td>alignment</td><td>distance</td><td>position</td><td>gene name</td><td>gene id</td></tr>
<tr><td>GU305914.1</td><td>4127</td><td>4149</td><td>-</td><td>0</td><td>ATGAGGATATCTACGTTTGA</td><td>CGG</td><td>||||||||[||||||||||||]PAM</td><td>NA</td><td>Intergenic</td><td>NA</td><td>NA</td></tr>
<tr><td>T34065</td><td>GGATATCTACGTTTGACGGACGG</td><td>1000</td><td></td><td></td><td></td><td></td><td></td><td></td><td></td><td></td><td></td></tr>
<tr><td>Oligo fwd</td><td>CACCGGATATCTACGTTTGACGGA</td><td></td><td></td><td></td><td></td><td></td><td></td><td></td><td></td><td></td><td></td></tr>
<tr><td>Oligo rev</td><td>AAACTCCGTCAAACGTAGATATCC</td><td></td><td></td><td></td><td></td><td></td><td></td><td></td><td></td><td></td><td></td></tr>
<tr><td>Chromosome</td><td>start</td><td>end</td><td>strand</td><td>MM</td><td>target_seq</td><td>PAM</td><td>alignment</td><td>distance</td><td>position</td><td>gene name</td><td>gene id</td></tr>
<tr><td>GU305914.1</td><td>4123</td><td>4145</td><td>-</td><td>0</td><td>GGATATCTACGTTTGACGGA</td><td>CGG</td><td>||||||||[||||||||||||]PAM</td><td>NA</td><td>Intergenic</td><td>NA</td><td>NA</td></tr>
<tr><td>T34066</td><td>GAAAGAAATACTCAGAACACCGG</td><td>1000</td><td></td><td></td><td></td><td></td><td></td><td></td><td></td><td></td><td></td></tr>
<tr><td>Oligo fwd</td><td>CACCGAAAGAAATACTCAGAACAC</td><td></td><td></td><td></td><td></td><td></td><td></td><td></td><td></td><td></td><td></td></tr>
<tr><td>Oligo rev</td><td>AAACGTGTTCTGAGTATTTCTTTC</td><td></td><td></td><td></td><td></td><td></td><td></td><td></td><td></td><td></td><td></td></tr>
<tr><td>Chromosome</td><td>start</td><td>end</td><td>strand</td><td>MM</td><td>target_seq</td><td>PAM</td><td>alignment</td><td>distance</td><td>position</td><td>gene name</td><td>gene id</td></tr>
<tr><td>GU305914.1</td><td>4101</td><td>4123</td><td>-</td><td>0</td><td>GAAAGAAATACTCAGAACAC</td><td>CGG</td><td>||||||||[||||||||||||]PAM</td><td>NA</td><td>Intergenic</td><td>NA</td><td>NA</td></tr>
<tr><td>T34067</td><td>ATACTCAGAACACCGGAAAATGG</td><td>1000</td><td></td><td></td><td></td><td></td><td></td><td></td><td></td><td></td><td></td></tr>
<tr><td>Oligo fwd</td><td>CACCATACTCAGAACACCGGAAAA</td><td></td><td></td><td></td><td></td><td></td><td></td><td></td><td></td><td></td><td></td></tr>
<tr><td>Oligo rev</td><td>AAACTTTTCCGGTGTTCTGAGTAT</td><td></td><td></td><td></td><td></td><td></td><td></td><td></td><td></td><td></td><td></td></tr>
<tr><td>Chromosome</td><td>start</td><td>end</td><td>strand</td><td>MM</td><td>target_seq</td><td>PAM</td><td>alignment</td><td>distance</td><td>position</td><td>gene name</td><td>gene id</td></tr>
<tr><td>GU305914.1</td><td>4094</td><td>4116</td><td>-</td><td>0</td><td>ATACTCAGAACACCGGAAAA</td><td>TGG</td><td>||||||||[||||||||||||]PAM</td><td>NA</td><td>Intergenic</td><td>NA</td><td>NA</td></tr>
<tr><td>T34068</td><td>ACCGGAAAATGGTTTACTCAAGG</td><td>1000</td><td></td><td></td><td></td><td></td><td></td><td></td><td></td><td></td><td></td></tr>
<tr><td>Oligo fwd</td><td>CACCACCGGAAAATGGTTTACTCA</td><td></td><td></td><td></td><td></td><td></td><td></td><td></td><td></td><td></td><td></td></tr>
<tr><td>Oligo rev</td><td>AAACTGAGTAAACCATTTTCCGGT</td><td></td><td></td><td></td><td></td><td></td><td></td><td></td><td></td><td></td><td></td></tr>
<tr><td>Chromosome</td><td>start</td><td>end</td><td>strand</td><td>MM</td><td>target_seq</td><td>PAM</td><td>alignment</td><td>distance</td><td>position</td><td>gene name</td><td>gene id</td></tr>
<tr><td>GU305914.1</td><td>4083</td><td>4105</td><td>-</td><td>0</td><td>ACCGGAAAATGGTTTACTCA</td><td>AGG</td><td>||||||||[||||||||||||]PAM</td><td>NA</td><td>Intergenic</td><td>NA</td><td>NA</td></tr>
<tr><td>T34069</td><td>CCGGAAAATGGTTTACTCAAGGG</td><td>1000</td><td></td><td></td><td></td><td></td><td></td><td></td><td></td><td></td><td></td></tr>
<tr><td>Oligo fwd</td><td>CACCCCGGAAAATGGTTTACTCAA</td><td></td><td></td><td></td><td></td><td></td><td></td><td></td><td></td><td></td><td></td></tr>
<tr><td>Oligo rev</td><td>AAACTTGAGTAAACCATTTTCCGG</td><td></td><td></td><td></td><td></td><td></td><td></td><td></td><td></td><td></td><td></td></tr>
<tr><td>Chromosome</td><td>start</td><td>end</td><td>strand</td><td>MM</td><td>target_seq</td><td>PAM</td><td>alignment</td><td>distance</td><td>position</td><td>gene name</td><td>gene id</td></tr>
<tr><td>GU305914.1</td><td>4082</td><td>4104</td><td>-</td><td>0</td><td>CCGGAAAATGGTTTACTCAA</td><td>GGG</td><td>||||||||[||||||||||||]PAM</td><td>NA</td><td>Intergenic</td><td>NA</td><td>NA</td></tr>
<tr><td>T34070</td><td>CGGAAAATGGTTTACTCAAGGGG</td><td>1000</td><td></td><td></td><td></td><td></td><td></td><td></td><td></td><td></td><td></td></tr>
<tr><td>Oligo fwd</td><td>CACCCGGAAAATGGTTTACTCAAG</td><td></td><td></td><td></td><td></td><td></td><td></td><td></td><td></td><td></td><td></td></tr>
<tr><td>Oligo rev</td><td>AAACCTTGAGTAAACCATTTTCCG</td><td></td><td></td><td></td><td></td><td></td><td></td><td></td><td></td><td></td><td></td></tr>
<tr><td>Chromosome</td><td>start</td><td>end</td><td>strand</td><td>MM</td><td>target_seq</td><td>PAM</td><td>alignment</td><td>distance</td><td>position</td><td>gene name</td><td>gene id</td></tr>
<tr><td>GU305914.1</td><td>4081</td><td>4103</td><td>-</td><td>0</td><td>CGGAAAATGGTTTACTCAAG</td><td>GGG</td><td>||||||||[||||||||||||]PAM</td><td>NA</td><td>Intergenic</td><td>NA</td><td>NA</td></tr>
<tr><td>T34071</td><td>ACTCAAGGGGATTTTTATTTAGG</td><td>1000</td><td></td><td></td><td></td><td></td><td></td><td></td><td></td><td></td><td></td></tr>
<tr><td>Oligo fwd</td><td>CACCACTCAAGGGGATTTTTATTT</td><td></td><td></td><td></td><td></td><td></td><td></td><td></td><td></td><td></td><td></td></tr>
<tr><td>Oligo rev</td><td>AAACAAATAAAAATCCCCTTGAGT</td><td></td><td></td><td></td><td></td><td></td><td></td><td></td><td></td><td></td><td></td></tr>
<tr><td>Chromosome</td><td>start</td><td>end</td><td>strand</td><td>MM</td><td>target_seq</td><td>PAM</td><td>alignment</td><td>distance</td><td>position</td><td>gene name</td><td>gene id</td></tr>
<tr><td>GU305914.1</td><td>4068</td><td>4090</td><td>-</td><td>0</td><td>ACTCAAGGGGATTTTTATTT</td><td>AGG</td><td>||||||||[||||||||||||]PAM</td><td>NA</td><td>Intergenic</td><td>NA</td><td>NA</td></tr>
<tr><td>T34072</td><td>CTCAAGGGGATTTTTATTTAGGG</td><td>1000</td><td></td><td></td><td></td><td></td><td></td><td></td><td></td><td></td><td></td></tr>
<tr><td>Oligo fwd</td><td>CACCCTCAAGGGGATTTTTATTTA</td><td></td><td></td><td></td><td></td><td></td><td></td><td></td><td></td><td></td><td></td></tr>
<tr><td>Oligo rev</td><td>AAACTAAATAAAAATCCCCTTGAG</td><td></td><td></td><td></td><td></td><td></td><td></td><td></td><td></td><td></td><td></td></tr>
<tr><td>Chromosome</td><td>start</td><td>end</td><td>strand</td><td>MM</td><td>target_seq</td><td>PAM</td><td>alignment</td><td>distance</td><td>position</td><td>gene name</td><td>gene id</td></tr>
<tr><td>GU305914.1</td><td>4067</td><td>4089</td><td>-</td><td>0</td><td>CTCAAGGGGATTTTTATTTA</td><td>GGG</td><td>||||||||[||||||||||||]PAM</td><td>NA</td><td>Intergenic</td><td>NA</td><td>NA</td></tr>
<tr><td>T34073</td><td>TCAAGGGGATTTTTATTTAGGGG</td><td>1000</td><td></td><td></td><td></td><td></td><td></td><td></td><td></td><td></td><td></td></tr>
<tr><td>Oligo fwd</td><td>CACCTCAAGGGGATTTTTATTTAG</td><td></td><td></td><td></td><td></td><td></td><td></td><td></td><td></td><td></td><td></td></tr>
<tr><td>Oligo rev</td><td>AAACCTAAATAAAAATCCCCTTGA</td><td></td><td></td><td></td><td></td><td></td><td></td><td></td><td></td><td></td><td></td></tr>
<tr><td>Chromosome</td><td>start</td><td>end</td><td>strand</td><td>MM</td><td>target_seq</td><td>PAM</td><td>alignment</td><td>distance</td><td>position</td><td>gene name</td><td>gene id</td></tr>
<tr><td>GU305914.1</td><td>4066</td><td>4088</td><td>-</td><td>0</td><td>TCAAGGGGATTTTTATTTAG</td><td>GGG</td><td>||||||||[||||||||||||]PAM</td><td>NA</td><td>Intergenic</td><td>NA</td><td>NA</td></tr>
<tr><td>T34074</td><td>CAAGGGGATTTTTATTTAGGGGG</td><td>1000</td><td></td><td></td><td></td><td></td><td></td><td></td><td></td><td></td><td></td></tr>
<tr><td>Oligo fwd</td><td>CACCCAAGGGGATTTTTATTTAGG</td><td></td><td></td><td></td><td></td><td></td><td></td><td></td><td></td><td></td><td></td></tr>
<tr><td>Oligo rev</td><td>AAACCCTAAATAAAAATCCCCTTG</td><td></td><td></td><td></td><td></td><td></td><td></td><td></td><td></td><td></td><td></td></tr>
<tr><td>Chromosome</td><td>start</td><td>end</td><td>strand</td><td>MM</td><td>target_seq</td><td>PAM</td><td>alignment</td><td>distance</td><td>position</td><td>gene name</td><td>gene id</td></tr>
<tr><td>GU305914.1</td><td>4065</td><td>4087</td><td>-</td><td>0</td><td>CAAGGGGATTTTTATTTAGG</td><td>GGG</td><td>||||||||[||||||||||||]PAM</td><td>NA</td><td>Intergenic</td><td>NA</td><td>NA</td></tr>
<tr><td>T34075</td><td>GGGGGTTAGTTACTCTTAAGCGG</td><td>1000</td><td></td><td></td><td></td><td></td><td></td><td></td><td></td><td></td><td></td></tr>
<tr><td>Oligo fwd</td><td>CACCGGGGGTTAGTTACTCTTAAG</td><td></td><td></td><td></td><td></td><td></td><td></td><td></td><td></td><td></td><td></td></tr>
<tr><td>Oligo rev</td><td>AAACCTTAAGAGTAACTAACCCCC</td><td></td><td></td><td></td><td></td><td></td><td></td><td></td><td></td><td></td><td></td></tr>
<tr><td>Chromosome</td><td>start</td><td>end</td><td>strand</td><td>MM</td><td>target_seq</td><td>PAM</td><td>alignment</td><td>distance</td><td>position</td><td>gene name</td><td>gene id</td></tr>
<tr><td>GU305914.1</td><td>4047</td><td>4069</td><td>-</td><td>0</td><td>GGGGGTTAGTTACTCTTAAG</td><td>CGG</td><td>||||||||[||||||||||||]PAM</td><td>NA</td><td>Intergenic</td><td>NA</td><td>NA</td></tr>
<tr><td>T34076</td><td>AATCATTACTGAAGAAACGACGG</td><td>1000</td><td></td><td></td><td></td><td></td><td></td><td></td><td></td><td></td><td></td></tr>
<tr><td>Oligo fwd</td><td>CACCAATCATTACTGAAGAAACGA</td><td></td><td></td><td></td><td></td><td></td><td></td><td></td><td></td><td></td><td></td></tr>
<tr><td>Oligo rev</td><td>AAACTCGTTTCTTCAGTAATGATT</td><td></td><td></td><td></td><td></td><td></td><td></td><td></td><td></td><td></td><td></td></tr>
<tr><td>Chromosome</td><td>start</td><td>end</td><td>strand</td><td>MM</td><td>target_seq</td><td>PAM</td><td>alignment</td><td>distance</td><td>position</td><td>gene name</td><td>gene id</td></tr>
<tr><td>GU305914.1</td><td>4007</td><td>4029</td><td>-</td><td>0</td><td>AATCATTACTGAAGAAACGA</td><td>CGG</td><td>||||||||[||||||||||||]PAM</td><td>NA</td><td>Intergenic</td><td>NA</td><td>NA</td></tr>
<tr><td>GU305914.1</td><td>216253</td><td>216275</td><td>-</td><td>4</td><td>GATCGTTACTGACGAATCGA</td><td>CGG</td><td>-|||-|||[||||-|||-|||]PAM</td><td>NA</td><td>Intergenic</td><td>NA</td><td>NA</td></tr>
<tr><td>T34077</td><td>GAAGAAACGACGGTATAATGAGG</td><td>1000</td><td></td><td></td><td></td><td></td><td></td><td></td><td></td><td></td><td></td></tr>
<tr><td>Oligo fwd</td><td>CACCGAAGAAACGACGGTATAATG</td><td></td><td></td><td></td><td></td><td></td><td></td><td></td><td></td><td></td><td></td></tr>
<tr><td>Oligo rev</td><td>AAACCATTATACCGTCGTTTCTTC</td><td></td><td></td><td></td><td></td><td></td><td></td><td></td><td></td><td></td><td></td></tr>
<tr><td>Chromosome</td><td>start</td><td>end</td><td>strand</td><td>MM</td><td>target_seq</td><td>PAM</td><td>alignment</td><td>distance</td><td>position</td><td>gene name</td><td>gene id</td></tr>
</table>

| | | | | | | | | | | | | |
|---|---|---|---|---|---|---|---|---|---|---|---|---|
| GU305914.1 | | 3997 | 4019 | - | 0 | GAAGAAACGACGGTATAATG | AGG | \|\|\|\|\|\|\|\|[\|\|\|\|\|\|\|\|\|\|\|\|]PAM | NA | Intergenic | NA | NA |
| T34078 | AAGAAACGACGGTATAATGAGGG | | 1000 | | | | | | | | | |
| Oligo fwd | CACCAAGAAACGACGGTATAATGA | | | | | | | | | | | |
| Oligo rev | AAACTCATTATACCGTCGTTTCTT | | | | | | | | | | | |
| Chromosome | start | | end | strand | MM | target_seq | PAM | alignment | distance | position | gene name | gene id |
| GU305914.1 | | 3996 | 4018 | - | 0 | AAGAAACGACGGTATAATGA | GGG | \|\|\|\|\|\|\|\|[\|\|\|\|\|\|\|\|\|\|\|\|]PAM | NA | Intergenic | NA | NA |
| T34079 | ACAAGTAAATCAGTACGAGAAGG | | 1000 | | | | | | | | | |
| Oligo fwd | CACCACAAGTAAATCAGTACGAGA | | | | | | | | | | | |
| Oligo rev | AAACTCTCGTACTGATTTACTTGT | | | | | | | | | | | |
| Chromosome | start | | end | strand | MM | target_seq | PAM | alignment | distance | position | gene name | gene id |
| GU305914.1 | | 3968 | 3990 | - | 0 | ACAAGTAAATCAGTACGAGA | AGG | \|\|\|\|\|\|\|\|[\|\|\|\|\|\|\|\|\|\|\|\|]PAM | NA | Intergenic | NA | NA |
| GU305914.1 | | 161268 | 161290 | + | 4 | CTAAGTAAAGCAGTACGAGG | CGG | --\|\|\|\|\|\|[\|-\|\|\|\|\|\|\|\|\|-]PAM | NA | Intergenic | NA | NA |
| T34080 | AAAAAGAAGAGTCACTATCACGG | | 1000 | | | | | | | | | |
| Oligo fwd | CACCAAAAAGAAGAGTCACTATCA | | | | | | | | | | | |
| Oligo rev | AAACTGATAGTGACTCTTCTTTTT | | | | | | | | | | | |
| Chromosome | start | | end | strand | MM | target_seq | PAM | alignment | distance | position | gene name | gene id |
| GU305914.1 | | 3937 | 3959 | - | 0 | AAAAAGAAGAGTCACTATCA | CGG | \|\|\|\|\|\|\|\|[\|\|\|\|\|\|\|\|\|\|\|\|]PAM | NA | Intergenic | NA | NA |
| T34081 | AGTCACTATCACGGTGAGCGTGG | | 1000 | | | | | | | | | |
| Oligo fwd | CACCAGTCACTATCACGGTGAGCG | | | | | | | | | | | |
| Oligo rev | AAACCGCTCACCGTGATAGTGACT | | | | | | | | | | | |
| Chromosome | start | | end | strand | MM | target_seq | PAM | alignment | distance | position | gene name | gene id |
| GU305914.1 | | 3928 | 3950 | - | 0 | AGTCACTATCACGGTGAGCG | TGG | \|\|\|\|\|\|\|\|[\|\|\|\|\|\|\|\|\|\|\|\|]PAM | NA | Intergenic | NA | NA |
| T34082 | AGCGTGGTCACGATCGTGTTTGG | | 1000 | | | | | | | | | |
| Oligo fwd | CACCAGCGTGGTCACGATCGTGTT | | | | | | | | | | | |
| Oligo rev | AAACAACACGATCGTGACCACGCT | | | | | | | | | | | |
| Chromosome | start | | end | strand | MM | target_seq | PAM | alignment | distance | position | gene name | gene id |
| GU305914.1 | | 3912 | 3934 | - | 0 | AGCGTGGTCACGATCGTGTT | TGG | \|\|\|\|\|\|\|\|[\|\|\|\|\|\|\|\|\|\|\|\|]PAM | NA | Intergenic | NA | NA |
| T34083 | CGTGTTTGGTGACTCATTCTTGG | | 1000 | | | | | | | | | |
| Oligo fwd | CACCCGTGTTTGGTGACTCATTCT | | | | | | | | | | | |
| Oligo rev | AAACAGAATGAGTCACCAAACACG | | | | | | | | | | | |
| Chromosome | start | | end | strand | MM | target_seq | PAM | alignment | distance | position | gene name | gene id |
| GU305914.1 | | 3898 | 3920 | - | 0 | CGTGTTTGGTGACTCATTCT | TGG | \|\|\|\|\|\|\|\|[\|\|\|\|\|\|\|\|\|\|\|\|]PAM | NA | Intergenic | NA | NA |
| T34084 | GTGTAAAATTCATCCCGTCTTGG | | 1000 | | | | | | | | | |
| Oligo fwd | CACCGTGTAAAATTCATCCCGTCT | | | | | | | | | | | |
| Oligo rev | AAACAGACGGGATGAATTTTACAC | | | | | | | | | | | |
| Chromosome | start | | end | strand | MM | target_seq | PAM | alignment | distance | position | gene name | gene id |
| GU305914.1 | | 3876 | 3898 | - | 0 | GTGTAAAATTCATCCCGTCT | TGG | \|\|\|\|\|\|\|\|[\|\|\|\|\|\|\|\|\|\|\|\|]PAM | NA | Intergenic | NA | NA |
| T34085 | TAAAATTCATCCCGTCTTGGAGG | | 1000 | | | | | | | | | |
| Oligo fwd | CACCTAAAATTCATCCCGTCTTGG | | | | | | | | | | | |
| Oligo rev | AAACCCAAGACGGGATGAATTTTA | | | | | | | | | | | |
| Chromosome | start | | end | strand | MM | target_seq | PAM | alignment | distance | position | gene name | gene id |
| GU305914.1 | | 3873 | 3895 | - | 0 | TAAAATTCATCCCGTCTTGG | AGG | \|\|\|\|\|\|\|\|[\|\|\|\|\|\|\|\|\|\|\|\|]PAM | NA | Intergenic | NA | NA |
| T34086 | AAAATTCATCCCGTCTTGGAGGG | | 1000 | | | | | | | | | |
| Oligo fwd | CACCAAAATTCATCCCGTCTTGGA | | | | | | | | | | | |
| Oligo rev | AAACTCCAAGACGGGATGAATTTT | | | | | | | | | | | |
| Chromosome | start | | end | strand | MM | target_seq | PAM | alignment | distance | position | gene name | gene id |
| GU305914.1 | | 3872 | 3894 | - | 0 | AAAATTCATCCCGTCTTGGA | GGG | \|\|\|\|\|\|\|\|[\|\|\|\|\|\|\|\|\|\|\|\|]PAM | NA | Intergenic | NA | NA |
| T34087 | AAATTCATCCCGTCTTGGAGGGG | | 1000 | | | | | | | | | |
| Oligo fwd | CACCAAATTCATCCCGTCTTGGAG | | | | | | | | | | | |
| Oligo rev | AAACCTCCAAGACGGGATGAATTT | | | | | | | | | | | |
| Chromosome | start | | end | strand | MM | target_seq | PAM | alignment | distance | position | gene name | gene id |
| GU305914.1 | | 3871 | 3893 | - | 0 | AAATTCATCCCGTCTTGGAG | GGG | \|\|\|\|\|\|\|\|[\|\|\|\|\|\|\|\|\|\|\|\|]PAM | NA | Intergenic | NA | NA |
| T34088 | CGTCTTGGAGGGGTCAGATCTGG | | 1000 | | | | | | | | | |
| Oligo fwd | CACCCGTCTTGGAGGGGTCAGATC | | | | | | | | | | | |
| Oligo rev | AAACGATCTGACCCCTCCAAGACG | | | | | | | | | | | |
| Chromosome | start | | end | strand | MM | target_seq | PAM | alignment | distance | position | gene name | gene id |
| GU305914.1 | | 3861 | 3883 | - | 0 | CGTCTTGGAGGGGTCAGATC | TGG | \|\|\|\|\|\|\|\|[\|\|\|\|\|\|\|\|\|\|\|\|]PAM | NA | Intergenic | NA | NA |
| T34089 | AGATCTGGAACACCGTAGTTTGG | | 1000 | | | | | | | | | |
| Oligo fwd | CACCAGATCTGGAACACCGTAGTT | | | | | | | | | | | |
| Oligo rev | AAACAACTACGGTGTTCCAGATCT | | | | | | | | | | | |
| Chromosome | start | | end | strand | MM | target_seq | PAM | alignment | distance | position | gene name | gene id |
| GU305914.1 | | 3846 | 3868 | - | 0 | AGATCTGGAACACCGTAGTT | TGG | \|\|\|\|\|\|\|\|[\|\|\|\|\|\|\|\|\|\|\|\|]PAM | NA | Intergenic | NA | NA |
| T34090 | GATCTGGAACACCGTAGTTTGGG | | 1000 | | | | | | | | | |
| Oligo fwd | CACCGATCTGGAACACCGTAGTTT | | | | | | | | | | | |
| Oligo rev | AAACAAACTACGGTGTTCCAGATC | | | | | | | | | | | |
| Chromosome | start | | end | strand | MM | target_seq | PAM | alignment | distance | position | gene name | gene id |
| GU305914.1 | | 3845 | 3867 | - | 0 | GATCTGGAACACCGTAGTTT | GGG | \|\|\|\|\|\|\|\|[\|\|\|\|\|\|\|\|\|\|\|\|]PAM | NA | Intergenic | NA | NA |
| T34091 | CTGGAACACCGTAGTTTGGGAGG | | 1000 | | | | | | | | | |
| Oligo fwd | CACCCTGGAACACCGTAGTTTGGG | | | | | | | | | | | |
| Oligo rev | AAACCCCAAACTACGGTGTTCCAG | | | | | | | | | | | |
| Chromosome | start | | end | strand | MM | target_seq | PAM | alignment | distance | position | gene name | gene id |
| GU305914.1 | | 3842 | 3864 | - | 0 | CTGGAACACCGTAGTTTGGG | AGG | \|\|\|\|\|\|\|\|[\|\|\|\|\|\|\|\|\|\|\|\|]PAM | NA | Intergenic | NA | NA |
| T34092 | TTCCGCAGCATCTCCTCATCCGG | | 1000 | | | | | | | | | |
| Oligo fwd | CACCTTCCGCAGCATCTCCTCATC | | | | | | | | | | | |

<table>
<tr><td>Oligo rev</td><td>AAACGATGAGGAGATGCTGCGGAA</td><td></td><td></td><td></td><td></td><td></td><td></td><td></td><td></td><td></td><td></td><td></td></tr>
<tr><td>Chromosome</td><td>start</td><td></td><td>end</td><td>strand</td><td>MM</td><td>target_seq</td><td>PAM</td><td>alignment</td><td>distance</td><td>position</td><td>gene name</td><td>gene id</td></tr>
<tr><td>GU305914.1</td><td></td><td>3801</td><td>3823</td><td>-</td><td>0</td><td>TTCCGCAGCATCTCCTCATC</td><td>CGG</td><td>||||||||[||||||||||||]PAM</td><td>NA</td><td>Intergenic</td><td>NA</td><td>NA</td></tr>
<tr><td>T34093</td><td>ATTTTGCTGTTAGATGTCCACGG</td><td></td><td>1000</td><td></td><td></td><td></td><td></td><td></td><td></td><td></td><td></td><td></td></tr>
<tr><td>Oligo fwd</td><td>CACCATTTTGCTGTTAGATGTCCA</td><td></td><td></td><td></td><td></td><td></td><td></td><td></td><td></td><td></td><td></td><td></td></tr>
<tr><td>Oligo rev</td><td>AAACTGGACATCTAACAGCAAAAT</td><td></td><td></td><td></td><td></td><td></td><td></td><td></td><td></td><td></td><td></td><td></td></tr>
<tr><td>Chromosome</td><td>start</td><td></td><td>end</td><td>strand</td><td>MM</td><td>target_seq</td><td>PAM</td><td>alignment</td><td>distance</td><td>position</td><td>gene name</td><td>gene id</td></tr>
<tr><td>GU305914.1</td><td></td><td>3771</td><td>3793</td><td>-</td><td>0</td><td>ATTTTGCTGTTAGATGTCCA</td><td>CGG</td><td>||||||||[||||||||||||]PAM</td><td>NA</td><td>Intergenic</td><td>NA</td><td>NA</td></tr>
<tr><td>T34094</td><td>ACTATCAGCGCTGCCTGTGCTGG</td><td></td><td>1000</td><td></td><td></td><td></td><td></td><td></td><td></td><td></td><td></td><td></td></tr>
<tr><td>Oligo fwd</td><td>CACCACTATCAGCGCTGCCTGTGC</td><td></td><td></td><td></td><td></td><td></td><td></td><td></td><td></td><td></td><td></td><td></td></tr>
<tr><td>Oligo rev</td><td>AAACGCACAGGCAGCGCTGATAGT</td><td></td><td></td><td></td><td></td><td></td><td></td><td></td><td></td><td></td><td></td><td></td></tr>
<tr><td>Chromosome</td><td>start</td><td></td><td>end</td><td>strand</td><td>MM</td><td>target_seq</td><td>PAM</td><td>alignment</td><td>distance</td><td>position</td><td>gene name</td><td>gene id</td></tr>
<tr><td>GU305914.1</td><td></td><td>3724</td><td>3746</td><td>-</td><td>0</td><td>ACTATCAGCGCTGCCTGTGC</td><td>TGG</td><td>||||||||[||||||||||||]PAM</td><td>NA</td><td>Intergenic</td><td>NA</td><td>NA</td></tr>
<tr><td>T34095</td><td>TATCAGTGCTGTTATCGACTTGG</td><td></td><td>1000</td><td></td><td></td><td></td><td></td><td></td><td></td><td></td><td></td><td></td></tr>
<tr><td>Oligo fwd</td><td>CACCTATCAGTGCTGTTATCGACT</td><td></td><td></td><td></td><td></td><td></td><td></td><td></td><td></td><td></td><td></td><td></td></tr>
<tr><td>Oligo rev</td><td>AAACAGTCGATAACAGCACTGATA</td><td></td><td></td><td></td><td></td><td></td><td></td><td></td><td></td><td></td><td></td><td></td></tr>
<tr><td>Chromosome</td><td>start</td><td></td><td>end</td><td>strand</td><td>MM</td><td>target_seq</td><td>PAM</td><td>alignment</td><td>distance</td><td>position</td><td>gene name</td><td>gene id</td></tr>
<tr><td>GU305914.1</td><td></td><td>3665</td><td>3687</td><td>-</td><td>0</td><td>TATCAGTGCTGTTATCGACT</td><td>TGG</td><td>||||||||[||||||||||||]PAM</td><td>NA</td><td>Intergenic</td><td>NA</td><td>NA</td></tr>
<tr><td>T34096</td><td>ATCCGTCCTTGACTGTCAACAGG</td><td></td><td>1000</td><td></td><td></td><td></td><td></td><td></td><td></td><td></td><td></td><td></td></tr>
<tr><td>Oligo fwd</td><td>CACCATCCGTCCTTGACTGTCAAC</td><td></td><td></td><td></td><td></td><td></td><td></td><td></td><td></td><td></td><td></td><td></td></tr>
<tr><td>Oligo rev</td><td>AAACGTTGACAGTCAAGGACGGAT</td><td></td><td></td><td></td><td></td><td></td><td></td><td></td><td></td><td></td><td></td><td></td></tr>
<tr><td>Chromosome</td><td>start</td><td></td><td>end</td><td>strand</td><td>MM</td><td>target_seq</td><td>PAM</td><td>alignment</td><td>distance</td><td>position</td><td>gene name</td><td>gene id</td></tr>
<tr><td>GU305914.1</td><td></td><td>3641</td><td>3663</td><td>-</td><td>0</td><td>ATCCGTCCTTGACTGTCAAC</td><td>AGG</td><td>||||||||[||||||||||||]PAM</td><td>NA</td><td>Intergenic</td><td>NA</td><td>NA</td></tr>
<tr><td>T34097</td><td>ATTCGTACGTAATCCGTGTATGG</td><td></td><td>1000</td><td></td><td></td><td></td><td></td><td></td><td></td><td></td><td></td><td></td></tr>
<tr><td>Oligo fwd</td><td>CACCATTCGTACGTAATCCGTGTA</td><td></td><td></td><td></td><td></td><td></td><td></td><td></td><td></td><td></td><td></td><td></td></tr>
<tr><td>Oligo rev</td><td>AAACTACACGGATTACGTACGAAT</td><td></td><td></td><td></td><td></td><td></td><td></td><td></td><td></td><td></td><td></td><td></td></tr>
<tr><td>Chromosome</td><td>start</td><td></td><td>end</td><td>strand</td><td>MM</td><td>target_seq</td><td>PAM</td><td>alignment</td><td>distance</td><td>position</td><td>gene name</td><td>gene id</td></tr>
<tr><td>GU305914.1</td><td></td><td>3616</td><td>3638</td><td>-</td><td>0</td><td>ATTCGTACGTAATCCGTGTA</td><td>TGG</td><td>||||||||[||||||||||||]PAM</td><td>NA</td><td>Intergenic</td><td>NA</td><td>NA</td></tr>
<tr><td>T34098</td><td>TTCGTACGTAATCCGTGTATGGG</td><td></td><td>1000</td><td></td><td></td><td></td><td></td><td></td><td></td><td></td><td></td><td></td></tr>
<tr><td>Oligo fwd</td><td>CACCTTCGTACGTAATCCGTGTAT</td><td></td><td></td><td></td><td></td><td></td><td></td><td></td><td></td><td></td><td></td><td></td></tr>
<tr><td>Oligo rev</td><td>AAACATACACGGATTACGTACGAA</td><td></td><td></td><td></td><td></td><td></td><td></td><td></td><td></td><td></td><td></td><td></td></tr>
<tr><td>Chromosome</td><td>start</td><td></td><td>end</td><td>strand</td><td>MM</td><td>target_seq</td><td>PAM</td><td>alignment</td><td>distance</td><td>position</td><td>gene name</td><td>gene id</td></tr>
<tr><td>GU305914.1</td><td></td><td>3615</td><td>3637</td><td>-</td><td>0</td><td>TTCGTACGTAATCCGTGTAT</td><td>GGG</td><td>||||||||[||||||||||||]PAM</td><td>NA</td><td>Intergenic</td><td>NA</td><td>NA</td></tr>
<tr><td>T34099</td><td>TGAAATTCAAAATAGCACGTAGG</td><td></td><td>1000</td><td></td><td></td><td></td><td></td><td></td><td></td><td></td><td></td><td></td></tr>
<tr><td>Oligo fwd</td><td>CACCTGAAATTCAAAATAGCACGT</td><td></td><td></td><td></td><td></td><td></td><td></td><td></td><td></td><td></td><td></td><td></td></tr>
<tr><td>Oligo rev</td><td>AAACACGTGCTATTTTGAATTTCA</td><td></td><td></td><td></td><td></td><td></td><td></td><td></td><td></td><td></td><td></td><td></td></tr>
<tr><td>Chromosome</td><td>start</td><td></td><td>end</td><td>strand</td><td>MM</td><td>target_seq</td><td>PAM</td><td>alignment</td><td>distance</td><td>position</td><td>gene name</td><td>gene id</td></tr>
<tr><td>GU305914.1</td><td></td><td>3590</td><td>3612</td><td>-</td><td>0</td><td>TGAAATTCAAAATAGCACGT</td><td>AGG</td><td>||||||||[||||||||||||]PAM</td><td>NA</td><td>Intergenic</td><td>NA</td><td>NA</td></tr>
<tr><td>T34100</td><td>AAAATAGCACGTAGGAATCTCGG</td><td></td><td>1000</td><td></td><td></td><td></td><td></td><td></td><td></td><td></td><td></td><td></td></tr>
<tr><td>Oligo fwd</td><td>CACCAAAATAGCACGTAGGAATCT</td><td></td><td></td><td></td><td></td><td></td><td></td><td></td><td></td><td></td><td></td><td></td></tr>
<tr><td>Oligo rev</td><td>AAACAGATTCCTACGTGCTATTTT</td><td></td><td></td><td></td><td></td><td></td><td></td><td></td><td></td><td></td><td></td><td></td></tr>
<tr><td>Chromosome</td><td>start</td><td></td><td>end</td><td>strand</td><td>MM</td><td>target_seq</td><td>PAM</td><td>alignment</td><td>distance</td><td>position</td><td>gene name</td><td>gene id</td></tr>
<tr><td>GU305914.1</td><td></td><td>3582</td><td>3604</td><td>-</td><td>0</td><td>AAAATAGCACGTAGGAATCT</td><td>CGG</td><td>||||||||[||||||||||||]PAM</td><td>NA</td><td>Intergenic</td><td>NA</td><td>NA</td></tr>
<tr><td>T34101</td><td>AGCACGTAGGAATCTCGGTTCGG</td><td></td><td>1000</td><td></td><td></td><td></td><td></td><td></td><td></td><td></td><td></td><td></td></tr>
<tr><td>Oligo fwd</td><td>CACCAGCACGTAGGAATCTCGGTT</td><td></td><td></td><td></td><td></td><td></td><td></td><td></td><td></td><td></td><td></td><td></td></tr>
<tr><td>Oligo rev</td><td>AAACAACCGAGATTCCTACGTGCT</td><td></td><td></td><td></td><td></td><td></td><td></td><td></td><td></td><td></td><td></td><td></td></tr>
<tr><td>Chromosome</td><td>start</td><td></td><td>end</td><td>strand</td><td>MM</td><td>target_seq</td><td>PAM</td><td>alignment</td><td>distance</td><td>position</td><td>gene name</td><td>gene id</td></tr>
<tr><td>GU305914.1</td><td></td><td>3577</td><td>3599</td><td>-</td><td>0</td><td>AGCACGTAGGAATCTCGGTT</td><td>CGG</td><td>||||||||[||||||||||||]PAM</td><td>NA</td><td>Intergenic</td><td>NA</td><td>NA</td></tr>
<tr><td>T34102</td><td>AGGAATCTCGGTTCGGAATTTGG</td><td></td><td>1000</td><td></td><td></td><td></td><td></td><td></td><td></td><td></td><td></td><td></td></tr>
<tr><td>Oligo fwd</td><td>CACCAGGAATCTCGGTTCGGAATT</td><td></td><td></td><td></td><td></td><td></td><td></td><td></td><td></td><td></td><td></td><td></td></tr>
<tr><td>Oligo rev</td><td>AAACAATTCCGAACCGAGATTCCT</td><td></td><td></td><td></td><td></td><td></td><td></td><td></td><td></td><td></td><td></td><td></td></tr>
<tr><td>Chromosome</td><td>start</td><td></td><td>end</td><td>strand</td><td>MM</td><td>target_seq</td><td>PAM</td><td>alignment</td><td>distance</td><td>position</td><td>gene name</td><td>gene id</td></tr>
<tr><td>GU305914.1</td><td></td><td>3570</td><td>3592</td><td>-</td><td>0</td><td>AGGAATCTCGGTTCGGAATT</td><td>TGG</td><td>||||||||[||||||||||||]PAM</td><td>NA</td><td>Intergenic</td><td>NA</td><td>NA</td></tr>
<tr><td>T34103</td><td>TCGTTACATCTACCGAATTCTGG</td><td></td><td>1000</td><td></td><td></td><td></td><td></td><td></td><td></td><td></td><td></td><td></td></tr>
<tr><td>Oligo fwd</td><td>CACCTCGTTACATCTACCGAATTC</td><td></td><td></td><td></td><td></td><td></td><td></td><td></td><td></td><td></td><td></td><td></td></tr>
<tr><td>Oligo rev</td><td>AAACGAATTCGGTAGATGTAACGA</td><td></td><td></td><td></td><td></td><td></td><td></td><td></td><td></td><td></td><td></td><td></td></tr>
<tr><td>Chromosome</td><td>start</td><td></td><td>end</td><td>strand</td><td>MM</td><td>target_seq</td><td>PAM</td><td>alignment</td><td>distance</td><td>position</td><td>gene name</td><td>gene id</td></tr>
<tr><td>GU305914.1</td><td></td><td>3530</td><td>3552</td><td>-</td><td>0</td><td>TCGTTACATCTACCGAATTC</td><td>TGG</td><td>||||||||[||||||||||||]PAM</td><td>NA</td><td>Intergenic</td><td>NA</td><td>NA</td></tr>
<tr><td>T34104</td><td>CTTGACGACGACCCATTCCAAGG</td><td></td><td>1000</td><td></td><td></td><td></td><td></td><td></td><td></td><td></td><td></td><td></td></tr>
<tr><td>Oligo fwd</td><td>CACCCTTGACGACGACCCATTCCA</td><td></td><td></td><td></td><td></td><td></td><td></td><td></td><td></td><td></td><td></td><td></td></tr>
<tr><td>Oligo rev</td><td>AAACTGGAATGGGTCGTCGTCAAG</td><td></td><td></td><td></td><td></td><td></td><td></td><td></td><td></td><td></td><td></td><td></td></tr>
<tr><td>Chromosome</td><td>start</td><td></td><td>end</td><td>strand</td><td>MM</td><td>target_seq</td><td>PAM</td><td>alignment</td><td>distance</td><td>position</td><td>gene name</td><td>gene id</td></tr>
<tr><td>GU305914.1</td><td></td><td>3505</td><td>3527</td><td>-</td><td>0</td><td>CTTGACGACGACCCATTCCA</td><td>AGG</td><td>||||||||[||||||||||||]PAM</td><td>NA</td><td>Intergenic</td><td>NA</td><td>NA</td></tr>
<tr><td>T34105</td><td>GAAGCATTCACTTTTACGTCAGG</td><td></td><td>1000</td><td></td><td></td><td></td><td></td><td></td><td></td><td></td><td></td><td></td></tr>
<tr><td>Oligo fwd</td><td>CACCGAAGCATTCACTTTTACGTC</td><td></td><td></td><td></td><td></td><td></td><td></td><td></td><td></td><td></td><td></td><td></td></tr>
<tr><td>Oligo rev</td><td>AAACGACGTAAAAGTGAATGCTTC</td><td></td><td></td><td></td><td></td><td></td><td></td><td></td><td></td><td></td><td></td><td></td></tr>
<tr><td>Chromosome</td><td>start</td><td></td><td>end</td><td>strand</td><td>MM</td><td>target_seq</td><td>PAM</td><td>alignment</td><td>distance</td><td>position</td><td>gene name</td><td>gene id</td></tr>
<tr><td>GU305914.1</td><td></td><td>3477</td><td>3499</td><td>-</td><td>0</td><td>GAAGCATTCACTTTTACGTC</td><td>AGG</td><td>||||||||[||||||||||||]PAM</td><td>NA</td><td>Intergenic</td><td>NA</td><td>NA</td></tr>
<tr><td>T34106</td><td>TTTTACGTCAGGAAGCTTACAGG</td><td></td><td>1000</td><td></td><td></td><td></td><td></td><td></td><td></td><td></td><td></td><td></td></tr>
<tr><td>Oligo fwd</td><td>CACCTTTTACGTCAGGAAGCTTAC</td><td></td><td></td><td></td><td></td><td></td><td></td><td></td><td></td><td></td><td></td><td></td></tr>
<tr><td>Oligo rev</td><td>AAACGTAAGCTTCCTGACGTAAAA</td><td></td><td></td><td></td><td></td><td></td><td></td><td></td><td></td><td></td><td></td><td></td></tr>
<tr><td>Chromosome</td><td>start</td><td></td><td>end</td><td>strand</td><td>MM</td><td>target_seq</td><td>PAM</td><td>alignment</td><td>distance</td><td>position</td><td>gene name</td><td>gene id</td></tr>
<tr><td>GU305914.1</td><td></td><td>3466</td><td>3488</td><td>-</td><td>0</td><td>TTTTACGTCAGGAAGCTTAC</td><td>AGG</td><td>||||||||[||||||||||||]PAM</td><td>NA</td><td>Intergenic</td><td>NA</td><td>NA</td></tr>
<tr><td>T34107</td><td>AATAACGTATCGCCCTTTGACGG</td><td></td><td>1000</td><td></td><td></td><td></td><td></td><td></td><td></td><td></td><td></td><td></td></tr>
</table>

Oligo fwd CACCAATAACGTATCGCCCTTTGA
Oligo rev AAACTCAAAGGGCGATACGTTATT
Chromosome start end strand MM target_seq PAM alignment distance position gene name gene id
GU305914.1 3441 3463 - 0 AATAACGTATCGCCCTTTGA CGG ||||||||[||||||||||||]PAM NA Intergenic NA NA

T34108 ATCGCCCTTTGACGGTATCTTGG 1000
Oligo fwd CACCATCGCCCTTTGACGGTATCT
Oligo rev AAACAGATACCGTCAAAGGGCGAT
Chromosome start end strand MM target_seq PAM alignment distance position gene name gene id
GU305914.1 3433 3455 - 0 ATCGCCCTTTGACGGTATCT TGG ||||||||[||||||||||||]PAM NA Intergenic NA NA

T34109 CCCTTTGACGGTATCTTGGTTGG 1000
Oligo fwd CACCCCCTTTGACGGTATCTTGGT
Oligo rev AAACACCAAGATACCGTCAAAGGG
Chromosome start end strand MM target_seq PAM alignment distance position gene name gene id
GU305914.1 3429 3451 - 0 CCCTTTGACGGTATCTTGGT TGG ||||||||[||||||||||||]PAM NA Intergenic NA NA

T34110 CCTTTGAAGTAGCAACCGAGAGG 1000
Oligo fwd CACCCCTTTGAAGTAGCAACCGAG
Oligo rev AAACCTCGGTTGCTACTTCAAAGG
Chromosome start end strand MM target_seq PAM alignment distance position gene name gene id
GU305914.1 3399 3421 - 0 CCTTTGAAGTAGCAACCGAG AGG ||||||||[||||||||||||]PAM NA Intergenic NA NA

T34111 ACAATGACTATCACCAACAACGG 1000
Oligo fwd CACCACAATGACTATCACCAACAA
Oligo rev AAACTTGTTGGTGATAGTCATTGT
Chromosome start end strand MM target_seq PAM alignment distance position gene name gene id
GU305914.1 3285 3307 - 0 ACAATGACTATCACCAACAA CGG ||||||||[||||||||||||]PAM NA Intergenic NA NA

T34112 TCTTGCTGCATCCGTCACTCTGG 1000
Oligo fwd CACCTCTTGCTGCATCCGTCACTC
Oligo rev AAACGAGTGACGGATGCAGCAAGA
Chromosome start end strand MM target_seq PAM alignment distance position gene name gene id
GU305914.1 3245 3267 - 0 TCTTGCTGCATCCGTCACTC TGG ||||||||[||||||||||||]PAM NA Intergenic NA NA

T34113 TTATATACGTGACTCAATAATGG 1000
Oligo fwd CACCTTATATACGTGACTCAATAA
Oligo rev AAACTTATTGAGTCACGTATATAA
Chromosome start end strand MM target_seq PAM alignment distance position gene name gene id
GU305914.1 3187 3209 - 0 TTATATACGTGACTCAATAA TGG ||||||||[||||||||||||]PAM NA Intergenic NA NA

T34114 TATATACGTGACTCAATAATGGG 1000
Oligo fwd CACCTATATACGTGACTCAATAAT
Oligo rev AAACATTATTGAGTCACGTATATA
Chromosome start end strand MM target_seq PAM alignment distance position gene name gene id
GU305914.1 3186 3208 - 0 TATATACGTGACTCAATAAT GGG ||||||||[||||||||||||]PAM NA Intergenic NA NA

T34115 ATACGTGACTCAATAATGGGTGG 1000
Oligo fwd CACCATACGTGACTCAATAATGGG
Oligo rev AAACCCCATTATTGAGTCACGTAT
Chromosome start end strand MM target_seq PAM alignment distance position gene name gene id
GU305914.1 3183 3205 - 0 ATACGTGACTCAATAATGGG TGG ||||||||[||||||||||||]PAM NA Intergenic NA NA

T34116 TACGTGACTCAATAATGGGTGGG 1000
Oligo fwd CACCTACGTGACTCAATAATGGGT
Oligo rev AAACACCCATTATTGAGTCACGTA
Chromosome start end strand MM target_seq PAM alignment distance position gene name gene id
GU305914.1 3182 3204 - 0 TACGTGACTCAATAATGGGT GGG ||||||||[||||||||||||]PAM NA Intergenic NA NA

T34117 TCAATAATGGGTGGGATGAATGG 1000
Oligo fwd CACCTCAATAATGGGTGGGATGAA
Oligo rev AAACTTCATCCCACCCATTATTGA
Chromosome start end strand MM target_seq PAM alignment distance position gene name gene id
GU305914.1 3174 3196 - 0 TCAATAATGGGTGGGATGAA TGG ||||||||[||||||||||||]PAM NA Intergenic NA NA

T34118 AATGGTACATACGTAATTATTGG 1000
Oligo fwd CACCAATGGTACATACGTAATTAT
Oligo rev AAACATAATTACGTATGTACCATT
Chromosome start end strand MM target_seq PAM alignment distance position gene name gene id
GU305914.1 3156 3178 - 0 AATGGTACATACGTAATTAT TGG ||||||||[||||||||||||]PAM NA Intergenic NA NA

T34119 ATGAAAAACAAAAAACACGACGG 1000
Oligo fwd CACCATGAAAAACAAAAAACACGA
Oligo rev AAACTCGTGTTTTTTGTTTTTCAT
Chromosome start end strand MM target_seq PAM alignment distance position gene name gene id
GU305914.1 3133 3155 - 0 ATGAAAAACAAAAAACACGA CGG ||||||||[||||||||||||]PAM NA Intergenic NA NA

T34120 ATCAGATATTTATTTATCTATGG 1000
Oligo fwd CACCATCAGATATTTATTTATCTA
Oligo rev AAACTAGATAAATAAATATCTGAT
Chromosome start end strand MM target_seq PAM alignment distance position gene name gene id
GU305914.1 3100 3122 - 0 ATCAGATATTTATTTATCTA TGG ||||||||[||||||||||||]PAM NA Intergenic NA NA

T34121 TCTATGGCTGTCGATCAGAATGG 1000
Oligo fwd CACCTCTATGGCTGTCGATCAGAA
Oligo rev AAACTTCTGATCGACAGCCATAGA
Chromosome start end strand MM target_seq PAM alignment distance position gene name gene id
GU305914.1 3084 3106 - 0 TCTATGGCTGTCGATCAGAA TGG ||||||||[||||||||||||]PAM NA Intergenic NA NA

T34122 CGATCAGAATGGAGAAACAACGG 1000
Oligo fwd CACCCGATCAGAATGGAGAAACAA
Oligo rev AAACTTGTTTCTCCATTCTGATCG
Chromosome start end strand MM target_seq PAM alignment distance position gene name gene id
GU305914.1 3073 3095 - 0 CGATCAGAATGGAGAAACAA CGG ||||||||[||||||||||||]PAM NA Intergenic NA NA

T34123 GGAGAAACAACGGAACAGTGTGG 1000
Oligo fwd CACCGGAGAAACAACGGAACAGTG
Oligo rev AAACCACTGTTCCGTTGTTTCTCC
Chromosome start end strand MM target_seq PAM alignment distance position gene name gene id
GU305914.1 3063 3085 - 0 GGAGAAACAACGGAACAGTG TGG ||||||||[||||||||||||]PAM NA Intergenic NA NA

T34124 GAGAAACAACGGAACAGTGTGGG 1000
Oligo fwd CACCGAGAAACAACGGAACAGTGT
Oligo rev AAACACACTGTTCCGTTGTTTCTC
Chromosome start end strand MM target_seq PAM alignment distance position gene name gene id
GU305914.1 3062 3084 - 0 GAGAAACAACGGAACAGTGT GGG ||||||||[||||||||||||]PAM NA Intergenic NA NA

T34125 AGAGTAACCCAACAAGAGCGCGG 1000
Oligo fwd CACCAGAGTAACCCAACAAGAGCG
Oligo rev AAACCGCTCTTGTTGGGTTACTCT
Chromosome start end strand MM target_seq PAM alignment distance position gene name gene id
GU305914.1 3033 3055 - 0 AGAGTAACCCAACAAGAGCG CGG ||||||||[||||||||||||]PAM NA Intergenic NA NA

T34126 GACCAGCGTCATTCGATCTCTGG 1000
Oligo fwd CACCGACCAGCGTCATTCGATCTC
Oligo rev AAACGAGATCGAATGACGCTGGTC
Chromosome start end strand MM target_seq PAM alignment distance position gene name gene id
GU305914.1 2968 2990 - 0 GACCAGCGTCATTCGATCTC TGG ||||||||[||||||||||||]PAM NA Intergenic NA NA

T34127 CTCTGGACGATACGTTTGATTGG 1000
Oligo fwd CACCCTCTGGACGATACGTTTGAT
Oligo rev AAACATCAAACGTATCGTCCAGAG
Chromosome start end strand MM target_seq PAM alignment distance position gene name gene id
GU305914.1 2951 2973 - 0 CTCTGGACGATACGTTTGAT TGG ||||||||[||||||||||||]PAM NA Intergenic NA NA

T34128 TCTGGACGATACGTTTGATTGGG 1000
Oligo fwd CACCTCTGGACGATACGTTTGATT
Oligo rev AAACAATCAAACGTATCGTCCAGA
Chromosome start end strand MM target_seq PAM alignment distance position gene name gene id
GU305914.1 2950 2972 - 0 TCTGGACGATACGTTTGATT GGG ||||||||[||||||||||||]PAM NA Intergenic NA NA

T34129 GGGATACTGTAGTTGTATTGAGG 1000
Oligo fwd CACCGGGATACTGTAGTTGTATTG
Oligo rev AAACCAATACAACTACAGTATCCC
Chromosome start end strand MM target_seq PAM alignment distance position gene name gene id
GU305914.1 2930 2952 - 0 GGGATACTGTAGTTGTATTG AGG ||||||||[||||||||||||]PAM NA Intergenic NA NA

T34130 CCACGAATCTCGTCGTATGTTGG 1000
Oligo fwd CACCCCACGAATCTCGTCGTATGT
Oligo rev AAACACATACGACGAGATTCGTGG
Chromosome start end strand MM target_seq PAM alignment distance position gene name gene id
GU305914.1 2891 2913 - 0 CCACGAATCTCGTCGTATGT TGG ||||||||[||||||||||||]PAM NA Intergenic NA NA

T34131 CGTCGTATGTTGGAAGCGAAAGG 1000
Oligo fwd CACCCGTCGTATGTTGGAAGCGAA
Oligo rev AAACTTCGCTTCCAACATACGACG
Chromosome start end strand MM target_seq PAM alignment distance position gene name gene id
GU305914.1 2881 2903 - 0 CGTCGTATGTTGGAAGCGAA AGG ||||||||[||||||||||||]PAM NA Intergenic NA NA

T34132 ACTCAAATTATTCTTGAGTCCGG 1000
Oligo fwd CACCACTCAAATTATTCTTGAGTC
Oligo rev AAACGACTCAAGAATAATTTGAGT
Chromosome start end strand MM target_seq PAM alignment distance position gene name gene id
GU305914.1 2853 2875 - 0 ACTCAAATTATTCTTGAGTC CGG ||||||||[||||||||||||]PAM NA Intergenic NA NA

T34133 ACACAGCCTTAGTACACATATGG 1000
Oligo fwd CACCACACAGCCTTAGTACACATA
Oligo rev AAACTATGTGTACTAAGGCTGTGT
Chromosome start end strand MM target_seq PAM alignment distance position gene name gene id
GU305914.1 2790 2812 - 0 ACACAGCCTTAGTACACATA TGG ||||||||[||||||||||||]PAM NA Intergenic NA NA

T34134 GTAAAGCTTGCCTGAATGATCGG 1000
Oligo fwd CACCGTAAAGCTTGCCTGAATGAT
Oligo rev AAACATCATTCAGGCAAGCTTTAC
Chromosome start end strand MM target_seq PAM alignment distance position gene name gene id
GU305914.1 2721 2743 - 0 GTAAAGCTTGCCTGAATGAT CGG ||||||||[||||||||||||]PAM NA Intergenic NA NA

T34135 GAAACCGTCTTTTAATCCCACGG 1000
Oligo fwd CACCGAAACCGTCTTTTAATCCCA
Oligo rev AAACTGGGATTAAAAGACGGTTTC
Chromosome start end strand MM target_seq PAM alignment distance position gene name gene id
GU305914.1 2694 2716 - 0 GAAACCGTCTTTTAATCCCA CGG ||||||||[||||||||||||]PAM NA Intergenic NA NA

T34136 CATATCATAGTAAGAAAAGATGG 1000
Oligo fwd CACCCATATCATAGTAAGAAAAGA
Oligo rev AAACTCTTTTCTTACTATGATATG
Chromosome start end strand MM target_seq PAM alignment distance position gene name gene id
GU305914.1 2649 2671 - 0 CATATCATAGTAAGAAAAGA TGG ||||||||[||||||||||||]PAM NA Intergenic NA NA

| | | | | | | | | | | | |
|---|---|---|---|---|---|---|---|---|---|---|---|
| T34137 | ATATCATAGTAAGAAAAGATGGG | 1000 | | | | | | | | | |
| Oligo fwd | CACCATATCATAGTAAGAAAAGAT | | | | | | | | | | |
| Oligo rev | AAACATCTTTTCTTACTATGATAT | | | | | | | | | | |
| Chromosome | start | end | strand | MM | target_seq | PAM | alignment | distance | position | gene name | gene id |
| GU305914.1 | 2648 | 2670 | - | 0 | ATATCATAGTAAGAAAAGAT | GGG | \|\|\|\|\|\|\|\|[\|\|\|\|\|\|\|\|\|\|\|\|]PAM | NA | Intergenic | NA | NA |
| T34138 | ATAGTAAGAAAAGATGGGTACGG | 1000 | | | | | | | | | |
| Oligo fwd | CACCATAGTAAGAAAAGATGGGTA | | | | | | | | | | |
| Oligo rev | AAACTACCCATCTTTTCTTACTAT | | | | | | | | | | |
| Chromosome | start | end | strand | MM | target_seq | PAM | alignment | distance | position | gene name | gene id |
| GU305914.1 | 2643 | 2665 | - | 0 | ATAGTAAGAAAAGATGGGTA | CGG | \|\|\|\|\|\|\|\|[\|\|\|\|\|\|\|\|\|\|\|\|]PAM | NA | Intergenic | NA | NA |
| T34139 | AAAGATGGGTACGGTATCTGTGG | 1000 | | | | | | | | | |
| Oligo fwd | CACCAAAGATGGGTACGGTATCTG | | | | | | | | | | |
| Oligo rev | AAACCAGATACCGTACCCATCTTT | | | | | | | | | | |
| Chromosome | start | end | strand | MM | target_seq | PAM | alignment | distance | position | gene name | gene id |
| GU305914.1 | 2634 | 2656 | - | 0 | AAAGATGGGTACGGTATCTG | TGG | \|\|\|\|\|\|\|\|[\|\|\|\|\|\|\|\|\|\|\|\|]PAM | NA | Intergenic | NA | NA |
| T34140 | GGCAGATTTAGCTGCCGAAGTGG | 1000 | | | | | | | | | |
| Oligo fwd | CACCGGCAGATTTAGCTGCCGAAG | | | | | | | | | | |
| Oligo rev | AAACCTTCGGCAGCTAAATCTGCC | | | | | | | | | | |
| Chromosome | start | end | strand | MM | target_seq | PAM | alignment | distance | position | gene name | gene id |
| GU305914.1 | 2613 | 2635 | - | 0 | GGCAGATTTAGCTGCCGAAG | TGG | \|\|\|\|\|\|\|\|[\|\|\|\|\|\|\|\|\|\|\|\|]PAM | NA | Intergenic | NA | NA |
| T34141 | AGCTGCCGAAGTGGTCGTCGAGG | 1000 | | | | | | | | | |
| Oligo fwd | CACCAGCTGCCGAAGTGGTCGTCG | | | | | | | | | | |
| Oligo rev | AAACCGACGACCACTTCGGCAGCT | | | | | | | | | | |
| Chromosome | start | end | strand | MM | target_seq | PAM | alignment | distance | position | gene name | gene id |
| GU305914.1 | 2604 | 2626 | - | 0 | AGCTGCCGAAGTGGTCGTCG | AGG | \|\|\|\|\|\|\|\|[\|\|\|\|\|\|\|\|\|\|\|\|]PAM | NA | Intergenic | NA | NA |
| T34142 | GGTCGTCGAGGTAAAATAAGAGG | 1000 | | | | | | | | | |
| Oligo fwd | CACCGGTCGTCGAGGTAAAATAAG | | | | | | | | | | |
| Oligo rev | AAACCTTATTTTACCTCGACGACC | | | | | | | | | | |
| Chromosome | start | end | strand | MM | target_seq | PAM | alignment | distance | position | gene name | gene id |
| GU305914.1 | 2592 | 2614 | - | 0 | GGTCGTCGAGGTAAAATAAG | AGG | \|\|\|\|\|\|\|\|[\|\|\|\|\|\|\|\|\|\|\|\|]PAM | NA | Intergenic | NA | NA |
| T34143 | ATAAGAGGCTGCGTCTACCGAGG | 1000 | | | | | | | | | |
| Oligo fwd | CACCATAAGAGGCTGCGTCTACCG | | | | | | | | | | |
| Oligo rev | AAACCGGTAGACGCAGCCTCTTAT | | | | | | | | | | |
| Chromosome | start | end | strand | MM | target_seq | PAM | alignment | distance | position | gene name | gene id |
| GU305914.1 | 2577 | 2599 | - | 0 | ATAAGAGGCTGCGTCTACCG | AGG | \|\|\|\|\|\|\|\|[\|\|\|\|\|\|\|\|\|\|\|\|]PAM | NA | Intergenic | NA | NA |
| T34144 | GCGTCTACCGAGGCGAAGAGCGG | 1000 | | | | | | | | | |
| Oligo fwd | CACCGCGTCTACCGAGGCGAAGAG | | | | | | | | | | |
| Oligo rev | AAACCTCTTCGCCTCGGTAGACGC | | | | | | | | | | |
| Chromosome | start | end | strand | MM | target_seq | PAM | alignment | distance | position | gene name | gene id |
| GU305914.1 | 2567 | 2589 | - | 0 | GCGTCTACCGAGGCGAAGAG | CGG | \|\|\|\|\|\|\|\|[\|\|\|\|\|\|\|\|\|\|\|\|]PAM | NA | Intergenic | NA | NA |
| T34145 | CGGTTTGATGAACCATTCAAAGG | 1000 | | | | | | | | | |
| Oligo fwd | CACCCGGTTTGATGAACCATTCAA | | | | | | | | | | |
| Oligo rev | AAACTTGAATGGTTCATCAAACCG | | | | | | | | | | |
| Chromosome | start | end | strand | MM | target_seq | PAM | alignment | distance | position | gene name | gene id |
| GU305914.1 | 2547 | 2569 | - | 0 | CGGTTTGATGAACCATTCAA | AGG | \|\|\|\|\|\|\|\|[\|\|\|\|\|\|\|\|\|\|\|\|]PAM | NA | Intergenic | NA | NA |
| T34146 | AGCGTTGACATAGACTGCTGCGG | 1000 | | | | | | | | | |
| Oligo fwd | CACCAGCGTTGACATAGACTGCTG | | | | | | | | | | |
| Oligo rev | AAACCAGCAGTCTATGTCAACGCT | | | | | | | | | | |
| Chromosome | start | end | strand | MM | target_seq | PAM | alignment | distance | position | gene name | gene id |
| GU305914.1 | 2517 | 2539 | - | 0 | AGCGTTGACATAGACTGCTG | CGG | \|\|\|\|\|\|\|\|[\|\|\|\|\|\|\|\|\|\|\|\|]PAM | NA | Intergenic | NA | NA |
| T34147 | GCGTTGACATAGACTGCTGCGGG | 1000 | | | | | | | | | |
| Oligo fwd | CACCGCGTTGACATAGACTGCTGC | | | | | | | | | | |
| Oligo rev | AAACGCAGCAGTCTATGTCAACGC | | | | | | | | | | |
| Chromosome | start | end | strand | MM | target_seq | PAM | alignment | distance | position | gene name | gene id |
| GU305914.1 | 2516 | 2538 | - | 0 | GCGTTGACATAGACTGCTGC | GGG | \|\|\|\|\|\|\|\|[\|\|\|\|\|\|\|\|\|\|\|\|]PAM | NA | Intergenic | NA | NA |
| T34148 | GACATAGACTGCTGCGGGACCGG | 1000 | | | | | | | | | |
| Oligo fwd | CACCGACATAGACTGCTGCGGGAC | | | | | | | | | | |
| Oligo rev | AAACGTCCCGCAGCAGTCTATGTC | | | | | | | | | | |
| Chromosome | start | end | strand | MM | target_seq | PAM | alignment | distance | position | gene name | gene id |
| GU305914.1 | 2511 | 2533 | - | 0 | GACATAGACTGCTGCGGGAC | CGG | \|\|\|\|\|\|\|\|[\|\|\|\|\|\|\|\|\|\|\|\|]PAM | NA | Intergenic | NA | NA |
| T34149 | ACATAGACTGCTGCGGGACCGGG | 1000 | | | | | | | | | |
| Oligo fwd | CACCACATAGACTGCTGCGGGACC | | | | | | | | | | |
| Oligo rev | AAACGGTCCCGCAGCAGTCTATGT | | | | | | | | | | |
| Chromosome | start | end | strand | MM | target_seq | PAM | alignment | distance | position | gene name | gene id |
| GU305914.1 | 2510 | 2532 | - | 0 | ACATAGACTGCTGCGGGACC | GGG | \|\|\|\|\|\|\|\|[\|\|\|\|\|\|\|\|\|\|\|\|]PAM | NA | Intergenic | NA | NA |
| T34150 | AATACCCTTATTTTCGTACAAGG | 1000 | | | | | | | | | |
| Oligo fwd | CACCAATACCCTTATTTTCGTACA | | | | | | | | | | |
| Oligo rev | AAACTGTACGAAAATAAGGGTATT | | | | | | | | | | |
| Chromosome | start | end | strand | MM | target_seq | PAM | alignment | distance | position | gene name | gene id |
| GU305914.1 | 2466 | 2488 | - | 0 | AATACCCTTATTTTCGTACA | AGG | \|\|\|\|\|\|\|\|[\|\|\|\|\|\|\|\|\|\|\|\|]PAM | NA | Intergenic | NA | NA |
| T34151 | TGCGTCTAAACAGACAACCGAGG | 1000 | | | | | | | | | |
| Oligo fwd | CACCTGCGTCTAAACAGACAACCG | | | | | | | | | | |
| Oligo rev | AAACCGGTTGTCTGTTTAGACGCA | | | | | | | | | | |
| Chromosome | start | end | strand | MM | target_seq | PAM | alignment | distance | position | gene name | gene id |

<table>
<tr><td>GU305914.1</td><td></td><td>2431</td><td>2453</td><td>-</td><td>0</td><td>TGCGTCTAAACAGACAACCG</td><td>AGG</td><td>||||||||[||||||||||||]PAM</td><td>NA</td><td>Intergenic</td><td>NA</td><td>NA</td></tr>
<tr><td>T34152</td><td>ACAGACAACCGAGGTCTTTACGG</td><td></td><td>1000</td><td></td><td></td><td></td><td></td><td></td><td></td><td></td><td></td><td></td></tr>
<tr><td>Oligo fwd</td><td>CACCACAGACAACCGAGGTCTTTA</td><td></td><td></td><td></td><td></td><td></td><td></td><td></td><td></td><td></td><td></td><td></td></tr>
<tr><td>Oligo rev</td><td>AAACTAAAGACCTCGGTTGTCTGT</td><td></td><td></td><td></td><td></td><td></td><td></td><td></td><td></td><td></td><td></td><td></td></tr>
<tr><td>Chromosome</td><td>start</td><td></td><td>end</td><td>strand</td><td>MM</td><td>target_seq</td><td>PAM</td><td>alignment</td><td>distance</td><td>position</td><td>gene name</td><td>gene id</td></tr>
<tr><td>GU305914.1</td><td></td><td>2422</td><td>2444</td><td>-</td><td>0</td><td>ACAGACAACCGAGGTCTTTA</td><td>CGG</td><td>||||||||[||||||||||||]PAM</td><td>NA</td><td>Intergenic</td><td>NA</td><td>NA</td></tr>
<tr><td>T34153</td><td>TGTCTTTTTTCCACCGTAATTGG</td><td></td><td>1000</td><td></td><td></td><td></td><td></td><td></td><td></td><td></td><td></td><td></td></tr>
<tr><td>Oligo fwd</td><td>CACCTGTCTTTTTTCCACCGTAAT</td><td></td><td></td><td></td><td></td><td></td><td></td><td></td><td></td><td></td><td></td><td></td></tr>
<tr><td>Oligo rev</td><td>AAACATTACGGTGGAAAAAAGACA</td><td></td><td></td><td></td><td></td><td></td><td></td><td></td><td></td><td></td><td></td><td></td></tr>
<tr><td>Chromosome</td><td>start</td><td></td><td>end</td><td>strand</td><td>MM</td><td>target_seq</td><td>PAM</td><td>alignment</td><td>distance</td><td>position</td><td>gene name</td><td>gene id</td></tr>
<tr><td>GU305914.1</td><td></td><td>2379</td><td>2401</td><td>-</td><td>0</td><td>TGTCTTTTTTCCACCGTAAT</td><td>TGG</td><td>||||||||[||||||||||||]PAM</td><td>NA</td><td>Intergenic</td><td>NA</td><td>NA</td></tr>
<tr><td>T34154</td><td>GTCTTTTTTCCACCGTAATTGGG</td><td></td><td>1000</td><td></td><td></td><td></td><td></td><td></td><td></td><td></td><td></td><td></td></tr>
<tr><td>Oligo fwd</td><td>CACCGTCTTTTTTCCACCGTAATT</td><td></td><td></td><td></td><td></td><td></td><td></td><td></td><td></td><td></td><td></td><td></td></tr>
<tr><td>Oligo rev</td><td>AAACAATTACGGTGGAAAAAAGAC</td><td></td><td></td><td></td><td></td><td></td><td></td><td></td><td></td><td></td><td></td><td></td></tr>
<tr><td>Chromosome</td><td>start</td><td></td><td>end</td><td>strand</td><td>MM</td><td>target_seq</td><td>PAM</td><td>alignment</td><td>distance</td><td>position</td><td>gene name</td><td>gene id</td></tr>
<tr><td>GU305914.1</td><td></td><td>2378</td><td>2400</td><td>-</td><td>0</td><td>GTCTTTTTTCCACCGTAATT</td><td>GGG</td><td>||||||||[||||||||||||]PAM</td><td>NA</td><td>Intergenic</td><td>NA</td><td>NA</td></tr>
<tr><td>T34155</td><td>ACAAAAGCAACCGTGTCAAACGG</td><td></td><td>1000</td><td></td><td></td><td></td><td></td><td></td><td></td><td></td><td></td><td></td></tr>
<tr><td>Oligo fwd</td><td>CACCACAAAAGCAACCGTGTCAAA</td><td></td><td></td><td></td><td></td><td></td><td></td><td></td><td></td><td></td><td></td><td></td></tr>
<tr><td>Oligo rev</td><td>AAACTTTGACACGGTTGCTTTTGT</td><td></td><td></td><td></td><td></td><td></td><td></td><td></td><td></td><td></td><td></td><td></td></tr>
<tr><td>Chromosome</td><td>start</td><td></td><td>end</td><td>strand</td><td>MM</td><td>target_seq</td><td>PAM</td><td>alignment</td><td>distance</td><td>position</td><td>gene name</td><td>gene id</td></tr>
<tr><td>GU305914.1</td><td></td><td>2314</td><td>2336</td><td>-</td><td>0</td><td>ACAAAAGCAACCGTGTCAAA</td><td>CGG</td><td>||||||||[||||||||||||]PAM</td><td>NA</td><td>Intergenic</td><td>NA</td><td>NA</td></tr>
<tr><td>T34156</td><td>CAAACGGAGACGTACATGTGCGG</td><td></td><td>1000</td><td></td><td></td><td></td><td></td><td></td><td></td><td></td><td></td><td></td></tr>
<tr><td>Oligo fwd</td><td>CACCCAAACGGAGACGTACATGTG</td><td></td><td></td><td></td><td></td><td></td><td></td><td></td><td></td><td></td><td></td><td></td></tr>
<tr><td>Oligo rev</td><td>AAACCACATGTACGTCTCCGTTTG</td><td></td><td></td><td></td><td></td><td></td><td></td><td></td><td></td><td></td><td></td><td></td></tr>
<tr><td>Chromosome</td><td>start</td><td></td><td>end</td><td>strand</td><td>MM</td><td>target_seq</td><td>PAM</td><td>alignment</td><td>distance</td><td>position</td><td>gene name</td><td>gene id</td></tr>
<tr><td>GU305914.1</td><td></td><td>2298</td><td>2320</td><td>-</td><td>0</td><td>CAAACGGAGACGTACATGTG</td><td>CGG</td><td>||||||||[||||||||||||]PAM</td><td>NA</td><td>Intergenic</td><td>NA</td><td>NA</td></tr>
<tr><td>T34157</td><td>CGGAGACGTACATGTGCGGACGG</td><td></td><td>1000</td><td></td><td></td><td></td><td></td><td></td><td></td><td></td><td></td><td></td></tr>
<tr><td>Oligo fwd</td><td>CACCCGGAGACGTACATGTGCGGA</td><td></td><td></td><td></td><td></td><td></td><td></td><td></td><td></td><td></td><td></td><td></td></tr>
<tr><td>Oligo rev</td><td>AAACTCCGCACATGTACGTCTCCG</td><td></td><td></td><td></td><td></td><td></td><td></td><td></td><td></td><td></td><td></td><td></td></tr>
<tr><td>Chromosome</td><td>start</td><td></td><td>end</td><td>strand</td><td>MM</td><td>target_seq</td><td>PAM</td><td>alignment</td><td>distance</td><td>position</td><td>gene name</td><td>gene id</td></tr>
<tr><td>GU305914.1</td><td></td><td>2294</td><td>2316</td><td>-</td><td>0</td><td>CGGAGACGTACATGTGCGGA</td><td>CGG</td><td>||||||||[||||||||||||]PAM</td><td>NA</td><td>Intergenic</td><td>NA</td><td>NA</td></tr>
<tr><td>T34158</td><td>TCATGTTTCTTCTCTATGACCGG</td><td></td><td>1000</td><td></td><td></td><td></td><td></td><td></td><td></td><td></td><td></td><td></td></tr>
<tr><td>Oligo fwd</td><td>CACCTCATGTTTCTTCTCTATGAC</td><td></td><td></td><td></td><td></td><td></td><td></td><td></td><td></td><td></td><td></td><td></td></tr>
<tr><td>Oligo rev</td><td>AAACGTCATAGAGAAGAAACATGA</td><td></td><td></td><td></td><td></td><td></td><td></td><td></td><td></td><td></td><td></td><td></td></tr>
<tr><td>Chromosome</td><td>start</td><td></td><td>end</td><td>strand</td><td>MM</td><td>target_seq</td><td>PAM</td><td>alignment</td><td>distance</td><td>position</td><td>gene name</td><td>gene id</td></tr>
<tr><td>GU305914.1</td><td></td><td>2251</td><td>2273</td><td>-</td><td>0</td><td>TCATGTTTCTTCTCTATGAC</td><td>CGG</td><td>||||||||[||||||||||||]PAM</td><td>NA</td><td>Intergenic</td><td>NA</td><td>NA</td></tr>
<tr><td>T34159</td><td>AGTGAATGAGTAGTATCATTTGG</td><td></td><td>1000</td><td></td><td></td><td></td><td></td><td></td><td></td><td></td><td></td><td></td></tr>
<tr><td>Oligo fwd</td><td>CACCAGTGAATGAGTAGTATCATT</td><td></td><td></td><td></td><td></td><td></td><td></td><td></td><td></td><td></td><td></td><td></td></tr>
<tr><td>Oligo rev</td><td>AAACAATGATACTACTCATTCACT</td><td></td><td></td><td></td><td></td><td></td><td></td><td></td><td></td><td></td><td></td><td></td></tr>
<tr><td>Chromosome</td><td>start</td><td></td><td>end</td><td>strand</td><td>MM</td><td>target_seq</td><td>PAM</td><td>alignment</td><td>distance</td><td>position</td><td>gene name</td><td>gene id</td></tr>
<tr><td>GU305914.1</td><td></td><td>2190</td><td>2212</td><td>-</td><td>0</td><td>AGTGAATGAGTAGTATCATT</td><td>TGG</td><td>||||||||[||||||||||||]PAM</td><td>NA</td><td>Intergenic</td><td>NA</td><td>NA</td></tr>
<tr><td>T34160</td><td>GGAGCGATGAGTCCCTATAAAGG</td><td></td><td>1000</td><td></td><td></td><td></td><td></td><td></td><td></td><td></td><td></td><td></td></tr>
<tr><td>Oligo fwd</td><td>CACCGGAGCGATGAGTCCCTATAA</td><td></td><td></td><td></td><td></td><td></td><td></td><td></td><td></td><td></td><td></td><td></td></tr>
<tr><td>Oligo rev</td><td>AAACTTATAGGGACTCATCGCTCC</td><td></td><td></td><td></td><td></td><td></td><td></td><td></td><td></td><td></td><td></td><td></td></tr>
<tr><td>Chromosome</td><td>start</td><td></td><td>end</td><td>strand</td><td>MM</td><td>target_seq</td><td>PAM</td><td>alignment</td><td>distance</td><td>position</td><td>gene name</td><td>gene id</td></tr>
<tr><td>GU305914.1</td><td></td><td>2169</td><td>2191</td><td>-</td><td>0</td><td>GGAGCGATGAGTCCCTATAA</td><td>AGG</td><td>||||||||[||||||||||||]PAM</td><td>NA</td><td>Intergenic</td><td>NA</td><td>NA</td></tr>
<tr><td>T34161</td><td>CGACATCATCTTCCCACGCTCGG</td><td></td><td>1000</td><td></td><td></td><td></td><td></td><td></td><td></td><td></td><td></td><td></td></tr>
<tr><td>Oligo fwd</td><td>CACCCGACATCATCTTCCCACGCT</td><td></td><td></td><td></td><td></td><td></td><td></td><td></td><td></td><td></td><td></td><td></td></tr>
<tr><td>Oligo rev</td><td>AAACAGCGTGGGAAGATGATGTCG</td><td></td><td></td><td></td><td></td><td></td><td></td><td></td><td></td><td></td><td></td><td></td></tr>
<tr><td>Chromosome</td><td>start</td><td></td><td>end</td><td>strand</td><td>MM</td><td>target_seq</td><td>PAM</td><td>alignment</td><td>distance</td><td>position</td><td>gene name</td><td>gene id</td></tr>
<tr><td>GU305914.1</td><td></td><td>2145</td><td>2167</td><td>-</td><td>0</td><td>CGACATCATCTTCCCACGCT</td><td>CGG</td><td>||||||||[||||||||||||]PAM</td><td>NA</td><td>Intergenic</td><td>NA</td><td>NA</td></tr>
<tr><td>T34162</td><td>GCGCAGCGCGTGAAACGAACAGG</td><td></td><td>1000</td><td></td><td></td><td></td><td></td><td></td><td></td><td></td><td></td><td></td></tr>
<tr><td>Oligo fwd</td><td>CACCGCGCAGCGCGTGAAACGAAC</td><td></td><td></td><td></td><td></td><td></td><td></td><td></td><td></td><td></td><td></td><td></td></tr>
<tr><td>Oligo rev</td><td>AAACGTTCGTTTCACGCGCTGCGC</td><td></td><td></td><td></td><td></td><td></td><td></td><td></td><td></td><td></td><td></td><td></td></tr>
<tr><td>Chromosome</td><td>start</td><td></td><td>end</td><td>strand</td><td>MM</td><td>target_seq</td><td>PAM</td><td>alignment</td><td>distance</td><td>position</td><td>gene name</td><td>gene id</td></tr>
<tr><td>GU305914.1</td><td></td><td>2115</td><td>2137</td><td>-</td><td>0</td><td>GCGCAGCGCGTGAAACGAAC</td><td>AGG</td><td>||||||||[||||||||||||]PAM</td><td>NA</td><td>Intergenic</td><td>NA</td><td>NA</td></tr>
<tr><td>T34163</td><td>AACAGGAGCTGCCTCGCAAGAGG</td><td></td><td>1000</td><td></td><td></td><td></td><td></td><td></td><td></td><td></td><td></td><td></td></tr>
<tr><td>Oligo fwd</td><td>CACCAACAGGAGCTGCCTCGCAAG</td><td></td><td></td><td></td><td></td><td></td><td></td><td></td><td></td><td></td><td></td><td></td></tr>
<tr><td>Oligo rev</td><td>AAACCTTGCGAGGCAGCTCCTGTT</td><td></td><td></td><td></td><td></td><td></td><td></td><td></td><td></td><td></td><td></td><td></td></tr>
<tr><td>Chromosome</td><td>start</td><td></td><td>end</td><td>strand</td><td>MM</td><td>target_seq</td><td>PAM</td><td>alignment</td><td>distance</td><td>position</td><td>gene name</td><td>gene id</td></tr>
<tr><td>GU305914.1</td><td></td><td>2098</td><td>2120</td><td>-</td><td>0</td><td>AACAGGAGCTGCCTCGCAAG</td><td>AGG</td><td>||||||||[||||||||||||]PAM</td><td>NA</td><td>Intergenic</td><td>NA</td><td>NA</td></tr>
<tr><td>T34164</td><td>AAGAGGAGAATAACGTCGAACGG</td><td></td><td>1000</td><td></td><td></td><td></td><td></td><td></td><td></td><td></td><td></td><td></td></tr>
<tr><td>Oligo fwd</td><td>CACCAAGAGGAGAATAACGTCGAA</td><td></td><td></td><td></td><td></td><td></td><td></td><td></td><td></td><td></td><td></td><td></td></tr>
<tr><td>Oligo rev</td><td>AAACTTCGACGTTATTCTCCTCTT</td><td></td><td></td><td></td><td></td><td></td><td></td><td></td><td></td><td></td><td></td><td></td></tr>
<tr><td>Chromosome</td><td>start</td><td></td><td>end</td><td>strand</td><td>MM</td><td>target_seq</td><td>PAM</td><td>alignment</td><td>distance</td><td>position</td><td>gene name</td><td>gene id</td></tr>
<tr><td>GU305914.1</td><td></td><td>2081</td><td>2103</td><td>-</td><td>0</td><td>AAGAGGAGAATAACGTCGAA</td><td>CGG</td><td>||||||||[||||||||||||]PAM</td><td>NA</td><td>Intergenic</td><td>NA</td><td>NA</td></tr>
<tr><td>T34165</td><td>GACTTTATTCTCTTCTCTCGAGG</td><td></td><td>1000</td><td></td><td></td><td></td><td></td><td></td><td></td><td></td><td></td><td></td></tr>
<tr><td>Oligo fwd</td><td>CACCGACTTTATTCTCTTCTCTCG</td><td></td><td></td><td></td><td></td><td></td><td></td><td></td><td></td><td></td><td></td><td></td></tr>
<tr><td>Oligo rev</td><td>AAACCGAGAGAAGAGAATAAAGTC</td><td></td><td></td><td></td><td></td><td></td><td></td><td></td><td></td><td></td><td></td><td></td></tr>
<tr><td>Chromosome</td><td>start</td><td></td><td>end</td><td>strand</td><td>MM</td><td>target_seq</td><td>PAM</td><td>alignment</td><td>distance</td><td>position</td><td>gene name</td><td>gene id</td></tr>
<tr><td>GU305914.1</td><td></td><td>2037</td><td>2059</td><td>-</td><td>0</td><td>GACTTTATTCTCTTCTCTCG</td><td>AGG</td><td>||||||||[||||||||||||]PAM</td><td>NA</td><td>Intergenic</td><td>NA</td><td>NA</td></tr>
<tr><td>T34166</td><td>TTATTCTCTTCTCTCGAGGTAGG</td><td></td><td>1000</td><td></td><td></td><td></td><td></td><td></td><td></td><td></td><td></td><td></td></tr>
<tr><td>Oligo fwd</td><td>CACCTTATTCTCTTCTCTCGAGGT</td><td></td><td></td><td></td><td></td><td></td><td></td><td></td><td></td><td></td><td></td><td></td></tr>
<tr><td>Oligo rev</td><td>AAACACCTCGAGAGAAGAGAATAA</td><td></td><td></td><td></td><td></td><td></td><td></td><td></td><td></td><td></td><td></td><td></td></tr>
</table>

<table>
<tr><td>Chromosome</td><td>start</td><td>end</td><td>strand</td><td>MM</td><td>target_seq</td><td>PAM</td><td>alignment</td><td>distance</td><td>position</td><td>gene name</td><td>gene id</td></tr>
<tr><td>GU305914.1</td><td>2033</td><td>2055</td><td>-</td><td>0</td><td>TTATTCTCTTCTCTCGAGGT</td><td>AGG</td><td>||||||||[||||||||||||]PAM</td><td>NA</td><td>Intergenic</td><td>NA</td><td>NA</td></tr>
<tr><td></td><td></td><td></td><td></td><td></td><td></td><td></td><td></td><td></td><td></td><td></td><td></td></tr>
<tr><td>T34167</td><td>TCTTCTCTCGAGGTAGGCTCAGG</td><td>1000</td><td></td><td></td><td></td><td></td><td></td><td></td><td></td><td></td><td></td></tr>
<tr><td>Oligo fwd</td><td>CACCTCTTCTCTCGAGGTAGGCTC</td><td></td><td></td><td></td><td></td><td></td><td></td><td></td><td></td><td></td><td></td></tr>
<tr><td>Oligo rev</td><td>AAACGAGCCTACCTCGAGAGAAGA</td><td></td><td></td><td></td><td></td><td></td><td></td><td></td><td></td><td></td><td></td></tr>
<tr><td>Chromosome</td><td>start</td><td>end</td><td>strand</td><td>MM</td><td>target_seq</td><td>PAM</td><td>alignment</td><td>distance</td><td>position</td><td>gene name</td><td>gene id</td></tr>
<tr><td>GU305914.1</td><td>2027</td><td>2049</td><td>-</td><td>0</td><td>TCTTCTCTCGAGGTAGGCTC</td><td>AGG</td><td>||||||||[||||||||||||]PAM</td><td>NA</td><td>Intergenic</td><td>NA</td><td>NA</td></tr>
<tr><td></td><td></td><td></td><td></td><td></td><td></td><td></td><td></td><td></td><td></td><td></td><td></td></tr>
<tr><td>T34168</td><td>CGTCTCGAGCGTAGTCTCCGCGG</td><td>1000</td><td></td><td></td><td></td><td></td><td></td><td></td><td></td><td></td><td></td></tr>
<tr><td>Oligo fwd</td><td>CACCCGTCTCGAGCGTAGTCTCCG</td><td></td><td></td><td></td><td></td><td></td><td></td><td></td><td></td><td></td><td></td></tr>
<tr><td>Oligo rev</td><td>AAACCGGAGACTACGCTCGAGACG</td><td></td><td></td><td></td><td></td><td></td><td></td><td></td><td></td><td></td><td></td></tr>
<tr><td>Chromosome</td><td>start</td><td>end</td><td>strand</td><td>MM</td><td>target_seq</td><td>PAM</td><td>alignment</td><td>distance</td><td>position</td><td>gene name</td><td>gene id</td></tr>
<tr><td>GU305914.1</td><td>1986</td><td>2008</td><td>-</td><td>0</td><td>CGTCTCGAGCGTAGTCTCCG</td><td>CGG</td><td>||||||||[||||||||||||]PAM</td><td>NA</td><td>Intergenic</td><td>NA</td><td>NA</td></tr>
<tr><td></td><td></td><td></td><td></td><td></td><td></td><td></td><td></td><td></td><td></td><td></td><td></td></tr>
<tr><td>T34169</td><td>CTCGAGCGTAGTCTCCGCGGTGG</td><td>1000</td><td></td><td></td><td></td><td></td><td></td><td></td><td></td><td></td><td></td></tr>
<tr><td>Oligo fwd</td><td>CACCCTCGAGCGTAGTCTCCGCGG</td><td></td><td></td><td></td><td></td><td></td><td></td><td></td><td></td><td></td><td></td></tr>
<tr><td>Oligo rev</td><td>AAACCCGCGGAGACTACGCTCGAG</td><td></td><td></td><td></td><td></td><td></td><td></td><td></td><td></td><td></td><td></td></tr>
<tr><td>Chromosome</td><td>start</td><td>end</td><td>strand</td><td>MM</td><td>target_seq</td><td>PAM</td><td>alignment</td><td>distance</td><td>position</td><td>gene name</td><td>gene id</td></tr>
<tr><td>GU305914.1</td><td>1983</td><td>2005</td><td>-</td><td>0</td><td>CTCGAGCGTAGTCTCCGCGG</td><td>TGG</td><td>||||||||[||||||||||||]PAM</td><td>NA</td><td>Intergenic</td><td>NA</td><td>NA</td></tr>
<tr><td></td><td></td><td></td><td></td><td></td><td></td><td></td><td></td><td></td><td></td><td></td><td></td></tr>
<tr><td>T34170</td><td>GTGGAGACGACGCCGCAGACTGG</td><td>1000</td><td></td><td></td><td></td><td></td><td></td><td></td><td></td><td></td><td></td></tr>
<tr><td>Oligo fwd</td><td>CACCGTGGAGACGACGCCGCAGAC</td><td></td><td></td><td></td><td></td><td></td><td></td><td></td><td></td><td></td><td></td></tr>
<tr><td>Oligo rev</td><td>AAACGTCTGCGGCGTCGTCTCCAC</td><td></td><td></td><td></td><td></td><td></td><td></td><td></td><td></td><td></td><td></td></tr>
<tr><td>Chromosome</td><td>start</td><td>end</td><td>strand</td><td>MM</td><td>target_seq</td><td>PAM</td><td>alignment</td><td>distance</td><td>position</td><td>gene name</td><td>gene id</td></tr>
<tr><td>GU305914.1</td><td>1964</td><td>1986</td><td>-</td><td>0</td><td>GTGGAGACGACGCCGCAGAC</td><td>TGG</td><td>||||||||[||||||||||||]PAM</td><td>NA</td><td>Intergenic</td><td>NA</td><td>NA</td></tr>
<tr><td></td><td></td><td></td><td></td><td></td><td></td><td></td><td></td><td></td><td></td><td></td><td></td></tr>
<tr><td>T34171</td><td>CGTCCGCTGCCGAGACTCCGTGG</td><td>1000</td><td></td><td></td><td></td><td></td><td></td><td></td><td></td><td></td><td></td></tr>
<tr><td>Oligo fwd</td><td>CACCCGTCCGCTGCCGAGACTCCG</td><td></td><td></td><td></td><td></td><td></td><td></td><td></td><td></td><td></td><td></td></tr>
<tr><td>Oligo rev</td><td>AAACCGGAGTCTCGGCAGCGGACG</td><td></td><td></td><td></td><td></td><td></td><td></td><td></td><td></td><td></td><td></td></tr>
<tr><td>Chromosome</td><td>start</td><td>end</td><td>strand</td><td>MM</td><td>target_seq</td><td>PAM</td><td>alignment</td><td>distance</td><td>position</td><td>gene name</td><td>gene id</td></tr>
<tr><td>GU305914.1</td><td>1916</td><td>1938</td><td>-</td><td>0</td><td>CGTCCGCTGCCGAGACTCCG</td><td>TGG</td><td>||||||||[||||||||||||]PAM</td><td>NA</td><td>Intergenic</td><td>NA</td><td>NA</td></tr>
<tr><td></td><td></td><td></td><td></td><td></td><td></td><td></td><td></td><td></td><td></td><td></td><td></td></tr>
<tr><td>T34172</td><td>GCTGCCGAGACTCCGTGGAACGG</td><td>1000</td><td></td><td></td><td></td><td></td><td></td><td></td><td></td><td></td><td></td></tr>
<tr><td>Oligo fwd</td><td>CACCGCTGCCGAGACTCCGTGGAA</td><td></td><td></td><td></td><td></td><td></td><td></td><td></td><td></td><td></td><td></td></tr>
<tr><td>Oligo rev</td><td>AAACTTCCACGGAGTCTCGGCAGC</td><td></td><td></td><td></td><td></td><td></td><td></td><td></td><td></td><td></td><td></td></tr>
<tr><td>Chromosome</td><td>start</td><td>end</td><td>strand</td><td>MM</td><td>target_seq</td><td>PAM</td><td>alignment</td><td>distance</td><td>position</td><td>gene name</td><td>gene id</td></tr>
<tr><td>GU305914.1</td><td>1911</td><td>1933</td><td>-</td><td>0</td><td>GCTGCCGAGACTCCGTGGAA</td><td>CGG</td><td>||||||||[||||||||||||]PAM</td><td>NA</td><td>Intergenic</td><td>NA</td><td>NA</td></tr>
<tr><td></td><td></td><td></td><td></td><td></td><td></td><td></td><td></td><td></td><td></td><td></td><td></td></tr>
<tr><td>T34173</td><td>ACGGCACAGATACCCAATTCCGG</td><td>1000</td><td></td><td></td><td></td><td></td><td></td><td></td><td></td><td></td><td></td></tr>
<tr><td>Oligo fwd</td><td>CACCACGGCACAGATACCCAATTC</td><td></td><td></td><td></td><td></td><td></td><td></td><td></td><td></td><td></td><td></td></tr>
<tr><td>Oligo rev</td><td>AAACGAATTGGGTATCTGTGCCGT</td><td></td><td></td><td></td><td></td><td></td><td></td><td></td><td></td><td></td><td></td></tr>
<tr><td>Chromosome</td><td>start</td><td>end</td><td>strand</td><td>MM</td><td>target_seq</td><td>PAM</td><td>alignment</td><td>distance</td><td>position</td><td>gene name</td><td>gene id</td></tr>
<tr><td>GU305914.1</td><td>1892</td><td>1914</td><td>-</td><td>0</td><td>ACGGCACAGATACCCAATTC</td><td>CGG</td><td>||||||||[||||||||||||]PAM</td><td>NA</td><td>Intergenic</td><td>NA</td><td>NA</td></tr>
<tr><td></td><td></td><td></td><td></td><td></td><td></td><td></td><td></td><td></td><td></td><td></td><td></td></tr>
<tr><td>T34174</td><td>CAACACGATCACCGTCACCACGG</td><td>1000</td><td></td><td></td><td></td><td></td><td></td><td></td><td></td><td></td><td></td></tr>
<tr><td>Oligo fwd</td><td>CACCCAACACGATCACCGTCACCA</td><td></td><td></td><td></td><td></td><td></td><td></td><td></td><td></td><td></td><td></td></tr>
<tr><td>Oligo rev</td><td>AAACTGGTGACGGTGATCGTGTTG</td><td></td><td></td><td></td><td></td><td></td><td></td><td></td><td></td><td></td><td></td></tr>
<tr><td>Chromosome</td><td>start</td><td>end</td><td>strand</td><td>MM</td><td>target_seq</td><td>PAM</td><td>alignment</td><td>distance</td><td>position</td><td>gene name</td><td>gene id</td></tr>
<tr><td>GU305914.1</td><td>1865</td><td>1887</td><td>-</td><td>0</td><td>CAACACGATCACCGTCACCA</td><td>CGG</td><td>||||||||[||||||||||||]PAM</td><td>NA</td><td>Intergenic</td><td>NA</td><td>NA</td></tr>
<tr><td></td><td></td><td></td><td></td><td></td><td></td><td></td><td></td><td></td><td></td><td></td><td></td></tr>
<tr><td>T34175</td><td>AACACGATCACCGTCACCACGGG</td><td>1000</td><td></td><td></td><td></td><td></td><td></td><td></td><td></td><td></td><td></td></tr>
<tr><td>Oligo fwd</td><td>CACCAACACGATCACCGTCACCAC</td><td></td><td></td><td></td><td></td><td></td><td></td><td></td><td></td><td></td><td></td></tr>
<tr><td>Oligo rev</td><td>AAACGTGGTGACGGTGATCGTGTT</td><td></td><td></td><td></td><td></td><td></td><td></td><td></td><td></td><td></td><td></td></tr>
<tr><td>Chromosome</td><td>start</td><td>end</td><td>strand</td><td>MM</td><td>target_seq</td><td>PAM</td><td>alignment</td><td>distance</td><td>position</td><td>gene name</td><td>gene id</td></tr>
<tr><td>GU305914.1</td><td>1864</td><td>1886</td><td>-</td><td>0</td><td>AACACGATCACCGTCACCAC</td><td>GGG</td><td>||||||||[||||||||||||]PAM</td><td>NA</td><td>Intergenic</td><td>NA</td><td>NA</td></tr>
<tr><td></td><td></td><td></td><td></td><td></td><td></td><td></td><td></td><td></td><td></td><td></td><td></td></tr>
<tr><td>T34176</td><td>GATCGCCGCCGCGTTCAGACCGG</td><td>1000</td><td></td><td></td><td></td><td></td><td></td><td></td><td></td><td></td><td></td></tr>
<tr><td>Oligo fwd</td><td>CACCGATCGCCGCCGCGTTCAGAC</td><td></td><td></td><td></td><td></td><td></td><td></td><td></td><td></td><td></td><td></td></tr>
<tr><td>Oligo rev</td><td>AAACGTCTGAACGCGGCGGCGATC</td><td></td><td></td><td></td><td></td><td></td><td></td><td></td><td></td><td></td><td></td></tr>
<tr><td>Chromosome</td><td>start</td><td>end</td><td>strand</td><td>MM</td><td>target_seq</td><td>PAM</td><td>alignment</td><td>distance</td><td>position</td><td>gene name</td><td>gene id</td></tr>
<tr><td>GU305914.1</td><td>1835</td><td>1857</td><td>-</td><td>0</td><td>GATCGCCGCCGCGTTCAGAC</td><td>CGG</td><td>||||||||[||||||||||||]PAM</td><td>NA</td><td>Intergenic</td><td>NA</td><td>NA</td></tr>
<tr><td></td><td></td><td></td><td></td><td></td><td></td><td></td><td></td><td></td><td></td><td></td><td></td></tr>
<tr><td>T34177</td><td>ATCGCCGCCGCGTTCAGACCGGG</td><td>1000</td><td></td><td></td><td></td><td></td><td></td><td></td><td></td><td></td><td></td></tr>
<tr><td>Oligo fwd</td><td>CACCATCGCCGCCGCGTTCAGACC</td><td></td><td></td><td></td><td></td><td></td><td></td><td></td><td></td><td></td><td></td></tr>
<tr><td>Oligo rev</td><td>AAACGGTCTGAACGCGGCGGCGAT</td><td></td><td></td><td></td><td></td><td></td><td></td><td></td><td></td><td></td><td></td></tr>
<tr><td>Chromosome</td><td>start</td><td>end</td><td>strand</td><td>MM</td><td>target_seq</td><td>PAM</td><td>alignment</td><td>distance</td><td>position</td><td>gene name</td><td>gene id</td></tr>
<tr><td>GU305914.1</td><td>1834</td><td>1856</td><td>-</td><td>0</td><td>ATCGCCGCCGCGTTCAGACC</td><td>GGG</td><td>||||||||[||||||||||||]PAM</td><td>NA</td><td>Intergenic</td><td>NA</td><td>NA</td></tr>
<tr><td></td><td></td><td></td><td></td><td></td><td></td><td></td><td></td><td></td><td></td><td></td><td></td></tr>
<tr><td>T34178</td><td>CATCCGATACACGACTCGTCTGG</td><td>1000</td><td></td><td></td><td></td><td></td><td></td><td></td><td></td><td></td><td></td></tr>
<tr><td>Oligo fwd</td><td>CACCCATCCGATACACGACTCGTC</td><td></td><td></td><td></td><td></td><td></td><td></td><td></td><td></td><td></td><td></td></tr>
<tr><td>Oligo rev</td><td>AAACGACGAGTCGTGTATCGGATG</td><td></td><td></td><td></td><td></td><td></td><td></td><td></td><td></td><td></td><td></td></tr>
<tr><td>Chromosome</td><td>start</td><td>end</td><td>strand</td><td>MM</td><td>target_seq</td><td>PAM</td><td>alignment</td><td>distance</td><td>position</td><td>gene name</td><td>gene id</td></tr>
<tr><td>GU305914.1</td><td>1804</td><td>1826</td><td>-</td><td>0</td><td>CATCCGATACACGACTCGTC</td><td>TGG</td><td>||||||||[||||||||||||]PAM</td><td>NA</td><td>Intergenic</td><td>NA</td><td>NA</td></tr>
<tr><td></td><td></td><td></td><td></td><td></td><td></td><td></td><td></td><td></td><td></td><td></td><td></td></tr>
<tr><td>T34179</td><td>CCGATACACGACTCGTCTGGCGG</td><td>1000</td><td></td><td></td><td></td><td></td><td></td><td></td><td></td><td></td><td></td></tr>
<tr><td>Oligo fwd</td><td>CACCCCGATACACGACTCGTCTGG</td><td></td><td></td><td></td><td></td><td></td><td></td><td></td><td></td><td></td><td></td></tr>
<tr><td>Oligo rev</td><td>AAACCCAGACGAGTCGTGTATCGG</td><td></td><td></td><td></td><td></td><td></td><td></td><td></td><td></td><td></td><td></td></tr>
<tr><td>Chromosome</td><td>start</td><td>end</td><td>strand</td><td>MM</td><td>target_seq</td><td>PAM</td><td>alignment</td><td>distance</td><td>position</td><td>gene name</td><td>gene id</td></tr>
<tr><td>GU305914.1</td><td>1801</td><td>1823</td><td>-</td><td>0</td><td>CCGATACACGACTCGTCTGG</td><td>CGG</td><td>||||||||[||||||||||||]PAM</td><td>NA</td><td>Intergenic</td><td>NA</td><td>NA</td></tr>
<tr><td></td><td></td><td></td><td></td><td></td><td></td><td></td><td></td><td></td><td></td><td></td><td></td></tr>
<tr><td>T34180</td><td>CTCGTCTGGCGGAGCGTCCTCGG</td><td>1000</td><td></td><td></td><td></td><td></td><td></td><td></td><td></td><td></td><td></td></tr>
<tr><td>Oligo fwd</td><td>CACCCTCGTCTGGCGGAGCGTCCT</td><td></td><td></td><td></td><td></td><td></td><td></td><td></td><td></td><td></td><td></td></tr>
<tr><td>Oligo rev</td><td>AAACAGGACGCTCCGCCAGACGAG</td><td></td><td></td><td></td><td></td><td></td><td></td><td></td><td></td><td></td><td></td></tr>
<tr><td>Chromosome</td><td>start</td><td>end</td><td>strand</td><td>MM</td><td>target_seq</td><td>PAM</td><td>alignment</td><td>distance</td><td>position</td><td>gene name</td><td>gene id</td></tr>
<tr><td>GU305914.1</td><td>1790</td><td>1812</td><td>-</td><td>0</td><td>CTCGTCTGGCGGAGCGTCCT</td><td>CGG</td><td>||||||||[||||||||||||]PAM</td><td>NA</td><td>Intergenic</td><td>NA</td><td>NA</td></tr>
<tr><td></td><td></td><td></td><td></td><td></td><td></td><td></td><td></td><td></td><td></td><td></td><td></td></tr>
<tr><td>T34181</td><td>GGAGCGTCCTCGGCCTTTCCAGG</td><td>1000</td><td></td><td></td><td></td><td></td><td></td><td></td><td></td><td></td><td></td></tr>
<tr><td>Oligo fwd</td><td>CACCGGAGCGTCCTCGGCCTTTCC</td><td></td><td></td><td></td><td></td><td></td><td></td><td></td><td></td><td></td><td></td></tr>
</table>

<table>
<tr><td>Oligo rev</td><td>AAACGGAAAGGCCGAGGACGCTCC</td><td></td><td></td><td></td><td></td><td></td><td></td><td></td><td></td><td></td><td></td><td></td></tr>
<tr><td>Chromosome</td><td>start</td><td></td><td>end</td><td>strand</td><td>MM</td><td>target_seq</td><td>PAM</td><td>alignment</td><td>distance</td><td>position</td><td>gene name</td><td>gene id</td></tr>
<tr><td>GU305914.1</td><td></td><td>1780</td><td>1802</td><td>-</td><td>0</td><td>GGAGCGTCCTCGGCCTTTCC</td><td>AGG</td><td>||||||||[||||||||||||]PAM</td><td>NA</td><td>Intergenic</td><td>NA</td><td>NA</td></tr>
<tr><td>T34182</td><td>CTCGGCCTTTCCAGGTACTTTGG</td><td></td><td>1000</td><td></td><td></td><td></td><td></td><td></td><td></td><td></td><td></td><td></td></tr>
<tr><td>Oligo fwd</td><td>CACCCTCGGCCTTTCCAGGTACTT</td><td></td><td></td><td></td><td></td><td></td><td></td><td></td><td></td><td></td><td></td><td></td></tr>
<tr><td>Oligo rev</td><td>AAACAAGTACCTGGAAAGGCCGAG</td><td></td><td></td><td></td><td></td><td></td><td></td><td></td><td></td><td></td><td></td><td></td></tr>
<tr><td>Chromosome</td><td>start</td><td></td><td>end</td><td>strand</td><td>MM</td><td>target_seq</td><td>PAM</td><td>alignment</td><td>distance</td><td>position</td><td>gene name</td><td>gene id</td></tr>
<tr><td>GU305914.1</td><td></td><td>1772</td><td>1794</td><td>-</td><td>0</td><td>CTCGGCCTTTCCAGGTACTT</td><td>TGG</td><td>||||||||[||||||||||||]PAM</td><td>NA</td><td>Intergenic</td><td>NA</td><td>NA</td></tr>
<tr><td>T34183</td><td>TCGGCCTTTCCAGGTACTTTGGG</td><td></td><td>1000</td><td></td><td></td><td></td><td></td><td></td><td></td><td></td><td></td><td></td></tr>
<tr><td>Oligo fwd</td><td>CACCTCGGCCTTTCCAGGTACTTT</td><td></td><td></td><td></td><td></td><td></td><td></td><td></td><td></td><td></td><td></td><td></td></tr>
<tr><td>Oligo rev</td><td>AAACAAAGTACCTGGAAAGGCCGA</td><td></td><td></td><td></td><td></td><td></td><td></td><td></td><td></td><td></td><td></td><td></td></tr>
<tr><td>Chromosome</td><td>start</td><td></td><td>end</td><td>strand</td><td>MM</td><td>target_seq</td><td>PAM</td><td>alignment</td><td>distance</td><td>position</td><td>gene name</td><td>gene id</td></tr>
<tr><td>GU305914.1</td><td></td><td>1771</td><td>1793</td><td>-</td><td>0</td><td>TCGGCCTTTCCAGGTACTTT</td><td>GGG</td><td>||||||||[||||||||||||]PAM</td><td>NA</td><td>Intergenic</td><td>NA</td><td>NA</td></tr>
<tr><td>T34184</td><td>GCCTTTCCAGGTACTTTGGGTGG</td><td></td><td>1000</td><td></td><td></td><td></td><td></td><td></td><td></td><td></td><td></td><td></td></tr>
<tr><td>Oligo fwd</td><td>CACCGCCTTTCCAGGTACTTTGGG</td><td></td><td></td><td></td><td></td><td></td><td></td><td></td><td></td><td></td><td></td><td></td></tr>
<tr><td>Oligo rev</td><td>AAACCCCAAAGTACCTGGAAAGGC</td><td></td><td></td><td></td><td></td><td></td><td></td><td></td><td></td><td></td><td></td><td></td></tr>
<tr><td>Chromosome</td><td>start</td><td></td><td>end</td><td>strand</td><td>MM</td><td>target_seq</td><td>PAM</td><td>alignment</td><td>distance</td><td>position</td><td>gene name</td><td>gene id</td></tr>
<tr><td>GU305914.1</td><td></td><td>1768</td><td>1790</td><td>-</td><td>0</td><td>GCCTTTCCAGGTACTTTGGG</td><td>TGG</td><td>||||||||[||||||||||||]PAM</td><td>NA</td><td>Intergenic</td><td>NA</td><td>NA</td></tr>
<tr><td>T34185</td><td>CCTTTCCAGGTACTTTGGGTGGG</td><td></td><td>1000</td><td></td><td></td><td></td><td></td><td></td><td></td><td></td><td></td><td></td></tr>
<tr><td>Oligo fwd</td><td>CACCCCTTTCCAGGTACTTTGGGT</td><td></td><td></td><td></td><td></td><td></td><td></td><td></td><td></td><td></td><td></td><td></td></tr>
<tr><td>Oligo rev</td><td>AAACACCCAAAGTACCTGGAAAGG</td><td></td><td></td><td></td><td></td><td></td><td></td><td></td><td></td><td></td><td></td><td></td></tr>
<tr><td>Chromosome</td><td>start</td><td></td><td>end</td><td>strand</td><td>MM</td><td>target_seq</td><td>PAM</td><td>alignment</td><td>distance</td><td>position</td><td>gene name</td><td>gene id</td></tr>
<tr><td>GU305914.1</td><td></td><td>1767</td><td>1789</td><td>-</td><td>0</td><td>CCTTTCCAGGTACTTTGGGT</td><td>GGG</td><td>||||||||[||||||||||||]PAM</td><td>NA</td><td>Intergenic</td><td>NA</td><td>NA</td></tr>
<tr><td>T34186</td><td>TCCAGGTACTTTGGGTGGGACGG</td><td></td><td>1000</td><td></td><td></td><td></td><td></td><td></td><td></td><td></td><td></td><td></td></tr>
<tr><td>Oligo fwd</td><td>CACCTCCAGGTACTTTGGGTGGGA</td><td></td><td></td><td></td><td></td><td></td><td></td><td></td><td></td><td></td><td></td><td></td></tr>
<tr><td>Oligo rev</td><td>AAACTCCCACCCAAAGTACCTGGA</td><td></td><td></td><td></td><td></td><td></td><td></td><td></td><td></td><td></td><td></td><td></td></tr>
<tr><td>Chromosome</td><td>start</td><td></td><td>end</td><td>strand</td><td>MM</td><td>target_seq</td><td>PAM</td><td>alignment</td><td>distance</td><td>position</td><td>gene name</td><td>gene id</td></tr>
<tr><td>GU305914.1</td><td></td><td>1763</td><td>1785</td><td>-</td><td>0</td><td>TCCAGGTACTTTGGGTGGGA</td><td>CGG</td><td>||||||||[||||||||||||]PAM</td><td>NA</td><td>Intergenic</td><td>NA</td><td>NA</td></tr>
<tr><td>T34187</td><td>AGGTACTTTGGGTGGGACGGCGG</td><td></td><td>1000</td><td></td><td></td><td></td><td></td><td></td><td></td><td></td><td></td><td></td></tr>
<tr><td>Oligo fwd</td><td>CACCAGGTACTTTGGGTGGGACGG</td><td></td><td></td><td></td><td></td><td></td><td></td><td></td><td></td><td></td><td></td><td></td></tr>
<tr><td>Oligo rev</td><td>AAACCCGTCCCACCCAAAGTACCT</td><td></td><td></td><td></td><td></td><td></td><td></td><td></td><td></td><td></td><td></td><td></td></tr>
<tr><td>Chromosome</td><td>start</td><td></td><td>end</td><td>strand</td><td>MM</td><td>target_seq</td><td>PAM</td><td>alignment</td><td>distance</td><td>position</td><td>gene name</td><td>gene id</td></tr>
<tr><td>GU305914.1</td><td></td><td>1760</td><td>1782</td><td>-</td><td>0</td><td>AGGTACTTTGGGTGGGACGG</td><td>CGG</td><td>||||||||[||||||||||||]PAM</td><td>NA</td><td>Intergenic</td><td>NA</td><td>NA</td></tr>
<tr><td>T34188</td><td>GGTACTTTGGGTGGGACGGCGGG</td><td></td><td>1000</td><td></td><td></td><td></td><td></td><td></td><td></td><td></td><td></td><td></td></tr>
<tr><td>Oligo fwd</td><td>CACCGGTACTTTGGGTGGGACGGC</td><td></td><td></td><td></td><td></td><td></td><td></td><td></td><td></td><td></td><td></td><td></td></tr>
<tr><td>Oligo rev</td><td>AAACGCCGTCCCACCCAAAGTACC</td><td></td><td></td><td></td><td></td><td></td><td></td><td></td><td></td><td></td><td></td><td></td></tr>
<tr><td>Chromosome</td><td>start</td><td></td><td>end</td><td>strand</td><td>MM</td><td>target_seq</td><td>PAM</td><td>alignment</td><td>distance</td><td>position</td><td>gene name</td><td>gene id</td></tr>
<tr><td>GU305914.1</td><td></td><td>1759</td><td>1781</td><td>-</td><td>0</td><td>GGTACTTTGGGTGGGACGGC</td><td>GGG</td><td>||||||||[||||||||||||]PAM</td><td>NA</td><td>Intergenic</td><td>NA</td><td>NA</td></tr>
<tr><td>T34189</td><td>GGGTGGGACGGCGGGTGCTTTGG</td><td></td><td>1000</td><td></td><td></td><td></td><td></td><td></td><td></td><td></td><td></td><td></td></tr>
<tr><td>Oligo fwd</td><td>CACCGGGTGGGACGGCGGGTGCTT</td><td></td><td></td><td></td><td></td><td></td><td></td><td></td><td></td><td></td><td></td><td></td></tr>
<tr><td>Oligo rev</td><td>AAACAAGCACCCGCCGTCCCACCC</td><td></td><td></td><td></td><td></td><td></td><td></td><td></td><td></td><td></td><td></td><td></td></tr>
<tr><td>Chromosome</td><td>start</td><td></td><td>end</td><td>strand</td><td>MM</td><td>target_seq</td><td>PAM</td><td>alignment</td><td>distance</td><td>position</td><td>gene name</td><td>gene id</td></tr>
<tr><td>GU305914.1</td><td></td><td>1751</td><td>1773</td><td>-</td><td>0</td><td>GGGTGGGACGGCGGGTGCTT</td><td>TGG</td><td>||||||||[||||||||||||]PAM</td><td>NA</td><td>Intergenic</td><td>NA</td><td>NA</td></tr>
<tr><td>T34190</td><td>GGTGGGACGGCGGGTGCTTTGGG</td><td></td><td>1000</td><td></td><td></td><td></td><td></td><td></td><td></td><td></td><td></td><td></td></tr>
<tr><td>Oligo fwd</td><td>CACCGGTGGGACGGCGGGTGCTTT</td><td></td><td></td><td></td><td></td><td></td><td></td><td></td><td></td><td></td><td></td><td></td></tr>
<tr><td>Oligo rev</td><td>AAACAAAGCACCCGCCGTCCCACC</td><td></td><td></td><td></td><td></td><td></td><td></td><td></td><td></td><td></td><td></td><td></td></tr>
<tr><td>Chromosome</td><td>start</td><td></td><td>end</td><td>strand</td><td>MM</td><td>target_seq</td><td>PAM</td><td>alignment</td><td>distance</td><td>position</td><td>gene name</td><td>gene id</td></tr>
<tr><td>GU305914.1</td><td></td><td>1750</td><td>1772</td><td>-</td><td>0</td><td>GGTGGGACGGCGGGTGCTTT</td><td>GGG</td><td>||||||||[||||||||||||]PAM</td><td>NA</td><td>Intergenic</td><td>NA</td><td>NA</td></tr>
<tr><td>T34191</td><td>GGGACGGCGGGTGCTTTGGGCGG</td><td></td><td>1000</td><td></td><td></td><td></td><td></td><td></td><td></td><td></td><td></td><td></td></tr>
<tr><td>Oligo fwd</td><td>CACCGGGACGGCGGGTGCTTTGGG</td><td></td><td></td><td></td><td></td><td></td><td></td><td></td><td></td><td></td><td></td><td></td></tr>
<tr><td>Oligo rev</td><td>AAACCCCAAAGCACCCGCCGTCCC</td><td></td><td></td><td></td><td></td><td></td><td></td><td></td><td></td><td></td><td></td><td></td></tr>
<tr><td>Chromosome</td><td>start</td><td></td><td>end</td><td>strand</td><td>MM</td><td>target_seq</td><td>PAM</td><td>alignment</td><td>distance</td><td>position</td><td>gene name</td><td>gene id</td></tr>
<tr><td>GU305914.1</td><td></td><td>1747</td><td>1769</td><td>-</td><td>0</td><td>GGGACGGCGGGTGCTTTGGG</td><td>CGG</td><td>||||||||[||||||||||||]PAM</td><td>NA</td><td>Intergenic</td><td>NA</td><td>NA</td></tr>
<tr><td>T34192</td><td>ACGGCGGGTGCTTTGGGCGGTGG</td><td></td><td>1000</td><td></td><td></td><td></td><td></td><td></td><td></td><td></td><td></td><td></td></tr>
<tr><td>Oligo fwd</td><td>CACCACGGCGGGTGCTTTGGGCGG</td><td></td><td></td><td></td><td></td><td></td><td></td><td></td><td></td><td></td><td></td><td></td></tr>
<tr><td>Oligo rev</td><td>AAACCCGCCCAAAGCACCCGCCGT</td><td></td><td></td><td></td><td></td><td></td><td></td><td></td><td></td><td></td><td></td><td></td></tr>
<tr><td>Chromosome</td><td>start</td><td></td><td>end</td><td>strand</td><td>MM</td><td>target_seq</td><td>PAM</td><td>alignment</td><td>distance</td><td>position</td><td>gene name</td><td>gene id</td></tr>
<tr><td>GU305914.1</td><td></td><td>1744</td><td>1766</td><td>-</td><td>0</td><td>ACGGCGGGTGCTTTGGGCGG</td><td>TGG</td><td>||||||||[||||||||||||]PAM</td><td>NA</td><td>Intergenic</td><td>NA</td><td>NA</td></tr>
<tr><td>T34193</td><td>CGGCGGGTGCTTTGGGCGGTGGG</td><td></td><td>1000</td><td></td><td></td><td></td><td></td><td></td><td></td><td></td><td></td><td></td></tr>
<tr><td>Oligo fwd</td><td>CACCCGGCGGGTGCTTTGGGCGGT</td><td></td><td></td><td></td><td></td><td></td><td></td><td></td><td></td><td></td><td></td><td></td></tr>
<tr><td>Oligo rev</td><td>AAACACCGCCCAAAGCACCCGCCG</td><td></td><td></td><td></td><td></td><td></td><td></td><td></td><td></td><td></td><td></td><td></td></tr>
<tr><td>Chromosome</td><td>start</td><td></td><td>end</td><td>strand</td><td>MM</td><td>target_seq</td><td>PAM</td><td>alignment</td><td>distance</td><td>position</td><td>gene name</td><td>gene id</td></tr>
<tr><td>GU305914.1</td><td></td><td>1743</td><td>1765</td><td>-</td><td>0</td><td>CGGCGGGTGCTTTGGGCGGT</td><td>GGG</td><td>||||||||[||||||||||||]PAM</td><td>NA</td><td>Intergenic</td><td>NA</td><td>NA</td></tr>
<tr><td>T34194</td><td>GCTTTGGGCGGTGGGACGTACGG</td><td></td><td>1000</td><td></td><td></td><td></td><td></td><td></td><td></td><td></td><td></td><td></td></tr>
<tr><td>Oligo fwd</td><td>CACCGCTTTGGGCGGTGGGACGTA</td><td></td><td></td><td></td><td></td><td></td><td></td><td></td><td></td><td></td><td></td><td></td></tr>
<tr><td>Oligo rev</td><td>AAACTACGTCCCACCGCCCAAAGC</td><td></td><td></td><td></td><td></td><td></td><td></td><td></td><td></td><td></td><td></td><td></td></tr>
<tr><td>Chromosome</td><td>start</td><td></td><td>end</td><td>strand</td><td>MM</td><td>target_seq</td><td>PAM</td><td>alignment</td><td>distance</td><td>position</td><td>gene name</td><td>gene id</td></tr>
<tr><td>GU305914.1</td><td></td><td>1735</td><td>1757</td><td>-</td><td>0</td><td>GCTTTGGGCGGTGGGACGTA</td><td>CGG</td><td>||||||||[||||||||||||]PAM</td><td>NA</td><td>Intergenic</td><td>NA</td><td>NA</td></tr>
<tr><td>T34195</td><td>TCGTATATGTTCTCGAAGTCCGG</td><td></td><td>1000</td><td></td><td></td><td></td><td></td><td></td><td></td><td></td><td></td><td></td></tr>
<tr><td>Oligo fwd</td><td>CACCTCGTATATGTTCTCGAAGTC</td><td></td><td></td><td></td><td></td><td></td><td></td><td></td><td></td><td></td><td></td><td></td></tr>
<tr><td>Oligo rev</td><td>AAACGACTTCGAGAACATATACGA</td><td></td><td></td><td></td><td></td><td></td><td></td><td></td><td></td><td></td><td></td><td></td></tr>
<tr><td>Chromosome</td><td>start</td><td></td><td>end</td><td>strand</td><td>MM</td><td>target_seq</td><td>PAM</td><td>alignment</td><td>distance</td><td>position</td><td>gene name</td><td>gene id</td></tr>
<tr><td>GU305914.1</td><td></td><td>1696</td><td>1718</td><td>-</td><td>0</td><td>TCGTATATGTTCTCGAAGTC</td><td>CGG</td><td>||||||||[||||||||||||]PAM</td><td>NA</td><td>Intergenic</td><td>NA</td><td>NA</td></tr>
<tr><td>T34196</td><td>TCGTATGTTGTGCGAGAATACGG</td><td></td><td>1000</td><td></td><td></td><td></td><td></td><td></td><td></td><td></td><td></td><td></td></tr>
</table>

Oligo fwd CACCTCGTATGTTGTGCGAGAATA
Oligo rev AAACTATTCTCGCACAACATACGA
Chromosome start end strand MM target_seq PAM alignment distance position gene name gene id
GU305914.1 1636 1658 - 0 TCGTATGTTGTGCGAGAATA CGG ||||||||[||||||||||||]PAM NA Intergenic NA NA

T34197 TGTGCGAGAATACGGCGAGACGG 1000
Oligo fwd CACCTGTGCGAGAATACGGCGAGA
Oligo rev AAACTCTCGCCGTATTCTCGCACA
Chromosome start end strand MM target_seq PAM alignment distance position gene name gene id
GU305914.1 1628 1650 - 0 TGTGCGAGAATACGGCGAGA CGG ||||||||[||||||||||||]PAM NA Intergenic NA NA

T34198 GCGAGAATACGGCGAGACGGAGG 1000
Oligo fwd CACCGCGAGAATACGGCGAGACGG
Oligo rev AAACCCGTCTCGCCGTATTCTCGC
Chromosome start end strand MM target_seq PAM alignment distance position gene name gene id
GU305914.1 1625 1647 - 0 GCGAGAATACGGCGAGACGG AGG ||||||||[||||||||||||]PAM NA Intergenic NA NA

T34199 GCGAGACGGAGGTCGCTCGCAGG 1000
Oligo fwd CACCGCGAGACGGAGGTCGCTCGC
Oligo rev AAACGCGAGCGACCTCCGTCTCGC
Chromosome start end strand MM target_seq PAM alignment distance position gene name gene id
GU305914.1 1614 1636 - 0 GCGAGACGGAGGTCGCTCGC AGG ||||||||[||||||||||||]PAM NA Intergenic NA NA

T34200 TTACACACCACGTCGCGTGTAGG 1000
Oligo fwd CACCTTACACACCACGTCGCGTGT
Oligo rev AAACACACGCGACGTGGTGTGTAA
Chromosome start end strand MM target_seq PAM alignment distance position gene name gene id
GU305914.1 1579 1601 - 0 TTACACACCACGTCGCGTGT AGG ||||||||[||||||||||||]PAM NA Intergenic NA NA

T34201 TTCTTTCCGTGTGTTGTTGTCGG 1000
Oligo fwd CACCTTCTTTCCGTGTGTTGTTGT
Oligo rev AAACACAACAACACACGGAAAGAA
Chromosome start end strand MM target_seq PAM alignment distance position gene name gene id
GU305914.1 1541 1563 - 0 TTCTTTCCGTGTGTTGTTGT CGG ||||||||[||||||||||||]PAM NA Intergenic NA NA

T34202 TGTTGTTGTCGGTGATGATGAGG 1000
Oligo fwd CACCTGTTGTTGTCGGTGATGATG
Oligo rev AAACCATCATCACCGACAACAACA
Chromosome start end strand MM target_seq PAM alignment distance position gene name gene id
GU305914.1 1530 1552 - 0 TGTTGTTGTCGGTGATGATG AGG ||||||||[||||||||||||]PAM NA Intergenic NA NA

T34203 GACGCAGAAACCTTGCCTACAGG 1000
Oligo fwd CACCGACGCAGAAACCTTGCCTAC
Oligo rev AAACGTAGGCAAGGTTTCTGCGTC
Chromosome start end strand MM target_seq PAM alignment distance position gene name gene id
GU305914.1 1490 1512 - 0 GACGCAGAAACCTTGCCTAC AGG ||||||||[||||||||||||]PAM NA Intergenic NA NA

T34204 TCCTTACCACGCTCGTACGACGG 1000
Oligo fwd CACCTCCTTACCACGCTCGTACGA
Oligo rev AAACTCGTACGAGCGTGGTAAGGA
Chromosome start end strand MM target_seq PAM alignment distance position gene name gene id
GU305914.1 1462 1484 - 0 TCCTTACCACGCTCGTACGA CGG ||||||||[||||||||||||]PAM NA Intergenic NA NA

T34205 GGTATGATGTACTGCATCGACGG 1000
Oligo fwd CACCGGTATGATGTACTGCATCGA
Oligo rev AAACTCGATGCAGTACATCATACC
Chromosome start end strand MM target_seq PAM alignment distance position gene name gene id
GU305914.1 1441 1463 - 0 GGTATGATGTACTGCATCGA CGG ||||||||[||||||||||||]PAM NA Intergenic NA NA

T34206 TCTGTGACTTAAAGAAGACCTGG 1000
Oligo fwd CACCTCTGTGACTTAAAGAAGACC
Oligo rev AAACGGTCTTCTTTAAGTCACAGA
Chromosome start end strand MM target_seq PAM alignment distance position gene name gene id
GU305914.1 1407 1429 - 0 TCTGTGACTTAAAGAAGACC TGG ||||||||[||||||||||||]PAM NA Intergenic NA NA

T34207 AAAGAAGACCTGGTTGATCACGG 1000
Oligo fwd CACCAAAGAAGACCTGGTTGATCA
Oligo rev AAACTGATCAACCAGGTCTTCTTT
Chromosome start end strand MM target_seq PAM alignment distance position gene name gene id
GU305914.1 1397 1419 - 0 AAAGAAGACCTGGTTGATCA CGG ||||||||[||||||||||||]PAM NA Intergenic NA NA

T34208 GGTTGATCACGGACGCCTGTAGG 1000
Oligo fwd CACCGGTTGATCACGGACGCCTGT
Oligo rev AAACACAGGCGTCCGTGATCAACC
Chromosome start end strand MM target_seq PAM alignment distance position gene name gene id
GU305914.1 1386 1408 - 0 GGTTGATCACGGACGCCTGT AGG ||||||||[||||||||||||]PAM NA Intergenic NA NA

T34209 GATCACGGACGCCTGTAGGCTGG 1000
Oligo fwd CACCGATCACGGACGCCTGTAGGC
Oligo rev AAACGCCTACAGGCGTCCGTGATC
Chromosome start end strand MM target_seq PAM alignment distance position gene name gene id
GU305914.1 1382 1404 - 0 GATCACGGACGCCTGTAGGC TGG ||||||||[||||||||||||]PAM NA Intergenic NA NA

T34210 ATCACGGACGCCTGTAGGCTGGG 1000
Oligo fwd CACCATCACGGACGCCTGTAGGCT
Oligo rev AAACAGCCTACAGGCGTCCGTGAT
Chromosome start end strand MM target_seq PAM alignment distance position gene name gene id
GU305914.1 1381 1403 - 0 ATCACGGACGCCTGTAGGCT GGG ||||||||[||||||||||||]PAM NA Intergenic NA NA

<table>
<tr><td>T34211</td><td>TCACGGACGCCTGTAGGCTGGGG</td><td></td><td>1000</td><td></td><td></td><td></td><td></td><td></td><td></td><td></td><td></td><td></td></tr>
<tr><td>Oligo fwd</td><td>CACCTCACGGACGCCTGTAGGCTG</td><td></td><td></td><td></td><td></td><td></td><td></td><td></td><td></td><td></td><td></td><td></td></tr>
<tr><td>Oligo rev</td><td>AAACCAGCCTACAGGCGTCCGTGA</td><td></td><td></td><td></td><td></td><td></td><td></td><td></td><td></td><td></td><td></td><td></td></tr>
<tr><td>Chromosome</td><td>start</td><td></td><td>end</td><td>strand</td><td>MM</td><td>target_seq</td><td>PAM</td><td>alignment</td><td>distance</td><td>position</td><td>gene name</td><td>gene id</td></tr>
<tr><td>GU305914.1</td><td></td><td>1380</td><td>1402</td><td>-</td><td>0</td><td>TCACGGACGCCTGTAGGCTG</td><td>GGG</td><td>||||||||[||||||||||||]PAM</td><td>NA</td><td>Intergenic</td><td>NA</td><td>NA</td></tr>
<tr><td>T34212</td><td>GCCCTCGATACACACTCCTTTGG</td><td></td><td>1000</td><td></td><td></td><td></td><td></td><td></td><td></td><td></td><td></td><td></td></tr>
<tr><td>Oligo fwd</td><td>CACCGCCCTCGATACACACTCCTT</td><td></td><td></td><td></td><td></td><td></td><td></td><td></td><td></td><td></td><td></td><td></td></tr>
<tr><td>Oligo rev</td><td>AAACAAGGAGTGTGTATCGAGGGC</td><td></td><td></td><td></td><td></td><td></td><td></td><td></td><td></td><td></td><td></td><td></td></tr>
<tr><td>Chromosome</td><td>start</td><td></td><td>end</td><td>strand</td><td>MM</td><td>target_seq</td><td>PAM</td><td>alignment</td><td>distance</td><td>position</td><td>gene name</td><td>gene id</td></tr>
<tr><td>GU305914.1</td><td></td><td>1334</td><td>1356</td><td>-</td><td>0</td><td>GCCCTCGATACACACTCCTT</td><td>TGG</td><td>||||||||[||||||||||||]PAM</td><td>NA</td><td>Intergenic</td><td>NA</td><td>NA</td></tr>
<tr><td>T34213</td><td>CCCTCGATACACACTCCTTTGGG</td><td></td><td>1000</td><td></td><td></td><td></td><td></td><td></td><td></td><td></td><td></td><td></td></tr>
<tr><td>Oligo fwd</td><td>CACCCCCTCGATACACACTCCTTT</td><td></td><td></td><td></td><td></td><td></td><td></td><td></td><td></td><td></td><td></td><td></td></tr>
<tr><td>Oligo rev</td><td>AAACAAAGGAGTGTGTATCGAGGG</td><td></td><td></td><td></td><td></td><td></td><td></td><td></td><td></td><td></td><td></td><td></td></tr>
<tr><td>Chromosome</td><td>start</td><td></td><td>end</td><td>strand</td><td>MM</td><td>target_seq</td><td>PAM</td><td>alignment</td><td>distance</td><td>position</td><td>gene name</td><td>gene id</td></tr>
<tr><td>GU305914.1</td><td></td><td>1333</td><td>1355</td><td>-</td><td>0</td><td>CCCTCGATACACACTCCTTT</td><td>GGG</td><td>||||||||[||||||||||||]PAM</td><td>NA</td><td>Intergenic</td><td>NA</td><td>NA</td></tr>
<tr><td>T34214</td><td>CAACCGAACACCGACACGCCCGG</td><td></td><td>1000</td><td></td><td></td><td></td><td></td><td></td><td></td><td></td><td></td><td></td></tr>
<tr><td>Oligo fwd</td><td>CACCCAACCGAACACCGACACGCC</td><td></td><td></td><td></td><td></td><td></td><td></td><td></td><td></td><td></td><td></td><td></td></tr>
<tr><td>Oligo rev</td><td>AAACGGCGTGTCGGTGTTCGGTTG</td><td></td><td></td><td></td><td></td><td></td><td></td><td></td><td></td><td></td><td></td><td></td></tr>
<tr><td>Chromosome</td><td>start</td><td></td><td>end</td><td>strand</td><td>MM</td><td>target_seq</td><td>PAM</td><td>alignment</td><td>distance</td><td>position</td><td>gene name</td><td>gene id</td></tr>
<tr><td>GU305914.1</td><td></td><td>1303</td><td>1325</td><td>-</td><td>0</td><td>CAACCGAACACCGACACGCC</td><td>CGG</td><td>||||||||[||||||||||||]PAM</td><td>NA</td><td>Intergenic</td><td>NA</td><td>NA</td></tr>
<tr><td>T34215</td><td>CAGTAATCGCGATCGATGTTAGG</td><td></td><td>1000</td><td></td><td></td><td></td><td></td><td></td><td></td><td></td><td></td><td></td></tr>
<tr><td>Oligo fwd</td><td>CACCCAGTAATCGCGATCGATGTT</td><td></td><td></td><td></td><td></td><td></td><td></td><td></td><td></td><td></td><td></td><td></td></tr>
<tr><td>Oligo rev</td><td>AAACAACATCGATCGCGATTACTG</td><td></td><td></td><td></td><td></td><td></td><td></td><td></td><td></td><td></td><td></td><td></td></tr>
<tr><td>Chromosome</td><td>start</td><td></td><td>end</td><td>strand</td><td>MM</td><td>target_seq</td><td>PAM</td><td>alignment</td><td>distance</td><td>position</td><td>gene name</td><td>gene id</td></tr>
<tr><td>GU305914.1</td><td></td><td>1276</td><td>1298</td><td>-</td><td>0</td><td>CAGTAATCGCGATCGATGTT</td><td>AGG</td><td>||||||||[||||||||||||]PAM</td><td>NA</td><td>Intergenic</td><td>NA</td><td>NA</td></tr>
<tr><td>T34216</td><td>TTCGCGTAGCAGTAGTACCCCGG</td><td></td><td>1000</td><td></td><td></td><td></td><td></td><td></td><td></td><td></td><td></td><td></td></tr>
<tr><td>Oligo fwd</td><td>CACCTTCGCGTAGCAGTAGTACCC</td><td></td><td></td><td></td><td></td><td></td><td></td><td></td><td></td><td></td><td></td><td></td></tr>
<tr><td>Oligo rev</td><td>AAACGGGTACTACTGCTACGCGAA</td><td></td><td></td><td></td><td></td><td></td><td></td><td></td><td></td><td></td><td></td><td></td></tr>
<tr><td>Chromosome</td><td>start</td><td></td><td>end</td><td>strand</td><td>MM</td><td>target_seq</td><td>PAM</td><td>alignment</td><td>distance</td><td>position</td><td>gene name</td><td>gene id</td></tr>
<tr><td>GU305914.1</td><td></td><td>1231</td><td>1253</td><td>-</td><td>0</td><td>TTCGCGTAGCAGTAGTACCC</td><td>CGG</td><td>||||||||[||||||||||||]PAM</td><td>NA</td><td>Intergenic</td><td>NA</td><td>NA</td></tr>
<tr><td>T34217</td><td>TAGCAGTAGTACCCCGGCAGCGG</td><td></td><td>1000</td><td></td><td></td><td></td><td></td><td></td><td></td><td></td><td></td><td></td></tr>
<tr><td>Oligo fwd</td><td>CACCTAGCAGTAGTACCCCGGCAG</td><td></td><td></td><td></td><td></td><td></td><td></td><td></td><td></td><td></td><td></td><td></td></tr>
<tr><td>Oligo rev</td><td>AAACCTGCCGGGGTACTACTGCTA</td><td></td><td></td><td></td><td></td><td></td><td></td><td></td><td></td><td></td><td></td><td></td></tr>
<tr><td>Chromosome</td><td>start</td><td></td><td>end</td><td>strand</td><td>MM</td><td>target_seq</td><td>PAM</td><td>alignment</td><td>distance</td><td>position</td><td>gene name</td><td>gene id</td></tr>
<tr><td>GU305914.1</td><td></td><td>1225</td><td>1247</td><td>-</td><td>0</td><td>TAGCAGTAGTACCCCGGCAG</td><td>CGG</td><td>||||||||[||||||||||||]PAM</td><td>NA</td><td>Intergenic</td><td>NA</td><td>NA</td></tr>
<tr><td>T34218</td><td>CAGCGGCGCCGACGACACAATGG</td><td></td><td>1000</td><td></td><td></td><td></td><td></td><td></td><td></td><td></td><td></td><td></td></tr>
<tr><td>Oligo fwd</td><td>CACCCAGCGGCGCCGACGACACAA</td><td></td><td></td><td></td><td></td><td></td><td></td><td></td><td></td><td></td><td></td><td></td></tr>
<tr><td>Oligo rev</td><td>AAACTTGTGTCGTCGGCGCCGCTG</td><td></td><td></td><td></td><td></td><td></td><td></td><td></td><td></td><td></td><td></td><td></td></tr>
<tr><td>Chromosome</td><td>start</td><td></td><td>end</td><td>strand</td><td>MM</td><td>target_seq</td><td>PAM</td><td>alignment</td><td>distance</td><td>position</td><td>gene name</td><td>gene id</td></tr>
<tr><td>GU305914.1</td><td></td><td>1208</td><td>1230</td><td>-</td><td>0</td><td>CAGCGGCGCCGACGACACAA</td><td>TGG</td><td>||||||||[||||||||||||]PAM</td><td>NA</td><td>Intergenic</td><td>NA</td><td>NA</td></tr>
<tr><td>T34219</td><td>AGCGGCGCCGACGACACAATGGG</td><td></td><td>1000</td><td></td><td></td><td></td><td></td><td></td><td></td><td></td><td></td><td></td></tr>
<tr><td>Oligo fwd</td><td>CACCAGCGGCGCCGACGACACAAT</td><td></td><td></td><td></td><td></td><td></td><td></td><td></td><td></td><td></td><td></td><td></td></tr>
<tr><td>Oligo rev</td><td>AAACATTGTGTCGTCGGCGCCGCT</td><td></td><td></td><td></td><td></td><td></td><td></td><td></td><td></td><td></td><td></td><td></td></tr>
<tr><td>Chromosome</td><td>start</td><td></td><td>end</td><td>strand</td><td>MM</td><td>target_seq</td><td>PAM</td><td>alignment</td><td>distance</td><td>position</td><td>gene name</td><td>gene id</td></tr>
<tr><td>GU305914.1</td><td></td><td>1207</td><td>1229</td><td>-</td><td>0</td><td>AGCGGCGCCGACGACACAAT</td><td>GGG</td><td>||||||||[||||||||||||]PAM</td><td>NA</td><td>Intergenic</td><td>NA</td><td>NA</td></tr>
<tr><td>T34220</td><td>GCGGCGCCGACGACACAATGGGG</td><td></td><td>1000</td><td></td><td></td><td></td><td></td><td></td><td></td><td></td><td></td><td></td></tr>
<tr><td>Oligo fwd</td><td>CACCGCGGCGCCGACGACACAATG</td><td></td><td></td><td></td><td></td><td></td><td></td><td></td><td></td><td></td><td></td><td></td></tr>
<tr><td>Oligo rev</td><td>AAACCATTGTGTCGTCGGCGCCGC</td><td></td><td></td><td></td><td></td><td></td><td></td><td></td><td></td><td></td><td></td><td></td></tr>
<tr><td>Chromosome</td><td>start</td><td></td><td>end</td><td>strand</td><td>MM</td><td>target_seq</td><td>PAM</td><td>alignment</td><td>distance</td><td>position</td><td>gene name</td><td>gene id</td></tr>
<tr><td>GU305914.1</td><td></td><td>1206</td><td>1228</td><td>-</td><td>0</td><td>GCGGCGCCGACGACACAATG</td><td>GGG</td><td>||||||||[||||||||||||]PAM</td><td>NA</td><td>Intergenic</td><td>NA</td><td>NA</td></tr>
<tr><td>T34221</td><td>CGACACAATGGGGACCACCGCGG</td><td></td><td>1000</td><td></td><td></td><td></td><td></td><td></td><td></td><td></td><td></td><td></td></tr>
<tr><td>Oligo fwd</td><td>CACCCGACACAATGGGGACCACCG</td><td></td><td></td><td></td><td></td><td></td><td></td><td></td><td></td><td></td><td></td><td></td></tr>
<tr><td>Oligo rev</td><td>AAACCGGTGGTCCCCATTGTGTCG</td><td></td><td></td><td></td><td></td><td></td><td></td><td></td><td></td><td></td><td></td><td></td></tr>
<tr><td>Chromosome</td><td>start</td><td></td><td>end</td><td>strand</td><td>MM</td><td>target_seq</td><td>PAM</td><td>alignment</td><td>distance</td><td>position</td><td>gene name</td><td>gene id</td></tr>
<tr><td>GU305914.1</td><td></td><td>1196</td><td>1218</td><td>-</td><td>0</td><td>CGACACAATGGGGACCACCG</td><td>CGG</td><td>||||||||[||||||||||||]PAM</td><td>NA</td><td>Intergenic</td><td>NA</td><td>NA</td></tr>
<tr><td>T34222</td><td>CTCGCAGCCGACAGACACCACGG</td><td></td><td>1000</td><td></td><td></td><td></td><td></td><td></td><td></td><td></td><td></td><td></td></tr>
<tr><td>Oligo fwd</td><td>CACCCTCGCAGCCGACAGACACCA</td><td></td><td></td><td></td><td></td><td></td><td></td><td></td><td></td><td></td><td></td><td></td></tr>
<tr><td>Oligo rev</td><td>AAACTGGTGTCTGTCGGCTGCGAG</td><td></td><td></td><td></td><td></td><td></td><td></td><td></td><td></td><td></td><td></td><td></td></tr>
<tr><td>Chromosome</td><td>start</td><td></td><td>end</td><td>strand</td><td>MM</td><td>target_seq</td><td>PAM</td><td>alignment</td><td>distance</td><td>position</td><td>gene name</td><td>gene id</td></tr>
<tr><td>GU305914.1</td><td></td><td>1155</td><td>1177</td><td>-</td><td>0</td><td>CTCGCAGCCGACAGACACCA</td><td>CGG</td><td>||||||||[||||||||||||]PAM</td><td>NA</td><td>Intergenic</td><td>NA</td><td>NA</td></tr>
<tr><td>T34223</td><td>TCGCAGCCGACAGACACCACGGG</td><td></td><td>1000</td><td></td><td></td><td></td><td></td><td></td><td></td><td></td><td></td><td></td></tr>
<tr><td>Oligo fwd</td><td>CACCTCGCAGCCGACAGACACCAC</td><td></td><td></td><td></td><td></td><td></td><td></td><td></td><td></td><td></td><td></td><td></td></tr>
<tr><td>Oligo rev</td><td>AAACGTGGTGTCTGTCGGCTGCGA</td><td></td><td></td><td></td><td></td><td></td><td></td><td></td><td></td><td></td><td></td><td></td></tr>
<tr><td>Chromosome</td><td>start</td><td></td><td>end</td><td>strand</td><td>MM</td><td>target_seq</td><td>PAM</td><td>alignment</td><td>distance</td><td>position</td><td>gene name</td><td>gene id</td></tr>
<tr><td>GU305914.1</td><td></td><td>1154</td><td>1176</td><td>-</td><td>0</td><td>TCGCAGCCGACAGACACCAC</td><td>GGG</td><td>||||||||[||||||||||||]PAM</td><td>NA</td><td>Intergenic</td><td>NA</td><td>NA</td></tr>
<tr><td>T34224</td><td>CGCAGCCGACAGACACCACGGGG</td><td></td><td>1000</td><td></td><td></td><td></td><td></td><td></td><td></td><td></td><td></td><td></td></tr>
<tr><td>Oligo fwd</td><td>CACCCGCAGCCGACAGACACCACG</td><td></td><td></td><td></td><td></td><td></td><td></td><td></td><td></td><td></td><td></td><td></td></tr>
<tr><td>Oligo rev</td><td>AAACCGTGGTGTCTGTCGGCTGCG</td><td></td><td></td><td></td><td></td><td></td><td></td><td></td><td></td><td></td><td></td><td></td></tr>
<tr><td>Chromosome</td><td>start</td><td></td><td>end</td><td>strand</td><td>MM</td><td>target_seq</td><td>PAM</td><td>alignment</td><td>distance</td><td>position</td><td>gene name</td><td>gene id</td></tr>
<tr><td>GU305914.1</td><td></td><td>1153</td><td>1175</td><td>-</td><td>0</td><td>CGCAGCCGACAGACACCACG</td><td>GGG</td><td>||||||||[||||||||||||]PAM</td><td>NA</td><td>Intergenic</td><td>NA</td><td>NA</td></tr>
<tr><td>T34225</td><td>TGCGTCGCTGCGCCTGCCTGCGG</td><td></td><td>1000</td><td></td><td></td><td></td><td></td><td></td><td></td><td></td><td></td><td></td></tr>
<tr><td>Oligo fwd</td><td>CACCTGCGTCGCTGCGCCTGCCTG</td><td></td><td></td><td></td><td></td><td></td><td></td><td></td><td></td><td></td><td></td><td></td></tr>
<tr><td>Oligo rev</td><td>AAACCAGGCAGGCGCAGCGACGCA</td><td></td><td></td><td></td><td></td><td></td><td></td><td></td><td></td><td></td><td></td><td></td></tr>
<tr><td>Chromosome</td><td>start</td><td></td><td>end</td><td>strand</td><td>MM</td><td>target_seq</td><td>PAM</td><td>alignment</td><td>distance</td><td>position</td><td>gene name</td><td>gene id</td></tr>
<tr><td>GU305914.1</td><td></td><td>1064</td><td>1086</td><td>-</td><td>0</td><td>TGCGTCGCTGCGCCTGCCTG</td><td>CGG</td><td>||||||||[||||||||||||]PAM</td><td>NA</td><td>Intergenic</td><td>NA</td><td>NA</td></tr>
</table>

T34226 GCGTCGCTGCGCCTGCCTGCGGG 1000
Oligo fwd CACCGCGTCGCTGCGCCTGCCTGC
Oligo rev AAACGCAGGCAGGCGCAGCGACGC
Chromosome start end strand MM target_seq PAM alignment distance position gene name gene id
GU305914.1 1063 1085 - 0 GCGTCGCTGCGCCTGCCTGC GGG ||||||||[||||||||||||]PAM NA Intergenic NA NA

T34227 CGTCGCTGCGCCTGCCTGCGGGG 1000
Oligo fwd CACCCGTCGCTGCGCCTGCCTGCG
Oligo rev AAACCGCAGGCAGGCGCAGCGACG
Chromosome start end strand MM target_seq PAM alignment distance position gene name gene id
GU305914.1 1062 1084 - 0 CGTCGCTGCGCCTGCCTGCG GGG ||||||||[||||||||||||]PAM NA Intergenic NA NA

T34228 GCACGCTCGTCTCCCGACAGCGG 1000
Oligo fwd CACCGCACGCTCGTCTCCCGACAG
Oligo rev AAACCTGTCGGGAGACGAGCGTGC
Chromosome start end strand MM target_seq PAM alignment distance position gene name gene id
GU305914.1 1040 1062 - 0 GCACGCTCGTCTCCCGACAG CGG ||||||||[||||||||||||]PAM NA Intergenic NA NA

T34229 TCTCCCGACAGCGGCTGCCATGG 1000
Oligo fwd CACCTCTCCCGACAGCGGCTGCCA
Oligo rev AAACTGGCAGCCGCTGTCGGGAGA
Chromosome start end strand MM target_seq PAM alignment distance position gene name gene id
GU305914.1 1031 1053 - 0 TCTCCCGACAGCGGCTGCCA TGG ||||||||[||||||||||||]PAM NA Intergenic NA NA

T34230 CAGCGGCTGCCATGGCACCGTGG 1000
Oligo fwd CACCCAGCGGCTGCCATGGCACCG
Oligo rev AAACCGGTGCCATGGCAGCCGCTG
Chromosome start end strand MM target_seq PAM alignment distance position gene name gene id
GU305914.1 1023 1045 - 0 CAGCGGCTGCCATGGCACCG TGG ||||||||[||||||||||||]PAM NA Intergenic NA NA

T34231 ATTTATCACCTCTGTCAGCTAGG 1000
Oligo fwd CACCATTTATCACCTCTGTCAGCT
Oligo rev AAACAGCTGACAGAGGTGATAAAT
Chromosome start end strand MM target_seq PAM alignment distance position gene name gene id
GU305914.1 996 1018 - 0 ATTTATCACCTCTGTCAGCT AGG ||||||||[||||||||||||]PAM NA Intergenic NA NA

T34232 TTTATCACCTCTGTCAGCTAGGG 1000
Oligo fwd CACCTTTATCACCTCTGTCAGCTA
Oligo rev AAACTAGCTGACAGAGGTGATAAA
Chromosome start end strand MM target_seq PAM alignment distance position gene name gene id
GU305914.1 995 1017 - 0 TTTATCACCTCTGTCAGCTA GGG ||||||||[||||||||||||]PAM NA Intergenic NA NA

T34233 TTATCACCTCTGTCAGCTAGGGG 1000
Oligo fwd CACCTTATCACCTCTGTCAGCTAG
Oligo rev AAACCTAGCTGACAGAGGTGATAA
Chromosome start end strand MM target_seq PAM alignment distance position gene name gene id
GU305914.1 994 1016 - 0 TTATCACCTCTGTCAGCTAG GGG ||||||||[||||||||||||]PAM NA Intergenic NA NA

T34234 TATCACCTCTGTCAGCTAGGGGG 1000
Oligo fwd CACCTATCACCTCTGTCAGCTAGG
Oligo rev AAACCCTAGCTGACAGAGGTGATA
Chromosome start end strand MM target_seq PAM alignment distance position gene name gene id
GU305914.1 993 1015 - 0 TATCACCTCTGTCAGCTAGG GGG ||||||||[||||||||||||]PAM NA Intergenic NA NA

T34235 GTCAGCTAGGGGGAATCAAGCGG 1000
Oligo fwd CACCGTCAGCTAGGGGGAATCAAG
Oligo rev AAACCTTGATTCCCCCTAGCTGAC
Chromosome start end strand MM target_seq PAM alignment distance position gene name gene id
GU305914.1 983 1005 - 0 GTCAGCTAGGGGGAATCAAG CGG ||||||||[||||||||||||]PAM NA Intergenic NA NA

T34236 CTAGGGGGAATCAAGCGGTAAGG 1000
Oligo fwd CACCCTAGGGGGAATCAAGCGGTA
Oligo rev AAACTACCGCTTGATTCCCCCTAG
Chromosome start end strand MM target_seq PAM alignment distance position gene name gene id
GU305914.1 978 1000 - 0 CTAGGGGGAATCAAGCGGTA AGG ||||||||[||||||||||||]PAM NA Intergenic NA NA

T34237 ATCAAGCGGTAAGGTCAGACCGG 1000
Oligo fwd CACCATCAAGCGGTAAGGTCAGAC
Oligo rev AAACGTCTGACCTTACCGCTTGAT
Chromosome start end strand MM target_seq PAM alignment distance position gene name gene id
GU305914.1 969 991 - 0 ATCAAGCGGTAAGGTCAGAC CGG ||||||||[||||||||||||]PAM NA Intergenic NA NA

T34238 AGCGGTAAGGTCAGACCGGACGG 1000
Oligo fwd CACCAGCGGTAAGGTCAGACCGGA
Oligo rev AAACTCCGGTCTGACCTTACCGCT
Chromosome start end strand MM target_seq PAM alignment distance position gene name gene id
GU305914.1 965 987 - 0 AGCGGTAAGGTCAGACCGGA CGG ||||||||[||||||||||||]PAM NA Intergenic NA NA

T34239 GGTAAGGTCAGACCGGACGGAGG 1000
Oligo fwd CACCGGTAAGGTCAGACCGGACGG
Oligo rev AAACCCGTCCGGTCTGACCTTACC
Chromosome start end strand MM target_seq PAM alignment distance position gene name gene id
GU305914.1 962 984 - 0 GGTAAGGTCAGACCGGACGG AGG ||||||||[||||||||||||]PAM NA Intergenic NA NA

T34240 GTAAGGTCAGACCGGACGGAGGG 1000
Oligo fwd CACCGTAAGGTCAGACCGGACGGA
Oligo rev AAACTCCGTCCGGTCTGACCTTAC
Chromosome start end strand MM target_seq PAM alignment distance position gene name gene id

<table>
<tr><td>GU305914.1</td><td></td><td>961</td><td>983</td><td>-</td><td>0</td><td>GTAAGGTCAGACCGGACGGA</td><td>GGG</td><td>||||||||[||||||||||||]PAM</td><td>NA</td><td>Intergenic</td><td>NA</td><td>NA</td></tr>
<tr><td>T34241</td><td>AGGTCAGACCGGACGGAGGGAGG</td><td></td><td>1000</td><td></td><td></td><td></td><td></td><td></td><td></td><td></td><td></td><td></td></tr>
<tr><td>Oligo fwd</td><td>CACCAGGTCAGACCGGACGGAGGG</td><td></td><td></td><td></td><td></td><td></td><td></td><td></td><td></td><td></td><td></td><td></td></tr>
<tr><td>Oligo rev</td><td>AAACCCCTCCGTCCGGTCTGACCT</td><td></td><td></td><td></td><td></td><td></td><td></td><td></td><td></td><td></td><td></td><td></td></tr>
<tr><td>Chromosome</td><td>start</td><td></td><td>end</td><td>strand</td><td>MM</td><td>target_seq</td><td>PAM</td><td>alignment</td><td>distance</td><td>position</td><td>gene name</td><td>gene id</td></tr>
<tr><td>GU305914.1</td><td></td><td>958</td><td>980</td><td>-</td><td>0</td><td>AGGTCAGACCGGACGGAGGG</td><td>AGG</td><td>||||||||[||||||||||||]PAM</td><td>NA</td><td>Intergenic</td><td>NA</td><td>NA</td></tr>
<tr><td>T34242</td><td>GGTCAGACCGGACGGAGGGAGGG</td><td></td><td>1000</td><td></td><td></td><td></td><td></td><td></td><td></td><td></td><td></td><td></td></tr>
<tr><td>Oligo fwd</td><td>CACCGGTCAGACCGGACGGAGGGA</td><td></td><td></td><td></td><td></td><td></td><td></td><td></td><td></td><td></td><td></td><td></td></tr>
<tr><td>Oligo rev</td><td>AAACTCCCTCCGTCCGGTCTGACC</td><td></td><td></td><td></td><td></td><td></td><td></td><td></td><td></td><td></td><td></td><td></td></tr>
<tr><td>Chromosome</td><td>start</td><td></td><td>end</td><td>strand</td><td>MM</td><td>target_seq</td><td>PAM</td><td>alignment</td><td>distance</td><td>position</td><td>gene name</td><td>gene id</td></tr>
<tr><td>GU305914.1</td><td></td><td>957</td><td>979</td><td>-</td><td>0</td><td>GGTCAGACCGGACGGAGGGA</td><td>GGG</td><td>||||||||[||||||||||||]PAM</td><td>NA</td><td>Intergenic</td><td>NA</td><td>NA</td></tr>
<tr><td>T34243</td><td>ACGAGTGACAACAGCCTACTTGG</td><td></td><td>1000</td><td></td><td></td><td></td><td></td><td></td><td></td><td></td><td></td><td></td></tr>
<tr><td>Oligo fwd</td><td>CACCACGAGTGACAACAGCCTACT</td><td></td><td></td><td></td><td></td><td></td><td></td><td></td><td></td><td></td><td></td><td></td></tr>
<tr><td>Oligo rev</td><td>AAACAGTAGGCTGTTGTCACTCGT</td><td></td><td></td><td></td><td></td><td></td><td></td><td></td><td></td><td></td><td></td><td></td></tr>
<tr><td>Chromosome</td><td>start</td><td></td><td>end</td><td>strand</td><td>MM</td><td>target_seq</td><td>PAM</td><td>alignment</td><td>distance</td><td>position</td><td>gene name</td><td>gene id</td></tr>
<tr><td>GU305914.1</td><td></td><td>930</td><td>952</td><td>-</td><td>0</td><td>ACGAGTGACAACAGCCTACT</td><td>TGG</td><td>||||||||[||||||||||||]PAM</td><td>NA</td><td>Intergenic</td><td>NA</td><td>NA</td></tr>
<tr><td>T34244</td><td>CGAGTGACAACAGCCTACTTGGG</td><td></td><td>1000</td><td></td><td></td><td></td><td></td><td></td><td></td><td></td><td></td><td></td></tr>
<tr><td>Oligo fwd</td><td>CACCCGAGTGACAACAGCCTACTT</td><td></td><td></td><td></td><td></td><td></td><td></td><td></td><td></td><td></td><td></td><td></td></tr>
<tr><td>Oligo rev</td><td>AAACAAGTAGGCTGTTGTCACTCG</td><td></td><td></td><td></td><td></td><td></td><td></td><td></td><td></td><td></td><td></td><td></td></tr>
<tr><td>Chromosome</td><td>start</td><td></td><td>end</td><td>strand</td><td>MM</td><td>target_seq</td><td>PAM</td><td>alignment</td><td>distance</td><td>position</td><td>gene name</td><td>gene id</td></tr>
<tr><td>GU305914.1</td><td></td><td>929</td><td>951</td><td>-</td><td>0</td><td>CGAGTGACAACAGCCTACTT</td><td>GGG</td><td>||||||||[||||||||||||]PAM</td><td>NA</td><td>Intergenic</td><td>NA</td><td>NA</td></tr>
<tr><td>T34245</td><td>GTGACAACAGCCTACTTGGGTGG</td><td></td><td>1000</td><td></td><td></td><td></td><td></td><td></td><td></td><td></td><td></td><td></td></tr>
<tr><td>Oligo fwd</td><td>CACCGTGACAACAGCCTACTTGGG</td><td></td><td></td><td></td><td></td><td></td><td></td><td></td><td></td><td></td><td></td><td></td></tr>
<tr><td>Oligo rev</td><td>AAACCCCAAGTAGGCTGTTGTCAC</td><td></td><td></td><td></td><td></td><td></td><td></td><td></td><td></td><td></td><td></td><td></td></tr>
<tr><td>Chromosome</td><td>start</td><td></td><td>end</td><td>strand</td><td>MM</td><td>target_seq</td><td>PAM</td><td>alignment</td><td>distance</td><td>position</td><td>gene name</td><td>gene id</td></tr>
<tr><td>GU305914.1</td><td></td><td>926</td><td>948</td><td>-</td><td>0</td><td>GTGACAACAGCCTACTTGGG</td><td>TGG</td><td>||||||||[||||||||||||]PAM</td><td>NA</td><td>Intergenic</td><td>NA</td><td>NA</td></tr>
<tr><td>T34246</td><td>TGACAACAGCCTACTTGGGTGGG</td><td></td><td>1000</td><td></td><td></td><td></td><td></td><td></td><td></td><td></td><td></td><td></td></tr>
<tr><td>Oligo fwd</td><td>CACCTGACAACAGCCTACTTGGGT</td><td></td><td></td><td></td><td></td><td></td><td></td><td></td><td></td><td></td><td></td><td></td></tr>
<tr><td>Oligo rev</td><td>AAACACCCAAGTAGGCTGTTGTCA</td><td></td><td></td><td></td><td></td><td></td><td></td><td></td><td></td><td></td><td></td><td></td></tr>
<tr><td>Chromosome</td><td>start</td><td></td><td>end</td><td>strand</td><td>MM</td><td>target_seq</td><td>PAM</td><td>alignment</td><td>distance</td><td>position</td><td>gene name</td><td>gene id</td></tr>
<tr><td>GU305914.1</td><td></td><td>925</td><td>947</td><td>-</td><td>0</td><td>TGACAACAGCCTACTTGGGT</td><td>GGG</td><td>||||||||[||||||||||||]PAM</td><td>NA</td><td>Intergenic</td><td>NA</td><td>NA</td></tr>
<tr><td>T34247</td><td>GCCTACTTGGGTGGGATCCTCGG</td><td></td><td>1000</td><td></td><td></td><td></td><td></td><td></td><td></td><td></td><td></td><td></td></tr>
<tr><td>Oligo fwd</td><td>CACCGCCTACTTGGGTGGGATCCT</td><td></td><td></td><td></td><td></td><td></td><td></td><td></td><td></td><td></td><td></td><td></td></tr>
<tr><td>Oligo rev</td><td>AAACAGGATCCCACCCAAGTAGGC</td><td></td><td></td><td></td><td></td><td></td><td></td><td></td><td></td><td></td><td></td><td></td></tr>
<tr><td>Chromosome</td><td>start</td><td></td><td>end</td><td>strand</td><td>MM</td><td>target_seq</td><td>PAM</td><td>alignment</td><td>distance</td><td>position</td><td>gene name</td><td>gene id</td></tr>
<tr><td>GU305914.1</td><td></td><td>917</td><td>939</td><td>-</td><td>0</td><td>GCCTACTTGGGTGGGATCCT</td><td>CGG</td><td>||||||||[||||||||||||]PAM</td><td>NA</td><td>Intergenic</td><td>NA</td><td>NA</td></tr>
<tr><td>T34248</td><td>CCTACTTGGGTGGGATCCTCGGG</td><td></td><td>1000</td><td></td><td></td><td></td><td></td><td></td><td></td><td></td><td></td><td></td></tr>
<tr><td>Oligo fwd</td><td>CACCCCTACTTGGGTGGGATCCTC</td><td></td><td></td><td></td><td></td><td></td><td></td><td></td><td></td><td></td><td></td><td></td></tr>
<tr><td>Oligo rev</td><td>AAACGAGGATCCCACCCAAGTAGG</td><td></td><td></td><td></td><td></td><td></td><td></td><td></td><td></td><td></td><td></td><td></td></tr>
<tr><td>Chromosome</td><td>start</td><td></td><td>end</td><td>strand</td><td>MM</td><td>target_seq</td><td>PAM</td><td>alignment</td><td>distance</td><td>position</td><td>gene name</td><td>gene id</td></tr>
<tr><td>GU305914.1</td><td></td><td>916</td><td>938</td><td>-</td><td>0</td><td>CCTACTTGGGTGGGATCCTC</td><td>GGG</td><td>||||||||[||||||||||||]PAM</td><td>NA</td><td>Intergenic</td><td>NA</td><td>NA</td></tr>
<tr><td>T34249</td><td>CTACTTGGGTGGGATCCTCGGGG</td><td></td><td>1000</td><td></td><td></td><td></td><td></td><td></td><td></td><td></td><td></td><td></td></tr>
<tr><td>Oligo fwd</td><td>CACCCTACTTGGGTGGGATCCTCG</td><td></td><td></td><td></td><td></td><td></td><td></td><td></td><td></td><td></td><td></td><td></td></tr>
<tr><td>Oligo rev</td><td>AAACCGAGGATCCCACCCAAGTAG</td><td></td><td></td><td></td><td></td><td></td><td></td><td></td><td></td><td></td><td></td><td></td></tr>
<tr><td>Chromosome</td><td>start</td><td></td><td>end</td><td>strand</td><td>MM</td><td>target_seq</td><td>PAM</td><td>alignment</td><td>distance</td><td>position</td><td>gene name</td><td>gene id</td></tr>
<tr><td>GU305914.1</td><td></td><td>915</td><td>937</td><td>-</td><td>0</td><td>CTACTTGGGTGGGATCCTCG</td><td>GGG</td><td>||||||||[||||||||||||]PAM</td><td>NA</td><td>Intergenic</td><td>NA</td><td>NA</td></tr>
<tr><td>T34250</td><td>CTTGGGTGGGATCCTCGGGGAGG</td><td></td><td>1000</td><td></td><td></td><td></td><td></td><td></td><td></td><td></td><td></td><td></td></tr>
<tr><td>Oligo fwd</td><td>CACCCTTGGGTGGGATCCTCGGGG</td><td></td><td></td><td></td><td></td><td></td><td></td><td></td><td></td><td></td><td></td><td></td></tr>
<tr><td>Oligo rev</td><td>AAACCCCCGAGGATCCCACCCAAG</td><td></td><td></td><td></td><td></td><td></td><td></td><td></td><td></td><td></td><td></td><td></td></tr>
<tr><td>Chromosome</td><td>start</td><td></td><td>end</td><td>strand</td><td>MM</td><td>target_seq</td><td>PAM</td><td>alignment</td><td>distance</td><td>position</td><td>gene name</td><td>gene id</td></tr>
<tr><td>GU305914.1</td><td></td><td>912</td><td>934</td><td>-</td><td>0</td><td>CTTGGGTGGGATCCTCGGGG</td><td>AGG</td><td>||||||||[||||||||||||]PAM</td><td>NA</td><td>Intergenic</td><td>NA</td><td>NA</td></tr>
<tr><td>T34251</td><td>ACACGCTCGAGATCGACTTGCGG</td><td></td><td>1000</td><td></td><td></td><td></td><td></td><td></td><td></td><td></td><td></td><td></td></tr>
<tr><td>Oligo fwd</td><td>CACCACACGCTCGAGATCGACTTG</td><td></td><td></td><td></td><td></td><td></td><td></td><td></td><td></td><td></td><td></td><td></td></tr>
<tr><td>Oligo rev</td><td>AAACCAAGTCGATCTCGAGCGTGT</td><td></td><td></td><td></td><td></td><td></td><td></td><td></td><td></td><td></td><td></td><td></td></tr>
<tr><td>Chromosome</td><td>start</td><td></td><td>end</td><td>strand</td><td>MM</td><td>target_seq</td><td>PAM</td><td>alignment</td><td>distance</td><td>position</td><td>gene name</td><td>gene id</td></tr>
<tr><td>GU305914.1</td><td></td><td>882</td><td>904</td><td>-</td><td>0</td><td>ACACGCTCGAGATCGACTTG</td><td>CGG</td><td>||||||||[||||||||||||]PAM</td><td>NA</td><td>Intergenic</td><td>NA</td><td>NA</td></tr>
<tr><td>T34252</td><td>CACGCTCGAGATCGACTTGCGGG</td><td></td><td>1000</td><td></td><td></td><td></td><td></td><td></td><td></td><td></td><td></td><td></td></tr>
<tr><td>Oligo fwd</td><td>CACCCACGCTCGAGATCGACTTGC</td><td></td><td></td><td></td><td></td><td></td><td></td><td></td><td></td><td></td><td></td><td></td></tr>
<tr><td>Oligo rev</td><td>AAACGCAAGTCGATCTCGAGCGTG</td><td></td><td></td><td></td><td></td><td></td><td></td><td></td><td></td><td></td><td></td><td></td></tr>
<tr><td>Chromosome</td><td>start</td><td></td><td>end</td><td>strand</td><td>MM</td><td>target_seq</td><td>PAM</td><td>alignment</td><td>distance</td><td>position</td><td>gene name</td><td>gene id</td></tr>
<tr><td>GU305914.1</td><td></td><td>881</td><td>903</td><td>-</td><td>0</td><td>CACGCTCGAGATCGACTTGC</td><td>GGG</td><td>||||||||[||||||||||||]PAM</td><td>NA</td><td>Intergenic</td><td>NA</td><td>NA</td></tr>
<tr><td>T34253</td><td>ATCGACTTGCGGGTACGTTGCGG</td><td></td><td>1000</td><td></td><td></td><td></td><td></td><td></td><td></td><td></td><td></td><td></td></tr>
<tr><td>Oligo fwd</td><td>CACCATCGACTTGCGGGTACGTTG</td><td></td><td></td><td></td><td></td><td></td><td></td><td></td><td></td><td></td><td></td><td></td></tr>
<tr><td>Oligo rev</td><td>AAACCAACGTACCCGCAAGTCGAT</td><td></td><td></td><td></td><td></td><td></td><td></td><td></td><td></td><td></td><td></td><td></td></tr>
<tr><td>Chromosome</td><td>start</td><td></td><td>end</td><td>strand</td><td>MM</td><td>target_seq</td><td>PAM</td><td>alignment</td><td>distance</td><td>position</td><td>gene name</td><td>gene id</td></tr>
<tr><td>GU305914.1</td><td></td><td>871</td><td>893</td><td>-</td><td>0</td><td>ATCGACTTGCGGGTACGTTG</td><td>CGG</td><td>||||||||[||||||||||||]PAM</td><td>NA</td><td>Intergenic</td><td>NA</td><td>NA</td></tr>
<tr><td>T34254</td><td>ACGTTGCGGATGAAGAGAATCGG</td><td></td><td>1000</td><td></td><td></td><td></td><td></td><td></td><td></td><td></td><td></td><td></td></tr>
<tr><td>Oligo fwd</td><td>CACCACGTTGCGGATGAAGAGAAT</td><td></td><td></td><td></td><td></td><td></td><td></td><td></td><td></td><td></td><td></td><td></td></tr>
<tr><td>Oligo rev</td><td>AAACATTCTCTTCATCCGCAACGT</td><td></td><td></td><td></td><td></td><td></td><td></td><td></td><td></td><td></td><td></td><td></td></tr>
<tr><td>Chromosome</td><td>start</td><td></td><td>end</td><td>strand</td><td>MM</td><td>target_seq</td><td>PAM</td><td>alignment</td><td>distance</td><td>position</td><td>gene name</td><td>gene id</td></tr>
<tr><td>GU305914.1</td><td></td><td>857</td><td>879</td><td>-</td><td>0</td><td>ACGTTGCGGATGAAGAGAAT</td><td>CGG</td><td>||||||||[||||||||||||]PAM</td><td>NA</td><td>Intergenic</td><td>NA</td><td>NA</td></tr>
<tr><td>T34255</td><td>CGTTGCGGATGAAGAGAATCGGG</td><td></td><td>1000</td><td></td><td></td><td></td><td></td><td></td><td></td><td></td><td></td><td></td></tr>
<tr><td>Oligo fwd</td><td>CACCCGTTGCGGATGAAGAGAATC</td><td></td><td></td><td></td><td></td><td></td><td></td><td></td><td></td><td></td><td></td><td></td></tr>
<tr><td>Oligo rev</td><td>AAACGATTCTCTTCATCCGCAACG</td><td></td><td></td><td></td><td></td><td></td><td></td><td></td><td></td><td></td><td></td><td></td></tr>
</table>

Chromosome start end strand MM target_seq PAM alignment distance position gene name gene id
GU305914.1 856 878 - 0 CGTTGCGGATGAAGAGAATC GGG ||||||||[||||||||||||]PAM NA Intergenic NA NA

T34256 GCGGATGAAGAGAATCGGGTTGG 1000
Oligo fwd CACCGCGGATGAAGAGAATCGGGT
Oligo rev AAACACCCGATTCTCTTCATCCGC
Chromosome start end strand MM target_seq PAM alignment distance position gene name gene id
GU305914.1 852 874 - 0 GCGGATGAAGAGAATCGGGT TGG ||||||||[||||||||||||]PAM NA Intergenic NA NA

T34257 TGAAGAGAATCGGGTTGGAACGG 1000
Oligo fwd CACCTGAAGAGAATCGGGTTGGAA
Oligo rev AAACTTCCAACCCGATTCTCTTCA
Chromosome start end strand MM target_seq PAM alignment distance position gene name gene id
GU305914.1 847 869 - 0 TGAAGAGAATCGGGTTGGAA CGG ||||||||[||||||||||||]PAM NA Intergenic NA NA

T34258 GTACGAGCTACCGTTCGACACGG 1000
Oligo fwd CACCGTACGAGCTACCGTTCGACA
Oligo rev AAACTGTCGAACGGTAGCTCGTAC
Chromosome start end strand MM target_seq PAM alignment distance position gene name gene id
GU305914.1 814 836 - 0 GTACGAGCTACCGTTCGACA CGG ||||||||[||||||||||||]PAM NA Intergenic NA NA

T34259 ACCCCCGACCCAAAACCCCGCGG 1000
Oligo fwd CACCACCCCCGACCCAAAACCCCG
Oligo rev AAACCGGGGTTTTGGGTCGGGGGT
Chromosome start end strand MM target_seq PAM alignment distance position gene name gene id
GU305914.1 737 759 - 0 ACCCCCGACCCAAAACCCCG CGG ||||||||[||||||||||||]PAM NA Intergenic NA NA

T34260 CCCCCGACCCAAAACCCCGCGGG 1000
Oligo fwd CACCCCCCCGACCCAAAACCCCGC
Oligo rev AAACGCGGGGTTTTGGGTCGGGGG
Chromosome start end strand MM target_seq PAM alignment distance position gene name gene id
GU305914.1 736 758 - 0 CCCCCGACCCAAAACCCCGC GGG ||||||||[||||||||||||]PAM NA Intergenic NA NA

T34261 CCCCGACCCAAAACCCCGCGGGG 1000
Oligo fwd CACCCCCCGACCCAAAACCCCGCG
Oligo rev AAACCGCGGGGTTTTGGGTCGGGG
Chromosome start end strand MM target_seq PAM alignment distance position gene name gene id
GU305914.1 735 757 - 0 CCCCGACCCAAAACCCCGCG GGG ||||||||[||||||||||||]PAM NA Intergenic NA NA

T34262 CCCGACCCAAAACCCCGCGGGGG 1000
Oligo fwd CACCCCCGACCCAAAACCCCGCGG
Oligo rev AAACCCGCGGGGTTTTGGGTCGGG
Chromosome start end strand MM target_seq PAM alignment distance position gene name gene id
GU305914.1 734 756 - 0 CCCGACCCAAAACCCCGCGG GGG ||||||||[||||||||||||]PAM NA Intergenic NA NA

T34263 CCGACCCAAAACCCCGCGGGGGG 1000
Oligo fwd CACCCCGACCCAAAACCCCGCGGG
Oligo rev AAACCCCGCGGGGTTTTGGGTCGG
Chromosome start end strand MM target_seq PAM alignment distance position gene name gene id
GU305914.1 733 755 - 0 CCGACCCAAAACCCCGCGGG GGG ||||||||[||||||||||||]PAM NA Intergenic NA NA

T34264 CGACCCAAAACCCCGCGGGGGGG 1000
Oligo fwd CACCCGACCCAAAACCCCGCGGGG
Oligo rev AAACCCCCGCGGGGTTTTGGGTCG
Chromosome start end strand MM target_seq PAM alignment distance position gene name gene id
GU305914.1 732 754 - 0 CGACCCAAAACCCCGCGGGG GGG ||||||||[||||||||||||]PAM NA Intergenic NA NA

T34265 GACCCAAAACCCCGCGGGGGGGG 1000
Oligo fwd CACCGACCCAAAACCCCGCGGGGG
Oligo rev AAACCCCCCGCGGGGTTTTGGGTC
Chromosome start end strand MM target_seq PAM alignment distance position gene name gene id
GU305914.1 731 753 - 0 GACCCAAAACCCCGCGGGGG GGG ||||||||[||||||||||||]PAM NA Intergenic NA NA

T34266 ACCCAAAACCCCGCGGGGGGGGG 1000
Oligo fwd CACCACCCAAAACCCCGCGGGGGG
Oligo rev AAACCCCCCCGCGGGGTTTTGGGT
Chromosome start end strand MM target_seq PAM alignment distance position gene name gene id
GU305914.1 730 752 - 0 ACCCAAAACCCCGCGGGGGG GGG ||||||||[||||||||||||]PAM NA Intergenic NA NA

T34267 CCCAAAACCCCGCGGGGGGGGGG 1000
Oligo fwd CACCCCCAAAACCCCGCGGGGGGG
Oligo rev AAACCCCCCCCGCGGGGTTTTGGG
Chromosome start end strand MM target_seq PAM alignment distance position gene name gene id
GU305914.1 729 751 - 0 CCCAAAACCCCGCGGGGGGG GGG ||||||||[||||||||||||]PAM NA Intergenic NA NA

T34268 GGTGCCCCGCCAATACTCCCCGG 1000
Oligo fwd CACCGGTGCCCCGCCAATACTCCC
Oligo rev AAACGGGAGTATTGGCGGGGCACC
Chromosome start end strand MM target_seq PAM alignment distance position gene name gene id
GU305914.1 708 730 - 0 GGTGCCCCGCCAATACTCCC CGG ||||||||[||||||||||||]PAM NA Intergenic NA NA

T34269 GTGCCCCGCCAATACTCCCCGGG 1000
Oligo fwd CACCGTGCCCCGCCAATACTCCCC
Oligo rev AAACGGGGAGTATTGGCGGGGCAC
Chromosome start end strand MM target_seq PAM alignment distance position gene name gene id
GU305914.1 707 729 - 0 GTGCCCCGCCAATACTCCCC GGG ||||||||[||||||||||||]PAM NA Intergenic NA NA

T34270 TTATACCCGCCCCCCCACCCCGG 1000
Oligo fwd CACCTTATACCCGCCCCCCCACCC

Oligo rev AAACGGGTGGGGGGGCGGGTATAA
Chromosome start end strand MM target_seq PAM alignment distance position gene name gene id
GU305914.1 682 704 - 0 TTATACCCGCCCCCCCACCC CGG ||||||||[||||||||||||]PAM NA Intergenic NA NA

T34271 TATACCCGCCCCCCCACCCCGGG 1000
Oligo fwd CACCTATACCCGCCCCCCCACCCC
Oligo rev AAACGGGGTGGGGGGGCGGGTATA
Chromosome start end strand MM target_seq PAM alignment distance position gene name gene id
GU305914.1 681 703 - 0 TATACCCGCCCCCCCACCCC GGG ||||||||[||||||||||||]PAM NA Intergenic NA NA

T34272 ATACCCGCCCCCCCACCCCGGGG 1000
Oligo fwd CACCATACCCGCCCCCCCACCCCG
Oligo rev AAACCGGGGTGGGGGGGCGGGTAT
Chromosome start end strand MM target_seq PAM alignment distance position gene name gene id
GU305914.1 680 702 - 0 ATACCCGCCCCCCCACCCCG GGG ||||||||[||||||||||||]PAM NA Intergenic NA NA

T34273 CCACCCCGGGGTCCGAAGAGCGG 1000
Oligo fwd CACCCCACCCCGGGGTCCGAAGAG
Oligo rev AAACCTCTTCGGACCCCGGGGTGG
Chromosome start end strand MM target_seq PAM alignment distance position gene name gene id
GU305914.1 668 690 - 0 CCACCCCGGGGTCCGAAGAG CGG ||||||||[||||||||||||]PAM NA Intergenic NA NA

T34274 GACTGTTGCCTCGAAACATACGG 1000
Oligo fwd CACCGACTGTTGCCTCGAAACATA
Oligo rev AAACTATGTTTCGAGGCAACAGTC
Chromosome start end strand MM target_seq PAM alignment distance position gene name gene id
GU305914.1 639 661 - 0 GACTGTTGCCTCGAAACATA CGG ||||||||[||||||||||||]PAM NA Intergenic NA NA

T34275 TGTTGCCTCGAAACATACGGCGG 1000
Oligo fwd CACCTGTTGCCTCGAAACATACGG
Oligo rev AAACCCGTATGTTTCGAGGCAACA
Chromosome start end strand MM target_seq PAM alignment distance position gene name gene id
GU305914.1 636 658 - 0 TGTTGCCTCGAAACATACGG CGG ||||||||[||||||||||||]PAM NA Intergenic NA NA

T34276 GAAACATACGGCGGCGCTCGAGG 1000
Oligo fwd CACCGAAACATACGGCGGCGCTCG
Oligo rev AAACCGAGCGCCGCCGTATGTTTC
Chromosome start end strand MM target_seq PAM alignment distance position gene name gene id
GU305914.1 627 649 - 0 GAAACATACGGCGGCGCTCG AGG ||||||||[||||||||||||]PAM NA Intergenic NA NA

T34277 CTCGAGGAACGCTCGCTTCACGG 1000
Oligo fwd CACCCTCGAGGAACGCTCGCTTCA
Oligo rev AAACTGAAGCGAGCGTTCCTCGAG
Chromosome start end strand MM target_seq PAM alignment distance position gene name gene id
GU305914.1 611 633 - 0 CTCGAGGAACGCTCGCTTCA CGG ||||||||[||||||||||||]PAM NA Intergenic NA NA

T34278 ACGCTCGCTTCACGGCTCTCCGG 1000
Oligo fwd CACCACGCTCGCTTCACGGCTCTC
Oligo rev AAACGAGAGCCGTGAAGCGAGCGT
Chromosome start end strand MM target_seq PAM alignment distance position gene name gene id
GU305914.1 603 625 - 0 ACGCTCGCTTCACGGCTCTC CGG ||||||||[||||||||||||]PAM NA Intergenic NA NA
GU305914.1 656 678 + 4 ACAGTCGCTTCACCGCTCTT CGG ||--||||[|||||-|||||-]PAM NA Intergenic NA NA

T34279 CGCTCGCTTCACGGCTCTCCGGG 1000
Oligo fwd CACCCGCTCGCTTCACGGCTCTCC
Oligo rev AAACGGAGAGCCGTGAAGCGAGCG
Chromosome start end strand MM target_seq PAM alignment distance position gene name gene id
GU305914.1 602 624 - 0 CGCTCGCTTCACGGCTCTCC GGG ||||||||[||||||||||||]PAM NA Intergenic NA NA

T34280 CGCTTCACGGCTCTCCGGGCCGG 1000
Oligo fwd CACCCGCTTCACGGCTCTCCGGGC
Oligo rev AAACGCCCGGAGAGCCGTGAAGCG
Chromosome start end strand MM target_seq PAM alignment distance position gene name gene id
GU305914.1 598 620 - 0 CGCTTCACGGCTCTCCGGGC CGG ||||||||[||||||||||||]PAM NA Intergenic NA NA

T34281 GCTTCACGGCTCTCCGGGCCGGG 1000
Oligo fwd CACCGCTTCACGGCTCTCCGGGCC
Oligo rev AAACGGCCCGGAGAGCCGTGAAGC
Chromosome start end strand MM target_seq PAM alignment distance position gene name gene id
GU305914.1 597 619 - 0 GCTTCACGGCTCTCCGGGCC GGG ||||||||[||||||||||||]PAM NA Intergenic NA NA

T34282 GGGCCGGGCGCCGCTCGTCGCGG 1000
Oligo fwd CACCGGGCCGGGCGCCGCTCGTCG
Oligo rev AAACCGACGAGCGGCGCCCGGCCC
Chromosome start end strand MM target_seq PAM alignment distance position gene name gene id
GU305914.1 582 604 - 0 GGGCCGGGCGCCGCTCGTCG CGG ||||||||[||||||||||||]PAM NA Intergenic NA NA

T34283 GTCGCGGAAGCTTCACACGCTGG 1000
Oligo fwd CACCGTCGCGGAAGCTTCACACGC
Oligo rev AAACGCGTGTGAAGCTTCCGCGAC
Chromosome start end strand MM target_seq PAM alignment distance position gene name gene id
GU305914.1 566 588 - 0 GTCGCGGAAGCTTCACACGC TGG ||||||||[||||||||||||]PAM NA Intergenic NA NA

T34284 GCGGAAGCTTCACACGCTGGCGG 1000
Oligo fwd CACCGCGGAAGCTTCACACGCTGG
Oligo rev AAACCCAGCGTGTGAAGCTTCCGC
Chromosome start end strand MM target_seq PAM alignment distance position gene name gene id
GU305914.1 563 585 - 0 GCGGAAGCTTCACACGCTGG CGG ||||||||[||||||||||||]PAM NA Intergenic NA NA

<table>
<tr><td>T34285</td><td>CGGAAGCTTCACACGCTGGCGGG</td><td>1000</td><td></td><td></td><td></td><td></td><td></td><td></td><td></td><td></td><td></td></tr>
<tr><td>Oligo fwd</td><td>CACCCGGAAGCTTCACACGCTGGC</td><td></td><td></td><td></td><td></td><td></td><td></td><td></td><td></td><td></td><td></td></tr>
<tr><td>Oligo rev</td><td>AAACGCCAGCGTGTGAAGCTTCCG</td><td></td><td></td><td></td><td></td><td></td><td></td><td></td><td></td><td></td><td></td></tr>
<tr><td>Chromosome</td><td>start</td><td>end</td><td>strand</td><td>MM</td><td>target_seq</td><td>PAM</td><td>alignment</td><td>distance</td><td>position</td><td>gene name</td><td>gene id</td></tr>
<tr><td>GU305914.1</td><td>562</td><td>584</td><td>-</td><td>0</td><td>CGGAAGCTTCACACGCTGGC</td><td>GGG</td><td>||||||||[||||||||||||]PAM</td><td>NA</td><td>Intergenic</td><td>NA</td><td>NA</td></tr>
<tr><td>T34286</td><td>GGAAGCTTCACACGCTGGCGGGG</td><td>1000</td><td></td><td></td><td></td><td></td><td></td><td></td><td></td><td></td><td></td></tr>
<tr><td>Oligo fwd</td><td>CACCGGAAGCTTCACACGCTGGCG</td><td></td><td></td><td></td><td></td><td></td><td></td><td></td><td></td><td></td><td></td></tr>
<tr><td>Oligo rev</td><td>AAACCGCCAGCGTGTGAAGCTTCC</td><td></td><td></td><td></td><td></td><td></td><td></td><td></td><td></td><td></td><td></td></tr>
<tr><td>Chromosome</td><td>start</td><td>end</td><td>strand</td><td>MM</td><td>target_seq</td><td>PAM</td><td>alignment</td><td>distance</td><td>position</td><td>gene name</td><td>gene id</td></tr>
<tr><td>GU305914.1</td><td>561</td><td>583</td><td>-</td><td>0</td><td>GGAAGCTTCACACGCTGGCG</td><td>GGG</td><td>||||||||[||||||||||||]PAM</td><td>NA</td><td>Intergenic</td><td>NA</td><td>NA</td></tr>
<tr><td>T34287</td><td>CGCCTCACCCCGCCTTCCCGCGG</td><td>1000</td><td></td><td></td><td></td><td></td><td></td><td></td><td></td><td></td><td></td></tr>
<tr><td>Oligo fwd</td><td>CACCCGCCTCACCCCGCCTTCCCG</td><td></td><td></td><td></td><td></td><td></td><td></td><td></td><td></td><td></td><td></td></tr>
<tr><td>Oligo rev</td><td>AAACCGGGAAGGCGGGGTGAGGCG</td><td></td><td></td><td></td><td></td><td></td><td></td><td></td><td></td><td></td><td></td></tr>
<tr><td>Chromosome</td><td>start</td><td>end</td><td>strand</td><td>MM</td><td>target_seq</td><td>PAM</td><td>alignment</td><td>distance</td><td>position</td><td>gene name</td><td>gene id</td></tr>
<tr><td>GU305914.1</td><td>538</td><td>560</td><td>-</td><td>0</td><td>CGCCTCACCCCGCCTTCCCG</td><td>CGG</td><td>||||||||[||||||||||||]PAM</td><td>NA</td><td>Intergenic</td><td>NA</td><td>NA</td></tr>
<tr><td>T34288</td><td>ACCCCGCCTTCCCGCGGACGCGG</td><td>1000</td><td></td><td></td><td></td><td></td><td></td><td></td><td></td><td></td><td></td></tr>
<tr><td>Oligo fwd</td><td>CACCACCCCGCCTTCCCGCGGACG</td><td></td><td></td><td></td><td></td><td></td><td></td><td></td><td></td><td></td><td></td></tr>
<tr><td>Oligo rev</td><td>AAACCGTCCGCGGGAAGGCGGGGT</td><td></td><td></td><td></td><td></td><td></td><td></td><td></td><td></td><td></td><td></td></tr>
<tr><td>Chromosome</td><td>start</td><td>end</td><td>strand</td><td>MM</td><td>target_seq</td><td>PAM</td><td>alignment</td><td>distance</td><td>position</td><td>gene name</td><td>gene id</td></tr>
<tr><td>GU305914.1</td><td>532</td><td>554</td><td>-</td><td>0</td><td>ACCCCGCCTTCCCGCGGACG</td><td>CGG</td><td>||||||||[||||||||||||]PAM</td><td>NA</td><td>Intergenic</td><td>NA</td><td>NA</td></tr>
<tr><td>T34289</td><td>CTTCCCGCGGACGCGGCGCGCGG</td><td>1000</td><td></td><td></td><td></td><td></td><td></td><td></td><td></td><td></td><td></td></tr>
<tr><td>Oligo fwd</td><td>CACCCTTCCCGCGGACGCGGCGCG</td><td></td><td></td><td></td><td></td><td></td><td></td><td></td><td></td><td></td><td></td></tr>
<tr><td>Oligo rev</td><td>AAACCGCGCCGCGTCCGCGGGAAG</td><td></td><td></td><td></td><td></td><td></td><td></td><td></td><td></td><td></td><td></td></tr>
<tr><td>Chromosome</td><td>start</td><td>end</td><td>strand</td><td>MM</td><td>target_seq</td><td>PAM</td><td>alignment</td><td>distance</td><td>position</td><td>gene name</td><td>gene id</td></tr>
<tr><td>GU305914.1</td><td>525</td><td>547</td><td>-</td><td>0</td><td>CTTCCCGCGGACGCGGCGCG</td><td>CGG</td><td>||||||||[||||||||||||]PAM</td><td>NA</td><td>Intergenic</td><td>NA</td><td>NA</td></tr>
<tr><td>T34290</td><td>CGCGGACGCGGCGCGCGGCCCGG</td><td>1000</td><td></td><td></td><td></td><td></td><td></td><td></td><td></td><td></td><td></td></tr>
<tr><td>Oligo fwd</td><td>CACCCGCGGACGCGGCGCGCGGCC</td><td></td><td></td><td></td><td></td><td></td><td></td><td></td><td></td><td></td><td></td></tr>
<tr><td>Oligo rev</td><td>AAACGGCCGCGCGCCGCGTCCGCG</td><td></td><td></td><td></td><td></td><td></td><td></td><td></td><td></td><td></td><td></td></tr>
<tr><td>Chromosome</td><td>start</td><td>end</td><td>strand</td><td>MM</td><td>target_seq</td><td>PAM</td><td>alignment</td><td>distance</td><td>position</td><td>gene name</td><td>gene id</td></tr>
<tr><td>GU305914.1</td><td>520</td><td>542</td><td>-</td><td>0</td><td>CGCGGACGCGGCGCGCGGCC</td><td>CGG</td><td>||||||||[||||||||||||]PAM</td><td>NA</td><td>Intergenic</td><td>NA</td><td>NA</td></tr>
<tr><td>T34291</td><td>GGACGCGGCGCGCGGCCCGGAGG</td><td>1000</td><td></td><td></td><td></td><td></td><td></td><td></td><td></td><td></td><td></td></tr>
<tr><td>Oligo fwd</td><td>CACCGGACGCGGCGCGCGGCCCGG</td><td></td><td></td><td></td><td></td><td></td><td></td><td></td><td></td><td></td><td></td></tr>
<tr><td>Oligo rev</td><td>AAACCCGGGCCGCGCGCCGCGTCC</td><td></td><td></td><td></td><td></td><td></td><td></td><td></td><td></td><td></td><td></td></tr>
<tr><td>Chromosome</td><td>start</td><td>end</td><td>strand</td><td>MM</td><td>target_seq</td><td>PAM</td><td>alignment</td><td>distance</td><td>position</td><td>gene name</td><td>gene id</td></tr>
<tr><td>GU305914.1</td><td>517</td><td>539</td><td>-</td><td>0</td><td>GGACGCGGCGCGCGGCCCGG</td><td>AGG</td><td>||||||||[||||||||||||]PAM</td><td>NA</td><td>Intergenic</td><td>NA</td><td>NA</td></tr>
<tr><td>T34292</td><td>CGCGGCCCGGAGGCTGACTCAGG</td><td>1000</td><td></td><td></td><td></td><td></td><td></td><td></td><td></td><td></td><td></td></tr>
<tr><td>Oligo fwd</td><td>CACCCGCGGCCCGGAGGCTGACTC</td><td></td><td></td><td></td><td></td><td></td><td></td><td></td><td></td><td></td><td></td></tr>
<tr><td>Oligo rev</td><td>AAACGAGTCAGCCTCCGGGCCGCG</td><td></td><td></td><td></td><td></td><td></td><td></td><td></td><td></td><td></td><td></td></tr>
<tr><td>Chromosome</td><td>start</td><td>end</td><td>strand</td><td>MM</td><td>target_seq</td><td>PAM</td><td>alignment</td><td>distance</td><td>position</td><td>gene name</td><td>gene id</td></tr>
<tr><td>GU305914.1</td><td>507</td><td>529</td><td>-</td><td>0</td><td>CGCGGCCCGGAGGCTGACTC</td><td>AGG</td><td>||||||||[||||||||||||]PAM</td><td>NA</td><td>Intergenic</td><td>NA</td><td>NA</td></tr>
<tr><td>T34293</td><td>GACTCAGGCGAGCGCGCGTCTGG</td><td>1000</td><td></td><td></td><td></td><td></td><td></td><td></td><td></td><td></td><td></td></tr>
<tr><td>Oligo fwd</td><td>CACCGACTCAGGCGAGCGCGCGTC</td><td></td><td></td><td></td><td></td><td></td><td></td><td></td><td></td><td></td><td></td></tr>
<tr><td>Oligo rev</td><td>AAACGACGCGCGCTCGCCTGAGTC</td><td></td><td></td><td></td><td></td><td></td><td></td><td></td><td></td><td></td><td></td></tr>
<tr><td>Chromosome</td><td>start</td><td>end</td><td>strand</td><td>MM</td><td>target_seq</td><td>PAM</td><td>alignment</td><td>distance</td><td>position</td><td>gene name</td><td>gene id</td></tr>
<tr><td>GU305914.1</td><td>492</td><td>514</td><td>-</td><td>0</td><td>GACTCAGGCGAGCGCGCGTC</td><td>TGG</td><td>||||||||[||||||||||||]PAM</td><td>NA</td><td>Intergenic</td><td>NA</td><td>NA</td></tr>
<tr><td>T34294</td><td>TACATGTCTCTCGCGCCGCGTGG</td><td>1000</td><td></td><td></td><td></td><td></td><td></td><td></td><td></td><td></td><td></td></tr>
<tr><td>Oligo fwd</td><td>CACCTACATGTCTCTCGCGCCGCG</td><td></td><td></td><td></td><td></td><td></td><td></td><td></td><td></td><td></td><td></td></tr>
<tr><td>Oligo rev</td><td>AAACCGCGGCGCGAGAGACATGTA</td><td></td><td></td><td></td><td></td><td></td><td></td><td></td><td></td><td></td><td></td></tr>
<tr><td>Chromosome</td><td>start</td><td>end</td><td>strand</td><td>MM</td><td>target_seq</td><td>PAM</td><td>alignment</td><td>distance</td><td>position</td><td>gene name</td><td>gene id</td></tr>
<tr><td>GU305914.1</td><td>440</td><td>462</td><td>-</td><td>0</td><td>TACATGTCTCTCGCGCCGCG</td><td>TGG</td><td>||||||||[||||||||||||]PAM</td><td>NA</td><td>Intergenic</td><td>NA</td><td>NA</td></tr>
<tr><td>T34295</td><td>CTCAGAAAACGAGCGTTCTACGG</td><td>1000</td><td></td><td></td><td></td><td></td><td></td><td></td><td></td><td></td><td></td></tr>
<tr><td>Oligo fwd</td><td>CACCCTCAGAAAACGAGCGTTCTA</td><td></td><td></td><td></td><td></td><td></td><td></td><td></td><td></td><td></td><td></td></tr>
<tr><td>Oligo rev</td><td>AAACTAGAACGCTCGTTTTCTGAG</td><td></td><td></td><td></td><td></td><td></td><td></td><td></td><td></td><td></td><td></td></tr>
<tr><td>Chromosome</td><td>start</td><td>end</td><td>strand</td><td>MM</td><td>target_seq</td><td>PAM</td><td>alignment</td><td>distance</td><td>position</td><td>gene name</td><td>gene id</td></tr>
<tr><td>GU305914.1</td><td>381</td><td>403</td><td>-</td><td>0</td><td>CTCAGAAAACGAGCGTTCTA</td><td>CGG</td><td>||||||||[||||||||||||]PAM</td><td>NA</td><td>Intergenic</td><td>NA</td><td>NA</td></tr>
<tr><td>T34296</td><td>ACGTTTTCGCGTCGCGCGCCCGG</td><td>1000</td><td></td><td></td><td></td><td></td><td></td><td></td><td></td><td></td><td></td></tr>
<tr><td>Oligo fwd</td><td>CACCACGTTTTCGCGTCGCGCGCC</td><td></td><td></td><td></td><td></td><td></td><td></td><td></td><td></td><td></td><td></td></tr>
<tr><td>Oligo rev</td><td>AAACGGCGCGCGACGCGAAAACGT</td><td></td><td></td><td></td><td></td><td></td><td></td><td></td><td></td><td></td><td></td></tr>
<tr><td>Chromosome</td><td>start</td><td>end</td><td>strand</td><td>MM</td><td>target_seq</td><td>PAM</td><td>alignment</td><td>distance</td><td>position</td><td>gene name</td><td>gene id</td></tr>
<tr><td>GU305914.1</td><td>316</td><td>338</td><td>-</td><td>0</td><td>ACGTTTTCGCGTCGCGCGCC</td><td>CGG</td><td>||||||||[||||||||||||]PAM</td><td>NA</td><td>Intergenic</td><td>NA</td><td>NA</td></tr>
<tr><td>T34297</td><td>CGTCGCGCGCCCGGCGCTTACGG</td><td>1000</td><td></td><td></td><td></td><td></td><td></td><td></td><td></td><td></td><td></td></tr>
<tr><td>Oligo fwd</td><td>CACCCGTCGCGCGCCCGGCGCTTA</td><td></td><td></td><td></td><td></td><td></td><td></td><td></td><td></td><td></td><td></td></tr>
<tr><td>Oligo rev</td><td>AAACTAAGCGCCGGGCGCGCGACG</td><td></td><td></td><td></td><td></td><td></td><td></td><td></td><td></td><td></td><td></td></tr>
<tr><td>Chromosome</td><td>start</td><td>end</td><td>strand</td><td>MM</td><td>target_seq</td><td>PAM</td><td>alignment</td><td>distance</td><td>position</td><td>gene name</td><td>gene id</td></tr>
<tr><td>GU305914.1</td><td>307</td><td>329</td><td>-</td><td>0</td><td>CGTCGCGCGCCCGGCGCTTA</td><td>CGG</td><td>||||||||[||||||||||||]PAM</td><td>NA</td><td>Intergenic</td><td>NA</td><td>NA</td></tr>
<tr><td>GU305914.1</td><td>97429</td><td>97451</td><td>+</td><td>4</td><td>CGTCCGGCGCACGGCGCTTC</td><td>TGG</td><td>||||--||[||-||||||||-]PAM</td><td>NA</td><td>Intergenic</td><td>NA</td><td>NA</td></tr>
<tr><td>T34298</td><td>GTCGCGCGCCCGGCGCTTACGGG</td><td>1000</td><td></td><td></td><td></td><td></td><td></td><td></td><td></td><td></td><td></td></tr>
<tr><td>Oligo fwd</td><td>CACCGTCGCGCGCCCGGCGCTTAC</td><td></td><td></td><td></td><td></td><td></td><td></td><td></td><td></td><td></td><td></td></tr>
<tr><td>Oligo rev</td><td>AAACGTAAGCGCCGGGCGCGCGAC</td><td></td><td></td><td></td><td></td><td></td><td></td><td></td><td></td><td></td><td></td></tr>
<tr><td>Chromosome</td><td>start</td><td>end</td><td>strand</td><td>MM</td><td>target_seq</td><td>PAM</td><td>alignment</td><td>distance</td><td>position</td><td>gene name</td><td>gene id</td></tr>
<tr><td>GU305914.1</td><td>306</td><td>328</td><td>-</td><td>0</td><td>GTCGCGCGCCCGGCGCTTAC</td><td>GGG</td><td>||||||||[||||||||||||]PAM</td><td>NA</td><td>Intergenic</td><td>NA</td><td>NA</td></tr>
<tr><td>T34299</td><td>TAGCAGAACGCGCTCCAAAAAGG</td><td>1000</td><td></td><td></td><td></td><td></td><td></td><td></td><td></td><td></td><td></td></tr>
<tr><td>Oligo fwd</td><td>CACCTAGCAGAACGCGCTCCAAAA</td><td></td><td></td><td></td><td></td><td></td><td></td><td></td><td></td><td></td><td></td></tr>
<tr><td>Oligo rev</td><td>AAACTTTTGGAGCGCGTTCTGCTA</td><td></td><td></td><td></td><td></td><td></td><td></td><td></td><td></td><td></td><td></td></tr>
<tr><td>Chromosome</td><td>start</td><td>end</td><td>strand</td><td>MM</td><td>target_seq</td><td>PAM</td><td>alignment</td><td>distance</td><td>position</td><td>gene name</td><td>gene id</td></tr>
</table>

| | | | | | | | | | | | | |
|---|---|---|---|---|---|---|---|---|---|---|---|---|
| GU305914.1 | | 268 | 290 | - | 0 | TAGCAGAACGCGCTCCAAAA | AGG | \|\|\|\|\|\|\|\|[\|\|\|\|\|\|\|\|\|\|\|\|]PAM | NA | Intergenic | NA | NA |
| T34300 | AGCAGAACGCGCTCCAAAAAGGG | | 1000 | | | | | | | | | |
| Oligo fwd | CACCAGCAGAACGCGCTCCAAAAA | | | | | | | | | | | |
| Oligo rev | AAACTTTTTGGAGCGCGTTCTGCT | | | | | | | | | | | |
| Chromosome | start | | end | strand | MM | target_seq | PAM | alignment | distance | position | gene name | gene id |
| GU305914.1 | | 267 | 289 | - | 0 | AGCAGAACGCGCTCCAAAAA | GGG | \|\|\|\|\|\|\|\|[\|\|\|\|\|\|\|\|\|\|\|\|]PAM | NA | Intergenic | NA | NA |
| T34301 | GCAGAACGCGCTCCAAAAAGGGG | | 1000 | | | | | | | | | |
| Oligo fwd | CACCGCAGAACGCGCTCCAAAAAG | | | | | | | | | | | |
| Oligo rev | AAACCTTTTTGGAGCGCGTTCTGC | | | | | | | | | | | |
| Chromosome | start | | end | strand | MM | target_seq | PAM | alignment | distance | position | gene name | gene id |
| GU305914.1 | | 266 | 288 | - | 0 | GCAGAACGCGCTCCAAAAAG | GGG | \|\|\|\|\|\|\|\|[\|\|\|\|\|\|\|\|\|\|\|\|]PAM | NA | Intergenic | NA | NA |
| T34302 | CGCCAGAATCTTCCCAGAAAAGG | | 1000 | | | | | | | | | |
| Oligo fwd | CACCCGCCAGAATCTTCCCAGAAA | | | | | | | | | | | |
| Oligo rev | AAACTTTCTGGGAAGATTCTGGCG | | | | | | | | | | | |
| Chromosome | start | | end | strand | MM | target_seq | PAM | alignment | distance | position | gene name | gene id |
| GU305914.1 | | 236 | 258 | - | 0 | CGCCAGAATCTTCCCAGAAA | AGG | \|\|\|\|\|\|\|\|[\|\|\|\|\|\|\|\|\|\|\|\|]PAM | NA | Intergenic | NA | NA |
| T34303 | GCCAGAATCTTCCCAGAAAAGGG | | 1000 | | | | | | | | | |
| Oligo fwd | CACCGCCAGAATCTTCCCAGAAAA | | | | | | | | | | | |
| Oligo rev | AAACTTTTCTGGGAAGATTCTGGC | | | | | | | | | | | |
| Chromosome | start | | end | strand | MM | target_seq | PAM | alignment | distance | position | gene name | gene id |
| GU305914.1 | | 235 | 257 | - | 0 | GCCAGAATCTTCCCAGAAAA | GGG | \|\|\|\|\|\|\|\|[\|\|\|\|\|\|\|\|\|\|\|\|]PAM | NA | Intergenic | NA | NA |
| T34304 | CCAGAATCTTCCCAGAAAAGGGG | | 1000 | | | | | | | | | |
| Oligo fwd | CACCCCAGAATCTTCCCAGAAAAG | | | | | | | | | | | |
| Oligo rev | AAACCTTTTCTGGGAAGATTCTGG | | | | | | | | | | | |
| Chromosome | start | | end | strand | MM | target_seq | PAM | alignment | distance | position | gene name | gene id |
| GU305914.1 | | 234 | 256 | - | 0 | CCAGAATCTTCCCAGAAAAG | GGG | \|\|\|\|\|\|\|\|[\|\|\|\|\|\|\|\|\|\|\|\|]PAM | NA | Intergenic | NA | NA |
| T34305 | CCCAGAAAAGGGGAACTTCTCGG | | 1000 | | | | | | | | | |
| Oligo fwd | CACCCCCAGAAAAGGGGAACTTCT | | | | | | | | | | | |
| Oligo rev | AAACAGAAGTTCCCCTTTTCTGGG | | | | | | | | | | | |
| Chromosome | start | | end | strand | MM | target_seq | PAM | alignment | distance | position | gene name | gene id |
| GU305914.1 | | 224 | 246 | - | 0 | CCCAGAAAAGGGGAACTTCT | CGG | \|\|\|\|\|\|\|\|[\|\|\|\|\|\|\|\|\|\|\|\|]PAM | NA | Intergenic | NA | NA |
| T34306 | GGGAACTTCTCGGAACGCTTTGG | | 1000 | | | | | | | | | |
| Oligo fwd | CACCGGGAACTTCTCGGAACGCTT | | | | | | | | | | | |
| Oligo rev | AAACAAGCGTTCCGAGAAGTTCCC | | | | | | | | | | | |
| Chromosome | start | | end | strand | MM | target_seq | PAM | alignment | distance | position | gene name | gene id |
| GU305914.1 | | 214 | 236 | - | 0 | GGGAACTTCTCGGAACGCTT | TGG | \|\|\|\|\|\|\|\|[\|\|\|\|\|\|\|\|\|\|\|\|]PAM | NA | Intergenic | NA | NA |
| T34307 | CTTTGGCACGAACATCTTCTTGG | | 1000 | | | | | | | | | |
| Oligo fwd | CACCCTTTGGCACGAACATCTTCT | | | | | | | | | | | |
| Oligo rev | AAACAGAAGATGTTCGTGCCAAAG | | | | | | | | | | | |
| Chromosome | start | | end | strand | MM | target_seq | PAM | alignment | distance | position | gene name | gene id |
| GU305914.1 | | 197 | 219 | - | 0 | CTTTGGCACGAACATCTTCT | TGG | \|\|\|\|\|\|\|\|[\|\|\|\|\|\|\|\|\|\|\|\|]PAM | NA | Intergenic | NA | NA |
| T34308 | GCACGAACATCTTCTTGGCACGG | | 1000 | | | | | | | | | |
| Oligo fwd | CACCGCACGAACATCTTCTTGGCA | | | | | | | | | | | |
| Oligo rev | AAACTGCCAAGAAGATGTTCGTGC | | | | | | | | | | | |
| Chromosome | start | | end | strand | MM | target_seq | PAM | alignment | distance | position | gene name | gene id |
| GU305914.1 | | 192 | 214 | - | 0 | GCACGAACATCTTCTTGGCA | CGG | \|\|\|\|\|\|\|\|[\|\|\|\|\|\|\|\|\|\|\|\|]PAM | NA | Intergenic | NA | NA |
| T34309 | TTGGCACGGTGCCACGCCCTCGG | | 1000 | | | | | | | | | |
| Oligo fwd | CACCTTGGCACGGTGCCACGCCCT | | | | | | | | | | | |
| Oligo rev | AAACAGGGCGTGGCACCGTGCCAA | | | | | | | | | | | |
| Chromosome | start | | end | strand | MM | target_seq | PAM | alignment | distance | position | gene name | gene id |
| GU305914.1 | | 178 | 200 | - | 0 | TTGGCACGGTGCCACGCCCT | CGG | \|\|\|\|\|\|\|\|[\|\|\|\|\|\|\|\|\|\|\|\|]PAM | NA | Intergenic | NA | NA |
| T34310 | AGCCACCTCATCGCACTCTCAGG | | 1000 | | | | | | | | | |
| Oligo fwd | CACCAGCCACCTCATCGCACTCTC | | | | | | | | | | | |
| Oligo rev | AAACGAGAGTGCGATGAGGTGGCT | | | | | | | | | | | |
| Chromosome | start | | end | strand | MM | target_seq | PAM | alignment | distance | position | gene name | gene id |
| GU305914.1 | | 143 | 165 | - | 0 | AGCCACCTCATCGCACTCTC | AGG | \|\|\|\|\|\|\|\|[\|\|\|\|\|\|\|\|\|\|\|\|]PAM | NA | Intergenic | NA | NA |
| T34311 | GATGCAACAGCGCCCTCTAGAGG | | 1000 | | | | | | | | | |
| Oligo fwd | CACCGATGCAACAGCGCCCTCTAG | | | | | | | | | | | |
| Oligo rev | AAACCTAGAGGGCGCTGTTGCATC | | | | | | | | | | | |
| Chromosome | start | | end | strand | MM | target_seq | PAM | alignment | distance | position | gene name | gene id |
| GU305914.1 | | 102 | 124 | - | 0 | GATGCAACAGCGCCCTCTAG | AGG | \|\|\|\|\|\|\|\|[\|\|\|\|\|\|\|\|\|\|\|\|]PAM | NA | Intergenic | NA | NA |
| T34312 | CCCCCATCAAAATACCCCCCCGG | | 1000 | | | | | | | | | |
| Oligo fwd | CACCCCCCCATCAAAATACCCCCC | | | | | | | | | | | |
| Oligo rev | AAACGGGGGGTATTTTGATGGGGG | | | | | | | | | | | |
| Chromosome | start | | end | strand | MM | target_seq | PAM | alignment | distance | position | gene name | gene id |
| GU305914.1 | | 65 | 87 | - | 0 | CCCCCATCAAAATACCCCCC | CGG | \|\|\|\|\|\|\|\|[\|\|\|\|\|\|\|\|\|\|\|\|]PAM | NA | Intergenic | NA | NA |
| T34313 | CCCCATCAAAATACCCCCCCGGG | | 1000 | | | | | | | | | |
| Oligo fwd | CACCCCCCATCAAAATACCCCCCC | | | | | | | | | | | |
| Oligo rev | AAACGGGGGGGTATTTTGATGGGG | | | | | | | | | | | |
| Chromosome | start | | end | strand | MM | target_seq | PAM | alignment | distance | position | gene name | gene id |
| GU305914.1 | | 64 | 86 | - | 0 | CCCCATCAAAATACCCCCCC | GGG | \|\|\|\|\|\|\|\|[\|\|\|\|\|\|\|\|\|\|\|\|]PAM | NA | Intergenic | NA | NA |
| T34314 | CCCATCAAAATACCCCCCCGGGG | | 1000 | | | | | | | | | |
| Oligo fwd | CACCCCCATCAAAATACCCCCCCG | | | | | | | | | | | |
| Oligo rev | AAACCGGGGGGGTATTTTGATGGG | | | | | | | | | | | |

<table>
<tr><td>Chromosome</td><td>start</td><td>end</td><td>strand</td><td>MM</td><td>target_seq</td><td>PAM</td><td>alignment</td><td>distance</td><td>position</td><td>gene name</td><td>gene id</td></tr>
<tr><td>GU305914.1</td><td>63</td><td>85</td><td>-</td><td>0</td><td>CCCATCAAAATACCCCCCCG</td><td>GGG</td><td>||||||||[||||||||||||]PAM</td><td>NA</td><td>Intergenic</td><td>NA</td><td>NA</td></tr>
<tr><td></td><td></td><td></td><td></td><td></td><td></td><td></td><td></td><td></td><td></td><td></td><td></td></tr>
<tr><td>T34315</td><td>CCATCAAAATACCCCCCCGGGGG</td><td>1000</td><td></td><td></td><td></td><td></td><td></td><td></td><td></td><td></td><td></td></tr>
<tr><td>Oligo fwd</td><td>CACCCCATCAAAATACCCCCCCGG</td><td></td><td></td><td></td><td></td><td></td><td></td><td></td><td></td><td></td><td></td></tr>
<tr><td>Oligo rev</td><td>AAACCCGGGGGGGTATTTTGATGG</td><td></td><td></td><td></td><td></td><td></td><td></td><td></td><td></td><td></td><td></td></tr>
<tr><td>Chromosome</td><td>start</td><td>end</td><td>strand</td><td>MM</td><td>target_seq</td><td>PAM</td><td>alignment</td><td>distance</td><td>position</td><td>gene name</td><td>gene id</td></tr>
<tr><td>GU305914.1</td><td>62</td><td>84</td><td>-</td><td>0</td><td>CCATCAAAATACCCCCCCGG</td><td>GGG</td><td>||||||||[||||||||||||]PAM</td><td>NA</td><td>Intergenic</td><td>NA</td><td>NA</td></tr>
<tr><td></td><td></td><td></td><td></td><td></td><td></td><td></td><td></td><td></td><td></td><td></td><td></td></tr>
<tr><td>T34316</td><td>GGGGCCCGCGCGCACTCAGACGG</td><td>1000</td><td></td><td></td><td></td><td></td><td></td><td></td><td></td><td></td><td></td></tr>
<tr><td>Oligo fwd</td><td>CACCGGGGCCCGCGCGCACTCAGA</td><td></td><td></td><td></td><td></td><td></td><td></td><td></td><td></td><td></td><td></td></tr>
<tr><td>Oligo rev</td><td>AAACTCTGAGTGCGCGCGGGCCCC</td><td></td><td></td><td></td><td></td><td></td><td></td><td></td><td></td><td></td><td></td></tr>
<tr><td>Chromosome</td><td>start</td><td>end</td><td>strand</td><td>MM</td><td>target_seq</td><td>PAM</td><td>alignment</td><td>distance</td><td>position</td><td>gene name</td><td>gene id</td></tr>
<tr><td>GU305914.1</td><td>43</td><td>65</td><td>-</td><td>0</td><td>GGGGCCCGCGCGCACTCAGA</td><td>CGG</td><td>||||||||[||||||||||||]PAM</td><td>NA</td><td>Intergenic</td><td>NA</td><td>NA</td></tr>
<tr><td>GU305914.1</td><td>230358</td><td>230380</td><td>-</td><td>1</td><td>TGGGCCCGCGCGCACTCAGA</td><td>CGG</td><td>-|||||||[||||||||||||]PAM</td><td>NA</td><td>Intergenic</td><td>NA</td><td>NA</td></tr>
<tr><td></td><td></td><td></td><td></td><td></td><td></td><td></td><td></td><td></td><td></td><td></td><td></td></tr>
<tr><td>T34317</td><td>CCCGCGCGCACTCAGACGGCCGG</td><td>1000</td><td></td><td></td><td></td><td></td><td></td><td></td><td></td><td></td><td></td></tr>
<tr><td>Oligo fwd</td><td>CACCCCCGCGCGCACTCAGACGGC</td><td></td><td></td><td></td><td></td><td></td><td></td><td></td><td></td><td></td><td></td></tr>
<tr><td>Oligo rev</td><td>AAACGCCGTCTGAGTGCGCGCGGG</td><td></td><td></td><td></td><td></td><td></td><td></td><td></td><td></td><td></td><td></td></tr>
<tr><td>Chromosome</td><td>start</td><td>end</td><td>strand</td><td>MM</td><td>target_seq</td><td>PAM</td><td>alignment</td><td>distance</td><td>position</td><td>gene name</td><td>gene id</td></tr>
<tr><td>GU305914.1</td><td>39</td><td>61</td><td>-</td><td>0</td><td>CCCGCGCGCACTCAGACGGC</td><td>CGG</td><td>||||||||[||||||||||||]PAM</td><td>NA</td><td>Intergenic</td><td>NA</td><td>NA</td></tr>
<tr><td>GU305914.1</td><td>230354</td><td>230376</td><td>-</td><td>0</td><td>CCCGCGCGCACTCAGACGGC</td><td>CGG</td><td>||||||||[||||||||||||]PAM</td><td>NA</td><td>Intergenic</td><td>NA</td><td>NA</td></tr>
<tr><td></td><td></td><td></td><td></td><td></td><td></td><td></td><td></td><td></td><td></td><td></td><td></td></tr>
<tr><td>T34318</td><td>CCGCGCGCACTCAGACGGCCGGG</td><td>1000</td><td></td><td></td><td></td><td></td><td></td><td></td><td></td><td></td><td></td></tr>
<tr><td>Oligo fwd</td><td>CACCCCGCGCGCACTCAGACGGCC</td><td></td><td></td><td></td><td></td><td></td><td></td><td></td><td></td><td></td><td></td></tr>
<tr><td>Oligo rev</td><td>AAACGGCCGTCTGAGTGCGCGCGG</td><td></td><td></td><td></td><td></td><td></td><td></td><td></td><td></td><td></td><td></td></tr>
<tr><td>Chromosome</td><td>start</td><td>end</td><td>strand</td><td>MM</td><td>target_seq</td><td>PAM</td><td>alignment</td><td>distance</td><td>position</td><td>gene name</td><td>gene id</td></tr>
<tr><td>GU305914.1</td><td>38</td><td>60</td><td>-</td><td>0</td><td>CCGCGCGCACTCAGACGGCC</td><td>GGG</td><td>||||||||[||||||||||||]PAM</td><td>NA</td><td>Intergenic</td><td>NA</td><td>NA</td></tr>
<tr><td>GU305914.1</td><td>230353</td><td>230375</td><td>-</td><td>0</td><td>CCGCGCGCACTCAGACGGCC</td><td>GGG</td><td>||||||||[||||||||||||]PAM</td><td>NA</td><td>Intergenic</td><td>NA</td><td>NA</td></tr>
<tr><td></td><td></td><td></td><td></td><td></td><td></td><td></td><td></td><td></td><td></td><td></td><td></td></tr>
<tr><td>T34319</td><td>CGCGCGCACTCAGACGGCCGGGG</td><td>1000</td><td></td><td></td><td></td><td></td><td></td><td></td><td></td><td></td><td></td></tr>
<tr><td>Oligo fwd</td><td>CACCCGCGCGCACTCAGACGGCCG</td><td></td><td></td><td></td><td></td><td></td><td></td><td></td><td></td><td></td><td></td></tr>
<tr><td>Oligo rev</td><td>AAACCGGCCGTCTGAGTGCGCGCG</td><td></td><td></td><td></td><td></td><td></td><td></td><td></td><td></td><td></td><td></td></tr>
<tr><td>Chromosome</td><td>start</td><td>end</td><td>strand</td><td>MM</td><td>target_seq</td><td>PAM</td><td>alignment</td><td>distance</td><td>position</td><td>gene name</td><td>gene id</td></tr>
<tr><td>GU305914.1</td><td>37</td><td>59</td><td>-</td><td>0</td><td>CGCGCGCACTCAGACGGCCG</td><td>GGG</td><td>||||||||[||||||||||||]PAM</td><td>NA</td><td>Intergenic</td><td>NA</td><td>NA</td></tr>
<tr><td>GU305914.1</td><td>230352</td><td>230374</td><td>-</td><td>0</td><td>CGCGCGCACTCAGACGGCCG</td><td>GGG</td><td>||||||||[||||||||||||]PAM</td><td>NA</td><td>Intergenic</td><td>NA</td><td>NA</td></tr>
<tr><td></td><td></td><td></td><td></td><td></td><td></td><td></td><td></td><td></td><td></td><td></td><td></td></tr>
<tr><td>T34320</td><td>GCGCGCACTCAGACGGCCGGGGG</td><td>1000</td><td></td><td></td><td></td><td></td><td></td><td></td><td></td><td></td><td></td></tr>
<tr><td>Oligo fwd</td><td>CACCGCGCGCACTCAGACGGCCGG</td><td></td><td></td><td></td><td></td><td></td><td></td><td></td><td></td><td></td><td></td></tr>
<tr><td>Oligo rev</td><td>AAACCCGGCCGTCTGAGTGCGCGC</td><td></td><td></td><td></td><td></td><td></td><td></td><td></td><td></td><td></td><td></td></tr>
<tr><td>Chromosome</td><td>start</td><td>end</td><td>strand</td><td>MM</td><td>target_seq</td><td>PAM</td><td>alignment</td><td>distance</td><td>position</td><td>gene name</td><td>gene id</td></tr>
<tr><td>GU305914.1</td><td>230351</td><td>230373</td><td>-</td><td>0</td><td>GCGCGCACTCAGACGGCCGG</td><td>GGG</td><td>||||||||[||||||||||||]PAM</td><td>NA</td><td>Intergenic</td><td>NA</td><td>NA</td></tr>
<tr><td>GU305914.1</td><td>36</td><td>58</td><td>-</td><td>0</td><td>GCGCGCACTCAGACGGCCGG</td><td>GGG</td><td>||||||||[||||||||||||]PAM</td><td>NA</td><td>Intergenic</td><td>NA</td><td>NA</td></tr>
<tr><td>GU305914.1</td><td>92153</td><td>92175</td><td>+</td><td>4</td><td>ACGCGCGCTCGGAGGGCCGG</td><td>CGG</td><td>-|||||-|[||-||-||||||]PAM</td><td>NA</td><td>Intergenic</td><td>NA</td><td>NA</td></tr>
<tr><td></td><td></td><td></td><td></td><td></td><td></td><td></td><td></td><td></td><td></td><td></td><td></td></tr>
<tr><td>T34321</td><td>CGCGCACTCAGACGGCCGGGGGG</td><td>1000</td><td></td><td></td><td></td><td></td><td></td><td></td><td></td><td></td><td></td></tr>
<tr><td>Oligo fwd</td><td>CACCCGCGCACTCAGACGGCCGGG</td><td></td><td></td><td></td><td></td><td></td><td></td><td></td><td></td><td></td><td></td></tr>
<tr><td>Oligo rev</td><td>AAACCCCGGCCGTCTGAGTGCGCG</td><td></td><td></td><td></td><td></td><td></td><td></td><td></td><td></td><td></td><td></td></tr>
<tr><td>Chromosome</td><td>start</td><td>end</td><td>strand</td><td>MM</td><td>target_seq</td><td>PAM</td><td>alignment</td><td>distance</td><td>position</td><td>gene name</td><td>gene id</td></tr>
<tr><td>GU305914.1</td><td>35</td><td>57</td><td>-</td><td>0</td><td>CGCGCACTCAGACGGCCGGG</td><td>GGG</td><td>||||||||[||||||||||||]PAM</td><td>NA</td><td>Intergenic</td><td>NA</td><td>NA</td></tr>
<tr><td>GU305914.1</td><td>230350</td><td>230372</td><td>-</td><td>0</td><td>CGCGCACTCAGACGGCCGGG</td><td>GGG</td><td>||||||||[||||||||||||]PAM</td><td>NA</td><td>Intergenic</td><td>NA</td><td>NA</td></tr>
<tr><td></td><td></td><td></td><td></td><td></td><td></td><td></td><td></td><td></td><td></td><td></td><td></td></tr>
<tr><td>T34322</td><td>GCGCACTCAGACGGCCGGGGGGG</td><td>1000</td><td></td><td></td><td></td><td></td><td></td><td></td><td></td><td></td><td></td></tr>
<tr><td>Oligo fwd</td><td>CACCGCGCACTCAGACGGCCGGGG</td><td></td><td></td><td></td><td></td><td></td><td></td><td></td><td></td><td></td><td></td></tr>
<tr><td>Oligo rev</td><td>AAACCCCCGGCCGTCTGAGTGCGC</td><td></td><td></td><td></td><td></td><td></td><td></td><td></td><td></td><td></td><td></td></tr>
<tr><td>Chromosome</td><td>start</td><td>end</td><td>strand</td><td>MM</td><td>target_seq</td><td>PAM</td><td>alignment</td><td>distance</td><td>position</td><td>gene name</td><td>gene id</td></tr>
<tr><td>GU305914.1</td><td>34</td><td>56</td><td>-</td><td>0</td><td>GCGCACTCAGACGGCCGGGG</td><td>GGG</td><td>||||||||[||||||||||||]PAM</td><td>NA</td><td>Intergenic</td><td>NA</td><td>NA</td></tr>
<tr><td>GU305914.1</td><td>230349</td><td>230371</td><td>-</td><td>0</td><td>GCGCACTCAGACGGCCGGGG</td><td>GGG</td><td>||||||||[||||||||||||]PAM</td><td>NA</td><td>Intergenic</td><td>NA</td><td>NA</td></tr>
<tr><td></td><td></td><td></td><td></td><td></td><td></td><td></td><td></td><td></td><td></td><td></td><td></td></tr>
<tr><td>T34323</td><td>CGCACTCAGACGGCCGGGGGGGG</td><td>1000</td><td></td><td></td><td></td><td></td><td></td><td></td><td></td><td></td><td></td></tr>
<tr><td>Oligo fwd</td><td>CACCCGCACTCAGACGGCCGGGGG</td><td></td><td></td><td></td><td></td><td></td><td></td><td></td><td></td><td></td><td></td></tr>
<tr><td>Oligo rev</td><td>AAACCCCCCGGCCGTCTGAGTGCG</td><td></td><td></td><td></td><td></td><td></td><td></td><td></td><td></td><td></td><td></td></tr>
<tr><td>Chromosome</td><td>start</td><td>end</td><td>strand</td><td>MM</td><td>target_seq</td><td>PAM</td><td>alignment</td><td>distance</td><td>position</td><td>gene name</td><td>gene id</td></tr>
<tr><td>GU305914.1</td><td>33</td><td>55</td><td>-</td><td>0</td><td>CGCACTCAGACGGCCGGGGG</td><td>GGG</td><td>||||||||[||||||||||||]PAM</td><td>NA</td><td>Intergenic</td><td>NA</td><td>NA</td></tr>
<tr><td>GU305914.1</td><td>28762</td><td>28784</td><td>-</td><td>4</td><td>GGCCCTCCGGCGGCCGGGGG</td><td>AGG</td><td>-||-|||-[|-||||||||||]PAM</td><td>NA</td><td>Intergenic</td><td>NA</td><td>NA</td></tr>
<tr><td></td><td></td><td></td><td></td><td></td><td></td><td></td><td></td><td></td><td></td><td></td><td></td></tr>
<tr><td>T34324</td><td>GCACTCAGACGGCCGGGGGGGGG</td><td>1000</td><td></td><td></td><td></td><td></td><td></td><td></td><td></td><td></td><td></td></tr>
<tr><td>Oligo fwd</td><td>CACCGCACTCAGACGGCCGGGGGG</td><td></td><td></td><td></td><td></td><td></td><td></td><td></td><td></td><td></td><td></td></tr>
<tr><td>Oligo rev</td><td>AAACCCCCCCGGCCGTCTGAGTGC</td><td></td><td></td><td></td><td></td><td></td><td></td><td></td><td></td><td></td><td></td></tr>
<tr><td>Chromosome</td><td>start</td><td>end</td><td>strand</td><td>MM</td><td>target_seq</td><td>PAM</td><td>alignment</td><td>distance</td><td>position</td><td>gene name</td><td>gene id</td></tr>
<tr><td>GU305914.1</td><td>32</td><td>54</td><td>-</td><td>0</td><td>GCACTCAGACGGCCGGGGGG</td><td>GGG</td><td>||||||||[||||||||||||]PAM</td><td>NA</td><td>Intergenic</td><td>NA</td><td>NA</td></tr>
</table>
